## Supplementary material for "Single-nucleotide Differences and Cell Type Decide the Subcellular Localization of miRNA Isoforms (isomiRs), tRNA-derived Fragments (tRFs) and rRNA-derived Fragments (rRFs)": Supp. Table S6

**Supplemental Table S6:** Data table of the differentially abundant (DA) short RNA for the cell line RNA between the three cell line fractions (Nucleus, Cytoplasm, Mitochondira, andMitoplast) with FDR < 0.05. The table is displayed as the output from DESeq2 where MedianFC (median fold change) is presented as raw reads . The table in decreasing order of FDR value. The annotation "am" is used to indicate abiguous short RNA, those that can be mapped to a noncoding RNA gene as well as somewhere else on the geneome. The annotation "multi" referrs to short RNA that map to locations in both the mitochondrial and nuclear genomes. The rRF are flanked by 50 nucleotdies upstream and downstream and their coordiantes reflect this. The coordiantes for the isomiR and rRF are from the hg38 human genome assembly while the coordintes for the tRF are from the hg19 human genome assembly.

<table>
<tr><th>RNA</th><th>Coordiantes</th><th>Start</th><th>End</th><th>Length</th><th>Sequence</th><th>MedianFC</th><th>log2FC</th><th>lfcSE</th><th>stat</th><th>p-value</th><th>adjp-value</th><th>Numerator/Denominator</th><th>Fraction</th></tr>
<tr><td>>hg38_21_+_8433172_8446622_RNA45SN1_45S_50ntFlanks@7979.8006.28|ACCTCAGATCAGACGTGGCGACCCGCTG</td><td>>hg38_21_+_8433172_8446622_RNA45SN1_45S_50ntFlanks</td><td>7979</td><td>8006</td><td>28</td><td>ACCTCAGATCAGACGTGGCGACCCGCTG</td><td>16501.8</td><td>-9.6</td><td>0.2</td><td>-38.7</td><td>0.0E+00</td><td>0.0E+00</td><td>BT20/MB468</td><td>NUC</td></tr>
<tr><td>>hg38_21_+_8433172_8446622_RNA45SN1_45S_50ntFlanks@9534.9553.20|AGGAAACTCTGGTGGAGGTC</td><td>>hg38_21_+_8433172_8446622_RNA45SN1_45S_50ntFlanks</td><td>9534</td><td>9553</td><td>20</td><td>AGGAAACTCTGGTGGAGGTC</td><td>200147.7</td><td>-13.7</td><td>0.4</td><td>-37.4</td><td>1.6E-306</td><td>4.4E-303</td><td>BT20/MB468</td><td>NUC</td></tr>
<tr><td>>hg38_21_+_8433172_8446622_RNA45SN1_45S_50ntFlanks@10350.10377.28|ACGAGAACTTTGAAGGCCGAAGTGGAGA</td><td>>hg38_21_+_8433172_8446622_RNA45SN1_45S_50ntFlanks</td><td>10350</td><td>10377</td><td>28</td><td>ACGAGAACTTTGAAGGCCGAAGTGGAGA</td><td>8600.7</td><td>-10.1</td><td>0.4</td><td>-26.9</td><td>5.8E-159</td><td>1.1E-155</td><td>BT20/MB468</td><td>NUC</td></tr>
<tr><td>>MI0000542|hsa-mir-320a&WithFlank&8|-|22244960|22245043@48.69.22&[MIMAT0000510&hsa-miR-320a-3p&offsets|0|0;m-60&8|-|22244975|22244996&offsets|0|0]|AAAAGCTGGGTTGAGAGGGCGA</td><td>>MI0000542|hsa-mir-320a&WithFlank&8|-|22244960|22245043</td><td>48</td><td>69</td><td>22</td><td>AAAAGCTGGGTTGAGAGGGCGA</td><td>83100.1</td><td>11.9</td><td>0.4</td><td>26.6</td><td>1.9E-155</td><td>1.1E-151</td><td>BT20/MB468</td><td>CYTO</td></tr>
<tr><td>>hg38_21_+_8433172_8446622_RNA45SN1_45S_50ntFlanks@7980.8006.27|CCTCAGATCAGACGTGGCGACCCGCTG</td><td>>hg38_21_+_8433172_8446622_RNA45SN1_45S_50ntFlanks</td><td>7980</td><td>8006</td><td>27</td><td>CCTCAGATCAGACGTGGCGACCCGCTG</td><td>13290.2</td><td>-9.9</td><td>0.4</td><td>-24.7</td><td>2.6E-134</td><td>3.5E-131</td><td>BT20/MB468</td><td>NUC</td></tr>
<tr><td>>hg38_21_+_8433172_8446622_RNA45SN1_45S_50ntFlanks@11783.11804.22|CATGAATGGATGAACGAGATTC</td><td>>hg38_21_+_8433172_8446622_RNA45SN1_45S_50ntFlanks</td><td>11783</td><td>11804</td><td>22</td><td>CATGAATGGATGAACGAGATTC</td><td>81892.4</td><td>-12.9</td><td>0.5</td><td>-24.1</td><td>1.3E-128</td><td>1.4E-125</td><td>BT20/MB468</td><td>NUC</td></tr>
<tr><td>>MI0000063|hsa-let-7b&WithFlank&22|+|46113680|46113774@12.33.22&[MIMAT0000063&hsa-let-7b-5p&offsets|0|0;m-14&22|+|46113691|46113712&offsets|0|0]|TGAGGTAGTAGGTTGTGTGGTT</td><td>>MI0000063|hsa-let-7b&WithFlank&22|+|46113680|46113774</td><td>12</td><td>33</td><td>22</td><td>TGAGGTAGTAGGTTGTGTGGTT</td><td>113044.9</td><td>8.8</td><td>0.4</td><td>23.9</td><td>9.4E-127</td><td>2.8E-123</td><td>BT20/MB468</td><td>CYTO</td></tr>
<tr><td>>hg38_21_+_8433172_8446622_RNA45SN1_45S_50ntFlanks@11783.11805.23|CATGAATGGATGAACGAGATTCC</td><td>>hg38_21_+_8433172_8446622_RNA45SN1_45S_50ntFlanks</td><td>11783</td><td>11805</td><td>23</td><td>CATGAATGGATGAACGAGATTCC</td><td>4086.9</td><td>-9.5</td><td>0.4</td><td>-23.7</td><td>7.6E-124</td><td>7.0E-121</td><td>BT20/MB468</td><td>NUC</td></tr>
<tr><td>>hg38_21_+_8433172_8446622_RNA45SN1_45S_50ntFlanks@9739.9758.20|AAACGATCTCAACCTATTCT</td><td>>hg38_21_+_8433172_8446622_RNA45SN1_45S_50ntFlanks</td><td>9739</td><td>9758</td><td>20</td><td>AAACGATCTCAACCTATTCT</td><td>4237.1</td><td>-7.9</td><td>0.3</td><td>-23.2</td><td>7.6E-119</td><td>6.0E-116</td><td>BT20/MB468</td><td>NUC</td></tr>
<tr><td>>MI0000542|hsa-mir-320a&WithFlank&8|-|22244960|22245043@48.70.23(+1U)&[MIMAT0000510&hsa-miR-320a-3p&offsets|0|+1(+1U);m-60&8|-|22244975|22244996&offsets|0|+1(+1U)]|AAAAGCTGGGTTGAGAGGGCGAAT</td><td>>MI0000542|hsa-mir-320a&WithFlank&8|-|22244960|22245043</td><td>48</td><td>70</td><td>23(+1U)</td><td>AAAAGCTGGGTTGAGAGGGCGAAT</td><td>35610.7</td><td>11.2</td><td>0.5</td><td>23.2</td><td>5.5E-119</td><td>1.1E-115</td><td>BT20/MB468</td><td>CYTO</td></tr>
<tr><td>>MI0000102|hsa-mir-100&WithFlank&11|-|122152223|122152314@19.39.21&[MIMAT0000098&hsa-miR-100-5p&offsets|0|-1;m-27&11|-|122152276|122152296&offsets|0|0]|AACCCGTAGATCCGAACTTGT</td><td>>MI0000102|hsa-mir-100&WithFlank&11|-|122152223|122152314</td><td>19</td><td>39</td><td>21</td><td>AACCCGTAGATCCGAACTTGT</td><td>209411.4</td><td>15.7</td><td>0.7</td><td>23.0</td><td>2.9E-117</td><td>1.7E-113</td><td>MB231/MB468</td><td>NUC</td></tr>
<tr><td>>am_trna10_ValCAC_5_-_180649395_180649467@1.29.29,trna12_ValAAC_5_-_180645270_180645342@1.29.29,trna132_ValAAC_6_-_27721179_27721251@1.29.29,trna136_ValAAC_6_-_27648885_27648957@1.29.29,trna139_ValAAC_6_-_27618707_27618779@1.29.29,trna18_ValCAC_5_-_180529253_180529325@1.29.29,trna2_ValAAC_3_+_169490018_169490090@1.29.29,trna2_ValCAC_5_+_180524070_180524142@1.29.29,trna4_ValAAC_5_+_180591154_180591226@1.29.29,trna5_ValAAC_5_+_180596610_180596682@1.29.29,trna6_ValCAC_5_+_180600650_180600722@1.29.29,trna85_ValCAC_1_-_161369490_161369562@1.29.29,trna90_ValCAC_1_-_149684088_149684161@1.29.29,trna98_ValCAC_1_-_149298555_149298627@1.29.29,trna9_ValCAC_6_+_26538282_26538354@1.29.29|GTTTCCGTAGTGTAGTGGTTATCACGTTC</td><td>>am_trna10_ValCAC_5_-_180649395_180649467</td><td>1</td><td>29</td><td>29</td><td>GTTTCCGTAGTGTAGTGGTTATCACGTTC</td><td>11806.4</td><td>-9.5</td><td>0.4</td><td>-22.3</td><td>3.3E-110</td><td>2.3E-107</td><td>BT20/MB468</td><td>NUC</td></tr>
<tr><td>>hg38_21_+_8433172_8446622_RNA45SN1_45S_50ntFlanks@10821.10840.20|GGAAGTCGGCAAGCCGGATC</td><td>>hg38_21_+_8433172_8446622_RNA45SN1_45S_50ntFlanks</td><td>10821</td><td>10840</td><td>20</td><td>GGAAGTCGGCAAGCCGGATC</td><td>3565.7</td><td>-8.5</td><td>0.4</td><td>-22.3</td><td>4.8E-110</td><td>3.0E-107</td><td>BT20/MB468</td><td>NUC</td></tr>
<tr><td>>MI0000542|hsa-mir-320a&WithFlank&8|-|22244960|22245043@48.69.22(+1U)&[MIMAT0000510&hsa-miR-320a-3p&offsets|0|0(+1U);m-60&8|-|22244975|22244996&offsets|0|0(+1U)]|AAAAGCTGGGTTGAGAGGGCGAT</td><td>>MI0000542|hsa-mir-320a&WithFlank&8|-|22244960|22245043</td><td>48</td><td>69</td><td>22(+1U)</td><td>AAAAGCTGGGTTGAGAGGGCGAT</td><td>49049.5</td><td>12.0</td><td>0.5</td><td>22.3</td><td>2.5E-110</td><td>3.7E-107</td><td>BT20/MB468</td><td>CYTO</td></tr>
<tr><td>>MI0000063|hsa-let-7b&WithFlank&22|+|46113680|46113774@12.34.23&[MIMAT0000063&hsa-let-7b-5p&offsets|0|+1;m-14&22|+|46113691|46113712&offsets|0|+1]|TGAGGTAGTAGGTTGTGTGGTTT</td><td>>MI0000063|hsa-let-7b&WithFlank&22|+|46113680|46113774</td><td>12</td><td>34</td><td>23</td><td>TGAGGTAGTAGGTTGTGTGGTTT</td><td>14220.2</td><td>8.6</td><td>0.4</td><td>22.1</td><td>3.9E-108</td><td>4.7E-105</td><td>BT20/MB468</td><td>CYTO</td></tr>
<tr><td>>hg38_21_+_8433172_8446622_RNA45SN1_45S_50ntFlanks@7983.8006.24|CAGATCAGACGTGGCGACCCGCTG</td><td>>hg38_21_+_8433172_8446622_RNA45SN1_45S_50ntFlanks</td><td>7983</td><td>8006</td><td>24</td><td>CAGATCAGACGTGGCGACCCGCTG</td><td>12332.2</td><td>-9.8</td><td>0.4</td><td>-22.0</td><td>5.4E-107</td><td>3.0E-104</td><td>BT20/MB468</td><td>NUC</td></tr>
<tr><td>>hg38_21_+_8433172_8446622_RNA45SN1_45S_50ntFlanks@12999.13027.29|CTGCGATCTATTGAAAGTCAGCCCTCGAC</td><td>>hg38_21_+_8433172_8446622_RNA45SN1_45S_50ntFlanks</td><td>12999</td><td>13027</td><td>29</td><td>CTGCGATCTATTGAAAGTCAGCCCTCGAC</td><td>2604.0</td><td>-8.4</td><td>0.4</td><td>-21.2</td><td>6.2E-100</td><td>3.1E-97</td><td>BT20/MB468</td><td>NUC</td></tr>
<tr><td>>hg38_21_+_8433172_8446622_RNA45SN1_45S_50ntFlanks@7982.8006.25|TCAGATCAGACGTGGCGACCCGCTG</td><td>>hg38_21_+_8433172_8446622_RNA45SN1_45S_50ntFlanks</td><td>7982</td><td>8006</td><td>25</td><td>TCAGATCAGACGTGGCGACCCGCTG</td><td>4041.1</td><td>-8.7</td><td>0.4</td><td>-21.2</td><td>1.6E-99</td><td>7.4E-97</td><td>BT20/MB468</td><td>NUC</td></tr>
<tr><td>>hg38_21_+_8433172_8446622_RNA45SN1_45S_50ntFlanks@9554.9575.22|CGTAGCGGTCCTGACGTGCAAA</td><td>>hg38_21_+_8433172_8446622_RNA45SN1_45S_50ntFlanks</td><td>9554</td><td>9575</td><td>22</td><td>CGTAGCGGTCCTGACGTGCAAA</td><td>1166.7</td><td>-5.8</td><td>0.3</td><td>-21.1</td><td>9.7E-99</td><td>4.1E-96</td><td>BT20/MB468</td><td>NUC</td></tr>
<tr><td>>hg38_21_+_8433172_8446622_RNA45SN1_45S_50ntFlanks@12936.12959.24|AAACCATTCGTAGACGACCTGCTT</td><td>>hg38_21_+_8433172_8446622_RNA45SN1_45S_50ntFlanks</td><td>12936</td><td>12959</td><td>24</td><td>AAACCATTCGTAGACGACCTGCTT</td><td>2181.7</td><td>-7.9</td><td>0.4</td><td>-21.0</td><td>4.4E-98</td><td>1.7E-95</td><td>BT20/MB468</td><td>NUC</td></tr>
<tr><td>>hg38_21_+_8433172_8446622_RNA45SN1_45S_50ntFlanks@12417.12441.25|TCGATGTCGGCTCTTCCTATCATTG</td><td>>hg38_21_+_8433172_8446622_RNA45SN1_45S_50ntFlanks</td><td>12417</td><td>12441</td><td>25</td><td>TCGATGTCGGCTCTTCCTATCATTG</td><td>2945.8</td><td>-8.3</td><td>0.4</td><td>-21.0</td><td>8.7E-98</td><td>3.2E-95</td><td>BT20/MB468</td><td>NUC</td></tr>
<tr><td>>hg38_21_+_8433172_8446622_RNA45SN1_45S_50ntFlanks@10473.10494.22|CTCGGCCGATCGAAAGGGAGTC</td><td>>hg38_21_+_8433172_8446622_RNA45SN1_45S_50ntFlanks</td><td>10473</td><td>10494</td><td>22</td><td>CTCGGCCGATCGAAAGGGAGTC</td><td>363.3</td><td>-5.2</td><td>0.2</td><td>-21.0</td><td>1.2E-97</td><td>4.3E-95</td><td>BT20/MB468</td><td>NUC</td></tr>
<tr><td>>am_MI0005763|hsa-mir-941-1&WithFlank&20|+|63919443|63919526@53.74.22(+1U);MI0005764|hsa-mir-941-2&WithFlank&20|+|63919499|63919582@53.74.22(+1U);MI0005765|hsa-mir-941-3&WithFlank&20|+|63919555|63919638@53.74.22(+1U);MI0005766|hsa-mir-941-4&WithFlank&20|+|63919750|63919833@53.74.22(+1U);MI0031520|hsa-mir-941-5&WithFlank&20|+|63919862|63919945@53.74.22(+1U)&[MIMAT0004984&hsa-miR-941&offsets|0|-1(+1U);m-263&20|+|63919495|63919517&offsets|0|-1(+1U)];[MIMAT0004984_1&hsa-miR-941&offsets|0|-1(+1U);m-263&20|+|63919551|63919573&offsets|0|-1(+1U)];[MIMAT0004984_2&hsa-miR-941&offsets|0|-1(+1U);m-263&20|+|63919607|63919629&offsets|0|-1(+1U)];[MIMAT0004984_3&hsa-miR-941&offsets|0|-1(+1U);m-263&20|+|63919802|63919824&offsets|0|-1(+1U)];[MIMAT0004984_4&hsa-miR-941&offsets|0|-1(+1U);m-263&20|+|63919914|63919936&offsets|0|-1(+1U)]|CACCCGGCTGTGTGCACATGTGT</td><td>>am_MI0005763|hsa-mir-941-1&WithFlank&20|+|63919443|63919526</td><td>53</td><td>74</td><td>22(+1U)</td><td>CACCCGGCTGTGTGCACATGTGT</td><td>8253.8</td><td>8.5</td><td>0.4</td><td>21.0</td><td>8.6E-98</td><td>8.6E-95</td><td>BT20/MB468</td><td>CYTO</td></tr>
<tr><td>>MI0001445|hsa-mir-423&WithFlank&17|+|30117073|30117178@59.81.23&[MIMAT0001340&hsa-miR-423-3p&offsets|0|0;m-100&17|+|30117131|30117153&offsets|0|0]|AGCTCGGTCTGAGGCCCCTCAGT</td><td>>MI0001445|hsa-mir-423&WithFlank&17|+|30117073|30117178</td><td>59</td><td>81</td><td>23</td><td>AGCTCGGTCTGAGGCCCCTCAGT</td><td>37483.9</td><td>8.5</td><td>0.4</td><td>20.9</td><td>8.5E-97</td><td>7.3E-94</td><td>BT20/MB468</td><td>CYTO</td></tr>
<tr><td>>hg38_21_+_8433172_8446622_RNA45SN1_45S_50ntFlanks@11784.11805.22|ATGAATGGATGAACGAGATTCC</td><td>>hg38_21_+_8433172_8446622_RNA45SN1_45S_50ntFlanks</td><td>11784</td><td>11805</td><td>22</td><td>ATGAATGGATGAACGAGATTCC</td><td>1053.5</td><td>-7.5</td><td>0.4</td><td>-20.8</td><td>1.0E-95</td><td>3.3E-93</td><td>BT20/MB468</td><td>NUC</td></tr>
<tr><td>>hg38_21_+_8433172_8446622_RNA45SN1_45S_50ntFlanks@12936.12960.25|AAACCATTCGTAGACGACCTGCTTC</td><td>>hg38_21_+_8433172_8446622_RNA45SN1_45S_50ntFlanks</td><td>12936</td><td>12960</td><td>25</td><td>AAACCATTCGTAGACGACCTGCTTC</td><td>1748.9</td><td>-6.2</td><td>0.3</td><td>-20.5</td><td>1.6E-93</td><td>4.9E-91</td><td>BT20/MB468</td><td>NUC</td></tr>
<tr><td>>hg38_21_+_8433172_8446622_RNA45SN1_45S_50ntFlanks@3769.3791.23|CGCACGGCCGGTACAGTGAAACT</td><td>>hg38_21_+_8433172_8446622_RNA45SN1_45S_50ntFlanks</td><td>3769</td><td>3791</td><td>23</td><td>CGCACGGCCGGTACAGTGAAACT</td><td>608.7</td><td>-6.5</td><td>0.3</td><td>-20.5</td><td>2.2E-93</td><td>6.2E-91</td><td>BT20/MB468</td><td>NUC</td></tr>
<tr><td>>hg38_21_+_8433172_8446622_RNA45SN1_45S_50ntFlanks@10350.10371.22|ACGAGAACTTTGAAGGCCGAAG</td><td>>hg38_21_+_8433172_8446622_RNA45SN1_45S_50ntFlanks</td><td>10350</td><td>10371</td><td>22</td><td>ACGAGAACTTTGAAGGCCGAAG</td><td>3190.3</td><td>-7.2</td><td>0.4</td><td>-20.5</td><td>2.8E-93</td><td>7.6E-91</td><td>BT20/MB468</td><td>NUC</td></tr>
<tr><td>>MI0000542|hsa-mir-320a&WithFlank&8|-|22244960|22245043@48.68.21(+1U)&[MIMAT0000510&hsa-miR-320a-3p&offsets|0|-1(+1U);m-60&8|-|22244975|22244996&offsets|0|-1(+1U)]|AAAAGCTGGGTTGAGAGGGCGT</td><td>>MI0000542|hsa-mir-320a&WithFlank&8|-|22244960|22245043</td><td>48</td><td>68</td><td>21(+1U)</td><td>AAAAGCTGGGTTGAGAGGGCGT</td><td>42922.8</td><td>11.8</td><td>0.6</td><td>20.4</td><td>1.1E-92</td><td>8.6E-90</td><td>BT20/MB468</td><td>CYTO</td></tr>
<tr><td>>MI0000734|hsa-mir-106b&WithFlank&7|-|100093987|100094080@58.78.21&[MIMAT0004672&hsa-miR-106b-3p&offsets|0|-1;m-87&7|-|100094003|100094023&offsets|0|0]|CCGCACTGTGGGTACTTGCTG</td><td>>MI0000734|hsa-mir-106b&WithFlank&7|-|100093987|100094080</td><td>58</td><td>78</td><td>21</td><td>CCGCACTGTGGGTACTTGCTG</td><td>7072.2</td><td>8.3</td><td>0.4</td><td>20.4</td><td>1.8E-92</td><td>1.2E-89</td><td>BT20/MB468</td><td>CYTO</td></tr>
<tr><td>>hg38_21_+_8433172_8446622_RNA45SN1_45S_50ntFlanks@4047.4066.20|ATTCGAACGTCTGCCCTATC</td><td>>hg38_21_+_8433172_8446622_RNA45SN1_45S_50ntFlanks</td><td>4047</td><td>4066</td><td>20</td><td>ATTCGAACGTCTGCCCTATC</td><td>7644.9</td><td>-9.0</td><td>0.4</td><td>-20.3</td><td>5.6E-92</td><td>1.5E-89</td><td>BT20/MB468</td><td>NUC</td></tr>
<tr><td>>hg38_21_+_8433172_8446622_RNA45SN1_45S_50ntFlanks@8176.8203.28|CTTCTGATCGAGGCCCAGCCCGTGGACG</td><td>>hg38_21_+_8433172_8446622_RNA45SN1_45S_50ntFlanks</td><td>8176</td><td>8203</td><td>28</td><td>CTTCTGATCGAGGCCCAGCCCGTGGACG</td><td>8167.6</td><td>-8.3</td><td>0.4</td><td>-20.2</td><td>9.6E-91</td><td>2.4E-88</td><td>BT20/MB468</td><td>NUC</td></tr>
<tr><td>>MI0000542|hsa-mir-320a&WithFlank&8|-|22244960|22245043@48.70.23&[MIMAT0000510&hsa-miR-320a-3p&offsets|0|+1;m-60&8|-|22244975|22244996&offsets|0|+1]|AAAAGCTGGGTTGAGAGGGCGAA</td><td>>MI0000542|hsa-mir-320a&WithFlank&8|-|22244960|22245043</td><td>48</td><td>70</td><td>23</td><td>AAAAGCTGGGTTGAGAGGGCGAA</td><td>41332.7</td><td>11.5</td><td>0.6</td><td>20.1</td><td>9.3E-90</td><td>5.6E-87</td><td>BT20/MB468</td><td>CYTO</td></tr>
<tr><td>>MI0000102|hsa-mir-100&WithFlank&11|-|122152223|122152314@19.40.22&[MIMAT0000098&hsa-miR-100-5p&offsets|0|0;m-27&11|-|122152276|122152296&offsets|0|+1]|AACCCGTAGATCCGAACTTGTG</td><td>>MI0000102|hsa-mir-100&WithFlank&11|-|122152223|122152314</td><td>19</td><td>40</td><td>22</td><td>AACCCGTAGATCCGAACTTGTG</td><td>862010.7</td><td>16.5</td><td>0.8</td><td>20.2</td><td>2.1E-90</td><td>6.3E-87</td><td>MB231/MB468</td><td>NUC</td></tr>
<tr><td>>hg38_21_+_8433172_8446622_RNA45SN1_45S_50ntFlanks@10351.10382.32|CGAGAACTTTGAAGGCCGAAGTGGAGAAGGGT</td><td>>hg38_21_+_8433172_8446622_RNA45SN1_45S_50ntFlanks</td><td>10351</td><td>10382</td><td>32</td><td>CGAGAACTTTGAAGGCCGAAGTGGAGAAGGGT</td><td>5025.8</td><td>-8.4</td><td>0.4</td><td>-19.9</td><td>2.1E-88</td><td>5.1E-86</td><td>BT20/MB468</td><td>NUC</td></tr>
<tr><td>>hg38_21_+_8433172_8446622_RNA45SN1_45S_50ntFlanks@10351.10383.33|CGAGAACTTTGAAGGCCGAAGTGGAGAAGGGTT</td><td>>hg38_21_+_8433172_8446622_RNA45SN1_45S_50ntFlanks</td><td>10351</td><td>10383</td><td>33</td><td>CGAGAACTTTGAAGGCCGAAGTGGAGAAGGGTT</td><td>104495.0</td><td>-12.1</td><td>0.6</td><td>-19.7</td><td>1.7E-86</td><td>4.0E-84</td><td>BT20/MB468</td><td>NUC</td></tr>
<tr><td>>am_MI0000809|hsa-mir-151a&WithFlank&8|-|140732558|140732659@53.72.20&[MIMAT0000757&hsa-miR-151a-3p&offsets|0|-1;m-30&8|-|140732587|140732607&offsets|0|-1]|CTAGACTGAAGCTCCTTGAG</td><td>>am_MI0000809|hsa-mir-151a&WithFlank&8|-|140732558|140732659</td><td>53</td><td>72</td><td>20</td><td>CTAGACTGAAGCTCCTTGAG</td><td>5606.0</td><td>9.2</td><td>0.5</td><td>19.6</td><td>1.4E-85</td><td>7.7E-83</td><td>BT20/MB468</td><td>CYTO</td></tr>
</table>

<table>
<tr><td>>MI0000255|hsa-mir-30d&WithFlank&8|-|134804870|134804951@12.33.22&[MIMAT0000245&hsa-miR-30d-5p&offsets|0|0;m-21&8|-|134804919|134804940&offsets|0|0]|TGTAAACATCCCCGACTGGAAG</td><td>>MI0000255|hsa-mir-30d&WithFlank&8|-|134804870|134804951</td><td>12</td><td>33</td><td>22</td><td>TGTAAACATCCCCGACTGGAAG</td><td>4767.3</td><td>8.3</td><td>0.4</td><td>19.6</td><td>1.3E-85</td><td>7.6E-82</td><td>BT20/MB468</td><td>MITO</td></tr>
<tr><td>>hg38_21_+_8433172_8446622_RNA45SN1_45S_50ntFlanks@7983.8004.22|CAGATCAGACGTGGCGACCCGC</td><td>>hg38_21_+_8433172_8446622_RNA45SN1_45S_50ntFlanks</td><td>7983</td><td>8004</td><td>22</td><td>CAGATCAGACGTGGCGACCCGC</td><td>1752.1</td><td>-7.6</td><td>0.4</td><td>-19.4</td><td>6.4E-84</td><td>1.4E-81</td><td>BT20/MB468</td><td>NUC</td></tr>
<tr><td>>MI0003560|hsa-mir-92b&WithFlank&1|+|155195171|155195278@67.88.22&[MIMAT0003218&hsa-miR-92b-3p&offsets|0|0;m-115&1|+|155195237|155195258&offsets|0|0]|TATTGCACTCGTCCCGGCCTCC</td><td>>MI0003560|hsa-mir-92b&WithFlank&1|+|155195171|155195278</td><td>67</td><td>88</td><td>22</td><td>TATTGCACTCGTCCCGGCCTCC</td><td>11517.3</td><td>9.4</td><td>0.5</td><td>19.3</td><td>2.6E-83</td><td>1.3E-80</td><td>BT20/MB468</td><td>CYTO</td></tr>
<tr><td>>hg38_21_+_8433172_8446622_RNA45SN1_45S_50ntFlanks@10679.10710.32|GGTTCCGGCGGCGTCCGGTGAGCTCTCGCTGG</td><td>>hg38_21_+_8433172_8446622_RNA45SN1_45S_50ntFlanks</td><td>10679</td><td>10710</td><td>32</td><td>GGTTCCGGCGGCGTCCGGTGAGCTCTCGCTGG</td><td>95923.2</td><td>-12.4</td><td>0.6</td><td>-19.5</td><td>2.3E-84</td><td>1.5E-80</td><td>MB231/MB468</td><td>MITO</td></tr>
<tr><td>>MI0000082|hsa-mir-25&WithFlank&7|-|100093554|100093649@58.79.22(+1A)&[MIMAT0000081&hsa-miR-25-3p&offsets|0|0(+1A);m-23&7|-|100093571|100093592&offsets|0|0(+1A)]|CATTGCACTTGTCTCGGTCTGAA</td><td>>MI0000082|hsa-mir-25&WithFlank&7|-|100093554|100093649</td><td>58</td><td>79</td><td>22(+1A)</td><td>CATTGCACTTGTCTCGGTCTGAA</td><td>3302.5</td><td>8.2</td><td>0.4</td><td>19.2</td><td>2.4E-82</td><td>1.1E-79</td><td>BT20/MB468</td><td>CYTO</td></tr>
<tr><td>>hg38_21_+_8433172_8446622_RNA45SN1_45S_50ntFlanks@6660.6689.30|GCGGTGGATCACTCGGCTCGTGCGTCGATG</td><td>>hg38_21_+_8433172_8446622_RNA45SN1_45S_50ntFlanks</td><td>6660</td><td>6689</td><td>30</td><td>GCGGTGGATCACTCGGCTCGTGCGTCGATG</td><td>1005.4</td><td>-4.2</td><td>0.2</td><td>-19.2</td><td>6.2E-82</td><td>1.3E-79</td><td>BT20/MB468</td><td>NUC</td></tr>
<tr><td>>hg38_21_+_8433172_8446622_RNA45SN1_45S_50ntFlanks@10678.10710.33|CGGTTCCGGCGGCGTCCGGTGAGCTCTCGCTGG</td><td>>hg38_21_+_8433172_8446622_RNA45SN1_45S_50ntFlanks</td><td>10678</td><td>10710</td><td>33</td><td>CGGTTCCGGCGGCGTCCGGTGAGCTCTCGCTGG</td><td>767552.3</td><td>-12.5</td><td>0.6</td><td>-19.3</td><td>4.3E-83</td><td>1.4E-79</td><td>MB231/MB468</td><td>MITO</td></tr>
<tr><td>>hg38_21_+_8433172_8446622_RNA45SN1_45S_50ntFlanks@12456.12487.32|CCAAGCGTTGGATTGTTCACCCACTAATAGGG</td><td>>hg38_21_+_8433172_8446622_RNA45SN1_45S_50ntFlanks</td><td>12456</td><td>12487</td><td>32</td><td>CCAAGCGTTGGATTGTTCACCCACTAATAGGG</td><td>72520.7</td><td>-11.4</td><td>0.6</td><td>-19.3</td><td>8.6E-83</td><td>1.7E-79</td><td>MB231/MB468</td><td>NUC</td></tr>
<tr><td>>hg38_21_+_8433172_8446622_RNA45SN1_45S_50ntFlanks@4590.4616.27|ATTTTGTTGGTTTTCGGAACTGAGGCC</td><td>>hg38_21_+_8433172_8446622_RNA45SN1_45S_50ntFlanks</td><td>4590</td><td>4616</td><td>27</td><td>ATTTTGTTGGTTTTCGGAACTGAGGCC</td><td>869.6</td><td>-6.8</td><td>0.4</td><td>-19.1</td><td>2.3E-81</td><td>4.8E-79</td><td>BT20/MB468</td><td>NUC</td></tr>
<tr><td>>hg38_21_+_8433172_8446622_RNA45SN1_45S_50ntFlanks@4724.4758.35|ATTAATCAAGAACGAAAGTCGGAGGTTCGAAGACG</td><td>>hg38_21_+_8433172_8446622_RNA45SN1_45S_50ntFlanks</td><td>4724</td><td>4758</td><td>35</td><td>ATTAATCAAGAACGAAAGTCGGAGGTTCGAAGACG</td><td>15286.2</td><td>-7.9</td><td>0.4</td><td>-19.0</td><td>7.5E-81</td><td>1.1E-77</td><td>MB231/MB468</td><td>NUC</td></tr>
<tr><td>>MI0000063|hsa-let-7b&WithFlank&22|+|46113680|46113774@12.33.22(+1A)&[MIMAT0000063&hsa-let-7b-5p&offsets|0|0(+1A);m-14&22|+|46113691|46113712&offsets|0|0(+1A)]|TGAGGTAGTAGGTTGTGTGGTTA</td><td>>MI0000063|hsa-let-7b&WithFlank&22|+|46113680|46113774</td><td>12</td><td>33</td><td>22(+1A)</td><td>TGAGGTAGTAGGTTGTGTGGTTA</td><td>5450.1</td><td>8.6</td><td>0.5</td><td>18.9</td><td>9.4E-80</td><td>4.0E-77</td><td>BT20/MB468</td><td>CYTO</td></tr>
<tr><td>>MI0000063|hsa-let-7b&WithFlank&22|+|46113680|46113774@12.34.23(+1U)&[MIMAT0000063&hsa-let-7b-5p&offsets|0|+1(+1U);m-14&22|+|46113691|46113712&offsets|0|+1(+1U)]|TGAGGTAGTAGGTTGTGTGGTTTT</td><td>>MI0000063|hsa-let-7b&WithFlank&22|+|46113680|46113774</td><td>12</td><td>34</td><td>23(+1U)</td><td>TGAGGTAGTAGGTTGTGTGGTTTT</td><td>1480.9</td><td>8.3</td><td>0.4</td><td>18.8</td><td>3.3E-79</td><td>1.3E-76</td><td>BT20/MB468</td><td>CYTO</td></tr>
<tr><td>>hg38_21_+_8433172_8446622_RNA45SN1_45S_50ntFlanks@10397.10424.28|AGTTGAACATGGGTCAGTCGGTCCTGAG</td><td>>hg38_21_+_8433172_8446622_RNA45SN1_45S_50ntFlanks</td><td>10397</td><td>10424</td><td>28</td><td>AGTTGAACATGGGTCAGTCGGTCCTGAG</td><td>4371.8</td><td>10.1</td><td>0.5</td><td>18.9</td><td>1.1E-79</td><td>3.2E-76</td><td>BT20/MB468</td><td>MITO</td></tr>
<tr><td>>MI0000063|hsa-let-7b&WithFlank&22|+|46113680|46113774@12.32.21(+1C)&[MIMAT0000063&hsa-let-7b-5p&offsets|0|-1(+1C);m-14&22|+|46113691|46113712&offsets|0|-1(+1C)]|TGAGGTAGTAGGTTGTGTGGTC</td><td>>MI0000063|hsa-let-7b&WithFlank&22|+|46113680|46113774</td><td>12</td><td>32</td><td>21(+1C)</td><td>TGAGGTAGTAGGTTGTGTGGTC</td><td>15547.3</td><td>9.1</td><td>0.5</td><td>18.8</td><td>1.3E-78</td><td>4.8E-76</td><td>BT20/MB468</td><td>CYTO</td></tr>
<tr><td>>MI0001445|hsa-mir-423&WithFlank&17|+|30117073|30117178@58.81.24&[MIMAT0001340&hsa-miR-423-3p&offsets|-1|0;m-100&17|+|30117131|30117153&offsets|-1|0]|AAGCTCGGTCTGAGGCCCCTCAGT</td><td>>MI0001445|hsa-mir-423&WithFlank&17|+|30117073|30117178</td><td>58</td><td>81</td><td>24</td><td>AAGCTCGGTCTGAGGCCCCTCAGT</td><td>5663.8</td><td>8.4</td><td>0.4</td><td>18.7</td><td>2.6E-78</td><td>9.1E-76</td><td>BT20/MB468</td><td>CYTO</td></tr>
<tr><td>>hg38_21_+_8433172_8446622_RNA45SN1_45S_50ntFlanks@12973.12991.19|TCGTACGTAGCAGAGCAGC</td><td>>hg38_21_+_8433172_8446622_RNA45SN1_45S_50ntFlanks</td><td>12973</td><td>12991</td><td>19</td><td>TCGTACGTAGCAGAGCAGC</td><td>13559.5</td><td>-9.4</td><td>0.5</td><td>-18.8</td><td>1.0E-78</td><td>1.2E-75</td><td>MB231/MB468</td><td>NUC</td></tr>
<tr><td>>am_MI0000809|hsa-mir-151a&WithFlank&8|-|140732558|140732659@53.72.20&[MIMAT0000757&hsa-miR-151a-3p&offsets|0|-1;m-30&8|-|140732587|140732607&offsets|0|-1]|CTAGACTGAAGCTCCTTGAG</td><td>>am_MI0000809|hsa-mir-151a&WithFlank&8|-|140732558|140732659</td><td>53</td><td>72</td><td>20</td><td>CTAGACTGAAGCTCCTTGAG</td><td>3886.1</td><td>8.8</td><td>0.5</td><td>18.8</td><td>6.2E-79</td><td>1.2E-75</td><td>BT20/MB468</td><td>MITO</td></tr>
<tr><td>>hg38_21_+_8433172_8446622_RNA45SN1_45S_50ntFlanks@10350.10383.34|ACGAGAACTTTGAAGGCCGAAGTGGAGAAGGGTT</td><td>>hg38_21_+_8433172_8446622_RNA45SN1_45S_50ntFlanks</td><td>10350</td><td>10383</td><td>34</td><td>ACGAGAACTTTGAAGGCCGAAGTGGAGAAGGGTT</td><td>48983.7</td><td>-11.1</td><td>0.6</td><td>-18.7</td><td>7.1E-78</td><td>1.4E-75</td><td>BT20/MB468</td><td>NUC</td></tr>
<tr><td>>MI0000650|hsa-mir-200c&WithFlank&12|+|6963693|6963772@50.72.23&[MIMAT0000617&hsa-miR-200c-3p&offsets|0|0;m-54&12|+|6963742|6963764&offsets|0|0]|TAATACTGCCGGGTAATGATGGA</td><td>>MI0000650|hsa-mir-200c&WithFlank&12|+|6963693|6963772</td><td>50</td><td>72</td><td>23</td><td>TAATACTGCCGGGTAATGATGGA</td><td>141359.9</td><td>11.1</td><td>0.6</td><td>18.8</td><td>7.4E-79</td><td>1.9E-75</td><td>BT20/MB231</td><td>NUC</td></tr>
<tr><td>>hg38_21_+_8433172_8446622_RNA45SN1_45S_50ntFlanks@7981.8006.26|CTCAGATCAGACGTGGCGACCCGCTG</td><td>>hg38_21_+_8433172_8446622_RNA45SN1_45S_50ntFlanks</td><td>7981</td><td>8006</td><td>26</td><td>CTCAGATCAGACGTGGCGACCCGCTG</td><td>10034.7</td><td>-9.4</td><td>0.5</td><td>-18.6</td><td>1.4E-77</td><td>2.7E-75</td><td>BT20/MB468</td><td>NUC</td></tr>
<tr><td>>hg38_21_+_8433172_8446622_RNA45SN1_45S_50ntFlanks@10350.10368.19|ACGAGAACTTTGAAGGCCG</td><td>>hg38_21_+_8433172_8446622_RNA45SN1_45S_50ntFlanks</td><td>10350</td><td>10368</td><td>19</td><td>ACGAGAACTTTGAAGGCCG</td><td>12589.1</td><td>-10.6</td><td>0.6</td><td>-18.6</td><td>1.9E-77</td><td>3.5E-75</td><td>BT20/MB468</td><td>NUC</td></tr>
<tr><td>>hg38_21_+_8433172_8446622_RNA45SN1_45S_50ntFlanks@12937.12960.24|AACCATTCGTAGACGACCTGCTTC</td><td>>hg38_21_+_8433172_8446622_RNA45SN1_45S_50ntFlanks</td><td>12937</td><td>12960</td><td>24</td><td>AACCATTCGTAGACGACCTGCTTC</td><td>1068.1</td><td>-5.1</td><td>0.3</td><td>-18.5</td><td>1.5E-76</td><td>2.7E-74</td><td>BT20/MB468</td><td>NUC</td></tr>
<tr><td>>trna11_GluTTC_15_-_26327381_26327452@1.33.33,trna3_GluTTC_13_-_45492062_45492133@1.33.33|TCCCACATGGTCTAGCGGTTAGGATTCCTGGTT</td><td>>trna11_GluTTC_15_-_26327381_26327452</td><td>1</td><td>33</td><td>33</td><td>TCCCACATGGTCTAGCGGTTAGGATTCCTGGTT</td><td>9513.3</td><td>9.1</td><td>0.5</td><td>18.5</td><td>9.2E-77</td><td>3.1E-74</td><td>BT20/MB468</td><td>CYTO</td></tr>
<tr><td>>MI0000542|hsa-mir-320a&WithFlank&8|-|22244960|22245043@48.69.22(+2U)&[MIMAT0000510&hsa-miR-320a-3p&offsets|0|0(+2U);m-60&8|-|22244975|22244996&offsets|0|0(+2U)]|AAAAGCTGGGTTGAGAGGGCGATT</td><td>>MI0000542|hsa-mir-320a&WithFlank&8|-|22244960|22245043</td><td>48</td><td>69</td><td>22(+2U)</td><td>AAAAGCTGGGTTGAGAGGGCGATT</td><td>27652.9</td><td>12.1</td><td>0.7</td><td>18.5</td><td>2.2E-76</td><td>7.0E-74</td><td>BT20/MB468</td><td>CYTO</td></tr>
<tr><td>>MI0000063|hsa-let-7b&WithFlank&22|+|46113680|46113774@12.33.22&[MIMAT0000063&hsa-let-7b-5p&offsets|0|0;m-14&22|+|46113691|46113712&offsets|0|0]|TGAGGTAGTAGGTTGTGTGGTT</td><td>>MI0000063|hsa-let-7b&WithFlank&22|+|46113680|46113774</td><td>12</td><td>33</td><td>22</td><td>TGAGGTAGTAGGTTGTGTGGTT</td><td>37454.9</td><td>8.4</td><td>0.5</td><td>18.5</td><td>1.4E-76</td><td>2.1E-73</td><td>BT20/MB468</td><td>MITO</td></tr>
<tr><td>>hg38_21_+_8433172_8446622_RNA45SN1_45S_50ntFlanks@10678.10711.34|CGGTTCCGGCGGCGTCCGGTGAGCTCTCGCTGGC</td><td>>hg38_21_+_8433172_8446622_RNA45SN1_45S_50ntFlanks</td><td>10678</td><td>10711</td><td>34</td><td>CGGTTCCGGCGGCGTCCGGTGAGCTCTCGCTGGC</td><td>187464.3</td><td>-11.7</td><td>0.6</td><td>-18.5</td><td>1.4E-76</td><td>3.1E-73</td><td>MB231/MB468</td><td>MITO</td></tr>
<tr><td>>hg38_21_+_8433172_8446622_RNA45SN1_45S_50ntFlanks@10351.10377.27|CGAGAACTTTGAAGGCCGAAGTGGAGA</td><td>>hg38_21_+_8433172_8446622_RNA45SN1_45S_50ntFlanks</td><td>10351</td><td>10377</td><td>27</td><td>CGAGAACTTTGAAGGCCGAAGTGGAGA</td><td>8798.5</td><td>-7.6</td><td>0.4</td><td>-18.5</td><td>4.7E-76</td><td>4.6E-73</td><td>MB231/MB468</td><td>NUC</td></tr>
<tr><td>>hg38_21_+_8433172_8446622_RNA45SN1_45S_50ntFlanks@6721.6743.23|AATTGCAGGACACATTGATCATC</td><td>>hg38_21_+_8433172_8446622_RNA45SN1_45S_50ntFlanks</td><td>6721</td><td>6743</td><td>23</td><td>AATTGCAGGACACATTGATCATC</td><td>20262.9</td><td>-10.8</td><td>0.6</td><td>-18.3</td><td>3.4E-75</td><td>5.8E-73</td><td>BT20/MB468</td><td>NUC</td></tr>
<tr><td>>hg38_21_+_8433172_8446622_RNA45SN1_45S_50ntFlanks@12457.12477.21|CAAGCGTTGGATTGTTCACCC</td><td>>hg38_21_+_8433172_8446622_RNA45SN1_45S_50ntFlanks</td><td>12457</td><td>12477</td><td>21</td><td>CAAGCGTTGGATTGTTCACCC</td><td>6290.2</td><td>-8.4</td><td>0.5</td><td>-18.3</td><td>5.1E-75</td><td>8.5E-73</td><td>BT20/MB468</td><td>NUC</td></tr>
<tr><td>>MI0000440|hsa-mir-27b&WithFlank&9|+|95085439|95085547@67.86.20&[MIMAT0000419&hsa-miR-27b-3p&offsets|0|-1;m-39&9|+|95085505|95085525&offsets|0|-1]|TTCACAGTGGCTAAGTTCTG</td><td>>MI0000440|hsa-mir-27b&WithFlank&9|+|95085439|95085547</td><td>67</td><td>86</td><td>20</td><td>TTCACAGTGGCTAAGTTCTG</td><td>4394.4</td><td>7.2</td><td>0.4</td><td>18.4</td><td>1.6E-75</td><td>1.9E-72</td><td>BT20/MB468</td><td>MITO</td></tr>
<tr><td>>hg38_21_+_8433172_8446622_RNA45SN1_45S_50ntFlanks@11572.11590.19|CCGACTTAGAACTGGTGCG</td><td>>hg38_21_+_8433172_8446622_RNA45SN1_45S_50ntFlanks</td><td>11572</td><td>11590</td><td>19</td><td>CCGACTTAGAACTGGTGCG</td><td>5881.1</td><td>-9.6</td><td>0.5</td><td>-18.3</td><td>1.5E-74</td><td>2.5E-72</td><td>BT20/MB468</td><td>NUC</td></tr>
<tr><td>>hg38_21_+_8433172_8446622_RNA45SN1_45S_50ntFlanks@6656.6681.26|CTTAGCGGTGGATCACTCGGCTCGTG</td><td>>hg38_21_+_8433172_8446622_RNA45SN1_45S_50ntFlanks</td><td>6656</td><td>6681</td><td>26</td><td>CTTAGCGGTGGATCACTCGGCTCGTG</td><td>3450.7</td><td>-6.7</td><td>0.4</td><td>-18.0</td><td>1.1E-72</td><td>1.7E-70</td><td>BT20/MB468</td><td>NUC</td></tr>
<tr><td>>hg38_21_+_8433172_8446622_RNA45SN1_45S_50ntFlanks@7979.8008.30|ACCTCAGATCAGACGTGGCGACCCGCTGAA</td><td>>hg38_21_+_8433172_8446622_RNA45SN1_45S_50ntFlanks</td><td>7979</td><td>8008</td><td>30</td><td>ACCTCAGATCAGACGTGGCGACCCGCTGAA</td><td>2954.5</td><td>-6.7</td><td>0.4</td><td>-18.0</td><td>1.1E-72</td><td>1.7E-70</td><td>BT20/MB468</td><td>NUC</td></tr>
<tr><td>>MI0000063|hsa-let-7b&WithFlank&22|+|46113680|46113774@12.32.21&[MIMAT0000063&hsa-let-7b-5p&offsets|0|-1;m-14&22|+|46113691|46113712&offsets|0|-1]|TGAGGTAGTAGGTTGTGTGGT</td><td>>MI0000063|hsa-let-7b&WithFlank&22|+|46113680|46113774</td><td>12</td><td>32</td><td>21</td><td>TGAGGTAGTAGGTTGTGTGGT</td><td>4256.9</td><td>8.3</td><td>0.5</td><td>18.1</td><td>2.6E-73</td><td>2.6E-70</td><td>BT20/MB468</td><td>MITO</td></tr>
<tr><td>>hg38_21_+_8433172_8446622_RNA45SN1_45S_50ntFlanks@12457.12476.20|CAAGCGTTGGATTGTTCACC</td><td>>hg38_21_+_8433172_8446622_RNA45SN1_45S_50ntFlanks</td><td>12457</td><td>12476</td><td>20</td><td>CAAGCGTTGGATTGTTCACC</td><td>8190.2</td><td>-7.4</td><td>0.4</td><td>-18.0</td><td>1.8E-72</td><td>2.6E-70</td><td>BT20/MB468</td><td>NUC</td></tr>
<tr><td>>hg38_21_+_8433172_8446622_RNA45SN1_45S_50ntFlanks@3835.3860.26|CTCCTCTCCTACTTGGATAACTGTGG</td><td>>hg38_21_+_8433172_8446622_RNA45SN1_45S_50ntFlanks</td><td>3835</td><td>3860</td><td>26</td><td>CTCCTCTCCTACTTGGATAACTGTGG</td><td>7685.3</td><td>-9.1</td><td>0.5</td><td>-18.0</td><td>1.9E-72</td><td>2.7E-70</td><td>BT20/MB468</td><td>NUC</td></tr>
<tr><td>>MI0000342|hsa-mir-200b&WithFlank&1|+|1167098|1167204@63.85.23&[MIMAT0000318&hsa-miR-200b-3p&offsets|0|+1;m-93&1|+|1167160|1167182&offsets|0|0]|TAATACTGCCTGGTAATGATGAC</td><td>>MI0000342|hsa-mir-200b&WithFlank&1|+|1167098|1167204</td><td>63</td><td>85</td><td>23</td><td>TAATACTGCCTGGTAATGATGAC</td><td>9242.0</td><td>-7.0</td><td>0.4</td><td>-18.0</td><td>3.2E-72</td><td>4.5E-70</td><td>BT20/MB468</td><td>NUC</td></tr>
<tr><td>>hg38_21_+_8433172_8446622_RNA45SN1_45S_50ntFlanks@11575.11594.20|ACTTAGAACTGGTGCGGACC</td><td>>hg38_21_+_8433172_8446622_RNA45SN1_45S_50ntFlanks</td><td>11575</td><td>11594</td><td>20</td><td>ACTTAGAACTGGTGCGGACC</td><td>5761.3</td><td>-8.3</td><td>0.5</td><td>-18.0</td><td>4.2E-72</td><td>5.8E-70</td><td>BT20/MB468</td><td>NUC</td></tr>
<tr><td>>MI0000734|hsa-mir-106b&WithFlank&7|-|100093987|100094080@58.78.21(+1U)&[MIMAT0004672&hsa-miR-106b-3p&offsets|0|-1(+1U);m-87&7|-|100094003|100094023&offsets|0|0(+1U)]|CCGCACTGTGGGTACTTGCTGT</td><td>>MI0000734|hsa-mir-106b&WithFlank&7|-|100093987|100094080</td><td>58</td><td>78</td><td>21(+1U)</td><td>CCGCACTGTGGGTACTTGCTGT</td><td>4181.9</td><td>7.8</td><td>0.4</td><td>17.9</td><td>9.4E-72</td><td>2.8E-69</td><td>BT20/MB468</td><td>CYTO</td></tr>
<tr><td>>MI0000283|hsa-mir-203a&WithFlank&14|+|104117399|104117520@72.92.21&[MIMAT0000264&hsa-miR-203a-3p&offsets|+1|0;m-79&14|+|104117469|104117490&offsets|+1|0]|TGAAATGTTTAGGACCACTAG</td><td>>MI0000283|hsa-mir-203a&WithFlank&14|+|104117399|104117520</td><td>72</td><td>92</td><td>21</td><td>TGAAATGTTTAGGACCACTAG</td><td>4114.8</td><td>9.3</td><td>0.5</td><td>17.9</td><td>1.0E-71</td><td>3.0E-69</td><td>BT20/MB468</td><td>CYTO</td></tr>
<tr><td>>MI0000299|hsa-mir-222&WithFlank&X|-|45747009|45747130@75.98.24(+1A)&[MIMAT0000279&hsa-miR-222-3p&offsets|0|+3(+1A);m-140&X|-|45747033|45747056&offsets|0|0(+1A)]|AGCTACATCTGGCTACTGGGTCTCA</td><td>>MI0000299|hsa-mir-222&WithFlank&X|-|45747009|45747130</td><td>75</td><td>98</td><td>24(+1A)</td><td>AGCTACATCTGGCTACTGGGTCTCA</td><td>3337.3</td><td>7.4</td><td>0.4</td><td>17.9</td><td>1.6E-71</td><td>4.4E-69</td><td>BT20/MB468</td><td>CYTO</td></tr>
<tr><td>>MI0001445|hsa-mir-423&WithFlank&17|+|30117073|30117178@59.80.22(+1C)&[MIMAT0001340&hsa-miR-423-3p&offsets|0|-1(+1C);m-100&17|+|30117131|30117153&offsets|0|-1(+1C)]|AGCTCGGTCTGAGGCCCCTCAGC</td><td>>MI0001445|hsa-mir-423&WithFlank&17|+|30117073|30117178</td><td>59</td><td>80</td><td>22(+1C)</td><td>AGCTCGGTCTGAGGCCCCTCAGC</td><td>4716.8</td><td>8.8</td><td>0.5</td><td>17.9</td><td>2.3E-71</td><td>5.7E-69</td><td>BT20/MB468</td><td>CYTO</td></tr>
<tr><td>>am_MI0000809|hsa-mir-151a&WithFlank&8|-|140732558|140732659@53.72.20(+1U)&[MIMAT0000757&hsa-miR-151a-3p&offsets|0|-1(+1U);m-30&8|-|140732587|140732607&offsets|0|-1(+1U)]|CTAGACTGAAGCTCCTTGAGT</td><td>>am_MI0000809|hsa-mir-151a&WithFlank&8|-|140732558|140732659</td><td>53</td><td>72</td><td>20(+1U)</td><td>CTAGACTGAAGCTCCTTGAGT</td><td>8251.8</td><td>9.5</td><td>0.5</td><td>17.9</td><td>2.2E-71</td><td>5.7E-69</td><td>BT20/MB468</td><td>CYTO</td></tr>
<tr><td>>MI0000063|hsa-let-7b&WithFlank&22|+|46113680|46113774@12.32.21&[MIMAT0000063&hsa-let-7b-5p&offsets|0|-1;m-14&22|+|46113691|46113712&offsets|0|-1]|TGAGGTAGTAGGTTGTGTGGT</td><td>>MI0000063|hsa-let-7b&WithFlank&22|+|46113680|46113774</td><td>12</td><td>32</td><td>21</td><td>TGAGGTAGTAGGTTGTGTGGT</td><td>13995.7</td><td>8.5</td><td>0.5</td><td>17.8</td><td>4.9E-71</td><td>1.2E-68</td><td>BT20/MB468</td><td>CYTO</td></tr>
<tr><td>>hg38_21_+_8433172_8446622_RNA45SN1_45S_50ntFlanks@10398.10424.27|GTTGAACATGGGTCAGTCGGTCCTGAG</td><td>>hg38_21_+_8433172_8446622_RNA45SN1_45S_50ntFlanks</td><td>10398</td><td>10424</td><td>27</td><td>GTTGAACATGGGTCAGTCGGTCCTGAG</td><td>18804.7</td><td>10.3</td><td>0.6</td><td>17.8</td><td>6.1E-71</td><td>5.2E-68</td><td>BT20/MB468</td><td>MITO</td></tr>
</table>

| | | | | | | | | | | | | | |
|---|---|---|---|---|---|---|---|---|---|---|---|---|---|
| >MI0003628\|hsa-mir-615&WithFlank&12\|+\|54033944\|54034051@67.88.22&[MIMAT0003283&hsa-miR-615-3p&offsets\|0\|0;m-495&12\|+\|54034010\|54034030&offsets\|0\|+1]\|TCCGAGCCTGGGTCTCCCTCTT | >MI0003628\|hsa-mir-615&WithFlank&12\|+\|54033944\|54034051 | 67 | 88 | 22 | TCCGAGCCTGGGTCTCCCTCTT | 2732.8 | 7.3 | 0.4 | 17.7 | 5.9E-70 | 1.4E-67 | BT20/MB468 | CYTO |
| >hg38_21_+_8433172_8446622_RNA45SN1_45S_50ntFlanks@11671.11702.32\|CCAGTGCTCTGAATGTCAAAGTGAAGAAATTC | >hg38_21_+_8433172_8446622_RNA45SN1_45S_50ntFlanks | 11671 | 11702 | 32 | CCAGTGCTCTGAATGTCAAAGTGAAGAAATTC | 7417.7 | -9.3 | 0.5 | -17.6 | 1.6E-69 | 2.2E-67 | BT20/MB468 | NUC |
| >MI0000279\|hsa-mir-196a-2&WithFlank&12\|+\|53991732\|53991853@31.52.22;MI0000238\|hsa-mir-196a-1&WithFlank&17\|-\|48632484\|48632565@13.34.22&[MIMAT0000226&hsa-miR-196a-5p&offsets\|0\|0;m-261&12\|+\|53991762\|53991783&offsets\|0\|0];[MIMAT0000226_1&hsa-miR-196a-5p&offsets\|0\|0;m-268&17\|-\|48632532\|48632553&offsets\|0\|0]\|TAGGTAGTTTCATGTTGTTGGG | >MI0000279\|hsa-mir-196a-2&WithFlank&12\|+\|53991732\|53991853 | 31 | 52 | 22 | TAGGTAGTTTCATGTTGTTGGG | 1758.4 | 7.8 | 0.4 | 17.7 | 3.0E-70 | 2.2E-67 | BT20/MB468 | MITO |
| >am_MI0005763\|hsa-mir-941-1&WithFlank&20\|+\|63919443\|63919526@53.74.22;MI0005764\|hsa-mir-941-2&WithFlank&20\|+\|63919499\|63919582@53.74.22;MI0005765\|hsa-mir-941-3&WithFlank&20\|+\|63919555\|63919638@53.74.22;MI0005766\|hsa-mir-941-4&WithFlank&20\|+\|63919750\|63919833@53.74.22;MI0031520\|hsa-mir-941-5&WithFlank&20\|+\|63919862\|63919945@53.74.22&[MIMAT0004984&hsa-miR-941&offsets\|0\|-1;m-263&20\|+\|63919495\|63919517&offsets\|0\|-1];[MIMAT0004984_1&hsa-miR-941&offsets\|0\|-1;m-263&20\|+\|63919551\|63919573&offsets\|0\|-1];[MIMAT0004984_2&hsa-miR-941&offsets\|0\|-1;m-263&20\|+\|63919607\|63919629&offsets\|0\|-1];[MIMAT0004984_3&hsa-miR-941&offsets\|0\|-1;m-263&20\|+\|63919802\|63919824&offsets\|0\|-1];[MIMAT0004984_4&hsa-miR-941&offsets\|0\|-1;m-263&20\|+\|63919914\|63919936&offsets\|0\|-1]\|CACCCGGCTGTGTGCACATGTG | >am_MI0005763\|hsa-mir-941-1&WithFlank&20\|+\|63919443\|63919526 | 53 | 74 | 22 | CACCCGGCTGTGTGCACATGTG | 1801.4 | 8.0 | 0.5 | 17.6 | 1.1E-69 | 2.5E-67 | BT20/MB468 | CYTO |
| >MI0000650\|hsa-mir-200c&WithFlank&12\|+\|6963693\|6963772@50.72.23(+1A)&[MIMAT0000617&hsa-miR-200c-3p&offsets\|0\|0(+1A);m-54&12\|+\|6963742\|6963764&offsets\|0\|0(+1A)]\|TAATACTGCCGGGTAATGATGGAA | >MI0000650\|hsa-mir-200c&WithFlank&12\|+\|6963693\|6963772 | 50 | 72 | 23(+1A) | TAATACTGCCGGGTAATGATGGAA | 27804.6 | 9.2 | 0.5 | 17.6 | 3.9E-69 | 8.3E-67 | BT20/MB468 | CYTO |
| >MI0000273\|hsa-mir-183&WithFlank&7\|-\|129774899\|129775020@34.55.22&[MIMAT0000261&hsa-miR-183-5p&offsets\|+1\|+1;m-41&7\|-\|129774966\|129774987&offsets\|0\|0]\|ATGGCACTGGTAGAATTCACTG | >MI0000273\|hsa-mir-183&WithFlank&7\|-\|129774899\|129775020 | 34 | 55 | 22 | ATGGCACTGGTAGAATTCACTG | 5263.7 | 8.1 | 0.5 | 17.6 | 1.8E-69 | 1.2E-66 | BT20/MB468 | MITO |
| >MI0000255\|hsa-mir-30d&WithFlank&8\|-\|134804870\|134804951@12.31.20&[MIMAT0000245&hsa-miR-30d-5p&offsets\|0\|-2;m-21&8\|-\|134804919\|134804940&offsets\|0\|-2]\|TGTAAACATCCCCGACTGGA | >MI0000255\|hsa-mir-30d&WithFlank&8\|-\|134804870\|134804951 | 12 | 31 | 20 | TGTAAACATCCCCGACTGGA | 3059.5 | 8.9 | 0.5 | 17.5 | 1.1E-68 | 2.2E-66 | BT20/MB468 | CYTO |
| >hg38_21_+_8433172_8446622_RNA45SN1_45S_50ntFlanks@11576.11594.19\|CTTAGAACTGGTGCGGACC | >hg38_21_+_8433172_8446622_RNA45SN1_45S_50ntFlanks | 11576 | 11594 | 19 | CTTAGAACTGGTGCGGACC | 3699.8 | -8.6 | 0.5 | -17.5 | 2.2E-68 | 2.8E-66 | BT20/MB468 | NUC |
| >hg38_21_+_8433172_8446622_RNA45SN1_45S_50ntFlanks@10680.10711.32\|GTTCCGGCGGCGTCCGGTGAGCTCTCGCTGGC | >hg38_21_+_8433172_8446622_RNA45SN1_45S_50ntFlanks | 10680 | 10711 | 32 | GTTCCGGCGGCGTCCGGTGAGCTCTCGCTGGC | 19473.6 | -9.6 | 0.5 | -17.5 | 1.4E-68 | 2.3E-65 | MB231/MB468 | MITO |
| >hg38_21_+_8433172_8446622_RNA45SN1_45S_50ntFlanks@4738.4758.21\|AAAGTCGGAGGTTCGAAGACG | >hg38_21_+_8433172_8446622_RNA45SN1_45S_50ntFlanks | 4738 | 4758 | 21 | AAAGTCGGAGGTTCGAAGACG | 6762.8 | -6.8 | 0.4 | -17.3 | 2.0E-67 | 2.6E-65 | BT20/MB468 | NUC |
| >MI0001445\|hsa-mir-423&WithFlank&17\|+\|30117073\|30117178@23.46.24&[MIMAT0004748&hsa-miR-423-5p&offsets\|0\|+1;m-141&17\|+\|30117095\|30117117&offsets\|0\|+1]\|TGAGGGGCAGAGAGCGAGACTTTT | >MI0001445\|hsa-mir-423&WithFlank&17\|+\|30117073\|30117178 | 23 | 46 | 24 | TGAGGGGCAGAGAGCGAGACTTTT | 3897.7 | 9.5 | 0.5 | 17.3 | 2.1E-67 | 4.2E-65 | BT20/MB468 | CYTO |
| >hg38_21_+_8433172_8446622_RNA45SN1_45S_50ntFlanks@10411.10429.19\|CAGTCGGTCCTGAGAGATG | >hg38_21_+_8433172_8446622_RNA45SN1_45S_50ntFlanks | 10411 | 10429 | 19 | CAGTCGGTCCTGAGAGATG | 18522.0 | -8.8 | 0.5 | -17.4 | 6.6E-68 | 5.6E-65 | MB231/MB468 | NUC |
| >hg38_21_+_8433172_8446622_RNA45SN1_45S_50ntFlanks@12419.12436.18\|GATGTCGGCTCTTCCTAT | >hg38_21_+_8433172_8446622_RNA45SN1_45S_50ntFlanks | 12419 | 12436 | 18 | GATGTCGGCTCTTCCTAT | 9794.6 | -8.0 | 0.5 | -17.3 | 5.1E-67 | 6.4E-65 | BT20/MB468 | NUC |
| >MI0000285\|hsa-mir-205&WithFlank&1\|+\|209432127\|209432248@40.60.21&[MIMAT0000266&hsa-miR-205-5p&offsets\|0\|-1;m-351&1\|+\|209432166\|209432187&offsets\|0\|-1]\|TCCTTCATTCCACCGGAGTCT | >MI0000285\|hsa-mir-205&WithFlank&1\|+\|209432127\|209432248 | 40 | 60 | 21 | TCCTTCATTCCACCGGAGTCT | 10662.6 | 7.4 | 0.4 | 17.3 | 3.7E-67 | 7.2E-65 | BT20/MB468 | CYTO |
| >MI0000650\|hsa-mir-200c&WithFlank&12\|+\|6963693\|6963772@50.72.23(+1U)&[MIMAT0000617&hsa-miR-200c-3p&offsets\|0\|0(+1U);m-54&12\|+\|6963742\|6963764&offsets\|0\|0(+1U)]\|TAATACTGCCGGGTAATGATGGAT | >MI0000650\|hsa-mir-200c&WithFlank&12\|+\|6963693\|6963772 | 50 | 72 | 23(+1U) | TAATACTGCCGGGTAATGATGGAT | 18018.2 | 8.6 | 0.5 | 17.3 | 4.4E-67 | 8.2E-65 | BT20/MB468 | CYTO |
| >hg38_21_+_8433172_8446622_RNA45SN1_45S_50ntFlanks@10678.10709.32\|CGGTTCCGGCGGCGTCCGGTGAGCTCTCGCTG | >hg38_21_+_8433172_8446622_RNA45SN1_45S_50ntFlanks | 10678 | 10709 | 32 | CGGTTCCGGCGGCGTCCGGTGAGCTCTCGCTG | 96670.0 | -11.2 | 0.6 | -17.4 | 7.9E-68 | 1.0E-64 | MB231/MB468 | MITO |
| >hg38_21_+_8433172_8446622_RNA45SN1_45S_50ntFlanks@12456.12477.22\|CCAAGCGTTGGATTGTTCACCC | >hg38_21_+_8433172_8446622_RNA45SN1_45S_50ntFlanks | 12456 | 12477 | 22 | CCAAGCGTTGGATTGTTCACCC | 10028.6 | -9.8 | 0.6 | -17.2 | 1.8E-66 | 2.2E-64 | BT20/MB468 | NUC |
| >am_MI0000809\|hsa-mir-151a&WithFlank&8\|-\|140732558\|140732659@53.73.21(+1U)&[MIMAT0000757&hsa-miR-151a-3p&offsets\|0\|0(+1U);m-30&8\|-\|140732587\|140732607&offsets\|0\|0(+1U)]\|CTAGACTGAAGCTCCTTGAGGT | >am_MI0000809\|hsa-mir-151a&WithFlank&8\|-\|140732558\|140732659 | 53 | 73 | 21(+1U) | CTAGACTGAAGCTCCTTGAGGT | 20140.2 | 9.4 | 0.5 | 17.2 | 3.1E-66 | 5.7E-64 | BT20/MB468 | CYTO |
| >MI0006273\|hsa-mir-1180&WithFlank&17\|-\|19344500\|19344580@47.68.22&[MIMAT0005825&hsa-miR-1180-3p&offsets\|0\|0;m-214&17\|-\|19344513\|19344534&offsets\|0\|0]\|TTTCCGGCTCGCGTGGGTGTGT | >MI0006273\|hsa-mir-1180&WithFlank&17\|-\|19344500\|19344580 | 47 | 68 | 22 | TTTCCGGCTCGCGTGGGTGTGT | 5506.2 | 9.3 | 0.5 | 17.2 | 3.4E-66 | 6.0E-64 | BT20/MB468 | CYTO |
| >MI0000266\|hsa-mir-10a&WithFlank&17\|-\|48579832\|48579953@28.49.22&[MIMAT0000253&hsa-miR-10a-5p&offsets\|0\|-1;m-18&17\|-\|48579905\|48579926&offsets\|0\|0]\|TACCCTGTAGATCCGAATTTGT | >MI0000266\|hsa-mir-10a&WithFlank&17\|-\|48579832\|48579953 | 28 | 49 | 22 | TACCCTGTAGATCCGAATTTGT | 8273.0 | 11.3 | 0.7 | 17.3 | 9.3E-67 | 6.8E-64 | MB231/MB468 | NUC |
| >MI0005559\|hsa-mir-744&WithFlank&17\|+\|12081893\|12082002@17.37.21&[MIMAT0004945&hsa-miR-744-5p&offsets\|0\|-1;m-172&17\|+\|12081909\|12081929&offsets\|0\|0]\|TGCGGGGCTAGGGCTAACAGC | >MI0005559\|hsa-mir-744&WithFlank&17\|+\|12081893\|12082002 | 17 | 37 | 21 | TGCGGGGCTAGGGCTAACAGC | 4919.8 | 8.5 | 0.5 | 17.1 | 1.5E-65 | 2.6E-63 | BT20/MB468 | CYTO |
| >MI0000239\|hsa-mir-197&WithFlank&1\|+\|109598887\|109598973@54.75.22&[MIMAT0000227&hsa-miR-197-3p&offsets\|0\|0;m-99&1\|+\|109598940\|109598961&offsets\|0\|0]\|TTCACCACCTTCTCCACCCAGC | >MI0000239\|hsa-mir-197&WithFlank&1\|+\|109598887\|109598973 | 54 | 75 | 22 | TTCACCACCTTCTCCACCCAGC | 11772.4 | 8.4 | 0.5 | 17.2 | 4.5E-66 | 2.7E-63 | BT20/MB468 | MITO |
| >MI0000746\|hsa-mir-99b&WithFlank&19\|+\|51692606\|51692687@51.73.23&[MIMAT0004678&hsa-miR-99b-3p&offsets\|0\|+1;m-151&19\|+\|51692656\|51692678&offsets\|0\|0]\|CAAGCTCGTGTCTGTGGGTCCGT | >MI0000746\|hsa-mir-99b&WithFlank&19\|+\|51692606\|51692687 | 51 | 73 | 23 | CAAGCTCGTGTCTGTGGGTCCGT | 3516.7 | 8.1 | 0.5 | 17.1 | 1.9E-65 | 3.2E-63 | BT20/MB468 | CYTO |
| >MI0000273\|hsa-mir-183&WithFlank&7\|-\|129774899\|129775020@33.55.23&[MIMAT0000261&hsa-miR-183-5p&offsets\|0\|+1;m-41&7\|-\|129774966\|129774987&offsets\|-1\|0]\|TATGGCACTGGTAGAATTCACTG | >MI0000273\|hsa-mir-183&WithFlank&7\|-\|129774899\|129775020 | 33 | 55 | 23 | TATGGCACTGGTAGAATTCACTG | 6934.3 | 7.5 | 0.4 | 17.0 | 6.1E-65 | 9.9E-63 | BT20/MB468 | CYTO |
| >MI0003560\|hsa-mir-92b&WithFlank&1\|+\|155195171\|155195278@67.87.21(+1U)&[MIMAT0003218&hsa-miR-92b-3p&offsets\|0\|-1(+1U);m-115&1\|+\|155195237\|155195258&offsets\|0\|-1(+1U)]\|TATTGCACTCGTCCCGGCCTCT | >MI0003560\|hsa-mir-92b&WithFlank&1\|+\|155195171\|155195278 | 67 | 87 | 21(+1U) | TATTGCACTCGTCCCGGCCTCT | 7391.2 | 10.1 | 0.6 | 17.0 | 7.5E-65 | 1.2E-62 | BT20/MB468 | CYTO |
| >hg38_21_+_8433172_8446622_RNA45SN1_45S_50ntFlanks@10351.10368.18\|CGAGAACTTTGAAGGCCG | >hg38_21_+_8433172_8446622_RNA45SN1_45S_50ntFlanks | 10351 | 10368 | 18 | CGAGAACTTTGAAGGCCG | 33114.6 | -11.8 | 0.7 | -17.0 | 1.0E-64 | 1.2E-62 | BT20/MB468 | NUC |
| >hg38_21_+_8433172_8446622_RNA45SN1_45S_50ntFlanks@10841.10864.24\|CGTAACTTCGGGATAAGGATTGGC | >hg38_21_+_8433172_8446622_RNA45SN1_45S_50ntFlanks | 10841 | 10864 | 24 | CGTAACTTCGGGATAAGGATTGGC | 4795.9 | -8.2 | 0.5 | -17.1 | 2.3E-65 | 1.5E-62 | MB231/MB468 | NUC |
| >MI0000086\|hsa-mir-28&WithFlank&3\|+\|188688775\|188688872@60.81.22&[MIMAT0004502&hsa-miR-28-3p&offsets\|0\|0;m-28&3\|+\|188688834\|188688855&offsets\|0\|0]\|CACTAGATTGTGAGCTCCTGGA | >MI0000086\|hsa-mir-28&WithFlank&3\|+\|188688775\|188688872 | 60 | 81 | 22 | CACTAGATTGTGAGCTCCTGGA | 42639.9 | 11.3 | 0.7 | 17.0 | 1.0E-64 | 1.5E-62 | BT20/MB468 | CYTO |
| >hg38_21_+_8433172_8446622_RNA45SN1_45S_50ntFlanks@12937.12959.23\|AACCATTCGTAGACGACCTGCTT | >hg38_21_+_8433172_8446622_RNA45SN1_45S_50ntFlanks | 12937 | 12959 | 23 | AACCATTCGTAGACGACCTGCTT | 1726.6 | -7.1 | 0.4 | -17.0 | 1.5E-64 | 1.8E-62 | BT20/MB468 | NUC |
| >hg38_21_+_8433172_8446622_RNA45SN1_45S_50ntFlanks@6700.6720.21\|CTAGCTGCGAGAATTAATGTG | >hg38_21_+_8433172_8446622_RNA45SN1_45S_50ntFlanks | 6700 | 6720 | 21 | CTAGCTGCGAGAATTAATGTG | 17699.9 | -10.0 | 0.6 | -17.0 | 1.5E-64 | 1.8E-62 | BT20/MB468 | NUC |
| >hg38_21_+_8433172_8446622_RNA45SN1_45S_50ntFlanks@11787.11804.18\|AATGGATGAACGAGATTC | >hg38_21_+_8433172_8446622_RNA45SN1_45S_50ntFlanks | 11787 | 11804 | 18 | AATGGATGAACGAGATTC | 16839.7 | -10.7 | 0.6 | -16.9 | 2.6E-64 | 3.0E-62 | BT20/MB468 | NUC |
| >am_MI0000809\|hsa-mir-151a&WithFlank&8\|-\|140732558\|140732659@53.72.20(+1A)&[MIMAT0000757&hsa-miR-151a-3p&offsets\|0\|-1(+1A);m-30&8\|-\|140732587\|140732607&offsets\|0\|-1(+1A)]\|CTAGACTGAAGCTCCTTGAGA | >am_MI0000809\|hsa-mir-151a&WithFlank&8\|-\|140732558\|140732659 | 53 | 72 | 20(+1A) | CTAGACTGAAGCTCCTTGAGA | 15502.1 | 9.5 | 0.6 | 17.0 | 5.5E-65 | 3.0E-62 | BT20/MB468 | MITO |
| >trna2_GlyGCC_21_-_18827107_18827177@1.31.31,trna35_GlyGCC_1_+_161413094_161413164@1.31.31,trna37_GlyGCC_1_+_161420467_161420537@1.31.31,trna39_GlyGCC_1_+_161427898_161427968@1.31.31,trna41_GlyGCC_1_+_161435258_161435328@1.31.31\|GCATGGGTGGTTCAGTGGTAGAATTCTCGCC | >trna2_GlyGCC_21_-_18827107_18827177 | 1 | 31 | 31 | GCATGGGTGGTTCAGTGGTAGAATTCTCGCC | 69846.7 | 6.8 | 0.4 | 16.9 | 2.6E-64 | 3.9E-62 | BT20/MB468 | CYTO |
| >hg38_21_+_8433172_8446622_RNA45SN1_45S_50ntFlanks@10391.10422.32\|AACAGCAGTTGAACATGGGTCAGTCGGTCCTG | >hg38_21_+_8433172_8446622_RNA45SN1_45S_50ntFlanks | 10391 | 10422 | 32 | AACAGCAGTTGAACATGGGTCAGTCGGTCCTG | 1016.9 | -7.0 | 0.4 | -16.9 | 3.9E-64 | 4.3E-62 | BT20/MB468 | NUC |
| >MI0005559\|hsa-mir-744&WithFlank&17\|+\|12081893\|12082002@17.38.22&[MIMAT0004945&hsa-miR-744-5p&offsets\|0\|0;m-172&17\|+\|12081909\|12081929&offsets\|0\|+1]\|TGCGGGGCTAGGGCTAACAGCA | >MI0005559\|hsa-mir-744&WithFlank&17\|+\|12081893\|12082002 | 17 | 38 | 22 | TGCGGGGCTAGGGCTAACAGCA | 1549.6 | 8.0 | 0.5 | 17.0 | 9.8E-65 | 4.9E-62 | BT20/MB468 | MITO |
| >MI0000650\|hsa-mir-200c&WithFlank&12\|+\|6963693\|6963772@50.72.23(+1A)&[MIMAT0000617&hsa-miR-200c-3p&offsets\|0\|0(+1A);m-54&12\|+\|6963742\|6963764&offsets\|0\|0(+1A)]\|TAATACTGCCGGGTAATGATGGAA | >MI0000650\|hsa-mir-200c&WithFlank&12\|+\|6963693\|6963772 | 50 | 72 | 23(+1A) | TAATACTGCCGGGTAATGATGGAA | 18156.1 | 9.2 | 0.5 | 17.0 | 1.5E-64 | 7.0E-62 | BT20/MB468 | MITO |
| >MI0000285\|hsa-mir-205&WithFlank&1\|+\|209432127\|209432248@40.59.20(+1C)&[MIMAT0000266&hsa-miR-205-5p&offsets\|0\|-2(+1C);m-351&1\|+\|209432166\|209432187&offsets\|0\|-2(+1C)]\|TCCTTCATTCCACCGGAGTCC | >MI0000285\|hsa-mir-205&WithFlank&1\|+\|209432127\|209432248 | 40 | 59 | 20(+1C) | TCCTTCATTCCACCGGAGTCC | 1542.5 | 7.6 | 0.5 | 16.9 | 4.8E-64 | 7.0E-62 | BT20/MB468 | CYTO |
| >MI0000746\|hsa-mir-99b&WithFlank&19\|+\|51692606\|51692687@13.32.20(+1U)&[MIMAT0000689&hsa-miR-99b-5p&offsets\|0\|-2(+1U);m-10&19\|+\|51692618\|51692639&offsets\|0\|-2(+1U)]\|CACCCGTAGAACCGACCTTGT | >MI0000746\|hsa-mir-99b&WithFlank&19\|+\|51692606\|51692687 | 13 | 32 | 20(+1U) | CACCCGTAGAACCGACCTTGT | 47999.3 | 11.6 | 0.7 | 16.9 | 7.2E-64 | 1.0E-61 | BT20/MB468 | CYTO |
| >hg38_21_+_8433172_8446622_RNA45SN1_45S_50ntFlanks@10678.10710.33\|CGGTTCCGGCGGCGTCCGGTGAGCTCTCGCTGG | >hg38_21_+_8433172_8446622_RNA45SN1_45S_50ntFlanks | 10678 | 10710 | 33 | CGGTTCCGGCGGCGTCCGGTGAGCTCTCGCTGG | 181639.9 | -12.4 | 0.7 | -17.1 | 1.0E-65 | 1.4E-61 | MB231/MB468 | MP |
| >hg38_21_+_8433172_8446622_RNA45SN1_45S_50ntFlanks@12003.12032.30\|GCGGGCCGCCGGTGAAATACCACTACTCTG | >hg38_21_+_8433172_8446622_RNA45SN1_45S_50ntFlanks | 12003 | 12032 | 30 | GCGGGCCGCCGGTGAAATACCACTACTCTG | 5762.7 | -9.1 | 0.5 | -16.8 | 2.1E-63 | 2.2E-61 | BT20/MB468 | NUC |
| >hg38_21_+_8433172_8446622_RNA45SN1_45S_50ntFlanks@12655.12674.20\|ATGACTGAACGCCTCTAAGT | >hg38_21_+_8433172_8446622_RNA45SN1_45S_50ntFlanks | 12655 | 12674 | 20 | ATGACTGAACGCCTCTAAGT | 20297.9 | -10.0 | 0.6 | -16.8 | 2.1E-63 | 2.2E-61 | BT20/MB468 | NUC |

<table>
<tr><th></th><th></th><th></th><th></th><th></th><th></th><th></th><th></th><th></th><th></th><th></th><th></th><th></th><th></th></tr>
<tr><td>>trna116_GluCTC_1_-_145399233_145399304@1.32.32,trna59_GluCTC_1_+_249168447_249168518@1.32.32,trna71_GluCTC_1_-_161439189_161439260@1.32.32,trna74_GluCTC_1_-_161431809_161431880@1.32.32,trna77_GluCTC_1_-_161424398_161424469@1.32.32,trna77_GluCTC_6_+_28949976_28950047@1.32.32,trna80_GluCTC_1_-_161417018_161417089@1.32.32,trna87_GluCTC_6_-_126101393_126101464@1.32.32|TCCCTGGTGGTCTAGTGGTTAGGATTCGGCGC</td><td>>trna116_GluCTC_1_-_145399233_145399304</td><td>1</td><td>32</td><td>32</td><td>TCCCTGGTGGTCTAGTGGTTAGGATTCGGCGC</td><td>8966.0</td><td>6.7</td><td>0.4</td><td>16.8</td><td>1.8E-63</td><td>2.6E-61</td><td>BT20/MB468</td><td>CYTO</td></tr>
<tr><td>>hg38_21_+_8433172_8446622_RNA45SN1_45S_50ntFlanks@6659.6689.31|AGCGGTGGATCACTCGGCTCGTGCGTCGATG</td><td>>hg38_21_+_8433172_8446622_RNA45SN1_45S_50ntFlanks</td><td>6659</td><td>6689</td><td>31</td><td>AGCGGTGGATCACTCGGCTCGTGCGTCGATG</td><td>519.8</td><td>-4.3</td><td>0.3</td><td>-16.7</td><td>7.3E-63</td><td>7.5E-61</td><td>BT20/MB468</td><td>NUC</td></tr>
<tr><td>>hg38_21_+_8433172_8446622_RNA45SN1_45S_50ntFlanks@11572.11601.30|CCGACTTAGAACTGGTGCGGACCAGGGGAA</td><td>>hg38_21_+_8433172_8446622_RNA45SN1_45S_50ntFlanks</td><td>11572</td><td>11601</td><td>30</td><td>CCGACTTAGAACTGGTGCGGACCAGGGGAA</td><td>22796.9</td><td>-10.4</td><td>0.6</td><td>-16.9</td><td>2.2E-64</td><td>1.4E-60</td><td>MB231/MB468</td><td>MP</td></tr>
<tr><td>>MI0000063|hsa-let-7b&WithFlank&22|+|46113680|46113774@12.33.22(+1C)&[MIMAT0000063&hsa-let-7b-5p&offsets|0|0(+1C);m-14&22|+|46113691|46113712&offsets|0|0(+1C)]|TGAGGTAGTAGGTTGTGTGGTTC</td><td>>MI0000063|hsa-let-7b&WithFlank&22|+|46113680|46113774</td><td>12</td><td>33</td><td>22(+1C)</td><td>TGAGGTAGTAGGTTGTGTGGTTC</td><td>1798.5</td><td>8.6</td><td>0.5</td><td>16.7</td><td>1.1E-62</td><td>1.6E-60</td><td>BT20/MB468</td><td>CYTO</td></tr>
<tr><td>>MI0000650|hsa-mir-200c&WithFlank&12|+|6963693|6963772@50.72.23(+1C)&[MIMAT0000617&hsa-miR-200c-3p&offsets|0|0(+1C);m-54&12|+|6963742|6963764&offsets|0|0(+1C)]|TAATACTGCCGGGTAATGATGGAC</td><td>>MI0000650|hsa-mir-200c&WithFlank&12|+|6963693|6963772</td><td>50</td><td>72</td><td>23(+1C)</td><td>TAATACTGCCGGGTAATGATGGAC</td><td>2738.0</td><td>8.8</td><td>0.5</td><td>16.7</td><td>1.5E-62</td><td>2.0E-60</td><td>BT20/MB468</td><td>CYTO</td></tr>
<tr><td>>hg38_21_+_8433172_8446622_RNA45SN1_45S_50ntFlanks@9534.9571.38|AGGAAACTCTGGTGGAGGTCCGTAGCGGTCCTGACGTG</td><td>>hg38_21_+_8433172_8446622_RNA45SN1_45S_50ntFlanks</td><td>9534</td><td>9571</td><td>38</td><td>AGGAAACTCTGGTGGAGGTCCGTAGCGGTCCTGACGTG</td><td>73220.7</td><td>-8.5</td><td>0.5</td><td>-16.8</td><td>4.1E-63</td><td>2.4E-60</td><td>MB231/MB468</td><td>NUC</td></tr>
<tr><td>>MI0000285|hsa-mir-205&WithFlank&1|+|209432127|209432248@40.59.20(+1A)&[MIMAT0000266&hsa-miR-205-5p&offsets|0|-2(+1A);m-351&1|+|209432166|209432187&offsets|0|-2(+1A)]|TCCTTCATTCCACCGGAGTCA</td><td>>MI0000285|hsa-mir-205&WithFlank&1|+|209432127|209432248</td><td>40</td><td>59</td><td>20(+1A)</td><td>TCCTTCATTCCACCGGAGTCA</td><td>1534.7</td><td>8.0</td><td>0.5</td><td>16.7</td><td>2.7E-62</td><td>3.5E-60</td><td>BT20/MB468</td><td>CYTO</td></tr>
<tr><td>>hg38_21_+_8433172_8446622_RNA45SN1_45S_50ntFlanks@11572.11603.32|CCGACTTAGAACTGGTGCGGACCAGGGGAATC</td><td>>hg38_21_+_8433172_8446622_RNA45SN1_45S_50ntFlanks</td><td>11572</td><td>11603</td><td>32</td><td>CCGACTTAGAACTGGTGCGGACCAGGGGAATC</td><td>22903.2</td><td>-10.6</td><td>0.6</td><td>-16.6</td><td>4.5E-62</td><td>4.6E-60</td><td>BT20/MB468</td><td>NUC</td></tr>
<tr><td>>hg38_21_+_8433172_8446622_RNA45SN1_45S_50ntFlanks@12935.12960.26|TAAACCATTCGTAGACGACCTGCTTC</td><td>>hg38_21_+_8433172_8446622_RNA45SN1_45S_50ntFlanks</td><td>12935</td><td>12960</td><td>26</td><td>TAAACCATTCGTAGACGACCTGCTTC</td><td>571.7</td><td>-5.7</td><td>0.3</td><td>-16.6</td><td>8.1E-62</td><td>8.1E-60</td><td>BT20/MB468</td><td>NUC</td></tr>
<tr><td>>MI0000440|hsa-mir-27b&WithFlank&9|+|95085439|95085547@67.88.22&[MIMAT0000419&hsa-miR-27b-3p&offsets|0|+1;m-39&9|+|95085505|95085525&offsets|0|+1]|TTCACAGTGGCTAAGTTCTGCA</td><td>>MI0000440|hsa-mir-27b&WithFlank&9|+|95085439|95085547</td><td>67</td><td>88</td><td>22</td><td>TTCACAGTGGCTAAGTTCTGCA</td><td>11144.5</td><td>7.5</td><td>0.5</td><td>16.6</td><td>6.8E-62</td><td>8.7E-60</td><td>BT20/MB468</td><td>CYTO</td></tr>
<tr><td>>MI0000063|hsa-let-7b&WithFlank&22|+|46113680|46113774@12.31.20&[MIMAT0000063&hsa-let-7b-5p&offsets|0|-2;m-14&22|+|46113691|46113712&offsets|0|-2]|TGAGGTAGTAGGTTGTGTGG</td><td>>MI0000063|hsa-let-7b&WithFlank&22|+|46113680|46113774</td><td>12</td><td>31</td><td>20</td><td>TGAGGTAGTAGGTTGTGTGG</td><td>1787.5</td><td>8.6</td><td>0.5</td><td>16.6</td><td>9.4E-62</td><td>1.2E-59</td><td>BT20/MB468</td><td>CYTO</td></tr>
<tr><td>>MI0000063|hsa-let-7b&WithFlank&22|+|46113680|46113774@12.32.21(+1C)&[MIMAT0000063&hsa-let-7b-5p&offsets|0|-1(+1C);m-14&22|+|46113691|46113712&offsets|0|-1(+1C)]|TGAGGTAGTAGGTTGTGTGGTC</td><td>>MI0000063|hsa-let-7b&WithFlank&22|+|46113680|46113774</td><td>12</td><td>32</td><td>21(+1C)</td><td>TGAGGTAGTAGGTTGTGTGGTC</td><td>5024.4</td><td>8.6</td><td>0.5</td><td>16.6</td><td>3.3E-62</td><td>1.4E-59</td><td>BT20/MB468</td><td>MITO</td></tr>
<tr><td>>hg38_21_+_8433172_8446622_RNA45SN1_45S_50ntFlanks@10350.10372.23|ACGAGAACTTTGAAGGCCGAAGT</td><td>>hg38_21_+_8433172_8446622_RNA45SN1_45S_50ntFlanks</td><td>10350</td><td>10372</td><td>23</td><td>ACGAGAACTTTGAAGGCCGAAGT</td><td>890.2</td><td>-4.8</td><td>0.3</td><td>-16.5</td><td>1.7E-61</td><td>1.7E-59</td><td>BT20/MB468</td><td>NUC</td></tr>
<tr><td>>hg38_21_+_8433172_8446622_RNA45SN1_45S_50ntFlanks@10351.10383.33|CGAGAACTTTGAAGGCCGAAGTGGAGAAGGGTT</td><td>>hg38_21_+_8433172_8446622_RNA45SN1_45S_50ntFlanks</td><td>10351</td><td>10383</td><td>33</td><td>CGAGAACTTTGAAGGCCGAAGTGGAGAAGGGTT</td><td>49410.1</td><td>-9.1</td><td>0.5</td><td>-16.6</td><td>4.5E-62</td><td>2.4E-59</td><td>MB231/MB468</td><td>NUC</td></tr>
<tr><td>>MI0006444|hsa-mir-1307&WithFlank&10|-|103394247|103394407@86.107.22(+1A)&[MIMAT0005951&hsa-miR-1307-3p&offsets|0|0(+1A);m-66&10|-|103394300|103394322&offsets|0|-1(+1A)]|ACTCGGCGTGGCGTCGGTCGTGA</td><td>>MI0006444|hsa-mir-1307&WithFlank&10|-|103394247|103394407</td><td>86</td><td>107</td><td>22(+1A)</td><td>ACTCGGCGTGGCGTCGGTCGTGA</td><td>1942.9</td><td>8.1</td><td>0.5</td><td>16.5</td><td>2.7E-61</td><td>3.3E-59</td><td>BT20/MB468</td><td>CYTO</td></tr>
<tr><td>>hg38_21_+_8433172_8446622_RNA45SN1_45S_50ntFlanks@9554.9574.21|CGTAGCGGTCCTGACGTGCAA</td><td>>hg38_21_+_8433172_8446622_RNA45SN1_45S_50ntFlanks</td><td>9554</td><td>9574</td><td>21</td><td>CGTAGCGGTCCTGACGTGCAA</td><td>4962.7</td><td>-8.0</td><td>0.5</td><td>-16.5</td><td>4.4E-61</td><td>4.3E-59</td><td>BT20/MB468</td><td>NUC</td></tr>
<tr><td>>hg38_21_+_8433172_8446622_RNA45SN1_45S_50ntFlanks@4723.4758.36|CATTAATCAAGAACGAAAGTCGGAGGTTCGAAGACG</td><td>>hg38_21_+_8433172_8446622_RNA45SN1_45S_50ntFlanks</td><td>4723</td><td>4758</td><td>36</td><td>CATTAATCAAGAACGAAAGTCGGAGGTTCGAAGACG</td><td>8774.8</td><td>-7.6</td><td>0.5</td><td>-16.6</td><td>9.0E-62</td><td>4.4E-59</td><td>MB231/MB468</td><td>NUC</td></tr>
<tr><td>>MI0000255|hsa-mir-30d&WithFlank&8|-|134804870|134804951@12.35.24(+1U)&[MIMAT0000245&hsa-miR-30d-5p&offsets|0|+2(+1U);m-21&8|-|134804919|134804940&offsets|0|+2(+1U)]|TGTAAACATCCCCGACTGGAAGCTT</td><td>>MI0000255|hsa-mir-30d&WithFlank&8|-|134804870|134804951</td><td>12</td><td>35</td><td>24(+1U)</td><td>TGTAAACATCCCCGACTGGAAGCTT</td><td>2332.4</td><td>8.0</td><td>0.5</td><td>16.4</td><td>9.4E-61</td><td>1.1E-58</td><td>BT20/MB468</td><td>CYTO</td></tr>
<tr><td>>MI0000082|hsa-mir-25&WithFlank&7|-|100093554|100093649@58.78.21(+1U)&[MIMAT0000081&hsa-miR-25-3p&offsets|0|-1(+1U);m-23&7|-|100093571|100093592&offsets|0|-1(+1U)]|CATTGCACTTGTCTCGGTCTGT</td><td>>MI0000082|hsa-mir-25&WithFlank&7|-|100093554|100093649</td><td>58</td><td>78</td><td>21(+1U)</td><td>CATTGCACTTGTCTCGGTCTGT</td><td>5925.4</td><td>8.5</td><td>0.5</td><td>16.4</td><td>1.1E-60</td><td>1.3E-58</td><td>BT20/MB468</td><td>CYTO</td></tr>
<tr><td>>MI0000266|hsa-mir-10a&WithFlank&17|-|48579832|48579953@28.49.22&[MIMAT0000253&hsa-miR-10a-5p&offsets|0|-1;m-18&17|-|48579905|48579926&offsets|0|0]|TACCCTGTAGATCCGAATTTGT</td><td>>MI0000266|hsa-mir-10a&WithFlank&17|-|48579832|48579953</td><td>28</td><td>49</td><td>22</td><td>TACCCTGTAGATCCGAATTTGT</td><td>2987.0</td><td>8.7</td><td>0.5</td><td>16.4</td><td>4.2E-60</td><td>4.0E-58</td><td>BT20/MB468</td><td>NUC</td></tr>
<tr><td>>trna111_HisGTG_1_-_147774845_147774916@-1T.31.32,trna118_HisGTG_1_-_145396881_145396952@-1T.31.32,trna16_HisGTG_1_+_146544773_146544844@-1T.31.32,trna1_HisGTG_15_+_45493349_45493420@-1T.31.32,trna21_HisGTG_1_+_147753471_147753542@-1T.31.32,trna33_HisGTG_6_+_27125906_27125977@-1T.31.32,trna7_HisGTG_9_-_14433938_14434009@-1T.31.32,trna8_HisGTG_15_-_45492611_45492682@-1T.31.32,trna9_HisGTG_15_-_45490804_45490875@-1T.31.32|TGCCGTGATCGTATAGTGGTTAGTACTCTGCG</td><td>>trna111_HisGTG_1_-_147774845_147774916</td><td>-1T</td><td>31</td><td>32</td><td>TGCCGTGATCGTATAGTGGTTAGTACTCTGCG</td><td>19758.2</td><td>-9.8</td><td>0.6</td><td>-16.3</td><td>4.8E-60</td><td>4.5E-58</td><td>BT20/MB468</td><td>NUC</td></tr>
<tr><td>>hg38_21_+_8433172_8446622_RNA45SN1_45S_50ntFlanks@12456.12478.23|CCAAGCGTTGGATTGTTCACCCA</td><td>>hg38_21_+_8433172_8446622_RNA45SN1_45S_50ntFlanks</td><td>12456</td><td>12478</td><td>23</td><td>CCAAGCGTTGGATTGTTCACCCA</td><td>10694.6</td><td>-8.2</td><td>0.5</td><td>-16.4</td><td>1.1E-60</td><td>5.0E-58</td><td>MB231/MB468</td><td>NUC</td></tr>
<tr><td>>MI0000811|hsa-mir-148b&WithFlank&12|+|54337210|54337320@69.90.22&[MIMAT0000759&hsa-miR-148b-3p&offsets|0|0;m-77&12|+|54337278|54337299&offsets|0|0]|TCAGTGCATCACAGAACTTTGT</td><td>>MI0000811|hsa-mir-148b&WithFlank&12|+|54337210|54337320</td><td>69</td><td>90</td><td>22</td><td>TCAGTGCATCACAGAACTTTGT</td><td>9269.4</td><td>6.8</td><td>0.4</td><td>16.3</td><td>4.9E-60</td><td>5.6E-58</td><td>BT20/MB468</td><td>CYTO</td></tr>
<tr><td>>hg38_21_+_8433172_8446622_RNA45SN1_45S_50ntFlanks@11999.12032.34|CCCTGCGGGCCGCCGGTGAAATACCACTACTCTG</td><td>>hg38_21_+_8433172_8446622_RNA45SN1_45S_50ntFlanks</td><td>11999</td><td>12032</td><td>34</td><td>CCCTGCGGGCCGCCGGTGAAATACCACTACTCTG</td><td>51312.1</td><td>-20.1</td><td>1.2</td><td>-16.3</td><td>7.2E-60</td><td>6.6E-58</td><td>BT20/MB468</td><td>NUC</td></tr>
<tr><td>>MI0000273|hsa-mir-183&WithFlank&7|-|129774899|129775020@34.55.22&[MIMAT0000261&hsa-miR-183-5p&offsets|+1|+1;m-41&7|-|129774966|129774987&offsets|0|0]|ATGGCACTGGTAGAATTCACTG</td><td>>MI0000273|hsa-mir-183&WithFlank&7|-|129774899|129775020</td><td>34</td><td>55</td><td>22</td><td>ATGGCACTGGTAGAATTCACTG</td><td>6297.1</td><td>7.8</td><td>0.5</td><td>16.3</td><td>1.0E-59</td><td>1.1E-57</td><td>BT20/MB468</td><td>CYTO</td></tr>
<tr><td>>MI0000093|hsa-mir-92a-1&WithFlank&13|+|91351308|91351397@55.75.21;MI0000094|hsa-mir-92a-2&WithFlank&X|-|134169532|134169618@55.75.21&[MIMAT0000092&hsa-miR-92a-3p&offsets|+1|0;m-9&13|+|91351361|91351382&offsets|+1|0];[MIMAT0000092_1&hsa-miR-92a-3p&offsets|+1|0;m-11&X|-|134169544|134169565&offsets|+1|0]|ATTGCACTTGTCCCGGCCTGT</td><td>>MI0000093|hsa-mir-92a-1&WithFlank&13|+|91351308|91351397</td><td>55</td><td>75</td><td>21</td><td>ATTGCACTTGTCCCGGCCTGT</td><td>997.9</td><td>6.2</td><td>0.4</td><td>16.3</td><td>1.1E-59</td><td>1.3E-57</td><td>BT20/MB468</td><td>CYTO</td></tr>
<tr><td>>MI0000809|hsa-mir-151a&WithFlank&8|-|140732558|140732659@53.74.22(+1A)&[MIMAT0000757&hsa-miR-151a-3p&offsets|0|+1(+1A);m-30&8|-|140732587|140732607&offsets|0|+1(+1A)]|CTAGACTGAAGCTCCTTGAGGAA</td><td>>MI0000809|hsa-mir-151a&WithFlank&8|-|140732558|140732659</td><td>53</td><td>74</td><td>22(+1A)</td><td>CTAGACTGAAGCTCCTTGAGGAA</td><td>9917.6</td><td>8.8</td><td>0.5</td><td>16.4</td><td>3.4E-60</td><td>1.4E-57</td><td>BT20/MB468</td><td>MITO</td></tr>
<tr><td>>hg38_21_+_8433172_8446622_RNA45SN1_45S_50ntFlanks@10678.10706.29|CGGTTCCGGCGGCGTCCGGTGAGCTCTCG</td><td>>hg38_21_+_8433172_8446622_RNA45SN1_45S_50ntFlanks</td><td>10678</td><td>10706</td><td>29</td><td>CGGTTCCGGCGGCGTCCGGTGAGCTCTCG</td><td>39279.0</td><td>-11.9</td><td>0.7</td><td>-16.4</td><td>1.3E-60</td><td>1.5E-57</td><td>MB231/MB468</td><td>MITO</td></tr>
<tr><td>>hg38_21_+_8433172_8446622_RNA45SN1_45S_50ntFlanks@6664.6686.23|TGGATCACTCGGCTCGTGCGTCG</td><td>>hg38_21_+_8433172_8446622_RNA45SN1_45S_50ntFlanks</td><td>6664</td><td>6686</td><td>23</td><td>TGGATCACTCGGCTCGTGCGTCG</td><td>369.5</td><td>-5.1</td><td>0.3</td><td>-16.2</td><td>2.2E-59</td><td>2.0E-57</td><td>BT20/MB468</td><td>NUC</td></tr>
<tr><td>>hg38_21_+_8433172_8446622_RNA45SN1_45S_50ntFlanks@9738.9758.21|GAAACGATCTCAACCTATTCT</td><td>>hg38_21_+_8433172_8446622_RNA45SN1_45S_50ntFlanks</td><td>9738</td><td>9758</td><td>21</td><td>GAAACGATCTCAACCTATTCT</td><td>3931.3</td><td>-8.9</td><td>0.6</td><td>-16.2</td><td>4.6E-59</td><td>4.1E-57</td><td>BT20/MB468</td><td>NUC</td></tr>
<tr><td>>hg38_21_+_8433172_8446622_RNA45SN1_45S_50ntFlanks@9554.9571.18|CGTAGCGGTCCTGACGTG</td><td>>hg38_21_+_8433172_8446622_RNA45SN1_45S_50ntFlanks</td><td>9554</td><td>9571</td><td>18</td><td>CGTAGCGGTCCTGACGTG</td><td>164859.8</td><td>-5.7</td><td>0.4</td><td>-16.2</td><td>4.3E-59</td><td>4.7E-57</td><td>BT20/MB468</td><td>CYTO</td></tr>
<tr><td>>MI0003591|hsa-mir-584&WithFlank&5|-|149062307|149062415@22.42.21&[MIMAT0003249&hsa-miR-584-5p&offsets|0|-1;m-167&5|-|149062374|149062394&offsets|0|0]|TTATGGTTTGCCTGGGACTGA</td><td>>MI0003591|hsa-mir-584&WithFlank&5|-|149062307|149062415</td><td>22</td><td>42</td><td>21</td><td>TTATGGTTTGCCTGGGACTGA</td><td>3957.5</td><td>9.2</td><td>0.6</td><td>16.3</td><td>1.9E-59</td><td>7.1E-57</td><td>BT20/MB468</td><td>MITO</td></tr>
<tr><td>>hg38_21_+_8433172_8446622_RNA45SN1_45S_50ntFlanks@7977.8006.30|CGACCTCAGATCAGACGTGGCGACCCGCTG</td><td>>hg38_21_+_8433172_8446622_RNA45SN1_45S_50ntFlanks</td><td>7977</td><td>8006</td><td>30</td><td>CGACCTCAGATCAGACGTGGCGACCCGCTG</td><td>4017.7</td><td>-7.2</td><td>0.4</td><td>-16.1</td><td>1.3E-58</td><td>1.1E-56</td><td>BT20/MB468</td><td>NUC</td></tr>
<tr><td>>hg38_21_+_8433172_8446622_RNA45SN1_45S_50ntFlanks@10400.10424.25|TGAACATGGGTCAGTCGGTCCTGAG</td><td>>hg38_21_+_8433172_8446622_RNA45SN1_45S_50ntFlanks</td><td>10400</td><td>10424</td><td>25</td><td>TGAACATGGGTCAGTCGGTCCTGAG</td><td>16808.5</td><td>10.0</td><td>0.6</td><td>16.2</td><td>3.3E-59</td><td>1.2E-56</td><td>BT20/MB468</td><td>MITO</td></tr>
<tr><td>>hg38_21_+_8433172_8446622_RNA45SN1_45S_50ntFlanks@12934.12956.23|CTAAACCATTCGTAGACGACCTG</td><td>>hg38_21_+_8433172_8446622_RNA45SN1_45S_50ntFlanks</td><td>12934</td><td>12956</td><td>23</td><td>CTAAACCATTCGTAGACGACCTG</td><td>16653.2</td><td>-9.3</td><td>0.6</td><td>-16.2</td><td>3.6E-59</td><td>1.5E-56</td><td>MB231/MB468</td><td>NUC</td></tr>
<tr><td>>MI0000255|hsa-mir-30d&WithFlank&8|-|134804870|134804951@12.34.23&[MIMAT0000245&hsa-miR-30d-5p&offsets|0|+1;m-21&8|-|134804919|134804940&offsets|0|+1]|TGTAAACATCCCCGACTGGAAGC</td><td>>MI0000255|hsa-mir-30d&WithFlank&8|-|134804870|134804951</td><td>12</td><td>34</td><td>23</td><td>TGTAAACATCCCCGACTGGAAGC</td><td>57000.2</td><td>8.8</td><td>0.5</td><td>16.1</td><td>2.0E-58</td><td>2.2E-56</td><td>BT20/MB468</td><td>CYTO</td></tr>
<tr><td>>hg38_21_+_8433172_8446622_RNA45SN1_45S_50ntFlanks@10678.10710.33|CGGTTCCGGCGGCGTCCGGTGAGCTCTCGCTGG</td><td>>hg38_21_+_8433172_8446622_RNA45SN1_45S_50ntFlanks</td><td>10678</td><td>10710</td><td>33</td><td>CGGTTCCGGCGGCGTCCGGTGAGCTCTCGCTGG</td><td>369560.2</td><td>-10.2</td><td>0.6</td><td>-16.2</td><td>5.6E-59</td><td>2.2E-56</td><td>MB231/MB468</td><td>NUC</td></tr>
<tr><td>>hg38_21_+_8433172_8446622_RNA45SN1_45S_50ntFlanks@475.510.36|CTTCGTGATCGATGTGGTGACGTCGTGCTCTCCCGG</td><td>>hg38_21_+_8433172_8446622_RNA45SN1_45S_50ntFlanks</td><td>475</td><td>510</td><td>36</td><td>CTTCGTGATCGATGTGGTGACGTCGTGCTCTCCCGG</td><td>2521.4</td><td>8.5</td><td>0.5</td><td>16.1</td><td>2.2E-58</td><td>2.3E-56</td><td>BT20/MB468</td><td>CYTO</td></tr>
<tr><td>>hg38_21_+_8433172_8446622_RNA45SN1_45S_50ntFlanks@10678.10712.35|CGGTTCCGGCGGCGTCCGGTGAGCTCTCGCTGGCC</td><td>>hg38_21_+_8433172_8446622_RNA45SN1_45S_50ntFlanks</td><td>10678</td><td>10712</td><td>35</td><td>CGGTTCCGGCGGCGTCCGGTGAGCTCTCGCTGGCC</td><td>18915.6</td><td>-9.4</td><td>0.6</td><td>-16.3</td><td>5.5E-60</td><td>2.4E-56</td><td>MB231/MB468</td><td>MP</td></tr>
<tr><td>>MI0003137|hsa-mir-193b&WithFlank&16|+|14303961|14304055@57.78.22&[MIMAT0002819&hsa-miR-193b-3p&offsets|0|0;m-166&16|+|14304017|14304038&offsets|0|0]|AACTGGCCCTCAAAGTCCCGCT</td><td>>MI0003137|hsa-mir-193b&WithFlank&16|+|14303961|14304055</td><td>57</td><td>78</td><td>22</td><td>AACTGGCCCTCAAAGTCCCGCT</td><td>3041.0</td><td>8.9</td><td>0.6</td><td>16.2</td><td>8.2E-59</td><td>2.7E-56</td><td>BT20/MB468</td><td>MITO</td></tr>
<tr><td>>hg38_21_+_8433172_8446622_RNA45SN1_45S_50ntFlanks@11593.11611.19|CCAGGGGAATCCGACTGTT</td><td>>hg38_21_+_8433172_8446622_RNA45SN1_45S_50ntFlanks</td><td>11593</td><td>11611</td><td>19</td><td>CCAGGGGAATCCGACTGTT</td><td>4975.8</td><td>-7.9</td><td>0.5</td><td>-16.1</td><td>3.8E-58</td><td>3.2E-56</td><td>BT20/MB468</td><td>NUC</td></tr>
</table>

<table>
<tr><td>>MI0000082|hsa-mir-25&WithFlank&7|-|100093554|100093649@58.79.22&[MIMAT0000081&hsa-miR-25-3p&offsets|0|0;m-23&7|-|100093571|100093592&offsets|0|0]|CATTGCACTTGTCTCGGTCTGA</td><td>>MI0000082|hsa-mir-25&WithFlank&7|-|100093554|100093649</td><td>58</td><td>79</td><td>22</td><td>CATTGCACTTGTCTCGGTCTGA</td><td>84951.8</td><td>7.8</td><td>0.5</td><td>16.1</td><td>4.3E-58</td><td>4.5E-56</td><td>BT20/MB468</td><td>CYTO</td></tr>
<tr><td>>hg38_21_+_8433172_8446622_RNA45SN1_45S_50ntFlanks@8191.8213.23|CAGCCCGTGGACGGTGTGAGGCC</td><td>>hg38_21_+_8433172_8446622_RNA45SN1_45S_50ntFlanks</td><td>8191</td><td>8213</td><td>23</td><td>CAGCCCGTGGACGGTGTGAGGCC</td><td>2280.9</td><td>-7.5</td><td>0.5</td><td>-16.1</td><td>5.6E-58</td><td>4.7E-56</td><td>BT20/MB468</td><td>NUC</td></tr>
<tr><td>>hg38_21_+_8433172_8446622_RNA45SN1_45S_50ntFlanks@12936.12957.22|AAACCATTCGTAGACGACCTGC</td><td>>hg38_21_+_8433172_8446622_RNA45SN1_45S_50ntFlanks</td><td>12936</td><td>12957</td><td>22</td><td>AAACCATTCGTAGACGACCTGC</td><td>1040.8</td><td>-6.1</td><td>0.4</td><td>-16.0</td><td>6.1E-58</td><td>5.1E-56</td><td>BT20/MB468</td><td>NUC</td></tr>
<tr><td>>MI0000065|hsa-let-7d&WithFlank&9|+|94178828|94178926@68.89.22&[MIMAT0004484&hsa-let-7d-3p&offsets|0|0;m-98&9|+|94178895|94178916&offsets|0|0]|CTATACGACCTGCTGCCTTTCT</td><td>>MI0000065|hsa-let-7d&WithFlank&9|+|94178828|94178926</td><td>68</td><td>89</td><td>22</td><td>CTATACGACCTGCTGCCTTTCT</td><td>5841.8</td><td>8.2</td><td>0.5</td><td>16.0</td><td>6.5E-58</td><td>6.6E-56</td><td>BT20/MB468</td><td>CYTO</td></tr>
<tr><td>>hg38_21_+_8433172_8446622_RNA45SN1_45S_50ntFlanks@12973.12990.18|TCGTACGTAGCAGAGCAG</td><td>>hg38_21_+_8433172_8446622_RNA45SN1_45S_50ntFlanks</td><td>12973</td><td>12990</td><td>18</td><td>TCGTACGTAGCAGAGCAG</td><td>24886.6</td><td>-11.3</td><td>0.7</td><td>-16.1</td><td>2.3E-58</td><td>8.4E-56</td><td>MB231/MB468</td><td>NUC</td></tr>
<tr><td>>MI0000255|hsa-mir-30d&WithFlank&8|-|134804870|134804951@12.34.23&[MIMAT0000245&hsa-miR-30d-5p&offsets|0|+1;m-21&8|-|134804919|134804940&offsets|0|+1]|TGTAAACATCCCCGACTGGAAGC</td><td>>MI0000255|hsa-mir-30d&WithFlank&8|-|134804870|134804951</td><td>12</td><td>34</td><td>23</td><td>TGTAAACATCCCCGACTGGAAGC</td><td>56472.7</td><td>8.5</td><td>0.5</td><td>16.1</td><td>2.9E-58</td><td>9.0E-56</td><td>BT20/MB468</td><td>MITO</td></tr>
<tr><td>>hg38_21_+_8433172_8446622_RNA45SN1_45S_50ntFlanks@6645.6681.37|CTCGTACGACTCTTAGCGGTGGATCACTCGGCTCGTG</td><td>>hg38_21_+_8433172_8446622_RNA45SN1_45S_50ntFlanks</td><td>6645</td><td>6681</td><td>37</td><td>CTCGTACGACTCTTAGCGGTGGATCACTCGGCTCGTG</td><td>5142.3</td><td>-5.0</td><td>0.3</td><td>-16.0</td><td>1.2E-57</td><td>9.8E-56</td><td>BT20/MB468</td><td>NUC</td></tr>
<tr><td>>hg38_21_+_8433172_8446622_RNA45SN1_45S_50ntFlanks@12405.12429.25|CTTTTTGATCCTTCGATGTCGGCTC</td><td>>hg38_21_+_8433172_8446622_RNA45SN1_45S_50ntFlanks</td><td>12405</td><td>12429</td><td>25</td><td>CTTTTTGATCCTTCGATGTCGGCTC</td><td>7281.5</td><td>-7.8</td><td>0.5</td><td>-16.1</td><td>4.0E-58</td><td>1.4E-55</td><td>MB231/MB468</td><td>NUC</td></tr>
<tr><td>>MI0000299|hsa-mir-222&WithFlank&X|-|45747009|45747130@75.99.25(+1U)&[MIMAT0000279&hsa-miR-222-3p&offsets|0|+4(+1U);m-140&X|-|45747033|45747056&offsets|0|+1(+1U)]|AGCTACATCTGGCTACTGGGTCTCTT</td><td>>MI0000299|hsa-mir-222&WithFlank&X|-|45747009|45747130</td><td>75</td><td>99</td><td>25(+1U)</td><td>AGCTACATCTGGCTACTGGGTCTCTT</td><td>2173.0</td><td>7.2</td><td>0.5</td><td>16.0</td><td>1.4E-57</td><td>1.4E-55</td><td>BT20/MB468</td><td>CYTO</td></tr>
<tr><td>>MI0000272|hsa-mir-182&WithFlank&7|-|129770377|129770498@29.50.22&[MIMAT0000259&hsa-miR-182-5p&offsets|0|-2;m-22&7|-|129770449|129770470&offsets|0|0]|TTTGGCAATGGTAGAACTCACA</td><td>>MI0000272|hsa-mir-182&WithFlank&7|-|129770377|129770498</td><td>29</td><td>50</td><td>22</td><td>TTTGGCAATGGTAGAACTCACA</td><td>2038.6</td><td>7.5</td><td>0.5</td><td>16.1</td><td>4.8E-58</td><td>1.4E-55</td><td>BT20/MB468</td><td>MITO</td></tr>
<tr><td>>MI0006273|hsa-mir-1180&WithFlank&17|-|19344500|19344580@47.68.22&[MIMAT0005825&hsa-miR-1180-3p&offsets|0|0;m-214&17|-|19344513|19344534&offsets|0|0]|TTTCCGGCTCGCGTGGGTGTGT</td><td>>MI0006273|hsa-mir-1180&WithFlank&17|-|19344500|19344580</td><td>47</td><td>68</td><td>22</td><td>TTTCCGGCTCGCGTGGGTGTGT</td><td>1786.0</td><td>7.5</td><td>0.5</td><td>16.0</td><td>8.6E-58</td><td>2.5E-55</td><td>BT20/MB468</td><td>MITO</td></tr>
<tr><td>>MI0000081|hsa-mir-24-2&WithFlank&19|-|13836281|13836365@56.77.22;MI0000080|hsa-mir-24-1&WithFlank&9|+|95086015|95086094@50.71.22&[MIMAT0000080&hsa-miR-24-3p&offsets|0|0;m-36&19|-|13836288|13836310&offsets|0|-1];[MIMAT0000080_1&hsa-miR-24-3p&offsets|0|0;m-35&9|+|95086064|95086086&offsets|0|-1]|TGGCTCAGTTCAGCAGGAACAG</td><td>>MI0000081|hsa-mir-24-2&WithFlank&19|-|13836281|13836365</td><td>56</td><td>77</td><td>22</td><td>TGGCTCAGTTCAGCAGGAACAG</td><td>14237.0</td><td>5.7</td><td>0.4</td><td>16.0</td><td>1.2E-57</td><td>3.4E-55</td><td>BT20/MB468</td><td>MITO</td></tr>
<tr><td>>MI0006444|hsa-mir-1307&WithFlank&10|-|103394247|103394407@86.108.23&[MIMAT0005951&hsa-miR-1307-3p&offsets|0|+1;m-66&10|-|103394300|103394322&offsets|0|0]|ACTCGGCGTGGCGTCGGTCGTGG</td><td>>MI0006444|hsa-mir-1307&WithFlank&10|-|103394247|103394407</td><td>86</td><td>108</td><td>23</td><td>ACTCGGCGTGGCGTCGGTCGTGG</td><td>7490.5</td><td>7.2</td><td>0.5</td><td>15.9</td><td>4.2E-57</td><td>4.1E-55</td><td>BT20/MB468</td><td>CYTO</td></tr>
<tr><td>>hg38_21_+_8433172_8446622_RNA45SN1_45S_50ntFlanks@12456.12487.32|CCAAGCGTTGGATTGTTCACCCACTAATAGGG</td><td>>hg38_21_+_8433172_8446622_RNA45SN1_45S_50ntFlanks</td><td>12456</td><td>12487</td><td>32</td><td>CCAAGCGTTGGATTGTTCACCCACTAATAGGG</td><td>157292.2</td><td>-21.7</td><td>1.4</td><td>-15.9</td><td>8.5E-57</td><td>6.9E-55</td><td>BT20/MB468</td><td>NUC</td></tr>
<tr><td>>MI0000255|hsa-mir-30d&WithFlank&8|-|134804870|134804951@12.33.22(+1U)&[MIMAT0000245&hsa-miR-30d-5p&offsets|0|0(+1U);m-21&8|-|134804919|134804940&offsets|0|0(+1U)]|TGTAAACATCCCCGACTGGAAGT</td><td>>MI0000255|hsa-mir-30d&WithFlank&8|-|134804870|134804951</td><td>12</td><td>33</td><td>22(+1U)</td><td>TGTAAACATCCCCGACTGGAAGT</td><td>14564.1</td><td>8.8</td><td>0.6</td><td>15.9</td><td>4.0E-57</td><td>1.0E-54</td><td>BT20/MB468</td><td>MITO</td></tr>
<tr><td>>hg38_21_+_8433172_8446622_RNA45SN1_45S_50ntFlanks@6654.6689.36|CTCTTAGCGGTGGATCACTCGGCTCGTGCGTCGATG</td><td>>hg38_21_+_8433172_8446622_RNA45SN1_45S_50ntFlanks</td><td>6654</td><td>6689</td><td>36</td><td>CTCTTAGCGGTGGATCACTCGGCTCGTGCGTCGATG</td><td>1552.2</td><td>-5.6</td><td>0.4</td><td>-15.9</td><td>1.3E-56</td><td>1.1E-54</td><td>BT20/MB468</td><td>NUC</td></tr>
<tr><td>>hg38_21_+_8433172_8446622_RNA45SN1_45S_50ntFlanks@12654.12675.22|TATGACTGAACGCCTCTAAGTC</td><td>>hg38_21_+_8433172_8446622_RNA45SN1_45S_50ntFlanks</td><td>12654</td><td>12675</td><td>22</td><td>TATGACTGAACGCCTCTAAGTC</td><td>5312.3</td><td>-7.9</td><td>0.5</td><td>-15.9</td><td>4.3E-57</td><td>1.4E-54</td><td>MB231/MB468</td><td>NUC</td></tr>
<tr><td>>hg38_21_+_8433172_8446622_RNA45SN1_45S_50ntFlanks@7984.8006.23|AGATCAGACGTGGCGACCCGCTG</td><td>>hg38_21_+_8433172_8446622_RNA45SN1_45S_50ntFlanks</td><td>7984</td><td>8006</td><td>23</td><td>AGATCAGACGTGGCGACCCGCTG</td><td>3617.6</td><td>-8.9</td><td>0.6</td><td>-15.8</td><td>2.6E-56</td><td>2.0E-54</td><td>BT20/MB468</td><td>NUC</td></tr>
<tr><td>>hg38_21_+_8433172_8446622_RNA45SN1_45S_50ntFlanks@12655.12675.21|ATGACTGAACGCCTCTAAGTC</td><td>>hg38_21_+_8433172_8446622_RNA45SN1_45S_50ntFlanks</td><td>12655</td><td>12675</td><td>21</td><td>ATGACTGAACGCCTCTAAGTC</td><td>9576.9</td><td>-8.2</td><td>0.5</td><td>-15.9</td><td>7.8E-57</td><td>2.4E-54</td><td>MB231/MB468</td><td>NUC</td></tr>
<tr><td>>MI0000086|hsa-mir-28&WithFlank&3|+|188688775|188688872@60.81.22&[MIMAT0004502&hsa-miR-28-3p&offsets|0|0;m-28&3|+|188688834|188688855&offsets|0|0]|CACTAGATTGTGAGCTCCTGGA</td><td>>MI0000086|hsa-mir-28&WithFlank&3|+|188688775|188688872</td><td>60</td><td>81</td><td>22</td><td>CACTAGATTGTGAGCTCCTGGA</td><td>6268.6</td><td>10.0</td><td>0.6</td><td>15.9</td><td>1.1E-56</td><td>2.7E-54</td><td>BT20/MB468</td><td>MITO</td></tr>
<tr><td>>MI0000542|hsa-mir-320a&WithFlank&8|-|22244960|22245043@48.68.21&[MIMAT0000510&hsa-miR-320a-3p&offsets|0|-1;m-60&8|-|22244975|22244996&offsets|0|-1]|AAAAGCTGGGTTGAGAGGGCG</td><td>>MI0000542|hsa-mir-320a&WithFlank&8|-|22244960|22245043</td><td>48</td><td>68</td><td>21</td><td>AAAAGCTGGGTTGAGAGGGCG</td><td>25203.4</td><td>10.5</td><td>0.7</td><td>15.8</td><td>6.3E-56</td><td>6.1E-54</td><td>BT20/MB468</td><td>CYTO</td></tr>
<tr><td>>hg38_21_+_8433172_8446622_RNA45SN1_45S_50ntFlanks@10410.10430.21|TCAGTCGGTCCTGAGAGATGG</td><td>>hg38_21_+_8433172_8446622_RNA45SN1_45S_50ntFlanks</td><td>10410</td><td>10430</td><td>21</td><td>TCAGTCGGTCCTGAGAGATGG</td><td>8368.6</td><td>-8.9</td><td>0.6</td><td>-15.8</td><td>2.5E-56</td><td>7.3E-54</td><td>MB231/MB468</td><td>NUC</td></tr>
<tr><td>>hg38_21_+_8433172_8446622_RNA45SN1_45S_50ntFlanks@9574.9597.24|AATCGGTCGTCCGACCTGGGTATA</td><td>>hg38_21_+_8433172_8446622_RNA45SN1_45S_50ntFlanks</td><td>9574</td><td>9597</td><td>24</td><td>AATCGGTCGTCCGACCTGGGTATA</td><td>1784.6</td><td>-7.8</td><td>0.5</td><td>-15.7</td><td>1.2E-55</td><td>9.4E-54</td><td>BT20/MB468</td><td>NUC</td></tr>
<tr><td>>hg38_21_+_8433172_8446622_RNA45SN1_45S_50ntFlanks@9534.9553.20|AGGAAACTCTGGTGGAGGTC</td><td>>hg38_21_+_8433172_8446622_RNA45SN1_45S_50ntFlanks</td><td>9534</td><td>9553</td><td>20</td><td>AGGAAACTCTGGTGGAGGTC</td><td>15088.6</td><td>-9.9</td><td>0.6</td><td>-15.9</td><td>1.1E-56</td><td>1.0E-53</td><td>MB231/MB468</td><td>MITO</td></tr>
<tr><td>>hg38_21_+_8433172_8446622_RNA45SN1_45S_50ntFlanks@12973.12991.19|TCGTACGTAGCAGAGCAGC</td><td>>hg38_21_+_8433172_8446622_RNA45SN1_45S_50ntFlanks</td><td>12973</td><td>12991</td><td>19</td><td>TCGTACGTAGCAGAGCAGC</td><td>28200.4</td><td>-19.2</td><td>1.2</td><td>-15.7</td><td>1.7E-55</td><td>1.3E-53</td><td>BT20/MB468</td><td>NUC</td></tr>
<tr><td>>hg38_21_+_8433172_8446622_RNA45SN1_45S_50ntFlanks@9534.9552.19|AGGAAACTCTGGTGGAGGT</td><td>>hg38_21_+_8433172_8446622_RNA45SN1_45S_50ntFlanks</td><td>9534</td><td>9552</td><td>19</td><td>AGGAAACTCTGGTGGAGGT</td><td>3798.9</td><td>-7.3</td><td>0.5</td><td>-15.8</td><td>5.3E-56</td><td>1.5E-53</td><td>MB231/MB468</td><td>NUC</td></tr>
<tr><td>>MI0005559|hsa-mir-744&WithFlank&17|+|12081893|12082002@17.38.22&[MIMAT0004945&hsa-miR-744-5p&offsets|0|0;m-172&17|+|12081909|12081929&offsets|0|+1]|TGCGGGGCTAGGGCTAACAGCA</td><td>>MI0005559|hsa-mir-744&WithFlank&17|+|12081893|12082002</td><td>17</td><td>38</td><td>22</td><td>TGCGGGGCTAGGGCTAACAGCA</td><td>1426.1</td><td>7.7</td><td>0.5</td><td>15.7</td><td>1.6E-55</td><td>1.6E-53</td><td>BT20/MB468</td><td>CYTO</td></tr>
<tr><td>>hg38_21_+_8433172_8446622_RNA45SN1_45S_50ntFlanks@10679.10710.32|GGTTCCGGCGGCGTCCGGTGAGCTCTCGCTGG</td><td>>hg38_21_+_8433172_8446622_RNA45SN1_45S_50ntFlanks</td><td>10679</td><td>10710</td><td>32</td><td>GGTTCCGGCGGCGTCCGGTGAGCTCTCGCTGG</td><td>41218.6</td><td>-10.0</td><td>0.6</td><td>-15.8</td><td>6.7E-56</td><td>1.8E-53</td><td>MB231/MB468</td><td>NUC</td></tr>
<tr><td>>hg38_21_+_8433172_8446622_RNA45SN1_45S_50ntFlanks@12418.12436.19|CGATGTCGGCTCTTCCTAT</td><td>>hg38_21_+_8433172_8446622_RNA45SN1_45S_50ntFlanks</td><td>12418</td><td>12436</td><td>19</td><td>CGATGTCGGCTCTTCCTAT</td><td>33579.9</td><td>-8.1</td><td>0.5</td><td>-15.7</td><td>7.6E-56</td><td>1.9E-53</td><td>MB231/MB468</td><td>NUC</td></tr>
<tr><td>>MI0000255|hsa-mir-30d&WithFlank&8|-|134804870|134804951@12.32.21&[MIMAT0000245&hsa-miR-30d-5p&offsets|0|-1;m-21&8|-|134804919|134804940&offsets|0|-1]|TGTAAACATCCCCGACTGGAA</td><td>>MI0000255|hsa-mir-30d&WithFlank&8|-|134804870|134804951</td><td>12</td><td>32</td><td>21</td><td>TGTAAACATCCCCGACTGGAA</td><td>1015.6</td><td>8.1</td><td>0.5</td><td>15.7</td><td>9.8E-56</td><td>2.3E-53</td><td>BT20/MB468</td><td>MITO</td></tr>
<tr><td>>MI0003628|hsa-mir-615&WithFlank&12|+|54033944|54034051@67.87.21&[MIMAT0003283&hsa-miR-615-3p&offsets|0|-1;m-495&12|+|54034010|54034030&offsets|0|0]|TCCGAGCCTGGGTCTCCCTCT</td><td>>MI0003628|hsa-mir-615&WithFlank&12|+|54033944|54034051</td><td>67</td><td>87</td><td>21</td><td>TCCGAGCCTGGGTCTCCCTCT</td><td>3208.5</td><td>7.1</td><td>0.5</td><td>15.7</td><td>2.8E-55</td><td>2.6E-53</td><td>BT20/MB468</td><td>CYTO</td></tr>
<tr><td>>hg38_21_+_8433172_8446622_RNA45SN1_45S_50ntFlanks@11582.11601.20|ACTGGTGCGGACCAGGGGAA</td><td>>hg38_21_+_8433172_8446622_RNA45SN1_45S_50ntFlanks</td><td>11582</td><td>11601</td><td>20</td><td>ACTGGTGCGGACCAGGGGAA</td><td>523.8</td><td>-5.7</td><td>0.4</td><td>-15.6</td><td>4.2E-55</td><td>3.2E-53</td><td>BT20/MB468</td><td>NUC</td></tr>
<tr><td>>hg38_21_+_8433172_8446622_RNA45SN1_45S_50ntFlanks@4046.4066.21|CATTCGAACGTCTGCCCTATC</td><td>>hg38_21_+_8433172_8446622_RNA45SN1_45S_50ntFlanks</td><td>4046</td><td>4066</td><td>21</td><td>CATTCGAACGTCTGCCCTATC</td><td>4491.8</td><td>-7.7</td><td>0.5</td><td>-15.6</td><td>4.5E-55</td><td>3.3E-53</td><td>BT20/MB468</td><td>NUC</td></tr>
<tr><td>>hg38_21_+_8433172_8446622_RNA45SN1_45S_50ntFlanks@4722.4758.37|TCATTAATCAAGAACGAAAGTCGGAGGTTCGAAGACG</td><td>>hg38_21_+_8433172_8446622_RNA45SN1_45S_50ntFlanks</td><td>4722</td><td>4758</td><td>37</td><td>TCATTAATCAAGAACGAAAGTCGGAGGTTCGAAGACG</td><td>11768.8</td><td>-7.9</td><td>0.5</td><td>-15.7</td><td>1.7E-55</td><td>4.2E-53</td><td>MB231/MB468</td><td>NUC</td></tr>
<tr><td>>hg38_21_+_8433172_8446622_RNA45SN1_45S_50ntFlanks@10678.10708.31|CGGTTCCGGCGGCGTCCGGTGAGCTCTCGCT</td><td>>hg38_21_+_8433172_8446622_RNA45SN1_45S_50ntFlanks</td><td>10678</td><td>10708</td><td>31</td><td>CGGTTCCGGCGGCGTCCGGTGAGCTCTCGCT</td><td>35379.3</td><td>-8.6</td><td>0.5</td><td>-15.7</td><td>1.8E-55</td><td>4.3E-53</td><td>MB231/MB468</td><td>NUC</td></tr>
<tr><td>>MI0000063|hsa-let-7b&WithFlank&22|+|46113680|46113774@12.34.23&[MIMAT0000063&hsa-let-7b-5p&offsets|0|+1;m-14&22|+|46113691|46113712&offsets|0|+1]|TGAGGTAGTAGGTTGTGTGGTTT</td><td>>MI0000063|hsa-let-7b&WithFlank&22|+|46113680|46113774</td><td>12</td><td>34</td><td>23</td><td>TGAGGTAGTAGGTTGTGTGGTTT</td><td>5843.2</td><td>8.4</td><td>0.5</td><td>15.7</td><td>2.0E-55</td><td>4.3E-53</td><td>BT20/MB468</td><td>MITO</td></tr>
<tr><td>>am_MI0000809|hsa-mir-151a&WithFlank&8|-|140732558|140732659@53.73.21&[MIMAT0000757&hsa-miR-151a-3p&offsets|0|0;m-30&8|-|140732587|140732607&offsets|0|0]|CTAGACTGAAGCTCCTTGAGG</td><td>>am_MI0000809|hsa-mir-151a&WithFlank&8|-|140732558|140732659</td><td>53</td><td>73</td><td>21</td><td>CTAGACTGAAGCTCCTTGAGG</td><td>121540.3</td><td>9.3</td><td>0.6</td><td>15.7</td><td>1.9E-55</td><td>4.3E-53</td><td>BT20/MB468</td><td>MITO</td></tr>
<tr><td>>MI0000440|hsa-mir-27b&WithFlank&9|+|95085439|95085547@67.87.21&[MIMAT0000419&hsa-miR-27b-3p&offsets|0|0;m-39&9|+|95085505|95085525&offsets|0|0]|TTCACAGTGGCTAAGTTCTGC</td><td>>MI0000440|hsa-mir-27b&WithFlank&9|+|95085439|95085547</td><td>67</td><td>87</td><td>21</td><td>TTCACAGTGGCTAAGTTCTGC</td><td>75993.0</td><td>7.7</td><td>0.5</td><td>15.6</td><td>5.3E-55</td><td>4.9E-53</td><td>BT20/MB468</td><td>CYTO</td></tr>
<tr><td>>MI0000440|hsa-mir-27b&WithFlank&9|+|95085439|95085547@67.88.22&[MIMAT0000419&hsa-miR-27b-3p&offsets|0|+1;m-39&9|+|95085505|95085525&offsets|0|+1]|TTCACAGTGGCTAAGTTCTGCA</td><td>>MI0000440|hsa-mir-27b&WithFlank&9|+|95085439|95085547</td><td>67</td><td>88</td><td>22</td><td>TTCACAGTGGCTAAGTTCTGCA</td><td>4860.9</td><td>7.2</td><td>0.5</td><td>15.7</td><td>2.3E-55</td><td>4.9E-53</td><td>BT20/MB468</td><td>MITO</td></tr>
<tr><td>>hg38_21_+_8433172_8446622_RNA45SN1_45S_50ntFlanks@10679.10708.30|GGTTCCGGCGGCGTCCGGTGAGCTCTCGCT</td><td>>hg38_21_+_8433172_8446622_RNA45SN1_45S_50ntFlanks</td><td>10679</td><td>10708</td><td>30</td><td>GGTTCCGGCGGCGTCCGGTGAGCTCTCGCT</td><td>9062.7</td><td>-8.5</td><td>0.5</td><td>-15.7</td><td>2.3E-55</td><td>5.1E-53</td><td>MB231/MB468</td><td>NUC</td></tr>
<tr><td>>hg38_21_+_8433172_8446622_RNA45SN1_45S_50ntFlanks@3987.4023.37|GCTTTGGTGACTCTAGATAACCTCGGGCCGATCGCAC</td><td>>hg38_21_+_8433172_8446622_RNA45SN1_45S_50ntFlanks</td><td>3987</td><td>4023</td><td>37</td><td>GCTTTGGTGACTCTAGATAACCTCGGGCCGATCGCAC</td><td>1315.2</td><td>7.3</td><td>0.5</td><td>15.6</td><td>6.8E-55</td><td>6.2E-53</td><td>BT20/MB468</td><td>CYTO</td></tr>
<tr><td>>hg38_21_+_8433172_8446622_RNA45SN1_45S_50ntFlanks@10679.10709.31|GGTTCCGGCGGCGTCCGGTGAGCTCTCGCTG</td><td>>hg38_21_+_8433172_8446622_RNA45SN1_45S_50ntFlanks</td><td>10679</td><td>10709</td><td>31</td><td>GGTTCCGGCGGCGTCCGGTGAGCTCTCGCTG</td><td>25541.9</td><td>-10.8</td><td>0.7</td><td>-15.7</td><td>1.3E-55</td><td>1.1E-52</td><td>MB231/MB468</td><td>MITO</td></tr>
<tr><td>>hg38_21_+_8433172_8446622_RNA45SN1_45S_50ntFlanks@12606.12625.20|ATGTGCTTGGCTGAGGAGCC</td><td>>hg38_21_+_8433172_8446622_RNA45SN1_45S_50ntFlanks</td><td>12606</td><td>12625</td><td>20</td><td>ATGTGCTTGGCTGAGGAGCC</td><td>1648.7</td><td>-7.6</td><td>0.5</td><td>-15.6</td><td>1.6E-54</td><td>1.2E-52</td><td>BT20/MB468</td><td>NUC</td></tr>
<tr><td>>MI0000102|hsa-mir-100&WithFlank&11|-|122152223|122152314@19.40.22(+1A)&[MIMAT0000098&hsa-miR-100-5p&offsets|0|0(+1A);m-27&11|-|122152276|122152296&offsets|0|+1(+1A)]|AACCCGTAGATCCGAACTTGTGA</td><td>>MI0000102|hsa-mir-100&WithFlank&11|-|122152223|122152314</td><td>19</td><td>40</td><td>22(+1A)</td><td>AACCCGTAGATCCGAACTTGTGA</td><td>53340.4</td><td>16.4</td><td>1.1</td><td>15.6</td><td>6.6E-55</td><td>1.4E-52</td><td>MB231/MB468</td><td>NUC</td></tr>
<tr><td>>MI0000255|hsa-mir-30d&WithFlank&8|-|134804870|134804951@12.33.22&[MIMAT0000245&hsa-miR-30d-5p&offsets|0|0;m-21&8|-|134804919|134804940&offsets|0|0]|TGTAAACATCCCCGACTGGAAG</td><td>>MI0000255|hsa-mir-30d&WithFlank&8|-|134804870|134804951</td><td>12</td><td>33</td><td>22</td><td>TGTAAACATCCCCGACTGGAAG</td><td>1464.6</td><td>4.2</td><td>0.3</td><td>15.5</td><td>2.2E-54</td><td>1.6E-52</td><td>BT20/MB468</td><td>NUC</td></tr>
<tr><td>>hg38_21_+_8433172_8446622_RNA45SN1_45S_50ntFlanks@9572.9605.34|CAAATCGGTCGTCCGACCTGGGTATAGGGGCGAA</td><td>>hg38_21_+_8433172_8446622_RNA45SN1_45S_50ntFlanks</td><td>9572</td><td>9605</td><td>34</td><td>CAAATCGGTCGTCCGACCTGGGTATAGGGGCGAA</td><td>676.1</td><td>-6.5</td><td>0.4</td><td>-15.5</td><td>2.2E-54</td><td>1.6E-52</td><td>BT20/MB468</td><td>NUC</td></tr>
<tr><td>>hg38_21_+_8433172_8446622_RNA45SN1_45S_50ntFlanks@12936.12958.23|AAACCATTCGTAGACGACCTGCT</td><td>>hg38_21_+_8433172_8446622_RNA45SN1_45S_50ntFlanks</td><td>12936</td><td>12958</td><td>23</td><td>AAACCATTCGTAGACGACCTGCT</td><td>1072.4</td><td>-6.1</td><td>0.4</td><td>-15.5</td><td>2.4E-54</td><td>1.7E-52</td><td>BT20/MB468</td><td>NUC</td></tr>
<tr><td>>hg38_21_+_8433172_8446622_RNA45SN1_45S_50ntFlanks@11641.11660.20|CGGCGGGTGTTGACGCGATG</td><td>>hg38_21_+_8433172_8446622_RNA45SN1_45S_50ntFlanks</td><td>11641</td><td>11660</td><td>20</td><td>CGGCGGGTGTTGACGCGATG</td><td>233672.2</td><td>-16.1</td><td>1.0</td><td>-15.7</td><td>8.3E-56</td><td>2.7E-52</td><td>MB231/MB468</td><td>MP</td></tr>
<tr><td>>hg38_21_+_8433172_8446622_RNA45SN1_45S_50ntFlanks@13002.13027.26|CGATCTATTGAAAGTCAGCCCTCGAC</td><td>>hg38_21_+_8433172_8446622_RNA45SN1_45S_50ntFlanks</td><td>13002</td><td>13027</td><td>26</td><td>CGATCTATTGAAAGTCAGCCCTCGAC</td><td>19196.2</td><td>-18.7</td><td>1.2</td><td>-15.5</td><td>4.3E-54</td><td>3.0E-52</td><td>BT20/MB468</td><td>NUC</td></tr>
<tr><td>>hg38_21_+_8433172_8446622_RNA45SN1_45S_50ntFlanks@12973.12990.18|TCGTACGTAGCAGAGCAG</td><td>>hg38_21_+_8433172_8446622_RNA45SN1_45S_50ntFlanks</td><td>12973</td><td>12990</td><td>18</td><td>TCGTACGTAGCAGAGCAG</td><td>52160.5</td><td>-20.1</td><td>1.3</td><td>-15.5</td><td>5.5E-54</td><td>3.8E-52</td><td>BT20/MB468</td><td>NUC</td></tr>
</table>

<table>
<tr><th></th><th></th><th></th><th></th><th></th><th></th><th></th><th></th><th></th><th></th><th></th><th></th><th></th><th></th></tr>
<tr><td>>MI0000650|hsa-mir-200c&WithFlank&12|+|6963693|6963772@50.70.21&[MIMAT0000617&hsa-miR-200c-3p&offsets|0|-2;m-54&12|+|6963742|6963764&offsets|0|-2]|TAATACTGCCGGGTAATGATG</td><td>>MI0000650|hsa-mir-200c&WithFlank&12|+|6963693|6963772</td><td>50</td><td>70</td><td>21</td><td>TAATACTGCCGGGTAATGATG</td><td>4881.0</td><td>9.2</td><td>0.6</td><td>15.5</td><td>2.6E-54</td><td>5.3E-52</td><td>BT20/MB468</td><td>MITO</td></tr>
<tr><td>>MI0000255|hsa-mir-30d&WithFlank&8|-|134804870|134804951@12.33.22(+1U)&[MIMAT0000245&hsa-miR-30d-5p&offsets|0|0(+1U);m-21&8|-|134804919|134804940&offsets|0|0(+1U)]|TGTAAACATCCCCGACTGGAAGT</td><td>>MI0000255|hsa-mir-30d&WithFlank&8|-|134804870|134804951</td><td>12</td><td>33</td><td>22(+1U)</td><td>TGTAAACATCCCCGACTGGAAGT</td><td>18062.7</td><td>9.8</td><td>0.6</td><td>15.5</td><td>5.9E-54</td><td>5.3E-52</td><td>BT20/MB468</td><td>CYTO</td></tr>
<tr><td>>hg38_21_+_8433172_8446622_RNA45SN1_45S_50ntFlanks@4731.4759.29|AAGAACGAAAGTCGGAGGTTCGAAGACGA</td><td>>hg38_21_+_8433172_8446622_RNA45SN1_45S_50ntFlanks</td><td>4731</td><td>4759</td><td>29</td><td>AAGAACGAAAGTCGGAGGTTCGAAGACGA</td><td>4598.7</td><td>-7.7</td><td>0.5</td><td>-15.4</td><td>1.0E-53</td><td>7.1E-52</td><td>BT20/MB468</td><td>NUC</td></tr>
<tr><td>>MI0000542|hsa-mir-320a&WithFlank&8|-|22244960|22245043@48.70.23&[MIMAT0000510&hsa-miR-320a-3p&offsets|0|+1;m-60&8|-|22244975|22244996&offsets|0|+1]|AAAAGCTGGGTTGAGAGGGCGAA</td><td>>MI0000542|hsa-mir-320a&WithFlank&8|-|22244960|22245043</td><td>48</td><td>70</td><td>23</td><td>AAAAGCTGGGTTGAGAGGGCGAA</td><td>5600.9</td><td>10.1</td><td>0.7</td><td>15.5</td><td>4.2E-54</td><td>8.3E-52</td><td>BT20/MB468</td><td>MITO</td></tr>
<tr><td>>hg38_21_+_8433172_8446622_RNA45SN1_45S_50ntFlanks@10402.10430.29|AACATGGGTCAGTCGGTCCTGAGAGATGG</td><td>>hg38_21_+_8433172_8446622_RNA45SN1_45S_50ntFlanks</td><td>10402</td><td>10430</td><td>29</td><td>AACATGGGTCAGTCGGTCCTGAGAGATGG</td><td>3519.1</td><td>-7.3</td><td>0.5</td><td>-15.5</td><td>4.5E-54</td><td>9.4E-52</td><td>MB231/MB468</td><td>NUC</td></tr>
<tr><td>>hg38_21_+_8433172_8446622_RNA45SN1_45S_50ntFlanks@10351.10378.28|CGAGAACTTTGAAGGCCGAAGTGGAGAA</td><td>>hg38_21_+_8433172_8446622_RNA45SN1_45S_50ntFlanks</td><td>10351</td><td>10378</td><td>28</td><td>CGAGAACTTTGAAGGCCGAAGTGGAGAA</td><td>2402.8</td><td>-6.7</td><td>0.4</td><td>-15.5</td><td>5.2E-54</td><td>1.0E-51</td><td>MB231/MB468</td><td>NUC</td></tr>
<tr><td>>hg38_21_+_8433172_8446622_RNA45SN1_45S_50ntFlanks@9534.9553.20|AGGAAACTCTGGTGGAGGTC</td><td>>hg38_21_+_8433172_8446622_RNA45SN1_45S_50ntFlanks</td><td>9534</td><td>9553</td><td>20</td><td>AGGAAACTCTGGTGGAGGTC</td><td>97472.0</td><td>-9.7</td><td>0.6</td><td>-15.5</td><td>5.2E-54</td><td>1.0E-51</td><td>MB231/MB468</td><td>NUC</td></tr>
<tr><td>>hg38_21_+_8433172_8446622_RNA45SN1_45S_50ntFlanks@11572.11600.29|CCGACTTAGAACTGGTGCGGACCAGGGGA</td><td>>hg38_21_+_8433172_8446622_RNA45SN1_45S_50ntFlanks</td><td>11572</td><td>11600</td><td>29</td><td>CCGACTTAGAACTGGTGCGGACCAGGGGA</td><td>13709.8</td><td>-9.4</td><td>0.6</td><td>-15.4</td><td>1.9E-53</td><td>1.3E-51</td><td>BT20/MB468</td><td>NUC</td></tr>
<tr><td>>hg38_21_+_8433172_8446622_RNA45SN1_45S_50ntFlanks@10350.10379.30|ACGAGAACTTTGAAGGCCGAAGTGGAGAAG</td><td>>hg38_21_+_8433172_8446622_RNA45SN1_45S_50ntFlanks</td><td>10350</td><td>10379</td><td>30</td><td>ACGAGAACTTTGAAGGCCGAAGTGGAGAAG</td><td>1450.1</td><td>-7.2</td><td>0.5</td><td>-15.4</td><td>2.1E-53</td><td>1.4E-51</td><td>BT20/MB468</td><td>NUC</td></tr>
<tr><td>>MI0000255|hsa-mir-30d&WithFlank&8|-|134804870|134804951@12.35.24&[MIMAT0000245&hsa-miR-30d-5p&offsets|0|+2;m-21&8|-|134804919|134804940&offsets|0|+2]|TGTAAACATCCCCGACTGGAAGCT</td><td>>MI0000255|hsa-mir-30d&WithFlank&8|-|134804870|134804951</td><td>12</td><td>35</td><td>24</td><td>TGTAAACATCCCCGACTGGAAGCT</td><td>96128.8</td><td>8.9</td><td>0.6</td><td>15.4</td><td>1.6E-53</td><td>1.4E-51</td><td>BT20/MB468</td><td>CYTO</td></tr>
<tr><td>>MI0000255|hsa-mir-30d&WithFlank&8|-|134804870|134804951@12.33.22&[MIMAT0000245&hsa-miR-30d-5p&offsets|0|0;m-21&8|-|134804919|134804940&offsets|0|0]|TGTAAACATCCCCGACTGGAAG</td><td>>MI0000255|hsa-mir-30d&WithFlank&8|-|134804870|134804951</td><td>12</td><td>33</td><td>22</td><td>TGTAAACATCCCCGACTGGAAG</td><td>7888.2</td><td>9.1</td><td>0.6</td><td>15.4</td><td>2.6E-53</td><td>2.2E-51</td><td>BT20/MB468</td><td>CYTO</td></tr>
<tr><td>>hg38_21_+_8433172_8446622_RNA45SN1_45S_50ntFlanks@12418.12443.26|CGATGTCGGCTCTTCCTATCATTGTG</td><td>>hg38_21_+_8433172_8446622_RNA45SN1_45S_50ntFlanks</td><td>12418</td><td>12443</td><td>26</td><td>CGATGTCGGCTCTTCCTATCATTGTG</td><td>32162.9</td><td>-10.1</td><td>0.7</td><td>-15.3</td><td>3.6E-53</td><td>2.4E-51</td><td>BT20/MB468</td><td>NUC</td></tr>
<tr><td>>hg38_21_+_8433172_8446622_RNA45SN1_45S_50ntFlanks@11768.11794.27|CTAATTAGTGACGCGCATGAATGGATG</td><td>>hg38_21_+_8433172_8446622_RNA45SN1_45S_50ntFlanks</td><td>11768</td><td>11794</td><td>27</td><td>CTAATTAGTGACGCGCATGAATGGATG</td><td>6951.5</td><td>-7.5</td><td>0.5</td><td>-15.4</td><td>2.0E-53</td><td>3.8E-51</td><td>MB231/MB468</td><td>NUC</td></tr>
<tr><td>>hg38_21_+_8433172_8446622_RNA45SN1_45S_50ntFlanks@11783.11804.22|CATGAATGGATGAACGAGATTC</td><td>>hg38_21_+_8433172_8446622_RNA45SN1_45S_50ntFlanks</td><td>11783</td><td>11804</td><td>22</td><td>CATGAATGGATGAACGAGATTC</td><td>31302.7</td><td>-10.0</td><td>0.6</td><td>-15.5</td><td>1.8E-54</td><td>4.8E-51</td><td>MB231/MB468</td><td>MP</td></tr>
<tr><td>>MI0000272|hsa-mir-182&WithFlank&7|-|129770377|129770498@29.53.25&[MIMAT0000259&hsa-miR-182-5p&offsets|0|+1;m-22&7|-|129770449|129770470&offsets|0|+3]|TTTGGCAATGGTAGAACTCACACTG</td><td>>MI0000272|hsa-mir-182&WithFlank&7|-|129770377|129770498</td><td>29</td><td>53</td><td>25</td><td>TTTGGCAATGGTAGAACTCACACTG</td><td>20804.0</td><td>7.9</td><td>0.5</td><td>15.3</td><td>5.9E-53</td><td>5.1E-51</td><td>BT20/MB468</td><td>CYTO</td></tr>
<tr><td>>MI0000102|hsa-mir-100&WithFlank&11|-|122152223|122152314@19.40.22&[MIMAT0000098&hsa-miR-100-5p&offsets|0|0;m-27&11|-|122152276|122152296&offsets|0|+1]|AACCCGTAGATCCGAACTTGTG</td><td>>MI0000102|hsa-mir-100&WithFlank&11|-|122152223|122152314</td><td>19</td><td>40</td><td>22</td><td>AACCCGTAGATCCGAACTTGTG</td><td>796867.6</td><td>15.6</td><td>1.0</td><td>15.6</td><td>8.7E-55</td><td>5.3E-51</td><td>MB231/MB468</td><td>CYTO</td></tr>
<tr><td>>hg38_21_+_8433172_8446622_RNA45SN1_45S_50ntFlanks@7975.8000.26|CGCGACCTCAGATCAGACGTGGCGAC</td><td>>hg38_21_+_8433172_8446622_RNA45SN1_45S_50ntFlanks</td><td>7975</td><td>8000</td><td>26</td><td>CGCGACCTCAGATCAGACGTGGCGAC</td><td>473.3</td><td>6.3</td><td>0.4</td><td>15.3</td><td>7.8E-53</td><td>6.6E-51</td><td>BT20/MB468</td><td>CYTO</td></tr>
<tr><td>>MI0000342|hsa-mir-200b&WithFlank&1|+|1167098|1167204@63.85.23(+1A)&[MIMAT0000318&hsa-miR-200b-3p&offsets|0|+1(+1A);m-93&1|+|1167160|1167182&offsets|0|0(+1A)]|TAATACTGCCTGGTAATGATGACA</td><td>>MI0000342|hsa-mir-200b&WithFlank&1|+|1167098|1167204</td><td>63</td><td>85</td><td>23(+1A)</td><td>TAATACTGCCTGGTAATGATGACA</td><td>1297.7</td><td>-6.8</td><td>0.4</td><td>-15.3</td><td>1.3E-52</td><td>8.4E-51</td><td>BT20/MB468</td><td>NUC</td></tr>
<tr><td>>MI0000088|hsa-mir-30a&WithFlank&6|-|71403545|71403627@54.74.21&[MIMAT0000088&hsa-miR-30a-3p&offsets|+1|0;m-38&6|-|71403554|71403575&offsets|+1|0]|TTTCAGTCGGATGTTTGCAGC</td><td>>MI0000088|hsa-mir-30a&WithFlank&6|-|71403545|71403627</td><td>54</td><td>74</td><td>21</td><td>TTTCAGTCGGATGTTTGCAGC</td><td>4013.4</td><td>7.2</td><td>0.5</td><td>15.3</td><td>1.0E-52</td><td>8.5E-51</td><td>BT20/MB468</td><td>CYTO</td></tr>
<tr><td>>hg38_21_+_8433172_8446622_RNA45SN1_45S_50ntFlanks@10678.10712.35|CGGTTCCGGCGGCGTCCGGTGAGCTCTCGCTGGCC</td><td>>hg38_21_+_8433172_8446622_RNA45SN1_45S_50ntFlanks</td><td>10678</td><td>10712</td><td>35</td><td>CGGTTCCGGCGGCGTCCGGTGAGCTCTCGCTGGCC</td><td>46769.0</td><td>-9.8</td><td>0.6</td><td>-15.4</td><td>1.2E-53</td><td>8.9E-51</td><td>MB231/MB468</td><td>MITO</td></tr>
<tr><td>>MI0001445|hsa-mir-423&WithFlank&17|+|30117073|30117178@23.43.21&[MIMAT0004748&hsa-miR-423-5p&offsets|0|-2;m-141&17|+|30117095|30117117&offsets|0|-2]|TGAGGGGCAGAGAGCGAGACT</td><td>>MI0001445|hsa-mir-423&WithFlank&17|+|30117073|30117178</td><td>23</td><td>43</td><td>21</td><td>TGAGGGGCAGAGAGCGAGACT</td><td>3887.4</td><td>9.4</td><td>0.6</td><td>15.3</td><td>1.6E-52</td><td>1.3E-50</td><td>BT20/MB468</td><td>CYTO</td></tr>
<tr><td>>hg38_21_+_8433172_8446622_RNA45SN1_45S_50ntFlanks@3988.4023.36|CTTTGGTGACTCTAGATAACCTCGGGCCGATCGCAC</td><td>>hg38_21_+_8433172_8446622_RNA45SN1_45S_50ntFlanks</td><td>3988</td><td>4023</td><td>36</td><td>CTTTGGTGACTCTAGATAACCTCGGGCCGATCGCAC</td><td>2778.0</td><td>6.7</td><td>0.4</td><td>15.2</td><td>1.9E-52</td><td>1.6E-50</td><td>BT20/MB468</td><td>CYTO</td></tr>
<tr><td>>hg38_21_+_8433172_8446622_RNA45SN1_45S_50ntFlanks@10680.10708.29|GTTCCGGCGGCGTCCGGTGAGCTCTCGCT</td><td>>hg38_21_+_8433172_8446622_RNA45SN1_45S_50ntFlanks</td><td>10680</td><td>10708</td><td>29</td><td>GTTCCGGCGGCGTCCGGTGAGCTCTCGCT</td><td>21907.6</td><td>-8.2</td><td>0.5</td><td>-15.3</td><td>1.0E-52</td><td>1.9E-50</td><td>MB231/MB468</td><td>NUC</td></tr>
<tr><td>>hg38_21_+_8433172_8446622_RNA45SN1_45S_50ntFlanks@10346.10372.27|TCAAACGAGAACTTTGAAGGCCGAAGT</td><td>>hg38_21_+_8433172_8446622_RNA45SN1_45S_50ntFlanks</td><td>10346</td><td>10372</td><td>27</td><td>TCAAACGAGAACTTTGAAGGCCGAAGT</td><td>337.7</td><td>-4.5</td><td>0.3</td><td>-15.2</td><td>3.3E-52</td><td>2.1E-50</td><td>BT20/MB468</td><td>NUC</td></tr>
<tr><td>>hg38_21_+_8433172_8446622_RNA45SN1_45S_50ntFlanks@12407.12432.26|TTTTGATCCTTCGATGTCGGCTCTTC</td><td>>hg38_21_+_8433172_8446622_RNA45SN1_45S_50ntFlanks</td><td>12407</td><td>12432</td><td>26</td><td>TTTTGATCCTTCGATGTCGGCTCTTC</td><td>856.2</td><td>-6.8</td><td>0.4</td><td>-15.2</td><td>3.3E-52</td><td>2.1E-50</td><td>BT20/MB468</td><td>NUC</td></tr>
<tr><td>>am_MI0005763|hsa-mir-941-1&WithFlank&20|+|63919443|63919526@53.75.23;MI0005764|hsa-mir-941-2&WithFlank&20|+|63919499|63919582@53.75.23;MI0005765|hsa-mir-941-3&WithFlank&20|+|63919555|63919638@53.75.23;MI0005766|hsa-mir-941-4&WithFlank&20|+|63919750|63919833@53.75.23;MI0031520|hsa-mir-941-5&WithFlank&20|+|63919862|63919945@53.75.23&[MIMAT0004984&hsa-miR-941&offsets|0|0;m-263&20|+|63919495|63919517&offsets|0|0];[MIMAT0004984_1&hsa-miR-941&offsets|0|0;m-263&20|+|63919551|63919573&offsets|0|0];[MIMAT0004984_2&hsa-miR-941&offsets|0|0;m-263&20|+|63919607|63919629&offsets|0|0];[MIMAT0004984_3&hsa-miR-941&offsets|0|0;m-263&20|+|63919802|63919824&offsets|0|0];[MIMAT0004984_4&hsa-miR-941&offsets|0|0;m-263&20|+|63919914|63919936&offsets|0|0]|CACCCGGCTGTGTGCACATGTGC</td><td>>am_MI0005763|hsa-mir-941-1&WithFlank&20|+|63919443|63919526</td><td>53</td><td>75</td><td>23</td><td>CACCCGGCTGTGTGCACATGTGC</td><td>17933.2</td><td>8.4</td><td>0.5</td><td>15.3</td><td>1.3E-52</td><td>2.6E-50</td><td>BT20/MB468</td><td>MITO</td></tr>
<tr><td>>hg38_21_+_8433172_8446622_RNA45SN1_45S_50ntFlanks@12476.12497.22|CCACTAATAGGGAACGTGAGCT</td><td>>hg38_21_+_8433172_8446622_RNA45SN1_45S_50ntFlanks</td><td>12476</td><td>12497</td><td>22</td><td>CCACTAATAGGGAACGTGAGCT</td><td>3485.9</td><td>-7.5</td><td>0.5</td><td>-15.2</td><td>4.2E-52</td><td>2.6E-50</td><td>BT20/MB468</td><td>NUC</td></tr>
<tr><td>>hg38_21_+_8433172_8446622_RNA45SN1_45S_50ntFlanks@6699.6720.22|GCTAGCTGCGAGAATTAATGTG</td><td>>hg38_21_+_8433172_8446622_RNA45SN1_45S_50ntFlanks</td><td>6699</td><td>6720</td><td>22</td><td>GCTAGCTGCGAGAATTAATGTG</td><td>119531.3</td><td>-11.0</td><td>0.7</td><td>-15.2</td><td>4.9E-52</td><td>3.0E-50</td><td>BT20/MB468</td><td>NUC</td></tr>
<tr><td>>hg38_21_+_8433172_8446622_RNA45SN1_45S_50ntFlanks@11661.11685.25|TGATTTCTGCCCAGTGCTCTGAATG</td><td>>hg38_21_+_8433172_8446622_RNA45SN1_45S_50ntFlanks</td><td>11661</td><td>11685</td><td>25</td><td>TGATTTCTGCCCAGTGCTCTGAATG</td><td>7198.1</td><td>-7.7</td><td>0.5</td><td>-15.2</td><td>1.8E-52</td><td>3.2E-50</td><td>MB231/MB468</td><td>NUC</td></tr>
<tr><td>>am_MI0000786|hsa-mir-378a&WithFlank&5|+|149732819|149732896@50.70.21&[MIMAT0000732&hsa-miR-378a-3p&offsets|+1|0;m-44&5|+|149732867|149732888&offsets|+1|0]|CTGGACTTGGAGTCAGAAGGC</td><td>>am_MI0000786|hsa-mir-378a&WithFlank&5|+|149732819|149732896</td><td>50</td><td>70</td><td>21</td><td>CTGGACTTGGAGTCAGAAGGC</td><td>3684.7</td><td>6.2</td><td>0.4</td><td>15.2</td><td>4.1E-52</td><td>3.3E-50</td><td>BT20/MB468</td><td>CYTO</td></tr>
<tr><td>>hg38_21_+_8433172_8446622_RNA45SN1_45S_50ntFlanks@7980.8008.29|CCTCAGATCAGACGTGGCGACCCGCTGAA</td><td>>hg38_21_+_8433172_8446622_RNA45SN1_45S_50ntFlanks</td><td>7980</td><td>8008</td><td>29</td><td>CCTCAGATCAGACGTGGCGACCCGCTGAA</td><td>2015.8</td><td>-7.0</td><td>0.5</td><td>-15.2</td><td>5.8E-52</td><td>3.6E-50</td><td>BT20/MB468</td><td>NUC</td></tr>
<tr><td>>MI0000273|hsa-mir-183&WithFlank&7|-|129774899|129775020@34.54.21(+1A)&[MIMAT0000261&hsa-miR-183-5p&offsets|+1|0(+1A);m-41&7|-|129774966|129774987&offsets|0|-1(+1A)]|ATGGCACTGGTAGAATTCACTA</td><td>>MI0000273|hsa-mir-183&WithFlank&7|-|129774899|129775020</td><td>34</td><td>54</td><td>21(+1A)</td><td>ATGGCACTGGTAGAATTCACTA</td><td>610.5</td><td>7.8</td><td>0.5</td><td>15.2</td><td>2.2E-52</td><td>4.1E-50</td><td>BT20/MB468</td><td>MITO</td></tr>
<tr><td>>MI0000447|hsa-mir-128-1&WithFlank&2|+|135665391|135665484@56.76.21;MI0000727|hsa-mir-128-2&WithFlank&3|+|35744470|35744565@58.78.21&[MIMAT0000424&hsa-miR-128-3p&offsets|0|0;m-108&2|+|135665446|135665466&offsets|0|0];[MIMAT0000424_1&hsa-miR-128-3p&offsets|0|0;m-117&3|+|35744527|35744547&offsets|0|0]|TCACAGTGAACCGGTCTCTTT</td><td>>MI0000447|hsa-mir-128-1&WithFlank&2|+|135665391|135665484</td><td>56</td><td>76</td><td>21</td><td>TCACAGTGAACCGGTCTCTTT</td><td>13705.8</td><td>6.1</td><td>0.4</td><td>15.2</td><td>5.3E-52</td><td>4.2E-50</td><td>BT20/MB468</td><td>CYTO</td></tr>
<tr><td>>hg38_21_+_8433172_8446622_RNA45SN1_45S_50ntFlanks@12624.12654.31|CCAATGGGGCGAAGCTACCATCTGTGGGATT</td><td>>hg38_21_+_8433172_8446622_RNA45SN1_45S_50ntFlanks</td><td>12624</td><td>12654</td><td>31</td><td>CCAATGGGGCGAAGCTACCATCTGTGGGATT</td><td>8752.3</td><td>-8.6</td><td>0.6</td><td>-15.2</td><td>2.4E-52</td><td>4.3E-50</td><td>BT20/MB468</td><td>MITO</td></tr>
<tr><td>>hg38_21_+_8433172_8446622_RNA45SN1_45S_50ntFlanks@10680.10710.31|GTTCCGGCGGCGTCCGGTGAGCTCTCGCTGG</td><td>>hg38_21_+_8433172_8446622_RNA45SN1_45S_50ntFlanks</td><td>10680</td><td>10710</td><td>31</td><td>GTTCCGGCGGCGTCCGGTGAGCTCTCGCTGG</td><td>37925.5</td><td>-10.0</td><td>0.7</td><td>-15.3</td><td>8.8E-53</td><td>5.2E-50</td><td>MB231/MB468</td><td>MITO</td></tr>
<tr><td>>hg38_21_+_8433172_8446622_RNA45SN1_45S_50ntFlanks@11209.11232.24|CGGCGACTCTGGACGCGAGCCGGG</td><td>>hg38_21_+_8433172_8446622_RNA45SN1_45S_50ntFlanks</td><td>11209</td><td>11232</td><td>24</td><td>CGGCGACTCTGGACGCGAGCCGGG</td><td>15652.1</td><td>-9.7</td><td>0.6</td><td>-15.3</td><td>8.1E-53</td><td>5.2E-50</td><td>MB231/MB468</td><td>MITO</td></tr>
<tr><td>>hg38_21_+_8433172_8446622_RNA45SN1_45S_50ntFlanks@6656.6689.34|CTTAGCGGTGGATCACTCGGCTCGTGCGTCGATG</td><td>>hg38_21_+_8433172_8446622_RNA45SN1_45S_50ntFlanks</td><td>6656</td><td>6689</td><td>34</td><td>CTTAGCGGTGGATCACTCGGCTCGTGCGTCGATG</td><td>1296.6</td><td>-5.9</td><td>0.4</td><td>-15.1</td><td>8.8E-52</td><td>5.3E-50</td><td>BT20/MB468</td><td>NUC</td></tr>
<tr><td>>hg38_21_+_8433172_8446622_RNA45SN1_45S_50ntFlanks@6655.6681.27|TCTTAGCGGTGGATCACTCGGCTCGTG</td><td>>hg38_21_+_8433172_8446622_RNA45SN1_45S_50ntFlanks</td><td>6655</td><td>6681</td><td>27</td><td>TCTTAGCGGTGGATCACTCGGCTCGTG</td><td>3305.2</td><td>-5.6</td><td>0.4</td><td>-15.1</td><td>9.1E-52</td><td>5.5E-50</td><td>BT20/MB468</td><td>NUC</td></tr>
<tr><td>>hg38_21_+_8433172_8446622_RNA45SN1_45S_50ntFlanks@10405.10424.20|ATGGGTCAGTCGGTCCTGAG</td><td>>hg38_21_+_8433172_8446622_RNA45SN1_45S_50ntFlanks</td><td>10405</td><td>10424</td><td>20</td><td>ATGGGTCAGTCGGTCCTGAG</td><td>15196.3</td><td>9.2</td><td>0.6</td><td>15.2</td><td>3.2E-52</td><td>5.6E-50</td><td>BT20/MB468</td><td>MITO</td></tr>
<tr><td>>MI0000469|hsa-mir-125a&WithFlank&19|+|51693248|51693345@21.42.22&[MIMAT0000443&hsa-miR-125a-5p&offsets|0|-2;m-63&19|+|51693268|51693289&offsets|0|0]|TCCCTGAGACCCTTTAACCTGT</td><td>>MI0000469|hsa-mir-125a&WithFlank&19|+|51693248|51693345</td><td>21</td><td>42</td><td>22</td><td>TCCCTGAGACCCTTTAACCTGT</td><td>11045.5</td><td>8.3</td><td>0.6</td><td>15.1</td><td>1.0E-51</td><td>8.1E-50</td><td>BT20/MB468</td><td>CYTO</td></tr>
<tr><td>>am_MI0000809|hsa-mir-151a&WithFlank&8|-|140732558|140732659@53.72.20(+1U)&[MIMAT0000757&hsa-miR-151a-3p&offsets|0|-1(+1U);m-30&8|-|140732587|140732607&offsets|0|-1(+1U)]|CTAGACTGAAGCTCCTTGAGT</td><td>>am_MI0000809|hsa-mir-151a&WithFlank&8|-|140732558|140732659</td><td>53</td><td>72</td><td>20(+1U)</td><td>CTAGACTGAAGCTCCTTGAGT</td><td>5949.2</td><td>9.4</td><td>0.6</td><td>15.2</td><td>4.8E-52</td><td>8.2E-50</td><td>BT20/MB468</td><td>MITO</td></tr>
<tr><td>>hg38_21_+_8433172_8446622_RNA45SN1_45S_50ntFlanks@10350.10369.20|ACGAGAACTTTGAAGGCCGA</td><td>>hg38_21_+_8433172_8446622_RNA45SN1_45S_50ntFlanks</td><td>10350</td><td>10369</td><td>20</td><td>ACGAGAACTTTGAAGGCCGA</td><td>932.2</td><td>-5.6</td><td>0.4</td><td>-15.1</td><td>1.5E-51</td><td>9.0E-50</td><td>BT20/MB468</td><td>NUC</td></tr>
<tr><td>>MI0000650|hsa-mir-200c&WithFlank&12|+|6963693|6963772@50.72.23&[MIMAT0000617&hsa-miR-200c-3p&offsets|0|0;m-54&12|+|6963742|6963764&offsets|0|0]|TAATACTGCCGGGTAATGATGGA</td><td>>MI0000650|hsa-mir-200c&WithFlank&12|+|6963693|6963772</td><td>50</td><td>72</td><td>23</td><td>TAATACTGCCGGGTAATGATGGA</td><td>217390.6</td><td>8.9</td><td>0.6</td><td>15.1</td><td>9.9E-52</td><td>1.7E-49</td><td>BT20/MB468</td><td>MITO</td></tr>
</table>

| | | | | | | | | | | | | | |
|---|---|---|---|---|---|---|---|---|---|---|---|---|---|
| >MI0000746\|hsa-mir-99b&WithFlank&19\|+\|51692606\|51692687@13.32.20(+1U)&[MIMAT0000689&hsa-miR-99b-5p&offsets\|0\|-2(+1U);m-10&19\|+\|51692618\|51692639&offsets\|0\|-2(+1U)]\|CACCCGTAGAACCGACCTTGT | >MI0000746\|hsa-mir-99b&WithFlank&19\|+\|51692606\|51692687 | 13 | 32 | 20(+1U) | CACCCGTAGAACCGACCTTGT | 7455.5 | 8.7 | 0.6 | 15.1 | 1.0E-51 | 1.7E-49 | BT20/MB468 | MITO |
| >MI0000542\|hsa-mir-320a&WithFlank&8\|-\|22244960\|22245043@48.69.22&[MIMAT0000510&hsa-miR-320a-3p&offsets\|0\|0;m-60&8\|-\|22244975\|22244996&offsets\|0\|0]\|AAAAGCTGGGTTGAGAGGGCGA | >MI0000542\|hsa-mir-320a&WithFlank&8\|-\|22244960\|22245043 | 48 | 69 | 22 | AAAAGCTGGGTTGAGAGGGCGA | 9518.7 | 10.0 | 0.7 | 15.1 | 1.2E-51 | 1.9E-49 | BT20/MB468 | MITO |
| >MI0000283\|hsa-mir-203a&WithFlank&14\|+\|104117399\|104117520@71.92.22&[MIMAT0000264&hsa-miR-203a-3p&offsets\|0\|0;m-79&14\|+\|104117469\|104117490&offsets\|0\|0]\|GTGAAATGTTTAGGACCACTAG | >MI0000283\|hsa-mir-203a&WithFlank&14\|+\|104117399\|104117520 | 71 | 92 | 22 | GTGAAATGTTTAGGACCACTAG | 3543.5 | 8.4 | 0.6 | 15.1 | 2.8E-51 | 2.1E-49 | BT20/MB468 | CYTO |
| >MI0001445\|hsa-mir-423&WithFlank&17\|+\|30117073\|30117178@59.81.23(+1U)&[MIMAT0001340&hsa-miR-423-3p&offsets\|0\|0(+1U);m-100&17\|+\|30117131\|30117153&offsets\|0\|0(+1U)]\|AGCTCGGTCTGAGGCCCCTCAGTT | >MI0001445\|hsa-mir-423&WithFlank&17\|+\|30117073\|30117178 | 59 | 81 | 23(+1U) | AGCTCGGTCTGAGGCCCCTCAGTT | 2473.2 | 8.1 | 0.5 | 15.1 | 3.2E-51 | 2.4E-49 | BT20/MB468 | CYTO |
| >hg38_21_+_8433172_8446622_RNA45SN1_45S_50ntFlanks@4565.4587.23\|AATAATGGAATAGGACCGCGGTT | >hg38_21_+_8433172_8446622_RNA45SN1_45S_50ntFlanks | 4565 | 4587 | 23 | AATAATGGAATAGGACCGCGGTT | 2247.3 | -7.5 | 0.5 | -15.0 | 4.7E-51 | 2.8E-49 | BT20/MB468 | NUC |
| >hg38_21_+_8433172_8446622_RNA45SN1_45S_50ntFlanks@10841.10865.25\|CGTAACTTCGGGATAAGGATTGGCT | >hg38_21_+_8433172_8446622_RNA45SN1_45S_50ntFlanks | 10841 | 10865 | 25 | CGTAACTTCGGGATAAGGATTGGCT | 53655.7 | -9.6 | 0.6 | -15.1 | 2.4E-51 | 4.2E-49 | MB231/MB468 | NUC |
| >hg38_21_+_8433172_8446622_RNA45SN1_45S_50ntFlanks@4592.4616.25\|TTTGTTGGTTTTCGGAACTGAGGCC | >hg38_21_+_8433172_8446622_RNA45SN1_45S_50ntFlanks | 4592 | 4616 | 25 | TTTGTTGGTTTTCGGAACTGAGGCC | 365.2 | -5.7 | 0.4 | -15.0 | 9.7E-51 | 5.6E-49 | BT20/MB468 | NUC |
| >am_MI0000809\|hsa-mir-151a&WithFlank&8\|-\|140732558\|140732659@53.72.20(+1U)&[MIMAT0000757&hsa-miR-151a-3p&offsets\|0\|-1(+1U);m-30&8\|-\|140732587\|140732607&offsets\|0\|-1(+1U)]\|CTAGACTGAAGCTCCTTGAGT | >am_MI0000809\|hsa-mir-151a&WithFlank&8\|-\|140732558\|140732659 | 53 | 72 | 20(+1U) | CTAGACTGAAGCTCCTTGAGT | 1070.5 | 3.9 | 0.3 | 15.0 | 1.1E-50 | 6.2E-49 | BT20/MB468 | NUC |
| >MI0000734\|hsa-mir-106b&WithFlank&7\|-\|100093987\|100094080@58.77.20&[MIMAT0004672&hsa-miR-106b-3p&offsets\|0\|-2;m-87&7\|-\|100094003\|100094023&offsets\|0\|-1]\|CCGCACTGTGGGTACTTGCT | >MI0000734\|hsa-mir-106b&WithFlank&7\|-\|100093987\|100094080 | 58 | 77 | 20 | CCGCACTGTGGGTACTTGCT | 2686.8 | 8.1 | 0.5 | 15.0 | 8.3E-51 | 6.2E-49 | BT20/MB468 | CYTO |
| >hg38_21_+_8433172_8446622_RNA45SN1_45S_50ntFlanks@12936.12956.21\|AAACCATTCGTAGACGACCTG | >hg38_21_+_8433172_8446622_RNA45SN1_45S_50ntFlanks | 12936 | 12956 | 21 | AAACCATTCGTAGACGACCTG | 13926.3 | -7.9 | 0.5 | -15.0 | 3.8E-51 | 6.4E-49 | MB231/MB468 | NUC |
| >hg38_21_+_8433172_8446622_RNA45SN1_45S_50ntFlanks@6655.6689.35\|TCTTAGCGGTGGATCACTCGGCTCGTGCGTCGATG | >hg38_21_+_8433172_8446622_RNA45SN1_45S_50ntFlanks | 6655 | 6689 | 35 | TCTTAGCGGTGGATCACTCGGCTCGTGCGTCGATG | 1921.9 | -4.5 | 0.3 | -15.0 | 1.1E-50 | 6.5E-49 | BT20/MB468 | NUC |
| >MI0000081\|hsa-mir-24-2&WithFlank&19\|-\|13836281\|13836365@56.76.21;MI0000080\|hsa-mir-24-1&WithFlank&9\|+\|95086015\|95086094@50.70.21&[MIMAT0000080&hsa-miR-24-3p&offsets\|0\|-1;m-36&19\|-\|13836288\|13836310&offsets\|0\|-2];[MIMAT0000080_1&hsa-miR-24-3p&offsets\|0\|-1;m-35&9\|+\|95086064\|95086086&offsets\|0\|-2]\|TGGCTCAGTTCAGCAGGAACA | >MI0000081\|hsa-mir-24-2&WithFlank&19\|-\|13836281\|13836365 | 56 | 76 | 21 | TGGCTCAGTTCAGCAGGAACA | 5186.5 | 5.7 | 0.4 | 15.0 | 5.4E-51 | 8.3E-49 | BT20/MB468 | MITO |
| >hg38_1_-_228634819_228635039_RNA5S12_5S_50ntFlanks@51.83.33\|GTCTACGGCCATACCACCCTGAACGCGCCCGAT | >hg38_1_-_228634819_228635039_RNA5S12_5S_50ntFlanks | 51 | 83 | 33 | GTCTACGGCCATACCACCCTGAACGCGCCCGAT | 24605.1 | -6.2 | 0.4 | -14.9 | 1.6E-50 | 9.1E-49 | BT20/MB468 | NUC |
| >hg38_21_+_8433172_8446622_RNA45SN1_45S_50ntFlanks@12420.12441.22\|ATGTCGGCTCTTCCTATCATTG | >hg38_21_+_8433172_8446622_RNA45SN1_45S_50ntFlanks | 12420 | 12441 | 22 | ATGTCGGCTCTTCCTATCATTG | 14915.4 | -9.7 | 0.6 | -14.9 | 1.6E-50 | 9.1E-49 | BT20/MB468 | NUC |
| >trna16_TyrGTA_14_-_21131351_21131444@59.76.18,trna17_TyrGTA_14_-_21128117_21128210@59.76.18,trna19_TyrGTA_14_-_21121258_21121351@59.76.18,trna2_TyrGTA_2_+_27273650_27273738@59.76.18,trna4_TyrGTA_8_+_67025602_67025694@59.76.18,trna5_TyrGTA_14_+_21151432_21151520@59.76.18,trna5_TyrGTA_8_+_67026223_67026311@59.76.18\|TTCCGGCTCGAAGGACCA | >trna16_TyrGTA_14_-_21131351_21131444 | 59 | 76 | 18 | TTCCGGCTCGAAGGACCA | 4383.7 | 7.9 | 0.5 | 15.0 | 1.3E-50 | 9.7E-49 | BT20/MB468 | CYTO |
| >hg38_21_+_8433172_8446622_RNA45SN1_45S_50ntFlanks@11579.11603.25\|AGAACTGGTGCGGACCAGGGGAATC | >hg38_21_+_8433172_8446622_RNA45SN1_45S_50ntFlanks | 11579 | 11603 | 25 | AGAACTGGTGCGGACCAGGGGAATC | 5234.5 | -8.5 | 0.6 | -14.9 | 2.9E-50 | 1.6E-48 | BT20/MB468 | NUC |
| >hg38_21_+_8433172_8446622_RNA45SN1_45S_50ntFlanks@12405.12433.29\|CTTTTTGATCCTTCGATGTCGGCTCTTCC | >hg38_21_+_8433172_8446622_RNA45SN1_45S_50ntFlanks | 12405 | 12433 | 29 | CTTTTTGATCCTTCGATGTCGGCTCTTCC | 27545.3 | -10.0 | 0.7 | -14.9 | 4.1E-50 | 2.2E-48 | BT20/MB468 | NUC |
| >hg38_21_+_8433172_8446622_RNA45SN1_45S_50ntFlanks@12405.12436.32\|CTTTTTGATCCTTCGATGTCGGCTCTTCCTAT | >hg38_21_+_8433172_8446622_RNA45SN1_45S_50ntFlanks | 12405 | 12436 | 32 | CTTTTTGATCCTTCGATGTCGGCTCTTCCTAT | 51446.5 | -11.1 | 0.7 | -14.9 | 2.1E-50 | 3.4E-48 | MB231/MB468 | NUC |
| >MI0000809\|hsa-mir-151a&WithFlank&8\|-\|140732558\|140732659@53.74.22&[MIMAT0000757&hsa-miR-151a-3p&offsets\|0\|+1;m-30&8\|-\|140732587\|140732607&offsets\|0\|+1]\|CTAGACTGAAGCTCCTTGAGGA | >MI0000809\|hsa-mir-151a&WithFlank&8\|-\|140732558\|140732659 | 53 | 74 | 22 | CTAGACTGAAGCTCCTTGAGGA | 25691.7 | 8.8 | 0.6 | 14.9 | 2.5E-50 | 3.8E-48 | BT20/MB468 | MITO |
| >hg38_21_+_8433172_8446622_RNA45SN1_45S_50ntFlanks@7979.8006.28\|ACCTCAGATCAGACGTGGCGACCCGCTG | >hg38_21_+_8433172_8446622_RNA45SN1_45S_50ntFlanks | 7979 | 8006 | 28 | ACCTCAGATCAGACGTGGCGACCCGCTG | 20486.3 | -9.2 | 0.6 | -15.0 | 7.5E-51 | 4.1E-48 | MB231/MB468 | MITO |
| >am_MI0000809\|hsa-mir-151a&WithFlank&8\|-\|140732558\|140732659@53.73.21(+1G)&[MIMAT0000757&hsa-miR-151a-3p&offsets\|0\|0(+1G);m-30&8\|-\|140732587\|140732607&offsets\|0\|0(+1G)]\|CTAGACTGAAGCTCCTTGAGGG | >am_MI0000809\|hsa-mir-151a&WithFlank&8\|-\|140732558\|140732659 | 53 | 73 | 21(+1G) | CTAGACTGAAGCTCCTTGAGGG | 1247.1 | 7.8 | 0.5 | 14.9 | 5.6E-50 | 4.1E-48 | BT20/MB468 | CYTO |
| >hg38_21_+_8433172_8446622_RNA45SN1_45S_50ntFlanks@10411.10430.20\|CAGTCGGTCCTGAGAGATGG | >hg38_21_+_8433172_8446622_RNA45SN1_45S_50ntFlanks | 10411 | 10430 | 20 | CAGTCGGTCCTGAGAGATGG | 20237.5 | -18.7 | 1.3 | -14.8 | 1.2E-49 | 6.5E-48 | BT20/MB468 | NUC |
| >hg38_21_+_8433172_8446622_RNA45SN1_45S_50ntFlanks@10351.10372.22\|CGAGAACTTTGAAGGCCGAAGT | >hg38_21_+_8433172_8446622_RNA45SN1_45S_50ntFlanks | 10351 | 10372 | 22 | CGAGAACTTTGAAGGCCGAAGT | 782.0 | -5.7 | 0.4 | -14.8 | 1.7E-49 | 9.2E-48 | BT20/MB468 | NUC |
| >hg38_21_+_8433172_8446622_RNA45SN1_45S_50ntFlanks@11572.11602.31\|CCGACTTAGAACTGGTGCGGACCAGGGGAAT | >hg38_21_+_8433172_8446622_RNA45SN1_45S_50ntFlanks | 11572 | 11602 | 31 | CCGACTTAGAACTGGTGCGGACCAGGGGAAT | 18740.7 | -9.8 | 0.7 | -14.8 | 2.2E-49 | 1.2E-47 | BT20/MB468 | NUC |
| >hg38_21_+_8433172_8446622_RNA45SN1_45S_50ntFlanks@9534.9553.20\|AGGAAACTCTGGTGGAGGTC | >hg38_21_+_8433172_8446622_RNA45SN1_45S_50ntFlanks | 9534 | 9553 | 20 | AGGAAACTCTGGTGGAGGTC | 54476.8 | -10.9 | 0.7 | -15.0 | 5.9E-51 | 1.3E-47 | MB231/MB468 | MP |
| >am_MI0005763\|hsa-mir-941-1&WithFlank&20\|+\|63919443\|63919526@53.74.22(+1A);MI0005764\|hsa-mir-941-2&WithFlank&20\|+\|63919499\|63919582@53.74.22(+1A);MI0005765\|hsa-mir-941-3&WithFlank&20\|+\|63919555\|63919638@53.74.22(+1A);MI0005766\|hsa-mir-941-4&WithFlank&20\|+\|63919750\|63919833@53.74.22(+1A);MI0031520\|hsa-mir-941-5&WithFlank&20\|+\|63919862\|63919945@53.74.22(+1A)&[MIMAT0004984&hsa-miR-941&offsets\|0\|-1(+1A);m-263&20\|+\|63919495\|63919517&offsets\|0\|-1(+1A)];[MIMAT0004984_1&hsa-miR-941&offsets\|0\|-1(+1A);m-263&20\|+\|63919551\|63919573&offsets\|0\|-1(+1A)];[MIMAT0004984_2&hsa-miR-941&offsets\|0\|-1(+1A);m-263&20\|+\|63919607\|63919629&offsets\|0\|-1(+1A)];[MIMAT0004984_3&hsa-miR-941&offsets\|0\|-1(+1A);m-263&20\|+\|63919802\|63919824&offsets\|0\|-1(+1A)];[MIMAT0004984_4&hsa-miR-941&offsets\|0\|-1(+1A);m-263&20\|+\|63919914\|63919936&offsets\|0\|-1(+1A)]\|CACCCGGCTGTGTGCACATGTGA | >am_MI0005763\|hsa-mir-941-1&WithFlank&20\|+\|63919443\|63919526 | 53 | 74 | 22(+1A) | CACCCGGCTGTGTGCACATGTGA | 2680.4 | 8.7 | 0.6 | 14.8 | 1.9E-49 | 1.4E-47 | BT20/MB468 | CYTO |
| >MI0000439\|hsa-mir-23b&WithFlank&9\|+\|95085202\|95085310@67.86.20(+1U)&[MIMAT0000418&hsa-miR-23b-3p&offsets\|+3\|0(+1U);m-42&9\|+\|95085265\|95085285&offsets\|+3\|+2(+1U)]\|ACATTGCCAGGGATTACCACT | >MI0000439\|hsa-mir-23b&WithFlank&9\|+\|95085202\|95085310 | 67 | 86 | 20(+1U) | ACATTGCCAGGGATTACCACT | 10933.4 | 17.7 | 1.2 | 14.7 | 4.1E-49 | 3.0E-47 | BT20/MB468 | CYTO |
| >MI0000253\|hsa-mir-148a&WithFlank&7\|-\|25949913\|25949992@12.33.22&[MIMAT0004549&hsa-miR-148a-5p&offsets\|0\|0;m-189&7\|-\|25949960\|25949981&offsets\|0\|0]\|AAAGTTCTGAGACACTCCGACT | >MI0000253\|hsa-mir-148a&WithFlank&7\|-\|25949913\|25949992 | 12 | 33 | 22 | AAAGTTCTGAGACACTCCGACT | 1136.6 | 7.4 | 0.5 | 14.7 | 4.8E-49 | 3.4E-47 | BT20/MB468 | CYTO |
| >MI0000283\|hsa-mir-203a&WithFlank&14\|+\|104117399\|104117520@72.93.22&[MIMAT0000264&hsa-miR-203a-3p&offsets\|+1\|+1;m-79&14\|+\|104117469\|104117490&offsets\|+1\|+1]\|TGAAATGTTTAGGACCACTAGA | >MI0000283\|hsa-mir-203a&WithFlank&14\|+\|104117399\|104117520 | 72 | 93 | 22 | TGAAATGTTTAGGACCACTAGA | 179.7 | 4.8 | 0.3 | 14.7 | 8.6E-49 | 4.5E-47 | BT20/MB468 | NUC |
| >hg38_21_+_8433172_8446622_RNA45SN1_45S_50ntFlanks@12456.12476.21\|CCAAGCGTTGGATTGTTCACC | >hg38_21_+_8433172_8446622_RNA45SN1_45S_50ntFlanks | 12456 | 12476 | 21 | CCAAGCGTTGGATTGTTCACC | 36662.4 | -10.7 | 0.7 | -14.7 | 1.1E-48 | 5.8E-47 | BT20/MB468 | NUC |
| >MI0000440\|hsa-mir-27b&WithFlank&9\|+\|95085439\|95085547@67.86.20(+1A)&[MIMAT0000419&hsa-miR-27b-3p&offsets\|0\|-1(+1A);m-39&9\|+\|95085505\|95085525&offsets\|0\|-1(+1A)]\|TTCACAGTGGCTAAGTTCTGA | >MI0000440\|hsa-mir-27b&WithFlank&9\|+\|95085439\|95085547 | 67 | 86 | 20(+1A) | TTCACAGTGGCTAAGTTCTGA | 772.0 | 4.2 | 0.3 | 14.7 | 1.1E-48 | 5.9E-47 | BT20/MB468 | NUC |
| >am_MI0000809\|hsa-mir-151a&WithFlank&8\|-\|140732558\|140732659@53.72.20(+1A)&[MIMAT0000757&hsa-miR-151a-3p&offsets\|0\|-1(+1A);m-30&8\|-\|140732587\|140732607&offsets\|0\|-1(+1A)]\|CTAGACTGAAGCTCCTTGAGA | >am_MI0000809\|hsa-mir-151a&WithFlank&8\|-\|140732558\|140732659 | 53 | 72 | 20(+1A) | CTAGACTGAAGCTCCTTGAGA | 32385.8 | 10.1 | 0.7 | 14.7 | 8.8E-49 | 6.2E-47 | BT20/MB468 | CYTO |
| >hg38_21_+_8433172_8446622_RNA45SN1_45S_50ntFlanks@7988.8008.21\|CAGACGTGGCGACCCGCTGAA | >hg38_21_+_8433172_8446622_RNA45SN1_45S_50ntFlanks | 7988 | 8008 | 21 | CAGACGTGGCGACCCGCTGAA | 15630.5 | -8.4 | 0.6 | -14.9 | 4.6E-50 | 8.6E-47 | MB231/MB468 | MP |
| >hg38_21_+_8433172_8446622_RNA45SN1_45S_50ntFlanks@10362.10383.22\|AAGGCCGAAGTGGAGAAGGGTT | >hg38_21_+_8433172_8446622_RNA45SN1_45S_50ntFlanks | 10362 | 10383 | 22 | AAGGCCGAAGTGGAGAAGGGTT | 4330.3 | -8.2 | 0.6 | -14.7 | 6.2E-49 | 9.8E-47 | MB231/MB468 | NUC |
| >MI0000266\|hsa-mir-10a&WithFlank&17\|-\|48579832\|48579953@29.50.22&[MIMAT0000253&hsa-miR-10a-5p&offsets\|+1\|0;m-18&17\|-\|48579905\|48579926&offsets\|+1\|+1]\|ACCCTGTAGATCCGAATTTGTG | >MI0000266\|hsa-mir-10a&WithFlank&17\|-\|48579832\|48579953 | 29 | 50 | 22 | ACCCTGTAGATCCGAATTTGTG | 18158.0 | 18.4 | 1.3 | 14.6 | 1.8E-48 | 1.3E-46 | BT20/MB468 | CYTO |
| >MI0000102\|hsa-mir-100&WithFlank&11\|-\|122152223\|122152314@19.40.22(+1U)&[MIMAT0000098&hsa-miR-100-5p&offsets\|0\|0(+1U);m-27&11\|-\|122152276\|122152296&offsets\|0\|+1(+1U)]\|AACCCGTAGATCCGAACTTGTGT | >MI0000102\|hsa-mir-100&WithFlank&11\|-\|122152223\|122152314 | 19 | 40 | 22(+1U) | AACCCGTAGATCCGAACTTGTGT | 65522.0 | 19.9 | 1.4 | 14.7 | 9.7E-49 | 1.5E-46 | MB231/MB468 | NUC |
| >MI0000066\|hsa-let-7e&WithFlank&19\|+\|51692780\|51692870@14.35.22&[MIMAT0000066&hsa-let-7e-5p&offsets\|0\|0;m-50&19\|+\|51692793\|51692814&offsets\|0\|0]\|TGAGGTAGGAGGTTGTATAGTT | >MI0000066\|hsa-let-7e&WithFlank&19\|+\|51692780\|51692870 | 14 | 35 | 22 | TGAGGTAGGAGGTTGTATAGTT | 42539.2 | 8.3 | 0.6 | 14.6 | 2.4E-48 | 1.7E-46 | BT20/MB468 | CYTO |
| >MI0000255\|hsa-mir-30d&WithFlank&8\|-\|134804870\|134804951@12.34.23(+1C)&[MIMAT0000245&hsa-miR-30d-5p&offsets\|0\|+1(+1C);m-21&8\|-\|134804919\|134804940&offsets\|0\|+1(+1C)]\|TGTAAACATCCCCGACTGGAAGCC | >MI0000255\|hsa-mir-30d&WithFlank&8\|-\|134804870\|134804951 | 12 | 34 | 23(+1C) | TGTAAACATCCCCGACTGGAAGCC | 14714.0 | 9.1 | 0.6 | 14.6 | 2.9E-48 | 1.9E-46 | BT20/MB468 | CYTO |

<table>
<tr><th></th><th></th><th></th><th></th><th></th><th></th><th></th><th></th><th></th><th></th><th></th><th></th><th></th><th></th></tr>
<tr><td>>MI0000650|hsa-mir-200c&WithFlank&12|+|6963693|6963772@50.70.21&[MIMAT0000617&hsa-miR-200c-3p&offsets|0|-2;m-54&12|+|6963742|6963764&offsets|0|-2]|TAATACTGCCGGGTAATGATG</td><td>>MI0000650|hsa-mir-200c&WithFlank&12|+|6963693|6963772</td><td>50</td><td>70</td><td>21</td><td>TAATACTGCCGGGTAATGATG</td><td>7233.5</td><td>9.2</td><td>0.6</td><td>14.6</td><td>2.8E-48</td><td>1.9E-46</td><td>BT20/MB468</td><td>CYTO</td></tr>
<tr><td>>MI0000273|hsa-mir-183&WithFlank&7|-|129774899|129775020@34.53.20(+1C)&[MIMAT0000261&hsa-miR-183-5p&offsets|+1|-1(+1C);m-41&7|-|129774966|129774987&offsets|0|-2(+1C)]|ATGGCACTGGTAGAATTCACC</td><td>>MI0000273|hsa-mir-183&WithFlank&7|-|129774899|129775020</td><td>34</td><td>53</td><td>20(+1C)</td><td>ATGGCACTGGTAGAATTCACC</td><td>798.0</td><td>8.3</td><td>0.6</td><td>14.7</td><td>1.3E-48</td><td>1.9E-46</td><td>BT20/MB468</td><td>MITO</td></tr>
<tr><td>>hg38_21_+_8433172_8446622_RNA45SN1_45S_50ntFlanks@10387.10424.38|TGTGAACAGCAGTTGAACATGGGTCAGTCGGTCCTGAG</td><td>>hg38_21_+_8433172_8446622_RNA45SN1_45S_50ntFlanks</td><td>10387</td><td>10424</td><td>38</td><td>TGTGAACAGCAGTTGAACATGGGTCAGTCGGTCCTGAG</td><td>18653.7</td><td>10.0</td><td>0.7</td><td>14.6</td><td>2.2E-48</td><td>3.1E-46</td><td>BT20/MB468</td><td>MITO</td></tr>
<tr><td>>MI0000440|hsa-mir-27b&WithFlank&9|+|95085439|95085547@67.85.19(+1A)&[MIMAT0000419&hsa-miR-27b-3p&offsets|0|-2(+1A);m-39&9|+|95085505|95085525&offsets|0|-2(+1A)]|TTCACAGTGGCTAAGTTCTA</td><td>>MI0000440|hsa-mir-27b&WithFlank&9|+|95085439|95085547</td><td>67</td><td>85</td><td>19(+1A)</td><td>TTCACAGTGGCTAAGTTCTA</td><td>431.2</td><td>7.2</td><td>0.5</td><td>14.6</td><td>2.3E-48</td><td>3.2E-46</td><td>BT20/MB468</td><td>MITO</td></tr>
<tr><td>>MI0000079|hsa-mir-23a&WithFlank&19|-|13836581|13836665@54.71.18&[MIMAT0000078&hsa-miR-23a-3p&offsets|+3|0;m-31&19|-|13836595|13836615&offsets|+3|0]|ACATTGCCAGGGATTTCC</td><td>>MI0000079|hsa-mir-23a&WithFlank&19|-|13836581|13836665</td><td>54</td><td>71</td><td>18</td><td>ACATTGCCAGGGATTTCC</td><td>13000.7</td><td>17.9</td><td>1.2</td><td>14.6</td><td>4.9E-48</td><td>3.2E-46</td><td>BT20/MB468</td><td>CYTO</td></tr>
<tr><td>>MI0000809|hsa-mir-151a&WithFlank&8|-|140732558|140732659@53.74.22&[MIMAT0000757&hsa-miR-151a-3p&offsets|0|+1;m-30&8|-|140732587|140732607&offsets|0|+1]|CTAGACTGAAGCTCCTTGAGGA</td><td>>MI0000809|hsa-mir-151a&WithFlank&8|-|140732558|140732659</td><td>53</td><td>74</td><td>22</td><td>CTAGACTGAAGCTCCTTGAGGA</td><td>33773.8</td><td>8.8</td><td>0.6</td><td>14.6</td><td>5.6E-48</td><td>3.6E-46</td><td>BT20/MB468</td><td>CYTO</td></tr>
<tr><td>>MI0000255|hsa-mir-30d&WithFlank&8|-|134804870|134804951@12.34.23&[MIMAT0000245&hsa-miR-30d-5p&offsets|0|+1;m-21&8|-|134804919|134804940&offsets|0|+1]|TGTAAACATCCCCGACTGGAAGC</td><td>>MI0000255|hsa-mir-30d&WithFlank&8|-|134804870|134804951</td><td>12</td><td>34</td><td>23</td><td>TGTAAACATCCCCGACTGGAAGC</td><td>10021.3</td><td>3.1</td><td>0.2</td><td>14.5</td><td>7.5E-48</td><td>3.9E-46</td><td>BT20/MB468</td><td>NUC</td></tr>
<tr><td>>MI0001445|hsa-mir-423&WithFlank&17|+|30117073|30117178@58.81.24&[MIMAT0001340&hsa-miR-423-3p&offsets|-1|0;m-100&17|+|30117131|30117153&offsets|-1|0]|AAGCTCGGTCTGAGGCCCCTCAGT</td><td>>MI0001445|hsa-mir-423&WithFlank&17|+|30117073|30117178</td><td>58</td><td>81</td><td>24</td><td>AAGCTCGGTCTGAGGCCCCTCAGT</td><td>8741.1</td><td>8.8</td><td>0.6</td><td>14.6</td><td>3.2E-48</td><td>4.3E-46</td><td>BT20/MB468</td><td>MITO</td></tr>
<tr><td>>MI0000255|hsa-mir-30d&WithFlank&8|-|134804870|134804951@12.33.22(+1A)&[MIMAT0000245&hsa-miR-30d-5p&offsets|0|0(+1A);m-21&8|-|134804919|134804940&offsets|0|0(+1A)]|TGTAAACATCCCCGACTGGAAGA</td><td>>MI0000255|hsa-mir-30d&WithFlank&8|-|134804870|134804951</td><td>12</td><td>33</td><td>22(+1A)</td><td>TGTAAACATCCCCGACTGGAAGA</td><td>2954.4</td><td>8.5</td><td>0.6</td><td>14.6</td><td>3.3E-48</td><td>4.4E-46</td><td>BT20/MB468</td><td>MITO</td></tr>
<tr><td>>MI0000266|hsa-mir-10a&WithFlank&17|-|48579832|48579953@28.50.23&[MIMAT0000253&hsa-miR-10a-5p&offsets|0|0;m-18&17|-|48579905|48579926&offsets|0|+1]|TACCCTGTAGATCCGAATTTGTG</td><td>>MI0000266|hsa-mir-10a&WithFlank&17|-|48579832|48579953</td><td>28</td><td>50</td><td>23</td><td>TACCCTGTAGATCCGAATTTGTG</td><td>27201.2</td><td>19.0</td><td>1.3</td><td>14.5</td><td>7.4E-48</td><td>4.8E-46</td><td>BT20/MB468</td><td>CYTO</td></tr>
<tr><td>>hg38_21_+_8433172_8446622_RNA45SN1_45S_50ntFlanks@12415.12441.27|CTTCGATGTCGGCTCTTCCTATCATTG</td><td>>hg38_21_+_8433172_8446622_RNA45SN1_45S_50ntFlanks</td><td>12415</td><td>12441</td><td>27</td><td>CTTCGATGTCGGCTCTTCCTATCATTG</td><td>5809.0</td><td>-7.2</td><td>0.5</td><td>-14.5</td><td>1.1E-47</td><td>5.4E-46</td><td>BT20/MB468</td><td>NUC</td></tr>
<tr><td>>MI0000283|hsa-mir-203a&WithFlank&14|+|104117399|104117520@71.92.22&[MIMAT0000264&hsa-miR-203a-3p&offsets|0|0;m-79&14|+|104117469|104117490&offsets|0|0]|GTGAAATGTTTAGGACCACTAG</td><td>>MI0000283|hsa-mir-203a&WithFlank&14|+|104117399|104117520</td><td>71</td><td>92</td><td>22</td><td>GTGAAATGTTTAGGACCACTAG</td><td>384.7</td><td>4.4</td><td>0.3</td><td>14.5</td><td>1.4E-47</td><td>7.1E-46</td><td>BT20/MB468</td><td>NUC</td></tr>
<tr><td>>MI0000650|hsa-mir-200c&WithFlank&12|+|6963693|6963772@50.71.22&[MIMAT0000617&hsa-miR-200c-3p&offsets|0|-1;m-54&12|+|6963742|6963764&offsets|0|-1]|TAATACTGCCGGGTAATGATGG</td><td>>MI0000650|hsa-mir-200c&WithFlank&12|+|6963693|6963772</td><td>50</td><td>71</td><td>22</td><td>TAATACTGCCGGGTAATGATGG</td><td>37598.4</td><td>8.7</td><td>0.6</td><td>14.5</td><td>1.1E-47</td><td>7.2E-46</td><td>BT20/MB468</td><td>CYTO</td></tr>
<tr><td>>hg38_21_+_8433172_8446622_RNA45SN1_45S_50ntFlanks@11669.11706.38|GCCCAGTGCTCTGAATGTCAAAGTGAAGAAATTCAATG</td><td>>hg38_21_+_8433172_8446622_RNA45SN1_45S_50ntFlanks</td><td>11669</td><td>11706</td><td>38</td><td>GCCCAGTGCTCTGAATGTCAAAGTGAAGAAATTCAATG</td><td>786.0</td><td>7.2</td><td>0.5</td><td>14.5</td><td>1.1E-47</td><td>7.2E-46</td><td>BT20/MB468</td><td>CYTO</td></tr>
<tr><td>>hg38_21_+_8433172_8446622_RNA45SN1_45S_50ntFlanks@11209.11232.24|CGGCGACTCTGGACGCGAGCCGGG</td><td>>hg38_21_+_8433172_8446622_RNA45SN1_45S_50ntFlanks</td><td>11209</td><td>11232</td><td>24</td><td>CGGCGACTCTGGACGCGAGCCGGG</td><td>20018.7</td><td>-12.7</td><td>0.9</td><td>-14.7</td><td>5.3E-49</td><td>8.7E-46</td><td>MB231/MB468</td><td>MP</td></tr>
<tr><td>>hg38_21_+_8433172_8446622_RNA45SN1_45S_50ntFlanks@10678.10706.29|CGGTTCCGGCGGCGTCCGGTGAGCTCTCG</td><td>>hg38_21_+_8433172_8446622_RNA45SN1_45S_50ntFlanks</td><td>10678</td><td>10706</td><td>29</td><td>CGGTTCCGGCGGCGTCCGGTGAGCTCTCG</td><td>40396.0</td><td>-13.5</td><td>0.9</td><td>-14.7</td><td>6.0E-49</td><td>8.7E-46</td><td>MB231/MB468</td><td>MP</td></tr>
<tr><td>>hg38_21_+_8433172_8446622_RNA45SN1_45S_50ntFlanks@4759.4780.22|ATCAGATACCGTCGTAGTTCCG</td><td>>hg38_21_+_8433172_8446622_RNA45SN1_45S_50ntFlanks</td><td>4759</td><td>4780</td><td>22</td><td>ATCAGATACCGTCGTAGTTCCG</td><td>19713.1</td><td>-18.7</td><td>1.3</td><td>-14.5</td><td>1.8E-47</td><td>8.9E-46</td><td>BT20/MB468</td><td>NUC</td></tr>
<tr><td>>MI0000079|hsa-mir-23a&WithFlank&19|-|13836581|13836665@54.72.19&[MIMAT0000078&hsa-miR-23a-3p&offsets|+3|+1;m-31&19|-|13836595|13836615&offsets|+3|+1]|ACATTGCCAGGGATTTCCA</td><td>>MI0000079|hsa-mir-23a&WithFlank&19|-|13836581|13836665</td><td>54</td><td>72</td><td>19</td><td>ACATTGCCAGGGATTTCCA</td><td>15102.0</td><td>18.1</td><td>1.3</td><td>14.5</td><td>1.8E-47</td><td>1.1E-45</td><td>BT20/MB468</td><td>CYTO</td></tr>
<tr><td>>hg38_21_+_8433172_8446622_RNA45SN1_45S_50ntFlanks@7976.8011.36|GCGACCTCAGATCAGACGTGGCGACCCGCTGAATTT</td><td>>hg38_21_+_8433172_8446622_RNA45SN1_45S_50ntFlanks</td><td>7976</td><td>8011</td><td>36</td><td>GCGACCTCAGATCAGACGTGGCGACCCGCTGAATTT</td><td>2681.9</td><td>6.7</td><td>0.5</td><td>14.5</td><td>8.9E-48</td><td>1.2E-45</td><td>BT20/MB468</td><td>MITO</td></tr>
<tr><td>>hg38_21_+_8433172_8446622_RNA45SN1_45S_50ntFlanks@4003.4024.22|ATAACCTCGGGCCGATCGCACG</td><td>>hg38_21_+_8433172_8446622_RNA45SN1_45S_50ntFlanks</td><td>4003</td><td>4024</td><td>22</td><td>ATAACCTCGGGCCGATCGCACG</td><td>3471.3</td><td>-6.0</td><td>0.4</td><td>-14.5</td><td>2.3E-47</td><td>1.4E-45</td><td>BT20/MB468</td><td>CYTO</td></tr>
<tr><td>>MI0001445|hsa-mir-423&WithFlank&17|+|30117073|30117178@59.77.19(+1U)&[MIMAT0001340&hsa-miR-423-3p&offsets|0|-4(+1U);m-100&17|+|30117131|30117153&offsets|0|-4(+1U)]|AGCTCGGTCTGAGGCCCCTT</td><td>>MI0001445|hsa-mir-423&WithFlank&17|+|30117073|30117178</td><td>59</td><td>77</td><td>19(+1U)</td><td>AGCTCGGTCTGAGGCCCCTT</td><td>1427.5</td><td>8.5</td><td>0.6</td><td>14.5</td><td>1.1E-47</td><td>1.5E-45</td><td>BT20/MB468</td><td>MITO</td></tr>
<tr><td>>hg38_21_+_8433172_8446622_RNA45SN1_45S_50ntFlanks@4731.4751.21|AAGAACGAAAGTCGGAGGTTC</td><td>>hg38_21_+_8433172_8446622_RNA45SN1_45S_50ntFlanks</td><td>4731</td><td>4751</td><td>21</td><td>AAGAACGAAAGTCGGAGGTTC</td><td>2396.5</td><td>-5.8</td><td>0.4</td><td>-14.4</td><td>3.4E-47</td><td>1.7E-45</td><td>BT20/MB468</td><td>NUC</td></tr>
<tr><td>>hg38_21_+_8433172_8446622_RNA45SN1_45S_50ntFlanks@11729.11751.23|TAACTATGACTCTCTTAAGGTAG</td><td>>hg38_21_+_8433172_8446622_RNA45SN1_45S_50ntFlanks</td><td>11729</td><td>11751</td><td>23</td><td>TAACTATGACTCTCTTAAGGTAG</td><td>83275.0</td><td>-9.4</td><td>0.7</td><td>-14.5</td><td>1.2E-47</td><td>1.9E-45</td><td>MB231/MB468</td><td>NUC</td></tr>
<tr><td>>hg38_21_+_8433172_8446622_RNA45SN1_45S_50ntFlanks@6664.6681.18|TGGATCACTCGGCTCGTG</td><td>>hg38_21_+_8433172_8446622_RNA45SN1_45S_50ntFlanks</td><td>6664</td><td>6681</td><td>18</td><td>TGGATCACTCGGCTCGTG</td><td>2755.9</td><td>-7.4</td><td>0.5</td><td>-14.4</td><td>4.1E-47</td><td>2.0E-45</td><td>BT20/MB468</td><td>NUC</td></tr>
<tr><td>>MI0000266|hsa-mir-10a&WithFlank&17|-|48579832|48579953@28.49.22&[MIMAT0000253&hsa-miR-10a-5p&offsets|0|-1;m-18&17|-|48579905|48579926&offsets|0|0]|TACCCTGTAGATCCGAATTTGT</td><td>>MI0000266|hsa-mir-10a&WithFlank&17|-|48579832|48579953</td><td>28</td><td>49</td><td>22</td><td>TACCCTGTAGATCCGAATTTGT</td><td>47387.3</td><td>14.8</td><td>1.0</td><td>14.4</td><td>4.2E-47</td><td>2.5E-45</td><td>BT20/MB468</td><td>CYTO</td></tr>
<tr><td>>MI0000273|hsa-mir-183&WithFlank&7|-|129774899|129775020@34.56.23&[MIMAT0000261&hsa-miR-183-5p&offsets|+1|+2;m-41&7|-|129774966|129774987&offsets|0|+1]|ATGGCACTGGTAGAATTCACTGT</td><td>>MI0000273|hsa-mir-183&WithFlank&7|-|129774899|129775020</td><td>34</td><td>56</td><td>23</td><td>ATGGCACTGGTAGAATTCACTGT</td><td>3653.1</td><td>7.7</td><td>0.5</td><td>14.4</td><td>4.2E-47</td><td>2.5E-45</td><td>BT20/MB468</td><td>CYTO</td></tr>
<tr><td>>hg38_21_+_8433172_8446622_RNA45SN1_45S_50ntFlanks@7975.8009.35|CGCGACCTCAGATCAGACGTGGCGACCCGCTGAAT</td><td>>hg38_21_+_8433172_8446622_RNA45SN1_45S_50ntFlanks</td><td>7975</td><td>8009</td><td>35</td><td>CGCGACCTCAGATCAGACGTGGCGACCCGCTGAAT</td><td>172155.0</td><td>8.4</td><td>0.6</td><td>14.4</td><td>4.3E-47</td><td>2.6E-45</td><td>BT20/MB468</td><td>CYTO</td></tr>
<tr><td>>hg38_21_+_8433172_8446622_RNA45SN1_45S_50ntFlanks@10411.10431.21|CAGTCGGTCCTGAGAGATGGG</td><td>>hg38_21_+_8433172_8446622_RNA45SN1_45S_50ntFlanks</td><td>10411</td><td>10431</td><td>21</td><td>CAGTCGGTCCTGAGAGATGGG</td><td>11850.4</td><td>-18.0</td><td>1.2</td><td>-14.4</td><td>5.7E-47</td><td>2.7E-45</td><td>BT20/MB468</td><td>NUC</td></tr>
<tr><td>>hg38_21_+_8433172_8446622_RNA45SN1_45S_50ntFlanks@10346.10368.23|TCAAACGAGAACTTTGAAGGCCG</td><td>>hg38_21_+_8433172_8446622_RNA45SN1_45S_50ntFlanks</td><td>10346</td><td>10368</td><td>23</td><td>TCAAACGAGAACTTTGAAGGCCG</td><td>5117.3</td><td>-7.9</td><td>0.5</td><td>-14.4</td><td>5.8E-47</td><td>2.8E-45</td><td>BT20/MB468</td><td>NUC</td></tr>
<tr><td>>MI0000086|hsa-mir-28&WithFlank&3|+|188688775|188688872@60.81.22&[MIMAT0004502&hsa-miR-28-3p&offsets|0|0;m-28&3|+|188688834|188688855&offsets|0|0]|CACTAGATTGTGAGCTCCTGGA</td><td>>MI0000086|hsa-mir-28&WithFlank&3|+|188688775|188688872</td><td>60</td><td>81</td><td>22</td><td>CACTAGATTGTGAGCTCCTGGA</td><td>1392.9</td><td>4.1</td><td>0.3</td><td>14.4</td><td>6.8E-47</td><td>3.2E-45</td><td>BT20/MB468</td><td>NUC</td></tr>
<tr><td>>hg38_21_+_8433172_8446622_RNA45SN1_45S_50ntFlanks@9554.9573.20|CGTAGCGGTCCTGACGTGCA</td><td>>hg38_21_+_8433172_8446622_RNA45SN1_45S_50ntFlanks</td><td>9554</td><td>9573</td><td>20</td><td>CGTAGCGGTCCTGACGTGCA</td><td>25211.8</td><td>-7.9</td><td>0.5</td><td>-14.4</td><td>7.4E-47</td><td>3.5E-45</td><td>BT20/MB468</td><td>NUC</td></tr>
<tr><td>>MI0000809|hsa-mir-151a&WithFlank&8|-|140732558|140732659@53.74.22(+1A)&[MIMAT0000757&hsa-miR-151a-3p&offsets|0|+1(+1A);m-30&8|-|140732587|140732607&offsets|0|+1(+1A)]|CTAGACTGAAGCTCCTTGAGGAA</td><td>>MI0000809|hsa-mir-151a&WithFlank&8|-|140732558|140732659</td><td>53</td><td>74</td><td>22(+1A)</td><td>CTAGACTGAAGCTCCTTGAGGAA</td><td>11004.3</td><td>8.6</td><td>0.6</td><td>14.4</td><td>6.1E-47</td><td>3.6E-45</td><td>BT20/MB468</td><td>CYTO</td></tr>
<tr><td>>MI0005756|hsa-mir-934&WithFlank&X|+|136550872|136550966@21.42.22(+1A)&[MIMAT0004977&hsa-miR-934&offsets|0|0(+1A);m-943&X|+|136550892|136550913&offsets|0|0(+1A)]|TGTCTACTACTGGAGACACTGGA</td><td>>MI0005756|hsa-mir-934&WithFlank&X|+|136550872|136550966</td><td>21</td><td>42</td><td>22(+1A)</td><td>TGTCTACTACTGGAGACACTGGA</td><td>2431.8</td><td>8.1</td><td>0.6</td><td>14.4</td><td>6.1E-47</td><td>3.6E-45</td><td>BT20/MB468</td><td>CYTO</td></tr>
<tr><td>>MI0000299|hsa-mir-222&WithFlank&X|-|45747009|45747130@75.99.25&[MIMAT0000279&hsa-miR-222-3p&offsets|0|+4;m-140&X|-|45747033|45747056&offsets|0|+1]|AGCTACATCTGGCTACTGGGTCTCT</td><td>>MI0000299|hsa-mir-222&WithFlank&X|-|45747009|45747130</td><td>75</td><td>99</td><td>25</td><td>AGCTACATCTGGCTACTGGGTCTCT</td><td>21811.2</td><td>7.3</td><td>0.5</td><td>14.4</td><td>6.3E-47</td><td>3.7E-45</td><td>BT20/MB468</td><td>CYTO</td></tr>
<tr><td>>hg38_21_+_8433172_8446622_RNA45SN1_45S_50ntFlanks@9564.9596.33|CTGACGTGCAAATCGGTCGTCCGACCTGGGTAT</td><td>>hg38_21_+_8433172_8446622_RNA45SN1_45S_50ntFlanks</td><td>9564</td><td>9596</td><td>33</td><td>CTGACGTGCAAATCGGTCGTCCGACCTGGGTAT</td><td>37510.1</td><td>-7.6</td><td>0.5</td><td>-14.4</td><td>3.3E-47</td><td>4.1E-45</td><td>BT20/MB468</td><td>MITO</td></tr>
<tr><td>>MI0000748|hsa-mir-130b&WithFlank&22|+|21653298|21653391@19.40.22&[MIMAT0004680&hsa-miR-130b-5p&offsets|0|+1;m-234&22|+|21653316|21653337&offsets|0|0]|ACTCTTTCCCTGTTGCACTACT</td><td>>MI0000748|hsa-mir-130b&WithFlank&22|+|21653298|21653391</td><td>19</td><td>40</td><td>22</td><td>ACTCTTTCCCTGTTGCACTACT</td><td>1912.6</td><td>8.0</td><td>0.6</td><td>14.4</td><td>7.6E-47</td><td>4.4E-45</td><td>BT20/MB468</td><td>CYTO</td></tr>
<tr><td>>hg38_21_+_8433172_8446622_RNA45SN1_45S_50ntFlanks@12456.12488.33|CCAAGCGTTGGATTGTTCACCCACTAATAGGGA</td><td>>hg38_21_+_8433172_8446622_RNA45SN1_45S_50ntFlanks</td><td>12456</td><td>12488</td><td>33</td><td>CCAAGCGTTGGATTGTTCACCCACTAATAGGGA</td><td>9924.8</td><td>-9.5</td><td>0.7</td><td>-14.4</td><td>4.6E-47</td><td>6.8E-45</td><td>MB231/MB468</td><td>NUC</td></tr>
<tr><td>>hg38_21_+_8433172_8446622_RNA45SN1_45S_50ntFlanks@6698.6720.23|AGCTAGCTGCGAGAATTAATGTG</td><td>>hg38_21_+_8433172_8446622_RNA45SN1_45S_50ntFlanks</td><td>6698</td><td>6720</td><td>23</td><td>AGCTAGCTGCGAGAATTAATGTG</td><td>68437.2</td><td>-10.1</td><td>0.7</td><td>-14.3</td><td>1.8E-46</td><td>8.3E-45</td><td>BT20/MB468</td><td>NUC</td></tr>
<tr><td>>MI0001445|hsa-mir-423&WithFlank&17|+|30117073|30117178@59.81.23&[MIMAT0001340&hsa-miR-423-3p&offsets|0|0;m-100&17|+|30117131|30117153&offsets|0|0]|AGCTCGGTCTGAGGCCCCTCAGT</td><td>>MI0001445|hsa-mir-423&WithFlank&17|+|30117073|30117178</td><td>59</td><td>81</td><td>23</td><td>AGCTCGGTCTGAGGCCCCTCAGT</td><td>52263.1</td><td>8.4</td><td>0.6</td><td>14.4</td><td>7.0E-47</td><td>8.6E-45</td><td>BT20/MB468</td><td>MITO</td></tr>
<tr><td>>hg38_21_+_8433172_8446622_RNA45SN1_45S_50ntFlanks@10403.10424.22|ACATGGGTCAGTCGGTCCTGAG</td><td>>hg38_21_+_8433172_8446622_RNA45SN1_45S_50ntFlanks</td><td>10403</td><td>10424</td><td>22</td><td>ACATGGGTCAGTCGGTCCTGAG</td><td>14008.4</td><td>7.7</td><td>0.5</td><td>14.4</td><td>7.4E-47</td><td>8.8E-45</td><td>BT20/MB468</td><td>MITO</td></tr>
<tr><td>>hg38_21_+_8433172_8446622_RNA45SN1_45S_50ntFlanks@4734.4759.26|AACGAAAGTCGGAGGTTCGAAGACGA</td><td>>hg38_21_+_8433172_8446622_RNA45SN1_45S_50ntFlanks</td><td>4734</td><td>4759</td><td>26</td><td>AACGAAAGTCGGAGGTTCGAAGACGA</td><td>1863.0</td><td>-7.1</td><td>0.5</td><td>-14.3</td><td>2.0E-46</td><td>9.2E-45</td><td>BT20/MB468</td><td>NUC</td></tr>
<tr><td>>hg38_21_+_8433172_8446622_RNA45SN1_45S_50ntFlanks@6663.6689.27|GTGGATCACTCGGCTCGTGCGTCGATG</td><td>>hg38_21_+_8433172_8446622_RNA45SN1_45S_50ntFlanks</td><td>6663</td><td>6689</td><td>27</td><td>GTGGATCACTCGGCTCGTGCGTCGATG</td><td>7456.6</td><td>-8.7</td><td>0.6</td><td>-14.5</td><td>2.1E-47</td><td>1.1E-44</td><td>MB231/MB468</td><td>MITO</td></tr>
<tr><td>>hg38_21_+_8433172_8446622_RNA45SN1_45S_50ntFlanks@12476.12496.21|CCACTAATAGGGAACGTGAGC</td><td>>hg38_21_+_8433172_8446622_RNA45SN1_45S_50ntFlanks</td><td>12476</td><td>12496</td><td>21</td><td>CCACTAATAGGGAACGTGAGC</td><td>4689.9</td><td>-8.3</td><td>0.6</td><td>-14.3</td><td>2.5E-46</td><td>1.1E-44</td><td>BT20/MB468</td><td>NUC</td></tr>
<tr><td>>hg38_21_+_8433172_8446622_RNA45SN1_45S_50ntFlanks@6719.6744.26|TGAATTGCAGGACACATTGATCATCG</td><td>>hg38_21_+_8433172_8446622_RNA45SN1_45S_50ntFlanks</td><td>6719</td><td>6744</td><td>26</td><td>TGAATTGCAGGACACATTGATCATCG</td><td>7235.8</td><td>-8.2</td><td>0.6</td><td>-14.4</td><td>9.8E-47</td><td>1.4E-44</td><td>MB231/MB468</td><td>NUC</td></tr>
<tr><td>>MI0000063|hsa-let-7b&WithFlank&22|+|46113680|46113774@12.31.20(+1C)&[MIMAT0000063&hsa-let-7b-5p&offsets|0|-2(+1C);m-14&22|+|46113691|46113712&offsets|0|-2(+1C)]|TGAGGTAGTAGGTTGTGTGGC</td><td>>MI0000063|hsa-let-7b&WithFlank&22|+|46113680|46113774</td><td>12</td><td>31</td><td>20(+1C)</td><td>TGAGGTAGTAGGTTGTGTGGC</td><td>1851.3</td><td>8.6</td><td>0.6</td><td>14.3</td><td>2.6E-46</td><td>1.5E-44</td><td>BT20/MB468</td><td>CYTO</td></tr>
<tr><td>>hg38_21_+_8433172_8446622_RNA45SN1_45S_50ntFlanks@4722.4758.37|TCATTAATCAAGAACGAAAGTCGGAGGTTCGAAGACG</td><td>>hg38_21_+_8433172_8446622_RNA45SN1_45S_50ntFlanks</td><td>4722</td><td>4758</td><td>37</td><td>TCATTAATCAAGAACGAAAGTCGGAGGTTCGAAGACG</td><td>6831.0</td><td>-7.6</td><td>0.5</td><td>-14.3</td><td>1.4E-46</td><td>1.6E-44</td><td>BT20/MB468</td><td>MITO</td></tr>
</table>

<table>
<tr><td>>MI0000093|hsa-mir-92a-1&WithFlank&13|+|91351308|91351397@54.75.22;MI0000094|hsa-mir-92a-2&WithFlank&X|-|134169532|134169618@54.75.22&[MIMAT0000092&hsa-miR-92a-3p&offsets|0|0;m-9&13|+|91351361|91351382&offsets|0|0];[MIMAT0000092_1&hsa-miR-92a-3p&offsets|0|0;m-11&X|-|134169544|134169565&offsets|0|0]|TATTGCACTTGTCCCGGCCTGT</td><td>>MI0000093|hsa-mir-92a-1&WithFlank&13|+|91351308|91351397</td><td>54</td><td>75</td><td>22</td><td>TATTGCACTTGTCCCGGCCTGT</td><td>68135.1</td><td>6.7</td><td>0.5</td><td>14.3</td><td>3.1E-46</td><td>1.8E-44</td><td>BT20/MB468</td><td>CYTO</td></tr>
<tr><td>>MI0000078|hsa-mir-22&WithFlank&17|-|1713897|1713993@59.80.22&[MIMAT0000077&hsa-miR-22-3p&offsets|0|0;m-2&17|-|1713914|1713935&offsets|0|0]|AAGCTGCCAGTTGAAGAACTGT</td><td>>MI0000078|hsa-mir-22&WithFlank&17|-|1713897|1713993</td><td>59</td><td>80</td><td>22</td><td>AAGCTGCCAGTTGAAGAACTGT</td><td>10021.5</td><td>7.8</td><td>0.5</td><td>14.3</td><td>1.6E-46</td><td>1.8E-44</td><td>BT20/MB468</td><td>MITO</td></tr>
<tr><td>>am_MI0000809|hsa-mir-151a&WithFlank&8|-|140732558|140732659@53.71.19&[MIMAT0000757&hsa-miR-151a-3p&offsets|0|-2;m-30&8|-|140732587|140732607&offsets|0|-2]|CTAGACTGAAGCTCCTTGA</td><td>>am_MI0000809|hsa-mir-151a&WithFlank&8|-|140732558|140732659</td><td>53</td><td>71</td><td>19</td><td>CTAGACTGAAGCTCCTTGA</td><td>277.9</td><td>4.2</td><td>0.3</td><td>14.2</td><td>5.9E-46</td><td>2.7E-44</td><td>BT20/MB468</td><td>NUC</td></tr>
<tr><td>>hg38_21_+_8433172_8446622_RNA45SN1_45S_50ntFlanks@10680.10710.31|GTTCCGGCGGCGTCCGGTGAGCTCTCGCTGG</td><td>>hg38_21_+_8433172_8446622_RNA45SN1_45S_50ntFlanks</td><td>10680</td><td>10710</td><td>31</td><td>GTTCCGGCGGCGTCCGGTGAGCTCTCGCTGG</td><td>16125.1</td><td>-8.7</td><td>0.6</td><td>-14.3</td><td>2.0E-46</td><td>2.9E-44</td><td>MB231/MB468</td><td>NUC</td></tr>
<tr><td>>am_MI0005763|hsa-mir-941-1&WithFlank&20|+|63919443|63919526@53.74.22(+1U);MI0005764|hsa-mir-941-2&WithFlank&20|+|63919499|63919582@53.74.22(+1U);MI0005765|hsa-mir-941-3&WithFlank&20|+|63919555|63919638@53.74.22(+1U);MI0005766|hsa-mir-941-4&WithFlank&20|+|63919750|63919833@53.74.22(+1U);MI0031520|hsa-mir-941-5&WithFlank&20|+|63919862|63919945@53.74.22(+1U)&[MIMAT0004984&hsa-miR-941&offsets|0|-1(+1U);m-263&20|+|63919495|63919517&offsets|0|-1(+1U)];[MIMAT0004984_1&hsa-miR-941&offsets|0|-1(+1U);m-263&20|+|63919551|63919573&offsets|0|-1(+1U)];[MIMAT0004984_2&hsa-miR-941&offsets|0|-1(+1U);m-263&20|+|63919607|63919629&offsets|0|-1(+1U)];[MIMAT0004984_3&hsa-miR-941&offsets|0|-1(+1U);m-263&20|+|63919802|63919824&offsets|0|-1(+1U)];[MIMAT0004984_4&hsa-miR-941&offsets|0|-1(+1U);m-263&20|+|63919914|63919936&offsets|0|-1(+1U)]|CACCCGGCTGTGTGCACATGTGT</td><td>>am_MI0005763|hsa-mir-941-1&WithFlank&20|+|63919443|63919526</td><td>53</td><td>74</td><td>22(+1U)</td><td>CACCCGGCTGTGTGCACATGTGT</td><td>4500.1</td><td>8.8</td><td>0.6</td><td>14.3</td><td>2.9E-46</td><td>3.3E-44</td><td>BT20/MB468</td><td>MITO</td></tr>
<tr><td>>hg38_21_+_8433172_8446622_RNA45SN1_45S_50ntFlanks@4734.4758.25|AACGAAAGTCGGAGGTTCGAAGACG</td><td>>hg38_21_+_8433172_8446622_RNA45SN1_45S_50ntFlanks</td><td>4734</td><td>4758</td><td>25</td><td>AACGAAAGTCGGAGGTTCGAAGACG</td><td>3823.0</td><td>-7.0</td><td>0.5</td><td>-14.2</td><td>9.5E-46</td><td>4.3E-44</td><td>BT20/MB468</td><td>NUC</td></tr>
<tr><td>>MI0000272|hsa-mir-182&WithFlank&7|-|129770377|129770498@29.52.24&[MIMAT0000259&hsa-miR-182-5p&offsets|0|0;m-22&7|-|129770449|129770470&offsets|0|+2]|TTTGGCAATGGTAGAACTCACACT</td><td>>MI0000272|hsa-mir-182&WithFlank&7|-|129770377|129770498</td><td>29</td><td>52</td><td>24</td><td>TTTGGCAATGGTAGAACTCACACT</td><td>163789.5</td><td>8.3</td><td>0.6</td><td>14.2</td><td>8.3E-46</td><td>4.7E-44</td><td>BT20/MB468</td><td>CYTO</td></tr>
<tr><td>>hg38_21_+_8433172_8446622_RNA45SN1_45S_50ntFlanks@4046.4063.18|CATTCGAACGTCTGCCCT</td><td>>hg38_21_+_8433172_8446622_RNA45SN1_45S_50ntFlanks</td><td>4046</td><td>4063</td><td>18</td><td>CATTCGAACGTCTGCCCT</td><td>6109.0</td><td>-8.7</td><td>0.6</td><td>-14.2</td><td>1.1E-45</td><td>4.9E-44</td><td>BT20/MB468</td><td>NUC</td></tr>
<tr><td>>trnaMT_PheGAA_MT_+_577_647@38.56.19|AATGTTTAGACGGGCTCAC</td><td>>trnaMT_PheGAA_MT_+_577_647</td><td>38</td><td>56</td><td>19</td><td>AATGTTTAGACGGGCTCAC</td><td>1475.0</td><td>9.0</td><td>0.6</td><td>14.2</td><td>5.2E-46</td><td>5.8E-44</td><td>BT20/MB468</td><td>MITO</td></tr>
<tr><td>>MI0000272|hsa-mir-182&WithFlank&7|-|129770377|129770498@29.51.23(+1C)&[MIMAT0000259&hsa-miR-182-5p&offsets|0|-1(+1C);m-22&7|-|129770449|129770470&offsets|0|+1(+1C)]|TTTGGCAATGGTAGAACTCACACC</td><td>>MI0000272|hsa-mir-182&WithFlank&7|-|129770377|129770498</td><td>29</td><td>51</td><td>23(+1C)</td><td>TTTGGCAATGGTAGAACTCACACC</td><td>2326.8</td><td>3.4</td><td>0.2</td><td>14.2</td><td>1.8E-45</td><td>7.9E-44</td><td>BT20/MB468</td><td>NUC</td></tr>
<tr><td>>hg38_21_+_8433172_8446622_RNA45SN1_45S_50ntFlanks@12405.12432.28|CTTTTTGATCCTTCGATGTCGGCTCTTC</td><td>>hg38_21_+_8433172_8446622_RNA45SN1_45S_50ntFlanks</td><td>12405</td><td>12432</td><td>28</td><td>CTTTTTGATCCTTCGATGTCGGCTCTTC</td><td>12537.2</td><td>-8.6</td><td>0.6</td><td>-14.2</td><td>5.8E-46</td><td>8.0E-44</td><td>MB231/MB468</td><td>NUC</td></tr>
<tr><td>>hg38_21_+_8433172_8446622_RNA45SN1_45S_50ntFlanks@12627.12654.28|ATGGGGCGAAGCTACCATCTGTGGGATT</td><td>>hg38_21_+_8433172_8446622_RNA45SN1_45S_50ntFlanks</td><td>12627</td><td>12654</td><td>28</td><td>ATGGGGCGAAGCTACCATCTGTGGGATT</td><td>7519.5</td><td>-8.7</td><td>0.6</td><td>-14.2</td><td>6.1E-46</td><td>8.2E-44</td><td>MB231/MB468</td><td>NUC</td></tr>
<tr><td>>MI0000272|hsa-mir-182&WithFlank&7|-|129770377|129770498@29.54.26&[MIMAT0000259&hsa-miR-182-5p&offsets|0|+2;m-22&7|-|129770449|129770470&offsets|0|+4]|TTTGGCAATGGTAGAACTCACACTGG</td><td>>MI0000272|hsa-mir-182&WithFlank&7|-|129770377|129770498</td><td>29</td><td>54</td><td>26</td><td>TTTGGCAATGGTAGAACTCACACTGG</td><td>8862.4</td><td>8.1</td><td>0.6</td><td>14.2</td><td>1.5E-45</td><td>8.3E-44</td><td>BT20/MB468</td><td>CYTO</td></tr>
<tr><td>>MI0000273|hsa-mir-183&WithFlank&7|-|129774899|129775020@33.54.22&[MIMAT0000261&hsa-miR-183-5p&offsets|0|0;m-41&7|-|129774966|129774987&offsets|-1|-1]|TATGGCACTGGTAGAATTCACT</td><td>>MI0000273|hsa-mir-183&WithFlank&7|-|129774899|129775020</td><td>33</td><td>54</td><td>22</td><td>TATGGCACTGGTAGAATTCACT</td><td>25373.1</td><td>7.3</td><td>0.5</td><td>14.2</td><td>1.6E-45</td><td>8.6E-44</td><td>BT20/MB468</td><td>CYTO</td></tr>
<tr><td>>MI0001445|hsa-mir-423&WithFlank&17|+|30117073|30117178@59.81.23(+1A)&[MIMAT0001340&hsa-miR-423-3p&offsets|0|0(+1A);m-100&17|+|30117131|30117153&offsets|0|0(+1A)]|AGCTCGGTCTGAGGCCCCTCAGTA</td><td>>MI0001445|hsa-mir-423&WithFlank&17|+|30117073|30117178</td><td>59</td><td>81</td><td>23(+1A)</td><td>AGCTCGGTCTGAGGCCCCTCAGTA</td><td>1964.7</td><td>8.2</td><td>0.6</td><td>14.2</td><td>1.6E-45</td><td>8.8E-44</td><td>BT20/MB468</td><td>CYTO</td></tr>
<tr><td>>MI0000273|hsa-mir-183&WithFlank&7|-|129774899|129775020@34.54.21&[MIMAT0000261&hsa-miR-183-5p&offsets|+1|0;m-41&7|-|129774966|129774987&offsets|0|-1]|ATGGCACTGGTAGAATTCACT</td><td>>MI0000273|hsa-mir-183&WithFlank&7|-|129774899|129775020</td><td>34</td><td>54</td><td>21</td><td>ATGGCACTGGTAGAATTCACT</td><td>7426.4</td><td>7.9</td><td>0.6</td><td>14.2</td><td>1.8E-45</td><td>9.7E-44</td><td>BT20/MB468</td><td>CYTO</td></tr>
<tr><td>>hg38_21_+_8433172_8446622_RNA45SN1_45S_50ntFlanks@11572.11605.34|CCGACTTAGAACTGGTGCGGACCAGGGGAATCCG</td><td>>hg38_21_+_8433172_8446622_RNA45SN1_45S_50ntFlanks</td><td>11572</td><td>11605</td><td>34</td><td>CCGACTTAGAACTGGTGCGGACCAGGGGAATCCG</td><td>5476.2</td><td>-7.4</td><td>0.5</td><td>-14.2</td><td>9.3E-46</td><td>1.2E-43</td><td>MB231/MB468</td><td>NUC</td></tr>
<tr><td>>MI0000767|hsa-mir-365a&WithFlank&16|+|14309279|14309377@62.83.22;MI0000769|hsa-mir-365b&WithFlank&17|+|31575405|31575527@74.95.22&[MIMAT0000710&hsa-miR-365a-3p&offsets|0|0;m-176&16|+|14309340|14309361&offsets|0|0];[MIMAT0022834&hsa-miR-365b-3p&offsets|0|0;m-175&17|+|31575478|31575499&offsets|0|0]|TAATGCCCCTAAAAATCCTTAT</td><td>>MI0000767|hsa-mir-365a&WithFlank&16|+|14309279|14309377</td><td>62</td><td>83</td><td>22</td><td>TAATGCCCCTAAAAATCCTTAT</td><td>3271.8</td><td>9.2</td><td>0.7</td><td>14.1</td><td>2.5E-45</td><td>1.3E-43</td><td>BT20/MB468</td><td>CYTO</td></tr>
<tr><td>>MI0000746|hsa-mir-99b&WithFlank&19|+|51692606|51692687@13.32.20&[MIMAT0000689&hsa-miR-99b-5p&offsets|0|-2;m-10&19|+|51692618|51692639&offsets|0|-2]|CACCCGTAGAACCGACCTTG</td><td>>MI0000746|hsa-mir-99b&WithFlank&19|+|51692606|51692687</td><td>13</td><td>32</td><td>20</td><td>CACCCGTAGAACCGACCTTG</td><td>6019.4</td><td>9.5</td><td>0.7</td><td>14.1</td><td>2.6E-45</td><td>1.4E-43</td><td>BT20/MB468</td><td>CYTO</td></tr>
<tr><td>>hg38_21_+_8433172_8446622_RNA45SN1_45S_50ntFlanks@12939.12956.18|CCATTCGTAGACGACCTG</td><td>>hg38_21_+_8433172_8446622_RNA45SN1_45S_50ntFlanks</td><td>12939</td><td>12956</td><td>18</td><td>CCATTCGTAGACGACCTG</td><td>1431.9</td><td>-5.5</td><td>0.4</td><td>-14.1</td><td>2.8E-45</td><td>1.5E-43</td><td>BT20/MB468</td><td>CYTO</td></tr>
<tr><td>>hg38_21_+_8433172_8446622_RNA45SN1_45S_50ntFlanks@12417.12443.27|TCGATGTCGGCTCTTCCTATCATTGTG</td><td>>hg38_21_+_8433172_8446622_RNA45SN1_45S_50ntFlanks</td><td>12417</td><td>12443</td><td>27</td><td>TCGATGTCGGCTCTTCCTATCATTGTG</td><td>6831.6</td><td>-8.0</td><td>0.6</td><td>-14.1</td><td>5.5E-45</td><td>2.4E-43</td><td>BT20/MB468</td><td>NUC</td></tr>
<tr><td>>MI0000088|hsa-mir-30a&WithFlank&6|-|71403545|71403627@54.73.20(+1U)&[MIMAT0000088&hsa-miR-30a-3p&offsets|+1|-1(+1U);m-38&6|-|71403554|71403575&offsets|+1|-1(+1U)]|TTTCAGTCGGATGTTTGCAGT</td><td>>MI0000088|hsa-mir-30a&WithFlank&6|-|71403545|71403627</td><td>54</td><td>73</td><td>20(+1U)</td><td>TTTCAGTCGGATGTTTGCAGT</td><td>3387.1</td><td>7.8</td><td>0.6</td><td>14.1</td><td>5.5E-45</td><td>2.9E-43</td><td>BT20/MB468</td><td>CYTO</td></tr>
<tr><td>>hg38_21_+_8433172_8446622_RNA45SN1_45S_50ntFlanks@13002.13028.27|CGATCTATTGAAAGTCAGCCCTCGACA</td><td>>hg38_21_+_8433172_8446622_RNA45SN1_45S_50ntFlanks</td><td>13002</td><td>13028</td><td>27</td><td>CGATCTATTGAAAGTCAGCCCTCGACA</td><td>8953.6</td><td>-17.6</td><td>1.2</td><td>-14.1</td><td>6.8E-45</td><td>3.0E-43</td><td>BT20/MB468</td><td>NUC</td></tr>
<tr><td>>MI0000272|hsa-mir-182&WithFlank&7|-|129770377|129770498@29.51.23&[MIMAT0000259&hsa-miR-182-5p&offsets|0|-1;m-22&7|-|129770449|129770470&offsets|0|+1]|TTTGGCAATGGTAGAACTCACAC</td><td>>MI0000272|hsa-mir-182&WithFlank&7|-|129770377|129770498</td><td>29</td><td>51</td><td>23</td><td>TTTGGCAATGGTAGAACTCACAC</td><td>6350.5</td><td>8.7</td><td>0.6</td><td>14.1</td><td>5.7E-45</td><td>3.0E-43</td><td>BT20/MB468</td><td>CYTO</td></tr>
<tr><td>>hg38_21_+_8433172_8446622_RNA45SN1_45S_50ntFlanks@12456.12484.29|CCAAGCGTTGGATTGTTCACCCACTAATA</td><td>>hg38_21_+_8433172_8446622_RNA45SN1_45S_50ntFlanks</td><td>12456</td><td>12484</td><td>29</td><td>CCAAGCGTTGGATTGTTCACCCACTAATA</td><td>27664.6</td><td>-19.2</td><td>1.4</td><td>-14.0</td><td>7.9E-45</td><td>3.4E-43</td><td>BT20/MB468</td><td>NUC</td></tr>
<tr><td>>hg38_21_+_8433172_8446622_RNA45SN1_45S_50ntFlanks@4728.4758.31|ATCAAGAACGAAAGTCGGAGGTTCGAAGACG</td><td>>hg38_21_+_8433172_8446622_RNA45SN1_45S_50ntFlanks</td><td>4728</td><td>4758</td><td>31</td><td>ATCAAGAACGAAAGTCGGAGGTTCGAAGACG</td><td>7666.0</td><td>-17.3</td><td>1.2</td><td>-14.0</td><td>9.2E-45</td><td>4.0E-43</td><td>BT20/MB468</td><td>NUC</td></tr>
<tr><td>>hg38_21_+_8433172_8446622_RNA45SN1_45S_50ntFlanks@11729.11749.21|TAACTATGACTCTCTTAAGGT</td><td>>hg38_21_+_8433172_8446622_RNA45SN1_45S_50ntFlanks</td><td>11729</td><td>11749</td><td>21</td><td>TAACTATGACTCTCTTAAGGT</td><td>7256.0</td><td>-6.4</td><td>0.5</td><td>-14.1</td><td>3.6E-45</td><td>4.6E-43</td><td>MB231/MB468</td><td>NUC</td></tr>
<tr><td>>MI0000299|hsa-mir-222&WithFlank&X|-|45747009|45747130@75.96.22&[MIMAT0000279&hsa-miR-222-3p&offsets|0|+1;m-140&X|-|45747033|45747056&offsets|0|-2]|AGCTACATCTGGCTACTGGGTC</td><td>>MI0000299|hsa-mir-222&WithFlank&X|-|45747009|45747130</td><td>75</td><td>96</td><td>22</td><td>AGCTACATCTGGCTACTGGGTC</td><td>1178.4</td><td>6.9</td><td>0.5</td><td>14.0</td><td>1.1E-44</td><td>5.5E-43</td><td>BT20/MB468</td><td>CYTO</td></tr>
<tr><td>>MI0000273|hsa-mir-183&WithFlank&7|-|129774899|129775020@34.54.21&[MIMAT0000261&hsa-miR-183-5p&offsets|+1|0;m-41&7|-|129774966|129774987&offsets|0|-1]|ATGGCACTGGTAGAATTCACT</td><td>>MI0000273|hsa-mir-183&WithFlank&7|-|129774899|129775020</td><td>34</td><td>54</td><td>21</td><td>ATGGCACTGGTAGAATTCACT</td><td>5721.7</td><td>8.0</td><td>0.6</td><td>14.1</td><td>6.0E-45</td><td>6.4E-43</td><td>BT20/MB468</td><td>MITO</td></tr>
<tr><td>>MI0001445|hsa-mir-423&WithFlank&17|+|30117073|30117178@59.80.22(+1A)&[MIMAT0001340&hsa-miR-423-3p&offsets|0|-1(+1A);m-100&17|+|30117131|30117153&offsets|0|-1(+1A)]|AGCTCGGTCTGAGGCCCCTCAGA</td><td>>MI0001445|hsa-mir-423&WithFlank&17|+|30117073|30117178</td><td>59</td><td>80</td><td>22(+1A)</td><td>AGCTCGGTCTGAGGCCCCTCAGA</td><td>1979.1</td><td>7.5</td><td>0.5</td><td>14.1</td><td>6.0E-45</td><td>6.4E-43</td><td>BT20/MB468</td><td>MITO</td></tr>
<tr><td>>MI0000734|hsa-mir-106b&WithFlank&7|-|100093987|100094080@58.78.21&[MIMAT0004672&hsa-miR-106b-3p&offsets|0|-1;m-87&7|-|100094003|100094023&offsets|0|0]|CCGCACTGTGGGTACTTGCTG</td><td>>MI0000734|hsa-mir-106b&WithFlank&7|-|100093987|100094080</td><td>58</td><td>78</td><td>21</td><td>CCGCACTGTGGGTACTTGCTG</td><td>2495.3</td><td>8.3</td><td>0.6</td><td>14.1</td><td>6.2E-45</td><td>6.5E-43</td><td>BT20/MB468</td><td>MITO</td></tr>
<tr><td>>hg38_21_+_8433172_8446622_RNA45SN1_45S_50ntFlanks@11572.11603.32|CCGACTTAGAACTGGTGCGGACCAGGGGAATC</td><td>>hg38_21_+_8433172_8446622_RNA45SN1_45S_50ntFlanks</td><td>11572</td><td>11603</td><td>32</td><td>CCGACTTAGAACTGGTGCGGACCAGGGGAATC</td><td>13253.2</td><td>-9.7</td><td>0.7</td><td>-14.2</td><td>6.1E-46</td><td>7.9E-43</td><td>MB231/MB468</td><td>MP</td></tr>
<tr><td>>MI0000299|hsa-mir-222&WithFlank&X|-|45747009|45747130@75.98.24(+1C)&[MIMAT0000279&hsa-miR-222-3p&offsets|0|+3(+1C);m-140&X|-|45747033|45747056&offsets|0|0(+1C)]|AGCTACATCTGGCTACTGGGTCTCC</td><td>>MI0000299|hsa-mir-222&WithFlank&X|-|45747009|45747130</td><td>75</td><td>98</td><td>24(+1C)</td><td>AGCTACATCTGGCTACTGGGTCTCC</td><td>4288.9</td><td>7.0</td><td>0.5</td><td>14.0</td><td>1.6E-44</td><td>8.1E-43</td><td>BT20/MB468</td><td>CYTO</td></tr>
<tr><td>>hg38_21_+_8433172_8446622_RNA45SN1_45S_50ntFlanks@7980.8004.25|CCTCAGATCAGACGTGGCGACCCGC</td><td>>hg38_21_+_8433172_8446622_RNA45SN1_45S_50ntFlanks</td><td>7980</td><td>8004</td><td>25</td><td>CCTCAGATCAGACGTGGCGACCCGC</td><td>470.7</td><td>-5.8</td><td>0.4</td><td>-14.0</td><td>2.1E-44</td><td>8.8E-43</td><td>BT20/MB468</td><td>NUC</td></tr>
<tr><td>>am_MI0000809|hsa-mir-151a&WithFlank&8|-|140732558|140732659@52.73.22&[MIMAT0000757&hsa-miR-151a-3p&offsets|-1|0;m-30&8|-|140732587|140732607&offsets|-1|0]|ACTAGACTGAAGCTCCTTGAGG</td><td>>am_MI0000809|hsa-mir-151a&WithFlank&8|-|140732558|140732659</td><td>52</td><td>73</td><td>22</td><td>ACTAGACTGAAGCTCCTTGAGG</td><td>3806.9</td><td>8.3</td><td>0.6</td><td>14.0</td><td>1.8E-44</td><td>9.0E-43</td><td>BT20/MB468</td><td>CYTO</td></tr>
<tr><td>>hg38_21_+_8433172_8446622_RNA45SN1_45S_50ntFlanks@12418.12444.27|CGATGTCGGCTCTTCCTATCATTGTGA</td><td>>hg38_21_+_8433172_8446622_RNA45SN1_45S_50ntFlanks</td><td>12418</td><td>12444</td><td>27</td><td>CGATGTCGGCTCTTCCTATCATTGTGA</td><td>7373.9</td><td>-8.1</td><td>0.6</td><td>-14.0</td><td>2.2E-44</td><td>9.2E-43</td><td>BT20/MB468</td><td>NUC</td></tr>
<tr><td>>trna116_GluCTC_1_-_145399233_145399304@1.34.34,trna59_GluCTC_1_+_249168447_249168518@1.34.34,trna71_GluCTC_1_-_161439189_161439260@1.34.34,trna74_GluCTC_1_-_161431809_161431880@1.34.34,trna77_GluCTC_1_-_161424398_161424469@1.34.34,trna77_GluCTC_6_+_28949976_28950047@1.34.34,trna80_GluCTC_1_-_161417018_161417089@1.34.34,trna87_GluCTC_6_-_126101393_126101464@1.34.34|TCCCTGGTGGTCTAGTGGTTAGGATTCGGCGCTC</td><td>>trna116_GluCTC_1_-_145399233_145399304</td><td>1</td><td>34</td><td>34</td><td>TCCCTGGTGGTCTAGTGGTTAGGATTCGGCGCTC</td><td>18510.8</td><td>8.9</td><td>0.6</td><td>14.0</td><td>2.1E-44</td><td>1.0E-42</td><td>BT20/MB468</td><td>CYTO</td></tr>
<tr><td>>hg38_21_+_8433172_8446622_RNA45SN1_45S_50ntFlanks@7982.8006.25|TCAGATCAGACGTGGCGACCCGCTG</td><td>>hg38_21_+_8433172_8446622_RNA45SN1_45S_50ntFlanks</td><td>7982</td><td>8006</td><td>25</td><td>TCAGATCAGACGTGGCGACCCGCTG</td><td>10707.3</td><td>-9.8</td><td>0.7</td><td>-14.1</td><td>2.2E-45</td><td>1.0E-42</td><td>MB231/MB468</td><td>MITO</td></tr>
<tr><td>>MI0001445|hsa-mir-423&WithFlank&17|+|30117073|30117178@58.78.21&[MIMAT0001340&hsa-miR-423-3p&offsets|-1|-3;m-100&17|+|30117131|30117153&offsets|-1|-3]|AAGCTCGGTCTGAGGCCCCTC</td><td>>MI0001445|hsa-mir-423&WithFlank&17|+|30117073|30117178</td><td>58</td><td>78</td><td>21</td><td>AAGCTCGGTCTGAGGCCCCTC</td><td>2550.6</td><td>8.3</td><td>0.6</td><td>14.0</td><td>1.0E-44</td><td>1.1E-42</td><td>BT20/MB468</td><td>MITO</td></tr>
</table>

<table>
<tr><th></th><th></th><th></th><th></th><th></th><th></th><th></th><th></th><th></th><th></th><th></th><th></th><th></th><th></th></tr>
<tr><td>>hg38_21_+_8433172_8446622_RNA45SN1_45S_50ntFlanks@4730.4758.29|CAAGAACGAAAGTCGGAGGTTCGAAGACG</td><td>>hg38_21_+_8433172_8446622_RNA45SN1_45S_50ntFlanks</td><td>4730</td><td>4758</td><td>29</td><td>CAAGAACGAAAGTCGGAGGTTCGAAGACG</td><td>6069.6</td><td>-8.2</td><td>0.6</td><td>-14.0</td><td>2.8E-44</td><td>1.2E-42</td><td>BT20/MB468</td><td>NUC</td></tr>
<tr><td>>hg38_21_+_8433172_8446622_RNA45SN1_45S_50ntFlanks@3985.4002.18|CGGCTTTGGTGACTCTAG</td><td>>hg38_21_+_8433172_8446622_RNA45SN1_45S_50ntFlanks</td><td>3985</td><td>4002</td><td>18</td><td>CGGCTTTGGTGACTCTAG</td><td>13594.7</td><td>-18.2</td><td>1.3</td><td>-13.9</td><td>3.2E-44</td><td>1.3E-42</td><td>BT20/MB468</td><td>NUC</td></tr>
<tr><td>>hg38_21_+_8433172_8446622_RNA45SN1_45S_50ntFlanks@12974.12991.18|CGTACGTAGCAGAGCAGC</td><td>>hg38_21_+_8433172_8446622_RNA45SN1_45S_50ntFlanks</td><td>12974</td><td>12991</td><td>18</td><td>CGTACGTAGCAGAGCAGC</td><td>10834.5</td><td>-8.3</td><td>0.6</td><td>-14.0</td><td>1.1E-44</td><td>1.4E-42</td><td>MB231/MB468</td><td>NUC</td></tr>
<tr><td>>hg38_21_+_8433172_8446622_RNA45SN1_45S_50ntFlanks@4724.4759.36|ATTAATCAAGAACGAAAGTCGGAGGTTCGAAGACGA</td><td>>hg38_21_+_8433172_8446622_RNA45SN1_45S_50ntFlanks</td><td>4724</td><td>4759</td><td>36</td><td>ATTAATCAAGAACGAAAGTCGGAGGTTCGAAGACGA</td><td>4745.2</td><td>-7.7</td><td>0.5</td><td>-14.0</td><td>1.1E-44</td><td>1.4E-42</td><td>MB231/MB468</td><td>NUC</td></tr>
<tr><td>>hg38_21_+_8433172_8446622_RNA45SN1_45S_50ntFlanks@10350.10377.28|ACGAGAACTTTGAAGGCCGAAGTGGAGA</td><td>>hg38_21_+_8433172_8446622_RNA45SN1_45S_50ntFlanks</td><td>10350</td><td>10377</td><td>28</td><td>ACGAGAACTTTGAAGGCCGAAGTGGAGA</td><td>4108.6</td><td>-6.1</td><td>0.4</td><td>-14.0</td><td>1.3E-44</td><td>1.6E-42</td><td>MB231/MB468</td><td>NUC</td></tr>
<tr><td>>MI0000269|hsa-mir-181a-2&WithFlank&9|+|124692436|124692557@83.104.22&[MIMAT0004558&hsa-miR-181a-2-3p&offsets|0|0;m-71&9|+|124692518|124692539&offsets|0|0]|ACCACTGACCGTTGACTGTACC</td><td>>MI0000269|hsa-mir-181a-2&WithFlank&9|+|124692436|124692557</td><td>83</td><td>104</td><td>22</td><td>ACCACTGACCGTTGACTGTACC</td><td>2657.5</td><td>8.2</td><td>0.6</td><td>13.9</td><td>3.4E-44</td><td>1.7E-42</td><td>BT20/MB468</td><td>CYTO</td></tr>
<tr><td>>hg38_21_+_8433172_8446622_RNA45SN1_45S_50ntFlanks@9568.9597.30|CGTGCAAATCGGTCGTCCGACCTGGGTATA</td><td>>hg38_21_+_8433172_8446622_RNA45SN1_45S_50ntFlanks</td><td>9568</td><td>9597</td><td>30</td><td>CGTGCAAATCGGTCGTCCGACCTGGGTATA</td><td>7546.9</td><td>-7.8</td><td>0.6</td><td>-14.0</td><td>2.4E-44</td><td>2.4E-42</td><td>BT20/MB468</td><td>MITO</td></tr>
<tr><td>>MI0000542|hsa-mir-320a&WithFlank&8|-|22244960|22245043@48.70.23(+1U)&[MIMAT0000510&hsa-miR-320a-3p&offsets|0|+1(+1U);m-60&8|-|22244975|22244996&offsets|0|+1(+1U)]|AAAAGCTGGGTTGAGAGGGCGAAT</td><td>>MI0000542|hsa-mir-320a&WithFlank&8|-|22244960|22245043</td><td>48</td><td>70</td><td>23(+1U)</td><td>AAAAGCTGGGTTGAGAGGGCGAAT</td><td>4874.5</td><td>9.9</td><td>0.7</td><td>14.0</td><td>2.4E-44</td><td>2.4E-42</td><td>BT20/MB468</td><td>MITO</td></tr>
<tr><td>>MI0000482|hsa-mir-185&WithFlank&22|+|20033133|20033226@21.42.22&[MIMAT0000455&hsa-miR-185-5p&offsets|0|0;m-145&22|+|20033153|20033174&offsets|0|0]|TGGAGAGAAAGGCAGTTCCTGA</td><td>>MI0000482|hsa-mir-185&WithFlank&22|+|20033133|20033226</td><td>21</td><td>42</td><td>22</td><td>TGGAGAGAAAGGCAGTTCCTGA</td><td>2188.8</td><td>7.3</td><td>0.5</td><td>14.0</td><td>2.6E-44</td><td>2.5E-42</td><td>BT20/MB468</td><td>MITO</td></tr>
<tr><td>>MI0000272|hsa-mir-182&WithFlank&7|-|129770377|129770498@29.50.22&[MIMAT0000259&hsa-miR-182-5p&offsets|0|-2;m-22&7|-|129770449|129770470&offsets|0|0]|TTTGGCAATGGTAGAACTCACA</td><td>>MI0000272|hsa-mir-182&WithFlank&7|-|129770377|129770498</td><td>29</td><td>50</td><td>22</td><td>TTTGGCAATGGTAGAACTCACA</td><td>9888.2</td><td>8.6</td><td>0.6</td><td>13.9</td><td>5.9E-44</td><td>2.9E-42</td><td>BT20/MB468</td><td>CYTO</td></tr>
<tr><td>>MI0001445|hsa-mir-423&WithFlank&17|+|30117073|30117178@23.45.23&[MIMAT0004748&hsa-miR-423-5p&offsets|0|0;m-141&17|+|30117095|30117117&offsets|0|0]|TGAGGGGCAGAGAGCGAGACTTT</td><td>>MI0001445|hsa-mir-423&WithFlank&17|+|30117073|30117178</td><td>23</td><td>45</td><td>23</td><td>TGAGGGGCAGAGAGCGAGACTTT</td><td>16829.4</td><td>9.5</td><td>0.7</td><td>13.9</td><td>6.3E-44</td><td>3.1E-42</td><td>BT20/MB468</td><td>CYTO</td></tr>
<tr><td>>hg38_21_+_8433172_8446622_RNA45SN1_45S_50ntFlanks@12008.12032.25|CCGCCGGTGAAATACCACTACTCTG</td><td>>hg38_21_+_8433172_8446622_RNA45SN1_45S_50ntFlanks</td><td>12008</td><td>12032</td><td>25</td><td>CCGCCGGTGAAATACCACTACTCTG</td><td>23463.8</td><td>-18.9</td><td>1.4</td><td>-13.9</td><td>7.7E-44</td><td>3.2E-42</td><td>BT20/MB468</td><td>NUC</td></tr>
<tr><td>>MI0000255|hsa-mir-30d&WithFlank&8|-|134804870|134804951@12.35.24&[MIMAT0000245&hsa-miR-30d-5p&offsets|0|+2;m-21&8|-|134804919|134804940&offsets|0|+2]|TGTAAACATCCCCGACTGGAAGCT</td><td>>MI0000255|hsa-mir-30d&WithFlank&8|-|134804870|134804951</td><td>12</td><td>35</td><td>24</td><td>TGTAAACATCCCCGACTGGAAGCT</td><td>70706.8</td><td>8.4</td><td>0.6</td><td>13.9</td><td>3.9E-44</td><td>3.8E-42</td><td>BT20/MB468</td><td>MITO</td></tr>
<tr><td>>hg38_21_+_8433172_8446622_RNA45SN1_45S_50ntFlanks@12456.12487.32|CCAAGCGTTGGATTGTTCACCCACTAATAGGG</td><td>>hg38_21_+_8433172_8446622_RNA45SN1_45S_50ntFlanks</td><td>12456</td><td>12487</td><td>32</td><td>CCAAGCGTTGGATTGTTCACCCACTAATAGGG</td><td>23134.2</td><td>-12.3</td><td>0.9</td><td>-14.0</td><td>9.0E-45</td><td>3.9E-42</td><td>MB231/MB468</td><td>MITO</td></tr>
<tr><td>>MI0000542|hsa-mir-320a&WithFlank&8|-|22244960|22245043@48.69.22(+1U)&[MIMAT0000510&hsa-miR-320a-3p&offsets|0|0(+1U);m-60&8|-|22244975|22244996&offsets|0|0(+1U)]|AAAAGCTGGGTTGAGAGGGCGAT</td><td>>MI0000542|hsa-mir-320a&WithFlank&8|-|22244960|22245043</td><td>48</td><td>69</td><td>22(+1U)</td><td>AAAAGCTGGGTTGAGAGGGCGAT</td><td>5528.6</td><td>10.1</td><td>0.7</td><td>13.9</td><td>4.5E-44</td><td>4.3E-42</td><td>BT20/MB468</td><td>MITO</td></tr>
<tr><td>>hg38_21_+_8433172_8446622_RNA45SN1_45S_50ntFlanks@4760.4780.21|TCAGATACCGTCGTAGTTCCG</td><td>>hg38_21_+_8433172_8446622_RNA45SN1_45S_50ntFlanks</td><td>4760</td><td>4780</td><td>21</td><td>TCAGATACCGTCGTAGTTCCG</td><td>8084.5</td><td>-17.4</td><td>1.3</td><td>-13.9</td><td>1.1E-43</td><td>4.4E-42</td><td>BT20/MB468</td><td>NUC</td></tr>
<tr><td>>MI0000272|hsa-mir-182&WithFlank&7|-|129770377|129770498@29.52.24(+1A)&[MIMAT0000259&hsa-miR-182-5p&offsets|0|0(+1A);m-22&7|-|129770449|129770470&offsets|0|+2(+1A)]|TTTGGCAATGGTAGAACTCACACTA</td><td>>MI0000272|hsa-mir-182&WithFlank&7|-|129770377|129770498</td><td>29</td><td>52</td><td>24(+1A)</td><td>TTTGGCAATGGTAGAACTCACACTA</td><td>4138.7</td><td>8.9</td><td>0.6</td><td>13.9</td><td>9.3E-44</td><td>4.5E-42</td><td>BT20/MB468</td><td>CYTO</td></tr>
<tr><td>>hg38_21_+_8433172_8446622_RNA45SN1_45S_50ntFlanks@12456.12484.29|CCAAGCGTTGGATTGTTCACCCACTAATA</td><td>>hg38_21_+_8433172_8446622_RNA45SN1_45S_50ntFlanks</td><td>12456</td><td>12484</td><td>29</td><td>CCAAGCGTTGGATTGTTCACCCACTAATA</td><td>12883.1</td><td>-9.2</td><td>0.7</td><td>-13.9</td><td>4.5E-44</td><td>5.3E-42</td><td>MB231/MB468</td><td>NUC</td></tr>
<tr><td>>hg38_21_+_8433172_8446622_RNA45SN1_45S_50ntFlanks@10402.10429.28|AACATGGGTCAGTCGGTCCTGAGAGATG</td><td>>hg38_21_+_8433172_8446622_RNA45SN1_45S_50ntFlanks</td><td>10402</td><td>10429</td><td>28</td><td>AACATGGGTCAGTCGGTCCTGAGAGATG</td><td>13515.3</td><td>-9.3</td><td>0.7</td><td>-13.8</td><td>1.3E-43</td><td>5.3E-42</td><td>BT20/MB468</td><td>NUC</td></tr>
<tr><td>>hg38_21_+_8433172_8446622_RNA45SN1_45S_50ntFlanks@8357.8379.23|AAAAGAACTTTGAAGAGAGAGTT</td><td>>hg38_21_+_8433172_8446622_RNA45SN1_45S_50ntFlanks</td><td>8357</td><td>8379</td><td>23</td><td>AAAAGAACTTTGAAGAGAGAGTT</td><td>91194.3</td><td>-11.5</td><td>0.8</td><td>-13.8</td><td>1.7E-43</td><td>6.8E-42</td><td>BT20/MB468</td><td>NUC</td></tr>
<tr><td>>MI0000811|hsa-mir-148b&WithFlank&12|+|54337210|54337320@69.90.22(+1U)&[MIMAT0000759&hsa-miR-148b-3p&offsets|0|0(+1U);m-77&12|+|54337278|54337299&offsets|0|0(+1U)]|TCAGTGCATCACAGAACTTTGTT</td><td>>MI0000811|hsa-mir-148b&WithFlank&12|+|54337210|54337320</td><td>69</td><td>90</td><td>22(+1U)</td><td>TCAGTGCATCACAGAACTTTGTT</td><td>6535.8</td><td>6.5</td><td>0.5</td><td>13.8</td><td>1.4E-43</td><td>6.9E-42</td><td>BT20/MB468</td><td>CYTO</td></tr>
<tr><td>>hg38_21_+_8433172_8446622_RNA45SN1_45S_50ntFlanks@7976.8010.35|GCGACCTCAGATCAGACGTGGCGACCCGCTGAATT</td><td>>hg38_21_+_8433172_8446622_RNA45SN1_45S_50ntFlanks</td><td>7976</td><td>8010</td><td>35</td><td>GCGACCTCAGATCAGACGTGGCGACCCGCTGAATT</td><td>2701.3</td><td>7.4</td><td>0.5</td><td>13.9</td><td>7.5E-44</td><td>7.0E-42</td><td>BT20/MB468</td><td>MITO</td></tr>
<tr><td>>MI0000650|hsa-mir-200c&WithFlank&12|+|6963693|6963772@50.72.23&[MIMAT0000617&hsa-miR-200c-3p&offsets|0|0;m-54&12|+|6963742|6963764&offsets|0|0]|TAATACTGCCGGGTAATGATGGA</td><td>>MI0000650|hsa-mir-200c&WithFlank&12|+|6963693|6963772</td><td>50</td><td>72</td><td>23</td><td>TAATACTGCCGGGTAATGATGGA</td><td>113585.7</td><td>9.8</td><td>0.7</td><td>14.1</td><td>2.3E-45</td><td>7.1E-42</td><td>BT20/MB231</td><td>MITO</td></tr>
<tr><td>>hg38_21_+_8433172_8446622_RNA45SN1_45S_50ntFlanks@11783.11803.21|CATGAATGGATGAACGAGATT</td><td>>hg38_21_+_8433172_8446622_RNA45SN1_45S_50ntFlanks</td><td>11783</td><td>11803</td><td>21</td><td>CATGAATGGATGAACGAGATT</td><td>17216.0</td><td>-7.5</td><td>0.5</td><td>-13.9</td><td>6.8E-44</td><td>7.8E-42</td><td>MB231/MB468</td><td>NUC</td></tr>
<tr><td>>hg38_21_+_8433172_8446622_RNA45SN1_45S_50ntFlanks@12627.12654.28|ATGGGGCGAAGCTACCATCTGTGGGATT</td><td>>hg38_21_+_8433172_8446622_RNA45SN1_45S_50ntFlanks</td><td>12627</td><td>12654</td><td>28</td><td>ATGGGGCGAAGCTACCATCTGTGGGATT</td><td>10539.6</td><td>-8.0</td><td>0.6</td><td>-13.9</td><td>9.3E-44</td><td>8.6E-42</td><td>BT20/MB468</td><td>MITO</td></tr>
<tr><td>>am_MI0000786|hsa-mir-378a&WithFlank&5|+|149732819|149732896@50.70.21(+1A)&[MIMAT0000732&hsa-miR-378a-3p&offsets|+1|0(+1A);m-44&5|+|149732867|149732888&offsets|+1|0(+1A)]|CTGGACTTGGAGTCAGAAGGCA</td><td>>am_MI0000786|hsa-mir-378a&WithFlank&5|+|149732819|149732896</td><td>50</td><td>70</td><td>21(+1A)</td><td>CTGGACTTGGAGTCAGAAGGCA</td><td>678.4</td><td>6.3</td><td>0.5</td><td>13.8</td><td>2.1E-43</td><td>1.0E-41</td><td>BT20/MB468</td><td>CYTO</td></tr>
<tr><td>>hg38_21_+_8433172_8446622_RNA45SN1_45S_50ntFlanks@4724.4758.35|ATTAATCAAGAACGAAAGTCGGAGGTTCGAAGACG</td><td>>hg38_21_+_8433172_8446622_RNA45SN1_45S_50ntFlanks</td><td>4724</td><td>4758</td><td>35</td><td>ATTAATCAAGAACGAAAGTCGGAGGTTCGAAGACG</td><td>36223.4</td><td>-11.2</td><td>0.8</td><td>-14.0</td><td>2.6E-44</td><td>1.1E-41</td><td>MB231/MB468</td><td>MITO</td></tr>
<tr><td>>hg38_21_+_8433172_8446622_RNA45SN1_45S_50ntFlanks@11783.11803.21|CATGAATGGATGAACGAGATT</td><td>>hg38_21_+_8433172_8446622_RNA45SN1_45S_50ntFlanks</td><td>11783</td><td>11803</td><td>21</td><td>CATGAATGGATGAACGAGATT</td><td>8644.3</td><td>-8.6</td><td>0.6</td><td>-14.0</td><td>1.0E-44</td><td>1.2E-41</td><td>MB231/MB468</td><td>MP</td></tr>
<tr><td>>MI0000081|hsa-mir-24-2&WithFlank&19|-|13836281|13836365@56.76.21(+1A);MI0000080|hsa-mir-24-1&WithFlank&9|+|95086015|95086094@50.70.21(+1A)&[MIMAT0000080&hsa-miR-24-3p&offsets|0|-1(+1A);m-36&19|-|13836288|13836310&offsets|0|-2(+1A)];[MIMAT0000080_1&hsa-miR-24-3p&offsets|0|-1(+1A);m-35&9|+|95086064|95086086&offsets|0|-2(+1A)]|TGGCTCAGTTCAGCAGGAACAA</td><td>>MI0000081|hsa-mir-24-2&WithFlank&19|-|13836281|13836365</td><td>56</td><td>76</td><td>21(+1A)</td><td>TGGCTCAGTTCAGCAGGAACAA</td><td>1513.3</td><td>5.9</td><td>0.4</td><td>13.8</td><td>2.4E-43</td><td>2.1E-41</td><td>BT20/MB468</td><td>MITO</td></tr>
<tr><td>>MI0003205|hsa-mir-532&WithFlank&X|+|50003142|50003244@26.46.21(+1C)&[MIMAT0002888&hsa-miR-532-5p&offsets|0|-1(+1C);m-65&X|+|50003167|50003188&offsets|0|-1(+1C)]|CATGCCTTGAGTGTAGGACCGC</td><td>>MI0003205|hsa-mir-532&WithFlank&X|+|50003142|50003244</td><td>26</td><td>46</td><td>21(+1C)</td><td>CATGCCTTGAGTGTAGGACCGC</td><td>849.6</td><td>6.1</td><td>0.4</td><td>13.8</td><td>4.6E-43</td><td>2.2E-41</td><td>BT20/MB468</td><td>CYTO</td></tr>
<tr><td>>MI0000438|hsa-mir-15b&WithFlank&3|+|160404582|160404691@26.47.22&[MIMAT0000417&hsa-miR-15b-5p&offsets|0|0;m-110&3|+|160404607|160404628&offsets|0|0]|TAGCAGCACATCATGGTTTACA</td><td>>MI0000438|hsa-mir-15b&WithFlank&3|+|160404582|160404691</td><td>26</td><td>47</td><td>22</td><td>TAGCAGCACATCATGGTTTACA</td><td>1739.5</td><td>6.7</td><td>0.5</td><td>13.8</td><td>5.1E-43</td><td>2.4E-41</td><td>BT20/MB468</td><td>CYTO</td></tr>
<tr><td>>am_MI0000809|hsa-mir-151a&WithFlank&8|-|140732558|140732659@53.73.21(+1U)&[MIMAT0000757&hsa-miR-151a-3p&offsets|0|0(+1U);m-30&8|-|140732587|140732607&offsets|0|0(+1U)]|CTAGACTGAAGCTCCTTGAGGT</td><td>>am_MI0000809|hsa-mir-151a&WithFlank&8|-|140732558|140732659</td><td>53</td><td>73</td><td>21(+1U)</td><td>CTAGACTGAAGCTCCTTGAGGT</td><td>2098.2</td><td>3.2</td><td>0.2</td><td>13.7</td><td>6.0E-43</td><td>2.4E-41</td><td>BT20/MB468</td><td>NUC</td></tr>
<tr><td>>hg38_21_+_8433172_8446622_RNA45SN1_45S_50ntFlanks@6661.6695.35|CGGTGGATCACTCGGCTCGTGCGTCGATGAAGAAC</td><td>>hg38_21_+_8433172_8446622_RNA45SN1_45S_50ntFlanks</td><td>6661</td><td>6695</td><td>35</td><td>CGGTGGATCACTCGGCTCGTGCGTCGATGAAGAAC</td><td>5607.1</td><td>7.0</td><td>0.5</td><td>13.8</td><td>2.8E-43</td><td>2.5E-41</td><td>BT20/MB468</td><td>MITO</td></tr>
<tr><td>>hg38_21_+_8433172_8446622_RNA45SN1_45S_50ntFlanks@11569.11603.35|CAGCCGACTTAGAACTGGTGCGGACCAGGGGAATC</td><td>>hg38_21_+_8433172_8446622_RNA45SN1_45S_50ntFlanks</td><td>11569</td><td>11603</td><td>35</td><td>CAGCCGACTTAGAACTGGTGCGGACCAGGGGAATC</td><td>19635.2</td><td>-10.0</td><td>0.7</td><td>-13.7</td><td>6.3E-43</td><td>2.5E-41</td><td>BT20/MB468</td><td>NUC</td></tr>
<tr><td>>hg38_21_+_8433172_8446622_RNA45SN1_45S_50ntFlanks@5472.5516.45|ACGGCCCTGGCGGAGCGCTGAGAAGACGGTCGAACTTGACTATCT</td><td>>hg38_21_+_8433172_8446622_RNA45SN1_45S_50ntFlanks</td><td>5472</td><td>5516</td><td>45</td><td>ACGGCCCTGGCGGAGCGCTGAGAAGACGGTCGAACTTGACTATCT</td><td>101253.5</td><td>11.0</td><td>0.8</td><td>14.0</td><td>8.4E-45</td><td>2.6E-41</td><td>MB231/MB468</td><td>CYTO</td></tr>
<tr><td>>MI0000440|hsa-mir-27b&WithFlank&9|+|95085439|95085547@67.87.21&[MIMAT0000419&hsa-miR-27b-3p&offsets|0|0;m-39&9|+|95085505|95085525&offsets|0|0]|TTCACAGTGGCTAAGTTCTGC</td><td>>MI0000440|hsa-mir-27b&WithFlank&9|+|95085439|95085547</td><td>67</td><td>87</td><td>21</td><td>TTCACAGTGGCTAAGTTCTGC</td><td>39383.0</td><td>7.3</td><td>0.5</td><td>13.8</td><td>2.9E-43</td><td>2.6E-41</td><td>BT20/MB468</td><td>MITO</td></tr>
<tr><td>>MI0000439|hsa-mir-23b&WithFlank&9|+|95085202|95085310@67.86.20&[MIMAT0000418&hsa-miR-23b-3p&offsets|+3|0;m-42&9|+|95085265|95085285&offsets|+3|+2]|ACATTGCCAGGGATTACCAC</td><td>>MI0000439|hsa-mir-23b&WithFlank&9|+|95085202|95085310</td><td>67</td><td>86</td><td>20</td><td>ACATTGCCAGGGATTACCAC</td><td>8529.9</td><td>17.3</td><td>1.3</td><td>13.7</td><td>6.3E-43</td><td>2.9E-41</td><td>BT20/MB468</td><td>CYTO</td></tr>
<tr><td>>MI0000102|hsa-mir-100&WithFlank&11|-|122152223|122152314@19.39.21&[MIMAT0000098&hsa-miR-100-5p&offsets|0|-1;m-27&11|-|122152276|122152296&offsets|0|0]|AACCCGTAGATCCGAACTTGT</td><td>>MI0000102|hsa-mir-100&WithFlank&11|-|122152223|122152314</td><td>19</td><td>39</td><td>21</td><td>AACCCGTAGATCCGAACTTGT</td><td>248700.7</td><td>16.4</td><td>1.2</td><td>14.0</td><td>1.5E-44</td><td>3.1E-41</td><td>MB231/MB468</td><td>CYTO</td></tr>
<tr><td>>trna111_HisGTG_1_-_147774845_147774916@-1G.31.32,trna118_HisGTG_1_-_145396881_145396952@-1G.31.32,trna16_HisGTG_1_+_146544773_146544844@-1G.31.32,trna1_HisGTG_15_+_45493349_45493420@-1G.31.32,trna21_HisGTG_1_+_147753471_147753542@-1G.31.32,trna33_HisGTG_6_+_27125906_27125977@-1G.31.32,trna7_HisGTG_9_-_14433938_14434009@-1G.31.32,trna8_HisGTG_15_-_45492611_45492682@-1G.31.32,trna9_HisGTG_15_-_45490804_45490875@-1G.31.32|GGCCGTGATCGTATAGTGGTTAGTACTCTGCG</td><td>>trna111_HisGTG_1_-_147774845_147774916</td><td>-1G</td><td>31</td><td>32</td><td>GGCCGTGATCGTATAGTGGTTAGTACTCTGCG</td><td>54269.8</td><td>-9.9</td><td>0.7</td><td>-13.7</td><td>8.0E-43</td><td>3.2E-41</td><td>BT20/MB468</td><td>NUC</td></tr>
<tr><td>>MI0003628|hsa-mir-615&WithFlank&12|+|54033944|54034051@67.86.20&[MIMAT0003283&hsa-miR-615-3p&offsets|0|-2;m-495&12|+|54034010|54034030&offsets|0|-1]|TCCGAGCCTGGGTCTCCCTC</td><td>>MI0003628|hsa-mir-615&WithFlank&12|+|54033944|54034051</td><td>67</td><td>86</td><td>20</td><td>TCCGAGCCTGGGTCTCCCTC</td><td>1508.5</td><td>8.3</td><td>0.6</td><td>13.8</td><td>4.1E-43</td><td>3.6E-41</td><td>BT20/MB468</td><td>MITO</td></tr>
<tr><td>>hg38_21_+_8433172_8446622_RNA45SN1_45S_50ntFlanks@6745.6771.27|ACACTTCGAACGCACTTGCGGCCCCGG</td><td>>hg38_21_+_8433172_8446622_RNA45SN1_45S_50ntFlanks</td><td>6745</td><td>6771</td><td>27</td><td>ACACTTCGAACGCACTTGCGGCCCCGG</td><td>11200.9</td><td>-9.8</td><td>0.7</td><td>-13.9</td><td>1.0E-43</td><td>3.9E-41</td><td>MB231/MB468</td><td>MITO</td></tr>
<tr><td>>MI0000272|hsa-mir-182&WithFlank&7|-|129770377|129770498@29.51.23(+1C)&[MIMAT0000259&hsa-miR-182-5p&offsets|0|-1(+1C);m-22&7|-|129770449|129770470&offsets|0|+1(+1C)]|TTTGGCAATGGTAGAACTCACACC</td><td>>MI0000272|hsa-mir-182&WithFlank&7|-|129770377|129770498</td><td>29</td><td>51</td><td>23(+1C)</td><td>TTTGGCAATGGTAGAACTCACACC</td><td>23898.1</td><td>8.5</td><td>0.6</td><td>13.7</td><td>9.6E-43</td><td>4.5E-41</td><td>BT20/MB468</td><td>CYTO</td></tr>
</table>

| | | | | | | | | | | | | | |
|---|---|---|---|---|---|---|---|---|---|---|---|---|---|
| >MI0000447\|hsa-mir-128-1&WithFlank&2\|+\|135665391\|135665484@56.77.22&[MIMAT0000424&hsa-miR-128-3p&offsets\|0\|+1;m-108&2\|+\|135665446\|135665466&offsets\|0\|+1]\|TCACAGTGAACCGGTCTCTTTT | >MI0000447\|hsa-mir-128-1&WithFlank&2\|+\|135665391\|135665484 | 56 | 77 | 22 | TCACAGTGAACCGGTCTCTTTT | 4063.5 | 5.8 | 0.4 | 13.7 | 9.8E-43 | 4.5E-41 | BT20/MB468 | CYTO |
| >hg38_21_+_8433172_8446622_RNA45SN1_45S_50ntFlanks@11727.11749.23\|AGTAACTATGACTCTCTTAAGGT | >hg38_21_+_8433172_8446622_RNA45SN1_45S_50ntFlanks | 11727 | 11749 | 23 | AGTAACTATGACTCTCTTAAGGT | 347.5 | -5.3 | 0.4 | -13.7 | 1.2E-42 | 4.8E-41 | BT20/MB468 | NUC |
| >MI0000066\|hsa-let-7e&WithFlank&19\|+\|51692780\|51692870@14.34.21&[MIMAT0000066&hsa-let-7e-5p&offsets\|0\|-1;m-50&19\|+\|51692793\|51692814&offsets\|0\|-1]\|TGAGGTAGGAGGTTGTATAGT | >MI0000066\|hsa-let-7e&WithFlank&19\|+\|51692780\|51692870 | 14 | 34 | 21 | TGAGGTAGGAGGTTGTATAGT | 6994.6 | 8.4 | 0.6 | 13.7 | 1.1E-42 | 4.9E-41 | BT20/MB468 | CYTO |
| >MI0000283\|hsa-mir-203a&WithFlank&14\|+\|104117399\|104117520@72.92.21&[MIMAT0000264&hsa-miR-203a-3p&offsets\|+1\|0;m-79&14\|+\|104117469\|104117490&offsets\|+1\|0]\|TGAAATGTTTAGGACCACTAG | >MI0000283\|hsa-mir-203a&WithFlank&14\|+\|104117399\|104117520 | 72 | 92 | 21 | TGAAATGTTTAGGACCACTAG | 448.9 | 4.2 | 0.3 | 13.7 | 1.3E-42 | 5.0E-41 | BT20/MB468 | NUC |
| >hg38_21_+_8433172_8446622_RNA45SN1_45S_50ntFlanks@8357.8378.22\|AAAAGAACTTTGAAGAGAGAGT | >hg38_21_+_8433172_8446622_RNA45SN1_45S_50ntFlanks | 8357 | 8378 | 22 | AAAAGAACTTTGAAGAGAGAGT | 31529.4 | -10.6 | 0.8 | -13.7 | 1.5E-42 | 6.0E-41 | BT20/MB468 | NUC |
| >hg38_21_+_8433172_8446622_RNA45SN1_45S_50ntFlanks@11786.11803.18\|GAATGGATGAACGAGATT | >hg38_21_+_8433172_8446622_RNA45SN1_45S_50ntFlanks | 11786 | 11803 | 18 | GAATGGATGAACGAGATT | 4010.3 | -9.1 | 0.7 | -13.7 | 1.6E-42 | 6.0E-41 | BT20/MB468 | NUC |
| >hg38_21_+_8433172_8446622_RNA45SN1_45S_50ntFlanks@7982.8004.23\|TCAGATCAGACGTGGCGACCCGC | >hg38_21_+_8433172_8446622_RNA45SN1_45S_50ntFlanks | 7982 | 8004 | 23 | TCAGATCAGACGTGGCGACCCGC | 462.2 | -6.1 | 0.4 | -13.7 | 1.6E-42 | 6.0E-41 | BT20/MB468 | NUC |
| >hg38_21_+_8433172_8446622_RNA45SN1_45S_50ntFlanks@8357.8377.21\|AAAAGAACTTTGAAGAGAGAG | >hg38_21_+_8433172_8446622_RNA45SN1_45S_50ntFlanks | 8357 | 8377 | 21 | AAAAGAACTTTGAAGAGAGAG | 10306.1 | -10.0 | 0.7 | -13.7 | 1.6E-42 | 6.1E-41 | BT20/MB468 | NUC |
| >am_trna128_GlyGCC_6_-_27870686_27870756@1.29.29,trna133_GlyCCC_1_-_16872434_16872504@1.29.29,trna18_GlyGCC_16_+_70822597_70822667@1.29.29,trna19_GlyGCC_16_+_70823410_70823480@1.29.29,trna19_GlyGCC_2_-_157257659_157257729@1.29.29,trna24_GlyGCC_16_-_70812942_70813012@1.29.29,trna25_GlyGCC_16_-_70812114_70812184@1.29.29,trna4_GlyCCC_1_+_17188416_17188486@1.29.29,trna5_GlyGCC_17_+_8029064_8029134@1.29.29,trna68_GlyGCC_1_-_161493637_161493707@1.29.29\|GCATTGGTGGTTCAGTGGTAGAATTCTCG | >am_trna128_GlyGCC_6_-_27870686_27870756 | 1 | 29 | 29 | GCATTGGTGGTTCAGTGGTAGAATTCTCG | 41059.8 | -11.0 | 0.8 | -13.7 | 1.6E-42 | 6.2E-41 | BT20/MB468 | NUC |
| >hg38_21_+_8433172_8446622_RNA45SN1_45S_50ntFlanks@11999.12031.33\|CCCTGCGGGCCGCCGGTGAAATACCACTACTCT | >hg38_21_+_8433172_8446622_RNA45SN1_45S_50ntFlanks | 11999 | 12031 | 33 | CCCTGCGGGCCGCCGGTGAAATACCACTACTCT | 7104.7 | -17.2 | 1.3 | -13.7 | 1.8E-42 | 6.9E-41 | BT20/MB468 | NUC |
| >MI0000063\|hsa-let-7b&WithFlank&22\|+\|46113680\|46113774@12.32.21(+1A)&[MIMAT0000063&hsa-let-7b-5p&offsets\|0\|-1(+1A);m-14&22\|+\|46113691\|46113712&offsets\|0\|-1(+1A)]\|TGAGGTAGTAGGTTGTGTGGTA | >MI0000063\|hsa-let-7b&WithFlank&22\|+\|46113680\|46113774 | 12 | 32 | 21(+1A) | TGAGGTAGTAGGTTGTGTGGTA | 1398.7 | 8.0 | 0.6 | 13.7 | 8.2E-43 | 7.0E-41 | BT20/MB468 | MITO |
| >hg38_21_+_8433172_8446622_RNA45SN1_45S_50ntFlanks@11572.11602.31\|CCGACTTAGAACTGGTGCGGACCAGGGGAAT | >hg38_21_+_8433172_8446622_RNA45SN1_45S_50ntFlanks | 11572 | 11602 | 31 | CCGACTTAGAACTGGTGCGGACCAGGGGAAT | 10176.9 | -8.3 | 0.6 | -13.9 | 8.0E-44 | 8.8E-41 | MB231/MB468 | MP |
| >MI0003628\|hsa-mir-615&WithFlank&12\|+\|54033944\|54034051@67.88.22&[MIMAT0003283&hsa-miR-615-3p&offsets\|0\|0;m-495&12\|+\|54034010\|54034030&offsets\|0\|+1]\|TCCGAGCCTGGGTCTCCCTCTT | >MI0003628\|hsa-mir-615&WithFlank&12\|+\|54033944\|54034051 | 67 | 88 | 22 | TCCGAGCCTGGGTCTCCCTCTT | 3489.8 | 7.4 | 0.5 | 13.7 | 1.1E-42 | 9.1E-41 | BT20/MB468 | MITO |
| >MI0000286\|hsa-mir-210&WithFlank&11\|-\|568083\|568204@72.93.22(+2A)&[MIMAT0000267&hsa-miR-210-3p&offsets\|0\|0(+2A);m-102&11\|-\|568112\|568133&offsets\|0\|0(+2A)]\|CTGTGCGTGTGACAGCGGCTGAAA | >MI0000286\|hsa-mir-210&WithFlank&11\|-\|568083\|568204 | 72 | 93 | 22(+2A) | CTGTGCGTGTGACAGCGGCTGAAA | 5755.6 | 9.8 | 0.7 | 13.7 | 1.2E-42 | 9.9E-41 | BT20/MB468 | MITO |
| >hg38_21_+_8433172_8446622_RNA45SN1_45S_50ntFlanks@12937.12956.20\|AACCATTCGTAGACGACCTG | >hg38_21_+_8433172_8446622_RNA45SN1_45S_50ntFlanks | 12937 | 12956 | 20 | AACCATTCGTAGACGACCTG | 9955.0 | -7.5 | 0.5 | -13.7 | 8.9E-43 | 1.0E-40 | MB231/MB468 | NUC |
| >MI0001445\|hsa-mir-423&WithFlank&17\|+\|30117073\|30117178@23.45.23&[MIMAT0004748&hsa-miR-423-5p&offsets\|0\|0;m-141&17\|+\|30117095\|30117117&offsets\|0\|0]\|TGAGGGGCAGAGAGCGAGACTTT | >MI0001445\|hsa-mir-423&WithFlank&17\|+\|30117073\|30117178 | 23 | 45 | 23 | TGAGGGGCAGAGAGCGAGACTTT | 3719.8 | 8.7 | 0.6 | 13.7 | 1.3E-42 | 1.0E-40 | BT20/MB468 | MITO |
| >hg38_21_+_8433172_8446622_RNA45SN1_45S_50ntFlanks@12163.12182.20\|ACTGGGGCGGTACACCTGTC | >hg38_21_+_8433172_8446622_RNA45SN1_45S_50ntFlanks | 12163 | 12182 | 20 | ACTGGGGCGGTACACCTGTC | 11938.2 | -11.7 | 0.8 | -13.9 | 1.1E-43 | 1.1E-40 | MB231/MB468 | MP |
| >MI0000255\|hsa-mir-30d&WithFlank&8\|-\|134804870\|134804951@12.34.23(+1A)&[MIMAT0000245&hsa-miR-30d-5p&offsets\|0\|+1(+1A);m-21&8\|-\|134804919\|134804940&offsets\|0\|+1(+1A)]\|TGTAAACATCCCCGACTGGAAGCA | >MI0000255\|hsa-mir-30d&WithFlank&8\|-\|134804870\|134804951 | 12 | 34 | 23(+1A) | TGTAAACATCCCCGACTGGAAGCA | 13126.0 | 9.7 | 0.7 | 13.6 | 2.5E-42 | 1.1E-40 | BT20/MB468 | CYTO |
| >hg38_21_+_8433172_8446622_RNA45SN1_45S_50ntFlanks@12633.12654.22\|CGAAGCTACCATCTGTGGGATT | >hg38_21_+_8433172_8446622_RNA45SN1_45S_50ntFlanks | 12633 | 12654 | 22 | CGAAGCTACCATCTGTGGGATT | 25938.4 | -18.5 | 1.3 | -13.8 | 1.3E-43 | 1.2E-40 | MB231/MB468 | MP |
| >hg38_21_+_8433172_8446622_RNA45SN1_45S_50ntFlanks@7975.8001.27\|CGCGACCTCAGATCAGACGTGGCGACC | >hg38_21_+_8433172_8446622_RNA45SN1_45S_50ntFlanks | 7975 | 8001 | 27 | CGCGACCTCAGATCAGACGTGGCGACC | 956.6 | 5.7 | 0.4 | 13.6 | 2.7E-42 | 1.2E-40 | BT20/MB468 | CYTO |
| >MI0000746\|hsa-mir-99b&WithFlank&19\|+\|51692606\|51692687@13.34.22(+1A)&[MIMAT0000689&hsa-miR-99b-5p&offsets\|0\|0(+1A);m-10&19\|+\|51692618\|51692639&offsets\|0\|0(+1A)]\|CACCCGTAGAACCGACCTTGCGA | >MI0000746\|hsa-mir-99b&WithFlank&19\|+\|51692606\|51692687 | 13 | 34 | 22(+1A) | CACCCGTAGAACCGACCTTGCGA | 5855.3 | 8.3 | 0.6 | 13.6 | 3.0E-42 | 1.3E-40 | BT20/MB468 | CYTO |
| >MI0000063\|hsa-let-7b&WithFlank&22\|+\|46113680\|46113774@12.32.21(+1A)&[MIMAT0000063&hsa-let-7b-5p&offsets\|0\|-1(+1A);m-14&22\|+\|46113691\|46113712&offsets\|0\|-1(+1A)]\|TGAGGTAGTAGGTTGTGTGGTA | >MI0000063\|hsa-let-7b&WithFlank&22\|+\|46113680\|46113774 | 12 | 32 | 21(+1A) | TGAGGTAGTAGGTTGTGTGGTA | 8193.8 | 9.9 | 0.7 | 13.6 | 3.1E-42 | 1.4E-40 | BT20/MB468 | CYTO |
| >MI0000272\|hsa-mir-182&WithFlank&7\|-\|129770377\|129770498@29.53.25(+1U)&[MIMAT0000259&hsa-miR-182-5p&offsets\|0\|+1(+1U);m-22&7\|-\|129770449\|129770470&offsets\|0\|+3(+1U)]\|TTTGGCAATGGTAGAACTCACACTGT | >MI0000272\|hsa-mir-182&WithFlank&7\|-\|129770377\|129770498 | 29 | 53 | 25(+1U) | TTTGGCAATGGTAGAACTCACACTGT | 4664.0 | 7.5 | 0.6 | 13.6 | 3.3E-42 | 1.4E-40 | BT20/MB468 | CYTO |
| >MI0000650\|hsa-mir-200c&WithFlank&12\|+\|6963693\|6963772@50.70.21(+1A)&[MIMAT0000617&hsa-miR-200c-3p&offsets\|0\|-2(+1A);m-54&12\|+\|6963742\|6963764&offsets\|0\|-2(+1A)]\|TAATACTGCCGGGTAATGATGA | >MI0000650\|hsa-mir-200c&WithFlank&12\|+\|6963693\|6963772 | 50 | 70 | 21(+1A) | TAATACTGCCGGGTAATGATGA | 6812.2 | 9.6 | 0.7 | 13.6 | 3.3E-42 | 1.4E-40 | BT20/MB468 | CYTO |
| >MI0000061\|hsa-let-7a-2&WithFlank&11\|-\|122146516\|122146599@11.33.23;MI0000062\|hsa-let-7a-3&WithFlank&22\|+\|46112743\|46112828@10.32.23;MI0000060\|hsa-let-7a-1&WithFlank&9\|+\|94175951\|94176042@12.34.23&[MIMAT0000062&hsa-let-7a-5p&offsets\|0\|+1;m-6&11\|-\|122146568\|122146589&offsets\|0\|+1];[MIMAT0000062_1&hsa-let-7a-5p&offsets\|0\|+1;m-5&22\|+\|46112752\|46112773&offsets\|0\|+1];[MIMAT0000062_2&hsa-let-7a-5p&offsets\|0\|+1;m-4&9\|+\|94175962\|94175983&offsets\|0\|+1]\|TGAGGTAGTAGGTTGTATAGTTT | >MI0000061\|hsa-let-7a-2&WithFlank&11\|-\|122146516\|122146599 | 11 | 33 | 23 | TGAGGTAGTAGGTTGTATAGTTT | 1571.2 | 7.3 | 0.5 | 13.7 | 2.0E-42 | 1.6E-40 | BT20/MB468 | MITO |
| >MI0000746\|hsa-mir-99b&WithFlank&19\|+\|51692606\|51692687@13.32.20&[MIMAT0000689&hsa-miR-99b-5p&offsets\|0\|-2;m-10&19\|+\|51692618\|51692639&offsets\|0\|-2]\|CACCCGTAGAACCGACCTTG | >MI0000746\|hsa-mir-99b&WithFlank&19\|+\|51692606\|51692687 | 13 | 32 | 20 | CACCCGTAGAACCGACCTTG | 2377.0 | 8.8 | 0.6 | 13.6 | 2.1E-42 | 1.7E-40 | BT20/MB468 | MITO |
| >hg38_21_+_8433172_8446622_RNA45SN1_45S_50ntFlanks@12935.12956.22\|TAAACCATTCGTAGACGACCTG | >hg38_21_+_8433172_8446622_RNA45SN1_45S_50ntFlanks | 12935 | 12956 | 22 | TAAACCATTCGTAGACGACCTG | 4264.3 | -7.4 | 0.5 | -13.7 | 1.6E-42 | 1.8E-40 | MB231/MB468 | NUC |
| >hg38_21_+_8433172_8446622_RNA45SN1_45S_50ntFlanks@7979.8006.28\|ACCTCAGATCAGACGTGGCGACCCGCTG | >hg38_21_+_8433172_8446622_RNA45SN1_45S_50ntFlanks | 7979 | 8006 | 28 | ACCTCAGATCAGACGTGGCGACCCGCTG | 8018.9 | -7.4 | 0.5 | -13.7 | 1.8E-42 | 1.9E-40 | MB231/MB468 | NUC |
| >MI0000093\|hsa-mir-92a-1&WithFlank&13\|+\|91351308\|91351397@54.74.21(+1C);MI0000094\|hsa-mir-92a-2&WithFlank&X\|-\|134169532\|134169618@54.74.21(+1C)&[MIMAT0000092&hsa-miR-92a-3p&offsets\|0\|-1(+1C);m-9&13\|+\|91351361\|91351382&offsets\|0\|-1(+1C)];[MIMAT0000092_1&hsa-miR-92a-3p&offsets\|0\|-1(+1C);m-11&X\|-\|134169544\|134169565&offsets\|0\|-1(+1C)]\|TATTGCACTTGTCCCGGCCTGC | >MI0000093\|hsa-mir-92a-1&WithFlank&13\|+\|91351308\|91351397 | 54 | 74 | 21(+1C) | TATTGCACTTGTCCCGGCCTGC | 9511.4 | 6.9 | 0.5 | 13.6 | 4.8E-42 | 2.1E-40 | BT20/MB468 | CYTO |
| >hg38_21_+_8433172_8446622_RNA45SN1_45S_50ntFlanks@12405.12434.30\|CTTTTTGATCCTTCGATGTCGGCTCTTCCT | >hg38_21_+_8433172_8446622_RNA45SN1_45S_50ntFlanks | 12405 | 12434 | 30 | CTTTTTGATCCTTCGATGTCGGCTCTTCCT | 47915.9 | -8.6 | 0.6 | -13.6 | 2.1E-42 | 2.2E-40 | MB231/MB468 | NUC |
| >hg38_21_+_8433172_8446622_RNA45SN1_45S_50ntFlanks@5493.5516.24\|GAAGACGGTCGAACTTGACTATCT | >hg38_21_+_8433172_8446622_RNA45SN1_45S_50ntFlanks | 5493 | 5516 | 24 | GAAGACGGTCGAACTTGACTATCT | 3147.3 | 8.2 | 0.6 | 13.6 | 5.3E-42 | 2.3E-40 | BT20/MB468 | CYTO |
| >hg38_21_+_8433172_8446622_RNA45SN1_45S_50ntFlanks@12405.12434.30\|CTTTTTGATCCTTCGATGTCGGCTCTTCCT | >hg38_21_+_8433172_8446622_RNA45SN1_45S_50ntFlanks | 12405 | 12434 | 30 | CTTTTTGATCCTTCGATGTCGGCTCTTCCT | 98894.9 | -10.9 | 0.8 | -13.6 | 6.1E-42 | 2.3E-40 | BT20/MB468 | NUC |
| >hg38_21_+_8433172_8446622_RNA45SN1_45S_50ntFlanks@12405.12434.30\|CTTTTTGATCCTTCGATGTCGGCTCTTCCT | >hg38_21_+_8433172_8446622_RNA45SN1_45S_50ntFlanks | 12405 | 12434 | 30 | CTTTTTGATCCTTCGATGTCGGCTCTTCCT | 15985.7 | -8.4 | 0.6 | -13.7 | 6.8E-43 | 2.5E-40 | MB231/MB468 | MITO |
| >hg38_21_+_8433172_8446622_RNA45SN1_45S_50ntFlanks@12476.12498.23\|CCACTAATAGGGAACGTGAGCTG | >hg38_21_+_8433172_8446622_RNA45SN1_45S_50ntFlanks | 12476 | 12498 | 23 | CCACTAATAGGGAACGTGAGCTG | 4881.1 | -16.7 | 1.2 | -13.6 | 6.8E-42 | 2.5E-40 | BT20/MB468 | NUC |
| >hg38_21_+_8433172_8446622_RNA45SN1_45S_50ntFlanks@10349.10377.29\|AACGAGAACTTTGAAGGCCGAAGTGGAGA | >hg38_21_+_8433172_8446622_RNA45SN1_45S_50ntFlanks | 10349 | 10377 | 29 | AACGAGAACTTTGAAGGCCGAAGTGGAGA | 3903.1 | -7.6 | 0.6 | -13.6 | 7.4E-42 | 2.7E-40 | BT20/MB468 | NUC |
| >hg38_21_+_8433172_8446622_RNA45SN1_45S_50ntFlanks@10410.10429.20\|TCAGTCGGTCCTGAGAGATG | >hg38_21_+_8433172_8446622_RNA45SN1_45S_50ntFlanks | 10410 | 10429 | 20 | TCAGTCGGTCCTGAGAGATG | 10668.6 | -7.2 | 0.5 | -13.6 | 2.7E-42 | 2.8E-40 | MB231/MB468 | NUC |
| >hg38_21_+_8433172_8446622_RNA45SN1_45S_50ntFlanks@10678.10711.34\|CGGTTCCGGCGGCGTCCGGTGAGCTCTCGCTGGC | >hg38_21_+_8433172_8446622_RNA45SN1_45S_50ntFlanks | 10678 | 10711 | 34 | CGGTTCCGGCGGCGTCCGGTGAGCTCTCGCTGGC | 92211.6 | -9.7 | 0.7 | -13.6 | 2.7E-42 | 2.8E-40 | MB231/MB468 | NUC |
| >trnaMT_PheGAA_MT_+_577_647@35.57.23\|GAAAATGTTTAGACGGGCTCACA | >trnaMT_PheGAA_MT_+_577_647 | 35 | 57 | 23 | GAAAATGTTTAGACGGGCTCACA | 4843.3 | 8.5 | 0.6 | 13.6 | 3.7E-42 | 2.9E-40 | BT20/MB468 | MITO |
| >hg38_21_+_8433172_8446622_RNA45SN1_45S_50ntFlanks@11570.11603.34\|AGCCGACTTAGAACTGGTGCGGACCAGGGGAATC | >hg38_21_+_8433172_8446622_RNA45SN1_45S_50ntFlanks | 11570 | 11603 | 34 | AGCCGACTTAGAACTGGTGCGGACCAGGGGAATC | 4191.2 | -7.7 | 0.6 | -13.5 | 8.7E-42 | 3.2E-40 | BT20/MB468 | NUC |
| >MI0000272\|hsa-mir-182&WithFlank&7\|-\|129770377\|129770498@29.51.23&[MIMAT0000259&hsa-miR-182-5p&offsets\|0\|-1;m-22&7\|-\|129770449\|129770470&offsets\|0\|+1]\|TTTGGCAATGGTAGAACTCACAC | >MI0000272\|hsa-mir-182&WithFlank&7\|-\|129770377\|129770498 | 29 | 51 | 23 | TTTGGCAATGGTAGAACTCACAC | 1896.9 | 8.0 | 0.6 | 13.6 | 4.2E-42 | 3.3E-40 | BT20/MB468 | MITO |

<table>
<tr><td></td><td></td><td></td><td></td><td></td><td></td><td></td><td></td><td></td><td></td><td></td><td></td><td></td><td></td></tr>
<tr><td>>MI0000746|hsa-mir-99b&WithFlank&19|+|51692606|51692687@13.34.22(+1U)&[MIMAT0000689&hsa-miR-99b-5p&offsets|0|0(+1U);m-10&19|+|51692618|51692639&offsets|0|0(+1U)]|CACCCGTAGAACCGACCTTGCGT</td><td>>MI0000746|hsa-mir-99b&WithFlank&19|+|51692606|51692687</td><td>13</td><td>34</td><td>22(+1U)</td><td>CACCCGTAGAACCGACCTTGCGT</td><td>6448.0</td><td>7.8</td><td>0.6</td><td>13.5</td><td>8.0E-42</td><td>3.4E-40</td><td>BT20/MB468</td><td>CYTO</td></tr>
<tr><td>>am_MI0000809|hsa-mir-151a&WithFlank&8|-|140732558|140732659@53.73.21(+1C)&[MIMAT0000757&hsa-miR-151a-3p&offsets|0|0(+1C);m-30&8|-|140732587|140732607&offsets|0|0(+1C)]|CTAGACTGAAGCTCCTTGAGGC</td><td>>am_MI0000809|hsa-mir-151a&WithFlank&8|-|140732558|140732659</td><td>53</td><td>73</td><td>21(+1C)</td><td>CTAGACTGAAGCTCCTTGAGGC</td><td>330.2</td><td>3.3</td><td>0.2</td><td>13.5</td><td>1.0E-41</td><td>3.7E-40</td><td>BT20/MB468</td><td>NUC</td></tr>
<tr><td>>hg38_21_+_8433172_8446622_RNA45SN1_45S_50ntFlanks@10347.10368.22|CAAACGAGAACTTTGAAGGCCG</td><td>>hg38_21_+_8433172_8446622_RNA45SN1_45S_50ntFlanks</td><td>10347</td><td>10368</td><td>22</td><td>CAAACGAGAACTTTGAAGGCCG</td><td>875.3</td><td>-7.0</td><td>0.5</td><td>-13.5</td><td>1.0E-41</td><td>3.7E-40</td><td>BT20/MB468</td><td>NUC</td></tr>
<tr><td>>hg38_21_+_8433172_8446622_RNA45SN1_45S_50ntFlanks@11729.11751.23|TAACTATGACTCTCTTAAGGTAG</td><td>>hg38_21_+_8433172_8446622_RNA45SN1_45S_50ntFlanks</td><td>11729</td><td>11751</td><td>23</td><td>TAACTATGACTCTCTTAAGGTAG</td><td>8704.1</td><td>-7.1</td><td>0.5</td><td>-13.6</td><td>4.9E-42</td><td>3.7E-40</td><td>BT20/MB468</td><td>MITO</td></tr>
<tr><td>>MI0000253|hsa-mir-148a&WithFlank&7|-|25949913|25949992@50.72.23&[MIMAT0000243&hsa-miR-148a-3p&offsets|0|+1;m-7&7|-|25949922|25949943&offsets|0|+1]|TCAGTGCACTACAGAACTTTGTC</td><td>>MI0000253|hsa-mir-148a&WithFlank&7|-|25949913|25949992</td><td>50</td><td>72</td><td>23</td><td>TCAGTGCACTACAGAACTTTGTC</td><td>3619.3</td><td>7.7</td><td>0.6</td><td>13.5</td><td>9.1E-42</td><td>3.8E-40</td><td>BT20/MB468</td><td>CYTO</td></tr>
<tr><td>>hg38_21_+_8433172_8446622_RNA45SN1_45S_50ntFlanks@3875.3898.24|ATACATGCCGACGGGCGCTGACCC</td><td>>hg38_21_+_8433172_8446622_RNA45SN1_45S_50ntFlanks</td><td>3875</td><td>3898</td><td>24</td><td>ATACATGCCGACGGGCGCTGACCC</td><td>5577.3</td><td>-6.8</td><td>0.5</td><td>-13.6</td><td>5.1E-42</td><td>3.9E-40</td><td>BT20/MB468</td><td>MITO</td></tr>
<tr><td>>hg38_21_+_8433172_8446622_RNA45SN1_45S_50ntFlanks@11768.11792.25|CTAATTAGTGACGCGCATGAATGGA</td><td>>hg38_21_+_8433172_8446622_RNA45SN1_45S_50ntFlanks</td><td>11768</td><td>11792</td><td>25</td><td>CTAATTAGTGACGCGCATGAATGGA</td><td>7699.8</td><td>-17.3</td><td>1.3</td><td>-13.5</td><td>1.1E-41</td><td>4.1E-40</td><td>BT20/MB468</td><td>NUC</td></tr>
<tr><td>>hg38_21_+_8433172_8446622_RNA45SN1_45S_50ntFlanks@9567.9596.30|ACGTGCAAATCGGTCGTCCGACCTGGGTAT</td><td>>hg38_21_+_8433172_8446622_RNA45SN1_45S_50ntFlanks</td><td>9567</td><td>9596</td><td>30</td><td>ACGTGCAAATCGGTCGTCCGACCTGGGTAT</td><td>7967.0</td><td>-17.4</td><td>1.3</td><td>-13.5</td><td>1.2E-41</td><td>4.2E-40</td><td>BT20/MB468</td><td>NUC</td></tr>
<tr><td>>hg38_21_+_8433172_8446622_RNA45SN1_45S_50ntFlanks@12456.12488.33|CCAAGCGTTGGATTGTTCACCCACTAATAGGGA</td><td>>hg38_21_+_8433172_8446622_RNA45SN1_45S_50ntFlanks</td><td>12456</td><td>12488</td><td>33</td><td>CCAAGCGTTGGATTGTTCACCCACTAATAGGGA</td><td>21561.8</td><td>-18.8</td><td>1.4</td><td>-13.5</td><td>1.2E-41</td><td>4.2E-40</td><td>BT20/MB468</td><td>NUC</td></tr>
<tr><td>>am_MI0000786|hsa-mir-378a&WithFlank&5|+|149732819|149732896@49.70.22(+1A)&[MIMAT0000732&hsa-miR-378a-3p&offsets|0|0(+1A);m-44&5|+|149732867|149732888&offsets|0|0(+1A)]|ACTGGACTTGGAGTCAGAAGGCA</td><td>>am_MI0000786|hsa-mir-378a&WithFlank&5|+|149732819|149732896</td><td>49</td><td>70</td><td>22(+1A)</td><td>ACTGGACTTGGAGTCAGAAGGCA</td><td>4694.7</td><td>5.8</td><td>0.4</td><td>13.5</td><td>1.1E-41</td><td>4.6E-40</td><td>BT20/MB468</td><td>CYTO</td></tr>
<tr><td>>hg38_21_+_8433172_8446622_RNA45SN1_45S_50ntFlanks@12995.13013.19|CTCGCTGCGATCTATTGAA</td><td>>hg38_21_+_8433172_8446622_RNA45SN1_45S_50ntFlanks</td><td>12995</td><td>13013</td><td>19</td><td>CTCGCTGCGATCTATTGAA</td><td>2256.2</td><td>-6.5</td><td>0.5</td><td>-13.5</td><td>1.4E-41</td><td>5.0E-40</td><td>BT20/MB468</td><td>NUC</td></tr>
<tr><td>>MI0000809|hsa-mir-151a&WithFlank&8|-|140732558|140732659@53.74.22(+1U)&[MIMAT0000757&hsa-miR-151a-3p&offsets|0|+1(+1U);m-30&8|-|140732587|140732607&offsets|0|+1(+1U)]|CTAGACTGAAGCTCCTTGAGGAT</td><td>>MI0000809|hsa-mir-151a&WithFlank&8|-|140732558|140732659</td><td>53</td><td>74</td><td>22(+1U)</td><td>CTAGACTGAAGCTCCTTGAGGAT</td><td>2631.7</td><td>8.6</td><td>0.6</td><td>13.6</td><td>6.8E-42</td><td>5.1E-40</td><td>BT20/MB468</td><td>MITO</td></tr>
<tr><td>>hg38_21_+_8433172_8446622_RNA45SN1_45S_50ntFlanks@11583.11611.29|CTGGTGCGGACCAGGGGAATCCGACTGTT</td><td>>hg38_21_+_8433172_8446622_RNA45SN1_45S_50ntFlanks</td><td>11583</td><td>11611</td><td>29</td><td>CTGGTGCGGACCAGGGGAATCCGACTGTT</td><td>4067.1</td><td>-7.1</td><td>0.5</td><td>-13.7</td><td>6.4E-43</td><td>5.5E-40</td><td>MB231/MB468</td><td>MP</td></tr>
<tr><td>>MI0006444|hsa-mir-1307&WithFlank&10|-|103394247|103394407@86.107.22(+1U)&[MIMAT0005951&hsa-miR-1307-3p&offsets|0|0(+1U);m-66&10|-|103394300|103394322&offsets|0|-1(+1U)]|ACTCGGCGTGGCGTCGGTCGTGT</td><td>>MI0006444|hsa-mir-1307&WithFlank&10|-|103394247|103394407</td><td>86</td><td>107</td><td>22(+1U)</td><td>ACTCGGCGTGGCGTCGGTCGTGT</td><td>1812.7</td><td>7.6</td><td>0.6</td><td>13.5</td><td>1.4E-41</td><td>5.9E-40</td><td>BT20/MB468</td><td>CYTO</td></tr>
<tr><td>>hg38_21_+_8433172_8446622_RNA45SN1_45S_50ntFlanks@12415.12432.18|CTTCGATGTCGGCTCTTC</td><td>>hg38_21_+_8433172_8446622_RNA45SN1_45S_50ntFlanks</td><td>12415</td><td>12432</td><td>18</td><td>CTTCGATGTCGGCTCTTC</td><td>11739.4</td><td>-8.1</td><td>0.6</td><td>-13.5</td><td>1.7E-41</td><td>6.1E-40</td><td>BT20/MB468</td><td>NUC</td></tr>
<tr><td>>hg38_21_+_8433172_8446622_RNA45SN1_45S_50ntFlanks@9504.9526.23|GTGAACTATGCCTGGGCAGGGCG</td><td>>hg38_21_+_8433172_8446622_RNA45SN1_45S_50ntFlanks</td><td>9504</td><td>9526</td><td>23</td><td>GTGAACTATGCCTGGGCAGGGCG</td><td>4095.8</td><td>-16.4</td><td>1.2</td><td>-13.5</td><td>2.2E-41</td><td>7.7E-40</td><td>BT20/MB468</td><td>NUC</td></tr>
<tr><td>>hg38_21_+_8433172_8446622_RNA45SN1_45S_50ntFlanks@10390.10424.35|GAACAGCAGTTGAACATGGGTCAGTCGGTCCTGAG</td><td>>hg38_21_+_8433172_8446622_RNA45SN1_45S_50ntFlanks</td><td>10390</td><td>10424</td><td>35</td><td>GAACAGCAGTTGAACATGGGTCAGTCGGTCCTGAG</td><td>7588.8</td><td>10.0</td><td>0.7</td><td>13.5</td><td>1.0E-41</td><td>7.7E-40</td><td>BT20/MB468</td><td>MITO</td></tr>
<tr><td>>hg38_21_+_8433172_8446622_RNA45SN1_45S_50ntFlanks@12459.12476.18|AGCGTTGGATTGTTCACC</td><td>>hg38_21_+_8433172_8446622_RNA45SN1_45S_50ntFlanks</td><td>12459</td><td>12476</td><td>18</td><td>AGCGTTGGATTGTTCACC</td><td>4359.4</td><td>-8.6</td><td>0.6</td><td>-13.5</td><td>2.3E-41</td><td>8.1E-40</td><td>BT20/MB468</td><td>NUC</td></tr>
<tr><td>>hg38_21_+_8433172_8446622_RNA45SN1_45S_50ntFlanks@6738.6771.34|ATCATCGACACTTCGAACGCACTTGCGGCCCCGG</td><td>>hg38_21_+_8433172_8446622_RNA45SN1_45S_50ntFlanks</td><td>6738</td><td>6771</td><td>34</td><td>ATCATCGACACTTCGAACGCACTTGCGGCCCCGG</td><td>6444.1</td><td>-17.1</td><td>1.3</td><td>-13.5</td><td>3.0E-41</td><td>1.0E-39</td><td>BT20/MB468</td><td>NUC</td></tr>
<tr><td>>hg38_21_+_8433172_8446622_RNA45SN1_45S_50ntFlanks@4003.4024.22|ATAACCTCGGGCCGATCGCACG</td><td>>hg38_21_+_8433172_8446622_RNA45SN1_45S_50ntFlanks</td><td>4003</td><td>4024</td><td>22</td><td>ATAACCTCGGGCCGATCGCACG</td><td>6653.8</td><td>-17.1</td><td>1.3</td><td>-13.4</td><td>3.2E-41</td><td>1.1E-39</td><td>BT20/MB468</td><td>NUC</td></tr>
<tr><td>>hg38_21_+_8433172_8446622_RNA45SN1_45S_50ntFlanks@6646.6670.25|TCGTACGACTCTTAGCGGTGGATCA</td><td>>hg38_21_+_8433172_8446622_RNA45SN1_45S_50ntFlanks</td><td>6646</td><td>6670</td><td>25</td><td>TCGTACGACTCTTAGCGGTGGATCA</td><td>4870.9</td><td>-7.3</td><td>0.5</td><td>-13.7</td><td>1.4E-42</td><td>1.1E-39</td><td>MB231/MB468</td><td>MP</td></tr>
<tr><td>>hg38_21_+_8433172_8446622_RNA45SN1_45S_50ntFlanks@10405.10424.20|ATGGGTCAGTCGGTCCTGAG</td><td>>hg38_21_+_8433172_8446622_RNA45SN1_45S_50ntFlanks</td><td>10405</td><td>10424</td><td>20</td><td>ATGGGTCAGTCGGTCCTGAG</td><td>1484.8</td><td>5.7</td><td>0.4</td><td>13.5</td><td>2.7E-41</td><td>1.1E-39</td><td>BT20/MB468</td><td>CYTO</td></tr>
<tr><td>>hg38_21_+_8433172_8446622_RNA45SN1_45S_50ntFlanks@9554.9571.18|CGTAGCGGTCCTGACGTG</td><td>>hg38_21_+_8433172_8446622_RNA45SN1_45S_50ntFlanks</td><td>9554</td><td>9571</td><td>18</td><td>CGTAGCGGTCCTGACGTG</td><td>321279.4</td><td>-10.5</td><td>0.8</td><td>-13.4</td><td>3.5E-41</td><td>1.2E-39</td><td>BT20/MB468</td><td>NUC</td></tr>
<tr><td>>MI0000098|hsa-mir-96&WithFlank&7|-|129774686|129774775@15.37.23&[MIMAT0000095&hsa-miR-96-5p&offsets|0|0;m-248&7|-|129774739|129774761&offsets|0|0]|TTTGGCACTAGCACATTTTTGCT</td><td>>MI0000098|hsa-mir-96&WithFlank&7|-|129774686|129774775</td><td>15</td><td>37</td><td>23</td><td>TTTGGCACTAGCACATTTTTGCT</td><td>4067.6</td><td>6.9</td><td>0.5</td><td>13.5</td><td>1.6E-41</td><td>1.2E-39</td><td>BT20/MB468</td><td>MITO</td></tr>
<tr><td>>hg38_21_+_8433172_8446622_RNA45SN1_45S_50ntFlanks@12437.12455.19|CATTGTGAAGCAGAATTCA</td><td>>hg38_21_+_8433172_8446622_RNA45SN1_45S_50ntFlanks</td><td>12437</td><td>12455</td><td>19</td><td>CATTGTGAAGCAGAATTCA</td><td>6827.8</td><td>-9.1</td><td>0.7</td><td>-13.5</td><td>1.2E-41</td><td>1.2E-39</td><td>MB231/MB468</td><td>NUC</td></tr>
<tr><td>>MI0003205|hsa-mir-532&WithFlank&X|+|50003142|50003244@26.47.22&[MIMAT0002888&hsa-miR-532-5p&offsets|0|0;m-65&X|+|50003167|50003188&offsets|0|0]|CATGCCTTGAGTGTAGGACCGT</td><td>>MI0003205|hsa-mir-532&WithFlank&X|+|50003142|50003244</td><td>26</td><td>47</td><td>22</td><td>CATGCCTTGAGTGTAGGACCGT</td><td>6410.0</td><td>6.0</td><td>0.4</td><td>13.4</td><td>3.2E-41</td><td>1.3E-39</td><td>BT20/MB468</td><td>CYTO</td></tr>
<tr><td>>am_MI0000786|hsa-mir-378a&WithFlank&5|+|149732819|149732896@49.70.22&[MIMAT0000732&hsa-miR-378a-3p&offsets|0|0;m-44&5|+|149732867|149732888&offsets|0|0]|ACTGGACTTGGAGTCAGAAGGC</td><td>>am_MI0000786|hsa-mir-378a&WithFlank&5|+|149732819|149732896</td><td>49</td><td>70</td><td>22</td><td>ACTGGACTTGGAGTCAGAAGGC</td><td>28763.2</td><td>5.5</td><td>0.4</td><td>13.4</td><td>3.3E-41</td><td>1.3E-39</td><td>BT20/MB468</td><td>CYTO</td></tr>
<tr><td>>MI0000542|hsa-mir-320a&WithFlank&8|-|22244960|22245043@48.71.24&[MIMAT0000510&hsa-miR-320a-3p&offsets|0|+2;m-60&8|-|22244975|22244996&offsets|0|+2]|AAAAGCTGGGTTGAGAGGGCGAAA</td><td>>MI0000542|hsa-mir-320a&WithFlank&8|-|22244960|22245043</td><td>48</td><td>71</td><td>24</td><td>AAAAGCTGGGTTGAGAGGGCGAAA</td><td>17406.1</td><td>12.3</td><td>0.9</td><td>13.4</td><td>3.6E-41</td><td>1.5E-39</td><td>BT20/MB468</td><td>CYTO</td></tr>
<tr><td>>hg38_21_+_8433172_8446622_RNA45SN1_45S_50ntFlanks@13002.13029.28|CGATCTATTGAAAGTCAGCCCTCGACAC</td><td>>hg38_21_+_8433172_8446622_RNA45SN1_45S_50ntFlanks</td><td>13002</td><td>13029</td><td>28</td><td>CGATCTATTGAAAGTCAGCCCTCGACAC</td><td>5975.1</td><td>-17.0</td><td>1.3</td><td>-13.4</td><td>4.5E-41</td><td>1.5E-39</td><td>BT20/MB468</td><td>NUC</td></tr>
<tr><td>>hg38_21_+_8433172_8446622_RNA45SN1_45S_50ntFlanks@8104.8143.40|CCGCGGCGGGGCGCGGGACATGTGGCGTACGGAAGACCCG</td><td>>hg38_21_+_8433172_8446622_RNA45SN1_45S_50ntFlanks</td><td>8104</td><td>8143</td><td>40</td><td>CCGCGGCGGGGCGCGGGACATGTGGCGTACGGAAGACCCG</td><td>38361.5</td><td>-13.5</td><td>1.0</td><td>-13.6</td><td>4.5E-42</td><td>1.5E-39</td><td>MB231/MB468</td><td>MITO</td></tr>
<tr><td>>MI0000087|hsa-mir-29a&WithFlank&7|-|130876741|130876816@48.68.21&[MIMAT0000086&hsa-miR-29a-3p&offsets|0|-1;m-26&7|-|130876749|130876769&offsets|0|0]|TAGCACCATCTGAAATCGGTT</td><td>>MI0000087|hsa-mir-29a&WithFlank&7|-|130876741|130876816</td><td>48</td><td>68</td><td>21</td><td>TAGCACCATCTGAAATCGGTT</td><td>6872.4</td><td>9.3</td><td>0.7</td><td>13.5</td><td>1.7E-41</td><td>1.7E-39</td><td>MB231/MB468</td><td>NUC</td></tr>
<tr><td>>hg38_21_+_8433172_8446622_RNA45SN1_45S_50ntFlanks@10687.10710.24|CGGCGTCCGGTGAGCTCTCGCTGG</td><td>>hg38_21_+_8433172_8446622_RNA45SN1_45S_50ntFlanks</td><td>10687</td><td>10710</td><td>24</td><td>CGGCGTCCGGTGAGCTCTCGCTGG</td><td>5155.8</td><td>-8.0</td><td>0.6</td><td>-13.5</td><td>1.8E-41</td><td>1.8E-39</td><td>MB231/MB468</td><td>NUC</td></tr>
<tr><td>>MI0000650|hsa-mir-200c&WithFlank&12|+|6963693|6963772@50.72.23(+1U)&[MIMAT0000617&hsa-miR-200c-3p&offsets|0|0(+1U);m-54&12|+|6963742|6963764&offsets|0|0(+1U)]|TAATACTGCCGGGTAATGATGGAT</td><td>>MI0000650|hsa-mir-200c&WithFlank&12|+|6963693|6963772</td><td>50</td><td>72</td><td>23(+1U)</td><td>TAATACTGCCGGGTAATGATGGAT</td><td>12286.4</td><td>8.9</td><td>0.7</td><td>13.5</td><td>2.6E-41</td><td>1.9E-39</td><td>BT20/MB468</td><td>MITO</td></tr>
<tr><td>>MI0000272|hsa-mir-182&WithFlank&7|-|129770377|129770498@29.52.24&[MIMAT0000259&hsa-miR-182-5p&offsets|0|0;m-22&7|-|129770449|129770470&offsets|0|+2]|TTTGGCAATGGTAGAACTCACACT</td><td>>MI0000272|hsa-mir-182&WithFlank&7|-|129770377|129770498</td><td>29</td><td>52</td><td>24</td><td>TTTGGCAATGGTAGAACTCACACT</td><td>33695.5</td><td>8.0</td><td>0.6</td><td>13.5</td><td>2.9E-41</td><td>2.1E-39</td><td>BT20/MB468</td><td>MITO</td></tr>
<tr><td>>hg38_21_+_8433172_8446622_RNA45SN1_45S_50ntFlanks@12420.12443.24|ATGTCGGCTCTTCCTATCATTGTG</td><td>>hg38_21_+_8433172_8446622_RNA45SN1_45S_50ntFlanks</td><td>12420</td><td>12443</td><td>24</td><td>ATGTCGGCTCTTCCTATCATTGTG</td><td>24356.8</td><td>-9.7</td><td>0.7</td><td>-13.4</td><td>6.6E-41</td><td>2.2E-39</td><td>BT20/MB468</td><td>NUC</td></tr>
<tr><td>>MI0001445|hsa-mir-423&WithFlank&17|+|30117073|30117178@59.80.22(+1C)&[MIMAT0001340&hsa-miR-423-3p&offsets|0|-1(+1C);m-100&17|+|30117131|30117153&offsets|0|-1(+1C)]|AGCTCGGTCTGAGGCCCCTCAGC</td><td>>MI0001445|hsa-mir-423&WithFlank&17|+|30117073|30117178</td><td>59</td><td>80</td><td>22(+1C)</td><td>AGCTCGGTCTGAGGCCCCTCAGC</td><td>6903.2</td><td>8.7</td><td>0.6</td><td>13.4</td><td>3.2E-41</td><td>2.2E-39</td><td>BT20/MB468</td><td>MITO</td></tr>
<tr><td>>MI0000102|hsa-mir-100&WithFlank&11|-|122152223|122152314@19.38.20(+1C)&[MIMAT0000098&hsa-miR-100-5p&offsets|0|-2(+1C);m-27&11|-|122152276|122152296&offsets|0|-1(+1C)]|AACCCGTAGATCCGAACTTGC</td><td>>MI0000102|hsa-mir-100&WithFlank&11|-|122152223|122152314</td><td>19</td><td>38</td><td>20(+1C)</td><td>AACCCGTAGATCCGAACTTGC</td><td>23463.5</td><td>18.4</td><td>1.4</td><td>13.5</td><td>2.3E-41</td><td>2.3E-39</td><td>MB231/MB468</td><td>NUC</td></tr>
<tr><td>>hg38_21_+_8433172_8446622_RNA45SN1_45S_50ntFlanks@475.505.31|CTTCGTGATCGATGTGGTGACGTCGTGCTCT</td><td>>hg38_21_+_8433172_8446622_RNA45SN1_45S_50ntFlanks</td><td>475</td><td>505</td><td>31</td><td>CTTCGTGATCGATGTGGTGACGTCGTGCTCT</td><td>10443.1</td><td>9.7</td><td>0.7</td><td>13.4</td><td>5.6E-41</td><td>2.3E-39</td><td>BT20/MB468</td><td>CYTO</td></tr>
<tr><td>>MI0000082|hsa-mir-25&WithFlank&7|-|100093554|100093649@58.79.22(+1U)&[MIMAT0000081&hsa-miR-25-3p&offsets|0|0(+1U);m-23&7|-|100093571|100093592&offsets|0|0(+1U)]|CATTGCACTTGTCTCGGTCTGAT</td><td>>MI0000082|hsa-mir-25&WithFlank&7|-|100093554|100093649</td><td>58</td><td>79</td><td>22(+1U)</td><td>CATTGCACTTGTCTCGGTCTGAT</td><td>2060.3</td><td>7.0</td><td>0.5</td><td>13.4</td><td>5.7E-41</td><td>2.3E-39</td><td>BT20/MB468</td><td>CYTO</td></tr>
<tr><td>>hg38_21_+_8433172_8446622_RNA45SN1_45S_50ntFlanks@10350.10378.29|ACGAGAACTTTGAAGGCCGAAGTGGAGAA</td><td>>hg38_21_+_8433172_8446622_RNA45SN1_45S_50ntFlanks</td><td>10350</td><td>10378</td><td>29</td><td>ACGAGAACTTTGAAGGCCGAAGTGGAGAA</td><td>1246.1</td><td>-5.4</td><td>0.4</td><td>-13.5</td><td>2.5E-41</td><td>2.4E-39</td><td>MB231/MB468</td><td>NUC</td></tr>
<tr><td>>hg38_21_+_8433172_8446622_RNA45SN1_45S_50ntFlanks@10687.10710.24|CGGCGTCCGGTGAGCTCTCGCTGG</td><td>>hg38_21_+_8433172_8446622_RNA45SN1_45S_50ntFlanks</td><td>10687</td><td>10710</td><td>24</td><td>CGGCGTCCGGTGAGCTCTCGCTGG</td><td>8453.1</td><td>-9.4</td><td>0.7</td><td>-13.6</td><td>7.5E-42</td><td>2.5E-39</td><td>MB231/MB468</td><td>MITO</td></tr>
<tr><td>>hg38_21_+_8433172_8446622_RNA45SN1_45S_50ntFlanks@9564.9597.34|CTGACGTGCAAATCGGTCGTCCGACCTGGGTATA</td><td>>hg38_21_+_8433172_8446622_RNA45SN1_45S_50ntFlanks</td><td>9564</td><td>9597</td><td>34</td><td>CTGACGTGCAAATCGGTCGTCCGACCTGGGTATA</td><td>17843.6</td><td>-6.9</td><td>0.5</td><td>-13.4</td><td>3.9E-41</td><td>2.7E-39</td><td>BT20/MB468</td><td>MITO</td></tr>
<tr><td>>MI0000650|hsa-mir-200c&WithFlank&12|+|6963693|6963772@50.72.23&[MIMAT0000617&hsa-miR-200c-3p&offsets|0|0;m-54&12|+|6963742|6963764&offsets|0|0]|TAATACTGCCGGGTAATGATGGA</td><td>>MI0000650|hsa-mir-200c&WithFlank&12|+|6963693|6963772</td><td>50</td><td>72</td><td>23</td><td>TAATACTGCCGGGTAATGATGGA</td><td>245214.0</td><td>8.8</td><td>0.7</td><td>13.4</td><td>6.8E-41</td><td>2.7E-39</td><td>BT20/MB468</td><td>CYTO</td></tr>
<tr><td>>hg38_21_+_8433172_8446622_RNA45SN1_45S_50ntFlanks@12418.12435.18|CGATGTCGGCTCTTCCTA</td><td>>hg38_21_+_8433172_8446622_RNA45SN1_45S_50ntFlanks</td><td>12418</td><td>12435</td><td>18</td><td>CGATGTCGGCTCTTCCTA</td><td>12421.1</td><td>-8.9</td><td>0.7</td><td>-13.5</td><td>1.1E-41</td><td>3.3E-39</td><td>MB231/MB468</td><td>MITO</td></tr>
<tr><td>>hg38_21_+_8433172_8446622_RNA45SN1_45S_50ntFlanks@11787.11804.18|AATGGATGAACGAGATTC</td><td>>hg38_21_+_8433172_8446622_RNA45SN1_45S_50ntFlanks</td><td>11787</td><td>11804</td><td>18</td><td>AATGGATGAACGAGATTC</td><td>14345.5</td><td>-8.8</td><td>0.6</td><td>-13.6</td><td>5.3E-42</td><td>3.8E-39</td><td>MB231/MB468</td><td>MP</td></tr>
<tr><td>>hg38_21_+_8433172_8446622_RNA45SN1_45S_50ntFlanks@12633.12653.21|CGAAGCTACCATCTGTGGGAT</td><td>>hg38_21_+_8433172_8446622_RNA45SN1_45S_50ntFlanks</td><td>12633</td><td>12653</td><td>21</td><td>CGAAGCTACCATCTGTGGGAT</td><td>19707.0</td><td>-18.1</td><td>1.3</td><td>-13.6</td><td>5.0E-42</td><td>3.8E-39</td><td>MB231/MB468</td><td>MP</td></tr>
<tr><td>>MI0000272|hsa-mir-182&WithFlank&7|-|129770377|129770498@29.53.25&[MIMAT0000259&hsa-miR-182-5p&offsets|0|+1;m-22&7|-|129770449|129770470&offsets|0|+3]|TTTGGCAATGGTAGAACTCACACTG</td><td>>MI0000272|hsa-mir-182&WithFlank&7|-|129770377|129770498</td><td>29</td><td>53</td><td>25</td><td>TTTGGCAATGGTAGAACTCACACTG</td><td>9142.5</td><td>8.0</td><td>0.6</td><td>13.4</td><td>5.6E-41</td><td>3.9E-39</td><td>BT20/MB468</td><td>MITO</td></tr>
<tr><td>>hg38_21_+_8433172_8446622_RNA45SN1_45S_50ntFlanks@4283.4317.35|CTTTAACGAGGATCCATTGGAGGGCAAGTCTGGTG</td><td>>hg38_21_+_8433172_8446622_RNA45SN1_45S_50ntFlanks</td><td>4283</td><td>4317</td><td>35</td><td>CTTTAACGAGGATCCATTGGAGGGCAAGTCTGGTG</td><td>11427.5</td><td>-17.9</td><td>1.3</td><td>-13.4</td><td>1.2E-40</td><td>3.9E-39</td><td>BT20/MB468</td><td>NUC</td></tr>
</table>

<table>
<tr><td>>hg38_21_+_8433172_8446622_RNA45SN1_45S_50ntFlanks@7973.8010.38|GACGCGACCTCAGATCAGACGTGGCGACCCGCTGAATT</td><td>>hg38_21_+_8433172_8446622_RNA45SN1_45S_50ntFlanks</td><td>7973</td><td>8010</td><td>38</td><td>GACGCGACCTCAGATCAGACGTGGCGACCCGCTGAATT</td><td>7884.9</td><td>6.6</td><td>0.5</td><td>13.4</td><td>6.0E-41</td><td>4.1E-39</td><td>BT20/MB468</td><td>MITO</td></tr>
<tr><td>>MI0001448|hsa-mir-425&WithFlank&3|-|49020142|49020240@20.43.24&[MIMAT0003393&hsa-miR-425-5p&offsets|0|+1;m-95&3|-|49020200|49020221&offsets|0|+2]|AATGACACGATCACTCCCGTTGAG</td><td>>MI0001448|hsa-mir-425&WithFlank&3|-|49020142|49020240</td><td>20</td><td>43</td><td>24</td><td>AATGACACGATCACTCCCGTTGAG</td><td>2568.0</td><td>7.2</td><td>0.5</td><td>13.4</td><td>1.1E-40</td><td>4.3E-39</td><td>BT20/MB468</td><td>CYTO</td></tr>
<tr><td>>trna152_ValCAC_6_-_27248049_27248121@1.32.32|GCTTCTGTAGTGTAGTGGTTATCACGTTCGCC</td><td>>trna152_ValCAC_6_-_27248049_27248121</td><td>1</td><td>32</td><td>32</td><td>GCTTCTGTAGTGTAGTGGTTATCACGTTCGCC</td><td>1112.4</td><td>6.4</td><td>0.5</td><td>13.4</td><td>1.1E-40</td><td>4.3E-39</td><td>BT20/MB468</td><td>CYTO</td></tr>
<tr><td>>MI0000767|hsa-mir-365a&WithFlank&16|+|14309279|14309377@62.84.23;MI0000769|hsa-mir-365b&WithFlank&17|+|31575405|31575527@74.96.23&[MIMAT0000710&hsa-miR-365a-3p&offsets|0|+1;m-176&16|+|14309340|14309361&offsets|0|+1];[MIMAT0022834&hsa-miR-365b-3p&offsets|0|+1;m-175&17|+|31575478|31575499&offsets|0|+1]|TAATGCCCCTAAAAATCCTTATT</td><td>>MI0000767|hsa-mir-365a&WithFlank&16|+|14309279|14309377</td><td>62</td><td>84</td><td>23</td><td>TAATGCCCCTAAAAATCCTTATT</td><td>182.0</td><td>3.1</td><td>0.2</td><td>13.3</td><td>1.3E-40</td><td>4.4E-39</td><td>BT20/MB468</td><td>NUC</td></tr>
<tr><td>>hg38_21_+_8433172_8446622_RNA45SN1_45S_50ntFlanks@10678.10706.29|CGGTTCCGGCGGCGTCCGGTGAGCTCTCG</td><td>>hg38_21_+_8433172_8446622_RNA45SN1_45S_50ntFlanks</td><td>10678</td><td>10706</td><td>29</td><td>CGGTTCCGGCGGCGTCCGGTGAGCTCTCG</td><td>17206.6</td><td>-9.1</td><td>0.7</td><td>-13.4</td><td>4.9E-41</td><td>4.6E-39</td><td>MB231/MB468</td><td>NUC</td></tr>
<tr><td>>MI0000266|hsa-mir-10a&WithFlank&17|-|48579832|48579953@28.49.22(+1A)&[MIMAT0000253&hsa-miR-10a-5p&offsets|0|-1(+1A);m-18&17|-|48579905|48579926&offsets|0|0(+1A)]|TACCCTGTAGATCCGAATTTGTA</td><td>>MI0000266|hsa-mir-10a&WithFlank&17|-|48579832|48579953</td><td>28</td><td>49</td><td>22(+1A)</td><td>TACCCTGTAGATCCGAATTTGTA</td><td>4782.3</td><td>16.5</td><td>1.2</td><td>13.4</td><td>1.2E-40</td><td>4.6E-39</td><td>BT20/MB468</td><td>CYTO</td></tr>
<tr><td>>hg38_21_+_8433172_8446622_RNA45SN1_45S_50ntFlanks@5487.5514.28|CGCTGAGAAGACGGTCGAACTTGACTAT</td><td>>hg38_21_+_8433172_8446622_RNA45SN1_45S_50ntFlanks</td><td>5487</td><td>5514</td><td>28</td><td>CGCTGAGAAGACGGTCGAACTTGACTAT</td><td>3424.3</td><td>-6.4</td><td>0.5</td><td>-13.4</td><td>7.2E-41</td><td>4.9E-39</td><td>BT20/MB468</td><td>MITO</td></tr>
<tr><td>>MI0000804|hsa-mir-328&WithFlank&16|-|67202315|67202401@54.75.22&[MIMAT0000752&hsa-miR-328-3p&offsets|0|0;m-160&16|-|67202327|67202348&offsets|0|0]|CTGGCCCTCTCTGCCCTTCCGT</td><td>>MI0000804|hsa-mir-328&WithFlank&16|-|67202315|67202401</td><td>54</td><td>75</td><td>22</td><td>CTGGCCCTCTCTGCCCTTCCGT</td><td>2359.6</td><td>10.1</td><td>0.8</td><td>13.3</td><td>1.3E-40</td><td>4.9E-39</td><td>BT20/MB468</td><td>CYTO</td></tr>
<tr><td>>MI0000342|hsa-mir-200b&WithFlank&1|+|1167098|1167204@63.84.22&[MIMAT0000318&hsa-miR-200b-3p&offsets|0|0;m-93&1|+|1167160|1167182&offsets|0|-1]|TAATACTGCCTGGTAATGATGA</td><td>>MI0000342|hsa-mir-200b&WithFlank&1|+|1167098|1167204</td><td>63</td><td>84</td><td>22</td><td>TAATACTGCCTGGTAATGATGA</td><td>1885.5</td><td>-6.1</td><td>0.5</td><td>-13.3</td><td>1.5E-40</td><td>5.0E-39</td><td>BT20/MB468</td><td>NUC</td></tr>
<tr><td>>am_trna146_GlnCTG_6_-_27515531_27515602@1.29.29,trna1_GlnCTG_6_+_18836402_18836473@1.29.29,trna3_GlnCTG_17_+_8023070_8023141@1.29.29,trna42_GlnCTG_6_+_27263212_27263283@1.29.29,trna49_GlnCTG_6_+_27487308_27487379@1.29.29,trna7_GlnCTG_15_-_66161400_66161471@1.29.29,trna99_GlnCTG_6_-_28909378_28909449@1.29.29|GGTTCCATGGTGTAATGGTTAGCACTCTG</td><td>>am_trna146_GlnCTG_6_-_27515531_27515602</td><td>1</td><td>29</td><td>29</td><td>GGTTCCATGGTGTAATGGTTAGCACTCTG</td><td>6532.5</td><td>-17.1</td><td>1.3</td><td>-13.3</td><td>1.6E-40</td><td>5.1E-39</td><td>BT20/MB468</td><td>NUC</td></tr>
<tr><td>>hg38_21_+_8433172_8446622_RNA45SN1_45S_50ntFlanks@10351.10369.19|CGAGAACTTTGAAGGCCGA</td><td>>hg38_21_+_8433172_8446622_RNA45SN1_45S_50ntFlanks</td><td>10351</td><td>10369</td><td>19</td><td>CGAGAACTTTGAAGGCCGA</td><td>1128.1</td><td>-6.5</td><td>0.5</td><td>-13.3</td><td>1.6E-40</td><td>5.3E-39</td><td>BT20/MB468</td><td>NUC</td></tr>
<tr><td>>hg38_21_+_8433172_8446622_RNA45SN1_45S_50ntFlanks@7974.8010.37|ACGCGACCTCAGATCAGACGTGGCGACCCGCTGAATT</td><td>>hg38_21_+_8433172_8446622_RNA45SN1_45S_50ntFlanks</td><td>7974</td><td>8010</td><td>37</td><td>ACGCGACCTCAGATCAGACGTGGCGACCCGCTGAATT</td><td>74351.0</td><td>6.7</td><td>0.5</td><td>13.4</td><td>8.2E-41</td><td>5.5E-39</td><td>BT20/MB468</td><td>MITO</td></tr>
<tr><td>>hg38_21_+_8433172_8446622_RNA45SN1_45S_50ntFlanks@11729.11748.20|TAACTATGACTCTCTTAAGG</td><td>>hg38_21_+_8433172_8446622_RNA45SN1_45S_50ntFlanks</td><td>11729</td><td>11748</td><td>20</td><td>TAACTATGACTCTCTTAAGG</td><td>15120.9</td><td>-9.7</td><td>0.7</td><td>-13.3</td><td>1.7E-40</td><td>5.5E-39</td><td>BT20/MB468</td><td>NUC</td></tr>
<tr><td>>hg38_21_+_8433172_8446622_RNA45SN1_45S_50ntFlanks@6734.6773.40|ATTGATCATCGACACTTCGAACGCACTTGCGGCCCCGGGT</td><td>>hg38_21_+_8433172_8446622_RNA45SN1_45S_50ntFlanks</td><td>6734</td><td>6773</td><td>40</td><td>ATTGATCATCGACACTTCGAACGCACTTGCGGCCCCGGGT</td><td>3801.7</td><td>7.8</td><td>0.6</td><td>13.4</td><td>8.9E-41</td><td>5.8E-39</td><td>BT20/MB468</td><td>MITO</td></tr>
<tr><td>>MI0000266|hsa-mir-10a&WithFlank&17|-|48579832|48579953@29.49.21&[MIMAT0000253&hsa-miR-10a-5p&offsets|+1|-1;m-18&17|-|48579905|48579926&offsets|+1|0]|ACCCTGTAGATCCGAATTTGT</td><td>>MI0000266|hsa-mir-10a&WithFlank&17|-|48579832|48579953</td><td>29</td><td>49</td><td>21</td><td>ACCCTGTAGATCCGAATTTGT</td><td>6160.7</td><td>16.8</td><td>1.3</td><td>13.3</td><td>1.8E-40</td><td>7.0E-39</td><td>BT20/MB468</td><td>CYTO</td></tr>
<tr><td>>hg38_21_+_8433172_8446622_RNA45SN1_45S_50ntFlanks@9554.9572.19|CGTAGCGGTCCTGACGTGC</td><td>>hg38_21_+_8433172_8446622_RNA45SN1_45S_50ntFlanks</td><td>9554</td><td>9572</td><td>19</td><td>CGTAGCGGTCCTGACGTGC</td><td>28506.4</td><td>-8.4</td><td>0.6</td><td>-13.3</td><td>2.2E-40</td><td>7.0E-39</td><td>BT20/MB468</td><td>NUC</td></tr>
<tr><td>>hg38_21_+_8433172_8446622_RNA45SN1_45S_50ntFlanks@10350.10380.31|ACGAGAACTTTGAAGGCCGAAGTGGAGAAGG</td><td>>hg38_21_+_8433172_8446622_RNA45SN1_45S_50ntFlanks</td><td>10350</td><td>10380</td><td>31</td><td>ACGAGAACTTTGAAGGCCGAAGTGGAGAAGG</td><td>714.4</td><td>-6.4</td><td>0.5</td><td>-13.3</td><td>2.3E-40</td><td>7.3E-39</td><td>BT20/MB468</td><td>NUC</td></tr>
<tr><td>>hg38_21_+_8433172_8446622_RNA45SN1_45S_50ntFlanks@11572.11603.32|CCGACTTAGAACTGGTGCGGACCAGGGGAATC</td><td>>hg38_21_+_8433172_8446622_RNA45SN1_45S_50ntFlanks</td><td>11572</td><td>11603</td><td>32</td><td>CCGACTTAGAACTGGTGCGGACCAGGGGAATC</td><td>11124.4</td><td>-6.9</td><td>0.5</td><td>-13.4</td><td>8.3E-41</td><td>7.7E-39</td><td>MB231/MB468</td><td>NUC</td></tr>
<tr><td>>hg38_21_+_8433172_8446622_RNA45SN1_45S_50ntFlanks@4002.4024.23|GATAACCTCGGGCCGATCGCACG</td><td>>hg38_21_+_8433172_8446622_RNA45SN1_45S_50ntFlanks</td><td>4002</td><td>4024</td><td>23</td><td>GATAACCTCGGGCCGATCGCACG</td><td>2386.7</td><td>-5.6</td><td>0.4</td><td>-13.3</td><td>2.0E-40</td><td>7.7E-39</td><td>BT20/MB468</td><td>CYTO</td></tr>
<tr><td>>hg38_21_+_8433172_8446622_RNA45SN1_45S_50ntFlanks@12004.12035.32|CGGGCCGCCGGTGAAATACCACTACTCTGATC</td><td>>hg38_21_+_8433172_8446622_RNA45SN1_45S_50ntFlanks</td><td>12004</td><td>12035</td><td>32</td><td>CGGGCCGCCGGTGAAATACCACTACTCTGATC</td><td>3728.8</td><td>8.6</td><td>0.6</td><td>13.3</td><td>1.2E-40</td><td>8.0E-39</td><td>BT20/MB468</td><td>MITO</td></tr>
<tr><td>>hg38_1_-_228634819_228635039_RNA5S12_5S_50ntFlanks@146.169.24|TGGGAATACCGGGTGCTGTAGGCT</td><td>>hg38_1_-_228634819_228635039_RNA5S12_5S_50ntFlanks</td><td>146</td><td>169</td><td>24</td><td>TGGGAATACCGGGTGCTGTAGGCT</td><td>2706.6</td><td>8.2</td><td>0.6</td><td>13.3</td><td>2.1E-40</td><td>8.1E-39</td><td>BT20/MB468</td><td>CYTO</td></tr>
<tr><td>>MI0001445|hsa-mir-423&WithFlank&17|+|30117073|30117178@59.81.23(+1U)&[MIMAT0001340&hsa-miR-423-3p&offsets|0|0(+1U);m-100&17|+|30117131|30117153&offsets|0|0(+1U)]|AGCTCGGTCTGAGGCCCCTCAGTT</td><td>>MI0001445|hsa-mir-423&WithFlank&17|+|30117073|30117178</td><td>59</td><td>81</td><td>23(+1U)</td><td>AGCTCGGTCTGAGGCCCCTCAGTT</td><td>4036.0</td><td>7.9</td><td>0.6</td><td>13.3</td><td>1.3E-40</td><td>8.6E-39</td><td>BT20/MB468</td><td>MITO</td></tr>
<tr><td>>hg38_21_+_8433172_8446622_RNA45SN1_45S_50ntFlanks@7975.8010.36|CGCGACCTCAGATCAGACGTGGCGACCCGCTGAATT</td><td>>hg38_21_+_8433172_8446622_RNA45SN1_45S_50ntFlanks</td><td>7975</td><td>8010</td><td>36</td><td>CGCGACCTCAGATCAGACGTGGCGACCCGCTGAATT</td><td>126164.3</td><td>8.5</td><td>0.6</td><td>13.3</td><td>2.3E-40</td><td>8.6E-39</td><td>BT20/MB468</td><td>CYTO</td></tr>
<tr><td>>MI0000061|hsa-let-7a-2&WithFlank&11|-|122146516|122146599@11.32.22;MI0000062|hsa-let-7a-3&WithFlank&22|+|46112743|46112828@10.31.22;MI0000060|hsa-let-7a-1&WithFlank&9|+|94175951|94176042@12.33.22&[MIMAT0000062&hsa-let-7a-5p&offsets|0|0;m-6&11|-|122146568|122146589&offsets|0|0];[MIMAT0000062_1&hsa-let-7a-5p&offsets|0|0;m-5&22|+|46112752|46112773&offsets|0|0];[MIMAT0000062_2&hsa-let-7a-5p&offsets|0|0;m-4&9|+|94175962|94175983&offsets|0|0]|TGAGGTAGTAGGTTGTATAGTT</td><td>>MI0000061|hsa-let-7a-2&WithFlank&11|-|122146516|122146599</td><td>11</td><td>32</td><td>22</td><td>TGAGGTAGTAGGTTGTATAGTT</td><td>64299.6</td><td>7.2</td><td>0.5</td><td>13.3</td><td>1.4E-40</td><td>9.0E-39</td><td>BT20/MB468</td><td>MITO</td></tr>
<tr><td>>hg38_21_+_8433172_8446622_RNA45SN1_45S_50ntFlanks@5444.5466.23|CTCGGATCGGCCCCGCCGGGGTC</td><td>>hg38_21_+_8433172_8446622_RNA45SN1_45S_50ntFlanks</td><td>5444</td><td>5466</td><td>23</td><td>CTCGGATCGGCCCCGCCGGGGTC</td><td>30600.4</td><td>-10.7</td><td>0.8</td><td>-13.5</td><td>3.1E-41</td><td>9.1E-39</td><td>MB231/MB468</td><td>MITO</td></tr>
<tr><td>>MI0000093|hsa-mir-92a-1&WithFlank&13|+|91351308|91351397@54.76.23&[MIMAT0000092&hsa-miR-92a-3p&offsets|0|+1;m-9&13|+|91351361|91351382&offsets|0|+1]|TATTGCACTTGTCCCGGCCTGTT</td><td>>MI0000093|hsa-mir-92a-1&WithFlank&13|+|91351308|91351397</td><td>54</td><td>76</td><td>23</td><td>TATTGCACTTGTCCCGGCCTGTT</td><td>4491.1</td><td>6.1</td><td>0.5</td><td>13.3</td><td>2.5E-40</td><td>9.2E-39</td><td>BT20/MB468</td><td>CYTO</td></tr>
<tr><td>>hg38_21_+_8433172_8446622_RNA45SN1_45S_50ntFlanks@12456.12476.21|CCAAGCGTTGGATTGTTCACC</td><td>>hg38_21_+_8433172_8446622_RNA45SN1_45S_50ntFlanks</td><td>12456</td><td>12476</td><td>21</td><td>CCAAGCGTTGGATTGTTCACC</td><td>50916.4</td><td>-10.5</td><td>0.8</td><td>-13.5</td><td>1.5E-41</td><td>1.0E-38</td><td>MB231/MB468</td><td>MP</td></tr>
<tr><td>>hg38_21_+_8433172_8446622_RNA45SN1_45S_50ntFlanks@11569.11603.35|CAGCCGACTTAGAACTGGTGCGGACCAGGGGAATC</td><td>>hg38_21_+_8433172_8446622_RNA45SN1_45S_50ntFlanks</td><td>11569</td><td>11603</td><td>35</td><td>CAGCCGACTTAGAACTGGTGCGGACCAGGGGAATC</td><td>18245.9</td><td>-9.9</td><td>0.7</td><td>-13.4</td><td>3.9E-41</td><td>1.1E-38</td><td>MB231/MB468</td><td>MITO</td></tr>
<tr><td>>hg38_21_+_8433172_8446622_RNA45SN1_45S_50ntFlanks@6721.6744.24|AATTGCAGGACACATTGATCATCG</td><td>>hg38_21_+_8433172_8446622_RNA45SN1_45S_50ntFlanks</td><td>6721</td><td>6744</td><td>24</td><td>AATTGCAGGACACATTGATCATCG</td><td>13090.3</td><td>-8.4</td><td>0.6</td><td>-13.3</td><td>1.2E-40</td><td>1.1E-38</td><td>MB231/MB468</td><td>NUC</td></tr>
<tr><td>>hg38_21_+_8433172_8446622_RNA45SN1_45S_50ntFlanks@9534.9563.30|AGGAAACTCTGGTGGAGGTCCGTAGCGGTC</td><td>>hg38_21_+_8433172_8446622_RNA45SN1_45S_50ntFlanks</td><td>9534</td><td>9563</td><td>30</td><td>AGGAAACTCTGGTGGAGGTCCGTAGCGGTC</td><td>32300.5</td><td>-9.7</td><td>0.7</td><td>-13.3</td><td>1.3E-40</td><td>1.1E-38</td><td>MB231/MB468</td><td>NUC</td></tr>
<tr><td>>hg38_21_+_8433172_8446622_RNA45SN1_45S_50ntFlanks@11575.11603.29|ACTTAGAACTGGTGCGGACCAGGGGAATC</td><td>>hg38_21_+_8433172_8446622_RNA45SN1_45S_50ntFlanks</td><td>11575</td><td>11603</td><td>29</td><td>ACTTAGAACTGGTGCGGACCAGGGGAATC</td><td>6254.9</td><td>-8.0</td><td>0.6</td><td>-13.3</td><td>1.4E-40</td><td>1.3E-38</td><td>MB231/MB468</td><td>NUC</td></tr>
<tr><td>>MI0000273|hsa-mir-183&WithFlank&7|-|129774899|129775020@34.53.20(+1C)&[MIMAT0000261&hsa-miR-183-5p&offsets|+1|-1(+1C);m-41&7|-|129774966|129774987&offsets|0|-2(+1C)]|ATGGCACTGGTAGAATTCACC</td><td>>MI0000273|hsa-mir-183&WithFlank&7|-|129774899|129775020</td><td>34</td><td>53</td><td>20(+1C)</td><td>ATGGCACTGGTAGAATTCACC</td><td>1054.5</td><td>8.1</td><td>0.6</td><td>13.3</td><td>3.5E-40</td><td>1.3E-38</td><td>BT20/MB468</td><td>CYTO</td></tr>
<tr><td>>MI0000078|hsa-mir-22&WithFlank&17|-|1713897|1713993@59.79.21(+1C)&[MIMAT0000077&hsa-miR-22-3p&offsets|0|-1(+1C);m-2&17|-|1713914|1713935&offsets|0|-1(+1C)]|AAGCTGCCAGTTGAAGAACTGC</td><td>>MI0000078|hsa-mir-22&WithFlank&17|-|1713897|1713993</td><td>59</td><td>79</td><td>21(+1C)</td><td>AAGCTGCCAGTTGAAGAACTGC</td><td>1337.5</td><td>7.8</td><td>0.6</td><td>13.3</td><td>2.1E-40</td><td>1.3E-38</td><td>BT20/MB468</td><td>MITO</td></tr>
<tr><td>>am_MI0000786|hsa-mir-378a&WithFlank&5|+|149732819|149732896@49.71.23&[MIMAT0000732&hsa-miR-378a-3p&offsets|0|+1;m-44&5|+|149732867|149732888&offsets|0|+1]|ACTGGACTTGGAGTCAGAAGGCC</td><td>>am_MI0000786|hsa-mir-378a&WithFlank&5|+|149732819|149732896</td><td>49</td><td>71</td><td>23</td><td>ACTGGACTTGGAGTCAGAAGGCC</td><td>449.9</td><td>5.6</td><td>0.4</td><td>13.3</td><td>3.7E-40</td><td>1.4E-38</td><td>BT20/MB468</td><td>CYTO</td></tr>
<tr><td>>MI0000465|hsa-mir-191&WithFlank&3|-|49020612|49020715@22.43.22(+1A)&[MIMAT0000440&hsa-miR-191-5p&offsets|0|-1(+1A);m-48&3|-|49020672|49020694&offsets|0|-1(+1A)]|CAACGGAATCCCAAAAGCAGCTA</td><td>>MI0000465|hsa-mir-191&WithFlank&3|-|49020612|49020715</td><td>22</td><td>43</td><td>22(+1A)</td><td>CAACGGAATCCCAAAAGCAGCTA</td><td>4189.1</td><td>8.3</td><td>0.6</td><td>13.3</td><td>3.7E-40</td><td>1.4E-38</td><td>BT20/MB468</td><td>CYTO</td></tr>
<tr><td>>hg38_21_+_8433172_8446622_RNA45SN1_45S_50ntFlanks@11874.11893.20|AGAAGACCCTGTTGAGCTTG</td><td>>hg38_21_+_8433172_8446622_RNA45SN1_45S_50ntFlanks</td><td>11874</td><td>11893</td><td>20</td><td>AGAAGACCCTGTTGAGCTTG</td><td>4792.9</td><td>-6.1</td><td>0.5</td><td>-13.3</td><td>3.7E-40</td><td>1.4E-38</td><td>BT20/MB468</td><td>CYTO</td></tr>
<tr><td>>hg38_21_+_8433172_8446622_RNA45SN1_45S_50ntFlanks@10411.10429.19|CAGTCGGTCCTGAGAGATG</td><td>>hg38_21_+_8433172_8446622_RNA45SN1_45S_50ntFlanks</td><td>10411</td><td>10429</td><td>19</td><td>CAGTCGGTCCTGAGAGATG</td><td>11198.7</td><td>-6.3</td><td>0.5</td><td>-13.3</td><td>4.5E-40</td><td>1.6E-38</td><td>BT20/MB468</td><td>CYTO</td></tr>
<tr><td>>hg38_21_+_8433172_8446622_RNA45SN1_45S_50ntFlanks@10351.10384.34|CGAGAACTTTGAAGGCCGAAGTGGAGAAGGGTTC</td><td>>hg38_21_+_8433172_8446622_RNA45SN1_45S_50ntFlanks</td><td>10351</td><td>10384</td><td>34</td><td>CGAGAACTTTGAAGGCCGAAGTGGAGAAGGGTTC</td><td>7036.3</td><td>-8.4</td><td>0.6</td><td>-13.3</td><td>2.0E-40</td><td>1.7E-38</td><td>MB231/MB468</td><td>NUC</td></tr>
<tr><td>>MI0000767|hsa-mir-365a&WithFlank&16|+|14309279|14309377@62.83.22;MI0000769|hsa-mir-365b&WithFlank&17|+|31575405|31575527@74.95.22&[MIMAT0000710&hsa-miR-365a-3p&offsets|0|0;m-176&16|+|14309340|14309361&offsets|0|0];[MIMAT0022834&hsa-miR-365b-3p&offsets|0|0;m-175&17|+|31575478|31575499&offsets|0|0]|TAATGCCCCTAAAAATCCTTAT</td><td>>MI0000767|hsa-mir-365a&WithFlank&16|+|14309279|14309377</td><td>62</td><td>83</td><td>22</td><td>TAATGCCCCTAAAAATCCTTAT</td><td>557.9</td><td>4.0</td><td>0.3</td><td>13.2</td><td>5.7E-40</td><td>1.8E-38</td><td>BT20/MB468</td><td>NUC</td></tr>
<tr><td>>MI0000440|hsa-mir-27b&WithFlank&9|+|95085439|95085547@67.86.20&[MIMAT0000419&hsa-miR-27b-3p&offsets|0|-1;m-39&9|+|95085505|95085525&offsets|0|-1]|TTCACAGTGGCTAAGTTCTG</td><td>>MI0000440|hsa-mir-27b&WithFlank&9|+|95085439|95085547</td><td>67</td><td>86</td><td>20</td><td>TTCACAGTGGCTAAGTTCTG</td><td>17052.9</td><td>8.2</td><td>0.6</td><td>13.2</td><td>5.5E-40</td><td>2.0E-38</td><td>BT20/MB468</td><td>CYTO</td></tr>
<tr><td>>hg38_21_+_8433172_8446622_RNA45SN1_45S_50ntFlanks@10350.10383.34|ACGAGAACTTTGAAGGCCGAAGTGGAGAAGGGTT</td><td>>hg38_21_+_8433172_8446622_RNA45SN1_45S_50ntFlanks</td><td>10350</td><td>10383</td><td>34</td><td>ACGAGAACTTTGAAGGCCGAAGTGGAGAAGGGTT</td><td>23181.3</td><td>-8.0</td><td>0.6</td><td>-13.3</td><td>2.5E-40</td><td>2.2E-38</td><td>MB231/MB468</td><td>NUC</td></tr>
<tr><td>>MI0000239|hsa-mir-197&WithFlank&1|+|109598887|109598973@54.74.21(+1U)&[MIMAT0000227&hsa-miR-197-3p&offsets|0|-1(+1U);m-99&1|+|109598940|109598961&offsets|0|-1(+1U)]|TTCACCACCTTCTCCACCCAGT</td><td>>MI0000239|hsa-mir-197&WithFlank&1|+|109598887|109598973</td><td>54</td><td>74</td><td>21(+1U)</td><td>TTCACCACCTTCTCCACCCAGT</td><td>2729.6</td><td>8.5</td><td>0.6</td><td>13.3</td><td>3.8E-40</td><td>2.4E-38</td><td>BT20/MB468</td><td>MITO</td></tr>
</table>

<table>
<tr><th></th><th></th><th></th><th></th><th></th><th></th><th></th><th></th><th></th><th></th><th></th><th></th><th></th><th></th></tr>
<tr><td>>hg38_21_+_8433172_8446622_RNA45SN1_45S_50ntFlanks@4723.4758.36|CATTAATCAAGAACGAAAGTCGGAGGTTCGAAGACG</td><td>>hg38_21_+_8433172_8446622_RNA45SN1_45S_50ntFlanks</td><td>4723</td><td>4758</td><td>36</td><td>CATTAATCAAGAACGAAAGTCGGAGGTTCGAAGACG</td><td>22750.2</td><td>-10.4</td><td>0.8</td><td>-13.4</td><td>8.8E-41</td><td>2.4E-38</td><td>MB231/MB468</td><td>MITO</td></tr>
<tr><td>>hg38_21_+_8433172_8446622_RNA45SN1_45S_50ntFlanks@9568.9597.30|CGTGCAAATCGGTCGTCCGACCTGGGTATA</td><td>>hg38_21_+_8433172_8446622_RNA45SN1_45S_50ntFlanks</td><td>9568</td><td>9597</td><td>30</td><td>CGTGCAAATCGGTCGTCCGACCTGGGTATA</td><td>5485.0</td><td>-16.8</td><td>1.3</td><td>-13.2</td><td>8.4E-40</td><td>2.6E-38</td><td>BT20/MB468</td><td>NUC</td></tr>
<tr><td>>hg38_21_+_8433172_8446622_RNA45SN1_45S_50ntFlanks@10349.10368.20|AACGAGAACTTTGAAGGCCG</td><td>>hg38_21_+_8433172_8446622_RNA45SN1_45S_50ntFlanks</td><td>10349</td><td>10368</td><td>20</td><td>AACGAGAACTTTGAAGGCCG</td><td>5894.8</td><td>-8.3</td><td>0.6</td><td>-13.2</td><td>8.5E-40</td><td>2.7E-38</td><td>BT20/MB468</td><td>NUC</td></tr>
<tr><td>>MI0003137|hsa-mir-193b&WithFlank&16|+|14303961|14304055@57.78.22&[MIMAT0002819&hsa-miR-193b-3p&offsets|0|0;m-166&16|+|14304017|14304038&offsets|0|0]|AACTGGCCCTCAAAGTCCCGCT</td><td>>MI0003137|hsa-mir-193b&WithFlank&16|+|14303961|14304055</td><td>57</td><td>78</td><td>22</td><td>AACTGGCCCTCAAAGTCCCGCT</td><td>4344.5</td><td>8.9</td><td>0.7</td><td>13.2</td><td>7.9E-40</td><td>2.8E-38</td><td>BT20/MB468</td><td>CYTO</td></tr>
<tr><td>>hg38_21_+_8433172_8446622_RNA45SN1_45S_50ntFlanks@11783.11806.24|CATGAATGGATGAACGAGATTCCC</td><td>>hg38_21_+_8433172_8446622_RNA45SN1_45S_50ntFlanks</td><td>11783</td><td>11806</td><td>24</td><td>CATGAATGGATGAACGAGATTCCC</td><td>3163.2</td><td>-16.1</td><td>1.2</td><td>-13.2</td><td>9.3E-40</td><td>2.9E-38</td><td>BT20/MB468</td><td>NUC</td></tr>
<tr><td>>MI0000088|hsa-mir-30a&WithFlank&6|-|71403545|71403627@12.34.23&[MIMAT0000087&hsa-miR-30a-5p&offsets|0|+1;m-15&6|-|71403595|71403616&offsets|0|+1]|TGTAAACATCCTCGACTGGAAGC</td><td>>MI0000088|hsa-mir-30a&WithFlank&6|-|71403545|71403627</td><td>12</td><td>34</td><td>23</td><td>TGTAAACATCCTCGACTGGAAGC</td><td>11321.1</td><td>6.4</td><td>0.5</td><td>13.2</td><td>5.1E-40</td><td>3.2E-38</td><td>BT20/MB468</td><td>MITO</td></tr>
<tr><td>>MI0000065|hsa-let-7d&WithFlank&9|+|94178828|94178926@14.34.21&[MIMAT0000065&hsa-let-7d-5p&offsets|0|-1;m-90&9|+|94178841|94178862&offsets|0|-1]|AGAGGTAGTAGGTTGCATAGT</td><td>>MI0000065|hsa-let-7d&WithFlank&9|+|94178828|94178926</td><td>14</td><td>34</td><td>21</td><td>AGAGGTAGTAGGTTGCATAGT</td><td>1289.1</td><td>7.7</td><td>0.6</td><td>13.2</td><td>5.8E-40</td><td>3.5E-38</td><td>BT20/MB468</td><td>MITO</td></tr>
<tr><td>>hg38_21_+_8433172_8446622_RNA45SN1_45S_50ntFlanks@11569.11602.34|CAGCCGACTTAGAACTGGTGCGGACCAGGGGAAT</td><td>>hg38_21_+_8433172_8446622_RNA45SN1_45S_50ntFlanks</td><td>11569</td><td>11602</td><td>34</td><td>CAGCCGACTTAGAACTGGTGCGGACCAGGGGAAT</td><td>8085.4</td><td>-7.2</td><td>0.5</td><td>-13.3</td><td>4.2E-40</td><td>3.6E-38</td><td>MB231/MB468</td><td>NUC</td></tr>
<tr><td>>hg38_21_+_8433172_8446622_RNA45SN1_45S_50ntFlanks@10841.10868.28|CGTAACTTCGGGATAAGGATTGGCTCTA</td><td>>hg38_21_+_8433172_8446622_RNA45SN1_45S_50ntFlanks</td><td>10841</td><td>10868</td><td>28</td><td>CGTAACTTCGGGATAAGGATTGGCTCTA</td><td>26946.8</td><td>-10.0</td><td>0.8</td><td>-13.2</td><td>1.2E-39</td><td>3.6E-38</td><td>BT20/MB468</td><td>NUC</td></tr>
<tr><td>>MI0000253|hsa-mir-148a&WithFlank&7|-|25949913|25949992@50.70.21(+1C)&[MIMAT0000243&hsa-miR-148a-3p&offsets|0|-1(+1C);m-7&7|-|25949922|25949943&offsets|0|-1(+1C)]|TCAGTGCACTACAGAACTTTGC</td><td>>MI0000253|hsa-mir-148a&WithFlank&7|-|25949913|25949992</td><td>50</td><td>70</td><td>21(+1C)</td><td>TCAGTGCACTACAGAACTTTGC</td><td>29097.5</td><td>7.5</td><td>0.6</td><td>13.2</td><td>1.2E-39</td><td>4.3E-38</td><td>BT20/MB468</td><td>CYTO</td></tr>
<tr><td>>hg38_21_+_8433172_8446622_RNA45SN1_45S_50ntFlanks@4726.4758.33|TAATCAAGAACGAAAGTCGGAGGTTCGAAGACG</td><td>>hg38_21_+_8433172_8446622_RNA45SN1_45S_50ntFlanks</td><td>4726</td><td>4758</td><td>33</td><td>TAATCAAGAACGAAAGTCGGAGGTTCGAAGACG</td><td>4933.4</td><td>-6.5</td><td>0.5</td><td>-13.2</td><td>5.7E-40</td><td>4.7E-38</td><td>MB231/MB468</td><td>NUC</td></tr>
<tr><td>>hg38_21_+_8433172_8446622_RNA45SN1_45S_50ntFlanks@10391.10409.19|AACAGCAGTTGAACATGGG</td><td>>hg38_21_+_8433172_8446622_RNA45SN1_45S_50ntFlanks</td><td>10391</td><td>10409</td><td>19</td><td>AACAGCAGTTGAACATGGG</td><td>4689.1</td><td>-16.6</td><td>1.3</td><td>-13.2</td><td>1.5E-39</td><td>4.8E-38</td><td>BT20/MB468</td><td>NUC</td></tr>
<tr><td>>hg38_21_+_8433172_8446622_RNA45SN1_45S_50ntFlanks@5473.5516.44|CGGCCCTGGCGGAGCGCTGAGAAGACGGTCGAACTTGACTATCT</td><td>>hg38_21_+_8433172_8446622_RNA45SN1_45S_50ntFlanks</td><td>5473</td><td>5516</td><td>44</td><td>CGGCCCTGGCGGAGCGCTGAGAAGACGGTCGAACTTGACTATCT</td><td>32197.0</td><td>10.3</td><td>0.8</td><td>13.4</td><td>3.2E-41</td><td>4.8E-38</td><td>MB231/MB468</td><td>CYTO</td></tr>
<tr><td>>MI0000061|hsa-let-7a-2&WithFlank&11|-|122146516|122146599@11.31.21;MI0000062|hsa-let-7a-3&WithFlank&22|+|46112743|46112828@10.30.21;MI0000060|hsa-let-7a-1&WithFlank&9|+|94175951|94176042@12.32.21&[MIMAT0000062&hsa-let-7a-5p&offsets|0|-1;m-6&11|-|122146568|122146589&offsets|0|-1];[MIMAT0000062_1&hsa-let-7a-5p&offsets|0|-1;m-5&22|+|46112752|46112773&offsets|0|-1];[MIMAT0000062_2&hsa-let-7a-5p&offsets|0|-1;m-4&9|+|94175962|94175983&offsets|0|-1]|TGAGGTAGTAGGTTGTATAGT</td><td>>MI0000061|hsa-let-7a-2&WithFlank&11|-|122146516|122146599</td><td>11</td><td>31</td><td>21</td><td>TGAGGTAGTAGGTTGTATAGT</td><td>5444.3</td><td>7.2</td><td>0.5</td><td>13.2</td><td>8.1E-40</td><td>4.9E-38</td><td>BT20/MB468</td><td>MITO</td></tr>
<tr><td>>hg38_21_+_8433172_8446622_RNA45SN1_45S_50ntFlanks@8357.8379.23|AAAAGAACTTTGAAGAGAGAGTT</td><td>>hg38_21_+_8433172_8446622_RNA45SN1_45S_50ntFlanks</td><td>8357</td><td>8379</td><td>23</td><td>AAAAGAACTTTGAAGAGAGAGTT</td><td>41335.7</td><td>-9.4</td><td>0.7</td><td>-13.2</td><td>6.3E-40</td><td>5.2E-38</td><td>MB231/MB468</td><td>NUC</td></tr>
<tr><td>>MI0000088|hsa-mir-30a&WithFlank&6|-|71403545|71403627@53.74.22&[MIMAT0000088&hsa-miR-30a-3p&offsets|0|0;m-38&6|-|71403554|71403575&offsets|0|0]|CTTTCAGTCGGATGTTTGCAGC</td><td>>MI0000088|hsa-mir-30a&WithFlank&6|-|71403545|71403627</td><td>53</td><td>74</td><td>22</td><td>CTTTCAGTCGGATGTTTGCAGC</td><td>11430.3</td><td>6.8</td><td>0.5</td><td>13.2</td><td>1.5E-39</td><td>5.4E-38</td><td>BT20/MB468</td><td>CYTO</td></tr>
<tr><td>>MI0000465|hsa-mir-191&WithFlank&3|-|49020612|49020715@22.43.22&[MIMAT0000440&hsa-miR-191-5p&offsets|0|-1;m-48&3|-|49020672|49020694&offsets|0|-1]|CAACGGAATCCCAAAAGCAGCT</td><td>>MI0000465|hsa-mir-191&WithFlank&3|-|49020612|49020715</td><td>22</td><td>43</td><td>22</td><td>CAACGGAATCCCAAAAGCAGCT</td><td>15680.9</td><td>7.5</td><td>0.6</td><td>13.2</td><td>1.5E-39</td><td>5.4E-38</td><td>BT20/MB468</td><td>CYTO</td></tr>
<tr><td>>MI0000086|hsa-mir-28&WithFlank&3|+|188688775|188688872@60.81.22(+1A)&[MIMAT0004502&hsa-miR-28-3p&offsets|0|0(+1A);m-28&3|+|188688834|188688855&offsets|0|0(+1A)]|CACTAGATTGTGAGCTCCTGGAA</td><td>>MI0000086|hsa-mir-28&WithFlank&3|+|188688775|188688872</td><td>60</td><td>81</td><td>22(+1A)</td><td>CACTAGATTGTGAGCTCCTGGAA</td><td>262.5</td><td>4.6</td><td>0.3</td><td>13.1</td><td>1.8E-39</td><td>5.6E-38</td><td>BT20/MB468</td><td>NUC</td></tr>
<tr><td>>hg38_21_+_8433172_8446622_RNA45SN1_45S_50ntFlanks@12457.12487.31|CAAGCGTTGGATTGTTCACCCACTAATAGGG</td><td>>hg38_21_+_8433172_8446622_RNA45SN1_45S_50ntFlanks</td><td>12457</td><td>12487</td><td>31</td><td>CAAGCGTTGGATTGTTCACCCACTAATAGGG</td><td>6261.9</td><td>-17.0</td><td>1.3</td><td>-13.1</td><td>1.8E-39</td><td>5.6E-38</td><td>BT20/MB468</td><td>NUC</td></tr>
<tr><td>>trna10_LysCTT_16_+_3241501_3241573@1.35.35,trna13_LysCTT_14_-_58706613_58706685@1.35.35,trna2_LysCTT_15_+_79152904_79152976@1.35.35|GCCCGGCTAGCTCAGTCGGTAGAGCATGGGACTCT</td><td>>trna10_LysCTT_16_+_3241501_3241573</td><td>1</td><td>35</td><td>35</td><td>GCCCGGCTAGCTCAGTCGGTAGAGCATGGGACTCT</td><td>352084.9</td><td>12.8</td><td>1.0</td><td>13.2</td><td>1.7E-39</td><td>5.9E-38</td><td>BT20/MB468</td><td>CYTO</td></tr>
<tr><td>>hg38_21_+_8433172_8446622_RNA45SN1_45S_50ntFlanks@12456.12473.18|CCAAGCGTTGGATTGTTC</td><td>>hg38_21_+_8433172_8446622_RNA45SN1_45S_50ntFlanks</td><td>12456</td><td>12473</td><td>18</td><td>CCAAGCGTTGGATTGTTC</td><td>46185.2</td><td>-10.4</td><td>0.8</td><td>-13.4</td><td>9.1E-41</td><td>5.9E-38</td><td>MB231/MB468</td><td>MP</td></tr>
<tr><td>>hg38_21_+_8433172_8446622_RNA45SN1_45S_50ntFlanks@4003.4024.22|ATAACCTCGGGCCGATCGCACG</td><td>>hg38_21_+_8433172_8446622_RNA45SN1_45S_50ntFlanks</td><td>4003</td><td>4024</td><td>22</td><td>ATAACCTCGGGCCGATCGCACG</td><td>3228.1</td><td>-7.2</td><td>0.5</td><td>-13.2</td><td>9.5E-40</td><td>7.7E-38</td><td>MB231/MB468</td><td>NUC</td></tr>
<tr><td>>MI0000065|hsa-let-7d&WithFlank&9|+|94178828|94178926@14.35.22&[MIMAT0000065&hsa-let-7d-5p&offsets|0|0;m-90&9|+|94178841|94178862&offsets|0|0]|AGAGGTAGTAGGTTGCATAGTT</td><td>>MI0000065|hsa-let-7d&WithFlank&9|+|94178828|94178926</td><td>14</td><td>35</td><td>22</td><td>AGAGGTAGTAGGTTGCATAGTT</td><td>8369.2</td><td>7.5</td><td>0.6</td><td>13.2</td><td>1.3E-39</td><td>7.7E-38</td><td>BT20/MB468</td><td>MITO</td></tr>
<tr><td>>hg38_21_+_8433172_8446622_RNA45SN1_45S_50ntFlanks@11316.11344.29|CACCCCACGTCTCGTCGCGCGCGCGTCCG</td><td>>hg38_21_+_8433172_8446622_RNA45SN1_45S_50ntFlanks</td><td>11316</td><td>11344</td><td>29</td><td>CACCCCACGTCTCGTCGCGCGCGCGTCCG</td><td>45556.1</td><td>-19.3</td><td>1.5</td><td>-13.3</td><td>3.1E-40</td><td>8.0E-38</td><td>MB231/MB468</td><td>MITO</td></tr>
<tr><td>>hg38_21_+_8433172_8446622_RNA45SN1_45S_50ntFlanks@12405.12432.28|CTTTTTGATCCTTCGATGTCGGCTCTTC</td><td>>hg38_21_+_8433172_8446622_RNA45SN1_45S_50ntFlanks</td><td>12405</td><td>12432</td><td>28</td><td>CTTTTTGATCCTTCGATGTCGGCTCTTC</td><td>5663.1</td><td>-8.5</td><td>0.6</td><td>-13.3</td><td>3.2E-40</td><td>8.0E-38</td><td>MB231/MB468</td><td>MITO</td></tr>
<tr><td>>hg38_21_+_8433172_8446622_RNA45SN1_45S_50ntFlanks@10351.10374.24|CGAGAACTTTGAAGGCCGAAGTGG</td><td>>hg38_21_+_8433172_8446622_RNA45SN1_45S_50ntFlanks</td><td>10351</td><td>10374</td><td>24</td><td>CGAGAACTTTGAAGGCCGAAGTGG</td><td>2868.0</td><td>-6.6</td><td>0.5</td><td>-13.2</td><td>1.0E-39</td><td>8.1E-38</td><td>MB231/MB468</td><td>NUC</td></tr>
<tr><td>>hg38_21_+_8433172_8446622_RNA45SN1_45S_50ntFlanks@12627.12653.27|ATGGGGCGAAGCTACCATCTGTGGGAT</td><td>>hg38_21_+_8433172_8446622_RNA45SN1_45S_50ntFlanks</td><td>12627</td><td>12653</td><td>27</td><td>ATGGGGCGAAGCTACCATCTGTGGGAT</td><td>4803.2</td><td>-16.7</td><td>1.3</td><td>-13.1</td><td>2.8E-39</td><td>8.4E-38</td><td>BT20/MB468</td><td>NUC</td></tr>
<tr><td>>hg38_21_+_8433172_8446622_RNA45SN1_45S_50ntFlanks@7984.8004.21|AGATCAGACGTGGCGACCCGC</td><td>>hg38_21_+_8433172_8446622_RNA45SN1_45S_50ntFlanks</td><td>7984</td><td>8004</td><td>21</td><td>AGATCAGACGTGGCGACCCGC</td><td>663.4</td><td>-6.1</td><td>0.5</td><td>-13.1</td><td>2.9E-39</td><td>8.8E-38</td><td>BT20/MB468</td><td>NUC</td></tr>
<tr><td>>hg38_21_+_8433172_8446622_RNA45SN1_45S_50ntFlanks@10351.10388.38|CGAGAACTTTGAAGGCCGAAGTGGAGAAGGGTTCCATG</td><td>>hg38_21_+_8433172_8446622_RNA45SN1_45S_50ntFlanks</td><td>10351</td><td>10388</td><td>38</td><td>CGAGAACTTTGAAGGCCGAAGTGGAGAAGGGTTCCATG</td><td>13214.0</td><td>-9.7</td><td>0.7</td><td>-13.3</td><td>3.8E-40</td><td>9.2E-38</td><td>MB231/MB468</td><td>MITO</td></tr>
<tr><td>>hg38_21_+_8433172_8446622_RNA45SN1_45S_50ntFlanks@12934.12959.26|CTAAACCATTCGTAGACGACCTGCTT</td><td>>hg38_21_+_8433172_8446622_RNA45SN1_45S_50ntFlanks</td><td>12934</td><td>12959</td><td>26</td><td>CTAAACCATTCGTAGACGACCTGCTT</td><td>1945.3</td><td>-6.6</td><td>0.5</td><td>-13.2</td><td>1.2E-39</td><td>9.3E-38</td><td>MB231/MB468</td><td>NUC</td></tr>
<tr><td>>MI0000542|hsa-mir-320a&WithFlank&8|-|22244960|22245043@48.68.21&[MIMAT0000510&hsa-miR-320a-3p&offsets|0|-1;m-60&8|-|22244975|22244996&offsets|0|-1]|AAAAGCTGGGTTGAGAGGGCG</td><td>>MI0000542|hsa-mir-320a&WithFlank&8|-|22244960|22245043</td><td>48</td><td>68</td><td>21</td><td>AAAAGCTGGGTTGAGAGGGCG</td><td>4400.1</td><td>9.3</td><td>0.7</td><td>13.1</td><td>1.8E-39</td><td>1.0E-37</td><td>BT20/MB468</td><td>MITO</td></tr>
<tr><td>>MI0003628|hsa-mir-615&WithFlank&12|+|54033944|54034051@67.87.21&[MIMAT0003283&hsa-miR-615-3p&offsets|0|-1;m-495&12|+|54034010|54034030&offsets|0|0]|TCCGAGCCTGGGTCTCCCTCT</td><td>>MI0003628|hsa-mir-615&WithFlank&12|+|54033944|54034051</td><td>67</td><td>87</td><td>21</td><td>TCCGAGCCTGGGTCTCCCTCT</td><td>5918.8</td><td>7.7</td><td>0.6</td><td>13.1</td><td>1.9E-39</td><td>1.1E-37</td><td>BT20/MB468</td><td>MITO</td></tr>
<tr><td>>hg38_21_+_8433172_8446622_RNA45SN1_45S_50ntFlanks@12607.12625.19|TGTGCTTGGCTGAGGAGCC</td><td>>hg38_21_+_8433172_8446622_RNA45SN1_45S_50ntFlanks</td><td>12607</td><td>12625</td><td>19</td><td>TGTGCTTGGCTGAGGAGCC</td><td>731.7</td><td>-5.8</td><td>0.4</td><td>-13.1</td><td>3.9E-39</td><td>1.2E-37</td><td>BT20/MB468</td><td>NUC</td></tr>
<tr><td>>MI0000081|hsa-mir-24-2&WithFlank&19|-|13836281|13836365@56.77.22(+1A);MI0000080|hsa-mir-24-1&WithFlank&9|+|95086015|95086094@50.71.22(+1A)&[MIMAT0000080&hsa-miR-24-3p&offsets|0|0(+1A);m-36&19|-|13836288|13836310&offsets|0|-1(+1A)];[MIMAT0000080_1&hsa-miR-24-3p&offsets|0|0(+1A);m-35&9|+|95086064|95086086&offsets|0|-1(+1A)]|TGGCTCAGTTCAGCAGGAACAGA</td><td>>MI0000081|hsa-mir-24-2&WithFlank&19|-|13836281|13836365</td><td>56</td><td>77</td><td>22(+1A)</td><td>TGGCTCAGTTCAGCAGGAACAGA</td><td>522.2</td><td>5.8</td><td>0.4</td><td>13.1</td><td>2.0E-39</td><td>1.2E-37</td><td>BT20/MB468</td><td>MITO</td></tr>
<tr><td>>hg38_21_+_8433172_8446622_RNA45SN1_45S_50ntFlanks@7980.8003.24|CCTCAGATCAGACGTGGCGACCCG</td><td>>hg38_21_+_8433172_8446622_RNA45SN1_45S_50ntFlanks</td><td>7980</td><td>8003</td><td>24</td><td>CCTCAGATCAGACGTGGCGACCCG</td><td>1598.6</td><td>-8.0</td><td>0.6</td><td>-13.1</td><td>4.4E-39</td><td>1.3E-37</td><td>BT20/MB468</td><td>NUC</td></tr>
<tr><td>>MI0000061|hsa-let-7a-2&WithFlank&11|-|122146516|122146599@11.31.21(+1C);MI0000062|hsa-let-7a-3&WithFlank&22|+|46112743|46112828@10.30.21(+1C);MI0000060|hsa-let-7a-1&WithFlank&9|+|94175951|94176042@12.32.21(+1C)&[MIMAT0000062&hsa-let-7a-5p&offsets|0|-1(+1C);m-6&11|-|122146568|122146589&offsets|0|-1(+1C)];[MIMAT0000062_1&hsa-let-7a-5p&offsets|0|-1(+1C);m-5&22|+|46112752|46112773&offsets|0|-1(+1C)];[MIMAT0000062_2&hsa-let-7a-5p&offsets|0|-1(+1C);m-4&9|+|94175962|94175983&offsets|0|-1(+1C)]|TGAGGTAGTAGGTTGTATAGTC</td><td>>MI0000061|hsa-let-7a-2&WithFlank&11|-|122146516|122146599</td><td>11</td><td>31</td><td>21(+1C)</td><td>TGAGGTAGTAGGTTGTATAGTC</td><td>8830.2</td><td>7.4</td><td>0.6</td><td>13.1</td><td>2.3E-39</td><td>1.4E-37</td><td>BT20/MB468</td><td>MITO</td></tr>
<tr><td>>hg38_21_+_8433172_8446622_RNA45SN1_45S_50ntFlanks@10411.10428.18|CAGTCGGTCCTGAGAGAT</td><td>>hg38_21_+_8433172_8446622_RNA45SN1_45S_50ntFlanks</td><td>10411</td><td>10428</td><td>18</td><td>CAGTCGGTCCTGAGAGAT</td><td>6013.1</td><td>-8.3</td><td>0.6</td><td>-13.1</td><td>1.8E-39</td><td>1.4E-37</td><td>MB231/MB468</td><td>NUC</td></tr>
<tr><td>>hg38_21_+_8433172_8446622_RNA45SN1_45S_50ntFlanks@10389.10407.19|TGAACAGCAGTTGAACATG</td><td>>hg38_21_+_8433172_8446622_RNA45SN1_45S_50ntFlanks</td><td>10389</td><td>10407</td><td>19</td><td>TGAACAGCAGTTGAACATG</td><td>4013.5</td><td>-16.4</td><td>1.3</td><td>-13.1</td><td>5.3E-39</td><td>1.6E-37</td><td>BT20/MB468</td><td>NUC</td></tr>
<tr><td>>MI0005559|hsa-mir-744&WithFlank&17|+|12081893|12082002@17.37.21&[MIMAT0004945&hsa-miR-744-5p&offsets|0|-1;m-172&17|+|12081909|12081929&offsets|0|0]|TGCGGGGCTAGGGCTAACAGC</td><td>>MI0005559|hsa-mir-744&WithFlank&17|+|12081893|12082002</td><td>17</td><td>37</td><td>21</td><td>TGCGGGGCTAGGGCTAACAGC</td><td>5514.6</td><td>9.5</td><td>0.7</td><td>13.1</td><td>2.8E-39</td><td>1.6E-37</td><td>BT20/MB468</td><td>MITO</td></tr>
<tr><td>>MI0000272|hsa-mir-182&WithFlank&7|-|129770377|129770498@29.53.25(+1A)&[MIMAT0000259&hsa-miR-182-5p&offsets|0|+1(+1A);m-22&7|-|129770449|129770470&offsets|0|+3(+1A)]|TTTGGCAATGGTAGAACTCACACTGA</td><td>>MI0000272|hsa-mir-182&WithFlank&7|-|129770377|129770498</td><td>29</td><td>53</td><td>25(+1A)</td><td>TTTGGCAATGGTAGAACTCACACTGA</td><td>2343.7</td><td>8.7</td><td>0.7</td><td>13.1</td><td>4.6E-39</td><td>1.6E-37</td><td>BT20/MB468</td><td>CYTO</td></tr>
<tr><td>>hg38_21_+_8433172_8446622_RNA45SN1_45S_50ntFlanks@11569.11602.34|CAGCCGACTTAGAACTGGTGCGGACCAGGGGAAT</td><td>>hg38_21_+_8433172_8446622_RNA45SN1_45S_50ntFlanks</td><td>11569</td><td>11602</td><td>34</td><td>CAGCCGACTTAGAACTGGTGCGGACCAGGGGAAT</td><td>16661.7</td><td>-9.2</td><td>0.7</td><td>-13.1</td><td>5.7E-39</td><td>1.7E-37</td><td>BT20/MB468</td><td>NUC</td></tr>
<tr><td>>MI0000255|hsa-mir-30d&WithFlank&8|-|134804870|134804951@12.31.20&[MIMAT0000245&hsa-miR-30d-5p&offsets|0|-2;m-21&8|-|134804919|134804940&offsets|0|-2]|TGTAAACATCCCCGACTGGA</td><td>>MI0000255|hsa-mir-30d&WithFlank&8|-|134804870|134804951</td><td>12</td><td>31</td><td>20</td><td>TGTAAACATCCCCGACTGGA</td><td>2109.1</td><td>7.6</td><td>0.6</td><td>13.1</td><td>3.3E-39</td><td>1.9E-37</td><td>BT20/MB468</td><td>MITO</td></tr>
<tr><td>>hg38_21_+_8433172_8446622_RNA45SN1_45S_50ntFlanks@11578.11601.24|TAGAACTGGTGCGGACCAGGGGAA</td><td>>hg38_21_+_8433172_8446622_RNA45SN1_45S_50ntFlanks</td><td>11578</td><td>11601</td><td>24</td><td>TAGAACTGGTGCGGACCAGGGGAA</td><td>660.9</td><td>-5.9</td><td>0.4</td><td>-13.0</td><td>6.5E-39</td><td>1.9E-37</td><td>BT20/MB468</td><td>NUC</td></tr>
<tr><td>>hg38_21_+_8433172_8446622_RNA45SN1_45S_50ntFlanks@12971.12990.20|TTTCGTACGTAGCAGAGCAG</td><td>>hg38_21_+_8433172_8446622_RNA45SN1_45S_50ntFlanks</td><td>12971</td><td>12990</td><td>20</td><td>TTTCGTACGTAGCAGAGCAG</td><td>3792.9</td><td>-16.3</td><td>1.3</td><td>-13.0</td><td>6.5E-39</td><td>1.9E-37</td><td>BT20/MB468</td><td>NUC</td></tr>
</table>

<table>
<tr><td>>hg38_21_+_8433172_8446622_RNA45SN1_45S_50ntFlanks@10690.10708.19|CGTCCGGTGAGCTCTCGCT</td><td>>hg38_21_+_8433172_8446622_RNA45SN1_45S_50ntFlanks</td><td>10690</td><td>10708</td><td>19</td><td>CGTCCGGTGAGCTCTCGCT</td><td>8419.0</td><td>-6.4</td><td>0.5</td><td>-13.1</td><td>5.5E-39</td><td>1.9E-37</td><td>BT20/MB468</td><td>CYTO</td></tr>
<tr><td>>hg38_21_+_8433172_8446622_RNA45SN1_45S_50ntFlanks@8381.8402.22|AAGAGGGCGTGAAACCGTTAAG</td><td>>hg38_21_+_8433172_8446622_RNA45SN1_45S_50ntFlanks</td><td>8381</td><td>8402</td><td>22</td><td>AAGAGGGCGTGAAACCGTTAAG</td><td>8550.0</td><td>-8.7</td><td>0.7</td><td>-13.3</td><td>3.3E-40</td><td>2.0E-37</td><td>MB231/MB468</td><td>MP</td></tr>
<tr><td>>hg38_21_+_8433172_8446622_RNA45SN1_45S_50ntFlanks@11769.11792.24|TAATTAGTGACGCGCATGAATGGA</td><td>>hg38_21_+_8433172_8446622_RNA45SN1_45S_50ntFlanks</td><td>11769</td><td>11792</td><td>24</td><td>TAATTAGTGACGCGCATGAATGGA</td><td>3050.2</td><td>-16.0</td><td>1.2</td><td>-13.0</td><td>7.7E-39</td><td>2.2E-37</td><td>BT20/MB468</td><td>NUC</td></tr>
<tr><td>>hg38_21_+_8433172_8446622_RNA45SN1_45S_50ntFlanks@12456.12478.23|CCAAGCGTTGGATTGTTCACCCA</td><td>>hg38_21_+_8433172_8446622_RNA45SN1_45S_50ntFlanks</td><td>12456</td><td>12478</td><td>23</td><td>CCAAGCGTTGGATTGTTCACCCA</td><td>11667.8</td><td>-9.4</td><td>0.7</td><td>-13.2</td><td>1.0E-39</td><td>2.3E-37</td><td>MB231/MB468</td><td>MITO</td></tr>
<tr><td>>MI0000239|hsa-mir-197&WithFlank&1|+|109598887|109598973@54.75.22&[MIMAT0000227&hsa-miR-197-3p&offsets|0|0;m-99&1|+|109598940|109598961&offsets|0|0]|TTCACCACCTTCTCCACCCAGC</td><td>>MI0000239|hsa-mir-197&WithFlank&1|+|109598887|109598973</td><td>54</td><td>75</td><td>22</td><td>TTCACCACCTTCTCCACCCAGC</td><td>25143.5</td><td>9.5</td><td>0.7</td><td>13.0</td><td>7.4E-39</td><td>2.6E-37</td><td>BT20/MB468</td><td>CYTO</td></tr>
<tr><td>>hg38_21_+_8433172_8446622_RNA45SN1_45S_50ntFlanks@11569.11586.18|CAGCCGACTTAGAACTGG</td><td>>hg38_21_+_8433172_8446622_RNA45SN1_45S_50ntFlanks</td><td>11569</td><td>11586</td><td>18</td><td>CAGCCGACTTAGAACTGG</td><td>743.9</td><td>-6.0</td><td>0.5</td><td>-13.0</td><td>8.9E-39</td><td>2.6E-37</td><td>BT20/MB468</td><td>NUC</td></tr>
<tr><td>>trnaMT_GluTTC_MT_-_14674_14742@40.58.19,trnalookalike8_GluTTC_5_-_93905172_93905240@40.58.19|TTGGTCGTGGTTGTAGTCC</td><td>>trnaMT_GluTTC_MT_-_14674_14742</td><td>40</td><td>58</td><td>19</td><td>TTGGTCGTGGTTGTAGTCC</td><td>1365.0</td><td>8.1</td><td>0.6</td><td>13.1</td><td>5.1E-39</td><td>2.8E-37</td><td>BT20/MB468</td><td>MITO</td></tr>
<tr><td>>hg38_21_+_8433172_8446622_RNA45SN1_45S_50ntFlanks@6663.6689.27|GTGGATCACTCGGCTCGTGCGTCGATG</td><td>>hg38_21_+_8433172_8446622_RNA45SN1_45S_50ntFlanks</td><td>6663</td><td>6689</td><td>27</td><td>GTGGATCACTCGGCTCGTGCGTCGATG</td><td>2196.7</td><td>-5.9</td><td>0.5</td><td>-13.0</td><td>1.1E-38</td><td>3.1E-37</td><td>BT20/MB468</td><td>NUC</td></tr>
<tr><td>>hg38_21_+_8433172_8446622_RNA45SN1_45S_50ntFlanks@10679.10709.31|GGTTCCGGCGGCGTCCGGTGAGCTCTCGCTG</td><td>>hg38_21_+_8433172_8446622_RNA45SN1_45S_50ntFlanks</td><td>10679</td><td>10709</td><td>31</td><td>GGTTCCGGCGGCGTCCGGTGAGCTCTCGCTG</td><td>10360.5</td><td>-9.2</td><td>0.7</td><td>-13.1</td><td>4.0E-39</td><td>3.1E-37</td><td>MB231/MB468</td><td>NUC</td></tr>
<tr><td>>MI0000650|hsa-mir-200c&WithFlank&12|+|6963693|6963772@50.71.22&[MIMAT0000617&hsa-miR-200c-3p&offsets|0|-1;m-54&12|+|6963742|6963764&offsets|0|-1]|TAATACTGCCGGGTAATGATGG</td><td>>MI0000650|hsa-mir-200c&WithFlank&12|+|6963693|6963772</td><td>50</td><td>71</td><td>22</td><td>TAATACTGCCGGGTAATGATGG</td><td>18928.4</td><td>8.3</td><td>0.6</td><td>13.1</td><td>5.8E-39</td><td>3.2E-37</td><td>BT20/MB468</td><td>MITO</td></tr>
<tr><td>>hg38_21_+_8433172_8446622_RNA45SN1_45S_50ntFlanks@4760.4780.21|TCAGATACCGTCGTAGTTCCG</td><td>>hg38_21_+_8433172_8446622_RNA45SN1_45S_50ntFlanks</td><td>4760</td><td>4780</td><td>21</td><td>TCAGATACCGTCGTAGTTCCG</td><td>3902.4</td><td>-7.2</td><td>0.5</td><td>-13.1</td><td>4.3E-39</td><td>3.3E-37</td><td>MB231/MB468</td><td>NUC</td></tr>
<tr><td>>MI0000063|hsa-let-7b&WithFlank&22|+|46113680|46113774@12.33.22(+1A)&[MIMAT0000063&hsa-let-7b-5p&offsets|0|0(+1A);m-14&22|+|46113691|46113712&offsets|0|0(+1A)]|TGAGGTAGTAGGTTGTGTGGTTA</td><td>>MI0000063|hsa-let-7b&WithFlank&22|+|46113680|46113774</td><td>12</td><td>33</td><td>22(+1A)</td><td>TGAGGTAGTAGGTTGTGTGGTTA</td><td>2243.4</td><td>7.8</td><td>0.6</td><td>13.0</td><td>6.5E-39</td><td>3.6E-37</td><td>BT20/MB468</td><td>MITO</td></tr>
<tr><td>>hg38_21_+_8433172_8446622_RNA45SN1_45S_50ntFlanks@11783.11804.22|CATGAATGGATGAACGAGATTC</td><td>>hg38_21_+_8433172_8446622_RNA45SN1_45S_50ntFlanks</td><td>11783</td><td>11804</td><td>22</td><td>CATGAATGGATGAACGAGATTC</td><td>12763.6</td><td>-8.8</td><td>0.7</td><td>-13.1</td><td>1.8E-39</td><td>4.0E-37</td><td>MB231/MB468</td><td>MITO</td></tr>
<tr><td>>MI0000266|hsa-mir-10a&WithFlank&17|-|48579832|48579953@28.48.21(+1C)&[MIMAT0000253&hsa-miR-10a-5p&offsets|0|-2(+1C);m-18&17|-|48579905|48579926&offsets|0|-1(+1C)]|TACCCTGTAGATCCGAATTTGC</td><td>>MI0000266|hsa-mir-10a&WithFlank&17|-|48579832|48579953</td><td>28</td><td>48</td><td>21(+1C)</td><td>TACCCTGTAGATCCGAATTTGC</td><td>6470.4</td><td>16.9</td><td>1.3</td><td>13.0</td><td>1.3E-38</td><td>4.3E-37</td><td>BT20/MB468</td><td>CYTO</td></tr>
<tr><td>>hg38_21_+_8433172_8446622_RNA45SN1_45S_50ntFlanks@12973.12992.20|TCGTACGTAGCAGAGCAGCT</td><td>>hg38_21_+_8433172_8446622_RNA45SN1_45S_50ntFlanks</td><td>12973</td><td>12992</td><td>20</td><td>TCGTACGTAGCAGAGCAGCT</td><td>1986.0</td><td>-5.9</td><td>0.5</td><td>-13.1</td><td>6.1E-39</td><td>4.6E-37</td><td>MB231/MB468</td><td>NUC</td></tr>
<tr><td>>hg38_21_+_8433172_8446622_RNA45SN1_45S_50ntFlanks@12418.12435.18|CGATGTCGGCTCTTCCTA</td><td>>hg38_21_+_8433172_8446622_RNA45SN1_45S_50ntFlanks</td><td>12418</td><td>12435</td><td>18</td><td>CGATGTCGGCTCTTCCTA</td><td>19436.5</td><td>-7.8</td><td>0.6</td><td>-13.0</td><td>6.4E-39</td><td>4.7E-37</td><td>MB231/MB468</td><td>NUC</td></tr>
<tr><td>>hg38_21_+_8433172_8446622_RNA45SN1_45S_50ntFlanks@11667.11702.36|CTGCCCAGTGCTCTGAATGTCAAAGTGAAGAAATTC</td><td>>hg38_21_+_8433172_8446622_RNA45SN1_45S_50ntFlanks</td><td>11667</td><td>11702</td><td>36</td><td>CTGCCCAGTGCTCTGAATGTCAAAGTGAAGAAATTC</td><td>524.0</td><td>4.5</td><td>0.3</td><td>13.0</td><td>1.5E-38</td><td>5.1E-37</td><td>BT20/MB468</td><td>CYTO</td></tr>
<tr><td>>am_trna128_GlyGCC_6_-_27870686_27870756@1.31.31,trna133_GlyCCC_1_-_16872434_16872504@1.31.31,trna18_GlyGCC_16_+_70822597_70822667@1.31.31,trna19_GlyGCC_16_+_70823410_70823480@1.31.31,trna19_GlyGCC_2_-_157257659_157257729@1.31.31,trna24_GlyGCC_16_-_70812942_70813012@1.31.31,trna25_GlyGCC_16_-_70812114_70812184@1.31.31,trna4_GlyCCC_1_+_17188416_17188486@1.31.31,trna5_GlyGCC_17_+_8029064_8029134@1.31.31,trna68_GlyGCC_1_-_161493637_161493707@1.31.31|GCATTGGTGGTTCAGTGGTAGAATTCTCGCC</td><td>>am_trna128_GlyGCC_6_-_27870686_27870756</td><td>1</td><td>31</td><td>31</td><td>GCATTGGTGGTTCAGTGGTAGAATTCTCGCC</td><td>31617.9</td><td>4.5</td><td>0.3</td><td>13.0</td><td>1.5E-38</td><td>5.2E-37</td><td>BT20/MB468</td><td>CYTO</td></tr>
<tr><td>>hg38_21_+_8433172_8446622_RNA45SN1_45S_50ntFlanks@9534.9553.20|AGGAAACTCTGGTGGAGGTC</td><td>>hg38_21_+_8433172_8446622_RNA45SN1_45S_50ntFlanks</td><td>9534</td><td>9553</td><td>20</td><td>AGGAAACTCTGGTGGAGGTC</td><td>6394.4</td><td>-5.9</td><td>0.5</td><td>-13.0</td><td>1.0E-38</td><td>5.5E-37</td><td>BT20/MB468</td><td>MITO</td></tr>
<tr><td>>hg38_21_+_8433172_8446622_RNA45SN1_45S_50ntFlanks@12008.12032.25|CCGCCGGTGAAATACCACTACTCTG</td><td>>hg38_21_+_8433172_8446622_RNA45SN1_45S_50ntFlanks</td><td>12008</td><td>12032</td><td>25</td><td>CCGCCGGTGAAATACCACTACTCTG</td><td>58930.9</td><td>-15.2</td><td>1.2</td><td>-13.1</td><td>2.6E-39</td><td>5.7E-37</td><td>MB231/MB468</td><td>MITO</td></tr>
<tr><td>>MI0000440|hsa-mir-27b&WithFlank&9|+|95085439|95085547@67.85.19(+1A)&[MIMAT0000419&hsa-miR-27b-3p&offsets|0|-2(+1A);m-39&9|+|95085505|95085525&offsets|0|-2(+1A)]|TTCACAGTGGCTAAGTTCTA</td><td>>MI0000440|hsa-mir-27b&WithFlank&9|+|95085439|95085547</td><td>67</td><td>85</td><td>19(+1A)</td><td>TTCACAGTGGCTAAGTTCTA</td><td>205.1</td><td>4.7</td><td>0.4</td><td>13.0</td><td>2.1E-38</td><td>6.0E-37</td><td>BT20/MB468</td><td>NUC</td></tr>
<tr><td>>trna10_ValCAC_5_-_180649395_180649467@59.76.18,trna12_ValAAC_5_-_180645270_180645342@59.76.18,trna132_ValAAC_6_-_27721179_27721251@59.76.18,trna136_ValAAC_6_-_27648885_27648957@59.76.18,trna15_ValAAC_5_-_180615416_180615488@59.76.18,trna18_ValCAC_5_-_180529253_180529325@59.76.18,trna2_ValAAC_3_+_169490018_169490090@59.76.18,trna2_ValCAC_5_+_180524070_180524142@59.76.18,trna4_ValAAC_5_+_180591154_180591226@59.76.18,trna5_ValAAC_5_+_180596610_180596682@59.76.18,trna6_ValCAC_5_+_180600650_180600722@59.76.18,trna85_ValCAC_1_-_161369490_161369562@59.76.18,trna90_ValCAC_1_-_149684088_149684161@60.77.18,trna9_ValCAC_6_+_26538282_26538354@59.76.18|AACCGGGCGGAAACACCA</td><td>>trna10_ValCAC_5_-_180649395_180649467</td><td>59</td><td>76</td><td>18</td><td>AACCGGGCGGAAACACCA</td><td>866.6</td><td>7.1</td><td>0.5</td><td>13.0</td><td>1.1E-38</td><td>6.1E-37</td><td>BT20/MB468</td><td>MITO</td></tr>
<tr><td>>hg38_21_+_8433172_8446622_RNA45SN1_45S_50ntFlanks@12633.12653.21|CGAAGCTACCATCTGTGGGAT</td><td>>hg38_21_+_8433172_8446622_RNA45SN1_45S_50ntFlanks</td><td>12633</td><td>12653</td><td>21</td><td>CGAAGCTACCATCTGTGGGAT</td><td>3144.3</td><td>-16.0</td><td>1.2</td><td>-12.9</td><td>2.5E-38</td><td>7.1E-37</td><td>BT20/MB468</td><td>NUC</td></tr>
<tr><td>>MI0000102|hsa-mir-100&WithFlank&11|-|122152223|122152314@19.40.22(+1U)&[MIMAT0000098&hsa-miR-100-5p&offsets|0|0(+1U);m-27&11|-|122152276|122152296&offsets|0|+1(+1U)]|AACCCGTAGATCCGAACTTGTGT</td><td>>MI0000102|hsa-mir-100&WithFlank&11|-|122152223|122152314</td><td>19</td><td>40</td><td>22(+1U)</td><td>AACCCGTAGATCCGAACTTGTGT</td><td>68447.4</td><td>20.1</td><td>1.5</td><td>13.2</td><td>6.2E-40</td><td>7.6E-37</td><td>MB231/MB468</td><td>CYTO</td></tr>
<tr><td>>hg38_21_+_8433172_8446622_RNA45SN1_45S_50ntFlanks@10690.10708.19|CGTCCGGTGAGCTCTCGCT</td><td>>hg38_21_+_8433172_8446622_RNA45SN1_45S_50ntFlanks</td><td>10690</td><td>10708</td><td>19</td><td>CGTCCGGTGAGCTCTCGCT</td><td>9914.8</td><td>-7.8</td><td>0.6</td><td>-13.2</td><td>7.6E-40</td><td>7.7E-37</td><td>MB231/MB468</td><td>CYTO</td></tr>
<tr><td>>hg38_21_+_8433172_8446622_RNA45SN1_45S_50ntFlanks@4657.4682.26|CTAGAGGTGAAATTCTTGGACCGGCG</td><td>>hg38_21_+_8433172_8446622_RNA45SN1_45S_50ntFlanks</td><td>4657</td><td>4682</td><td>26</td><td>CTAGAGGTGAAATTCTTGGACCGGCG</td><td>3015.0</td><td>-16.0</td><td>1.2</td><td>-12.9</td><td>2.7E-38</td><td>7.8E-37</td><td>BT20/MB468</td><td>NUC</td></tr>
<tr><td>>hg38_21_+_8433172_8446622_RNA45SN1_45S_50ntFlanks@3705.3725.21|TACCTGGTTGATCCTGCCAGT</td><td>>hg38_21_+_8433172_8446622_RNA45SN1_45S_50ntFlanks</td><td>3705</td><td>3725</td><td>21</td><td>TACCTGGTTGATCCTGCCAGT</td><td>36820.1</td><td>9.6</td><td>0.7</td><td>12.9</td><td>2.8E-38</td><td>9.5E-37</td><td>BT20/MB468</td><td>CYTO</td></tr>
<tr><td>>MI0000433|hsa-let-7g&WithFlank&3|-|52268272|52268367@11.31.21(+1C)&[MIMAT0000414&hsa-let-7g-5p&offsets|0|-1(+1C);m-58&3|-|52268336|52268357&offsets|0|-1(+1C)]|TGAGGTAGTAGTTTGTACAGTC</td><td>>MI0000433|hsa-let-7g&WithFlank&3|-|52268272|52268367</td><td>11</td><td>31</td><td>21(+1C)</td><td>TGAGGTAGTAGTTTGTACAGTC</td><td>11640.0</td><td>7.7</td><td>0.6</td><td>13.0</td><td>1.8E-38</td><td>9.5E-37</td><td>BT20/MB468</td><td>MITO</td></tr>
<tr><td>>MI0000469|hsa-mir-125a&WithFlank&19|+|51693248|51693345@21.41.21(+1C)&[MIMAT0000443&hsa-miR-125a-5p&offsets|0|-3(+1C);m-63&19|+|51693268|51693289&offsets|0|-1(+1C)]|TCCCTGAGACCCTTTAACCTGC</td><td>>MI0000469|hsa-mir-125a&WithFlank&19|+|51693248|51693345</td><td>21</td><td>41</td><td>21(+1C)</td><td>TCCCTGAGACCCTTTAACCTGC</td><td>1414.6</td><td>8.5</td><td>0.7</td><td>12.9</td><td>3.2E-38</td><td>1.1E-36</td><td>BT20/MB468</td><td>CYTO</td></tr>
<tr><td>>hg38_21_+_8433172_8446622_RNA45SN1_45S_50ntFlanks@10772.10796.25|AGGTCTCCAAGGTGAACAGCCTCTG</td><td>>hg38_21_+_8433172_8446622_RNA45SN1_45S_50ntFlanks</td><td>10772</td><td>10796</td><td>25</td><td>AGGTCTCCAAGGTGAACAGCCTCTG</td><td>1321.5</td><td>-7.3</td><td>0.6</td><td>-12.9</td><td>3.9E-38</td><td>1.1E-36</td><td>BT20/MB468</td><td>NUC</td></tr>
<tr><td>>hg38_21_+_8433172_8446622_RNA45SN1_45S_50ntFlanks@11641.11669.29|CGGCGGGTGTTGACGCGATGTGATTTCTG</td><td>>hg38_21_+_8433172_8446622_RNA45SN1_45S_50ntFlanks</td><td>11641</td><td>11669</td><td>29</td><td>CGGCGGGTGTTGACGCGATGTGATTTCTG</td><td>4325.1</td><td>-16.5</td><td>1.3</td><td>-12.9</td><td>4.2E-38</td><td>1.2E-36</td><td>BT20/MB468</td><td>NUC</td></tr>
<tr><td>>hg38_21_+_8433172_8446622_RNA45SN1_45S_50ntFlanks@12991.13015.25|CTCCCTCGCTGCGATCTATTGAAAG</td><td>>hg38_21_+_8433172_8446622_RNA45SN1_45S_50ntFlanks</td><td>12991</td><td>13015</td><td>25</td><td>CTCCCTCGCTGCGATCTATTGAAAG</td><td>18409.3</td><td>-9.3</td><td>0.7</td><td>-13.0</td><td>1.6E-38</td><td>1.2E-36</td><td>MB231/MB468</td><td>NUC</td></tr>
<tr><td>>MI0000440|hsa-mir-27b&WithFlank&9|+|95085439|95085547@67.88.22(+1A)&[MIMAT0000419&hsa-miR-27b-3p&offsets|0|+1(+1A);m-39&9|+|95085505|95085525&offsets|0|+1(+1A)]|TTCACAGTGGCTAAGTTCTGCAA</td><td>>MI0000440|hsa-mir-27b&WithFlank&9|+|95085439|95085547</td><td>67</td><td>88</td><td>22(+1A)</td><td>TTCACAGTGGCTAAGTTCTGCAA</td><td>925.7</td><td>7.0</td><td>0.5</td><td>12.9</td><td>3.7E-38</td><td>1.2E-36</td><td>BT20/MB468</td><td>CYTO</td></tr>
<tr><td>>hg38_21_+_8433172_8446622_RNA45SN1_45S_50ntFlanks@6719.6740.22|TGAATTGCAGGACACATTGATC</td><td>>hg38_21_+_8433172_8446622_RNA45SN1_45S_50ntFlanks</td><td>6719</td><td>6740</td><td>22</td><td>TGAATTGCAGGACACATTGATC</td><td>6666.2</td><td>-7.2</td><td>0.6</td><td>-13.0</td><td>1.9E-38</td><td>1.3E-36</td><td>MB231/MB468</td><td>NUC</td></tr>
<tr><td>>am_MI0000809|hsa-mir-151a&WithFlank&8|-|140732558|140732659@53.73.21(+1U)&[MIMAT0000757&hsa-miR-151a-3p&offsets|0|0(+1U);m-30&8|-|140732587|140732607&offsets|0|0(+1U)]|CTAGACTGAAGCTCCTTGAGGT</td><td>>am_MI0000809|hsa-mir-151a&WithFlank&8|-|140732558|140732659</td><td>53</td><td>73</td><td>21(+1U)</td><td>CTAGACTGAAGCTCCTTGAGGT</td><td>15708.5</td><td>9.5</td><td>0.7</td><td>12.9</td><td>2.6E-38</td><td>1.4E-36</td><td>BT20/MB468</td><td>MITO</td></tr>
<tr><td>>hg38_21_+_8433172_8446622_RNA45SN1_45S_50ntFlanks@10410.10429.20|TCAGTCGGTCCTGAGAGATG</td><td>>hg38_21_+_8433172_8446622_RNA45SN1_45S_50ntFlanks</td><td>10410</td><td>10429</td><td>20</td><td>TCAGTCGGTCCTGAGAGATG</td><td>6473.1</td><td>-5.4</td><td>0.4</td><td>-12.9</td><td>4.3E-38</td><td>1.4E-36</td><td>BT20/MB468</td><td>CYTO</td></tr>
<tr><td>>hg38_21_+_8433172_8446622_RNA45SN1_45S_50ntFlanks@11569.11601.33|CAGCCGACTTAGAACTGGTGCGGACCAGGGGAA</td><td>>hg38_21_+_8433172_8446622_RNA45SN1_45S_50ntFlanks</td><td>11569</td><td>11601</td><td>33</td><td>CAGCCGACTTAGAACTGGTGCGGACCAGGGGAA</td><td>9447.3</td><td>-9.6</td><td>0.7</td><td>-13.1</td><td>2.7E-39</td><td>1.6E-36</td><td>MB231/MB468</td><td>MP</td></tr>
<tr><td>>MI0000079|hsa-mir-23a&WithFlank&19|-|13836581|13836665@54.73.20&[MIMAT0000078&hsa-miR-23a-3p&offsets|+3|+2;m-31&19|-|13836595|13836615&offsets|+3|+2]|ACATTGCCAGGGATTTCCAA</td><td>>MI0000079|hsa-mir-23a&WithFlank&19|-|13836581|13836665</td><td>54</td><td>73</td><td>20</td><td>ACATTGCCAGGGATTTCCAA</td><td>3784.9</td><td>16.1</td><td>1.3</td><td>12.9</td><td>5.5E-38</td><td>1.8E-36</td><td>BT20/MB468</td><td>CYTO</td></tr>
<tr><td>>hg38_21_+_8433172_8446622_RNA45SN1_45S_50ntFlanks@10408.10430.23|GGTCAGTCGGTCCTGAGAGATGG</td><td>>hg38_21_+_8433172_8446622_RNA45SN1_45S_50ntFlanks</td><td>10408</td><td>10430</td><td>23</td><td>GGTCAGTCGGTCCTGAGAGATGG</td><td>7087.9</td><td>-17.2</td><td>1.3</td><td>-12.9</td><td>6.6E-38</td><td>1.8E-36</td><td>BT20/MB468</td><td>NUC</td></tr>
<tr><td>>MI0000478|hsa-mir-149&WithFlank&2|+|240455995|240456095@21.43.23&[MIMAT0000450&hsa-miR-149-5p&offsets|0|0;m-195&2|+|240456015|240456037&offsets|0|0]|TCTGGCTCCGTGTCTTCACTCCC</td><td>>MI0000478|hsa-mir-149&WithFlank&2|+|240455995|240456095</td><td>21</td><td>43</td><td>23</td><td>TCTGGCTCCGTGTCTTCACTCCC</td><td>2811.3</td><td>15.9</td><td>1.2</td><td>12.9</td><td>4.2E-38</td><td>2.2E-36</td><td>BT20/MB468</td><td>MITO</td></tr>
<tr><td>>MI0000272|hsa-mir-182&WithFlank&7|-|129770377|129770498@29.52.24&[MIMAT0000259&hsa-miR-182-5p&offsets|0|0;m-22&7|-|129770449|129770470&offsets|0|+2]|TTTGGCAATGGTAGAACTCACACT</td><td>>MI0000272|hsa-mir-182&WithFlank&7|-|129770377|129770498</td><td>29</td><td>52</td><td>24</td><td>TTTGGCAATGGTAGAACTCACACT</td><td>16529.0</td><td>2.9</td><td>0.2</td><td>12.9</td><td>8.1E-38</td><td>2.2E-36</td><td>BT20/MB468</td><td>NUC</td></tr>
<tr><td>>MI0000746|hsa-mir-99b&WithFlank&19|+|51692606|51692687@13.34.22&[MIMAT0000689&hsa-miR-99b-5p&offsets|0|0;m-10&19|+|51692618|51692639&offsets|0|0]|CACCCGTAGAACCGACCTTGCG</td><td>>MI0000746|hsa-mir-99b&WithFlank&19|+|51692606|51692687</td><td>13</td><td>34</td><td>22</td><td>CACCCGTAGAACCGACCTTGCG</td><td>23902.1</td><td>6.6</td><td>0.5</td><td>12.9</td><td>4.5E-38</td><td>2.3E-36</td><td>BT20/MB468</td><td>MITO</td></tr>
<tr><td>>MI0003591|hsa-mir-584&WithFlank&5|-|149062307|149062415@22.42.21&[MIMAT0003249&hsa-miR-584-5p&offsets|0|-1;m-167&5|-|149062374|149062394&offsets|0|0]|TTATGGTTTGCCTGGGACTGA</td><td>>MI0003591|hsa-mir-584&WithFlank&5|-|149062307|149062415</td><td>22</td><td>42</td><td>21</td><td>TTATGGTTTGCCTGGGACTGA</td><td>480.9</td><td>3.4</td><td>0.3</td><td>12.8</td><td>9.3E-38</td><td>2.6E-36</td><td>BT20/MB468</td><td>NUC</td></tr>
</table>

<table>
<tr><th></th><th></th><th></th><th></th><th></th><th></th><th></th><th></th><th></th><th></th><th></th><th></th><th></th><th></th></tr>
<tr><td>>hg38_21_+_8433172_8446622_RNA45SN1_45S_50ntFlanks@12436.12455.20|TCATTGTGAAGCAGAATTCA</td><td>>hg38_21_+_8433172_8446622_RNA45SN1_45S_50ntFlanks</td><td>12436</td><td>12455</td><td>20</td><td>TCATTGTGAAGCAGAATTCA</td><td>3520.3</td><td>-16.2</td><td>1.3</td><td>-12.8</td><td>9.9E-38</td><td>2.7E-36</td><td>BT20/MB468</td><td>NUC</td></tr>
<tr><td>>MI0000086|hsa-mir-28&WithFlank&3|+|188688775|188688872@60.81.22(+1A)&[MIMAT0004502&hsa-miR-28-3p&offsets|0|0(+1A);m-28&3|+|188688834|188688855&offsets|0|0(+1A)]|CACTAGATTGTGAGCTCCTGGAA</td><td>>MI0000086|hsa-mir-28&WithFlank&3|+|188688775|188688872</td><td>60</td><td>81</td><td>22(+1A)</td><td>CACTAGATTGTGAGCTCCTGGAA</td><td>6907.7</td><td>11.5</td><td>0.9</td><td>12.8</td><td>9.2E-38</td><td>3.0E-36</td><td>BT20/MB468</td><td>CYTO</td></tr>
<tr><td>>hg38_21_+_8433172_8446622_RNA45SN1_45S_50ntFlanks@11323.11344.22|CGTCTCGTCGCGCGCGCGTCCG</td><td>>hg38_21_+_8433172_8446622_RNA45SN1_45S_50ntFlanks</td><td>11323</td><td>11344</td><td>22</td><td>CGTCTCGTCGCGCGCGCGTCCG</td><td>32900.1</td><td>-18.8</td><td>1.4</td><td>-13.0</td><td>1.5E-38</td><td>3.1E-36</td><td>MB231/MB468</td><td>MITO</td></tr>
<tr><td>>hg38_21_+_8433172_8446622_RNA45SN1_45S_50ntFlanks@11569.11601.33|CAGCCGACTTAGAACTGGTGCGGACCAGGGGAA</td><td>>hg38_21_+_8433172_8446622_RNA45SN1_45S_50ntFlanks</td><td>11569</td><td>11601</td><td>33</td><td>CAGCCGACTTAGAACTGGTGCGGACCAGGGGAA</td><td>16013.7</td><td>-8.5</td><td>0.7</td><td>-12.9</td><td>4.6E-38</td><td>3.2E-36</td><td>MB231/MB468</td><td>NUC</td></tr>
<tr><td>>hg38_21_+_8433172_8446622_RNA45SN1_45S_50ntFlanks@10350.10373.24|ACGAGAACTTTGAAGGCCGAAGTG</td><td>>hg38_21_+_8433172_8446622_RNA45SN1_45S_50ntFlanks</td><td>10350</td><td>10373</td><td>24</td><td>ACGAGAACTTTGAAGGCCGAAGTG</td><td>1045.7</td><td>-6.6</td><td>0.5</td><td>-12.8</td><td>1.2E-37</td><td>3.3E-36</td><td>BT20/MB468</td><td>NUC</td></tr>
<tr><td>>MI0000095|hsa-mir-93&WithFlank&7|-|100093762|100093853@17.39.23(+1A)&[MIMAT0000093&hsa-miR-93-5p&offsets|0|0(+1A);m-29&7|-|100093815|100093837&offsets|0|0(+1A)]|CAAAGTGCTGTTCGTGCAGGTAGA</td><td>>MI0000095|hsa-mir-93&WithFlank&7|-|100093762|100093853</td><td>17</td><td>39</td><td>23(+1A)</td><td>CAAAGTGCTGTTCGTGCAGGTAGA</td><td>4362.7</td><td>6.8</td><td>0.5</td><td>12.8</td><td>1.0E-37</td><td>3.3E-36</td><td>BT20/MB468</td><td>CYTO</td></tr>
<tr><td>>hg38_21_+_8433172_8446622_RNA45SN1_45S_50ntFlanks@12995.13017.23|CTCGCTGCGATCTATTGAAAGTC</td><td>>hg38_21_+_8433172_8446622_RNA45SN1_45S_50ntFlanks</td><td>12995</td><td>13017</td><td>23</td><td>CTCGCTGCGATCTATTGAAAGTC</td><td>1433.4</td><td>6.7</td><td>0.5</td><td>13.1</td><td>4.0E-39</td><td>3.5E-36</td><td>MB231/MB468</td><td>CYTO</td></tr>
<tr><td>>MI0000469|hsa-mir-125a&WithFlank&19|+|51693248|51693345@21.42.22(+1A)&[MIMAT0000443&hsa-miR-125a-5p&offsets|0|-2(+1A);m-63&19|+|51693268|51693289&offsets|0|0(+1A)]|TCCCTGAGACCCTTTAACCTGTA</td><td>>MI0000469|hsa-mir-125a&WithFlank&19|+|51693248|51693345</td><td>21</td><td>42</td><td>22(+1A)</td><td>TCCCTGAGACCCTTTAACCTGTA</td><td>7755.1</td><td>8.7</td><td>0.7</td><td>12.8</td><td>1.1E-37</td><td>3.5E-36</td><td>BT20/MB468</td><td>CYTO</td></tr>
<tr><td>>hg38_21_+_8433172_8446622_RNA45SN1_45S_50ntFlanks@8114.8143.30|GCGCGGGACATGTGGCGTACGGAAGACCCG</td><td>>hg38_21_+_8433172_8446622_RNA45SN1_45S_50ntFlanks</td><td>8114</td><td>8143</td><td>30</td><td>GCGCGGGACATGTGGCGTACGGAAGACCCG</td><td>20990.6</td><td>-18.2</td><td>1.4</td><td>-13.0</td><td>6.5E-39</td><td>3.6E-36</td><td>MB231/MB468</td><td>MP</td></tr>
<tr><td>>hg38_21_+_8433172_8446622_RNA45SN1_45S_50ntFlanks@9572.9597.26|CAAATCGGTCGTCCGACCTGGGTATA</td><td>>hg38_21_+_8433172_8446622_RNA45SN1_45S_50ntFlanks</td><td>9572</td><td>9597</td><td>26</td><td>CAAATCGGTCGTCCGACCTGGGTATA</td><td>16481.2</td><td>-12.6</td><td>1.0</td><td>-13.0</td><td>6.6E-39</td><td>3.6E-36</td><td>MB231/MB468</td><td>MP</td></tr>
<tr><td>>MI0000279|hsa-mir-196a-2&WithFlank&12|+|53991732|53991853@31.53.23&[MIMAT0000226&hsa-miR-196a-5p&offsets|0|+1;m-261&12|+|53991762|53991783&offsets|0|+1]|TAGGTAGTTTCATGTTGTTGGGA</td><td>>MI0000279|hsa-mir-196a-2&WithFlank&12|+|53991732|53991853</td><td>31</td><td>53</td><td>23</td><td>TAGGTAGTTTCATGTTGTTGGGA</td><td>3906.4</td><td>8.5</td><td>0.7</td><td>12.8</td><td>1.1E-37</td><td>3.7E-36</td><td>BT20/MB468</td><td>CYTO</td></tr>
<tr><td>>hg38_21_+_8433172_8446622_RNA45SN1_45S_50ntFlanks@12624.12654.31|CCAATGGGGCGAAGCTACCATCTGTGGGATT</td><td>>hg38_21_+_8433172_8446622_RNA45SN1_45S_50ntFlanks</td><td>12624</td><td>12654</td><td>31</td><td>CCAATGGGGCGAAGCTACCATCTGTGGGATT</td><td>20314.2</td><td>-18.1</td><td>1.4</td><td>-13.0</td><td>1.8E-38</td><td>3.8E-36</td><td>MB231/MB468</td><td>MITO</td></tr>
<tr><td>>MI0000266|hsa-mir-10a&WithFlank&17|-|48579832|48579953@29.49.21(+1A)&[MIMAT0000253&hsa-miR-10a-5p&offsets|+1|-1(+1A);m-18&17|-|48579905|48579926&offsets|+1|0(+1A)]|ACCCTGTAGATCCGAATTTGTA</td><td>>MI0000266|hsa-mir-10a&WithFlank&17|-|48579832|48579953</td><td>29</td><td>49</td><td>21(+1A)</td><td>ACCCTGTAGATCCGAATTTGTA</td><td>3383.2</td><td>16.0</td><td>1.2</td><td>12.8</td><td>1.2E-37</td><td>3.9E-36</td><td>BT20/MB468</td><td>CYTO</td></tr>
<tr><td>>hg38_21_+_8433172_8446622_RNA45SN1_45S_50ntFlanks@11729.11751.23|TAACTATGACTCTCTTAAGGTAG</td><td>>hg38_21_+_8433172_8446622_RNA45SN1_45S_50ntFlanks</td><td>11729</td><td>11751</td><td>23</td><td>TAACTATGACTCTCTTAAGGTAG</td><td>174675.8</td><td>-17.4</td><td>1.4</td><td>-12.8</td><td>1.4E-37</td><td>3.9E-36</td><td>BT20/MB468</td><td>NUC</td></tr>
<tr><td>>hg38_21_+_8433172_8446622_RNA45SN1_45S_50ntFlanks@11672.11700.29|CAGTGCTCTGAATGTCAAAGTGAAGAAAT</td><td>>hg38_21_+_8433172_8446622_RNA45SN1_45S_50ntFlanks</td><td>11672</td><td>11700</td><td>29</td><td>CAGTGCTCTGAATGTCAAAGTGAAGAAAT</td><td>516.9</td><td>-5.7</td><td>0.4</td><td>-12.8</td><td>1.5E-37</td><td>3.9E-36</td><td>BT20/MB468</td><td>NUC</td></tr>
<tr><td>>MI0000650|hsa-mir-200c&WithFlank&12|+|6963693|6963772@50.72.23(+1A)&[MIMAT0000617&hsa-miR-200c-3p&offsets|0|0(+1A);m-54&12|+|6963742|6963764&offsets|0|0(+1A)]|TAATACTGCCGGGTAATGATGGAA</td><td>>MI0000650|hsa-mir-200c&WithFlank&12|+|6963693|6963772</td><td>50</td><td>72</td><td>23(+1A)</td><td>TAATACTGCCGGGTAATGATGGAA</td><td>12098.5</td><td>12.1</td><td>0.9</td><td>13.1</td><td>3.0E-39</td><td>4.0E-36</td><td>BT20/MB231</td><td>NUC</td></tr>
<tr><td>>hg38_21_+_8433172_8446622_RNA45SN1_45S_50ntFlanks@3705.3728.24|TACCTGGTTGATCCTGCCAGTAGC</td><td>>hg38_21_+_8433172_8446622_RNA45SN1_45S_50ntFlanks</td><td>3705</td><td>3728</td><td>24</td><td>TACCTGGTTGATCCTGCCAGTAGC</td><td>12140.0</td><td>8.4</td><td>0.7</td><td>12.8</td><td>1.3E-37</td><td>4.1E-36</td><td>BT20/MB468</td><td>CYTO</td></tr>
<tr><td>>hg38_21_+_8433172_8446622_RNA45SN1_45S_50ntFlanks@12241.12272.32|CAGAAGGGCAAAAGCTCGCTTGATCTTGATTT</td><td>>hg38_21_+_8433172_8446622_RNA45SN1_45S_50ntFlanks</td><td>12241</td><td>12272</td><td>32</td><td>CAGAAGGGCAAAAGCTCGCTTGATCTTGATTT</td><td>616.9</td><td>-5.9</td><td>0.5</td><td>-12.8</td><td>1.6E-37</td><td>4.2E-36</td><td>BT20/MB468</td><td>NUC</td></tr>
<tr><td>>hg38_21_+_8433172_8446622_RNA45SN1_45S_50ntFlanks@10771.10805.35|CAGGTCTCCAAGGTGAACAGCCTCTGGCATGTTGG</td><td>>hg38_21_+_8433172_8446622_RNA45SN1_45S_50ntFlanks</td><td>10771</td><td>10805</td><td>35</td><td>CAGGTCTCCAAGGTGAACAGCCTCTGGCATGTTGG</td><td>2628.4</td><td>-15.8</td><td>1.2</td><td>-12.8</td><td>1.7E-37</td><td>4.6E-36</td><td>BT20/MB468</td><td>NUC</td></tr>
<tr><td>>MI0000273|hsa-mir-183&WithFlank&7|-|129774899|129775020@33.55.23&[MIMAT0000261&hsa-miR-183-5p&offsets|0|+1;m-41&7|-|129774966|129774987&offsets|-1|0]|TATGGCACTGGTAGAATTCACTG</td><td>>MI0000273|hsa-mir-183&WithFlank&7|-|129774899|129775020</td><td>33</td><td>55</td><td>23</td><td>TATGGCACTGGTAGAATTCACTG</td><td>4483.5</td><td>7.8</td><td>0.6</td><td>12.8</td><td>9.1E-38</td><td>4.7E-36</td><td>BT20/MB468</td><td>MITO</td></tr>
<tr><td>>MI0000273|hsa-mir-183&WithFlank&7|-|129774899|129775020@33.53.21(+1C)&[MIMAT0000261&hsa-miR-183-5p&offsets|0|-1(+1C);m-41&7|-|129774966|129774987&offsets|-1|-2(+1C)]|TATGGCACTGGTAGAATTCACC</td><td>>MI0000273|hsa-mir-183&WithFlank&7|-|129774899|129775020</td><td>33</td><td>53</td><td>21(+1C)</td><td>TATGGCACTGGTAGAATTCACC</td><td>3394.6</td><td>7.6</td><td>0.6</td><td>12.8</td><td>1.6E-37</td><td>5.2E-36</td><td>BT20/MB468</td><td>CYTO</td></tr>
<tr><td>>MI0016005|hsa-mir-3615&WithFlank&17|+|74748607|74748705@57.78.22&[MIMAT0017994&hsa-miR-3615&offsets|0|+1;m-271&17|+|74748663|74748684&offsets|0|0]|TCTCTCGGCTCCTCGCGGCTCG</td><td>>MI0016005|hsa-mir-3615&WithFlank&17|+|74748607|74748705</td><td>57</td><td>78</td><td>22</td><td>TCTCTCGGCTCCTCGCGGCTCG</td><td>1520.1</td><td>5.6</td><td>0.4</td><td>12.8</td><td>1.7E-37</td><td>5.3E-36</td><td>BT20/MB468</td><td>CYTO</td></tr>
<tr><td>>hg38_21_+_8433172_8446622_RNA45SN1_45S_50ntFlanks@11235.11254.20|CTTCCCGTGGATCGCCCCAG</td><td>>hg38_21_+_8433172_8446622_RNA45SN1_45S_50ntFlanks</td><td>11235</td><td>11254</td><td>20</td><td>CTTCCCGTGGATCGCCCCAG</td><td>3652.1</td><td>-16.3</td><td>1.3</td><td>-12.8</td><td>2.0E-37</td><td>5.3E-36</td><td>BT20/MB468</td><td>NUC</td></tr>
<tr><td>>hg38_21_+_8433172_8446622_RNA45SN1_45S_50ntFlanks@11576.11605.30|CTTAGAACTGGTGCGGACCAGGGGAATCCG</td><td>>hg38_21_+_8433172_8446622_RNA45SN1_45S_50ntFlanks</td><td>11576</td><td>11605</td><td>30</td><td>CTTAGAACTGGTGCGGACCAGGGGAATCCG</td><td>3210.4</td><td>-16.1</td><td>1.3</td><td>-12.8</td><td>2.0E-37</td><td>5.4E-36</td><td>BT20/MB468</td><td>NUC</td></tr>
<tr><td>>hg38_21_+_8433172_8446622_RNA45SN1_45S_50ntFlanks@11669.11706.38|GCCCAGTGCTCTGAATGTCAAAGTGAAGAAATTCAATG</td><td>>hg38_21_+_8433172_8446622_RNA45SN1_45S_50ntFlanks</td><td>11669</td><td>11706</td><td>38</td><td>GCCCAGTGCTCTGAATGTCAAAGTGAAGAAATTCAATG</td><td>3071.4</td><td>8.9</td><td>0.7</td><td>13.0</td><td>7.1E-39</td><td>5.4E-36</td><td>MB231/MB468</td><td>CYTO</td></tr>
<tr><td>>MI0000815|hsa-mir-339&WithFlank&7|-|1022927|1023032@21.43.23&[MIMAT0000764&hsa-miR-339-5p&offsets|0|0;m-163&7|-|1022990|1023012&offsets|0|0]|TCCCTGTCCTCCAGGAGCTCACG</td><td>>MI0000815|hsa-mir-339&WithFlank&7|-|1022927|1023032</td><td>21</td><td>43</td><td>23</td><td>TCCCTGTCCTCCAGGAGCTCACG</td><td>2323.2</td><td>7.3</td><td>0.6</td><td>12.8</td><td>1.8E-37</td><td>5.5E-36</td><td>BT20/MB468</td><td>CYTO</td></tr>
<tr><td>>hg38_21_+_8433172_8446622_RNA45SN1_45S_50ntFlanks@7981.8003.23|CTCAGATCAGACGTGGCGACCCG</td><td>>hg38_21_+_8433172_8446622_RNA45SN1_45S_50ntFlanks</td><td>7981</td><td>8003</td><td>23</td><td>CTCAGATCAGACGTGGCGACCCG</td><td>725.5</td><td>-7.0</td><td>0.5</td><td>-12.8</td><td>2.1E-37</td><td>5.6E-36</td><td>BT20/MB468</td><td>NUC</td></tr>
<tr><td>>hg38_21_+_8433172_8446622_RNA45SN1_45S_50ntFlanks@10679.10708.30|GGTTCCGGCGGCGTCCGGTGAGCTCTCGCT</td><td>>hg38_21_+_8433172_8446622_RNA45SN1_45S_50ntFlanks</td><td>10679</td><td>10708</td><td>30</td><td>GGTTCCGGCGGCGTCCGGTGAGCTCTCGCT</td><td>16872.6</td><td>-8.7</td><td>0.7</td><td>-12.9</td><td>3.2E-38</td><td>6.3E-36</td><td>MB231/MB468</td><td>MITO</td></tr>
<tr><td>>hg38_21_+_8433172_8446622_RNA45SN1_45S_50ntFlanks@6738.6770.33|ATCATCGACACTTCGAACGCACTTGCGGCCCCG</td><td>>hg38_21_+_8433172_8446622_RNA45SN1_45S_50ntFlanks</td><td>6738</td><td>6770</td><td>33</td><td>ATCATCGACACTTCGAACGCACTTGCGGCCCCG</td><td>5784.8</td><td>-16.9</td><td>1.3</td><td>-12.8</td><td>2.5E-37</td><td>6.6E-36</td><td>BT20/MB468</td><td>NUC</td></tr>
<tr><td>>MI0000650|hsa-mir-200c&WithFlank&12|+|6963693|6963772@49.72.24&[MIMAT0000617&hsa-miR-200c-3p&offsets|-1|0;m-54&12|+|6963742|6963764&offsets|-1|0]|CTAATACTGCCGGGTAATGATGGA</td><td>>MI0000650|hsa-mir-200c&WithFlank&12|+|6963693|6963772</td><td>49</td><td>72</td><td>24</td><td>CTAATACTGCCGGGTAATGATGGA</td><td>679.6</td><td>7.7</td><td>0.6</td><td>12.8</td><td>1.3E-37</td><td>6.6E-36</td><td>BT20/MB468</td><td>MITO</td></tr>
<tr><td>>hg38_21_+_8433172_8446622_RNA45SN1_45S_50ntFlanks@10350.10376.27|ACGAGAACTTTGAAGGCCGAAGTGGAG</td><td>>hg38_21_+_8433172_8446622_RNA45SN1_45S_50ntFlanks</td><td>10350</td><td>10376</td><td>27</td><td>ACGAGAACTTTGAAGGCCGAAGTGGAG</td><td>2320.5</td><td>-6.3</td><td>0.5</td><td>-12.8</td><td>9.7E-38</td><td>6.8E-36</td><td>MB231/MB468</td><td>NUC</td></tr>
<tr><td>>hg38_21_+_8433172_8446622_RNA45SN1_45S_50ntFlanks@10679.10711.33|GGTTCCGGCGGCGTCCGGTGAGCTCTCGCTGGC</td><td>>hg38_21_+_8433172_8446622_RNA45SN1_45S_50ntFlanks</td><td>10679</td><td>10711</td><td>33</td><td>GGTTCCGGCGGCGTCCGGTGAGCTCTCGCTGGC</td><td>22603.3</td><td>-12.0</td><td>0.9</td><td>-12.9</td><td>3.6E-38</td><td>6.9E-36</td><td>MB231/MB468</td><td>MITO</td></tr>
<tr><td>>hg38_21_+_8433172_8446622_RNA45SN1_45S_50ntFlanks@11572.11603.32|CCGACTTAGAACTGGTGCGGACCAGGGGAATC</td><td>>hg38_21_+_8433172_8446622_RNA45SN1_45S_50ntFlanks</td><td>11572</td><td>11603</td><td>32</td><td>CCGACTTAGAACTGGTGCGGACCAGGGGAATC</td><td>20369.6</td><td>-9.5</td><td>0.7</td><td>-12.9</td><td>3.7E-38</td><td>6.9E-36</td><td>MB231/MB468</td><td>MITO</td></tr>
<tr><td>>hg38_21_+_8433172_8446622_RNA45SN1_45S_50ntFlanks@11576.11609.34|CTTAGAACTGGTGCGGACCAGGGGAATCCGACTG</td><td>>hg38_21_+_8433172_8446622_RNA45SN1_45S_50ntFlanks</td><td>11576</td><td>11609</td><td>34</td><td>CTTAGAACTGGTGCGGACCAGGGGAATCCGACTG</td><td>1085.1</td><td>-6.5</td><td>0.5</td><td>-12.8</td><td>2.8E-37</td><td>7.3E-36</td><td>BT20/MB468</td><td>NUC</td></tr>
<tr><td>>hg38_21_+_8433172_8446622_RNA45SN1_45S_50ntFlanks@5446.5466.21|CGGATCGGCCCCGCCGGGGTC</td><td>>hg38_21_+_8433172_8446622_RNA45SN1_45S_50ntFlanks</td><td>5446</td><td>5466</td><td>21</td><td>CGGATCGGCCCCGCCGGGGTC</td><td>54301.7</td><td>-19.6</td><td>1.5</td><td>-13.0</td><td>1.4E-38</td><td>7.3E-36</td><td>MB231/MB468</td><td>MP</td></tr>
<tr><td>>hg38_21_+_8433172_8446622_RNA45SN1_45S_50ntFlanks@12420.12445.26|ATGTCGGCTCTTCCTATCATTGTGAA</td><td>>hg38_21_+_8433172_8446622_RNA45SN1_45S_50ntFlanks</td><td>12420</td><td>12445</td><td>26</td><td>ATGTCGGCTCTTCCTATCATTGTGAA</td><td>6190.5</td><td>-8.6</td><td>0.7</td><td>-12.8</td><td>3.1E-37</td><td>8.0E-36</td><td>BT20/MB468</td><td>NUC</td></tr>
<tr><td>>hg38_21_+_8433172_8446622_RNA45SN1_45S_50ntFlanks@11572.11600.29|CCGACTTAGAACTGGTGCGGACCAGGGGA</td><td>>hg38_21_+_8433172_8446622_RNA45SN1_45S_50ntFlanks</td><td>11572</td><td>11600</td><td>29</td><td>CCGACTTAGAACTGGTGCGGACCAGGGGA</td><td>9564.6</td><td>-8.4</td><td>0.6</td><td>-13.0</td><td>1.7E-38</td><td>8.8E-36</td><td>MB231/MB468</td><td>MP</td></tr>
<tr><td>>hg38_21_+_8433172_8446622_RNA45SN1_45S_50ntFlanks@12996.13017.22|TCGCTGCGATCTATTGAAAGTC</td><td>>hg38_21_+_8433172_8446622_RNA45SN1_45S_50ntFlanks</td><td>12996</td><td>13017</td><td>22</td><td>TCGCTGCGATCTATTGAAAGTC</td><td>2342.7</td><td>7.1</td><td>0.5</td><td>13.0</td><td>1.3E-38</td><td>9.1E-36</td><td>MB231/MB468</td><td>CYTO</td></tr>
<tr><td>>MI0000285|hsa-mir-205&WithFlank&1|+|209432127|209432248@40.62.23&[MIMAT0000266&hsa-miR-205-5p&offsets|0|+1;m-351&1|+|209432166|209432187&offsets|0|+1]|TCCTTCATTCCACCGGAGTCTGT</td><td>>MI0000285|hsa-mir-205&WithFlank&1|+|209432127|209432248</td><td>40</td><td>62</td><td>23</td><td>TCCTTCATTCCACCGGAGTCTGT</td><td>125601.1</td><td>7.7</td><td>0.6</td><td>12.8</td><td>2.9E-37</td><td>9.2E-36</td><td>BT20/MB468</td><td>CYTO</td></tr>
<tr><td>>hg38_21_+_8433172_8446622_RNA45SN1_45S_50ntFlanks@7978.8006.29|GACCTCAGATCAGACGTGGCGACCCGCTG</td><td>>hg38_21_+_8433172_8446622_RNA45SN1_45S_50ntFlanks</td><td>7978</td><td>8006</td><td>29</td><td>GACCTCAGATCAGACGTGGCGACCCGCTG</td><td>857.1</td><td>-5.7</td><td>0.4</td><td>-12.7</td><td>4.0E-37</td><td>1.0E-35</td><td>BT20/MB468</td><td>NUC</td></tr>
<tr><td>>hg38_21_+_8433172_8446622_RNA45SN1_45S_50ntFlanks@11669.11692.24|GCCCAGTGCTCTGAATGTCAAAGT</td><td>>hg38_21_+_8433172_8446622_RNA45SN1_45S_50ntFlanks</td><td>11669</td><td>11692</td><td>24</td><td>GCCCAGTGCTCTGAATGTCAAAGT</td><td>433.9</td><td>7.1</td><td>0.6</td><td>12.8</td><td>2.0E-37</td><td>1.0E-35</td><td>BT20/MB468</td><td>MITO</td></tr>
<tr><td>>hg38_21_+_8433172_8446622_RNA45SN1_45S_50ntFlanks@9568.9597.30|CGTGCAAATCGGTCGTCCGACCTGGGTATA</td><td>>hg38_21_+_8433172_8446622_RNA45SN1_45S_50ntFlanks</td><td>9568</td><td>9597</td><td>30</td><td>CGTGCAAATCGGTCGTCCGACCTGGGTATA</td><td>17765.1</td><td>-11.6</td><td>0.9</td><td>-12.9</td><td>5.9E-38</td><td>1.1E-35</td><td>MB231/MB468</td><td>MITO</td></tr>
<tr><td>>hg38_21_+_8433172_8446622_RNA45SN1_45S_50ntFlanks@6721.6745.25|AATTGCAGGACACATTGATCATCGA</td><td>>hg38_21_+_8433172_8446622_RNA45SN1_45S_50ntFlanks</td><td>6721</td><td>6745</td><td>25</td><td>AATTGCAGGACACATTGATCATCGA</td><td>3103.5</td><td>-16.0</td><td>1.3</td><td>-12.7</td><td>4.3E-37</td><td>1.1E-35</td><td>BT20/MB468</td><td>NUC</td></tr>
<tr><td>>hg38_21_+_8433172_8446622_RNA45SN1_45S_50ntFlanks@12163.12180.18|ACTGGGGCGGTACACCTG</td><td>>hg38_21_+_8433172_8446622_RNA45SN1_45S_50ntFlanks</td><td>12163</td><td>12180</td><td>18</td><td>ACTGGGGCGGTACACCTG</td><td>10032.3</td><td>-17.2</td><td>1.3</td><td>-13.0</td><td>2.3E-38</td><td>1.1E-35</td><td>MB231/MB468</td><td>MP</td></tr>
<tr><td>>hg38_21_+_8433172_8446622_RNA45SN1_45S_50ntFlanks@12456.12476.21|CCAAGCGTTGGATTGTTCACC</td><td>>hg38_21_+_8433172_8446622_RNA45SN1_45S_50ntFlanks</td><td>12456</td><td>12476</td><td>21</td><td>CCAAGCGTTGGATTGTTCACC</td><td>23063.4</td><td>-9.1</td><td>0.7</td><td>-12.8</td><td>8.7E-38</td><td>1.5E-35</td><td>MB231/MB468</td><td>MITO</td></tr>
<tr><td>>hg38_21_+_8433172_8446622_RNA45SN1_45S_50ntFlanks@10819.10840.22|AGGGAAGTCGGCAAGCCGGATC</td><td>>hg38_21_+_8433172_8446622_RNA45SN1_45S_50ntFlanks</td><td>10819</td><td>10840</td><td>22</td><td>AGGGAAGTCGGCAAGCCGGATC</td><td>6414.7</td><td>-7.6</td><td>0.6</td><td>-12.8</td><td>2.2E-37</td><td>1.5E-35</td><td>MB231/MB468</td><td>NUC</td></tr>
</table>

<table>
<tr><td>>MI0000750|hsa-mir-26a-2&WithFlank&12|-|57824603|57824698@20.40.21;MI0000083|hsa-mir-26a-1&WithFlank&3|+|37969398|37969486@16.36.21&[MIMAT0000082&hsa-miR-26a-5p&offsets|0|-1;m-33&12|-|57824658|57824679&offsets|0|-1];[MIMAT0000082_1&hsa-miR-26a-5p&offsets|0|-1;m-34&3|+|37969413|37969434&offsets|0|-1]|TTCAAGTAATCCAGGATAGGC</td><td>>MI0000750|hsa-mir-26a-2&WithFlank&12|-|57824603|57824698</td><td>20</td><td>40</td><td>21</td><td>TTCAAGTAATCCAGGATAGGC</td><td>2014.4</td><td>6.9</td><td>0.5</td><td>12.7</td><td>5.1E-37</td><td>1.6E-35</td><td>BT20/MB468</td><td>CYTO</td></tr>
<tr><td>>hg38_21_+_8433172_8446622_RNA45SN1_45S_50ntFlanks@7983.8006.24|CAGATCAGACGTGGCGACCCGCTG</td><td>>hg38_21_+_8433172_8446622_RNA45SN1_45S_50ntFlanks</td><td>7983</td><td>8006</td><td>24</td><td>CAGATCAGACGTGGCGACCCGCTG</td><td>5844.8</td><td>-8.0</td><td>0.6</td><td>-12.8</td><td>2.5E-37</td><td>1.7E-35</td><td>MB231/MB468</td><td>NUC</td></tr>
<tr><td>>MI0000081|hsa-mir-24-2&WithFlank&19|-|13836281|13836365@56.75.20;MI0000080|hsa-mir-24-1&WithFlank&9|+|95086015|95086094@50.69.20&[MIMAT0000080&hsa-miR-24-3p&offsets|0|-2;m-36&19|-|13836288|13836310&offsets|0|-3];[MIMAT0000080_1&hsa-miR-24-3p&offsets|0|-2;m-35&9|+|95086064|95086086&offsets|0|-3]|TGGCTCAGTTCAGCAGGAAC</td><td>>MI0000081|hsa-mir-24-2&WithFlank&19|-|13836281|13836365</td><td>56</td><td>75</td><td>20</td><td>TGGCTCAGTTCAGCAGGAAC</td><td>17236.4</td><td>6.5</td><td>0.5</td><td>12.7</td><td>3.9E-37</td><td>2.0E-35</td><td>BT20/MB468</td><td>MITO</td></tr>
<tr><td>>hg38_21_+_8433172_8446622_RNA45SN1_45S_50ntFlanks@11567.11588.22|AGCAGCCGACTTAGAACTGGTG</td><td>>hg38_21_+_8433172_8446622_RNA45SN1_45S_50ntFlanks</td><td>11567</td><td>11588</td><td>22</td><td>AGCAGCCGACTTAGAACTGGTG</td><td>496.0</td><td>-6.0</td><td>0.5</td><td>-12.7</td><td>7.9E-37</td><td>2.0E-35</td><td>BT20/MB468</td><td>NUC</td></tr>
<tr><td>>MI0000066|hsa-let-7e&WithFlank&19|+|51692780|51692870@14.35.22(+1U)&[MIMAT0000066&hsa-let-7e-5p&offsets|0|0(+1U);m-50&19|+|51692793|51692814&offsets|0|0(+1U)]|TGAGGTAGGAGGTTGTATAGTTT</td><td>>MI0000066|hsa-let-7e&WithFlank&19|+|51692780|51692870</td><td>14</td><td>35</td><td>22(+1U)</td><td>TGAGGTAGGAGGTTGTATAGTTT</td><td>2195.7</td><td>7.7</td><td>0.6</td><td>12.7</td><td>6.6E-37</td><td>2.0E-35</td><td>BT20/MB468</td><td>CYTO</td></tr>
<tr><td>>hg38_21_+_8433172_8446622_RNA45SN1_45S_50ntFlanks@6697.6718.22|CAGCTAGCTGCGAGAATTAATG</td><td>>hg38_21_+_8433172_8446622_RNA45SN1_45S_50ntFlanks</td><td>6697</td><td>6718</td><td>22</td><td>CAGCTAGCTGCGAGAATTAATG</td><td>2017.7</td><td>-5.6</td><td>0.4</td><td>-12.7</td><td>4.1E-37</td><td>2.1E-35</td><td>BT20/MB468</td><td>MITO</td></tr>
<tr><td>>MI0005756|hsa-mir-934&WithFlank&X|+|136550872|136550966@21.42.22&[MIMAT0004977&hsa-miR-934&offsets|0|0;m-943&X|+|136550892|136550913&offsets|0|0]|TGTCTACTACTGGAGACACTGG</td><td>>MI0005756|hsa-mir-934&WithFlank&X|+|136550872|136550966</td><td>21</td><td>42</td><td>22</td><td>TGTCTACTACTGGAGACACTGG</td><td>1772.3</td><td>7.3</td><td>0.6</td><td>12.7</td><td>6.8E-37</td><td>2.1E-35</td><td>BT20/MB468</td><td>CYTO</td></tr>
<tr><td>>hg38_21_+_8433172_8446622_RNA45SN1_45S_50ntFlanks@9554.9577.24|CGTAGCGGTCCTGACGTGCAAATC</td><td>>hg38_21_+_8433172_8446622_RNA45SN1_45S_50ntFlanks</td><td>9554</td><td>9577</td><td>24</td><td>CGTAGCGGTCCTGACGTGCAAATC</td><td>1952.6</td><td>-4.9</td><td>0.4</td><td>-12.7</td><td>8.8E-37</td><td>2.2E-35</td><td>BT20/MB468</td><td>NUC</td></tr>
<tr><td>>hg38_21_+_8433172_8446622_RNA45SN1_45S_50ntFlanks@11572.11591.20|CCGACTTAGAACTGGTGCGG</td><td>>hg38_21_+_8433172_8446622_RNA45SN1_45S_50ntFlanks</td><td>11572</td><td>11591</td><td>20</td><td>CCGACTTAGAACTGGTGCGG</td><td>14372.9</td><td>-12.0</td><td>0.9</td><td>-12.9</td><td>4.9E-38</td><td>2.3E-35</td><td>MB231/MB468</td><td>MP</td></tr>
<tr><td>>hg38_21_+_8433172_8446622_RNA45SN1_45S_50ntFlanks@5485.5514.30|AGCGCTGAGAAGACGGTCGAACTTGACTAT</td><td>>hg38_21_+_8433172_8446622_RNA45SN1_45S_50ntFlanks</td><td>5485</td><td>5514</td><td>30</td><td>AGCGCTGAGAAGACGGTCGAACTTGACTAT</td><td>17968.6</td><td>-18.0</td><td>1.4</td><td>-12.9</td><td>5.3E-38</td><td>2.4E-35</td><td>MB231/MB468</td><td>MP</td></tr>
<tr><td>>MI0003643|hsa-mir-629&WithFlank&15|-|70079366|70079474@28.49.22&[MIMAT0004810&hsa-miR-629-5p&offsets|0|+1;m-150&15|-|70079427|70079447&offsets|0|+1]|TGGGTTTACGTTGGGAGAACTT</td><td>>MI0003643|hsa-mir-629&WithFlank&15|-|70079366|70079474</td><td>28</td><td>49</td><td>22</td><td>TGGGTTTACGTTGGGAGAACTT</td><td>4949.8</td><td>16.5</td><td>1.3</td><td>12.7</td><td>8.1E-37</td><td>2.5E-35</td><td>BT20/MB468</td><td>CYTO</td></tr>
<tr><td>>MI0000809|hsa-mir-151a&WithFlank&8|-|140732558|140732659@51.73.23&[MIMAT0000757&hsa-miR-151a-3p&offsets|-2|0;m-30&8|-|140732587|140732607&offsets|-2|0]|TACTAGACTGAAGCTCCTTGAGG</td><td>>MI0000809|hsa-mir-151a&WithFlank&8|-|140732558|140732659</td><td>51</td><td>73</td><td>23</td><td>TACTAGACTGAAGCTCCTTGAGG</td><td>10432.7</td><td>8.3</td><td>0.7</td><td>12.7</td><td>8.2E-37</td><td>2.5E-35</td><td>BT20/MB468</td><td>CYTO</td></tr>
<tr><td>>MI0000279|hsa-mir-196a-2&WithFlank&12|+|53991732|53991853@31.52.22;MI0000238|hsa-mir-196a-1&WithFlank&17|-|48632484|48632565@13.34.22&[MIMAT0000226&hsa-miR-196a-5p&offsets|0|0;m-261&12|+|53991762|53991783&offsets|0|0];[MIMAT0000226_1&hsa-miR-196a-5p&offsets|0|0;m-268&17|-|48632532|48632553&offsets|0|0]|TAGGTAGTTTCATGTTGTTGGG</td><td>>MI0000279|hsa-mir-196a-2&WithFlank&12|+|53991732|53991853</td><td>31</td><td>52</td><td>22</td><td>TAGGTAGTTTCATGTTGTTGGG</td><td>9349.5</td><td>8.1</td><td>0.6</td><td>12.7</td><td>8.4E-37</td><td>2.6E-35</td><td>BT20/MB468</td><td>CYTO</td></tr>
<tr><td>>hg38_21_+_8433172_8446622_RNA45SN1_45S_50ntFlanks@12476.12500.25|CCACTAATAGGGAACGTGAGCTGGG</td><td>>hg38_21_+_8433172_8446622_RNA45SN1_45S_50ntFlanks</td><td>12476</td><td>12500</td><td>25</td><td>CCACTAATAGGGAACGTGAGCTGGG</td><td>2087.8</td><td>-15.5</td><td>1.2</td><td>-12.6</td><td>1.1E-36</td><td>2.9E-35</td><td>BT20/MB468</td><td>NUC</td></tr>
<tr><td>>hg38_21_+_8433172_8446622_RNA45SN1_45S_50ntFlanks@12324.12346.23|TTGGGTTTTAAGCAGGAGGTGTC</td><td>>hg38_21_+_8433172_8446622_RNA45SN1_45S_50ntFlanks</td><td>12324</td><td>12346</td><td>23</td><td>TTGGGTTTTAAGCAGGAGGTGTC</td><td>4690.8</td><td>-16.6</td><td>1.3</td><td>-12.6</td><td>1.2E-36</td><td>2.9E-35</td><td>BT20/MB468</td><td>NUC</td></tr>
<tr><td>>hg38_21_+_8433172_8446622_RNA45SN1_45S_50ntFlanks@12416.12436.21|TTCGATGTCGGCTCTTCCTAT</td><td>>hg38_21_+_8433172_8446622_RNA45SN1_45S_50ntFlanks</td><td>12416</td><td>12436</td><td>21</td><td>TTCGATGTCGGCTCTTCCTAT</td><td>3350.2</td><td>-7.2</td><td>0.6</td><td>-12.6</td><td>1.2E-36</td><td>3.0E-35</td><td>BT20/MB468</td><td>NUC</td></tr>
<tr><td>>hg38_21_+_8433172_8446622_RNA45SN1_45S_50ntFlanks@11209.11232.24|CGGCGACTCTGGACGCGAGCCGGG</td><td>>hg38_21_+_8433172_8446622_RNA45SN1_45S_50ntFlanks</td><td>11209</td><td>11232</td><td>24</td><td>CGGCGACTCTGGACGCGAGCCGGG</td><td>17408.3</td><td>-8.3</td><td>0.6</td><td>-12.7</td><td>4.4E-37</td><td>3.0E-35</td><td>MB231/MB468</td><td>NUC</td></tr>
<tr><td>>MI0000809|hsa-mir-151a&WithFlank&8|-|140732558|140732659@51.72.22&[MIMAT0000757&hsa-miR-151a-3p&offsets|-2|-1;m-30&8|-|140732587|140732607&offsets|-2|-1]|TACTAGACTGAAGCTCCTTGAG</td><td>>MI0000809|hsa-mir-151a&WithFlank&8|-|140732558|140732659</td><td>51</td><td>72</td><td>22</td><td>TACTAGACTGAAGCTCCTTGAG</td><td>833.2</td><td>7.7</td><td>0.6</td><td>12.7</td><td>6.1E-37</td><td>3.0E-35</td><td>BT20/MB468</td><td>MITO</td></tr>
<tr><td>>MI0000255|hsa-mir-30d&WithFlank&8|-|134804870|134804951@12.34.23(+1C)&[MIMAT0000245&hsa-miR-30d-5p&offsets|0|+1(+1C);m-21&8|-|134804919|134804940&offsets|0|+1(+1C)]|TGTAAACATCCCCGACTGGAAGCC</td><td>>MI0000255|hsa-mir-30d&WithFlank&8|-|134804870|134804951</td><td>12</td><td>34</td><td>23(+1C)</td><td>TGTAAACATCCCCGACTGGAAGCC</td><td>11596.1</td><td>8.5</td><td>0.7</td><td>12.7</td><td>6.9E-37</td><td>3.4E-35</td><td>BT20/MB468</td><td>MITO</td></tr>
<tr><td>>hg38_21_+_8433172_8446622_RNA45SN1_45S_50ntFlanks@11561.11585.25|GCGCCTAGCAGCCGACTTAGAACTG</td><td>>hg38_21_+_8433172_8446622_RNA45SN1_45S_50ntFlanks</td><td>11561</td><td>11585</td><td>25</td><td>GCGCCTAGCAGCCGACTTAGAACTG</td><td>3505.8</td><td>-16.2</td><td>1.3</td><td>-12.6</td><td>1.4E-36</td><td>3.4E-35</td><td>BT20/MB468</td><td>NUC</td></tr>
<tr><td>>MI0000471|hsa-mir-126&WithFlank&9|+|136670596|136670692@58.79.22&[MIMAT0000445&hsa-miR-126-3p&offsets|0|0;m-40&9|+|136670653|136670674&offsets|0|0]|TCGTACCGTGAGTAATAATGCG</td><td>>MI0000471|hsa-mir-126&WithFlank&9|+|136670596|136670692</td><td>58</td><td>79</td><td>22</td><td>TCGTACCGTGAGTAATAATGCG</td><td>2836.5</td><td>9.4</td><td>0.7</td><td>12.7</td><td>5.2E-37</td><td>3.5E-35</td><td>MB231/MB468</td><td>NUC</td></tr>
<tr><td>>hg38_21_+_8433172_8446622_RNA45SN1_45S_50ntFlanks@11569.11600.32|CAGCCGACTTAGAACTGGTGCGGACCAGGGGA</td><td>>hg38_21_+_8433172_8446622_RNA45SN1_45S_50ntFlanks</td><td>11569</td><td>11600</td><td>32</td><td>CAGCCGACTTAGAACTGGTGCGGACCAGGGGA</td><td>6728.4</td><td>-7.0</td><td>0.6</td><td>-12.7</td><td>5.4E-37</td><td>3.6E-35</td><td>MB231/MB468</td><td>NUC</td></tr>
<tr><td>>MI0000087|hsa-mir-29a&WithFlank&7|-|130876741|130876816@47.68.22&[MIMAT0000086&hsa-miR-29a-3p&offsets|-1|-1;m-26&7|-|130876749|130876769&offsets|-1|0]|CTAGCACCATCTGAAATCGGTT</td><td>>MI0000087|hsa-mir-29a&WithFlank&7|-|130876741|130876816</td><td>47</td><td>68</td><td>22</td><td>CTAGCACCATCTGAAATCGGTT</td><td>14260.4</td><td>8.3</td><td>0.7</td><td>12.7</td><td>5.4E-37</td><td>3.6E-35</td><td>MB231/MB468</td><td>NUC</td></tr>
<tr><td>>am_MI0005763|hsa-mir-941-1&WithFlank&20|+|63919443|63919526@53.75.23;MI0005764|hsa-mir-941-2&WithFlank&20|+|63919499|63919582@53.75.23;MI0005765|hsa-mir-941-3&WithFlank&20|+|63919555|63919638@53.75.23;MI0005766|hsa-mir-941-4&WithFlank&20|+|63919750|63919833@53.75.23;MI0031520|hsa-mir-941-5&WithFlank&20|+|63919862|63919945@53.75.23&[MIMAT0004984&hsa-miR-941&offsets|0|0;m-263&20|+|63919495|63919517&offsets|0|0];[MIMAT0004984_1&hsa-miR-941&offsets|0|0;m-263&20|+|63919551|63919573&offsets|0|0];[MIMAT0004984_2&hsa-miR-941&offsets|0|0;m-263&20|+|63919607|63919629&offsets|0|0];[MIMAT0004984_3&hsa-miR-941&offsets|0|0;m-263&20|+|63919802|63919824&offsets|0|0];[MIMAT0004984_4&hsa-miR-941&offsets|0|0;m-263&20|+|63919914|63919936&offsets|0|0]|CACCCGGCTGTGTGCACATGTGC</td><td>>am_MI0005763|hsa-mir-941-1&WithFlank&20|+|63919443|63919526</td><td>53</td><td>75</td><td>23</td><td>CACCCGGCTGTGTGCACATGTGC</td><td>25213.6</td><td>7.6</td><td>0.6</td><td>12.6</td><td>1.2E-36</td><td>3.6E-35</td><td>BT20/MB468</td><td>CYTO</td></tr>
<tr><td>>MI0000272|hsa-mir-182&WithFlank&7|-|129770377|129770498@29.51.23(+1C)&[MIMAT0000259&hsa-miR-182-5p&offsets|0|-1(+1C);m-22&7|-|129770449|129770470&offsets|0|+1(+1C)]|TTTGGCAATGGTAGAACTCACACC</td><td>>MI0000272|hsa-mir-182&WithFlank&7|-|129770377|129770498</td><td>29</td><td>51</td><td>23(+1C)</td><td>TTTGGCAATGGTAGAACTCACACC</td><td>4912.9</td><td>8.3</td><td>0.7</td><td>12.7</td><td>7.6E-37</td><td>3.7E-35</td><td>BT20/MB468</td><td>MITO</td></tr>
<tr><td>>hg38_21_+_8433172_8446622_RNA45SN1_45S_50ntFlanks@8357.8375.19|AAAAGAACTTTGAAGAGAG</td><td>>hg38_21_+_8433172_8446622_RNA45SN1_45S_50ntFlanks</td><td>8357</td><td>8375</td><td>19</td><td>AAAAGAACTTTGAAGAGAG</td><td>5468.7</td><td>-8.4</td><td>0.7</td><td>-12.6</td><td>1.5E-36</td><td>3.7E-35</td><td>BT20/MB468</td><td>NUC</td></tr>
<tr><td>>hg38_21_+_8433172_8446622_RNA45SN1_45S_50ntFlanks@12324.12344.21|TTGGGTTTTAAGCAGGAGGTG</td><td>>hg38_21_+_8433172_8446622_RNA45SN1_45S_50ntFlanks</td><td>12324</td><td>12344</td><td>21</td><td>TTGGGTTTTAAGCAGGAGGTG</td><td>3094.0</td><td>-16.0</td><td>1.3</td><td>-12.6</td><td>1.5E-36</td><td>3.8E-35</td><td>BT20/MB468</td><td>NUC</td></tr>
<tr><td>>MI0000266|hsa-mir-10a&WithFlank&17|-|48579832|48579953@29.51.23&[MIMAT0000253&hsa-miR-10a-5p&offsets|+1|+1;m-18&17|-|48579905|48579926&offsets|+1|+2]|ACCCTGTAGATCCGAATTTGTGT</td><td>>MI0000266|hsa-mir-10a&WithFlank&17|-|48579832|48579953</td><td>29</td><td>51</td><td>23</td><td>ACCCTGTAGATCCGAATTTGTGT</td><td>2560.0</td><td>15.6</td><td>1.2</td><td>12.6</td><td>1.4E-36</td><td>4.2E-35</td><td>BT20/MB468</td><td>CYTO</td></tr>
<tr><td>>MI0000439|hsa-mir-23b&WithFlank&9|+|95085202|95085310@67.84.18&[MIMAT0000418&hsa-miR-23b-3p&offsets|+3|-2;m-42&9|+|95085265|95085285&offsets|+3|0]|ACATTGCCAGGGATTACC</td><td>>MI0000439|hsa-mir-23b&WithFlank&9|+|95085202|95085310</td><td>67</td><td>84</td><td>18</td><td>ACATTGCCAGGGATTACC</td><td>3299.2</td><td>15.9</td><td>1.3</td><td>12.6</td><td>1.6E-36</td><td>4.8E-35</td><td>BT20/MB468</td><td>CYTO</td></tr>
<tr><td>>MI0000433|hsa-let-7g&WithFlank&3|-|52268272|52268367@11.32.22(+1A)&[MIMAT0000414&hsa-let-7g-5p&offsets|0|0(+1A);m-58&3|-|52268336|52268357&offsets|0|0(+1A)]|TGAGGTAGTAGTTTGTACAGTTA</td><td>>MI0000433|hsa-let-7g&WithFlank&3|-|52268272|52268367</td><td>11</td><td>32</td><td>22(+1A)</td><td>TGAGGTAGTAGTTTGTACAGTTA</td><td>1451.8</td><td>7.2</td><td>0.6</td><td>12.7</td><td>1.0E-36</td><td>4.9E-35</td><td>BT20/MB468</td><td>MITO</td></tr>
<tr><td>>MI0003643|hsa-mir-629&WithFlank&15|-|70079366|70079474@28.49.22&[MIMAT0004810&hsa-miR-629-5p&offsets|0|+1;m-150&15|-|70079427|70079447&offsets|0|+1]|TGGGTTTACGTTGGGAGAACTT</td><td>>MI0003643|hsa-mir-629&WithFlank&15|-|70079366|70079474</td><td>28</td><td>49</td><td>22</td><td>TGGGTTTACGTTGGGAGAACTT</td><td>3159.8</td><td>16.1</td><td>1.3</td><td>12.7</td><td>1.1E-36</td><td>5.2E-35</td><td>BT20/MB468</td><td>MITO</td></tr>
<tr><td>>hg38_21_+_8433172_8446622_RNA45SN1_45S_50ntFlanks@12937.12961.25|AACCATTCGTAGACGACCTGCTTCT</td><td>>hg38_21_+_8433172_8446622_RNA45SN1_45S_50ntFlanks</td><td>12937</td><td>12961</td><td>25</td><td>AACCATTCGTAGACGACCTGCTTCT</td><td>1735.1</td><td>-15.2</td><td>1.2</td><td>-12.6</td><td>2.2E-36</td><td>5.5E-35</td><td>BT20/MB468</td><td>NUC</td></tr>
<tr><td>>hg38_21_+_8433172_8446622_RNA45SN1_45S_50ntFlanks@3875.3898.24|ATACATGCCGACGGGCGCTGACCC</td><td>>hg38_21_+_8433172_8446622_RNA45SN1_45S_50ntFlanks</td><td>3875</td><td>3898</td><td>24</td><td>ATACATGCCGACGGGCGCTGACCC</td><td>13132.0</td><td>-11.5</td><td>0.9</td><td>-12.7</td><td>3.3E-37</td><td>5.7E-35</td><td>MB231/MB468</td><td>MITO</td></tr>
<tr><td>>MI0000286|hsa-mir-210&WithFlank&11|-|568083|568204@72.93.22&[MIMAT0000267&hsa-miR-210-3p&offsets|0|0;m-102&11|-|568112|568133&offsets|0|0]|CTGTGCGTGTGACAGCGGCTGA</td><td>>MI0000286|hsa-mir-210&WithFlank&11|-|568083|568204</td><td>72</td><td>93</td><td>22</td><td>CTGTGCGTGTGACAGCGGCTGA</td><td>834.7</td><td>8.4</td><td>0.7</td><td>12.6</td><td>1.2E-36</td><td>5.9E-35</td><td>BT20/MB468</td><td>MITO</td></tr>
<tr><td>>MI0000650|hsa-mir-200c&WithFlank&12|+|6963693|6963772@50.71.22(+1U)&[MIMAT0000617&hsa-miR-200c-3p&offsets|0|-1(+1U);m-54&12|+|6963742|6963764&offsets|0|-1(+1U)]|TAATACTGCCGGGTAATGATGGT</td><td>>MI0000650|hsa-mir-200c&WithFlank&12|+|6963693|6963772</td><td>50</td><td>71</td><td>22(+1U)</td><td>TAATACTGCCGGGTAATGATGGT</td><td>3335.6</td><td>8.4</td><td>0.7</td><td>12.6</td><td>1.3E-36</td><td>6.3E-35</td><td>BT20/MB468</td><td>MITO</td></tr>
<tr><td>>trna111_HisGTG_1_-_147774845_147774916@-1T.23.24,trna118_HisGTG_1_-_145396881_145396952@-1T.23.24,trna16_HisGTG_1_+_146544773_146544844@-1T.23.24,trna1_HisGTG_15_+_45493349_45493420@-1T.23.24,trna21_HisGTG_1_+_147753471_147753542@-1T.23.24,trna33_HisGTG_6_+_27125906_27125977@-1T.23.24,trna7_HisGTG_9_-_14433938_14434009@-1T.23.24,trna8_HisGTG_15_-_45492611_45492682@-1T.23.24,trna9_HisGTG_15_-_45490804_45490875@-1T.23.24|TGCCGTGATCGTATAGTGGTTAGT</td><td>>trna111_HisGTG_1_-_147774845_147774916</td><td>-1T</td><td>23</td><td>24</td><td>TGCCGTGATCGTATAGTGGTTAGT</td><td>6295.5</td><td>9.5</td><td>0.8</td><td>12.7</td><td>1.1E-36</td><td>6.8E-35</td><td>MB231/MB468</td><td>NUC</td></tr>
<tr><td>>hg38_21_+_8433172_8446622_RNA45SN1_45S_50ntFlanks@10411.10429.19|CAGTCGGTCCTGAGAGATG</td><td>>hg38_21_+_8433172_8446622_RNA45SN1_45S_50ntFlanks</td><td>10411</td><td>10429</td><td>19</td><td>CAGTCGGTCCTGAGAGATG</td><td>11100.6</td><td>-9.2</td><td>0.7</td><td>-12.7</td><td>4.1E-37</td><td>6.9E-35</td><td>MB231/MB468</td><td>MITO</td></tr>
<tr><td>>hg38_21_+_8433172_8446622_RNA45SN1_45S_50ntFlanks@10349.10383.35|AACGAGAACTTTGAAGGCCGAAGTGGAGAAGGGTT</td><td>>hg38_21_+_8433172_8446622_RNA45SN1_45S_50ntFlanks</td><td>10349</td><td>10383</td><td>35</td><td>AACGAGAACTTTGAAGGCCGAAGTGGAGAAGGGTT</td><td>22359.4</td><td>-8.7</td><td>0.7</td><td>-12.6</td><td>2.9E-36</td><td>7.1E-35</td><td>BT20/MB468</td><td>NUC</td></tr>
</table>

<table>
<tr><td>>hg38_21_+_8433172_8446622_RNA45SN1_45S_50ntFlanks@9567.9596.30|ACGTGCAAATCGGTCGTCCGACCTGGGTAT</td><td>>hg38_21_+_8433172_8446622_RNA45SN1_45S_50ntFlanks</td><td>9567</td><td>9596</td><td>30</td><td>ACGTGCAAATCGGTCGTCCGACCTGGGTAT</td><td>16997.4</td><td>-11.6</td><td>0.9</td><td>-12.7</td><td>4.4E-37</td><td>7.2E-35</td><td>MB231/MB468</td><td>MITO</td></tr>
<tr><td>>hg38_21_+_8433172_8446622_RNA45SN1_45S_50ntFlanks@6743.6771.29|CGACACTTCGAACGCACTTGCGGCCCCGG</td><td>>hg38_21_+_8433172_8446622_RNA45SN1_45S_50ntFlanks</td><td>6743</td><td>6771</td><td>29</td><td>CGACACTTCGAACGCACTTGCGGCCCCGG</td><td>1791.9</td><td>-4.6</td><td>0.4</td><td>-12.6</td><td>1.6E-36</td><td>7.7E-35</td><td>BT20/MB468</td><td>MITO</td></tr>
<tr><td>>MI0000478|hsa-mir-149&WithFlank&2|+|240455995|240456095@21.42.22&[MIMAT0000450&hsa-miR-149-5p&offsets|0|-1;m-195&2|+|240456015|240456037&offsets|0|-1]|TCTGGCTCCGTGTCTTCACTCC</td><td>>MI0000478|hsa-mir-149&WithFlank&2|+|240455995|240456095</td><td>21</td><td>42</td><td>22</td><td>TCTGGCTCCGTGTCTTCACTCC</td><td>1685.0</td><td>15.2</td><td>1.2</td><td>12.6</td><td>1.7E-36</td><td>7.7E-35</td><td>BT20/MB468</td><td>MITO</td></tr>
<tr><td>>hg38_21_+_8433172_8446622_RNA45SN1_45S_50ntFlanks@12934.12955.22|CTAAACCATTCGTAGACGACCT</td><td>>hg38_21_+_8433172_8446622_RNA45SN1_45S_50ntFlanks</td><td>12934</td><td>12955</td><td>22</td><td>CTAAACCATTCGTAGACGACCT</td><td>1889.9</td><td>-15.3</td><td>1.2</td><td>-12.6</td><td>3.2E-36</td><td>7.8E-35</td><td>BT20/MB468</td><td>NUC</td></tr>
<tr><td>>hg38_21_+_8433172_8446622_RNA45SN1_45S_50ntFlanks@6699.6717.19|GCTAGCTGCGAGAATTAAT</td><td>>hg38_21_+_8433172_8446622_RNA45SN1_45S_50ntFlanks</td><td>6699</td><td>6717</td><td>19</td><td>GCTAGCTGCGAGAATTAAT</td><td>1990.8</td><td>-15.4</td><td>1.2</td><td>-12.6</td><td>3.3E-36</td><td>8.1E-35</td><td>BT20/MB468</td><td>NUC</td></tr>
<tr><td>>hg38_21_+_8433172_8446622_RNA45SN1_45S_50ntFlanks@7983.8003.21|CAGATCAGACGTGGCGACCCG</td><td>>hg38_21_+_8433172_8446622_RNA45SN1_45S_50ntFlanks</td><td>7983</td><td>8003</td><td>21</td><td>CAGATCAGACGTGGCGACCCG</td><td>1675.1</td><td>-7.7</td><td>0.6</td><td>-12.6</td><td>3.4E-36</td><td>8.2E-35</td><td>BT20/MB468</td><td>NUC</td></tr>
<tr><td>>hg38_21_+_8433172_8446622_RNA45SN1_45S_50ntFlanks@12159.12180.22|TTTGACTGGGGCGGTACACCTG</td><td>>hg38_21_+_8433172_8446622_RNA45SN1_45S_50ntFlanks</td><td>12159</td><td>12180</td><td>22</td><td>TTTGACTGGGGCGGTACACCTG</td><td>8572.0</td><td>-16.9</td><td>1.3</td><td>-12.8</td><td>2.0E-37</td><td>8.5E-35</td><td>MB231/MB468</td><td>MP</td></tr>
<tr><td>>MI0000285|hsa-mir-205&WithFlank&1|+|209432127|209432248@40.61.22(+1A)&[MIMAT0000266&hsa-miR-205-5p&offsets|0|0(+1A);m-351&1|+|209432166|209432187&offsets|0|0(+1A)]|TCCTTCATTCCACCGGAGTCTGA</td><td>>MI0000285|hsa-mir-205&WithFlank&1|+|209432127|209432248</td><td>40</td><td>61</td><td>22(+1A)</td><td>TCCTTCATTCCACCGGAGTCTGA</td><td>33164.8</td><td>18.6</td><td>1.4</td><td>12.9</td><td>5.6E-38</td><td>8.6E-35</td><td>BT20/MB231</td><td>MITO</td></tr>
<tr><td>>hg38_21_+_8433172_8446622_RNA45SN1_45S_50ntFlanks@12435.12455.21|ATCATTGTGAAGCAGAATTCA</td><td>>hg38_21_+_8433172_8446622_RNA45SN1_45S_50ntFlanks</td><td>12435</td><td>12455</td><td>21</td><td>ATCATTGTGAAGCAGAATTCA</td><td>2348.3</td><td>-15.6</td><td>1.2</td><td>-12.6</td><td>3.6E-36</td><td>8.7E-35</td><td>BT20/MB468</td><td>NUC</td></tr>
<tr><td>>hg38_21_+_8433172_8446622_RNA45SN1_45S_50ntFlanks@12997.13036.40|CGCTGCGATCTATTGAAAGTCAGCCCTCGACACAAGGGTT</td><td>>hg38_21_+_8433172_8446622_RNA45SN1_45S_50ntFlanks</td><td>12997</td><td>13036</td><td>40</td><td>CGCTGCGATCTATTGAAAGTCAGCCCTCGACACAAGGGTT</td><td>3516.2</td><td>7.5</td><td>0.6</td><td>12.8</td><td>1.5E-37</td><td>8.9E-35</td><td>MB231/MB468</td><td>CYTO</td></tr>
<tr><td>>hg38_21_+_8433172_8446622_RNA45SN1_45S_50ntFlanks@11785.11804.20|TGAATGGATGAACGAGATTC</td><td>>hg38_21_+_8433172_8446622_RNA45SN1_45S_50ntFlanks</td><td>11785</td><td>11804</td><td>20</td><td>TGAATGGATGAACGAGATTC</td><td>9957.7</td><td>-8.7</td><td>0.7</td><td>-12.8</td><td>2.1E-37</td><td>9.0E-35</td><td>MB231/MB468</td><td>MP</td></tr>
<tr><td>>hg38_21_+_8433172_8446622_RNA45SN1_45S_50ntFlanks@6661.6685.25|CGGTGGATCACTCGGCTCGTGCGTC</td><td>>hg38_21_+_8433172_8446622_RNA45SN1_45S_50ntFlanks</td><td>6661</td><td>6685</td><td>25</td><td>CGGTGGATCACTCGGCTCGTGCGTC</td><td>8443.0</td><td>6.4</td><td>0.5</td><td>12.6</td><td>2.1E-36</td><td>9.6E-35</td><td>BT20/MB468</td><td>MITO</td></tr>
<tr><td>>MI0001445|hsa-mir-423&WithFlank&17|+|30117073|30117178@58.80.23(+1C)&[MIMAT0001340&hsa-miR-423-3p&offsets|-1|-1(+1C);m-100&17|+|30117131|30117153&offsets|-1|-1(+1C)]|AAGCTCGGTCTGAGGCCCCTCAGC</td><td>>MI0001445|hsa-mir-423&WithFlank&17|+|30117073|30117178</td><td>58</td><td>80</td><td>23(+1C)</td><td>AAGCTCGGTCTGAGGCCCCTCAGC</td><td>1156.7</td><td>8.9</td><td>0.7</td><td>12.6</td><td>2.1E-36</td><td>9.8E-35</td><td>BT20/MB468</td><td>MITO</td></tr>
<tr><td>>hg38_21_+_8433172_8446622_RNA45SN1_45S_50ntFlanks@8356.8379.24|GAAAAGAACTTTGAAGAGAGAGTT</td><td>>hg38_21_+_8433172_8446622_RNA45SN1_45S_50ntFlanks</td><td>8356</td><td>8379</td><td>24</td><td>GAAAAGAACTTTGAAGAGAGAGTT</td><td>28013.9</td><td>-10.3</td><td>0.8</td><td>-12.5</td><td>4.4E-36</td><td>1.1E-34</td><td>BT20/MB468</td><td>NUC</td></tr>
<tr><td>>hg38_21_+_8433172_8446622_RNA45SN1_45S_50ntFlanks@12627.12653.27|ATGGGGCGAAGCTACCATCTGTGGGAT</td><td>>hg38_21_+_8433172_8446622_RNA45SN1_45S_50ntFlanks</td><td>12627</td><td>12653</td><td>27</td><td>ATGGGGCGAAGCTACCATCTGTGGGAT</td><td>10637.0</td><td>-17.2</td><td>1.4</td><td>-12.8</td><td>2.6E-37</td><td>1.1E-34</td><td>MB231/MB468</td><td>MP</td></tr>
<tr><td>>trna111_HisGTG_1_-_147774845_147774916@-1G.33.34,trna118_HisGTG_1_-_145396881_145396952@-1G.33.34,trna16_HisGTG_1_+_146544773_146544844@-1G.33.34,trna1_HisGTG_15_+_45493349_45493420@-1G.33.34,trna21_HisGTG_1_+_147753471_147753542@-1G.33.34,trna33_HisGTG_6_+_27125906_27125977@-1G.33.34,trna7_HisGTG_9_-_14433938_14434009@-1G.33.34,trna8_HisGTG_15_-_45492611_45492682@-1G.33.34,trna9_HisGTG_15_-_45490804_45490875@-1G.33.34|GGCCGTGATCGTATAGTGGTTAGTACTCTGCGTT</td><td>>trna111_HisGTG_1_-_147774845_147774916</td><td>-1G</td><td>33</td><td>34</td><td>GGCCGTGATCGTATAGTGGTTAGTACTCTGCGTT</td><td>94099.4</td><td>11.8</td><td>0.9</td><td>12.8</td><td>2.0E-37</td><td>1.1E-34</td><td>MB231/MB468</td><td>CYTO</td></tr>
<tr><td>>hg38_21_+_8433172_8446622_RNA45SN1_45S_50ntFlanks@12605.12626.22|TATGTGCTTGGCTGAGGAGCCA</td><td>>hg38_21_+_8433172_8446622_RNA45SN1_45S_50ntFlanks</td><td>12605</td><td>12626</td><td>22</td><td>TATGTGCTTGGCTGAGGAGCCA</td><td>8973.8</td><td>-17.6</td><td>1.4</td><td>-12.5</td><td>4.8E-36</td><td>1.2E-34</td><td>BT20/MB468</td><td>NUC</td></tr>
<tr><td>>hg38_21_+_8433172_8446622_RNA45SN1_45S_50ntFlanks@3775.3792.18|GCCGGTACAGTGAAACTG</td><td>>hg38_21_+_8433172_8446622_RNA45SN1_45S_50ntFlanks</td><td>3775</td><td>3792</td><td>18</td><td>GCCGGTACAGTGAAACTG</td><td>5741.1</td><td>-16.9</td><td>1.3</td><td>-12.5</td><td>4.9E-36</td><td>1.2E-34</td><td>BT20/MB468</td><td>NUC</td></tr>
<tr><td>>hg38_21_+_8433172_8446622_RNA45SN1_45S_50ntFlanks@9562.9596.35|TCCTGACGTGCAAATCGGTCGTCCGACCTGGGTAT</td><td>>hg38_21_+_8433172_8446622_RNA45SN1_45S_50ntFlanks</td><td>9562</td><td>9596</td><td>35</td><td>TCCTGACGTGCAAATCGGTCGTCCGACCTGGGTAT</td><td>2530.2</td><td>-6.6</td><td>0.5</td><td>-12.6</td><td>2.7E-36</td><td>1.2E-34</td><td>BT20/MB468</td><td>MITO</td></tr>
<tr><td>>MI0000812|hsa-mir-331&WithFlank&12|+|95308414|95308519@67.87.21(+1U)&[MIMAT0000760&hsa-miR-331-3p&offsets|0|0(+1U);m-192&12|+|95308480|95308501&offsets|0|-1(+1U)]|GCCCCTGGGCCTATCCTAGAAT</td><td>>MI0000812|hsa-mir-331&WithFlank&12|+|95308414|95308519</td><td>67</td><td>87</td><td>21(+1U)</td><td>GCCCCTGGGCCTATCCTAGAAT</td><td>987.9</td><td>7.5</td><td>0.6</td><td>12.6</td><td>2.7E-36</td><td>1.2E-34</td><td>BT20/MB468</td><td>MITO</td></tr>
<tr><td>>hg38_21_+_8433172_8446622_RNA45SN1_45S_50ntFlanks@9567.9596.30|ACGTGCAAATCGGTCGTCCGACCTGGGTAT</td><td>>hg38_21_+_8433172_8446622_RNA45SN1_45S_50ntFlanks</td><td>9567</td><td>9596</td><td>30</td><td>ACGTGCAAATCGGTCGTCCGACCTGGGTAT</td><td>7306.3</td><td>-7.4</td><td>0.6</td><td>-12.6</td><td>2.8E-36</td><td>1.3E-34</td><td>BT20/MB468</td><td>MITO</td></tr>
<tr><td>>MI0000301|hsa-mir-224&WithFlank&X|-|151958572|151958664@14.35.22&[MIMAT0000281&hsa-miR-224-5p&offsets|+1|-2;m-238&X|-|151958630|151958651&offsets|0|0]|CAAGTCACTAGTGGTTCCGTTT</td><td>>MI0000301|hsa-mir-224&WithFlank&X|-|151958572|151958664</td><td>14</td><td>35</td><td>22</td><td>CAAGTCACTAGTGGTTCCGTTT</td><td>2432.5</td><td>7.5</td><td>0.6</td><td>12.5</td><td>5.7E-36</td><td>1.3E-34</td><td>BT20/MB468</td><td>NUC</td></tr>
<tr><td>>hg38_21_+_8433172_8446622_RNA45SN1_45S_50ntFlanks@4759.4779.21|ATCAGATACCGTCGTAGTTCC</td><td>>hg38_21_+_8433172_8446622_RNA45SN1_45S_50ntFlanks</td><td>4759</td><td>4779</td><td>21</td><td>ATCAGATACCGTCGTAGTTCC</td><td>2117.2</td><td>-15.5</td><td>1.2</td><td>-12.5</td><td>5.6E-36</td><td>1.3E-34</td><td>BT20/MB468</td><td>NUC</td></tr>
<tr><td>>hg38_21_+_8433172_8446622_RNA45SN1_45S_50ntFlanks@9738.9767.30|GAAACGATCTCAACCTATTCTCAAACTTTA</td><td>>hg38_21_+_8433172_8446622_RNA45SN1_45S_50ntFlanks</td><td>9738</td><td>9767</td><td>30</td><td>GAAACGATCTCAACCTATTCTCAAACTTTA</td><td>10820.1</td><td>-8.4</td><td>0.7</td><td>-12.7</td><td>8.5E-37</td><td>1.3E-34</td><td>MB231/MB468</td><td>MITO</td></tr>
<tr><td>>am_MI0003581|hsa-mir-574&WithFlank&4|+|38868026|38868133@31.53.23&[MIMAT0004795&hsa-miR-574-5p&offsets|0|0;m-213&4|+|38868056|38868078&offsets|0|0]|TGAGTGTGTGTGTGTGAGTGTGT</td><td>>am_MI0003581|hsa-mir-574&WithFlank&4|+|38868026|38868133</td><td>31</td><td>53</td><td>23</td><td>TGAGTGTGTGTGTGTGAGTGTGT</td><td>434.7</td><td>7.3</td><td>0.6</td><td>12.6</td><td>3.5E-36</td><td>1.5E-34</td><td>BT20/MB468</td><td>MITO</td></tr>
<tr><td>>MI0000065|hsa-let-7d&WithFlank&9|+|94178828|94178926@14.34.21(+1C)&[MIMAT0000065&hsa-let-7d-5p&offsets|0|-1(+1C);m-90&9|+|94178841|94178862&offsets|0|-1(+1C)]|AGAGGTAGTAGGTTGCATAGTC</td><td>>MI0000065|hsa-let-7d&WithFlank&9|+|94178828|94178926</td><td>14</td><td>34</td><td>21(+1C)</td><td>AGAGGTAGTAGGTTGCATAGTC</td><td>1128.1</td><td>7.8</td><td>0.6</td><td>12.6</td><td>3.5E-36</td><td>1.5E-34</td><td>BT20/MB468</td><td>MITO</td></tr>
<tr><td>>hg38_21_+_8433172_8446622_RNA45SN1_45S_50ntFlanks@3840.3861.22|CTCCTACTTGGATAACTGTGGT</td><td>>hg38_21_+_8433172_8446622_RNA45SN1_45S_50ntFlanks</td><td>3840</td><td>3861</td><td>22</td><td>CTCCTACTTGGATAACTGTGGT</td><td>1517.1</td><td>5.0</td><td>0.4</td><td>12.5</td><td>5.9E-36</td><td>1.8E-34</td><td>BT20/MB468</td><td>CYTO</td></tr>
<tr><td>>hg38_21_+_8433172_8446622_RNA45SN1_45S_50ntFlanks@6657.6681.25|TTAGCGGTGGATCACTCGGCTCGTG</td><td>>hg38_21_+_8433172_8446622_RNA45SN1_45S_50ntFlanks</td><td>6657</td><td>6681</td><td>25</td><td>TTAGCGGTGGATCACTCGGCTCGTG</td><td>897.2</td><td>-5.6</td><td>0.4</td><td>-12.5</td><td>8.9E-36</td><td>2.1E-34</td><td>BT20/MB468</td><td>NUC</td></tr>
<tr><td>>MI0000066|hsa-let-7e&WithFlank&19|+|51692780|51692870@14.36.23&[MIMAT0000066&hsa-let-7e-5p&offsets|0|+1;m-50&19|+|51692793|51692814&offsets|0|+1]|TGAGGTAGGAGGTTGTATAGTTG</td><td>>MI0000066|hsa-let-7e&WithFlank&19|+|51692780|51692870</td><td>14</td><td>36</td><td>23</td><td>TGAGGTAGGAGGTTGTATAGTTG</td><td>472.7</td><td>7.3</td><td>0.6</td><td>12.5</td><td>4.8E-36</td><td>2.1E-34</td><td>BT20/MB468</td><td>MITO</td></tr>
<tr><td>>MI0000272|hsa-mir-182&WithFlank&7|-|129770377|129770498@29.54.26&[MIMAT0000259&hsa-miR-182-5p&offsets|0|+2;m-22&7|-|129770449|129770470&offsets|0|+4]|TTTGGCAATGGTAGAACTCACACTGG</td><td>>MI0000272|hsa-mir-182&WithFlank&7|-|129770377|129770498</td><td>29</td><td>54</td><td>26</td><td>TTTGGCAATGGTAGAACTCACACTGG</td><td>4878.4</td><td>8.0</td><td>0.6</td><td>12.5</td><td>5.1E-36</td><td>2.2E-34</td><td>BT20/MB468</td><td>MITO</td></tr>
<tr><td>>MI0000066|hsa-let-7e&WithFlank&19|+|51692780|51692870@14.35.22&[MIMAT0000066&hsa-let-7e-5p&offsets|0|0;m-50&19|+|51692793|51692814&offsets|0|0]|TGAGGTAGGAGGTTGTATAGTT</td><td>>MI0000066|hsa-let-7e&WithFlank&19|+|51692780|51692870</td><td>14</td><td>35</td><td>22</td><td>TGAGGTAGGAGGTTGTATAGTT</td><td>10972.2</td><td>7.1</td><td>0.6</td><td>12.5</td><td>5.1E-36</td><td>2.2E-34</td><td>BT20/MB468</td><td>MITO</td></tr>
<tr><td>>hg38_21_+_8433172_8446622_RNA45SN1_45S_50ntFlanks@7980.8006.27|CCTCAGATCAGACGTGGCGACCCGCTG</td><td>>hg38_21_+_8433172_8446622_RNA45SN1_45S_50ntFlanks</td><td>7980</td><td>8006</td><td>27</td><td>CCTCAGATCAGACGTGGCGACCCGCTG</td><td>6291.9</td><td>-8.0</td><td>0.6</td><td>-12.6</td><td>3.6E-36</td><td>2.3E-34</td><td>MB231/MB468</td><td>NUC</td></tr>
<tr><td>>hg38_21_+_8433172_8446622_RNA45SN1_45S_50ntFlanks@6658.6689.32|TAGCGGTGGATCACTCGGCTCGTGCGTCGATG</td><td>>hg38_21_+_8433172_8446622_RNA45SN1_45S_50ntFlanks</td><td>6658</td><td>6689</td><td>32</td><td>TAGCGGTGGATCACTCGGCTCGTGCGTCGATG</td><td>361.9</td><td>-4.2</td><td>0.3</td><td>-12.5</td><td>1.0E-35</td><td>2.4E-34</td><td>BT20/MB468</td><td>NUC</td></tr>
<tr><td>>hg38_21_+_8433172_8446622_RNA45SN1_45S_50ntFlanks@12457.12488.32|CAAGCGTTGGATTGTTCACCCACTAATAGGGA</td><td>>hg38_21_+_8433172_8446622_RNA45SN1_45S_50ntFlanks</td><td>12457</td><td>12488</td><td>32</td><td>CAAGCGTTGGATTGTTCACCCACTAATAGGGA</td><td>2081.1</td><td>-15.4</td><td>1.2</td><td>-12.5</td><td>1.1E-35</td><td>2.5E-34</td><td>BT20/MB468</td><td>NUC</td></tr>
<tr><td>>MI0000465|hsa-mir-191&WithFlank&3|-|49020612|49020715@22.42.21(+1C)&[MIMAT0000440&hsa-miR-191-5p&offsets|0|-2(+1C);m-48&3|-|49020672|49020694&offsets|0|-2(+1C)]|CAACGGAATCCCAAAAGCAGCC</td><td>>MI0000465|hsa-mir-191&WithFlank&3|-|49020612|49020715</td><td>22</td><td>42</td><td>21(+1C)</td><td>CAACGGAATCCCAAAAGCAGCC</td><td>2293.7</td><td>7.6</td><td>0.6</td><td>12.5</td><td>8.3E-36</td><td>2.5E-34</td><td>BT20/MB468</td><td>CYTO</td></tr>
<tr><td>>hg38_21_+_8433172_8446622_RNA45SN1_45S_50ntFlanks@12008.12032.25|CCGCCGGTGAAATACCACTACTCTG</td><td>>hg38_21_+_8433172_8446622_RNA45SN1_45S_50ntFlanks</td><td>12008</td><td>12032</td><td>25</td><td>CCGCCGGTGAAATACCACTACTCTG</td><td>11541.9</td><td>-9.6</td><td>0.8</td><td>-12.6</td><td>4.0E-36</td><td>2.5E-34</td><td>MB231/MB468</td><td>NUC</td></tr>
<tr><td>>MI0000087|hsa-mir-29a&WithFlank&7|-|130876741|130876816@48.69.22&[MIMAT0000086&hsa-miR-29a-3p&offsets|0|0;m-26&7|-|130876749|130876769&offsets|0|+1]|TAGCACCATCTGAAATCGGTTA</td><td>>MI0000087|hsa-mir-29a&WithFlank&7|-|130876741|130876816</td><td>48</td><td>69</td><td>22</td><td>TAGCACCATCTGAAATCGGTTA</td><td>5831.7</td><td>9.5</td><td>0.8</td><td>12.5</td><td>4.4E-36</td><td>2.7E-34</td><td>MB231/MB468</td><td>NUC</td></tr>
<tr><td>>hg38_21_+_8433172_8446622_RNA45SN1_45S_50ntFlanks@6721.6746.26|AATTGCAGGACACATTGATCATCGAC</td><td>>hg38_21_+_8433172_8446622_RNA45SN1_45S_50ntFlanks</td><td>6721</td><td>6746</td><td>26</td><td>AATTGCAGGACACATTGATCATCGAC</td><td>1750.9</td><td>-7.3</td><td>0.6</td><td>-12.5</td><td>4.4E-36</td><td>2.7E-34</td><td>MB231/MB468</td><td>NUC</td></tr>
<tr><td>>MI0000439|hsa-mir-23b&WithFlank&9|+|95085202|95085310@67.86.20(+1C)&[MIMAT0000418&hsa-miR-23b-3p&offsets|+3|0(+1C);m-42&9|+|95085265|95085285&offsets|+3|+2(+1C)]|ACATTGCCAGGGATTACCACC</td><td>>MI0000439|hsa-mir-23b&WithFlank&9|+|95085202|95085310</td><td>67</td><td>86</td><td>20(+1C)</td><td>ACATTGCCAGGGATTACCACC</td><td>1676.7</td><td>15.0</td><td>1.2</td><td>12.5</td><td>9.3E-36</td><td>2.7E-34</td><td>BT20/MB468</td><td>CYTO</td></tr>
<tr><td>>hg38_21_+_8433172_8446622_RNA45SN1_45S_50ntFlanks@11572.11601.30|CCGACTTAGAACTGGTGCGGACCAGGGGAA</td><td>>hg38_21_+_8433172_8446622_RNA45SN1_45S_50ntFlanks</td><td>11572</td><td>11601</td><td>30</td><td>CCGACTTAGAACTGGTGCGGACCAGGGGAA</td><td>9899.0</td><td>-8.6</td><td>0.7</td><td>-12.6</td><td>1.9E-36</td><td>2.9E-34</td><td>MB231/MB468</td><td>MITO</td></tr>
<tr><td>>hg38_21_+_8433172_8446622_RNA45SN1_45S_50ntFlanks@12627.12659.33|ATGGGGCGAAGCTACCATCTGTGGGATTATGAC</td><td>>hg38_21_+_8433172_8446622_RNA45SN1_45S_50ntFlanks</td><td>12627</td><td>12659</td><td>33</td><td>ATGGGGCGAAGCTACCATCTGTGGGATTATGAC</td><td>4085.1</td><td>-16.4</td><td>1.3</td><td>-12.5</td><td>1.3E-35</td><td>3.1E-34</td><td>BT20/MB468</td><td>NUC</td></tr>
<tr><td>>hg38_21_+_8433172_8446622_RNA45SN1_45S_50ntFlanks@12627.12654.28|ATGGGGCGAAGCTACCATCTGTGGGATT</td><td>>hg38_21_+_8433172_8446622_RNA45SN1_45S_50ntFlanks</td><td>12627</td><td>12654</td><td>28</td><td>ATGGGGCGAAGCTACCATCTGTGGGATT</td><td>14925.3</td><td>-17.7</td><td>1.4</td><td>-12.7</td><td>8.2E-37</td><td>3.3E-34</td><td>MB231/MB468</td><td>MP</td></tr>
<tr><td>>hg38_21_+_8433172_8446622_RNA45SN1_45S_50ntFlanks@9567.9597.31|ACGTGCAAATCGGTCGTCCGACCTGGGTATA</td><td>>hg38_21_+_8433172_8446622_RNA45SN1_45S_50ntFlanks</td><td>9567</td><td>9597</td><td>31</td><td>ACGTGCAAATCGGTCGTCCGACCTGGGTATA</td><td>7084.2</td><td>-6.7</td><td>0.5</td><td>-12.5</td><td>8.1E-36</td><td>3.5E-34</td><td>BT20/MB468</td><td>MITO</td></tr>
<tr><td>>hg38_21_+_8433172_8446622_RNA45SN1_45S_50ntFlanks@12626.12653.28|AATGGGGCGAAGCTACCATCTGTGGGAT</td><td>>hg38_21_+_8433172_8446622_RNA45SN1_45S_50ntFlanks</td><td>12626</td><td>12653</td><td>28</td><td>AATGGGGCGAAGCTACCATCTGTGGGAT</td><td>11291.4</td><td>-17.3</td><td>1.4</td><td>-12.7</td><td>9.1E-37</td><td>3.5E-34</td><td>MB231/MB468</td><td>MP</td></tr>
<tr><td>>hg38_21_+_8433172_8446622_RNA45SN1_45S_50ntFlanks@12003.12032.30|GCGGGCCGCCGGTGAAATACCACTACTCTG</td><td>>hg38_21_+_8433172_8446622_RNA45SN1_45S_50ntFlanks</td><td>12003</td><td>12032</td><td>30</td><td>GCGGGCCGCCGGTGAAATACCACTACTCTG</td><td>6771.8</td><td>-8.3</td><td>0.7</td><td>-12.7</td><td>9.9E-37</td><td>3.7E-34</td><td>MB231/MB468</td><td>MP</td></tr>
</table>

| | | | | | | | | | | | | | |
|---|---|---|---|---|---|---|---|---|---|---|---|---|---|
| >hg38_21_+_8433172_8446622_RNA45SN1_45S_50ntFlanks@12991.13011.21\|CTCCCTCGCTGCGATCTATTG | >hg38_21_+_8433172_8446622_RNA45SN1_45S_50ntFlanks | 12991 | 13011 | 21 | CTCCCTCGCTGCGATCTATTG | 10935.2 | -17.8 | 1.4 | -12.4 | 1.7E-35 | 3.9E-34 | BT20/MB468 | NUC |
| >MI0000299\|hsa-mir-222&WithFlank&X\|-\|45747009\|45747130@75.97.23(+1U)&[MIMAT0000279&hsa-miR-222-3p&offsets\|0\|+2(+1U);m-140&X\|-\|45747033\|45747056&offsets\|0\|-1(+1U)]\|AGCTACATCTGGCTACTGGGTCTT | >MI0000299\|hsa-mir-222&WithFlank&X\|-\|45747009\|45747130 | 75 | 97 | 23(+1U) | AGCTACATCTGGCTACTGGGTCTT | 2274.8 | 7.6 | 0.6 | 12.5 | 1.4E-35 | 4.0E-34 | BT20/MB468 | CYTO |
| >MI0000264\|hsa-mir-7-2&WithFlank&15\|+\|88611819\|88611940@38.60.23(+1A);MI0000265\|hsa-mir-7-3&WithFlank&19\|+\|4770664\|4770785@37.59.23(+1A);MI0000263\|hsa-mir-7-1&WithFlank&9\|-\|83969742\|83969863@30.52.23(+1A)&[MIMAT0000252&hsa-miR-7-5p&offsets\|0\|-1(+1A);m-400&15\|+\|88611856\|88611879&offsets\|0\|-1(+1A)];[MIMAT0000252_1&hsa-miR-7-5p&offsets\|0\|-1(+1A);m-402&19\|+\|4770700\|4770723&offsets\|0\|-1(+1A)];[MIMAT0000252_2&hsa-miR-7-5p&offsets\|0\|-1(+1A);m-401&9\|-\|83969811\|83969834&offsets\|0\|-1(+1A)]\|TGGAAGACTAGTGATTTTGTTGTA | >MI0000264\|hsa-mir-7-2&WithFlank&15\|+\|88611819\|88611940 | 38 | 60 | 23(+1A) | TGGAAGACTAGTGATTTTGTTGTA | 858.2 | 5.6 | 0.4 | 12.5 | 7.9E-36 | 4.8E-34 | MB231/MB468 | NUC |
| >hg38_21_+_8433172_8446622_RNA45SN1_45S_50ntFlanks@10458.10477.20\|ATGGCCTCCGTTGCCCTCGG | >hg38_21_+_8433172_8446622_RNA45SN1_45S_50ntFlanks | 10458 | 10477 | 20 | ATGGCCTCCGTTGCCCTCGG | 3190.3 | -16.1 | 1.3 | -12.4 | 2.2E-35 | 5.0E-34 | BT20/MB468 | NUC |
| >hg38_21_+_8433172_8446622_RNA45SN1_45S_50ntFlanks@11781.11806.26\|CGCATGAATGGATGAACGAGATTCCC | >hg38_21_+_8433172_8446622_RNA45SN1_45S_50ntFlanks | 11781 | 11806 | 26 | CGCATGAATGGATGAACGAGATTCCC | 1780.3 | -15.2 | 1.2 | -12.4 | 2.5E-35 | 5.6E-34 | BT20/MB468 | NUC |
| >MI0000746\|hsa-mir-99b&WithFlank&19\|+\|51692606\|51692687@13.34.22(+1A)&[MIMAT0000689&hsa-miR-99b-5p&offsets\|0\|0(+1A);m-10&19\|+\|51692618\|51692639&offsets\|0\|0(+1A)]\|CACCCGTAGAACCGACCTTGCGA | >MI0000746\|hsa-mir-99b&WithFlank&19\|+\|51692606\|51692687 | 13 | 34 | 22(+1A) | CACCCGTAGAACCGACCTTGCGA | 1364.5 | 7.3 | 0.6 | 12.5 | 1.3E-35 | 5.7E-34 | BT20/MB468 | MITO |
| >hg38_21_+_8433172_8446622_RNA45SN1_45S_50ntFlanks@11894.11912.19\|ACTCTAGTCTGGCACGGTG | >hg38_21_+_8433172_8446622_RNA45SN1_45S_50ntFlanks | 11894 | 11912 | 19 | ACTCTAGTCTGGCACGGTG | 2643.9 | -15.8 | 1.3 | -12.4 | 2.5E-35 | 5.7E-34 | BT20/MB468 | NUC |
| >hg38_21_+_8433172_8446622_RNA45SN1_45S_50ntFlanks@12635.12653.19\|AAGCTACCATCTGTGGGAT | >hg38_21_+_8433172_8446622_RNA45SN1_45S_50ntFlanks | 12635 | 12653 | 19 | AAGCTACCATCTGTGGGAT | 10167.4 | -17.2 | 1.4 | -12.6 | 1.6E-36 | 5.8E-34 | MB231/MB468 | MP |
| >hg38_21_+_8433172_8446622_RNA45SN1_45S_50ntFlanks@12991.13019.29\|CTCCCTCGCTGCGATCTATTGAAAGTCAG | >hg38_21_+_8433172_8446622_RNA45SN1_45S_50ntFlanks | 12991 | 13019 | 29 | CTCCCTCGCTGCGATCTATTGAAAGTCAG | 4134.2 | -16.4 | 1.3 | -12.4 | 2.6E-35 | 6.0E-34 | BT20/MB468 | NUC |
| >hg38_21_+_8433172_8446622_RNA45SN1_45S_50ntFlanks@12995.13036.42\|CTCGCTGCGATCTATTGAAAGTCAGCCCTCGACACAAGGGTT | >hg38_21_+_8433172_8446622_RNA45SN1_45S_50ntFlanks | 12995 | 13036 | 42 | CTCGCTGCGATCTATTGAAAGTCAGCCCTCGACACAAGGGTT | 6424.9 | 8.8 | 0.7 | 12.6 | 1.2E-36 | 6.0E-34 | MB231/MB468 | CYTO |
| >hg38_21_+_8433172_8446622_RNA45SN1_45S_50ntFlanks@12412.12443.32\|ATCCTTCGATGTCGGCTCTTCCTATCATTGTG | >hg38_21_+_8433172_8446622_RNA45SN1_45S_50ntFlanks | 12412 | 12443 | 32 | ATCCTTCGATGTCGGCTCTTCCTATCATTGTG | 4044.5 | -16.4 | 1.3 | -12.4 | 2.9E-35 | 6.5E-34 | BT20/MB468 | NUC |
| >MI0000650\|hsa-mir-200c&WithFlank&12\|+\|6963693\|6963772@50.71.22&[MIMAT0000617&hsa-miR-200c-3p&offsets\|0\|-1;m-54&12\|+\|6963742\|6963764&offsets\|0\|-1]\|TAATACTGCCGGGTAATGATGG | >MI0000650\|hsa-mir-200c&WithFlank&12\|+\|6963693\|6963772 | 50 | 71 | 22 | TAATACTGCCGGGTAATGATGG | 9552.3 | 9.2 | 0.7 | 12.7 | 6.6E-37 | 6.7E-34 | BT20/MB231 | MITO |
| >trna111_HisGTG_1_-_147774845_147774916@1.34.34,trna118_HisGTG_1_-_145396881_145396952@1.34.34,trna16_HisGTG_1_+_146544773_146544844@1.34.34,trna1_HisGTG_15_+_45493349_45493420@1.34.34,trna21_HisGTG_1_+_147753471_147753542@1.34.34,trna33_HisGTG_6_+_27125906_27125977@1.34.34,trna7_HisGTG_9_-_14433938_14434009@1.34.34,trna8_HisGTG_15_-_45492611_45492682@1.34.34,trna9_HisGTG_15_-_45490804_45490875@1.34.34\|GCCGTGATCGTATAGTGGTTAGTACTCTGCGTTG | >trna111_HisGTG_1_-_147774845_147774916 | 1 | 34 | 34 | GCCGTGATCGTATAGTGGTTAGTACTCTGCGTTG | 4695.5 | 8.0 | 0.6 | 12.4 | 2.4E-35 | 6.9E-34 | BT20/MB468 | CYTO |
| >hg38_21_+_8433172_8446622_RNA45SN1_45S_50ntFlanks@12626.12654.29\|AATGGGGCGAAGCTACCATCTGTGGGATT | >hg38_21_+_8433172_8446622_RNA45SN1_45S_50ntFlanks | 12626 | 12654 | 29 | AATGGGGCGAAGCTACCATCTGTGGGATT | 5276.9 | -4.7 | 0.4 | -12.4 | 2.5E-35 | 7.2E-34 | BT20/MB468 | CYTO |
| >hg38_21_+_8433172_8446622_RNA45SN1_45S_50ntFlanks@4163.4184.22\|CATCCAAGGAAGGCAGCAGGCG | >hg38_21_+_8433172_8446622_RNA45SN1_45S_50ntFlanks | 4163 | 4184 | 22 | CATCCAAGGAAGGCAGCAGGCG | 3177.6 | -16.1 | 1.3 | -12.4 | 3.2E-35 | 7.2E-34 | BT20/MB468 | NUC |
| >hg38_21_+_8433172_8446622_RNA45SN1_45S_50ntFlanks@12936.12955.20\|AAACCATTCGTAGACGACCT | >hg38_21_+_8433172_8446622_RNA45SN1_45S_50ntFlanks | 12936 | 12955 | 20 | AAACCATTCGTAGACGACCT | 1496.5 | -15.0 | 1.2 | -12.4 | 3.2E-35 | 7.3E-34 | BT20/MB468 | NUC |
| >hg38_21_+_8433172_8446622_RNA45SN1_45S_50ntFlanks@4730.4751.22\|CAAGAACGAAAGTCGGAGGTTC | >hg38_21_+_8433172_8446622_RNA45SN1_45S_50ntFlanks | 4730 | 4751 | 22 | CAAGAACGAAAGTCGGAGGTTC | 938.9 | -6.3 | 0.5 | -12.4 | 3.3E-35 | 7.5E-34 | BT20/MB468 | NUC |
| >am_trna128_GlyGCC_6_-_27870686_27870756@1.30.30,trna133_GlyCCC_1_-_16872434_16872504@1.30.30,trna18_GlyGCC_16_+_70822597_70822667@1.30.30,trna19_GlyGCC_16_+_70823410_70823480@1.30.30,trna19_GlyGCC_2_-_157257659_157257729@1.30.30,trna24_GlyGCC_16_-_70812942_70813012@1.30.30,trna25_GlyGCC_16_-_70812114_70812184@1.30.30,trna4_GlyCCC_1_+_17188416_17188486@1.30.30,trna5_GlyGCC_17_+_8029064_8029134@1.30.30,trna68_GlyGCC_1_-_161493637_161493707@1.30.30\|GCATTGGTGGTTCAGTGGTAGAATTCTCGC | >am_trna128_GlyGCC_6_-_27870686_27870756 | 1 | 30 | 30 | GCATTGGTGGTTCAGTGGTAGAATTCTCGC | 14157.9 | -8.0 | 0.6 | -12.4 | 3.5E-35 | 7.9E-34 | BT20/MB468 | NUC |
| >MI0001445\|hsa-mir-423&WithFlank&17\|+\|30117073\|30117178@59.81.23(+1A)&[MIMAT0001340&hsa-miR-423-3p&offsets\|0\|0(+1A);m-100&17\|+\|30117131\|30117153&offsets\|0\|0(+1A)]\|AGCTCGGTCTGAGGCCCCTCAGTA | >MI0001445\|hsa-mir-423&WithFlank&17\|+\|30117073\|30117178 | 59 | 81 | 23(+1A) | AGCTCGGTCTGAGGCCCCTCAGTA | 3183.2 | 7.9 | 0.6 | 12.4 | 1.9E-35 | 8.0E-34 | BT20/MB468 | MITO |
| >hg38_21_+_8433172_8446622_RNA45SN1_45S_50ntFlanks@12002.12032.31\|TGCGGGCCGCCGGTGAAATACCACTACTCTG | >hg38_21_+_8433172_8446622_RNA45SN1_45S_50ntFlanks | 12002 | 12032 | 31 | TGCGGGCCGCCGGTGAAATACCACTACTCTG | 8049.0 | -8.8 | 0.7 | -12.5 | 5.5E-36 | 8.4E-34 | MB231/MB468 | MITO |
| >hg38_21_+_8433172_8446622_RNA45SN1_45S_50ntFlanks@10350.10382.33\|ACGAGAACTTTGAAGGCCGAAGTGGAGAAGGGT | >hg38_21_+_8433172_8446622_RNA45SN1_45S_50ntFlanks | 10350 | 10382 | 33 | ACGAGAACTTTGAAGGCCGAAGTGGAGAAGGGT | 2711.3 | -7.0 | 0.6 | -12.4 | 4.0E-35 | 8.8E-34 | BT20/MB468 | NUC |
| >hg38_21_+_8433172_8446622_RNA45SN1_45S_50ntFlanks@12456.12485.30\|CCAAGCGTTGGATTGTTCACCCACTAATAG | >hg38_21_+_8433172_8446622_RNA45SN1_45S_50ntFlanks | 12456 | 12485 | 30 | CCAAGCGTTGGATTGTTCACCCACTAATAG | 11896.5 | -18.0 | 1.5 | -12.4 | 4.0E-35 | 8.8E-34 | BT20/MB468 | NUC |
| >hg38_21_+_8433172_8446622_RNA45SN1_45S_50ntFlanks@10466.10497.32\|CGTTGCCCTCGGCCGATCGAAAGGGAGTCGGG | >hg38_21_+_8433172_8446622_RNA45SN1_45S_50ntFlanks | 10466 | 10497 | 32 | CGTTGCCCTCGGCCGATCGAAAGGGAGTCGGG | 1922.9 | -15.3 | 1.2 | -12.4 | 4.1E-35 | 9.1E-34 | BT20/MB468 | NUC |
| >hg38_21_+_8433172_8446622_RNA45SN1_45S_50ntFlanks@12627.12654.28\|ATGGGGCGAAGCTACCATCTGTGGGATT | >hg38_21_+_8433172_8446622_RNA45SN1_45S_50ntFlanks | 12627 | 12654 | 28 | ATGGGGCGAAGCTACCATCTGTGGGATT | 25562.7 | -14.4 | 1.2 | -12.5 | 6.2E-36 | 9.1E-34 | MB231/MB468 | MITO |
| >hg38_21_+_8433172_8446622_RNA45SN1_45S_50ntFlanks@10679.10706.28\|GGTTCCGGCGGCGTCCGGTGAGCTCTCG | >hg38_21_+_8433172_8446622_RNA45SN1_45S_50ntFlanks | 10679 | 10706 | 28 | GGTTCCGGCGGCGTCCGGTGAGCTCTCG | 12072.1 | -17.4 | 1.4 | -12.6 | 2.7E-36 | 9.5E-34 | MB231/MB468 | MP |
| >MI0000734\|hsa-mir-106b&WithFlank&7\|-\|100093987\|100094080@58.79.22&[MIMAT0004672&hsa-miR-106b-3p&offsets\|0\|0;m-87&7\|-\|100094003\|100094023&offsets\|0\|+1]\|CCGCACTGTGGGTACTTGCTGC | >MI0000734\|hsa-mir-106b&WithFlank&7\|-\|100093987\|100094080 | 58 | 79 | 22 | CCGCACTGTGGGTACTTGCTGC | 13356.7 | 7.0 | 0.6 | 12.4 | 3.4E-35 | 9.8E-34 | BT20/MB468 | CYTO |
| >hg38_21_+_8433172_8446622_RNA45SN1_45S_50ntFlanks@7980.8006.27\|CCTCAGATCAGACGTGGCGACCCGCTG | >hg38_21_+_8433172_8446622_RNA45SN1_45S_50ntFlanks | 7980 | 8006 | 27 | CCTCAGATCAGACGTGGCGACCCGCTG | 8380.5 | -16.9 | 1.3 | -12.6 | 3.2E-36 | 1.1E-33 | MB231/MB468 | MP |
| >hg38_21_+_8433172_8446622_RNA45SN1_45S_50ntFlanks@12992.13014.23\|TCCCTCGCTGCGATCTATTGAAA | >hg38_21_+_8433172_8446622_RNA45SN1_45S_50ntFlanks | 12992 | 13014 | 23 | TCCCTCGCTGCGATCTATTGAAA | 3017.0 | -16.0 | 1.3 | -12.3 | 5.1E-35 | 1.1E-33 | BT20/MB468 | NUC |
| >hg38_21_+_8433172_8446622_RNA45SN1_45S_50ntFlanks@5446.5466.21\|CGGATCGGCCCCGCCGGGGTC | >hg38_21_+_8433172_8446622_RNA45SN1_45S_50ntFlanks | 5446 | 5466 | 21 | CGGATCGGCCCCGCCGGGGTC | 24779.8 | -18.4 | 1.5 | -12.5 | 8.2E-36 | 1.2E-33 | MB231/MB468 | MITO |
| >MI0002468\|hsa-mir-484&WithFlank&16\|+\|15643288\|15643378@14.35.22(+1U)&[MIMAT0002174&hsa-miR-484&offsets\|0\|0(+1U);m-127&16\|+\|15643301\|15643321&offsets\|0\|+1(+1U)]\|TCAGGCTCAGTCCCCTCCCGATT | >MI0002468\|hsa-mir-484&WithFlank&16\|+\|15643288\|15643378 | 14 | 35 | 22(+1U) | TCAGGCTCAGTCCCCTCCCGATT | 823.3 | 6.5 | 0.5 | 12.4 | 4.1E-35 | 1.2E-33 | BT20/MB468 | CYTO |
| >hg38_21_+_8433172_8446622_RNA45SN1_45S_50ntFlanks@11317.11344.28\|ACCCCACGTCTCGTCGCGCGCGCGTCCG | >hg38_21_+_8433172_8446622_RNA45SN1_45S_50ntFlanks | 11317 | 11344 | 28 | ACCCCACGTCTCGTCGCGCGCGCGTCCG | 23321.5 | -18.3 | 1.5 | -12.5 | 8.4E-36 | 1.2E-33 | MB231/MB468 | MITO |
| >hg38_21_+_8433172_8446622_RNA45SN1_45S_50ntFlanks@11999.12030.32\|CCCTGCGGGCCGCCGGTGAAATACCACTACTC | >hg38_21_+_8433172_8446622_RNA45SN1_45S_50ntFlanks | 11999 | 12030 | 32 | CCCTGCGGGCCGCCGGTGAAATACCACTACTC | 3463.9 | -16.2 | 1.3 | -12.3 | 5.5E-35 | 1.2E-33 | BT20/MB468 | NUC |
| >hg38_21_+_8433172_8446622_RNA45SN1_45S_50ntFlanks@11999.12032.34\|CCCTGCGGGCCGCCGGTGAAATACCACTACTCTG | >hg38_21_+_8433172_8446622_RNA45SN1_45S_50ntFlanks | 11999 | 12032 | 34 | CCCTGCGGGCCGCCGGTGAAATACCACTACTCTG | 22028.5 | -18.2 | 1.5 | -12.5 | 8.9E-36 | 1.2E-33 | MB231/MB468 | MITO |
| >hg38_21_+_8433172_8446622_RNA45SN1_45S_50ntFlanks@12973.13001.29\|TCGTACGTAGCAGAGCAGCTCCCTCGCTG | >hg38_21_+_8433172_8446622_RNA45SN1_45S_50ntFlanks | 12973 | 13001 | 29 | TCGTACGTAGCAGAGCAGCTCCCTCGCTG | 1346.7 | -14.8 | 1.2 | -12.3 | 5.9E-35 | 1.3E-33 | BT20/MB468 | NUC |
| >hg38_21_+_8433172_8446622_RNA45SN1_45S_50ntFlanks@11641.11665.25\|CGGCGGGTGTTGACGCGATGTGATT | >hg38_21_+_8433172_8446622_RNA45SN1_45S_50ntFlanks | 11641 | 11665 | 25 | CGGCGGGTGTTGACGCGATGTGATT | 6805.8 | -8.0 | 0.6 | -12.4 | 3.1E-35 | 1.3E-33 | BT20/MB468 | MITO |
| >MI0000285\|hsa-mir-205&WithFlank&1\|+\|209432127\|209432248@40.62.23&[MIMAT0000266&hsa-miR-205-5p&offsets\|0\|+1;m-351&1\|+\|209432166\|209432187&offsets\|0\|+1]\|TCCTTCATTCCACCGGAGTCTGT | >MI0000285\|hsa-mir-205&WithFlank&1\|+\|209432127\|209432248 | 40 | 62 | 23 | TCCTTCATTCCACCGGAGTCTGT | 41271.6 | 15.9 | 1.3 | 12.6 | 1.6E-36 | 1.4E-33 | BT20/MB231 | NUC |
| >MI0000102\|hsa-mir-100&WithFlank&11\|-\|122152223\|122152314@19.40.22(+1C)&[MIMAT0000098&hsa-miR-100-5p&offsets\|0\|0(+1C);m-27&11\|-\|122152276\|122152296&offsets\|0\|+1(+1C)]\|AACCCGTAGATCCGAACTTGTGC | >MI0000102\|hsa-mir-100&WithFlank&11\|-\|122152223\|122152314 | 19 | 40 | 22(+1C) | AACCCGTAGATCCGAACTTGTGC | 7402.1 | 16.7 | 1.3 | 12.4 | 2.2E-35 | 1.4E-33 | MB231/MB468 | NUC |
| >hg38_21_+_8433172_8446622_RNA45SN1_45S_50ntFlanks@10840.10865.26\|CCGTAACTTCGGGATAAGGATTGGCT | >hg38_21_+_8433172_8446622_RNA45SN1_45S_50ntFlanks | 10840 | 10865 | 26 | CCGTAACTTCGGGATAAGGATTGGCT | 2285.1 | -6.2 | 0.5 | -12.4 | 2.3E-35 | 1.4E-33 | MB231/MB468 | NUC |
| >MI0000253\|hsa-mir-148a&WithFlank&7\|-\|25949913\|25949992@50.71.22&[MIMAT0000243&hsa-miR-148a-3p&offsets\|0\|0;m-7&7\|-\|25949922\|25949943&offsets\|0\|0]\|TCAGTGCACTACAGAACTTTGT | >MI0000253\|hsa-mir-148a&WithFlank&7\|-\|25949913\|25949992 | 50 | 71 | 22 | TCAGTGCACTACAGAACTTTGT | 228851.8 | 7.3 | 0.6 | 12.3 | 4.9E-35 | 1.4E-33 | BT20/MB468 | CYTO |
| >MI0000266\|hsa-mir-10a&WithFlank&17\|-\|48579832\|48579953@28.51.24&[MIMAT0000253&hsa-miR-10a-5p&offsets\|0\|+1;m-18&17\|-\|48579905\|48579926&offsets\|0\|+2]\|TACCCTGTAGATCCGAATTTGTGT | >MI0000266\|hsa-mir-10a&WithFlank&17\|-\|48579832\|48579953 | 28 | 51 | 24 | TACCCTGTAGATCCGAATTTGTGT | 2617.3 | 15.6 | 1.3 | 12.3 | 5.0E-35 | 1.4E-33 | BT20/MB468 | CYTO |
| >hg38_21_+_8433172_8446622_RNA45SN1_45S_50ntFlanks@3775.3792.18\|GCCGGTACAGTGAAACTG | >hg38_21_+_8433172_8446622_RNA45SN1_45S_50ntFlanks | 3775 | 3792 | 18 | GCCGGTACAGTGAAACTG | 2014.2 | -6.2 | 0.5 | -12.3 | 5.0E-35 | 1.4E-33 | BT20/MB468 | CYTO |

| | | | | | | | | | | | | | |
|---|---|---|---|---|---|---|---|---|---|---|---|---|---|
| >MI0000102\|hsa-mir-100&WithFlank&11\|-\|122152223\|122152314@19.39.21(+1A)&[MIMAT0000098&hsa-miR-100-5p&offsets\|0\|-1(+1A);m-27&11\|-\|122152276\|122152296&offsets\|0\|0(+1A)]\|AACCCGTAGATCCGAACTTGTA | >MI0000102\|hsa-mir-100&WithFlank&11\|-\|122152223\|122152314 | 19 | 39 | 21(+1A) | AACCCGTAGATCCGAACTTGTA | 66875.5 | 16.8 | 1.4 | 12.4 | 2.4E-35 | 1.4E-33 | MB231/MB468 | NUC |
| >hg38_21_+_8433172_8446622_RNA45SN1_45S_50ntFlanks@3984.4002.19\|GCGGCTTTGGTGACTCTAG | >hg38_21_+_8433172_8446622_RNA45SN1_45S_50ntFlanks | 3984 | 4002 | 19 | GCGGCTTTGGTGACTCTAG | 8498.5 | -17.5 | 1.4 | -12.3 | 6.8E-35 | 1.5E-33 | BT20/MB468 | NUC |
| >MI0000301\|hsa-mir-224&WithFlank&X\|-\|151958572\|151958664@14.36.23&[MIMAT0000281&hsa-miR-224-5p&offsets\|+1\|-1;m-238&X\|-\|151958630\|151958651&offsets\|0\|+1]\|CAAGTCACTAGTGGTTCCGTTTA | >MI0000301\|hsa-mir-224&WithFlank&X\|-\|151958572\|151958664 | 14 | 36 | 23 | CAAGTCACTAGTGGTTCCGTTTA | 3558.8 | 16.1 | 1.3 | 12.3 | 5.3E-35 | 1.5E-33 | BT20/MB468 | CYTO |
| >hg38_21_+_8433172_8446622_RNA45SN1_45S_50ntFlanks@10690.10709.20\|CGTCCGGTGAGCTCTCGCTG | >hg38_21_+_8433172_8446622_RNA45SN1_45S_50ntFlanks | 10690 | 10709 | 20 | CGTCCGGTGAGCTCTCGCTG | 2466.9 | -9.0 | 0.7 | -12.6 | 3.2E-36 | 1.5E-33 | MB231/MB468 | CYTO |
| >hg38_21_+_8433172_8446622_RNA45SN1_45S_50ntFlanks@12438.12455.18\|ATTGTGAAGCAGAATTCA | >hg38_21_+_8433172_8446622_RNA45SN1_45S_50ntFlanks | 12438 | 12455 | 18 | ATTGTGAAGCAGAATTCA | 2298.5 | -15.6 | 1.3 | -12.3 | 7.0E-35 | 1.5E-33 | BT20/MB468 | NUC |
| >hg38_21_+_8433172_8446622_RNA45SN1_45S_50ntFlanks@8380.8405.26\|CAAGAGGGCGTGAAACCGTTAAGAGG | >hg38_21_+_8433172_8446622_RNA45SN1_45S_50ntFlanks | 8380 | 8405 | 26 | CAAGAGGGCGTGAAACCGTTAAGAGG | 6362.6 | -9.5 | 0.8 | -12.5 | 4.7E-36 | 1.6E-33 | MB231/MB468 | MP |
| >MI0000285\|hsa-mir-205&WithFlank&1\|+\|209432127\|209432248@40.61.22(+1A)&[MIMAT0000266&hsa-miR-205-5p&offsets\|0\|0(+1A);m-351&1\|+\|209432166\|209432187&offsets\|0\|0(+1A)]\|TCCTTCATTCCACCGGAGTCTGA | >MI0000285\|hsa-mir-205&WithFlank&1\|+\|209432127\|209432248 | 40 | 61 | 22(+1A) | TCCTTCATTCCACCGGAGTCTGA | 31842.9 | 18.7 | 1.5 | 12.6 | 2.5E-36 | 1.6E-33 | BT20/MB231 | NUC |
| >MI0001445\|hsa-mir-423&WithFlank&17\|+\|30117073\|30117178@59.80.22&[MIMAT0001340&hsa-miR-423-3p&offsets\|0\|-1;m-100&17\|+\|30117131\|30117153&offsets\|0\|-1]\|AGCTCGGTCTGAGGCCCCTCAG | >MI0001445\|hsa-mir-423&WithFlank&17\|+\|30117073\|30117178 | 59 | 80 | 22 | AGCTCGGTCTGAGGCCCCTCAG | 2359.8 | 7.7 | 0.6 | 12.3 | 5.8E-35 | 1.6E-33 | BT20/MB468 | CYTO |
| >hg38_21_+_8433172_8446622_RNA45SN1_45S_50ntFlanks@11672.11709.38\|CAGTGCTCTGAATGTCAAAGTGAAGAAATTCAATGAAG | >hg38_21_+_8433172_8446622_RNA45SN1_45S_50ntFlanks | 11672 | 11709 | 38 | CAGTGCTCTGAATGTCAAAGTGAAGAAATTCAATGAAG | 780.6 | -5.6 | 0.5 | -12.3 | 8.4E-35 | 1.8E-33 | BT20/MB468 | NUC |
| >hg38_21_+_8433172_8446622_RNA45SN1_45S_50ntFlanks@12458.12487.30\|AAGCGTTGGATTGTTCACCCACTAATAGGG | >hg38_21_+_8433172_8446622_RNA45SN1_45S_50ntFlanks | 12458 | 12487 | 30 | AAGCGTTGGATTGTTCACCCACTAATAGGG | 2533.7 | -15.7 | 1.3 | -12.3 | 8.7E-35 | 1.9E-33 | BT20/MB468 | NUC |
| >MI0000102\|hsa-mir-100&WithFlank&11\|-\|122152223\|122152314@19.39.21(+1U)&[MIMAT0000098&hsa-miR-100-5p&offsets\|0\|-1(+1U);m-27&11\|-\|122152276\|122152296&offsets\|0\|0(+1U)]\|AACCCGTAGATCCGAACTTGTT | >MI0000102\|hsa-mir-100&WithFlank&11\|-\|122152223\|122152314 | 19 | 39 | 21(+1U) | AACCCGTAGATCCGAACTTGTT | 6196.1 | 16.5 | 1.3 | 12.4 | 3.3E-35 | 2.0E-33 | MB231/MB468 | NUC |
| >hg38_21_+_8433172_8446622_RNA45SN1_45S_50ntFlanks@12412.12434.23\|ATCCTTCGATGTCGGCTCTTCCT | >hg38_21_+_8433172_8446622_RNA45SN1_45S_50ntFlanks | 12412 | 12434 | 23 | ATCCTTCGATGTCGGCTCTTCCT | 3168.7 | -16.1 | 1.3 | -12.3 | 9.3E-35 | 2.0E-33 | BT20/MB468 | NUC |
| >hg38_1_-_228634819_228635039_RNA5S12_5S_50ntFlanks@84.112.29\|CTCGTCTGATCTCGGAAGCTAAGCAGGGT | >hg38_1_-_228634819_228635039_RNA5S12_5S_50ntFlanks | 84 | 112 | 29 | CTCGTCTGATCTCGGAAGCTAAGCAGGGT | 1882.5 | -15.3 | 1.2 | -12.3 | 9.4E-35 | 2.0E-33 | BT20/MB468 | NUC |
| >hg38_21_+_8433172_8446622_RNA45SN1_45S_50ntFlanks@11783.11804.22\|CATGAATGGATGAACGAGATTC | >hg38_21_+_8433172_8446622_RNA45SN1_45S_50ntFlanks | 11783 | 11804 | 22 | CATGAATGGATGAACGAGATTC | 40556.5 | -8.5 | 0.7 | -12.4 | 3.5E-35 | 2.1E-33 | MB231/MB468 | NUC |
| >hg38_21_+_8433172_8446622_RNA45SN1_45S_50ntFlanks@10351.10383.33\|CGAGAACTTTGAAGGCCGAAGTGGAGAAGGGTT | >hg38_21_+_8433172_8446622_RNA45SN1_45S_50ntFlanks | 10351 | 10383 | 33 | CGAGAACTTTGAAGGCCGAAGTGGAGAAGGGTT | 50233.8 | -10.4 | 0.8 | -12.5 | 6.4E-36 | 2.1E-33 | MB231/MB468 | MP |
| >hg38_21_+_8433172_8446622_RNA45SN1_45S_50ntFlanks@11781.11805.25\|CGCATGAATGGATGAACGAGATTCC | >hg38_21_+_8433172_8446622_RNA45SN1_45S_50ntFlanks | 11781 | 11805 | 25 | CGCATGAATGGATGAACGAGATTCC | 1773.2 | -15.2 | 1.2 | -12.3 | 1.0E-34 | 2.2E-33 | BT20/MB468 | NUC |
| >am_MI0000809\|hsa-mir-151a&WithFlank&8\|-\|140732558\|140732659@53.73.21(+2U)&[MIMAT0000757&hsa-miR-151a-3p&offsets\|0\|0(+2U);m-30&8\|-\|140732587\|140732607&offsets\|0\|0(+2U)]\|CTAGACTGAAGCTCCTTGAGGTT | >am_MI0000809\|hsa-mir-151a&WithFlank&8\|-\|140732558\|140732659 | 53 | 73 | 21(+2U) | CTAGACTGAAGCTCCTTGAGGTT | 1137.2 | 9.2 | 0.7 | 12.3 | 5.4E-35 | 2.3E-33 | BT20/MB468 | MITO |
| >MI0000061\|hsa-let-7a-2&WithFlank&11\|-\|122146516\|122146599@11.31.21;MI0000062\|hsa-let-7a-3&WithFlank&22\|+\|46112743\|46112828@10.30.21;MI0000060\|hsa-let-7a-1&WithFlank&9\|+\|94175951\|94176042@12.32.21&[MIMAT0000062&hsa-let-7a-5p&offsets\|0\|-1;m-6&11\|-\|122146568\|122146589&offsets\|0\|-1];[MIMAT0000062_1&hsa-let-7a-5p&offsets\|0\|-1;m-5&22\|+\|46112752\|46112773&offsets\|0\|-1];[MIMAT0000062_2&hsa-let-7a-5p&offsets\|0\|-1;m-4&9\|+\|94175962\|94175983&offsets\|0\|-1]\|TGAGGTAGTAGGTTGTATAGT | >MI0000061\|hsa-let-7a-2&WithFlank&11\|-\|122146516\|122146599 | 11 | 31 | 21 | TGAGGTAGTAGGTTGTATAGT | 14650.4 | 7.7 | 0.6 | 12.3 | 8.3E-35 | 2.3E-33 | BT20/MB468 | CYTO |
| >MI0002468\|hsa-mir-484&WithFlank&16\|+\|15643288\|15643378@14.35.22&[MIMAT0002174&hsa-miR-484&offsets\|0\|0;m-127&16\|+\|15643301\|15643321&offsets\|0\|+1]\|TCAGGCTCAGTCCCCTCCCGAT | >MI0002468\|hsa-mir-484&WithFlank&16\|+\|15643288\|15643378 | 14 | 35 | 22 | TCAGGCTCAGTCCCCTCCCGAT | 2016.6 | 7.2 | 0.6 | 12.3 | 8.8E-35 | 2.4E-33 | BT20/MB468 | CYTO |
| >hg38_21_+_8433172_8446622_RNA45SN1_45S_50ntFlanks@10392.10409.18\|ACAGCAGTTGAACATGGG | >hg38_21_+_8433172_8446622_RNA45SN1_45S_50ntFlanks | 10392 | 10409 | 18 | ACAGCAGTTGAACATGGG | 4939.7 | -16.7 | 1.4 | -12.3 | 1.2E-34 | 2.5E-33 | BT20/MB468 | NUC |
| >MI0000077\|hsa-mir-21&WithFlank&17\|+\|59841260\|59841343@14.36.23(+1C)&[MIMAT0000076&hsa-miR-21-5p&offsets\|0\|+1(+1C);m-1&17\|+\|59841273\|59841294&offsets\|0\|+1(+1C)]\|TAGCTTATCAGACTGATGTTGACC | >MI0000077\|hsa-mir-21&WithFlank&17\|+\|59841260\|59841343 | 14 | 36 | 23(+1C) | TAGCTTATCAGACTGATGTTGACC | 42711.8 | 7.3 | 0.6 | 12.4 | 4.4E-35 | 2.6E-33 | MB231/MB468 | NUC |
| >hg38_21_+_8433172_8446622_RNA45SN1_45S_50ntFlanks@10678.10709.32\|CGGTTCCGGCGGCGTCCGGTGAGCTCTCGCTG | >hg38_21_+_8433172_8446622_RNA45SN1_45S_50ntFlanks | 10678 | 10709 | 32 | CGGTTCCGGCGGCGTCCGGTGAGCTCTCGCTG | 18672.5 | -12.1 | 1.0 | -12.5 | 8.1E-36 | 2.6E-33 | MB231/MB468 | MP |
| >hg38_21_+_8433172_8446622_RNA45SN1_45S_50ntFlanks@12418.12436.19\|CGATGTCGGCTCTTCCTAT | >hg38_21_+_8433172_8446622_RNA45SN1_45S_50ntFlanks | 12418 | 12436 | 19 | CGATGTCGGCTCTTCCTAT | 30570.7 | -9.1 | 0.7 | -12.4 | 1.9E-35 | 2.6E-33 | MB231/MB468 | MITO |
| >hg38_21_+_8433172_8446622_RNA45SN1_45S_50ntFlanks@10678.10712.35\|CGGTTCCGGCGGCGTCCGGTGAGCTCTCGCTGGCC | >hg38_21_+_8433172_8446622_RNA45SN1_45S_50ntFlanks | 10678 | 10712 | 35 | CGGTTCCGGCGGCGTCCGGTGAGCTCTCGCTGGCC | 19476.9 | -8.5 | 0.7 | -12.3 | 5.0E-35 | 2.9E-33 | MB231/MB468 | NUC |
| >am_MI0000809\|hsa-mir-151a&WithFlank&8\|-\|140732558\|140732659@53.73.21(+1C)&[MIMAT0000757&hsa-miR-151a-3p&offsets\|0\|0(+1C);m-30&8\|-\|140732587\|140732607&offsets\|0\|0(+1C)]\|CTAGACTGAAGCTCCTTGAGGC | >am_MI0000809\|hsa-mir-151a&WithFlank&8\|-\|140732558\|140732659 | 53 | 73 | 21(+1C) | CTAGACTGAAGCTCCTTGAGGC | 2534.1 | 9.5 | 0.8 | 12.3 | 7.1E-35 | 3.0E-33 | BT20/MB468 | MITO |
| >hg38_21_+_8433172_8446622_RNA45SN1_45S_50ntFlanks@3771.3792.22\|CACGGCCGGTACAGTGAAACTG | >hg38_21_+_8433172_8446622_RNA45SN1_45S_50ntFlanks | 3771 | 3792 | 22 | CACGGCCGGTACAGTGAAACTG | 2171.9 | -15.5 | 1.3 | -12.3 | 1.5E-34 | 3.1E-33 | BT20/MB468 | NUC |
| >MI0001448\|hsa-mir-425&WithFlank&3\|-\|49020142\|49020240@20.44.25&[MIMAT0003393&hsa-miR-425-5p&offsets\|0\|+2;m-95&3\|-\|49020200\|49020221&offsets\|0\|+3]\|AATGACACGATCACTCCCGTTGAGT | >MI0001448\|hsa-mir-425&WithFlank&3\|-\|49020142\|49020240 | 20 | 44 | 25 | AATGACACGATCACTCCCGTTGAGT | 10831.5 | 8.1 | 0.7 | 12.3 | 1.1E-34 | 3.1E-33 | BT20/MB468 | CYTO |
| >MI0000727\|hsa-mir-128-2&WithFlank&3\|+\|35744470\|35744565@58.79.22&[MIMAT0000424_1&hsa-miR-128-3p&offsets\|0\|+1;m-117&3\|+\|35744527\|35744547&offsets\|0\|+1]\|TCACAGTGAACCGGTCTCTTTC | >MI0000727\|hsa-mir-128-2&WithFlank&3\|+\|35744470\|35744565 | 58 | 79 | 22 | TCACAGTGAACCGGTCTCTTTC | 439.2 | 6.2 | 0.5 | 12.3 | 1.2E-34 | 3.2E-33 | BT20/MB468 | CYTO |
| >hg38_21_+_8433172_8446622_RNA45SN1_45S_50ntFlanks@4722.4758.37\|TCATTAATCAAGAACGAAAGTCGGAGGTTCGAAGACG | >hg38_21_+_8433172_8446622_RNA45SN1_45S_50ntFlanks | 4722 | 4758 | 37 | TCATTAATCAAGAACGAAAGTCGGAGGTTCGAAGACG | 24532.1 | -13.0 | 1.1 | -12.3 | 1.5E-34 | 3.2E-33 | BT20/MB468 | NUC |
| >MI0000809\|hsa-mir-151a&WithFlank&8\|-\|140732558\|140732659@51.73.23&[MIMAT0000757&hsa-miR-151a-3p&offsets\|-2\|0;m-30&8\|-\|140732587\|140732607&offsets\|-2\|0]\|TACTAGACTGAAGCTCCTTGAGG | >MI0000809\|hsa-mir-151a&WithFlank&8\|-\|140732558\|140732659 | 51 | 73 | 23 | TACTAGACTGAAGCTCCTTGAGG | 5256.2 | 8.3 | 0.7 | 12.3 | 7.9E-35 | 3.2E-33 | BT20/MB468 | MITO |
| >hg38_21_+_8433172_8446622_RNA45SN1_45S_50ntFlanks@10361.10383.23\|GAAGGCCGAAGTGGAGAAGGGTT | >hg38_21_+_8433172_8446622_RNA45SN1_45S_50ntFlanks | 10361 | 10383 | 23 | GAAGGCCGAAGTGGAGAAGGGTT | 1785.6 | -15.2 | 1.2 | -12.3 | 1.6E-34 | 3.3E-33 | BT20/MB468 | NUC |
| >MI0000446\|hsa-mir-125b-1&WithFlank&11\|-\|122099751\|122099850@21.43.23&[MIMAT0000423&hsa-miR-125b-5p&offsets\|0\|+1;m-67&11\|-\|122099809\|122099830&offsets\|0\|+1]\|TCCCTGAGACCCTAACTTGTGAT | >MI0000446\|hsa-mir-125b-1&WithFlank&11\|-\|122099751\|122099850 | 21 | 43 | 23 | TCCCTGAGACCCTAACTTGTGAT | 1858.4 | 9.3 | 0.8 | 12.3 | 5.9E-35 | 3.4E-33 | MB231/MB468 | NUC |
| >hg38_21_+_8433172_8446622_RNA45SN1_45S_50ntFlanks@12008.12043.36\|CCGCCGGTGAAATACCACTACTCTGATCGTTTTTTC | >hg38_21_+_8433172_8446622_RNA45SN1_45S_50ntFlanks | 12008 | 12043 | 36 | CCGCCGGTGAAATACCACTACTCTGATCGTTTTTTC | 27459.6 | -18.5 | 1.5 | -12.4 | 2.6E-35 | 3.5E-33 | MB231/MB468 | MITO |
| >trna134_GluTTC_1_-_16861774_16861845@1.33.33,trna5_GluTTC_1_+_17199078_17199149@1.33.33,trna84_GluTTC_1_-_161391883_161391954@1.33.33,trna94_GluTTC_1_-_149664355_149664427@1.33.33\|TCCCTGGTGGTCTAGTGGCTAGGATTCGGCGCT | >trna134_GluTTC_1_-_16861774_16861845 | 1 | 33 | 33 | TCCCTGGTGGTCTAGTGGCTAGGATTCGGCGCT | 9944.4 | 9.5 | 0.8 | 12.3 | 1.3E-34 | 3.5E-33 | BT20/MB468 | CYTO |
| >hg38_21_+_8433172_8446622_RNA45SN1_45S_50ntFlanks@4759.4778.20\|ATCAGATACCGTCGTAGTTC | >hg38_21_+_8433172_8446622_RNA45SN1_45S_50ntFlanks | 4759 | 4778 | 20 | ATCAGATACCGTCGTAGTTC | 3102.1 | -16.0 | 1.3 | -12.2 | 1.9E-34 | 3.9E-33 | BT20/MB468 | NUC |
| >hg38_21_+_8433172_8446622_RNA45SN1_45S_50ntFlanks@3844.3866.23\|TACTTGGATAACTGTGGTAATTC | >hg38_21_+_8433172_8446622_RNA45SN1_45S_50ntFlanks | 3844 | 3866 | 23 | TACTTGGATAACTGTGGTAATTC | 383.3 | 5.5 | 0.4 | 12.3 | 1.4E-34 | 3.9E-33 | BT20/MB468 | CYTO |
| >MI0000272\|hsa-mir-182&WithFlank&7\|-\|129770377\|129770498@29.52.24(+1A)&[MIMAT0000259&hsa-miR-182-5p&offsets\|0\|0(+1A);m-22&7\|-\|129770449\|129770470&offsets\|0\|+2(+1A)]\|TTTGGCAATGGTAGAACTCACACTA | >MI0000272\|hsa-mir-182&WithFlank&7\|-\|129770377\|129770498 | 29 | 52 | 24(+1A) | TTTGGCAATGGTAGAACTCACACTA | 1108.1 | 8.3 | 0.7 | 12.3 | 1.0E-34 | 4.1E-33 | BT20/MB468 | MITO |
| >hg38_21_+_8433172_8446622_RNA45SN1_45S_50ntFlanks@10404.10430.27\|CATGGGTCAGTCGGTCCTGAGAGATGG | >hg38_21_+_8433172_8446622_RNA45SN1_45S_50ntFlanks | 10404 | 10430 | 27 | CATGGGTCAGTCGGTCCTGAGAGATGG | 2123.1 | -15.5 | 1.3 | -12.2 | 2.0E-34 | 4.2E-33 | BT20/MB468 | NUC |
| >hg38_21_+_8433172_8446622_RNA45SN1_45S_50ntFlanks@12405.12433.29\|CTTTTTGATCCTTCGATGTCGGCTCTTCC | >hg38_21_+_8433172_8446622_RNA45SN1_45S_50ntFlanks | 12405 | 12433 | 29 | CTTTTTGATCCTTCGATGTCGGCTCTTCC | 13332.4 | -6.5 | 0.5 | -12.3 | 7.6E-35 | 4.3E-33 | MB231/MB468 | NUC |
| >hg38_21_+_8433172_8446622_RNA45SN1_45S_50ntFlanks@12420.12444.25\|ATGTCGGCTCTTCCTATCATTGTGA | >hg38_21_+_8433172_8446622_RNA45SN1_45S_50ntFlanks | 12420 | 12444 | 25 | ATGTCGGCTCTTCCTATCATTGTGA | 9436.8 | -8.3 | 0.7 | -12.2 | 2.2E-34 | 4.5E-33 | BT20/MB468 | NUC |
| >hg38_21_+_8433172_8446622_RNA45SN1_45S_50ntFlanks@10690.10708.19\|CGTCCGGTGAGCTCTCGCT | >hg38_21_+_8433172_8446622_RNA45SN1_45S_50ntFlanks | 10690 | 10708 | 19 | CGTCCGGTGAGCTCTCGCT | 8282.0 | -9.0 | 0.7 | -12.4 | 3.5E-35 | 4.6E-33 | MB231/MB468 | MITO |
| >hg38_21_+_8433172_8446622_RNA45SN1_45S_50ntFlanks@10841.10866.26\|CGTAACTTCGGGATAAGGATTGGCTC | >hg38_21_+_8433172_8446622_RNA45SN1_45S_50ntFlanks | 10841 | 10866 | 26 | CGTAACTTCGGGATAAGGATTGGCTC | 3836.9 | -8.1 | 0.7 | -12.2 | 2.3E-34 | 4.8E-33 | BT20/MB468 | NUC |
| >hg38_21_+_8433172_8446622_RNA45SN1_45S_50ntFlanks@4759.4783.25\|ATCAGATACCGTCGTAGTTCCGACC | >hg38_21_+_8433172_8446622_RNA45SN1_45S_50ntFlanks | 4759 | 4783 | 25 | ATCAGATACCGTCGTAGTTCCGACC | 1504.2 | -15.0 | 1.2 | -12.2 | 2.5E-34 | 5.1E-33 | BT20/MB468 | NUC |
| >MI0000285\|hsa-mir-205&WithFlank&1\|+\|209432127\|209432248@40.61.22&[MIMAT0000266&hsa-miR-205-5p&offsets\|0\|0;m-351&1\|+\|209432166\|209432187&offsets\|0\|0]\|TCCTTCATTCCACCGGAGTCTG | >MI0000285\|hsa-mir-205&WithFlank&1\|+\|209432127\|209432248 | 40 | 61 | 22 | TCCTTCATTCCACCGGAGTCTG | 27780.5 | 16.1 | 1.3 | 12.5 | 1.0E-35 | 5.3E-33 | BT20/MB231 | NUC |
| >hg38_21_+_8433172_8446622_RNA45SN1_45S_50ntFlanks@6660.6695.36\|GCGGTGGATCACTCGGCTCGTGCGTCGATGAAGAAC | >hg38_21_+_8433172_8446622_RNA45SN1_45S_50ntFlanks | 6660 | 6695 | 36 | GCGGTGGATCACTCGGCTCGTGCGTCGATGAAGAAC | 649.0 | 6.6 | 0.5 | 12.3 | 1.3E-34 | 5.4E-33 | BT20/MB468 | MITO |

<table>
<tr><td>>hg38_21_+_8433172_8446622_RNA45SN1_45S_50ntFlanks@10843.10868.26|TAACTTCGGGATAAGGATTGGCTCTA</td><td>>hg38_21_+_8433172_8446622_RNA45SN1_45S_50ntFlanks</td><td>10843</td><td>10868</td><td>26</td><td>TAACTTCGGGATAAGGATTGGCTCTA</td><td>2501.4</td><td>-15.7</td><td>1.3</td><td>-12.2</td><td>2.8E-34</td><td>5.7E-33</td><td>BT20/MB468</td><td>NUC</td></tr>
<tr><td>>hg38_21_+_8433172_8446622_RNA45SN1_45S_50ntFlanks@12008.12031.24|CCGCCGGTGAAATACCACTACTCT</td><td>>hg38_21_+_8433172_8446622_RNA45SN1_45S_50ntFlanks</td><td>12008</td><td>12031</td><td>24</td><td>CCGCCGGTGAAATACCACTACTCT</td><td>4556.5</td><td>-6.2</td><td>0.5</td><td>-12.3</td><td>1.4E-34</td><td>5.8E-33</td><td>BT20/MB468</td><td>MITO</td></tr>
<tr><td>>MI0000087|hsa-mir-29a&WithFlank&7|-|130876741|130876816@47.68.22&[MIMAT0000086&hsa-miR-29a-3p&offsets|-1|-1;m-26&7|-|130876749|130876769&offsets|-1|0]|CTAGCACCATCTGAAATCGGTT</td><td>>MI0000087|hsa-mir-29a&WithFlank&7|-|130876741|130876816</td><td>47</td><td>68</td><td>22</td><td>CTAGCACCATCTGAAATCGGTT</td><td>15445.2</td><td>8.9</td><td>0.7</td><td>12.3</td><td>1.4E-34</td><td>5.8E-33</td><td>BT20/MB468</td><td>MITO</td></tr>
<tr><td>>hg38_21_+_8433172_8446622_RNA45SN1_45S_50ntFlanks@11784.11804.21|ATGAATGGATGAACGAGATTC</td><td>>hg38_21_+_8433172_8446622_RNA45SN1_45S_50ntFlanks</td><td>11784</td><td>11804</td><td>21</td><td>ATGAATGGATGAACGAGATTC</td><td>10717.9</td><td>-7.0</td><td>0.6</td><td>-12.3</td><td>1.1E-34</td><td>6.1E-33</td><td>MB231/MB468</td><td>NUC</td></tr>
<tr><td>>hg38_21_+_8433172_8446622_RNA45SN1_45S_50ntFlanks@7980.8006.27|CCTCAGATCAGACGTGGCGACCCGCTG</td><td>>hg38_21_+_8433172_8446622_RNA45SN1_45S_50ntFlanks</td><td>7980</td><td>8006</td><td>27</td><td>CCTCAGATCAGACGTGGCGACCCGCTG</td><td>19128.1</td><td>-11.5</td><td>0.9</td><td>-12.3</td><td>5.0E-35</td><td>6.3E-33</td><td>MB231/MB468</td><td>MITO</td></tr>
<tr><td>>MI0000079|hsa-mir-23a&WithFlank&19|-|13836581|13836665@54.72.19&[MIMAT0000078&hsa-miR-23a-3p&offsets|+3|+1;m-31&19|-|13836595|13836615&offsets|+3|+1]|ACATTGCCAGGGATTTCCA</td><td>>MI0000079|hsa-mir-23a&WithFlank&19|-|13836581|13836665</td><td>54</td><td>72</td><td>19</td><td>ACATTGCCAGGGATTTCCA</td><td>3602.2</td><td>16.3</td><td>1.3</td><td>12.3</td><td>1.6E-34</td><td>6.4E-33</td><td>BT20/MB468</td><td>MITO</td></tr>
<tr><td>>trnaMT_PheGAA_MT_+_577_647@39.56.18|ATGTTTAGACGGGCTCAC</td><td>>trnaMT_PheGAA_MT_+_577_647</td><td>39</td><td>56</td><td>18</td><td>ATGTTTAGACGGGCTCAC</td><td>14184.8</td><td>7.6</td><td>0.6</td><td>12.3</td><td>1.7E-34</td><td>6.5E-33</td><td>BT20/MB468</td><td>MITO</td></tr>
<tr><td>>hg38_21_+_8433172_8446622_RNA45SN1_45S_50ntFlanks@6661.6681.21|CGGTGGATCACTCGGCTCGTG</td><td>>hg38_21_+_8433172_8446622_RNA45SN1_45S_50ntFlanks</td><td>6661</td><td>6681</td><td>21</td><td>CGGTGGATCACTCGGCTCGTG</td><td>3845.3</td><td>-3.8</td><td>0.3</td><td>-12.2</td><td>3.2E-34</td><td>6.5E-33</td><td>BT20/MB468</td><td>NUC</td></tr>
<tr><td>>hg38_21_+_8433172_8446622_RNA45SN1_45S_50ntFlanks@12625.12654.30|CAATGGGGCGAAGCTACCATCTGTGGGATT</td><td>>hg38_21_+_8433172_8446622_RNA45SN1_45S_50ntFlanks</td><td>12625</td><td>12654</td><td>30</td><td>CAATGGGGCGAAGCTACCATCTGTGGGATT</td><td>4896.8</td><td>-6.8</td><td>0.6</td><td>-12.3</td><td>1.2E-34</td><td>6.5E-33</td><td>MB231/MB468</td><td>NUC</td></tr>
<tr><td>>am_MI0000809|hsa-mir-151a&WithFlank&8|-|140732558|140732659@53.73.21(+2U)&[MIMAT0000757&hsa-miR-151a-3p&offsets|0|0(+2U);m-30&8|-|140732587|140732607&offsets|0|0(+2U)]|CTAGACTGAAGCTCCTTGAGGTT</td><td>>am_MI0000809|hsa-mir-151a&WithFlank&8|-|140732558|140732659</td><td>53</td><td>73</td><td>21(+2U)</td><td>CTAGACTGAAGCTCCTTGAGGTT</td><td>187.5</td><td>4.0</td><td>0.3</td><td>12.2</td><td>3.2E-34</td><td>6.6E-33</td><td>BT20/MB468</td><td>NUC</td></tr>
<tr><td>>hg38_21_+_8433172_8446622_RNA45SN1_45S_50ntFlanks@11208.11232.25|GCGGCGACTCTGGACGCGAGCCGGG</td><td>>hg38_21_+_8433172_8446622_RNA45SN1_45S_50ntFlanks</td><td>11208</td><td>11232</td><td>25</td><td>GCGGCGACTCTGGACGCGAGCCGGG</td><td>12859.6</td><td>-17.5</td><td>1.4</td><td>-12.4</td><td>2.2E-35</td><td>6.7E-33</td><td>MB231/MB468</td><td>MP</td></tr>
<tr><td>>MI0000108|hsa-mir-103a-2&WithFlank&20|+|3917488|3917577@54.76.23;MI0000109|hsa-mir-103a-1&WithFlank&5|-|168560890|168560979@54.76.23&[MIMAT0000101&hsa-miR-103a-3p&offsets|0|0;m-12&20|+|3917541|3917563&offsets|0|0];[MIMAT0000101_1&hsa-miR-103a-3p&offsets|0|0;m-13&5|-|168560904|168560926&offsets|0|0;m-44220&5|-|168560908|168560926&offsets|0|+4]|AGCAGCATTGTACAGGGCTATGA</td><td>>MI0000108|hsa-mir-103a-2&WithFlank&20|+|3917488|3917577</td><td>54</td><td>76</td><td>23</td><td>AGCAGCATTGTACAGGGCTATGA</td><td>35914.3</td><td>7.4</td><td>0.6</td><td>12.2</td><td>2.5E-34</td><td>6.9E-33</td><td>BT20/MB468</td><td>CYTO</td></tr>
<tr><td>>MI0000746|hsa-mir-99b&WithFlank&19|+|51692606|51692687@13.33.21&[MIMAT0000689&hsa-miR-99b-5p&offsets|0|-1;m-10&19|+|51692618|51692639&offsets|0|-1]|CACCCGTAGAACCGACCTTGC</td><td>>MI0000746|hsa-mir-99b&WithFlank&19|+|51692606|51692687</td><td>13</td><td>33</td><td>21</td><td>CACCCGTAGAACCGACCTTGC</td><td>28514.7</td><td>7.9</td><td>0.6</td><td>12.2</td><td>2.5E-34</td><td>6.9E-33</td><td>BT20/MB468</td><td>CYTO</td></tr>
<tr><td>>hg38_21_+_8433172_8446622_RNA45SN1_45S_50ntFlanks@12636.12654.19|AGCTACCATCTGTGGGATT</td><td>>hg38_21_+_8433172_8446622_RNA45SN1_45S_50ntFlanks</td><td>12636</td><td>12654</td><td>19</td><td>AGCTACCATCTGTGGGATT</td><td>6337.3</td><td>-16.5</td><td>1.3</td><td>-12.4</td><td>2.5E-35</td><td>7.4E-33</td><td>MB231/MB468</td><td>MP</td></tr>
<tr><td>>hg38_21_+_8433172_8446622_RNA45SN1_45S_50ntFlanks@8114.8140.27|GCGCGGGACATGTGGCGTACGGAAGAC</td><td>>hg38_21_+_8433172_8446622_RNA45SN1_45S_50ntFlanks</td><td>8114</td><td>8140</td><td>27</td><td>GCGCGGGACATGTGGCGTACGGAAGAC</td><td>14596.4</td><td>-17.7</td><td>1.4</td><td>-12.4</td><td>2.5E-35</td><td>7.4E-33</td><td>MB231/MB468</td><td>MP</td></tr>
<tr><td>>hg38_21_+_8433172_8446622_RNA45SN1_45S_50ntFlanks@7974.8009.36|ACGCGACCTCAGATCAGACGTGGCGACCCGCTGAAT</td><td>>hg38_21_+_8433172_8446622_RNA45SN1_45S_50ntFlanks</td><td>7974</td><td>8009</td><td>36</td><td>ACGCGACCTCAGATCAGACGTGGCGACCCGCTGAAT</td><td>27264.7</td><td>8.5</td><td>0.7</td><td>12.2</td><td>2.8E-34</td><td>7.5E-33</td><td>BT20/MB468</td><td>CYTO</td></tr>
<tr><td>>hg38_21_+_8433172_8446622_RNA45SN1_45S_50ntFlanks@5445.5466.22|TCGGATCGGCCCCGCCGGGGTC</td><td>>hg38_21_+_8433172_8446622_RNA45SN1_45S_50ntFlanks</td><td>5445</td><td>5466</td><td>22</td><td>TCGGATCGGCCCCGCCGGGGTC</td><td>3782.4</td><td>-7.2</td><td>0.6</td><td>-12.3</td><td>1.4E-34</td><td>7.5E-33</td><td>MB231/MB468</td><td>NUC</td></tr>
<tr><td>>hg38_21_+_8433172_8446622_RNA45SN1_45S_50ntFlanks@10350.10368.19|ACGAGAACTTTGAAGGCCG</td><td>>hg38_21_+_8433172_8446622_RNA45SN1_45S_50ntFlanks</td><td>10350</td><td>10368</td><td>19</td><td>ACGAGAACTTTGAAGGCCG</td><td>5173.1</td><td>-7.9</td><td>0.6</td><td>-12.3</td><td>6.5E-35</td><td>8.2E-33</td><td>MB231/MB468</td><td>MITO</td></tr>
<tr><td>>hg38_21_+_8433172_8446622_RNA45SN1_45S_50ntFlanks@4560.4587.28|CTAGGAATAATGGAATAGGACCGCGGTT</td><td>>hg38_21_+_8433172_8446622_RNA45SN1_45S_50ntFlanks</td><td>4560</td><td>4587</td><td>28</td><td>CTAGGAATAATGGAATAGGACCGCGGTT</td><td>1483.1</td><td>-15.0</td><td>1.2</td><td>-12.2</td><td>4.2E-34</td><td>8.4E-33</td><td>BT20/MB468</td><td>NUC</td></tr>
<tr><td>>hg38_21_+_8433172_8446622_RNA45SN1_45S_50ntFlanks@10678.10711.34|CGGTTCCGGCGGCGTCCGGTGAGCTCTCGCTGGC</td><td>>hg38_21_+_8433172_8446622_RNA45SN1_45S_50ntFlanks</td><td>10678</td><td>10711</td><td>34</td><td>CGGTTCCGGCGGCGTCCGGTGAGCTCTCGCTGGC</td><td>21291.2</td><td>-12.4</td><td>1.0</td><td>-12.4</td><td>2.9E-35</td><td>8.4E-33</td><td>MB231/MB468</td><td>MP</td></tr>
<tr><td>>hg38_21_+_8433172_8446622_RNA45SN1_45S_50ntFlanks@7981.8006.26|CTCAGATCAGACGTGGCGACCCGCTG</td><td>>hg38_21_+_8433172_8446622_RNA45SN1_45S_50ntFlanks</td><td>7981</td><td>8006</td><td>26</td><td>CTCAGATCAGACGTGGCGACCCGCTG</td><td>7678.8</td><td>-16.8</td><td>1.4</td><td>-12.4</td><td>3.1E-35</td><td>8.7E-33</td><td>MB231/MB468</td><td>MP</td></tr>
<tr><td>>MI0000061|hsa-let-7a-2&WithFlank&11|-|122146516|122146599@11.32.22;MI0000062|hsa-let-7a-3&WithFlank&22|+|46112743|46112828@10.31.22;MI0000060|hsa-let-7a-1&WithFlank&9|+|94175951|94176042@12.33.22&[MIMAT0000062&hsa-let-7a-5p&offsets|0|0;m-6&11|-|122146568|122146589&offsets|0|0];[MIMAT0000062_1&hsa-let-7a-5p&offsets|0|0;m-5&22|+|46112752|46112773&offsets|0|0];[MIMAT0000062_2&hsa-let-7a-5p&offsets|0|0;m-4&9|+|94175962|94175983&offsets|0|0]|TGAGGTAGTAGGTTGTATAGTT</td><td>>MI0000061|hsa-let-7a-2&WithFlank&11|-|122146516|122146599</td><td>11</td><td>32</td><td>22</td><td>TGAGGTAGTAGGTTGTATAGTT</td><td>157493.0</td><td>7.4</td><td>0.6</td><td>12.2</td><td>3.5E-34</td><td>9.3E-33</td><td>BT20/MB468</td><td>CYTO</td></tr>
<tr><td>>hg38_21_+_8433172_8446622_RNA45SN1_45S_50ntFlanks@10362.10384.23|AAGGCCGAAGTGGAGAAGGGTTC</td><td>>hg38_21_+_8433172_8446622_RNA45SN1_45S_50ntFlanks</td><td>10362</td><td>10384</td><td>23</td><td>AAGGCCGAAGTGGAGAAGGGTTC</td><td>2177.8</td><td>-15.5</td><td>1.3</td><td>-12.2</td><td>5.0E-34</td><td>1.0E-32</td><td>BT20/MB468</td><td>NUC</td></tr>
<tr><td>>hg38_21_+_8433172_8446622_RNA45SN1_45S_50ntFlanks@10679.10710.32|GGTTCCGGCGGCGTCCGGTGAGCTCTCGCTGG</td><td>>hg38_21_+_8433172_8446622_RNA45SN1_45S_50ntFlanks</td><td>10679</td><td>10710</td><td>32</td><td>GGTTCCGGCGGCGTCCGGTGAGCTCTCGCTGG</td><td>29995.9</td><td>-12.6</td><td>1.0</td><td>-12.4</td><td>3.7E-35</td><td>1.0E-32</td><td>MB231/MB468</td><td>MP</td></tr>
<tr><td>>hg38_21_+_8433172_8446622_RNA45SN1_45S_50ntFlanks@12991.13013.23|CTCCCTCGCTGCGATCTATTGAA</td><td>>hg38_21_+_8433172_8446622_RNA45SN1_45S_50ntFlanks</td><td>12991</td><td>13013</td><td>23</td><td>CTCCCTCGCTGCGATCTATTGAA</td><td>45156.3</td><td>-9.7</td><td>0.8</td><td>-12.2</td><td>1.9E-34</td><td>1.0E-32</td><td>MB231/MB468</td><td>NUC</td></tr>
<tr><td>>hg38_21_+_8433172_8446622_RNA45SN1_45S_50ntFlanks@5444.5466.23|CTCGGATCGGCCCCGCCGGGGTC</td><td>>hg38_21_+_8433172_8446622_RNA45SN1_45S_50ntFlanks</td><td>5444</td><td>5466</td><td>23</td><td>CTCGGATCGGCCCCGCCGGGGTC</td><td>18409.6</td><td>-8.9</td><td>0.7</td><td>-12.2</td><td>2.0E-34</td><td>1.0E-32</td><td>MB231/MB468</td><td>NUC</td></tr>
<tr><td>>MI0000433|hsa-let-7g&WithFlank&3|-|52268272|52268367@11.32.22&[MIMAT0000414&hsa-let-7g-5p&offsets|0|0;m-58&3|-|52268336|52268357&offsets|0|0]|TGAGGTAGTAGTTTGTACAGTT</td><td>>MI0000433|hsa-let-7g&WithFlank&3|-|52268272|52268367</td><td>11</td><td>32</td><td>22</td><td>TGAGGTAGTAGTTTGTACAGTT</td><td>84569.4</td><td>7.5</td><td>0.6</td><td>12.2</td><td>2.7E-34</td><td>1.1E-32</td><td>BT20/MB468</td><td>MITO</td></tr>
<tr><td>>hg38_21_+_8433172_8446622_RNA45SN1_45S_50ntFlanks@9534.9570.37|AGGAAACTCTGGTGGAGGTCCGTAGCGGTCCTGACGT</td><td>>hg38_21_+_8433172_8446622_RNA45SN1_45S_50ntFlanks</td><td>9534</td><td>9570</td><td>37</td><td>AGGAAACTCTGGTGGAGGTCCGTAGCGGTCCTGACGT</td><td>11874.5</td><td>-9.0</td><td>0.7</td><td>-12.2</td><td>5.3E-34</td><td>1.1E-32</td><td>BT20/MB468</td><td>NUC</td></tr>
<tr><td>>hg38_21_+_8433172_8446622_RNA45SN1_45S_50ntFlanks@10680.10708.29|GTTCCGGCGGCGTCCGGTGAGCTCTCGCT</td><td>>hg38_21_+_8433172_8446622_RNA45SN1_45S_50ntFlanks</td><td>10680</td><td>10708</td><td>29</td><td>GTTCCGGCGGCGTCCGGTGAGCTCTCGCT</td><td>38330.1</td><td>-8.6</td><td>0.7</td><td>-12.3</td><td>9.0E-35</td><td>1.1E-32</td><td>MB231/MB468</td><td>MITO</td></tr>
<tr><td>>hg38_21_+_8433172_8446622_RNA45SN1_45S_50ntFlanks@7985.8006.22|GATCAGACGTGGCGACCCGCTG</td><td>>hg38_21_+_8433172_8446622_RNA45SN1_45S_50ntFlanks</td><td>7985</td><td>8006</td><td>22</td><td>GATCAGACGTGGCGACCCGCTG</td><td>2303.9</td><td>-15.6</td><td>1.3</td><td>-12.1</td><td>5.8E-34</td><td>1.2E-32</td><td>BT20/MB468</td><td>NUC</td></tr>
<tr><td>>MI0000077|hsa-mir-21&WithFlank&17|+|59841260|59841343@15.36.22&[MIMAT0000076&hsa-miR-21-5p&offsets|+1|+1;m-1&17|+|59841273|59841294&offsets|+1|+1]|AGCTTATCAGACTGATGTTGAC</td><td>>MI0000077|hsa-mir-21&WithFlank&17|+|59841260|59841343</td><td>15</td><td>36</td><td>22</td><td>AGCTTATCAGACTGATGTTGAC</td><td>1335.6</td><td>7.0</td><td>0.6</td><td>12.2</td><td>3.0E-34</td><td>1.2E-32</td><td>BT20/MB468</td><td>MITO</td></tr>
<tr><td>>MI0000065|hsa-let-7d&WithFlank&9|+|94178828|94178926@68.89.22&[MIMAT0004484&hsa-let-7d-3p&offsets|0|0;m-98&9|+|94178895|94178916&offsets|0|0]|CTATACGACCTGCTGCCTTTCT</td><td>>MI0000065|hsa-let-7d&WithFlank&9|+|94178828|94178926</td><td>68</td><td>89</td><td>22</td><td>CTATACGACCTGCTGCCTTTCT</td><td>4500.9</td><td>7.8</td><td>0.6</td><td>12.2</td><td>3.0E-34</td><td>1.2E-32</td><td>BT20/MB468</td><td>MITO</td></tr>
<tr><td>>hg38_21_+_8433172_8446622_RNA45SN1_45S_50ntFlanks@11579.11602.24|AGAACTGGTGCGGACCAGGGGAAT</td><td>>hg38_21_+_8433172_8446622_RNA45SN1_45S_50ntFlanks</td><td>11579</td><td>11602</td><td>24</td><td>AGAACTGGTGCGGACCAGGGGAAT</td><td>667.6</td><td>-6.4</td><td>0.5</td><td>-12.1</td><td>6.2E-34</td><td>1.2E-32</td><td>BT20/MB468</td><td>NUC</td></tr>
<tr><td>>hg38_21_+_8433172_8446622_RNA45SN1_45S_50ntFlanks@10353.10383.31|AGAACTTTGAAGGCCGAAGTGGAGAAGGGTT</td><td>>hg38_21_+_8433172_8446622_RNA45SN1_45S_50ntFlanks</td><td>10353</td><td>10383</td><td>31</td><td>AGAACTTTGAAGGCCGAAGTGGAGAAGGGTT</td><td>1167.2</td><td>-6.7</td><td>0.6</td><td>-12.1</td><td>6.4E-34</td><td>1.3E-32</td><td>BT20/MB468</td><td>NUC</td></tr>
<tr><td>>trna119_LysCTT_1_-_145395522_145395594@1.29.29,trna11_LysCTT_5_-_180648979_180649051@1.29.29,trna13_LysCTT_6_+_26556774_26556846@1.29.29,trna32_LysCTT_16_-_3207406_3207478@1.29.29,trna7_LysCTT_16_+_3225692_3225764@1.29.29,trna9_LysCTT_5_+_180634755_180634827@1.29.29|GCCCGGCTAGCTCAGTCGGTAGAGCATGA</td><td>>trna119_LysCTT_1_-_145395522_145395594</td><td>1</td><td>29</td><td>29</td><td>GCCCGGCTAGCTCAGTCGGTAGAGCATGA</td><td>2016.2</td><td>-15.4</td><td>1.3</td><td>-12.1</td><td>6.4E-34</td><td>1.3E-32</td><td>BT20/MB468</td><td>NUC</td></tr>
<tr><td>>MI0000264|hsa-mir-7-2&WithFlank&15|+|88611819|88611940@38.60.23(+1A);MI0000265|hsa-mir-7-3&WithFlank&19|+|4770664|4770785@37.59.23(+1A);MI0000263|hsa-mir-7-1&WithFlank&9|-|83969742|83969863@30.52.23(+1A)&[MIMAT0000252&hsa-miR-7-5p&offsets|0|-1(+1A);m-400&15|+|88611856|88611879&offsets|0|-1(+1A)];[MIMAT0000252_1&hsa-miR-7-5p&offsets|0|-1(+1A);m-402&19|+|4770700|4770723&offsets|0|-1(+1A)];[MIMAT0000252_2&hsa-miR-7-5p&offsets|0|-1(+1A);m-401&9|-|83969811|83969834&offsets|0|-1(+1A)]|TGGAAGACTAGTGATTTTGTTGTA</td><td>>MI0000264|hsa-mir-7-2&WithFlank&15|+|88611819|88611940</td><td>38</td><td>60</td><td>23(+1A)</td><td>TGGAAGACTAGTGATTTTGTTGTA</td><td>2033.3</td><td>7.7</td><td>0.6</td><td>12.2</td><td>3.5E-34</td><td>1.3E-32</td><td>BT20/MB468</td><td>MITO</td></tr>
<tr><td>>MI0001733|hsa-mir-452&WithFlank&X|-|151959622|151959718@20.42.23&[MIMAT0001635&hsa-miR-452-5p&offsets|0|+1;m-188&X|-|151959677|151959699&offsets|0|0]|AACTGTTTGCAGAGGAAACTGAG</td><td>>MI0001733|hsa-mir-452&WithFlank&X|-|151959622|151959718</td><td>20</td><td>42</td><td>23</td><td>AACTGTTTGCAGAGGAAACTGAG</td><td>1817.6</td><td>15.1</td><td>1.2</td><td>12.2</td><td>5.0E-34</td><td>1.3E-32</td><td>BT20/MB468</td><td>CYTO</td></tr>
<tr><td>>hg38_21_+_8433172_8446622_RNA45SN1_45S_50ntFlanks@10680.10709.30|GTTCCGGCGGCGTCCGGTGAGCTCTCGCTG</td><td>>hg38_21_+_8433172_8446622_RNA45SN1_45S_50ntFlanks</td><td>10680</td><td>10709</td><td>30</td><td>GTTCCGGCGGCGTCCGGTGAGCTCTCGCTG</td><td>9910.6</td><td>-8.4</td><td>0.7</td><td>-12.3</td><td>1.1E-34</td><td>1.3E-32</td><td>MB231/MB468</td><td>MITO</td></tr>
<tr><td>>hg38_21_+_8433172_8446622_RNA45SN1_45S_50ntFlanks@4724.4752.29|ATTAATCAAGAACGAAAGTCGGAGGTTCG</td><td>>hg38_21_+_8433172_8446622_RNA45SN1_45S_50ntFlanks</td><td>4724</td><td>4752</td><td>29</td><td>ATTAATCAAGAACGAAAGTCGGAGGTTCG</td><td>1482.4</td><td>-15.0</td><td>1.2</td><td>-12.1</td><td>7.1E-34</td><td>1.4E-32</td><td>BT20/MB468</td><td>NUC</td></tr>
<tr><td>>hg38_21_+_8433172_8446622_RNA45SN1_45S_50ntFlanks@12159.12179.21|TTTGACTGGGGCGGTACACCT</td><td>>hg38_21_+_8433172_8446622_RNA45SN1_45S_50ntFlanks</td><td>12159</td><td>12179</td><td>21</td><td>TTTGACTGGGGCGGTACACCT</td><td>8465.0</td><td>-16.9</td><td>1.4</td><td>-12.3</td><td>5.5E-35</td><td>1.5E-32</td><td>MB231/MB468</td><td>MP</td></tr>
<tr><td>>am_MI0000786|hsa-mir-378a&WithFlank&5|+|149732819|149732896@49.69.21(+1U)&[MIMAT0000732&hsa-miR-378a-3p&offsets|0|-1(+1U);m-44&5|+|149732867|149732888&offsets|0|-1(+1U)]|ACTGGACTTGGAGTCAGAAGGT</td><td>>am_MI0000786|hsa-mir-378a&WithFlank&5|+|149732819|149732896</td><td>49</td><td>69</td><td>21(+1U)</td><td>ACTGGACTTGGAGTCAGAAGGT</td><td>9431.7</td><td>6.4</td><td>0.5</td><td>12.2</td><td>5.6E-34</td><td>1.5E-32</td><td>BT20/MB468</td><td>CYTO</td></tr>
<tr><td>>MI0001641|hsa-mir-429&WithFlank&1|+|1168999|1169093@57.78.22&[MIMAT0001536&hsa-miR-429&offsets|0|0;m-207&1|+|1169055|1169076&offsets|0|0]|TAATACTGTCTGGTAAAACCGT</td><td>>MI0001641|hsa-mir-429&WithFlank&1|+|1168999|1169093</td><td>57</td><td>78</td><td>22</td><td>TAATACTGTCTGGTAAAACCGT</td><td>499.1</td><td>-5.6</td><td>0.5</td><td>-12.1</td><td>7.8E-34</td><td>1.5E-32</td><td>BT20/MB468</td><td>NUC</td></tr>
<tr><td>>hg38_21_+_8433172_8446622_RNA45SN1_45S_50ntFlanks@10680.10712.33|GTTCCGGCGGCGTCCGGTGAGCTCTCGCTGGCC</td><td>>hg38_21_+_8433172_8446622_RNA45SN1_45S_50ntFlanks</td><td>10680</td><td>10712</td><td>33</td><td>GTTCCGGCGGCGTCCGGTGAGCTCTCGCTGGCC</td><td>2906.1</td><td>-7.4</td><td>0.6</td><td>-12.3</td><td>5.8E-35</td><td>1.5E-32</td><td>MB231/MB468</td><td>MP</td></tr>
</table>

<table>
<tr><th></th><th></th><th></th><th></th><th></th><th></th><th></th><th></th><th></th><th></th><th></th><th></th><th></th><th></th></tr>
<tr><td>>hg38_21_+_8433172_8446622_RNA45SN1_45S_50ntFlanks@8114.8138.25|GCGCGGGACATGTGGCGTACGGAAG</td><td>>hg38_21_+_8433172_8446622_RNA45SN1_45S_50ntFlanks</td><td>8114</td><td>8138</td><td>25</td><td>GCGCGGGACATGTGGCGTACGGAAG</td><td>22354.6</td><td>-18.3</td><td>1.5</td><td>-12.3</td><td>5.9E-35</td><td>1.5E-32</td><td>MB231/MB468</td><td>MP</td></tr>
<tr><td>>hg38_21_+_8433172_8446622_RNA45SN1_45S_50ntFlanks@12412.12432.21|ATCCTTCGATGTCGGCTCTTC</td><td>>hg38_21_+_8433172_8446622_RNA45SN1_45S_50ntFlanks</td><td>12412</td><td>12432</td><td>21</td><td>ATCCTTCGATGTCGGCTCTTC</td><td>3819.8</td><td>-16.3</td><td>1.3</td><td>-12.1</td><td>7.9E-34</td><td>1.5E-32</td><td>BT20/MB468</td><td>NUC</td></tr>
<tr><td>>hg38_21_+_8433172_8446622_RNA45SN1_45S_50ntFlanks@11564.11591.28|CCTAGCAGCCGACTTAGAACTGGTGCGG</td><td>>hg38_21_+_8433172_8446622_RNA45SN1_45S_50ntFlanks</td><td>11564</td><td>11591</td><td>28</td><td>CCTAGCAGCCGACTTAGAACTGGTGCGG</td><td>1574.9</td><td>-15.0</td><td>1.2</td><td>-12.1</td><td>8.6E-34</td><td>1.7E-32</td><td>BT20/MB468</td><td>NUC</td></tr>
<tr><td>>hg38_21_+_8433172_8446622_RNA45SN1_45S_50ntFlanks@7975.8011.37|CGCGACCTCAGATCAGACGTGGCGACCCGCTGAATTT</td><td>>hg38_21_+_8433172_8446622_RNA45SN1_45S_50ntFlanks</td><td>7975</td><td>8011</td><td>37</td><td>CGCGACCTCAGATCAGACGTGGCGACCCGCTGAATTT</td><td>74550.9</td><td>7.2</td><td>0.6</td><td>12.1</td><td>6.9E-34</td><td>1.8E-32</td><td>BT20/MB468</td><td>CYTO</td></tr>
<tr><td>>hg38_21_+_8433172_8446622_RNA45SN1_45S_50ntFlanks@12633.12654.22|CGAAGCTACCATCTGTGGGATT</td><td>>hg38_21_+_8433172_8446622_RNA45SN1_45S_50ntFlanks</td><td>12633</td><td>12654</td><td>22</td><td>CGAAGCTACCATCTGTGGGATT</td><td>29575.7</td><td>-12.3</td><td>1.0</td><td>-12.3</td><td>1.5E-34</td><td>1.8E-32</td><td>MB231/MB468</td><td>MITO</td></tr>
<tr><td>>am_MI0000809|hsa-mir-151a&WithFlank&8|-|140732558|140732659@53.73.21&[MIMAT0000757&hsa-miR-151a-3p&offsets|0|0;m-30&8|-|140732587|140732607&offsets|0|0]|CTAGACTGAAGCTCCTTGAGG</td><td>>am_MI0000809|hsa-mir-151a&WithFlank&8|-|140732558|140732659</td><td>53</td><td>73</td><td>21</td><td>CTAGACTGAAGCTCCTTGAGG</td><td>178685.8</td><td>9.1</td><td>0.7</td><td>12.1</td><td>7.0E-34</td><td>1.8E-32</td><td>BT20/MB468</td><td>CYTO</td></tr>
<tr><td>>MI0000264|hsa-mir-7-2&WithFlank&15|+|88611819|88611940@38.60.23(+1C);MI0000265|hsa-mir-7-3&WithFlank&19|+|4770664|4770785@37.59.23(+1C);MI0000263|hsa-mir-7-1&WithFlank&9|-|83969742|83969863@30.52.23(+1C)&[MIMAT0000252&hsa-miR-7-5p&offsets|0|-1(+1C);m-400&15|+|88611856|88611879&offsets|0|-1(+1C)];[MIMAT0000252_1&hsa-miR-7-5p&offsets|0|-1(+1C);m-402&19|+|4770700|4770723&offsets|0|-1(+1C)];[MIMAT0000252_2&hsa-miR-7-5p&offsets|0|-1(+1C);m-401&9|-|83969811|83969834&offsets|0|-1(+1C)]|TGGAAGACTAGTGATTTTGTTGTC</td><td>>MI0000264|hsa-mir-7-2&WithFlank&15|+|88611819|88611940</td><td>38</td><td>60</td><td>23(+1C)</td><td>TGGAAGACTAGTGATTTTGTTGTC</td><td>11138.7</td><td>7.8</td><td>0.6</td><td>12.2</td><td>4.9E-34</td><td>1.9E-32</td><td>BT20/MB468</td><td>MITO</td></tr>
<tr><td>>MI0000266|hsa-mir-10a&WithFlank&17|-|48579832|48579953@29.50.22&[MIMAT0000253&hsa-miR-10a-5p&offsets|+1|0;m-18&17|-|48579905|48579926&offsets|+1|+1]|ACCCTGTAGATCCGAATTTGTG</td><td>>MI0000266|hsa-mir-10a&WithFlank&17|-|48579832|48579953</td><td>29</td><td>50</td><td>22</td><td>ACCCTGTAGATCCGAATTTGTG</td><td>7133.9</td><td>17.3</td><td>1.4</td><td>12.2</td><td>5.3E-34</td><td>2.0E-32</td><td>BT20/MB468</td><td>MITO</td></tr>
<tr><td>>hg38_21_+_8433172_8446622_RNA45SN1_45S_50ntFlanks@13004.13027.24|ATCTATTGAAAGTCAGCCCTCGAC</td><td>>hg38_21_+_8433172_8446622_RNA45SN1_45S_50ntFlanks</td><td>13004</td><td>13027</td><td>24</td><td>ATCTATTGAAAGTCAGCCCTCGAC</td><td>4849.5</td><td>-16.7</td><td>1.4</td><td>-12.1</td><td>1.0E-33</td><td>2.0E-32</td><td>BT20/MB468</td><td>NUC</td></tr>
<tr><td>>hg38_21_+_8433172_8446622_RNA45SN1_45S_50ntFlanks@10351.10383.33|CGAGAACTTTGAAGGCCGAAGTGGAGAAGGGTT</td><td>>hg38_21_+_8433172_8446622_RNA45SN1_45S_50ntFlanks</td><td>10351</td><td>10383</td><td>33</td><td>CGAGAACTTTGAAGGCCGAAGTGGAGAAGGGTT</td><td>25967.2</td><td>-8.5</td><td>0.7</td><td>-12.2</td><td>1.7E-34</td><td>2.0E-32</td><td>MB231/MB468</td><td>MITO</td></tr>
<tr><td>>hg38_21_+_8433172_8446622_RNA45SN1_45S_50ntFlanks@5488.5515.28|GCTGAGAAGACGGTCGAACTTGACTATC</td><td>>hg38_21_+_8433172_8446622_RNA45SN1_45S_50ntFlanks</td><td>5488</td><td>5515</td><td>28</td><td>GCTGAGAAGACGGTCGAACTTGACTATC</td><td>565.9</td><td>6.6</td><td>0.5</td><td>12.1</td><td>7.7E-34</td><td>2.0E-32</td><td>BT20/MB468</td><td>CYTO</td></tr>
<tr><td>>hg38_21_+_8433172_8446622_RNA45SN1_45S_50ntFlanks@7986.8006.21|ATCAGACGTGGCGACCCGCTG</td><td>>hg38_21_+_8433172_8446622_RNA45SN1_45S_50ntFlanks</td><td>7986</td><td>8006</td><td>21</td><td>ATCAGACGTGGCGACCCGCTG</td><td>23232.8</td><td>-18.4</td><td>1.5</td><td>-12.3</td><td>7.9E-35</td><td>2.0E-32</td><td>MB231/MB468</td><td>MP</td></tr>
<tr><td>>hg38_21_+_8433172_8446622_RNA45SN1_45S_50ntFlanks@7974.8011.38|ACGCGACCTCAGATCAGACGTGGCGACCCGCTGAATTT</td><td>>hg38_21_+_8433172_8446622_RNA45SN1_45S_50ntFlanks</td><td>7974</td><td>8011</td><td>38</td><td>ACGCGACCTCAGATCAGACGTGGCGACCCGCTGAATTT</td><td>16361.1</td><td>7.6</td><td>0.6</td><td>12.4</td><td>4.7E-35</td><td>2.1E-32</td><td>MB231/MB468</td><td>CYTO</td></tr>
<tr><td>>MI0001445|hsa-mir-423&WithFlank&17|+|30117073|30117178@59.80.22&[MIMAT0001340&hsa-miR-423-3p&offsets|0|-1;m-100&17|+|30117131|30117153&offsets|0|-1]|AGCTCGGTCTGAGGCCCCTCAG</td><td>>MI0001445|hsa-mir-423&WithFlank&17|+|30117073|30117178</td><td>59</td><td>80</td><td>22</td><td>AGCTCGGTCTGAGGCCCCTCAG</td><td>2755.6</td><td>7.6</td><td>0.6</td><td>12.2</td><td>5.6E-34</td><td>2.1E-32</td><td>BT20/MB468</td><td>MITO</td></tr>
<tr><td>>hg38_21_+_8433172_8446622_RNA45SN1_45S_50ntFlanks@4723.4752.30|CATTAATCAAGAACGAAAGTCGGAGGTTCG</td><td>>hg38_21_+_8433172_8446622_RNA45SN1_45S_50ntFlanks</td><td>4723</td><td>4752</td><td>30</td><td>CATTAATCAAGAACGAAAGTCGGAGGTTCG</td><td>1269.8</td><td>-14.7</td><td>1.2</td><td>-12.1</td><td>1.2E-33</td><td>2.3E-32</td><td>BT20/MB468</td><td>NUC</td></tr>
<tr><td>>hg38_21_+_8433172_8446622_RNA45SN1_45S_50ntFlanks@12626.12654.29|AATGGGGCGAAGCTACCATCTGTGGGATT</td><td>>hg38_21_+_8433172_8446622_RNA45SN1_45S_50ntFlanks</td><td>12626</td><td>12654</td><td>29</td><td>AATGGGGCGAAGCTACCATCTGTGGGATT</td><td>9438.7</td><td>-7.1</td><td>0.6</td><td>-12.2</td><td>4.6E-34</td><td>2.4E-32</td><td>MB231/MB468</td><td>NUC</td></tr>
<tr><td>>hg38_21_+_8433172_8446622_RNA45SN1_45S_50ntFlanks@12456.12486.31|CCAAGCGTTGGATTGTTCACCCACTAATAGG</td><td>>hg38_21_+_8433172_8446622_RNA45SN1_45S_50ntFlanks</td><td>12456</td><td>12486</td><td>31</td><td>CCAAGCGTTGGATTGTTCACCCACTAATAGG</td><td>14056.0</td><td>-8.7</td><td>0.7</td><td>-12.2</td><td>4.9E-34</td><td>2.6E-32</td><td>MB231/MB468</td><td>NUC</td></tr>
<tr><td>>hg38_21_+_8433172_8446622_RNA45SN1_45S_50ntFlanks@4722.4759.38|TCATTAATCAAGAACGAAAGTCGGAGGTTCGAAGACGA</td><td>>hg38_21_+_8433172_8446622_RNA45SN1_45S_50ntFlanks</td><td>4722</td><td>4759</td><td>38</td><td>TCATTAATCAAGAACGAAAGTCGGAGGTTCGAAGACGA</td><td>2318.2</td><td>-6.0</td><td>0.5</td><td>-12.1</td><td>6.8E-34</td><td>2.6E-32</td><td>BT20/MB468</td><td>MITO</td></tr>
<tr><td>>MI0000266|hsa-mir-10a&WithFlank&17|-|48579832|48579953@28.50.23&[MIMAT0000253&hsa-miR-10a-5p&offsets|0|0;m-18&17|-|48579905|48579926&offsets|0|+1]|TACCCTGTAGATCCGAATTTGTG</td><td>>MI0000266|hsa-mir-10a&WithFlank&17|-|48579832|48579953</td><td>28</td><td>50</td><td>23</td><td>TACCCTGTAGATCCGAATTTGTG</td><td>10651.2</td><td>17.8</td><td>1.5</td><td>12.1</td><td>6.9E-34</td><td>2.6E-32</td><td>BT20/MB468</td><td>MITO</td></tr>
<tr><td>>MI0000650|hsa-mir-200c&WithFlank&12|+|6963693|6963772@50.71.22(+1U)&[MIMAT0000617&hsa-miR-200c-3p&offsets|0|-1(+1U);m-54&12|+|6963742|6963764&offsets|0|-1(+1U)]|TAATACTGCCGGGTAATGATGGT</td><td>>MI0000650|hsa-mir-200c&WithFlank&12|+|6963693|6963772</td><td>50</td><td>71</td><td>22(+1U)</td><td>TAATACTGCCGGGTAATGATGGT</td><td>5142.3</td><td>9.0</td><td>0.7</td><td>12.1</td><td>1.1E-33</td><td>2.8E-32</td><td>BT20/MB468</td><td>CYTO</td></tr>
<tr><td>>MI0006444|hsa-mir-1307&WithFlank&10|-|103394247|103394407@87.107.21(+1U)&[MIMAT0005951&hsa-miR-1307-3p&offsets|+1|0(+1U);m-66&10|-|103394300|103394322&offsets|+1|-1(+1U)]|CTCGGCGTGGCGTCGGTCGTGT</td><td>>MI0006444|hsa-mir-1307&WithFlank&10|-|103394247|103394407</td><td>87</td><td>107</td><td>21(+1U)</td><td>CTCGGCGTGGCGTCGGTCGTGT</td><td>509.3</td><td>6.9</td><td>0.6</td><td>12.1</td><td>1.1E-33</td><td>2.9E-32</td><td>BT20/MB468</td><td>CYTO</td></tr>
<tr><td>>hg38_21_+_8433172_8446622_RNA45SN1_45S_50ntFlanks@11318.11344.27|CCCCACGTCTCGTCGCGCGCGCGTCCG</td><td>>hg38_21_+_8433172_8446622_RNA45SN1_45S_50ntFlanks</td><td>11318</td><td>11344</td><td>27</td><td>CCCCACGTCTCGTCGCGCGCGCGTCCG</td><td>17501.0</td><td>-17.9</td><td>1.5</td><td>-12.2</td><td>2.5E-34</td><td>2.9E-32</td><td>MB231/MB468</td><td>MITO</td></tr>
<tr><td>>hg38_21_+_8433172_8446622_RNA45SN1_45S_50ntFlanks@11233.11254.22|CCCTTCCCGTGGATCGCCCCAG</td><td>>hg38_21_+_8433172_8446622_RNA45SN1_45S_50ntFlanks</td><td>11233</td><td>11254</td><td>22</td><td>CCCTTCCCGTGGATCGCCCCAG</td><td>3670.5</td><td>-16.3</td><td>1.3</td><td>-12.1</td><td>1.5E-33</td><td>2.9E-32</td><td>BT20/MB468</td><td>NUC</td></tr>
<tr><td>>hg38_21_+_8433172_8446622_RNA45SN1_45S_50ntFlanks@7980.8007.28|CCTCAGATCAGACGTGGCGACCCGCTGA</td><td>>hg38_21_+_8433172_8446622_RNA45SN1_45S_50ntFlanks</td><td>7980</td><td>8007</td><td>28</td><td>CCTCAGATCAGACGTGGCGACCCGCTGA</td><td>332.6</td><td>-5.4</td><td>0.4</td><td>-12.1</td><td>1.6E-33</td><td>3.1E-32</td><td>BT20/MB468</td><td>NUC</td></tr>
<tr><td>>hg38_21_+_8433172_8446622_RNA45SN1_45S_50ntFlanks@12073.12095.23|CCGAGGGGCTCTCGCTTCTGGCG</td><td>>hg38_21_+_8433172_8446622_RNA45SN1_45S_50ntFlanks</td><td>12073</td><td>12095</td><td>23</td><td>CCGAGGGGCTCTCGCTTCTGGCG</td><td>2349.3</td><td>-15.6</td><td>1.3</td><td>-12.1</td><td>1.6E-33</td><td>3.1E-32</td><td>BT20/MB468</td><td>NUC</td></tr>
<tr><td>>MI0000299|hsa-mir-222&WithFlank&X|-|45747009|45747130@75.97.23&[MIMAT0000279&hsa-miR-222-3p&offsets|0|+2;m-140&X|-|45747033|45747056&offsets|0|-1]|AGCTACATCTGGCTACTGGGTCT</td><td>>MI0000299|hsa-mir-222&WithFlank&X|-|45747009|45747130</td><td>75</td><td>97</td><td>23</td><td>AGCTACATCTGGCTACTGGGTCT</td><td>3963.9</td><td>7.1</td><td>0.6</td><td>12.1</td><td>1.2E-33</td><td>3.2E-32</td><td>BT20/MB468</td><td>CYTO</td></tr>
<tr><td>>hg38_21_+_8433172_8446622_RNA45SN1_45S_50ntFlanks@11874.11900.27|AGAAGACCCTGTTGAGCTTGACTCTAG</td><td>>hg38_21_+_8433172_8446622_RNA45SN1_45S_50ntFlanks</td><td>11874</td><td>11900</td><td>27</td><td>AGAAGACCCTGTTGAGCTTGACTCTAG</td><td>1973.6</td><td>-15.4</td><td>1.3</td><td>-12.1</td><td>1.7E-33</td><td>3.2E-32</td><td>BT20/MB468</td><td>NUC</td></tr>
<tr><td>>hg38_21_+_8433172_8446622_RNA45SN1_45S_50ntFlanks@7986.8006.21|ATCAGACGTGGCGACCCGCTG</td><td>>hg38_21_+_8433172_8446622_RNA45SN1_45S_50ntFlanks</td><td>7986</td><td>8006</td><td>21</td><td>ATCAGACGTGGCGACCCGCTG</td><td>2183.5</td><td>-15.5</td><td>1.3</td><td>-12.1</td><td>1.7E-33</td><td>3.3E-32</td><td>BT20/MB468</td><td>NUC</td></tr>
<tr><td>>MI0000285|hsa-mir-205&WithFlank&1|+|209432127|209432248@40.61.22(+1C)&[MIMAT0000266&hsa-miR-205-5p&offsets|0|0(+1C);m-351&1|+|209432166|209432187&offsets|0|0(+1C)]|TCCTTCATTCCACCGGAGTCTGC</td><td>>MI0000285|hsa-mir-205&WithFlank&1|+|209432127|209432248</td><td>40</td><td>61</td><td>22(+1C)</td><td>TCCTTCATTCCACCGGAGTCTGC</td><td>16459.5</td><td>7.7</td><td>0.6</td><td>12.1</td><td>1.4E-33</td><td>3.5E-32</td><td>BT20/MB468</td><td>CYTO</td></tr>
<tr><td>>hg38_21_+_8433172_8446622_RNA45SN1_45S_50ntFlanks@10678.10709.32|CGGTTCCGGCGGCGTCCGGTGAGCTCTCGCTG</td><td>>hg38_21_+_8433172_8446622_RNA45SN1_45S_50ntFlanks</td><td>10678</td><td>10709</td><td>32</td><td>CGGTTCCGGCGGCGTCCGGTGAGCTCTCGCTG</td><td>60198.7</td><td>-9.4</td><td>0.8</td><td>-12.1</td><td>7.2E-34</td><td>3.8E-32</td><td>MB231/MB468</td><td>NUC</td></tr>
<tr><td>>hg38_21_+_8433172_8446622_RNA45SN1_45S_50ntFlanks@11594.11611.18|CAGGGGAATCCGACTGTT</td><td>>hg38_21_+_8433172_8446622_RNA45SN1_45S_50ntFlanks</td><td>11594</td><td>11611</td><td>18</td><td>CAGGGGAATCCGACTGTT</td><td>3713.1</td><td>-6.7</td><td>0.6</td><td>-12.0</td><td>2.1E-33</td><td>3.9E-32</td><td>BT20/MB468</td><td>NUC</td></tr>
<tr><td>>MI0000108|hsa-mir-103a-2&WithFlank&20|+|3917488|3917577@54.76.23(+1U);MI0000109|hsa-mir-103a-1&WithFlank&5|-|168560890|168560979@54.76.23(+1U)&[MIMAT0000101&hsa-miR-103a-3p&offsets|0|0(+1U);m-12&20|+|3917541|3917563&offsets|0|0(+1U)];[MIMAT0000101_1&hsa-miR-103a-3p&offsets|0|0(+1U);m-13&5|-|168560904|168560926&offsets|0|0(+1U);m-44220&5|-|168560908|168560926&offsets|0|+4(+1U)]|AGCAGCATTGTACAGGGCTATGAT</td><td>>MI0000108|hsa-mir-103a-2&WithFlank&20|+|3917488|3917577</td><td>54</td><td>76</td><td>23(+1U)</td><td>AGCAGCATTGTACAGGGCTATGAT</td><td>3008.2</td><td>7.0</td><td>0.6</td><td>12.1</td><td>1.5E-33</td><td>3.9E-32</td><td>BT20/MB468</td><td>CYTO</td></tr>
<tr><td>>hg38_21_+_8433172_8446622_RNA45SN1_45S_50ntFlanks@7974.8010.37|ACGCGACCTCAGATCAGACGTGGCGACCCGCTGAATT</td><td>>hg38_21_+_8433172_8446622_RNA45SN1_45S_50ntFlanks</td><td>7974</td><td>8010</td><td>37</td><td>ACGCGACCTCAGATCAGACGTGGCGACCCGCTGAATT</td><td>16227.5</td><td>8.5</td><td>0.7</td><td>12.1</td><td>1.6E-33</td><td>4.0E-32</td><td>BT20/MB468</td><td>CYTO</td></tr>
<tr><td>>hg38_21_+_8433172_8446622_RNA45SN1_45S_50ntFlanks@11552.11569.18|CCTCGGCCGGCGCCTAGC</td><td>>hg38_21_+_8433172_8446622_RNA45SN1_45S_50ntFlanks</td><td>11552</td><td>11569</td><td>18</td><td>CCTCGGCCGGCGCCTAGC</td><td>1395.1</td><td>-6.8</td><td>0.6</td><td>-12.0</td><td>2.2E-33</td><td>4.2E-32</td><td>BT20/MB468</td><td>NUC</td></tr>
<tr><td>>hg38_21_+_8433172_8446622_RNA45SN1_45S_50ntFlanks@7984.8006.23|AGATCAGACGTGGCGACCCGCTG</td><td>>hg38_21_+_8433172_8446622_RNA45SN1_45S_50ntFlanks</td><td>7984</td><td>8006</td><td>23</td><td>AGATCAGACGTGGCGACCCGCTG</td><td>13048.7</td><td>-17.5</td><td>1.4</td><td>-12.2</td><td>1.8E-34</td><td>4.6E-32</td><td>MB231/MB468</td><td>MP</td></tr>
<tr><td>>hg38_21_+_8433172_8446622_RNA45SN1_45S_50ntFlanks@10574.10608.35|AAGCCGGCGGGAGCCCCGGGGAGAGTTCTCTTTTC</td><td>>hg38_21_+_8433172_8446622_RNA45SN1_45S_50ntFlanks</td><td>10574</td><td>10608</td><td>35</td><td>AAGCCGGCGGGAGCCCCGGGGAGAGTTCTCTTTTC</td><td>2523.7</td><td>-15.7</td><td>1.3</td><td>-12.0</td><td>2.5E-33</td><td>4.7E-32</td><td>BT20/MB468</td><td>NUC</td></tr>
<tr><td>>hg38_21_+_8433172_8446622_RNA45SN1_45S_50ntFlanks@12627.12653.27|ATGGGGCGAAGCTACCATCTGTGGGAT</td><td>>hg38_21_+_8433172_8446622_RNA45SN1_45S_50ntFlanks</td><td>12627</td><td>12653</td><td>27</td><td>ATGGGGCGAAGCTACCATCTGTGGGAT</td><td>11523.0</td><td>-17.3</td><td>1.4</td><td>-12.2</td><td>4.2E-34</td><td>4.7E-32</td><td>MB231/MB468</td><td>MITO</td></tr>
<tr><td>>hg38_1_-_228634819_228635039_RNA5S12_5S_50ntFlanks@146.170.25|TGGGAATACCGGGTGCTGTAGGCTT</td><td>>hg38_1_-_228634819_228635039_RNA5S12_5S_50ntFlanks</td><td>146</td><td>170</td><td>25</td><td>TGGGAATACCGGGTGCTGTAGGCTT</td><td>7945.1</td><td>7.8</td><td>0.6</td><td>12.1</td><td>1.9E-33</td><td>4.8E-32</td><td>BT20/MB468</td><td>CYTO</td></tr>
<tr><td>>hg38_21_+_8433172_8446622_RNA45SN1_45S_50ntFlanks@12405.12424.20|CTTTTTGATCCTTCGATGTC</td><td>>hg38_21_+_8433172_8446622_RNA45SN1_45S_50ntFlanks</td><td>12405</td><td>12424</td><td>20</td><td>CTTTTTGATCCTTCGATGTC</td><td>1265.2</td><td>-14.7</td><td>1.2</td><td>-12.0</td><td>2.6E-33</td><td>4.9E-32</td><td>BT20/MB468</td><td>NUC</td></tr>
<tr><td>>hg38_21_+_8433172_8446622_RNA45SN1_45S_50ntFlanks@7983.8006.24|CAGATCAGACGTGGCGACCCGCTG</td><td>>hg38_21_+_8433172_8446622_RNA45SN1_45S_50ntFlanks</td><td>7983</td><td>8006</td><td>24</td><td>CAGATCAGACGTGGCGACCCGCTG</td><td>36523.1</td><td>-12.1</td><td>1.0</td><td>-12.2</td><td>4.5E-34</td><td>5.0E-32</td><td>MB231/MB468</td><td>MITO</td></tr>
<tr><td>>MI0000542|hsa-mir-320a&WithFlank&8|-|22244960|22245043@48.69.22(+1G)&[MIMAT0000510&hsa-miR-320a-3p&offsets|0|0(+1G);m-60&8|-|22244975|22244996&offsets|0|0(+1G)]|AAAAGCTGGGTTGAGAGGGCGAG</td><td>>MI0000542|hsa-mir-320a&WithFlank&8|-|22244960|22245043</td><td>48</td><td>69</td><td>22(+1G)</td><td>AAAAGCTGGGTTGAGAGGGCGAG</td><td>3252.6</td><td>15.9</td><td>1.3</td><td>12.0</td><td>2.0E-33</td><td>5.2E-32</td><td>BT20/MB468</td><td>CYTO</td></tr>
<tr><td>>hg38_21_+_8433172_8446622_RNA45SN1_45S_50ntFlanks@485.505.21|GATGTGGTGACGTCGTGCTCT</td><td>>hg38_21_+_8433172_8446622_RNA45SN1_45S_50ntFlanks</td><td>485</td><td>505</td><td>21</td><td>GATGTGGTGACGTCGTGCTCT</td><td>4706.0</td><td>16.7</td><td>1.4</td><td>12.1</td><td>1.4E-33</td><td>5.2E-32</td><td>BT20/MB468</td><td>MITO</td></tr>
<tr><td>>MI0000439|hsa-mir-23b&WithFlank&9|+|95085202|95085310@67.86.20(+1U)&[MIMAT0000418&hsa-miR-23b-3p&offsets|+3|0(+1U);m-42&9|+|95085265|95085285&offsets|+3|+2(+1U)]|ACATTGCCAGGGATTACCACT</td><td>>MI0000439|hsa-mir-23b&WithFlank&9|+|95085202|95085310</td><td>67</td><td>86</td><td>20(+1U)</td><td>ACATTGCCAGGGATTACCACT</td><td>7632.0</td><td>17.4</td><td>1.4</td><td>12.1</td><td>1.5E-33</td><td>5.3E-32</td><td>BT20/MB468</td><td>MITO</td></tr>
<tr><td>>hg38_21_+_8433172_8446622_RNA45SN1_45S_50ntFlanks@10399.10424.26|TTGAACATGGGTCAGTCGGTCCTGAG</td><td>>hg38_21_+_8433172_8446622_RNA45SN1_45S_50ntFlanks</td><td>10399</td><td>10424</td><td>26</td><td>TTGAACATGGGTCAGTCGGTCCTGAG</td><td>3595.6</td><td>8.0</td><td>0.7</td><td>12.1</td><td>1.5E-33</td><td>5.4E-32</td><td>BT20/MB468</td><td>MITO</td></tr>
<tr><td>>MI0000542|hsa-mir-320a&WithFlank&8|-|22244960|22245043@49.69.21(+1U)&[MIMAT0000510&hsa-miR-320a-3p&offsets|+1|0(+1U);m-60&8|-|22244975|22244996&offsets|+1|0(+1U)]|AAAGCTGGGTTGAGAGGGCGAT</td><td>>MI0000542|hsa-mir-320a&WithFlank&8|-|22244960|22245043</td><td>49</td><td>69</td><td>21(+1U)</td><td>AAAGCTGGGTTGAGAGGGCGAT</td><td>1555.4</td><td>14.9</td><td>1.2</td><td>12.0</td><td>2.2E-33</td><td>5.5E-32</td><td>BT20/MB468</td><td>CYTO</td></tr>
</table>

<table>
<tr><th></th><th></th><th></th><th></th><th></th><th></th><th></th><th></th><th></th><th></th><th></th><th></th><th></th><th></th></tr>
<tr><td>>hg38_21_+_8433172_8446622_RNA45SN1_45S_50ntFlanks@10351.10373.23|CGAGAACTTTGAAGGCCGAAGTG</td><td>>hg38_21_+_8433172_8446622_RNA45SN1_45S_50ntFlanks</td><td>10351</td><td>10373</td><td>23</td><td>CGAGAACTTTGAAGGCCGAAGTG</td><td>1255.1</td><td>-7.2</td><td>0.6</td><td>-12.0</td><td>3.0E-33</td><td>5.5E-32</td><td>BT20/MB468</td><td>NUC</td></tr>
<tr><td>>hg38_21_+_8433172_8446622_RNA45SN1_45S_50ntFlanks@10406.10424.19|TGGGTCAGTCGGTCCTGAG</td><td>>hg38_21_+_8433172_8446622_RNA45SN1_45S_50ntFlanks</td><td>10406</td><td>10424</td><td>19</td><td>TGGGTCAGTCGGTCCTGAG</td><td>4078.3</td><td>8.0</td><td>0.7</td><td>12.1</td><td>1.6E-33</td><td>5.8E-32</td><td>BT20/MB468</td><td>MITO</td></tr>
<tr><td>>MI0000082|hsa-mir-25&WithFlank&7|-|100093554|100093649@58.77.20(+1A)&[MIMAT0000081&hsa-miR-25-3p&offsets|0|-2(+1A);m-23&7|-|100093571|100093592&offsets|0|-2(+1A)]|CATTGCACTTGTCTCGGTCTA</td><td>>MI0000082|hsa-mir-25&WithFlank&7|-|100093554|100093649</td><td>58</td><td>77</td><td>20(+1A)</td><td>CATTGCACTTGTCTCGGTCTA</td><td>1454.0</td><td>7.9</td><td>0.7</td><td>12.0</td><td>2.4E-33</td><td>6.1E-32</td><td>BT20/MB468</td><td>CYTO</td></tr>
<tr><td>>MI0000465|hsa-mir-191&WithFlank&3|-|49020612|49020715@22.42.21(+1A)&[MIMAT0000440&hsa-miR-191-5p&offsets|0|-2(+1A);m-48&3|-|49020672|49020694&offsets|0|-2(+1A)]|CAACGGAATCCCAAAAGCAGCA</td><td>>MI0000465|hsa-mir-191&WithFlank&3|-|49020612|49020715</td><td>22</td><td>42</td><td>21(+1A)</td><td>CAACGGAATCCCAAAAGCAGCA</td><td>1982.0</td><td>8.4</td><td>0.7</td><td>12.0</td><td>2.6E-33</td><td>6.4E-32</td><td>BT20/MB468</td><td>CYTO</td></tr>
<tr><td>>hg38_21_+_8433172_8446622_RNA45SN1_45S_50ntFlanks@12456.12483.28|CCAAGCGTTGGATTGTTCACCCACTAAT</td><td>>hg38_21_+_8433172_8446622_RNA45SN1_45S_50ntFlanks</td><td>12456</td><td>12483</td><td>28</td><td>CCAAGCGTTGGATTGTTCACCCACTAAT</td><td>5625.6</td><td>-16.9</td><td>1.4</td><td>-12.0</td><td>3.5E-33</td><td>6.4E-32</td><td>BT20/MB468</td><td>NUC</td></tr>
<tr><td>>hg38_21_+_8433172_8446622_RNA45SN1_45S_50ntFlanks@12163.12181.19|ACTGGGGCGGTACACCTGT</td><td>>hg38_21_+_8433172_8446622_RNA45SN1_45S_50ntFlanks</td><td>12163</td><td>12181</td><td>19</td><td>ACTGGGGCGGTACACCTGT</td><td>6121.8</td><td>-16.4</td><td>1.3</td><td>-12.2</td><td>2.6E-34</td><td>6.5E-32</td><td>MB231/MB468</td><td>MP</td></tr>
<tr><td>>hg38_21_+_8433172_8446622_RNA45SN1_45S_50ntFlanks@12626.12654.29|AATGGGGCGAAGCTACCATCTGTGGGATT</td><td>>hg38_21_+_8433172_8446622_RNA45SN1_45S_50ntFlanks</td><td>12626</td><td>12654</td><td>29</td><td>AATGGGGCGAAGCTACCATCTGTGGGATT</td><td>15989.8</td><td>-11.7</td><td>1.0</td><td>-12.2</td><td>2.8E-34</td><td>6.7E-32</td><td>MB231/MB468</td><td>MP</td></tr>
<tr><td>>MI0000650|hsa-mir-200c&WithFlank&12|+|6963693|6963772@50.70.21(+1A)&[MIMAT0000617&hsa-miR-200c-3p&offsets|0|-2(+1A);m-54&12|+|6963742|6963764&offsets|0|-2(+1A)]|TAATACTGCCGGGTAATGATGA</td><td>>MI0000650|hsa-mir-200c&WithFlank&12|+|6963693|6963772</td><td>50</td><td>70</td><td>21(+1A)</td><td>TAATACTGCCGGGTAATGATGA</td><td>2536.0</td><td>8.7</td><td>0.7</td><td>12.1</td><td>1.9E-33</td><td>6.8E-32</td><td>BT20/MB468</td><td>MITO</td></tr>
<tr><td>>hg38_21_+_8433172_8446622_RNA45SN1_45S_50ntFlanks@4722.4752.31|TCATTAATCAAGAACGAAAGTCGGAGGTTCG</td><td>>hg38_21_+_8433172_8446622_RNA45SN1_45S_50ntFlanks</td><td>4722</td><td>4752</td><td>31</td><td>TCATTAATCAAGAACGAAAGTCGGAGGTTCG</td><td>1527.2</td><td>-15.0</td><td>1.3</td><td>-12.0</td><td>3.8E-33</td><td>7.1E-32</td><td>BT20/MB468</td><td>NUC</td></tr>
<tr><td>>hg38_21_+_8433172_8446622_RNA45SN1_45S_50ntFlanks@11569.11603.35|CAGCCGACTTAGAACTGGTGCGGACCAGGGGAATC</td><td>>hg38_21_+_8433172_8446622_RNA45SN1_45S_50ntFlanks</td><td>11569</td><td>11603</td><td>35</td><td>CAGCCGACTTAGAACTGGTGCGGACCAGGGGAATC</td><td>9457.6</td><td>-8.0</td><td>0.7</td><td>-12.1</td><td>1.4E-33</td><td>7.5E-32</td><td>MB231/MB468</td><td>NUC</td></tr>
<tr><td>>hg38_21_+_8433172_8446622_RNA45SN1_45S_50ntFlanks@10840.10865.26|CCGTAACTTCGGGATAAGGATTGGCT</td><td>>hg38_21_+_8433172_8446622_RNA45SN1_45S_50ntFlanks</td><td>10840</td><td>10865</td><td>26</td><td>CCGTAACTTCGGGATAAGGATTGGCT</td><td>3296.3</td><td>-5.5</td><td>0.5</td><td>-12.0</td><td>3.1E-33</td><td>7.7E-32</td><td>BT20/MB468</td><td>CYTO</td></tr>
<tr><td>>hg38_21_+_8433172_8446622_RNA45SN1_45S_50ntFlanks@12991.13016.26|CTCCCTCGCTGCGATCTATTGAAAGT</td><td>>hg38_21_+_8433172_8446622_RNA45SN1_45S_50ntFlanks</td><td>12991</td><td>13016</td><td>26</td><td>CTCCCTCGCTGCGATCTATTGAAAGT</td><td>2433.1</td><td>-5.7</td><td>0.5</td><td>-12.1</td><td>1.5E-33</td><td>7.7E-32</td><td>MB231/MB468</td><td>NUC</td></tr>
<tr><td>>MI0000087|hsa-mir-29a&WithFlank&7|-|130876741|130876816@47.68.22&[MIMAT0000086&hsa-miR-29a-3p&offsets|-1|-1;m-26&7|-|130876749|130876769&offsets|-1|0]|CTAGCACCATCTGAAATCGGTT</td><td>>MI0000087|hsa-mir-29a&WithFlank&7|-|130876741|130876816</td><td>47</td><td>68</td><td>22</td><td>CTAGCACCATCTGAAATCGGTT</td><td>14595.6</td><td>9.3</td><td>0.8</td><td>12.0</td><td>3.3E-33</td><td>8.1E-32</td><td>BT20/MB468</td><td>CYTO</td></tr>
<tr><td>>hg38_21_+_8433172_8446622_RNA45SN1_45S_50ntFlanks@11661.11679.19|TGATTTCTGCCCAGTGCTC</td><td>>hg38_21_+_8433172_8446622_RNA45SN1_45S_50ntFlanks</td><td>11661</td><td>11679</td><td>19</td><td>TGATTTCTGCCCAGTGCTC</td><td>4564.7</td><td>-16.6</td><td>1.4</td><td>-12.0</td><td>4.4E-33</td><td>8.2E-32</td><td>BT20/MB468</td><td>NUC</td></tr>
<tr><td>>hg38_21_+_8433172_8446622_RNA45SN1_45S_50ntFlanks@3705.3735.31|TACCTGGTTGATCCTGCCAGTAGCATATGCT</td><td>>hg38_21_+_8433172_8446622_RNA45SN1_45S_50ntFlanks</td><td>3705</td><td>3735</td><td>31</td><td>TACCTGGTTGATCCTGCCAGTAGCATATGCT</td><td>3177.5</td><td>8.5</td><td>0.7</td><td>12.0</td><td>3.3E-33</td><td>8.3E-32</td><td>BT20/MB468</td><td>CYTO</td></tr>
<tr><td>>trna10_AspGTC_12_-_125424193_125424264@1.30.30,trna12_AspGTC_12_-_125411891_125411962@1.30.30,trna38_AspGTC_17_-_8125556_8125627@1.30.30,trna45_AspGTC_6_+_27447453_27447524@1.30.30,trna48_AspGTC_6_+_27471523_27471594@1.30.30,trna4_AspGTC_12_+_96429799_96429870@1.30.30,trna69_AspGTC_1_-_161440205_161440276@1.30.30,trna72_AspGTC_1_-_161432824_161432895@1.30.30,trna75_AspGTC_1_-_161425414_161425485@1.30.30,trna78_AspGTC_1_-_161418033_161418104@1.30.30,trna81_AspGTC_1_-_161410615_161410686@1.30.30|TCCTCGTTAGTATAGTGGTGAGTATCCCCG</td><td>>trna10_AspGTC_12_-_125424193_125424264</td><td>1</td><td>30</td><td>30</td><td>TCCTCGTTAGTATAGTGGTGAGTATCCCCG</td><td>1245.8</td><td>-14.7</td><td>1.2</td><td>-12.0</td><td>4.6E-33</td><td>8.5E-32</td><td>BT20/MB468</td><td>NUC</td></tr>
<tr><td>>hg38_21_+_8433172_8446622_RNA45SN1_45S_50ntFlanks@7991.8008.18|ACGTGGCGACCCGCTGAA</td><td>>hg38_21_+_8433172_8446622_RNA45SN1_45S_50ntFlanks</td><td>7991</td><td>8008</td><td>18</td><td>ACGTGGCGACCCGCTGAA</td><td>45690.2</td><td>-9.8</td><td>0.8</td><td>-12.2</td><td>3.7E-34</td><td>8.7E-32</td><td>MB231/MB468</td><td>MP</td></tr>
<tr><td>>hg38_21_+_8433172_8446622_RNA45SN1_45S_50ntFlanks@8114.8143.30|GCGCGGGACATGTGGCGTACGGAAGACCCG</td><td>>hg38_21_+_8433172_8446622_RNA45SN1_45S_50ntFlanks</td><td>8114</td><td>8143</td><td>30</td><td>GCGCGGGACATGTGGCGTACGGAAGACCCG</td><td>14544.0</td><td>-9.5</td><td>0.8</td><td>-12.5</td><td>6.7E-36</td><td>9.1E-32</td><td>BT20/MB468</td><td>MP</td></tr>
<tr><td>>hg38_21_+_8433172_8446622_RNA45SN1_45S_50ntFlanks@10410.10429.20|TCAGTCGGTCCTGAGAGATG</td><td>>hg38_21_+_8433172_8446622_RNA45SN1_45S_50ntFlanks</td><td>10410</td><td>10429</td><td>20</td><td>TCAGTCGGTCCTGAGAGATG</td><td>3860.2</td><td>-7.7</td><td>0.6</td><td>-12.1</td><td>8.5E-34</td><td>9.3E-32</td><td>MB231/MB468</td><td>MITO</td></tr>
<tr><td>>hg38_21_+_8433172_8446622_RNA45SN1_45S_50ntFlanks@12405.12425.21|CTTTTTGATCCTTCGATGTCG</td><td>>hg38_21_+_8433172_8446622_RNA45SN1_45S_50ntFlanks</td><td>12405</td><td>12425</td><td>21</td><td>CTTTTTGATCCTTCGATGTCG</td><td>999.6</td><td>-14.4</td><td>1.2</td><td>-12.0</td><td>5.1E-33</td><td>9.4E-32</td><td>BT20/MB468</td><td>NUC</td></tr>
<tr><td>>hg38_21_+_8433172_8446622_RNA45SN1_45S_50ntFlanks@11573.11590.18|CGACTTAGAACTGGTGCG</td><td>>hg38_21_+_8433172_8446622_RNA45SN1_45S_50ntFlanks</td><td>11573</td><td>11590</td><td>18</td><td>CGACTTAGAACTGGTGCG</td><td>392.5</td><td>-5.6</td><td>0.5</td><td>-12.0</td><td>5.1E-33</td><td>9.4E-32</td><td>BT20/MB468</td><td>NUC</td></tr>
<tr><td>>MI0000087|hsa-mir-29a&WithFlank&7|-|130876741|130876816@47.67.21(+1C)&[MIMAT0000086&hsa-miR-29a-3p&offsets|-1|-2(+1C);m-26&7|-|130876749|130876769&offsets|-1|-1(+1C)]|CTAGCACCATCTGAAATCGGTC</td><td>>MI0000087|hsa-mir-29a&WithFlank&7|-|130876741|130876816</td><td>47</td><td>67</td><td>21(+1C)</td><td>CTAGCACCATCTGAAATCGGTC</td><td>1997.6</td><td>9.4</td><td>0.8</td><td>12.0</td><td>2.6E-33</td><td>9.5E-32</td><td>BT20/MB468</td><td>MITO</td></tr>
<tr><td>>hg38_21_+_8433172_8446622_RNA45SN1_45S_50ntFlanks@3838.3871.34|CTCTCCTACTTGGATAACTGTGGTAATTCTAGAG</td><td>>hg38_21_+_8433172_8446622_RNA45SN1_45S_50ntFlanks</td><td>3838</td><td>3871</td><td>34</td><td>CTCTCCTACTTGGATAACTGTGGTAATTCTAGAG</td><td>1083.4</td><td>-6.4</td><td>0.5</td><td>-12.0</td><td>5.4E-33</td><td>9.8E-32</td><td>BT20/MB468</td><td>NUC</td></tr>
<tr><td>>hg38_21_+_8433172_8446622_RNA45SN1_45S_50ntFlanks@6719.6746.28|TGAATTGCAGGACACATTGATCATCGAC</td><td>>hg38_21_+_8433172_8446622_RNA45SN1_45S_50ntFlanks</td><td>6719</td><td>6746</td><td>28</td><td>TGAATTGCAGGACACATTGATCATCGAC</td><td>1632.3</td><td>-15.1</td><td>1.3</td><td>-12.0</td><td>5.6E-33</td><td>1.0E-31</td><td>BT20/MB468</td><td>NUC</td></tr>
<tr><td>>MI0000077|hsa-mir-21&WithFlank&17|+|59841260|59841343@14.35.22&[MIMAT0000076&hsa-miR-21-5p&offsets|0|0;m-1&17|+|59841273|59841294&offsets|0|0]|TAGCTTATCAGACTGATGTTGA</td><td>>MI0000077|hsa-mir-21&WithFlank&17|+|59841260|59841343</td><td>14</td><td>35</td><td>22</td><td>TAGCTTATCAGACTGATGTTGA</td><td>64099.2</td><td>6.5</td><td>0.5</td><td>12.0</td><td>2.2E-33</td><td>1.1E-31</td><td>MB231/MB468</td><td>NUC</td></tr>
<tr><td>>MI0000285|hsa-mir-205&WithFlank&1|+|209432127|209432248@40.62.23&[MIMAT0000266&hsa-miR-205-5p&offsets|0|+1;m-351&1|+|209432166|209432187&offsets|0|+1]|TCCTTCATTCCACCGGAGTCTGT</td><td>>MI0000285|hsa-mir-205&WithFlank&1|+|209432127|209432248</td><td>40</td><td>62</td><td>23</td><td>TCCTTCATTCCACCGGAGTCTGT</td><td>28557.7</td><td>18.4</td><td>1.5</td><td>12.3</td><td>1.5E-34</td><td>1.1E-31</td><td>BT20/MB231</td><td>MITO</td></tr>
<tr><td>>hg38_21_+_8433172_8446622_RNA45SN1_45S_50ntFlanks@4738.4759.22|AAAGTCGGAGGTTCGAAGACGA</td><td>>hg38_21_+_8433172_8446622_RNA45SN1_45S_50ntFlanks</td><td>4738</td><td>4759</td><td>22</td><td>AAAGTCGGAGGTTCGAAGACGA</td><td>621.3</td><td>-6.5</td><td>0.5</td><td>-12.0</td><td>6.3E-33</td><td>1.1E-31</td><td>BT20/MB468</td><td>NUC</td></tr>
<tr><td>>hg38_21_+_8433172_8446622_RNA45SN1_45S_50ntFlanks@10389.10409.21|TGAACAGCAGTTGAACATGGG</td><td>>hg38_21_+_8433172_8446622_RNA45SN1_45S_50ntFlanks</td><td>10389</td><td>10409</td><td>21</td><td>TGAACAGCAGTTGAACATGGG</td><td>2612.4</td><td>-15.8</td><td>1.3</td><td>-12.0</td><td>6.4E-33</td><td>1.2E-31</td><td>BT20/MB468</td><td>NUC</td></tr>
<tr><td>>hg38_21_+_8433172_8446622_RNA45SN1_45S_50ntFlanks@3705.3743.39|TACCTGGTTGATCCTGCCAGTAGCATATGCTTGTCTCAA</td><td>>hg38_21_+_8433172_8446622_RNA45SN1_45S_50ntFlanks</td><td>3705</td><td>3743</td><td>39</td><td>TACCTGGTTGATCCTGCCAGTAGCATATGCTTGTCTCAA</td><td>6680.9</td><td>8.5</td><td>0.7</td><td>12.0</td><td>4.7E-33</td><td>1.2E-31</td><td>BT20/MB468</td><td>CYTO</td></tr>
<tr><td>>MI0000762|hsa-mir-362&WithFlank&X|+|50008958|50009034@11.34.24&[MIMAT0000705&hsa-miR-362-5p&offsets|0|0;m-267&X|+|50008968|50008991&offsets|0|0]|AATCCTTGGAACCTAGGTGTGAGT</td><td>>MI0000762|hsa-mir-362&WithFlank&X|+|50008958|50009034</td><td>11</td><td>34</td><td>24</td><td>AATCCTTGGAACCTAGGTGTGAGT</td><td>521.3</td><td>7.0</td><td>0.6</td><td>12.0</td><td>4.8E-33</td><td>1.2E-31</td><td>BT20/MB468</td><td>CYTO</td></tr>
<tr><td>>hg38_21_+_8433172_8446622_RNA45SN1_45S_50ntFlanks@10731.10755.25|AGGGTGTAAATCTCGCGCCGGGCCG</td><td>>hg38_21_+_8433172_8446622_RNA45SN1_45S_50ntFlanks</td><td>10731</td><td>10755</td><td>25</td><td>AGGGTGTAAATCTCGCGCCGGGCCG</td><td>1979.8</td><td>-15.4</td><td>1.3</td><td>-11.9</td><td>6.6E-33</td><td>1.2E-31</td><td>BT20/MB468</td><td>NUC</td></tr>
<tr><td>>am_MI0000809|hsa-mir-151a&WithFlank&8|-|140732558|140732659@52.73.22&[MIMAT0000757&hsa-miR-151a-3p&offsets|-1|0;m-30&8|-|140732587|140732607&offsets|-1|0]|ACTAGACTGAAGCTCCTTGAGG</td><td>>am_MI0000809|hsa-mir-151a&WithFlank&8|-|140732558|140732659</td><td>52</td><td>73</td><td>22</td><td>ACTAGACTGAAGCTCCTTGAGG</td><td>1902.0</td><td>8.0</td><td>0.7</td><td>12.0</td><td>3.5E-33</td><td>1.2E-31</td><td>BT20/MB468</td><td>MITO</td></tr>
<tr><td>>hg38_21_+_8433172_8446622_RNA45SN1_45S_50ntFlanks@10680.10712.33|GTTCCGGCGGCGTCCGGTGAGCTCTCGCTGGCC</td><td>>hg38_21_+_8433172_8446622_RNA45SN1_45S_50ntFlanks</td><td>10680</td><td>10712</td><td>33</td><td>GTTCCGGCGGCGTCCGGTGAGCTCTCGCTGGCC</td><td>9690.7</td><td>-7.0</td><td>0.6</td><td>-12.1</td><td>1.2E-33</td><td>1.2E-31</td><td>MB231/MB468</td><td>MITO</td></tr>
<tr><td>>hg38_21_+_8433172_8446622_RNA45SN1_45S_50ntFlanks@7973.8011.39|GACGCGACCTCAGATCAGACGTGGCGACCCGCTGAATTT</td><td>>hg38_21_+_8433172_8446622_RNA45SN1_45S_50ntFlanks</td><td>7973</td><td>8011</td><td>39</td><td>GACGCGACCTCAGATCAGACGTGGCGACCCGCTGAATTT</td><td>2239.3</td><td>7.8</td><td>0.6</td><td>12.2</td><td>3.1E-34</td><td>1.3E-31</td><td>MB231/MB468</td><td>CYTO</td></tr>
<tr><td>>am_MI0000786|hsa-mir-378a&WithFlank&5|+|149732819|149732896@49.69.21(+1A)&[MIMAT0000732&hsa-miR-378a-3p&offsets|0|-1(+1A);m-44&5|+|149732867|149732888&offsets|0|-1(+1A)]|ACTGGACTTGGAGTCAGAAGGA</td><td>>am_MI0000786|hsa-mir-378a&WithFlank&5|+|149732819|149732896</td><td>49</td><td>69</td><td>21(+1A)</td><td>ACTGGACTTGGAGTCAGAAGGA</td><td>3641.2</td><td>6.5</td><td>0.5</td><td>12.0</td><td>5.3E-33</td><td>1.3E-31</td><td>BT20/MB468</td><td>CYTO</td></tr>
<tr><td>>hg38_21_+_8433172_8446622_RNA45SN1_45S_50ntFlanks@11671.11706.36|CCAGTGCTCTGAATGTCAAAGTGAAGAAATTCAATG</td><td>>hg38_21_+_8433172_8446622_RNA45SN1_45S_50ntFlanks</td><td>11671</td><td>11706</td><td>36</td><td>CCAGTGCTCTGAATGTCAAAGTGAAGAAATTCAATG</td><td>2051.2</td><td>-7.7</td><td>0.6</td><td>-11.9</td><td>7.3E-33</td><td>1.3E-31</td><td>BT20/MB468</td><td>NUC</td></tr>
<tr><td>>hg38_21_+_8433172_8446622_RNA45SN1_45S_50ntFlanks@9714.9737.24|CGAATGATTAGAGGTCTTGGGGCC</td><td>>hg38_21_+_8433172_8446622_RNA45SN1_45S_50ntFlanks</td><td>9714</td><td>9737</td><td>24</td><td>CGAATGATTAGAGGTCTTGGGGCC</td><td>1815.2</td><td>-15.3</td><td>1.3</td><td>-11.9</td><td>7.7E-33</td><td>1.4E-31</td><td>BT20/MB468</td><td>NUC</td></tr>
<tr><td>>hg38_21_+_8433172_8446622_RNA45SN1_45S_50ntFlanks@12934.12956.23|CTAAACCATTCGTAGACGACCTG</td><td>>hg38_21_+_8433172_8446622_RNA45SN1_45S_50ntFlanks</td><td>12934</td><td>12956</td><td>23</td><td>CTAAACCATTCGTAGACGACCTG</td><td>6411.6</td><td>-10.5</td><td>0.9</td><td>-12.1</td><td>1.3E-33</td><td>1.4E-31</td><td>MB231/MB468</td><td>MITO</td></tr>
<tr><td>>trna119_LysCTT_1_-_145395522_145395594@1.35.35,trna11_LysCTT_5_-_180648979_180649051@1.35.35,trna13_LysCTT_6_+_26556774_26556846@1.35.35,trna7_LysCTT_16_+_3225692_3225764@1.35.35,trna9_LysCTT_5_+_180634755_180634827@1.35.35|GCCCGGCTAGCTCAGTCGGTAGAGCATGAGACTCT</td><td>>trna119_LysCTT_1_-_145395522_145395594</td><td>1</td><td>35</td><td>35</td><td>GCCCGGCTAGCTCAGTCGGTAGAGCATGAGACTCT</td><td>50737.1</td><td>10.8</td><td>0.9</td><td>12.0</td><td>5.9E-33</td><td>1.4E-31</td><td>BT20/MB468</td><td>CYTO</td></tr>
<tr><td>>hg38_21_+_8433172_8446622_RNA45SN1_45S_50ntFlanks@7982.8006.25|TCAGATCAGACGTGGCGACCCGCTG</td><td>>hg38_21_+_8433172_8446622_RNA45SN1_45S_50ntFlanks</td><td>7982</td><td>8006</td><td>25</td><td>TCAGATCAGACGTGGCGACCCGCTG</td><td>4885.5</td><td>-16.1</td><td>1.3</td><td>-12.1</td><td>6.2E-34</td><td>1.5E-31</td><td>MB231/MB468</td><td>MP</td></tr>
<tr><td>>MI0000746|hsa-mir-99b&WithFlank&19|+|51692606|51692687@13.33.21(+1A)&[MIMAT0000689&hsa-miR-99b-5p&offsets|0|-1(+1A);m-10&19|+|51692618|51692639&offsets|0|-1(+1A)]|CACCCGTAGAACCGACCTTGCA</td><td>>MI0000746|hsa-mir-99b&WithFlank&19|+|51692606|51692687</td><td>13</td><td>33</td><td>21(+1A)</td><td>CACCCGTAGAACCGACCTTGCA</td><td>4270.2</td><td>6.8</td><td>0.6</td><td>12.0</td><td>4.1E-33</td><td>1.5E-31</td><td>BT20/MB468</td><td>MITO</td></tr>
<tr><td>>hg38_21_+_8433172_8446622_RNA45SN1_45S_50ntFlanks@12008.12030.23|CCGCCGGTGAAATACCACTACTC</td><td>>hg38_21_+_8433172_8446622_RNA45SN1_45S_50ntFlanks</td><td>12008</td><td>12030</td><td>23</td><td>CCGCCGGTGAAATACCACTACTC</td><td>8114.9</td><td>-16.8</td><td>1.4</td><td>-12.1</td><td>1.4E-33</td><td>1.5E-31</td><td>MB231/MB468</td><td>MITO</td></tr>
<tr><td>>MI0000301|hsa-mir-224&WithFlank&X|-|151958572|151958664@14.34.21(+1C)&[MIMAT0000281&hsa-miR-224-5p&offsets|+1|-3(+1C);m-238&X|-|151958630|151958651&offsets|0|-1(+1C)]|CAAGTCACTAGTGGTTCCGTTC</td><td>>MI0000301|hsa-mir-224&WithFlank&X|-|151958572|151958664</td><td>14</td><td>34</td><td>21(+1C)</td><td>CAAGTCACTAGTGGTTCCGTTC</td><td>2247.9</td><td>15.4</td><td>1.3</td><td>12.0</td><td>6.1E-33</td><td>1.5E-31</td><td>BT20/MB468</td><td>CYTO</td></tr>
</table>

| | | | | | | | | | | | | | |
|---|---|---|---|---|---|---|---|---|---|---|---|---|---|
| >hg38_21_+_8433172_8446622_RNA45SN1_45S_50ntFlanks@11561.11603.43\|GCGCCTAGCAGCCGACTTAGAACTGGTGCGGACCAGGGGAATC | >hg38_21_+_8433172_8446622_RNA45SN1_45S_50ntFlanks | 11561 | 11603 | 43 | GCGCCTAGCAGCCGACTTAGAACTGGTGCGGACCAGGGGAATC | 7103.9 | -17.2 | 1.4 | -11.9 | 8.5E-33 | 1.5E-31 | BT20/MB468 | NUC |
| >MI0000102\|hsa-mir-100&WithFlank&11\|-\|122152223\|122152314@19.38.20(+1A)&[MIMAT0000098&hsa-miR-100-5p&offsets\|0\|-2(+1A);m-27&11\|-\|122152276\|122152296&offsets\|0\|-1(+1A)]\|AACCCGTAGATCCGAACTTGA | >MI0000102\|hsa-mir-100&WithFlank&11\|-\|122152223\|122152314 | 19 | 38 | 20(+1A) | AACCCGTAGATCCGAACTTGA | 3758.4 | 15.8 | 1.3 | 12.0 | 3.1E-33 | 1.5E-31 | MB231/MB468 | NUC |
| >hg38_21_+_8433172_8446622_RNA45SN1_45S_50ntFlanks@12159.12181.23\|TTTGACTGGGGCGGTACACCTGT | >hg38_21_+_8433172_8446622_RNA45SN1_45S_50ntFlanks | 12159 | 12181 | 23 | TTTGACTGGGGCGGTACACCTGT | 5669.1 | -16.3 | 1.3 | -12.1 | 6.8E-34 | 1.6E-31 | MB231/MB468 | MP |
| >MI0000273\|hsa-mir-183&WithFlank&7\|-\|129774899\|129775020@33.54.22(+1A)&[MIMAT0000261&hsa-miR-183-5p&offsets\|0\|0(+1A);m-41&7\|-\|129774966\|129774987&offsets\|-1\|-1(+1A)]\|TATGGCACTGGTAGAATTCACTA | >MI0000273\|hsa-mir-183&WithFlank&7\|-\|129774899\|129775020 | 33 | 54 | 22(+1A) | TATGGCACTGGTAGAATTCACTA | 539.6 | 7.6 | 0.6 | 12.0 | 4.5E-33 | 1.6E-31 | BT20/MB468 | MITO |
| >hg38_21_+_8433172_8446622_RNA45SN1_45S_50ntFlanks@10361.10383.23\|GAAGGCCGAAGTGGAGAAGGGTT | >hg38_21_+_8433172_8446622_RNA45SN1_45S_50ntFlanks | 10361 | 10383 | 23 | GAAGGCCGAAGTGGAGAAGGGTT | 863.8 | -5.6 | 0.5 | -12.0 | 3.3E-33 | 1.6E-31 | MB231/MB468 | NUC |
| >hg38_21_+_8433172_8446622_RNA45SN1_45S_50ntFlanks@10410.10431.22\|TCAGTCGGTCCTGAGAGATGGG | >hg38_21_+_8433172_8446622_RNA45SN1_45S_50ntFlanks | 10410 | 10431 | 22 | TCAGTCGGTCCTGAGAGATGGG | 4351.4 | -8.3 | 0.7 | -12.0 | 3.2E-33 | 1.6E-31 | MB231/MB468 | NUC |
| >hg38_21_+_8433172_8446622_RNA45SN1_45S_50ntFlanks@11641.11665.25\|CGGCGGGTGTTGACGCGATGTGATT | >hg38_21_+_8433172_8446622_RNA45SN1_45S_50ntFlanks | 11641 | 11665 | 25 | CGGCGGGTGTTGACGCGATGTGATT | 3394.4 | -8.0 | 0.7 | -12.0 | 3.2E-33 | 1.6E-31 | MB231/MB468 | NUC |
| >hg38_21_+_8433172_8446622_RNA45SN1_45S_50ntFlanks@9570.9595.26\|TGCAAATCGGTCGTCCGACCTGGGTA | >hg38_21_+_8433172_8446622_RNA45SN1_45S_50ntFlanks | 9570 | 9595 | 26 | TGCAAATCGGTCGTCCGACCTGGGTA | 2902.2 | -6.3 | 0.5 | -12.2 | 4.3E-34 | 1.7E-31 | MB231/MB468 | CYTO |
| >hg38_21_+_8433172_8446622_RNA45SN1_45S_50ntFlanks@9545.9564.20\|GTGGAGGTCCGTAGCGGTCC | >hg38_21_+_8433172_8446622_RNA45SN1_45S_50ntFlanks | 9545 | 9564 | 20 | GTGGAGGTCCGTAGCGGTCC | 2110.6 | -15.5 | 1.3 | -11.9 | 9.5E-33 | 1.7E-31 | BT20/MB468 | NUC |
| >MI0003591\|hsa-mir-584&WithFlank&5\|-\|149062307\|149062415@22.41.20(+1U)&[MIMAT0003249&hsa-miR-584-5p&offsets\|0\|-2(+1U);m-167&5\|-\|149062374\|149062394&offsets\|0\|-1(+1U)]\|TTATGGTTTGCCTGGGACTGT | >MI0003591\|hsa-mir-584&WithFlank&5\|-\|149062307\|149062415 | 22 | 41 | 20(+1U) | TTATGGTTTGCCTGGGACTGT | 140.4 | 4.1 | 0.3 | 11.9 | 9.8E-33 | 1.7E-31 | BT20/MB468 | NUC |
| >hg38_21_+_8433172_8446622_RNA45SN1_45S_50ntFlanks@7983.8006.24\|CAGATCAGACGTGGCGACCCGCTG | >hg38_21_+_8433172_8446622_RNA45SN1_45S_50ntFlanks | 7983 | 8006 | 24 | CAGATCAGACGTGGCGACCCGCTG | 19029.8 | -13.8 | 1.1 | -12.1 | 7.8E-34 | 1.7E-31 | MB231/MB468 | MP |
| >MI0000077\|hsa-mir-21&WithFlank&17\|+\|59841260\|59841343@14.37.24&[MIMAT0000076&hsa-miR-21-5p&offsets\|0\|+2;m-1&17\|+\|59841273\|59841294&offsets\|0\|+2]\|TAGCTTATCAGACTGATGTTGACT | >MI0000077\|hsa-mir-21&WithFlank&17\|+\|59841260\|59841343 | 14 | 37 | 24 | TAGCTTATCAGACTGATGTTGACT | 244423.8 | 7.3 | 0.6 | 12.0 | 3.7E-33 | 1.8E-31 | MB231/MB468 | NUC |
| >hg38_21_+_8433172_8446622_RNA45SN1_45S_50ntFlanks@5527.5550.24\|AAAGTCGTAACAAGGTTTCCGTAG | >hg38_21_+_8433172_8446622_RNA45SN1_45S_50ntFlanks | 5527 | 5550 | 24 | AAAGTCGTAACAAGGTTTCCGTAG | 4178.2 | -16.5 | 1.4 | -11.9 | 1.1E-32 | 1.9E-31 | BT20/MB468 | NUC |
| >trna111_HisGTG_1_-_147774845_147774916@-1A.29.30,trna118_HisGTG_1_-_145396881_145396952@-1A.29.30,trna16_HisGTG_1_+_146544773_146544844@-1A.29.30,trna1_HisGTG_15_+_45493349_45493420@-1A.29.30,trna21_HisGTG_1_+_147753471_147753542@-1A.29.30,trna33_HisGTG_6_+_27125906_27125977@-1A.29.30,trna7_HisGTG_9_-_14433938_14434009@-1A.29.30,trna8_HisGTG_15_-_45492611_45492682@-1A.29.30,trna9_HisGTG_15_-_45490804_45490875@-1A.29.30\|AGCCGTGATCGTATAGTGGTTAGTACTCTG | >trna111_HisGTG_1_-_147774845_147774916 | -1A | 29 | 30 | AGCCGTGATCGTATAGTGGTTAGTACTCTG | 1763.3 | -15.2 | 1.3 | -11.9 | 1.1E-32 | 2.0E-31 | BT20/MB468 | NUC |
| >hg38_21_+_8433172_8446622_RNA45SN1_45S_50ntFlanks@10680.10711.32\|GTTCCGGCGGCGTCCGGTGAGCTCTCGCTGGC | >hg38_21_+_8433172_8446622_RNA45SN1_45S_50ntFlanks | 10680 | 10711 | 32 | GTTCCGGCGGCGTCCGGTGAGCTCTCGCTGGC | 10308.2 | -8.6 | 0.7 | -12.0 | 4.2E-33 | 2.0E-31 | MB231/MB468 | NUC |
| >MI0003560\|hsa-mir-92b&WithFlank&1\|+\|155195171\|155195278@67.88.22&[MIMAT0003218&hsa-miR-92b-3p&offsets\|0\|0;m-115&1\|+\|155195237\|155195258&offsets\|0\|0]\|TATTGCACTCGTCCCGGCCTCC | >MI0003560\|hsa-mir-92b&WithFlank&1\|+\|155195171\|155195278 | 67 | 88 | 22 | TATTGCACTCGTCCCGGCCTCC | 6690.7 | 9.0 | 0.8 | 12.0 | 5.8E-33 | 2.0E-31 | BT20/MB468 | MITO |
| >hg38_21_+_8433172_8446622_RNA45SN1_45S_50ntFlanks@11874.11899.26\|AGAAGACCCTGTTGAGCTTGACTCTA | >hg38_21_+_8433172_8446622_RNA45SN1_45S_50ntFlanks | 11874 | 11899 | 26 | AGAAGACCCTGTTGAGCTTGACTCTA | 1173.1 | -14.6 | 1.2 | -11.9 | 1.2E-32 | 2.1E-31 | BT20/MB468 | NUC |
| >hg38_21_+_8433172_8446622_RNA45SN1_45S_50ntFlanks@10841.10865.25\|CGTAACTTCGGGATAAGGATTGGCT | >hg38_21_+_8433172_8446622_RNA45SN1_45S_50ntFlanks | 10841 | 10865 | 25 | CGTAACTTCGGGATAAGGATTGGCT | 37381.9 | -10.1 | 0.8 | -12.1 | 9.4E-34 | 2.1E-31 | MB231/MB468 | MP |
| >MI0000342\|hsa-mir-200b&WithFlank&1\|+\|1167098\|1167204@63.85.23(+1U)&[MIMAT0000318&hsa-miR-200b-3p&offsets\|0\|+1(+1U);m-93&1\|+\|1167160\|1167182&offsets\|0\|0(+1U)]\|TAATACTGCCTGGTAATGATGACT | >MI0000342\|hsa-mir-200b&WithFlank&1\|+\|1167098\|1167204 | 63 | 85 | 23(+1U) | TAATACTGCCTGGTAATGATGACT | 3449.6 | -7.4 | 0.6 | -11.9 | 1.2E-32 | 2.1E-31 | BT20/MB468 | NUC |
| >MI0000077\|hsa-mir-21&WithFlank&17\|+\|59841260\|59841343@14.35.22(+1A)&[MIMAT0000076&hsa-miR-21-5p&offsets\|0\|0(+1A);m-1&17\|+\|59841273\|59841294&offsets\|0\|0(+1A)]\|TAGCTTATCAGACTGATGTTGAA | >MI0000077\|hsa-mir-21&WithFlank&17\|+\|59841260\|59841343 | 14 | 35 | 22(+1A) | TAGCTTATCAGACTGATGTTGAA | 12149.9 | 6.8 | 0.6 | 12.0 | 4.5E-33 | 2.2E-31 | MB231/MB468 | NUC |
| >hg38_21_+_8433172_8446622_RNA45SN1_45S_50ntFlanks@12365.12382.18\|ATAACTGGCTTGTGGCGG | >hg38_21_+_8433172_8446622_RNA45SN1_45S_50ntFlanks | 12365 | 12382 | 18 | ATAACTGGCTTGTGGCGG | 938.7 | -14.3 | 1.2 | -11.9 | 1.3E-32 | 2.2E-31 | BT20/MB468 | NUC |
| >MI0000650\|hsa-mir-200c&WithFlank&12\|+\|6963693\|6963772@49.72.24&[MIMAT0000617&hsa-miR-200c-3p&offsets\|-1\|0;m-54&12\|+\|6963742\|6963764&offsets\|-1\|0]\|CTAATACTGCCGGGTAATGATGGA | >MI0000650\|hsa-mir-200c&WithFlank&12\|+\|6963693\|6963772 | 49 | 72 | 24 | CTAATACTGCCGGGTAATGATGGA | 1019.0 | 7.9 | 0.7 | 11.9 | 9.5E-33 | 2.3E-31 | BT20/MB468 | CYTO |
| >hg38_21_+_8433172_8446622_RNA45SN1_45S_50ntFlanks@12456.12484.29\|CCAAGCGTTGGATTGTTCACCCACTAATA | >hg38_21_+_8433172_8446622_RNA45SN1_45S_50ntFlanks | 12456 | 12484 | 29 | CCAAGCGTTGGATTGTTCACCCACTAATA | 12622.6 | -11.4 | 0.9 | -12.0 | 2.3E-33 | 2.3E-31 | MB231/MB468 | MITO |
| >MI0000082\|hsa-mir-25&WithFlank&7\|-\|100093554\|100093649@58.78.21&[MIMAT0000081&hsa-miR-25-3p&offsets\|0\|-1;m-23&7\|-\|100093571\|100093592&offsets\|0\|-1]\|CATTGCACTTGTCTCGGTCTG | >MI0000082\|hsa-mir-25&WithFlank&7\|-\|100093554\|100093649 | 58 | 78 | 21 | CATTGCACTTGTCTCGGTCTG | 7215.8 | 7.2 | 0.6 | 11.9 | 7.0E-33 | 2.4E-31 | BT20/MB468 | MITO |
| >trnaMT_GluTTC_MT_-_14674_14742@40.66.27,trnalookalike8_GluTTC_5_-_93905172_93905240@40.66.27\|TTGGTCGTGGTTGTAGTCCGTGCGAGA | >trnaMT_GluTTC_MT_-_14674_14742 | 40 | 66 | 27 | TTGGTCGTGGTTGTAGTCCGTGCGAGA | 1642.6 | 8.0 | 0.7 | 11.9 | 7.0E-33 | 2.4E-31 | BT20/MB468 | MITO |
| >MI0000456\|hsa-mir-140&WithFlank&16\|+\|69933075\|69933186@69.89.21&[MIMAT0004597&hsa-miR-140-3p&offsets\|+1\|+1;m-45&16\|+\|69933143\|69933164&offsets\|0\|-1]\|ACCACAGGGTAGAACCACGGA | >MI0000456\|hsa-mir-140&WithFlank&16\|+\|69933075\|69933186 | 69 | 89 | 21 | ACCACAGGGTAGAACCACGGA | 2237.6 | 15.4 | 1.3 | 11.9 | 1.0E-32 | 2.4E-31 | BT20/MB468 | CYTO |
| >trna2_GlyGCC_21_-_18827107_18827177@1.28.28,trna35_GlyGCC_1_+_161413094_161413164@1.28.28,trna37_GlyGCC_1_+_161420467_161420537@1.28.28,trna39_GlyGCC_1_+_161427898_161427968@1.28.28,trna41_GlyGCC_1_+_161435258_161435328@1.28.28\|GCATGGGTGGTTCAGTGGTAGAATTCTC | >trna2_GlyGCC_21_-_18827107_18827177 | 1 | 28 | 28 | GCATGGGTGGTTCAGTGGTAGAATTCTC | 22745.5 | -10.2 | 0.8 | -12.1 | 1.1E-33 | 2.4E-31 | MB231/MB468 | MP |
| >hg38_21_+_8433172_8446622_RNA45SN1_45S_50ntFlanks@4853.4883.31\|AAGTCTTTGGGTTCCGGGGGGAGTATGGTTG | >hg38_21_+_8433172_8446622_RNA45SN1_45S_50ntFlanks | 4853 | 4883 | 31 | AAGTCTTTGGGTTCCGGGGGGAGTATGGTTG | 1306.2 | -14.8 | 1.2 | -11.9 | 1.4E-32 | 2.5E-31 | BT20/MB468 | NUC |
| >MI0000102\|hsa-mir-100&WithFlank&11\|-\|122152223\|122152314@19.38.20&[MIMAT0000098&hsa-miR-100-5p&offsets\|0\|-2;m-27&11\|-\|122152276\|122152296&offsets\|0\|-1]\|AACCCGTAGATCCGAACTTG | >MI0000102\|hsa-mir-100&WithFlank&11\|-\|122152223\|122152314 | 19 | 38 | 20 | AACCCGTAGATCCGAACTTG | 35762.3 | 19.0 | 1.6 | 12.0 | 5.2E-33 | 2.5E-31 | MB231/MB468 | NUC |
| >hg38_21_+_8433172_8446622_RNA45SN1_45S_50ntFlanks@10841.10865.25\|CGTAACTTCGGGATAAGGATTGGCT | >hg38_21_+_8433172_8446622_RNA45SN1_45S_50ntFlanks | 10841 | 10865 | 25 | CGTAACTTCGGGATAAGGATTGGCT | 16397.6 | -9.3 | 0.8 | -12.0 | 2.6E-33 | 2.6E-31 | MB231/MB468 | MITO |
| >hg38_21_+_8433172_8446622_RNA45SN1_45S_50ntFlanks@10572.10608.37\|AGAAGCCGGCGGGAGCCCCGGGGAGAGTTCTCTTTTC | >hg38_21_+_8433172_8446622_RNA45SN1_45S_50ntFlanks | 10572 | 10608 | 37 | AGAAGCCGGCGGGAGCCCCGGGGAGAGTTCTCTTTTC | 2681.6 | -15.8 | 1.3 | -11.9 | 1.5E-32 | 2.6E-31 | BT20/MB468 | NUC |
| >hg38_21_+_8433172_8446622_RNA45SN1_45S_50ntFlanks@12163.12182.20\|ACTGGGGCGGTACACCTGTC | >hg38_21_+_8433172_8446622_RNA45SN1_45S_50ntFlanks | 12163 | 12182 | 20 | ACTGGGGCGGTACACCTGTC | 10442.0 | -10.2 | 0.8 | -12.0 | 2.7E-33 | 2.7E-31 | MB231/MB468 | MITO |
| >hg38_21_+_8433172_8446622_RNA45SN1_45S_50ntFlanks@7986.8006.21\|ATCAGACGTGGCGACCCGCTG | >hg38_21_+_8433172_8446622_RNA45SN1_45S_50ntFlanks | 7986 | 8006 | 21 | ATCAGACGTGGCGACCCGCTG | 15017.5 | -17.7 | 1.5 | -12.0 | 2.8E-33 | 2.7E-31 | MB231/MB468 | MITO |
| >hg38_21_+_8433172_8446622_RNA45SN1_45S_50ntFlanks@11161.11179.19\|CACGGTCCCCCGCGAGGGG | >hg38_21_+_8433172_8446622_RNA45SN1_45S_50ntFlanks | 11161 | 11179 | 19 | CACGGTCCCCCGCGAGGGG | 1970.4 | -15.4 | 1.3 | -11.9 | 1.6E-32 | 2.8E-31 | BT20/MB468 | NUC |
| >hg38_21_+_8433172_8446622_RNA45SN1_45S_50ntFlanks@12937.12955.19\|AACCATTCGTAGACGACCT | >hg38_21_+_8433172_8446622_RNA45SN1_45S_50ntFlanks | 12937 | 12955 | 19 | AACCATTCGTAGACGACCT | 1075.0 | -14.5 | 1.2 | -11.9 | 1.7E-32 | 2.9E-31 | BT20/MB468 | NUC |
| >hg38_21_+_8433172_8446622_RNA45SN1_45S_50ntFlanks@11641.11670.30\|CGGCGGGTGTTGACGCGATGTGATTTCTGC | >hg38_21_+_8433172_8446622_RNA45SN1_45S_50ntFlanks | 11641 | 11670 | 30 | CGGCGGGTGTTGACGCGATGTGATTTCTGC | 8600.5 | -17.5 | 1.5 | -11.9 | 1.8E-32 | 3.0E-31 | BT20/MB468 | NUC |
| >hg38_21_+_8433172_8446622_RNA45SN1_45S_50ntFlanks@11672.11692.21\|CAGTGCTCTGAATGTCAAAGT | >hg38_21_+_8433172_8446622_RNA45SN1_45S_50ntFlanks | 11672 | 11692 | 21 | CAGTGCTCTGAATGTCAAAGT | 601.3 | 6.1 | 0.5 | 11.9 | 9.0E-33 | 3.1E-31 | BT20/MB468 | MITO |
| >hg38_21_+_8433172_8446622_RNA45SN1_45S_50ntFlanks@6719.6749.31\|TGAATTGCAGGACACATTGATCATCGACACT | >hg38_21_+_8433172_8446622_RNA45SN1_45S_50ntFlanks | 6719 | 6749 | 31 | TGAATTGCAGGACACATTGATCATCGACACT | 1374.3 | -14.9 | 1.3 | -11.9 | 1.8E-32 | 3.1E-31 | BT20/MB468 | NUC |
| >hg38_21_+_8433172_8446622_RNA45SN1_45S_50ntFlanks@7984.8003.20\|AGATCAGACGTGGCGACCCG | >hg38_21_+_8433172_8446622_RNA45SN1_45S_50ntFlanks | 7984 | 8003 | 20 | AGATCAGACGTGGCGACCCG | 1096.5 | -7.3 | 0.6 | -11.9 | 1.8E-32 | 3.1E-31 | BT20/MB468 | NUC |
| >hg38_21_+_8433172_8446622_RNA45SN1_45S_50ntFlanks@3930.3959.30\|ATCAAAACCAACCCGGTCAGCCCCTCTCCG | >hg38_21_+_8433172_8446622_RNA45SN1_45S_50ntFlanks | 3930 | 3959 | 30 | ATCAAAACCAACCCGGTCAGCCCCTCTCCG | 1688.5 | -15.1 | 1.3 | -11.9 | 1.9E-32 | 3.2E-31 | BT20/MB468 | NUC |
| >hg38_21_+_8433172_8446622_RNA45SN1_45S_50ntFlanks@12008.12031.24\|CCGCCGGTGAAATACCACTACTCT | >hg38_21_+_8433172_8446622_RNA45SN1_45S_50ntFlanks | 12008 | 12031 | 24 | CCGCCGGTGAAATACCACTACTCT | 9648.7 | -17.1 | 1.4 | -12.1 | 1.5E-33 | 3.2E-31 | MB231/MB468 | MP |
| >hg38_21_+_8433172_8446622_RNA45SN1_45S_50ntFlanks@11641.11665.25\|CGGCGGGTGTTGACGCGATGTGATT | >hg38_21_+_8433172_8446622_RNA45SN1_45S_50ntFlanks | 11641 | 11665 | 25 | CGGCGGGTGTTGACGCGATGTGATT | 14211.0 | -17.7 | 1.5 | -12.1 | 1.6E-33 | 3.3E-31 | MB231/MB468 | MP |
| >MI0000088\|hsa-mir-30a&WithFlank&6\|-\|71403545\|71403627@53.73.21(+1U)&[MIMAT0000088&hsa-miR-30a-3p&offsets\|0\|-1(+1U);m-38&6\|-\|71403554\|71403575&offsets\|0\|-1(+1U)]\|CTTTCAGTCGGATGTTTGCAGT | >MI0000088\|hsa-mir-30a&WithFlank&6\|-\|71403545\|71403627 | 53 | 73 | 21(+1U) | CTTTCAGTCGGATGTTTGCAGT | 8089.3 | 7.9 | 0.7 | 11.9 | 1.4E-32 | 3.3E-31 | BT20/MB468 | CYTO |
| >hg38_21_+_8433172_8446622_RNA45SN1_45S_50ntFlanks@12459.12487.29\|AGCGTTGGATTGTTCACCCACTAATAGGG | >hg38_21_+_8433172_8446622_RNA45SN1_45S_50ntFlanks | 12459 | 12487 | 29 | AGCGTTGGATTGTTCACCCACTAATAGGG | 2636.6 | -15.8 | 1.3 | -11.9 | 2.0E-32 | 3.4E-31 | BT20/MB468 | NUC |
| >MI0000440\|hsa-mir-27b&WithFlank&9\|+\|95085439\|95085547@67.86.20(+1U)&[MIMAT0000419&hsa-miR-27b-3p&offsets\|0\|-1(+1U);m-39&9\|+\|95085505\|95085525&offsets\|0\|-1(+1U)]\|TTCACAGTGGCTAAGTTCTGT | >MI0000440\|hsa-mir-27b&WithFlank&9\|+\|95085439\|95085547 | 67 | 86 | 20(+1U) | TTCACAGTGGCTAAGTTCTGT | 20741.4 | 8.6 | 0.7 | 11.9 | 1.4E-32 | 3.4E-31 | BT20/MB468 | CYTO |
| >hg38_21_+_8433172_8446622_RNA45SN1_45S_50ntFlanks@6721.6740.20\|AATTGCAGGACACATTGATC | >hg38_21_+_8433172_8446622_RNA45SN1_45S_50ntFlanks | 6721 | 6740 | 20 | AATTGCAGGACACATTGATC | 9494.9 | -7.8 | 0.6 | -11.9 | 7.3E-33 | 3.5E-31 | MB231/MB468 | NUC |

<table>
<tr><td>>hg38_21_+_8433172_8446622_RNA45SN1_45S_50ntFlanks@9533.9553.21|GAGGAAACTCTGGTGGAGGTC</td><td>>hg38_21_+_8433172_8446622_RNA45SN1_45S_50ntFlanks</td><td>9533</td><td>9553</td><td>21</td><td>GAGGAAACTCTGGTGGAGGTC</td><td>27798.7</td><td>-10.1</td><td>0.9</td><td>-11.9</td><td>2.0E-32</td><td>3.5E-31</td><td>BT20/MB468</td><td>NUC</td></tr>
<tr><td>>MI0000102|hsa-mir-100&WithFlank&11|-|122152223|122152314@19.40.22(+1A)&[MIMAT0000098&hsa-miR-100-5p&offsets|0|0(+1A);m-27&11|-|122152276|122152296&offsets|0|+1(+1A)]|AACCCGTAGATCCGAACTTGTGA</td><td>>MI0000102|hsa-mir-100&WithFlank&11|-|122152223|122152314</td><td>19</td><td>40</td><td>22(+1A)</td><td>AACCCGTAGATCCGAACTTGTGA</td><td>30980.2</td><td>16.3</td><td>1.3</td><td>12.1</td><td>9.8E-34</td><td>3.5E-31</td><td>MB231/MB468</td><td>CYTO</td></tr>
<tr><td>>hg38_21_+_8433172_8446622_RNA45SN1_45S_50ntFlanks@12419.12441.23|GATGTCGGCTCTTCCTATCATTG</td><td>>hg38_21_+_8433172_8446622_RNA45SN1_45S_50ntFlanks</td><td>12419</td><td>12441</td><td>23</td><td>GATGTCGGCTCTTCCTATCATTG</td><td>1691.4</td><td>-5.3</td><td>0.4</td><td>-11.9</td><td>2.2E-32</td><td>3.6E-31</td><td>BT20/MB468</td><td>NUC</td></tr>
<tr><td>>hg38_21_+_8433172_8446622_RNA45SN1_45S_50ntFlanks@4733.4767.35|GAACGAAAGTCGGAGGTTCGAAGACGATCAGATAC</td><td>>hg38_21_+_8433172_8446622_RNA45SN1_45S_50ntFlanks</td><td>4733</td><td>4767</td><td>35</td><td>GAACGAAAGTCGGAGGTTCGAAGACGATCAGATAC</td><td>1645.2</td><td>10.1</td><td>0.8</td><td>11.9</td><td>1.5E-32</td><td>3.7E-31</td><td>BT20/MB468</td><td>CYTO</td></tr>
<tr><td>>hg38_21_+_8433172_8446622_RNA45SN1_45S_50ntFlanks@11320.11344.25|CCACGTCTCGTCGCGCGCGCGTCCG</td><td>>hg38_21_+_8433172_8446622_RNA45SN1_45S_50ntFlanks</td><td>11320</td><td>11344</td><td>25</td><td>CCACGTCTCGTCGCGCGCGCGTCCG</td><td>9367.7</td><td>-17.0</td><td>1.4</td><td>-12.0</td><td>3.9E-33</td><td>3.7E-31</td><td>MB231/MB468</td><td>MITO</td></tr>
<tr><td>>MI0000746|hsa-mir-99b&WithFlank&19|+|51692606|51692687@51.72.22&[MIMAT0004678&hsa-miR-99b-3p&offsets|0|0;m-151&19|+|51692656|51692678&offsets|0|-1]|CAAGCTCGTGTCTGTGGGTCCG</td><td>>MI0000746|hsa-mir-99b&WithFlank&19|+|51692606|51692687</td><td>51</td><td>72</td><td>22</td><td>CAAGCTCGTGTCTGTGGGTCCG</td><td>3722.9</td><td>7.0</td><td>0.6</td><td>11.9</td><td>1.6E-32</td><td>3.8E-31</td><td>BT20/MB468</td><td>CYTO</td></tr>
<tr><td>>hg38_1_-_228634819_228635039_RNA5S12_5S_50ntFlanks@145.169.25|CTGGGAATACCGGGTGCTGTAGGCT</td><td>>hg38_1_-_228634819_228635039_RNA5S12_5S_50ntFlanks</td><td>145</td><td>169</td><td>25</td><td>CTGGGAATACCGGGTGCTGTAGGCT</td><td>1370.5</td><td>7.0</td><td>0.6</td><td>11.9</td><td>1.6E-32</td><td>3.9E-31</td><td>BT20/MB468</td><td>CYTO</td></tr>
<tr><td>>hg38_21_+_8433172_8446622_RNA45SN1_45S_50ntFlanks@4759.4793.35|ATCAGATACCGTCGTAGTTCCGACCATAAACGATG</td><td>>hg38_21_+_8433172_8446622_RNA45SN1_45S_50ntFlanks</td><td>4759</td><td>4793</td><td>35</td><td>ATCAGATACCGTCGTAGTTCCGACCATAAACGATG</td><td>1200.9</td><td>-14.7</td><td>1.2</td><td>-11.8</td><td>2.5E-32</td><td>4.2E-31</td><td>BT20/MB468</td><td>NUC</td></tr>
<tr><td>>MI0000266|hsa-mir-10a&WithFlank&17|-|48579832|48579953@29.50.22(+1A)&[MIMAT0000253&hsa-miR-10a-5p&offsets|+1|0(+1A);m-18&17|-|48579905|48579926&offsets|+1|+1(+1A)]|ACCCTGTAGATCCGAATTTGTGA</td><td>>MI0000266|hsa-mir-10a&WithFlank&17|-|48579832|48579953</td><td>29</td><td>50</td><td>22(+1A)</td><td>ACCCTGTAGATCCGAATTTGTGA</td><td>2149.1</td><td>15.3</td><td>1.3</td><td>11.9</td><td>1.8E-32</td><td>4.2E-31</td><td>BT20/MB468</td><td>CYTO</td></tr>
<tr><td>>hg38_21_+_8433172_8446622_RNA45SN1_45S_50ntFlanks@9545.9571.27|GTGGAGGTCCGTAGCGGTCCTGACGTG</td><td>>hg38_21_+_8433172_8446622_RNA45SN1_45S_50ntFlanks</td><td>9545</td><td>9571</td><td>27</td><td>GTGGAGGTCCGTAGCGGTCCTGACGTG</td><td>8294.2</td><td>-7.6</td><td>0.6</td><td>-12.0</td><td>4.5E-33</td><td>4.2E-31</td><td>MB231/MB468</td><td>MITO</td></tr>
<tr><td>>hg38_21_+_8433172_8446622_RNA45SN1_45S_50ntFlanks@9572.9597.26|CAAATCGGTCGTCCGACCTGGGTATA</td><td>>hg38_21_+_8433172_8446622_RNA45SN1_45S_50ntFlanks</td><td>9572</td><td>9597</td><td>26</td><td>CAAATCGGTCGTCCGACCTGGGTATA</td><td>2461.2</td><td>-15.7</td><td>1.3</td><td>-11.8</td><td>2.6E-32</td><td>4.3E-31</td><td>BT20/MB468</td><td>NUC</td></tr>
<tr><td>>hg38_21_+_8433172_8446622_RNA45SN1_45S_50ntFlanks@12456.12484.29|CCAAGCGTTGGATTGTTCACCCACTAATA</td><td>>hg38_21_+_8433172_8446622_RNA45SN1_45S_50ntFlanks</td><td>12456</td><td>12484</td><td>29</td><td>CCAAGCGTTGGATTGTTCACCCACTAATA</td><td>5628.1</td><td>-16.3</td><td>1.4</td><td>-12.0</td><td>2.1E-33</td><td>4.3E-31</td><td>MB231/MB468</td><td>MP</td></tr>
<tr><td>>hg38_21_+_8433172_8446622_RNA45SN1_45S_50ntFlanks@12934.12961.28|CTAAACCATTCGTAGACGACCTGCTTCT</td><td>>hg38_21_+_8433172_8446622_RNA45SN1_45S_50ntFlanks</td><td>12934</td><td>12961</td><td>28</td><td>CTAAACCATTCGTAGACGACCTGCTTCT</td><td>1722.0</td><td>-6.6</td><td>0.6</td><td>-11.9</td><td>9.5E-33</td><td>4.5E-31</td><td>MB231/MB468</td><td>NUC</td></tr>
<tr><td>>hg38_21_+_8433172_8446622_RNA45SN1_45S_50ntFlanks@11728.11751.24|GTAACTATGACTCTCTTAAGGTAG</td><td>>hg38_21_+_8433172_8446622_RNA45SN1_45S_50ntFlanks</td><td>11728</td><td>11751</td><td>24</td><td>GTAACTATGACTCTCTTAAGGTAG</td><td>4888.8</td><td>-16.7</td><td>1.4</td><td>-11.8</td><td>2.7E-32</td><td>4.6E-31</td><td>BT20/MB468</td><td>NUC</td></tr>
<tr><td>>hg38_21_+_8433172_8446622_RNA45SN1_45S_50ntFlanks@4367.4398.32|CAGTTAAAAAGCTCGTAGTTGGATCTTGGGAG</td><td>>hg38_21_+_8433172_8446622_RNA45SN1_45S_50ntFlanks</td><td>4367</td><td>4398</td><td>32</td><td>CAGTTAAAAAGCTCGTAGTTGGATCTTGGGAG</td><td>2599.8</td><td>-15.8</td><td>1.3</td><td>-11.8</td><td>2.8E-32</td><td>4.6E-31</td><td>BT20/MB468</td><td>NUC</td></tr>
<tr><td>>hg38_21_+_8433172_8446622_RNA45SN1_45S_50ntFlanks@12628.12653.26|TGGGGCGAAGCTACCATCTGTGGGAT</td><td>>hg38_21_+_8433172_8446622_RNA45SN1_45S_50ntFlanks</td><td>12628</td><td>12653</td><td>26</td><td>TGGGGCGAAGCTACCATCTGTGGGAT</td><td>1609.0</td><td>-15.1</td><td>1.3</td><td>-11.8</td><td>2.8E-32</td><td>4.7E-31</td><td>BT20/MB468</td><td>NUC</td></tr>
<tr><td>>hg38_21_+_8433172_8446622_RNA45SN1_45S_50ntFlanks@11594.11611.18|CAGGGGAATCCGACTGTT</td><td>>hg38_21_+_8433172_8446622_RNA45SN1_45S_50ntFlanks</td><td>11594</td><td>11611</td><td>18</td><td>CAGGGGAATCCGACTGTT</td><td>1798.2</td><td>-5.0</td><td>0.4</td><td>-11.9</td><td>1.1E-32</td><td>4.9E-31</td><td>MB231/MB468</td><td>NUC</td></tr>
<tr><td>>am_MI0000809|hsa-mir-151a&WithFlank&8|-|140732558|140732659@53.72.20(+1C)&[MIMAT0000757&hsa-miR-151a-3p&offsets|0|-1(+1C);m-30&8|-|140732587|140732607&offsets|0|-1(+1C)]|CTAGACTGAAGCTCCTTGAGC</td><td>>am_MI0000809|hsa-mir-151a&WithFlank&8|-|140732558|140732659</td><td>53</td><td>72</td><td>20(+1C)</td><td>CTAGACTGAAGCTCCTTGAGC</td><td>1613.5</td><td>11.3</td><td>1.0</td><td>11.9</td><td>2.1E-32</td><td>5.0E-31</td><td>BT20/MB468</td><td>CYTO</td></tr>
<tr><td>>MI0000734|hsa-mir-106b&WithFlank&7|-|100093987|100094080@58.77.20&[MIMAT0004672&hsa-miR-106b-3p&offsets|0|-2;m-87&7|-|100094003|100094023&offsets|0|-1]|CCGCACTGTGGGTACTTGCT</td><td>>MI0000734|hsa-mir-106b&WithFlank&7|-|100093987|100094080</td><td>58</td><td>77</td><td>20</td><td>CCGCACTGTGGGTACTTGCT</td><td>1361.7</td><td>8.8</td><td>0.7</td><td>11.9</td><td>1.5E-32</td><td>5.1E-31</td><td>BT20/MB468</td><td>MITO</td></tr>
<tr><td>>hg38_21_+_8433172_8446622_RNA45SN1_45S_50ntFlanks@8380.8405.26|CAAGAGGGCGTGAAACCGTTAAGAGG</td><td>>hg38_21_+_8433172_8446622_RNA45SN1_45S_50ntFlanks</td><td>8380</td><td>8405</td><td>26</td><td>CAAGAGGGCGTGAAACCGTTAAGAGG</td><td>2440.2</td><td>-15.7</td><td>1.3</td><td>-11.8</td><td>3.1E-32</td><td>5.1E-31</td><td>BT20/MB468</td><td>NUC</td></tr>
<tr><td>>hg38_21_+_8433172_8446622_RNA45SN1_45S_50ntFlanks@9545.9566.22|GTGGAGGTCCGTAGCGGTCCTG</td><td>>hg38_21_+_8433172_8446622_RNA45SN1_45S_50ntFlanks</td><td>9545</td><td>9566</td><td>22</td><td>GTGGAGGTCCGTAGCGGTCCTG</td><td>2133.4</td><td>-15.5</td><td>1.3</td><td>-11.8</td><td>3.1E-32</td><td>5.2E-31</td><td>BT20/MB468</td><td>NUC</td></tr>
<tr><td>>hg38_21_+_8433172_8446622_RNA45SN1_45S_50ntFlanks@11874.11906.33|AGAAGACCCTGTTGAGCTTGACTCTAGTCTGGC</td><td>>hg38_21_+_8433172_8446622_RNA45SN1_45S_50ntFlanks</td><td>11874</td><td>11906</td><td>33</td><td>AGAAGACCCTGTTGAGCTTGACTCTAGTCTGGC</td><td>1418.0</td><td>-14.9</td><td>1.3</td><td>-11.8</td><td>3.2E-32</td><td>5.2E-31</td><td>BT20/MB468</td><td>NUC</td></tr>
<tr><td>>hg38_21_+_8433172_8446622_RNA45SN1_45S_50ntFlanks@5485.5509.25|AGCGCTGAGAAGACGGTCGAACTTG</td><td>>hg38_21_+_8433172_8446622_RNA45SN1_45S_50ntFlanks</td><td>5485</td><td>5509</td><td>25</td><td>AGCGCTGAGAAGACGGTCGAACTTG</td><td>3752.4</td><td>-10.4</td><td>0.9</td><td>-12.0</td><td>2.6E-33</td><td>5.2E-31</td><td>MB231/MB468</td><td>MP</td></tr>
<tr><td>>hg38_21_+_8433172_8446622_RNA45SN1_45S_50ntFlanks@11570.11602.33|AGCCGACTTAGAACTGGTGCGGACCAGGGGAAT</td><td>>hg38_21_+_8433172_8446622_RNA45SN1_45S_50ntFlanks</td><td>11570</td><td>11602</td><td>33</td><td>AGCCGACTTAGAACTGGTGCGGACCAGGGGAAT</td><td>3125.9</td><td>-7.1</td><td>0.6</td><td>-11.8</td><td>3.3E-32</td><td>5.4E-31</td><td>BT20/MB468</td><td>NUC</td></tr>
<tr><td>>MI0000071|hsa-mir-17&WithFlank&13|+|91350599|91350694@20.42.23(+1A)&[MIMAT0000070&hsa-miR-17-5p&offsets|0|0(+1A);m-88&13|+|91350618|91350640&offsets|0|0(+1A)]|CAAAGTGCTTACAGTGCAGGTAGA</td><td>>MI0000071|hsa-mir-17&WithFlank&13|+|91350599|91350694</td><td>20</td><td>42</td><td>23(+1A)</td><td>CAAAGTGCTTACAGTGCAGGTAGA</td><td>3496.2</td><td>5.3</td><td>0.4</td><td>11.8</td><td>2.4E-32</td><td>5.5E-31</td><td>BT20/MB468</td><td>CYTO</td></tr>
<tr><td>>MI0000301|hsa-mir-224&WithFlank&X|-|151958572|151958664@14.35.22&[MIMAT0000281&hsa-miR-224-5p&offsets|+1|-2;m-238&X|-|151958630|151958651&offsets|0|0]|CAAGTCACTAGTGGTTCCGTTT</td><td>>MI0000301|hsa-mir-224&WithFlank&X|-|151958572|151958664</td><td>14</td><td>35</td><td>22</td><td>CAAGTCACTAGTGGTTCCGTTT</td><td>21058.0</td><td>14.1</td><td>1.2</td><td>11.8</td><td>2.4E-32</td><td>5.5E-31</td><td>BT20/MB468</td><td>CYTO</td></tr>
<tr><td>>MI0000465|hsa-mir-191&WithFlank&3|-|49020612|49020715@22.42.21&[MIMAT0000440&hsa-miR-191-5p&offsets|0|-2;m-48&3|-|49020672|49020694&offsets|0|-2]|CAACGGAATCCCAAAAGCAGC</td><td>>MI0000465|hsa-mir-191&WithFlank&3|-|49020612|49020715</td><td>22</td><td>42</td><td>21</td><td>CAACGGAATCCCAAAAGCAGC</td><td>1360.6</td><td>7.6</td><td>0.6</td><td>11.8</td><td>2.4E-32</td><td>5.5E-31</td><td>BT20/MB468</td><td>CYTO</td></tr>
<tr><td>>hg38_21_+_8433172_8446622_RNA45SN1_45S_50ntFlanks@8168.8186.19|CCCAAGTCCTTCTGATCGA</td><td>>hg38_21_+_8433172_8446622_RNA45SN1_45S_50ntFlanks</td><td>8168</td><td>8186</td><td>19</td><td>CCCAAGTCCTTCTGATCGA</td><td>3066.2</td><td>-16.0</td><td>1.4</td><td>-11.8</td><td>3.4E-32</td><td>5.6E-31</td><td>BT20/MB468</td><td>NUC</td></tr>
<tr><td>>hg38_21_+_8433172_8446622_RNA45SN1_45S_50ntFlanks@11323.11343.21|CGTCTCGTCGCGCGCGCGTCC</td><td>>hg38_21_+_8433172_8446622_RNA45SN1_45S_50ntFlanks</td><td>11323</td><td>11343</td><td>21</td><td>CGTCTCGTCGCGCGCGCGTCC</td><td>11451.4</td><td>-9.3</td><td>0.8</td><td>-12.0</td><td>6.0E-33</td><td>5.6E-31</td><td>MB231/MB468</td><td>MITO</td></tr>
<tr><td>>hg38_21_+_8433172_8446622_RNA45SN1_45S_50ntFlanks@12000.12032.33|CCTGCGGGCCGCCGGTGAAATACCACTACTCTG</td><td>>hg38_21_+_8433172_8446622_RNA45SN1_45S_50ntFlanks</td><td>12000</td><td>12032</td><td>33</td><td>CCTGCGGGCCGCCGGTGAAATACCACTACTCTG</td><td>2743.8</td><td>-15.8</td><td>1.3</td><td>-11.8</td><td>3.5E-32</td><td>5.7E-31</td><td>BT20/MB468</td><td>NUC</td></tr>
<tr><td>>MI0000447|hsa-mir-128-1&WithFlank&2|+|135665391|135665484@56.75.20;MI0000727|hsa-mir-128-2&WithFlank&3|+|35744470|35744565@58.77.20&[MIMAT0000424&hsa-miR-128-3p&offsets|0|-1;m-108&2|+|135665446|135665466&offsets|0|-1];[MIMAT0000424_1&hsa-miR-128-3p&offsets|0|-1;m-117&3|+|35744527|35744547&offsets|0|-1]|TCACAGTGAACCGGTCTCTT</td><td>>MI0000447|hsa-mir-128-1&WithFlank&2|+|135665391|135665484</td><td>56</td><td>75</td><td>20</td><td>TCACAGTGAACCGGTCTCTT</td><td>1455.1</td><td>6.1</td><td>0.5</td><td>11.8</td><td>2.5E-32</td><td>5.7E-31</td><td>BT20/MB468</td><td>CYTO</td></tr>
<tr><td>>MI0000465|hsa-mir-191&WithFlank&3|-|49020612|49020715@22.44.23&[MIMAT0000440&hsa-miR-191-5p&offsets|0|0;m-48&3|-|49020672|49020694&offsets|0|0]|CAACGGAATCCCAAAAGCAGCTG</td><td>>MI0000465|hsa-mir-191&WithFlank&3|-|49020612|49020715</td><td>22</td><td>44</td><td>23</td><td>CAACGGAATCCCAAAAGCAGCTG</td><td>26522.0</td><td>7.4</td><td>0.6</td><td>11.8</td><td>2.5E-32</td><td>5.8E-31</td><td>BT20/MB468</td><td>CYTO</td></tr>
<tr><td>>hg38_21_+_8433172_8446622_RNA45SN1_45S_50ntFlanks@4130.4162.33|ATTCCGGAGAGGGAGCCTGAGAAACGGCTACCA</td><td>>hg38_21_+_8433172_8446622_RNA45SN1_45S_50ntFlanks</td><td>4130</td><td>4162</td><td>33</td><td>ATTCCGGAGAGGGAGCCTGAGAAACGGCTACCA</td><td>1604.6</td><td>-15.1</td><td>1.3</td><td>-11.8</td><td>3.6E-32</td><td>5.9E-31</td><td>BT20/MB468</td><td>NUC</td></tr>
<tr><td>>hg38_21_+_8433172_8446622_RNA45SN1_45S_50ntFlanks@12436.12454.19|TCATTGTGAAGCAGAATTC</td><td>>hg38_21_+_8433172_8446622_RNA45SN1_45S_50ntFlanks</td><td>12436</td><td>12454</td><td>19</td><td>TCATTGTGAAGCAGAATTC</td><td>2308.5</td><td>-15.6</td><td>1.3</td><td>-11.8</td><td>3.6E-32</td><td>5.9E-31</td><td>BT20/MB468</td><td>NUC</td></tr>
<tr><td>>MI0000261|hsa-mir-139&WithFlank&11|-|72615057|72615136@13.35.23&[MIMAT0000250&hsa-miR-139-5p&offsets|0|0;m-186&11|-|72615102|72615124&offsets|0|0]|TCTACAGTGCACGTGTCTCCAGT</td><td>>MI0000261|hsa-mir-139&WithFlank&11|-|72615057|72615136</td><td>13</td><td>35</td><td>23</td><td>TCTACAGTGCACGTGTCTCCAGT</td><td>1188.9</td><td>14.5</td><td>1.2</td><td>11.8</td><td>2.6E-32</td><td>5.9E-31</td><td>BT20/MB468</td><td>CYTO</td></tr>
<tr><td>>hg38_21_+_8433172_8446622_RNA45SN1_45S_50ntFlanks@9504.9526.23|GTGAACTATGCCTGGGCAGGGCG</td><td>>hg38_21_+_8433172_8446622_RNA45SN1_45S_50ntFlanks</td><td>9504</td><td>9526</td><td>23</td><td>GTGAACTATGCCTGGGCAGGGCG</td><td>31536.0</td><td>-18.8</td><td>1.6</td><td>-12.0</td><td>3.0E-33</td><td>6.1E-31</td><td>MB231/MB468</td><td>MP</td></tr>
<tr><td>>MI0000273|hsa-mir-183&WithFlank&7|-|129774899|129775020@33.56.24&[MIMAT0000261&hsa-miR-183-5p&offsets|0|+2;m-41&7|-|129774966|129774987&offsets|-1|+1]|TATGGCACTGGTAGAATTCACTGT</td><td>>MI0000273|hsa-mir-183&WithFlank&7|-|129774899|129775020</td><td>33</td><td>56</td><td>24</td><td>TATGGCACTGGTAGAATTCACTGT</td><td>2276.5</td><td>7.4</td><td>0.6</td><td>11.8</td><td>2.8E-32</td><td>6.4E-31</td><td>BT20/MB468</td><td>CYTO</td></tr>
<tr><td>>hg38_21_+_8433172_8446622_RNA45SN1_45S_50ntFlanks@7974.8013.40|ACGCGACCTCAGATCAGACGTGGCGACCCGCTGAATTTAA</td><td>>hg38_21_+_8433172_8446622_RNA45SN1_45S_50ntFlanks</td><td>7974</td><td>8013</td><td>40</td><td>ACGCGACCTCAGATCAGACGTGGCGACCCGCTGAATTTAA</td><td>1658.1</td><td>7.7</td><td>0.6</td><td>12.1</td><td>1.9E-33</td><td>6.5E-31</td><td>MB231/MB468</td><td>CYTO</td></tr>
<tr><td>>trna130_GlnTTG_6_-_27763640_27763711@1.29.29,trna173_GlnTTG_6_-_26311975_26312046@1.29.29,trna174_GlnTTG_6_-_26311424_26311495@1.29.29|GGCCCCATGGTGTAATGGTTAGCACTCTG</td><td>>trna130_GlnTTG_6_-_27763640_27763711</td><td>1</td><td>29</td><td>29</td><td>GGCCCCATGGTGTAATGGTTAGCACTCTG</td><td>1788.6</td><td>-15.2</td><td>1.3</td><td>-11.8</td><td>4.1E-32</td><td>6.6E-31</td><td>BT20/MB468</td><td>NUC</td></tr>
<tr><td>>MI0005756|hsa-mir-934&WithFlank&X|+|136550872|136550966@21.41.21&[MIMAT0004977&hsa-miR-934&offsets|0|-1;m-943&X|+|136550892|136550913&offsets|0|-1]|TGTCTACTACTGGAGACACTG</td><td>>MI0005756|hsa-mir-934&WithFlank&X|+|136550872|136550966</td><td>21</td><td>41</td><td>21</td><td>TGTCTACTACTGGAGACACTG</td><td>3153.9</td><td>7.1</td><td>0.6</td><td>11.9</td><td>1.9E-32</td><td>6.6E-31</td><td>BT20/MB468</td><td>MITO</td></tr>
<tr><td>>hg38_21_+_8433172_8446622_RNA45SN1_45S_50ntFlanks@8380.8402.23|CAAGAGGGCGTGAAACCGTTAAG</td><td>>hg38_21_+_8433172_8446622_RNA45SN1_45S_50ntFlanks</td><td>8380</td><td>8402</td><td>23</td><td>CAAGAGGGCGTGAAACCGTTAAG</td><td>1280.7</td><td>-14.7</td><td>1.3</td><td>-11.8</td><td>4.1E-32</td><td>6.6E-31</td><td>BT20/MB468</td><td>NUC</td></tr>
<tr><td>>hg38_21_+_8433172_8446622_RNA45SN1_45S_50ntFlanks@10409.10430.22|GTCAGTCGGTCCTGAGAGATGG</td><td>>hg38_21_+_8433172_8446622_RNA45SN1_45S_50ntFlanks</td><td>10409</td><td>10430</td><td>22</td><td>GTCAGTCGGTCCTGAGAGATGG</td><td>1216.8</td><td>-6.1</td><td>0.5</td><td>-11.9</td><td>1.4E-32</td><td>6.6E-31</td><td>MB231/MB468</td><td>NUC</td></tr>
<tr><td>>hg38_21_+_8433172_8446622_RNA45SN1_45S_50ntFlanks@10683.10706.24|CCGGCGGCGTCCGGTGAGCTCTCG</td><td>>hg38_21_+_8433172_8446622_RNA45SN1_45S_50ntFlanks</td><td>10683</td><td>10706</td><td>24</td><td>CCGGCGGCGTCCGGTGAGCTCTCG</td><td>14252.0</td><td>-8.3</td><td>0.7</td><td>-11.9</td><td>1.5E-32</td><td>6.6E-31</td><td>MB231/MB468</td><td>NUC</td></tr>
<tr><td>>hg38_21_+_8433172_8446622_RNA45SN1_45S_50ntFlanks@11583.11603.21|CTGGTGCGGACCAGGGGAATC</td><td>>hg38_21_+_8433172_8446622_RNA45SN1_45S_50ntFlanks</td><td>11583</td><td>11603</td><td>21</td><td>CTGGTGCGGACCAGGGGAATC</td><td>1920.7</td><td>-5.6</td><td>0.5</td><td>-11.9</td><td>1.5E-32</td><td>6.6E-31</td><td>MB231/MB468</td><td>NUC</td></tr>
<tr><td>>MI0000439|hsa-mir-23b&WithFlank&9|+|95085202|95085310@67.86.20(+1A)&[MIMAT0000418&hsa-miR-23b-3p&offsets|+3|0(+1A);m-42&9|+|95085265|95085285&offsets|+3|+2(+1A)]|ACATTGCCAGGGATTACCACA</td><td>>MI0000439|hsa-mir-23b&WithFlank&9|+|95085202|95085310</td><td>67</td><td>86</td><td>20(+1A)</td><td>ACATTGCCAGGGATTACCACA</td><td>1476.3</td><td>14.8</td><td>1.3</td><td>11.8</td><td>2.9E-32</td><td>6.6E-31</td><td>BT20/MB468</td><td>CYTO</td></tr>
<tr><td>>hg38_21_+_8433172_8446622_RNA45SN1_45S_50ntFlanks@4724.4759.36|ATTAATCAAGAACGAAAGTCGGAGGTTCGAAGACGA</td><td>>hg38_21_+_8433172_8446622_RNA45SN1_45S_50ntFlanks</td><td>4724</td><td>4759</td><td>36</td><td>ATTAATCAAGAACGAAAGTCGGAGGTTCGAAGACGA</td><td>9390.9</td><td>-11.5</td><td>1.0</td><td>-11.9</td><td>7.3E-33</td><td>6.7E-31</td><td>MB231/MB468</td><td>MITO</td></tr>
<tr><td>>hg38_21_+_8433172_8446622_RNA45SN1_45S_50ntFlanks@11577.11605.29|TTAGAACTGGTGCGGACCAGGGGAATCCG</td><td>>hg38_21_+_8433172_8446622_RNA45SN1_45S_50ntFlanks</td><td>11577</td><td>11605</td><td>29</td><td>TTAGAACTGGTGCGGACCAGGGGAATCCG</td><td>1362.4</td><td>-14.8</td><td>1.3</td><td>-11.8</td><td>4.2E-32</td><td>6.8E-31</td><td>BT20/MB468</td><td>NUC</td></tr>
<tr><td>>hg38_21_+_8433172_8446622_RNA45SN1_45S_50ntFlanks@11668.11706.39|TGCCCAGTGCTCTGAATGTCAAAGTGAAGAAATTCAATG</td><td>>hg38_21_+_8433172_8446622_RNA45SN1_45S_50ntFlanks</td><td>11668</td><td>11706</td><td>39</td><td>TGCCCAGTGCTCTGAATGTCAAAGTGAAGAAATTCAATG</td><td>3212.8</td><td>8.3</td><td>0.7</td><td>12.0</td><td>2.1E-33</td><td>6.9E-31</td><td>MB231/MB468</td><td>CYTO</td></tr>
</table>

<table>
<tr><th></th><th></th><th></th><th></th><th></th><th></th><th></th><th></th><th></th><th></th><th></th><th></th><th></th><th></th></tr>
<tr><td>>hg38_21_+_8433172_8446622_RNA45SN1_45S_50ntFlanks@10841.10874.34|CGTAACTTCGGGATAAGGATTGGCTCTAAGGGCT</td><td>>hg38_21_+_8433172_8446622_RNA45SN1_45S_50ntFlanks</td><td>10841</td><td>10874</td><td>34</td><td>CGTAACTTCGGGATAAGGATTGGCTCTAAGGGCT</td><td>1283.8</td><td>-5.9</td><td>0.5</td><td>-11.9</td><td>1.7E-32</td><td>7.5E-31</td><td>MB231/MB468</td><td>NUC</td></tr>
<tr><td>>MI0000255|hsa-mir-30d&WithFlank&8|-|134804870|134804951@12.32.21&[MIMAT0000245&hsa-miR-30d-5p&offsets|0|-1;m-21&8|-|134804919|134804940&offsets|0|-1]|TGTAAACATCCCCGACTGGAA</td><td>>MI0000255|hsa-mir-30d&WithFlank&8|-|134804870|134804951</td><td>12</td><td>32</td><td>21</td><td>TGTAAACATCCCCGACTGGAA</td><td>1823.3</td><td>9.6</td><td>0.8</td><td>11.8</td><td>3.4E-32</td><td>7.7E-31</td><td>BT20/MB468</td><td>CYTO</td></tr>
<tr><td>>hg38_21_+_8433172_8446622_RNA45SN1_45S_50ntFlanks@12893.12932.40|CCTCGCCCGTCACGCACCGCACGTTCGTGGGGAACCTGGC</td><td>>hg38_21_+_8433172_8446622_RNA45SN1_45S_50ntFlanks</td><td>12893</td><td>12932</td><td>40</td><td>CCTCGCCCGTCACGCACCGCACGTTCGTGGGGAACCTGGC</td><td>6507.2</td><td>-8.6</td><td>0.7</td><td>-11.9</td><td>8.8E-33</td><td>8.0E-31</td><td>MB231/MB468</td><td>MITO</td></tr>
<tr><td>>hg38_21_+_8433172_8446622_RNA45SN1_45S_50ntFlanks@9539.9571.33|ACTCTGGTGGAGGTCCGTAGCGGTCCTGACGTG</td><td>>hg38_21_+_8433172_8446622_RNA45SN1_45S_50ntFlanks</td><td>9539</td><td>9571</td><td>33</td><td>ACTCTGGTGGAGGTCCGTAGCGGTCCTGACGTG</td><td>3589.6</td><td>-5.7</td><td>0.5</td><td>-11.9</td><td>1.8E-32</td><td>8.0E-31</td><td>MB231/MB468</td><td>NUC</td></tr>
<tr><td>>MI0000478|hsa-mir-149&WithFlank&2|+|240455995|240456095@21.42.22(+1U)&[MIMAT0000450&hsa-miR-149-5p&offsets|0|-1(+1U);m-195&2|+|240456015|240456037&offsets|0|-1(+1U)]|TCTGGCTCCGTGTCTTCACTCCT</td><td>>MI0000478|hsa-mir-149&WithFlank&2|+|240455995|240456095</td><td>21</td><td>42</td><td>22(+1U)</td><td>TCTGGCTCCGTGTCTTCACTCCT</td><td>1368.7</td><td>14.9</td><td>1.3</td><td>11.8</td><td>2.4E-32</td><td>8.0E-31</td><td>BT20/MB468</td><td>MITO</td></tr>
<tr><td>>hg38_21_+_8433172_8446622_RNA45SN1_45S_50ntFlanks@12405.12435.31|CTTTTTGATCCTTCGATGTCGGCTCTTCCTA</td><td>>hg38_21_+_8433172_8446622_RNA45SN1_45S_50ntFlanks</td><td>12405</td><td>12435</td><td>31</td><td>CTTTTTGATCCTTCGATGTCGGCTCTTCCTA</td><td>8479.7</td><td>-10.8</td><td>0.9</td><td>-11.9</td><td>9.1E-33</td><td>8.1E-31</td><td>MB231/MB468</td><td>MITO</td></tr>
<tr><td>>MI0000469|hsa-mir-125a&WithFlank&19|+|51693248|51693345@21.44.24&[MIMAT0000443&hsa-miR-125a-5p&offsets|0|0;m-63&19|+|51693268|51693289&offsets|0|+2]|TCCCTGAGACCCTTTAACCTGTGA</td><td>>MI0000469|hsa-mir-125a&WithFlank&19|+|51693248|51693345</td><td>21</td><td>44</td><td>24</td><td>TCCCTGAGACCCTTTAACCTGTGA</td><td>9753.8</td><td>7.5</td><td>0.6</td><td>11.8</td><td>3.6E-32</td><td>8.2E-31</td><td>BT20/MB468</td><td>CYTO</td></tr>
<tr><td>>hg38_21_+_8433172_8446622_RNA45SN1_45S_50ntFlanks@7981.8004.24|CTCAGATCAGACGTGGCGACCCGC</td><td>>hg38_21_+_8433172_8446622_RNA45SN1_45S_50ntFlanks</td><td>7981</td><td>8004</td><td>24</td><td>CTCAGATCAGACGTGGCGACCCGC</td><td>419.2</td><td>-5.7</td><td>0.5</td><td>-11.8</td><td>5.2E-32</td><td>8.4E-31</td><td>BT20/MB468</td><td>NUC</td></tr>
<tr><td>>MI0000086|hsa-mir-28&WithFlank&3|+|188688775|188688872@60.81.22&[MIMAT0004502&hsa-miR-28-3p&offsets|0|0;m-28&3|+|188688834|188688855&offsets|0|0]|CACTAGATTGTGAGCTCCTGGA</td><td>>MI0000086|hsa-mir-28&WithFlank&3|+|188688775|188688872</td><td>60</td><td>81</td><td>22</td><td>CACTAGATTGTGAGCTCCTGGA</td><td>3814.3</td><td>6.7</td><td>0.6</td><td>11.9</td><td>2.0E-32</td><td>8.7E-31</td><td>MB231/MB468</td><td>NUC</td></tr>
<tr><td>>hg38_21_+_8433172_8446622_RNA45SN1_45S_50ntFlanks@11882.11904.23|CTGTTGAGCTTGACTCTAGTCTG</td><td>>hg38_21_+_8433172_8446622_RNA45SN1_45S_50ntFlanks</td><td>11882</td><td>11904</td><td>23</td><td>CTGTTGAGCTTGACTCTAGTCTG</td><td>1030.0</td><td>-14.4</td><td>1.2</td><td>-11.8</td><td>5.6E-32</td><td>9.0E-31</td><td>BT20/MB468</td><td>NUC</td></tr>
<tr><td>>hg38_21_+_8433172_8446622_RNA45SN1_45S_50ntFlanks@9504.9525.22|GTGAACTATGCCTGGGCAGGGC</td><td>>hg38_21_+_8433172_8446622_RNA45SN1_45S_50ntFlanks</td><td>9504</td><td>9525</td><td>22</td><td>GTGAACTATGCCTGGGCAGGGC</td><td>993.1</td><td>-14.4</td><td>1.2</td><td>-11.8</td><td>5.7E-32</td><td>9.1E-31</td><td>BT20/MB468</td><td>NUC</td></tr>
<tr><td>>hg38_21_+_8433172_8446622_RNA45SN1_45S_50ntFlanks@12324.12348.25|TTGGGTTTTAAGCAGGAGGTGTCAG</td><td>>hg38_21_+_8433172_8446622_RNA45SN1_45S_50ntFlanks</td><td>12324</td><td>12348</td><td>25</td><td>TTGGGTTTTAAGCAGGAGGTGTCAG</td><td>1292.9</td><td>-14.8</td><td>1.3</td><td>-11.8</td><td>5.8E-32</td><td>9.1E-31</td><td>BT20/MB468</td><td>NUC</td></tr>
<tr><td>>hg38_21_+_8433172_8446622_RNA45SN1_45S_50ntFlanks@4722.4751.30|TCATTAATCAAGAACGAAAGTCGGAGGTTC</td><td>>hg38_21_+_8433172_8446622_RNA45SN1_45S_50ntFlanks</td><td>4722</td><td>4751</td><td>30</td><td>TCATTAATCAAGAACGAAAGTCGGAGGTTC</td><td>1296.7</td><td>-14.8</td><td>1.3</td><td>-11.8</td><td>6.0E-32</td><td>9.5E-31</td><td>BT20/MB468</td><td>NUC</td></tr>
<tr><td>>hg38_21_+_8433172_8446622_RNA45SN1_45S_50ntFlanks@10382.10424.43|TTCCATGTGAACAGCAGTTGAACATGGGTCAGTCGGTCCTGAG</td><td>>hg38_21_+_8433172_8446622_RNA45SN1_45S_50ntFlanks</td><td>10382</td><td>10424</td><td>43</td><td>TTCCATGTGAACAGCAGTTGAACATGGGTCAGTCGGTCCTGAG</td><td>9917.5</td><td>10.5</td><td>0.9</td><td>11.8</td><td>2.9E-32</td><td>9.8E-31</td><td>BT20/MB468</td><td>MITO</td></tr>
<tr><td>>MI0000077|hsa-mir-21&WithFlank&17|+|59841260|59841343@14.36.23&[MIMAT0000076&hsa-miR-21-5p&offsets|0|+1;m-1&17|+|59841273|59841294&offsets|0|+1]|TAGCTTATCAGACTGATGTTGAC</td><td>>MI0000077|hsa-mir-21&WithFlank&17|+|59841260|59841343</td><td>14</td><td>36</td><td>23</td><td>TAGCTTATCAGACTGATGTTGAC</td><td>895680.7</td><td>6.9</td><td>0.6</td><td>11.8</td><td>2.3E-32</td><td>9.9E-31</td><td>MB231/MB468</td><td>NUC</td></tr>
<tr><td>>hg38_21_+_8433172_8446622_RNA45SN1_45S_50ntFlanks@12405.12437.33|CTTTTTGATCCTTCGATGTCGGCTCTTCCTATC</td><td>>hg38_21_+_8433172_8446622_RNA45SN1_45S_50ntFlanks</td><td>12405</td><td>12437</td><td>33</td><td>CTTTTTGATCCTTCGATGTCGGCTCTTCCTATC</td><td>7920.9</td><td>-7.7</td><td>0.6</td><td>-11.9</td><td>1.1E-32</td><td>9.9E-31</td><td>MB231/MB468</td><td>MITO</td></tr>
<tr><td>>hg38_21_+_8433172_8446622_RNA45SN1_45S_50ntFlanks@3769.3792.24|CGCACGGCCGGTACAGTGAAACTG</td><td>>hg38_21_+_8433172_8446622_RNA45SN1_45S_50ntFlanks</td><td>3769</td><td>3792</td><td>24</td><td>CGCACGGCCGGTACAGTGAAACTG</td><td>1635.9</td><td>-15.1</td><td>1.3</td><td>-11.8</td><td>6.3E-32</td><td>1.0E-30</td><td>BT20/MB468</td><td>NUC</td></tr>
<tr><td>>MI0000299|hsa-mir-222&WithFlank&X|-|45747009|45747130@75.99.25(+2U)&[MIMAT0000279&hsa-miR-222-3p&offsets|0|+4(+2U);m-140&X|-|45747033|45747056&offsets|0|+1(+2U)]|AGCTACATCTGGCTACTGGGTCTCTTT</td><td>>MI0000299|hsa-mir-222&WithFlank&X|-|45747009|45747130</td><td>75</td><td>99</td><td>25(+2U)</td><td>AGCTACATCTGGCTACTGGGTCTCTTT</td><td>764.8</td><td>6.9</td><td>0.6</td><td>11.8</td><td>4.6E-32</td><td>1.0E-30</td><td>BT20/MB468</td><td>CYTO</td></tr>
<tr><td>>hg38_21_+_8433172_8446622_RNA45SN1_45S_50ntFlanks@7986.8003.18|ATCAGACGTGGCGACCCG</td><td>>hg38_21_+_8433172_8446622_RNA45SN1_45S_50ntFlanks</td><td>7986</td><td>8003</td><td>18</td><td>ATCAGACGTGGCGACCCG</td><td>5256.9</td><td>-16.8</td><td>1.4</td><td>-11.8</td><td>7.1E-32</td><td>1.1E-30</td><td>BT20/MB468</td><td>NUC</td></tr>
<tr><td>>hg38_21_+_8433172_8446622_RNA45SN1_45S_50ntFlanks@7991.8012.22|ACGTGGCGACCCGCTGAATTTA</td><td>>hg38_21_+_8433172_8446622_RNA45SN1_45S_50ntFlanks</td><td>7991</td><td>8012</td><td>22</td><td>ACGTGGCGACCCGCTGAATTTA</td><td>6066.6</td><td>-8.3</td><td>0.7</td><td>-12.0</td><td>5.7E-33</td><td>1.1E-30</td><td>MB231/MB468</td><td>MP</td></tr>
<tr><td>>hg38_21_+_8433172_8446622_RNA45SN1_45S_50ntFlanks@12159.12179.21|TTTGACTGGGGCGGTACACCT</td><td>>hg38_21_+_8433172_8446622_RNA45SN1_45S_50ntFlanks</td><td>12159</td><td>12179</td><td>21</td><td>TTTGACTGGGGCGGTACACCT</td><td>1672.5</td><td>-15.1</td><td>1.3</td><td>-11.7</td><td>7.1E-32</td><td>1.1E-30</td><td>BT20/MB468</td><td>NUC</td></tr>
<tr><td>>hg38_21_+_8433172_8446622_RNA45SN1_45S_50ntFlanks@6664.6681.18|TGGATCACTCGGCTCGTG</td><td>>hg38_21_+_8433172_8446622_RNA45SN1_45S_50ntFlanks</td><td>6664</td><td>6681</td><td>18</td><td>TGGATCACTCGGCTCGTG</td><td>6623.3</td><td>-16.6</td><td>1.4</td><td>-12.0</td><td>5.9E-33</td><td>1.2E-30</td><td>MB231/MB468</td><td>MP</td></tr>
<tr><td>>MI0016005|hsa-mir-3615&WithFlank&17|+|74748607|74748705@57.79.23&[MIMAT0017994&hsa-miR-3615&offsets|0|+2;m-271&17|+|74748663|74748684&offsets|0|+1]|TCTCTCGGCTCCTCGCGGCTCGC</td><td>>MI0016005|hsa-mir-3615&WithFlank&17|+|74748607|74748705</td><td>57</td><td>79</td><td>23</td><td>TCTCTCGGCTCCTCGCGGCTCGC</td><td>432.3</td><td>6.0</td><td>0.5</td><td>11.8</td><td>5.3E-32</td><td>1.2E-30</td><td>BT20/MB468</td><td>CYTO</td></tr>
<tr><td>>hg38_21_+_8433172_8446622_RNA45SN1_45S_50ntFlanks@9534.9563.30|AGGAAACTCTGGTGGAGGTCCGTAGCGGTC</td><td>>hg38_21_+_8433172_8446622_RNA45SN1_45S_50ntFlanks</td><td>9534</td><td>9563</td><td>30</td><td>AGGAAACTCTGGTGGAGGTCCGTAGCGGTC</td><td>20374.2</td><td>-9.9</td><td>0.8</td><td>-12.0</td><td>6.2E-33</td><td>1.2E-30</td><td>MB231/MB468</td><td>MP</td></tr>
<tr><td>>hg38_21_+_8433172_8446622_RNA45SN1_45S_50ntFlanks@6743.6770.28|CGACACTTCGAACGCACTTGCGGCCCCG</td><td>>hg38_21_+_8433172_8446622_RNA45SN1_45S_50ntFlanks</td><td>6743</td><td>6770</td><td>28</td><td>CGACACTTCGAACGCACTTGCGGCCCCG</td><td>1339.3</td><td>-4.9</td><td>0.4</td><td>-11.8</td><td>3.7E-32</td><td>1.2E-30</td><td>BT20/MB468</td><td>MITO</td></tr>
<tr><td>>hg38_21_+_8433172_8446622_RNA45SN1_45S_50ntFlanks@11729.11752.24|TAACTATGACTCTCTTAAGGTAGC</td><td>>hg38_21_+_8433172_8446622_RNA45SN1_45S_50ntFlanks</td><td>11729</td><td>11752</td><td>24</td><td>TAACTATGACTCTCTTAAGGTAGC</td><td>1114.4</td><td>-14.5</td><td>1.2</td><td>-11.7</td><td>8.0E-32</td><td>1.3E-30</td><td>BT20/MB468</td><td>NUC</td></tr>
<tr><td>>hg38_21_+_8433172_8446622_RNA45SN1_45S_50ntFlanks@9534.9571.38|AGGAAACTCTGGTGGAGGTCCGTAGCGGTCCTGACGTG</td><td>>hg38_21_+_8433172_8446622_RNA45SN1_45S_50ntFlanks</td><td>9534</td><td>9571</td><td>38</td><td>AGGAAACTCTGGTGGAGGTCCGTAGCGGTCCTGACGTG</td><td>153179.2</td><td>-11.8</td><td>1.0</td><td>-11.7</td><td>8.1E-32</td><td>1.3E-30</td><td>BT20/MB468</td><td>NUC</td></tr>
<tr><td>>MI0000077|hsa-mir-21&WithFlank&17|+|59841260|59841343@15.36.22(+1A)&[MIMAT0000076&hsa-miR-21-5p&offsets|+1|+1(+1A);m-1&17|+|59841273|59841294&offsets|+1|+1(+1A)]|AGCTTATCAGACTGATGTTGACA</td><td>>MI0000077|hsa-mir-21&WithFlank&17|+|59841260|59841343</td><td>15</td><td>36</td><td>22(+1A)</td><td>AGCTTATCAGACTGATGTTGACA</td><td>515.4</td><td>7.3</td><td>0.6</td><td>11.8</td><td>3.9E-32</td><td>1.3E-30</td><td>BT20/MB468</td><td>MITO</td></tr>
<tr><td>>hg38_21_+_8433172_8446622_RNA45SN1_45S_50ntFlanks@6745.6771.27|ACACTTCGAACGCACTTGCGGCCCCGG</td><td>>hg38_21_+_8433172_8446622_RNA45SN1_45S_50ntFlanks</td><td>6745</td><td>6771</td><td>27</td><td>ACACTTCGAACGCACTTGCGGCCCCGG</td><td>6840.1</td><td>-16.6</td><td>1.4</td><td>-11.9</td><td>6.9E-33</td><td>1.3E-30</td><td>MB231/MB468</td><td>MP</td></tr>
<tr><td>>hg38_21_+_8433172_8446622_RNA45SN1_45S_50ntFlanks@12637.12654.18|GCTACCATCTGTGGGATT</td><td>>hg38_21_+_8433172_8446622_RNA45SN1_45S_50ntFlanks</td><td>12637</td><td>12654</td><td>18</td><td>GCTACCATCTGTGGGATT</td><td>3881.2</td><td>-15.8</td><td>1.3</td><td>-11.9</td><td>7.2E-33</td><td>1.4E-30</td><td>MB231/MB468</td><td>MP</td></tr>
<tr><td>>hg38_21_+_8433172_8446622_RNA45SN1_45S_50ntFlanks@11873.11893.21|AAGAAGACCCTGTTGAGCTTG</td><td>>hg38_21_+_8433172_8446622_RNA45SN1_45S_50ntFlanks</td><td>11873</td><td>11893</td><td>21</td><td>AAGAAGACCCTGTTGAGCTTG</td><td>8900.3</td><td>-17.0</td><td>1.4</td><td>-11.9</td><td>7.4E-33</td><td>1.4E-30</td><td>MB231/MB468</td><td>MP</td></tr>
<tr><td>>hg38_21_+_8433172_8446622_RNA45SN1_45S_50ntFlanks@12008.12043.36|CCGCCGGTGAAATACCACTACTCTGATCGTTTTTTC</td><td>>hg38_21_+_8433172_8446622_RNA45SN1_45S_50ntFlanks</td><td>12008</td><td>12043</td><td>36</td><td>CCGCCGGTGAAATACCACTACTCTGATCGTTTTTTC</td><td>22830.2</td><td>-18.9</td><td>1.6</td><td>-11.7</td><td>8.8E-32</td><td>1.4E-30</td><td>BT20/MB468</td><td>NUC</td></tr>
<tr><td>>hg38_21_+_8433172_8446622_RNA45SN1_45S_50ntFlanks@10411.10430.20|CAGTCGGTCCTGAGAGATGG</td><td>>hg38_21_+_8433172_8446622_RNA45SN1_45S_50ntFlanks</td><td>10411</td><td>10430</td><td>20</td><td>CAGTCGGTCCTGAGAGATGG</td><td>6706.7</td><td>-16.6</td><td>1.4</td><td>-11.9</td><td>7.6E-33</td><td>1.4E-30</td><td>MB231/MB468</td><td>MP</td></tr>
<tr><td>>hg38_21_+_8433172_8446622_RNA45SN1_45S_50ntFlanks@9568.9596.29|CGTGCAAATCGGTCGTCCGACCTGGGTAT</td><td>>hg38_21_+_8433172_8446622_RNA45SN1_45S_50ntFlanks</td><td>9568</td><td>9596</td><td>29</td><td>CGTGCAAATCGGTCGTCCGACCTGGGTAT</td><td>5311.1</td><td>-16.2</td><td>1.4</td><td>-11.9</td><td>7.8E-33</td><td>1.4E-30</td><td>MB231/MB468</td><td>MP</td></tr>
<tr><td>>hg38_21_+_8433172_8446622_RNA45SN1_45S_50ntFlanks@11670.11701.32|CCCAGTGCTCTGAATGTCAAAGTGAAGAAATT</td><td>>hg38_21_+_8433172_8446622_RNA45SN1_45S_50ntFlanks</td><td>11670</td><td>11701</td><td>32</td><td>CCCAGTGCTCTGAATGTCAAAGTGAAGAAATT</td><td>2897.6</td><td>-6.3</td><td>0.5</td><td>-11.8</td><td>3.3E-32</td><td>1.4E-30</td><td>MB231/MB468</td><td>NUC</td></tr>
<tr><td>>hg38_21_+_8433172_8446622_RNA45SN1_45S_50ntFlanks@12405.12441.37|CTTTTTGATCCTTCGATGTCGGCTCTTCCTATCATTG</td><td>>hg38_21_+_8433172_8446622_RNA45SN1_45S_50ntFlanks</td><td>12405</td><td>12441</td><td>37</td><td>CTTTTTGATCCTTCGATGTCGGCTCTTCCTATCATTG</td><td>19224.2</td><td>-8.5</td><td>0.7</td><td>-11.8</td><td>3.3E-32</td><td>1.4E-30</td><td>MB231/MB468</td><td>NUC</td></tr>
<tr><td>>hg38_21_+_8433172_8446622_RNA45SN1_45S_50ntFlanks@10411.10443.33|CAGTCGGTCCTGAGAGATGGGCGAGCGCCGTTC</td><td>>hg38_21_+_8433172_8446622_RNA45SN1_45S_50ntFlanks</td><td>10411</td><td>10443</td><td>33</td><td>CAGTCGGTCCTGAGAGATGGGCGAGCGCCGTTC</td><td>1118.9</td><td>-5.8</td><td>0.5</td><td>-11.8</td><td>7.0E-32</td><td>1.6E-30</td><td>BT20/MB468</td><td>CYTO</td></tr>
<tr><td>>MI0000285|hsa-mir-205&WithFlank&1|+|209432127|209432248@40.60.21(+1A)&[MIMAT0000266&hsa-miR-205-5p&offsets|0|-1(+1A);m-351&1|+|209432166|209432187&offsets|0|-1(+1A)]|TCCTTCATTCCACCGGAGTCTA</td><td>>MI0000285|hsa-mir-205&WithFlank&1|+|209432127|209432248</td><td>40</td><td>60</td><td>21(+1A)</td><td>TCCTTCATTCCACCGGAGTCTA</td><td>14165.7</td><td>8.0</td><td>0.7</td><td>11.7</td><td>7.1E-32</td><td>1.6E-30</td><td>BT20/MB468</td><td>CYTO</td></tr>
<tr><td>>hg38_21_+_8433172_8446622_RNA45SN1_45S_50ntFlanks@4557.4587.31|CAGCTAGGAATAATGGAATAGGACCGCGGTT</td><td>>hg38_21_+_8433172_8446622_RNA45SN1_45S_50ntFlanks</td><td>4557</td><td>4587</td><td>31</td><td>CAGCTAGGAATAATGGAATAGGACCGCGGTT</td><td>1112.9</td><td>-14.5</td><td>1.2</td><td>-11.7</td><td>1.0E-31</td><td>1.6E-30</td><td>BT20/MB468</td><td>NUC</td></tr>
<tr><td>>hg38_21_+_8433172_8446622_RNA45SN1_45S_50ntFlanks@7989.8012.24|AGACGTGGCGACCCGCTGAATTTA</td><td>>hg38_21_+_8433172_8446622_RNA45SN1_45S_50ntFlanks</td><td>7989</td><td>8012</td><td>24</td><td>AGACGTGGCGACCCGCTGAATTTA</td><td>6747.9</td><td>-7.7</td><td>0.6</td><td>-11.9</td><td>9.6E-33</td><td>1.7E-30</td><td>MB231/MB468</td><td>MP</td></tr>
<tr><td>>hg38_21_+_8433172_8446622_RNA45SN1_45S_50ntFlanks@10512.10533.22|TCCGGAGTGGCGGAGATGGGCG</td><td>>hg38_21_+_8433172_8446622_RNA45SN1_45S_50ntFlanks</td><td>10512</td><td>10533</td><td>22</td><td>TCCGGAGTGGCGGAGATGGGCG</td><td>17216.6</td><td>-17.9</td><td>1.5</td><td>-11.9</td><td>9.9E-33</td><td>1.7E-30</td><td>MB231/MB468</td><td>MP</td></tr>
<tr><td>>hg38_21_+_8433172_8446622_RNA45SN1_45S_50ntFlanks@12633.12653.21|CGAAGCTACCATCTGTGGGAT</td><td>>hg38_21_+_8433172_8446622_RNA45SN1_45S_50ntFlanks</td><td>12633</td><td>12653</td><td>21</td><td>CGAAGCTACCATCTGTGGGAT</td><td>16225.7</td><td>-17.8</td><td>1.5</td><td>-11.9</td><td>2.0E-32</td><td>1.7E-30</td><td>MB231/MB468</td><td>MITO</td></tr>
<tr><td>>MI0000254|hsa-mir-30c-2&WithFlank&6|-|71376954|71377037@53.74.22&[MIMAT0004550&hsa-miR-30c-2-3p&offsets|0|0;m-206&6|-|71376964|71376985&offsets|0|0]|CTGGGAGAAGGCTGTTTACTCT</td><td>>MI0000254|hsa-mir-30c-2&WithFlank&6|-|71376954|71377037</td><td>53</td><td>74</td><td>22</td><td>CTGGGAGAAGGCTGTTTACTCT</td><td>3031.2</td><td>7.1</td><td>0.6</td><td>11.7</td><td>8.0E-32</td><td>1.8E-30</td><td>BT20/MB468</td><td>CYTO</td></tr>
<tr><td>>hg38_21_+_8433172_8446622_RNA45SN1_45S_50ntFlanks@12478.12497.20|ACTAATAGGGAACGTGAGCT</td><td>>hg38_21_+_8433172_8446622_RNA45SN1_45S_50ntFlanks</td><td>12478</td><td>12497</td><td>20</td><td>ACTAATAGGGAACGTGAGCT</td><td>607.3</td><td>4.2</td><td>0.4</td><td>11.8</td><td>5.5E-32</td><td>1.8E-30</td><td>BT20/MB468</td><td>MITO</td></tr>
<tr><td>>MI0000272|hsa-mir-182&WithFlank&7|-|129770377|129770498@29.51.23(+1A)&[MIMAT0000259&hsa-miR-182-5p&offsets|0|-1(+1A);m-22&7|-|129770449|129770470&offsets|0|+1(+1A)]|TTTGGCAATGGTAGAACTCACACA</td><td>>MI0000272|hsa-mir-182&WithFlank&7|-|129770377|129770498</td><td>29</td><td>51</td><td>23(+1A)</td><td>TTTGGCAATGGTAGAACTCACACA</td><td>18440.1</td><td>9.5</td><td>0.8</td><td>11.7</td><td>8.4E-32</td><td>1.8E-30</td><td>BT20/MB468</td><td>CYTO</td></tr>
<tr><td>>hg38_21_+_8433172_8446622_RNA45SN1_45S_50ntFlanks@12414.12448.35|CCTTCGATGTCGGCTCTTCCTATCATTGTGAAGCA</td><td>>hg38_21_+_8433172_8446622_RNA45SN1_45S_50ntFlanks</td><td>12414</td><td>12448</td><td>35</td><td>CCTTCGATGTCGGCTCTTCCTATCATTGTGAAGCA</td><td>778.3</td><td>6.8</td><td>0.6</td><td>11.8</td><td>5.8E-32</td><td>1.9E-30</td><td>BT20/MB468</td><td>MITO</td></tr>
<tr><td>>hg38_21_+_8433172_8446622_RNA45SN1_45S_50ntFlanks@4657.4689.33|CTAGAGGTGAAATTCTTGGACCGGCGCAAGACG</td><td>>hg38_21_+_8433172_8446622_RNA45SN1_45S_50ntFlanks</td><td>4657</td><td>4689</td><td>33</td><td>CTAGAGGTGAAATTCTTGGACCGGCGCAAGACG</td><td>25947.4</td><td>-18.4</td><td>1.6</td><td>-11.8</td><td>2.4E-32</td><td>2.0E-30</td><td>MB231/MB468</td><td>MITO</td></tr>
<tr><td>>hg38_21_+_8433172_8446622_RNA45SN1_45S_50ntFlanks@12001.12031.31|CTGCGGGCCGCCGGTGAAATACCACTACTCT</td><td>>hg38_21_+_8433172_8446622_RNA45SN1_45S_50ntFlanks</td><td>12001</td><td>12031</td><td>31</td><td>CTGCGGGCCGCCGGTGAAATACCACTACTCT</td><td>1406.2</td><td>-14.9</td><td>1.3</td><td>-11.7</td><td>1.3E-31</td><td>2.1E-30</td><td>BT20/MB468</td><td>NUC</td></tr>
<tr><td>>hg38_21_+_8433172_8446622_RNA45SN1_45S_50ntFlanks@12366.12383.18|TAACTGGCTTGTGGCGGC</td><td>>hg38_21_+_8433172_8446622_RNA45SN1_45S_50ntFlanks</td><td>12366</td><td>12383</td><td>18</td><td>TAACTGGCTTGTGGCGGC</td><td>951.2</td><td>-14.3</td><td>1.2</td><td>-11.7</td><td>1.4E-31</td><td>2.1E-30</td><td>BT20/MB468</td><td>NUC</td></tr>
</table>

| | | | | | | | | | | | | | |
|---|---|---|---|---|---|---|---|---|---|---|---|---|---|
| >hg38_21_+_8433172_8446622_RNA45SN1_45S_50ntFlanks@6740.6771.32\|CATCGACACTTCGAACGCACTTGCGGCCCCGG | >hg38_21_+_8433172_8446622_RNA45SN1_45S_50ntFlanks | 6740 | 6771 | 32 | CATCGACACTTCGAACGCACTTGCGGCCCCGG | 3311.7 | -5.4 | 0.5 | -11.8 | 6.7E-32 | 2.2E-30 | BT20/MB468 | MITO |
| >hg38_21_+_8433172_8446622_RNA45SN1_45S_50ntFlanks@11768.11804.37\|CTAATTAGTGACGCGCATGAATGGATGAACGAGATTC | >hg38_21_+_8433172_8446622_RNA45SN1_45S_50ntFlanks | 11768 | 11804 | 37 | CTAATTAGTGACGCGCATGAATGGATGAACGAGATTC | 1796.1 | -15.2 | 1.3 | -11.7 | 1.4E-31 | 2.2E-30 | BT20/MB468 | NUC |
| >hg38_21_+_8433172_8446622_RNA45SN1_45S_50ntFlanks@8381.8402.22\|AAGAGGGCGTGAAACCGTTAAG | >hg38_21_+_8433172_8446622_RNA45SN1_45S_50ntFlanks | 8381 | 8402 | 22 | AAGAGGGCGTGAAACCGTTAAG | 1170.1 | -14.6 | 1.3 | -11.7 | 1.5E-31 | 2.2E-30 | BT20/MB468 | NUC |
| >hg38_21_+_8433172_8446622_RNA45SN1_45S_50ntFlanks@4760.4778.19\|TCAGATACCGTCGTAGTTC | >hg38_21_+_8433172_8446622_RNA45SN1_45S_50ntFlanks | 4760 | 4778 | 19 | TCAGATACCGTCGTAGTTC | 1133.8 | -14.6 | 1.2 | -11.7 | 1.5E-31 | 2.3E-30 | BT20/MB468 | NUC |
| >MI0000255\|hsa-mir-30d&WithFlank&8\|-\|134804870\|134804951@12.34.23(+1A)&[MIMAT0000245&hsa-miR-30d-5p&offsets\|0\|+1(+1A);m-21&8\|-\|134804919\|134804940&offsets\|0\|+1(+1A)]\|TGTAAACATCCCCGACTGGAAGCA | >MI0000255\|hsa-mir-30d&WithFlank&8\|-\|134804870\|134804951 | 12 | 34 | 23(+1A) | TGTAAACATCCCCGACTGGAAGCA | 6408.4 | 8.3 | 0.7 | 11.7 | 7.3E-32 | 2.4E-30 | BT20/MB468 | MITO |
| >hg38_21_+_8433172_8446622_RNA45SN1_45S_50ntFlanks@7989.8008.20\|AGACGTGGCGACCCGCTGAA | >hg38_21_+_8433172_8446622_RNA45SN1_45S_50ntFlanks | 7989 | 8008 | 20 | AGACGTGGCGACCCGCTGAA | 33942.3 | -7.7 | 0.6 | -11.9 | 1.4E-32 | 2.4E-30 | MB231/MB468 | MP |
| >hg38_21_+_8433172_8446622_RNA45SN1_45S_50ntFlanks@4723.4751.29\|CATTAATCAAGAACGAAAGTCGGAGGTTC | >hg38_21_+_8433172_8446622_RNA45SN1_45S_50ntFlanks | 4723 | 4751 | 29 | CATTAATCAAGAACGAAAGTCGGAGGTTC | 1092.7 | -14.5 | 1.2 | -11.7 | 1.6E-31 | 2.4E-30 | BT20/MB468 | NUC |
| >hg38_21_+_8433172_8446622_RNA45SN1_45S_50ntFlanks@6719.6745.27\|TGAATTGCAGGACACATTGATCATCGA | >hg38_21_+_8433172_8446622_RNA45SN1_45S_50ntFlanks | 6719 | 6745 | 27 | TGAATTGCAGGACACATTGATCATCGA | 1213.0 | -14.7 | 1.3 | -11.7 | 1.7E-31 | 2.5E-30 | BT20/MB468 | NUC |
| >MI0000299\|hsa-mir-222&WithFlank&X\|-\|45747009\|45747130@75.95.21&[MIMAT0000279&hsa-miR-222-3p&offsets\|0\|0;m-140&X\|-\|45747033\|45747056&offsets\|0\|-3]\|AGCTACATCTGGCTACTGGGT | >MI0000299\|hsa-mir-222&WithFlank&X\|-\|45747009\|45747130 | 75 | 95 | 21 | AGCTACATCTGGCTACTGGGT | 1513.2 | 6.8 | 0.6 | 11.7 | 1.2E-31 | 2.6E-30 | BT20/MB468 | CYTO |
| >MI0000077\|hsa-mir-21&WithFlank&17\|+\|59841260\|59841343@14.37.24&[MIMAT0000076&hsa-miR-21-5p&offsets\|0\|+2;m-1&17\|+\|59841273\|59841294&offsets\|0\|+2]\|TAGCTTATCAGACTGATGTTGACT | >MI0000077\|hsa-mir-21&WithFlank&17\|+\|59841260\|59841343 | 14 | 37 | 24 | TAGCTTATCAGACTGATGTTGACT | 108894.5 | 7.4 | 0.6 | 11.7 | 7.9E-32 | 2.6E-30 | BT20/MB468 | MITO |
| >hg38_21_+_8433172_8446622_RNA45SN1_45S_50ntFlanks@10846.10868.23\|CTTCGGGATAAGGATTGGCTCTA | >hg38_21_+_8433172_8446622_RNA45SN1_45S_50ntFlanks | 10846 | 10868 | 23 | CTTCGGGATAAGGATTGGCTCTA | 2663.9 | -15.8 | 1.4 | -11.7 | 1.7E-31 | 2.6E-30 | BT20/MB468 | NUC |
| >hg38_21_+_8433172_8446622_RNA45SN1_45S_50ntFlanks@6698.6718.21\|AGCTAGCTGCGAGAATTAATG | >hg38_21_+_8433172_8446622_RNA45SN1_45S_50ntFlanks | 6698 | 6718 | 21 | AGCTAGCTGCGAGAATTAATG | 8220.1 | -8.1 | 0.7 | -11.8 | 3.1E-32 | 2.6E-30 | MB231/MB468 | MITO |
| >MI0000446\|hsa-mir-125b-1&WithFlank&11\|-\|122099751\|122099850@21.42.22(+1A);MI0000470\|hsa-mir-125b-2&WithFlank&21\|+\|16590231\|16590331@23.44.22(+1A)&[MIMAT0000423&hsa-miR-125b-5p&offsets\|0\|0(+1A);m-67&11\|-\|122099809\|122099830&offsets\|0\|0(+1A)];[MIMAT0000423_1&hsa-miR-125b-5p&offsets\|0\|0(+1A);m-68&21\|+\|16590253\|16590274&offsets\|0\|0(+1A)]\|TCCCTGAGACCCTAACTTGTGAA | >MI0000446\|hsa-mir-125b-1&WithFlank&11\|-\|122099751\|122099850 | 21 | 42 | 22(+1A) | TCCCTGAGACCCTAACTTGTGAA | 4761.5 | 7.7 | 0.7 | 11.8 | 6.4E-32 | 2.7E-30 | MB231/MB468 | NUC |
| >MI0000253\|hsa-mir-148a&WithFlank&7\|-\|25949913\|25949992@50.71.22&[MIMAT0000243&hsa-miR-148a-3p&offsets\|0\|0;m-7&7\|-\|25949922\|25949943&offsets\|0\|0]\|TCAGTGCACTACAGAACTTTGT | >MI0000253\|hsa-mir-148a&WithFlank&7\|-\|25949913\|25949992 | 50 | 71 | 22 | TCAGTGCACTACAGAACTTTGT | 172764.8 | 7.5 | 0.6 | 11.7 | 8.5E-32 | 2.7E-30 | BT20/MB468 | MITO |
| >hg38_21_+_8433172_8446622_RNA45SN1_45S_50ntFlanks@11885.11905.21\|TTGAGCTTGACTCTAGTCTGG | >hg38_21_+_8433172_8446622_RNA45SN1_45S_50ntFlanks | 11885 | 11905 | 21 | TTGAGCTTGACTCTAGTCTGG | 949.1 | -14.3 | 1.2 | -11.7 | 1.8E-31 | 2.8E-30 | BT20/MB468 | NUC |
| >MI0000441\|hsa-mir-30b&WithFlank&8\|-\|134800514\|134800613@23.44.22&[MIMAT0000420&hsa-miR-30b-5p&offsets\|0\|0;m-76&8\|-\|134800570\|134800591&offsets\|0\|0]\|TGTAAACATCCTACACTCAGCT | >MI0000441\|hsa-mir-30b&WithFlank&8\|-\|134800514\|134800613 | 23 | 44 | 22 | TGTAAACATCCTACACTCAGCT | 4158.6 | 8.0 | 0.7 | 11.7 | 1.3E-31 | 2.8E-30 | BT20/MB468 | CYTO |
| >hg38_21_+_8433172_8446622_RNA45SN1_45S_50ntFlanks@485.506.22\|GATGTGGTGACGTCGTGCTCTC | >hg38_21_+_8433172_8446622_RNA45SN1_45S_50ntFlanks | 485 | 506 | 22 | GATGTGGTGACGTCGTGCTCTC | 3103.6 | 16.1 | 1.4 | 11.7 | 9.1E-32 | 2.9E-30 | BT20/MB468 | MITO |
| >MI0000079\|hsa-mir-23a&WithFlank&19\|-\|13836581\|13836665@54.72.19&[MIMAT0000078&hsa-miR-23a-3p&offsets\|+3\|+1;m-31&19\|-\|13836595\|13836615&offsets\|+3\|+1]\|ACATTGCCAGGGATTTCCA | >MI0000079\|hsa-mir-23a&WithFlank&19\|-\|13836581\|13836665 | 54 | 72 | 19 | ACATTGCCAGGGATTTCCA | 1523.6 | 13.4 | 1.1 | 11.7 | 2.0E-31 | 3.0E-30 | BT20/MB468 | NUC |
| >MI0000102\|hsa-mir-100&WithFlank&11\|-\|122152223\|122152314@19.38.20(+1C)&[MIMAT0000098&hsa-miR-100-5p&offsets\|0\|-2(+1C);m-27&11\|-\|122152276\|122152296&offsets\|0\|-1(+1C)]\|AACCCGTAGATCCGAACTTGC | >MI0000102\|hsa-mir-100&WithFlank&11\|-\|122152223\|122152314 | 19 | 38 | 20(+1C) | AACCCGTAGATCCGAACTTGC | 27599.9 | 18.8 | 1.6 | 11.9 | 9.9E-33 | 3.0E-30 | MB231/MB468 | CYTO |
| >hg38_21_+_8433172_8446622_RNA45SN1_45S_50ntFlanks@10408.10431.24\|GGTCAGTCGGTCCTGAGAGATGGG | >hg38_21_+_8433172_8446622_RNA45SN1_45S_50ntFlanks | 10408 | 10431 | 24 | GGTCAGTCGGTCCTGAGAGATGGG | 3447.2 | -16.2 | 1.4 | -11.7 | 2.0E-31 | 3.1E-30 | BT20/MB468 | NUC |
| >MI0000469\|hsa-mir-125a&WithFlank&19\|+\|51693248\|51693345@21.43.23&[MIMAT0000443&hsa-miR-125a-5p&offsets\|0\|-1;m-63&19\|+\|51693268\|51693289&offsets\|0\|+1]\|TCCCTGAGACCCTTTAACCTGTG | >MI0000469\|hsa-mir-125a&WithFlank&19\|+\|51693248\|51693345 | 21 | 43 | 23 | TCCCTGAGACCCTTTAACCTGTG | 52823.1 | 7.9 | 0.7 | 11.7 | 1.0E-31 | 3.2E-30 | BT20/MB468 | MITO |
| >hg38_21_+_8433172_8446622_RNA45SN1_45S_50ntFlanks@9562.9596.35\|TCCTGACGTGCAAATCGGTCGTCCGACCTGGGTAT | >hg38_21_+_8433172_8446622_RNA45SN1_45S_50ntFlanks | 9562 | 9596 | 35 | TCCTGACGTGCAAATCGGTCGTCCGACCTGGGTAT | 3261.4 | -16.1 | 1.4 | -11.7 | 2.1E-31 | 3.2E-30 | BT20/MB468 | NUC |
| >MI0000439\|hsa-mir-23b&WithFlank&9\|+\|95085202\|95085310@67.86.20(+1U)&[MIMAT0000418&hsa-miR-23b-3p&offsets\|+3\|0(+1U);m-42&9\|+\|95085265\|95085285&offsets\|+3\|+2(+1U)]\|ACATTGCCAGGGATTACCACT | >MI0000439\|hsa-mir-23b&WithFlank&9\|+\|95085202\|95085310 | 67 | 86 | 20(+1U) | ACATTGCCAGGGATTACCACT | 1267.2 | 13.1 | 1.1 | 11.7 | 2.2E-31 | 3.3E-30 | BT20/MB468 | NUC |
| >hg38_21_+_8433172_8446622_RNA45SN1_45S_50ntFlanks@11641.11660.20\|CGGCGGGTGTTGACGCGATG | >hg38_21_+_8433172_8446622_RNA45SN1_45S_50ntFlanks | 11641 | 11660 | 20 | CGGCGGGTGTTGACGCGATG | 77978.8 | -15.3 | 1.3 | -11.8 | 3.9E-32 | 3.3E-30 | MB231/MB468 | MITO |
| >MI0000082\|hsa-mir-25&WithFlank&7\|-\|100093554\|100093649@58.79.22(+1A)&[MIMAT0000081&hsa-miR-25-3p&offsets\|0\|0(+1A);m-23&7\|-\|100093571\|100093592&offsets\|0\|0(+1A)]\|CATTGCACTTGTCTCGGTCTGAA | >MI0000082\|hsa-mir-25&WithFlank&7\|-\|100093554\|100093649 | 58 | 79 | 22(+1A) | CATTGCACTTGTCTCGGTCTGAA | 1111.8 | 7.7 | 0.7 | 11.7 | 1.0E-31 | 3.3E-30 | BT20/MB468 | MITO |
| >hg38_21_+_8433172_8446622_RNA45SN1_45S_50ntFlanks@12001.12030.30\|CTGCGGGCCGCCGGTGAAATACCACTACTC | >hg38_21_+_8433172_8446622_RNA45SN1_45S_50ntFlanks | 12001 | 12030 | 30 | CTGCGGGCCGCCGGTGAAATACCACTACTC | 1299.4 | -14.8 | 1.3 | -11.7 | 2.3E-31 | 3.4E-30 | BT20/MB468 | NUC |
| >MI0000285\|hsa-mir-205&WithFlank&1\|+\|209432127\|209432248@40.61.22&[MIMAT0000266&hsa-miR-205-5p&offsets\|0\|0;m-351&1\|+\|209432166\|209432187&offsets\|0\|0]\|TCCTTCATTCCACCGGAGTCTG | >MI0000285\|hsa-mir-205&WithFlank&1\|+\|209432127\|209432248 | 40 | 61 | 22 | TCCTTCATTCCACCGGAGTCTG | 19718.1 | 17.9 | 1.5 | 12.0 | 5.7E-33 | 3.5E-30 | BT20/MB231 | MITO |
| >hg38_21_+_8433172_8446622_RNA45SN1_45S_50ntFlanks@13002.13027.26\|CGATCTATTGAAAGTCAGCCCTCGAC | >hg38_21_+_8433172_8446622_RNA45SN1_45S_50ntFlanks | 13002 | 13027 | 26 | CGATCTATTGAAAGTCAGCCCTCGAC | 11096.4 | -15.7 | 1.3 | -11.7 | 1.6E-31 | 3.5E-30 | BT20/MB468 | CYTO |
| >MI0000065\|hsa-let-7d&WithFlank&9\|+\|94178828\|94178926@14.35.22&[MIMAT0000065&hsa-let-7d-5p&offsets\|0\|0;m-90&9\|+\|94178841\|94178862&offsets\|0\|0]\|AGAGGTAGTAGGTTGCATAGTT | >MI0000065\|hsa-let-7d&WithFlank&9\|+\|94178828\|94178926 | 14 | 35 | 22 | AGAGGTAGTAGGTTGCATAGTT | 23933.3 | 8.2 | 0.7 | 11.7 | 1.6E-31 | 3.5E-30 | BT20/MB468 | CYTO |
| >MI0003776\|hsa-mir-320b-1&WithFlank&1\|+\|116671740\|116671823@48.68.21;MI0003839\|hsa-mir-320b-2&WithFlank&1\|-\|224257034\|224257116@47.67.21&[MIMAT0005792&hsa-miR-320b&offsets\|0\|-1;m-183&1\|+\|116671787\|116671806&offsets\|0\|+1];[MIMAT0005792_1&hsa-miR-320b&offsets\|0\|-1;m-183&1\|-\|224257051\|224257070&offsets\|0\|+1]\|AAAAGCTGGGTTGAGAGGGCA | >MI0003776\|hsa-mir-320b-1&WithFlank&1\|+\|116671740\|116671823 | 48 | 68 | 21 | AAAAGCTGGGTTGAGAGGGCA | 8192.3 | 11.7 | 1.0 | 11.7 | 1.7E-31 | 3.6E-30 | BT20/MB468 | CYTO |
| >hg38_21_+_8433172_8446622_RNA45SN1_45S_50ntFlanks@12934.12953.20\|CTAAACCATTCGTAGACGAC | >hg38_21_+_8433172_8446622_RNA45SN1_45S_50ntFlanks | 12934 | 12953 | 20 | CTAAACCATTCGTAGACGAC | 1696.0 | -15.2 | 1.3 | -11.6 | 2.5E-31 | 3.7E-30 | BT20/MB468 | NUC |
| >hg38_21_+_8433172_8446622_RNA45SN1_45S_50ntFlanks@10687.10706.20\|CGGCGTCCGGTGAGCTCTCG | >hg38_21_+_8433172_8446622_RNA45SN1_45S_50ntFlanks | 10687 | 10706 | 20 | CGGCGTCCGGTGAGCTCTCG | 5911.5 | -16.3 | 1.4 | -11.8 | 4.5E-32 | 3.7E-30 | MB231/MB468 | MITO |
| >hg38_21_+_8433172_8446622_RNA45SN1_45S_50ntFlanks@8198.8215.18\|TGGACGGTGTGAGGCCGG | >hg38_21_+_8433172_8446622_RNA45SN1_45S_50ntFlanks | 8198 | 8215 | 18 | TGGACGGTGTGAGGCCGG | 1219.4 | -14.7 | 1.3 | -11.6 | 2.6E-31 | 3.8E-30 | BT20/MB468 | NUC |
| >hg38_21_+_8433172_8446622_RNA45SN1_45S_50ntFlanks@12412.12436.25\|ATCCTTCGATGTCGGCTCTTCCTAT | >hg38_21_+_8433172_8446622_RNA45SN1_45S_50ntFlanks | 12412 | 12436 | 25 | ATCCTTCGATGTCGGCTCTTCCTAT | 4397.4 | -16.5 | 1.4 | -11.6 | 2.8E-31 | 4.1E-30 | BT20/MB468 | NUC |
| >hg38_21_+_8433172_8446622_RNA45SN1_45S_50ntFlanks@4759.4780.22\|ATCAGATACCGTCGTAGTTCCG | >hg38_21_+_8433172_8446622_RNA45SN1_45S_50ntFlanks | 4759 | 4780 | 22 | ATCAGATACCGTCGTAGTTCCG | 8157.1 | -15.3 | 1.3 | -11.7 | 1.9E-31 | 4.1E-30 | BT20/MB468 | CYTO |
| >hg38_21_+_8433172_8446622_RNA45SN1_45S_50ntFlanks@12417.12436.20\|TCGATGTCGGCTCTTCCTAT | >hg38_21_+_8433172_8446622_RNA45SN1_45S_50ntFlanks | 12417 | 12436 | 20 | TCGATGTCGGCTCTTCCTAT | 11823.1 | -6.3 | 0.5 | -11.7 | 9.9E-32 | 4.2E-30 | MB231/MB468 | NUC |
| >hg38_21_+_8433172_8446622_RNA45SN1_45S_50ntFlanks@6698.6717.20\|AGCTAGCTGCGAGAATTAAT | >hg38_21_+_8433172_8446622_RNA45SN1_45S_50ntFlanks | 6698 | 6717 | 20 | AGCTAGCTGCGAGAATTAAT | 878.5 | -14.2 | 1.2 | -11.6 | 3.0E-31 | 4.4E-30 | BT20/MB468 | NUC |
| >trna10_ValCAC_5_-_180649395_180649467@59.76.18,trna12_ValAAC_5_-_180645270_180645342@59.76.18,trna132_ValAAC_6_-_27721179_27721251@59.76.18,trna136_ValAAC_6_-_27648885_27648957@59.76.18,trna15_ValAAC_5_-_180615416_180615488@59.76.18,trna18_ValCAC_5_-_180529253_180529325@59.76.18,trna2_ValAAC_3_+_169490018_169490090@59.76.18,trna2_ValCAC_5_+_180524070_180524142@59.76.18,trna4_ValAAC_5_+_180591154_180591226@59.76.18,trna5_ValAAC_5_+_180596610_180596682@59.76.18,trna6_ValCAC_5_+_180600650_180600722@59.76.18,trna85_ValCAC_1_-_161369490_161369562@59.76.18,trna90_ValCAC_1_-_149684088_149684161@60.77.18,trna9_ValCAC_6_+_26538282_26538354@59.76.18\|AACCGGGCGGAAACACCA | >trna10_ValCAC_5_-_180649395_180649467 | 59 | 76 | 18 | AACCGGGCGGAAACACCA | 622.9 | 6.1 | 0.5 | 11.7 | 2.0E-31 | 4.4E-30 | BT20/MB468 | CYTO |
| >hg38_21_+_8433172_8446622_RNA45SN1_45S_50ntFlanks@9504.9524.21\|GTGAACTATGCCTGGGCAGGG | >hg38_21_+_8433172_8446622_RNA45SN1_45S_50ntFlanks | 9504 | 9524 | 21 | GTGAACTATGCCTGGGCAGGG | 1096.3 | -14.5 | 1.2 | -11.6 | 3.1E-31 | 4.5E-30 | BT20/MB468 | NUC |
| >hg38_21_+_8433172_8446622_RNA45SN1_45S_50ntFlanks@4047.4067.21\|ATTCGAACGTCTGCCCTATCA | >hg38_21_+_8433172_8446622_RNA45SN1_45S_50ntFlanks | 4047 | 4067 | 21 | ATTCGAACGTCTGCCCTATCA | 1483.6 | -15.0 | 1.3 | -11.6 | 3.1E-31 | 4.5E-30 | BT20/MB468 | NUC |
| >hg38_21_+_8433172_8446622_RNA45SN1_45S_50ntFlanks@482.507.26\|ATCGATGTGGTGACGTCGTGCTCTCC | >hg38_21_+_8433172_8446622_RNA45SN1_45S_50ntFlanks | 482 | 507 | 26 | ATCGATGTGGTGACGTCGTGCTCTCC | 1844.9 | -15.3 | 1.3 | -11.6 | 3.2E-31 | 4.6E-30 | BT20/MB468 | NUC |
| >MI0000273\|hsa-mir-183&WithFlank&7\|-\|129774899\|129775020@33.53.21(+1A)&[MIMAT0000261&hsa-miR-183-5p&offsets\|0\|-1(+1A);m-41&7\|-\|129774966\|129774987&offsets\|-1\|-2(+1A)]\|TATGGCACTGGTAGAATTCACA | >MI0000273\|hsa-mir-183&WithFlank&7\|-\|129774899\|129775020 | 33 | 53 | 21(+1A) | TATGGCACTGGTAGAATTCACA | 619.2 | 7.1 | 0.6 | 11.7 | 1.5E-31 | 4.7E-30 | BT20/MB468 | MITO |
| >hg38_21_+_8433172_8446622_RNA45SN1_45S_50ntFlanks@10391.10420.30\|AACAGCAGTTGAACATGGGTCAGTCGGTCC | >hg38_21_+_8433172_8446622_RNA45SN1_45S_50ntFlanks | 10391 | 10420 | 30 | AACAGCAGTTGAACATGGGTCAGTCGGTCC | 1732.2 | -15.2 | 1.3 | -11.6 | 3.3E-31 | 4.8E-30 | BT20/MB468 | NUC |
| >MI0000439\|hsa-mir-23b&WithFlank&9\|+\|95085202\|95085310@67.85.19&[MIMAT0000418&hsa-miR-23b-3p&offsets\|+3\|-1;m-42&9\|+\|95085265\|95085285&offsets\|+3\|+1]\|ACATTGCCAGGGATTACCA | >MI0000439\|hsa-mir-23b&WithFlank&9\|+\|95085202\|95085310 | 67 | 85 | 19 | ACATTGCCAGGGATTACCA | 1632.7 | 14.9 | 1.3 | 11.6 | 2.3E-31 | 5.0E-30 | BT20/MB468 | CYTO |
| >hg38_21_+_8433172_8446622_RNA45SN1_45S_50ntFlanks@11873.11906.34\|AAGAAGACCCTGTTGAGCTTGACTCTAGTCTGGC | >hg38_21_+_8433172_8446622_RNA45SN1_45S_50ntFlanks | 11873 | 11906 | 34 | AAGAAGACCCTGTTGAGCTTGACTCTAGTCTGGC | 1361.6 | -14.8 | 1.3 | -11.6 | 3.5E-31 | 5.1E-30 | BT20/MB468 | NUC |

| | | | | | | | | | | | | | |
|---|---|---|---|---|---|---|---|---|---|---|---|---|---|
| >MI0000447\|hsa-mir-128-1&WithFlank&2\|+\|135665391\|135665484@56.76.21(+1A);MI0000727\|hsa-mir-128-2&WithFlank&3\|+\|35744470\|35744565@58.78.21(+1A)&[MIMAT0000424&hsa-miR-128-3p&offsets\|0\|0(+1A);m-108&2\|+\|135665446\|135665466&offsets\|0\|0(+1A)];[MIMAT0000424_1&hsa-miR-128-3p&offsets\|0\|0(+1A);m-117&3\|+\|35744527\|35744547&offsets\|0\|0(+1A)]\|TCACAGTGAACCGGTCTCTTTA | >MI0000447\|hsa-mir-128-1&WithFlank&2\|+\|135665391\|135665484 | 56 | 76 | 21(+1A) | TCACAGTGAACCGGTCTCTTTA | 2248.4 | 5.5 | 0.5 | 11.6 | 2.4E-31 | 5.2E-30 | BT20/MB468 | CYTO |
| >hg38_21_+_8433172_8446622_RNA45SN1_45S_50ntFlanks@10381.10424.44\|GTTCCATGTGAACAGCAGTTGAACATGGGTCAGTCGGTCCTGAG | >hg38_21_+_8433172_8446622_RNA45SN1_45S_50ntFlanks | 10381 | 10424 | 44 | GTTCCATGTGAACAGCAGTTGAACATGGGTCAGTCGGTCCTGAG | 7339.9 | 9.9 | 0.8 | 11.7 | 1.7E-31 | 5.2E-30 | BT20/MB468 | MITO |
| >hg38_21_+_8433172_8446622_RNA45SN1_45S_50ntFlanks@12640.12667.28\|ACCATCTGTGGGATTATGACTGAACGCC | >hg38_21_+_8433172_8446622_RNA45SN1_45S_50ntFlanks | 12640 | 12667 | 28 | ACCATCTGTGGGATTATGACTGAACGCC | 332.5 | 6.0 | 0.5 | 11.7 | 1.8E-31 | 5.5E-30 | BT20/MB468 | MITO |
| >hg38_21_+_8433172_8446622_RNA45SN1_45S_50ntFlanks@12324.12346.23\|TTGGGTTTTAAGCAGGAGGTGTC | >hg38_21_+_8433172_8446622_RNA45SN1_45S_50ntFlanks | 12324 | 12346 | 23 | TTGGGTTTTAAGCAGGAGGTGTC | 4714.8 | -16.1 | 1.4 | -11.8 | 3.3E-32 | 5.5E-30 | MB231/MB468 | MP |
| >hg38_21_+_8433172_8446622_RNA45SN1_45S_50ntFlanks@12412.12441.30\|ATCCTTCGATGTCGGCTCTTCCTATCATTG | >hg38_21_+_8433172_8446622_RNA45SN1_45S_50ntFlanks | 12412 | 12441 | 30 | ATCCTTCGATGTCGGCTCTTCCTATCATTG | 1686.6 | -15.1 | 1.3 | -11.6 | 3.8E-31 | 5.6E-30 | BT20/MB468 | NUC |
| >hg38_21_+_8433172_8446622_RNA45SN1_45S_50ntFlanks@4726.4758.33\|TAATCAAGAACGAAAGTCGGAGGTTCGAAGACG | >hg38_21_+_8433172_8446622_RNA45SN1_45S_50ntFlanks | 4726 | 4758 | 33 | TAATCAAGAACGAAAGTCGGAGGTTCGAAGACG | 5852.7 | -9.1 | 0.8 | -11.8 | 7.0E-32 | 5.7E-30 | MB231/MB468 | MITO |
| >hg38_21_+_8433172_8446622_RNA45SN1_45S_50ntFlanks@8114.8143.30\|GCGCGGGACATGTGGCGTACGGAAGACCCG | >hg38_21_+_8433172_8446622_RNA45SN1_45S_50ntFlanks | 8114 | 8143 | 30 | GCGCGGGACATGTGGCGTACGGAAGACCCG | 7564.2 | -16.7 | 1.4 | -11.8 | 7.0E-32 | 5.7E-30 | MB231/MB468 | MITO |
| >am_trnaMT_ValTAC_MT_+_1602_1670@51.68.18\|CTTAACTTGACCGCTCTG | >am_trnaMT_ValTAC_MT_+_1602_1670 | 51 | 68 | 18 | CTTAACTTGACCGCTCTG | 113866.0 | -13.0 | 1.1 | -11.6 | 4.1E-31 | 6.0E-30 | BT20/MB468 | NUC |
| >hg38_21_+_8433172_8446622_RNA45SN1_45S_50ntFlanks@10403.10430.28\|ACATGGGTCAGTCGGTCCTGAGAGATGG | >hg38_21_+_8433172_8446622_RNA45SN1_45S_50ntFlanks | 10403 | 10430 | 28 | ACATGGGTCAGTCGGTCCTGAGAGATGG | 881.1 | -14.2 | 1.2 | -11.6 | 4.1E-31 | 6.0E-30 | BT20/MB468 | NUC |
| >MI0000088\|hsa-mir-30a&WithFlank&6\|-\|71403545\|71403627@54.73.20(+2U)&[MIMAT0000088&hsa-miR-30a-3p&offsets\|+1\|-1(+2U);m-38&6\|-\|71403554\|71403575&offsets\|+1\|-1(+2U)]\|TTTCAGTCGGATGTTTGCAGTT | >MI0000088\|hsa-mir-30a&WithFlank&6\|-\|71403545\|71403627 | 54 | 73 | 20(+2U) | TTTCAGTCGGATGTTTGCAGTT | 593.8 | 6.9 | 0.6 | 11.6 | 2.9E-31 | 6.1E-30 | BT20/MB468 | CYTO |
| >hg38_21_+_8433172_8446622_RNA45SN1_45S_50ntFlanks@10351.10368.18\|CGAGAACTTTGAAGGCCG | >hg38_21_+_8433172_8446622_RNA45SN1_45S_50ntFlanks | 10351 | 10368 | 18 | CGAGAACTTTGAAGGCCG | 15589.1 | -7.6 | 0.7 | -11.7 | 1.4E-31 | 6.1E-30 | MB231/MB468 | NUC |
| >MI0000299\|hsa-mir-222&WithFlank&X\|-\|45747009\|45747130@75.96.22(+1C)&[MIMAT0000279&hsa-miR-222-3p&offsets\|0\|+1(+1C);m-140&X\|-\|45747033\|45747056&offsets\|0\|-2(+1C)]\|AGCTACATCTGGCTACTGGGTCC | >MI0000299\|hsa-mir-222&WithFlank&X\|-\|45747009\|45747130 | 75 | 96 | 22(+1C) | AGCTACATCTGGCTACTGGGTCC | 597.3 | 7.0 | 0.6 | 11.6 | 2.9E-31 | 6.2E-30 | BT20/MB468 | CYTO |
| >hg38_21_+_8433172_8446622_RNA45SN1_45S_50ntFlanks@12992.13012.21\|TCCCTCGCTGCGATCTATTGA | >hg38_21_+_8433172_8446622_RNA45SN1_45S_50ntFlanks | 12992 | 13012 | 21 | TCCCTCGCTGCGATCTATTGA | 2542.0 | -15.7 | 1.4 | -11.6 | 4.4E-31 | 6.3E-30 | BT20/MB468 | NUC |
| >hg38_21_+_8433172_8446622_RNA45SN1_45S_50ntFlanks@9736.9765.30\|CCGAAACGATCTCAACCTATTCTCAAACTT | >hg38_21_+_8433172_8446622_RNA45SN1_45S_50ntFlanks | 9736 | 9765 | 30 | CCGAAACGATCTCAACCTATTCTCAAACTT | 2886.6 | -15.9 | 1.4 | -11.6 | 4.4E-31 | 6.3E-30 | BT20/MB468 | NUC |
| >hg38_21_+_8433172_8446622_RNA45SN1_45S_50ntFlanks@10396.10424.29\|CAGTTGAACATGGGTCAGTCGGTCCTGAG | >hg38_21_+_8433172_8446622_RNA45SN1_45S_50ntFlanks | 10396 | 10424 | 29 | CAGTTGAACATGGGTCAGTCGGTCCTGAG | 28428.9 | 7.7 | 0.7 | 11.7 | 2.1E-31 | 6.4E-30 | BT20/MB468 | MITO |
| >hg38_21_+_8433172_8446622_RNA45SN1_45S_50ntFlanks@9564.9596.33\|CTGACGTGCAAATCGGTCGTCCGACCTGGGTAT | >hg38_21_+_8433172_8446622_RNA45SN1_45S_50ntFlanks | 9564 | 9596 | 33 | CTGACGTGCAAATCGGTCGTCCGACCTGGGTAT | 90395.8 | -10.1 | 0.9 | -11.7 | 8.1E-32 | 6.4E-30 | MB231/MB468 | MITO |
| >hg38_21_+_8433172_8446622_RNA45SN1_45S_50ntFlanks@6746.6770.25\|CACTTCGAACGCACTTGCGGCCCCG | >hg38_21_+_8433172_8446622_RNA45SN1_45S_50ntFlanks | 6746 | 6770 | 25 | CACTTCGAACGCACTTGCGGCCCCG | 1696.9 | -8.7 | 0.7 | -11.8 | 2.3E-32 | 6.6E-30 | MB231/MB468 | CYTO |
| >hg38_21_+_8433172_8446622_RNA45SN1_45S_50ntFlanks@12633.12651.19\|CGAAGCTACCATCTGTGGG | >hg38_21_+_8433172_8446622_RNA45SN1_45S_50ntFlanks | 12633 | 12651 | 19 | CGAAGCTACCATCTGTGGG | 2137.8 | -8.0 | 0.7 | -11.8 | 4.0E-32 | 6.7E-30 | MB231/MB468 | MP |
| >hg38_21_+_8433172_8446622_RNA45SN1_45S_50ntFlanks@10786.10818.33\|AACAGCCTCTGGCATGTTGGAACAATGTAGGTA | >hg38_21_+_8433172_8446622_RNA45SN1_45S_50ntFlanks | 10786 | 10818 | 33 | AACAGCCTCTGGCATGTTGGAACAATGTAGGTA | 7260.3 | -10.9 | 0.9 | -11.8 | 4.1E-32 | 6.8E-30 | MB231/MB468 | MP |
| >hg38_21_+_8433172_8446622_RNA45SN1_45S_50ntFlanks@4753.4782.30\|AAGACGATCAGATACCGTCGTAGTTCCGAC | >hg38_21_+_8433172_8446622_RNA45SN1_45S_50ntFlanks | 4753 | 4782 | 30 | AAGACGATCAGATACCGTCGTAGTTCCGAC | 921.2 | -14.3 | 1.2 | -11.6 | 4.7E-31 | 6.8E-30 | BT20/MB468 | NUC |
| >trna10_LysCTT_16_+_3241501_3241573@14.31.18,trna13_LysCTT_14_-_58706613_58706685@14.31.18,trna2_LysCTT_15_+_79152904_79152976@14.31.18,trna4_LysCTT_18_-_43669271_43669343@14.31.18\|AGTCGGTAGAGCATGGGA | >trna10_LysCTT_16_+_3241501_3241573 | 14 | 31 | 18 | AGTCGGTAGAGCATGGGA | 2642.8 | -15.8 | 1.4 | -11.6 | 4.8E-31 | 6.9E-30 | BT20/MB468 | NUC |
| >hg38_21_+_8433172_8446622_RNA45SN1_45S_50ntFlanks@10408.10429.22\|GGTCAGTCGGTCCTGAGAGATG | >hg38_21_+_8433172_8446622_RNA45SN1_45S_50ntFlanks | 10408 | 10429 | 22 | GGTCAGTCGGTCCTGAGAGATG | 6929.2 | -8.3 | 0.7 | -11.7 | 1.7E-31 | 7.0E-30 | MB231/MB468 | NUC |
| >hg38_21_+_8433172_8446622_RNA45SN1_45S_50ntFlanks@6663.6681.19\|GTGGATCACTCGGCTCGTG | >hg38_21_+_8433172_8446622_RNA45SN1_45S_50ntFlanks | 6663 | 6681 | 19 | GTGGATCACTCGGCTCGTG | 1727.4 | -6.5 | 0.6 | -11.7 | 1.7E-31 | 7.0E-30 | MB231/MB468 | NUC |
| >hg38_21_+_8433172_8446622_RNA45SN1_45S_50ntFlanks@10683.10706.24\|CCGGCGGCGTCCGGTGAGCTCTCG | >hg38_21_+_8433172_8446622_RNA45SN1_45S_50ntFlanks | 10683 | 10706 | 24 | CCGGCGGCGTCCGGTGAGCTCTCG | 8636.8 | -16.9 | 1.4 | -11.8 | 4.3E-32 | 7.1E-30 | MB231/MB468 | MP |
| >hg38_21_+_8433172_8446622_RNA45SN1_45S_50ntFlanks@12626.12654.29\|AATGGGGCGAAGCTACCATCTGTGGGATT | >hg38_21_+_8433172_8446622_RNA45SN1_45S_50ntFlanks | 12626 | 12654 | 29 | AATGGGGCGAAGCTACCATCTGTGGGATT | 10222.5 | -7.1 | 0.6 | -11.6 | 2.3E-31 | 7.1E-30 | BT20/MB468 | MITO |
| >hg38_21_+_8433172_8446622_RNA45SN1_45S_50ntFlanks@9504.9526.23\|GTGAACTATGCCTGGGCAGGGCG | >hg38_21_+_8433172_8446622_RNA45SN1_45S_50ntFlanks | 9504 | 9526 | 23 | GTGAACTATGCCTGGGCAGGGCG | 7637.2 | -16.7 | 1.4 | -11.7 | 9.2E-32 | 7.2E-30 | MB231/MB468 | MITO |
| >hg38_21_+_8433172_8446622_RNA45SN1_45S_50ntFlanks@8170.8197.28\|CAAGTCCTTCTGATCGAGGCCCAGCCCG | >hg38_21_+_8433172_8446622_RNA45SN1_45S_50ntFlanks | 8170 | 8197 | 28 | CAAGTCCTTCTGATCGAGGCCCAGCCCG | 7887.7 | -17.4 | 1.5 | -11.6 | 5.2E-31 | 7.4E-30 | BT20/MB468 | NUC |
| >hg38_21_+_8433172_8446622_RNA45SN1_45S_50ntFlanks@11661.11681.21\|TGATTTCTGCCCAGTGCTCTG | >hg38_21_+_8433172_8446622_RNA45SN1_45S_50ntFlanks | 11661 | 11681 | 21 | TGATTTCTGCCCAGTGCTCTG | 7861.5 | -7.7 | 0.7 | -11.7 | 1.8E-31 | 7.4E-30 | MB231/MB468 | NUC |
| >hg38_21_+_8433172_8446622_RNA45SN1_45S_50ntFlanks@7981.8006.26\|CTCAGATCAGACGTGGCGACCCGCTG | >hg38_21_+_8433172_8446622_RNA45SN1_45S_50ntFlanks | 7981 | 8006 | 26 | CTCAGATCAGACGTGGCGACCCGCTG | 4673.2 | -7.8 | 0.7 | -11.7 | 1.8E-31 | 7.4E-30 | MB231/MB468 | NUC |
| >hg38_21_+_8433172_8446622_RNA45SN1_45S_50ntFlanks@12008.12031.24\|CCGCCGGTGAAATACCACTACTCT | >hg38_21_+_8433172_8446622_RNA45SN1_45S_50ntFlanks | 12008 | 12031 | 24 | CCGCCGGTGAAATACCACTACTCT | 10631.7 | -13.0 | 1.1 | -11.7 | 9.6E-32 | 7.4E-30 | MB231/MB468 | MITO |
| >hg38_1_-_228634819_228635039_RNA5S12_5S_50ntFlanks@130.170.41\|TGGATGGGAGACCGCCTGGGAATACCGGGTGCTGTAGGCTT | >hg38_1_-_228634819_228635039_RNA5S12_5S_50ntFlanks | 130 | 170 | 41 | TGGATGGGAGACCGCCTGGGAATACCGGGTGCTGTAGGCTT | 6880.2 | 9.4 | 0.8 | 11.8 | 2.8E-32 | 7.7E-30 | MB231/MB468 | CYTO |
| >hg38_21_+_8433172_8446622_RNA45SN1_45S_50ntFlanks@12078.12095.18\|GGGCTCTCGCTTCTGGCG | >hg38_21_+_8433172_8446622_RNA45SN1_45S_50ntFlanks | 12078 | 12095 | 18 | GGGCTCTCGCTTCTGGCG | 1698.5 | -15.2 | 1.3 | -11.6 | 5.5E-31 | 7.8E-30 | BT20/MB468 | NUC |
| >hg38_21_+_8433172_8446622_RNA45SN1_45S_50ntFlanks@12633.12654.22\|CGAAGCTACCATCTGTGGGATT | >hg38_21_+_8433172_8446622_RNA45SN1_45S_50ntFlanks | 12633 | 12654 | 22 | CGAAGCTACCATCTGTGGGATT | 2756.1 | -6.7 | 0.6 | -11.7 | 2.0E-31 | 8.1E-30 | MB231/MB468 | NUC |
| >hg38_21_+_8433172_8446622_RNA45SN1_45S_50ntFlanks@6661.6678.18\|CGGTGGATCACTCGGCTC | >hg38_21_+_8433172_8446622_RNA45SN1_45S_50ntFlanks | 6661 | 6678 | 18 | CGGTGGATCACTCGGCTC | 963.8 | 5.7 | 0.5 | 11.6 | 4.0E-31 | 8.5E-30 | BT20/MB468 | CYTO |
| >hg38_21_+_8433172_8446622_RNA45SN1_45S_50ntFlanks@11729.11749.21\|TAACTATGACTCTCTTAAGGT | >hg38_21_+_8433172_8446622_RNA45SN1_45S_50ntFlanks | 11729 | 11749 | 21 | TAACTATGACTCTCTTAAGGT | 4890.3 | -5.8 | 0.5 | -11.6 | 4.0E-31 | 8.5E-30 | BT20/MB468 | CYTO |
| >MI0000272\|hsa-mir-182&WithFlank&7\|-\|129770377\|129770498@29.51.23&[MIMAT0000259&hsa-miR-182-5p&offsets\|0\|-1;m-22&7\|-\|129770449\|129770470&offsets\|0\|+1]\|TTTGGCAATGGTAGAACTCACAC | >MI0000272\|hsa-mir-182&WithFlank&7\|-\|129770377\|129770498 | 29 | 51 | 23 | TTTGGCAATGGTAGAACTCACAC | 983.7 | 3.2 | 0.3 | 11.6 | 6.2E-31 | 8.8E-30 | BT20/MB468 | NUC |
| >hg38_21_+_8433172_8446622_RNA45SN1_45S_50ntFlanks@12420.12437.18\|ATGTCGGCTCTTCCTATC | >hg38_21_+_8433172_8446622_RNA45SN1_45S_50ntFlanks | 12420 | 12437 | 18 | ATGTCGGCTCTTCCTATC | 24026.8 | -8.0 | 0.7 | -11.6 | 6.3E-31 | 8.9E-30 | BT20/MB468 | NUC |
| >MI0003778\|hsa-mir-320c-1&WithFlank&18\|+\|21683512\|21683595@48.67.20;MI0008191\|hsa-mir-320c-2&WithFlank&18\|+\|24321669\|24321752@48.67.20&[MIMAT0005793&hsa-miR-320c&offsets\|0\|0;m-222&18\|+\|21683559\|21683578&offsets\|0\|0];[MIMAT0005793_1&hsa-miR-320c&offsets\|0\|0;m-219&18\|+\|24321716\|24321735&offsets\|0\|0]\|AAAAGCTGGGTTGAGAGGGT | >MI0003778\|hsa-mir-320c-1&WithFlank&18\|+\|21683512\|21683595 | 48 | 67 | 20 | AAAAGCTGGGTTGAGAGGGT | 2970.7 | 15.8 | 1.4 | 11.6 | 4.3E-31 | 9.0E-30 | BT20/MB468 | CYTO |
| >trna16_GlnTTG_17_+_47269890_47269961@1.29.29,trna64_GlnTTG_6_+_28557156_28557227@1.29.29,trna84_GlnTTG_6_+_145503859_145503930@1.29.29\|GGTCCCATGGTGTAATGGTTAGCACTCTG | >trna16_GlnTTG_17_+_47269890_47269961 | 1 | 29 | 29 | GGTCCCATGGTGTAATGGTTAGCACTCTG | 2111.4 | -15.5 | 1.3 | -11.6 | 6.5E-31 | 9.1E-30 | BT20/MB468 | NUC |
| >hg38_21_+_8433172_8446622_RNA45SN1_45S_50ntFlanks@11316.11342.27\|CACCCCACGTCTCGTCGCGCGCGCGTC | >hg38_21_+_8433172_8446622_RNA45SN1_45S_50ntFlanks | 11316 | 11342 | 27 | CACCCCACGTCTCGTCGCGCGCGCGTC | 11284.2 | -17.2 | 1.5 | -11.7 | 1.2E-31 | 9.4E-30 | MB231/MB468 | MITO |
| >hg38_21_+_8433172_8446622_RNA45SN1_45S_50ntFlanks@13001.13027.27\|GCGATCTATTGAAAGTCAGCCCTCGAC | >hg38_21_+_8433172_8446622_RNA45SN1_45S_50ntFlanks | 13001 | 13027 | 27 | GCGATCTATTGAAAGTCAGCCCTCGAC | 852.5 | -14.2 | 1.2 | -11.6 | 6.7E-31 | 9.5E-30 | BT20/MB468 | NUC |
| >hg38_21_+_8433172_8446622_RNA45SN1_45S_50ntFlanks@10679.10706.28\|GGTTCCGGCGGCGTCCGGTGAGCTCTCG | >hg38_21_+_8433172_8446622_RNA45SN1_45S_50ntFlanks | 10679 | 10706 | 28 | GGTTCCGGCGGCGTCCGGTGAGCTCTCG | 7692.7 | -16.7 | 1.4 | -11.7 | 1.3E-31 | 9.9E-30 | MB231/MB468 | MITO |
| >hg38_21_+_8433172_8446622_RNA45SN1_45S_50ntFlanks@4759.4782.24\|ATCAGATACCGTCGTAGTTCCGAC | >hg38_21_+_8433172_8446622_RNA45SN1_45S_50ntFlanks | 4759 | 4782 | 24 | ATCAGATACCGTCGTAGTTCCGAC | 1129.0 | -14.6 | 1.3 | -11.6 | 7.1E-31 | 1.0E-29 | BT20/MB468 | NUC |
| >hg38_21_+_8433172_8446622_RNA45SN1_45S_50ntFlanks@12655.12675.21\|ATGACTGAACGCCTCTAAGTC | >hg38_21_+_8433172_8446622_RNA45SN1_45S_50ntFlanks | 12655 | 12675 | 21 | ATGACTGAACGCCTCTAAGTC | 8914.2 | -9.9 | 0.8 | -11.7 | 1.3E-31 | 1.0E-29 | MB231/MB468 | MITO |
| >hg38_21_+_8433172_8446622_RNA45SN1_45S_50ntFlanks@7984.8006.23\|AGATCAGACGTGGCGACCCGCTG | >hg38_21_+_8433172_8446622_RNA45SN1_45S_50ntFlanks | 7984 | 8006 | 23 | AGATCAGACGTGGCGACCCGCTG | 12777.7 | -17.4 | 1.5 | -11.7 | 1.4E-31 | 1.0E-29 | MB231/MB468 | MITO |
| >MI0000440\|hsa-mir-27b&WithFlank&9\|+\|95085439\|95085547@67.86.20(+1U)&[MIMAT0000419&hsa-miR-27b-3p&offsets\|0\|-1(+1U);m-39&9\|+\|95085505\|95085525&offsets\|0\|-1(+1U)]\|TTCACAGTGGCTAAGTTCTGT | >MI0000440\|hsa-mir-27b&WithFlank&9\|+\|95085439\|95085547 | 67 | 86 | 20(+1U) | TTCACAGTGGCTAAGTTCTGT | 7539.5 | 7.3 | 0.6 | 11.6 | 3.4E-31 | 1.0E-29 | BT20/MB468 | MITO |
| >hg38_21_+_8433172_8446622_RNA45SN1_45S_50ntFlanks@12625.12653.29\|CAATGGGGCGAAGCTACCATCTGTGGGAT | >hg38_21_+_8433172_8446622_RNA45SN1_45S_50ntFlanks | 12625 | 12653 | 29 | CAATGGGGCGAAGCTACCATCTGTGGGAT | 1958.7 | -5.7 | 0.5 | -11.6 | 3.4E-31 | 1.0E-29 | BT20/MB468 | MITO |
| >hg38_21_+_8433172_8446622_RNA45SN1_45S_50ntFlanks@10727.10755.29\|GGAGAGGGTGTAAATCTCGCGCCGGGCCG | >hg38_21_+_8433172_8446622_RNA45SN1_45S_50ntFlanks | 10727 | 10755 | 29 | GGAGAGGGTGTAAATCTCGCGCCGGGCCG | 3818.5 | -16.3 | 1.4 | -11.5 | 7.8E-31 | 1.1E-29 | BT20/MB468 | NUC |
| >hg38_21_+_8433172_8446622_RNA45SN1_45S_50ntFlanks@12157.12180.24\|AGTTTGACTGGGGCGGTACACCTG | >hg38_21_+_8433172_8446622_RNA45SN1_45S_50ntFlanks | 12157 | 12180 | 24 | AGTTTGACTGGGGCGGTACACCTG | 4654.2 | -16.0 | 1.4 | -11.8 | 6.8E-32 | 1.1E-29 | MB231/MB468 | MP |

<table>
<tr><td>>MI0000746|hsa-mir-99b&WithFlank&19|+|51692606|51692687@51.73.23&[MIMAT0004678&hsa-miR-99b-3p&offsets|0|+1;m-151&19|+|51692656|51692678&offsets|0|0]|CAAGCTCGTGTCTGTGGGTCCGT</td><td>>MI0000746|hsa-mir-99b&WithFlank&19|+|51692606|51692687</td><td>51</td><td>73</td><td>23</td><td>CAAGCTCGTGTCTGTGGGTCCGT</td><td>1024.4</td><td>7.7</td><td>0.7</td><td>11.6</td><td>3.6E-31</td><td>1.1E-29</td><td>BT20/MB468</td><td>MITO</td></tr>
<tr><td>>hg38_21_+_8433172_8446622_RNA45SN1_45S_50ntFlanks@11845.11872.28|GGAACGGGCTTGGCGGAATCAGCGGGGA</td><td>>hg38_21_+_8433172_8446622_RNA45SN1_45S_50ntFlanks</td><td>11845</td><td>11872</td><td>28</td><td>GGAACGGGCTTGGCGGAATCAGCGGGGA</td><td>936.7</td><td>-14.3</td><td>1.2</td><td>-11.5</td><td>8.2E-31</td><td>1.1E-29</td><td>BT20/MB468</td><td>NUC</td></tr>
<tr><td>>hg38_21_+_8433172_8446622_RNA45SN1_45S_50ntFlanks@4297.4321.25|CATTGGAGGGCAAGTCTGGTGCCAG</td><td>>hg38_21_+_8433172_8446622_RNA45SN1_45S_50ntFlanks</td><td>4297</td><td>4321</td><td>25</td><td>CATTGGAGGGCAAGTCTGGTGCCAG</td><td>692.4</td><td>-13.9</td><td>1.2</td><td>-11.5</td><td>8.5E-31</td><td>1.2E-29</td><td>BT20/MB468</td><td>NUC</td></tr>
<tr><td>>hg38_21_+_8433172_8446622_RNA45SN1_45S_50ntFlanks@12002.12032.31|TGCGGGCCGCCGGTGAAATACCACTACTCTG</td><td>>hg38_21_+_8433172_8446622_RNA45SN1_45S_50ntFlanks</td><td>12002</td><td>12032</td><td>31</td><td>TGCGGGCCGCCGGTGAAATACCACTACTCTG</td><td>4691.0</td><td>-16.1</td><td>1.4</td><td>-11.7</td><td>7.5E-32</td><td>1.2E-29</td><td>MB231/MB468</td><td>MP</td></tr>
<tr><td>>hg38_21_+_8433172_8446622_RNA45SN1_45S_50ntFlanks@10408.10426.19|GGTCAGTCGGTCCTGAGAG</td><td>>hg38_21_+_8433172_8446622_RNA45SN1_45S_50ntFlanks</td><td>10408</td><td>10426</td><td>19</td><td>GGTCAGTCGGTCCTGAGAG</td><td>2693.9</td><td>-15.8</td><td>1.4</td><td>-11.5</td><td>8.8E-31</td><td>1.2E-29</td><td>BT20/MB468</td><td>NUC</td></tr>
<tr><td>>MI0000273|hsa-mir-183&WithFlank&7|-|129774899|129775020@33.53.21(+1C)&[MIMAT0000261&hsa-miR-183-5p&offsets|0|-1(+1C);m-41&7|-|129774966|129774987&offsets|-1|-2(+1C)]|TATGGCACTGGTAGAATTCACC</td><td>>MI0000273|hsa-mir-183&WithFlank&7|-|129774899|129775020</td><td>33</td><td>53</td><td>21(+1C)</td><td>TATGGCACTGGTAGAATTCACC</td><td>1890.9</td><td>7.5</td><td>0.6</td><td>11.6</td><td>4.2E-31</td><td>1.3E-29</td><td>BT20/MB468</td><td>MITO</td></tr>
<tr><td>>hg38_21_+_8433172_8446622_RNA45SN1_45S_50ntFlanks@11670.11687.18|CCCAGTGCTCTGAATGTC</td><td>>hg38_21_+_8433172_8446622_RNA45SN1_45S_50ntFlanks</td><td>11670</td><td>11687</td><td>18</td><td>CCCAGTGCTCTGAATGTC</td><td>2137.9</td><td>-6.5</td><td>0.6</td><td>-11.5</td><td>9.5E-31</td><td>1.3E-29</td><td>BT20/MB468</td><td>NUC</td></tr>
<tr><td>>hg38_21_+_8433172_8446622_RNA45SN1_45S_50ntFlanks@10420.10443.24|CTGAGAGATGGGCGAGCGCCGTTC</td><td>>hg38_21_+_8433172_8446622_RNA45SN1_45S_50ntFlanks</td><td>10420</td><td>10443</td><td>24</td><td>CTGAGAGATGGGCGAGCGCCGTTC</td><td>1082.0</td><td>-14.5</td><td>1.3</td><td>-11.5</td><td>9.6E-31</td><td>1.3E-29</td><td>BT20/MB468</td><td>NUC</td></tr>
<tr><td>>hg38_21_+_8433172_8446622_RNA45SN1_45S_50ntFlanks@4555.4587.33|CGCAGCTAGGAATAATGGAATAGGACCGCGGTT</td><td>>hg38_21_+_8433172_8446622_RNA45SN1_45S_50ntFlanks</td><td>4555</td><td>4587</td><td>33</td><td>CGCAGCTAGGAATAATGGAATAGGACCGCGGTT</td><td>693.3</td><td>-13.9</td><td>1.2</td><td>-11.5</td><td>9.6E-31</td><td>1.3E-29</td><td>BT20/MB468</td><td>NUC</td></tr>
<tr><td>>hg38_21_+_8433172_8446622_RNA45SN1_45S_50ntFlanks@3771.3799.29|CACGGCCGGTACAGTGAAACTGCGAATGG</td><td>>hg38_21_+_8433172_8446622_RNA45SN1_45S_50ntFlanks</td><td>3771</td><td>3799</td><td>29</td><td>CACGGCCGGTACAGTGAAACTGCGAATGG</td><td>734.7</td><td>-13.9</td><td>1.2</td><td>-11.5</td><td>9.8E-31</td><td>1.3E-29</td><td>BT20/MB468</td><td>NUC</td></tr>
<tr><td>>MI0000746|hsa-mir-99b&WithFlank&19|+|51692606|51692687@51.71.21(+1A)&[MIMAT0004678&hsa-miR-99b-3p&offsets|0|-1(+1A);m-151&19|+|51692656|51692678&offsets|0|-2(+1A)]|CAAGCTCGTGTCTGTGGGTCCA</td><td>>MI0000746|hsa-mir-99b&WithFlank&19|+|51692606|51692687</td><td>51</td><td>71</td><td>21(+1A)</td><td>CAAGCTCGTGTCTGTGGGTCCA</td><td>1284.1</td><td>7.8</td><td>0.7</td><td>11.6</td><td>6.5E-31</td><td>1.4E-29</td><td>BT20/MB468</td><td>CYTO</td></tr>
<tr><td>>hg38_21_+_8433172_8446622_RNA45SN1_45S_50ntFlanks@7975.8002.28|CGCGACCTCAGATCAGACGTGGCGACCC</td><td>>hg38_21_+_8433172_8446622_RNA45SN1_45S_50ntFlanks</td><td>7975</td><td>8002</td><td>28</td><td>CGCGACCTCAGATCAGACGTGGCGACCC</td><td>1761.4</td><td>5.0</td><td>0.4</td><td>11.6</td><td>6.5E-31</td><td>1.4E-29</td><td>BT20/MB468</td><td>CYTO</td></tr>
<tr><td>>hg38_21_+_8433172_8446622_RNA45SN1_45S_50ntFlanks@8176.8202.27|CTTCTGATCGAGGCCCAGCCCGTGGAC</td><td>>hg38_21_+_8433172_8446622_RNA45SN1_45S_50ntFlanks</td><td>8176</td><td>8202</td><td>27</td><td>CTTCTGATCGAGGCCCAGCCCGTGGAC</td><td>1557.2</td><td>-3.5</td><td>0.3</td><td>-11.5</td><td>9.9E-31</td><td>1.4E-29</td><td>BT20/MB468</td><td>NUC</td></tr>
<tr><td>>hg38_21_+_8433172_8446622_RNA45SN1_45S_50ntFlanks@6738.6771.34|ATCATCGACACTTCGAACGCACTTGCGGCCCCGG</td><td>>hg38_21_+_8433172_8446622_RNA45SN1_45S_50ntFlanks</td><td>6738</td><td>6771</td><td>34</td><td>ATCATCGACACTTCGAACGCACTTGCGGCCCCGG</td><td>6672.7</td><td>-14.8</td><td>1.3</td><td>-11.6</td><td>4.6E-31</td><td>1.4E-29</td><td>BT20/MB468</td><td>MITO</td></tr>
<tr><td>>hg38_21_+_8433172_8446622_RNA45SN1_45S_50ntFlanks@11115.11133.19|CGGCAGGGGCCGGCGGCGG</td><td>>hg38_21_+_8433172_8446622_RNA45SN1_45S_50ntFlanks</td><td>11115</td><td>11133</td><td>19</td><td>CGGCAGGGGCCGGCGGCGG</td><td>1799.2</td><td>-15.2</td><td>1.3</td><td>-11.5</td><td>1.1E-30</td><td>1.4E-29</td><td>BT20/MB468</td><td>NUC</td></tr>
<tr><td>>hg38_21_+_8433172_8446622_RNA45SN1_45S_50ntFlanks@12044.12071.28|ACTGACCCGGTGAGGCGGGGGGGCGAGC</td><td>>hg38_21_+_8433172_8446622_RNA45SN1_45S_50ntFlanks</td><td>12044</td><td>12071</td><td>28</td><td>ACTGACCCGGTGAGGCGGGGGGGCGAGC</td><td>1283.4</td><td>-14.8</td><td>1.3</td><td>-11.5</td><td>1.1E-30</td><td>1.4E-29</td><td>BT20/MB468</td><td>NUC</td></tr>
<tr><td>>hg38_21_+_8433172_8446622_RNA45SN1_45S_50ntFlanks@11885.11904.20|TTGAGCTTGACTCTAGTCTG</td><td>>hg38_21_+_8433172_8446622_RNA45SN1_45S_50ntFlanks</td><td>11885</td><td>11904</td><td>20</td><td>TTGAGCTTGACTCTAGTCTG</td><td>832.1</td><td>-14.1</td><td>1.2</td><td>-11.5</td><td>1.1E-30</td><td>1.5E-29</td><td>BT20/MB468</td><td>NUC</td></tr>
<tr><td>>MI0000469|hsa-mir-125a&WithFlank&19|+|51693248|51693345@21.43.23&[MIMAT0000443&hsa-miR-125a-5p&offsets|0|-1;m-63&19|+|51693268|51693289&offsets|0|+1]|TCCCTGAGACCCTTTAACCTGTG</td><td>>MI0000469|hsa-mir-125a&WithFlank&19|+|51693248|51693345</td><td>21</td><td>43</td><td>23</td><td>TCCCTGAGACCCTTTAACCTGTG</td><td>53581.0</td><td>8.1</td><td>0.7</td><td>11.6</td><td>7.0E-31</td><td>1.5E-29</td><td>BT20/MB468</td><td>CYTO</td></tr>
<tr><td>>MI0000469|hsa-mir-125a&WithFlank&19|+|51693248|51693345@21.42.22&[MIMAT0000443&hsa-miR-125a-5p&offsets|0|-2;m-63&19|+|51693268|51693289&offsets|0|0]|TCCCTGAGACCCTTTAACCTGT</td><td>>MI0000469|hsa-mir-125a&WithFlank&19|+|51693248|51693345</td><td>21</td><td>42</td><td>22</td><td>TCCCTGAGACCCTTTAACCTGT</td><td>7864.1</td><td>7.2</td><td>0.6</td><td>11.6</td><td>5.0E-31</td><td>1.5E-29</td><td>BT20/MB468</td><td>MITO</td></tr>
<tr><td>>MI0000098|hsa-mir-96&WithFlank&7|-|129774686|129774775@15.36.22(+1C)&[MIMAT0000095&hsa-miR-96-5p&offsets|0|-1(+1C);m-248&7|-|129774739|129774761&offsets|0|-1(+1C)]|TTTGGCACTAGCACATTTTTGCC</td><td>>MI0000098|hsa-mir-96&WithFlank&7|-|129774686|129774775</td><td>15</td><td>36</td><td>22(+1C)</td><td>TTTGGCACTAGCACATTTTTGCC</td><td>624.1</td><td>7.3</td><td>0.6</td><td>11.6</td><td>5.2E-31</td><td>1.6E-29</td><td>BT20/MB468</td><td>MITO</td></tr>
<tr><td>>hg38_21_+_8433172_8446622_RNA45SN1_45S_50ntFlanks@9567.9597.31|ACGTGCAAATCGGTCGTCCGACCTGGGTATA</td><td>>hg38_21_+_8433172_8446622_RNA45SN1_45S_50ntFlanks</td><td>9567</td><td>9597</td><td>31</td><td>ACGTGCAAATCGGTCGTCCGACCTGGGTATA</td><td>2803.7</td><td>-7.2</td><td>0.6</td><td>-11.6</td><td>3.8E-31</td><td>1.6E-29</td><td>MB231/MB468</td><td>NUC</td></tr>
<tr><td>>hg38_21_+_8433172_8446622_RNA45SN1_45S_50ntFlanks@12991.13013.23|CTCCCTCGCTGCGATCTATTGAA</td><td>>hg38_21_+_8433172_8446622_RNA45SN1_45S_50ntFlanks</td><td>12991</td><td>13013</td><td>23</td><td>CTCCCTCGCTGCGATCTATTGAA</td><td>89741.1</td><td>-16.4</td><td>1.4</td><td>-11.5</td><td>1.2E-30</td><td>1.6E-29</td><td>BT20/MB468</td><td>NUC</td></tr>
<tr><td>>MI0000299|hsa-mir-222&WithFlank&X|-|45747009|45747130@75.99.25(+3U)&[MIMAT0000279&hsa-miR-222-3p&offsets|0|+4(+3U);m-140&X|-|45747033|45747056&offsets|0|+1(+3U)]|AGCTACATCTGGCTACTGGGTCTCTTTT</td><td>>MI0000299|hsa-mir-222&WithFlank&X|-|45747009|45747130</td><td>75</td><td>99</td><td>25(+3U)</td><td>AGCTACATCTGGCTACTGGGTCTCTTTT</td><td>2874.4</td><td>8.9</td><td>0.8</td><td>11.8</td><td>6.1E-32</td><td>1.6E-29</td><td>MB231/MB468</td><td>CYTO</td></tr>
<tr><td>>hg38_21_+_8433172_8446622_RNA45SN1_45S_50ntFlanks@6723.6744.22|TTGCAGGACACATTGATCATCG</td><td>>hg38_21_+_8433172_8446622_RNA45SN1_45S_50ntFlanks</td><td>6723</td><td>6744</td><td>22</td><td>TTGCAGGACACATTGATCATCG</td><td>1375.4</td><td>-14.9</td><td>1.3</td><td>-11.5</td><td>1.2E-30</td><td>1.6E-29</td><td>BT20/MB468</td><td>NUC</td></tr>
<tr><td>>hg38_21_+_8433172_8446622_RNA45SN1_45S_50ntFlanks@10679.10711.33|GGTTCCGGCGGCGTCCGGTGAGCTCTCGCTGGC</td><td>>hg38_21_+_8433172_8446622_RNA45SN1_45S_50ntFlanks</td><td>10679</td><td>10711</td><td>33</td><td>GGTTCCGGCGGCGTCCGGTGAGCTCTCGCTGGC</td><td>8825.4</td><td>-9.5</td><td>0.8</td><td>-11.6</td><td>4.1E-31</td><td>1.7E-29</td><td>MB231/MB468</td><td>NUC</td></tr>
<tr><td>>hg38_21_+_8433172_8446622_RNA45SN1_45S_50ntFlanks@12934.12956.23|CTAAACCATTCGTAGACGACCTG</td><td>>hg38_21_+_8433172_8446622_RNA45SN1_45S_50ntFlanks</td><td>12934</td><td>12956</td><td>23</td><td>CTAAACCATTCGTAGACGACCTG</td><td>33999.9</td><td>-15.2</td><td>1.3</td><td>-11.5</td><td>1.2E-30</td><td>1.7E-29</td><td>BT20/MB468</td><td>NUC</td></tr>
<tr><td>>hg38_21_+_8433172_8446622_RNA45SN1_45S_50ntFlanks@8168.8187.20|CCCAAGTCCTTCTGATCGAG</td><td>>hg38_21_+_8433172_8446622_RNA45SN1_45S_50ntFlanks</td><td>8168</td><td>8187</td><td>20</td><td>CCCAAGTCCTTCTGATCGAG</td><td>2196.5</td><td>-15.5</td><td>1.3</td><td>-11.5</td><td>1.2E-30</td><td>1.7E-29</td><td>BT20/MB468</td><td>NUC</td></tr>
<tr><td>>hg38_21_+_8433172_8446622_RNA45SN1_45S_50ntFlanks@7978.8009.32|GACCTCAGATCAGACGTGGCGACCCGCTGAAT</td><td>>hg38_21_+_8433172_8446622_RNA45SN1_45S_50ntFlanks</td><td>7978</td><td>8009</td><td>32</td><td>GACCTCAGATCAGACGTGGCGACCCGCTGAAT</td><td>618.4</td><td>5.3</td><td>0.5</td><td>11.6</td><td>5.7E-31</td><td>1.7E-29</td><td>BT20/MB468</td><td>MITO</td></tr>
<tr><td>>MI0000433|hsa-let-7g&WithFlank&3|-|52268272|52268367@11.31.21(+1A)&[MIMAT0000414&hsa-let-7g-5p&offsets|0|-1(+1A);m-58&3|-|52268336|52268357&offsets|0|-1(+1A)]|TGAGGTAGTAGTTTGTACAGTA</td><td>>MI0000433|hsa-let-7g&WithFlank&3|-|52268272|52268367</td><td>11</td><td>31</td><td>21(+1A)</td><td>TGAGGTAGTAGTTTGTACAGTA</td><td>2281.1</td><td>7.5</td><td>0.6</td><td>11.6</td><td>5.8E-31</td><td>1.7E-29</td><td>BT20/MB468</td><td>MITO</td></tr>
<tr><td>>hg38_21_+_8433172_8446622_RNA45SN1_45S_50ntFlanks@11561.11586.26|GCGCCTAGCAGCCGACTTAGAACTGG</td><td>>hg38_21_+_8433172_8446622_RNA45SN1_45S_50ntFlanks</td><td>11561</td><td>11586</td><td>26</td><td>GCGCCTAGCAGCCGACTTAGAACTGG</td><td>1515.3</td><td>-15.0</td><td>1.3</td><td>-11.5</td><td>1.3E-30</td><td>1.7E-29</td><td>BT20/MB468</td><td>NUC</td></tr>
<tr><td>>MI0000082|hsa-mir-25&WithFlank&7|-|100093554|100093649@58.79.22(+2A)&[MIMAT0000081&hsa-miR-25-3p&offsets|0|0(+2A);m-23&7|-|100093571|100093592&offsets|0|0(+2A)]|CATTGCACTTGTCTCGGTCTGAAA</td><td>>MI0000082|hsa-mir-25&WithFlank&7|-|100093554|100093649</td><td>58</td><td>79</td><td>22(+2A)</td><td>CATTGCACTTGTCTCGGTCTGAAA</td><td>628.8</td><td>6.7</td><td>0.6</td><td>11.6</td><td>6.0E-31</td><td>1.8E-29</td><td>BT20/MB468</td><td>MITO</td></tr>
<tr><td>>hg38_21_+_8433172_8446622_RNA45SN1_45S_50ntFlanks@9567.9596.30|ACGTGCAAATCGGTCGTCCGACCTGGGTAT</td><td>>hg38_21_+_8433172_8446622_RNA45SN1_45S_50ntFlanks</td><td>9567</td><td>9596</td><td>30</td><td>ACGTGCAAATCGGTCGTCCGACCTGGGTAT</td><td>3828.2</td><td>-7.9</td><td>0.7</td><td>-11.6</td><td>4.4E-31</td><td>1.8E-29</td><td>MB231/MB468</td><td>NUC</td></tr>
<tr><td>>hg38_21_+_8433172_8446622_RNA45SN1_45S_50ntFlanks@6660.6681.22|GCGGTGGATCACTCGGCTCGTG</td><td>>hg38_21_+_8433172_8446622_RNA45SN1_45S_50ntFlanks</td><td>6660</td><td>6681</td><td>22</td><td>GCGGTGGATCACTCGGCTCGTG</td><td>1096.5</td><td>-5.5</td><td>0.5</td><td>-11.5</td><td>1.3E-30</td><td>1.8E-29</td><td>BT20/MB468</td><td>NUC</td></tr>
<tr><td>>hg38_21_+_8433172_8446622_RNA45SN1_45S_50ntFlanks@6746.6770.25|CACTTCGAACGCACTTGCGGCCCCG</td><td>>hg38_21_+_8433172_8446622_RNA45SN1_45S_50ntFlanks</td><td>6746</td><td>6770</td><td>25</td><td>CACTTCGAACGCACTTGCGGCCCCG</td><td>1377.7</td><td>-14.9</td><td>1.3</td><td>-11.5</td><td>1.4E-30</td><td>1.8E-29</td><td>BT20/MB468</td><td>NUC</td></tr>
<tr><td>>MI0000285|hsa-mir-205&WithFlank&1|+|209432127|209432248@40.62.23(+1A)&[MIMAT0000266&hsa-miR-205-5p&offsets|0|+1(+1A);m-351&1|+|209432166|209432187&offsets|0|+1(+1A)]|TCCTTCATTCCACCGGAGTCTGTA</td><td>>MI0000285|hsa-mir-205&WithFlank&1|+|209432127|209432248</td><td>40</td><td>62</td><td>23(+1A)</td><td>TCCTTCATTCCACCGGAGTCTGTA</td><td>6033.4</td><td>7.2</td><td>0.6</td><td>11.5</td><td>9.3E-31</td><td>1.9E-29</td><td>BT20/MB468</td><td>CYTO</td></tr>
<tr><td>>hg38_21_+_8433172_8446622_RNA45SN1_45S_50ntFlanks@11718.11738.21|AACGGCGGGAGTAACTATGAC</td><td>>hg38_21_+_8433172_8446622_RNA45SN1_45S_50ntFlanks</td><td>11718</td><td>11738</td><td>21</td><td>AACGGCGGGAGTAACTATGAC</td><td>2904.2</td><td>-15.9</td><td>1.4</td><td>-11.5</td><td>1.4E-30</td><td>1.9E-29</td><td>BT20/MB468</td><td>NUC</td></tr>
<tr><td>>MI0000650|hsa-mir-200c&WithFlank&12|+|6963693|6963772@49.70.22&[MIMAT0000617&hsa-miR-200c-3p&offsets|-1|-2;m-54&12|+|6963742|6963764&offsets|-1|-2]|CTAATACTGCCGGGTAATGATG</td><td>>MI0000650|hsa-mir-200c&WithFlank&12|+|6963693|6963772</td><td>49</td><td>70</td><td>22</td><td>CTAATACTGCCGGGTAATGATG</td><td>115.3</td><td>3.8</td><td>0.3</td><td>11.5</td><td>1.4E-30</td><td>1.9E-29</td><td>BT20/MB468</td><td>NUC</td></tr>
<tr><td>>hg38_21_+_8433172_8446622_RNA45SN1_45S_50ntFlanks@11784.11806.23|ATGAATGGATGAACGAGATTCCC</td><td>>hg38_21_+_8433172_8446622_RNA45SN1_45S_50ntFlanks</td><td>11784</td><td>11806</td><td>23</td><td>ATGAATGGATGAACGAGATTCCC</td><td>835.0</td><td>-14.1</td><td>1.2</td><td>-11.5</td><td>1.5E-30</td><td>2.0E-29</td><td>BT20/MB468</td><td>NUC</td></tr>
<tr><td>>hg38_21_+_8433172_8446622_RNA45SN1_45S_50ntFlanks@8182.8204.23|ATCGAGGCCCAGCCCGTGGACGG</td><td>>hg38_21_+_8433172_8446622_RNA45SN1_45S_50ntFlanks</td><td>8182</td><td>8204</td><td>23</td><td>ATCGAGGCCCAGCCCGTGGACGG</td><td>798.7</td><td>-14.1</td><td>1.2</td><td>-11.5</td><td>1.6E-30</td><td>2.1E-29</td><td>BT20/MB468</td><td>NUC</td></tr>
<tr><td>>MI0000746|hsa-mir-99b&WithFlank&19|+|51692606|51692687@13.32.20(+1A)&[MIMAT0000689&hsa-miR-99b-5p&offsets|0|-2(+1A);m-10&19|+|51692618|51692639&offsets|0|-2(+1A)]|CACCCGTAGAACCGACCTTGA</td><td>>MI0000746|hsa-mir-99b&WithFlank&19|+|51692606|51692687</td><td>13</td><td>32</td><td>20(+1A)</td><td>CACCCGTAGAACCGACCTTGA</td><td>511.1</td><td>7.2</td><td>0.6</td><td>11.6</td><td>7.3E-31</td><td>2.1E-29</td><td>BT20/MB468</td><td>MITO</td></tr>
<tr><td>>MI0000439|hsa-mir-23b&WithFlank&9|+|95085202|95085310@67.85.19(+1U)&[MIMAT0000418&hsa-miR-23b-3p&offsets|+3|-1(+1U);m-42&9|+|95085265|95085285&offsets|+3|+1(+1U)]|ACATTGCCAGGGATTACCAT</td><td>>MI0000439|hsa-mir-23b&WithFlank&9|+|95085202|95085310</td><td>67</td><td>85</td><td>19(+1U)</td><td>ACATTGCCAGGGATTACCAT</td><td>2576.5</td><td>15.6</td><td>1.4</td><td>11.5</td><td>1.0E-30</td><td>2.1E-29</td><td>BT20/MB468</td><td>CYTO</td></tr>
<tr><td>>trna116_GluCTC_1_-_145399233_145399304@32.71.40,trna71_GluCTC_1_-_161439189_161439260@32.71.40,trna74_GluCTC_1_-_161431809_161431880@32.71.40,trna77_GluCTC_1_-_161424398_161424469@32.71.40,trna77_GluCTC_6_+_28949976_28950047@32.71.40,trna80_GluCTC_1_-_161417018_161417089@32.71.40,trna87_GluCTC_6_-_126101393_126101464@32.71.40|CTCTCACCGCCGCGGCCCGGGTTCGATTCCCGGTCAGGGA</td><td>>trna116_GluCTC_1_-_145399233_145399304</td><td>32</td><td>71</td><td>40</td><td>CTCTCACCGCCGCGGCCCGGGTTCGATTCCCGGTCAGGGA</td><td>1813.3</td><td>-15.3</td><td>1.3</td><td>-11.5</td><td>1.7E-30</td><td>2.2E-29</td><td>BT20/MB468</td><td>NUC</td></tr>
<tr><td>>hg38_21_+_8433172_8446622_RNA45SN1_45S_50ntFlanks@11315.11344.30|CCACCCCACGTCTCGTCGCGCGCGCGTCCG</td><td>>hg38_21_+_8433172_8446622_RNA45SN1_45S_50ntFlanks</td><td>11315</td><td>11344</td><td>30</td><td>CCACCCCACGTCTCGTCGCGCGCGCGTCCG</td><td>9356.7</td><td>-16.2</td><td>1.4</td><td>-11.6</td><td>5.7E-31</td><td>2.3E-29</td><td>MB231/MB468</td><td>NUC</td></tr>
<tr><td>>hg38_21_+_8433172_8446622_RNA45SN1_45S_50ntFlanks@10395.10424.30|GCAGTTGAACATGGGTCAGTCGGTCCTGAG</td><td>>hg38_21_+_8433172_8446622_RNA45SN1_45S_50ntFlanks</td><td>10395</td><td>10424</td><td>30</td><td>GCAGTTGAACATGGGTCAGTCGGTCCTGAG</td><td>15566.7</td><td>8.1</td><td>0.7</td><td>11.5</td><td>8.2E-31</td><td>2.4E-29</td><td>BT20/MB468</td><td>MITO</td></tr>
<tr><td>>hg38_21_+_8433172_8446622_RNA45SN1_45S_50ntFlanks@12934.12956.23|CTAAACCATTCGTAGACGACCTG</td><td>>hg38_21_+_8433172_8446622_RNA45SN1_45S_50ntFlanks</td><td>12934</td><td>12956</td><td>23</td><td>CTAAACCATTCGTAGACGACCTG</td><td>4580.3</td><td>-16.0</td><td>1.4</td><td>-11.7</td><td>1.5E-31</td><td>2.4E-29</td><td>MB231/MB468</td><td>MP</td></tr>
</table>

<table>
<tr><td>>MI0000285|hsa-mir-205&WithFlank&1|+|209432127|209432248@40.61.22(+1A)&[MIMAT0000266&hsa-miR-205-5p&offsets|0|0(+1A);m-351&1|+|209432166|209432187&offsets|0|0(+1A)]|TCCTTCATTCCACCGGAGTCTGA</td><td>>MI0000285|hsa-mir-205&WithFlank&1|+|209432127|209432248</td><td>40</td><td>61</td><td>22(+1A)</td><td>TCCTTCATTCCACCGGAGTCTGA</td><td>3270.9</td><td>-15.5</td><td>1.3</td><td>-11.7</td><td>1.6E-31</td><td>2.5E-29</td><td>MB231/MB468</td><td>MP</td></tr>
<tr><td>>hg38_21_+_8433172_8446622_RNA45SN1_45S_50ntFlanks@6663.6681.19|GTGGATCACTCGGCTCGTG</td><td>>hg38_21_+_8433172_8446622_RNA45SN1_45S_50ntFlanks</td><td>6663</td><td>6681</td><td>19</td><td>GTGGATCACTCGGCTCGTG</td><td>6678.5</td><td>-16.6</td><td>1.4</td><td>-11.7</td><td>1.7E-31</td><td>2.6E-29</td><td>MB231/MB468</td><td>MP</td></tr>
<tr><td>>MI0003834|hsa-mir-769&WithFlank&19|+|46018926|46019055@36.57.22&[MIMAT0003886&hsa-miR-769-5p&offsets|0|0;m-216&19|+|46018961|46018982&offsets|0|0]|TGAGACCTCTGGGTTCTGAGCT</td><td>>MI0003834|hsa-mir-769&WithFlank&19|+|46018926|46019055</td><td>36</td><td>57</td><td>22</td><td>TGAGACCTCTGGGTTCTGAGCT</td><td>803.1</td><td>6.2</td><td>0.5</td><td>11.5</td><td>9.0E-31</td><td>2.6E-29</td><td>BT20/MB468</td><td>MITO</td></tr>
<tr><td>>hg38_21_+_8433172_8446622_RNA45SN1_45S_50ntFlanks@3837.3868.32|CCTCTCCTACTTGGATAACTGTGGTAATTCTA</td><td>>hg38_21_+_8433172_8446622_RNA45SN1_45S_50ntFlanks</td><td>3837</td><td>3868</td><td>32</td><td>CCTCTCCTACTTGGATAACTGTGGTAATTCTA</td><td>969.0</td><td>-14.3</td><td>1.3</td><td>-11.5</td><td>2.0E-30</td><td>2.6E-29</td><td>BT20/MB468</td><td>NUC</td></tr>
<tr><td>>hg38_21_+_8433172_8446622_RNA45SN1_45S_50ntFlanks@12893.12933.41|CCTCGCCCGTCACGCACCGCACGTTCGTGGGGAACCTGGCG</td><td>>hg38_21_+_8433172_8446622_RNA45SN1_45S_50ntFlanks</td><td>12893</td><td>12933</td><td>41</td><td>CCTCGCCCGTCACGCACCGCACGTTCGTGGGGAACCTGGCG</td><td>6916.4</td><td>-16.5</td><td>1.4</td><td>-11.6</td><td>3.7E-31</td><td>2.7E-29</td><td>MB231/MB468</td><td>MITO</td></tr>
<tr><td>>hg38_21_+_8433172_8446622_RNA45SN1_45S_50ntFlanks@6733.6770.38|CATTGATCATCGACACTTCGAACGCACTTGCGGCCCCG</td><td>>hg38_21_+_8433172_8446622_RNA45SN1_45S_50ntFlanks</td><td>6733</td><td>6770</td><td>38</td><td>CATTGATCATCGACACTTCGAACGCACTTGCGGCCCCG</td><td>6057.1</td><td>-7.4</td><td>0.6</td><td>-11.6</td><td>3.7E-31</td><td>2.7E-29</td><td>MB231/MB468</td><td>MITO</td></tr>
<tr><td>>hg38_21_+_8433172_8446622_RNA45SN1_45S_50ntFlanks@12406.12434.29|TTTTTGATCCTTCGATGTCGGCTCTTCCT</td><td>>hg38_21_+_8433172_8446622_RNA45SN1_45S_50ntFlanks</td><td>12406</td><td>12434</td><td>29</td><td>TTTTTGATCCTTCGATGTCGGCTCTTCCT</td><td>4419.7</td><td>-6.7</td><td>0.6</td><td>-11.5</td><td>2.0E-30</td><td>2.7E-29</td><td>BT20/MB468</td><td>NUC</td></tr>
<tr><td>>hg38_21_+_8433172_8446622_RNA45SN1_45S_50ntFlanks@3985.4013.29|CGGCTTTGGTGACTCTAGATAACCTCGGG</td><td>>hg38_21_+_8433172_8446622_RNA45SN1_45S_50ntFlanks</td><td>3985</td><td>4013</td><td>29</td><td>CGGCTTTGGTGACTCTAGATAACCTCGGG</td><td>1141.3</td><td>-14.6</td><td>1.3</td><td>-11.5</td><td>2.1E-30</td><td>2.7E-29</td><td>BT20/MB468</td><td>NUC</td></tr>
<tr><td>>hg38_21_+_8433172_8446622_RNA45SN1_45S_50ntFlanks@4283.4307.25|CTTTAACGAGGATCCATTGGAGGGC</td><td>>hg38_21_+_8433172_8446622_RNA45SN1_45S_50ntFlanks</td><td>4283</td><td>4307</td><td>25</td><td>CTTTAACGAGGATCCATTGGAGGGC</td><td>938.3</td><td>-14.3</td><td>1.2</td><td>-11.5</td><td>2.1E-30</td><td>2.7E-29</td><td>BT20/MB468</td><td>NUC</td></tr>
<tr><td>>hg38_21_+_8433172_8446622_RNA45SN1_45S_50ntFlanks@9564.9596.33|CTGACGTGCAAATCGGTCGTCCGACCTGGGTAT</td><td>>hg38_21_+_8433172_8446622_RNA45SN1_45S_50ntFlanks</td><td>9564</td><td>9596</td><td>33</td><td>CTGACGTGCAAATCGGTCGTCCGACCTGGGTAT</td><td>34895.6</td><td>-8.0</td><td>0.7</td><td>-11.6</td><td>7.0E-31</td><td>2.8E-29</td><td>MB231/MB468</td><td>NUC</td></tr>
<tr><td>>MI0001733|hsa-mir-452&WithFlank&X|-|151959622|151959718@20.42.23&[MIMAT0001635&hsa-miR-452-5p&offsets|0|+1;m-188&X|-|151959677|151959699&offsets|0|0]|AACTGTTTGCAGAGGAAACTGAG</td><td>>MI0001733|hsa-mir-452&WithFlank&X|-|151959622|151959718</td><td>20</td><td>42</td><td>23</td><td>AACTGTTTGCAGAGGAAACTGAG</td><td>1110.2</td><td>14.6</td><td>1.3</td><td>11.5</td><td>9.6E-31</td><td>2.8E-29</td><td>BT20/MB468</td><td>MITO</td></tr>
<tr><td>>hg38_21_+_8433172_8446622_RNA45SN1_45S_50ntFlanks@6738.6772.35|ATCATCGACACTTCGAACGCACTTGCGGCCCCGGG</td><td>>hg38_21_+_8433172_8446622_RNA45SN1_45S_50ntFlanks</td><td>6738</td><td>6772</td><td>35</td><td>ATCATCGACACTTCGAACGCACTTGCGGCCCCGGG</td><td>1732.3</td><td>-15.2</td><td>1.3</td><td>-11.5</td><td>2.1E-30</td><td>2.8E-29</td><td>BT20/MB468</td><td>NUC</td></tr>
<tr><td>>hg38_21_+_8433172_8446622_RNA45SN1_45S_50ntFlanks@11766.11803.38|ATCTAATTAGTGACGCGCATGAATGGATGAACGAGATT</td><td>>hg38_21_+_8433172_8446622_RNA45SN1_45S_50ntFlanks</td><td>11766</td><td>11803</td><td>38</td><td>ATCTAATTAGTGACGCGCATGAATGGATGAACGAGATT</td><td>930.0</td><td>-14.3</td><td>1.2</td><td>-11.5</td><td>2.2E-30</td><td>2.8E-29</td><td>BT20/MB468</td><td>NUC</td></tr>
<tr><td>>hg38_21_+_8433172_8446622_RNA45SN1_45S_50ntFlanks@9568.9597.30|CGTGCAAATCGGTCGTCCGACCTGGGTATA</td><td>>hg38_21_+_8433172_8446622_RNA45SN1_45S_50ntFlanks</td><td>9568</td><td>9597</td><td>30</td><td>CGTGCAAATCGGTCGTCCGACCTGGGTATA</td><td>4392.0</td><td>-16.0</td><td>1.4</td><td>-11.7</td><td>1.9E-31</td><td>2.9E-29</td><td>MB231/MB468</td><td>MP</td></tr>
<tr><td>>hg38_21_+_8433172_8446622_RNA45SN1_45S_50ntFlanks@3831.3860.30|CTCGCTCCTCTCCTACTTGGATAACTGTGG</td><td>>hg38_21_+_8433172_8446622_RNA45SN1_45S_50ntFlanks</td><td>3831</td><td>3860</td><td>30</td><td>CTCGCTCCTCTCCTACTTGGATAACTGTGG</td><td>1881.8</td><td>-5.6</td><td>0.5</td><td>-11.5</td><td>7.6E-31</td><td>3.0E-29</td><td>MB231/MB468</td><td>NUC</td></tr>
<tr><td>>hg38_21_+_8433172_8446622_RNA45SN1_45S_50ntFlanks@4367.4396.30|CAGTTAAAAAGCTCGTAGTTGGATCTTGGG</td><td>>hg38_21_+_8433172_8446622_RNA45SN1_45S_50ntFlanks</td><td>4367</td><td>4396</td><td>30</td><td>CAGTTAAAAAGCTCGTAGTTGGATCTTGGG</td><td>871.9</td><td>-14.2</td><td>1.2</td><td>-11.4</td><td>2.4E-30</td><td>3.1E-29</td><td>BT20/MB468</td><td>NUC</td></tr>
<tr><td>>hg38_21_+_8433172_8446622_RNA45SN1_45S_50ntFlanks@12456.12477.22|CCAAGCGTTGGATTGTTCACCC</td><td>>hg38_21_+_8433172_8446622_RNA45SN1_45S_50ntFlanks</td><td>12456</td><td>12477</td><td>22</td><td>CCAAGCGTTGGATTGTTCACCC</td><td>6211.5</td><td>-8.1</td><td>0.7</td><td>-11.6</td><td>4.6E-31</td><td>3.2E-29</td><td>MB231/MB468</td><td>MITO</td></tr>
<tr><td>>hg38_21_+_8433172_8446622_RNA45SN1_45S_50ntFlanks@5484.5514.31|GAGCGCTGAGAAGACGGTCGAACTTGACTAT</td><td>>hg38_21_+_8433172_8446622_RNA45SN1_45S_50ntFlanks</td><td>5484</td><td>5514</td><td>31</td><td>GAGCGCTGAGAAGACGGTCGAACTTGACTAT</td><td>9143.9</td><td>-11.2</td><td>1.0</td><td>-11.6</td><td>4.6E-31</td><td>3.2E-29</td><td>MB231/MB468</td><td>MITO</td></tr>
<tr><td>>hg38_21_+_8433172_8446622_RNA45SN1_45S_50ntFlanks@9570.9596.27|TGCAAATCGGTCGTCCGACCTGGGTAT</td><td>>hg38_21_+_8433172_8446622_RNA45SN1_45S_50ntFlanks</td><td>9570</td><td>9596</td><td>27</td><td>TGCAAATCGGTCGTCCGACCTGGGTAT</td><td>1752.2</td><td>-15.2</td><td>1.3</td><td>-11.4</td><td>2.5E-30</td><td>3.3E-29</td><td>BT20/MB468</td><td>NUC</td></tr>
<tr><td>>hg38_21_+_8433172_8446622_RNA45SN1_45S_50ntFlanks@11578.11603.26|TAGAACTGGTGCGGACCAGGGGAATC</td><td>>hg38_21_+_8433172_8446622_RNA45SN1_45S_50ntFlanks</td><td>11578</td><td>11603</td><td>26</td><td>TAGAACTGGTGCGGACCAGGGGAATC</td><td>1628.9</td><td>-7.2</td><td>0.6</td><td>-11.4</td><td>2.5E-30</td><td>3.3E-29</td><td>BT20/MB468</td><td>NUC</td></tr>
<tr><td>>hg38_21_+_8433172_8446622_RNA45SN1_45S_50ntFlanks@12007.12032.26|GCCGCCGGTGAAATACCACTACTCTG</td><td>>hg38_21_+_8433172_8446622_RNA45SN1_45S_50ntFlanks</td><td>12007</td><td>12032</td><td>26</td><td>GCCGCCGGTGAAATACCACTACTCTG</td><td>23449.8</td><td>-8.9</td><td>0.8</td><td>-11.6</td><td>4.8E-31</td><td>3.3E-29</td><td>MB231/MB468</td><td>MITO</td></tr>
<tr><td>>hg38_21_+_8433172_8446622_RNA45SN1_45S_50ntFlanks@12007.12043.37|GCCGCCGGTGAAATACCACTACTCTGATCGTTTTTTC</td><td>>hg38_21_+_8433172_8446622_RNA45SN1_45S_50ntFlanks</td><td>12007</td><td>12043</td><td>37</td><td>GCCGCCGGTGAAATACCACTACTCTGATCGTTTTTTC</td><td>10541.2</td><td>-9.2</td><td>0.8</td><td>-11.6</td><td>4.7E-31</td><td>3.3E-29</td><td>MB231/MB468</td><td>MITO</td></tr>
<tr><td>>hg38_21_+_8433172_8446622_RNA45SN1_45S_50ntFlanks@8105.8143.39|CGCGGCGGGGCGCGGGACATGTGGCGTACGGAAGACCCG</td><td>>hg38_21_+_8433172_8446622_RNA45SN1_45S_50ntFlanks</td><td>8105</td><td>8143</td><td>39</td><td>CGCGGCGGGGCGCGGGACATGTGGCGTACGGAAGACCCG</td><td>47183.3</td><td>-15.2</td><td>1.3</td><td>-11.6</td><td>4.8E-31</td><td>3.3E-29</td><td>MB231/MB468</td><td>MITO</td></tr>
<tr><td>>hg38_21_+_8433172_8446622_RNA45SN1_45S_50ntFlanks@4722.4758.37|TCATTAATCAAGAACGAAAGTCGGAGGTTCGAAGACG</td><td>>hg38_21_+_8433172_8446622_RNA45SN1_45S_50ntFlanks</td><td>4722</td><td>4758</td><td>37</td><td>TCATTAATCAAGAACGAAAGTCGGAGGTTCGAAGACG</td><td>16654.7</td><td>-11.3</td><td>1.0</td><td>-11.6</td><td>5.1E-31</td><td>3.5E-29</td><td>MB231/MB468</td><td>MITO</td></tr>
<tr><td>>hg38_21_+_8433172_8446622_RNA45SN1_45S_50ntFlanks@482.505.24|ATCGATGTGGTGACGTCGTGCTCT</td><td>>hg38_21_+_8433172_8446622_RNA45SN1_45S_50ntFlanks</td><td>482</td><td>505</td><td>24</td><td>ATCGATGTGGTGACGTCGTGCTCT</td><td>1481.0</td><td>-15.0</td><td>1.3</td><td>-11.4</td><td>2.7E-30</td><td>3.5E-29</td><td>BT20/MB468</td><td>NUC</td></tr>
<tr><td>>hg38_21_+_8433172_8446622_RNA45SN1_45S_50ntFlanks@10680.10709.30|GTTCCGGCGGCGTCCGGTGAGCTCTCGCTG</td><td>>hg38_21_+_8433172_8446622_RNA45SN1_45S_50ntFlanks</td><td>10680</td><td>10709</td><td>30</td><td>GTTCCGGCGGCGTCCGGTGAGCTCTCGCTG</td><td>3029.3</td><td>-7.1</td><td>0.6</td><td>-11.5</td><td>9.4E-31</td><td>3.7E-29</td><td>MB231/MB468</td><td>NUC</td></tr>
<tr><td>>trnaMT_PheGAA_MT_+_577_647@35.54.20|GAAAATGTTTAGACGGGCTC</td><td>>trnaMT_PheGAA_MT_+_577_647</td><td>35</td><td>54</td><td>20</td><td>GAAAATGTTTAGACGGGCTC</td><td>1588.5</td><td>8.2</td><td>0.7</td><td>11.5</td><td>1.3E-30</td><td>3.9E-29</td><td>BT20/MB468</td><td>MITO</td></tr>
<tr><td>>MI0000253|hsa-mir-148a&WithFlank&7|-|25949913|25949992@50.71.22(+1U)&[MIMAT0000243&hsa-miR-148a-3p&offsets|0|0(+1U);m-7&7|-|25949922|25949943&offsets|0|0(+1U)]|TCAGTGCACTACAGAACTTTGTT</td><td>>MI0000253|hsa-mir-148a&WithFlank&7|-|25949913|25949992</td><td>50</td><td>71</td><td>22(+1U)</td><td>TCAGTGCACTACAGAACTTTGTT</td><td>1128.9</td><td>8.1</td><td>0.7</td><td>11.5</td><td>1.9E-30</td><td>4.0E-29</td><td>BT20/MB468</td><td>CYTO</td></tr>
<tr><td>>hg38_21_+_8433172_8446622_RNA45SN1_45S_50ntFlanks@12406.12436.31|TTTTTGATCCTTCGATGTCGGCTCTTCCTAT</td><td>>hg38_21_+_8433172_8446622_RNA45SN1_45S_50ntFlanks</td><td>12406</td><td>12436</td><td>31</td><td>TTTTTGATCCTTCGATGTCGGCTCTTCCTAT</td><td>4140.1</td><td>-16.4</td><td>1.4</td><td>-11.4</td><td>3.1E-30</td><td>4.0E-29</td><td>BT20/MB468</td><td>NUC</td></tr>
<tr><td>>hg38_21_+_8433172_8446622_RNA45SN1_45S_50ntFlanks@12008.12030.23|CCGCCGGTGAAATACCACTACTC</td><td>>hg38_21_+_8433172_8446622_RNA45SN1_45S_50ntFlanks</td><td>12008</td><td>12030</td><td>23</td><td>CCGCCGGTGAAATACCACTACTC</td><td>6777.0</td><td>-16.6</td><td>1.4</td><td>-11.6</td><td>2.7E-31</td><td>4.0E-29</td><td>MB231/MB468</td><td>MP</td></tr>
<tr><td>>hg38_21_+_8433172_8446622_RNA45SN1_45S_50ntFlanks@12044.12072.29|ACTGACCCGGTGAGGCGGGGGGGCGAGCC</td><td>>hg38_21_+_8433172_8446622_RNA45SN1_45S_50ntFlanks</td><td>12044</td><td>12072</td><td>29</td><td>ACTGACCCGGTGAGGCGGGGGGGCGAGCC</td><td>1829.1</td><td>-15.3</td><td>1.3</td><td>-11.4</td><td>3.1E-30</td><td>4.0E-29</td><td>BT20/MB468</td><td>NUC</td></tr>
<tr><td>>hg38_21_+_8433172_8446622_RNA45SN1_45S_50ntFlanks@11890.11912.23|CTTGACTCTAGTCTGGCACGGTG</td><td>>hg38_21_+_8433172_8446622_RNA45SN1_45S_50ntFlanks</td><td>11890</td><td>11912</td><td>23</td><td>CTTGACTCTAGTCTGGCACGGTG</td><td>711.5</td><td>-13.9</td><td>1.2</td><td>-11.4</td><td>3.2E-30</td><td>4.1E-29</td><td>BT20/MB468</td><td>NUC</td></tr>
<tr><td>>MI0000061|hsa-let-7a-2&WithFlank&11|-|122146516|122146599@11.33.23;MI0000062|hsa-let-7a-3&WithFlank&22|+|46112743|46112828@10.32.23;MI0000060|hsa-let-7a-1&WithFlank&9|+|94175951|94176042@12.34.23&[MIMAT0000062&hsa-let-7a-5p&offsets|0|+1;m-6&11|-|122146568|122146589&offsets|0|+1];[MIMAT0000062_1&hsa-let-7a-5p&offsets|0|+1;m-5&22|+|46112752|46112773&offsets|0|+1];[MIMAT0000062_2&hsa-let-7a-5p&offsets|0|+1;m-4&9|+|94175962|94175983&offsets|0|+1]|TGAGGTAGTAGGTTGTATAGTTT</td><td>>MI0000061|hsa-let-7a-2&WithFlank&11|-|122146516|122146599</td><td>11</td><td>33</td><td>23</td><td>TGAGGTAGTAGGTTGTATAGTTT</td><td>3278.8</td><td>7.3</td><td>0.6</td><td>11.5</td><td>2.1E-30</td><td>4.2E-29</td><td>BT20/MB468</td><td>CYTO</td></tr>
<tr><td>>MI0000439|hsa-mir-23b&WithFlank&9|+|95085202|95085310@67.85.19(+2U)&[MIMAT0000418&hsa-miR-23b-3p&offsets|+3|-1(+2U);m-42&9|+|95085265|95085285&offsets|+3|+1(+2U)]|ACATTGCCAGGGATTACCATT</td><td>>MI0000439|hsa-mir-23b&WithFlank&9|+|95085202|95085310</td><td>67</td><td>85</td><td>19(+2U)</td><td>ACATTGCCAGGGATTACCATT</td><td>879.3</td><td>14.0</td><td>1.2</td><td>11.5</td><td>2.1E-30</td><td>4.4E-29</td><td>BT20/MB468</td><td>CYTO</td></tr>
<tr><td>>MI0000542|hsa-mir-320a&WithFlank&8|-|22244960|22245043@48.68.21(+1C)&[MIMAT0000510&hsa-miR-320a-3p&offsets|0|-1(+1C);m-60&8|-|22244975|22244996&offsets|0|-1(+1C)]|AAAAGCTGGGTTGAGAGGGCGC</td><td>>MI0000542|hsa-mir-320a&WithFlank&8|-|22244960|22245043</td><td>48</td><td>68</td><td>21(+1C)</td><td>AAAAGCTGGGTTGAGAGGGCGC</td><td>5822.2</td><td>12.6</td><td>1.1</td><td>11.5</td><td>2.1E-30</td><td>4.4E-29</td><td>BT20/MB468</td><td>CYTO</td></tr>
<tr><td>>MI0000102|hsa-mir-100&WithFlank&11|-|122152223|122152314@19.41.23&[MIMAT0000098&hsa-miR-100-5p&offsets|0|+1;m-27&11|-|122152276|122152296&offsets|0|+2]|AACCCGTAGATCCGAACTTGTGG</td><td>>MI0000102|hsa-mir-100&WithFlank&11|-|122152223|122152314</td><td>19</td><td>41</td><td>23</td><td>AACCCGTAGATCCGAACTTGTGG</td><td>3330.4</td><td>15.6</td><td>1.4</td><td>11.5</td><td>1.1E-30</td><td>4.4E-29</td><td>MB231/MB468</td><td>NUC</td></tr>
<tr><td>>hg38_21_+_8433172_8446622_RNA45SN1_45S_50ntFlanks@12412.12433.22|ATCCTTCGATGTCGGCTCTTCC</td><td>>hg38_21_+_8433172_8446622_RNA45SN1_45S_50ntFlanks</td><td>12412</td><td>12433</td><td>22</td><td>ATCCTTCGATGTCGGCTCTTCC</td><td>1107.4</td><td>-14.5</td><td>1.3</td><td>-11.4</td><td>3.5E-30</td><td>4.5E-29</td><td>BT20/MB468</td><td>NUC</td></tr>
<tr><td>>hg38_21_+_8433172_8446622_RNA45SN1_45S_50ntFlanks@3872.3898.27|CTAATACATGCCGACGGGCGCTGACCC</td><td>>hg38_21_+_8433172_8446622_RNA45SN1_45S_50ntFlanks</td><td>3872</td><td>3898</td><td>27</td><td>CTAATACATGCCGACGGGCGCTGACCC</td><td>2597.3</td><td>-15.8</td><td>1.4</td><td>-11.4</td><td>3.6E-30</td><td>4.6E-29</td><td>BT20/MB468</td><td>NUC</td></tr>
<tr><td>>hg38_21_+_8433172_8446622_RNA45SN1_45S_50ntFlanks@6719.6742.24|TGAATTGCAGGACACATTGATCAT</td><td>>hg38_21_+_8433172_8446622_RNA45SN1_45S_50ntFlanks</td><td>6719</td><td>6742</td><td>24</td><td>TGAATTGCAGGACACATTGATCAT</td><td>1613.5</td><td>-15.1</td><td>1.3</td><td>-11.4</td><td>3.6E-30</td><td>4.6E-29</td><td>BT20/MB468</td><td>NUC</td></tr>
<tr><td>>MI0000483|hsa-mir-186&WithFlank&1|-|71067625|71067722@21.42.22&[MIMAT0000456&hsa-miR-186-5p&offsets|0|0;m-84&1|-|71067681|71067702&offsets|0|0]|CAAAGAATTCTCCTTTTGGGCT</td><td>>MI0000483|hsa-mir-186&WithFlank&1|-|71067625|71067722</td><td>21</td><td>42</td><td>22</td><td>CAAAGAATTCTCCTTTTGGGCT</td><td>1549.8</td><td>6.0</td><td>0.5</td><td>11.5</td><td>1.6E-30</td><td>4.7E-29</td><td>BT20/MB468</td><td>MITO</td></tr>
<tr><td>>hg38_21_+_8433172_8446622_RNA45SN1_45S_50ntFlanks@12957.12980.24|CTTCTGGGTCGGGGTTTCGTACGT</td><td>>hg38_21_+_8433172_8446622_RNA45SN1_45S_50ntFlanks</td><td>12957</td><td>12980</td><td>24</td><td>CTTCTGGGTCGGGGTTTCGTACGT</td><td>1321.6</td><td>-14.8</td><td>1.3</td><td>-11.4</td><td>3.7E-30</td><td>4.7E-29</td><td>BT20/MB468</td><td>NUC</td></tr>
<tr><td>>hg38_21_+_8433172_8446622_RNA45SN1_45S_50ntFlanks@11270.11295.26|CGGCCGCCCCCGGGGAGCCCGGCGGG</td><td>>hg38_21_+_8433172_8446622_RNA45SN1_45S_50ntFlanks</td><td>11270</td><td>11295</td><td>26</td><td>CGGCCGCCCCCGGGGAGCCCGGCGGG</td><td>6195.2</td><td>-17.0</td><td>1.5</td><td>-11.4</td><td>3.9E-30</td><td>4.9E-29</td><td>BT20/MB468</td><td>NUC</td></tr>
<tr><td>>trna111_HisGTG_1_-_147774845_147774916@-1T.33.34,trna118_HisGTG_1_-_145396881_145396952@-1T.33.34,trna16_HisGTG_1_+_146544773_146544844@-1T.33.34,trna1_HisGTG_15_+_45493349_45493420@-1T.33.34,trna21_HisGTG_1_+_147753471_147753542@-1T.33.34,trna33_HisGTG_6_+_27125906_27125977@-1T.33.34,trna7_HisGTG_9_-_14433938_14434009@-1T.33.34,trna8_HisGTG_15_-_45492611_45492682@-1T.33.34,trna9_HisGTG_15_-_45490804_45490875@-1T.33.34|TGCCGTGATCGTATAGTGGTTAGTACTCTGCGTT</td><td>>trna111_HisGTG_1_-_147774845_147774916</td><td>-1T</td><td>33</td><td>34</td><td>TGCCGTGATCGTATAGTGGTTAGTACTCTGCGTT</td><td>2674.9</td><td>7.9</td><td>0.7</td><td>11.4</td><td>2.4E-30</td><td>4.9E-29</td><td>BT20/MB468</td><td>CYTO</td></tr>
<tr><td>>hg38_21_+_8433172_8446622_RNA45SN1_45S_50ntFlanks@10386.10424.39|ATGTGAACAGCAGTTGAACATGGGTCAGTCGGTCCTGAG</td><td>>hg38_21_+_8433172_8446622_RNA45SN1_45S_50ntFlanks</td><td>10386</td><td>10424</td><td>39</td><td>ATGTGAACAGCAGTTGAACATGGGTCAGTCGGTCCTGAG</td><td>18699.8</td><td>9.7</td><td>0.8</td><td>11.5</td><td>1.7E-30</td><td>5.0E-29</td><td>BT20/MB468</td><td>MITO</td></tr>
</table>

<table>
<tr><td>>hg38_21_+_8433172_8446622_RNA45SN1_45S_50ntFlanks@9568.9597.30|CGTGCAAATCGGTCGTCCGACCTGGGTATA</td><td>>hg38_21_+_8433172_8446622_RNA45SN1_45S_50ntFlanks</td><td>9568</td><td>9597</td><td>30</td><td>CGTGCAAATCGGTCGTCCGACCTGGGTATA</td><td>2667.9</td><td>-7.6</td><td>0.7</td><td>-11.5</td><td>1.3E-30</td><td>5.0E-29</td><td>MB231/MB468</td><td>NUC</td></tr>
<tr><td>>hg38_21_+_8433172_8446622_RNA45SN1_45S_50ntFlanks@3705.3736.32|TACCTGGTTGATCCTGCCAGTAGCATATGCTT</td><td>>hg38_21_+_8433172_8446622_RNA45SN1_45S_50ntFlanks</td><td>3705</td><td>3736</td><td>32</td><td>TACCTGGTTGATCCTGCCAGTAGCATATGCTT</td><td>3208.2</td><td>7.4</td><td>0.6</td><td>11.4</td><td>2.7E-30</td><td>5.4E-29</td><td>BT20/MB468</td><td>CYTO</td></tr>
<tr><td>>hg38_21_+_8433172_8446622_RNA45SN1_45S_50ntFlanks@10346.10374.29|TCAAACGAGAACTTTGAAGGCCGAAGTGG</td><td>>hg38_21_+_8433172_8446622_RNA45SN1_45S_50ntFlanks</td><td>10346</td><td>10374</td><td>29</td><td>TCAAACGAGAACTTTGAAGGCCGAAGTGG</td><td>3223.6</td><td>-7.8</td><td>0.7</td><td>-11.6</td><td>3.7E-31</td><td>5.5E-29</td><td>MB231/MB468</td><td>MP</td></tr>
<tr><td>>hg38_21_+_8433172_8446622_RNA45SN1_45S_50ntFlanks@12406.12433.28|TTTTTGATCCTTCGATGTCGGCTCTTCC</td><td>>hg38_21_+_8433172_8446622_RNA45SN1_45S_50ntFlanks</td><td>12406</td><td>12433</td><td>28</td><td>TTTTTGATCCTTCGATGTCGGCTCTTCC</td><td>1018.8</td><td>-5.4</td><td>0.5</td><td>-11.4</td><td>4.4E-30</td><td>5.5E-29</td><td>BT20/MB468</td><td>NUC</td></tr>
<tr><td>>am_MI0000809|hsa-mir-151a&WithFlank&8|-|140732558|140732659@53.72.20&[MIMAT0000757&hsa-miR-151a-3p&offsets|0|-1;m-30&8|-|140732587|140732607&offsets|0|-1]|CTAGACTGAAGCTCCTTGAG</td><td>>am_MI0000809|hsa-mir-151a&WithFlank&8|-|140732558|140732659</td><td>53</td><td>72</td><td>20</td><td>CTAGACTGAAGCTCCTTGAG</td><td>810.9</td><td>3.9</td><td>0.3</td><td>11.4</td><td>4.4E-30</td><td>5.6E-29</td><td>BT20/MB468</td><td>NUC</td></tr>
<tr><td>>hg38_21_+_8433172_8446622_RNA45SN1_45S_50ntFlanks@8182.8206.25|ATCGAGGCCCAGCCCGTGGACGGTG</td><td>>hg38_21_+_8433172_8446622_RNA45SN1_45S_50ntFlanks</td><td>8182</td><td>8206</td><td>25</td><td>ATCGAGGCCCAGCCCGTGGACGGTG</td><td>1052.2</td><td>-14.5</td><td>1.3</td><td>-11.4</td><td>4.4E-30</td><td>5.6E-29</td><td>BT20/MB468</td><td>NUC</td></tr>
<tr><td>>hg38_21_+_8433172_8446622_RNA45SN1_45S_50ntFlanks@8357.8379.23|AAAAGAACTTTGAAGAGAGAGTT</td><td>>hg38_21_+_8433172_8446622_RNA45SN1_45S_50ntFlanks</td><td>8357</td><td>8379</td><td>23</td><td>AAAAGAACTTTGAAGAGAGAGTT</td><td>90380.4</td><td>-13.2</td><td>1.1</td><td>-11.6</td><td>3.9E-31</td><td>5.8E-29</td><td>MB231/MB468</td><td>MP</td></tr>
<tr><td>>MI0000273|hsa-mir-183&WithFlank&7|-|129774899|129775020@34.56.23&[MIMAT0000261&hsa-miR-183-5p&offsets|+1|+2;m-41&7|-|129774966|129774987&offsets|0|+1]|ATGGCACTGGTAGAATTCACTGT</td><td>>MI0000273|hsa-mir-183&WithFlank&7|-|129774899|129775020</td><td>34</td><td>56</td><td>23</td><td>ATGGCACTGGTAGAATTCACTGT</td><td>952.4</td><td>7.5</td><td>0.7</td><td>11.5</td><td>2.1E-30</td><td>6.0E-29</td><td>BT20/MB468</td><td>MITO</td></tr>
<tr><td>>hg38_21_+_8433172_8446622_RNA45SN1_45S_50ntFlanks@10404.10429.26|CATGGGTCAGTCGGTCCTGAGAGATG</td><td>>hg38_21_+_8433172_8446622_RNA45SN1_45S_50ntFlanks</td><td>10404</td><td>10429</td><td>26</td><td>CATGGGTCAGTCGGTCCTGAGAGATG</td><td>4523.9</td><td>-8.1</td><td>0.7</td><td>-11.4</td><td>4.9E-30</td><td>6.1E-29</td><td>BT20/MB468</td><td>NUC</td></tr>
<tr><td>>hg38_21_+_8433172_8446622_RNA45SN1_45S_50ntFlanks@8380.8402.23|CAAGAGGGCGTGAAACCGTTAAG</td><td>>hg38_21_+_8433172_8446622_RNA45SN1_45S_50ntFlanks</td><td>8380</td><td>8402</td><td>23</td><td>CAAGAGGGCGTGAAACCGTTAAG</td><td>9970.7</td><td>-11.2</td><td>1.0</td><td>-11.6</td><td>4.2E-31</td><td>6.1E-29</td><td>MB231/MB468</td><td>MP</td></tr>
<tr><td>>hg38_21_+_8433172_8446622_RNA45SN1_45S_50ntFlanks@12567.12587.21|CTCAGTACGAGAGGAACCGCA</td><td>>hg38_21_+_8433172_8446622_RNA45SN1_45S_50ntFlanks</td><td>12567</td><td>12587</td><td>21</td><td>CTCAGTACGAGAGGAACCGCA</td><td>847.5</td><td>-14.2</td><td>1.2</td><td>-11.4</td><td>4.9E-30</td><td>6.2E-29</td><td>BT20/MB468</td><td>NUC</td></tr>
<tr><td>>am_trnaMT_ValTAC_MT_+_1602_1670@51.68.18|CTTAACTTGACCGCTCTG</td><td>>am_trnaMT_ValTAC_MT_+_1602_1670</td><td>51</td><td>68</td><td>18</td><td>CTTAACTTGACCGCTCTG</td><td>13660.8</td><td>-11.9</td><td>1.0</td><td>-11.6</td><td>4.3E-31</td><td>6.2E-29</td><td>MB231/MB468</td><td>MP</td></tr>
<tr><td>>hg38_21_+_8433172_8446622_RNA45SN1_45S_50ntFlanks@12936.12956.21|AAACCATTCGTAGACGACCTG</td><td>>hg38_21_+_8433172_8446622_RNA45SN1_45S_50ntFlanks</td><td>12936</td><td>12956</td><td>21</td><td>AAACCATTCGTAGACGACCTG</td><td>4354.8</td><td>-16.0</td><td>1.4</td><td>-11.6</td><td>4.5E-31</td><td>6.4E-29</td><td>MB231/MB468</td><td>MP</td></tr>
<tr><td>>MI0000301|hsa-mir-224&WithFlank&X|-|151958572|151958664@14.34.21&[MIMAT0000281&hsa-miR-224-5p&offsets|+1|-3;m-238&X|-|151958630|151958651&offsets|0|-1]|CAAGTCACTAGTGGTTCCGTT</td><td>>MI0000301|hsa-mir-224&WithFlank&X|-|151958572|151958664</td><td>14</td><td>34</td><td>21</td><td>CAAGTCACTAGTGGTTCCGTT</td><td>2229.7</td><td>15.4</td><td>1.3</td><td>11.4</td><td>3.3E-30</td><td>6.7E-29</td><td>BT20/MB468</td><td>CYTO</td></tr>
<tr><td>>hg38_21_+_8433172_8446622_RNA45SN1_45S_50ntFlanks@10375.10406.32|AGAAGGGTTCCATGTGAACAGCAGTTGAACAT</td><td>>hg38_21_+_8433172_8446622_RNA45SN1_45S_50ntFlanks</td><td>10375</td><td>10406</td><td>32</td><td>AGAAGGGTTCCATGTGAACAGCAGTTGAACAT</td><td>1091.0</td><td>-14.5</td><td>1.3</td><td>-11.4</td><td>5.4E-30</td><td>6.8E-29</td><td>BT20/MB468</td><td>NUC</td></tr>
<tr><td>>hg38_21_+_8433172_8446622_RNA45SN1_45S_50ntFlanks@8196.8215.20|CGTGGACGGTGTGAGGCCGG</td><td>>hg38_21_+_8433172_8446622_RNA45SN1_45S_50ntFlanks</td><td>8196</td><td>8215</td><td>20</td><td>CGTGGACGGTGTGAGGCCGG</td><td>5047.1</td><td>-8.1</td><td>0.7</td><td>-11.6</td><td>4.8E-31</td><td>6.8E-29</td><td>MB231/MB468</td><td>MP</td></tr>
<tr><td>>MI0000079|hsa-mir-23a&WithFlank&19|-|13836581|13836665@54.73.20(+1U)&[MIMAT0000078&hsa-miR-23a-3p&offsets|+3|+2(+1U);m-31&19|-|13836595|13836615&offsets|+3|+2(+1U)]|ACATTGCCAGGGATTTCCAAT</td><td>>MI0000079|hsa-mir-23a&WithFlank&19|-|13836581|13836665</td><td>54</td><td>73</td><td>20(+1U)</td><td>ACATTGCCAGGGATTTCCAAT</td><td>1757.0</td><td>15.0</td><td>1.3</td><td>11.4</td><td>3.4E-30</td><td>6.8E-29</td><td>BT20/MB468</td><td>CYTO</td></tr>
<tr><td>>hg38_21_+_8433172_8446622_RNA45SN1_45S_50ntFlanks@10817.10840.24|TAAGGGAAGTCGGCAAGCCGGATC</td><td>>hg38_21_+_8433172_8446622_RNA45SN1_45S_50ntFlanks</td><td>10817</td><td>10840</td><td>24</td><td>TAAGGGAAGTCGGCAAGCCGGATC</td><td>790.1</td><td>-14.1</td><td>1.2</td><td>-11.4</td><td>5.5E-30</td><td>6.9E-29</td><td>BT20/MB468</td><td>NUC</td></tr>
<tr><td>>hg38_21_+_8433172_8446622_RNA45SN1_45S_50ntFlanks@10410.10432.23|TCAGTCGGTCCTGAGAGATGGGC</td><td>>hg38_21_+_8433172_8446622_RNA45SN1_45S_50ntFlanks</td><td>10410</td><td>10432</td><td>23</td><td>TCAGTCGGTCCTGAGAGATGGGC</td><td>868.3</td><td>-4.8</td><td>0.4</td><td>-11.4</td><td>3.5E-30</td><td>6.9E-29</td><td>BT20/MB468</td><td>CYTO</td></tr>
<tr><td>>MI0000266|hsa-mir-10a&WithFlank&17|-|48579832|48579953@28.50.23&[MIMAT0000253&hsa-miR-10a-5p&offsets|0|0;m-18&17|-|48579905|48579926&offsets|0|+1]|TACCCTGTAGATCCGAATTTGTG</td><td>>MI0000266|hsa-mir-10a&WithFlank&17|-|48579832|48579953</td><td>28</td><td>50</td><td>23</td><td>TACCCTGTAGATCCGAATTTGTG</td><td>3987.0</td><td>16.0</td><td>1.4</td><td>11.6</td><td>2.8E-31</td><td>7.0E-29</td><td>MB231/MB468</td><td>CYTO</td></tr>
<tr><td>>MI0000082|hsa-mir-25&WithFlank&7|-|100093554|100093649@58.79.22&[MIMAT0000081&hsa-miR-25-3p&offsets|0|0;m-23&7|-|100093571|100093592&offsets|0|0]|CATTGCACTTGTCTCGGTCTGA</td><td>>MI0000082|hsa-mir-25&WithFlank&7|-|100093554|100093649</td><td>58</td><td>79</td><td>22</td><td>CATTGCACTTGTCTCGGTCTGA</td><td>55739.8</td><td>7.8</td><td>0.7</td><td>11.4</td><td>2.5E-30</td><td>7.1E-29</td><td>BT20/MB468</td><td>MITO</td></tr>
<tr><td>>hg38_21_+_8433172_8446622_RNA45SN1_45S_50ntFlanks@12456.12476.21|CCAAGCGTTGGATTGTTCACC</td><td>>hg38_21_+_8433172_8446622_RNA45SN1_45S_50ntFlanks</td><td>12456</td><td>12476</td><td>21</td><td>CCAAGCGTTGGATTGTTCACC</td><td>30716.9</td><td>-7.1</td><td>0.6</td><td>-11.4</td><td>3.7E-30</td><td>7.3E-29</td><td>BT20/MB468</td><td>CYTO</td></tr>
<tr><td>>hg38_21_+_8433172_8446622_RNA45SN1_45S_50ntFlanks@11768.11791.24|CTAATTAGTGACGCGCATGAATGG</td><td>>hg38_21_+_8433172_8446622_RNA45SN1_45S_50ntFlanks</td><td>11768</td><td>11791</td><td>24</td><td>CTAATTAGTGACGCGCATGAATGG</td><td>2614.0</td><td>-15.8</td><td>1.4</td><td>-11.4</td><td>5.9E-30</td><td>7.4E-29</td><td>BT20/MB468</td><td>NUC</td></tr>
<tr><td>>MI0000746|hsa-mir-99b&WithFlank&19|+|51692606|51692687@13.34.22(+1U)&[MIMAT0000689&hsa-miR-99b-5p&offsets|0|0(+1U);m-10&19|+|51692618|51692639&offsets|0|0(+1U)]|CACCCGTAGAACCGACCTTGCGT</td><td>>MI0000746|hsa-mir-99b&WithFlank&19|+|51692606|51692687</td><td>13</td><td>34</td><td>22(+1U)</td><td>CACCCGTAGAACCGACCTTGCGT</td><td>2033.9</td><td>7.1</td><td>0.6</td><td>11.4</td><td>2.6E-30</td><td>7.4E-29</td><td>BT20/MB468</td><td>MITO</td></tr>
<tr><td>>hg38_21_+_8433172_8446622_RNA45SN1_45S_50ntFlanks@8174.8203.30|TCCTTCTGATCGAGGCCCAGCCCGTGGACG</td><td>>hg38_21_+_8433172_8446622_RNA45SN1_45S_50ntFlanks</td><td>8174</td><td>8203</td><td>30</td><td>TCCTTCTGATCGAGGCCCAGCCCGTGGACG</td><td>660.6</td><td>-5.2</td><td>0.5</td><td>-11.4</td><td>6.0E-30</td><td>7.4E-29</td><td>BT20/MB468</td><td>NUC</td></tr>
<tr><td>>MI0000447|hsa-mir-128-1&WithFlank&2|+|135665391|135665484@56.75.20(+1C);MI0000727|hsa-mir-128-2&WithFlank&3|+|35744470|35744565@58.77.20(+1C)&[MIMAT0000424&hsa-miR-128-3p&offsets|0|-1(+1C);m-108&2|+|135665446|135665466&offsets|0|-1(+1C)];[MIMAT0000424_1&hsa-miR-128-3p&offsets|0|-1(+1C);m-117&3|+|35744527|35744547&offsets|0|-1(+1C)]|TCACAGTGAACCGGTCTCTTC</td><td>>MI0000447|hsa-mir-128-1&WithFlank&2|+|135665391|135665484</td><td>56</td><td>75</td><td>20(+1C)</td><td>TCACAGTGAACCGGTCTCTTC</td><td>1579.2</td><td>6.2</td><td>0.5</td><td>11.4</td><td>3.8E-30</td><td>7.5E-29</td><td>BT20/MB468</td><td>CYTO</td></tr>
<tr><td>>hg38_21_+_8433172_8446622_RNA45SN1_45S_50ntFlanks@7988.8012.25|CAGACGTGGCGACCCGCTGAATTTA</td><td>>hg38_21_+_8433172_8446622_RNA45SN1_45S_50ntFlanks</td><td>7988</td><td>8012</td><td>25</td><td>CAGACGTGGCGACCCGCTGAATTTA</td><td>12311.4</td><td>-7.8</td><td>0.7</td><td>-11.5</td><td>1.1E-30</td><td>7.5E-29</td><td>MB231/MB468</td><td>MITO</td></tr>
<tr><td>>hg38_21_+_8433172_8446622_RNA45SN1_45S_50ntFlanks@9504.9526.23|GTGAACTATGCCTGGGCAGGGCG</td><td>>hg38_21_+_8433172_8446622_RNA45SN1_45S_50ntFlanks</td><td>9504</td><td>9526</td><td>23</td><td>GTGAACTATGCCTGGGCAGGGCG</td><td>5114.4</td><td>-11.6</td><td>1.0</td><td>-11.6</td><td>3.2E-31</td><td>7.7E-29</td><td>MB231/MB468</td><td>CYTO</td></tr>
<tr><td>>am_trna111_HisGTG_1_-_147774845_147774916@1.28.28,trna118_HisGTG_1_-_145396881_145396952@1.28.28,trna16_HisGTG_1_+_146544773_146544844@1.28.28,trna1_HisGTG_15_+_45493349_45493420@1.28.28,trna21_HisGTG_1_+_147753471_147753542@1.28.28,trna33_HisGTG_6_+_27125906_27125977@1.28.28,trna7_HisGTG_9_-_14433938_14434009@1.28.28,trna8_HisGTG_15_-_45492611_45492682@1.28.28,trna9_HisGTG_15_-_45490804_45490875@1.28.28|GCCGTGATCGTATAGTGGTTAGTACTCT</td><td>>am_trna111_HisGTG_1_-_147774845_147774916</td><td>1</td><td>28</td><td>28</td><td>GCCGTGATCGTATAGTGGTTAGTACTCT</td><td>1806.5</td><td>-6.4</td><td>0.6</td><td>-11.4</td><td>6.3E-30</td><td>7.8E-29</td><td>BT20/MB468</td><td>NUC</td></tr>
<tr><td>>hg38_21_+_8433172_8446622_RNA45SN1_45S_50ntFlanks@12997.13017.21|CGCTGCGATCTATTGAAAGTC</td><td>>hg38_21_+_8433172_8446622_RNA45SN1_45S_50ntFlanks</td><td>12997</td><td>13017</td><td>21</td><td>CGCTGCGATCTATTGAAAGTC</td><td>2676.9</td><td>6.7</td><td>0.6</td><td>11.6</td><td>3.3E-31</td><td>7.8E-29</td><td>MB231/MB468</td><td>CYTO</td></tr>
<tr><td>>MI0000087|hsa-mir-29a&WithFlank&7|-|130876741|130876816@48.69.22&[MIMAT0000086&hsa-miR-29a-3p&offsets|0|0;m-26&7|-|130876749|130876769&offsets|0|+1]|TAGCACCATCTGAAATCGGTTA</td><td>>MI0000087|hsa-mir-29a&WithFlank&7|-|130876741|130876816</td><td>48</td><td>69</td><td>22</td><td>TAGCACCATCTGAAATCGGTTA</td><td>5729.5</td><td>9.4</td><td>0.8</td><td>11.4</td><td>4.0E-30</td><td>7.9E-29</td><td>BT20/MB468</td><td>CYTO</td></tr>
<tr><td>>MI0001445|hsa-mir-423&WithFlank&17|+|30117073|30117178@59.80.22(+1G)&[MIMAT0001340&hsa-miR-423-3p&offsets|0|-1(+1G);m-100&17|+|30117131|30117153&offsets|0|-1(+1G)]|AGCTCGGTCTGAGGCCCCTCAGG</td><td>>MI0001445|hsa-mir-423&WithFlank&17|+|30117073|30117178</td><td>59</td><td>80</td><td>22(+1G)</td><td>AGCTCGGTCTGAGGCCCCTCAGG</td><td>885.0</td><td>7.3</td><td>0.6</td><td>11.4</td><td>4.1E-30</td><td>8.2E-29</td><td>BT20/MB468</td><td>CYTO</td></tr>
<tr><td>>MI0000077|hsa-mir-21&WithFlank&17|+|59841260|59841343@14.36.23&[MIMAT0000076&hsa-miR-21-5p&offsets|0|+1;m-1&17|+|59841273|59841294&offsets|0|+1]|TAGCTTATCAGACTGATGTTGAC</td><td>>MI0000077|hsa-mir-21&WithFlank&17|+|59841260|59841343</td><td>14</td><td>36</td><td>23</td><td>TAGCTTATCAGACTGATGTTGAC</td><td>1595637.4</td><td>7.9</td><td>0.7</td><td>11.4</td><td>4.2E-30</td><td>8.2E-29</td><td>BT20/MB468</td><td>CYTO</td></tr>
<tr><td>>hg38_21_+_8433172_8446622_RNA45SN1_45S_50ntFlanks@11566.11584.19|TAGCAGCCGACTTAGAACT</td><td>>hg38_21_+_8433172_8446622_RNA45SN1_45S_50ntFlanks</td><td>11566</td><td>11584</td><td>19</td><td>TAGCAGCCGACTTAGAACT</td><td>400.9</td><td>6.6</td><td>0.6</td><td>11.4</td><td>2.9E-30</td><td>8.2E-29</td><td>BT20/MB468</td><td>MITO</td></tr>
<tr><td>>hg38_21_+_8433172_8446622_RNA45SN1_45S_50ntFlanks@10388.10424.37|GTGAACAGCAGTTGAACATGGGTCAGTCGGTCCTGAG</td><td>>hg38_21_+_8433172_8446622_RNA45SN1_45S_50ntFlanks</td><td>10388</td><td>10424</td><td>37</td><td>GTGAACAGCAGTTGAACATGGGTCAGTCGGTCCTGAG</td><td>27502.5</td><td>8.5</td><td>0.7</td><td>11.4</td><td>3.0E-30</td><td>8.3E-29</td><td>BT20/MB468</td><td>MITO</td></tr>
<tr><td>>hg38_21_+_8433172_8446622_RNA45SN1_45S_50ntFlanks@8177.8195.19|TTCTGATCGAGGCCCAGCC</td><td>>hg38_21_+_8433172_8446622_RNA45SN1_45S_50ntFlanks</td><td>8177</td><td>8195</td><td>19</td><td>TTCTGATCGAGGCCCAGCC</td><td>794.5</td><td>4.9</td><td>0.4</td><td>11.4</td><td>4.2E-30</td><td>8.3E-29</td><td>BT20/MB468</td><td>CYTO</td></tr>
<tr><td>>trna119_LysCTT_1_-_145395522_145395594@1.30.30,trna11_LysCTT_5_-_180648979_180649051@1.30.30,trna13_LysCTT_6_+_26556774_26556846@1.30.30,trna32_LysCTT_16_-_3207406_3207478@1.30.30,trna7_LysCTT_16_+_3225692_3225764@1.30.30,trna9_LysCTT_5_+_180634755_180634827@1.30.30|GCCCGGCTAGCTCAGTCGGTAGAGCATGAG</td><td>>trna119_LysCTT_1_-_145395522_145395594</td><td>1</td><td>30</td><td>30</td><td>GCCCGGCTAGCTCAGTCGGTAGAGCATGAG</td><td>20722.7</td><td>-12.4</td><td>1.1</td><td>-11.4</td><td>6.7E-30</td><td>8.3E-29</td><td>BT20/MB468</td><td>NUC</td></tr>
<tr><td>>hg38_21_+_8433172_8446622_RNA45SN1_45S_50ntFlanks@12605.12625.21|TATGTGCTTGGCTGAGGAGCC</td><td>>hg38_21_+_8433172_8446622_RNA45SN1_45S_50ntFlanks</td><td>12605</td><td>12625</td><td>21</td><td>TATGTGCTTGGCTGAGGAGCC</td><td>9594.7</td><td>-7.3</td><td>0.6</td><td>-11.5</td><td>2.3E-30</td><td>8.7E-29</td><td>MB231/MB468</td><td>NUC</td></tr>
<tr><td>>MI0000061|hsa-let-7a-2&WithFlank&11|-|122146516|122146599@11.30.20(+1C);MI0000062|hsa-let-7a-3&WithFlank&22|+|46112743|46112828@10.29.20(+1C);MI0000060|hsa-let-7a-1&WithFlank&9|+|94175951|94176042@12.31.20(+1C)&[MIMAT0000062&hsa-let-7a-5p&offsets|0|-2(+1C);m-6&11|-|122146568|122146589&offsets|0|-2(+1C)];[MIMAT0000062_1&hsa-let-7a-5p&offsets|0|-2(+1C);m-5&22|+|46112752|46112773&offsets|0|-2(+1C)];[MIMAT0000062_2&hsa-let-7a-5p&offsets|0|-2(+1C);m-4&9|+|94175962|94175983&offsets|0|-2(+1C)]|TGAGGTAGTAGGTTGTATAGC</td><td>>MI0000061|hsa-let-7a-2&WithFlank&11|-|122146516|122146599</td><td>11</td><td>30</td><td>20(+1C)</td><td>TGAGGTAGTAGGTTGTATAGC</td><td>705.8</td><td>7.8</td><td>0.7</td><td>11.4</td><td>3.2E-30</td><td>8.7E-29</td><td>BT20/MB468</td><td>MITO</td></tr>
<tr><td>>hg38_21_+_8433172_8446622_RNA45SN1_45S_50ntFlanks@6700.6720.21|CTAGCTGCGAGAATTAATGTG</td><td>>hg38_21_+_8433172_8446622_RNA45SN1_45S_50ntFlanks</td><td>6700</td><td>6720</td><td>21</td><td>CTAGCTGCGAGAATTAATGTG</td><td>15214.8</td><td>-4.9</td><td>0.4</td><td>-11.4</td><td>4.5E-30</td><td>8.8E-29</td><td>BT20/MB468</td><td>CYTO</td></tr>
<tr><td>>hg38_21_+_8433172_8446622_RNA45SN1_45S_50ntFlanks@10408.10435.28|GGTCAGTCGGTCCTGAGAGATGGGCGAG</td><td>>hg38_21_+_8433172_8446622_RNA45SN1_45S_50ntFlanks</td><td>10408</td><td>10435</td><td>28</td><td>GGTCAGTCGGTCCTGAGAGATGGGCGAG</td><td>5091.1</td><td>-16.7</td><td>1.5</td><td>-11.4</td><td>7.3E-30</td><td>9.0E-29</td><td>BT20/MB468</td><td>NUC</td></tr>
<tr><td>>hg38_21_+_8433172_8446622_RNA45SN1_45S_50ntFlanks@11577.11601.25|TTAGAACTGGTGCGGACCAGGGGAA</td><td>>hg38_21_+_8433172_8446622_RNA45SN1_45S_50ntFlanks</td><td>11577</td><td>11601</td><td>25</td><td>TTAGAACTGGTGCGGACCAGGGGAA</td><td>769.6</td><td>-14.0</td><td>1.2</td><td>-11.4</td><td>7.3E-30</td><td>9.1E-29</td><td>BT20/MB468</td><td>NUC</td></tr>
<tr><td>>MI0000283|hsa-mir-203a&WithFlank&14|+|104117399|104117520@72.93.22&[MIMAT0000264&hsa-miR-203a-3p&offsets|+1|+1;m-79&14|+|104117469|104117490&offsets|+1|+1]|TGAAATGTTTAGGACCACTAGA</td><td>>MI0000283|hsa-mir-203a&WithFlank&14|+|104117399|104117520</td><td>72</td><td>93</td><td>22</td><td>TGAAATGTTTAGGACCACTAGA</td><td>1270.7</td><td>14.6</td><td>1.3</td><td>11.4</td><td>4.7E-30</td><td>9.2E-29</td><td>BT20/MB468</td><td>CYTO</td></tr>
</table>

| | | | | | | | | | | | | | |
|---|---|---|---|---|---|---|---|---|---|---|---|---|---|
| >MI0000279\|hsa-mir-196a-2&WithFlank&12\|+\|53991732\|53991853@31.51.21(+1A);MI0000238\|hsa-mir-196a-1&WithFlank&17\|-\|48632484\|48632565@13.33.21(+1A)&[MIMAT0000226&hsa-miR-196a-5p&offsets\|0\|-1(+1A);m-261&12\|+\|53991762\|53991783&offsets\|0\|-1(+1A)];[MIMAT0000226_1&hsa-miR-196a-5p&offsets\|0\|-1(+1A);m-268&17\|-\|48632532\|48632553&offsets\|0\|-1(+1A)]\|TAGGTAGTTTCATGTTGTTGGA | >MI0000279\|hsa-mir-196a-2&WithFlank&12\|+\|53991732\|53991853 | 31 | 51 | 21(+1A) | TAGGTAGTTTCATGTTGTTGGA | 1960.5 | 8.9 | 0.8 | 11.4 | 4.8E-30 | 9.3E-29 | BT20/MB468 | CYTO |
| >hg38_21_+_8433172_8446622_RNA45SN1_45S_50ntFlanks@4567.4587.21\|TAATGGAATAGGACCGCGGTT | >hg38_21_+_8433172_8446622_RNA45SN1_45S_50ntFlanks | 4567 | 4587 | 21 | TAATGGAATAGGACCGCGGTT | 1459.0 | -14.9 | 1.3 | -11.3 | 7.6E-30 | 9.3E-29 | BT20/MB468 | NUC |
| >hg38_21_+_8433172_8446622_RNA45SN1_45S_50ntFlanks@9572.9596.25\|CAAATCGGTCGTCCGACCTGGGTAT | >hg38_21_+_8433172_8446622_RNA45SN1_45S_50ntFlanks | 9572 | 9596 | 25 | CAAATCGGTCGTCCGACCTGGGTAT | 23710.9 | -12.2 | 1.1 | -11.5 | 1.4E-30 | 9.4E-29 | MB231/MB468 | MITO |
| >hg38_21_+_8433172_8446622_RNA45SN1_45S_50ntFlanks@10680.10710.31\|GTTCCGGCGGCGTCCGGTGAGCTCTCGCTGG | >hg38_21_+_8433172_8446622_RNA45SN1_45S_50ntFlanks | 10680 | 10710 | 31 | GTTCCGGCGGCGTCCGGTGAGCTCTCGCTGG | 16510.2 | -11.4 | 1.0 | -11.6 | 6.8E-31 | 9.4E-29 | MB231/MB468 | MP |
| >hg38_21_+_8433172_8446622_RNA45SN1_45S_50ntFlanks@8115.8140.26\|CGCGGGACATGTGGCGTACGGAAGAC | >hg38_21_+_8433172_8446622_RNA45SN1_45S_50ntFlanks | 8115 | 8140 | 26 | CGCGGGACATGTGGCGTACGGAAGAC | 6751.9 | -16.6 | 1.4 | -11.6 | 7.0E-31 | 9.7E-29 | MB231/MB468 | MP |
| >hg38_21_+_8433172_8446622_RNA45SN1_45S_50ntFlanks@11891.11912.22\|TTGACTCTAGTCTGGCACGGTG | >hg38_21_+_8433172_8446622_RNA45SN1_45S_50ntFlanks | 11891 | 11912 | 22 | TTGACTCTAGTCTGGCACGGTG | 1925.2 | -4.7 | 0.4 | -11.4 | 5.0E-30 | 9.7E-29 | BT20/MB468 | CYTO |
| >MI0000434\|hsa-let-7i&WithFlank&12\|+\|62603680\|62603775@12.33.22&[MIMAT0000415&hsa-let-7i-5p&offsets\|0\|0;m-49&12\|+\|62603691\|62603712&offsets\|0\|0]\|TGAGGTAGTAGTTTGTGCTGTT | >MI0000434\|hsa-let-7i&WithFlank&12\|+\|62603680\|62603775 | 12 | 33 | 22 | TGAGGTAGTAGTTTGTGCTGTT | 117731.4 | 6.4 | 0.6 | 11.4 | 5.1E-30 | 9.9E-29 | BT20/MB468 | CYTO |
| >hg38_21_+_8433172_8446622_RNA45SN1_45S_50ntFlanks@12626.12653.28\|AATGGGGCGAAGCTACCATCTGTGGGAT | >hg38_21_+_8433172_8446622_RNA45SN1_45S_50ntFlanks | 12626 | 12653 | 28 | AATGGGGCGAAGCTACCATCTGTGGGAT | 3287.9 | -5.8 | 0.5 | -11.6 | 4.4E-31 | 1.0E-28 | MB231/MB468 | CYTO |
| >hg38_21_+_8433172_8446622_RNA45SN1_45S_50ntFlanks@12628.12654.27\|TGGGGCGAAGCTACCATCTGTGGGATT | >hg38_21_+_8433172_8446622_RNA45SN1_45S_50ntFlanks | 12628 | 12654 | 27 | TGGGGCGAAGCTACCATCTGTGGGATT | 9152.1 | -16.9 | 1.5 | -11.5 | 1.6E-30 | 1.0E-28 | MB231/MB468 | MITO |
| >hg38_21_+_8433172_8446622_RNA45SN1_45S_50ntFlanks@4722.4759.38\|TCATTAATCAAGAACGAAAGTCGGAGGTTCGAAGACGA | >hg38_21_+_8433172_8446622_RNA45SN1_45S_50ntFlanks | 4722 | 4759 | 38 | TCATTAATCAAGAACGAAAGTCGGAGGTTCGAAGACGA | 5453.3 | -8.3 | 0.7 | -11.5 | 1.5E-30 | 1.0E-28 | MB231/MB468 | MITO |
| >hg38_21_+_8433172_8446622_RNA45SN1_45S_50ntFlanks@9352.9370.19\|CCGTGGCGCAATGAAGGTG | >hg38_21_+_8433172_8446622_RNA45SN1_45S_50ntFlanks | 9352 | 9370 | 19 | CCGTGGCGCAATGAAGGTG | 1259.3 | -14.7 | 1.3 | -11.3 | 8.2E-30 | 1.0E-28 | BT20/MB468 | NUC |
| >hg38_21_+_8433172_8446622_RNA45SN1_45S_50ntFlanks@12396.12416.21\|GCGACGTCGCTTTTTGATCCT | >hg38_21_+_8433172_8446622_RNA45SN1_45S_50ntFlanks | 12396 | 12416 | 21 | GCGACGTCGCTTTTTGATCCT | 1039.7 | -6.3 | 0.6 | -11.3 | 8.3E-30 | 1.0E-28 | BT20/MB468 | NUC |
| >MI0000433\|hsa-let-7g&WithFlank&3\|-\|52268272\|52268367@11.31.21&[MIMAT0000414&hsa-let-7g-5p&offsets\|0\|-1;m-58&3\|-\|52268336\|52268357&offsets\|0\|-1]\|TGAGGTAGTAGTTTGTACAGT | >MI0000433\|hsa-let-7g&WithFlank&3\|-\|52268272\|52268367 | 11 | 31 | 21 | TGAGGTAGTAGTTTGTACAGT | 3673.9 | 7.7 | 0.7 | 11.4 | 3.9E-30 | 1.1E-28 | BT20/MB468 | MITO |
| >MI0000440\|hsa-mir-27b&WithFlank&9\|+\|95085439\|95085547@70.87.18&[MIMAT0000419&hsa-miR-27b-3p&offsets\|+3\|0;m-39&9\|+\|95085505\|95085525&offsets\|+3\|0]\|ACAGTGGCTAAGTTCTGC | >MI0000440\|hsa-mir-27b&WithFlank&9\|+\|95085439\|95085547 | 70 | 87 | 18 | ACAGTGGCTAAGTTCTGC | 1314.5 | 14.6 | 1.3 | 11.4 | 5.6E-30 | 1.1E-28 | BT20/MB468 | CYTO |
| >am_trnaMT_PheGAA_MT_+_577_647@40.57.18\|TGTTTAGACGGGCTCACA | >am_trnaMT_PheGAA_MT_+_577_647 | 40 | 57 | 18 | TGTTTAGACGGGCTCACA | 2122.3 | 6.7 | 0.6 | 11.4 | 4.0E-30 | 1.1E-28 | BT20/MB468 | MITO |
| >hg38_21_+_8433172_8446622_RNA45SN1_45S_50ntFlanks@12617.12653.37\|TGAGGAGCCAATGGGGCGAAGCTACCATCTGTGGGAT | >hg38_21_+_8433172_8446622_RNA45SN1_45S_50ntFlanks | 12617 | 12653 | 37 | TGAGGAGCCAATGGGGCGAAGCTACCATCTGTGGGAT | 1208.8 | -14.7 | 1.3 | -11.3 | 9.7E-30 | 1.2E-28 | BT20/MB468 | NUC |
| >MI0000066\|hsa-let-7e&WithFlank&19\|+\|51692780\|51692870@14.34.21&[MIMAT0000066&hsa-let-7e-5p&offsets\|0\|-1;m-50&19\|+\|51692793\|51692814&offsets\|0\|-1]\|TGAGGTAGGAGGTTGTATAGT | >MI0000066\|hsa-let-7e&WithFlank&19\|+\|51692780\|51692870 | 14 | 34 | 21 | TGAGGTAGGAGGTTGTATAGT | 1409.1 | 7.0 | 0.6 | 11.4 | 4.4E-30 | 1.2E-28 | BT20/MB468 | MITO |
| >hg38_21_+_8433172_8446622_RNA45SN1_45S_50ntFlanks@7979.8004.26\|ACCTCAGATCAGACGTGGCGACCCGC | >hg38_21_+_8433172_8446622_RNA45SN1_45S_50ntFlanks | 7979 | 8004 | 26 | ACCTCAGATCAGACGTGGCGACCCGC | 463.7 | -5.3 | 0.5 | -11.3 | 9.9E-30 | 1.2E-28 | BT20/MB468 | NUC |
| >hg38_21_+_8433172_8446622_RNA45SN1_45S_50ntFlanks@7986.8003.18\|ATCAGACGTGGCGACCCG | >hg38_21_+_8433172_8446622_RNA45SN1_45S_50ntFlanks | 7986 | 8003 | 18 | ATCAGACGTGGCGACCCG | 3477.0 | -15.6 | 1.4 | -11.5 | 8.9E-31 | 1.2E-28 | MB231/MB468 | MP |
| >hg38_21_+_8433172_8446622_RNA45SN1_45S_50ntFlanks@10866.10884.19\|CTAAGGGCTGGGTCGGTCG | >hg38_21_+_8433172_8446622_RNA45SN1_45S_50ntFlanks | 10866 | 10884 | 19 | CTAAGGGCTGGGTCGGTCG | 1863.4 | -15.3 | 1.4 | -11.3 | 9.9E-30 | 1.2E-28 | BT20/MB468 | NUC |
| >MI0000440\|hsa-mir-27b&WithFlank&9\|+\|95085439\|95085547@67.88.22(+1A)&[MIMAT0000419&hsa-miR-27b-3p&offsets\|0\|+1(+1A);m-39&9\|+\|95085505\|95085525&offsets\|0\|+1(+1A)]\|TTCACAGTGGCTAAGTTCTGCAA | >MI0000440\|hsa-mir-27b&WithFlank&9\|+\|95085439\|95085547 | 67 | 88 | 22(+1A) | TTCACAGTGGCTAAGTTCTGCAA | 645.0 | 6.6 | 0.6 | 11.4 | 4.5E-30 | 1.2E-28 | BT20/MB468 | MITO |
| >hg38_21_+_8433172_8446622_RNA45SN1_45S_50ntFlanks@11668.11692.25\|TGCCCAGTGCTCTGAATGTCAAAGT | >hg38_21_+_8433172_8446622_RNA45SN1_45S_50ntFlanks | 11668 | 11692 | 25 | TGCCCAGTGCTCTGAATGTCAAAGT | 346.6 | 6.6 | 0.6 | 11.4 | 4.5E-30 | 1.2E-28 | BT20/MB468 | MITO |
| >am_MI0000809\|hsa-mir-151a&WithFlank&8\|-\|140732558\|140732659@17.37.21&[MIMAT0004697&hsa-miR-151a-5p&offsets\|0\|0;m-75&8\|-\|140732623\|140732643&offsets\|0\|0]\|TCGAGGAGCTCACAGTCTAGT | >am_MI0000809\|hsa-mir-151a&WithFlank&8\|-\|140732558\|140732659 | 17 | 37 | 21 | TCGAGGAGCTCACAGTCTAGT | 4584.3 | 10.4 | 0.9 | 11.4 | 6.5E-30 | 1.2E-28 | BT20/MB468 | CYTO |
| >hg38_21_+_8433172_8446622_RNA45SN1_45S_50ntFlanks@8114.8143.30\|GCGCGGGACATGTGGCGTACGGAAGACCCG | >hg38_21_+_8433172_8446622_RNA45SN1_45S_50ntFlanks | 8114 | 8143 | 30 | GCGCGGGACATGTGGCGTACGGAAGACCCG | 1011.6 | -14.4 | 1.3 | -11.3 | 1.0E-29 | 1.2E-28 | BT20/MB468 | NUC |
| >hg38_21_+_8433172_8446622_RNA45SN1_45S_50ntFlanks@10411.10430.20\|CAGTCGGTCCTGAGAGATGG | >hg38_21_+_8433172_8446622_RNA45SN1_45S_50ntFlanks | 10411 | 10430 | 20 | CAGTCGGTCCTGAGAGATGG | 6364.7 | -16.4 | 1.4 | -11.5 | 2.0E-30 | 1.3E-28 | MB231/MB468 | MITO |
| >hg38_21_+_8433172_8446622_RNA45SN1_45S_50ntFlanks@6741.6770.30\|ATCGACACTTCGAACGCACTTGCGGCCCCG | >hg38_21_+_8433172_8446622_RNA45SN1_45S_50ntFlanks | 6741 | 6770 | 30 | ATCGACACTTCGAACGCACTTGCGGCCCCG | 2865.0 | -15.3 | 1.3 | -11.5 | 9.7E-31 | 1.3E-28 | MB231/MB468 | MP |
| >hg38_21_+_8433172_8446622_RNA45SN1_45S_50ntFlanks@4472.4494.23\|CTTAGCTGAGTGTCCCGCGGGGC | >hg38_21_+_8433172_8446622_RNA45SN1_45S_50ntFlanks | 4472 | 4494 | 23 | CTTAGCTGAGTGTCCCGCGGGGC | 1171.0 | -14.6 | 1.3 | -11.3 | 1.1E-29 | 1.3E-28 | BT20/MB468 | NUC |
| >hg38_21_+_8433172_8446622_RNA45SN1_45S_50ntFlanks@3985.4026.42\|CGGCTTTGGTGACTCTAGATAACCTCGGGCCGATCGCACGCC | >hg38_21_+_8433172_8446622_RNA45SN1_45S_50ntFlanks | 3985 | 4026 | 42 | CGGCTTTGGTGACTCTAGATAACCTCGGGCCGATCGCACGCC | 1853.9 | -15.3 | 1.4 | -11.3 | 1.1E-29 | 1.4E-28 | BT20/MB468 | NUC |
| >hg38_21_+_8433172_8446622_RNA45SN1_45S_50ntFlanks@12157.12181.25\|AGTTTGACTGGGGCGGTACACCTGT | >hg38_21_+_8433172_8446622_RNA45SN1_45S_50ntFlanks | 12157 | 12181 | 25 | AGTTTGACTGGGGCGGTACACCTGT | 3027.2 | -15.4 | 1.3 | -11.5 | 1.0E-30 | 1.4E-28 | MB231/MB468 | MP |
| >hg38_21_+_8433172_8446622_RNA45SN1_45S_50ntFlanks@8356.8378.23\|GAAAAGAACTTTGAAGAGAGAGT | >hg38_21_+_8433172_8446622_RNA45SN1_45S_50ntFlanks | 8356 | 8378 | 23 | GAAAAGAACTTTGAAGAGAGAGT | 5485.3 | -8.3 | 0.7 | -11.3 | 1.1E-29 | 1.4E-28 | BT20/MB468 | NUC |
| >hg38_21_+_8433172_8446622_RNA45SN1_45S_50ntFlanks@6731.6770.40\|CACATTGATCATCGACACTTCGAACGCACTTGCGGCCCCG | >hg38_21_+_8433172_8446622_RNA45SN1_45S_50ntFlanks | 6731 | 6770 | 40 | CACATTGATCATCGACACTTCGAACGCACTTGCGGCCCCG | 4044.4 | -16.4 | 1.5 | -11.3 | 1.1E-29 | 1.4E-28 | BT20/MB468 | NUC |
| >hg38_21_+_8433172_8446622_RNA45SN1_45S_50ntFlanks@8113.8140.28\|GGCGCGGGACATGTGGCGTACGGAAGAC | >hg38_21_+_8433172_8446622_RNA45SN1_45S_50ntFlanks | 8113 | 8140 | 28 | GGCGCGGGACATGTGGCGTACGGAAGAC | 5497.6 | -16.3 | 1.4 | -11.5 | 1.0E-30 | 1.4E-28 | MB231/MB468 | MP |
| >hg38_21_+_8433172_8446622_RNA45SN1_45S_50ntFlanks@11572.11591.20\|CCGACTTAGAACTGGTGCGG | >hg38_21_+_8433172_8446622_RNA45SN1_45S_50ntFlanks | 11572 | 11591 | 20 | CCGACTTAGAACTGGTGCGG | 4361.3 | -8.2 | 0.7 | -11.5 | 2.2E-30 | 1.4E-28 | MB231/MB468 | MITO |
| >hg38_21_+_8433172_8446622_RNA45SN1_45S_50ntFlanks@11874.11906.33\|AGAAGACCCTGTTGAGCTTGACTCTAGTCTGGC | >hg38_21_+_8433172_8446622_RNA45SN1_45S_50ntFlanks | 11874 | 11906 | 33 | AGAAGACCCTGTTGAGCTTGACTCTAGTCTGGC | 4497.5 | -15.9 | 1.4 | -11.5 | 2.2E-30 | 1.4E-28 | MB231/MB468 | MITO |
| >hg38_21_+_8433172_8446622_RNA45SN1_45S_50ntFlanks@10099.10117.19\|CGCGCGCGCGTGTGGTGTG | >hg38_21_+_8433172_8446622_RNA45SN1_45S_50ntFlanks | 10099 | 10117 | 19 | CGCGCGCGCGTGTGGTGTG | 4125.8 | -15.9 | 1.4 | -11.5 | 1.1E-30 | 1.5E-28 | MB231/MB468 | MP |
| >hg38_21_+_8433172_8446622_RNA45SN1_45S_50ntFlanks@8113.8143.31\|GGCGCGGGACATGTGGCGTACGGAAGACCCG | >hg38_21_+_8433172_8446622_RNA45SN1_45S_50ntFlanks | 8113 | 8143 | 31 | GGCGCGGGACATGTGGCGTACGGAAGACCCG | 4562.3 | -16.0 | 1.4 | -11.5 | 1.1E-30 | 1.5E-28 | MB231/MB468 | MP |
| >hg38_21_+_8433172_8446622_RNA45SN1_45S_50ntFlanks@4570.4587.18\|TGGAATAGGACCGCGGTT | >hg38_21_+_8433172_8446622_RNA45SN1_45S_50ntFlanks | 4570 | 4587 | 18 | TGGAATAGGACCGCGGTT | 1450.8 | -14.9 | 1.3 | -11.3 | 1.2E-29 | 1.5E-28 | BT20/MB468 | NUC |
| >hg38_21_+_8433172_8446622_RNA45SN1_45S_50ntFlanks@8115.8138.24\|CGCGGGACATGTGGCGTACGGAAG | >hg38_21_+_8433172_8446622_RNA45SN1_45S_50ntFlanks | 8115 | 8138 | 24 | CGCGGGACATGTGGCGTACGGAAG | 9155.0 | -17.0 | 1.5 | -11.5 | 1.2E-30 | 1.5E-28 | MB231/MB468 | MP |
| >hg38_21_+_8433172_8446622_RNA45SN1_45S_50ntFlanks@13002.13025.24\|CGATCTATTGAAAGTCAGCCCTCG | >hg38_21_+_8433172_8446622_RNA45SN1_45S_50ntFlanks | 13002 | 13025 | 24 | CGATCTATTGAAAGTCAGCCCTCG | 622.8 | -13.7 | 1.2 | -11.3 | 1.3E-29 | 1.5E-28 | BT20/MB468 | NUC |
| >MI0000434\|hsa-let-7i&WithFlank&12\|+\|62603680\|62603775@12.32.21&[MIMAT0000415&hsa-let-7i-5p&offsets\|0\|-1;m-49&12\|+\|62603691\|62603712&offsets\|0\|-1]\|TGAGGTAGTAGTTTGTGCTGT | >MI0000434\|hsa-let-7i&WithFlank&12\|+\|62603680\|62603775 | 12 | 32 | 21 | TGAGGTAGTAGTTTGTGCTGT | 5592.2 | 7.0 | 0.6 | 11.3 | 8.1E-30 | 1.5E-28 | BT20/MB468 | CYTO |
| >MI0000440\|hsa-mir-27b&WithFlank&9\|+\|95085439\|95085547@67.86.20(+1A)&[MIMAT0000419&hsa-miR-27b-3p&offsets\|0\|-1(+1A);m-39&9\|+\|95085505\|95085525&offsets\|0\|-1(+1A)]\|TTCACAGTGGCTAAGTTCTGA | >MI0000440\|hsa-mir-27b&WithFlank&9\|+\|95085439\|95085547 | 67 | 86 | 20(+1A) | TTCACAGTGGCTAAGTTCTGA | 8191.4 | 8.8 | 0.8 | 11.3 | 8.1E-30 | 1.5E-28 | BT20/MB468 | CYTO |
| >hg38_21_+_8433172_8446622_RNA45SN1_45S_50ntFlanks@12786.12805.20\|CCCGCGCGCGCGGGAGGGCG | >hg38_21_+_8433172_8446622_RNA45SN1_45S_50ntFlanks | 12786 | 12805 | 20 | CCCGCGCGCGCGGGAGGGCG | 2235.4 | -15.6 | 1.4 | -11.3 | 1.3E-29 | 1.5E-28 | BT20/MB468 | NUC |
| >hg38_21_+_8433172_8446622_RNA45SN1_45S_50ntFlanks@4728.4759.32\|ATCAAGAACGAAAGTCGGAGGTTCGAAGACGA | >hg38_21_+_8433172_8446622_RNA45SN1_45S_50ntFlanks | 4728 | 4759 | 32 | ATCAAGAACGAAAGTCGGAGGTTCGAAGACGA | 929.1 | -14.3 | 1.3 | -11.3 | 1.3E-29 | 1.6E-28 | BT20/MB468 | NUC |
| >hg38_21_+_8433172_8446622_RNA45SN1_45S_50ntFlanks@4047.4065.19\|ATTCGAACGTCTGCCCTAT | >hg38_21_+_8433172_8446622_RNA45SN1_45S_50ntFlanks | 4047 | 4065 | 19 | ATTCGAACGTCTGCCCTAT | 938.4 | -14.3 | 1.3 | -11.3 | 1.3E-29 | 1.6E-28 | BT20/MB468 | NUC |
| >hg38_21_+_8433172_8446622_RNA45SN1_45S_50ntFlanks@6721.6742.22\|AATTGCAGGACACATTGATCAT | >hg38_21_+_8433172_8446622_RNA45SN1_45S_50ntFlanks | 6721 | 6742 | 22 | AATTGCAGGACACATTGATCAT | 1511.9 | -15.0 | 1.3 | -11.3 | 1.3E-29 | 1.6E-28 | BT20/MB468 | NUC |
| >hg38_21_+_8433172_8446622_RNA45SN1_45S_50ntFlanks@11208.11232.25\|GCGGCGACTCTGGACGCGAGCCGGG | >hg38_21_+_8433172_8446622_RNA45SN1_45S_50ntFlanks | 11208 | 11232 | 25 | GCGGCGACTCTGGACGCGAGCCGGG | 8622.0 | -13.0 | 1.1 | -11.4 | 2.5E-30 | 1.6E-28 | MB231/MB468 | MITO |
| >hg38_21_+_8433172_8446622_RNA45SN1_45S_50ntFlanks@11766.11790.25\|ATCTAATTAGTGACGCGCATGAATG | >hg38_21_+_8433172_8446622_RNA45SN1_45S_50ntFlanks | 11766 | 11790 | 25 | ATCTAATTAGTGACGCGCATGAATG | 966.6 | -14.3 | 1.3 | -11.3 | 1.3E-29 | 1.6E-28 | BT20/MB468 | NUC |
| >hg38_21_+_8433172_8446622_RNA45SN1_45S_50ntFlanks@12456.12489.34\|CCAAGCGTTGGATTGTTCACCCACTAATAGGGAA | >hg38_21_+_8433172_8446622_RNA45SN1_45S_50ntFlanks | 12456 | 12489 | 34 | CCAAGCGTTGGATTGTTCACCCACTAATAGGGAA | 5154.9 | -16.8 | 1.5 | -11.3 | 1.4E-29 | 1.6E-28 | BT20/MB468 | NUC |
| >hg38_21_+_8433172_8446622_RNA45SN1_45S_50ntFlanks@3985.4014.30\|CGGCTTTGGTGACTCTAGATAACCTCGGGC | >hg38_21_+_8433172_8446622_RNA45SN1_45S_50ntFlanks | 3985 | 4014 | 30 | CGGCTTTGGTGACTCTAGATAACCTCGGGC | 795.8 | -14.1 | 1.2 | -11.3 | 1.4E-29 | 1.6E-28 | BT20/MB468 | NUC |

<table>
<tr><td>>MI0000811|hsa-mir-148b&WithFlank&12|+|54337210|54337320@69.89.21(+1C)&[MIMAT0000759&hsa-miR-148b-3p&offsets|0|-1(+1C);m-77&12|+|54337278|54337299&offsets|0|-1(+1C)]|TCAGTGCATCACAGAACTTTGC</td><td>>MI0000811|hsa-mir-148b&WithFlank&12|+|54337210|54337320</td><td>69</td><td>89</td><td>21(+1C)</td><td>TCAGTGCATCACAGAACTTTGC</td><td>1191.8</td><td>6.8</td><td>0.6</td><td>11.3</td><td>8.6E-30</td><td>1.6E-28</td><td>BT20/MB468</td><td>CYTO</td></tr>
<tr><td>>hg38_21_+_8433172_8446622_RNA45SN1_45S_50ntFlanks@8174.8202.29|TCCTTCTGATCGAGGCCCAGCCCGTGGAC</td><td>>hg38_21_+_8433172_8446622_RNA45SN1_45S_50ntFlanks</td><td>8174</td><td>8202</td><td>29</td><td>TCCTTCTGATCGAGGCCCAGCCCGTGGAC</td><td>561.8</td><td>7.1</td><td>0.6</td><td>11.3</td><td>8.7E-30</td><td>1.6E-28</td><td>BT20/MB468</td><td>CYTO</td></tr>
<tr><td>>MI0000066|hsa-let-7e&WithFlank&19|+|51692780|51692870@14.34.21(+1C)&[MIMAT0000066&hsa-let-7e-5p&offsets|0|-1(+1C);m-50&19|+|51692793|51692814&offsets|0|-1(+1C)]|TGAGGTAGGAGGTTGTATAGTC</td><td>>MI0000066|hsa-let-7e&WithFlank&19|+|51692780|51692870</td><td>14</td><td>34</td><td>21(+1C)</td><td>TGAGGTAGGAGGTTGTATAGTC</td><td>1430.2</td><td>7.2</td><td>0.6</td><td>11.4</td><td>6.5E-30</td><td>1.8E-28</td><td>BT20/MB468</td><td>MITO</td></tr>
<tr><td>>hg38_21_+_8433172_8446622_RNA45SN1_45S_50ntFlanks@10346.10373.28|TCAAACGAGAACTTTGAAGGCCGAAGTG</td><td>>hg38_21_+_8433172_8446622_RNA45SN1_45S_50ntFlanks</td><td>10346</td><td>10373</td><td>28</td><td>TCAAACGAGAACTTTGAAGGCCGAAGTG</td><td>705.4</td><td>-5.5</td><td>0.5</td><td>-11.3</td><td>1.5E-29</td><td>1.8E-28</td><td>BT20/MB468</td><td>NUC</td></tr>
<tr><td>>MI0000061|hsa-let-7a-2&WithFlank&11|-|122146516|122146599@11.32.22(+1A);MI0000062|hsa-let-7a-3&WithFlank&22|+|46112743|46112828@10.31.22(+1A);MI0000060|hsa-let-7a-1&WithFlank&9|+|94175951|94176042@12.33.22(+1A)&[MIMAT0000062&hsa-let-7a-5p&offsets|0|0(+1A);m-6&11|-|122146568|122146589&offsets|0|0(+1A)];[MIMAT0000062_1&hsa-let-7a-5p&offsets|0|0(+1A);m-5&22|+|46112752|46112773&offsets|0|0(+1A)];[MIMAT0000062_2&hsa-let-7a-5p&offsets|0|0(+1A);m-4&9|+|94175962|94175983&offsets|0|0(+1A)]|TGAGGTAGTAGGTTGTATAGTTA</td><td>>MI0000061|hsa-let-7a-2&WithFlank&11|-|122146516|122146599</td><td>11</td><td>32</td><td>22(+1A)</td><td>TGAGGTAGTAGGTTGTATAGTTA</td><td>2751.5</td><td>7.1</td><td>0.6</td><td>11.3</td><td>9.4E-30</td><td>1.8E-28</td><td>BT20/MB468</td><td>CYTO</td></tr>
<tr><td>>hg38_21_+_8433172_8446622_RNA45SN1_45S_50ntFlanks@10786.10818.33|AACAGCCTCTGGCATGTTGGAACAATGTAGGTA</td><td>>hg38_21_+_8433172_8446622_RNA45SN1_45S_50ntFlanks</td><td>10786</td><td>10818</td><td>33</td><td>AACAGCCTCTGGCATGTTGGAACAATGTAGGTA</td><td>1416.5</td><td>-5.5</td><td>0.5</td><td>-11.4</td><td>4.7E-30</td><td>1.8E-28</td><td>MB231/MB468</td><td>NUC</td></tr>
<tr><td>>hg38_21_+_8433172_8446622_RNA45SN1_45S_50ntFlanks@11315.11344.30|CCACCCCACGTCTCGTCGCGCGCGCGTCCG</td><td>>hg38_21_+_8433172_8446622_RNA45SN1_45S_50ntFlanks</td><td>11315</td><td>11344</td><td>30</td><td>CCACCCCACGTCTCGTCGCGCGCGCGTCCG</td><td>7724.3</td><td>-16.7</td><td>1.5</td><td>-11.4</td><td>2.9E-30</td><td>1.8E-28</td><td>MB231/MB468</td><td>MITO</td></tr>
<tr><td>>hg38_21_+_8433172_8446622_RNA45SN1_45S_50ntFlanks@10408.10428.21|GGTCAGTCGGTCCTGAGAGAT</td><td>>hg38_21_+_8433172_8446622_RNA45SN1_45S_50ntFlanks</td><td>10408</td><td>10428</td><td>21</td><td>GGTCAGTCGGTCCTGAGAGAT</td><td>1580.1</td><td>-15.1</td><td>1.3</td><td>-11.3</td><td>1.6E-29</td><td>1.9E-28</td><td>BT20/MB468</td><td>NUC</td></tr>
<tr><td>>hg38_21_+_8433172_8446622_RNA45SN1_45S_50ntFlanks@9305.9338.34|CCAAGGAGTCTAACACGTGCGCGAGTCGGGGGCT</td><td>>hg38_21_+_8433172_8446622_RNA45SN1_45S_50ntFlanks</td><td>9305</td><td>9338</td><td>34</td><td>CCAAGGAGTCTAACACGTGCGCGAGTCGGGGGCT</td><td>3446.0</td><td>-15.6</td><td>1.4</td><td>-11.5</td><td>1.5E-30</td><td>1.9E-28</td><td>MB231/MB468</td><td>MP</td></tr>
<tr><td>>MI0000439|hsa-mir-23b&WithFlank&9|+|95085202|95085310@67.86.20&[MIMAT0000418&hsa-miR-23b-3p&offsets|+3|0;m-42&9|+|95085265|95085285&offsets|+3|+2]|ACATTGCCAGGGATTACCAC</td><td>>MI0000439|hsa-mir-23b&WithFlank&9|+|95085202|95085310</td><td>67</td><td>86</td><td>20</td><td>ACATTGCCAGGGATTACCAC</td><td>1377.5</td><td>14.9</td><td>1.3</td><td>11.4</td><td>7.4E-30</td><td>2.0E-28</td><td>BT20/MB468</td><td>MITO</td></tr>
<tr><td>>hg38_21_+_8433172_8446622_RNA45SN1_45S_50ntFlanks@12474.12498.25|ACCCACTAATAGGGAACGTGAGCTG</td><td>>hg38_21_+_8433172_8446622_RNA45SN1_45S_50ntFlanks</td><td>12474</td><td>12498</td><td>25</td><td>ACCCACTAATAGGGAACGTGAGCTG</td><td>613.7</td><td>-13.7</td><td>1.2</td><td>-11.3</td><td>1.7E-29</td><td>2.0E-28</td><td>BT20/MB468</td><td>NUC</td></tr>
<tr><td>>MI0000066|hsa-let-7e&WithFlank&19|+|51692780|51692870@14.34.21(+1C)&[MIMAT0000066&hsa-let-7e-5p&offsets|0|-1(+1C);m-50&19|+|51692793|51692814&offsets|0|-1(+1C)]|TGAGGTAGGAGGTTGTATAGTC</td><td>>MI0000066|hsa-let-7e&WithFlank&19|+|51692780|51692870</td><td>14</td><td>34</td><td>21(+1C)</td><td>TGAGGTAGGAGGTTGTATAGTC</td><td>5991.6</td><td>8.5</td><td>0.7</td><td>11.3</td><td>1.1E-29</td><td>2.0E-28</td><td>BT20/MB468</td><td>CYTO</td></tr>
<tr><td>>hg38_21_+_8433172_8446622_RNA45SN1_45S_50ntFlanks@9568.9596.29|CGTGCAAATCGGTCGTCCGACCTGGGTAT</td><td>>hg38_21_+_8433172_8446622_RNA45SN1_45S_50ntFlanks</td><td>9568</td><td>9596</td><td>29</td><td>CGTGCAAATCGGTCGTCCGACCTGGGTAT</td><td>14712.1</td><td>-13.1</td><td>1.2</td><td>-11.4</td><td>3.3E-30</td><td>2.0E-28</td><td>MB231/MB468</td><td>MITO</td></tr>
<tr><td>>hg38_21_+_8433172_8446622_RNA45SN1_45S_50ntFlanks@7975.8010.36|CGCGACCTCAGATCAGACGTGGCGACCCGCTGAATT</td><td>>hg38_21_+_8433172_8446622_RNA45SN1_45S_50ntFlanks</td><td>7975</td><td>8010</td><td>36</td><td>CGCGACCTCAGATCAGACGTGGCGACCCGCTGAATT</td><td>744986.8</td><td>6.7</td><td>0.6</td><td>11.3</td><td>7.8E-30</td><td>2.1E-28</td><td>BT20/MB468</td><td>MITO</td></tr>
<tr><td>>hg38_21_+_8433172_8446622_RNA45SN1_45S_50ntFlanks@10362.10388.27|AAGGCCGAAGTGGAGAAGGGTTCCATG</td><td>>hg38_21_+_8433172_8446622_RNA45SN1_45S_50ntFlanks</td><td>10362</td><td>10388</td><td>27</td><td>AAGGCCGAAGTGGAGAAGGGTTCCATG</td><td>8241.8</td><td>-17.4</td><td>1.5</td><td>-11.3</td><td>1.8E-29</td><td>2.1E-28</td><td>BT20/MB468</td><td>NUC</td></tr>
<tr><td>>MI0000089|hsa-mir-31&WithFlank&9|-|21512109|21512191@14.36.23&[MIMAT0000089&hsa-miR-31-5p&offsets|0|+2;m-360&9|-|21512157|21512178&offsets|0|+1]|AGGCAAGATGCTGGCATAGCTGT</td><td>>MI0000089|hsa-mir-31&WithFlank&9|-|21512109|21512191</td><td>14</td><td>36</td><td>23</td><td>AGGCAAGATGCTGGCATAGCTGT</td><td>2396.3</td><td>8.2</td><td>0.7</td><td>11.3</td><td>1.2E-29</td><td>2.2E-28</td><td>BT20/MB468</td><td>CYTO</td></tr>
<tr><td>>hg38_21_+_8433172_8446622_RNA45SN1_45S_50ntFlanks@13002.13028.27|CGATCTATTGAAAGTCAGCCCTCGACA</td><td>>hg38_21_+_8433172_8446622_RNA45SN1_45S_50ntFlanks</td><td>13002</td><td>13028</td><td>27</td><td>CGATCTATTGAAAGTCAGCCCTCGACA</td><td>4565.0</td><td>-15.9</td><td>1.4</td><td>-11.4</td><td>3.6E-30</td><td>2.2E-28</td><td>MB231/MB468</td><td>MITO</td></tr>
<tr><td>>hg38_21_+_8433172_8446622_RNA45SN1_45S_50ntFlanks@12405.12443.39|CTTTTTGATCCTTCGATGTCGGCTCTTCCTATCATTGTG</td><td>>hg38_21_+_8433172_8446622_RNA45SN1_45S_50ntFlanks</td><td>12405</td><td>12443</td><td>39</td><td>CTTTTTGATCCTTCGATGTCGGCTCTTCCTATCATTGTG</td><td>46977.6</td><td>-8.9</td><td>0.8</td><td>-11.4</td><td>6.0E-30</td><td>2.3E-28</td><td>MB231/MB468</td><td>NUC</td></tr>
<tr><td>>MI0000082|hsa-mir-25&WithFlank&7|-|100093554|100093649@58.78.21(+1U)&[MIMAT0000081&hsa-miR-25-3p&offsets|0|-1(+1U);m-23&7|-|100093571|100093592&offsets|0|-1(+1U)]|CATTGCACTTGTCTCGGTCTGT</td><td>>MI0000082|hsa-mir-25&WithFlank&7|-|100093554|100093649</td><td>58</td><td>78</td><td>21(+1U)</td><td>CATTGCACTTGTCTCGGTCTGT</td><td>3460.5</td><td>8.4</td><td>0.7</td><td>11.3</td><td>8.7E-30</td><td>2.3E-28</td><td>BT20/MB468</td><td>MITO</td></tr>
<tr><td>>hg38_21_+_8433172_8446622_RNA45SN1_45S_50ntFlanks@12008.12032.25|CCGCCGGTGAAATACCACTACTCTG</td><td>>hg38_21_+_8433172_8446622_RNA45SN1_45S_50ntFlanks</td><td>12008</td><td>12032</td><td>25</td><td>CCGCCGGTGAAATACCACTACTCTG</td><td>68425.1</td><td>-15.0</td><td>1.3</td><td>-11.5</td><td>1.9E-30</td><td>2.4E-28</td><td>MB231/MB468</td><td>MP</td></tr>
<tr><td>>hg38_21_+_8433172_8446622_RNA45SN1_45S_50ntFlanks@13004.13028.25|ATCTATTGAAAGTCAGCCCTCGACA</td><td>>hg38_21_+_8433172_8446622_RNA45SN1_45S_50ntFlanks</td><td>13004</td><td>13028</td><td>25</td><td>ATCTATTGAAAGTCAGCCCTCGACA</td><td>1223.9</td><td>-14.7</td><td>1.3</td><td>-11.3</td><td>2.1E-29</td><td>2.5E-28</td><td>BT20/MB468</td><td>NUC</td></tr>
<tr><td>>MI0000440|hsa-mir-27b&WithFlank&9|+|95085439|95085547@67.86.20(+1A)&[MIMAT0000419&hsa-miR-27b-3p&offsets|0|-1(+1A);m-39&9|+|95085505|95085525&offsets|0|-1(+1A)]|TTCACAGTGGCTAAGTTCTGA</td><td>>MI0000440|hsa-mir-27b&WithFlank&9|+|95085439|95085547</td><td>67</td><td>86</td><td>20(+1A)</td><td>TTCACAGTGGCTAAGTTCTGA</td><td>1418.3</td><td>6.3</td><td>0.6</td><td>11.4</td><td>6.6E-30</td><td>2.5E-28</td><td>MB231/MB468</td><td>NUC</td></tr>
<tr><td>>hg38_21_+_8433172_8446622_RNA45SN1_45S_50ntFlanks@11613.11640.28|AATTAAAACAAAGCATCGCGAAGGCCCG</td><td>>hg38_21_+_8433172_8446622_RNA45SN1_45S_50ntFlanks</td><td>11613</td><td>11640</td><td>28</td><td>AATTAAAACAAAGCATCGCGAAGGCCCG</td><td>3357.3</td><td>-15.6</td><td>1.4</td><td>-11.5</td><td>2.0E-30</td><td>2.5E-28</td><td>MB231/MB468</td><td>MP</td></tr>
<tr><td>>MI0001733|hsa-mir-452&WithFlank&X|-|151959622|151959718@20.43.24&[MIMAT0001635&hsa-miR-452-5p&offsets|0|+2;m-188&X|-|151959677|151959699&offsets|0|+1]|AACTGTTTGCAGAGGAAACTGAGA</td><td>>MI0001733|hsa-mir-452&WithFlank&X|-|151959622|151959718</td><td>20</td><td>43</td><td>24</td><td>AACTGTTTGCAGAGGAAACTGAGA</td><td>1821.9</td><td>15.1</td><td>1.3</td><td>11.3</td><td>1.4E-29</td><td>2.7E-28</td><td>BT20/MB468</td><td>CYTO</td></tr>
<tr><td>>MI0001445|hsa-mir-423&WithFlank&17|+|30117073|30117178@59.78.20&[MIMAT0001340&hsa-miR-423-3p&offsets|0|-3;m-100&17|+|30117131|30117153&offsets|0|-3]|AGCTCGGTCTGAGGCCCCTC</td><td>>MI0001445|hsa-mir-423&WithFlank&17|+|30117073|30117178</td><td>59</td><td>78</td><td>20</td><td>AGCTCGGTCTGAGGCCCCTC</td><td>5726.1</td><td>8.2</td><td>0.7</td><td>11.3</td><td>1.4E-29</td><td>2.7E-28</td><td>BT20/MB468</td><td>CYTO</td></tr>
<tr><td>>hg38_21_+_8433172_8446622_RNA45SN1_45S_50ntFlanks@10349.10378.30|AACGAGAACTTTGAAGGCCGAAGTGGAGAA</td><td>>hg38_21_+_8433172_8446622_RNA45SN1_45S_50ntFlanks</td><td>10349</td><td>10378</td><td>30</td><td>AACGAGAACTTTGAAGGCCGAAGTGGAGAA</td><td>1135.0</td><td>-6.6</td><td>0.6</td><td>-11.2</td><td>2.3E-29</td><td>2.7E-28</td><td>BT20/MB468</td><td>NUC</td></tr>
<tr><td>>MI0000079|hsa-mir-23a&WithFlank&19|-|13836581|13836665@54.71.18&[MIMAT0000078&hsa-miR-23a-3p&offsets|+3|0;m-31&19|-|13836595|13836615&offsets|+3|0]|ACATTGCCAGGGATTTCC</td><td>>MI0000079|hsa-mir-23a&WithFlank&19|-|13836581|13836665</td><td>54</td><td>71</td><td>18</td><td>ACATTGCCAGGGATTTCC</td><td>2555.8</td><td>15.8</td><td>1.4</td><td>11.3</td><td>1.1E-29</td><td>2.8E-28</td><td>BT20/MB468</td><td>MITO</td></tr>
<tr><td>>hg38_21_+_8433172_8446622_RNA45SN1_45S_50ntFlanks@10729.10755.27|AGAGGGTGTAAATCTCGCGCCGGGCCG</td><td>>hg38_21_+_8433172_8446622_RNA45SN1_45S_50ntFlanks</td><td>10729</td><td>10755</td><td>27</td><td>AGAGGGTGTAAATCTCGCGCCGGGCCG</td><td>1725.8</td><td>-15.2</td><td>1.4</td><td>-11.2</td><td>2.5E-29</td><td>2.9E-28</td><td>BT20/MB468</td><td>NUC</td></tr>
<tr><td>>hg38_21_+_8433172_8446622_RNA45SN1_45S_50ntFlanks@482.516.35|ATCGATGTGGTGACGTCGTGCTCTCCCGGGCCGGG</td><td>>hg38_21_+_8433172_8446622_RNA45SN1_45S_50ntFlanks</td><td>482</td><td>516</td><td>35</td><td>ATCGATGTGGTGACGTCGTGCTCTCCCGGGCCGGG</td><td>1247.9</td><td>-14.7</td><td>1.3</td><td>-11.2</td><td>2.5E-29</td><td>2.9E-28</td><td>BT20/MB468</td><td>NUC</td></tr>
<tr><td>>hg38_21_+_8433172_8446622_RNA45SN1_45S_50ntFlanks@9570.9597.28|TGCAAATCGGTCGTCCGACCTGGGTATA</td><td>>hg38_21_+_8433172_8446622_RNA45SN1_45S_50ntFlanks</td><td>9570</td><td>9597</td><td>28</td><td>TGCAAATCGGTCGTCCGACCTGGGTATA</td><td>1363.8</td><td>-14.8</td><td>1.3</td><td>-11.2</td><td>2.5E-29</td><td>2.9E-28</td><td>BT20/MB468</td><td>NUC</td></tr>
<tr><td>>hg38_21_+_8433172_8446622_RNA45SN1_45S_50ntFlanks@3838.3862.25|CTCTCCTACTTGGATAACTGTGGTA</td><td>>hg38_21_+_8433172_8446622_RNA45SN1_45S_50ntFlanks</td><td>3838</td><td>3862</td><td>25</td><td>CTCTCCTACTTGGATAACTGTGGTA</td><td>348.7</td><td>-4.7</td><td>0.4</td><td>-11.2</td><td>2.6E-29</td><td>3.0E-28</td><td>BT20/MB468</td><td>NUC</td></tr>
<tr><td>>hg38_21_+_8433172_8446622_RNA45SN1_45S_50ntFlanks@12934.12954.21|CTAAACCATTCGTAGACGACC</td><td>>hg38_21_+_8433172_8446622_RNA45SN1_45S_50ntFlanks</td><td>12934</td><td>12954</td><td>21</td><td>CTAAACCATTCGTAGACGACC</td><td>701.2</td><td>-13.9</td><td>1.2</td><td>-11.2</td><td>2.6E-29</td><td>3.1E-28</td><td>BT20/MB468</td><td>NUC</td></tr>
<tr><td>>hg38_21_+_8433172_8446622_RNA45SN1_45S_50ntFlanks@5491.5512.22|GAGAAGACGGTCGAACTTGACT</td><td>>hg38_21_+_8433172_8446622_RNA45SN1_45S_50ntFlanks</td><td>5491</td><td>5512</td><td>22</td><td>GAGAAGACGGTCGAACTTGACT</td><td>342.5</td><td>5.7</td><td>0.5</td><td>11.5</td><td>1.5E-30</td><td>3.1E-28</td><td>MB231/MB468</td><td>CYTO</td></tr>
<tr><td>>hg38_21_+_8433172_8446622_RNA45SN1_45S_50ntFlanks@10684.10710.27|CGGCGGCGTCCGGTGAGCTCTCGCTGG</td><td>>hg38_21_+_8433172_8446622_RNA45SN1_45S_50ntFlanks</td><td>10684</td><td>10710</td><td>27</td><td>CGGCGGCGTCCGGTGAGCTCTCGCTGG</td><td>9275.4</td><td>-6.6</td><td>0.6</td><td>-11.5</td><td>1.5E-30</td><td>3.1E-28</td><td>MB231/MB468</td><td>CYTO</td></tr>
<tr><td>>hg38_21_+_8433172_8446622_RNA45SN1_45S_50ntFlanks@6721.6739.19|AATTGCAGGACACATTGAT</td><td>>hg38_21_+_8433172_8446622_RNA45SN1_45S_50ntFlanks</td><td>6721</td><td>6739</td><td>19</td><td>AATTGCAGGACACATTGAT</td><td>3664.5</td><td>-16.3</td><td>1.4</td><td>-11.2</td><td>2.7E-29</td><td>3.1E-28</td><td>BT20/MB468</td><td>NUC</td></tr>
<tr><td>>hg38_21_+_8433172_8446622_RNA45SN1_45S_50ntFlanks@7989.8006.18|AGACGTGGCGACCCGCTG</td><td>>hg38_21_+_8433172_8446622_RNA45SN1_45S_50ntFlanks</td><td>7989</td><td>8006</td><td>18</td><td>AGACGTGGCGACCCGCTG</td><td>10790.4</td><td>-9.3</td><td>0.8</td><td>-11.4</td><td>5.2E-30</td><td>3.1E-28</td><td>MB231/MB468</td><td>MITO</td></tr>
<tr><td>>hg38_21_+_8433172_8446622_RNA45SN1_45S_50ntFlanks@8170.8195.26|CAAGTCCTTCTGATCGAGGCCCAGCC</td><td>>hg38_21_+_8433172_8446622_RNA45SN1_45S_50ntFlanks</td><td>8170</td><td>8195</td><td>26</td><td>CAAGTCCTTCTGATCGAGGCCCAGCC</td><td>1366.7</td><td>-14.8</td><td>1.3</td><td>-11.2</td><td>2.7E-29</td><td>3.2E-28</td><td>BT20/MB468</td><td>NUC</td></tr>
<tr><td>>hg38_21_+_8433172_8446622_RNA45SN1_45S_50ntFlanks@3874.3898.25|AATACATGCCGACGGGCGCTGACCC</td><td>>hg38_21_+_8433172_8446622_RNA45SN1_45S_50ntFlanks</td><td>3874</td><td>3898</td><td>25</td><td>AATACATGCCGACGGGCGCTGACCC</td><td>2675.1</td><td>-5.9</td><td>0.5</td><td>-11.3</td><td>1.2E-29</td><td>3.2E-28</td><td>BT20/MB468</td><td>MITO</td></tr>
<tr><td>>MI0000070|hsa-mir-16-1&WithFlank&13|-|50048967|50049067@20.41.22;MI0000115|hsa-mir-16-2&WithFlank&3|+|160404739|160404831@16.37.22&[MIMAT0000069&hsa-miR-16-5p&offsets|0|0;m-62&13|-|50049027|50049048&offsets|0|0];[MIMAT0000069_1&hsa-miR-16-5p&offsets|0|0;m-61&3|+|160404754|160404775&offsets|0|0]|TAGCAGCACGTAAATATTGGCG</td><td>>MI0000070|hsa-mir-16-1&WithFlank&13|-|50048967|50049067</td><td>20</td><td>41</td><td>22</td><td>TAGCAGCACGTAAATATTGGCG</td><td>7574.2</td><td>7.9</td><td>0.7</td><td>11.3</td><td>1.8E-29</td><td>3.3E-28</td><td>BT20/MB468</td><td>CYTO</td></tr>
<tr><td>>MI0000266|hsa-mir-10a&WithFlank&17|-|48579832|48579953@28.50.23(+1A)&[MIMAT0000253&hsa-miR-10a-5p&offsets|0|0(+1A);m-18&17|-|48579905|48579926&offsets|0|+1(+1A)]|TACCCTGTAGATCCGAATTTGTGA</td><td>>MI0000266|hsa-mir-10a&WithFlank&17|-|48579832|48579953</td><td>28</td><td>50</td><td>23(+1A)</td><td>TACCCTGTAGATCCGAATTTGTGA</td><td>2333.6</td><td>15.4</td><td>1.4</td><td>11.3</td><td>1.8E-29</td><td>3.3E-28</td><td>BT20/MB468</td><td>CYTO</td></tr>
<tr><td>>MI0000542|hsa-mir-320a&WithFlank&8|-|22244960|22245043@48.68.21(+4U)&[MIMAT0000510&hsa-miR-320a-3p&offsets|0|-1(+4U);m-60&8|-|22244975|22244996&offsets|0|-1(+4U)]|AAAAGCTGGGTTGAGAGGGCGTTTT</td><td>>MI0000542|hsa-mir-320a&WithFlank&8|-|22244960|22245043</td><td>48</td><td>68</td><td>21(+4U)</td><td>AAAAGCTGGGTTGAGAGGGCGTTTT</td><td>900.4</td><td>14.1</td><td>1.2</td><td>11.3</td><td>1.8E-29</td><td>3.4E-28</td><td>BT20/MB468</td><td>CYTO</td></tr>
<tr><td>>am_MI0000809|hsa-mir-151a&WithFlank&8|-|140732558|140732659@53.71.19(+1A)&[MIMAT0000757&hsa-miR-151a-3p&offsets|0|-2(+1A);m-30&8|-|140732587|140732607&offsets|0|-2(+1A)]|CTAGACTGAAGCTCCTTGAA</td><td>>am_MI0000809|hsa-mir-151a&WithFlank&8|-|140732558|140732659</td><td>53</td><td>71</td><td>19(+1A)</td><td>CTAGACTGAAGCTCCTTGAA</td><td>856.3</td><td>14.0</td><td>1.2</td><td>11.3</td><td>1.9E-29</td><td>3.4E-28</td><td>BT20/MB468</td><td>CYTO</td></tr>
<tr><td>>hg38_21_+_8433172_8446622_RNA45SN1_45S_50ntFlanks@3837.3860.24|CCTCTCCTACTTGGATAACTGTGG</td><td>>hg38_21_+_8433172_8446622_RNA45SN1_45S_50ntFlanks</td><td>3837</td><td>3860</td><td>24</td><td>CCTCTCCTACTTGGATAACTGTGG</td><td>713.1</td><td>-13.9</td><td>1.2</td><td>-11.2</td><td>3.0E-29</td><td>3.5E-28</td><td>BT20/MB468</td><td>NUC</td></tr>
<tr><td>>hg38_21_+_8433172_8446622_RNA45SN1_45S_50ntFlanks@12973.12990.18|TCGTACGTAGCAGAGCAG</td><td>>hg38_21_+_8433172_8446622_RNA45SN1_45S_50ntFlanks</td><td>12973</td><td>12990</td><td>18</td><td>TCGTACGTAGCAGAGCAG</td><td>3829.3</td><td>-15.8</td><td>1.4</td><td>-11.4</td><td>2.9E-30</td><td>3.5E-28</td><td>MB231/MB468</td><td>MP</td></tr>
</table>

<table>
<tr><td>>hg38_21_+_8433172_8446622_RNA45SN1_45S_50ntFlanks@4588.4614.27|CTATTTTGTTGGTTTTCGGAACTGAGG</td><td>>hg38_21_+_8433172_8446622_RNA45SN1_45S_50ntFlanks</td><td>4588</td><td>4614</td><td>27</td><td>CTATTTTGTTGGTTTTCGGAACTGAGG</td><td>1362.6</td><td>-14.8</td><td>1.3</td><td>-11.2</td><td>3.1E-29</td><td>3.6E-28</td><td>BT20/MB468</td><td>NUC</td></tr>
<tr><td>>hg38_21_+_8433172_8446622_RNA45SN1_45S_50ntFlanks@11208.11225.18|GCGGCGACTCTGGACGCG</td><td>>hg38_21_+_8433172_8446622_RNA45SN1_45S_50ntFlanks</td><td>11208</td><td>11225</td><td>18</td><td>GCGGCGACTCTGGACGCG</td><td>17509.1</td><td>-13.7</td><td>1.2</td><td>-11.4</td><td>2.9E-30</td><td>3.6E-28</td><td>MB231/MB468</td><td>MP</td></tr>
<tr><td>>hg38_21_+_8433172_8446622_RNA45SN1_45S_50ntFlanks@12412.12444.33|ATCCTTCGATGTCGGCTCTTCCTATCATTGTGA</td><td>>hg38_21_+_8433172_8446622_RNA45SN1_45S_50ntFlanks</td><td>12412</td><td>12444</td><td>33</td><td>ATCCTTCGATGTCGGCTCTTCCTATCATTGTGA</td><td>980.0</td><td>-14.4</td><td>1.3</td><td>-11.2</td><td>3.2E-29</td><td>3.6E-28</td><td>BT20/MB468</td><td>NUC</td></tr>
<tr><td>>MI0000266|hsa-mir-10a&WithFlank&17|-|48579832|48579953@29.50.22&[MIMAT0000253&hsa-miR-10a-5p&offsets|+1|0;m-18&17|-|48579905|48579926&offsets|+1|+1]|ACCCTGTAGATCCGAATTTGTG</td><td>>MI0000266|hsa-mir-10a&WithFlank&17|-|48579832|48579953</td><td>29</td><td>50</td><td>22</td><td>ACCCTGTAGATCCGAATTTGTG</td><td>3481.9</td><td>15.8</td><td>1.4</td><td>11.5</td><td>1.8E-30</td><td>3.6E-28</td><td>MB231/MB468</td><td>CYTO</td></tr>
<tr><td>>hg38_21_+_8433172_8446622_RNA45SN1_45S_50ntFlanks@11566.11583.18|TAGCAGCCGACTTAGAAC</td><td>>hg38_21_+_8433172_8446622_RNA45SN1_45S_50ntFlanks</td><td>11566</td><td>11583</td><td>18</td><td>TAGCAGCCGACTTAGAAC</td><td>681.7</td><td>6.7</td><td>0.6</td><td>11.5</td><td>1.9E-30</td><td>3.6E-28</td><td>MB231/MB468</td><td>CYTO</td></tr>
<tr><td>>hg38_21_+_8433172_8446622_RNA45SN1_45S_50ntFlanks@6778.6806.29|CCCGGGGCTACGCCTGTCTGAGCGTCGCT</td><td>>hg38_21_+_8433172_8446622_RNA45SN1_45S_50ntFlanks</td><td>6778</td><td>6806</td><td>29</td><td>CCCGGGGCTACGCCTGTCTGAGCGTCGCT</td><td>1042.3</td><td>5.5</td><td>0.5</td><td>11.5</td><td>1.9E-30</td><td>3.6E-28</td><td>MB231/MB468</td><td>CYTO</td></tr>
<tr><td>>MI0003643|hsa-mir-629&WithFlank&15|-|70079366|70079474@28.48.21&[MIMAT0004810&hsa-miR-629-5p&offsets|0|0;m-150&15|-|70079427|70079447&offsets|0|0]|TGGGTTTACGTTGGGAGAACT</td><td>>MI0003643|hsa-mir-629&WithFlank&15|-|70079366|70079474</td><td>28</td><td>48</td><td>21</td><td>TGGGTTTACGTTGGGAGAACT</td><td>756.5</td><td>14.0</td><td>1.2</td><td>11.3</td><td>1.4E-29</td><td>3.7E-28</td><td>BT20/MB468</td><td>MITO</td></tr>
<tr><td>>MI0000079|hsa-mir-23a&WithFlank&19|-|13836581|13836665@54.71.18&[MIMAT0000078&hsa-miR-23a-3p&offsets|+3|0;m-31&19|-|13836595|13836615&offsets|+3|0]|ACATTGCCAGGGATTTCC</td><td>>MI0000079|hsa-mir-23a&WithFlank&19|-|13836581|13836665</td><td>54</td><td>71</td><td>18</td><td>ACATTGCCAGGGATTTCC</td><td>1057.3</td><td>12.9</td><td>1.1</td><td>11.2</td><td>3.2E-29</td><td>3.7E-28</td><td>BT20/MB468</td><td>NUC</td></tr>
<tr><td>>MI0000102|hsa-mir-100&WithFlank&11|-|122152223|122152314@19.38.20(+1G)&[MIMAT0000098&hsa-miR-100-5p&offsets|0|-2(+1G);m-27&11|-|122152276|122152296&offsets|0|-1(+1G)]|AACCCGTAGATCCGAACTTGG</td><td>>MI0000102|hsa-mir-100&WithFlank&11|-|122152223|122152314</td><td>19</td><td>38</td><td>20(+1G)</td><td>AACCCGTAGATCCGAACTTGG</td><td>3476.8</td><td>15.7</td><td>1.4</td><td>11.3</td><td>9.9E-30</td><td>3.7E-28</td><td>MB231/MB468</td><td>NUC</td></tr>
<tr><td>>hg38_21_+_8433172_8446622_RNA45SN1_45S_50ntFlanks@10346.10374.29|TCAAACGAGAACTTTGAAGGCCGAAGTGG</td><td>>hg38_21_+_8433172_8446622_RNA45SN1_45S_50ntFlanks</td><td>10346</td><td>10374</td><td>29</td><td>TCAAACGAGAACTTTGAAGGCCGAAGTGG</td><td>1152.0</td><td>-4.9</td><td>0.4</td><td>-11.3</td><td>1.0E-29</td><td>3.8E-28</td><td>MB231/MB468</td><td>NUC</td></tr>
<tr><td>>hg38_21_+_8433172_8446622_RNA45SN1_45S_50ntFlanks@12957.12989.33|CTTCTGGGTCGGGGTTTCGTACGTAGCAGAGCA</td><td>>hg38_21_+_8433172_8446622_RNA45SN1_45S_50ntFlanks</td><td>12957</td><td>12989</td><td>33</td><td>CTTCTGGGTCGGGGTTTCGTACGTAGCAGAGCA</td><td>1334.6</td><td>-14.8</td><td>1.3</td><td>-11.2</td><td>3.5E-29</td><td>4.0E-28</td><td>BT20/MB468</td><td>NUC</td></tr>
<tr><td>>MI0006444|hsa-mir-1307&WithFlank&10|-|103394247|103394407@87.108.22&[MIMAT0005951&hsa-miR-1307-3p&offsets|+1|+1;m-66&10|-|103394300|103394322&offsets|+1|0]|CTCGGCGTGGCGTCGGTCGTGG</td><td>>MI0006444|hsa-mir-1307&WithFlank&10|-|103394247|103394407</td><td>87</td><td>108</td><td>22</td><td>CTCGGCGTGGCGTCGGTCGTGG</td><td>2320.5</td><td>6.6</td><td>0.6</td><td>11.3</td><td>2.2E-29</td><td>4.0E-28</td><td>BT20/MB468</td><td>CYTO</td></tr>
<tr><td>>MI0000734|hsa-mir-106b&WithFlank&7|-|100093987|100094080@58.78.21(+1U)&[MIMAT0004672&hsa-miR-106b-3p&offsets|0|-1(+1U);m-87&7|-|100094003|100094023&offsets|0|0(+1U)]|CCGCACTGTGGGTACTTGCTGT</td><td>>MI0000734|hsa-mir-106b&WithFlank&7|-|100093987|100094080</td><td>58</td><td>78</td><td>21(+1U)</td><td>CCGCACTGTGGGTACTTGCTGT</td><td>1650.0</td><td>7.7</td><td>0.7</td><td>11.3</td><td>1.6E-29</td><td>4.2E-28</td><td>BT20/MB468</td><td>MITO</td></tr>
<tr><td>>hg38_21_+_8433172_8446622_RNA45SN1_45S_50ntFlanks@12397.12417.21|CGACGTCGCTTTTTGATCCTT</td><td>>hg38_21_+_8433172_8446622_RNA45SN1_45S_50ntFlanks</td><td>12397</td><td>12417</td><td>21</td><td>CGACGTCGCTTTTTGATCCTT</td><td>2056.2</td><td>-7.1</td><td>0.6</td><td>-11.2</td><td>3.7E-29</td><td>4.2E-28</td><td>BT20/MB468</td><td>NUC</td></tr>
<tr><td>>MI0000088|hsa-mir-30a&WithFlank&6|-|71403545|71403627@54.73.20(+1U)&[MIMAT0000088&hsa-miR-30a-3p&offsets|+1|-1(+1U);m-38&6|-|71403554|71403575&offsets|+1|-1(+1U)]|TTTCAGTCGGATGTTTGCAGT</td><td>>MI0000088|hsa-mir-30a&WithFlank&6|-|71403545|71403627</td><td>54</td><td>73</td><td>20(+1U)</td><td>TTTCAGTCGGATGTTTGCAGT</td><td>1221.8</td><td>7.5</td><td>0.7</td><td>11.3</td><td>1.6E-29</td><td>4.2E-28</td><td>BT20/MB468</td><td>MITO</td></tr>
<tr><td>>MI0000478|hsa-mir-149&WithFlank&2|+|240455995|240456095@21.41.21(+1U)&[MIMAT0000450&hsa-miR-149-5p&offsets|0|-2(+1U);m-195&2|+|240456015|240456037&offsets|0|-2(+1U)]|TCTGGCTCCGTGTCTTCACTCT</td><td>>MI0000478|hsa-mir-149&WithFlank&2|+|240455995|240456095</td><td>21</td><td>41</td><td>21(+1U)</td><td>TCTGGCTCCGTGTCTTCACTCT</td><td>632.9</td><td>13.8</td><td>1.2</td><td>11.3</td><td>1.6E-29</td><td>4.2E-28</td><td>BT20/MB468</td><td>MITO</td></tr>
<tr><td>>hg38_21_+_8433172_8446622_RNA45SN1_45S_50ntFlanks@12417.12435.19|TCGATGTCGGCTCTTCCTA</td><td>>hg38_21_+_8433172_8446622_RNA45SN1_45S_50ntFlanks</td><td>12417</td><td>12435</td><td>19</td><td>TCGATGTCGGCTCTTCCTA</td><td>3701.4</td><td>-7.2</td><td>0.6</td><td>-11.4</td><td>7.1E-30</td><td>4.2E-28</td><td>MB231/MB468</td><td>MITO</td></tr>
<tr><td>>MI0000082|hsa-mir-25&WithFlank&7|-|100093554|100093649@58.77.20(+1A)&[MIMAT0000081&hsa-miR-25-3p&offsets|0|-2(+1A);m-23&7|-|100093571|100093592&offsets|0|-2(+1A)]|CATTGCACTTGTCTCGGTCTA</td><td>>MI0000082|hsa-mir-25&WithFlank&7|-|100093554|100093649</td><td>58</td><td>77</td><td>20(+1A)</td><td>CATTGCACTTGTCTCGGTCTA</td><td>895.6</td><td>7.4</td><td>0.7</td><td>11.3</td><td>1.7E-29</td><td>4.3E-28</td><td>BT20/MB468</td><td>MITO</td></tr>
<tr><td>>hg38_21_+_8433172_8446622_RNA45SN1_45S_50ntFlanks@7975.8011.37|CGCGACCTCAGATCAGACGTGGCGACCCGCTGAATTT</td><td>>hg38_21_+_8433172_8446622_RNA45SN1_45S_50ntFlanks</td><td>7975</td><td>8011</td><td>37</td><td>CGCGACCTCAGATCAGACGTGGCGACCCGCTGAATTT</td><td>132026.5</td><td>7.7</td><td>0.7</td><td>11.5</td><td>2.3E-30</td><td>4.3E-28</td><td>MB231/MB468</td><td>CYTO</td></tr>
<tr><td>>am_MI0000809|hsa-mir-151a&WithFlank&8|-|140732558|140732659@17.37.21&[MIMAT0004697&hsa-miR-151a-5p&offsets|0|0;m-75&8|-|140732623|140732643&offsets|0|0]|TCGAGGAGCTCACAGTCTAGT</td><td>>am_MI0000809|hsa-mir-151a&WithFlank&8|-|140732558|140732659</td><td>17</td><td>37</td><td>21</td><td>TCGAGGAGCTCACAGTCTAGT</td><td>911.2</td><td>5.8</td><td>0.5</td><td>11.2</td><td>3.9E-29</td><td>4.4E-28</td><td>BT20/MB468</td><td>NUC</td></tr>
<tr><td>>hg38_21_+_8433172_8446622_RNA45SN1_45S_50ntFlanks@9504.9533.30|GTGAACTATGCCTGGGCAGGGCGAAGCCAG</td><td>>hg38_21_+_8433172_8446622_RNA45SN1_45S_50ntFlanks</td><td>9504</td><td>9533</td><td>30</td><td>GTGAACTATGCCTGGGCAGGGCGAAGCCAG</td><td>1776.1</td><td>-15.2</td><td>1.4</td><td>-11.2</td><td>3.9E-29</td><td>4.4E-28</td><td>BT20/MB468</td><td>NUC</td></tr>
<tr><td>>hg38_21_+_8433172_8446622_RNA45SN1_45S_50ntFlanks@10391.10419.29|AACAGCAGTTGAACATGGGTCAGTCGGTC</td><td>>hg38_21_+_8433172_8446622_RNA45SN1_45S_50ntFlanks</td><td>10391</td><td>10419</td><td>29</td><td>AACAGCAGTTGAACATGGGTCAGTCGGTC</td><td>1662.3</td><td>-15.1</td><td>1.4</td><td>-11.2</td><td>4.0E-29</td><td>4.5E-28</td><td>BT20/MB468</td><td>NUC</td></tr>
<tr><td>>hg38_21_+_8433172_8446622_RNA45SN1_45S_50ntFlanks@10679.10711.33|GGTTCCGGCGGCGTCCGGTGAGCTCTCGCTGGC</td><td>>hg38_21_+_8433172_8446622_RNA45SN1_45S_50ntFlanks</td><td>10679</td><td>10711</td><td>33</td><td>GGTTCCGGCGGCGTCCGGTGAGCTCTCGCTGGC</td><td>3825.5</td><td>-15.8</td><td>1.4</td><td>-11.4</td><td>3.7E-30</td><td>4.5E-28</td><td>MB231/MB468</td><td>MP</td></tr>
<tr><td>>hg38_21_+_8433172_8446622_RNA45SN1_45S_50ntFlanks@3834.3860.27|GCTCCTCTCCTACTTGGATAACTGTGG</td><td>>hg38_21_+_8433172_8446622_RNA45SN1_45S_50ntFlanks</td><td>3834</td><td>3860</td><td>27</td><td>GCTCCTCTCCTACTTGGATAACTGTGG</td><td>567.3</td><td>-13.6</td><td>1.2</td><td>-11.2</td><td>4.1E-29</td><td>4.6E-28</td><td>BT20/MB468</td><td>NUC</td></tr>
<tr><td>>hg38_21_+_8433172_8446622_RNA45SN1_45S_50ntFlanks@5491.5516.26|GAGAAGACGGTCGAACTTGACTATCT</td><td>>hg38_21_+_8433172_8446622_RNA45SN1_45S_50ntFlanks</td><td>5491</td><td>5516</td><td>26</td><td>GAGAAGACGGTCGAACTTGACTATCT</td><td>9012.7</td><td>7.7</td><td>0.7</td><td>11.4</td><td>2.6E-30</td><td>4.6E-28</td><td>MB231/MB468</td><td>CYTO</td></tr>
<tr><td>>trna116_GluCTC_1_-_145399233_145399304@32.73.42,trna71_GluCTC_1_-_161439189_161439260@32.73.42,trna74_GluCTC_1_-_161431809_161431880@32.73.42,trna77_GluCTC_1_-_161424398_161424469@32.73.42,trna77_GluCTC_6_+_28949976_28950047@32.73.42,trna80_GluCTC_1_-_161417018_161417089@32.73.42,trna87_GluCTC_6_-_126101393_126101464@32.73.42|CTCTCACCGCCGCGGCCCGGGTTCGATTCCCGGTCAGGGAAC</td><td>>trna116_GluCTC_1_-_145399233_145399304</td><td>32</td><td>73</td><td>42</td><td>CTCTCACCGCCGCGGCCCGGGTTCGATTCCCGGTCAGGGAAC</td><td>1675.2</td><td>-15.1</td><td>1.4</td><td>-11.2</td><td>4.1E-29</td><td>4.6E-28</td><td>BT20/MB468</td><td>NUC</td></tr>
<tr><td>>hg38_21_+_8433172_8446622_RNA45SN1_45S_50ntFlanks@6661.6701.41|CGGTGGATCACTCGGCTCGTGCGTCGATGAAGAACGCAGCT</td><td>>hg38_21_+_8433172_8446622_RNA45SN1_45S_50ntFlanks</td><td>6661</td><td>6701</td><td>41</td><td>CGGTGGATCACTCGGCTCGTGCGTCGATGAAGAACGCAGCT</td><td>6931.5</td><td>7.4</td><td>0.7</td><td>11.3</td><td>1.8E-29</td><td>4.7E-28</td><td>BT20/MB468</td><td>MITO</td></tr>
<tr><td>>hg38_21_+_8433172_8446622_RNA45SN1_45S_50ntFlanks@7991.8008.18|ACGTGGCGACCCGCTGAA</td><td>>hg38_21_+_8433172_8446622_RNA45SN1_45S_50ntFlanks</td><td>7991</td><td>8008</td><td>18</td><td>ACGTGGCGACCCGCTGAA</td><td>20419.6</td><td>-6.1</td><td>0.5</td><td>-11.3</td><td>1.8E-29</td><td>4.7E-28</td><td>BT20/MB468</td><td>MITO</td></tr>
<tr><td>>hg38_21_+_8433172_8446622_RNA45SN1_45S_50ntFlanks@6746.6770.25|CACTTCGAACGCACTTGCGGCCCCG</td><td>>hg38_21_+_8433172_8446622_RNA45SN1_45S_50ntFlanks</td><td>6746</td><td>6770</td><td>25</td><td>CACTTCGAACGCACTTGCGGCCCCG</td><td>2990.3</td><td>-15.4</td><td>1.4</td><td>-11.4</td><td>4.0E-30</td><td>4.8E-28</td><td>MB231/MB468</td><td>MP</td></tr>
<tr><td>>hg38_21_+_8433172_8446622_RNA45SN1_45S_50ntFlanks@11781.11804.24|CGCATGAATGGATGAACGAGATTC</td><td>>hg38_21_+_8433172_8446622_RNA45SN1_45S_50ntFlanks</td><td>11781</td><td>11804</td><td>24</td><td>CGCATGAATGGATGAACGAGATTC</td><td>6852.6</td><td>-16.6</td><td>1.5</td><td>-11.4</td><td>4.1E-30</td><td>4.9E-28</td><td>MB231/MB468</td><td>MP</td></tr>
<tr><td>>hg38_21_+_8433172_8446622_RNA45SN1_45S_50ntFlanks@4728.4751.24|ATCAAGAACGAAAGTCGGAGGTTC</td><td>>hg38_21_+_8433172_8446622_RNA45SN1_45S_50ntFlanks</td><td>4728</td><td>4751</td><td>24</td><td>ATCAAGAACGAAAGTCGGAGGTTC</td><td>1466.9</td><td>-14.9</td><td>1.3</td><td>-11.2</td><td>4.4E-29</td><td>5.0E-28</td><td>BT20/MB468</td><td>NUC</td></tr>
<tr><td>>MI0000299|hsa-mir-222&WithFlank&X|-|45747009|45747130@75.99.25(+2U)&[MIMAT0000279&hsa-miR-222-3p&offsets|0|+4(+2U);m-140&X|-|45747033|45747056&offsets|0|+1(+2U)]|AGCTACATCTGGCTACTGGGTCTCTTT</td><td>>MI0000299|hsa-mir-222&WithFlank&X|-|45747009|45747130</td><td>75</td><td>99</td><td>25(+2U)</td><td>AGCTACATCTGGCTACTGGGTCTCTTT</td><td>4093.4</td><td>9.1</td><td>0.8</td><td>11.4</td><td>3.2E-30</td><td>5.5E-28</td><td>MB231/MB468</td><td>CYTO</td></tr>
<tr><td>>hg38_21_+_8433172_8446622_RNA45SN1_45S_50ntFlanks@11783.11800.18|CATGAATGGATGAACGAG</td><td>>hg38_21_+_8433172_8446622_RNA45SN1_45S_50ntFlanks</td><td>11783</td><td>11800</td><td>18</td><td>CATGAATGGATGAACGAG</td><td>1998.0</td><td>-15.4</td><td>1.4</td><td>-11.2</td><td>4.9E-29</td><td>5.5E-28</td><td>BT20/MB468</td><td>NUC</td></tr>
<tr><td>>hg38_21_+_8433172_8446622_RNA45SN1_45S_50ntFlanks@12412.12431.20|ATCCTTCGATGTCGGCTCTT</td><td>>hg38_21_+_8433172_8446622_RNA45SN1_45S_50ntFlanks</td><td>12412</td><td>12431</td><td>20</td><td>ATCCTTCGATGTCGGCTCTT</td><td>774.3</td><td>-14.0</td><td>1.3</td><td>-11.2</td><td>4.9E-29</td><td>5.5E-28</td><td>BT20/MB468</td><td>NUC</td></tr>
<tr><td>>MI0000093|hsa-mir-92a-1&WithFlank&13|+|91351308|91351397@54.75.22(+1C);MI0000094|hsa-mir-92a-2&WithFlank&X|-|134169532|134169618@54.75.22(+1C)&[MIMAT0000092&hsa-miR-92a-3p&offsets|0|0(+1C);m-9&13|+|91351361|91351382&offsets|0|0(+1C)];[MIMAT0000092_1&hsa-miR-92a-3p&offsets|0|0(+1C);m-11&X|-|134169544|134169565&offsets|0|0(+1C)]|TATTGCACTTGTCCCGGCCTGTC</td><td>>MI0000093|hsa-mir-92a-1&WithFlank&13|+|91351308|91351397</td><td>54</td><td>75</td><td>22(+1C)</td><td>TATTGCACTTGTCCCGGCCTGTC</td><td>581.4</td><td>6.2</td><td>0.6</td><td>11.2</td><td>3.1E-29</td><td>5.6E-28</td><td>BT20/MB468</td><td>CYTO</td></tr>
<tr><td>>hg38_21_+_8433172_8446622_RNA45SN1_45S_50ntFlanks@8357.8378.22|AAAAGAACTTTGAAGAGAGAGT</td><td>>hg38_21_+_8433172_8446622_RNA45SN1_45S_50ntFlanks</td><td>8357</td><td>8378</td><td>22</td><td>AAAAGAACTTTGAAGAGAGAGT</td><td>44711.5</td><td>-10.3</td><td>0.9</td><td>-11.4</td><td>4.8E-30</td><td>5.6E-28</td><td>MB231/MB468</td><td>MP</td></tr>
<tr><td>>hg38_21_+_8433172_8446622_RNA45SN1_45S_50ntFlanks@11581.11609.29|AACTGGTGCGGACCAGGGGAATCCGACTG</td><td>>hg38_21_+_8433172_8446622_RNA45SN1_45S_50ntFlanks</td><td>11581</td><td>11609</td><td>29</td><td>AACTGGTGCGGACCAGGGGAATCCGACTG</td><td>3341.7</td><td>-7.0</td><td>0.6</td><td>-11.4</td><td>4.8E-30</td><td>5.6E-28</td><td>MB231/MB468</td><td>MP</td></tr>
<tr><td>>hg38_21_+_8433172_8446622_RNA45SN1_45S_50ntFlanks@12412.12437.26|ATCCTTCGATGTCGGCTCTTCCTATC</td><td>>hg38_21_+_8433172_8446622_RNA45SN1_45S_50ntFlanks</td><td>12412</td><td>12437</td><td>26</td><td>ATCCTTCGATGTCGGCTCTTCCTATC</td><td>1235.6</td><td>-14.7</td><td>1.3</td><td>-11.2</td><td>5.1E-29</td><td>5.7E-28</td><td>BT20/MB468</td><td>NUC</td></tr>
<tr><td>>hg38_21_+_8433172_8446622_RNA45SN1_45S_50ntFlanks@10687.10706.20|CGGCGTCCGGTGAGCTCTCG</td><td>>hg38_21_+_8433172_8446622_RNA45SN1_45S_50ntFlanks</td><td>10687</td><td>10706</td><td>20</td><td>CGGCGTCCGGTGAGCTCTCG</td><td>4083.0</td><td>-15.9</td><td>1.4</td><td>-11.4</td><td>5.0E-30</td><td>5.8E-28</td><td>MB231/MB468</td><td>MP</td></tr>
<tr><td>>hg38_21_+_8433172_8446622_RNA45SN1_45S_50ntFlanks@12635.12657.23|AAGCTACCATCTGTGGGATTATG</td><td>>hg38_21_+_8433172_8446622_RNA45SN1_45S_50ntFlanks</td><td>12635</td><td>12657</td><td>23</td><td>AAGCTACCATCTGTGGGATTATG</td><td>2952.0</td><td>-15.4</td><td>1.4</td><td>-11.4</td><td>5.1E-30</td><td>5.8E-28</td><td>MB231/MB468</td><td>MP</td></tr>
<tr><td>>hg38_21_+_8433172_8446622_RNA45SN1_45S_50ntFlanks@12628.12654.27|TGGGGCGAAGCTACCATCTGTGGGATT</td><td>>hg38_21_+_8433172_8446622_RNA45SN1_45S_50ntFlanks</td><td>12628</td><td>12654</td><td>27</td><td>TGGGGCGAAGCTACCATCTGTGGGATT</td><td>3727.0</td><td>-6.9</td><td>0.6</td><td>-11.3</td><td>2.3E-29</td><td>5.8E-28</td><td>BT20/MB468</td><td>MITO</td></tr>
<tr><td>>hg38_21_+_8433172_8446622_RNA45SN1_45S_50ntFlanks@10350.10367.18|ACGAGAACTTTGAAGGCC</td><td>>hg38_21_+_8433172_8446622_RNA45SN1_45S_50ntFlanks</td><td>10350</td><td>10367</td><td>18</td><td>ACGAGAACTTTGAAGGCC</td><td>1923.5</td><td>-7.3</td><td>0.7</td><td>-11.2</td><td>5.3E-29</td><td>5.9E-28</td><td>BT20/MB468</td><td>NUC</td></tr>
<tr><td>>hg38_21_+_8433172_8446622_RNA45SN1_45S_50ntFlanks@10690.10708.19|CGTCCGGTGAGCTCTCGCT</td><td>>hg38_21_+_8433172_8446622_RNA45SN1_45S_50ntFlanks</td><td>10690</td><td>10708</td><td>19</td><td>CGTCCGGTGAGCTCTCGCT</td><td>6192.9</td><td>-7.0</td><td>0.6</td><td>-11.3</td><td>1.7E-29</td><td>6.1E-28</td><td>MB231/MB468</td><td>NUC</td></tr>
<tr><td>>hg38_21_+_8433172_8446622_RNA45SN1_45S_50ntFlanks@4129.4162.34|GATTCCGGAGAGGGAGCCTGAGAAACGGCTACCA</td><td>>hg38_21_+_8433172_8446622_RNA45SN1_45S_50ntFlanks</td><td>4129</td><td>4162</td><td>34</td><td>GATTCCGGAGAGGGAGCCTGAGAAACGGCTACCA</td><td>964.0</td><td>-14.3</td><td>1.3</td><td>-11.2</td><td>5.6E-29</td><td>6.2E-28</td><td>BT20/MB468</td><td>NUC</td></tr>
<tr><td>>hg38_21_+_8433172_8446622_RNA45SN1_45S_50ntFlanks@11999.12032.34|CCCTGCGGGCCGCCGGTGAAATACCACTACTCTG</td><td>>hg38_21_+_8433172_8446622_RNA45SN1_45S_50ntFlanks</td><td>11999</td><td>12032</td><td>34</td><td>CCCTGCGGGCCGCCGGTGAAATACCACTACTCTG</td><td>9609.5</td><td>-8.3</td><td>0.7</td><td>-11.2</td><td>2.5E-29</td><td>6.3E-28</td><td>BT20/MB468</td><td>MITO</td></tr>
<tr><td>>hg38_21_+_8433172_8446622_RNA45SN1_45S_50ntFlanks@9568.9596.29|CGTGCAAATCGGTCGTCCGACCTGGGTAT</td><td>>hg38_21_+_8433172_8446622_RNA45SN1_45S_50ntFlanks</td><td>9568</td><td>9596</td><td>29</td><td>CGTGCAAATCGGTCGTCCGACCTGGGTAT</td><td>6276.7</td><td>-14.7</td><td>1.3</td><td>-11.2</td><td>2.5E-29</td><td>6.3E-28</td><td>BT20/MB468</td><td>MITO</td></tr>
</table>

<table>
<tr><td></td><td></td><td></td><td></td><td></td><td></td><td></td><td></td><td></td><td></td><td></td><td></td><td></td><td></td></tr>
<tr><td>>MI0000542|hsa-mir-320a&WithFlank&8|-|22244960|22245043@48.68.21(+2U)&[MIMAT0000510&hsa-miR-320a-3p&offsets|0|-1(+2U);m-60&8|-|22244975|22244996&offsets|0|-1(+2U)]|AAAAGCTGGGTTGAGAGGGCGTT</td><td>>MI0000542|hsa-mir-320a&WithFlank&8|-|22244960|22245043</td><td>48</td><td>68</td><td>21(+2U)</td><td>AAAAGCTGGGTTGAGAGGGCGTT</td><td>623.4</td><td>4.5</td><td>0.4</td><td>11.2</td><td>5.7E-29</td><td>6.3E-28</td><td>BT20/MB468</td><td>NUC</td></tr>
<tr><td>>hg38_21_+_8433172_8446622_RNA45SN1_45S_50ntFlanks@11076.11103.28|CGCCGCGTGGGGGCGGCGGCGGGGGGAG</td><td>>hg38_21_+_8433172_8446622_RNA45SN1_45S_50ntFlanks</td><td>11076</td><td>11103</td><td>28</td><td>CGCCGCGTGGGGGCGGCGGCGGGGGGAG</td><td>1928.1</td><td>-15.3</td><td>1.4</td><td>-11.2</td><td>5.9E-29</td><td>6.5E-28</td><td>BT20/MB468</td><td>NUC</td></tr>
<tr><td>>hg38_21_+_8433172_8446622_RNA45SN1_45S_50ntFlanks@13008.13027.20|ATTGAAAGTCAGCCCTCGAC</td><td>>hg38_21_+_8433172_8446622_RNA45SN1_45S_50ntFlanks</td><td>13008</td><td>13027</td><td>20</td><td>ATTGAAAGTCAGCCCTCGAC</td><td>622.4</td><td>-13.7</td><td>1.2</td><td>-11.2</td><td>5.9E-29</td><td>6.5E-28</td><td>BT20/MB468</td><td>NUC</td></tr>
<tr><td>>hg38_21_+_8433172_8446622_RNA45SN1_45S_50ntFlanks@10786.10817.32|AACAGCCTCTGGCATGTTGGAACAATGTAGGT</td><td>>hg38_21_+_8433172_8446622_RNA45SN1_45S_50ntFlanks</td><td>10786</td><td>10817</td><td>32</td><td>AACAGCCTCTGGCATGTTGGAACAATGTAGGT</td><td>1807.8</td><td>-7.1</td><td>0.6</td><td>-11.4</td><td>5.8E-30</td><td>6.6E-28</td><td>MB231/MB468</td><td>MP</td></tr>
<tr><td>>MI0000285|hsa-mir-205&WithFlank&1|+|209432127|209432248@40.61.22(+1A)&[MIMAT0000266&hsa-miR-205-5p&offsets|0|0(+1A);m-351&1|+|209432166|209432187&offsets|0|0(+1A)]|TCCTTCATTCCACCGGAGTCTGA</td><td>>MI0000285|hsa-mir-205&WithFlank&1|+|209432127|209432248</td><td>40</td><td>61</td><td>22(+1A)</td><td>TCCTTCATTCCACCGGAGTCTGA</td><td>76361.2</td><td>10.5</td><td>0.9</td><td>11.7</td><td>1.8E-31</td><td>6.6E-28</td><td>BT20/MB231</td><td>CYTO</td></tr>
<tr><td>>hg38_21_+_8433172_8446622_RNA45SN1_45S_50ntFlanks@12488.12521.34|AACGTGAGCTGGGTTTAGACCGTCGTGAGACAGG</td><td>>hg38_21_+_8433172_8446622_RNA45SN1_45S_50ntFlanks</td><td>12488</td><td>12521</td><td>34</td><td>AACGTGAGCTGGGTTTAGACCGTCGTGAGACAGG</td><td>1113.1</td><td>-14.5</td><td>1.3</td><td>-11.2</td><td>6.0E-29</td><td>6.6E-28</td><td>BT20/MB468</td><td>NUC</td></tr>
<tr><td>>hg38_21_+_8433172_8446622_RNA45SN1_45S_50ntFlanks@12476.12499.24|CCACTAATAGGGAACGTGAGCTGG</td><td>>hg38_21_+_8433172_8446622_RNA45SN1_45S_50ntFlanks</td><td>12476</td><td>12499</td><td>24</td><td>CCACTAATAGGGAACGTGAGCTGG</td><td>780.0</td><td>-14.0</td><td>1.3</td><td>-11.2</td><td>6.1E-29</td><td>6.7E-28</td><td>BT20/MB468</td><td>NUC</td></tr>
<tr><td>>hg38_21_+_8433172_8446622_RNA45SN1_45S_50ntFlanks@10781.10807.27|AGGTGAACAGCCTCTGGCATGTTGGAA</td><td>>hg38_21_+_8433172_8446622_RNA45SN1_45S_50ntFlanks</td><td>10781</td><td>10807</td><td>27</td><td>AGGTGAACAGCCTCTGGCATGTTGGAA</td><td>843.7</td><td>-14.1</td><td>1.3</td><td>-11.2</td><td>6.2E-29</td><td>6.8E-28</td><td>BT20/MB468</td><td>NUC</td></tr>
<tr><td>>MI0000299|hsa-mir-222&WithFlank&X|-|45747009|45747130@75.99.25(+1U)&[MIMAT0000279&hsa-miR-222-3p&offsets|0|+4(+1U);m-140&X|-|45747033|45747056&offsets|0|+1(+1U)]|AGCTACATCTGGCTACTGGGTCTCTT</td><td>>MI0000299|hsa-mir-222&WithFlank&X|-|45747009|45747130</td><td>75</td><td>99</td><td>25(+1U)</td><td>AGCTACATCTGGCTACTGGGTCTCTT</td><td>6286.0</td><td>8.5</td><td>0.7</td><td>11.4</td><td>4.2E-30</td><td>6.9E-28</td><td>MB231/MB468</td><td>CYTO</td></tr>
<tr><td>>hg38_21_+_8433172_8446622_RNA45SN1_45S_50ntFlanks@11581.11605.25|AACTGGTGCGGACCAGGGGAATCCG</td><td>>hg38_21_+_8433172_8446622_RNA45SN1_45S_50ntFlanks</td><td>11581</td><td>11605</td><td>25</td><td>AACTGGTGCGGACCAGGGGAATCCG</td><td>2925.8</td><td>-6.1</td><td>0.5</td><td>-11.4</td><td>4.1E-30</td><td>6.9E-28</td><td>MB231/MB468</td><td>CYTO</td></tr>
<tr><td>>MI0000542|hsa-mir-320a&WithFlank&8|-|22244960|22245043@49.68.20(+2U)&[MIMAT0000510&hsa-miR-320a-3p&offsets|+1|-1(+2U);m-60&8|-|22244975|22244996&offsets|+1|-1(+2U)]|AAAGCTGGGTTGAGAGGGCGTT</td><td>>MI0000542|hsa-mir-320a&WithFlank&8|-|22244960|22245043</td><td>49</td><td>68</td><td>20(+2U)</td><td>AAAGCTGGGTTGAGAGGGCGTT</td><td>752.0</td><td>13.8</td><td>1.2</td><td>11.2</td><td>3.9E-29</td><td>7.1E-28</td><td>BT20/MB468</td><td>CYTO</td></tr>
<tr><td>>hg38_21_+_8433172_8446622_RNA45SN1_45S_50ntFlanks@11845.11872.28|GGAACGGGCTTGGCGGAATCAGCGGGGA</td><td>>hg38_21_+_8433172_8446622_RNA45SN1_45S_50ntFlanks</td><td>11845</td><td>11872</td><td>28</td><td>GGAACGGGCTTGGCGGAATCAGCGGGGA</td><td>3411.2</td><td>-15.6</td><td>1.4</td><td>-11.4</td><td>6.3E-30</td><td>7.1E-28</td><td>MB231/MB468</td><td>MP</td></tr>
<tr><td>>MI0000434|hsa-let-7i&WithFlank&12|+|62603680|62603775@12.32.21&[MIMAT0000415&hsa-let-7i-5p&offsets|0|-1;m-49&12|+|62603691|62603712&offsets|0|-1]|TGAGGTAGTAGTTTGTGCTGT</td><td>>MI0000434|hsa-let-7i&WithFlank&12|+|62603680|62603775</td><td>12</td><td>32</td><td>21</td><td>TGAGGTAGTAGTTTGTGCTGT</td><td>2706.7</td><td>7.4</td><td>0.7</td><td>11.2</td><td>3.0E-29</td><td>7.5E-28</td><td>BT20/MB468</td><td>MITO</td></tr>
<tr><td>>MI0003560|hsa-mir-92b&WithFlank&1|+|155195171|155195278@67.87.21(+1U)&[MIMAT0003218&hsa-miR-92b-3p&offsets|0|-1(+1U);m-115&1|+|155195237|155195258&offsets|0|-1(+1U)]|TATTGCACTCGTCCCGGCCTCT</td><td>>MI0003560|hsa-mir-92b&WithFlank&1|+|155195171|155195278</td><td>67</td><td>87</td><td>21(+1U)</td><td>TATTGCACTCGTCCCGGCCTCT</td><td>3510.0</td><td>9.4</td><td>0.8</td><td>11.2</td><td>3.0E-29</td><td>7.5E-28</td><td>BT20/MB468</td><td>MITO</td></tr>
<tr><td>>hg38_21_+_8433172_8446622_RNA45SN1_45S_50ntFlanks@12434.12455.22|TATCATTGTGAAGCAGAATTCA</td><td>>hg38_21_+_8433172_8446622_RNA45SN1_45S_50ntFlanks</td><td>12434</td><td>12455</td><td>22</td><td>TATCATTGTGAAGCAGAATTCA</td><td>678.9</td><td>-13.8</td><td>1.2</td><td>-11.2</td><td>6.9E-29</td><td>7.5E-28</td><td>BT20/MB468</td><td>NUC</td></tr>
<tr><td>>hg38_21_+_8433172_8446622_RNA45SN1_45S_50ntFlanks@11581.11603.23|AACTGGTGCGGACCAGGGGAATC</td><td>>hg38_21_+_8433172_8446622_RNA45SN1_45S_50ntFlanks</td><td>11581</td><td>11603</td><td>23</td><td>AACTGGTGCGGACCAGGGGAATC</td><td>7486.3</td><td>-5.2</td><td>0.5</td><td>-11.2</td><td>4.2E-29</td><td>7.7E-28</td><td>BT20/MB468</td><td>CYTO</td></tr>
<tr><td>>hg38_21_+_8433172_8446622_RNA45SN1_45S_50ntFlanks@10841.10861.21|CGTAACTTCGGGATAAGGATT</td><td>>hg38_21_+_8433172_8446622_RNA45SN1_45S_50ntFlanks</td><td>10841</td><td>10861</td><td>21</td><td>CGTAACTTCGGGATAAGGATT</td><td>1739.2</td><td>-6.0</td><td>0.5</td><td>-11.2</td><td>7.1E-29</td><td>7.7E-28</td><td>BT20/MB468</td><td>NUC</td></tr>
<tr><td>>hg38_21_+_8433172_8446622_RNA45SN1_45S_50ntFlanks@12476.12496.21|CCACTAATAGGGAACGTGAGC</td><td>>hg38_21_+_8433172_8446622_RNA45SN1_45S_50ntFlanks</td><td>12476</td><td>12496</td><td>21</td><td>CCACTAATAGGGAACGTGAGC</td><td>2802.6</td><td>-6.7</td><td>0.6</td><td>-11.4</td><td>7.0E-30</td><td>7.8E-28</td><td>MB231/MB468</td><td>MP</td></tr>
<tr><td>>hg38_21_+_8433172_8446622_RNA45SN1_45S_50ntFlanks@10687.10711.25|CGGCGTCCGGTGAGCTCTCGCTGGC</td><td>>hg38_21_+_8433172_8446622_RNA45SN1_45S_50ntFlanks</td><td>10687</td><td>10711</td><td>25</td><td>CGGCGTCCGGTGAGCTCTCGCTGGC</td><td>2651.8</td><td>-15.2</td><td>1.3</td><td>-11.4</td><td>7.2E-30</td><td>8.0E-28</td><td>MB231/MB468</td><td>MP</td></tr>
<tr><td>>MI0000279|hsa-mir-196a-2&WithFlank&12|+|53991732|53991853@31.52.22(+1U);MI0000238|hsa-mir-196a-1&WithFlank&17|-|48632484|48632565@13.34.22(+1U)&[MIMAT0000226&hsa-miR-196a-5p&offsets|0|0(+1U);m-261&12|+|53991762|53991783&offsets|0|0(+1U)];[MIMAT0000226_1&hsa-miR-196a-5p&offsets|0|0(+1U);m-268&17|-|48632532|48632553&offsets|0|0(+1U)]|TAGGTAGTTTCATGTTGTTGGGT</td><td>>MI0000279|hsa-mir-196a-2&WithFlank&12|+|53991732|53991853</td><td>31</td><td>52</td><td>22(+1U)</td><td>TAGGTAGTTTCATGTTGTTGGGT</td><td>859.3</td><td>7.4</td><td>0.7</td><td>11.2</td><td>4.4E-29</td><td>8.0E-28</td><td>BT20/MB468</td><td>CYTO</td></tr>
<tr><td>>hg38_21_+_8433172_8446622_RNA45SN1_45S_50ntFlanks@12935.12956.22|TAAACCATTCGTAGACGACCTG</td><td>>hg38_21_+_8433172_8446622_RNA45SN1_45S_50ntFlanks</td><td>12935</td><td>12956</td><td>22</td><td>TAAACCATTCGTAGACGACCTG</td><td>2837.3</td><td>-5.6</td><td>0.5</td><td>-11.2</td><td>4.5E-29</td><td>8.2E-28</td><td>BT20/MB468</td><td>CYTO</td></tr>
<tr><td>>hg38_21_+_8433172_8446622_RNA45SN1_45S_50ntFlanks@8380.8400.21|CAAGAGGGCGTGAAACCGTTA</td><td>>hg38_21_+_8433172_8446622_RNA45SN1_45S_50ntFlanks</td><td>8380</td><td>8400</td><td>21</td><td>CAAGAGGGCGTGAAACCGTTA</td><td>1186.4</td><td>-14.6</td><td>1.3</td><td>-11.1</td><td>7.7E-29</td><td>8.4E-28</td><td>BT20/MB468</td><td>NUC</td></tr>
<tr><td>>hg38_21_+_8433172_8446622_RNA45SN1_45S_50ntFlanks@11641.11669.29|CGGCGGGTGTTGACGCGATGTGATTTCTG</td><td>>hg38_21_+_8433172_8446622_RNA45SN1_45S_50ntFlanks</td><td>11641</td><td>11669</td><td>29</td><td>CGGCGGGTGTTGACGCGATGTGATTTCTG</td><td>9269.0</td><td>-17.0</td><td>1.5</td><td>-11.3</td><td>7.9E-30</td><td>8.6E-28</td><td>MB231/MB468</td><td>MP</td></tr>
<tr><td>>hg38_21_+_8433172_8446622_RNA45SN1_45S_50ntFlanks@4724.4757.34|ATTAATCAAGAACGAAAGTCGGAGGTTCGAAGAC</td><td>>hg38_21_+_8433172_8446622_RNA45SN1_45S_50ntFlanks</td><td>4724</td><td>4757</td><td>34</td><td>ATTAATCAAGAACGAAAGTCGGAGGTTCGAAGAC</td><td>587.9</td><td>-13.6</td><td>1.2</td><td>-11.1</td><td>8.0E-29</td><td>8.7E-28</td><td>BT20/MB468</td><td>NUC</td></tr>
<tr><td>>hg38_21_+_8433172_8446622_RNA45SN1_45S_50ntFlanks@10362.10382.21|AAGGCCGAAGTGGAGAAGGGT</td><td>>hg38_21_+_8433172_8446622_RNA45SN1_45S_50ntFlanks</td><td>10362</td><td>10382</td><td>21</td><td>AAGGCCGAAGTGGAGAAGGGT</td><td>706.2</td><td>-13.9</td><td>1.2</td><td>-11.1</td><td>8.0E-29</td><td>8.7E-28</td><td>BT20/MB468</td><td>NUC</td></tr>
<tr><td>>hg38_21_+_8433172_8446622_RNA45SN1_45S_50ntFlanks@11613.11640.28|AATTAAAACAAAGCATCGCGAAGGCCCG</td><td>>hg38_21_+_8433172_8446622_RNA45SN1_45S_50ntFlanks</td><td>11613</td><td>11640</td><td>28</td><td>AATTAAAACAAAGCATCGCGAAGGCCCG</td><td>821.2</td><td>-14.1</td><td>1.3</td><td>-11.1</td><td>8.0E-29</td><td>8.7E-28</td><td>BT20/MB468</td><td>NUC</td></tr>
<tr><td>>hg38_21_+_8433172_8446622_RNA45SN1_45S_50ntFlanks@4726.4752.27|TAATCAAGAACGAAAGTCGGAGGTTCG</td><td>>hg38_21_+_8433172_8446622_RNA45SN1_45S_50ntFlanks</td><td>4726</td><td>4752</td><td>27</td><td>TAATCAAGAACGAAAGTCGGAGGTTCG</td><td>638.9</td><td>-13.7</td><td>1.2</td><td>-11.1</td><td>8.1E-29</td><td>8.8E-28</td><td>BT20/MB468</td><td>NUC</td></tr>
<tr><td>>hg38_21_+_8433172_8446622_RNA45SN1_45S_50ntFlanks@6719.6741.23|TGAATTGCAGGACACATTGATCA</td><td>>hg38_21_+_8433172_8446622_RNA45SN1_45S_50ntFlanks</td><td>6719</td><td>6741</td><td>23</td><td>TGAATTGCAGGACACATTGATCA</td><td>1063.4</td><td>-14.5</td><td>1.3</td><td>-11.1</td><td>8.2E-29</td><td>8.8E-28</td><td>BT20/MB468</td><td>NUC</td></tr>
<tr><td>>hg38_21_+_8433172_8446622_RNA45SN1_45S_50ntFlanks@7971.7988.18|GAGACGCGACCTCAGATC</td><td>>hg38_21_+_8433172_8446622_RNA45SN1_45S_50ntFlanks</td><td>7971</td><td>7988</td><td>18</td><td>GAGACGCGACCTCAGATC</td><td>684.7</td><td>7.6</td><td>0.7</td><td>11.2</td><td>3.6E-29</td><td>8.9E-28</td><td>BT20/MB468</td><td>MITO</td></tr>
<tr><td>>hg38_21_+_8433172_8446622_RNA45SN1_45S_50ntFlanks@12417.12444.28|TCGATGTCGGCTCTTCCTATCATTGTGA</td><td>>hg38_21_+_8433172_8446622_RNA45SN1_45S_50ntFlanks</td><td>12417</td><td>12444</td><td>28</td><td>TCGATGTCGGCTCTTCCTATCATTGTGA</td><td>2294.8</td><td>-6.4</td><td>0.6</td><td>-11.1</td><td>8.4E-29</td><td>9.1E-28</td><td>BT20/MB468</td><td>NUC</td></tr>
<tr><td>>hg38_21_+_8433172_8446622_RNA45SN1_45S_50ntFlanks@13006.13027.22|CTATTGAAAGTCAGCCCTCGAC</td><td>>hg38_21_+_8433172_8446622_RNA45SN1_45S_50ntFlanks</td><td>13006</td><td>13027</td><td>22</td><td>CTATTGAAAGTCAGCCCTCGAC</td><td>954.0</td><td>-14.3</td><td>1.3</td><td>-11.1</td><td>8.6E-29</td><td>9.3E-28</td><td>BT20/MB468</td><td>NUC</td></tr>
<tr><td>>hg38_21_+_8433172_8446622_RNA45SN1_45S_50ntFlanks@5523.5550.28|AGTAAAAGTCGTAACAAGGTTTCCGTAG</td><td>>hg38_21_+_8433172_8446622_RNA45SN1_45S_50ntFlanks</td><td>5523</td><td>5550</td><td>28</td><td>AGTAAAAGTCGTAACAAGGTTTCCGTAG</td><td>2218.5</td><td>-15.5</td><td>1.4</td><td>-11.1</td><td>8.6E-29</td><td>9.3E-28</td><td>BT20/MB468</td><td>NUC</td></tr>
<tr><td>>hg38_21_+_8433172_8446622_RNA45SN1_45S_50ntFlanks@8182.8208.27|ATCGAGGCCCAGCCCGTGGACGGTGTG</td><td>>hg38_21_+_8433172_8446622_RNA45SN1_45S_50ntFlanks</td><td>8182</td><td>8208</td><td>27</td><td>ATCGAGGCCCAGCCCGTGGACGGTGTG</td><td>1364.1</td><td>-14.8</td><td>1.3</td><td>-11.1</td><td>8.9E-29</td><td>9.5E-28</td><td>BT20/MB468</td><td>NUC</td></tr>
<tr><td>>hg38_21_+_8433172_8446622_RNA45SN1_45S_50ntFlanks@3848.3866.19|TGGATAACTGTGGTAATTC</td><td>>hg38_21_+_8433172_8446622_RNA45SN1_45S_50ntFlanks</td><td>3848</td><td>3866</td><td>19</td><td>TGGATAACTGTGGTAATTC</td><td>509.6</td><td>6.2</td><td>0.6</td><td>11.2</td><td>5.6E-29</td><td>1.0E-27</td><td>BT20/MB468</td><td>CYTO</td></tr>
<tr><td>>MI0000266|hsa-mir-10a&WithFlank&17|-|48579832|48579953@29.49.21&[MIMAT0000253&hsa-miR-10a-5p&offsets|+1|-1;m-18&17|-|48579905|48579926&offsets|+1|0]|ACCCTGTAGATCCGAATTTGT</td><td>>MI0000266|hsa-mir-10a&WithFlank&17|-|48579832|48579953</td><td>29</td><td>49</td><td>21</td><td>ACCCTGTAGATCCGAATTTGT</td><td>1608.6</td><td>15.1</td><td>1.3</td><td>11.2</td><td>4.3E-29</td><td>1.1E-27</td><td>BT20/MB468</td><td>MITO</td></tr>
<tr><td>>hg38_21_+_8433172_8446622_RNA45SN1_45S_50ntFlanks@12570.12587.18|AGTACGAGAGGAACCGCA</td><td>>hg38_21_+_8433172_8446622_RNA45SN1_45S_50ntFlanks</td><td>12570</td><td>12587</td><td>18</td><td>AGTACGAGAGGAACCGCA</td><td>1280.9</td><td>-14.7</td><td>1.3</td><td>-11.1</td><td>1.0E-28</td><td>1.1E-27</td><td>BT20/MB468</td><td>NUC</td></tr>
<tr><td>>MI0000746|hsa-mir-99b&WithFlank&19|+|51692606|51692687@51.72.22&[MIMAT0004678&hsa-miR-99b-3p&offsets|0|0;m-151&19|+|51692656|51692678&offsets|0|-1]|CAAGCTCGTGTCTGTGGGTCCG</td><td>>MI0000746|hsa-mir-99b&WithFlank&19|+|51692606|51692687</td><td>51</td><td>72</td><td>22</td><td>CAAGCTCGTGTCTGTGGGTCCG</td><td>2330.7</td><td>7.1</td><td>0.6</td><td>11.2</td><td>4.5E-29</td><td>1.1E-27</td><td>BT20/MB468</td><td>MITO</td></tr>
<tr><td>>hg38_21_+_8433172_8446622_RNA45SN1_45S_50ntFlanks@5476.5516.41|CCCTGGCGGAGCGCTGAGAAGACGGTCGAACTTGACTATCT</td><td>>hg38_21_+_8433172_8446622_RNA45SN1_45S_50ntFlanks</td><td>5476</td><td>5516</td><td>41</td><td>CCCTGGCGGAGCGCTGAGAAGACGGTCGAACTTGACTATCT</td><td>3692.4</td><td>11.6</td><td>1.0</td><td>11.4</td><td>7.1E-30</td><td>1.1E-27</td><td>MB231/MB468</td><td>CYTO</td></tr>
<tr><td>>hg38_21_+_8433172_8446622_RNA45SN1_45S_50ntFlanks@10684.10706.23|CGGCGGCGTCCGGTGAGCTCTCG</td><td>>hg38_21_+_8433172_8446622_RNA45SN1_45S_50ntFlanks</td><td>10684</td><td>10706</td><td>23</td><td>CGGCGGCGTCCGGTGAGCTCTCG</td><td>5396.1</td><td>-16.3</td><td>1.4</td><td>-11.3</td><td>1.1E-29</td><td>1.2E-27</td><td>MB231/MB468</td><td>MP</td></tr>
<tr><td>>hg38_21_+_8433172_8446622_RNA45SN1_45S_50ntFlanks@3711.3737.27|GTTGATCCTGCCAGTAGCATATGCTTG</td><td>>hg38_21_+_8433172_8446622_RNA45SN1_45S_50ntFlanks</td><td>3711</td><td>3737</td><td>27</td><td>GTTGATCCTGCCAGTAGCATATGCTTG</td><td>3168.9</td><td>-16.1</td><td>1.4</td><td>-11.1</td><td>1.1E-28</td><td>1.2E-27</td><td>BT20/MB468</td><td>NUC</td></tr>
<tr><td>>hg38_21_+_8433172_8446622_RNA45SN1_45S_50ntFlanks@13002.13028.27|CGATCTATTGAAAGTCAGCCCTCGACA</td><td>>hg38_21_+_8433172_8446622_RNA45SN1_45S_50ntFlanks</td><td>13002</td><td>13028</td><td>27</td><td>CGATCTATTGAAAGTCAGCCCTCGACA</td><td>4335.2</td><td>-15.0</td><td>1.3</td><td>-11.2</td><td>3.3E-29</td><td>1.2E-27</td><td>MB231/MB468</td><td>NUC</td></tr>
<tr><td>>hg38_21_+_8433172_8446622_RNA45SN1_45S_50ntFlanks@10680.10712.33|GTTCCGGCGGCGTCCGGTGAGCTCTCGCTGGCC</td><td>>hg38_21_+_8433172_8446622_RNA45SN1_45S_50ntFlanks</td><td>10680</td><td>10712</td><td>33</td><td>GTTCCGGCGGCGTCCGGTGAGCTCTCGCTGGCC</td><td>6877.8</td><td>-6.6</td><td>0.6</td><td>-11.2</td><td>3.3E-29</td><td>1.2E-27</td><td>MB231/MB468</td><td>NUC</td></tr>
<tr><td>>hg38_21_+_8433172_8446622_RNA45SN1_45S_50ntFlanks@5494.5516.23|AAGACGGTCGAACTTGACTATCT</td><td>>hg38_21_+_8433172_8446622_RNA45SN1_45S_50ntFlanks</td><td>5494</td><td>5516</td><td>23</td><td>AAGACGGTCGAACTTGACTATCT</td><td>6197.2</td><td>7.3</td><td>0.7</td><td>11.2</td><td>6.8E-29</td><td>1.2E-27</td><td>BT20/MB468</td><td>CYTO</td></tr>
<tr><td>>MI0000253|hsa-mir-148a&WithFlank&7|-|25949913|25949992@50.70.21(+1C)&[MIMAT0000243&hsa-miR-148a-3p&offsets|0|-1(+1C);m-7&7|-|25949922|25949943&offsets|0|-1(+1C)]|TCAGTGCACTACAGAACTTTGC</td><td>>MI0000253|hsa-mir-148a&WithFlank&7|-|25949913|25949992</td><td>50</td><td>70</td><td>21(+1C)</td><td>TCAGTGCACTACAGAACTTTGC</td><td>22785.8</td><td>7.6</td><td>0.7</td><td>11.2</td><td>5.0E-29</td><td>1.2E-27</td><td>BT20/MB468</td><td>MITO</td></tr>
<tr><td>>hg38_21_+_8433172_8446622_RNA45SN1_45S_50ntFlanks@10839.10865.27|TCCGTAACTTCGGGATAAGGATTGGCT</td><td>>hg38_21_+_8433172_8446622_RNA45SN1_45S_50ntFlanks</td><td>10839</td><td>10865</td><td>27</td><td>TCCGTAACTTCGGGATAAGGATTGGCT</td><td>918.0</td><td>-14.3</td><td>1.3</td><td>-11.1</td><td>1.2E-28</td><td>1.3E-27</td><td>BT20/MB468</td><td>NUC</td></tr>
<tr><td>>hg38_21_+_8433172_8446622_RNA45SN1_45S_50ntFlanks@4297.4320.24|CATTGGAGGGCAAGTCTGGTGCCA</td><td>>hg38_21_+_8433172_8446622_RNA45SN1_45S_50ntFlanks</td><td>4297</td><td>4320</td><td>24</td><td>CATTGGAGGGCAAGTCTGGTGCCA</td><td>478.6</td><td>-13.3</td><td>1.2</td><td>-11.1</td><td>1.2E-28</td><td>1.3E-27</td><td>BT20/MB468</td><td>NUC</td></tr>
<tr><td>>hg38_21_+_8433172_8446622_RNA45SN1_45S_50ntFlanks@12995.13038.44|CTCGCTGCGATCTATTGAAAGTCAGCCCTCGACACAAGGGTTTG</td><td>>hg38_21_+_8433172_8446622_RNA45SN1_45S_50ntFlanks</td><td>12995</td><td>13038</td><td>44</td><td>CTCGCTGCGATCTATTGAAAGTCAGCCCTCGACACAAGGGTTTG</td><td>9482.7</td><td>10.2</td><td>0.9</td><td>11.3</td><td>8.3E-30</td><td>1.3E-27</td><td>MB231/MB468</td><td>CYTO</td></tr>
</table>

| | | | | | | | | | | | | | |
|---|---|---|---|---|---|---|---|---|---|---|---|---|---|
| >am_trna134_GluTTC_1_-_16861774_16861845@1.29.29,trna5_GluTTC_1_+_17199078_17199149@1.29.29,trna84_GluTTC_1_-_161391883_161391954@1.29.29,trna94_GluTTC_1_-_149664355_149664427@1.29.29\|TCCCTGGTGGTCTAGTGGCTAGGATTCGG | >am_trna134_GluTTC_1_-_16861774_16861845 | 1 | 29 | 29 | TCCCTGGTGGTCTAGTGGCTAGGATTCGG | 4344.5 | -7.0 | 0.6 | -11.3 | 8.5E-30 | 1.3E-27 | MB231/MB468 | CYTO |
| >hg38_21_+_8433172_8446622_RNA45SN1_45S_50ntFlanks@11561.11592.32\|GCGCCTAGCAGCCGACTTAGAACTGGTGCGGA | >hg38_21_+_8433172_8446622_RNA45SN1_45S_50ntFlanks | 11561 | 11592 | 32 | GCGCCTAGCAGCCGACTTAGAACTGGTGCGGA | 949.7 | -14.3 | 1.3 | -11.1 | 1.2E-28 | 1.3E-27 | BT20/MB468 | NUC |
| >hg38_21_+_8433172_8446622_RNA45SN1_45S_50ntFlanks@13002.13029.28\|CGATCTATTGAAAGTCAGCCCTCGACAC | >hg38_21_+_8433172_8446622_RNA45SN1_45S_50ntFlanks | 13002 | 13029 | 28 | CGATCTATTGAAAGTCAGCCCTCGACAC | 3674.6 | -15.6 | 1.4 | -11.3 | 2.2E-29 | 1.3E-27 | MB231/MB468 | MITO |
| >hg38_21_+_8433172_8446622_RNA45SN1_45S_50ntFlanks@8182.8210.29\|ATCGAGGCCCAGCCCGTGGACGGTGTGAG | >hg38_21_+_8433172_8446622_RNA45SN1_45S_50ntFlanks | 8182 | 8210 | 29 | ATCGAGGCCCAGCCCGTGGACGGTGTGAG | 1963.3 | -15.4 | 1.4 | -11.1 | 1.3E-28 | 1.3E-27 | BT20/MB468 | NUC |
| >MI0000650\|hsa-mir-200c&WithFlank&12\|+\|6963693\|6963772@50.71.22(+1G)&[MIMAT0000617&hsa-miR-200c-3p&offsets\|0\|-1(+1G);m-54&12\|+\|6963742\|6963764&offsets\|0\|-1(+1G)]\|TAATACTGCCGGGTAATGATGGG | >MI0000650\|hsa-mir-200c&WithFlank&12\|+\|6963693\|6963772 | 50 | 71 | 22(+1G) | TAATACTGCCGGGTAATGATGGG | 5799.2 | 7.4 | 0.7 | 11.1 | 7.5E-29 | 1.3E-27 | BT20/MB468 | CYTO |
| >trnaMT_ValTAC_MT_+_1602_1670@55.72.18\|ACTTGACCGCTCTGACCA | >trnaMT_ValTAC_MT_+_1602_1670 | 55 | 72 | 18 | ACTTGACCGCTCTGACCA | 3499.9 | 8.5 | 0.8 | 11.1 | 7.7E-29 | 1.4E-27 | BT20/MB468 | CYTO |
| >hg38_21_+_8433172_8446622_RNA45SN1_45S_50ntFlanks@4443.4471.29\|CCCTTGCCTCTCGGCGCCCCCTCGATGCT | >hg38_21_+_8433172_8446622_RNA45SN1_45S_50ntFlanks | 4443 | 4471 | 29 | CCCTTGCCTCTCGGCGCCCCCTCGATGCT | 3687.4 | -15.6 | 1.4 | -11.2 | 2.4E-29 | 1.4E-27 | MB231/MB468 | MITO |
| >MI0001733\|hsa-mir-452&WithFlank&X\|-\|151959622\|151959718@20.43.24&[MIMAT0001635&hsa-miR-452-5p&offsets\|0\|+2;m-188&X\|-\|151959677\|151959699&offsets\|0\|+1]\|AACTGTTTGCAGAGGAAACTGAGA | >MI0001733\|hsa-mir-452&WithFlank&X\|-\|151959622\|151959718 | 20 | 43 | 24 | AACTGTTTGCAGAGGAAACTGAGA | 827.8 | 14.2 | 1.3 | 11.2 | 5.9E-29 | 1.4E-27 | BT20/MB468 | MITO |
| >hg38_21_+_8433172_8446622_RNA45SN1_45S_50ntFlanks@8380.8397.18\|CAAGAGGGCGTGAAACCG | >hg38_21_+_8433172_8446622_RNA45SN1_45S_50ntFlanks | 8380 | 8397 | 18 | CAAGAGGGCGTGAAACCG | 1135.0 | -14.6 | 1.3 | -11.1 | 1.4E-28 | 1.5E-27 | BT20/MB468 | NUC |
| >MI0000102\|hsa-mir-100&WithFlank&11\|-\|122152223\|122152314@19.38.20&[MIMAT0000098&hsa-miR-100-5p&offsets\|0\|-2;m-27&11\|-\|122152276\|122152296&offsets\|0\|-1]\|AACCCGTAGATCCGAACTTG | >MI0000102\|hsa-mir-100&WithFlank&11\|-\|122152223\|122152314 | 19 | 38 | 20 | AACCCGTAGATCCGAACTTG | 34563.7 | 19.1 | 1.7 | 11.3 | 9.9E-30 | 1.5E-27 | MB231/MB468 | CYTO |
| >MI0000102\|hsa-mir-100&WithFlank&11\|-\|122152223\|122152314@19.40.22(+2A)&[MIMAT0000098&hsa-miR-100-5p&offsets\|0\|0(+2A);m-27&11\|-\|122152276\|122152296&offsets\|0\|+1(+2A)]\|AACCCGTAGATCCGAACTTGTGAA | >MI0000102\|hsa-mir-100&WithFlank&11\|-\|122152223\|122152314 | 19 | 40 | 22(+2A) | AACCCGTAGATCCGAACTTGTGAA | 2390.1 | 15.1 | 1.4 | 11.2 | 4.2E-29 | 1.5E-27 | MB231/MB468 | NUC |
| >MI0000102\|hsa-mir-100&WithFlank&11\|-\|122152223\|122152314@19.40.22(+1C)&[MIMAT0000098&hsa-miR-100-5p&offsets\|0\|0(+1C);m-27&11\|-\|122152276\|122152296&offsets\|0\|+1(+1C)]\|AACCCGTAGATCCGAACTTGTGC | >MI0000102\|hsa-mir-100&WithFlank&11\|-\|122152223\|122152314 | 19 | 40 | 22(+1C) | AACCCGTAGATCCGAACTTGTGC | 7787.1 | 16.9 | 1.5 | 11.3 | 1.1E-29 | 1.5E-27 | MB231/MB468 | CYTO |
| >hg38_21_+_8433172_8446622_RNA45SN1_45S_50ntFlanks@7975.8013.39\|CGCGACCTCAGATCAGACGTGGCGACCCGCTGAATTTAA | >hg38_21_+_8433172_8446622_RNA45SN1_45S_50ntFlanks | 7975 | 8013 | 39 | CGCGACCTCAGATCAGACGTGGCGACCCGCTGAATTTAA | 12413.3 | 8.1 | 0.7 | 11.3 | 1.1E-29 | 1.5E-27 | MB231/MB468 | CYTO |
| >hg38_21_+_8433172_8446622_RNA45SN1_45S_50ntFlanks@9352.9370.19\|CCGTGGCGCAATGAAGGTG | >hg38_21_+_8433172_8446622_RNA45SN1_45S_50ntFlanks | 9352 | 9370 | 19 | CCGTGGCGCAATGAAGGTG | 5050.1 | -16.1 | 1.4 | -11.2 | 2.7E-29 | 1.6E-27 | MB231/MB468 | MITO |
| >hg38_21_+_8433172_8446622_RNA45SN1_45S_50ntFlanks@12646.12669.24\|TGTGGGATTATGACTGAACGCCTC | >hg38_21_+_8433172_8446622_RNA45SN1_45S_50ntFlanks | 12646 | 12669 | 24 | TGTGGGATTATGACTGAACGCCTC | 928.2 | 7.3 | 0.7 | 11.2 | 6.6E-29 | 1.6E-27 | BT20/MB468 | MITO |
| >hg38_21_+_8433172_8446622_RNA45SN1_45S_50ntFlanks@10771.10796.26\|CAGGTCTCCAAGGTGAACAGCCTCTG | >hg38_21_+_8433172_8446622_RNA45SN1_45S_50ntFlanks | 10771 | 10796 | 26 | CAGGTCTCCAAGGTGAACAGCCTCTG | 1102.5 | -14.5 | 1.3 | -11.1 | 1.5E-28 | 1.6E-27 | BT20/MB468 | NUC |
| >hg38_21_+_8433172_8446622_RNA45SN1_45S_50ntFlanks@4760.4779.20\|TCAGATACCGTCGTAGTTCC | >hg38_21_+_8433172_8446622_RNA45SN1_45S_50ntFlanks | 4760 | 4779 | 20 | TCAGATACCGTCGTAGTTCC | 950.7 | -14.3 | 1.3 | -11.1 | 1.6E-28 | 1.6E-27 | BT20/MB468 | NUC |
| >MI0000299\|hsa-mir-222&WithFlank&X\|-\|45747009\|45747130@75.98.24&[MIMAT0000279&hsa-miR-222-3p&offsets\|0\|+3;m-140&X\|-\|45747033\|45747056&offsets\|0\|0]\|AGCTACATCTGGCTACTGGGTCTC | >MI0000299\|hsa-mir-222&WithFlank&X\|-\|45747009\|45747130 | 75 | 98 | 24 | AGCTACATCTGGCTACTGGGTCTC | 10907.5 | 6.8 | 0.6 | 11.1 | 9.2E-29 | 1.6E-27 | BT20/MB468 | CYTO |
| >hg38_21_+_8433172_8446622_RNA45SN1_45S_50ntFlanks@485.507.23\|GATGTGGTGACGTCGTGCTCTCC | >hg38_21_+_8433172_8446622_RNA45SN1_45S_50ntFlanks | 485 | 507 | 23 | GATGTGGTGACGTCGTGCTCTCC | 778.1 | 13.9 | 1.2 | 11.1 | 9.3E-29 | 1.6E-27 | BT20/MB468 | CYTO |
| >hg38_21_+_8433172_8446622_RNA45SN1_45S_50ntFlanks@11582.11603.22\|ACTGGTGCGGACCAGGGGAATC | >hg38_21_+_8433172_8446622_RNA45SN1_45S_50ntFlanks | 11582 | 11603 | 22 | ACTGGTGCGGACCAGGGGAATC | 1440.3 | -5.5 | 0.5 | -11.1 | 1.6E-28 | 1.6E-27 | BT20/MB468 | NUC |
| >hg38_21_+_8433172_8446622_RNA45SN1_45S_50ntFlanks@4064.4101.38\|ATCAACTTTCGATGGTAGTCGCCGTGCCTACCATGGTG | >hg38_21_+_8433172_8446622_RNA45SN1_45S_50ntFlanks | 4064 | 4101 | 38 | ATCAACTTTCGATGGTAGTCGCCGTGCCTACCATGGTG | 549.7 | -13.5 | 1.2 | -11.1 | 1.6E-28 | 1.7E-27 | BT20/MB468 | NUC |
| >hg38_21_+_8433172_8446622_RNA45SN1_45S_50ntFlanks@12934.12965.32\|CTAAACCATTCGTAGACGACCTGCTTCTGGGT | >hg38_21_+_8433172_8446622_RNA45SN1_45S_50ntFlanks | 12934 | 12965 | 32 | CTAAACCATTCGTAGACGACCTGCTTCTGGGT | 1364.1 | -14.8 | 1.3 | -11.1 | 1.6E-28 | 1.7E-27 | BT20/MB468 | NUC |
| >hg38_21_+_8433172_8446622_RNA45SN1_45S_50ntFlanks@3773.3792.20\|CGGCCGGTACAGTGAAACTG | >hg38_21_+_8433172_8446622_RNA45SN1_45S_50ntFlanks | 3773 | 3792 | 20 | CGGCCGGTACAGTGAAACTG | 968.1 | -14.3 | 1.3 | -11.1 | 1.6E-28 | 1.7E-27 | BT20/MB468 | NUC |
| >hg38_21_+_8433172_8446622_RNA45SN1_45S_50ntFlanks@12329.12346.18\|TTTTAAGCAGGAGGTGTC | >hg38_21_+_8433172_8446622_RNA45SN1_45S_50ntFlanks | 12329 | 12346 | 18 | TTTTAAGCAGGAGGTGTC | 936.6 | -14.3 | 1.3 | -11.1 | 1.6E-28 | 1.7E-27 | BT20/MB468 | NUC |
| >hg38_21_+_8433172_8446622_RNA45SN1_45S_50ntFlanks@5487.5514.28\|CGCTGAGAAGACGGTCGAACTTGACTAT | >hg38_21_+_8433172_8446622_RNA45SN1_45S_50ntFlanks | 5487 | 5514 | 28 | CGCTGAGAAGACGGTCGAACTTGACTAT | 8251.1 | -12.8 | 1.1 | -11.2 | 3.0E-29 | 1.7E-27 | MB231/MB468 | MITO |
| >hg38_21_+_8433172_8446622_RNA45SN1_45S_50ntFlanks@12002.12032.31\|TGCGGGCCGCCGGTGAAATACCACTACTCTG | >hg38_21_+_8433172_8446622_RNA45SN1_45S_50ntFlanks | 12002 | 12032 | 31 | TGCGGGCCGCCGGTGAAATACCACTACTCTG | 3419.4 | -5.3 | 0.5 | -11.2 | 7.1E-29 | 1.7E-27 | BT20/MB468 | MITO |
| >hg38_21_+_8433172_8446622_RNA45SN1_45S_50ntFlanks@11918.11951.34\|ACATGAGAGGTGTAGAATAAGTGGGAGGCCCCCG | >hg38_21_+_8433172_8446622_RNA45SN1_45S_50ntFlanks | 11918 | 11951 | 34 | ACATGAGAGGTGTAGAATAAGTGGGAGGCCCCCG | 624.1 | -13.7 | 1.2 | -11.1 | 1.7E-28 | 1.8E-27 | BT20/MB468 | NUC |
| >hg38_21_+_8433172_8446622_RNA45SN1_45S_50ntFlanks@10386.10410.25\|ATGTGAACAGCAGTTGAACATGGGT | >hg38_21_+_8433172_8446622_RNA45SN1_45S_50ntFlanks | 10386 | 10410 | 25 | ATGTGAACAGCAGTTGAACATGGGT | 589.4 | -13.6 | 1.2 | -11.1 | 1.7E-28 | 1.8E-27 | BT20/MB468 | NUC |
| >MI0000342\|hsa-mir-200b&WithFlank&1\|+\|1167098\|1167204@63.83.21&[MIMAT0000318&hsa-miR-200b-3p&offsets\|0\|-1;m-93&1\|+\|1167160\|1167182&offsets\|0\|-2]\|TAATACTGCCTGGTAATGATG | >MI0000342\|hsa-mir-200b&WithFlank&1\|+\|1167098\|1167204 | 63 | 83 | 21 | TAATACTGCCTGGTAATGATG | 611.2 | -6.0 | 0.5 | -11.1 | 1.7E-28 | 1.8E-27 | BT20/MB468 | NUC |
| >MI0000462\|hsa-mir-152&WithFlank&17\|-\|48037155\|48037253@60.81.22&[MIMAT0000438&hsa-miR-152-3p&offsets\|0\|+1;m-97&17\|-\|48037174\|48037194&offsets\|0\|+1]\|TCAGTGCATGACAGAACTTGGG | >MI0000462\|hsa-mir-152&WithFlank&17\|-\|48037155\|48037253 | 60 | 81 | 22 | TCAGTGCATGACAGAACTTGGG | 780.9 | 14.1 | 1.3 | 11.1 | 7.5E-29 | 1.8E-27 | BT20/MB468 | MITO |
| >trna10_LysCTT_16_+_3241501_3241573@1.36.36,trna13_LysCTT_14_-_58706613_58706685@1.36.36,trna2_LysCTT_15_+_79152904_79152976@1.36.36\|GCCCGGCTAGCTCAGTCGGTAGAGCATGGGACTCTT | >trna10_LysCTT_16_+_3241501_3241573 | 1 | 36 | 36 | GCCCGGCTAGCTCAGTCGGTAGAGCATGGGACTCTT | 11948.4 | 9.1 | 0.8 | 11.1 | 1.0E-28 | 1.8E-27 | BT20/MB468 | CYTO |
| >MI0000082\|hsa-mir-25&WithFlank&7\|-\|100093554\|100093649@58.78.21&[MIMAT0000081&hsa-miR-25-3p&offsets\|0\|-1;m-23&7\|-\|100093571\|100093592&offsets\|0\|-1]\|CATTGCACTTGTCTCGGTCTG | >MI0000082\|hsa-mir-25&WithFlank&7\|-\|100093554\|100093649 | 58 | 78 | 21 | CATTGCACTTGTCTCGGTCTG | 8037.6 | 6.8 | 0.6 | 11.1 | 1.0E-28 | 1.8E-27 | BT20/MB468 | CYTO |
| >hg38_21_+_8433172_8446622_RNA45SN1_45S_50ntFlanks@4279.4314.36\|AATCCTTTAACGAGGATCCATTGGAGGGCAAGTCTG | >hg38_21_+_8433172_8446622_RNA45SN1_45S_50ntFlanks | 4279 | 4314 | 36 | AATCCTTTAACGAGGATCCATTGGAGGGCAAGTCTG | 836.3 | -14.1 | 1.3 | -11.1 | 1.8E-28 | 1.8E-27 | BT20/MB468 | NUC |
| >hg38_21_+_8433172_8446622_RNA45SN1_45S_50ntFlanks@6719.6744.26\|TGAATTGCAGGACACATTGATCATCG | >hg38_21_+_8433172_8446622_RNA45SN1_45S_50ntFlanks | 6719 | 6744 | 26 | TGAATTGCAGGACACATTGATCATCG | 4746.0 | -10.6 | 0.9 | -11.3 | 1.7E-29 | 1.9E-27 | MB231/MB468 | MP |
| >hg38_21_+_8433172_8446622_RNA45SN1_45S_50ntFlanks@6738.6770.33\|ATCATCGACACTTCGAACGCACTTGCGGCCCCG | >hg38_21_+_8433172_8446622_RNA45SN1_45S_50ntFlanks | 6738 | 6770 | 33 | ATCATCGACACTTCGAACGCACTTGCGGCCCCG | 5512.7 | -16.3 | 1.4 | -11.3 | 1.7E-29 | 1.9E-27 | MB231/MB468 | MP |
| >hg38_21_+_8433172_8446622_RNA45SN1_45S_50ntFlanks@8182.8223.42\|ATCGAGGCCCAGCCCGTGGACGGTGTGAGGCCGGTAGCGGCC | >hg38_21_+_8433172_8446622_RNA45SN1_45S_50ntFlanks | 8182 | 8223 | 42 | ATCGAGGCCCAGCCCGTGGACGGTGTGAGGCCGGTAGCGGCC | 1581.5 | -15.1 | 1.4 | -11.1 | 1.9E-28 | 1.9E-27 | BT20/MB468 | NUC |
| >hg38_21_+_8433172_8446622_RNA45SN1_45S_50ntFlanks@13006.13025.20\|CTATTGAAAGTCAGCCCTCG | >hg38_21_+_8433172_8446622_RNA45SN1_45S_50ntFlanks | 13006 | 13025 | 20 | CTATTGAAAGTCAGCCCTCG | 1738.9 | -15.2 | 1.4 | -11.1 | 1.9E-28 | 1.9E-27 | BT20/MB468 | NUC |
| >hg38_21_+_8433172_8446622_RNA45SN1_45S_50ntFlanks@3873.3898.26\|TAATACATGCCGACGGGCGCTGACCC | >hg38_21_+_8433172_8446622_RNA45SN1_45S_50ntFlanks | 3873 | 3898 | 26 | TAATACATGCCGACGGGCGCTGACCC | 4495.9 | -16.6 | 1.5 | -11.1 | 1.9E-28 | 2.0E-27 | BT20/MB468 | NUC |
| >hg38_21_+_8433172_8446622_RNA45SN1_45S_50ntFlanks@9352.9370.19\|CCGTGGCGCAATGAAGGTG | >hg38_21_+_8433172_8446622_RNA45SN1_45S_50ntFlanks | 9352 | 9370 | 19 | CCGTGGCGCAATGAAGGTG | 6048.5 | -16.4 | 1.5 | -11.3 | 1.9E-29 | 2.0E-27 | MB231/MB468 | MP |
| >hg38_21_+_8433172_8446622_RNA45SN1_45S_50ntFlanks@12974.13001.28\|CGTACGTAGCAGAGCAGCTCCCTCGCTG | >hg38_21_+_8433172_8446622_RNA45SN1_45S_50ntFlanks | 12974 | 13001 | 28 | CGTACGTAGCAGAGCAGCTCCCTCGCTG | 567.1 | -13.6 | 1.2 | -11.1 | 2.1E-28 | 2.1E-27 | BT20/MB468 | NUC |
| >hg38_21_+_8433172_8446622_RNA45SN1_45S_50ntFlanks@12126.12150.25\|CGACCCGCTCCGGGGACAGTGCCAG | >hg38_21_+_8433172_8446622_RNA45SN1_45S_50ntFlanks | 12126 | 12150 | 25 | CGACCCGCTCCGGGGACAGTGCCAG | 1063.5 | -14.5 | 1.3 | -11.1 | 2.1E-28 | 2.2E-27 | BT20/MB468 | NUC |
| >hg38_21_+_8433172_8446622_RNA45SN1_45S_50ntFlanks@12328.12346.19\|GTTTTAAGCAGGAGGTGTC | >hg38_21_+_8433172_8446622_RNA45SN1_45S_50ntFlanks | 12328 | 12346 | 19 | GTTTTAAGCAGGAGGTGTC | 555.4 | -13.5 | 1.2 | -11.1 | 2.1E-28 | 2.2E-27 | BT20/MB468 | NUC |
| >MI0000301\|hsa-mir-224&WithFlank&X\|-\|151958572\|151958664@14.37.24&[MIMAT0000281&hsa-miR-224-5p&offsets\|+1\|0;m-238&X\|-\|151958630\|151958651&offsets\|0\|+2]\|CAAGTCACTAGTGGTTCCGTTTAG | >MI0000301\|hsa-mir-224&WithFlank&X\|-\|151958572\|151958664 | 14 | 37 | 24 | CAAGTCACTAGTGGTTCCGTTTAG | 6329.3 | 14.5 | 1.3 | 11.1 | 1.3E-28 | 2.2E-27 | BT20/MB468 | CYTO |
| >hg38_21_+_8433172_8446622_RNA45SN1_45S_50ntFlanks@4160.4182.23\|CCACATCCAAGGAAGGCAGCAGG | >hg38_21_+_8433172_8446622_RNA45SN1_45S_50ntFlanks | 4160 | 4182 | 23 | CCACATCCAAGGAAGGCAGCAGG | 820.8 | -14.1 | 1.3 | -11.1 | 2.2E-28 | 2.2E-27 | BT20/MB468 | NUC |
| >MI0000102\|hsa-mir-100&WithFlank&11\|-\|122152223\|122152314@19.40.22(+1C)&[MIMAT0000098&hsa-miR-100-5p&offsets\|0\|0(+1C);m-27&11\|-\|122152276\|122152296&offsets\|0\|+1(+1C)]\|AACCCGTAGATCCGAACTTGTGC | >MI0000102\|hsa-mir-100&WithFlank&11\|-\|122152223\|122152314 | 19 | 40 | 22(+1C) | AACCCGTAGATCCGAACTTGTGC | 5610.7 | 15.7 | 1.4 | 11.3 | 2.1E-29 | 2.2E-27 | MB231/MB468 | MP |
| >MI0000272\|hsa-mir-182&WithFlank&7\|-\|129770377\|129770498@29.50.22&[MIMAT0000259&hsa-miR-182-5p&offsets\|0\|-2;m-22&7\|-\|129770449\|129770470&offsets\|0\|0]\|TTTGGCAATGGTAGAACTCACA | >MI0000272\|hsa-mir-182&WithFlank&7\|-\|129770377\|129770498 | 29 | 50 | 22 | TTTGGCAATGGTAGAACTCACA | 1028.9 | 2.9 | 0.3 | 11.0 | 2.2E-28 | 2.3E-27 | BT20/MB468 | NUC |
| >hg38_21_+_8433172_8446622_RNA45SN1_45S_50ntFlanks@4283.4314.32\|CTTTAACGAGGATCCATTGGAGGGCAAGTCTG | >hg38_21_+_8433172_8446622_RNA45SN1_45S_50ntFlanks | 4283 | 4314 | 32 | CTTTAACGAGGATCCATTGGAGGGCAAGTCTG | 1788.6 | -15.2 | 1.4 | -11.0 | 2.3E-28 | 2.3E-27 | BT20/MB468 | NUC |
| >hg38_21_+_8433172_8446622_RNA45SN1_45S_50ntFlanks@11583.11612.30\|CTGGTGCGGACCAGGGGAATCCGACTGTTT | >hg38_21_+_8433172_8446622_RNA45SN1_45S_50ntFlanks | 11583 | 11612 | 30 | CTGGTGCGGACCAGGGGAATCCGACTGTTT | 1486.7 | -6.2 | 0.6 | -11.3 | 2.2E-29 | 2.3E-27 | MB231/MB468 | MP |
| >hg38_21_+_8433172_8446622_RNA45SN1_45S_50ntFlanks@10392.10410.19\|ACAGCAGTTGAACATGGGT | >hg38_21_+_8433172_8446622_RNA45SN1_45S_50ntFlanks | 10392 | 10410 | 19 | ACAGCAGTTGAACATGGGT | 5897.6 | -10.3 | 0.9 | -11.2 | 6.6E-29 | 2.4E-27 | MB231/MB468 | NUC |

<table>
<tr><th></th><th></th><th></th><th></th><th></th><th></th><th></th><th></th><th></th><th></th><th></th><th></th><th></th><th></th></tr>
<tr><td>>MI0000446|hsa-mir-125b-1&WithFlank&11|-|122099751|122099850@21.42.22;MI0000470|hsa-mir-125b-2&WithFlank&21|+|16590231|16590331@23.44.22&[MIMAT0000423&hsa-miR-125b-5p&offsets|0|0;m-67&11|-|122099809|122099830&offsets|0|0];[MIMAT0000423_1&hsa-miR-125b-5p&offsets|0|0;m-68&21|+|16590253|16590274&offsets|0|0]|TCCCTGAGACCCTAACTTGTGA</td><td>>MI0000446|hsa-mir-125b-1&WithFlank&11|-|122099751|122099850</td><td>21</td><td>42</td><td>22</td><td>TCCCTGAGACCCTAACTTGTGA</td><td>21622.9</td><td>7.9</td><td>0.7</td><td>11.2</td><td>6.7E-29</td><td>2.4E-27</td><td>MB231/MB468</td><td>NUC</td></tr>
<tr><td>>am_MI0000809|hsa-mir-151a&WithFlank&8|-|140732558|140732659@53.72.20(+1C)&[MIMAT0000757&hsa-miR-151a-3p&offsets|0|-1(+1C);m-30&8|-|140732587|140732607&offsets|0|-1(+1C)]|CTAGACTGAAGCTCCTTGAGC</td><td>>am_MI0000809|hsa-mir-151a&WithFlank&8|-|140732558|140732659</td><td>53</td><td>72</td><td>20(+1C)</td><td>CTAGACTGAAGCTCCTTGAGC</td><td>209.4</td><td>3.9</td><td>0.4</td><td>11.0</td><td>2.4E-28</td><td>2.4E-27</td><td>BT20/MB468</td><td>NUC</td></tr>
<tr><td>>hg38_21_+_8433172_8446622_RNA45SN1_45S_50ntFlanks@11729.11767.39|TAACTATGACTCTCTTAAGGTAGCCAAATGCCTCGTCAT</td><td>>hg38_21_+_8433172_8446622_RNA45SN1_45S_50ntFlanks</td><td>11729</td><td>11767</td><td>39</td><td>TAACTATGACTCTCTTAAGGTAGCCAAATGCCTCGTCAT</td><td>3980.6</td><td>-16.4</td><td>1.5</td><td>-11.0</td><td>2.4E-28</td><td>2.4E-27</td><td>BT20/MB468</td><td>NUC</td></tr>
<tr><td>>hg38_21_+_8433172_8446622_RNA45SN1_45S_50ntFlanks@11668.11707.40|TGCCCAGTGCTCTGAATGTCAAAGTGAAGAAATTCAATGA</td><td>>hg38_21_+_8433172_8446622_RNA45SN1_45S_50ntFlanks</td><td>11668</td><td>11707</td><td>40</td><td>TGCCCAGTGCTCTGAATGTCAAAGTGAAGAAATTCAATGA</td><td>1948.0</td><td>9.5</td><td>0.8</td><td>11.3</td><td>1.8E-29</td><td>2.5E-27</td><td>MB231/MB468</td><td>CYTO</td></tr>
<tr><td>>hg38_21_+_8433172_8446622_RNA45SN1_45S_50ntFlanks@10587.10607.21|CCCCGGGGAGAGTTCTCTTTT</td><td>>hg38_21_+_8433172_8446622_RNA45SN1_45S_50ntFlanks</td><td>10587</td><td>10607</td><td>21</td><td>CCCCGGGGAGAGTTCTCTTTT</td><td>595.3</td><td>-13.6</td><td>1.2</td><td>-11.0</td><td>2.5E-28</td><td>2.6E-27</td><td>BT20/MB468</td><td>NUC</td></tr>
<tr><td>>trna116_GluCTC_1_-_145399233_145399304@32.70.39,trna71_GluCTC_1_-_161439189_161439260@32.70.39,trna74_GluCTC_1_-_161431809_161431880@32.70.39,trna77_GluCTC_1_-_161424398_161424469@32.70.39,trna77_GluCTC_6_+_28949976_28950047@32.70.39,trna80_GluCTC_1_-_161417018_161417089@32.70.39,trna87_GluCTC_6_-_126101393_126101464@32.70.39|CTCTCACCGCCGCGGCCCGGGTTCGATTCCCGGTCAGGG</td><td>>trna116_GluCTC_1_-_145399233_145399304</td><td>32</td><td>70</td><td>39</td><td>CTCTCACCGCCGCGGCCCGGGTTCGATTCCCGGTCAGGG</td><td>5265.0</td><td>-15.3</td><td>1.4</td><td>-11.1</td><td>7.3E-29</td><td>2.6E-27</td><td>MB231/MB468</td><td>NUC</td></tr>
<tr><td>>hg38_21_+_8433172_8446622_RNA45SN1_45S_50ntFlanks@10391.10408.18|AACAGCAGTTGAACATGG</td><td>>hg38_21_+_8433172_8446622_RNA45SN1_45S_50ntFlanks</td><td>10391</td><td>10408</td><td>18</td><td>AACAGCAGTTGAACATGG</td><td>917.2</td><td>-14.3</td><td>1.3</td><td>-11.0</td><td>2.6E-28</td><td>2.6E-27</td><td>BT20/MB468</td><td>NUC</td></tr>
<tr><td>>hg38_21_+_8433172_8446622_RNA45SN1_45S_50ntFlanks@9572.9595.24|CAAATCGGTCGTCCGACCTGGGTA</td><td>>hg38_21_+_8433172_8446622_RNA45SN1_45S_50ntFlanks</td><td>9572</td><td>9595</td><td>24</td><td>CAAATCGGTCGTCCGACCTGGGTA</td><td>4416.8</td><td>-6.3</td><td>0.6</td><td>-11.1</td><td>1.5E-28</td><td>2.6E-27</td><td>BT20/MB468</td><td>CYTO</td></tr>
<tr><td>>hg38_21_+_8433172_8446622_RNA45SN1_45S_50ntFlanks@5486.5514.29|GCGCTGAGAAGACGGTCGAACTTGACTAT</td><td>>hg38_21_+_8433172_8446622_RNA45SN1_45S_50ntFlanks</td><td>5486</td><td>5514</td><td>29</td><td>GCGCTGAGAAGACGGTCGAACTTGACTAT</td><td>4566.0</td><td>-16.0</td><td>1.4</td><td>-11.2</td><td>2.6E-29</td><td>2.7E-27</td><td>MB231/MB468</td><td>MP</td></tr>
<tr><td>>hg38_21_+_8433172_8446622_RNA45SN1_45S_50ntFlanks@6651.6668.18|CGACTCTTAGCGGTGGAT</td><td>>hg38_21_+_8433172_8446622_RNA45SN1_45S_50ntFlanks</td><td>6651</td><td>6668</td><td>18</td><td>CGACTCTTAGCGGTGGAT</td><td>14332.2</td><td>4.7</td><td>0.4</td><td>11.1</td><td>1.5E-28</td><td>2.7E-27</td><td>BT20/MB468</td><td>CYTO</td></tr>
<tr><td>>MI0000299|hsa-mir-222&WithFlank&X|-|45747009|45747130@75.98.24(+1G)&[MIMAT0000279&hsa-miR-222-3p&offsets|0|+3(+1G);m-140&X|-|45747033|45747056&offsets|0|0(+1G)]|AGCTACATCTGGCTACTGGGTCTCG</td><td>>MI0000299|hsa-mir-222&WithFlank&X|-|45747009|45747130</td><td>75</td><td>98</td><td>24(+1G)</td><td>AGCTACATCTGGCTACTGGGTCTCG</td><td>890.1</td><td>6.4</td><td>0.6</td><td>11.1</td><td>1.5E-28</td><td>2.7E-27</td><td>BT20/MB468</td><td>CYTO</td></tr>
<tr><td>>hg38_21_+_8433172_8446622_RNA45SN1_45S_50ntFlanks@10350.10374.25|ACGAGAACTTTGAAGGCCGAAGTGG</td><td>>hg38_21_+_8433172_8446622_RNA45SN1_45S_50ntFlanks</td><td>10350</td><td>10374</td><td>25</td><td>ACGAGAACTTTGAAGGCCGAAGTGG</td><td>2602.2</td><td>-5.7</td><td>0.5</td><td>-11.1</td><td>7.8E-29</td><td>2.8E-27</td><td>MB231/MB468</td><td>NUC</td></tr>
<tr><td>>hg38_21_+_8433172_8446622_RNA45SN1_45S_50ntFlanks@10786.10815.30|AACAGCCTCTGGCATGTTGGAACAATGTAG</td><td>>hg38_21_+_8433172_8446622_RNA45SN1_45S_50ntFlanks</td><td>10786</td><td>10815</td><td>30</td><td>AACAGCCTCTGGCATGTTGGAACAATGTAG</td><td>743.2</td><td>-14.0</td><td>1.3</td><td>-11.0</td><td>2.9E-28</td><td>2.9E-27</td><td>BT20/MB468</td><td>NUC</td></tr>
<tr><td>>hg38_21_+_8433172_8446622_RNA45SN1_45S_50ntFlanks@7981.8006.26|CTCAGATCAGACGTGGCGACCCGCTG</td><td>>hg38_21_+_8433172_8446622_RNA45SN1_45S_50ntFlanks</td><td>7981</td><td>8006</td><td>26</td><td>CTCAGATCAGACGTGGCGACCCGCTG</td><td>14839.3</td><td>-10.9</td><td>1.0</td><td>-11.2</td><td>5.2E-29</td><td>2.9E-27</td><td>MB231/MB468</td><td>MITO</td></tr>
<tr><td>>am_MI0000786|hsa-mir-378a&WithFlank&5|+|149732819|149732896@50.69.20(+1U)&[MIMAT0000732&hsa-miR-378a-3p&offsets|+1|-1(+1U);m-44&5|+|149732867|149732888&offsets|+1|-1(+1U)]|CTGGACTTGGAGTCAGAAGGT</td><td>>am_MI0000786|hsa-mir-378a&WithFlank&5|+|149732819|149732896</td><td>50</td><td>69</td><td>20(+1U)</td><td>CTGGACTTGGAGTCAGAAGGT</td><td>1212.5</td><td>7.4</td><td>0.7</td><td>11.1</td><td>1.7E-28</td><td>3.0E-27</td><td>BT20/MB468</td><td>CYTO</td></tr>
<tr><td>>hg38_21_+_8433172_8446622_RNA45SN1_45S_50ntFlanks@12405.12427.23|CTTTTTGATCCTTCGATGTCGGC</td><td>>hg38_21_+_8433172_8446622_RNA45SN1_45S_50ntFlanks</td><td>12405</td><td>12427</td><td>23</td><td>CTTTTTGATCCTTCGATGTCGGC</td><td>486.5</td><td>-13.4</td><td>1.2</td><td>-11.0</td><td>3.0E-28</td><td>3.1E-27</td><td>BT20/MB468</td><td>NUC</td></tr>
<tr><td>>hg38_21_+_8433172_8446622_RNA45SN1_45S_50ntFlanks@11581.11605.25|AACTGGTGCGGACCAGGGGAATCCG</td><td>>hg38_21_+_8433172_8446622_RNA45SN1_45S_50ntFlanks</td><td>11581</td><td>11605</td><td>25</td><td>AACTGGTGCGGACCAGGGGAATCCG</td><td>1649.1</td><td>-6.8</td><td>0.6</td><td>-11.1</td><td>8.8E-29</td><td>3.1E-27</td><td>MB231/MB468</td><td>NUC</td></tr>
<tr><td>>hg38_21_+_8433172_8446622_RNA45SN1_45S_50ntFlanks@6700.6717.18|CTAGCTGCGAGAATTAAT</td><td>>hg38_21_+_8433172_8446622_RNA45SN1_45S_50ntFlanks</td><td>6700</td><td>6717</td><td>18</td><td>CTAGCTGCGAGAATTAAT</td><td>808.2</td><td>-14.1</td><td>1.3</td><td>-11.0</td><td>3.1E-28</td><td>3.1E-27</td><td>BT20/MB468</td><td>NUC</td></tr>
<tr><td>>MI0000088|hsa-mir-30a&WithFlank&6|-|71403545|71403627@13.35.23&[MIMAT0000087&hsa-miR-30a-5p&offsets|+1|+2;m-15&6|-|71403595|71403616&offsets|+1|+2]|GTAAACATCCTCGACTGGAAGCT</td><td>>MI0000088|hsa-mir-30a&WithFlank&6|-|71403545|71403627</td><td>13</td><td>35</td><td>23</td><td>GTAAACATCCTCGACTGGAAGCT</td><td>552.0</td><td>6.8</td><td>0.6</td><td>11.1</td><td>1.3E-28</td><td>3.2E-27</td><td>BT20/MB468</td><td>MITO</td></tr>
<tr><td>>hg38_21_+_8433172_8446622_RNA45SN1_45S_50ntFlanks@4160.4184.25|CCACATCCAAGGAAGGCAGCAGGCG</td><td>>hg38_21_+_8433172_8446622_RNA45SN1_45S_50ntFlanks</td><td>4160</td><td>4184</td><td>25</td><td>CCACATCCAAGGAAGGCAGCAGGCG</td><td>775.6</td><td>-14.0</td><td>1.3</td><td>-11.0</td><td>3.2E-28</td><td>3.2E-27</td><td>BT20/MB468</td><td>NUC</td></tr>
<tr><td>>MI0000746|hsa-mir-99b&WithFlank&19|+|51692606|51692687@13.34.22&[MIMAT0000689&hsa-miR-99b-5p&offsets|0|0;m-10&19|+|51692618|51692639&offsets|0|0]|CACCCGTAGAACCGACCTTGCG</td><td>>MI0000746|hsa-mir-99b&WithFlank&19|+|51692606|51692687</td><td>13</td><td>34</td><td>22</td><td>CACCCGTAGAACCGACCTTGCG</td><td>128484.8</td><td>7.7</td><td>0.7</td><td>11.1</td><td>1.8E-28</td><td>3.2E-27</td><td>BT20/MB468</td><td>CYTO</td></tr>
<tr><td>>hg38_21_+_8433172_8446622_RNA45SN1_45S_50ntFlanks@10843.10862.20|TAACTTCGGGATAAGGATTG</td><td>>hg38_21_+_8433172_8446622_RNA45SN1_45S_50ntFlanks</td><td>10843</td><td>10862</td><td>20</td><td>TAACTTCGGGATAAGGATTG</td><td>621.6</td><td>-13.7</td><td>1.2</td><td>-11.0</td><td>3.2E-28</td><td>3.2E-27</td><td>BT20/MB468</td><td>NUC</td></tr>
<tr><td>>hg38_21_+_8433172_8446622_RNA45SN1_45S_50ntFlanks@9534.9574.41|AGGAAACTCTGGTGGAGGTCCGTAGCGGTCCTGACGTGCAA</td><td>>hg38_21_+_8433172_8446622_RNA45SN1_45S_50ntFlanks</td><td>9534</td><td>9574</td><td>41</td><td>AGGAAACTCTGGTGGAGGTCCGTAGCGGTCCTGACGTGCAA</td><td>4552.0</td><td>-16.6</td><td>1.5</td><td>-11.0</td><td>3.2E-28</td><td>3.2E-27</td><td>BT20/MB468</td><td>NUC</td></tr>
<tr><td>>hg38_21_+_8433172_8446622_RNA45SN1_45S_50ntFlanks@11212.11232.21|CGACTCTGGACGCGAGCCGGG</td><td>>hg38_21_+_8433172_8446622_RNA45SN1_45S_50ntFlanks</td><td>11212</td><td>11232</td><td>21</td><td>CGACTCTGGACGCGAGCCGGG</td><td>2544.5</td><td>-15.2</td><td>1.4</td><td>-11.2</td><td>3.2E-29</td><td>3.3E-27</td><td>MB231/MB468</td><td>MP</td></tr>
<tr><td>>MI0000434|hsa-let-7i&WithFlank&12|+|62603680|62603775@12.32.21(+1A)&[MIMAT0000415&hsa-let-7i-5p&offsets|0|-1(+1A);m-49&12|+|62603691|62603712&offsets|0|-1(+1A)]|TGAGGTAGTAGTTTGTGCTGTA</td><td>>MI0000434|hsa-let-7i&WithFlank&12|+|62603680|62603775</td><td>12</td><td>32</td><td>21(+1A)</td><td>TGAGGTAGTAGTTTGTGCTGTA</td><td>2795.1</td><td>7.8</td><td>0.7</td><td>11.1</td><td>1.4E-28</td><td>3.3E-27</td><td>BT20/MB468</td><td>MITO</td></tr>
<tr><td>>trna15_ValAAC_5_-_180615416_180615488@1.29.29|GTTTCCGTAGTGTAGTGGTCATCACGTTC</td><td>>trna15_ValAAC_5_-_180615416_180615488</td><td>1</td><td>29</td><td>29</td><td>GTTTCCGTAGTGTAGTGGTCATCACGTTC</td><td>1139.2</td><td>-14.6</td><td>1.3</td><td>-11.0</td><td>3.3E-28</td><td>3.3E-27</td><td>BT20/MB468</td><td>NUC</td></tr>
<tr><td>>hg38_21_+_8433172_8446622_RNA45SN1_45S_50ntFlanks@9564.9592.29|CTGACGTGCAAATCGGTCGTCCGACCTGG</td><td>>hg38_21_+_8433172_8446622_RNA45SN1_45S_50ntFlanks</td><td>9564</td><td>9592</td><td>29</td><td>CTGACGTGCAAATCGGTCGTCCGACCTGG</td><td>2434.8</td><td>-6.1</td><td>0.6</td><td>-11.1</td><td>2.0E-28</td><td>3.4E-27</td><td>BT20/MB468</td><td>CYTO</td></tr>
<tr><td>>hg38_21_+_8433172_8446622_RNA45SN1_45S_50ntFlanks@485.507.23|GATGTGGTGACGTCGTGCTCTCC</td><td>>hg38_21_+_8433172_8446622_RNA45SN1_45S_50ntFlanks</td><td>485</td><td>507</td><td>23</td><td>GATGTGGTGACGTCGTGCTCTCC</td><td>1267.5</td><td>13.1</td><td>1.2</td><td>11.0</td><td>3.4E-28</td><td>3.4E-27</td><td>BT20/MB468</td><td>NUC</td></tr>
<tr><td>>hg38_21_+_8433172_8446622_RNA45SN1_45S_50ntFlanks@6646.6668.23|TCGTACGACTCTTAGCGGTGGAT</td><td>>hg38_21_+_8433172_8446622_RNA45SN1_45S_50ntFlanks</td><td>6646</td><td>6668</td><td>23</td><td>TCGTACGACTCTTAGCGGTGGAT</td><td>13684.0</td><td>5.7</td><td>0.5</td><td>11.1</td><td>2.0E-28</td><td>3.5E-27</td><td>BT20/MB468</td><td>CYTO</td></tr>
<tr><td>>hg38_21_+_8433172_8446622_RNA45SN1_45S_50ntFlanks@11323.11342.20|CGTCTCGTCGCGCGCGCGTC</td><td>>hg38_21_+_8433172_8446622_RNA45SN1_45S_50ntFlanks</td><td>11323</td><td>11342</td><td>20</td><td>CGTCTCGTCGCGCGCGCGTC</td><td>5131.8</td><td>-10.5</td><td>0.9</td><td>-11.2</td><td>3.4E-29</td><td>3.5E-27</td><td>MB231/MB468</td><td>MP</td></tr>
<tr><td>>hg38_21_+_8433172_8446622_RNA45SN1_45S_50ntFlanks@12605.12626.22|TATGTGCTTGGCTGAGGAGCCA</td><td>>hg38_21_+_8433172_8446622_RNA45SN1_45S_50ntFlanks</td><td>12605</td><td>12626</td><td>22</td><td>TATGTGCTTGGCTGAGGAGCCA</td><td>2814.0</td><td>-15.3</td><td>1.4</td><td>-11.2</td><td>3.5E-29</td><td>3.5E-27</td><td>MB231/MB468</td><td>MP</td></tr>
<tr><td>>hg38_21_+_8433172_8446622_RNA45SN1_45S_50ntFlanks@6721.6741.21|AATTGCAGGACACATTGATCA</td><td>>hg38_21_+_8433172_8446622_RNA45SN1_45S_50ntFlanks</td><td>6721</td><td>6741</td><td>21</td><td>AATTGCAGGACACATTGATCA</td><td>1399.5</td><td>-14.9</td><td>1.4</td><td>-11.0</td><td>3.6E-28</td><td>3.5E-27</td><td>BT20/MB468</td><td>NUC</td></tr>
<tr><td>>hg38_21_+_8433172_8446622_RNA45SN1_45S_50ntFlanks@12934.12956.23|CTAAACCATTCGTAGACGACCTG</td><td>>hg38_21_+_8433172_8446622_RNA45SN1_45S_50ntFlanks</td><td>12934</td><td>12956</td><td>23</td><td>CTAAACCATTCGTAGACGACCTG</td><td>8705.0</td><td>-7.5</td><td>0.7</td><td>-11.1</td><td>2.1E-28</td><td>3.6E-27</td><td>BT20/MB468</td><td>CYTO</td></tr>
<tr><td>>hg38_21_+_8433172_8446622_RNA45SN1_45S_50ntFlanks@6652.6693.42|GACTCTTAGCGGTGGATCACTCGGCTCGTGCGTCGATGAAGA</td><td>>hg38_21_+_8433172_8446622_RNA45SN1_45S_50ntFlanks</td><td>6652</td><td>6693</td><td>42</td><td>GACTCTTAGCGGTGGATCACTCGGCTCGTGCGTCGATGAAGA</td><td>4703.3</td><td>5.4</td><td>0.5</td><td>11.1</td><td>1.5E-28</td><td>3.6E-27</td><td>BT20/MB468</td><td>MITO</td></tr>
<tr><td>>hg38_21_+_8433172_8446622_RNA45SN1_45S_50ntFlanks@10571.10596.26|GAGAAGCCGGCGGGAGCCCCGGGGAG</td><td>>hg38_21_+_8433172_8446622_RNA45SN1_45S_50ntFlanks</td><td>10571</td><td>10596</td><td>26</td><td>GAGAAGCCGGCGGGAGCCCCGGGGAG</td><td>815.2</td><td>-14.1</td><td>1.3</td><td>-11.0</td><td>3.6E-28</td><td>3.6E-27</td><td>BT20/MB468</td><td>NUC</td></tr>
<tr><td>>hg38_21_+_8433172_8446622_RNA45SN1_45S_50ntFlanks@12006.12035.30|GGCCGCCGGTGAAATACCACTACTCTGATC</td><td>>hg38_21_+_8433172_8446622_RNA45SN1_45S_50ntFlanks</td><td>12006</td><td>12035</td><td>30</td><td>GGCCGCCGGTGAAATACCACTACTCTGATC</td><td>1190.1</td><td>6.5</td><td>0.6</td><td>11.1</td><td>1.5E-28</td><td>3.6E-27</td><td>BT20/MB468</td><td>MITO</td></tr>
<tr><td>>hg38_21_+_8433172_8446622_RNA45SN1_45S_50ntFlanks@12418.12435.18|CGATGTCGGCTCTTCCTA</td><td>>hg38_21_+_8433172_8446622_RNA45SN1_45S_50ntFlanks</td><td>12418</td><td>12435</td><td>18</td><td>CGATGTCGGCTCTTCCTA</td><td>2890.3</td><td>-15.4</td><td>1.4</td><td>-11.2</td><td>3.7E-29</td><td>3.7E-27</td><td>MB231/MB468</td><td>MP</td></tr>
<tr><td>>hg38_21_+_8433172_8446622_RNA45SN1_45S_50ntFlanks@12456.12476.21|CCAAGCGTTGGATTGTTCACC</td><td>>hg38_21_+_8433172_8446622_RNA45SN1_45S_50ntFlanks</td><td>12456</td><td>12476</td><td>21</td><td>CCAAGCGTTGGATTGTTCACC</td><td>9831.5</td><td>-6.1</td><td>0.5</td><td>-11.1</td><td>1.6E-28</td><td>3.7E-27</td><td>BT20/MB468</td><td>MITO</td></tr>
<tr><td>>MI0000767|hsa-mir-365a&WithFlank&16|+|14309279|14309377@62.83.22;MI0000769|hsa-mir-365b&WithFlank&17|+|31575405|31575527@74.95.22&[MIMAT0000710&hsa-miR-365a-3p&offsets|0|0;m-176&16|+|14309340|14309361&offsets|0|0];[MIMAT0022834&hsa-miR-365b-3p&offsets|0|0;m-175&17|+|31575478|31575499&offsets|0|0]|TAATGCCCCTAAAAATCCTTAT</td><td>>MI0000767|hsa-mir-365a&WithFlank&16|+|14309279|14309377</td><td>62</td><td>83</td><td>22</td><td>TAATGCCCCTAAAAATCCTTAT</td><td>2135.8</td><td>8.7</td><td>0.8</td><td>11.1</td><td>1.6E-28</td><td>3.8E-27</td><td>BT20/MB468</td><td>MITO</td></tr>
<tr><td>>hg38_21_+_8433172_8446622_RNA45SN1_45S_50ntFlanks@11581.11603.23|AACTGGTGCGGACCAGGGGAATC</td><td>>hg38_21_+_8433172_8446622_RNA45SN1_45S_50ntFlanks</td><td>11581</td><td>11603</td><td>23</td><td>AACTGGTGCGGACCAGGGGAATC</td><td>4462.8</td><td>-8.3</td><td>0.7</td><td>-11.2</td><td>6.8E-29</td><td>3.8E-27</td><td>MB231/MB468</td><td>MITO</td></tr>
<tr><td>>MI0000746|hsa-mir-99b&WithFlank&19|+|51692606|51692687@13.32.20(+2U)&[MIMAT0000689&hsa-miR-99b-5p&offsets|0|-2(+2U);m-10&19|+|51692618|51692639&offsets|0|-2(+2U)]|CACCCGTAGAACCGACCTTGTT</td><td>>MI0000746|hsa-mir-99b&WithFlank&19|+|51692606|51692687</td><td>13</td><td>32</td><td>20(+2U)</td><td>CACCCGTAGAACCGACCTTGTT</td><td>557.6</td><td>13.6</td><td>1.2</td><td>11.1</td><td>1.6E-28</td><td>3.8E-27</td><td>BT20/MB468</td><td>MITO</td></tr>
<tr><td>>hg38_21_+_8433172_8446622_RNA45SN1_45S_50ntFlanks@12991.13010.20|CTCCCTCGCTGCGATCTATT</td><td>>hg38_21_+_8433172_8446622_RNA45SN1_45S_50ntFlanks</td><td>12991</td><td>13010</td><td>20</td><td>CTCCCTCGCTGCGATCTATT</td><td>1829.7</td><td>-15.3</td><td>1.4</td><td>-11.0</td><td>3.9E-28</td><td>3.9E-27</td><td>BT20/MB468</td><td>NUC</td></tr>
<tr><td>>hg38_21_+_8433172_8446622_RNA45SN1_45S_50ntFlanks@9575.9603.29|ATCGGTCGTCCGACCTGGGTATAGGGGCG</td><td>>hg38_21_+_8433172_8446622_RNA45SN1_45S_50ntFlanks</td><td>9575</td><td>9603</td><td>29</td><td>ATCGGTCGTCCGACCTGGGTATAGGGGCG</td><td>1105.2</td><td>-14.5</td><td>1.3</td><td>-11.0</td><td>3.9E-28</td><td>3.9E-27</td><td>BT20/MB468</td><td>NUC</td></tr>
<tr><td>>hg38_21_+_8433172_8446622_RNA45SN1_45S_50ntFlanks@12405.12431.27|CTTTTTGATCCTTCGATGTCGGCTCTT</td><td>>hg38_21_+_8433172_8446622_RNA45SN1_45S_50ntFlanks</td><td>12405</td><td>12431</td><td>27</td><td>CTTTTTGATCCTTCGATGTCGGCTCTT</td><td>4852.6</td><td>-16.1</td><td>1.4</td><td>-11.2</td><td>4.0E-29</td><td>3.9E-27</td><td>MB231/MB468</td><td>MP</td></tr>
<tr><td>>hg38_21_+_8433172_8446622_RNA45SN1_45S_50ntFlanks@4657.4684.28|CTAGAGGTGAAATTCTTGGACCGGCGCA</td><td>>hg38_21_+_8433172_8446622_RNA45SN1_45S_50ntFlanks</td><td>4657</td><td>4684</td><td>28</td><td>CTAGAGGTGAAATTCTTGGACCGGCGCA</td><td>617.0</td><td>-13.7</td><td>1.2</td><td>-11.0</td><td>4.1E-28</td><td>4.0E-27</td><td>BT20/MB468</td><td>NUC</td></tr>
<tr><td>>hg38_21_+_8433172_8446622_RNA45SN1_45S_50ntFlanks@4003.4026.24|ATAACCTCGGGCCGATCGCACGCC</td><td>>hg38_21_+_8433172_8446622_RNA45SN1_45S_50ntFlanks</td><td>4003</td><td>4026</td><td>24</td><td>ATAACCTCGGGCCGATCGCACGCC</td><td>2064.4</td><td>-15.4</td><td>1.4</td><td>-11.0</td><td>4.1E-28</td><td>4.0E-27</td><td>BT20/MB468</td><td>NUC</td></tr>
<tr><td>>hg38_21_+_8433172_8446622_RNA45SN1_45S_50ntFlanks@4728.4752.25|ATCAAGAACGAAAGTCGGAGGTTCG</td><td>>hg38_21_+_8433172_8446622_RNA45SN1_45S_50ntFlanks</td><td>4728</td><td>4752</td><td>25</td><td>ATCAAGAACGAAAGTCGGAGGTTCG</td><td>957.1</td><td>-14.3</td><td>1.3</td><td>-11.0</td><td>4.1E-28</td><td>4.0E-27</td><td>BT20/MB468</td><td>NUC</td></tr>
</table>

<table>
<tr><th></th><th></th><th></th><th></th><th></th><th></th><th></th><th></th><th></th><th></th><th></th><th></th><th></th><th></th></tr>
<tr><td>>MI0000783|hsa-mir-375&WithFlank&2|-|219001639|219001714@46.67.22&[MIMAT0000728&hsa-miR-375-3p&offsets|0|0;m-92&2|-|219001648|219001669&offsets|0|0]|TTTGTTCGTTCGGCTCGCGTGA</td><td>>MI0000783|hsa-mir-375&WithFlank&2|-|219001639|219001714</td><td>46</td><td>67</td><td>22</td><td>TTTGTTCGTTCGGCTCGCGTGA</td><td>398.3</td><td>4.7</td><td>0.4</td><td>11.0</td><td>2.4E-28</td><td>4.1E-27</td><td>BT20/MB468</td><td>CYTO</td></tr>
<tr><td>>hg38_21_+_8433172_8446622_RNA45SN1_45S_50ntFlanks@13004.13025.22|ATCTATTGAAAGTCAGCCCTCG</td><td>>hg38_21_+_8433172_8446622_RNA45SN1_45S_50ntFlanks</td><td>13004</td><td>13025</td><td>22</td><td>ATCTATTGAAAGTCAGCCCTCG</td><td>2268.7</td><td>-15.6</td><td>1.4</td><td>-11.0</td><td>4.1E-28</td><td>4.1E-27</td><td>BT20/MB468</td><td>NUC</td></tr>
<tr><td>>hg38_21_+_8433172_8446622_RNA45SN1_45S_50ntFlanks@7973.8009.37|GACGCGACCTCAGATCAGACGTGGCGACCCGCTGAAT</td><td>>hg38_21_+_8433172_8446622_RNA45SN1_45S_50ntFlanks</td><td>7973</td><td>8009</td><td>37</td><td>GACGCGACCTCAGATCAGACGTGGCGACCCGCTGAAT</td><td>2482.8</td><td>7.7</td><td>0.7</td><td>11.2</td><td>3.0E-29</td><td>4.1E-27</td><td>MB231/MB468</td><td>CYTO</td></tr>
<tr><td>>hg38_21_+_8433172_8446622_RNA45SN1_45S_50ntFlanks@10375.10407.33|AGAAGGGTTCCATGTGAACAGCAGTTGAACATG</td><td>>hg38_21_+_8433172_8446622_RNA45SN1_45S_50ntFlanks</td><td>10375</td><td>10407</td><td>33</td><td>AGAAGGGTTCCATGTGAACAGCAGTTGAACATG</td><td>916.1</td><td>-14.3</td><td>1.3</td><td>-11.0</td><td>4.3E-28</td><td>4.2E-27</td><td>BT20/MB468</td><td>NUC</td></tr>
<tr><td>>MI0000279|hsa-mir-196a-2&WithFlank&12|+|53991732|53991853@31.52.22;MI0000238|hsa-mir-196a-1&WithFlank&17|-|48632484|48632565@13.34.22&[MIMAT0000226&hsa-miR-196a-5p&offsets|0|0;m-261&12|+|53991762|53991783&offsets|0|0];[MIMAT0000226_1&hsa-miR-196a-5p&offsets|0|0;m-268&17|-|48632532|48632553&offsets|0|0]|TAGGTAGTTTCATGTTGTTGGG</td><td>>MI0000279|hsa-mir-196a-2&WithFlank&12|+|53991732|53991853</td><td>31</td><td>52</td><td>22</td><td>TAGGTAGTTTCATGTTGTTGGG</td><td>3731.1</td><td>15.6</td><td>1.4</td><td>11.3</td><td>9.6E-30</td><td>4.2E-27</td><td>BT20/MB231</td><td>NUC</td></tr>
<tr><td>>hg38_21_+_8433172_8446622_RNA45SN1_45S_50ntFlanks@12625.12659.35|CAATGGGGCGAAGCTACCATCTGTGGGATTATGAC</td><td>>hg38_21_+_8433172_8446622_RNA45SN1_45S_50ntFlanks</td><td>12625</td><td>12659</td><td>35</td><td>CAATGGGGCGAAGCTACCATCTGTGGGATTATGAC</td><td>580.3</td><td>-13.6</td><td>1.2</td><td>-11.0</td><td>4.3E-28</td><td>4.2E-27</td><td>BT20/MB468</td><td>NUC</td></tr>
<tr><td>>hg38_21_+_8433172_8446622_RNA45SN1_45S_50ntFlanks@4512.4537.26|AAAAAATTAGAGTGTTCAAAGCAGGC</td><td>>hg38_21_+_8433172_8446622_RNA45SN1_45S_50ntFlanks</td><td>4512</td><td>4537</td><td>26</td><td>AAAAAATTAGAGTGTTCAAAGCAGGC</td><td>500.1</td><td>-13.4</td><td>1.2</td><td>-11.0</td><td>4.3E-28</td><td>4.2E-27</td><td>BT20/MB468</td><td>NUC</td></tr>
<tr><td>>hg38_21_+_8433172_8446622_RNA45SN1_45S_50ntFlanks@6699.6718.20|GCTAGCTGCGAGAATTAATG</td><td>>hg38_21_+_8433172_8446622_RNA45SN1_45S_50ntFlanks</td><td>6699</td><td>6718</td><td>20</td><td>GCTAGCTGCGAGAATTAATG</td><td>8400.7</td><td>-6.1</td><td>0.5</td><td>-11.1</td><td>1.8E-28</td><td>4.3E-27</td><td>BT20/MB468</td><td>MITO</td></tr>
<tr><td>>hg38_21_+_8433172_8446622_RNA45SN1_45S_50ntFlanks@9567.9592.26|ACGTGCAAATCGGTCGTCCGACCTGG</td><td>>hg38_21_+_8433172_8446622_RNA45SN1_45S_50ntFlanks</td><td>9567</td><td>9592</td><td>26</td><td>ACGTGCAAATCGGTCGTCCGACCTGG</td><td>666.1</td><td>-13.8</td><td>1.3</td><td>-11.0</td><td>4.4E-28</td><td>4.3E-27</td><td>BT20/MB468</td><td>NUC</td></tr>
<tr><td>>hg38_21_+_8433172_8446622_RNA45SN1_45S_50ntFlanks@11873.11906.34|AAGAAGACCCTGTTGAGCTTGACTCTAGTCTGGC</td><td>>hg38_21_+_8433172_8446622_RNA45SN1_45S_50ntFlanks</td><td>11873</td><td>11906</td><td>34</td><td>AAGAAGACCCTGTTGAGCTTGACTCTAGTCTGGC</td><td>3134.3</td><td>-15.4</td><td>1.4</td><td>-11.1</td><td>7.6E-29</td><td>4.3E-27</td><td>MB231/MB468</td><td>MITO</td></tr>
<tr><td>>hg38_21_+_8433172_8446622_RNA45SN1_45S_50ntFlanks@12712.12742.31|CGGAGCCTCGGTTGGCCTCGGATAGCCGGTC</td><td>>hg38_21_+_8433172_8446622_RNA45SN1_45S_50ntFlanks</td><td>12712</td><td>12742</td><td>31</td><td>CGGAGCCTCGGTTGGCCTCGGATAGCCGGTC</td><td>5105.5</td><td>-16.2</td><td>1.4</td><td>-11.2</td><td>4.4E-29</td><td>4.3E-27</td><td>MB231/MB468</td><td>MP</td></tr>
<tr><td>>hg38_21_+_8433172_8446622_RNA45SN1_45S_50ntFlanks@6740.6770.31|CATCGACACTTCGAACGCACTTGCGGCCCCG</td><td>>hg38_21_+_8433172_8446622_RNA45SN1_45S_50ntFlanks</td><td>6740</td><td>6770</td><td>31</td><td>CATCGACACTTCGAACGCACTTGCGGCCCCG</td><td>2734.1</td><td>-15.3</td><td>1.4</td><td>-11.2</td><td>4.4E-29</td><td>4.3E-27</td><td>MB231/MB468</td><td>MP</td></tr>
<tr><td>>MI0000087|hsa-mir-29a&WithFlank&7|-|130876741|130876816@48.68.21&[MIMAT0000086&hsa-miR-29a-3p&offsets|0|-1;m-26&7|-|130876749|130876769&offsets|0|0]|TAGCACCATCTGAAATCGGTT</td><td>>MI0000087|hsa-mir-29a&WithFlank&7|-|130876741|130876816</td><td>48</td><td>68</td><td>21</td><td>TAGCACCATCTGAAATCGGTT</td><td>8943.2</td><td>9.1</td><td>0.8</td><td>11.0</td><td>2.5E-28</td><td>4.4E-27</td><td>BT20/MB468</td><td>CYTO</td></tr>
<tr><td>>hg38_21_+_8433172_8446622_RNA45SN1_45S_50ntFlanks@8380.8404.25|CAAGAGGGCGTGAAACCGTTAAGAG</td><td>>hg38_21_+_8433172_8446622_RNA45SN1_45S_50ntFlanks</td><td>8380</td><td>8404</td><td>25</td><td>CAAGAGGGCGTGAAACCGTTAAGAG</td><td>641.4</td><td>-13.8</td><td>1.3</td><td>-11.0</td><td>4.5E-28</td><td>4.4E-27</td><td>BT20/MB468</td><td>NUC</td></tr>
<tr><td>>hg38_21_+_8433172_8446622_RNA45SN1_45S_50ntFlanks@12676.12708.33|AGAATCCCGCCCAGGCGGAACGATACGGCAGCG</td><td>>hg38_21_+_8433172_8446622_RNA45SN1_45S_50ntFlanks</td><td>12676</td><td>12708</td><td>33</td><td>AGAATCCCGCCCAGGCGGAACGATACGGCAGCG</td><td>522.1</td><td>-13.5</td><td>1.2</td><td>-11.0</td><td>4.5E-28</td><td>4.4E-27</td><td>BT20/MB468</td><td>NUC</td></tr>
<tr><td>>hg38_21_+_8433172_8446622_RNA45SN1_45S_50ntFlanks@3992.4023.32|GGTGACTCTAGATAACCTCGGGCCGATCGCAC</td><td>>hg38_21_+_8433172_8446622_RNA45SN1_45S_50ntFlanks</td><td>3992</td><td>4023</td><td>32</td><td>GGTGACTCTAGATAACCTCGGGCCGATCGCAC</td><td>732.6</td><td>6.0</td><td>0.5</td><td>11.0</td><td>2.7E-28</td><td>4.5E-27</td><td>BT20/MB468</td><td>CYTO</td></tr>
<tr><td>>am_MI0000809|hsa-mir-151a&WithFlank&8|-|140732558|140732659@53.72.20(+1U)&[MIMAT0000757&hsa-miR-151a-3p&offsets|0|-1(+1U);m-30&8|-|140732587|140732607&offsets|0|-1(+1U)]|CTAGACTGAAGCTCCTTGAGT</td><td>>am_MI0000809|hsa-mir-151a&WithFlank&8|-|140732558|140732659</td><td>53</td><td>72</td><td>20(+1U)</td><td>CTAGACTGAAGCTCCTTGAGT</td><td>3241.8</td><td>6.7</td><td>0.6</td><td>11.1</td><td>1.3E-28</td><td>4.5E-27</td><td>MB231/MB468</td><td>NUC</td></tr>
<tr><td>>MI0000298|hsa-mir-221&WithFlank&X|-|45746151|45746272@71.92.22(+1U)&[MIMAT0000278&hsa-miR-221-3p&offsets|0|-1(+1U);m-91&X|-|45746181|45746202&offsets|0|0(+1U)]|AGCTACATTGTCTGCTGGGTTTT</td><td>>MI0000298|hsa-mir-221&WithFlank&X|-|45746151|45746272</td><td>71</td><td>92</td><td>22(+1U)</td><td>AGCTACATTGTCTGCTGGGTTTT</td><td>3753.1</td><td>6.9</td><td>0.6</td><td>11.1</td><td>1.3E-28</td><td>4.5E-27</td><td>MB231/MB468</td><td>NUC</td></tr>
<tr><td>>hg38_21_+_8433172_8446622_RNA45SN1_45S_50ntFlanks@6745.6771.27|ACACTTCGAACGCACTTGCGGCCCCGG</td><td>>hg38_21_+_8433172_8446622_RNA45SN1_45S_50ntFlanks</td><td>6745</td><td>6771</td><td>27</td><td>ACACTTCGAACGCACTTGCGGCCCCGG</td><td>1542.5</td><td>-6.7</td><td>0.6</td><td>-11.1</td><td>1.3E-28</td><td>4.6E-27</td><td>MB231/MB468</td><td>NUC</td></tr>
<tr><td>>hg38_21_+_8433172_8446622_RNA45SN1_45S_50ntFlanks@3705.3743.39|TACCTGGTTGATCCTGCCAGTAGCATATGCTTGTCTCAA</td><td>>hg38_21_+_8433172_8446622_RNA45SN1_45S_50ntFlanks</td><td>3705</td><td>3743</td><td>39</td><td>TACCTGGTTGATCCTGCCAGTAGCATATGCTTGTCTCAA</td><td>3620.5</td><td>7.3</td><td>0.7</td><td>11.2</td><td>3.5E-29</td><td>4.7E-27</td><td>MB231/MB468</td><td>CYTO</td></tr>
<tr><td>>MI0006444|hsa-mir-1307&WithFlank&10|-|103394247|103394407@86.107.22&[MIMAT0005951&hsa-miR-1307-3p&offsets|0|0;m-66&10|-|103394300|103394322&offsets|0|-1]|ACTCGGCGTGGCGTCGGTCGTG</td><td>>MI0006444|hsa-mir-1307&WithFlank&10|-|103394247|103394407</td><td>86</td><td>107</td><td>22</td><td>ACTCGGCGTGGCGTCGGTCGTG</td><td>1130.6</td><td>6.6</td><td>0.6</td><td>11.0</td><td>2.8E-28</td><td>4.7E-27</td><td>BT20/MB468</td><td>CYTO</td></tr>
<tr><td>>hg38_21_+_8433172_8446622_RNA45SN1_45S_50ntFlanks@11208.11227.20|GCGGCGACTCTGGACGCGAG</td><td>>hg38_21_+_8433172_8446622_RNA45SN1_45S_50ntFlanks</td><td>11208</td><td>11227</td><td>20</td><td>GCGGCGACTCTGGACGCGAG</td><td>3704.6</td><td>-15.7</td><td>1.4</td><td>-11.2</td><td>4.9E-29</td><td>4.7E-27</td><td>MB231/MB468</td><td>MP</td></tr>
<tr><td>>hg38_21_+_8433172_8446622_RNA45SN1_45S_50ntFlanks@12628.12654.27|TGGGGCGAAGCTACCATCTGTGGGATT</td><td>>hg38_21_+_8433172_8446622_RNA45SN1_45S_50ntFlanks</td><td>12628</td><td>12654</td><td>27</td><td>TGGGGCGAAGCTACCATCTGTGGGATT</td><td>3864.7</td><td>-15.8</td><td>1.4</td><td>-11.2</td><td>4.9E-29</td><td>4.7E-27</td><td>MB231/MB468</td><td>MP</td></tr>
<tr><td>>hg38_21_+_8433172_8446622_RNA45SN1_45S_50ntFlanks@7986.8005.20|ATCAGACGTGGCGACCCGCT</td><td>>hg38_21_+_8433172_8446622_RNA45SN1_45S_50ntFlanks</td><td>7986</td><td>8005</td><td>20</td><td>ATCAGACGTGGCGACCCGCT</td><td>596.5</td><td>-13.6</td><td>1.2</td><td>-11.0</td><td>5.0E-28</td><td>4.8E-27</td><td>BT20/MB468</td><td>NUC</td></tr>
<tr><td>>hg38_21_+_8433172_8446622_RNA45SN1_45S_50ntFlanks@6661.6700.40|CGGTGGATCACTCGGCTCGTGCGTCGATGAAGAACGCAGC</td><td>>hg38_21_+_8433172_8446622_RNA45SN1_45S_50ntFlanks</td><td>6661</td><td>6700</td><td>40</td><td>CGGTGGATCACTCGGCTCGTGCGTCGATGAAGAACGCAGC</td><td>4218.5</td><td>8.2</td><td>0.7</td><td>11.1</td><td>2.1E-28</td><td>4.8E-27</td><td>BT20/MB468</td><td>MITO</td></tr>
<tr><td>>hg38_21_+_8433172_8446622_RNA45SN1_45S_50ntFlanks@9563.9596.34|CCTGACGTGCAAATCGGTCGTCCGACCTGGGTAT</td><td>>hg38_21_+_8433172_8446622_RNA45SN1_45S_50ntFlanks</td><td>9563</td><td>9596</td><td>34</td><td>CCTGACGTGCAAATCGGTCGTCCGACCTGGGTAT</td><td>3315.4</td><td>-5.7</td><td>0.5</td><td>-11.1</td><td>2.1E-28</td><td>4.8E-27</td><td>BT20/MB468</td><td>MITO</td></tr>
<tr><td>>hg38_21_+_8433172_8446622_RNA45SN1_45S_50ntFlanks@12008.12034.27|CCGCCGGTGAAATACCACTACTCTGAT</td><td>>hg38_21_+_8433172_8446622_RNA45SN1_45S_50ntFlanks</td><td>12008</td><td>12034</td><td>27</td><td>CCGCCGGTGAAATACCACTACTCTGAT</td><td>528.5</td><td>-13.5</td><td>1.2</td><td>-11.0</td><td>5.0E-28</td><td>4.9E-27</td><td>BT20/MB468</td><td>NUC</td></tr>
<tr><td>>hg38_21_+_8433172_8446622_RNA45SN1_45S_50ntFlanks@2959.2979.21|CGACGTGTGGCGTGGGTCGAC</td><td>>hg38_21_+_8433172_8446622_RNA45SN1_45S_50ntFlanks</td><td>2959</td><td>2979</td><td>21</td><td>CGACGTGTGGCGTGGGTCGAC</td><td>843.0</td><td>-14.1</td><td>1.3</td><td>-11.0</td><td>5.1E-28</td><td>4.9E-27</td><td>BT20/MB468</td><td>NUC</td></tr>
<tr><td>>hg38_21_+_8433172_8446622_RNA45SN1_45S_50ntFlanks@12008.12030.23|CCGCCGGTGAAATACCACTACTC</td><td>>hg38_21_+_8433172_8446622_RNA45SN1_45S_50ntFlanks</td><td>12008</td><td>12030</td><td>23</td><td>CCGCCGGTGAAATACCACTACTC</td><td>1172.9</td><td>-14.6</td><td>1.3</td><td>-11.0</td><td>5.2E-28</td><td>5.0E-27</td><td>BT20/MB468</td><td>NUC</td></tr>
<tr><td>>hg38_21_+_8433172_8446622_RNA45SN1_45S_50ntFlanks@10393.10420.28|CAGCAGTTGAACATGGGTCAGTCGGTCC</td><td>>hg38_21_+_8433172_8446622_RNA45SN1_45S_50ntFlanks</td><td>10393</td><td>10420</td><td>28</td><td>CAGCAGTTGAACATGGGTCAGTCGGTCC</td><td>888.1</td><td>-14.2</td><td>1.3</td><td>-11.0</td><td>5.3E-28</td><td>5.1E-27</td><td>BT20/MB468</td><td>NUC</td></tr>
<tr><td>>hg38_21_+_8433172_8446622_RNA45SN1_45S_50ntFlanks@6737.6770.34|GATCATCGACACTTCGAACGCACTTGCGGCCCCG</td><td>>hg38_21_+_8433172_8446622_RNA45SN1_45S_50ntFlanks</td><td>6737</td><td>6770</td><td>34</td><td>GATCATCGACACTTCGAACGCACTTGCGGCCCCG</td><td>1345.5</td><td>-14.8</td><td>1.4</td><td>-11.0</td><td>5.3E-28</td><td>5.1E-27</td><td>BT20/MB468</td><td>NUC</td></tr>
<tr><td>>hg38_21_+_8433172_8446622_RNA45SN1_45S_50ntFlanks@4472.4493.22|CTTAGCTGAGTGTCCCGCGGGG</td><td>>hg38_21_+_8433172_8446622_RNA45SN1_45S_50ntFlanks</td><td>4472</td><td>4493</td><td>22</td><td>CTTAGCTGAGTGTCCCGCGGGG</td><td>531.5</td><td>-13.5</td><td>1.2</td><td>-11.0</td><td>5.3E-28</td><td>5.1E-27</td><td>BT20/MB468</td><td>NUC</td></tr>
<tr><td>>hg38_21_+_8433172_8446622_RNA45SN1_45S_50ntFlanks@12432.12455.24|CCTATCATTGTGAAGCAGAATTCA</td><td>>hg38_21_+_8433172_8446622_RNA45SN1_45S_50ntFlanks</td><td>12432</td><td>12455</td><td>24</td><td>CCTATCATTGTGAAGCAGAATTCA</td><td>935.0</td><td>-14.3</td><td>1.3</td><td>-11.0</td><td>5.4E-28</td><td>5.2E-27</td><td>BT20/MB468</td><td>NUC</td></tr>
<tr><td>>MI0006444|hsa-mir-1307&WithFlank&10|-|103394247|103394407@87.108.22&[MIMAT0005951&hsa-miR-1307-3p&offsets|+1|+1;m-66&10|-|103394300|103394322&offsets|+1|0]|CTCGGCGTGGCGTCGGTCGTGG</td><td>>MI0006444|hsa-mir-1307&WithFlank&10|-|103394247|103394407</td><td>87</td><td>108</td><td>22</td><td>CTCGGCGTGGCGTCGGTCGTGG</td><td>1817.8</td><td>6.8</td><td>0.6</td><td>11.0</td><td>2.2E-28</td><td>5.2E-27</td><td>BT20/MB468</td><td>MITO</td></tr>
<tr><td>>hg38_21_+_8433172_8446622_RNA45SN1_45S_50ntFlanks@10572.10596.25|AGAAGCCGGCGGGAGCCCCGGGGAG</td><td>>hg38_21_+_8433172_8446622_RNA45SN1_45S_50ntFlanks</td><td>10572</td><td>10596</td><td>25</td><td>AGAAGCCGGCGGGAGCCCCGGGGAG</td><td>672.9</td><td>-13.8</td><td>1.3</td><td>-11.0</td><td>5.4E-28</td><td>5.2E-27</td><td>BT20/MB468</td><td>NUC</td></tr>
<tr><td>>hg38_21_+_8433172_8446622_RNA45SN1_45S_50ntFlanks@485.521.37|GATGTGGTGACGTCGTGCTCTCCCGGGCCGGGTCCGA</td><td>>hg38_21_+_8433172_8446622_RNA45SN1_45S_50ntFlanks</td><td>485</td><td>521</td><td>37</td><td>GATGTGGTGACGTCGTGCTCTCCCGGGCCGGGTCCGA</td><td>973.9</td><td>14.4</td><td>1.3</td><td>11.0</td><td>2.3E-28</td><td>5.2E-27</td><td>BT20/MB468</td><td>MITO</td></tr>
<tr><td>>hg38_21_+_8433172_8446622_RNA45SN1_45S_50ntFlanks@11585.11623.39|GGTGCGGACCAGGGGAATCCGACTGTTTAATTAAAACAA</td><td>>hg38_21_+_8433172_8446622_RNA45SN1_45S_50ntFlanks</td><td>11585</td><td>11623</td><td>39</td><td>GGTGCGGACCAGGGGAATCCGACTGTTTAATTAAAACAA</td><td>1404.1</td><td>14.9</td><td>1.4</td><td>11.0</td><td>2.3E-28</td><td>5.2E-27</td><td>BT20/MB468</td><td>MITO</td></tr>
<tr><td>>hg38_21_+_8433172_8446622_RNA45SN1_45S_50ntFlanks@10392.10420.29|ACAGCAGTTGAACATGGGTCAGTCGGTCC</td><td>>hg38_21_+_8433172_8446622_RNA45SN1_45S_50ntFlanks</td><td>10392</td><td>10420</td><td>29</td><td>ACAGCAGTTGAACATGGGTCAGTCGGTCC</td><td>1927.2</td><td>-15.3</td><td>1.4</td><td>-11.0</td><td>5.5E-28</td><td>5.3E-27</td><td>BT20/MB468</td><td>NUC</td></tr>
<tr><td>>MI0000471|hsa-mir-126&WithFlank&9|+|136670596|136670692@57.79.23&[MIMAT0000445&hsa-miR-126-3p&offsets|-1|0;m-40&9|+|136670653|136670674&offsets|-1|0]|CTCGTACCGTGAGTAATAATGCG</td><td>>MI0000471|hsa-mir-126&WithFlank&9|+|136670596|136670692</td><td>57</td><td>79</td><td>23</td><td>CTCGTACCGTGAGTAATAATGCG</td><td>1163.2</td><td>14.6</td><td>1.3</td><td>11.0</td><td>2.3E-28</td><td>5.3E-27</td><td>BT20/MB468</td><td>MITO</td></tr>
<tr><td>>hg38_21_+_8433172_8446622_RNA45SN1_45S_50ntFlanks@4099.4117.19|GTGACCACGGGTGACGGGG</td><td>>hg38_21_+_8433172_8446622_RNA45SN1_45S_50ntFlanks</td><td>4099</td><td>4117</td><td>19</td><td>GTGACCACGGGTGACGGGG</td><td>432.8</td><td>-13.2</td><td>1.2</td><td>-11.0</td><td>5.6E-28</td><td>5.4E-27</td><td>BT20/MB468</td><td>NUC</td></tr>
<tr><td>>hg38_21_+_8433172_8446622_RNA45SN1_45S_50ntFlanks@12934.12972.39|CTAAACCATTCGTAGACGACCTGCTTCTGGGTCGGGGTT</td><td>>hg38_21_+_8433172_8446622_RNA45SN1_45S_50ntFlanks</td><td>12934</td><td>12972</td><td>39</td><td>CTAAACCATTCGTAGACGACCTGCTTCTGGGTCGGGGTT</td><td>2622.7</td><td>-15.8</td><td>1.4</td><td>-11.0</td><td>5.8E-28</td><td>5.5E-27</td><td>BT20/MB468</td><td>NUC</td></tr>
<tr><td>>MI0000061|hsa-let-7a-2&WithFlank&11|-|122146516|122146599@11.28.18;MI0000064|hsa-let-7c&WithFlank&21|+|16539822|16539917@17.34.18;MI0000062|hsa-let-7a-3&WithFlank&22|+|46112743|46112828@10.27.18;MI0000060|hsa-let-7a-1&WithFlank&9|+|94175951|94176042@12.29.18&[MIMAT0000062&hsa-let-7a-5p&offsets|0|-4;m-6&11|-|122146568|122146589&offsets|0|-4];[MIMAT0000064&hsa-let-7c-5p&offsets|0|-4;m-37&21|+|16539838|16539859&offsets|0|-4];[MIMAT0000062_1&hsa-let-7a-5p&offsets|0|-4;m-5&22|+|46112752|46112773&offsets|0|-4];[MIMAT0000062_2&hsa-let-7a-5p&offsets|0|-4;m-4&9|+|94175962|94175983&offsets|0|-4]|TGAGGTAGTAGGTTGTAT</td><td>>MI0000061|hsa-let-7a-2&WithFlank&11|-|122146516|122146599</td><td>11</td><td>28</td><td>18</td><td>TGAGGTAGTAGGTTGTAT</td><td>2125.4</td><td>7.6</td><td>0.7</td><td>11.0</td><td>2.4E-28</td><td>5.5E-27</td><td>BT20/MB468</td><td>MITO</td></tr>
<tr><td>>hg38_21_+_8433172_8446622_RNA45SN1_45S_50ntFlanks@9534.9571.38|AGGAAACTCTGGTGGAGGTCCGTAGCGGTCCTGACGTG</td><td>>hg38_21_+_8433172_8446622_RNA45SN1_45S_50ntFlanks</td><td>9534</td><td>9571</td><td>38</td><td>AGGAAACTCTGGTGGAGGTCCGTAGCGGTCCTGACGTG</td><td>24699.9</td><td>-5.4</td><td>0.5</td><td>-11.0</td><td>3.3E-28</td><td>5.6E-27</td><td>BT20/MB468</td><td>CYTO</td></tr>
<tr><td>>hg38_21_+_8433172_8446622_RNA45SN1_45S_50ntFlanks@5525.5550.26|TAAAAGTCGTAACAAGGTTTCCGTAG</td><td>>hg38_21_+_8433172_8446622_RNA45SN1_45S_50ntFlanks</td><td>5525</td><td>5550</td><td>26</td><td>TAAAAGTCGTAACAAGGTTTCCGTAG</td><td>1436.5</td><td>-14.9</td><td>1.4</td><td>-11.0</td><td>5.9E-28</td><td>5.6E-27</td><td>BT20/MB468</td><td>NUC</td></tr>
<tr><td>>hg38_21_+_8433172_8446622_RNA45SN1_45S_50ntFlanks@9736.9755.20|CCGAAACGATCTCAACCTAT</td><td>>hg38_21_+_8433172_8446622_RNA45SN1_45S_50ntFlanks</td><td>9736</td><td>9755</td><td>20</td><td>CCGAAACGATCTCAACCTAT</td><td>701.4</td><td>-13.9</td><td>1.3</td><td>-11.0</td><td>6.0E-28</td><td>5.7E-27</td><td>BT20/MB468</td><td>NUC</td></tr>
<tr><td>>MI0000299|hsa-mir-222&WithFlank&X|-|45747009|45747130@75.99.25(+3U)&[MIMAT0000279&hsa-miR-222-3p&offsets|0|+4(+3U);m-140&X|-|45747033|45747056&offsets|0|+1(+3U)]|AGCTACATCTGGCTACTGGGTCTCTTTT</td><td>>MI0000299|hsa-mir-222&WithFlank&X|-|45747009|45747130</td><td>75</td><td>99</td><td>25(+3U)</td><td>AGCTACATCTGGCTACTGGGTCTCTTTT</td><td>468.5</td><td>6.5</td><td>0.6</td><td>11.0</td><td>3.4E-28</td><td>5.7E-27</td><td>BT20/MB468</td><td>CYTO</td></tr>
</table>

<table>
<tr><td>>hg38_21_+_8433172_8446622_RNA45SN1_45S_50ntFlanks@485.517.33|GATGTGGTGACGTCGTGCTCTCCCGGGCCGGGT</td><td>>hg38_21_+_8433172_8446622_RNA45SN1_45S_50ntFlanks</td><td>485</td><td>517</td><td>33</td><td>GATGTGGTGACGTCGTGCTCTCCCGGGCCGGGT</td><td>1426.4</td><td>14.9</td><td>1.4</td><td>11.0</td><td>2.6E-28</td><td>5.8E-27</td><td>BT20/MB468</td><td>MITO</td></tr>
<tr><td>>hg38_21_+_8433172_8446622_RNA45SN1_45S_50ntFlanks@11316.11338.23|CACCCCACGTCTCGTCGCGCGCG</td><td>>hg38_21_+_8433172_8446622_RNA45SN1_45S_50ntFlanks</td><td>11316</td><td>11338</td><td>23</td><td>CACCCCACGTCTCGTCGCGCGCG</td><td>7363.3</td><td>-16.6</td><td>1.5</td><td>-11.1</td><td>1.1E-28</td><td>5.9E-27</td><td>MB231/MB468</td><td>MITO</td></tr>
<tr><td>>hg38_21_+_8433172_8446622_RNA45SN1_45S_50ntFlanks@9738.9755.18|GAAACGATCTCAACCTAT</td><td>>hg38_21_+_8433172_8446622_RNA45SN1_45S_50ntFlanks</td><td>9738</td><td>9755</td><td>18</td><td>GAAACGATCTCAACCTAT</td><td>1260.6</td><td>-14.7</td><td>1.3</td><td>-11.0</td><td>6.4E-28</td><td>6.0E-27</td><td>BT20/MB468</td><td>NUC</td></tr>
<tr><td>>hg38_21_+_8433172_8446622_RNA45SN1_45S_50ntFlanks@10349.10384.36|AACGAGAACTTTGAAGGCCGAAGTGGAGAAGGGTTC</td><td>>hg38_21_+_8433172_8446622_RNA45SN1_45S_50ntFlanks</td><td>10349</td><td>10384</td><td>36</td><td>AACGAGAACTTTGAAGGCCGAAGTGGAGAAGGGTTC</td><td>4039.5</td><td>-7.6</td><td>0.7</td><td>-11.0</td><td>6.5E-28</td><td>6.1E-27</td><td>BT20/MB468</td><td>NUC</td></tr>
<tr><td>>hg38_21_+_8433172_8446622_RNA45SN1_45S_50ntFlanks@4727.4758.32|AATCAAGAACGAAAGTCGGAGGTTCGAAGACG</td><td>>hg38_21_+_8433172_8446622_RNA45SN1_45S_50ntFlanks</td><td>4727</td><td>4758</td><td>32</td><td>AATCAAGAACGAAAGTCGGAGGTTCGAAGACG</td><td>1277.5</td><td>-5.3</td><td>0.5</td><td>-11.0</td><td>6.6E-28</td><td>6.2E-27</td><td>BT20/MB468</td><td>NUC</td></tr>
<tr><td>>hg38_21_+_8433172_8446622_RNA45SN1_45S_50ntFlanks@7974.8010.37|ACGCGACCTCAGATCAGACGTGGCGACCCGCTGAATT</td><td>>hg38_21_+_8433172_8446622_RNA45SN1_45S_50ntFlanks</td><td>7974</td><td>8010</td><td>37</td><td>ACGCGACCTCAGATCAGACGTGGCGACCCGCTGAATT</td><td>12964.9</td><td>7.9</td><td>0.7</td><td>11.2</td><td>4.8E-29</td><td>6.3E-27</td><td>MB231/MB468</td><td>CYTO</td></tr>
<tr><td>>hg38_21_+_8433172_8446622_RNA45SN1_45S_50ntFlanks@12632.12653.22|GCGAAGCTACCATCTGTGGGAT</td><td>>hg38_21_+_8433172_8446622_RNA45SN1_45S_50ntFlanks</td><td>12632</td><td>12653</td><td>22</td><td>GCGAAGCTACCATCTGTGGGAT</td><td>2794.3</td><td>-15.3</td><td>1.4</td><td>-11.2</td><td>6.8E-29</td><td>6.5E-27</td><td>MB231/MB468</td><td>MP</td></tr>
<tr><td>>trna116_GluCTC_1_-_145399233_145399304@1.36.36,trna59_GluCTC_1_+_249168447_249168518@1.36.36,trna71_GluCTC_1_-_161439189_161439260@1.36.36,trna74_GluCTC_1_-_161431809_161431880@1.36.36,trna77_GluCTC_1_-_161424398_161424469@1.36.36,trna77_GluCTC_6_+_28949976_28950047@1.36.36,trna80_GluCTC_1_-_161417018_161417089@1.36.36,trna87_GluCTC_6_-_126101393_126101464@1.36.36|TCCCTGGTGGTCTAGTGGTTAGGATTCGGCGCTCTC</td><td>>trna116_GluCTC_1_-_145399233_145399304</td><td>1</td><td>36</td><td>36</td><td>TCCCTGGTGGTCTAGTGGTTAGGATTCGGCGCTCTC</td><td>10110.7</td><td>8.8</td><td>0.8</td><td>11.0</td><td>3.9E-28</td><td>6.6E-27</td><td>BT20/MB468</td><td>CYTO</td></tr>
<tr><td>>MI0000061|hsa-let-7a-2&WithFlank&11|-|122146516|122146599@11.31.21(+1C);MI0000062|hsa-let-7a-3&WithFlank&22|+|46112743|46112828@10.30.21(+1C);MI0000060|hsa-let-7a-1&WithFlank&9|+|94175951|94176042@12.32.21(+1C)&[MIMAT0000062&hsa-let-7a-5p&offsets|0|-1(+1C);m-6&11|-|122146568|122146589&offsets|0|-1(+1C)];[MIMAT0000062_1&hsa-let-7a-5p&offsets|0|-1(+1C);m-5&22|+|46112752|46112773&offsets|0|-1(+1C)];[MIMAT0000062_2&hsa-let-7a-5p&offsets|0|-1(+1C);m-4&9|+|94175962|94175983&offsets|0|-1(+1C)]|TGAGGTAGTAGGTTGTATAGTC</td><td>>MI0000061|hsa-let-7a-2&WithFlank&11|-|122146516|122146599</td><td>11</td><td>31</td><td>21(+1C)</td><td>TGAGGTAGTAGGTTGTATAGTC</td><td>21023.2</td><td>7.6</td><td>0.7</td><td>11.0</td><td>4.0E-28</td><td>6.7E-27</td><td>BT20/MB468</td><td>CYTO</td></tr>
<tr><td>>hg38_21_+_8433172_8446622_RNA45SN1_45S_50ntFlanks@11316.11344.29|CACCCCACGTCTCGTCGCGCGCGCGTCCG</td><td>>hg38_21_+_8433172_8446622_RNA45SN1_45S_50ntFlanks</td><td>11316</td><td>11344</td><td>29</td><td>CACCCCACGTCTCGTCGCGCGCGCGTCCG</td><td>38330.5</td><td>-12.0</td><td>1.1</td><td>-11.1</td><td>2.0E-28</td><td>6.7E-27</td><td>MB231/MB468</td><td>NUC</td></tr>
<tr><td>>MI0000750|hsa-mir-26a-2&WithFlank&12|-|57824603|57824698@20.39.20;MI0000083|hsa-mir-26a-1&WithFlank&3|+|37969398|37969486@16.35.20&[MIMAT0000082&hsa-miR-26a-5p&offsets|0|-2;m-33&12|-|57824658|57824679&offsets|0|-2];[MIMAT0000082_1&hsa-miR-26a-5p&offsets|0|-2;m-34&3|+|37969413|37969434&offsets|0|-2]|TTCAAGTAATCCAGGATAGG</td><td>>MI0000750|hsa-mir-26a-2&WithFlank&12|-|57824603|57824698</td><td>20</td><td>39</td><td>20</td><td>TTCAAGTAATCCAGGATAGG</td><td>1285.6</td><td>7.2</td><td>0.7</td><td>11.0</td><td>4.1E-28</td><td>6.8E-27</td><td>BT20/MB468</td><td>CYTO</td></tr>
<tr><td>>hg38_21_+_8433172_8446622_RNA45SN1_45S_50ntFlanks@11890.11912.23|CTTGACTCTAGTCTGGCACGGTG</td><td>>hg38_21_+_8433172_8446622_RNA45SN1_45S_50ntFlanks</td><td>11890</td><td>11912</td><td>23</td><td>CTTGACTCTAGTCTGGCACGGTG</td><td>1428.4</td><td>-4.9</td><td>0.4</td><td>-11.0</td><td>4.1E-28</td><td>6.8E-27</td><td>BT20/MB468</td><td>CYTO</td></tr>
<tr><td>>hg38_21_+_8433172_8446622_RNA45SN1_45S_50ntFlanks@12405.12443.39|CTTTTTGATCCTTCGATGTCGGCTCTTCCTATCATTGTG</td><td>>hg38_21_+_8433172_8446622_RNA45SN1_45S_50ntFlanks</td><td>12405</td><td>12443</td><td>39</td><td>CTTTTTGATCCTTCGATGTCGGCTCTTCCTATCATTGTG</td><td>11904.8</td><td>-8.1</td><td>0.7</td><td>-11.1</td><td>1.2E-28</td><td>6.9E-27</td><td>MB231/MB468</td><td>MITO</td></tr>
<tr><td>>am_trnaMT_ValTAC_MT_+_1602_1670@51.68.18|CTTAACTTGACCGCTCTG</td><td>>am_trnaMT_ValTAC_MT_+_1602_1670</td><td>51</td><td>68</td><td>18</td><td>CTTAACTTGACCGCTCTG</td><td>21600.4</td><td>-9.1</td><td>0.8</td><td>-11.1</td><td>1.3E-28</td><td>6.9E-27</td><td>MB231/MB468</td><td>MITO</td></tr>
<tr><td>>hg38_21_+_8433172_8446622_RNA45SN1_45S_50ntFlanks@12995.13014.20|CTCGCTGCGATCTATTGAAA</td><td>>hg38_21_+_8433172_8446622_RNA45SN1_45S_50ntFlanks</td><td>12995</td><td>13014</td><td>20</td><td>CTCGCTGCGATCTATTGAAA</td><td>850.2</td><td>-5.5</td><td>0.5</td><td>-10.9</td><td>7.4E-28</td><td>7.0E-27</td><td>BT20/MB468</td><td>NUC</td></tr>
<tr><td>>hg38_21_+_8433172_8446622_RNA45SN1_45S_50ntFlanks@12435.12454.20|ATCATTGTGAAGCAGAATTC</td><td>>hg38_21_+_8433172_8446622_RNA45SN1_45S_50ntFlanks</td><td>12435</td><td>12454</td><td>20</td><td>ATCATTGTGAAGCAGAATTC</td><td>987.2</td><td>-14.4</td><td>1.3</td><td>-10.9</td><td>7.6E-28</td><td>7.1E-27</td><td>BT20/MB468</td><td>NUC</td></tr>
<tr><td>>hg38_21_+_8433172_8446622_RNA45SN1_45S_50ntFlanks@7968.7995.28|TCCGAGACGCGACCTCAGATCAGACGTG</td><td>>hg38_21_+_8433172_8446622_RNA45SN1_45S_50ntFlanks</td><td>7968</td><td>7995</td><td>28</td><td>TCCGAGACGCGACCTCAGATCAGACGTG</td><td>341.3</td><td>4.8</td><td>0.4</td><td>11.1</td><td>2.1E-28</td><td>7.2E-27</td><td>MB231/MB468</td><td>NUC</td></tr>
<tr><td>>am_trna111_HisGTG_1_-_147774845_147774916@-1G.29.30,trna118_HisGTG_1_-_145396881_145396952@-1G.29.30,trna16_HisGTG_1_+_146544773_146544844@-1G.29.30,trna1_HisGTG_15_+_45493349_45493420@-1G.29.30,trna21_HisGTG_1_+_147753471_147753542@-1G.29.30,trna33_HisGTG_6_+_27125906_27125977@-1G.29.30,trna7_HisGTG_9_-_14433938_14434009@-1G.29.30,trna8_HisGTG_15_-_45492611_45492682@-1G.29.30,trna9_HisGTG_15_-_45490804_45490875@-1G.29.30|GGCCGTGATCGTATAGTGGTTAGTACTCTG</td><td>>am_trna111_HisGTG_1_-_147774845_147774916</td><td>-1G</td><td>29</td><td>30</td><td>GGCCGTGATCGTATAGTGGTTAGTACTCTG</td><td>29539.2</td><td>-12.6</td><td>1.2</td><td>-10.9</td><td>7.8E-28</td><td>7.3E-27</td><td>BT20/MB468</td><td>NUC</td></tr>
<tr><td>>trnaMT_PheGAA_MT_+_577_647@38.57.20|AATGTTTAGACGGGCTCACA</td><td>>trnaMT_PheGAA_MT_+_577_647</td><td>38</td><td>57</td><td>20</td><td>AATGTTTAGACGGGCTCACA</td><td>648.3</td><td>6.8</td><td>0.6</td><td>11.0</td><td>3.3E-28</td><td>7.4E-27</td><td>BT20/MB468</td><td>MITO</td></tr>
<tr><td>>hg38_21_+_8433172_8446622_RNA45SN1_45S_50ntFlanks@4297.4319.23|CATTGGAGGGCAAGTCTGGTGCC</td><td>>hg38_21_+_8433172_8446622_RNA45SN1_45S_50ntFlanks</td><td>4297</td><td>4319</td><td>23</td><td>CATTGGAGGGCAAGTCTGGTGCC</td><td>498.9</td><td>-13.4</td><td>1.2</td><td>-10.9</td><td>7.9E-28</td><td>7.4E-27</td><td>BT20/MB468</td><td>NUC</td></tr>
<tr><td>>hg38_21_+_8433172_8446622_RNA45SN1_45S_50ntFlanks@11641.11669.29|CGGCGGGTGTTGACGCGATGTGATTTCTG</td><td>>hg38_21_+_8433172_8446622_RNA45SN1_45S_50ntFlanks</td><td>11641</td><td>11669</td><td>29</td><td>CGGCGGGTGTTGACGCGATGTGATTTCTG</td><td>7262.4</td><td>-16.6</td><td>1.5</td><td>-11.1</td><td>1.4E-28</td><td>7.4E-27</td><td>MB231/MB468</td><td>MITO</td></tr>
<tr><td>>hg38_21_+_8433172_8446622_RNA45SN1_45S_50ntFlanks@7975.8011.37|CGCGACCTCAGATCAGACGTGGCGACCCGCTGAATTT</td><td>>hg38_21_+_8433172_8446622_RNA45SN1_45S_50ntFlanks</td><td>7975</td><td>8011</td><td>37</td><td>CGCGACCTCAGATCAGACGTGGCGACCCGCTGAATTT</td><td>709767.5</td><td>6.2</td><td>0.6</td><td>11.0</td><td>3.4E-28</td><td>7.6E-27</td><td>BT20/MB468</td><td>MITO</td></tr>
<tr><td>>hg38_21_+_8433172_8446622_RNA45SN1_45S_50ntFlanks@11672.11703.32|CAGTGCTCTGAATGTCAAAGTGAAGAAATTCA</td><td>>hg38_21_+_8433172_8446622_RNA45SN1_45S_50ntFlanks</td><td>11672</td><td>11703</td><td>32</td><td>CAGTGCTCTGAATGTCAAAGTGAAGAAATTCA</td><td>1172.4</td><td>-6.2</td><td>0.6</td><td>-10.9</td><td>8.2E-28</td><td>7.7E-27</td><td>BT20/MB468</td><td>NUC</td></tr>
<tr><td>>MI0000767|hsa-mir-365a&WithFlank&16|+|14309279|14309377@62.82.21;MI0000769|hsa-mir-365b&WithFlank&17|+|31575405|31575527@74.94.21&[MIMAT0000710&hsa-miR-365a-3p&offsets|0|-1;m-176&16|+|14309340|14309361&offsets|0|-1];[MIMAT0022834&hsa-miR-365b-3p&offsets|0|-1;m-175&17|+|31575478|31575499&offsets|0|-1]|TAATGCCCCTAAAAATCCTTA</td><td>>MI0000767|hsa-mir-365a&WithFlank&16|+|14309279|14309377</td><td>62</td><td>82</td><td>21</td><td>TAATGCCCCTAAAAATCCTTA</td><td>596.8</td><td>13.7</td><td>1.2</td><td>11.0</td><td>3.4E-28</td><td>7.7E-27</td><td>BT20/MB468</td><td>MITO</td></tr>
<tr><td>>hg38_21_+_8433172_8446622_RNA45SN1_45S_50ntFlanks@12655.12675.21|ATGACTGAACGCCTCTAAGTC</td><td>>hg38_21_+_8433172_8446622_RNA45SN1_45S_50ntFlanks</td><td>12655</td><td>12675</td><td>21</td><td>ATGACTGAACGCCTCTAAGTC</td><td>19649.2</td><td>-12.1</td><td>1.1</td><td>-10.9</td><td>8.3E-28</td><td>7.8E-27</td><td>BT20/MB468</td><td>NUC</td></tr>
<tr><td>>hg38_21_+_8433172_8446622_RNA45SN1_45S_50ntFlanks@12935.12955.21|TAAACCATTCGTAGACGACCT</td><td>>hg38_21_+_8433172_8446622_RNA45SN1_45S_50ntFlanks</td><td>12935</td><td>12955</td><td>21</td><td>TAAACCATTCGTAGACGACCT</td><td>499.8</td><td>-13.4</td><td>1.2</td><td>-10.9</td><td>8.3E-28</td><td>7.8E-27</td><td>BT20/MB468</td><td>NUC</td></tr>
<tr><td>>MI0000077|hsa-mir-21&WithFlank&17|+|59841260|59841343@14.36.23(+1C)&[MIMAT0000076&hsa-miR-21-5p&offsets|0|+1(+1C);m-1&17|+|59841273|59841294&offsets|0|+1(+1C)]|TAGCTTATCAGACTGATGTTGACC</td><td>>MI0000077|hsa-mir-21&WithFlank&17|+|59841260|59841343</td><td>14</td><td>36</td><td>23(+1C)</td><td>TAGCTTATCAGACTGATGTTGACC</td><td>22948.7</td><td>7.6</td><td>0.7</td><td>11.0</td><td>3.5E-28</td><td>7.8E-27</td><td>BT20/MB468</td><td>MITO</td></tr>
<tr><td>>MI0001445|hsa-mir-423&WithFlank&17|+|30117073|30117178@59.79.21&[MIMAT0001340&hsa-miR-423-3p&offsets|0|-2;m-100&17|+|30117131|30117153&offsets|0|-2]|AGCTCGGTCTGAGGCCCCTCA</td><td>>MI0001445|hsa-mir-423&WithFlank&17|+|30117073|30117178</td><td>59</td><td>79</td><td>21</td><td>AGCTCGGTCTGAGGCCCCTCA</td><td>1573.3</td><td>7.6</td><td>0.7</td><td>11.0</td><td>3.5E-28</td><td>7.9E-27</td><td>BT20/MB468</td><td>MITO</td></tr>
<tr><td>>hg38_21_+_8433172_8446622_RNA45SN1_45S_50ntFlanks@13002.13023.22|CGATCTATTGAAAGTCAGCCCT</td><td>>hg38_21_+_8433172_8446622_RNA45SN1_45S_50ntFlanks</td><td>13002</td><td>13023</td><td>22</td><td>CGATCTATTGAAAGTCAGCCCT</td><td>1798.9</td><td>-15.2</td><td>1.4</td><td>-10.9</td><td>8.5E-28</td><td>7.9E-27</td><td>BT20/MB468</td><td>NUC</td></tr>
<tr><td>>hg38_21_+_8433172_8446622_RNA45SN1_45S_50ntFlanks@3939.3959.21|AACCCGGTCAGCCCCTCTCCG</td><td>>hg38_21_+_8433172_8446622_RNA45SN1_45S_50ntFlanks</td><td>3939</td><td>3959</td><td>21</td><td>AACCCGGTCAGCCCCTCTCCG</td><td>3239.9</td><td>-15.4</td><td>1.4</td><td>-11.1</td><td>1.5E-28</td><td>8.0E-27</td><td>MB231/MB468</td><td>MITO</td></tr>
<tr><td>>am_MI0000809|hsa-mir-151a&WithFlank&8|-|140732558|140732659@53.73.21(+1C)&[MIMAT0000757&hsa-miR-151a-3p&offsets|0|0(+1C);m-30&8|-|140732587|140732607&offsets|0|0(+1C)]|CTAGACTGAAGCTCCTTGAGGC</td><td>>am_MI0000809|hsa-mir-151a&WithFlank&8|-|140732558|140732659</td><td>53</td><td>73</td><td>21(+1C)</td><td>CTAGACTGAAGCTCCTTGAGGC</td><td>3075.4</td><td>9.8</td><td>0.9</td><td>11.0</td><td>4.8E-28</td><td>8.0E-27</td><td>BT20/MB468</td><td>CYTO</td></tr>
<tr><td>>hg38_21_+_8433172_8446622_RNA45SN1_45S_50ntFlanks@12159.12180.22|TTTGACTGGGGCGGTACACCTG</td><td>>hg38_21_+_8433172_8446622_RNA45SN1_45S_50ntFlanks</td><td>12159</td><td>12180</td><td>22</td><td>TTTGACTGGGGCGGTACACCTG</td><td>1421.3</td><td>-14.9</td><td>1.4</td><td>-10.9</td><td>8.7E-28</td><td>8.1E-27</td><td>BT20/MB468</td><td>NUC</td></tr>
<tr><td>>hg38_21_+_8433172_8446622_RNA45SN1_45S_50ntFlanks@6663.6688.26|GTGGATCACTCGGCTCGTGCGTCGAT</td><td>>hg38_21_+_8433172_8446622_RNA45SN1_45S_50ntFlanks</td><td>6663</td><td>6688</td><td>26</td><td>GTGGATCACTCGGCTCGTGCGTCGAT</td><td>296.0</td><td>-3.4</td><td>0.3</td><td>-10.9</td><td>8.8E-28</td><td>8.1E-27</td><td>BT20/MB468</td><td>NUC</td></tr>
<tr><td>>MI0000542|hsa-mir-320a&WithFlank&8|-|22244960|22245043@49.69.21(+2U)&[MIMAT0000510&hsa-miR-320a-3p&offsets|+1|0(+2U);m-60&8|-|22244975|22244996&offsets|+1|0(+2U)]|AAAGCTGGGTTGAGAGGGCGATT</td><td>>MI0000542|hsa-mir-320a&WithFlank&8|-|22244960|22245043</td><td>49</td><td>69</td><td>21(+2U)</td><td>AAAGCTGGGTTGAGAGGGCGATT</td><td>719.6</td><td>13.7</td><td>1.3</td><td>11.0</td><td>4.9E-28</td><td>8.2E-27</td><td>BT20/MB468</td><td>CYTO</td></tr>
<tr><td>>hg38_21_+_8433172_8446622_RNA45SN1_45S_50ntFlanks@4723.4759.37|CATTAATCAAGAACGAAAGTCGGAGGTTCGAAGACGA</td><td>>hg38_21_+_8433172_8446622_RNA45SN1_45S_50ntFlanks</td><td>4723</td><td>4759</td><td>37</td><td>CATTAATCAAGAACGAAAGTCGGAGGTTCGAAGACGA</td><td>4886.1</td><td>-10.2</td><td>0.9</td><td>-11.1</td><td>1.5E-28</td><td>8.2E-27</td><td>MB231/MB468</td><td>MITO</td></tr>
<tr><td>>hg38_21_+_8433172_8446622_RNA45SN1_45S_50ntFlanks@11234.11254.21|CCTTCCCGTGGATCGCCCCAG</td><td>>hg38_21_+_8433172_8446622_RNA45SN1_45S_50ntFlanks</td><td>11234</td><td>11254</td><td>21</td><td>CCTTCCCGTGGATCGCCCCAG</td><td>555.1</td><td>-13.5</td><td>1.2</td><td>-10.9</td><td>8.9E-28</td><td>8.2E-27</td><td>BT20/MB468</td><td>NUC</td></tr>
<tr><td>>hg38_21_+_8433172_8446622_RNA45SN1_45S_50ntFlanks@7989.8006.18|AGACGTGGCGACCCGCTG</td><td>>hg38_21_+_8433172_8446622_RNA45SN1_45S_50ntFlanks</td><td>7989</td><td>8006</td><td>18</td><td>AGACGTGGCGACCCGCTG</td><td>1199.8</td><td>-5.5</td><td>0.5</td><td>-10.9</td><td>8.9E-28</td><td>8.2E-27</td><td>BT20/MB468</td><td>NUC</td></tr>
<tr><td>>hg38_21_+_8433172_8446622_RNA45SN1_45S_50ntFlanks@6737.6771.35|GATCATCGACACTTCGAACGCACTTGCGGCCCCGG</td><td>>hg38_21_+_8433172_8446622_RNA45SN1_45S_50ntFlanks</td><td>6737</td><td>6771</td><td>35</td><td>GATCATCGACACTTCGAACGCACTTGCGGCCCCGG</td><td>1096.3</td><td>-14.5</td><td>1.3</td><td>-10.9</td><td>9.2E-28</td><td>8.5E-27</td><td>BT20/MB468</td><td>NUC</td></tr>
<tr><td>>hg38_21_+_8433172_8446622_RNA45SN1_45S_50ntFlanks@8182.8203.22|ATCGAGGCCCAGCCCGTGGACG</td><td>>hg38_21_+_8433172_8446622_RNA45SN1_45S_50ntFlanks</td><td>8182</td><td>8203</td><td>22</td><td>ATCGAGGCCCAGCCCGTGGACG</td><td>666.1</td><td>-13.8</td><td>1.3</td><td>-10.9</td><td>9.6E-28</td><td>8.9E-27</td><td>BT20/MB468</td><td>NUC</td></tr>
<tr><td>>hg38_21_+_8433172_8446622_RNA45SN1_45S_50ntFlanks@11766.11791.26|ATCTAATTAGTGACGCGCATGAATGG</td><td>>hg38_21_+_8433172_8446622_RNA45SN1_45S_50ntFlanks</td><td>11766</td><td>11791</td><td>26</td><td>ATCTAATTAGTGACGCGCATGAATGG</td><td>1251.6</td><td>-14.7</td><td>1.3</td><td>-10.9</td><td>9.7E-28</td><td>8.9E-27</td><td>BT20/MB468</td><td>NUC</td></tr>
<tr><td>>hg38_21_+_8433172_8446622_RNA45SN1_45S_50ntFlanks@5524.5550.27|GTAAAAGTCGTAACAAGGTTTCCGTAG</td><td>>hg38_21_+_8433172_8446622_RNA45SN1_45S_50ntFlanks</td><td>5524</td><td>5550</td><td>27</td><td>GTAAAAGTCGTAACAAGGTTTCCGTAG</td><td>1264.7</td><td>-14.7</td><td>1.3</td><td>-10.9</td><td>9.7E-28</td><td>8.9E-27</td><td>BT20/MB468</td><td>NUC</td></tr>
</table>

<table>
<tr><td>>trna111_HisGTG_1_-_147774845_147774916@-1G.32.33,trna118_HisGTG_1_-_145396881_145396952@-1G.32.33,trna16_HisGTG_1_+_146544773_146544844@-1G.32.33,trna1_HisGTG_15_+_45493349_45493420@-1G.32.33,trna21_HisGTG_1_+_147753471_147753542@-1G.32.33,trna33_HisGTG_6_+_27125906_27125977@-1G.32.33,trna7_HisGTG_9_-_14433938_14434009@-1G.32.33,trna8_HisGTG_15_-_45492611_45492682@-1G.32.33,trna9_HisGTG_15_-_45490804_45490875@-1G.32.33|GGCCGTGATCGTATAGTGGTTAGTACTCTGCGT</td><td>>trna111_HisGTG_1_-_147774845_147774916</td><td>-1G</td><td>32</td><td>33</td><td>GGCCGTGATCGTATAGTGGTTAGTACTCTGCGT</td><td>7246.7</td><td>13.2</td><td>1.2</td><td>11.2</td><td>7.1E-29</td><td>9.1E-27</td><td>MB231/MB468</td><td>CYTO</td></tr>
<tr><td>>hg38_21_+_8433172_8446622_RNA45SN1_45S_50ntFlanks@12157.12180.24|AGTTTGACTGGGGCGGTACACCTG</td><td>>hg38_21_+_8433172_8446622_RNA45SN1_45S_50ntFlanks</td><td>12157</td><td>12180</td><td>24</td><td>AGTTTGACTGGGGCGGTACACCTG</td><td>1158.3</td><td>-14.6</td><td>1.3</td><td>-10.9</td><td>9.9E-28</td><td>9.1E-27</td><td>BT20/MB468</td><td>NUC</td></tr>
<tr><td>>MI0000298|hsa-mir-221&WithFlank&X|-|45746151|45746272@71.91.21&[MIMAT0000278&hsa-miR-221-3p&offsets|0|-2;m-91&X|-|45746181|45746202&offsets|0|-1]|AGCTACATTGTCTGCTGGGTT</td><td>>MI0000298|hsa-mir-221&WithFlank&X|-|45746151|45746272</td><td>71</td><td>91</td><td>21</td><td>AGCTACATTGTCTGCTGGGTT</td><td>1030.9</td><td>6.8</td><td>0.6</td><td>11.0</td><td>5.5E-28</td><td>9.2E-27</td><td>BT20/MB468</td><td>CYTO</td></tr>
<tr><td>>hg38_21_+_8433172_8446622_RNA45SN1_45S_50ntFlanks@12008.12033.26|CCGCCGGTGAAATACCACTACTCTGA</td><td>>hg38_21_+_8433172_8446622_RNA45SN1_45S_50ntFlanks</td><td>12008</td><td>12033</td><td>26</td><td>CCGCCGGTGAAATACCACTACTCTGA</td><td>902.6</td><td>-14.2</td><td>1.3</td><td>-10.9</td><td>1.0E-27</td><td>9.2E-27</td><td>BT20/MB468</td><td>NUC</td></tr>
<tr><td>>hg38_1_-_228634819_228635039_RNA5S12_5S_50ntFlanks@131.170.40|GGATGGGAGACCGCCTGGGAATACCGGGTGCTGTAGGCTT</td><td>>hg38_1_-_228634819_228635039_RNA5S12_5S_50ntFlanks</td><td>131</td><td>170</td><td>40</td><td>GGATGGGAGACCGCCTGGGAATACCGGGTGCTGTAGGCTT</td><td>3668.5</td><td>8.9</td><td>0.8</td><td>11.1</td><td>7.6E-29</td><td>9.4E-27</td><td>MB231/MB468</td><td>CYTO</td></tr>
<tr><td>>hg38_21_+_8433172_8446622_RNA45SN1_45S_50ntFlanks@7977.8011.35|CGACCTCAGATCAGACGTGGCGACCCGCTGAATTT</td><td>>hg38_21_+_8433172_8446622_RNA45SN1_45S_50ntFlanks</td><td>7977</td><td>8011</td><td>35</td><td>CGACCTCAGATCAGACGTGGCGACCCGCTGAATTT</td><td>495.3</td><td>5.7</td><td>0.5</td><td>11.0</td><td>5.8E-28</td><td>9.5E-27</td><td>BT20/MB468</td><td>CYTO</td></tr>
<tr><td>>hg38_21_+_8433172_8446622_RNA45SN1_45S_50ntFlanks@8183.8223.41|TCGAGGCCCAGCCCGTGGACGGTGTGAGGCCGGTAGCGGCC</td><td>>hg38_21_+_8433172_8446622_RNA45SN1_45S_50ntFlanks</td><td>8183</td><td>8223</td><td>41</td><td>TCGAGGCCCAGCCCGTGGACGGTGTGAGGCCGGTAGCGGCC</td><td>1606.1</td><td>-15.1</td><td>1.4</td><td>-10.9</td><td>1.1E-27</td><td>9.6E-27</td><td>BT20/MB468</td><td>NUC</td></tr>
<tr><td>>hg38_21_+_8433172_8446622_RNA45SN1_45S_50ntFlanks@4723.4758.36|CATTAATCAAGAACGAAAGTCGGAGGTTCGAAGACG</td><td>>hg38_21_+_8433172_8446622_RNA45SN1_45S_50ntFlanks</td><td>4723</td><td>4758</td><td>36</td><td>CATTAATCAAGAACGAAAGTCGGAGGTTCGAAGACG</td><td>9203.5</td><td>-7.2</td><td>0.7</td><td>-11.0</td><td>4.4E-28</td><td>9.7E-27</td><td>BT20/MB468</td><td>MITO</td></tr>
<tr><td>>MI0000477|hsa-mir-146a&WithFlank&5|+|160485346|160485456@27.48.22&[MIMAT0000449&hsa-miR-146a-5p&offsets|0|0;m-129&5|+|160485372|160485393&offsets|0|0]|TGAGAACTGAATTCCATGGGTT</td><td>>MI0000477|hsa-mir-146a&WithFlank&5|+|160485346|160485456</td><td>27</td><td>48</td><td>22</td><td>TGAGAACTGAATTCCATGGGTT</td><td>25575.3</td><td>14.8</td><td>1.3</td><td>11.0</td><td>2.9E-28</td><td>9.7E-27</td><td>MB231/MB468</td><td>NUC</td></tr>
<tr><td>>hg38_21_+_8433172_8446622_RNA45SN1_45S_50ntFlanks@5492.5526.35|AGAAGACGGTCGAACTTGACTATCTAGAGGAAGTA</td><td>>hg38_21_+_8433172_8446622_RNA45SN1_45S_50ntFlanks</td><td>5492</td><td>5526</td><td>35</td><td>AGAAGACGGTCGAACTTGACTATCTAGAGGAAGTA</td><td>3015.8</td><td>-7.9</td><td>0.7</td><td>-11.1</td><td>1.9E-28</td><td>1.0E-26</td><td>MB231/MB468</td><td>MITO</td></tr>
<tr><td>>hg38_21_+_8433172_8446622_RNA45SN1_45S_50ntFlanks@6647.6665.19|CGTACGACTCTTAGCGGTG</td><td>>hg38_21_+_8433172_8446622_RNA45SN1_45S_50ntFlanks</td><td>6647</td><td>6665</td><td>19</td><td>CGTACGACTCTTAGCGGTG</td><td>31725.1</td><td>6.6</td><td>0.6</td><td>11.0</td><td>4.6E-28</td><td>1.0E-26</td><td>BT20/MB468</td><td>MITO</td></tr>
<tr><td>>hg38_21_+_8433172_8446622_RNA45SN1_45S_50ntFlanks@11209.11254.46|CGGCGACTCTGGACGCGAGCCGGGCCCTTCCCGTGGATCGCCCCAG</td><td>>hg38_21_+_8433172_8446622_RNA45SN1_45S_50ntFlanks</td><td>11209</td><td>11254</td><td>46</td><td>CGGCGACTCTGGACGCGAGCCGGGCCCTTCCCGTGGATCGCCCCAG</td><td>19925.1</td><td>-10.1</td><td>0.9</td><td>-11.1</td><td>1.9E-28</td><td>1.0E-26</td><td>MB231/MB468</td><td>MITO</td></tr>
<tr><td>>hg38_21_+_8433172_8446622_RNA45SN1_45S_50ntFlanks@12942.12965.24|TTCGTAGACGACCTGCTTCTGGGT</td><td>>hg38_21_+_8433172_8446622_RNA45SN1_45S_50ntFlanks</td><td>12942</td><td>12965</td><td>24</td><td>TTCGTAGACGACCTGCTTCTGGGT</td><td>612.5</td><td>-13.7</td><td>1.3</td><td>-10.9</td><td>1.1E-27</td><td>1.0E-26</td><td>BT20/MB468</td><td>NUC</td></tr>
<tr><td>>hg38_21_+_8433172_8446622_RNA45SN1_45S_50ntFlanks@10734.10755.22|GTGTAAATCTCGCGCCGGGCCG</td><td>>hg38_21_+_8433172_8446622_RNA45SN1_45S_50ntFlanks</td><td>10734</td><td>10755</td><td>22</td><td>GTGTAAATCTCGCGCCGGGCCG</td><td>1040.3</td><td>-14.4</td><td>1.3</td><td>-10.9</td><td>1.1E-27</td><td>1.0E-26</td><td>BT20/MB468</td><td>NUC</td></tr>
<tr><td>>hg38_21_+_8433172_8446622_RNA45SN1_45S_50ntFlanks@12456.12475.20|CCAAGCGTTGGATTGTTCAC</td><td>>hg38_21_+_8433172_8446622_RNA45SN1_45S_50ntFlanks</td><td>12456</td><td>12475</td><td>20</td><td>CCAAGCGTTGGATTGTTCAC</td><td>7966.0</td><td>-9.2</td><td>0.8</td><td>-11.0</td><td>3.0E-28</td><td>1.0E-26</td><td>MB231/MB468</td><td>NUC</td></tr>
<tr><td>>MI0000542|hsa-mir-320a&WithFlank&8|-|22244960|22245043@48.70.23(+2U)&[MIMAT0000510&hsa-miR-320a-3p&offsets|0|+1(+2U);m-60&8|-|22244975|22244996&offsets|0|+1(+2U)]|AAAAGCTGGGTTGAGAGGGCGAATT</td><td>>MI0000542|hsa-mir-320a&WithFlank&8|-|22244960|22245043</td><td>48</td><td>70</td><td>23(+2U)</td><td>AAAAGCTGGGTTGAGAGGGCGAATT</td><td>2350.8</td><td>11.5</td><td>1.0</td><td>11.0</td><td>6.3E-28</td><td>1.0E-26</td><td>BT20/MB468</td><td>CYTO</td></tr>
<tr><td>>MI0000734|hsa-mir-106b&WithFlank&7|-|100093987|100094080@58.79.22&[MIMAT0004672&hsa-miR-106b-3p&offsets|0|0;m-87&7|-|100094003|100094023&offsets|0|+1]|CCGCACTGTGGGTACTTGCTGC</td><td>>MI0000734|hsa-mir-106b&WithFlank&7|-|100093987|100094080</td><td>58</td><td>79</td><td>22</td><td>CCGCACTGTGGGTACTTGCTGC</td><td>5975.5</td><td>7.4</td><td>0.7</td><td>11.0</td><td>4.8E-28</td><td>1.1E-26</td><td>BT20/MB468</td><td>MITO</td></tr>
<tr><td>>hg38_21_+_8433172_8446622_RNA45SN1_45S_50ntFlanks@6721.6744.24|AATTGCAGGACACATTGATCATCG</td><td>>hg38_21_+_8433172_8446622_RNA45SN1_45S_50ntFlanks</td><td>6721</td><td>6744</td><td>24</td><td>AATTGCAGGACACATTGATCATCG</td><td>2799.2</td><td>-6.0</td><td>0.5</td><td>-11.0</td><td>4.8E-28</td><td>1.1E-26</td><td>BT20/MB468</td><td>MITO</td></tr>
<tr><td>>hg38_21_+_8433172_8446622_RNA45SN1_45S_50ntFlanks@6721.6744.24|AATTGCAGGACACATTGATCATCG</td><td>>hg38_21_+_8433172_8446622_RNA45SN1_45S_50ntFlanks</td><td>6721</td><td>6744</td><td>24</td><td>AATTGCAGGACACATTGATCATCG</td><td>27226.8</td><td>-12.4</td><td>1.1</td><td>-10.9</td><td>1.2E-27</td><td>1.1E-26</td><td>BT20/MB468</td><td>NUC</td></tr>
<tr><td>>hg38_21_+_8433172_8446622_RNA45SN1_45S_50ntFlanks@9544.9566.23|GGTGGAGGTCCGTAGCGGTCCTG</td><td>>hg38_21_+_8433172_8446622_RNA45SN1_45S_50ntFlanks</td><td>9544</td><td>9566</td><td>23</td><td>GGTGGAGGTCCGTAGCGGTCCTG</td><td>444.9</td><td>-13.2</td><td>1.2</td><td>-10.9</td><td>1.2E-27</td><td>1.1E-26</td><td>BT20/MB468</td><td>NUC</td></tr>
<tr><td>>hg38_21_+_8433172_8446622_RNA45SN1_45S_50ntFlanks@12456.12488.33|CCAAGCGTTGGATTGTTCACCCACTAATAGGGA</td><td>>hg38_21_+_8433172_8446622_RNA45SN1_45S_50ntFlanks</td><td>12456</td><td>12488</td><td>33</td><td>CCAAGCGTTGGATTGTTCACCCACTAATAGGGA</td><td>7476.4</td><td>-16.7</td><td>1.5</td><td>-11.1</td><td>1.1E-28</td><td>1.1E-26</td><td>MB231/MB468</td><td>MP</td></tr>
<tr><td>>hg38_21_+_8433172_8446622_RNA45SN1_45S_50ntFlanks@10356.10383.28|ACTTTGAAGGCCGAAGTGGAGAAGGGTT</td><td>>hg38_21_+_8433172_8446622_RNA45SN1_45S_50ntFlanks</td><td>10356</td><td>10383</td><td>28</td><td>ACTTTGAAGGCCGAAGTGGAGAAGGGTT</td><td>536.2</td><td>-13.5</td><td>1.2</td><td>-10.9</td><td>1.2E-27</td><td>1.1E-26</td><td>BT20/MB468</td><td>NUC</td></tr>
<tr><td>>MI0000102|hsa-mir-100&WithFlank&11|-|122152223|122152314@19.39.21(+1U)&[MIMAT0000098&hsa-miR-100-5p&offsets|0|-1(+1U);m-27&11|-|122152276|122152296&offsets|0|0(+1U)]|AACCCGTAGATCCGAACTTGTT</td><td>>MI0000102|hsa-mir-100&WithFlank&11|-|122152223|122152314</td><td>19</td><td>39</td><td>21(+1U)</td><td>AACCCGTAGATCCGAACTTGTT</td><td>6738.1</td><td>16.7</td><td>1.5</td><td>11.1</td><td>9.0E-29</td><td>1.1E-26</td><td>MB231/MB468</td><td>CYTO</td></tr>
<tr><td>>hg38_21_+_8433172_8446622_RNA45SN1_45S_50ntFlanks@8357.8377.21|AAAAGAACTTTGAAGAGAGAG</td><td>>hg38_21_+_8433172_8446622_RNA45SN1_45S_50ntFlanks</td><td>8357</td><td>8377</td><td>21</td><td>AAAAGAACTTTGAAGAGAGAG</td><td>3913.1</td><td>-7.6</td><td>0.7</td><td>-11.1</td><td>2.1E-28</td><td>1.1E-26</td><td>MB231/MB468</td><td>MITO</td></tr>
<tr><td>>hg38_21_+_8433172_8446622_RNA45SN1_45S_50ntFlanks@10410.10433.24|TCAGTCGGTCCTGAGAGATGGGCG</td><td>>hg38_21_+_8433172_8446622_RNA45SN1_45S_50ntFlanks</td><td>10410</td><td>10433</td><td>24</td><td>TCAGTCGGTCCTGAGAGATGGGCG</td><td>803.1</td><td>-14.1</td><td>1.3</td><td>-10.9</td><td>1.2E-27</td><td>1.1E-26</td><td>BT20/MB468</td><td>NUC</td></tr>
<tr><td>>hg38_21_+_8433172_8446622_RNA45SN1_45S_50ntFlanks@9534.9564.31|AGGAAACTCTGGTGGAGGTCCGTAGCGGTCC</td><td>>hg38_21_+_8433172_8446622_RNA45SN1_45S_50ntFlanks</td><td>9534</td><td>9564</td><td>31</td><td>AGGAAACTCTGGTGGAGGTCCGTAGCGGTCC</td><td>5600.4</td><td>-7.7</td><td>0.7</td><td>-11.0</td><td>3.4E-28</td><td>1.1E-26</td><td>MB231/MB468</td><td>NUC</td></tr>
<tr><td>>hg38_21_+_8433172_8446622_RNA45SN1_45S_50ntFlanks@11874.11898.25|AGAAGACCCTGTTGAGCTTGACTCT</td><td>>hg38_21_+_8433172_8446622_RNA45SN1_45S_50ntFlanks</td><td>11874</td><td>11898</td><td>25</td><td>AGAAGACCCTGTTGAGCTTGACTCT</td><td>1836.8</td><td>-4.0</td><td>0.4</td><td>-10.9</td><td>6.9E-28</td><td>1.1E-26</td><td>BT20/MB468</td><td>CYTO</td></tr>
<tr><td>>hg38_21_+_8433172_8446622_RNA45SN1_45S_50ntFlanks@6738.6770.33|ATCATCGACACTTCGAACGCACTTGCGGCCCCG</td><td>>hg38_21_+_8433172_8446622_RNA45SN1_45S_50ntFlanks</td><td>6738</td><td>6770</td><td>33</td><td>ATCATCGACACTTCGAACGCACTTGCGGCCCCG</td><td>4777.7</td><td>-14.3</td><td>1.3</td><td>-11.0</td><td>5.2E-28</td><td>1.1E-26</td><td>BT20/MB468</td><td>MITO</td></tr>
<tr><td>>hg38_21_+_8433172_8446622_RNA45SN1_45S_50ntFlanks@10401.10424.24|GAACATGGGTCAGTCGGTCCTGAG</td><td>>hg38_21_+_8433172_8446622_RNA45SN1_45S_50ntFlanks</td><td>10401</td><td>10424</td><td>24</td><td>GAACATGGGTCAGTCGGTCCTGAG</td><td>20842.2</td><td>7.2</td><td>0.7</td><td>11.0</td><td>5.2E-28</td><td>1.1E-26</td><td>BT20/MB468</td><td>MITO</td></tr>
<tr><td>>hg38_21_+_8433172_8446622_RNA45SN1_45S_50ntFlanks@5417.5445.29|CTACCGATTGGATGGTTTAGTGAGGCCCT</td><td>>hg38_21_+_8433172_8446622_RNA45SN1_45S_50ntFlanks</td><td>5417</td><td>5445</td><td>29</td><td>CTACCGATTGGATGGTTTAGTGAGGCCCT</td><td>1148.0</td><td>-14.6</td><td>1.3</td><td>-10.9</td><td>1.3E-27</td><td>1.1E-26</td><td>BT20/MB468</td><td>NUC</td></tr>
<tr><td>>MI0000804|hsa-mir-328&WithFlank&16|-|67202315|67202401@54.75.22&[MIMAT0000752&hsa-miR-328-3p&offsets|0|0;m-160&16|-|67202327|67202348&offsets|0|0]|CTGGCCCTCTCTGCCCTTCCGT</td><td>>MI0000804|hsa-mir-328&WithFlank&16|-|67202315|67202401</td><td>54</td><td>75</td><td>22</td><td>CTGGCCCTCTCTGCCCTTCCGT</td><td>1265.4</td><td>8.4</td><td>0.8</td><td>11.0</td><td>5.3E-28</td><td>1.1E-26</td><td>BT20/MB468</td><td>MITO</td></tr>
<tr><td>>hg38_21_+_8433172_8446622_RNA45SN1_45S_50ntFlanks@11999.12032.34|CCCTGCGGGCCGCCGGTGAAATACCACTACTCTG</td><td>>hg38_21_+_8433172_8446622_RNA45SN1_45S_50ntFlanks</td><td>11999</td><td>12032</td><td>34</td><td>CCCTGCGGGCCGCCGGTGAAATACCACTACTCTG</td><td>3486.9</td><td>-15.6</td><td>1.4</td><td>-11.1</td><td>1.2E-28</td><td>1.2E-26</td><td>MB231/MB468</td><td>MP</td></tr>
<tr><td>>hg38_21_+_8433172_8446622_RNA45SN1_45S_50ntFlanks@12417.12437.21|TCGATGTCGGCTCTTCCTATC</td><td>>hg38_21_+_8433172_8446622_RNA45SN1_45S_50ntFlanks</td><td>12417</td><td>12437</td><td>21</td><td>TCGATGTCGGCTCTTCCTATC</td><td>7226.3</td><td>-7.1</td><td>0.6</td><td>-10.9</td><td>1.3E-27</td><td>1.2E-26</td><td>BT20/MB468</td><td>NUC</td></tr>
<tr><td>>hg38_21_+_8433172_8446622_RNA45SN1_45S_50ntFlanks@6740.6771.32|CATCGACACTTCGAACGCACTTGCGGCCCCGG</td><td>>hg38_21_+_8433172_8446622_RNA45SN1_45S_50ntFlanks</td><td>6740</td><td>6771</td><td>32</td><td>CATCGACACTTCGAACGCACTTGCGGCCCCGG</td><td>7604.7</td><td>-10.2</td><td>0.9</td><td>-11.0</td><td>2.3E-28</td><td>1.2E-26</td><td>MB231/MB468</td><td>MITO</td></tr>
<tr><td>>hg38_21_+_8433172_8446622_RNA45SN1_45S_50ntFlanks@13004.13023.20|ATCTATTGAAAGTCAGCCCT</td><td>>hg38_21_+_8433172_8446622_RNA45SN1_45S_50ntFlanks</td><td>13004</td><td>13023</td><td>20</td><td>ATCTATTGAAAGTCAGCCCT</td><td>417.7</td><td>-13.1</td><td>1.2</td><td>-10.9</td><td>1.3E-27</td><td>1.2E-26</td><td>BT20/MB468</td><td>NUC</td></tr>
<tr><td>>hg38_21_+_8433172_8446622_RNA45SN1_45S_50ntFlanks@482.517.36|ATCGATGTGGTGACGTCGTGCTCTCCCGGGCCGGGT</td><td>>hg38_21_+_8433172_8446622_RNA45SN1_45S_50ntFlanks</td><td>482</td><td>517</td><td>36</td><td>ATCGATGTGGTGACGTCGTGCTCTCCCGGGCCGGGT</td><td>1209.4</td><td>-14.7</td><td>1.3</td><td>-10.9</td><td>1.3E-27</td><td>1.2E-26</td><td>BT20/MB468</td><td>NUC</td></tr>
<tr><td>>MI0000283|hsa-mir-203a&WithFlank&14|+|104117399|104117520@71.92.22&[MIMAT0000264&hsa-miR-203a-3p&offsets|0|0;m-79&14|+|104117469|104117490&offsets|0|0]|GTGAAATGTTTAGGACCACTAG</td><td>>MI0000283|hsa-mir-203a&WithFlank&14|+|104117399|104117520</td><td>71</td><td>92</td><td>22</td><td>GTGAAATGTTTAGGACCACTAG</td><td>815.5</td><td>7.8</td><td>0.7</td><td>11.0</td><td>5.5E-28</td><td>1.2E-26</td><td>BT20/MB468</td><td>MITO</td></tr>
<tr><td>>MI0000342|hsa-mir-200b&WithFlank&1|+|1167098|1167204@63.84.22(+1U)&[MIMAT0000318&hsa-miR-200b-3p&offsets|0|0(+1U);m-93&1|+|1167160|1167182&offsets|0|-1(+1U)]|TAATACTGCCTGGTAATGATGAT</td><td>>MI0000342|hsa-mir-200b&WithFlank&1|+|1167098|1167204</td><td>63</td><td>84</td><td>22(+1U)</td><td>TAATACTGCCTGGTAATGATGAT</td><td>738.3</td><td>-6.2</td><td>0.6</td><td>-10.9</td><td>1.3E-27</td><td>1.2E-26</td><td>BT20/MB468</td><td>NUC</td></tr>
<tr><td>>hg38_21_+_8433172_8446622_RNA45SN1_45S_50ntFlanks@8114.8139.26|GCGCGGGACATGTGGCGTACGGAAGA</td><td>>hg38_21_+_8433172_8446622_RNA45SN1_45S_50ntFlanks</td><td>8114</td><td>8139</td><td>26</td><td>GCGCGGGACATGTGGCGTACGGAAGA</td><td>4074.2</td><td>-15.9</td><td>1.4</td><td>-11.1</td><td>1.3E-28</td><td>1.2E-26</td><td>MB231/MB468</td><td>MP</td></tr>
<tr><td>>MI0000086|hsa-mir-28&WithFlank&3|+|188688775|188688872@20.40.21&[MIMAT0000085&hsa-miR-28-5p&offsets|0|-1;m-133&3|+|188688794|188688815&offsets|0|-1]|AAGGAGCTCACAGTCTATTGA</td><td>>MI0000086|hsa-mir-28&WithFlank&3|+|188688775|188688872</td><td>20</td><td>40</td><td>21</td><td>AAGGAGCTCACAGTCTATTGA</td><td>93.0</td><td>3.7</td><td>0.3</td><td>10.9</td><td>1.4E-27</td><td>1.2E-26</td><td>BT20/MB468</td><td>NUC</td></tr>
<tr><td>>hg38_21_+_8433172_8446622_RNA45SN1_45S_50ntFlanks@12417.12435.19|TCGATGTCGGCTCTTCCTA</td><td>>hg38_21_+_8433172_8446622_RNA45SN1_45S_50ntFlanks</td><td>12417</td><td>12435</td><td>19</td><td>TCGATGTCGGCTCTTCCTA</td><td>6568.0</td><td>-6.3</td><td>0.6</td><td>-11.0</td><td>3.7E-28</td><td>1.2E-26</td><td>MB231/MB468</td><td>NUC</td></tr>
<tr><td>>hg38_21_+_8433172_8446622_RNA45SN1_45S_50ntFlanks@11515.11539.25|GTCCGCCCCCGGGGCCGCGGTTCCG</td><td>>hg38_21_+_8433172_8446622_RNA45SN1_45S_50ntFlanks</td><td>11515</td><td>11539</td><td>25</td><td>GTCCGCCCCCGGGGCCGCGGTTCCG</td><td>869.2</td><td>-14.2</td><td>1.3</td><td>-10.9</td><td>1.4E-27</td><td>1.3E-26</td><td>BT20/MB468</td><td>NUC</td></tr>
<tr><td>>MI0000471|hsa-mir-126&WithFlank&9|+|136670596|136670692@58.79.22&[MIMAT0000445&hsa-miR-126-3p&offsets|0|0;m-40&9|+|136670653|136670674&offsets|0|0]|TCGTACCGTGAGTAATAATGCG</td><td>>MI0000471|hsa-mir-126&WithFlank&9|+|136670596|136670692</td><td>58</td><td>79</td><td>22</td><td>TCGTACCGTGAGTAATAATGCG</td><td>7686.5</td><td>11.2</td><td>1.0</td><td>10.9</td><td>7.9E-28</td><td>1.3E-26</td><td>BT20/MB468</td><td>CYTO</td></tr>
<tr><td>>MI0000439|hsa-mir-23b&WithFlank&9|+|95085202|95085310@67.86.20&[MIMAT0000418&hsa-miR-23b-3p&offsets|+3|0;m-42&9|+|95085265|95085285&offsets|+3|+2]|ACATTGCCAGGGATTACCAC</td><td>>MI0000439|hsa-mir-23b&WithFlank&9|+|95085202|95085310</td><td>67</td><td>86</td><td>20</td><td>ACATTGCCAGGGATTACCAC</td><td>556.1</td><td>12.0</td><td>1.1</td><td>10.9</td><td>1.5E-27</td><td>1.3E-26</td><td>BT20/MB468</td><td>NUC</td></tr>
</table>

<table>
<tr><td>>MI0000108|hsa-mir-103a-2&WithFlank&20|+|3917488|3917577@54.77.24;MI0000109|hsa-mir-103a-1&WithFlank&5|-|168560890|168560979@54.77.24&[MIMAT0000101&hsa-miR-103a-3p&offsets|0|+1;m-12&20|+|3917541|3917563&offsets|0|+1];[MIMAT0000101_1&hsa-miR-103a-3p&offsets|0|+1;m-13&5|-|168560904|168560926&offsets|0|+1;m-44220&5|-|168560908|168560926&offsets|0|+5]|AGCAGCATTGTACAGGGCTATGAA</td><td>>MI0000108|hsa-mir-103a-2&WithFlank&20|+|3917488|3917577</td><td>54</td><td>77</td><td>24</td><td>AGCAGCATTGTACAGGGCTATGAA</td><td>1345.1</td><td>7.2</td><td>0.7</td><td>10.9</td><td>8.0E-28</td><td>1.3E-26</td><td>BT20/MB468</td><td>CYTO</td></tr>
<tr><td>>hg38_21_+_8433172_8446622_RNA45SN1_45S_50ntFlanks@10737.10767.31|TAAATCTCGCGCCGGGCCGTACCCATATCCG</td><td>>hg38_21_+_8433172_8446622_RNA45SN1_45S_50ntFlanks</td><td>10737</td><td>10767</td><td>31</td><td>TAAATCTCGCGCCGGGCCGTACCCATATCCG</td><td>1146.9</td><td>-14.6</td><td>1.3</td><td>-10.9</td><td>1.5E-27</td><td>1.3E-26</td><td>BT20/MB468</td><td>NUC</td></tr>
<tr><td>>MI0000285|hsa-mir-205&WithFlank&1|+|209432127|209432248@40.61.22(+2A)&[MIMAT0000266&hsa-miR-205-5p&offsets|0|0(+2A);m-351&1|+|209432166|209432187&offsets|0|0(+2A)]|TCCTTCATTCCACCGGAGTCTGAA</td><td>>MI0000285|hsa-mir-205&WithFlank&1|+|209432127|209432248</td><td>40</td><td>61</td><td>22(+2A)</td><td>TCCTTCATTCCACCGGAGTCTGAA</td><td>5414.4</td><td>7.6</td><td>0.7</td><td>10.9</td><td>8.3E-28</td><td>1.4E-26</td><td>BT20/MB468</td><td>CYTO</td></tr>
<tr><td>>hg38_21_+_8433172_8446622_RNA45SN1_45S_50ntFlanks@10411.10431.21|CAGTCGGTCCTGAGAGATGGG</td><td>>hg38_21_+_8433172_8446622_RNA45SN1_45S_50ntFlanks</td><td>10411</td><td>10431</td><td>21</td><td>CAGTCGGTCCTGAGAGATGGG</td><td>3364.4</td><td>-15.6</td><td>1.4</td><td>-11.1</td><td>1.5E-28</td><td>1.4E-26</td><td>MB231/MB468</td><td>MP</td></tr>
<tr><td>>hg38_21_+_8433172_8446622_RNA45SN1_45S_50ntFlanks@12004.12034.31|CGGGCCGCCGGTGAAATACCACTACTCTGAT</td><td>>hg38_21_+_8433172_8446622_RNA45SN1_45S_50ntFlanks</td><td>12004</td><td>12034</td><td>31</td><td>CGGGCCGCCGGTGAAATACCACTACTCTGAT</td><td>761.6</td><td>7.5</td><td>0.7</td><td>11.0</td><td>6.3E-28</td><td>1.4E-26</td><td>BT20/MB468</td><td>MITO</td></tr>
<tr><td>>MI0001448|hsa-mir-425&WithFlank&3|-|49020142|49020240@20.43.24(+1C)&[MIMAT0003393&hsa-miR-425-5p&offsets|0|+1(+1C);m-95&3|-|49020200|49020221&offsets|0|+2(+1C)]|AATGACACGATCACTCCCGTTGAGC</td><td>>MI0001448|hsa-mir-425&WithFlank&3|-|49020142|49020240</td><td>20</td><td>43</td><td>24(+1C)</td><td>AATGACACGATCACTCCCGTTGAGC</td><td>1461.6</td><td>8.4</td><td>0.8</td><td>10.9</td><td>8.5E-28</td><td>1.4E-26</td><td>BT20/MB468</td><td>CYTO</td></tr>
<tr><td>>hg38_21_+_8433172_8446622_RNA45SN1_45S_50ntFlanks@6743.6771.29|CGACACTTCGAACGCACTTGCGGCCCCGG</td><td>>hg38_21_+_8433172_8446622_RNA45SN1_45S_50ntFlanks</td><td>6743</td><td>6771</td><td>29</td><td>CGACACTTCGAACGCACTTGCGGCCCCGG</td><td>4158.7</td><td>-9.7</td><td>0.9</td><td>-11.0</td><td>2.7E-28</td><td>1.4E-26</td><td>MB231/MB468</td><td>MITO</td></tr>
<tr><td>>hg38_21_+_8433172_8446622_RNA45SN1_45S_50ntFlanks@10771.10806.36|CAGGTCTCCAAGGTGAACAGCCTCTGGCATGTTGGA</td><td>>hg38_21_+_8433172_8446622_RNA45SN1_45S_50ntFlanks</td><td>10771</td><td>10806</td><td>36</td><td>CAGGTCTCCAAGGTGAACAGCCTCTGGCATGTTGGA</td><td>1951.1</td><td>-15.4</td><td>1.4</td><td>-10.9</td><td>1.6E-27</td><td>1.4E-26</td><td>BT20/MB468</td><td>NUC</td></tr>
<tr><td>>hg38_21_+_8433172_8446622_RNA45SN1_45S_50ntFlanks@9305.9338.34|CCAAGGAGTCTAACACGTGCGCGAGTCGGGGGCT</td><td>>hg38_21_+_8433172_8446622_RNA45SN1_45S_50ntFlanks</td><td>9305</td><td>9338</td><td>34</td><td>CCAAGGAGTCTAACACGTGCGCGAGTCGGGGGCT</td><td>745.4</td><td>-14.0</td><td>1.3</td><td>-10.9</td><td>1.7E-27</td><td>1.5E-26</td><td>BT20/MB468</td><td>NUC</td></tr>
<tr><td>>hg38_21_+_8433172_8446622_RNA45SN1_45S_50ntFlanks@3899.3919.21|CCTTCGCGGGGGGGATGCGTG</td><td>>hg38_21_+_8433172_8446622_RNA45SN1_45S_50ntFlanks</td><td>3899</td><td>3919</td><td>21</td><td>CCTTCGCGGGGGGGATGCGTG</td><td>8332.0</td><td>-16.8</td><td>1.5</td><td>-11.0</td><td>2.9E-28</td><td>1.5E-26</td><td>MB231/MB468</td><td>MITO</td></tr>
<tr><td>>MI0000266|hsa-mir-10a&WithFlank&17|-|48579832|48579953@28.48.21(+1C)&[MIMAT0000253&hsa-miR-10a-5p&offsets|0|-2(+1C);m-18&17|-|48579905|48579926&offsets|0|-1(+1C)]|TACCCTGTAGATCCGAATTTGC</td><td>>MI0000266|hsa-mir-10a&WithFlank&17|-|48579832|48579953</td><td>28</td><td>48</td><td>21(+1C)</td><td>TACCCTGTAGATCCGAATTTGC</td><td>1429.2</td><td>14.9</td><td>1.4</td><td>10.9</td><td>7.0E-28</td><td>1.5E-26</td><td>BT20/MB468</td><td>MITO</td></tr>
<tr><td>>hg38_21_+_8433172_8446622_RNA45SN1_45S_50ntFlanks@6740.6772.33|CATCGACACTTCGAACGCACTTGCGGCCCCGGG</td><td>>hg38_21_+_8433172_8446622_RNA45SN1_45S_50ntFlanks</td><td>6740</td><td>6772</td><td>33</td><td>CATCGACACTTCGAACGCACTTGCGGCCCCGGG</td><td>1023.7</td><td>-14.4</td><td>1.3</td><td>-10.9</td><td>1.7E-27</td><td>1.5E-26</td><td>BT20/MB468</td><td>NUC</td></tr>
<tr><td>>hg38_21_+_8433172_8446622_RNA45SN1_45S_50ntFlanks@10678.10708.31|CGGTTCCGGCGGCGTCCGGTGAGCTCTCGCT</td><td>>hg38_21_+_8433172_8446622_RNA45SN1_45S_50ntFlanks</td><td>10678</td><td>10708</td><td>31</td><td>CGGTTCCGGCGGCGTCCGGTGAGCTCTCGCT</td><td>51056.1</td><td>-8.6</td><td>0.8</td><td>-11.0</td><td>3.0E-28</td><td>1.5E-26</td><td>MB231/MB468</td><td>MITO</td></tr>
<tr><td>>hg38_21_+_8433172_8446622_RNA45SN1_45S_50ntFlanks@11874.11908.35|AGAAGACCCTGTTGAGCTTGACTCTAGTCTGGCAC</td><td>>hg38_21_+_8433172_8446622_RNA45SN1_45S_50ntFlanks</td><td>11874</td><td>11908</td><td>35</td><td>AGAAGACCCTGTTGAGCTTGACTCTAGTCTGGCAC</td><td>759.0</td><td>-14.0</td><td>1.3</td><td>-10.9</td><td>1.7E-27</td><td>1.5E-26</td><td>BT20/MB468</td><td>NUC</td></tr>
<tr><td>>hg38_21_+_8433172_8446622_RNA45SN1_45S_50ntFlanks@11319.11344.26|CCCACGTCTCGTCGCGCGCGCGTCCG</td><td>>hg38_21_+_8433172_8446622_RNA45SN1_45S_50ntFlanks</td><td>11319</td><td>11344</td><td>26</td><td>CCCACGTCTCGTCGCGCGCGCGTCCG</td><td>5108.5</td><td>-16.1</td><td>1.5</td><td>-11.0</td><td>3.1E-28</td><td>1.5E-26</td><td>MB231/MB468</td><td>MITO</td></tr>
<tr><td>>MI0000078|hsa-mir-22&WithFlank&17|-|1713897|1713993@59.80.22&[MIMAT0000077&hsa-miR-22-3p&offsets|0|0;m-2&17|-|1713914|1713935&offsets|0|0]|AAGCTGCCAGTTGAAGAACTGT</td><td>>MI0000078|hsa-mir-22&WithFlank&17|-|1713897|1713993</td><td>59</td><td>80</td><td>22</td><td>AAGCTGCCAGTTGAAGAACTGT</td><td>22491.6</td><td>9.2</td><td>0.8</td><td>10.9</td><td>9.5E-28</td><td>1.5E-26</td><td>BT20/MB468</td><td>CYTO</td></tr>
<tr><td>>hg38_21_+_8433172_8446622_RNA45SN1_45S_50ntFlanks@13002.13028.27|CGATCTATTGAAAGTCAGCCCTCGACA</td><td>>hg38_21_+_8433172_8446622_RNA45SN1_45S_50ntFlanks</td><td>13002</td><td>13028</td><td>27</td><td>CGATCTATTGAAAGTCAGCCCTCGACA</td><td>5031.9</td><td>-14.6</td><td>1.3</td><td>-10.9</td><td>9.8E-28</td><td>1.6E-26</td><td>BT20/MB468</td><td>CYTO</td></tr>
<tr><td>>hg38_21_+_8433172_8446622_RNA45SN1_45S_50ntFlanks@11572.11602.31|CCGACTTAGAACTGGTGCGGACCAGGGGAAT</td><td>>hg38_21_+_8433172_8446622_RNA45SN1_45S_50ntFlanks</td><td>11572</td><td>11602</td><td>31</td><td>CCGACTTAGAACTGGTGCGGACCAGGGGAAT</td><td>6845.3</td><td>-7.5</td><td>0.7</td><td>-11.0</td><td>3.2E-28</td><td>1.6E-26</td><td>MB231/MB468</td><td>MITO</td></tr>
<tr><td>>hg38_21_+_8433172_8446622_RNA45SN1_45S_50ntFlanks@10360.10383.24|TGAAGGCCGAAGTGGAGAAGGGTT</td><td>>hg38_21_+_8433172_8446622_RNA45SN1_45S_50ntFlanks</td><td>10360</td><td>10383</td><td>24</td><td>TGAAGGCCGAAGTGGAGAAGGGTT</td><td>506.9</td><td>-13.4</td><td>1.2</td><td>-10.9</td><td>1.8E-27</td><td>1.6E-26</td><td>BT20/MB468</td><td>NUC</td></tr>
<tr><td>>hg38_21_+_8433172_8446622_RNA45SN1_45S_50ntFlanks@12939.12961.23|CCATTCGTAGACGACCTGCTTCT</td><td>>hg38_21_+_8433172_8446622_RNA45SN1_45S_50ntFlanks</td><td>12939</td><td>12961</td><td>23</td><td>CCATTCGTAGACGACCTGCTTCT</td><td>422.2</td><td>-13.1</td><td>1.2</td><td>-10.9</td><td>1.9E-27</td><td>1.6E-26</td><td>BT20/MB468</td><td>NUC</td></tr>
<tr><td>>hg38_21_+_8433172_8446622_RNA45SN1_45S_50ntFlanks@10728.10755.28|GAGAGGGTGTAAATCTCGCGCCGGGCCG</td><td>>hg38_21_+_8433172_8446622_RNA45SN1_45S_50ntFlanks</td><td>10728</td><td>10755</td><td>28</td><td>GAGAGGGTGTAAATCTCGCGCCGGGCCG</td><td>1375.3</td><td>-14.9</td><td>1.4</td><td>-10.9</td><td>1.9E-27</td><td>1.7E-26</td><td>BT20/MB468</td><td>NUC</td></tr>
<tr><td>>hg38_21_+_8433172_8446622_RNA45SN1_45S_50ntFlanks@6721.6746.26|AATTGCAGGACACATTGATCATCGAC</td><td>>hg38_21_+_8433172_8446622_RNA45SN1_45S_50ntFlanks</td><td>6721</td><td>6746</td><td>26</td><td>AATTGCAGGACACATTGATCATCGAC</td><td>3401.3</td><td>-15.6</td><td>1.4</td><td>-11.1</td><td>1.8E-28</td><td>1.7E-26</td><td>MB231/MB468</td><td>MP</td></tr>
<tr><td>>hg38_21_+_8433172_8446622_RNA45SN1_45S_50ntFlanks@3705.3742.38|TACCTGGTTGATCCTGCCAGTAGCATATGCTTGTCTCA</td><td>>hg38_21_+_8433172_8446622_RNA45SN1_45S_50ntFlanks</td><td>3705</td><td>3742</td><td>38</td><td>TACCTGGTTGATCCTGCCAGTAGCATATGCTTGTCTCA</td><td>10177.6</td><td>8.3</td><td>0.8</td><td>10.9</td><td>1.1E-27</td><td>1.7E-26</td><td>BT20/MB468</td><td>CYTO</td></tr>
<tr><td>>hg38_21_+_8433172_8446622_RNA45SN1_45S_50ntFlanks@12347.12365.19|AGAAAAGTTACCACAGGGA</td><td>>hg38_21_+_8433172_8446622_RNA45SN1_45S_50ntFlanks</td><td>12347</td><td>12365</td><td>19</td><td>AGAAAAGTTACCACAGGGA</td><td>576.4</td><td>-13.6</td><td>1.3</td><td>-10.9</td><td>1.9E-27</td><td>1.7E-26</td><td>BT20/MB468</td><td>NUC</td></tr>
<tr><td>>hg38_21_+_8433172_8446622_RNA45SN1_45S_50ntFlanks@6648.6681.34|GTACGACTCTTAGCGGTGGATCACTCGGCTCGTG</td><td>>hg38_21_+_8433172_8446622_RNA45SN1_45S_50ntFlanks</td><td>6648</td><td>6681</td><td>34</td><td>GTACGACTCTTAGCGGTGGATCACTCGGCTCGTG</td><td>4427.9</td><td>-3.9</td><td>0.4</td><td>-10.9</td><td>2.0E-27</td><td>1.7E-26</td><td>BT20/MB468</td><td>NUC</td></tr>
<tr><td>>MI0000079|hsa-mir-23a&WithFlank&19|-|13836581|13836665@54.72.19(+1U)&[MIMAT0000078&hsa-miR-23a-3p&offsets|+3|+1(+1U);m-31&19|-|13836595|13836615&offsets|+3|+1(+1U)]|ACATTGCCAGGGATTTCCAT</td><td>>MI0000079|hsa-mir-23a&WithFlank&19|-|13836581|13836665</td><td>54</td><td>72</td><td>19(+1U)</td><td>ACATTGCCAGGGATTTCCAT</td><td>1002.9</td><td>14.2</td><td>1.3</td><td>10.9</td><td>1.1E-27</td><td>1.8E-26</td><td>BT20/MB468</td><td>CYTO</td></tr>
<tr><td>>MI0000650|hsa-mir-200c&WithFlank&12|+|6963693|6963772@50.69.20(+1A)&[MIMAT0000617&hsa-miR-200c-3p&offsets|0|-3(+1A);m-54&12|+|6963742|6963764&offsets|0|-3(+1A)]|TAATACTGCCGGGTAATGATA</td><td>>MI0000650|hsa-mir-200c&WithFlank&12|+|6963693|6963772</td><td>50</td><td>69</td><td>20(+1A)</td><td>TAATACTGCCGGGTAATGATA</td><td>1183.0</td><td>14.5</td><td>1.3</td><td>10.9</td><td>1.1E-27</td><td>1.8E-26</td><td>BT20/MB468</td><td>CYTO</td></tr>
<tr><td>>hg38_21_+_8433172_8446622_RNA45SN1_45S_50ntFlanks@4283.4316.34|CTTTAACGAGGATCCATTGGAGGGCAAGTCTGGT</td><td>>hg38_21_+_8433172_8446622_RNA45SN1_45S_50ntFlanks</td><td>4283</td><td>4316</td><td>34</td><td>CTTTAACGAGGATCCATTGGAGGGCAAGTCTGGT</td><td>534.4</td><td>-13.5</td><td>1.2</td><td>-10.8</td><td>2.0E-27</td><td>1.8E-26</td><td>BT20/MB468</td><td>NUC</td></tr>
<tr><td>>hg38_21_+_8433172_8446622_RNA45SN1_45S_50ntFlanks@12636.12653.18|AGCTACCATCTGTGGGAT</td><td>>hg38_21_+_8433172_8446622_RNA45SN1_45S_50ntFlanks</td><td>12636</td><td>12653</td><td>18</td><td>AGCTACCATCTGTGGGAT</td><td>2476.4</td><td>-15.1</td><td>1.4</td><td>-11.1</td><td>1.9E-28</td><td>1.8E-26</td><td>MB231/MB468</td><td>MP</td></tr>
<tr><td>>hg38_21_+_8433172_8446622_RNA45SN1_45S_50ntFlanks@12405.12428.24|CTTTTTGATCCTTCGATGTCGGCT</td><td>>hg38_21_+_8433172_8446622_RNA45SN1_45S_50ntFlanks</td><td>12405</td><td>12428</td><td>24</td><td>CTTTTTGATCCTTCGATGTCGGCT</td><td>407.4</td><td>-13.1</td><td>1.2</td><td>-10.8</td><td>2.1E-27</td><td>1.8E-26</td><td>BT20/MB468</td><td>NUC</td></tr>
<tr><td>>MI0000088|hsa-mir-30a&WithFlank&6|-|71403545|71403627@54.74.21&[MIMAT0000088&hsa-miR-30a-3p&offsets|+1|0;m-38&6|-|71403554|71403575&offsets|+1|0]|TTTCAGTCGGATGTTTGCAGC</td><td>>MI0000088|hsa-mir-30a&WithFlank&6|-|71403545|71403627</td><td>54</td><td>74</td><td>21</td><td>TTTCAGTCGGATGTTTGCAGC</td><td>1318.0</td><td>7.1</td><td>0.7</td><td>10.9</td><td>8.6E-28</td><td>1.8E-26</td><td>BT20/MB468</td><td>MITO</td></tr>
<tr><td>>hg38_21_+_8433172_8446622_RNA45SN1_45S_50ntFlanks@4854.4883.30|AGTCTTTGGGTTCCGGGGGGAGTATGGTTG</td><td>>hg38_21_+_8433172_8446622_RNA45SN1_45S_50ntFlanks</td><td>4854</td><td>4883</td><td>30</td><td>AGTCTTTGGGTTCCGGGGGGAGTATGGTTG</td><td>561.4</td><td>-13.6</td><td>1.3</td><td>-10.8</td><td>2.2E-27</td><td>1.9E-26</td><td>BT20/MB468</td><td>NUC</td></tr>
<tr><td>>hg38_21_+_8433172_8446622_RNA45SN1_45S_50ntFlanks@3705.3722.18|TACCTGGTTGATCCTGCC</td><td>>hg38_21_+_8433172_8446622_RNA45SN1_45S_50ntFlanks</td><td>3705</td><td>3722</td><td>18</td><td>TACCTGGTTGATCCTGCC</td><td>37679.7</td><td>5.8</td><td>0.5</td><td>10.9</td><td>1.2E-27</td><td>1.9E-26</td><td>BT20/MB468</td><td>CYTO</td></tr>
<tr><td>>hg38_21_+_8433172_8446622_RNA45SN1_45S_50ntFlanks@5472.5512.41|ACGGCCCTGGCGGAGCGCTGAGAAGACGGTCGAACTTGACT</td><td>>hg38_21_+_8433172_8446622_RNA45SN1_45S_50ntFlanks</td><td>5472</td><td>5512</td><td>41</td><td>ACGGCCCTGGCGGAGCGCTGAGAAGACGGTCGAACTTGACT</td><td>9907.1</td><td>7.9</td><td>0.7</td><td>11.1</td><td>1.6E-28</td><td>2.0E-26</td><td>MB231/MB468</td><td>CYTO</td></tr>
<tr><td>>hg38_21_+_8433172_8446622_RNA45SN1_45S_50ntFlanks@9554.9571.18|CGTAGCGGTCCTGACGTG</td><td>>hg38_21_+_8433172_8446622_RNA45SN1_45S_50ntFlanks</td><td>9554</td><td>9571</td><td>18</td><td>CGTAGCGGTCCTGACGTG</td><td>85407.2</td><td>-7.5</td><td>0.7</td><td>-11.0</td><td>4.1E-28</td><td>2.0E-26</td><td>MB231/MB468</td><td>MITO</td></tr>
<tr><td>>hg38_21_+_8433172_8446622_RNA45SN1_45S_50ntFlanks@5488.5516.29|GCTGAGAAGACGGTCGAACTTGACTATCT</td><td>>hg38_21_+_8433172_8446622_RNA45SN1_45S_50ntFlanks</td><td>5488</td><td>5516</td><td>29</td><td>GCTGAGAAGACGGTCGAACTTGACTATCT</td><td>8739.1</td><td>8.4</td><td>0.8</td><td>10.9</td><td>1.3E-27</td><td>2.0E-26</td><td>BT20/MB468</td><td>CYTO</td></tr>
<tr><td>>hg38_21_+_8433172_8446622_RNA45SN1_45S_50ntFlanks@11894.11916.23|ACTCTAGTCTGGCACGGTGAAGA</td><td>>hg38_21_+_8433172_8446622_RNA45SN1_45S_50ntFlanks</td><td>11894</td><td>11916</td><td>23</td><td>ACTCTAGTCTGGCACGGTGAAGA</td><td>1406.0</td><td>-14.9</td><td>1.4</td><td>-10.8</td><td>2.4E-27</td><td>2.1E-26</td><td>BT20/MB468</td><td>NUC</td></tr>
<tr><td>>hg38_21_+_8433172_8446622_RNA45SN1_45S_50ntFlanks@12044.12068.25|ACTGACCCGGTGAGGCGGGGGGGCG</td><td>>hg38_21_+_8433172_8446622_RNA45SN1_45S_50ntFlanks</td><td>12044</td><td>12068</td><td>25</td><td>ACTGACCCGGTGAGGCGGGGGGGCG</td><td>1604.9</td><td>-15.1</td><td>1.4</td><td>-10.8</td><td>2.4E-27</td><td>2.1E-26</td><td>BT20/MB468</td><td>NUC</td></tr>
<tr><td>>hg38_21_+_8433172_8446622_RNA45SN1_45S_50ntFlanks@6745.6772.28|ACACTTCGAACGCACTTGCGGCCCCGGG</td><td>>hg38_21_+_8433172_8446622_RNA45SN1_45S_50ntFlanks</td><td>6745</td><td>6772</td><td>28</td><td>ACACTTCGAACGCACTTGCGGCCCCGGG</td><td>509.2</td><td>-13.4</td><td>1.2</td><td>-10.8</td><td>2.4E-27</td><td>2.1E-26</td><td>BT20/MB468</td><td>NUC</td></tr>
<tr><td>>hg38_21_+_8433172_8446622_RNA45SN1_45S_50ntFlanks@9736.9754.19|CCGAAACGATCTCAACCTA</td><td>>hg38_21_+_8433172_8446622_RNA45SN1_45S_50ntFlanks</td><td>9736</td><td>9754</td><td>19</td><td>CCGAAACGATCTCAACCTA</td><td>957.1</td><td>-14.3</td><td>1.3</td><td>-10.8</td><td>2.4E-27</td><td>2.1E-26</td><td>BT20/MB468</td><td>NUC</td></tr>
<tr><td>>hg38_21_+_8433172_8446622_RNA45SN1_45S_50ntFlanks@6719.6737.19|TGAATTGCAGGACACATTG</td><td>>hg38_21_+_8433172_8446622_RNA45SN1_45S_50ntFlanks</td><td>6719</td><td>6737</td><td>19</td><td>TGAATTGCAGGACACATTG</td><td>2020.5</td><td>-14.8</td><td>1.3</td><td>-11.0</td><td>2.3E-28</td><td>2.1E-26</td><td>MB231/MB468</td><td>MP</td></tr>
<tr><td>>MI0000266|hsa-mir-10a&WithFlank&17|-|48579832|48579953@28.48.21(+1A)&[MIMAT0000253&hsa-miR-10a-5p&offsets|0|-2(+1A);m-18&17|-|48579905|48579926&offsets|0|-1(+1A)]|TACCCTGTAGATCCGAATTTGA</td><td>>MI0000266|hsa-mir-10a&WithFlank&17|-|48579832|48579953</td><td>28</td><td>48</td><td>21(+1A)</td><td>TACCCTGTAGATCCGAATTTGA</td><td>2406.6</td><td>15.5</td><td>1.4</td><td>10.9</td><td>1.3E-27</td><td>2.1E-26</td><td>BT20/MB468</td><td>CYTO</td></tr>
<tr><td>>MI0000102|hsa-mir-100&WithFlank&11|-|122152223|122152314@19.39.21(+1C)&[MIMAT0000098&hsa-miR-100-5p&offsets|0|-1(+1C);m-27&11|-|122152276|122152296&offsets|0|0(+1C)]|AACCCGTAGATCCGAACTTGTC</td><td>>MI0000102|hsa-mir-100&WithFlank&11|-|122152223|122152314</td><td>19</td><td>39</td><td>21(+1C)</td><td>AACCCGTAGATCCGAACTTGTC</td><td>2107.5</td><td>14.9</td><td>1.4</td><td>11.0</td><td>6.5E-28</td><td>2.2E-26</td><td>MB231/MB468</td><td>NUC</td></tr>
<tr><td>>hg38_21_+_8433172_8446622_RNA45SN1_45S_50ntFlanks@12628.12659.32|TGGGGCGAAGCTACCATCTGTGGGATTATGAC</td><td>>hg38_21_+_8433172_8446622_RNA45SN1_45S_50ntFlanks</td><td>12628</td><td>12659</td><td>32</td><td>TGGGGCGAAGCTACCATCTGTGGGATTATGAC</td><td>1081.3</td><td>-14.5</td><td>1.3</td><td>-10.8</td><td>2.5E-27</td><td>2.2E-26</td><td>BT20/MB468</td><td>NUC</td></tr>
<tr><td>>hg38_21_+_8433172_8446622_RNA45SN1_45S_50ntFlanks@7974.8011.38|ACGCGACCTCAGATCAGACGTGGCGACCCGCTGAATTT</td><td>>hg38_21_+_8433172_8446622_RNA45SN1_45S_50ntFlanks</td><td>7974</td><td>8011</td><td>38</td><td>ACGCGACCTCAGATCAGACGTGGCGACCCGCTGAATTT</td><td>65269.5</td><td>5.9</td><td>0.5</td><td>10.9</td><td>1.0E-27</td><td>2.2E-26</td><td>BT20/MB468</td><td>MITO</td></tr>
<tr><td>>MI0000746|hsa-mir-99b&WithFlank&19|+|51692606|51692687@13.33.21&[MIMAT0000689&hsa-miR-99b-5p&offsets|0|-1;m-10&19|+|51692618|51692639&offsets|0|-1]|CACCCGTAGAACCGACCTTGC</td><td>>MI0000746|hsa-mir-99b&WithFlank&19|+|51692606|51692687</td><td>13</td><td>33</td><td>21</td><td>CACCCGTAGAACCGACCTTGC</td><td>8684.4</td><td>6.6</td><td>0.6</td><td>10.9</td><td>1.0E-27</td><td>2.2E-26</td><td>BT20/MB468</td><td>MITO</td></tr>
<tr><td>>hg38_21_+_8433172_8446622_RNA45SN1_45S_50ntFlanks@12405.12431.27|CTTTTTGATCCTTCGATGTCGGCTCTT</td><td>>hg38_21_+_8433172_8446622_RNA45SN1_45S_50ntFlanks</td><td>12405</td><td>12431</td><td>27</td><td>CTTTTTGATCCTTCGATGTCGGCTCTT</td><td>2023.0</td><td>-5.2</td><td>0.5</td><td>-10.9</td><td>1.0E-27</td><td>2.2E-26</td><td>BT20/MB468</td><td>MITO</td></tr>
</table>

<table>
<tr><th></th><th></th><th></th><th></th><th></th><th></th><th></th><th></th><th></th><th></th><th></th><th></th><th></th><th></th></tr>
<tr><td>>MI0000299|hsa-mir-222&WithFlank&X|-|45747009|45747130@75.96.22(+1A)&[MIMAT0000279&hsa-miR-222-3p&offsets|0|+1(+1A);m-140&X|-|45747033|45747056&offsets|0|-2(+1A)]|AGCTACATCTGGCTACTGGGTCA</td><td>>MI0000299|hsa-mir-222&WithFlank&X|-|45747009|45747130</td><td>75</td><td>96</td><td>22(+1A)</td><td>AGCTACATCTGGCTACTGGGTCA</td><td>646.6</td><td>7.7</td><td>0.7</td><td>10.9</td><td>1.4E-27</td><td>2.2E-26</td><td>BT20/MB468</td><td>CYTO</td></tr>
<tr><td>>trna59_GluCTC_1_+_249168447_249168518@32.70.39|CTCTCACCGCCGCGGCCCGGGTTCGATTCCCGGTCAGGA</td><td>>trna59_GluCTC_1_+_249168447_249168518</td><td>32</td><td>70</td><td>39</td><td>CTCTCACCGCCGCGGCCCGGGTTCGATTCCCGGTCAGGA</td><td>856.7</td><td>-14.2</td><td>1.3</td><td>-10.8</td><td>2.6E-27</td><td>2.3E-26</td><td>BT20/MB468</td><td>NUC</td></tr>
<tr><td>>hg38_21_+_8433172_8446622_RNA45SN1_45S_50ntFlanks@5445.5466.22|TCGGATCGGCCCCGCCGGGGTC</td><td>>hg38_21_+_8433172_8446622_RNA45SN1_45S_50ntFlanks</td><td>5445</td><td>5466</td><td>22</td><td>TCGGATCGGCCCCGCCGGGGTC</td><td>18984.0</td><td>-11.7</td><td>1.1</td><td>-11.0</td><td>2.5E-28</td><td>2.3E-26</td><td>MB231/MB468</td><td>MP</td></tr>
<tr><td>>hg38_21_+_8433172_8446622_RNA45SN1_45S_50ntFlanks@3985.4002.18|CGGCTTTGGTGACTCTAG</td><td>>hg38_21_+_8433172_8446622_RNA45SN1_45S_50ntFlanks</td><td>3985</td><td>4002</td><td>18</td><td>CGGCTTTGGTGACTCTAG</td><td>8784.1</td><td>-16.9</td><td>1.5</td><td>-11.0</td><td>4.7E-28</td><td>2.3E-26</td><td>MB231/MB468</td><td>MITO</td></tr>
<tr><td>>MI0003137|hsa-mir-193b&WithFlank&16|+|14303961|14304055@57.78.22&[MIMAT0002819&hsa-miR-193b-3p&offsets|0|0;m-166&16|+|14304017|14304038&offsets|0|0]|AACTGGCCCTCAAAGTCCCGCT</td><td>>MI0003137|hsa-mir-193b&WithFlank&16|+|14303961|14304055</td><td>57</td><td>78</td><td>22</td><td>AACTGGCCCTCAAAGTCCCGCT</td><td>1493.9</td><td>5.8</td><td>0.5</td><td>10.8</td><td>2.7E-27</td><td>2.4E-26</td><td>BT20/MB468</td><td>NUC</td></tr>
<tr><td>>hg38_21_+_8433172_8446622_RNA45SN1_45S_50ntFlanks@12418.12437.20|CGATGTCGGCTCTTCCTATC</td><td>>hg38_21_+_8433172_8446622_RNA45SN1_45S_50ntFlanks</td><td>12418</td><td>12437</td><td>20</td><td>CGATGTCGGCTCTTCCTATC</td><td>18535.2</td><td>-7.8</td><td>0.7</td><td>-10.8</td><td>2.8E-27</td><td>2.4E-26</td><td>BT20/MB468</td><td>NUC</td></tr>
<tr><td>>MI0003763|hsa-mir-767&WithFlank&X|-|152393415|152393535@33.55.23&[MIMAT0003882&hsa-miR-767-5p&offsets|0|0;m-788&X|-|152393481|152393503&offsets|0|0]|TGCACCATGGTTGTCTGAGCATG</td><td>>MI0003763|hsa-mir-767&WithFlank&X|-|152393415|152393535</td><td>33</td><td>55</td><td>23</td><td>TGCACCATGGTTGTCTGAGCATG</td><td>581.1</td><td>5.3</td><td>0.5</td><td>10.9</td><td>1.5E-27</td><td>2.4E-26</td><td>BT20/MB468</td><td>CYTO</td></tr>
<tr><td>>MI0000542|hsa-mir-320a&WithFlank&8|-|22244960|22245043@48.68.21(+3U)&[MIMAT0000510&hsa-miR-320a-3p&offsets|0|-1(+3U);m-60&8|-|22244975|22244996&offsets|0|-1(+3U)]|AAAAGCTGGGTTGAGAGGGCGTTT</td><td>>MI0000542|hsa-mir-320a&WithFlank&8|-|22244960|22245043</td><td>48</td><td>68</td><td>21(+3U)</td><td>AAAAGCTGGGTTGAGAGGGCGTTT</td><td>7814.9</td><td>14.3</td><td>1.3</td><td>10.9</td><td>1.5E-27</td><td>2.4E-26</td><td>BT20/MB468</td><td>CYTO</td></tr>
<tr><td>>hg38_21_+_8433172_8446622_RNA45SN1_45S_50ntFlanks@8105.8140.36|CGCGGCGGGGCGCGGGACATGTGGCGTACGGAAGAC</td><td>>hg38_21_+_8433172_8446622_RNA45SN1_45S_50ntFlanks</td><td>8105</td><td>8140</td><td>36</td><td>CGCGGCGGGGCGCGGGACATGTGGCGTACGGAAGAC</td><td>6317.7</td><td>-16.4</td><td>1.5</td><td>-11.0</td><td>5.0E-28</td><td>2.4E-26</td><td>MB231/MB468</td><td>MITO</td></tr>
<tr><td>>hg38_21_+_8433172_8446622_RNA45SN1_45S_50ntFlanks@11885.11904.20|TTGAGCTTGACTCTAGTCTG</td><td>>hg38_21_+_8433172_8446622_RNA45SN1_45S_50ntFlanks</td><td>11885</td><td>11904</td><td>20</td><td>TTGAGCTTGACTCTAGTCTG</td><td>2672.1</td><td>-15.2</td><td>1.4</td><td>-11.0</td><td>2.7E-28</td><td>2.5E-26</td><td>MB231/MB468</td><td>MP</td></tr>
<tr><td>>hg38_21_+_8433172_8446622_RNA45SN1_45S_50ntFlanks@12633.12657.25|CGAAGCTACCATCTGTGGGATTATG</td><td>>hg38_21_+_8433172_8446622_RNA45SN1_45S_50ntFlanks</td><td>12633</td><td>12657</td><td>25</td><td>CGAAGCTACCATCTGTGGGATTATG</td><td>2534.6</td><td>-15.2</td><td>1.4</td><td>-11.0</td><td>2.8E-28</td><td>2.5E-26</td><td>MB231/MB468</td><td>MP</td></tr>
<tr><td>>hg38_21_+_8433172_8446622_RNA45SN1_45S_50ntFlanks@8472.8489.18|CGGCCCGGCGGATCTTTC</td><td>>hg38_21_+_8433172_8446622_RNA45SN1_45S_50ntFlanks</td><td>8472</td><td>8489</td><td>18</td><td>CGGCCCGGCGGATCTTTC</td><td>620.3</td><td>-13.7</td><td>1.3</td><td>-10.8</td><td>3.0E-27</td><td>2.6E-26</td><td>BT20/MB468</td><td>NUC</td></tr>
<tr><td>>hg38_21_+_8433172_8446622_RNA45SN1_45S_50ntFlanks@12151.12180.30|GTGGGGAGTTTGACTGGGGCGGTACACCTG</td><td>>hg38_21_+_8433172_8446622_RNA45SN1_45S_50ntFlanks</td><td>12151</td><td>12180</td><td>30</td><td>GTGGGGAGTTTGACTGGGGCGGTACACCTG</td><td>874.9</td><td>-14.2</td><td>1.3</td><td>-10.8</td><td>3.0E-27</td><td>2.6E-26</td><td>BT20/MB468</td><td>NUC</td></tr>
<tr><td>>MI0003776|hsa-mir-320b-1&WithFlank&1|+|116671740|116671823@48.67.20;MI0003839|hsa-mir-320b-2&WithFlank&1|-|224257034|224257116@47.66.20;MI0000542|hsa-mir-320a&WithFlank&8|-|22244960|22245043@48.67.20&[MIMAT0005792&hsa-miR-320b&offsets|0|-2;m-183&1|+|116671787|116671806&offsets|0|0];[MIMAT0005792_1&hsa-miR-320b&offsets|0|-2;m-183&1|-|224257051|224257070&offsets|0|0];[MIMAT0000510&hsa-miR-320a-3p&offsets|0|-2;m-60&8|-|22244975|22244996&offsets|0|-2]|AAAAGCTGGGTTGAGAGGGC</td><td>>MI0003776|hsa-mir-320b-1&WithFlank&1|+|116671740|116671823</td><td>48</td><td>67</td><td>20</td><td>AAAAGCTGGGTTGAGAGGGC</td><td>1446.4</td><td>9.9</td><td>0.9</td><td>10.9</td><td>1.2E-27</td><td>2.6E-26</td><td>BT20/MB468</td><td>MITO</td></tr>
<tr><td>>MI0000264|hsa-mir-7-2&WithFlank&15|+|88611819|88611940@38.60.23;MI0000265|hsa-mir-7-3&WithFlank&19|+|4770664|4770785@37.59.23;MI0000263|hsa-mir-7-1&WithFlank&9|-|83969742|83969863@30.52.23&[MIMAT0000252&hsa-miR-7-5p&offsets|0|-1;m-400&15|+|88611856|88611879&offsets|0|-1];[MIMAT0000252_1&hsa-miR-7-5p&offsets|0|-1;m-402&19|+|4770700|4770723&offsets|0|-1];[MIMAT0000252_2&hsa-miR-7-5p&offsets|0|-1;m-401&9|-|83969811|83969834&offsets|0|-1]|TGGAAGACTAGTGATTTTGTTGT</td><td>>MI0000264|hsa-mir-7-2&WithFlank&15|+|88611819|88611940</td><td>38</td><td>60</td><td>23</td><td>TGGAAGACTAGTGATTTTGTTGT</td><td>6089.6</td><td>7.4</td><td>0.7</td><td>10.9</td><td>1.3E-27</td><td>2.6E-26</td><td>BT20/MB468</td><td>MITO</td></tr>
<tr><td>>MI0000266|hsa-mir-10a&WithFlank&17|-|48579832|48579953@29.49.21&[MIMAT0000253&hsa-miR-10a-5p&offsets|+1|-1;m-18&17|-|48579905|48579926&offsets|+1|0]|ACCCTGTAGATCCGAATTTGT</td><td>>MI0000266|hsa-mir-10a&WithFlank&17|-|48579832|48579953</td><td>29</td><td>49</td><td>21</td><td>ACCCTGTAGATCCGAATTTGT</td><td>1376.5</td><td>14.3</td><td>1.3</td><td>10.9</td><td>8.0E-28</td><td>2.6E-26</td><td>MB231/MB468</td><td>NUC</td></tr>
<tr><td>>hg38_21_+_8433172_8446622_RNA45SN1_45S_50ntFlanks@12456.12478.23|CCAAGCGTTGGATTGTTCACCCA</td><td>>hg38_21_+_8433172_8446622_RNA45SN1_45S_50ntFlanks</td><td>12456</td><td>12478</td><td>23</td><td>CCAAGCGTTGGATTGTTCACCCA</td><td>4945.2</td><td>-6.7</td><td>0.6</td><td>-10.9</td><td>1.3E-27</td><td>2.6E-26</td><td>BT20/MB468</td><td>MITO</td></tr>
<tr><td>>MI0000272|hsa-mir-182&WithFlank&7|-|129770377|129770498@29.51.23(+1G)&[MIMAT0000259&hsa-miR-182-5p&offsets|0|-1(+1G);m-22&7|-|129770449|129770470&offsets|0|+1(+1G)]|TTTGGCAATGGTAGAACTCACACG</td><td>>MI0000272|hsa-mir-182&WithFlank&7|-|129770377|129770498</td><td>29</td><td>51</td><td>23(+1G)</td><td>TTTGGCAATGGTAGAACTCACACG</td><td>443.8</td><td>6.7</td><td>0.6</td><td>10.9</td><td>1.3E-27</td><td>2.7E-26</td><td>BT20/MB468</td><td>MITO</td></tr>
<tr><td>>hg38_21_+_8433172_8446622_RNA45SN1_45S_50ntFlanks@8170.8215.46|CAAGTCCTTCTGATCGAGGCCCAGCCCGTGGACGGTGTGAGGCCGG</td><td>>hg38_21_+_8433172_8446622_RNA45SN1_45S_50ntFlanks</td><td>8170</td><td>8215</td><td>46</td><td>CAAGTCCTTCTGATCGAGGCCCAGCCCGTGGACGGTGTGAGGCCGG</td><td>4469.0</td><td>-16.6</td><td>1.5</td><td>-10.8</td><td>3.2E-27</td><td>2.7E-26</td><td>BT20/MB468</td><td>NUC</td></tr>
<tr><td>>hg38_21_+_8433172_8446622_RNA45SN1_45S_50ntFlanks@12628.12653.26|TGGGGCGAAGCTACCATCTGTGGGAT</td><td>>hg38_21_+_8433172_8446622_RNA45SN1_45S_50ntFlanks</td><td>12628</td><td>12653</td><td>26</td><td>TGGGGCGAAGCTACCATCTGTGGGAT</td><td>2399.7</td><td>-15.1</td><td>1.4</td><td>-11.0</td><td>3.2E-28</td><td>2.8E-26</td><td>MB231/MB468</td><td>MP</td></tr>
<tr><td>>hg38_21_+_8433172_8446622_RNA45SN1_45S_50ntFlanks@13002.13027.26|CGATCTATTGAAAGTCAGCCCTCGAC</td><td>>hg38_21_+_8433172_8446622_RNA45SN1_45S_50ntFlanks</td><td>13002</td><td>13027</td><td>26</td><td>CGATCTATTGAAAGTCAGCCCTCGAC</td><td>2007.1</td><td>-14.8</td><td>1.3</td><td>-11.0</td><td>3.2E-28</td><td>2.8E-26</td><td>MB231/MB468</td><td>MP</td></tr>
<tr><td>>hg38_21_+_8433172_8446622_RNA45SN1_45S_50ntFlanks@12159.12181.23|TTTGACTGGGGCGGTACACCTGT</td><td>>hg38_21_+_8433172_8446622_RNA45SN1_45S_50ntFlanks</td><td>12159</td><td>12181</td><td>23</td><td>TTTGACTGGGGCGGTACACCTGT</td><td>850.8</td><td>-14.2</td><td>1.3</td><td>-10.8</td><td>3.4E-27</td><td>2.9E-26</td><td>BT20/MB468</td><td>NUC</td></tr>
<tr><td>>hg38_21_+_8433172_8446622_RNA45SN1_45S_50ntFlanks@10389.10408.20|TGAACAGCAGTTGAACATGG</td><td>>hg38_21_+_8433172_8446622_RNA45SN1_45S_50ntFlanks</td><td>10389</td><td>10408</td><td>20</td><td>TGAACAGCAGTTGAACATGG</td><td>581.6</td><td>-13.6</td><td>1.3</td><td>-10.8</td><td>3.4E-27</td><td>2.9E-26</td><td>BT20/MB468</td><td>NUC</td></tr>
<tr><td>>hg38_21_+_8433172_8446622_RNA45SN1_45S_50ntFlanks@12033.12053.21|ATCGTTTTTTCACTGACCCGG</td><td>>hg38_21_+_8433172_8446622_RNA45SN1_45S_50ntFlanks</td><td>12033</td><td>12053</td><td>21</td><td>ATCGTTTTTTCACTGACCCGG</td><td>496.6</td><td>-13.4</td><td>1.2</td><td>-10.8</td><td>3.4E-27</td><td>2.9E-26</td><td>BT20/MB468</td><td>NUC</td></tr>
<tr><td>>hg38_21_+_8433172_8446622_RNA45SN1_45S_50ntFlanks@12431.12455.25|TCCTATCATTGTGAAGCAGAATTCA</td><td>>hg38_21_+_8433172_8446622_RNA45SN1_45S_50ntFlanks</td><td>12431</td><td>12455</td><td>25</td><td>TCCTATCATTGTGAAGCAGAATTCA</td><td>641.8</td><td>-13.8</td><td>1.3</td><td>-10.8</td><td>3.4E-27</td><td>2.9E-26</td><td>BT20/MB468</td><td>NUC</td></tr>
<tr><td>>hg38_21_+_8433172_8446622_RNA45SN1_45S_50ntFlanks@4044.4063.20|CCCATTCGAACGTCTGCCCT</td><td>>hg38_21_+_8433172_8446622_RNA45SN1_45S_50ntFlanks</td><td>4044</td><td>4063</td><td>20</td><td>CCCATTCGAACGTCTGCCCT</td><td>620.0</td><td>-13.7</td><td>1.3</td><td>-10.8</td><td>3.4E-27</td><td>2.9E-26</td><td>BT20/MB468</td><td>NUC</td></tr>
<tr><td>>hg38_21_+_8433172_8446622_RNA45SN1_45S_50ntFlanks@12626.12654.29|AATGGGGCGAAGCTACCATCTGTGGGATT</td><td>>hg38_21_+_8433172_8446622_RNA45SN1_45S_50ntFlanks</td><td>12626</td><td>12654</td><td>29</td><td>AATGGGGCGAAGCTACCATCTGTGGGATT</td><td>25106.2</td><td>-11.5</td><td>1.0</td><td>-11.0</td><td>6.0E-28</td><td>2.9E-26</td><td>MB231/MB468</td><td>MITO</td></tr>
<tr><td>>am_MI0000809|hsa-mir-151a&WithFlank&8|-|140732558|140732659@53.73.21(+2U)&[MIMAT0000757&hsa-miR-151a-3p&offsets|0|0(+2U);m-30&8|-|140732587|140732607&offsets|0|0(+2U)]|CTAGACTGAAGCTCCTTGAGGTT</td><td>>am_MI0000809|hsa-mir-151a&WithFlank&8|-|140732558|140732659</td><td>53</td><td>73</td><td>21(+2U)</td><td>CTAGACTGAAGCTCCTTGAGGTT</td><td>999.0</td><td>7.6</td><td>0.7</td><td>10.9</td><td>8.9E-28</td><td>2.9E-26</td><td>MB231/MB468</td><td>NUC</td></tr>
<tr><td>>hg38_21_+_8433172_8446622_RNA45SN1_45S_50ntFlanks@9545.9567.23|GTGGAGGTCCGTAGCGGTCCTGA</td><td>>hg38_21_+_8433172_8446622_RNA45SN1_45S_50ntFlanks</td><td>9545</td><td>9567</td><td>23</td><td>GTGGAGGTCCGTAGCGGTCCTGA</td><td>566.9</td><td>-13.6</td><td>1.3</td><td>-10.8</td><td>3.5E-27</td><td>3.0E-26</td><td>BT20/MB468</td><td>NUC</td></tr>
<tr><td>>MI0000093|hsa-mir-92a-1&WithFlank&13|+|91351308|91351397@54.75.22(+2A);MI0000094|hsa-mir-92a-2&WithFlank&X|-|134169532|134169618@54.75.22(+2A)&[MIMAT0000092&hsa-miR-92a-3p&offsets|0|0(+2A);m-9&13|+|91351361|91351382&offsets|0|0(+2A)];[MIMAT0000092_1&hsa-miR-92a-3p&offsets|0|0(+2A);m-11&X|-|134169544|134169565&offsets|0|0(+2A)]|TATTGCACTTGTCCCGGCCTGTAA</td><td>>MI0000093|hsa-mir-92a-1&WithFlank&13|+|91351308|91351397</td><td>54</td><td>75</td><td>22(+2A)</td><td>TATTGCACTTGTCCCGGCCTGTAA</td><td>845.8</td><td>6.3</td><td>0.6</td><td>10.9</td><td>1.9E-27</td><td>3.0E-26</td><td>BT20/MB468</td><td>CYTO</td></tr>
<tr><td>>hg38_21_+_8433172_8446622_RNA45SN1_45S_50ntFlanks@5472.5490.19|ACGGCCCTGGCGGAGCGCT</td><td>>hg38_21_+_8433172_8446622_RNA45SN1_45S_50ntFlanks</td><td>5472</td><td>5490</td><td>19</td><td>ACGGCCCTGGCGGAGCGCT</td><td>1517.0</td><td>7.2</td><td>0.7</td><td>11.0</td><td>2.6E-28</td><td>3.0E-26</td><td>MB231/MB468</td><td>CYTO</td></tr>
<tr><td>>hg38_21_+_8433172_8446622_RNA45SN1_45S_50ntFlanks@8114.8140.27|GCGCGGGACATGTGGCGTACGGAAGAC</td><td>>hg38_21_+_8433172_8446622_RNA45SN1_45S_50ntFlanks</td><td>8114</td><td>8140</td><td>27</td><td>GCGCGGGACATGTGGCGTACGGAAGAC</td><td>3683.9</td><td>-15.6</td><td>1.4</td><td>-11.0</td><td>6.4E-28</td><td>3.1E-26</td><td>MB231/MB468</td><td>MITO</td></tr>
<tr><td>>hg38_21_+_8433172_8446622_RNA45SN1_45S_50ntFlanks@11733.11751.19|TATGACTCTCTTAAGGTAG</td><td>>hg38_21_+_8433172_8446622_RNA45SN1_45S_50ntFlanks</td><td>11733</td><td>11751</td><td>19</td><td>TATGACTCTCTTAAGGTAG</td><td>693.0</td><td>-13.9</td><td>1.3</td><td>-10.8</td><td>3.6E-27</td><td>3.1E-26</td><td>BT20/MB468</td><td>NUC</td></tr>
<tr><td>>hg38_21_+_8433172_8446622_RNA45SN1_45S_50ntFlanks@12973.13000.28|TCGTACGTAGCAGAGCAGCTCCCTCGCT</td><td>>hg38_21_+_8433172_8446622_RNA45SN1_45S_50ntFlanks</td><td>12973</td><td>13000</td><td>28</td><td>TCGTACGTAGCAGAGCAGCTCCCTCGCT</td><td>546.6</td><td>-13.5</td><td>1.3</td><td>-10.8</td><td>3.7E-27</td><td>3.1E-26</td><td>BT20/MB468</td><td>NUC</td></tr>
<tr><td>>hg38_21_+_8433172_8446622_RNA45SN1_45S_50ntFlanks@9739.9757.19|AAACGATCTCAACCTATTC</td><td>>hg38_21_+_8433172_8446622_RNA45SN1_45S_50ntFlanks</td><td>9739</td><td>9757</td><td>19</td><td>AAACGATCTCAACCTATTC</td><td>1247.9</td><td>-14.7</td><td>1.4</td><td>-10.8</td><td>3.7E-27</td><td>3.1E-26</td><td>BT20/MB468</td><td>NUC</td></tr>
<tr><td>>hg38_21_+_8433172_8446622_RNA45SN1_45S_50ntFlanks@5491.5516.26|GAGAAGACGGTCGAACTTGACTATCT</td><td>>hg38_21_+_8433172_8446622_RNA45SN1_45S_50ntFlanks</td><td>5491</td><td>5516</td><td>26</td><td>GAGAAGACGGTCGAACTTGACTATCT</td><td>70788.4</td><td>10.9</td><td>1.0</td><td>10.9</td><td>2.0E-27</td><td>3.1E-26</td><td>BT20/MB468</td><td>CYTO</td></tr>
<tr><td>>hg38_21_+_8433172_8446622_RNA45SN1_45S_50ntFlanks@12459.12488.30|AGCGTTGGATTGTTCACCCACTAATAGGGA</td><td>>hg38_21_+_8433172_8446622_RNA45SN1_45S_50ntFlanks</td><td>12459</td><td>12488</td><td>30</td><td>AGCGTTGGATTGTTCACCCACTAATAGGGA</td><td>554.2</td><td>-13.5</td><td>1.3</td><td>-10.8</td><td>3.8E-27</td><td>3.2E-26</td><td>BT20/MB468</td><td>NUC</td></tr>
<tr><td>>hg38_21_+_8433172_8446622_RNA45SN1_45S_50ntFlanks@11581.11603.23|AACTGGTGCGGACCAGGGGAATC</td><td>>hg38_21_+_8433172_8446622_RNA45SN1_45S_50ntFlanks</td><td>11581</td><td>11603</td><td>23</td><td>AACTGGTGCGGACCAGGGGAATC</td><td>5833.5</td><td>-8.6</td><td>0.8</td><td>-10.8</td><td>3.9E-27</td><td>3.3E-26</td><td>BT20/MB468</td><td>NUC</td></tr>
<tr><td>>hg38_21_+_8433172_8446622_RNA45SN1_45S_50ntFlanks@4657.4689.33|CTAGAGGTGAAATTCTTGGACCGGCGCAAGACG</td><td>>hg38_21_+_8433172_8446622_RNA45SN1_45S_50ntFlanks</td><td>4657</td><td>4689</td><td>33</td><td>CTAGAGGTGAAATTCTTGGACCGGCGCAAGACG</td><td>5484.2</td><td>-16.3</td><td>1.5</td><td>-11.0</td><td>3.8E-28</td><td>3.3E-26</td><td>MB231/MB468</td><td>MP</td></tr>
<tr><td>>hg38_21_+_8433172_8446622_RNA45SN1_45S_50ntFlanks@12488.12518.31|AACGTGAGCTGGGTTTAGACCGTCGTGAGAC</td><td>>hg38_21_+_8433172_8446622_RNA45SN1_45S_50ntFlanks</td><td>12488</td><td>12518</td><td>31</td><td>AACGTGAGCTGGGTTTAGACCGTCGTGAGAC</td><td>440.1</td><td>-13.2</td><td>1.2</td><td>-10.8</td><td>4.0E-27</td><td>3.3E-26</td><td>BT20/MB468</td><td>NUC</td></tr>
<tr><td>>hg38_21_+_8433172_8446622_RNA45SN1_45S_50ntFlanks@12161.12180.20|TGACTGGGGCGGTACACCTG</td><td>>hg38_21_+_8433172_8446622_RNA45SN1_45S_50ntFlanks</td><td>12161</td><td>12180</td><td>20</td><td>TGACTGGGGCGGTACACCTG</td><td>1755.7</td><td>-14.6</td><td>1.3</td><td>-11.0</td><td>3.8E-28</td><td>3.3E-26</td><td>MB231/MB468</td><td>MP</td></tr>
<tr><td>>hg38_21_+_8433172_8446622_RNA45SN1_45S_50ntFlanks@12974.12991.18|CGTACGTAGCAGAGCAGC</td><td>>hg38_21_+_8433172_8446622_RNA45SN1_45S_50ntFlanks</td><td>12974</td><td>12991</td><td>18</td><td>CGTACGTAGCAGAGCAGC</td><td>22132.3</td><td>-14.6</td><td>1.4</td><td>-10.8</td><td>4.0E-27</td><td>3.3E-26</td><td>BT20/MB468</td><td>NUC</td></tr>
<tr><td>>MI0003776|hsa-mir-320b-1&WithFlank&1|+|116671740|116671823@48.66.19;MI0003839|hsa-mir-320b-2&WithFlank&1|-|224257034|224257116@47.65.19;MI0003778|hsa-mir-320c-1&WithFlank&18|+|21683512|21683595@48.66.19;MI0008191|hsa-mir-320c-2&WithFlank&18|+|24321669|24321752@48.66.19;MI0000542|hsa-mir-320a&WithFlank&8|-|22244960|22245043@48.66.19&[MIMAT0005792&hsa-miR-320b&offsets|0|-3;m-183&1|+|116671787|116671806&offsets|0|-1];[MIMAT0005792_1&hsa-miR-320b&offsets|0|-3;m-183&1|-|224257051|224257070&offsets|0|-1];[MIMAT0005793&hsa-miR-320c&offsets|0|-1;m-222&18|+|21683559|21683578&offsets|0|-1];[MIMAT0005793_1&hsa-miR-320c&offsets|0|-1;m-219&18|+|24321716|24321735&offsets|0|-1];[MIMAT0000510&hsa-miR-320a-3p&offsets|0|-3;m-60&8|-|22244975|22244996&offsets|0|-3]|AAAAGCTGGGTTGAGAGGG</td><td>>MI0003776|hsa-mir-320b-1&WithFlank&1|+|116671740|116671823</td><td>48</td><td>66</td><td>19</td><td>AAAAGCTGGGTTGAGAGGG</td><td>949.8</td><td>14.1</td><td>1.3</td><td>10.8</td><td>2.1E-27</td><td>3.4E-26</td><td>BT20/MB468</td><td>CYTO</td></tr>
</table>

| | | | | | | | | | | | | | |
|---|---|---|---|---|---|---|---|---|---|---|---|---|---|
| >hg38_21_+_8433172_8446622_RNA45SN1_45S_50ntFlanks@12162.12180.19\|GACTGGGGCGGTACACCTG | >hg38_21_+_8433172_8446622_RNA45SN1_45S_50ntFlanks | 12162 | 12180 | 19 | GACTGGGGCGGTACACCTG | 2203.8 | -15.0 | 1.4 | -11.0 | 3.9E-28 | 3.4E-26 | MB231/MB468 | MP |
| >hg38_21_+_8433172_8446622_RNA45SN1_45S_50ntFlanks@11612.11640.29\|TAATTAAAACAAAGCATCGCGAAGGCCCG | >hg38_21_+_8433172_8446622_RNA45SN1_45S_50ntFlanks | 11612 | 11640 | 29 | TAATTAAAACAAAGCATCGCGAAGGCCCG | 701.9 | -13.9 | 1.3 | -10.8 | 4.1E-27 | 3.4E-26 | BT20/MB468 | NUC |
| >hg38_21_+_8433172_8446622_RNA45SN1_45S_50ntFlanks@8105.8143.39\|CGCGGCGGGGCGCGGGACATGTGGCGTACGGAAGACCCG | >hg38_21_+_8433172_8446622_RNA45SN1_45S_50ntFlanks | 8105 | 8143 | 39 | CGCGGCGGGGCGCGGGACATGTGGCGTACGGAAGACCCG | 71564.2 | -12.8 | 1.2 | -11.0 | 4.0E-28 | 3.5E-26 | MB231/MB468 | MP |
| >hg38_21_+_8433172_8446622_RNA45SN1_45S_50ntFlanks@6738.6772.35\|ATCATCGACACTTCGAACGCACTTGCGGCCCCGGG | >hg38_21_+_8433172_8446622_RNA45SN1_45S_50ntFlanks | 6738 | 6772 | 35 | ATCATCGACACTTCGAACGCACTTGCGGCCCCGGG | 4937.0 | -16.1 | 1.5 | -10.9 | 7.3E-28 | 3.5E-26 | MB231/MB468 | MITO |
| >hg38_21_+_8433172_8446622_RNA45SN1_45S_50ntFlanks@4723.4748.26\|CATTAATCAAGAACGAAAGTCGGAGG | >hg38_21_+_8433172_8446622_RNA45SN1_45S_50ntFlanks | 4723 | 4748 | 26 | CATTAATCAAGAACGAAAGTCGGAGG | 615.5 | -13.7 | 1.3 | -10.8 | 4.2E-27 | 3.5E-26 | BT20/MB468 | NUC |
| >MI0000102\|hsa-mir-100&WithFlank&11\|-\|122152223\|122152314@19.39.21(+1A)&[MIMAT0000098&hsa-miR-100-5p&offsets\|0\|-1(+1A);m-27&11\|-\|122152276\|122152296&offsets\|0\|0(+1A)]\|AACCCGTAGATCCGAACTTGTA | >MI0000102\|hsa-mir-100&WithFlank&11\|-\|122152223\|122152314 | 19 | 39 | 21(+1A) | AACCCGTAGATCCGAACTTGTA | 98713.4 | 16.7 | 1.5 | 11.0 | 3.0E-28 | 3.5E-26 | MB231/MB468 | CYTO |
| >hg38_21_+_8433172_8446622_RNA45SN1_45S_50ntFlanks@12996.13036.41\|TCGCTGCGATCTATTGAAAGTCAGCCCTCGACACAAGGGTT | >hg38_21_+_8433172_8446622_RNA45SN1_45S_50ntFlanks | 12996 | 13036 | 41 | TCGCTGCGATCTATTGAAAGTCAGCCCTCGACACAAGGGTT | 3748.6 | 7.6 | 0.7 | 11.0 | 3.1E-28 | 3.6E-26 | MB231/MB468 | CYTO |
| >hg38_21_+_8433172_8446622_RNA45SN1_45S_50ntFlanks@11569.11602.34\|CAGCCGACTTAGAACTGGTGCGGACCAGGGGAAT | >hg38_21_+_8433172_8446622_RNA45SN1_45S_50ntFlanks | 11569 | 11602 | 34 | CAGCCGACTTAGAACTGGTGCGGACCAGGGGAAT | 4613.8 | -8.2 | 0.7 | -11.0 | 4.2E-28 | 3.6E-26 | MB231/MB468 | MP |
| >hg38_21_+_8433172_8446622_RNA45SN1_45S_50ntFlanks@12934.12951.18\|CTAAACCATTCGTAGACG | >hg38_21_+_8433172_8446622_RNA45SN1_45S_50ntFlanks | 12934 | 12951 | 18 | CTAAACCATTCGTAGACG | 1042.6 | -14.5 | 1.3 | -10.8 | 4.3E-27 | 3.6E-26 | BT20/MB468 | NUC |
| >MI0000273\|hsa-mir-183&WithFlank&7\|-\|129774899\|129775020@33.54.22&[MIMAT0000261&hsa-miR-183-5p&offsets\|0\|0;m-41&7\|-\|129774966\|129774987&offsets\|-1\|-1]\|TATGGCACTGGTAGAATTCACT | >MI0000273\|hsa-mir-183&WithFlank&7\|-\|129774899\|129775020 | 33 | 54 | 22 | TATGGCACTGGTAGAATTCACT | 13441.3 | 7.1 | 0.7 | 10.9 | 1.7E-27 | 3.6E-26 | BT20/MB468 | MITO |
| >MI0000285\|hsa-mir-205&WithFlank&1\|+\|209432127\|209432248@40.61.22(+1A)&[MIMAT0000266&hsa-miR-205-5p&offsets\|0\|0(+1A);m-351&1\|+\|209432166\|209432187&offsets\|0\|0(+1A)]\|TCCTTCATTCCACCGGAGTCTGA | >MI0000285\|hsa-mir-205&WithFlank&1\|+\|209432127\|209432248 | 40 | 61 | 22(+1A) | TCCTTCATTCCACCGGAGTCTGA | 2911.8 | -15.3 | 1.4 | -10.9 | 7.6E-28 | 3.6E-26 | MB231/MB468 | MITO |
| >hg38_21_+_8433172_8446622_RNA45SN1_45S_50ntFlanks@12627.12661.35\|ATGGGGCGAAGCTACCATCTGTGGGATTATGACTG | >hg38_21_+_8433172_8446622_RNA45SN1_45S_50ntFlanks | 12627 | 12661 | 35 | ATGGGGCGAAGCTACCATCTGTGGGATTATGACTG | 3370.0 | -15.6 | 1.4 | -11.0 | 4.3E-28 | 3.6E-26 | MB231/MB468 | MP |
| >hg38_21_+_8433172_8446622_RNA45SN1_45S_50ntFlanks@7978.8010.33\|GACCTCAGATCAGACGTGGCGACCCGCTGAATT | >hg38_21_+_8433172_8446622_RNA45SN1_45S_50ntFlanks | 7978 | 8010 | 33 | GACCTCAGATCAGACGTGGCGACCCGCTGAATT | 832.5 | 6.7 | 0.6 | 10.9 | 1.8E-27 | 3.7E-26 | BT20/MB468 | MITO |
| >hg38_21_+_8433172_8446622_RNA45SN1_45S_50ntFlanks@7983.8007.25\|CAGATCAGACGTGGCGACCCGCTGA | >hg38_21_+_8433172_8446622_RNA45SN1_45S_50ntFlanks | 7983 | 8007 | 25 | CAGATCAGACGTGGCGACCCGCTGA | 446.7 | -6.5 | 0.6 | -10.8 | 4.5E-27 | 3.8E-26 | BT20/MB468 | NUC |
| >hg38_21_+_8433172_8446622_RNA45SN1_45S_50ntFlanks@7982.8006.25\|TCAGATCAGACGTGGCGACCCGCTG | >hg38_21_+_8433172_8446622_RNA45SN1_45S_50ntFlanks | 7982 | 8006 | 25 | TCAGATCAGACGTGGCGACCCGCTG | 1929.5 | -6.7 | 0.6 | -10.9 | 1.2E-27 | 3.8E-26 | MB231/MB468 | NUC |
| >hg38_21_+_8433172_8446622_RNA45SN1_45S_50ntFlanks@4046.4065.20\|CATTCGAACGTCTGCCCTAT | >hg38_21_+_8433172_8446622_RNA45SN1_45S_50ntFlanks | 4046 | 4065 | 20 | CATTCGAACGTCTGCCCTAT | 445.2 | -13.2 | 1.2 | -10.8 | 4.6E-27 | 3.9E-26 | BT20/MB468 | NUC |
| >hg38_21_+_8433172_8446622_RNA45SN1_45S_50ntFlanks@12635.12654.20\|AAGCTACCATCTGTGGGATT | >hg38_21_+_8433172_8446622_RNA45SN1_45S_50ntFlanks | 12635 | 12654 | 20 | AAGCTACCATCTGTGGGATT | 24166.3 | -13.6 | 1.2 | -11.0 | 4.7E-28 | 4.0E-26 | MB231/MB468 | MP |
| >hg38_21_+_8433172_8446622_RNA45SN1_45S_50ntFlanks@8472.8491.20\|CGGCCCGGCGGATCTTTCCC | >hg38_21_+_8433172_8446622_RNA45SN1_45S_50ntFlanks | 8472 | 8491 | 20 | CGGCCCGGCGGATCTTTCCC | 1525.6 | -15.0 | 1.4 | -10.8 | 4.8E-27 | 4.0E-26 | BT20/MB468 | NUC |
| >hg38_21_+_8433172_8446622_RNA45SN1_45S_50ntFlanks@3986.4010.25\|GGCTTTGGTGACTCTAGATAACCTC | >hg38_21_+_8433172_8446622_RNA45SN1_45S_50ntFlanks | 3986 | 4010 | 25 | GGCTTTGGTGACTCTAGATAACCTC | 758.7 | 6.8 | 0.6 | 10.9 | 1.9E-27 | 4.0E-26 | BT20/MB468 | MITO |
| >MI0000283\|hsa-mir-203a&WithFlank&14\|+\|104117399\|104117520@71.93.23&[MIMAT0000264&hsa-miR-203a-3p&offsets\|0\|+1;m-79&14\|+\|104117469\|104117490&offsets\|0\|+1]\|GTGAAATGTTTAGGACCACTAGA | >MI0000283\|hsa-mir-203a&WithFlank&14\|+\|104117399\|104117520 | 71 | 93 | 23 | GTGAAATGTTTAGGACCACTAGA | 753.1 | 13.8 | 1.3 | 10.8 | 2.6E-27 | 4.0E-26 | BT20/MB468 | CYTO |
| >hg38_21_+_8433172_8446622_RNA45SN1_45S_50ntFlanks@482.520.39\|ATCGATGTGGTGACGTCGTGCTCTCCCGGGCCGGGTCCG | >hg38_21_+_8433172_8446622_RNA45SN1_45S_50ntFlanks | 482 | 520 | 39 | ATCGATGTGGTGACGTCGTGCTCTCCCGGGCCGGGTCCG | 1171.0 | -14.6 | 1.4 | -10.8 | 4.9E-27 | 4.1E-26 | BT20/MB468 | NUC |
| >hg38_21_+_8433172_8446622_RNA45SN1_45S_50ntFlanks@11672.11702.31\|CAGTGCTCTGAATGTCAAAGTGAAGAAATTC | >hg38_21_+_8433172_8446622_RNA45SN1_45S_50ntFlanks | 11672 | 11702 | 31 | CAGTGCTCTGAATGTCAAAGTGAAGAAATTC | 3872.0 | -7.0 | 0.6 | -10.8 | 4.9E-27 | 4.1E-26 | BT20/MB468 | NUC |
| >hg38_21_+_8433172_8446622_RNA45SN1_45S_50ntFlanks@11561.11588.28\|GCGCCTAGCAGCCGACTTAGAACTGGTG | >hg38_21_+_8433172_8446622_RNA45SN1_45S_50ntFlanks | 11561 | 11588 | 28 | GCGCCTAGCAGCCGACTTAGAACTGGTG | 827.3 | -14.1 | 1.3 | -10.8 | 5.0E-27 | 4.1E-26 | BT20/MB468 | NUC |
| >MI0000650\|hsa-mir-200c&WithFlank&12\|+\|6963693\|6963772@50.71.22(+1G)&[MIMAT0000617&hsa-miR-200c-3p&offsets\|0\|-1(+1G);m-54&12\|+\|6963742\|6963764&offsets\|0\|-1(+1G)]\|TAATACTGCCGGGTAATGATGGG | >MI0000650\|hsa-mir-200c&WithFlank&12\|+\|6963693\|6963772 | 50 | 71 | 22(+1G) | TAATACTGCCGGGTAATGATGGG | 4258.7 | 15.8 | 1.4 | 11.1 | 1.1E-28 | 4.2E-26 | BT20/MB231 | NUC |
| >hg38_21_+_8433172_8446622_RNA45SN1_45S_50ntFlanks@11270.11293.24\|CGGCCGCCCCCGGGGAGCCCGGCG | >hg38_21_+_8433172_8446622_RNA45SN1_45S_50ntFlanks | 11270 | 11293 | 24 | CGGCCGCCCCCGGGGAGCCCGGCG | 1266.6 | -14.7 | 1.4 | -10.8 | 5.1E-27 | 4.2E-26 | BT20/MB468 | NUC |
| >hg38_21_+_8433172_8446622_RNA45SN1_45S_50ntFlanks@4122.4145.24\|AGGGTTCGATTCCGGAGAGGGAGC | >hg38_21_+_8433172_8446622_RNA45SN1_45S_50ntFlanks | 4122 | 4145 | 24 | AGGGTTCGATTCCGGAGAGGGAGC | 653.7 | -13.8 | 1.3 | -10.8 | 5.1E-27 | 4.2E-26 | BT20/MB468 | NUC |
| >MI0003137\|hsa-mir-193b&WithFlank&16\|+\|14303961\|14304055@57.77.21(+1C)&[MIMAT0002819&hsa-miR-193b-3p&offsets\|0\|-1(+1C);m-166&16\|+\|14304017\|14304038&offsets\|0\|-1(+1C)]\|AACTGGCCCTCAAAGTCCCGCC | >MI0003137\|hsa-mir-193b&WithFlank&16\|+\|14303961\|14304055 | 57 | 77 | 21(+1C) | AACTGGCCCTCAAAGTCCCGCC | 506.0 | 13.4 | 1.2 | 10.8 | 2.1E-27 | 4.3E-26 | BT20/MB468 | MITO |
| >hg38_21_+_8433172_8446622_RNA45SN1_45S_50ntFlanks@4724.4758.35\|ATTAATCAAGAACGAAAGTCGGAGGTTCGAAGACG | >hg38_21_+_8433172_8446622_RNA45SN1_45S_50ntFlanks | 4724 | 4758 | 35 | ATTAATCAAGAACGAAAGTCGGAGGTTCGAAGACG | 16921.0 | -9.8 | 0.9 | -11.0 | 5.2E-28 | 4.4E-26 | MB231/MB468 | MP |
| >hg38_21_+_8433172_8446622_RNA45SN1_45S_50ntFlanks@4759.4786.28\|ATCAGATACCGTCGTAGTTCCGACCATA | >hg38_21_+_8433172_8446622_RNA45SN1_45S_50ntFlanks | 4759 | 4786 | 28 | ATCAGATACCGTCGTAGTTCCGACCATA | 470.5 | -13.3 | 1.2 | -10.8 | 5.3E-27 | 4.4E-26 | BT20/MB468 | NUC |
| >am_MI0000809\|hsa-mir-151a&WithFlank&8\|-\|140732558\|140732659@53.73.21(+1U)&[MIMAT0000757&hsa-miR-151a-3p&offsets\|0\|0(+1U);m-30&8\|-\|140732587\|140732607&offsets\|0\|0(+1U)]\|CTAGACTGAAGCTCCTTGAGGT | >am_MI0000809\|hsa-mir-151a&WithFlank&8\|-\|140732558\|140732659 | 53 | 73 | 21(+1U) | CTAGACTGAAGCTCCTTGAGGT | 8098.9 | 6.4 | 0.6 | 10.9 | 1.4E-27 | 4.5E-26 | MB231/MB468 | NUC |
| >hg38_21_+_8433172_8446622_RNA45SN1_45S_50ntFlanks@12436.12455.20\|TCATTGTGAAGCAGAATTCA | >hg38_21_+_8433172_8446622_RNA45SN1_45S_50ntFlanks | 12436 | 12455 | 20 | TCATTGTGAAGCAGAATTCA | 5742.3 | -10.7 | 1.0 | -11.0 | 5.4E-28 | 4.5E-26 | MB231/MB468 | MP |
| >trna101_AlaAGC_6_-_28831462_28831533@58.75.18,trna102_AlaAGC_6_-_28806221_28806292@58.75.18,trna110_AlaTGC_6_-_28757547_28757618@58.75.18,trna113_AlaTGC_6_-_28726141_28726212@58.75.18,trna65_AlaAGC_6_+_28574933_28575004@58.75.18\|TCCCCGGCACCTCCACCA | >trna101_AlaAGC_6_-_28831462_28831533 | 58 | 75 | 18 | TCCCCGGCACCTCCACCA | 20844.0 | 8.3 | 0.8 | 10.8 | 3.0E-27 | 4.6E-26 | BT20/MB468 | CYTO |
| >hg38_21_+_8433172_8446622_RNA45SN1_45S_50ntFlanks@6772.6797.26\|GTTCCTCCCGGGGCTACGCCTGTCTG | >hg38_21_+_8433172_8446622_RNA45SN1_45S_50ntFlanks | 6772 | 6797 | 26 | GTTCCTCCCGGGGCTACGCCTGTCTG | 510.4 | -13.4 | 1.2 | -10.8 | 5.6E-27 | 4.6E-26 | BT20/MB468 | NUC |
| >hg38_21_+_8433172_8446622_RNA45SN1_45S_50ntFlanks@3984.4005.22\|GCGGCTTTGGTGACTCTAGATA | >hg38_21_+_8433172_8446622_RNA45SN1_45S_50ntFlanks | 3984 | 4005 | 22 | GCGGCTTTGGTGACTCTAGATA | 1416.1 | -14.9 | 1.4 | -10.8 | 5.6E-27 | 4.6E-26 | BT20/MB468 | NUC |
| >MI0000471\|hsa-mir-126&WithFlank&9\|+\|136670596\|136670692@58.78.21&[MIMAT0000445&hsa-miR-126-3p&offsets\|0\|-1;m-40&9\|+\|136670653\|136670674&offsets\|0\|-1]\|TCGTACCGTGAGTAATAATGC | >MI0000471\|hsa-mir-126&WithFlank&9\|+\|136670596\|136670692 | 58 | 78 | 21 | TCGTACCGTGAGTAATAATGC | 2667.9 | 10.8 | 1.0 | 10.8 | 2.3E-27 | 4.7E-26 | BT20/MB468 | MITO |
| >MI0000650\|hsa-mir-200c&WithFlank&12\|+\|6963693\|6963772@50.72.23(+2A)&[MIMAT0000617&hsa-miR-200c-3p&offsets\|0\|0(+2A);m-54&12\|+\|6963742\|6963764&offsets\|0\|0(+2A)]\|TAATACTGCCGGGTAATGATGGAAA | >MI0000650\|hsa-mir-200c&WithFlank&12\|+\|6963693\|6963772 | 50 | 72 | 23(+2A) | TAATACTGCCGGGTAATGATGGAAA | 1190.0 | 9.2 | 0.8 | 10.8 | 2.3E-27 | 4.7E-26 | BT20/MB468 | MITO |
| >MI0000086\|hsa-mir-28&WithFlank&3\|+\|188688775\|188688872@60.82.23&[MIMAT0004502&hsa-miR-28-3p&offsets\|0\|+1;m-28&3\|+\|188688834\|188688855&offsets\|0\|+1]\|CACTAGATTGTGAGCTCCTGGAG | >MI0000086\|hsa-mir-28&WithFlank&3\|+\|188688775\|188688872 | 60 | 82 | 23 | CACTAGATTGTGAGCTCCTGGAG | 561.4 | 13.6 | 1.3 | 10.8 | 2.3E-27 | 4.7E-26 | BT20/MB468 | MITO |
| >MI0000067\|hsa-let-7f-1&WithFlank&9\|+\|94176341\|94176439@13.34.22;MI0000068\|hsa-let-7f-2&WithFlank&X\|-\|53557186\|53557280@14.35.22&[MIMAT0000067&hsa-let-7f-5p&offsets\|0\|0;m-20&9\|+\|94176353\|94176374&offsets\|0\|0];[MIMAT0000067_1&hsa-let-7f-5p&offsets\|0\|0;m-19&X\|-\|53557246\|53557267&offsets\|0\|0]\|TGAGGTAGTAGATTGTATAGTT | >MI0000067\|hsa-let-7f-1&WithFlank&9\|+\|94176341\|94176439 | 13 | 34 | 22 | TGAGGTAGTAGATTGTATAGTT | 90726.3 | 7.1 | 0.7 | 10.8 | 2.4E-27 | 4.9E-26 | BT20/MB468 | MITO |
| >hg38_21_+_8433172_8446622_RNA45SN1_45S_50ntFlanks@6732.6758.27\|ACATTGATCATCGACACTTCGAACGCA | >hg38_21_+_8433172_8446622_RNA45SN1_45S_50ntFlanks | 6732 | 6758 | 27 | ACATTGATCATCGACACTTCGAACGCA | 406.6 | -4.8 | 0.4 | -10.7 | 5.9E-27 | 4.9E-26 | BT20/MB468 | NUC |
| >MI0000266\|hsa-mir-10a&WithFlank&17\|-\|48579832\|48579953@28.49.22&[MIMAT0000253&hsa-miR-10a-5p&offsets\|0\|-1;m-18&17\|-\|48579905\|48579926&offsets\|0\|0]\|TACCCTGTAGATCCGAATTTGT | >MI0000266\|hsa-mir-10a&WithFlank&17\|-\|48579832\|48579953 | 28 | 49 | 22 | TACCCTGTAGATCCGAATTTGT | 8790.2 | 12.1 | 1.1 | 11.0 | 4.5E-28 | 4.9E-26 | MB231/MB468 | CYTO |
| >hg38_21_+_8433172_8446622_RNA45SN1_45S_50ntFlanks@5493.5516.24\|GAAGACGGTCGAACTTGACTATCT | >hg38_21_+_8433172_8446622_RNA45SN1_45S_50ntFlanks | 5493 | 5516 | 24 | GAAGACGGTCGAACTTGACTATCT | 2665.8 | 7.7 | 0.7 | 11.0 | 4.5E-28 | 4.9E-26 | MB231/MB468 | CYTO |
| >hg38_21_+_8433172_8446622_RNA45SN1_45S_50ntFlanks@11998.12032.35\|GCCCTGCGGGCCGCCGGTGAAATACCACTACTCTG | >hg38_21_+_8433172_8446622_RNA45SN1_45S_50ntFlanks | 11998 | 12032 | 35 | GCCCTGCGGGCCGCCGGTGAAATACCACTACTCTG | 894.3 | -14.2 | 1.3 | -10.7 | 6.0E-27 | 5.0E-26 | BT20/MB468 | NUC |
| >hg38_21_+_8433172_8446622_RNA45SN1_45S_50ntFlanks@9556.9582.27\|TAGCGGTCCTGACGTGCAAATCGGTCG | >hg38_21_+_8433172_8446622_RNA45SN1_45S_50ntFlanks | 9556 | 9582 | 27 | TAGCGGTCCTGACGTGCAAATCGGTCG | 1786.9 | -15.2 | 1.4 | -10.7 | 6.1E-27 | 5.0E-26 | BT20/MB468 | NUC |
| >hg38_21_+_8433172_8446622_RNA45SN1_45S_50ntFlanks@5444.5466.23\|CTCGGATCGGCCCCGCCGGGGTC | >hg38_21_+_8433172_8446622_RNA45SN1_45S_50ntFlanks | 5444 | 5466 | 23 | CTCGGATCGGCCCCGCCGGGGTC | 41214.1 | -12.2 | 1.1 | -11.0 | 6.1E-28 | 5.1E-26 | MB231/MB468 | MP |
| >hg38_21_+_8433172_8446622_RNA45SN1_45S_50ntFlanks@8423.8442.20\|CGCAGTCCGCCCGGAGGATT | >hg38_21_+_8433172_8446622_RNA45SN1_45S_50ntFlanks | 8423 | 8442 | 20 | CGCAGTCCGCCCGGAGGATT | 2504.8 | -15.2 | 1.4 | -11.0 | 6.2E-28 | 5.1E-26 | MB231/MB468 | MP |
| >hg38_21_+_8433172_8446622_RNA45SN1_45S_50ntFlanks@10411.10432.22\|CAGTCGGTCCTGAGAGATGGGC | >hg38_21_+_8433172_8446622_RNA45SN1_45S_50ntFlanks | 10411 | 10432 | 22 | CAGTCGGTCCTGAGAGATGGGC | 2233.1 | -15.0 | 1.4 | -11.0 | 6.2E-28 | 5.1E-26 | MB231/MB468 | MP |

<table>
<tr><td>>hg38_21_+_8433172_8446622_RNA45SN1_45S_50ntFlanks@4047.4064.18|ATTCGAACGTCTGCCCTA</td><td>>hg38_21_+_8433172_8446622_RNA45SN1_45S_50ntFlanks</td><td>4047</td><td>4064</td><td>18</td><td>ATTCGAACGTCTGCCCTA</td><td>608.8</td><td>-13.7</td><td>1.3</td><td>-10.7</td><td>6.3E-27</td><td>5.1E-26</td><td>BT20/MB468</td><td>NUC</td></tr>
<tr><td>>hg38_21_+_8433172_8446622_RNA45SN1_45S_50ntFlanks@9033.9058.26|CTCGTGGGGGGGCCGGGCCACCCCTC</td><td>>hg38_21_+_8433172_8446622_RNA45SN1_45S_50ntFlanks</td><td>9033</td><td>9058</td><td>26</td><td>CTCGTGGGGGGGCCGGGCCACCCCTC</td><td>3114.9</td><td>-15.5</td><td>1.4</td><td>-11.0</td><td>6.3E-28</td><td>5.1E-26</td><td>MB231/MB468</td><td>MP</td></tr>
<tr><td>>hg38_21_+_8433172_8446622_RNA45SN1_45S_50ntFlanks@12991.13027.37|CTCCCTCGCTGCGATCTATTGAAAGTCAGCCCTCGAC</td><td>>hg38_21_+_8433172_8446622_RNA45SN1_45S_50ntFlanks</td><td>12991</td><td>13027</td><td>37</td><td>CTCCCTCGCTGCGATCTATTGAAAGTCAGCCCTCGAC</td><td>1996.6</td><td>-15.4</td><td>1.4</td><td>-10.7</td><td>6.3E-27</td><td>5.1E-26</td><td>BT20/MB468</td><td>NUC</td></tr>
<tr><td>>hg38_21_+_8433172_8446622_RNA45SN1_45S_50ntFlanks@9534.9565.32|AGGAAACTCTGGTGGAGGTCCGTAGCGGTCCT</td><td>>hg38_21_+_8433172_8446622_RNA45SN1_45S_50ntFlanks</td><td>9534</td><td>9565</td><td>32</td><td>AGGAAACTCTGGTGGAGGTCCGTAGCGGTCCT</td><td>2942.6</td><td>-6.2</td><td>0.6</td><td>-10.9</td><td>1.6E-27</td><td>5.2E-26</td><td>MB231/MB468</td><td>NUC</td></tr>
<tr><td>>hg38_21_+_8433172_8446622_RNA45SN1_45S_50ntFlanks@6647.6664.18|CGTACGACTCTTAGCGGT</td><td>>hg38_21_+_8433172_8446622_RNA45SN1_45S_50ntFlanks</td><td>6647</td><td>6664</td><td>18</td><td>CGTACGACTCTTAGCGGT</td><td>22561.0</td><td>6.1</td><td>0.6</td><td>10.8</td><td>2.6E-27</td><td>5.3E-26</td><td>BT20/MB468</td><td>MITO</td></tr>
<tr><td>>hg38_21_+_8433172_8446622_RNA45SN1_45S_50ntFlanks@12991.13012.22|CTCCCTCGCTGCGATCTATTGA</td><td>>hg38_21_+_8433172_8446622_RNA45SN1_45S_50ntFlanks</td><td>12991</td><td>13012</td><td>22</td><td>CTCCCTCGCTGCGATCTATTGA</td><td>12191.9</td><td>-8.4</td><td>0.8</td><td>-10.9</td><td>1.7E-27</td><td>5.3E-26</td><td>MB231/MB468</td><td>NUC</td></tr>
<tr><td>>hg38_21_+_8433172_8446622_RNA45SN1_45S_50ntFlanks@8105.8143.39|CGCGGCGGGGCGCGGGACATGTGGCGTACGGAAGACCCG</td><td>>hg38_21_+_8433172_8446622_RNA45SN1_45S_50ntFlanks</td><td>8105</td><td>8143</td><td>39</td><td>CGCGGCGGGGCGCGGGACATGTGGCGTACGGAAGACCCG</td><td>9107.7</td><td>-12.6</td><td>1.2</td><td>-10.9</td><td>1.7E-27</td><td>5.4E-26</td><td>MB231/MB468</td><td>NUC</td></tr>
<tr><td>>hg38_21_+_8433172_8446622_RNA45SN1_45S_50ntFlanks@12407.12426.20|TTTTGATCCTTCGATGTCGG</td><td>>hg38_21_+_8433172_8446622_RNA45SN1_45S_50ntFlanks</td><td>12407</td><td>12426</td><td>20</td><td>TTTTGATCCTTCGATGTCGG</td><td>351.2</td><td>-12.9</td><td>1.2</td><td>-10.7</td><td>6.8E-27</td><td>5.6E-26</td><td>BT20/MB468</td><td>NUC</td></tr>
<tr><td>>hg38_21_+_8433172_8446622_RNA45SN1_45S_50ntFlanks@11588.11623.36|GCGGACCAGGGGAATCCGACTGTTTAATTAAAACAA</td><td>>hg38_21_+_8433172_8446622_RNA45SN1_45S_50ntFlanks</td><td>11588</td><td>11623</td><td>36</td><td>GCGGACCAGGGGAATCCGACTGTTTAATTAAAACAA</td><td>705.1</td><td>13.9</td><td>1.3</td><td>10.8</td><td>2.8E-27</td><td>5.7E-26</td><td>BT20/MB468</td><td>MITO</td></tr>
<tr><td>>hg38_21_+_8433172_8446622_RNA45SN1_45S_50ntFlanks@12008.12029.22|CCGCCGGTGAAATACCACTACT</td><td>>hg38_21_+_8433172_8446622_RNA45SN1_45S_50ntFlanks</td><td>12008</td><td>12029</td><td>22</td><td>CCGCCGGTGAAATACCACTACT</td><td>2274.2</td><td>-15.0</td><td>1.4</td><td>-10.9</td><td>7.0E-28</td><td>5.7E-26</td><td>MB231/MB468</td><td>MP</td></tr>
<tr><td>>hg38_21_+_8433172_8446622_RNA45SN1_45S_50ntFlanks@10798.10818.21|CATGTTGGAACAATGTAGGTA</td><td>>hg38_21_+_8433172_8446622_RNA45SN1_45S_50ntFlanks</td><td>10798</td><td>10818</td><td>21</td><td>CATGTTGGAACAATGTAGGTA</td><td>2118.8</td><td>-14.9</td><td>1.4</td><td>-10.9</td><td>7.1E-28</td><td>5.7E-26</td><td>MB231/MB468</td><td>MP</td></tr>
<tr><td>>hg38_21_+_8433172_8446622_RNA45SN1_45S_50ntFlanks@3899.3919.21|CCTTCGCGGGGGGGATGCGTG</td><td>>hg38_21_+_8433172_8446622_RNA45SN1_45S_50ntFlanks</td><td>3899</td><td>3919</td><td>21</td><td>CCTTCGCGGGGGGGATGCGTG</td><td>7600.9</td><td>-16.8</td><td>1.5</td><td>-10.9</td><td>7.1E-28</td><td>5.7E-26</td><td>MB231/MB468</td><td>MP</td></tr>
<tr><td>>hg38_21_+_8433172_8446622_RNA45SN1_45S_50ntFlanks@8342.8375.34|CGTAAGGGAAAGTTGAAAAGAACTTTGAAGAGAG</td><td>>hg38_21_+_8433172_8446622_RNA45SN1_45S_50ntFlanks</td><td>8342</td><td>8375</td><td>34</td><td>CGTAAGGGAAAGTTGAAAAGAACTTTGAAGAGAG</td><td>671.8</td><td>13.9</td><td>1.3</td><td>10.8</td><td>2.8E-27</td><td>5.7E-26</td><td>BT20/MB468</td><td>MITO</td></tr>
<tr><td>>hg38_21_+_8433172_8446622_RNA45SN1_45S_50ntFlanks@5487.5526.40|CGCTGAGAAGACGGTCGAACTTGACTATCTAGAGGAAGTA</td><td>>hg38_21_+_8433172_8446622_RNA45SN1_45S_50ntFlanks</td><td>5487</td><td>5526</td><td>40</td><td>CGCTGAGAAGACGGTCGAACTTGACTATCTAGAGGAAGTA</td><td>2768.1</td><td>-15.2</td><td>1.4</td><td>-10.9</td><td>1.2E-27</td><td>5.7E-26</td><td>MB231/MB468</td><td>MITO</td></tr>
<tr><td>>hg38_21_+_8433172_8446622_RNA45SN1_45S_50ntFlanks@8357.8379.23|AAAAGAACTTTGAAGAGAGAGTT</td><td>>hg38_21_+_8433172_8446622_RNA45SN1_45S_50ntFlanks</td><td>8357</td><td>8379</td><td>23</td><td>AAAAGAACTTTGAAGAGAGAGTT</td><td>48407.9</td><td>-10.0</td><td>0.9</td><td>-10.9</td><td>1.2E-27</td><td>5.7E-26</td><td>MB231/MB468</td><td>MITO</td></tr>
<tr><td>>hg38_21_+_8433172_8446622_RNA45SN1_45S_50ntFlanks@11534.11560.27|GTTCCGCGCGGCGCCTCGCCTCGGCCG</td><td>>hg38_21_+_8433172_8446622_RNA45SN1_45S_50ntFlanks</td><td>11534</td><td>11560</td><td>27</td><td>GTTCCGCGCGGCGCCTCGCCTCGGCCG</td><td>1147.4</td><td>-14.6</td><td>1.4</td><td>-10.7</td><td>7.1E-27</td><td>5.8E-26</td><td>BT20/MB468</td><td>NUC</td></tr>
<tr><td>>hg38_21_+_8433172_8446622_RNA45SN1_45S_50ntFlanks@12991.13013.23|CTCCCTCGCTGCGATCTATTGAA</td><td>>hg38_21_+_8433172_8446622_RNA45SN1_45S_50ntFlanks</td><td>12991</td><td>13013</td><td>23</td><td>CTCCCTCGCTGCGATCTATTGAA</td><td>5451.9</td><td>-7.5</td><td>0.7</td><td>-10.8</td><td>2.9E-27</td><td>5.8E-26</td><td>BT20/MB468</td><td>MITO</td></tr>
<tr><td>>hg38_21_+_8433172_8446622_RNA45SN1_45S_50ntFlanks@12157.12181.25|AGTTTGACTGGGGCGGTACACCTGT</td><td>>hg38_21_+_8433172_8446622_RNA45SN1_45S_50ntFlanks</td><td>12157</td><td>12181</td><td>25</td><td>AGTTTGACTGGGGCGGTACACCTGT</td><td>725.0</td><td>-13.9</td><td>1.3</td><td>-10.7</td><td>7.2E-27</td><td>5.9E-26</td><td>BT20/MB468</td><td>NUC</td></tr>
<tr><td>>MI0000079|hsa-mir-23a&WithFlank&19|-|13836581|13836665@51.72.22&[MIMAT0000078&hsa-miR-23a-3p&offsets|0|+1;m-31&19|-|13836595|13836615&offsets|0|+1]|ATCACATTGCCAGGGATTTCCA</td><td>>MI0000079|hsa-mir-23a&WithFlank&19|-|13836581|13836665</td><td>51</td><td>72</td><td>22</td><td>ATCACATTGCCAGGGATTTCCA</td><td>487.2</td><td>-13.4</td><td>1.2</td><td>-10.7</td><td>7.3E-27</td><td>5.9E-26</td><td>BT20/MB468</td><td>NUC</td></tr>
<tr><td>>hg38_21_+_8433172_8446622_RNA45SN1_45S_50ntFlanks@9305.9336.32|CCAAGGAGTCTAACACGTGCGCGAGTCGGGGG</td><td>>hg38_21_+_8433172_8446622_RNA45SN1_45S_50ntFlanks</td><td>9305</td><td>9336</td><td>32</td><td>CCAAGGAGTCTAACACGTGCGCGAGTCGGGGG</td><td>1928.1</td><td>-14.8</td><td>1.4</td><td>-10.9</td><td>7.6E-28</td><td>6.0E-26</td><td>MB231/MB468</td><td>MP</td></tr>
<tr><td>>MI0000542|hsa-mir-320a&WithFlank&8|-|22244960|22245043@49.70.22(+1U)&[MIMAT0000510&hsa-miR-320a-3p&offsets|+1|+1(+1U);m-60&8|-|22244975|22244996&offsets|+1|+1(+1U)]|AAAGCTGGGTTGAGAGGGCGAAT</td><td>>MI0000542|hsa-mir-320a&WithFlank&8|-|22244960|22245043</td><td>49</td><td>70</td><td>22(+1U)</td><td>AAAGCTGGGTTGAGAGGGCGAAT</td><td>1718.2</td><td>11.0</td><td>1.0</td><td>10.8</td><td>3.9E-27</td><td>6.1E-26</td><td>BT20/MB468</td><td>CYTO</td></tr>
<tr><td>>hg38_21_+_8433172_8446622_RNA45SN1_45S_50ntFlanks@12411.12443.33|GATCCTTCGATGTCGGCTCTTCCTATCATTGTG</td><td>>hg38_21_+_8433172_8446622_RNA45SN1_45S_50ntFlanks</td><td>12411</td><td>12443</td><td>33</td><td>GATCCTTCGATGTCGGCTCTTCCTATCATTGTG</td><td>845.4</td><td>-14.1</td><td>1.3</td><td>-10.7</td><td>7.5E-27</td><td>6.1E-26</td><td>BT20/MB468</td><td>NUC</td></tr>
<tr><td>>hg38_21_+_8433172_8446622_RNA45SN1_45S_50ntFlanks@51.72.22|GCTGACACGCTGTCCTCTGGCG</td><td>>hg38_21_+_8433172_8446622_RNA45SN1_45S_50ntFlanks</td><td>51</td><td>72</td><td>22</td><td>GCTGACACGCTGTCCTCTGGCG</td><td>1895.8</td><td>-6.8</td><td>0.6</td><td>-10.7</td><td>7.5E-27</td><td>6.1E-26</td><td>BT20/MB468</td><td>NUC</td></tr>
<tr><td>>hg38_21_+_8433172_8446622_RNA45SN1_45S_50ntFlanks@12412.12435.24|ATCCTTCGATGTCGGCTCTTCCTA</td><td>>hg38_21_+_8433172_8446622_RNA45SN1_45S_50ntFlanks</td><td>12412</td><td>12435</td><td>24</td><td>ATCCTTCGATGTCGGCTCTTCCTA</td><td>2099.3</td><td>-15.5</td><td>1.4</td><td>-10.7</td><td>7.5E-27</td><td>6.1E-26</td><td>BT20/MB468</td><td>NUC</td></tr>
<tr><td>>hg38_21_+_8433172_8446622_RNA45SN1_45S_50ntFlanks@10574.10607.34|AAGCCGGCGGGAGCCCCGGGGAGAGTTCTCTTTT</td><td>>hg38_21_+_8433172_8446622_RNA45SN1_45S_50ntFlanks</td><td>10574</td><td>10607</td><td>34</td><td>AAGCCGGCGGGAGCCCCGGGGAGAGTTCTCTTTT</td><td>717.7</td><td>-13.9</td><td>1.3</td><td>-10.7</td><td>7.6E-27</td><td>6.1E-26</td><td>BT20/MB468</td><td>NUC</td></tr>
<tr><td>>hg38_21_+_8433172_8446622_RNA45SN1_45S_50ntFlanks@12141.12158.18|ACAGTGCCAGGTGGGGAG</td><td>>hg38_21_+_8433172_8446622_RNA45SN1_45S_50ntFlanks</td><td>12141</td><td>12158</td><td>18</td><td>ACAGTGCCAGGTGGGGAG</td><td>451.6</td><td>-13.2</td><td>1.2</td><td>-10.7</td><td>7.6E-27</td><td>6.1E-26</td><td>BT20/MB468</td><td>NUC</td></tr>
<tr><td>>hg38_21_+_8433172_8446622_RNA45SN1_45S_50ntFlanks@11316.11342.27|CACCCCACGTCTCGTCGCGCGCGCGTC</td><td>>hg38_21_+_8433172_8446622_RNA45SN1_45S_50ntFlanks</td><td>11316</td><td>11342</td><td>27</td><td>CACCCCACGTCTCGTCGCGCGCGCGTC</td><td>8749.7</td><td>-12.9</td><td>1.2</td><td>-10.9</td><td>1.9E-27</td><td>6.1E-26</td><td>MB231/MB468</td><td>NUC</td></tr>
<tr><td>>hg38_21_+_8433172_8446622_RNA45SN1_45S_50ntFlanks@12001.12032.32|CTGCGGGCCGCCGGTGAAATACCACTACTCTG</td><td>>hg38_21_+_8433172_8446622_RNA45SN1_45S_50ntFlanks</td><td>12001</td><td>12032</td><td>32</td><td>CTGCGGGCCGCCGGTGAAATACCACTACTCTG</td><td>2645.2</td><td>-15.2</td><td>1.4</td><td>-10.9</td><td>8.0E-28</td><td>6.3E-26</td><td>MB231/MB468</td><td>MP</td></tr>
<tr><td>>hg38_21_+_8433172_8446622_RNA45SN1_45S_50ntFlanks@11027.11060.34|CTCTCTCTCTCTCTCCCCCGCTCCCCGTCCTCCC</td><td>>hg38_21_+_8433172_8446622_RNA45SN1_45S_50ntFlanks</td><td>11027</td><td>11060</td><td>34</td><td>CTCTCTCTCTCTCTCCCCCGCTCCCCGTCCTCCC</td><td>2730.4</td><td>-15.2</td><td>1.4</td><td>-10.9</td><td>1.4E-27</td><td>6.3E-26</td><td>MB231/MB468</td><td>MITO</td></tr>
<tr><td>>hg38_21_+_8433172_8446622_RNA45SN1_45S_50ntFlanks@12785.12805.21|CCCCGCGCGCGCGGGAGGGCG</td><td>>hg38_21_+_8433172_8446622_RNA45SN1_45S_50ntFlanks</td><td>12785</td><td>12805</td><td>21</td><td>CCCCGCGCGCGCGGGAGGGCG</td><td>5668.3</td><td>-16.3</td><td>1.5</td><td>-10.9</td><td>1.4E-27</td><td>6.3E-26</td><td>MB231/MB468</td><td>MITO</td></tr>
<tr><td>>hg38_21_+_8433172_8446622_RNA45SN1_45S_50ntFlanks@12457.12486.30|CAAGCGTTGGATTGTTCACCCACTAATAGG</td><td>>hg38_21_+_8433172_8446622_RNA45SN1_45S_50ntFlanks</td><td>12457</td><td>12486</td><td>30</td><td>CAAGCGTTGGATTGTTCACCCACTAATAGG</td><td>1047.2</td><td>-14.5</td><td>1.3</td><td>-10.7</td><td>7.9E-27</td><td>6.4E-26</td><td>BT20/MB468</td><td>NUC</td></tr>
<tr><td>>hg38_21_+_8433172_8446622_RNA45SN1_45S_50ntFlanks@10348.10368.21|AAACGAGAACTTTGAAGGCCG</td><td>>hg38_21_+_8433172_8446622_RNA45SN1_45S_50ntFlanks</td><td>10348</td><td>10368</td><td>21</td><td>AAACGAGAACTTTGAAGGCCG</td><td>2839.0</td><td>-7.5</td><td>0.7</td><td>-10.7</td><td>8.0E-27</td><td>6.4E-26</td><td>BT20/MB468</td><td>NUC</td></tr>
<tr><td>>hg38_21_+_8433172_8446622_RNA45SN1_45S_50ntFlanks@12624.12657.34|CCAATGGGGCGAAGCTACCATCTGTGGGATTATG</td><td>>hg38_21_+_8433172_8446622_RNA45SN1_45S_50ntFlanks</td><td>12624</td><td>12657</td><td>34</td><td>CCAATGGGGCGAAGCTACCATCTGTGGGATTATG</td><td>1215.4</td><td>-14.7</td><td>1.4</td><td>-10.7</td><td>8.0E-27</td><td>6.4E-26</td><td>BT20/MB468</td><td>NUC</td></tr>
<tr><td>>MI0000239|hsa-mir-197&WithFlank&1|+|109598887|109598973@54.72.19&[MIMAT0000227&hsa-miR-197-3p&offsets|0|-3;m-99&1|+|109598940|109598961&offsets|0|-3]|TTCACCACCTTCTCCACCC</td><td>>MI0000239|hsa-mir-197&WithFlank&1|+|109598887|109598973</td><td>54</td><td>72</td><td>19</td><td>TTCACCACCTTCTCCACCC</td><td>490.8</td><td>13.4</td><td>1.2</td><td>10.8</td><td>3.2E-27</td><td>6.4E-26</td><td>BT20/MB468</td><td>MITO</td></tr>
<tr><td>>hg38_21_+_8433172_8446622_RNA45SN1_45S_50ntFlanks@12417.12441.25|TCGATGTCGGCTCTTCCTATCATTG</td><td>>hg38_21_+_8433172_8446622_RNA45SN1_45S_50ntFlanks</td><td>12417</td><td>12441</td><td>25</td><td>TCGATGTCGGCTCTTCCTATCATTG</td><td>1615.2</td><td>-3.4</td><td>0.3</td><td>-10.8</td><td>4.2E-27</td><td>6.5E-26</td><td>BT20/MB468</td><td>CYTO</td></tr>
<tr><td>>hg38_21_+_8433172_8446622_RNA45SN1_45S_50ntFlanks@3985.4009.25|CGGCTTTGGTGACTCTAGATAACCT</td><td>>hg38_21_+_8433172_8446622_RNA45SN1_45S_50ntFlanks</td><td>3985</td><td>4009</td><td>25</td><td>CGGCTTTGGTGACTCTAGATAACCT</td><td>478.0</td><td>-13.3</td><td>1.2</td><td>-10.7</td><td>8.1E-27</td><td>6.5E-26</td><td>BT20/MB468</td><td>NUC</td></tr>
<tr><td>>MI0000299|hsa-mir-222&WithFlank&X|-|45747009|45747130@75.98.24(+1C)&[MIMAT0000279&hsa-miR-222-3p&offsets|0|+3(+1C);m-140&X|-|45747033|45747056&offsets|0|0(+1C)]|AGCTACATCTGGCTACTGGGTCTCC</td><td>>MI0000299|hsa-mir-222&WithFlank&X|-|45747009|45747130</td><td>75</td><td>98</td><td>24(+1C)</td><td>AGCTACATCTGGCTACTGGGTCTCC</td><td>10348.5</td><td>8.0</td><td>0.7</td><td>11.0</td><td>6.1E-28</td><td>6.5E-26</td><td>MB231/MB468</td><td>CYTO</td></tr>
<tr><td>>hg38_21_+_8433172_8446622_RNA45SN1_45S_50ntFlanks@10806.10826.21|AACAATGTAGGTAAGGGAAGT</td><td>>hg38_21_+_8433172_8446622_RNA45SN1_45S_50ntFlanks</td><td>10806</td><td>10826</td><td>21</td><td>AACAATGTAGGTAAGGGAAGT</td><td>1132.1</td><td>-14.6</td><td>1.4</td><td>-10.7</td><td>8.2E-27</td><td>6.5E-26</td><td>BT20/MB468</td><td>NUC</td></tr>
<tr><td>>hg38_21_+_8433172_8446622_RNA45SN1_45S_50ntFlanks@4385.4423.39|TTGGATCTTGGGAGCGGGCGGGCGGTCCGCCGCGAGGCG</td><td>>hg38_21_+_8433172_8446622_RNA45SN1_45S_50ntFlanks</td><td>4385</td><td>4423</td><td>39</td><td>TTGGATCTTGGGAGCGGGCGGGCGGTCCGCCGCGAGGCG</td><td>605.6</td><td>-13.7</td><td>1.3</td><td>-10.7</td><td>8.2E-27</td><td>6.5E-26</td><td>BT20/MB468</td><td>NUC</td></tr>
<tr><td>>hg38_21_+_8433172_8446622_RNA45SN1_45S_50ntFlanks@11613.11640.28|AATTAAAACAAAGCATCGCGAAGGCCCG</td><td>>hg38_21_+_8433172_8446622_RNA45SN1_45S_50ntFlanks</td><td>11613</td><td>11640</td><td>28</td><td>AATTAAAACAAAGCATCGCGAAGGCCCG</td><td>4383.5</td><td>-15.9</td><td>1.5</td><td>-10.9</td><td>1.4E-27</td><td>6.5E-26</td><td>MB231/MB468</td><td>MITO</td></tr>
<tr><td>>MI0000264|hsa-mir-7-2&WithFlank&15|+|88611819|88611940@38.59.22(+1C);MI0000265|hsa-mir-7-3&WithFlank&19|+|4770664|4770785@37.58.22(+1C);MI0000263|hsa-mir-7-1&WithFlank&9|-|83969742|83969863@30.51.22(+1C)&[MIMAT0000252&hsa-miR-7-5p&offsets|0|-2(+1C);m-400&15|+|88611856|88611879&offsets|0|-2(+1C)];[MIMAT0000252_1&hsa-miR-7-5p&offsets|0|-2(+1C);m-402&19|+|4770700|4770723&offsets|0|-2(+1C)];[MIMAT0000252_2&hsa-miR-7-5p&offsets|0|-2(+1C);m-401&9|-|83969811|83969834&offsets|0|-2(+1C)]|TGGAAGACTAGTGATTTTGTTGC</td><td>>MI0000264|hsa-mir-7-2&WithFlank&15|+|88611819|88611940</td><td>38</td><td>59</td><td>22(+1C)</td><td>TGGAAGACTAGTGATTTTGTTGC</td><td>837.8</td><td>7.8</td><td>0.7</td><td>10.8</td><td>3.3E-27</td><td>6.6E-26</td><td>BT20/MB468</td><td>MITO</td></tr>
<tr><td>>hg38_21_+_8433172_8446622_RNA45SN1_45S_50ntFlanks@11208.11225.18|GCGGCGACTCTGGACGCG</td><td>>hg38_21_+_8433172_8446622_RNA45SN1_45S_50ntFlanks</td><td>11208</td><td>11225</td><td>18</td><td>GCGGCGACTCTGGACGCG</td><td>2735.6</td><td>-15.2</td><td>1.4</td><td>-10.9</td><td>1.5E-27</td><td>6.7E-26</td><td>MB231/MB468</td><td>MITO</td></tr>
<tr><td>>hg38_21_+_8433172_8446622_RNA45SN1_45S_50ntFlanks@482.506.25|ATCGATGTGGTGACGTCGTGCTCTC</td><td>>hg38_21_+_8433172_8446622_RNA45SN1_45S_50ntFlanks</td><td>482</td><td>506</td><td>25</td><td>ATCGATGTGGTGACGTCGTGCTCTC</td><td>1650.5</td><td>-15.1</td><td>1.4</td><td>-10.7</td><td>8.3E-27</td><td>6.7E-26</td><td>BT20/MB468</td><td>NUC</td></tr>
<tr><td>>hg38_21_+_8433172_8446622_RNA45SN1_45S_50ntFlanks@10391.10426.36|AACAGCAGTTGAACATGGGTCAGTCGGTCCTGAGAG</td><td>>hg38_21_+_8433172_8446622_RNA45SN1_45S_50ntFlanks</td><td>10391</td><td>10426</td><td>36</td><td>AACAGCAGTTGAACATGGGTCAGTCGGTCCTGAGAG</td><td>1060.4</td><td>-14.5</td><td>1.4</td><td>-10.7</td><td>8.3E-27</td><td>6.7E-26</td><td>BT20/MB468</td><td>NUC</td></tr>
<tr><td>>hg38_21_+_8433172_8446622_RNA45SN1_45S_50ntFlanks@8357.8379.23|AAAAGAACTTTGAAGAGAGAGTT</td><td>>hg38_21_+_8433172_8446622_RNA45SN1_45S_50ntFlanks</td><td>8357</td><td>8379</td><td>23</td><td>AAAAGAACTTTGAAGAGAGAGTT</td><td>29118.1</td><td>-9.5</td><td>0.9</td><td>-11.0</td><td>6.4E-28</td><td>6.7E-26</td><td>MB231/MB468</td><td>CYTO</td></tr>
<tr><td>>hg38_21_+_8433172_8446622_RNA45SN1_45S_50ntFlanks@485.506.22|GATGTGGTGACGTCGTGCTCTC</td><td>>hg38_21_+_8433172_8446622_RNA45SN1_45S_50ntFlanks</td><td>485</td><td>506</td><td>22</td><td>GATGTGGTGACGTCGTGCTCTC</td><td>2263.3</td><td>15.4</td><td>1.4</td><td>10.8</td><td>4.3E-27</td><td>6.7E-26</td><td>BT20/MB468</td><td>CYTO</td></tr>
<tr><td>>hg38_21_+_8433172_8446622_RNA45SN1_45S_50ntFlanks@12477.12496.20|CACTAATAGGGAACGTGAGC</td><td>>hg38_21_+_8433172_8446622_RNA45SN1_45S_50ntFlanks</td><td>12477</td><td>12496</td><td>20</td><td>CACTAATAGGGAACGTGAGC</td><td>767.0</td><td>-4.3</td><td>0.4</td><td>-10.7</td><td>8.5E-27</td><td>6.8E-26</td><td>BT20/MB468</td><td>NUC</td></tr>
<tr><td>>hg38_21_+_8433172_8446622_RNA45SN1_45S_50ntFlanks@3999.4024.26|CTAGATAACCTCGGGCCGATCGCACG</td><td>>hg38_21_+_8433172_8446622_RNA45SN1_45S_50ntFlanks</td><td>3999</td><td>4024</td><td>26</td><td>CTAGATAACCTCGGGCCGATCGCACG</td><td>725.5</td><td>-13.9</td><td>1.3</td><td>-10.7</td><td>8.7E-27</td><td>6.9E-26</td><td>BT20/MB468</td><td>NUC</td></tr>
<tr><td>>hg38_21_+_8433172_8446622_RNA45SN1_45S_50ntFlanks@5488.5516.29|GCTGAGAAGACGGTCGAACTTGACTATCT</td><td>>hg38_21_+_8433172_8446622_RNA45SN1_45S_50ntFlanks</td><td>5488</td><td>5516</td><td>29</td><td>GCTGAGAAGACGGTCGAACTTGACTATCT</td><td>1259.4</td><td>7.7</td><td>0.7</td><td>10.8</td><td>3.5E-27</td><td>6.9E-26</td><td>BT20/MB468</td><td>MITO</td></tr>
</table>

| | | | | | | | | | | | | | |
|---|---|---|---|---|---|---|---|---|---|---|---|---|---|
| >hg38_21_+_8433172_8446622_RNA45SN1_45S_50ntFlanks@4122.4144.23\|AGGGTTCGATTCCGGAGAGGGAG | >hg38_21_+_8433172_8446622_RNA45SN1_45S_50ntFlanks | 4122 | 4144 | 23 | AGGGTTCGATTCCGGAGAGGGAG | 537.5 | -13.5 | 1.3 | -10.7 | 8.7E-27 | 6.9E-26 | BT20/MB468 | NUC |
| >MI0000266\|hsa-mir-10a&WithFlank&17\|-\|48579832\|48579953@28.48.21(+1C)&[MIMAT0000253&hsa-miR-10a-5p&offsets\|0\|-2(+1C);m-18&17\|-\|48579905\|48579926&offsets\|0\|-1(+1C)]\|TACCCTGTAGATCCGAATTTGC | >MI0000266\|hsa-mir-10a&WithFlank&17\|-\|48579832\|48579953 | 28 | 48 | 21(+1C) | TACCCTGTAGATCCGAATTTGC | 908.3 | 13.7 | 1.3 | 10.8 | 2.2E-27 | 7.0E-26 | MB231/MB468 | NUC |
| >am_trna119_LysCTT_1_-_145395522_145395594@14.31.18,trna11_LysCTT_5_-_180648979_180649051@14.31.18,trna128_LysCTT_1_-_55423542_55423614@14.31.18,trna13_LysCTT_6_+_26556774_26556846@14.31.18,trna32_LysCTT_16_-_3207406_3207478@14.31.18,trna6_LysCTT_19_-_52425396_52425468@14.31.18,trna7_LysCTT_16_+_3225692_3225764@14.31.18,trna9_LysCTT_5_+_180634755_180634827@14.31.18\|AGTCGGTAGAGCATGAGA | >am_trna119_LysCTT_1_-_145395522_145395594 | 14 | 31 | 18 | AGTCGGTAGAGCATGAGA | 1245.2 | -14.7 | 1.4 | -10.7 | 9.0E-27 | 7.2E-26 | BT20/MB468 | NUC |
| >MI0006444\|hsa-mir-1307&WithFlank&10\|-\|103394247\|103394407@86.107.22(+1A)&[MIMAT0005951&hsa-miR-1307-3p&offsets\|0\|0(+1A);m-66&10\|-\|103394300\|103394322&offsets\|0\|-1(+1A)]\|ACTCGGCGTGGCGTCGGTCGTGA | >MI0006444\|hsa-mir-1307&WithFlank&10\|-\|103394247\|103394407 | 86 | 107 | 22(+1A) | ACTCGGCGTGGCGTCGGTCGTGA | 931.9 | 7.4 | 0.7 | 10.8 | 3.7E-27 | 7.3E-26 | BT20/MB468 | MITO |
| >MI0000077\|hsa-mir-21&WithFlank&17\|+\|59841260\|59841343@14.35.22(+1U)&[MIMAT0000076&hsa-miR-21-5p&offsets\|0\|0(+1U);m-1&17\|+\|59841273\|59841294&offsets\|0\|0(+1U)]\|TAGCTTATCAGACTGATGTTGAT | >MI0000077\|hsa-mir-21&WithFlank&17\|+\|59841260\|59841343 | 14 | 35 | 22(+1U) | TAGCTTATCAGACTGATGTTGAT | 67996.8 | 7.0 | 0.6 | 10.8 | 2.3E-27 | 7.4E-26 | MB231/MB468 | NUC |
| >MI0000087\|hsa-mir-29a&WithFlank&7\|-\|130876741\|130876816@47.67.21(+1C)&[MIMAT0000086&hsa-miR-29a-3p&offsets\|-1\|-2(+1C);m-26&7\|-\|130876749\|130876769&offsets\|-1\|-1(+1C)]\|CTAGCACCATCTGAAATCGGTC | >MI0000087\|hsa-mir-29a&WithFlank&7\|-\|130876741\|130876816 | 47 | 67 | 21(+1C) | CTAGCACCATCTGAAATCGGTC | 1372.8 | 8.3 | 0.8 | 10.8 | 2.4E-27 | 7.4E-26 | MB231/MB468 | NUC |
| >hg38_21_+_8433172_8446622_RNA45SN1_45S_50ntFlanks@8115.8143.29\|CGCGGGACATGTGGCGTACGGAAGACCCG | >hg38_21_+_8433172_8446622_RNA45SN1_45S_50ntFlanks | 8115 | 8143 | 29 | CGCGGGACATGTGGCGTACGGAAGACCCG | 19884.7 | -13.7 | 1.3 | -10.9 | 1.0E-27 | 7.9E-26 | MB231/MB468 | MP |
| >hg38_21_+_8433172_8446622_RNA45SN1_45S_50ntFlanks@7980.8008.29\|CCTCAGATCAGACGTGGCGACCCGCTGAA | >hg38_21_+_8433172_8446622_RNA45SN1_45S_50ntFlanks | 7980 | 8008 | 29 | CCTCAGATCAGACGTGGCGACCCGCTGAA | 2180.0 | -9.4 | 0.9 | -10.9 | 1.0E-27 | 8.0E-26 | MB231/MB468 | MP |
| >MI0000299\|hsa-mir-222&WithFlank&X\|-\|45747009\|45747130@75.97.23(+1U)&[MIMAT0000279&hsa-miR-222-3p&offsets\|0\|+2(+1U);m-140&X\|-\|45747033\|45747056&offsets\|0\|-1(+1U)]\|AGCTACATCTGGCTACTGGGTCTT | >MI0000299\|hsa-mir-222&WithFlank&X\|-\|45747009\|45747130 | 75 | 97 | 23(+1U) | AGCTACATCTGGCTACTGGGTCTT | 3856.9 | 8.1 | 0.7 | 10.9 | 7.7E-28 | 8.0E-26 | MB231/MB468 | CYTO |
| >MI0000746\|hsa-mir-99b&WithFlank&19\|+\|51692606\|51692687@51.73.23(+1U)&[MIMAT0004678&hsa-miR-99b-3p&offsets\|0\|+1(+1U);m-151&19\|+\|51692656\|51692678&offsets\|0\|0(+1U)]\|CAAGCTCGTGTCTGTGGGTCCGTT | >MI0000746\|hsa-mir-99b&WithFlank&19\|+\|51692606\|51692687 | 51 | 73 | 23(+1U) | CAAGCTCGTGTCTGTGGGTCCGTT | 674.1 | 8.1 | 0.7 | 10.8 | 5.2E-27 | 8.0E-26 | BT20/MB468 | CYTO |
| >MI0000446\|hsa-mir-125b-1&WithFlank&11\|-\|122099751\|122099850@21.41.21(+1U);MI0000470\|hsa-mir-125b-2&WithFlank&21\|+\|16590231\|16590331@23.43.21(+1U)&[MIMAT0000423&hsa-miR-125b-5p&offsets\|0\|-1(+1U);m-67&11\|-\|122099809\|122099830&offsets\|0\|-1(+1U)];[MIMAT0000423_1&hsa-miR-125b-5p&offsets\|0\|-1(+1U);m-68&21\|+\|16590253\|16590274&offsets\|0\|-1(+1U)]\|TCCCTGAGACCCTAACTTGTGT | >MI0000446\|hsa-mir-125b-1&WithFlank&11\|-\|122099751\|122099850 | 21 | 41 | 21(+1U) | TCCCTGAGACCCTAACTTGTGT | 968.4 | 7.7 | 0.7 | 10.8 | 2.6E-27 | 8.1E-26 | MB231/MB468 | NUC |
| >MI0000440\|hsa-mir-27b&WithFlank&9\|+\|95085439\|95085547@67.87.21(+2U)&[MIMAT0000419&hsa-miR-27b-3p&offsets\|0\|0(+2U);m-39&9\|+\|95085505\|95085525&offsets\|0\|0(+2U)]\|TTCACAGTGGCTAAGTTCTGCTT | >MI0000440\|hsa-mir-27b&WithFlank&9\|+\|95085439\|95085547 | 67 | 87 | 21(+2U) | TTCACAGTGGCTAAGTTCTGCTT | 357.4 | 6.6 | 0.6 | 10.8 | 2.6E-27 | 8.2E-26 | MB231/MB468 | NUC |
| >hg38_21_+_8433172_8446622_RNA45SN1_45S_50ntFlanks@11621.11640.20\|CAAAGCATCGCGAAGGCCCG | >hg38_21_+_8433172_8446622_RNA45SN1_45S_50ntFlanks | 11621 | 11640 | 20 | CAAAGCATCGCGAAGGCCCG | 3565.8 | -15.7 | 1.4 | -10.9 | 1.1E-27 | 8.2E-26 | MB231/MB468 | MP |
| >MI0000440\|hsa-mir-27b&WithFlank&9\|+\|95085439\|95085547@67.86.20(+1A)&[MIMAT0000419&hsa-miR-27b-3p&offsets\|0\|-1(+1A);m-39&9\|+\|95085505\|95085525&offsets\|0\|-1(+1A)]\|TTCACAGTGGCTAAGTTCTGA | >MI0000440\|hsa-mir-27b&WithFlank&9\|+\|95085439\|95085547 | 67 | 86 | 20(+1A) | TTCACAGTGGCTAAGTTCTGA | 1881.7 | 7.1 | 0.7 | 10.8 | 4.3E-27 | 8.5E-26 | BT20/MB468 | MITO |
| >hg38_21_+_8433172_8446622_RNA45SN1_45S_50ntFlanks@10845.10862.18\|ACTTCGGGATAAGGATTG | >hg38_21_+_8433172_8446622_RNA45SN1_45S_50ntFlanks | 10845 | 10862 | 18 | ACTTCGGGATAAGGATTG | 1109.8 | -14.5 | 1.4 | -10.7 | 1.1E-26 | 8.5E-26 | BT20/MB468 | NUC |
| >hg38_21_+_8433172_8446622_RNA45SN1_45S_50ntFlanks@11873.11893.21\|AAGAAGACCCTGTTGAGCTTG | >hg38_21_+_8433172_8446622_RNA45SN1_45S_50ntFlanks | 11873 | 11893 | 21 | AAGAAGACCCTGTTGAGCTTG | 2991.5 | -15.3 | 1.4 | -10.9 | 1.9E-27 | 8.6E-26 | MB231/MB468 | MITO |
| >hg38_21_+_8433172_8446622_RNA45SN1_45S_50ntFlanks@12457.12475.19\|CAAGCGTTGGATTGTTCAC | >hg38_21_+_8433172_8446622_RNA45SN1_45S_50ntFlanks | 12457 | 12475 | 19 | CAAGCGTTGGATTGTTCAC | 816.0 | -14.1 | 1.3 | -10.7 | 1.1E-26 | 8.6E-26 | BT20/MB468 | NUC |
| >MI0000065\|hsa-let-7d&WithFlank&9\|+\|94178828\|94178926@14.34.21(+1C)&[MIMAT0000065&hsa-let-7d-5p&offsets\|0\|-1(+1C);m-90&9\|+\|94178841\|94178862&offsets\|0\|-1(+1C)]\|AGAGGTAGTAGGTTGCATAGTC | >MI0000065\|hsa-let-7d&WithFlank&9\|+\|94178828\|94178926 | 14 | 34 | 21(+1C) | AGAGGTAGTAGGTTGCATAGTC | 3150.9 | 8.2 | 0.8 | 10.8 | 5.7E-27 | 8.7E-26 | BT20/MB468 | CYTO |
| >trna10_LysCTT_16_+_3241501_3241573@1.33.33,trna13_LysCTT_14_-_58706613_58706685@1.33.33,trna2_LysCTT_15_+_79152904_79152976@1.33.33\|GCCCGGCTAGCTCAGTCGGTAGAGCATGGGACT | >trna10_LysCTT_16_+_3241501_3241573 | 1 | 33 | 33 | GCCCGGCTAGCTCAGTCGGTAGAGCATGGGACT | 1205.9 | 7.5 | 0.7 | 10.8 | 5.7E-27 | 8.8E-26 | BT20/MB468 | CYTO |
| >hg38_21_+_8433172_8446622_RNA45SN1_45S_50ntFlanks@11581.11611.31\|AACTGGTGCGGACCAGGGGAATCCGACTGTT | >hg38_21_+_8433172_8446622_RNA45SN1_45S_50ntFlanks | 11581 | 11611 | 31 | AACTGGTGCGGACCAGGGGAATCCGACTGTT | 4071.3 | -6.6 | 0.6 | -10.9 | 2.0E-27 | 8.8E-26 | MB231/MB468 | MITO |
| >hg38_21_+_8433172_8446622_RNA45SN1_45S_50ntFlanks@12157.12179.23\|AGTTTGACTGGGGCGGTACACCT | >hg38_21_+_8433172_8446622_RNA45SN1_45S_50ntFlanks | 12157 | 12179 | 23 | AGTTTGACTGGGGCGGTACACCT | 2793.3 | -15.3 | 1.4 | -10.9 | 1.1E-27 | 8.8E-26 | MB231/MB468 | MP |
| >hg38_21_+_8433172_8446622_RNA45SN1_45S_50ntFlanks@5485.5510.26\|AGCGCTGAGAAGACGGTCGAACTTGA | >hg38_21_+_8433172_8446622_RNA45SN1_45S_50ntFlanks | 5485 | 5510 | 26 | AGCGCTGAGAAGACGGTCGAACTTGA | 1904.5 | -14.8 | 1.4 | -10.9 | 1.2E-27 | 8.8E-26 | MB231/MB468 | MP |
| >hg38_21_+_8433172_8446622_RNA45SN1_45S_50ntFlanks@10806.10823.18\|AACAATGTAGGTAAGGGA | >hg38_21_+_8433172_8446622_RNA45SN1_45S_50ntFlanks | 10806 | 10823 | 18 | AACAATGTAGGTAAGGGA | 1142.9 | -14.6 | 1.4 | -10.7 | 1.1E-26 | 8.9E-26 | BT20/MB468 | NUC |
| >hg38_21_+_8433172_8446622_RNA45SN1_45S_50ntFlanks@12892.12933.42\|CCCTCGCCCGTCACGCACCGCACGTTCGTGGGGAACCTGGCG | >hg38_21_+_8433172_8446622_RNA45SN1_45S_50ntFlanks | 12892 | 12933 | 42 | CCCTCGCCCGTCACGCACCGCACGTTCGTGGGGAACCTGGCG | 3346.0 | -15.5 | 1.4 | -10.8 | 2.0E-27 | 8.9E-26 | MB231/MB468 | MITO |
| >MI0006444\|hsa-mir-1307&WithFlank&10\|-\|103394247\|103394407@86.108.23&[MIMAT0005951&hsa-miR-1307-3p&offsets\|0\|+1;m-66&10\|-\|103394300\|103394322&offsets\|0\|0]\|ACTCGGCGTGGCGTCGGTCGTGG | >MI0006444\|hsa-mir-1307&WithFlank&10\|-\|103394247\|103394407 | 86 | 108 | 23 | ACTCGGCGTGGCGTCGGTCGTGG | 4230.7 | 7.2 | 0.7 | 10.8 | 4.6E-27 | 9.0E-26 | BT20/MB468 | MITO |
| >MI0000264\|hsa-mir-7-2&WithFlank&15\|+\|88611819\|88611940@38.61.24;MI0000265\|hsa-mir-7-3&WithFlank&19\|+\|4770664\|4770785@37.60.24;MI0000263\|hsa-mir-7-1&WithFlank&9\|-\|83969742\|83969863@30.53.24&[MIMAT0000252&hsa-miR-7-5p&offsets\|0\|0;m-400&15\|+\|88611856\|88611879&offsets\|0\|0];[MIMAT0000252_1&hsa-miR-7-5p&offsets\|0\|0;m-402&19\|+\|4770700\|4770723&offsets\|0\|0];[MIMAT0000252_2&hsa-miR-7-5p&offsets\|0\|0;m-401&9\|-\|83969811\|83969834&offsets\|0\|0]\|TGGAAGACTAGTGATTTTGTTGTT | >MI0000264\|hsa-mir-7-2&WithFlank&15\|+\|88611819\|88611940 | 38 | 61 | 24 | TGGAAGACTAGTGATTTTGTTGTT | 84616.2 | 7.7 | 0.7 | 10.8 | 4.7E-27 | 9.2E-26 | BT20/MB468 | MITO |
| >MI0000266\|hsa-mir-10a&WithFlank&17\|-\|48579832\|48579953@29.50.22(+1A)&[MIMAT0000253&hsa-miR-10a-5p&offsets\|+1\|0(+1A);m-18&17\|-\|48579905\|48579926&offsets\|+1\|+1(+1A)]\|ACCCTGTAGATCCGAATTTGTGA | >MI0000266\|hsa-mir-10a&WithFlank&17\|-\|48579832\|48579953 | 29 | 50 | 22(+1A) | ACCCTGTAGATCCGAATTTGTGA | 1344.3 | 14.9 | 1.4 | 10.8 | 4.7E-27 | 9.2E-26 | BT20/MB468 | MITO |
| >hg38_21_+_8433172_8446622_RNA45SN1_45S_50ntFlanks@11575.11592.18\|ACTTAGAACTGGTGCGGA | >hg38_21_+_8433172_8446622_RNA45SN1_45S_50ntFlanks | 11575 | 11592 | 18 | ACTTAGAACTGGTGCGGA | 4844.4 | -16.1 | 1.5 | -10.9 | 1.2E-27 | 9.3E-26 | MB231/MB468 | MP |
| >hg38_21_+_8433172_8446622_RNA45SN1_45S_50ntFlanks@10687.10709.23\|CGGCGTCCGGTGAGCTCTCGCTG | >hg38_21_+_8433172_8446622_RNA45SN1_45S_50ntFlanks | 10687 | 10709 | 23 | CGGCGTCCGGTGAGCTCTCGCTG | 1938.8 | -14.8 | 1.4 | -10.9 | 1.2E-27 | 9.3E-26 | MB231/MB468 | MP |
| >hg38_21_+_8433172_8446622_RNA45SN1_45S_50ntFlanks@10736.10755.20\|GTAAATCTCGCGCCGGGCCG | >hg38_21_+_8433172_8446622_RNA45SN1_45S_50ntFlanks | 10736 | 10755 | 20 | GTAAATCTCGCGCCGGGCCG | 1329.7 | -14.8 | 1.4 | -10.7 | 1.2E-26 | 9.4E-26 | BT20/MB468 | NUC |
| >hg38_21_+_8433172_8446622_RNA45SN1_45S_50ntFlanks@12635.12659.25\|AAGCTACCATCTGTGGGATTATGAC | >hg38_21_+_8433172_8446622_RNA45SN1_45S_50ntFlanks | 12635 | 12659 | 25 | AAGCTACCATCTGTGGGATTATGAC | 6762.0 | -16.6 | 1.5 | -10.9 | 1.3E-27 | 9.5E-26 | MB231/MB468 | MP |
| >trna119_LysCTT_1_-_145395522_145395594@1.30.30,trna11_LysCTT_5_-_180648979_180649051@1.30.30,trna13_LysCTT_6_+_26556774_26556846@1.30.30,trna32_LysCTT_16_-_3207406_3207478@1.30.30,trna7_LysCTT_16_+_3225692_3225764@1.30.30,trna9_LysCTT_5_+_180634755_180634827@1.30.30\|GCCCGGCTAGCTCAGTCGGTAGAGCATGAG | >trna119_LysCTT_1_-_145395522_145395594 | 1 | 30 | 30 | GCCCGGCTAGCTCAGTCGGTAGAGCATGAG | 4260.0 | -7.3 | 0.7 | -10.9 | 1.3E-27 | 9.6E-26 | MB231/MB468 | MP |
| >hg38_21_+_8433172_8446622_RNA45SN1_45S_50ntFlanks@11641.11659.19\|CGGCGGGTGTTGACGCGAT | >hg38_21_+_8433172_8446622_RNA45SN1_45S_50ntFlanks | 11641 | 11659 | 19 | CGGCGGGTGTTGACGCGAT | 3673.3 | -15.7 | 1.4 | -10.9 | 1.3E-27 | 9.6E-26 | MB231/MB468 | MP |
| >hg38_21_+_8433172_8446622_RNA45SN1_45S_50ntFlanks@12159.12182.24\|TTTGACTGGGGCGGTACACCTGTC | >hg38_21_+_8433172_8446622_RNA45SN1_45S_50ntFlanks | 12159 | 12182 | 24 | TTTGACTGGGGCGGTACACCTGTC | 7017.4 | -16.6 | 1.5 | -10.9 | 1.3E-27 | 9.6E-26 | MB231/MB468 | MP |
| >hg38_21_+_8433172_8446622_RNA45SN1_45S_50ntFlanks@12456.12484.29\|CCAAGCGTTGGATTGTTCACCCACTAATA | >hg38_21_+_8433172_8446622_RNA45SN1_45S_50ntFlanks | 12456 | 12484 | 29 | CCAAGCGTTGGATTGTTCACCCACTAATA | 5313.1 | -7.3 | 0.7 | -10.8 | 4.9E-27 | 9.6E-26 | BT20/MB468 | MITO |
| >hg38_21_+_8433172_8446622_RNA45SN1_45S_50ntFlanks@11517.11539.23\|CCGCCCCCGGGGCCGCGGTTCCG | >hg38_21_+_8433172_8446622_RNA45SN1_45S_50ntFlanks | 11517 | 11539 | 23 | CCGCCCCCGGGGCCGCGGTTCCG | 1213.9 | -14.7 | 1.4 | -10.7 | 1.3E-26 | 9.9E-26 | BT20/MB468 | NUC |
| >hg38_21_+_8433172_8446622_RNA45SN1_45S_50ntFlanks@8117.8145.29\|CGGGACATGTGGCGTACGGAAGACCCGCT | >hg38_21_+_8433172_8446622_RNA45SN1_45S_50ntFlanks | 8117 | 8145 | 29 | CGGGACATGTGGCGTACGGAAGACCCGCT | 439.2 | 7.1 | 0.7 | 10.8 | 5.3E-27 | 1.0E-25 | BT20/MB468 | MITO |
| >MI0000285\|hsa-mir-205&WithFlank&1\|+\|209432127\|209432248@40.60.21(+1A)&[MIMAT0000266&hsa-miR-205-5p&offsets\|0\|-1(+1A);m-351&1\|+\|209432166\|209432187&offsets\|0\|-1(+1A)]\|TCCTTCATTCCACCGGAGTCTA | >MI0000285\|hsa-mir-205&WithFlank&1\|+\|209432127\|209432248 | 40 | 60 | 21(+1A) | TCCTTCATTCCACCGGAGTCTA | 8267.4 | 12.4 | 1.1 | 11.2 | 5.7E-29 | 1.0E-25 | BT20/MB231 | CYTO |
| >hg38_21_+_8433172_8446622_RNA45SN1_45S_50ntFlanks@12405.12426.22\|CTTTTTGATCCTTCGATGTCGG | >hg38_21_+_8433172_8446622_RNA45SN1_45S_50ntFlanks | 12405 | 12426 | 22 | CTTTTTGATCCTTCGATGTCGG | 3809.3 | -15.8 | 1.4 | -10.9 | 1.4E-27 | 1.1E-25 | MB231/MB468 | MP |

| | | | | | | | | | | | | | |
|---|---|---|---|---|---|---|---|---|---|---|---|---|---|
| >hg38_21_+_8433172_8446622_RNA45SN1_45S_50ntFlanks@12991.13013.23\|CTCCCTCGCTGCGATCTATTGAA | >hg38_21_+_8433172_8446622_RNA45SN1_45S_50ntFlanks | 12991 | 13013 | 23 | CTCCCTCGCTGCGATCTATTGAA | 13194.2 | -11.0 | 1.0 | -10.8 | 2.4E-27 | 1.1E-25 | MB231/MB468 | MITO |
| >hg38_21_+_8433172_8446622_RNA45SN1_45S_50ntFlanks@4731.4758.28\|AAGAACGAAAGTCGGAGGTTCGAAGACG | >hg38_21_+_8433172_8446622_RNA45SN1_45S_50ntFlanks | 4731 | 4758 | 28 | AAGAACGAAAGTCGGAGGTTCGAAGACG | 16585.1 | -7.4 | 0.7 | -10.7 | 1.4E-26 | 1.1E-25 | BT20/MB468 | NUC |
| >MI0000061\|hsa-let-7a-2&WithFlank&11\|-\|122146516\|122146599@11.32.22(+1A);MI0000062\|hsa-let-7a-3&WithFlank&22\|+\|46112743\|46112828@10.31.22(+1A);MI0000060\|hsa-let-7a-1&WithFlank&9\|+\|94175951\|94176042@12.33.22(+1A)&[MIMAT0000062&hsa-let-7a-5p&offsets\|0\|0(+1A);m-6&11\|-\|122146568\|122146589&offsets\|0\|0(+1A)];[MIMAT0000062_1&hsa-let-7a-5p&offsets\|0\|0(+1A);m-5&22\|+\|46112752\|46112773&offsets\|0\|0(+1A)];[MIMAT0000062_2&hsa-let-7a-5p&offsets\|0\|0(+1A);m-4&9\|+\|94175962\|94175983&offsets\|0\|0(+1A)]\|TGAGGTAGTAGGTTGTATAGTTA | >MI0000061\|hsa-let-7a-2&WithFlank&11\|-\|122146516\|122146599 | 11 | 32 | 22(+1A) | TGAGGTAGTAGGTTGTATAGTTA | 893.9 | 6.7 | 0.6 | 10.8 | 5.6E-27 | 1.1E-25 | BT20/MB468 | MITO |
| >hg38_21_+_8433172_8446622_RNA45SN1_45S_50ntFlanks@12957.12991.35\|CTTCTGGGTCGGGGTTTCGTACGTAGCAGAGCAGC | >hg38_21_+_8433172_8446622_RNA45SN1_45S_50ntFlanks | 12957 | 12991 | 35 | CTTCTGGGTCGGGGTTTCGTACGTAGCAGAGCAGC | 811.6 | -14.1 | 1.3 | -10.7 | 1.4E-26 | 1.1E-25 | BT20/MB468 | NUC |
| >hg38_21_+_8433172_8446622_RNA45SN1_45S_50ntFlanks@4760.4782.23\|TCAGATACCGTCGTAGTTCCGAC | >hg38_21_+_8433172_8446622_RNA45SN1_45S_50ntFlanks | 4760 | 4782 | 23 | TCAGATACCGTCGTAGTTCCGAC | 995.3 | -14.4 | 1.3 | -10.7 | 1.4E-26 | 1.1E-25 | BT20/MB468 | NUC |
| >hg38_21_+_8433172_8446622_RNA45SN1_45S_50ntFlanks@12159.12179.21\|TTTGACTGGGGCGGTACACCT | >hg38_21_+_8433172_8446622_RNA45SN1_45S_50ntFlanks | 12159 | 12179 | 21 | TTTGACTGGGGCGGTACACCT | 4603.5 | -16.0 | 1.5 | -10.8 | 2.5E-27 | 1.1E-25 | MB231/MB468 | MITO |
| >hg38_21_+_8433172_8446622_RNA45SN1_45S_50ntFlanks@10351.10390.40\|CGAGAACTTTGAAGGCCGAAGTGGAGAAGGGTTCCATGTG | >hg38_21_+_8433172_8446622_RNA45SN1_45S_50ntFlanks | 10351 | 10390 | 40 | CGAGAACTTTGAAGGCCGAAGTGGAGAAGGGTTCCATGTG | 10153.6 | -9.1 | 0.8 | -10.9 | 1.5E-27 | 1.1E-25 | MB231/MB468 | MP |
| >hg38_21_+_8433172_8446622_RNA45SN1_45S_50ntFlanks@13007.13028.22\|TATTGAAAGTCAGCCCTCGACA | >hg38_21_+_8433172_8446622_RNA45SN1_45S_50ntFlanks | 13007 | 13028 | 22 | TATTGAAAGTCAGCCCTCGACA | 663.5 | -13.8 | 1.3 | -10.7 | 1.4E-26 | 1.1E-25 | BT20/MB468 | NUC |
| >hg38_21_+_8433172_8446622_RNA45SN1_45S_50ntFlanks@11586.11605.20\|GTGCGGACCAGGGGAATCCG | >hg38_21_+_8433172_8446622_RNA45SN1_45S_50ntFlanks | 11586 | 11605 | 20 | GTGCGGACCAGGGGAATCCG | 559.7 | -13.6 | 1.3 | -10.7 | 1.4E-26 | 1.1E-25 | BT20/MB468 | NUC |
| >MI0000488\|hsa-mir-194-1&WithFlank&1\|-\|220118151\|220118247@21.42.22;MI0000732\|hsa-mir-194-2&WithFlank&11\|-\|64891349\|64891445@21.42.22&[MIMAT0000460&hsa-miR-194-5p&offsets\|0\|0;m-131&1\|-\|220118206\|220118227&offsets\|0\|0];[MIMAT0000460_1&hsa-miR-194-5p&offsets\|0\|0;m-126&11\|-\|64891404\|64891425&offsets\|0\|0]\|TGTAACAGCAACTCCATGTGGA | >MI0000488\|hsa-mir-194-1&WithFlank&1\|-\|220118151\|220118247 | 21 | 42 | 22 | TGTAACAGCAACTCCATGTGGA | 677.1 | 6.3 | 0.6 | 10.7 | 7.5E-27 | 1.1E-25 | BT20/MB468 | CYTO |
| >MI0000102\|hsa-mir-100&WithFlank&11\|-\|122152223\|122152314@19.38.20(+1A)&[MIMAT0000098&hsa-miR-100-5p&offsets\|0\|-2(+1A);m-27&11\|-\|122152276\|122152296&offsets\|0\|-1(+1A)]\|AACCCGTAGATCCGAACTTGA | >MI0000102\|hsa-mir-100&WithFlank&11\|-\|122152223\|122152314 | 19 | 38 | 20(+1A) | AACCCGTAGATCCGAACTTGA | 6631.5 | 15.9 | 1.5 | 10.9 | 1.6E-27 | 1.1E-25 | MB231/MB468 | MP |
| >hg38_21_+_8433172_8446622_RNA45SN1_45S_50ntFlanks@974.992.19\|CGCGTGGGTCCTGAGGGAG | >hg38_21_+_8433172_8446622_RNA45SN1_45S_50ntFlanks | 974 | 992 | 19 | CGCGTGGGTCCTGAGGGAG | 615.6 | -13.7 | 1.3 | -10.7 | 1.5E-26 | 1.2E-25 | BT20/MB468 | NUC |
| >hg38_21_+_8433172_8446622_RNA45SN1_45S_50ntFlanks@11575.11609.35\|ACTTAGAACTGGTGCGGACCAGGGGAATCCGACTG | >hg38_21_+_8433172_8446622_RNA45SN1_45S_50ntFlanks | 11575 | 11609 | 35 | ACTTAGAACTGGTGCGGACCAGGGGAATCCGACTG | 793.4 | -5.8 | 0.5 | -10.7 | 1.5E-26 | 1.2E-25 | BT20/MB468 | NUC |
| >MI0000650\|hsa-mir-200c&WithFlank&12\|+\|6963693\|6963772@50.71.22(+1U)&[MIMAT0000617&hsa-miR-200c-3p&offsets\|0\|-1(+1U);m-54&12\|+\|6963742\|6963764&offsets\|0\|-1(+1U)]\|TAATACTGCCGGGTAATGATGGT | >MI0000650\|hsa-mir-200c&WithFlank&12\|+\|6963693\|6963772 | 50 | 71 | 22(+1U) | TAATACTGCCGGGTAATGATGGT | 3421.6 | 13.1 | 1.2 | 11.0 | 3.7E-28 | 1.2E-25 | BT20/MB231 | NUC |
| >MI0000471\|hsa-mir-126&WithFlank&9\|+\|136670596\|136670692@58.79.22&[MIMAT0000445&hsa-miR-126-3p&offsets\|0\|0;m-40&9\|+\|136670653\|136670674&offsets\|0\|0]\|TCGTACCGTGAGTAATAATGCG | >MI0000471\|hsa-mir-126&WithFlank&9\|+\|136670596\|136670692 | 58 | 79 | 22 | TCGTACCGTGAGTAATAATGCG | 4879.7 | 10.9 | 1.0 | 10.7 | 6.3E-27 | 1.2E-25 | BT20/MB468 | MITO |
| >MI0000650\|hsa-mir-200c&WithFlank&12\|+\|6963693\|6963772@50.71.22(+1G)&[MIMAT0000617&hsa-miR-200c-3p&offsets\|0\|-1(+1G);m-54&12\|+\|6963742\|6963764&offsets\|0\|-1(+1G)]\|TAATACTGCCGGGTAATGATGGG | >MI0000650\|hsa-mir-200c&WithFlank&12\|+\|6963693\|6963772 | 50 | 71 | 22(+1G) | TAATACTGCCGGGTAATGATGGG | 2804.8 | 7.9 | 0.7 | 10.7 | 6.3E-27 | 1.2E-25 | BT20/MB468 | MITO |
| >MI0000077\|hsa-mir-21&WithFlank&17\|+\|59841260\|59841343@15.36.22&[MIMAT0000076&hsa-miR-21-5p&offsets\|+1\|+1;m-1&17\|+\|59841273\|59841294&offsets\|+1\|+1]\|AGCTTATCAGACTGATGTTGAC | >MI0000077\|hsa-mir-21&WithFlank&17\|+\|59841260\|59841343 | 15 | 36 | 22 | AGCTTATCAGACTGATGTTGAC | 1485.9 | 7.8 | 0.7 | 10.7 | 7.9E-27 | 1.2E-25 | BT20/MB468 | CYTO |
| >MI0000067\|hsa-let-7f-1&WithFlank&9\|+\|94176341\|94176439@13.33.21;MI0000068\|hsa-let-7f-2&WithFlank&X\|-\|53557186\|53557280@14.34.21&[MIMAT0000067&hsa-let-7f-5p&offsets\|0\|-1;m-20&9\|+\|94176353\|94176374&offsets\|0\|-1];[MIMAT0000067_1&hsa-let-7f-5p&offsets\|0\|-1;m-19&X\|-\|53557246\|53557267&offsets\|0\|-1]\|TGAGGTAGTAGATTGTATAGT | >MI0000067\|hsa-let-7f-1&WithFlank&9\|+\|94176341\|94176439 | 13 | 33 | 21 | TGAGGTAGTAGATTGTATAGT | 6348.6 | 7.0 | 0.7 | 10.7 | 6.3E-27 | 1.2E-25 | BT20/MB468 | MITO |
| >hg38_21_+_8433172_8446622_RNA45SN1_45S_50ntFlanks@10343.10362.20\|TATTCAAACGAGAACTTTGA | >hg38_21_+_8433172_8446622_RNA45SN1_45S_50ntFlanks | 10343 | 10362 | 20 | TATTCAAACGAGAACTTTGA | 626.4 | 7.2 | 0.7 | 10.7 | 6.4E-27 | 1.2E-25 | BT20/MB468 | MITO |
| >hg38_21_+_8433172_8446622_RNA45SN1_45S_50ntFlanks@11669.11687.19\|GCCCAGTGCTCTGAATGTC | >hg38_21_+_8433172_8446622_RNA45SN1_45S_50ntFlanks | 11669 | 11687 | 19 | GCCCAGTGCTCTGAATGTC | 382.4 | 5.0 | 0.5 | 10.7 | 6.4E-27 | 1.2E-25 | BT20/MB468 | MITO |
| >hg38_21_+_8433172_8446622_RNA45SN1_45S_50ntFlanks@11578.11605.28\|TAGAACTGGTGCGGACCAGGGGAATCCG | >hg38_21_+_8433172_8446622_RNA45SN1_45S_50ntFlanks | 11578 | 11605 | 28 | TAGAACTGGTGCGGACCAGGGGAATCCG | 556.8 | -4.1 | 0.4 | -10.7 | 8.1E-27 | 1.2E-25 | BT20/MB468 | CYTO |
| >hg38_21_+_8433172_8446622_RNA45SN1_45S_50ntFlanks@4045.4065.21\|CCATTCGAACGTCTGCCCTAT | >hg38_21_+_8433172_8446622_RNA45SN1_45S_50ntFlanks | 4045 | 4065 | 21 | CCATTCGAACGTCTGCCCTAT | 431.2 | -13.2 | 1.2 | -10.7 | 1.6E-26 | 1.3E-25 | BT20/MB468 | NUC |
| >MI0000456\|hsa-mir-140&WithFlank&16\|+\|69933075\|69933186@68.88.21&[MIMAT0004597&hsa-miR-140-3p&offsets\|0\|0;m-45&16\|+\|69933143\|69933164&offsets\|-1\|-2]\|TACCACAGGGTAGAACCACGG | >MI0000456\|hsa-mir-140&WithFlank&16\|+\|69933075\|69933186 | 68 | 88 | 21 | TACCACAGGGTAGAACCACGG | 565.2 | 13.4 | 1.2 | 10.7 | 8.3E-27 | 1.3E-25 | BT20/MB468 | CYTO |
| >hg38_21_+_8433172_8446622_RNA45SN1_45S_50ntFlanks@6646.6669.24\|TCGTACGACTCTTAGCGGTGGATC | >hg38_21_+_8433172_8446622_RNA45SN1_45S_50ntFlanks | 6646 | 6669 | 24 | TCGTACGACTCTTAGCGGTGGATC | 39541.8 | 4.9 | 0.5 | 10.7 | 8.4E-27 | 1.3E-25 | BT20/MB468 | CYTO |
| >hg38_21_+_8433172_8446622_RNA45SN1_45S_50ntFlanks@9339.9370.32\|CGCACGAAAGCCGCCGTGGCGCAATGAAGGTG | >hg38_21_+_8433172_8446622_RNA45SN1_45S_50ntFlanks | 9339 | 9370 | 32 | CGCACGAAAGCCGCCGTGGCGCAATGAAGGTG | 9731.5 | -17.1 | 1.6 | -10.9 | 1.8E-27 | 1.3E-25 | MB231/MB468 | MP |
| >hg38_21_+_8433172_8446622_RNA45SN1_45S_50ntFlanks@9545.9563.19\|GTGGAGGTCCGTAGCGGTC | >hg38_21_+_8433172_8446622_RNA45SN1_45S_50ntFlanks | 9545 | 9563 | 19 | GTGGAGGTCCGTAGCGGTC | 9882.6 | -8.8 | 0.8 | -10.8 | 4.2E-27 | 1.3E-25 | MB231/MB468 | NUC |
| >hg38_21_+_8433172_8446622_RNA45SN1_45S_50ntFlanks@10761.10785.25\|ATATCCGCAGCAGGTCTCCAAGGTG | >hg38_21_+_8433172_8446622_RNA45SN1_45S_50ntFlanks | 10761 | 10785 | 25 | ATATCCGCAGCAGGTCTCCAAGGTG | 433.5 | -13.2 | 1.2 | -10.7 | 1.7E-26 | 1.3E-25 | BT20/MB468 | NUC |
| >MI0000809\|hsa-mir-151a&WithFlank&8\|-\|140732558\|140732659@53.75.23&[MIMAT0000757&hsa-miR-151a-3p&offsets\|0\|+2;m-30&8\|-\|140732587\|140732607&offsets\|0\|+2]\|CTAGACTGAAGCTCCTTGAGGAC | >MI0000809\|hsa-mir-151a&WithFlank&8\|-\|140732558\|140732659 | 53 | 75 | 23 | CTAGACTGAAGCTCCTTGAGGAC | 431.9 | 13.0 | 1.2 | 10.7 | 8.7E-27 | 1.3E-25 | BT20/MB468 | CYTO |
| >hg38_21_+_8433172_8446622_RNA45SN1_45S_50ntFlanks@12416.12438.23\|TTCGATGTCGGCTCTTCCTATCA | >hg38_21_+_8433172_8446622_RNA45SN1_45S_50ntFlanks | 12416 | 12438 | 23 | TTCGATGTCGGCTCTTCCTATCA | 1127.1 | 5.6 | 0.5 | 10.7 | 8.7E-27 | 1.3E-25 | BT20/MB468 | CYTO |
| >hg38_21_+_8433172_8446622_RNA45SN1_45S_50ntFlanks@3837.3869.33\|CCTCTCCTACTTGGATAACTGTGGTAATTCTAG | >hg38_21_+_8433172_8446622_RNA45SN1_45S_50ntFlanks | 3837 | 3869 | 33 | CCTCTCCTACTTGGATAACTGTGGTAATTCTAG | 320.1 | -12.7 | 1.2 | -10.7 | 1.7E-26 | 1.3E-25 | BT20/MB468 | NUC |
| >hg38_21_+_8433172_8446622_RNA45SN1_45S_50ntFlanks@10794.10818.25\|CTGGCATGTTGGAACAATGTAGGTA | >hg38_21_+_8433172_8446622_RNA45SN1_45S_50ntFlanks | 10794 | 10818 | 25 | CTGGCATGTTGGAACAATGTAGGTA | 2894.0 | -15.4 | 1.4 | -10.9 | 1.9E-27 | 1.4E-25 | MB231/MB468 | MP |
| >hg38_21_+_8433172_8446622_RNA45SN1_45S_50ntFlanks@11847.11872.26\|AACGGGCTTGGCGGAATCAGCGGGGA | >hg38_21_+_8433172_8446622_RNA45SN1_45S_50ntFlanks | 11847 | 11872 | 26 | AACGGGCTTGGCGGAATCAGCGGGGA | 2324.3 | -15.0 | 1.4 | -10.9 | 1.9E-27 | 1.4E-25 | MB231/MB468 | MP |
| >hg38_21_+_8433172_8446622_RNA45SN1_45S_50ntFlanks@9572.9595.24\|CAAATCGGTCGTCCGACCTGGGTA | >hg38_21_+_8433172_8446622_RNA45SN1_45S_50ntFlanks | 9572 | 9595 | 24 | CAAATCGGTCGTCCGACCTGGGTA | 11474.3 | -17.4 | 1.6 | -10.9 | 1.9E-27 | 1.4E-25 | MB231/MB468 | MP |
| >hg38_21_+_8433172_8446622_RNA45SN1_45S_50ntFlanks@4558.4587.30\|AGCTAGGAATAATGGAATAGGACCGCGGTT | >hg38_21_+_8433172_8446622_RNA45SN1_45S_50ntFlanks | 4558 | 4587 | 30 | AGCTAGGAATAATGGAATAGGACCGCGGTT | 566.5 | -13.6 | 1.3 | -10.6 | 1.8E-26 | 1.4E-25 | BT20/MB468 | NUC |
| >hg38_21_+_8433172_8446622_RNA45SN1_45S_50ntFlanks@12992.13011.20\|TCCCTCGCTGCGATCTATTG | >hg38_21_+_8433172_8446622_RNA45SN1_45S_50ntFlanks | 12992 | 13011 | 20 | TCCCTCGCTGCGATCTATTG | 1273.2 | -14.7 | 1.4 | -10.6 | 1.8E-26 | 1.4E-25 | BT20/MB468 | NUC |
| >hg38_21_+_8433172_8446622_RNA45SN1_45S_50ntFlanks@12455.12487.33\|ACCAAGCGTTGGATTGTTCACCCACTAATAGGG | >hg38_21_+_8433172_8446622_RNA45SN1_45S_50ntFlanks | 12455 | 12487 | 33 | ACCAAGCGTTGGATTGTTCACCCACTAATAGGG | 649.1 | -13.8 | 1.3 | -10.6 | 1.8E-26 | 1.4E-25 | BT20/MB468 | NUC |
| >MI0005756\|hsa-mir-934&WithFlank&X\|+\|136550872\|136550966@21.41.21&[MIMAT0004977&hsa-miR-934&offsets\|0\|-1;m-943&X\|+\|136550892\|136550913&offsets\|0\|-1]\|TGTCTACTACTGGAGACACTG | >MI0005756\|hsa-mir-934&WithFlank&X\|+\|136550872\|136550966 | 21 | 41 | 21 | TGTCTACTACTGGAGACACTG | 502.3 | 4.7 | 0.4 | 10.6 | 1.9E-26 | 1.4E-25 | BT20/MB468 | NUC |
| >hg38_21_+_8433172_8446622_RNA45SN1_45S_50ntFlanks@12957.12982.26\|CTTCTGGGTCGGGGTTTCGTACGTAG | >hg38_21_+_8433172_8446622_RNA45SN1_45S_50ntFlanks | 12957 | 12982 | 26 | CTTCTGGGTCGGGGTTTCGTACGTAG | 3123.6 | -15.4 | 1.4 | -10.8 | 3.3E-27 | 1.5E-25 | MB231/MB468 | MITO |
| >MI0000285\|hsa-mir-205&WithFlank&1\|+\|209432127\|209432248@40.60.21&[MIMAT0000266&hsa-miR-205-5p&offsets\|0\|-1;m-351&1\|+\|209432166\|209432187&offsets\|0\|-1]\|TCCTTCATTCCACCGGAGTCT | >MI0000285\|hsa-mir-205&WithFlank&1\|+\|209432127\|209432248 | 40 | 60 | 21 | TCCTTCATTCCACCGGAGTCT | 6003.5 | 16.3 | 1.5 | 11.0 | 5.0E-28 | 1.5E-25 | BT20/MB231 | NUC |
| >MI0000266\|hsa-mir-10a&WithFlank&17\|-\|48579832\|48579953@28.48.21(+1G)&[MIMAT0000253&hsa-miR-10a-5p&offsets\|0\|-2(+1G);m-18&17\|-\|48579905\|48579926&offsets\|0\|-1(+1G)]\|TACCCTGTAGATCCGAATTTGG | >MI0000266\|hsa-mir-10a&WithFlank&17\|-\|48579832\|48579953 | 28 | 48 | 21(+1G) | TACCCTGTAGATCCGAATTTGG | 1250.4 | 14.5 | 1.4 | 10.7 | 9.7E-27 | 1.5E-25 | BT20/MB468 | CYTO |
| >hg38_21_+_8433172_8446622_RNA45SN1_45S_50ntFlanks@11795.11820.26\|AACGAGATTCCCACTGTCCCTACCTA | >hg38_21_+_8433172_8446622_RNA45SN1_45S_50ntFlanks | 11795 | 11820 | 26 | AACGAGATTCCCACTGTCCCTACCTA | 647.3 | -13.8 | 1.3 | -10.6 | 1.9E-26 | 1.5E-25 | BT20/MB468 | NUC |
| >MI0000542\|hsa-mir-320a&WithFlank&8\|-\|22244960\|22245043@49.69.21&[MIMAT0000510&hsa-miR-320a-3p&offsets\|+1\|0;m-60&8\|-\|22244975\|22244996&offsets\|+1\|0]\|AAAGCTGGGTTGAGAGGGCGA | >MI0000542\|hsa-mir-320a&WithFlank&8\|-\|22244960\|22245043 | 49 | 69 | 21 | AAAGCTGGGTTGAGAGGGCGA | 2097.0 | 11.3 | 1.1 | 10.7 | 9.9E-27 | 1.5E-25 | BT20/MB468 | CYTO |
| >hg38_21_+_8433172_8446622_RNA45SN1_45S_50ntFlanks@11739.11768.30\|TCTCTTAAGGTAGCCAAATGCCTCGTCATC | >hg38_21_+_8433172_8446622_RNA45SN1_45S_50ntFlanks | 11739 | 11768 | 30 | TCTCTTAAGGTAGCCAAATGCCTCGTCATC | 519.3 | -13.4 | 1.3 | -10.6 | 1.9E-26 | 1.5E-25 | BT20/MB468 | NUC |
| >hg38_21_+_8433172_8446622_RNA45SN1_45S_50ntFlanks@10402.10426.25\|AACATGGGTCAGTCGGTCCTGAGAG | >hg38_21_+_8433172_8446622_RNA45SN1_45S_50ntFlanks | 10402 | 10426 | 25 | AACATGGGTCAGTCGGTCCTGAGAG | 846.2 | -14.2 | 1.3 | -10.6 | 2.0E-26 | 1.5E-25 | BT20/MB468 | NUC |
| >MI0000077\|hsa-mir-21&WithFlank&17\|+\|59841260\|59841343@14.36.23&[MIMAT0000076&hsa-miR-21-5p&offsets\|0\|+1;m-1&17\|+\|59841273\|59841294&offsets\|0\|+1]\|TAGCTTATCAGACTGATGTTGAC | >MI0000077\|hsa-mir-21&WithFlank&17\|+\|59841260\|59841343 | 14 | 36 | 23 | TAGCTTATCAGACTGATGTTGAC | 378957.4 | 6.6 | 0.6 | 10.7 | 7.9E-27 | 1.5E-25 | BT20/MB468 | MITO |
| >hg38_21_+_8433172_8446622_RNA45SN1_45S_50ntFlanks@12159.12180.22\|TTTGACTGGGGCGGTACACCTG | >hg38_21_+_8433172_8446622_RNA45SN1_45S_50ntFlanks | 12159 | 12180 | 22 | TTTGACTGGGGCGGTACACCTG | 3934.2 | -15.7 | 1.5 | -10.8 | 3.5E-27 | 1.5E-25 | MB231/MB468 | MITO |

<table>
<tr><td>>MI0000478|hsa-mir-149&WithFlank&2|+|240455995|240456095@21.42.22(+1U)&[MIMAT0000450&hsa-miR-149-5p&offsets|0|-1(+1U);m-195&2|+|240456015|240456037&offsets|0|-1(+1U)]|TCTGGCTCCGTGTCTTCACTCCT</td><td>>MI0000478|hsa-mir-149&WithFlank&2|+|240455995|240456095</td><td>21</td><td>42</td><td>22(+1U)</td><td>TCTGGCTCCGTGTCTTCACTCCT</td><td>2703.1</td><td>15.7</td><td>1.5</td><td>10.7</td><td>1.0E-26</td><td>1.5E-25</td><td>BT20/MB468</td><td>CYTO</td></tr>
<tr><td>>hg38_21_+_8433172_8446622_RNA45SN1_45S_50ntFlanks@11233.11254.22|CCCTTCCCGTGGATCGCCCCAG</td><td>>hg38_21_+_8433172_8446622_RNA45SN1_45S_50ntFlanks</td><td>11233</td><td>11254</td><td>22</td><td>CCCTTCCCGTGGATCGCCCCAG</td><td>3282.2</td><td>-15.5</td><td>1.4</td><td>-10.8</td><td>3.6E-27</td><td>1.5E-25</td><td>MB231/MB468</td><td>MITO</td></tr>
<tr><td>>hg38_21_+_8433172_8446622_RNA45SN1_45S_50ntFlanks@11317.11342.26|ACCCCACGTCTCGTCGCGCGCGCGTC</td><td>>hg38_21_+_8433172_8446622_RNA45SN1_45S_50ntFlanks</td><td>11317</td><td>11342</td><td>26</td><td>ACCCCACGTCTCGTCGCGCGCGCGTC</td><td>4272.4</td><td>-15.8</td><td>1.5</td><td>-10.8</td><td>3.6E-27</td><td>1.5E-25</td><td>MB231/MB468</td><td>MITO</td></tr>
<tr><td>>hg38_21_+_8433172_8446622_RNA45SN1_45S_50ntFlanks@11586.11603.18|GTGCGGACCAGGGGAATC</td><td>>hg38_21_+_8433172_8446622_RNA45SN1_45S_50ntFlanks</td><td>11586</td><td>11603</td><td>18</td><td>GTGCGGACCAGGGGAATC</td><td>3174.6</td><td>-9.8</td><td>0.9</td><td>-10.8</td><td>5.0E-27</td><td>1.5E-25</td><td>MB231/MB468</td><td>NUC</td></tr>
<tr><td>>MI0000301|hsa-mir-224&WithFlank&X|-|151958572|151958664@14.34.21(+1A)&[MIMAT0000281&hsa-miR-224-5p&offsets|+1|-3(+1A);m-238&X|-|151958630|151958651&offsets|0|-1(+1A)]|CAAGTCACTAGTGGTTCCGTTA</td><td>>MI0000301|hsa-mir-224&WithFlank&X|-|151958572|151958664</td><td>14</td><td>34</td><td>21(+1A)</td><td>CAAGTCACTAGTGGTTCCGTTA</td><td>770.2</td><td>13.8</td><td>1.3</td><td>10.7</td><td>1.0E-26</td><td>1.5E-25</td><td>BT20/MB468</td><td>CYTO</td></tr>
<tr><td>>hg38_21_+_8433172_8446622_RNA45SN1_45S_50ntFlanks@4555.4573.19|CGCAGCTAGGAATAATGGA</td><td>>hg38_21_+_8433172_8446622_RNA45SN1_45S_50ntFlanks</td><td>4555</td><td>4573</td><td>19</td><td>CGCAGCTAGGAATAATGGA</td><td>1070.4</td><td>-14.5</td><td>1.4</td><td>-10.6</td><td>2.1E-26</td><td>1.6E-25</td><td>BT20/MB468</td><td>NUC</td></tr>
<tr><td>>MI0000101|hsa-mir-99a&WithFlank&21|+|16539083|16539175@19.38.20(+1C)&[MIMAT0000097&hsa-miR-99a-5p&offsets|0|-2(+1C);m-64&21|+|16539101|16539121&offsets|0|-1(+1C)]|AACCCGTAGATCCGATCTTGC</td><td>>MI0000101|hsa-mir-99a&WithFlank&21|+|16539083|16539175</td><td>19</td><td>38</td><td>20(+1C)</td><td>AACCCGTAGATCCGATCTTGC</td><td>452.4</td><td>-13.2</td><td>1.2</td><td>-10.6</td><td>2.1E-26</td><td>1.6E-25</td><td>BT20/MB468</td><td>NUC</td></tr>
<tr><td>>hg38_21_+_8433172_8446622_RNA45SN1_45S_50ntFlanks@10357.10388.32|CTTTGAAGGCCGAAGTGGAGAAGGGTTCCATG</td><td>>hg38_21_+_8433172_8446622_RNA45SN1_45S_50ntFlanks</td><td>10357</td><td>10388</td><td>32</td><td>CTTTGAAGGCCGAAGTGGAGAAGGGTTCCATG</td><td>1700.5</td><td>-15.2</td><td>1.4</td><td>-10.6</td><td>2.1E-26</td><td>1.6E-25</td><td>BT20/MB468</td><td>NUC</td></tr>
<tr><td>>hg38_21_+_8433172_8446622_RNA45SN1_45S_50ntFlanks@11821.11844.24|CTATCCAGCGAAACCACAGCCAAG</td><td>>hg38_21_+_8433172_8446622_RNA45SN1_45S_50ntFlanks</td><td>11821</td><td>11844</td><td>24</td><td>CTATCCAGCGAAACCACAGCCAAG</td><td>676.0</td><td>-13.8</td><td>1.3</td><td>-10.6</td><td>2.2E-26</td><td>1.7E-25</td><td>BT20/MB468</td><td>NUC</td></tr>
<tr><td>>hg38_21_+_8433172_8446622_RNA45SN1_45S_50ntFlanks@12417.12436.20|TCGATGTCGGCTCTTCCTAT</td><td>>hg38_21_+_8433172_8446622_RNA45SN1_45S_50ntFlanks</td><td>12417</td><td>12436</td><td>20</td><td>TCGATGTCGGCTCTTCCTAT</td><td>10162.6</td><td>-7.6</td><td>0.7</td><td>-10.8</td><td>3.9E-27</td><td>1.7E-25</td><td>MB231/MB468</td><td>MITO</td></tr>
<tr><td>>MI0000088|hsa-mir-30a&WithFlank&6|-|71403545|71403627@12.35.24(+1U)&[MIMAT0000087&hsa-miR-30a-5p&offsets|0|+2(+1U);m-15&6|-|71403595|71403616&offsets|0|+2(+1U)]|TGTAAACATCCTCGACTGGAAGCTT</td><td>>MI0000088|hsa-mir-30a&WithFlank&6|-|71403545|71403627</td><td>12</td><td>35</td><td>24(+1U)</td><td>TGTAAACATCCTCGACTGGAAGCTT</td><td>4811.3</td><td>8.5</td><td>0.8</td><td>10.8</td><td>5.5E-27</td><td>1.7E-25</td><td>MB231/MB468</td><td>NUC</td></tr>
<tr><td>>hg38_21_+_8433172_8446622_RNA45SN1_45S_50ntFlanks@12126.12146.21|CGACCCGCTCCGGGGACAGTG</td><td>>hg38_21_+_8433172_8446622_RNA45SN1_45S_50ntFlanks</td><td>12126</td><td>12146</td><td>21</td><td>CGACCCGCTCCGGGGACAGTG</td><td>3503.5</td><td>-15.6</td><td>1.4</td><td>-10.8</td><td>2.4E-27</td><td>1.7E-25</td><td>MB231/MB468</td><td>MP</td></tr>
<tr><td>>MI0000811|hsa-mir-148b&WithFlank&12|+|54337210|54337320@69.91.23&[MIMAT0000759&hsa-miR-148b-3p&offsets|0|+1;m-77&12|+|54337278|54337299&offsets|0|+1]|TCAGTGCATCACAGAACTTTGTC</td><td>>MI0000811|hsa-mir-148b&WithFlank&12|+|54337210|54337320</td><td>69</td><td>91</td><td>23</td><td>TCAGTGCATCACAGAACTTTGTC</td><td>875.3</td><td>6.5</td><td>0.6</td><td>10.7</td><td>1.1E-26</td><td>1.7E-25</td><td>BT20/MB468</td><td>CYTO</td></tr>
<tr><td>>hg38_21_+_8433172_8446622_RNA45SN1_45S_50ntFlanks@12686.12708.23|CCAGGCGGAACGATACGGCAGCG</td><td>>hg38_21_+_8433172_8446622_RNA45SN1_45S_50ntFlanks</td><td>12686</td><td>12708</td><td>23</td><td>CCAGGCGGAACGATACGGCAGCG</td><td>317.3</td><td>-12.7</td><td>1.2</td><td>-10.6</td><td>2.2E-26</td><td>1.7E-25</td><td>BT20/MB468</td><td>NUC</td></tr>
<tr><td>>am_MI0003776|hsa-mir-320b-1&WithFlank&1|+|116671740|116671823@48.65.18;MI0003839|hsa-mir-320b-2&WithFlank&1|-|224257034|224257116@47.64.18;MI0008190|hsa-mir-320d-1&WithFlank&13|-|40727810|40727893@48.65.18;MI0003778|hsa-mir-320c-1&WithFlank&18|+|21683512|21683595@48.65.18;MI0008191|hsa-mir-320c-2&WithFlank&18|+|24321669|24321752@48.65.18;MI0000542|hsa-mir-320a&WithFlank&8|-|22244960|22245043@48.65.18;MI0008192|hsa-mir-320d-2&WithFlank&X|-|140926154|140926237@48.65.18&[MIMAT0005792&hsa-miR-320b&offsets|0|-4;m-183&1|+|116671787|116671806&offsets|0|-2];[MIMAT0005792_1&hsa-miR-320b&offsets|0|-4;m-183&1|-|224257051|224257070&offsets|0|-2];[MIMAT0006764&hsa-miR-320d&offsets|0|-1;m-246&13|-|40727829|40727846&offsets|0|0];[MIMAT0005793&hsa-miR-320c&offsets|0|-2;m-222&18|+|21683559|21683578&offsets|0|-2];[MIMAT0005793_1&hsa-miR-320c&offsets|0|-2;m-219&18|+|24321716|24321735&offsets|0|-2];[MIMAT0000510&hsa-miR-320a-3p&offsets|0|-4;m-60&8|-|22244975|22244996&offsets|0|-4];[MIMAT0006764_1&hsa-miR-320d&offsets|0|-1;m-253&X|-|140926173|140926190&offsets|0|0]|AAAAGCTGGGTTGAGAGG</td><td>>am_MI0003776|hsa-mir-320b-1&WithFlank&1|+|116671740|116671823</td><td>48</td><td>65</td><td>18</td><td>AAAAGCTGGGTTGAGAGG</td><td>1693.9</td><td>15.0</td><td>1.4</td><td>10.7</td><td>1.2E-26</td><td>1.8E-25</td><td>BT20/MB468</td><td>CYTO</td></tr>
<tr><td>>hg38_21_+_8433172_8446622_RNA45SN1_45S_50ntFlanks@12655.12675.21|ATGACTGAACGCCTCTAAGTC</td><td>>hg38_21_+_8433172_8446622_RNA45SN1_45S_50ntFlanks</td><td>12655</td><td>12675</td><td>21</td><td>ATGACTGAACGCCTCTAAGTC</td><td>1534.2</td><td>-14.4</td><td>1.3</td><td>-10.8</td><td>2.6E-27</td><td>1.8E-25</td><td>MB231/MB468</td><td>MP</td></tr>
<tr><td>>hg38_21_+_8433172_8446622_RNA45SN1_45S_50ntFlanks@8380.8400.21|CAAGAGGGCGTGAAACCGTTA</td><td>>hg38_21_+_8433172_8446622_RNA45SN1_45S_50ntFlanks</td><td>8380</td><td>8400</td><td>21</td><td>CAAGAGGGCGTGAAACCGTTA</td><td>3662.4</td><td>-15.6</td><td>1.4</td><td>-10.8</td><td>4.4E-27</td><td>1.8E-25</td><td>MB231/MB468</td><td>MITO</td></tr>
<tr><td>>hg38_21_+_8433172_8446622_RNA45SN1_45S_50ntFlanks@4045.4067.23|CCATTCGAACGTCTGCCCTATCA</td><td>>hg38_21_+_8433172_8446622_RNA45SN1_45S_50ntFlanks</td><td>4045</td><td>4067</td><td>23</td><td>CCATTCGAACGTCTGCCCTATCA</td><td>554.5</td><td>-13.5</td><td>1.3</td><td>-10.6</td><td>2.6E-26</td><td>1.9E-25</td><td>BT20/MB468</td><td>NUC</td></tr>
<tr><td>>hg38_21_+_8433172_8446622_RNA45SN1_45S_50ntFlanks@12539.12569.31|ATGATGTGTTGTTGCCATGGTAATCCTGCTC</td><td>>hg38_21_+_8433172_8446622_RNA45SN1_45S_50ntFlanks</td><td>12539</td><td>12569</td><td>31</td><td>ATGATGTGTTGTTGCCATGGTAATCCTGCTC</td><td>4125.5</td><td>-15.8</td><td>1.5</td><td>-10.8</td><td>4.8E-27</td><td>2.0E-25</td><td>MB231/MB468</td><td>MITO</td></tr>
<tr><td>>hg38_21_+_8433172_8446622_RNA45SN1_45S_50ntFlanks@10680.10706.27|GTTCCGGCGGCGTCCGGTGAGCTCTCG</td><td>>hg38_21_+_8433172_8446622_RNA45SN1_45S_50ntFlanks</td><td>10680</td><td>10706</td><td>27</td><td>GTTCCGGCGGCGTCCGGTGAGCTCTCG</td><td>15685.0</td><td>-8.3</td><td>0.8</td><td>-10.7</td><td>6.6E-27</td><td>2.0E-25</td><td>MB231/MB468</td><td>NUC</td></tr>
<tr><td>>hg38_21_+_8433172_8446622_RNA45SN1_45S_50ntFlanks@11572.11591.20|CCGACTTAGAACTGGTGCGG</td><td>>hg38_21_+_8433172_8446622_RNA45SN1_45S_50ntFlanks</td><td>11572</td><td>11591</td><td>20</td><td>CCGACTTAGAACTGGTGCGG</td><td>1822.3</td><td>-5.5</td><td>0.5</td><td>-10.7</td><td>6.6E-27</td><td>2.0E-25</td><td>MB231/MB468</td><td>NUC</td></tr>
<tr><td>>MI0000266|hsa-mir-10a&WithFlank&17|-|48579832|48579953@29.48.20(+1C)&[MIMAT0000253&hsa-miR-10a-5p&offsets|+1|-2(+1C);m-18&17|-|48579905|48579926&offsets|+1|-1(+1C)]|ACCCTGTAGATCCGAATTTGC</td><td>>MI0000266|hsa-mir-10a&WithFlank&17|-|48579832|48579953</td><td>29</td><td>48</td><td>20(+1C)</td><td>ACCCTGTAGATCCGAATTTGC</td><td>862.5</td><td>14.0</td><td>1.3</td><td>10.7</td><td>1.4E-26</td><td>2.0E-25</td><td>BT20/MB468</td><td>CYTO</td></tr>
<tr><td>>hg38_21_+_8433172_8446622_RNA45SN1_45S_50ntFlanks@3831.3850.20|CTCGCTCCTCTCCTACTTGG</td><td>>hg38_21_+_8433172_8446622_RNA45SN1_45S_50ntFlanks</td><td>3831</td><td>3850</td><td>20</td><td>CTCGCTCCTCTCCTACTTGG</td><td>361.1</td><td>-12.9</td><td>1.2</td><td>-10.6</td><td>2.7E-26</td><td>2.0E-25</td><td>BT20/MB468</td><td>NUC</td></tr>
<tr><td>>hg38_21_+_8433172_8446622_RNA45SN1_45S_50ntFlanks@9562.9597.36|TCCTGACGTGCAAATCGGTCGTCCGACCTGGGTATA</td><td>>hg38_21_+_8433172_8446622_RNA45SN1_45S_50ntFlanks</td><td>9562</td><td>9597</td><td>36</td><td>TCCTGACGTGCAAATCGGTCGTCCGACCTGGGTATA</td><td>1695.1</td><td>-15.2</td><td>1.4</td><td>-10.6</td><td>2.7E-26</td><td>2.0E-25</td><td>BT20/MB468</td><td>NUC</td></tr>
<tr><td>>hg38_21_+_8433172_8446622_RNA45SN1_45S_50ntFlanks@4731.4759.29|AAGAACGAAAGTCGGAGGTTCGAAGACGA</td><td>>hg38_21_+_8433172_8446622_RNA45SN1_45S_50ntFlanks</td><td>4731</td><td>4759</td><td>29</td><td>AAGAACGAAAGTCGGAGGTTCGAAGACGA</td><td>6122.2</td><td>-6.5</td><td>0.6</td><td>-10.8</td><td>2.9E-27</td><td>2.1E-25</td><td>MB231/MB468</td><td>MP</td></tr>
<tr><td>>hg38_21_+_8433172_8446622_RNA45SN1_45S_50ntFlanks@12419.12462.44|GATGTCGGCTCTTCCTATCATTGTGAAGCAGAATTCACCAAGCG</td><td>>hg38_21_+_8433172_8446622_RNA45SN1_45S_50ntFlanks</td><td>12419</td><td>12462</td><td>44</td><td>GATGTCGGCTCTTCCTATCATTGTGAAGCAGAATTCACCAAGCG</td><td>1590.6</td><td>14.6</td><td>1.3</td><td>10.8</td><td>2.1E-27</td><td>2.1E-25</td><td>MB231/MB468</td><td>CYTO</td></tr>
<tr><td>>MI0003643|hsa-mir-629&WithFlank&15|-|70079366|70079474@28.48.21&[MIMAT0004810&hsa-miR-629-5p&offsets|0|0;m-150&15|-|70079427|70079447&offsets|0|0]|TGGGTTTACGTTGGGAGAACT</td><td>>MI0003643|hsa-mir-629&WithFlank&15|-|70079366|70079474</td><td>28</td><td>48</td><td>21</td><td>TGGGTTTACGTTGGGAGAACT</td><td>1274.8</td><td>14.6</td><td>1.4</td><td>10.7</td><td>1.4E-26</td><td>2.1E-25</td><td>BT20/MB468</td><td>CYTO</td></tr>
<tr><td>>MI0000650|hsa-mir-200c&WithFlank&12|+|6963693|6963772@50.71.22&[MIMAT0000617&hsa-miR-200c-3p&offsets|0|-1;m-54&12|+|6963742|6963764&offsets|0|-1]|TAATACTGCCGGGTAATGATGG</td><td>>MI0000650|hsa-mir-200c&WithFlank&12|+|6963693|6963772</td><td>50</td><td>71</td><td>22</td><td>TAATACTGCCGGGTAATGATGG</td><td>18002.3</td><td>11.6</td><td>1.1</td><td>10.9</td><td>8.1E-28</td><td>2.1E-25</td><td>BT20/MB231</td><td>NUC</td></tr>
<tr><td>>hg38_21_+_8433172_8446622_RNA45SN1_45S_50ntFlanks@11873.11893.21|AAGAAGACCCTGTTGAGCTTG</td><td>>hg38_21_+_8433172_8446622_RNA45SN1_45S_50ntFlanks</td><td>11873</td><td>11893</td><td>21</td><td>AAGAAGACCCTGTTGAGCTTG</td><td>3882.3</td><td>-5.6</td><td>0.5</td><td>-10.7</td><td>1.4E-26</td><td>2.1E-25</td><td>BT20/MB468</td><td>CYTO</td></tr>
<tr><td>>hg38_21_+_8433172_8446622_RNA45SN1_45S_50ntFlanks@11534.11553.20|GTTCCGCGCGGCGCCTCGCC</td><td>>hg38_21_+_8433172_8446622_RNA45SN1_45S_50ntFlanks</td><td>11534</td><td>11553</td><td>20</td><td>GTTCCGCGCGGCGCCTCGCC</td><td>1908.6</td><td>-15.3</td><td>1.4</td><td>-10.6</td><td>2.8E-26</td><td>2.1E-25</td><td>BT20/MB468</td><td>NUC</td></tr>
<tr><td>>hg38_21_+_8433172_8446622_RNA45SN1_45S_50ntFlanks@12539.12560.22|ATGATGTGTTGTTGCCATGGTA</td><td>>hg38_21_+_8433172_8446622_RNA45SN1_45S_50ntFlanks</td><td>12539</td><td>12560</td><td>22</td><td>ATGATGTGTTGTTGCCATGGTA</td><td>566.0</td><td>-13.6</td><td>1.3</td><td>-10.6</td><td>2.8E-26</td><td>2.1E-25</td><td>BT20/MB468</td><td>NUC</td></tr>
<tr><td>>hg38_21_+_8433172_8446622_RNA45SN1_45S_50ntFlanks@6746.6771.26|CACTTCGAACGCACTTGCGGCCCCGG</td><td>>hg38_21_+_8433172_8446622_RNA45SN1_45S_50ntFlanks</td><td>6746</td><td>6771</td><td>26</td><td>CACTTCGAACGCACTTGCGGCCCCGG</td><td>1820.8</td><td>-14.7</td><td>1.4</td><td>-10.8</td><td>3.1E-27</td><td>2.1E-25</td><td>MB231/MB468</td><td>MP</td></tr>
<tr><td>>MI0000446|hsa-mir-125b-1&WithFlank&11|-|122099751|122099850@21.41.21;MI0000470|hsa-mir-125b-2&WithFlank&21|+|16590231|16590331@23.43.21&[MIMAT0000423&hsa-miR-125b-5p&offsets|0|-1;m-67&11|-|122099809|122099830&offsets|0|-1];[MIMAT0000423_1&hsa-miR-125b-5p&offsets|0|-1;m-68&21|+|16590253|16590274&offsets|0|-1]|TCCCTGAGACCCTAACTTGTG</td><td>>MI0000446|hsa-mir-125b-1&WithFlank&11|-|122099751|122099850</td><td>21</td><td>41</td><td>21</td><td>TCCCTGAGACCCTAACTTGTG</td><td>10608.4</td><td>7.1</td><td>0.7</td><td>10.7</td><td>7.1E-27</td><td>2.1E-25</td><td>MB231/MB468</td><td>NUC</td></tr>
<tr><td>>hg38_21_+_8433172_8446622_RNA45SN1_45S_50ntFlanks@10351.10390.40|CGAGAACTTTGAAGGCCGAAGTGGAGAAGGGTTCCATGTG</td><td>>hg38_21_+_8433172_8446622_RNA45SN1_45S_50ntFlanks</td><td>10351</td><td>10390</td><td>40</td><td>CGAGAACTTTGAAGGCCGAAGTGGAGAAGGGTTCCATGTG</td><td>23604.3</td><td>-9.0</td><td>0.8</td><td>-10.8</td><td>5.2E-27</td><td>2.1E-25</td><td>MB231/MB468</td><td>MITO</td></tr>
<tr><td>>hg38_21_+_8433172_8446622_RNA45SN1_45S_50ntFlanks@8169.8186.18|CCAAGTCCTTCTGATCGA</td><td>>hg38_21_+_8433172_8446622_RNA45SN1_45S_50ntFlanks</td><td>8169</td><td>8186</td><td>18</td><td>CCAAGTCCTTCTGATCGA</td><td>656.2</td><td>-13.8</td><td>1.3</td><td>-10.6</td><td>2.9E-26</td><td>2.2E-25</td><td>BT20/MB468</td><td>NUC</td></tr>
<tr><td>>hg38_21_+_8433172_8446622_RNA45SN1_45S_50ntFlanks@11564.11585.22|CCTAGCAGCCGACTTAGAACTG</td><td>>hg38_21_+_8433172_8446622_RNA45SN1_45S_50ntFlanks</td><td>11564</td><td>11585</td><td>22</td><td>CCTAGCAGCCGACTTAGAACTG</td><td>2726.5</td><td>-5.9</td><td>0.5</td><td>-10.7</td><td>7.3E-27</td><td>2.2E-25</td><td>MB231/MB468</td><td>NUC</td></tr>
<tr><td>>hg38_21_+_8433172_8446622_RNA45SN1_45S_50ntFlanks@482.515.34|ATCGATGTGGTGACGTCGTGCTCTCCCGGGCCGG</td><td>>hg38_21_+_8433172_8446622_RNA45SN1_45S_50ntFlanks</td><td>482</td><td>515</td><td>34</td><td>ATCGATGTGGTGACGTCGTGCTCTCCCGGGCCGG</td><td>952.4</td><td>-14.3</td><td>1.4</td><td>-10.6</td><td>3.0E-26</td><td>2.2E-25</td><td>BT20/MB468</td><td>NUC</td></tr>
<tr><td>>hg38_21_+_8433172_8446622_RNA45SN1_45S_50ntFlanks@10411.10432.22|CAGTCGGTCCTGAGAGATGGGC</td><td>>hg38_21_+_8433172_8446622_RNA45SN1_45S_50ntFlanks</td><td>10411</td><td>10432</td><td>22</td><td>CAGTCGGTCCTGAGAGATGGGC</td><td>1538.3</td><td>-5.7</td><td>0.5</td><td>-10.7</td><td>1.5E-26</td><td>2.3E-25</td><td>BT20/MB468</td><td>CYTO</td></tr>
<tr><td>>MI0000439|hsa-mir-23b&WithFlank&9|+|95085202|95085310@67.84.18&[MIMAT0000418&hsa-miR-23b-3p&offsets|+3|-2;m-42&9|+|95085265|95085285&offsets|+3|0]|ACATTGCCAGGGATTACC</td><td>>MI0000439|hsa-mir-23b&WithFlank&9|+|95085202|95085310</td><td>67</td><td>84</td><td>18</td><td>ACATTGCCAGGGATTACC</td><td>871.7</td><td>14.2</td><td>1.3</td><td>10.7</td><td>1.2E-26</td><td>2.3E-25</td><td>BT20/MB468</td><td>MITO</td></tr>
<tr><td>>hg38_21_+_8433172_8446622_RNA45SN1_45S_50ntFlanks@12970.12990.21|GTTTCGTACGTAGCAGAGCAG</td><td>>hg38_21_+_8433172_8446622_RNA45SN1_45S_50ntFlanks</td><td>12970</td><td>12990</td><td>21</td><td>GTTTCGTACGTAGCAGAGCAG</td><td>559.2</td><td>-13.6</td><td>1.3</td><td>-10.6</td><td>3.0E-26</td><td>2.3E-25</td><td>BT20/MB468</td><td>NUC</td></tr>
<tr><td>>hg38_21_+_8433172_8446622_RNA45SN1_45S_50ntFlanks@10829.10858.30|GCAAGCCGGATCCGTAACTTCGGGATAAGG</td><td>>hg38_21_+_8433172_8446622_RNA45SN1_45S_50ntFlanks</td><td>10829</td><td>10858</td><td>30</td><td>GCAAGCCGGATCCGTAACTTCGGGATAAGG</td><td>609.3</td><td>-13.7</td><td>1.3</td><td>-10.6</td><td>3.0E-26</td><td>2.3E-25</td><td>BT20/MB468</td><td>NUC</td></tr>
<tr><td>>hg38_21_+_8433172_8446622_RNA45SN1_45S_50ntFlanks@11702.11728.27|CAATGAAGCGCGGGTAAACGGCGGGAG</td><td>>hg38_21_+_8433172_8446622_RNA45SN1_45S_50ntFlanks</td><td>11702</td><td>11728</td><td>27</td><td>CAATGAAGCGCGGGTAAACGGCGGGAG</td><td>561.4</td><td>-13.6</td><td>1.3</td><td>-10.6</td><td>3.1E-26</td><td>2.3E-25</td><td>BT20/MB468</td><td>NUC</td></tr>
<tr><td>>hg38_21_+_8433172_8446622_RNA45SN1_45S_50ntFlanks@6745.6770.26|ACACTTCGAACGCACTTGCGGCCCCG</td><td>>hg38_21_+_8433172_8446622_RNA45SN1_45S_50ntFlanks</td><td>6745</td><td>6770</td><td>26</td><td>ACACTTCGAACGCACTTGCGGCCCCG</td><td>2499.8</td><td>-7.1</td><td>0.7</td><td>-10.7</td><td>8.0E-27</td><td>2.4E-25</td><td>MB231/MB468</td><td>NUC</td></tr>
</table>

| | | | | | | | | | | | | | |
|---|---|---|---|---|---|---|---|---|---|---|---|---|---|
| >MI0000469\|hsa-mir-125a&WithFlank&19\|+\|51693248\|51693345@21.43.23(+1U)&[MIMAT0000443&hsa-miR-125a-5p&offsets\|0\|-1(+1U);m-63&19\|+\|51693268\|51693289&offsets\|0\|+1(+1U)]\|TCCCTGAGACCCTTTAACCTGTGT | >MI0000469\|hsa-mir-125a&WithFlank&19\|+\|51693248\|51693345 | 21 | 43 | 23(+1U) | TCCCTGAGACCCTTTAACCTGTGT | 2113.4 | 7.4 | 0.7 | 10.7 | 1.6E-26 | 2.4E-25 | BT20/MB468 | CYTO |
| >hg38_21_+_8433172_8446622_RNA45SN1_45S_50ntFlanks@12419.12438.20\|GATGTCGGCTCTTCCTATCA | >hg38_21_+_8433172_8446622_RNA45SN1_45S_50ntFlanks | 12419 | 12438 | 20 | GATGTCGGCTCTTCCTATCA | 1190.9 | 4.8 | 0.5 | 10.7 | 1.7E-26 | 2.4E-25 | BT20/MB468 | CYTO |
| >trna111_HisGTG_1_-_147774845_147774916@-1T.30.31,trna118_HisGTG_1_-_145396881_145396952@-1T.30.31,trna16_HisGTG_1_+_146544773_146544844@-1T.30.31,trna1_HisGTG_15_+_45493349_45493420@-1T.30.31,trna21_HisGTG_1_+_147753471_147753542@-1T.30.31,trna33_HisGTG_6_+_27125906_27125977@-1T.30.31,trna7_HisGTG_9_-_14433938_14434009@-1T.30.31,trna8_HisGTG_15_-_45492611_45492682@-1T.30.31,trna9_HisGTG_15_-_45490804_45490875@-1T.30.31\|TGCCGTGATCGTATAGTGGTTAGTACTCTGC | >trna111_HisGTG_1_-_147774845_147774916 | -1T | 30 | 31 | TGCCGTGATCGTATAGTGGTTAGTACTCTGC | 1323.8 | -6.6 | 0.6 | -10.6 | 3.3E-26 | 2.5E-25 | BT20/MB468 | NUC |
| >hg38_21_+_8433172_8446622_RNA45SN1_45S_50ntFlanks@10398.10424.27\|GTTGAACATGGGTCAGTCGGTCCTGAG | >hg38_21_+_8433172_8446622_RNA45SN1_45S_50ntFlanks | 10398 | 10424 | 27 | GTTGAACATGGGTCAGTCGGTCCTGAG | 1851.8 | 7.8 | 0.7 | 10.6 | 1.8E-26 | 2.6E-25 | BT20/MB468 | CYTO |
| >MI0000299\|hsa-mir-222&WithFlank&X\|-\|45747009\|45747130@75.99.25&[MIMAT0000279&hsa-miR-222-3p&offsets\|0\|+4;m-140&X\|-\|45747033\|45747056&offsets\|0\|+1]\|AGCTACATCTGGCTACTGGGTCTCT | >MI0000299\|hsa-mir-222&WithFlank&X\|-\|45747009\|45747130 | 75 | 99 | 25 | AGCTACATCTGGCTACTGGGTCTCT | 57600.1 | 8.4 | 0.8 | 10.8 | 2.6E-27 | 2.6E-25 | MB231/MB468 | CYTO |
| >MI0000253\|hsa-mir-148a&WithFlank&7\|-\|25949913\|25949992@50.72.23&[MIMAT0000243&hsa-miR-148a-3p&offsets\|0\|+1;m-7&7\|-\|25949922\|25949943&offsets\|0\|+1]\|TCAGTGCACTACAGAACTTTGTC | >MI0000253\|hsa-mir-148a&WithFlank&7\|-\|25949913\|25949992 | 50 | 72 | 23 | TCAGTGCACTACAGAACTTTGTC | 1997.9 | 7.2 | 0.7 | 10.7 | 1.4E-26 | 2.6E-25 | BT20/MB468 | MITO |
| >hg38_21_+_8433172_8446622_RNA45SN1_45S_50ntFlanks@12539.12566.28\|ATGATGTGTTGTTGCCATGGTAATCCTG | >hg38_21_+_8433172_8446622_RNA45SN1_45S_50ntFlanks | 12539 | 12566 | 28 | ATGATGTGTTGTTGCCATGGTAATCCTG | 2165.7 | -14.9 | 1.4 | -10.8 | 3.8E-27 | 2.6E-25 | MB231/MB468 | MP |
| >hg38_21_+_8433172_8446622_RNA45SN1_45S_50ntFlanks@12638.12661.24\|CTACCATCTGTGGGATTATGACTG | >hg38_21_+_8433172_8446622_RNA45SN1_45S_50ntFlanks | 12638 | 12661 | 24 | CTACCATCTGTGGGATTATGACTG | 6483.6 | -16.5 | 1.5 | -10.8 | 3.9E-27 | 2.7E-25 | MB231/MB468 | MP |
| >hg38_21_+_8433172_8446622_RNA45SN1_45S_50ntFlanks@10410.10429.20\|TCAGTCGGTCCTGAGAGATG | >hg38_21_+_8433172_8446622_RNA45SN1_45S_50ntFlanks | 10410 | 10429 | 20 | TCAGTCGGTCCTGAGAGATG | 4506.5 | -10.6 | 1.0 | -10.8 | 3.9E-27 | 2.7E-25 | MB231/MB468 | MP |
| >hg38_21_+_8433172_8446622_RNA45SN1_45S_50ntFlanks@11561.11603.43\|GCGCCTAGCAGCCGACTTAGAACTGGTGCGGACCAGGGGAATC | >hg38_21_+_8433172_8446622_RNA45SN1_45S_50ntFlanks | 11561 | 11603 | 43 | GCGCCTAGCAGCCGACTTAGAACTGGTGCGGACCAGGGGAATC | 10071.5 | -13.2 | 1.2 | -10.7 | 6.5E-27 | 2.7E-25 | MB231/MB468 | MITO |
| >MI0000465\|hsa-mir-191&WithFlank&3\|-\|49020612\|49020715@22.44.23&[MIMAT0000440&hsa-miR-191-5p&offsets\|0\|0;m-48&3\|-\|49020672\|49020694&offsets\|0\|0]\|CAACGGAATCCCAAAAGCAGCTG | >MI0000465\|hsa-mir-191&WithFlank&3\|-\|49020612\|49020715 | 22 | 44 | 23 | CAACGGAATCCCAAAAGCAGCTG | 22904.5 | 7.6 | 0.7 | 10.7 | 1.4E-26 | 2.7E-25 | BT20/MB468 | MITO |
| >hg38_21_+_8433172_8446622_RNA45SN1_45S_50ntFlanks@4283.4321.39\|CTTTAACGAGGATCCATTGGAGGGCAAGTCTGGTGCCAG | >hg38_21_+_8433172_8446622_RNA45SN1_45S_50ntFlanks | 4283 | 4321 | 39 | CTTTAACGAGGATCCATTGGAGGGCAAGTCTGGTGCCAG | 1094.3 | -14.5 | 1.4 | -10.6 | 3.6E-26 | 2.7E-25 | BT20/MB468 | NUC |
| >hg38_21_+_8433172_8446622_RNA45SN1_45S_50ntFlanks@10808.10840.33\|CAATGTAGGTAAGGGAAGTCGGCAAGCCGGATC | >hg38_21_+_8433172_8446622_RNA45SN1_45S_50ntFlanks | 10808 | 10840 | 33 | CAATGTAGGTAAGGGAAGTCGGCAAGCCGGATC | 2143.5 | -6.4 | 0.6 | -10.7 | 1.5E-26 | 2.7E-25 | BT20/MB468 | MITO |
| >hg38_21_+_8433172_8446622_RNA45SN1_45S_50ntFlanks@12456.12485.30\|CCAAGCGTTGGATTGTTCACCCACTAATAG | >hg38_21_+_8433172_8446622_RNA45SN1_45S_50ntFlanks | 12456 | 12485 | 30 | CCAAGCGTTGGATTGTTCACCCACTAATAG | 2180.4 | -15.0 | 1.4 | -10.8 | 4.1E-27 | 2.8E-25 | MB231/MB468 | MP |
| >hg38_21_+_8433172_8446622_RNA45SN1_45S_50ntFlanks@11619.11640.22\|AACAAAGCATCGCGAAGGCCCG | >hg38_21_+_8433172_8446622_RNA45SN1_45S_50ntFlanks | 11619 | 11640 | 22 | AACAAAGCATCGCGAAGGCCCG | 2579.0 | -15.2 | 1.4 | -10.8 | 4.1E-27 | 2.8E-25 | MB231/MB468 | MP |
| >hg38_21_+_8433172_8446622_RNA45SN1_45S_50ntFlanks@6671.6699.29\|CTCGGCTCGTGCGTCGATGAAGAACGCAG | >hg38_21_+_8433172_8446622_RNA45SN1_45S_50ntFlanks | 6671 | 6699 | 29 | CTCGGCTCGTGCGTCGATGAAGAACGCAG | 7185.6 | -10.4 | 1.0 | -10.8 | 4.1E-27 | 2.8E-25 | MB231/MB468 | MP |
| >hg38_21_+_8433172_8446622_RNA45SN1_45S_50ntFlanks@12616.12653.38\|CTGAGGAGCCAATGGGGCGAAGCTACCATCTGTGGGAT | >hg38_21_+_8433172_8446622_RNA45SN1_45S_50ntFlanks | 12616 | 12653 | 38 | CTGAGGAGCCAATGGGGCGAAGCTACCATCTGTGGGAT | 1130.3 | -14.6 | 1.4 | -10.6 | 3.8E-26 | 2.9E-25 | BT20/MB468 | NUC |
| >hg38_21_+_8433172_8446622_RNA45SN1_45S_50ntFlanks@12638.12657.20\|CTACCATCTGTGGGATTATG | >hg38_21_+_8433172_8446622_RNA45SN1_45S_50ntFlanks | 12638 | 12657 | 20 | CTACCATCTGTGGGATTATG | 1540.2 | -14.5 | 1.3 | -10.8 | 4.3E-27 | 2.9E-25 | MB231/MB468 | MP |
| >hg38_21_+_8433172_8446622_RNA45SN1_45S_50ntFlanks@11886.11905.20\|TGAGCTTGACTCTAGTCTGG | >hg38_21_+_8433172_8446622_RNA45SN1_45S_50ntFlanks | 11886 | 11905 | 20 | TGAGCTTGACTCTAGTCTGG | 378.6 | -13.0 | 1.2 | -10.6 | 3.8E-26 | 2.9E-25 | BT20/MB468 | NUC |
| >hg38_21_+_8433172_8446622_RNA45SN1_45S_50ntFlanks@12045.12070.26\|CTGACCCGGTGAGGCGGGGGGGCGAG | >hg38_21_+_8433172_8446622_RNA45SN1_45S_50ntFlanks | 12045 | 12070 | 26 | CTGACCCGGTGAGGCGGGGGGGCGAG | 1017.8 | -14.4 | 1.4 | -10.6 | 3.9E-26 | 2.9E-25 | BT20/MB468 | NUC |
| >MI0000273\|hsa-mir-183&WithFlank&7\|-\|129774899\|129775020@34.54.21(+1A)&[MIMAT0000261&hsa-miR-183-5p&offsets\|+1\|0(+1A);m-41&7\|-\|129774966\|129774987&offsets\|0\|-1(+1A)]\|ATGGCACTGGTAGAATTCACTA | >MI0000273\|hsa-mir-183&WithFlank&7\|-\|129774899\|129775020 | 34 | 54 | 21(+1A) | ATGGCACTGGTAGAATTCACTA | 134.3 | 3.6 | 0.3 | 10.6 | 3.9E-26 | 2.9E-25 | BT20/MB468 | NUC |
| >hg38_21_+_8433172_8446622_RNA45SN1_45S_50ntFlanks@12324.12344.21\|TTGGGTTTTAAGCAGGAGGTG | >hg38_21_+_8433172_8446622_RNA45SN1_45S_50ntFlanks | 12324 | 12344 | 21 | TTGGGTTTTAAGCAGGAGGTG | 2649.0 | -15.2 | 1.4 | -10.8 | 4.4E-27 | 2.9E-25 | MB231/MB468 | MP |
| >hg38_21_+_8433172_8446622_RNA45SN1_45S_50ntFlanks@12418.12436.19\|CGATGTCGGCTCTTCCTAT | >hg38_21_+_8433172_8446622_RNA45SN1_45S_50ntFlanks | 12418 | 12436 | 19 | CGATGTCGGCTCTTCCTAT | 12877.6 | -6.6 | 0.6 | -10.7 | 1.6E-26 | 2.9E-25 | BT20/MB468 | MITO |
| >hg38_21_+_8433172_8446622_RNA45SN1_45S_50ntFlanks@11317.11338.22\|ACCCCACGTCTCGTCGCGCGCG | >hg38_21_+_8433172_8446622_RNA45SN1_45S_50ntFlanks | 11317 | 11338 | 22 | ACCCCACGTCTCGTCGCGCGCG | 1245.6 | -8.8 | 0.8 | -10.8 | 3.0E-27 | 2.9E-25 | MB231/MB468 | CYTO |
| >hg38_21_+_8433172_8446622_RNA45SN1_45S_50ntFlanks@11999.12031.33\|CCCTGCGGGCCGCCGGTGAAATACCACTACTCT | >hg38_21_+_8433172_8446622_RNA45SN1_45S_50ntFlanks | 11999 | 12031 | 33 | CCCTGCGGGCCGCCGGTGAAATACCACTACTCT | 2661.1 | -15.2 | 1.4 | -10.7 | 7.2E-27 | 2.9E-25 | MB231/MB468 | MITO |
| >hg38_21_+_8433172_8446622_RNA45SN1_45S_50ntFlanks@11209.11234.26\|CGGCGACTCTGGACGCGAGCCGGGCC | >hg38_21_+_8433172_8446622_RNA45SN1_45S_50ntFlanks | 11209 | 11234 | 26 | CGGCGACTCTGGACGCGAGCCGGGCC | 2246.5 | -15.0 | 1.4 | -10.8 | 4.4E-27 | 2.9E-25 | MB231/MB468 | MP |
| >hg38_21_+_8433172_8446622_RNA45SN1_45S_50ntFlanks@6773.6797.25\|TTCCTCCCGGGGCTACGCCTGTCTG | >hg38_21_+_8433172_8446622_RNA45SN1_45S_50ntFlanks | 6773 | 6797 | 25 | TTCCTCCCGGGGCTACGCCTGTCTG | 458.1 | -13.3 | 1.3 | -10.6 | 4.0E-26 | 3.0E-25 | BT20/MB468 | NUC |
| >hg38_21_+_8433172_8446622_RNA45SN1_45S_50ntFlanks@9736.9763.28\|CCGAAACGATCTCAACCTATTCTCAAAC | >hg38_21_+_8433172_8446622_RNA45SN1_45S_50ntFlanks | 9736 | 9763 | 28 | CCGAAACGATCTCAACCTATTCTCAAAC | 400.4 | -13.1 | 1.2 | -10.6 | 4.0E-26 | 3.0E-25 | BT20/MB468 | NUC |
| >hg38_21_+_8433172_8446622_RNA45SN1_45S_50ntFlanks@11849.11872.24\|CGGGCTTGGCGGAATCAGCGGGGA | >hg38_21_+_8433172_8446622_RNA45SN1_45S_50ntFlanks | 11849 | 11872 | 24 | CGGGCTTGGCGGAATCAGCGGGGA | 2226.9 | -15.0 | 1.4 | -10.8 | 4.6E-27 | 3.0E-25 | MB231/MB468 | MP |
| >hg38_21_+_8433172_8446622_RNA45SN1_45S_50ntFlanks@3773.3799.27\|CGGCCGGTACAGTGAAACTGCGAATGG | >hg38_21_+_8433172_8446622_RNA45SN1_45S_50ntFlanks | 3773 | 3799 | 27 | CGGCCGGTACAGTGAAACTGCGAATGG | 311.6 | -12.7 | 1.2 | -10.6 | 4.1E-26 | 3.0E-25 | BT20/MB468 | NUC |
| >hg38_21_+_8433172_8446622_RNA45SN1_45S_50ntFlanks@4736.4759.24\|CGAAAGTCGGAGGTTCGAAGACGA | >hg38_21_+_8433172_8446622_RNA45SN1_45S_50ntFlanks | 4736 | 4759 | 24 | CGAAAGTCGGAGGTTCGAAGACGA | 372.3 | -13.0 | 1.2 | -10.6 | 4.1E-26 | 3.0E-25 | BT20/MB468 | NUC |
| >hg38_21_+_8433172_8446622_RNA45SN1_45S_50ntFlanks@11781.11803.23\|CGCATGAATGGATGAACGAGATT | >hg38_21_+_8433172_8446622_RNA45SN1_45S_50ntFlanks | 11781 | 11803 | 23 | CGCATGAATGGATGAACGAGATT | 3148.8 | -6.6 | 0.6 | -10.7 | 1.0E-26 | 3.0E-25 | MB231/MB468 | NUC |
| >hg38_21_+_8433172_8446622_RNA45SN1_45S_50ntFlanks@11873.11904.32\|AAGAAGACCCTGTTGAGCTTGACTCTAGTCTG | >hg38_21_+_8433172_8446622_RNA45SN1_45S_50ntFlanks | 11873 | 11904 | 32 | AAGAAGACCCTGTTGAGCTTGACTCTAGTCTG | 3226.3 | -15.5 | 1.4 | -10.8 | 4.8E-27 | 3.1E-25 | MB231/MB468 | MP |
| >MI0000750\|hsa-mir-26a-2&WithFlank&12\|-\|57824603\|57824698@20.41.22(+1A);MI0000083\|hsa-mir-26a-1&WithFlank&3\|+\|37969398\|37969486@16.37.22(+1A)&[MIMAT0000082&hsa-miR-26a-5p&offsets\|0\|0(+1A);m-33&12\|-\|57824658\|57824679&offsets\|0\|0(+1A)];[MIMAT0000082_1&hsa-miR-26a-5p&offsets\|0\|0(+1A);m-34&3\|+\|37969413\|37969434&offsets\|0\|0(+1A)]\|TTCAAGTAATCCAGGATAGGCTA | >MI0000750\|hsa-mir-26a-2&WithFlank&12\|-\|57824603\|57824698 | 20 | 41 | 22(+1A) | TTCAAGTAATCCAGGATAGGCTA | 1996.3 | 7.2 | 0.7 | 10.6 | 2.1E-26 | 3.1E-25 | BT20/MB468 | CYTO |
| >hg38_21_+_8433172_8446622_RNA45SN1_45S_50ntFlanks@12628.12653.26\|TGGGGCGAAGCTACCATCTGTGGGAT | >hg38_21_+_8433172_8446622_RNA45SN1_45S_50ntFlanks | 12628 | 12653 | 26 | TGGGGCGAAGCTACCATCTGTGGGAT | 3404.1 | -15.5 | 1.4 | -10.7 | 7.7E-27 | 3.1E-25 | MB231/MB468 | MITO |
| >hg38_21_+_8433172_8446622_RNA45SN1_45S_50ntFlanks@10781.10818.38\|AGGTGAACAGCCTCTGGCATGTTGGAACAATGTAGGTA | >hg38_21_+_8433172_8446622_RNA45SN1_45S_50ntFlanks | 10781 | 10818 | 38 | AGGTGAACAGCCTCTGGCATGTTGGAACAATGTAGGTA | 999.4 | -5.9 | 0.6 | -10.7 | 1.1E-26 | 3.2E-25 | MB231/MB468 | NUC |
| >hg38_21_+_8433172_8446622_RNA45SN1_45S_50ntFlanks@6773.6793.21\|TTCCTCCCGGGGCTACGCCTG | >hg38_21_+_8433172_8446622_RNA45SN1_45S_50ntFlanks | 6773 | 6793 | 21 | TTCCTCCCGGGGCTACGCCTG | 1284.7 | -14.8 | 1.4 | -10.6 | 4.3E-26 | 3.2E-25 | BT20/MB468 | NUC |
| >MI0000061\|hsa-let-7a-2&WithFlank&11\|-\|122146516\|122146599@11.30.20(+1C);MI0000062\|hsa-let-7a-3&WithFlank&22\|+\|46112743\|46112828@10.29.20(+1C);MI0000060\|hsa-let-7a-1&WithFlank&9\|+\|94175951\|94176042@12.31.20(+1C)&[MIMAT0000062&hsa-let-7a-5p&offsets\|0\|-2(+1C);m-6&11\|-\|122146568\|122146589&offsets\|0\|-2(+1C)];[MIMAT0000062_1&hsa-let-7a-5p&offsets\|0\|-2(+1C);m-5&22\|+\|46112752\|46112773&offsets\|0\|-2(+1C)];[MIMAT0000062_2&hsa-let-7a-5p&offsets\|0\|-2(+1C);m-4&9\|+\|94175962\|94175983&offsets\|0\|-2(+1C)]\|TGAGGTAGTAGGTTGTATAGC | >MI0000061\|hsa-let-7a-2&WithFlank&11\|-\|122146516\|122146599 | 11 | 30 | 20(+1C) | TGAGGTAGTAGGTTGTATAGC | 1785.3 | 7.8 | 0.7 | 10.6 | 2.3E-26 | 3.3E-25 | BT20/MB468 | CYTO |
| >trna11_GluTTC_15_-_26327381_26327452@1.33.33,trna3_GluTTC_13_-_45492062_45492133@1.33.33\|TCCCACATGGTCTAGCGGTTAGGATTCCTGGTT | >trna11_GluTTC_15_-_26327381_26327452 | 1 | 33 | 33 | TCCCACATGGTCTAGCGGTTAGGATTCCTGGTT | 10057.4 | 8.9 | 0.8 | 10.8 | 3.5E-27 | 3.4E-25 | MB231/MB468 | CYTO |
| >hg38_21_+_8433172_8446622_RNA45SN1_45S_50ntFlanks@11581.11603.23\|AACTGGTGCGGACCAGGGGAATC | >hg38_21_+_8433172_8446622_RNA45SN1_45S_50ntFlanks | 11581 | 11603 | 23 | AACTGGTGCGGACCAGGGGAATC | 2704.5 | -6.3 | 0.6 | -10.7 | 1.2E-26 | 3.4E-25 | MB231/MB468 | NUC |
| >hg38_21_+_8433172_8446622_RNA45SN1_45S_50ntFlanks@3872.3898.27\|CTAATACATGCCGACGGGCGCTGACCC | >hg38_21_+_8433172_8446622_RNA45SN1_45S_50ntFlanks | 3872 | 3898 | 27 | CTAATACATGCCGACGGGCGCTGACCC | 3818.8 | -15.7 | 1.5 | -10.7 | 8.8E-27 | 3.5E-25 | MB231/MB468 | MITO |
| >hg38_21_+_8433172_8446622_RNA45SN1_45S_50ntFlanks@12934.12952.19\|CTAAACCATTCGTAGACGA | >hg38_21_+_8433172_8446622_RNA45SN1_45S_50ntFlanks | 12934 | 12952 | 19 | CTAAACCATTCGTAGACGA | 621.1 | -13.7 | 1.3 | -10.6 | 4.8E-26 | 3.6E-25 | BT20/MB468 | NUC |
| >hg38_21_+_8433172_8446622_RNA45SN1_45S_50ntFlanks@9454.9503.50\|CCGCGCCGGGGAGGTGGAGCACGAGCGCACGTGTTAGGACCCGAAAGATG | >hg38_21_+_8433172_8446622_RNA45SN1_45S_50ntFlanks | 9454 | 9503 | 50 | CCGCGCCGGGGAGGTGGAGCACGAGCGCACGTGTTAGGACCCGAAAGATG | 9983.6 | -17.1 | 1.6 | -10.7 | 8.9E-27 | 3.6E-25 | MB231/MB468 | MITO |

| | | | | | | | | | | | | | |
|---|---|---|---|---|---|---|---|---|---|---|---|---|---|
| >MI0000077\|hsa-mir-21&WithFlank&17\|+\|59841260\|59841343@14.35.22(+2A)&[MIMAT0000076&hsa-miR-21-5p&offsets\|0\|0(+2A);m-1&17\|+\|59841273\|59841294&offsets\|0\|0(+2A)]\|TAGCTTATCAGACTGATGTTGAAA | >MI0000077\|hsa-mir-21&WithFlank&17\|+\|59841260\|59841343 | 14 | 35 | 22(+2A) | TAGCTTATCAGACTGATGTTGAAA | 575.4 | 6.4 | 0.6 | 10.6 | 1.9E-26 | 3.6E-25 | BT20/MB468 | MITO |
| >hg38_21_+_8433172_8446622_RNA45SN1_45S_50ntFlanks@485.510.26\|GATGTGGTGACGTCGTGCTCTCCCGG | >hg38_21_+_8433172_8446622_RNA45SN1_45S_50ntFlanks | 485 | 510 | 26 | GATGTGGTGACGTCGTGCTCTCCCGG | 877.6 | 14.2 | 1.3 | 10.6 | 1.9E-26 | 3.6E-25 | BT20/MB468 | MITO |
| >hg38_21_+_8433172_8446622_RNA45SN1_45S_50ntFlanks@4733.4768.36\|GAACGAAAGTCGGAGGTTCGAAGACGATCAGATACC | >hg38_21_+_8433172_8446622_RNA45SN1_45S_50ntFlanks | 4733 | 4768 | 36 | GAACGAAAGTCGGAGGTTCGAAGACGATCAGATACC | 590.6 | 13.5 | 1.3 | 10.6 | 2.5E-26 | 3.6E-25 | BT20/MB468 | CYTO |
| >hg38_21_+_8433172_8446622_RNA45SN1_45S_50ntFlanks@10845.10872.28\|ACTTCGGGATAAGGATTGGCTCTAAGGG | >hg38_21_+_8433172_8446622_RNA45SN1_45S_50ntFlanks | 10845 | 10872 | 28 | ACTTCGGGATAAGGATTGGCTCTAAGGG | 508.8 | -13.4 | 1.3 | -10.6 | 4.9E-26 | 3.6E-25 | BT20/MB468 | NUC |
| >hg38_21_+_8433172_8446622_RNA45SN1_45S_50ntFlanks@12973.12990.18\|TCGTACGTAGCAGAGCAG | >hg38_21_+_8433172_8446622_RNA45SN1_45S_50ntFlanks | 12973 | 12990 | 18 | TCGTACGTAGCAGAGCAG | 4538.6 | -11.9 | 1.1 | -10.7 | 9.3E-27 | 3.7E-25 | MB231/MB468 | MITO |
| >hg38_21_+_8433172_8446622_RNA45SN1_45S_50ntFlanks@11821.11842.22\|CTATCCAGCGAAACCACAGCCA | >hg38_21_+_8433172_8446622_RNA45SN1_45S_50ntFlanks | 11821 | 11842 | 22 | CTATCCAGCGAAACCACAGCCA | 507.2 | -13.4 | 1.3 | -10.6 | 5.1E-26 | 3.8E-25 | BT20/MB468 | NUC |
| >hg38_21_+_8433172_8446622_RNA45SN1_45S_50ntFlanks@10838.10868.31\|ATCCGTAACTTCGGGATAAGGATTGGCTCTA | >hg38_21_+_8433172_8446622_RNA45SN1_45S_50ntFlanks | 10838 | 10868 | 31 | ATCCGTAACTTCGGGATAAGGATTGGCTCTA | 435.4 | -13.2 | 1.3 | -10.5 | 5.1E-26 | 3.8E-25 | BT20/MB468 | NUC |
| >hg38_21_+_8433172_8446622_RNA45SN1_45S_50ntFlanks@6721.6743.23\|AATTGCAGGACACATTGATCATC | >hg38_21_+_8433172_8446622_RNA45SN1_45S_50ntFlanks | 6721 | 6743 | 23 | AATTGCAGGACACATTGATCATC | 3326.1 | -6.7 | 0.6 | -10.7 | 9.5E-27 | 3.8E-25 | MB231/MB468 | MITO |
| >trna129_MetCAT_6_-_27870271_27870342@31.51.21,trna142_MetCAT_6_-_27560600_27560671@31.51.21,trna150_MetCAT_6_-_27300764_27300835@31.51.21,trna169_MetCAT_6_-_26330529_26330600@31.51.21,trna171_MetCAT_6_-_26313352_26313423@31.51.21,trna20_MetCAT_17_-_80452597_80452668@31.51.21,trna2_MetCAT_6_+_26286754_26286825@31.51.21,trna32_MetCAT_1_+_153643726_153643797@31.51.21,trna61_MetCAT_6_+_27745664_27745735@31.51.21\|CCCATAACCCAGAGGTCGATG | >trna129_MetCAT_6_-_27870271_27870342 | 31 | 51 | 21 | CCCATAACCCAGAGGTCGATG | 1223.7 | -14.7 | 1.4 | -10.5 | 5.1E-26 | 3.8E-25 | BT20/MB468 | NUC |
| >MI0000471\|hsa-mir-126&WithFlank&9\|+\|136670596\|136670692@58.79.22&[MIMAT0000445&hsa-miR-126-3p&offsets\|0\|0;m-40&9\|+\|136670653\|136670674&offsets\|0\|0]\|TCGTACCGTGAGTAATAATGCG | >MI0000471\|hsa-mir-126&WithFlank&9\|+\|136670596\|136670692 | 58 | 79 | 22 | TCGTACCGTGAGTAATAATGCG | 646.5 | 7.4 | 0.7 | 10.8 | 4.0E-27 | 3.8E-25 | MB231/MB468 | CYTO |
| >hg38_21_+_8433172_8446622_RNA45SN1_45S_50ntFlanks@9570.9596.27\|TGCAAATCGGTCGTCCGACCTGGGTAT | >hg38_21_+_8433172_8446622_RNA45SN1_45S_50ntFlanks | 9570 | 9596 | 27 | TGCAAATCGGTCGTCCGACCTGGGTAT | 4818.0 | -16.1 | 1.5 | -10.8 | 5.9E-27 | 3.9E-25 | MB231/MB468 | MP |
| >MI0000067\|hsa-let-7f-1&WithFlank&9\|+\|94176341\|94176439@13.33.21(+1C);MI0000068\|hsa-let-7f-2&WithFlank&X\|-\|53557186\|53557280@14.34.21(+1C)&[MIMAT0000067&hsa-let-7f-5p&offsets\|0\|-1(+1C);m-20&9\|+\|94176353\|94176374&offsets\|0\|-1(+1C)];[MIMAT0000067_1&hsa-let-7f-5p&offsets\|0\|-1(+1C);m-19&X\|-\|53557246\|53557267&offsets\|0\|-1(+1C)]\|TGAGGTAGTAGATTGTATAGTC | >MI0000067\|hsa-let-7f-1&WithFlank&9\|+\|94176341\|94176439 | 13 | 33 | 21(+1C) | TGAGGTAGTAGATTGTATAGTC | 12089.4 | 7.2 | 0.7 | 10.6 | 2.1E-26 | 3.9E-25 | BT20/MB468 | MITO |
| >MI0000447\|hsa-mir-128-1&WithFlank&2\|+\|135665391\|135665484@56.76.21(+2A);MI0000727\|hsa-mir-128-2&WithFlank&3\|+\|35744470\|35744565@58.78.21(+2A)&[MIMAT0000424&hsa-miR-128-3p&offsets\|0\|0(+2A);m-108&2\|+\|135665446\|135665466&offsets\|0\|0(+2A)];[MIMAT0000424_1&hsa-miR-128-3p&offsets\|0\|0(+2A);m-117&3\|+\|35744527\|35744547&offsets\|0\|0(+2A)]\|TCACAGTGAACCGGTCTCTTTAA | >MI0000447\|hsa-mir-128-1&WithFlank&2\|+\|135665391\|135665484 | 56 | 76 | 21(+2A) | TCACAGTGAACCGGTCTCTTTAA | 1198.9 | 5.3 | 0.5 | 10.6 | 2.7E-26 | 3.9E-25 | BT20/MB468 | CYTO |
| >hg38_21_+_8433172_8446622_RNA45SN1_45S_50ntFlanks@11199.11225.27\|CCGGCGGCGGCGGCGACTCTGGACGCG | >hg38_21_+_8433172_8446622_RNA45SN1_45S_50ntFlanks | 11199 | 11225 | 27 | CCGGCGGCGGCGGCGACTCTGGACGCG | 2392.1 | -15.0 | 1.4 | -10.7 | 9.9E-27 | 3.9E-25 | MB231/MB468 | MITO |
| >hg38_21_+_8433172_8446622_RNA45SN1_45S_50ntFlanks@9545.9571.27\|GTGGAGGTCCGTAGCGGTCCTGACGTG | >hg38_21_+_8433172_8446622_RNA45SN1_45S_50ntFlanks | 9545 | 9571 | 27 | GTGGAGGTCCGTAGCGGTCCTGACGTG | 22571.7 | -5.3 | 0.5 | -10.6 | 2.7E-26 | 3.9E-25 | BT20/MB468 | CYTO |
| >MI0000088\|hsa-mir-30a&WithFlank&6\|-\|71403545\|71403627@53.73.21&[MIMAT0000088&hsa-miR-30a-3p&offsets\|0\|-1;m-38&6\|-\|71403554\|71403575&offsets\|0\|-1]\|CTTTCAGTCGGATGTTTGCAG | >MI0000088\|hsa-mir-30a&WithFlank&6\|-\|71403545\|71403627 | 53 | 73 | 21 | CTTTCAGTCGGATGTTTGCAG | 1102.2 | 7.4 | 0.7 | 10.6 | 2.8E-26 | 4.0E-25 | BT20/MB468 | CYTO |
| >hg38_21_+_8433172_8446622_RNA45SN1_45S_50ntFlanks@11564.11602.39\|CCTAGCAGCCGACTTAGAACTGGTGCGGACCAGGGGAAT | >hg38_21_+_8433172_8446622_RNA45SN1_45S_50ntFlanks | 11564 | 11602 | 39 | CCTAGCAGCCGACTTAGAACTGGTGCGGACCAGGGGAAT | 7374.7 | -8.2 | 0.8 | -10.7 | 1.0E-26 | 4.1E-25 | MB231/MB468 | MITO |
| >hg38_21_+_8433172_8446622_RNA45SN1_45S_50ntFlanks@10458.10477.20\|ATGGCCTCCGTTGCCCTCGG | >hg38_21_+_8433172_8446622_RNA45SN1_45S_50ntFlanks | 10458 | 10477 | 20 | ATGGCCTCCGTTGCCCTCGG | 2109.6 | -14.8 | 1.4 | -10.7 | 1.1E-26 | 4.1E-25 | MB231/MB468 | MITO |
| >hg38_21_+_8433172_8446622_RNA45SN1_45S_50ntFlanks@7988.8009.22\|CAGACGTGGCGACCCGCTGAAT | >hg38_21_+_8433172_8446622_RNA45SN1_45S_50ntFlanks | 7988 | 8009 | 22 | CAGACGTGGCGACCCGCTGAAT | 3811.2 | -6.3 | 0.6 | -10.7 | 6.4E-27 | 4.1E-25 | MB231/MB468 | MP |
| >MI0000298\|hsa-mir-221&WithFlank&X\|-\|45746151\|45746272@71.91.21(+1C)&[MIMAT0000278&hsa-miR-221-3p&offsets\|0\|-2(+1C);m-91&X\|-\|45746181\|45746202&offsets\|0\|-1(+1C)]\|AGCTACATTGTCTGCTGGGTTC | >MI0000298\|hsa-mir-221&WithFlank&X\|-\|45746151\|45746272 | 71 | 91 | 21(+1C) | AGCTACATTGTCTGCTGGGTTC | 2193.2 | 7.5 | 0.7 | 10.7 | 1.4E-26 | 4.2E-25 | MB231/MB468 | NUC |
| >MI0000809\|hsa-mir-151a&WithFlank&8\|-\|140732558\|140732659@53.74.22&[MIMAT0000757&hsa-miR-151a-3p&offsets\|0\|+1;m-30&8\|-\|140732587\|140732607&offsets\|0\|+1]\|CTAGACTGAAGCTCCTTGAGGA | >MI0000809\|hsa-mir-151a&WithFlank&8\|-\|140732558\|140732659 | 53 | 74 | 22 | CTAGACTGAAGCTCCTTGAGGA | 9366.9 | 5.1 | 0.5 | 10.7 | 1.4E-26 | 4.2E-25 | MB231/MB468 | NUC |
| >hg38_21_+_8433172_8446622_RNA45SN1_45S_50ntFlanks@10404.10429.26\|CATGGGTCAGTCGGTCCTGAGAGATG | >hg38_21_+_8433172_8446622_RNA45SN1_45S_50ntFlanks | 10404 | 10429 | 26 | CATGGGTCAGTCGGTCCTGAGAGATG | 2240.4 | -4.7 | 0.4 | -10.7 | 1.5E-26 | 4.2E-25 | MB231/MB468 | NUC |
| >hg38_21_+_8433172_8446622_RNA45SN1_45S_50ntFlanks@12655.12675.21\|ATGACTGAACGCCTCTAAGTC | >hg38_21_+_8433172_8446622_RNA45SN1_45S_50ntFlanks | 12655 | 12675 | 21 | ATGACTGAACGCCTCTAAGTC | 14559.0 | -7.8 | 0.7 | -10.6 | 2.9E-26 | 4.2E-25 | BT20/MB468 | CYTO |
| >hg38_1_-_228634819_228635039_RNA5S12_5S_50ntFlanks@130.171.42\|TGGATGGGAGACCGCCTGGGAATACCGGGTGCTGTAGGCTTT | >hg38_1_-_228634819_228635039_RNA5S12_5S_50ntFlanks | 130 | 171 | 42 | TGGATGGGAGACCGCCTGGGAATACCGGGTGCTGTAGGCTTT | 5157.3 | 9.5 | 0.9 | 10.8 | 4.5E-27 | 4.3E-25 | MB231/MB468 | CYTO |
| >hg38_21_+_8433172_8446622_RNA45SN1_45S_50ntFlanks@3985.4002.18\|CGGCTTTGGTGACTCTAG | >hg38_21_+_8433172_8446622_RNA45SN1_45S_50ntFlanks | 3985 | 4002 | 18 | CGGCTTTGGTGACTCTAG | 7548.4 | -16.7 | 1.6 | -10.7 | 6.7E-27 | 4.3E-25 | MB231/MB468 | MP |
| >hg38_21_+_8433172_8446622_RNA45SN1_45S_50ntFlanks@3835.3869.35\|CTCCTCTCCTACTTGGATAACTGTGGTAATTCTAG | >hg38_21_+_8433172_8446622_RNA45SN1_45S_50ntFlanks | 3835 | 3869 | 35 | CTCCTCTCCTACTTGGATAACTGTGGTAATTCTAG | 583.9 | -13.6 | 1.3 | -10.5 | 5.9E-26 | 4.3E-25 | BT20/MB468 | NUC |
| >hg38_21_+_8433172_8446622_RNA45SN1_45S_50ntFlanks@9033.9059.27\|CTCGTGGGGGGGCCGGGCCACCCCTCC | >hg38_21_+_8433172_8446622_RNA45SN1_45S_50ntFlanks | 9033 | 9059 | 27 | CTCGTGGGGGGGCCGGGCCACCCCTCC | 2494.6 | -15.1 | 1.4 | -10.7 | 6.7E-27 | 4.3E-25 | MB231/MB468 | MP |
| >hg38_21_+_8433172_8446622_RNA45SN1_45S_50ntFlanks@6746.6771.26\|CACTTCGAACGCACTTGCGGCCCCGG | >hg38_21_+_8433172_8446622_RNA45SN1_45S_50ntFlanks | 6746 | 6771 | 26 | CACTTCGAACGCACTTGCGGCCCCGG | 2333.5 | -15.0 | 1.4 | -10.7 | 1.1E-26 | 4.4E-25 | MB231/MB468 | MITO |
| >MI0003667\|hsa-mir-652&WithFlank&X\|+\|110055323\|110055432@67.87.21(+1U)&[MIMAT0003322&hsa-miR-652-3p&offsets\|0\|0(+1U);m-148&X\|+\|110055389\|110055410&offsets\|0\|-1(+1U)]\|AATGGCGCCACTAGGGTTGTGT | >MI0003667\|hsa-mir-652&WithFlank&X\|+\|110055323\|110055432 | 67 | 87 | 21(+1U) | AATGGCGCCACTAGGGTTGTGT | 164.4 | 4.0 | 0.4 | 10.5 | 5.9E-26 | 4.4E-25 | BT20/MB468 | NUC |
| >MI0000077\|hsa-mir-21&WithFlank&17\|+\|59841260\|59841343@14.36.23(+1A)&[MIMAT0000076&hsa-miR-21-5p&offsets\|0\|+1(+1A);m-1&17\|+\|59841273\|59841294&offsets\|0\|+1(+1A)]\|TAGCTTATCAGACTGATGTTGACA | >MI0000077\|hsa-mir-21&WithFlank&17\|+\|59841260\|59841343 | 14 | 36 | 23(+1A) | TAGCTTATCAGACTGATGTTGACA | 75238.5 | 6.7 | 0.6 | 10.6 | 2.4E-26 | 4.4E-25 | BT20/MB468 | MITO |
| >hg38_21_+_8433172_8446622_RNA45SN1_45S_50ntFlanks@9534.9570.37\|AGGAAACTCTGGTGGAGGTCCGTAGCGGTCCTGACGT | >hg38_21_+_8433172_8446622_RNA45SN1_45S_50ntFlanks | 9534 | 9570 | 37 | AGGAAACTCTGGTGGAGGTCCGTAGCGGTCCTGACGT | 5649.9 | -5.6 | 0.5 | -10.7 | 1.5E-26 | 4.4E-25 | MB231/MB468 | NUC |
| >MI0000477\|hsa-mir-146a&WithFlank&5\|+\|160485346\|160485456@27.48.22&[MIMAT0000449&hsa-miR-146a-5p&offsets\|0\|0;m-129&5\|+\|160485372\|160485393&offsets\|0\|0]\|TGAGAACTGAATTCCATGGGTT | >MI0000477\|hsa-mir-146a&WithFlank&5\|+\|160485346\|160485456 | 27 | 48 | 22 | TGAGAACTGAATTCCATGGGTT | 20807.7 | 17.6 | 1.6 | 10.7 | 1.2E-26 | 4.5E-25 | MB231/MB468 | MITO |
| >hg38_21_+_8433172_8446622_RNA45SN1_45S_50ntFlanks@11569.11601.33\|CAGCCGACTTAGAACTGGTGCGGACCAGGGGAA | >hg38_21_+_8433172_8446622_RNA45SN1_45S_50ntFlanks | 11569 | 11601 | 33 | CAGCCGACTTAGAACTGGTGCGGACCAGGGGAA | 8542.1 | -9.0 | 0.8 | -10.7 | 1.2E-26 | 4.5E-25 | MB231/MB468 | MITO |
| >MI0000093\|hsa-mir-92a-1&WithFlank&13\|+\|91351308\|91351397@54.74.21(+1A);MI0000094\|hsa-mir-92a-2&WithFlank&X\|-\|134169532\|134169618@54.74.21(+1A)&[MIMAT0000092&hsa-miR-92a-3p&offsets\|0\|-1(+1A);m-9&13\|+\|91351361\|91351382&offsets\|0\|-1(+1A)];[MIMAT0000092_1&hsa-miR-92a-3p&offsets\|0\|-1(+1A);m-11&X\|-\|134169544\|134169565&offsets\|0\|-1(+1A)]\|TATTGCACTTGTCCCGGCCTGA | >MI0000093\|hsa-mir-92a-1&WithFlank&13\|+\|91351308\|91351397 | 54 | 74 | 21(+1A) | TATTGCACTTGTCCCGGCCTGA | 4045.2 | 7.5 | 0.7 | 10.6 | 3.2E-26 | 4.6E-25 | BT20/MB468 | CYTO |
| >hg38_21_+_8433172_8446622_RNA45SN1_45S_50ntFlanks@12420.12440.21\|ATGTCGGCTCTTCCTATCATT | >hg38_21_+_8433172_8446622_RNA45SN1_45S_50ntFlanks | 12420 | 12440 | 21 | ATGTCGGCTCTTCCTATCATT | 2046.3 | -5.8 | 0.6 | -10.5 | 6.3E-26 | 4.6E-25 | BT20/MB468 | NUC |
| >MI0000272\|hsa-mir-182&WithFlank&7\|-\|129770377\|129770498@29.51.23(+1A)&[MIMAT0000259&hsa-miR-182-5p&offsets\|0\|-1(+1A);m-22&7\|-\|129770449\|129770470&offsets\|0\|+1(+1A)]\|TTTGGCAATGGTAGAACTCACACA | >MI0000272\|hsa-mir-182&WithFlank&7\|-\|129770377\|129770498 | 29 | 51 | 23(+1A) | TTTGGCAATGGTAGAACTCACACA | 1022.1 | 3.8 | 0.4 | 10.5 | 6.3E-26 | 4.6E-25 | BT20/MB468 | NUC |
| >hg38_21_+_8433172_8446622_RNA45SN1_45S_50ntFlanks@6745.6770.26\|ACACTTCGAACGCACTTGCGGCCCCG | >hg38_21_+_8433172_8446622_RNA45SN1_45S_50ntFlanks | 6745 | 6770 | 26 | ACACTTCGAACGCACTTGCGGCCCCG | 12520.0 | -10.4 | 1.0 | -10.7 | 1.2E-26 | 4.6E-25 | MB231/MB468 | MITO |
| >hg38_21_+_8433172_8446622_RNA45SN1_45S_50ntFlanks@12539.12569.31\|ATGATGTGTTGTTGCCATGGTAATCCTGCTC | >hg38_21_+_8433172_8446622_RNA45SN1_45S_50ntFlanks | 12539 | 12569 | 31 | ATGATGTGTTGTTGCCATGGTAATCCTGCTC | 2396.2 | -15.1 | 1.4 | -10.7 | 7.2E-27 | 4.6E-25 | MB231/MB468 | MP |
| >MI0001733\|hsa-mir-452&WithFlank&X\|-\|151959622\|151959718@23.43.21&[MIMAT0001635&hsa-miR-452-5p&offsets\|+3\|+2;m-188&X\|-\|151959677\|151959699&offsets\|+3\|+1]\|TGTTTGCAGAGGAAACTGAGA | >MI0001733\|hsa-mir-452&WithFlank&X\|-\|151959622\|151959718 | 23 | 43 | 21 | TGTTTGCAGAGGAAACTGAGA | 1318.2 | 14.6 | 1.4 | 10.6 | 3.2E-26 | 4.6E-25 | BT20/MB468 | CYTO |

<table>
<tr><td>>hg38_21_+_8433172_8446622_RNA45SN1_45S_50ntFlanks@10349.10371.23|AACGAGAACTTTGAAGGCCGAAG</td><td>>hg38_21_+_8433172_8446622_RNA45SN1_45S_50ntFlanks</td><td>10349</td><td>10371</td><td>23</td><td>AACGAGAACTTTGAAGGCCGAAG</td><td>1793.6</td><td>-4.6</td><td>0.4</td><td>-10.5</td><td>6.3E-26</td><td>4.6E-25</td><td>BT20/MB468</td><td>NUC</td></tr>
<tr><td>>hg38_21_+_8433172_8446622_RNA45SN1_45S_50ntFlanks@10684.10711.28|CGGCGGCGTCCGGTGAGCTCTCGCTGGC</td><td>>hg38_21_+_8433172_8446622_RNA45SN1_45S_50ntFlanks</td><td>10684</td><td>10711</td><td>28</td><td>CGGCGGCGTCCGGTGAGCTCTCGCTGGC</td><td>13131.3</td><td>-9.1</td><td>0.9</td><td>-10.5</td><td>6.4E-26</td><td>4.7E-25</td><td>BT20/MB468</td><td>NUC</td></tr>
<tr><td>>hg38_21_+_8433172_8446622_RNA45SN1_45S_50ntFlanks@4240.4258.19|ACTCTTTCGAGGCCCTGTA</td><td>>hg38_21_+_8433172_8446622_RNA45SN1_45S_50ntFlanks</td><td>4240</td><td>4258</td><td>19</td><td>ACTCTTTCGAGGCCCTGTA</td><td>363.8</td><td>-12.9</td><td>1.2</td><td>-10.5</td><td>6.4E-26</td><td>4.7E-25</td><td>BT20/MB468</td><td>NUC</td></tr>
<tr><td>>hg38_21_+_8433172_8446622_RNA45SN1_45S_50ntFlanks@12457.12484.28|CAAGCGTTGGATTGTTCACCCACTAATA</td><td>>hg38_21_+_8433172_8446622_RNA45SN1_45S_50ntFlanks</td><td>12457</td><td>12484</td><td>28</td><td>CAAGCGTTGGATTGTTCACCCACTAATA</td><td>1063.5</td><td>-14.5</td><td>1.4</td><td>-10.5</td><td>6.4E-26</td><td>4.7E-25</td><td>BT20/MB468</td><td>NUC</td></tr>
<tr><td>>hg38_21_+_8433172_8446622_RNA45SN1_45S_50ntFlanks@12954.12982.29|CTGCTTCTGGGTCGGGGTTTCGTACGTAG</td><td>>hg38_21_+_8433172_8446622_RNA45SN1_45S_50ntFlanks</td><td>12954</td><td>12982</td><td>29</td><td>CTGCTTCTGGGTCGGGGTTTCGTACGTAG</td><td>953.1</td><td>-14.3</td><td>1.4</td><td>-10.5</td><td>6.5E-26</td><td>4.8E-25</td><td>BT20/MB468</td><td>NUC</td></tr>
<tr><td>>MI0000272|hsa-mir-182&WithFlank&7|-|129770377|129770498@29.53.25(+1U)&[MIMAT0000259&hsa-miR-182-5p&offsets|0|+1(+1U);m-22&7|-|129770449|129770470&offsets|0|+3(+1U)]|TTTGGCAATGGTAGAACTCACACTGT</td><td>>MI0000272|hsa-mir-182&WithFlank&7|-|129770377|129770498</td><td>29</td><td>53</td><td>25(+1U)</td><td>TTTGGCAATGGTAGAACTCACACTGT</td><td>1513.9</td><td>7.7</td><td>0.7</td><td>10.6</td><td>2.6E-26</td><td>4.8E-25</td><td>BT20/MB468</td><td>MITO</td></tr>
<tr><td>>MI0000088|hsa-mir-30a&WithFlank&6|-|71403545|71403627@13.35.23&[MIMAT0000087&hsa-miR-30a-5p&offsets|+1|+2;m-15&6|-|71403595|71403616&offsets|+1|+2]|GTAAACATCCTCGACTGGAAGCT</td><td>>MI0000088|hsa-mir-30a&WithFlank&6|-|71403545|71403627</td><td>13</td><td>35</td><td>23</td><td>GTAAACATCCTCGACTGGAAGCT</td><td>1360.3</td><td>7.9</td><td>0.7</td><td>10.7</td><td>1.7E-26</td><td>4.8E-25</td><td>MB231/MB468</td><td>NUC</td></tr>
<tr><td>>hg38_21_+_8433172_8446622_RNA45SN1_45S_50ntFlanks@12991.13014.24|CTCCCTCGCTGCGATCTATTGAAA</td><td>>hg38_21_+_8433172_8446622_RNA45SN1_45S_50ntFlanks</td><td>12991</td><td>13014</td><td>24</td><td>CTCCCTCGCTGCGATCTATTGAAA</td><td>14716.1</td><td>-8.6</td><td>0.8</td><td>-10.7</td><td>1.7E-26</td><td>4.9E-25</td><td>MB231/MB468</td><td>NUC</td></tr>
<tr><td>>hg38_21_+_8433172_8446622_RNA45SN1_45S_50ntFlanks@9027.9065.39|CGTCTCCTCGTGGGGGGGCCGGGCCACCCCTCCCACGGC</td><td>>hg38_21_+_8433172_8446622_RNA45SN1_45S_50ntFlanks</td><td>9027</td><td>9065</td><td>39</td><td>CGTCTCCTCGTGGGGGGGCCGGGCCACCCCTCCCACGGC</td><td>362.7</td><td>-12.9</td><td>1.2</td><td>-10.5</td><td>6.7E-26</td><td>4.9E-25</td><td>BT20/MB468</td><td>NUC</td></tr>
<tr><td>>hg38_21_+_8433172_8446622_RNA45SN1_45S_50ntFlanks@9564.9596.33|CTGACGTGCAAATCGGTCGTCCGACCTGGGTAT</td><td>>hg38_21_+_8433172_8446622_RNA45SN1_45S_50ntFlanks</td><td>9564</td><td>9596</td><td>33</td><td>CTGACGTGCAAATCGGTCGTCCGACCTGGGTAT</td><td>72039.4</td><td>-13.1</td><td>1.2</td><td>-10.5</td><td>6.8E-26</td><td>4.9E-25</td><td>BT20/MB468</td><td>NUC</td></tr>
<tr><td>>hg38_21_+_8433172_8446622_RNA45SN1_45S_50ntFlanks@12626.12653.28|AATGGGGCGAAGCTACCATCTGTGGGAT</td><td>>hg38_21_+_8433172_8446622_RNA45SN1_45S_50ntFlanks</td><td>12626</td><td>12653</td><td>28</td><td>AATGGGGCGAAGCTACCATCTGTGGGAT</td><td>10914.9</td><td>-12.7</td><td>1.2</td><td>-10.7</td><td>1.3E-26</td><td>4.9E-25</td><td>MB231/MB468</td><td>MITO</td></tr>
<tr><td>>MI0000088|hsa-mir-30a&WithFlank&6|-|71403545|71403627@12.34.23&[MIMAT0000087&hsa-miR-30a-5p&offsets|0|+1;m-15&6|-|71403595|71403616&offsets|0|+1]|TGTAAACATCCTCGACTGGAAGC</td><td>>MI0000088|hsa-mir-30a&WithFlank&6|-|71403545|71403627</td><td>12</td><td>34</td><td>23</td><td>TGTAAACATCCTCGACTGGAAGC</td><td>13646.8</td><td>6.8</td><td>0.6</td><td>10.6</td><td>3.5E-26</td><td>5.0E-25</td><td>BT20/MB468</td><td>CYTO</td></tr>
<tr><td>>hg38_21_+_8433172_8446622_RNA45SN1_45S_50ntFlanks@6778.6806.29|CCCGGGGCTACGCCTGTCTGAGCGTCGCT</td><td>>hg38_21_+_8433172_8446622_RNA45SN1_45S_50ntFlanks</td><td>6778</td><td>6806</td><td>29</td><td>CCCGGGGCTACGCCTGTCTGAGCGTCGCT</td><td>1280.4</td><td>6.1</td><td>0.6</td><td>10.6</td><td>3.5E-26</td><td>5.0E-25</td><td>BT20/MB468</td><td>CYTO</td></tr>
<tr><td>>trna2_GlyGCC_21_-_18827107_18827177@1.30.30,trna35_GlyGCC_1_+_161413094_161413164@1.30.30,trna37_GlyGCC_1_+_161420467_161420537@1.30.30,trna39_GlyGCC_1_+_161427898_161427968@1.30.30,trna41_GlyGCC_1_+_161435258_161435328@1.30.30|GCATGGGTGGTTCAGTGGTAGAATTCTCGC</td><td>>trna2_GlyGCC_21_-_18827107_18827177</td><td>1</td><td>30</td><td>30</td><td>GCATGGGTGGTTCAGTGGTAGAATTCTCGC</td><td>8308.3</td><td>-6.4</td><td>0.6</td><td>-10.5</td><td>6.9E-26</td><td>5.0E-25</td><td>BT20/MB468</td><td>NUC</td></tr>
<tr><td>>hg38_21_+_8433172_8446622_RNA45SN1_45S_50ntFlanks@10408.10425.18|GGTCAGTCGGTCCTGAGA</td><td>>hg38_21_+_8433172_8446622_RNA45SN1_45S_50ntFlanks</td><td>10408</td><td>10425</td><td>18</td><td>GGTCAGTCGGTCCTGAGA</td><td>722.4</td><td>-13.9</td><td>1.3</td><td>-10.5</td><td>7.0E-26</td><td>5.1E-25</td><td>BT20/MB468</td><td>NUC</td></tr>
<tr><td>>hg38_21_+_8433172_8446622_RNA45SN1_45S_50ntFlanks@12958.12982.25|TTCTGGGTCGGGGTTTCGTACGTAG</td><td>>hg38_21_+_8433172_8446622_RNA45SN1_45S_50ntFlanks</td><td>12958</td><td>12982</td><td>25</td><td>TTCTGGGTCGGGGTTTCGTACGTAG</td><td>521.5</td><td>-13.5</td><td>1.3</td><td>-10.5</td><td>7.0E-26</td><td>5.1E-25</td><td>BT20/MB468</td><td>NUC</td></tr>
<tr><td>>hg38_21_+_8433172_8446622_RNA45SN1_45S_50ntFlanks@10366.10383.18|CCGAAGTGGAGAAGGGTT</td><td>>hg38_21_+_8433172_8446622_RNA45SN1_45S_50ntFlanks</td><td>10366</td><td>10383</td><td>18</td><td>CCGAAGTGGAGAAGGGTT</td><td>302.4</td><td>-12.7</td><td>1.2</td><td>-10.5</td><td>7.0E-26</td><td>5.1E-25</td><td>BT20/MB468</td><td>NUC</td></tr>
<tr><td>>MI0003560|hsa-mir-92b&WithFlank&1|+|155195171|155195278@67.86.20(+1U)&[MIMAT0003218&hsa-miR-92b-3p&offsets|0|-2(+1U);m-115&1|+|155195237|155195258&offsets|0|-2(+1U)]|TATTGCACTCGTCCCGGCCTT</td><td>>MI0003560|hsa-mir-92b&WithFlank&1|+|155195171|155195278</td><td>67</td><td>86</td><td>20(+1U)</td><td>TATTGCACTCGTCCCGGCCTT</td><td>737.0</td><td>13.8</td><td>1.3</td><td>10.6</td><td>3.6E-26</td><td>5.1E-25</td><td>BT20/MB468</td><td>CYTO</td></tr>
<tr><td>>hg38_21_+_8433172_8446622_RNA45SN1_45S_50ntFlanks@11586.11611.26|GTGCGGACCAGGGGAATCCGACTGTT</td><td>>hg38_21_+_8433172_8446622_RNA45SN1_45S_50ntFlanks</td><td>11586</td><td>11611</td><td>26</td><td>GTGCGGACCAGGGGAATCCGACTGTT</td><td>4195.9</td><td>-5.9</td><td>0.6</td><td>-10.5</td><td>7.1E-26</td><td>5.1E-25</td><td>BT20/MB468</td><td>NUC</td></tr>
<tr><td>>hg38_21_+_8433172_8446622_RNA45SN1_45S_50ntFlanks@10412.10430.19|AGTCGGTCCTGAGAGATGG</td><td>>hg38_21_+_8433172_8446622_RNA45SN1_45S_50ntFlanks</td><td>10412</td><td>10430</td><td>19</td><td>AGTCGGTCCTGAGAGATGG</td><td>369.9</td><td>-13.0</td><td>1.2</td><td>-10.5</td><td>7.2E-26</td><td>5.2E-25</td><td>BT20/MB468</td><td>NUC</td></tr>
<tr><td>>hg38_21_+_8433172_8446622_RNA45SN1_45S_50ntFlanks@13002.13029.28|CGATCTATTGAAAGTCAGCCCTCGACAC</td><td>>hg38_21_+_8433172_8446622_RNA45SN1_45S_50ntFlanks</td><td>13002</td><td>13029</td><td>28</td><td>CGATCTATTGAAAGTCAGCCCTCGACAC</td><td>2911.7</td><td>-14.5</td><td>1.4</td><td>-10.6</td><td>1.8E-26</td><td>5.2E-25</td><td>MB231/MB468</td><td>NUC</td></tr>
<tr><td>>hg38_21_+_8433172_8446622_RNA45SN1_45S_50ntFlanks@11729.11753.25|TAACTATGACTCTCTTAAGGTAGCC</td><td>>hg38_21_+_8433172_8446622_RNA45SN1_45S_50ntFlanks</td><td>11729</td><td>11753</td><td>25</td><td>TAACTATGACTCTCTTAAGGTAGCC</td><td>659.3</td><td>-13.8</td><td>1.3</td><td>-10.5</td><td>7.3E-26</td><td>5.2E-25</td><td>BT20/MB468</td><td>NUC</td></tr>
<tr><td>>MI0000440|hsa-mir-27b&WithFlank&9|+|95085439|95085547@67.85.19(+1A)&[MIMAT0000419&hsa-miR-27b-3p&offsets|0|-2(+1A);m-39&9|+|95085505|95085525&offsets|0|-2(+1A)]|TTCACAGTGGCTAAGTTCTA</td><td>>MI0000440|hsa-mir-27b&WithFlank&9|+|95085439|95085547</td><td>67</td><td>85</td><td>19(+1A)</td><td>TTCACAGTGGCTAAGTTCTA</td><td>369.3</td><td>6.7</td><td>0.6</td><td>10.6</td><td>1.9E-26</td><td>5.3E-25</td><td>MB231/MB468</td><td>NUC</td></tr>
<tr><td>>MI0000285|hsa-mir-205&WithFlank&1|+|209432127|209432248@40.59.20(+1G)&[MIMAT0000266&hsa-miR-205-5p&offsets|0|-2(+1G);m-351&1|+|209432166|209432187&offsets|0|-2(+1G)]|TCCTTCATTCCACCGGAGTCG</td><td>>MI0000285|hsa-mir-205&WithFlank&1|+|209432127|209432248</td><td>40</td><td>59</td><td>20(+1G)</td><td>TCCTTCATTCCACCGGAGTCG</td><td>393.5</td><td>6.3</td><td>0.6</td><td>10.6</td><td>3.7E-26</td><td>5.3E-25</td><td>BT20/MB468</td><td>CYTO</td></tr>
<tr><td>>hg38_21_+_8433172_8446622_RNA45SN1_45S_50ntFlanks@3769.3799.31|CGCACGGCCGGTACAGTGAAACTGCGAATGG</td><td>>hg38_21_+_8433172_8446622_RNA45SN1_45S_50ntFlanks</td><td>3769</td><td>3799</td><td>31</td><td>CGCACGGCCGGTACAGTGAAACTGCGAATGG</td><td>391.4</td><td>-13.0</td><td>1.2</td><td>-10.5</td><td>7.4E-26</td><td>5.3E-25</td><td>BT20/MB468</td><td>NUC</td></tr>
<tr><td>>hg38_21_+_8433172_8446622_RNA45SN1_45S_50ntFlanks@12937.12961.25|AACCATTCGTAGACGACCTGCTTCT</td><td>>hg38_21_+_8433172_8446622_RNA45SN1_45S_50ntFlanks</td><td>12937</td><td>12961</td><td>25</td><td>AACCATTCGTAGACGACCTGCTTCT</td><td>855.3</td><td>-5.3</td><td>0.5</td><td>-10.6</td><td>2.0E-26</td><td>5.5E-25</td><td>MB231/MB468</td><td>NUC</td></tr>
<tr><td>>hg38_21_+_8433172_8446622_RNA45SN1_45S_50ntFlanks@10346.10367.22|TCAAACGAGAACTTTGAAGGCC</td><td>>hg38_21_+_8433172_8446622_RNA45SN1_45S_50ntFlanks</td><td>10346</td><td>10367</td><td>22</td><td>TCAAACGAGAACTTTGAAGGCC</td><td>324.9</td><td>-4.6</td><td>0.4</td><td>-10.5</td><td>7.6E-26</td><td>5.5E-25</td><td>BT20/MB468</td><td>NUC</td></tr>
<tr><td>>hg38_21_+_8433172_8446622_RNA45SN1_45S_50ntFlanks@10690.10710.21|CGTCCGGTGAGCTCTCGCTGG</td><td>>hg38_21_+_8433172_8446622_RNA45SN1_45S_50ntFlanks</td><td>10690</td><td>10710</td><td>21</td><td>CGTCCGGTGAGCTCTCGCTGG</td><td>800.7</td><td>-14.1</td><td>1.3</td><td>-10.5</td><td>7.6E-26</td><td>5.5E-25</td><td>BT20/MB468</td><td>NUC</td></tr>
<tr><td>>hg38_21_+_8433172_8446622_RNA45SN1_45S_50ntFlanks@9738.9762.25|GAAACGATCTCAACCTATTCTCAAA</td><td>>hg38_21_+_8433172_8446622_RNA45SN1_45S_50ntFlanks</td><td>9738</td><td>9762</td><td>25</td><td>GAAACGATCTCAACCTATTCTCAAA</td><td>461.3</td><td>-13.3</td><td>1.3</td><td>-10.5</td><td>7.7E-26</td><td>5.5E-25</td><td>BT20/MB468</td><td>NUC</td></tr>
<tr><td>>hg38_21_+_8433172_8446622_RNA45SN1_45S_50ntFlanks@11318.11342.25|CCCCACGTCTCGTCGCGCGCGCGTC</td><td>>hg38_21_+_8433172_8446622_RNA45SN1_45S_50ntFlanks</td><td>11318</td><td>11342</td><td>25</td><td>CCCCACGTCTCGTCGCGCGCGCGTC</td><td>2880.3</td><td>-14.5</td><td>1.4</td><td>-10.6</td><td>2.0E-26</td><td>5.5E-25</td><td>MB231/MB468</td><td>NUC</td></tr>
<tr><td>>hg38_21_+_8433172_8446622_RNA45SN1_45S_50ntFlanks@10840.10865.26|CCGTAACTTCGGGATAAGGATTGGCT</td><td>>hg38_21_+_8433172_8446622_RNA45SN1_45S_50ntFlanks</td><td>10840</td><td>10865</td><td>26</td><td>CCGTAACTTCGGGATAAGGATTGGCT</td><td>2031.4</td><td>-7.4</td><td>0.7</td><td>-10.7</td><td>1.5E-26</td><td>5.5E-25</td><td>MB231/MB468</td><td>MITO</td></tr>
<tr><td>>MI0000285|hsa-mir-205&WithFlank&1|+|209432127|209432248@40.62.23&[MIMAT0000266&hsa-miR-205-5p&offsets|0|+1;m-351&1|+|209432166|209432187&offsets|0|+1]|TCCTTCATTCCACCGGAGTCTGT</td><td>>MI0000285|hsa-mir-205&WithFlank&1|+|209432127|209432248</td><td>40</td><td>62</td><td>23</td><td>TCCTTCATTCCACCGGAGTCTGT</td><td>75198.9</td><td>10.0</td><td>0.9</td><td>11.0</td><td>4.6E-28</td><td>5.6E-25</td><td>BT20/MB231</td><td>CYTO</td></tr>
<tr><td>>hg38_21_+_8433172_8446622_RNA45SN1_45S_50ntFlanks@7975.8005.31|CGCGACCTCAGATCAGACGTGGCGACCCGCT</td><td>>hg38_21_+_8433172_8446622_RNA45SN1_45S_50ntFlanks</td><td>7975</td><td>8005</td><td>31</td><td>CGCGACCTCAGATCAGACGTGGCGACCCGCT</td><td>830759.4</td><td>7.5</td><td>0.7</td><td>10.6</td><td>3.0E-26</td><td>5.6E-25</td><td>BT20/MB468</td><td>MITO</td></tr>
<tr><td>>hg38_21_+_8433172_8446622_RNA45SN1_45S_50ntFlanks@11918.11951.34|ACATGAGAGGTGTAGAATAAGTGGGAGGCCCCCG</td><td>>hg38_21_+_8433172_8446622_RNA45SN1_45S_50ntFlanks</td><td>11918</td><td>11951</td><td>34</td><td>ACATGAGAGGTGTAGAATAAGTGGGAGGCCCCCG</td><td>1651.1</td><td>-8.6</td><td>0.8</td><td>-10.7</td><td>6.0E-27</td><td>5.6E-25</td><td>MB231/MB468</td><td>CYTO</td></tr>
<tr><td>>hg38_21_+_8433172_8446622_RNA45SN1_45S_50ntFlanks@3815.3845.31|ATGGTTCCTTTGGTCGCTCGCTCCTCTCCTA</td><td>>hg38_21_+_8433172_8446622_RNA45SN1_45S_50ntFlanks</td><td>3815</td><td>3845</td><td>31</td><td>ATGGTTCCTTTGGTCGCTCGCTCCTCTCCTA</td><td>488.3</td><td>-13.4</td><td>1.3</td><td>-10.5</td><td>7.9E-26</td><td>5.6E-25</td><td>BT20/MB468</td><td>NUC</td></tr>
<tr><td>>hg38_21_+_8433172_8446622_RNA45SN1_45S_50ntFlanks@6731.6771.41|CACATTGATCATCGACACTTCGAACGCACTTGCGGCCCCGG</td><td>>hg38_21_+_8433172_8446622_RNA45SN1_45S_50ntFlanks</td><td>6731</td><td>6771</td><td>41</td><td>CACATTGATCATCGACACTTCGAACGCACTTGCGGCCCCGG</td><td>3273.7</td><td>-16.1</td><td>1.5</td><td>-10.5</td><td>7.9E-26</td><td>5.7E-25</td><td>BT20/MB468</td><td>NUC</td></tr>
<tr><td>>MI0000086|hsa-mir-28&WithFlank&3|+|188688775|188688872@60.81.22(+1U)&[MIMAT0004502&hsa-miR-28-3p&offsets|0|0(+1U);m-28&3|+|188688834|188688855&offsets|0|0(+1U)]|CACTAGATTGTGAGCTCCTGGAT</td><td>>MI0000086|hsa-mir-28&WithFlank&3|+|188688775|188688872</td><td>60</td><td>81</td><td>22(+1U)</td><td>CACTAGATTGTGAGCTCCTGGAT</td><td>663.8</td><td>13.8</td><td>1.3</td><td>10.6</td><td>3.1E-26</td><td>5.7E-25</td><td>BT20/MB468</td><td>MITO</td></tr>
<tr><td>>hg38_21_+_8433172_8446622_RNA45SN1_45S_50ntFlanks@3872.3899.28|CTAATACATGCCGACGGGCGCTGACCCC</td><td>>hg38_21_+_8433172_8446622_RNA45SN1_45S_50ntFlanks</td><td>3872</td><td>3899</td><td>28</td><td>CTAATACATGCCGACGGGCGCTGACCCC</td><td>443.3</td><td>-13.2</td><td>1.3</td><td>-10.5</td><td>8.1E-26</td><td>5.7E-25</td><td>BT20/MB468</td><td>NUC</td></tr>
<tr><td>>MI0000077|hsa-mir-21&WithFlank&17|+|59841260|59841343@13.35.23&[MIMAT0000076&hsa-miR-21-5p&offsets|-1|0;m-1&17|+|59841273|59841294&offsets|-1|0]|GTAGCTTATCAGACTGATGTTGA</td><td>>MI0000077|hsa-mir-21&WithFlank&17|+|59841260|59841343</td><td>13</td><td>35</td><td>23</td><td>GTAGCTTATCAGACTGATGTTGA</td><td>466.9</td><td>5.6</td><td>0.5</td><td>10.6</td><td>2.1E-26</td><td>5.8E-25</td><td>MB231/MB468</td><td>NUC</td></tr>
<tr><td>>hg38_21_+_8433172_8446622_RNA45SN1_45S_50ntFlanks@10794.10818.25|CTGGCATGTTGGAACAATGTAGGTA</td><td>>hg38_21_+_8433172_8446622_RNA45SN1_45S_50ntFlanks</td><td>10794</td><td>10818</td><td>25</td><td>CTGGCATGTTGGAACAATGTAGGTA</td><td>3497.8</td><td>-9.8</td><td>0.9</td><td>-10.7</td><td>1.5E-26</td><td>5.8E-25</td><td>MB231/MB468</td><td>MITO</td></tr>
<tr><td>>MI0000266|hsa-mir-10a&WithFlank&17|-|48579832|48579953@28.48.21&[MIMAT0000253&hsa-miR-10a-5p&offsets|0|-2;m-18&17|-|48579905|48579926&offsets|0|-1]|TACCCTGTAGATCCGAATTTG</td><td>>MI0000266|hsa-mir-10a&WithFlank&17|-|48579832|48579953</td><td>28</td><td>48</td><td>21</td><td>TACCCTGTAGATCCGAATTTG</td><td>1386.7</td><td>14.7</td><td>1.4</td><td>10.6</td><td>4.1E-26</td><td>5.9E-25</td><td>BT20/MB468</td><td>CYTO</td></tr>
<tr><td>>hg38_21_+_8433172_8446622_RNA45SN1_45S_50ntFlanks@8114.8138.25|GCGCGGGACATGTGGCGTACGGAAG</td><td>>hg38_21_+_8433172_8446622_RNA45SN1_45S_50ntFlanks</td><td>8114</td><td>8138</td><td>25</td><td>GCGCGGGACATGTGGCGTACGGAAG</td><td>3815.5</td><td>-15.7</td><td>1.5</td><td>-10.7</td><td>1.6E-26</td><td>5.9E-25</td><td>MB231/MB468</td><td>MITO</td></tr>
<tr><td>>hg38_21_+_8433172_8446622_RNA45SN1_45S_50ntFlanks@3803.3829.27|ATTAAATCAGTTATGGTTCCTTTGGTC</td><td>>hg38_21_+_8433172_8446622_RNA45SN1_45S_50ntFlanks</td><td>3803</td><td>3829</td><td>27</td><td>ATTAAATCAGTTATGGTTCCTTTGGTC</td><td>355.8</td><td>-12.9</td><td>1.2</td><td>-10.5</td><td>8.3E-26</td><td>5.9E-25</td><td>BT20/MB468</td><td>NUC</td></tr>
<tr><td>>hg38_21_+_8433172_8446622_RNA45SN1_45S_50ntFlanks@11621.11640.20|CAAAGCATCGCGAAGGCCCG</td><td>>hg38_21_+_8433172_8446622_RNA45SN1_45S_50ntFlanks</td><td>11621</td><td>11640</td><td>20</td><td>CAAAGCATCGCGAAGGCCCG</td><td>3910.3</td><td>-15.7</td><td>1.5</td><td>-10.7</td><td>1.6E-26</td><td>6.1E-25</td><td>MB231/MB468</td><td>MITO</td></tr>
<tr><td>>hg38_21_+_8433172_8446622_RNA45SN1_45S_50ntFlanks@4161.4182.22|CACATCCAAGGAAGGCAGCAGG</td><td>>hg38_21_+_8433172_8446622_RNA45SN1_45S_50ntFlanks</td><td>4161</td><td>4182</td><td>22</td><td>CACATCCAAGGAAGGCAGCAGG</td><td>392.0</td><td>-13.0</td><td>1.2</td><td>-10.5</td><td>8.6E-26</td><td>6.1E-25</td><td>BT20/MB468</td><td>NUC</td></tr>
<tr><td>>hg38_21_+_8433172_8446622_RNA45SN1_45S_50ntFlanks@11233.11250.18|CCCTTCCCGTGGATCGCC</td><td>>hg38_21_+_8433172_8446622_RNA45SN1_45S_50ntFlanks</td><td>11233</td><td>11250</td><td>18</td><td>CCCTTCCCGTGGATCGCC</td><td>1127.6</td><td>-14.6</td><td>1.4</td><td>-10.5</td><td>8.7E-26</td><td>6.2E-25</td><td>BT20/MB468</td><td>NUC</td></tr>
<tr><td>>MI0000111|hsa-mir-105-1&WithFlank&X|-|152392213|152392305@19.41.23;MI0000112|hsa-mir-105-2&WithFlank&X|-|152394406|152394498@19.41.23&[MIMAT0000102&hsa-miR-105-5p&offsets|0|0;m-735&X|-|152392266|152392287&offsets|0|+1];[MIMAT0000102_1&hsa-miR-105-5p&offsets|0|0;m-735&X|-|152394459|152394480&offsets|0|+1]|TCAAATGCTCAGACTCCTGTGGT</td><td>>MI0000111|hsa-mir-105-1&WithFlank&X|-|152392213|152392305</td><td>19</td><td>41</td><td>23</td><td>TCAAATGCTCAGACTCCTGTGGT</td><td>340.0</td><td>5.9</td><td>0.6</td><td>10.6</td><td>4.4E-26</td><td>6.2E-25</td><td>BT20/MB468</td><td>CYTO</td></tr>
</table>

<table>
<tr><td>>hg38_21_+_8433172_8446622_RNA45SN1_45S_50ntFlanks@11821.11856.36|CTATCCAGCGAAACCACAGCCAAGGGAACGGGCTTG</td><td>>hg38_21_+_8433172_8446622_RNA45SN1_45S_50ntFlanks</td><td>11821</td><td>11856</td><td>36</td><td>CTATCCAGCGAAACCACAGCCAAGGGAACGGGCTTG</td><td>397.6</td><td>-13.1</td><td>1.2</td><td>-10.5</td><td>8.8E-26</td><td>6.2E-25</td><td>BT20/MB468</td><td>NUC</td></tr>
<tr><td>>hg38_21_+_8433172_8446622_RNA45SN1_45S_50ntFlanks@4555.4577.23|CGCAGCTAGGAATAATGGAATAG</td><td>>hg38_21_+_8433172_8446622_RNA45SN1_45S_50ntFlanks</td><td>4555</td><td>4577</td><td>23</td><td>CGCAGCTAGGAATAATGGAATAG</td><td>429.2</td><td>-13.2</td><td>1.3</td><td>-10.5</td><td>8.8E-26</td><td>6.3E-25</td><td>BT20/MB468</td><td>NUC</td></tr>
<tr><td>>hg38_21_+_8433172_8446622_RNA45SN1_45S_50ntFlanks@4002.4024.23|GATAACCTCGGGCCGATCGCACG</td><td>>hg38_21_+_8433172_8446622_RNA45SN1_45S_50ntFlanks</td><td>4002</td><td>4024</td><td>23</td><td>GATAACCTCGGGCCGATCGCACG</td><td>2664.9</td><td>-6.5</td><td>0.6</td><td>-10.6</td><td>2.3E-26</td><td>6.3E-25</td><td>MB231/MB468</td><td>NUC</td></tr>
<tr><td>>hg38_21_+_8433172_8446622_RNA45SN1_45S_50ntFlanks@10351.10375.25|CGAGAACTTTGAAGGCCGAAGTGGA</td><td>>hg38_21_+_8433172_8446622_RNA45SN1_45S_50ntFlanks</td><td>10351</td><td>10375</td><td>25</td><td>CGAGAACTTTGAAGGCCGAAGTGGA</td><td>481.9</td><td>-13.3</td><td>1.3</td><td>-10.5</td><td>8.9E-26</td><td>6.3E-25</td><td>BT20/MB468</td><td>NUC</td></tr>
<tr><td>>MI0000082|hsa-mir-25&WithFlank&7|-|100093554|100093649@58.78.21(+1C)&[MIMAT0000081&hsa-miR-25-3p&offsets|0|-1(+1C);m-23&7|-|100093571|100093592&offsets|0|-1(+1C)]|CATTGCACTTGTCTCGGTCTGC</td><td>>MI0000082|hsa-mir-25&WithFlank&7|-|100093554|100093649</td><td>58</td><td>78</td><td>21(+1C)</td><td>CATTGCACTTGTCTCGGTCTGC</td><td>729.2</td><td>8.5</td><td>0.8</td><td>10.6</td><td>4.5E-26</td><td>6.4E-25</td><td>BT20/MB468</td><td>CYTO</td></tr>
<tr><td>>hg38_21_+_8433172_8446622_RNA45SN1_45S_50ntFlanks@12363.12382.20|GGATAACTGGCTTGTGGCGG</td><td>>hg38_21_+_8433172_8446622_RNA45SN1_45S_50ntFlanks</td><td>12363</td><td>12382</td><td>20</td><td>GGATAACTGGCTTGTGGCGG</td><td>1067.8</td><td>-14.5</td><td>1.4</td><td>-10.5</td><td>9.0E-26</td><td>6.4E-25</td><td>BT20/MB468</td><td>NUC</td></tr>
<tr><td>>hg38_21_+_8433172_8446622_RNA45SN1_45S_50ntFlanks@9539.9563.25|ACTCTGGTGGAGGTCCGTAGCGGTC</td><td>>hg38_21_+_8433172_8446622_RNA45SN1_45S_50ntFlanks</td><td>9539</td><td>9563</td><td>25</td><td>ACTCTGGTGGAGGTCCGTAGCGGTC</td><td>988.7</td><td>-14.4</td><td>1.4</td><td>-10.5</td><td>9.1E-26</td><td>6.4E-25</td><td>BT20/MB468</td><td>NUC</td></tr>
<tr><td>>hg38_21_+_8433172_8446622_RNA45SN1_45S_50ntFlanks@10421.10443.23|TGAGAGATGGGCGAGCGCCGTTC</td><td>>hg38_21_+_8433172_8446622_RNA45SN1_45S_50ntFlanks</td><td>10421</td><td>10443</td><td>23</td><td>TGAGAGATGGGCGAGCGCCGTTC</td><td>305.3</td><td>-12.7</td><td>1.2</td><td>-10.5</td><td>9.2E-26</td><td>6.5E-25</td><td>BT20/MB468</td><td>NUC</td></tr>
<tr><td>>hg38_21_+_8433172_8446622_RNA45SN1_45S_50ntFlanks@5485.5526.42|AGCGCTGAGAAGACGGTCGAACTTGACTATCTAGAGGAAGTA</td><td>>hg38_21_+_8433172_8446622_RNA45SN1_45S_50ntFlanks</td><td>5485</td><td>5526</td><td>42</td><td>AGCGCTGAGAAGACGGTCGAACTTGACTATCTAGAGGAAGTA</td><td>2768.8</td><td>-15.2</td><td>1.4</td><td>-10.6</td><td>1.8E-26</td><td>6.5E-25</td><td>MB231/MB468</td><td>MITO</td></tr>
<tr><td>>hg38_21_+_8433172_8446622_RNA45SN1_45S_50ntFlanks@11564.11588.25|CCTAGCAGCCGACTTAGAACTGGTG</td><td>>hg38_21_+_8433172_8446622_RNA45SN1_45S_50ntFlanks</td><td>11564</td><td>11588</td><td>25</td><td>CCTAGCAGCCGACTTAGAACTGGTG</td><td>880.4</td><td>-5.1</td><td>0.5</td><td>-10.6</td><td>2.4E-26</td><td>6.5E-25</td><td>MB231/MB468</td><td>NUC</td></tr>
<tr><td>>MI0000736|hsa-mir-30c-1&WithFlank&1|+|40757278|40757378@23.44.22(+1A);MI0000254|hsa-mir-30c-2&WithFlank&6|-|71376954|71377037@13.34.22(+1A)&[MIMAT0000244&hsa-miR-30c-5p&offsets|0|-1(+1A);m-57&1|+|40757300|40757323&offsets|0|-2(+1A)];[MIMAT0000244_1&hsa-miR-30c-5p&offsets|0|-1(+1A);m-56&6|-|71377002|71377025&offsets|0|-2(+1A)]|TGTAAACATCCTACACTCTCAGA</td><td>>MI0000736|hsa-mir-30c-1&WithFlank&1|+|40757278|40757378</td><td>23</td><td>44</td><td>22(+1A)</td><td>TGTAAACATCCTACACTCTCAGA</td><td>1248.8</td><td>6.9</td><td>0.7</td><td>10.6</td><td>4.7E-26</td><td>6.6E-25</td><td>BT20/MB468</td><td>CYTO</td></tr>
<tr><td>>hg38_21_+_8433172_8446622_RNA45SN1_45S_50ntFlanks@11668.11706.39|TGCCCAGTGCTCTGAATGTCAAAGTGAAGAAATTCAATG</td><td>>hg38_21_+_8433172_8446622_RNA45SN1_45S_50ntFlanks</td><td>11668</td><td>11706</td><td>39</td><td>TGCCCAGTGCTCTGAATGTCAAAGTGAAGAAATTCAATG</td><td>692.0</td><td>6.3</td><td>0.6</td><td>10.6</td><td>4.7E-26</td><td>6.6E-25</td><td>BT20/MB468</td><td>CYTO</td></tr>
<tr><td>>hg38_21_+_8433172_8446622_RNA45SN1_45S_50ntFlanks@4555.4576.22|CGCAGCTAGGAATAATGGAATA</td><td>>hg38_21_+_8433172_8446622_RNA45SN1_45S_50ntFlanks</td><td>4555</td><td>4576</td><td>22</td><td>CGCAGCTAGGAATAATGGAATA</td><td>453.9</td><td>-13.3</td><td>1.3</td><td>-10.5</td><td>9.3E-26</td><td>6.6E-25</td><td>BT20/MB468</td><td>NUC</td></tr>
<tr><td>>hg38_21_+_8433172_8446622_RNA45SN1_45S_50ntFlanks@6659.6681.23|AGCGGTGGATCACTCGGCTCGTG</td><td>>hg38_21_+_8433172_8446622_RNA45SN1_45S_50ntFlanks</td><td>6659</td><td>6681</td><td>23</td><td>AGCGGTGGATCACTCGGCTCGTG</td><td>924.2</td><td>-5.0</td><td>0.5</td><td>-10.5</td><td>9.6E-26</td><td>6.7E-25</td><td>BT20/MB468</td><td>NUC</td></tr>
<tr><td>>hg38_21_+_8433172_8446622_RNA45SN1_45S_50ntFlanks@11785.11806.22|TGAATGGATGAACGAGATTCCC</td><td>>hg38_21_+_8433172_8446622_RNA45SN1_45S_50ntFlanks</td><td>11785</td><td>11806</td><td>22</td><td>TGAATGGATGAACGAGATTCCC</td><td>575.2</td><td>-13.6</td><td>1.3</td><td>-10.5</td><td>9.6E-26</td><td>6.8E-25</td><td>BT20/MB468</td><td>NUC</td></tr>
<tr><td>>hg38_21_+_8433172_8446622_RNA45SN1_45S_50ntFlanks@12437.12455.19|CATTGTGAAGCAGAATTCA</td><td>>hg38_21_+_8433172_8446622_RNA45SN1_45S_50ntFlanks</td><td>12437</td><td>12455</td><td>19</td><td>CATTGTGAAGCAGAATTCA</td><td>14736.2</td><td>-13.2</td><td>1.2</td><td>-10.6</td><td>1.9E-26</td><td>6.8E-25</td><td>MB231/MB468</td><td>MITO</td></tr>
<tr><td>>MI0000261|hsa-mir-139&WithFlank&11|-|72615057|72615136@13.35.23&[MIMAT0000250&hsa-miR-139-5p&offsets|0|0;m-186&11|-|72615102|72615124&offsets|0|0]|TCTACAGTGCACGTGTCTCCAGT</td><td>>MI0000261|hsa-mir-139&WithFlank&11|-|72615057|72615136</td><td>13</td><td>35</td><td>23</td><td>TCTACAGTGCACGTGTCTCCAGT</td><td>1736.1</td><td>14.7</td><td>1.4</td><td>10.6</td><td>2.5E-26</td><td>6.9E-25</td><td>MB231/MB468</td><td>NUC</td></tr>
<tr><td>>hg38_21_+_8433172_8446622_RNA45SN1_45S_50ntFlanks@12133.12150.18|CTCCGGGGACAGTGCCAG</td><td>>hg38_21_+_8433172_8446622_RNA45SN1_45S_50ntFlanks</td><td>12133</td><td>12150</td><td>18</td><td>CTCCGGGGACAGTGCCAG</td><td>4055.7</td><td>-16.4</td><td>1.6</td><td>-10.5</td><td>9.8E-26</td><td>6.9E-25</td><td>BT20/MB468</td><td>NUC</td></tr>
<tr><td>>MI0000285|hsa-mir-205&WithFlank&1|+|209432127|209432248@40.61.22&[MIMAT0000266&hsa-miR-205-5p&offsets|0|0;m-351&1|+|209432166|209432187&offsets|0|0]|TCCTTCATTCCACCGGAGTCTG</td><td>>MI0000285|hsa-mir-205&WithFlank&1|+|209432127|209432248</td><td>40</td><td>61</td><td>22</td><td>TCCTTCATTCCACCGGAGTCTG</td><td>49687.5</td><td>10.4</td><td>1.0</td><td>10.9</td><td>9.6E-28</td><td>7.0E-25</td><td>BT20/MB231</td><td>CYTO</td></tr>
<tr><td>>MI0000650|hsa-mir-200c&WithFlank&12|+|6963693|6963772@50.72.23(+1U)&[MIMAT0000617&hsa-miR-200c-3p&offsets|0|0(+1U);m-54&12|+|6963742|6963764&offsets|0|0(+1U)]|TAATACTGCCGGGTAATGATGGAT</td><td>>MI0000650|hsa-mir-200c&WithFlank&12|+|6963693|6963772</td><td>50</td><td>72</td><td>23(+1U)</td><td>TAATACTGCCGGGTAATGATGGAT</td><td>10588.6</td><td>9.1</td><td>0.8</td><td>10.9</td><td>8.3E-28</td><td>7.0E-25</td><td>BT20/MB231</td><td>CYTO</td></tr>
<tr><td>>hg38_21_+_8433172_8446622_RNA45SN1_45S_50ntFlanks@6656.6681.26|CTTAGCGGTGGATCACTCGGCTCGTG</td><td>>hg38_21_+_8433172_8446622_RNA45SN1_45S_50ntFlanks</td><td>6656</td><td>6681</td><td>26</td><td>CTTAGCGGTGGATCACTCGGCTCGTG</td><td>2059.2</td><td>-6.2</td><td>0.6</td><td>-10.7</td><td>7.7E-27</td><td>7.0E-25</td><td>MB231/MB468</td><td>CYTO</td></tr>
<tr><td>>hg38_1_-_228634819_228635039_RNA5S12_5S_50ntFlanks@51.91.41|GTCTACGGCCATACCACCCTGAACGCGCCCGATCTCGTCTG</td><td>>hg38_1_-_228634819_228635039_RNA5S12_5S_50ntFlanks</td><td>51</td><td>91</td><td>41</td><td>GTCTACGGCCATACCACCCTGAACGCGCCCGATCTCGTCTG</td><td>940.6</td><td>6.2</td><td>0.6</td><td>10.6</td><td>3.9E-26</td><td>7.1E-25</td><td>BT20/MB468</td><td>MITO</td></tr>
<tr><td>>hg38_21_+_8433172_8446622_RNA45SN1_45S_50ntFlanks@8115.8143.29|CGCGGGACATGTGGCGTACGGAAGACCCG</td><td>>hg38_21_+_8433172_8446622_RNA45SN1_45S_50ntFlanks</td><td>8115</td><td>8143</td><td>29</td><td>CGCGGGACATGTGGCGTACGGAAGACCCG</td><td>13968.2</td><td>-12.8</td><td>1.2</td><td>-10.6</td><td>2.0E-26</td><td>7.1E-25</td><td>MB231/MB468</td><td>MITO</td></tr>
<tr><td>>MI0000650|hsa-mir-200c&WithFlank&12|+|6963693|6963772@50.70.21&[MIMAT0000617&hsa-miR-200c-3p&offsets|0|-2;m-54&12|+|6963742|6963764&offsets|0|-2]|TAATACTGCCGGGTAATGATG</td><td>>MI0000650|hsa-mir-200c&WithFlank&12|+|6963693|6963772</td><td>50</td><td>70</td><td>21</td><td>TAATACTGCCGGGTAATGATG</td><td>2687.5</td><td>15.0</td><td>1.4</td><td>10.9</td><td>1.4E-27</td><td>7.1E-25</td><td>BT20/MB231</td><td>MITO</td></tr>
<tr><td>>MI0000809|hsa-mir-151a&WithFlank&8|-|140732558|140732659@53.74.22(+1U)&[MIMAT0000757&hsa-miR-151a-3p&offsets|0|+1(+1U);m-30&8|-|140732587|140732607&offsets|0|+1(+1U)]|CTAGACTGAAGCTCCTTGAGGAT</td><td>>MI0000809|hsa-mir-151a&WithFlank&8|-|140732558|140732659</td><td>53</td><td>74</td><td>22(+1U)</td><td>CTAGACTGAAGCTCCTTGAGGAT</td><td>345.4</td><td>2.6</td><td>0.3</td><td>10.5</td><td>1.0E-25</td><td>7.2E-25</td><td>BT20/MB468</td><td>NUC</td></tr>
<tr><td>>hg38_21_+_8433172_8446622_RNA45SN1_45S_50ntFlanks@12159.12182.24|TTTGACTGGGGCGGTACACCTGTC</td><td>>hg38_21_+_8433172_8446622_RNA45SN1_45S_50ntFlanks</td><td>12159</td><td>12182</td><td>24</td><td>TTTGACTGGGGCGGTACACCTGTC</td><td>301.8</td><td>-12.7</td><td>1.2</td><td>-10.5</td><td>1.0E-25</td><td>7.2E-25</td><td>BT20/MB468</td><td>NUC</td></tr>
<tr><td>>MI0000061|hsa-let-7a-2&WithFlank&11|-|122146516|122146599@11.28.18;MI0000064|hsa-let-7c&WithFlank&21|+|16539822|16539917@17.34.18;MI0000062|hsa-let-7a-3&WithFlank&22|+|46112743|46112828@10.27.18;MI0000060|hsa-let-7a-1&WithFlank&9|+|94175951|94176042@12.29.18&[MIMAT0000062&hsa-let-7a-5p&offsets|0|-4;m-6&11|-|122146568|122146589&offsets|0|-4];[MIMAT0000064&hsa-let-7c-5p&offsets|0|-4;m-37&21|+|16539838|16539859&offsets|0|-4];[MIMAT0000062_1&hsa-let-7a-5p&offsets|0|-4;m-5&22|+|46112752|46112773&offsets|0|-4];[MIMAT0000062_2&hsa-let-7a-5p&offsets|0|-4;m-4&9|+|94175962|94175983&offsets|0|-4]|TGAGGTAGTAGGTTGTAT</td><td>>MI0000061|hsa-let-7a-2&WithFlank&11|-|122146516|122146599</td><td>11</td><td>28</td><td>18</td><td>TGAGGTAGTAGGTTGTAT</td><td>6962.0</td><td>8.1</td><td>0.8</td><td>10.5</td><td>5.3E-26</td><td>7.4E-25</td><td>BT20/MB468</td><td>CYTO</td></tr>
<tr><td>>MI0000465|hsa-mir-191&WithFlank&3|-|49020612|49020715@22.44.23(+1A)&[MIMAT0000440&hsa-miR-191-5p&offsets|0|0(+1A);m-48&3|-|49020672|49020694&offsets|0|0(+1A)]|CAACGGAATCCCAAAAGCAGCTGA</td><td>>MI0000465|hsa-mir-191&WithFlank&3|-|49020612|49020715</td><td>22</td><td>44</td><td>23(+1A)</td><td>CAACGGAATCCCAAAAGCAGCTGA</td><td>744.5</td><td>7.1</td><td>0.7</td><td>10.5</td><td>5.3E-26</td><td>7.4E-25</td><td>BT20/MB468</td><td>CYTO</td></tr>
<tr><td>>hg38_21_+_8433172_8446622_RNA45SN1_45S_50ntFlanks@11885.11903.19|TTGAGCTTGACTCTAGTCT</td><td>>hg38_21_+_8433172_8446622_RNA45SN1_45S_50ntFlanks</td><td>11885</td><td>11903</td><td>19</td><td>TTGAGCTTGACTCTAGTCT</td><td>321.4</td><td>-12.8</td><td>1.2</td><td>-10.5</td><td>1.1E-25</td><td>7.4E-25</td><td>BT20/MB468</td><td>NUC</td></tr>
<tr><td>>MI0000088|hsa-mir-30a&WithFlank&6|-|71403545|71403627@12.35.24(+2U)&[MIMAT0000087&hsa-miR-30a-5p&offsets|0|+2(+2U);m-15&6|-|71403595|71403616&offsets|0|+2(+2U)]|TGTAAACATCCTCGACTGGAAGCTTT</td><td>>MI0000088|hsa-mir-30a&WithFlank&6|-|71403545|71403627</td><td>12</td><td>35</td><td>24(+2U)</td><td>TGTAAACATCCTCGACTGGAAGCTTT</td><td>1863.8</td><td>9.0</td><td>0.8</td><td>10.6</td><td>2.8E-26</td><td>7.5E-25</td><td>MB231/MB468</td><td>NUC</td></tr>
<tr><td>>hg38_21_+_8433172_8446622_RNA45SN1_45S_50ntFlanks@12624.12657.34|CCAATGGGGCGAAGCTACCATCTGTGGGATTATG</td><td>>hg38_21_+_8433172_8446622_RNA45SN1_45S_50ntFlanks</td><td>12624</td><td>12657</td><td>34</td><td>CCAATGGGGCGAAGCTACCATCTGTGGGATTATG</td><td>5022.4</td><td>-10.0</td><td>0.9</td><td>-10.6</td><td>2.1E-26</td><td>7.5E-25</td><td>MB231/MB468</td><td>MITO</td></tr>
<tr><td>>hg38_21_+_8433172_8446622_RNA45SN1_45S_50ntFlanks@4657.4686.30|CTAGAGGTGAAATTCTTGGACCGGCGCAAG</td><td>>hg38_21_+_8433172_8446622_RNA45SN1_45S_50ntFlanks</td><td>4657</td><td>4686</td><td>30</td><td>CTAGAGGTGAAATTCTTGGACCGGCGCAAG</td><td>494.8</td><td>-13.4</td><td>1.3</td><td>-10.5</td><td>1.1E-25</td><td>7.5E-25</td><td>BT20/MB468</td><td>NUC</td></tr>
<tr><td>>MI0000440|hsa-mir-27b&WithFlank&9|+|95085439|95085547@67.85.19&[MIMAT0000419&hsa-miR-27b-3p&offsets|0|-2;m-39&9|+|95085505|95085525&offsets|0|-2]|TTCACAGTGGCTAAGTTCT</td><td>>MI0000440|hsa-mir-27b&WithFlank&9|+|95085439|95085547</td><td>67</td><td>85</td><td>19</td><td>TTCACAGTGGCTAAGTTCT</td><td>3127.1</td><td>7.7</td><td>0.7</td><td>10.5</td><td>5.5E-26</td><td>7.6E-25</td><td>BT20/MB468</td><td>CYTO</td></tr>
<tr><td>>hg38_21_+_8433172_8446622_RNA45SN1_45S_50ntFlanks@3875.3898.24|ATACATGCCGACGGGCGCTGACCC</td><td>>hg38_21_+_8433172_8446622_RNA45SN1_45S_50ntFlanks</td><td>3875</td><td>3898</td><td>24</td><td>ATACATGCCGACGGGCGCTGACCC</td><td>3368.0</td><td>-15.6</td><td>1.5</td><td>-10.7</td><td>1.2E-26</td><td>7.7E-25</td><td>MB231/MB468</td><td>MP</td></tr>
<tr><td>>hg38_21_+_8433172_8446622_RNA45SN1_45S_50ntFlanks@4449.4471.23|CCTCTCGGCGCCCCCTCGATGCT</td><td>>hg38_21_+_8433172_8446622_RNA45SN1_45S_50ntFlanks</td><td>4449</td><td>4471</td><td>23</td><td>CCTCTCGGCGCCCCCTCGATGCT</td><td>3826.0</td><td>-15.7</td><td>1.5</td><td>-10.6</td><td>2.1E-26</td><td>7.7E-25</td><td>MB231/MB468</td><td>MITO</td></tr>
<tr><td>>hg38_21_+_8433172_8446622_RNA45SN1_45S_50ntFlanks@13002.13029.28|CGATCTATTGAAAGTCAGCCCTCGACAC</td><td>>hg38_21_+_8433172_8446622_RNA45SN1_45S_50ntFlanks</td><td>13002</td><td>13029</td><td>28</td><td>CGATCTATTGAAAGTCAGCCCTCGACAC</td><td>2783.0</td><td>-13.8</td><td>1.3</td><td>-10.5</td><td>5.7E-26</td><td>7.8E-25</td><td>BT20/MB468</td><td>CYTO</td></tr>
<tr><td>>hg38_21_+_8433172_8446622_RNA45SN1_45S_50ntFlanks@12416.12440.25|TTCGATGTCGGCTCTTCCTATCATT</td><td>>hg38_21_+_8433172_8446622_RNA45SN1_45S_50ntFlanks</td><td>12416</td><td>12440</td><td>25</td><td>TTCGATGTCGGCTCTTCCTATCATT</td><td>1192.6</td><td>5.0</td><td>0.5</td><td>10.5</td><td>5.7E-26</td><td>7.9E-25</td><td>BT20/MB468</td><td>CYTO</td></tr>
<tr><td>>hg38_21_+_8433172_8446622_RNA45SN1_45S_50ntFlanks@10421.10457.37|TGAGAGATGGGCGAGCGCCGTTCCGAAGGGACGGGCG</td><td>>hg38_21_+_8433172_8446622_RNA45SN1_45S_50ntFlanks</td><td>10421</td><td>10457</td><td>37</td><td>TGAGAGATGGGCGAGCGCCGTTCCGAAGGGACGGGCG</td><td>1085.8</td><td>-14.5</td><td>1.4</td><td>-10.5</td><td>1.1E-25</td><td>7.9E-25</td><td>BT20/MB468</td><td>NUC</td></tr>
<tr><td>>hg38_21_+_8433172_8446622_RNA45SN1_45S_50ntFlanks@10350.10388.39|ACGAGAACTTTGAAGGCCGAAGTGGAGAAGGGTTCCATG</td><td>>hg38_21_+_8433172_8446622_RNA45SN1_45S_50ntFlanks</td><td>10350</td><td>10388</td><td>39</td><td>ACGAGAACTTTGAAGGCCGAAGTGGAGAAGGGTTCCATG</td><td>4984.7</td><td>-7.9</td><td>0.7</td><td>-10.6</td><td>2.2E-26</td><td>7.9E-25</td><td>MB231/MB468</td><td>MITO</td></tr>
<tr><td>>hg38_21_+_8433172_8446622_RNA45SN1_45S_50ntFlanks@5489.5514.26|CTGAGAAGACGGTCGAACTTGACTAT</td><td>>hg38_21_+_8433172_8446622_RNA45SN1_45S_50ntFlanks</td><td>5489</td><td>5514</td><td>26</td><td>CTGAGAAGACGGTCGAACTTGACTAT</td><td>7471.7</td><td>-10.3</td><td>1.0</td><td>-10.7</td><td>1.3E-26</td><td>8.0E-25</td><td>MB231/MB468</td><td>MP</td></tr>
<tr><td>>hg38_21_+_8433172_8446622_RNA45SN1_45S_50ntFlanks@4046.4063.18|CATTCGAACGTCTGCCCT</td><td>>hg38_21_+_8433172_8446622_RNA45SN1_45S_50ntFlanks</td><td>4046</td><td>4063</td><td>18</td><td>CATTCGAACGTCTGCCCT</td><td>1623.7</td><td>-14.5</td><td>1.4</td><td>-10.7</td><td>1.3E-26</td><td>8.0E-25</td><td>MB231/MB468</td><td>MP</td></tr>
<tr><td>>hg38_21_+_8433172_8446622_RNA45SN1_45S_50ntFlanks@3872.3898.27|CTAATACATGCCGACGGGCGCTGACCC</td><td>>hg38_21_+_8433172_8446622_RNA45SN1_45S_50ntFlanks</td><td>3872</td><td>3898</td><td>27</td><td>CTAATACATGCCGACGGGCGCTGACCC</td><td>1233.4</td><td>-5.2</td><td>0.5</td><td>-10.5</td><td>5.8E-26</td><td>8.0E-25</td><td>BT20/MB468</td><td>CYTO</td></tr>
<tr><td>>MI0000466|hsa-mir-9-1&WithFlank&1|-|156420335|156420435@22.44.23;MI0000468|hsa-mir-9-3&WithFlank&15|+|89368011|89368112@22.44.23;MI0000467|hsa-mir-9-2&WithFlank&5|-|88666847|88666945@22.44.23&[MIMAT0000441&hsa-miR-9-5p&offsets|0|0;m-104&1|-|156420392|156420414&offsets|0|0];[MIMAT0000441_1&hsa-miR-9-5p&offsets|0|0;m-105&15|+|89368032|89368054&offsets|0|0];[MIMAT0000441_2&hsa-miR-9-5p&offsets|0|0;m-104&5|-|88666902|88666924&offsets|0|0]|TCTTTGGTTATCTAGCTGTATGA</td><td>>MI0000466|hsa-mir-9-1&WithFlank&1|-|156420335|156420435</td><td>22</td><td>44</td><td>23</td><td>TCTTTGGTTATCTAGCTGTATGA</td><td>1817.6</td><td>11.5</td><td>1.1</td><td>10.5</td><td>6.0E-26</td><td>8.2E-25</td><td>BT20/MB468</td><td>CYTO</td></tr>
</table>

| | | | | | | | | | | | | | |
|---|---|---|---|---|---|---|---|---|---|---|---|---|---|
| >hg38_21_+_8433172_8446622_RNA45SN1_45S_50ntFlanks@4722.4757.36\|TCATTAATCAAGAACGAAAGTCGGAGGTTCGAAGAC | >hg38_21_+_8433172_8446622_RNA45SN1_45S_50ntFlanks | 4722 | 4757 | 36 | TCATTAATCAAGAACGAAAGTCGGAGGTTCGAAGAC | 473.8 | -13.3 | 1.3 | -10.5 | 1.2E-25 | 8.3E-25 | BT20/MB468 | NUC |
| >hg38_21_+_8433172_8446622_RNA45SN1_45S_50ntFlanks@3802.3832.31\|CATTAAATCAGTTATGGTTCCTTTGGTCGCT | >hg38_21_+_8433172_8446622_RNA45SN1_45S_50ntFlanks | 3802 | 3832 | 31 | CATTAAATCAGTTATGGTTCCTTTGGTCGCT | 316.4 | -12.7 | 1.2 | -10.5 | 1.2E-25 | 8.4E-25 | BT20/MB468 | NUC |
| >MI0003188\|hsa-mir-503&WithFlank&X\|-\|134546322\|134546404@12.31.20&[MIMAT0002874&hsa-miR-503-5p&offsets\|0\|-3;m-297&X\|-\|134546371\|134546393&offsets\|0\|-3]\|TAGCAGCGGGAACAGTTCTG | >MI0003188\|hsa-mir-503&WithFlank&X\|-\|134546322\|134546404 | 12 | 31 | 20 | TAGCAGCGGGAACAGTTCTG | 603.4 | 13.7 | 1.3 | 10.6 | 4.7E-26 | 8.5E-25 | BT20/MB468 | MITO |
| >MI0000809\|hsa-mir-151a&WithFlank&8\|-\|140732558\|140732659@53.74.22(+1U)&[MIMAT0000757&hsa-miR-151a-3p&offsets\|0\|+1(+1U);m-30&8\|-\|140732587\|140732607&offsets\|0\|+1(+1U)]\|CTAGACTGAAGCTCCTTGAGGAT | >MI0000809\|hsa-mir-151a&WithFlank&8\|-\|140732558\|140732659 | 53 | 74 | 22(+1U) | CTAGACTGAAGCTCCTTGAGGAT | 1383.2 | 5.9 | 0.6 | 10.6 | 3.1E-26 | 8.5E-25 | MB231/MB468 | NUC |
| >MI0000650\|hsa-mir-200c&WithFlank&12\|+\|6963693\|6963772@50.72.23(+1U)&[MIMAT0000617&hsa-miR-200c-3p&offsets\|0\|0(+1U);m-54&12\|+\|6963742\|6963764&offsets\|0\|0(+1U)]\|TAATACTGCCGGGTAATGATGGAT | >MI0000650\|hsa-mir-200c&WithFlank&12\|+\|6963693\|6963772 | 50 | 72 | 23(+1U) | TAATACTGCCGGGTAATGATGGAT | 6199.3 | 8.5 | 0.8 | 10.9 | 1.9E-27 | 8.6E-25 | BT20/MB231 | MITO |
| >hg38_21_+_8433172_8446622_RNA45SN1_45S_50ntFlanks@4390.4408.19\|TCTTGGGAGCGGGCGGGCG | >hg38_21_+_8433172_8446622_RNA45SN1_45S_50ntFlanks | 4390 | 4408 | 19 | TCTTGGGAGCGGGCGGGCG | 392.9 | -13.0 | 1.2 | -10.5 | 1.2E-25 | 8.6E-25 | BT20/MB468 | NUC |
| >hg38_21_+_8433172_8446622_RNA45SN1_45S_50ntFlanks@9535.9553.19\|GGAAACTCTGGTGGAGGTC | >hg38_21_+_8433172_8446622_RNA45SN1_45S_50ntFlanks | 9535 | 9553 | 19 | GGAAACTCTGGTGGAGGTC | 303.3 | -12.7 | 1.2 | -10.5 | 1.2E-25 | 8.7E-25 | BT20/MB468 | NUC |
| >MI0000650\|hsa-mir-200c&WithFlank&12\|+\|6963693\|6963772@50.70.21(+1A)&[MIMAT0000617&hsa-miR-200c-3p&offsets\|0\|-2(+1A);m-54&12\|+\|6963742\|6963764&offsets\|0\|-2(+1A)]\|TAATACTGCCGGGTAATGATGA | >MI0000650\|hsa-mir-200c&WithFlank&12\|+\|6963693\|6963772 | 50 | 70 | 21(+1A) | TAATACTGCCGGGTAATGATGA | 2912.4 | 12.8 | 1.2 | 10.8 | 3.7E-27 | 8.7E-25 | BT20/MB231 | NUC |
| >hg38_21_+_8433172_8446622_RNA45SN1_45S_50ntFlanks@3802.3829.28\|CATTAAATCAGTTATGGTTCCTTTGGTC | >hg38_21_+_8433172_8446622_RNA45SN1_45S_50ntFlanks | 3802 | 3829 | 28 | CATTAAATCAGTTATGGTTCCTTTGGTC | 490.8 | -13.4 | 1.3 | -10.5 | 1.3E-25 | 8.7E-25 | BT20/MB468 | NUC |
| >hg38_21_+_8433172_8446622_RNA45SN1_45S_50ntFlanks@10458.10477.20\|ATGGCCTCCGTTGCCCTCGG | >hg38_21_+_8433172_8446622_RNA45SN1_45S_50ntFlanks | 10458 | 10477 | 20 | ATGGCCTCCGTTGCCCTCGG | 2518.7 | -15.2 | 1.4 | -10.7 | 1.4E-26 | 9.0E-25 | MB231/MB468 | MP |
| >hg38_21_+_8433172_8446622_RNA45SN1_45S_50ntFlanks@10844.10865.22\|AACTTCGGGATAAGGATTGGCT | >hg38_21_+_8433172_8446622_RNA45SN1_45S_50ntFlanks | 10844 | 10865 | 22 | AACTTCGGGATAAGGATTGGCT | 997.9 | -14.4 | 1.4 | -10.5 | 1.3E-25 | 9.1E-25 | BT20/MB468 | NUC |
| >hg38_21_+_8433172_8446622_RNA45SN1_45S_50ntFlanks@12456.12487.32\|CCAAGCGTTGGATTGTTCACCCACTAATAGGG | >hg38_21_+_8433172_8446622_RNA45SN1_45S_50ntFlanks | 12456 | 12487 | 32 | CCAAGCGTTGGATTGTTCACCCACTAATAGGG | 23434.7 | -13.5 | 1.3 | -10.7 | 1.5E-26 | 9.1E-25 | MB231/MB468 | MP |
| >MI0000434\|hsa-let-7i&WithFlank&12\|+\|62603680\|62603775@12.33.22(+1A)&[MIMAT0000415&hsa-let-7i-5p&offsets\|0\|0(+1A);m-49&12\|+\|62603691\|62603712&offsets\|0\|0(+1A)]\|TGAGGTAGTAGTTTGTGCTGTTA | >MI0000434\|hsa-let-7i&WithFlank&12\|+\|62603680\|62603775 | 12 | 33 | 22(+1A) | TGAGGTAGTAGTTTGTGCTGTTA | 4604.5 | 7.1 | 0.7 | 10.5 | 5.3E-26 | 9.6E-25 | BT20/MB468 | MITO |
| >MI0000746\|hsa-mir-99b&WithFlank&19\|+\|51692606\|51692687@13.32.20(+2U)&[MIMAT0000689&hsa-miR-99b-5p&offsets\|0\|-2(+2U);m-10&19\|+\|51692618\|51692639&offsets\|0\|-2(+2U)]\|CACCCGTAGAACCGACCTTGTT | >MI0000746\|hsa-mir-99b&WithFlank&19\|+\|51692606\|51692687 | 13 | 32 | 20(+2U) | CACCCGTAGAACCGACCTTGTT | 4534.3 | 16.4 | 1.6 | 10.5 | 7.0E-26 | 9.6E-25 | BT20/MB468 | CYTO |
| >hg38_21_+_8433172_8446622_RNA45SN1_45S_50ntFlanks@9504.9525.22\|GTGAACTATGCCTGGGCAGGGC | >hg38_21_+_8433172_8446622_RNA45SN1_45S_50ntFlanks | 9504 | 9525 | 22 | GTGAACTATGCCTGGGCAGGGC | 1158.6 | -8.1 | 0.8 | -10.7 | 1.1E-26 | 9.6E-25 | MB231/MB468 | CYTO |
| >hg38_21_+_8433172_8446622_RNA45SN1_45S_50ntFlanks@12000.12031.32\|CCTGCGGGCCGCCGGTGAAATACCACTACTCT | >hg38_21_+_8433172_8446622_RNA45SN1_45S_50ntFlanks | 12000 | 12031 | 32 | CCTGCGGGCCGCCGGTGAAATACCACTACTCT | 306.1 | -12.7 | 1.2 | -10.5 | 1.4E-25 | 9.8E-25 | BT20/MB468 | NUC |
| >hg38_21_+_8433172_8446622_RNA45SN1_45S_50ntFlanks@10351.10383.33\|CGAGAACTTTGAAGGCCGAAGTGGAGAAGGGTT | >hg38_21_+_8433172_8446622_RNA45SN1_45S_50ntFlanks | 10351 | 10383 | 33 | CGAGAACTTTGAAGGCCGAAGTGGAGAAGGGTT | 18125.7 | -5.6 | 0.5 | -10.7 | 1.1E-26 | 9.8E-25 | MB231/MB468 | CYTO |
| >hg38_1_-_228634819_228635039_RNA5S12_5S_50ntFlanks@147.169.23\|GGGAATACCGGGTGCTGTAGGCT | >hg38_1_-_228634819_228635039_RNA5S12_5S_50ntFlanks | 147 | 169 | 23 | GGGAATACCGGGTGCTGTAGGCT | 2527.9 | 6.1 | 0.6 | 10.5 | 7.2E-26 | 9.9E-25 | BT20/MB468 | CYTO |
| >hg38_21_+_8433172_8446622_RNA45SN1_45S_50ntFlanks@8182.8211.30\|ATCGAGGCCCAGCCCGTGGACGGTGTGAGG | >hg38_21_+_8433172_8446622_RNA45SN1_45S_50ntFlanks | 8182 | 8211 | 30 | ATCGAGGCCCAGCCCGTGGACGGTGTGAGG | 1146.1 | -14.6 | 1.4 | -10.5 | 1.4E-25 | 1.0E-24 | BT20/MB468 | NUC |
| >hg38_21_+_8433172_8446622_RNA45SN1_45S_50ntFlanks@11575.11607.33\|ACTTAGAACTGGTGCGGACCAGGGGAATCCGAC | >hg38_21_+_8433172_8446622_RNA45SN1_45S_50ntFlanks | 11575 | 11607 | 33 | ACTTAGAACTGGTGCGGACCAGGGGAATCCGAC | 873.5 | -5.8 | 0.6 | -10.5 | 1.4E-25 | 1.0E-24 | BT20/MB468 | NUC |
| >MI0000102\|hsa-mir-100&WithFlank&11\|-\|122152223\|122152314@20.40.21&[MIMAT0000098&hsa-miR-100-5p&offsets\|+1\|0;m-27&11\|-\|122152276\|122152296&offsets\|+1\|+1]\|ACCCGTAGATCCGAACTTGTG | >MI0000102\|hsa-mir-100&WithFlank&11\|-\|122152223\|122152314 | 20 | 40 | 21 | ACCCGTAGATCCGAACTTGTG | 1687.6 | 14.6 | 1.4 | 10.6 | 3.8E-26 | 1.0E-24 | MB231/MB468 | NUC |
| >hg38_21_+_8433172_8446622_RNA45SN1_45S_50ntFlanks@10808.10826.19\|CAATGTAGGTAAGGGAAGT | >hg38_21_+_8433172_8446622_RNA45SN1_45S_50ntFlanks | 10808 | 10826 | 19 | CAATGTAGGTAAGGGAAGT | 3329.9 | -15.5 | 1.5 | -10.6 | 2.9E-26 | 1.0E-24 | MB231/MB468 | MITO |
| >hg38_21_+_8433172_8446622_RNA45SN1_45S_50ntFlanks@11576.11603.28\|CTTAGAACTGGTGCGGACCAGGGGAATC | >hg38_21_+_8433172_8446622_RNA45SN1_45S_50ntFlanks | 11576 | 11603 | 28 | CTTAGAACTGGTGCGGACCAGGGGAATC | 2284.8 | -14.9 | 1.4 | -10.6 | 3.0E-26 | 1.0E-24 | MB231/MB468 | MITO |
| >hg38_21_+_8433172_8446622_RNA45SN1_45S_50ntFlanks@3854.3882.29\|ACTGTGGTAATTCTAGAGCTAATACATGC | >hg38_21_+_8433172_8446622_RNA45SN1_45S_50ntFlanks | 3854 | 3882 | 29 | ACTGTGGTAATTCTAGAGCTAATACATGC | 459.9 | -13.3 | 1.3 | -10.4 | 1.5E-25 | 1.0E-24 | BT20/MB468 | NUC |
| >hg38_21_+_8433172_8446622_RNA45SN1_45S_50ntFlanks@10866.10902.37\|CTAAGGGCTGGGTCGGTCGGGCTGGGGCGCGAAGCGG | >hg38_21_+_8433172_8446622_RNA45SN1_45S_50ntFlanks | 10866 | 10902 | 37 | CTAAGGGCTGGGTCGGTCGGGCTGGGGCGCGAAGCGG | 2796.3 | -15.2 | 1.4 | -10.6 | 3.0E-26 | 1.0E-24 | MB231/MB468 | MITO |
| >trna116_GluCTC_1_-_145399233_145399304@1.34.34,trna59_GluCTC_1_+_249168447_249168518@1.34.34,trna71_GluCTC_1_-_161439189_161439260@1.34.34,trna74_GluCTC_1_-_161431809_161431880@1.34.34,trna77_GluCTC_1_-_161424398_161424469@1.34.34,trna77_GluCTC_6_+_28949976_28950047@1.34.34,trna80_GluCTC_1_-_161417018_161417089@1.34.34,trna87_GluCTC_6_-_126101393_126101464@1.34.34\|TCCCTGGTGGTCTAGTGGTTAGGATTCGGCGCTC | >trna116_GluCTC_1_-_145399233_145399304 | 1 | 34 | 34 | TCCCTGGTGGTCTAGTGGTTAGGATTCGGCGCTC | 14561.5 | 8.3 | 0.8 | 10.7 | 1.2E-26 | 1.1E-24 | MB231/MB468 | CYTO |
| >hg38_21_+_8433172_8446622_RNA45SN1_45S_50ntFlanks@12994.13013.20\|CCTCGCTGCGATCTATTGAA | >hg38_21_+_8433172_8446622_RNA45SN1_45S_50ntFlanks | 12994 | 13013 | 20 | CCTCGCTGCGATCTATTGAA | 2831.3 | -5.2 | 0.5 | -10.6 | 4.0E-26 | 1.1E-24 | MB231/MB468 | NUC |
| >hg38_21_+_8433172_8446622_RNA45SN1_45S_50ntFlanks@10391.10430.40\|AACAGCAGTTGAACATGGGTCAGTCGGTCCTGAGAGATGG | >hg38_21_+_8433172_8446622_RNA45SN1_45S_50ntFlanks | 10391 | 10430 | 40 | AACAGCAGTTGAACATGGGTCAGTCGGTCCTGAGAGATGG | 2342.8 | -15.6 | 1.5 | -10.4 | 1.6E-25 | 1.1E-24 | BT20/MB468 | NUC |
| >hg38_21_+_8433172_8446622_RNA45SN1_45S_50ntFlanks@12141.12162.22\|ACAGTGCCAGGTGGGGAGTTTG | >hg38_21_+_8433172_8446622_RNA45SN1_45S_50ntFlanks | 12141 | 12162 | 22 | ACAGTGCCAGGTGGGGAGTTTG | 425.0 | -13.2 | 1.3 | -10.4 | 1.6E-25 | 1.1E-24 | BT20/MB468 | NUC |
| >hg38_21_+_8433172_8446622_RNA45SN1_45S_50ntFlanks@9454.9494.41\|CCGCGCCGGGGAGGTGGAGCACGAGCGCACGTGTTAGGACC | >hg38_21_+_8433172_8446622_RNA45SN1_45S_50ntFlanks | 9454 | 9494 | 41 | CCGCGCCGGGGAGGTGGAGCACGAGCGCACGTGTTAGGACC | 334.4 | -12.8 | 1.2 | -10.4 | 1.6E-25 | 1.1E-24 | BT20/MB468 | NUC |
| >hg38_21_+_8433172_8446622_RNA45SN1_45S_50ntFlanks@12456.12478.23\|CCAAGCGTTGGATTGTTCACCCA | >hg38_21_+_8433172_8446622_RNA45SN1_45S_50ntFlanks | 12456 | 12478 | 23 | CCAAGCGTTGGATTGTTCACCCA | 9960.0 | -6.5 | 0.6 | -10.5 | 8.0E-26 | 1.1E-24 | BT20/MB468 | CYTO |
| >hg38_21_+_8433172_8446622_RNA45SN1_45S_50ntFlanks@11322.11344.23\|ACGTCTCGTCGCGCGCGCGTCCG | >hg38_21_+_8433172_8446622_RNA45SN1_45S_50ntFlanks | 11322 | 11344 | 23 | ACGTCTCGTCGCGCGCGCGTCCG | 16184.5 | -8.9 | 0.8 | -10.6 | 3.1E-26 | 1.1E-24 | MB231/MB468 | MITO |
| >hg38_21_+_8433172_8446622_RNA45SN1_45S_50ntFlanks@11572.11601.30\|CCGACTTAGAACTGGTGCGGACCAGGGGAA | >hg38_21_+_8433172_8446622_RNA45SN1_45S_50ntFlanks | 11572 | 11601 | 30 | CCGACTTAGAACTGGTGCGGACCAGGGGAA | 20237.4 | -7.7 | 0.7 | -10.6 | 4.1E-26 | 1.1E-24 | MB231/MB468 | NUC |
| >am_trna117_GlyTCC_1_-_145397864_145397935@1.29.29,trna45_GlyTCC_1_+_161500903_161500974@1.29.29,trna70_GlyTCC_1_-_161439547_161439618@1.29.29,trna73_GlyTCC_1_-_161432166_161432237@1.29.29,trna76_GlyTCC_1_-_161424756_161424827@1.29.29,trna79_GlyTCC_1_-_161417375_161417446@1.29.29\|GCGTTGGTGGTATAGTGGTGAGCATAGCT | >am_trna117_GlyTCC_1_-_145397864_145397935 | 1 | 29 | 29 | GCGTTGGTGGTATAGTGGTGAGCATAGCT | 3948.5 | -7.4 | 0.7 | -10.7 | 1.3E-26 | 1.1E-24 | MB231/MB468 | CYTO |
| >hg38_21_+_8433172_8446622_RNA45SN1_45S_50ntFlanks@12157.12179.23\|AGTTTGACTGGGGCGGTACACCT | >hg38_21_+_8433172_8446622_RNA45SN1_45S_50ntFlanks | 12157 | 12179 | 23 | AGTTTGACTGGGGCGGTACACCT | 548.4 | -13.5 | 1.3 | -10.4 | 1.7E-25 | 1.1E-24 | BT20/MB468 | NUC |
| >hg38_21_+_8433172_8446622_RNA45SN1_45S_50ntFlanks@8380.8405.26\|CAAGAGGGCGTGAAACCGTTAAGAGG | >hg38_21_+_8433172_8446622_RNA45SN1_45S_50ntFlanks | 8380 | 8405 | 26 | CAAGAGGGCGTGAAACCGTTAAGAGG | 1653.9 | -5.8 | 0.5 | -10.7 | 1.4E-26 | 1.1E-24 | MB231/MB468 | CYTO |
| >MI0000093\|hsa-mir-92a-1&WithFlank&13\|+\|91351308\|91351397@54.74.21;MI0000094\|hsa-mir-92a-2&WithFlank&X\|-\|134169532\|134169618@54.74.21&[MIMAT0000092&hsa-miR-92a-3p&offsets\|0\|-1;m-9&13\|+\|91351361\|91351382&offsets\|0\|-1];[MIMAT0000092_1&hsa-miR-92a-3p&offsets\|0\|-1;m-11&X\|-\|134169544\|134169565&offsets\|0\|-1]\|TATTGCACTTGTCCCGGCCTG | >MI0000093\|hsa-mir-92a-1&WithFlank&13\|+\|91351308\|91351397 | 54 | 74 | 21 | TATTGCACTTGTCCCGGCCTG | 825.4 | 5.9 | 0.6 | 10.5 | 8.5E-26 | 1.2E-24 | BT20/MB468 | CYTO |
| >hg38_21_+_8433172_8446622_RNA45SN1_45S_50ntFlanks@11209.11227.19\|CGGCGACTCTGGACGCGAG | >hg38_21_+_8433172_8446622_RNA45SN1_45S_50ntFlanks | 11209 | 11227 | 19 | CGGCGACTCTGGACGCGAG | 2805.1 | -15.3 | 1.4 | -10.6 | 1.9E-26 | 1.2E-24 | MB231/MB468 | MP |
| >trna111_HisGTG_1_-_147774845_147774916@15.33.19,trna118_HisGTG_1_-_145396881_145396952@15.33.19,trna16_HisGTG_1_+_146544773_146544844@15.33.19,trna1_HisGTG_15_+_45493349_45493420@15.33.19,trna21_HisGTG_1_+_147753471_147753542@15.33.19,trna33_HisGTG_6_+_27125906_27125977@15.33.19,trna7_HisGTG_9_-_14433938_14434009@15.33.19,trna8_HisGTG_15_-_45492611_45492682@15.33.19,trna9_HisGTG_15_-_45490804_45490875@15.33.19\|GTGGTTAGTACTCTGCGTT | >trna111_HisGTG_1_-_147774845_147774916 | 15 | 33 | 19 | GTGGTTAGTACTCTGCGTT | 301.0 | 5.6 | 0.5 | 10.5 | 8.5E-26 | 1.2E-24 | BT20/MB468 | CYTO |
| >hg38_21_+_8433172_8446622_RNA45SN1_45S_50ntFlanks@4722.4752.31\|TCATTAATCAAGAACGAAAGTCGGAGGTTCG | >hg38_21_+_8433172_8446622_RNA45SN1_45S_50ntFlanks | 4722 | 4752 | 31 | TCATTAATCAAGAACGAAAGTCGGAGGTTCG | 1916.9 | -14.7 | 1.4 | -10.6 | 3.3E-26 | 1.2E-24 | MB231/MB468 | MITO |
| >MI0001446\|hsa-mir-424&WithFlank&X\|-\|134546608\|134546717@54.75.22&[MIMAT0004749&hsa-miR-424-3p&offsets\|0\|+1;m-305&X\|-\|134546644\|134546664&offsets\|0\|+1]\|CAAAACGTGAGGCGCTGCTATA | >MI0001446\|hsa-mir-424&WithFlank&X\|-\|134546608\|134546717 | 54 | 75 | 22 | CAAAACGTGAGGCGCTGCTATA | 776.1 | 13.9 | 1.3 | 10.5 | 8.6E-26 | 1.2E-24 | BT20/MB468 | CYTO |

<table>
<tr><th></th><th></th><th></th><th></th><th></th><th></th><th></th><th></th><th></th><th></th><th></th><th></th><th></th><th></th></tr>
<tr><td>>MI0000650|hsa-mir-200c&WithFlank&12|+|6963693|6963772@50.72.23(+1A)&[MIMAT0000617&hsa-miR-200c-3p&offsets|0|0(+1A);m-54&12|+|6963742|6963764&offsets|0|0(+1A)]|TAATACTGCCGGGTAATGATGGAA</td><td>>MI0000650|hsa-mir-200c&WithFlank&12|+|6963693|6963772</td><td>50</td><td>72</td><td>23(+1A)</td><td>TAATACTGCCGGGTAATGATGGAA</td><td>10425.6</td><td>10.8</td><td>1.0</td><td>10.8</td><td>3.0E-27</td><td>1.2E-24</td><td>BT20/MB231</td><td>MITO</td></tr>
<tr><td>>hg38_21_+_8433172_8446622_RNA45SN1_45S_50ntFlanks@6646.6665.20|TCGTACGACTCTTAGCGGTG</td><td>>hg38_21_+_8433172_8446622_RNA45SN1_45S_50ntFlanks</td><td>6646</td><td>6665</td><td>20</td><td>TCGTACGACTCTTAGCGGTG</td><td>197336.0</td><td>7.1</td><td>0.7</td><td>10.5</td><td>6.7E-26</td><td>1.2E-24</td><td>BT20/MB468</td><td>MITO</td></tr>
<tr><td>>hg38_21_+_8433172_8446622_RNA45SN1_45S_50ntFlanks@12405.12422.18|CTTTTTGATCCTTCGATG</td><td>>hg38_21_+_8433172_8446622_RNA45SN1_45S_50ntFlanks</td><td>12405</td><td>12422</td><td>18</td><td>CTTTTTGATCCTTCGATG</td><td>405.1</td><td>-13.1</td><td>1.3</td><td>-10.4</td><td>1.8E-25</td><td>1.2E-24</td><td>BT20/MB468</td><td>NUC</td></tr>
<tr><td>>hg38_21_+_8433172_8446622_RNA45SN1_45S_50ntFlanks@5446.5467.22|CGGATCGGCCCCGCCGGGGTCG</td><td>>hg38_21_+_8433172_8446622_RNA45SN1_45S_50ntFlanks</td><td>5446</td><td>5467</td><td>22</td><td>CGGATCGGCCCCGCCGGGGTCG</td><td>434.6</td><td>-13.2</td><td>1.3</td><td>-10.4</td><td>1.8E-25</td><td>1.3E-24</td><td>BT20/MB468</td><td>NUC</td></tr>
<tr><td>>hg38_21_+_8433172_8446622_RNA45SN1_45S_50ntFlanks@12411.12434.24|GATCCTTCGATGTCGGCTCTTCCT</td><td>>hg38_21_+_8433172_8446622_RNA45SN1_45S_50ntFlanks</td><td>12411</td><td>12434</td><td>24</td><td>GATCCTTCGATGTCGGCTCTTCCT</td><td>758.8</td><td>-14.0</td><td>1.3</td><td>-10.4</td><td>1.9E-25</td><td>1.3E-24</td><td>BT20/MB468</td><td>NUC</td></tr>
<tr><td>>MI0000078|hsa-mir-22&WithFlank&17|-|1713897|1713993@59.79.21(+1C)&[MIMAT0000077&hsa-miR-22-3p&offsets|0|-1(+1C);m-2&17|-|1713914|1713935&offsets|0|-1(+1C)]|AAGCTGCCAGTTGAAGAACTGC</td><td>>MI0000078|hsa-mir-22&WithFlank&17|-|1713897|1713993</td><td>59</td><td>79</td><td>21(+1C)</td><td>AAGCTGCCAGTTGAAGAACTGC</td><td>1124.7</td><td>7.6</td><td>0.7</td><td>10.6</td><td>4.8E-26</td><td>1.3E-24</td><td>MB231/MB468</td><td>NUC</td></tr>
<tr><td>>hg38_21_+_8433172_8446622_RNA45SN1_45S_50ntFlanks@12456.12473.18|CCAAGCGTTGGATTGTTC</td><td>>hg38_21_+_8433172_8446622_RNA45SN1_45S_50ntFlanks</td><td>12456</td><td>12473</td><td>18</td><td>CCAAGCGTTGGATTGTTC</td><td>11728.7</td><td>-8.0</td><td>0.7</td><td>-10.7</td><td>1.5E-26</td><td>1.3E-24</td><td>MB231/MB468</td><td>CYTO</td></tr>
<tr><td>>hg38_21_+_8433172_8446622_RNA45SN1_45S_50ntFlanks@3775.3792.18|GCCGGTACAGTGAAACTG</td><td>>hg38_21_+_8433172_8446622_RNA45SN1_45S_50ntFlanks</td><td>3775</td><td>3792</td><td>18</td><td>GCCGGTACAGTGAAACTG</td><td>2754.1</td><td>-6.7</td><td>0.6</td><td>-10.6</td><td>4.9E-26</td><td>1.3E-24</td><td>MB231/MB468</td><td>NUC</td></tr>
<tr><td>>hg38_21_+_8433172_8446622_RNA45SN1_45S_50ntFlanks@11874.11910.37|AGAAGACCCTGTTGAGCTTGACTCTAGTCTGGCACGG</td><td>>hg38_21_+_8433172_8446622_RNA45SN1_45S_50ntFlanks</td><td>11874</td><td>11910</td><td>37</td><td>AGAAGACCCTGTTGAGCTTGACTCTAGTCTGGCACGG</td><td>1941.5</td><td>-14.7</td><td>1.4</td><td>-10.6</td><td>3.8E-26</td><td>1.3E-24</td><td>MB231/MB468</td><td>MITO</td></tr>
<tr><td>>hg38_21_+_8433172_8446622_RNA45SN1_45S_50ntFlanks@11999.12031.33|CCCTGCGGGCCGCCGGTGAAATACCACTACTCT</td><td>>hg38_21_+_8433172_8446622_RNA45SN1_45S_50ntFlanks</td><td>11999</td><td>12031</td><td>33</td><td>CCCTGCGGGCCGCCGGTGAAATACCACTACTCT</td><td>3482.8</td><td>-12.1</td><td>1.1</td><td>-10.6</td><td>4.9E-26</td><td>1.3E-24</td><td>MB231/MB468</td><td>NUC</td></tr>
<tr><td>>hg38_21_+_8433172_8446622_RNA45SN1_45S_50ntFlanks@10786.10817.32|AACAGCCTCTGGCATGTTGGAACAATGTAGGT</td><td>>hg38_21_+_8433172_8446622_RNA45SN1_45S_50ntFlanks</td><td>10786</td><td>10817</td><td>32</td><td>AACAGCCTCTGGCATGTTGGAACAATGTAGGT</td><td>1387.4</td><td>-6.5</td><td>0.6</td><td>-10.6</td><td>3.8E-26</td><td>1.3E-24</td><td>MB231/MB468</td><td>MITO</td></tr>
<tr><td>>MI0000462|hsa-mir-152&WithFlank&17|-|48037155|48037253@60.81.22(+1U)&[MIMAT0000438&hsa-miR-152-3p&offsets|0|+1(+1U);m-97&17|-|48037174|48037194&offsets|0|+1(+1U)]|TCAGTGCATGACAGAACTTGGGT</td><td>>MI0000462|hsa-mir-152&WithFlank&17|-|48037155|48037253</td><td>60</td><td>81</td><td>22(+1U)</td><td>TCAGTGCATGACAGAACTTGGGT</td><td>1047.4</td><td>14.3</td><td>1.4</td><td>10.5</td><td>9.7E-26</td><td>1.3E-24</td><td>BT20/MB468</td><td>CYTO</td></tr>
<tr><td>>hg38_21_+_8433172_8446622_RNA45SN1_45S_50ntFlanks@4006.4025.20|ACCTCGGGCCGATCGCACGC</td><td>>hg38_21_+_8433172_8446622_RNA45SN1_45S_50ntFlanks</td><td>4006</td><td>4025</td><td>20</td><td>ACCTCGGGCCGATCGCACGC</td><td>1304.3</td><td>-14.8</td><td>1.4</td><td>-10.4</td><td>1.9E-25</td><td>1.3E-24</td><td>BT20/MB468</td><td>NUC</td></tr>
<tr><td>>hg38_21_+_8433172_8446622_RNA45SN1_45S_50ntFlanks@9572.9597.26|CAAATCGGTCGTCCGACCTGGGTATA</td><td>>hg38_21_+_8433172_8446622_RNA45SN1_45S_50ntFlanks</td><td>9572</td><td>9597</td><td>26</td><td>CAAATCGGTCGTCCGACCTGGGTATA</td><td>27943.7</td><td>-11.6</td><td>1.1</td><td>-10.6</td><td>3.9E-26</td><td>1.3E-24</td><td>MB231/MB468</td><td>MITO</td></tr>
<tr><td>>hg38_21_+_8433172_8446622_RNA45SN1_45S_50ntFlanks@482.514.33|ATCGATGTGGTGACGTCGTGCTCTCCCGGGCCG</td><td>>hg38_21_+_8433172_8446622_RNA45SN1_45S_50ntFlanks</td><td>482</td><td>514</td><td>33</td><td>ATCGATGTGGTGACGTCGTGCTCTCCCGGGCCG</td><td>1093.0</td><td>-14.5</td><td>1.4</td><td>-10.4</td><td>1.9E-25</td><td>1.3E-24</td><td>BT20/MB468</td><td>NUC</td></tr>
<tr><td>>hg38_21_+_8433172_8446622_RNA45SN1_45S_50ntFlanks@12157.12180.24|AGTTTGACTGGGGCGGTACACCTG</td><td>>hg38_21_+_8433172_8446622_RNA45SN1_45S_50ntFlanks</td><td>12157</td><td>12180</td><td>24</td><td>AGTTTGACTGGGGCGGTACACCTG</td><td>2680.9</td><td>-15.2</td><td>1.4</td><td>-10.6</td><td>3.9E-26</td><td>1.3E-24</td><td>MB231/MB468</td><td>MITO</td></tr>
<tr><td>>hg38_21_+_8433172_8446622_RNA45SN1_45S_50ntFlanks@10411.10430.20|CAGTCGGTCCTGAGAGATGG</td><td>>hg38_21_+_8433172_8446622_RNA45SN1_45S_50ntFlanks</td><td>10411</td><td>10430</td><td>20</td><td>CAGTCGGTCCTGAGAGATGG</td><td>9581.7</td><td>-10.4</td><td>1.0</td><td>-10.5</td><td>5.1E-26</td><td>1.3E-24</td><td>MB231/MB468</td><td>NUC</td></tr>
<tr><td>>hg38_21_+_8433172_8446622_RNA45SN1_45S_50ntFlanks@10727.10759.33|GGAGAGGGTGTAAATCTCGCGCCGGGCCGTACC</td><td>>hg38_21_+_8433172_8446622_RNA45SN1_45S_50ntFlanks</td><td>10727</td><td>10759</td><td>33</td><td>GGAGAGGGTGTAAATCTCGCGCCGGGCCGTACC</td><td>504.5</td><td>-13.4</td><td>1.3</td><td>-10.4</td><td>2.0E-25</td><td>1.3E-24</td><td>BT20/MB468</td><td>NUC</td></tr>
<tr><td>>am_trna2_GlyGCC_21_-_18827107_18827177@57.74.18,trna35_GlyGCC_1_+_161413094_161413164@57.74.18,trna37_GlyGCC_1_+_161420467_161420537@57.74.18,trna39_GlyGCC_1_+_161427898_161427968@57.74.18,trna41_GlyGCC_1_+_161435258_161435328@57.74.18|TTCCCGGCCCATGCACCA</td><td>>am_trna2_GlyGCC_21_-_18827107_18827177</td><td>57</td><td>74</td><td>18</td><td>TTCCCGGCCCATGCACCA</td><td>2078.8</td><td>11.2</td><td>1.1</td><td>10.5</td><td>9.9E-26</td><td>1.3E-24</td><td>BT20/MB468</td><td>CYTO</td></tr>
<tr><td>>hg38_21_+_8433172_8446622_RNA45SN1_45S_50ntFlanks@12633.12659.27|CGAAGCTACCATCTGTGGGATTATGAC</td><td>>hg38_21_+_8433172_8446622_RNA45SN1_45S_50ntFlanks</td><td>12633</td><td>12659</td><td>27</td><td>CGAAGCTACCATCTGTGGGATTATGAC</td><td>3604.8</td><td>-15.7</td><td>1.5</td><td>-10.6</td><td>2.2E-26</td><td>1.3E-24</td><td>MB231/MB468</td><td>MP</td></tr>
<tr><td>>hg38_21_+_8433172_8446622_RNA45SN1_45S_50ntFlanks@12936.12956.21|AAACCATTCGTAGACGACCTG</td><td>>hg38_21_+_8433172_8446622_RNA45SN1_45S_50ntFlanks</td><td>12936</td><td>12956</td><td>21</td><td>AAACCATTCGTAGACGACCTG</td><td>28588.9</td><td>-12.1</td><td>1.2</td><td>-10.4</td><td>2.0E-25</td><td>1.4E-24</td><td>BT20/MB468</td><td>NUC</td></tr>
<tr><td>>MI0000477|hsa-mir-146a&WithFlank&5|+|160485346|160485456@27.48.22&[MIMAT0000449&hsa-miR-146a-5p&offsets|0|0;m-129&5|+|160485372|160485393&offsets|0|0]|TGAGAACTGAATTCCATGGGTT</td><td>>MI0000477|hsa-mir-146a&WithFlank&5|+|160485346|160485456</td><td>27</td><td>48</td><td>22</td><td>TGAGAACTGAATTCCATGGGTT</td><td>12919.0</td><td>17.7</td><td>1.7</td><td>10.7</td><td>1.6E-26</td><td>1.4E-24</td><td>MB231/MB468</td><td>CYTO</td></tr>
<tr><td>>hg38_21_+_8433172_8446622_RNA45SN1_45S_50ntFlanks@12685.12708.24|CCCAGGCGGAACGATACGGCAGCG</td><td>>hg38_21_+_8433172_8446622_RNA45SN1_45S_50ntFlanks</td><td>12685</td><td>12708</td><td>24</td><td>CCCAGGCGGAACGATACGGCAGCG</td><td>412.3</td><td>-13.1</td><td>1.3</td><td>-10.4</td><td>2.0E-25</td><td>1.4E-24</td><td>BT20/MB468</td><td>NUC</td></tr>
<tr><td>>MI0003834|hsa-mir-769&WithFlank&19|+|46018926|46019055@36.57.22&[MIMAT0003886&hsa-miR-769-5p&offsets|0|0;m-216&19|+|46018961|46018982&offsets|0|0]|TGAGACCTCTGGGTTCTGAGCT</td><td>>MI0003834|hsa-mir-769&WithFlank&19|+|46018926|46019055</td><td>36</td><td>57</td><td>22</td><td>TGAGACCTCTGGGTTCTGAGCT</td><td>2156.6</td><td>7.6</td><td>0.7</td><td>10.5</td><td>1.0E-25</td><td>1.4E-24</td><td>BT20/MB468</td><td>CYTO</td></tr>
<tr><td>>hg38_21_+_8433172_8446622_RNA45SN1_45S_50ntFlanks@12476.12498.23|CCACTAATAGGGAACGTGAGCTG</td><td>>hg38_21_+_8433172_8446622_RNA45SN1_45S_50ntFlanks</td><td>12476</td><td>12498</td><td>23</td><td>CCACTAATAGGGAACGTGAGCTG</td><td>2667.8</td><td>-15.2</td><td>1.4</td><td>-10.6</td><td>2.2E-26</td><td>1.4E-24</td><td>MB231/MB468</td><td>MP</td></tr>
<tr><td>>hg38_21_+_8433172_8446622_RNA45SN1_45S_50ntFlanks@8104.8140.37|CCGCGGCGGGGCGCGGGACATGTGGCGTACGGAAGAC</td><td>>hg38_21_+_8433172_8446622_RNA45SN1_45S_50ntFlanks</td><td>8104</td><td>8140</td><td>37</td><td>CCGCGGCGGGGCGCGGGACATGTGGCGTACGGAAGAC</td><td>4221.7</td><td>-15.8</td><td>1.5</td><td>-10.6</td><td>4.1E-26</td><td>1.4E-24</td><td>MB231/MB468</td><td>MITO</td></tr>
<tr><td>>trna10_LysCTT_16_+_3241501_3241573@1.29.29,trna13_LysCTT_14_-_58706613_58706685@1.29.29,trna2_LysCTT_15_+_79152904_79152976@1.29.29|GCCCGGCTAGCTCAGTCGGTAGAGCATGG</td><td>>trna10_LysCTT_16_+_3241501_3241573</td><td>1</td><td>29</td><td>29</td><td>GCCCGGCTAGCTCAGTCGGTAGAGCATGG</td><td>635.0</td><td>-13.7</td><td>1.3</td><td>-10.4</td><td>2.1E-25</td><td>1.4E-24</td><td>BT20/MB468</td><td>NUC</td></tr>
<tr><td>>hg38_21_+_8433172_8446622_RNA45SN1_45S_50ntFlanks@12159.12181.23|TTTGACTGGGGCGGTACACCTGT</td><td>>hg38_21_+_8433172_8446622_RNA45SN1_45S_50ntFlanks</td><td>12159</td><td>12181</td><td>23</td><td>TTTGACTGGGGCGGTACACCTGT</td><td>2989.7</td><td>-15.3</td><td>1.5</td><td>-10.6</td><td>4.2E-26</td><td>1.4E-24</td><td>MB231/MB468</td><td>MITO</td></tr>
<tr><td>>hg38_21_+_8433172_8446622_RNA45SN1_45S_50ntFlanks@6739.6770.32|TCATCGACACTTCGAACGCACTTGCGGCCCCG</td><td>>hg38_21_+_8433172_8446622_RNA45SN1_45S_50ntFlanks</td><td>6739</td><td>6770</td><td>32</td><td>TCATCGACACTTCGAACGCACTTGCGGCCCCG</td><td>672.9</td><td>-13.8</td><td>1.3</td><td>-10.4</td><td>2.1E-25</td><td>1.4E-24</td><td>BT20/MB468</td><td>NUC</td></tr>
<tr><td>>hg38_21_+_8433172_8446622_RNA45SN1_45S_50ntFlanks@10423.10457.35|AGAGATGGGCGAGCGCCGTTCCGAAGGGACGGGCG</td><td>>hg38_21_+_8433172_8446622_RNA45SN1_45S_50ntFlanks</td><td>10423</td><td>10457</td><td>35</td><td>AGAGATGGGCGAGCGCCGTTCCGAAGGGACGGGCG</td><td>1414.1</td><td>-14.9</td><td>1.4</td><td>-10.4</td><td>2.1E-25</td><td>1.4E-24</td><td>BT20/MB468</td><td>NUC</td></tr>
<tr><td>>hg38_21_+_8433172_8446622_RNA45SN1_45S_50ntFlanks@3988.4013.26|CTTTGGTGACTCTAGATAACCTCGGG</td><td>>hg38_21_+_8433172_8446622_RNA45SN1_45S_50ntFlanks</td><td>3988</td><td>4013</td><td>26</td><td>CTTTGGTGACTCTAGATAACCTCGGG</td><td>364.5</td><td>-12.9</td><td>1.2</td><td>-10.4</td><td>2.1E-25</td><td>1.4E-24</td><td>BT20/MB468</td><td>NUC</td></tr>
<tr><td>>hg38_21_+_8433172_8446622_RNA45SN1_45S_50ntFlanks@10770.10805.36|GCAGGTCTCCAAGGTGAACAGCCTCTGGCATGTTGG</td><td>>hg38_21_+_8433172_8446622_RNA45SN1_45S_50ntFlanks</td><td>10770</td><td>10805</td><td>36</td><td>GCAGGTCTCCAAGGTGAACAGCCTCTGGCATGTTGG</td><td>372.5</td><td>-13.0</td><td>1.2</td><td>-10.4</td><td>2.2E-25</td><td>1.5E-24</td><td>BT20/MB468</td><td>NUC</td></tr>
<tr><td>>hg38_21_+_8433172_8446622_RNA45SN1_45S_50ntFlanks@8472.8492.21|CGGCCCGGCGGATCTTTCCCG</td><td>>hg38_21_+_8433172_8446622_RNA45SN1_45S_50ntFlanks</td><td>8472</td><td>8492</td><td>21</td><td>CGGCCCGGCGGATCTTTCCCG</td><td>1144.1</td><td>-14.6</td><td>1.4</td><td>-10.4</td><td>2.2E-25</td><td>1.5E-24</td><td>BT20/MB468</td><td>NUC</td></tr>
<tr><td>>hg38_21_+_8433172_8446622_RNA45SN1_45S_50ntFlanks@9004.9026.23|TCTCCCCCGCGGGGGCGCGCCGG</td><td>>hg38_21_+_8433172_8446622_RNA45SN1_45S_50ntFlanks</td><td>9004</td><td>9026</td><td>23</td><td>TCTCCCCCGCGGGGGCGCGCCGG</td><td>8200.4</td><td>-16.8</td><td>1.6</td><td>-10.6</td><td>4.4E-26</td><td>1.5E-24</td><td>MB231/MB468</td><td>MITO</td></tr>
<tr><td>>hg38_21_+_8433172_8446622_RNA45SN1_45S_50ntFlanks@10727.10755.29|GGAGAGGGTGTAAATCTCGCGCCGGGCCG</td><td>>hg38_21_+_8433172_8446622_RNA45SN1_45S_50ntFlanks</td><td>10727</td><td>10755</td><td>29</td><td>GGAGAGGGTGTAAATCTCGCGCCGGGCCG</td><td>1703.7</td><td>-14.6</td><td>1.4</td><td>-10.6</td><td>2.5E-26</td><td>1.5E-24</td><td>MB231/MB468</td><td>MP</td></tr>
<tr><td>>hg38_21_+_8433172_8446622_RNA45SN1_45S_50ntFlanks@11208.11254.47|GCGGCGACTCTGGACGCGAGCCGGGCCCTTCCCGTGGATCGCCCCAG</td><td>>hg38_21_+_8433172_8446622_RNA45SN1_45S_50ntFlanks</td><td>11208</td><td>11254</td><td>47</td><td>GCGGCGACTCTGGACGCGAGCCGGGCCCTTCCCGTGGATCGCCCCAG</td><td>12825.0</td><td>-11.8</td><td>1.1</td><td>-10.6</td><td>4.5E-26</td><td>1.5E-24</td><td>MB231/MB468</td><td>MITO</td></tr>
<tr><td>>hg38_21_+_8433172_8446622_RNA45SN1_45S_50ntFlanks@11564.11591.28|CCTAGCAGCCGACTTAGAACTGGTGCGG</td><td>>hg38_21_+_8433172_8446622_RNA45SN1_45S_50ntFlanks</td><td>11564</td><td>11591</td><td>28</td><td>CCTAGCAGCCGACTTAGAACTGGTGCGG</td><td>3248.0</td><td>-15.5</td><td>1.5</td><td>-10.6</td><td>2.5E-26</td><td>1.5E-24</td><td>MB231/MB468</td><td>MP</td></tr>
<tr><td>>trna2_GlyGCC_21_-_18827107_18827177@1.28.28,trna35_GlyGCC_1_+_161413094_161413164@1.28.28,trna37_GlyGCC_1_+_161420467_161420537@1.28.28,trna39_GlyGCC_1_+_161427898_161427968@1.28.28,trna41_GlyGCC_1_+_161435258_161435328@1.28.28|GCATGGGTGGTTCAGTGGTAGAATTCTC</td><td>>trna2_GlyGCC_21_-_18827107_18827177</td><td>1</td><td>28</td><td>28</td><td>GCATGGGTGGTTCAGTGGTAGAATTCTC</td><td>4511.7</td><td>-15.9</td><td>1.5</td><td>-10.6</td><td>4.6E-26</td><td>1.5E-24</td><td>MB231/MB468</td><td>MITO</td></tr>
<tr><td>>hg38_21_+_8433172_8446622_RNA45SN1_45S_50ntFlanks@10806.10829.24|AACAATGTAGGTAAGGGAAGTCGG</td><td>>hg38_21_+_8433172_8446622_RNA45SN1_45S_50ntFlanks</td><td>10806</td><td>10829</td><td>24</td><td>AACAATGTAGGTAAGGGAAGTCGG</td><td>647.3</td><td>-13.8</td><td>1.3</td><td>-10.4</td><td>2.3E-25</td><td>1.5E-24</td><td>BT20/MB468</td><td>NUC</td></tr>
<tr><td>>MI0000088|hsa-mir-30a&WithFlank&6|-|71403545|71403627@12.33.22&[MIMAT0000087&hsa-miR-30a-5p&offsets|0|0;m-15&6|-|71403595|71403616&offsets|0|0]|TGTAAACATCCTCGACTGGAAG</td><td>>MI0000088|hsa-mir-30a&WithFlank&6|-|71403545|71403627</td><td>12</td><td>33</td><td>22</td><td>TGTAAACATCCTCGACTGGAAG</td><td>1540.9</td><td>6.3</td><td>0.6</td><td>10.5</td><td>8.5E-26</td><td>1.5E-24</td><td>BT20/MB468</td><td>MITO</td></tr>
<tr><td>>hg38_21_+_8433172_8446622_RNA45SN1_45S_50ntFlanks@13002.13027.26|CGATCTATTGAAAGTCAGCCCTCGAC</td><td>>hg38_21_+_8433172_8446622_RNA45SN1_45S_50ntFlanks</td><td>13002</td><td>13027</td><td>26</td><td>CGATCTATTGAAAGTCAGCCCTCGAC</td><td>3325.9</td><td>-13.8</td><td>1.3</td><td>-10.5</td><td>8.6E-26</td><td>1.5E-24</td><td>BT20/MB468</td><td>MITO</td></tr>
<tr><td>>hg38_21_+_8433172_8446622_RNA45SN1_45S_50ntFlanks@6776.6803.28|CTCCCGGGGCTACGCCTGTCTGAGCGTC</td><td>>hg38_21_+_8433172_8446622_RNA45SN1_45S_50ntFlanks</td><td>6776</td><td>6803</td><td>28</td><td>CTCCCGGGGCTACGCCTGTCTGAGCGTC</td><td>1325.4</td><td>-6.0</td><td>0.6</td><td>-10.4</td><td>2.3E-25</td><td>1.6E-24</td><td>BT20/MB468</td><td>NUC</td></tr>
<tr><td>>hg38_21_+_8433172_8446622_RNA45SN1_45S_50ntFlanks@11881.11904.24|CCTGTTGAGCTTGACTCTAGTCTG</td><td>>hg38_21_+_8433172_8446622_RNA45SN1_45S_50ntFlanks</td><td>11881</td><td>11904</td><td>24</td><td>CCTGTTGAGCTTGACTCTAGTCTG</td><td>2836.6</td><td>-15.3</td><td>1.4</td><td>-10.6</td><td>2.6E-26</td><td>1.6E-24</td><td>MB231/MB468</td><td>MP</td></tr>
<tr><td>>hg38_21_+_8433172_8446622_RNA45SN1_45S_50ntFlanks@9568.9595.28|CGTGCAAATCGGTCGTCCGACCTGGGTA</td><td>>hg38_21_+_8433172_8446622_RNA45SN1_45S_50ntFlanks</td><td>9568</td><td>9595</td><td>28</td><td>CGTGCAAATCGGTCGTCCGACCTGGGTA</td><td>2253.7</td><td>-15.0</td><td>1.4</td><td>-10.6</td><td>2.6E-26</td><td>1.6E-24</td><td>MB231/MB468</td><td>MP</td></tr>
<tr><td>>hg38_21_+_8433172_8446622_RNA45SN1_45S_50ntFlanks@12606.12624.19|ATGTGCTTGGCTGAGGAGC</td><td>>hg38_21_+_8433172_8446622_RNA45SN1_45S_50ntFlanks</td><td>12606</td><td>12624</td><td>19</td><td>ATGTGCTTGGCTGAGGAGC</td><td>1406.5</td><td>-6.1</td><td>0.6</td><td>-10.4</td><td>2.4E-25</td><td>1.6E-24</td><td>BT20/MB468</td><td>NUC</td></tr>
<tr><td>>hg38_21_+_8433172_8446622_RNA45SN1_45S_50ntFlanks@7989.8007.19|AGACGTGGCGACCCGCTGA</td><td>>hg38_21_+_8433172_8446622_RNA45SN1_45S_50ntFlanks</td><td>7989</td><td>8007</td><td>19</td><td>AGACGTGGCGACCCGCTGA</td><td>8037.7</td><td>-7.6</td><td>0.7</td><td>-10.6</td><td>2.7E-26</td><td>1.6E-24</td><td>MB231/MB468</td><td>MP</td></tr>
<tr><td>>hg38_21_+_8433172_8446622_RNA45SN1_45S_50ntFlanks@11235.11269.35|CTTCCCGTGGATCGCCCCAGCTGCGGCGGGCGTCG</td><td>>hg38_21_+_8433172_8446622_RNA45SN1_45S_50ntFlanks</td><td>11235</td><td>11269</td><td>35</td><td>CTTCCCGTGGATCGCCCCAGCTGCGGCGGGCGTCG</td><td>2397.0</td><td>-15.0</td><td>1.4</td><td>-10.6</td><td>4.9E-26</td><td>1.6E-24</td><td>MB231/MB468</td><td>MITO</td></tr>
<tr><td>>hg38_21_+_8433172_8446622_RNA45SN1_45S_50ntFlanks@9714.9737.24|CGAATGATTAGAGGTCTTGGGGCC</td><td>>hg38_21_+_8433172_8446622_RNA45SN1_45S_50ntFlanks</td><td>9714</td><td>9737</td><td>24</td><td>CGAATGATTAGAGGTCTTGGGGCC</td><td>6678.3</td><td>-16.5</td><td>1.6</td><td>-10.6</td><td>4.9E-26</td><td>1.6E-24</td><td>MB231/MB468</td><td>MITO</td></tr>
<tr><td>>hg38_21_+_8433172_8446622_RNA45SN1_45S_50ntFlanks@10382.10424.43|TTCCATGTGAACAGCAGTTGAACATGGGTCAGTCGGTCCTGAG</td><td>>hg38_21_+_8433172_8446622_RNA45SN1_45S_50ntFlanks</td><td>10382</td><td>10424</td><td>43</td><td>TTCCATGTGAACAGCAGTTGAACATGGGTCAGTCGGTCCTGAG</td><td>2164.5</td><td>9.5</td><td>0.9</td><td>10.6</td><td>2.0E-26</td><td>1.6E-24</td><td>MB231/MB468</td><td>CYTO</td></tr>
<tr><td>>hg38_21_+_8433172_8446622_RNA45SN1_45S_50ntFlanks@10369.10407.39|AAGTGGAGAAGGGTTCCATGTGAACAGCAGTTGAACATG</td><td>>hg38_21_+_8433172_8446622_RNA45SN1_45S_50ntFlanks</td><td>10369</td><td>10407</td><td>39</td><td>AAGTGGAGAAGGGTTCCATGTGAACAGCAGTTGAACATG</td><td>2321.3</td><td>-15.6</td><td>1.5</td><td>-10.4</td><td>2.4E-25</td><td>1.6E-24</td><td>BT20/MB468</td><td>NUC</td></tr>
</table>

<table>
<tr><td>>hg38_21_+_8433172_8446622_RNA45SN1_45S_50ntFlanks@11318.11342.25|CCCCACGTCTCGTCGCGCGCGCGTC</td><td>>hg38_21_+_8433172_8446622_RNA45SN1_45S_50ntFlanks</td><td>11318</td><td>11342</td><td>25</td><td>CCCCACGTCTCGTCGCGCGCGCGTC</td><td>3029.3</td><td>-15.3</td><td>1.5</td><td>-10.6</td><td>5.0E-26</td><td>1.6E-24</td><td>MB231/MB468</td><td>MITO</td></tr>
<tr><td>>MI0000098|hsa-mir-96&WithFlank&7|-|129774686|129774775@15.37.23&[MIMAT0000095&hsa-miR-96-5p&offsets|0|0;m-248&7|-|129774739|129774761&offsets|0|0]|TTTGGCACTAGCACATTTTTGCT</td><td>>MI0000098|hsa-mir-96&WithFlank&7|-|129774686|129774775</td><td>15</td><td>37</td><td>23</td><td>TTTGGCACTAGCACATTTTTGCT</td><td>4791.8</td><td>6.8</td><td>0.7</td><td>10.5</td><td>1.2E-25</td><td>1.6E-24</td><td>BT20/MB468</td><td>CYTO</td></tr>
<tr><td>>MI0000478|hsa-mir-149&WithFlank&2|+|240455995|240456095@21.41.21&[MIMAT0000450&hsa-miR-149-5p&offsets|0|-2;m-195&2|+|240456015|240456037&offsets|0|-2]|TCTGGCTCCGTGTCTTCACTC</td><td>>MI0000478|hsa-mir-149&WithFlank&2|+|240455995|240456095</td><td>21</td><td>41</td><td>21</td><td>TCTGGCTCCGTGTCTTCACTC</td><td>745.7</td><td>13.8</td><td>1.3</td><td>10.5</td><td>1.2E-25</td><td>1.6E-24</td><td>BT20/MB468</td><td>CYTO</td></tr>
<tr><td>>hg38_21_+_8433172_8446622_RNA45SN1_45S_50ntFlanks@11322.11344.23|ACGTCTCGTCGCGCGCGCGTCCG</td><td>>hg38_21_+_8433172_8446622_RNA45SN1_45S_50ntFlanks</td><td>11322</td><td>11344</td><td>23</td><td>ACGTCTCGTCGCGCGCGCGTCCG</td><td>3379.6</td><td>-15.6</td><td>1.5</td><td>-10.6</td><td>2.7E-26</td><td>1.6E-24</td><td>MB231/MB468</td><td>MP</td></tr>
<tr><td>>hg38_1_-_228634819_228635039_RNA5S12_5S_50ntFlanks@84.111.28|CTCGTCTGATCTCGGAAGCTAAGCAGGG</td><td>>hg38_1_-_228634819_228635039_RNA5S12_5S_50ntFlanks</td><td>84</td><td>111</td><td>28</td><td>CTCGTCTGATCTCGGAAGCTAAGCAGGG</td><td>596.3</td><td>-13.6</td><td>1.3</td><td>-10.4</td><td>2.5E-25</td><td>1.7E-24</td><td>BT20/MB468</td><td>NUC</td></tr>
<tr><td>>hg38_21_+_8433172_8446622_RNA45SN1_45S_50ntFlanks@11661.11685.25|TGATTTCTGCCCAGTGCTCTGAATG</td><td>>hg38_21_+_8433172_8446622_RNA45SN1_45S_50ntFlanks</td><td>11661</td><td>11685</td><td>25</td><td>TGATTTCTGCCCAGTGCTCTGAATG</td><td>6477.9</td><td>-10.3</td><td>1.0</td><td>-10.6</td><td>2.8E-26</td><td>1.7E-24</td><td>MB231/MB468</td><td>MP</td></tr>
<tr><td>>hg38_21_+_8433172_8446622_RNA45SN1_45S_50ntFlanks@11580.11603.24|GAACTGGTGCGGACCAGGGGAATC</td><td>>hg38_21_+_8433172_8446622_RNA45SN1_45S_50ntFlanks</td><td>11580</td><td>11603</td><td>24</td><td>GAACTGGTGCGGACCAGGGGAATC</td><td>6612.7</td><td>-4.7</td><td>0.4</td><td>-10.5</td><td>1.2E-25</td><td>1.7E-24</td><td>BT20/MB468</td><td>CYTO</td></tr>
<tr><td>>trnaMT_GluTTC_MT_-_14674_14742@39.61.23,trnalookalike8_GluTTC_5_-_93905172_93905240@39.61.23|ATTGGTCGTGGTTGTAGTCCGTG</td><td>>trnaMT_GluTTC_MT_-_14674_14742</td><td>39</td><td>61</td><td>23</td><td>ATTGGTCGTGGTTGTAGTCCGTG</td><td>1318.8</td><td>7.8</td><td>0.7</td><td>10.5</td><td>9.4E-26</td><td>1.7E-24</td><td>BT20/MB468</td><td>MITO</td></tr>
<tr><td>>hg38_21_+_8433172_8446622_RNA45SN1_45S_50ntFlanks@5487.5514.28|CGCTGAGAAGACGGTCGAACTTGACTAT</td><td>>hg38_21_+_8433172_8446622_RNA45SN1_45S_50ntFlanks</td><td>5487</td><td>5514</td><td>28</td><td>CGCTGAGAAGACGGTCGAACTTGACTAT</td><td>14356.5</td><td>-13.1</td><td>1.2</td><td>-10.6</td><td>2.9E-26</td><td>1.7E-24</td><td>MB231/MB468</td><td>MP</td></tr>
<tr><td>>MI0000272|hsa-mir-182&WithFlank&7|-|129770377|129770498@29.54.26&[MIMAT0000259&hsa-miR-182-5p&offsets|0|+2;m-22&7|-|129770449|129770470&offsets|0|+4]|TTTGGCAATGGTAGAACTCACACTGG</td><td>>MI0000272|hsa-mir-182&WithFlank&7|-|129770377|129770498</td><td>29</td><td>54</td><td>26</td><td>TTTGGCAATGGTAGAACTCACACTGG</td><td>1588.0</td><td>2.5</td><td>0.2</td><td>10.4</td><td>2.6E-25</td><td>1.7E-24</td><td>BT20/MB468</td><td>NUC</td></tr>
<tr><td>>MI0000088|hsa-mir-30a&WithFlank&6|-|71403545|71403627@12.35.24&[MIMAT0000087&hsa-miR-30a-5p&offsets|0|+2;m-15&6|-|71403595|71403616&offsets|0|+2]|TGTAAACATCCTCGACTGGAAGCT</td><td>>MI0000088|hsa-mir-30a&WithFlank&6|-|71403545|71403627</td><td>12</td><td>35</td><td>24</td><td>TGTAAACATCCTCGACTGGAAGCT</td><td>26593.0</td><td>6.1</td><td>0.6</td><td>10.5</td><td>9.6E-26</td><td>1.7E-24</td><td>BT20/MB468</td><td>MITO</td></tr>
<tr><td>>am_trnaMT_PheGAA_MT_+_577_647@40.58.19|TGTTTAGACGGGCTCACAT</td><td>>am_trnaMT_PheGAA_MT_+_577_647</td><td>40</td><td>58</td><td>19</td><td>TGTTTAGACGGGCTCACAT</td><td>984.5</td><td>7.3</td><td>0.7</td><td>10.5</td><td>9.7E-26</td><td>1.7E-24</td><td>BT20/MB468</td><td>MITO</td></tr>
<tr><td>>MI0000456|hsa-mir-140&WithFlank&16|+|69933075|69933186@69.90.22&[MIMAT0004597&hsa-miR-140-3p&offsets|+1|+2;m-45&16|+|69933143|69933164&offsets|0|0]|ACCACAGGGTAGAACCACGGAC</td><td>>MI0000456|hsa-mir-140&WithFlank&16|+|69933075|69933186</td><td>69</td><td>90</td><td>22</td><td>ACCACAGGGTAGAACCACGGAC</td><td>281.3</td><td>5.1</td><td>0.5</td><td>10.4</td><td>2.6E-25</td><td>1.7E-24</td><td>BT20/MB468</td><td>NUC</td></tr>
<tr><td>>MI0000462|hsa-mir-152&WithFlank&17|-|48037155|48037253@60.81.22(+1A)&[MIMAT0000438&hsa-miR-152-3p&offsets|0|+1(+1A);m-97&17|-|48037174|48037194&offsets|0|+1(+1A)]|TCAGTGCATGACAGAACTTGGGA</td><td>>MI0000462|hsa-mir-152&WithFlank&17|-|48037155|48037253</td><td>60</td><td>81</td><td>22(+1A)</td><td>TCAGTGCATGACAGAACTTGGGA</td><td>769.2</td><td>13.8</td><td>1.3</td><td>10.5</td><td>1.3E-25</td><td>1.8E-24</td><td>BT20/MB468</td><td>CYTO</td></tr>
<tr><td>>hg38_21_+_8433172_8446622_RNA45SN1_45S_50ntFlanks@10781.10805.25|AGGTGAACAGCCTCTGGCATGTTGG</td><td>>hg38_21_+_8433172_8446622_RNA45SN1_45S_50ntFlanks</td><td>10781</td><td>10805</td><td>25</td><td>AGGTGAACAGCCTCTGGCATGTTGG</td><td>321.8</td><td>-12.8</td><td>1.2</td><td>-10.4</td><td>2.6E-25</td><td>1.8E-24</td><td>BT20/MB468</td><td>NUC</td></tr>
<tr><td>>trna10_MetCAT_8_-_124169470_124169542@31.51.21,trna22_MetCAT_16_-_87417628_87417700@31.51.21|TCTCATAATCTGAAGGTCGTG</td><td>>trna10_MetCAT_8_-_124169470_124169542</td><td>31</td><td>51</td><td>21</td><td>TCTCATAATCTGAAGGTCGTG</td><td>830.7</td><td>-14.1</td><td>1.4</td><td>-10.4</td><td>2.7E-25</td><td>1.8E-24</td><td>BT20/MB468</td><td>NUC</td></tr>
<tr><td>>hg38_21_+_8433172_8446622_RNA45SN1_45S_50ntFlanks@11767.11804.38|TCTAATTAGTGACGCGCATGAATGGATGAACGAGATTC</td><td>>hg38_21_+_8433172_8446622_RNA45SN1_45S_50ntFlanks</td><td>11767</td><td>11804</td><td>38</td><td>TCTAATTAGTGACGCGCATGAATGGATGAACGAGATTC</td><td>440.6</td><td>-13.2</td><td>1.3</td><td>-10.4</td><td>2.7E-25</td><td>1.8E-24</td><td>BT20/MB468</td><td>NUC</td></tr>
<tr><td>>hg38_21_+_8433172_8446622_RNA45SN1_45S_50ntFlanks@12444.12484.41|AAGCAGAATTCACCAAGCGTTGGATTGTTCACCCACTAATA</td><td>>hg38_21_+_8433172_8446622_RNA45SN1_45S_50ntFlanks</td><td>12444</td><td>12484</td><td>41</td><td>AAGCAGAATTCACCAAGCGTTGGATTGTTCACCCACTAATA</td><td>2146.1</td><td>-15.5</td><td>1.5</td><td>-10.4</td><td>2.7E-25</td><td>1.8E-24</td><td>BT20/MB468</td><td>NUC</td></tr>
<tr><td>>am_MI0000809|hsa-mir-151a&WithFlank&8|-|140732558|140732659@53.73.21(+3U)&[MIMAT0000757&hsa-miR-151a-3p&offsets|0|0(+3U);m-30&8|-|140732587|140732607&offsets|0|0(+3U)]|CTAGACTGAAGCTCCTTGAGGTTT</td><td>>am_MI0000809|hsa-mir-151a&WithFlank&8|-|140732558|140732659</td><td>53</td><td>73</td><td>21(+3U)</td><td>CTAGACTGAAGCTCCTTGAGGTTT</td><td>452.2</td><td>13.1</td><td>1.3</td><td>10.5</td><td>1.4E-25</td><td>1.8E-24</td><td>BT20/MB468</td><td>CYTO</td></tr>
<tr><td>>MI0000478|hsa-mir-149&WithFlank&2|+|240455995|240456095@21.43.23&[MIMAT0000450&hsa-miR-149-5p&offsets|0|0;m-195&2|+|240456015|240456037&offsets|0|0]|TCTGGCTCCGTGTCTTCACTCCC</td><td>>MI0000478|hsa-mir-149&WithFlank&2|+|240455995|240456095</td><td>21</td><td>43</td><td>23</td><td>TCTGGCTCCGTGTCTTCACTCCC</td><td>4249.0</td><td>12.1</td><td>1.2</td><td>10.5</td><td>1.4E-25</td><td>1.8E-24</td><td>BT20/MB468</td><td>CYTO</td></tr>
<tr><td>>hg38_21_+_8433172_8446622_RNA45SN1_45S_50ntFlanks@12935.12957.23|TAAACCATTCGTAGACGACCTGC</td><td>>hg38_21_+_8433172_8446622_RNA45SN1_45S_50ntFlanks</td><td>12935</td><td>12957</td><td>23</td><td>TAAACCATTCGTAGACGACCTGC</td><td>289.0</td><td>-12.6</td><td>1.2</td><td>-10.4</td><td>2.8E-25</td><td>1.8E-24</td><td>BT20/MB468</td><td>NUC</td></tr>
<tr><td>>hg38_21_+_8433172_8446622_RNA45SN1_45S_50ntFlanks@3985.4004.20|CGGCTTTGGTGACTCTAGAT</td><td>>hg38_21_+_8433172_8446622_RNA45SN1_45S_50ntFlanks</td><td>3985</td><td>4004</td><td>20</td><td>CGGCTTTGGTGACTCTAGAT</td><td>896.2</td><td>-14.2</td><td>1.4</td><td>-10.4</td><td>2.8E-25</td><td>1.9E-24</td><td>BT20/MB468</td><td>NUC</td></tr>
<tr><td>>hg38_21_+_8433172_8446622_RNA45SN1_45S_50ntFlanks@4723.4758.36|CATTAATCAAGAACGAAAGTCGGAGGTTCGAAGACG</td><td>>hg38_21_+_8433172_8446622_RNA45SN1_45S_50ntFlanks</td><td>4723</td><td>4758</td><td>36</td><td>CATTAATCAAGAACGAAAGTCGGAGGTTCGAAGACG</td><td>10008.0</td><td>-9.4</td><td>0.9</td><td>-10.6</td><td>3.2E-26</td><td>1.9E-24</td><td>MB231/MB468</td><td>MP</td></tr>
<tr><td>>hg38_21_+_8433172_8446622_RNA45SN1_45S_50ntFlanks@9739.9765.27|AAACGATCTCAACCTATTCTCAAACTT</td><td>>hg38_21_+_8433172_8446622_RNA45SN1_45S_50ntFlanks</td><td>9739</td><td>9765</td><td>27</td><td>AAACGATCTCAACCTATTCTCAAACTT</td><td>29961.1</td><td>-9.1</td><td>0.9</td><td>-10.5</td><td>5.7E-26</td><td>1.9E-24</td><td>MB231/MB468</td><td>MITO</td></tr>
<tr><td>>hg38_21_+_8433172_8446622_RNA45SN1_45S_50ntFlanks@4547.4565.19|CTGGATACCGCAGCTAGGA</td><td>>hg38_21_+_8433172_8446622_RNA45SN1_45S_50ntFlanks</td><td>4547</td><td>4565</td><td>19</td><td>CTGGATACCGCAGCTAGGA</td><td>295.6</td><td>-12.6</td><td>1.2</td><td>-10.4</td><td>2.8E-25</td><td>1.9E-24</td><td>BT20/MB468</td><td>NUC</td></tr>
<tr><td>>hg38_21_+_8433172_8446622_RNA45SN1_45S_50ntFlanks@11737.11768.32|ACTCTCTTAAGGTAGCCAAATGCCTCGTCATC</td><td>>hg38_21_+_8433172_8446622_RNA45SN1_45S_50ntFlanks</td><td>11737</td><td>11768</td><td>32</td><td>ACTCTCTTAAGGTAGCCAAATGCCTCGTCATC</td><td>291.5</td><td>-12.6</td><td>1.2</td><td>-10.4</td><td>2.9E-25</td><td>1.9E-24</td><td>BT20/MB468</td><td>NUC</td></tr>
<tr><td>>hg38_21_+_8433172_8446622_RNA45SN1_45S_50ntFlanks@6772.6803.32|GTTCCTCCCGGGGCTACGCCTGTCTGAGCGTC</td><td>>hg38_21_+_8433172_8446622_RNA45SN1_45S_50ntFlanks</td><td>6772</td><td>6803</td><td>32</td><td>GTTCCTCCCGGGGCTACGCCTGTCTGAGCGTC</td><td>402.0</td><td>-13.1</td><td>1.3</td><td>-10.4</td><td>2.9E-25</td><td>1.9E-24</td><td>BT20/MB468</td><td>NUC</td></tr>
<tr><td>>hg38_21_+_8433172_8446622_RNA45SN1_45S_50ntFlanks@11561.11592.32|GCGCCTAGCAGCCGACTTAGAACTGGTGCGGA</td><td>>hg38_21_+_8433172_8446622_RNA45SN1_45S_50ntFlanks</td><td>11561</td><td>11592</td><td>32</td><td>GCGCCTAGCAGCCGACTTAGAACTGGTGCGGA</td><td>2860.4</td><td>-9.3</td><td>0.9</td><td>-10.5</td><td>6.0E-26</td><td>2.0E-24</td><td>MB231/MB468</td><td>MITO</td></tr>
<tr><td>>hg38_21_+_8433172_8446622_RNA45SN1_45S_50ntFlanks@12001.12032.32|CTGCGGGCCGCCGGTGAAATACCACTACTCTG</td><td>>hg38_21_+_8433172_8446622_RNA45SN1_45S_50ntFlanks</td><td>12001</td><td>12032</td><td>32</td><td>CTGCGGGCCGCCGGTGAAATACCACTACTCTG</td><td>2319.1</td><td>-5.3</td><td>0.5</td><td>-10.5</td><td>1.1E-25</td><td>2.0E-24</td><td>BT20/MB468</td><td>MITO</td></tr>
<tr><td>>hg38_21_+_8433172_8446622_RNA45SN1_45S_50ntFlanks@485.515.31|GATGTGGTGACGTCGTGCTCTCCCGGGCCGG</td><td>>hg38_21_+_8433172_8446622_RNA45SN1_45S_50ntFlanks</td><td>485</td><td>515</td><td>31</td><td>GATGTGGTGACGTCGTGCTCTCCCGGGCCGG</td><td>520.3</td><td>11.9</td><td>1.1</td><td>10.4</td><td>3.0E-25</td><td>2.0E-24</td><td>BT20/MB468</td><td>NUC</td></tr>
<tr><td>>MI0000439|hsa-mir-23b&WithFlank&9|+|95085202|95085310@67.84.18&[MIMAT0000418&hsa-miR-23b-3p&offsets|+3|-2;m-42&9|+|95085265|95085285&offsets|+3|0]|ACATTGCCAGGGATTACC</td><td>>MI0000439|hsa-mir-23b&WithFlank&9|+|95085202|95085310</td><td>67</td><td>84</td><td>18</td><td>ACATTGCCAGGGATTACC</td><td>297.6</td><td>11.1</td><td>1.1</td><td>10.4</td><td>3.0E-25</td><td>2.0E-24</td><td>BT20/MB468</td><td>NUC</td></tr>
<tr><td>>MI0000760|hsa-mir-361&WithFlank&X|-|85903630|85903713@52.75.24&[MIMAT0004682&hsa-miR-361-3p&offsets|+1|+2;m-124&X|-|85903642|85903663&offsets|+1|+3]|CCCCCAGGTGTGATTCTGATTTGC</td><td>>MI0000760|hsa-mir-361&WithFlank&X|-|85903630|85903713</td><td>52</td><td>75</td><td>24</td><td>CCCCCAGGTGTGATTCTGATTTGC</td><td>789.2</td><td>6.4</td><td>0.6</td><td>10.4</td><td>1.5E-25</td><td>2.0E-24</td><td>BT20/MB468</td><td>CYTO</td></tr>
<tr><td>>MI0000469|hsa-mir-125a&WithFlank&19|+|51693248|51693345@21.41.21(+1C)&[MIMAT0000443&hsa-miR-125a-5p&offsets|0|-3(+1C);m-63&19|+|51693268|51693289&offsets|0|-1(+1C)]|TCCCTGAGACCCTTTAACCTGC</td><td>>MI0000469|hsa-mir-125a&WithFlank&19|+|51693248|51693345</td><td>21</td><td>41</td><td>21(+1C)</td><td>TCCCTGAGACCCTTTAACCTGC</td><td>1055.3</td><td>7.3</td><td>0.7</td><td>10.5</td><td>1.1E-25</td><td>2.0E-24</td><td>BT20/MB468</td><td>MITO</td></tr>
<tr><td>>hg38_21_+_8433172_8446622_RNA45SN1_45S_50ntFlanks@9539.9566.28|ACTCTGGTGGAGGTCCGTAGCGGTCCTG</td><td>>hg38_21_+_8433172_8446622_RNA45SN1_45S_50ntFlanks</td><td>9539</td><td>9566</td><td>28</td><td>ACTCTGGTGGAGGTCCGTAGCGGTCCTG</td><td>384.7</td><td>-13.0</td><td>1.3</td><td>-10.4</td><td>3.1E-25</td><td>2.0E-24</td><td>BT20/MB468</td><td>NUC</td></tr>
<tr><td>>hg38_21_+_8433172_8446622_RNA45SN1_45S_50ntFlanks@12635.12653.19|AAGCTACCATCTGTGGGAT</td><td>>hg38_21_+_8433172_8446622_RNA45SN1_45S_50ntFlanks</td><td>12635</td><td>12653</td><td>19</td><td>AAGCTACCATCTGTGGGAT</td><td>458.7</td><td>-13.3</td><td>1.3</td><td>-10.4</td><td>3.1E-25</td><td>2.0E-24</td><td>BT20/MB468</td><td>NUC</td></tr>
<tr><td>>hg38_21_+_8433172_8446622_RNA45SN1_45S_50ntFlanks@10392.10410.19|ACAGCAGTTGAACATGGGT</td><td>>hg38_21_+_8433172_8446622_RNA45SN1_45S_50ntFlanks</td><td>10392</td><td>10410</td><td>19</td><td>ACAGCAGTTGAACATGGGT</td><td>4084.8</td><td>-10.1</td><td>1.0</td><td>-10.6</td><td>3.5E-26</td><td>2.0E-24</td><td>MB231/MB468</td><td>MP</td></tr>
<tr><td>>hg38_21_+_8433172_8446622_RNA45SN1_45S_50ntFlanks@10764.10785.22|TCCGCAGCAGGTCTCCAAGGTG</td><td>>hg38_21_+_8433172_8446622_RNA45SN1_45S_50ntFlanks</td><td>10764</td><td>10785</td><td>22</td><td>TCCGCAGCAGGTCTCCAAGGTG</td><td>367.9</td><td>-12.9</td><td>1.2</td><td>-10.4</td><td>3.1E-25</td><td>2.0E-24</td><td>BT20/MB468</td><td>NUC</td></tr>
<tr><td>>hg38_21_+_8433172_8446622_RNA45SN1_45S_50ntFlanks@10735.10752.18|TGTAAATCTCGCGCCGGG</td><td>>hg38_21_+_8433172_8446622_RNA45SN1_45S_50ntFlanks</td><td>10735</td><td>10752</td><td>18</td><td>TGTAAATCTCGCGCCGGG</td><td>471.5</td><td>-13.3</td><td>1.3</td><td>-10.4</td><td>3.1E-25</td><td>2.1E-24</td><td>BT20/MB468</td><td>NUC</td></tr>
<tr><td>>hg38_21_+_8433172_8446622_RNA45SN1_45S_50ntFlanks@8356.8379.24|GAAAAGAACTTTGAAGAGAGAGTT</td><td>>hg38_21_+_8433172_8446622_RNA45SN1_45S_50ntFlanks</td><td>8356</td><td>8379</td><td>24</td><td>GAAAAGAACTTTGAAGAGAGAGTT</td><td>12781.5</td><td>-7.9</td><td>0.8</td><td>-10.5</td><td>7.9E-26</td><td>2.1E-24</td><td>MB231/MB468</td><td>NUC</td></tr>
<tr><td>>trnaMT_GluTTC_MT_-_14674_14742@35.71.37,trnalookalike8_GluTTC_5_-_93905172_93905240@35.71.37|TATCATTGGTCGTGGTTGTAGTCCGTGCGAGAATACC</td><td>>trnaMT_GluTTC_MT_-_14674_14742</td><td>35</td><td>71</td><td>37</td><td>TATCATTGGTCGTGGTTGTAGTCCGTGCGAGAATACC</td><td>2122.2</td><td>15.1</td><td>1.4</td><td>10.6</td><td>2.6E-26</td><td>2.1E-24</td><td>MB231/MB468</td><td>CYTO</td></tr>
<tr><td>>MI0003513|hsa-mir-455&WithFlank&9|+|114209428|114209535@22.44.23&[MIMAT0003150&hsa-miR-455-5p&offsets|0|+1;m-232&9|+|114209449|114209470&offsets|0|+1]|TATGTGCCTTTGGACTACATCGT</td><td>>MI0003513|hsa-mir-455&WithFlank&9|+|114209428|114209535</td><td>22</td><td>44</td><td>23</td><td>TATGTGCCTTTGGACTACATCGT</td><td>936.4</td><td>13.8</td><td>1.3</td><td>10.5</td><td>8.1E-26</td><td>2.1E-24</td><td>MB231/MB468</td><td>NUC</td></tr>
<tr><td>>hg38_21_+_8433172_8446622_RNA45SN1_45S_50ntFlanks@12151.12179.29|GTGGGGAGTTTGACTGGGGCGGTACACCT</td><td>>hg38_21_+_8433172_8446622_RNA45SN1_45S_50ntFlanks</td><td>12151</td><td>12179</td><td>29</td><td>GTGGGGAGTTTGACTGGGGCGGTACACCT</td><td>529.4</td><td>-13.5</td><td>1.3</td><td>-10.4</td><td>3.2E-25</td><td>2.1E-24</td><td>BT20/MB468</td><td>NUC</td></tr>
<tr><td>>hg38_21_+_8433172_8446622_RNA45SN1_45S_50ntFlanks@11873.11905.33|AAGAAGACCCTGTTGAGCTTGACTCTAGTCTGG</td><td>>hg38_21_+_8433172_8446622_RNA45SN1_45S_50ntFlanks</td><td>11873</td><td>11905</td><td>33</td><td>AAGAAGACCCTGTTGAGCTTGACTCTAGTCTGG</td><td>453.1</td><td>-13.2</td><td>1.3</td><td>-10.4</td><td>3.3E-25</td><td>2.1E-24</td><td>BT20/MB468</td><td>NUC</td></tr>
<tr><td>>hg38_21_+_8433172_8446622_RNA45SN1_45S_50ntFlanks@7986.8008.23|ATCAGACGTGGCGACCCGCTGAA</td><td>>hg38_21_+_8433172_8446622_RNA45SN1_45S_50ntFlanks</td><td>7986</td><td>8008</td><td>23</td><td>ATCAGACGTGGCGACCCGCTGAA</td><td>4652.0</td><td>-16.0</td><td>1.5</td><td>-10.6</td><td>3.6E-26</td><td>2.1E-24</td><td>MB231/MB468</td><td>MP</td></tr>
<tr><td>>hg38_21_+_8433172_8446622_RNA45SN1_45S_50ntFlanks@12995.13016.22|CTCGCTGCGATCTATTGAAAGT</td><td>>hg38_21_+_8433172_8446622_RNA45SN1_45S_50ntFlanks</td><td>12995</td><td>13016</td><td>22</td><td>CTCGCTGCGATCTATTGAAAGT</td><td>1168.1</td><td>5.9</td><td>0.6</td><td>10.4</td><td>1.6E-25</td><td>2.1E-24</td><td>BT20/MB468</td><td>CYTO</td></tr>
<tr><td>>hg38_21_+_8433172_8446622_RNA45SN1_45S_50ntFlanks@11918.11956.39|ACATGAGAGGTGTAGAATAAGTGGGAGGCCCCCGGCGCC</td><td>>hg38_21_+_8433172_8446622_RNA45SN1_45S_50ntFlanks</td><td>11918</td><td>11956</td><td>39</td><td>ACATGAGAGGTGTAGAATAAGTGGGAGGCCCCCGGCGCC</td><td>488.9</td><td>-13.4</td><td>1.3</td><td>-10.4</td><td>3.3E-25</td><td>2.1E-24</td><td>BT20/MB468</td><td>NUC</td></tr>
<tr><td>>hg38_21_+_8433172_8446622_RNA45SN1_45S_50ntFlanks@11618.11640.23|AAACAAAGCATCGCGAAGGCCCG</td><td>>hg38_21_+_8433172_8446622_RNA45SN1_45S_50ntFlanks</td><td>11618</td><td>11640</td><td>23</td><td>AAACAAAGCATCGCGAAGGCCCG</td><td>2108.2</td><td>-14.9</td><td>1.4</td><td>-10.6</td><td>3.7E-26</td><td>2.2E-24</td><td>MB231/MB468</td><td>MP</td></tr>
<tr><td>>hg38_21_+_8433172_8446622_RNA45SN1_45S_50ntFlanks@11115.11132.18|CGGCAGGGGCCGGCGGCG</td><td>>hg38_21_+_8433172_8446622_RNA45SN1_45S_50ntFlanks</td><td>11115</td><td>11132</td><td>18</td><td>CGGCAGGGGCCGGCGGCG</td><td>893.9</td><td>-14.2</td><td>1.4</td><td>-10.4</td><td>3.3E-25</td><td>2.2E-24</td><td>BT20/MB468</td><td>NUC</td></tr>
<tr><td>>hg38_21_+_8433172_8446622_RNA45SN1_45S_50ntFlanks@11323.11344.22|CGTCTCGTCGCGCGCGCGTCCG</td><td>>hg38_21_+_8433172_8446622_RNA45SN1_45S_50ntFlanks</td><td>11323</td><td>11344</td><td>22</td><td>CGTCTCGTCGCGCGCGCGTCCG</td><td>11400.5</td><td>-13.0</td><td>1.2</td><td>-10.6</td><td>3.7E-26</td><td>2.2E-24</td><td>MB231/MB468</td><td>MP</td></tr>
<tr><td>>hg38_21_+_8433172_8446622_RNA45SN1_45S_50ntFlanks@3840.3867.28|CTCCTACTTGGATAACTGTGGTAATTCT</td><td>>hg38_21_+_8433172_8446622_RNA45SN1_45S_50ntFlanks</td><td>3840</td><td>3867</td><td>28</td><td>CTCCTACTTGGATAACTGTGGTAATTCT</td><td>618.3</td><td>4.8</td><td>0.5</td><td>10.4</td><td>1.7E-25</td><td>2.2E-24</td><td>BT20/MB468</td><td>CYTO</td></tr>
</table>

<table>
<tr><th></th><th></th><th></th><th></th><th></th><th></th><th></th><th></th><th></th><th></th><th></th><th></th><th></th><th></th></tr>
<tr><td>>hg38_21_+_8433172_8446622_RNA45SN1_45S_50ntFlanks@4726.4751.26|TAATCAAGAACGAAAGTCGGAGGTTC</td><td>>hg38_21_+_8433172_8446622_RNA45SN1_45S_50ntFlanks</td><td>4726</td><td>4751</td><td>26</td><td>TAATCAAGAACGAAAGTCGGAGGTTC</td><td>427.5</td><td>-13.2</td><td>1.3</td><td>-10.4</td><td>3.4E-25</td><td>2.2E-24</td><td>BT20/MB468</td><td>NUC</td></tr>
<tr><td>>hg38_21_+_8433172_8446622_RNA45SN1_45S_50ntFlanks@12957.12982.26|CTTCTGGGTCGGGGTTTCGTACGTAG</td><td>>hg38_21_+_8433172_8446622_RNA45SN1_45S_50ntFlanks</td><td>12957</td><td>12982</td><td>26</td><td>CTTCTGGGTCGGGGTTTCGTACGTAG</td><td>6048.2</td><td>-12.4</td><td>1.2</td><td>-10.5</td><td>8.7E-26</td><td>2.2E-24</td><td>MB231/MB468</td><td>NUC</td></tr>
<tr><td>>hg38_21_+_8433172_8446622_RNA45SN1_45S_50ntFlanks@11781.11803.23|CGCATGAATGGATGAACGAGATT</td><td>>hg38_21_+_8433172_8446622_RNA45SN1_45S_50ntFlanks</td><td>11781</td><td>11803</td><td>23</td><td>CGCATGAATGGATGAACGAGATT</td><td>2551.1</td><td>-15.2</td><td>1.4</td><td>-10.6</td><td>3.9E-26</td><td>2.2E-24</td><td>MB231/MB468</td><td>MP</td></tr>
<tr><td>>hg38_21_+_8433172_8446622_RNA45SN1_45S_50ntFlanks@6698.6718.21|AGCTAGCTGCGAGAATTAATG</td><td>>hg38_21_+_8433172_8446622_RNA45SN1_45S_50ntFlanks</td><td>6698</td><td>6718</td><td>21</td><td>AGCTAGCTGCGAGAATTAATG</td><td>3438.4</td><td>-5.5</td><td>0.5</td><td>-10.5</td><td>1.3E-25</td><td>2.3E-24</td><td>BT20/MB468</td><td>MITO</td></tr>
<tr><td>>hg38_21_+_8433172_8446622_RNA45SN1_45S_50ntFlanks@11316.11344.29|CACCCCACGTCTCGTCGCGCGCGCGTCCG</td><td>>hg38_21_+_8433172_8446622_RNA45SN1_45S_50ntFlanks</td><td>11316</td><td>11344</td><td>29</td><td>CACCCCACGTCTCGTCGCGCGCGCGTCCG</td><td>1563.4</td><td>-14.5</td><td>1.4</td><td>-10.6</td><td>3.9E-26</td><td>2.3E-24</td><td>MB231/MB468</td><td>MP</td></tr>
<tr><td>>trnaMT_PheGAA_MT_+_577_647@1.19.19|GTTTATGTAGCTTACCTCC</td><td>>trnaMT_PheGAA_MT_+_577_647</td><td>1</td><td>19</td><td>19</td><td>GTTTATGTAGCTTACCTCC</td><td>614.0</td><td>7.1</td><td>0.7</td><td>10.5</td><td>9.0E-26</td><td>2.3E-24</td><td>MB231/MB468</td><td>NUC</td></tr>
<tr><td>>hg38_21_+_8433172_8446622_RNA45SN1_45S_50ntFlanks@5485.5514.30|AGCGCTGAGAAGACGGTCGAACTTGACTAT</td><td>>hg38_21_+_8433172_8446622_RNA45SN1_45S_50ntFlanks</td><td>5485</td><td>5514</td><td>30</td><td>AGCGCTGAGAAGACGGTCGAACTTGACTAT</td><td>9884.8</td><td>-12.4</td><td>1.2</td><td>-10.5</td><td>7.4E-26</td><td>2.4E-24</td><td>MB231/MB468</td><td>MITO</td></tr>
<tr><td>>trnaMT_SerGCT_MT_+_12207_12265@8.38.31|CTCACAAGAACTGCTAACTCATGCCCCCATG</td><td>>trnaMT_SerGCT_MT_+_12207_12265</td><td>8</td><td>38</td><td>31</td><td>CTCACAAGAACTGCTAACTCATGCCCCCATG</td><td>4131.2</td><td>-16.4</td><td>1.6</td><td>-10.4</td><td>3.7E-25</td><td>2.4E-24</td><td>BT20/MB468</td><td>NUC</td></tr>
<tr><td>>hg38_21_+_8433172_8446622_RNA45SN1_45S_50ntFlanks@5431.5451.21|GTTTAGTGAGGCCCTCGGATC</td><td>>hg38_21_+_8433172_8446622_RNA45SN1_45S_50ntFlanks</td><td>5431</td><td>5451</td><td>21</td><td>GTTTAGTGAGGCCCTCGGATC</td><td>484.0</td><td>-13.3</td><td>1.3</td><td>-10.4</td><td>3.7E-25</td><td>2.4E-24</td><td>BT20/MB468</td><td>NUC</td></tr>
<tr><td>>trna116_GluCTC_1_-_145399233_145399304@32.70.39,trna71_GluCTC_1_-_161439189_161439260@32.70.39,trna74_GluCTC_1_-_161431809_161431880@32.70.39,trna77_GluCTC_1_-_161424398_161424469@32.70.39,trna77_GluCTC_6_+_28949976_28950047@32.70.39,trna80_GluCTC_1_-_161417018_161417089@32.70.39,trna87_GluCTC_6_-_126101393_126101464@32.70.39|CTCTCACCGCCGCGGCCCGGGTTCGATTCCCGGTCAGGG</td><td>>trna116_GluCTC_1_-_145399233_145399304</td><td>32</td><td>70</td><td>39</td><td>CTCTCACCGCCGCGGCCCGGGTTCGATTCCCGGTCAGGG</td><td>10745.0</td><td>-11.4</td><td>1.1</td><td>-10.4</td><td>3.7E-25</td><td>2.4E-24</td><td>BT20/MB468</td><td>NUC</td></tr>
<tr><td>>hg38_21_+_8433172_8446622_RNA45SN1_45S_50ntFlanks@12008.12029.22|CCGCCGGTGAAATACCACTACT</td><td>>hg38_21_+_8433172_8446622_RNA45SN1_45S_50ntFlanks</td><td>12008</td><td>12029</td><td>22</td><td>CCGCCGGTGAAATACCACTACT</td><td>2095.8</td><td>-14.8</td><td>1.4</td><td>-10.5</td><td>7.5E-26</td><td>2.4E-24</td><td>MB231/MB468</td><td>MITO</td></tr>
<tr><td>>hg38_21_+_8433172_8446622_RNA45SN1_45S_50ntFlanks@5477.5516.40|CCTGGCGGAGCGCTGAGAAGACGGTCGAACTTGACTATCT</td><td>>hg38_21_+_8433172_8446622_RNA45SN1_45S_50ntFlanks</td><td>5477</td><td>5516</td><td>40</td><td>CCTGGCGGAGCGCTGAGAAGACGGTCGAACTTGACTATCT</td><td>2536.2</td><td>11.0</td><td>1.0</td><td>10.6</td><td>3.1E-26</td><td>2.4E-24</td><td>MB231/MB468</td><td>CYTO</td></tr>
<tr><td>>MI0000462|hsa-mir-152&WithFlank&17|-|48037155|48037253@60.81.22&[MIMAT0000438&hsa-miR-152-3p&offsets|0|+1;m-97&17|-|48037174|48037194&offsets|0|+1]|TCAGTGCATGACAGAACTTGGG</td><td>>MI0000462|hsa-mir-152&WithFlank&17|-|48037155|48037253</td><td>60</td><td>81</td><td>22</td><td>TCAGTGCATGACAGAACTTGGG</td><td>1393.4</td><td>14.7</td><td>1.4</td><td>10.4</td><td>1.9E-25</td><td>2.5E-24</td><td>BT20/MB468</td><td>CYTO</td></tr>
<tr><td>>MI0000301|hsa-mir-224&WithFlank&X|-|151958572|151958664@14.34.21&[MIMAT0000281&hsa-miR-224-5p&offsets|+1|-3;m-238&X|-|151958630|151958651&offsets|0|-1]|CAAGTCACTAGTGGTTCCGTT</td><td>>MI0000301|hsa-mir-224&WithFlank&X|-|151958572|151958664</td><td>14</td><td>34</td><td>21</td><td>CAAGTCACTAGTGGTTCCGTT</td><td>1556.5</td><td>15.1</td><td>1.4</td><td>10.5</td><td>1.4E-25</td><td>2.5E-24</td><td>BT20/MB468</td><td>MITO</td></tr>
<tr><td>>hg38_21_+_8433172_8446622_RNA45SN1_45S_50ntFlanks@12992.13017.26|TCCCTCGCTGCGATCTATTGAAAGTC</td><td>>hg38_21_+_8433172_8446622_RNA45SN1_45S_50ntFlanks</td><td>12992</td><td>13017</td><td>26</td><td>TCCCTCGCTGCGATCTATTGAAAGTC</td><td>309.1</td><td>-12.7</td><td>1.2</td><td>-10.4</td><td>3.8E-25</td><td>2.5E-24</td><td>BT20/MB468</td><td>NUC</td></tr>
<tr><td>>MI0000298|hsa-mir-221&WithFlank&X|-|45746151|45746272@71.92.22&[MIMAT0000278&hsa-miR-221-3p&offsets|0|-1;m-91&X|-|45746181|45746202&offsets|0|0]|AGCTACATTGTCTGCTGGGTTT</td><td>>MI0000298|hsa-mir-221&WithFlank&X|-|45746151|45746272</td><td>71</td><td>92</td><td>22</td><td>AGCTACATTGTCTGCTGGGTTT</td><td>20461.9</td><td>7.4</td><td>0.7</td><td>10.5</td><td>9.7E-26</td><td>2.5E-24</td><td>MB231/MB468</td><td>NUC</td></tr>
<tr><td>>hg38_21_+_8433172_8446622_RNA45SN1_45S_50ntFlanks@9504.9526.23|GTGAACTATGCCTGGGCAGGGCG</td><td>>hg38_21_+_8433172_8446622_RNA45SN1_45S_50ntFlanks</td><td>9504</td><td>9526</td><td>23</td><td>GTGAACTATGCCTGGGCAGGGCG</td><td>1988.6</td><td>-13.9</td><td>1.3</td><td>-10.5</td><td>9.8E-26</td><td>2.5E-24</td><td>MB231/MB468</td><td>NUC</td></tr>
<tr><td>>hg38_21_+_8433172_8446622_RNA45SN1_45S_50ntFlanks@3840.3868.29|CTCCTACTTGGATAACTGTGGTAATTCTA</td><td>>hg38_21_+_8433172_8446622_RNA45SN1_45S_50ntFlanks</td><td>3840</td><td>3868</td><td>29</td><td>CTCCTACTTGGATAACTGTGGTAATTCTA</td><td>1916.4</td><td>4.0</td><td>0.4</td><td>10.4</td><td>1.9E-25</td><td>2.5E-24</td><td>BT20/MB468</td><td>CYTO</td></tr>
<tr><td>>MI0001733|hsa-mir-452&WithFlank&X|-|151959622|151959718@23.43.21&[MIMAT0001635&hsa-miR-452-5p&offsets|+3|+2;m-188&X|-|151959677|151959699&offsets|+3|+1]|TGTTTGCAGAGGAAACTGAGA</td><td>>MI0001733|hsa-mir-452&WithFlank&X|-|151959622|151959718</td><td>23</td><td>43</td><td>21</td><td>TGTTTGCAGAGGAAACTGAGA</td><td>649.5</td><td>13.8</td><td>1.3</td><td>10.5</td><td>1.5E-25</td><td>2.6E-24</td><td>BT20/MB468</td><td>MITO</td></tr>
<tr><td>>hg38_21_+_8433172_8446622_RNA45SN1_45S_50ntFlanks@13009.13027.19|TTGAAAGTCAGCCCTCGAC</td><td>>hg38_21_+_8433172_8446622_RNA45SN1_45S_50ntFlanks</td><td>13009</td><td>13027</td><td>19</td><td>TTGAAAGTCAGCCCTCGAC</td><td>345.4</td><td>-12.9</td><td>1.2</td><td>-10.4</td><td>4.0E-25</td><td>2.6E-24</td><td>BT20/MB468</td><td>NUC</td></tr>
<tr><td>>MI0003591|hsa-mir-584&WithFlank&5|-|149062307|149062415@22.42.21&[MIMAT0003249&hsa-miR-584-5p&offsets|0|-1;m-167&5|-|149062374|149062394&offsets|0|0]|TTATGGTTTGCCTGGGACTGA</td><td>>MI0003591|hsa-mir-584&WithFlank&5|-|149062307|149062415</td><td>22</td><td>42</td><td>21</td><td>TTATGGTTTGCCTGGGACTGA</td><td>2046.6</td><td>9.3</td><td>0.9</td><td>10.4</td><td>2.0E-25</td><td>2.6E-24</td><td>BT20/MB468</td><td>CYTO</td></tr>
<tr><td>>MI0000102|hsa-mir-100&WithFlank&11|-|122152223|122152314@19.38.20(+1A)&[MIMAT0000098&hsa-miR-100-5p&offsets|0|-2(+1A);m-27&11|-|122152276|122152296&offsets|0|-1(+1A)]|AACCCGTAGATCCGAACTTGA</td><td>>MI0000102|hsa-mir-100&WithFlank&11|-|122152223|122152314</td><td>19</td><td>38</td><td>20(+1A)</td><td>AACCCGTAGATCCGAACTTGA</td><td>7438.4</td><td>16.9</td><td>1.6</td><td>10.6</td><td>3.4E-26</td><td>2.6E-24</td><td>MB231/MB468</td><td>CYTO</td></tr>
<tr><td>>hg38_21_+_8433172_8446622_RNA45SN1_45S_50ntFlanks@9534.9553.20|AGGAAACTCTGGTGGAGGTC</td><td>>hg38_21_+_8433172_8446622_RNA45SN1_45S_50ntFlanks</td><td>9534</td><td>9553</td><td>20</td><td>AGGAAACTCTGGTGGAGGTC</td><td>124313.7</td><td>-8.2</td><td>0.8</td><td>-10.6</td><td>3.4E-26</td><td>2.6E-24</td><td>MB231/MB468</td><td>CYTO</td></tr>
<tr><td>>hg38_21_+_8433172_8446622_RNA45SN1_45S_50ntFlanks@11161.11178.18|CACGGTCCCCCGCGAGGG</td><td>>hg38_21_+_8433172_8446622_RNA45SN1_45S_50ntFlanks</td><td>11161</td><td>11178</td><td>18</td><td>CACGGTCCCCCGCGAGGG</td><td>2148.8</td><td>-14.9</td><td>1.4</td><td>-10.5</td><td>8.3E-26</td><td>2.6E-24</td><td>MB231/MB468</td><td>MITO</td></tr>
<tr><td>>hg38_21_+_8433172_8446622_RNA45SN1_45S_50ntFlanks@10410.10430.21|TCAGTCGGTCCTGAGAGATGG</td><td>>hg38_21_+_8433172_8446622_RNA45SN1_45S_50ntFlanks</td><td>10410</td><td>10430</td><td>21</td><td>TCAGTCGGTCCTGAGAGATGG</td><td>2861.3</td><td>-15.3</td><td>1.5</td><td>-10.6</td><td>4.7E-26</td><td>2.7E-24</td><td>MB231/MB468</td><td>MP</td></tr>
<tr><td>>hg38_21_+_8433172_8446622_RNA45SN1_45S_50ntFlanks@6719.6736.18|TGAATTGCAGGACACATT</td><td>>hg38_21_+_8433172_8446622_RNA45SN1_45S_50ntFlanks</td><td>6719</td><td>6736</td><td>18</td><td>TGAATTGCAGGACACATT</td><td>1532.0</td><td>-9.0</td><td>0.9</td><td>-10.6</td><td>4.7E-26</td><td>2.7E-24</td><td>MB231/MB468</td><td>MP</td></tr>
<tr><td>>MI0000285|hsa-mir-205&WithFlank&1|+|209432127|209432248@40.61.22(+1C)&[MIMAT0000266&hsa-miR-205-5p&offsets|0|0(+1C);m-351&1|+|209432166|209432187&offsets|0|0(+1C)]|TCCTTCATTCCACCGGAGTCTGC</td><td>>MI0000285|hsa-mir-205&WithFlank&1|+|209432127|209432248</td><td>40</td><td>61</td><td>22(+1C)</td><td>TCCTTCATTCCACCGGAGTCTGC</td><td>6515.8</td><td>16.4</td><td>1.5</td><td>10.7</td><td>1.2E-26</td><td>2.7E-24</td><td>BT20/MB231</td><td>NUC</td></tr>
<tr><td>>MI0000440|hsa-mir-27b&WithFlank&9|+|95085439|95085547@70.87.18&[MIMAT0000419&hsa-miR-27b-3p&offsets|+3|0;m-39&9|+|95085505|95085525&offsets|+3|0]|ACAGTGGCTAAGTTCTGC</td><td>>MI0000440|hsa-mir-27b&WithFlank&9|+|95085439|95085547</td><td>70</td><td>87</td><td>18</td><td>ACAGTGGCTAAGTTCTGC</td><td>747.6</td><td>14.0</td><td>1.3</td><td>10.4</td><td>1.6E-25</td><td>2.7E-24</td><td>BT20/MB468</td><td>MITO</td></tr>
<tr><td>>hg38_21_+_8433172_8446622_RNA45SN1_45S_50ntFlanks@9454.9503.50|CCGCGCCGGGGAGGTGGAGCACGAGCGCACGTGTTAGGACCCGAAAGATG</td><td>>hg38_21_+_8433172_8446622_RNA45SN1_45S_50ntFlanks</td><td>9454</td><td>9503</td><td>50</td><td>CCGCGCCGGGGAGGTGGAGCACGAGCGCACGTGTTAGGACCCGAAAGATG</td><td>6519.8</td><td>-17.1</td><td>1.7</td><td>-10.3</td><td>4.2E-25</td><td>2.7E-24</td><td>BT20/MB468</td><td>NUC</td></tr>
<tr><td>>MI0000298|hsa-mir-221&WithFlank&X|-|45746151|45746272@71.91.21(+1A)&[MIMAT0000278&hsa-miR-221-3p&offsets|0|-2(+1A);m-91&X|-|45746181|45746202&offsets|0|-1(+1A)]|AGCTACATTGTCTGCTGGGTTA</td><td>>MI0000298|hsa-mir-221&WithFlank&X|-|45746151|45746272</td><td>71</td><td>91</td><td>21(+1A)</td><td>AGCTACATTGTCTGCTGGGTTA</td><td>503.4</td><td>7.4</td><td>0.7</td><td>10.5</td><td>1.1E-25</td><td>2.8E-24</td><td>MB231/MB468</td><td>NUC</td></tr>
<tr><td>>MI0000272|hsa-mir-182&WithFlank&7|-|129770377|129770498@29.50.22(+1U)&[MIMAT0000259&hsa-miR-182-5p&offsets|0|-2(+1U);m-22&7|-|129770449|129770470&offsets|0|0(+1U)]|TTTGGCAATGGTAGAACTCACAT</td><td>>MI0000272|hsa-mir-182&WithFlank&7|-|129770377|129770498</td><td>29</td><td>50</td><td>22(+1U)</td><td>TTTGGCAATGGTAGAACTCACAT</td><td>180.6</td><td>4.0</td><td>0.4</td><td>10.3</td><td>4.3E-25</td><td>2.8E-24</td><td>BT20/MB468</td><td>NUC</td></tr>
<tr><td>>hg38_21_+_8433172_8446622_RNA45SN1_45S_50ntFlanks@12991.13014.24|CTCCCTCGCTGCGATCTATTGAAA</td><td>>hg38_21_+_8433172_8446622_RNA45SN1_45S_50ntFlanks</td><td>12991</td><td>13014</td><td>24</td><td>CTCCCTCGCTGCGATCTATTGAAA</td><td>29319.8</td><td>-14.8</td><td>1.4</td><td>-10.3</td><td>4.3E-25</td><td>2.8E-24</td><td>BT20/MB468</td><td>NUC</td></tr>
<tr><td>>hg38_21_+_8433172_8446622_RNA45SN1_45S_50ntFlanks@7986.8012.27|ATCAGACGTGGCGACCCGCTGAATTTA</td><td>>hg38_21_+_8433172_8446622_RNA45SN1_45S_50ntFlanks</td><td>7986</td><td>8012</td><td>27</td><td>ATCAGACGTGGCGACCCGCTGAATTTA</td><td>2445.7</td><td>-15.1</td><td>1.4</td><td>-10.6</td><td>4.9E-26</td><td>2.8E-24</td><td>MB231/MB468</td><td>MP</td></tr>
<tr><td>>hg38_21_+_8433172_8446622_RNA45SN1_45S_50ntFlanks@7981.8008.28|CTCAGATCAGACGTGGCGACCCGCTGAA</td><td>>hg38_21_+_8433172_8446622_RNA45SN1_45S_50ntFlanks</td><td>7981</td><td>8008</td><td>28</td><td>CTCAGATCAGACGTGGCGACCCGCTGAA</td><td>3092.4</td><td>-6.3</td><td>0.6</td><td>-10.5</td><td>8.9E-26</td><td>2.8E-24</td><td>MB231/MB468</td><td>MITO</td></tr>
<tr><td>>hg38_21_+_8433172_8446622_RNA45SN1_45S_50ntFlanks@7973.8009.37|GACGCGACCTCAGATCAGACGTGGCGACCCGCTGAAT</td><td>>hg38_21_+_8433172_8446622_RNA45SN1_45S_50ntFlanks</td><td>7973</td><td>8009</td><td>37</td><td>GACGCGACCTCAGATCAGACGTGGCGACCCGCTGAAT</td><td>2417.8</td><td>7.9</td><td>0.8</td><td>10.4</td><td>2.2E-25</td><td>2.9E-24</td><td>BT20/MB468</td><td>CYTO</td></tr>
<tr><td>>hg38_21_+_8433172_8446622_RNA45SN1_45S_50ntFlanks@9574.9595.22|AATCGGTCGTCCGACCTGGGTA</td><td>>hg38_21_+_8433172_8446622_RNA45SN1_45S_50ntFlanks</td><td>9574</td><td>9595</td><td>22</td><td>AATCGGTCGTCCGACCTGGGTA</td><td>2812.1</td><td>-4.5</td><td>0.4</td><td>-10.4</td><td>2.2E-25</td><td>2.9E-24</td><td>BT20/MB468</td><td>CYTO</td></tr>
<tr><td>>trna111_HisGTG_1_-_147774845_147774916@-1G.34.35,trna118_HisGTG_1_-_145396881_145396952@-1G.34.35,trna16_HisGTG_1_+_146544773_146544844@-1G.34.35,trna1_HisGTG_15_+_45493349_45493420@-1G.34.35,trna21_HisGTG_1_+_147753471_147753542@-1G.34.35,trna33_HisGTG_6_+_27125906_27125977@-1G.34.35,trna7_HisGTG_9_-_14433938_14434009@-1G.34.35,trna8_HisGTG_15_-_45492611_45492682@-1G.34.35,trna9_HisGTG_15_-_45490804_45490875@-1G.34.35|GGCCGTGATCGTATAGTGGTTAGTACTCTGCGTTG</td><td>>trna111_HisGTG_1_-_147774845_147774916</td><td>-1G</td><td>34</td><td>35</td><td>GGCCGTGATCGTATAGTGGTTAGTACTCTGCGTTG</td><td>58247.1</td><td>12.7</td><td>1.2</td><td>10.6</td><td>3.8E-26</td><td>2.9E-24</td><td>MB231/MB468</td><td>CYTO</td></tr>
<tr><td>>MI0000446|hsa-mir-125b-1&WithFlank&11|-|122099751|122099850@61.82.22&[MIMAT0004592&hsa-miR-125b-1-3p&offsets|0|0;m-485&11|-|122099769|122099790&offsets|0|0]|ACGGGTTAGGCTCTTGGGAGCT</td><td>>MI0000446|hsa-mir-125b-1&WithFlank&11|-|122099751|122099850</td><td>61</td><td>82</td><td>22</td><td>ACGGGTTAGGCTCTTGGGAGCT</td><td>1285.4</td><td>14.3</td><td>1.4</td><td>10.6</td><td>3.9E-26</td><td>2.9E-24</td><td>MB231/MB468</td><td>CYTO</td></tr>
<tr><td>>hg38_21_+_8433172_8446622_RNA45SN1_45S_50ntFlanks@11799.11820.22|AGATTCCCACTGTCCCTACCTA</td><td>>hg38_21_+_8433172_8446622_RNA45SN1_45S_50ntFlanks</td><td>11799</td><td>11820</td><td>22</td><td>AGATTCCCACTGTCCCTACCTA</td><td>406.8</td><td>-13.1</td><td>1.3</td><td>-10.3</td><td>4.5E-25</td><td>2.9E-24</td><td>BT20/MB468</td><td>NUC</td></tr>
<tr><td>>MI0003643|hsa-mir-629&WithFlank&15|-|70079366|70079474@28.49.22&[MIMAT0004810&hsa-miR-629-5p&offsets|0|+1;m-150&15|-|70079427|70079447&offsets|0|+1]|TGGGTTTACGTTGGGAGAACTT</td><td>>MI0003643|hsa-mir-629&WithFlank&15|-|70079366|70079474</td><td>28</td><td>49</td><td>22</td><td>TGGGTTTACGTTGGGAGAACTT</td><td>2404.8</td><td>14.5</td><td>1.4</td><td>10.5</td><td>5.2E-26</td><td>2.9E-24</td><td>MB231/MB468</td><td>MP</td></tr>
<tr><td>>hg38_21_+_8433172_8446622_RNA45SN1_45S_50ntFlanks@5489.5509.21|CTGAGAAGACGGTCGAACTTG</td><td>>hg38_21_+_8433172_8446622_RNA45SN1_45S_50ntFlanks</td><td>5489</td><td>5509</td><td>21</td><td>CTGAGAAGACGGTCGAACTTG</td><td>1488.8</td><td>-14.4</td><td>1.4</td><td>-10.5</td><td>5.2E-26</td><td>2.9E-24</td><td>MB231/MB468</td><td>MP</td></tr>
<tr><td>>MI0000805|hsa-mir-342&WithFlank&14|+|100109649|100109759@67.90.24&[MIMAT0000753&hsa-miR-342-3p&offsets|0|+1;m-128&14|+|100109715|100109737&offsets|0|+1]|TCTCACACAGAAATCGCACCCGTC</td><td>>MI0000805|hsa-mir-342&WithFlank&14|+|100109649|100109759</td><td>67</td><td>90</td><td>24</td><td>TCTCACACAGAAATCGCACCCGTC</td><td>1568.4</td><td>7.2</td><td>0.7</td><td>10.4</td><td>2.3E-25</td><td>2.9E-24</td><td>BT20/MB468</td><td>CYTO</td></tr>
<tr><td>>hg38_21_+_8433172_8446622_RNA45SN1_45S_50ntFlanks@11792.11820.29|ATGAACGAGATTCCCACTGTCCCTACCTA</td><td>>hg38_21_+_8433172_8446622_RNA45SN1_45S_50ntFlanks</td><td>11792</td><td>11820</td><td>29</td><td>ATGAACGAGATTCCCACTGTCCCTACCTA</td><td>480.3</td><td>-13.3</td><td>1.3</td><td>-10.3</td><td>4.6E-25</td><td>3.0E-24</td><td>BT20/MB468</td><td>NUC</td></tr>
<tr><td>>hg38_21_+_8433172_8446622_RNA45SN1_45S_50ntFlanks@6784.6803.20|GCTACGCCTGTCTGAGCGTC</td><td>>hg38_21_+_8433172_8446622_RNA45SN1_45S_50ntFlanks</td><td>6784</td><td>6803</td><td>20</td><td>GCTACGCCTGTCTGAGCGTC</td><td>3266.9</td><td>-15.5</td><td>1.5</td><td>-10.5</td><td>5.3E-26</td><td>3.0E-24</td><td>MB231/MB468</td><td>MP</td></tr>
<tr><td>>hg38_21_+_8433172_8446622_RNA45SN1_45S_50ntFlanks@5420.5445.26|CCGATTGGATGGTTTAGTGAGGCCCT</td><td>>hg38_21_+_8433172_8446622_RNA45SN1_45S_50ntFlanks</td><td>5420</td><td>5445</td><td>26</td><td>CCGATTGGATGGTTTAGTGAGGCCCT</td><td>2728.7</td><td>-15.8</td><td>1.5</td><td>-10.3</td><td>4.7E-25</td><td>3.0E-24</td><td>BT20/MB468</td><td>NUC</td></tr>
<tr><td>>trna15_ValAAC_5_-_180615416_180615488@1.29.29|GTTTCCGTAGTGTAGTGGTCATCACGTTC</td><td>>trna15_ValAAC_5_-_180615416_180615488</td><td>1</td><td>29</td><td>29</td><td>GTTTCCGTAGTGTAGTGGTCATCACGTTC</td><td>2219.0</td><td>-9.0</td><td>0.8</td><td>-10.6</td><td>4.0E-26</td><td>3.0E-24</td><td>MB231/MB468</td><td>CYTO</td></tr>
</table>

| | | | | | | | | | | | | | |
|---|---|---|---|---|---|---|---|---|---|---|---|---|---|
| >hg38_21_+_8433172_8446622_RNA45SN1_45S_50ntFlanks@7991.8012.22\|ACGTGGCGACCCGCTGAATTTA | >hg38_21_+_8433172_8446622_RNA45SN1_45S_50ntFlanks | 7991 | 8012 | 22 | ACGTGGCGACCCGCTGAATTTA | 25043.0 | -7.8 | 0.7 | -10.5 | 9.5E-26 | 3.0E-24 | MB231/MB468 | MITO |
| >hg38_21_+_8433172_8446622_RNA45SN1_45S_50ntFlanks@9575.9597.23\|ATCGGTCGTCCGACCTGGGTATA | >hg38_21_+_8433172_8446622_RNA45SN1_45S_50ntFlanks | 9575 | 9597 | 23 | ATCGGTCGTCCGACCTGGGTATA | 584.2 | -13.6 | 1.3 | -10.3 | 4.7E-25 | 3.0E-24 | BT20/MB468 | NUC |
| >hg38_21_+_8433172_8446622_RNA45SN1_45S_50ntFlanks@11316.11343.28\|CACCCCACGTCTCGTCGCGCGCGCGTCC | >hg38_21_+_8433172_8446622_RNA45SN1_45S_50ntFlanks | 11316 | 11343 | 28 | CACCCCACGTCTCGTCGCGCGCGCGTCC | 2485.6 | -15.1 | 1.4 | -10.5 | 9.8E-26 | 3.1E-24 | MB231/MB468 | MITO |
| >hg38_21_+_8433172_8446622_RNA45SN1_45S_50ntFlanks@12151.12182.32\|GTGGGGAGTTTGACTGGGGCGGTACACCTGTC | >hg38_21_+_8433172_8446622_RNA45SN1_45S_50ntFlanks | 12151 | 12182 | 32 | GTGGGGAGTTTGACTGGGGCGGTACACCTGTC | 776.9 | -14.0 | 1.4 | -10.3 | 4.8E-25 | 3.1E-24 | BT20/MB468 | NUC |
| >hg38_21_+_8433172_8446622_RNA45SN1_45S_50ntFlanks@11197.11225.29\|GGCCGGCGGCGGCGGCGACTCTGGACGCG | >hg38_21_+_8433172_8446622_RNA45SN1_45S_50ntFlanks | 11197 | 11225 | 29 | GGCCGGCGGCGGCGGCGACTCTGGACGCG | 334.5 | -12.8 | 1.2 | -10.3 | 4.8E-25 | 3.1E-24 | BT20/MB468 | NUC |
| >hg38_21_+_8433172_8446622_RNA45SN1_45S_50ntFlanks@10686.10710.25\|GCGGCGTCCGGTGAGCTCTCGCTGG | >hg38_21_+_8433172_8446622_RNA45SN1_45S_50ntFlanks | 10686 | 10710 | 25 | GCGGCGTCCGGTGAGCTCTCGCTGG | 2678.7 | -5.6 | 0.5 | -10.6 | 4.2E-26 | 3.1E-24 | MB231/MB468 | CYTO |
| >hg38_21_+_8433172_8446622_RNA45SN1_45S_50ntFlanks@4559.4587.29\|GCTAGGAATAATGGAATAGGACCGCGGTT | >hg38_21_+_8433172_8446622_RNA45SN1_45S_50ntFlanks | 4559 | 4587 | 29 | GCTAGGAATAATGGAATAGGACCGCGGTT | 411.5 | -13.1 | 1.3 | -10.3 | 4.9E-25 | 3.1E-24 | BT20/MB468 | NUC |
| >MI0000264\|hsa-mir-7-2&WithFlank&15\|+\|88611819\|88611940@38.61.24(+1A);MI0000265\|hsa-mir-7-3&WithFlank&19\|+\|4770664\|4770785@37.60.24(+1A);MI0000263\|hsa-mir-7-1&WithFlank&9\|-\|83969742\|83969863@30.53.24(+1A)&[MIMAT0000252&hsa-miR-7-5p&offsets\|0\|0(+1A);m-400&15\|+\|88611856\|88611879&offsets\|0\|0(+1A)];[MIMAT0000252_1&hsa-miR-7-5p&offsets\|0\|0(+1A);m-402&19\|+\|4770700\|4770723&offsets\|0\|0(+1A)];[MIMAT0000252_2&hsa-miR-7-5p&offsets\|0\|0(+1A);m-401&9\|-\|83969811\|83969834&offsets\|0\|0(+1A)]\|TGGAAGACTAGTGATTTTGTTGTTA | >MI0000264\|hsa-mir-7-2&WithFlank&15\|+\|88611819\|88611940 | 38 | 61 | 24(+1A) | TGGAAGACTAGTGATTTTGTTGTTA | 610.0 | 7.2 | 0.7 | 10.4 | 1.8E-25 | 3.1E-24 | BT20/MB468 | MITO |
| >hg38_21_+_8433172_8446622_RNA45SN1_45S_50ntFlanks@11599.11623.25\|GAATCCGACTGTTTAATTAAAACAA | >hg38_21_+_8433172_8446622_RNA45SN1_45S_50ntFlanks | 11599 | 11623 | 25 | GAATCCGACTGTTTAATTAAAACAA | 512.2 | 13.5 | 1.3 | 10.4 | 1.8E-25 | 3.2E-24 | BT20/MB468 | MITO |
| >MI0000750\|hsa-mir-26a-2&WithFlank&12\|-\|57824603\|57824698@23.41.19;MI0000083\|hsa-mir-26a-1&WithFlank&3\|+\|37969398\|37969486@19.37.19&[MIMAT0000082&hsa-miR-26a-5p&offsets\|+3\|0;m-33&12\|-\|57824658\|57824679&offsets\|+3\|0];[MIMAT0000082_1&hsa-miR-26a-5p&offsets\|+3\|0;m-34&3\|+\|37969413\|37969434&offsets\|+3\|0]\|AAGTAATCCAGGATAGGCT | >MI0000750\|hsa-mir-26a-2&WithFlank&12\|-\|57824603\|57824698 | 23 | 41 | 19 | AAGTAATCCAGGATAGGCT | 1561.2 | 14.9 | 1.4 | 10.4 | 2.4E-25 | 3.2E-24 | BT20/MB468 | CYTO |
| >hg38_21_+_8433172_8446622_RNA45SN1_45S_50ntFlanks@485.507.23\|GATGTGGTGACGTCGTGCTCTCC | >hg38_21_+_8433172_8446622_RNA45SN1_45S_50ntFlanks | 485 | 507 | 23 | GATGTGGTGACGTCGTGCTCTCC | 1559.1 | 15.1 | 1.4 | 10.4 | 1.8E-25 | 3.2E-24 | BT20/MB468 | MITO |
| >hg38_21_+_8433172_8446622_RNA45SN1_45S_50ntFlanks@11784.11803.20\|ATGAATGGATGAACGAGATT | >hg38_21_+_8433172_8446622_RNA45SN1_45S_50ntFlanks | 11784 | 11803 | 20 | ATGAATGGATGAACGAGATT | 5070.9 | -6.7 | 0.6 | -10.5 | 1.3E-25 | 3.3E-24 | MB231/MB468 | NUC |
| >hg38_21_+_8433172_8446622_RNA45SN1_45S_50ntFlanks@11767.11791.25\|TCTAATTAGTGACGCGCATGAATGG | >hg38_21_+_8433172_8446622_RNA45SN1_45S_50ntFlanks | 11767 | 11791 | 25 | TCTAATTAGTGACGCGCATGAATGG | 830.5 | -14.1 | 1.4 | -10.3 | 5.2E-25 | 3.3E-24 | BT20/MB468 | NUC |
| >hg38_21_+_8433172_8446622_RNA45SN1_45S_50ntFlanks@10411.10433.23\|CAGTCGGTCCTGAGAGATGGGCG | >hg38_21_+_8433172_8446622_RNA45SN1_45S_50ntFlanks | 10411 | 10433 | 23 | CAGTCGGTCCTGAGAGATGGGCG | 3624.8 | -15.7 | 1.5 | -10.5 | 6.0E-26 | 3.3E-24 | MB231/MB468 | MP |
| >hg38_21_+_8433172_8446622_RNA45SN1_45S_50ntFlanks@11581.11605.25\|AACTGGTGCGGACCAGGGGAATCCG | >hg38_21_+_8433172_8446622_RNA45SN1_45S_50ntFlanks | 11581 | 11605 | 25 | AACTGGTGCGGACCAGGGGAATCCG | 3564.4 | -8.1 | 0.8 | -10.5 | 6.0E-26 | 3.3E-24 | MB231/MB468 | MP |
| >hg38_21_+_8433172_8446622_RNA45SN1_45S_50ntFlanks@12151.12181.31\|GTGGGGAGTTTGACTGGGGCGGTACACCTGT | >hg38_21_+_8433172_8446622_RNA45SN1_45S_50ntFlanks | 12151 | 12181 | 31 | GTGGGGAGTTTGACTGGGGCGGTACACCTGT | 476.7 | -13.3 | 1.3 | -10.3 | 5.4E-25 | 3.5E-24 | BT20/MB468 | NUC |
| >hg38_21_+_8433172_8446622_RNA45SN1_45S_50ntFlanks@482.522.41\|ATCGATGTGGTGACGTCGTGCTCTCCCGGGCCGGGTCCGAG | >hg38_21_+_8433172_8446622_RNA45SN1_45S_50ntFlanks | 482 | 522 | 41 | ATCGATGTGGTGACGTCGTGCTCTCCCGGGCCGGGTCCGAG | 1915.1 | -15.3 | 1.5 | -10.3 | 5.4E-25 | 3.5E-24 | BT20/MB468 | NUC |
| >hg38_21_+_8433172_8446622_RNA45SN1_45S_50ntFlanks@12458.12477.20\|AAGCGTTGGATTGTTCACCC | >hg38_21_+_8433172_8446622_RNA45SN1_45S_50ntFlanks | 12458 | 12477 | 20 | AAGCGTTGGATTGTTCACCC | 1751.5 | -4.9 | 0.5 | -10.3 | 5.5E-25 | 3.5E-24 | BT20/MB468 | NUC |
| >MI0000749\|hsa-mir-30e&WithFlank&1\|+\|40754349\|40754452@65.85.21(+1U)&[MIMAT0000693&hsa-miR-30e-3p&offsets\|0\|-1(+1U);m-25&1\|+\|40754413\|40754434&offsets\|0\|-1(+1U)]\|CTTTCAGTCGGATGTTTACAGT | >MI0000749\|hsa-mir-30e&WithFlank&1\|+\|40754349\|40754452 | 65 | 85 | 21(+1U) | CTTTCAGTCGGATGTTTACAGT | 2840.7 | 6.0 | 0.6 | 10.4 | 2.7E-25 | 3.5E-24 | BT20/MB468 | CYTO |
| >hg38_21_+_8433172_8446622_RNA45SN1_45S_50ntFlanks@3828.3860.33\|TCGCTCGCTCCTCTCCTACTTGGATAACTGTGG | >hg38_21_+_8433172_8446622_RNA45SN1_45S_50ntFlanks | 3828 | 3860 | 33 | TCGCTCGCTCCTCTCCTACTTGGATAACTGTGG | 381.5 | -13.0 | 1.3 | -10.3 | 5.6E-25 | 3.6E-24 | BT20/MB468 | NUC |
| >hg38_21_+_8433172_8446622_RNA45SN1_45S_50ntFlanks@5487.5515.29\|CGCTGAGAAGACGGTCGAACTTGACTATC | >hg38_21_+_8433172_8446622_RNA45SN1_45S_50ntFlanks | 5487 | 5515 | 29 | CGCTGAGAAGACGGTCGAACTTGACTATC | 1591.9 | -14.5 | 1.4 | -10.5 | 6.5E-26 | 3.6E-24 | MB231/MB468 | MP |
| >hg38_21_+_8433172_8446622_RNA45SN1_45S_50ntFlanks@12624.12653.30\|CCAATGGGGCGAAGCTACCATCTGTGGGAT | >hg38_21_+_8433172_8446622_RNA45SN1_45S_50ntFlanks | 12624 | 12653 | 30 | CCAATGGGGCGAAGCTACCATCTGTGGGAT | 2112.6 | -14.8 | 1.4 | -10.5 | 1.1E-25 | 3.6E-24 | MB231/MB468 | MITO |
| >MI0000088\|hsa-mir-30a&WithFlank&6\|-\|71403545\|71403627@53.73.21(+1U)&[MIMAT0000088&hsa-miR-30a-3p&offsets\|0\|-1(+1U);m-38&6\|-\|71403554\|71403575&offsets\|0\|-1(+1U)]\|CTTTCAGTCGGATGTTTGCAGT | >MI0000088\|hsa-mir-30a&WithFlank&6\|-\|71403545\|71403627 | 53 | 73 | 21(+1U) | CTTTCAGTCGGATGTTTGCAGT | 10266.0 | 5.7 | 0.5 | 10.5 | 1.4E-25 | 3.6E-24 | MB231/MB468 | NUC |
| >hg38_21_+_8433172_8446622_RNA45SN1_45S_50ntFlanks@4657.4680.24\|CTAGAGGTGAAATTCTTGGACCGG | >hg38_21_+_8433172_8446622_RNA45SN1_45S_50ntFlanks | 4657 | 4680 | 24 | CTAGAGGTGAAATTCTTGGACCGG | 324.7 | -12.8 | 1.2 | -10.3 | 5.7E-25 | 3.6E-24 | BT20/MB468 | NUC |
| >hg38_21_+_8433172_8446622_RNA45SN1_45S_50ntFlanks@11516.11539.24\|TCCGCCCCCGGGGCCGCGGTTCCG | >hg38_21_+_8433172_8446622_RNA45SN1_45S_50ntFlanks | 11516 | 11539 | 24 | TCCGCCCCCGGGGCCGCGGTTCCG | 470.8 | -13.3 | 1.3 | -10.3 | 5.7E-25 | 3.6E-24 | BT20/MB468 | NUC |
| >hg38_21_+_8433172_8446622_RNA45SN1_45S_50ntFlanks@6745.6771.27\|ACACTTCGAACGCACTTGCGGCCCCGG | >hg38_21_+_8433172_8446622_RNA45SN1_45S_50ntFlanks | 6745 | 6771 | 27 | ACACTTCGAACGCACTTGCGGCCCCGG | 4959.5 | -6.7 | 0.6 | -10.4 | 2.1E-25 | 3.7E-24 | BT20/MB468 | MITO |
| >hg38_21_+_8433172_8446622_RNA45SN1_45S_50ntFlanks@6747.6770.24\|ACTTCGAACGCACTTGCGGCCCCG | >hg38_21_+_8433172_8446622_RNA45SN1_45S_50ntFlanks | 6747 | 6770 | 24 | ACTTCGAACGCACTTGCGGCCCCG | 1751.9 | -14.6 | 1.4 | -10.5 | 6.7E-26 | 3.7E-24 | MB231/MB468 | MP |
| >hg38_21_+_8433172_8446622_RNA45SN1_45S_50ntFlanks@8355.8375.21\|TGAAAAGAACTTTGAAGAGAG | >hg38_21_+_8433172_8446622_RNA45SN1_45S_50ntFlanks | 8355 | 8375 | 21 | TGAAAAGAACTTTGAAGAGAG | 1665.9 | 15.2 | 1.5 | 10.4 | 2.2E-25 | 3.7E-24 | BT20/MB468 | MITO |
| >MI0001445\|hsa-mir-423&WithFlank&17\|+\|30117073\|30117178@23.41.19&[MIMAT0004748&hsa-miR-423-5p&offsets\|0\|-4;m-141&17\|+\|30117095\|30117117&offsets\|0\|-4]\|TGAGGGGCAGAGAGCGAGA | >MI0001445\|hsa-mir-423&WithFlank&17\|+\|30117073\|30117178 | 23 | 41 | 19 | TGAGGGGCAGAGAGCGAGA | 517.9 | 13.3 | 1.3 | 10.4 | 2.9E-25 | 3.7E-24 | BT20/MB468 | CYTO |
| >MI0003599\|hsa-mir-589&WithFlank&7\|-\|5495813\|5495923@30.50.21&[MIMAT0004799&hsa-miR-589-5p&offsets\|0\|-1;m-162&7\|-\|5495874\|5495894&offsets\|0\|0]\|TGAGAACCACGTCTGCTCTGA | >MI0003599\|hsa-mir-589&WithFlank&7\|-\|5495813\|5495923 | 30 | 50 | 21 | TGAGAACCACGTCTGCTCTGA | 87.8 | 3.8 | 0.4 | 10.3 | 5.9E-25 | 3.8E-24 | BT20/MB468 | NUC |
| >hg38_21_+_8433172_8446622_RNA45SN1_45S_50ntFlanks@11572.11591.20\|CCGACTTAGAACTGGTGCGG | >hg38_21_+_8433172_8446622_RNA45SN1_45S_50ntFlanks | 11572 | 11591 | 20 | CCGACTTAGAACTGGTGCGG | 9605.2 | -7.7 | 0.7 | -11.0 | 5.6E-28 | 3.8E-24 | BT20/MB468 | MP |
| >hg38_21_+_8433172_8446622_RNA45SN1_45S_50ntFlanks@10845.10865.21\|ACTTCGGGATAAGGATTGGCT | >hg38_21_+_8433172_8446622_RNA45SN1_45S_50ntFlanks | 10845 | 10865 | 21 | ACTTCGGGATAAGGATTGGCT | 2203.2 | -14.9 | 1.4 | -10.5 | 1.2E-25 | 3.8E-24 | MB231/MB468 | MITO |
| >trna11_GluTTC_15_-_26327381_26327452@1.33.33,trna3_GluTTC_13_-_45492062_45492133@1.33.33\|TCCCACATGGTCTAGCGGTTAGGATTCCTGGTT | >trna11_GluTTC_15_-_26327381_26327452 | 1 | 33 | 33 | TCCCACATGGTCTAGCGGTTAGGATTCCTGGTT | 3707.6 | 8.3 | 0.8 | 10.5 | 1.2E-25 | 3.8E-24 | MB231/MB468 | MITO |
| >hg38_21_+_8433172_8446622_RNA45SN1_45S_50ntFlanks@6722.6745.24\|ATTGCAGGACACATTGATCATCGA | >hg38_21_+_8433172_8446622_RNA45SN1_45S_50ntFlanks | 6722 | 6745 | 24 | ATTGCAGGACACATTGATCATCGA | 382.9 | -13.0 | 1.3 | -10.3 | 6.1E-25 | 3.9E-24 | BT20/MB468 | NUC |
| >MI0000086\|hsa-mir-28&WithFlank&3\|+\|188688775\|188688872@61.82.22&[MIMAT0004502&hsa-miR-28-3p&offsets\|+1\|+1;m-28&3\|+\|188688834\|188688855&offsets\|+1\|+1]\|ACTAGATTGTGAGCTCCTGGAG | >MI0000086\|hsa-mir-28&WithFlank&3\|+\|188688775\|188688872 | 61 | 82 | 22 | ACTAGATTGTGAGCTCCTGGAG | 351.1 | 12.9 | 1.2 | 10.4 | 2.2E-25 | 3.9E-24 | BT20/MB468 | MITO |
| >hg38_21_+_8433172_8446622_RNA45SN1_45S_50ntFlanks@13003.13027.25\|GATCTATTGAAAGTCAGCCCTCGAC | >hg38_21_+_8433172_8446622_RNA45SN1_45S_50ntFlanks | 13003 | 13027 | 25 | GATCTATTGAAAGTCAGCCCTCGAC | 503.7 | -13.4 | 1.3 | -10.3 | 6.1E-25 | 3.9E-24 | BT20/MB468 | NUC |
| >hg38_21_+_8433172_8446622_RNA45SN1_45S_50ntFlanks@11918.11954.37\|ACATGAGAGGTGTAGAATAAGTGGGAGGCCCCCGGCG | >hg38_21_+_8433172_8446622_RNA45SN1_45S_50ntFlanks | 11918 | 11954 | 37 | ACATGAGAGGTGTAGAATAAGTGGGAGGCCCCCGGCG | 451.3 | -13.2 | 1.3 | -10.3 | 6.1E-25 | 3.9E-24 | BT20/MB468 | NUC |
| >hg38_21_+_8433172_8446622_RNA45SN1_45S_50ntFlanks@10361.10384.24\|GAAGGCCGAAGTGGAGAAGGGTTC | >hg38_21_+_8433172_8446622_RNA45SN1_45S_50ntFlanks | 10361 | 10384 | 24 | GAAGGCCGAAGTGGAGAAGGGTTC | 311.6 | -12.7 | 1.2 | -10.3 | 6.2E-25 | 3.9E-24 | BT20/MB468 | NUC |
| >hg38_21_+_8433172_8446622_RNA45SN1_45S_50ntFlanks@4550.4573.24\|GATACCGCAGCTAGGAATAATGGA | >hg38_21_+_8433172_8446622_RNA45SN1_45S_50ntFlanks | 4550 | 4573 | 24 | GATACCGCAGCTAGGAATAATGGA | 326.3 | -12.8 | 1.2 | -10.3 | 6.2E-25 | 3.9E-24 | BT20/MB468 | NUC |
| >MI0000301\|hsa-mir-224&WithFlank&X\|-\|151958572\|151958664@14.35.22(+1U)&[MIMAT0000281&hsa-miR-224-5p&offsets\|+1\|-2(+1U);m-238&X\|-\|151958630\|151958651&offsets\|0\|0(+1U)]\|CAAGTCACTAGTGGTTCCGTTTT | >MI0000301\|hsa-mir-224&WithFlank&X\|-\|151958572\|151958664 | 14 | 35 | 22(+1U) | CAAGTCACTAGTGGTTCCGTTTT | 401.7 | 12.9 | 1.2 | 10.4 | 3.1E-25 | 4.0E-24 | BT20/MB468 | CYTO |
| >MI0003560\|hsa-mir-92b&WithFlank&1\|+\|155195171\|155195278@67.88.22(+2U)&[MIMAT0003218&hsa-miR-92b-3p&offsets\|0\|0(+2U);m-115&1\|+\|155195237\|155195258&offsets\|0\|0(+2U)]\|TATTGCACTCGTCCCGGCCTCCTT | >MI0003560\|hsa-mir-92b&WithFlank&1\|+\|155195171\|155195278 | 67 | 88 | 22(+2U) | TATTGCACTCGTCCCGGCCTCCTT | 424.1 | 13.0 | 1.3 | 10.4 | 3.1E-25 | 4.0E-24 | BT20/MB468 | CYTO |
| >MI0000285\|hsa-mir-205&WithFlank&1\|+\|209432127\|209432248@40.61.22(+1G)&[MIMAT0000266&hsa-miR-205-5p&offsets\|0\|0(+1G);m-351&1\|+\|209432166\|209432187&offsets\|0\|0(+1G)]\|TCCTTCATTCCACCGGAGTCTGG | >MI0000285\|hsa-mir-205&WithFlank&1\|+\|209432127\|209432248 | 40 | 61 | 22(+1G) | TCCTTCATTCCACCGGAGTCTGG | 3219.4 | 11.6 | 1.1 | 10.7 | 6.6E-27 | 4.0E-24 | BT20/MB231 | CYTO |
| >hg38_21_+_8433172_8446622_RNA45SN1_45S_50ntFlanks@6732.6773.42\|ACATTGATCATCGACACTTCGAACGCACTTGCGGCCCCGGGT | >hg38_21_+_8433172_8446622_RNA45SN1_45S_50ntFlanks | 6732 | 6773 | 42 | ACATTGATCATCGACACTTCGAACGCACTTGCGGCCCCGGGT | 2881.3 | 7.2 | 0.7 | 10.4 | 2.4E-25 | 4.0E-24 | BT20/MB468 | MITO |
| >hg38_21_+_8433172_8446622_RNA45SN1_45S_50ntFlanks@5527.5549.23\|AAAGTCGTAACAAGGTTTCCGTA | >hg38_21_+_8433172_8446622_RNA45SN1_45S_50ntFlanks | 5527 | 5549 | 23 | AAAGTCGTAACAAGGTTTCCGTA | 403.7 | -13.1 | 1.3 | -10.3 | 6.4E-25 | 4.1E-24 | BT20/MB468 | NUC |

<table>
<tr><td>>hg38_21_+_8433172_8446622_RNA45SN1_45S_50ntFlanks@10781.10806.26|AGGTGAACAGCCTCTGGCATGTTGGA</td><td>>hg38_21_+_8433172_8446622_RNA45SN1_45S_50ntFlanks</td><td>10781</td><td>10806</td><td>26</td><td>AGGTGAACAGCCTCTGGCATGTTGGA</td><td>464.9</td><td>-13.3</td><td>1.3</td><td>-10.3</td><td>6.5E-25</td><td>4.1E-24</td><td>BT20/MB468</td><td>NUC</td></tr>
<tr><td>>hg38_21_+_8433172_8446622_RNA45SN1_45S_50ntFlanks@7976.7994.19|GCGACCTCAGATCAGACGT</td><td>>hg38_21_+_8433172_8446622_RNA45SN1_45S_50ntFlanks</td><td>7976</td><td>7994</td><td>19</td><td>GCGACCTCAGATCAGACGT</td><td>763.1</td><td>6.1</td><td>0.6</td><td>10.4</td><td>2.4E-25</td><td>4.1E-24</td><td>BT20/MB468</td><td>MITO</td></tr>
<tr><td>>hg38_21_+_8433172_8446622_RNA45SN1_45S_50ntFlanks@12476.12497.22|CCACTAATAGGGAACGTGAGCT</td><td>>hg38_21_+_8433172_8446622_RNA45SN1_45S_50ntFlanks</td><td>12476</td><td>12497</td><td>22</td><td>CCACTAATAGGGAACGTGAGCT</td><td>1793.2</td><td>-6.3</td><td>0.6</td><td>-10.5</td><td>7.6E-26</td><td>4.2E-24</td><td>MB231/MB468</td><td>MP</td></tr>
<tr><td>>hg38_21_+_8433172_8446622_RNA45SN1_45S_50ntFlanks@12957.12990.34|CTTCTGGGTCGGGGTTTCGTACGTAGCAGAGCAG</td><td>>hg38_21_+_8433172_8446622_RNA45SN1_45S_50ntFlanks</td><td>12957</td><td>12990</td><td>34</td><td>CTTCTGGGTCGGGGTTTCGTACGTAGCAGAGCAG</td><td>2157.1</td><td>-14.0</td><td>1.3</td><td>-10.4</td><td>1.7E-25</td><td>4.3E-24</td><td>MB231/MB468</td><td>NUC</td></tr>
<tr><td>>trna10_LysCTT_16_+_3241501_3241573@14.32.19,trna13_LysCTT_14_-_58706613_58706685@14.32.19,trna2_LysCTT_15_+_79152904_79152976@14.32.19,trna4_LysCTT_18_-_43669271_43669343@14.32.19|AGTCGGTAGAGCATGGGAC</td><td>>trna10_LysCTT_16_+_3241501_3241573</td><td>14</td><td>32</td><td>19</td><td>AGTCGGTAGAGCATGGGAC</td><td>329.7</td><td>-12.8</td><td>1.2</td><td>-10.3</td><td>6.8E-25</td><td>4.3E-24</td><td>BT20/MB468</td><td>NUC</td></tr>
<tr><td>>MI0000469|hsa-mir-125a&WithFlank&19|+|51693248|51693345@21.42.22(+1A)&[MIMAT0000443&hsa-miR-125a-5p&offsets|0|-2(+1A);m-63&19|+|51693268|51693289&offsets|0|0(+1A)]|TCCCTGAGACCCTTTAACCTGTA</td><td>>MI0000469|hsa-mir-125a&WithFlank&19|+|51693248|51693345</td><td>21</td><td>42</td><td>22(+1A)</td><td>TCCCTGAGACCCTTTAACCTGTA</td><td>5179.6</td><td>7.8</td><td>0.8</td><td>10.4</td><td>2.6E-25</td><td>4.3E-24</td><td>BT20/MB468</td><td>MITO</td></tr>
<tr><td>>MI0000266|hsa-mir-10a&WithFlank&17|-|48579832|48579953@28.50.23&[MIMAT0000253&hsa-miR-10a-5p&offsets|0|0;m-18&17|-|48579905|48579926&offsets|0|+1]|TACCCTGTAGATCCGAATTTGTG</td><td>>MI0000266|hsa-mir-10a&WithFlank&17|-|48579832|48579953</td><td>28</td><td>50</td><td>23</td><td>TACCCTGTAGATCCGAATTTGTG</td><td>4142.1</td><td>13.1</td><td>1.3</td><td>10.4</td><td>1.8E-25</td><td>4.4E-24</td><td>MB231/MB468</td><td>NUC</td></tr>
<tr><td>>hg38_21_+_8433172_8446622_RNA45SN1_45S_50ntFlanks@6664.6681.18|TGGATCACTCGGCTCGTG</td><td>>hg38_21_+_8433172_8446622_RNA45SN1_45S_50ntFlanks</td><td>6664</td><td>6681</td><td>18</td><td>TGGATCACTCGGCTCGTG</td><td>1321.7</td><td>-5.8</td><td>0.6</td><td>-10.4</td><td>1.8E-25</td><td>4.4E-24</td><td>MB231/MB468</td><td>NUC</td></tr>
<tr><td>>hg38_21_+_8433172_8446622_RNA45SN1_45S_50ntFlanks@12405.12437.33|CTTTTTGATCCTTCGATGTCGGCTCTTCCTATC</td><td>>hg38_21_+_8433172_8446622_RNA45SN1_45S_50ntFlanks</td><td>12405</td><td>12437</td><td>33</td><td>CTTTTTGATCCTTCGATGTCGGCTCTTCCTATC</td><td>17555.0</td><td>-7.6</td><td>0.7</td><td>-10.4</td><td>1.8E-25</td><td>4.4E-24</td><td>MB231/MB468</td><td>NUC</td></tr>
<tr><td>>hg38_21_+_8433172_8446622_RNA45SN1_45S_50ntFlanks@12624.12661.38|CCAATGGGGCGAAGCTACCATCTGTGGGATTATGACTG</td><td>>hg38_21_+_8433172_8446622_RNA45SN1_45S_50ntFlanks</td><td>12624</td><td>12661</td><td>38</td><td>CCAATGGGGCGAAGCTACCATCTGTGGGATTATGACTG</td><td>1995.3</td><td>-5.3</td><td>0.5</td><td>-10.4</td><td>1.8E-25</td><td>4.4E-24</td><td>MB231/MB468</td><td>NUC</td></tr>
<tr><td>>trna10_GlyTCC_17_+_8124866_8124937@1.30.30|GCGTTGGTGGTATAGTGGTAAGCATAGCTG</td><td>>trna10_GlyTCC_17_+_8124866_8124937</td><td>1</td><td>30</td><td>30</td><td>GCGTTGGTGGTATAGTGGTAAGCATAGCTG</td><td>1043.2</td><td>-8.3</td><td>0.8</td><td>-10.5</td><td>6.1E-26</td><td>4.4E-24</td><td>MB231/MB468</td><td>CYTO</td></tr>
<tr><td>>MI0000477|hsa-mir-146a&WithFlank&5|+|160485346|160485456@27.48.22(+1U)&[MIMAT0000449&hsa-miR-146a-5p&offsets|0|0(+1U);m-129&5|+|160485372|160485393&offsets|0|0(+1U)]|TGAGAACTGAATTCCATGGGTTT</td><td>>MI0000477|hsa-mir-146a&WithFlank&5|+|160485346|160485456</td><td>27</td><td>48</td><td>22(+1U)</td><td>TGAGAACTGAATTCCATGGGTTT</td><td>7025.9</td><td>16.0</td><td>1.5</td><td>10.5</td><td>8.2E-26</td><td>4.5E-24</td><td>MB231/MB468</td><td>MP</td></tr>
<tr><td>>MI0000078|hsa-mir-22&WithFlank&17|-|1713897|1713993@59.79.21(+1C)&[MIMAT0000077&hsa-miR-22-3p&offsets|0|-1(+1C);m-2&17|-|1713914|1713935&offsets|0|-1(+1C)]|AAGCTGCCAGTTGAAGAACTGC</td><td>>MI0000078|hsa-mir-22&WithFlank&17|-|1713897|1713993</td><td>59</td><td>79</td><td>21(+1C)</td><td>AAGCTGCCAGTTGAAGAACTGC</td><td>3150.3</td><td>9.3</td><td>0.9</td><td>10.4</td><td>3.5E-25</td><td>4.5E-24</td><td>BT20/MB468</td><td>CYTO</td></tr>
<tr><td>>hg38_21_+_8433172_8446622_RNA45SN1_45S_50ntFlanks@10841.10862.22|CGTAACTTCGGGATAAGGATTG</td><td>>hg38_21_+_8433172_8446622_RNA45SN1_45S_50ntFlanks</td><td>10841</td><td>10862</td><td>22</td><td>CGTAACTTCGGGATAAGGATTG</td><td>3703.5</td><td>-15.7</td><td>1.5</td><td>-10.5</td><td>8.3E-26</td><td>4.5E-24</td><td>MB231/MB468</td><td>MP</td></tr>
<tr><td>>hg38_21_+_8433172_8446622_RNA45SN1_45S_50ntFlanks@12405.12435.31|CTTTTTGATCCTTCGATGTCGGCTCTTCCTA</td><td>>hg38_21_+_8433172_8446622_RNA45SN1_45S_50ntFlanks</td><td>12405</td><td>12435</td><td>31</td><td>CTTTTTGATCCTTCGATGTCGGCTCTTCCTA</td><td>1878.5</td><td>-14.7</td><td>1.4</td><td>-10.5</td><td>8.3E-26</td><td>4.5E-24</td><td>MB231/MB468</td><td>MP</td></tr>
<tr><td>>hg38_21_+_8433172_8446622_RNA45SN1_45S_50ntFlanks@4724.4759.36|ATTAATCAAGAACGAAAGTCGGAGGTTCGAAGACGA</td><td>>hg38_21_+_8433172_8446622_RNA45SN1_45S_50ntFlanks</td><td>4724</td><td>4759</td><td>36</td><td>ATTAATCAAGAACGAAAGTCGGAGGTTCGAAGACGA</td><td>3844.8</td><td>-6.2</td><td>0.6</td><td>-10.4</td><td>2.7E-25</td><td>4.5E-24</td><td>BT20/MB468</td><td>MITO</td></tr>
<tr><td>>hg38_21_+_8433172_8446622_RNA45SN1_45S_50ntFlanks@475.505.31|CTTCGTGATCGATGTGGTGACGTCGTGCTCT</td><td>>hg38_21_+_8433172_8446622_RNA45SN1_45S_50ntFlanks</td><td>475</td><td>505</td><td>31</td><td>CTTCGTGATCGATGTGGTGACGTCGTGCTCT</td><td>4935.9</td><td>8.3</td><td>0.8</td><td>10.5</td><td>6.4E-26</td><td>4.6E-24</td><td>MB231/MB468</td><td>CYTO</td></tr>
<tr><td>>hg38_21_+_8433172_8446622_RNA45SN1_45S_50ntFlanks@8381.8400.20|AAGAGGGCGTGAAACCGTTA</td><td>>hg38_21_+_8433172_8446622_RNA45SN1_45S_50ntFlanks</td><td>8381</td><td>8400</td><td>20</td><td>AAGAGGGCGTGAAACCGTTA</td><td>506.4</td><td>-13.4</td><td>1.3</td><td>-10.3</td><td>7.4E-25</td><td>4.6E-24</td><td>BT20/MB468</td><td>NUC</td></tr>
<tr><td>>hg38_21_+_8433172_8446622_RNA45SN1_45S_50ntFlanks@9568.9595.28|CGTGCAAATCGGTCGTCCGACCTGGGTA</td><td>>hg38_21_+_8433172_8446622_RNA45SN1_45S_50ntFlanks</td><td>9568</td><td>9595</td><td>28</td><td>CGTGCAAATCGGTCGTCCGACCTGGGTA</td><td>4611.8</td><td>-10.0</td><td>1.0</td><td>-10.4</td><td>1.5E-25</td><td>4.6E-24</td><td>MB231/MB468</td><td>MITO</td></tr>
<tr><td>>hg38_21_+_8433172_8446622_RNA45SN1_45S_50ntFlanks@10351.10385.35|CGAGAACTTTGAAGGCCGAAGTGGAGAAGGGTTCC</td><td>>hg38_21_+_8433172_8446622_RNA45SN1_45S_50ntFlanks</td><td>10351</td><td>10385</td><td>35</td><td>CGAGAACTTTGAAGGCCGAAGTGGAGAAGGGTTCC</td><td>725.1</td><td>-13.9</td><td>1.4</td><td>-10.3</td><td>7.4E-25</td><td>4.7E-24</td><td>BT20/MB468</td><td>NUC</td></tr>
<tr><td>>hg38_21_+_8433172_8446622_RNA45SN1_45S_50ntFlanks@9504.9533.30|GTGAACTATGCCTGGGCAGGGCGAAGCCAG</td><td>>hg38_21_+_8433172_8446622_RNA45SN1_45S_50ntFlanks</td><td>9504</td><td>9533</td><td>30</td><td>GTGAACTATGCCTGGGCAGGGCGAAGCCAG</td><td>2408.5</td><td>-15.0</td><td>1.4</td><td>-10.4</td><td>1.5E-25</td><td>4.7E-24</td><td>MB231/MB468</td><td>MITO</td></tr>
<tr><td>>hg38_21_+_8433172_8446622_RNA45SN1_45S_50ntFlanks@11874.11909.36|AGAAGACCCTGTTGAGCTTGACTCTAGTCTGGCACG</td><td>>hg38_21_+_8433172_8446622_RNA45SN1_45S_50ntFlanks</td><td>11874</td><td>11909</td><td>36</td><td>AGAAGACCCTGTTGAGCTTGACTCTAGTCTGGCACG</td><td>517.7</td><td>-13.4</td><td>1.3</td><td>-10.3</td><td>7.5E-25</td><td>4.7E-24</td><td>BT20/MB468</td><td>NUC</td></tr>
<tr><td>>hg38_21_+_8433172_8446622_RNA45SN1_45S_50ntFlanks@10683.10711.29|CCGGCGGCGTCCGGTGAGCTCTCGCTGGC</td><td>>hg38_21_+_8433172_8446622_RNA45SN1_45S_50ntFlanks</td><td>10683</td><td>10711</td><td>29</td><td>CCGGCGGCGTCCGGTGAGCTCTCGCTGGC</td><td>3296.2</td><td>-7.5</td><td>0.7</td><td>-10.4</td><td>1.5E-25</td><td>4.7E-24</td><td>MB231/MB468</td><td>MITO</td></tr>
<tr><td>>hg38_21_+_8433172_8446622_RNA45SN1_45S_50ntFlanks@10869.10902.34|AGGGCTGGGTCGGTCGGGCTGGGGCGCGAAGCGG</td><td>>hg38_21_+_8433172_8446622_RNA45SN1_45S_50ntFlanks</td><td>10869</td><td>10902</td><td>34</td><td>AGGGCTGGGTCGGTCGGGCTGGGGCGCGAAGCGG</td><td>2847.7</td><td>-15.3</td><td>1.5</td><td>-10.5</td><td>8.8E-26</td><td>4.7E-24</td><td>MB231/MB468</td><td>MP</td></tr>
<tr><td>>hg38_21_+_8433172_8446622_RNA45SN1_45S_50ntFlanks@10786.10825.40|AACAGCCTCTGGCATGTTGGAACAATGTAGGTAAGGGAAG</td><td>>hg38_21_+_8433172_8446622_RNA45SN1_45S_50ntFlanks</td><td>10786</td><td>10825</td><td>40</td><td>AACAGCCTCTGGCATGTTGGAACAATGTAGGTAAGGGAAG</td><td>1069.2</td><td>-14.5</td><td>1.4</td><td>-10.3</td><td>7.6E-25</td><td>4.7E-24</td><td>BT20/MB468</td><td>NUC</td></tr>
<tr><td>>hg38_21_+_8433172_8446622_RNA45SN1_45S_50ntFlanks@10362.10381.20|AAGGCCGAAGTGGAGAAGGG</td><td>>hg38_21_+_8433172_8446622_RNA45SN1_45S_50ntFlanks</td><td>10362</td><td>10381</td><td>20</td><td>AAGGCCGAAGTGGAGAAGGG</td><td>389.6</td><td>-13.0</td><td>1.3</td><td>-10.3</td><td>7.6E-25</td><td>4.8E-24</td><td>BT20/MB468</td><td>NUC</td></tr>
<tr><td>>MI0000077|hsa-mir-21&WithFlank&17|+|59841260|59841343@14.34.21(+2C)&[MIMAT0000076&hsa-miR-21-5p&offsets|0|-1(+2C);m-1&17|+|59841273|59841294&offsets|0|-1(+2C)]|TAGCTTATCAGACTGATGTTGCC</td><td>>MI0000077|hsa-mir-21&WithFlank&17|+|59841260|59841343</td><td>14</td><td>34</td><td>21(+2C)</td><td>TAGCTTATCAGACTGATGTTGCC</td><td>2426.3</td><td>8.1</td><td>0.8</td><td>10.4</td><td>2.0E-25</td><td>4.9E-24</td><td>MB231/MB468</td><td>NUC</td></tr>
<tr><td>>hg38_21_+_8433172_8446622_RNA45SN1_45S_50ntFlanks@6745.6769.25|ACACTTCGAACGCACTTGCGGCCCC</td><td>>hg38_21_+_8433172_8446622_RNA45SN1_45S_50ntFlanks</td><td>6745</td><td>6769</td><td>25</td><td>ACACTTCGAACGCACTTGCGGCCCC</td><td>354.9</td><td>-12.9</td><td>1.3</td><td>-10.3</td><td>8.0E-25</td><td>5.0E-24</td><td>BT20/MB468</td><td>NUC</td></tr>
<tr><td>>hg38_21_+_8433172_8446622_RNA45SN1_45S_50ntFlanks@12456.12474.19|CCAAGCGTTGGATTGTTCA</td><td>>hg38_21_+_8433172_8446622_RNA45SN1_45S_50ntFlanks</td><td>12456</td><td>12474</td><td>19</td><td>CCAAGCGTTGGATTGTTCA</td><td>12568.0</td><td>-8.3</td><td>0.8</td><td>-10.5</td><td>7.0E-26</td><td>5.0E-24</td><td>MB231/MB468</td><td>CYTO</td></tr>
<tr><td>>hg38_21_+_8433172_8446622_RNA45SN1_45S_50ntFlanks@9545.9566.22|GTGGAGGTCCGTAGCGGTCCTG</td><td>>hg38_21_+_8433172_8446622_RNA45SN1_45S_50ntFlanks</td><td>9545</td><td>9566</td><td>22</td><td>GTGGAGGTCCGTAGCGGTCCTG</td><td>2699.4</td><td>-8.4</td><td>0.8</td><td>-10.5</td><td>7.1E-26</td><td>5.0E-24</td><td>MB231/MB468</td><td>CYTO</td></tr>
<tr><td>>hg38_21_+_8433172_8446622_RNA45SN1_45S_50ntFlanks@11729.11751.23|TAACTATGACTCTCTTAAGGTAG</td><td>>hg38_21_+_8433172_8446622_RNA45SN1_45S_50ntFlanks</td><td>11729</td><td>11751</td><td>23</td><td>TAACTATGACTCTCTTAAGGTAG</td><td>45012.0</td><td>-8.0</td><td>0.8</td><td>-10.5</td><td>7.2E-26</td><td>5.0E-24</td><td>MB231/MB468</td><td>CYTO</td></tr>
<tr><td>>hg38_21_+_8433172_8446622_RNA45SN1_45S_50ntFlanks@12324.12348.25|TTGGGTTTTAAGCAGGAGGTGTCAG</td><td>>hg38_21_+_8433172_8446622_RNA45SN1_45S_50ntFlanks</td><td>12324</td><td>12348</td><td>25</td><td>TTGGGTTTTAAGCAGGAGGTGTCAG</td><td>1144.1</td><td>-14.0</td><td>1.3</td><td>-10.5</td><td>9.3E-26</td><td>5.0E-24</td><td>MB231/MB468</td><td>MP</td></tr>
<tr><td>>MI0000101|hsa-mir-99a&WithFlank&21|+|16539083|16539175@18.39.22&[MIMAT0000097&hsa-miR-99a-5p&offsets|-1|-1;m-64&21|+|16539101|16539121&offsets|-1|0]|AAACCCGTAGATCCGATCTTGT</td><td>>MI0000101|hsa-mir-99a&WithFlank&21|+|16539083|16539175</td><td>18</td><td>39</td><td>22</td><td>AAACCCGTAGATCCGATCTTGT</td><td>427.7</td><td>-13.2</td><td>1.3</td><td>-10.3</td><td>8.0E-25</td><td>5.0E-24</td><td>BT20/MB468</td><td>NUC</td></tr>
<tr><td>>hg38_21_+_8433172_8446622_RNA45SN1_45S_50ntFlanks@3835.3860.26|CTCCTCTCCTACTTGGATAACTGTGG</td><td>>hg38_21_+_8433172_8446622_RNA45SN1_45S_50ntFlanks</td><td>3835</td><td>3860</td><td>26</td><td>CTCCTCTCCTACTTGGATAACTGTGG</td><td>3676.5</td><td>-5.3</td><td>0.5</td><td>-10.4</td><td>2.1E-25</td><td>5.0E-24</td><td>MB231/MB468</td><td>NUC</td></tr>
<tr><td>>hg38_21_+_8433172_8446622_RNA45SN1_45S_50ntFlanks@10760.10785.26|CATATCCGCAGCAGGTCTCCAAGGTG</td><td>>hg38_21_+_8433172_8446622_RNA45SN1_45S_50ntFlanks</td><td>10760</td><td>10785</td><td>26</td><td>CATATCCGCAGCAGGTCTCCAAGGTG</td><td>323.0</td><td>-12.8</td><td>1.2</td><td>-10.3</td><td>8.1E-25</td><td>5.1E-24</td><td>BT20/MB468</td><td>NUC</td></tr>
<tr><td>>hg38_21_+_8433172_8446622_RNA45SN1_45S_50ntFlanks@12395.12414.20|AGCGACGTCGCTTTTTGATC</td><td>>hg38_21_+_8433172_8446622_RNA45SN1_45S_50ntFlanks</td><td>12395</td><td>12414</td><td>20</td><td>AGCGACGTCGCTTTTTGATC</td><td>1018.6</td><td>5.0</td><td>0.5</td><td>10.5</td><td>7.5E-26</td><td>5.1E-24</td><td>MB231/MB468</td><td>CYTO</td></tr>
<tr><td>>hg38_21_+_8433172_8446622_RNA45SN1_45S_50ntFlanks@5477.5516.40|CCTGGCGGAGCGCTGAGAAGACGGTCGAACTTGACTATCT</td><td>>hg38_21_+_8433172_8446622_RNA45SN1_45S_50ntFlanks</td><td>5477</td><td>5516</td><td>40</td><td>CCTGGCGGAGCGCTGAGAAGACGGTCGAACTTGACTATCT</td><td>1567.8</td><td>15.1</td><td>1.5</td><td>10.4</td><td>3.0E-25</td><td>5.1E-24</td><td>BT20/MB468</td><td>MITO</td></tr>
<tr><td>>hg38_21_+_8433172_8446622_RNA45SN1_45S_50ntFlanks@11534.11552.19|GTTCCGCGCGGCGCCTCGC</td><td>>hg38_21_+_8433172_8446622_RNA45SN1_45S_50ntFlanks</td><td>11534</td><td>11552</td><td>19</td><td>GTTCCGCGCGGCGCCTCGC</td><td>4088.1</td><td>-16.4</td><td>1.6</td><td>-10.3</td><td>8.3E-25</td><td>5.2E-24</td><td>BT20/MB468</td><td>NUC</td></tr>
<tr><td>>MI0003560|hsa-mir-92b&WithFlank&1|+|155195171|155195278@67.88.22(+1A)&[MIMAT0003218&hsa-miR-92b-3p&offsets|0|0(+1A);m-115&1|+|155195237|155195258&offsets|0|0(+1A)]|TATTGCACTCGTCCCGGCCTCCA</td><td>>MI0003560|hsa-mir-92b&WithFlank&1|+|155195171|155195278</td><td>67</td><td>88</td><td>22(+1A)</td><td>TATTGCACTCGTCCCGGCCTCCA</td><td>953.1</td><td>10.4</td><td>1.0</td><td>10.4</td><td>4.0E-25</td><td>5.2E-24</td><td>BT20/MB468</td><td>CYTO</td></tr>
<tr><td>>hg38_21_+_8433172_8446622_RNA45SN1_45S_50ntFlanks@4760.4782.23|TCAGATACCGTCGTAGTTCCGAC</td><td>>hg38_21_+_8433172_8446622_RNA45SN1_45S_50ntFlanks</td><td>4760</td><td>4782</td><td>23</td><td>TCAGATACCGTCGTAGTTCCGAC</td><td>1713.4</td><td>-14.6</td><td>1.4</td><td>-10.5</td><td>9.7E-26</td><td>5.2E-24</td><td>MB231/MB468</td><td>MP</td></tr>
<tr><td>>hg38_21_+_8433172_8446622_RNA45SN1_45S_50ntFlanks@10681.10710.30|TTCCGGCGGCGTCCGGTGAGCTCTCGCTGG</td><td>>hg38_21_+_8433172_8446622_RNA45SN1_45S_50ntFlanks</td><td>10681</td><td>10710</td><td>30</td><td>TTCCGGCGGCGTCCGGTGAGCTCTCGCTGG</td><td>1698.5</td><td>-14.6</td><td>1.4</td><td>-10.5</td><td>9.7E-26</td><td>5.2E-24</td><td>MB231/MB468</td><td>MP</td></tr>
<tr><td>>hg38_21_+_8433172_8446622_RNA45SN1_45S_50ntFlanks@7981.7999.19|CTCAGATCAGACGTGGCGA</td><td>>hg38_21_+_8433172_8446622_RNA45SN1_45S_50ntFlanks</td><td>7981</td><td>7999</td><td>19</td><td>CTCAGATCAGACGTGGCGA</td><td>341.0</td><td>-12.8</td><td>1.2</td><td>-10.3</td><td>8.4E-25</td><td>5.2E-24</td><td>BT20/MB468</td><td>NUC</td></tr>
<tr><td>>hg38_21_+_8433172_8446622_RNA45SN1_45S_50ntFlanks@12435.12455.21|ATCATTGTGAAGCAGAATTCA</td><td>>hg38_21_+_8433172_8446622_RNA45SN1_45S_50ntFlanks</td><td>12435</td><td>12455</td><td>21</td><td>ATCATTGTGAAGCAGAATTCA</td><td>3004.3</td><td>-15.4</td><td>1.5</td><td>-10.5</td><td>9.9E-26</td><td>5.3E-24</td><td>MB231/MB468</td><td>MP</td></tr>
<tr><td>>hg38_21_+_8433172_8446622_RNA45SN1_45S_50ntFlanks@9534.9558.25|AGGAAACTCTGGTGGAGGTCCGTAG</td><td>>hg38_21_+_8433172_8446622_RNA45SN1_45S_50ntFlanks</td><td>9534</td><td>9558</td><td>25</td><td>AGGAAACTCTGGTGGAGGTCCGTAG</td><td>427.4</td><td>-13.2</td><td>1.3</td><td>-10.3</td><td>8.5E-25</td><td>5.3E-24</td><td>BT20/MB468</td><td>NUC</td></tr>
<tr><td>>hg38_21_+_8433172_8446622_RNA45SN1_45S_50ntFlanks@11843.11872.30|AGGGAACGGGCTTGGCGGAATCAGCGGGGA</td><td>>hg38_21_+_8433172_8446622_RNA45SN1_45S_50ntFlanks</td><td>11843</td><td>11872</td><td>30</td><td>AGGGAACGGGCTTGGCGGAATCAGCGGGGA</td><td>1598.6</td><td>-14.5</td><td>1.4</td><td>-10.5</td><td>1.0E-25</td><td>5.3E-24</td><td>MB231/MB468</td><td>MP</td></tr>
<tr><td>>hg38_21_+_8433172_8446622_RNA45SN1_45S_50ntFlanks@12157.12182.26|AGTTTGACTGGGGCGGTACACCTGTC</td><td>>hg38_21_+_8433172_8446622_RNA45SN1_45S_50ntFlanks</td><td>12157</td><td>12182</td><td>26</td><td>AGTTTGACTGGGGCGGTACACCTGTC</td><td>4867.3</td><td>-16.1</td><td>1.5</td><td>-10.5</td><td>1.0E-25</td><td>5.3E-24</td><td>MB231/MB468</td><td>MP</td></tr>
<tr><td>>hg38_21_+_8433172_8446622_RNA45SN1_45S_50ntFlanks@12995.13016.22|CTCGCTGCGATCTATTGAAAGT</td><td>>hg38_21_+_8433172_8446622_RNA45SN1_45S_50ntFlanks</td><td>12995</td><td>13016</td><td>22</td><td>CTCGCTGCGATCTATTGAAAGT</td><td>1158.0</td><td>5.6</td><td>0.5</td><td>10.5</td><td>7.9E-26</td><td>5.4E-24</td><td>MB231/MB468</td><td>CYTO</td></tr>
<tr><td>>hg38_21_+_8433172_8446622_RNA45SN1_45S_50ntFlanks@10410.10433.24|TCAGTCGGTCCTGAGAGATGGGCG</td><td>>hg38_21_+_8433172_8446622_RNA45SN1_45S_50ntFlanks</td><td>10410</td><td>10433</td><td>24</td><td>TCAGTCGGTCCTGAGAGATGGGCG</td><td>2140.3</td><td>-14.9</td><td>1.4</td><td>-10.5</td><td>1.0E-25</td><td>5.4E-24</td><td>MB231/MB468</td><td>MP</td></tr>
<tr><td>>MI0000434|hsa-let-7i&WithFlank&12|+|62603680|62603775@12.33.22(+1A)&[MIMAT0000415&hsa-let-7i-5p&offsets|0|0(+1A);m-49&12|+|62603691|62603712&offsets|0|0(+1A)]|TGAGGTAGTAGTTTGTGCTGTTA</td><td>>MI0000434|hsa-let-7i&WithFlank&12|+|62603680|62603775</td><td>12</td><td>33</td><td>22(+1A)</td><td>TGAGGTAGTAGTTTGTGCTGTTA</td><td>5344.3</td><td>6.5</td><td>0.6</td><td>10.3</td><td>4.2E-25</td><td>5.4E-24</td><td>BT20/MB468</td><td>CYTO</td></tr>
</table>

| | | | | | | | | | | | | | |
|---|---|---|---|---|---|---|---|---|---|---|---|---|---|
| >hg38_21_+_8433172_8446622_RNA45SN1_45S_50ntFlanks@11208.11234.27\|GCGGCGACTCTGGACGCGAGCCGGGCC | >hg38_21_+_8433172_8446622_RNA45SN1_45S_50ntFlanks | 11208 | 11234 | 27 | GCGGCGACTCTGGACGCGAGCCGGGCC | 2720.6 | -15.3 | 1.5 | -10.5 | 1.1E-25 | 5.5E-24 | MB231/MB468 | MP |
| >trna111_HisGTG_1_-_147774845_147774916@1.33.33,trna118_HisGTG_1_-_145396881_145396952@1.33.33,trna16_HisGTG_1_+_146544773_146544844@1.33.33,trna1_HisGTG_15_+_45493349_45493420@1.33.33,trna21_HisGTG_1_+_147753471_147753542@1.33.33,trna33_HisGTG_6_+_27125906_27125977@1.33.33,trna7_HisGTG_9_-_14433938_14434009@1.33.33,trna8_HisGTG_15_-_45492611_45492682@1.33.33,trna9_HisGTG_15_-_45490804_45490875@1.33.33\|GCCGTGATCGTATAGTGGTTAGTACTCTGCGTT | >trna111_HisGTG_1_-_147774845_147774916 | 1 | 33 | 33 | GCCGTGATCGTATAGTGGTTAGTACTCTGCGTT | 18224.6 | 8.6 | 0.8 | 10.3 | 4.4E-25 | 5.6E-24 | BT20/MB468 | CYTO |
| >hg38_21_+_8433172_8446622_RNA45SN1_45S_50ntFlanks@4161.4184.24\|CACATCCAAGGAAGGCAGCAGGCG | >hg38_21_+_8433172_8446622_RNA45SN1_45S_50ntFlanks | 4161 | 4184 | 24 | CACATCCAAGGAAGGCAGCAGGCG | 387.4 | -13.0 | 1.3 | -10.3 | 9.0E-25 | 5.6E-24 | BT20/MB468 | NUC |
| >hg38_21_+_8433172_8446622_RNA45SN1_45S_50ntFlanks@6773.6803.31\|TTCCTCCCGGGGCTACGCCTGTCTGAGCGTC | >hg38_21_+_8433172_8446622_RNA45SN1_45S_50ntFlanks | 6773 | 6803 | 31 | TTCCTCCCGGGGCTACGCCTGTCTGAGCGTC | 1274.0 | -14.2 | 1.4 | -10.5 | 1.1E-25 | 5.6E-24 | MB231/MB468 | MP |
| >hg38_21_+_8433172_8446622_RNA45SN1_45S_50ntFlanks@10513.10534.22\|CCGGAGTGGCGGAGATGGGCGC | >hg38_21_+_8433172_8446622_RNA45SN1_45S_50ntFlanks | 10513 | 10534 | 22 | CCGGAGTGGCGGAGATGGGCGC | 3294.3 | -15.5 | 1.5 | -10.5 | 1.1E-25 | 5.7E-24 | MB231/MB468 | MP |
| >hg38_21_+_8433172_8446622_RNA45SN1_45S_50ntFlanks@9575.9596.22\|ATCGGTCGTCCGACCTGGGTAT | >hg38_21_+_8433172_8446622_RNA45SN1_45S_50ntFlanks | 9575 | 9596 | 22 | ATCGGTCGTCCGACCTGGGTAT | 556.0 | -13.5 | 1.3 | -10.3 | 9.2E-25 | 5.7E-24 | BT20/MB468 | NUC |
| >hg38_21_+_8433172_8446622_RNA45SN1_45S_50ntFlanks@11534.11553.20\|GTTCCGCGCGGCGCCTCGCC | >hg38_21_+_8433172_8446622_RNA45SN1_45S_50ntFlanks | 11534 | 11553 | 20 | GTTCCGCGCGGCGCCTCGCC | 1233.8 | -9.0 | 0.9 | -10.5 | 8.6E-26 | 5.7E-24 | MB231/MB468 | CYTO |
| >hg38_21_+_8433172_8446622_RNA45SN1_45S_50ntFlanks@10727.10756.30\|GGAGAGGGTGTAAATCTCGCGCCGGGCCGT | >hg38_21_+_8433172_8446622_RNA45SN1_45S_50ntFlanks | 10727 | 10756 | 30 | GGAGAGGGTGTAAATCTCGCGCCGGGCCGT | 670.0 | -13.8 | 1.3 | -10.3 | 9.4E-25 | 5.8E-24 | BT20/MB468 | NUC |
| >hg38_21_+_8433172_8446622_RNA45SN1_45S_50ntFlanks@11874.11904.31\|AGAAGACCCTGTTGAGCTTGACTCTAGTCTG | >hg38_21_+_8433172_8446622_RNA45SN1_45S_50ntFlanks | 11874 | 11904 | 31 | AGAAGACCCTGTTGAGCTTGACTCTAGTCTG | 3183.5 | -15.5 | 1.5 | -10.5 | 1.1E-25 | 5.9E-24 | MB231/MB468 | MP |
| >hg38_21_+_8433172_8446622_RNA45SN1_45S_50ntFlanks@9343.9370.28\|CGAAAGCCGCCGTGGCGCAATGAAGGTG | >hg38_21_+_8433172_8446622_RNA45SN1_45S_50ntFlanks | 9343 | 9370 | 28 | CGAAAGCCGCCGTGGCGCAATGAAGGTG | 2331.5 | -15.1 | 1.4 | -10.5 | 1.1E-25 | 5.9E-24 | MB231/MB468 | MP |
| >hg38_21_+_8433172_8446622_RNA45SN1_45S_50ntFlanks@11561.11591.31\|GCGCCTAGCAGCCGACTTAGAACTGGTGCGG | >hg38_21_+_8433172_8446622_RNA45SN1_45S_50ntFlanks | 11561 | 11591 | 31 | GCGCCTAGCAGCCGACTTAGAACTGGTGCGG | 904.0 | -4.7 | 0.5 | -10.4 | 3.5E-25 | 5.9E-24 | BT20/MB468 | MITO |
| >hg38_21_+_8433172_8446622_RNA45SN1_45S_50ntFlanks@10689.10706.18\|GCGTCCGGTGAGCTCTCG | >hg38_21_+_8433172_8446622_RNA45SN1_45S_50ntFlanks | 10689 | 10706 | 18 | GCGTCCGGTGAGCTCTCG | 1911.9 | -14.7 | 1.4 | -10.4 | 2.0E-25 | 5.9E-24 | MB231/MB468 | MITO |
| >MI0000301\|hsa-mir-224&WithFlank&X\|-\|151958572\|151958664@14.35.22&[MIMAT0000281&hsa-miR-224-5p&offsets\|+1\|-2;m-238&X\|-\|151958630\|151958651&offsets\|0\|0]\|CAAGTCACTAGTGGTTCCGTTT | >MI0000301\|hsa-mir-224&WithFlank&X\|-\|151958572\|151958664 | 14 | 35 | 22 | CAAGTCACTAGTGGTTCCGTTT | 1754.7 | 8.1 | 0.8 | 10.4 | 2.4E-25 | 5.9E-24 | MB231/MB468 | NUC |
| >MI0000299\|hsa-mir-222&WithFlank&X\|-\|45747009\|45747130@75.98.24(+1A)&[MIMAT0000279&hsa-miR-222-3p&offsets\|0\|+3(+1A);m-140&X\|-\|45747033\|45747056&offsets\|0\|0(+1A)]\|AGCTACATCTGGCTACTGGGTCTCA | >MI0000299\|hsa-mir-222&WithFlank&X\|-\|45747009\|45747130 | 75 | 98 | 24(+1A) | AGCTACATCTGGCTACTGGGTCTCA | 6348.8 | 8.1 | 0.8 | 10.5 | 9.1E-26 | 6.0E-24 | MB231/MB468 | CYTO |
| >hg38_21_+_8433172_8446622_RNA45SN1_45S_50ntFlanks@10786.10818.33\|AACAGCCTCTGGCATGTTGGAACAATGTAGGTA | >hg38_21_+_8433172_8446622_RNA45SN1_45S_50ntFlanks | 10786 | 10818 | 33 | AACAGCCTCTGGCATGTTGGAACAATGTAGGTA | 5084.2 | -6.6 | 0.6 | -10.5 | 9.1E-26 | 6.0E-24 | MB231/MB468 | CYTO |
| >hg38_21_+_8433172_8446622_RNA45SN1_45S_50ntFlanks@3769.3794.26\|CGCACGGCCGGTACAGTGAAACTGCG | >hg38_21_+_8433172_8446622_RNA45SN1_45S_50ntFlanks | 3769 | 3794 | 26 | CGCACGGCCGGTACAGTGAAACTGCG | 760.2 | -14.0 | 1.4 | -10.3 | 9.7E-25 | 6.0E-24 | BT20/MB468 | NUC |
| >hg38_21_+_8433172_8446622_RNA45SN1_45S_50ntFlanks@6652.6695.44\|GACTCTTAGCGGTGGATCACTCGGCTCGTGCGTCGATGAAGAAC | >hg38_21_+_8433172_8446622_RNA45SN1_45S_50ntFlanks | 6652 | 6695 | 44 | GACTCTTAGCGGTGGATCACTCGGCTCGTGCGTCGATGAAGAAC | 9102.9 | 6.9 | 0.7 | 10.4 | 3.6E-25 | 6.1E-24 | BT20/MB468 | MITO |
| >hg38_21_+_8433172_8446622_RNA45SN1_45S_50ntFlanks@7986.8005.20\|ATCAGACGTGGCGACCCGCT | >hg38_21_+_8433172_8446622_RNA45SN1_45S_50ntFlanks | 7986 | 8005 | 20 | ATCAGACGTGGCGACCCGCT | 2797.4 | -15.2 | 1.5 | -10.4 | 2.0E-25 | 6.1E-24 | MB231/MB468 | MITO |
| >hg38_21_+_8433172_8446622_RNA45SN1_45S_50ntFlanks@12717.12747.31\|CCTCGGTTGGCCTCGGATAGCCGGTCCCCCG | >hg38_21_+_8433172_8446622_RNA45SN1_45S_50ntFlanks | 12717 | 12747 | 31 | CCTCGGTTGGCCTCGGATAGCCGGTCCCCCG | 1216.3 | -14.1 | 1.3 | -10.5 | 1.2E-25 | 6.1E-24 | MB231/MB468 | MP |
| >MI0000079\|hsa-mir-23a&WithFlank&19\|-\|13836581\|13836665@51.71.21&[MIMAT0000078&hsa-miR-23a-3p&offsets\|0\|0;m-31&19\|-\|13836595\|13836615&offsets\|0\|0]\|ATCACATTGCCAGGGATTTCC | >MI0000079\|hsa-mir-23a&WithFlank&19\|-\|13836581\|13836665 | 51 | 71 | 21 | ATCACATTGCCAGGGATTTCC | 327.0 | -12.8 | 1.2 | -10.3 | 1.0E-24 | 6.2E-24 | BT20/MB468 | NUC |
| >hg38_21_+_8433172_8446622_RNA45SN1_45S_50ntFlanks@4046.4068.23\|CATTCGAACGTCTGCCCTATCAA | >hg38_21_+_8433172_8446622_RNA45SN1_45S_50ntFlanks | 4046 | 4068 | 23 | CATTCGAACGTCTGCCCTATCAA | 348.8 | -12.9 | 1.3 | -10.3 | 1.0E-24 | 6.2E-24 | BT20/MB468 | NUC |
| >hg38_21_+_8433172_8446622_RNA45SN1_45S_50ntFlanks@4590.4614.25\|ATTTTGTTGGTTTTCGGAACTGAGG | >hg38_21_+_8433172_8446622_RNA45SN1_45S_50ntFlanks | 4590 | 4614 | 25 | ATTTTGTTGGTTTTCGGAACTGAGG | 362.2 | -12.9 | 1.3 | -10.3 | 1.0E-24 | 6.2E-24 | BT20/MB468 | NUC |
| >hg38_21_+_8433172_8446622_RNA45SN1_45S_50ntFlanks@4724.4752.29\|ATTAATCAAGAACGAAAGTCGGAGGTTCG | >hg38_21_+_8433172_8446622_RNA45SN1_45S_50ntFlanks | 4724 | 4752 | 29 | ATTAATCAAGAACGAAAGTCGGAGGTTCG | 1345.4 | -14.3 | 1.4 | -10.5 | 1.2E-25 | 6.3E-24 | MB231/MB468 | MP |
| >hg38_21_+_8433172_8446622_RNA45SN1_45S_50ntFlanks@5485.5513.29\|AGCGCTGAGAAGACGGTCGAACTTGACTA | >hg38_21_+_8433172_8446622_RNA45SN1_45S_50ntFlanks | 5485 | 5513 | 29 | AGCGCTGAGAAGACGGTCGAACTTGACTA | 1789.4 | -14.7 | 1.4 | -10.5 | 1.3E-25 | 6.4E-24 | MB231/MB468 | MP |
| >hg38_21_+_8433172_8446622_RNA45SN1_45S_50ntFlanks@8380.8400.21\|CAAGAGGGCGTGAAACCGTTA | >hg38_21_+_8433172_8446622_RNA45SN1_45S_50ntFlanks | 8380 | 8400 | 21 | CAAGAGGGCGTGAAACCGTTA | 2662.7 | -9.2 | 0.9 | -10.5 | 9.8E-26 | 6.4E-24 | MB231/MB468 | CYTO |
| >hg38_21_+_8433172_8446622_RNA45SN1_45S_50ntFlanks@3705.3723.19\|TACCTGGTTGATCCTGCCA | >hg38_21_+_8433172_8446622_RNA45SN1_45S_50ntFlanks | 3705 | 3723 | 19 | TACCTGGTTGATCCTGCCA | 1934.3 | 5.2 | 0.5 | 10.3 | 5.0E-25 | 6.4E-24 | BT20/MB468 | CYTO |
| >hg38_21_+_8433172_8446622_RNA45SN1_45S_50ntFlanks@4283.4317.35\|CTTTAACGAGGATCCATTGGAGGGCAAGTCTGGTG | >hg38_21_+_8433172_8446622_RNA45SN1_45S_50ntFlanks | 4283 | 4317 | 35 | CTTTAACGAGGATCCATTGGAGGGCAAGTCTGGTG | 1335.5 | -5.5 | 0.5 | -10.3 | 5.0E-25 | 6.4E-24 | BT20/MB468 | CYTO |
| >MI0003205\|hsa-mir-532&WithFlank&X\|+\|50003142\|50003244@26.47.22(+1A)&[MIMAT0002888&hsa-miR-532-5p&offsets\|0\|0(+1A);m-65&X\|+\|50003167\|50003188&offsets\|0\|0(+1A)]\|CATGCCTTGAGTGTAGGACCGTA | >MI0003205\|hsa-mir-532&WithFlank&X\|+\|50003142\|50003244 | 26 | 47 | 22(+1A) | CATGCCTTGAGTGTAGGACCGTA | 247.6 | 5.5 | 0.5 | 10.3 | 5.1E-25 | 6.4E-24 | BT20/MB468 | CYTO |
| >MI0000542\|hsa-mir-320a&WithFlank&8\|-\|22244960\|22245043@47.69.23&[MIMAT0000510&hsa-miR-320a-3p&offsets\|-1\|0;m-60&8\|-\|22244975\|22244996&offsets\|-1\|0]\|GAAAAGCTGGGTTGAGAGGGCGA | >MI0000542\|hsa-mir-320a&WithFlank&8\|-\|22244960\|22245043 | 47 | 69 | 23 | GAAAAGCTGGGTTGAGAGGGCGA | 523.8 | 13.3 | 1.3 | 10.3 | 5.2E-25 | 6.5E-24 | BT20/MB468 | CYTO |
| >hg38_21_+_8433172_8446622_RNA45SN1_45S_50ntFlanks@12630.12654.25\|GGGCGAAGCTACCATCTGTGGGATT | >hg38_21_+_8433172_8446622_RNA45SN1_45S_50ntFlanks | 12630 | 12654 | 25 | GGGCGAAGCTACCATCTGTGGGATT | 420.4 | -13.1 | 1.3 | -10.3 | 1.1E-24 | 6.6E-24 | BT20/MB468 | NUC |
| >hg38_21_+_8433172_8446622_RNA45SN1_45S_50ntFlanks@12634.12654.21\|GAAGCTACCATCTGTGGGATT | >hg38_21_+_8433172_8446622_RNA45SN1_45S_50ntFlanks | 12634 | 12654 | 21 | GAAGCTACCATCTGTGGGATT | 1184.7 | -14.1 | 1.3 | -10.5 | 1.3E-25 | 6.7E-24 | MB231/MB468 | MP |
| >hg38_21_+_8433172_8446622_RNA45SN1_45S_50ntFlanks@10350.10375.26\|ACGAGAACTTTGAAGGCCGAAGTGGA | >hg38_21_+_8433172_8446622_RNA45SN1_45S_50ntFlanks | 10350 | 10375 | 26 | ACGAGAACTTTGAAGGCCGAAGTGGA | 289.3 | -12.6 | 1.2 | -10.3 | 1.1E-24 | 6.7E-24 | BT20/MB468 | NUC |
| >hg38_21_+_8433172_8446622_RNA45SN1_45S_50ntFlanks@9204.9222.19\|TCGGGGCCACGCGCGCGTC | >hg38_21_+_8433172_8446622_RNA45SN1_45S_50ntFlanks | 9204 | 9222 | 19 | TCGGGGCCACGCGCGCGTC | 2161.6 | -14.9 | 1.4 | -10.5 | 1.3E-25 | 6.7E-24 | MB231/MB468 | MP |
| >hg38_21_+_8433172_8446622_RNA45SN1_45S_50ntFlanks@4641.4663.23\|CATTCGTATTGCGCCGCTAGAGG | >hg38_21_+_8433172_8446622_RNA45SN1_45S_50ntFlanks | 4641 | 4663 | 23 | CATTCGTATTGCGCCGCTAGAGG | 723.6 | -6.3 | 0.6 | -10.5 | 1.3E-25 | 6.7E-24 | MB231/MB468 | MP |
| >hg38_21_+_8433172_8446622_RNA45SN1_45S_50ntFlanks@9534.9566.33\|AGGAAACTCTGGTGGAGGTCCGTAGCGGTCCTG | >hg38_21_+_8433172_8446622_RNA45SN1_45S_50ntFlanks | 9534 | 9566 | 33 | AGGAAACTCTGGTGGAGGTCCGTAGCGGTCCTG | 2087.6 | -14.9 | 1.4 | -10.5 | 1.3E-25 | 6.7E-24 | MB231/MB468 | MP |
| >hg38_21_+_8433172_8446622_RNA45SN1_45S_50ntFlanks@10512.10534.23\|TCCGGAGTGGCGGAGATGGGCGC | >hg38_21_+_8433172_8446622_RNA45SN1_45S_50ntFlanks | 10512 | 10534 | 23 | TCCGGAGTGGCGGAGATGGGCGC | 2193.5 | -15.0 | 1.4 | -10.5 | 1.3E-25 | 6.7E-24 | MB231/MB468 | MP |
| >hg38_21_+_8433172_8446622_RNA45SN1_45S_50ntFlanks@11561.11591.31\|GCGCCTAGCAGCCGACTTAGAACTGGTGCGG | >hg38_21_+_8433172_8446622_RNA45SN1_45S_50ntFlanks | 11561 | 11591 | 31 | GCGCCTAGCAGCCGACTTAGAACTGGTGCGG | 598.9 | -13.7 | 1.3 | -10.3 | 1.1E-24 | 6.8E-24 | BT20/MB468 | NUC |
| >hg38_MT_+_1621_3279_RNR2_16S_50ntFlanks@51.72.22\|GCTAAACCTAGCCCCAAACCCA | >hg38_MT_+_1621_3279_RNR2_16S_50ntFlanks | 51 | 72 | 22 | GCTAAACCTAGCCCCAAACCCA | 149.8 | 3.3 | 0.3 | 10.3 | 1.1E-24 | 6.8E-24 | BT20/MB468 | NUC |
| >hg38_21_+_8433172_8446622_RNA45SN1_45S_50ntFlanks@6741.6771.31\|ATCGACACTTCGAACGCACTTGCGGCCCCGG | >hg38_21_+_8433172_8446622_RNA45SN1_45S_50ntFlanks | 6741 | 6771 | 31 | ATCGACACTTCGAACGCACTTGCGGCCCCGG | 1906.8 | -14.8 | 1.4 | -10.5 | 1.4E-25 | 6.8E-24 | MB231/MB468 | MP |
| >hg38_21_+_8433172_8446622_RNA45SN1_45S_50ntFlanks@5449.5466.18\|ATCGGCCCCGCCGGGGTC | >hg38_21_+_8433172_8446622_RNA45SN1_45S_50ntFlanks | 5449 | 5466 | 18 | ATCGGCCCCGCCGGGGTC | 3770.8 | -15.7 | 1.5 | -10.5 | 1.4E-25 | 6.8E-24 | MB231/MB468 | MP |
| >hg38_21_+_8433172_8446622_RNA45SN1_45S_50ntFlanks@6719.6744.26\|TGAATTGCAGGACACATTGATCATCG | >hg38_21_+_8433172_8446622_RNA45SN1_45S_50ntFlanks | 6719 | 6744 | 26 | TGAATTGCAGGACACATTGATCATCG | 4700.1 | -9.6 | 0.9 | -10.4 | 2.3E-25 | 6.8E-24 | MB231/MB468 | MITO |
| >hg38_21_+_8433172_8446622_RNA45SN1_45S_50ntFlanks@4370.4398.29\|TTAAAAAGCTCGTAGTTGGATCTTGGGAG | >hg38_21_+_8433172_8446622_RNA45SN1_45S_50ntFlanks | 4370 | 4398 | 29 | TTAAAAAGCTCGTAGTTGGATCTTGGGAG | 788.5 | -14.0 | 1.4 | -10.3 | 1.1E-24 | 6.9E-24 | BT20/MB468 | NUC |
| >hg38_21_+_8433172_8446622_RNA45SN1_45S_50ntFlanks@6721.6753.33\|AATTGCAGGACACATTGATCATCGACACTTCGA | >hg38_21_+_8433172_8446622_RNA45SN1_45S_50ntFlanks | 6721 | 6753 | 33 | AATTGCAGGACACATTGATCATCGACACTTCGA | 479.5 | -13.3 | 1.3 | -10.3 | 1.1E-24 | 6.9E-24 | BT20/MB468 | NUC |
| >hg38_21_+_8433172_8446622_RNA45SN1_45S_50ntFlanks@12416.12439.24\|TTCGATGTCGGCTCTTCCTATCAT | >hg38_21_+_8433172_8446622_RNA45SN1_45S_50ntFlanks | 12416 | 12439 | 24 | TTCGATGTCGGCTCTTCCTATCAT | 971.3 | 5.2 | 0.5 | 10.3 | 5.5E-25 | 6.9E-24 | BT20/MB468 | CYTO |
| >MI0003560\|hsa-mir-92b&WithFlank&1\|+\|155195171\|155195278@67.87.21(+1U)&[MIMAT0003218&hsa-miR-92b-3p&offsets\|0\|-1(+1U);m-115&1\|+\|155195237\|155195258&offsets\|0\|-1(+1U)]\|TATTGCACTCGTCCCGGCCTCT | >MI0003560\|hsa-mir-92b&WithFlank&1\|+\|155195171\|155195278 | 67 | 87 | 21(+1U) | TATTGCACTCGTCCCGGCCTCT | 2812.5 | 8.5 | 0.8 | 10.5 | 1.1E-25 | 7.0E-24 | MB231/MB468 | CYTO |
| >hg38_21_+_8433172_8446622_RNA45SN1_45S_50ntFlanks@9308.9336.29\|AGGAGTCTAACACGTGCGCGAGTCGGGGG | >hg38_21_+_8433172_8446622_RNA45SN1_45S_50ntFlanks | 9308 | 9336 | 29 | AGGAGTCTAACACGTGCGCGAGTCGGGGG | 3004.0 | -15.4 | 1.5 | -10.5 | 1.4E-25 | 7.0E-24 | MB231/MB468 | MP |
| >MI0000077\|hsa-mir-21&WithFlank&17\|+\|59841260\|59841343@16.36.21&[MIMAT0000076&hsa-miR-21-5p&offsets\|+2\|+1;m-1&17\|+\|59841273\|59841294&offsets\|+2\|+1]\|GCTTATCAGACTGATGTTGAC | >MI0000077\|hsa-mir-21&WithFlank&17\|+\|59841260\|59841343 | 16 | 36 | 21 | GCTTATCAGACTGATGTTGAC | 724.6 | 13.8 | 1.3 | 10.3 | 5.6E-25 | 7.1E-24 | BT20/MB468 | CYTO |
| >hg38_21_+_8433172_8446622_RNA45SN1_45S_50ntFlanks@6733.6771.39\|CATTGATCATCGACACTTCGAACGCACTTGCGGCCCCGG | >hg38_21_+_8433172_8446622_RNA45SN1_45S_50ntFlanks | 6733 | 6771 | 39 | CATTGATCATCGACACTTCGAACGCACTTGCGGCCCCGG | 8265.8 | -7.5 | 0.7 | -10.4 | 2.4E-25 | 7.1E-24 | MB231/MB468 | MITO |
| >MI0003643\|hsa-mir-629&WithFlank&15\|-\|70079366\|70079474@28.49.22&[MIMAT0004810&hsa-miR-629-5p&offsets\|0\|+1;m-150&15\|-\|70079427\|70079447&offsets\|0\|+1]\|TGGGTTTACGTTGGGAGAACTT | >MI0003643\|hsa-mir-629&WithFlank&15\|-\|70079366\|70079474 | 28 | 49 | 22 | TGGGTTTACGTTGGGAGAACTT | 468.9 | 11.7 | 1.1 | 10.3 | 1.2E-24 | 7.1E-24 | BT20/MB468 | NUC |

<table>
<tr><td>>hg38_21_+_8433172_8446622_RNA45SN1_45S_50ntFlanks@10571.10591.21|GAGAAGCCGGCGGGAGCCCCG</td><td>>hg38_21_+_8433172_8446622_RNA45SN1_45S_50ntFlanks</td><td>10571</td><td>10591</td><td>21</td><td>GAGAAGCCGGCGGGAGCCCCG</td><td>1748.3</td><td>-15.2</td><td>1.5</td><td>-10.3</td><td>1.2E-24</td><td>7.2E-24</td><td>BT20/MB468</td><td>NUC</td></tr>
<tr><td>>hg38_21_+_8433172_8446622_RNA45SN1_45S_50ntFlanks@9534.9572.39|AGGAAACTCTGGTGGAGGTCCGTAGCGGTCCTGACGTGC</td><td>>hg38_21_+_8433172_8446622_RNA45SN1_45S_50ntFlanks</td><td>9534</td><td>9572</td><td>39</td><td>AGGAAACTCTGGTGGAGGTCCGTAGCGGTCCTGACGTGC</td><td>19588.7</td><td>-9.9</td><td>1.0</td><td>-10.3</td><td>1.2E-24</td><td>7.2E-24</td><td>BT20/MB468</td><td>NUC</td></tr>
<tr><td>>hg38_21_+_8433172_8446622_RNA45SN1_45S_50ntFlanks@10846.10865.20|CTTCGGGATAAGGATTGGCT</td><td>>hg38_21_+_8433172_8446622_RNA45SN1_45S_50ntFlanks</td><td>10846</td><td>10865</td><td>20</td><td>CTTCGGGATAAGGATTGGCT</td><td>2694.7</td><td>-15.3</td><td>1.5</td><td>-10.4</td><td>1.5E-25</td><td>7.2E-24</td><td>MB231/MB468</td><td>MP</td></tr>
<tr><td>>hg38_21_+_8433172_8446622_RNA45SN1_45S_50ntFlanks@12676.12708.33|AGAATCCCGCCCAGGCGGAACGATACGGCAGCG</td><td>>hg38_21_+_8433172_8446622_RNA45SN1_45S_50ntFlanks</td><td>12676</td><td>12708</td><td>33</td><td>AGAATCCCGCCCAGGCGGAACGATACGGCAGCG</td><td>1145.4</td><td>-14.0</td><td>1.3</td><td>-10.4</td><td>1.5E-25</td><td>7.2E-24</td><td>MB231/MB468</td><td>MP</td></tr>
<tr><td>>hg38_21_+_8433172_8446622_RNA45SN1_45S_50ntFlanks@4160.4186.27|CCACATCCAAGGAAGGCAGCAGGCGCG</td><td>>hg38_21_+_8433172_8446622_RNA45SN1_45S_50ntFlanks</td><td>4160</td><td>4186</td><td>27</td><td>CCACATCCAAGGAAGGCAGCAGGCGCG</td><td>355.7</td><td>-12.9</td><td>1.3</td><td>-10.2</td><td>1.2E-24</td><td>7.3E-24</td><td>BT20/MB468</td><td>NUC</td></tr>
<tr><td>>hg38_21_+_8433172_8446622_RNA45SN1_45S_50ntFlanks@3835.3859.25|CTCCTCTCCTACTTGGATAACTGTG</td><td>>hg38_21_+_8433172_8446622_RNA45SN1_45S_50ntFlanks</td><td>3835</td><td>3859</td><td>25</td><td>CTCCTCTCCTACTTGGATAACTGTG</td><td>359.3</td><td>-12.9</td><td>1.3</td><td>-10.2</td><td>1.2E-24</td><td>7.3E-24</td><td>BT20/MB468</td><td>NUC</td></tr>
<tr><td>>hg38_21_+_8433172_8446622_RNA45SN1_45S_50ntFlanks@6732.6752.21|ACATTGATCATCGACACTTCG</td><td>>hg38_21_+_8433172_8446622_RNA45SN1_45S_50ntFlanks</td><td>6732</td><td>6752</td><td>21</td><td>ACATTGATCATCGACACTTCG</td><td>591.1</td><td>-4.3</td><td>0.4</td><td>-10.2</td><td>1.2E-24</td><td>7.3E-24</td><td>BT20/MB468</td><td>NUC</td></tr>
<tr><td>>hg38_21_+_8433172_8446622_RNA45SN1_45S_50ntFlanks@8107.8143.37|CGGCGGGGCGCGGGACATGTGGCGTACGGAAGACCCG</td><td>>hg38_21_+_8433172_8446622_RNA45SN1_45S_50ntFlanks</td><td>8107</td><td>8143</td><td>37</td><td>CGGCGGGGCGCGGGACATGTGGCGTACGGAAGACCCG</td><td>2208.5</td><td>-15.0</td><td>1.4</td><td>-10.4</td><td>1.5E-25</td><td>7.3E-24</td><td>MB231/MB468</td><td>MP</td></tr>
<tr><td>>hg38_21_+_8433172_8446622_RNA45SN1_45S_50ntFlanks@11853.11872.20|CTTGGCGGAATCAGCGGGGA</td><td>>hg38_21_+_8433172_8446622_RNA45SN1_45S_50ntFlanks</td><td>11853</td><td>11872</td><td>20</td><td>CTTGGCGGAATCAGCGGGGA</td><td>431.9</td><td>-13.2</td><td>1.3</td><td>-10.2</td><td>1.2E-24</td><td>7.3E-24</td><td>BT20/MB468</td><td>NUC</td></tr>
<tr><td>>hg38_21_+_8433172_8446622_RNA45SN1_45S_50ntFlanks@10684.10711.28|CGGCGGCGTCCGGTGAGCTCTCGCTGGC</td><td>>hg38_21_+_8433172_8446622_RNA45SN1_45S_50ntFlanks</td><td>10684</td><td>10711</td><td>28</td><td>CGGCGGCGTCCGGTGAGCTCTCGCTGGC</td><td>7905.3</td><td>-6.7</td><td>0.6</td><td>-10.4</td><td>2.5E-25</td><td>7.4E-24</td><td>MB231/MB468</td><td>MITO</td></tr>
<tr><td>>MI0000264|hsa-mir-7-2&WithFlank&15|+|88611819|88611940@38.61.24(+1A);MI0000265|hsa-mir-7-3&WithFlank&19|+|4770664|4770785@37.60.24(+1A);MI0000263|hsa-mir-7-1&WithFlank&9|-|83969742|83969863@30.53.24(+1A)&[MIMAT0000252&hsa-miR-7-5p&offsets|0|0(+1A);m-400&15|+|88611856|88611879&offsets|0|0(+1A)];[MIMAT0000252_1&hsa-miR-7-5p&offsets|0|0(+1A);m-402&19|+|4770700|4770723&offsets|0|0(+1A)];[MIMAT0000252_2&hsa-miR-7-5p&offsets|0|0(+1A);m-401&9|-|83969811|83969834&offsets|0|0(+1A)]|TGGAAGACTAGTGATTTTGTTGTTA</td><td>>MI0000264|hsa-mir-7-2&WithFlank&15|+|88611819|88611940</td><td>38</td><td>61</td><td>24(+1A)</td><td>TGGAAGACTAGTGATTTTGTTGTTA</td><td>410.3</td><td>5.6</td><td>0.5</td><td>10.4</td><td>3.0E-25</td><td>7.4E-24</td><td>MB231/MB468</td><td>NUC</td></tr>
<tr><td>>hg38_21_+_8433172_8446622_RNA45SN1_45S_50ntFlanks@12218.12236.19|GAGGACAGAAACCTCCCGT</td><td>>hg38_21_+_8433172_8446622_RNA45SN1_45S_50ntFlanks</td><td>12218</td><td>12236</td><td>19</td><td>GAGGACAGAAACCTCCCGT</td><td>600.0</td><td>13.7</td><td>1.3</td><td>10.3</td><td>4.4E-25</td><td>7.4E-24</td><td>BT20/MB468</td><td>MITO</td></tr>
<tr><td>>MI0000239|hsa-mir-197&WithFlank&1|+|109598887|109598973@54.76.23&[MIMAT0000227&hsa-miR-197-3p&offsets|0|+1;m-99&1|+|109598940|109598961&offsets|0|+1]|TTCACCACCTTCTCCACCCAGCA</td><td>>MI0000239|hsa-mir-197&WithFlank&1|+|109598887|109598973</td><td>54</td><td>76</td><td>23</td><td>TTCACCACCTTCTCCACCCAGCA</td><td>72.7</td><td>3.6</td><td>0.4</td><td>10.2</td><td>1.2E-24</td><td>7.4E-24</td><td>BT20/MB468</td><td>NUC</td></tr>
<tr><td>>am_trna162_MetCAT_6_-_26758550_26758622@31.51.21,trna164_MetCAT_6_-_26735574_26735646@31.51.21,trna20_MetCAT_16_+_71460396_71460468@31.51.21,trna21_MetCAT_6_+_26701712_26701784@31.51.21,trna27_MetCAT_6_+_26766444_26766516@31.51.21,trna75_MetCAT_6_+_28912352_28912424@31.51.21,trna97_MetCAT_6_-_28921042_28921114@31.51.21|TCTCATAATCTGAAGGTCCTG</td><td>>am_trna162_MetCAT_6_-_26758550_26758622</td><td>31</td><td>51</td><td>21</td><td>TCTCATAATCTGAAGGTCCTG</td><td>464.7</td><td>-13.3</td><td>1.3</td><td>-10.2</td><td>1.2E-24</td><td>7.4E-24</td><td>BT20/MB468</td><td>NUC</td></tr>
<tr><td>>hg38_21_+_8433172_8446622_RNA45SN1_45S_50ntFlanks@11209.11230.22|CGGCGACTCTGGACGCGAGCCG</td><td>>hg38_21_+_8433172_8446622_RNA45SN1_45S_50ntFlanks</td><td>11209</td><td>11230</td><td>22</td><td>CGGCGACTCTGGACGCGAGCCG</td><td>1543.9</td><td>-14.5</td><td>1.4</td><td>-10.4</td><td>1.5E-25</td><td>7.4E-24</td><td>MB231/MB468</td><td>MP</td></tr>
<tr><td>>hg38_21_+_8433172_8446622_RNA45SN1_45S_50ntFlanks@11704.11728.25|ATGAAGCGCGGGTAAACGGCGGGAG</td><td>>hg38_21_+_8433172_8446622_RNA45SN1_45S_50ntFlanks</td><td>11704</td><td>11728</td><td>25</td><td>ATGAAGCGCGGGTAAACGGCGGGAG</td><td>1976.4</td><td>-14.7</td><td>1.4</td><td>-10.4</td><td>2.5E-25</td><td>7.5E-24</td><td>MB231/MB468</td><td>MITO</td></tr>
<tr><td>>hg38_21_+_8433172_8446622_RNA45SN1_45S_50ntFlanks@10808.10840.33|CAATGTAGGTAAGGGAAGTCGGCAAGCCGGATC</td><td>>hg38_21_+_8433172_8446622_RNA45SN1_45S_50ntFlanks</td><td>10808</td><td>10840</td><td>33</td><td>CAATGTAGGTAAGGGAAGTCGGCAAGCCGGATC</td><td>2154.6</td><td>-14.9</td><td>1.4</td><td>-10.4</td><td>1.5E-25</td><td>7.5E-24</td><td>MB231/MB468</td><td>MP</td></tr>
<tr><td>>hg38_21_+_8433172_8446622_RNA45SN1_45S_50ntFlanks@10128.10149.22|CGGCGGCGGCGGCGGCGGCGGG</td><td>>hg38_21_+_8433172_8446622_RNA45SN1_45S_50ntFlanks</td><td>10128</td><td>10149</td><td>22</td><td>CGGCGGCGGCGGCGGCGGCGGG</td><td>317.8</td><td>-12.7</td><td>1.2</td><td>-10.2</td><td>1.3E-24</td><td>7.6E-24</td><td>BT20/MB468</td><td>NUC</td></tr>
<tr><td>>hg38_21_+_8433172_8446622_RNA45SN1_45S_50ntFlanks@4722.4759.38|TCATTAATCAAGAACGAAAGTCGGAGGTTCGAAGACGA</td><td>>hg38_21_+_8433172_8446622_RNA45SN1_45S_50ntFlanks</td><td>4722</td><td>4759</td><td>38</td><td>TCATTAATCAAGAACGAAAGTCGGAGGTTCGAAGACGA</td><td>10516.5</td><td>-12.0</td><td>1.2</td><td>-10.2</td><td>1.2E-24</td><td>7.6E-24</td><td>BT20/MB468</td><td>NUC</td></tr>
<tr><td>>hg38_21_+_8433172_8446622_RNA45SN1_45S_50ntFlanks@8845.8866.22|CGGCGGCGGCGGCGGCGGCGGG</td><td>>hg38_21_+_8433172_8446622_RNA45SN1_45S_50ntFlanks</td><td>8845</td><td>8866</td><td>22</td><td>CGGCGGCGGCGGCGGCGGCGGG</td><td>317.8</td><td>-12.7</td><td>1.2</td><td>-10.2</td><td>1.3E-24</td><td>7.6E-24</td><td>BT20/MB468</td><td>NUC</td></tr>
<tr><td>>MI0001733|hsa-mir-452&WithFlank&X|-|151959622|151959718@20.42.23(+1U)&[MIMAT0001635&hsa-miR-452-5p&offsets|0|+1(+1U);m-188&X|-|151959677|151959699&offsets|0|0(+1U)]|AACTGTTTGCAGAGGAAACTGAGT</td><td>>MI0001733|hsa-mir-452&WithFlank&X|-|151959622|151959718</td><td>20</td><td>42</td><td>23(+1U)</td><td>AACTGTTTGCAGAGGAAACTGAGT</td><td>457.2</td><td>13.1</td><td>1.3</td><td>10.3</td><td>6.0E-25</td><td>7.6E-24</td><td>BT20/MB468</td><td>CYTO</td></tr>
<tr><td>>hg38_1_-_228634819_228635039_RNA5S12_5S_50ntFlanks@146.171.26|TGGGAATACCGGGTGCTGTAGGCTTT</td><td>>hg38_1_-_228634819_228635039_RNA5S12_5S_50ntFlanks</td><td>146</td><td>171</td><td>26</td><td>TGGGAATACCGGGTGCTGTAGGCTTT</td><td>4387.8</td><td>7.8</td><td>0.8</td><td>10.3</td><td>6.1E-25</td><td>7.7E-24</td><td>BT20/MB468</td><td>CYTO</td></tr>
<tr><td>>MI0000750|hsa-mir-26a-2&WithFlank&12|-|57824603|57824698@23.41.19;MI0000083|hsa-mir-26a-1&WithFlank&3|+|37969398|37969486@19.37.19&[MIMAT0000082&hsa-miR-26a-5p&offsets|+3|0;m-33&12|-|57824658|57824679&offsets|+3|0];[MIMAT0000082_1&hsa-miR-26a-5p&offsets|+3|0;m-34&3|+|37969413|37969434&offsets|+3|0]|AAGTAATCCAGGATAGGCT</td><td>>MI0000750|hsa-mir-26a-2&WithFlank&12|-|57824603|57824698</td><td>23</td><td>41</td><td>19</td><td>AAGTAATCCAGGATAGGCT</td><td>1118.0</td><td>14.6</td><td>1.4</td><td>10.3</td><td>4.6E-25</td><td>7.7E-24</td><td>BT20/MB468</td><td>MITO</td></tr>
<tr><td>>MI0000477|hsa-mir-146a&WithFlank&5|+|160485346|160485456@27.49.23&[MIMAT0000449&hsa-miR-146a-5p&offsets|0|+1;m-129&5|+|160485372|160485393&offsets|0|+1]|TGAGAACTGAATTCCATGGGTTG</td><td>>MI0000477|hsa-mir-146a&WithFlank&5|+|160485346|160485456</td><td>27</td><td>49</td><td>23</td><td>TGAGAACTGAATTCCATGGGTTG</td><td>4454.1</td><td>15.3</td><td>1.5</td><td>10.4</td><td>1.6E-25</td><td>7.7E-24</td><td>MB231/MB468</td><td>MP</td></tr>
<tr><td>>hg38_21_+_8433172_8446622_RNA45SN1_45S_50ntFlanks@475.510.36|CTTCGTGATCGATGTGGTGACGTCGTGCTCTCCCGG</td><td>>hg38_21_+_8433172_8446622_RNA45SN1_45S_50ntFlanks</td><td>475</td><td>510</td><td>36</td><td>CTTCGTGATCGATGTGGTGACGTCGTGCTCTCCCGG</td><td>1671.8</td><td>7.7</td><td>0.7</td><td>10.5</td><td>1.2E-25</td><td>7.8E-24</td><td>MB231/MB468</td><td>CYTO</td></tr>
<tr><td>>hg38_21_+_8433172_8446622_RNA45SN1_45S_50ntFlanks@12635.12661.27|AAGCTACCATCTGTGGGATTATGACTG</td><td>>hg38_21_+_8433172_8446622_RNA45SN1_45S_50ntFlanks</td><td>12635</td><td>12661</td><td>27</td><td>AAGCTACCATCTGTGGGATTATGACTG</td><td>24013.8</td><td>-8.8</td><td>0.8</td><td>-10.4</td><td>2.7E-25</td><td>7.9E-24</td><td>MB231/MB468</td><td>MITO</td></tr>
<tr><td>>MI0000077|hsa-mir-21&WithFlank&17|+|59841260|59841343@14.34.21&[MIMAT0000076&hsa-miR-21-5p&offsets|0|-1;m-1&17|+|59841273|59841294&offsets|0|-1]|TAGCTTATCAGACTGATGTTG</td><td>>MI0000077|hsa-mir-21&WithFlank&17|+|59841260|59841343</td><td>14</td><td>34</td><td>21</td><td>TAGCTTATCAGACTGATGTTG</td><td>7057.7</td><td>6.4</td><td>0.6</td><td>10.3</td><td>4.8E-25</td><td>7.9E-24</td><td>BT20/MB468</td><td>MITO</td></tr>
<tr><td>>hg38_21_+_8433172_8446622_RNA45SN1_45S_50ntFlanks@10479.10497.19|CGATCGAAAGGGAGTCGGG</td><td>>hg38_21_+_8433172_8446622_RNA45SN1_45S_50ntFlanks</td><td>10479</td><td>10497</td><td>19</td><td>CGATCGAAAGGGAGTCGGG</td><td>444.0</td><td>-13.2</td><td>1.3</td><td>-10.2</td><td>1.3E-24</td><td>7.9E-24</td><td>BT20/MB468</td><td>NUC</td></tr>
<tr><td>>MI0001445|hsa-mir-423&WithFlank&17|+|30117073|30117178@59.78.20&[MIMAT0001340&hsa-miR-423-3p&offsets|0|-3;m-100&17|+|30117131|30117153&offsets|0|-3]|AGCTCGGTCTGAGGCCCCTC</td><td>>MI0001445|hsa-mir-423&WithFlank&17|+|30117073|30117178</td><td>59</td><td>78</td><td>20</td><td>AGCTCGGTCTGAGGCCCCTC</td><td>11003.4</td><td>8.1</td><td>0.8</td><td>10.3</td><td>4.8E-25</td><td>8.0E-24</td><td>BT20/MB468</td><td>MITO</td></tr>
<tr><td>>MI0000301|hsa-mir-224&WithFlank&X|-|151958572|151958664@14.36.23(+1A)&[MIMAT0000281&hsa-miR-224-5p&offsets|+1|-1(+1A);m-238&X|-|151958630|151958651&offsets|0|+1(+1A)]|CAAGTCACTAGTGGTTCCGTTTAA</td><td>>MI0000301|hsa-mir-224&WithFlank&X|-|151958572|151958664</td><td>14</td><td>36</td><td>23(+1A)</td><td>CAAGTCACTAGTGGTTCCGTTTAA</td><td>687.4</td><td>13.7</td><td>1.3</td><td>10.3</td><td>6.4E-25</td><td>8.0E-24</td><td>BT20/MB468</td><td>CYTO</td></tr>
<tr><td>>hg38_21_+_8433172_8446622_RNA45SN1_45S_50ntFlanks@4043.4063.21|ACCCATTCGAACGTCTGCCCT</td><td>>hg38_21_+_8433172_8446622_RNA45SN1_45S_50ntFlanks</td><td>4043</td><td>4063</td><td>21</td><td>ACCCATTCGAACGTCTGCCCT</td><td>1124.6</td><td>-14.6</td><td>1.4</td><td>-10.2</td><td>1.3E-24</td><td>8.0E-24</td><td>BT20/MB468</td><td>NUC</td></tr>
<tr><td>>hg38_21_+_8433172_8446622_RNA45SN1_45S_50ntFlanks@4399.4423.25|CGGGCGGGCGGTCCGCCGCGAGGCG</td><td>>hg38_21_+_8433172_8446622_RNA45SN1_45S_50ntFlanks</td><td>4399</td><td>4423</td><td>25</td><td>CGGGCGGGCGGTCCGCCGCGAGGCG</td><td>645.8</td><td>-13.8</td><td>1.3</td><td>-10.2</td><td>1.3E-24</td><td>8.1E-24</td><td>BT20/MB468</td><td>NUC</td></tr>
<tr><td>>hg38_21_+_8433172_8446622_RNA45SN1_45S_50ntFlanks@12126.12150.25|CGACCCGCTCCGGGGACAGTGCCAG</td><td>>hg38_21_+_8433172_8446622_RNA45SN1_45S_50ntFlanks</td><td>12126</td><td>12150</td><td>25</td><td>CGACCCGCTCCGGGGACAGTGCCAG</td><td>3127.1</td><td>-15.4</td><td>1.5</td><td>-10.4</td><td>2.8E-25</td><td>8.2E-24</td><td>MB231/MB468</td><td>MITO</td></tr>
<tr><td>>hg38_21_+_8433172_8446622_RNA45SN1_45S_50ntFlanks@11270.11295.26|CGGCCGCCCCCGGGGAGCCCGGCGGG</td><td>>hg38_21_+_8433172_8446622_RNA45SN1_45S_50ntFlanks</td><td>11270</td><td>11295</td><td>26</td><td>CGGCCGCCCCCGGGGAGCCCGGCGGG</td><td>4287.3</td><td>-15.8</td><td>1.5</td><td>-10.4</td><td>2.8E-25</td><td>8.2E-24</td><td>MB231/MB468</td><td>MITO</td></tr>
<tr><td>>hg38_21_+_8433172_8446622_RNA45SN1_45S_50ntFlanks@485.505.21|GATGTGGTGACGTCGTGCTCT</td><td>>hg38_21_+_8433172_8446622_RNA45SN1_45S_50ntFlanks</td><td>485</td><td>505</td><td>21</td><td>GATGTGGTGACGTCGTGCTCT</td><td>1968.3</td><td>15.2</td><td>1.5</td><td>10.3</td><td>6.7E-25</td><td>8.3E-24</td><td>BT20/MB468</td><td>CYTO</td></tr>
<tr><td>>hg38_21_+_8433172_8446622_RNA45SN1_45S_50ntFlanks@10513.10533.21|CCGGAGTGGCGGAGATGGGCG</td><td>>hg38_21_+_8433172_8446622_RNA45SN1_45S_50ntFlanks</td><td>10513</td><td>10533</td><td>21</td><td>CCGGAGTGGCGGAGATGGGCG</td><td>6478.1</td><td>-16.4</td><td>1.6</td><td>-10.4</td><td>2.9E-25</td><td>8.4E-24</td><td>MB231/MB468</td><td>MITO</td></tr>
<tr><td>>hg38_21_+_8433172_8446622_RNA45SN1_45S_50ntFlanks@10786.10818.33|AACAGCCTCTGGCATGTTGGAACAATGTAGGTA</td><td>>hg38_21_+_8433172_8446622_RNA45SN1_45S_50ntFlanks</td><td>10786</td><td>10818</td><td>33</td><td>AACAGCCTCTGGCATGTTGGAACAATGTAGGTA</td><td>3761.0</td><td>-5.0</td><td>0.5</td><td>-10.3</td><td>5.1E-25</td><td>8.5E-24</td><td>BT20/MB468</td><td>MITO</td></tr>
<tr><td>>hg38_21_+_8433172_8446622_RNA45SN1_45S_50ntFlanks@485.514.30|GATGTGGTGACGTCGTGCTCTCCCGGGCCG</td><td>>hg38_21_+_8433172_8446622_RNA45SN1_45S_50ntFlanks</td><td>485</td><td>514</td><td>30</td><td>GATGTGGTGACGTCGTGCTCTCCCGGGCCG</td><td>771.6</td><td>14.1</td><td>1.4</td><td>10.3</td><td>5.2E-25</td><td>8.6E-24</td><td>BT20/MB468</td><td>MITO</td></tr>
<tr><td>>hg38_21_+_8433172_8446622_RNA45SN1_45S_50ntFlanks@4760.4793.34|TCAGATACCGTCGTAGTTCCGACCATAAACGATG</td><td>>hg38_21_+_8433172_8446622_RNA45SN1_45S_50ntFlanks</td><td>4760</td><td>4793</td><td>34</td><td>TCAGATACCGTCGTAGTTCCGACCATAAACGATG</td><td>941.1</td><td>-14.3</td><td>1.4</td><td>-10.2</td><td>1.4E-24</td><td>8.6E-24</td><td>BT20/MB468</td><td>NUC</td></tr>
<tr><td>>hg38_21_+_8433172_8446622_RNA45SN1_45S_50ntFlanks@12633.12661.29|CGAAGCTACCATCTGTGGGATTATGACTG</td><td>>hg38_21_+_8433172_8446622_RNA45SN1_45S_50ntFlanks</td><td>12633</td><td>12661</td><td>29</td><td>CGAAGCTACCATCTGTGGGATTATGACTG</td><td>8857.8</td><td>-12.8</td><td>1.2</td><td>-10.4</td><td>1.8E-25</td><td>8.7E-24</td><td>MB231/MB468</td><td>MP</td></tr>
<tr><td>>MI0000086|hsa-mir-28&WithFlank&3|+|188688775|188688872@62.82.21&[MIMAT0004502&hsa-miR-28-3p&offsets|+2|+1;m-28&3|+|188688834|188688855&offsets|+2|+1]|CTAGATTGTGAGCTCCTGGAG</td><td>>MI0000086|hsa-mir-28&WithFlank&3|+|188688775|188688872</td><td>62</td><td>82</td><td>21</td><td>CTAGATTGTGAGCTCCTGGAG</td><td>697.4</td><td>13.7</td><td>1.3</td><td>10.3</td><td>6.9E-25</td><td>8.7E-24</td><td>BT20/MB468</td><td>CYTO</td></tr>
<tr><td>>hg38_21_+_8433172_8446622_RNA45SN1_45S_50ntFlanks@11593.11611.19|CCAGGGGAATCCGACTGTT</td><td>>hg38_21_+_8433172_8446622_RNA45SN1_45S_50ntFlanks</td><td>11593</td><td>11611</td><td>19</td><td>CCAGGGGAATCCGACTGTT</td><td>3536.2</td><td>-8.4</td><td>0.8</td><td>-10.4</td><td>1.8E-25</td><td>8.7E-24</td><td>MB231/MB468</td><td>MP</td></tr>
<tr><td>>hg38_21_+_8433172_8446622_RNA45SN1_45S_50ntFlanks@5485.5518.34|AGCGCTGAGAAGACGGTCGAACTTGACTATCTAG</td><td>>hg38_21_+_8433172_8446622_RNA45SN1_45S_50ntFlanks</td><td>5485</td><td>5518</td><td>34</td><td>AGCGCTGAGAAGACGGTCGAACTTGACTATCTAG</td><td>1952.7</td><td>-14.8</td><td>1.4</td><td>-10.4</td><td>1.8E-25</td><td>8.8E-24</td><td>MB231/MB468</td><td>MP</td></tr>
<tr><td>>hg38_21_+_8433172_8446622_RNA45SN1_45S_50ntFlanks@5472.5511.40|ACGGCCCTGGCGGAGCGCTGAGAAGACGGTCGAACTTGAC</td><td>>hg38_21_+_8433172_8446622_RNA45SN1_45S_50ntFlanks</td><td>5472</td><td>5511</td><td>40</td><td>ACGGCCCTGGCGGAGCGCTGAGAAGACGGTCGAACTTGAC</td><td>1761.0</td><td>8.6</td><td>0.8</td><td>10.5</td><td>1.4E-25</td><td>8.8E-24</td><td>MB231/MB468</td><td>CYTO</td></tr>
<tr><td>>MI0000283|hsa-mir-203a&WithFlank&14|+|104117399|104117520@72.92.21&[MIMAT0000264&hsa-miR-203a-3p&offsets|+1|0;m-79&14|+|104117469|104117490&offsets|+1|0]|TGAAATGTTTAGGACCACTAG</td><td>>MI0000283|hsa-mir-203a&WithFlank&14|+|104117399|104117520</td><td>72</td><td>92</td><td>21</td><td>TGAAATGTTTAGGACCACTAG</td><td>2345.6</td><td>11.9</td><td>1.1</td><td>10.7</td><td>1.7E-26</td><td>8.8E-24</td><td>BT20/MB231</td><td>CYTO</td></tr>
</table>

<table>
<tr><td>>trna12_TrpCCA_17_+_19411494_19411565@30.52.23,trna168_TrpCCA_6_-_26331672_26331743@30.52.23,trna170_TrpCCA_6_-_26319330_26319401@30.52.23,trna1_TrpCCA_7_+_99067307_99067378@30.52.23,trna39_TrpCCA_17_-_8124187_8124258@30.52.23,trna6_TrpCCA_17_+_8089676_8089747@30.52.23|ACTCCAGATCAGAAGGTTGCGTG</td><td>>trna12_TrpCCA_17_+_19411494_19411565</td><td>30</td><td>52</td><td>23</td><td>ACTCCAGATCAGAAGGTTGCGTG</td><td>1039.6</td><td>-14.4</td><td>1.4</td><td>-10.2</td><td>1.5E-24</td><td>8.8E-24</td><td>BT20/MB468</td><td>NUC</td></tr>
<tr><td>>hg38_21_+_8433172_8446622_RNA45SN1_45S_50ntFlanks@12456.12477.22|CCAAGCGTTGGATTGTTCACCC</td><td>>hg38_21_+_8433172_8446622_RNA45SN1_45S_50ntFlanks</td><td>12456</td><td>12477</td><td>22</td><td>CCAAGCGTTGGATTGTTCACCC</td><td>7019.4</td><td>-5.5</td><td>0.5</td><td>-10.3</td><td>7.1E-25</td><td>8.9E-24</td><td>BT20/MB468</td><td>CYTO</td></tr>
<tr><td>>hg38_21_+_8433172_8446622_RNA45SN1_45S_50ntFlanks@8179.8211.33|CTGATCGAGGCCCAGCCCGTGGACGGTGTGAGG</td><td>>hg38_21_+_8433172_8446622_RNA45SN1_45S_50ntFlanks</td><td>8179</td><td>8211</td><td>33</td><td>CTGATCGAGGCCCAGCCCGTGGACGGTGTGAGG</td><td>493.0</td><td>-13.4</td><td>1.3</td><td>-10.2</td><td>1.5E-24</td><td>8.9E-24</td><td>BT20/MB468</td><td>NUC</td></tr>
<tr><td>>MI0000264|hsa-mir-7-2&WithFlank&15|+|88611819|88611940@38.58.21;MI0000265|hsa-mir-7-3&WithFlank&19|+|4770664|4770785@37.57.21;MI0000263|hsa-mir-7-1&WithFlank&9|-|83969742|83969863@30.50.21&[MIMAT0000252&hsa-miR-7-5p&offsets|0|-3;m-400&15|+|88611856|88611879&offsets|0|-3];[MIMAT0000252_1&hsa-miR-7-5p&offsets|0|-3;m-402&19|+|4770700|4770723&offsets|0|-3];[MIMAT0000252_2&hsa-miR-7-5p&offsets|0|-3;m-401&9|-|83969811|83969834&offsets|0|-3]|TGGAAGACTAGTGATTTTGTT</td><td>>MI0000264|hsa-mir-7-2&WithFlank&15|+|88611819|88611940</td><td>38</td><td>58</td><td>21</td><td>TGGAAGACTAGTGATTTTGTT</td><td>752.7</td><td>7.1</td><td>0.7</td><td>10.3</td><td>5.5E-25</td><td>9.1E-24</td><td>BT20/MB468</td><td>MITO</td></tr>
<tr><td>>trna10_AspGTC_12_-_125424193_125424264@1.31.31,trna12_AspGTC_12_-_125411891_125411962@1.31.31,trna38_AspGTC_17_-_8125556_8125627@1.31.31,trna45_AspGTC_6_+_27447453_27447524@1.31.31,trna48_AspGTC_6_+_27471523_27471594@1.31.31,trna4_AspGTC_12_+_96429799_96429870@1.31.31,trna69_AspGTC_1_-_161440205_161440276@1.31.31,trna72_AspGTC_1_-_161432824_161432895@1.31.31,trna75_AspGTC_1_-_161425414_161425485@1.31.31,trna78_AspGTC_1_-_161418033_161418104@1.31.31,trna81_AspGTC_1_-_161410615_161410686@1.31.31|TCCTCGTTAGTATAGTGGTGAGTATCCCCGC</td><td>>trna10_AspGTC_12_-_125424193_125424264</td><td>1</td><td>31</td><td>31</td><td>TCCTCGTTAGTATAGTGGTGAGTATCCCCGC</td><td>387.2</td><td>-13.0</td><td>1.3</td><td>-10.2</td><td>1.5E-24</td><td>9.1E-24</td><td>BT20/MB468</td><td>NUC</td></tr>
<tr><td>>hg38_21_+_8433172_8446622_RNA45SN1_45S_50ntFlanks@12033.12053.21|ATCGTTTTTTCACTGACCCGG</td><td>>hg38_21_+_8433172_8446622_RNA45SN1_45S_50ntFlanks</td><td>12033</td><td>12053</td><td>21</td><td>ATCGTTTTTTCACTGACCCGG</td><td>1867.1</td><td>-14.6</td><td>1.4</td><td>-10.4</td><td>3.2E-25</td><td>9.2E-24</td><td>MB231/MB468</td><td>MITO</td></tr>
<tr><td>>hg38_21_+_8433172_8446622_RNA45SN1_45S_50ntFlanks@4730.4752.23|CAAGAACGAAAGTCGGAGGTTCG</td><td>>hg38_21_+_8433172_8446622_RNA45SN1_45S_50ntFlanks</td><td>4730</td><td>4752</td><td>23</td><td>CAAGAACGAAAGTCGGAGGTTCG</td><td>358.1</td><td>-12.9</td><td>1.3</td><td>-10.2</td><td>1.5E-24</td><td>9.2E-24</td><td>BT20/MB468</td><td>NUC</td></tr>
<tr><td>>hg38_21_+_8433172_8446622_RNA45SN1_45S_50ntFlanks@9454.9493.40|CCGCGCCGGGGAGGTGGAGCACGAGCGCACGTGTTAGGAC</td><td>>hg38_21_+_8433172_8446622_RNA45SN1_45S_50ntFlanks</td><td>9454</td><td>9493</td><td>40</td><td>CCGCGCCGGGGAGGTGGAGCACGAGCGCACGTGTTAGGAC</td><td>312.2</td><td>-12.7</td><td>1.2</td><td>-10.2</td><td>1.6E-24</td><td>9.4E-24</td><td>BT20/MB468</td><td>NUC</td></tr>
<tr><td>>MI0003599|hsa-mir-589&WithFlank&7|-|5495813|5495923@30.50.21&[MIMAT0004799&hsa-miR-589-5p&offsets|0|-1;m-162&7|-|5495874|5495894&offsets|0|0]|TGAGAACCACGTCTGCTCTGA</td><td>>MI0003599|hsa-mir-589&WithFlank&7|-|5495813|5495923</td><td>30</td><td>50</td><td>21</td><td>TGAGAACCACGTCTGCTCTGA</td><td>513.0</td><td>7.5</td><td>0.7</td><td>10.4</td><td>3.9E-25</td><td>9.5E-24</td><td>MB231/MB468</td><td>NUC</td></tr>
<tr><td>>am_MI0000786|hsa-mir-378a&WithFlank&5|+|149732819|149732896@50.70.21(+1U)&[MIMAT0000732&hsa-miR-378a-3p&offsets|+1|0(+1U);m-44&5|+|149732867|149732888&offsets|+1|0(+1U)]|CTGGACTTGGAGTCAGAAGGCT</td><td>>am_MI0000786|hsa-mir-378a&WithFlank&5|+|149732819|149732896</td><td>50</td><td>70</td><td>21(+1U)</td><td>CTGGACTTGGAGTCAGAAGGCT</td><td>1271.9</td><td>5.9</td><td>0.6</td><td>10.3</td><td>7.6E-25</td><td>9.5E-24</td><td>BT20/MB468</td><td>CYTO</td></tr>
<tr><td>>hg38_21_+_8433172_8446622_RNA45SN1_45S_50ntFlanks@6645.6668.24|CTCGTACGACTCTTAGCGGTGGAT</td><td>>hg38_21_+_8433172_8446622_RNA45SN1_45S_50ntFlanks</td><td>6645</td><td>6668</td><td>24</td><td>CTCGTACGACTCTTAGCGGTGGAT</td><td>2797.4</td><td>5.7</td><td>0.6</td><td>10.3</td><td>7.8E-25</td><td>9.7E-24</td><td>BT20/MB468</td><td>CYTO</td></tr>
<tr><td>>hg38_21_+_8433172_8446622_RNA45SN1_45S_50ntFlanks@4047.4072.26|ATTCGAACGTCTGCCCTATCAACTTT</td><td>>hg38_21_+_8433172_8446622_RNA45SN1_45S_50ntFlanks</td><td>4047</td><td>4072</td><td>26</td><td>ATTCGAACGTCTGCCCTATCAACTTT</td><td>361.8</td><td>-12.9</td><td>1.3</td><td>-10.2</td><td>1.6E-24</td><td>9.8E-24</td><td>BT20/MB468</td><td>NUC</td></tr>
<tr><td>>MI0000465|hsa-mir-191&WithFlank&3|-|49020612|49020715@22.45.24&[MIMAT0000440&hsa-miR-191-5p&offsets|0|+1;m-48&3|-|49020672|49020694&offsets|0|+1]|CAACGGAATCCCAAAAGCAGCTGT</td><td>>MI0000465|hsa-mir-191&WithFlank&3|-|49020612|49020715</td><td>22</td><td>45</td><td>24</td><td>CAACGGAATCCCAAAAGCAGCTGT</td><td>3770.7</td><td>6.9</td><td>0.7</td><td>10.3</td><td>7.9E-25</td><td>9.8E-24</td><td>BT20/MB468</td><td>CYTO</td></tr>
<tr><td>>hg38_21_+_8433172_8446622_RNA45SN1_45S_50ntFlanks@10914.10931.18|CGCCGCGGCTGGACGAGG</td><td>>hg38_21_+_8433172_8446622_RNA45SN1_45S_50ntFlanks</td><td>10914</td><td>10931</td><td>18</td><td>CGCCGCGGCTGGACGAGG</td><td>1694.7</td><td>-14.6</td><td>1.4</td><td>-10.4</td><td>2.1E-25</td><td>9.8E-24</td><td>MB231/MB468</td><td>MP</td></tr>
<tr><td>>hg38_21_+_8433172_8446622_RNA45SN1_45S_50ntFlanks@12540.12569.30|TGATGTGTTGTTGCCATGGTAATCCTGCTC</td><td>>hg38_21_+_8433172_8446622_RNA45SN1_45S_50ntFlanks</td><td>12540</td><td>12569</td><td>30</td><td>TGATGTGTTGTTGCCATGGTAATCCTGCTC</td><td>571.7</td><td>-13.6</td><td>1.3</td><td>-10.2</td><td>1.7E-24</td><td>9.9E-24</td><td>BT20/MB468</td><td>NUC</td></tr>
<tr><td>>trna10_LysCTT_16_+_3241501_3241573@1.34.34,trna13_LysCTT_14_-_58706613_58706685@1.34.34,trna2_LysCTT_15_+_79152904_79152976@1.34.34|GCCCGGCTAGCTCAGTCGGTAGAGCATGGGACTC</td><td>>trna10_LysCTT_16_+_3241501_3241573</td><td>1</td><td>34</td><td>34</td><td>GCCCGGCTAGCTCAGTCGGTAGAGCATGGGACTC</td><td>2441.6</td><td>7.6</td><td>0.7</td><td>10.3</td><td>8.0E-25</td><td>1.0E-23</td><td>BT20/MB468</td><td>CYTO</td></tr>
<tr><td>>hg38_21_+_8433172_8446622_RNA45SN1_45S_50ntFlanks@12419.12448.30|GATGTCGGCTCTTCCTATCATTGTGAAGCA</td><td>>hg38_21_+_8433172_8446622_RNA45SN1_45S_50ntFlanks</td><td>12419</td><td>12448</td><td>30</td><td>GATGTCGGCTCTTCCTATCATTGTGAAGCA</td><td>660.2</td><td>5.0</td><td>0.5</td><td>10.3</td><td>6.1E-25</td><td>1.0E-23</td><td>BT20/MB468</td><td>MITO</td></tr>
<tr><td>>hg38_21_+_8433172_8446622_RNA45SN1_45S_50ntFlanks@11641.11663.23|CGGCGGGTGTTGACGCGATGTGA</td><td>>hg38_21_+_8433172_8446622_RNA45SN1_45S_50ntFlanks</td><td>11641</td><td>11663</td><td>23</td><td>CGGCGGGTGTTGACGCGATGTGA</td><td>2137.9</td><td>-14.9</td><td>1.4</td><td>-10.4</td><td>2.1E-25</td><td>1.0E-23</td><td>MB231/MB468</td><td>MP</td></tr>
<tr><td>>hg38_21_+_8433172_8446622_RNA45SN1_45S_50ntFlanks@12991.13008.18|CTCCCTCGCTGCGATCTA</td><td>>hg38_21_+_8433172_8446622_RNA45SN1_45S_50ntFlanks</td><td>12991</td><td>13008</td><td>18</td><td>CTCCCTCGCTGCGATCTA</td><td>1294.4</td><td>-14.8</td><td>1.4</td><td>-10.2</td><td>1.7E-24</td><td>1.0E-23</td><td>BT20/MB468</td><td>NUC</td></tr>
<tr><td>>hg38_21_+_8433172_8446622_RNA45SN1_45S_50ntFlanks@11767.11794.28|TCTAATTAGTGACGCGCATGAATGGATG</td><td>>hg38_21_+_8433172_8446622_RNA45SN1_45S_50ntFlanks</td><td>11767</td><td>11794</td><td>28</td><td>TCTAATTAGTGACGCGCATGAATGGATG</td><td>2878.7</td><td>-5.6</td><td>0.5</td><td>-10.3</td><td>4.3E-25</td><td>1.0E-23</td><td>MB231/MB468</td><td>NUC</td></tr>
<tr><td>>hg38_21_+_8433172_8446622_RNA45SN1_45S_50ntFlanks@10869.10902.34|AGGGCTGGGTCGGTCGGGCTGGGGCGCGAAGCGG</td><td>>hg38_21_+_8433172_8446622_RNA45SN1_45S_50ntFlanks</td><td>10869</td><td>10902</td><td>34</td><td>AGGGCTGGGTCGGTCGGGCTGGGGCGCGAAGCGG</td><td>1026.9</td><td>-8.6</td><td>0.8</td><td>-10.4</td><td>1.6E-25</td><td>1.0E-23</td><td>MB231/MB468</td><td>CYTO</td></tr>
<tr><td>>hg38_21_+_8433172_8446622_RNA45SN1_45S_50ntFlanks@12633.12652.20|CGAAGCTACCATCTGTGGGA</td><td>>hg38_21_+_8433172_8446622_RNA45SN1_45S_50ntFlanks</td><td>12633</td><td>12652</td><td>20</td><td>CGAAGCTACCATCTGTGGGA</td><td>3060.0</td><td>-15.4</td><td>1.5</td><td>-10.4</td><td>2.2E-25</td><td>1.0E-23</td><td>MB231/MB468</td><td>MP</td></tr>
<tr><td>>hg38_21_+_8433172_8446622_RNA45SN1_45S_50ntFlanks@4724.4758.35|ATTAATCAAGAACGAAAGTCGGAGGTTCGAAGACG</td><td>>hg38_21_+_8433172_8446622_RNA45SN1_45S_50ntFlanks</td><td>4724</td><td>4758</td><td>35</td><td>ATTAATCAAGAACGAAAGTCGGAGGTTCGAAGACG</td><td>14663.9</td><td>-7.2</td><td>0.7</td><td>-10.3</td><td>6.3E-25</td><td>1.0E-23</td><td>BT20/MB468</td><td>MITO</td></tr>
<tr><td>>hg38_21_+_8433172_8446622_RNA45SN1_45S_50ntFlanks@3900.3919.20|CTTCGCGGGGGGGATGCGTG</td><td>>hg38_21_+_8433172_8446622_RNA45SN1_45S_50ntFlanks</td><td>3900</td><td>3919</td><td>20</td><td>CTTCGCGGGGGGGATGCGTG</td><td>3013.2</td><td>-15.4</td><td>1.5</td><td>-10.4</td><td>2.2E-25</td><td>1.0E-23</td><td>MB231/MB468</td><td>MP</td></tr>
<tr><td>>hg38_21_+_8433172_8446622_RNA45SN1_45S_50ntFlanks@9537.9563.27|AAACTCTGGTGGAGGTCCGTAGCGGTC</td><td>>hg38_21_+_8433172_8446622_RNA45SN1_45S_50ntFlanks</td><td>9537</td><td>9563</td><td>27</td><td>AAACTCTGGTGGAGGTCCGTAGCGGTC</td><td>512.8</td><td>-13.4</td><td>1.3</td><td>-10.2</td><td>1.8E-24</td><td>1.0E-23</td><td>BT20/MB468</td><td>NUC</td></tr>
<tr><td>>hg38_21_+_8433172_8446622_RNA45SN1_45S_50ntFlanks@12159.12178.20|TTTGACTGGGGCGGTACACC</td><td>>hg38_21_+_8433172_8446622_RNA45SN1_45S_50ntFlanks</td><td>12159</td><td>12178</td><td>20</td><td>TTTGACTGGGGCGGTACACC</td><td>1554.0</td><td>-14.5</td><td>1.4</td><td>-10.4</td><td>2.2E-25</td><td>1.0E-23</td><td>MB231/MB468</td><td>MP</td></tr>
<tr><td>>hg38_21_+_8433172_8446622_RNA45SN1_45S_50ntFlanks@10805.10826.22|GAACAATGTAGGTAAGGGAAGT</td><td>>hg38_21_+_8433172_8446622_RNA45SN1_45S_50ntFlanks</td><td>10805</td><td>10826</td><td>22</td><td>GAACAATGTAGGTAAGGGAAGT</td><td>528.6</td><td>-13.5</td><td>1.3</td><td>-10.2</td><td>1.8E-24</td><td>1.1E-23</td><td>BT20/MB468</td><td>NUC</td></tr>
<tr><td>>hg38_21_+_8433172_8446622_RNA45SN1_45S_50ntFlanks@11233.11253.21|CCCTTCCCGTGGATCGCCCCA</td><td>>hg38_21_+_8433172_8446622_RNA45SN1_45S_50ntFlanks</td><td>11233</td><td>11253</td><td>21</td><td>CCCTTCCCGTGGATCGCCCCA</td><td>804.3</td><td>-14.1</td><td>1.4</td><td>-10.2</td><td>1.8E-24</td><td>1.1E-23</td><td>BT20/MB468</td><td>NUC</td></tr>
<tr><td>>hg38_21_+_8433172_8446622_RNA45SN1_45S_50ntFlanks@12081.12110.30|CTCTCGCTTCTGGCGCCAAGCGCCCGGCCG</td><td>>hg38_21_+_8433172_8446622_RNA45SN1_45S_50ntFlanks</td><td>12081</td><td>12110</td><td>30</td><td>CTCTCGCTTCTGGCGCCAAGCGCCCGGCCG</td><td>979.1</td><td>-8.5</td><td>0.8</td><td>-10.4</td><td>1.7E-25</td><td>1.1E-23</td><td>MB231/MB468</td><td>CYTO</td></tr>
<tr><td>>hg38_21_+_8433172_8446622_RNA45SN1_45S_50ntFlanks@11907.11929.23|ACGGTGAAGAGACATGAGAGGTG</td><td>>hg38_21_+_8433172_8446622_RNA45SN1_45S_50ntFlanks</td><td>11907</td><td>11929</td><td>23</td><td>ACGGTGAAGAGACATGAGAGGTG</td><td>335.7</td><td>-12.8</td><td>1.3</td><td>-10.2</td><td>1.8E-24</td><td>1.1E-23</td><td>BT20/MB468</td><td>NUC</td></tr>
<tr><td>>hg38_21_+_8433172_8446622_RNA45SN1_45S_50ntFlanks@10772.10805.34|AGGTCTCCAAGGTGAACAGCCTCTGGCATGTTGG</td><td>>hg38_21_+_8433172_8446622_RNA45SN1_45S_50ntFlanks</td><td>10772</td><td>10805</td><td>34</td><td>AGGTCTCCAAGGTGAACAGCCTCTGGCATGTTGG</td><td>320.5</td><td>-12.7</td><td>1.2</td><td>-10.2</td><td>1.8E-24</td><td>1.1E-23</td><td>BT20/MB468</td><td>NUC</td></tr>
<tr><td>>MI0000650|hsa-mir-200c&WithFlank&12|+|6963693|6963772@50.72.23(+2U)&[MIMAT0000617&hsa-miR-200c-3p&offsets|0|0(+2U);m-54&12|+|6963742|6963764&offsets|0|0(+2U)]|TAATACTGCCGGGTAATGATGGATT</td><td>>MI0000650|hsa-mir-200c&WithFlank&12|+|6963693|6963772</td><td>50</td><td>72</td><td>23(+2U)</td><td>TAATACTGCCGGGTAATGATGGATT</td><td>954.2</td><td>7.6</td><td>0.7</td><td>10.3</td><td>8.9E-25</td><td>1.1E-23</td><td>BT20/MB468</td><td>CYTO</td></tr>
<tr><td>>hg38_21_+_8433172_8446622_RNA45SN1_45S_50ntFlanks@12242.12272.31|AGAAGGGCAAAAGCTCGCTTGATCTTGATTT</td><td>>hg38_21_+_8433172_8446622_RNA45SN1_45S_50ntFlanks</td><td>12242</td><td>12272</td><td>31</td><td>AGAAGGGCAAAAGCTCGCTTGATCTTGATTT</td><td>570.4</td><td>-5.6</td><td>0.5</td><td>-10.2</td><td>1.9E-24</td><td>1.1E-23</td><td>BT20/MB468</td><td>NUC</td></tr>
<tr><td>>MI0000255|hsa-mir-30d&WithFlank&8|-|134804870|134804951@12.33.22(+1U)&[MIMAT0000245&hsa-miR-30d-5p&offsets|0|0(+1U);m-21&8|-|134804919|134804940&offsets|0|0(+1U)]|TGTAAACATCCCCGACTGGAAGT</td><td>>MI0000255|hsa-mir-30d&WithFlank&8|-|134804870|134804951</td><td>12</td><td>33</td><td>22(+1U)</td><td>TGTAAACATCCCCGACTGGAAGT</td><td>2807.9</td><td>4.1</td><td>0.4</td><td>10.2</td><td>1.9E-24</td><td>1.1E-23</td><td>BT20/MB468</td><td>NUC</td></tr>
<tr><td>>hg38_21_+_8433172_8446622_RNA45SN1_45S_50ntFlanks@6748.6770.23|CTTCGAACGCACTTGCGGCCCCG</td><td>>hg38_21_+_8433172_8446622_RNA45SN1_45S_50ntFlanks</td><td>6748</td><td>6770</td><td>23</td><td>CTTCGAACGCACTTGCGGCCCCG</td><td>1370.6</td><td>-14.3</td><td>1.4</td><td>-10.4</td><td>2.4E-25</td><td>1.1E-23</td><td>MB231/MB468</td><td>MP</td></tr>
<tr><td>>hg38_21_+_8433172_8446622_RNA45SN1_45S_50ntFlanks@11612.11640.29|TAATTAAAACAAAGCATCGCGAAGGCCCG</td><td>>hg38_21_+_8433172_8446622_RNA45SN1_45S_50ntFlanks</td><td>11612</td><td>11640</td><td>29</td><td>TAATTAAAACAAAGCATCGCGAAGGCCCG</td><td>3320.1</td><td>-15.5</td><td>1.5</td><td>-10.4</td><td>3.9E-25</td><td>1.1E-23</td><td>MB231/MB468</td><td>MITO</td></tr>
<tr><td>>hg38_21_+_8433172_8446622_RNA45SN1_45S_50ntFlanks@12405.12441.37|CTTTTTGATCCTTCGATGTCGGCTCTTCCTATCATTG</td><td>>hg38_21_+_8433172_8446622_RNA45SN1_45S_50ntFlanks</td><td>12405</td><td>12441</td><td>37</td><td>CTTTTTGATCCTTCGATGTCGGCTCTTCCTATCATTG</td><td>2186.8</td><td>-4.8</td><td>0.5</td><td>-10.3</td><td>7.0E-25</td><td>1.1E-23</td><td>BT20/MB468</td><td>MITO</td></tr>
<tr><td>>hg38_21_+_8433172_8446622_RNA45SN1_45S_50ntFlanks@12685.12703.19|CCCAGGCGGAACGATACGG</td><td>>hg38_21_+_8433172_8446622_RNA45SN1_45S_50ntFlanks</td><td>12685</td><td>12703</td><td>19</td><td>CCCAGGCGGAACGATACGG</td><td>319.1</td><td>-12.7</td><td>1.2</td><td>-10.2</td><td>1.9E-24</td><td>1.1E-23</td><td>BT20/MB468</td><td>NUC</td></tr>
<tr><td>>hg38_21_+_8433172_8446622_RNA45SN1_45S_50ntFlanks@4550.4572.23|GATACCGCAGCTAGGAATAATGG</td><td>>hg38_21_+_8433172_8446622_RNA45SN1_45S_50ntFlanks</td><td>4550</td><td>4572</td><td>23</td><td>GATACCGCAGCTAGGAATAATGG</td><td>467.2</td><td>-13.3</td><td>1.3</td><td>-10.2</td><td>2.0E-24</td><td>1.2E-23</td><td>BT20/MB468</td><td>NUC</td></tr>
<tr><td>>MI0000061|hsa-let-7a-2&WithFlank&11|-|122146516|122146599@11.30.20(+1C);MI0000062|hsa-let-7a-3&WithFlank&22|+|46112743|46112828@10.29.20(+1C);MI0000060|hsa-let-7a-1&WithFlank&9|+|94175951|94176042@12.31.20(+1C)&[MIMAT0000062&hsa-let-7a-5p&offsets|0|-2(+1C);m-6&11|-|122146568|122146589&offsets|0|-2(+1C)];[MIMAT0000062_1&hsa-let-7a-5p&offsets|0|-2(+1C);m-5&22|+|46112752|46112773&offsets|0|-2(+1C)];[MIMAT0000062_2&hsa-let-7a-5p&offsets|0|-2(+1C);m-4&9|+|94175962|94175983&offsets|0|-2(+1C)]|TGAGGTAGTAGGTTGTATAGC</td><td>>MI0000061|hsa-let-7a-2&WithFlank&11|-|122146516|122146599</td><td>11</td><td>30</td><td>20(+1C)</td><td>TGAGGTAGTAGGTTGTATAGC</td><td>1183.6</td><td>5.8</td><td>0.6</td><td>10.3</td><td>4.9E-25</td><td>1.2E-23</td><td>MB231/MB468</td><td>NUC</td></tr>
</table>

<table>
<tr><td>>hg38_21_+_8433172_8446622_RNA45SN1_45S_50ntFlanks@9531.9571.41|CAGAGGAAACTCTGGTGGAGGTCCGTAGCGGTCCTGACGTG</td><td>>hg38_21_+_8433172_8446622_RNA45SN1_45S_50ntFlanks</td><td>9531</td><td>9571</td><td>41</td><td>CAGAGGAAACTCTGGTGGAGGTCCGTAGCGGTCCTGACGTG</td><td>467.2</td><td>12.9</td><td>1.2</td><td>10.4</td><td>1.9E-25</td><td>1.2E-23</td><td>MB231/MB468</td><td>CYTO</td></tr>
<tr><td>>hg38_21_+_8433172_8446622_RNA45SN1_45S_50ntFlanks@7990.8008.19|GACGTGGCGACCCGCTGAA</td><td>>hg38_21_+_8433172_8446622_RNA45SN1_45S_50ntFlanks</td><td>7990</td><td>8008</td><td>19</td><td>GACGTGGCGACCCGCTGAA</td><td>5801.7</td><td>-7.4</td><td>0.7</td><td>-10.4</td><td>2.6E-25</td><td>1.2E-23</td><td>MB231/MB468</td><td>MP</td></tr>
<tr><td>>hg38_21_+_8433172_8446622_RNA45SN1_45S_50ntFlanks@12997.13017.21|CGCTGCGATCTATTGAAAGTC</td><td>>hg38_21_+_8433172_8446622_RNA45SN1_45S_50ntFlanks</td><td>12997</td><td>13017</td><td>21</td><td>CGCTGCGATCTATTGAAAGTC</td><td>1532.9</td><td>6.1</td><td>0.6</td><td>10.3</td><td>5.0E-25</td><td>1.2E-23</td><td>MB231/MB468</td><td>NUC</td></tr>
<tr><td>>am_trna2_ArgCCT_16_+_3202901_3202973@14.31.18,trna3_ArgCCT_7_+_139025446_139025518@14.31.18|AATGGATAAGGCATTGGC</td><td>>am_trna2_ArgCCT_16_+_3202901_3202973</td><td>14</td><td>31</td><td>18</td><td>AATGGATAAGGCATTGGC</td><td>1281.0</td><td>-14.7</td><td>1.4</td><td>-10.2</td><td>2.0E-24</td><td>1.2E-23</td><td>BT20/MB468</td><td>NUC</td></tr>
<tr><td>>hg38_21_+_8433172_8446622_RNA45SN1_45S_50ntFlanks@3840.3865.26|CTCCTACTTGGATAACTGTGGTAATT</td><td>>hg38_21_+_8433172_8446622_RNA45SN1_45S_50ntFlanks</td><td>3840</td><td>3865</td><td>26</td><td>CTCCTACTTGGATAACTGTGGTAATT</td><td>1053.9</td><td>5.4</td><td>0.5</td><td>10.3</td><td>9.8E-25</td><td>1.2E-23</td><td>BT20/MB468</td><td>CYTO</td></tr>
<tr><td>>hg38_21_+_8433172_8446622_RNA45SN1_45S_50ntFlanks@4734.4768.35|AACGAAAGTCGGAGGTTCGAAGACGATCAGATACC</td><td>>hg38_21_+_8433172_8446622_RNA45SN1_45S_50ntFlanks</td><td>4734</td><td>4768</td><td>35</td><td>AACGAAAGTCGGAGGTTCGAAGACGATCAGATACC</td><td>629.3</td><td>13.6</td><td>1.3</td><td>10.3</td><td>9.8E-25</td><td>1.2E-23</td><td>BT20/MB468</td><td>CYTO</td></tr>
<tr><td>>MI0000298|hsa-mir-221&WithFlank&X|-|45746151|45746272@71.93.23(+1U)&[MIMAT0000278&hsa-miR-221-3p&offsets|0|0(+1U);m-91&X|-|45746181|45746202&offsets|0|+1(+1U)]|AGCTACATTGTCTGCTGGGTTTCT</td><td>>MI0000298|hsa-mir-221&WithFlank&X|-|45746151|45746272</td><td>71</td><td>93</td><td>23(+1U)</td><td>AGCTACATTGTCTGCTGGGTTTCT</td><td>9058.3</td><td>6.0</td><td>0.6</td><td>10.3</td><td>9.9E-25</td><td>1.2E-23</td><td>BT20/MB468</td><td>CYTO</td></tr>
<tr><td>>hg38_21_+_8433172_8446622_RNA45SN1_45S_50ntFlanks@12971.12990.20|TTTCGTACGTAGCAGAGCAG</td><td>>hg38_21_+_8433172_8446622_RNA45SN1_45S_50ntFlanks</td><td>12971</td><td>12990</td><td>20</td><td>TTTCGTACGTAGCAGAGCAG</td><td>1824.6</td><td>-13.8</td><td>1.3</td><td>-10.3</td><td>5.1E-25</td><td>1.2E-23</td><td>MB231/MB468</td><td>NUC</td></tr>
<tr><td>>hg38_21_+_8433172_8446622_RNA45SN1_45S_50ntFlanks@9534.9572.39|AGGAAACTCTGGTGGAGGTCCGTAGCGGTCCTGACGTGC</td><td>>hg38_21_+_8433172_8446622_RNA45SN1_45S_50ntFlanks</td><td>9534</td><td>9572</td><td>39</td><td>AGGAAACTCTGGTGGAGGTCCGTAGCGGTCCTGACGTGC</td><td>9696.2</td><td>-6.8</td><td>0.7</td><td>-10.3</td><td>5.1E-25</td><td>1.2E-23</td><td>MB231/MB468</td><td>NUC</td></tr>
<tr><td>>hg38_21_+_8433172_8446622_RNA45SN1_45S_50ntFlanks@11766.11789.24|ATCTAATTAGTGACGCGCATGAAT</td><td>>hg38_21_+_8433172_8446622_RNA45SN1_45S_50ntFlanks</td><td>11766</td><td>11789</td><td>24</td><td>ATCTAATTAGTGACGCGCATGAAT</td><td>588.9</td><td>-13.6</td><td>1.3</td><td>-10.2</td><td>2.1E-24</td><td>1.2E-23</td><td>BT20/MB468</td><td>NUC</td></tr>
<tr><td>>hg38_21_+_8433172_8446622_RNA45SN1_45S_50ntFlanks@5487.5509.23|CGCTGAGAAGACGGTCGAACTTG</td><td>>hg38_21_+_8433172_8446622_RNA45SN1_45S_50ntFlanks</td><td>5487</td><td>5509</td><td>23</td><td>CGCTGAGAAGACGGTCGAACTTG</td><td>1946.7</td><td>-14.8</td><td>1.4</td><td>-10.4</td><td>2.6E-25</td><td>1.2E-23</td><td>MB231/MB468</td><td>MP</td></tr>
<tr><td>>hg38_21_+_8433172_8446622_RNA45SN1_45S_50ntFlanks@10420.10457.38|CTGAGAGATGGGCGAGCGCCGTTCCGAAGGGACGGGCG</td><td>>hg38_21_+_8433172_8446622_RNA45SN1_45S_50ntFlanks</td><td>10420</td><td>10457</td><td>38</td><td>CTGAGAGATGGGCGAGCGCCGTTCCGAAGGGACGGGCG</td><td>1660.4</td><td>-15.1</td><td>1.5</td><td>-10.2</td><td>2.1E-24</td><td>1.2E-23</td><td>BT20/MB468</td><td>NUC</td></tr>
<tr><td>>MI0000266|hsa-mir-10a&WithFlank&17|-|48579832|48579953@28.49.22(+1U)&[MIMAT0000253&hsa-miR-10a-5p&offsets|0|-1(+1U);m-18&17|-|48579905|48579926&offsets|0|0(+1U)]|TACCCTGTAGATCCGAATTTGTT</td><td>>MI0000266|hsa-mir-10a&WithFlank&17|-|48579832|48579953</td><td>28</td><td>49</td><td>22(+1U)</td><td>TACCCTGTAGATCCGAATTTGTT</td><td>516.0</td><td>13.3</td><td>1.3</td><td>10.3</td><td>1.0E-24</td><td>1.2E-23</td><td>BT20/MB468</td><td>CYTO</td></tr>
<tr><td>>MI0000301|hsa-mir-224&WithFlank&X|-|151958572|151958664@14.36.23&[MIMAT0000281&hsa-miR-224-5p&offsets|+1|-1;m-238&X|-|151958630|151958651&offsets|0|+1]|CAAGTCACTAGTGGTTCCGTTTA</td><td>>MI0000301|hsa-mir-224&WithFlank&X|-|151958572|151958664</td><td>14</td><td>36</td><td>23</td><td>CAAGTCACTAGTGGTTCCGTTTA</td><td>1539.3</td><td>15.1</td><td>1.5</td><td>10.3</td><td>7.5E-25</td><td>1.2E-23</td><td>BT20/MB468</td><td>MITO</td></tr>
<tr><td>>hg38_21_+_8433172_8446622_RNA45SN1_45S_50ntFlanks@11572.11602.31|CCGACTTAGAACTGGTGCGGACCAGGGGAAT</td><td>>hg38_21_+_8433172_8446622_RNA45SN1_45S_50ntFlanks</td><td>11572</td><td>11602</td><td>31</td><td>CCGACTTAGAACTGGTGCGGACCAGGGGAAT</td><td>9182.0</td><td>-6.0</td><td>0.6</td><td>-10.3</td><td>5.2E-25</td><td>1.2E-23</td><td>MB231/MB468</td><td>NUC</td></tr>
<tr><td>>hg38_21_+_8433172_8446622_RNA45SN1_45S_50ntFlanks@10587.10608.22|CCCCGGGGAGAGTTCTCTTTTC</td><td>>hg38_21_+_8433172_8446622_RNA45SN1_45S_50ntFlanks</td><td>10587</td><td>10608</td><td>22</td><td>CCCCGGGGAGAGTTCTCTTTTC</td><td>378.4</td><td>-13.0</td><td>1.3</td><td>-10.2</td><td>2.1E-24</td><td>1.2E-23</td><td>BT20/MB468</td><td>NUC</td></tr>
<tr><td>>hg38_1_-_228634819_228635039_RNA5S12_5S_50ntFlanks@149.170.22|GAATACCGGGTGCTGTAGGCTT</td><td>>hg38_1_-_228634819_228635039_RNA5S12_5S_50ntFlanks</td><td>149</td><td>170</td><td>22</td><td>GAATACCGGGTGCTGTAGGCTT</td><td>5301.3</td><td>6.2</td><td>0.6</td><td>10.3</td><td>1.0E-24</td><td>1.2E-23</td><td>BT20/MB468</td><td>CYTO</td></tr>
<tr><td>>MI0000286|hsa-mir-210&WithFlank&11|-|568083|568204@72.94.23&[MIMAT0000267&hsa-miR-210-3p&offsets|0|+1;m-102&11|-|568112|568133&offsets|0|+1]|CTGTGCGTGTGACAGCGGCTGAT</td><td>>MI0000286|hsa-mir-210&WithFlank&11|-|568083|568204</td><td>72</td><td>94</td><td>23</td><td>CTGTGCGTGTGACAGCGGCTGAT</td><td>466.6</td><td>13.3</td><td>1.3</td><td>10.3</td><td>7.7E-25</td><td>1.3E-23</td><td>BT20/MB468</td><td>MITO</td></tr>
<tr><td>>hg38_21_+_8433172_8446622_RNA45SN1_45S_50ntFlanks@9306.9336.31|CAAGGAGTCTAACACGTGCGCGAGTCGGGGG</td><td>>hg38_21_+_8433172_8446622_RNA45SN1_45S_50ntFlanks</td><td>9306</td><td>9336</td><td>31</td><td>CAAGGAGTCTAACACGTGCGCGAGTCGGGGG</td><td>1314.4</td><td>-14.2</td><td>1.4</td><td>-10.4</td><td>2.7E-25</td><td>1.3E-23</td><td>MB231/MB468</td><td>MP</td></tr>
<tr><td>>MI0000542|hsa-mir-320a&WithFlank&8|-|22244960|22245043@48.70.23(+1C)&[MIMAT0000510&hsa-miR-320a-3p&offsets|0|+1(+1C);m-60&8|-|22244975|22244996&offsets|0|+1(+1C)]|AAAAGCTGGGTTGAGAGGGCGAAC</td><td>>MI0000542|hsa-mir-320a&WithFlank&8|-|22244960|22245043</td><td>48</td><td>70</td><td>23(+1C)</td><td>AAAAGCTGGGTTGAGAGGGCGAAC</td><td>685.8</td><td>13.9</td><td>1.3</td><td>10.3</td><td>7.8E-25</td><td>1.3E-23</td><td>BT20/MB468</td><td>MITO</td></tr>
<tr><td>>MI0000266|hsa-mir-10a&WithFlank&17|-|48579832|48579953@28.50.23(+1A)&[MIMAT0000253&hsa-miR-10a-5p&offsets|0|0(+1A);m-18&17|-|48579905|48579926&offsets|0|+1(+1A)]|TACCCTGTAGATCCGAATTTGTGA</td><td>>MI0000266|hsa-mir-10a&WithFlank&17|-|48579832|48579953</td><td>28</td><td>50</td><td>23(+1A)</td><td>TACCCTGTAGATCCGAATTTGTGA</td><td>1313.9</td><td>14.8</td><td>1.4</td><td>10.3</td><td>7.9E-25</td><td>1.3E-23</td><td>BT20/MB468</td><td>MITO</td></tr>
<tr><td>>hg38_21_+_8433172_8446622_RNA45SN1_45S_50ntFlanks@9129.9152.24|CGGGTCGGGGGGCGGGGCGGACTG</td><td>>hg38_21_+_8433172_8446622_RNA45SN1_45S_50ntFlanks</td><td>9129</td><td>9152</td><td>24</td><td>CGGGTCGGGGGGCGGGGCGGACTG</td><td>3665.6</td><td>-15.6</td><td>1.5</td><td>-10.3</td><td>4.4E-25</td><td>1.3E-23</td><td>MB231/MB468</td><td>MITO</td></tr>
<tr><td>>MI0000465|hsa-mir-191&WithFlank&3|-|49020612|49020715@22.43.22(+1A)&[MIMAT0000440&hsa-miR-191-5p&offsets|0|-1(+1A);m-48&3|-|49020672|49020694&offsets|0|-1(+1A)]|CAACGGAATCCCAAAAGCAGCTA</td><td>>MI0000465|hsa-mir-191&WithFlank&3|-|49020612|49020715</td><td>22</td><td>43</td><td>22(+1A)</td><td>CAACGGAATCCCAAAAGCAGCTA</td><td>2381.6</td><td>7.7</td><td>0.7</td><td>10.3</td><td>8.0E-25</td><td>1.3E-23</td><td>BT20/MB468</td><td>MITO</td></tr>
<tr><td>>hg38_21_+_8433172_8446622_RNA45SN1_45S_50ntFlanks@12991.13018.28|CTCCCTCGCTGCGATCTATTGAAAGTCA</td><td>>hg38_21_+_8433172_8446622_RNA45SN1_45S_50ntFlanks</td><td>12991</td><td>13018</td><td>28</td><td>CTCCCTCGCTGCGATCTATTGAAAGTCA</td><td>752.9</td><td>-14.0</td><td>1.4</td><td>-10.2</td><td>2.2E-24</td><td>1.3E-23</td><td>BT20/MB468</td><td>NUC</td></tr>
<tr><td>>MI0000266|hsa-mir-10a&WithFlank&17|-|48579832|48579953@29.49.21&[MIMAT0000253&hsa-miR-10a-5p&offsets|+1|-1;m-18&17|-|48579905|48579926&offsets|+1|0]|ACCCTGTAGATCCGAATTTGT</td><td>>MI0000266|hsa-mir-10a&WithFlank&17|-|48579832|48579953</td><td>29</td><td>49</td><td>21</td><td>ACCCTGTAGATCCGAATTTGT</td><td>389.4</td><td>11.4</td><td>1.1</td><td>10.2</td><td>2.2E-24</td><td>1.3E-23</td><td>BT20/MB468</td><td>NUC</td></tr>
<tr><td>>hg38_21_+_8433172_8446622_RNA45SN1_45S_50ntFlanks@10346.10368.23|TCAAACGAGAACTTTGAAGGCCG</td><td>>hg38_21_+_8433172_8446622_RNA45SN1_45S_50ntFlanks</td><td>10346</td><td>10368</td><td>23</td><td>TCAAACGAGAACTTTGAAGGCCG</td><td>5724.1</td><td>-7.2</td><td>0.7</td><td>-10.3</td><td>4.6E-25</td><td>1.3E-23</td><td>MB231/MB468</td><td>MITO</td></tr>
<tr><td>>hg38_21_+_8433172_8446622_RNA45SN1_45S_50ntFlanks@4724.4755.32|ATTAATCAAGAACGAAAGTCGGAGGTTCGAAG</td><td>>hg38_21_+_8433172_8446622_RNA45SN1_45S_50ntFlanks</td><td>4724</td><td>4755</td><td>32</td><td>ATTAATCAAGAACGAAAGTCGGAGGTTCGAAG</td><td>345.6</td><td>-12.9</td><td>1.3</td><td>-10.2</td><td>2.2E-24</td><td>1.3E-23</td><td>BT20/MB468</td><td>NUC</td></tr>
<tr><td>>hg38_21_+_8433172_8446622_RNA45SN1_45S_50ntFlanks@7984.8008.25|AGATCAGACGTGGCGACCCGCTGAA</td><td>>hg38_21_+_8433172_8446622_RNA45SN1_45S_50ntFlanks</td><td>7984</td><td>8008</td><td>25</td><td>AGATCAGACGTGGCGACCCGCTGAA</td><td>5806.3</td><td>-10.2</td><td>1.0</td><td>-10.4</td><td>2.9E-25</td><td>1.3E-23</td><td>MB231/MB468</td><td>MP</td></tr>
<tr><td>>hg38_21_+_8433172_8446622_RNA45SN1_45S_50ntFlanks@6647.6670.24|CGTACGACTCTTAGCGGTGGATCA</td><td>>hg38_21_+_8433172_8446622_RNA45SN1_45S_50ntFlanks</td><td>6647</td><td>6670</td><td>24</td><td>CGTACGACTCTTAGCGGTGGATCA</td><td>777.7</td><td>-6.4</td><td>0.6</td><td>-10.4</td><td>2.9E-25</td><td>1.3E-23</td><td>MB231/MB468</td><td>MP</td></tr>
<tr><td>>MI0001445|hsa-mir-423&WithFlank&17|+|30117073|30117178@59.77.19(+1A)&[MIMAT0001340&hsa-miR-423-3p&offsets|0|-4(+1A);m-100&17|+|30117131|30117153&offsets|0|-4(+1A)]|AGCTCGGTCTGAGGCCCCTA</td><td>>MI0001445|hsa-mir-423&WithFlank&17|+|30117073|30117178</td><td>59</td><td>77</td><td>19(+1A)</td><td>AGCTCGGTCTGAGGCCCCTA</td><td>487.8</td><td>13.4</td><td>1.3</td><td>10.3</td><td>8.3E-25</td><td>1.3E-23</td><td>BT20/MB468</td><td>MITO</td></tr>
<tr><td>>hg38_21_+_8433172_8446622_RNA45SN1_45S_50ntFlanks@11323.11343.21|CGTCTCGTCGCGCGCGCGTCC</td><td>>hg38_21_+_8433172_8446622_RNA45SN1_45S_50ntFlanks</td><td>11323</td><td>11343</td><td>21</td><td>CGTCTCGTCGCGCGCGCGTCC</td><td>14303.1</td><td>-8.3</td><td>0.8</td><td>-10.3</td><td>5.7E-25</td><td>1.3E-23</td><td>MB231/MB468</td><td>NUC</td></tr>
<tr><td>>hg38_21_+_8433172_8446622_RNA45SN1_45S_50ntFlanks@485.525.41|GATGTGGTGACGTCGTGCTCTCCCGGGCCGGGTCCGAGCCG</td><td>>hg38_21_+_8433172_8446622_RNA45SN1_45S_50ntFlanks</td><td>485</td><td>525</td><td>41</td><td>GATGTGGTGACGTCGTGCTCTCCCGGGCCGGGTCCGAGCCG</td><td>456.5</td><td>13.3</td><td>1.3</td><td>10.3</td><td>8.4E-25</td><td>1.4E-23</td><td>BT20/MB468</td><td>MITO</td></tr>
<tr><td>>hg38_21_+_8433172_8446622_RNA45SN1_45S_50ntFlanks@10690.10709.20|CGTCCGGTGAGCTCTCGCTG</td><td>>hg38_21_+_8433172_8446622_RNA45SN1_45S_50ntFlanks</td><td>10690</td><td>10709</td><td>20</td><td>CGTCCGGTGAGCTCTCGCTG</td><td>1184.2</td><td>-14.1</td><td>1.4</td><td>-10.4</td><td>3.0E-25</td><td>1.4E-23</td><td>MB231/MB468</td><td>MP</td></tr>
<tr><td>>hg38_21_+_8433172_8446622_RNA45SN1_45S_50ntFlanks@12159.12182.24|TTTGACTGGGGCGGTACACCTGTC</td><td>>hg38_21_+_8433172_8446622_RNA45SN1_45S_50ntFlanks</td><td>12159</td><td>12182</td><td>24</td><td>TTTGACTGGGGCGGTACACCTGTC</td><td>2977.0</td><td>-15.3</td><td>1.5</td><td>-10.3</td><td>4.7E-25</td><td>1.4E-23</td><td>MB231/MB468</td><td>MITO</td></tr>
<tr><td>>hg38_21_+_8433172_8446622_RNA45SN1_45S_50ntFlanks@3840.3862.23|CTCCTACTTGGATAACTGTGGTA</td><td>>hg38_21_+_8433172_8446622_RNA45SN1_45S_50ntFlanks</td><td>3840</td><td>3862</td><td>23</td><td>CTCCTACTTGGATAACTGTGGTA</td><td>3217.9</td><td>6.0</td><td>0.6</td><td>10.3</td><td>1.1E-24</td><td>1.4E-23</td><td>BT20/MB468</td><td>CYTO</td></tr>
<tr><td>>trna129_MetCAT_6_-_27870271_27870342@14.31.18,trna142_MetCAT_6_-_27560600_27560671@14.31.18,trna150_MetCAT_6_-_27300764_27300835@14.31.18,trna151_ThrCGT_6_-_27271568_27271639@14.31.18,trna169_MetCAT_6_-_26330529_26330600@14.31.18,trna171_MetCAT_6_-_26313352_26313423@14.31.18,trna20_MetCAT_17_-_80452597_80452668@14.31.18,trna2_MetCAT_6_+_26286754_26286825@14.31.18,trna32_MetCAT_1_+_153643726_153643797@14.31.18,trna61_MetCAT_6_+_27745664_27745735@14.31.18|AGCGGAAGCGTGCTGGGC</td><td>>trna129_MetCAT_6_-_27870271_27870342</td><td>14</td><td>31</td><td>18</td><td>AGCGGAAGCGTGCTGGGC</td><td>774.0</td><td>-14.0</td><td>1.4</td><td>-10.2</td><td>2.4E-24</td><td>1.4E-23</td><td>BT20/MB468</td><td>NUC</td></tr>
<tr><td>>hg38_21_+_8433172_8446622_RNA45SN1_45S_50ntFlanks@12163.12180.18|ACTGGGGCGGTACACCTG</td><td>>hg38_21_+_8433172_8446622_RNA45SN1_45S_50ntFlanks</td><td>12163</td><td>12180</td><td>18</td><td>ACTGGGGCGGTACACCTG</td><td>5269.3</td><td>-16.1</td><td>1.6</td><td>-10.3</td><td>4.9E-25</td><td>1.4E-23</td><td>MB231/MB468</td><td>MITO</td></tr>
<tr><td>>hg38_21_+_8433172_8446622_RNA45SN1_45S_50ntFlanks@11874.11906.33|AGAAGACCCTGTTGAGCTTGACTCTAGTCTGGC</td><td>>hg38_21_+_8433172_8446622_RNA45SN1_45S_50ntFlanks</td><td>11874</td><td>11906</td><td>33</td><td>AGAAGACCCTGTTGAGCTTGACTCTAGTCTGGC</td><td>1468.6</td><td>-14.4</td><td>1.4</td><td>-10.4</td><td>3.1E-25</td><td>1.4E-23</td><td>MB231/MB468</td><td>MP</td></tr>
<tr><td>>am_trna21_ThrTGT_14_-_21081949_21082021@59.76.18|TTCTCGCTGGGGCCTCCA</td><td>>am_trna21_ThrTGT_14_-_21081949_21082021</td><td>59</td><td>76</td><td>18</td><td>TTCTCGCTGGGGCCTCCA</td><td>454.4</td><td>9.6</td><td>0.9</td><td>10.3</td><td>1.1E-24</td><td>1.4E-23</td><td>BT20/MB468</td><td>CYTO</td></tr>
<tr><td>>hg38_21_+_8433172_8446622_RNA45SN1_45S_50ntFlanks@6734.6774.41|ATTGATCATCGACACTTCGAACGCACTTGCGGCCCCGGGTT</td><td>>hg38_21_+_8433172_8446622_RNA45SN1_45S_50ntFlanks</td><td>6734</td><td>6774</td><td>41</td><td>ATTGATCATCGACACTTCGAACGCACTTGCGGCCCCGGGTT</td><td>2706.4</td><td>8.1</td><td>0.8</td><td>10.3</td><td>8.7E-25</td><td>1.4E-23</td><td>BT20/MB468</td><td>MITO</td></tr>
<tr><td>>hg38_21_+_8433172_8446622_RNA45SN1_45S_50ntFlanks@12433.12454.22|CTATCATTGTGAAGCAGAATTC</td><td>>hg38_21_+_8433172_8446622_RNA45SN1_45S_50ntFlanks</td><td>12433</td><td>12454</td><td>22</td><td>CTATCATTGTGAAGCAGAATTC</td><td>582.7</td><td>-13.6</td><td>1.3</td><td>-10.2</td><td>2.4E-24</td><td>1.4E-23</td><td>BT20/MB468</td><td>NUC</td></tr>
<tr><td>>MI0000298|hsa-mir-221&WithFlank&X|-|45746151|45746272@71.92.22&[MIMAT0000278&hsa-miR-221-3p&offsets|0|-1;m-91&X|-|45746181|45746202&offsets|0|0]|AGCTACATTGTCTGCTGGGTTT</td><td>>MI0000298|hsa-mir-221&WithFlank&X|-|45746151|45746272</td><td>71</td><td>92</td><td>22</td><td>AGCTACATTGTCTGCTGGGTTT</td><td>11062.7</td><td>6.5</td><td>0.6</td><td>10.3</td><td>1.2E-24</td><td>1.4E-23</td><td>BT20/MB468</td><td>CYTO</td></tr>
<tr><td>>hg38_21_+_8433172_8446622_RNA45SN1_45S_50ntFlanks@4103.4127.25|CCACGGGTGACGGGGAATCAGGGTT</td><td>>hg38_21_+_8433172_8446622_RNA45SN1_45S_50ntFlanks</td><td>4103</td><td>4127</td><td>25</td><td>CCACGGGTGACGGGGAATCAGGGTT</td><td>579.5</td><td>-13.6</td><td>1.3</td><td>-10.2</td><td>2.4E-24</td><td>1.4E-23</td><td>BT20/MB468</td><td>NUC</td></tr>
<tr><td>>hg38_21_+_8433172_8446622_RNA45SN1_45S_50ntFlanks@9532.9572.41|AGAGGAAACTCTGGTGGAGGTCCGTAGCGGTCCTGACGTGC</td><td>>hg38_21_+_8433172_8446622_RNA45SN1_45S_50ntFlanks</td><td>9532</td><td>9572</td><td>41</td><td>AGAGGAAACTCTGGTGGAGGTCCGTAGCGGTCCTGACGTGC</td><td>853.9</td><td>14.2</td><td>1.4</td><td>10.3</td><td>9.0E-25</td><td>1.4E-23</td><td>BT20/MB468</td><td>MITO</td></tr>
<tr><td>>hg38_21_+_8433172_8446622_RNA45SN1_45S_50ntFlanks@8381.8404.24|AAGAGGGCGTGAAACCGTTAAGAG</td><td>>hg38_21_+_8433172_8446622_RNA45SN1_45S_50ntFlanks</td><td>8381</td><td>8404</td><td>24</td><td>AAGAGGGCGTGAAACCGTTAAGAG</td><td>335.2</td><td>-12.8</td><td>1.3</td><td>-10.2</td><td>2.5E-24</td><td>1.5E-23</td><td>BT20/MB468</td><td>NUC</td></tr>
</table>

| | | | | | | | | | | | | | |
|---|---|---|---|---|---|---|---|---|---|---|---|---|---|
| >hg38_21_+_8433172_8446622_RNA45SN1_45S_50ntFlanks@11208.11234.27\|GCGGCGACTCTGGACGCGAGCCGGGCC | >hg38_21_+_8433172_8446622_RNA45SN1_45S_50ntFlanks | 11208 | 11234 | 27 | GCGGCGACTCTGGACGCGAGCCGGGCC | 2773.8 | -9.8 | 1.0 | -10.3 | 5.2E-25 | 1.5E-23 | MB231/MB468 | MITO |
| >hg38_21_+_8433172_8446622_RNA45SN1_45S_50ntFlanks@12416.12433.18\|TTCGATGTCGGCTCTTCC | >hg38_21_+_8433172_8446622_RNA45SN1_45S_50ntFlanks | 12416 | 12433 | 18 | TTCGATGTCGGCTCTTCC | 380.4 | 4.1 | 0.4 | 10.3 | 9.3E-25 | 1.5E-23 | BT20/MB468 | MITO |
| >hg38_21_+_8433172_8446622_RNA45SN1_45S_50ntFlanks@7983.8005.23\|CAGATCAGACGTGGCGACCCGCT | >hg38_21_+_8433172_8446622_RNA45SN1_45S_50ntFlanks | 7983 | 8005 | 23 | CAGATCAGACGTGGCGACCCGCT | 2557.0 | -7.0 | 0.7 | -10.3 | 5.3E-25 | 1.5E-23 | MB231/MB468 | MITO |
| >hg38_21_+_8433172_8446622_RNA45SN1_45S_50ntFlanks@9180.9197.18\|CCGTCGGGCCCGGGGGAG | >hg38_21_+_8433172_8446622_RNA45SN1_45S_50ntFlanks | 9180 | 9197 | 18 | CCGTCGGGCCCGGGGGAG | 4955.9 | -16.1 | 1.6 | -10.3 | 5.3E-25 | 1.5E-23 | MB231/MB468 | MITO |
| >MI0000086\|hsa-mir-28&WithFlank&3\|+\|188688775\|188688872@60.80.21(+1U)&[MIMAT0004502&hsa-miR-28-3p&offsets\|0\|-1(+1U);m-28&3\|+\|188688834\|188688855&offsets\|0\|-1(+1U)]\|CACTAGATTGTGAGCTCCTGGT | >MI0000086\|hsa-mir-28&WithFlank&3\|+\|188688775\|188688872 | 60 | 80 | 21(+1U) | CACTAGATTGTGAGCTCCTGGT | 1733.9 | 15.0 | 1.5 | 10.2 | 1.2E-24 | 1.5E-23 | BT20/MB468 | CYTO |
| >MI0000542\|hsa-mir-320a&WithFlank&8\|-\|22244960\|22245043@47.68.22(+1U)&[MIMAT0000510&hsa-miR-320a-3p&offsets\|-1\|-1(+1U);m-60&8\|-\|22244975\|22244996&offsets\|-1\|-1(+1U)]\|GAAAAGCTGGGTTGAGAGGGCGT | >MI0000542\|hsa-mir-320a&WithFlank&8\|-\|22244960\|22245043 | 47 | 68 | 22(+1U) | GAAAAGCTGGGTTGAGAGGGCGT | 486.5 | 13.2 | 1.3 | 10.2 | 1.3E-24 | 1.5E-23 | BT20/MB468 | CYTO |
| >MI0000088\|hsa-mir-30a&WithFlank&6\|-\|71403545\|71403627@53.73.21&[MIMAT0000088&hsa-miR-30a-3p&offsets\|0\|-1;m-38&6\|-\|71403554\|71403575&offsets\|0\|-1]\|CTTTCAGTCGGATGTTTGCAG | >MI0000088\|hsa-mir-30a&WithFlank&6\|-\|71403545\|71403627 | 53 | 73 | 21 | CTTTCAGTCGGATGTTTGCAG | 397.3 | 6.4 | 0.6 | 10.3 | 9.6E-25 | 1.5E-23 | BT20/MB468 | MITO |
| >hg38_21_+_8433172_8446622_RNA45SN1_45S_50ntFlanks@11876.11893.18\|AAGACCCTGTTGAGCTTG | >hg38_21_+_8433172_8446622_RNA45SN1_45S_50ntFlanks | 11876 | 11893 | 18 | AAGACCCTGTTGAGCTTG | 4567.5 | -10.2 | 1.0 | -10.4 | 3.4E-25 | 1.5E-23 | MB231/MB468 | MP |
| >hg38_21_+_8433172_8446622_RNA45SN1_45S_50ntFlanks@10808.10829.22\|CAATGTAGGTAAGGGAAGTCGG | >hg38_21_+_8433172_8446622_RNA45SN1_45S_50ntFlanks | 10808 | 10829 | 22 | CAATGTAGGTAAGGGAAGTCGG | 1197.7 | -14.7 | 1.4 | -10.2 | 2.7E-24 | 1.5E-23 | BT20/MB468 | NUC |
| >hg38_21_+_8433172_8446622_RNA45SN1_45S_50ntFlanks@12628.12654.27\|TGGGGCGAAGCTACCATCTGTGGGATT | >hg38_21_+_8433172_8446622_RNA45SN1_45S_50ntFlanks | 12628 | 12654 | 27 | TGGGGCGAAGCTACCATCTGTGGGATT | 3411.8 | -11.6 | 1.1 | -10.3 | 6.6E-25 | 1.6E-23 | MB231/MB468 | NUC |
| >hg38_21_+_8433172_8446622_RNA45SN1_45S_50ntFlanks@10393.10429.37\|CAGCAGTTGAACATGGGTCAGTCGGTCCTGAGAGATG | >hg38_21_+_8433172_8446622_RNA45SN1_45S_50ntFlanks | 10393 | 10429 | 37 | CAGCAGTTGAACATGGGTCAGTCGGTCCTGAGAGATG | 4833.8 | -8.1 | 0.8 | -10.2 | 2.7E-24 | 1.6E-23 | BT20/MB468 | NUC |
| >hg38_21_+_8433172_8446622_RNA45SN1_45S_50ntFlanks@4564.4587.24\|GAATAATGGAATAGGACCGCGGTT | >hg38_21_+_8433172_8446622_RNA45SN1_45S_50ntFlanks | 4564 | 4587 | 24 | GAATAATGGAATAGGACCGCGGTT | 804.8 | -14.1 | 1.4 | -10.2 | 2.7E-24 | 1.6E-23 | BT20/MB468 | NUC |
| >am_MI0000809\|hsa-mir-151a&WithFlank&8\|-\|140732558\|140732659@53.70.18(+3G)&[MIMAT0000757&hsa-miR-151a-3p&offsets\|0\|-3(+3G);m-30&8\|-\|140732587\|140732607&offsets\|0\|-3(+3G)]\|CTAGACTGAAGCTCCTTGGGG | >am_MI0000809\|hsa-mir-151a&WithFlank&8\|-\|140732558\|140732659 | 53 | 70 | 18(+3G) | CTAGACTGAAGCTCCTTGGGG | 504.1 | 13.4 | 1.3 | 10.3 | 1.0E-24 | 1.6E-23 | BT20/MB468 | MITO |
| >hg38_21_+_8433172_8446622_RNA45SN1_45S_50ntFlanks@6740.6770.31\|CATCGACACTTCGAACGCACTTGCGGCCCCG | >hg38_21_+_8433172_8446622_RNA45SN1_45S_50ntFlanks | 6740 | 6770 | 31 | CATCGACACTTCGAACGCACTTGCGGCCCCG | 2162.8 | -5.7 | 0.6 | -10.3 | 1.0E-24 | 1.6E-23 | BT20/MB468 | MITO |
| >hg38_21_+_8433172_8446622_RNA45SN1_45S_50ntFlanks@13002.13028.27\|CGATCTATTGAAAGTCAGCCCTCGACA | >hg38_21_+_8433172_8446622_RNA45SN1_45S_50ntFlanks | 13002 | 13028 | 27 | CGATCTATTGAAAGTCAGCCCTCGACA | 1494.4 | -14.4 | 1.4 | -10.4 | 3.5E-25 | 1.6E-23 | MB231/MB468 | MP |
| >hg38_21_+_8433172_8446622_RNA45SN1_45S_50ntFlanks@3776.3810.35\|CCGGTACAGTGAAACTGCGAATGGCTCATTAAATC | >hg38_21_+_8433172_8446622_RNA45SN1_45S_50ntFlanks | 3776 | 3810 | 35 | CCGGTACAGTGAAACTGCGAATGGCTCATTAAATC | 370.3 | -13.0 | 1.3 | -10.2 | 2.8E-24 | 1.6E-23 | BT20/MB468 | NUC |
| >MI0000440\|hsa-mir-27b&WithFlank&9\|+\|95085439\|95085547@67.85.19(+1A)&[MIMAT0000419&hsa-miR-27b-3p&offsets\|0\|-2(+1A);m-39&9\|+\|95085505\|95085525&offsets\|0\|-2(+1A)]\|TTCACAGTGGCTAAGTTCTA | >MI0000440\|hsa-mir-27b&WithFlank&9\|+\|95085439\|95085547 | 67 | 85 | 19(+1A) | TTCACAGTGGCTAAGTTCTA | 3250.8 | 9.1 | 0.9 | 10.2 | 1.3E-24 | 1.6E-23 | BT20/MB468 | CYTO |
| >hg38_21_+_8433172_8446622_RNA45SN1_45S_50ntFlanks@4733.4758.26\|GAACGAAAGTCGGAGGTTCGAAGACG | >hg38_21_+_8433172_8446622_RNA45SN1_45S_50ntFlanks | 4733 | 4758 | 26 | GAACGAAAGTCGGAGGTTCGAAGACG | 1960.0 | -6.2 | 0.6 | -10.2 | 2.8E-24 | 1.6E-23 | BT20/MB468 | NUC |
| >trnaMT_PheGAA_MT_+_577_647@36.56.21\|AAAATGTTTAGACGGGCTCAC | >trnaMT_PheGAA_MT_+_577_647 | 36 | 56 | 21 | AAAATGTTTAGACGGGCTCAC | 1581.8 | 8.4 | 0.8 | 10.3 | 1.0E-24 | 1.6E-23 | BT20/MB468 | MITO |
| >hg38_21_+_8433172_8446622_RNA45SN1_45S_50ntFlanks@11578.11613.36\|TAGAACTGGTGCGGACCAGGGGAATCCGACTGTTTA | >hg38_21_+_8433172_8446622_RNA45SN1_45S_50ntFlanks | 11578 | 11613 | 36 | TAGAACTGGTGCGGACCAGGGGAATCCGACTGTTTA | 2231.0 | 6.5 | 0.6 | 10.3 | 1.0E-24 | 1.6E-23 | BT20/MB468 | MITO |
| >MI0000102\|hsa-mir-100&WithFlank&11\|-\|122152223\|122152314@19.41.23&[MIMAT0000098&hsa-miR-100-5p&offsets\|0\|+1;m-27&11\|-\|122152276\|122152296&offsets\|0\|+2]\|AACCCGTAGATCCGAACTTGTGG | >MI0000102\|hsa-mir-100&WithFlank&11\|-\|122152223\|122152314 | 19 | 41 | 23 | AACCCGTAGATCCGAACTTGTGG | 1785.1 | 14.0 | 1.4 | 10.4 | 3.7E-25 | 1.6E-23 | MB231/MB468 | MP |
| >hg38_21_+_8433172_8446622_RNA45SN1_45S_50ntFlanks@13002.13030.29\|CGATCTATTGAAAGTCAGCCCTCGACACA | >hg38_21_+_8433172_8446622_RNA45SN1_45S_50ntFlanks | 13002 | 13030 | 29 | CGATCTATTGAAAGTCAGCCCTCGACACA | 505.5 | -13.4 | 1.3 | -10.2 | 2.9E-24 | 1.7E-23 | BT20/MB468 | NUC |
| >hg38_21_+_8433172_8446622_RNA45SN1_45S_50ntFlanks@3984.4004.21\|GCGGCTTTGGTGACTCTAGAT | >hg38_21_+_8433172_8446622_RNA45SN1_45S_50ntFlanks | 3984 | 4004 | 21 | GCGGCTTTGGTGACTCTAGAT | 717.0 | -13.9 | 1.4 | -10.2 | 2.9E-24 | 1.7E-23 | BT20/MB468 | NUC |
| >hg38_21_+_8433172_8446622_RNA45SN1_45S_50ntFlanks@8107.8140.34\|CGGCGGGGCGCGGGACATGTGGCGTACGGAAGAC | >hg38_21_+_8433172_8446622_RNA45SN1_45S_50ntFlanks | 8107 | 8140 | 34 | CGGCGGGGCGCGGGACATGTGGCGTACGGAAGAC | 1176.3 | -14.1 | 1.4 | -10.4 | 3.7E-25 | 1.7E-23 | MB231/MB468 | MP |
| >hg38_21_+_8433172_8446622_RNA45SN1_45S_50ntFlanks@6700.6718.19\|CTAGCTGCGAGAATTAATG | >hg38_21_+_8433172_8446622_RNA45SN1_45S_50ntFlanks | 6700 | 6718 | 19 | CTAGCTGCGAGAATTAATG | 5134.9 | -6.7 | 0.7 | -10.3 | 1.1E-24 | 1.7E-23 | BT20/MB468 | MITO |
| >hg38_21_+_8433172_8446622_RNA45SN1_45S_50ntFlanks@12632.12654.23\|GCGAAGCTACCATCTGTGGGATT | >hg38_21_+_8433172_8446622_RNA45SN1_45S_50ntFlanks | 12632 | 12654 | 23 | GCGAAGCTACCATCTGTGGGATT | 6380.3 | -10.0 | 1.0 | -10.3 | 6.1E-25 | 1.7E-23 | MB231/MB468 | MITO |
| >MI0000478\|hsa-mir-149&WithFlank&2\|+\|240455995\|240456095@21.42.22(+1A)&[MIMAT0000450&hsa-miR-149-5p&offsets\|0\|-1(+1A);m-195&2\|+\|240456015\|240456037&offsets\|0\|-1(+1A)]\|TCTGGCTCCGTGTCTTCACTCCA | >MI0000478\|hsa-mir-149&WithFlank&2\|+\|240455995\|240456095 | 21 | 42 | 22(+1A) | TCTGGCTCCGTGTCTTCACTCCA | 445.7 | 13.3 | 1.3 | 10.3 | 1.1E-24 | 1.7E-23 | BT20/MB468 | MITO |
| >hg38_21_+_8433172_8446622_RNA45SN1_45S_50ntFlanks@10808.10825.18\|CAATGTAGGTAAGGGAAG | >hg38_21_+_8433172_8446622_RNA45SN1_45S_50ntFlanks | 10808 | 10825 | 18 | CAATGTAGGTAAGGGAAG | 3289.7 | -15.5 | 1.5 | -10.3 | 6.3E-25 | 1.8E-23 | MB231/MB468 | MITO |
| >hg38_21_+_8433172_8446622_RNA45SN1_45S_50ntFlanks@10393.10410.18\|CAGCAGTTGAACATGGGT | >hg38_21_+_8433172_8446622_RNA45SN1_45S_50ntFlanks | 10393 | 10410 | 18 | CAGCAGTTGAACATGGGT | 2578.9 | -5.7 | 0.5 | -10.4 | 2.9E-25 | 1.8E-23 | MB231/MB468 | CYTO |
| >MI0001445\|hsa-mir-423&WithFlank&17\|+\|30117073\|30117178@23.44.22&[MIMAT0004748&hsa-miR-423-5p&offsets\|0\|-1;m-141&17\|+\|30117095\|30117117&offsets\|0\|-1]\|TGAGGGGCAGAGAGCGAGACTT | >MI0001445\|hsa-mir-423&WithFlank&17\|+\|30117073\|30117178 | 23 | 44 | 22 | TGAGGGGCAGAGAGCGAGACTT | 441.1 | 13.2 | 1.3 | 10.3 | 1.1E-24 | 1.8E-23 | BT20/MB468 | MITO |
| >hg38_21_+_8433172_8446622_RNA45SN1_45S_50ntFlanks@11674.11692.19\|GTGCTCTGAATGTCAAAGT | >hg38_21_+_8433172_8446622_RNA45SN1_45S_50ntFlanks | 11674 | 11692 | 19 | GTGCTCTGAATGTCAAAGT | 410.2 | 13.1 | 1.3 | 10.3 | 1.2E-24 | 1.8E-23 | BT20/MB468 | MITO |
| >MI0000067\|hsa-let-7f-1&WithFlank&9\|+\|94176341\|94176439@13.30.18;MI0000068\|hsa-let-7f-2&WithFlank&X\|-\|53557186\|53557280@14.31.18&[MIMAT0000067&hsa-let-7f-5p&offsets\|0\|-4;m-20&9\|+\|94176353\|94176374&offsets\|0\|-4];[MIMAT0000067_1&hsa-let-7f-5p&offsets\|0\|-4;m-19&X\|-\|53557246\|53557267&offsets\|0\|-4]\|TGAGGTAGTAGATTGTAT | >MI0000067\|hsa-let-7f-1&WithFlank&9\|+\|94176341\|94176439 | 13 | 30 | 18 | TGAGGTAGTAGATTGTAT | 3324.4 | 7.3 | 0.7 | 10.3 | 1.2E-24 | 1.8E-23 | BT20/MB468 | MITO |
| >hg38_21_+_8433172_8446622_RNA45SN1_45S_50ntFlanks@10402.10430.29\|AACATGGGTCAGTCGGTCCTGAGAGATGG | >hg38_21_+_8433172_8446622_RNA45SN1_45S_50ntFlanks | 10402 | 10430 | 29 | AACATGGGTCAGTCGGTCCTGAGAGATGG | 2211.5 | -9.4 | 0.9 | -10.4 | 4.1E-25 | 1.8E-23 | MB231/MB468 | MP |
| >hg38_21_+_8433172_8446622_RNA45SN1_45S_50ntFlanks@11161.11178.18\|CACGGTCCCCCGCGAGGG | >hg38_21_+_8433172_8446622_RNA45SN1_45S_50ntFlanks | 11161 | 11178 | 18 | CACGGTCCCCCGCGAGGG | 1673.0 | -13.7 | 1.3 | -10.3 | 7.9E-25 | 1.8E-23 | MB231/MB468 | NUC |
| >MI0001445\|hsa-mir-423&WithFlank&17\|+\|30117073\|30117178@59.80.22(+1G)&[MIMAT0001340&hsa-miR-423-3p&offsets\|0\|-1(+1G);m-100&17\|+\|30117131\|30117153&offsets\|0\|-1(+1G)]\|AGCTCGGTCTGAGGCCCCTCAGG | >MI0001445\|hsa-mir-423&WithFlank&17\|+\|30117073\|30117178 | 59 | 80 | 22(+1G) | AGCTCGGTCTGAGGCCCCTCAGG | 464.0 | 7.0 | 0.7 | 10.2 | 1.2E-24 | 1.8E-23 | BT20/MB468 | MITO |
| >MI0000746\|hsa-mir-99b&WithFlank&19\|+\|51692606\|51692687@13.33.21(+1U)&[MIMAT0000689&hsa-miR-99b-5p&offsets\|0\|-1(+1U);m-10&19\|+\|51692618\|51692639&offsets\|0\|-1(+1U)]\|CACCCGTAGAACCGACCTTGCT | >MI0000746\|hsa-mir-99b&WithFlank&19\|+\|51692606\|51692687 | 13 | 33 | 21(+1U) | CACCCGTAGAACCGACCTTGCT | 497.6 | 6.5 | 0.6 | 10.2 | 1.2E-24 | 1.9E-23 | BT20/MB468 | MITO |
| >MI0000087\|hsa-mir-29a&WithFlank&7\|-\|130876741\|130876816@48.68.21&[MIMAT0000086&hsa-miR-29a-3p&offsets\|0\|-1;m-26&7\|-\|130876749\|130876769&offsets\|0\|0]\|TAGCACCATCTGAAATCGGTT | >MI0000087\|hsa-mir-29a&WithFlank&7\|-\|130876741\|130876816 | 48 | 68 | 21 | TAGCACCATCTGAAATCGGTT | 4687.4 | 8.2 | 0.8 | 10.2 | 1.2E-24 | 1.9E-23 | BT20/MB468 | MITO |
| >hg38_21_+_8433172_8446622_RNA45SN1_45S_50ntFlanks@12615.12654.40\|GCTGAGGAGCCAATGGGGCGAAGCTACCATCTGTGGGATT | >hg38_21_+_8433172_8446622_RNA45SN1_45S_50ntFlanks | 12615 | 12654 | 40 | GCTGAGGAGCCAATGGGGCGAAGCTACCATCTGTGGGATT | 17002.0 | -11.1 | 1.1 | -10.3 | 6.8E-25 | 1.9E-23 | MB231/MB468 | MITO |
| >hg38_21_+_8433172_8446622_RNA45SN1_45S_50ntFlanks@11594.11611.18\|CAGGGGAATCCGACTGTT | >hg38_21_+_8433172_8446622_RNA45SN1_45S_50ntFlanks | 11594 | 11611 | 18 | CAGGGGAATCCGACTGTT | 2025.4 | -7.0 | 0.7 | -10.3 | 4.2E-25 | 1.9E-23 | MB231/MB468 | MP |
| >hg38_21_+_8433172_8446622_RNA45SN1_45S_50ntFlanks@4730.4759.30\|CAAGAACGAAAGTCGGAGGTTCGAAGACGA | >hg38_21_+_8433172_8446622_RNA45SN1_45S_50ntFlanks | 4730 | 4759 | 30 | CAAGAACGAAAGTCGGAGGTTCGAAGACGA | 1710.3 | -6.7 | 0.6 | -10.3 | 4.2E-25 | 1.9E-23 | MB231/MB468 | MP |
| >MI0000301\|hsa-mir-224&WithFlank&X\|-\|151958572\|151958664@14.34.21(+1C)&[MIMAT0000281&hsa-miR-224-5p&offsets\|+1\|-3(+1C);m-238&X\|-\|151958630\|151958651&offsets\|0\|-1(+1C)]\|CAAGTCACTAGTGGTTCCGTTC | >MI0000301\|hsa-mir-224&WithFlank&X\|-\|151958572\|151958664 | 14 | 34 | 21(+1C) | CAAGTCACTAGTGGTTCCGTTC | 1088.1 | 14.5 | 1.4 | 10.2 | 1.2E-24 | 1.9E-23 | BT20/MB468 | MITO |
| >hg38_1_-_228634819_228635039_RNA5S12_5S_50ntFlanks@145.170.26\|CTGGGAATACCGGGTGCTGTAGGCTT | >hg38_1_-_228634819_228635039_RNA5S12_5S_50ntFlanks | 145 | 170 | 26 | CTGGGAATACCGGGTGCTGTAGGCTT | 3084.6 | 6.9 | 0.7 | 10.2 | 1.6E-24 | 1.9E-23 | BT20/MB468 | CYTO |
| >hg38_21_+_8433172_8446622_RNA45SN1_45S_50ntFlanks@10841.10858.18\|CGTAACTTCGGGATAAGG | >hg38_21_+_8433172_8446622_RNA45SN1_45S_50ntFlanks | 10841 | 10858 | 18 | CGTAACTTCGGGATAAGG | 608.1 | -13.7 | 1.3 | -10.1 | 3.3E-24 | 1.9E-23 | BT20/MB468 | NUC |
| >MI0000542\|hsa-mir-320a&WithFlank&8\|-\|22244960\|22245043@48.69.22(+3U)&[MIMAT0000510&hsa-miR-320a-3p&offsets\|0\|0(+3U);m-60&8\|-\|22244975\|22244996&offsets\|0\|0(+3U)]\|AAAAGCTGGGTTGAGAGGGCGATTT | >MI0000542\|hsa-mir-320a&WithFlank&8\|-\|22244960\|22245043 | 48 | 69 | 22(+3U) | AAAAGCTGGGTTGAGAGGGCGATTT | 1114.0 | 12.1 | 1.2 | 10.2 | 1.6E-24 | 1.9E-23 | BT20/MB468 | CYTO |
| >MI0000471\|hsa-mir-126&WithFlank&9\|+\|136670596\|136670692@58.77.20(+1U)&[MIMAT0000445&hsa-miR-126-3p&offsets\|0\|-2(+1U);m-40&9\|+\|136670653\|136670674&offsets\|0\|-2(+1U)]\|TCGTACCGTGAGTAATAATGT | >MI0000471\|hsa-mir-126&WithFlank&9\|+\|136670596\|136670692 | 58 | 77 | 20(+1U) | TCGTACCGTGAGTAATAATGT | 495.1 | 12.8 | 1.2 | 10.3 | 8.4E-25 | 1.9E-23 | MB231/MB468 | NUC |
| >hg38_21_+_8433172_8446622_RNA45SN1_45S_50ntFlanks@7986.8006.21\|ATCAGACGTGGCGACCCGCTG | >hg38_21_+_8433172_8446622_RNA45SN1_45S_50ntFlanks | 7986 | 8006 | 21 | ATCAGACGTGGCGACCCGCTG | 6600.9 | -14.8 | 1.4 | -10.2 | 1.3E-24 | 2.0E-23 | BT20/MB468 | MITO |
| >hg38_21_+_8433172_8446622_RNA45SN1_45S_50ntFlanks@11848.11872.25\|ACGGGCTTGGCGGAATCAGCGGGGA | >hg38_21_+_8433172_8446622_RNA45SN1_45S_50ntFlanks | 11848 | 11872 | 25 | ACGGGCTTGGCGGAATCAGCGGGGA | 1350.3 | -14.3 | 1.4 | -10.3 | 4.4E-25 | 2.0E-23 | MB231/MB468 | MP |

<table>
<tr><td>>MI0000093|hsa-mir-92a-1&WithFlank&13|+|91351308|91351397@54.75.22(+1A);MI0000094|hsa-mir-92a-2&WithFlank&X|-|134169532|134169618@54.75.22(+1A)&[MIMAT0000092&hsa-miR-92a-3p&offsets|0|0(+1A);m-9&13|+|91351361|91351382&offsets|0|0(+1A)];[MIMAT0000092_1&hsa-miR-92a-3p&offsets|0|0(+1A);m-11&X|-|134169544|134169565&offsets|0|0(+1A)]|TATTGCACTTGTCCCGGCCTGTA</td><td>>MI0000093|hsa-mir-92a-1&WithFlank&13|+|91351308|91351397</td><td>54</td><td>75</td><td>22(+1A)</td><td>TATTGCACTTGTCCCGGCCTGTA</td><td>14645.8</td><td>6.3</td><td>0.6</td><td>10.2</td><td>1.6E-24</td><td>2.0E-23</td><td>BT20/MB468</td><td>CYTO</td></tr>
<tr><td>>hg38_21_+_8433172_8446622_RNA45SN1_45S_50ntFlanks@12357.12376.20|CCACAGGGATAACTGGCTTG</td><td>>hg38_21_+_8433172_8446622_RNA45SN1_45S_50ntFlanks</td><td>12357</td><td>12376</td><td>20</td><td>CCACAGGGATAACTGGCTTG</td><td>1311.6</td><td>-14.2</td><td>1.4</td><td>-10.3</td><td>4.5E-25</td><td>2.0E-23</td><td>MB231/MB468</td><td>MP</td></tr>
<tr><td>>hg38_21_+_8433172_8446622_RNA45SN1_45S_50ntFlanks@7974.8009.36|ACGCGACCTCAGATCAGACGTGGCGACCCGCTGAAT</td><td>>hg38_21_+_8433172_8446622_RNA45SN1_45S_50ntFlanks</td><td>7974</td><td>8009</td><td>36</td><td>ACGCGACCTCAGATCAGACGTGGCGACCCGCTGAAT</td><td>20543.7</td><td>7.9</td><td>0.8</td><td>10.4</td><td>3.3E-25</td><td>2.0E-23</td><td>MB231/MB468</td><td>CYTO</td></tr>
<tr><td>>hg38_21_+_8433172_8446622_RNA45SN1_45S_50ntFlanks@11873.11908.36|AAGAAGACCCTGTTGAGCTTGACTCTAGTCTGGCAC</td><td>>hg38_21_+_8433172_8446622_RNA45SN1_45S_50ntFlanks</td><td>11873</td><td>11908</td><td>36</td><td>AAGAAGACCCTGTTGAGCTTGACTCTAGTCTGGCAC</td><td>596.1</td><td>-13.6</td><td>1.3</td><td>-10.1</td><td>3.5E-24</td><td>2.0E-23</td><td>BT20/MB468</td><td>NUC</td></tr>
<tr><td>>MI0000542|hsa-mir-320a&WithFlank&8|-|22244960|22245043@48.69.22(+1C)&[MIMAT0000510&hsa-miR-320a-3p&offsets|0|0(+1C);m-60&8|-|22244975|22244996&offsets|0|0(+1C)]|AAAAGCTGGGTTGAGAGGGCGAC</td><td>>MI0000542|hsa-mir-320a&WithFlank&8|-|22244960|22245043</td><td>48</td><td>69</td><td>22(+1C)</td><td>AAAAGCTGGGTTGAGAGGGCGAC</td><td>6342.8</td><td>13.8</td><td>1.4</td><td>10.2</td><td>1.7E-24</td><td>2.0E-23</td><td>BT20/MB468</td><td>CYTO</td></tr>
<tr><td>>MI0000456|hsa-mir-140&WithFlank&16|+|69933075|69933186@69.91.23&[MIMAT0004597&hsa-miR-140-3p&offsets|+1|+3;m-45&16|+|69933143|69933164&offsets|0|+1]|ACCACAGGGTAGAACCACGGACA</td><td>>MI0000456|hsa-mir-140&WithFlank&16|+|69933075|69933186</td><td>69</td><td>91</td><td>23</td><td>ACCACAGGGTAGAACCACGGACA</td><td>150.6</td><td>5.0</td><td>0.5</td><td>10.1</td><td>3.5E-24</td><td>2.0E-23</td><td>BT20/MB468</td><td>NUC</td></tr>
<tr><td>>hg38_21_+_8433172_8446622_RNA45SN1_45S_50ntFlanks@10808.10825.18|CAATGTAGGTAAGGGAAG</td><td>>hg38_21_+_8433172_8446622_RNA45SN1_45S_50ntFlanks</td><td>10808</td><td>10825</td><td>18</td><td>CAATGTAGGTAAGGGAAG</td><td>3105.0</td><td>-16.0</td><td>1.6</td><td>-10.1</td><td>3.5E-24</td><td>2.0E-23</td><td>BT20/MB468</td><td>NUC</td></tr>
<tr><td>>hg38_21_+_8433172_8446622_RNA45SN1_45S_50ntFlanks@11849.11873.25|CGGGCTTGGCGGAATCAGCGGGGAA</td><td>>hg38_21_+_8433172_8446622_RNA45SN1_45S_50ntFlanks</td><td>11849</td><td>11873</td><td>25</td><td>CGGGCTTGGCGGAATCAGCGGGGAA</td><td>327.6</td><td>-12.8</td><td>1.3</td><td>-10.1</td><td>3.6E-24</td><td>2.1E-23</td><td>BT20/MB468</td><td>NUC</td></tr>
<tr><td>>hg38_21_+_8433172_8446622_RNA45SN1_45S_50ntFlanks@11612.11640.29|TAATTAAAACAAAGCATCGCGAAGGCCCG</td><td>>hg38_21_+_8433172_8446622_RNA45SN1_45S_50ntFlanks</td><td>11612</td><td>11640</td><td>29</td><td>TAATTAAAACAAAGCATCGCGAAGGCCCG</td><td>1266.2</td><td>-14.2</td><td>1.4</td><td>-10.3</td><td>4.7E-25</td><td>2.1E-23</td><td>MB231/MB468</td><td>MP</td></tr>
<tr><td>>MI0000734|hsa-mir-106b&WithFlank&7|-|100093987|100094080@56.77.22&[MIMAT0004672&hsa-miR-106b-3p&offsets|-2|-2;m-87&7|-|100094003|100094023&offsets|-2|-1]|TACCGCACTGTGGGTACTTGCT</td><td>>MI0000734|hsa-mir-106b&WithFlank&7|-|100093987|100094080</td><td>56</td><td>77</td><td>22</td><td>TACCGCACTGTGGGTACTTGCT</td><td>2163.1</td><td>9.5</td><td>0.9</td><td>10.2</td><td>1.7E-24</td><td>2.1E-23</td><td>BT20/MB468</td><td>CYTO</td></tr>
<tr><td>>MI0000261|hsa-mir-139&WithFlank&11|-|72615057|72615136@13.35.23&[MIMAT0000250&hsa-miR-139-5p&offsets|0|0;m-186&11|-|72615102|72615124&offsets|0|0]|TCTACAGTGCACGTGTCTCCAGT</td><td>>MI0000261|hsa-mir-139&WithFlank&11|-|72615057|72615136</td><td>13</td><td>35</td><td>23</td><td>TCTACAGTGCACGTGTCTCCAGT</td><td>2083.3</td><td>15.0</td><td>1.4</td><td>10.4</td><td>3.5E-25</td><td>2.1E-23</td><td>MB231/MB468</td><td>CYTO</td></tr>
<tr><td>>hg38_21_+_8433172_8446622_RNA45SN1_45S_50ntFlanks@12407.12426.20|TTTTGATCCTTCGATGTCGG</td><td>>hg38_21_+_8433172_8446622_RNA45SN1_45S_50ntFlanks</td><td>12407</td><td>12426</td><td>20</td><td>TTTTGATCCTTCGATGTCGG</td><td>581.1</td><td>-8.2</td><td>0.8</td><td>-10.4</td><td>3.5E-25</td><td>2.1E-23</td><td>MB231/MB468</td><td>CYTO</td></tr>
<tr><td>>trnaMT_SerGCT_MT_+_12207_12265@10.38.29|CACAAGAACTGCTAACTCATGCCCCCATG</td><td>>trnaMT_SerGCT_MT_+_12207_12265</td><td>10</td><td>38</td><td>29</td><td>CACAAGAACTGCTAACTCATGCCCCCATG</td><td>3101.5</td><td>-16.0</td><td>1.6</td><td>-10.1</td><td>3.7E-24</td><td>2.1E-23</td><td>BT20/MB468</td><td>NUC</td></tr>
<tr><td>>hg38_21_+_8433172_8446622_RNA45SN1_45S_50ntFlanks@12936.12956.21|AAACCATTCGTAGACGACCTG</td><td>>hg38_21_+_8433172_8446622_RNA45SN1_45S_50ntFlanks</td><td>12936</td><td>12956</td><td>21</td><td>AAACCATTCGTAGACGACCTG</td><td>5449.9</td><td>-9.8</td><td>1.0</td><td>-10.3</td><td>7.8E-25</td><td>2.1E-23</td><td>MB231/MB468</td><td>MITO</td></tr>
<tr><td>>hg38_21_+_8433172_8446622_RNA45SN1_45S_50ntFlanks@11641.11685.45|CGGCGGGTGTTGACGCGATGTGATTTCTGCCCAGTGCTCTGAATG</td><td>>hg38_21_+_8433172_8446622_RNA45SN1_45S_50ntFlanks</td><td>11641</td><td>11685</td><td>45</td><td>CGGCGGGTGTTGACGCGATGTGATTTCTGCCCAGTGCTCTGAATG</td><td>3515.9</td><td>-16.2</td><td>1.6</td><td>-10.1</td><td>3.7E-24</td><td>2.2E-23</td><td>BT20/MB468</td><td>NUC</td></tr>
<tr><td>>hg38_21_+_8433172_8446622_RNA45SN1_45S_50ntFlanks@7986.8004.19|ATCAGACGTGGCGACCCGC</td><td>>hg38_21_+_8433172_8446622_RNA45SN1_45S_50ntFlanks</td><td>7986</td><td>8004</td><td>19</td><td>ATCAGACGTGGCGACCCGC</td><td>696.2</td><td>-13.9</td><td>1.4</td><td>-10.1</td><td>3.8E-24</td><td>2.2E-23</td><td>BT20/MB468</td><td>NUC</td></tr>
<tr><td>>hg38_21_+_8433172_8446622_RNA45SN1_45S_50ntFlanks@3813.3842.30|TTATGGTTCCTTTGGTCGCTCGCTCCTCTC</td><td>>hg38_21_+_8433172_8446622_RNA45SN1_45S_50ntFlanks</td><td>3813</td><td>3842</td><td>30</td><td>TTATGGTTCCTTTGGTCGCTCGCTCCTCTC</td><td>718.2</td><td>-13.9</td><td>1.4</td><td>-10.1</td><td>3.8E-24</td><td>2.2E-23</td><td>BT20/MB468</td><td>NUC</td></tr>
<tr><td>>hg38_21_+_8433172_8446622_RNA45SN1_45S_50ntFlanks@11666.11707.42|TCTGCCCAGTGCTCTGAATGTCAAAGTGAAGAAATTCAATGA</td><td>>hg38_21_+_8433172_8446622_RNA45SN1_45S_50ntFlanks</td><td>11666</td><td>11707</td><td>42</td><td>TCTGCCCAGTGCTCTGAATGTCAAAGTGAAGAAATTCAATGA</td><td>585.7</td><td>13.2</td><td>1.3</td><td>10.4</td><td>3.8E-25</td><td>2.2E-23</td><td>MB231/MB468</td><td>CYTO</td></tr>
<tr><td>>hg38_21_+_8433172_8446622_RNA45SN1_45S_50ntFlanks@4102.4129.28|ACCACGGGTGACGGGGAATCAGGGTTCG</td><td>>hg38_21_+_8433172_8446622_RNA45SN1_45S_50ntFlanks</td><td>4102</td><td>4129</td><td>28</td><td>ACCACGGGTGACGGGGAATCAGGGTTCG</td><td>475.5</td><td>-13.3</td><td>1.3</td><td>-10.1</td><td>3.9E-24</td><td>2.2E-23</td><td>BT20/MB468</td><td>NUC</td></tr>
<tr><td>>hg38_21_+_8433172_8446622_RNA45SN1_45S_50ntFlanks@6740.6771.32|CATCGACACTTCGAACGCACTTGCGGCCCCGG</td><td>>hg38_21_+_8433172_8446622_RNA45SN1_45S_50ntFlanks</td><td>6740</td><td>6771</td><td>32</td><td>CATCGACACTTCGAACGCACTTGCGGCCCCGG</td><td>1400.7</td><td>-6.4</td><td>0.6</td><td>-10.3</td><td>9.6E-25</td><td>2.2E-23</td><td>MB231/MB468</td><td>NUC</td></tr>
<tr><td>>hg38_21_+_8433172_8446622_RNA45SN1_45S_50ntFlanks@10781.10801.21|AGGTGAACAGCCTCTGGCATG</td><td>>hg38_21_+_8433172_8446622_RNA45SN1_45S_50ntFlanks</td><td>10781</td><td>10801</td><td>21</td><td>AGGTGAACAGCCTCTGGCATG</td><td>1392.5</td><td>-14.3</td><td>1.4</td><td>-10.3</td><td>5.1E-25</td><td>2.2E-23</td><td>MB231/MB468</td><td>MP</td></tr>
<tr><td>>hg38_21_+_8433172_8446622_RNA45SN1_45S_50ntFlanks@7986.8008.23|ATCAGACGTGGCGACCCGCTGAA</td><td>>hg38_21_+_8433172_8446622_RNA45SN1_45S_50ntFlanks</td><td>7986</td><td>8008</td><td>23</td><td>ATCAGACGTGGCGACCCGCTGAA</td><td>2855.3</td><td>-15.3</td><td>1.5</td><td>-10.3</td><td>8.1E-25</td><td>2.2E-23</td><td>MB231/MB468</td><td>MITO</td></tr>
<tr><td>>hg38_21_+_8433172_8446622_RNA45SN1_45S_50ntFlanks@10350.10383.34|ACGAGAACTTTGAAGGCCGAAGTGGAGAAGGGTT</td><td>>hg38_21_+_8433172_8446622_RNA45SN1_45S_50ntFlanks</td><td>10350</td><td>10383</td><td>34</td><td>ACGAGAACTTTGAAGGCCGAAGTGGAGAAGGGTT</td><td>17234.9</td><td>-8.5</td><td>0.8</td><td>-10.3</td><td>5.2E-25</td><td>2.3E-23</td><td>MB231/MB468</td><td>MP</td></tr>
<tr><td>>MI0000478|hsa-mir-149&WithFlank&2|+|240455995|240456095@21.41.21(+1U)&[MIMAT0000450&hsa-miR-149-5p&offsets|0|-2(+1U);m-195&2|+|240456015|240456037&offsets|0|-2(+1U)]|TCTGGCTCCGTGTCTTCACTCT</td><td>>MI0000478|hsa-mir-149&WithFlank&2|+|240455995|240456095</td><td>21</td><td>41</td><td>21(+1U)</td><td>TCTGGCTCCGTGTCTTCACTCT</td><td>1853.0</td><td>15.1</td><td>1.5</td><td>10.2</td><td>1.9E-24</td><td>2.3E-23</td><td>BT20/MB468</td><td>CYTO</td></tr>
<tr><td>>hg38_21_+_8433172_8446622_RNA45SN1_45S_50ntFlanks@9939.9968.30|GCAGGACGGTGGCCATGGAAGTCGGAATCC</td><td>>hg38_21_+_8433172_8446622_RNA45SN1_45S_50ntFlanks</td><td>9939</td><td>9968</td><td>30</td><td>GCAGGACGGTGGCCATGGAAGTCGGAATCC</td><td>296.5</td><td>6.5</td><td>0.6</td><td>10.2</td><td>1.5E-24</td><td>2.3E-23</td><td>BT20/MB468</td><td>MITO</td></tr>
<tr><td>>hg38_21_+_8433172_8446622_RNA45SN1_45S_50ntFlanks@8169.8187.19|CCAAGTCCTTCTGATCGAG</td><td>>hg38_21_+_8433172_8446622_RNA45SN1_45S_50ntFlanks</td><td>8169</td><td>8187</td><td>19</td><td>CCAAGTCCTTCTGATCGAG</td><td>548.4</td><td>-13.5</td><td>1.3</td><td>-10.1</td><td>4.1E-24</td><td>2.4E-23</td><td>BT20/MB468</td><td>NUC</td></tr>
<tr><td>>trna32_LysCTT_16_-_3207406_3207478@1.35.35|GCCCGGCTAGCTCAGTCGGTAGAGCATGAGACCCT</td><td>>trna32_LysCTT_16_-_3207406_3207478</td><td>1</td><td>35</td><td>35</td><td>GCCCGGCTAGCTCAGTCGGTAGAGCATGAGACCCT</td><td>453.6</td><td>13.1</td><td>1.3</td><td>10.2</td><td>2.0E-24</td><td>2.4E-23</td><td>BT20/MB468</td><td>CYTO</td></tr>
<tr><td>>MI0000239|hsa-mir-197&WithFlank&1|+|109598887|109598973@54.75.22(+1U)&[MIMAT0000227&hsa-miR-197-3p&offsets|0|0(+1U);m-99&1|+|109598940|109598961&offsets|0|0(+1U)]|TTCACCACCTTCTCCACCCAGCT</td><td>>MI0000239|hsa-mir-197&WithFlank&1|+|109598887|109598973</td><td>54</td><td>75</td><td>22(+1U)</td><td>TTCACCACCTTCTCCACCCAGCT</td><td>524.5</td><td>13.3</td><td>1.3</td><td>10.2</td><td>2.0E-24</td><td>2.4E-23</td><td>BT20/MB468</td><td>CYTO</td></tr>
<tr><td>>hg38_1_-_228634819_228635039_RNA5S12_5S_50ntFlanks@84.101.18|CTCGTCTGATCTCGGAAG</td><td>>hg38_1_-_228634819_228635039_RNA5S12_5S_50ntFlanks</td><td>84</td><td>101</td><td>18</td><td>CTCGTCTGATCTCGGAAG</td><td>1530.4</td><td>-14.4</td><td>1.4</td><td>-10.3</td><td>5.6E-25</td><td>2.4E-23</td><td>MB231/MB468</td><td>MP</td></tr>
<tr><td>>hg38_21_+_8433172_8446622_RNA45SN1_45S_50ntFlanks@9739.9757.19|AAACGATCTCAACCTATTC</td><td>>hg38_21_+_8433172_8446622_RNA45SN1_45S_50ntFlanks</td><td>9739</td><td>9757</td><td>19</td><td>AAACGATCTCAACCTATTC</td><td>1360.4</td><td>-14.3</td><td>1.4</td><td>-10.3</td><td>5.6E-25</td><td>2.4E-23</td><td>MB231/MB468</td><td>MP</td></tr>
<tr><td>>hg38_21_+_8433172_8446622_RNA45SN1_45S_50ntFlanks@6690.6720.31|AAGAACGCAGCTAGCTGCGAGAATTAATGTG</td><td>>hg38_21_+_8433172_8446622_RNA45SN1_45S_50ntFlanks</td><td>6690</td><td>6720</td><td>31</td><td>AAGAACGCAGCTAGCTGCGAGAATTAATGTG</td><td>1091.2</td><td>-14.0</td><td>1.4</td><td>-10.3</td><td>5.7E-25</td><td>2.4E-23</td><td>MB231/MB468</td><td>MP</td></tr>
<tr><td>>hg38_21_+_8433172_8446622_RNA45SN1_45S_50ntFlanks@3987.4024.38|GCTTTGGTGACTCTAGATAACCTCGGGCCGATCGCACG</td><td>>hg38_21_+_8433172_8446622_RNA45SN1_45S_50ntFlanks</td><td>3987</td><td>4024</td><td>38</td><td>GCTTTGGTGACTCTAGATAACCTCGGGCCGATCGCACG</td><td>1361.9</td><td>-6.8</td><td>0.7</td><td>-10.3</td><td>9.0E-25</td><td>2.5E-23</td><td>MB231/MB468</td><td>MITO</td></tr>
<tr><td>>hg38_21_+_8433172_8446622_RNA45SN1_45S_50ntFlanks@12157.12181.25|AGTTTGACTGGGGCGGTACACCTGT</td><td>>hg38_21_+_8433172_8446622_RNA45SN1_45S_50ntFlanks</td><td>12157</td><td>12181</td><td>25</td><td>AGTTTGACTGGGGCGGTACACCTGT</td><td>1925.9</td><td>-14.7</td><td>1.4</td><td>-10.3</td><td>9.1E-25</td><td>2.5E-23</td><td>MB231/MB468</td><td>MITO</td></tr>
<tr><td>>am_MI0000809|hsa-mir-151a&WithFlank&8|-|140732558|140732659@53.73.21(+1C)&[MIMAT0000757&hsa-miR-151a-3p&offsets|0|0(+1C);m-30&8|-|140732587|140732607&offsets|0|0(+1C)]|CTAGACTGAAGCTCCTTGAGGC</td><td>>am_MI0000809|hsa-mir-151a&WithFlank&8|-|140732558|140732659</td><td>53</td><td>73</td><td>21(+1C)</td><td>CTAGACTGAAGCTCCTTGAGGC</td><td>1019.3</td><td>6.2</td><td>0.6</td><td>10.3</td><td>1.1E-24</td><td>2.5E-23</td><td>MB231/MB468</td><td>NUC</td></tr>
<tr><td>>am_MI0031829|hsa-mir-9901&WithFlank&16|+|34163572|34163679@25.50.26&[MIMAT0039321&hsa-miR-9901&offsets|-5|0;TJU_CMC_MD2.ID01155.5p-miR&offsets|0|+3;m-2778&16|+|34163596|34163618&offsets|0|+3]|CGTCCCGGTCGCCGCGGTTCGCCGCC</td><td>>am_MI0031829|hsa-mir-9901&WithFlank&16|+|34163572|34163679</td><td>25</td><td>50</td><td>26</td><td>CGTCCCGGTCGCCGCGGTTCGCCGCC</td><td>500.6</td><td>-13.4</td><td>1.3</td><td>-10.1</td><td>4.4E-24</td><td>2.5E-23</td><td>BT20/MB468</td><td>NUC</td></tr>
<tr><td>>hg38_21_+_8433172_8446622_RNA45SN1_45S_50ntFlanks@7340.7365.26|CGTCCCGGTCGCCGCGGTTCGCCGCC</td><td>>hg38_21_+_8433172_8446622_RNA45SN1_45S_50ntFlanks</td><td>7340</td><td>7365</td><td>26</td><td>CGTCCCGGTCGCCGCGGTTCGCCGCC</td><td>500.6</td><td>-13.4</td><td>1.3</td><td>-10.1</td><td>4.4E-24</td><td>2.5E-23</td><td>BT20/MB468</td><td>NUC</td></tr>
<tr><td>>hg38_21_+_8433172_8446622_RNA45SN1_45S_50ntFlanks@12578.12604.27|AGGAACCGCAGGTTCAGACATTTGGTG</td><td>>hg38_21_+_8433172_8446622_RNA45SN1_45S_50ntFlanks</td><td>12578</td><td>12604</td><td>27</td><td>AGGAACCGCAGGTTCAGACATTTGGTG</td><td>1016.2</td><td>-13.9</td><td>1.3</td><td>-10.3</td><td>5.8E-25</td><td>2.5E-23</td><td>MB231/MB468</td><td>MP</td></tr>
<tr><td>>hg38_21_+_8433172_8446622_RNA45SN1_45S_50ntFlanks@8114.8141.28|GCGCGGGACATGTGGCGTACGGAAGACC</td><td>>hg38_21_+_8433172_8446622_RNA45SN1_45S_50ntFlanks</td><td>8114</td><td>8141</td><td>28</td><td>GCGCGGGACATGTGGCGTACGGAAGACC</td><td>1973.8</td><td>-14.7</td><td>1.4</td><td>-10.3</td><td>9.5E-25</td><td>2.6E-23</td><td>MB231/MB468</td><td>MITO</td></tr>
<tr><td>>hg38_21_+_8433172_8446622_RNA45SN1_45S_50ntFlanks@4370.4396.27|TTAAAAAGCTCGTAGTTGGATCTTGGG</td><td>>hg38_21_+_8433172_8446622_RNA45SN1_45S_50ntFlanks</td><td>4370</td><td>4396</td><td>27</td><td>TTAAAAAGCTCGTAGTTGGATCTTGGG</td><td>293.8</td><td>-12.6</td><td>1.2</td><td>-10.1</td><td>4.5E-24</td><td>2.6E-23</td><td>BT20/MB468</td><td>NUC</td></tr>
<tr><td>>hg38_21_+_8433172_8446622_RNA45SN1_45S_50ntFlanks@12935.12953.19|TAAACCATTCGTAGACGAC</td><td>>hg38_21_+_8433172_8446622_RNA45SN1_45S_50ntFlanks</td><td>12935</td><td>12953</td><td>19</td><td>TAAACCATTCGTAGACGAC</td><td>359.8</td><td>-12.9</td><td>1.3</td><td>-10.1</td><td>4.5E-24</td><td>2.6E-23</td><td>BT20/MB468</td><td>NUC</td></tr>
<tr><td>>hg38_21_+_8433172_8446622_RNA45SN1_45S_50ntFlanks@8184.8223.40|CGAGGCCCAGCCCGTGGACGGTGTGAGGCCGGTAGCGGCC</td><td>>hg38_21_+_8433172_8446622_RNA45SN1_45S_50ntFlanks</td><td>8184</td><td>8223</td><td>40</td><td>CGAGGCCCAGCCCGTGGACGGTGTGAGGCCGGTAGCGGCC</td><td>853.6</td><td>-14.2</td><td>1.4</td><td>-10.1</td><td>4.6E-24</td><td>2.6E-23</td><td>BT20/MB468</td><td>NUC</td></tr>
<tr><td>>hg38_21_+_8433172_8446622_RNA45SN1_45S_50ntFlanks@12949.12982.34|ACGACCTGCTTCTGGGTCGGGGTTTCGTACGTAG</td><td>>hg38_21_+_8433172_8446622_RNA45SN1_45S_50ntFlanks</td><td>12949</td><td>12982</td><td>34</td><td>ACGACCTGCTTCTGGGTCGGGGTTTCGTACGTAG</td><td>415.6</td><td>-13.1</td><td>1.3</td><td>-10.1</td><td>4.6E-24</td><td>2.6E-23</td><td>BT20/MB468</td><td>NUC</td></tr>
<tr><td>>hg38_21_+_8433172_8446622_RNA45SN1_45S_50ntFlanks@11918.11950.33|ACATGAGAGGTGTAGAATAAGTGGGAGGCCCCC</td><td>>hg38_21_+_8433172_8446622_RNA45SN1_45S_50ntFlanks</td><td>11918</td><td>11950</td><td>33</td><td>ACATGAGAGGTGTAGAATAAGTGGGAGGCCCCC</td><td>736.0</td><td>-13.9</td><td>1.4</td><td>-10.1</td><td>4.7E-24</td><td>2.7E-23</td><td>BT20/MB468</td><td>NUC</td></tr>
<tr><td>>hg38_21_+_8433172_8446622_RNA45SN1_45S_50ntFlanks@8168.8213.46|CCCAAGTCCTTCTGATCGAGGCCCAGCCCGTGGACGGTGTGAGGCC</td><td>>hg38_21_+_8433172_8446622_RNA45SN1_45S_50ntFlanks</td><td>8168</td><td>8213</td><td>46</td><td>CCCAAGTCCTTCTGATCGAGGCCCAGCCCGTGGACGGTGTGAGGCC</td><td>996.0</td><td>-14.4</td><td>1.4</td><td>-10.1</td><td>4.7E-24</td><td>2.7E-23</td><td>BT20/MB468</td><td>NUC</td></tr>
<tr><td>>MI0000114|hsa-mir-107&WithFlank&10|-|89592741|89592833@56.76.21;MI0000108|hsa-mir-103a-2&WithFlank&20|+|3917488|3917577@54.74.21;MI0000109|hsa-mir-103a-1&WithFlank&5|-|168560890|168560979@54.74.21&[MIMAT0000104&hsa-miR-107&offsets|0|-2;m-78&10|-|89592758|89592778&offsets|0|0];[MIMAT0000101&hsa-miR-103a-3p&offsets|0|-2;m-12&20|+|3917541|3917563&offsets|0|-2];[MIMAT0000101_1&hsa-miR-103a-3p&offsets|0|-2;m-13&5|-|168560904|168560926&offsets|0|-2;m-44220&5|-|168560908|168560926&offsets|0|+2]|AGCAGCATTGTACAGGGCTAT</td><td>>MI0000114|hsa-mir-107&WithFlank&10|-|89592741|89592833</td><td>56</td><td>76</td><td>21</td><td>AGCAGCATTGTACAGGGCTAT</td><td>799.2</td><td>7.5</td><td>0.7</td><td>10.2</td><td>2.3E-24</td><td>2.7E-23</td><td>BT20/MB468</td><td>CYTO</td></tr>
<tr><td>>MI0000650|hsa-mir-200c&WithFlank&12|+|6963693|6963772@50.72.23(+1A)&[MIMAT0000617&hsa-miR-200c-3p&offsets|0|0(+1A);m-54&12|+|6963742|6963764&offsets|0|0(+1A)]|TAATACTGCCGGGTAATGATGGAA</td><td>>MI0000650|hsa-mir-200c&WithFlank&12|+|6963693|6963772</td><td>50</td><td>72</td><td>23(+1A)</td><td>TAATACTGCCGGGTAATGATGGAA</td><td>16095.2</td><td>9.3</td><td>0.9</td><td>10.5</td><td>5.9E-26</td><td>2.7E-23</td><td>BT20/MB231</td><td>CYTO</td></tr>
</table>

<table>
<tr><th></th><th></th><th></th><th></th><th></th><th></th><th></th><th></th><th></th><th></th><th></th><th></th><th></th><th></th></tr>
<tr><td>>MI0000471|hsa-mir-126&WithFlank&9|+|136670596|136670692@58.77.20&[MIMAT0000445&hsa-miR-126-3p&offsets|0|-2;m-40&9|+|136670653|136670674&offsets|0|-2]|TCGTACCGTGAGTAATAATG</td><td>>MI0000471|hsa-mir-126&WithFlank&9|+|136670596|136670692</td><td>58</td><td>77</td><td>20</td><td>TCGTACCGTGAGTAATAATG</td><td>604.9</td><td>13.7</td><td>1.3</td><td>10.2</td><td>1.8E-24</td><td>2.7E-23</td><td>BT20/MB468</td><td>MITO</td></tr>
<tr><td>>hg38_21_+_8433172_8446622_RNA45SN1_45S_50ntFlanks@12942.12961.20|TTCGTAGACGACCTGCTTCT</td><td>>hg38_21_+_8433172_8446622_RNA45SN1_45S_50ntFlanks</td><td>12942</td><td>12961</td><td>20</td><td>TTCGTAGACGACCTGCTTCT</td><td>654.9</td><td>-5.1</td><td>0.5</td><td>-10.1</td><td>4.8E-24</td><td>2.7E-23</td><td>BT20/MB468</td><td>NUC</td></tr>
<tr><td>>hg38_21_+_8433172_8446622_RNA45SN1_45S_50ntFlanks@4443.4471.29|CCCTTGCCTCTCGGCGCCCCCTCGATGCT</td><td>>hg38_21_+_8433172_8446622_RNA45SN1_45S_50ntFlanks</td><td>4443</td><td>4471</td><td>29</td><td>CCCTTGCCTCTCGGCGCCCCCTCGATGCT</td><td>4105.1</td><td>-16.4</td><td>1.6</td><td>-10.1</td><td>4.9E-24</td><td>2.8E-23</td><td>BT20/MB468</td><td>NUC</td></tr>
<tr><td>>MI0000285|hsa-mir-205&WithFlank&1|+|209432127|209432248@40.61.22&[MIMAT0000266&hsa-miR-205-5p&offsets|0|0;m-351&1|+|209432166|209432187&offsets|0|0]|TCCTTCATTCCACCGGAGTCTG</td><td>>MI0000285|hsa-mir-205&WithFlank&1|+|209432127|209432248</td><td>40</td><td>61</td><td>22</td><td>TCCTTCATTCCACCGGAGTCTG</td><td>84187.3</td><td>7.4</td><td>0.7</td><td>10.2</td><td>2.3E-24</td><td>2.8E-23</td><td>BT20/MB468</td><td>CYTO</td></tr>
<tr><td>>hg38_21_+_8433172_8446622_RNA45SN1_45S_50ntFlanks@13004.13029.26|ATCTATTGAAAGTCAGCCCTCGACAC</td><td>>hg38_21_+_8433172_8446622_RNA45SN1_45S_50ntFlanks</td><td>13004</td><td>13029</td><td>26</td><td>ATCTATTGAAAGTCAGCCCTCGACAC</td><td>352.3</td><td>-12.9</td><td>1.3</td><td>-10.1</td><td>4.9E-24</td><td>2.8E-23</td><td>BT20/MB468</td><td>NUC</td></tr>
<tr><td>>trna14_LysTTT_11_-_59327808_59327880@1.24.24,trna2_LysTTT_17_+_8022473_8022545@1.24.24,trna54_LysTTT_1_+_204475655_204475727@1.24.24,trna5_LysTTT_11_+_59323902_59323974@1.24.24,trna62_LysTTT_1_-_204476158_204476230@1.24.24,trna76_LysTTT_6_+_28918806_28918878@1.24.24|GCCCGGATAGCTCAGTCGGTAGAG</td><td>>trna14_LysTTT_11_-_59327808_59327880</td><td>1</td><td>24</td><td>24</td><td>GCCCGGATAGCTCAGTCGGTAGAG</td><td>585.6</td><td>-13.6</td><td>1.3</td><td>-10.1</td><td>4.9E-24</td><td>2.8E-23</td><td>BT20/MB468</td><td>NUC</td></tr>
<tr><td>>hg38_21_+_8433172_8446622_RNA45SN1_45S_50ntFlanks@12456.12474.19|CCAAGCGTTGGATTGTTCA</td><td>>hg38_21_+_8433172_8446622_RNA45SN1_45S_50ntFlanks</td><td>12456</td><td>12474</td><td>19</td><td>CCAAGCGTTGGATTGTTCA</td><td>3468.4</td><td>-16.2</td><td>1.6</td><td>-10.1</td><td>4.9E-24</td><td>2.8E-23</td><td>BT20/MB468</td><td>NUC</td></tr>
<tr><td>>hg38_21_+_8433172_8446622_RNA45SN1_45S_50ntFlanks@10771.10797.27|CAGGTCTCCAAGGTGAACAGCCTCTGG</td><td>>hg38_21_+_8433172_8446622_RNA45SN1_45S_50ntFlanks</td><td>10771</td><td>10797</td><td>27</td><td>CAGGTCTCCAAGGTGAACAGCCTCTGG</td><td>668.8</td><td>-13.8</td><td>1.4</td><td>-10.1</td><td>5.0E-24</td><td>2.8E-23</td><td>BT20/MB468</td><td>NUC</td></tr>
<tr><td>>hg38_21_+_8433172_8446622_RNA45SN1_45S_50ntFlanks@6731.6758.28|CACATTGATCATCGACACTTCGAACGCA</td><td>>hg38_21_+_8433172_8446622_RNA45SN1_45S_50ntFlanks</td><td>6731</td><td>6758</td><td>28</td><td>CACATTGATCATCGACACTTCGAACGCA</td><td>456.2</td><td>-13.3</td><td>1.3</td><td>-10.1</td><td>5.0E-24</td><td>2.8E-23</td><td>BT20/MB468</td><td>NUC</td></tr>
<tr><td>>hg38_21_+_8433172_8446622_RNA45SN1_45S_50ntFlanks@11999.12043.45|CCCTGCGGGCCGCCGGTGAAATACCACTACTCTGATCGTTTTTTC</td><td>>hg38_21_+_8433172_8446622_RNA45SN1_45S_50ntFlanks</td><td>11999</td><td>12043</td><td>45</td><td>CCCTGCGGGCCGCCGGTGAAATACCACTACTCTGATCGTTTTTTC</td><td>4126.7</td><td>-15.8</td><td>1.5</td><td>-10.3</td><td>1.0E-24</td><td>2.8E-23</td><td>MB231/MB468</td><td>MITO</td></tr>
<tr><td>>hg38_21_+_8433172_8446622_RNA45SN1_45S_50ntFlanks@12619.12654.36|AGGAGCCAATGGGGCGAAGCTACCATCTGTGGGATT</td><td>>hg38_21_+_8433172_8446622_RNA45SN1_45S_50ntFlanks</td><td>12619</td><td>12654</td><td>36</td><td>AGGAGCCAATGGGGCGAAGCTACCATCTGTGGGATT</td><td>4360.7</td><td>-10.1</td><td>1.0</td><td>-10.3</td><td>1.0E-24</td><td>2.8E-23</td><td>MB231/MB468</td><td>MITO</td></tr>
<tr><td>>hg38_21_+_8433172_8446622_RNA45SN1_45S_50ntFlanks@11209.11231.23|CGGCGACTCTGGACGCGAGCCGG</td><td>>hg38_21_+_8433172_8446622_RNA45SN1_45S_50ntFlanks</td><td>11209</td><td>11231</td><td>23</td><td>CGGCGACTCTGGACGCGAGCCGG</td><td>1182.9</td><td>-5.7</td><td>0.6</td><td>-10.2</td><td>1.2E-24</td><td>2.8E-23</td><td>MB231/MB468</td><td>NUC</td></tr>
<tr><td>>hg38_21_+_8433172_8446622_RNA45SN1_45S_50ntFlanks@12633.12652.20|CGAAGCTACCATCTGTGGGA</td><td>>hg38_21_+_8433172_8446622_RNA45SN1_45S_50ntFlanks</td><td>12633</td><td>12652</td><td>20</td><td>CGAAGCTACCATCTGTGGGA</td><td>409.2</td><td>-13.1</td><td>1.3</td><td>-10.1</td><td>5.0E-24</td><td>2.8E-23</td><td>BT20/MB468</td><td>NUC</td></tr>
<tr><td>>hg38_21_+_8433172_8446622_RNA45SN1_45S_50ntFlanks@4002.4039.38|GATAACCTCGGGCCGATCGCACGCCCCCCGTGGCGGCG</td><td>>hg38_21_+_8433172_8446622_RNA45SN1_45S_50ntFlanks</td><td>4002</td><td>4039</td><td>38</td><td>GATAACCTCGGGCCGATCGCACGCCCCCCGTGGCGGCG</td><td>522.8</td><td>-13.5</td><td>1.3</td><td>-10.1</td><td>5.0E-24</td><td>2.8E-23</td><td>BT20/MB468</td><td>NUC</td></tr>
<tr><td>>hg38_21_+_8433172_8446622_RNA45SN1_45S_50ntFlanks@10362.10390.29|AAGGCCGAAGTGGAGAAGGGTTCCATGTG</td><td>>hg38_21_+_8433172_8446622_RNA45SN1_45S_50ntFlanks</td><td>10362</td><td>10390</td><td>29</td><td>AAGGCCGAAGTGGAGAAGGGTTCCATGTG</td><td>3065.7</td><td>-15.4</td><td>1.5</td><td>-10.3</td><td>6.6E-25</td><td>2.8E-23</td><td>MB231/MB468</td><td>MP</td></tr>
<tr><td>>hg38_21_+_8433172_8446622_RNA45SN1_45S_50ntFlanks@6647.6676.30|CGTACGACTCTTAGCGGTGGATCACTCGGC</td><td>>hg38_21_+_8433172_8446622_RNA45SN1_45S_50ntFlanks</td><td>6647</td><td>6676</td><td>30</td><td>CGTACGACTCTTAGCGGTGGATCACTCGGC</td><td>4185.8</td><td>6.1</td><td>0.6</td><td>10.2</td><td>1.9E-24</td><td>2.9E-23</td><td>BT20/MB468</td><td>MITO</td></tr>
<tr><td>>hg38_21_+_8433172_8446622_RNA45SN1_45S_50ntFlanks@6651.6670.20|CGACTCTTAGCGGTGGATCA</td><td>>hg38_21_+_8433172_8446622_RNA45SN1_45S_50ntFlanks</td><td>6651</td><td>6670</td><td>20</td><td>CGACTCTTAGCGGTGGATCA</td><td>2048.8</td><td>-7.2</td><td>0.7</td><td>-10.3</td><td>6.7E-25</td><td>2.9E-23</td><td>MB231/MB468</td><td>MP</td></tr>
<tr><td>>hg38_21_+_8433172_8446622_RNA45SN1_45S_50ntFlanks@4103.4121.19|CCACGGGTGACGGGGAATC</td><td>>hg38_21_+_8433172_8446622_RNA45SN1_45S_50ntFlanks</td><td>4103</td><td>4121</td><td>19</td><td>CCACGGGTGACGGGGAATC</td><td>1947.3</td><td>-15.4</td><td>1.5</td><td>-10.1</td><td>5.2E-24</td><td>2.9E-23</td><td>BT20/MB468</td><td>NUC</td></tr>
<tr><td>>hg38_21_+_8433172_8446622_RNA45SN1_45S_50ntFlanks@12638.12659.22|CTACCATCTGTGGGATTATGAC</td><td>>hg38_21_+_8433172_8446622_RNA45SN1_45S_50ntFlanks</td><td>12638</td><td>12659</td><td>22</td><td>CTACCATCTGTGGGATTATGAC</td><td>2744.2</td><td>-15.3</td><td>1.5</td><td>-10.3</td><td>6.9E-25</td><td>2.9E-23</td><td>MB231/MB468</td><td>MP</td></tr>
<tr><td>>hg38_21_+_8433172_8446622_RNA45SN1_45S_50ntFlanks@10510.10533.24|AATCCGGAGTGGCGGAGATGGGCG</td><td>>hg38_21_+_8433172_8446622_RNA45SN1_45S_50ntFlanks</td><td>10510</td><td>10533</td><td>24</td><td>AATCCGGAGTGGCGGAGATGGGCG</td><td>3446.1</td><td>-15.5</td><td>1.5</td><td>-10.3</td><td>1.1E-24</td><td>2.9E-23</td><td>MB231/MB468</td><td>MITO</td></tr>
<tr><td>>MI0000266|hsa-mir-10a&WithFlank&17|-|48579832|48579953@28.48.21(+1C)&[MIMAT0000253&hsa-miR-10a-5p&offsets|0|-2(+1C);m-18&17|-|48579905|48579926&offsets|0|-1(+1C)]|TACCCTGTAGATCCGAATTTGC</td><td>>MI0000266|hsa-mir-10a&WithFlank&17|-|48579832|48579953</td><td>28</td><td>48</td><td>21(+1C)</td><td>TACCCTGTAGATCCGAATTTGC</td><td>446.2</td><td>11.6</td><td>1.2</td><td>10.1</td><td>5.3E-24</td><td>3.0E-23</td><td>BT20/MB468</td><td>NUC</td></tr>
<tr><td>>hg38_21_+_8433172_8446622_RNA45SN1_45S_50ntFlanks@10351.10374.24|CGAGAACTTTGAAGGCCGAAGTGG</td><td>>hg38_21_+_8433172_8446622_RNA45SN1_45S_50ntFlanks</td><td>10351</td><td>10374</td><td>24</td><td>CGAGAACTTTGAAGGCCGAAGTGG</td><td>4753.4</td><td>-9.6</td><td>0.9</td><td>-10.3</td><td>1.1E-24</td><td>3.0E-23</td><td>MB231/MB468</td><td>MITO</td></tr>
<tr><td>>hg38_21_+_8433172_8446622_RNA45SN1_45S_50ntFlanks@8113.8139.27|GGCGCGGGACATGTGGCGTACGGAAGA</td><td>>hg38_21_+_8433172_8446622_RNA45SN1_45S_50ntFlanks</td><td>8113</td><td>8139</td><td>27</td><td>GGCGCGGGACATGTGGCGTACGGAAGA</td><td>1638.1</td><td>-14.5</td><td>1.4</td><td>-10.3</td><td>7.1E-25</td><td>3.0E-23</td><td>MB231/MB468</td><td>MP</td></tr>
<tr><td>>MI0000746|hsa-mir-99b&WithFlank&19|+|51692606|51692687@13.31.19(+1A)&[MIMAT0000689&hsa-miR-99b-5p&offsets|0|-3(+1A);m-10&19|+|51692618|51692639&offsets|0|-3(+1A)]|CACCCGTAGAACCGACCTTA</td><td>>MI0000746|hsa-mir-99b&WithFlank&19|+|51692606|51692687</td><td>13</td><td>31</td><td>19(+1A)</td><td>CACCCGTAGAACCGACCTTA</td><td>977.8</td><td>14.2</td><td>1.4</td><td>10.2</td><td>2.6E-24</td><td>3.1E-23</td><td>BT20/MB468</td><td>CYTO</td></tr>
<tr><td>>hg38_1_-_228634819_228635039_RNA5S12_5S_50ntFlanks@134.170.37|TGGGAGACCGCCTGGGAATACCGGGTGCTGTAGGCTT</td><td>>hg38_1_-_228634819_228635039_RNA5S12_5S_50ntFlanks</td><td>134</td><td>170</td><td>37</td><td>TGGGAGACCGCCTGGGAATACCGGGTGCTGTAGGCTT</td><td>47574.3</td><td>6.5</td><td>0.6</td><td>10.2</td><td>2.6E-24</td><td>3.1E-23</td><td>BT20/MB468</td><td>CYTO</td></tr>
<tr><td>>MI0000065|hsa-let-7d&WithFlank&9|+|94178828|94178926@14.34.21&[MIMAT0000065&hsa-let-7d-5p&offsets|0|-1;m-90&9|+|94178841|94178862&offsets|0|-1]|AGAGGTAGTAGGTTGCATAGT</td><td>>MI0000065|hsa-let-7d&WithFlank&9|+|94178828|94178926</td><td>14</td><td>34</td><td>21</td><td>AGAGGTAGTAGGTTGCATAGT</td><td>3417.0</td><td>7.9</td><td>0.8</td><td>10.2</td><td>2.6E-24</td><td>3.1E-23</td><td>BT20/MB468</td><td>CYTO</td></tr>
<tr><td>>hg38_21_+_8433172_8446622_RNA45SN1_45S_50ntFlanks@12636.12661.26|AGCTACCATCTGTGGGATTATGACTG</td><td>>hg38_21_+_8433172_8446622_RNA45SN1_45S_50ntFlanks</td><td>12636</td><td>12661</td><td>26</td><td>AGCTACCATCTGTGGGATTATGACTG</td><td>4541.4</td><td>-16.0</td><td>1.6</td><td>-10.3</td><td>7.3E-25</td><td>3.1E-23</td><td>MB231/MB468</td><td>MP</td></tr>
<tr><td>>hg38_21_+_8433172_8446622_RNA45SN1_45S_50ntFlanks@11769.11786.18|TAATTAGTGACGCGCATG</td><td>>hg38_21_+_8433172_8446622_RNA45SN1_45S_50ntFlanks</td><td>11769</td><td>11786</td><td>18</td><td>TAATTAGTGACGCGCATG</td><td>630.0</td><td>-13.7</td><td>1.4</td><td>-10.1</td><td>5.5E-24</td><td>3.1E-23</td><td>BT20/MB468</td><td>NUC</td></tr>
<tr><td>>MI0000767|hsa-mir-365a&WithFlank&16|+|14309279|14309377@62.81.20;MI0000769|hsa-mir-365b&WithFlank&17|+|31575405|31575527@74.93.20&[MIMAT0000710&hsa-miR-365a-3p&offsets|0|-2;m-176&16|+|14309340|14309361&offsets|0|-2];[MIMAT0022834&hsa-miR-365b-3p&offsets|0|-2;m-175&17|+|31575478|31575499&offsets|0|-2]|TAATGCCCCTAAAAATCCTT</td><td>>MI0000767|hsa-mir-365a&WithFlank&16|+|14309279|14309377</td><td>62</td><td>81</td><td>20</td><td>TAATGCCCCTAAAAATCCTT</td><td>407.8</td><td>13.1</td><td>1.3</td><td>10.2</td><td>2.1E-24</td><td>3.1E-23</td><td>BT20/MB468</td><td>MITO</td></tr>
<tr><td>>MI0000286|hsa-mir-210&WithFlank&11|-|568083|568204@72.93.22(+2A)&[MIMAT0000267&hsa-miR-210-3p&offsets|0|0(+2A);m-102&11|-|568112|568133&offsets|0|0(+2A)]|CTGTGCGTGTGACAGCGGCTGAAA</td><td>>MI0000286|hsa-mir-210&WithFlank&11|-|568083|568204</td><td>72</td><td>93</td><td>22(+2A)</td><td>CTGTGCGTGTGACAGCGGCTGAAA</td><td>3840.7</td><td>9.5</td><td>0.9</td><td>10.2</td><td>2.7E-24</td><td>3.1E-23</td><td>BT20/MB468</td><td>CYTO</td></tr>
<tr><td>>am_MI0000809|hsa-mir-151a&WithFlank&8|-|140732558|140732659@54.73.20&[MIMAT0000757&hsa-miR-151a-3p&offsets|+1|0;m-30&8|-|140732587|140732607&offsets|+1|0]|TAGACTGAAGCTCCTTGAGG</td><td>>am_MI0000809|hsa-mir-151a&WithFlank&8|-|140732558|140732659</td><td>54</td><td>73</td><td>20</td><td>TAGACTGAAGCTCCTTGAGG</td><td>284.9</td><td>12.4</td><td>1.2</td><td>10.2</td><td>2.7E-24</td><td>3.1E-23</td><td>BT20/MB468</td><td>CYTO</td></tr>
<tr><td>>hg38_21_+_8433172_8446622_RNA45SN1_45S_50ntFlanks@10687.10711.25|CGGCGTCCGGTGAGCTCTCGCTGGC</td><td>>hg38_21_+_8433172_8446622_RNA45SN1_45S_50ntFlanks</td><td>10687</td><td>10711</td><td>25</td><td>CGGCGTCCGGTGAGCTCTCGCTGGC</td><td>6467.0</td><td>-7.7</td><td>0.8</td><td>-10.3</td><td>1.2E-24</td><td>3.1E-23</td><td>MB231/MB468</td><td>MITO</td></tr>
<tr><td>>hg38_21_+_8433172_8446622_RNA45SN1_45S_50ntFlanks@6647.6681.35|CGTACGACTCTTAGCGGTGGATCACTCGGCTCGTG</td><td>>hg38_21_+_8433172_8446622_RNA45SN1_45S_50ntFlanks</td><td>6647</td><td>6681</td><td>35</td><td>CGTACGACTCTTAGCGGTGGATCACTCGGCTCGTG</td><td>6194.3</td><td>-3.2</td><td>0.3</td><td>-10.1</td><td>5.6E-24</td><td>3.1E-23</td><td>BT20/MB468</td><td>NUC</td></tr>
<tr><td>>hg38_21_+_8433172_8446622_RNA45SN1_45S_50ntFlanks@10407.10430.24|GGGTCAGTCGGTCCTGAGAGATGG</td><td>>hg38_21_+_8433172_8446622_RNA45SN1_45S_50ntFlanks</td><td>10407</td><td>10430</td><td>24</td><td>GGGTCAGTCGGTCCTGAGAGATGG</td><td>372.8</td><td>-13.0</td><td>1.3</td><td>-10.1</td><td>5.7E-24</td><td>3.2E-23</td><td>BT20/MB468</td><td>NUC</td></tr>
<tr><td>>hg38_21_+_8433172_8446622_RNA45SN1_45S_50ntFlanks@6645.6667.23|CTCGTACGACTCTTAGCGGTGGA</td><td>>hg38_21_+_8433172_8446622_RNA45SN1_45S_50ntFlanks</td><td>6645</td><td>6667</td><td>23</td><td>CTCGTACGACTCTTAGCGGTGGA</td><td>12979.0</td><td>6.6</td><td>0.6</td><td>10.2</td><td>2.1E-24</td><td>3.2E-23</td><td>BT20/MB468</td><td>MITO</td></tr>
<tr><td>>hg38_21_+_8433172_8446622_RNA45SN1_45S_50ntFlanks@12151.12180.30|GTGGGGAGTTTGACTGGGGCGGTACACCTG</td><td>>hg38_21_+_8433172_8446622_RNA45SN1_45S_50ntFlanks</td><td>12151</td><td>12180</td><td>30</td><td>GTGGGGAGTTTGACTGGGGCGGTACACCTG</td><td>1116.2</td><td>-14.0</td><td>1.4</td><td>-10.3</td><td>7.5E-25</td><td>3.2E-23</td><td>MB231/MB468</td><td>MP</td></tr>
<tr><td>>hg38_21_+_8433172_8446622_RNA45SN1_45S_50ntFlanks@10572.10589.18|AGAAGCCGGCGGGAGCCC</td><td>>hg38_21_+_8433172_8446622_RNA45SN1_45S_50ntFlanks</td><td>10572</td><td>10589</td><td>18</td><td>AGAAGCCGGCGGGAGCCC</td><td>386.5</td><td>-13.0</td><td>1.3</td><td>-10.1</td><td>5.8E-24</td><td>3.3E-23</td><td>BT20/MB468</td><td>NUC</td></tr>
<tr><td>>hg38_21_+_8433172_8446622_RNA45SN1_45S_50ntFlanks@10805.10823.19|GAACAATGTAGGTAAGGGA</td><td>>hg38_21_+_8433172_8446622_RNA45SN1_45S_50ntFlanks</td><td>10805</td><td>10823</td><td>19</td><td>GAACAATGTAGGTAAGGGA</td><td>475.4</td><td>-13.3</td><td>1.3</td><td>-10.1</td><td>5.8E-24</td><td>3.3E-23</td><td>BT20/MB468</td><td>NUC</td></tr>
<tr><td>>hg38_21_+_8433172_8446622_RNA45SN1_45S_50ntFlanks@10128.10147.20|CGGCGGCGGCGGCGGCGGCG</td><td>>hg38_21_+_8433172_8446622_RNA45SN1_45S_50ntFlanks</td><td>10128</td><td>10147</td><td>20</td><td>CGGCGGCGGCGGCGGCGGCG</td><td>3758.3</td><td>-15.7</td><td>1.5</td><td>-10.2</td><td>1.2E-24</td><td>3.3E-23</td><td>MB231/MB468</td><td>MITO</td></tr>
<tr><td>>hg38_21_+_8433172_8446622_RNA45SN1_45S_50ntFlanks@8845.8864.20|CGGCGGCGGCGGCGGCGGCG</td><td>>hg38_21_+_8433172_8446622_RNA45SN1_45S_50ntFlanks</td><td>8845</td><td>8864</td><td>20</td><td>CGGCGGCGGCGGCGGCGGCG</td><td>3758.3</td><td>-15.7</td><td>1.5</td><td>-10.2</td><td>1.2E-24</td><td>3.3E-23</td><td>MB231/MB468</td><td>MITO</td></tr>
<tr><td>>hg38_21_+_8433172_8446622_RNA45SN1_45S_50ntFlanks@11572.11611.40|CCGACTTAGAACTGGTGCGGACCAGGGGAATCCGACTGTT</td><td>>hg38_21_+_8433172_8446622_RNA45SN1_45S_50ntFlanks</td><td>11572</td><td>11611</td><td>40</td><td>CCGACTTAGAACTGGTGCGGACCAGGGGAATCCGACTGTT</td><td>2689.4</td><td>-6.8</td><td>0.7</td><td>-10.1</td><td>6.0E-24</td><td>3.3E-23</td><td>BT20/MB468</td><td>NUC</td></tr>
<tr><td>>hg38_21_+_8433172_8446622_RNA45SN1_45S_50ntFlanks@7973.8009.37|GACGCGACCTCAGATCAGACGTGGCGACCCGCTGAAT</td><td>>hg38_21_+_8433172_8446622_RNA45SN1_45S_50ntFlanks</td><td>7973</td><td>8009</td><td>37</td><td>GACGCGACCTCAGATCAGACGTGGCGACCCGCTGAAT</td><td>10323.6</td><td>6.1</td><td>0.6</td><td>10.2</td><td>2.2E-24</td><td>3.3E-23</td><td>BT20/MB468</td><td>MITO</td></tr>
<tr><td>>hg38_21_+_8433172_8446622_RNA45SN1_45S_50ntFlanks@8168.8214.47|CCCAAGTCCTTCTGATCGAGGCCCAGCCCGTGGACGGTGTGAGGCCG</td><td>>hg38_21_+_8433172_8446622_RNA45SN1_45S_50ntFlanks</td><td>8168</td><td>8214</td><td>47</td><td>CCCAAGTCCTTCTGATCGAGGCCCAGCCCGTGGACGGTGTGAGGCCG</td><td>2244.3</td><td>-15.6</td><td>1.5</td><td>-10.1</td><td>6.0E-24</td><td>3.4E-23</td><td>BT20/MB468</td><td>NUC</td></tr>
<tr><td>>MI0000264|hsa-mir-7-2&WithFlank&15|+|88611819|88611940@38.59.22;MI0000265|hsa-mir-7-3&WithFlank&19|+|4770664|4770785@37.58.22;MI0000263|hsa-mir-7-1&WithFlank&9|-|83969742|83969863@30.51.22&[MIMAT0000252&hsa-miR-7-5p&offsets|0|-2;m-400&15|+|88611856|88611879&offsets|0|-2];[MIMAT0000252_1&hsa-miR-7-5p&offsets|0|-2;m-402&19|+|4770700|4770723&offsets|0|-2];[MIMAT0000252_2&hsa-miR-7-5p&offsets|0|-2;m-401&9|-|83969811|83969834&offsets|0|-2]|TGGAAGACTAGTGATTTTGTTG</td><td>>MI0000264|hsa-mir-7-2&WithFlank&15|+|88611819|88611940</td><td>38</td><td>59</td><td>22</td><td>TGGAAGACTAGTGATTTTGTTG</td><td>928.5</td><td>6.0</td><td>0.6</td><td>10.2</td><td>1.5E-24</td><td>3.4E-23</td><td>MB231/MB468</td><td>NUC</td></tr>
<tr><td>>hg38_21_+_8433172_8446622_RNA45SN1_45S_50ntFlanks@4661.4682.22|AGGTGAAATTCTTGGACCGGCG</td><td>>hg38_21_+_8433172_8446622_RNA45SN1_45S_50ntFlanks</td><td>4661</td><td>4682</td><td>22</td><td>AGGTGAAATTCTTGGACCGGCG</td><td>285.9</td><td>-12.6</td><td>1.2</td><td>-10.1</td><td>6.0E-24</td><td>3.4E-23</td><td>BT20/MB468</td><td>NUC</td></tr>
<tr><td>>hg38_21_+_8433172_8446622_RNA45SN1_45S_50ntFlanks@10408.10429.22|GGTCAGTCGGTCCTGAGAGATG</td><td>>hg38_21_+_8433172_8446622_RNA45SN1_45S_50ntFlanks</td><td>10408</td><td>10429</td><td>22</td><td>GGTCAGTCGGTCCTGAGAGATG</td><td>2812.8</td><td>-6.3</td><td>0.6</td><td>-10.2</td><td>2.9E-24</td><td>3.4E-23</td><td>BT20/MB468</td><td>CYTO</td></tr>
</table>

| | | | | | | | | | | | | | |
|---|---|---|---|---|---|---|---|---|---|---|---|---|---|
| >hg38_21_+_8433172_8446622_RNA45SN1_45S_50ntFlanks@10479.10497.19\|CGATCGAAAGGGAGTCGGG | >hg38_21_+_8433172_8446622_RNA45SN1_45S_50ntFlanks | 10479 | 10497 | 19 | CGATCGAAAGGGAGTCGGG | 1869.0 | -14.7 | 1.4 | -10.3 | 8.1E-25 | 3.4E-23 | MB231/MB468 | MP |
| >am_MI0003639\|hsa-mir-625&WithFlank&14\|+\|65471096\|65471192@58.78.21(+1U)&[MIMAT0004808&hsa-miR-625-3p&offsets\|0\|-1(+1U);m-137&14\|+\|65471153\|65471174&offsets\|0\|-1(+1U)]\|GACTATAGAACTTTCCCCCTCT | >am_MI0003639\|hsa-mir-625&WithFlank&14\|+\|65471096\|65471192 | 58 | 78 | 21(+1U) | GACTATAGAACTTTCCCCCTCT | 266.8 | 6.2 | 0.6 | 10.2 | 1.5E-24 | 3.4E-23 | MB231/MB468 | NUC |
| >MI0000746\|hsa-mir-99b&WithFlank&19\|+\|51692606\|51692687@13.33.21(+1A)&[MIMAT0000689&hsa-miR-99b-5p&offsets\|0\|-1(+1A);m-10&19\|+\|51692618\|51692639&offsets\|0\|-1(+1A)]\|CACCCGTAGAACCGACCTTGCA | >MI0000746\|hsa-mir-99b&WithFlank&19\|+\|51692606\|51692687 | 13 | 33 | 21(+1A) | CACCCGTAGAACCGACCTTGCA | 40351.2 | 8.8 | 0.9 | 10.2 | 2.9E-24 | 3.4E-23 | BT20/MB468 | CYTO |
| >hg38_21_+_8433172_8446622_RNA45SN1_45S_50ntFlanks@13006.13028.23\|CTATTGAAAGTCAGCCCTCGACA | >hg38_21_+_8433172_8446622_RNA45SN1_45S_50ntFlanks | 13006 | 13028 | 23 | CTATTGAAAGTCAGCCCTCGACA | 475.2 | -13.3 | 1.3 | -10.1 | 6.3E-24 | 3.5E-23 | BT20/MB468 | NUC |
| >trna143_LysTTT_6_-_27559593_27559665@13.30.18,trna149_LysTTT_6_-_27302769_27302841@13.30.18,trna14_LysTTT_11_-_59327808_59327880@13.30.18,trna2_LysTTT_17_+_8022473_8022545@13.30.18,trna54_LysTTT_1_+_204475655_204475727@13.30.18,trna5_LysTTT_11_+_59323902_59323974@13.30.18,trna62_LysTTT_1_-_204476158_204476230@13.30.18,trna76_LysTTT_6_+_28918806_28918878@13.30.18\|CAGTCGGTAGAGCATCAG | >trna143_LysTTT_6_-_27559593_27559665 | 13 | 30 | 18 | CAGTCGGTAGAGCATCAG | 665.8 | -13.8 | 1.4 | -10.1 | 6.3E-24 | 3.5E-23 | BT20/MB468 | NUC |
| >hg38_21_+_8433172_8446622_RNA45SN1_45S_50ntFlanks@7985.8006.22\|GATCAGACGTGGCGACCCGCTG | >hg38_21_+_8433172_8446622_RNA45SN1_45S_50ntFlanks | 7985 | 8006 | 22 | GATCAGACGTGGCGACCCGCTG | 9500.7 | -12.6 | 1.2 | -10.3 | 8.4E-25 | 3.5E-23 | MB231/MB468 | MP |
| >hg38_21_+_8433172_8446622_RNA45SN1_45S_50ntFlanks@12456.12488.33\|CCAAGCGTTGGATTGTTCACCCACTAATAGGGA | >hg38_21_+_8433172_8446622_RNA45SN1_45S_50ntFlanks | 12456 | 12488 | 33 | CCAAGCGTTGGATTGTTCACCCACTAATAGGGA | 5548.9 | -10.0 | 1.0 | -10.2 | 1.4E-24 | 3.6E-23 | MB231/MB468 | MITO |
| >hg38_21_+_8433172_8446622_RNA45SN1_45S_50ntFlanks@12476.12493.18\|CCACTAATAGGGAACGTG | >hg38_21_+_8433172_8446622_RNA45SN1_45S_50ntFlanks | 12476 | 12493 | 18 | CCACTAATAGGGAACGTG | 553.2 | -13.5 | 1.3 | -10.1 | 6.4E-24 | 3.6E-23 | BT20/MB468 | NUC |
| >hg38_21_+_8433172_8446622_RNA45SN1_45S_50ntFlanks@11703.11728.26\|AATGAAGCGCGGGTAAACGGCGGGAG | >hg38_21_+_8433172_8446622_RNA45SN1_45S_50ntFlanks | 11703 | 11728 | 26 | AATGAAGCGCGGGTAAACGGCGGGAG | 8152.5 | -11.1 | 1.1 | -10.1 | 6.4E-24 | 3.6E-23 | BT20/MB468 | NUC |
| >hg38_21_+_8433172_8446622_RNA45SN1_45S_50ntFlanks@12054.12071.18\|TGAGGCGGGGGGGCGAGC | >hg38_21_+_8433172_8446622_RNA45SN1_45S_50ntFlanks | 12054 | 12071 | 18 | TGAGGCGGGGGGGCGAGC | 523.2 | -13.5 | 1.3 | -10.1 | 6.5E-24 | 3.6E-23 | BT20/MB468 | NUC |
| >hg38_21_+_8433172_8446622_RNA45SN1_45S_50ntFlanks@993.1027.35\|CTCGTCGGTGTGGGGTTCGAGGCGGTTTGAGTGAG | >hg38_21_+_8433172_8446622_RNA45SN1_45S_50ntFlanks | 993 | 1027 | 35 | CTCGTCGGTGTGGGGTTCGAGGCGGTTTGAGTGAG | 308.5 | -12.7 | 1.3 | -10.1 | 6.6E-24 | 3.6E-23 | BT20/MB468 | NUC |
| >hg38_21_+_8433172_8446622_RNA45SN1_45S_50ntFlanks@6748.6770.23\|CTTCGAACGCACTTGCGGCCCCG | >hg38_21_+_8433172_8446622_RNA45SN1_45S_50ntFlanks | 6748 | 6770 | 23 | CTTCGAACGCACTTGCGGCCCCG | 306.7 | -12.7 | 1.3 | -10.1 | 6.6E-24 | 3.7E-23 | BT20/MB468 | NUC |
| >MI0000737\|hsa-mir-200a&WithFlank&1\|+\|1167857\|1167958@22.43.22&[MIMAT0001620&hsa-miR-200a-5p&offsets\|0\|0;m-154&1\|+\|1167878\|1167899&offsets\|0\|0]\|CATCTTACCGGACAGTGCTGGA | >MI0000737\|hsa-mir-200a&WithFlank&1\|+\|1167857\|1167958 | 22 | 43 | 22 | CATCTTACCGGACAGTGCTGGA | 462.9 | -13.3 | 1.3 | -10.1 | 6.6E-24 | 3.7E-23 | BT20/MB468 | NUC |
| >hg38_21_+_8433172_8446622_RNA45SN1_45S_50ntFlanks@11661.11687.27\|TGATTTCTGCCCAGTGCTCTGAATGTC | >hg38_21_+_8433172_8446622_RNA45SN1_45S_50ntFlanks | 11661 | 11687 | 27 | TGATTTCTGCCCAGTGCTCTGAATGTC | 405.3 | -4.9 | 0.5 | -10.1 | 6.7E-24 | 3.7E-23 | BT20/MB468 | NUC |
| >hg38_21_+_8433172_8446622_RNA45SN1_45S_50ntFlanks@12996.13016.21\|TCGCTGCGATCTATTGAAAGT | >hg38_21_+_8433172_8446622_RNA45SN1_45S_50ntFlanks | 12996 | 13016 | 21 | TCGCTGCGATCTATTGAAAGT | 454.1 | 5.4 | 0.5 | 10.2 | 3.2E-24 | 3.7E-23 | BT20/MB468 | CYTO |
| >MI0000446\|hsa-mir-125b-1&WithFlank&11\|-\|122099751\|122099850@61.82.22&[MIMAT0004592&hsa-miR-125b-1-3p&offsets\|0\|0;m-485&11\|-\|122099769\|122099790&offsets\|0\|0]\|ACGGGTTAGGCTCTTGGGAGCT | >MI0000446\|hsa-mir-125b-1&WithFlank&11\|-\|122099751\|122099850 | 61 | 82 | 22 | ACGGGTTAGGCTCTTGGGAGCT | 595.7 | 13.1 | 1.3 | 10.2 | 1.7E-24 | 3.8E-23 | MB231/MB468 | NUC |
| >hg38_21_+_8433172_8446622_RNA45SN1_45S_50ntFlanks@12456.12486.31\|CCAAGCGTTGGATTGTTCACCCACTAATAGG | >hg38_21_+_8433172_8446622_RNA45SN1_45S_50ntFlanks | 12456 | 12486 | 31 | CCAAGCGTTGGATTGTTCACCCACTAATAGG | 3115.9 | -5.6 | 0.5 | -10.2 | 2.5E-24 | 3.8E-23 | BT20/MB468 | MITO |
| >hg38_21_+_8433172_8446622_RNA45SN1_45S_50ntFlanks@11092.11114.23\|CGGCGGGGGGAGAAGGGTCGGGG | >hg38_21_+_8433172_8446622_RNA45SN1_45S_50ntFlanks | 11092 | 11114 | 23 | CGGCGGGGGGAGAAGGGTCGGGG | 359.2 | -12.9 | 1.3 | -10.1 | 6.8E-24 | 3.8E-23 | BT20/MB468 | NUC |
| >hg38_21_+_8433172_8446622_RNA45SN1_45S_50ntFlanks@12488.12509.22\|AACGTGAGCTGGGTTTAGACCG | >hg38_21_+_8433172_8446622_RNA45SN1_45S_50ntFlanks | 12488 | 12509 | 22 | AACGTGAGCTGGGTTTAGACCG | 3244.4 | -15.5 | 1.5 | -10.3 | 9.0E-25 | 3.8E-23 | MB231/MB468 | MP |
| >MI0000734\|hsa-mir-106b&WithFlank&7\|-\|100093987\|100094080@56.76.21(+1C)&[MIMAT0004672&hsa-miR-106b-3p&offsets\|-2\|-3(+1C);m-87&7\|-\|100094003\|100094023&offsets\|-2\|-2(+1C)]\|TACCGCACTGTGGGTACTTGCC | >MI0000734\|hsa-mir-106b&WithFlank&7\|-\|100093987\|100094080 | 56 | 76 | 21(+1C) | TACCGCACTGTGGGTACTTGCC | 284.0 | 12.4 | 1.2 | 10.2 | 3.3E-24 | 3.8E-23 | BT20/MB468 | CYTO |
| >MI0003628\|hsa-mir-615&WithFlank&12\|+\|54033944\|54034051@67.86.20(+1C)&[MIMAT0003283&hsa-miR-615-3p&offsets\|0\|-2(+1C);m-495&12\|+\|54034010\|54034030&offsets\|0\|-1(+1C)]\|TCCGAGCCTGGGTCTCCCTCC | >MI0003628\|hsa-mir-615&WithFlank&12\|+\|54033944\|54034051 | 67 | 86 | 20(+1C) | TCCGAGCCTGGGTCTCCCTCC | 452.0 | 7.6 | 0.8 | 10.2 | 3.3E-24 | 3.8E-23 | BT20/MB468 | CYTO |
| >hg38_1_-_228634819_228635039_RNA5S12_5S_50ntFlanks@151.169.19\|ATACCGGGTGCTGTAGGCT | >hg38_1_-_228634819_228635039_RNA5S12_5S_50ntFlanks | 151 | 169 | 19 | ATACCGGGTGCTGTAGGCT | 1677.2 | 6.1 | 0.6 | 10.2 | 3.3E-24 | 3.8E-23 | BT20/MB468 | CYTO |
| >hg38_21_+_8433172_8446622_RNA45SN1_45S_50ntFlanks@12126.12146.21\|CGACCCGCTCCGGGGACAGTG | >hg38_21_+_8433172_8446622_RNA45SN1_45S_50ntFlanks | 12126 | 12146 | 21 | CGACCCGCTCCGGGGACAGTG | 314.4 | -12.7 | 1.3 | -10.1 | 7.0E-24 | 3.9E-23 | BT20/MB468 | NUC |
| >hg38_21_+_8433172_8446622_RNA45SN1_45S_50ntFlanks@8168.8212.45\|CCCAAGTCCTTCTGATCGAGGCCCAGCCCGTGGACGGTGTGAGGC | >hg38_21_+_8433172_8446622_RNA45SN1_45S_50ntFlanks | 8168 | 8212 | 45 | CCCAAGTCCTTCTGATCGAGGCCCAGCCCGTGGACGGTGTGAGGC | 2418.2 | -15.7 | 1.6 | -10.1 | 7.0E-24 | 3.9E-23 | BT20/MB468 | NUC |
| >MI0000065\|hsa-let-7d&WithFlank&9\|+\|94178828\|94178926@68.88.21(+1C)&[MIMAT0004484&hsa-let-7d-3p&offsets\|0\|-1(+1C);m-98&9\|+\|94178895\|94178916&offsets\|0\|-1(+1C)]\|CTATACGACCTGCTGCCTTTCC | >MI0000065\|hsa-let-7d&WithFlank&9\|+\|94178828\|94178926 | 68 | 88 | 21(+1C) | CTATACGACCTGCTGCCTTTCC | 722.4 | 7.9 | 0.8 | 10.2 | 2.6E-24 | 3.9E-23 | BT20/MB468 | MITO |
| >am_trnaMT_GluTTC_MT_-_14674_14742@27.52.26,trnalookalike8_GluTTC_5_-_93905172_93905240@27.52.26\|GTTTTTCATATCATTGGTCGTGGTTG | >am_trnaMT_GluTTC_MT_-_14674_14742 | 27 | 52 | 26 | GTTTTTCATATCATTGGTCGTGGTTG | 455.8 | -13.3 | 1.3 | -10.1 | 7.2E-24 | 3.9E-23 | BT20/MB468 | NUC |
| >hg38_21_+_8433172_8446622_RNA45SN1_45S_50ntFlanks@11601.11620.20\|ATCCGACTGTTTAATTAAAA | >hg38_21_+_8433172_8446622_RNA45SN1_45S_50ntFlanks | 11601 | 11620 | 20 | ATCCGACTGTTTAATTAAAA | 688.8 | -13.9 | 1.4 | -10.1 | 7.2E-24 | 3.9E-23 | BT20/MB468 | NUC |
| >hg38_21_+_8433172_8446622_RNA45SN1_45S_50ntFlanks@12656.12675.20\|TGACTGAACGCCTCTAAGTC | >hg38_21_+_8433172_8446622_RNA45SN1_45S_50ntFlanks | 12656 | 12675 | 20 | TGACTGAACGCCTCTAAGTC | 544.9 | -13.5 | 1.3 | -10.1 | 7.2E-24 | 4.0E-23 | BT20/MB468 | NUC |
| >hg38_21_+_8433172_8446622_RNA45SN1_45S_50ntFlanks@3985.4026.42\|CGGCTTTGGTGACTCTAGATAACCTCGGGCCGATCGCACGCC | >hg38_21_+_8433172_8446622_RNA45SN1_45S_50ntFlanks | 3985 | 4026 | 42 | CGGCTTTGGTGACTCTAGATAACCTCGGGCCGATCGCACGCC | 5224.4 | -16.1 | 1.6 | -10.2 | 1.5E-24 | 4.0E-23 | MB231/MB468 | MITO |
| >hg38_21_+_8433172_8446622_RNA45SN1_45S_50ntFlanks@11586.11603.18\|GTGCGGACCAGGGGAATC | >hg38_21_+_8433172_8446622_RNA45SN1_45S_50ntFlanks | 11586 | 11603 | 18 | GTGCGGACCAGGGGAATC | 3112.1 | -15.5 | 1.5 | -10.3 | 9.5E-25 | 4.0E-23 | MB231/MB468 | MP |
| >hg38_21_+_8433172_8446622_RNA45SN1_45S_50ntFlanks@4003.4042.40\|ATAACCTCGGGCCGATCGCACGCCCCCCGTGGCGGCGACG | >hg38_21_+_8433172_8446622_RNA45SN1_45S_50ntFlanks | 4003 | 4042 | 40 | ATAACCTCGGGCCGATCGCACGCCCCCCGTGGCGGCGACG | 1678.9 | -14.5 | 1.4 | -10.2 | 1.5E-24 | 4.0E-23 | MB231/MB468 | MITO |
| >hg38_21_+_8433172_8446622_RNA45SN1_45S_50ntFlanks@6721.6745.25\|AATTGCAGGACACATTGATCATCGA | >hg38_21_+_8433172_8446622_RNA45SN1_45S_50ntFlanks | 6721 | 6745 | 25 | AATTGCAGGACACATTGATCATCGA | 1467.4 | -13.5 | 1.3 | -10.2 | 1.8E-24 | 4.0E-23 | MB231/MB468 | NUC |
| >hg38_21_+_8433172_8446622_RNA45SN1_45S_50ntFlanks@11890.11907.18\|CTTGACTCTAGTCTGGCA | >hg38_21_+_8433172_8446622_RNA45SN1_45S_50ntFlanks | 11890 | 11907 | 18 | CTTGACTCTAGTCTGGCA | 1711.5 | -14.6 | 1.4 | -10.3 | 9.7E-25 | 4.0E-23 | MB231/MB468 | MP |
| >am_MI0000809\|hsa-mir-151a&WithFlank&8\|-\|140732558\|140732659@53.72.20(+1A)&[MIMAT0000757&hsa-miR-151a-3p&offsets\|0\|-1(+1A);m-30&8\|-\|140732587\|140732607&offsets\|0\|-1(+1A)]\|CTAGACTGAAGCTCCTTGAGA | >am_MI0000809\|hsa-mir-151a&WithFlank&8\|-\|140732558\|140732659 | 53 | 72 | 20(+1A) | CTAGACTGAAGCTCCTTGAGA | 4152.7 | 5.6 | 0.5 | 10.2 | 1.8E-24 | 4.1E-23 | MB231/MB468 | NUC |
| >hg38_21_+_8433172_8446622_RNA45SN1_45S_50ntFlanks@12044.12070.27\|ACTGACCCGGTGAGGCGGGGGGGCGAG | >hg38_21_+_8433172_8446622_RNA45SN1_45S_50ntFlanks | 12044 | 12070 | 27 | ACTGACCCGGTGAGGCGGGGGGGCGAG | 3978.4 | -15.7 | 1.5 | -10.2 | 1.6E-24 | 4.1E-23 | MB231/MB468 | MITO |
| >hg38_21_+_8433172_8446622_RNA45SN1_45S_50ntFlanks@12627.12659.33\|ATGGGGCGAAGCTACCATCTGTGGGATTATGAC | >hg38_21_+_8433172_8446622_RNA45SN1_45S_50ntFlanks | 12627 | 12659 | 33 | ATGGGGCGAAGCTACCATCTGTGGGATTATGAC | 2045.3 | -7.7 | 0.8 | -10.2 | 1.8E-24 | 4.1E-23 | MB231/MB468 | NUC |
| >hg38_21_+_8433172_8446622_RNA45SN1_45S_50ntFlanks@11210.11232.23\|GGCGACTCTGGACGCGAGCCGGG | >hg38_21_+_8433172_8446622_RNA45SN1_45S_50ntFlanks | 11210 | 11232 | 23 | GGCGACTCTGGACGCGAGCCGGG | 2171.8 | -14.9 | 1.5 | -10.3 | 9.9E-25 | 4.1E-23 | MB231/MB468 | MP |
| >hg38_21_+_8433172_8446622_RNA45SN1_45S_50ntFlanks@12405.12432.28\|CTTTTTGATCCTTCGATGTCGGCTCTTC | >hg38_21_+_8433172_8446622_RNA45SN1_45S_50ntFlanks | 12405 | 12432 | 28 | CTTTTTGATCCTTCGATGTCGGCTCTTC | 6644.6 | -12.4 | 1.2 | -10.3 | 9.9E-25 | 4.1E-23 | MB231/MB468 | MP |
| >hg38_21_+_8433172_8446622_RNA45SN1_45S_50ntFlanks@12421.12443.23\|TGTCGGCTCTTCCTATCATTGTG | >hg38_21_+_8433172_8446622_RNA45SN1_45S_50ntFlanks | 12421 | 12443 | 23 | TGTCGGCTCTTCCTATCATTGTG | 339.2 | -12.8 | 1.3 | -10.1 | 7.6E-24 | 4.1E-23 | BT20/MB468 | NUC |
| >hg38_21_+_8433172_8446622_RNA45SN1_45S_50ntFlanks@11729.11751.23\|TAACTATGACTCTCTTAAGGTAG | >hg38_21_+_8433172_8446622_RNA45SN1_45S_50ntFlanks | 11729 | 11751 | 23 | TAACTATGACTCTCTTAAGGTAG | 20519.1 | -10.5 | 1.0 | -10.2 | 1.6E-24 | 4.2E-23 | MB231/MB468 | MITO |
| >hg38_21_+_8433172_8446622_RNA45SN1_45S_50ntFlanks@11579.11605.27\|AGAACTGGTGCGGACCAGGGGAATCCG | >hg38_21_+_8433172_8446622_RNA45SN1_45S_50ntFlanks | 11579 | 11605 | 27 | AGAACTGGTGCGGACCAGGGGAATCCG | 2338.4 | -15.1 | 1.5 | -10.3 | 1.0E-24 | 4.2E-23 | MB231/MB468 | MP |
| >hg38_21_+_8433172_8446622_RNA45SN1_45S_50ntFlanks@10914.10936.23\|CGCCGCGGCTGGACGAGGCGCCG | >hg38_21_+_8433172_8446622_RNA45SN1_45S_50ntFlanks | 10914 | 10936 | 23 | CGCCGCGGCTGGACGAGGCGCCG | 1253.9 | -14.2 | 1.4 | -10.3 | 1.0E-24 | 4.2E-23 | MB231/MB468 | MP |
| >hg38_1_-_228634819_228635039_RNA5S12_5S_50ntFlanks@130.170.41\|TGGATGGGAGACCGCCTGGGAATACCGGGTGCTGTAGGCTT | >hg38_1_-_228634819_228635039_RNA5S12_5S_50ntFlanks | 130 | 170 | 41 | TGGATGGGAGACCGCCTGGGAATACCGGGTGCTGTAGGCTT | 7068.9 | 9.7 | 1.0 | 10.1 | 3.7E-24 | 4.3E-23 | BT20/MB468 | CYTO |
| >hg38_21_+_8433172_8446622_RNA45SN1_45S_50ntFlanks@11890.11912.23\|CTTGACTCTAGTCTGGCACGGTG | >hg38_21_+_8433172_8446622_RNA45SN1_45S_50ntFlanks | 11890 | 11912 | 23 | CTTGACTCTAGTCTGGCACGGTG | 1478.4 | -14.4 | 1.4 | -10.3 | 1.0E-24 | 4.3E-23 | MB231/MB468 | MP |
| >hg38_21_+_8433172_8446622_RNA45SN1_45S_50ntFlanks@11667.11707.41\|CTGCCCAGTGCTCTGAATGTCAAAGTGAAGAAATTCAATGA | >hg38_21_+_8433172_8446622_RNA45SN1_45S_50ntFlanks | 11667 | 11707 | 41 | CTGCCCAGTGCTCTGAATGTCAAAGTGAAGAAATTCAATGA | 468.3 | 12.9 | 1.3 | 10.3 | 7.5E-25 | 4.3E-23 | MB231/MB468 | CYTO |
| >hg38_21_+_8433172_8446622_RNA45SN1_45S_50ntFlanks@8552.8569.18\|CTCCCCGGAGGGGGCGGG | >hg38_21_+_8433172_8446622_RNA45SN1_45S_50ntFlanks | 8552 | 8569 | 18 | CTCCCCGGAGGGGGCGGG | 542.7 | -13.5 | 1.3 | -10.1 | 7.9E-24 | 4.3E-23 | BT20/MB468 | NUC |
| >hg38_21_+_8433172_8446622_RNA45SN1_45S_50ntFlanks@11759.11777.19\|CCTCGTCATCTAATTAGTG | >hg38_21_+_8433172_8446622_RNA45SN1_45S_50ntFlanks | 11759 | 11777 | 19 | CCTCGTCATCTAATTAGTG | 353.4 | -12.9 | 1.3 | -10.1 | 7.9E-24 | 4.3E-23 | BT20/MB468 | NUC |

<table>
<tr><th></th><th></th><th></th><th></th><th></th><th></th><th></th><th></th><th></th><th></th><th></th><th></th><th></th><th></th></tr>
<tr><td>>hg38_21_+_8433172_8446622_RNA45SN1_45S_50ntFlanks@3782.3802.21|CAGTGAAACTGCGAATGGCTC</td><td>>hg38_21_+_8433172_8446622_RNA45SN1_45S_50ntFlanks</td><td>3782</td><td>3802</td><td>21</td><td>CAGTGAAACTGCGAATGGCTC</td><td>798.5</td><td>-14.1</td><td>1.4</td><td>-10.1</td><td>7.9E-24</td><td>4.3E-23</td><td>BT20/MB468</td><td>NUC</td></tr>
<tr><td>>MI0000299|hsa-mir-222&WithFlank&X|-|45747009|45747130@75.98.24&[MIMAT0000279&hsa-miR-222-3p&offsets|0|+3;m-140&X|-|45747033|45747056&offsets|0|0]|AGCTACATCTGGCTACTGGGTCTC</td><td>>MI0000299|hsa-mir-222&WithFlank&X|-|45747009|45747130</td><td>75</td><td>98</td><td>24</td><td>AGCTACATCTGGCTACTGGGTCTC</td><td>30061.0</td><td>8.0</td><td>0.8</td><td>10.3</td><td>7.6E-25</td><td>4.3E-23</td><td>MB231/MB468</td><td>CYTO</td></tr>
<tr><td>>hg38_21_+_8433172_8446622_RNA45SN1_45S_50ntFlanks@12628.12657.30|TGGGGCGAAGCTACCATCTGTGGGATTATG</td><td>>hg38_21_+_8433172_8446622_RNA45SN1_45S_50ntFlanks</td><td>12628</td><td>12657</td><td>30</td><td>TGGGGCGAAGCTACCATCTGTGGGATTATG</td><td>525.1</td><td>-13.5</td><td>1.3</td><td>-10.1</td><td>8.0E-24</td><td>4.4E-23</td><td>BT20/MB468</td><td>NUC</td></tr>
<tr><td>>hg38_21_+_8433172_8446622_RNA45SN1_45S_50ntFlanks@8169.8212.44|CCAAGTCCTTCTGATCGAGGCCCAGCCCGTGGACGGTGTGAGGC</td><td>>hg38_21_+_8433172_8446622_RNA45SN1_45S_50ntFlanks</td><td>8169</td><td>8212</td><td>44</td><td>CCAAGTCCTTCTGATCGAGGCCCAGCCCGTGGACGGTGTGAGGC</td><td>1608.6</td><td>-15.1</td><td>1.5</td><td>-10.1</td><td>8.0E-24</td><td>4.4E-23</td><td>BT20/MB468</td><td>NUC</td></tr>
<tr><td>>hg38_21_+_8433172_8446622_RNA45SN1_45S_50ntFlanks@12937.12957.21|AACCATTCGTAGACGACCTGC</td><td>>hg38_21_+_8433172_8446622_RNA45SN1_45S_50ntFlanks</td><td>12937</td><td>12957</td><td>21</td><td>AACCATTCGTAGACGACCTGC</td><td>794.6</td><td>-5.6</td><td>0.6</td><td>-10.1</td><td>8.0E-24</td><td>4.4E-23</td><td>BT20/MB468</td><td>NUC</td></tr>
<tr><td>>hg38_21_+_8433172_8446622_RNA45SN1_45S_50ntFlanks@12644.12661.18|TCTGTGGGATTATGACTG</td><td>>hg38_21_+_8433172_8446622_RNA45SN1_45S_50ntFlanks</td><td>12644</td><td>12661</td><td>18</td><td>TCTGTGGGATTATGACTG</td><td>2547.1</td><td>-15.2</td><td>1.5</td><td>-10.3</td><td>1.1E-24</td><td>4.4E-23</td><td>MB231/MB468</td><td>MP</td></tr>
<tr><td>>hg38_21_+_8433172_8446622_RNA45SN1_45S_50ntFlanks@12366.12383.18|TAACTGGCTTGTGGCGGC</td><td>>hg38_21_+_8433172_8446622_RNA45SN1_45S_50ntFlanks</td><td>12366</td><td>12383</td><td>18</td><td>TAACTGGCTTGTGGCGGC</td><td>978.5</td><td>-13.8</td><td>1.3</td><td>-10.3</td><td>1.1E-24</td><td>4.4E-23</td><td>MB231/MB468</td><td>MP</td></tr>
<tr><td>>MI0000266|hsa-mir-10a&WithFlank&17|-|48579832|48579953@28.51.24&[MIMAT0000253&hsa-miR-10a-5p&offsets|0|+1;m-18&17|-|48579905|48579926&offsets|0|+2]|TACCCTGTAGATCCGAATTTGTGT</td><td>>MI0000266|hsa-mir-10a&WithFlank&17|-|48579832|48579953</td><td>28</td><td>51</td><td>24</td><td>TACCCTGTAGATCCGAATTTGTGT</td><td>641.7</td><td>13.2</td><td>1.3</td><td>10.2</td><td>2.0E-24</td><td>4.4E-23</td><td>MB231/MB468</td><td>NUC</td></tr>
<tr><td>>hg38_21_+_8433172_8446622_RNA45SN1_45S_50ntFlanks@12456.12483.28|CCAAGCGTTGGATTGTTCACCCACTAAT</td><td>>hg38_21_+_8433172_8446622_RNA45SN1_45S_50ntFlanks</td><td>12456</td><td>12483</td><td>28</td><td>CCAAGCGTTGGATTGTTCACCCACTAAT</td><td>1928.5</td><td>-7.1</td><td>0.7</td><td>-10.2</td><td>1.7E-24</td><td>4.4E-23</td><td>MB231/MB468</td><td>MITO</td></tr>
<tr><td>>hg38_21_+_8433172_8446622_RNA45SN1_45S_50ntFlanks@12627.12654.28|ATGGGGCGAAGCTACCATCTGTGGGATT</td><td>>hg38_21_+_8433172_8446622_RNA45SN1_45S_50ntFlanks</td><td>12627</td><td>12654</td><td>28</td><td>ATGGGGCGAAGCTACCATCTGTGGGATT</td><td>4059.5</td><td>-5.7</td><td>0.6</td><td>-10.1</td><td>3.9E-24</td><td>4.5E-23</td><td>BT20/MB468</td><td>CYTO</td></tr>
<tr><td>>hg38_21_+_8433172_8446622_RNA45SN1_45S_50ntFlanks@10830.10858.29|CAAGCCGGATCCGTAACTTCGGGATAAGG</td><td>>hg38_21_+_8433172_8446622_RNA45SN1_45S_50ntFlanks</td><td>10830</td><td>10858</td><td>29</td><td>CAAGCCGGATCCGTAACTTCGGGATAAGG</td><td>390.1</td><td>-13.0</td><td>1.3</td><td>-10.1</td><td>8.2E-24</td><td>4.5E-23</td><td>BT20/MB468</td><td>NUC</td></tr>
<tr><td>>hg38_21_+_8433172_8446622_RNA45SN1_45S_50ntFlanks@10574.10591.18|AAGCCGGCGGGAGCCCCG</td><td>>hg38_21_+_8433172_8446622_RNA45SN1_45S_50ntFlanks</td><td>10574</td><td>10591</td><td>18</td><td>AAGCCGGCGGGAGCCCCG</td><td>416.9</td><td>-13.1</td><td>1.3</td><td>-10.1</td><td>8.3E-24</td><td>4.5E-23</td><td>BT20/MB468</td><td>NUC</td></tr>
<tr><td>>MI0000266|hsa-mir-10a&WithFlank&17|-|48579832|48579953@29.49.21&[MIMAT0000253&hsa-miR-10a-5p&offsets|+1|-1;m-18&17|-|48579905|48579926&offsets|+1|0]|ACCCTGTAGATCCGAATTTGT</td><td>>MI0000266|hsa-mir-10a&WithFlank&17|-|48579832|48579953</td><td>29</td><td>49</td><td>21</td><td>ACCCTGTAGATCCGAATTTGT</td><td>1422.3</td><td>14.5</td><td>1.4</td><td>10.3</td><td>8.0E-25</td><td>4.5E-23</td><td>MB231/MB468</td><td>CYTO</td></tr>
<tr><td>>MI0000067|hsa-let-7f-1&WithFlank&9|+|94176341|94176439@13.34.22;MI0000068|hsa-let-7f-2&WithFlank&X|-|53557186|53557280@14.35.22&[MIMAT0000067&hsa-let-7f-5p&offsets|0|0;m-20&9|+|94176353|94176374&offsets|0|0];[MIMAT0000067_1&hsa-let-7f-5p&offsets|0|0;m-19&X|-|53557246|53557267&offsets|0|0]|TGAGGTAGTAGATTGTATAGTT</td><td>>MI0000067|hsa-let-7f-1&WithFlank&9|+|94176341|94176439</td><td>13</td><td>34</td><td>22</td><td>TGAGGTAGTAGATTGTATAGTT</td><td>200839.1</td><td>6.6</td><td>0.7</td><td>10.1</td><td>4.0E-24</td><td>4.6E-23</td><td>BT20/MB468</td><td>CYTO</td></tr>
<tr><td>>MI0000465|hsa-mir-191&WithFlank&3|-|49020612|49020715@22.43.22&[MIMAT0000440&hsa-miR-191-5p&offsets|0|-1;m-48&3|-|49020672|49020694&offsets|0|-1]|CAACGGAATCCCAAAAGCAGCT</td><td>>MI0000465|hsa-mir-191&WithFlank&3|-|49020612|49020715</td><td>22</td><td>43</td><td>22</td><td>CAACGGAATCCCAAAAGCAGCT</td><td>10380.0</td><td>7.3</td><td>0.7</td><td>10.2</td><td>3.1E-24</td><td>4.6E-23</td><td>BT20/MB468</td><td>MITO</td></tr>
<tr><td>>hg38_21_+_8433172_8446622_RNA45SN1_45S_50ntFlanks@12457.12485.29|CAAGCGTTGGATTGTTCACCCACTAATAG</td><td>>hg38_21_+_8433172_8446622_RNA45SN1_45S_50ntFlanks</td><td>12457</td><td>12485</td><td>29</td><td>CAAGCGTTGGATTGTTCACCCACTAATAG</td><td>525.6</td><td>-13.5</td><td>1.3</td><td>-10.1</td><td>8.4E-24</td><td>4.6E-23</td><td>BT20/MB468</td><td>NUC</td></tr>
<tr><td>>MI0000266|hsa-mir-10a&WithFlank&17|-|48579832|48579953@28.50.23(+1C)&[MIMAT0000253&hsa-miR-10a-5p&offsets|0|0(+1C);m-18&17|-|48579905|48579926&offsets|0|+1(+1C)]|TACCCTGTAGATCCGAATTTGTGC</td><td>>MI0000266|hsa-mir-10a&WithFlank&17|-|48579832|48579953</td><td>28</td><td>50</td><td>23(+1C)</td><td>TACCCTGTAGATCCGAATTTGTGC</td><td>386.5</td><td>12.8</td><td>1.3</td><td>10.1</td><td>4.0E-24</td><td>4.6E-23</td><td>BT20/MB468</td><td>CYTO</td></tr>
<tr><td>>MI0000253|hsa-mir-148a&WithFlank&7|-|25949913|25949992@50.70.21(+1A)&[MIMAT0000243&hsa-miR-148a-3p&offsets|0|-1(+1A);m-7&7|-|25949922|25949943&offsets|0|-1(+1A)]|TCAGTGCACTACAGAACTTTGA</td><td>>MI0000253|hsa-mir-148a&WithFlank&7|-|25949913|25949992</td><td>50</td><td>70</td><td>21(+1A)</td><td>TCAGTGCACTACAGAACTTTGA</td><td>9828.4</td><td>8.3</td><td>0.8</td><td>10.1</td><td>4.0E-24</td><td>4.6E-23</td><td>BT20/MB468</td><td>CYTO</td></tr>
<tr><td>>MI0001733|hsa-mir-452&WithFlank&X|-|151959622|151959718@20.41.22&[MIMAT0001635&hsa-miR-452-5p&offsets|0|0;m-188&X|-|151959677|151959699&offsets|0|-1]|AACTGTTTGCAGAGGAAACTGA</td><td>>MI0001733|hsa-mir-452&WithFlank&X|-|151959622|151959718</td><td>20</td><td>41</td><td>22</td><td>AACTGTTTGCAGAGGAAACTGA</td><td>349.5</td><td>12.7</td><td>1.3</td><td>10.1</td><td>4.2E-24</td><td>4.8E-23</td><td>BT20/MB468</td><td>CYTO</td></tr>
<tr><td>>hg38_21_+_8433172_8446622_RNA45SN1_45S_50ntFlanks@6719.6746.28|TGAATTGCAGGACACATTGATCATCGAC</td><td>>hg38_21_+_8433172_8446622_RNA45SN1_45S_50ntFlanks</td><td>6719</td><td>6746</td><td>28</td><td>TGAATTGCAGGACACATTGATCATCGAC</td><td>1475.5</td><td>-14.4</td><td>1.4</td><td>-10.3</td><td>1.2E-24</td><td>4.8E-23</td><td>MB231/MB468</td><td>MP</td></tr>
<tr><td>>hg38_21_+_8433172_8446622_RNA45SN1_45S_50ntFlanks@3989.4023.35|TTTGGTGACTCTAGATAACCTCGGGCCGATCGCAC</td><td>>hg38_21_+_8433172_8446622_RNA45SN1_45S_50ntFlanks</td><td>3989</td><td>4023</td><td>35</td><td>TTTGGTGACTCTAGATAACCTCGGGCCGATCGCAC</td><td>559.3</td><td>5.4</td><td>0.5</td><td>10.1</td><td>4.2E-24</td><td>4.8E-23</td><td>BT20/MB468</td><td>CYTO</td></tr>
<tr><td>>hg38_21_+_8433172_8446622_RNA45SN1_45S_50ntFlanks@12002.12043.42|TGCGGGCCGCCGGTGAAATACCACTACTCTGATCGTTTTTTC</td><td>>hg38_21_+_8433172_8446622_RNA45SN1_45S_50ntFlanks</td><td>12002</td><td>12043</td><td>42</td><td>TGCGGGCCGCCGGTGAAATACCACTACTCTGATCGTTTTTTC</td><td>4446.3</td><td>-16.5</td><td>1.6</td><td>-10.1</td><td>8.8E-24</td><td>4.8E-23</td><td>BT20/MB468</td><td>NUC</td></tr>
<tr><td>>am_MI0000239|hsa-mir-197&WithFlank&1|+|109598887|109598973@54.71.18(+1U)&[MIMAT0000227&hsa-miR-197-3p&offsets|0|-4(+1U);m-99&1|+|109598940|109598961&offsets|0|-4(+1U)]|TTCACCACCTTCTCCACCT</td><td>>am_MI0000239|hsa-mir-197&WithFlank&1|+|109598887|109598973</td><td>54</td><td>71</td><td>18(+1U)</td><td>TTCACCACCTTCTCCACCT</td><td>556.8</td><td>13.4</td><td>1.3</td><td>10.1</td><td>4.2E-24</td><td>4.8E-23</td><td>BT20/MB468</td><td>CYTO</td></tr>
<tr><td>>MI0000255|hsa-mir-30d&WithFlank&8|-|134804870|134804951@12.35.24(+1U)&[MIMAT0000245&hsa-miR-30d-5p&offsets|0|+2(+1U);m-21&8|-|134804919|134804940&offsets|0|+2(+1U)]|TGTAAACATCCCCGACTGGAAGCTT</td><td>>MI0000255|hsa-mir-30d&WithFlank&8|-|134804870|134804951</td><td>12</td><td>35</td><td>24(+1U)</td><td>TGTAAACATCCCCGACTGGAAGCTT</td><td>2680.5</td><td>6.8</td><td>0.7</td><td>10.2</td><td>2.2E-24</td><td>4.9E-23</td><td>MB231/MB468</td><td>NUC</td></tr>
<tr><td>>hg38_21_+_8433172_8446622_RNA45SN1_45S_50ntFlanks@8113.8138.26|GGCGCGGGACATGTGGCGTACGGAAG</td><td>>hg38_21_+_8433172_8446622_RNA45SN1_45S_50ntFlanks</td><td>8113</td><td>8138</td><td>26</td><td>GGCGCGGGACATGTGGCGTACGGAAG</td><td>9023.5</td><td>-12.9</td><td>1.3</td><td>-10.2</td><td>1.2E-24</td><td>4.9E-23</td><td>MB231/MB468</td><td>MP</td></tr>
<tr><td>>hg38_21_+_8433172_8446622_RNA45SN1_45S_50ntFlanks@11873.11905.33|AAGAAGACCCTGTTGAGCTTGACTCTAGTCTGG</td><td>>hg38_21_+_8433172_8446622_RNA45SN1_45S_50ntFlanks</td><td>11873</td><td>11905</td><td>33</td><td>AAGAAGACCCTGTTGAGCTTGACTCTAGTCTGG</td><td>1562.8</td><td>-14.5</td><td>1.4</td><td>-10.2</td><td>1.2E-24</td><td>4.9E-23</td><td>MB231/MB468</td><td>MP</td></tr>
<tr><td>>hg38_21_+_8433172_8446622_RNA45SN1_45S_50ntFlanks@12627.12652.26|ATGGGGCGAAGCTACCATCTGTGGGA</td><td>>hg38_21_+_8433172_8446622_RNA45SN1_45S_50ntFlanks</td><td>12627</td><td>12652</td><td>26</td><td>ATGGGGCGAAGCTACCATCTGTGGGA</td><td>427.7</td><td>-13.2</td><td>1.3</td><td>-10.1</td><td>9.1E-24</td><td>5.0E-23</td><td>BT20/MB468</td><td>NUC</td></tr>
<tr><td>>hg38_21_+_8433172_8446622_RNA45SN1_45S_50ntFlanks@6700.6718.19|CTAGCTGCGAGAATTAATG</td><td>>hg38_21_+_8433172_8446622_RNA45SN1_45S_50ntFlanks</td><td>6700</td><td>6718</td><td>19</td><td>CTAGCTGCGAGAATTAATG</td><td>8943.8</td><td>-6.8</td><td>0.7</td><td>-10.1</td><td>4.3E-24</td><td>5.0E-23</td><td>BT20/MB468</td><td>CYTO</td></tr>
<tr><td>>hg38_21_+_8433172_8446622_RNA45SN1_45S_50ntFlanks@6646.6681.36|TCGTACGACTCTTAGCGGTGGATCACTCGGCTCGTG</td><td>>hg38_21_+_8433172_8446622_RNA45SN1_45S_50ntFlanks</td><td>6646</td><td>6681</td><td>36</td><td>TCGTACGACTCTTAGCGGTGGATCACTCGGCTCGTG</td><td>32841.6</td><td>-4.2</td><td>0.4</td><td>-10.1</td><td>9.2E-24</td><td>5.0E-23</td><td>BT20/MB468</td><td>NUC</td></tr>
<tr><td>>hg38_21_+_8433172_8446622_RNA45SN1_45S_50ntFlanks@7983.8006.24|CAGATCAGACGTGGCGACCCGCTG</td><td>>hg38_21_+_8433172_8446622_RNA45SN1_45S_50ntFlanks</td><td>7983</td><td>8006</td><td>24</td><td>CAGATCAGACGTGGCGACCCGCTG</td><td>16105.3</td><td>-6.7</td><td>0.7</td><td>-10.1</td><td>3.3E-24</td><td>5.0E-23</td><td>BT20/MB468</td><td>MITO</td></tr>
<tr><td>>hg38_21_+_8433172_8446622_RNA45SN1_45S_50ntFlanks@4730.4768.39|CAAGAACGAAAGTCGGAGGTTCGAAGACGATCAGATACC</td><td>>hg38_21_+_8433172_8446622_RNA45SN1_45S_50ntFlanks</td><td>4730</td><td>4768</td><td>39</td><td>CAAGAACGAAAGTCGGAGGTTCGAAGACGATCAGATACC</td><td>774.8</td><td>13.9</td><td>1.4</td><td>10.1</td><td>4.4E-24</td><td>5.0E-23</td><td>BT20/MB468</td><td>CYTO</td></tr>
<tr><td>>hg38_21_+_8433172_8446622_RNA45SN1_45S_50ntFlanks@12416.12439.24|TTCGATGTCGGCTCTTCCTATCAT</td><td>>hg38_21_+_8433172_8446622_RNA45SN1_45S_50ntFlanks</td><td>12416</td><td>12439</td><td>24</td><td>TTCGATGTCGGCTCTTCCTATCAT</td><td>2709.2</td><td>6.5</td><td>0.6</td><td>10.3</td><td>9.0E-25</td><td>5.0E-23</td><td>MB231/MB468</td><td>CYTO</td></tr>
<tr><td>>hg38_21_+_8433172_8446622_RNA45SN1_45S_50ntFlanks@10689.10709.21|GCGTCCGGTGAGCTCTCGCTG</td><td>>hg38_21_+_8433172_8446622_RNA45SN1_45S_50ntFlanks</td><td>10689</td><td>10709</td><td>21</td><td>GCGTCCGGTGAGCTCTCGCTG</td><td>843.9</td><td>-8.1</td><td>0.8</td><td>-10.3</td><td>9.1E-25</td><td>5.0E-23</td><td>MB231/MB468</td><td>CYTO</td></tr>
<tr><td>>hg38_21_+_8433172_8446622_RNA45SN1_45S_50ntFlanks@9027.9059.33|CGTCTCCTCGTGGGGGGGCCGGCCACCCCTCC</td><td>>hg38_21_+_8433172_8446622_RNA45SN1_45S_50ntFlanks</td><td>9027</td><td>9059</td><td>33</td><td>CGTCTCCTCGTGGGGGGGCCGGCCACCCCTCC</td><td>1864.2</td><td>-14.6</td><td>1.4</td><td>-10.2</td><td>2.0E-24</td><td>5.1E-23</td><td>MB231/MB468</td><td>MITO</td></tr>
<tr><td>>hg38_21_+_8433172_8446622_RNA45SN1_45S_50ntFlanks@12044.12072.29|ACTGACCCGGTGAGGCGGGGGGGCGAGCC</td><td>>hg38_21_+_8433172_8446622_RNA45SN1_45S_50ntFlanks</td><td>12044</td><td>12072</td><td>29</td><td>ACTGACCCGGTGAGGCGGGGGGGCGAGCC</td><td>3376.9</td><td>-15.5</td><td>1.5</td><td>-10.2</td><td>2.0E-24</td><td>5.1E-23</td><td>MB231/MB468</td><td>MITO</td></tr>
<tr><td>>hg38_21_+_8433172_8446622_RNA45SN1_45S_50ntFlanks@4220.4242.23|ACGAAAAATAACAATACAGGACT</td><td>>hg38_21_+_8433172_8446622_RNA45SN1_45S_50ntFlanks</td><td>4220</td><td>4242</td><td>23</td><td>ACGAAAAATAACAATACAGGACT</td><td>1710.2</td><td>-6.9</td><td>0.7</td><td>-10.2</td><td>2.0E-24</td><td>5.1E-23</td><td>MB231/MB468</td><td>MITO</td></tr>
<tr><td>>hg38_21_+_8433172_8446622_RNA45SN1_45S_50ntFlanks@11886.11904.19|TGAGCTTGACTCTAGTCTG</td><td>>hg38_21_+_8433172_8446622_RNA45SN1_45S_50ntFlanks</td><td>11886</td><td>11904</td><td>19</td><td>TGAGCTTGACTCTAGTCTG</td><td>1371.0</td><td>-14.3</td><td>1.4</td><td>-10.2</td><td>1.3E-24</td><td>5.1E-23</td><td>MB231/MB468</td><td>MP</td></tr>
<tr><td>>hg38_21_+_8433172_8446622_RNA45SN1_45S_50ntFlanks@482.508.27|ATCGATGTGGTGACGTCGTGCTCTCCC</td><td>>hg38_21_+_8433172_8446622_RNA45SN1_45S_50ntFlanks</td><td>482</td><td>508</td><td>27</td><td>ATCGATGTGGTGACGTCGTGCTCTCCC</td><td>423.0</td><td>-13.2</td><td>1.3</td><td>-10.0</td><td>9.5E-24</td><td>5.1E-23</td><td>BT20/MB468</td><td>NUC</td></tr>
<tr><td>>hg38_21_+_8433172_8446622_RNA45SN1_45S_50ntFlanks@8115.8142.28|CGCGGGACATGTGGCGTACGGAAGACCC</td><td>>hg38_21_+_8433172_8446622_RNA45SN1_45S_50ntFlanks</td><td>8115</td><td>8142</td><td>28</td><td>CGCGGGACATGTGGCGTACGGAAGACCC</td><td>1522.4</td><td>-14.4</td><td>1.4</td><td>-10.2</td><td>1.3E-24</td><td>5.1E-23</td><td>MB231/MB468</td><td>MP</td></tr>
<tr><td>>MI0000446|hsa-mir-125b-1&WithFlank&11|-|122099751|122099850@61.81.21&[MIMAT0004592&hsa-miR-125b-1-3p&offsets|0|-1;m-485&11|-|122099769|122099790&offsets|0|-1]|ACGGGTTAGGCTCTTGGGAGC</td><td>>MI0000446|hsa-mir-125b-1&WithFlank&11|-|122099751|122099850</td><td>61</td><td>81</td><td>21</td><td>ACGGGTTAGGCTCTTGGGAGC</td><td>1425.5</td><td>14.5</td><td>1.4</td><td>10.3</td><td>9.4E-25</td><td>5.2E-23</td><td>MB231/MB468</td><td>CYTO</td></tr>
<tr><td>>hg38_21_+_8433172_8446622_RNA45SN1_45S_50ntFlanks@12712.12747.36|CGGAGCCTCGGTTGGCCTCGGATAGCCGGTCCCCCG</td><td>>hg38_21_+_8433172_8446622_RNA45SN1_45S_50ntFlanks</td><td>12712</td><td>12747</td><td>36</td><td>CGGAGCCTCGGTTGGCCTCGGATAGCCGGTCCCCCG</td><td>1854.1</td><td>-14.7</td><td>1.4</td><td>-10.2</td><td>1.3E-24</td><td>5.2E-23</td><td>MB231/MB468</td><td>MP</td></tr>
<tr><td>>hg38_21_+_8433172_8446622_RNA45SN1_45S_50ntFlanks@8380.8399.20|CAAGAGGGCGTGAAACCGTT</td><td>>hg38_21_+_8433172_8446622_RNA45SN1_45S_50ntFlanks</td><td>8380</td><td>8399</td><td>20</td><td>CAAGAGGGCGTGAAACCGTT</td><td>333.6</td><td>-12.8</td><td>1.3</td><td>-10.0</td><td>9.7E-24</td><td>5.2E-23</td><td>BT20/MB468</td><td>NUC</td></tr>
<tr><td>>hg38_21_+_8433172_8446622_RNA45SN1_45S_50ntFlanks@11873.11905.33|AAGAAGACCCTGTTGAGCTTGACTCTAGTCTGG</td><td>>hg38_21_+_8433172_8446622_RNA45SN1_45S_50ntFlanks</td><td>11873</td><td>11905</td><td>33</td><td>AAGAAGACCCTGTTGAGCTTGACTCTAGTCTGG</td><td>1622.2</td><td>-14.4</td><td>1.4</td><td>-10.2</td><td>2.1E-24</td><td>5.3E-23</td><td>MB231/MB468</td><td>MITO</td></tr>
<tr><td>>am_trnaMT_ValTAC_MT_+_1602_1670@51.68.18|CTTAACTTGACCGCTCTG</td><td>>am_trnaMT_ValTAC_MT_+_1602_1670</td><td>51</td><td>68</td><td>18</td><td>CTTAACTTGACCGCTCTG</td><td>49091.7</td><td>-11.1</td><td>1.1</td><td>-10.2</td><td>2.4E-24</td><td>5.3E-23</td><td>MB231/MB468</td><td>NUC</td></tr>
<tr><td>>MI0000299|hsa-mir-222&WithFlank&X|-|45747009|45747130@75.95.21&[MIMAT0000279&hsa-miR-222-3p&offsets|0|0;m-140&X|-|45747033|45747056&offsets|0|-3]|AGCTACATCTGGCTACTGGGT</td><td>>MI0000299|hsa-mir-222&WithFlank&X|-|45747009|45747130</td><td>75</td><td>95</td><td>21</td><td>AGCTACATCTGGCTACTGGGT</td><td>1562.5</td><td>6.3</td><td>0.6</td><td>10.1</td><td>3.6E-24</td><td>5.3E-23</td><td>BT20/MB468</td><td>MITO</td></tr>
<tr><td>>hg38_21_+_8433172_8446622_RNA45SN1_45S_50ntFlanks@11572.11592.21|CCGACTTAGAACTGGTGCGGA</td><td>>hg38_21_+_8433172_8446622_RNA45SN1_45S_50ntFlanks</td><td>11572</td><td>11592</td><td>21</td><td>CCGACTTAGAACTGGTGCGGA</td><td>6116.4</td><td>-16.4</td><td>1.6</td><td>-10.2</td><td>1.3E-24</td><td>5.3E-23</td><td>MB231/MB468</td><td>MP</td></tr>
<tr><td>>hg38_21_+_8433172_8446622_RNA45SN1_45S_50ntFlanks@485.508.24|GATGTGGTGACGTCGTGCTCTCCC</td><td>>hg38_21_+_8433172_8446622_RNA45SN1_45S_50ntFlanks</td><td>485</td><td>508</td><td>24</td><td>GATGTGGTGACGTCGTGCTCTCCC</td><td>363.9</td><td>11.3</td><td>1.1</td><td>10.0</td><td>9.9E-24</td><td>5.3E-23</td><td>BT20/MB468</td><td>NUC</td></tr>
<tr><td>>hg38_21_+_8433172_8446622_RNA45SN1_45S_50ntFlanks@11535.11552.18|TTCCGCGCGGCGCCTCGC</td><td>>hg38_21_+_8433172_8446622_RNA45SN1_45S_50ntFlanks</td><td>11535</td><td>11552</td><td>18</td><td>TTCCGCGCGGCGCCTCGC</td><td>1148.7</td><td>-14.6</td><td>1.5</td><td>-10.0</td><td>9.9E-24</td><td>5.3E-23</td><td>BT20/MB468</td><td>NUC</td></tr>
<tr><td>>MI0000283|hsa-mir-203a&WithFlank&14|+|104117399|104117520@71.91.21(+1A)&[MIMAT0000264&hsa-miR-203a-3p&offsets|0|-1(+1A);m-79&14|+|104117469|104117490&offsets|0|-1(+1A)]|GTGAAATGTTTAGGACCACTAA</td><td>>MI0000283|hsa-mir-203a&WithFlank&14|+|104117399|104117520</td><td>71</td><td>91</td><td>21(+1A)</td><td>GTGAAATGTTTAGGACCACTAA</td><td>588.2</td><td>13.5</td><td>1.3</td><td>10.1</td><td>4.7E-24</td><td>5.3E-23</td><td>BT20/MB468</td><td>CYTO</td></tr>
<tr><td>>MI0000433|hsa-let-7g&WithFlank&3|-|52268272|52268367@11.33.23&[MIMAT0000414&hsa-let-7g-5p&offsets|0|+1;m-58&3|-|52268336|52268357&offsets|0|+1]|TGAGGTAGTAGTTTGTACAGTTT</td><td>>MI0000433|hsa-let-7g&WithFlank&3|-|52268272|52268367</td><td>11</td><td>33</td><td>23</td><td>TGAGGTAGTAGTTTGTACAGTTT</td><td>1132.0</td><td>7.2</td><td>0.7</td><td>10.1</td><td>3.6E-24</td><td>5.3E-23</td><td>BT20/MB468</td><td>MITO</td></tr>
</table>

<table>
<tr><td>>MI0001733|hsa-mir-452&WithFlank&X|-|151959622|151959718@23.44.22&[MIMAT0001635&hsa-miR-452-5p&offsets|+3|+3;m-188&X|-|151959677|151959699&offsets|+3|+2]|TGTTTGCAGAGGAAACTGAGAC</td><td>>MI0001733|hsa-mir-452&WithFlank&X|-|151959622|151959718</td><td>23</td><td>44</td><td>22</td><td>TGTTTGCAGAGGAAACTGAGAC</td><td>573.0</td><td>13.6</td><td>1.3</td><td>10.1</td><td>3.6E-24</td><td>5.3E-23</td><td>BT20/MB468</td><td>MITO</td></tr>
<tr><td>>MI0000266|hsa-mir-10a&WithFlank&17|-|48579832|48579953@29.51.23&[MIMAT0000253&hsa-miR-10a-5p&offsets|+1|+1;m-18&17|-|48579905|48579926&offsets|+1|+2]|ACCCTGTAGATCCGAATTTGTGT</td><td>>MI0000266|hsa-mir-10a&WithFlank&17|-|48579832|48579953</td><td>29</td><td>51</td><td>23</td><td>ACCCTGTAGATCCGAATTTGTGT</td><td>719.8</td><td>13.4</td><td>1.3</td><td>10.2</td><td>2.4E-24</td><td>5.4E-23</td><td>MB231/MB468</td><td>NUC</td></tr>
<tr><td>>hg38_21_+_8433172_8446622_RNA45SN1_45S_50ntFlanks@12539.12565.27|ATGATGTGTTGTTGCCATGGTAATCCT</td><td>>hg38_21_+_8433172_8446622_RNA45SN1_45S_50ntFlanks</td><td>12539</td><td>12565</td><td>27</td><td>ATGATGTGTTGTTGCCATGGTAATCCT</td><td>918.9</td><td>-14.3</td><td>1.4</td><td>-10.0</td><td>1.0E-23</td><td>5.4E-23</td><td>BT20/MB468</td><td>NUC</td></tr>
<tr><td>>hg38_21_+_8433172_8446622_RNA45SN1_45S_50ntFlanks@9538.9563.26|AACTCTGGTGGAGGTCCGTAGCGGTC</td><td>>hg38_21_+_8433172_8446622_RNA45SN1_45S_50ntFlanks</td><td>9538</td><td>9563</td><td>26</td><td>AACTCTGGTGGAGGTCCGTAGCGGTC</td><td>543.0</td><td>-13.5</td><td>1.3</td><td>-10.0</td><td>1.0E-23</td><td>5.4E-23</td><td>BT20/MB468</td><td>NUC</td></tr>
<tr><td>>hg38_21_+_8433172_8446622_RNA45SN1_45S_50ntFlanks@4147.4175.29|TGAGAAACGGCTACCACATCCAAGGAAGG</td><td>>hg38_21_+_8433172_8446622_RNA45SN1_45S_50ntFlanks</td><td>4147</td><td>4175</td><td>29</td><td>TGAGAAACGGCTACCACATCCAAGGAAGG</td><td>472.6</td><td>-13.3</td><td>1.3</td><td>-10.0</td><td>1.0E-23</td><td>5.5E-23</td><td>BT20/MB468</td><td>NUC</td></tr>
<tr><td>>hg38_21_+_8433172_8446622_RNA45SN1_45S_50ntFlanks@6646.6667.22|TCGTACGACTCTTAGCGGTGGA</td><td>>hg38_21_+_8433172_8446622_RNA45SN1_45S_50ntFlanks</td><td>6646</td><td>6667</td><td>22</td><td>TCGTACGACTCTTAGCGGTGGA</td><td>18662.7</td><td>6.7</td><td>0.7</td><td>10.1</td><td>4.8E-24</td><td>5.5E-23</td><td>BT20/MB468</td><td>CYTO</td></tr>
<tr><td>>am_MI0000786|hsa-mir-378a&WithFlank&5|+|149732819|149732896@49.69.21(+1U)&[MIMAT0000732&hsa-miR-378a-3p&offsets|0|-1(+1U);m-44&5|+|149732867|149732888&offsets|0|-1(+1U)]|ACTGGACTTGGAGTCAGAAGGT</td><td>>am_MI0000786|hsa-mir-378a&WithFlank&5|+|149732819|149732896</td><td>49</td><td>69</td><td>21(+1U)</td><td>ACTGGACTTGGAGTCAGAAGGT</td><td>3606.7</td><td>5.8</td><td>0.6</td><td>10.1</td><td>3.7E-24</td><td>5.5E-23</td><td>BT20/MB468</td><td>MITO</td></tr>
<tr><td>>MI0000102|hsa-mir-100&WithFlank&11|-|122152223|122152314@20.40.21&[MIMAT0000098&hsa-miR-100-5p&offsets|+1|0;m-27&11|-|122152276|122152296&offsets|+1|+1]|ACCCGTAGATCCGAACTTGTG</td><td>>MI0000102|hsa-mir-100&WithFlank&11|-|122152223|122152314</td><td>20</td><td>40</td><td>21</td><td>ACCCGTAGATCCGAACTTGTG</td><td>2196.6</td><td>15.1</td><td>1.5</td><td>10.3</td><td>1.0E-24</td><td>5.5E-23</td><td>MB231/MB468</td><td>CYTO</td></tr>
<tr><td>>hg38_21_+_8433172_8446622_RNA45SN1_45S_50ntFlanks@5482.5514.33|CGGAGCGCTGAGAAGACGGTCGAACTTGACTAT</td><td>>hg38_21_+_8433172_8446622_RNA45SN1_45S_50ntFlanks</td><td>5482</td><td>5514</td><td>33</td><td>CGGAGCGCTGAGAAGACGGTCGAACTTGACTAT</td><td>772.1</td><td>-14.0</td><td>1.4</td><td>-10.0</td><td>1.0E-23</td><td>5.5E-23</td><td>BT20/MB468</td><td>NUC</td></tr>
<tr><td>>hg38_21_+_8433172_8446622_RNA45SN1_45S_50ntFlanks@10355.10388.34|AACTTTGAAGGCCGAAGTGGAGAAGGGTTCCATG</td><td>>hg38_21_+_8433172_8446622_RNA45SN1_45S_50ntFlanks</td><td>10355</td><td>10388</td><td>34</td><td>AACTTTGAAGGCCGAAGTGGAGAAGGGTTCCATG</td><td>1904.3</td><td>-15.3</td><td>1.5</td><td>-10.0</td><td>1.0E-23</td><td>5.5E-23</td><td>BT20/MB468</td><td>NUC</td></tr>
<tr><td>>hg38_21_+_8433172_8446622_RNA45SN1_45S_50ntFlanks@4047.4081.35|ATTCGAACGTCTGCCCTATCAACTTTCGATGGTAG</td><td>>hg38_21_+_8433172_8446622_RNA45SN1_45S_50ntFlanks</td><td>4047</td><td>4081</td><td>35</td><td>ATTCGAACGTCTGCCCTATCAACTTTCGATGGTAG</td><td>620.8</td><td>-13.7</td><td>1.4</td><td>-10.0</td><td>1.0E-23</td><td>5.5E-23</td><td>BT20/MB468</td><td>NUC</td></tr>
<tr><td>>hg38_21_+_8433172_8446622_RNA45SN1_45S_50ntFlanks@12407.12443.37|TTTTGATCCTTCGATGTCGGCTCTTCCTATCATTGTG</td><td>>hg38_21_+_8433172_8446622_RNA45SN1_45S_50ntFlanks</td><td>12407</td><td>12443</td><td>37</td><td>TTTTGATCCTTCGATGTCGGCTCTTCCTATCATTGTG</td><td>11244.2</td><td>-8.2</td><td>0.8</td><td>-10.0</td><td>1.0E-23</td><td>5.6E-23</td><td>BT20/MB468</td><td>NUC</td></tr>
<tr><td>>hg38_21_+_8433172_8446622_RNA45SN1_45S_50ntFlanks@12636.12657.22|AGCTACCATCTGTGGGATTATG</td><td>>hg38_21_+_8433172_8446622_RNA45SN1_45S_50ntFlanks</td><td>12636</td><td>12657</td><td>22</td><td>AGCTACCATCTGTGGGATTATG</td><td>1081.4</td><td>-13.9</td><td>1.4</td><td>-10.2</td><td>1.4E-24</td><td>5.6E-23</td><td>MB231/MB468</td><td>MP</td></tr>
<tr><td>>hg38_21_+_8433172_8446622_RNA45SN1_45S_50ntFlanks@6721.6738.18|AATTGCAGGACACATTGA</td><td>>hg38_21_+_8433172_8446622_RNA45SN1_45S_50ntFlanks</td><td>6721</td><td>6738</td><td>18</td><td>AATTGCAGGACACATTGA</td><td>610.7</td><td>-13.7</td><td>1.4</td><td>-10.0</td><td>1.0E-23</td><td>5.6E-23</td><td>BT20/MB468</td><td>NUC</td></tr>
<tr><td>>hg38_21_+_8433172_8446622_RNA45SN1_45S_50ntFlanks@13005.13027.23|TCTATTGAAAGTCAGCCCTCGAC</td><td>>hg38_21_+_8433172_8446622_RNA45SN1_45S_50ntFlanks</td><td>13005</td><td>13027</td><td>23</td><td>TCTATTGAAAGTCAGCCCTCGAC</td><td>390.2</td><td>-13.0</td><td>1.3</td><td>-10.0</td><td>1.1E-23</td><td>5.7E-23</td><td>BT20/MB468</td><td>NUC</td></tr>
<tr><td>>hg38_21_+_8433172_8446622_RNA45SN1_45S_50ntFlanks@9203.9222.20|CTCGGGGCCACGCGCGCGTC</td><td>>hg38_21_+_8433172_8446622_RNA45SN1_45S_50ntFlanks</td><td>9203</td><td>9222</td><td>20</td><td>CTCGGGGCCACGCGCGCGTC</td><td>1900.5</td><td>-14.8</td><td>1.4</td><td>-10.2</td><td>1.4E-24</td><td>5.7E-23</td><td>MB231/MB468</td><td>MP</td></tr>
<tr><td>>hg38_21_+_8433172_8446622_RNA45SN1_45S_50ntFlanks@12574.12604.31|CGAGAGGAACCGCAGGTTCAGACATTTGGTG</td><td>>hg38_21_+_8433172_8446622_RNA45SN1_45S_50ntFlanks</td><td>12574</td><td>12604</td><td>31</td><td>CGAGAGGAACCGCAGGTTCAGACATTTGGTG</td><td>952.6</td><td>-13.8</td><td>1.3</td><td>-10.2</td><td>1.4E-24</td><td>5.7E-23</td><td>MB231/MB468</td><td>MP</td></tr>
<tr><td>>hg38_21_+_8433172_8446622_RNA45SN1_45S_50ntFlanks@12324.12346.23|TTGGGTTTTAAGCAGGAGGTGTC</td><td>>hg38_21_+_8433172_8446622_RNA45SN1_45S_50ntFlanks</td><td>12324</td><td>12346</td><td>23</td><td>TTGGGTTTTAAGCAGGAGGTGTC</td><td>2268.8</td><td>-14.1</td><td>1.4</td><td>-10.2</td><td>2.6E-24</td><td>5.7E-23</td><td>MB231/MB468</td><td>NUC</td></tr>
<tr><td>>hg38_21_+_8433172_8446622_RNA45SN1_45S_50ntFlanks@12934.12966.33|CTAAACCATTCGTAGACGACCTGCTTCTGGGTC</td><td>>hg38_21_+_8433172_8446622_RNA45SN1_45S_50ntFlanks</td><td>12934</td><td>12966</td><td>33</td><td>CTAAACCATTCGTAGACGACCTGCTTCTGGGTC</td><td>689.5</td><td>-13.9</td><td>1.4</td><td>-10.0</td><td>1.1E-23</td><td>5.7E-23</td><td>BT20/MB468</td><td>NUC</td></tr>
<tr><td>>trna6_TrpCCA_12_+_98898030_98898101@30.52.23|ACTCCAGATCAGAAGGCTGCGTG</td><td>>trna6_TrpCCA_12_+_98898030_98898101</td><td>30</td><td>52</td><td>23</td><td>ACTCCAGATCAGAAGGCTGCGTG</td><td>855.1</td><td>-14.2</td><td>1.4</td><td>-10.0</td><td>1.1E-23</td><td>5.8E-23</td><td>BT20/MB468</td><td>NUC</td></tr>
<tr><td>>hg38_21_+_8433172_8446622_RNA45SN1_45S_50ntFlanks@5478.5516.39|CTGGCGGAGCGCTGAGAAGACGGTCGAACTTGACTATCT</td><td>>hg38_21_+_8433172_8446622_RNA45SN1_45S_50ntFlanks</td><td>5478</td><td>5516</td><td>39</td><td>CTGGCGGAGCGCTGAGAAGACGGTCGAACTTGACTATCT</td><td>347.5</td><td>12.7</td><td>1.3</td><td>10.1</td><td>5.1E-24</td><td>5.8E-23</td><td>BT20/MB468</td><td>CYTO</td></tr>
<tr><td>>MI0000440|hsa-mir-27b&WithFlank&9|+|95085439|95085547@67.87.21(+1U)&[MIMAT0000419&hsa-miR-27b-3p&offsets|0|0(+1U);m-39&9|+|95085505|95085525&offsets|0|0(+1U)]|TTCACAGTGGCTAAGTTCTGCT</td><td>>MI0000440|hsa-mir-27b&WithFlank&9|+|95085439|95085547</td><td>67</td><td>87</td><td>21(+1U)</td><td>TTCACAGTGGCTAAGTTCTGCT</td><td>886.6</td><td>6.7</td><td>0.7</td><td>10.2</td><td>2.6E-24</td><td>5.8E-23</td><td>MB231/MB468</td><td>NUC</td></tr>
<tr><td>>hg38_21_+_8433172_8446622_RNA45SN1_45S_50ntFlanks@10571.10608.38|GAGAAGCCGGCGGGAGCCCCGGGGAGAGTTCTCTTTTC</td><td>>hg38_21_+_8433172_8446622_RNA45SN1_45S_50ntFlanks</td><td>10571</td><td>10608</td><td>38</td><td>GAGAAGCCGGCGGGAGCCCCGGGGAGAGTTCTCTTTTC</td><td>772.3</td><td>-14.0</td><td>1.4</td><td>-10.0</td><td>1.1E-23</td><td>5.9E-23</td><td>BT20/MB468</td><td>NUC</td></tr>
<tr><td>>hg38_21_+_8433172_8446622_RNA45SN1_45S_50ntFlanks@12161.12181.21|TGACTGGGGCGGTACACCTGT</td><td>>hg38_21_+_8433172_8446622_RNA45SN1_45S_50ntFlanks</td><td>12161</td><td>12181</td><td>21</td><td>TGACTGGGGCGGTACACCTGT</td><td>882.7</td><td>-13.6</td><td>1.3</td><td>-10.2</td><td>1.5E-24</td><td>5.9E-23</td><td>MB231/MB468</td><td>MP</td></tr>
<tr><td>>hg38_21_+_8433172_8446622_RNA45SN1_45S_50ntFlanks@12712.12745.34|CGGAGCCTCGGTTGGCCTCGGATAGCCGGTCCCC</td><td>>hg38_21_+_8433172_8446622_RNA45SN1_45S_50ntFlanks</td><td>12712</td><td>12745</td><td>34</td><td>CGGAGCCTCGGTTGGCCTCGGATAGCCGGTCCCC</td><td>2265.7</td><td>-15.0</td><td>1.5</td><td>-10.2</td><td>1.5E-24</td><td>5.9E-23</td><td>MB231/MB468</td><td>MP</td></tr>
<tr><td>>hg38_21_+_8433172_8446622_RNA45SN1_45S_50ntFlanks@9198.9222.25|GTTCTCTCGGGGCCACGCGCGCGTC</td><td>>hg38_21_+_8433172_8446622_RNA45SN1_45S_50ntFlanks</td><td>9198</td><td>9222</td><td>25</td><td>GTTCTCTCGGGGCCACGCGCGCGTC</td><td>2141.0</td><td>-14.9</td><td>1.5</td><td>-10.2</td><td>1.5E-24</td><td>5.9E-23</td><td>MB231/MB468</td><td>MP</td></tr>
<tr><td>>hg38_21_+_8433172_8446622_RNA45SN1_45S_50ntFlanks@9572.9596.25|CAAATCGGTCGTCCGACCTGGGTAT</td><td>>hg38_21_+_8433172_8446622_RNA45SN1_45S_50ntFlanks</td><td>9572</td><td>9596</td><td>25</td><td>CAAATCGGTCGTCCGACCTGGGTAT</td><td>24994.1</td><td>-13.7</td><td>1.3</td><td>-10.2</td><td>1.5E-24</td><td>5.9E-23</td><td>MB231/MB468</td><td>MP</td></tr>
<tr><td>>hg38_21_+_8433172_8446622_RNA45SN1_45S_50ntFlanks@10683.10709.27|CCGGCGGCGTCCGGTGAGCTCTCGCTG</td><td>>hg38_21_+_8433172_8446622_RNA45SN1_45S_50ntFlanks</td><td>10683</td><td>10709</td><td>27</td><td>CCGGCGGCGTCCGGTGAGCTCTCGCTG</td><td>1097.8</td><td>-14.0</td><td>1.4</td><td>-10.2</td><td>1.5E-24</td><td>5.9E-23</td><td>MB231/MB468</td><td>MP</td></tr>
<tr><td>>MI0001445|hsa-mir-423&WithFlank&17|+|30117073|30117178@59.78.20(+1U)&[MIMAT0001340&hsa-miR-423-3p&offsets|0|-3(+1U);m-100&17|+|30117131|30117153&offsets|0|-3(+1U)]|AGCTCGGTCTGAGGCCCCTCT</td><td>>MI0001445|hsa-mir-423&WithFlank&17|+|30117073|30117178</td><td>59</td><td>78</td><td>20(+1U)</td><td>AGCTCGGTCTGAGGCCCCTCT</td><td>290.0</td><td>12.4</td><td>1.2</td><td>10.1</td><td>5.3E-24</td><td>5.9E-23</td><td>BT20/MB468</td><td>CYTO</td></tr>
<tr><td>>hg38_21_+_8433172_8446622_RNA45SN1_45S_50ntFlanks@6747.6770.24|ACTTCGAACGCACTTGCGGCCCCG</td><td>>hg38_21_+_8433172_8446622_RNA45SN1_45S_50ntFlanks</td><td>6747</td><td>6770</td><td>24</td><td>ACTTCGAACGCACTTGCGGCCCCG</td><td>796.9</td><td>-8.2</td><td>0.8</td><td>-10.3</td><td>1.1E-24</td><td>5.9E-23</td><td>MB231/MB468</td><td>CYTO</td></tr>
<tr><td>>hg38_21_+_8433172_8446622_RNA45SN1_45S_50ntFlanks@6738.6770.33|ATCATCGACACTTCGAACGCACTTGCGGCCCCG</td><td>>hg38_21_+_8433172_8446622_RNA45SN1_45S_50ntFlanks</td><td>6738</td><td>6770</td><td>33</td><td>ATCATCGACACTTCGAACGCACTTGCGGCCCCG</td><td>2884.2</td><td>-14.5</td><td>1.4</td><td>-10.2</td><td>2.7E-24</td><td>6.0E-23</td><td>MB231/MB468</td><td>NUC</td></tr>
<tr><td>>hg38_21_+_8433172_8446622_RNA45SN1_45S_50ntFlanks@11873.11890.18|AAGAAGACCCTGTTGAGC</td><td>>hg38_21_+_8433172_8446622_RNA45SN1_45S_50ntFlanks</td><td>11873</td><td>11890</td><td>18</td><td>AAGAAGACCCTGTTGAGC</td><td>1929.8</td><td>-14.8</td><td>1.4</td><td>-10.2</td><td>1.5E-24</td><td>6.0E-23</td><td>MB231/MB468</td><td>MP</td></tr>
<tr><td>>hg38_21_+_8433172_8446622_RNA45SN1_45S_50ntFlanks@12405.12436.32|CTTTTTGATCCTTCGATGTCGGCTCTTCCTAT</td><td>>hg38_21_+_8433172_8446622_RNA45SN1_45S_50ntFlanks</td><td>12405</td><td>12436</td><td>32</td><td>CTTTTTGATCCTTCGATGTCGGCTCTTCCTAT</td><td>1738.0</td><td>-14.6</td><td>1.4</td><td>-10.2</td><td>1.6E-24</td><td>6.0E-23</td><td>MB231/MB468</td><td>MP</td></tr>
<tr><td>>hg38_21_+_8433172_8446622_RNA45SN1_45S_50ntFlanks@6747.6771.25|ACTTCGAACGCACTTGCGGCCCCGG</td><td>>hg38_21_+_8433172_8446622_RNA45SN1_45S_50ntFlanks</td><td>6747</td><td>6771</td><td>25</td><td>ACTTCGAACGCACTTGCGGCCCCGG</td><td>1191.4</td><td>-14.1</td><td>1.4</td><td>-10.2</td><td>1.6E-24</td><td>6.0E-23</td><td>MB231/MB468</td><td>MP</td></tr>
<tr><td>>hg38_21_+_8433172_8446622_RNA45SN1_45S_50ntFlanks@9564.9592.29|CTGACGTGCAAATCGGTCGTCCGACCTGG</td><td>>hg38_21_+_8433172_8446622_RNA45SN1_45S_50ntFlanks</td><td>9564</td><td>9592</td><td>29</td><td>CTGACGTGCAAATCGGTCGTCCGACCTGG</td><td>1200.6</td><td>-14.1</td><td>1.4</td><td>-10.2</td><td>1.6E-24</td><td>6.0E-23</td><td>MB231/MB468</td><td>MP</td></tr>
<tr><td>>hg38_21_+_8433172_8446622_RNA45SN1_45S_50ntFlanks@9568.9592.25|CGTGCAAATCGGTCGTCCGACCTGG</td><td>>hg38_21_+_8433172_8446622_RNA45SN1_45S_50ntFlanks</td><td>9568</td><td>9592</td><td>25</td><td>CGTGCAAATCGGTCGTCCGACCTGG</td><td>1014.6</td><td>-13.9</td><td>1.4</td><td>-10.2</td><td>1.6E-24</td><td>6.0E-23</td><td>MB231/MB468</td><td>MP</td></tr>
<tr><td>>hg38_21_+_8433172_8446622_RNA45SN1_45S_50ntFlanks@12076.12095.20|AGGGGCTCTCGCTTCTGGCG</td><td>>hg38_21_+_8433172_8446622_RNA45SN1_45S_50ntFlanks</td><td>12076</td><td>12095</td><td>20</td><td>AGGGGCTCTCGCTTCTGGCG</td><td>4171.7</td><td>-5.8</td><td>0.6</td><td>-10.2</td><td>2.8E-24</td><td>6.1E-23</td><td>MB231/MB468</td><td>NUC</td></tr>
<tr><td>>hg38_21_+_8433172_8446622_RNA45SN1_45S_50ntFlanks@4728.4755.28|ATCAAGAACGAAAGTCGGAGGTTCGAAG</td><td>>hg38_21_+_8433172_8446622_RNA45SN1_45S_50ntFlanks</td><td>4728</td><td>4755</td><td>28</td><td>ATCAAGAACGAAAGTCGGAGGTTCGAAG</td><td>287.5</td><td>-12.6</td><td>1.3</td><td>-10.0</td><td>1.1E-23</td><td>6.1E-23</td><td>BT20/MB468</td><td>NUC</td></tr>
<tr><td>>MI0000102|hsa-mir-100&WithFlank&11|-|122152223|122152314@19.40.22(+2U)&[MIMAT0000098&hsa-miR-100-5p&offsets|0|0(+2U);m-27&11|-|122152276|122152296&offsets|0|+1(+2U)]|AACCCGTAGATCCGAACTTGTGTT</td><td>>MI0000102|hsa-mir-100&WithFlank&11|-|122152223|122152314</td><td>19</td><td>40</td><td>22(+2U)</td><td>AACCCGTAGATCCGAACTTGTGTT</td><td>833.0</td><td>13.6</td><td>1.3</td><td>10.2</td><td>2.8E-24</td><td>6.1E-23</td><td>MB231/MB468</td><td>NUC</td></tr>
<tr><td>>MI0000283|hsa-mir-203a&WithFlank&14|+|104117399|104117520@72.93.22&[MIMAT0000264&hsa-miR-203a-3p&offsets|+1|+1;m-79&14|+|104117469|104117490&offsets|+1|+1]|TGAAATGTTTAGGACCACTAGA</td><td>>MI0000283|hsa-mir-203a&WithFlank&14|+|104117399|104117520</td><td>72</td><td>93</td><td>22</td><td>TGAAATGTTTAGGACCACTAGA</td><td>347.4</td><td>12.9</td><td>1.3</td><td>10.1</td><td>4.2E-24</td><td>6.1E-23</td><td>BT20/MB468</td><td>MITO</td></tr>
<tr><td>>hg38_21_+_8433172_8446622_RNA45SN1_45S_50ntFlanks@11665.11692.28|TTCTGCCCAGTGCTCTGAATGTCAAAGT</td><td>>hg38_21_+_8433172_8446622_RNA45SN1_45S_50ntFlanks</td><td>11665</td><td>11692</td><td>28</td><td>TTCTGCCCAGTGCTCTGAATGTCAAAGT</td><td>332.3</td><td>12.8</td><td>1.3</td><td>10.1</td><td>4.2E-24</td><td>6.1E-23</td><td>BT20/MB468</td><td>MITO</td></tr>
<tr><td>>hg38_21_+_8433172_8446622_RNA45SN1_45S_50ntFlanks@9532.9572.41|AGAGGAAACTCTGGTGGAGGTCCGTAGCGGTCCTGACGTGC</td><td>>hg38_21_+_8433172_8446622_RNA45SN1_45S_50ntFlanks</td><td>9532</td><td>9572</td><td>41</td><td>AGAGGAAACTCTGGTGGAGGTCCGTAGCGGTCCTGACGTGC</td><td>414.4</td><td>12.7</td><td>1.2</td><td>10.3</td><td>1.2E-24</td><td>6.2E-23</td><td>MB231/MB468</td><td>CYTO</td></tr>
<tr><td>>hg38_21_+_8433172_8446622_RNA45SN1_45S_50ntFlanks@11999.12032.34|CCCTGCGGGCCGCCGGTGAAATACCACTACTCTG</td><td>>hg38_21_+_8433172_8446622_RNA45SN1_45S_50ntFlanks</td><td>11999</td><td>12032</td><td>34</td><td>CCCTGCGGGCCGCCGGTGAAATACCACTACTCTG</td><td>24807.6</td><td>-11.7</td><td>1.1</td><td>-10.2</td><td>2.9E-24</td><td>6.2E-23</td><td>MB231/MB468</td><td>NUC</td></tr>
<tr><td>>hg38_21_+_8433172_8446622_RNA45SN1_45S_50ntFlanks@5522.5550.29|AAGTAAAAGTCGTAACAAGGTTTCCGTAG</td><td>>hg38_21_+_8433172_8446622_RNA45SN1_45S_50ntFlanks</td><td>5522</td><td>5550</td><td>29</td><td>AAGTAAAAGTCGTAACAAGGTTTCCGTAG</td><td>473.7</td><td>-13.3</td><td>1.3</td><td>-10.0</td><td>1.2E-23</td><td>6.2E-23</td><td>BT20/MB468</td><td>NUC</td></tr>
<tr><td>>hg38_21_+_8433172_8446622_RNA45SN1_45S_50ntFlanks@4000.4025.26|TAGATAACCTCGGGCCGATCGCACGC</td><td>>hg38_21_+_8433172_8446622_RNA45SN1_45S_50ntFlanks</td><td>4000</td><td>4025</td><td>26</td><td>TAGATAACCTCGGGCCGATCGCACGC</td><td>582.4</td><td>-13.6</td><td>1.4</td><td>-10.0</td><td>1.2E-23</td><td>6.2E-23</td><td>BT20/MB468</td><td>NUC</td></tr>
<tr><td>>hg38_21_+_8433172_8446622_RNA45SN1_45S_50ntFlanks@11901.11920.20|TCTGGCACGGTGAAGAGACA</td><td>>hg38_21_+_8433172_8446622_RNA45SN1_45S_50ntFlanks</td><td>11901</td><td>11920</td><td>20</td><td>TCTGGCACGGTGAAGAGACA</td><td>410.6</td><td>-13.1</td><td>1.3</td><td>-10.0</td><td>1.2E-23</td><td>6.3E-23</td><td>BT20/MB468</td><td>NUC</td></tr>
<tr><td>>hg38_21_+_8433172_8446622_RNA45SN1_45S_50ntFlanks@12488.12520.33|AACGTGAGCTGGGTTTAGACCGTCGTGAGACAG</td><td>>hg38_21_+_8433172_8446622_RNA45SN1_45S_50ntFlanks</td><td>12488</td><td>12520</td><td>33</td><td>AACGTGAGCTGGGTTTAGACCGTCGTGAGACAG</td><td>488.4</td><td>-13.4</td><td>1.3</td><td>-10.0</td><td>1.2E-23</td><td>6.3E-23</td><td>BT20/MB468</td><td>NUC</td></tr>
</table>

<table>
<tr><td>>MI0000093|hsa-mir-92a-1&WithFlank&13|+|91351308|91351397@54.75.22(+1A);MI0000094|hsa-mir-92a-2&WithFlank&X|-|134169532|134169618@54.75.22(+1A)&[MIMAT0000092&hsa-miR-92a-3p&offsets|0|0(+1A);m-9&13|+|91351361|91351382&offsets|0|0(+1A)];[MIMAT0000092_1&hsa-miR-92a-3p&offsets|0|0(+1A);m-11&X|-|134169544|134169565&offsets|0|0(+1A)]|TATTGCACTTGTCCCGGCCTGTA</td><td>>MI0000093|hsa-mir-92a-1&WithFlank&13|+|91351308|91351397</td><td>54</td><td>75</td><td>22(+1A)</td><td>TATTGCACTTGTCCCGGCCTGTA</td><td>10003.8</td><td>5.8</td><td>0.6</td><td>10.1</td><td>4.3E-24</td><td>6.3E-23</td><td>BT20/MB468</td><td>MITO</td></tr>
<tr><td>>hg38_21_+_8433172_8446622_RNA45SN1_45S_50ntFlanks@5515.5535.21|CTAGAGGAAGTAAAAGTCGTA</td><td>>hg38_21_+_8433172_8446622_RNA45SN1_45S_50ntFlanks</td><td>5515</td><td>5535</td><td>21</td><td>CTAGAGGAAGTAAAAGTCGTA</td><td>329.4</td><td>-12.8</td><td>1.3</td><td>-10.0</td><td>1.2E-23</td><td>6.3E-23</td><td>BT20/MB468</td><td>NUC</td></tr>
<tr><td>>hg38_21_+_8433172_8446622_RNA45SN1_45S_50ntFlanks@9621.9650.30|TCTAGTAGCTGGTTCCCTCCGAAGTTTCCC</td><td>>hg38_21_+_8433172_8446622_RNA45SN1_45S_50ntFlanks</td><td>9621</td><td>9650</td><td>30</td><td>TCTAGTAGCTGGTTCCCTCCGAAGTTTCCC</td><td>335.2</td><td>-12.8</td><td>1.3</td><td>-10.0</td><td>1.2E-23</td><td>6.3E-23</td><td>BT20/MB468</td><td>NUC</td></tr>
<tr><td>>hg38_21_+_8433172_8446622_RNA45SN1_45S_50ntFlanks@11317.11338.22|ACCCCACGTCTCGTCGCGCGCG</td><td>>hg38_21_+_8433172_8446622_RNA45SN1_45S_50ntFlanks</td><td>11317</td><td>11338</td><td>22</td><td>ACCCCACGTCTCGTCGCGCGCG</td><td>3550.3</td><td>-15.6</td><td>1.5</td><td>-10.2</td><td>2.5E-24</td><td>6.3E-23</td><td>MB231/MB468</td><td>MITO</td></tr>
<tr><td>>hg38_21_+_8433172_8446622_RNA45SN1_45S_50ntFlanks@4003.4024.22|ATAACCTCGGGCCGATCGCACG</td><td>>hg38_21_+_8433172_8446622_RNA45SN1_45S_50ntFlanks</td><td>4003</td><td>4024</td><td>22</td><td>ATAACCTCGGGCCGATCGCACG</td><td>4656.8</td><td>-9.8</td><td>1.0</td><td>-10.2</td><td>2.5E-24</td><td>6.4E-23</td><td>MB231/MB468</td><td>MITO</td></tr>
<tr><td>>hg38_21_+_8433172_8446622_RNA45SN1_45S_50ntFlanks@3986.4023.38|GGCTTTGGTGACTCTAGATAACCTCGGGCCGATCGCAC</td><td>>hg38_21_+_8433172_8446622_RNA45SN1_45S_50ntFlanks</td><td>3986</td><td>4023</td><td>38</td><td>GGCTTTGGTGACTCTAGATAACCTCGGGCCGATCGCAC</td><td>1746.9</td><td>7.1</td><td>0.7</td><td>10.1</td><td>5.7E-24</td><td>6.4E-23</td><td>BT20/MB468</td><td>CYTO</td></tr>
<tr><td>>MI0000285|hsa-mir-205&WithFlank&1|+|209432127|209432248@40.61.22(+1A)&[MIMAT0000266&hsa-miR-205-5p&offsets|0|0(+1A);m-351&1|+|209432166|209432187&offsets|0|0(+1A)]|TCCTTCATTCCACCGGAGTCTGA</td><td>>MI0000285|hsa-mir-205&WithFlank&1|+|209432127|209432248</td><td>40</td><td>61</td><td>22(+1A)</td><td>TCCTTCATTCCACCGGAGTCTGA</td><td>129550.2</td><td>7.2</td><td>0.7</td><td>10.1</td><td>5.7E-24</td><td>6.4E-23</td><td>BT20/MB468</td><td>CYTO</td></tr>
<tr><td>>hg38_21_+_8433172_8446622_RNA45SN1_45S_50ntFlanks@9504.9521.18|GTGAACTATGCCTGGGCA</td><td>>hg38_21_+_8433172_8446622_RNA45SN1_45S_50ntFlanks</td><td>9504</td><td>9521</td><td>18</td><td>GTGAACTATGCCTGGGCA</td><td>1708.1</td><td>-6.0</td><td>0.6</td><td>-10.2</td><td>3.0E-24</td><td>6.5E-23</td><td>MB231/MB468</td><td>NUC</td></tr>
<tr><td>>hg38_21_+_8433172_8446622_RNA45SN1_45S_50ntFlanks@12405.12436.32|CTTTTTGATCCTTCGATGTCGGCTCTTCCTAT</td><td>>hg38_21_+_8433172_8446622_RNA45SN1_45S_50ntFlanks</td><td>12405</td><td>12436</td><td>32</td><td>CTTTTTGATCCTTCGATGTCGGCTCTTCCTAT</td><td>24344.4</td><td>-10.7</td><td>1.1</td><td>-10.2</td><td>2.6E-24</td><td>6.5E-23</td><td>MB231/MB468</td><td>MITO</td></tr>
<tr><td>>hg38_21_+_8433172_8446622_RNA45SN1_45S_50ntFlanks@10781.10808.28|AGGTGAACAGCCTCTGGCATGTTGGAAC</td><td>>hg38_21_+_8433172_8446622_RNA45SN1_45S_50ntFlanks</td><td>10781</td><td>10808</td><td>28</td><td>AGGTGAACAGCCTCTGGCATGTTGGAAC</td><td>329.2</td><td>-12.8</td><td>1.3</td><td>-10.0</td><td>1.2E-23</td><td>6.5E-23</td><td>BT20/MB468</td><td>NUC</td></tr>
<tr><td>>hg38_21_+_8433172_8446622_RNA45SN1_45S_50ntFlanks@4047.4068.22|ATTCGAACGTCTGCCCTATCAA</td><td>>hg38_21_+_8433172_8446622_RNA45SN1_45S_50ntFlanks</td><td>4047</td><td>4068</td><td>22</td><td>ATTCGAACGTCTGCCCTATCAA</td><td>507.0</td><td>-13.4</td><td>1.3</td><td>-10.0</td><td>1.2E-23</td><td>6.6E-23</td><td>BT20/MB468</td><td>NUC</td></tr>
<tr><td>>hg38_21_+_8433172_8446622_RNA45SN1_45S_50ntFlanks@11661.11685.25|TGATTTCTGCCCAGTGCTCTGAATG</td><td>>hg38_21_+_8433172_8446622_RNA45SN1_45S_50ntFlanks</td><td>11661</td><td>11685</td><td>25</td><td>TGATTTCTGCCCAGTGCTCTGAATG</td><td>8614.3</td><td>-10.1</td><td>1.0</td><td>-10.2</td><td>2.7E-24</td><td>6.6E-23</td><td>MB231/MB468</td><td>MITO</td></tr>
<tr><td>>hg38_21_+_8433172_8446622_RNA45SN1_45S_50ntFlanks@6737.6771.35|GATCATCGACACTTCGAACGCACTTGCGGCCCCGG</td><td>>hg38_21_+_8433172_8446622_RNA45SN1_45S_50ntFlanks</td><td>6737</td><td>6771</td><td>35</td><td>GATCATCGACACTTCGAACGCACTTGCGGCCCCGG</td><td>2026.4</td><td>-14.8</td><td>1.5</td><td>-10.2</td><td>2.7E-24</td><td>6.6E-23</td><td>MB231/MB468</td><td>MITO</td></tr>
<tr><td>>hg38_21_+_8433172_8446622_RNA45SN1_45S_50ntFlanks@9534.9566.33|AGGAAACTCTGGTGGAGGTCCGTAGCGGTCCTG</td><td>>hg38_21_+_8433172_8446622_RNA45SN1_45S_50ntFlanks</td><td>9534</td><td>9566</td><td>33</td><td>AGGAAACTCTGGTGGAGGTCCGTAGCGGTCCTG</td><td>1654.6</td><td>-14.5</td><td>1.4</td><td>-10.2</td><td>2.7E-24</td><td>6.6E-23</td><td>MB231/MB468</td><td>MITO</td></tr>
<tr><td>>MI0003560|hsa-mir-92b&WithFlank&1|+|155195171|155195278@67.88.22(+3U)&[MIMAT0003218&hsa-miR-92b-3p&offsets|0|0(+3U);m-115&1|+|155195237|155195258&offsets|0|0(+3U)]|TATTGCACTCGTCCCGGCCTCCTTT</td><td>>MI0003560|hsa-mir-92b&WithFlank&1|+|155195171|155195278</td><td>67</td><td>88</td><td>22(+3U)</td><td>TATTGCACTCGTCCCGGCCTCCTTT</td><td>376.1</td><td>12.8</td><td>1.3</td><td>10.1</td><td>5.9E-24</td><td>6.6E-23</td><td>BT20/MB468</td><td>CYTO</td></tr>
<tr><td>>hg38_21_+_8433172_8446622_RNA45SN1_45S_50ntFlanks@8113.8141.29|GGCGCGGGACATGTGGCGTACGGAAGACC</td><td>>hg38_21_+_8433172_8446622_RNA45SN1_45S_50ntFlanks</td><td>8113</td><td>8141</td><td>29</td><td>GGCGCGGGACATGTGGCGTACGGAAGACC</td><td>2025.1</td><td>-14.8</td><td>1.5</td><td>-10.2</td><td>1.7E-24</td><td>6.7E-23</td><td>MB231/MB468</td><td>MP</td></tr>
<tr><td>>trna116_GluCTC_1_-_145399233_145399304@1.39.39,trna59_GluCTC_1_+_249168447_249168518@1.39.39,trna71_GluCTC_1_-_161439189_161439260@1.39.39,trna74_GluCTC_1_-_161431809_161431880@1.39.39,trna77_GluCTC_1_-_161424398_161424469@1.39.39,trna77_GluCTC_6_+_28949976_28950047@1.39.39,trna80_GluCTC_1_-_161417018_161417089@1.39.39,trna87_GluCTC_6_-_126101393_126101464@1.39.39|TCCCTGGTGGTCTAGTGGTTAGGATTCGGCGCTCTCACC</td><td>>trna116_GluCTC_1_-_145399233_145399304</td><td>1</td><td>39</td><td>39</td><td>TCCCTGGTGGTCTAGTGGTTAGGATTCGGCGCTCTCACC</td><td>9995.4</td><td>8.4</td><td>0.8</td><td>10.1</td><td>4.6E-24</td><td>6.7E-23</td><td>BT20/MB468</td><td>MITO</td></tr>
<tr><td>>MI0000255|hsa-mir-30d&WithFlank&8|-|134804870|134804951@12.35.24(+1U)&[MIMAT0000245&hsa-miR-30d-5p&offsets|0|+2(+1U);m-21&8|-|134804919|134804940&offsets|0|+2(+1U)]|TGTAAACATCCCCGACTGGAAGCTT</td><td>>MI0000255|hsa-mir-30d&WithFlank&8|-|134804870|134804951</td><td>12</td><td>35</td><td>24(+1U)</td><td>TGTAAACATCCCCGACTGGAAGCTT</td><td>1943.6</td><td>7.8</td><td>0.8</td><td>10.1</td><td>4.6E-24</td><td>6.7E-23</td><td>BT20/MB468</td><td>MITO</td></tr>
<tr><td>>hg38_21_+_8433172_8446622_RNA45SN1_45S_50ntFlanks@10914.10932.19|CGCCGCGGCTGGACGAGGC</td><td>>hg38_21_+_8433172_8446622_RNA45SN1_45S_50ntFlanks</td><td>10914</td><td>10932</td><td>19</td><td>CGCCGCGGCTGGACGAGGC</td><td>1276.3</td><td>-14.2</td><td>1.4</td><td>-10.2</td><td>1.8E-24</td><td>6.8E-23</td><td>MB231/MB468</td><td>MP</td></tr>
<tr><td>>hg38_21_+_8433172_8446622_RNA45SN1_45S_50ntFlanks@8170.8215.46|CAAGTCCTTCTGATCGAGGCCCAGCCCGTGGACGGTGTGAGGCCGG</td><td>>hg38_21_+_8433172_8446622_RNA45SN1_45S_50ntFlanks</td><td>8170</td><td>8215</td><td>46</td><td>CAAGTCCTTCTGATCGAGGCCCAGCCCGTGGACGGTGTGAGGCCGG</td><td>3829.9</td><td>-15.7</td><td>1.5</td><td>-10.2</td><td>2.7E-24</td><td>6.8E-23</td><td>MB231/MB468</td><td>MITO</td></tr>
<tr><td>>hg38_21_+_8433172_8446622_RNA45SN1_45S_50ntFlanks@12053.12070.18|GTGAGGCGGGGGGGCGAG</td><td>>hg38_21_+_8433172_8446622_RNA45SN1_45S_50ntFlanks</td><td>12053</td><td>12070</td><td>18</td><td>GTGAGGCGGGGGGGCGAG</td><td>761.8</td><td>-14.0</td><td>1.4</td><td>-10.0</td><td>1.3E-23</td><td>6.8E-23</td><td>BT20/MB468</td><td>NUC</td></tr>
<tr><td>>hg38_21_+_8433172_8446622_RNA45SN1_45S_50ntFlanks@11567.11584.18|AGCAGCCGACTTAGAACT</td><td>>hg38_21_+_8433172_8446622_RNA45SN1_45S_50ntFlanks</td><td>11567</td><td>11584</td><td>18</td><td>AGCAGCCGACTTAGAACT</td><td>2351.6</td><td>6.4</td><td>0.6</td><td>10.1</td><td>4.8E-24</td><td>7.0E-23</td><td>BT20/MB468</td><td>MITO</td></tr>
<tr><td>>hg38_21_+_8433172_8446622_RNA45SN1_45S_50ntFlanks@12405.12424.20|CTTTTTGATCCTTCGATGTC</td><td>>hg38_21_+_8433172_8446622_RNA45SN1_45S_50ntFlanks</td><td>12405</td><td>12424</td><td>20</td><td>CTTTTTGATCCTTCGATGTC</td><td>1448.8</td><td>-14.4</td><td>1.4</td><td>-10.2</td><td>1.8E-24</td><td>7.0E-23</td><td>MB231/MB468</td><td>MP</td></tr>
<tr><td>>hg38_21_+_8433172_8446622_RNA45SN1_45S_50ntFlanks@9578.9603.26|GGTCGTCCGACCTGGGTATAGGGGCG</td><td>>hg38_21_+_8433172_8446622_RNA45SN1_45S_50ntFlanks</td><td>9578</td><td>9603</td><td>26</td><td>GGTCGTCCGACCTGGGTATAGGGGCG</td><td>466.3</td><td>13.3</td><td>1.3</td><td>10.1</td><td>4.9E-24</td><td>7.1E-23</td><td>BT20/MB468</td><td>MITO</td></tr>
<tr><td>>trna116_GluCTC_1_-_145399233_145399304@32.72.41,trna71_GluCTC_1_-_161439189_161439260@32.72.41,trna74_GluCTC_1_-_161431809_161431880@32.72.41,trna77_GluCTC_1_-_161424398_161424469@32.72.41,trna77_GluCTC_6_+_28949976_28950047@32.72.41,trna80_GluCTC_1_-_161417018_161417089@32.72.41,trna87_GluCTC_6_-_126101393_126101464@32.72.41|CTCTCACCGCCGCGGCCCGGGTTCGATTCCCGGTCAGGGAA</td><td>>trna116_GluCTC_1_-_145399233_145399304</td><td>32</td><td>72</td><td>41</td><td>CTCTCACCGCCGCGGCCCGGGTTCGATTCCCGGTCAGGGAA</td><td>633.9</td><td>-13.7</td><td>1.4</td><td>-10.0</td><td>1.4E-23</td><td>7.2E-23</td><td>BT20/MB468</td><td>NUC</td></tr>
<tr><td>>hg38_21_+_8433172_8446622_RNA45SN1_45S_50ntFlanks@10351.10384.34|CGAGAACTTTGAAGGCCGAAGTGGAGAAGGGTTC</td><td>>hg38_21_+_8433172_8446622_RNA45SN1_45S_50ntFlanks</td><td>10351</td><td>10384</td><td>34</td><td>CGAGAACTTTGAAGGCCGAAGTGGAGAAGGGTTC</td><td>14921.6</td><td>-11.8</td><td>1.2</td><td>-10.0</td><td>1.4E-23</td><td>7.2E-23</td><td>BT20/MB468</td><td>NUC</td></tr>
<tr><td>>hg38_21_+_8433172_8446622_RNA45SN1_45S_50ntFlanks@11614.11640.27|ATTAAAACAAAGCATCGCGAAGGCCCG</td><td>>hg38_21_+_8433172_8446622_RNA45SN1_45S_50ntFlanks</td><td>11614</td><td>11640</td><td>27</td><td>ATTAAAACAAAGCATCGCGAAGGCCCG</td><td>1105.0</td><td>-14.0</td><td>1.4</td><td>-10.2</td><td>1.9E-24</td><td>7.2E-23</td><td>MB231/MB468</td><td>MP</td></tr>
<tr><td>>hg38_21_+_8433172_8446622_RNA45SN1_45S_50ntFlanks@4760.4780.21|TCAGATACCGTCGTAGTTCCG</td><td>>hg38_21_+_8433172_8446622_RNA45SN1_45S_50ntFlanks</td><td>4760</td><td>4780</td><td>21</td><td>TCAGATACCGTCGTAGTTCCG</td><td>2707.7</td><td>-15.3</td><td>1.5</td><td>-10.2</td><td>1.9E-24</td><td>7.2E-23</td><td>MB231/MB468</td><td>MP</td></tr>
<tr><td>>hg38_21_+_8433172_8446622_RNA45SN1_45S_50ntFlanks@12635.12653.19|AAGCTACCATCTGTGGGAT</td><td>>hg38_21_+_8433172_8446622_RNA45SN1_45S_50ntFlanks</td><td>12635</td><td>12653</td><td>19</td><td>AAGCTACCATCTGTGGGAT</td><td>4631.2</td><td>-16.0</td><td>1.6</td><td>-10.2</td><td>2.9E-24</td><td>7.2E-23</td><td>MB231/MB468</td><td>MITO</td></tr>
<tr><td>>MI0000102|hsa-mir-100&WithFlank&11|-|122152223|122152314@19.41.23&[MIMAT0000098&hsa-miR-100-5p&offsets|0|+1;m-27&11|-|122152276|122152296&offsets|0|+2]|AACCCGTAGATCCGAACTTGTGG</td><td>>MI0000102|hsa-mir-100&WithFlank&11|-|122152223|122152314</td><td>19</td><td>41</td><td>23</td><td>AACCCGTAGATCCGAACTTGTGG</td><td>2069.5</td><td>15.0</td><td>1.5</td><td>10.2</td><td>1.4E-24</td><td>7.3E-23</td><td>MB231/MB468</td><td>CYTO</td></tr>
<tr><td>>MI0000077|hsa-mir-21&WithFlank&17|+|59841260|59841343@14.36.23(+1A)&[MIMAT0000076&hsa-miR-21-5p&offsets|0|+1(+1A);m-1&17|+|59841273|59841294&offsets|0|+1(+1A)]|TAGCTTATCAGACTGATGTTGACA</td><td>>MI0000077|hsa-mir-21&WithFlank&17|+|59841260|59841343</td><td>14</td><td>36</td><td>23(+1A)</td><td>TAGCTTATCAGACTGATGTTGACA</td><td>93592.7</td><td>5.7</td><td>0.6</td><td>10.1</td><td>3.4E-24</td><td>7.3E-23</td><td>MB231/MB468</td><td>NUC</td></tr>
<tr><td>>hg38_21_+_8433172_8446622_RNA45SN1_45S_50ntFlanks@11235.11254.20|CTTCCCGTGGATCGCCCCAG</td><td>>hg38_21_+_8433172_8446622_RNA45SN1_45S_50ntFlanks</td><td>11235</td><td>11254</td><td>20</td><td>CTTCCCGTGGATCGCCCCAG</td><td>2248.7</td><td>-15.0</td><td>1.5</td><td>-10.2</td><td>2.0E-24</td><td>7.4E-23</td><td>MB231/MB468</td><td>MP</td></tr>
<tr><td>>hg38_21_+_8433172_8446622_RNA45SN1_45S_50ntFlanks@10687.10711.25|CGGCGTCCGGTGAGCTCTCGCTGGC</td><td>>hg38_21_+_8433172_8446622_RNA45SN1_45S_50ntFlanks</td><td>10687</td><td>10711</td><td>25</td><td>CGGCGTCCGGTGAGCTCTCGCTGGC</td><td>5199.0</td><td>-6.8</td><td>0.7</td><td>-10.1</td><td>3.5E-24</td><td>7.5E-23</td><td>MB231/MB468</td><td>NUC</td></tr>
<tr><td>>hg38_21_+_8433172_8446622_RNA45SN1_45S_50ntFlanks@9567.9592.26|ACGTGCAAATCGGTCGTCCGACCTGG</td><td>>hg38_21_+_8433172_8446622_RNA45SN1_45S_50ntFlanks</td><td>9567</td><td>9592</td><td>26</td><td>ACGTGCAAATCGGTCGTCCGACCTGG</td><td>1016.8</td><td>-13.9</td><td>1.4</td><td>-10.2</td><td>2.0E-24</td><td>7.6E-23</td><td>MB231/MB468</td><td>MP</td></tr>
<tr><td>>MI0000650|hsa-mir-200c&WithFlank&12|+|6963693|6963772@50.72.23&[MIMAT0000617&hsa-miR-200c-3p&offsets|0|0;m-54&12|+|6963742|6963764&offsets|0|0]|TAATACTGCCGGGTAATGATGGA</td><td>>MI0000650|hsa-mir-200c&WithFlank&12|+|6963693|6963772</td><td>50</td><td>72</td><td>23</td><td>TAATACTGCCGGGTAATGATGGA</td><td>141451.6</td><td>9.6</td><td>0.9</td><td>10.4</td><td>1.9E-25</td><td>7.7E-23</td><td>BT20/MB231</td><td>CYTO</td></tr>
<tr><td>>hg38_21_+_8433172_8446622_RNA45SN1_45S_50ntFlanks@6653.6689.37|ACTCTTAGCGGTGGATCACTCGGCTCGTGCGTCGATG</td><td>>hg38_21_+_8433172_8446622_RNA45SN1_45S_50ntFlanks</td><td>6653</td><td>6689</td><td>37</td><td>ACTCTTAGCGGTGGATCACTCGGCTCGTGCGTCGATG</td><td>3098.9</td><td>-4.0</td><td>0.4</td><td>-10.0</td><td>1.5E-23</td><td>7.8E-23</td><td>BT20/MB468</td><td>NUC</td></tr>
<tr><td>>MI0000070|hsa-mir-16-1&WithFlank&13|-|50048967|50049067@20.40.21;MI0000115|hsa-mir-16-2&WithFlank&3|+|160404739|160404831@16.36.21&[MIMAT0000069&hsa-miR-16-5p&offsets|0|-1;m-62&13|-|50049027|50049048&offsets|0|-1];[MIMAT0000069_1&hsa-miR-16-5p&offsets|0|-1;m-61&3|+|160404754|160404775&offsets|0|-1]|TAGCAGCACGTAAATATTGGC</td><td>>MI0000070|hsa-mir-16-1&WithFlank&13|-|50048967|50049067</td><td>20</td><td>40</td><td>21</td><td>TAGCAGCACGTAAATATTGGC</td><td>2992.1</td><td>7.3</td><td>0.7</td><td>10.1</td><td>6.9E-24</td><td>7.8E-23</td><td>BT20/MB468</td><td>CYTO</td></tr>
<tr><td>>MI0000102|hsa-mir-100&WithFlank&11|-|122152223|122152314@19.40.22(+2A)&[MIMAT0000098&hsa-miR-100-5p&offsets|0|0(+2A);m-27&11|-|122152276|122152296&offsets|0|+1(+2A)]|AACCCGTAGATCCGAACTTGTGAA</td><td>>MI0000102|hsa-mir-100&WithFlank&11|-|122152223|122152314</td><td>19</td><td>40</td><td>22(+2A)</td><td>AACCCGTAGATCCGAACTTGTGAA</td><td>1347.1</td><td>14.4</td><td>1.4</td><td>10.2</td><td>1.5E-24</td><td>7.8E-23</td><td>MB231/MB468</td><td>CYTO</td></tr>
<tr><td>>hg38_21_+_8433172_8446622_RNA45SN1_45S_50ntFlanks@8115.8140.26|CGCGGGACATGTGGCGTACGGAAGAC</td><td>>hg38_21_+_8433172_8446622_RNA45SN1_45S_50ntFlanks</td><td>8115</td><td>8140</td><td>26</td><td>CGCGGGACATGTGGCGTACGGAAGAC</td><td>1860.8</td><td>-14.6</td><td>1.4</td><td>-10.2</td><td>3.2E-24</td><td>7.9E-23</td><td>MB231/MB468</td><td>MITO</td></tr>
<tr><td>>hg38_21_+_8433172_8446622_RNA45SN1_45S_50ntFlanks@9739.9758.20|AAACGATCTCAACCTATTCT</td><td>>hg38_21_+_8433172_8446622_RNA45SN1_45S_50ntFlanks</td><td>9739</td><td>9758</td><td>20</td><td>AAACGATCTCAACCTATTCT</td><td>10751.6</td><td>-8.3</td><td>0.8</td><td>-10.2</td><td>3.2E-24</td><td>7.9E-23</td><td>MB231/MB468</td><td>MITO</td></tr>
<tr><td>>hg38_21_+_8433172_8446622_RNA45SN1_45S_50ntFlanks@10362.10383.22|AAGGCCGAAGTGGAGAAGGGTT</td><td>>hg38_21_+_8433172_8446622_RNA45SN1_45S_50ntFlanks</td><td>10362</td><td>10383</td><td>22</td><td>AAGGCCGAAGTGGAGAAGGGTT</td><td>3772.6</td><td>-9.9</td><td>1.0</td><td>-10.2</td><td>2.1E-24</td><td>7.9E-23</td><td>MB231/MB468</td><td>MP</td></tr>
<tr><td>>hg38_21_+_8433172_8446622_RNA45SN1_45S_50ntFlanks@11792.11810.19|ATGAACGAGATTCCCACTG</td><td>>hg38_21_+_8433172_8446622_RNA45SN1_45S_50ntFlanks</td><td>11792</td><td>11810</td><td>19</td><td>ATGAACGAGATTCCCACTG</td><td>333.1</td><td>-12.8</td><td>1.3</td><td>-10.0</td><td>1.5E-23</td><td>8.0E-23</td><td>BT20/MB468</td><td>NUC</td></tr>
<tr><td>>hg38_21_+_8433172_8446622_RNA45SN1_45S_50ntFlanks@11561.11588.28|GCGCCTAGCAGCCGACTTAGAACTGGTG</td><td>>hg38_21_+_8433172_8446622_RNA45SN1_45S_50ntFlanks</td><td>11561</td><td>11588</td><td>28</td><td>GCGCCTAGCAGCCGACTTAGAACTGGTG</td><td>1252.9</td><td>-14.2</td><td>1.4</td><td>-10.2</td><td>2.2E-24</td><td>8.1E-23</td><td>MB231/MB468</td><td>MP</td></tr>
</table>

<table>
<tr><td>>hg38_21_+_8433172_8446622_RNA45SN1_45S_50ntFlanks@10388.10409.22|GTGAACAGCAGTTGAACATGGG</td><td>>hg38_21_+_8433172_8446622_RNA45SN1_45S_50ntFlanks</td><td>10388</td><td>10409</td><td>22</td><td>GTGAACAGCAGTTGAACATGGG</td><td>371.9</td><td>-13.0</td><td>1.3</td><td>-10.0</td><td>1.5E-23</td><td>8.1E-23</td><td>BT20/MB468</td><td>NUC</td></tr>
<tr><td>>hg38_21_+_8433172_8446622_RNA45SN1_45S_50ntFlanks@6737.6770.34|GATCATCGACACTTCGAACGCACTTGCGGCCCCG</td><td>>hg38_21_+_8433172_8446622_RNA45SN1_45S_50ntFlanks</td><td>6737</td><td>6770</td><td>34</td><td>GATCATCGACACTTCGAACGCACTTGCGGCCCCG</td><td>1848.4</td><td>-14.6</td><td>1.4</td><td>-10.1</td><td>3.3E-24</td><td>8.1E-23</td><td>MB231/MB468</td><td>MITO</td></tr>
<tr><td>>hg38_21_+_8433172_8446622_RNA45SN1_45S_50ntFlanks@11844.11872.29|GGGAACGGGCTTGGCGGAATCAGCGGGGA</td><td>>hg38_21_+_8433172_8446622_RNA45SN1_45S_50ntFlanks</td><td>11844</td><td>11872</td><td>29</td><td>GGGAACGGGCTTGGCGGAATCAGCGGGGA</td><td>1280.2</td><td>-14.2</td><td>1.4</td><td>-10.2</td><td>2.2E-24</td><td>8.1E-23</td><td>MB231/MB468</td><td>MP</td></tr>
<tr><td>>hg38_21_+_8433172_8446622_RNA45SN1_45S_50ntFlanks@9545.9571.27|GTGGAGGTCCGTAGCGGTCCTGACGTG</td><td>>hg38_21_+_8433172_8446622_RNA45SN1_45S_50ntFlanks</td><td>9545</td><td>9571</td><td>27</td><td>GTGGAGGTCCGTAGCGGTCCTGACGTG</td><td>9871.6</td><td>-7.8</td><td>0.8</td><td>-10.2</td><td>2.2E-24</td><td>8.2E-23</td><td>MB231/MB468</td><td>MP</td></tr>
<tr><td>>hg38_21_+_8433172_8446622_RNA45SN1_45S_50ntFlanks@12419.12440.22|GATGTCGGCTCTTCCTATCATT</td><td>>hg38_21_+_8433172_8446622_RNA45SN1_45S_50ntFlanks</td><td>12419</td><td>12440</td><td>22</td><td>GATGTCGGCTCTTCCTATCATT</td><td>772.3</td><td>4.2</td><td>0.4</td><td>10.1</td><td>7.3E-24</td><td>8.2E-23</td><td>BT20/MB468</td><td>CYTO</td></tr>
<tr><td>>MI0003560|hsa-mir-92b&WithFlank&1|+|155195171|155195278@67.88.22&[MIMAT0003218&hsa-miR-92b-3p&offsets|0|0;m-115&1|+|155195237|155195258&offsets|0|0]|TATTGCACTCGTCCCGGCCTCC</td><td>>MI0003560|hsa-mir-92b&WithFlank&1|+|155195171|155195278</td><td>67</td><td>88</td><td>22</td><td>TATTGCACTCGTCCCGGCCTCC</td><td>4939.3</td><td>7.9</td><td>0.8</td><td>10.2</td><td>1.6E-24</td><td>8.2E-23</td><td>MB231/MB468</td><td>CYTO</td></tr>
<tr><td>>hg38_21_+_8433172_8446622_RNA45SN1_45S_50ntFlanks@11577.11605.29|TTAGAACTGGTGCGGACCAGGGGAATCCG</td><td>>hg38_21_+_8433172_8446622_RNA45SN1_45S_50ntFlanks</td><td>11577</td><td>11605</td><td>29</td><td>TTAGAACTGGTGCGGACCAGGGGAATCCG</td><td>693.2</td><td>-4.7</td><td>0.5</td><td>-10.1</td><td>7.4E-24</td><td>8.3E-23</td><td>BT20/MB468</td><td>CYTO</td></tr>
<tr><td>>hg38_21_+_8433172_8446622_RNA45SN1_45S_50ntFlanks@11535.11553.19|TTCCGCGCGGCGCCTCGCC</td><td>>hg38_21_+_8433172_8446622_RNA45SN1_45S_50ntFlanks</td><td>11535</td><td>11553</td><td>19</td><td>TTCCGCGCGGCGCCTCGCC</td><td>497.4</td><td>-13.4</td><td>1.3</td><td>-10.0</td><td>1.6E-23</td><td>8.3E-23</td><td>BT20/MB468</td><td>NUC</td></tr>
<tr><td>>am_trna162_MetCAT_6_-_26758550_26758622@31.53.23,trna164_MetCAT_6_-_26735574_26735646@31.53.23,trna20_MetCAT_16_+_71460396_71460468@31.53.23,trna21_MetCAT_6_+_26701712_26701784@31.53.23,trna27_MetCAT_6_+_26766444_26766516@31.53.23,trna75_MetCAT_6_+_28912352_28912424@31.53.23,trna97_MetCAT_6_-_28921042_28921114@31.53.23|TCTCATAATCTGAAGGTCCTGAG</td><td>>am_trna162_MetCAT_6_-_26758550_26758622</td><td>31</td><td>53</td><td>23</td><td>TCTCATAATCTGAAGGTCCTGAG</td><td>354.1</td><td>-12.9</td><td>1.3</td><td>-10.0</td><td>1.6E-23</td><td>8.3E-23</td><td>BT20/MB468</td><td>NUC</td></tr>
<tr><td>>hg38_21_+_8433172_8446622_RNA45SN1_45S_50ntFlanks@12997.13039.43|CGCTGCGATCTATTGAAAGTCAGCCCTCGACACAAGGGTTTGT</td><td>>hg38_21_+_8433172_8446622_RNA45SN1_45S_50ntFlanks</td><td>12997</td><td>13039</td><td>43</td><td>CGCTGCGATCTATTGAAAGTCAGCCCTCGACACAAGGGTTTGT</td><td>437667.0</td><td>9.1</td><td>0.9</td><td>10.2</td><td>1.6E-24</td><td>8.4E-23</td><td>MB231/MB468</td><td>CYTO</td></tr>
<tr><td>>hg38_21_+_8433172_8446622_RNA45SN1_45S_50ntFlanks@12624.12659.36|CCAATGGGGCGAAGCTACCATCTGTGGGATTATGAC</td><td>>hg38_21_+_8433172_8446622_RNA45SN1_45S_50ntFlanks</td><td>12624</td><td>12659</td><td>36</td><td>CCAATGGGGCGAAGCTACCATCTGTGGGATTATGAC</td><td>674.3</td><td>-13.8</td><td>1.4</td><td>-10.0</td><td>1.6E-23</td><td>8.4E-23</td><td>BT20/MB468</td><td>NUC</td></tr>
<tr><td>>hg38_21_+_8433172_8446622_RNA45SN1_45S_50ntFlanks@10351.10374.24|CGAGAACTTTGAAGGCCGAAGTGG</td><td>>hg38_21_+_8433172_8446622_RNA45SN1_45S_50ntFlanks</td><td>10351</td><td>10374</td><td>24</td><td>CGAGAACTTTGAAGGCCGAAGTGG</td><td>6099.8</td><td>-10.9</td><td>1.1</td><td>-10.0</td><td>1.6E-23</td><td>8.5E-23</td><td>BT20/MB468</td><td>NUC</td></tr>
<tr><td>>hg38_21_+_8433172_8446622_RNA45SN1_45S_50ntFlanks@4002.4026.25|GATAACCTCGGGCCGATCGCACGCC</td><td>>hg38_21_+_8433172_8446622_RNA45SN1_45S_50ntFlanks</td><td>4002</td><td>4026</td><td>25</td><td>GATAACCTCGGGCCGATCGCACGCC</td><td>788.6</td><td>-14.0</td><td>1.4</td><td>-10.0</td><td>1.6E-23</td><td>8.5E-23</td><td>BT20/MB468</td><td>NUC</td></tr>
<tr><td>>hg38_21_+_8433172_8446622_RNA45SN1_45S_50ntFlanks@10683.10706.24|CCGGCGGCGTCCGGTGAGCTCTCG</td><td>>hg38_21_+_8433172_8446622_RNA45SN1_45S_50ntFlanks</td><td>10683</td><td>10706</td><td>24</td><td>CCGGCGGCGTCCGGTGAGCTCTCG</td><td>28751.4</td><td>-10.0</td><td>1.0</td><td>-10.0</td><td>1.6E-23</td><td>8.6E-23</td><td>BT20/MB468</td><td>NUC</td></tr>
<tr><td>>hg38_21_+_8433172_8446622_RNA45SN1_45S_50ntFlanks@12054.12072.19|TGAGGCGGGGGGGCGAGCC</td><td>>hg38_21_+_8433172_8446622_RNA45SN1_45S_50ntFlanks</td><td>12054</td><td>12072</td><td>19</td><td>TGAGGCGGGGGGGCGAGCC</td><td>730.1</td><td>-13.9</td><td>1.4</td><td>-10.0</td><td>1.6E-23</td><td>8.6E-23</td><td>BT20/MB468</td><td>NUC</td></tr>
<tr><td>>am_trna2_GlyGCC_21_-_18827107_18827177@1.18.18,trna35_GlyGCC_1_+_161413094_161413164@1.18.18,trna37_GlyGCC_1_+_161420467_161420537@1.18.18,trna39_GlyGCC_1_+_161427898_161427968@1.18.18,trna41_GlyGCC_1_+_161435258_161435328@1.18.18|GCATGGGTGGTTCAGTGG</td><td>>am_trna2_GlyGCC_21_-_18827107_18827177</td><td>1</td><td>18</td><td>18</td><td>GCATGGGTGGTTCAGTGG</td><td>553.4</td><td>7.3</td><td>0.7</td><td>10.1</td><td>7.7E-24</td><td>8.6E-23</td><td>BT20/MB468</td><td>CYTO</td></tr>
<tr><td>>hg38_21_+_8433172_8446622_RNA45SN1_45S_50ntFlanks@12163.12183.21|ACTGGGGCGGTACACCTGTCA</td><td>>hg38_21_+_8433172_8446622_RNA45SN1_45S_50ntFlanks</td><td>12163</td><td>12183</td><td>21</td><td>ACTGGGGCGGTACACCTGTCA</td><td>1149.7</td><td>-14.0</td><td>1.4</td><td>-10.2</td><td>2.3E-24</td><td>8.6E-23</td><td>MB231/MB468</td><td>MP</td></tr>
<tr><td>>hg38_21_+_8433172_8446622_RNA45SN1_45S_50ntFlanks@3874.3898.25|AATACATGCCGACGGGCGCTGACCC</td><td>>hg38_21_+_8433172_8446622_RNA45SN1_45S_50ntFlanks</td><td>3874</td><td>3898</td><td>25</td><td>AATACATGCCGACGGGCGCTGACCC</td><td>2255.2</td><td>-15.0</td><td>1.5</td><td>-10.2</td><td>2.3E-24</td><td>8.6E-23</td><td>MB231/MB468</td><td>MP</td></tr>
<tr><td>>hg38_21_+_8433172_8446622_RNA45SN1_45S_50ntFlanks@12641.12661.21|CCATCTGTGGGATTATGACTG</td><td>>hg38_21_+_8433172_8446622_RNA45SN1_45S_50ntFlanks</td><td>12641</td><td>12661</td><td>21</td><td>CCATCTGTGGGATTATGACTG</td><td>2943.4</td><td>-15.4</td><td>1.5</td><td>-10.2</td><td>2.4E-24</td><td>8.7E-23</td><td>MB231/MB468</td><td>MP</td></tr>
<tr><td>>MI0000255|hsa-mir-30d&WithFlank&8|-|134804870|134804951@12.33.22(+1A)&[MIMAT0000245&hsa-miR-30d-5p&offsets|0|0(+1A);m-21&8|-|134804919|134804940&offsets|0|0(+1A)]|TGTAAACATCCCCGACTGGAAGA</td><td>>MI0000255|hsa-mir-30d&WithFlank&8|-|134804870|134804951</td><td>12</td><td>33</td><td>22(+1A)</td><td>TGTAAACATCCCCGACTGGAAGA</td><td>597.3</td><td>3.9</td><td>0.4</td><td>10.0</td><td>1.7E-23</td><td>8.7E-23</td><td>BT20/MB468</td><td>NUC</td></tr>
<tr><td>>hg38_21_+_8433172_8446622_RNA45SN1_45S_50ntFlanks@485.515.31|GATGTGGTGACGTCGTGCTCTCCCGGGCCGG</td><td>>hg38_21_+_8433172_8446622_RNA45SN1_45S_50ntFlanks</td><td>485</td><td>515</td><td>31</td><td>GATGTGGTGACGTCGTGCTCTCCCGGGCCGG</td><td>781.3</td><td>14.1</td><td>1.4</td><td>10.1</td><td>6.0E-24</td><td>8.7E-23</td><td>BT20/MB468</td><td>MITO</td></tr>
<tr><td>>MI0000266|hsa-mir-10a&WithFlank&17|-|48579832|48579953@28.50.23(+1A)&[MIMAT0000253&hsa-miR-10a-5p&offsets|0|0(+1A);m-18&17|-|48579905|48579926&offsets|0|+1(+1A)]|TACCCTGTAGATCCGAATTTGTGA</td><td>>MI0000266|hsa-mir-10a&WithFlank&17|-|48579832|48579953</td><td>28</td><td>50</td><td>23(+1A)</td><td>TACCCTGTAGATCCGAATTTGTGA</td><td>571.2</td><td>13.1</td><td>1.3</td><td>10.1</td><td>4.1E-24</td><td>8.8E-23</td><td>MB231/MB468</td><td>NUC</td></tr>
<tr><td>>hg38_21_+_8433172_8446622_RNA45SN1_45S_50ntFlanks@11316.11336.21|CACCCCACGTCTCGTCGCGCG</td><td>>hg38_21_+_8433172_8446622_RNA45SN1_45S_50ntFlanks</td><td>11316</td><td>11336</td><td>21</td><td>CACCCCACGTCTCGTCGCGCG</td><td>5994.6</td><td>-12.6</td><td>1.2</td><td>-10.1</td><td>3.6E-24</td><td>8.8E-23</td><td>MB231/MB468</td><td>MITO</td></tr>
<tr><td>>hg38_21_+_8433172_8446622_RNA45SN1_45S_50ntFlanks@4725.4767.43|TTAATCAAGAACGAAAGTCGGAGGTTCGAAGACGATCAGATAC</td><td>>hg38_21_+_8433172_8446622_RNA45SN1_45S_50ntFlanks</td><td>4725</td><td>4767</td><td>43</td><td>TTAATCAAGAACGAAAGTCGGAGGTTCGAAGACGATCAGATAC</td><td>1044.5</td><td>14.0</td><td>1.4</td><td>10.2</td><td>1.7E-24</td><td>8.8E-23</td><td>MB231/MB468</td><td>CYTO</td></tr>
<tr><td>>MI0000816|hsa-mir-335&WithFlank&7|+|130496105|130496210@58.79.22&[MIMAT0004703&hsa-miR-335-3p&offsets|0|0;m-159&7|+|130496162|130496183&offsets|0|0]|TTTTTCATTATTGCTCCTGACC</td><td>>MI0000816|hsa-mir-335&WithFlank&7|+|130496105|130496210</td><td>58</td><td>79</td><td>22</td><td>TTTTTCATTATTGCTCCTGACC</td><td>3136.7</td><td>7.1</td><td>0.7</td><td>10.1</td><td>7.9E-24</td><td>8.8E-23</td><td>BT20/MB468</td><td>CYTO</td></tr>
<tr><td>>MI0000285|hsa-mir-205&WithFlank&1|+|209432127|209432248@40.61.22(+1C)&[MIMAT0000266&hsa-miR-205-5p&offsets|0|0(+1C);m-351&1|+|209432166|209432187&offsets|0|0(+1C)]|TCCTTCATTCCACCGGAGTCTGC</td><td>>MI0000285|hsa-mir-205&WithFlank&1|+|209432127|209432248</td><td>40</td><td>61</td><td>22(+1C)</td><td>TCCTTCATTCCACCGGAGTCTGC</td><td>4208.8</td><td>15.7</td><td>1.5</td><td>10.4</td><td>2.6E-25</td><td>8.9E-23</td><td>BT20/MB231</td><td>MITO</td></tr>
<tr><td>>MI0000439|hsa-mir-23b&WithFlank&9|+|95085202|95085310@67.85.19&[MIMAT0000418&hsa-miR-23b-3p&offsets|+3|-1;m-42&9|+|95085265|95085285&offsets|+3|+1]|ACATTGCCAGGGATTACCA</td><td>>MI0000439|hsa-mir-23b&WithFlank&9|+|95085202|95085310</td><td>67</td><td>85</td><td>19</td><td>ACATTGCCAGGGATTACCA</td><td>487.7</td><td>13.4</td><td>1.3</td><td>10.1</td><td>6.1E-24</td><td>8.9E-23</td><td>BT20/MB468</td><td>MITO</td></tr>
<tr><td>>MI0000067|hsa-let-7f-1&WithFlank&9|+|94176341|94176439@13.33.21;MI0000068|hsa-let-7f-2&WithFlank&X|-|53557186|53557280@14.34.21&[MIMAT0000067&hsa-let-7f-5p&offsets|0|-1;m-20&9|+|94176353|94176374&offsets|0|-1];[MIMAT0000067_1&hsa-let-7f-5p&offsets|0|-1;m-19&X|-|53557246|53557267&offsets|0|-1]|TGAGGTAGTAGATTGTATAGT</td><td>>MI0000067|hsa-let-7f-1&WithFlank&9|+|94176341|94176439</td><td>13</td><td>33</td><td>21</td><td>TGAGGTAGTAGATTGTATAGT</td><td>16637.5</td><td>6.9</td><td>0.7</td><td>10.1</td><td>8.1E-24</td><td>9.0E-23</td><td>BT20/MB468</td><td>CYTO</td></tr>
<tr><td>>hg38_21_+_8433172_8446622_RNA45SN1_45S_50ntFlanks@12405.12443.39|CTTTTTGATCCTTCGATGTCGGCTCTTCCTATCATTGTG</td><td>>hg38_21_+_8433172_8446622_RNA45SN1_45S_50ntFlanks</td><td>12405</td><td>12443</td><td>39</td><td>CTTTTTGATCCTTCGATGTCGGCTCTTCCTATCATTGTG</td><td>99246.9</td><td>-13.6</td><td>1.4</td><td>-10.0</td><td>1.7E-23</td><td>9.0E-23</td><td>BT20/MB468</td><td>NUC</td></tr>
<tr><td>>MI0000439|hsa-mir-23b&WithFlank&9|+|95085202|95085310@67.87.21&[MIMAT0000418&hsa-miR-23b-3p&offsets|+3|+1;m-42&9|+|95085265|95085285&offsets|+3|+3]|ACATTGCCAGGGATTACCACG</td><td>>MI0000439|hsa-mir-23b&WithFlank&9|+|95085202|95085310</td><td>67</td><td>87</td><td>21</td><td>ACATTGCCAGGGATTACCACG</td><td>418.6</td><td>13.0</td><td>1.3</td><td>10.1</td><td>8.2E-24</td><td>9.0E-23</td><td>BT20/MB468</td><td>CYTO</td></tr>
<tr><td>>hg38_21_+_8433172_8446622_RNA45SN1_45S_50ntFlanks@12539.12566.28|ATGATGTGTTGTTGCCATGGTAATCCTG</td><td>>hg38_21_+_8433172_8446622_RNA45SN1_45S_50ntFlanks</td><td>12539</td><td>12566</td><td>28</td><td>ATGATGTGTTGTTGCCATGGTAATCCTG</td><td>1628.1</td><td>-14.5</td><td>1.4</td><td>-10.1</td><td>3.8E-24</td><td>9.1E-23</td><td>MB231/MB468</td><td>MITO</td></tr>
<tr><td>>hg38_21_+_8433172_8446622_RNA45SN1_45S_50ntFlanks@10517.10534.18|AGTGGCGGAGATGGGCGC</td><td>>hg38_21_+_8433172_8446622_RNA45SN1_45S_50ntFlanks</td><td>10517</td><td>10534</td><td>18</td><td>AGTGGCGGAGATGGGCGC</td><td>1875.5</td><td>-14.7</td><td>1.4</td><td>-10.2</td><td>2.5E-24</td><td>9.1E-23</td><td>MB231/MB468</td><td>MP</td></tr>
<tr><td>>hg38_21_+_8433172_8446622_RNA45SN1_45S_50ntFlanks@13002.13026.25|CGATCTATTGAAAGTCAGCCCTCGA</td><td>>hg38_21_+_8433172_8446622_RNA45SN1_45S_50ntFlanks</td><td>13002</td><td>13026</td><td>25</td><td>CGATCTATTGAAAGTCAGCCCTCGA</td><td>320.9</td><td>-12.8</td><td>1.3</td><td>-10.0</td><td>1.8E-23</td><td>9.2E-23</td><td>BT20/MB468</td><td>NUC</td></tr>
<tr><td>>hg38_21_+_8433172_8446622_RNA45SN1_45S_50ntFlanks@11641.11669.29|CGGCGGGTGTTGACGCGATGTGATTTCTG</td><td>>hg38_21_+_8433172_8446622_RNA45SN1_45S_50ntFlanks</td><td>11641</td><td>11669</td><td>29</td><td>CGGCGGGTGTTGACGCGATGTGATTTCTG</td><td>2142.1</td><td>-14.0</td><td>1.4</td><td>-10.1</td><td>4.3E-24</td><td>9.2E-23</td><td>MB231/MB468</td><td>NUC</td></tr>
<tr><td>>hg38_21_+_8433172_8446622_RNA45SN1_45S_50ntFlanks@7976.8009.34|GCGACCTCAGATCAGACGTGGCGACCCGCTGAAT</td><td>>hg38_21_+_8433172_8446622_RNA45SN1_45S_50ntFlanks</td><td>7976</td><td>8009</td><td>34</td><td>GCGACCTCAGATCAGACGTGGCGACCCGCTGAAT</td><td>2797.6</td><td>6.5</td><td>0.6</td><td>10.1</td><td>6.5E-24</td><td>9.4E-23</td><td>BT20/MB468</td><td>MITO</td></tr>
<tr><td>>hg38_21_+_8433172_8446622_RNA45SN1_45S_50ntFlanks@10808.10826.19|CAATGTAGGTAAGGGAAGT</td><td>>hg38_21_+_8433172_8446622_RNA45SN1_45S_50ntFlanks</td><td>10808</td><td>10826</td><td>19</td><td>CAATGTAGGTAAGGGAAGT</td><td>1382.0</td><td>-8.0</td><td>0.8</td><td>-10.2</td><td>1.8E-24</td><td>9.4E-23</td><td>MB231/MB468</td><td>CYTO</td></tr>
<tr><td>>hg38_21_+_8433172_8446622_RNA45SN1_45S_50ntFlanks@5420.5443.24|CCGATTGGATGGTTTAGTGAGGCC</td><td>>hg38_21_+_8433172_8446622_RNA45SN1_45S_50ntFlanks</td><td>5420</td><td>5443</td><td>24</td><td>CCGATTGGATGGTTTAGTGAGGCC</td><td>2596.7</td><td>-15.2</td><td>1.5</td><td>-10.2</td><td>2.6E-24</td><td>9.5E-23</td><td>MB231/MB468</td><td>MP</td></tr>
<tr><td>>hg38_21_+_8433172_8446622_RNA45SN1_45S_50ntFlanks@12405.12445.41|CTTTTTGATCCTTCGATGTCGGCTCTTCCTATCATTGTGAA</td><td>>hg38_21_+_8433172_8446622_RNA45SN1_45S_50ntFlanks</td><td>12405</td><td>12445</td><td>41</td><td>CTTTTTGATCCTTCGATGTCGGCTCTTCCTATCATTGTGAA</td><td>1658.9</td><td>-15.1</td><td>1.5</td><td>-10.0</td><td>1.8E-23</td><td>9.5E-23</td><td>BT20/MB468</td><td>NUC</td></tr>
<tr><td>>hg38_21_+_8433172_8446622_RNA45SN1_45S_50ntFlanks@9738.9758.21|GAAACGATCTCAACCTATTCT</td><td>>hg38_21_+_8433172_8446622_RNA45SN1_45S_50ntFlanks</td><td>9738</td><td>9758</td><td>21</td><td>GAAACGATCTCAACCTATTCT</td><td>10225.6</td><td>-9.2</td><td>0.9</td><td>-10.1</td><td>4.0E-24</td><td>9.6E-23</td><td>MB231/MB468</td><td>MITO</td></tr>
<tr><td>>MI0000285|hsa-mir-205&WithFlank&1|+|209432127|209432248@40.62.23(+1U)&[MIMAT0000266&hsa-miR-205-5p&offsets|0|+1(+1U);m-351&1|+|209432166|209432187&offsets|0|+1(+1U)]|TCCTTCATTCCACCGGAGTCTGTT</td><td>>MI0000285|hsa-mir-205&WithFlank&1|+|209432127|209432248</td><td>40</td><td>62</td><td>23(+1U)</td><td>TCCTTCATTCCACCGGAGTCTGTT</td><td>3245.7</td><td>7.2</td><td>0.7</td><td>10.1</td><td>8.8E-24</td><td>9.7E-23</td><td>BT20/MB468</td><td>CYTO</td></tr>
<tr><td>>hg38_21_+_8433172_8446622_RNA45SN1_45S_50ntFlanks@11235.11254.20|CTTCCCGTGGATCGCCCCAG</td><td>>hg38_21_+_8433172_8446622_RNA45SN1_45S_50ntFlanks</td><td>11235</td><td>11254</td><td>20</td><td>CTTCCCGTGGATCGCCCCAG</td><td>4320.0</td><td>-11.7</td><td>1.2</td><td>-10.1</td><td>4.1E-24</td><td>9.7E-23</td><td>MB231/MB468</td><td>MITO</td></tr>
<tr><td>>hg38_21_+_8433172_8446622_RNA45SN1_45S_50ntFlanks@3984.4001.18|GCGGCTTTGGTGACTCTA</td><td>>hg38_21_+_8433172_8446622_RNA45SN1_45S_50ntFlanks</td><td>3984</td><td>4001</td><td>18</td><td>GCGGCTTTGGTGACTCTA</td><td>3583.1</td><td>-15.6</td><td>1.5</td><td>-10.1</td><td>4.1E-24</td><td>9.7E-23</td><td>MB231/MB468</td><td>MITO</td></tr>
<tr><td>>hg38_21_+_8433172_8446622_RNA45SN1_45S_50ntFlanks@11320.11342.23|CCACGTCTCGTCGCGCGCGCGTC</td><td>>hg38_21_+_8433172_8446622_RNA45SN1_45S_50ntFlanks</td><td>11320</td><td>11342</td><td>23</td><td>CCACGTCTCGTCGCGCGCGCGTC</td><td>2260.1</td><td>-14.9</td><td>1.5</td><td>-10.1</td><td>4.1E-24</td><td>9.7E-23</td><td>MB231/MB468</td><td>MITO</td></tr>
<tr><td>>MI0000086|hsa-mir-28&WithFlank&3|+|188688775|188688872@60.81.22(+2A)&[MIMAT0004502&hsa-miR-28-3p&offsets|0|0(+2A);m-28&3|+|188688834|188688855&offsets|0|0(+2A)]|CACTAGATTGTGAGCTCCTGGAAA</td><td>>MI0000086|hsa-mir-28&WithFlank&3|+|188688775|188688872</td><td>60</td><td>81</td><td>22(+2A)</td><td>CACTAGATTGTGAGCTCCTGGAAA</td><td>753.5</td><td>13.8</td><td>1.4</td><td>10.1</td><td>8.8E-24</td><td>9.7E-23</td><td>BT20/MB468</td><td>CYTO</td></tr>
<tr><td>>hg38_1_-_228634819_228635039_RNA5S12_5S_50ntFlanks@137.171.35|GAGACCGCCTGGGAATACCGGGTGCTGTAGGCTTT</td><td>>hg38_1_-_228634819_228635039_RNA5S12_5S_50ntFlanks</td><td>137</td><td>171</td><td>35</td><td>GAGACCGCCTGGGAATACCGGGTGCTGTAGGCTTT</td><td>1776.2</td><td>7.5</td><td>0.8</td><td>10.1</td><td>8.9E-24</td><td>9.8E-23</td><td>BT20/MB468</td><td>CYTO</td></tr>
</table>

<table>
<tr><td>>hg38_21_+_8433172_8446622_RNA45SN1_45S_50ntFlanks@4731.4768.38|AAGAACGAAAGTCGGAGGTTCGAAGACGATCAGATACC</td><td>>hg38_21_+_8433172_8446622_RNA45SN1_45S_50ntFlanks</td><td>4731</td><td>4768</td><td>38</td><td>AAGAACGAAAGTCGGAGGTTCGAAGACGATCAGATACC</td><td>3235.2</td><td>10.5</td><td>1.0</td><td>10.1</td><td>8.9E-24</td><td>9.8E-23</td><td>BT20/MB468</td><td>CYTO</td></tr>
<tr><td>>hg38_21_+_8433172_8446622_RNA45SN1_45S_50ntFlanks@11552.11574.23|CCTCGGCCGGCGCCTAGCAGCCG</td><td>>hg38_21_+_8433172_8446622_RNA45SN1_45S_50ntFlanks</td><td>11552</td><td>11574</td><td>23</td><td>CCTCGGCCGGCGCCTAGCAGCCG</td><td>354.0</td><td>-12.9</td><td>1.3</td><td>-10.0</td><td>1.9E-23</td><td>9.8E-23</td><td>BT20/MB468</td><td>NUC</td></tr>
<tr><td>>hg38_21_+_8433172_8446622_RNA45SN1_45S_50ntFlanks@12974.12992.19|CGTACGTAGCAGAGCAGCT</td><td>>hg38_21_+_8433172_8446622_RNA45SN1_45S_50ntFlanks</td><td>12974</td><td>12992</td><td>19</td><td>CGTACGTAGCAGAGCAGCT</td><td>632.5</td><td>-4.5</td><td>0.4</td><td>-10.1</td><td>4.6E-24</td><td>9.8E-23</td><td>MB231/MB468</td><td>NUC</td></tr>
<tr><td>>hg38_21_+_8433172_8446622_RNA45SN1_45S_50ntFlanks@5527.5550.24|AAAGTCGTAACAAGGTTTCCGTAG</td><td>>hg38_21_+_8433172_8446622_RNA45SN1_45S_50ntFlanks</td><td>5527</td><td>5550</td><td>24</td><td>AAAGTCGTAACAAGGTTTCCGTAG</td><td>1503.6</td><td>-14.4</td><td>1.4</td><td>-10.2</td><td>2.7E-24</td><td>9.9E-23</td><td>MB231/MB468</td><td>MP</td></tr>
<tr><td>>hg38_21_+_8433172_8446622_RNA45SN1_45S_50ntFlanks@5071.5110.40|TTAATTCCGATAACGAACGAGACTCTGGCATGCTAACTAG</td><td>>hg38_21_+_8433172_8446622_RNA45SN1_45S_50ntFlanks</td><td>5071</td><td>5110</td><td>40</td><td>TTAATTCCGATAACGAACGAGACTCTGGCATGCTAACTAG</td><td>451.4</td><td>-13.2</td><td>1.3</td><td>-10.0</td><td>1.9E-23</td><td>9.9E-23</td><td>BT20/MB468</td><td>NUC</td></tr>
<tr><td>>hg38_21_+_8433172_8446622_RNA45SN1_45S_50ntFlanks@11778.11795.18|ACGCGCATGAATGGATGA</td><td>>hg38_21_+_8433172_8446622_RNA45SN1_45S_50ntFlanks</td><td>11778</td><td>11795</td><td>18</td><td>ACGCGCATGAATGGATGA</td><td>317.1</td><td>-12.7</td><td>1.3</td><td>-10.0</td><td>1.9E-23</td><td>9.9E-23</td><td>BT20/MB468</td><td>NUC</td></tr>
<tr><td>>MI0005757|hsa-mir-935&WithFlank&19|+|53982301|53982403@63.84.22&[MIMAT0004978&hsa-miR-935&offsets|+1|0;m-549&19|+|53982363|53982384&offsets|0|0]|CAGTTACCGCTTCCGCTACCGC</td><td>>MI0005757|hsa-mir-935&WithFlank&19|+|53982301|53982403</td><td>63</td><td>84</td><td>22</td><td>CAGTTACCGCTTCCGCTACCGC</td><td>412.4</td><td>13.1</td><td>1.3</td><td>10.1</td><td>6.9E-24</td><td>9.9E-23</td><td>BT20/MB468</td><td>MITO</td></tr>
<tr><td>>am_MI0003639|hsa-mir-625&WithFlank&14|+|65471096|65471192@58.80.23&[MIMAT0004808&hsa-miR-625-3p&offsets|0|+1;m-137&14|+|65471153|65471174&offsets|0|+1]|GACTATAGAACTTTCCCCCTCAT</td><td>>am_MI0003639|hsa-mir-625&WithFlank&14|+|65471096|65471192</td><td>58</td><td>80</td><td>23</td><td>GACTATAGAACTTTCCCCCTCAT</td><td>542.5</td><td>6.6</td><td>0.6</td><td>10.1</td><td>4.7E-24</td><td>9.9E-23</td><td>MB231/MB468</td><td>NUC</td></tr>
<tr><td>>hg38_21_+_8433172_8446622_RNA45SN1_45S_50ntFlanks@11579.11603.25|AGAACTGGTGCGGACCAGGGGAATC</td><td>>hg38_21_+_8433172_8446622_RNA45SN1_45S_50ntFlanks</td><td>11579</td><td>11603</td><td>25</td><td>AGAACTGGTGCGGACCAGGGGAATC</td><td>3917.8</td><td>-9.4</td><td>0.9</td><td>-10.1</td><td>4.2E-24</td><td>1.0E-22</td><td>MB231/MB468</td><td>MITO</td></tr>
<tr><td>>hg38_21_+_8433172_8446622_RNA45SN1_45S_50ntFlanks@4657.4682.26|CTAGAGGTGAAATTCTTGGACCGGCG</td><td>>hg38_21_+_8433172_8446622_RNA45SN1_45S_50ntFlanks</td><td>4657</td><td>4682</td><td>26</td><td>CTAGAGGTGAAATTCTTGGACCGGCG</td><td>1457.6</td><td>-13.5</td><td>1.3</td><td>-10.1</td><td>4.8E-24</td><td>1.0E-22</td><td>MB231/MB468</td><td>NUC</td></tr>
<tr><td>>MI0005756|hsa-mir-934&WithFlank&X|+|136550872|136550966@21.41.21&[MIMAT0004977&hsa-miR-934&offsets|0|-1;m-943&X|+|136550892|136550913&offsets|0|-1]|TGTCTACTACTGGAGACACTG</td><td>>MI0005756|hsa-mir-934&WithFlank&X|+|136550872|136550966</td><td>21</td><td>41</td><td>21</td><td>TGTCTACTACTGGAGACACTG</td><td>1657.4</td><td>14.3</td><td>1.4</td><td>10.4</td><td>3.2E-25</td><td>1.0E-22</td><td>BT20/MB231</td><td>MITO</td></tr>
<tr><td>>hg38_21_+_8433172_8446622_RNA45SN1_45S_50ntFlanks@4099.4121.23|GTGACCACGGGTGACGGGGAATC</td><td>>hg38_21_+_8433172_8446622_RNA45SN1_45S_50ntFlanks</td><td>4099</td><td>4121</td><td>23</td><td>GTGACCACGGGTGACGGGGAATC</td><td>815.4</td><td>-14.1</td><td>1.4</td><td>-10.0</td><td>1.9E-23</td><td>1.0E-22</td><td>BT20/MB468</td><td>NUC</td></tr>
<tr><td>>hg38_21_+_8433172_8446622_RNA45SN1_45S_50ntFlanks@7983.8005.23|CAGATCAGACGTGGCGACCCGCT</td><td>>hg38_21_+_8433172_8446622_RNA45SN1_45S_50ntFlanks</td><td>7983</td><td>8005</td><td>23</td><td>CAGATCAGACGTGGCGACCCGCT</td><td>1451.9</td><td>-5.7</td><td>0.6</td><td>-10.0</td><td>1.9E-23</td><td>1.0E-22</td><td>BT20/MB468</td><td>NUC</td></tr>
<tr><td>>hg38_21_+_8433172_8446622_RNA45SN1_45S_50ntFlanks@5485.5515.31|AGCGCTGAGAAGACGGTCGAACTTGACTATC</td><td>>hg38_21_+_8433172_8446622_RNA45SN1_45S_50ntFlanks</td><td>5485</td><td>5515</td><td>31</td><td>AGCGCTGAGAAGACGGTCGAACTTGACTATC</td><td>1459.6</td><td>-14.4</td><td>1.4</td><td>-10.2</td><td>2.8E-24</td><td>1.0E-22</td><td>MB231/MB468</td><td>MP</td></tr>
<tr><td>>hg38_21_+_8433172_8446622_RNA45SN1_45S_50ntFlanks@9738.9765.28|GAAACGATCTCAACCTATTCTCAAACTT</td><td>>hg38_21_+_8433172_8446622_RNA45SN1_45S_50ntFlanks</td><td>9738</td><td>9765</td><td>28</td><td>GAAACGATCTCAACCTATTCTCAAACTT</td><td>1518.6</td><td>-14.4</td><td>1.4</td><td>-10.2</td><td>2.8E-24</td><td>1.0E-22</td><td>MB231/MB468</td><td>MP</td></tr>
<tr><td>>MI0003820|hsa-mir-454&WithFlank&17|-|59137752|59137878@70.93.24&[MIMAT0003885&hsa-miR-454-3p&offsets|0|+1;m-272&17|-|59137787|59137809&offsets|0|+1]|TAGTGCAATATTGCTTATAGGGTT</td><td>>MI0003820|hsa-mir-454&WithFlank&17|-|59137752|59137878</td><td>70</td><td>93</td><td>24</td><td>TAGTGCAATATTGCTTATAGGGTT</td><td>547.7</td><td>6.8</td><td>0.7</td><td>10.0</td><td>9.2E-24</td><td>1.0E-22</td><td>BT20/MB468</td><td>CYTO</td></tr>
<tr><td>>hg38_21_+_8433172_8446622_RNA45SN1_45S_50ntFlanks@10381.10424.44|GTTCCATGTGAACAGCAGTTGAACATGGGTCAGTCGGTCCTGAG</td><td>>hg38_21_+_8433172_8446622_RNA45SN1_45S_50ntFlanks</td><td>10381</td><td>10424</td><td>44</td><td>GTTCCATGTGAACAGCAGTTGAACATGGGTCAGTCGGTCCTGAG</td><td>1254.2</td><td>8.9</td><td>0.9</td><td>10.2</td><td>2.0E-24</td><td>1.0E-22</td><td>MB231/MB468</td><td>CYTO</td></tr>
<tr><td>>hg38_21_+_8433172_8446622_RNA45SN1_45S_50ntFlanks@12161.12179.19|TGACTGGGGCGGTACACCT</td><td>>hg38_21_+_8433172_8446622_RNA45SN1_45S_50ntFlanks</td><td>12161</td><td>12179</td><td>19</td><td>TGACTGGGGCGGTACACCT</td><td>1678.1</td><td>-14.6</td><td>1.4</td><td>-10.2</td><td>2.8E-24</td><td>1.0E-22</td><td>MB231/MB468</td><td>MP</td></tr>
<tr><td>>hg38_21_+_8433172_8446622_RNA45SN1_45S_50ntFlanks@4742.4759.18|TCGGAGGTTCGAAGACGA</td><td>>hg38_21_+_8433172_8446622_RNA45SN1_45S_50ntFlanks</td><td>4742</td><td>4759</td><td>18</td><td>TCGGAGGTTCGAAGACGA</td><td>820.3</td><td>-4.4</td><td>0.4</td><td>-10.0</td><td>2.0E-23</td><td>1.0E-22</td><td>BT20/MB468</td><td>NUC</td></tr>
<tr><td>>hg38_21_+_8433172_8446622_RNA45SN1_45S_50ntFlanks@12324.12345.22|TTGGGTTTTAAGCAGGAGGTGT</td><td>>hg38_21_+_8433172_8446622_RNA45SN1_45S_50ntFlanks</td><td>12324</td><td>12345</td><td>22</td><td>TTGGGTTTTAAGCAGGAGGTGT</td><td>704.0</td><td>-13.9</td><td>1.4</td><td>-10.0</td><td>2.0E-23</td><td>1.0E-22</td><td>BT20/MB468</td><td>NUC</td></tr>
<tr><td>>hg38_MT_+_1621_3279_RNR2_16S_50ntFlanks@1216.1262.47|CAACCTCCGAGCAGTACATGCTAAGACTTCACCAGTCAAAGCGAACT</td><td>>hg38_MT_+_1621_3279_RNR2_16S_50ntFlanks</td><td>1216</td><td>1262</td><td>47</td><td>CAACCTCCGAGCAGTACATGCTAAGACTTCACCAGTCAAAGCGAACT</td><td>568.7</td><td>13.2</td><td>1.3</td><td>10.2</td><td>2.1E-24</td><td>1.1E-22</td><td>MB231/MB468</td><td>CYTO</td></tr>
<tr><td>>MI0000266|hsa-mir-10a&WithFlank&17|-|48579832|48579953@29.50.22(+1C)&[MIMAT0000253&hsa-miR-10a-5p&offsets|+1|0(+1C);m-18&17|-|48579905|48579926&offsets|+1|+1(+1C)]|ACCCTGTAGATCCGAATTTGTGC</td><td>>MI0000266|hsa-mir-10a&WithFlank&17|-|48579832|48579953</td><td>29</td><td>50</td><td>22(+1C)</td><td>ACCCTGTAGATCCGAATTTGTGC</td><td>340.4</td><td>12.7</td><td>1.3</td><td>10.0</td><td>9.7E-24</td><td>1.1E-22</td><td>BT20/MB468</td><td>CYTO</td></tr>
<tr><td>>hg38_21_+_8433172_8446622_RNA45SN1_45S_50ntFlanks@10411.10428.18|CAGTCGGTCCTGAGAGAT</td><td>>hg38_21_+_8433172_8446622_RNA45SN1_45S_50ntFlanks</td><td>10411</td><td>10428</td><td>18</td><td>CAGTCGGTCCTGAGAGAT</td><td>3171.1</td><td>-6.1</td><td>0.6</td><td>-10.0</td><td>9.8E-24</td><td>1.1E-22</td><td>BT20/MB468</td><td>CYTO</td></tr>
<tr><td>>hg38_21_+_8433172_8446622_RNA45SN1_45S_50ntFlanks@11873.11906.34|AAGAAGACCCTGTTGAGCTTGACTCTAGTCTGGC</td><td>>hg38_21_+_8433172_8446622_RNA45SN1_45S_50ntFlanks</td><td>11873</td><td>11906</td><td>34</td><td>AAGAAGACCCTGTTGAGCTTGACTCTAGTCTGGC</td><td>1392.5</td><td>-14.3</td><td>1.4</td><td>-10.2</td><td>3.0E-24</td><td>1.1E-22</td><td>MB231/MB468</td><td>MP</td></tr>
<tr><td>>hg38_21_+_8433172_8446622_RNA45SN1_45S_50ntFlanks@6646.6664.19|TCGTACGACTCTTAGCGGT</td><td>>hg38_21_+_8433172_8446622_RNA45SN1_45S_50ntFlanks</td><td>6646</td><td>6664</td><td>19</td><td>TCGTACGACTCTTAGCGGT</td><td>196677.0</td><td>6.4</td><td>0.6</td><td>10.1</td><td>7.5E-24</td><td>1.1E-22</td><td>BT20/MB468</td><td>MITO</td></tr>
<tr><td>>hg38_21_+_8433172_8446622_RNA45SN1_45S_50ntFlanks@11874.11905.32|AGAAGACCCTGTTGAGCTTGACTCTAGTCTGG</td><td>>hg38_21_+_8433172_8446622_RNA45SN1_45S_50ntFlanks</td><td>11874</td><td>11905</td><td>32</td><td>AGAAGACCCTGTTGAGCTTGACTCTAGTCTGG</td><td>1706.9</td><td>-14.5</td><td>1.4</td><td>-10.1</td><td>4.5E-24</td><td>1.1E-22</td><td>MB231/MB468</td><td>MITO</td></tr>
<tr><td>>hg38_21_+_8433172_8446622_RNA45SN1_45S_50ntFlanks@12709.12745.37|CCGCGGAGCCTCGGTTGGCCTCGGATAGCCGGTCCCC</td><td>>hg38_21_+_8433172_8446622_RNA45SN1_45S_50ntFlanks</td><td>12709</td><td>12745</td><td>37</td><td>CCGCGGAGCCTCGGTTGGCCTCGGATAGCCGGTCCCC</td><td>1570.8</td><td>-14.4</td><td>1.4</td><td>-10.1</td><td>4.6E-24</td><td>1.1E-22</td><td>MB231/MB468</td><td>MITO</td></tr>
<tr><td>>hg38_21_+_8433172_8446622_RNA45SN1_45S_50ntFlanks@10680.10708.29|GTTCCGGCGGCGTCCGGTGAGCTCTCGCT</td><td>>hg38_21_+_8433172_8446622_RNA45SN1_45S_50ntFlanks</td><td>10680</td><td>10708</td><td>29</td><td>GTTCCGGCGGCGTCCGGTGAGCTCTCGCT</td><td>9984.5</td><td>-8.5</td><td>0.8</td><td>-10.2</td><td>3.0E-24</td><td>1.1E-22</td><td>MB231/MB468</td><td>MP</td></tr>
<tr><td>>hg38_21_+_8433172_8446622_RNA45SN1_45S_50ntFlanks@11882.11904.23|CTGTTGAGCTTGACTCTAGTCTG</td><td>>hg38_21_+_8433172_8446622_RNA45SN1_45S_50ntFlanks</td><td>11882</td><td>11904</td><td>23</td><td>CTGTTGAGCTTGACTCTAGTCTG</td><td>1498.0</td><td>-14.4</td><td>1.4</td><td>-10.2</td><td>3.0E-24</td><td>1.1E-22</td><td>MB231/MB468</td><td>MP</td></tr>
<tr><td>>hg38_21_+_8433172_8446622_RNA45SN1_45S_50ntFlanks@12638.12661.24|CTACCATCTGTGGGATTATGACTG</td><td>>hg38_21_+_8433172_8446622_RNA45SN1_45S_50ntFlanks</td><td>12638</td><td>12661</td><td>24</td><td>CTACCATCTGTGGGATTATGACTG</td><td>9198.5</td><td>-9.0</td><td>0.9</td><td>-10.1</td><td>4.6E-24</td><td>1.1E-22</td><td>MB231/MB468</td><td>MITO</td></tr>
<tr><td>>hg38_21_+_8433172_8446622_RNA45SN1_45S_50ntFlanks@6740.6771.32|CATCGACACTTCGAACGCACTTGCGGCCCCGG</td><td>>hg38_21_+_8433172_8446622_RNA45SN1_45S_50ntFlanks</td><td>6740</td><td>6771</td><td>32</td><td>CATCGACACTTCGAACGCACTTGCGGCCCCGG</td><td>1936.4</td><td>-14.8</td><td>1.5</td><td>-10.2</td><td>3.0E-24</td><td>1.1E-22</td><td>MB231/MB468</td><td>MP</td></tr>
<tr><td>>MI0000650|hsa-mir-200c&WithFlank&12|+|6963693|6963772@50.69.20(+1A)&[MIMAT0000617&hsa-miR-200c-3p&offsets|0|-3(+1A);m-54&12|+|6963742|6963764&offsets|0|-3(+1A)]|TAATACTGCCGGGTAATGATA</td><td>>MI0000650|hsa-mir-200c&WithFlank&12|+|6963693|6963772</td><td>50</td><td>69</td><td>20(+1A)</td><td>TAATACTGCCGGGTAATGATA</td><td>520.9</td><td>13.5</td><td>1.3</td><td>10.1</td><td>7.6E-24</td><td>1.1E-22</td><td>BT20/MB468</td><td>MITO</td></tr>
<tr><td>>MI0000809|hsa-mir-151a&WithFlank&8|-|140732558|140732659@53.74.22(+2A)&[MIMAT0000757&hsa-miR-151a-3p&offsets|0|+1(+2A);m-30&8|-|140732587|140732607&offsets|0|+1(+2A)]|CTAGACTGAAGCTCCTTGAGGAAA</td><td>>MI0000809|hsa-mir-151a&WithFlank&8|-|140732558|140732659</td><td>53</td><td>74</td><td>22(+2A)</td><td>CTAGACTGAAGCTCCTTGAGGAAA</td><td>1533.2</td><td>9.0</td><td>0.9</td><td>10.1</td><td>7.7E-24</td><td>1.1E-22</td><td>BT20/MB468</td><td>MITO</td></tr>
<tr><td>>hg38_21_+_8433172_8446622_RNA45SN1_45S_50ntFlanks@12619.12654.36|AGGAGCCAATGGGGCGAAGCTACCATCTGTGGGATT</td><td>>hg38_21_+_8433172_8446622_RNA45SN1_45S_50ntFlanks</td><td>12619</td><td>12654</td><td>36</td><td>AGGAGCCAATGGGGCGAAGCTACCATCTGTGGGATT</td><td>1480.2</td><td>-14.4</td><td>1.4</td><td>-10.2</td><td>3.1E-24</td><td>1.1E-22</td><td>MB231/MB468</td><td>MP</td></tr>
<tr><td>>MI0000255|hsa-mir-30d&WithFlank&8|-|134804870|134804951@12.32.21&[MIMAT0000245&hsa-miR-30d-5p&offsets|0|-1;m-21&8|-|134804919|134804940&offsets|0|-1]|TGTAAACATCCCCGACTGGAA</td><td>>MI0000255|hsa-mir-30d&WithFlank&8|-|134804870|134804951</td><td>12</td><td>32</td><td>21</td><td>TGTAAACATCCCCGACTGGAA</td><td>284.2</td><td>4.0</td><td>0.4</td><td>10.0</td><td>2.1E-23</td><td>1.1E-22</td><td>BT20/MB468</td><td>NUC</td></tr>
<tr><td>>hg38_21_+_8433172_8446622_RNA45SN1_45S_50ntFlanks@13002.13031.30|CGATCTATTGAAAGTCAGCCCTCGACACAA</td><td>>hg38_21_+_8433172_8446622_RNA45SN1_45S_50ntFlanks</td><td>13002</td><td>13031</td><td>30</td><td>CGATCTATTGAAAGTCAGCCCTCGACACAA</td><td>395.7</td><td>-13.1</td><td>1.3</td><td>-10.0</td><td>2.1E-23</td><td>1.1E-22</td><td>BT20/MB468</td><td>NUC</td></tr>
<tr><td>>hg38_21_+_8433172_8446622_RNA45SN1_45S_50ntFlanks@4726.4767.42|TAATCAAGAACGAAAGTCGGAGGTTCGAAGACGATCAGATAC</td><td>>hg38_21_+_8433172_8446622_RNA45SN1_45S_50ntFlanks</td><td>4726</td><td>4767</td><td>42</td><td>TAATCAAGAACGAAAGTCGGAGGTTCGAAGACGATCAGATAC</td><td>794.7</td><td>13.6</td><td>1.3</td><td>10.2</td><td>2.2E-24</td><td>1.1E-22</td><td>MB231/MB468</td><td>CYTO</td></tr>
<tr><td>>hg38_21_+_8433172_8446622_RNA45SN1_45S_50ntFlanks@10805.10822.18|GAACAATGTAGGTAAGGG</td><td>>hg38_21_+_8433172_8446622_RNA45SN1_45S_50ntFlanks</td><td>10805</td><td>10822</td><td>18</td><td>GAACAATGTAGGTAAGGG</td><td>453.3</td><td>-13.3</td><td>1.3</td><td>-10.0</td><td>2.2E-23</td><td>1.1E-22</td><td>BT20/MB468</td><td>NUC</td></tr>
<tr><td>>hg38_21_+_8433172_8446622_RNA45SN1_45S_50ntFlanks@12405.12431.27|CTTTTTGATCCTTCGATGTCGGCTCTT</td><td>>hg38_21_+_8433172_8446622_RNA45SN1_45S_50ntFlanks</td><td>12405</td><td>12431</td><td>27</td><td>CTTTTTGATCCTTCGATGTCGGCTCTT</td><td>4689.1</td><td>-9.5</td><td>0.9</td><td>-10.1</td><td>4.8E-24</td><td>1.1E-22</td><td>MB231/MB468</td><td>MITO</td></tr>
<tr><td>>hg38_1_-_228634819_228635039_RNA5S12_5S_50ntFlanks@51.89.39|GTCTACGGCCATACCACCCTGAACGCGCCCGATCTCGTC</td><td>>hg38_1_-_228634819_228635039_RNA5S12_5S_50ntFlanks</td><td>51</td><td>89</td><td>39</td><td>GTCTACGGCCATACCACCCTGAACGCGCCCGATCTCGTC</td><td>76254.4</td><td>10.3</td><td>1.0</td><td>10.0</td><td>1.0E-23</td><td>1.1E-22</td><td>BT20/MB468</td><td>CYTO</td></tr>
<tr><td>>hg38_21_+_8433172_8446622_RNA45SN1_45S_50ntFlanks@12605.12625.21|TATGTGCTTGGCTGAGGAGCC</td><td>>hg38_21_+_8433172_8446622_RNA45SN1_45S_50ntFlanks</td><td>12605</td><td>12625</td><td>21</td><td>TATGTGCTTGGCTGAGGAGCC</td><td>2742.8</td><td>-7.3</td><td>0.7</td><td>-10.1</td><td>4.9E-24</td><td>1.1E-22</td><td>MB231/MB468</td><td>MITO</td></tr>
<tr><td>>hg38_21_+_8433172_8446622_RNA45SN1_45S_50ntFlanks@12417.12435.19|TCGATGTCGGCTCTTCCTA</td><td>>hg38_21_+_8433172_8446622_RNA45SN1_45S_50ntFlanks</td><td>12417</td><td>12435</td><td>19</td><td>TCGATGTCGGCTCTTCCTA</td><td>1143.7</td><td>-14.0</td><td>1.4</td><td>-10.2</td><td>3.2E-24</td><td>1.1E-22</td><td>MB231/MB468</td><td>MP</td></tr>
<tr><td>>MI0000447|hsa-mir-128-1&WithFlank&2|+|135665391|135665484@56.75.20(+2A);MI0000727|hsa-mir-128-2&WithFlank&3|+|35744470|35744565@58.77.20(+2A)&[MIMAT0000424&hsa-miR-128-3p&offsets|0|-1(+2A);m-108&2|+|135665446|135665466&offsets|0|-1(+2A)];[MIMAT0000424_1&hsa-miR-128-3p&offsets|0|-1(+2A);m-117&3|+|35744527|35744547&offsets|0|-1(+2A)]|TCACAGTGAACCGGTCTCTTAA</td><td>>MI0000447|hsa-mir-128-1&WithFlank&2|+|135665391|135665484</td><td>56</td><td>75</td><td>20(+2A)</td><td>TCACAGTGAACCGGTCTCTTAA</td><td>349.3</td><td>5.7</td><td>0.6</td><td>10.0</td><td>1.0E-23</td><td>1.1E-22</td><td>BT20/MB468</td><td>CYTO</td></tr>
<tr><td>>MI0000239|hsa-mir-197&WithFlank&1|+|109598887|109598973@57.75.19&[MIMAT0000227&hsa-miR-197-3p&offsets|+3|0;m-99&1|+|109598940|109598961&offsets|+3|0]|ACCACCTTCTCCACCCAGC</td><td>>MI0000239|hsa-mir-197&WithFlank&1|+|109598887|109598973</td><td>57</td><td>75</td><td>19</td><td>ACCACCTTCTCCACCCAGC</td><td>416.7</td><td>13.0</td><td>1.3</td><td>10.0</td><td>1.1E-23</td><td>1.1E-22</td><td>BT20/MB468</td><td>CYTO</td></tr>
<tr><td>>hg38_21_+_8433172_8446622_RNA45SN1_45S_50ntFlanks@8105.8138.34|CGCGGCGGGGCGCGGGACATGTGGCGTACGGAAG</td><td>>hg38_21_+_8433172_8446622_RNA45SN1_45S_50ntFlanks</td><td>8105</td><td>8138</td><td>34</td><td>CGCGGCGGGGCGCGGGACATGTGGCGTACGGAAG</td><td>4614.7</td><td>-8.9</td><td>0.9</td><td>-10.1</td><td>5.0E-24</td><td>1.1E-22</td><td>MB231/MB468</td><td>MITO</td></tr>
<tr><td>>hg38_21_+_8433172_8446622_RNA45SN1_45S_50ntFlanks@9198.9225.28|GTTCTCTCGGGGCCACGCGCGCGTCCCC</td><td>>hg38_21_+_8433172_8446622_RNA45SN1_45S_50ntFlanks</td><td>9198</td><td>9225</td><td>28</td><td>GTTCTCTCGGGGCCACGCGCGCGTCCCC</td><td>1484.3</td><td>-14.3</td><td>1.4</td><td>-10.1</td><td>5.0E-24</td><td>1.1E-22</td><td>MB231/MB468</td><td>MITO</td></tr>
<tr><td>>hg38_21_+_8433172_8446622_RNA45SN1_45S_50ntFlanks@5472.5511.40|ACGGCCCTGGCGGAGCGCTGAGAAGACGGTCGAACTTGAC</td><td>>hg38_21_+_8433172_8446622_RNA45SN1_45S_50ntFlanks</td><td>5472</td><td>5511</td><td>40</td><td>ACGGCCCTGGCGGAGCGCTGAGAAGACGGTCGAACTTGAC</td><td>593.7</td><td>7.2</td><td>0.7</td><td>10.0</td><td>1.1E-23</td><td>1.1E-22</td><td>BT20/MB468</td><td>CYTO</td></tr>
<tr><td>>hg38_21_+_8433172_8446622_RNA45SN1_45S_50ntFlanks@9941.9968.28|AGGACGGTGGCCATGGAAGTCGGAATCC</td><td>>hg38_21_+_8433172_8446622_RNA45SN1_45S_50ntFlanks</td><td>9941</td><td>9968</td><td>28</td><td>AGGACGGTGGCCATGGAAGTCGGAATCC</td><td>480.2</td><td>13.4</td><td>1.3</td><td>10.1</td><td>8.1E-24</td><td>1.2E-22</td><td>BT20/MB468</td><td>MITO</td></tr>
<tr><td>>hg38_21_+_8433172_8446622_RNA45SN1_45S_50ntFlanks@11873.11903.31|AAGAAGACCCTGTTGAGCTTGACTCTAGTCT</td><td>>hg38_21_+_8433172_8446622_RNA45SN1_45S_50ntFlanks</td><td>11873</td><td>11903</td><td>31</td><td>AAGAAGACCCTGTTGAGCTTGACTCTAGTCT</td><td>354.8</td><td>-12.9</td><td>1.3</td><td>-10.0</td><td>2.3E-23</td><td>1.2E-22</td><td>BT20/MB468</td><td>NUC</td></tr>
</table>

<table>
<tr><th></th><th></th><th></th><th></th><th></th><th></th><th></th><th></th><th></th><th></th><th></th><th></th><th></th><th></th></tr>
<tr><td>>MI0000750|hsa-mir-26a-2&WithFlank&12|-|57824603|57824698@20.40.21(+1A);MI0000083|hsa-mir-26a-1&WithFlank&3|+|37969398|37969486@16.36.21(+1A)&[MIMAT0000082&hsa-miR-26a-5p&offsets|0|-1(+1A);m-33&12|-|57824658|57824679&offsets|0|-1(+1A)];[MIMAT0000082_1&hsa-miR-26a-5p&offsets|0|-1(+1A);m-34&3|+|37969413|37969434&offsets|0|-1(+1A)]|TTCAAGTAATCCAGGATAGGCA</td><td>>MI0000750|hsa-mir-26a-2&WithFlank&12|-|57824603|57824698</td><td>20</td><td>40</td><td>21(+1A)</td><td>TTCAAGTAATCCAGGATAGGCA</td><td>12255.8</td><td>6.9</td><td>0.7</td><td>10.0</td><td>1.1E-23</td><td>1.2E-22</td><td>BT20/MB468</td><td>CYTO</td></tr>
<tr><td>>hg38_21_+_8433172_8446622_RNA45SN1_45S_50ntFlanks@10369.10401.33|AAGTGGAGAAGGGTTCCATGTGAACAGCAGTTG</td><td>>hg38_21_+_8433172_8446622_RNA45SN1_45S_50ntFlanks</td><td>10369</td><td>10401</td><td>33</td><td>AAGTGGAGAAGGGTTCCATGTGAACAGCAGTTG</td><td>415.2</td><td>-13.1</td><td>1.3</td><td>-10.0</td><td>2.3E-23</td><td>1.2E-22</td><td>BT20/MB468</td><td>NUC</td></tr>
<tr><td>>hg38_21_+_8433172_8446622_RNA45SN1_45S_50ntFlanks@475.516.42|CTTCGTGATCGATGTGGTGACGTCGTGCTCTCCCGGGCCGGG</td><td>>hg38_21_+_8433172_8446622_RNA45SN1_45S_50ntFlanks</td><td>475</td><td>516</td><td>42</td><td>CTTCGTGATCGATGTGGTGACGTCGTGCTCTCCCGGGCCGGG</td><td>1368.2</td><td>9.2</td><td>0.9</td><td>10.2</td><td>2.4E-24</td><td>1.2E-22</td><td>MB231/MB468</td><td>CYTO</td></tr>
<tr><td>>hg38_21_+_8433172_8446622_RNA45SN1_45S_50ntFlanks@12995.13017.23|CTCGCTGCGATCTATTGAAAGTC</td><td>>hg38_21_+_8433172_8446622_RNA45SN1_45S_50ntFlanks</td><td>12995</td><td>13017</td><td>23</td><td>CTCGCTGCGATCTATTGAAAGTC</td><td>819.9</td><td>6.2</td><td>0.6</td><td>10.0</td><td>1.1E-23</td><td>1.2E-22</td><td>BT20/MB468</td><td>CYTO</td></tr>
<tr><td>>trna112_GlnCTG_1_-_147737382_147737453@30.50.21,trna131_GlnCTG_6_-_27759135_27759206@30.50.21,trna146_GlnCTG_6_-_27515531_27515602@30.50.21,trna15_GlnCTG_1_+_145963304_145963375@30.50.21,trna19_GlnCTG_1_+_147505038_147505109@30.50.21,trna1_GlnCTG_6_+_18836402_18836473@30.50.21,trna28_GlnCTG_1_+_149186125_149186196@30.50.21,trna3_GlnCTG_17_+_8023070_8023141@30.50.21,trna49_GlnCTG_6_+_27487308_27487379@30.50.21,trna7_GlnCTG_15_-_66161400_66161471@30.50.21,trna99_GlnCTG_6_-_28909378_28909449@30.50.21|GACTCTGAATCCAGCGATCCG</td><td>>trna112_GlnCTG_1_-_147737382_147737453</td><td>30</td><td>50</td><td>21</td><td>GACTCTGAATCCAGCGATCCG</td><td>814.7</td><td>-14.1</td><td>1.4</td><td>-10.0</td><td>2.3E-23</td><td>1.2E-22</td><td>BT20/MB468</td><td>NUC</td></tr>
<tr><td>>hg38_21_+_8433172_8446622_RNA45SN1_45S_50ntFlanks@11669.11708.40|GCCCAGTGCTCTGAATGTCAAAGTGAAGAAATTCAATGAA</td><td>>hg38_21_+_8433172_8446622_RNA45SN1_45S_50ntFlanks</td><td>11669</td><td>11708</td><td>40</td><td>GCCCAGTGCTCTGAATGTCAAAGTGAAGAAATTCAATGAA</td><td>438.6</td><td>12.8</td><td>1.3</td><td>10.2</td><td>2.5E-24</td><td>1.2E-22</td><td>MB231/MB468</td><td>CYTO</td></tr>
<tr><td>>hg38_21_+_8433172_8446622_RNA45SN1_45S_50ntFlanks@11783.11803.21|CATGAATGGATGAACGAGATT</td><td>>hg38_21_+_8433172_8446622_RNA45SN1_45S_50ntFlanks</td><td>11783</td><td>11803</td><td>21</td><td>CATGAATGGATGAACGAGATT</td><td>2352.9</td><td>-6.7</td><td>0.7</td><td>-10.1</td><td>5.3E-24</td><td>1.2E-22</td><td>MB231/MB468</td><td>MITO</td></tr>
<tr><td>>hg38_21_+_8433172_8446622_RNA45SN1_45S_50ntFlanks@8472.8497.26|CGGCCCGGCGGATCTTTCCCGCCCCC</td><td>>hg38_21_+_8433172_8446622_RNA45SN1_45S_50ntFlanks</td><td>8472</td><td>8497</td><td>26</td><td>CGGCCCGGCGGATCTTTCCCGCCCCC</td><td>1653.8</td><td>-14.5</td><td>1.4</td><td>-10.1</td><td>5.3E-24</td><td>1.2E-22</td><td>MB231/MB468</td><td>MITO</td></tr>
<tr><td>>MI0000285|hsa-mir-205&WithFlank&1|+|209432127|209432248@40.60.21&[MIMAT0000266&hsa-miR-205-5p&offsets|0|-1;m-351&1|+|209432166|209432187&offsets|0|-1]|TCCTTCATTCCACCGGAGTCT</td><td>>MI0000285|hsa-mir-205&WithFlank&1|+|209432127|209432248</td><td>40</td><td>60</td><td>21</td><td>TCCTTCATTCCACCGGAGTCT</td><td>2721.0</td><td>15.0</td><td>1.5</td><td>10.3</td><td>4.4E-25</td><td>1.2E-22</td><td>BT20/MB231</td><td>MITO</td></tr>
<tr><td>>hg38_21_+_8433172_8446622_RNA45SN1_45S_50ntFlanks@10922.10939.18|CTGGACGAGGCGCCGCCG</td><td>>hg38_21_+_8433172_8446622_RNA45SN1_45S_50ntFlanks</td><td>10922</td><td>10939</td><td>18</td><td>CTGGACGAGGCGCCGCCG</td><td>2282.1</td><td>-15.0</td><td>1.5</td><td>-10.1</td><td>3.5E-24</td><td>1.2E-22</td><td>MB231/MB468</td><td>MP</td></tr>
<tr><td>>hg38_21_+_8433172_8446622_RNA45SN1_45S_50ntFlanks@8356.8379.24|GAAAAGAACTTTGAAGAGAGAGTT</td><td>>hg38_21_+_8433172_8446622_RNA45SN1_45S_50ntFlanks</td><td>8356</td><td>8379</td><td>24</td><td>GAAAAGAACTTTGAAGAGAGAGTT</td><td>20344.7</td><td>-11.4</td><td>1.1</td><td>-10.1</td><td>3.5E-24</td><td>1.2E-22</td><td>MB231/MB468</td><td>MP</td></tr>
<tr><td>>hg38_21_+_8433172_8446622_RNA45SN1_45S_50ntFlanks@12717.12742.26|CCTCGGTTGGCCTCGGATAGCCGGTC</td><td>>hg38_21_+_8433172_8446622_RNA45SN1_45S_50ntFlanks</td><td>12717</td><td>12742</td><td>26</td><td>CCTCGGTTGGCCTCGGATAGCCGGTC</td><td>1567.4</td><td>-14.5</td><td>1.4</td><td>-10.1</td><td>3.5E-24</td><td>1.2E-22</td><td>MB231/MB468</td><td>MP</td></tr>
<tr><td>>hg38_21_+_8433172_8446622_RNA45SN1_45S_50ntFlanks@12933.12953.21|GCTAAACCATTCGTAGACGAC</td><td>>hg38_21_+_8433172_8446622_RNA45SN1_45S_50ntFlanks</td><td>12933</td><td>12953</td><td>21</td><td>GCTAAACCATTCGTAGACGAC</td><td>681.3</td><td>-13.8</td><td>1.4</td><td>-10.0</td><td>2.5E-23</td><td>1.3E-22</td><td>BT20/MB468</td><td>NUC</td></tr>
<tr><td>>hg38_21_+_8433172_8446622_RNA45SN1_45S_50ntFlanks@10458.10478.21|ATGGCCTCCGTTGCCCTCGGC</td><td>>hg38_21_+_8433172_8446622_RNA45SN1_45S_50ntFlanks</td><td>10458</td><td>10478</td><td>21</td><td>ATGGCCTCCGTTGCCCTCGGC</td><td>1181.7</td><td>-4.3</td><td>0.4</td><td>-10.1</td><td>8.9E-24</td><td>1.3E-22</td><td>BT20/MB468</td><td>MITO</td></tr>
<tr><td>>hg38_21_+_8433172_8446622_RNA45SN1_45S_50ntFlanks@11583.11605.23|CTGGTGCGGACCAGGGGAATCCG</td><td>>hg38_21_+_8433172_8446622_RNA45SN1_45S_50ntFlanks</td><td>11583</td><td>11605</td><td>23</td><td>CTGGTGCGGACCAGGGGAATCCG</td><td>1258.9</td><td>-14.2</td><td>1.4</td><td>-10.1</td><td>3.6E-24</td><td>1.3E-22</td><td>MB231/MB468</td><td>MP</td></tr>
<tr><td>>hg38_21_+_8433172_8446622_RNA45SN1_45S_50ntFlanks@9738.9765.28|GAAACGATCTCAACCTATTCTCAAACTT</td><td>>hg38_21_+_8433172_8446622_RNA45SN1_45S_50ntFlanks</td><td>9738</td><td>9765</td><td>28</td><td>GAAACGATCTCAACCTATTCTCAAACTT</td><td>18338.3</td><td>-7.7</td><td>0.8</td><td>-10.1</td><td>9.0E-24</td><td>1.3E-22</td><td>BT20/MB468</td><td>MITO</td></tr>
<tr><td>>MI0000466|hsa-mir-9-1&WithFlank&1|-|156420335|156420435@22.43.22;MI0000468|hsa-mir-9-3&WithFlank&15|+|89368011|89368112@22.43.22;MI0000467|hsa-mir-9-2&WithFlank&5|-|88666847|88666945@22.43.22&[MIMAT0000441&hsa-miR-9-5p&offsets|0|-1;m-104&1|-|156420392|156420414&offsets|0|-1];[MIMAT0000441_1&hsa-miR-9-5p&offsets|0|-1;m-105&15|+|89368032|89368054&offsets|0|-1];[MIMAT0000441_2&hsa-miR-9-5p&offsets|0|-1;m-104&5|-|88666902|88666924&offsets|0|-1]|TCTTTGGTTATCTAGCTGTATG</td><td>>MI0000466|hsa-mir-9-1&WithFlank&1|-|156420335|156420435</td><td>22</td><td>43</td><td>22</td><td>TCTTTGGTTATCTAGCTGTATG</td><td>311.7</td><td>5.2</td><td>0.5</td><td>10.0</td><td>2.5E-23</td><td>1.3E-22</td><td>BT20/MB468</td><td>NUC</td></tr>
<tr><td>>hg38_21_+_8433172_8446622_RNA45SN1_45S_50ntFlanks@3787.3812.26|AAACTGCGAATGGCTCATTAAATCAG</td><td>>hg38_21_+_8433172_8446622_RNA45SN1_45S_50ntFlanks</td><td>3787</td><td>3812</td><td>26</td><td>AAACTGCGAATGGCTCATTAAATCAG</td><td>1128.2</td><td>-14.6</td><td>1.5</td><td>-9.9</td><td>2.5E-23</td><td>1.3E-22</td><td>BT20/MB468</td><td>NUC</td></tr>
<tr><td>>am_MI0000086|hsa-mir-28&WithFlank&3|+|188688775|188688872@60.78.19&[MIMAT0004502&hsa-miR-28-3p&offsets|0|-3;m-28&3|+|188688834|188688855&offsets|0|-3]|CACTAGATTGTGAGCTCCT</td><td>>am_MI0000086|hsa-mir-28&WithFlank&3|+|188688775|188688872</td><td>60</td><td>78</td><td>19</td><td>CACTAGATTGTGAGCTCCT</td><td>652.0</td><td>13.6</td><td>1.4</td><td>10.0</td><td>1.2E-23</td><td>1.3E-22</td><td>BT20/MB468</td><td>CYTO</td></tr>
<tr><td>>hg38_21_+_8433172_8446622_RNA45SN1_45S_50ntFlanks@12957.12982.26|CTTCTGGGTCGGGGTTTCGTACGTAG</td><td>>hg38_21_+_8433172_8446622_RNA45SN1_45S_50ntFlanks</td><td>12957</td><td>12982</td><td>26</td><td>CTTCTGGGTCGGGGTTTCGTACGTAG</td><td>1280.8</td><td>-14.2</td><td>1.4</td><td>-10.1</td><td>3.8E-24</td><td>1.3E-22</td><td>MB231/MB468</td><td>MP</td></tr>
<tr><td>>hg38_21_+_8433172_8446622_RNA45SN1_45S_50ntFlanks@13001.13028.28|GCGATCTATTGAAAGTCAGCCCTCGACA</td><td>>hg38_21_+_8433172_8446622_RNA45SN1_45S_50ntFlanks</td><td>13001</td><td>13028</td><td>28</td><td>GCGATCTATTGAAAGTCAGCCCTCGACA</td><td>329.1</td><td>-12.8</td><td>1.3</td><td>-9.9</td><td>2.6E-23</td><td>1.3E-22</td><td>BT20/MB468</td><td>NUC</td></tr>
<tr><td>>hg38_21_+_8433172_8446622_RNA45SN1_45S_50ntFlanks@11211.11232.22|GCGACTCTGGACGCGAGCCGGG</td><td>>hg38_21_+_8433172_8446622_RNA45SN1_45S_50ntFlanks</td><td>11211</td><td>11232</td><td>22</td><td>GCGACTCTGGACGCGAGCCGGG</td><td>1523.5</td><td>-14.4</td><td>1.4</td><td>-10.1</td><td>3.8E-24</td><td>1.3E-22</td><td>MB231/MB468</td><td>MP</td></tr>
<tr><td>>hg38_21_+_8433172_8446622_RNA45SN1_45S_50ntFlanks@12939.12956.18|CCATTCGTAGACGACCTG</td><td>>hg38_21_+_8433172_8446622_RNA45SN1_45S_50ntFlanks</td><td>12939</td><td>12956</td><td>18</td><td>CCATTCGTAGACGACCTG</td><td>1111.0</td><td>-14.0</td><td>1.4</td><td>-10.1</td><td>3.9E-24</td><td>1.3E-22</td><td>MB231/MB468</td><td>MP</td></tr>
<tr><td>>hg38_21_+_8433172_8446622_RNA45SN1_45S_50ntFlanks@7989.8014.26|AGACGTGGCGACCCGCTGAATTTAAG</td><td>>hg38_21_+_8433172_8446622_RNA45SN1_45S_50ntFlanks</td><td>7989</td><td>8014</td><td>26</td><td>AGACGTGGCGACCCGCTGAATTTAAG</td><td>1382.8</td><td>-6.9</td><td>0.7</td><td>-10.1</td><td>3.9E-24</td><td>1.3E-22</td><td>MB231/MB468</td><td>MP</td></tr>
<tr><td>>MI0006273|hsa-mir-1180&WithFlank&17|-|19344500|19344580@47.67.21(+1C)&[MIMAT0005825&hsa-miR-1180-3p&offsets|0|-1(+1C);m-214&17|-|19344513|19344534&offsets|0|-1(+1C)]|TTTCCGGCTCGCGTGGGTGTGC</td><td>>MI0006273|hsa-mir-1180&WithFlank&17|-|19344500|19344580</td><td>47</td><td>67</td><td>21(+1C)</td><td>TTTCCGGCTCGCGTGGGTGTGC</td><td>819.6</td><td>10.6</td><td>1.1</td><td>10.0</td><td>1.3E-23</td><td>1.4E-22</td><td>BT20/MB468</td><td>CYTO</td></tr>
<tr><td>>MI0000456|hsa-mir-140&WithFlank&16|+|69933075|69933186@69.90.22&[MIMAT0004597&hsa-miR-140-3p&offsets|+1|+2;m-45&16|+|69933143|69933164&offsets|0|0]|ACCACAGGGTAGAACCACGGAC</td><td>>MI0000456|hsa-mir-140&WithFlank&16|+|69933075|69933186</td><td>69</td><td>90</td><td>22</td><td>ACCACAGGGTAGAACCACGGAC</td><td>569.0</td><td>13.6</td><td>1.4</td><td>10.0</td><td>9.7E-24</td><td>1.4E-22</td><td>BT20/MB468</td><td>MITO</td></tr>
<tr><td>>hg38_21_+_8433172_8446622_RNA45SN1_45S_50ntFlanks@6653.6680.28|ACTCTTAGCGGTGGATCACTCGGCTCGT</td><td>>hg38_21_+_8433172_8446622_RNA45SN1_45S_50ntFlanks</td><td>6653</td><td>6680</td><td>28</td><td>ACTCTTAGCGGTGGATCACTCGGCTCGT</td><td>419.3</td><td>-3.0</td><td>0.3</td><td>-9.9</td><td>2.7E-23</td><td>1.4E-22</td><td>BT20/MB468</td><td>NUC</td></tr>
<tr><td>>MI0000063|hsa-let-7b&WithFlank&22|+|46113680|46113774@12.32.21(+1G)&[MIMAT0000063&hsa-let-7b-5p&offsets|0|-1(+1G);m-14&22|+|46113691|46113712&offsets|0|-1(+1G)]|TGAGGTAGTAGGTTGTGTGGTG</td><td>>MI0000063|hsa-let-7b&WithFlank&22|+|46113680|46113774</td><td>12</td><td>32</td><td>21(+1G)</td><td>TGAGGTAGTAGGTTGTGTGGTG</td><td>2541.8</td><td>7.9</td><td>0.8</td><td>10.0</td><td>1.3E-23</td><td>1.4E-22</td><td>BT20/MB468</td><td>CYTO</td></tr>
<tr><td>>hg38_21_+_8433172_8446622_RNA45SN1_45S_50ntFlanks@8115.8138.24|CGCGGGACATGTGGCGTACGGAAG</td><td>>hg38_21_+_8433172_8446622_RNA45SN1_45S_50ntFlanks</td><td>8115</td><td>8138</td><td>24</td><td>CGCGGGACATGTGGCGTACGGAAG</td><td>1881.4</td><td>-14.7</td><td>1.5</td><td>-10.1</td><td>6.1E-24</td><td>1.4E-22</td><td>MB231/MB468</td><td>MITO</td></tr>
<tr><td>>hg38_21_+_8433172_8446622_RNA45SN1_45S_50ntFlanks@12008.12041.34|CCGCCGGTGAAATACCACTACTCTGATCGTTTTT</td><td>>hg38_21_+_8433172_8446622_RNA45SN1_45S_50ntFlanks</td><td>12008</td><td>12041</td><td>34</td><td>CCGCCGGTGAAATACCACTACTCTGATCGTTTTT</td><td>1285.4</td><td>-14.8</td><td>1.5</td><td>-9.9</td><td>2.7E-23</td><td>1.4E-22</td><td>BT20/MB468</td><td>NUC</td></tr>
<tr><td>>MI0000760|hsa-mir-361&WithFlank&X|-|85903630|85903713@12.33.22&[MIMAT0000703&hsa-miR-361-5p&offsets|0|0;m-89&X|-|85903681|85903702&offsets|0|0]|TTATCAGAATCTCCAGGGGTAC</td><td>>MI0000760|hsa-mir-361&WithFlank&X|-|85903630|85903713</td><td>12</td><td>33</td><td>22</td><td>TTATCAGAATCTCCAGGGGTAC</td><td>517.3</td><td>6.8</td><td>0.7</td><td>10.0</td><td>1.3E-23</td><td>1.4E-22</td><td>BT20/MB468</td><td>CYTO</td></tr>
<tr><td>>MI0000102|hsa-mir-100&WithFlank&11|-|122152223|122152314@19.38.20(+2G)&[MIMAT0000098&hsa-miR-100-5p&offsets|0|-2(+2G);m-27&11|-|122152276|122152296&offsets|0|-1(+2G)]|AACCCGTAGATCCGAACTTGGG</td><td>>MI0000102|hsa-mir-100&WithFlank&11|-|122152223|122152314</td><td>19</td><td>38</td><td>20(+2G)</td><td>AACCCGTAGATCCGAACTTGGG</td><td>1299.5</td><td>14.2</td><td>1.4</td><td>10.1</td><td>6.7E-24</td><td>1.4E-22</td><td>MB231/MB468</td><td>NUC</td></tr>
<tr><td>>hg38_21_+_8433172_8446622_RNA45SN1_45S_50ntFlanks@11561.11591.31|GCGCCTAGCAGCCGACTTAGAACTGGTGCGG</td><td>>hg38_21_+_8433172_8446622_RNA45SN1_45S_50ntFlanks</td><td>11561</td><td>11591</td><td>31</td><td>GCGCCTAGCAGCCGACTTAGAACTGGTGCGG</td><td>2067.5</td><td>-11.0</td><td>1.1</td><td>-10.1</td><td>6.2E-24</td><td>1.4E-22</td><td>MB231/MB468</td><td>MITO</td></tr>
<tr><td>>hg38_21_+_8433172_8446622_RNA45SN1_45S_50ntFlanks@12567.12587.21|CTCAGTACGAGAGGAACCGCA</td><td>>hg38_21_+_8433172_8446622_RNA45SN1_45S_50ntFlanks</td><td>12567</td><td>12587</td><td>21</td><td>CTCAGTACGAGAGGAACCGCA</td><td>1391.1</td><td>-14.2</td><td>1.4</td><td>-10.1</td><td>6.2E-24</td><td>1.4E-22</td><td>MB231/MB468</td><td>MITO</td></tr>
<tr><td>>hg38_21_+_8433172_8446622_RNA45SN1_45S_50ntFlanks@6746.6771.26|CACTTCGAACGCACTTGCGGCCCCGG</td><td>>hg38_21_+_8433172_8446622_RNA45SN1_45S_50ntFlanks</td><td>6746</td><td>6771</td><td>26</td><td>CACTTCGAACGCACTTGCGGCCCCGG</td><td>649.4</td><td>-7.4</td><td>0.7</td><td>-10.2</td><td>2.9E-24</td><td>1.4E-22</td><td>MB231/MB468</td><td>CYTO</td></tr>
<tr><td>>hg38_21_+_8433172_8446622_RNA45SN1_45S_50ntFlanks@12456.12477.22|CCAAGCGTTGGATTGTTCACCC</td><td>>hg38_21_+_8433172_8446622_RNA45SN1_45S_50ntFlanks</td><td>12456</td><td>12477</td><td>22</td><td>CCAAGCGTTGGATTGTTCACCC</td><td>4740.1</td><td>-6.4</td><td>0.6</td><td>-10.1</td><td>6.8E-24</td><td>1.4E-22</td><td>MB231/MB468</td><td>NUC</td></tr>
<tr><td>>hg38_21_+_8433172_8446622_RNA45SN1_45S_50ntFlanks@12932.12956.25|CGCTAAACCATTCGTAGACGACCTG</td><td>>hg38_21_+_8433172_8446622_RNA45SN1_45S_50ntFlanks</td><td>12932</td><td>12956</td><td>25</td><td>CGCTAAACCATTCGTAGACGACCTG</td><td>990.5</td><td>-13.8</td><td>1.4</td><td>-10.1</td><td>4.1E-24</td><td>1.4E-22</td><td>MB231/MB468</td><td>MP</td></tr>
<tr><td>>MI0003643|hsa-mir-629&WithFlank&15|-|70079366|70079474@28.48.21(+1C)&[MIMAT0004810&hsa-miR-629-5p&offsets|0|0(+1C);m-150&15|-|70079427|70079447&offsets|0|0(+1C)]|TGGGTTTACGTTGGGAGAACTC</td><td>>MI0003643|hsa-mir-629&WithFlank&15|-|70079366|70079474</td><td>28</td><td>48</td><td>21(+1C)</td><td>TGGGTTTACGTTGGGAGAACTC</td><td>404.9</td><td>13.1</td><td>1.3</td><td>10.0</td><td>1.0E-23</td><td>1.4E-22</td><td>BT20/MB468</td><td>MITO</td></tr>
<tr><td>>hg38_21_+_8433172_8446622_RNA45SN1_45S_50ntFlanks@11601.11619.19|ATCCGACTGTTTAATTAAA</td><td>>hg38_21_+_8433172_8446622_RNA45SN1_45S_50ntFlanks</td><td>11601</td><td>11619</td><td>19</td><td>ATCCGACTGTTTAATTAAA</td><td>611.7</td><td>-13.7</td><td>1.4</td><td>-9.9</td><td>2.8E-23</td><td>1.4E-22</td><td>BT20/MB468</td><td>NUC</td></tr>
<tr><td>>MI0000063|hsa-let-7b&WithFlank&22|+|46113680|46113774@12.33.22(+1G)&[MIMAT0000063&hsa-let-7b-5p&offsets|0|0(+1G);m-14&22|+|46113691|46113712&offsets|0|0(+1G)]|TGAGGTAGTAGGTTGTGTGGTTG</td><td>>MI0000063|hsa-let-7b&WithFlank&22|+|46113680|46113774</td><td>12</td><td>33</td><td>22(+1G)</td><td>TGAGGTAGTAGGTTGTGTGGTTG</td><td>756.3</td><td>8.7</td><td>0.9</td><td>10.0</td><td>1.4E-23</td><td>1.5E-22</td><td>BT20/MB468</td><td>CYTO</td></tr>
<tr><td>>MI0003643|hsa-mir-629&WithFlank&15|-|70079366|70079474@28.48.21(+1C)&[MIMAT0004810&hsa-miR-629-5p&offsets|0|0(+1C);m-150&15|-|70079427|70079447&offsets|0|0(+1C)]|TGGGTTTACGTTGGGAGAACTC</td><td>>MI0003643|hsa-mir-629&WithFlank&15|-|70079366|70079474</td><td>28</td><td>48</td><td>21(+1C)</td><td>TGGGTTTACGTTGGGAGAACTC</td><td>675.5</td><td>13.7</td><td>1.4</td><td>10.0</td><td>1.4E-23</td><td>1.5E-22</td><td>BT20/MB468</td><td>CYTO</td></tr>
<tr><td>>MI0000750|hsa-mir-26a-2&WithFlank&12|-|57824603|57824698@20.41.22;MI0000083|hsa-mir-26a-1&WithFlank&3|+|37969398|37969486@16.37.22&[MIMAT0000082&hsa-miR-26a-5p&offsets|0|0;m-33&12|-|57824658|57824679&offsets|0|0];[MIMAT0000082_1&hsa-miR-26a-5p&offsets|0|0;m-34&3|+|37969413|37969434&offsets|0|0]|TTCAAGTAATCCAGGATAGGCT</td><td>>MI0000750|hsa-mir-26a-2&WithFlank&12|-|57824603|57824698</td><td>20</td><td>41</td><td>22</td><td>TTCAAGTAATCCAGGATAGGCT</td><td>102158.0</td><td>6.3</td><td>0.6</td><td>10.0</td><td>1.4E-23</td><td>1.5E-22</td><td>BT20/MB468</td><td>CYTO</td></tr>
</table>

| | | | | | | | | | | | | | |
|---|---|---|---|---|---|---|---|---|---|---|---|---|---|
| >hg38_21_+_8433172_8446622_RNA45SN1_45S_50ntFlanks@12008.12038.31\|CCGCCGGTGAAATACCACTACTCTGATCGTT | >hg38_21_+_8433172_8446622_RNA45SN1_45S_50ntFlanks | 12008 | 12038 | 31 | CCGCCGGTGAAATACCACTACTCTGATCGTT | 511.6 | -13.4 | 1.4 | -9.9 | 2.9E-23 | 1.5E-22 | BT20/MB468 | NUC |
| >hg38_21_+_8433172_8446622_RNA45SN1_45S_50ntFlanks@12411.12436.26\|GATCCTTCGATGTCGGCTCTTCCTAT | >hg38_21_+_8433172_8446622_RNA45SN1_45S_50ntFlanks | 12411 | 12436 | 26 | GATCCTTCGATGTCGGCTCTTCCTAT | 846.2 | -14.2 | 1.4 | -9.9 | 2.9E-23 | 1.5E-22 | BT20/MB468 | NUC |
| >hg38_21_+_8433172_8446622_RNA45SN1_45S_50ntFlanks@3785.3812.28\|TGAAACTGCGAATGGCTCATTAAATCAG | >hg38_21_+_8433172_8446622_RNA45SN1_45S_50ntFlanks | 3785 | 3812 | 28 | TGAAACTGCGAATGGCTCATTAAATCAG | 1619.2 | -15.1 | 1.5 | -9.9 | 3.0E-23 | 1.5E-22 | BT20/MB468 | NUC |
| >hg38_21_+_8433172_8446622_RNA45SN1_45S_50ntFlanks@11314.11344.31\|CCCACCCCACGTCTCGTCGCGCGCGCGTCCG | >hg38_21_+_8433172_8446622_RNA45SN1_45S_50ntFlanks | 11314 | 11344 | 31 | CCCACCCCACGTCTCGTCGCGCGCGCGTCCG | 2428.3 | -11.6 | 1.1 | -10.1 | 7.3E-24 | 1.5E-22 | MB231/MB468 | NUC |
| >am_trnaMT_ValTAC_MT_+_1602_1670@50.68.19\|ACTTAACTTGACCGCTCTG | >am_trnaMT_ValTAC_MT_+_1602_1670 | 50 | 68 | 19 | ACTTAACTTGACCGCTCTG | 52217.6 | -10.4 | 1.0 | -9.9 | 3.0E-23 | 1.5E-22 | BT20/MB468 | NUC |
| >hg38_21_+_8433172_8446622_RNA45SN1_45S_50ntFlanks@8182.8209.28\|ATCGAGGCCCAGCCCGTGGACGGTGTGA | >hg38_21_+_8433172_8446622_RNA45SN1_45S_50ntFlanks | 8182 | 8209 | 28 | ATCGAGGCCCAGCCCGTGGACGGTGTGA | 748.7 | -14.0 | 1.4 | -9.9 | 3.0E-23 | 1.5E-22 | BT20/MB468 | NUC |
| >hg38_21_+_8433172_8446622_RNA45SN1_45S_50ntFlanks@4163.4182.20\|CATCCAAGGAAGGCAGCAGG | >hg38_21_+_8433172_8446622_RNA45SN1_45S_50ntFlanks | 4163 | 4182 | 20 | CATCCAAGGAAGGCAGCAGG | 1490.4 | -13.5 | 1.3 | -10.1 | 7.5E-24 | 1.5E-22 | MB231/MB468 | NUC |
| >hg38_21_+_8433172_8446622_RNA45SN1_45S_50ntFlanks@7991.8012.22\|ACGTGGCGACCCGCTGAATTTA | >hg38_21_+_8433172_8446622_RNA45SN1_45S_50ntFlanks | 7991 | 8012 | 22 | ACGTGGCGACCCGCTGAATTTA | 2580.9 | -6.5 | 0.7 | -9.9 | 3.1E-23 | 1.6E-22 | BT20/MB468 | NUC |
| >hg38_21_+_8433172_8446622_RNA45SN1_45S_50ntFlanks@485.527.43\|GATGTGGTGACGTCGTGCTCTCCCGGGCCGGGTCCGAGCCGCG | >hg38_21_+_8433172_8446622_RNA45SN1_45S_50ntFlanks | 485 | 527 | 43 | GATGTGGTGACGTCGTGCTCTCCCGGGCCGGGTCCGAGCCGCG | 412.0 | 13.1 | 1.3 | 10.0 | 1.1E-23 | 1.6E-22 | BT20/MB468 | MITO |
| >hg38_21_+_8433172_8446622_RNA45SN1_45S_50ntFlanks@12405.12436.32\|CTTTTTGATCCTTCGATGTCGGCTCTTCCTAT | >hg38_21_+_8433172_8446622_RNA45SN1_45S_50ntFlanks | 12405 | 12436 | 32 | CTTTTTGATCCTTCGATGTCGGCTCTTCCTAT | 106180.3 | -15.9 | 1.6 | -9.9 | 3.1E-23 | 1.6E-22 | BT20/MB468 | NUC |
| >hg38_21_+_8433172_8446622_RNA45SN1_45S_50ntFlanks@11575.11593.19\|ACTTAGAACTGGTGCGGAC | >hg38_21_+_8433172_8446622_RNA45SN1_45S_50ntFlanks | 11575 | 11593 | 19 | ACTTAGAACTGGTGCGGAC | 1690.1 | -6.2 | 0.6 | -10.1 | 7.7E-24 | 1.6E-22 | MB231/MB468 | NUC |
| >hg38_21_+_8433172_8446622_RNA45SN1_45S_50ntFlanks@12991.13015.25\|CTCCCTCGCTGCGATCTATTGAAAG | >hg38_21_+_8433172_8446622_RNA45SN1_45S_50ntFlanks | 12991 | 13015 | 25 | CTCCCTCGCTGCGATCTATTGAAAG | 36830.5 | -14.8 | 1.5 | -9.9 | 3.1E-23 | 1.6E-22 | BT20/MB468 | NUC |
| >trna116_GluCTC_1_-_145399233_145399304@1.31.31,trna59_GluCTC_1_+_249168447_249168518@1.31.31,trna71_GluCTC_1_-_161439189_161439260@1.31.31,trna74_GluCTC_1_-_161431809_161431880@1.31.31,trna77_GluCTC_1_-_161424398_161424469@1.31.31,trna77_GluCTC_6_+_28949976_28950047@1.31.31,trna80_GluCTC_1_-_161417018_161417089@1.31.31,trna87_GluCTC_6_-_126101393_126101464@1.31.31\|TCCCTGGTGGTCTAGTGGTTAGGATTCGGCG | >trna116_GluCTC_1_-_145399233_145399304 | 1 | 31 | 31 | TCCCTGGTGGTCTAGTGGTTAGGATTCGGCG | 277886.7 | -9.8 | 1.0 | -9.9 | 3.2E-23 | 1.6E-22 | BT20/MB468 | NUC |
| >hg38_21_+_8433172_8446622_RNA45SN1_45S_50ntFlanks@11161.11178.18\|CACGGTCCCCCGCGAGGG | >hg38_21_+_8433172_8446622_RNA45SN1_45S_50ntFlanks | 11161 | 11178 | 18 | CACGGTCCCCCGCGAGGG | 1905.0 | -14.8 | 1.5 | -10.1 | 4.6E-24 | 1.6E-22 | MB231/MB468 | MP |
| >MI0000446\|hsa-mir-125b-1&WithFlank&11\|-\|122099751\|122099850@61.80.20(+1U)&[MIMAT0004592&hsa-miR-125b-1-3p&offsets\|0\|-2(+1U);m-485&11\|-\|122099769\|122099790&offsets\|0\|-2(+1U)]\|ACGGGTTAGGCTCTTGGGAGT | >MI0000446\|hsa-mir-125b-1&WithFlank&11\|-\|122099751\|122099850 | 61 | 80 | 20(+1U) | ACGGGTTAGGCTCTTGGGAGT | 1342.6 | 14.4 | 1.4 | 10.1 | 3.4E-24 | 1.6E-22 | MB231/MB468 | CYTO |
| >hg38_21_+_8433172_8446622_RNA45SN1_45S_50ntFlanks@11768.11806.39\|CTAATTAGTGACGCGCATGAATGGATGAACGAGATTCCC | >hg38_21_+_8433172_8446622_RNA45SN1_45S_50ntFlanks | 11768 | 11806 | 39 | CTAATTAGTGACGCGCATGAATGGATGAACGAGATTCCC | 855.8 | -14.2 | 1.4 | -9.9 | 3.2E-23 | 1.6E-22 | BT20/MB468 | NUC |
| >MI0000542\|hsa-mir-320a&WithFlank&8\|-\|22244960\|22245043@48.68.21(+2U)&[MIMAT0000510&hsa-miR-320a-3p&offsets\|0\|-1(+2U);m-60&8\|-\|22244975\|22244996&offsets\|0\|-1(+2U)]\|AAAAGCTGGGTTGAGAGGGCGTT | >MI0000542\|hsa-mir-320a&WithFlank&8\|-\|22244960\|22245043 | 48 | 68 | 21(+2U) | AAAAGCTGGGTTGAGAGGGCGTT | 18057.6 | 13.1 | 1.3 | 10.0 | 1.5E-23 | 1.6E-22 | BT20/MB468 | CYTO |
| >trnaMT_SerGCT_MT_+_12207_12265@7.38.32\|GCTCACAAGAACTGCTAACTCATGCCCCCATG | >trnaMT_SerGCT_MT_+_12207_12265 | 7 | 38 | 32 | GCTCACAAGAACTGCTAACTCATGCCCCCATG | 1016.1 | -14.4 | 1.5 | -9.9 | 3.2E-23 | 1.6E-22 | BT20/MB468 | NUC |
| >MI0000088\|hsa-mir-30a&WithFlank&6\|-\|71403545\|71403627@12.34.23(+1C)&[MIMAT0000087&hsa-miR-30a-5p&offsets\|0\|+1(+1C);m-15&6\|-\|71403595\|71403616&offsets\|0\|+1(+1C)]\|TGTAAACATCCTCGACTGGAAGCC | >MI0000088\|hsa-mir-30a&WithFlank&6\|-\|71403545\|71403627 | 12 | 34 | 23(+1C) | TGTAAACATCCTCGACTGGAAGCC | 4231.8 | 6.3 | 0.6 | 10.0 | 1.2E-23 | 1.6E-22 | BT20/MB468 | MITO |
| >hg38_21_+_8433172_8446622_RNA45SN1_45S_50ntFlanks@10735.10767.33\|TGTAAATCTCGCGCCGGGCCGTACCCATATCCG | >hg38_21_+_8433172_8446622_RNA45SN1_45S_50ntFlanks | 10735 | 10767 | 33 | TGTAAATCTCGCGCCGGGCCGTACCCATATCCG | 967.7 | -14.3 | 1.4 | -9.9 | 3.3E-23 | 1.6E-22 | BT20/MB468 | NUC |
| >hg38_21_+_8433172_8446622_RNA45SN1_45S_50ntFlanks@10843.10863.21\|TAACTTCGGGATAAGGATTGG | >hg38_21_+_8433172_8446622_RNA45SN1_45S_50ntFlanks | 10843 | 10863 | 21 | TAACTTCGGGATAAGGATTGG | 379.3 | -13.0 | 1.3 | -9.9 | 3.3E-23 | 1.6E-22 | BT20/MB468 | NUC |
| >MI0000102\|hsa-mir-100&WithFlank&11\|-\|122152223\|122152314@19.40.22(+1C)&[MIMAT0000098&hsa-miR-100-5p&offsets\|0\|0(+1C);m-27&11\|-\|122152276\|122152296&offsets\|0\|+1(+1C)]\|AACCCGTAGATCCGAACTTGTGC | >MI0000102\|hsa-mir-100&WithFlank&11\|-\|122152223\|122152314 | 19 | 40 | 22(+1C) | AACCCGTAGATCCGAACTTGTGC | 7990.1 | 16.2 | 1.6 | 10.1 | 7.3E-24 | 1.7E-22 | MB231/MB468 | MITO |
| >MI0003188\|hsa-mir-503&WithFlank&X\|-\|134546322\|134546404@12.31.20&[MIMAT0002874&hsa-miR-503-5p&offsets\|0\|-3;m-297&X\|-\|134546371\|134546393&offsets\|0\|-3]\|TAGCAGCGGGAACAGTTCTG | >MI0003188\|hsa-mir-503&WithFlank&X\|-\|134546322\|134546404 | 12 | 31 | 20 | TAGCAGCGGGAACAGTTCTG | 754.8 | 13.8 | 1.4 | 10.0 | 1.6E-23 | 1.7E-22 | BT20/MB468 | CYTO |
| >hg38_21_+_8433172_8446622_RNA45SN1_45S_50ntFlanks@10099.10117.19\|CGCGCGCGCGTGTGGTGTG | >hg38_21_+_8433172_8446622_RNA45SN1_45S_50ntFlanks | 10099 | 10117 | 19 | CGCGCGCGCGTGTGGTGTG | 2097.1 | -14.8 | 1.5 | -10.1 | 7.5E-24 | 1.7E-22 | MB231/MB468 | MITO |
| >hg38_21_+_8433172_8446622_RNA45SN1_45S_50ntFlanks@12712.12751.40\|CGGAGCCTCGGTTGGCCTCGGATAGCCGGTCCCCCGCCTG | >hg38_21_+_8433172_8446622_RNA45SN1_45S_50ntFlanks | 12712 | 12751 | 40 | CGGAGCCTCGGTTGGCCTCGGATAGCCGGTCCCCCGCCTG | 2358.1 | -15.0 | 1.5 | -10.1 | 7.5E-24 | 1.7E-22 | MB231/MB468 | MITO |
| >MI0000465\|hsa-mir-191&WithFlank&3\|-\|49020612\|49020715@23.44.22&[MIMAT0000440&hsa-miR-191-5p&offsets\|+1\|0;m-48&3\|-\|49020672\|49020694&offsets\|+1\|0]\|AACGGAATCCCAAAAGCAGCTG | >MI0000465\|hsa-mir-191&WithFlank&3\|-\|49020612\|49020715 | 23 | 44 | 22 | AACGGAATCCCAAAAGCAGCTG | 657.3 | 7.1 | 0.7 | 10.0 | 1.6E-23 | 1.7E-22 | BT20/MB468 | CYTO |
| >trna116_GluCTC_1_-_145399233_145399304@32.75.44,trna71_GluCTC_1_-_161439189_161439260@32.75.44,trna74_GluCTC_1_-_161431809_161431880@32.75.44,trna77_GluCTC_1_-_161424398_161424469@32.75.44,trna77_GluCTC_6_+_28949976_28950047@32.75.44,trna80_GluCTC_1_-_161417018_161417089@32.75.44,trna87_GluCTC_6_-_126101393_126101464@32.75.44\|CTCTCACCGCCGCGGCCCGGGTTCGATTCCCGGTCAGGGAACCA | >trna116_GluCTC_1_-_145399233_145399304 | 32 | 75 | 44 | CTCTCACCGCCGCGGCCCGGGTTCGATTCCCGGTCAGGGAACCA | 2878.4 | -15.9 | 1.6 | -9.9 | 3.4E-23 | 1.7E-22 | BT20/MB468 | NUC |
| >MI0001733\|hsa-mir-452&WithFlank&X\|-\|151959622\|151959718@20.43.24&[MIMAT0001635&hsa-miR-452-5p&offsets\|0\|+2;m-188&X\|-\|151959677\|151959699&offsets\|0\|+1]\|AACTGTTTGCAGAGGAAACTGAGA | >MI0001733\|hsa-mir-452&WithFlank&X\|-\|151959622\|151959718 | 20 | 43 | 24 | AACTGTTTGCAGAGGAAACTGAGA | 301.0 | 9.1 | 0.9 | 9.9 | 3.4E-23 | 1.7E-22 | BT20/MB468 | NUC |
| >MI0000102\|hsa-mir-100&WithFlank&11\|-\|122152223\|122152314@19.39.21(+1C)&[MIMAT0000098&hsa-miR-100-5p&offsets\|0\|-1(+1C);m-27&11\|-\|122152276\|122152296&offsets\|0\|0(+1C)]\|AACCCGTAGATCCGAACTTGTC | >MI0000102\|hsa-mir-100&WithFlank&11\|-\|122152223\|122152314 | 19 | 39 | 21(+1C) | AACCCGTAGATCCGAACTTGTC | 2100.1 | 15.0 | 1.5 | 10.1 | 3.6E-24 | 1.7E-22 | MB231/MB468 | CYTO |
| >hg38_21_+_8433172_8446622_RNA45SN1_45S_50ntFlanks@10513.10534.22\|CCGGAGTGGCGGAGATGGGCGC | >hg38_21_+_8433172_8446622_RNA45SN1_45S_50ntFlanks | 10513 | 10534 | 22 | CCGGAGTGGCGGAGATGGGCGC | 730.0 | -13.9 | 1.4 | -9.9 | 3.4E-23 | 1.7E-22 | BT20/MB468 | NUC |
| >hg38_21_+_8433172_8446622_RNA45SN1_45S_50ntFlanks@12405.12435.31\|CTTTTTGATCCTTCGATGTCGGCTCTTCCTA | >hg38_21_+_8433172_8446622_RNA45SN1_45S_50ntFlanks | 12405 | 12435 | 31 | CTTTTTGATCCTTCGATGTCGGCTCTTCCTA | 44361.7 | -15.2 | 1.5 | -9.9 | 3.5E-23 | 1.7E-22 | BT20/MB468 | NUC |
| >hg38_21_+_8433172_8446622_RNA45SN1_45S_50ntFlanks@12436.12473.38\|TCATTGTGAAGCAGAATTCACCAAGCGTTGGATTGTTC | >hg38_21_+_8433172_8446622_RNA45SN1_45S_50ntFlanks | 12436 | 12473 | 38 | TCATTGTGAAGCAGAATTCACCAAGCGTTGGATTGTTC | 974.6 | -14.4 | 1.4 | -9.9 | 3.5E-23 | 1.7E-22 | BT20/MB468 | NUC |
| >hg38_21_+_8433172_8446622_RNA45SN1_45S_50ntFlanks@12624.12654.31\|CCAATGGGGCGAAGCTACCATCTGTGGGATT | >hg38_21_+_8433172_8446622_RNA45SN1_45S_50ntFlanks | 12624 | 12654 | 31 | CCAATGGGGCGAAGCTACCATCTGTGGGATT | 10009.3 | -12.6 | 1.2 | -10.1 | 5.1E-24 | 1.7E-22 | MB231/MB468 | MP |
| >hg38_21_+_8433172_8446622_RNA45SN1_45S_50ntFlanks@3984.4001.18\|GCGGCTTTGGTGACTCTA | >hg38_21_+_8433172_8446622_RNA45SN1_45S_50ntFlanks | 3984 | 4001 | 18 | GCGGCTTTGGTGACTCTA | 9144.0 | -11.3 | 1.1 | -9.9 | 3.5E-23 | 1.7E-22 | BT20/MB468 | NUC |
| >hg38_21_+_8433172_8446622_RNA45SN1_45S_50ntFlanks@4353.4391.39\|TATTAAAGTTGCTGCAGTTAAAAAGCTCGTAGTTGGATC | >hg38_21_+_8433172_8446622_RNA45SN1_45S_50ntFlanks | 4353 | 4391 | 39 | TATTAAAGTTGCTGCAGTTAAAAAGCTCGTAGTTGGATC | 452.1 | -13.2 | 1.3 | -9.9 | 3.5E-23 | 1.8E-22 | BT20/MB468 | NUC |
| >hg38_21_+_8433172_8446622_RNA45SN1_45S_50ntFlanks@9535.9563.29\|GGAAACTCTGGTGGAGGTCCGTAGCGGTC | >hg38_21_+_8433172_8446622_RNA45SN1_45S_50ntFlanks | 9535 | 9563 | 29 | GGAAACTCTGGTGGAGGTCCGTAGCGGTC | 419.7 | -13.1 | 1.3 | -9.9 | 3.5E-23 | 1.8E-22 | BT20/MB468 | NUC |
| >hg38_21_+_8433172_8446622_RNA45SN1_45S_50ntFlanks@11874.11893.20\|AGAAGACCCTGTTGAGCTTG | >hg38_21_+_8433172_8446622_RNA45SN1_45S_50ntFlanks | 11874 | 11893 | 20 | AGAAGACCCTGTTGAGCTTG | 8334.6 | -12.5 | 1.2 | -10.1 | 5.1E-24 | 1.8E-22 | MB231/MB468 | MP |
| >hg38_21_+_8433172_8446622_RNA45SN1_45S_50ntFlanks@7989.8006.18\|AGACGTGGCGACCCGCTG | >hg38_21_+_8433172_8446622_RNA45SN1_45S_50ntFlanks | 7989 | 8006 | 18 | AGACGTGGCGACCCGCTG | 24837.0 | -11.5 | 1.1 | -10.1 | 5.1E-24 | 1.8E-22 | MB231/MB468 | MP |
| >hg38_21_+_8433172_8446622_RNA45SN1_45S_50ntFlanks@12933.12955.23\|GCTAAACCATTCGTAGACGACCT | >hg38_21_+_8433172_8446622_RNA45SN1_45S_50ntFlanks | 12933 | 12955 | 23 | GCTAAACCATTCGTAGACGACCT | 380.8 | -13.0 | 1.3 | -9.9 | 3.5E-23 | 1.8E-22 | BT20/MB468 | NUC |
| >hg38_21_+_8433172_8446622_RNA45SN1_45S_50ntFlanks@4739.4758.20\|AAGTCGGAGGTTCGAAGACG | >hg38_21_+_8433172_8446622_RNA45SN1_45S_50ntFlanks | 4739 | 4758 | 20 | AAGTCGGAGGTTCGAAGACG | 318.2 | -12.7 | 1.3 | -9.9 | 3.5E-23 | 1.8E-22 | BT20/MB468 | NUC |
| >MI0000481\|hsa-mir-184&WithFlank&15\|+\|79209782\|79209877@59.80.22&[MIMAT0000454&hsa-miR-184&offsets\|0\|0;m-460&15\|+\|79209840\|79209861&offsets\|0\|0]\|TGGACGGAGAACTGATAAGGGT | >MI0000481\|hsa-mir-184&WithFlank&15\|+\|79209782\|79209877 | 59 | 80 | 22 | TGGACGGAGAACTGATAAGGGT | 464.4 | 13.1 | 1.3 | 10.0 | 1.7E-23 | 1.8E-22 | BT20/MB468 | CYTO |
| >hg38_21_+_8433172_8446622_RNA45SN1_45S_50ntFlanks@11701.11728.28\|TCAATGAAGCGCGGGTAAACGGCGGGAG | >hg38_21_+_8433172_8446622_RNA45SN1_45S_50ntFlanks | 11701 | 11728 | 28 | TCAATGAAGCGCGGGTAAACGGCGGGAG | 370.2 | -13.0 | 1.3 | -9.9 | 3.5E-23 | 1.8E-22 | BT20/MB468 | NUC |
| >hg38_21_+_8433172_8446622_RNA45SN1_45S_50ntFlanks@10680.10706.27\|GTTCCGGCGGCGTCCGGTGAGCTCTCG | >hg38_21_+_8433172_8446622_RNA45SN1_45S_50ntFlanks | 10680 | 10706 | 27 | GTTCCGGCGGCGTCCGGTGAGCTCTCG | 10446.7 | -12.8 | 1.3 | -10.1 | 5.2E-24 | 1.8E-22 | MB231/MB468 | MP |
| >hg38_21_+_8433172_8446622_RNA45SN1_45S_50ntFlanks@11641.11670.30\|CGGCGGGTGTTGACGCGATGTGATTTCTGC | >hg38_21_+_8433172_8446622_RNA45SN1_45S_50ntFlanks | 11641 | 11670 | 30 | CGGCGGGTGTTGACGCGATGTGATTTCTGC | 19706.1 | -14.3 | 1.4 | -10.1 | 8.0E-24 | 1.8E-22 | MB231/MB468 | MITO |
| >hg38_21_+_8433172_8446622_RNA45SN1_45S_50ntFlanks@8079.8104.26\|AAGAGCCCAGCGCCGAATCCCCGCCC | >hg38_21_+_8433172_8446622_RNA45SN1_45S_50ntFlanks | 8079 | 8104 | 26 | AAGAGCCCAGCGCCGAATCCCCGCCC | 2025.2 | -14.8 | 1.5 | -10.1 | 8.0E-24 | 1.8E-22 | MB231/MB468 | MITO |
| >hg38_21_+_8433172_8446622_RNA45SN1_45S_50ntFlanks@8472.8490.19\|CGGCCCGGCGGATCTTTCC | >hg38_21_+_8433172_8446622_RNA45SN1_45S_50ntFlanks | 8472 | 8490 | 19 | CGGCCCGGCGGATCTTTCC | 1189.1 | -14.6 | 1.5 | -9.9 | 3.6E-23 | 1.8E-22 | BT20/MB468 | NUC |

<table>
<tr><th></th><th></th><th></th><th></th><th></th><th></th><th></th><th></th><th></th><th></th><th></th><th></th><th></th><th></th></tr>
<tr><td>>MI0000285|hsa-mir-205&WithFlank&1|+|209432127|209432248@40.61.22(+3A)&[MIMAT0000266&hsa-miR-205-5p&offsets|0|0(+3A);m-351&1|+|209432166|209432187&offsets|0|0(+3A)]|TCCTTCATTCCACCGGAGTCTGAAA</td><td>>MI0000285|hsa-mir-205&WithFlank&1|+|209432127|209432248</td><td>40</td><td>61</td><td>22(+3A)</td><td>TCCTTCATTCCACCGGAGTCTGAAA</td><td>874.8</td><td>7.4</td><td>0.7</td><td>10.0</td><td>1.7E-23</td><td>1.8E-22</td><td>BT20/MB468</td><td>CYTO</td></tr>
<tr><td>>MI0000542|hsa-mir-320a&WithFlank&8|-|22244960|22245043@50.70.21&[MIMAT0000510&hsa-miR-320a-3p&offsets|+2|+1;m-60&8|-|22244975|22244996&offsets|+2|+1]|AAGCTGGGTTGAGAGGGCGAA</td><td>>MI0000542|hsa-mir-320a&WithFlank&8|-|22244960|22245043</td><td>50</td><td>70</td><td>21</td><td>AAGCTGGGTTGAGAGGGCGAA</td><td>416.7</td><td>13.0</td><td>1.3</td><td>10.0</td><td>1.7E-23</td><td>1.8E-22</td><td>BT20/MB468</td><td>CYTO</td></tr>
<tr><td>>hg38_21_+_8433172_8446622_RNA45SN1_45S_50ntFlanks@6732.6770.39|ACATTGATCATCGACACTTCGAACGCACTTGCGGCCCCG</td><td>>hg38_21_+_8433172_8446622_RNA45SN1_45S_50ntFlanks</td><td>6732</td><td>6770</td><td>39</td><td>ACATTGATCATCGACACTTCGAACGCACTTGCGGCCCCG</td><td>3130.4</td><td>-7.9</td><td>0.8</td><td>-9.9</td><td>3.6E-23</td><td>1.8E-22</td><td>BT20/MB468</td><td>NUC</td></tr>
<tr><td>>hg38_21_+_8433172_8446622_RNA45SN1_45S_50ntFlanks@13002.13039.38|CGATCTATTGAAAGTCAGCCCTCGACACAAGGGTTTGT</td><td>>hg38_21_+_8433172_8446622_RNA45SN1_45S_50ntFlanks</td><td>13002</td><td>13039</td><td>38</td><td>CGATCTATTGAAAGTCAGCCCTCGACACAAGGGTTTGT</td><td>3051.7</td><td>-16.0</td><td>1.6</td><td>-9.9</td><td>3.7E-23</td><td>1.8E-22</td><td>BT20/MB468</td><td>NUC</td></tr>
<tr><td>>hg38_21_+_8433172_8446622_RNA45SN1_45S_50ntFlanks@6772.6793.22|GTTCCTCCCGGGGCTACGCCTG</td><td>>hg38_21_+_8433172_8446622_RNA45SN1_45S_50ntFlanks</td><td>6772</td><td>6793</td><td>22</td><td>GTTCCTCCCGGGGCTACGCCTG</td><td>582.1</td><td>-13.6</td><td>1.4</td><td>-9.9</td><td>3.7E-23</td><td>1.8E-22</td><td>BT20/MB468</td><td>NUC</td></tr>
<tr><td>>hg38_21_+_8433172_8446622_RNA45SN1_45S_50ntFlanks@11874.11907.34|AGAAGACCCTGTTGAGCTTGACTCTAGTCTGGCA</td><td>>hg38_21_+_8433172_8446622_RNA45SN1_45S_50ntFlanks</td><td>11874</td><td>11907</td><td>34</td><td>AGAAGACCCTGTTGAGCTTGACTCTAGTCTGGCA</td><td>327.8</td><td>-12.8</td><td>1.3</td><td>-9.9</td><td>3.7E-23</td><td>1.8E-22</td><td>BT20/MB468</td><td>NUC</td></tr>
<tr><td>>hg38_21_+_8433172_8446622_RNA45SN1_45S_50ntFlanks@11641.11670.30|CGGCGGGTGTTGACGCGATGTGATTTCTGC</td><td>>hg38_21_+_8433172_8446622_RNA45SN1_45S_50ntFlanks</td><td>11641</td><td>11670</td><td>30</td><td>CGGCGGGTGTTGACGCGATGTGATTTCTGC</td><td>10565.2</td><td>-17.2</td><td>1.7</td><td>-10.1</td><td>5.3E-24</td><td>1.8E-22</td><td>MB231/MB468</td><td>MP</td></tr>
<tr><td>>hg38_21_+_8433172_8446622_RNA45SN1_45S_50ntFlanks@11564.11591.28|CCTAGCAGCCGACTTAGAACTGGTGCGG</td><td>>hg38_21_+_8433172_8446622_RNA45SN1_45S_50ntFlanks</td><td>11564</td><td>11591</td><td>28</td><td>CCTAGCAGCCGACTTAGAACTGGTGCGG</td><td>3414.0</td><td>-9.4</td><td>0.9</td><td>-10.1</td><td>8.2E-24</td><td>1.8E-22</td><td>MB231/MB468</td><td>MITO</td></tr>
<tr><td>>hg38_21_+_8433172_8446622_RNA45SN1_45S_50ntFlanks@8107.8138.32|CGGCGGGGCGCGGGACATGTGGCGTACGGAAG</td><td>>hg38_21_+_8433172_8446622_RNA45SN1_45S_50ntFlanks</td><td>8107</td><td>8138</td><td>32</td><td>CGGCGGGGCGCGGGACATGTGGCGTACGGAAG</td><td>1441.7</td><td>-14.4</td><td>1.4</td><td>-10.1</td><td>5.4E-24</td><td>1.9E-22</td><td>MB231/MB468</td><td>MP</td></tr>
<tr><td>>hg38_21_+_8433172_8446622_RNA45SN1_45S_50ntFlanks@8472.8492.21|CGGCCCGGCGGATCTTTCCCG</td><td>>hg38_21_+_8433172_8446622_RNA45SN1_45S_50ntFlanks</td><td>8472</td><td>8492</td><td>21</td><td>CGGCCCGGCGGATCTTTCCCG</td><td>3307.3</td><td>-15.5</td><td>1.5</td><td>-10.1</td><td>8.5E-24</td><td>1.9E-22</td><td>MB231/MB468</td><td>MITO</td></tr>
<tr><td>>hg38_21_+_8433172_8446622_RNA45SN1_45S_50ntFlanks@11874.11905.32|AGAAGACCCTGTTGAGCTTGACTCTAGTCTGG</td><td>>hg38_21_+_8433172_8446622_RNA45SN1_45S_50ntFlanks</td><td>11874</td><td>11905</td><td>32</td><td>AGAAGACCCTGTTGAGCTTGACTCTAGTCTGG</td><td>1623.2</td><td>-14.5</td><td>1.4</td><td>-10.1</td><td>5.5E-24</td><td>1.9E-22</td><td>MB231/MB468</td><td>MP</td></tr>
<tr><td>>MI0000279|hsa-mir-196a-2&WithFlank&12|+|53991732|53991853@31.53.23&[MIMAT0000226&hsa-miR-196a-5p&offsets|0|+1;m-261&12|+|53991762|53991783&offsets|0|+1]|TAGGTAGTTTCATGTTGTTGGGA</td><td>>MI0000279|hsa-mir-196a-2&WithFlank&12|+|53991732|53991853</td><td>31</td><td>53</td><td>23</td><td>TAGGTAGTTTCATGTTGTTGGGA</td><td>1433.9</td><td>14.3</td><td>1.4</td><td>10.3</td><td>9.4E-25</td><td>1.9E-22</td><td>BT20/MB231</td><td>NUC</td></tr>
<tr><td>>hg38_21_+_8433172_8446622_RNA45SN1_45S_50ntFlanks@10806.10825.20|AACAATGTAGGTAAGGGAAG</td><td>>hg38_21_+_8433172_8446622_RNA45SN1_45S_50ntFlanks</td><td>10806</td><td>10825</td><td>20</td><td>AACAATGTAGGTAAGGGAAG</td><td>1230.7</td><td>-14.7</td><td>1.5</td><td>-9.9</td><td>3.8E-23</td><td>1.9E-22</td><td>BT20/MB468</td><td>NUC</td></tr>
<tr><td>>hg38_21_+_8433172_8446622_RNA45SN1_45S_50ntFlanks@485.514.30|GATGTGGTGACGTCGTGCTCTCCCGGGCCG</td><td>>hg38_21_+_8433172_8446622_RNA45SN1_45S_50ntFlanks</td><td>485</td><td>514</td><td>30</td><td>GATGTGGTGACGTCGTGCTCTCCCGGGCCG</td><td>459.7</td><td>13.1</td><td>1.3</td><td>10.0</td><td>1.8E-23</td><td>1.9E-22</td><td>BT20/MB468</td><td>CYTO</td></tr>
<tr><td>>hg38_21_+_8433172_8446622_RNA45SN1_45S_50ntFlanks@10350.10368.19|ACGAGAACTTTGAAGGCCG</td><td>>hg38_21_+_8433172_8446622_RNA45SN1_45S_50ntFlanks</td><td>10350</td><td>10368</td><td>19</td><td>ACGAGAACTTTGAAGGCCG</td><td>5352.4</td><td>-6.1</td><td>0.6</td><td>-10.1</td><td>4.0E-24</td><td>1.9E-22</td><td>MB231/MB468</td><td>CYTO</td></tr>
<tr><td>>trna139_ValAAC_6_-_27618707_27618779@36.56.21|CACGCGAAAGGTCCCTGGATC</td><td>>trna139_ValAAC_6_-_27618707_27618779</td><td>36</td><td>56</td><td>21</td><td>CACGCGAAAGGTCCCTGGATC</td><td>674.1</td><td>13.7</td><td>1.4</td><td>10.0</td><td>1.8E-23</td><td>1.9E-22</td><td>BT20/MB468</td><td>CYTO</td></tr>
<tr><td>>MI0000449|hsa-mir-132&WithFlank&17|-|2049902|2050014@65.86.22&[MIMAT0000426&hsa-miR-132-3p&offsets|0|0;m-138&17|-|2049929|2049950&offsets|0|0]|TAACAGTCTACAGCCATGGTCG</td><td>>MI0000449|hsa-mir-132&WithFlank&17|-|2049902|2050014</td><td>65</td><td>86</td><td>22</td><td>TAACAGTCTACAGCCATGGTCG</td><td>1138.0</td><td>10.8</td><td>1.1</td><td>10.0</td><td>1.8E-23</td><td>1.9E-22</td><td>BT20/MB468</td><td>CYTO</td></tr>
<tr><td>>hg38_21_+_8433172_8446622_RNA45SN1_45S_50ntFlanks@12636.12654.19|AGCTACCATCTGTGGGATT</td><td>>hg38_21_+_8433172_8446622_RNA45SN1_45S_50ntFlanks</td><td>12636</td><td>12654</td><td>19</td><td>AGCTACCATCTGTGGGATT</td><td>4566.3</td><td>-7.4</td><td>0.7</td><td>-10.6</td><td>4.2E-26</td><td>1.9E-22</td><td>BT20/MB468</td><td>MP</td></tr>
<tr><td>>hg38_21_+_8433172_8446622_RNA45SN1_45S_50ntFlanks@6773.6799.27|TTCCTCCCGGGGCTACGCCTGTCTGAG</td><td>>hg38_21_+_8433172_8446622_RNA45SN1_45S_50ntFlanks</td><td>6773</td><td>6799</td><td>27</td><td>TTCCTCCCGGGGCTACGCCTGTCTGAG</td><td>1554.8</td><td>-14.4</td><td>1.4</td><td>-10.1</td><td>8.7E-24</td><td>1.9E-22</td><td>MB231/MB468</td><td>MITO</td></tr>
<tr><td>>hg38_21_+_8433172_8446622_RNA45SN1_45S_50ntFlanks@7988.8007.20|CAGACGTGGCGACCCGCTGA</td><td>>hg38_21_+_8433172_8446622_RNA45SN1_45S_50ntFlanks</td><td>7988</td><td>8007</td><td>20</td><td>CAGACGTGGCGACCCGCTGA</td><td>3566.1</td><td>-7.6</td><td>0.8</td><td>-10.1</td><td>8.7E-24</td><td>1.9E-22</td><td>MB231/MB468</td><td>MITO</td></tr>
<tr><td>>hg38_21_+_8433172_8446622_RNA45SN1_45S_50ntFlanks@5485.5518.34|AGCGCTGAGAAGACGGTCGAACTTGACTATCTAG</td><td>>hg38_21_+_8433172_8446622_RNA45SN1_45S_50ntFlanks</td><td>5485</td><td>5518</td><td>34</td><td>AGCGCTGAGAAGACGGTCGAACTTGACTATCTAG</td><td>1537.1</td><td>-14.4</td><td>1.4</td><td>-10.1</td><td>8.8E-24</td><td>1.9E-22</td><td>MB231/MB468</td><td>MITO</td></tr>
<tr><td>>hg38_21_+_8433172_8446622_RNA45SN1_45S_50ntFlanks@9621.9648.28|TCTAGTAGCTGGTTCCCTCCGAAGTTTC</td><td>>hg38_21_+_8433172_8446622_RNA45SN1_45S_50ntFlanks</td><td>9621</td><td>9648</td><td>28</td><td>TCTAGTAGCTGGTTCCCTCCGAAGTTTC</td><td>349.7</td><td>-12.9</td><td>1.3</td><td>-9.9</td><td>3.9E-23</td><td>1.9E-22</td><td>BT20/MB468</td><td>NUC</td></tr>
<tr><td>>MI0000737|hsa-mir-200a&WithFlank&1|+|1167857|1167958@22.43.22&[MIMAT0001620&hsa-miR-200a-5p&offsets|0|0;m-154&1|+|1167878|1167899&offsets|0|0]|CATCTTACCGGACAGTGCTGGA</td><td>>MI0000737|hsa-mir-200a&WithFlank&1|+|1167857|1167958</td><td>22</td><td>43</td><td>22</td><td>CATCTTACCGGACAGTGCTGGA</td><td>1263.6</td><td>-6.2</td><td>0.6</td><td>-10.1</td><td>5.7E-24</td><td>2.0E-22</td><td>MB231/MB468</td><td>MP</td></tr>
<tr><td>>trna116_GluCTC_1_-_145399233_145399304@1.36.36,trna59_GluCTC_1_+_249168447_249168518@1.36.36,trna71_GluCTC_1_-_161439189_161439260@1.36.36,trna74_GluCTC_1_-_161431809_161431880@1.36.36,trna77_GluCTC_1_-_161424398_161424469@1.36.36,trna77_GluCTC_6_+_28949976_28950047@1.36.36,trna80_GluCTC_1_-_161417018_161417089@1.36.36,trna87_GluCTC_6_-_126101393_126101464@1.36.36|TCCCTGGTGGTCTAGTGGTTAGGATTCGGCGCTCTC</td><td>>trna116_GluCTC_1_-_145399233_145399304</td><td>1</td><td>36</td><td>36</td><td>TCCCTGGTGGTCTAGTGGTTAGGATTCGGCGCTCTC</td><td>30672.2</td><td>10.2</td><td>1.0</td><td>10.1</td><td>4.2E-24</td><td>2.0E-22</td><td>MB231/MB468</td><td>CYTO</td></tr>
<tr><td>>hg38_21_+_8433172_8446622_RNA45SN1_45S_50ntFlanks@4069.4101.33|CTTTCGATGGTAGTCGCCGTGCCTACCATGGTG</td><td>>hg38_21_+_8433172_8446622_RNA45SN1_45S_50ntFlanks</td><td>4069</td><td>4101</td><td>33</td><td>CTTTCGATGGTAGTCGCCGTGCCTACCATGGTG</td><td>435.1</td><td>-13.2</td><td>1.3</td><td>-9.9</td><td>4.0E-23</td><td>2.0E-22</td><td>BT20/MB468</td><td>NUC</td></tr>
<tr><td>>hg38_21_+_8433172_8446622_RNA45SN1_45S_50ntFlanks@11891.11912.22|TTGACTCTAGTCTGGCACGGTG</td><td>>hg38_21_+_8433172_8446622_RNA45SN1_45S_50ntFlanks</td><td>11891</td><td>11912</td><td>22</td><td>TTGACTCTAGTCTGGCACGGTG</td><td>1021.2</td><td>-13.9</td><td>1.4</td><td>-10.1</td><td>6.0E-24</td><td>2.0E-22</td><td>MB231/MB468</td><td>MP</td></tr>
<tr><td>>MI0000465|hsa-mir-191&WithFlank&3|-|49020612|49020715@22.42.21(+1G)&[MIMAT0000440&hsa-miR-191-5p&offsets|0|-2(+1G);m-48&3|-|49020672|49020694&offsets|0|-2(+1G)]|CAACGGAATCCCAAAAGCAGCG</td><td>>MI0000465|hsa-mir-191&WithFlank&3|-|49020612|49020715</td><td>22</td><td>42</td><td>21(+1G)</td><td>CAACGGAATCCCAAAAGCAGCG</td><td>568.6</td><td>6.5</td><td>0.7</td><td>10.0</td><td>1.9E-23</td><td>2.0E-22</td><td>BT20/MB468</td><td>CYTO</td></tr>
<tr><td>>hg38_21_+_8433172_8446622_RNA45SN1_45S_50ntFlanks@8423.8469.47|CGCAGTCCGCCCGGAGGATTCAACCCGGCGGCGGGTCCGGCCGTGTC</td><td>>hg38_21_+_8433172_8446622_RNA45SN1_45S_50ntFlanks</td><td>8423</td><td>8469</td><td>47</td><td>CGCAGTCCGCCCGGAGGATTCAACCCGGCGGCGGGTCCGGCCGTGTC</td><td>3040.3</td><td>-15.4</td><td>1.5</td><td>-10.0</td><td>9.3E-24</td><td>2.0E-22</td><td>MB231/MB468</td><td>MITO</td></tr>
<tr><td>>MI0015866|hsa-mir-4326&WithFlank&20|+|63286802|63286872@17.40.24&[MIMAT0016888&hsa-miR-4326&offsets|0|+4;m-339&20|+|63286818|63286841&offsets|0|0]|TGTTCCTCTGTCTCCCAGACTCTG</td><td>>MI0015866|hsa-mir-4326&WithFlank&20|+|63286802|63286872</td><td>17</td><td>40</td><td>24</td><td>TGTTCCTCTGTCTCCCAGACTCTG</td><td>1116.5</td><td>10.4</td><td>1.0</td><td>10.0</td><td>1.9E-23</td><td>2.0E-22</td><td>BT20/MB468</td><td>CYTO</td></tr>
<tr><td>>hg38_21_+_8433172_8446622_RNA45SN1_45S_50ntFlanks@11874.11905.32|AGAAGACCCTGTTGAGCTTGACTCTAGTCTGG</td><td>>hg38_21_+_8433172_8446622_RNA45SN1_45S_50ntFlanks</td><td>11874</td><td>11905</td><td>32</td><td>AGAAGACCCTGTTGAGCTTGACTCTAGTCTGG</td><td>356.9</td><td>-12.9</td><td>1.3</td><td>-9.9</td><td>4.1E-23</td><td>2.0E-22</td><td>BT20/MB468</td><td>NUC</td></tr>
<tr><td>>hg38_21_+_8433172_8446622_RNA45SN1_45S_50ntFlanks@12415.12443.29|CTTCGATGTCGGCTCTTCCTATCATTGTG</td><td>>hg38_21_+_8433172_8446622_RNA45SN1_45S_50ntFlanks</td><td>12415</td><td>12443</td><td>29</td><td>CTTCGATGTCGGCTCTTCCTATCATTGTG</td><td>7126.7</td><td>-6.5</td><td>0.7</td><td>-9.9</td><td>4.1E-23</td><td>2.0E-22</td><td>BT20/MB468</td><td>NUC</td></tr>
<tr><td>>MI0006371|hsa-mir-1304&WithFlank&11|-|93733668|93733770@23.44.22&[MIMAT0005892&hsa-miR-1304-5p&offsets|-3|-3;m-632&11|-|93733727|93733748&offsets|0|0]|CGGTTTGAGGCTACAGTGAGAT</td><td>>MI0006371|hsa-mir-1304&WithFlank&11|-|93733668|93733770</td><td>23</td><td>44</td><td>22</td><td>CGGTTTGAGGCTACAGTGAGAT</td><td>288.7</td><td>12.4</td><td>1.2</td><td>10.0</td><td>2.0E-23</td><td>2.1E-22</td><td>BT20/MB468</td><td>CYTO</td></tr>
<tr><td>>hg38_21_+_8433172_8446622_RNA45SN1_45S_50ntFlanks@7979.8008.30|ACCTCAGATCAGACGTGGCGACCCGCTGAA</td><td>>hg38_21_+_8433172_8446622_RNA45SN1_45S_50ntFlanks</td><td>7979</td><td>8008</td><td>30</td><td>ACCTCAGATCAGACGTGGCGACCCGCTGAA</td><td>2490.0</td><td>-7.0</td><td>0.7</td><td>-10.1</td><td>6.1E-24</td><td>2.1E-22</td><td>MB231/MB468</td><td>MP</td></tr>
<tr><td>>hg38_21_+_8433172_8446622_RNA45SN1_45S_50ntFlanks@12636.12659.24|AGCTACCATCTGTGGGATTATGAC</td><td>>hg38_21_+_8433172_8446622_RNA45SN1_45S_50ntFlanks</td><td>12636</td><td>12659</td><td>24</td><td>AGCTACCATCTGTGGGATTATGAC</td><td>2808.3</td><td>-15.3</td><td>1.5</td><td>-10.1</td><td>6.1E-24</td><td>2.1E-22</td><td>MB231/MB468</td><td>MP</td></tr>
<tr><td>>MI0000478|hsa-mir-149&WithFlank&2|+|240455995|240456095@21.42.22&[MIMAT0000450&hsa-miR-149-5p&offsets|0|-1;m-195&2|+|240456015|240456037&offsets|0|-1]|TCTGGCTCCGTGTCTTCACTCC</td><td>>MI0000478|hsa-mir-149&WithFlank&2|+|240455995|240456095</td><td>21</td><td>42</td><td>22</td><td>TCTGGCTCCGTGTCTTCACTCC</td><td>3220.6</td><td>11.6</td><td>1.2</td><td>10.0</td><td>2.0E-23</td><td>2.1E-22</td><td>BT20/MB468</td><td>CYTO</td></tr>
<tr><td>>hg38_21_+_8433172_8446622_RNA45SN1_45S_50ntFlanks@485.522.38|GATGTGGTGACGTCGTGCTCTCCCGGGCCGGGTCCGAG</td><td>>hg38_21_+_8433172_8446622_RNA45SN1_45S_50ntFlanks</td><td>485</td><td>522</td><td>38</td><td>GATGTGGTGACGTCGTGCTCTCCCGGGCCGGGTCCGAG</td><td>782.6</td><td>14.1</td><td>1.4</td><td>10.0</td><td>1.5E-23</td><td>2.1E-22</td><td>BT20/MB468</td><td>MITO</td></tr>
<tr><td>>MI0000650|hsa-mir-200c&WithFlank&12|+|6963693|6963772@50.69.20&[MIMAT0000617&hsa-miR-200c-3p&offsets|0|-3;m-54&12|+|6963742|6963764&offsets|0|-3]|TAATACTGCCGGGTAATGAT</td><td>>MI0000650|hsa-mir-200c&WithFlank&12|+|6963693|6963772</td><td>50</td><td>69</td><td>20</td><td>TAATACTGCCGGGTAATGAT</td><td>1502.3</td><td>14.2</td><td>1.4</td><td>10.3</td><td>8.1E-25</td><td>2.1E-22</td><td>BT20/MB231</td><td>MITO</td></tr>
<tr><td>>trna37_ValAAC_6_+_27203288_27203360@1.29.29|GTTTCCGTAGTGTAGTGGTTATCACGTTT</td><td>>trna37_ValAAC_6_+_27203288_27203360</td><td>1</td><td>29</td><td>29</td><td>GTTTCCGTAGTGTAGTGGTTATCACGTTT</td><td>548.4</td><td>-13.5</td><td>1.4</td><td>-9.9</td><td>4.3E-23</td><td>2.1E-22</td><td>BT20/MB468</td><td>NUC</td></tr>
<tr><td>>MI0000650|hsa-mir-200c&WithFlank&12|+|6963693|6963772@50.71.22&[MIMAT0000617&hsa-miR-200c-3p&offsets|0|-1;m-54&12|+|6963742|6963764&offsets|0|-1]|TAATACTGCCGGGTAATGATGG</td><td>>MI0000650|hsa-mir-200c&WithFlank&12|+|6963693|6963772</td><td>50</td><td>71</td><td>22</td><td>TAATACTGCCGGGTAATGATGG</td><td>21683.1</td><td>9.2</td><td>0.9</td><td>10.3</td><td>5.8E-25</td><td>2.1E-22</td><td>BT20/MB231</td><td>CYTO</td></tr>
<tr><td>>trna129_MetCAT_6_-_27870271_27870342@31.51.21,trna142_MetCAT_6_-_27560600_27560671@31.51.21,trna150_MetCAT_6_-_27300764_27300835@31.51.21,trna169_MetCAT_6_-_26330529_26330600@31.51.21,trna171_MetCAT_6_-_26313352_26313423@31.51.21,trna20_MetCAT_17_-_80452597_80452668@31.51.21,trna2_MetCAT_6_+_26286754_26286825@31.51.21,trna32_MetCAT_1_+_153643726_153643797@31.51.21,trna61_MetCAT_6_+_27745664_27745735@31.51.21|CCCATAACCCAGAGGTCGATG</td><td>>trna129_MetCAT_6_-_27870271_27870342</td><td>31</td><td>51</td><td>21</td><td>CCCATAACCCAGAGGTCGATG</td><td>1967.0</td><td>-14.8</td><td>1.5</td><td>-10.1</td><td>6.3E-24</td><td>2.1E-22</td><td>MB231/MB468</td><td>MP</td></tr>
<tr><td>>hg38_21_+_8433172_8446622_RNA45SN1_45S_50ntFlanks@12157.12182.26|AGTTTGACTGGGGCGGTACACCTGTC</td><td>>hg38_21_+_8433172_8446622_RNA45SN1_45S_50ntFlanks</td><td>12157</td><td>12182</td><td>26</td><td>AGTTTGACTGGGGCGGTACACCTGTC</td><td>1865.4</td><td>-14.6</td><td>1.5</td><td>-10.0</td><td>9.7E-24</td><td>2.1E-22</td><td>MB231/MB468</td><td>MITO</td></tr>
<tr><td>>hg38_21_+_8433172_8446622_RNA45SN1_45S_50ntFlanks@9567.9597.31|ACGTGCAAATCGGTCGTCCGACCTGGGTATA</td><td>>hg38_21_+_8433172_8446622_RNA45SN1_45S_50ntFlanks</td><td>9567</td><td>9597</td><td>31</td><td>ACGTGCAAATCGGTCGTCCGACCTGGGTATA</td><td>16345.1</td><td>-10.4</td><td>1.0</td><td>-10.0</td><td>9.9E-24</td><td>2.1E-22</td><td>MB231/MB468</td><td>MITO</td></tr>
<tr><td>>hg38_21_+_8433172_8446622_RNA45SN1_45S_50ntFlanks@3738.3756.19|TCTCAAAGATTAAGCCATG</td><td>>hg38_21_+_8433172_8446622_RNA45SN1_45S_50ntFlanks</td><td>3738</td><td>3756</td><td>19</td><td>TCTCAAAGATTAAGCCATG</td><td>1633.9</td><td>-14.5</td><td>1.4</td><td>-10.1</td><td>6.4E-24</td><td>2.2E-22</td><td>MB231/MB468</td><td>MP</td></tr>
<tr><td>>hg38_21_+_8433172_8446622_RNA45SN1_45S_50ntFlanks@11208.11230.23|GCGGCGACTCTGGACGCGAGCCG</td><td>>hg38_21_+_8433172_8446622_RNA45SN1_45S_50ntFlanks</td><td>11208</td><td>11230</td><td>23</td><td>GCGGCGACTCTGGACGCGAGCCG</td><td>1482.1</td><td>-14.4</td><td>1.4</td><td>-10.1</td><td>6.5E-24</td><td>2.2E-22</td><td>MB231/MB468</td><td>MP</td></tr>
<tr><td>>hg38_21_+_8433172_8446622_RNA45SN1_45S_50ntFlanks@10346.10383.38|TCAAACGAGAACTTTGAAGGCCGAAGTGGAGAAGGGTT</td><td>>hg38_21_+_8433172_8446622_RNA45SN1_45S_50ntFlanks</td><td>10346</td><td>10383</td><td>38</td><td>TCAAACGAGAACTTTGAAGGCCGAAGTGGAGAAGGGTT</td><td>1083.8</td><td>-6.7</td><td>0.7</td><td>-9.9</td><td>4.4E-23</td><td>2.2E-22</td><td>BT20/MB468</td><td>NUC</td></tr>
<tr><td>>hg38_21_+_8433172_8446622_RNA45SN1_45S_50ntFlanks@8198.8215.18|TGGACGGTGTGAGGCCGG</td><td>>hg38_21_+_8433172_8446622_RNA45SN1_45S_50ntFlanks</td><td>8198</td><td>8215</td><td>18</td><td>TGGACGGTGTGAGGCCGG</td><td>588.8</td><td>-5.3</td><td>0.5</td><td>-10.0</td><td>1.1E-23</td><td>2.2E-22</td><td>MB231/MB468</td><td>NUC</td></tr>
</table>

| | | | | | | | | | | | | | |
|---|---|---|---|---|---|---|---|---|---|---|---|---|---|
| >hg38_21_+_8433172_8446622_RNA45SN1_45S_50ntFlanks@9570.9595.26\|TGCAAATCGGTCGTCCGACCTGGGTA | >hg38_21_+_8433172_8446622_RNA45SN1_45S_50ntFlanks | 9570 | 9595 | 26 | TGCAAATCGGTCGTCCGACCTGGGTA | 2444.2 | -5.6 | 0.6 | -10.0 | 2.1E-23 | 2.2E-22 | BT20/MB468 | CYTO |
| >MI0000266\|hsa-mir-10a&WithFlank&17\|-\|48579832\|48579953@27.49.23&[MIMAT0000253&hsa-miR-10a-5p&offsets\|-1\|-1;m-18&17\|-\|48579905\|48579926&offsets\|-1\|0]\|ATACCCTGTAGATCCGAATTTGT | >MI0000266\|hsa-mir-10a&WithFlank&17\|-\|48579832\|48579953 | 27 | 49 | 23 | ATACCCTGTAGATCCGAATTTGT | 369.3 | 12.8 | 1.3 | 10.0 | 2.2E-23 | 2.2E-22 | BT20/MB468 | CYTO |
| >MI0000269\|hsa-mir-181a-2&WithFlank&9\|+\|124692436\|124692557@83.103.21(+1A)&[MIMAT0004558&hsa-miR-181a-2-3p&offsets\|0\|-1(+1A);m-71&9\|+\|124692518\|124692539&offsets\|0\|-1(+1A)]\|ACCACTGACCGTTGACTGTACA | >MI0000269\|hsa-mir-181a-2&WithFlank&9\|+\|124692436\|124692557 | 83 | 103 | 21(+1A) | ACCACTGACCGTTGACTGTACA | 568.5 | 13.4 | 1.3 | 10.0 | 2.2E-23 | 2.2E-22 | BT20/MB468 | CYTO |
| >hg38_21_+_8433172_8446622_RNA45SN1_45S_50ntFlanks@9738.9755.18\|GAAACGATCTCAACCTAT | >hg38_21_+_8433172_8446622_RNA45SN1_45S_50ntFlanks | 9738 | 9755 | 18 | GAAACGATCTCAACCTAT | 908.6 | -13.7 | 1.4 | -10.1 | 6.7E-24 | 2.2E-22 | MB231/MB468 | MP |
| >hg38_21_+_8433172_8446622_RNA45SN1_45S_50ntFlanks@6743.6771.29\|CGACACTTCGAACGCACTTGCGGCCCCGG | >hg38_21_+_8433172_8446622_RNA45SN1_45S_50ntFlanks | 6743 | 6771 | 29 | CGACACTTCGAACGCACTTGCGGCCCCGG | 2574.8 | -15.2 | 1.5 | -10.1 | 6.8E-24 | 2.2E-22 | MB231/MB468 | MP |
| >hg38_21_+_8433172_8446622_RNA45SN1_45S_50ntFlanks@9304.9338.35\|ACCAAGGAGTCTAACACGTGCGCGAGTCGGGGGCT | >hg38_21_+_8433172_8446622_RNA45SN1_45S_50ntFlanks | 9304 | 9338 | 35 | ACCAAGGAGTCTAACACGTGCGCGAGTCGGGGGCT | 1454.1 | -14.3 | 1.4 | -10.0 | 1.0E-23 | 2.2E-22 | MB231/MB468 | MITO |
| >hg38_21_+_8433172_8446622_RNA45SN1_45S_50ntFlanks@3705.3739.35\|TACCTGGTTGATCCTGCCAGTAGCATATGCTTGTC | >hg38_21_+_8433172_8446622_RNA45SN1_45S_50ntFlanks | 3705 | 3739 | 35 | TACCTGGTTGATCCTGCCAGTAGCATATGCTTGTC | 555.7 | 2.3 | 0.2 | 9.9 | 4.6E-23 | 2.3E-22 | BT20/MB468 | NUC |
| >MI0000264\|hsa-mir-7-2&WithFlank&15\|+\|88611819\|88611940@38.60.23(+1C);MI0000265\|hsa-mir-7-3&WithFlank&19\|+\|4770664\|4770785@37.59.23(+1C);MI0000263\|hsa-mir-7-1&WithFlank&9\|-\|83969742\|83969863@30.52.23(+1C)&[MIMAT0000252&hsa-miR-7-5p&offsets\|0\|-1(+1C);m-400&15\|+\|88611856\|88611879&offsets\|0\|-1(+1C)];[MIMAT0000252_1&hsa-miR-7-5p&offsets\|0\|-1(+1C);m-402&19\|+\|4770700\|4770723&offsets\|0\|-1(+1C)];[MIMAT0000252_2&hsa-miR-7-5p&offsets\|0\|-1(+1C);m-401&9\|-\|83969811\|83969834&offsets\|0\|-1(+1C)]\|TGGAAGACTAGTGATTTTGTTGTC | >MI0000264\|hsa-mir-7-2&WithFlank&15\|+\|88611819\|88611940 | 38 | 60 | 23(+1C) | TGGAAGACTAGTGATTTTGTTGTC | 4641.0 | 5.4 | 0.5 | 10.0 | 1.1E-23 | 2.3E-22 | MB231/MB468 | NUC |
| >hg38_21_+_8433172_8446622_RNA45SN1_45S_50ntFlanks@12991.13019.29\|CTCCCTCGCTGCGATCTATTGAAAGTCAG | >hg38_21_+_8433172_8446622_RNA45SN1_45S_50ntFlanks | 12991 | 13019 | 29 | CTCCCTCGCTGCGATCTATTGAAAGTCAG | 1982.4 | -13.9 | 1.4 | -10.0 | 1.1E-23 | 2.3E-22 | MB231/MB468 | NUC |
| >hg38_21_+_8433172_8446622_RNA45SN1_45S_50ntFlanks@10771.10792.22\|CAGGTCTCCAAGGTGAACAGCC | >hg38_21_+_8433172_8446622_RNA45SN1_45S_50ntFlanks | 10771 | 10792 | 22 | CAGGTCTCCAAGGTGAACAGCC | 369.5 | -13.0 | 1.3 | -9.9 | 4.7E-23 | 2.3E-22 | BT20/MB468 | NUC |
| >hg38_21_+_8433172_8446622_RNA45SN1_45S_50ntFlanks@485.511.27\|GATGTGGTGACGTCGTGCTCTCCCGGG | >hg38_21_+_8433172_8446622_RNA45SN1_45S_50ntFlanks | 485 | 511 | 27 | GATGTGGTGACGTCGTGCTCTCCCGGG | 523.7 | 13.5 | 1.4 | 10.0 | 1.7E-23 | 2.3E-22 | BT20/MB468 | MITO |
| >hg38_21_+_8433172_8446622_RNA45SN1_45S_50ntFlanks@485.506.22\|GATGTGGTGACGTCGTGCTCTC | >hg38_21_+_8433172_8446622_RNA45SN1_45S_50ntFlanks | 485 | 506 | 22 | GATGTGGTGACGTCGTGCTCTC | 2312.5 | 11.7 | 1.2 | 9.9 | 4.8E-23 | 2.4E-22 | BT20/MB468 | NUC |
| >hg38_21_+_8433172_8446622_RNA45SN1_45S_50ntFlanks@5487.5510.24\|CGCTGAGAAGACGGTCGAACTTGA | >hg38_21_+_8433172_8446622_RNA45SN1_45S_50ntFlanks | 5487 | 5510 | 24 | CGCTGAGAAGACGGTCGAACTTGA | 1096.6 | -14.0 | 1.4 | -10.1 | 7.1E-24 | 2.4E-22 | MB231/MB468 | MP |
| >hg38_21_+_8433172_8446622_RNA45SN1_45S_50ntFlanks@12539.12567.29\|ATGATGTGTTGTTGCCATGGTAATCCTGC | >hg38_21_+_8433172_8446622_RNA45SN1_45S_50ntFlanks | 12539 | 12567 | 29 | ATGATGTGTTGTTGCCATGGTAATCCTGC | 1175.5 | -14.1 | 1.4 | -10.1 | 7.2E-24 | 2.4E-22 | MB231/MB468 | MP |
| >hg38_21_+_8433172_8446622_RNA45SN1_45S_50ntFlanks@12044.12068.25\|ACTGACCCGGTGAGGCGGGGGGGCG | >hg38_21_+_8433172_8446622_RNA45SN1_45S_50ntFlanks | 12044 | 12068 | 25 | ACTGACCCGGTGAGGCGGGGGGGCG | 2890.0 | -15.3 | 1.5 | -10.0 | 1.1E-23 | 2.4E-22 | MB231/MB468 | MITO |
| >hg38_21_+_8433172_8446622_RNA45SN1_45S_50ntFlanks@9562.9596.35\|TCCTGACGTGCAAATCGGTCGTCCGACCTGGGTAT | >hg38_21_+_8433172_8446622_RNA45SN1_45S_50ntFlanks | 9562 | 9596 | 35 | TCCTGACGTGCAAATCGGTCGTCCGACCTGGGTAT | 5996.2 | -11.8 | 1.2 | -10.0 | 1.1E-23 | 2.4E-22 | MB231/MB468 | MITO |
| >hg38_21_+_8433172_8446622_RNA45SN1_45S_50ntFlanks@11314.11344.31\|CCCACCCCACGTCTCGTCGCGCGCGCGTCCG | >hg38_21_+_8433172_8446622_RNA45SN1_45S_50ntFlanks | 11314 | 11344 | 31 | CCCACCCCACGTCTCGTCGCGCGCGCGTCCG | 1732.2 | -14.5 | 1.4 | -10.0 | 1.1E-23 | 2.4E-22 | MB231/MB468 | MITO |
| >hg38_21_+_8433172_8446622_RNA45SN1_45S_50ntFlanks@11586.11603.18\|GTGCGGACCAGGGGAATC | >hg38_21_+_8433172_8446622_RNA45SN1_45S_50ntFlanks | 11586 | 11603 | 18 | GTGCGGACCAGGGGAATC | 1195.1 | -14.0 | 1.4 | -10.0 | 1.1E-23 | 2.4E-22 | MB231/MB468 | MITO |
| >hg38_21_+_8433172_8446622_RNA45SN1_45S_50ntFlanks@11273.11293.21\|CCGCCCCCGGGGAGCCCGGCG | >hg38_21_+_8433172_8446622_RNA45SN1_45S_50ntFlanks | 11273 | 11293 | 21 | CCGCCCCCGGGGAGCCCGGCG | 548.6 | -13.5 | 1.4 | -9.9 | 4.9E-23 | 2.4E-22 | BT20/MB468 | NUC |
| >MI0000440\|hsa-mir-27b&WithFlank&9\|+\|95085439\|95085547@67.86.20(+1U)&[MIMAT0000419&hsa-miR-27b-3p&offsets\|0\|-1(+1U);m-39&9\|+\|95085505\|95085525&offsets\|0\|-1(+1U)]\|TTCACAGTGGCTAAGTTCTGT | >MI0000440\|hsa-mir-27b&WithFlank&9\|+\|95085439\|95085547 | 67 | 86 | 20(+1U) | TTCACAGTGGCTAAGTTCTGT | 5063.5 | 6.5 | 0.6 | 10.0 | 1.2E-23 | 2.4E-22 | MB231/MB468 | NUC |
| >hg38_21_+_8433172_8446622_RNA45SN1_45S_50ntFlanks@10512.10533.22\|TCCGGAGTGGCGGAGATGGGCG | >hg38_21_+_8433172_8446622_RNA45SN1_45S_50ntFlanks | 10512 | 10533 | 22 | TCCGGAGTGGCGGAGATGGGCG | 4239.3 | -15.8 | 1.6 | -10.0 | 1.2E-23 | 2.5E-22 | MB231/MB468 | MITO |
| >MI0000067\|hsa-let-7f-1&WithFlank&9\|+\|94176341\|94176439@13.32.20(+1C);MI0000068\|hsa-let-7f-2&WithFlank&X\|-\|53557186\|53557280@14.33.20(+1C)&[MIMAT0000067&hsa-let-7f-5p&offsets\|0\|-2(+1C);m-20&9\|+\|94176353\|94176374&offsets\|0\|-2(+1C)];[MIMAT0000067_1&hsa-let-7f-5p&offsets\|0\|-2(+1C);m-19&X\|-\|53557246\|53557267&offsets\|0\|-2(+1C)]\|TGAGGTAGTAGATTGTATAGC | >MI0000067\|hsa-let-7f-1&WithFlank&9\|+\|94176341\|94176439 | 13 | 32 | 20(+1C) | TGAGGTAGTAGATTGTATAGC | 774.8 | 6.8 | 0.7 | 10.0 | 1.8E-23 | 2.5E-22 | BT20/MB468 | MITO |
| >MI0000298\|hsa-mir-221&WithFlank&X\|-\|45746151\|45746272@71.93.23(+1C)&[MIMAT0000278&hsa-miR-221-3p&offsets\|0\|0(+1C);m-91&X\|-\|45746181\|45746202&offsets\|0\|+1(+1C)]\|AGCTACATTGTCTGCTGGGTTTCC | >MI0000298\|hsa-mir-221&WithFlank&X\|-\|45746151\|45746272 | 71 | 93 | 23(+1C) | AGCTACATTGTCTGCTGGGTTTCC | 1314.5 | 5.9 | 0.6 | 10.0 | 1.2E-23 | 2.5E-22 | MB231/MB468 | NUC |
| >hg38_21_+_8433172_8446622_RNA45SN1_45S_50ntFlanks@6738.6771.34\|ATCATCGACACTTCGAACGCACTTGCGGCCCCGG | >hg38_21_+_8433172_8446622_RNA45SN1_45S_50ntFlanks | 6738 | 6771 | 34 | ATCATCGACACTTCGAACGCACTTGCGGCCCCGG | 3914.9 | -15.8 | 1.6 | -10.1 | 7.6E-24 | 2.5E-22 | MB231/MB468 | MP |
| >hg38_21_+_8433172_8446622_RNA45SN1_45S_50ntFlanks@3769.3786.18\|CGCACGGCCGGTACAGTG | >hg38_21_+_8433172_8446622_RNA45SN1_45S_50ntFlanks | 3769 | 3786 | 18 | CGCACGGCCGGTACAGTG | 384.5 | -13.0 | 1.3 | -9.9 | 5.1E-23 | 2.5E-22 | BT20/MB468 | NUC |
| >hg38_21_+_8433172_8446622_RNA45SN1_45S_50ntFlanks@12456.12479.24\|CCAAGCGTTGGATTGTTCACCCAC | >hg38_21_+_8433172_8446622_RNA45SN1_45S_50ntFlanks | 12456 | 12479 | 24 | CCAAGCGTTGGATTGTTCACCCAC | 1639.2 | -14.5 | 1.4 | -10.1 | 7.6E-24 | 2.5E-22 | MB231/MB468 | MP |
| >MI0001446\|hsa-mir-424&WithFlank&X\|-\|134546608\|134546717@54.74.21&[MIMAT0004749&hsa-miR-424-3p&offsets\|0\|0;m-305&X\|-\|134546644\|134546664&offsets\|0\|0]\|CAAAACGTGAGGCGCTGCTAT | >MI0001446\|hsa-mir-424&WithFlank&X\|-\|134546608\|134546717 | 54 | 74 | 21 | CAAAACGTGAGGCGCTGCTAT | 1774.7 | 12.7 | 1.3 | 10.0 | 2.4E-23 | 2.5E-22 | BT20/MB468 | CYTO |
| >hg38_21_+_8433172_8446622_RNA45SN1_45S_50ntFlanks@12456.12473.18\|CCAAGCGTTGGATTGTTC | >hg38_21_+_8433172_8446622_RNA45SN1_45S_50ntFlanks | 12456 | 12473 | 18 | CCAAGCGTTGGATTGTTC | 9915.6 | -7.5 | 0.8 | -10.0 | 2.4E-23 | 2.5E-22 | BT20/MB468 | CYTO |
| >hg38_21_+_8433172_8446622_RNA45SN1_45S_50ntFlanks@11641.11664.24\|CGGCGGGTGTTGACGCGATGTGAT | >hg38_21_+_8433172_8446622_RNA45SN1_45S_50ntFlanks | 11641 | 11664 | 24 | CGGCGGGTGTTGACGCGATGTGAT | 4443.4 | -16.0 | 1.6 | -10.1 | 7.7E-24 | 2.5E-22 | MB231/MB468 | MP |
| >hg38_21_+_8433172_8446622_RNA45SN1_45S_50ntFlanks@5484.5514.31\|GAGCGCTGAGAAGACGGTCGAACTTGACTAT | >hg38_21_+_8433172_8446622_RNA45SN1_45S_50ntFlanks | 5484 | 5514 | 31 | GAGCGCTGAGAAGACGGTCGAACTTGACTAT | 4835.8 | -8.1 | 0.8 | -10.0 | 1.3E-23 | 2.6E-22 | MB231/MB468 | NUC |
| >hg38_21_+_8433172_8446622_RNA45SN1_45S_50ntFlanks@6719.6738.20\|TGAATTGCAGGACACATTGA | >hg38_21_+_8433172_8446622_RNA45SN1_45S_50ntFlanks | 6719 | 6738 | 20 | TGAATTGCAGGACACATTGA | 443.8 | -13.2 | 1.3 | -9.9 | 5.2E-23 | 2.6E-22 | BT20/MB468 | NUC |
| >hg38_21_+_8433172_8446622_RNA45SN1_45S_50ntFlanks@10769.10796.28\|AGCAGGTCTCCAAGGTGAACAGCCTCTG | >hg38_21_+_8433172_8446622_RNA45SN1_45S_50ntFlanks | 10769 | 10796 | 28 | AGCAGGTCTCCAAGGTGAACAGCCTCTG | 439.3 | -4.6 | 0.5 | -9.9 | 5.3E-23 | 2.6E-22 | BT20/MB468 | NUC |
| >hg38_21_+_8433172_8446622_RNA45SN1_45S_50ntFlanks@6661.6704.44\|CGGTGGATCACTCGGCTCGTGCGTCGATGAAGAACGCAGCTAGC | >hg38_21_+_8433172_8446622_RNA45SN1_45S_50ntFlanks | 6661 | 6704 | 44 | CGGTGGATCACTCGGCTCGTGCGTCGATGAAGAACGCAGCTAGC | 962.5 | 7.3 | 0.7 | 10.0 | 1.9E-23 | 2.6E-22 | BT20/MB468 | MITO |
| >MI0000088\|hsa-mir-30a&WithFlank&6\|-\|71403545\|71403627@12.34.23(+1A)&[MIMAT0000087&hsa-miR-30a-5p&offsets\|0\|+1(+1A);m-15&6\|-\|71403595\|71403616&offsets\|0\|+1(+1A)]\|TGTAAACATCCTCGACTGGAAGCA | >MI0000088\|hsa-mir-30a&WithFlank&6\|-\|71403545\|71403627 | 12 | 34 | 23(+1A) | TGTAAACATCCTCGACTGGAAGCA | 6124.5 | 7.7 | 0.8 | 10.0 | 1.3E-23 | 2.6E-22 | MB231/MB468 | NUC |
| >hg38_21_+_8433172_8446622_RNA45SN1_45S_50ntFlanks@11321.11344.24\|CACGTCTCGTCGCGCGCGCGTCCG | >hg38_21_+_8433172_8446622_RNA45SN1_45S_50ntFlanks | 11321 | 11344 | 24 | CACGTCTCGTCGCGCGCGCGTCCG | 14999.5 | -11.5 | 1.1 | -10.0 | 1.2E-23 | 2.6E-22 | MB231/MB468 | MITO |
| >hg38_21_+_8433172_8446622_RNA45SN1_45S_50ntFlanks@9534.9562.29\|AGGAAACTCTGGTGGAGGTCCGTAGCGGT | >hg38_21_+_8433172_8446622_RNA45SN1_45S_50ntFlanks | 9534 | 9562 | 29 | AGGAAACTCTGGTGGAGGTCCGTAGCGGT | 3764.1 | -7.3 | 0.7 | -10.0 | 1.3E-23 | 2.7E-22 | MB231/MB468 | NUC |
| >trnaMT_PheGAA_MT_+_577_647@35.54.20\|GAAAATGTTTAGACGGGCTC | >trnaMT_PheGAA_MT_+_577_647 | 35 | 54 | 20 | GAAAATGTTTAGACGGGCTC | 307.2 | 12.5 | 1.3 | 9.9 | 2.6E-23 | 2.7E-22 | BT20/MB468 | CYTO |
| >hg38_21_+_8433172_8446622_RNA45SN1_45S_50ntFlanks@12632.12653.22\|GCGAAGCTACCATCTGTGGGAT | >hg38_21_+_8433172_8446622_RNA45SN1_45S_50ntFlanks | 12632 | 12653 | 22 | GCGAAGCTACCATCTGTGGGAT | 575.9 | -13.6 | 1.4 | -9.9 | 5.5E-23 | 2.7E-22 | BT20/MB468 | NUC |
| >hg38_21_+_8433172_8446622_RNA45SN1_45S_50ntFlanks@10866.10902.37\|CTAAGGGCTGGGTCGGTCGGGCTGGGGCGCGAAGCGG | >hg38_21_+_8433172_8446622_RNA45SN1_45S_50ntFlanks | 10866 | 10902 | 37 | CTAAGGGCTGGGTCGGTCGGGCTGGGGCGCGAAGCGG | 1236.0 | -14.1 | 1.4 | -10.1 | 8.3E-24 | 2.7E-22 | MB231/MB468 | MP |
| >hg38_21_+_8433172_8446622_RNA45SN1_45S_50ntFlanks@4588.4616.29\|CTATTTTGTTGGTTTTCGGAACTGAGGCC | >hg38_21_+_8433172_8446622_RNA45SN1_45S_50ntFlanks | 4588 | 4616 | 29 | CTATTTTGTTGGTTTTCGGAACTGAGGCC | 2387.5 | -15.1 | 1.5 | -10.1 | 8.2E-24 | 2.7E-22 | MB231/MB468 | MP |
| >hg38_21_+_8433172_8446622_RNA45SN1_45S_50ntFlanks@11874.11907.34\|AGAAGACCCTGTTGAGCTTGACTCTAGTCTGGCA | >hg38_21_+_8433172_8446622_RNA45SN1_45S_50ntFlanks | 11874 | 11907 | 34 | AGAAGACCCTGTTGAGCTTGACTCTAGTCTGGCA | 1240.0 | -14.1 | 1.4 | -10.0 | 1.3E-23 | 2.7E-22 | MB231/MB468 | MITO |
| >MI0000285\|hsa-mir-205&WithFlank&1\|+\|209432127\|209432248@40.60.21(+1A)&[MIMAT0000266&hsa-miR-205-5p&offsets\|0\|-1(+1A);m-351&1\|+\|209432166\|209432187&offsets\|0\|-1(+1A)]\|TCCTTCATTCCACCGGAGTCTA | >MI0000285\|hsa-mir-205&WithFlank&1\|+\|209432127\|209432248 | 40 | 60 | 21(+1A) | TCCTTCATTCCACCGGAGTCTA | 4484.8 | 15.9 | 1.6 | 10.2 | 1.5E-24 | 2.7E-22 | BT20/MB231 | NUC |
| >hg38_21_+_8433172_8446622_RNA45SN1_45S_50ntFlanks@12163.12181.19\|ACTGGGGCGGTACACCTGT | >hg38_21_+_8433172_8446622_RNA45SN1_45S_50ntFlanks | 12163 | 12181 | 19 | ACTGGGGCGGTACACCTGT | 4857.3 | -16.0 | 1.6 | -10.0 | 1.3E-23 | 2.7E-22 | MB231/MB468 | MITO |
| >hg38_21_+_8433172_8446622_RNA45SN1_45S_50ntFlanks@10351.10384.34\|CGAGAACTTTGAAGGCCGAAGTGGAGAAGGGTTC | >hg38_21_+_8433172_8446622_RNA45SN1_45S_50ntFlanks | 10351 | 10384 | 34 | CGAGAACTTTGAAGGCCGAAGTGGAGAAGGGTTC | 2799.0 | -5.5 | 0.5 | -10.1 | 5.9E-24 | 2.7E-22 | MB231/MB468 | CYTO |
| >MI0003591\|hsa-mir-584&WithFlank&5\|-\|149062307\|149062415@22.42.21(+2A)&[MIMAT0003249&hsa-miR-584-5p&offsets\|0\|-1(+2A);m-167&5\|-\|149062374\|149062394&offsets\|0\|0(+2A)]\|TTATGGTTTGCCTGGGACTGAAA | >MI0003591\|hsa-mir-584&WithFlank&5\|-\|149062307\|149062415 | 22 | 42 | 21(+2A) | TTATGGTTTGCCTGGGACTGAAA | 428.0 | 13.2 | 1.3 | 10.0 | 2.0E-23 | 2.7E-22 | BT20/MB468 | MITO |
| >hg38_21_+_8433172_8446622_RNA45SN1_45S_50ntFlanks@12942.12972.31\|TTCGTAGACGACCTGCTTCTGGGTCGGGGTT | >hg38_21_+_8433172_8446622_RNA45SN1_45S_50ntFlanks | 12942 | 12972 | 31 | TTCGTAGACGACCTGCTTCTGGGTCGGGGTT | 1493.1 | -14.4 | 1.4 | -10.1 | 8.4E-24 | 2.7E-22 | MB231/MB468 | MP |

| | | | | | | | | | | | | | |
|---|---|---|---|---|---|---|---|---|---|---|---|---|---|
| >MI0001445\|hsa-mir-423&WithFlank&17\|+\|30117073\|30117178@23.44.22(+1A)&[MIMAT0004748&hsa-miR-423-5p&offsets\|0\|-1(+1A);m-141&17\|+\|30117095\|30117117&offsets\|0\|-1(+1A)]\|TGAGGGGCAGAGAGCGAGACTTA | >MI0001445\|hsa-mir-423&WithFlank&17\|+\|30117073\|30117178 | 23 | 44 | 22(+1A) | TGAGGGGCAGAGAGCGAGACTTA | 497.1 | 13.2 | 1.3 | 9.9 | 2.7E-23 | 2.8E-22 | BT20/MB468 | CYTO |
| >hg38_21_+_8433172_8446622_RNA45SN1_45S_50ntFlanks@11541.11584.44\|GCGGCGCCTCGCCTCGGCCGGCGCCTAGCAGCCGACTTAGAACT | >hg38_21_+_8433172_8446622_RNA45SN1_45S_50ntFlanks | 11541 | 11584 | 44 | GCGGCGCCTCGCCTCGGCCGGCGCCTAGCAGCCGACTTAGAACT | 403.2 | 12.7 | 1.3 | 10.1 | 6.0E-24 | 2.8E-22 | MB231/MB468 | CYTO |
| >hg38_21_+_8433172_8446622_RNA45SN1_45S_50ntFlanks@5490.5532.43\|TGAGAAGACGGTCGAACTTGACTATCTAGAGGAAGTAAAAGTC | >hg38_21_+_8433172_8446622_RNA45SN1_45S_50ntFlanks | 5490 | 5532 | 43 | TGAGAAGACGGTCGAACTTGACTATCTAGAGGAAGTAAAAGTC | 2123.6 | 15.1 | 1.5 | 10.1 | 6.0E-24 | 2.8E-22 | MB231/MB468 | CYTO |
| >hg38_21_+_8433172_8446622_RNA45SN1_45S_50ntFlanks@9574.9596.23\|AATCGGTCGTCCGACCTGGGTAT | >hg38_21_+_8433172_8446622_RNA45SN1_45S_50ntFlanks | 9574 | 9596 | 23 | AATCGGTCGTCCGACCTGGGTAT | 13655.9 | -8.1 | 0.8 | -10.0 | 1.3E-23 | 2.8E-22 | MB231/MB468 | MITO |
| >hg38_21_+_8433172_8446622_RNA45SN1_45S_50ntFlanks@12936.12961.26\|AAACCATTCGTAGACGACCTGCTTCT | >hg38_21_+_8433172_8446622_RNA45SN1_45S_50ntFlanks | 12936 | 12961 | 26 | AAACCATTCGTAGACGACCTGCTTCT | 1137.7 | -4.9 | 0.5 | -10.0 | 1.4E-23 | 2.9E-22 | MB231/MB468 | NUC |
| >hg38_21_+_8433172_8446622_RNA45SN1_45S_50ntFlanks@6772.6799.28\|GTTCCTCCCGGGGCTACGCCTGTCTGAG | >hg38_21_+_8433172_8446622_RNA45SN1_45S_50ntFlanks | 6772 | 6799 | 28 | GTTCCTCCCGGGGCTACGCCTGTCTGAG | 4253.9 | -11.6 | 1.2 | -10.0 | 1.4E-23 | 2.9E-22 | MB231/MB468 | MITO |
| >hg38_21_+_8433172_8446622_RNA45SN1_45S_50ntFlanks@11235.11254.20\|CTTCCCGTGGATCGCCCCAG | >hg38_21_+_8433172_8446622_RNA45SN1_45S_50ntFlanks | 11235 | 11254 | 20 | CTTCCCGTGGATCGCCCCAG | 1800.9 | -13.8 | 1.4 | -10.0 | 1.5E-23 | 2.9E-22 | MB231/MB468 | NUC |
| >MI0000266\|hsa-mir-10a&WithFlank&17\|-\|48579832\|48579953@29.51.23&[MIMAT0000253&hsa-miR-10a-5p&offsets\|+1\|+1;m-18&17\|-\|48579905\|48579926&offsets\|+1\|+2]\|ACCCTGTAGATCCGAATTTGTGT | >MI0000266\|hsa-mir-10a&WithFlank&17\|-\|48579832\|48579953 | 29 | 51 | 23 | ACCCTGTAGATCCGAATTTGTGT | 858.1 | 14.2 | 1.4 | 10.0 | 2.1E-23 | 2.9E-22 | BT20/MB468 | MITO |
| >hg38_21_+_8433172_8446622_RNA45SN1_45S_50ntFlanks@13004.13027.24\|ATCTATTGAAAGTCAGCCCTCGAC | >hg38_21_+_8433172_8446622_RNA45SN1_45S_50ntFlanks | 13004 | 13027 | 24 | ATCTATTGAAAGTCAGCCCTCGAC | 1368.2 | -14.2 | 1.4 | -10.0 | 1.4E-23 | 2.9E-22 | MB231/MB468 | MITO |
| >hg38_21_+_8433172_8446622_RNA45SN1_45S_50ntFlanks@11804.11821.18\|CCCACTGTCCCTACCTAC | >hg38_21_+_8433172_8446622_RNA45SN1_45S_50ntFlanks | 11804 | 11821 | 18 | CCCACTGTCCCTACCTAC | 1089.5 | -14.5 | 1.5 | -9.9 | 6.0E-23 | 2.9E-22 | BT20/MB468 | NUC |
| >hg38_21_+_8433172_8446622_RNA45SN1_45S_50ntFlanks@4554.4572.19\|CCGCAGCTAGGAATAATGG | >hg38_21_+_8433172_8446622_RNA45SN1_45S_50ntFlanks | 4554 | 4572 | 19 | CCGCAGCTAGGAATAATGG | 364.6 | -12.9 | 1.3 | -9.9 | 6.1E-23 | 3.0E-22 | BT20/MB468 | NUC |
| >hg38_21_+_8433172_8446622_RNA45SN1_45S_50ntFlanks@12602.12626.25\|GTGTATGTGCTTGGCTGAGGAGCCA | >hg38_21_+_8433172_8446622_RNA45SN1_45S_50ntFlanks | 12602 | 12626 | 25 | GTGTATGTGCTTGGCTGAGGAGCCA | 362.6 | -12.9 | 1.3 | -9.9 | 6.3E-23 | 3.1E-22 | BT20/MB468 | NUC |
| >MI0000266\|hsa-mir-10a&WithFlank&17\|-\|48579832\|48579953@29.50.22(+1A)&[MIMAT0000253&hsa-miR-10a-5p&offsets\|+1\|0(+1A);m-18&17\|-\|48579905\|48579926&offsets\|+1\|+1(+1A)]\|ACCCTGTAGATCCGAATTTGTGA | >MI0000266\|hsa-mir-10a&WithFlank&17\|-\|48579832\|48579953 | 29 | 50 | 22(+1A) | ACCCTGTAGATCCGAATTTGTGA | 187.0 | 10.4 | 1.1 | 9.9 | 6.3E-23 | 3.1E-22 | BT20/MB468 | NUC |
| >MI0000650\|hsa-mir-200c&WithFlank&12\|+\|6963693\|6963772@50.71.22(+1G)&[MIMAT0000617&hsa-miR-200c-3p&offsets\|0\|-1(+1G);m-54&12\|+\|6963742\|6963764&offsets\|0\|-1(+1G)]\|TAATACTGCCGGGTAATGATGGG | >MI0000650\|hsa-mir-200c&WithFlank&12\|+\|6963693\|6963772 | 50 | 71 | 22(+1G) | TAATACTGCCGGGTAATGATGGG | 1473.0 | 14.1 | 1.4 | 10.2 | 1.3E-24 | 3.1E-22 | BT20/MB231 | MITO |
| >hg38_21_+_8433172_8446622_RNA45SN1_45S_50ntFlanks@12044.12071.28\|ACTGACCCGGTGAGGCGGGGGGGCGAGC | >hg38_21_+_8433172_8446622_RNA45SN1_45S_50ntFlanks | 12044 | 12071 | 28 | ACTGACCCGGTGAGGCGGGGGGGCGAGC | 2478.0 | -15.1 | 1.5 | -10.0 | 1.5E-23 | 3.1E-22 | MB231/MB468 | MITO |
| >hg38_21_+_8433172_8446622_RNA45SN1_45S_50ntFlanks@8570.8593.24\|CTCCGGCGGGTGCGGGGGTGGGCG | >hg38_21_+_8433172_8446622_RNA45SN1_45S_50ntFlanks | 8570 | 8593 | 24 | CTCCGGCGGGTGCGGGGGTGGGCG | 803.5 | -14.1 | 1.4 | -9.9 | 6.4E-23 | 3.1E-22 | BT20/MB468 | NUC |
| >MI0000816\|hsa-mir-335&WithFlank&7\|+\|130496105\|130496210@58.78.21(+1U)&[MIMAT0004703&hsa-miR-335-3p&offsets\|0\|-1(+1U);m-159&7\|+\|130496162\|130496183&offsets\|0\|-1(+1U)]\|TTTTTCATTATTGCTCCTGACT | >MI0000816\|hsa-mir-335&WithFlank&7\|+\|130496105\|130496210 | 58 | 78 | 21(+1U) | TTTTTCATTATTGCTCCTGACT | 603.0 | 5.1 | 0.5 | 10.0 | 1.6E-23 | 3.2E-22 | MB231/MB468 | NUC |
| >MI0000439\|hsa-mir-23b&WithFlank&9\|+\|95085202\|95085310@67.86.20(+1C)&[MIMAT0000418&hsa-miR-23b-3p&offsets\|+3\|0(+1C);m-42&9\|+\|95085265\|95085285&offsets\|+3\|+2(+1C)]\|ACATTGCCAGGGATTACCACC | >MI0000439\|hsa-mir-23b&WithFlank&9\|+\|95085202\|95085310 | 67 | 86 | 20(+1C) | ACATTGCCAGGGATTACCACC | 1215.4 | 14.7 | 1.5 | 10.0 | 2.3E-23 | 3.2E-22 | BT20/MB468 | MITO |
| >hg38_21_+_8433172_8446622_RNA45SN1_45S_50ntFlanks@12625.12654.30\|CAATGGGGCGAAGCTACCATCTGTGGGATT | >hg38_21_+_8433172_8446622_RNA45SN1_45S_50ntFlanks | 12625 | 12654 | 30 | CAATGGGGCGAAGCTACCATCTGTGGGATT | 14530.3 | -10.6 | 1.1 | -10.0 | 1.6E-23 | 3.2E-22 | MB231/MB468 | MITO |
| >hg38_21_+_8433172_8446622_RNA45SN1_45S_50ntFlanks@6719.6743.25\|TGAATTGCAGGACACATTGATCATC | >hg38_21_+_8433172_8446622_RNA45SN1_45S_50ntFlanks | 6719 | 6743 | 25 | TGAATTGCAGGACACATTGATCATC | 23210.3 | -12.0 | 1.2 | -9.9 | 6.6E-23 | 3.2E-22 | BT20/MB468 | NUC |
| >MI0001445\|hsa-mir-423&WithFlank&17\|+\|30117073\|30117178@59.82.24&[MIMAT0001340&hsa-miR-423-3p&offsets\|0\|+1;m-100&17\|+\|30117131\|30117153&offsets\|0\|+1]\|AGCTCGGTCTGAGGCCCCTCAGTC | >MI0001445\|hsa-mir-423&WithFlank&17\|+\|30117073\|30117178 | 59 | 82 | 24 | AGCTCGGTCTGAGGCCCCTCAGTC | 341.1 | 12.7 | 1.3 | 9.9 | 3.1E-23 | 3.2E-22 | BT20/MB468 | CYTO |
| >hg38_21_+_8433172_8446622_RNA45SN1_45S_50ntFlanks@7985.8005.21\|GATCAGACGTGGCGACCCGCT | >hg38_21_+_8433172_8446622_RNA45SN1_45S_50ntFlanks | 7985 | 8005 | 21 | GATCAGACGTGGCGACCCGCT | 316.5 | -12.7 | 1.3 | -9.9 | 6.7E-23 | 3.3E-22 | BT20/MB468 | NUC |
| >hg38_21_+_8433172_8446622_RNA45SN1_45S_50ntFlanks@12073.12097.25\|CCGAGGGGCTCTCGCTTCTGGCGCC | >hg38_21_+_8433172_8446622_RNA45SN1_45S_50ntFlanks | 12073 | 12097 | 25 | CCGAGGGGCTCTCGCTTCTGGCGCC | 2129.4 | -5.9 | 0.6 | -10.0 | 1.6E-23 | 3.3E-22 | MB231/MB468 | NUC |
| >hg38_21_+_8433172_8446622_RNA45SN1_45S_50ntFlanks@12456.12478.23\|CCAAGCGTTGGATTGTTCACCCA | >hg38_21_+_8433172_8446622_RNA45SN1_45S_50ntFlanks | 12456 | 12478 | 23 | CCAAGCGTTGGATTGTTCACCCA | 8714.4 | -12.5 | 1.2 | -10.0 | 1.0E-23 | 3.3E-22 | MB231/MB468 | MP |
| >hg38_21_+_8433172_8446622_RNA45SN1_45S_50ntFlanks@10840.10865.26\|CCGTAACTTCGGGATAAGGATTGGCT | >hg38_21_+_8433172_8446622_RNA45SN1_45S_50ntFlanks | 10840 | 10865 | 26 | CCGTAACTTCGGGATAAGGATTGGCT | 3910.8 | -5.3 | 0.5 | -10.1 | 7.3E-24 | 3.3E-22 | MB231/MB468 | CYTO |
| >MI0005756\|hsa-mir-934&WithFlank&X\|+\|136550872\|136550966@21.41.21&[MIMAT0004977&hsa-miR-934&offsets\|0\|-1;m-943&X\|+\|136550892\|136550913&offsets\|0\|-1]\|TGTCTACTACTGGAGACACTG | >MI0005756\|hsa-mir-934&WithFlank&X\|+\|136550872\|136550966 | 21 | 41 | 21 | TGTCTACTACTGGAGACACTG | 1616.3 | 14.4 | 1.4 | 10.2 | 1.9E-24 | 3.3E-22 | BT20/MB231 | NUC |
| >hg38_21_+_8433172_8446622_RNA45SN1_45S_50ntFlanks@4000.4042.43\|TAGATAACCTCGGGCCGATCGCACGCCCCCCGTGGCGGCGACG | >hg38_21_+_8433172_8446622_RNA45SN1_45S_50ntFlanks | 4000 | 4042 | 43 | TAGATAACCTCGGGCCGATCGCACGCCCCCCGTGGCGGCGACG | 880.3 | -14.2 | 1.4 | -9.8 | 6.9E-23 | 3.3E-22 | BT20/MB468 | NUC |
| >MI0000471\|hsa-mir-126&WithFlank&9\|+\|136670596\|136670692@58.77.20(+1U)&[MIMAT0000445&hsa-miR-126-3p&offsets\|0\|-2(+1U);m-40&9\|+\|136670653\|136670674&offsets\|0\|-2(+1U)]\|TCGTACCGTGAGTAATAATGT | >MI0000471\|hsa-mir-126&WithFlank&9\|+\|136670596\|136670692 | 58 | 77 | 20(+1U) | TCGTACCGTGAGTAATAATGT | 625.1 | 13.8 | 1.4 | 10.0 | 2.4E-23 | 3.4E-22 | BT20/MB468 | MITO |
| >hg38_21_+_8433172_8446622_RNA45SN1_45S_50ntFlanks@12005.12035.31\|GGGCCGCCGGTGAAATACCACTACTCTGATC | >hg38_21_+_8433172_8446622_RNA45SN1_45S_50ntFlanks | 12005 | 12035 | 31 | GGGCCGCCGGTGAAATACCACTACTCTGATC | 820.3 | 7.7 | 0.8 | 10.0 | 2.4E-23 | 3.4E-22 | BT20/MB468 | MITO |
| >hg38_21_+_8433172_8446622_RNA45SN1_45S_50ntFlanks@12008.12033.26\|CCGCCGGTGAAATACCACTACTCTGA | >hg38_21_+_8433172_8446622_RNA45SN1_45S_50ntFlanks | 12008 | 12033 | 26 | CCGCCGGTGAAATACCACTACTCTGA | 1297.1 | -14.1 | 1.4 | -10.0 | 1.6E-23 | 3.4E-22 | MB231/MB468 | MITO |
| >hg38_21_+_8433172_8446622_RNA45SN1_45S_50ntFlanks@11584.11613.30\|TGGTGCGGACCAGGGGAATCCGACTGTTTA | >hg38_21_+_8433172_8446622_RNA45SN1_45S_50ntFlanks | 11584 | 11613 | 30 | TGGTGCGGACCAGGGGAATCCGACTGTTTA | 313.3 | 6.2 | 0.6 | 10.0 | 2.5E-23 | 3.4E-22 | BT20/MB468 | MITO |
| >hg38_21_+_8433172_8446622_RNA45SN1_45S_50ntFlanks@10350.10383.34\|ACGAGAACTTTGAAGGCCGAAGTGGAGAAGGGTT | >hg38_21_+_8433172_8446622_RNA45SN1_45S_50ntFlanks | 10350 | 10383 | 34 | ACGAGAACTTTGAAGGCCGAAGTGGAGAAGGGTT | 10035.8 | -7.2 | 0.7 | -10.0 | 1.6E-23 | 3.4E-22 | MB231/MB468 | MITO |
| >hg38_21_+_8433172_8446622_RNA45SN1_45S_50ntFlanks@7986.8006.21\|ATCAGACGTGGCGACCCGCTG | >hg38_21_+_8433172_8446622_RNA45SN1_45S_50ntFlanks | 7986 | 8006 | 21 | ATCAGACGTGGCGACCCGCTG | 16481.0 | -17.1 | 1.6 | -10.5 | 9.9E-26 | 3.4E-22 | BT20/MB468 | MP |
| >hg38_21_+_8433172_8446622_RNA45SN1_45S_50ntFlanks@11321.11342.22\|CACGTCTCGTCGCGCGCGCGTC | >hg38_21_+_8433172_8446622_RNA45SN1_45S_50ntFlanks | 11321 | 11342 | 22 | CACGTCTCGTCGCGCGCGCGTC | 2782.6 | -8.4 | 0.8 | -10.0 | 1.7E-23 | 3.4E-22 | MB231/MB468 | MITO |
| >MI0000086\|hsa-mir-28&WithFlank&3\|+\|188688775\|188688872@61.82.22&[MIMAT0004502&hsa-miR-28-3p&offsets\|+1\|+1;m-28&3\|+\|188688834\|188688855&offsets\|+1\|+1]\|ACTAGATTGTGAGCTCCTGGAG | >MI0000086\|hsa-mir-28&WithFlank&3\|+\|188688775\|188688872 | 61 | 82 | 22 | ACTAGATTGTGAGCTCCTGGAG | 1889.7 | 11.3 | 1.1 | 9.9 | 3.3E-23 | 3.4E-22 | BT20/MB468 | CYTO |
| >MI0000465\|hsa-mir-191&WithFlank&3\|-\|49020612\|49020715@22.42.21(+1C)&[MIMAT0000440&hsa-miR-191-5p&offsets\|0\|-2(+1C);m-48&3\|-\|49020672\|49020694&offsets\|0\|-2(+1C)]\|CAACGGAATCCCAAAAGCAGCC | >MI0000465\|hsa-mir-191&WithFlank&3\|-\|49020612\|49020715 | 22 | 42 | 21(+1C) | CAACGGAATCCCAAAAGCAGCC | 1612.6 | 7.7 | 0.8 | 10.0 | 2.5E-23 | 3.5E-22 | BT20/MB468 | MITO |
| >hg38_21_+_8433172_8446622_RNA45SN1_45S_50ntFlanks@6661.6678.18\|CGGTGGATCACTCGGCTC | >hg38_21_+_8433172_8446622_RNA45SN1_45S_50ntFlanks | 6661 | 6678 | 18 | CGGTGGATCACTCGGCTC | 3784.9 | 8.0 | 0.8 | 9.9 | 2.5E-23 | 3.5E-22 | BT20/MB468 | MITO |
| >hg38_21_+_8433172_8446622_RNA45SN1_45S_50ntFlanks@10689.10706.18\|GCGTCCGGTGAGCTCTCG | >hg38_21_+_8433172_8446622_RNA45SN1_45S_50ntFlanks | 10689 | 10706 | 18 | GCGTCCGGTGAGCTCTCG | 2186.6 | -15.0 | 1.5 | -10.0 | 1.1E-23 | 3.5E-22 | MB231/MB468 | MP |
| >trnaMT_MetCAT_MT_+_4402_4469@37.71.35\|CCGAAAATGTTGGTTATACCCTTCCCGTACTACCA | >trnaMT_MetCAT_MT_+_4402_4469 | 37 | 71 | 35 | CCGAAAATGTTGGTTATACCCTTCCCGTACTACCA | 257.3 | 12.3 | 1.2 | 9.9 | 3.4E-23 | 3.5E-22 | BT20/MB468 | CYTO |
| >hg38_21_+_8433172_8446622_RNA45SN1_45S_50ntFlanks@11729.11767.39\|TAACTATGACTCTCTTAAGGTAGCCAAATGCCTCGTCAT | >hg38_21_+_8433172_8446622_RNA45SN1_45S_50ntFlanks | 11729 | 11767 | 39 | TAACTATGACTCTCTTAAGGTAGCCAAATGCCTCGTCAT | 1917.1 | -14.7 | 1.5 | -10.0 | 1.7E-23 | 3.5E-22 | MB231/MB468 | MITO |
| >MI0000285\|hsa-mir-205&WithFlank&1\|+\|209432127\|209432248@40.61.22&[MIMAT0000266&hsa-miR-205-5p&offsets\|0\|0;m-351&1\|+\|209432166\|209432187&offsets\|0\|0]\|TCCTTCATTCCACCGGAGTCTG | >MI0000285\|hsa-mir-205&WithFlank&1\|+\|209432127\|209432248 | 40 | 61 | 22 | TCCTTCATTCCACCGGAGTCTG | 1511.2 | -14.3 | 1.4 | -10.0 | 1.7E-23 | 3.5E-22 | MB231/MB468 | MITO |
| >hg38_21_+_8433172_8446622_RNA45SN1_45S_50ntFlanks@12456.12486.31\|CCAAGCGTTGGATTGTTCACCCACTAATAGG | >hg38_21_+_8433172_8446622_RNA45SN1_45S_50ntFlanks | 12456 | 12486 | 31 | CCAAGCGTTGGATTGTTCACCCACTAATAGG | 7136.3 | -9.7 | 1.0 | -10.0 | 1.7E-23 | 3.5E-22 | MB231/MB468 | MITO |
| >hg38_21_+_8433172_8446622_RNA45SN1_45S_50ntFlanks@3873.3898.26\|TAATACATGCCGACGGGCGCTGACCC | >hg38_21_+_8433172_8446622_RNA45SN1_45S_50ntFlanks | 3873 | 3898 | 26 | TAATACATGCCGACGGGCGCTGACCC | 4524.7 | -11.7 | 1.2 | -10.0 | 1.7E-23 | 3.5E-22 | MB231/MB468 | MITO |
| >hg38_21_+_8433172_8446622_RNA45SN1_45S_50ntFlanks@7973.8013.41\|GACGCGACCTCAGATCAGACGTGGCGACCCGCTGAATTTAA | >hg38_21_+_8433172_8446622_RNA45SN1_45S_50ntFlanks | 7973 | 8013 | 41 | GACGCGACCTCAGATCAGACGTGGCGACCCGCTGAATTTAA | 311.1 | 5.0 | 0.5 | 9.9 | 2.6E-23 | 3.5E-22 | BT20/MB468 | MITO |
| >hg38_21_+_8433172_8446622_RNA45SN1_45S_50ntFlanks@12456.12480.25\|CCAAGCGTTGGATTGTTCACCCACT | >hg38_21_+_8433172_8446622_RNA45SN1_45S_50ntFlanks | 12456 | 12480 | 25 | CCAAGCGTTGGATTGTTCACCCACT | 4982.5 | -8.3 | 0.8 | -10.0 | 1.1E-23 | 3.6E-22 | MB231/MB468 | MP |
| >hg38_21_+_8433172_8446622_RNA45SN1_45S_50ntFlanks@485.517.33\|GATGTGGTGACGTCGTGCTCTCCCGGGCCGGGT | >hg38_21_+_8433172_8446622_RNA45SN1_45S_50ntFlanks | 485 | 517 | 33 | GATGTGGTGACGTCGTGCTCTCCCGGGCCGGGT | 751.2 | 12.4 | 1.3 | 9.8 | 7.4E-23 | 3.6E-22 | BT20/MB468 | NUC |
| >hg38_21_+_8433172_8446622_RNA45SN1_45S_50ntFlanks@11653.11691.39\|ACGCGATGTGATTTCTGCCCAGTGCTCTGAATGTCAAAG | >hg38_21_+_8433172_8446622_RNA45SN1_45S_50ntFlanks | 11653 | 11691 | 39 | ACGCGATGTGATTTCTGCCCAGTGCTCTGAATGTCAAAG | 348.6 | 6.9 | 0.7 | 9.9 | 2.7E-23 | 3.6E-22 | BT20/MB468 | MITO |
| >MI0003763\|hsa-mir-767&WithFlank&X\|-\|152393415\|152393535@33.56.24&[MIMAT0003882&hsa-miR-767-5p&offsets\|0\|+1;m-788&X\|-\|152393481\|152393503&offsets\|0\|+1]\|TGCACCATGGTTGTCTGAGCATGC | >MI0003763\|hsa-mir-767&WithFlank&X\|-\|152393415\|152393535 | 33 | 56 | 24 | TGCACCATGGTTGTCTGAGCATGC | 1181.0 | 6.1 | 0.6 | 9.9 | 3.5E-23 | 3.6E-22 | BT20/MB468 | CYTO |

<table>
<tr><th></th><th></th><th></th><th></th><th></th><th></th><th></th><th></th><th></th><th></th><th></th><th></th><th></th><th></th></tr>
<tr><td>>MI0000086|hsa-mir-28&WithFlank&3|+|188688775|188688872@60.82.23(+1U)&[MIMAT0004502&hsa-miR-28-3p&offsets|0|+1(+1U);m-28&3|+|188688834|188688855&offsets|0|+1(+1U)]|CACTAGATTGTGAGCTCCTGGAGT</td><td>>MI0000086|hsa-mir-28&WithFlank&3|+|188688775|188688872</td><td>60</td><td>82</td><td>23(+1U)</td><td>CACTAGATTGTGAGCTCCTGGAGT</td><td>539.3</td><td>13.3</td><td>1.3</td><td>9.9</td><td>3.6E-23</td><td>3.6E-22</td><td>BT20/MB468</td><td>CYTO</td></tr>
<tr><td>>hg38_21_+_8433172_8446622_RNA45SN1_45S_50ntFlanks@12624.12657.34|CCAATGGGGCGAAGCTACCATCTGTGGGATTATG</td><td>>hg38_21_+_8433172_8446622_RNA45SN1_45S_50ntFlanks</td><td>12624</td><td>12657</td><td>34</td><td>CCAATGGGGCGAAGCTACCATCTGTGGGATTATG</td><td>2128.4</td><td>-5.8</td><td>0.6</td><td>-9.9</td><td>2.7E-23</td><td>3.7E-22</td><td>BT20/MB468</td><td>MITO</td></tr>
<tr><td>>MI0000088|hsa-mir-30a&WithFlank&6|-|71403545|71403627@54.73.20(+1U)&[MIMAT0000088&hsa-miR-30a-3p&offsets|+1|-1(+1U);m-38&6|-|71403554|71403575&offsets|+1|-1(+1U)]|TTTCAGTCGGATGTTTGCAGT</td><td>>MI0000088|hsa-mir-30a&WithFlank&6|-|71403545|71403627</td><td>54</td><td>73</td><td>20(+1U)</td><td>TTTCAGTCGGATGTTTGCAGT</td><td>4372.6</td><td>7.0</td><td>0.7</td><td>10.0</td><td>1.9E-23</td><td>3.7E-22</td><td>MB231/MB468</td><td>NUC</td></tr>
<tr><td>>hg38_21_+_8433172_8446622_RNA45SN1_45S_50ntFlanks@8154.8181.28|CCGCTCGTGGGGGGCCCAAGTCCTTCTG</td><td>>hg38_21_+_8433172_8446622_RNA45SN1_45S_50ntFlanks</td><td>8154</td><td>8181</td><td>28</td><td>CCGCTCGTGGGGGGCCCAAGTCCTTCTG</td><td>1759.9</td><td>-5.6</td><td>0.6</td><td>-10.0</td><td>1.9E-23</td><td>3.7E-22</td><td>MB231/MB468</td><td>NUC</td></tr>
<tr><td>>hg38_21_+_8433172_8446622_RNA45SN1_45S_50ntFlanks@10869.10902.34|AGGGCTGGGTCGGTCGGGCTGGGGCGCGAAGCGG</td><td>>hg38_21_+_8433172_8446622_RNA45SN1_45S_50ntFlanks</td><td>10869</td><td>10902</td><td>34</td><td>AGGGCTGGGTCGGTCGGGCTGGGGCGCGAAGCGG</td><td>2408.6</td><td>-15.0</td><td>1.5</td><td>-10.0</td><td>1.8E-23</td><td>3.7E-22</td><td>MB231/MB468</td><td>MITO</td></tr>
<tr><td>>hg38_21_+_8433172_8446622_RNA45SN1_45S_50ntFlanks@4163.4181.19|CATCCAAGGAAGGCAGCAG</td><td>>hg38_21_+_8433172_8446622_RNA45SN1_45S_50ntFlanks</td><td>4163</td><td>4181</td><td>19</td><td>CATCCAAGGAAGGCAGCAG</td><td>316.4</td><td>-12.7</td><td>1.3</td><td>-9.8</td><td>7.6E-23</td><td>3.7E-22</td><td>BT20/MB468</td><td>NUC</td></tr>
<tr><td>>MI0000299|hsa-mir-222&WithFlank&X|-|45747009|45747130@75.96.22(+1A)&[MIMAT0000279&hsa-miR-222-3p&offsets|0|+1(+1A);m-140&X|-|45747033|45747056&offsets|0|-2(+1A)]|AGCTACATCTGGCTACTGGGTCA</td><td>>MI0000299|hsa-mir-222&WithFlank&X|-|45747009|45747130</td><td>75</td><td>96</td><td>22(+1A)</td><td>AGCTACATCTGGCTACTGGGTCA</td><td>583.4</td><td>6.9</td><td>0.7</td><td>9.9</td><td>2.7E-23</td><td>3.7E-22</td><td>BT20/MB468</td><td>MITO</td></tr>
<tr><td>>hg38_21_+_8433172_8446622_RNA45SN1_45S_50ntFlanks@11037.11075.39|CTCTCCCCCGCTCCCCGTCCTCCCCCCTCCCCGGGGGAG</td><td>>hg38_21_+_8433172_8446622_RNA45SN1_45S_50ntFlanks</td><td>11037</td><td>11075</td><td>39</td><td>CTCTCCCCCGCTCCCCGTCCTCCCCCCTCCCCGGGGGAG</td><td>871.9</td><td>-14.2</td><td>1.4</td><td>-9.8</td><td>7.6E-23</td><td>3.7E-22</td><td>BT20/MB468</td><td>NUC</td></tr>
<tr><td>>hg38_21_+_8433172_8446622_RNA45SN1_45S_50ntFlanks@11561.11591.31|GCGCCTAGCAGCCGACTTAGAACTGGTGCGG</td><td>>hg38_21_+_8433172_8446622_RNA45SN1_45S_50ntFlanks</td><td>11561</td><td>11591</td><td>31</td><td>GCGCCTAGCAGCCGACTTAGAACTGGTGCGG</td><td>1377.5</td><td>-14.3</td><td>1.4</td><td>-10.0</td><td>1.1E-23</td><td>3.7E-22</td><td>MB231/MB468</td><td>MP</td></tr>
<tr><td>>hg38_21_+_8433172_8446622_RNA45SN1_45S_50ntFlanks@12959.12993.35|TCTGGGTCGGGGTTTCGTACGTAGCAGAGCAGCTC</td><td>>hg38_21_+_8433172_8446622_RNA45SN1_45S_50ntFlanks</td><td>12959</td><td>12993</td><td>35</td><td>TCTGGGTCGGGGTTTCGTACGTAGCAGAGCAGCTC</td><td>2050.0</td><td>8.2</td><td>0.8</td><td>10.1</td><td>8.3E-24</td><td>3.7E-22</td><td>MB231/MB468</td><td>CYTO</td></tr>
<tr><td>>hg38_21_+_8433172_8446622_RNA45SN1_45S_50ntFlanks@7974.8016.43|ACGCGACCTCAGATCAGACGTGGCGACCCGCTGAATTTAAGCA</td><td>>hg38_21_+_8433172_8446622_RNA45SN1_45S_50ntFlanks</td><td>7974</td><td>8016</td><td>43</td><td>ACGCGACCTCAGATCAGACGTGGCGACCCGCTGAATTTAAGCA</td><td>1575.6</td><td>6.9</td><td>0.7</td><td>10.1</td><td>8.3E-24</td><td>3.7E-22</td><td>MB231/MB468</td><td>CYTO</td></tr>
<tr><td>>MI0000440|hsa-mir-27b&WithFlank&9|+|95085439|95085547@67.87.21&[MIMAT0000419&hsa-miR-27b-3p&offsets|0|0;m-39&9|+|95085505|95085525&offsets|0|0]|TTCACAGTGGCTAAGTTCTGC</td><td>>MI0000440|hsa-mir-27b&WithFlank&9|+|95085439|95085547</td><td>67</td><td>87</td><td>21</td><td>TTCACAGTGGCTAAGTTCTGC</td><td>34935.5</td><td>6.3</td><td>0.6</td><td>10.0</td><td>1.9E-23</td><td>3.8E-22</td><td>MB231/MB468</td><td>NUC</td></tr>
<tr><td>>MI0000283|hsa-mir-203a&WithFlank&14|+|104117399|104117520@72.91.20(+1A)&[MIMAT0000264&hsa-miR-203a-3p&offsets|+1|-1(+1A);m-79&14|+|104117469|104117490&offsets|+1|-1(+1A)]|TGAAATGTTTAGGACCACTAA</td><td>>MI0000283|hsa-mir-203a&WithFlank&14|+|104117399|104117520</td><td>72</td><td>91</td><td>20(+1A)</td><td>TGAAATGTTTAGGACCACTAA</td><td>711.1</td><td>13.7</td><td>1.4</td><td>9.9</td><td>3.7E-23</td><td>3.8E-22</td><td>BT20/MB468</td><td>CYTO</td></tr>
<tr><td>>hg38_21_+_8433172_8446622_RNA45SN1_45S_50ntFlanks@9203.9220.18|CTCGGGGCCACGCGCGCG</td><td>>hg38_21_+_8433172_8446622_RNA45SN1_45S_50ntFlanks</td><td>9203</td><td>9220</td><td>18</td><td>CTCGGGGCCACGCGCGCG</td><td>1564.6</td><td>-14.5</td><td>1.4</td><td>-10.0</td><td>1.2E-23</td><td>3.8E-22</td><td>MB231/MB468</td><td>MP</td></tr>
<tr><td>>hg38_21_+_8433172_8446622_RNA45SN1_45S_50ntFlanks@11783.11800.18|CATGAATGGATGAACGAG</td><td>>hg38_21_+_8433172_8446622_RNA45SN1_45S_50ntFlanks</td><td>11783</td><td>11800</td><td>18</td><td>CATGAATGGATGAACGAG</td><td>1032.0</td><td>-13.9</td><td>1.4</td><td>-10.0</td><td>1.2E-23</td><td>3.8E-22</td><td>MB231/MB468</td><td>MP</td></tr>
<tr><td>>hg38_21_+_8433172_8446622_RNA45SN1_45S_50ntFlanks@4722.4751.30|TCATTAATCAAGAACGAAAGTCGGAGGTTC</td><td>>hg38_21_+_8433172_8446622_RNA45SN1_45S_50ntFlanks</td><td>4722</td><td>4751</td><td>30</td><td>TCATTAATCAAGAACGAAAGTCGGAGGTTC</td><td>1114.1</td><td>-14.0</td><td>1.4</td><td>-10.0</td><td>1.2E-23</td><td>3.8E-22</td><td>MB231/MB468</td><td>MP</td></tr>
<tr><td>>hg38_21_+_8433172_8446622_RNA45SN1_45S_50ntFlanks@10821.10840.20|GGAAGTCGGCAAGCCGGATC</td><td>>hg38_21_+_8433172_8446622_RNA45SN1_45S_50ntFlanks</td><td>10821</td><td>10840</td><td>20</td><td>GGAAGTCGGCAAGCCGGATC</td><td>1743.6</td><td>-5.2</td><td>0.5</td><td>-10.0</td><td>2.0E-23</td><td>4.0E-22</td><td>MB231/MB468</td><td>NUC</td></tr>
<tr><td>>MI0000108|hsa-mir-103a-2&WithFlank&20|+|3917488|3917577@54.75.22;MI0000109|hsa-mir-103a-1&WithFlank&5|-|168560890|168560979@54.75.22&[MIMAT0000101&hsa-miR-103a-3p&offsets|0|-1;m-12&20|+|3917541|3917563&offsets|0|-1];[MIMAT0000101_1&hsa-miR-103a-3p&offsets|0|-1;m-13&5|-|168560904|168560926&offsets|0|-1;m-44220&5|-|168560908|168560926&offsets|0|+3]|AGCAGCATTGTACAGGGCTATG</td><td>>MI0000108|hsa-mir-103a-2&WithFlank&20|+|3917488|3917577</td><td>54</td><td>75</td><td>22</td><td>AGCAGCATTGTACAGGGCTATG</td><td>7155.5</td><td>7.0</td><td>0.7</td><td>9.9</td><td>3.9E-23</td><td>4.0E-22</td><td>BT20/MB468</td><td>CYTO</td></tr>
<tr><td>>hg38_21_+_8433172_8446622_RNA45SN1_45S_50ntFlanks@12045.12072.28|CTGACCCGGTGAGGCGGGGGGGCGAGCC</td><td>>hg38_21_+_8433172_8446622_RNA45SN1_45S_50ntFlanks</td><td>12045</td><td>12072</td><td>28</td><td>CTGACCCGGTGAGGCGGGGGGGCGAGCC</td><td>547.8</td><td>-13.5</td><td>1.4</td><td>-9.8</td><td>8.3E-23</td><td>4.0E-22</td><td>BT20/MB468</td><td>NUC</td></tr>
<tr><td>>hg38_21_+_8433172_8446622_RNA45SN1_45S_50ntFlanks@12419.12447.29|GATGTCGGCTCTTCCTATCATTGTGAAGC</td><td>>hg38_21_+_8433172_8446622_RNA45SN1_45S_50ntFlanks</td><td>12419</td><td>12447</td><td>29</td><td>GATGTCGGCTCTTCCTATCATTGTGAAGC</td><td>3449.4</td><td>7.2</td><td>0.7</td><td>10.1</td><td>9.1E-24</td><td>4.0E-22</td><td>MB231/MB468</td><td>CYTO</td></tr>
<tr><td>>hg38_21_+_8433172_8446622_RNA45SN1_45S_50ntFlanks@5216.5233.18|CCGGGGCTGCACGCGCGC</td><td>>hg38_21_+_8433172_8446622_RNA45SN1_45S_50ntFlanks</td><td>5216</td><td>5233</td><td>18</td><td>CCGGGGCTGCACGCGCGC</td><td>1327.8</td><td>11.1</td><td>1.1</td><td>10.1</td><td>9.2E-24</td><td>4.0E-22</td><td>MB231/MB468</td><td>CYTO</td></tr>
<tr><td>>MI0006273|hsa-mir-1180&WithFlank&17|-|19344500|19344580@47.68.22(+1U)&[MIMAT0005825&hsa-miR-1180-3p&offsets|0|0(+1U);m-214&17|-|19344513|19344534&offsets|0|0(+1U)]|TTTCCGGCTCGCGTGGGTGTGTT</td><td>>MI0006273|hsa-mir-1180&WithFlank&17|-|19344500|19344580</td><td>47</td><td>68</td><td>22(+1U)</td><td>TTTCCGGCTCGCGTGGGTGTGTT</td><td>305.3</td><td>12.5</td><td>1.3</td><td>9.9</td><td>4.0E-23</td><td>4.0E-22</td><td>BT20/MB468</td><td>CYTO</td></tr>
<tr><td>>MI0000266|hsa-mir-10a&WithFlank&17|-|48579832|48579953@28.50.23&[MIMAT0000253&hsa-miR-10a-5p&offsets|0|0;m-18&17|-|48579905|48579926&offsets|0|+1]|TACCCTGTAGATCCGAATTTGTG</td><td>>MI0000266|hsa-mir-10a&WithFlank&17|-|48579832|48579953</td><td>28</td><td>50</td><td>23</td><td>TACCCTGTAGATCCGAATTTGTG</td><td>3381.2</td><td>14.9</td><td>1.5</td><td>10.0</td><td>2.0E-23</td><td>4.1E-22</td><td>MB231/MB468</td><td>MITO</td></tr>
<tr><td>>hg38_21_+_8433172_8446622_RNA45SN1_45S_50ntFlanks@11321.11344.24|CACGTCTCGTCGCGCGCGCGTCCG</td><td>>hg38_21_+_8433172_8446622_RNA45SN1_45S_50ntFlanks</td><td>11321</td><td>11344</td><td>24</td><td>CACGTCTCGTCGCGCGCGCGTCCG</td><td>1837.0</td><td>-14.7</td><td>1.5</td><td>-10.0</td><td>1.3E-23</td><td>4.1E-22</td><td>MB231/MB468</td><td>MP</td></tr>
<tr><td>>hg38_21_+_8433172_8446622_RNA45SN1_45S_50ntFlanks@9565.9596.32|TGACGTGCAAATCGGTCGTCCGACCTGGGTAT</td><td>>hg38_21_+_8433172_8446622_RNA45SN1_45S_50ntFlanks</td><td>9565</td><td>9596</td><td>32</td><td>TGACGTGCAAATCGGTCGTCCGACCTGGGTAT</td><td>5782.7</td><td>-6.1</td><td>0.6</td><td>-9.9</td><td>4.0E-23</td><td>4.1E-22</td><td>BT20/MB468</td><td>CYTO</td></tr>
<tr><td>>MI0000298|hsa-mir-221&WithFlank&X|-|45746151|45746272@31.55.25(+1C)&[MIMAT0004568&hsa-miR-221-5p&offsets|0|+3(+1C);m-405&X|-|45746220|45746242&offsets|0|+2(+1C)]|ACCTGGCATACAATGTAGATTTCTGC</td><td>>MI0000298|hsa-mir-221&WithFlank&X|-|45746151|45746272</td><td>31</td><td>55</td><td>25(+1C)</td><td>ACCTGGCATACAATGTAGATTTCTGC</td><td>204.6</td><td>5.8</td><td>0.6</td><td>10.0</td><td>2.1E-23</td><td>4.1E-22</td><td>MB231/MB468</td><td>NUC</td></tr>
<tr><td>>hg38_21_+_8433172_8446622_RNA45SN1_45S_50ntFlanks@11704.11728.25|ATGAAGCGCGGGTAAACGGCGGGAG</td><td>>hg38_21_+_8433172_8446622_RNA45SN1_45S_50ntFlanks</td><td>11704</td><td>11728</td><td>25</td><td>ATGAAGCGCGGGTAAACGGCGGGAG</td><td>1722.9</td><td>-14.6</td><td>1.5</td><td>-10.0</td><td>1.3E-23</td><td>4.1E-22</td><td>MB231/MB468</td><td>MP</td></tr>
<tr><td>>hg38_21_+_8433172_8446622_RNA45SN1_45S_50ntFlanks@9459.9503.45|CCGGGGAGGTGGAGCACGAGCGCACGTGTTAGGACCCGAAAGATG</td><td>>hg38_21_+_8433172_8446622_RNA45SN1_45S_50ntFlanks</td><td>9459</td><td>9503</td><td>45</td><td>CCGGGGAGGTGGAGCACGAGCGCACGTGTTAGGACCCGAAAGATG</td><td>2860.5</td><td>-15.3</td><td>1.5</td><td>-10.0</td><td>2.1E-23</td><td>4.2E-22</td><td>MB231/MB468</td><td>MITO</td></tr>
<tr><td>>MI0000440|hsa-mir-27b&WithFlank&9|+|95085439|95085547@67.85.19&[MIMAT0000419&hsa-miR-27b-3p&offsets|0|-2;m-39&9|+|95085505|95085525&offsets|0|-2]|TTCACAGTGGCTAAGTTCT</td><td>>MI0000440|hsa-mir-27b&WithFlank&9|+|95085439|95085547</td><td>67</td><td>85</td><td>19</td><td>TTCACAGTGGCTAAGTTCT</td><td>1307.1</td><td>6.1</td><td>0.6</td><td>9.9</td><td>3.1E-23</td><td>4.2E-22</td><td>BT20/MB468</td><td>MITO</td></tr>
<tr><td>>MI0003628|hsa-mir-615&WithFlank&12|+|54033944|54034051@67.87.21(+1A)&[MIMAT0003283&hsa-miR-615-3p&offsets|0|-1(+1A);m-495&12|+|54034010|54034030&offsets|0|0(+1A)]|TCCGAGCCTGGGTCTCCCTCTA</td><td>>MI0003628|hsa-mir-615&WithFlank&12|+|54033944|54034051</td><td>67</td><td>87</td><td>21(+1A)</td><td>TCCGAGCCTGGGTCTCCCTCTA</td><td>333.5</td><td>12.6</td><td>1.3</td><td>9.9</td><td>4.2E-23</td><td>4.2E-22</td><td>BT20/MB468</td><td>CYTO</td></tr>
<tr><td>>MI0000061|hsa-let-7a-2&WithFlank&11|-|122146516|122146599@11.31.21(+1A);MI0000062|hsa-let-7a-3&WithFlank&22|+|46112743|46112828@10.30.21(+1A);MI0000060|hsa-let-7a-1&WithFlank&9|+|94175951|94176042@12.32.21(+1A)&[MIMAT0000062&hsa-let-7a-5p&offsets|0|-1(+1A);m-6&11|-|122146568|122146589&offsets|0|-1(+1A)];[MIMAT0000062_1&hsa-let-7a-5p&offsets|0|-1(+1A);m-5&22|+|46112752|46112773&offsets|0|-1(+1A)];[MIMAT0000062_2&hsa-let-7a-5p&offsets|0|-1(+1A);m-4&9|+|94175962|94175983&offsets|0|-1(+1A)]|TGAGGTAGTAGGTTGTATAGTA</td><td>>MI0000061|hsa-let-7a-2&WithFlank&11|-|122146516|122146599</td><td>11</td><td>31</td><td>21(+1A)</td><td>TGAGGTAGTAGGTTGTATAGTA</td><td>1325.5</td><td>6.9</td><td>0.7</td><td>9.9</td><td>3.2E-23</td><td>4.3E-22</td><td>BT20/MB468</td><td>MITO</td></tr>
<tr><td>>hg38_21_+_8433172_8446622_RNA45SN1_45S_50ntFlanks@10411.10433.23|CAGTCGGTCCTGAGAGATGGGCG</td><td>>hg38_21_+_8433172_8446622_RNA45SN1_45S_50ntFlanks</td><td>10411</td><td>10433</td><td>23</td><td>CAGTCGGTCCTGAGAGATGGGCG</td><td>560.4</td><td>-13.6</td><td>1.4</td><td>-9.8</td><td>8.8E-23</td><td>4.3E-22</td><td>BT20/MB468</td><td>NUC</td></tr>
<tr><td>>MI0005756|hsa-mir-934&WithFlank&X|+|136550872|136550966@21.42.22(+1A)&[MIMAT0004977&hsa-miR-934&offsets|0|0(+1A);m-943&X|+|136550892|136550913&offsets|0|0(+1A)]|TGTCTACTACTGGAGACACTGGA</td><td>>MI0005756|hsa-mir-934&WithFlank&X|+|136550872|136550966</td><td>21</td><td>42</td><td>22(+1A)</td><td>TGTCTACTACTGGAGACACTGGA</td><td>1747.2</td><td>14.5</td><td>1.4</td><td>10.2</td><td>2.7E-24</td><td>4.3E-22</td><td>BT20/MB231</td><td>NUC</td></tr>
<tr><td>>hg38_21_+_8433172_8446622_RNA45SN1_45S_50ntFlanks@5474.5516.43|GGCCCTGGCGGAGCGCTGAGAAGACGGTCGAACTTGACTATCT</td><td>>hg38_21_+_8433172_8446622_RNA45SN1_45S_50ntFlanks</td><td>5474</td><td>5516</td><td>43</td><td>GGCCCTGGCGGAGCGCTGAGAAGACGGTCGAACTTGACTATCT</td><td>273.7</td><td>12.1</td><td>1.2</td><td>10.0</td><td>9.9E-24</td><td>4.3E-22</td><td>MB231/MB468</td><td>CYTO</td></tr>
<tr><td>>MI0000067|hsa-let-7f-1&WithFlank&9|+|94176341|94176439@13.34.22(+1A);MI0000068|hsa-let-7f-2&WithFlank&X|-|53557186|53557280@14.35.22(+1A)&[MIMAT0000067&hsa-let-7f-5p&offsets|0|0(+1A);m-20&9|+|94176353|94176374&offsets|0|0(+1A)];[MIMAT0000067_1&hsa-let-7f-5p&offsets|0|0(+1A);m-19&X|-|53557246|53557267&offsets|0|0(+1A)]|TGAGGTAGTAGATTGTATAGTTA</td><td>>MI0000067|hsa-let-7f-1&WithFlank&9|+|94176341|94176439</td><td>13</td><td>34</td><td>22(+1A)</td><td>TGAGGTAGTAGATTGTATAGTTA</td><td>1476.7</td><td>7.1</td><td>0.7</td><td>9.9</td><td>3.2E-23</td><td>4.4E-22</td><td>BT20/MB468</td><td>MITO</td></tr>
<tr><td>>hg38_21_+_8433172_8446622_RNA45SN1_45S_50ntFlanks@7973.8011.39|GACGCGACCTCAGATCAGACGTGGCGACCCGCTGAATTT</td><td>>hg38_21_+_8433172_8446622_RNA45SN1_45S_50ntFlanks</td><td>7973</td><td>8011</td><td>39</td><td>GACGCGACCTCAGATCAGACGTGGCGACCCGCTGAATTT</td><td>6655.6</td><td>5.7</td><td>0.6</td><td>9.9</td><td>3.2E-23</td><td>4.4E-22</td><td>BT20/MB468</td><td>MITO</td></tr>
<tr><td>>hg38_21_+_8433172_8446622_RNA45SN1_45S_50ntFlanks@10389.10407.19|TGAACAGCAGTTGAACATG</td><td>>hg38_21_+_8433172_8446622_RNA45SN1_45S_50ntFlanks</td><td>10389</td><td>10407</td><td>19</td><td>TGAACAGCAGTTGAACATG</td><td>1934.3</td><td>-6.0</td><td>0.6</td><td>-10.0</td><td>2.2E-23</td><td>4.4E-22</td><td>MB231/MB468</td><td>NUC</td></tr>
<tr><td>>MI0000650|hsa-mir-200c&WithFlank&12|+|6963693|6963772@50.69.20&[MIMAT0000617&hsa-miR-200c-3p&offsets|0|-3;m-54&12|+|6963742|6963764&offsets|0|-3]|TAATACTGCCGGGTAATGAT</td><td>>MI0000650|hsa-mir-200c&WithFlank&12|+|6963693|6963772</td><td>50</td><td>69</td><td>20</td><td>TAATACTGCCGGGTAATGAT</td><td>2797.8</td><td>10.1</td><td>1.0</td><td>9.9</td><td>3.3E-23</td><td>4.4E-22</td><td>BT20/MB468</td><td>MITO</td></tr>
<tr><td>>MI0000102|hsa-mir-100&WithFlank&11|-|122152223|122152314@19.38.20(+1G)&[MIMAT0000098&hsa-miR-100-5p&offsets|0|-2(+1G);m-27&11|-|122152276|122152296&offsets|0|-1(+1G)]|AACCCGTAGATCCGAACTTGG</td><td>>MI0000102|hsa-mir-100&WithFlank&11|-|122152223|122152314</td><td>19</td><td>38</td><td>20(+1G)</td><td>AACCCGTAGATCCGAACTTGG</td><td>2302.3</td><td>14.4</td><td>1.4</td><td>10.0</td><td>1.4E-23</td><td>4.4E-22</td><td>MB231/MB468</td><td>MP</td></tr>
<tr><td>>hg38_21_+_8433172_8446622_RNA45SN1_45S_50ntFlanks@8380.8404.25|CAAGAGGGCGTGAAACCGTTAAGAG</td><td>>hg38_21_+_8433172_8446622_RNA45SN1_45S_50ntFlanks</td><td>8380</td><td>8404</td><td>25</td><td>CAAGAGGGCGTGAAACCGTTAAGAG</td><td>952.7</td><td>-13.8</td><td>1.4</td><td>-10.0</td><td>1.4E-23</td><td>4.5E-22</td><td>MB231/MB468</td><td>MP</td></tr>
</table>

| | | | | | | | | | | | | | |
|---|---|---|---|---|---|---|---|---|---|---|---|---|---|
| >MI0000108\|hsa-mir-103a-2&WithFlank&20\|+\|3917488\|3917577@54.75.22(+1U);MI0000109\|hsa-mir-103a-1&WithFlank&5\|-\|168560890\|168560979@54.75.22(+1U)&[MIMAT0000101&hsa-miR-103a-3p&offsets\|0\|-1(+1U);m-12&20\|+\|3917541\|3917563&offsets\|0\|-1(+1U)];[MIMAT0000101_1&hsa-miR-103a-3p&offsets\|0\|-1(+1U);m-13&5\|-\|168560904\|168560926&offsets\|0\|-1(+1U);m-44220&5\|-\|168560908\|168560926&offsets\|0\|+3(+1U)]\|AGCAGCATTGTACAGGGCTATGT | >MI0000108\|hsa-mir-103a-2&WithFlank&20\|+\|3917488\|3917577 | 54 | 75 | 22(+1U) | AGCAGCATTGTACAGGGCTATGT | 1537.0 | 7.4 | 0.8 | 9.9 | 4.5E-23 | 4.5E-22 | BT20/MB468 | CYTO |
| >hg38_21_+_8433172_8446622_RNA45SN1_45S_50ntFlanks@3985.4024.40\|CGGCTTTGGTGACTCTAGATAACCTCGGGCCGATCGCACG | >hg38_21_+_8433172_8446622_RNA45SN1_45S_50ntFlanks | 3985 | 4024 | 40 | CGGCTTTGGTGACTCTAGATAACCTCGGGCCGATCGCACG | 4771.2 | -16.1 | 1.6 | -10.0 | 1.4E-23 | 4.5E-22 | MB231/MB468 | MP |
| >hg38_21_+_8433172_8446622_RNA45SN1_45S_50ntFlanks@12405.12434.30\|CTTTTTGATCCTTCGATGTCGGCTCTTCCT | >hg38_21_+_8433172_8446622_RNA45SN1_45S_50ntFlanks | 12405 | 12434 | 30 | CTTTTTGATCCTTCGATGTCGGCTCTTCCT | 6872.5 | -4.7 | 0.5 | -9.9 | 3.4E-23 | 4.5E-22 | BT20/MB468 | MITO |
| >hg38_1_-_228634819_228635039_RNA5S12_5S_50ntFlanks@146.170.25\|TGGGAATACCGGGTGCTGTAGGCTT | >hg38_1_-_228634819_228635039_RNA5S12_5S_50ntFlanks | 146 | 170 | 25 | TGGGAATACCGGGTGCTGTAGGCTT | 5672.2 | 7.1 | 0.7 | 10.0 | 1.0E-23 | 4.5E-22 | MB231/MB468 | CYTO |
| >hg38_21_+_8433172_8446622_RNA45SN1_45S_50ntFlanks@4734.4767.34\|AACGAAAGTCGGAGGTTCGAAGACGATCAGATAC | >hg38_21_+_8433172_8446622_RNA45SN1_45S_50ntFlanks | 4734 | 4767 | 34 | AACGAAAGTCGGAGGTTCGAAGACGATCAGATAC | 1170.0 | 12.1 | 1.2 | 9.9 | 4.6E-23 | 4.6E-22 | BT20/MB468 | CYTO |
| >hg38_21_+_8433172_8446622_RNA45SN1_45S_50ntFlanks@10681.10706.26\|TTCCGGCGGCGTCCGGTGAGCTCTCG | >hg38_21_+_8433172_8446622_RNA45SN1_45S_50ntFlanks | 10681 | 10706 | 26 | TTCCGGCGGCGTCCGGTGAGCTCTCG | 1467.1 | -14.4 | 1.4 | -10.0 | 1.5E-23 | 4.6E-22 | MB231/MB468 | MP |
| >hg38_21_+_8433172_8446622_RNA45SN1_45S_50ntFlanks@10128.10147.20\|CGGCGGCGGCGGCGGCGGCG | >hg38_21_+_8433172_8446622_RNA45SN1_45S_50ntFlanks | 10128 | 10147 | 20 | CGGCGGCGGCGGCGGCGGCG | 1868.4 | -14.7 | 1.5 | -10.0 | 1.5E-23 | 4.6E-22 | MB231/MB468 | MP |
| >hg38_21_+_8433172_8446622_RNA45SN1_45S_50ntFlanks@8845.8864.20\|CGGCGGCGGCGGCGGCGGCG | >hg38_21_+_8433172_8446622_RNA45SN1_45S_50ntFlanks | 8845 | 8864 | 20 | CGGCGGCGGCGGCGGCGGCG | 1868.4 | -14.7 | 1.5 | -10.0 | 1.5E-23 | 4.6E-22 | MB231/MB468 | MP |
| >hg38_21_+_8433172_8446622_RNA45SN1_45S_50ntFlanks@10408.10435.28\|GGTCAGTCGGTCCTGAGAGATGGGCGAG | >hg38_21_+_8433172_8446622_RNA45SN1_45S_50ntFlanks | 10408 | 10435 | 28 | GGTCAGTCGGTCCTGAGAGATGGGCGAG | 2156.8 | -14.9 | 1.5 | -10.0 | 1.5E-23 | 4.6E-22 | MB231/MB468 | MP |
| >hg38_21_+_8433172_8446622_RNA45SN1_45S_50ntFlanks@4657.4688.32\|CTAGAGGTGAAATTCTTGGACCGGCGCAAGAC | >hg38_21_+_8433172_8446622_RNA45SN1_45S_50ntFlanks | 4657 | 4688 | 32 | CTAGAGGTGAAATTCTTGGACCGGCGCAAGAC | 1121.2 | -14.6 | 1.5 | -9.8 | 9.6E-23 | 4.7E-22 | BT20/MB468 | NUC |
| >hg38_21_+_8433172_8446622_RNA45SN1_45S_50ntFlanks@10574.10612.39\|AAGCCGGCGGGAGCCCCGGGGAGAGTTCTCTTTTCTTTG | >hg38_21_+_8433172_8446622_RNA45SN1_45S_50ntFlanks | 10574 | 10612 | 39 | AAGCCGGCGGGAGCCCCGGGGAGAGTTCTCTTTTCTTTG | 548.1 | -13.5 | 1.4 | -9.8 | 9.7E-23 | 4.7E-22 | BT20/MB468 | NUC |
| >MI0000079\|hsa-mir-23a&WithFlank&19\|-\|13836581\|13836665@54.73.20&[MIMAT0000078&hsa-miR-23a-3p&offsets\|+3\|+2;m-31&19\|-\|13836595\|13836615&offsets\|+3\|+2]\|ACATTGCCAGGGATTTCCAA | >MI0000079\|hsa-mir-23a&WithFlank&19\|-\|13836581\|13836665 | 54 | 73 | 20 | ACATTGCCAGGGATTTCCAA | 658.4 | 13.8 | 1.4 | 9.9 | 3.5E-23 | 4.7E-22 | BT20/MB468 | MITO |
| >hg38_21_+_8433172_8446622_RNA45SN1_45S_50ntFlanks@6654.6681.28\|CTCTTAGCGGTGGATCACTCGGCTCGTG | >hg38_21_+_8433172_8446622_RNA45SN1_45S_50ntFlanks | 6654 | 6681 | 28 | CTCTTAGCGGTGGATCACTCGGCTCGTG | 2173.3 | -7.0 | 0.7 | -10.0 | 1.5E-23 | 4.7E-22 | MB231/MB468 | MP |
| >hg38_21_+_8433172_8446622_RNA45SN1_45S_50ntFlanks@9306.9335.30\|CAAGGAGTCTAACACGTGCGCGAGTCGGGG | >hg38_21_+_8433172_8446622_RNA45SN1_45S_50ntFlanks | 9306 | 9335 | 30 | CAAGGAGTCTAACACGTGCGCGAGTCGGGG | 1218.6 | -14.1 | 1.4 | -10.0 | 1.5E-23 | 4.7E-22 | MB231/MB468 | MP |
| >hg38_21_+_8433172_8446622_RNA45SN1_45S_50ntFlanks@12990.13019.30\|GCTCCCTCGCTGCGATCTATTGAAAGTCAG | >hg38_21_+_8433172_8446622_RNA45SN1_45S_50ntFlanks | 12990 | 13019 | 30 | GCTCCCTCGCTGCGATCTATTGAAAGTCAG | 318.5 | -12.7 | 1.3 | -9.8 | 9.8E-23 | 4.8E-22 | BT20/MB468 | NUC |
| >hg38_21_+_8433172_8446622_RNA45SN1_45S_50ntFlanks@9567.9586.20\|ACGTGCAAATCGGTCGTCCG | >hg38_21_+_8433172_8446622_RNA45SN1_45S_50ntFlanks | 9567 | 9586 | 20 | ACGTGCAAATCGGTCGTCCG | 412.1 | -13.1 | 1.3 | -9.8 | 9.9E-23 | 4.8E-22 | BT20/MB468 | NUC |
| >hg38_21_+_8433172_8446622_RNA45SN1_45S_50ntFlanks@8168.8203.36\|CCCAAGTCCTTCTGATCGAGGCCCAGCCCGTGGACG | >hg38_21_+_8433172_8446622_RNA45SN1_45S_50ntFlanks | 8168 | 8203 | 36 | CCCAAGTCCTTCTGATCGAGGCCCAGCCCGTGGACG | 451.1 | -13.2 | 1.3 | -9.8 | 9.9E-23 | 4.8E-22 | BT20/MB468 | NUC |
| >am_MI0000786\|hsa-mir-378a&WithFlank&5\|+\|149732819\|149732896@49.70.22(+1U)&[MIMAT0000732&hsa-miR-378a-3p&offsets\|0\|0(+1U);m-44&5\|+\|149732867\|149732888&offsets\|0\|0(+1U)]\|ACTGGACTTGGAGTCAGAAGGCT | >am_MI0000786\|hsa-mir-378a&WithFlank&5\|+\|149732819\|149732896 | 49 | 70 | 22(+1U) | ACTGGACTTGGAGTCAGAAGGCT | 2869.6 | 5.2 | 0.5 | 9.9 | 4.7E-23 | 4.8E-22 | BT20/MB468 | CYTO |
| >hg38_21_+_8433172_8446622_RNA45SN1_45S_50ntFlanks@11580.11605.26\|GAACTGGTGCGGACCAGGGGAATCCG | >hg38_21_+_8433172_8446622_RNA45SN1_45S_50ntFlanks | 11580 | 11605 | 26 | GAACTGGTGCGGACCAGGGGAATCCG | 2107.3 | -5.4 | 0.5 | -10.0 | 1.1E-23 | 4.8E-22 | MB231/MB468 | CYTO |
| >hg38_21_+_8433172_8446622_RNA45SN1_45S_50ntFlanks@7985.8004.20\|GATCAGACGTGGCGACCCGC | >hg38_21_+_8433172_8446622_RNA45SN1_45S_50ntFlanks | 7985 | 8004 | 20 | GATCAGACGTGGCGACCCGC | 361.2 | -12.9 | 1.3 | -9.8 | 1.0E-22 | 4.8E-22 | BT20/MB468 | NUC |
| >hg38_21_+_8433172_8446622_RNA45SN1_45S_50ntFlanks@12405.12429.25\|CTTTTTGATCCTTCGATGTCGGCTC | >hg38_21_+_8433172_8446622_RNA45SN1_45S_50ntFlanks | 12405 | 12429 | 25 | CTTTTTGATCCTTCGATGTCGGCTC | 1599.0 | -14.5 | 1.5 | -10.0 | 1.5E-23 | 4.8E-22 | MB231/MB468 | MP |
| >hg38_21_+_8433172_8446622_RNA45SN1_45S_50ntFlanks@12366.12383.18\|TAACTGGCTTGTGGCGGC | >hg38_21_+_8433172_8446622_RNA45SN1_45S_50ntFlanks | 12366 | 12383 | 18 | TAACTGGCTTGTGGCGGC | 2382.0 | -8.9 | 0.9 | -10.0 | 1.1E-23 | 4.8E-22 | MB231/MB468 | CYTO |
| >hg38_21_+_8433172_8446622_RNA45SN1_45S_50ntFlanks@3775.3792.18\|GCCGGTACAGTGAAACTG | >hg38_21_+_8433172_8446622_RNA45SN1_45S_50ntFlanks | 3775 | 3792 | 18 | GCCGGTACAGTGAAACTG | 3473.5 | -15.6 | 1.6 | -10.0 | 1.6E-23 | 4.8E-22 | MB231/MB468 | MP |
| >hg38_21_+_8433172_8446622_RNA45SN1_45S_50ntFlanks@9027.9058.32\|CGTCTCCTCGTGGGGGGGCCGGGCCACCCCTC | >hg38_21_+_8433172_8446622_RNA45SN1_45S_50ntFlanks | 9027 | 9058 | 32 | CGTCTCCTCGTGGGGGGGCCGGGCCACCCCTC | 1445.4 | -14.3 | 1.4 | -10.0 | 2.4E-23 | 4.8E-22 | MB231/MB468 | MITO |
| >hg38_21_+_8433172_8446622_RNA45SN1_45S_50ntFlanks@12570.12587.18\|AGTACGAGAGGAACCGCA | >hg38_21_+_8433172_8446622_RNA45SN1_45S_50ntFlanks | 12570 | 12587 | 18 | AGTACGAGAGGAACCGCA | 1320.8 | -14.2 | 1.4 | -10.0 | 2.5E-23 | 4.9E-22 | MB231/MB468 | MITO |
| >hg38_21_+_8433172_8446622_RNA45SN1_45S_50ntFlanks@3985.4024.40\|CGGCTTTGGTGACTCTAGATAACCTCGGGCCGATCGCACG | >hg38_21_+_8433172_8446622_RNA45SN1_45S_50ntFlanks | 3985 | 4024 | 40 | CGGCTTTGGTGACTCTAGATAACCTCGGGCCGATCGCACG | 2909.4 | -15.3 | 1.5 | -10.0 | 2.5E-23 | 4.9E-22 | MB231/MB468 | MITO |
| >hg38_21_+_8433172_8446622_RNA45SN1_45S_50ntFlanks@11783.11806.24\|CATGAATGGATGAACGAGATTCCC | >hg38_21_+_8433172_8446622_RNA45SN1_45S_50ntFlanks | 11783 | 11806 | 24 | CATGAATGGATGAACGAGATTCCC | 1564.3 | -5.5 | 0.6 | -9.9 | 2.6E-23 | 4.9E-22 | MB231/MB468 | NUC |
| >hg38_21_+_8433172_8446622_RNA45SN1_45S_50ntFlanks@12709.12742.34\|CCGCGGAGCCTCGGTTGGCCTCGGATAGCCGGTC | >hg38_21_+_8433172_8446622_RNA45SN1_45S_50ntFlanks | 12709 | 12742 | 34 | CCGCGGAGCCTCGGTTGGCCTCGGATAGCCGGTC | 1425.3 | -14.3 | 1.4 | -10.0 | 2.5E-23 | 4.9E-22 | MB231/MB468 | MITO |
| >hg38_21_+_8433172_8446622_RNA45SN1_45S_50ntFlanks@4723.4752.30\|CATTAATCAAGAACGAAAGTCGGAGGTTCG | >hg38_21_+_8433172_8446622_RNA45SN1_45S_50ntFlanks | 4723 | 4752 | 30 | CATTAATCAAGAACGAAAGTCGGAGGTTCG | 2441.4 | -15.0 | 1.5 | -10.0 | 2.5E-23 | 4.9E-22 | MB231/MB468 | MITO |
| >hg38_21_+_8433172_8446622_RNA45SN1_45S_50ntFlanks@10388.10424.37\|GTGAACAGCAGTTGAACATGGGTCAGTCGGTCCTGAG | >hg38_21_+_8433172_8446622_RNA45SN1_45S_50ntFlanks | 10388 | 10424 | 37 | GTGAACAGCAGTTGAACATGGGTCAGTCGGTCCTGAG | 2532.1 | 5.2 | 0.5 | 9.9 | 4.9E-23 | 5.0E-22 | BT20/MB468 | CYTO |
| >MI0000433\|hsa-let-7g&WithFlank&3\|-\|52268272\|52268367@11.31.21&[MIMAT0000414&hsa-let-7g-5p&offsets\|0\|-1;m-58&3\|-\|52268336\|52268357&offsets\|0\|-1]\|TGAGGTAGTAGTTTGTACAGT | >MI0000433\|hsa-let-7g&WithFlank&3\|-\|52268272\|52268367 | 11 | 31 | 21 | TGAGGTAGTAGTTTGTACAGT | 7091.7 | 7.4 | 0.7 | 9.9 | 4.9E-23 | 5.0E-22 | BT20/MB468 | CYTO |
| >hg38_21_+_8433172_8446622_RNA45SN1_45S_50ntFlanks@10678.10708.31\|CGGTTCCGGCGGCGTCCGGTGAGCTCTCGCT | >hg38_21_+_8433172_8446622_RNA45SN1_45S_50ntFlanks | 10678 | 10708 | 31 | CGGTTCCGGCGGCGTCCGGTGAGCTCTCGCT | 10752.9 | -8.8 | 0.9 | -10.0 | 1.6E-23 | 5.0E-22 | MB231/MB468 | MP |
| >hg38_21_+_8433172_8446622_RNA45SN1_45S_50ntFlanks@6748.6771.24\|CTTCGAACGCACTTGCGGCCCCGG | >hg38_21_+_8433172_8446622_RNA45SN1_45S_50ntFlanks | 6748 | 6771 | 24 | CTTCGAACGCACTTGCGGCCCCGG | 884.9 | -13.7 | 1.4 | -10.0 | 1.6E-23 | 5.0E-22 | MB231/MB468 | MP |
| >hg38_21_+_8433172_8446622_RNA45SN1_45S_50ntFlanks@12476.12498.23\|CCACTAATAGGGAACGTGAGCTG | >hg38_21_+_8433172_8446622_RNA45SN1_45S_50ntFlanks | 12476 | 12498 | 23 | CCACTAATAGGGAACGTGAGCTG | 2365.1 | -9.3 | 0.9 | -9.9 | 2.6E-23 | 5.0E-22 | MB231/MB468 | NUC |
| >hg38_21_+_8433172_8446622_RNA45SN1_45S_50ntFlanks@4283.4317.35\|CTTTAACGAGGATCCATTGGAGGGCAAGTCTGGTG | >hg38_21_+_8433172_8446622_RNA45SN1_45S_50ntFlanks | 4283 | 4317 | 35 | CTTTAACGAGGATCCATTGGAGGGCAAGTCTGGTG | 5718.1 | -12.3 | 1.2 | -9.9 | 2.6E-23 | 5.0E-22 | MB231/MB468 | NUC |
| >hg38_21_+_8433172_8446622_RNA45SN1_45S_50ntFlanks@3984.4024.41\|GCGGCTTTGGTGACTCTAGATAACCTCGGGCCGATCGCACG | >hg38_21_+_8433172_8446622_RNA45SN1_45S_50ntFlanks | 3984 | 4024 | 41 | GCGGCTTTGGTGACTCTAGATAACCTCGGGCCGATCGCACG | 3064.9 | -15.4 | 1.5 | -9.9 | 2.5E-23 | 5.0E-22 | MB231/MB468 | MITO |
| >hg38_21_+_8433172_8446622_RNA45SN1_45S_50ntFlanks@11668.11708.41\|TGCCCAGTGCTCTGAATGTCAAAGTGAAGAAATTCAATGAA | >hg38_21_+_8433172_8446622_RNA45SN1_45S_50ntFlanks | 11668 | 11708 | 41 | TGCCCAGTGCTCTGAATGTCAAAGTGAAGAAATTCAATGAA | 364.2 | 12.5 | 1.2 | 10.0 | 1.2E-23 | 5.1E-22 | MB231/MB468 | CYTO |
| >hg38_21_+_8433172_8446622_RNA45SN1_45S_50ntFlanks@12435.12455.21\|ATCATTGTGAAGCAGAATTCA | >hg38_21_+_8433172_8446622_RNA45SN1_45S_50ntFlanks | 12435 | 12455 | 21 | ATCATTGTGAAGCAGAATTCA | 1116.3 | -13.1 | 1.3 | -9.9 | 2.7E-23 | 5.1E-22 | MB231/MB468 | NUC |
| >hg38_21_+_8433172_8446622_RNA45SN1_45S_50ntFlanks@12324.12346.23\|TTGGGTTTTAAGCAGGAGGTGTC | >hg38_21_+_8433172_8446622_RNA45SN1_45S_50ntFlanks | 12324 | 12346 | 23 | TTGGGTTTTAAGCAGGAGGTGTC | 1986.9 | -14.7 | 1.5 | -9.9 | 2.6E-23 | 5.1E-22 | MB231/MB468 | MITO |
| >hg38_21_+_8433172_8446622_RNA45SN1_45S_50ntFlanks@6742.6770.29\|TCGACACTTCGAACGCACTTGCGGCCCCG | >hg38_21_+_8433172_8446622_RNA45SN1_45S_50ntFlanks | 6742 | 6770 | 29 | TCGACACTTCGAACGCACTTGCGGCCCCG | 753.0 | -13.4 | 1.3 | -10.0 | 1.7E-23 | 5.2E-22 | MB231/MB468 | MP |
| >MI0000285\|hsa-mir-205&WithFlank&1\|+\|209432127\|209432248@40.61.22(+2A)&[MIMAT0000266&hsa-miR-205-5p&offsets\|0\|0(+2A);m-351&1\|+\|209432166\|209432187&offsets\|0\|0(+2A)]\|TCCTTCATTCCACCGGAGTCTGAA | >MI0000285\|hsa-mir-205&WithFlank&1\|+\|209432127\|209432248 | 40 | 61 | 22(+2A) | TCCTTCATTCCACCGGAGTCTGAA | 3103.8 | 12.3 | 1.2 | 10.2 | 1.6E-24 | 5.2E-22 | BT20/MB231 | CYTO |
| >hg38_21_+_8433172_8446622_RNA45SN1_45S_50ntFlanks@12151.12182.32\|GTGGGGAGTTTGACTGGGGCGGTACACCTGTC | >hg38_21_+_8433172_8446622_RNA45SN1_45S_50ntFlanks | 12151 | 12182 | 32 | GTGGGGAGTTTGACTGGGGCGGTACACCTGTC | 1152.2 | -14.0 | 1.4 | -10.0 | 1.7E-23 | 5.2E-22 | MB231/MB468 | MP |
| >hg38_21_+_8433172_8446622_RNA45SN1_45S_50ntFlanks@10873.10892.20\|CTGGGTCGGTCGGGCTGGGG | >hg38_21_+_8433172_8446622_RNA45SN1_45S_50ntFlanks | 10873 | 10892 | 20 | CTGGGTCGGTCGGGCTGGGG | 1620.8 | -14.5 | 1.5 | -10.0 | 1.7E-23 | 5.2E-22 | MB231/MB468 | MP |
| >MI0000542\|hsa-mir-320a&WithFlank&8\|-\|22244960\|22245043@48.70.23(+1G)&[MIMAT0000510&hsa-miR-320a-3p&offsets\|0\|+1(+1G);m-60&8\|-\|22244975\|22244996&offsets\|0\|+1(+1G)]\|AAAAGCTGGGTTGAGAGGGCGAAG | >MI0000542\|hsa-mir-320a&WithFlank&8\|-\|22244960\|22245043 | 48 | 70 | 23(+1G) | AAAAGCTGGGTTGAGAGGGCGAAG | 402.1 | 13.1 | 1.3 | 9.9 | 3.9E-23 | 5.2E-22 | BT20/MB468 | MITO |
| >hg38_21_+_8433172_8446622_RNA45SN1_45S_50ntFlanks@9027.9064.38\|CGTCTCCTCGTGGGGGGGCCGGGCCACCCCTCCCACGG | >hg38_21_+_8433172_8446622_RNA45SN1_45S_50ntFlanks | 9027 | 9064 | 38 | CGTCTCCTCGTGGGGGGGCCGGGCCACCCCTCCCACGG | 2798.5 | -15.2 | 1.5 | -9.9 | 2.7E-23 | 5.2E-22 | MB231/MB468 | MITO |
| >hg38_21_+_8433172_8446622_RNA45SN1_45S_50ntFlanks@12463.12496.34\|TTGGATTGTTCACCCACTAATAGGGAACGTGAGC | >hg38_21_+_8433172_8446622_RNA45SN1_45S_50ntFlanks | 12463 | 12496 | 34 | TTGGATTGTTCACCCACTAATAGGGAACGTGAGC | 544.9 | 5.3 | 0.5 | 9.9 | 5.2E-23 | 5.2E-22 | BT20/MB468 | CYTO |
| >hg38_21_+_8433172_8446622_RNA45SN1_45S_50ntFlanks@12004.12035.32\|CGGGCCGCCGGTGAAATACCACTACTCTGATC | >hg38_21_+_8433172_8446622_RNA45SN1_45S_50ntFlanks | 12004 | 12035 | 32 | CGGGCCGCCGGTGAAATACCACTACTCTGATC | 797.1 | 7.7 | 0.8 | 9.9 | 5.2E-23 | 5.2E-22 | BT20/MB468 | CYTO |
| >MI0000456\|hsa-mir-140&WithFlank&16\|+\|69933075\|69933186@69.88.20&[MIMAT0004597&hsa-miR-140-3p&offsets\|+1\|0;m-45&16\|+\|69933143\|69933164&offsets\|0\|-2]\|ACCACAGGGTAGAACCACGG | >MI0000456\|hsa-mir-140&WithFlank&16\|+\|69933075\|69933186 | 69 | 88 | 20 | ACCACAGGGTAGAACCACGG | 364.3 | 12.8 | 1.3 | 9.9 | 5.2E-23 | 5.3E-22 | BT20/MB468 | CYTO |
| >hg38_21_+_8433172_8446622_RNA45SN1_45S_50ntFlanks@12637.12654.18\|GCTACCATCTGTGGGATT | >hg38_21_+_8433172_8446622_RNA45SN1_45S_50ntFlanks | 12637 | 12654 | 18 | GCTACCATCTGTGGGATT | 2751.9 | -6.9 | 0.7 | -10.4 | 1.9E-25 | 5.3E-22 | BT20/MB468 | MP |

<table>
<tr><th></th><th></th><th></th><th></th><th></th><th></th><th></th><th></th><th></th><th></th><th></th><th></th><th></th><th></th><th></th></tr>
<tr><td>>hg38_21_+_8433172_8446622_RNA45SN1_45S_50ntFlanks@485.509.25|GATGTGGTGACGTCGTGCTCTCCCG</td><td>>hg38_21_+_8433172_8446622_RNA45SN1_45S_50ntFlanks</td><td>485</td><td>509</td><td>25</td><td>GATGTGGTGACGTCGTGCTCTCCCG</td><td>671.4</td><td>13.9</td><td>1.4</td><td>9.9</td><td>4.0E-23</td><td>5.3E-22</td><td>BT20/MB468</td><td>MITO</td></tr>
<tr><td>>hg38_21_+_8433172_8446622_RNA45SN1_45S_50ntFlanks@8357.8391.35|AAAAGAACTTTGAAGAGAGAGTTCAAGAGGGCGTG</td><td>>hg38_21_+_8433172_8446622_RNA45SN1_45S_50ntFlanks</td><td>8357</td><td>8391</td><td>35</td><td>AAAAGAACTTTGAAGAGAGAGTTCAAGAGGGCGTG</td><td>2661.0</td><td>-6.8</td><td>0.7</td><td>-9.8</td><td>1.1E-22</td><td>5.4E-22</td><td>BT20/MB468</td><td>NUC</td></tr>
<tr><td>>hg38_21_+_8433172_8446622_RNA45SN1_45S_50ntFlanks@482.518.37|ATCGATGTGGTGACGTCGTGCTCTCCCGGGCCGGGTC</td><td>>hg38_21_+_8433172_8446622_RNA45SN1_45S_50ntFlanks</td><td>482</td><td>518</td><td>37</td><td>ATCGATGTGGTGACGTCGTGCTCTCCCGGGCCGGGTC</td><td>443.5</td><td>-13.2</td><td>1.3</td><td>-9.8</td><td>1.1E-22</td><td>5.4E-22</td><td>BT20/MB468</td><td>NUC</td></tr>
<tr><td>>MI0000434|hsa-let-7i&WithFlank&12|+|62603680|62603775@12.32.21(+1C)&[MIMAT0000415&hsa-let-7i-5p&offsets|0|-1(+1C);m-49&12|+|62603691|62603712&offsets|0|-1(+1C)]|TGAGGTAGTAGTTTGTGCTGTC</td><td>>MI0000434|hsa-let-7i&WithFlank&12|+|62603680|62603775</td><td>12</td><td>32</td><td>21(+1C)</td><td>TGAGGTAGTAGTTTGTGCTGTC</td><td>14808.3</td><td>6.6</td><td>0.7</td><td>9.9</td><td>5.4E-23</td><td>5.4E-22</td><td>BT20/MB468</td><td>CYTO</td></tr>
<tr><td>>hg38_21_+_8433172_8446622_RNA45SN1_45S_50ntFlanks@6786.6806.21|TACGCCTGTCTGAGCGTCGCT</td><td>>hg38_21_+_8433172_8446622_RNA45SN1_45S_50ntFlanks</td><td>6786</td><td>6806</td><td>21</td><td>TACGCCTGTCTGAGCGTCGCT</td><td>2074.4</td><td>6.4</td><td>0.6</td><td>9.9</td><td>5.4E-23</td><td>5.4E-22</td><td>BT20/MB468</td><td>CYTO</td></tr>
<tr><td>>hg38_21_+_8433172_8446622_RNA45SN1_45S_50ntFlanks@3831.3852.22|CTCGCTCCTCTCCTACTTGGAT</td><td>>hg38_21_+_8433172_8446622_RNA45SN1_45S_50ntFlanks</td><td>3831</td><td>3852</td><td>22</td><td>CTCGCTCCTCTCCTACTTGGAT</td><td>563.2</td><td>-13.6</td><td>1.4</td><td>-9.8</td><td>1.1E-22</td><td>5.4E-22</td><td>BT20/MB468</td><td>NUC</td></tr>
<tr><td>>hg38_21_+_8433172_8446622_RNA45SN1_45S_50ntFlanks@8154.8181.28|CCGCTCGTGGGGGGCCCAAGTCCTTCTG</td><td>>hg38_21_+_8433172_8446622_RNA45SN1_45S_50ntFlanks</td><td>8154</td><td>8181</td><td>28</td><td>CCGCTCGTGGGGGGCCCAAGTCCTTCTG</td><td>5673.6</td><td>-5.5</td><td>0.6</td><td>-9.9</td><td>4.1E-23</td><td>5.4E-22</td><td>BT20/MB468</td><td>MITO</td></tr>
<tr><td>>hg38_21_+_8433172_8446622_RNA45SN1_45S_50ntFlanks@8353.8375.23|GTTGAAAAGAACTTTGAAGAGAG</td><td>>hg38_21_+_8433172_8446622_RNA45SN1_45S_50ntFlanks</td><td>8353</td><td>8375</td><td>23</td><td>GTTGAAAAGAACTTTGAAGAGAG</td><td>1246.5</td><td>14.7</td><td>1.5</td><td>9.9</td><td>4.1E-23</td><td>5.5E-22</td><td>BT20/MB468</td><td>MITO</td></tr>
<tr><td>>hg38_21_+_8433172_8446622_RNA45SN1_45S_50ntFlanks@10869.10892.24|AGGGCTGGGTCGGTCGGGCTGGGG</td><td>>hg38_21_+_8433172_8446622_RNA45SN1_45S_50ntFlanks</td><td>10869</td><td>10892</td><td>24</td><td>AGGGCTGGGTCGGTCGGGCTGGGG</td><td>2261.0</td><td>-15.0</td><td>1.5</td><td>-10.0</td><td>1.8E-23</td><td>5.5E-22</td><td>MB231/MB468</td><td>MP</td></tr>
<tr><td>>hg38_21_+_8433172_8446622_RNA45SN1_45S_50ntFlanks@4557.4577.21|CAGCTAGGAATAATGGAATAG</td><td>>hg38_21_+_8433172_8446622_RNA45SN1_45S_50ntFlanks</td><td>4557</td><td>4577</td><td>21</td><td>CAGCTAGGAATAATGGAATAG</td><td>265.4</td><td>-12.5</td><td>1.3</td><td>-9.8</td><td>1.2E-22</td><td>5.5E-22</td><td>BT20/MB468</td><td>NUC</td></tr>
<tr><td>>hg38_21_+_8433172_8446622_RNA45SN1_45S_50ntFlanks@6646.6665.20|TCGTACGACTCTTAGCGGTG</td><td>>hg38_21_+_8433172_8446622_RNA45SN1_45S_50ntFlanks</td><td>6646</td><td>6665</td><td>20</td><td>TCGTACGACTCTTAGCGGTG</td><td>64127.7</td><td>6.2</td><td>0.6</td><td>9.9</td><td>5.6E-23</td><td>5.6E-22</td><td>BT20/MB468</td><td>CYTO</td></tr>
<tr><td>>hg38_21_+_8433172_8446622_RNA45SN1_45S_50ntFlanks@11200.11225.26|CGGCGGCGGCGGCGACTCTGGACGCG</td><td>>hg38_21_+_8433172_8446622_RNA45SN1_45S_50ntFlanks</td><td>11200</td><td>11225</td><td>26</td><td>CGGCGGCGGCGGCGACTCTGGACGCG</td><td>1298.7</td><td>-14.2</td><td>1.4</td><td>-10.0</td><td>1.8E-23</td><td>5.6E-22</td><td>MB231/MB468</td><td>MP</td></tr>
<tr><td>>MI0000266|hsa-mir-10a&WithFlank&17|-|48579832|48579953@29.49.21(+1A)&[MIMAT0000253&hsa-miR-10a-5p&offsets|+1|-1(+1A);m-18&17|-|48579905|48579926&offsets|+1|0(+1A)]|ACCCTGTAGATCCGAATTTGTA</td><td>>MI0000266|hsa-mir-10a&WithFlank&17|-|48579832|48579953</td><td>29</td><td>49</td><td>21(+1A)</td><td>ACCCTGTAGATCCGAATTTGTA</td><td>195.5</td><td>10.4</td><td>1.1</td><td>9.8</td><td>1.2E-22</td><td>5.6E-22</td><td>BT20/MB468</td><td>NUC</td></tr>
<tr><td>>trnaMT_GluTTC_MT_-_14674_14742@38.72.35,trnalookalike8_GluTTC_5_-_93905172_93905240@38.72.35|CATTGGTCGTGGTTGTAGTCCGTGCGAGAATACCA</td><td>>trnaMT_GluTTC_MT_-_14674_14742</td><td>38</td><td>72</td><td>35</td><td>CATTGGTCGTGGTTGTAGTCCGTGCGAGAATACCA</td><td>15975.6</td><td>6.8</td><td>0.7</td><td>9.9</td><td>4.3E-23</td><td>5.7E-22</td><td>BT20/MB468</td><td>MITO</td></tr>
<tr><td>>hg38_21_+_8433172_8446622_RNA45SN1_45S_50ntFlanks@12417.12445.29|TCGATGTCGGCTCTTCCTATCATTGTGAA</td><td>>hg38_21_+_8433172_8446622_RNA45SN1_45S_50ntFlanks</td><td>12417</td><td>12445</td><td>29</td><td>TCGATGTCGGCTCTTCCTATCATTGTGAA</td><td>906.7</td><td>-5.9</td><td>0.6</td><td>-9.8</td><td>1.2E-22</td><td>5.8E-22</td><td>BT20/MB468</td><td>NUC</td></tr>
<tr><td>>MI0000269|hsa-mir-181a-2&WithFlank&9|+|124692436|124692557@83.104.22&[MIMAT0004558&hsa-miR-181a-2-3p&offsets|0|0;m-71&9|+|124692518|124692539&offsets|0|0]|ACCACTGACCGTTGACTGTACC</td><td>>MI0000269|hsa-mir-181a-2&WithFlank&9|+|124692436|124692557</td><td>83</td><td>104</td><td>22</td><td>ACCACTGACCGTTGACTGTACC</td><td>2699.0</td><td>8.0</td><td>0.8</td><td>10.0</td><td>1.4E-23</td><td>5.8E-22</td><td>MB231/MB468</td><td>CYTO</td></tr>
<tr><td>>hg38_21_+_8433172_8446622_RNA45SN1_45S_50ntFlanks@12655.12674.20|ATGACTGAACGCCTCTAAGT</td><td>>hg38_21_+_8433172_8446622_RNA45SN1_45S_50ntFlanks</td><td>12655</td><td>12674</td><td>20</td><td>ATGACTGAACGCCTCTAAGT</td><td>1172.1</td><td>-14.1</td><td>1.4</td><td>-10.0</td><td>1.9E-23</td><td>5.8E-22</td><td>MB231/MB468</td><td>MP</td></tr>
<tr><td>>hg38_21_+_8433172_8446622_RNA45SN1_45S_50ntFlanks@9545.9566.22|GTGGAGGTCCGTAGCGGTCCTG</td><td>>hg38_21_+_8433172_8446622_RNA45SN1_45S_50ntFlanks</td><td>9545</td><td>9566</td><td>22</td><td>GTGGAGGTCCGTAGCGGTCCTG</td><td>1253.3</td><td>-14.2</td><td>1.4</td><td>-10.0</td><td>1.9E-23</td><td>5.8E-22</td><td>MB231/MB468</td><td>MP</td></tr>
<tr><td>>hg38_21_+_8433172_8446622_RNA45SN1_45S_50ntFlanks@4733.4767.35|GAACGAAAGTCGGAGGTTCGAAGACGATCAGATAC</td><td>>hg38_21_+_8433172_8446622_RNA45SN1_45S_50ntFlanks</td><td>4733</td><td>4767</td><td>35</td><td>GAACGAAAGTCGGAGGTTCGAAGACGATCAGATAC</td><td>1767.8</td><td>9.9</td><td>1.0</td><td>10.0</td><td>1.4E-23</td><td>5.8E-22</td><td>MB231/MB468</td><td>CYTO</td></tr>
<tr><td>>MI0000456|hsa-mir-140&WithFlank&16|+|69933075|69933186@68.89.22(+1A)&[MIMAT0004597&hsa-miR-140-3p&offsets|0|+1(+1A);m-45&16|+|69933143|69933164&offsets|-1|-1(+1A)]|TACCACAGGGTAGAACCACGGAA</td><td>>MI0000456|hsa-mir-140&WithFlank&16|+|69933075|69933186</td><td>68</td><td>89</td><td>22(+1A)</td><td>TACCACAGGGTAGAACCACGGAA</td><td>353.6</td><td>12.7</td><td>1.3</td><td>9.9</td><td>5.9E-23</td><td>5.9E-22</td><td>BT20/MB468</td><td>CYTO</td></tr>
<tr><td>>hg38_21_+_8433172_8446622_RNA45SN1_45S_50ntFlanks@12717.12757.41|CCTCGGTTGGCCTCGGATAGCCGGTCCCCCGCCTGTCCCCG</td><td>>hg38_21_+_8433172_8446622_RNA45SN1_45S_50ntFlanks</td><td>12717</td><td>12757</td><td>41</td><td>CCTCGGTTGGCCTCGGATAGCCGGTCCCCCGCCTGTCCCCG</td><td>668.3</td><td>-13.8</td><td>1.4</td><td>-9.8</td><td>1.2E-22</td><td>5.9E-22</td><td>BT20/MB468</td><td>NUC</td></tr>
<tr><td>>hg38_1_-_228634819_228635039_RNA5S12_5S_50ntFlanks@134.169.36|TGGGAGACCGCCTGGGAATACCGGGTGCTGTAGGCT</td><td>>hg38_1_-_228634819_228635039_RNA5S12_5S_50ntFlanks</td><td>134</td><td>169</td><td>36</td><td>TGGGAGACCGCCTGGGAATACCGGGTGCTGTAGGCT</td><td>17589.7</td><td>7.5</td><td>0.8</td><td>9.9</td><td>6.0E-23</td><td>5.9E-22</td><td>BT20/MB468</td><td>CYTO</td></tr>
<tr><td>>hg38_21_+_8433172_8446622_RNA45SN1_45S_50ntFlanks@11641.11659.19|CGGCGGGTGTTGACGCGAT</td><td>>hg38_21_+_8433172_8446622_RNA45SN1_45S_50ntFlanks</td><td>11641</td><td>11659</td><td>19</td><td>CGGCGGGTGTTGACGCGAT</td><td>2224.1</td><td>-14.9</td><td>1.5</td><td>-9.9</td><td>3.0E-23</td><td>5.9E-22</td><td>MB231/MB468</td><td>MITO</td></tr>
<tr><td>>hg38_21_+_8433172_8446622_RNA45SN1_45S_50ntFlanks@10411.10430.20|CAGTCGGTCCTGAGAGATGG</td><td>>hg38_21_+_8433172_8446622_RNA45SN1_45S_50ntFlanks</td><td>10411</td><td>10430</td><td>20</td><td>CAGTCGGTCCTGAGAGATGG</td><td>2680.2</td><td>-6.1</td><td>0.6</td><td>-9.9</td><td>4.5E-23</td><td>5.9E-22</td><td>BT20/MB468</td><td>MITO</td></tr>
<tr><td>>hg38_21_+_8433172_8446622_RNA45SN1_45S_50ntFlanks@10534.10567.34|CCGCGAGGCGTCCAGTGCGGTAACGCGACCGATC</td><td>>hg38_21_+_8433172_8446622_RNA45SN1_45S_50ntFlanks</td><td>10534</td><td>10567</td><td>34</td><td>CCGCGAGGCGTCCAGTGCGGTAACGCGACCGATC</td><td>1029.7</td><td>-13.9</td><td>1.4</td><td>-10.0</td><td>2.0E-23</td><td>6.0E-22</td><td>MB231/MB468</td><td>MP</td></tr>
<tr><td>>hg38_21_+_8433172_8446622_RNA45SN1_45S_50ntFlanks@10534.10570.37|CCGCGAGGCGTCCAGTGCGGTAACGCGACCGATCCCG</td><td>>hg38_21_+_8433172_8446622_RNA45SN1_45S_50ntFlanks</td><td>10534</td><td>10570</td><td>37</td><td>CCGCGAGGCGTCCAGTGCGGTAACGCGACCGATCCCG</td><td>332.2</td><td>-12.8</td><td>1.3</td><td>-9.8</td><td>1.2E-22</td><td>6.0E-22</td><td>BT20/MB468</td><td>NUC</td></tr>
<tr><td>>MI0000456|hsa-mir-140&WithFlank&16|+|69933075|69933186@69.89.21(+1U)&[MIMAT0004597&hsa-miR-140-3p&offsets|+1|+1(+1U);m-45&16|+|69933143|69933164&offsets|0|-1(+1U)]|ACCACAGGGTAGAACCACGGAT</td><td>>MI0000456|hsa-mir-140&WithFlank&16|+|69933075|69933186</td><td>69</td><td>89</td><td>21(+1U)</td><td>ACCACAGGGTAGAACCACGGAT</td><td>663.8</td><td>13.6</td><td>1.4</td><td>9.9</td><td>6.1E-23</td><td>6.0E-22</td><td>BT20/MB468</td><td>CYTO</td></tr>
<tr><td>>hg38_21_+_8433172_8446622_RNA45SN1_45S_50ntFlanks@12627.12657.31|ATGGGGCGAAGCTACCATCTGTGGGATTATG</td><td>>hg38_21_+_8433172_8446622_RNA45SN1_45S_50ntFlanks</td><td>12627</td><td>12657</td><td>31</td><td>ATGGGGCGAAGCTACCATCTGTGGGATTATG</td><td>1416.8</td><td>-14.3</td><td>1.4</td><td>-10.0</td><td>2.0E-23</td><td>6.1E-22</td><td>MB231/MB468</td><td>MP</td></tr>
<tr><td>>hg38_21_+_8433172_8446622_RNA45SN1_45S_50ntFlanks@4735.4767.33|ACGAAAGTCGGAGGTTCGAAGACGATCAGATAC</td><td>>hg38_21_+_8433172_8446622_RNA45SN1_45S_50ntFlanks</td><td>4735</td><td>4767</td><td>33</td><td>ACGAAAGTCGGAGGTTCGAAGACGATCAGATAC</td><td>397.1</td><td>12.9</td><td>1.3</td><td>9.9</td><td>6.2E-23</td><td>6.1E-22</td><td>BT20/MB468</td><td>CYTO</td></tr>
<tr><td>>hg38_21_+_8433172_8446622_RNA45SN1_45S_50ntFlanks@12162.12181.20|GACTGGGGCGGTACACCTGT</td><td>>hg38_21_+_8433172_8446622_RNA45SN1_45S_50ntFlanks</td><td>12162</td><td>12181</td><td>20</td><td>GACTGGGGCGGTACACCTGT</td><td>856.9</td><td>-13.6</td><td>1.4</td><td>-10.0</td><td>2.0E-23</td><td>6.2E-22</td><td>MB231/MB468</td><td>MP</td></tr>
<tr><td>>MI0000749|hsa-mir-30e&WithFlank&1|+|40754349|40754452@65.86.22&[MIMAT0000693&hsa-miR-30e-3p&offsets|0|0;m-25&1|+|40754413|40754434&offsets|0|0]|CTTTCAGTCGGATGTTTACAGC</td><td>>MI0000749|hsa-mir-30e&WithFlank&1|+|40754349|40754452</td><td>65</td><td>86</td><td>22</td><td>CTTTCAGTCGGATGTTTACAGC</td><td>2742.0</td><td>4.7</td><td>0.5</td><td>9.9</td><td>6.3E-23</td><td>6.2E-22</td><td>BT20/MB468</td><td>CYTO</td></tr>
<tr><td>>am_trnaMT_MetCAT_MT_+_4402_4469@1.49.49|AGTAAGGTCAGCTAAATAAGCTATCGGGCCCATACCCCGAAAATGTTGG</td><td>>am_trnaMT_MetCAT_MT_+_4402_4469</td><td>1</td><td>49</td><td>49</td><td>AGTAAGGTCAGCTAAATAAGCTATCGGGCCCATACCCCGAAAATGTTGG</td><td>625.4</td><td>27.2</td><td>2.6</td><td>10.4</td><td>2.7E-25</td><td>6.2E-22</td><td>BT20/MB468</td><td>MP</td></tr>
<tr><td>>hg38_21_+_8433172_8446622_RNA45SN1_45S_50ntFlanks@4283.4319.37|CTTTAACGAGGATCCATTGGAGGGCAAGTCTGGTGCC</td><td>>hg38_21_+_8433172_8446622_RNA45SN1_45S_50ntFlanks</td><td>4283</td><td>4319</td><td>37</td><td>CTTTAACGAGGATCCATTGGAGGGCAAGTCTGGTGCC</td><td>378.7</td><td>-13.0</td><td>1.3</td><td>-9.8</td><td>1.3E-22</td><td>6.2E-22</td><td>BT20/MB468</td><td>NUC</td></tr>
<tr><td>>hg38_21_+_8433172_8446622_RNA45SN1_45S_50ntFlanks@4732.4758.27|AGAACGAAAGTCGGAGGTTCGAAGACG</td><td>>hg38_21_+_8433172_8446622_RNA45SN1_45S_50ntFlanks</td><td>4732</td><td>4758</td><td>27</td><td>AGAACGAAAGTCGGAGGTTCGAAGACG</td><td>3802.6</td><td>-6.7</td><td>0.7</td><td>-9.8</td><td>1.3E-22</td><td>6.3E-22</td><td>BT20/MB468</td><td>NUC</td></tr>
<tr><td>>MI0005756|hsa-mir-934&WithFlank&X|+|136550872|136550966@21.42.22&[MIMAT0004977&hsa-miR-934&offsets|0|0;m-943&X|+|136550892|136550913&offsets|0|0]|TGTCTACTACTGGAGACACTGG</td><td>>MI0005756|hsa-mir-934&WithFlank&X|+|136550872|136550966</td><td>21</td><td>42</td><td>22</td><td>TGTCTACTACTGGAGACACTGG</td><td>2281.1</td><td>14.9</td><td>1.5</td><td>10.1</td><td>4.1E-24</td><td>6.3E-22</td><td>BT20/MB231</td><td>NUC</td></tr>
<tr><td>>hg38_21_+_8433172_8446622_RNA45SN1_45S_50ntFlanks@11845.11873.29|GGAACGGGCTTGGCGGAATCAGCGGGGAA</td><td>>hg38_21_+_8433172_8446622_RNA45SN1_45S_50ntFlanks</td><td>11845</td><td>11873</td><td>29</td><td>GGAACGGGCTTGGCGGAATCAGCGGGGAA</td><td>1037.9</td><td>-13.9</td><td>1.4</td><td>-10.0</td><td>2.1E-23</td><td>6.3E-22</td><td>MB231/MB468</td><td>MP</td></tr>
<tr><td>>hg38_21_+_8433172_8446622_RNA45SN1_45S_50ntFlanks@12406.12437.32|TTTTTGATCCTTCGATGTCGGCTCTTCCTATC</td><td>>hg38_21_+_8433172_8446622_RNA45SN1_45S_50ntFlanks</td><td>12406</td><td>12437</td><td>32</td><td>TTTTTGATCCTTCGATGTCGGCTCTTCCTATC</td><td>1573.0</td><td>-7.2</td><td>0.7</td><td>-9.8</td><td>1.3E-22</td><td>6.3E-22</td><td>BT20/MB468</td><td>NUC</td></tr>
<tr><td>>hg38_21_+_8433172_8446622_RNA45SN1_45S_50ntFlanks@10362.10388.27|AAGGCCGAAGTGGAGAAGGGTTCCATG</td><td>>hg38_21_+_8433172_8446622_RNA45SN1_45S_50ntFlanks</td><td>10362</td><td>10388</td><td>27</td><td>AAGGCCGAAGTGGAGAAGGGTTCCATG</td><td>4411.4</td><td>-16.0</td><td>1.6</td><td>-10.0</td><td>2.1E-23</td><td>6.3E-22</td><td>MB231/MB468</td><td>MP</td></tr>
<tr><td>>MI0000263|hsa-mir-7-1&WithFlank&9|-|83969742|83969863@30.54.25&[MIMAT0000252_2&hsa-miR-7-5p&offsets|0|+1;m-401&9|-|83969811|83969834&offsets|0|+1]|TGGAAGACTAGTGATTTTGTTGTTT</td><td>>MI0000263|hsa-mir-7-1&WithFlank&9|-|83969742|83969863</td><td>30</td><td>54</td><td>25</td><td>TGGAAGACTAGTGATTTTGTTGTTT</td><td>1377.6</td><td>6.1</td><td>0.6</td><td>9.9</td><td>3.4E-23</td><td>6.4E-22</td><td>MB231/MB468</td><td>NUC</td></tr>
<tr><td>>MI0000477|hsa-mir-146a&WithFlank&5|+|160485346|160485456@27.47.21&[MIMAT0000449&hsa-miR-146a-5p&offsets|0|-1;m-129&5|+|160485372|160485393&offsets|0|-1]|TGAGAACTGAATTCCATGGGT</td><td>>MI0000477|hsa-mir-146a&WithFlank&5|+|160485346|160485456</td><td>27</td><td>47</td><td>21</td><td>TGAGAACTGAATTCCATGGGT</td><td>2643.1</td><td>15.3</td><td>1.5</td><td>9.9</td><td>3.3E-23</td><td>6.4E-22</td><td>MB231/MB468</td><td>NUC</td></tr>
<tr><td>>hg38_21_+_8433172_8446622_RNA45SN1_45S_50ntFlanks@4723.4752.30|CATTAATCAAGAACGAAAGTCGGAGGTTCG</td><td>>hg38_21_+_8433172_8446622_RNA45SN1_45S_50ntFlanks</td><td>4723</td><td>4752</td><td>30</td><td>CATTAATCAAGAACGAAAGTCGGAGGTTCG</td><td>1314.0</td><td>-14.2</td><td>1.4</td><td>-10.0</td><td>2.1E-23</td><td>6.4E-22</td><td>MB231/MB468</td><td>MP</td></tr>
<tr><td>>hg38_21_+_8433172_8446622_RNA45SN1_45S_50ntFlanks@10389.10419.31|TGAACAGCAGTTGAACATGGGTCAGTCGGTC</td><td>>hg38_21_+_8433172_8446622_RNA45SN1_45S_50ntFlanks</td><td>10389</td><td>10419</td><td>31</td><td>TGAACAGCAGTTGAACATGGGTCAGTCGGTC</td><td>345.0</td><td>-12.9</td><td>1.3</td><td>-9.8</td><td>1.3E-22</td><td>6.4E-22</td><td>BT20/MB468</td><td>NUC</td></tr>
<tr><td>>MI0000542|hsa-mir-320a&WithFlank&8|-|22244960|22245043@47.68.22&[MIMAT0000510&hsa-miR-320a-3p&offsets|-1|-1;m-60&8|-|22244975|22244996&offsets|-1|-1]|GAAAAGCTGGGTTGAGAGGGCG</td><td>>MI0000542|hsa-mir-320a&WithFlank&8|-|22244960|22245043</td><td>47</td><td>68</td><td>22</td><td>GAAAAGCTGGGTTGAGAGGGCG</td><td>454.8</td><td>13.1</td><td>1.3</td><td>9.9</td><td>6.5E-23</td><td>6.4E-22</td><td>BT20/MB468</td><td>CYTO</td></tr>
<tr><td>>hg38_21_+_8433172_8446622_RNA45SN1_45S_50ntFlanks@6733.6773.41|CATTGATCATCGACACTTCGAACGCACTTGCGGCCCCGGGT</td><td>>hg38_21_+_8433172_8446622_RNA45SN1_45S_50ntFlanks</td><td>6733</td><td>6773</td><td>41</td><td>CATTGATCATCGACACTTCGAACGCACTTGCGGCCCCGGGT</td><td>1415.0</td><td>6.2</td><td>0.6</td><td>9.9</td><td>4.9E-23</td><td>6.4E-22</td><td>BT20/MB468</td><td>MITO</td></tr>
<tr><td>>MI0005756|hsa-mir-934&WithFlank&X|+|136550872|136550966@21.41.21&[MIMAT0004977&hsa-miR-934&offsets|0|-1;m-943&X|+|136550892|136550913&offsets|0|-1]|TGTCTACTACTGGAGACACTG</td><td>>MI0005756|hsa-mir-934&WithFlank&X|+|136550872|136550966</td><td>21</td><td>41</td><td>21</td><td>TGTCTACTACTGGAGACACTG</td><td>1520.3</td><td>6.9</td><td>0.7</td><td>9.9</td><td>6.5E-23</td><td>6.4E-22</td><td>BT20/MB468</td><td>CYTO</td></tr>
<tr><td>>hg38_21_+_8433172_8446622_RNA45SN1_45S_50ntFlanks@9027.9065.39|CGTCTCCTCGTGGGGGGGCCGGGCCACCCCTCCCACGGC</td><td>>hg38_21_+_8433172_8446622_RNA45SN1_45S_50ntFlanks</td><td>9027</td><td>9065</td><td>39</td><td>CGTCTCCTCGTGGGGGGGCCGGGCCACCCCTCCCACGGC</td><td>2341.6</td><td>-15.0</td><td>1.5</td><td>-9.9</td><td>3.3E-23</td><td>6.5E-22</td><td>MB231/MB468</td><td>MITO</td></tr>
<tr><td>>trnaMT_GluTTC_MT_-_14674_14742@37.71.35,trnalookalike8_GluTTC_5_-_93905172_93905240@37.71.35|TCATTGGTCGTGGTTGTAGTCCGTGCGAGAATACC</td><td>>trnaMT_GluTTC_MT_-_14674_14742</td><td>37</td><td>71</td><td>35</td><td>TCATTGGTCGTGGTTGTAGTCCGTGCGAGAATACC</td><td>1454.2</td><td>14.5</td><td>1.5</td><td>10.0</td><td>1.6E-23</td><td>6.5E-22</td><td>MB231/MB468</td><td>CYTO</td></tr>
<tr><td>>MI0000734|hsa-mir-106b&WithFlank&7|-|100093987|100094080@56.76.21(+1A)&[MIMAT0004672&hsa-miR-106b-3p&offsets|-2|-3(+1A);m-87&7|-|100094003|100094023&offsets|-2|-2(+1A)]|TACCGCACTGTGGGTACTTGCA</td><td>>MI0000734|hsa-mir-106b&WithFlank&7|-|100093987|100094080</td><td>56</td><td>76</td><td>21(+1A)</td><td>TACCGCACTGTGGGTACTTGCA</td><td>270.1</td><td>12.3</td><td>1.3</td><td>9.9</td><td>6.6E-23</td><td>6.5E-22</td><td>BT20/MB468</td><td>CYTO</td></tr>
</table>

<table>
<tr><td>>hg38_21_+_8433172_8446622_RNA45SN1_45S_50ntFlanks@11617.11640.24|AAAACAAAGCATCGCGAAGGCCCG</td><td>>hg38_21_+_8433172_8446622_RNA45SN1_45S_50ntFlanks</td><td>11617</td><td>11640</td><td>24</td><td>AAAACAAAGCATCGCGAAGGCCCG</td><td>1212.9</td><td>-14.1</td><td>1.4</td><td>-10.0</td><td>2.2E-23</td><td>6.6E-22</td><td>MB231/MB468</td><td>MP</td></tr>
<tr><td>>hg38_21_+_8433172_8446622_RNA45SN1_45S_50ntFlanks@4443.4472.30|CCCTTGCCTCTCGGCGCCCCCTCGATGCTC</td><td>>hg38_21_+_8433172_8446622_RNA45SN1_45S_50ntFlanks</td><td>4443</td><td>4472</td><td>30</td><td>CCCTTGCCTCTCGGCGCCCCCTCGATGCTC</td><td>1266.7</td><td>-14.1</td><td>1.4</td><td>-9.9</td><td>3.4E-23</td><td>6.6E-22</td><td>MB231/MB468</td><td>MITO</td></tr>
<tr><td>>hg38_21_+_8433172_8446622_RNA45SN1_45S_50ntFlanks@5527.5550.24|AAAGTCGTAACAAGGTTTCCGTAG</td><td>>hg38_21_+_8433172_8446622_RNA45SN1_45S_50ntFlanks</td><td>5527</td><td>5550</td><td>24</td><td>AAAGTCGTAACAAGGTTTCCGTAG</td><td>757.1</td><td>-8.3</td><td>0.8</td><td>-10.0</td><td>1.6E-23</td><td>6.6E-22</td><td>MB231/MB468</td><td>CYTO</td></tr>
<tr><td>>hg38_21_+_8433172_8446622_RNA45SN1_45S_50ntFlanks@6647.6668.22|CGTACGACTCTTAGCGGTGGAT</td><td>>hg38_21_+_8433172_8446622_RNA45SN1_45S_50ntFlanks</td><td>6647</td><td>6668</td><td>22</td><td>CGTACGACTCTTAGCGGTGGAT</td><td>2088.9</td><td>5.4</td><td>0.5</td><td>9.9</td><td>6.8E-23</td><td>6.6E-22</td><td>BT20/MB468</td><td>CYTO</td></tr>
<tr><td>>hg38_21_+_8433172_8446622_RNA45SN1_45S_50ntFlanks@11583.11611.29|CTGGTGCGGACCAGGGGAATCCGACTGTT</td><td>>hg38_21_+_8433172_8446622_RNA45SN1_45S_50ntFlanks</td><td>11583</td><td>11611</td><td>29</td><td>CTGGTGCGGACCAGGGGAATCCGACTGTT</td><td>6799.3</td><td>-5.7</td><td>0.6</td><td>-9.9</td><td>3.4E-23</td><td>6.7E-22</td><td>MB231/MB468</td><td>MITO</td></tr>
<tr><td>>MI0000103|hsa-mir-101-1&WithFlank&1|-|65058428|65058514@53.74.22;MI0000739|hsa-mir-101-2&WithFlank&9|+|4850291|4850381@55.76.22&[MIMAT0000099&hsa-miR-101-3p&offsets|0|+1;m-17&1|-|65058443|65058462&offsets|0|+2];[MIMAT0000099_1&hsa-miR-101-3p&offsets|0|+1;m-16&9|+|4850345|4850364&offsets|0|+2]|TACAGTACTGTGATAACTGAAG</td><td>>MI0000103|hsa-mir-101-1&WithFlank&1|-|65058428|65058514</td><td>53</td><td>74</td><td>22</td><td>TACAGTACTGTGATAACTGAAG</td><td>1276.7</td><td>6.9</td><td>0.7</td><td>9.9</td><td>6.8E-23</td><td>6.7E-22</td><td>BT20/MB468</td><td>CYTO</td></tr>
<tr><td>>hg38_21_+_8433172_8446622_RNA45SN1_45S_50ntFlanks@8115.8139.25|CGCGGGACATGTGGCGTACGGAAGA</td><td>>hg38_21_+_8433172_8446622_RNA45SN1_45S_50ntFlanks</td><td>8115</td><td>8139</td><td>25</td><td>CGCGGGACATGTGGCGTACGGAAGA</td><td>1773.1</td><td>-14.7</td><td>1.5</td><td>-10.0</td><td>2.2E-23</td><td>6.7E-22</td><td>MB231/MB468</td><td>MP</td></tr>
<tr><td>>MI0000067|hsa-let-7f-1&WithFlank&9|+|94176341|94176439@69.89.21(+1U)&[MIMAT0004486&hsa-let-7f-1-3p&offsets|0|-1(+1U);m-275&9|+|94176409|94176430&offsets|0|-1(+1U)]|CTATACAATCTATTGCCTTCCT</td><td>>MI0000067|hsa-let-7f-1&WithFlank&9|+|94176341|94176439</td><td>69</td><td>89</td><td>21(+1U)</td><td>CTATACAATCTATTGCCTTCCT</td><td>363.5</td><td>5.4</td><td>0.5</td><td>9.9</td><td>3.6E-23</td><td>6.7E-22</td><td>MB231/MB468</td><td>NUC</td></tr>
<tr><td>>MI0000266|hsa-mir-10a&WithFlank&17|-|48579832|48579953@29.51.23&[MIMAT0000253&hsa-miR-10a-5p&offsets|+1|+1;m-18&17|-|48579905|48579926&offsets|+1|+2]|ACCCTGTAGATCCGAATTTGTGT</td><td>>MI0000266|hsa-mir-10a&WithFlank&17|-|48579832|48579953</td><td>29</td><td>51</td><td>23</td><td>ACCCTGTAGATCCGAATTTGTGT</td><td>188.5</td><td>10.4</td><td>1.1</td><td>9.8</td><td>1.4E-22</td><td>6.8E-22</td><td>BT20/MB468</td><td>NUC</td></tr>
<tr><td>>hg38_21_+_8433172_8446622_RNA45SN1_45S_50ntFlanks@3990.4023.34|TTGGTGACTCTAGATAACCTCGGGCCGATCGCAC</td><td>>hg38_21_+_8433172_8446622_RNA45SN1_45S_50ntFlanks</td><td>3990</td><td>4023</td><td>34</td><td>TTGGTGACTCTAGATAACCTCGGGCCGATCGCAC</td><td>1289.8</td><td>5.1</td><td>0.5</td><td>9.8</td><td>6.9E-23</td><td>6.8E-22</td><td>BT20/MB468</td><td>CYTO</td></tr>
<tr><td>>hg38_21_+_8433172_8446622_RNA45SN1_45S_50ntFlanks@6719.6744.26|TGAATTGCAGGACACATTGATCATCG</td><td>>hg38_21_+_8433172_8446622_RNA45SN1_45S_50ntFlanks</td><td>6719</td><td>6744</td><td>26</td><td>TGAATTGCAGGACACATTGATCATCG</td><td>2012.7</td><td>-5.7</td><td>0.6</td><td>-9.9</td><td>5.2E-23</td><td>6.8E-22</td><td>BT20/MB468</td><td>MITO</td></tr>
<tr><td>>hg38_21_+_8433172_8446622_RNA45SN1_45S_50ntFlanks@12635.12653.19|AAGCTACCATCTGTGGGAT</td><td>>hg38_21_+_8433172_8446622_RNA45SN1_45S_50ntFlanks</td><td>12635</td><td>12653</td><td>19</td><td>AAGCTACCATCTGTGGGAT</td><td>7415.2</td><td>-15.9</td><td>1.5</td><td>-10.4</td><td>3.5E-25</td><td>6.9E-22</td><td>BT20/MB468</td><td>MP</td></tr>
<tr><td>>hg38_21_+_8433172_8446622_RNA45SN1_45S_50ntFlanks@12388.12433.46|CGTTCATAGCGACGTCGCTTTTTGATCCTTCGATGTCGGCTCTTCC</td><td>>hg38_21_+_8433172_8446622_RNA45SN1_45S_50ntFlanks</td><td>12388</td><td>12433</td><td>46</td><td>CGTTCATAGCGACGTCGCTTTTTGATCCTTCGATGTCGGCTCTTCC</td><td>2037.0</td><td>8.0</td><td>0.8</td><td>10.0</td><td>1.7E-23</td><td>6.9E-22</td><td>MB231/MB468</td><td>CYTO</td></tr>
<tr><td>>hg38_21_+_8433172_8446622_RNA45SN1_45S_50ntFlanks@11561.11592.32|GCGCCTAGCAGCCGACTTAGAACTGGTGCGGA</td><td>>hg38_21_+_8433172_8446622_RNA45SN1_45S_50ntFlanks</td><td>11561</td><td>11592</td><td>32</td><td>GCGCCTAGCAGCCGACTTAGAACTGGTGCGGA</td><td>2525.3</td><td>-15.2</td><td>1.5</td><td>-10.0</td><td>2.3E-23</td><td>6.9E-22</td><td>MB231/MB468</td><td>MP</td></tr>
<tr><td>>am_MI0000809|hsa-mir-151a&WithFlank&8|-|140732558|140732659@53.73.21(+3U)&[MIMAT0000757&hsa-miR-151a-3p&offsets|0|0(+3U);m-30&8|-|140732587|140732607&offsets|0|0(+3U)]|CTAGACTGAAGCTCCTTGAGGTTT</td><td>>am_MI0000809|hsa-mir-151a&WithFlank&8|-|140732558|140732659</td><td>53</td><td>73</td><td>21(+3U)</td><td>CTAGACTGAAGCTCCTTGAGGTTT</td><td>457.3</td><td>6.9</td><td>0.7</td><td>9.9</td><td>3.7E-23</td><td>7.0E-22</td><td>MB231/MB468</td><td>NUC</td></tr>
<tr><td>>MI0000266|hsa-mir-10a&WithFlank&17|-|48579832|48579953@28.49.22(+1A)&[MIMAT0000253&hsa-miR-10a-5p&offsets|0|-1(+1A);m-18&17|-|48579905|48579926&offsets|0|0(+1A)]|TACCCTGTAGATCCGAATTTGTA</td><td>>MI0000266|hsa-mir-10a&WithFlank&17|-|48579832|48579953</td><td>28</td><td>49</td><td>22(+1A)</td><td>TACCCTGTAGATCCGAATTTGTA</td><td>1210.4</td><td>14.7</td><td>1.5</td><td>9.9</td><td>5.4E-23</td><td>7.1E-22</td><td>BT20/MB468</td><td>MITO</td></tr>
<tr><td>>hg38_21_+_8433172_8446622_RNA45SN1_45S_50ntFlanks@10361.10390.30|GAAGGCCGAAGTGGAGAAGGGTTCCATGTG</td><td>>hg38_21_+_8433172_8446622_RNA45SN1_45S_50ntFlanks</td><td>10361</td><td>10390</td><td>30</td><td>GAAGGCCGAAGTGGAGAAGGGTTCCATGTG</td><td>830.7</td><td>-14.1</td><td>1.4</td><td>-9.8</td><td>1.5E-22</td><td>7.1E-22</td><td>BT20/MB468</td><td>NUC</td></tr>
<tr><td>>hg38_21_+_8433172_8446622_RNA45SN1_45S_50ntFlanks@8106.8140.35|GCGGCGGGGCGCGGGACATGTGGCGTACGGAAGAC</td><td>>hg38_21_+_8433172_8446622_RNA45SN1_45S_50ntFlanks</td><td>8106</td><td>8140</td><td>35</td><td>GCGGCGGGGCGCGGGACATGTGGCGTACGGAAGAC</td><td>1202.4</td><td>-14.1</td><td>1.4</td><td>-10.0</td><td>2.4E-23</td><td>7.1E-22</td><td>MB231/MB468</td><td>MP</td></tr>
<tr><td>>hg38_21_+_8433172_8446622_RNA45SN1_45S_50ntFlanks@4723.4759.37|CATTAATCAAGAACGAAAGTCGGAGGTTCGAAGACGA</td><td>>hg38_21_+_8433172_8446622_RNA45SN1_45S_50ntFlanks</td><td>4723</td><td>4759</td><td>37</td><td>CATTAATCAAGAACGAAAGTCGGAGGTTCGAAGACGA</td><td>4799.2</td><td>-10.8</td><td>1.1</td><td>-9.8</td><td>1.5E-22</td><td>7.1E-22</td><td>BT20/MB468</td><td>NUC</td></tr>
<tr><td>>hg38_21_+_8433172_8446622_RNA45SN1_45S_50ntFlanks@9343.9370.28|CGAAAGCCGCCGTGGCGCAATGAAGGTG</td><td>>hg38_21_+_8433172_8446622_RNA45SN1_45S_50ntFlanks</td><td>9343</td><td>9370</td><td>28</td><td>CGAAAGCCGCCGTGGCGCAATGAAGGTG</td><td>1394.1</td><td>-14.2</td><td>1.4</td><td>-9.9</td><td>3.7E-23</td><td>7.1E-22</td><td>MB231/MB468</td><td>MITO</td></tr>
<tr><td>>hg38_21_+_8433172_8446622_RNA45SN1_45S_50ntFlanks@12627.12661.35|ATGGGGCGAAGCTACCATCTGTGGGATTATGACTG</td><td>>hg38_21_+_8433172_8446622_RNA45SN1_45S_50ntFlanks</td><td>12627</td><td>12661</td><td>35</td><td>ATGGGGCGAAGCTACCATCTGTGGGATTATGACTG</td><td>8315.6</td><td>-10.2</td><td>1.0</td><td>-9.9</td><td>3.7E-23</td><td>7.2E-22</td><td>MB231/MB468</td><td>MITO</td></tr>
<tr><td>>MI0000086|hsa-mir-28&WithFlank&3|+|188688775|188688872@60.79.20(+1A)&[MIMAT0004502&hsa-miR-28-3p&offsets|0|-2(+1A);m-28&3|+|188688834|188688855&offsets|0|-2(+1A)]|CACTAGATTGTGAGCTCCTGA</td><td>>MI0000086|hsa-mir-28&WithFlank&3|+|188688775|188688872</td><td>60</td><td>79</td><td>20(+1A)</td><td>CACTAGATTGTGAGCTCCTGA</td><td>1227.6</td><td>14.5</td><td>1.5</td><td>9.8</td><td>7.4E-23</td><td>7.2E-22</td><td>BT20/MB468</td><td>CYTO</td></tr>
<tr><td>>hg38_21_+_8433172_8446622_RNA45SN1_45S_50ntFlanks@10466.10497.32|CGTTGCCCTCGGCCGATCGAAAGGGAGTCGGG</td><td>>hg38_21_+_8433172_8446622_RNA45SN1_45S_50ntFlanks</td><td>10466</td><td>10497</td><td>32</td><td>CGTTGCCCTCGGCCGATCGAAAGGGAGTCGGG</td><td>895.2</td><td>-13.7</td><td>1.4</td><td>-10.0</td><td>2.5E-23</td><td>7.3E-22</td><td>MB231/MB468</td><td>MP</td></tr>
<tr><td>>hg38_21_+_8433172_8446622_RNA45SN1_45S_50ntFlanks@11769.11806.38|TAATTAGTGACGCGCATGAATGGATGAACGAGATTCCC</td><td>>hg38_21_+_8433172_8446622_RNA45SN1_45S_50ntFlanks</td><td>11769</td><td>11806</td><td>38</td><td>TAATTAGTGACGCGCATGAATGGATGAACGAGATTCCC</td><td>747.4</td><td>-14.0</td><td>1.4</td><td>-9.8</td><td>1.5E-22</td><td>7.4E-22</td><td>BT20/MB468</td><td>NUC</td></tr>
<tr><td>>trna111_HisGTG_1_-_147774845_147774916@-1T.34.35,trna118_HisGTG_1_-_145396881_145396952@-1T.34.35,trna16_HisGTG_1_+_146544773_146544844@-1T.34.35,trna1_HisGTG_15_+_45493349_45493420@-1T.34.35,trna21_HisGTG_1_+_147753471_147753542@-1T.34.35,trna33_HisGTG_6_+_27125906_27125977@-1T.34.35,trna7_HisGTG_9_-_14433938_14434009@-1T.34.35,trna8_HisGTG_15_-_45492611_45492682@-1T.34.35,trna9_HisGTG_15_-_45490804_45490875@-1T.34.35|TGCCGTGATCGTATAGTGGTTAGTACTCTGCGTTG</td><td>>trna111_HisGTG_1_-_147774845_147774916</td><td>-1T</td><td>34</td><td>35</td><td>TGCCGTGATCGTATAGTGGTTAGTACTCTGCGTTG</td><td>6944.2</td><td>9.4</td><td>0.9</td><td>10.0</td><td>1.8E-23</td><td>7.4E-22</td><td>MB231/MB468</td><td>CYTO</td></tr>
<tr><td>>hg38_21_+_8433172_8446622_RNA45SN1_45S_50ntFlanks@4823.4859.37|ATGACCCGCCGGGCAGCTTCCGGGAAACCAAAGTCTT</td><td>>hg38_21_+_8433172_8446622_RNA45SN1_45S_50ntFlanks</td><td>4823</td><td>4859</td><td>37</td><td>ATGACCCGCCGGGCAGCTTCCGGGAAACCAAAGTCTT</td><td>781.1</td><td>13.9</td><td>1.4</td><td>9.8</td><td>7.6E-23</td><td>7.4E-22</td><td>BT20/MB468</td><td>CYTO</td></tr>
<tr><td>>hg38_21_+_8433172_8446622_RNA45SN1_45S_50ntFlanks@3984.4002.19|GCGGCTTTGGTGACTCTAG</td><td>>hg38_21_+_8433172_8446622_RNA45SN1_45S_50ntFlanks</td><td>3984</td><td>4002</td><td>19</td><td>GCGGCTTTGGTGACTCTAG</td><td>5247.4</td><td>-16.1</td><td>1.6</td><td>-9.9</td><td>3.9E-23</td><td>7.4E-22</td><td>MB231/MB468</td><td>MITO</td></tr>
<tr><td>>MI0001445|hsa-mir-423&WithFlank&17|+|30117073|30117178@23.42.20(+1A)&[MIMAT0004748&hsa-miR-423-5p&offsets|0|-3(+1A);m-141&17|+|30117095|30117117&offsets|0|-3(+1A)]|TGAGGGGCAGAGAGCGAGACA</td><td>>MI0001445|hsa-mir-423&WithFlank&17|+|30117073|30117178</td><td>23</td><td>42</td><td>20(+1A)</td><td>TGAGGGGCAGAGAGCGAGACA</td><td>587.1</td><td>13.5</td><td>1.4</td><td>9.8</td><td>7.7E-23</td><td>7.5E-22</td><td>BT20/MB468</td><td>CYTO</td></tr>
<tr><td>>hg38_21_+_8433172_8446622_RNA45SN1_45S_50ntFlanks@4730.4768.39|CAAGAACGAAAGTCGGAGGTTCGAAGACGATCAGATACC</td><td>>hg38_21_+_8433172_8446622_RNA45SN1_45S_50ntFlanks</td><td>4730</td><td>4768</td><td>39</td><td>CAAGAACGAAAGTCGGAGGTTCGAAGACGATCAGATACC</td><td>890.8</td><td>13.8</td><td>1.4</td><td>10.0</td><td>1.9E-23</td><td>7.6E-22</td><td>MB231/MB468</td><td>CYTO</td></tr>
<tr><td>>hg38_21_+_8433172_8446622_RNA45SN1_45S_50ntFlanks@9622.9648.27|CTAGTAGCTGGTTCCCTCCGAAGTTTC</td><td>>hg38_21_+_8433172_8446622_RNA45SN1_45S_50ntFlanks</td><td>9622</td><td>9648</td><td>27</td><td>CTAGTAGCTGGTTCCCTCCGAAGTTTC</td><td>1261.4</td><td>-14.1</td><td>1.4</td><td>-9.9</td><td>3.9E-23</td><td>7.6E-22</td><td>MB231/MB468</td><td>MITO</td></tr>
<tr><td>>MI0000067|hsa-let-7f-1&WithFlank&9|+|94176341|94176439@13.35.23&[MIMAT0000067&hsa-let-7f-5p&offsets|0|+1;m-20&9|+|94176353|94176374&offsets|0|+1]|TGAGGTAGTAGATTGTATAGTTG</td><td>>MI0000067|hsa-let-7f-1&WithFlank&9|+|94176341|94176439</td><td>13</td><td>35</td><td>23</td><td>TGAGGTAGTAGATTGTATAGTTG</td><td>661.3</td><td>7.0</td><td>0.7</td><td>9.8</td><td>7.8E-23</td><td>7.6E-22</td><td>BT20/MB468</td><td>CYTO</td></tr>
<tr><td>>am_MI0005763|hsa-mir-941-1&WithFlank&20|+|63919443|63919526@53.73.21;MI0005764|hsa-mir-941-2&WithFlank&20|+|63919499|63919582@53.73.21;MI0005765|hsa-mir-941-3&WithFlank&20|+|63919555|63919638@53.73.21;MI0005766|hsa-mir-941-4&WithFlank&20|+|63919750|63919833@53.73.21;MI0031520|hsa-mir-941-5&WithFlank&20|+|63919862|63919945@53.73.21&[MIMAT0004984&hsa-miR-941&offsets|0|-2;m-263&20|+|63919495|63919517&offsets|0|-2];[MIMAT0004984_1&hsa-miR-941&offsets|0|-2;m-263&20|+|63919551|63919573&offsets|0|-2];[MIMAT0004984_2&hsa-miR-941&offsets|0|-2;m-263&20|+|63919607|63919629&offsets|0|-2];[MIMAT0004984_3&hsa-miR-941&offsets|0|-2;m-263&20|+|63919802|63919824&offsets|0|-2];[MIMAT0004984_4&hsa-miR-941&offsets|0|-2;m-263&20|+|63919914|63919936&offsets|0|-2]|CACCCGGCTGTGTGCACATGT</td><td>>am_MI0005763|hsa-mir-941-1&WithFlank&20|+|63919443|63919526</td><td>53</td><td>73</td><td>21</td><td>CACCCGGCTGTGTGCACATGT</td><td>249.6</td><td>12.2</td><td>1.2</td><td>9.8</td><td>7.8E-23</td><td>7.6E-22</td><td>BT20/MB468</td><td>CYTO</td></tr>
<tr><td>>MI0000804|hsa-mir-328&WithFlank&16|-|67202315|67202401@54.75.22(+1U)&[MIMAT0000752&hsa-miR-328-3p&offsets|0|0(+1U);m-160&16|-|67202327|67202348&offsets|0|0(+1U)]|CTGGCCCTCTCTGCCCTTCCGTT</td><td>>MI0000804|hsa-mir-328&WithFlank&16|-|67202315|67202401</td><td>54</td><td>75</td><td>22(+1U)</td><td>CTGGCCCTCTCTGCCCTTCCGTT</td><td>270.4</td><td>12.3</td><td>1.3</td><td>9.8</td><td>7.9E-23</td><td>7.6E-22</td><td>BT20/MB468</td><td>CYTO</td></tr>
<tr><td>>hg38_21_+_8433172_8446622_RNA45SN1_45S_50ntFlanks@5420.5443.24|CCGATTGGATGGTTTAGTGAGGCC</td><td>>hg38_21_+_8433172_8446622_RNA45SN1_45S_50ntFlanks</td><td>5420</td><td>5443</td><td>24</td><td>CCGATTGGATGGTTTAGTGAGGCC</td><td>1533.6</td><td>-14.4</td><td>1.5</td><td>-9.9</td><td>4.0E-23</td><td>7.6E-22</td><td>MB231/MB468</td><td>MITO</td></tr>
<tr><td>>hg38_21_+_8433172_8446622_RNA45SN1_45S_50ntFlanks@8380.8402.23|CAAGAGGGCGTGAAACCGTTAAG</td><td>>hg38_21_+_8433172_8446622_RNA45SN1_45S_50ntFlanks</td><td>8380</td><td>8402</td><td>23</td><td>CAAGAGGGCGTGAAACCGTTAAG</td><td>7124.4</td><td>-12.3</td><td>1.2</td><td>-9.9</td><td>4.0E-23</td><td>7.7E-22</td><td>MB231/MB468</td><td>MITO</td></tr>
<tr><td>>MI0000441|hsa-mir-30b&WithFlank&8|-|134800514|134800613@23.44.22&[MIMAT0000420&hsa-miR-30b-5p&offsets|0|0;m-76&8|-|134800570|134800591&offsets|0|0]|TGTAAACATCCTACACTCAGCT</td><td>>MI0000441|hsa-mir-30b&WithFlank&8|-|134800514|134800613</td><td>23</td><td>44</td><td>22</td><td>TGTAAACATCCTACACTCAGCT</td><td>3277.7</td><td>6.6</td><td>0.7</td><td>9.9</td><td>4.1E-23</td><td>7.7E-22</td><td>MB231/MB468</td><td>NUC</td></tr>
<tr><td>>MI0003776|hsa-mir-320b-1&WithFlank&1|+|116671740|116671823@48.66.19(+1A);MI0003839|hsa-mir-320b-2&WithFlank&1|-|224257034|224257116@47.65.19(+1A);MI0003778|hsa-mir-320c-1&WithFlank&18|+|21683512|21683595@48.66.19(+1A);MI0008191|hsa-mir-320c-2&WithFlank&18|+|24321669|24321752@48.66.19(+1A);MI0000542|hsa-mir-320a&WithFlank&8|-|22244960|22245043@48.66.19(+1A)&[MIMAT0005792&hsa-miR-320b&offsets|0|-3(+1A);m-183&1|+|116671787|116671806&offsets|0|-1(+1A)];[MIMAT0005792_1&hsa-miR-320b&offsets|0|-3(+1A);m-183&1|-|224257051|224257070&offsets|0|-1(+1A)];[MIMAT0005793&hsa-miR-320c&offsets|0|-1(+1A);m-222&18|+|21683559|21683578&offsets|0|-1(+1A)];[MIMAT0005793_1&hsa-miR-320c&offsets|0|-1(+1A);m-219&18|+|24321716|24321735&offsets|0|-1(+1A)];[MIMAT0000510&hsa-miR-320a-3p&offsets|0|-3(+1A);m-60&8|-|22244975|22244996&offsets|0|-3(+1A)]|AAAAGCTGGGTTGAGAGGGA</td><td>>MI0003776|hsa-mir-320b-1&WithFlank&1|+|116671740|116671823</td><td>48</td><td>66</td><td>19(+1A)</td><td>AAAAGCTGGGTTGAGAGGGA</td><td>1522.1</td><td>11.1</td><td>1.1</td><td>9.8</td><td>8.1E-23</td><td>7.8E-22</td><td>BT20/MB468</td><td>CYTO</td></tr>
</table>

<table>
<tr><th></th><th></th><th></th><th></th><th></th><th></th><th></th><th></th><th></th><th></th><th></th><th></th><th></th><th></th></tr>
<tr><td>>MI0000070|hsa-mir-16-1&WithFlank&13|-|50048967|50049067@20.40.21(+1A);MI0000115|hsa-mir-16-2&WithFlank&3|+|160404739|160404831@16.36.21(+1A)&[MIMAT0000069&hsa-miR-16-5p&offsets|0|-1(+1A);m-62&13|-|50049027|50049048&offsets|0|-1(+1A)];[MIMAT0000069_1&hsa-miR-16-5p&offsets|0|-1(+1A);m-61&3|+|160404754|160404775&offsets|0|-1(+1A)]|TAGCAGCACGTAAATATTGGCA</td><td>>MI0000070|hsa-mir-16-1&WithFlank&13|-|50048967|50049067</td><td>20</td><td>40</td><td>21(+1A)</td><td>TAGCAGCACGTAAATATTGGCA</td><td>2637.9</td><td>8.4</td><td>0.9</td><td>9.8</td><td>8.2E-23</td><td>7.9E-22</td><td>BT20/MB468</td><td>CYTO</td></tr>
<tr><td>>MI0000065|hsa-let-7d&WithFlank&9|+|94178828|94178926@68.88.21&[MIMAT0004484&hsa-let-7d-3p&offsets|0|-1;m-98&9|+|94178895|94178916&offsets|0|-1]|CTATACGACCTGCTGCCTTTC</td><td>>MI0000065|hsa-let-7d&WithFlank&9|+|94178828|94178926</td><td>68</td><td>88</td><td>21</td><td>CTATACGACCTGCTGCCTTTC</td><td>505.0</td><td>6.9</td><td>0.7</td><td>9.9</td><td>6.0E-23</td><td>7.9E-22</td><td>BT20/MB468</td><td>MITO</td></tr>
<tr><td>>hg38_21_+_8433172_8446622_RNA45SN1_45S_50ntFlanks@7975.8010.36|CGCGACCTCAGATCAGACGTGGCGACCCGCTGAATT</td><td>>hg38_21_+_8433172_8446622_RNA45SN1_45S_50ntFlanks</td><td>7975</td><td>8010</td><td>36</td><td>CGCGACCTCAGATCAGACGTGGCGACCCGCTGAATT</td><td>106695.2</td><td>8.0</td><td>0.8</td><td>10.0</td><td>2.0E-23</td><td>7.9E-22</td><td>MB231/MB468</td><td>CYTO</td></tr>
<tr><td>>MI0000736|hsa-mir-30c-1&WithFlank&1|+|40757278|40757378@23.45.23;MI0000254|hsa-mir-30c-2&WithFlank&6|-|71376954|71377037@13.35.23&[MIMAT0000244&hsa-miR-30c-5p&offsets|0|0;m-57&1|+|40757300|40757323&offsets|0|-1];[MIMAT0000244_1&hsa-miR-30c-5p&offsets|0|0;m-56&6|-|71377002|71377025&offsets|0|-1]|TGTAAACATCCTACACTCTCAGC</td><td>>MI0000736|hsa-mir-30c-1&WithFlank&1|+|40757278|40757378</td><td>23</td><td>45</td><td>23</td><td>TGTAAACATCCTACACTCTCAGC</td><td>15125.8</td><td>6.2</td><td>0.6</td><td>9.8</td><td>8.3E-23</td><td>8.0E-22</td><td>BT20/MB468</td><td>CYTO</td></tr>
<tr><td>>hg38_21_+_8433172_8446622_RNA45SN1_45S_50ntFlanks@10101.10127.27|CGCGCGCGTGTGGTGTGCGTCGGAGGG</td><td>>hg38_21_+_8433172_8446622_RNA45SN1_45S_50ntFlanks</td><td>10101</td><td>10127</td><td>27</td><td>CGCGCGCGTGTGGTGTGCGTCGGAGGG</td><td>2461.7</td><td>-15.0</td><td>1.5</td><td>-9.9</td><td>4.2E-23</td><td>8.0E-22</td><td>MB231/MB468</td><td>MITO</td></tr>
<tr><td>>MI0000299|hsa-mir-222&WithFlank&X|-|45747009|45747130@75.97.23&[MIMAT0000279&hsa-miR-222-3p&offsets|0|+2;m-140&X|-|45747033|45747056&offsets|0|-1]|AGCTACATCTGGCTACTGGGTCT</td><td>>MI0000299|hsa-mir-222&WithFlank&X|-|45747009|45747130</td><td>75</td><td>97</td><td>23</td><td>AGCTACATCTGGCTACTGGGTCT</td><td>5124.9</td><td>6.8</td><td>0.7</td><td>9.9</td><td>6.2E-23</td><td>8.0E-22</td><td>BT20/MB468</td><td>MITO</td></tr>
<tr><td>>hg38_21_+_8433172_8446622_RNA45SN1_45S_50ntFlanks@3984.4026.43|GCGGCTTTGGTGACTCTAGATAACCTCGGGCCGATCGCACGCC</td><td>>hg38_21_+_8433172_8446622_RNA45SN1_45S_50ntFlanks</td><td>3984</td><td>4026</td><td>43</td><td>GCGGCTTTGGTGACTCTAGATAACCTCGGGCCGATCGCACGCC</td><td>403.7</td><td>-13.1</td><td>1.3</td><td>-9.8</td><td>1.7E-22</td><td>8.0E-22</td><td>BT20/MB468</td><td>NUC</td></tr>
<tr><td>>hg38_21_+_8433172_8446622_RNA45SN1_45S_50ntFlanks@9739.9760.22|AAACGATCTCAACCTATTCTCA</td><td>>hg38_21_+_8433172_8446622_RNA45SN1_45S_50ntFlanks</td><td>9739</td><td>9760</td><td>22</td><td>AAACGATCTCAACCTATTCTCA</td><td>353.7</td><td>-12.9</td><td>1.3</td><td>-9.8</td><td>1.7E-22</td><td>8.0E-22</td><td>BT20/MB468</td><td>NUC</td></tr>
<tr><td>>hg38_21_+_8433172_8446622_RNA45SN1_45S_50ntFlanks@11208.11233.26|GCGGCGACTCTGGACGCGAGCCGGGC</td><td>>hg38_21_+_8433172_8446622_RNA45SN1_45S_50ntFlanks</td><td>11208</td><td>11233</td><td>26</td><td>GCGGCGACTCTGGACGCGAGCCGGGC</td><td>4590.8</td><td>-7.1</td><td>0.7</td><td>-9.9</td><td>4.3E-23</td><td>8.1E-22</td><td>MB231/MB468</td><td>NUC</td></tr>
<tr><td>>hg38_21_+_8433172_8446622_RNA45SN1_45S_50ntFlanks@12416.12440.25|TTCGATGTCGGCTCTTCCTATCATT</td><td>>hg38_21_+_8433172_8446622_RNA45SN1_45S_50ntFlanks</td><td>12416</td><td>12440</td><td>25</td><td>TTCGATGTCGGCTCTTCCTATCATT</td><td>501.5</td><td>5.1</td><td>0.5</td><td>9.9</td><td>6.3E-23</td><td>8.1E-22</td><td>BT20/MB468</td><td>MITO</td></tr>
<tr><td>>MI0003513|hsa-mir-455&WithFlank&9|+|114209428|114209535@22.44.23&[MIMAT0003150&hsa-miR-455-5p&offsets|0|+1;m-232&9|+|114209449|114209470&offsets|0|+1]|TATGTGCCTTTGGACTACATCGT</td><td>>MI0003513|hsa-mir-455&WithFlank&9|+|114209428|114209535</td><td>22</td><td>44</td><td>23</td><td>TATGTGCCTTTGGACTACATCGT</td><td>1005.4</td><td>13.2</td><td>1.3</td><td>9.9</td><td>2.8E-23</td><td>8.2E-22</td><td>MB231/MB468</td><td>MP</td></tr>
<tr><td>>MI0000285|hsa-mir-205&WithFlank&1|+|209432127|209432248@40.60.21(+1U)&[MIMAT0000266&hsa-miR-205-5p&offsets|0|-1(+1U);m-351&1|+|209432166|209432187&offsets|0|-1(+1U)]|TCCTTCATTCCACCGGAGTCTT</td><td>>MI0000285|hsa-mir-205&WithFlank&1|+|209432127|209432248</td><td>40</td><td>60</td><td>21(+1U)</td><td>TCCTTCATTCCACCGGAGTCTT</td><td>2397.7</td><td>8.0</td><td>0.8</td><td>9.8</td><td>8.5E-23</td><td>8.2E-22</td><td>BT20/MB468</td><td>CYTO</td></tr>
<tr><td>>hg38_21_+_8433172_8446622_RNA45SN1_45S_50ntFlanks@10348.10383.36|AAACGAGAACTTTGAAGGCCGAAGTGGAGAAGGGTT</td><td>>hg38_21_+_8433172_8446622_RNA45SN1_45S_50ntFlanks</td><td>10348</td><td>10383</td><td>36</td><td>AAACGAGAACTTTGAAGGCCGAAGTGGAGAAGGGTT</td><td>3675.5</td><td>-6.7</td><td>0.7</td><td>-9.8</td><td>1.7E-22</td><td>8.2E-22</td><td>BT20/MB468</td><td>NUC</td></tr>
<tr><td>>hg38_1_-_228634819_228635039_RNA5S12_5S_50ntFlanks@146.171.26|TGGGAATACCGGGTGCTGTAGGCTTT</td><td>>hg38_1_-_228634819_228635039_RNA5S12_5S_50ntFlanks</td><td>146</td><td>171</td><td>26</td><td>TGGGAATACCGGGTGCTGTAGGCTTT</td><td>4819.1</td><td>7.6</td><td>0.8</td><td>10.0</td><td>2.1E-23</td><td>8.3E-22</td><td>MB231/MB468</td><td>CYTO</td></tr>
<tr><td>>hg38_21_+_8433172_8446622_RNA45SN1_45S_50ntFlanks@10922.10946.25|CTGGACGAGGCGCCGCCGCCCCCCC</td><td>>hg38_21_+_8433172_8446622_RNA45SN1_45S_50ntFlanks</td><td>10922</td><td>10946</td><td>25</td><td>CTGGACGAGGCGCCGCCGCCCCCCC</td><td>1923.7</td><td>-13.6</td><td>1.4</td><td>-10.0</td><td>2.1E-23</td><td>8.3E-22</td><td>MB231/MB468</td><td>CYTO</td></tr>
<tr><td>>hg38_21_+_8433172_8446622_RNA45SN1_45S_50ntFlanks@10392.10431.40|ACAGCAGTTGAACATGGGTCAGTCGGTCCTGAGAGATGGG</td><td>>hg38_21_+_8433172_8446622_RNA45SN1_45S_50ntFlanks</td><td>10392</td><td>10431</td><td>40</td><td>ACAGCAGTTGAACATGGGTCAGTCGGTCCTGAGAGATGGG</td><td>1106.5</td><td>-14.5</td><td>1.5</td><td>-9.8</td><td>1.7E-22</td><td>8.3E-22</td><td>BT20/MB468</td><td>NUC</td></tr>
<tr><td>>hg38_21_+_8433172_8446622_RNA45SN1_45S_50ntFlanks@6746.6770.25|CACTTCGAACGCACTTGCGGCCCCG</td><td>>hg38_21_+_8433172_8446622_RNA45SN1_45S_50ntFlanks</td><td>6746</td><td>6770</td><td>25</td><td>CACTTCGAACGCACTTGCGGCCCCG</td><td>3141.9</td><td>-11.3</td><td>1.1</td><td>-9.9</td><td>4.4E-23</td><td>8.4E-22</td><td>MB231/MB468</td><td>MITO</td></tr>
<tr><td>>MI0000434|hsa-let-7i&WithFlank&12|+|62603680|62603775@12.31.20(+1C)&[MIMAT0000415&hsa-let-7i-5p&offsets|0|-2(+1C);m-49&12|+|62603691|62603712&offsets|0|-2(+1C)]|TGAGGTAGTAGTTTGTGCTGC</td><td>>MI0000434|hsa-let-7i&WithFlank&12|+|62603680|62603775</td><td>12</td><td>31</td><td>20(+1C)</td><td>TGAGGTAGTAGTTTGTGCTGC</td><td>728.0</td><td>7.3</td><td>0.7</td><td>9.8</td><td>8.7E-23</td><td>8.4E-22</td><td>BT20/MB468</td><td>CYTO</td></tr>
<tr><td>>hg38_21_+_8433172_8446622_RNA45SN1_45S_50ntFlanks@11452.11478.27|CTCCTCGGCGCGCGGCGGCGGCGGCGG</td><td>>hg38_21_+_8433172_8446622_RNA45SN1_45S_50ntFlanks</td><td>11452</td><td>11478</td><td>27</td><td>CTCCTCGGCGCGCGGCGGCGGCGGCGG</td><td>1155.3</td><td>-14.0</td><td>1.4</td><td>-9.9</td><td>2.8E-23</td><td>8.4E-22</td><td>MB231/MB468</td><td>MP</td></tr>
<tr><td>>MI0000070|hsa-mir-16-1&WithFlank&13|-|50048967|50049067@20.42.23;MI0000115|hsa-mir-16-2&WithFlank&3|+|160404739|160404831@16.38.23&[MIMAT0000069&hsa-miR-16-5p&offsets|0|+1;m-62&13|-|50049027|50049048&offsets|0|+1];[MIMAT0000069_1&hsa-miR-16-5p&offsets|0|+1;m-61&3|+|160404754|160404775&offsets|0|+1]|TAGCAGCACGTAAATATTGGCGT</td><td>>MI0000070|hsa-mir-16-1&WithFlank&13|-|50048967|50049067</td><td>20</td><td>42</td><td>23</td><td>TAGCAGCACGTAAATATTGGCGT</td><td>3620.3</td><td>7.5</td><td>0.8</td><td>9.8</td><td>8.8E-23</td><td>8.4E-22</td><td>BT20/MB468</td><td>CYTO</td></tr>
<tr><td>>hg38_21_+_8433172_8446622_RNA45SN1_45S_50ntFlanks@4554.4573.20|CCGCAGCTAGGAATAATGGA</td><td>>hg38_21_+_8433172_8446622_RNA45SN1_45S_50ntFlanks</td><td>4554</td><td>4573</td><td>20</td><td>CCGCAGCTAGGAATAATGGA</td><td>479.4</td><td>-13.3</td><td>1.4</td><td>-9.8</td><td>1.8E-22</td><td>8.5E-22</td><td>BT20/MB468</td><td>NUC</td></tr>
<tr><td>>MI0003778|hsa-mir-320c-1&WithFlank&18|+|21683512|21683595@48.67.20;MI0008191|hsa-mir-320c-2&WithFlank&18|+|24321669|24321752@48.67.20&[MIMAT0005793&hsa-miR-320c&offsets|0|0;m-222&18|+|21683559|21683578&offsets|0|0];[MIMAT0005793_1&hsa-miR-320c&offsets|0|0;m-219&18|+|24321716|24321735&offsets|0|0]|AAAAGCTGGGTTGAGAGGGT</td><td>>MI0003778|hsa-mir-320c-1&WithFlank&18|+|21683512|21683595</td><td>48</td><td>67</td><td>20</td><td>AAAAGCTGGGTTGAGAGGGT</td><td>431.8</td><td>13.2</td><td>1.3</td><td>9.9</td><td>6.5E-23</td><td>8.5E-22</td><td>BT20/MB468</td><td>MITO</td></tr>
<tr><td>>hg38_21_+_8433172_8446622_RNA45SN1_45S_50ntFlanks@11316.11340.25|CACCCCACGTCTCGTCGCGCGCGCG</td><td>>hg38_21_+_8433172_8446622_RNA45SN1_45S_50ntFlanks</td><td>11316</td><td>11340</td><td>25</td><td>CACCCCACGTCTCGTCGCGCGCGCG</td><td>3290.5</td><td>-15.5</td><td>1.6</td><td>-9.9</td><td>4.5E-23</td><td>8.5E-22</td><td>MB231/MB468</td><td>MITO</td></tr>
<tr><td>>hg38_21_+_8433172_8446622_RNA45SN1_45S_50ntFlanks@9960.9994.35|TCGGAATCCGCTAAGGAGTGTGTAACAACTCACCT</td><td>>hg38_21_+_8433172_8446622_RNA45SN1_45S_50ntFlanks</td><td>9960</td><td>9994</td><td>35</td><td>TCGGAATCCGCTAAGGAGTGTGTAACAACTCACCT</td><td>418.5</td><td>13.2</td><td>1.3</td><td>9.9</td><td>6.6E-23</td><td>8.5E-22</td><td>BT20/MB468</td><td>MITO</td></tr>
<tr><td>>hg38_21_+_8433172_8446622_RNA45SN1_45S_50ntFlanks@6777.6806.30|TCCCGGGGCTACGCCTGTCTGAGCGTCGCT</td><td>>hg38_21_+_8433172_8446622_RNA45SN1_45S_50ntFlanks</td><td>6777</td><td>6806</td><td>30</td><td>TCCCGGGGCTACGCCTGTCTGAGCGTCGCT</td><td>489.2</td><td>5.4</td><td>0.5</td><td>10.0</td><td>2.2E-23</td><td>8.6E-22</td><td>MB231/MB468</td><td>CYTO</td></tr>
<tr><td>>MI0000102|hsa-mir-100&WithFlank&11|-|122152223|122152314@19.38.20(+1G)&[MIMAT0000098&hsa-miR-100-5p&offsets|0|-2(+1G);m-27&11|-|122152276|122152296&offsets|0|-1(+1G)]|AACCCGTAGATCCGAACTTGG</td><td>>MI0000102|hsa-mir-100&WithFlank&11|-|122152223|122152314</td><td>19</td><td>38</td><td>20(+1G)</td><td>AACCCGTAGATCCGAACTTGG</td><td>4324.1</td><td>16.1</td><td>1.6</td><td>10.0</td><td>2.2E-23</td><td>8.7E-22</td><td>MB231/MB468</td><td>CYTO</td></tr>
<tr><td>>MI0000748|hsa-mir-130b&WithFlank&22|+|21653298|21653391@19.40.22&[MIMAT0004680&hsa-miR-130b-5p&offsets|0|+1;m-234&22|+|21653316|21653337&offsets|0|0]|ACTCTTTCCCTGTTGCACTACT</td><td>>MI0000748|hsa-mir-130b&WithFlank&22|+|21653298|21653391</td><td>19</td><td>40</td><td>22</td><td>ACTCTTTCCCTGTTGCACTACT</td><td>1935.2</td><td>7.8</td><td>0.8</td><td>10.0</td><td>2.2E-23</td><td>8.7E-22</td><td>MB231/MB468</td><td>CYTO</td></tr>
<tr><td>>am_MI0005763|hsa-mir-941-1&WithFlank&20|+|63919443|63919526@53.74.22(+1U);MI0005764|hsa-mir-941-2&WithFlank&20|+|63919499|63919582@53.74.22(+1U);MI0005765|hsa-mir-941-3&WithFlank&20|+|63919555|63919638@53.74.22(+1U);MI0005766|hsa-mir-941-4&WithFlank&20|+|63919750|63919833@53.74.22(+1U);MI0031520|hsa-mir-941-5&WithFlank&20|+|63919862|63919945@53.74.22(+1U)&[MIMAT0004984&hsa-miR-941&offsets|0|-1(+1U);m-263&20|+|63919495|63919517&offsets|0|-1(+1U)];[MIMAT0004984_1&hsa-miR-941&offsets|0|-1(+1U);m-263&20|+|63919551|63919573&offsets|0|-1(+1U)];[MIMAT0004984_2&hsa-miR-941&offsets|0|-1(+1U);m-263&20|+|63919607|63919629&offsets|0|-1(+1U)];[MIMAT0004984_3&hsa-miR-941&offsets|0|-1(+1U);m-263&20|+|63919802|63919824&offsets|0|-1(+1U)];[MIMAT0004984_4&hsa-miR-941&offsets|0|-1(+1U);m-263&20|+|63919914|63919936&offsets|0|-1(+1U)]|CACCCGGCTGTGTGCACATGTGT</td><td>>am_MI0005763|hsa-mir-941-1&WithFlank&20|+|63919443|63919526</td><td>53</td><td>74</td><td>22(+1U)</td><td>CACCCGGCTGTGTGCACATGTGT</td><td>3737.4</td><td>7.1</td><td>0.7</td><td>10.0</td><td>2.2E-23</td><td>8.7E-22</td><td>MB231/MB468</td><td>CYTO</td></tr>
<tr><td>>trnaMT_PheGAA_MT_+_577_647@35.56.22|GAAAATGTTTAGACGGGCTCAC</td><td>>trnaMT_PheGAA_MT_+_577_647</td><td>35</td><td>56</td><td>22</td><td>GAAAATGTTTAGACGGGCTCAC</td><td>397.8</td><td>12.9</td><td>1.3</td><td>9.8</td><td>9.1E-23</td><td>8.7E-22</td><td>BT20/MB468</td><td>CYTO</td></tr>
<tr><td>>MI0000272|hsa-mir-182&WithFlank&7|-|129770377|129770498@29.54.26(+1U)&[MIMAT0000259&hsa-miR-182-5p&offsets|0|+2(+1U);m-22&7|-|129770449|129770470&offsets|0|+4(+1U)]|TTTGGCAATGGTAGAACTCACACTGGT</td><td>>MI0000272|hsa-mir-182&WithFlank&7|-|129770377|129770498</td><td>29</td><td>54</td><td>26(+1U)</td><td>TTTGGCAATGGTAGAACTCACACTGGT</td><td>718.7</td><td>7.3</td><td>0.7</td><td>9.9</td><td>6.8E-23</td><td>8.8E-22</td><td>BT20/MB468</td><td>MITO</td></tr>
<tr><td>>MI0000266|hsa-mir-10a&WithFlank&17|-|48579832|48579953@29.50.22(+1A)&[MIMAT0000253&hsa-miR-10a-5p&offsets|+1|0(+1A);m-18&17|-|48579905|48579926&offsets|+1|+1(+1A)]|ACCCTGTAGATCCGAATTTGTGA</td><td>>MI0000266|hsa-mir-10a&WithFlank&17|-|48579832|48579953</td><td>29</td><td>50</td><td>22(+1A)</td><td>ACCCTGTAGATCCGAATTTGTGA</td><td>534.8</td><td>13.0</td><td>1.3</td><td>9.9</td><td>4.7E-23</td><td>8.8E-22</td><td>MB231/MB468</td><td>NUC</td></tr>
<tr><td>>hg38_21_+_8433172_8446622_RNA45SN1_45S_50ntFlanks@3845.3866.22|ACTTGGATAACTGTGGTAATTC</td><td>>hg38_21_+_8433172_8446622_RNA45SN1_45S_50ntFlanks</td><td>3845</td><td>3866</td><td>22</td><td>ACTTGGATAACTGTGGTAATTC</td><td>409.0</td><td>5.0</td><td>0.5</td><td>9.8</td><td>9.3E-23</td><td>8.9E-22</td><td>BT20/MB468</td><td>CYTO</td></tr>
<tr><td>>hg38_21_+_8433172_8446622_RNA45SN1_45S_50ntFlanks@3835.3872.38|CTCCTCTCCTACTTGGATAACTGTGGTAATTCTAGAGC</td><td>>hg38_21_+_8433172_8446622_RNA45SN1_45S_50ntFlanks</td><td>3835</td><td>3872</td><td>38</td><td>CTCCTCTCCTACTTGGATAACTGTGGTAATTCTAGAGC</td><td>1042.7</td><td>-14.5</td><td>1.5</td><td>-9.7</td><td>1.9E-22</td><td>8.9E-22</td><td>BT20/MB468</td><td>NUC</td></tr>
<tr><td>>hg38_21_+_8433172_8446622_RNA45SN1_45S_50ntFlanks@7986.8004.19|ATCAGACGTGGCGACCCGC</td><td>>hg38_21_+_8433172_8446622_RNA45SN1_45S_50ntFlanks</td><td>7986</td><td>8004</td><td>19</td><td>ATCAGACGTGGCGACCCGC</td><td>1246.6</td><td>-14.1</td><td>1.4</td><td>-9.9</td><td>3.0E-23</td><td>9.0E-22</td><td>MB231/MB468</td><td>MP</td></tr>
<tr><td>>hg38_21_+_8433172_8446622_RNA45SN1_45S_50ntFlanks@7975.8013.39|CGCGACCTCAGATCAGACGTGGCGACCCGCTGAATTTAA</td><td>>hg38_21_+_8433172_8446622_RNA45SN1_45S_50ntFlanks</td><td>7975</td><td>8013</td><td>39</td><td>CGCGACCTCAGATCAGACGTGGCGACCCGCTGAATTTAA</td><td>4083.7</td><td>6.7</td><td>0.7</td><td>9.8</td><td>9.4E-23</td><td>9.0E-22</td><td>BT20/MB468</td><td>CYTO</td></tr>
<tr><td>>hg38_21_+_8433172_8446622_RNA45SN1_45S_50ntFlanks@10845.10865.21|ACTTCGGGATAAGGATTGGCT</td><td>>hg38_21_+_8433172_8446622_RNA45SN1_45S_50ntFlanks</td><td>10845</td><td>10865</td><td>21</td><td>ACTTCGGGATAAGGATTGGCT</td><td>7343.0</td><td>-12.2</td><td>1.2</td><td>-9.9</td><td>4.9E-23</td><td>9.1E-22</td><td>MB231/MB468</td><td>NUC</td></tr>
<tr><td>>hg38_21_+_8433172_8446622_RNA45SN1_45S_50ntFlanks@12405.12437.33|CTTTTTGATCCTTCGATGTCGGCTCTTCCTATC</td><td>>hg38_21_+_8433172_8446622_RNA45SN1_45S_50ntFlanks</td><td>12405</td><td>12437</td><td>33</td><td>CTTTTTGATCCTTCGATGTCGGCTCTTCCTATC</td><td>931.6</td><td>-13.7</td><td>1.4</td><td>-9.9</td><td>3.1E-23</td><td>9.1E-22</td><td>MB231/MB468</td><td>MP</td></tr>
</table>

<table>
<tr><th></th><th></th><th></th><th></th><th></th><th></th><th></th><th></th><th></th><th></th><th></th><th></th><th></th><th></th></tr>
<tr><td>>MI0003776|hsa-mir-320b-1&WithFlank&1|+|116671740|116671823@48.66.19;MI0003839|hsa-mir-320b-2&WithFlank&1|-|224257034|224257116@47.65.19;MI0003778|hsa-mir-320c-1&WithFlank&18|+|21683512|21683595@48.66.19;MI0008191|hsa-mir-320c-2&WithFlank&18|+|24321669|24321752@48.66.19;MI0000542|hsa-mir-320a&WithFlank&8|-|22244960|22245043@48.66.19&[MIMAT0005792&hsa-miR-320b&offsets|0|-3;m-183&1|+|116671787|116671806&offsets|0|-1];[MIMAT0005792_1&hsa-miR-320b&offsets|0|-3;m-183&1|-|224257051|224257070&offsets|0|-1];[MIMAT0005793&hsa-miR-320c&offsets|0|-1;m-222&18|+|21683559|21683578&offsets|0|-1];[MIMAT0005793_1&hsa-miR-320c&offsets|0|-1;m-219&18|+|24321716|24321735&offsets|0|-1];[MIMAT0000510&hsa-miR-320a-3p&offsets|0|-3;m-60&8|-|22244975|22244996&offsets|0|-3]|AAAAGCTGGGTTGAGAGGG</td><td>>MI0003776|hsa-mir-320b-1&WithFlank&1|+|116671740|116671823</td><td>48</td><td>66</td><td>19</td><td>AAAAGCTGGGTTGAGAGGG</td><td>322.5</td><td>12.8</td><td>1.3</td><td>9.8</td><td>7.1E-23</td><td>9.1E-22</td><td>BT20/MB468</td><td>MITO</td></tr>
<tr><td>>hg38_21_+_8433172_8446622_RNA45SN1_45S_50ntFlanks@9534.9552.19|AGGAAACTCTGGTGGAGGT</td><td>>hg38_21_+_8433172_8446622_RNA45SN1_45S_50ntFlanks</td><td>9534</td><td>9552</td><td>19</td><td>AGGAAACTCTGGTGGAGGT</td><td>3411.8</td><td>-5.6</td><td>0.6</td><td>-10.0</td><td>2.4E-23</td><td>9.1E-22</td><td>MB231/MB468</td><td>CYTO</td></tr>
<tr><td>>hg38_21_+_8433172_8446622_RNA45SN1_45S_50ntFlanks@11569.11602.34|CAGCCGACTTAGAACTGGTGCGGACCAGGGGAAT</td><td>>hg38_21_+_8433172_8446622_RNA45SN1_45S_50ntFlanks</td><td>11569</td><td>11602</td><td>34</td><td>CAGCCGACTTAGAACTGGTGCGGACCAGGGGAAT</td><td>6557.8</td><td>-7.8</td><td>0.8</td><td>-9.9</td><td>4.9E-23</td><td>9.1E-22</td><td>MB231/MB468</td><td>MITO</td></tr>
<tr><td>>hg38_21_+_8433172_8446622_RNA45SN1_45S_50ntFlanks@11784.11804.21|ATGAATGGATGAACGAGATTC</td><td>>hg38_21_+_8433172_8446622_RNA45SN1_45S_50ntFlanks</td><td>11784</td><td>11804</td><td>21</td><td>ATGAATGGATGAACGAGATTC</td><td>2258.2</td><td>-6.7</td><td>0.7</td><td>-9.9</td><td>4.9E-23</td><td>9.2E-22</td><td>MB231/MB468</td><td>MITO</td></tr>
<tr><td>>hg38_21_+_8433172_8446622_RNA45SN1_45S_50ntFlanks@8079.8103.25|AAGAGCCCAGCGCCGAATCCCCGCC</td><td>>hg38_21_+_8433172_8446622_RNA45SN1_45S_50ntFlanks</td><td>8079</td><td>8103</td><td>25</td><td>AAGAGCCCAGCGCCGAATCCCCGCC</td><td>1457.4</td><td>-14.3</td><td>1.4</td><td>-9.9</td><td>4.9E-23</td><td>9.2E-22</td><td>MB231/MB468</td><td>MITO</td></tr>
<tr><td>>MI0000086|hsa-mir-28&WithFlank&3|+|188688775|188688872@60.81.22(+1A)&[MIMAT0004502&hsa-miR-28-3p&offsets|0|0(+1A);m-28&3|+|188688834|188688855&offsets|0|0(+1A)]|CACTAGATTGTGAGCTCCTGGAA</td><td>>MI0000086|hsa-mir-28&WithFlank&3|+|188688775|188688872</td><td>60</td><td>81</td><td>22(+1A)</td><td>CACTAGATTGTGAGCTCCTGGAA</td><td>520.4</td><td>6.7</td><td>0.7</td><td>9.9</td><td>4.9E-23</td><td>9.2E-22</td><td>MB231/MB468</td><td>NUC</td></tr>
<tr><td>>hg38_21_+_8433172_8446622_RNA45SN1_45S_50ntFlanks@10408.10430.23|GGTCAGTCGGTCCTGAGAGATGG</td><td>>hg38_21_+_8433172_8446622_RNA45SN1_45S_50ntFlanks</td><td>10408</td><td>10430</td><td>23</td><td>GGTCAGTCGGTCCTGAGAGATGG</td><td>3527.7</td><td>-12.1</td><td>1.2</td><td>-9.9</td><td>4.9E-23</td><td>9.2E-22</td><td>MB231/MB468</td><td>NUC</td></tr>
<tr><td>>MI0001445|hsa-mir-423&WithFlank&17|+|30117073|30117178@58.77.20(+1U)&[MIMAT0001340&hsa-miR-423-3p&offsets|-1|-4(+1U);m-100&17|+|30117131|30117153&offsets|-1|-4(+1U)]|AAGCTCGGTCTGAGGCCCCTT</td><td>>MI0001445|hsa-mir-423&WithFlank&17|+|30117073|30117178</td><td>58</td><td>77</td><td>20(+1U)</td><td>AAGCTCGGTCTGAGGCCCCTT</td><td>321.7</td><td>12.8</td><td>1.3</td><td>9.8</td><td>7.2E-23</td><td>9.2E-22</td><td>BT20/MB468</td><td>MITO</td></tr>
<tr><td>>MI0000471|hsa-mir-126&WithFlank&9|+|136670596|136670692@58.78.21&[MIMAT0000445&hsa-miR-126-3p&offsets|0|-1;m-40&9|+|136670653|136670674&offsets|0|-1]|TCGTACCGTGAGTAATAATGC</td><td>>MI0000471|hsa-mir-126&WithFlank&9|+|136670596|136670692</td><td>58</td><td>78</td><td>21</td><td>TCGTACCGTGAGTAATAATGC</td><td>2802.7</td><td>9.6</td><td>1.0</td><td>9.9</td><td>5.0E-23</td><td>9.2E-22</td><td>MB231/MB468</td><td>NUC</td></tr>
<tr><td>>hg38_21_+_8433172_8446622_RNA45SN1_45S_50ntFlanks@10351.10380.30|CGAGAACTTTGAAGGCCGAAGTGGAGAAGG</td><td>>hg38_21_+_8433172_8446622_RNA45SN1_45S_50ntFlanks</td><td>10351</td><td>10380</td><td>30</td><td>CGAGAACTTTGAAGGCCGAAGTGGAGAAGG</td><td>806.2</td><td>-4.4</td><td>0.4</td><td>-9.9</td><td>5.0E-23</td><td>9.3E-22</td><td>MB231/MB468</td><td>NUC</td></tr>
<tr><td>>hg38_21_+_8433172_8446622_RNA45SN1_45S_50ntFlanks@9454.9488.35|CCGCGCCGGGGAGGTGGAGCACGAGCGCACGTGTT</td><td>>hg38_21_+_8433172_8446622_RNA45SN1_45S_50ntFlanks</td><td>9454</td><td>9488</td><td>35</td><td>CCGCGCCGGGGAGGTGGAGCACGAGCGCACGTGTT</td><td>315.8</td><td>-12.7</td><td>1.3</td><td>-9.7</td><td>2.0E-22</td><td>9.3E-22</td><td>BT20/MB468</td><td>NUC</td></tr>
<tr><td>>hg38_21_+_8433172_8446622_RNA45SN1_45S_50ntFlanks@8168.8197.30|CCCAAGTCCTTCTGATCGAGGCCCAGCCCG</td><td>>hg38_21_+_8433172_8446622_RNA45SN1_45S_50ntFlanks</td><td>8168</td><td>8197</td><td>30</td><td>CCCAAGTCCTTCTGATCGAGGCCCAGCCCG</td><td>2918.9</td><td>-15.9</td><td>1.6</td><td>-9.7</td><td>2.0E-22</td><td>9.6E-22</td><td>BT20/MB468</td><td>NUC</td></tr>
<tr><td>>MI0000462|hsa-mir-152&WithFlank&17|-|48037155|48037253@60.80.21&[MIMAT0000438&hsa-miR-152-3p&offsets|0|0;m-97&17|-|48037174|48037194&offsets|0|0]|TCAGTGCATGACAGAACTTGG</td><td>>MI0000462|hsa-mir-152&WithFlank&17|-|48037155|48037253</td><td>60</td><td>80</td><td>21</td><td>TCAGTGCATGACAGAACTTGG</td><td>1446.2</td><td>14.8</td><td>1.5</td><td>9.8</td><td>1.0E-22</td><td>9.6E-22</td><td>BT20/MB468</td><td>CYTO</td></tr>
<tr><td>>MI0000279|hsa-mir-196a-2&WithFlank&12|+|53991732|53991853@31.53.23&[MIMAT0000226&hsa-miR-196a-5p&offsets|0|+1;m-261&12|+|53991762|53991783&offsets|0|+1]|TAGGTAGTTTCATGTTGTTGGGA</td><td>>MI0000279|hsa-mir-196a-2&WithFlank&12|+|53991732|53991853</td><td>31</td><td>53</td><td>23</td><td>TAGGTAGTTTCATGTTGTTGGGA</td><td>2159.5</td><td>11.2</td><td>1.1</td><td>10.2</td><td>3.2E-24</td><td>9.7E-22</td><td>BT20/MB231</td><td>CYTO</td></tr>
<tr><td>>hg38_21_+_8433172_8446622_RNA45SN1_45S_50ntFlanks@10808.10831.24|CAATGTAGGTAAGGGAAGTCGGCA</td><td>>hg38_21_+_8433172_8446622_RNA45SN1_45S_50ntFlanks</td><td>10808</td><td>10831</td><td>24</td><td>CAATGTAGGTAAGGGAAGTCGGCA</td><td>399.9</td><td>-13.1</td><td>1.3</td><td>-9.7</td><td>2.1E-22</td><td>9.7E-22</td><td>BT20/MB468</td><td>NUC</td></tr>
<tr><td>>MI0000542|hsa-mir-320a&WithFlank&8|-|22244960|22245043@50.69.20&[MIMAT0000510&hsa-miR-320a-3p&offsets|+2|0;m-60&8|-|22244975|22244996&offsets|+2|0]|AAGCTGGGTTGAGAGGGCGA</td><td>>MI0000542|hsa-mir-320a&WithFlank&8|-|22244960|22245043</td><td>50</td><td>69</td><td>20</td><td>AAGCTGGGTTGAGAGGGCGA</td><td>334.4</td><td>12.6</td><td>1.3</td><td>9.8</td><td>1.0E-22</td><td>9.8E-22</td><td>BT20/MB468</td><td>CYTO</td></tr>
<tr><td>>hg38_21_+_8433172_8446622_RNA45SN1_45S_50ntFlanks@10534.10570.37|CCGCGAGGCGTCCAGTGCGGTAACGCGACCGATCCCG</td><td>>hg38_21_+_8433172_8446622_RNA45SN1_45S_50ntFlanks</td><td>10534</td><td>10570</td><td>37</td><td>CCGCGAGGCGTCCAGTGCGGTAACGCGACCGATCCCG</td><td>1753.3</td><td>-14.6</td><td>1.5</td><td>-9.9</td><td>5.2E-23</td><td>9.8E-22</td><td>MB231/MB468</td><td>MITO</td></tr>
<tr><td>>hg38_21_+_8433172_8446622_RNA45SN1_45S_50ntFlanks@10690.10709.20|CGTCCGGTGAGCTCTCGCTG</td><td>>hg38_21_+_8433172_8446622_RNA45SN1_45S_50ntFlanks</td><td>10690</td><td>10709</td><td>20</td><td>CGTCCGGTGAGCTCTCGCTG</td><td>2389.2</td><td>-11.0</td><td>1.1</td><td>-9.9</td><td>5.3E-23</td><td>9.8E-22</td><td>MB231/MB468</td><td>MITO</td></tr>
<tr><td>>hg38_21_+_8433172_8446622_RNA45SN1_45S_50ntFlanks@11874.11900.27|AGAAGACCCTGTTGAGCTTGACTCTAG</td><td>>hg38_21_+_8433172_8446622_RNA45SN1_45S_50ntFlanks</td><td>11874</td><td>11900</td><td>27</td><td>AGAAGACCCTGTTGAGCTTGACTCTAG</td><td>1064.9</td><td>-13.8</td><td>1.4</td><td>-9.9</td><td>5.3E-23</td><td>9.8E-22</td><td>MB231/MB468</td><td>MITO</td></tr>
<tr><td>>hg38_21_+_8433172_8446622_RNA45SN1_45S_50ntFlanks@10737.10769.33|TAAATCTCGCGCCGGGCCGTACCCATATCCGCA</td><td>>hg38_21_+_8433172_8446622_RNA45SN1_45S_50ntFlanks</td><td>10737</td><td>10769</td><td>33</td><td>TAAATCTCGCGCCGGGCCGTACCCATATCCGCA</td><td>525.1</td><td>-13.5</td><td>1.4</td><td>-9.7</td><td>2.1E-22</td><td>9.8E-22</td><td>BT20/MB468</td><td>NUC</td></tr>
<tr><td>>MI0005559|hsa-mir-744&WithFlank&17|+|12081893|12082002@17.36.20(+1A)&[MIMAT0004945&hsa-miR-744-5p&offsets|0|-2(+1A);m-172&17|+|12081909|12081929&offsets|0|-1(+1A)]|TGCGGGGCTAGGGCTAACAGA</td><td>>MI0005559|hsa-mir-744&WithFlank&17|+|12081893|12082002</td><td>17</td><td>36</td><td>20(+1A)</td><td>TGCGGGGCTAGGGCTAACAGA</td><td>330.1</td><td>12.6</td><td>1.3</td><td>9.8</td><td>1.0E-22</td><td>9.9E-22</td><td>BT20/MB468</td><td>CYTO</td></tr>
<tr><td>>hg38_21_+_8433172_8446622_RNA45SN1_45S_50ntFlanks@6646.6676.31|TCGTACGACTCTTAGCGGTGGATCACTCGGC</td><td>>hg38_21_+_8433172_8446622_RNA45SN1_45S_50ntFlanks</td><td>6646</td><td>6676</td><td>31</td><td>TCGTACGACTCTTAGCGGTGGATCACTCGGC</td><td>20439.6</td><td>6.1</td><td>0.6</td><td>9.8</td><td>7.8E-23</td><td>9.9E-22</td><td>BT20/MB468</td><td>MITO</td></tr>
<tr><td>>hg38_21_+_8433172_8446622_RNA45SN1_45S_50ntFlanks@11539.11583.45|GCGCGGCGCCTCGCCTCGGCCGGCGCCTAGCAGCCGACTTAGAAC</td><td>>hg38_21_+_8433172_8446622_RNA45SN1_45S_50ntFlanks</td><td>11539</td><td>11583</td><td>45</td><td>GCGCGGCGCCTCGCCTCGGCCGGCGCCTAGCAGCCGACTTAGAAC</td><td>460.5</td><td>12.9</td><td>1.3</td><td>9.9</td><td>2.7E-23</td><td>1.0E-21</td><td>MB231/MB468</td><td>CYTO</td></tr>
<tr><td>>am_MI0000809|hsa-mir-151a&WithFlank&8|-|140732558|140732659@53.73.21(+1U)&[MIMAT0000757&hsa-miR-151a-3p&offsets|0|0(+1U);m-30&8|-|140732587|140732607&offsets|0|0(+1U)]|CTAGACTGAAGCTCCTTGAGGT</td><td>>am_MI0000809|hsa-mir-151a&WithFlank&8|-|140732558|140732659</td><td>53</td><td>73</td><td>21(+1U)</td><td>CTAGACTGAAGCTCCTTGAGGT</td><td>6057.9</td><td>7.4</td><td>0.7</td><td>9.9</td><td>2.7E-23</td><td>1.0E-21</td><td>MB231/MB468</td><td>CYTO</td></tr>
<tr><td>>hg38_21_+_8433172_8446622_RNA45SN1_45S_50ntFlanks@4725.4768.44|TTAATCAAGAACGAAAGTCGGAGGTTCGAAGACGATCAGATACC</td><td>>hg38_21_+_8433172_8446622_RNA45SN1_45S_50ntFlanks</td><td>4725</td><td>4768</td><td>44</td><td>TTAATCAAGAACGAAAGTCGGAGGTTCGAAGACGATCAGATACC</td><td>696.1</td><td>13.4</td><td>1.4</td><td>9.9</td><td>2.7E-23</td><td>1.0E-21</td><td>MB231/MB468</td><td>CYTO</td></tr>
<tr><td>>MI0000079|hsa-mir-23a&WithFlank&19|-|13836581|13836665@54.73.20&[MIMAT0000078&hsa-miR-23a-3p&offsets|+3|+2;m-31&19|-|13836595|13836615&offsets|+3|+2]|ACATTGCCAGGGATTTCCAA</td><td>>MI0000079|hsa-mir-23a&WithFlank&19|-|13836581|13836665</td><td>54</td><td>73</td><td>20</td><td>ACATTGCCAGGGATTTCCAA</td><td>428.3</td><td>11.6</td><td>1.2</td><td>9.7</td><td>2.2E-22</td><td>1.0E-21</td><td>BT20/MB468</td><td>NUC</td></tr>
<tr><td>>hg38_21_+_8433172_8446622_RNA45SN1_45S_50ntFlanks@4006.4026.21|ACCTCGGGCCGATCGCACGCC</td><td>>hg38_21_+_8433172_8446622_RNA45SN1_45S_50ntFlanks</td><td>4006</td><td>4026</td><td>21</td><td>ACCTCGGGCCGATCGCACGCC</td><td>689.7</td><td>-13.9</td><td>1.4</td><td>-9.7</td><td>2.2E-22</td><td>1.0E-21</td><td>BT20/MB468</td><td>NUC</td></tr>
<tr><td>>hg38_21_+_8433172_8446622_RNA45SN1_45S_50ntFlanks@485.516.32|GATGTGGTGACGTCGTGCTCTCCCGGGCCGGG</td><td>>hg38_21_+_8433172_8446622_RNA45SN1_45S_50ntFlanks</td><td>485</td><td>516</td><td>32</td><td>GATGTGGTGACGTCGTGCTCTCCCGGGCCGGG</td><td>826.3</td><td>14.2</td><td>1.4</td><td>9.8</td><td>8.0E-23</td><td>1.0E-21</td><td>BT20/MB468</td><td>MITO</td></tr>
<tr><td>>MI0000088|hsa-mir-30a&WithFlank&6|-|71403545|71403627@12.33.22&[MIMAT0000087&hsa-miR-30a-5p&offsets|0|0;m-15&6|-|71403595|71403616&offsets|0|0]|TGTAAACATCCTCGACTGGAAG</td><td>>MI0000088|hsa-mir-30a&WithFlank&6|-|71403545|71403627</td><td>12</td><td>33</td><td>22</td><td>TGTAAACATCCTCGACTGGAAG</td><td>2491.5</td><td>7.0</td><td>0.7</td><td>9.8</td><td>1.1E-22</td><td>1.0E-21</td><td>BT20/MB468</td><td>CYTO</td></tr>
<tr><td>>trna134_GluTTC_1_-_16861774_16861845@1.35.35,trna5_GluTTC_1_+_17199078_17199149@1.35.35,trna84_GluTTC_1_-_161391883_161391954@1.35.35,trna94_GluTTC_1_-_149664355_149664427@1.35.35|TCCCTGGTGGTCTAGTGGCTAGGATTCGGCGCTTT</td><td>>trna134_GluTTC_1_-_16861774_16861845</td><td>1</td><td>35</td><td>35</td><td>TCCCTGGTGGTCTAGTGGCTAGGATTCGGCGCTTT</td><td>313.5</td><td>12.5</td><td>1.3</td><td>9.8</td><td>1.1E-22</td><td>1.0E-21</td><td>BT20/MB468</td><td>CYTO</td></tr>
<tr><td>>MI0000266|hsa-mir-10a&WithFlank&17|-|48579832|48579953@29.49.21(+1A)&[MIMAT0000253&hsa-miR-10a-5p&offsets|+1|-1(+1A);m-18&17|-|48579905|48579926&offsets|+1|0(+1A)]|ACCCTGTAGATCCGAATTTGTA</td><td>>MI0000266|hsa-mir-10a&WithFlank&17|-|48579832|48579953</td><td>29</td><td>49</td><td>21(+1A)</td><td>ACCCTGTAGATCCGAATTTGTA</td><td>834.6</td><td>14.2</td><td>1.4</td><td>9.8</td><td>8.2E-23</td><td>1.0E-21</td><td>BT20/MB468</td><td>MITO</td></tr>
<tr><td>>MI0003591|hsa-mir-584&WithFlank&5|-|149062307|149062415@22.43.22&[MIMAT0003249&hsa-miR-584-5p&offsets|0|0;m-167&5|-|149062374|149062394&offsets|0|+1]|TTATGGTTTGCCTGGGACTGAG</td><td>>MI0003591|hsa-mir-584&WithFlank&5|-|149062307|149062415</td><td>22</td><td>43</td><td>22</td><td>TTATGGTTTGCCTGGGACTGAG</td><td>422.5</td><td>13.2</td><td>1.3</td><td>9.8</td><td>8.2E-23</td><td>1.0E-21</td><td>BT20/MB468</td><td>MITO</td></tr>
<tr><td>>hg38_21_+_8433172_8446622_RNA45SN1_45S_50ntFlanks@12991.13027.37|CTCCCTCGCTGCGATCTATTGAAAGTCAGCCCTCGAC</td><td>>hg38_21_+_8433172_8446622_RNA45SN1_45S_50ntFlanks</td><td>12991</td><td>13027</td><td>37</td><td>CTCCCTCGCTGCGATCTATTGAAAGTCAGCCCTCGAC</td><td>1525.8</td><td>-14.4</td><td>1.5</td><td>-9.9</td><td>5.8E-23</td><td>1.1E-21</td><td>MB231/MB468</td><td>MITO</td></tr>
<tr><td>>hg38_21_+_8433172_8446622_RNA45SN1_45S_50ntFlanks@10571.10589.19|GAGAAGCCGGCGGGAGCCC</td><td>>hg38_21_+_8433172_8446622_RNA45SN1_45S_50ntFlanks</td><td>10571</td><td>10589</td><td>19</td><td>GAGAAGCCGGCGGGAGCCC</td><td>368.2</td><td>-12.9</td><td>1.3</td><td>-9.7</td><td>2.3E-22</td><td>1.1E-21</td><td>BT20/MB468</td><td>NUC</td></tr>
<tr><td>>hg38_21_+_8433172_8446622_RNA45SN1_45S_50ntFlanks@10771.10807.37|CAGGTCTCCAAGGTGAACAGCCTCTGGCATGTTGGAA</td><td>>hg38_21_+_8433172_8446622_RNA45SN1_45S_50ntFlanks</td><td>10771</td><td>10807</td><td>37</td><td>CAGGTCTCCAAGGTGAACAGCCTCTGGCATGTTGGAA</td><td>364.0</td><td>-12.9</td><td>1.3</td><td>-9.7</td><td>2.3E-22</td><td>1.1E-21</td><td>BT20/MB468</td><td>NUC</td></tr>
<tr><td>>hg38_21_+_8433172_8446622_RNA45SN1_45S_50ntFlanks@13002.13032.31|CGATCTATTGAAAGTCAGCCCTCGACACAAG</td><td>>hg38_21_+_8433172_8446622_RNA45SN1_45S_50ntFlanks</td><td>13002</td><td>13032</td><td>31</td><td>CGATCTATTGAAAGTCAGCCCTCGACACAAG</td><td>432.6</td><td>-13.2</td><td>1.4</td><td>-9.7</td><td>2.3E-22</td><td>1.1E-21</td><td>BT20/MB468</td><td>NUC</td></tr>
<tr><td>>hg38_21_+_8433172_8446622_RNA45SN1_45S_50ntFlanks@9198.9222.25|GTTCTCTCGGGGCCACGCGCGCGTC</td><td>>hg38_21_+_8433172_8446622_RNA45SN1_45S_50ntFlanks</td><td>9198</td><td>9222</td><td>25</td><td>GTTCTCTCGGGGCCACGCGCGCGTC</td><td>6063.1</td><td>-12.6</td><td>1.3</td><td>-9.9</td><td>6.0E-23</td><td>1.1E-21</td><td>MB231/MB468</td><td>MITO</td></tr>
<tr><td>>hg38_21_+_8433172_8446622_RNA45SN1_45S_50ntFlanks@6721.6758.38|AATTGCAGGACACATTGATCATCGACACTTCGAACGCA</td><td>>hg38_21_+_8433172_8446622_RNA45SN1_45S_50ntFlanks</td><td>6721</td><td>6758</td><td>38</td><td>AATTGCAGGACACATTGATCATCGACACTTCGAACGCA</td><td>1633.3</td><td>-4.9</td><td>0.5</td><td>-9.8</td><td>1.2E-22</td><td>1.1E-21</td><td>BT20/MB468</td><td>CYTO</td></tr>
<tr><td>>MI0000298|hsa-mir-221&WithFlank&X|-|45746151|45746272@71.90.20&[MIMAT0000278&hsa-miR-221-3p&offsets|0|-3;m-91&X|-|45746181|45746202&offsets|0|-2]|AGCTACATTGTCTGCTGGGT</td><td>>MI0000298|hsa-mir-221&WithFlank&X|-|45746151|45746272</td><td>71</td><td>90</td><td>20</td><td>AGCTACATTGTCTGCTGGGT</td><td>533.9</td><td>6.9</td><td>0.7</td><td>9.9</td><td>6.1E-23</td><td>1.1E-21</td><td>MB231/MB468</td><td>NUC</td></tr>
<tr><td>>hg38_21_+_8433172_8446622_RNA45SN1_45S_50ntFlanks@12285.12323.39|CAGACCGTGAAAGCGGGGCCTCACGATCCTTCTGACCTT</td><td>>hg38_21_+_8433172_8446622_RNA45SN1_45S_50ntFlanks</td><td>12285</td><td>12323</td><td>39</td><td>CAGACCGTGAAAGCGGGGCCTCACGATCCTTCTGACCTT</td><td>2487.9</td><td>-15.1</td><td>1.5</td><td>-9.9</td><td>6.1E-23</td><td>1.1E-21</td><td>MB231/MB468</td><td>MITO</td></tr>
<tr><td>>hg38_21_+_8433172_8446622_RNA45SN1_45S_50ntFlanks@6719.6746.28|TGAATTGCAGGACACATTGATCATCGAC</td><td>>hg38_21_+_8433172_8446622_RNA45SN1_45S_50ntFlanks</td><td>6719</td><td>6746</td><td>28</td><td>TGAATTGCAGGACACATTGATCATCGAC</td><td>1390.3</td><td>-8.5</td><td>0.9</td><td>-9.9</td><td>6.1E-23</td><td>1.1E-21</td><td>MB231/MB468</td><td>MITO</td></tr>
<tr><td>>hg38_21_+_8433172_8446622_RNA45SN1_45S_50ntFlanks@9457.9503.47|CGCCGGGGAGGTGGAGCACGAGCGCACGTGTTAGGACCCGAAAGATG</td><td>>hg38_21_+_8433172_8446622_RNA45SN1_45S_50ntFlanks</td><td>9457</td><td>9503</td><td>47</td><td>CGCCGGGGAGGTGGAGCACGAGCGCACGTGTTAGGACCCGAAAGATG</td><td>2259.0</td><td>-14.9</td><td>1.5</td><td>-9.9</td><td>6.1E-23</td><td>1.1E-21</td><td>MB231/MB468</td><td>MITO</td></tr>
<tr><td>>hg38_21_+_8433172_8446622_RNA45SN1_45S_50ntFlanks@11564.11600.37|CCTAGCAGCCGACTTAGAACTGGTGCGGACCAGGGGA</td><td>>hg38_21_+_8433172_8446622_RNA45SN1_45S_50ntFlanks</td><td>11564</td><td>11600</td><td>37</td><td>CCTAGCAGCCGACTTAGAACTGGTGCGGACCAGGGGA</td><td>2240.8</td><td>-6.9</td><td>0.7</td><td>-9.9</td><td>6.2E-23</td><td>1.1E-21</td><td>MB231/MB468</td><td>MITO</td></tr>
<tr><td>>hg38_21_+_8433172_8446622_RNA45SN1_45S_50ntFlanks@11161.11188.28|CACGGTCCCCCGCGAGGGGGGCCCGGGC</td><td>>hg38_21_+_8433172_8446622_RNA45SN1_45S_50ntFlanks</td><td>11161</td><td>11188</td><td>28</td><td>CACGGTCCCCCGCGAGGGGGGCCCGGGC</td><td>2154.8</td><td>-15.5</td><td>1.6</td><td>-9.7</td><td>2.4E-22</td><td>1.1E-21</td><td>BT20/MB468</td><td>NUC</td></tr>
</table>

<table>
<tr><td>>hg38_21_+_8433172_8446622_RNA45SN1_45S_50ntFlanks@10511.10533.23|ATCCGGAGTGGCGGAGATGGGCG</td><td>>hg38_21_+_8433172_8446622_RNA45SN1_45S_50ntFlanks</td><td>10511</td><td>10533</td><td>23</td><td>ATCCGGAGTGGCGGAGATGGGCG</td><td>2667.5</td><td>-15.2</td><td>1.5</td><td>-9.9</td><td>3.9E-23</td><td>1.1E-21</td><td>MB231/MB468</td><td>MP</td></tr>
<tr><td>>MI0000239|hsa-mir-197&WithFlank&1|+|109598887|109598973@54.74.21(+1U)&[MIMAT0000227&hsa-miR-197-3p&offsets|0|-1(+1U);m-99&1|+|109598940|109598961&offsets|0|-1(+1U)]|TTCACCACCTTCTCCACCCAGT</td><td>>MI0000239|hsa-mir-197&WithFlank&1|+|109598887|109598973</td><td>54</td><td>74</td><td>21(+1U)</td><td>TTCACCACCTTCTCCACCCAGT</td><td>8309.1</td><td>10.9</td><td>1.1</td><td>9.8</td><td>1.2E-22</td><td>1.1E-21</td><td>BT20/MB468</td><td>CYTO</td></tr>
<tr><td>>MI0000767|hsa-mir-365a&WithFlank&16|+|14309279|14309377@62.82.21;MI0000769|hsa-mir-365b&WithFlank&17|+|31575405|31575527@74.94.21&[MIMAT0000710&hsa-miR-365a-3p&offsets|0|-1;m-176&16|+|14309340|14309361&offsets|0|-1];[MIMAT0022834&hsa-miR-365b-3p&offsets|0|-1;m-175&17|+|31575478|31575499&offsets|0|-1]|TAATGCCCCTAAAAATCCTTA</td><td>>MI0000767|hsa-mir-365a&WithFlank&16|+|14309279|14309377</td><td>62</td><td>82</td><td>21</td><td>TAATGCCCCTAAAAATCCTTA</td><td>130.9</td><td>4.3</td><td>0.4</td><td>9.7</td><td>2.4E-22</td><td>1.1E-21</td><td>BT20/MB468</td><td>NUC</td></tr>
<tr><td>>hg38_21_+_8433172_8446622_RNA45SN1_45S_50ntFlanks@6671.6699.29|CTCGGCTCGTGCGTCGATGAAGAACGCAG</td><td>>hg38_21_+_8433172_8446622_RNA45SN1_45S_50ntFlanks</td><td>6671</td><td>6699</td><td>29</td><td>CTCGGCTCGTGCGTCGATGAAGAACGCAG</td><td>38221.6</td><td>-10.0</td><td>1.0</td><td>-9.9</td><td>6.3E-23</td><td>1.1E-21</td><td>MB231/MB468</td><td>MITO</td></tr>
<tr><td>>trnaMT_GluTTC_MT_-_14674_14742@38.72.35,trnalookalike8_GluTTC_5_-_93905172_93905240@38.72.35|CATTGGTCGTGGTTGTAGTCCGTGCGAGAATACCA</td><td>>trnaMT_GluTTC_MT_-_14674_14742</td><td>38</td><td>72</td><td>35</td><td>CATTGGTCGTGGTTGTAGTCCGTGCGAGAATACCA</td><td>3569.8</td><td>9.9</td><td>1.0</td><td>9.9</td><td>3.0E-23</td><td>1.1E-21</td><td>MB231/MB468</td><td>CYTO</td></tr>
<tr><td>>hg38_21_+_8433172_8446622_RNA45SN1_45S_50ntFlanks@11785.11804.20|TGAATGGATGAACGAGATTC</td><td>>hg38_21_+_8433172_8446622_RNA45SN1_45S_50ntFlanks</td><td>11785</td><td>11804</td><td>20</td><td>TGAATGGATGAACGAGATTC</td><td>29610.0</td><td>-12.2</td><td>1.3</td><td>-9.7</td><td>2.4E-22</td><td>1.1E-21</td><td>BT20/MB468</td><td>NUC</td></tr>
<tr><td>>hg38_21_+_8433172_8446622_RNA45SN1_45S_50ntFlanks@11534.11560.27|GTTCCGCGCGGCGCCTCGCCTCGGCCG</td><td>>hg38_21_+_8433172_8446622_RNA45SN1_45S_50ntFlanks</td><td>11534</td><td>11560</td><td>27</td><td>GTTCCGCGCGGCGCCTCGCCTCGGCCG</td><td>1177.3</td><td>-14.0</td><td>1.4</td><td>-9.9</td><td>6.3E-23</td><td>1.2E-21</td><td>MB231/MB468</td><td>MITO</td></tr>
<tr><td>>hg38_21_+_8433172_8446622_RNA45SN1_45S_50ntFlanks@11873.11908.36|AAGAAGACCCTGTTGAGCTTGACTCTAGTCTGGCAC</td><td>>hg38_21_+_8433172_8446622_RNA45SN1_45S_50ntFlanks</td><td>11873</td><td>11908</td><td>36</td><td>AAGAAGACCCTGTTGAGCTTGACTCTAGTCTGGCAC</td><td>1101.3</td><td>-13.9</td><td>1.4</td><td>-9.9</td><td>6.4E-23</td><td>1.2E-21</td><td>MB231/MB468</td><td>MITO</td></tr>
<tr><td>>hg38_21_+_8433172_8446622_RNA45SN1_45S_50ntFlanks@9305.9335.31|CCAAGGAGTCTAACACGTGCGCGAGTCGGGG</td><td>>hg38_21_+_8433172_8446622_RNA45SN1_45S_50ntFlanks</td><td>9305</td><td>9335</td><td>31</td><td>CCAAGGAGTCTAACACGTGCGCGAGTCGGGG</td><td>1008.8</td><td>-13.8</td><td>1.4</td><td>-9.9</td><td>4.0E-23</td><td>1.2E-21</td><td>MB231/MB468</td><td>MP</td></tr>
<tr><td>>MI0003560|hsa-mir-92b&WithFlank&1|+|155195171|155195278@67.86.20&[MIMAT0003218&hsa-miR-92b-3p&offsets|0|-2;m-115&1|+|155195237|155195258&offsets|0|-2]|TATTGCACTCGTCCCGGCCT</td><td>>MI0003560|hsa-mir-92b&WithFlank&1|+|155195171|155195278</td><td>67</td><td>86</td><td>20</td><td>TATTGCACTCGTCCCGGCCT</td><td>407.3</td><td>6.6</td><td>0.7</td><td>9.9</td><td>6.4E-23</td><td>1.2E-21</td><td>MB231/MB468</td><td>NUC</td></tr>
<tr><td>>hg38_21_+_8433172_8446622_RNA45SN1_45S_50ntFlanks@11667.11705.39|CTGCCCAGTGCTCTGAATGTCAAAGTGAAGAAATTCAAT</td><td>>hg38_21_+_8433172_8446622_RNA45SN1_45S_50ntFlanks</td><td>11667</td><td>11705</td><td>39</td><td>CTGCCCAGTGCTCTGAATGTCAAAGTGAAGAAATTCAAT</td><td>238.7</td><td>11.9</td><td>1.2</td><td>9.9</td><td>3.1E-23</td><td>1.2E-21</td><td>MB231/MB468</td><td>CYTO</td></tr>
<tr><td>>MI0000254|hsa-mir-30c-2&WithFlank&6|-|71376954|71377037@53.74.22&[MIMAT0004550&hsa-miR-30c-2-3p&offsets|0|0;m-206&6|-|71376964|71376985&offsets|0|0]|CTGGGAGAAGGCTGTTTACTCT</td><td>>MI0000254|hsa-mir-30c-2&WithFlank&6|-|71376954|71377037</td><td>53</td><td>74</td><td>22</td><td>CTGGGAGAAGGCTGTTTACTCT</td><td>777.1</td><td>7.5</td><td>0.8</td><td>9.8</td><td>9.3E-23</td><td>1.2E-21</td><td>BT20/MB468</td><td>MITO</td></tr>
<tr><td>>hg38_21_+_8433172_8446622_RNA45SN1_45S_50ntFlanks@10389.10410.22|TGAACAGCAGTTGAACATGGGT</td><td>>hg38_21_+_8433172_8446622_RNA45SN1_45S_50ntFlanks</td><td>10389</td><td>10410</td><td>22</td><td>TGAACAGCAGTTGAACATGGGT</td><td>1657.4</td><td>-14.6</td><td>1.5</td><td>-9.9</td><td>4.1E-23</td><td>1.2E-21</td><td>MB231/MB468</td><td>MP</td></tr>
<tr><td>>hg38_21_+_8433172_8446622_RNA45SN1_45S_50ntFlanks@11317.11343.27|ACCCCACGTCTCGTCGCGCGCGCGTCC</td><td>>hg38_21_+_8433172_8446622_RNA45SN1_45S_50ntFlanks</td><td>11317</td><td>11343</td><td>27</td><td>ACCCCACGTCTCGTCGCGCGCGCGTCC</td><td>1494.6</td><td>-14.3</td><td>1.5</td><td>-9.9</td><td>6.6E-23</td><td>1.2E-21</td><td>MB231/MB468</td><td>MITO</td></tr>
<tr><td>>hg38_21_+_8433172_8446622_RNA45SN1_45S_50ntFlanks@9532.9573.42|AGAGGAAACTCTGGTGGAGGTCCGTAGCGGTCCTGACGTGCA</td><td>>hg38_21_+_8433172_8446622_RNA45SN1_45S_50ntFlanks</td><td>9532</td><td>9573</td><td>42</td><td>AGAGGAAACTCTGGTGGAGGTCCGTAGCGGTCCTGACGTGCA</td><td>336.6</td><td>12.4</td><td>1.2</td><td>9.9</td><td>3.2E-23</td><td>1.2E-21</td><td>MB231/MB468</td><td>CYTO</td></tr>
<tr><td>>hg38_21_+_8433172_8446622_RNA45SN1_45S_50ntFlanks@3802.3830.29|CATTAAATCAGTTATGGTTCCTTTGGTCG</td><td>>hg38_21_+_8433172_8446622_RNA45SN1_45S_50ntFlanks</td><td>3802</td><td>3830</td><td>29</td><td>CATTAAATCAGTTATGGTTCCTTTGGTCG</td><td>265.8</td><td>-12.5</td><td>1.3</td><td>-9.7</td><td>2.6E-22</td><td>1.2E-21</td><td>BT20/MB468</td><td>NUC</td></tr>
<tr><td>>hg38_21_+_8433172_8446622_RNA45SN1_45S_50ntFlanks@4367.4389.23|CAGTTAAAAAGCTCGTAGTTGGA</td><td>>hg38_21_+_8433172_8446622_RNA45SN1_45S_50ntFlanks</td><td>4367</td><td>4389</td><td>23</td><td>CAGTTAAAAAGCTCGTAGTTGGA</td><td>1860.5</td><td>-15.3</td><td>1.6</td><td>-9.7</td><td>2.6E-22</td><td>1.2E-21</td><td>BT20/MB468</td><td>NUC</td></tr>
<tr><td>>MI0000077|hsa-mir-21&WithFlank&17|+|59841260|59841343@14.36.23(+1C)&[MIMAT0000076&hsa-miR-21-5p&offsets|0|+1(+1C);m-1&17|+|59841273|59841294&offsets|0|+1(+1C)]|TAGCTTATCAGACTGATGTTGACC</td><td>>MI0000077|hsa-mir-21&WithFlank&17|+|59841260|59841343</td><td>14</td><td>36</td><td>23(+1C)</td><td>TAGCTTATCAGACTGATGTTGACC</td><td>59522.5</td><td>8.1</td><td>0.8</td><td>9.8</td><td>1.3E-22</td><td>1.2E-21</td><td>BT20/MB468</td><td>CYTO</td></tr>
<tr><td>>MI0000283|hsa-mir-203a&WithFlank&14|+|104117399|104117520@71.92.22&[MIMAT0000264&hsa-miR-203a-3p&offsets|0|0;m-79&14|+|104117469|104117490&offsets|0|0]|GTGAAATGTTTAGGACCACTAG</td><td>>MI0000283|hsa-mir-203a&WithFlank&14|+|104117399|104117520</td><td>71</td><td>92</td><td>22</td><td>GTGAAATGTTTAGGACCACTAG</td><td>2039.8</td><td>10.6</td><td>1.0</td><td>10.1</td><td>4.3E-24</td><td>1.2E-21</td><td>BT20/MB231</td><td>CYTO</td></tr>
<tr><td>>hg38_21_+_8433172_8446622_RNA45SN1_45S_50ntFlanks@12892.12930.39|CCCTCGCCCGTCACGCACCGCACGTTCGTGGGGAACCTG</td><td>>hg38_21_+_8433172_8446622_RNA45SN1_45S_50ntFlanks</td><td>12892</td><td>12930</td><td>39</td><td>CCCTCGCCCGTCACGCACCGCACGTTCGTGGGGAACCTG</td><td>1352.6</td><td>-14.2</td><td>1.4</td><td>-9.9</td><td>6.8E-23</td><td>1.2E-21</td><td>MB231/MB468</td><td>MITO</td></tr>
<tr><td>>hg38_21_+_8433172_8446622_RNA45SN1_45S_50ntFlanks@11538.11583.46|CGCGCGGCGCCTCGCCTCGGCCGGCGCCTAGCAGCCGACTTAGAAC</td><td>>hg38_21_+_8433172_8446622_RNA45SN1_45S_50ntFlanks</td><td>11538</td><td>11583</td><td>46</td><td>CGCGCGGCGCCTCGCCTCGGCCGGCGCCTAGCAGCCGACTTAGAAC</td><td>269.8</td><td>12.1</td><td>1.2</td><td>9.9</td><td>3.3E-23</td><td>1.2E-21</td><td>MB231/MB468</td><td>CYTO</td></tr>
<tr><td>>MI0000650|hsa-mir-200c&WithFlank&12|+|6963693|6963772@50.72.23(+1C)&[MIMAT0000617&hsa-miR-200c-3p&offsets|0|0(+1C);m-54&12|+|6963742|6963764&offsets|0|0(+1C)]|TAATACTGCCGGGTAATGATGGAC</td><td>>MI0000650|hsa-mir-200c&WithFlank&12|+|6963693|6963772</td><td>50</td><td>72</td><td>23(+1C)</td><td>TAATACTGCCGGGTAATGATGGAC</td><td>1155.2</td><td>11.1</td><td>1.1</td><td>10.1</td><td>8.5E-24</td><td>1.2E-21</td><td>BT20/MB231</td><td>NUC</td></tr>
<tr><td>>hg38_21_+_8433172_8446622_RNA45SN1_45S_50ntFlanks@7983.8007.25|CAGATCAGACGTGGCGACCCGCTGA</td><td>>hg38_21_+_8433172_8446622_RNA45SN1_45S_50ntFlanks</td><td>7983</td><td>8007</td><td>25</td><td>CAGATCAGACGTGGCGACCCGCTGA</td><td>1057.9</td><td>-13.9</td><td>1.4</td><td>-9.9</td><td>4.3E-23</td><td>1.2E-21</td><td>MB231/MB468</td><td>MP</td></tr>
<tr><td>>hg38_21_+_8433172_8446622_RNA45SN1_45S_50ntFlanks@485.511.27|GATGTGGTGACGTCGTGCTCTCCCGGG</td><td>>hg38_21_+_8433172_8446622_RNA45SN1_45S_50ntFlanks</td><td>485</td><td>511</td><td>27</td><td>GATGTGGTGACGTCGTGCTCTCCCGGG</td><td>320.0</td><td>11.2</td><td>1.1</td><td>9.7</td><td>2.7E-22</td><td>1.2E-21</td><td>BT20/MB468</td><td>NUC</td></tr>
<tr><td>>trna14_LysTTT_11_-_59327808_59327880@1.33.33,trna2_LysTTT_17_+_8022473_8022545@1.33.33,trna54_LysTTT_1_+_204475655_204475727@1.33.33,trna5_LysTTT_11_+_59323902_59323974@1.33.33,trna62_LysTTT_1_-_204476158_204476230@1.33.33,trna76_LysTTT_6_+_28918806_28918878@1.33.33|GCCCGGATAGCTCAGTCGGTAGAGCATCAGACT</td><td>>trna14_LysTTT_11_-_59327808_59327880</td><td>1</td><td>33</td><td>33</td><td>GCCCGGATAGCTCAGTCGGTAGAGCATCAGACT</td><td>4059.1</td><td>7.3</td><td>0.8</td><td>9.8</td><td>1.3E-22</td><td>1.2E-21</td><td>BT20/MB468</td><td>CYTO</td></tr>
<tr><td>>MI0000456|hsa-mir-140&WithFlank&16|+|69933075|69933186@69.89.21(+1A)&[MIMAT0004597&hsa-miR-140-3p&offsets|+1|+1(+1A);m-45&16|+|69933143|69933164&offsets|0|-1(+1A)]|ACCACAGGGTAGAACCACGGAA</td><td>>MI0000456|hsa-mir-140&WithFlank&16|+|69933075|69933186</td><td>69</td><td>89</td><td>21(+1A)</td><td>ACCACAGGGTAGAACCACGGAA</td><td>400.7</td><td>12.9</td><td>1.3</td><td>9.8</td><td>1.3E-22</td><td>1.3E-21</td><td>BT20/MB468</td><td>CYTO</td></tr>
<tr><td>>hg38_21_+_8433172_8446622_RNA45SN1_45S_50ntFlanks@10362.10388.27|AAGGCCGAAGTGGAGAAGGGTTCCATG</td><td>>hg38_21_+_8433172_8446622_RNA45SN1_45S_50ntFlanks</td><td>10362</td><td>10388</td><td>27</td><td>AAGGCCGAAGTGGAGAAGGGTTCCATG</td><td>1993.4</td><td>-14.7</td><td>1.5</td><td>-9.8</td><td>7.0E-23</td><td>1.3E-21</td><td>MB231/MB468</td><td>MITO</td></tr>
<tr><td>>hg38_21_+_8433172_8446622_RNA45SN1_45S_50ntFlanks@11641.11679.39|CGGCGGGTGTTGACGCGATGTGATTTCTGCCCAGTGCTC</td><td>>hg38_21_+_8433172_8446622_RNA45SN1_45S_50ntFlanks</td><td>11641</td><td>11679</td><td>39</td><td>CGGCGGGTGTTGACGCGATGTGATTTCTGCCCAGTGCTC</td><td>1887.2</td><td>-14.7</td><td>1.5</td><td>-9.8</td><td>7.0E-23</td><td>1.3E-21</td><td>MB231/MB468</td><td>MITO</td></tr>
<tr><td>>hg38_21_+_8433172_8446622_RNA45SN1_45S_50ntFlanks@12461.12487.27|CGTTGGATTGTTCACCCACTAATAGGG</td><td>>hg38_21_+_8433172_8446622_RNA45SN1_45S_50ntFlanks</td><td>12461</td><td>12487</td><td>27</td><td>CGTTGGATTGTTCACCCACTAATAGGG</td><td>326.5</td><td>-12.8</td><td>1.3</td><td>-9.7</td><td>2.7E-22</td><td>1.3E-21</td><td>BT20/MB468</td><td>NUC</td></tr>
<tr><td>>hg38_21_+_8433172_8446622_RNA45SN1_45S_50ntFlanks@11575.11592.18|ACTTAGAACTGGTGCGGA</td><td>>hg38_21_+_8433172_8446622_RNA45SN1_45S_50ntFlanks</td><td>11575</td><td>11592</td><td>18</td><td>ACTTAGAACTGGTGCGGA</td><td>2690.3</td><td>-9.4</td><td>1.0</td><td>-9.8</td><td>6.9E-23</td><td>1.3E-21</td><td>MB231/MB468</td><td>NUC</td></tr>
<tr><td>>hg38_21_+_8433172_8446622_RNA45SN1_45S_50ntFlanks@11669.11691.23|GCCCAGTGCTCTGAATGTCAAAG</td><td>>hg38_21_+_8433172_8446622_RNA45SN1_45S_50ntFlanks</td><td>11669</td><td>11691</td><td>23</td><td>GCCCAGTGCTCTGAATGTCAAAG</td><td>464.4</td><td>6.0</td><td>0.6</td><td>9.9</td><td>3.5E-23</td><td>1.3E-21</td><td>MB231/MB468</td><td>CYTO</td></tr>
<tr><td>>hg38_21_+_8433172_8446622_RNA45SN1_45S_50ntFlanks@12002.12043.42|TGCGGGCCGCCGGTGAAATACCACTACTCTGATCGTTTTTTC</td><td>>hg38_21_+_8433172_8446622_RNA45SN1_45S_50ntFlanks</td><td>12002</td><td>12043</td><td>42</td><td>TGCGGGCCGCCGGTGAAATACCACTACTCTGATCGTTTTTTC</td><td>3180.9</td><td>-15.4</td><td>1.6</td><td>-9.8</td><td>7.2E-23</td><td>1.3E-21</td><td>MB231/MB468</td><td>MITO</td></tr>
<tr><td>>MI0000079|hsa-mir-23a&WithFlank&19|-|13836581|13836665@54.71.18(+1G)&[MIMAT0000078&hsa-miR-23a-3p&offsets|+3|0(+1G);m-31&19|-|13836595|13836615&offsets|+3|0(+1G)]|ACATTGCCAGGGATTTCCG</td><td>>MI0000079|hsa-mir-23a&WithFlank&19|-|13836581|13836665</td><td>54</td><td>71</td><td>18(+1G)</td><td>ACATTGCCAGGGATTTCCG</td><td>587.3</td><td>13.5</td><td>1.4</td><td>9.8</td><td>1.4E-22</td><td>1.3E-21</td><td>BT20/MB468</td><td>CYTO</td></tr>
<tr><td>>hg38_21_+_8433172_8446622_RNA45SN1_45S_50ntFlanks@482.521.40|ATCGATGTGGTGACGTCGTGCTCTCCCGGGCCGGGTCCGA</td><td>>hg38_21_+_8433172_8446622_RNA45SN1_45S_50ntFlanks</td><td>482</td><td>521</td><td>40</td><td>ATCGATGTGGTGACGTCGTGCTCTCCCGGGCCGGGTCCGA</td><td>1878.1</td><td>-15.3</td><td>1.6</td><td>-9.7</td><td>2.8E-22</td><td>1.3E-21</td><td>BT20/MB468</td><td>NUC</td></tr>
<tr><td>>hg38_21_+_8433172_8446622_RNA45SN1_45S_50ntFlanks@475.495.21|CTTCGTGATCGATGTGGTGAC</td><td>>hg38_21_+_8433172_8446622_RNA45SN1_45S_50ntFlanks</td><td>475</td><td>495</td><td>21</td><td>CTTCGTGATCGATGTGGTGAC</td><td>397.0</td><td>12.9</td><td>1.3</td><td>9.8</td><td>1.4E-22</td><td>1.3E-21</td><td>BT20/MB468</td><td>CYTO</td></tr>
<tr><td>>MI0000809|hsa-mir-151a&WithFlank&8|-|140732558|140732659@51.72.22(+1A)&[MIMAT0000757&hsa-miR-151a-3p&offsets|-2|-1(+1A);m-30&8|-|140732587|140732607&offsets|-2|-1(+1A)]|TACTAGACTGAAGCTCCTTGAGA</td><td>>MI0000809|hsa-mir-151a&WithFlank&8|-|140732558|140732659</td><td>51</td><td>72</td><td>22(+1A)</td><td>TACTAGACTGAAGCTCCTTGAGA</td><td>2064.4</td><td>9.1</td><td>0.9</td><td>9.8</td><td>1.4E-22</td><td>1.3E-21</td><td>BT20/MB468</td><td>CYTO</td></tr>
<tr><td>>hg38_21_+_8433172_8446622_RNA45SN1_45S_50ntFlanks@11873.11892.20|AAGAAGACCCTGTTGAGCTT</td><td>>hg38_21_+_8433172_8446622_RNA45SN1_45S_50ntFlanks</td><td>11873</td><td>11892</td><td>20</td><td>AAGAAGACCCTGTTGAGCTT</td><td>1108.0</td><td>-14.0</td><td>1.4</td><td>-9.9</td><td>4.5E-23</td><td>1.3E-21</td><td>MB231/MB468</td><td>MP</td></tr>
<tr><td>>hg38_21_+_8433172_8446622_RNA45SN1_45S_50ntFlanks@4728.4758.31|ATCAAGAACGAAAGTCGGAGGTTCGAAGACG</td><td>>hg38_21_+_8433172_8446622_RNA45SN1_45S_50ntFlanks</td><td>4728</td><td>4758</td><td>31</td><td>ATCAAGAACGAAAGTCGGAGGTTCGAAGACG</td><td>4458.0</td><td>-15.2</td><td>1.5</td><td>-10.3</td><td>8.2E-25</td><td>1.3E-21</td><td>BT20/MB468</td><td>MP</td></tr>
<tr><td>>hg38_21_+_8433172_8446622_RNA45SN1_45S_50ntFlanks@7985.8006.22|GATCAGACGTGGCGACCCGCTG</td><td>>hg38_21_+_8433172_8446622_RNA45SN1_45S_50ntFlanks</td><td>7985</td><td>8006</td><td>22</td><td>GATCAGACGTGGCGACCCGCTG</td><td>6693.1</td><td>-15.8</td><td>1.5</td><td>-10.3</td><td>8.7E-25</td><td>1.3E-21</td><td>BT20/MB468</td><td>MP</td></tr>
<tr><td>>hg38_21_+_8433172_8446622_RNA45SN1_45S_50ntFlanks@11669.11687.19|GCCCAGTGCTCTGAATGTC</td><td>>hg38_21_+_8433172_8446622_RNA45SN1_45S_50ntFlanks</td><td>11669</td><td>11687</td><td>19</td><td>GCCCAGTGCTCTGAATGTC</td><td>2828.0</td><td>6.3</td><td>0.6</td><td>9.9</td><td>3.6E-23</td><td>1.3E-21</td><td>MB231/MB468</td><td>CYTO</td></tr>
<tr><td>>hg38_21_+_8433172_8446622_RNA45SN1_45S_50ntFlanks@12995.13039.45|CTCGCTGCGATCTATTGAAAGTCAGCCCTCGACACAAGGGTTTGT</td><td>>hg38_21_+_8433172_8446622_RNA45SN1_45S_50ntFlanks</td><td>12995</td><td>13039</td><td>45</td><td>CTCGCTGCGATCTATTGAAAGTCAGCCCTCGACACAAGGGTTTGT</td><td>1183582.6</td><td>9.6</td><td>1.0</td><td>9.9</td><td>3.6E-23</td><td>1.3E-21</td><td>MB231/MB468</td><td>CYTO</td></tr>
<tr><td>>hg38_21_+_8433172_8446622_RNA45SN1_45S_50ntFlanks@10410.10432.23|TCAGTCGGTCCTGAGAGATGGGC</td><td>>hg38_21_+_8433172_8446622_RNA45SN1_45S_50ntFlanks</td><td>10410</td><td>10432</td><td>23</td><td>TCAGTCGGTCCTGAGAGATGGGC</td><td>1044.5</td><td>-13.9</td><td>1.4</td><td>-9.9</td><td>4.6E-23</td><td>1.3E-21</td><td>MB231/MB468</td><td>MP</td></tr>
<tr><td>>hg38_21_+_8433172_8446622_RNA45SN1_45S_50ntFlanks@11873.11910.38|AAGAAGACCCTGTTGAGCTTGACTCTAGTCTGGCACGG</td><td>>hg38_21_+_8433172_8446622_RNA45SN1_45S_50ntFlanks</td><td>11873</td><td>11910</td><td>38</td><td>AAGAAGACCCTGTTGAGCTTGACTCTAGTCTGGCACGG</td><td>1086.1</td><td>-13.9</td><td>1.4</td><td>-9.8</td><td>7.5E-23</td><td>1.3E-21</td><td>MB231/MB468</td><td>MITO</td></tr>
<tr><td>>MI0000471|hsa-mir-126&WithFlank&9|+|136670596|136670692@58.79.22&[MIMAT0000445&hsa-miR-126-3p&offsets|0|0;m-40&9|+|136670653|136670674&offsets|0|0]|TCGTACCGTGAGTAATAATGCG</td><td>>MI0000471|hsa-mir-126&WithFlank&9|+|136670596|136670692</td><td>58</td><td>79</td><td>22</td><td>TCGTACCGTGAGTAATAATGCG</td><td>1040.9</td><td>7.0</td><td>0.7</td><td>9.7</td><td>2.9E-22</td><td>1.3E-21</td><td>BT20/MB468</td><td>NUC</td></tr>
<tr><td>>hg38_21_+_8433172_8446622_RNA45SN1_45S_50ntFlanks@12324.12344.21|TTGGGTTTTAAGCAGGAGGTG</td><td>>hg38_21_+_8433172_8446622_RNA45SN1_45S_50ntFlanks</td><td>12324</td><td>12344</td><td>21</td><td>TTGGGTTTTAAGCAGGAGGTG</td><td>1525.7</td><td>-13.5</td><td>1.4</td><td>-9.8</td><td>7.5E-23</td><td>1.4E-21</td><td>MB231/MB468</td><td>NUC</td></tr>
<tr><td>>hg38_21_+_8433172_8446622_RNA45SN1_45S_50ntFlanks@12619.12653.35|AGGAGCCAATGGGGCGAAGCTACCATCTGTGGGAT</td><td>>hg38_21_+_8433172_8446622_RNA45SN1_45S_50ntFlanks</td><td>12619</td><td>12653</td><td>35</td><td>AGGAGCCAATGGGGCGAAGCTACCATCTGTGGGAT</td><td>1037.0</td><td>-13.9</td><td>1.4</td><td>-9.9</td><td>4.7E-23</td><td>1.4E-21</td><td>MB231/MB468</td><td>MP</td></tr>
</table>

<table>
<tr><td>>hg38_21_+_8433172_8446622_RNA45SN1_45S_50ntFlanks@9033.9056.24|CTCGTGGGGGGGCCGGGCCACCCC</td><td>>hg38_21_+_8433172_8446622_RNA45SN1_45S_50ntFlanks</td><td>9033</td><td>9056</td><td>24</td><td>CTCGTGGGGGGGCCGGGCCACCCC</td><td>1279.7</td><td>-14.2</td><td>1.4</td><td>-9.9</td><td>4.7E-23</td><td>1.4E-21</td><td>MB231/MB468</td><td>MP</td></tr>
<tr><td>>hg38_21_+_8433172_8446622_RNA45SN1_45S_50ntFlanks@12456.12474.19|CCAAGCGTTGGATTGTTCA</td><td>>hg38_21_+_8433172_8446622_RNA45SN1_45S_50ntFlanks</td><td>12456</td><td>12474</td><td>19</td><td>CCAAGCGTTGGATTGTTCA</td><td>11023.6</td><td>-11.0</td><td>1.1</td><td>-9.9</td><td>4.8E-23</td><td>1.4E-21</td><td>MB231/MB468</td><td>MP</td></tr>
<tr><td>>trna116_GluCTC_1_-_145399233_145399304@1.35.35,trna59_GluCTC_1_+_249168447_249168518@1.35.35,trna71_GluCTC_1_-_161439189_161439260@1.35.35,trna74_GluCTC_1_-_161431809_161431880@1.35.35,trna77_GluCTC_1_-_161424398_161424469@1.35.35,trna77_GluCTC_6_+_28949976_28950047@1.35.35,trna80_GluCTC_1_-_161417018_161417089@1.35.35,trna87_GluCTC_6_-_126101393_126101464@1.35.35|TCCCTGGTGGTCTAGTGGTTAGGATTCGGCGCTCT</td><td>>trna116_GluCTC_1_-_145399233_145399304</td><td>1</td><td>35</td><td>35</td><td>TCCCTGGTGGTCTAGTGGTTAGGATTCGGCGCTCT</td><td>49057.4</td><td>9.7</td><td>1.0</td><td>9.8</td><td>1.5E-22</td><td>1.4E-21</td><td>BT20/MB468</td><td>CYTO</td></tr>
<tr><td>>hg38_1_-_228634819_228635039_RNA5S12_5S_50ntFlanks@138.170.33|AGACCGCCTGGGAATACCGGGTGCTGTAGGCTT</td><td>>hg38_1_-_228634819_228635039_RNA5S12_5S_50ntFlanks</td><td>138</td><td>170</td><td>33</td><td>AGACCGCCTGGGAATACCGGGTGCTGTAGGCTT</td><td>1803.7</td><td>7.1</td><td>0.7</td><td>9.8</td><td>1.5E-22</td><td>1.4E-21</td><td>BT20/MB468</td><td>CYTO</td></tr>
<tr><td>>hg38_1_-_228634819_228635039_RNA5S12_5S_50ntFlanks@152.169.18|TACCGGGTGCTGTAGGCT</td><td>>hg38_1_-_228634819_228635039_RNA5S12_5S_50ntFlanks</td><td>152</td><td>169</td><td>18</td><td>TACCGGGTGCTGTAGGCT</td><td>1602.0</td><td>6.2</td><td>0.6</td><td>9.8</td><td>1.5E-22</td><td>1.4E-21</td><td>BT20/MB468</td><td>CYTO</td></tr>
<tr><td>>hg38_21_+_8433172_8446622_RNA45SN1_45S_50ntFlanks@10807.10826.20|ACAATGTAGGTAAGGGAAGT</td><td>>hg38_21_+_8433172_8446622_RNA45SN1_45S_50ntFlanks</td><td>10807</td><td>10826</td><td>20</td><td>ACAATGTAGGTAAGGGAAGT</td><td>554.7</td><td>-13.5</td><td>1.4</td><td>-9.7</td><td>3.0E-22</td><td>1.4E-21</td><td>BT20/MB468</td><td>NUC</td></tr>
<tr><td>>hg38_21_+_8433172_8446622_RNA45SN1_45S_50ntFlanks@6738.6770.33|ATCATCGACACTTCGAACGCACTTGCGGCCCCG</td><td>>hg38_21_+_8433172_8446622_RNA45SN1_45S_50ntFlanks</td><td>6738</td><td>6770</td><td>33</td><td>ATCATCGACACTTCGAACGCACTTGCGGCCCCG</td><td>1742.2</td><td>-13.1</td><td>1.3</td><td>-9.8</td><td>1.5E-22</td><td>1.4E-21</td><td>BT20/MB468</td><td>CYTO</td></tr>
<tr><td>>hg38_21_+_8433172_8446622_RNA45SN1_45S_50ntFlanks@11092.11116.25|CGGCGGGGGGAGAAGGGTCGGGGCG</td><td>>hg38_21_+_8433172_8446622_RNA45SN1_45S_50ntFlanks</td><td>11092</td><td>11116</td><td>25</td><td>CGGCGGGGGGAGAAGGGTCGGGGCG</td><td>427.7</td><td>-13.2</td><td>1.4</td><td>-9.7</td><td>3.0E-22</td><td>1.4E-21</td><td>BT20/MB468</td><td>NUC</td></tr>
<tr><td>>am_trnaMT_MetCAT_MT_+_4402_4469@1.50.50|AGTAAGGTCAGCTAAATAAGCTATCGGGCCCATACCCCGAAAATGTTGGT</td><td>>am_trnaMT_MetCAT_MT_+_4402_4469</td><td>1</td><td>50</td><td>50</td><td>AGTAAGGTCAGCTAAATAAGCTATCGGGCCCATACCCCGAAAATGTTGGT</td><td>500.6</td><td>26.9</td><td>2.6</td><td>10.3</td><td>1.0E-24</td><td>1.4E-21</td><td>BT20/MB468</td><td>MP</td></tr>
<tr><td>>MI0000439|hsa-mir-23b&WithFlank&9|+|95085202|95085310@67.85.19&[MIMAT0000418&hsa-miR-23b-3p&offsets|+3|-1;m-42&9|+|95085265|95085285&offsets|+3|+1]|ACATTGCCAGGGATTACCA</td><td>>MI0000439|hsa-mir-23b&WithFlank&9|+|95085202|95085310</td><td>67</td><td>85</td><td>19</td><td>ACATTGCCAGGGATTACCA</td><td>175.8</td><td>10.3</td><td>1.1</td><td>9.7</td><td>3.1E-22</td><td>1.4E-21</td><td>BT20/MB468</td><td>NUC</td></tr>
<tr><td>>hg38_21_+_8433172_8446622_RNA45SN1_45S_50ntFlanks@8157.8178.22|CTCGTGGGGGGCCCAAGTCCTT</td><td>>hg38_21_+_8433172_8446622_RNA45SN1_45S_50ntFlanks</td><td>8157</td><td>8178</td><td>22</td><td>CTCGTGGGGGGCCCAAGTCCTT</td><td>875.9</td><td>-13.6</td><td>1.4</td><td>-9.9</td><td>5.0E-23</td><td>1.4E-21</td><td>MB231/MB468</td><td>MP</td></tr>
<tr><td>>trna111_HisGTG_1_-_147774845_147774916@-1G.33.34,trna118_HisGTG_1_-_145396881_145396952@-1G.33.34,trna16_HisGTG_1_+_146544773_146544844@-1G.33.34,trna1_HisGTG_15_+_45493349_45493420@-1G.33.34,trna21_HisGTG_1_+_147753471_147753542@-1G.33.34,trna33_HisGTG_6_+_27125906_27125977@-1G.33.34,trna7_HisGTG_9_-_14433938_14434009@-1G.33.34,trna8_HisGTG_15_-_45492611_45492682@-1G.33.34,trna9_HisGTG_15_-_45490804_45490875@-1G.33.34|GGCCGTGATCGTATAGTGGTTAGTACTCTGCGTT</td><td>>trna111_HisGTG_1_-_147774845_147774916</td><td>-1G</td><td>33</td><td>34</td><td>GGCCGTGATCGTATAGTGGTTAGTACTCTGCGTT</td><td>15873.1</td><td>8.5</td><td>0.9</td><td>9.9</td><td>5.1E-23</td><td>1.5E-21</td><td>MB231/MB468</td><td>MP</td></tr>
<tr><td>>hg38_21_+_8433172_8446622_RNA45SN1_45S_50ntFlanks@6657.6689.33|TTAGCGGTGGATCACTCGGCTCGTGCGTCGATG</td><td>>hg38_21_+_8433172_8446622_RNA45SN1_45S_50ntFlanks</td><td>6657</td><td>6689</td><td>33</td><td>TTAGCGGTGGATCACTCGGCTCGTGCGTCGATG</td><td>318.4</td><td>-4.3</td><td>0.4</td><td>-9.7</td><td>3.2E-22</td><td>1.5E-21</td><td>BT20/MB468</td><td>NUC</td></tr>
<tr><td>>am_trnaMT_SerGCT_MT_+_12207_12265@16.38.23|AACTGCTAACTCATGCCCCCATG</td><td>>am_trnaMT_SerGCT_MT_+_12207_12265</td><td>16</td><td>38</td><td>23</td><td>AACTGCTAACTCATGCCCCCATG</td><td>1677.8</td><td>-15.1</td><td>1.6</td><td>-9.7</td><td>3.2E-22</td><td>1.5E-21</td><td>BT20/MB468</td><td>NUC</td></tr>
<tr><td>>hg38_21_+_8433172_8446622_RNA45SN1_45S_50ntFlanks@10421.10455.35|TGAGAGATGGGCGAGCGCCGTTCCGAAGGGACGGG</td><td>>hg38_21_+_8433172_8446622_RNA45SN1_45S_50ntFlanks</td><td>10421</td><td>10455</td><td>35</td><td>TGAGAGATGGGCGAGCGCCGTTCCGAAGGGACGGG</td><td>511.0</td><td>-13.4</td><td>1.4</td><td>-9.7</td><td>3.2E-22</td><td>1.5E-21</td><td>BT20/MB468</td><td>NUC</td></tr>
<tr><td>>hg38_21_+_8433172_8446622_RNA45SN1_45S_50ntFlanks@12126.12151.26|CGACCCGCTCCGGGGACAGTGCCAGG</td><td>>hg38_21_+_8433172_8446622_RNA45SN1_45S_50ntFlanks</td><td>12126</td><td>12151</td><td>26</td><td>CGACCCGCTCCGGGGACAGTGCCAGG</td><td>513.2</td><td>-13.4</td><td>1.4</td><td>-9.7</td><td>3.2E-22</td><td>1.5E-21</td><td>BT20/MB468</td><td>NUC</td></tr>
<tr><td>>hg38_1_-_228634819_228635039_RNA5S12_5S_50ntFlanks@149.169.21|GAATACCGGGTGCTGTAGGCT</td><td>>hg38_1_-_228634819_228635039_RNA5S12_5S_50ntFlanks</td><td>149</td><td>169</td><td>21</td><td>GAATACCGGGTGCTGTAGGCT</td><td>2027.4</td><td>6.6</td><td>0.7</td><td>9.8</td><td>1.6E-22</td><td>1.5E-21</td><td>BT20/MB468</td><td>CYTO</td></tr>
<tr><td>>MI0000264|hsa-mir-7-2&WithFlank&15|+|88611819|88611940@38.59.22(+1C);MI0000265|hsa-mir-7-3&WithFlank&19|+|4770664|4770785@37.58.22(+1C);MI0000263|hsa-mir-7-1&WithFlank&9|-|83969742|83969863@30.51.22(+1C)&[MIMAT0000252&hsa-miR-7-5p&offsets|0|-2(+1C);m-400&15|+|88611856|88611879&offsets|0|-2(+1C)];[MIMAT0000252_1&hsa-miR-7-5p&offsets|0|-2(+1C);m-402&19|+|4770700|4770723&offsets|0|-2(+1C)];[MIMAT0000252_2&hsa-miR-7-5p&offsets|0|-2(+1C);m-401&9|-|83969811|83969834&offsets|0|-2(+1C)]|TGGAAGACTAGTGATTTTGTTGC</td><td>>MI0000264|hsa-mir-7-2&WithFlank&15|+|88611819|88611940</td><td>38</td><td>59</td><td>22(+1C)</td><td>TGGAAGACTAGTGATTTTGTTGC</td><td>594.3</td><td>5.7</td><td>0.6</td><td>9.8</td><td>8.3E-23</td><td>1.5E-21</td><td>MB231/MB468</td><td>NUC</td></tr>
<tr><td>>hg38_21_+_8433172_8446622_RNA45SN1_45S_50ntFlanks@7977.8008.32|CGACCTCAGATCAGACGTGGCGACCCGCTGAA</td><td>>hg38_21_+_8433172_8446622_RNA45SN1_45S_50ntFlanks</td><td>7977</td><td>8008</td><td>32</td><td>CGACCTCAGATCAGACGTGGCGACCCGCTGAA</td><td>1114.6</td><td>-4.7</td><td>0.5</td><td>-9.7</td><td>3.3E-22</td><td>1.5E-21</td><td>BT20/MB468</td><td>NUC</td></tr>
<tr><td>>MI0001445|hsa-mir-423&WithFlank&17|+|30117073|30117178@23.43.21&[MIMAT0004748&hsa-miR-423-5p&offsets|0|-2;m-141&17|+|30117095|30117117&offsets|0|-2]|TGAGGGGCAGAGAGCGAGACT</td><td>>MI0001445|hsa-mir-423&WithFlank&17|+|30117073|30117178</td><td>23</td><td>43</td><td>21</td><td>TGAGGGGCAGAGAGCGAGACT</td><td>2207.2</td><td>8.4</td><td>0.8</td><td>9.9</td><td>4.2E-23</td><td>1.5E-21</td><td>MB231/MB468</td><td>CYTO</td></tr>
<tr><td>>hg38_21_+_8433172_8446622_RNA45SN1_45S_50ntFlanks@12935.12956.22|TAAACCATTCGTAGACGACCTG</td><td>>hg38_21_+_8433172_8446622_RNA45SN1_45S_50ntFlanks</td><td>12935</td><td>12956</td><td>22</td><td>TAAACCATTCGTAGACGACCTG</td><td>3389.2</td><td>-5.3</td><td>0.5</td><td>-9.9</td><td>4.3E-23</td><td>1.5E-21</td><td>MB231/MB468</td><td>CYTO</td></tr>
<tr><td>>hg38_21_+_8433172_8446622_RNA45SN1_45S_50ntFlanks@11874.11910.37|AGAAGACCCTGTTGAGCTTGACTCTAGTCTGGCACGG</td><td>>hg38_21_+_8433172_8446622_RNA45SN1_45S_50ntFlanks</td><td>11874</td><td>11910</td><td>37</td><td>AGAAGACCCTGTTGAGCTTGACTCTAGTCTGGCACGG</td><td>483.1</td><td>-13.3</td><td>1.4</td><td>-9.7</td><td>3.3E-22</td><td>1.5E-21</td><td>BT20/MB468</td><td>NUC</td></tr>
<tr><td>>hg38_21_+_8433172_8446622_RNA45SN1_45S_50ntFlanks@12405.12435.31|CTTTTTGATCCTTCGATGTCGGCTCTTCCTA</td><td>>hg38_21_+_8433172_8446622_RNA45SN1_45S_50ntFlanks</td><td>12405</td><td>12435</td><td>31</td><td>CTTTTTGATCCTTCGATGTCGGCTCTTCCTA</td><td>3627.5</td><td>-6.3</td><td>0.6</td><td>-9.8</td><td>1.2E-22</td><td>1.5E-21</td><td>BT20/MB468</td><td>MITO</td></tr>
<tr><td>>hg38_21_+_8433172_8446622_RNA45SN1_45S_50ntFlanks@13002.13028.27|CGATCTATTGAAAGTCAGCCCTCGACA</td><td>>hg38_21_+_8433172_8446622_RNA45SN1_45S_50ntFlanks</td><td>13002</td><td>13028</td><td>27</td><td>CGATCTATTGAAAGTCAGCCCTCGACA</td><td>1972.4</td><td>-13.0</td><td>1.3</td><td>-9.8</td><td>1.2E-22</td><td>1.5E-21</td><td>BT20/MB468</td><td>MITO</td></tr>
<tr><td>>MI0000440|hsa-mir-27b&WithFlank&9|+|95085439|95085547@67.86.20&[MIMAT0000419&hsa-miR-27b-3p&offsets|0|-1;m-39&9|+|95085505|95085525&offsets|0|-1]|TTCACAGTGGCTAAGTTCTG</td><td>>MI0000440|hsa-mir-27b&WithFlank&9|+|95085439|95085547</td><td>67</td><td>86</td><td>20</td><td>TTCACAGTGGCTAAGTTCTG</td><td>5354.6</td><td>6.2</td><td>0.6</td><td>9.8</td><td>8.5E-23</td><td>1.5E-21</td><td>MB231/MB468</td><td>NUC</td></tr>
<tr><td>>hg38_21_+_8433172_8446622_RNA45SN1_45S_50ntFlanks@11118.11145.28|CAGGGGCCGGCGGCGGCCCGCCGCGGGG</td><td>>hg38_21_+_8433172_8446622_RNA45SN1_45S_50ntFlanks</td><td>11118</td><td>11145</td><td>28</td><td>CAGGGGCCGGCGGCGGCCCGCCGCGGGG</td><td>834.1</td><td>-14.1</td><td>1.5</td><td>-9.7</td><td>3.3E-22</td><td>1.5E-21</td><td>BT20/MB468</td><td>NUC</td></tr>
<tr><td>>hg38_21_+_8433172_8446622_RNA45SN1_45S_50ntFlanks@5476.5516.41|CCCTGGCGGAGCGCTGAGAAGACGGTCGAACTTGACTATCT</td><td>>hg38_21_+_8433172_8446622_RNA45SN1_45S_50ntFlanks</td><td>5476</td><td>5516</td><td>41</td><td>CCCTGGCGGAGCGCTGAGAAGACGGTCGAACTTGACTATCT</td><td>2937.6</td><td>16.0</td><td>1.6</td><td>9.8</td><td>1.2E-22</td><td>1.6E-21</td><td>BT20/MB468</td><td>MITO</td></tr>
<tr><td>>hg38_21_+_8433172_8446622_RNA45SN1_45S_50ntFlanks@10391.10410.20|AACAGCAGTTGAACATGGGT</td><td>>hg38_21_+_8433172_8446622_RNA45SN1_45S_50ntFlanks</td><td>10391</td><td>10410</td><td>20</td><td>AACAGCAGTTGAACATGGGT</td><td>5589.8</td><td>-9.6</td><td>1.0</td><td>-9.8</td><td>8.7E-23</td><td>1.6E-21</td><td>MB231/MB468</td><td>NUC</td></tr>
<tr><td>>hg38_21_+_8433172_8446622_RNA45SN1_45S_50ntFlanks@12709.12742.34|CCGCGGAGCCTCGGTTGGCCTCGGATAGCCGGTC</td><td>>hg38_21_+_8433172_8446622_RNA45SN1_45S_50ntFlanks</td><td>12709</td><td>12742</td><td>34</td><td>CCGCGGAGCCTCGGTTGGCCTCGGATAGCCGGTC</td><td>819.4</td><td>-13.5</td><td>1.4</td><td>-9.9</td><td>5.5E-23</td><td>1.6E-21</td><td>MB231/MB468</td><td>MP</td></tr>
<tr><td>>hg38_21_+_8433172_8446622_RNA45SN1_45S_50ntFlanks@9201.9218.18|CTCTCGGGGCCACGCGCG</td><td>>hg38_21_+_8433172_8446622_RNA45SN1_45S_50ntFlanks</td><td>9201</td><td>9218</td><td>18</td><td>CTCTCGGGGCCACGCGCG</td><td>1057.7</td><td>-13.9</td><td>1.4</td><td>-9.9</td><td>5.5E-23</td><td>1.6E-21</td><td>MB231/MB468</td><td>MP</td></tr>
<tr><td>>hg38_21_+_8433172_8446622_RNA45SN1_45S_50ntFlanks@12432.12454.23|CCTATCATTGTGAAGCAGAATTC</td><td>>hg38_21_+_8433172_8446622_RNA45SN1_45S_50ntFlanks</td><td>12432</td><td>12454</td><td>23</td><td>CCTATCATTGTGAAGCAGAATTC</td><td>291.9</td><td>-12.6</td><td>1.3</td><td>-9.7</td><td>3.4E-22</td><td>1.6E-21</td><td>BT20/MB468</td><td>NUC</td></tr>
<tr><td>>hg38_21_+_8433172_8446622_RNA45SN1_45S_50ntFlanks@11873.11904.32|AAGAAGACCCTGTTGAGCTTGACTCTAGTCTG</td><td>>hg38_21_+_8433172_8446622_RNA45SN1_45S_50ntFlanks</td><td>11873</td><td>11904</td><td>32</td><td>AAGAAGACCCTGTTGAGCTTGACTCTAGTCTG</td><td>1257.1</td><td>-14.1</td><td>1.4</td><td>-9.8</td><td>9.0E-23</td><td>1.6E-21</td><td>MB231/MB468</td><td>MITO</td></tr>
<tr><td>>hg38_21_+_8433172_8446622_RNA45SN1_45S_50ntFlanks@12619.12653.35|AGGAGCCAATGGGGCGAAGCTACCATCTGTGGGAT</td><td>>hg38_21_+_8433172_8446622_RNA45SN1_45S_50ntFlanks</td><td>12619</td><td>12653</td><td>35</td><td>AGGAGCCAATGGGGCGAAGCTACCATCTGTGGGAT</td><td>1562.3</td><td>-14.4</td><td>1.5</td><td>-9.8</td><td>9.1E-23</td><td>1.6E-21</td><td>MB231/MB468</td><td>MITO</td></tr>
<tr><td>>MI0000061|hsa-let-7a-2&WithFlank&11|-|122146516|122146599@11.32.22(+1C);MI0000062|hsa-let-7a-3&WithFlank&22|+|46112743|46112828@10.31.22(+1C);MI0000060|hsa-let-7a-1&WithFlank&9|+|94175951|94176042@12.33.22(+1C)&[MIMAT0000062&hsa-let-7a-5p&offsets|0|0(+1C);m-6&11|-|122146568|122146589&offsets|0|0(+1C)];[MIMAT0000062_1&hsa-let-7a-5p&offsets|0|0(+1C);m-5&22|+|46112752|46112773&offsets|0|0(+1C)];[MIMAT0000062_2&hsa-let-7a-5p&offsets|0|0(+1C);m-4&9|+|94175962|94175983&offsets|0|0(+1C)]|TGAGGTAGTAGGTTGTATAGTTC</td><td>>MI0000061|hsa-let-7a-2&WithFlank&11|-|122146516|122146599</td><td>11</td><td>32</td><td>22(+1C)</td><td>TGAGGTAGTAGGTTGTATAGTTC</td><td>302.3</td><td>5.7</td><td>0.6</td><td>9.8</td><td>8.9E-23</td><td>1.6E-21</td><td>MB231/MB468</td><td>NUC</td></tr>
<tr><td>>hg38_21_+_8433172_8446622_RNA45SN1_45S_50ntFlanks@12154.12180.27|GGGAGTTTGACTGGGGCGGTACACCTG</td><td>>hg38_21_+_8433172_8446622_RNA45SN1_45S_50ntFlanks</td><td>12154</td><td>12180</td><td>27</td><td>GGGAGTTTGACTGGGGCGGTACACCTG</td><td>959.8</td><td>-13.8</td><td>1.4</td><td>-9.9</td><td>5.6E-23</td><td>1.6E-21</td><td>MB231/MB468</td><td>MP</td></tr>
<tr><td>>hg38_21_+_8433172_8446622_RNA45SN1_45S_50ntFlanks@4283.4320.38|CTTTAACGAGGATCCATTGGAGGGCAAGTCTGGTGCCA</td><td>>hg38_21_+_8433172_8446622_RNA45SN1_45S_50ntFlanks</td><td>4283</td><td>4320</td><td>38</td><td>CTTTAACGAGGATCCATTGGAGGGCAAGTCTGGTGCCA</td><td>397.3</td><td>-13.1</td><td>1.3</td><td>-9.7</td><td>3.5E-22</td><td>1.6E-21</td><td>BT20/MB468</td><td>NUC</td></tr>
<tr><td>>hg38_21_+_8433172_8446622_RNA45SN1_45S_50ntFlanks@8180.8211.32|TGATCGAGGCCCAGCCCGTGGACGGTGTGAGG</td><td>>hg38_21_+_8433172_8446622_RNA45SN1_45S_50ntFlanks</td><td>8180</td><td>8211</td><td>32</td><td>TGATCGAGGCCCAGCCCGTGGACGGTGTGAGG</td><td>674.7</td><td>-13.8</td><td>1.4</td><td>-9.7</td><td>3.5E-22</td><td>1.6E-21</td><td>BT20/MB468</td><td>NUC</td></tr>
<tr><td>>hg38_21_+_8433172_8446622_RNA45SN1_45S_50ntFlanks@12008.12042.35|CCGCCGGTGAAATACCACTACTCTGATCGTTTTTT</td><td>>hg38_21_+_8433172_8446622_RNA45SN1_45S_50ntFlanks</td><td>12008</td><td>12042</td><td>35</td><td>CCGCCGGTGAAATACCACTACTCTGATCGTTTTTT</td><td>2117.6</td><td>-15.5</td><td>1.6</td><td>-9.7</td><td>3.5E-22</td><td>1.6E-21</td><td>BT20/MB468</td><td>NUC</td></tr>
<tr><td>>hg38_21_+_8433172_8446622_RNA45SN1_45S_50ntFlanks@4472.4494.23|CTTAGCTGAGTGTCCCGCGGGGC</td><td>>hg38_21_+_8433172_8446622_RNA45SN1_45S_50ntFlanks</td><td>4472</td><td>4494</td><td>23</td><td>CTTAGCTGAGTGTCCCGCGGGGC</td><td>1219.8</td><td>-14.0</td><td>1.4</td><td>-9.8</td><td>9.2E-23</td><td>1.6E-21</td><td>MB231/MB468</td><td>MITO</td></tr>
<tr><td>>hg38_21_+_8433172_8446622_RNA45SN1_45S_50ntFlanks@10410.10430.21|TCAGTCGGTCCTGAGAGATGG</td><td>>hg38_21_+_8433172_8446622_RNA45SN1_45S_50ntFlanks</td><td>10410</td><td>10430</td><td>21</td><td>TCAGTCGGTCCTGAGAGATGG</td><td>5797.0</td><td>-7.3</td><td>0.8</td><td>-9.8</td><td>1.8E-22</td><td>1.6E-21</td><td>BT20/MB468</td><td>CYTO</td></tr>
<tr><td>>hg38_21_+_8433172_8446622_RNA45SN1_45S_50ntFlanks@11564.11592.29|CCTAGCAGCCGACTTAGAACTGGTGCGGA</td><td>>hg38_21_+_8433172_8446622_RNA45SN1_45S_50ntFlanks</td><td>11564</td><td>11592</td><td>29</td><td>CCTAGCAGCCGACTTAGAACTGGTGCGGA</td><td>324.0</td><td>-12.8</td><td>1.3</td><td>-9.7</td><td>3.6E-22</td><td>1.7E-21</td><td>BT20/MB468</td><td>NUC</td></tr>
<tr><td>>hg38_21_+_8433172_8446622_RNA45SN1_45S_50ntFlanks@9572.9595.24|CAAATCGGTCGTCCGACCTGGGTA</td><td>>hg38_21_+_8433172_8446622_RNA45SN1_45S_50ntFlanks</td><td>9572</td><td>9595</td><td>24</td><td>CAAATCGGTCGTCCGACCTGGGTA</td><td>5260.2</td><td>-7.0</td><td>0.7</td><td>-9.9</td><td>4.7E-23</td><td>1.7E-21</td><td>MB231/MB468</td><td>CYTO</td></tr>
</table>

<table>
<tr><td>>MI0000089|hsa-mir-31&WithFlank&9|-|21512109|21512191@14.35.22&[MIMAT0000089&hsa-miR-31-5p&offsets|0|+1;m-360&9|-|21512157|21512178&offsets|0|0]|AGGCAAGATGCTGGCATAGCTG</td><td>>MI0000089|hsa-mir-31&WithFlank&9|-|21512109|21512191</td><td>14</td><td>35</td><td>22</td><td>AGGCAAGATGCTGGCATAGCTG</td><td>799.7</td><td>7.8</td><td>0.8</td><td>9.8</td><td>1.3E-22</td><td>1.7E-21</td><td>BT20/MB468</td><td>MITO</td></tr>
<tr><td>>hg38_21_+_8433172_8446622_RNA45SN1_45S_50ntFlanks@5490.5516.27|TGAGAAGACGGTCGAACTTGACTATCT</td><td>>hg38_21_+_8433172_8446622_RNA45SN1_45S_50ntFlanks</td><td>5490</td><td>5516</td><td>27</td><td>TGAGAAGACGGTCGAACTTGACTATCT</td><td>331.1</td><td>4.9</td><td>0.5</td><td>9.8</td><td>9.3E-23</td><td>1.7E-21</td><td>MB231/MB468</td><td>NUC</td></tr>
<tr><td>>hg38_21_+_8433172_8446622_RNA45SN1_45S_50ntFlanks@10922.10946.25|CTGGACGAGGCGCCGCCGCCCCCCC</td><td>>hg38_21_+_8433172_8446622_RNA45SN1_45S_50ntFlanks</td><td>10922</td><td>10946</td><td>25</td><td>CTGGACGAGGCGCCGCCGCCCCCCC</td><td>1309.7</td><td>-14.1</td><td>1.4</td><td>-9.8</td><td>9.5E-23</td><td>1.7E-21</td><td>MB231/MB468</td><td>MITO</td></tr>
<tr><td>>MI0003591|hsa-mir-584&WithFlank&5|-|149062307|149062415@22.42.21&[MIMAT0003249&hsa-miR-584-5p&offsets|0|-1;m-167&5|-|149062374|149062394&offsets|0|0]|TTATGGTTTGCCTGGGACTGA</td><td>>MI0003591|hsa-mir-584&WithFlank&5|-|149062307|149062415</td><td>22</td><td>42</td><td>21</td><td>TTATGGTTTGCCTGGGACTGA</td><td>1579.7</td><td>6.3</td><td>0.6</td><td>9.8</td><td>9.4E-23</td><td>1.7E-21</td><td>MB231/MB468</td><td>NUC</td></tr>
<tr><td>>hg38_21_+_8433172_8446622_RNA45SN1_45S_50ntFlanks@8176.8196.21|CTTCTGATCGAGGCCCAGCCC</td><td>>hg38_21_+_8433172_8446622_RNA45SN1_45S_50ntFlanks</td><td>8176</td><td>8196</td><td>21</td><td>CTTCTGATCGAGGCCCAGCCC</td><td>1151.7</td><td>-6.1</td><td>0.6</td><td>-9.7</td><td>3.7E-22</td><td>1.7E-21</td><td>BT20/MB468</td><td>NUC</td></tr>
<tr><td>>hg38_21_+_8433172_8446622_RNA45SN1_45S_50ntFlanks@10806.10823.18|AACAATGTAGGTAAGGGA</td><td>>hg38_21_+_8433172_8446622_RNA45SN1_45S_50ntFlanks</td><td>10806</td><td>10823</td><td>18</td><td>AACAATGTAGGTAAGGGA</td><td>1141.2</td><td>-13.9</td><td>1.4</td><td>-9.8</td><td>9.7E-23</td><td>1.7E-21</td><td>MB231/MB468</td><td>MITO</td></tr>
<tr><td>>hg38_21_+_8433172_8446622_RNA45SN1_45S_50ntFlanks@10684.10709.26|CGGCGGCGTCCGGTGAGCTCTCGCTG</td><td>>hg38_21_+_8433172_8446622_RNA45SN1_45S_50ntFlanks</td><td>10684</td><td>10709</td><td>26</td><td>CGGCGGCGTCCGGTGAGCTCTCGCTG</td><td>5053.1</td><td>-6.9</td><td>0.7</td><td>-9.8</td><td>9.8E-23</td><td>1.7E-21</td><td>MB231/MB468</td><td>MITO</td></tr>
<tr><td>>MI0001445|hsa-mir-423&WithFlank&17|+|30117073|30117178@23.45.23(+1C)&[MIMAT0004748&hsa-miR-423-5p&offsets|0|0(+1C);m-141&17|+|30117095|30117117&offsets|0|0(+1C)]|TGAGGGGCAGAGAGCGAGACTTTC</td><td>>MI0001445|hsa-mir-423&WithFlank&17|+|30117073|30117178</td><td>23</td><td>45</td><td>23(+1C)</td><td>TGAGGGGCAGAGAGCGAGACTTTC</td><td>308.2</td><td>12.5</td><td>1.3</td><td>9.7</td><td>1.9E-22</td><td>1.7E-21</td><td>BT20/MB468</td><td>CYTO</td></tr>
<tr><td>>hg38_21_+_8433172_8446622_RNA45SN1_45S_50ntFlanks@11752.11777.26|CCAAATGCCTCGTCATCTAATTAGTG</td><td>>hg38_21_+_8433172_8446622_RNA45SN1_45S_50ntFlanks</td><td>11752</td><td>11777</td><td>26</td><td>CCAAATGCCTCGTCATCTAATTAGTG</td><td>306.6</td><td>-12.7</td><td>1.3</td><td>-9.7</td><td>3.7E-22</td><td>1.7E-21</td><td>BT20/MB468</td><td>NUC</td></tr>
<tr><td>>hg38_21_+_8433172_8446622_RNA45SN1_45S_50ntFlanks@4163.4184.22|CATCCAAGGAAGGCAGCAGGCG</td><td>>hg38_21_+_8433172_8446622_RNA45SN1_45S_50ntFlanks</td><td>4163</td><td>4184</td><td>22</td><td>CATCCAAGGAAGGCAGCAGGCG</td><td>1546.7</td><td>-13.6</td><td>1.4</td><td>-9.8</td><td>9.7E-23</td><td>1.7E-21</td><td>MB231/MB468</td><td>NUC</td></tr>
<tr><td>>hg38_21_+_8433172_8446622_RNA45SN1_45S_50ntFlanks@7975.8009.35|CGCGACCTCAGATCAGACGTGGCGACCCGCTGAAT</td><td>>hg38_21_+_8433172_8446622_RNA45SN1_45S_50ntFlanks</td><td>7975</td><td>8009</td><td>35</td><td>CGCGACCTCAGATCAGACGTGGCGACCCGCTGAAT</td><td>136455.3</td><td>7.8</td><td>0.8</td><td>9.9</td><td>4.9E-23</td><td>1.7E-21</td><td>MB231/MB468</td><td>CYTO</td></tr>
<tr><td>>hg38_21_+_8433172_8446622_RNA45SN1_45S_50ntFlanks@6721.6744.24|AATTGCAGGACACATTGATCATCG</td><td>>hg38_21_+_8433172_8446622_RNA45SN1_45S_50ntFlanks</td><td>6721</td><td>6744</td><td>24</td><td>AATTGCAGGACACATTGATCATCG</td><td>6514.1</td><td>-9.6</td><td>1.0</td><td>-9.8</td><td>1.0E-22</td><td>1.7E-21</td><td>MB231/MB468</td><td>MITO</td></tr>
<tr><td>>hg38_21_+_8433172_8446622_RNA45SN1_45S_50ntFlanks@8177.8202.26|TTCTGATCGAGGCCCAGCCCGTGGAC</td><td>>hg38_21_+_8433172_8446622_RNA45SN1_45S_50ntFlanks</td><td>8177</td><td>8202</td><td>26</td><td>TTCTGATCGAGGCCCAGCCCGTGGAC</td><td>343.0</td><td>5.1</td><td>0.5</td><td>9.7</td><td>1.9E-22</td><td>1.7E-21</td><td>BT20/MB468</td><td>CYTO</td></tr>
<tr><td>>hg38_21_+_8433172_8446622_RNA45SN1_45S_50ntFlanks@9129.9152.24|CGGGTCGGGGGGCGGGGCGGACTG</td><td>>hg38_21_+_8433172_8446622_RNA45SN1_45S_50ntFlanks</td><td>9129</td><td>9152</td><td>24</td><td>CGGGTCGGGGGGCGGGGCGGACTG</td><td>2248.9</td><td>-15.0</td><td>1.5</td><td>-9.9</td><td>6.1E-23</td><td>1.7E-21</td><td>MB231/MB468</td><td>MP</td></tr>
<tr><td>>MI0000542|hsa-mir-320a&WithFlank&8|-|22244960|22245043@50.71.22&[MIMAT0000510&hsa-miR-320a-3p&offsets|+2|+2;m-60&8|-|22244975|22244996&offsets|+2|+2]|AAGCTGGGTTGAGAGGGCGAAA</td><td>>MI0000542|hsa-mir-320a&WithFlank&8|-|22244960|22245043</td><td>50</td><td>71</td><td>22</td><td>AAGCTGGGTTGAGAGGGCGAAA</td><td>466.1</td><td>13.1</td><td>1.3</td><td>9.7</td><td>1.9E-22</td><td>1.8E-21</td><td>BT20/MB468</td><td>CYTO</td></tr>
<tr><td>>hg38_21_+_8433172_8446622_RNA45SN1_45S_50ntFlanks@12935.12956.22|TAAACCATTCGTAGACGACCTG</td><td>>hg38_21_+_8433172_8446622_RNA45SN1_45S_50ntFlanks</td><td>12935</td><td>12956</td><td>22</td><td>TAAACCATTCGTAGACGACCTG</td><td>893.1</td><td>-13.7</td><td>1.4</td><td>-9.9</td><td>6.2E-23</td><td>1.8E-21</td><td>MB231/MB468</td><td>MP</td></tr>
<tr><td>>hg38_21_+_8433172_8446622_RNA45SN1_45S_50ntFlanks@12624.12653.30|CCAATGGGGCGAAGCTACCATCTGTGGGAT</td><td>>hg38_21_+_8433172_8446622_RNA45SN1_45S_50ntFlanks</td><td>12624</td><td>12653</td><td>30</td><td>CCAATGGGGCGAAGCTACCATCTGTGGGAT</td><td>1890.9</td><td>-14.7</td><td>1.5</td><td>-9.9</td><td>6.3E-23</td><td>1.8E-21</td><td>MB231/MB468</td><td>MP</td></tr>
<tr><td>>hg38_21_+_8433172_8446622_RNA45SN1_45S_50ntFlanks@8169.8214.46|CCAAGTCCTTCTGATCGAGGCCCAGCCCGTGGACGGTGTGAGGCCG</td><td>>hg38_21_+_8433172_8446622_RNA45SN1_45S_50ntFlanks</td><td>8169</td><td>8214</td><td>46</td><td>CCAAGTCCTTCTGATCGAGGCCCAGCCCGTGGACGGTGTGAGGCCG</td><td>1035.1</td><td>-14.4</td><td>1.5</td><td>-9.7</td><td>3.9E-22</td><td>1.8E-21</td><td>BT20/MB468</td><td>NUC</td></tr>
<tr><td>>hg38_21_+_8433172_8446622_RNA45SN1_45S_50ntFlanks@4726.4768.43|TAATCAAGAACGAAAGTCGGAGGTTCGAAGACGATCAGATACC</td><td>>hg38_21_+_8433172_8446622_RNA45SN1_45S_50ntFlanks</td><td>4726</td><td>4768</td><td>43</td><td>TAATCAAGAACGAAAGTCGGAGGTTCGAAGACGATCAGATACC</td><td>490.3</td><td>12.9</td><td>1.3</td><td>9.9</td><td>5.2E-23</td><td>1.8E-21</td><td>MB231/MB468</td><td>CYTO</td></tr>
<tr><td>>hg38_21_+_8433172_8446622_RNA45SN1_45S_50ntFlanks@11873.11898.26|AAGAAGACCCTGTTGAGCTTGACTCT</td><td>>hg38_21_+_8433172_8446622_RNA45SN1_45S_50ntFlanks</td><td>11873</td><td>11898</td><td>26</td><td>AAGAAGACCCTGTTGAGCTTGACTCT</td><td>1447.7</td><td>-6.5</td><td>0.7</td><td>-9.8</td><td>1.0E-22</td><td>1.8E-21</td><td>MB231/MB468</td><td>MITO</td></tr>
<tr><td>>hg38_21_+_8433172_8446622_RNA45SN1_45S_50ntFlanks@12405.12426.22|CTTTTTGATCCTTCGATGTCGG</td><td>>hg38_21_+_8433172_8446622_RNA45SN1_45S_50ntFlanks</td><td>12405</td><td>12426</td><td>22</td><td>CTTTTTGATCCTTCGATGTCGG</td><td>6149.9</td><td>-12.8</td><td>1.3</td><td>-9.7</td><td>4.0E-22</td><td>1.8E-21</td><td>BT20/MB468</td><td>NUC</td></tr>
<tr><td>>hg38_21_+_8433172_8446622_RNA45SN1_45S_50ntFlanks@11576.11603.28|CTTAGAACTGGTGCGGACCAGGGGAATC</td><td>>hg38_21_+_8433172_8446622_RNA45SN1_45S_50ntFlanks</td><td>11576</td><td>11603</td><td>28</td><td>CTTAGAACTGGTGCGGACCAGGGGAATC</td><td>8988.2</td><td>-10.9</td><td>1.1</td><td>-9.7</td><td>4.0E-22</td><td>1.8E-21</td><td>BT20/MB468</td><td>NUC</td></tr>
<tr><td>>MI0000266|hsa-mir-10a&WithFlank&17|-|48579832|48579953@29.50.22&[MIMAT0000253&hsa-miR-10a-5p&offsets|+1|0;m-18&17|-|48579905|48579926&offsets|+1|+1]|ACCCTGTAGATCCGAATTTGTG</td><td>>MI0000266|hsa-mir-10a&WithFlank&17|-|48579832|48579953</td><td>29</td><td>50</td><td>22</td><td>ACCCTGTAGATCCGAATTTGTG</td><td>2535.6</td><td>14.5</td><td>1.5</td><td>9.8</td><td>1.1E-22</td><td>1.8E-21</td><td>MB231/MB468</td><td>MITO</td></tr>
<tr><td>>MI0000068|hsa-let-7f-2&WithFlank&X|-|53557186|53557280@14.36.23&[MIMAT0000067_1&hsa-let-7f-5p&offsets|0|+1;m-19&X|-|53557246|53557267&offsets|0|+1]|TGAGGTAGTAGATTGTATAGTTT</td><td>>MI0000068|hsa-let-7f-2&WithFlank&X|-|53557186|53557280</td><td>14</td><td>36</td><td>23</td><td>TGAGGTAGTAGATTGTATAGTTT</td><td>1931.7</td><td>6.8</td><td>0.7</td><td>9.8</td><td>1.5E-22</td><td>1.8E-21</td><td>BT20/MB468</td><td>MITO</td></tr>
<tr><td>>hg38_21_+_8433172_8446622_RNA45SN1_45S_50ntFlanks@8170.8212.43|CAAGTCCTTCTGATCGAGGCCCAGCCCGTGGACGGTGTGAGGC</td><td>>hg38_21_+_8433172_8446622_RNA45SN1_45S_50ntFlanks</td><td>8170</td><td>8212</td><td>43</td><td>CAAGTCCTTCTGATCGAGGCCCAGCCCGTGGACGGTGTGAGGC</td><td>13859.8</td><td>-11.8</td><td>1.2</td><td>-9.7</td><td>4.1E-22</td><td>1.9E-21</td><td>BT20/MB468</td><td>NUC</td></tr>
<tr><td>>MI0000077|hsa-mir-21&WithFlank&17|+|59841260|59841343@14.34.21(+1U)&[MIMAT0000076&hsa-miR-21-5p&offsets|0|-1(+1U);m-1&17|+|59841273|59841294&offsets|0|-1(+1U)]|TAGCTTATCAGACTGATGTTGT</td><td>>MI0000077|hsa-mir-21&WithFlank&17|+|59841260|59841343</td><td>14</td><td>34</td><td>21(+1U)</td><td>TAGCTTATCAGACTGATGTTGT</td><td>831.8</td><td>7.0</td><td>0.7</td><td>9.8</td><td>1.1E-22</td><td>1.9E-21</td><td>MB231/MB468</td><td>NUC</td></tr>
<tr><td>>hg38_21_+_8433172_8446622_RNA45SN1_45S_50ntFlanks@7985.8006.22|GATCAGACGTGGCGACCCGCTG</td><td>>hg38_21_+_8433172_8446622_RNA45SN1_45S_50ntFlanks</td><td>7985</td><td>8006</td><td>22</td><td>GATCAGACGTGGCGACCCGCTG</td><td>3743.8</td><td>-14.0</td><td>1.4</td><td>-9.8</td><td>1.5E-22</td><td>1.9E-21</td><td>BT20/MB468</td><td>MITO</td></tr>
<tr><td>>hg38_1_-_228634819_228635039_RNA5S12_5S_50ntFlanks@128.170.43|CTTGGATGGGAGACCGCCTGGGAATACCGGGTGCTGTAGGCTT</td><td>>hg38_1_-_228634819_228635039_RNA5S12_5S_50ntFlanks</td><td>128</td><td>170</td><td>43</td><td>CTTGGATGGGAGACCGCCTGGGAATACCGGGTGCTGTAGGCTT</td><td>3161.8</td><td>8.4</td><td>0.8</td><td>9.9</td><td>5.4E-23</td><td>1.9E-21</td><td>MB231/MB468</td><td>CYTO</td></tr>
<tr><td>>hg38_21_+_8433172_8446622_RNA45SN1_45S_50ntFlanks@12633.12654.22|CGAAGCTACCATCTGTGGGATT</td><td>>hg38_21_+_8433172_8446622_RNA45SN1_45S_50ntFlanks</td><td>12633</td><td>12654</td><td>22</td><td>CGAAGCTACCATCTGTGGGATT</td><td>2856.7</td><td>-6.4</td><td>0.7</td><td>-9.7</td><td>2.1E-22</td><td>1.9E-21</td><td>BT20/MB468</td><td>CYTO</td></tr>
<tr><td>>hg38_21_+_8433172_8446622_RNA45SN1_45S_50ntFlanks@10351.10374.24|CGAGAACTTTGAAGGCCGAAGTGG</td><td>>hg38_21_+_8433172_8446622_RNA45SN1_45S_50ntFlanks</td><td>10351</td><td>10374</td><td>24</td><td>CGAGAACTTTGAAGGCCGAAGTGG</td><td>10190.2</td><td>-11.2</td><td>1.1</td><td>-9.9</td><td>6.8E-23</td><td>1.9E-21</td><td>MB231/MB468</td><td>MP</td></tr>
<tr><td>>hg38_21_+_8433172_8446622_RNA45SN1_45S_50ntFlanks@12289.12323.35|CCGTGAAAGCGGGGCCTCACGATCCTTCTGACCTT</td><td>>hg38_21_+_8433172_8446622_RNA45SN1_45S_50ntFlanks</td><td>12289</td><td>12323</td><td>35</td><td>CCGTGAAAGCGGGGCCTCACGATCCTTCTGACCTT</td><td>5538.5</td><td>-16.2</td><td>1.7</td><td>-9.8</td><td>1.1E-22</td><td>1.9E-21</td><td>MB231/MB468</td><td>MITO</td></tr>
<tr><td>>MI0000736|hsa-mir-30c-1&WithFlank&1|+|40757278|40757378@23.44.22(+1U);MI0000254|hsa-mir-30c-2&WithFlank&6|-|71376954|71377037@13.34.22(+1U)&[MIMAT0000244&hsa-miR-30c-5p&offsets|0|-1(+1U);m-57&1|+|40757300|40757323&offsets|0|-2(+1U)];[MIMAT0000244_1&hsa-miR-30c-5p&offsets|0|-1(+1U);m-56&6|-|71377002|71377025&offsets|0|-2(+1U)]|TGTAAACATCCTACACTCTCAGT</td><td>>MI0000736|hsa-mir-30c-1&WithFlank&1|+|40757278|40757378</td><td>23</td><td>44</td><td>22(+1U)</td><td>TGTAAACATCCTACACTCTCAGT</td><td>4285.7</td><td>6.8</td><td>0.7</td><td>9.7</td><td>2.1E-22</td><td>1.9E-21</td><td>BT20/MB468</td><td>CYTO</td></tr>
<tr><td>>MI0000298|hsa-mir-221&WithFlank&X|-|45746151|45746272@31.56.26&[MIMAT0004568&hsa-miR-221-5p&offsets|0|+4;m-405&X|-|45746220|45746242&offsets|0|+3]|ACCTGGCATACAATGTAGATTTCTGT</td><td>>MI0000298|hsa-mir-221&WithFlank&X|-|45746151|45746272</td><td>31</td><td>56</td><td>26</td><td>ACCTGGCATACAATGTAGATTTCTGT</td><td>3500.6</td><td>6.8</td><td>0.7</td><td>9.9</td><td>5.5E-23</td><td>1.9E-21</td><td>MB231/MB468</td><td>CYTO</td></tr>
<tr><td>>hg38_21_+_8433172_8446622_RNA45SN1_45S_50ntFlanks@5419.5445.27|ACCGATTGGATGGTTTAGTGAGGCCCT</td><td>>hg38_21_+_8433172_8446622_RNA45SN1_45S_50ntFlanks</td><td>5419</td><td>5445</td><td>27</td><td>ACCGATTGGATGGTTTAGTGAGGCCCT</td><td>563.8</td><td>-13.6</td><td>1.4</td><td>-9.7</td><td>4.2E-22</td><td>1.9E-21</td><td>BT20/MB468</td><td>NUC</td></tr>
<tr><td>>hg38_21_+_8433172_8446622_RNA45SN1_45S_50ntFlanks@8115.8141.27|CGCGGGACATGTGGCGTACGGAAGACC</td><td>>hg38_21_+_8433172_8446622_RNA45SN1_45S_50ntFlanks</td><td>8115</td><td>8141</td><td>27</td><td>CGCGGGACATGTGGCGTACGGAAGACC</td><td>1396.1</td><td>-14.2</td><td>1.5</td><td>-9.8</td><td>1.1E-22</td><td>2.0E-21</td><td>MB231/MB468</td><td>MITO</td></tr>
<tr><td>>MI0000266|hsa-mir-10a&WithFlank&17|-|48579832|48579953@28.49.22(+1A)&[MIMAT0000253&hsa-miR-10a-5p&offsets|0|-1(+1A);m-18&17|-|48579905|48579926&offsets|0|0(+1A)]|TACCCTGTAGATCCGAATTTGTA</td><td>>MI0000266|hsa-mir-10a&WithFlank&17|-|48579832|48579953</td><td>28</td><td>49</td><td>22(+1A)</td><td>TACCCTGTAGATCCGAATTTGTA</td><td>352.4</td><td>12.4</td><td>1.3</td><td>9.8</td><td>1.1E-22</td><td>2.0E-21</td><td>MB231/MB468</td><td>NUC</td></tr>
<tr><td>>MI0000542|hsa-mir-320a&WithFlank&8|-|22244960|22245043@48.69.22(+1C)&[MIMAT0000510&hsa-miR-320a-3p&offsets|0|0(+1C);m-60&8|-|22244975|22244996&offsets|0|0(+1C)]|AAAAGCTGGGTTGAGAGGGCGAC</td><td>>MI0000542|hsa-mir-320a&WithFlank&8|-|22244960|22245043</td><td>48</td><td>69</td><td>22(+1C)</td><td>AAAAGCTGGGTTGAGAGGGCGAC</td><td>702.6</td><td>13.9</td><td>1.4</td><td>9.8</td><td>1.6E-22</td><td>2.0E-21</td><td>BT20/MB468</td><td>MITO</td></tr>
<tr><td>>hg38_21_+_8433172_8446622_RNA45SN1_45S_50ntFlanks@8352.8375.24|AGTTGAAAAGAACTTTGAAGAGAG</td><td>>hg38_21_+_8433172_8446622_RNA45SN1_45S_50ntFlanks</td><td>8352</td><td>8375</td><td>24</td><td>AGTTGAAAAGAACTTTGAAGAGAG</td><td>1208.5</td><td>14.7</td><td>1.5</td><td>9.8</td><td>1.6E-22</td><td>2.0E-21</td><td>BT20/MB468</td><td>MITO</td></tr>
<tr><td>>hg38_21_+_8433172_8446622_RNA45SN1_45S_50ntFlanks@10771.10816.46|CAGGTCTCCAAGGTGAACAGCCTCTGGCATGTTGGAACAATGTAGG</td><td>>hg38_21_+_8433172_8446622_RNA45SN1_45S_50ntFlanks</td><td>10771</td><td>10816</td><td>46</td><td>CAGGTCTCCAAGGTGAACAGCCTCTGGCATGTTGGAACAATGTAGG</td><td>1344.2</td><td>-14.8</td><td>1.5</td><td>-9.7</td><td>4.3E-22</td><td>2.0E-21</td><td>BT20/MB468</td><td>NUC</td></tr>
<tr><td>>hg38_21_+_8433172_8446622_RNA45SN1_45S_50ntFlanks@12407.12441.35|TTTTGATCCTTCGATGTCGGCTCTTCCTATCATTG</td><td>>hg38_21_+_8433172_8446622_RNA45SN1_45S_50ntFlanks</td><td>12407</td><td>12441</td><td>35</td><td>TTTTGATCCTTCGATGTCGGCTCTTCCTATCATTG</td><td>4435.4</td><td>-8.0</td><td>0.8</td><td>-9.7</td><td>4.3E-22</td><td>2.0E-21</td><td>BT20/MB468</td><td>NUC</td></tr>
<tr><td>>hg38_21_+_8433172_8446622_RNA45SN1_45S_50ntFlanks@6778.6806.29|CCCGGGGCTACGCCTGTCTGAGCGTCGCT</td><td>>hg38_21_+_8433172_8446622_RNA45SN1_45S_50ntFlanks</td><td>6778</td><td>6806</td><td>29</td><td>CCCGGGGCTACGCCTGTCTGAGCGTCGCT</td><td>1000.1</td><td>4.7</td><td>0.5</td><td>9.8</td><td>1.1E-22</td><td>2.0E-21</td><td>MB231/MB468</td><td>NUC</td></tr>
<tr><td>>MI0000650|hsa-mir-200c&WithFlank&12|+|6963693|6963772@50.70.21(+1A)&[MIMAT0000617&hsa-miR-200c-3p&offsets|0|-2(+1A);m-54&12|+|6963742|6963764&offsets|0|-2(+1A)]|TAATACTGCCGGGTAATGATGA</td><td>>MI0000650|hsa-mir-200c&WithFlank&12|+|6963693|6963772</td><td>50</td><td>70</td><td>21(+1A)</td><td>TAATACTGCCGGGTAATGATGA</td><td>1257.6</td><td>13.9</td><td>1.4</td><td>10.1</td><td>9.0E-24</td><td>2.0E-21</td><td>BT20/MB231</td><td>MITO</td></tr>
<tr><td>>hg38_21_+_8433172_8446622_RNA45SN1_45S_50ntFlanks@12003.12043.41|GCGGGCCGCCGGTGAAATACCACTACTCTGATCGTTTTTTC</td><td>>hg38_21_+_8433172_8446622_RNA45SN1_45S_50ntFlanks</td><td>12003</td><td>12043</td><td>41</td><td>GCGGGCCGCCGGTGAAATACCACTACTCTGATCGTTTTTTC</td><td>2135.4</td><td>-15.5</td><td>1.6</td><td>-9.7</td><td>4.4E-22</td><td>2.0E-21</td><td>BT20/MB468</td><td>NUC</td></tr>
<tr><td>>hg38_21_+_8433172_8446622_RNA45SN1_45S_50ntFlanks@10107.10127.21|CGTGTGGTGTGCGTCGGAGGG</td><td>>hg38_21_+_8433172_8446622_RNA45SN1_45S_50ntFlanks</td><td>10107</td><td>10127</td><td>21</td><td>CGTGTGGTGTGCGTCGGAGGG</td><td>1013.4</td><td>-13.8</td><td>1.4</td><td>-9.8</td><td>7.1E-23</td><td>2.0E-21</td><td>MB231/MB468</td><td>MP</td></tr>
<tr><td>>hg38_21_+_8433172_8446622_RNA45SN1_45S_50ntFlanks@11849.11872.24|CGGGCTTGGCGGAATCAGCGGGGA</td><td>>hg38_21_+_8433172_8446622_RNA45SN1_45S_50ntFlanks</td><td>11849</td><td>11872</td><td>24</td><td>CGGGCTTGGCGGAATCAGCGGGGA</td><td>1321.7</td><td>-14.2</td><td>1.4</td><td>-9.8</td><td>1.2E-22</td><td>2.0E-21</td><td>MB231/MB468</td><td>MITO</td></tr>
<tr><td>>MI0000469|hsa-mir-125a&WithFlank&19|+|51693248|51693345@21.44.24&[MIMAT0000443&hsa-miR-125a-5p&offsets|0|0;m-63&19|+|51693268|51693289&offsets|0|+2]|TCCCTGAGACCCTTTAACCTGTGA</td><td>>MI0000469|hsa-mir-125a&WithFlank&19|+|51693248|51693345</td><td>21</td><td>44</td><td>24</td><td>TCCCTGAGACCCTTTAACCTGTGA</td><td>7345.6</td><td>6.9</td><td>0.7</td><td>9.8</td><td>1.6E-22</td><td>2.0E-21</td><td>BT20/MB468</td><td>MITO</td></tr>
</table>

<table>
<tr><td>>hg38_21_+_8433172_8446622_RNA45SN1_45S_50ntFlanks@13002.13029.28|CGATCTATTGAAAGTCAGCCCTCGACAC</td><td>>hg38_21_+_8433172_8446622_RNA45SN1_45S_50ntFlanks</td><td>13002</td><td>13029</td><td>28</td><td>CGATCTATTGAAAGTCAGCCCTCGACAC</td><td>1566.3</td><td>-12.7</td><td>1.3</td><td>-9.8</td><td>1.6E-22</td><td>2.0E-21</td><td>BT20/MB468</td><td>MITO</td></tr>
<tr><td>>MI0003591|hsa-mir-584&WithFlank&5|-|149062307|149062415@22.40.19&[MIMAT0003249&hsa-miR-584-5p&offsets|0|-3;m-167&5|-|149062374|149062394&offsets|0|-2]|TTATGGTTTGCCTGGGACT</td><td>>MI0003591|hsa-mir-584&WithFlank&5|-|149062307|149062415</td><td>22</td><td>40</td><td>19</td><td>TTATGGTTTGCCTGGGACT</td><td>556.8</td><td>6.5</td><td>0.7</td><td>9.8</td><td>1.2E-22</td><td>2.0E-21</td><td>MB231/MB468</td><td>NUC</td></tr>
<tr><td>>hg38_21_+_8433172_8446622_RNA45SN1_45S_50ntFlanks@11572.11612.41|CCGACTTAGAACTGGTGCGGACCAGGGGAATCCGACTGTTT</td><td>>hg38_21_+_8433172_8446622_RNA45SN1_45S_50ntFlanks</td><td>11572</td><td>11612</td><td>41</td><td>CCGACTTAGAACTGGTGCGGACCAGGGGAATCCGACTGTTT</td><td>7206.7</td><td>-8.0</td><td>0.8</td><td>-9.7</td><td>4.5E-22</td><td>2.0E-21</td><td>BT20/MB468</td><td>NUC</td></tr>
<tr><td>>hg38_21_+_8433172_8446622_RNA45SN1_45S_50ntFlanks@9531.9572.42|CAGAGGAAACTCTGGTGGAGGTCCGTAGCGGTCCTGACGTGC</td><td>>hg38_21_+_8433172_8446622_RNA45SN1_45S_50ntFlanks</td><td>9531</td><td>9572</td><td>42</td><td>CAGAGGAAACTCTGGTGGAGGTCCGTAGCGGTCCTGACGTGC</td><td>668.7</td><td>13.8</td><td>1.4</td><td>9.8</td><td>1.7E-22</td><td>2.1E-21</td><td>BT20/MB468</td><td>MITO</td></tr>
<tr><td>>MI0000750|hsa-mir-26a-2&WithFlank&12|-|57824603|57824698@20.41.22(+2U);MI0000083|hsa-mir-26a-1&WithFlank&3|+|37969398|37969486@16.37.22(+2U)&[MIMAT0000082&hsa-miR-26a-5p&offsets|0|0(+2U);m-33&12|-|57824658|57824679&offsets|0|0(+2U)];[MIMAT0000082_1&hsa-miR-26a-5p&offsets|0|0(+2U);m-34&3|+|37969413|37969434&offsets|0|0(+2U)]|TTCAAGTAATCCAGGATAGGCTTT</td><td>>MI0000750|hsa-mir-26a-2&WithFlank&12|-|57824603|57824698</td><td>20</td><td>41</td><td>22(+2U)</td><td>TTCAAGTAATCCAGGATAGGCTTT</td><td>2492.2</td><td>6.9</td><td>0.7</td><td>9.7</td><td>2.3E-22</td><td>2.1E-21</td><td>BT20/MB468</td><td>CYTO</td></tr>
<tr><td>>hg38_21_+_8433172_8446622_RNA45SN1_45S_50ntFlanks@11874.11904.31|AGAAGACCCTGTTGAGCTTGACTCTAGTCTG</td><td>>hg38_21_+_8433172_8446622_RNA45SN1_45S_50ntFlanks</td><td>11874</td><td>11904</td><td>31</td><td>AGAAGACCCTGTTGAGCTTGACTCTAGTCTG</td><td>1350.4</td><td>-14.2</td><td>1.4</td><td>-9.8</td><td>1.2E-22</td><td>2.1E-21</td><td>MB231/MB468</td><td>MITO</td></tr>
<tr><td>>hg38_21_+_8433172_8446622_RNA45SN1_45S_50ntFlanks@10794.10815.22|CTGGCATGTTGGAACAATGTAG</td><td>>hg38_21_+_8433172_8446622_RNA45SN1_45S_50ntFlanks</td><td>10794</td><td>10815</td><td>22</td><td>CTGGCATGTTGGAACAATGTAG</td><td>276.1</td><td>-12.5</td><td>1.3</td><td>-9.7</td><td>4.6E-22</td><td>2.1E-21</td><td>BT20/MB468</td><td>NUC</td></tr>
<tr><td>>MI0000285|hsa-mir-205&WithFlank&1|+|209432127|209432248@40.60.21(+1A)&[MIMAT0000266&hsa-miR-205-5p&offsets|0|-1(+1A);m-351&1|+|209432166|209432187&offsets|0|-1(+1A)]|TCCTTCATTCCACCGGAGTCTA</td><td>>MI0000285|hsa-mir-205&WithFlank&1|+|209432127|209432248</td><td>40</td><td>60</td><td>21(+1A)</td><td>TCCTTCATTCCACCGGAGTCTA</td><td>2540.4</td><td>14.9</td><td>1.5</td><td>10.0</td><td>1.0E-23</td><td>2.1E-21</td><td>BT20/MB231</td><td>MITO</td></tr>
<tr><td>>hg38_21_+_8433172_8446622_RNA45SN1_45S_50ntFlanks@11580.11603.24|GAACTGGTGCGGACCAGGGGAATC</td><td>>hg38_21_+_8433172_8446622_RNA45SN1_45S_50ntFlanks</td><td>11580</td><td>11603</td><td>24</td><td>GAACTGGTGCGGACCAGGGGAATC</td><td>2628.7</td><td>-5.8</td><td>0.6</td><td>-9.8</td><td>1.2E-22</td><td>2.1E-21</td><td>MB231/MB468</td><td>NUC</td></tr>
<tr><td>>hg38_21_+_8433172_8446622_RNA45SN1_45S_50ntFlanks@482.510.29|ATCGATGTGGTGACGTCGTGCTCTCCCGG</td><td>>hg38_21_+_8433172_8446622_RNA45SN1_45S_50ntFlanks</td><td>482</td><td>510</td><td>29</td><td>ATCGATGTGGTGACGTCGTGCTCTCCCGG</td><td>460.8</td><td>-13.3</td><td>1.4</td><td>-9.7</td><td>4.6E-22</td><td>2.1E-21</td><td>BT20/MB468</td><td>NUC</td></tr>
<tr><td>>hg38_21_+_8433172_8446622_RNA45SN1_45S_50ntFlanks@12635.12661.27|AAGCTACCATCTGTGGGATTATGACTG</td><td>>hg38_21_+_8433172_8446622_RNA45SN1_45S_50ntFlanks</td><td>12635</td><td>12661</td><td>27</td><td>AAGCTACCATCTGTGGGATTATGACTG</td><td>20978.7</td><td>-11.3</td><td>1.2</td><td>-9.8</td><td>7.5E-23</td><td>2.1E-21</td><td>MB231/MB468</td><td>MP</td></tr>
<tr><td>>MI0000456|hsa-mir-140&WithFlank&16|+|69933075|69933186@68.90.23&[MIMAT0004597&hsa-miR-140-3p&offsets|0|+2;m-45&16|+|69933143|69933164&offsets|-1|0]|TACCACAGGGTAGAACCACGGAC</td><td>>MI0000456|hsa-mir-140&WithFlank&16|+|69933075|69933186</td><td>68</td><td>90</td><td>23</td><td>TACCACAGGGTAGAACCACGGAC</td><td>1768.2</td><td>10.8</td><td>1.1</td><td>9.7</td><td>2.3E-22</td><td>2.1E-21</td><td>BT20/MB468</td><td>CYTO</td></tr>
<tr><td>>hg38_21_+_8433172_8446622_RNA45SN1_45S_50ntFlanks@8845.8867.23|CGGCGGCGGCGGCGGCGGCGGGA</td><td>>hg38_21_+_8433172_8446622_RNA45SN1_45S_50ntFlanks</td><td>8845</td><td>8867</td><td>23</td><td>CGGCGGCGGCGGCGGCGGCGGGA</td><td>521.0</td><td>-13.5</td><td>1.4</td><td>-9.7</td><td>4.7E-22</td><td>2.2E-21</td><td>BT20/MB468</td><td>NUC</td></tr>
<tr><td>>hg38_21_+_8433172_8446622_RNA45SN1_45S_50ntFlanks@7975.8013.39|CGCGACCTCAGATCAGACGTGGCGACCCGCTGAATTTAA</td><td>>hg38_21_+_8433172_8446622_RNA45SN1_45S_50ntFlanks</td><td>7975</td><td>8013</td><td>39</td><td>CGCGACCTCAGATCAGACGTGGCGACCCGCTGAATTTAA</td><td>27481.6</td><td>5.3</td><td>0.5</td><td>9.8</td><td>1.7E-22</td><td>2.2E-21</td><td>BT20/MB468</td><td>MITO</td></tr>
<tr><td>>hg38_21_+_8433172_8446622_RNA45SN1_45S_50ntFlanks@6647.6672.26|CGTACGACTCTTAGCGGTGGATCACT</td><td>>hg38_21_+_8433172_8446622_RNA45SN1_45S_50ntFlanks</td><td>6647</td><td>6672</td><td>26</td><td>CGTACGACTCTTAGCGGTGGATCACT</td><td>1297.6</td><td>6.8</td><td>0.7</td><td>9.7</td><td>2.4E-22</td><td>2.2E-21</td><td>BT20/MB468</td><td>CYTO</td></tr>
<tr><td>>MI0000102|hsa-mir-100&WithFlank&11|-|122152223|122152314@19.39.21(+1C)&[MIMAT0000098&hsa-miR-100-5p&offsets|0|-1(+1C);m-27&11|-|122152276|122152296&offsets|0|0(+1C)]|AACCCGTAGATCCGAACTTGTC</td><td>>MI0000102|hsa-mir-100&WithFlank&11|-|122152223|122152314</td><td>19</td><td>39</td><td>21(+1C)</td><td>AACCCGTAGATCCGAACTTGTC</td><td>1945.4</td><td>14.1</td><td>1.4</td><td>9.8</td><td>7.8E-23</td><td>2.2E-21</td><td>MB231/MB468</td><td>MP</td></tr>
<tr><td>>trna20_GluTTC_2_-_131094701_131094772@1.33.33,trna5_GluTTC_13_-_41634874_41634945@1.33.33|TCCCATATGGTCTAGCGGTTAGGATTCCTGGTT</td><td>>trna20_GluTTC_2_-_131094701_131094772</td><td>1</td><td>33</td><td>33</td><td>TCCCATATGGTCTAGCGGTTAGGATTCCTGGTT</td><td>3770.1</td><td>11.7</td><td>1.2</td><td>9.9</td><td>6.3E-23</td><td>2.2E-21</td><td>MB231/MB468</td><td>CYTO</td></tr>
<tr><td>>trna10_GlyTCC_17_+_8124866_8124937@1.30.30|GCGTTGGTGGTATAGTGGTAAGCATAGCTG</td><td>>trna10_GlyTCC_17_+_8124866_8124937</td><td>1</td><td>30</td><td>30</td><td>GCGTTGGTGGTATAGTGGTAAGCATAGCTG</td><td>571.7</td><td>-13.6</td><td>1.4</td><td>-9.7</td><td>4.8E-22</td><td>2.2E-21</td><td>BT20/MB468</td><td>NUC</td></tr>
<tr><td>>hg38_21_+_8433172_8446622_RNA45SN1_45S_50ntFlanks@10116.10136.21|TGCGTCGGAGGGCGGCGGCGG</td><td>>hg38_21_+_8433172_8446622_RNA45SN1_45S_50ntFlanks</td><td>10116</td><td>10136</td><td>21</td><td>TGCGTCGGAGGGCGGCGGCGG</td><td>1038.4</td><td>-13.8</td><td>1.4</td><td>-9.8</td><td>1.3E-22</td><td>2.2E-21</td><td>MB231/MB468</td><td>MITO</td></tr>
<tr><td>>hg38_21_+_8433172_8446622_RNA45SN1_45S_50ntFlanks@11315.11342.28|CCACCCCACGTCTCGTCGCGCGCGCGTC</td><td>>hg38_21_+_8433172_8446622_RNA45SN1_45S_50ntFlanks</td><td>11315</td><td>11342</td><td>28</td><td>CCACCCCACGTCTCGTCGCGCGCGCGTC</td><td>1297.6</td><td>-14.1</td><td>1.4</td><td>-9.8</td><td>1.3E-22</td><td>2.2E-21</td><td>MB231/MB468</td><td>MITO</td></tr>
<tr><td>>hg38_21_+_8433172_8446622_RNA45SN1_45S_50ntFlanks@4657.4689.33|CTAGAGGTGAAATTCTTGGACCGGCGCAAGACG</td><td>>hg38_21_+_8433172_8446622_RNA45SN1_45S_50ntFlanks</td><td>4657</td><td>4689</td><td>33</td><td>CTAGAGGTGAAATTCTTGGACCGGCGCAAGACG</td><td>4174.1</td><td>-11.8</td><td>1.2</td><td>-9.8</td><td>1.3E-22</td><td>2.2E-21</td><td>MB231/MB468</td><td>NUC</td></tr>
<tr><td>>hg38_21_+_8433172_8446622_RNA45SN1_45S_50ntFlanks@4733.4768.36|GAACGAAAGTCGGAGGTTCGAAGACGATCAGATACC</td><td>>hg38_21_+_8433172_8446622_RNA45SN1_45S_50ntFlanks</td><td>4733</td><td>4768</td><td>36</td><td>GAACGAAAGTCGGAGGTTCGAAGACGATCAGATACC</td><td>622.4</td><td>13.3</td><td>1.3</td><td>9.9</td><td>6.4E-23</td><td>2.2E-21</td><td>MB231/MB468</td><td>CYTO</td></tr>
<tr><td>>hg38_21_+_8433172_8446622_RNA45SN1_45S_50ntFlanks@11614.11640.27|ATTAAAACAAAGCATCGCGAAGGCCCG</td><td>>hg38_21_+_8433172_8446622_RNA45SN1_45S_50ntFlanks</td><td>11614</td><td>11640</td><td>27</td><td>ATTAAAACAAAGCATCGCGAAGGCCCG</td><td>1344.1</td><td>-14.2</td><td>1.4</td><td>-9.8</td><td>1.3E-22</td><td>2.2E-21</td><td>MB231/MB468</td><td>MITO</td></tr>
<tr><td>>hg38_21_+_8433172_8446622_RNA45SN1_45S_50ntFlanks@12893.12935.43|CCTCGCCCGTCACGCACCGCACGTTCGTGGGGAACCTGGCGCT</td><td>>hg38_21_+_8433172_8446622_RNA45SN1_45S_50ntFlanks</td><td>12893</td><td>12935</td><td>43</td><td>CCTCGCCCGTCACGCACCGCACGTTCGTGGGGAACCTGGCGCT</td><td>1529.7</td><td>-14.4</td><td>1.5</td><td>-9.8</td><td>1.3E-22</td><td>2.2E-21</td><td>MB231/MB468</td><td>MITO</td></tr>
<tr><td>>hg38_21_+_8433172_8446622_RNA45SN1_45S_50ntFlanks@3866.3898.33|CTAGAGCTAATACATGCCGACGGGCGCTGACCC</td><td>>hg38_21_+_8433172_8446622_RNA45SN1_45S_50ntFlanks</td><td>3866</td><td>3898</td><td>33</td><td>CTAGAGCTAATACATGCCGACGGGCGCTGACCC</td><td>1204.6</td><td>-14.0</td><td>1.4</td><td>-9.8</td><td>1.3E-22</td><td>2.2E-21</td><td>MB231/MB468</td><td>MITO</td></tr>
<tr><td>>hg38_21_+_8433172_8446622_RNA45SN1_45S_50ntFlanks@10727.10752.26|GGAGAGGGTGTAAATCTCGCGCCGGG</td><td>>hg38_21_+_8433172_8446622_RNA45SN1_45S_50ntFlanks</td><td>10727</td><td>10752</td><td>26</td><td>GGAGAGGGTGTAAATCTCGCGCCGGG</td><td>984.0</td><td>-13.8</td><td>1.4</td><td>-9.8</td><td>8.0E-23</td><td>2.2E-21</td><td>MB231/MB468</td><td>MP</td></tr>
<tr><td>>hg38_21_+_8433172_8446622_RNA45SN1_45S_50ntFlanks@9930.9959.30|ATATAGACAGCAGGACGGTGGCCATGGAAG</td><td>>hg38_21_+_8433172_8446622_RNA45SN1_45S_50ntFlanks</td><td>9930</td><td>9959</td><td>30</td><td>ATATAGACAGCAGGACGGTGGCCATGGAAG</td><td>4709.0</td><td>-16.0</td><td>1.6</td><td>-9.8</td><td>1.3E-22</td><td>2.2E-21</td><td>MB231/MB468</td><td>MITO</td></tr>
<tr><td>>hg38_21_+_8433172_8446622_RNA45SN1_45S_50ntFlanks@11536.11553.18|TCCGCGCGGCGCCTCGCC</td><td>>hg38_21_+_8433172_8446622_RNA45SN1_45S_50ntFlanks</td><td>11536</td><td>11553</td><td>18</td><td>TCCGCGCGGCGCCTCGCC</td><td>423.2</td><td>-13.2</td><td>1.4</td><td>-9.6</td><td>5.0E-22</td><td>2.2E-21</td><td>BT20/MB468</td><td>NUC</td></tr>
<tr><td>>hg38_21_+_8433172_8446622_RNA45SN1_45S_50ntFlanks@7985.8008.24|GATCAGACGTGGCGACCCGCTGAA</td><td>>hg38_21_+_8433172_8446622_RNA45SN1_45S_50ntFlanks</td><td>7985</td><td>8008</td><td>24</td><td>GATCAGACGTGGCGACCCGCTGAA</td><td>357.5</td><td>-12.9</td><td>1.3</td><td>-9.6</td><td>5.0E-22</td><td>2.2E-21</td><td>BT20/MB468</td><td>NUC</td></tr>
<tr><td>>MI0000446|hsa-mir-125b-1&WithFlank&11|-|122099751|122099850@21.40.20;MI0000470|hsa-mir-125b-2&WithFlank&21|+|16590231|16590331@23.42.20&[MIMAT0000423&hsa-miR-125b-5p&offsets|0|-2;m-67&11|-|122099809|122099830&offsets|0|-2];[MIMAT0000423_1&hsa-miR-125b-5p&offsets|0|-2;m-68&21|+|16590253|16590274&offsets|0|-2]|TCCCTGAGACCCTAACTTGT</td><td>>MI0000446|hsa-mir-125b-1&WithFlank&11|-|122099751|122099850</td><td>21</td><td>40</td><td>20</td><td>TCCCTGAGACCCTAACTTGT</td><td>2432.0</td><td>7.0</td><td>0.7</td><td>9.8</td><td>1.3E-22</td><td>2.3E-21</td><td>MB231/MB468</td><td>NUC</td></tr>
<tr><td>>hg38_21_+_8433172_8446622_RNA45SN1_45S_50ntFlanks@6739.6771.33|TCATCGACACTTCGAACGCACTTGCGGCCCCGG</td><td>>hg38_21_+_8433172_8446622_RNA45SN1_45S_50ntFlanks</td><td>6739</td><td>6771</td><td>33</td><td>TCATCGACACTTCGAACGCACTTGCGGCCCCGG</td><td>1900.1</td><td>-8.7</td><td>0.9</td><td>-9.8</td><td>1.4E-22</td><td>2.3E-21</td><td>MB231/MB468</td><td>MITO</td></tr>
<tr><td>>hg38_21_+_8433172_8446622_RNA45SN1_45S_50ntFlanks@5484.5518.35|GAGCGCTGAGAAGACGGTCGAACTTGACTATCTAG</td><td>>hg38_21_+_8433172_8446622_RNA45SN1_45S_50ntFlanks</td><td>5484</td><td>5518</td><td>35</td><td>GAGCGCTGAGAAGACGGTCGAACTTGACTATCTAG</td><td>554.4</td><td>-13.5</td><td>1.4</td><td>-9.6</td><td>5.0E-22</td><td>2.3E-21</td><td>BT20/MB468</td><td>NUC</td></tr>
<tr><td>>hg38_21_+_8433172_8446622_RNA45SN1_45S_50ntFlanks@9033.9059.27|CTCGTGGGGGGGCCGGGCCACCCCTCC</td><td>>hg38_21_+_8433172_8446622_RNA45SN1_45S_50ntFlanks</td><td>9033</td><td>9059</td><td>27</td><td>CTCGTGGGGGGGCCGGGCCACCCCTCC</td><td>2649.5</td><td>-15.2</td><td>1.5</td><td>-9.8</td><td>1.4E-22</td><td>2.3E-21</td><td>MB231/MB468</td><td>MITO</td></tr>
<tr><td>>MI0000085|hsa-mir-27a&WithFlank&19|-|13836434|13836523@57.76.20(+3U)&[MIMAT0000084&hsa-miR-27a-3p&offsets|0|-1(+3U);m-43&19|-|13836447|13836467&offsets|0|-1(+3U)]|TTCACAGTGGCTAAGTTCCGTTT</td><td>>MI0000085|hsa-mir-27a&WithFlank&19|-|13836434|13836523</td><td>57</td><td>76</td><td>20(+3U)</td><td>TTCACAGTGGCTAAGTTCCGTTT</td><td>858.1</td><td>6.8</td><td>0.7</td><td>9.8</td><td>1.3E-22</td><td>2.3E-21</td><td>MB231/MB468</td><td>NUC</td></tr>
<tr><td>>hg38_21_+_8433172_8446622_RNA45SN1_45S_50ntFlanks@8196.8224.29|CGTGGACGGTGTGAGGCCGGTAGCGGCCC</td><td>>hg38_21_+_8433172_8446622_RNA45SN1_45S_50ntFlanks</td><td>8196</td><td>8224</td><td>29</td><td>CGTGGACGGTGTGAGGCCGGTAGCGGCCC</td><td>1316.2</td><td>-5.2</td><td>0.5</td><td>-9.8</td><td>1.3E-22</td><td>2.3E-21</td><td>MB231/MB468</td><td>NUC</td></tr>
<tr><td>>hg38_21_+_8433172_8446622_RNA45SN1_45S_50ntFlanks@10841.10861.21|CGTAACTTCGGGATAAGGATT</td><td>>hg38_21_+_8433172_8446622_RNA45SN1_45S_50ntFlanks</td><td>10841</td><td>10861</td><td>21</td><td>CGTAACTTCGGGATAAGGATT</td><td>1820.4</td><td>-6.2</td><td>0.6</td><td>-9.8</td><td>8.2E-23</td><td>2.3E-21</td><td>MB231/MB468</td><td>MP</td></tr>
<tr><td>>hg38_21_+_8433172_8446622_RNA45SN1_45S_50ntFlanks@3984.4024.41|GCGGCTTTGGTGACTCTAGATAACCTCGGGCCGATCGCACG</td><td>>hg38_21_+_8433172_8446622_RNA45SN1_45S_50ntFlanks</td><td>3984</td><td>4024</td><td>41</td><td>GCGGCTTTGGTGACTCTAGATAACCTCGGGCCGATCGCACG</td><td>4263.9</td><td>-15.9</td><td>1.6</td><td>-9.8</td><td>8.3E-23</td><td>2.3E-21</td><td>MB231/MB468</td><td>MP</td></tr>
<tr><td>>MI0003591|hsa-mir-584&WithFlank&5|-|149062307|149062415@22.42.21(+1U)&[MIMAT0003249&hsa-miR-584-5p&offsets|0|-1(+1U);m-167&5|-|149062374|149062394&offsets|0|0(+1U)]|TTATGGTTTGCCTGGGACTGAT</td><td>>MI0003591|hsa-mir-584&WithFlank&5|-|149062307|149062415</td><td>22</td><td>42</td><td>21(+1U)</td><td>TTATGGTTTGCCTGGGACTGAT</td><td>632.3</td><td>13.8</td><td>1.4</td><td>9.7</td><td>1.9E-22</td><td>2.3E-21</td><td>BT20/MB468</td><td>MITO</td></tr>
<tr><td>>hg38_21_+_8433172_8446622_RNA45SN1_45S_50ntFlanks@12937.12958.22|AACCATTCGTAGACGACCTGCT</td><td>>hg38_21_+_8433172_8446622_RNA45SN1_45S_50ntFlanks</td><td>12937</td><td>12958</td><td>22</td><td>AACCATTCGTAGACGACCTGCT</td><td>746.0</td><td>-5.4</td><td>0.6</td><td>-9.6</td><td>5.1E-22</td><td>2.3E-21</td><td>BT20/MB468</td><td>NUC</td></tr>
<tr><td>>hg38_21_+_8433172_8446622_RNA45SN1_45S_50ntFlanks@11920.11948.29|ATGAGAGGTGTAGAATAAGTGGGAGGCCC</td><td>>hg38_21_+_8433172_8446622_RNA45SN1_45S_50ntFlanks</td><td>11920</td><td>11948</td><td>29</td><td>ATGAGAGGTGTAGAATAAGTGGGAGGCCC</td><td>1903.1</td><td>-14.8</td><td>1.5</td><td>-9.8</td><td>8.3E-23</td><td>2.3E-21</td><td>MB231/MB468</td><td>MP</td></tr>
<tr><td>>hg38_21_+_8433172_8446622_RNA45SN1_45S_50ntFlanks@11729.11751.23|TAACTATGACTCTCTTAAGGTAG</td><td>>hg38_21_+_8433172_8446622_RNA45SN1_45S_50ntFlanks</td><td>11729</td><td>11751</td><td>23</td><td>TAACTATGACTCTCTTAAGGTAG</td><td>8237.8</td><td>-10.6</td><td>1.1</td><td>-9.8</td><td>8.4E-23</td><td>2.3E-21</td><td>MB231/MB468</td><td>MP</td></tr>
<tr><td>>hg38_21_+_8433172_8446622_RNA45SN1_45S_50ntFlanks@8154.8181.28|CCGCTCGTGGGGGGCCCAAGTCCTTCTG</td><td>>hg38_21_+_8433172_8446622_RNA45SN1_45S_50ntFlanks</td><td>8154</td><td>8181</td><td>28</td><td>CCGCTCGTGGGGGGCCCAAGTCCTTCTG</td><td>13471.5</td><td>-10.2</td><td>1.0</td><td>-9.8</td><td>1.4E-22</td><td>2.3E-21</td><td>MB231/MB468</td><td>MITO</td></tr>
<tr><td>>hg38_21_+_8433172_8446622_RNA45SN1_45S_50ntFlanks@10731.10752.22|AGGGTGTAAATCTCGCGCCGGG</td><td>>hg38_21_+_8433172_8446622_RNA45SN1_45S_50ntFlanks</td><td>10731</td><td>10752</td><td>22</td><td>AGGGTGTAAATCTCGCGCCGGG</td><td>700.1</td><td>-13.3</td><td>1.4</td><td>-9.8</td><td>8.5E-23</td><td>2.4E-21</td><td>MB231/MB468</td><td>MP</td></tr>
<tr><td>>hg38_21_+_8433172_8446622_RNA45SN1_45S_50ntFlanks@12405.12425.21|CTTTTTGATCCTTCGATGTCG</td><td>>hg38_21_+_8433172_8446622_RNA45SN1_45S_50ntFlanks</td><td>12405</td><td>12425</td><td>21</td><td>CTTTTTGATCCTTCGATGTCG</td><td>842.9</td><td>-13.6</td><td>1.4</td><td>-9.8</td><td>8.6E-23</td><td>2.4E-21</td><td>MB231/MB468</td><td>MP</td></tr>
<tr><td>>hg38_21_+_8433172_8446622_RNA45SN1_45S_50ntFlanks@8112.8140.29|GGGCGCGGGACATGTGGCGTACGGAAGAC</td><td>>hg38_21_+_8433172_8446622_RNA45SN1_45S_50ntFlanks</td><td>8112</td><td>8140</td><td>29</td><td>GGGCGCGGGACATGTGGCGTACGGAAGAC</td><td>1866.3</td><td>-14.7</td><td>1.5</td><td>-9.8</td><td>8.7E-23</td><td>2.4E-21</td><td>MB231/MB468</td><td>MP</td></tr>
<tr><td>>hg38_21_+_8433172_8446622_RNA45SN1_45S_50ntFlanks@10914.10936.23|CGCCGCGGCTGGACGAGGCGCCG</td><td>>hg38_21_+_8433172_8446622_RNA45SN1_45S_50ntFlanks</td><td>10914</td><td>10936</td><td>23</td><td>CGCCGCGGCTGGACGAGGCGCCG</td><td>1748.9</td><td>-14.6</td><td>1.5</td><td>-9.8</td><td>1.4E-22</td><td>2.4E-21</td><td>MB231/MB468</td><td>MITO</td></tr>
</table>

| | | | | | | | | | | | | | |
|---|---|---|---|---|---|---|---|---|---|---|---|---|---|
| >MI0000285\|hsa-mir-205&WithFlank&1\|+\|209432127\|209432248@40.61.22(+1A)&[MIMAT0000266&hsa-miR-205-5p&offsets\|0\|0(+1A);m-351&1\|+\|209432166\|209432187&offsets\|0\|0(+1A)]\|TCCTTCATTCCACCGGAGTCTGA | >MI0000285\|hsa-mir-205&WithFlank&1\|+\|209432127\|209432248 | 40 | 61 | 22(+1A) | TCCTTCATTCCACCGGAGTCTGA | 2720.3 | -14.4 | 1.5 | -9.8 | 1.4E-22 | 2.5E-21 | MB231/MB468 | NUC |
| >hg38_21_+_8433172_8446622_RNA45SN1_45S_50ntFlanks@5474.5512.39\|GGCCCTGGCGGAGCGCTGAGAAGACGGTCGAACTTGACT | >hg38_21_+_8433172_8446622_RNA45SN1_45S_50ntFlanks | 5474 | 5512 | 39 | GGCCCTGGCGGAGCGCTGAGAAGACGGTCGAACTTGACT | 210.7 | 11.7 | 1.2 | 9.8 | 7.2E-23 | 2.5E-21 | MB231/MB468 | CYTO |
| >MI0000734\|hsa-mir-106b&WithFlank&7\|-\|100093987\|100094080@58.76.19(+1A)&[MIMAT0004672&hsa-miR-106b-3p&offsets\|0\|-3(+1A);m-87&7\|-\|100094003\|100094023&offsets\|0\|-2(+1A)]\|CCGCACTGTGGGTACTTGCA | >MI0000734\|hsa-mir-106b&WithFlank&7\|-\|100093987\|100094080 | 58 | 76 | 19(+1A) | CCGCACTGTGGGTACTTGCA | 348.1 | 12.7 | 1.3 | 9.7 | 2.7E-22 | 2.5E-21 | BT20/MB468 | CYTO |
| >hg38_21_+_8433172_8446622_RNA45SN1_45S_50ntFlanks@10342.10361.20\|ATATTCAAACGAGAACTTTG | >hg38_21_+_8433172_8446622_RNA45SN1_45S_50ntFlanks | 10342 | 10361 | 20 | ATATTCAAACGAGAACTTTG | 1730.7 | -8.9 | 0.9 | -9.8 | 9.0E-23 | 2.5E-21 | MB231/MB468 | MP |
| >MI0000285\|hsa-mir-205&WithFlank&1\|+\|209432127\|209432248@40.62.23&[MIMAT0000266&hsa-miR-205-5p&offsets\|0\|+1;m-351&1\|+\|209432166\|209432187&offsets\|0\|+1]\|TCCTTCATTCCACCGGAGTCTGT | >MI0000285\|hsa-mir-205&WithFlank&1\|+\|209432127\|209432248 | 40 | 62 | 23 | TCCTTCATTCCACCGGAGTCTGT | 1251.9 | -14.1 | 1.4 | -9.8 | 1.5E-22 | 2.5E-21 | MB231/MB468 | MITO |
| >hg38_21_+_8433172_8446622_RNA45SN1_45S_50ntFlanks@8845.8867.23\|CGGCGGCGGCGGCGGCGGCGGGA | >hg38_21_+_8433172_8446622_RNA45SN1_45S_50ntFlanks | 8845 | 8867 | 23 | CGGCGGCGGCGGCGGCGGCGGGA | 2427.3 | -15.0 | 1.5 | -9.8 | 1.5E-22 | 2.5E-21 | MB231/MB468 | MITO |
| >MI0000078\|hsa-mir-22&WithFlank&17\|-\|1713897\|1713993@59.80.22&[MIMAT0000077&hsa-miR-22-3p&offsets\|0\|0;m-2&17\|-\|1713914\|1713935&offsets\|0\|0]\|AAGCTGCCAGTTGAAGAACTGT | >MI0000078\|hsa-mir-22&WithFlank&17\|-\|1713897\|1713993 | 59 | 80 | 22 | AAGCTGCCAGTTGAAGAACTGT | 9869.1 | 7.7 | 0.8 | 9.8 | 1.4E-22 | 2.5E-21 | MB231/MB468 | NUC |
| >hg38_21_+_8433172_8446622_RNA45SN1_45S_50ntFlanks@4449.4472.24\|CCTCTCGGCGCCCCCTCGATGCTC | >hg38_21_+_8433172_8446622_RNA45SN1_45S_50ntFlanks | 4449 | 4472 | 24 | CCTCTCGGCGCCCCCTCGATGCTC | 2811.8 | -15.2 | 1.6 | -9.8 | 1.5E-22 | 2.5E-21 | MB231/MB468 | MITO |
| >hg38_21_+_8433172_8446622_RNA45SN1_45S_50ntFlanks@11323.11340.18\|CGTCTCGTCGCGCGCGCG | >hg38_21_+_8433172_8446622_RNA45SN1_45S_50ntFlanks | 11323 | 11340 | 18 | CGTCTCGTCGCGCGCGCG | 2273.8 | -14.9 | 1.5 | -9.8 | 1.5E-22 | 2.5E-21 | MB231/MB468 | MITO |
| >hg38_21_+_8433172_8446622_RNA45SN1_45S_50ntFlanks@11586.11613.28\|GTGCGGACCAGGGGAATCCGACTGTTTA | >hg38_21_+_8433172_8446622_RNA45SN1_45S_50ntFlanks | 11586 | 11613 | 28 | GTGCGGACCAGGGGAATCCGACTGTTTA | 425.4 | -4.9 | 0.5 | -9.6 | 5.6E-22 | 2.5E-21 | BT20/MB468 | NUC |
| >hg38_21_+_8433172_8446622_RNA45SN1_45S_50ntFlanks@11620.11640.21\|ACAAAGCATCGCGAAGGCCCG | >hg38_21_+_8433172_8446622_RNA45SN1_45S_50ntFlanks | 11620 | 11640 | 21 | ACAAAGCATCGCGAAGGCCCG | 4174.1 | -11.7 | 1.2 | -9.8 | 9.2E-23 | 2.5E-21 | MB231/MB468 | MP |
| >hg38_21_+_8433172_8446622_RNA45SN1_45S_50ntFlanks@4588.4613.26\|CTATTTTGTTGGTTTTCGGAACTGAG | >hg38_21_+_8433172_8446622_RNA45SN1_45S_50ntFlanks | 4588 | 4613 | 26 | CTATTTTGTTGGTTTTCGGAACTGAG | 951.4 | -13.8 | 1.4 | -9.8 | 9.2E-23 | 2.5E-21 | MB231/MB468 | MP |
| >hg38_21_+_8433172_8446622_RNA45SN1_45S_50ntFlanks@11235.11252.18\|CTTCCCGTGGATCGCCCC | >hg38_21_+_8433172_8446622_RNA45SN1_45S_50ntFlanks | 11235 | 11252 | 18 | CTTCCCGTGGATCGCCCC | 808.2 | -13.5 | 1.4 | -9.8 | 9.3E-23 | 2.5E-21 | MB231/MB468 | MP |
| >hg38_21_+_8433172_8446622_RNA45SN1_45S_50ntFlanks@4380.4423.44\|CGTAGTTGGATCTTGGGAGCGGGCGGGCGGTCCGCCGCGAGGCG | >hg38_21_+_8433172_8446622_RNA45SN1_45S_50ntFlanks | 4380 | 4423 | 44 | CGTAGTTGGATCTTGGGAGCGGGCGGGCGGTCCGCCGCGAGGCG | 620.8 | -13.7 | 1.4 | -9.6 | 5.6E-22 | 2.5E-21 | BT20/MB468 | NUC |
| >hg38_21_+_8433172_8446622_RNA45SN1_45S_50ntFlanks@9166.9197.32\|CCGGGCGGGTCGCGCCGTCGGGCCCGGGGGAG | >hg38_21_+_8433172_8446622_RNA45SN1_45S_50ntFlanks | 9166 | 9197 | 32 | CCGGGCGGGTCGCGCCGTCGGGCCCGGGGGAG | 1106.3 | -14.5 | 1.5 | -9.6 | 5.6E-22 | 2.5E-21 | BT20/MB468 | NUC |
| >MI0000746\|hsa-mir-99b&WithFlank&19\|+\|51692606\|51692687@13.30.18&[MIMAT0000689&hsa-miR-99b-5p&offsets\|0\|-4;m-10&19\|+\|51692618\|51692639&offsets\|0\|-4]\|CACCCGTAGAACCGACCT | >MI0000746\|hsa-mir-99b&WithFlank&19\|+\|51692606\|51692687 | 13 | 30 | 18 | CACCCGTAGAACCGACCT | 310.5 | 12.5 | 1.3 | 9.7 | 2.8E-22 | 2.5E-21 | BT20/MB468 | CYTO |
| >hg38_21_+_8433172_8446622_RNA45SN1_45S_50ntFlanks@10411.10435.25\|CAGTCGGTCCTGAGAGATGGGCGAG | >hg38_21_+_8433172_8446622_RNA45SN1_45S_50ntFlanks | 10411 | 10435 | 25 | CAGTCGGTCCTGAGAGATGGGCGAG | 2993.8 | -15.4 | 1.6 | -9.8 | 9.4E-23 | 2.6E-21 | MB231/MB468 | MP |
| >hg38_21_+_8433172_8446622_RNA45SN1_45S_50ntFlanks@12008.12035.28\|CCGCCGGTGAAATACCACTACTCTGATC | >hg38_21_+_8433172_8446622_RNA45SN1_45S_50ntFlanks | 12008 | 12035 | 28 | CCGCCGGTGAAATACCACTACTCTGATC | 898.0 | -13.7 | 1.4 | -9.8 | 9.5E-23 | 2.6E-21 | MB231/MB468 | MP |
| >hg38_21_+_8433172_8446622_RNA45SN1_45S_50ntFlanks@4289.4317.29\|CGAGGATCCATTGGAGGGCAAGTCTGGTG | >hg38_21_+_8433172_8446622_RNA45SN1_45S_50ntFlanks | 4289 | 4317 | 29 | CGAGGATCCATTGGAGGGCAAGTCTGGTG | 372.7 | -13.0 | 1.3 | -9.6 | 5.7E-22 | 2.6E-21 | BT20/MB468 | NUC |
| >hg38_21_+_8433172_8446622_RNA45SN1_45S_50ntFlanks@9202.9222.21\|TCTCGGGGCCACGCGCGCGTC | >hg38_21_+_8433172_8446622_RNA45SN1_45S_50ntFlanks | 9202 | 9222 | 21 | TCTCGGGGCCACGCGCGCGTC | 1362.5 | -14.3 | 1.5 | -9.8 | 9.5E-23 | 2.6E-21 | MB231/MB468 | MP |
| >hg38_21_+_8433172_8446622_RNA45SN1_45S_50ntFlanks@7974.8005.32\|ACGCGACCTCAGATCAGACGTGGCGACCCGCT | >hg38_21_+_8433172_8446622_RNA45SN1_45S_50ntFlanks | 7974 | 8005 | 32 | ACGCGACCTCAGATCAGACGTGGCGACCCGCT | 82039.7 | 7.4 | 0.8 | 9.7 | 2.1E-22 | 2.6E-21 | BT20/MB468 | MITO |
| >hg38_21_+_8433172_8446622_RNA45SN1_45S_50ntFlanks@3813.3850.38\|TTATGGTTCCTTTGGTCGCTCGCTCCTCTCCTACTTGG | >hg38_21_+_8433172_8446622_RNA45SN1_45S_50ntFlanks | 3813 | 3850 | 38 | TTATGGTTCCTTTGGTCGCTCGCTCCTCTCCTACTTGG | 650.6 | -13.8 | 1.4 | -9.6 | 5.7E-22 | 2.6E-21 | BT20/MB468 | NUC |
| >hg38_21_+_8433172_8446622_RNA45SN1_45S_50ntFlanks@12447.12484.38\|CAGAATTCACCAAGCGTTGGATTGTTCACCCACTAATA | >hg38_21_+_8433172_8446622_RNA45SN1_45S_50ntFlanks | 12447 | 12484 | 38 | CAGAATTCACCAAGCGTTGGATTGTTCACCCACTAATA | 511.9 | -13.4 | 1.4 | -9.6 | 5.8E-22 | 2.6E-21 | BT20/MB468 | NUC |
| >am_trna18_ArgCCT_17_+_73030001_73030073@14.31.18,trna21_ArgCCT_17_-_73030526_73030598@14.31.18\|AATGGATAAGGCACTGGC | >am_trna18_ArgCCT_17_+_73030001_73030073 | 14 | 31 | 18 | AATGGATAAGGCACTGGC | 350.1 | -12.9 | 1.3 | -9.6 | 5.8E-22 | 2.6E-21 | BT20/MB468 | NUC |
| >hg38_21_+_8433172_8446622_RNA45SN1_45S_50ntFlanks@11315.11342.28\|CCACCCCACGTCTCGTCGCGCGCGCGTC | >hg38_21_+_8433172_8446622_RNA45SN1_45S_50ntFlanks | 11315 | 11342 | 28 | CCACCCCACGTCTCGTCGCGCGCGCGTC | 1600.4 | -13.6 | 1.4 | -9.8 | 1.5E-22 | 2.6E-21 | MB231/MB468 | NUC |
| >trnaMT_ValTAC_MT_+_1602_1670@49.70.22\|AACTTAACTTGACCGCTCTGAC | >trnaMT_ValTAC_MT_+_1602_1670 | 49 | 70 | 22 | AACTTAACTTGACCGCTCTGAC | 315.9 | 12.6 | 1.3 | 9.7 | 2.9E-22 | 2.6E-21 | BT20/MB468 | CYTO |
| >hg38_21_+_8433172_8446622_RNA45SN1_45S_50ntFlanks@10458.10497.40\|ATGGCCTCCGTTGCCCTCGGCCGATCGAAAGGGAGTCGGG | >hg38_21_+_8433172_8446622_RNA45SN1_45S_50ntFlanks | 10458 | 10497 | 40 | ATGGCCTCCGTTGCCCTCGGCCGATCGAAAGGGAGTCGGG | 2010.9 | -14.8 | 1.5 | -9.8 | 1.6E-22 | 2.6E-21 | MB231/MB468 | MITO |
| >hg38_21_+_8433172_8446622_RNA45SN1_45S_50ntFlanks@11562.11603.42\|CGCCTAGCAGCCGACTTAGAACTGGTGCGGACCAGGGGAATC | >hg38_21_+_8433172_8446622_RNA45SN1_45S_50ntFlanks | 11562 | 11603 | 42 | CGCCTAGCAGCCGACTTAGAACTGGTGCGGACCAGGGGAATC | 675.1 | -13.8 | 1.4 | -9.6 | 5.9E-22 | 2.6E-21 | BT20/MB468 | NUC |
| >MI0000102\|hsa-mir-100&WithFlank&11\|-\|122152223\|122152314@19.37.19(+1A)&[MIMAT0000098&hsa-miR-100-5p&offsets\|0\|-3(+1A);m-27&11\|-\|122152276\|122152296&offsets\|0\|-2(+1A)]\|AACCCGTAGATCCGAACTTA | >MI0000102\|hsa-mir-100&WithFlank&11\|-\|122152223\|122152314 | 19 | 37 | 19(+1A) | AACCCGTAGATCCGAACTTA | 2310.3 | 15.1 | 1.5 | 9.8 | 1.5E-22 | 2.6E-21 | MB231/MB468 | NUC |
| >hg38_21_+_8433172_8446622_RNA45SN1_45S_50ntFlanks@9304.9338.35\|ACCAAGGAGTCTAACACGTGCGCGAGTCGGGGGCT | >hg38_21_+_8433172_8446622_RNA45SN1_45S_50ntFlanks | 9304 | 9338 | 35 | ACCAAGGAGTCTAACACGTGCGCGAGTCGGGGGCT | 1573.1 | -14.5 | 1.5 | -9.8 | 9.8E-23 | 2.7E-21 | MB231/MB468 | MP |
| >hg38_21_+_8433172_8446622_RNA45SN1_45S_50ntFlanks@10403.10429.27\|ACATGGGTCAGTCGGTCCTGAGAGATG | >hg38_21_+_8433172_8446622_RNA45SN1_45S_50ntFlanks | 10403 | 10429 | 27 | ACATGGGTCAGTCGGTCCTGAGAGATG | 1853.6 | -6.9 | 0.7 | -9.6 | 6.0E-22 | 2.7E-21 | BT20/MB468 | NUC |
| >hg38_1_-_228634819_228635039_RNA5S12_5S_50ntFlanks@138.169.32\|AGACCGCCTGGGAATACCGGGTGCTGTAGGCT | >hg38_1_-_228634819_228635039_RNA5S12_5S_50ntFlanks | 138 | 169 | 32 | AGACCGCCTGGGAATACCGGGTGCTGTAGGCT | 1026.5 | 6.9 | 0.7 | 9.7 | 3.0E-22 | 2.7E-21 | BT20/MB468 | CYTO |
| >MI0000456\|hsa-mir-140&WithFlank&16\|+\|69933075\|69933186@69.90.22&[MIMAT0004597&hsa-miR-140-3p&offsets\|+1\|+2;m-45&16\|+\|69933143\|69933164&offsets\|0\|0]\|ACCACAGGGTAGAACCACGGAC | >MI0000456\|hsa-mir-140&WithFlank&16\|+\|69933075\|69933186 | 69 | 90 | 22 | ACCACAGGGTAGAACCACGGAC | 961.7 | 7.9 | 0.8 | 9.8 | 1.6E-22 | 2.7E-21 | MB231/MB468 | NUC |
| >hg38_21_+_8433172_8446622_RNA45SN1_45S_50ntFlanks@6656.6681.26\|CTTAGCGGTGGATCACTCGGCTCGTG | >hg38_21_+_8433172_8446622_RNA45SN1_45S_50ntFlanks | 6656 | 6681 | 26 | CTTAGCGGTGGATCACTCGGCTCGTG | 5814.5 | -10.7 | 1.1 | -9.8 | 1.0E-22 | 2.7E-21 | MB231/MB468 | MP |
| >hg38_21_+_8433172_8446622_RNA45SN1_45S_50ntFlanks@11641.11661.21\|CGGCGGGTGTTGACGCGATGT | >hg38_21_+_8433172_8446622_RNA45SN1_45S_50ntFlanks | 11641 | 11661 | 21 | CGGCGGGTGTTGACGCGATGT | 1027.8 | -13.9 | 1.4 | -9.8 | 1.0E-22 | 2.7E-21 | MB231/MB468 | MP |
| >hg38_21_+_8433172_8446622_RNA45SN1_45S_50ntFlanks@7986.8006.21\|ATCAGACGTGGCGACCCGCTG | >hg38_21_+_8433172_8446622_RNA45SN1_45S_50ntFlanks | 7986 | 8006 | 21 | ATCAGACGTGGCGACCCGCTG | 1282.5 | -12.6 | 1.3 | -9.7 | 3.0E-22 | 2.7E-21 | BT20/MB468 | CYTO |
| >hg38_21_+_8433172_8446622_RNA45SN1_45S_50ntFlanks@10411.10429.19\|CAGTCGGTCCTGAGAGATG | >hg38_21_+_8433172_8446622_RNA45SN1_45S_50ntFlanks | 10411 | 10429 | 19 | CAGTCGGTCCTGAGAGATG | 38635.7 | -12.0 | 1.2 | -9.6 | 6.1E-22 | 2.7E-21 | BT20/MB468 | NUC |
| >hg38_21_+_8433172_8446622_RNA45SN1_45S_50ntFlanks@12008.12033.26\|CCGCCGGTGAAATACCACTACTCTGA | >hg38_21_+_8433172_8446622_RNA45SN1_45S_50ntFlanks | 12008 | 12033 | 26 | CCGCCGGTGAAATACCACTACTCTGA | 954.1 | -13.8 | 1.4 | -9.8 | 1.0E-22 | 2.7E-21 | MB231/MB468 | MP |
| >hg38_21_+_8433172_8446622_RNA45SN1_45S_50ntFlanks@10873.10902.30\|CTGGGTCGGTCGGGCTGGGGCGCGAAGCGG | >hg38_21_+_8433172_8446622_RNA45SN1_45S_50ntFlanks | 10873 | 10902 | 30 | CTGGGTCGGTCGGGCTGGGGCGCGAAGCGG | 1367.9 | -14.3 | 1.5 | -9.8 | 1.0E-22 | 2.8E-21 | MB231/MB468 | MP |
| >hg38_21_+_8433172_8446622_RNA45SN1_45S_50ntFlanks@10351.10368.18\|CGAGAACTTTGAAGGCCG | >hg38_21_+_8433172_8446622_RNA45SN1_45S_50ntFlanks | 10351 | 10368 | 18 | CGAGAACTTTGAAGGCCG | 28840.6 | -10.9 | 1.1 | -9.8 | 1.0E-22 | 2.8E-21 | MB231/MB468 | MP |
| >hg38_21_+_8433172_8446622_RNA45SN1_45S_50ntFlanks@11766.11794.29\|ATCTAATTAGTGACGCGCATGAATGGATG | >hg38_21_+_8433172_8446622_RNA45SN1_45S_50ntFlanks | 11766 | 11794 | 29 | ATCTAATTAGTGACGCGCATGAATGGATG | 317.4 | -12.7 | 1.3 | -9.6 | 6.2E-22 | 2.8E-21 | BT20/MB468 | NUC |
| >MI0000299\|hsa-mir-222&WithFlank&X\|-\|45747009\|45747130@75.96.22(+1C)&[MIMAT0000279&hsa-miR-222-3p&offsets\|0\|+1(+1C);m-140&X\|-\|45747033\|45747056&offsets\|0\|-2(+1C)]\|AGCTACATCTGGCTACTGGGTCC | >MI0000299\|hsa-mir-222&WithFlank&X\|-\|45747009\|45747130 | 75 | 96 | 22(+1C) | AGCTACATCTGGCTACTGGGTCC | 796.3 | 7.2 | 0.7 | 9.7 | 2.3E-22 | 2.8E-21 | BT20/MB468 | MITO |
| >hg38_21_+_8433172_8446622_RNA45SN1_45S_50ntFlanks@5446.5467.22\|CGGATCGGCCCCGCCGGGGTCG | >hg38_21_+_8433172_8446622_RNA45SN1_45S_50ntFlanks | 5446 | 5467 | 22 | CGGATCGGCCCCGCCGGGGTCG | 1928.4 | -14.8 | 1.5 | -9.8 | 1.1E-22 | 2.8E-21 | MB231/MB468 | MP |
| >MI0000079\|hsa-mir-23a&WithFlank&19\|-\|13836581\|13836665@54.72.19(+2U)&[MIMAT0000078&hsa-miR-23a-3p&offsets\|+3\|+1(+2U);m-31&19\|-\|13836595\|13836615&offsets\|+3\|+1(+2U)]\|ACATTGCCAGGGATTTCCATT | >MI0000079\|hsa-mir-23a&WithFlank&19\|-\|13836581\|13836665 | 54 | 72 | 19(+2U) | ACATTGCCAGGGATTTCCATT | 397.2 | 12.9 | 1.3 | 9.7 | 3.1E-22 | 2.8E-21 | BT20/MB468 | CYTO |
| >MI0000476\|hsa-mir-138-1&WithFlank&3\|+\|44114206\|44114316@29.52.24&[MIMAT0000430_1&hsa-miR-138-5p&offsets\|0\|+1;m-497&3\|+\|44114234\|44114256&offsets\|0\|+1]\|AGCTGGTGTTGTGAATCAGGCCGT | >MI0000476\|hsa-mir-138-1&WithFlank&3\|+\|44114206\|44114316 | 29 | 52 | 24 | AGCTGGTGTTGTGAATCAGGCCGT | 834.7 | 12.9 | 1.3 | 9.8 | 1.1E-22 | 2.8E-21 | MB231/MB468 | MP |
| >MI0000434\|hsa-let-7i&WithFlank&12\|+\|62603680\|62603775@12.33.22(+1U)&[MIMAT0000415&hsa-let-7i-5p&offsets\|0\|0(+1U);m-49&12\|+\|62603691\|62603712&offsets\|0\|0(+1U)]\|TGAGGTAGTAGTTTGTGCTGTTT | >MI0000434\|hsa-let-7i&WithFlank&12\|+\|62603680\|62603775 | 12 | 33 | 22(+1U) | TGAGGTAGTAGTTTGTGCTGTTT | 3436.5 | 6.3 | 0.7 | 9.7 | 3.2E-22 | 2.9E-21 | BT20/MB468 | CYTO |
| >hg38_21_+_8433172_8446622_RNA45SN1_45S_50ntFlanks@4759.4780.22\|ATCAGATACCGTCGTAGTTCCG | >hg38_21_+_8433172_8446622_RNA45SN1_45S_50ntFlanks | 4759 | 4780 | 22 | ATCAGATACCGTCGTAGTTCCG | 5223.7 | -14.4 | 1.5 | -9.7 | 2.4E-22 | 2.9E-21 | BT20/MB468 | MITO |
| >hg38_21_+_8433172_8446622_RNA45SN1_45S_50ntFlanks@9736.9765.30\|CCGAAACGATCTCAACCTATTCTCAAACTT | >hg38_21_+_8433172_8446622_RNA45SN1_45S_50ntFlanks | 9736 | 9765 | 30 | CCGAAACGATCTCAACCTATTCTCAAACTT | 5400.6 | -12.4 | 1.3 | -9.8 | 1.8E-22 | 2.9E-21 | MB231/MB468 | MITO |

| | | | | | | | | | | | | | |
|---|---|---|---|---|---|---|---|---|---|---|---|---|---|
| >hg38_21_+_8433172_8446622_RNA45SN1_45S_50ntFlanks@9738.9765.28\|GAAACGATCTCAACCTATTCTCAAACTT | >hg38_21_+_8433172_8446622_RNA45SN1_45S_50ntFlanks | 9738 | 9765 | 28 | GAAACGATCTCAACCTATTCTCAAACTT | 43363.1 | -9.6 | 1.0 | -9.8 | 1.8E-22 | 2.9E-21 | MB231/MB468 | MITO |
| >hg38_21_+_8433172_8446622_RNA45SN1_45S_50ntFlanks@11233.11255.23\|CCCTTCCCGTGGATCGCCCCAGC | >hg38_21_+_8433172_8446622_RNA45SN1_45S_50ntFlanks | 11233 | 11255 | 23 | CCCTTCCCGTGGATCGCCCCAGC | 310.1 | -12.7 | 1.3 | -9.6 | 6.5E-22 | 2.9E-21 | BT20/MB468 | NUC |
| >trna116_GluCTC_1_-_145399233_145399304@32.74.43,trna71_GluCTC_1_-_161439189_161439260@32.74.43,trna74_GluCTC_1_-_161431809_161431880@32.74.43,trna77_GluCTC_1_-_161424398_161424469@32.74.43,trna77_GluCTC_6_+_28949976_28950047@32.74.43,trna80_GluCTC_1_-_161417018_161417089@32.74.43,trna87_GluCTC_6_-_126101393_126101464@32.74.43\|CTCTCACCGCCGCGGCCCGGGTTCGATTCCCGGTCAGGGAACC | >trna116_GluCTC_1_-_145399233_145399304 | 32 | 74 | 43 | CTCTCACCGCCGCGGCCCGGGTTCGATTCCCGGTCAGGGAACC | 515.9 | -13.4 | 1.4 | -9.6 | 6.5E-22 | 2.9E-21 | BT20/MB468 | NUC |
| >hg38_21_+_8433172_8446622_RNA45SN1_45S_50ntFlanks@12634.12653.20\|GAAGCTACCATCTGTGGGAT | >hg38_21_+_8433172_8446622_RNA45SN1_45S_50ntFlanks | 12634 | 12653 | 20 | GAAGCTACCATCTGTGGGAT | 669.5 | -13.3 | 1.4 | -9.8 | 1.1E-22 | 2.9E-21 | MB231/MB468 | MP |
| >hg38_21_+_8433172_8446622_RNA45SN1_45S_50ntFlanks@11561.11585.25\|GCGCCTAGCAGCCGACTTAGAACTG | >hg38_21_+_8433172_8446622_RNA45SN1_45S_50ntFlanks | 11561 | 11585 | 25 | GCGCCTAGCAGCCGACTTAGAACTG | 1747.4 | -6.0 | 0.6 | -9.8 | 1.7E-22 | 2.9E-21 | MB231/MB468 | NUC |
| >hg38_21_+_8433172_8446622_RNA45SN1_45S_50ntFlanks@4003.4026.24\|ATAACCTCGGGCCGATCGCACGCC | >hg38_21_+_8433172_8446622_RNA45SN1_45S_50ntFlanks | 4003 | 4026 | 24 | ATAACCTCGGGCCGATCGCACGCC | 2035.7 | -14.8 | 1.5 | -9.8 | 1.8E-22 | 2.9E-21 | MB231/MB468 | MITO |
| >MI0000082\|hsa-mir-25&WithFlank&7\|-\|100093554\|100093649@58.77.20(+1A)&[MIMAT0000081&hsa-miR-25-3p&offsets\|0\|-2(+1A);m-23&7\|-\|100093571\|100093592&offsets\|0\|-2(+1A)]\|CATTGCACTTGTCTCGGTCTA | >MI0000082\|hsa-mir-25&WithFlank&7\|-\|100093554\|100093649 | 58 | 77 | 20(+1A) | CATTGCACTTGTCTCGGTCTA | 522.2 | 4.9 | 0.5 | 9.8 | 1.7E-22 | 2.9E-21 | MB231/MB468 | NUC |
| >hg38_21_+_8433172_8446622_RNA45SN1_45S_50ntFlanks@12423.12457.35\|TCGGCTCTTCCTATCATTGTGAAGCAGAATTCACC | >hg38_21_+_8433172_8446622_RNA45SN1_45S_50ntFlanks | 12423 | 12457 | 35 | TCGGCTCTTCCTATCATTGTGAAGCAGAATTCACC | 725.3 | 5.8 | 0.6 | 9.8 | 8.7E-23 | 2.9E-21 | MB231/MB468 | CYTO |
| >hg38_21_+_8433172_8446622_RNA45SN1_45S_50ntFlanks@10766.10797.32\|CGCAGCAGGTCTCCAAGGTGAACAGCCTCTGG | >hg38_21_+_8433172_8446622_RNA45SN1_45S_50ntFlanks | 10766 | 10797 | 32 | CGCAGCAGGTCTCCAAGGTGAACAGCCTCTGG | 331.1 | -12.8 | 1.3 | -9.6 | 6.6E-22 | 3.0E-21 | BT20/MB468 | NUC |
| >hg38_21_+_8433172_8446622_RNA45SN1_45S_50ntFlanks@12044.12070.27\|ACTGACCCGGTGAGGCGGGGGGGCGAG | >hg38_21_+_8433172_8446622_RNA45SN1_45S_50ntFlanks | 12044 | 12070 | 27 | ACTGACCCGGTGAGGCGGGGGGGCGAG | 1449.0 | -14.4 | 1.5 | -9.8 | 1.1E-22 | 3.0E-21 | MB231/MB468 | MP |
| >hg38_21_+_8433172_8446622_RNA45SN1_45S_50ntFlanks@10361.10388.28\|GAAGGCCGAAGTGGAGAAGGGTTCCATG | >hg38_21_+_8433172_8446622_RNA45SN1_45S_50ntFlanks | 10361 | 10388 | 28 | GAAGGCCGAAGTGGAGAAGGGTTCCATG | 1311.5 | -14.8 | 1.5 | -9.6 | 6.7E-22 | 3.0E-21 | BT20/MB468 | NUC |
| >hg38_21_+_8433172_8446622_RNA45SN1_45S_50ntFlanks@485.524.40\|GATGTGGTGACGTCGTGCTCTCCCGGGCCGGGTCCGAGCC | >hg38_21_+_8433172_8446622_RNA45SN1_45S_50ntFlanks | 485 | 524 | 40 | GATGTGGTGACGTCGTGCTCTCCCGGGCCGGGTCCGAGCC | 365.2 | 13.0 | 1.3 | 9.7 | 2.4E-22 | 3.0E-21 | BT20/MB468 | MITO |
| >hg38_21_+_8433172_8446622_RNA45SN1_45S_50ntFlanks@4759.4779.21\|ATCAGATACCGTCGTAGTTCC | >hg38_21_+_8433172_8446622_RNA45SN1_45S_50ntFlanks | 4759 | 4779 | 21 | ATCAGATACCGTCGTAGTTCC | 1020.0 | -13.0 | 1.3 | -9.8 | 1.8E-22 | 3.0E-21 | MB231/MB468 | NUC |
| >hg38_21_+_8433172_8446622_RNA45SN1_45S_50ntFlanks@4045.4063.19\|CCATTCGAACGTCTGCCCT | >hg38_21_+_8433172_8446622_RNA45SN1_45S_50ntFlanks | 4045 | 4063 | 19 | CCATTCGAACGTCTGCCCT | 772.4 | -13.5 | 1.4 | -9.8 | 1.1E-22 | 3.0E-21 | MB231/MB468 | MP |
| >hg38_21_+_8433172_8446622_RNA45SN1_45S_50ntFlanks@9714.9737.24\|CGAATGATTAGAGGTCTTGGGGCC | >hg38_21_+_8433172_8446622_RNA45SN1_45S_50ntFlanks | 9714 | 9737 | 24 | CGAATGATTAGAGGTCTTGGGGCC | 1377.6 | -14.3 | 1.5 | -9.8 | 1.1E-22 | 3.0E-21 | MB231/MB468 | MP |
| >MI0000764\|hsa-mir-363&WithFlank&X\|-\|134169372\|134169458@56.77.22&[MIMAT0000707&hsa-miR-363-3p&offsets\|0\|0;m-178&X\|-\|134169381\|134169403&offsets\|0\|-1]\|AATTGCACGGTATCCATCTGTA | >MI0000764\|hsa-mir-363&WithFlank&X\|-\|134169372\|134169458 | 56 | 77 | 22 | AATTGCACGGTATCCATCTGTA | 1193.3 | 13.8 | 1.4 | 10.0 | 1.6E-23 | 3.0E-21 | BT20/MB231 | MITO |
| >hg38_21_+_8433172_8446622_RNA45SN1_45S_50ntFlanks@12714.12751.38\|GAGCCTCGGTTGGCCTCGGATAGCCGGTCCCCCGCCTG | >hg38_21_+_8433172_8446622_RNA45SN1_45S_50ntFlanks | 12714 | 12751 | 38 | GAGCCTCGGTTGGCCTCGGATAGCCGGTCCCCCGCCTG | 1196.3 | -14.0 | 1.4 | -9.7 | 1.9E-22 | 3.0E-21 | MB231/MB468 | MITO |
| >MI0000440\|hsa-mir-27b&WithFlank&9\|+\|95085439\|95085547@67.88.22(+1U)&[MIMAT0000419&hsa-miR-27b-3p&offsets\|0\|+1(+1U);m-39&9\|+\|95085505\|95085525&offsets\|0\|+1(+1U)]\|TTCACAGTGGCTAAGTTCTGCAT | >MI0000440\|hsa-mir-27b&WithFlank&9\|+\|95085439\|95085547 | 67 | 88 | 22(+1U) | TTCACAGTGGCTAAGTTCTGCAT | 588.3 | 8.1 | 0.8 | 9.7 | 3.4E-22 | 3.1E-21 | BT20/MB468 | CYTO |
| >trnaMT_GluTTC_MT_-_14674_14742@20.58.39,trnalookalike8_GluTTC_5_-_93905172_93905240@20.58.39\|AACGATGGTTTTTCATATCATTGGTCGTGGTTGTAGTCC | >trnaMT_GluTTC_MT_-_14674_14742 | 20 | 58 | 39 | AACGATGGTTTTTCATATCATTGGTCGTGGTTGTAGTCC | 393.0 | 13.1 | 1.3 | 9.7 | 2.5E-22 | 3.1E-21 | BT20/MB468 | MITO |
| >hg38_21_+_8433172_8446622_RNA45SN1_45S_50ntFlanks@11876.11906.31\|AAGACCCTGTTGAGCTTGACTCTAGTCTGGC | >hg38_21_+_8433172_8446622_RNA45SN1_45S_50ntFlanks | 11876 | 11906 | 31 | AAGACCCTGTTGAGCTTGACTCTAGTCTGGC | 1379.3 | -14.2 | 1.5 | -9.7 | 1.9E-22 | 3.1E-21 | MB231/MB468 | MITO |
| >hg38_21_+_8433172_8446622_RNA45SN1_45S_50ntFlanks@9568.9594.27\|CGTGCAAATCGGTCGTCCGACCTGGGT | >hg38_21_+_8433172_8446622_RNA45SN1_45S_50ntFlanks | 9568 | 9594 | 27 | CGTGCAAATCGGTCGTCCGACCTGGGT | 888.9 | -13.7 | 1.4 | -9.8 | 1.2E-22 | 3.2E-21 | MB231/MB468 | MP |
| >MI0001445\|hsa-mir-423&WithFlank&17\|+\|30117073\|30117178@23.44.22(+1C)&[MIMAT0004748&hsa-miR-423-5p&offsets\|0\|-1(+1C);m-141&17\|+\|30117095\|30117117&offsets\|0\|-1(+1C)]\|TGAGGGGCAGAGAGCGAGACTTC | >MI0001445\|hsa-mir-423&WithFlank&17\|+\|30117073\|30117178 | 23 | 44 | 22(+1C) | TGAGGGGCAGAGAGCGAGACTTC | 396.9 | 13.1 | 1.3 | 9.7 | 2.6E-22 | 3.2E-21 | BT20/MB468 | MITO |
| >hg38_1_-_228634819_228635039_RNA5S12_5S_50ntFlanks@145.171.27\|CTGGGAATACCGGGTGCTGTAGGCTTT | >hg38_1_-_228634819_228635039_RNA5S12_5S_50ntFlanks | 145 | 171 | 27 | CTGGGAATACCGGGTGCTGTAGGCTTT | 2368.2 | 6.3 | 0.7 | 9.7 | 3.6E-22 | 3.2E-21 | BT20/MB468 | CYTO |
| >trna146_GlnCTG_6_-_27515531_27515602@17.54.38,trna1_GlnCTG_6_+_18836402_18836473@17.54.38,trna3_GlnCTG_17_+_8023070_8023141@17.54.38,trna49_GlnCTG_6_+_27487308_27487379@17.54.38,trna7_GlnCTG_15_-_66161400_66161471@17.54.38,trna99_GlnCTG_6_-_28909378_28909449@17.54.38\|GGTTAGCACTCTGGACTCTGAATCCAGCGATCCGAGTT | >trna146_GlnCTG_6_-_27515531_27515602 | 17 | 54 | 38 | GGTTAGCACTCTGGACTCTGAATCCAGCGATCCGAGTT | 1114.3 | 14.4 | 1.5 | 9.7 | 3.6E-22 | 3.2E-21 | BT20/MB468 | CYTO |
| >hg38_21_+_8433172_8446622_RNA45SN1_45S_50ntFlanks@11027.11062.36\|CTCTCTCTCTCTCTCCCCCGCTCCCCGTCCTCCCCC | >hg38_21_+_8433172_8446622_RNA45SN1_45S_50ntFlanks | 11027 | 11062 | 36 | CTCTCTCTCTCTCTCCCCCGCTCCCCGTCCTCCCCC | 425.7 | -13.2 | 1.4 | -9.6 | 7.3E-22 | 3.2E-21 | BT20/MB468 | NUC |
| >hg38_21_+_8433172_8446622_RNA45SN1_45S_50ntFlanks@10410.10435.26\|TCAGTCGGTCCTGAGAGATGGGCGAG | >hg38_21_+_8433172_8446622_RNA45SN1_45S_50ntFlanks | 10410 | 10435 | 26 | TCAGTCGGTCCTGAGAGATGGGCGAG | 1915.2 | -14.8 | 1.5 | -9.8 | 1.2E-22 | 3.3E-21 | MB231/MB468 | MP |
| >MI0000542\|hsa-mir-320a&WithFlank&8\|-\|22244960\|22245043@48.71.24(+1C)&[MIMAT0000510&hsa-miR-320a-3p&offsets\|0\|+2(+1C);m-60&8\|-\|22244975\|22244996&offsets\|0\|+2(+1C)]\|AAAAGCTGGGTTGAGAGGGCGAAAC | >MI0000542\|hsa-mir-320a&WithFlank&8\|-\|22244960\|22245043 | 48 | 71 | 24(+1C) | AAAAGCTGGGTTGAGAGGGCGAAAC | 413.6 | 12.9 | 1.3 | 9.7 | 3.6E-22 | 3.3E-21 | BT20/MB468 | CYTO |
| >hg38_21_+_8433172_8446622_RNA45SN1_45S_50ntFlanks@12444.12485.42\|AAGCAGAATTCACCAAGCGTTGGATTGTTCACCCACTAATAG | >hg38_21_+_8433172_8446622_RNA45SN1_45S_50ntFlanks | 12444 | 12485 | 42 | AAGCAGAATTCACCAAGCGTTGGATTGTTCACCCACTAATAG | 1438.4 | -14.9 | 1.6 | -9.6 | 7.3E-22 | 3.3E-21 | BT20/MB468 | NUC |
| >hg38_21_+_8433172_8446622_RNA45SN1_45S_50ntFlanks@11781.11806.26\|CGCATGAATGGATGAACGAGATTCCC | >hg38_21_+_8433172_8446622_RNA45SN1_45S_50ntFlanks | 11781 | 11806 | 26 | CGCATGAATGGATGAACGAGATTCCC | 1138.4 | -14.0 | 1.4 | -9.8 | 1.2E-22 | 3.3E-21 | MB231/MB468 | MP |
| >am_MI0015825\|hsa-mir-378c&WithFlank&10\|-\|130962582\|130962674@17.34.18;MI0016749\|hsa-mir-378d-1&WithFlank&4\|-\|5923269\|5923334@37.54.18;MI0000786\|hsa-mir-378a&WithFlank&5\|+\|149732819\|149732896@49.66.18;MI0003840\|hsa-mir-378d-2&WithFlank&8\|-\|93916016\|93916125@28.45.18&[MIMAT0016847&hsa-miR-378c&offsets\|0\|-7;m-130&10\|-\|130962639\|130962658&offsets\|0\|-2];[MIMAT0018926&hsa-miR-378d&offsets\|0\|-2;m-228&4\|-\|5923281\|5923298&offsets\|0\|0];[MIMAT0000732&hsa-miR-378a-3p&offsets\|0\|-4;m-44&5\|+\|149732867\|149732888&offsets\|0\|-4];[MIMAT0018926_1&hsa-miR-378d&offsets\|0\|-2;m-227&8\|-\|93916081\|93916098&offsets\|0\|0]\|ACTGGACTTGGAGTCAGA | >am_MI0015825\|hsa-mir-378c&WithFlank&10\|-\|130962582\|130962674 | 17 | 34 | 18 | ACTGGACTTGGAGTCAGA | 551.2 | 6.0 | 0.6 | 9.7 | 3.7E-22 | 3.3E-21 | BT20/MB468 | CYTO |
| >hg38_21_+_8433172_8446622_RNA45SN1_45S_50ntFlanks@10409.10429.21\|GTCAGTCGGTCCTGAGAGATG | >hg38_21_+_8433172_8446622_RNA45SN1_45S_50ntFlanks | 10409 | 10429 | 21 | GTCAGTCGGTCCTGAGAGATG | 4205.7 | -7.4 | 0.8 | -9.6 | 7.4E-22 | 3.3E-21 | BT20/MB468 | NUC |
| >hg38_21_+_8433172_8446622_RNA45SN1_45S_50ntFlanks@7974.8011.38\|ACGCGACCTCAGATCAGACGTGGCGACCCGCTGAATTT | >hg38_21_+_8433172_8446622_RNA45SN1_45S_50ntFlanks | 7974 | 8011 | 38 | ACGCGACCTCAGATCAGACGTGGCGACCCGCTGAATTT | 7835.0 | 6.8 | 0.7 | 9.7 | 3.7E-22 | 3.3E-21 | BT20/MB468 | CYTO |
| >MI0000439\|hsa-mir-23b&WithFlank&9\|+\|95085202\|95085310@67.86.20(+1A)&[MIMAT0000418&hsa-miR-23b-3p&offsets\|+3\|0(+1A);m-42&9\|+\|95085265\|95085285&offsets\|+3\|+2(+1A)]\|ACATTGCCAGGGATTACCACA | >MI0000439\|hsa-mir-23b&WithFlank&9\|+\|95085202\|95085310 | 67 | 86 | 20(+1A) | ACATTGCCAGGGATTACCACA | 661.9 | 13.8 | 1.4 | 9.7 | 2.7E-22 | 3.3E-21 | BT20/MB468 | MITO |
| >hg38_21_+_8433172_8446622_RNA45SN1_45S_50ntFlanks@9308.9338.31\|AGGAGTCTAACACGTGCGCGAGTCGGGGGCT | >hg38_21_+_8433172_8446622_RNA45SN1_45S_50ntFlanks | 9308 | 9338 | 31 | AGGAGTCTAACACGTGCGCGAGTCGGGGGCT | 1325.9 | -14.2 | 1.5 | -9.7 | 2.1E-22 | 3.4E-21 | MB231/MB468 | MITO |
| >hg38_21_+_8433172_8446622_RNA45SN1_45S_50ntFlanks@11781.11798.18\|CGCATGAATGGATGAACG | >hg38_21_+_8433172_8446622_RNA45SN1_45S_50ntFlanks | 11781 | 11798 | 18 | CGCATGAATGGATGAACG | 937.8 | -13.7 | 1.4 | -9.8 | 1.3E-22 | 3.4E-21 | MB231/MB468 | MP |
| >hg38_21_+_8433172_8446622_RNA45SN1_45S_50ntFlanks@10420.10455.36\|CTGAGAGATGGGCGAGCGCCGTTCCGAAGGGACGGG | >hg38_21_+_8433172_8446622_RNA45SN1_45S_50ntFlanks | 10420 | 10455 | 36 | CTGAGAGATGGGCGAGCGCCGTTCCGAAGGGACGGG | 922.0 | -14.3 | 1.5 | -9.6 | 7.5E-22 | 3.4E-21 | BT20/MB468 | NUC |
| >hg38_21_+_8433172_8446622_RNA45SN1_45S_50ntFlanks@10795.10818.24\|TGGCATGTTGGAACAATGTAGGTA | >hg38_21_+_8433172_8446622_RNA45SN1_45S_50ntFlanks | 10795 | 10818 | 24 | TGGCATGTTGGAACAATGTAGGTA | 725.7 | -13.4 | 1.4 | -9.8 | 1.3E-22 | 3.4E-21 | MB231/MB468 | MP |
| >hg38_21_+_8433172_8446622_RNA45SN1_45S_50ntFlanks@11323.11340.18\|CGTCTCGTCGCGCGCGCG | >hg38_21_+_8433172_8446622_RNA45SN1_45S_50ntFlanks | 11323 | 11340 | 18 | CGTCTCGTCGCGCGCGCG | 764.7 | -13.4 | 1.4 | -9.8 | 1.3E-22 | 3.4E-21 | MB231/MB468 | MP |
| >hg38_21_+_8433172_8446622_RNA45SN1_45S_50ntFlanks@11703.11728.26\|AATGAAGCGCGGGTAAACGGCGGGAG | >hg38_21_+_8433172_8446622_RNA45SN1_45S_50ntFlanks | 11703 | 11728 | 26 | AATGAAGCGCGGGTAAACGGCGGGAG | 3908.7 | -9.4 | 1.0 | -9.7 | 2.0E-22 | 3.4E-21 | MB231/MB468 | NUC |
| >hg38_21_+_8433172_8446622_RNA45SN1_45S_50ntFlanks@9574.9597.24\|AATCGGTCGTCCGACCTGGGTATA | >hg38_21_+_8433172_8446622_RNA45SN1_45S_50ntFlanks | 9574 | 9597 | 24 | AATCGGTCGTCCGACCTGGGTATA | 18116.5 | -7.7 | 0.8 | -9.7 | 2.1E-22 | 3.4E-21 | MB231/MB468 | MITO |
| >MI0000434\|hsa-let-7i&WithFlank&12\|+\|62603680\|62603775@12.33.22(+1U)&[MIMAT0000415&hsa-let-7i-5p&offsets\|0\|0(+1U);m-49&12\|+\|62603691\|62603712&offsets\|0\|0(+1U)]\|TGAGGTAGTAGTTTGTGCTGTTT | >MI0000434\|hsa-let-7i&WithFlank&12\|+\|62603680\|62603775 | 12 | 33 | 22(+1U) | TGAGGTAGTAGTTTGTGCTGTTT | 3138.5 | 7.4 | 0.8 | 9.7 | 2.8E-22 | 3.4E-21 | BT20/MB468 | MITO |
| >hg38_21_+_8433172_8446622_RNA45SN1_45S_50ntFlanks@6645.6665.21\|CTCGTACGACTCTTAGCGGTG | >hg38_21_+_8433172_8446622_RNA45SN1_45S_50ntFlanks | 6645 | 6665 | 21 | CTCGTACGACTCTTAGCGGTG | 38307.4 | 6.4 | 0.7 | 9.7 | 2.8E-22 | 3.4E-21 | BT20/MB468 | MITO |
| >hg38_21_+_8433172_8446622_RNA45SN1_45S_50ntFlanks@485.520.36\|GATGTGGTGACGTCGTGCTCTCCCGGGCCGGGTCCG | >hg38_21_+_8433172_8446622_RNA45SN1_45S_50ntFlanks | 485 | 520 | 36 | GATGTGGTGACGTCGTGCTCTCCCGGGCCGGGTCCG | 749.1 | 14.0 | 1.4 | 9.7 | 2.8E-22 | 3.4E-21 | BT20/MB468 | MITO |
| >hg38_21_+_8433172_8446622_RNA45SN1_45S_50ntFlanks@11876.11905.30\|AAGACCCTGTTGAGCTTGACTCTAGTCTGG | >hg38_21_+_8433172_8446622_RNA45SN1_45S_50ntFlanks | 11876 | 11905 | 30 | AAGACCCTGTTGAGCTTGACTCTAGTCTGG | 763.5 | -13.4 | 1.4 | -9.8 | 1.3E-22 | 3.5E-21 | MB231/MB468 | MP |

| | | | | | | | | | | | | | |
|---|---|---|---|---|---|---|---|---|---|---|---|---|---|
| >hg38_21_+_8433172_8446622_RNA45SN1_45S_50ntFlanks@8114.8142.29\|GCGCGGGACATGTGGCGTACGGAAGACCC | >hg38_21_+_8433172_8446622_RNA45SN1_45S_50ntFlanks | 8114 | 8142 | 29 | GCGCGGGACATGTGGCGTACGGAAGACCC | 1480.2 | -14.4 | 1.5 | -9.8 | 1.3E-22 | 3.5E-21 | MB231/MB468 | MP |
| >hg38_21_+_8433172_8446622_RNA45SN1_45S_50ntFlanks@11999.12030.32\|CCCTGCGGGCCGCCGGTGAAATACCACTACTC | >hg38_21_+_8433172_8446622_RNA45SN1_45S_50ntFlanks | 11999 | 12030 | 32 | CCCTGCGGGCCGCCGGTGAAATACCACTACTC | 1716.2 | -13.7 | 1.4 | -9.7 | 2.1E-22 | 3.5E-21 | MB231/MB468 | NUC |
| >hg38_21_+_8433172_8446622_RNA45SN1_45S_50ntFlanks@10807.10828.22\|ACAATGTAGGTAAGGGAAGTCG | >hg38_21_+_8433172_8446622_RNA45SN1_45S_50ntFlanks | 10807 | 10828 | 22 | ACAATGTAGGTAAGGGAAGTCG | 460.8 | -13.3 | 1.4 | -9.6 | 7.9E-22 | 3.5E-21 | BT20/MB468 | NUC |
| >hg38_21_+_8433172_8446622_RNA45SN1_45S_50ntFlanks@7986.8004.19\|ATCAGACGTGGCGACCCGC | >hg38_21_+_8433172_8446622_RNA45SN1_45S_50ntFlanks | 7986 | 8004 | 19 | ATCAGACGTGGCGACCCGC | 1423.2 | -14.3 | 1.5 | -9.7 | 2.2E-22 | 3.5E-21 | MB231/MB468 | MITO |
| >hg38_21_+_8433172_8446622_RNA45SN1_45S_50ntFlanks@9166.9197.32\|CCGGGCGGGTCGCGCCGTCGGGCCCGGGGGAG | >hg38_21_+_8433172_8446622_RNA45SN1_45S_50ntFlanks | 9166 | 9197 | 32 | CCGGGCGGGTCGCGCCGTCGGGCCCGGGGGAG | 5373.2 | -16.2 | 1.7 | -9.7 | 2.2E-22 | 3.5E-21 | MB231/MB468 | MITO |
| >hg38_21_+_8433172_8446622_RNA45SN1_45S_50ntFlanks@3930.3959.30\|ATCAAAACCAACCCGGTCAGCCCCTCTCCG | >hg38_21_+_8433172_8446622_RNA45SN1_45S_50ntFlanks | 3930 | 3959 | 30 | ATCAAAACCAACCCGGTCAGCCCCTCTCCG | 1109.4 | -13.9 | 1.4 | -9.7 | 2.2E-22 | 3.6E-21 | MB231/MB468 | MITO |
| >hg38_21_+_8433172_8446622_RNA45SN1_45S_50ntFlanks@10512.10534.23\|TCCGGAGTGGCGGAGATGGGCGC | >hg38_21_+_8433172_8446622_RNA45SN1_45S_50ntFlanks | 10512 | 10534 | 23 | TCCGGAGTGGCGGAGATGGGCGC | 610.1 | -13.7 | 1.4 | -9.6 | 8.0E-22 | 3.6E-21 | BT20/MB468 | NUC |
| >hg38_21_+_8433172_8446622_RNA45SN1_45S_50ntFlanks@12003.12035.33\|GCGGGCCGCCGGTGAAATACCACTACTCTGATC | >hg38_21_+_8433172_8446622_RNA45SN1_45S_50ntFlanks | 12003 | 12035 | 33 | GCGGGCCGCCGGTGAAATACCACTACTCTGATC | 1575.8 | 5.4 | 0.6 | 9.7 | 3.0E-22 | 3.6E-21 | BT20/MB468 | MITO |
| >hg38_21_+_8433172_8446622_RNA45SN1_45S_50ntFlanks@10771.10797.27\|CAGGTCTCCAAGGTGAACAGCCTCTGG | >hg38_21_+_8433172_8446622_RNA45SN1_45S_50ntFlanks | 10771 | 10797 | 27 | CAGGTCTCCAAGGTGAACAGCCTCTGG | 926.7 | -13.7 | 1.4 | -9.8 | 1.4E-22 | 3.6E-21 | MB231/MB468 | MP |
| >hg38_21_+_8433172_8446622_RNA45SN1_45S_50ntFlanks@12615.12653.39\|GCTGAGGAGCCAATGGGGCGAAGCTACCATCTGTGGGAT | >hg38_21_+_8433172_8446622_RNA45SN1_45S_50ntFlanks | 12615 | 12653 | 39 | GCTGAGGAGCCAATGGGGCGAAGCTACCATCTGTGGGAT | 5130.1 | -11.9 | 1.2 | -9.7 | 2.3E-22 | 3.6E-21 | MB231/MB468 | MITO |
| >hg38_21_+_8433172_8446622_RNA45SN1_45S_50ntFlanks@10362.10379.18\|AAGGCCGAAGTGGAGAAG | >hg38_21_+_8433172_8446622_RNA45SN1_45S_50ntFlanks | 10362 | 10379 | 18 | AAGGCCGAAGTGGAGAAG | 474.0 | -13.3 | 1.4 | -9.6 | 8.2E-22 | 3.6E-21 | BT20/MB468 | NUC |
| >MI0000061\|hsa-let-7a-2&WithFlank&11\|-\|122146516\|122146599@11.31.21(+1C);MI0000062\|hsa-let-7a-3&WithFlank&22\|+\|46112743\|46112828@10.30.21(+1C);MI0000060\|hsa-let-7a-1&WithFlank&9\|+\|94175951\|94176042@12.32.21(+1C)&[MIMAT0000062&hsa-let-7a-5p&offsets\|0\|-1(+1C);m-6&11\|-\|122146568\|122146589&offsets\|0\|-1(+1C)];[MIMAT0000062_1&hsa-let-7a-5p&offsets\|0\|-1(+1C);m-5&22\|+\|46112752\|46112773&offsets\|0\|-1(+1C)];[MIMAT0000062_2&hsa-let-7a-5p&offsets\|0\|-1(+1C);m-4&9\|+\|94175962\|94175983&offsets\|0\|-1(+1C)]\|TGAGGTAGTAGGTTGTATAGTC | >MI0000061\|hsa-let-7a-2&WithFlank&11\|-\|122146516\|122146599 | 11 | 31 | 21(+1C) | TGAGGTAGTAGGTTGTATAGTC | 11934.5 | 5.2 | 0.5 | 9.7 | 2.2E-22 | 3.7E-21 | MB231/MB468 | NUC |
| >hg38_21_+_8433172_8446622_RNA45SN1_45S_50ntFlanks@9504.9533.30\|GTGAACTATGCCTGGGCAGGGCGAAGCCAG | >hg38_21_+_8433172_8446622_RNA45SN1_45S_50ntFlanks | 9504 | 9533 | 30 | GTGAACTATGCCTGGGCAGGGCGAAGCCAG | 2032.7 | -14.9 | 1.5 | -9.8 | 1.4E-22 | 3.7E-21 | MB231/MB468 | MP |
| >trnaMT_GluTTC_MT_-_14674_14742@39.71.33,trnalookalike8_GluTTC_5_-_93905172_93905240@39.71.33\|ATTGGTCGTGGTTGTAGTCCGTGCGAGAATACC | >trnaMT_GluTTC_MT_-_14674_14742 | 39 | 71 | 33 | ATTGGTCGTGGTTGTAGTCCGTGCGAGAATACC | 3503.8 | 7.4 | 0.8 | 9.7 | 3.1E-22 | 3.8E-21 | BT20/MB468 | MITO |
| >am_MI0000809\|hsa-mir-151a&WithFlank&8\|-\|140732558\|140732659@53.72.20&[MIMAT0000757&hsa-miR-151a-3p&offsets\|0\|-1;m-30&8\|-\|140732587\|140732607&offsets\|0\|-1]\|CTAGACTGAAGCTCCTTGAG | >am_MI0000809\|hsa-mir-151a&WithFlank&8\|-\|140732558\|140732659 | 53 | 72 | 20 | CTAGACTGAAGCTCCTTGAG | 1707.3 | 6.2 | 0.6 | 9.7 | 2.2E-22 | 3.8E-21 | MB231/MB468 | NUC |
| >hg38_21_+_8433172_8446622_RNA45SN1_45S_50ntFlanks@12418.12441.24\|CGATGTCGGCTCTTCCTATCATTG | >hg38_21_+_8433172_8446622_RNA45SN1_45S_50ntFlanks | 12418 | 12441 | 24 | CGATGTCGGCTCTTCCTATCATTG | 3052.9 | -9.8 | 1.0 | -9.6 | 8.5E-22 | 3.8E-21 | BT20/MB468 | NUC |
| >MI0003628\|hsa-mir-615&WithFlank&12\|+\|54033944\|54034051@67.86.20(+1A)&[MIMAT0003283&hsa-miR-615-3p&offsets\|0\|-2(+1A);m-495&12\|+\|54034010\|54034030&offsets\|0\|-1(+1A)]\|TCCGAGCCTGGGTCTCCCTCA | >MI0003628\|hsa-mir-615&WithFlank&12\|+\|54033944\|54034051 | 67 | 86 | 20(+1A) | TCCGAGCCTGGGTCTCCCTCA | 527.2 | 7.8 | 0.8 | 9.7 | 4.2E-22 | 3.8E-21 | BT20/MB468 | CYTO |
| >trna116_GluCTC_1_-_145399233_145399304@1.35.35,trna59_GluCTC_1_+_249168447_249168518@1.35.35,trna71_GluCTC_1_-_161439189_161439260@1.35.35,trna74_GluCTC_1_-_161431809_161431880@1.35.35,trna77_GluCTC_1_-_161424398_161424469@1.35.35,trna77_GluCTC_6_+_28949976_28950047@1.35.35,trna80_GluCTC_1_-_161417018_161417089@1.35.35,trna87_GluCTC_6_-_126101393_126101464@1.35.35\|TCCCTGGTGGTCTAGTGGTTAGGATTCGGCGCTCT | >trna116_GluCTC_1_-_145399233_145399304 | 1 | 35 | 35 | TCCCTGGTGGTCTAGTGGTTAGGATTCGGCGCTCT | 11491.4 | 8.3 | 0.9 | 9.7 | 3.2E-22 | 3.8E-21 | BT20/MB468 | MITO |
| >hg38_21_+_8433172_8446622_RNA45SN1_45S_50ntFlanks@4724.4752.29\|ATTAATCAAGAACGAAAGTCGGAGGTTCG | >hg38_21_+_8433172_8446622_RNA45SN1_45S_50ntFlanks | 4724 | 4752 | 29 | ATTAATCAAGAACGAAAGTCGGAGGTTCG | 2238.2 | -14.9 | 1.5 | -9.7 | 2.4E-22 | 3.8E-21 | MB231/MB468 | MITO |
| >hg38_21_+_8433172_8446622_RNA45SN1_45S_50ntFlanks@11768.11792.25\|CTAATTAGTGACGCGCATGAATGGA | >hg38_21_+_8433172_8446622_RNA45SN1_45S_50ntFlanks | 11768 | 11792 | 25 | CTAATTAGTGACGCGCATGAATGGA | 1244.0 | -14.1 | 1.4 | -9.8 | 1.5E-22 | 3.9E-21 | MB231/MB468 | MP |
| >hg38_21_+_8433172_8446622_RNA45SN1_45S_50ntFlanks@11577.11603.27\|TTAGAACTGGTGCGGACCAGGGGAATC | >hg38_21_+_8433172_8446622_RNA45SN1_45S_50ntFlanks | 11577 | 11603 | 27 | TTAGAACTGGTGCGGACCAGGGGAATC | 2122.4 | -5.6 | 0.6 | -9.8 | 1.1E-22 | 3.9E-21 | MB231/MB468 | CYTO |
| >MI0000063\|hsa-let-7b&WithFlank&22\|+\|46113680\|46113774@12.29.18&[MIMAT0000063&hsa-let-7b-5p&offsets\|0\|-4;m-14&22\|+\|46113691\|46113712&offsets\|0\|-4]\|TGAGGTAGTAGGTTGTGT | >MI0000063\|hsa-let-7b&WithFlank&22\|+\|46113680\|46113774 | 12 | 29 | 18 | TGAGGTAGTAGGTTGTGT | 715.4 | 10.1 | 1.0 | 9.7 | 4.3E-22 | 3.9E-21 | BT20/MB468 | CYTO |
| >hg38_21_+_8433172_8446622_RNA45SN1_45S_50ntFlanks@11208.11231.24\|GCGGCGACTCTGGACGCGAGCCGG | >hg38_21_+_8433172_8446622_RNA45SN1_45S_50ntFlanks | 11208 | 11231 | 24 | GCGGCGACTCTGGACGCGAGCCGG | 1171.9 | -14.1 | 1.4 | -9.8 | 1.5E-22 | 3.9E-21 | MB231/MB468 | MP |
| >hg38_21_+_8433172_8446622_RNA45SN1_45S_50ntFlanks@11323.11343.21\|CGTCTCGTCGCGCGCGCGTCC | >hg38_21_+_8433172_8446622_RNA45SN1_45S_50ntFlanks | 11323 | 11343 | 21 | CGTCTCGTCGCGCGCGCGTCC | 1221.5 | -14.1 | 1.4 | -9.8 | 1.5E-22 | 3.9E-21 | MB231/MB468 | MP |
| >hg38_21_+_8433172_8446622_RNA45SN1_45S_50ntFlanks@13004.13027.24\|ATCTATTGAAAGTCAGCCCTCGAC | >hg38_21_+_8433172_8446622_RNA45SN1_45S_50ntFlanks | 13004 | 13027 | 24 | ATCTATTGAAAGTCAGCCCTCGAC | 2432.0 | -14.2 | 1.5 | -9.7 | 2.3E-22 | 3.9E-21 | MB231/MB468 | NUC |
| >hg38_21_+_8433172_8446622_RNA45SN1_45S_50ntFlanks@4730.4758.29\|CAAGAACGAAAGTCGGAGGTTCGAAGACG | >hg38_21_+_8433172_8446622_RNA45SN1_45S_50ntFlanks | 4730 | 4758 | 29 | CAAGAACGAAAGTCGGAGGTTCGAAGACG | 3022.3 | -5.0 | 0.5 | -9.7 | 2.3E-22 | 3.9E-21 | MB231/MB468 | NUC |
| >hg38_21_+_8433172_8446622_RNA45SN1_45S_50ntFlanks@12654.12675.22\|TATGACTGAACGCCTCTAAGTC | >hg38_21_+_8433172_8446622_RNA45SN1_45S_50ntFlanks | 12654 | 12675 | 22 | TATGACTGAACGCCTCTAAGTC | 3526.4 | -9.5 | 1.0 | -9.7 | 2.4E-22 | 3.9E-21 | MB231/MB468 | MITO |
| >hg38_21_+_8433172_8446622_RNA45SN1_45S_50ntFlanks@4473.4493.21\|TTAGCTGAGTGTCCCGCGGGG | >hg38_21_+_8433172_8446622_RNA45SN1_45S_50ntFlanks | 4473 | 4493 | 21 | TTAGCTGAGTGTCCCGCGGGG | 321.5 | -12.8 | 1.3 | -9.6 | 8.8E-22 | 3.9E-21 | BT20/MB468 | NUC |
| >hg38_21_+_8433172_8446622_RNA45SN1_45S_50ntFlanks@12456.12473.18\|CCAAGCGTTGGATTGTTC | >hg38_21_+_8433172_8446622_RNA45SN1_45S_50ntFlanks | 12456 | 12473 | 18 | CCAAGCGTTGGATTGTTC | 31758.8 | -9.4 | 0.9 | -10.2 | 3.2E-24 | 3.9E-21 | BT20/MB468 | MP |
| >hg38_21_+_8433172_8446622_RNA45SN1_45S_50ntFlanks@4005.4026.22\|AACCTCGGGCCGATCGCACGCC | >hg38_21_+_8433172_8446622_RNA45SN1_45S_50ntFlanks | 4005 | 4026 | 22 | AACCTCGGGCCGATCGCACGCC | 774.4 | -14.0 | 1.5 | -9.6 | 8.8E-22 | 3.9E-21 | BT20/MB468 | NUC |
| >hg38_21_+_8433172_8446622_RNA45SN1_45S_50ntFlanks@3822.3853.32\|CTTTGGTCGCTCGCTCCTCTCCTACTTGGATA | >hg38_21_+_8433172_8446622_RNA45SN1_45S_50ntFlanks | 3822 | 3853 | 32 | CTTTGGTCGCTCGCTCCTCTCCTACTTGGATA | 429.9 | -13.2 | 1.4 | -9.6 | 8.9E-22 | 3.9E-21 | BT20/MB468 | NUC |
| >hg38_21_+_8433172_8446622_RNA45SN1_45S_50ntFlanks@4724.4759.36\|ATTAATCAAGAACGAAAGTCGGAGGTTCGAAGACGA | >hg38_21_+_8433172_8446622_RNA45SN1_45S_50ntFlanks | 4724 | 4759 | 36 | ATTAATCAAGAACGAAAGTCGGAGGTTCGAAGACGA | 9998.5 | -11.1 | 1.2 | -9.6 | 9.0E-22 | 4.0E-21 | BT20/MB468 | NUC |
| >trna116_GluCTC_1_-_145399233_145399304@3.36.34,trna59_GluCTC_1_+_249168447_249168518@3.36.34,trna71_GluCTC_1_-_161439189_161439260@3.36.34,trna74_GluCTC_1_-_161431809_161431880@3.36.34,trna77_GluCTC_1_-_161424398_161424469@3.36.34,trna77_GluCTC_6_+_28949976_28950047@3.36.34,trna80_GluCTC_1_-_161417018_161417089@3.36.34,trna87_GluCTC_6_-_126101393_126101464@3.36.34\|CCTGGTGGTCTAGTGGTTAGGATTCGGCGCTCTC | >trna116_GluCTC_1_-_145399233_145399304 | 3 | 36 | 34 | CCTGGTGGTCTAGTGGTTAGGATTCGGCGCTCTC | 266.8 | 12.1 | 1.2 | 9.8 | 1.2E-22 | 4.0E-21 | MB231/MB468 | CYTO |
| >hg38_21_+_8433172_8446622_RNA45SN1_45S_50ntFlanks@10678.10695.18\|CGGTTCCGGCGGCGTCCG | >hg38_21_+_8433172_8446622_RNA45SN1_45S_50ntFlanks | 10678 | 10695 | 18 | CGGTTCCGGCGGCGTCCG | 296.7 | -12.6 | 1.3 | -9.6 | 9.0E-22 | 4.0E-21 | BT20/MB468 | NUC |
| >hg38_21_+_8433172_8446622_RNA45SN1_45S_50ntFlanks@12632.12654.23\|GCGAAGCTACCATCTGTGGGATT | >hg38_21_+_8433172_8446622_RNA45SN1_45S_50ntFlanks | 12632 | 12654 | 23 | GCGAAGCTACCATCTGTGGGATT | 4600.2 | -11.7 | 1.2 | -9.8 | 1.6E-22 | 4.0E-21 | MB231/MB468 | MP |
| >hg38_21_+_8433172_8446622_RNA45SN1_45S_50ntFlanks@4367.4394.28\|CAGTTAAAAAGCTCGTAGTTGGATCTTG | >hg38_21_+_8433172_8446622_RNA45SN1_45S_50ntFlanks | 4367 | 4394 | 28 | CAGTTAAAAAGCTCGTAGTTGGATCTTG | 504.8 | -13.4 | 1.4 | -9.6 | 9.1E-22 | 4.0E-21 | BT20/MB468 | NUC |
| >MI0000264\|hsa-mir-7-2&WithFlank&15\|+\|88611819\|88611940@38.60.23(+1G);MI0000265\|hsa-mir-7-3&WithFlank&19\|+\|4770664\|4770785@37.59.23(+1G);MI0000263\|hsa-mir-7-1&WithFlank&9\|-\|83969742\|83969863@30.52.23(+1G)&[MIMAT0000252&hsa-miR-7-5p&offsets\|0\|-1(+1G);m-400&15\|+\|88611856\|88611879&offsets\|0\|-1(+1G)];[MIMAT0000252_1&hsa-miR-7-5p&offsets\|0\|-1(+1G);m-402&19\|+\|4770700\|4770723&offsets\|0\|-1(+1G)];[MIMAT0000252_2&hsa-miR-7-5p&offsets\|0\|-1(+1G);m-401&9\|-\|83969811\|83969834&offsets\|0\|-1(+1G)]\|TGGAAGACTAGTGATTTTGTTGTG | >MI0000264\|hsa-mir-7-2&WithFlank&15\|+\|88611819\|88611940 | 38 | 60 | 23(+1G) | TGGAAGACTAGTGATTTTGTTGTG | 527.4 | 5.5 | 0.6 | 9.7 | 2.4E-22 | 4.0E-21 | MB231/MB468 | NUC |
| >hg38_21_+_8433172_8446622_RNA45SN1_45S_50ntFlanks@8552.8569.18\|CTCCCCGGAGGGGGCGGG | >hg38_21_+_8433172_8446622_RNA45SN1_45S_50ntFlanks | 8552 | 8569 | 18 | CTCCCCGGAGGGGGCGGG | 2713.6 | -15.3 | 1.6 | -9.8 | 1.6E-22 | 4.1E-21 | MB231/MB468 | MP |
| >am_trna111_HisGTG_1_-_147774845_147774916@-1G.28.29,trna118_HisGTG_1_-_145396881_145396952@-1G.28.29,trna16_HisGTG_1_+_146544773_146544844@-1G.28.29,trna1_HisGTG_15_+_45493349_45493420@-1G.28.29,trna21_HisGTG_1_+_147753471_147753542@-1G.28.29,trna33_HisGTG_6_+_27125906_27125977@-1G.28.29,trna7_HisGTG_9_-_14433938_14434009@-1G.28.29,trna8_HisGTG_15_-_45492611_45492682@-1G.28.29,trna9_HisGTG_15_-_45490804_45490875@-1G.28.29\|GGCCGTGATCGTATAGTGGTTAGTACTCT | >am_trna111_HisGTG_1_-_147774845_147774916 | -1G | 28 | 29 | GGCCGTGATCGTATAGTGGTTAGTACTCT | 4530.5 | -7.2 | 0.7 | -9.6 | 9.2E-22 | 4.1E-21 | BT20/MB468 | NUC |

| | | | | | | | | | | | | | |
|---|---|---|---|---|---|---|---|---|---|---|---|---|---|
| >MI0003591\|hsa-mir-584&WithFlank&5\|-\|149062307\|149062415@22.42.21(+1A)&[MIMAT0003249&hsa-miR-584-5p&offsets\|0\|-1(+1A);m-167&5\|-\|149062374\|149062394&offsets\|0\|0(+1A)]\|TTATGGTTTGCCTGGGACTGAA | >MI0003591\|hsa-mir-584&WithFlank&5\|-\|149062307\|149062415 | 22 | 42 | 21(+1A) | TTATGGTTTGCCTGGGACTGAA | 62.6 | 3.8 | 0.4 | 9.6 | 9.2E-22 | 4.1E-21 | BT20/MB468 | NUC |
| >hg38_21_+_8433172_8446622_RNA45SN1_45S_50ntFlanks@4051.4076.26\|GAACGTCTGCCCTATCAACTTTCGAT | >hg38_21_+_8433172_8446622_RNA45SN1_45S_50ntFlanks | 4051 | 4076 | 26 | GAACGTCTGCCCTATCAACTTTCGAT | 200.8 | 11.7 | 1.2 | 9.8 | 1.2E-22 | 4.1E-21 | MB231/MB468 | CYTO |
| >hg38_21_+_8433172_8446622_RNA45SN1_45S_50ntFlanks@11564.11601.38\|CCTAGCAGCCGACTTAGAACTGGTGCGGACCAGGGGAA | >hg38_21_+_8433172_8446622_RNA45SN1_45S_50ntFlanks | 11564 | 11601 | 38 | CCTAGCAGCCGACTTAGAACTGGTGCGGACCAGGGGAA | 10736.3 | -10.2 | 1.0 | -9.7 | 2.5E-22 | 4.1E-21 | MB231/MB468 | MITO |
| >MI0000266\|hsa-mir-10a&WithFlank&17\|-\|48579832\|48579953@28.48.21(+1C)&[MIMAT0000253&hsa-miR-10a-5p&offsets\|0\|-2(+1C);m-18&17\|-\|48579905\|48579926&offsets\|0\|-1(+1C)]\|TACCCTGTAGATCCGAATTTGC | >MI0000266\|hsa-mir-10a&WithFlank&17\|-\|48579832\|48579953 | 28 | 48 | 21(+1C) | TACCCTGTAGATCCGAATTTGC | 972.0 | 13.9 | 1.4 | 9.8 | 1.2E-22 | 4.1E-21 | MB231/MB468 | CYTO |
| >hg38_1_-_228634819_228635039_RNA5S12_5S_50ntFlanks@137.171.35\|GAGACCGCCTGGGAATACCGGGTGCTGTAGGCTTT | >hg38_1_-_228634819_228635039_RNA5S12_5S_50ntFlanks | 137 | 171 | 35 | GAGACCGCCTGGGAATACCGGGTGCTGTAGGCTTT | 2537.2 | 7.8 | 0.8 | 9.8 | 1.2E-22 | 4.1E-21 | MB231/MB468 | CYTO |
| >hg38_21_+_8433172_8446622_RNA45SN1_45S_50ntFlanks@12637.12657.21\|GCTACCATCTGTGGGATTATG | >hg38_21_+_8433172_8446622_RNA45SN1_45S_50ntFlanks | 12637 | 12657 | 21 | GCTACCATCTGTGGGATTATG | 763.3 | -13.4 | 1.4 | -9.8 | 1.6E-22 | 4.1E-21 | MB231/MB468 | MP |
| >MI0000095\|hsa-mir-93&WithFlank&7\|-\|100093762\|100093853@17.39.23&[MIMAT0000093&hsa-miR-93-5p&offsets\|0\|0;m-29&7\|-\|100093815\|100093837&offsets\|0\|0]\|CAAAGTGCTGTTCGTGCAGGTAG | >MI0000095\|hsa-mir-93&WithFlank&7\|-\|100093762\|100093853 | 17 | 39 | 23 | CAAAGTGCTGTTCGTGCAGGTAG | 15354.3 | 6.3 | 0.7 | 9.7 | 4.6E-22 | 4.1E-21 | BT20/MB468 | CYTO |
| >hg38_21_+_8433172_8446622_RNA45SN1_45S_50ntFlanks@10866.10892.27\|CTAAGGGCTGGGTCGGTCGGGCTGGGG | >hg38_21_+_8433172_8446622_RNA45SN1_45S_50ntFlanks | 10866 | 10892 | 27 | CTAAGGGCTGGGTCGGTCGGGCTGGGG | 1583.2 | -14.5 | 1.5 | -9.8 | 1.6E-22 | 4.2E-21 | MB231/MB468 | MP |
| >hg38_21_+_8433172_8446622_RNA45SN1_45S_50ntFlanks@3774.3792.19\|GGCCGGTACAGTGAAACTG | >hg38_21_+_8433172_8446622_RNA45SN1_45S_50ntFlanks | 3774 | 3792 | 19 | GGCCGGTACAGTGAAACTG | 356.2 | -12.9 | 1.3 | -9.6 | 9.4E-22 | 4.2E-21 | BT20/MB468 | NUC |
| >hg38_21_+_8433172_8446622_RNA45SN1_45S_50ntFlanks@12288.12323.36\|ACCGTGAAAGCGGGGCCTCACGATCCTTCTGACCTT | >hg38_21_+_8433172_8446622_RNA45SN1_45S_50ntFlanks | 12288 | 12323 | 36 | ACCGTGAAAGCGGGGCCTCACGATCCTTCTGACCTT | 614.0 | -13.7 | 1.4 | -9.6 | 9.4E-22 | 4.2E-21 | BT20/MB468 | NUC |
| >hg38_21_+_8433172_8446622_RNA45SN1_45S_50ntFlanks@4443.4471.29\|CCCTTGCCTCTCGGCGCCCCCTCGATGCT | >hg38_21_+_8433172_8446622_RNA45SN1_45S_50ntFlanks | 4443 | 4471 | 29 | CCCTTGCCTCTCGGCGCCCCCTCGATGCT | 1583.7 | -12.7 | 1.3 | -9.7 | 3.5E-22 | 4.2E-21 | BT20/MB468 | MITO |
| >hg38_21_+_8433172_8446622_RNA45SN1_45S_50ntFlanks@11575.11605.31\|ACTTAGAACTGGTGCGGACCAGGGGAATCCG | >hg38_21_+_8433172_8446622_RNA45SN1_45S_50ntFlanks | 11575 | 11605 | 31 | ACTTAGAACTGGTGCGGACCAGGGGAATCCG | 1501.7 | -14.4 | 1.5 | -9.8 | 1.6E-22 | 4.2E-21 | MB231/MB468 | MP |
| >hg38_21_+_8433172_8446622_RNA45SN1_45S_50ntFlanks@11572.11600.29\|CCGACTTAGAACTGGTGCGGACCAGGGGA | >hg38_21_+_8433172_8446622_RNA45SN1_45S_50ntFlanks | 11572 | 11600 | 29 | CCGACTTAGAACTGGTGCGGACCAGGGGA | 2778.4 | -6.4 | 0.7 | -9.7 | 2.6E-22 | 4.2E-21 | MB231/MB468 | MITO |
| >hg38_21_+_8433172_8446622_RNA45SN1_45S_50ntFlanks@4445.4471.27\|CTTGCCTCTCGGCGCCCCCTCGATGCT | >hg38_21_+_8433172_8446622_RNA45SN1_45S_50ntFlanks | 4445 | 4471 | 27 | CTTGCCTCTCGGCGCCCCCTCGATGCT | 1163.9 | -14.0 | 1.4 | -9.7 | 2.7E-22 | 4.2E-21 | MB231/MB468 | MITO |
| >hg38_21_+_8433172_8446622_RNA45SN1_45S_50ntFlanks@10394.10424.31\|AGCAGTTGAACATGGGTCAGTCGGTCCTGAG | >hg38_21_+_8433172_8446622_RNA45SN1_45S_50ntFlanks | 10394 | 10424 | 31 | AGCAGTTGAACATGGGTCAGTCGGTCCTGAG | 12871.8 | 6.9 | 0.7 | 9.7 | 3.5E-22 | 4.2E-21 | BT20/MB468 | MITO |
| >hg38_21_+_8433172_8446622_RNA45SN1_45S_50ntFlanks@12456.12475.20\|CCAAGCGTTGGATTGTTCAC | >hg38_21_+_8433172_8446622_RNA45SN1_45S_50ntFlanks | 12456 | 12475 | 20 | CCAAGCGTTGGATTGTTCAC | 8867.9 | -12.1 | 1.2 | -9.7 | 2.7E-22 | 4.2E-21 | MB231/MB468 | MITO |
| >hg38_21_+_8433172_8446622_RNA45SN1_45S_50ntFlanks@12453.12496.44\|TCACCAAGCGTTGGATTGTTCACCCACTAATAGGGAACGTGAGC | >hg38_21_+_8433172_8446622_RNA45SN1_45S_50ntFlanks | 12453 | 12496 | 44 | TCACCAAGCGTTGGATTGTTCACCCACTAATAGGGAACGTGAGC | 248.2 | 12.0 | 1.2 | 9.8 | 1.3E-22 | 4.3E-21 | MB231/MB468 | CYTO |
| >hg38_21_+_8433172_8446622_RNA45SN1_45S_50ntFlanks@11161.11179.19\|CACGGTCCCCCGCGAGGGG | >hg38_21_+_8433172_8446622_RNA45SN1_45S_50ntFlanks | 11161 | 11179 | 19 | CACGGTCCCCCGCGAGGGG | 1591.4 | -13.4 | 1.4 | -9.8 | 1.3E-22 | 4.3E-21 | MB231/MB468 | CYTO |
| >trnaMT_LeuTAA_MT_+_3230_3304@16.45.30\|CCCGGTAATCGCATAAAACTTAAAACTTTA | >trnaMT_LeuTAA_MT_+_3230_3304 | 16 | 45 | 30 | CCCGGTAATCGCATAAAACTTAAAACTTTA | 597.8 | -13.6 | 1.4 | -9.6 | 9.7E-22 | 4.3E-21 | BT20/MB468 | NUC |
| >MI0000298\|hsa-mir-221&WithFlank&X\|-\|45746151\|45746272@31.56.26&[MIMAT0004568&hsa-miR-221-5p&offsets\|0\|+4;m-405&X\|-\|45746220\|45746242&offsets\|0\|+3]\|ACCTGGCATACAATGTAGATTTCTGT | >MI0000298\|hsa-mir-221&WithFlank&X\|-\|45746151\|45746272 | 31 | 56 | 26 | ACCTGGCATACAATGTAGATTTCTGT | 1917.6 | 5.7 | 0.6 | 9.7 | 2.6E-22 | 4.3E-21 | MB231/MB468 | NUC |
| >hg38_21_+_8433172_8446622_RNA45SN1_45S_50ntFlanks@12633.12657.25\|CGAAGCTACCATCTGTGGGATTATG | >hg38_21_+_8433172_8446622_RNA45SN1_45S_50ntFlanks | 12633 | 12657 | 25 | CGAAGCTACCATCTGTGGGATTATG | 2918.8 | -9.5 | 1.0 | -9.7 | 2.8E-22 | 4.4E-21 | MB231/MB468 | MITO |
| >hg38_21_+_8433172_8446622_RNA45SN1_45S_50ntFlanks@7986.8006.21\|ATCAGACGTGGCGACCCGCTG | >hg38_21_+_8433172_8446622_RNA45SN1_45S_50ntFlanks | 7986 | 8006 | 21 | ATCAGACGTGGCGACCCGCTG | 1024.6 | -13.0 | 1.3 | -9.7 | 2.6E-22 | 4.4E-21 | MB231/MB468 | NUC |
| >MI0000439\|hsa-mir-23b&WithFlank&9\|+\|95085202\|95085310@67.86.20(+1C)&[MIMAT0000418&hsa-miR-23b-3p&offsets\|+3\|0(+1C);m-42&9\|+\|95085265\|95085285&offsets\|+3\|+2(+1C)]\|ACATTGCCAGGGATTACCACC | >MI0000439\|hsa-mir-23b&WithFlank&9\|+\|95085202\|95085310 | 67 | 86 | 20(+1C) | ACATTGCCAGGGATTACCACC | 208.0 | 10.5 | 1.1 | 9.6 | 9.9E-22 | 4.4E-21 | BT20/MB468 | NUC |
| >hg38_21_+_8433172_8446622_RNA45SN1_45S_50ntFlanks@5491.5511.21\|GAGAAGACGGTCGAACTTGAC | >hg38_21_+_8433172_8446622_RNA45SN1_45S_50ntFlanks | 5491 | 5511 | 21 | GAGAAGACGGTCGAACTTGAC | 238.5 | 11.9 | 1.2 | 9.8 | 1.3E-22 | 4.4E-21 | MB231/MB468 | CYTO |
| >hg38_21_+_8433172_8446622_RNA45SN1_45S_50ntFlanks@12892.12933.42\|CCCTCGCCCGTCACGCACCGCACGTTCGTGGGGAACCTGGCG | >hg38_21_+_8433172_8446622_RNA45SN1_45S_50ntFlanks | 12892 | 12933 | 42 | CCCTCGCCCGTCACGCACCGCACGTTCGTGGGGAACCTGGCG | 462.4 | -13.3 | 1.4 | -9.6 | 9.9E-22 | 4.4E-21 | BT20/MB468 | NUC |
| >MI0000465\|hsa-mir-191&WithFlank&3\|-\|49020612\|49020715@22.42.21&[MIMAT0000440&hsa-miR-191-5p&offsets\|0\|-2;m-48&3\|-\|49020672\|49020694&offsets\|0\|-2]\|CAACGGAATCCCAAAAGCAGC | >MI0000465\|hsa-mir-191&WithFlank&3\|-\|49020612\|49020715 | 22 | 42 | 21 | CAACGGAATCCCAAAAGCAGC | 654.2 | 7.6 | 0.8 | 9.7 | 3.7E-22 | 4.4E-21 | BT20/MB468 | MITO |
| >hg38_21_+_8433172_8446622_RNA45SN1_45S_50ntFlanks@10679.10705.27\|GGTTCCGGCGGCGTCCGGTGAGCTCTC | >hg38_21_+_8433172_8446622_RNA45SN1_45S_50ntFlanks | 10679 | 10705 | 27 | GGTTCCGGCGGCGTCCGGTGAGCTCTC | 863.6 | -13.6 | 1.4 | -9.8 | 1.7E-22 | 4.4E-21 | MB231/MB468 | MP |
| >hg38_21_+_8433172_8446622_RNA45SN1_45S_50ntFlanks@485.505.21\|GATGTGGTGACGTCGTGCTCT | >hg38_21_+_8433172_8446622_RNA45SN1_45S_50ntFlanks | 485 | 505 | 21 | GATGTGGTGACGTCGTGCTCT | 4457.1 | 12.4 | 1.3 | 9.6 | 1.0E-21 | 4.4E-21 | BT20/MB468 | NUC |
| >hg38_21_+_8433172_8446622_RNA45SN1_45S_50ntFlanks@10846.10865.20\|CTTCGGGATAAGGATTGGCT | >hg38_21_+_8433172_8446622_RNA45SN1_45S_50ntFlanks | 10846 | 10865 | 20 | CTTCGGGATAAGGATTGGCT | 1208.7 | -14.0 | 1.4 | -9.7 | 2.8E-22 | 4.4E-21 | MB231/MB468 | MITO |
| >hg38_21_+_8433172_8446622_RNA45SN1_45S_50ntFlanks@12623.12654.32\|GCCAATGGGGCGAAGCTACCATCTGTGGGATT | >hg38_21_+_8433172_8446622_RNA45SN1_45S_50ntFlanks | 12623 | 12654 | 32 | GCCAATGGGGCGAAGCTACCATCTGTGGGATT | 935.1 | -13.7 | 1.4 | -9.7 | 2.8E-22 | 4.4E-21 | MB231/MB468 | MITO |
| >MI0000298\|hsa-mir-221&WithFlank&X\|-\|45746151\|45746272@71.93.23(+1C)&[MIMAT0000278&hsa-miR-221-3p&offsets\|0\|0(+1C);m-91&X\|-\|45746181\|45746202&offsets\|0\|+1(+1C)]\|AGCTACATTGTCTGCTGGGTTTCC | >MI0000298\|hsa-mir-221&WithFlank&X\|-\|45746151\|45746272 | 71 | 93 | 23(+1C) | AGCTACATTGTCTGCTGGGTTTCC | 1350.9 | 6.1 | 0.6 | 9.6 | 5.0E-22 | 4.5E-21 | BT20/MB468 | CYTO |
| >hg38_21_+_8433172_8446622_RNA45SN1_45S_50ntFlanks@12627.12654.28\|ATGGGGCGAAGCTACCATCTGTGGGATT | >hg38_21_+_8433172_8446622_RNA45SN1_45S_50ntFlanks | 12627 | 12654 | 28 | ATGGGGCGAAGCTACCATCTGTGGGATT | 4852.2 | -5.8 | 0.6 | -9.8 | 1.4E-22 | 4.5E-21 | MB231/MB468 | CYTO |
| >hg38_21_+_8433172_8446622_RNA45SN1_45S_50ntFlanks@10819.10838.20\|AGGGAAGTCGGCAAGCCGGA | >hg38_21_+_8433172_8446622_RNA45SN1_45S_50ntFlanks | 10819 | 10838 | 20 | AGGGAAGTCGGCAAGCCGGA | 456.0 | -13.3 | 1.4 | -9.6 | 1.0E-21 | 4.5E-21 | BT20/MB468 | NUC |
| >MI0000088\|hsa-mir-30a&WithFlank&6\|-\|71403545\|71403627@53.73.21(+2U)&[MIMAT0000088&hsa-miR-30a-3p&offsets\|0\|-1(+2U);m-38&6\|-\|71403554\|71403575&offsets\|0\|-1(+2U)]\|CTTTCAGTCGGATGTTTGCAGTT | >MI0000088\|hsa-mir-30a&WithFlank&6\|-\|71403545\|71403627 | 53 | 73 | 21(+2U) | CTTTCAGTCGGATGTTTGCAGTT | 808.5 | 6.1 | 0.6 | 9.7 | 2.7E-22 | 4.5E-21 | MB231/MB468 | NUC |
| >hg38_21_+_8433172_8446622_RNA45SN1_45S_50ntFlanks@6666.6689.24\|GATCACTCGGCTCGTGCGTCGATG | >hg38_21_+_8433172_8446622_RNA45SN1_45S_50ntFlanks | 6666 | 6689 | 24 | GATCACTCGGCTCGTGCGTCGATG | 1203.9 | -14.1 | 1.4 | -9.8 | 1.8E-22 | 4.5E-21 | MB231/MB468 | MP |
| >MI0001733\|hsa-mir-452&WithFlank&X\|-\|151959622\|151959718@23.43.21&[MIMAT0001635&hsa-miR-452-5p&offsets\|+3\|+2;m-188&X\|-\|151959677\|151959699&offsets\|+3\|+1]\|TGTTTGCAGAGGAAACTGAGA | >MI0001733\|hsa-mir-452&WithFlank&X\|-\|151959622\|151959718 | 23 | 43 | 21 | TGTTTGCAGAGGAAACTGAGA | 180.7 | 10.3 | 1.1 | 9.6 | 1.0E-21 | 4.5E-21 | BT20/MB468 | NUC |
| >MI0000088\|hsa-mir-30a&WithFlank&6\|-\|71403545\|71403627@53.73.21(+1U)&[MIMAT0000088&hsa-miR-30a-3p&offsets\|0\|-1(+1U);m-38&6\|-\|71403554\|71403575&offsets\|0\|-1(+1U)]\|CTTTCAGTCGGATGTTTGCAGT | >MI0000088\|hsa-mir-30a&WithFlank&6\|-\|71403545\|71403627 | 53 | 73 | 21(+1U) | CTTTCAGTCGGATGTTTGCAGT | 2523.8 | 6.4 | 0.7 | 9.7 | 3.8E-22 | 4.5E-21 | BT20/MB468 | MITO |
| >hg38_21_+_8433172_8446622_RNA45SN1_45S_50ntFlanks@4201.4219.19\|CCGACCCGGGGAGGTAGTG | >hg38_21_+_8433172_8446622_RNA45SN1_45S_50ntFlanks | 4201 | 4219 | 19 | CCGACCCGGGGAGGTAGTG | 712.5 | -13.3 | 1.4 | -9.8 | 1.8E-22 | 4.6E-21 | MB231/MB468 | MP |
| >MI0001448\|hsa-mir-425&WithFlank&3\|-\|49020142\|49020240@20.42.23&[MIMAT0003393&hsa-miR-425-5p&offsets\|0\|0;m-95&3\|-\|49020200\|49020221&offsets\|0\|+1]\|AATGACACGATCACTCCCGTTGA | >MI0001448\|hsa-mir-425&WithFlank&3\|-\|49020142\|49020240 | 20 | 42 | 23 | AATGACACGATCACTCCCGTTGA | 810.7 | 8.5 | 0.9 | 9.6 | 5.1E-22 | 4.6E-21 | BT20/MB468 | CYTO |
| >hg38_21_+_8433172_8446622_RNA45SN1_45S_50ntFlanks@8021.8046.26\|AGTCAGCGGAGGAGAAGAAACTAACC | >hg38_21_+_8433172_8446622_RNA45SN1_45S_50ntFlanks | 8021 | 8046 | 26 | AGTCAGCGGAGGAGAAGAAACTAACC | 182.5 | 11.5 | 1.2 | 9.8 | 1.4E-22 | 4.6E-21 | MB231/MB468 | CYTO |
| >hg38_21_+_8433172_8446622_RNA45SN1_45S_50ntFlanks@8380.8406.27\|CAAGAGGGCGTGAAACCGTTAAGAGGT | >hg38_21_+_8433172_8446622_RNA45SN1_45S_50ntFlanks | 8380 | 8406 | 27 | CAAGAGGGCGTGAAACCGTTAAGAGGT | 2856.3 | -6.6 | 0.7 | -9.8 | 1.8E-22 | 4.6E-21 | MB231/MB468 | MP |
| >hg38_21_+_8433172_8446622_RNA45SN1_45S_50ntFlanks@9454.9488.35\|CCGCGCCGGGGAGGTGGAGCACGAGCGCACGTGTT | >hg38_21_+_8433172_8446622_RNA45SN1_45S_50ntFlanks | 9454 | 9488 | 35 | CCGCGCCGGGGAGGTGGAGCACGAGCGCACGTGTT | 1094.8 | -14.0 | 1.4 | -9.8 | 1.8E-22 | 4.6E-21 | MB231/MB468 | MP |
| >hg38_21_+_8433172_8446622_RNA45SN1_45S_50ntFlanks@10663.10706.44\|CCTTGGAAAGCGTCGCGGTTCCGGCGGCGTCCGGTGAGCTCTCG | >hg38_21_+_8433172_8446622_RNA45SN1_45S_50ntFlanks | 10663 | 10706 | 44 | CCTTGGAAAGCGTCGCGGTTCCGGCGGCGTCCGGTGAGCTCTCG | 1582.1 | -15.1 | 1.6 | -9.6 | 1.0E-21 | 4.6E-21 | BT20/MB468 | NUC |
| >MI0000061\|hsa-let-7a-2&WithFlank&11\|-\|122146516\|122146599@11.31.21;MI0000062\|hsa-let-7a-3&WithFlank&22\|+\|46112743\|46112828@10.30.21;MI0000060\|hsa-let-7a-1&WithFlank&9\|+\|94175951\|94176042@12.32.21&[MIMAT0000062&hsa-let-7a-5p&offsets\|0\|-1;m-6&11\|-\|122146568\|122146589&offsets\|0\|-1];[MIMAT0000062_1&hsa-let-7a-5p&offsets\|0\|-1;m-5&22\|+\|46112752\|46112773&offsets\|0\|-1];[MIMAT0000062_2&hsa-let-7a-5p&offsets\|0\|-1;m-4&9\|+\|94175962\|94175983&offsets\|0\|-1]\|TGAGGTAGTAGGTTGTATAGT | >MI0000061\|hsa-let-7a-2&WithFlank&11\|-\|122146516\|122146599 | 11 | 31 | 21 | TGAGGTAGTAGGTTGTATAGT | 12390.1 | 5.7 | 0.6 | 9.7 | 2.8E-22 | 4.6E-21 | MB231/MB468 | NUC |
| >hg38_21_+_8433172_8446622_RNA45SN1_45S_50ntFlanks@5467.5514.48\|GGCCCACGGCCCTGGCGGAGCGCTGAGAAGACGGTCGAACTTGACTAT | >hg38_21_+_8433172_8446622_RNA45SN1_45S_50ntFlanks | 5467 | 5514 | 48 | GGCCCACGGCCCTGGCGGAGCGCTGAGAAGACGGTCGAACTTGACTAT | 4333.3 | -16.5 | 1.7 | -9.6 | 1.1E-21 | 4.6E-21 | BT20/MB468 | NUC |
| >hg38_21_+_8433172_8446622_RNA45SN1_45S_50ntFlanks@12991.13012.22\|CTCCCTCGCTGCGATCTATTGA | >hg38_21_+_8433172_8446622_RNA45SN1_45S_50ntFlanks | 12991 | 13012 | 22 | CTCCCTCGCTGCGATCTATTGA | 24126.2 | -14.3 | 1.5 | -9.6 | 1.1E-21 | 4.7E-21 | BT20/MB468 | NUC |

| | | | | | | | | | | | | | |
|---|---|---|---|---|---|---|---|---|---|---|---|---|---|
| >trnaMT_GluTTC_MT_-_14674_14742@38.71.34,trnalookalike8_GluTTC_5_-_93905172_93905240@38.71.34\|CATTGGTCGTGGTTGTAGTCCGTGCGAGAATACC | >trnaMT_GluTTC_MT_-_14674_14742 | 38 | 71 | 34 | CATTGGTCGTGGTTGTAGTCCGTGCGAGAATACC | 1311.5 | 7.3 | 0.8 | 9.7 | 3.9E-22 | 4.7E-21 | BT20/MB468 | MITO |
| >hg38_21_+_8433172_8446622_RNA45SN1_45S_50ntFlanks@9739.9765.27\|AAACGATCTCAACCTATTCTCAAACTT | >hg38_21_+_8433172_8446622_RNA45SN1_45S_50ntFlanks | 9739 | 9765 | 27 | AAACGATCTCAACCTATTCTCAAACTT | 1490.9 | -14.4 | 1.5 | -9.8 | 1.8E-22 | 4.7E-21 | MB231/MB468 | MP |
| >hg38_21_+_8433172_8446622_RNA45SN1_45S_50ntFlanks@11885.11905.21\|TTGAGCTTGACTCTAGTCTGG | >hg38_21_+_8433172_8446622_RNA45SN1_45S_50ntFlanks | 11885 | 11905 | 21 | TTGAGCTTGACTCTAGTCTGG | 864.8 | -13.6 | 1.4 | -9.7 | 1.8E-22 | 4.7E-21 | MB231/MB468 | MP |
| >MI0000456\|hsa-mir-140&WithFlank&16\|+\|69933075\|69933186@68.89.22&[MIMAT0004597&hsa-miR-140-3p&offsets\|0\|+1;m-45&16\|+\|69933143\|69933164&offsets\|-1\|-1]\|TACCACAGGGTAGAACCACGGA | >MI0000456\|hsa-mir-140&WithFlank&16\|+\|69933075\|69933186 | 68 | 89 | 22 | TACCACAGGGTAGAACCACGGA | 2655.8 | 11.3 | 1.2 | 9.6 | 5.3E-22 | 4.7E-21 | BT20/MB468 | CYTO |
| >MI0000102\|hsa-mir-100&WithFlank&11\|-\|122152223\|122152314@19.40.22&[MIMAT0000098&hsa-miR-100-5p&offsets\|0\|0;m-27&11\|-\|122152276\|122152296&offsets\|0\|+1]\|AACCCGTAGATCCGAACTTGTG | >MI0000102\|hsa-mir-100&WithFlank&11\|-\|122152223\|122152314 | 19 | 40 | 22 | AACCCGTAGATCCGAACTTGTG | 96928.0 | 12.8 | 1.3 | 9.6 | 5.3E-22 | 4.7E-21 | BT20/MB468 | CYTO |
| >am_MI0039740\|hsa-mir-12136&WithFlank&1\|-\|632609\|632691@7.31.25&[MIMAT0049032&hsa-miR-12136&offsets\|0\|+7]\|GAAAAAGTCATGGAGGCCATGGGGT | >am_MI0039740\|hsa-mir-12136&WithFlank&1\|-\|632609\|632691 | 7 | 31 | 25 | GAAAAAGTCATGGAGGCCATGGGGT | 113.6 | 3.5 | 0.4 | 9.6 | 1.1E-21 | 4.7E-21 | BT20/MB468 | NUC |
| >trnaMT_SerTGA_MT_-_7446_7514@1.25.25,trnalookalike6_SerTGA_1_-_567997_568065@1.25.25\|GAAAAAGTCATGGAGGCCATGGGGT | >trnaMT_SerTGA_MT_-_7446_7514 | 1 | 25 | 25 | GAAAAAGTCATGGAGGCCATGGGGT | 113.6 | 3.5 | 0.4 | 9.6 | 1.1E-21 | 4.7E-21 | BT20/MB468 | NUC |
| >hg38_21_+_8433172_8446622_RNA45SN1_45S_50ntFlanks@10369.10406.38\|AAGTGGAGAAGGGTTCCATGTGAACAGCAGTTGAACAT | >hg38_21_+_8433172_8446622_RNA45SN1_45S_50ntFlanks | 10369 | 10406 | 38 | AAGTGGAGAAGGGTTCCATGTGAACAGCAGTTGAACAT | 681.9 | -13.8 | 1.4 | -9.6 | 1.1E-21 | 4.7E-21 | BT20/MB468 | NUC |
| >hg38_21_+_8433172_8446622_RNA45SN1_45S_50ntFlanks@10391.10429.39\|AACAGCAGTTGAACATGGGTCAGTCGGTCCTGAGAGATG | >hg38_21_+_8433172_8446622_RNA45SN1_45S_50ntFlanks | 10391 | 10429 | 39 | AACAGCAGTTGAACATGGGTCAGTCGGTCCTGAGAGATG | 5040.2 | -6.6 | 0.7 | -9.6 | 1.1E-21 | 4.8E-21 | BT20/MB468 | NUC |
| >MI0000456\|hsa-mir-140&WithFlank&16\|+\|69933075\|69933186@68.91.24&[MIMAT0004597&hsa-miR-140-3p&offsets\|0\|+3;m-45&16\|+\|69933143\|69933164&offsets\|-1\|+1]\|TACCACAGGGTAGAACCACGGACA | >MI0000456\|hsa-mir-140&WithFlank&16\|+\|69933075\|69933186 | 68 | 91 | 24 | TACCACAGGGTAGAACCACGGACA | 976.0 | 10.2 | 1.1 | 9.6 | 5.5E-22 | 4.9E-21 | BT20/MB468 | CYTO |
| >hg38_21_+_8433172_8446622_RNA45SN1_45S_50ntFlanks@9720.9737.18\|ATTAGAGGTCTTGGGGCC | >hg38_21_+_8433172_8446622_RNA45SN1_45S_50ntFlanks | 9720 | 9737 | 18 | ATTAGAGGTCTTGGGGCC | 1291.0 | -14.2 | 1.5 | -9.7 | 1.9E-22 | 4.9E-21 | MB231/MB468 | MP |
| >MI0000266\|hsa-mir-10a&WithFlank&17\|-\|48579832\|48579953@29.49.21(+1U)&[MIMAT0000253&hsa-miR-10a-5p&offsets\|+1\|-1(+1U);m-18&17\|-\|48579905\|48579926&offsets\|+1\|0(+1U)]\|ACCCTGTAGATCCGAATTTGTT | >MI0000266\|hsa-mir-10a&WithFlank&17\|-\|48579832\|48579953 | 29 | 49 | 21(+1U) | ACCCTGTAGATCCGAATTTGTT | 329.6 | 12.6 | 1.3 | 9.6 | 5.5E-22 | 4.9E-21 | BT20/MB468 | CYTO |
| >hg38_21_+_8433172_8446622_RNA45SN1_45S_50ntFlanks@9554.9571.18\|CGTAGCGGTCCTGACGTG | >hg38_21_+_8433172_8446622_RNA45SN1_45S_50ntFlanks | 9554 | 9571 | 18 | CGTAGCGGTCCTGACGTG | 152171.0 | -6.3 | 0.6 | -9.7 | 3.0E-22 | 4.9E-21 | MB231/MB468 | NUC |
| >hg38_21_+_8433172_8446622_RNA45SN1_45S_50ntFlanks@12710.12742.33\|CGCGGAGCCTCGGTTGGCCTCGGATAGCCGGTC | >hg38_21_+_8433172_8446622_RNA45SN1_45S_50ntFlanks | 12710 | 12742 | 33 | CGCGGAGCCTCGGTTGGCCTCGGATAGCCGGTC | 777.8 | -13.5 | 1.4 | -9.7 | 2.0E-22 | 4.9E-21 | MB231/MB468 | MP |
| >hg38_21_+_8433172_8446622_RNA45SN1_45S_50ntFlanks@7985.8003.19\|GATCAGACGTGGCGACCCG | >hg38_21_+_8433172_8446622_RNA45SN1_45S_50ntFlanks | 7985 | 8003 | 19 | GATCAGACGTGGCGACCCG | 636.0 | -13.7 | 1.4 | -9.6 | 1.1E-21 | 4.9E-21 | BT20/MB468 | NUC |
| >hg38_21_+_8433172_8446622_RNA45SN1_45S_50ntFlanks@10458.10497.40\|ATGGCCTCCGTTGCCCTCGGCCGATCGAAAGGGAGTCGGG | >hg38_21_+_8433172_8446622_RNA45SN1_45S_50ntFlanks | 10458 | 10497 | 40 | ATGGCCTCCGTTGCCCTCGGCCGATCGAAAGGGAGTCGGG | 904.8 | -14.2 | 1.5 | -9.6 | 1.1E-21 | 4.9E-21 | BT20/MB468 | NUC |
| >hg38_21_+_8433172_8446622_RNA45SN1_45S_50ntFlanks@8176.8223.48\|CTTCTGATCGAGGCCCAGCCCGTGGACGGTGTGAGGCCGGTAGCGGCC | >hg38_21_+_8433172_8446622_RNA45SN1_45S_50ntFlanks | 8176 | 8223 | 48 | CTTCTGATCGAGGCCCAGCCCGTGGACGGTGTGAGGCCGGTAGCGGCC | 2209.6 | -15.5 | 1.6 | -9.6 | 1.1E-21 | 5.0E-21 | BT20/MB468 | NUC |
| >hg38_21_+_8433172_8446622_RNA45SN1_45S_50ntFlanks@11572.11603.32\|CCGACTTAGAACTGGTGCGGACCAGGGGAATC | >hg38_21_+_8433172_8446622_RNA45SN1_45S_50ntFlanks | 11572 | 11603 | 32 | CCGACTTAGAACTGGTGCGGACCAGGGGAATC | 5054.6 | -4.9 | 0.5 | -9.6 | 5.6E-22 | 5.0E-21 | BT20/MB468 | CYTO |
| >hg38_21_+_8433172_8446622_RNA45SN1_45S_50ntFlanks@11561.11601.41\|GCGCCTAGCAGCCGACTTAGAACTGGTGCGGACCAGGGGAA | >hg38_21_+_8433172_8446622_RNA45SN1_45S_50ntFlanks | 11561 | 11601 | 41 | GCGCCTAGCAGCCGACTTAGAACTGGTGCGGACCAGGGGAA | 6314.5 | -12.2 | 1.3 | -9.7 | 3.2E-22 | 5.0E-21 | MB231/MB468 | MITO |
| >hg38_21_+_8433172_8446622_RNA45SN1_45S_50ntFlanks@9806.9838.33\|CGTGGAATGCGAGTGCCTAGTGGGCCACTTTTG | >hg38_21_+_8433172_8446622_RNA45SN1_45S_50ntFlanks | 9806 | 9838 | 33 | CGTGGAATGCGAGTGCCTAGTGGGCCACTTTTG | 1155.0 | -14.0 | 1.4 | -9.7 | 3.2E-22 | 5.0E-21 | MB231/MB468 | MITO |
| >hg38_21_+_8433172_8446622_RNA45SN1_45S_50ntFlanks@12157.12179.23\|AGTTTGACTGGGGCGGTACACCT | >hg38_21_+_8433172_8446622_RNA45SN1_45S_50ntFlanks | 12157 | 12179 | 23 | AGTTTGACTGGGGCGGTACACCT | 1348.6 | -14.2 | 1.5 | -9.7 | 3.2E-22 | 5.0E-21 | MB231/MB468 | MITO |
| >hg38_21_+_8433172_8446622_RNA45SN1_45S_50ntFlanks@12405.12433.29\|CTTTTTGATCCTTCGATGTCGGCTCTTCC | >hg38_21_+_8433172_8446622_RNA45SN1_45S_50ntFlanks | 12405 | 12433 | 29 | CTTTTTGATCCTTCGATGTCGGCTCTTCC | 3435.1 | -6.0 | 0.6 | -9.7 | 3.2E-22 | 5.0E-21 | MB231/MB468 | MITO |
| >hg38_21_+_8433172_8446622_RNA45SN1_45S_50ntFlanks@10348.10379.32\|AAACGAGAACTTTGAAGGCCGAAGTGGAGAAG | >hg38_21_+_8433172_8446622_RNA45SN1_45S_50ntFlanks | 10348 | 10379 | 32 | AAACGAGAACTTTGAAGGCCGAAGTGGAGAAG | 288.1 | -4.0 | 0.4 | -9.6 | 1.2E-21 | 5.0E-21 | BT20/MB468 | NUC |
| >hg38_21_+_8433172_8446622_RNA45SN1_45S_50ntFlanks@6647.6667.21\|CGTACGACTCTTAGCGGTGGA | >hg38_21_+_8433172_8446622_RNA45SN1_45S_50ntFlanks | 6647 | 6667 | 21 | CGTACGACTCTTAGCGGTGGA | 2531.9 | 6.3 | 0.7 | 9.6 | 5.7E-22 | 5.1E-21 | BT20/MB468 | CYTO |
| >trna32_LysCTT_16_-_3207406_3207478@1.33.33\|GCCCGGCTAGCTCAGTCGGTAGAGCATGAGACC | >trna32_LysCTT_16_-_3207406_3207478 | 1 | 33 | 33 | GCCCGGCTAGCTCAGTCGGTAGAGCATGAGACC | 897.6 | 5.7 | 0.6 | 9.6 | 5.8E-22 | 5.1E-21 | BT20/MB468 | CYTO |
| >hg38_21_+_8433172_8446622_RNA45SN1_45S_50ntFlanks@7986.8012.27\|ATCAGACGTGGCGACCCGCTGAATTTA | >hg38_21_+_8433172_8446622_RNA45SN1_45S_50ntFlanks | 7986 | 8012 | 27 | ATCAGACGTGGCGACCCGCTGAATTTA | 288.5 | -12.6 | 1.3 | -9.6 | 1.2E-21 | 5.1E-21 | BT20/MB468 | NUC |
| >hg38_21_+_8433172_8446622_RNA45SN1_45S_50ntFlanks@12717.12757.41\|CCTCGGTTGGCCTCGGATAGCCGGTCCCCCGCCTGTCCCCG | >hg38_21_+_8433172_8446622_RNA45SN1_45S_50ntFlanks | 12717 | 12757 | 41 | CCTCGGTTGGCCTCGGATAGCCGGTCCCCCGCCTGTCCCCG | 1253.3 | -14.1 | 1.5 | -9.7 | 3.3E-22 | 5.1E-21 | MB231/MB468 | MITO |
| >MI0000086\|hsa-mir-28&WithFlank&3\|+\|188688775\|188688872@60.80.21&[MIMAT0004502&hsa-miR-28-3p&offsets\|0\|-1;m-28&3\|+\|188688834\|188688855&offsets\|0\|-1]\|CACTAGATTGTGAGCTCCTGG | >MI0000086\|hsa-mir-28&WithFlank&3\|+\|188688775\|188688872 | 60 | 80 | 21 | CACTAGATTGTGAGCTCCTGG | 5553.9 | 10.7 | 1.1 | 9.6 | 5.8E-22 | 5.1E-21 | BT20/MB468 | CYTO |
| >hg38_21_+_8433172_8446622_RNA45SN1_45S_50ntFlanks@11209.11231.23\|CGGCGACTCTGGACGCGAGCCGG | >hg38_21_+_8433172_8446622_RNA45SN1_45S_50ntFlanks | 11209 | 11231 | 23 | CGGCGACTCTGGACGCGAGCCGG | 1148.9 | -14.0 | 1.4 | -9.7 | 2.0E-22 | 5.1E-21 | MB231/MB468 | MP |
| >hg38_21_+_8433172_8446622_RNA45SN1_45S_50ntFlanks@11580.11609.30\|GAACTGGTGCGGACCAGGGGAATCCGACTG | >hg38_21_+_8433172_8446622_RNA45SN1_45S_50ntFlanks | 11580 | 11609 | 30 | GAACTGGTGCGGACCAGGGGAATCCGACTG | 2059.0 | -6.4 | 0.7 | -9.7 | 2.0E-22 | 5.1E-21 | MB231/MB468 | MP |
| >hg38_21_+_8433172_8446622_RNA45SN1_45S_50ntFlanks@11583.11603.21\|CTGGTGCGGACCAGGGGAATC | >hg38_21_+_8433172_8446622_RNA45SN1_45S_50ntFlanks | 11583 | 11603 | 21 | CTGGTGCGGACCAGGGGAATC | 3997.7 | -7.4 | 0.8 | -9.6 | 1.2E-21 | 5.1E-21 | BT20/MB468 | NUC |
| >hg38_21_+_8433172_8446622_RNA45SN1_45S_50ntFlanks@12288.12318.31\|ACCGTGAAAGCGGGGCCTCACGATCCTTCTG | >hg38_21_+_8433172_8446622_RNA45SN1_45S_50ntFlanks | 12288 | 12318 | 31 | ACCGTGAAAGCGGGGCCTCACGATCCTTCTG | 532.7 | -13.5 | 1.4 | -9.6 | 1.2E-21 | 5.1E-21 | BT20/MB468 | NUC |
| >hg38_21_+_8433172_8446622_RNA45SN1_45S_50ntFlanks@8356.8375.20\|GAAAAGAACTTTGAAGAGAG | >hg38_21_+_8433172_8446622_RNA45SN1_45S_50ntFlanks | 8356 | 8375 | 20 | GAAAAGAACTTTGAAGAGAG | 719.3 | -6.2 | 0.6 | -9.6 | 1.2E-21 | 5.1E-21 | BT20/MB468 | NUC |
| >hg38_21_+_8433172_8446622_RNA45SN1_45S_50ntFlanks@4731.4765.35\|AAGAACGAAAGTCGGAGGTTCGAAGACGATCAGAT | >hg38_21_+_8433172_8446622_RNA45SN1_45S_50ntFlanks | 4731 | 4765 | 35 | AAGAACGAAAGTCGGAGGTTCGAAGACGATCAGAT | 470.8 | 4.9 | 0.5 | 9.7 | 4.3E-22 | 5.1E-21 | BT20/MB468 | MITO |
| >trna17_ValTAC_11_-_59318102_59318174@1.30.30,trna4_ValTAC_X_-_18693029_18693101@1.30.30\|GGTTCCATAGTGTAGTGGTTATCACGTCTG | >trna17_ValTAC_11_-_59318102_59318174 | 1 | 30 | 30 | GGTTCCATAGTGTAGTGGTTATCACGTCTG | 474.3 | -13.3 | 1.4 | -9.6 | 1.2E-21 | 5.2E-21 | BT20/MB468 | NUC |
| >hg38_21_+_8433172_8446622_RNA45SN1_45S_50ntFlanks@10411.10428.18\|CAGTCGGTCCTGAGAGAT | >hg38_21_+_8433172_8446622_RNA45SN1_45S_50ntFlanks | 10411 | 10428 | 18 | CAGTCGGTCCTGAGAGAT | 12401.6 | -13.6 | 1.4 | -9.6 | 1.2E-21 | 5.2E-21 | BT20/MB468 | NUC |
| >hg38_21_+_8433172_8446622_RNA45SN1_45S_50ntFlanks@9567.9597.31\|ACGTGCAAATCGGTCGTCCGACCTGGGTATA | >hg38_21_+_8433172_8446622_RNA45SN1_45S_50ntFlanks | 9567 | 9597 | 31 | ACGTGCAAATCGGTCGTCCGACCTGGGTATA | 4403.1 | -8.3 | 0.8 | -9.8 | 1.6E-22 | 5.2E-21 | MB231/MB468 | CYTO |
| >hg38_21_+_8433172_8446622_RNA45SN1_45S_50ntFlanks@11579.11609.31\|AGAACTGGTGCGGACCAGGGGAATCCGACTG | >hg38_21_+_8433172_8446622_RNA45SN1_45S_50ntFlanks | 11579 | 11609 | 31 | AGAACTGGTGCGGACCAGGGGAATCCGACTG | 1428.0 | -6.6 | 0.7 | -9.7 | 2.1E-22 | 5.3E-21 | MB231/MB468 | MP |
| >MI0000439\|hsa-mir-23b&WithFlank&9\|+\|95085202\|95085310@67.85.19(+1U)&[MIMAT0000418&hsa-miR-23b-3p&offsets\|+3\|-1(+1U);m-42&9\|+\|95085265\|95085285&offsets\|+3\|+1(+1U)]\|ACATTGCCAGGGATTACCAT | >MI0000439\|hsa-mir-23b&WithFlank&9\|+\|95085202\|95085310 | 67 | 85 | 19(+1U) | ACATTGCCAGGGATTACCAT | 164.6 | 10.2 | 1.1 | 9.6 | 1.2E-21 | 5.3E-21 | BT20/MB468 | NUC |
| >hg38_21_+_8433172_8446622_RNA45SN1_45S_50ntFlanks@10351.10374.24\|CGAGAACTTTGAAGGCCGAAGTGG | >hg38_21_+_8433172_8446622_RNA45SN1_45S_50ntFlanks | 10351 | 10374 | 24 | CGAGAACTTTGAAGGCCGAAGTGG | 2038.3 | -5.4 | 0.6 | -9.7 | 4.4E-22 | 5.3E-21 | BT20/MB468 | MITO |
| >hg38_21_+_8433172_8446622_RNA45SN1_45S_50ntFlanks@11570.11603.34\|AGCCGACTTAGAACTGGTGCGGACCAGGGGAATC | >hg38_21_+_8433172_8446622_RNA45SN1_45S_50ntFlanks | 11570 | 11603 | 34 | AGCCGACTTAGAACTGGTGCGGACCAGGGGAATC | 2021.6 | -5.8 | 0.6 | -9.7 | 3.3E-22 | 5.4E-21 | MB231/MB468 | NUC |
| >hg38_21_+_8433172_8446622_RNA45SN1_45S_50ntFlanks@7983.8005.23\|CAGATCAGACGTGGCGACCCGCT | >hg38_21_+_8433172_8446622_RNA45SN1_45S_50ntFlanks | 7983 | 8005 | 23 | CAGATCAGACGTGGCGACCCGCT | 694.2 | -5.1 | 0.5 | -9.7 | 3.3E-22 | 5.4E-21 | MB231/MB468 | NUC |
| >trna116_GluCTC_1_-_145399233_145399304@1.33.33,trna59_GluCTC_1_+_249168447_249168518@1.33.33,trna71_GluCTC_1_-_161439189_161439260@1.33.33,trna74_GluCTC_1_-_161431809_161431880@1.33.33,trna77_GluCTC_1_-_161424398_161424469@1.33.33,trna77_GluCTC_6_+_28949976_28950047@1.33.33,trna80_GluCTC_1_-_161417018_161417089@1.33.33,trna87_GluCTC_6_-_126101393_126101464@1.33.33\|TCCCTGGTGGTCTAGTGGTTAGGATTCGGCGCT | >trna116_GluCTC_1_-_145399233_145399304 | 1 | 33 | 33 | TCCCTGGTGGTCTAGTGGTTAGGATTCGGCGCT | 163647.8 | 8.3 | 0.9 | 9.6 | 6.2E-22 | 5.5E-21 | BT20/MB468 | CYTO |
| >hg38_21_+_8433172_8446622_RNA45SN1_45S_50ntFlanks@11564.11582.19\|CCTAGCAGCCGACTTAGAA | >hg38_21_+_8433172_8446622_RNA45SN1_45S_50ntFlanks | 11564 | 11582 | 19 | CCTAGCAGCCGACTTAGAA | 640.4 | -13.2 | 1.4 | -9.7 | 2.2E-22 | 5.5E-21 | MB231/MB468 | MP |
| >MI0000816\|hsa-mir-335&WithFlank&7\|+\|130496105\|130496210@58.78.21(+1A)&[MIMAT0004703&hsa-miR-335-3p&offsets\|0\|-1(+1A);m-159&7\|+\|130496162\|130496183&offsets\|0\|-1(+1A)]\|TTTTTCATTATTGCTCCTGACA | >MI0000816\|hsa-mir-335&WithFlank&7\|+\|130496105\|130496210 | 58 | 78 | 21(+1A) | TTTTTCATTATTGCTCCTGACA | 263.9 | 5.3 | 0.5 | 9.7 | 3.4E-22 | 5.5E-21 | MB231/MB468 | NUC |
| >trna5_AspGTC_12_+_98897281_98897352@35.74.40\|TCACGCGGGAGACCGGGGTTCAATTCCCCGACGGGGAGCC | >trna5_AspGTC_12_+_98897281_98897352 | 35 | 74 | 40 | TCACGCGGGAGACCGGGGTTCAATTCCCCGACGGGGAGCC | 3782.3 | 15.9 | 1.6 | 9.8 | 1.7E-22 | 5.5E-21 | MB231/MB468 | CYTO |
| >MI0000266\|hsa-mir-10a&WithFlank&17\|-\|48579832\|48579953@28.49.22(+1A)&[MIMAT0000253&hsa-miR-10a-5p&offsets\|0\|-1(+1A);m-18&17\|-\|48579905\|48579926&offsets\|0\|0(+1A)]\|TACCCTGTAGATCCGAATTTGTA | >MI0000266\|hsa-mir-10a&WithFlank&17\|-\|48579832\|48579953 | 28 | 49 | 22(+1A) | TACCCTGTAGATCCGAATTTGTA | 280.8 | 11.0 | 1.1 | 9.6 | 1.3E-21 | 5.6E-21 | BT20/MB468 | NUC |
| >hg38_21_+_8433172_8446622_RNA45SN1_45S_50ntFlanks@4283.4317.35\|CTTTAACGAGGATCCATTGGAGGGCAAGTCTGGTG | >hg38_21_+_8433172_8446622_RNA45SN1_45S_50ntFlanks | 4283 | 4317 | 35 | CTTTAACGAGGATCCATTGGAGGGCAAGTCTGGTG | 4841.0 | -16.1 | 1.7 | -9.7 | 2.3E-22 | 5.6E-21 | MB231/MB468 | MP |

| | | | | | | | | | | | | | |
|---|---|---|---|---|---|---|---|---|---|---|---|---|---|
| >hg38_21_+_8433172_8446622_RNA45SN1_45S_50ntFlanks@9567.9596.30\|ACGTGCAAATCGGTCGTCCGACCTGGGTAT | >hg38_21_+_8433172_8446622_RNA45SN1_45S_50ntFlanks | 9567 | 9596 | 30 | ACGTGCAAATCGGTCGTCCGACCTGGGTAT | 5764.9 | -11.9 | 1.2 | -9.7 | 2.3E-22 | 5.7E-21 | MB231/MB468 | MP |
| >hg38_1_-_228634819_228635039_RNA5S12_5S_50ntFlanks@139.169.31\|GACCGCCTGGGAATACCGGGTGCTGTAGGCT | >hg38_1_-_228634819_228635039_RNA5S12_5S_50ntFlanks | 139 | 169 | 31 | GACCGCCTGGGAATACCGGGTGCTGTAGGCT | 2114.2 | 7.0 | 0.7 | 9.6 | 6.5E-22 | 5.7E-21 | BT20/MB468 | CYTO |
| >hg38_21_+_8433172_8446622_RNA45SN1_45S_50ntFlanks@10788.10818.31\|CAGCCTCTGGCATGTTGGAACAATGTAGGTA | >hg38_21_+_8433172_8446622_RNA45SN1_45S_50ntFlanks | 10788 | 10818 | 31 | CAGCCTCTGGCATGTTGGAACAATGTAGGTA | 3378.4 | -9.3 | 1.0 | -9.7 | 2.3E-22 | 5.7E-21 | MB231/MB468 | MP |
| >hg38_21_+_8433172_8446622_RNA45SN1_45S_50ntFlanks@12635.12652.18\|AAGCTACCATCTGTGGGA | >hg38_21_+_8433172_8446622_RNA45SN1_45S_50ntFlanks | 12635 | 12652 | 18 | AAGCTACCATCTGTGGGA | 1694.6 | -14.6 | 1.5 | -9.7 | 2.3E-22 | 5.7E-21 | MB231/MB468 | MP |
| >hg38_21_+_8433172_8446622_RNA45SN1_45S_50ntFlanks@12454.12496.43\|CACCAAGCGTTGGATTGTTCACCCACTAATAGGGAACGTGAGC | >hg38_21_+_8433172_8446622_RNA45SN1_45S_50ntFlanks | 12454 | 12496 | 43 | CACCAAGCGTTGGATTGTTCACCCACTAATAGGGAACGTGAGC | 235.1 | 11.9 | 1.2 | 9.8 | 1.8E-22 | 5.7E-21 | MB231/MB468 | CYTO |
| >hg38_21_+_8433172_8446622_RNA45SN1_45S_50ntFlanks@11270.11295.26\|CGGCCGCCCCCGGGGAGCCCGGCGGG | >hg38_21_+_8433172_8446622_RNA45SN1_45S_50ntFlanks | 11270 | 11295 | 26 | CGGCCGCCCCCGGGGAGCCCGGCGGG | 1121.4 | -12.8 | 1.3 | -9.8 | 1.8E-22 | 5.7E-21 | MB231/MB468 | CYTO |
| >hg38_21_+_8433172_8446622_RNA45SN1_45S_50ntFlanks@11575.11602.28\|ACTTAGAACTGGTGCGGACCAGGGGAAT | >hg38_21_+_8433172_8446622_RNA45SN1_45S_50ntFlanks | 11575 | 11602 | 28 | ACTTAGAACTGGTGCGGACCAGGGGAAT | 1816.9 | -5.5 | 0.6 | -9.8 | 1.8E-22 | 5.7E-21 | MB231/MB468 | CYTO |
| >hg38_21_+_8433172_8446622_RNA45SN1_45S_50ntFlanks@11660.11685.26\|GTGATTTCTGCCCAGTGCTCTGAATG | >hg38_21_+_8433172_8446622_RNA45SN1_45S_50ntFlanks | 11660 | 11685 | 26 | GTGATTTCTGCCCAGTGCTCTGAATG | 825.2 | -5.0 | 0.5 | -9.7 | 3.5E-22 | 5.7E-21 | MB231/MB468 | NUC |
| >hg38_21_+_8433172_8446622_RNA45SN1_45S_50ntFlanks@4588.4616.29\|CTATTTTGTTGGTTTTCGGAACTGAGGCC | >hg38_21_+_8433172_8446622_RNA45SN1_45S_50ntFlanks | 4588 | 4616 | 29 | CTATTTTGTTGGTTTTCGGAACTGAGGCC | 1592.4 | -10.6 | 1.1 | -9.7 | 3.7E-22 | 5.8E-21 | MB231/MB468 | MITO |
| >hg38_21_+_8433172_8446622_RNA45SN1_45S_50ntFlanks@10912.10932.21\|CGCGCCGCGGCTGGACGAGGC | >hg38_21_+_8433172_8446622_RNA45SN1_45S_50ntFlanks | 10912 | 10932 | 21 | CGCGCCGCGGCTGGACGAGGC | 794.9 | -13.5 | 1.4 | -9.7 | 2.3E-22 | 5.8E-21 | MB231/MB468 | MP |
| >hg38_21_+_8433172_8446622_RNA45SN1_45S_50ntFlanks@10866.10894.29\|CTAAGGGCTGGGTCGGTCGGGCTGGGGCG | >hg38_21_+_8433172_8446622_RNA45SN1_45S_50ntFlanks | 10866 | 10894 | 29 | CTAAGGGCTGGGTCGGTCGGGCTGGGGCG | 1063.4 | -13.8 | 1.4 | -9.7 | 3.7E-22 | 5.8E-21 | MB231/MB468 | MITO |
| >hg38_21_+_8433172_8446622_RNA45SN1_45S_50ntFlanks@12007.12030.24\|GCCGCCGGTGAAATACCACTACTC | >hg38_21_+_8433172_8446622_RNA45SN1_45S_50ntFlanks | 12007 | 12030 | 24 | GCCGCCGGTGAAATACCACTACTC | 4065.1 | -7.5 | 0.8 | -9.7 | 3.7E-22 | 5.8E-21 | MB231/MB468 | MITO |
| >hg38_21_+_8433172_8446622_RNA45SN1_45S_50ntFlanks@5421.5445.25\|CGATTGGATGGTTTAGTGAGGCCCT | >hg38_21_+_8433172_8446622_RNA45SN1_45S_50ntFlanks | 5421 | 5445 | 25 | CGATTGGATGGTTTAGTGAGGCCCT | 1271.4 | -14.7 | 1.5 | -9.5 | 1.3E-21 | 5.8E-21 | BT20/MB468 | NUC |
| >hg38_21_+_8433172_8446622_RNA45SN1_45S_50ntFlanks@12654.12675.22\|TATGACTGAACGCCTCTAAGTC | >hg38_21_+_8433172_8446622_RNA45SN1_45S_50ntFlanks | 12654 | 12675 | 22 | TATGACTGAACGCCTCTAAGTC | 840.2 | -13.6 | 1.4 | -9.7 | 2.4E-22 | 5.8E-21 | MB231/MB468 | MP |
| >hg38_21_+_8433172_8446622_RNA45SN1_45S_50ntFlanks@7986.8014.29\|ATCAGACGTGGCGACCCGCTGAATTTAAG | >hg38_21_+_8433172_8446622_RNA45SN1_45S_50ntFlanks | 7986 | 8014 | 29 | ATCAGACGTGGCGACCCGCTGAATTTAAG | 679.6 | -13.3 | 1.4 | -9.7 | 2.4E-22 | 5.9E-21 | MB231/MB468 | MP |
| >hg38_21_+_8433172_8446622_RNA45SN1_45S_50ntFlanks@11575.11601.27\|ACTTAGAACTGGTGCGGACCAGGGGAA | >hg38_21_+_8433172_8446622_RNA45SN1_45S_50ntFlanks | 11575 | 11601 | 27 | ACTTAGAACTGGTGCGGACCAGGGGAA | 1454.3 | -14.4 | 1.5 | -9.7 | 2.4E-22 | 5.9E-21 | MB231/MB468 | MP |
| >hg38_21_+_8433172_8446622_RNA45SN1_45S_50ntFlanks@8381.8400.20\|AAGAGGGCGTGAAACCGTTA | >hg38_21_+_8433172_8446622_RNA45SN1_45S_50ntFlanks | 8381 | 8400 | 20 | AAGAGGGCGTGAAACCGTTA | 876.1 | -7.8 | 0.8 | -9.7 | 1.9E-22 | 5.9E-21 | MB231/MB468 | CYTO |
| >MI0000061\|hsa-let-7a-2&WithFlank&11\|-\|122146516\|122146599@11.31.21(+1A);MI0000062\|hsa-let-7a-3&WithFlank&22\|+\|46112743\|46112828@10.30.21(+1A);MI0000060\|hsa-let-7a-1&WithFlank&9\|+\|94175951\|94176042@12.32.21(+1A)&[MIMAT0000062&hsa-let-7a-5p&offsets\|0\|-1(+1A);m-6&11\|-\|122146568\|122146589&offsets\|0\|-1(+1A)];[MIMAT0000062_1&hsa-let-7a-5p&offsets\|0\|-1(+1A);m-5&22\|+\|46112752\|46112773&offsets\|0\|-1(+1A)];[MIMAT0000062_2&hsa-let-7a-5p&offsets\|0\|-1(+1A);m-4&9\|+\|94175962\|94175983&offsets\|0\|-1(+1A)]\|TGAGGTAGTAGGTTGTATAGTA | >MI0000061\|hsa-let-7a-2&WithFlank&11\|-\|122146516\|122146599 | 11 | 31 | 21(+1A) | TGAGGTAGTAGGTTGTATAGTA | 6614.0 | 8.2 | 0.9 | 9.6 | 6.8E-22 | 5.9E-21 | BT20/MB468 | CYTO |
| >MI0000809\|hsa-mir-151a&WithFlank&8\|-\|140732558\|140732659@51.72.22(+1U)&[MIMAT0000757&hsa-miR-151a-3p&offsets\|-2\|-1(+1U);m-30&8\|-\|140732587\|140732607&offsets\|-2\|-1(+1U)]\|TACTAGACTGAAGCTCCTTGAGT | >MI0000809\|hsa-mir-151a&WithFlank&8\|-\|140732558\|140732659 | 51 | 72 | 22(+1U) | TACTAGACTGAAGCTCCTTGAGT | 799.0 | 8.2 | 0.9 | 9.6 | 6.8E-22 | 6.0E-21 | BT20/MB468 | CYTO |
| >MI0001445\|hsa-mir-423&WithFlank&17\|+\|30117073\|30117178@23.46.24&[MIMAT0004748&hsa-miR-423-5p&offsets\|0\|+1;m-141&17\|+\|30117095\|30117117&offsets\|0\|+1]\|TGAGGGGCAGAGAGCGAGACTTTT | >MI0001445\|hsa-mir-423&WithFlank&17\|+\|30117073\|30117178 | 23 | 46 | 24 | TGAGGGGCAGAGAGCGAGACTTTT | 1642.0 | 8.0 | 0.8 | 9.7 | 1.9E-22 | 6.0E-21 | MB231/MB468 | CYTO |
| >hg38_21_+_8433172_8446622_RNA45SN1_45S_50ntFlanks@9564.9597.34\|CTGACGTGCAAATCGGTCGTCCGACCTGGGTATA | >hg38_21_+_8433172_8446622_RNA45SN1_45S_50ntFlanks | 9564 | 9597 | 34 | CTGACGTGCAAATCGGTCGTCCGACCTGGGTATA | 13007.6 | -6.7 | 0.7 | -9.7 | 3.7E-22 | 6.0E-21 | MB231/MB468 | NUC |
| >hg38_21_+_8433172_8446622_RNA45SN1_45S_50ntFlanks@10689.10709.21\|GCGTCCGGTGAGCTCTCGCTG | >hg38_21_+_8433172_8446622_RNA45SN1_45S_50ntFlanks | 10689 | 10709 | 21 | GCGTCCGGTGAGCTCTCGCTG | 887.4 | -13.6 | 1.4 | -9.7 | 3.9E-22 | 6.0E-21 | MB231/MB468 | MITO |
| >hg38_21_+_8433172_8446622_RNA45SN1_45S_50ntFlanks@4760.4780.21\|TCAGATACCGTCGTAGTTCCG | >hg38_21_+_8433172_8446622_RNA45SN1_45S_50ntFlanks | 4760 | 4780 | 21 | TCAGATACCGTCGTAGTTCCG | 4755.9 | -11.8 | 1.2 | -9.7 | 3.9E-22 | 6.1E-21 | MB231/MB468 | MITO |
| >hg38_21_+_8433172_8446622_RNA45SN1_45S_50ntFlanks@11729.11749.21\|TAACTATGACTCTCTTAAGGT | >hg38_21_+_8433172_8446622_RNA45SN1_45S_50ntFlanks | 11729 | 11749 | 21 | TAACTATGACTCTCTTAAGGT | 5823.6 | -5.4 | 0.6 | -9.7 | 2.0E-22 | 6.1E-21 | MB231/MB468 | CYTO |
| >hg38_21_+_8433172_8446622_RNA45SN1_45S_50ntFlanks@4723.4758.36\|CATTAATCAAGAACGAAAGTCGGAGGTTCGAAGACG | >hg38_21_+_8433172_8446622_RNA45SN1_45S_50ntFlanks | 4723 | 4758 | 36 | CATTAATCAAGAACGAAAGTCGGAGGTTCGAAGACG | 18122.1 | -11.3 | 1.2 | -9.5 | 1.4E-21 | 6.2E-21 | BT20/MB468 | NUC |
| >hg38_21_+_8433172_8446622_RNA45SN1_45S_50ntFlanks@10128.10149.22\|CGGCGGCGGCGGCGGCGGCGGG | >hg38_21_+_8433172_8446622_RNA45SN1_45S_50ntFlanks | 10128 | 10149 | 22 | CGGCGGCGGCGGCGGCGGCGGG | 2882.3 | -15.3 | 1.6 | -9.7 | 4.1E-22 | 6.2E-21 | MB231/MB468 | MITO |
| >hg38_21_+_8433172_8446622_RNA45SN1_45S_50ntFlanks@8845.8866.22\|CGGCGGCGGCGGCGGCGGCGGG | >hg38_21_+_8433172_8446622_RNA45SN1_45S_50ntFlanks | 8845 | 8866 | 22 | CGGCGGCGGCGGCGGCGGCGGG | 2882.3 | -15.3 | 1.6 | -9.7 | 4.1E-22 | 6.2E-21 | MB231/MB468 | MITO |
| >hg38_21_+_8433172_8446622_RNA45SN1_45S_50ntFlanks@6671.6698.28\|CTCGGCTCGTGCGTCGATGAAGAACGCA | >hg38_21_+_8433172_8446622_RNA45SN1_45S_50ntFlanks | 6671 | 6698 | 28 | CTCGGCTCGTGCGTCGATGAAGAACGCA | 1547.6 | -14.5 | 1.5 | -9.7 | 2.5E-22 | 6.2E-21 | MB231/MB468 | MP |
| >hg38_21_+_8433172_8446622_RNA45SN1_45S_50ntFlanks@11027.11051.25\|CTCTCTCTCTCTCTCCCCCGCTCCC | >hg38_21_+_8433172_8446622_RNA45SN1_45S_50ntFlanks | 11027 | 11051 | 25 | CTCTCTCTCTCTCTCCCCCGCTCCC | 1316.6 | -14.1 | 1.5 | -9.7 | 4.1E-22 | 6.3E-21 | MB231/MB468 | MITO |
| >hg38_21_+_8433172_8446622_RNA45SN1_45S_50ntFlanks@10411.10428.18\|CAGTCGGTCCTGAGAGAT | >hg38_21_+_8433172_8446622_RNA45SN1_45S_50ntFlanks | 10411 | 10428 | 18 | CAGTCGGTCCTGAGAGAT | 1739.1 | -14.6 | 1.5 | -9.7 | 2.6E-22 | 6.3E-21 | MB231/MB468 | MP |
| >hg38_21_+_8433172_8446622_RNA45SN1_45S_50ntFlanks@6731.6770.40\|CACATTGATCATCGACACTTCGAACGCACTTGCGGCCCCG | >hg38_21_+_8433172_8446622_RNA45SN1_45S_50ntFlanks | 6731 | 6770 | 40 | CACATTGATCATCGACACTTCGAACGCACTTGCGGCCCCG | 4216.2 | -9.2 | 0.9 | -9.7 | 4.1E-22 | 6.3E-21 | MB231/MB468 | MITO |
| >hg38_21_+_8433172_8446622_RNA45SN1_45S_50ntFlanks@485.510.26\|GATGTGGTGACGTCGTGCTCTCCCGG | >hg38_21_+_8433172_8446622_RNA45SN1_45S_50ntFlanks | 485 | 510 | 26 | GATGTGGTGACGTCGTGCTCTCCCGG | 428.6 | 11.6 | 1.2 | 9.5 | 1.5E-21 | 6.3E-21 | BT20/MB468 | NUC |
| >MI0000809\|hsa-mir-151a&WithFlank&8\|-\|140732558\|140732659@53.74.22(+2U)&[MIMAT0000757&hsa-miR-151a-3p&offsets\|0\|+1(+2U);m-30&8\|-\|140732587\|140732607&offsets\|0\|+1(+2U)]\|CTAGACTGAAGCTCCTTGAGGATT | >MI0000809\|hsa-mir-151a&WithFlank&8\|-\|140732558\|140732659 | 53 | 74 | 22(+2U) | CTAGACTGAAGCTCCTTGAGGATT | 693.0 | 6.4 | 0.7 | 9.7 | 3.9E-22 | 6.3E-21 | MB231/MB468 | NUC |
| >hg38_21_+_8433172_8446622_RNA45SN1_45S_50ntFlanks@4588.4616.29\|CTATTTTGTTGGTTTTCGGAACTGAGGCC | >hg38_21_+_8433172_8446622_RNA45SN1_45S_50ntFlanks | 4588 | 4616 | 29 | CTATTTTGTTGGTTTTCGGAACTGAGGCC | 692.9 | -4.5 | 0.5 | -9.6 | 5.4E-22 | 6.4E-21 | BT20/MB468 | MITO |
| >MI0000086\|hsa-mir-28&WithFlank&3\|+\|188688775\|188688872@60.82.23(+1A)&[MIMAT0004502&hsa-miR-28-3p&offsets\|0\|+1(+1A);m-28&3\|+\|188688834\|188688855&offsets\|0\|+1(+1A)]\|CACTAGATTGTGAGCTCCTGGAGA | >MI0000086\|hsa-mir-28&WithFlank&3\|+\|188688775\|188688872 | 60 | 82 | 23(+1A) | CACTAGATTGTGAGCTCCTGGAGA | 326.6 | 12.6 | 1.3 | 9.6 | 7.4E-22 | 6.5E-21 | BT20/MB468 | CYTO |
| >MI0000816\|hsa-mir-335&WithFlank&7\|+\|130496105\|130496210@58.78.21(+1U)&[MIMAT0004703&hsa-miR-335-3p&offsets\|0\|-1(+1U);m-159&7\|+\|130496162\|130496183&offsets\|0\|-1(+1U)]\|TTTTTCATTATTGCTCCTGACT | >MI0000816\|hsa-mir-335&WithFlank&7\|+\|130496105\|130496210 | 58 | 78 | 21(+1U) | TTTTTCATTATTGCTCCTGACT | 1490.6 | 7.6 | 0.8 | 9.6 | 7.4E-22 | 6.5E-21 | BT20/MB468 | CYTO |
| >hg38_21_+_8433172_8446622_RNA45SN1_45S_50ntFlanks@5472.5520.49\|ACGGCCCTGGCGGAGCGCTGAGAAGACGGTCGAACTTGACTATCTAGAG | >hg38_21_+_8433172_8446622_RNA45SN1_45S_50ntFlanks | 5472 | 5520 | 49 | ACGGCCCTGGCGGAGCGCTGAGAAGACGGTCGAACTTGACTATCTAGAG | 250.9 | 12.0 | 1.2 | 9.7 | 2.1E-22 | 6.5E-21 | MB231/MB468 | CYTO |
| >hg38_21_+_8433172_8446622_RNA45SN1_45S_50ntFlanks@11665.11706.42\|TTCTGCCCAGTGCTCTGAATGTCAAAGTGAAGAAATTCAATG | >hg38_21_+_8433172_8446622_RNA45SN1_45S_50ntFlanks | 11665 | 11706 | 42 | TTCTGCCCAGTGCTCTGAATGTCAAAGTGAAGAAATTCAATG | 683.5 | 13.4 | 1.4 | 9.7 | 2.1E-22 | 6.5E-21 | MB231/MB468 | CYTO |
| >hg38_21_+_8433172_8446622_RNA45SN1_45S_50ntFlanks@11317.11336.20\|ACCCCACGTCTCGTCGCGCG | >hg38_21_+_8433172_8446622_RNA45SN1_45S_50ntFlanks | 11317 | 11336 | 20 | ACCCCACGTCTCGTCGCGCG | 2800.4 | -15.2 | 1.6 | -9.7 | 4.3E-22 | 6.5E-21 | MB231/MB468 | MITO |
| >hg38_21_+_8433172_8446622_RNA45SN1_45S_50ntFlanks@12936.12966.31\|AAACCATTCGTAGACGACCTGCTTCTGGGTC | >hg38_21_+_8433172_8446622_RNA45SN1_45S_50ntFlanks | 12936 | 12966 | 31 | AAACCATTCGTAGACGACCTGCTTCTGGGTC | 383.3 | -13.0 | 1.4 | -9.5 | 1.5E-21 | 6.7E-21 | BT20/MB468 | NUC |
| >MI0000478\|hsa-mir-149&WithFlank&2\|+\|240455995\|240456095@21.42.22(+1A)&[MIMAT0000450&hsa-miR-149-5p&offsets\|0\|-1(+1A);m-195&2\|+\|240456015\|240456037&offsets\|0\|-1(+1A)]\|TCTGGCTCCGTGTCTTCACTCCA | >MI0000478\|hsa-mir-149&WithFlank&2\|+\|240455995\|240456095 | 21 | 42 | 22(+1A) | TCTGGCTCCGTGTCTTCACTCCA | 1390.0 | 14.7 | 1.5 | 9.6 | 7.6E-22 | 6.7E-21 | BT20/MB468 | CYTO |
| >hg38_21_+_8433172_8446622_RNA45SN1_45S_50ntFlanks@4560.4579.20\|CTAGGAATAATGGAATAGGA | >hg38_21_+_8433172_8446622_RNA45SN1_45S_50ntFlanks | 4560 | 4579 | 20 | CTAGGAATAATGGAATAGGA | 458.0 | -13.3 | 1.4 | -9.5 | 1.6E-21 | 6.7E-21 | BT20/MB468 | NUC |
| >hg38_21_+_8433172_8446622_RNA45SN1_45S_50ntFlanks@9531.9573.43\|CAGAGGAAACTCTGGTGGAGGTCCGTAGCGGTCCTGACGTGCA | >hg38_21_+_8433172_8446622_RNA45SN1_45S_50ntFlanks | 9531 | 9573 | 43 | CAGAGGAAACTCTGGTGGAGGTCCGTAGCGGTCCTGACGTGCA | 178.9 | 11.5 | 1.2 | 9.7 | 2.2E-22 | 6.7E-21 | MB231/MB468 | CYTO |
| >hg38_21_+_8433172_8446622_RNA45SN1_45S_50ntFlanks@10783.10816.34\|GTGAACAGCCTCTGGCATGTTGGAACAATGTAGG | >hg38_21_+_8433172_8446622_RNA45SN1_45S_50ntFlanks | 10783 | 10816 | 34 | GTGAACAGCCTCTGGCATGTTGGAACAATGTAGG | 392.9 | -13.0 | 1.4 | -9.5 | 1.6E-21 | 6.7E-21 | BT20/MB468 | NUC |
| >hg38_21_+_8433172_8446622_RNA45SN1_45S_50ntFlanks@12539.12569.31\|ATGATGTGTTGTTGCCATGGTAATCCTGCTC | >hg38_21_+_8433172_8446622_RNA45SN1_45S_50ntFlanks | 12539 | 12569 | 31 | ATGATGTGTTGTTGCCATGGTAATCCTGCTC | 8208.0 | -13.2 | 1.4 | -9.5 | 1.6E-21 | 6.7E-21 | BT20/MB468 | NUC |
| >hg38_21_+_8433172_8446622_RNA45SN1_45S_50ntFlanks@8356.8379.24\|GAAAAGAACTTTGAAGAGAGAGTT | >hg38_21_+_8433172_8446622_RNA45SN1_45S_50ntFlanks | 8356 | 8379 | 24 | GAAAAGAACTTTGAAGAGAGAGTT | 7037.6 | -7.7 | 0.8 | -9.7 | 4.5E-22 | 6.8E-21 | MB231/MB468 | MITO |

| | | | | | | | | | | | | | |
|---|---|---|---|---|---|---|---|---|---|---|---|---|---|
| >am_trna13_LysCTT_14_-_58706613_58706685@31.51.21,trna2_LysCTT_15_+_79152904_79152976@31.51.21,trna4_LysCTT_18_-_43669271_43669343@31.51.21\|ACTCTTAATCCCAGGGTCGTG | >am_trna13_LysCTT_14_-_58706613_58706685 | 31 | 51 | 21 | ACTCTTAATCCCAGGGTCGTG | 353.1 | -12.9 | 1.4 | -9.5 | 1.6E-21 | 6.8E-21 | BT20/MB468 | NUC |
| >hg38_21_+_8433172_8446622_RNA45SN1_45S_50ntFlanks@13005.13025.21\|TCTATTGAAAGTCAGCCCTCG | >hg38_21_+_8433172_8446622_RNA45SN1_45S_50ntFlanks | 13005 | 13025 | 21 | TCTATTGAAAGTCAGCCCTCG | 414.0 | -13.1 | 1.4 | -9.5 | 1.6E-21 | 6.8E-21 | BT20/MB468 | NUC |
| >hg38_21_+_8433172_8446622_RNA45SN1_45S_50ntFlanks@12435.12455.21\|ATCATTGTGAAGCAGAATTCA | >hg38_21_+_8433172_8446622_RNA45SN1_45S_50ntFlanks | 12435 | 12455 | 21 | ATCATTGTGAAGCAGAATTCA | 1178.5 | -14.0 | 1.4 | -9.7 | 4.6E-22 | 6.9E-21 | MB231/MB468 | MITO |
| >hg38_21_+_8433172_8446622_RNA45SN1_45S_50ntFlanks@11766.11786.21\|ATCTAATTAGTGACGCGCATG | >hg38_21_+_8433172_8446622_RNA45SN1_45S_50ntFlanks | 11766 | 11786 | 21 | ATCTAATTAGTGACGCGCATG | 379.1 | -13.0 | 1.4 | -9.5 | 1.6E-21 | 6.9E-21 | BT20/MB468 | NUC |
| >hg38_21_+_8433172_8446622_RNA45SN1_45S_50ntFlanks@10731.10755.25\|AGGGTGTAAATCTCGCGCCGGGCCG | >hg38_21_+_8433172_8446622_RNA45SN1_45S_50ntFlanks | 10731 | 10755 | 25 | AGGGTGTAAATCTCGCGCCGGGCCG | 697.4 | -13.3 | 1.4 | -9.7 | 2.8E-22 | 6.9E-21 | MB231/MB468 | MP |
| >hg38_21_+_8433172_8446622_RNA45SN1_45S_50ntFlanks@11669.11705.37\|GCCCAGTGCTCTGAATGTCAAAGTGAAGAAATTCAAT | >hg38_21_+_8433172_8446622_RNA45SN1_45S_50ntFlanks | 11669 | 11705 | 37 | GCCCAGTGCTCTGAATGTCAAAGTGAAGAAATTCAAT | 384.5 | 12.6 | 1.3 | 9.7 | 2.3E-22 | 6.9E-21 | MB231/MB468 | CYTO |
| >hg38_21_+_8433172_8446622_RNA45SN1_45S_50ntFlanks@10761.10793.33\|ATATCCGCAGCAGGTCTCCAAGGTGAACAGCCT | >hg38_21_+_8433172_8446622_RNA45SN1_45S_50ntFlanks | 10761 | 10793 | 33 | ATATCCGCAGCAGGTCTCCAAGGTGAACAGCCT | 387.8 | -13.0 | 1.4 | -9.5 | 1.6E-21 | 6.9E-21 | BT20/MB468 | NUC |
| >hg38_21_+_8433172_8446622_RNA45SN1_45S_50ntFlanks@11593.11612.20\|CCAGGGGAATCCGACTGTTT | >hg38_21_+_8433172_8446622_RNA45SN1_45S_50ntFlanks | 11593 | 11612 | 20 | CCAGGGGAATCCGACTGTTT | 1889.7 | -7.2 | 0.7 | -9.7 | 4.6E-22 | 6.9E-21 | MB231/MB468 | MITO |
| >hg38_21_+_8433172_8446622_RNA45SN1_45S_50ntFlanks@11874.11900.27\|AGAAGACCCTGTTGAGCTTGACTCTAG | >hg38_21_+_8433172_8446622_RNA45SN1_45S_50ntFlanks | 11874 | 11900 | 27 | AGAAGACCCTGTTGAGCTTGACTCTAG | 816.2 | -13.5 | 1.4 | -9.7 | 2.9E-22 | 7.0E-21 | MB231/MB468 | MP |
| >hg38_21_+_8433172_8446622_RNA45SN1_45S_50ntFlanks@12456.12481.26\|CCAAGCGTTGGATTGTTCACCCACTA | >hg38_21_+_8433172_8446622_RNA45SN1_45S_50ntFlanks | 12456 | 12481 | 26 | CCAAGCGTTGGATTGTTCACCCACTA | 1640.7 | -15.1 | 1.6 | -9.5 | 1.6E-21 | 7.0E-21 | BT20/MB468 | NUC |
| >hg38_21_+_8433172_8446622_RNA45SN1_45S_50ntFlanks@3882.3899.18\|CCGACGGGCGCTGACCCC | >hg38_21_+_8433172_8446622_RNA45SN1_45S_50ntFlanks | 3882 | 3899 | 18 | CCGACGGGCGCTGACCCC | 5327.6 | -12.4 | 1.3 | -9.7 | 4.7E-22 | 7.0E-21 | MB231/MB468 | MITO |
| >hg38_21_+_8433172_8446622_RNA45SN1_45S_50ntFlanks@9621.9650.30\|TCTAGTAGCTGGTTCCCTCCGAAGTTTCCC | >hg38_21_+_8433172_8446622_RNA45SN1_45S_50ntFlanks | 9621 | 9650 | 30 | TCTAGTAGCTGGTTCCCTCCGAAGTTTCCC | 901.5 | -13.6 | 1.4 | -9.7 | 4.7E-22 | 7.0E-21 | MB231/MB468 | MITO |
| >MI0000102\|hsa-mir-100&WithFlank&11\|-\|122152223\|122152314@19.36.18&[MIMAT0000098&hsa-miR-100-5p&offsets\|0\|-4;m-27&11\|-\|122152276\|122152296&offsets\|0\|-3]\|AACCCGTAGATCCGAACT | >MI0000102\|hsa-mir-100&WithFlank&11\|-\|122152223\|122152314 | 19 | 36 | 18 | AACCCGTAGATCCGAACT | 375.3 | 12.4 | 1.3 | 9.7 | 4.4E-22 | 7.0E-21 | MB231/MB468 | NUC |
| >hg38_21_+_8433172_8446622_RNA45SN1_45S_50ntFlanks@6698.6718.21\|AGCTAGCTGCGAGAATTAATG | >hg38_21_+_8433172_8446622_RNA45SN1_45S_50ntFlanks | 6698 | 6718 | 21 | AGCTAGCTGCGAGAATTAATG | 12740.7 | -6.3 | 0.7 | -9.7 | 4.4E-22 | 7.0E-21 | MB231/MB468 | NUC |
| >MI0000068\|hsa-let-7f-2&WithFlank&X\|-\|53557186\|53557280@14.36.23&[MIMAT0000067_1&hsa-let-7f-5p&offsets\|0\|+1;m-19&X\|-\|53557246\|53557267&offsets\|0\|+1]\|TGAGGTAGTAGATTGTATAGTTT | >MI0000068\|hsa-let-7f-2&WithFlank&X\|-\|53557186\|53557280 | 14 | 36 | 23 | TGAGGTAGTAGATTGTATAGTTT | 3018.9 | 4.4 | 0.5 | 9.7 | 4.4E-22 | 7.0E-21 | MB231/MB468 | NUC |
| >hg38_21_+_8433172_8446622_RNA45SN1_45S_50ntFlanks@4721.4765.45\|TTCATTAATCAAGAACGAAAGTCGGAGGTTCGAAGACGATCAGAT | >hg38_21_+_8433172_8446622_RNA45SN1_45S_50ntFlanks | 4721 | 4765 | 45 | TTCATTAATCAAGAACGAAAGTCGGAGGTTCGAAGACGATCAGAT | 228.4 | 11.8 | 1.2 | 9.7 | 2.3E-22 | 7.1E-21 | MB231/MB468 | CYTO |
| >MI0000298\|hsa-mir-221&WithFlank&X\|-\|45746151\|45746272@71.92.22(+2U)&[MIMAT0000278&hsa-miR-221-3p&offsets\|0\|-1(+2U);m-91&X\|-\|45746181\|45746202&offsets\|0\|0(+2U)]\|AGCTACATTGTCTGCTGGGTTTTT | >MI0000298\|hsa-mir-221&WithFlank&X\|-\|45746151\|45746272 | 71 | 92 | 22(+2U) | AGCTACATTGTCTGCTGGGTTTTT | 394.7 | 6.9 | 0.7 | 9.7 | 4.4E-22 | 7.1E-21 | MB231/MB468 | NUC |
| >hg38_21_+_8433172_8446622_RNA45SN1_45S_50ntFlanks@6721.6744.24\|AATTGCAGGACACATTGATCATCG | >hg38_21_+_8433172_8446622_RNA45SN1_45S_50ntFlanks | 6721 | 6744 | 24 | AATTGCAGGACACATTGATCATCG | 6571.1 | -10.0 | 1.0 | -9.7 | 2.9E-22 | 7.1E-21 | MB231/MB468 | MP |
| >hg38_21_+_8433172_8446622_RNA45SN1_45S_50ntFlanks@6699.6718.20\|GCTAGCTGCGAGAATTAATG | >hg38_21_+_8433172_8446622_RNA45SN1_45S_50ntFlanks | 6699 | 6718 | 20 | GCTAGCTGCGAGAATTAATG | 26040.1 | -6.9 | 0.7 | -9.7 | 4.5E-22 | 7.1E-21 | MB231/MB468 | NUC |
| >hg38_21_+_8433172_8446622_RNA45SN1_45S_50ntFlanks@12458.12476.19\|AAGCGTTGGATTGTTCACC | >hg38_21_+_8433172_8446622_RNA45SN1_45S_50ntFlanks | 12458 | 12476 | 19 | AAGCGTTGGATTGTTCACC | 3990.8 | -5.0 | 0.5 | -9.5 | 1.7E-21 | 7.1E-21 | BT20/MB468 | NUC |
| >MI0000086\|hsa-mir-28&WithFlank&3\|+\|188688775\|188688872@60.80.21&[MIMAT0004502&hsa-miR-28-3p&offsets\|0\|-1;m-28&3\|+\|188688834\|188688855&offsets\|0\|-1]\|CACTAGATTGTGAGCTCCTGG | >MI0000086\|hsa-mir-28&WithFlank&3\|+\|188688775\|188688872 | 60 | 80 | 21 | CACTAGATTGTGAGCTCCTGG | 892.9 | 6.6 | 0.7 | 9.7 | 4.5E-22 | 7.2E-21 | MB231/MB468 | NUC |
| >am_trna146_GlnCTG_6_-_27515531_27515602@1.29.29,trna1_GlnCTG_6_+_18836402_18836473@1.29.29,trna3_GlnCTG_17_+_8023070_8023141@1.29.29,trna42_GlnCTG_6_+_27263212_27263283@1.29.29,trna49_GlnCTG_6_+_27487308_27487379@1.29.29,trna7_GlnCTG_15_-_66161400_66161471@1.29.29,trna99_GlnCTG_6_-_28909378_28909449@1.29.29\|GGTTCCATGGTGTAATGGTTAGCACTCTG | >am_trna146_GlnCTG_6_-_27515531_27515602 | 1 | 29 | 29 | GGTTCCATGGTGTAATGGTTAGCACTCTG | 19227.8 | -6.6 | 0.7 | -9.7 | 2.4E-22 | 7.2E-21 | MB231/MB468 | CYTO |
| >hg38_21_+_8433172_8446622_RNA45SN1_45S_50ntFlanks@10396.10435.40\|CAGTTGAACATGGGTCAGTCGGTCCTGAGAGATGGGCGAG | >hg38_21_+_8433172_8446622_RNA45SN1_45S_50ntFlanks | 10396 | 10435 | 40 | CAGTTGAACATGGGTCAGTCGGTCCTGAGAGATGGGCGAG | 705.1 | -13.9 | 1.5 | -9.5 | 1.7E-21 | 7.2E-21 | BT20/MB468 | NUC |
| >hg38_21_+_8433172_8446622_RNA45SN1_45S_50ntFlanks@9546.9572.27\|TGGAGGTCCGTAGCGGTCCTGACGTGC | >hg38_21_+_8433172_8446622_RNA45SN1_45S_50ntFlanks | 9546 | 9572 | 27 | TGGAGGTCCGTAGCGGTCCTGACGTGC | 1085.7 | 5.6 | 0.6 | 9.6 | 6.2E-22 | 7.3E-21 | BT20/MB468 | MITO |
| >hg38_21_+_8433172_8446622_RNA45SN1_45S_50ntFlanks@6719.6743.25\|TGAATTGCAGGACACATTGATCATC | >hg38_21_+_8433172_8446622_RNA45SN1_45S_50ntFlanks | 6719 | 6743 | 25 | TGAATTGCAGGACACATTGATCATC | 11027.0 | -7.1 | 0.7 | -9.7 | 4.6E-22 | 7.3E-21 | MB231/MB468 | NUC |
| >trnaMT_GluTTC_MT_-_14674_14742@36.72.37,trnalookalike8_GluTTC_5_-_93905172_93905240@36.72.37\|ATCATTGGTCGTGGTTGTAGTCCGTGCGAGAATACCA | >trnaMT_GluTTC_MT_-_14674_14742 | 36 | 72 | 37 | ATCATTGGTCGTGGTTGTAGTCCGTGCGAGAATACCA | 386.4 | -13.0 | 1.4 | -9.5 | 1.7E-21 | 7.4E-21 | BT20/MB468 | NUC |
| >hg38_21_+_8433172_8446622_RNA45SN1_45S_50ntFlanks@8114.8141.28\|GCGCGGGACATGTGGCGTACGGAAGACC | >hg38_21_+_8433172_8446622_RNA45SN1_45S_50ntFlanks | 8114 | 8141 | 28 | GCGCGGGACATGTGGCGTACGGAAGACC | 7131.5 | -12.3 | 1.3 | -9.7 | 3.1E-22 | 7.5E-21 | MB231/MB468 | MP |
| >hg38_21_+_8433172_8446622_RNA45SN1_45S_50ntFlanks@9738.9760.23\|GAAACGATCTCAACCTATTCTCA | >hg38_21_+_8433172_8446622_RNA45SN1_45S_50ntFlanks | 9738 | 9760 | 23 | GAAACGATCTCAACCTATTCTCA | 326.2 | -12.8 | 1.3 | -9.5 | 1.8E-21 | 7.5E-21 | BT20/MB468 | NUC |
| >trna10_AlaCGC_6_+_26553731_26553802@58.75.18,trna119_AlaCGC_6_-_28641613_28641684@58.75.18,trna13_AlaCGC_2_+_157257281_157257352@58.75.18,trna13_AlaTGC_12_-_125406301_125406372@58.75.18,trna66_AlaTGC_6_+_28611222_28611293@58.75.18,trna8_AlaTGC_12_+_125424512_125424583@58.75.18,trna8_AlaTGC_5_+_180633868_180633939@58.75.18\|TCCCCGGCATCTCCACCA | >trna10_AlaCGC_6_+_26553731_26553802 | 58 | 75 | 18 | TCCCCGGCATCTCCACCA | 26313.3 | 8.4 | 0.9 | 9.6 | 8.7E-22 | 7.6E-21 | BT20/MB468 | CYTO |
| >hg38_21_+_8433172_8446622_RNA45SN1_45S_50ntFlanks@11710.11728.19\|CGCGGGTAAACGGCGGGAG | >hg38_21_+_8433172_8446622_RNA45SN1_45S_50ntFlanks | 11710 | 11728 | 19 | CGCGGGTAAACGGCGGGAG | 940.3 | -13.7 | 1.4 | -9.7 | 3.1E-22 | 7.6E-21 | MB231/MB468 | MP |
| >hg38_21_+_8433172_8446622_RNA45SN1_45S_50ntFlanks@10128.10149.22\|CGGCGGCGGCGGCGGCGGCGGG | >hg38_21_+_8433172_8446622_RNA45SN1_45S_50ntFlanks | 10128 | 10149 | 22 | CGGCGGCGGCGGCGGCGGCGGG | 1722.0 | -14.6 | 1.5 | -9.7 | 3.2E-22 | 7.7E-21 | MB231/MB468 | MP |
| >hg38_21_+_8433172_8446622_RNA45SN1_45S_50ntFlanks@8845.8866.22\|CGGCGGCGGCGGCGGCGGCGGG | >hg38_21_+_8433172_8446622_RNA45SN1_45S_50ntFlanks | 8845 | 8866 | 22 | CGGCGGCGGCGGCGGCGGCGGG | 1722.0 | -14.6 | 1.5 | -9.7 | 3.2E-22 | 7.7E-21 | MB231/MB468 | MP |
| >hg38_21_+_8433172_8446622_RNA45SN1_45S_50ntFlanks@7980.8008.29\|CCTCAGATCAGACGTGGCGACCCGCTGAA | >hg38_21_+_8433172_8446622_RNA45SN1_45S_50ntFlanks | 7980 | 8008 | 29 | CCTCAGATCAGACGTGGCGACCCGCTGAA | 4072.9 | -6.6 | 0.7 | -9.6 | 5.2E-22 | 7.8E-21 | MB231/MB468 | MITO |
| >hg38_21_+_8433172_8446622_RNA45SN1_45S_50ntFlanks@3939.3958.20\|AACCCGGTCAGCCCCTCTCC | >hg38_21_+_8433172_8446622_RNA45SN1_45S_50ntFlanks | 3939 | 3958 | 20 | AACCCGGTCAGCCCCTCTCC | 856.5 | -13.5 | 1.4 | -9.6 | 5.3E-22 | 7.9E-21 | MB231/MB468 | MITO |
| >hg38_21_+_8433172_8446622_RNA45SN1_45S_50ntFlanks@10866.10894.29\|CTAAGGGCTGGGTCGGTCGGGCTGGGGCG | >hg38_21_+_8433172_8446622_RNA45SN1_45S_50ntFlanks | 10866 | 10894 | 29 | CTAAGGGCTGGGTCGGTCGGGCTGGGGCG | 1126.1 | -14.0 | 1.4 | -9.7 | 3.3E-22 | 7.9E-21 | MB231/MB468 | MP |
| >hg38_21_+_8433172_8446622_RNA45SN1_45S_50ntFlanks@12589.12614.26\|GTTCAGACATTTGGTGTATGTGCTTG | >hg38_21_+_8433172_8446622_RNA45SN1_45S_50ntFlanks | 12589 | 12614 | 26 | GTTCAGACATTTGGTGTATGTGCTTG | 948.7 | -13.8 | 1.4 | -9.7 | 3.3E-22 | 7.9E-21 | MB231/MB468 | MP |
| >hg38_21_+_8433172_8446622_RNA45SN1_45S_50ntFlanks@3854.3871.18\|ACTGTGGTAATTCTAGAG | >hg38_21_+_8433172_8446622_RNA45SN1_45S_50ntFlanks | 3854 | 3871 | 18 | ACTGTGGTAATTCTAGAG | 728.9 | -13.4 | 1.4 | -9.7 | 3.3E-22 | 8.0E-21 | MB231/MB468 | MP |
| >hg38_21_+_8433172_8446622_RNA45SN1_45S_50ntFlanks@12160.12180.21\|TTGACTGGGGCGGTACACCTG | >hg38_21_+_8433172_8446622_RNA45SN1_45S_50ntFlanks | 12160 | 12180 | 21 | TTGACTGGGGCGGTACACCTG | 570.0 | -13.0 | 1.3 | -9.7 | 3.3E-22 | 8.0E-21 | MB231/MB468 | MP |
| >hg38_21_+_8433172_8446622_RNA45SN1_45S_50ntFlanks@12416.12435.20\|TTCGATGTCGGCTCTTCCTA | >hg38_21_+_8433172_8446622_RNA45SN1_45S_50ntFlanks | 12416 | 12435 | 20 | TTCGATGTCGGCTCTTCCTA | 2123.8 | -6.8 | 0.7 | -9.5 | 1.9E-21 | 8.0E-21 | BT20/MB468 | NUC |
| >hg38_21_+_8433172_8446622_RNA45SN1_45S_50ntFlanks@11564.11603.40\|CCTAGCAGCCGACTTAGAACTGGTGCGGACCAGGGGAATC | >hg38_21_+_8433172_8446622_RNA45SN1_45S_50ntFlanks | 11564 | 11603 | 40 | CCTAGCAGCCGACTTAGAACTGGTGCGGACCAGGGGAATC | 16388.0 | -10.7 | 1.1 | -9.6 | 5.4E-22 | 8.0E-21 | MB231/MB468 | MITO |
| >hg38_21_+_8433172_8446622_RNA45SN1_45S_50ntFlanks@11703.11728.26\|AATGAAGCGCGGGTAAACGGCGGGAG | >hg38_21_+_8433172_8446622_RNA45SN1_45S_50ntFlanks | 11703 | 11728 | 26 | AATGAAGCGCGGGTAAACGGCGGGAG | 3834.4 | -11.3 | 1.2 | -9.6 | 5.4E-22 | 8.0E-21 | MB231/MB468 | MITO |
| >hg38_21_+_8433172_8446622_RNA45SN1_45S_50ntFlanks@10794.10822.29\|CTGGCATGTTGGAACAATGTAGGTAAGGG | >hg38_21_+_8433172_8446622_RNA45SN1_45S_50ntFlanks | 10794 | 10822 | 29 | CTGGCATGTTGGAACAATGTAGGTAAGGG | 746.2 | -13.4 | 1.4 | -9.7 | 3.4E-22 | 8.1E-21 | MB231/MB468 | MP |
| >hg38_21_+_8433172_8446622_RNA45SN1_45S_50ntFlanks@4726.4758.33\|TAATCAAGAACGAAAGTCGGAGGTTCGAAGACG | >hg38_21_+_8433172_8446622_RNA45SN1_45S_50ntFlanks | 4726 | 4758 | 33 | TAATCAAGAACGAAAGTCGGAGGTTCGAAGACG | 4338.4 | -7.7 | 0.8 | -9.7 | 3.4E-22 | 8.1E-21 | MB231/MB468 | MP |
| >hg38_21_+_8433172_8446622_RNA45SN1_45S_50ntFlanks@10794.10817.24\|CTGGCATGTTGGAACAATGTAGGT | >hg38_21_+_8433172_8446622_RNA45SN1_45S_50ntFlanks | 10794 | 10817 | 24 | CTGGCATGTTGGAACAATGTAGGT | 997.2 | -13.8 | 1.4 | -9.7 | 3.4E-22 | 8.1E-21 | MB231/MB468 | MP |
| >MI0000102\|hsa-mir-100&WithFlank&11\|-\|122152223\|122152314@19.38.20(+1A)&[MIMAT0000098&hsa-miR-100-5p&offsets\|0\|-2(+1A);m-27&11\|-\|122152276\|122152296&offsets\|0\|-1(+1A)]\|AACCCGTAGATCCGAACTTGA | >MI0000102\|hsa-mir-100&WithFlank&11\|-\|122152223\|122152314 | 19 | 38 | 20(+1A) | AACCCGTAGATCCGAACTTGA | 4068.5 | 15.2 | 1.6 | 9.6 | 5.5E-22 | 8.1E-21 | MB231/MB468 | MITO |
| >hg38_21_+_8433172_8446622_RNA45SN1_45S_50ntFlanks@11999.12043.45\|CCCTGCGGGCCGCCGGTGAAATACCACTACTCTGATCGTTTTTTC | >hg38_21_+_8433172_8446622_RNA45SN1_45S_50ntFlanks | 11999 | 12043 | 45 | CCCTGCGGGCCGCCGGTGAAATACCACTACTCTGATCGTTTTTTC | 4987.3 | -16.7 | 1.8 | -9.5 | 1.9E-21 | 8.2E-21 | BT20/MB468 | NUC |
| >hg38_21_+_8433172_8446622_RNA45SN1_45S_50ntFlanks@12416.12438.23\|TTCGATGTCGGCTCTTCCTATCA | >hg38_21_+_8433172_8446622_RNA45SN1_45S_50ntFlanks | 12416 | 12438 | 23 | TTCGATGTCGGCTCTTCCTATCA | 3936.5 | 7.1 | 0.7 | 9.7 | 2.8E-22 | 8.3E-21 | MB231/MB468 | CYTO |
| >hg38_21_+_8433172_8446622_RNA45SN1_45S_50ntFlanks@12044.12068.25\|ACTGACCCGGTGAGGCGGGGGGGCG | >hg38_21_+_8433172_8446622_RNA45SN1_45S_50ntFlanks | 12044 | 12068 | 25 | ACTGACCCGGTGAGGCGGGGGGGCG | 1308.3 | -14.2 | 1.5 | -9.7 | 3.5E-22 | 8.3E-21 | MB231/MB468 | MP |

| | | | | | | | | | | | | | |
|---|---|---|---|---|---|---|---|---|---|---|---|---|---|
| >hg38_21_+_8433172_8446622_RNA45SN1_45S_50ntFlanks@9574.9596.23\|AATCGGTCGTCCGACCTGGGTAT | >hg38_21_+_8433172_8446622_RNA45SN1_45S_50ntFlanks | 9574 | 9596 | 23 | AATCGGTCGTCCGACCTGGGTAT | 3322.4 | -6.8 | 0.7 | -9.6 | 9.5E-22 | 8.3E-21 | BT20/MB468 | CYTO |
| >hg38_21_+_8433172_8446622_RNA45SN1_45S_50ntFlanks@4728.4748.21\|ATCAAGAACGAAAGTCGGAGG | >hg38_21_+_8433172_8446622_RNA45SN1_45S_50ntFlanks | 4728 | 4748 | 21 | ATCAAGAACGAAAGTCGGAGG | 267.1 | -12.5 | 1.3 | -9.5 | 1.9E-21 | 8.3E-21 | BT20/MB468 | NUC |
| >MI0000440\|hsa-mir-27b&WithFlank&9\|+\|95085439\|95085547@67.87.21(+1U)&[MIMAT0000419&hsa-miR-27b-3p&offsets\|0\|0(+1U);m-39&9\|+\|95085505\|95085525&offsets\|0\|0(+1U)]\|TTCACAGTGGCTAAGTTCTGCT | >MI0000440\|hsa-mir-27b&WithFlank&9\|+\|95085439\|95085547 | 67 | 87 | 21(+1U) | TTCACAGTGGCTAAGTTCTGCT | 951.6 | 7.3 | 0.8 | 9.6 | 9.6E-22 | 8.3E-21 | BT20/MB468 | CYTO |
| >hg38_21_+_8433172_8446622_RNA45SN1_45S_50ntFlanks@9564.9596.33\|CTGACGTGCAAATCGGTCGTCCGACCTGGGTAT | >hg38_21_+_8433172_8446622_RNA45SN1_45S_50ntFlanks | 9564 | 9596 | 33 | CTGACGTGCAAATCGGTCGTCCGACCTGGGTAT | 7605.5 | -10.6 | 1.1 | -9.7 | 3.5E-22 | 8.3E-21 | MB231/MB468 | MP |
| >hg38_21_+_8433172_8446622_RNA45SN1_45S_50ntFlanks@6657.6681.25\|TTAGCGGTGGATCACTCGGCTCGTG | >hg38_21_+_8433172_8446622_RNA45SN1_45S_50ntFlanks | 6657 | 6681 | 25 | TTAGCGGTGGATCACTCGGCTCGTG | 1852.0 | -14.7 | 1.5 | -9.7 | 3.5E-22 | 8.4E-21 | MB231/MB468 | MP |
| >hg38_21_+_8433172_8446622_RNA45SN1_45S_50ntFlanks@10735.10769.35\|TGTAAATCTCGCGCCGGGCCGTACCCATATCCGCA | >hg38_21_+_8433172_8446622_RNA45SN1_45S_50ntFlanks | 10735 | 10769 | 35 | TGTAAATCTCGCGCCGGGCCGTACCCATATCCGCA | 432.7 | -13.2 | 1.4 | -9.5 | 2.0E-21 | 8.4E-21 | BT20/MB468 | NUC |
| >hg38_21_+_8433172_8446622_RNA45SN1_45S_50ntFlanks@11705.11728.24\|TGAAGCGCGGGTAAACGGCGGGAG | >hg38_21_+_8433172_8446622_RNA45SN1_45S_50ntFlanks | 11705 | 11728 | 24 | TGAAGCGCGGGTAAACGGCGGGAG | 1062.6 | -13.8 | 1.4 | -9.6 | 5.7E-22 | 8.4E-21 | MB231/MB468 | MITO |
| >hg38_21_+_8433172_8446622_RNA45SN1_45S_50ntFlanks@12934.12953.20\|CTAAACCATTCGTAGACGAC | >hg38_21_+_8433172_8446622_RNA45SN1_45S_50ntFlanks | 12934 | 12953 | 20 | CTAAACCATTCGTAGACGAC | 990.9 | -13.8 | 1.4 | -9.7 | 3.6E-22 | 8.4E-21 | MB231/MB468 | MP |
| >am_trna11_ProAGG_16_+_3241989_3242060@1.27.27,trna12_ProAGG_6_+_26555498_26555569@1.27.27,trna12_ProTGG_11_-_75946869_75946940@1.27.27,trna14_ProTGG_5_-_180615854_180615925@1.27.27,trna22_ProAGG_14_-_21081560_21081631@1.27.27,trna23_ProAGG_14_-_21077495_21077566@1.27.27,trna28_ProTGG_16_-_3234133_3234204@1.27.27,trna29_ProAGG_16_-_3232635_3232706@1.27.27,trna2_ProAGG_7_+_128423504_128423575@1.27.27,trna30_ProCGG_6_+_27059521_27059592@1.27.27,trna37_ProCGG_17_-_8126151_8126222@1.27.27,trna3_ProTGG_16_+_3208923_3208994@1.27.27,trna52_ProCGG_1_+_167683962_167684033@1.27.27,trna65_ProAGG_1_-_167684725_167684796@1.27.27,trna6_ProCGG_16_+_3222049_3222120@1.27.27,trna6_ProTGG_14_+_21152175_21152246@1.27.27,trna8_ProTGG_16_+_3238094_3238165@1.27.27,trna9_ProAGG_11_+_75946557_75946628@1.27.27,trna9_ProAGG_16_+_3239634_3239705@1.27.27\|GGCTCGTTGGTCTAGGGGTATGATTCT | >am_trna11_ProAGG_16_+_3241989_3242060 | 1 | 27 | 27 | GGCTCGTTGGTCTAGGGGTATGATTCT | 1372.1 | -14.8 | 1.6 | -9.5 | 2.0E-21 | 8.4E-21 | BT20/MB468 | NUC |
| >hg38_21_+_8433172_8446622_RNA45SN1_45S_50ntFlanks@12949.12990.42\|ACGACCTGCTTCTGGGTCGGGGTTTCGTACGTAGCAGAGCAG | >hg38_21_+_8433172_8446622_RNA45SN1_45S_50ntFlanks | 12949 | 12990 | 42 | ACGACCTGCTTCTGGGTCGGGGTTTCGTACGTAGCAGAGCAG | 1779.8 | -15.2 | 1.6 | -9.5 | 2.0E-21 | 8.4E-21 | BT20/MB468 | NUC |
| >hg38_21_+_8433172_8446622_RNA45SN1_45S_50ntFlanks@12455.12476.22\|ACCAAGCGTTGGATTGTTCACC | >hg38_21_+_8433172_8446622_RNA45SN1_45S_50ntFlanks | 12455 | 12476 | 22 | ACCAAGCGTTGGATTGTTCACC | 573.7 | -4.5 | 0.5 | -9.5 | 2.0E-21 | 8.4E-21 | BT20/MB468 | NUC |
| >MI0000077\|hsa-mir-21&WithFlank&17\|+\|59841260\|59841343@14.31.18&[MIMAT0000076&hsa-miR-21-5p&offsets\|0\|-4;m-1&17\|+\|59841273\|59841294&offsets\|0\|-4]\|TAGCTTATCAGACTGATG | >MI0000077\|hsa-mir-21&WithFlank&17\|+\|59841260\|59841343 | 14 | 31 | 18 | TAGCTTATCAGACTGATG | 1865.5 | 7.4 | 0.8 | 9.6 | 9.9E-22 | 8.6E-21 | BT20/MB468 | CYTO |
| >hg38_21_+_8433172_8446622_RNA45SN1_45S_50ntFlanks@10404.10429.26\|CATGGGTCAGTCGGTCCTGAGAGATG | >hg38_21_+_8433172_8446622_RNA45SN1_45S_50ntFlanks | 10404 | 10429 | 26 | CATGGGTCAGTCGGTCCTGAGAGATG | 1850.5 | -5.9 | 0.6 | -9.7 | 3.7E-22 | 8.6E-21 | MB231/MB468 | MP |
| >MI0000077\|hsa-mir-21&WithFlank&17\|+\|59841260\|59841343@14.37.24&[MIMAT0000076&hsa-miR-21-5p&offsets\|0\|+2;m-1&17\|+\|59841273\|59841294&offsets\|0\|+2]\|TAGCTTATCAGACTGATGTTGACT | >MI0000077\|hsa-mir-21&WithFlank&17\|+\|59841260\|59841343 | 14 | 37 | 24 | TAGCTTATCAGACTGATGTTGACT | 261332.8 | 7.8 | 0.8 | 9.6 | 1.0E-21 | 8.6E-21 | BT20/MB468 | CYTO |
| >hg38_21_+_8433172_8446622_RNA45SN1_45S_50ntFlanks@7986.8003.18\|ATCAGACGTGGCGACCCG | >hg38_21_+_8433172_8446622_RNA45SN1_45S_50ntFlanks | 7986 | 8003 | 18 | ATCAGACGTGGCGACCCG | 1343.0 | -12.5 | 1.3 | -9.6 | 7.3E-22 | 8.7E-21 | BT20/MB468 | MITO |
| >hg38_21_+_8433172_8446622_RNA45SN1_45S_50ntFlanks@485.516.32\|GATGTGGTGACGTCGTGCTCTCCCGGGCCGGG | >hg38_21_+_8433172_8446622_RNA45SN1_45S_50ntFlanks | 485 | 516 | 32 | GATGTGGTGACGTCGTGCTCTCCCGGGCCGGG | 562.4 | 8.6 | 0.9 | 9.5 | 2.1E-21 | 8.7E-21 | BT20/MB468 | NUC |
| >hg38_21_+_8433172_8446622_RNA45SN1_45S_50ntFlanks@4729.4758.30\|TCAAGAACGAAAGTCGGAGGTTCGAAGACG | >hg38_21_+_8433172_8446622_RNA45SN1_45S_50ntFlanks | 4729 | 4758 | 30 | TCAAGAACGAAAGTCGGAGGTTCGAAGACG | 2295.1 | -5.3 | 0.6 | -9.6 | 5.5E-22 | 8.7E-21 | MB231/MB468 | NUC |
| >MI0000061\|hsa-let-7a-2&WithFlank&11\|-\|122146516\|122146599@11.29.19;MI0000062\|hsa-let-7a-3&WithFlank&22\|+\|46112743\|46112828@10.28.19;MI0000060\|hsa-let-7a-1&WithFlank&9\|+\|94175951\|94176042@12.30.19&[MIMAT0000062&hsa-let-7a-5p&offsets\|0\|-3;m-6&11\|-\|122146568\|122146589&offsets\|0\|-3];[MIMAT0000062_1&hsa-let-7a-5p&offsets\|0\|-3;m-5&22\|+\|46112752\|46112773&offsets\|0\|-3];[MIMAT0000062_2&hsa-let-7a-5p&offsets\|0\|-3;m-4&9\|+\|94175962\|94175983&offsets\|0\|-3]\|TGAGGTAGTAGGTTGTATA | >MI0000061\|hsa-let-7a-2&WithFlank&11\|-\|122146516\|122146599 | 11 | 29 | 19 | TGAGGTAGTAGGTTGTATA | 5225.7 | 8.1 | 0.8 | 9.6 | 1.0E-21 | 8.7E-21 | BT20/MB468 | CYTO |
| >hg38_21_+_8433172_8446622_RNA45SN1_45S_50ntFlanks@4367.4389.23\|CAGTTAAAAAGCTCGTAGTTGGA | >hg38_21_+_8433172_8446622_RNA45SN1_45S_50ntFlanks | 4367 | 4389 | 23 | CAGTTAAAAAGCTCGTAGTTGGA | 1277.4 | -14.2 | 1.5 | -9.7 | 3.7E-22 | 8.8E-21 | MB231/MB468 | MP |
| >hg38_21_+_8433172_8446622_RNA45SN1_45S_50ntFlanks@3870.3898.29\|AGCTAATACATGCCGACGGGCGCTGACCC | >hg38_21_+_8433172_8446622_RNA45SN1_45S_50ntFlanks | 3870 | 3898 | 29 | AGCTAATACATGCCGACGGGCGCTGACCC | 1239.1 | -14.1 | 1.5 | -9.6 | 6.0E-22 | 8.8E-21 | MB231/MB468 | MITO |
| >hg38_21_+_8433172_8446622_RNA45SN1_45S_50ntFlanks@4449.4470.22\|CCTCTCGGCGCCCCCTCGATGC | >hg38_21_+_8433172_8446622_RNA45SN1_45S_50ntFlanks | 4449 | 4470 | 22 | CCTCTCGGCGCCCCCTCGATGC | 459.1 | -13.3 | 1.4 | -9.5 | 2.1E-21 | 8.9E-21 | BT20/MB468 | NUC |
| >hg38_21_+_8433172_8446622_RNA45SN1_45S_50ntFlanks@10805.10825.21\|GAACAATGTAGGTAAGGGAAG | >hg38_21_+_8433172_8446622_RNA45SN1_45S_50ntFlanks | 10805 | 10825 | 21 | GAACAATGTAGGTAAGGGAAG | 521.6 | -13.5 | 1.4 | -9.5 | 2.1E-21 | 8.9E-21 | BT20/MB468 | NUC |
| >trna10_AspGTC_12_-_125424193_125424264@36.75.40,trna12_AspGTC_12_-_125411891_125411962@36.75.40,trna144_AspGTC_6_-_27551236_27551307@36.75.40,trna38_AspGTC_17_-_8125556_8125627@36.75.40,trna45_AspGTC_6_+_27447453_27447524@36.75.40,trna48_AspGTC_6_+_27471523_27471594@36.75.40,trna4_AspGTC_12_+_96429799_96429870@36.75.40,trna69_AspGTC_1_-_161440205_161440276@36.75.40,trna72_AspGTC_1_-_161432824_161432895@36.75.40,trna75_AspGTC_1_-_161425414_161425485@36.75.40,trna78_AspGTC_1_-_161418033_161418104@36.75.40,trna81_AspGTC_1_-_161410615_161410686@36.75.40\|CACGCGGGAGACCGGGGTTCGATTCCCCGACGGGGAGCCA | >trna10_AspGTC_12_-_125424193_125424264 | 36 | 75 | 40 | CACGCGGGAGACCGGGGTTCGATTCCCCGACGGGGAGCCA | 2768.9 | 7.1 | 0.7 | 9.6 | 1.0E-21 | 8.9E-21 | BT20/MB468 | CYTO |
| >hg38_21_+_8433172_8446622_RNA45SN1_45S_50ntFlanks@10351.10378.28\|CGAGAACTTTGAAGGCCGAAGTGGAGAA | >hg38_21_+_8433172_8446622_RNA45SN1_45S_50ntFlanks | 10351 | 10378 | 28 | CGAGAACTTTGAAGGCCGAAGTGGAGAA | 5057.2 | -10.3 | 1.1 | -9.5 | 2.1E-21 | 9.0E-21 | BT20/MB468 | NUC |
| >hg38_21_+_8433172_8446622_RNA45SN1_45S_50ntFlanks@482.519.38\|ATCGATGTGGTGACGTCGTGCTCTCCCGGGCCGGGTCC | >hg38_21_+_8433172_8446622_RNA45SN1_45S_50ntFlanks | 482 | 519 | 38 | ATCGATGTGGTGACGTCGTGCTCTCCCGGGCCGGGTCC | 345.5 | -12.9 | 1.4 | -9.5 | 2.1E-21 | 9.0E-21 | BT20/MB468 | NUC |
| >MI0000077\|hsa-mir-21&WithFlank&17\|+\|59841260\|59841343@14.31.18&[MIMAT0000076&hsa-miR-21-5p&offsets\|0\|-4;m-1&17\|+\|59841273\|59841294&offsets\|0\|-4]\|TAGCTTATCAGACTGATG | >MI0000077\|hsa-mir-21&WithFlank&17\|+\|59841260\|59841343 | 14 | 31 | 18 | TAGCTTATCAGACTGATG | 419.7 | 6.0 | 0.6 | 9.6 | 7.6E-22 | 9.0E-21 | BT20/MB468 | MITO |
| >hg38_21_+_8433172_8446622_RNA45SN1_45S_50ntFlanks@475.502.28\|CTTCGTGATCGATGTGGTGACGTCGTGC | >hg38_21_+_8433172_8446622_RNA45SN1_45S_50ntFlanks | 475 | 502 | 28 | CTTCGTGATCGATGTGGTGACGTCGTGC | 359.6 | 12.7 | 1.3 | 9.6 | 1.0E-21 | 9.0E-21 | BT20/MB468 | CYTO |
| >hg38_21_+_8433172_8446622_RNA45SN1_45S_50ntFlanks@10352.10383.32\|GAGAACTTTGAAGGCCGAAGTGGAGAAGGGTT | >hg38_21_+_8433172_8446622_RNA45SN1_45S_50ntFlanks | 10352 | 10383 | 32 | GAGAACTTTGAAGGCCGAAGTGGAGAAGGGTT | 534.1 | -5.3 | 0.6 | -9.5 | 2.1E-21 | 9.1E-21 | BT20/MB468 | NUC |
| >hg38_21_+_8433172_8446622_RNA45SN1_45S_50ntFlanks@11575.11593.19\|ACTTAGAACTGGTGCGGAC | >hg38_21_+_8433172_8446622_RNA45SN1_45S_50ntFlanks | 11575 | 11593 | 19 | ACTTAGAACTGGTGCGGAC | 2536.7 | -5.4 | 0.6 | -9.7 | 3.0E-22 | 9.1E-21 | MB231/MB468 | CYTO |
| >MI0000298\|hsa-mir-221&WithFlank&X\|-\|45746151\|45746272@71.93.23(+1U)&[MIMAT0000278&hsa-miR-221-3p&offsets\|0\|0(+1U);m-91&X\|-\|45746181\|45746202&offsets\|0\|+1(+1U)]\|AGCTACATTGTCTGCTGGGTTTCT | >MI0000298\|hsa-mir-221&WithFlank&X\|-\|45746151\|45746272 | 71 | 93 | 23(+1U) | AGCTACATTGTCTGCTGGGTTTCT | 11565.4 | 5.9 | 0.6 | 9.6 | 5.8E-22 | 9.1E-21 | MB231/MB468 | NUC |
| >hg38_21_+_8433172_8446622_RNA45SN1_45S_50ntFlanks@11566.11584.19\|TAGCAGCCGACTTAGAACT | >hg38_21_+_8433172_8446622_RNA45SN1_45S_50ntFlanks | 11566 | 11584 | 19 | TAGCAGCCGACTTAGAACT | 2121.3 | 7.1 | 0.7 | 9.7 | 3.1E-22 | 9.1E-21 | MB231/MB468 | CYTO |
| >MI0000456\|hsa-mir-140&WithFlank&16\|+\|69933075\|69933186@68.90.23(+1U)&[MIMAT0004597&hsa-miR-140-3p&offsets\|0\|+2(+1U);m-45&16\|+\|69933143\|69933164&offsets\|-1\|0(+1U)]\|TACCACAGGGTAGAACCACGGACT | >MI0000456\|hsa-mir-140&WithFlank&16\|+\|69933075\|69933186 | 68 | 90 | 23(+1U) | TACCACAGGGTAGAACCACGGACT | 500.8 | 10.9 | 1.1 | 9.6 | 1.1E-21 | 9.1E-21 | BT20/MB468 | CYTO |
| >MI0000279\|hsa-mir-196a-2&WithFlank&12\|+\|53991732\|53991853@31.51.21;MI0000238\|hsa-mir-196a-1&WithFlank&17\|-\|48632484\|48632565@13.33.21&[MIMAT0000226&hsa-miR-196a-5p&offsets\|0\|-1;m-261&12\|+\|53991762\|53991783&offsets\|0\|-1];[MIMAT0000226_1&hsa-miR-196a-5p&offsets\|0\|-1;m-268&17\|-\|48632532\|48632553&offsets\|0\|-1]\|TAGGTAGTTTCATGTTGTTGG | >MI0000279\|hsa-mir-196a-2&WithFlank&12\|+\|53991732\|53991853 | 31 | 51 | 21 | TAGGTAGTTTCATGTTGTTGG | 903.6 | 13.6 | 1.4 | 9.9 | 6.7E-23 | 9.2E-21 | BT20/MB231 | NUC |
| >MI0000477\|hsa-mir-146a&WithFlank&5\|+\|160485346\|160485456@27.47.21(+1C)&[MIMAT0000449&hsa-miR-146a-5p&offsets\|0\|-1(+1C);m-129&5\|+\|160485372\|160485393&offsets\|0\|-1(+1C)]\|TGAGAACTGAATTCCATGGGTC | >MI0000477\|hsa-mir-146a&WithFlank&5\|+\|160485346\|160485456 | 27 | 47 | 21(+1C) | TGAGAACTGAATTCCATGGGTC | 2841.9 | 15.4 | 1.6 | 9.6 | 5.9E-22 | 9.3E-21 | MB231/MB468 | NUC |
| >hg38_21_+_8433172_8446622_RNA45SN1_45S_50ntFlanks@4759.4778.20\|ATCAGATACCGTCGTAGTTC | >hg38_21_+_8433172_8446622_RNA45SN1_45S_50ntFlanks | 4759 | 4778 | 20 | ATCAGATACCGTCGTAGTTC | 1538.2 | -13.6 | 1.4 | -9.6 | 6.0E-22 | 9.3E-21 | MB231/MB468 | NUC |
| >hg38_21_+_8433172_8446622_RNA45SN1_45S_50ntFlanks@12456.12483.28\|CCAAGCGTTGGATTGTTCACCCACTAAT | >hg38_21_+_8433172_8446622_RNA45SN1_45S_50ntFlanks | 12456 | 12483 | 28 | CCAAGCGTTGGATTGTTCACCCACTAAT | 1127.4 | -5.6 | 0.6 | -9.6 | 1.1E-21 | 9.3E-21 | BT20/MB468 | CYTO |
| >hg38_21_+_8433172_8446622_RNA45SN1_45S_50ntFlanks@10873.10894.22\|CTGGGTCGGTCGGGCTGGGGCG | >hg38_21_+_8433172_8446622_RNA45SN1_45S_50ntFlanks | 10873 | 10894 | 22 | CTGGGTCGGTCGGGCTGGGGCG | 1086.0 | -13.9 | 1.4 | -9.7 | 4.0E-22 | 9.4E-21 | MB231/MB468 | MP |

| | | | | | | | | | | | | | |
|---|---|---|---|---|---|---|---|---|---|---|---|---|---|
| >hg38_21_+_8433172_8446622_RNA45SN1_45S_50ntFlanks@4163.4184.22\|CATCCAAGGAAGGCAGCAGGCG | >hg38_21_+_8433172_8446622_RNA45SN1_45S_50ntFlanks | 4163 | 4184 | 22 | CATCCAAGGAAGGCAGCAGGCG | 1287.4 | -14.2 | 1.5 | -9.7 | 4.0E-22 | 9.4E-21 | MB231/MB468 | MP |
| >MI0000767\|hsa-mir-365a&WithFlank&16\|+\|14309279\|14309377@62.82.21(+1C);MI0000769\|hsa-mir-365b&WithFlank&17\|+\|31575405\|31575527@74.94.21(+1C)&[MIMAT0000710&hsa-miR-365a-3p&offsets\|0\|-1(+1C);m-176&16\|+\|14309340\|14309361&offsets\|0\|-1(+1C)];[MIMAT0022834&hsa-miR-365b-3p&offsets\|0\|-1(+1C);m-175&17\|+\|31575478\|31575499&offsets\|0\|-1(+1C)]\|TAATGCCCCTAAAAATCCTTAC | >MI0000767\|hsa-mir-365a&WithFlank&16\|+\|14309279\|14309377 | 62 | 82 | 21(+1C) | TAATGCCCCTAAAAATCCTTAC | 413.7 | 12.9 | 1.4 | 9.6 | 1.1E-21 | 9.4E-21 | BT20/MB468 | CYTO |
| >am_trna128_GlyGCC_6_-_27870686_27870756@1.32.32,trna133_GlyCCC_1_-_16872434_16872504@1.32.32,trna18_GlyGCC_16_+_70822597_70822667@1.32.32,trna19_GlyGCC_16_+_70823410_70823480@1.32.32,trna19_GlyGCC_2_-_157257659_157257729@1.32.32,trna24_GlyGCC_16_-_70812942_70813012@1.32.32,trna25_GlyGCC_16_-_70812114_70812184@1.32.32,trna4_GlyCCC_1_+_17188416_17188486@1.32.32,trna5_GlyGCC_17_+_8029064_8029134@1.32.32,trna68_GlyGCC_1_-_161493637_161493707@1.32.32\|GCATTGGTGGTTCAGTGGTAGAATTCTCGCCT | >am_trna128_GlyGCC_6_-_27870686_27870756 | 1 | 32 | 32 | GCATTGGTGGTTCAGTGGTAGAATTCTCGCCT | 9826.2 | 6.4 | 0.7 | 9.6 | 1.1E-21 | 9.5E-21 | BT20/MB468 | CYTO |
| >hg38_21_+_8433172_8446622_RNA45SN1_45S_50ntFlanks@12456.12474.19\|CCAAGCGTTGGATTGTTCA | >hg38_21_+_8433172_8446622_RNA45SN1_45S_50ntFlanks | 12456 | 12474 | 19 | CCAAGCGTTGGATTGTTCA | 10626.1 | -7.6 | 0.8 | -9.6 | 1.1E-21 | 9.5E-21 | BT20/MB468 | CYTO |
| >hg38_21_+_8433172_8446622_RNA45SN1_45S_50ntFlanks@6772.6799.28\|GTTCCTCCCGGGGCTACGCCTGTCTGAG | >hg38_21_+_8433172_8446622_RNA45SN1_45S_50ntFlanks | 6772 | 6799 | 28 | GTTCCTCCCGGGGCTACGCCTGTCTGAG | 792.7 | -13.5 | 1.4 | -9.7 | 4.0E-22 | 9.5E-21 | MB231/MB468 | MP |
| >hg38_21_+_8433172_8446622_RNA45SN1_45S_50ntFlanks@8106.8143.38\|GCGGCGGGGCGCGGGACATGTGGCGTACGGAAGACCCG | >hg38_21_+_8433172_8446622_RNA45SN1_45S_50ntFlanks | 8106 | 8143 | 38 | GCGGCGGGGCGCGGGACATGTGGCGTACGGAAGACCCG | 1437.4 | -14.3 | 1.5 | -9.6 | 6.5E-22 | 9.5E-21 | MB231/MB468 | MITO |
| >MI0000077\|hsa-mir-21&WithFlank&17\|+\|59841260\|59841343@14.36.23(+1A)&[MIMAT0000076&hsa-miR-21-5p&offsets\|0\|+1(+1A);m-1&17\|+\|59841273\|59841294&offsets\|0\|+1(+1A)]\|TAGCTTATCAGACTGATGTTGACA | >MI0000077\|hsa-mir-21&WithFlank&17\|+\|59841260\|59841343 | 14 | 36 | 23(+1A) | TAGCTTATCAGACTGATGTTGACA | 285137.8 | 7.4 | 0.8 | 9.6 | 1.1E-21 | 9.6E-21 | BT20/MB468 | CYTO |
| >hg38_21_+_8433172_8446622_RNA45SN1_45S_50ntFlanks@10351.10377.27\|CGAGAACTTTGAAGGCCGAAGTGGAGA | >hg38_21_+_8433172_8446622_RNA45SN1_45S_50ntFlanks | 10351 | 10377 | 27 | CGAGAACTTTGAAGGCCGAAGTGGAGA | 4544.2 | -5.1 | 0.5 | -9.7 | 3.3E-22 | 9.6E-21 | MB231/MB468 | CYTO |
| >hg38_21_+_8433172_8446622_RNA45SN1_45S_50ntFlanks@9033.9058.26\|CTCGTGGGGGGGCCGGGCCACCCCTC | >hg38_21_+_8433172_8446622_RNA45SN1_45S_50ntFlanks | 9033 | 9058 | 26 | CTCGTGGGGGGGCCGGGCCACCCCTC | 1651.8 | -14.5 | 1.5 | -9.6 | 6.6E-22 | 9.6E-21 | MB231/MB468 | MITO |
| >hg38_21_+_8433172_8446622_RNA45SN1_45S_50ntFlanks@12415.12435.21\|CTTCGATGTCGGCTCTTCCTA | >hg38_21_+_8433172_8446622_RNA45SN1_45S_50ntFlanks | 12415 | 12435 | 21 | CTTCGATGTCGGCTCTTCCTA | 684.4 | -13.3 | 1.4 | -9.7 | 4.1E-22 | 9.7E-21 | MB231/MB468 | MP |
| >MI0000750\|hsa-mir-26a-2&WithFlank&12\|-\|57824603\|57824698@20.41.22(+1U);MI0000083\|hsa-mir-26a-1&WithFlank&3\|+\|37969398\|37969486@16.37.22(+1U)&[MIMAT0000082&hsa-miR-26a-5p&offsets\|0\|0(+1U);m-33&12\|-\|57824658\|57824679&offsets\|0\|0(+1U)];[MIMAT0000082_1&hsa-miR-26a-5p&offsets\|0\|0(+1U);m-34&3\|+\|37969413\|37969434&offsets\|0\|0(+1U)]\|TTCAAGTAATCCAGGATAGGCTT | >MI0000750\|hsa-mir-26a-2&WithFlank&12\|-\|57824603\|57824698 | 20 | 41 | 22(+1U) | TTCAAGTAATCCAGGATAGGCTT | 5451.0 | 7.2 | 0.8 | 9.6 | 1.1E-21 | 9.7E-21 | BT20/MB468 | CYTO |
| >MI0000809\|hsa-mir-151a&WithFlank&8\|-\|140732558\|140732659@53.75.23&[MIMAT0000757&hsa-miR-151a-3p&offsets\|0\|+2;m-30&8\|-\|140732587\|140732607&offsets\|0\|+2]\|CTAGACTGAAGCTCCTTGAGGAC | >MI0000809\|hsa-mir-151a&WithFlank&8\|-\|140732558\|140732659 | 53 | 75 | 23 | CTAGACTGAAGCTCCTTGAGGAC | 433.5 | 13.2 | 1.4 | 9.6 | 8.3E-22 | 9.7E-21 | BT20/MB468 | MITO |
| >hg38_21_+_8433172_8446622_RNA45SN1_45S_50ntFlanks@11873.11900.28\|AAGAAGACCCTGTTGAGCTTGACTCTAG | >hg38_21_+_8433172_8446622_RNA45SN1_45S_50ntFlanks | 11873 | 11900 | 28 | AAGAAGACCCTGTTGAGCTTGACTCTAG | 909.1 | -13.7 | 1.4 | -9.7 | 4.2E-22 | 9.8E-21 | MB231/MB468 | MP |
| >hg38_21_+_8433172_8446622_RNA45SN1_45S_50ntFlanks@9504.9522.19\|GTGAACTATGCCTGGGCAG | >hg38_21_+_8433172_8446622_RNA45SN1_45S_50ntFlanks | 9504 | 9522 | 19 | GTGAACTATGCCTGGGCAG | 1211.0 | -14.1 | 1.5 | -9.7 | 4.2E-22 | 9.8E-21 | MB231/MB468 | MP |
| >hg38_21_+_8433172_8446622_RNA45SN1_45S_50ntFlanks@12991.13025.35\|CTCCCTCGCTGCGATCTATTGAAAGTCAGCCCTCG | >hg38_21_+_8433172_8446622_RNA45SN1_45S_50ntFlanks | 12991 | 13025 | 35 | CTCCCTCGCTGCGATCTATTGAAAGTCAGCCCTCG | 388.0 | -13.0 | 1.4 | -9.5 | 2.3E-21 | 9.8E-21 | BT20/MB468 | NUC |
| >hg38_21_+_8433172_8446622_RNA45SN1_45S_50ntFlanks@10840.10867.28\|CCGTAACTTCGGGATAAGGATTGGCTCT | >hg38_21_+_8433172_8446622_RNA45SN1_45S_50ntFlanks | 10840 | 10867 | 28 | CCGTAACTTCGGGATAAGGATTGGCTCT | 821.1 | -5.6 | 0.6 | -9.5 | 2.3E-21 | 9.8E-21 | BT20/MB468 | NUC |
| >hg38_21_+_8433172_8446622_RNA45SN1_45S_50ntFlanks@12991.13009.19\|CTCCCTCGCTGCGATCTAT | >hg38_21_+_8433172_8446622_RNA45SN1_45S_50ntFlanks | 12991 | 13009 | 19 | CTCCCTCGCTGCGATCTAT | 1429.9 | -14.9 | 1.6 | -9.5 | 2.3E-21 | 9.9E-21 | BT20/MB468 | NUC |
| >hg38_21_+_8433172_8446622_RNA45SN1_45S_50ntFlanks@8170.8188.19\|CAAGTCCTTCTGATCGAGG | >hg38_21_+_8433172_8446622_RNA45SN1_45S_50ntFlanks | 8170 | 8188 | 19 | CAAGTCCTTCTGATCGAGG | 944.4 | -13.7 | 1.4 | -9.7 | 4.3E-22 | 9.9E-21 | MB231/MB468 | MP |
| >hg38_21_+_8433172_8446622_RNA45SN1_45S_50ntFlanks@11581.11611.31\|AACTGGTGCGGACCAGGGGAATCCGACTGTT | >hg38_21_+_8433172_8446622_RNA45SN1_45S_50ntFlanks | 11581 | 11611 | 31 | AACTGGTGCGGACCAGGGGAATCCGACTGTT | 2711.9 | -7.0 | 0.7 | -9.7 | 4.3E-22 | 1.0E-20 | MB231/MB468 | MP |
| >hg38_21_+_8433172_8446622_RNA45SN1_45S_50ntFlanks@3899.3927.29\|CCTTCGCGGGGGGGATGCGTGCATTTATC | >hg38_21_+_8433172_8446622_RNA45SN1_45S_50ntFlanks | 3899 | 3927 | 29 | CCTTCGCGGGGGGGATGCGTGCATTTATC | 1567.7 | -14.4 | 1.5 | -9.6 | 6.8E-22 | 1.0E-20 | MB231/MB468 | MITO |
| >hg38_21_+_8433172_8446622_RNA45SN1_45S_50ntFlanks@5446.5466.21\|CGGATCGGCCCCGCCGGGGTC | >hg38_21_+_8433172_8446622_RNA45SN1_45S_50ntFlanks | 5446 | 5466 | 21 | CGGATCGGCCCCGCCGGGGTC | 5856.2 | -11.6 | 1.2 | -9.6 | 6.4E-22 | 1.0E-20 | MB231/MB468 | NUC |
| >am_MI0000809\|hsa-mir-151a&WithFlank&8\|-\|140732558\|140732659@53.72.20(+1C)&[MIMAT0000757&hsa-miR-151a-3p&offsets\|0\|-1(+1C);m-30&8\|-\|140732587\|140732607&offsets\|0\|-1(+1C)]\|CTAGACTGAAGCTCCTTGAGC | >am_MI0000809\|hsa-mir-151a&WithFlank&8\|-\|140732558\|140732659 | 53 | 72 | 20(+1C) | CTAGACTGAAGCTCCTTGAGC | 468.4 | 6.3 | 0.7 | 9.6 | 6.5E-22 | 1.0E-20 | MB231/MB468 | NUC |
| >am_trna106_PheGAA_6_-_28775610_28775682@31.52.22,trna109_PheGAA_6_-_28758499_28758571@31.52.22,trna11_PheGAA_12_-_125412389_125412461@31.52.22,trna13_PheGAA_11_-_59333853_59333925@31.52.22,trna14_PheGAA_19_-_1383361_1383433@31.52.22,trna15_PheGAA_11_-_59324970_59325042@31.52.22,trna1_PheGAA_13_-_95201904_95201976@31.52.22,trna72_PheGAA_6_+_28732164_28732236@31.52.22,trna96_PheGAA_6_-_28949449_28949521@31.52.22\|ACTGAAGATCTAAAGGTCCCTG | >am_trna106_PheGAA_6_-_28775610_28775682 | 31 | 52 | 22 | ACTGAAGATCTAAAGGTCCCTG | 465.1 | -13.3 | 1.4 | -9.5 | 2.4E-21 | 1.0E-20 | BT20/MB468 | NUC |
| >hg38_21_+_8433172_8446622_RNA45SN1_45S_50ntFlanks@4975.4994.20\|CCCGGACACGGACAGGATTG | >hg38_21_+_8433172_8446622_RNA45SN1_45S_50ntFlanks | 4975 | 4994 | 20 | CCCGGACACGGACAGGATTG | 1094.7 | -14.0 | 1.4 | -9.7 | 4.4E-22 | 1.0E-20 | MB231/MB468 | MP |
| >hg38_21_+_8433172_8446622_RNA45SN1_45S_50ntFlanks@12785.12805.21\|CCCCGCGCGCGCGGGAGGGCG | >hg38_21_+_8433172_8446622_RNA45SN1_45S_50ntFlanks | 12785 | 12805 | 21 | CCCCGCGCGCGCGGGAGGGCG | 1728.7 | -14.6 | 1.5 | -9.7 | 4.4E-22 | 1.0E-20 | MB231/MB468 | MP |
| >hg38_21_+_8433172_8446622_RNA45SN1_45S_50ntFlanks@6655.6676.22\|TCTTAGCGGTGGATCACTCGGC | >hg38_21_+_8433172_8446622_RNA45SN1_45S_50ntFlanks | 6655 | 6676 | 22 | TCTTAGCGGTGGATCACTCGGC | 431.8 | 6.7 | 0.7 | 9.6 | 8.8E-22 | 1.0E-20 | BT20/MB468 | MITO |
| >hg38_21_+_8433172_8446622_RNA45SN1_45S_50ntFlanks@9739.9759.21\|AAACGATCTCAACCTATTCTC | >hg38_21_+_8433172_8446622_RNA45SN1_45S_50ntFlanks | 9739 | 9759 | 21 | AAACGATCTCAACCTATTCTC | 6373.8 | -8.3 | 0.9 | -9.6 | 7.1E-22 | 1.0E-20 | MB231/MB468 | MITO |
| >hg38_21_+_8433172_8446622_RNA45SN1_45S_50ntFlanks@9736.9757.22\|CCGAAACGATCTCAACCTATTC | >hg38_21_+_8433172_8446622_RNA45SN1_45S_50ntFlanks | 9736 | 9757 | 22 | CCGAAACGATCTCAACCTATTC | 310.2 | -12.7 | 1.3 | -9.5 | 2.5E-21 | 1.0E-20 | BT20/MB468 | NUC |
| >hg38_21_+_8433172_8446622_RNA45SN1_45S_50ntFlanks@4287.4317.31\|AACGAGGATCCATTGGAGGGCAAGTCTGGTG | >hg38_21_+_8433172_8446622_RNA45SN1_45S_50ntFlanks | 4287 | 4317 | 31 | AACGAGGATCCATTGGAGGGCAAGTCTGGTG | 384.6 | -13.0 | 1.4 | -9.5 | 2.5E-21 | 1.0E-20 | BT20/MB468 | NUC |
| >hg38_21_+_8433172_8446622_RNA45SN1_45S_50ntFlanks@6743.6772.30\|CGACACTTCGAACGCACTTGCGGCCCCGGG | >hg38_21_+_8433172_8446622_RNA45SN1_45S_50ntFlanks | 6743 | 6772 | 30 | CGACACTTCGAACGCACTTGCGGCCCCGGG | 603.8 | -13.7 | 1.4 | -9.5 | 2.5E-21 | 1.1E-20 | BT20/MB468 | NUC |
| >hg38_21_+_8433172_8446622_RNA45SN1_45S_50ntFlanks@7968.7988.21\|TCCGAGACGCGACCTCAGATC | >hg38_21_+_8433172_8446622_RNA45SN1_45S_50ntFlanks | 7968 | 7988 | 21 | TCCGAGACGCGACCTCAGATC | 321.4 | 12.8 | 1.3 | 9.6 | 9.0E-22 | 1.1E-20 | BT20/MB468 | MITO |
| >hg38_21_+_8433172_8446622_RNA45SN1_45S_50ntFlanks@11716.11740.25\|TAAACGGCGGGAGTAACTATGACTC | >hg38_21_+_8433172_8446622_RNA45SN1_45S_50ntFlanks | 11716 | 11740 | 25 | TAAACGGCGGGAGTAACTATGACTC | 497.1 | -13.4 | 1.4 | -9.5 | 2.5E-21 | 1.1E-20 | BT20/MB468 | NUC |
| >MI0000746\|hsa-mir-99b&WithFlank&19\|+\|51692606\|51692687@13.33.21(+1U)&[MIMAT0000689&hsa-miR-99b-5p&offsets\|0\|-1(+1U);m-10&19\|+\|51692618\|51692639&offsets\|0\|-1(+1U)]\|CACCCGTAGAACCGACCTTGCT | >MI0000746\|hsa-mir-99b&WithFlank&19\|+\|51692606\|51692687 | 13 | 33 | 21(+1U) | CACCCGTAGAACCGACCTTGCT | 4258.1 | 9.0 | 0.9 | 9.6 | 1.3E-21 | 1.1E-20 | BT20/MB468 | CYTO |
| >MI0000285\|hsa-mir-205&WithFlank&1\|+\|209432127\|209432248@40.61.22(+1G)&[MIMAT0000266&hsa-miR-205-5p&offsets\|0\|0(+1G);m-351&1\|+\|209432166\|209432187&offsets\|0\|0(+1G)]\|TCCTTCATTCCACCGGAGTCTGG | >MI0000285\|hsa-mir-205&WithFlank&1\|+\|209432127\|209432248 | 40 | 61 | 22(+1G) | TCCTTCATTCCACCGGAGTCTGG | 5479.7 | 6.1 | 0.6 | 9.6 | 1.3E-21 | 1.1E-20 | BT20/MB468 | CYTO |
| >hg38_21_+_8433172_8446622_RNA45SN1_45S_50ntFlanks@10516.10533.18\|GAGTGGCGGAGATGGGCG | >hg38_21_+_8433172_8446622_RNA45SN1_45S_50ntFlanks | 10516 | 10533 | 18 | GAGTGGCGGAGATGGGCG | 1432.9 | -14.3 | 1.5 | -9.6 | 7.5E-22 | 1.1E-20 | MB231/MB468 | MITO |
| >hg38_21_+_8433172_8446622_RNA45SN1_45S_50ntFlanks@9573.9596.24\|AAATCGGTCGTCCGACCTGGGTAT | >hg38_21_+_8433172_8446622_RNA45SN1_45S_50ntFlanks | 9573 | 9596 | 24 | AAATCGGTCGTCCGACCTGGGTAT | 10632.8 | -10.9 | 1.1 | -9.7 | 4.8E-22 | 1.1E-20 | MB231/MB468 | MP |
| >hg38_21_+_8433172_8446622_RNA45SN1_45S_50ntFlanks@10687.10710.24\|CGGCGTCCGGTGAGCTCTCGCTGG | >hg38_21_+_8433172_8446622_RNA45SN1_45S_50ntFlanks | 10687 | 10710 | 24 | CGGCGTCCGGTGAGCTCTCGCTGG | 7404.0 | -7.7 | 0.8 | -9.7 | 3.8E-22 | 1.1E-20 | MB231/MB468 | CYTO |
| >hg38_21_+_8433172_8446622_RNA45SN1_45S_50ntFlanks@12539.12573.35\|ATGATGTGTTGTTGCCATGGTAATCCTGCTCAGTA | >hg38_21_+_8433172_8446622_RNA45SN1_45S_50ntFlanks | 12539 | 12573 | 35 | ATGATGTGTTGTTGCCATGGTAATCCTGCTCAGTA | 1078.8 | -13.9 | 1.4 | -9.6 | 7.6E-22 | 1.1E-20 | MB231/MB468 | MITO |
| >hg38_21_+_8433172_8446622_RNA45SN1_45S_50ntFlanks@6644.6683.40\|CCTCGTACGACTCTTAGCGGTGGATCACTCGGCTCGTGCG | >hg38_21_+_8433172_8446622_RNA45SN1_45S_50ntFlanks | 6644 | 6683 | 40 | CCTCGTACGACTCTTAGCGGTGGATCACTCGGCTCGTGCG | 1203.4 | 7.7 | 0.8 | 9.6 | 7.2E-22 | 1.1E-20 | MB231/MB468 | NUC |
| >hg38_21_+_8433172_8446622_RNA45SN1_45S_50ntFlanks@12932.12956.25\|CGCTAAACCATTCGTAGACGACCTG | >hg38_21_+_8433172_8446622_RNA45SN1_45S_50ntFlanks | 12932 | 12956 | 25 | CGCTAAACCATTCGTAGACGACCTG | 1095.5 | -5.2 | 0.5 | -9.6 | 7.2E-22 | 1.1E-20 | MB231/MB468 | NUC |
| >hg38_21_+_8433172_8446622_RNA45SN1_45S_50ntFlanks@8380.8400.21\|CAAGAGGGCGTGAAACCGTTA | >hg38_21_+_8433172_8446622_RNA45SN1_45S_50ntFlanks | 8380 | 8400 | 21 | CAAGAGGGCGTGAAACCGTTA | 5894.1 | -12.2 | 1.3 | -9.7 | 4.8E-22 | 1.1E-20 | MB231/MB468 | MP |
| >hg38_21_+_8433172_8446622_RNA45SN1_45S_50ntFlanks@12658.12675.18\|ACTGAACGCCTCTAAGTC | >hg38_21_+_8433172_8446622_RNA45SN1_45S_50ntFlanks | 12658 | 12675 | 18 | ACTGAACGCCTCTAAGTC | 1526.4 | -6.1 | 0.6 | -9.6 | 7.3E-22 | 1.1E-20 | MB231/MB468 | NUC |
| >hg38_21_+_8433172_8446622_RNA45SN1_45S_50ntFlanks@10107.10125.19\|CGTGTGGTGTGCGTCGGAG | >hg38_21_+_8433172_8446622_RNA45SN1_45S_50ntFlanks | 10107 | 10125 | 19 | CGTGTGGTGTGCGTCGGAG | 1036.2 | -13.9 | 1.4 | -9.6 | 4.9E-22 | 1.1E-20 | MB231/MB468 | MP |

<table>
<tr><th></th><th></th><th></th><th></th><th></th><th></th><th></th><th></th><th></th><th></th><th></th><th></th><th></th><th></th></tr>
<tr><td>>hg38_21_+_8433172_8446622_RNA45SN1_45S_50ntFlanks@12289.12323.35|CCGTGAAAGCGGGGCCTCACGATCCTTCTGACCTT</td><td>>hg38_21_+_8433172_8446622_RNA45SN1_45S_50ntFlanks</td><td>12289</td><td>12323</td><td>35</td><td>CCGTGAAAGCGGGGCCTCACGATCCTTCTGACCTT</td><td>396.7</td><td>-13.1</td><td>1.4</td><td>-9.5</td><td>2.7E-21</td><td>1.1E-20</td><td>BT20/MB468</td><td>NUC</td></tr>
<tr><td>>hg38_21_+_8433172_8446622_RNA45SN1_45S_50ntFlanks@11999.12033.35|CCCTGCGGGCCGCCGGTGAAATACCACTACTCTGA</td><td>>hg38_21_+_8433172_8446622_RNA45SN1_45S_50ntFlanks</td><td>11999</td><td>12033</td><td>35</td><td>CCCTGCGGGCCGCCGGTGAAATACCACTACTCTGA</td><td>1172.0</td><td>-14.0</td><td>1.5</td><td>-9.6</td><td>7.8E-22</td><td>1.1E-20</td><td>MB231/MB468</td><td>MITO</td></tr>
<tr><td>>hg38_21_+_8433172_8446622_RNA45SN1_45S_50ntFlanks@5445.5467.23|TCGGATCGGCCCCGCCGGGGTCG</td><td>>hg38_21_+_8433172_8446622_RNA45SN1_45S_50ntFlanks</td><td>5445</td><td>5467</td><td>23</td><td>TCGGATCGGCCCCGCCGGGGTCG</td><td>1059.5</td><td>-13.9</td><td>1.4</td><td>-9.6</td><td>5.0E-22</td><td>1.1E-20</td><td>MB231/MB468</td><td>MP</td></tr>
<tr><td>>hg38_21_+_8433172_8446622_RNA45SN1_45S_50ntFlanks@10404.10424.21|CATGGGTCAGTCGGTCCTGAG</td><td>>hg38_21_+_8433172_8446622_RNA45SN1_45S_50ntFlanks</td><td>10404</td><td>10424</td><td>21</td><td>CATGGGTCAGTCGGTCCTGAG</td><td>13636.6</td><td>5.6</td><td>0.6</td><td>9.6</td><td>9.8E-22</td><td>1.1E-20</td><td>BT20/MB468</td><td>MITO</td></tr>
<tr><td>>hg38_21_+_8433172_8446622_RNA45SN1_45S_50ntFlanks@7975.8016.42|CGCGACCTCAGATCAGACGTGGCGACCCGCTGAATTTAAGCA</td><td>>hg38_21_+_8433172_8446622_RNA45SN1_45S_50ntFlanks</td><td>7975</td><td>8016</td><td>42</td><td>CGCGACCTCAGATCAGACGTGGCGACCCGCTGAATTTAAGCA</td><td>11676.8</td><td>6.8</td><td>0.7</td><td>9.7</td><td>4.0E-22</td><td>1.2E-20</td><td>MB231/MB468</td><td>CYTO</td></tr>
<tr><td>>hg38_21_+_8433172_8446622_RNA45SN1_45S_50ntFlanks@11669.11710.42|GCCCAGTGCTCTGAATGTCAAAGTGAAGAAATTCAATGAAGC</td><td>>hg38_21_+_8433172_8446622_RNA45SN1_45S_50ntFlanks</td><td>11669</td><td>11710</td><td>42</td><td>GCCCAGTGCTCTGAATGTCAAAGTGAAGAAATTCAATGAAGC</td><td>928.3</td><td>13.9</td><td>1.4</td><td>9.7</td><td>4.0E-22</td><td>1.2E-20</td><td>MB231/MB468</td><td>CYTO</td></tr>
<tr><td>>hg38_21_+_8433172_8446622_RNA45SN1_45S_50ntFlanks@12407.12436.30|TTTTGATCCTTCGATGTCGGCTCTTCCTAT</td><td>>hg38_21_+_8433172_8446622_RNA45SN1_45S_50ntFlanks</td><td>12407</td><td>12436</td><td>30</td><td>TTTTGATCCTTCGATGTCGGCTCTTCCTAT</td><td>6078.1</td><td>-6.5</td><td>0.7</td><td>-9.6</td><td>7.6E-22</td><td>1.2E-20</td><td>MB231/MB468</td><td>NUC</td></tr>
<tr><td>>hg38_21_+_8433172_8446622_RNA45SN1_45S_50ntFlanks@8380.8405.26|CAAGAGGGCGTGAAACCGTTAAGAGG</td><td>>hg38_21_+_8433172_8446622_RNA45SN1_45S_50ntFlanks</td><td>8380</td><td>8405</td><td>26</td><td>CAAGAGGGCGTGAAACCGTTAAGAGG</td><td>5373.6</td><td>-12.4</td><td>1.3</td><td>-9.6</td><td>8.1E-22</td><td>1.2E-20</td><td>MB231/MB468</td><td>MITO</td></tr>
<tr><td>>hg38_21_+_8433172_8446622_RNA45SN1_45S_50ntFlanks@3782.3812.31|CAGTGAAACTGCGAATGGCTCATTAAATCAG</td><td>>hg38_21_+_8433172_8446622_RNA45SN1_45S_50ntFlanks</td><td>3782</td><td>3812</td><td>31</td><td>CAGTGAAACTGCGAATGGCTCATTAAATCAG</td><td>1108.8</td><td>-14.5</td><td>1.5</td><td>-9.5</td><td>2.8E-21</td><td>1.2E-20</td><td>BT20/MB468</td><td>NUC</td></tr>
<tr><td>>hg38_21_+_8433172_8446622_RNA45SN1_45S_50ntFlanks@3899.3920.22|CCTTCGCGGGGGGGATGCGTGC</td><td>>hg38_21_+_8433172_8446622_RNA45SN1_45S_50ntFlanks</td><td>3899</td><td>3920</td><td>22</td><td>CCTTCGCGGGGGGGATGCGTGC</td><td>1875.1</td><td>-15.3</td><td>1.6</td><td>-9.5</td><td>2.8E-21</td><td>1.2E-20</td><td>BT20/MB468</td><td>NUC</td></tr>
<tr><td>>hg38_21_+_8433172_8446622_RNA45SN1_45S_50ntFlanks@11564.11605.42|CCTAGCAGCCGACTTAGAACTGGTGCGGACCAGGGGAATCCG</td><td>>hg38_21_+_8433172_8446622_RNA45SN1_45S_50ntFlanks</td><td>11564</td><td>11605</td><td>42</td><td>CCTAGCAGCCGACTTAGAACTGGTGCGGACCAGGGGAATCCG</td><td>1212.4</td><td>-14.7</td><td>1.5</td><td>-9.5</td><td>2.8E-21</td><td>1.2E-20</td><td>BT20/MB468</td><td>NUC</td></tr>
<tr><td>>hg38_21_+_8433172_8446622_RNA45SN1_45S_50ntFlanks@11665.11707.43|TTCTGCCCAGTGCTCTGAATGTCAAAGTGAAGAAATTCAATGA</td><td>>hg38_21_+_8433172_8446622_RNA45SN1_45S_50ntFlanks</td><td>11665</td><td>11707</td><td>43</td><td>TTCTGCCCAGTGCTCTGAATGTCAAAGTGAAGAAATTCAATGA</td><td>250.2</td><td>12.0</td><td>1.2</td><td>9.7</td><td>4.1E-22</td><td>1.2E-20</td><td>MB231/MB468</td><td>CYTO</td></tr>
<tr><td>>trna111_HisGTG_1_-_147774845_147774916@-1T.23.24,trna118_HisGTG_1_-_145396881_145396952@-1T.23.24,trna16_HisGTG_1_+_146544773_146544844@-1T.23.24,trna1_HisGTG_15_+_45493349_45493420@-1T.23.24,trna21_HisGTG_1_+_147753471_147753542@-1T.23.24,trna33_HisGTG_6_+_27125906_27125977@-1T.23.24,trna7_HisGTG_9_-_14433938_14434009@-1T.23.24,trna8_HisGTG_15_-_45492611_45492682@-1T.23.24,trna9_HisGTG_15_-_45490804_45490875@-1T.23.24|TGCCGTGATCGTATAGTGGTTAGT</td><td>>trna111_HisGTG_1_-_147774845_147774916</td><td>-1T</td><td>23</td><td>24</td><td>TGCCGTGATCGTATAGTGGTTAGT</td><td>2263.9</td><td>8.3</td><td>0.9</td><td>9.7</td><td>4.2E-22</td><td>1.2E-20</td><td>MB231/MB468</td><td>CYTO</td></tr>
<tr><td>>hg38_21_+_8433172_8446622_RNA45SN1_45S_50ntFlanks@10402.10431.30|AACATGGGTCAGTCGGTCCTGAGAGATGGG</td><td>>hg38_21_+_8433172_8446622_RNA45SN1_45S_50ntFlanks</td><td>10402</td><td>10431</td><td>30</td><td>AACATGGGTCAGTCGGTCCTGAGAGATGGG</td><td>1362.8</td><td>-4.9</td><td>0.5</td><td>-9.7</td><td>4.2E-22</td><td>1.2E-20</td><td>MB231/MB468</td><td>CYTO</td></tr>
<tr><td>>hg38_21_+_8433172_8446622_RNA45SN1_45S_50ntFlanks@11598.11623.26|GGAATCCGACTGTTTAATTAAAACAA</td><td>>hg38_21_+_8433172_8446622_RNA45SN1_45S_50ntFlanks</td><td>11598</td><td>11623</td><td>26</td><td>GGAATCCGACTGTTTAATTAAAACAA</td><td>305.0</td><td>12.7</td><td>1.3</td><td>9.6</td><td>1.0E-21</td><td>1.2E-20</td><td>BT20/MB468</td><td>MITO</td></tr>
<tr><td>>hg38_21_+_8433172_8446622_RNA45SN1_45S_50ntFlanks@7974.8005.32|ACGCGACCTCAGATCAGACGTGGCGACCCGCT</td><td>>hg38_21_+_8433172_8446622_RNA45SN1_45S_50ntFlanks</td><td>7974</td><td>8005</td><td>32</td><td>ACGCGACCTCAGATCAGACGTGGCGACCCGCT</td><td>5004.2</td><td>5.7</td><td>0.6</td><td>9.5</td><td>1.4E-21</td><td>1.2E-20</td><td>BT20/MB468</td><td>CYTO</td></tr>
<tr><td>>hg38_21_+_8433172_8446622_RNA45SN1_45S_50ntFlanks@10690.10710.21|CGTCCGGTGAGCTCTCGCTGG</td><td>>hg38_21_+_8433172_8446622_RNA45SN1_45S_50ntFlanks</td><td>10690</td><td>10710</td><td>21</td><td>CGTCCGGTGAGCTCTCGCTGG</td><td>1720.1</td><td>-14.6</td><td>1.5</td><td>-9.6</td><td>5.2E-22</td><td>1.2E-20</td><td>MB231/MB468</td><td>MP</td></tr>
<tr><td>>hg38_21_+_8433172_8446622_RNA45SN1_45S_50ntFlanks@12456.12481.26|CCAAGCGTTGGATTGTTCACCCACTA</td><td>>hg38_21_+_8433172_8446622_RNA45SN1_45S_50ntFlanks</td><td>12456</td><td>12481</td><td>26</td><td>CCAAGCGTTGGATTGTTCACCCACTA</td><td>1341.6</td><td>-14.3</td><td>1.5</td><td>-9.6</td><td>5.3E-22</td><td>1.2E-20</td><td>MB231/MB468</td><td>MP</td></tr>
<tr><td>>hg38_21_+_8433172_8446622_RNA45SN1_45S_50ntFlanks@13002.13023.22|CGATCTATTGAAAGTCAGCCCT</td><td>>hg38_21_+_8433172_8446622_RNA45SN1_45S_50ntFlanks</td><td>13002</td><td>13023</td><td>22</td><td>CGATCTATTGAAAGTCAGCCCT</td><td>1072.0</td><td>-13.9</td><td>1.4</td><td>-9.6</td><td>5.3E-22</td><td>1.2E-20</td><td>MB231/MB468</td><td>MP</td></tr>
<tr><td>>hg38_21_+_8433172_8446622_RNA45SN1_45S_50ntFlanks@12366.12383.18|TAACTGGCTTGTGGCGGC</td><td>>hg38_21_+_8433172_8446622_RNA45SN1_45S_50ntFlanks</td><td>12366</td><td>12383</td><td>18</td><td>TAACTGGCTTGTGGCGGC</td><td>1194.8</td><td>-14.0</td><td>1.5</td><td>-9.6</td><td>8.3E-22</td><td>1.2E-20</td><td>MB231/MB468</td><td>MITO</td></tr>
<tr><td>>hg38_21_+_8433172_8446622_RNA45SN1_45S_50ntFlanks@4723.4767.45|CATTAATCAAGAACGAAAGTCGGAGGTTCGAAGACGATCAGATAC</td><td>>hg38_21_+_8433172_8446622_RNA45SN1_45S_50ntFlanks</td><td>4723</td><td>4767</td><td>45</td><td>CATTAATCAAGAACGAAAGTCGGAGGTTCGAAGACGATCAGATAC</td><td>659.8</td><td>13.4</td><td>1.4</td><td>9.7</td><td>4.3E-22</td><td>1.2E-20</td><td>MB231/MB468</td><td>CYTO</td></tr>
<tr><td>>hg38_21_+_8433172_8446622_RNA45SN1_45S_50ntFlanks@12405.12437.33|CTTTTTGATCCTTCGATGTCGGCTCTTCCTATC</td><td>>hg38_21_+_8433172_8446622_RNA45SN1_45S_50ntFlanks</td><td>12405</td><td>12437</td><td>33</td><td>CTTTTTGATCCTTCGATGTCGGCTCTTCCTATC</td><td>3462.7</td><td>-4.6</td><td>0.5</td><td>-9.6</td><td>1.0E-21</td><td>1.2E-20</td><td>BT20/MB468</td><td>MITO</td></tr>
<tr><td>>hg38_21_+_8433172_8446622_RNA45SN1_45S_50ntFlanks@11235.11254.20|CTTCCCGTGGATCGCCCCAG</td><td>>hg38_21_+_8433172_8446622_RNA45SN1_45S_50ntFlanks</td><td>11235</td><td>11254</td><td>20</td><td>CTTCCCGTGGATCGCCCCAG</td><td>1833.3</td><td>-12.9</td><td>1.4</td><td>-9.6</td><td>1.0E-21</td><td>1.2E-20</td><td>BT20/MB468</td><td>MITO</td></tr>
<tr><td>>trna119_LysCTT_1_-_145395522_145395594@1.36.36,trna11_LysCTT_5_-_180648979_180649051@1.36.36,trna13_LysCTT_6_+_26556774_26556846@1.36.36,trna7_LysCTT_16_+_3225692_3225764@1.36.36,trna9_LysCTT_5_+_180634755_180634827@1.36.36|GCCCGGCTAGCTCAGTCGGTAGAGCATGAGACTCTT</td><td>>trna119_LysCTT_1_-_145395522_145395594</td><td>1</td><td>36</td><td>36</td><td>GCCCGGCTAGCTCAGTCGGTAGAGCATGAGACTCTT</td><td>6703.8</td><td>8.5</td><td>0.9</td><td>9.5</td><td>1.4E-21</td><td>1.2E-20</td><td>BT20/MB468</td><td>CYTO</td></tr>
<tr><td>>hg38_21_+_8433172_8446622_RNA45SN1_45S_50ntFlanks@12635.12654.20|AAGCTACCATCTGTGGGATT</td><td>>hg38_21_+_8433172_8446622_RNA45SN1_45S_50ntFlanks</td><td>12635</td><td>12654</td><td>20</td><td>AAGCTACCATCTGTGGGATT</td><td>19809.8</td><td>-13.3</td><td>1.4</td><td>-9.6</td><td>8.5E-22</td><td>1.2E-20</td><td>MB231/MB468</td><td>MITO</td></tr>
<tr><td>>hg38_21_+_8433172_8446622_RNA45SN1_45S_50ntFlanks@11918.11948.31|ACATGAGAGGTGTAGAATAAGTGGGAGGCCC</td><td>>hg38_21_+_8433172_8446622_RNA45SN1_45S_50ntFlanks</td><td>11918</td><td>11948</td><td>31</td><td>ACATGAGAGGTGTAGAATAAGTGGGAGGCCC</td><td>2044.0</td><td>-14.9</td><td>1.5</td><td>-9.6</td><td>5.4E-22</td><td>1.2E-20</td><td>MB231/MB468</td><td>MP</td></tr>
<tr><td>>am_trnaMT_GluTTC_MT_-_14674_14742@27.49.23,trnalookalike8_GluTTC_5_-_93905172_93905240@27.49.23|GTTTTTCATATCATTGGTCGTGG</td><td>>am_trnaMT_GluTTC_MT_-_14674_14742</td><td>27</td><td>49</td><td>23</td><td>GTTTTTCATATCATTGGTCGTGG</td><td>255.1</td><td>-12.4</td><td>1.3</td><td>-9.5</td><td>3.0E-21</td><td>1.2E-20</td><td>BT20/MB468</td><td>NUC</td></tr>
<tr><td>>hg38_21_+_8433172_8446622_RNA45SN1_45S_50ntFlanks@10808.10826.19|CAATGTAGGTAAGGGAAGT</td><td>>hg38_21_+_8433172_8446622_RNA45SN1_45S_50ntFlanks</td><td>10808</td><td>10826</td><td>19</td><td>CAATGTAGGTAAGGGAAGT</td><td>807.3</td><td>-13.5</td><td>1.4</td><td>-9.6</td><td>5.5E-22</td><td>1.2E-20</td><td>MB231/MB468</td><td>MP</td></tr>
<tr><td>>hg38_21_+_8433172_8446622_RNA45SN1_45S_50ntFlanks@6743.6770.28|CGACACTTCGAACGCACTTGCGGCCCCG</td><td>>hg38_21_+_8433172_8446622_RNA45SN1_45S_50ntFlanks</td><td>6743</td><td>6770</td><td>28</td><td>CGACACTTCGAACGCACTTGCGGCCCCG</td><td>3217.8</td><td>-11.2</td><td>1.2</td><td>-9.6</td><td>5.5E-22</td><td>1.2E-20</td><td>MB231/MB468</td><td>MP</td></tr>
<tr><td>>MI0000650|hsa-mir-200c&WithFlank&12|+|6963693|6963772@50.69.20&[MIMAT0000617&hsa-miR-200c-3p&offsets|0|-3;m-54&12|+|6963742|6963764&offsets|0|-3]|TAATACTGCCGGGTAATGAT</td><td>>MI0000650|hsa-mir-200c&WithFlank&12|+|6963693|6963772</td><td>50</td><td>69</td><td>20</td><td>TAATACTGCCGGGTAATGAT</td><td>1833.2</td><td>14.6</td><td>1.5</td><td>9.8</td><td>9.6E-23</td><td>1.2E-20</td><td>BT20/MB231</td><td>NUC</td></tr>
<tr><td>>hg38_21_+_8433172_8446622_RNA45SN1_45S_50ntFlanks@10240.10259.20|CTACGCCGCGACGAGTAGGA</td><td>>hg38_21_+_8433172_8446622_RNA45SN1_45S_50ntFlanks</td><td>10240</td><td>10259</td><td>20</td><td>CTACGCCGCGACGAGTAGGA</td><td>1851.2</td><td>6.0</td><td>0.6</td><td>9.6</td><td>1.1E-21</td><td>1.2E-20</td><td>BT20/MB468</td><td>MITO</td></tr>
<tr><td>>MI0001733|hsa-mir-452&WithFlank&X|-|151959622|151959718@20.42.23&[MIMAT0001635&hsa-miR-452-5p&offsets|0|+1;m-188&X|-|151959677|151959699&offsets|0|0]|AACTGTTTGCAGAGGAAACTGAG</td><td>>MI0001733|hsa-mir-452&WithFlank&X|-|151959622|151959718</td><td>20</td><td>42</td><td>23</td><td>AACTGTTTGCAGAGGAAACTGAG</td><td>334.6</td><td>9.2</td><td>1.0</td><td>9.5</td><td>3.0E-21</td><td>1.3E-20</td><td>BT20/MB468</td><td>NUC</td></tr>
<tr><td>>hg38_21_+_8433172_8446622_RNA45SN1_45S_50ntFlanks@12633.12657.25|CGAAGCTACCATCTGTGGGATTATG</td><td>>hg38_21_+_8433172_8446622_RNA45SN1_45S_50ntFlanks</td><td>12633</td><td>12657</td><td>25</td><td>CGAAGCTACCATCTGTGGGATTATG</td><td>497.9</td><td>-13.4</td><td>1.4</td><td>-9.5</td><td>3.0E-21</td><td>1.3E-20</td><td>BT20/MB468</td><td>NUC</td></tr>
<tr><td>>hg38_21_+_8433172_8446622_RNA45SN1_45S_50ntFlanks@12934.12952.19|CTAAACCATTCGTAGACGA</td><td>>hg38_21_+_8433172_8446622_RNA45SN1_45S_50ntFlanks</td><td>12934</td><td>12952</td><td>19</td><td>CTAAACCATTCGTAGACGA</td><td>681.5</td><td>-13.3</td><td>1.4</td><td>-9.6</td><td>5.6E-22</td><td>1.3E-20</td><td>MB231/MB468</td><td>MP</td></tr>
<tr><td>>MI0000086|hsa-mir-28&WithFlank&3|+|188688775|188688872@61.81.21&[MIMAT0004502&hsa-miR-28-3p&offsets|+1|0;m-28&3|+|188688834|188688855&offsets|+1|0]|ACTAGATTGTGAGCTCCTGGA</td><td>>MI0000086|hsa-mir-28&WithFlank&3|+|188688775|188688872</td><td>61</td><td>81</td><td>21</td><td>ACTAGATTGTGAGCTCCTGGA</td><td>484.3</td><td>13.2</td><td>1.4</td><td>9.5</td><td>1.5E-21</td><td>1.3E-20</td><td>BT20/MB468</td><td>CYTO</td></tr>
<tr><td>>hg38_21_+_8433172_8446622_RNA45SN1_45S_50ntFlanks@11318.11343.26|CCCCACGTCTCGTCGCGCGCGCGTCC</td><td>>hg38_21_+_8433172_8446622_RNA45SN1_45S_50ntFlanks</td><td>11318</td><td>11343</td><td>26</td><td>CCCCACGTCTCGTCGCGCGCGCGTCC</td><td>1141.7</td><td>-13.9</td><td>1.5</td><td>-9.6</td><td>8.9E-22</td><td>1.3E-20</td><td>MB231/MB468</td><td>MITO</td></tr>
<tr><td>>MI0000299|hsa-mir-222&WithFlank&X|-|45747009|45747130@75.96.22&[MIMAT0000279&hsa-miR-222-3p&offsets|0|+1;m-140&X|-|45747033|45747056&offsets|0|-2]|AGCTACATCTGGCTACTGGGTC</td><td>>MI0000299|hsa-mir-222&WithFlank&X|-|45747009|45747130</td><td>75</td><td>96</td><td>22</td><td>AGCTACATCTGGCTACTGGGTC</td><td>1206.3</td><td>6.5</td><td>0.7</td><td>9.6</td><td>1.1E-21</td><td>1.3E-20</td><td>BT20/MB468</td><td>MITO</td></tr>
<tr><td>>hg38_21_+_8433172_8446622_RNA45SN1_45S_50ntFlanks@11669.11704.36|GCCCAGTGCTCTGAATGTCAAAGTGAAGAAATTCAA</td><td>>hg38_21_+_8433172_8446622_RNA45SN1_45S_50ntFlanks</td><td>11669</td><td>11704</td><td>36</td><td>GCCCAGTGCTCTGAATGTCAAAGTGAAGAAATTCAA</td><td>481.1</td><td>6.1</td><td>0.6</td><td>9.7</td><td>4.6E-22</td><td>1.3E-20</td><td>MB231/MB468</td><td>CYTO</td></tr>
<tr><td>>hg38_21_+_8433172_8446622_RNA45SN1_45S_50ntFlanks@8552.8581.30|CTCCCCGGAGGGGGCGGGCTCCGGCGGGTG</td><td>>hg38_21_+_8433172_8446622_RNA45SN1_45S_50ntFlanks</td><td>8552</td><td>8581</td><td>30</td><td>CTCCCCGGAGGGGGCGGGCTCCGGCGGGTG</td><td>562.9</td><td>-13.6</td><td>1.4</td><td>-9.5</td><td>3.1E-21</td><td>1.3E-20</td><td>BT20/MB468</td><td>NUC</td></tr>
<tr><td>>hg38_21_+_8433172_8446622_RNA45SN1_45S_50ntFlanks@11212.11231.20|CGACTCTGGACGCGAGCCGG</td><td>>hg38_21_+_8433172_8446622_RNA45SN1_45S_50ntFlanks</td><td>11212</td><td>11231</td><td>20</td><td>CGACTCTGGACGCGAGCCGG</td><td>425.8</td><td>-7.4</td><td>0.8</td><td>-9.7</td><td>4.7E-22</td><td>1.3E-20</td><td>MB231/MB468</td><td>CYTO</td></tr>
<tr><td>>hg38_21_+_8433172_8446622_RNA45SN1_45S_50ntFlanks@11580.11611.32|GAACTGGTGCGGACCAGGGGAATCCGACTGTT</td><td>>hg38_21_+_8433172_8446622_RNA45SN1_45S_50ntFlanks</td><td>11580</td><td>11611</td><td>32</td><td>GAACTGGTGCGGACCAGGGGAATCCGACTGTT</td><td>5196.1</td><td>-5.4</td><td>0.6</td><td>-9.6</td><td>9.2E-22</td><td>1.3E-20</td><td>MB231/MB468</td><td>MITO</td></tr>
<tr><td>>hg38_21_+_8433172_8446622_RNA45SN1_45S_50ntFlanks@6738.6771.34|ATCATCGACACTTCGAACGCACTTGCGGCCCCGG</td><td>>hg38_21_+_8433172_8446622_RNA45SN1_45S_50ntFlanks</td><td>6738</td><td>6771</td><td>34</td><td>ATCATCGACACTTCGAACGCACTTGCGGCCCCGG</td><td>15865.5</td><td>-12.3</td><td>1.3</td><td>-9.6</td><td>9.2E-22</td><td>1.3E-20</td><td>MB231/MB468</td><td>MITO</td></tr>
<tr><td>>hg38_21_+_8433172_8446622_RNA45SN1_45S_50ntFlanks@6738.6771.34|ATCATCGACACTTCGAACGCACTTGCGGCCCCGG</td><td>>hg38_21_+_8433172_8446622_RNA45SN1_45S_50ntFlanks</td><td>6738</td><td>6771</td><td>34</td><td>ATCATCGACACTTCGAACGCACTTGCGGCCCCGG</td><td>1018.8</td><td>-12.3</td><td>1.3</td><td>-9.5</td><td>1.6E-21</td><td>1.3E-20</td><td>BT20/MB468</td><td>CYTO</td></tr>
<tr><td>>hg38_21_+_8433172_8446622_RNA45SN1_45S_50ntFlanks@4773.4793.21|TAGTTCCGACCATAAACGATG</td><td>>hg38_21_+_8433172_8446622_RNA45SN1_45S_50ntFlanks</td><td>4773</td><td>4793</td><td>21</td><td>TAGTTCCGACCATAAACGATG</td><td>1149.6</td><td>-14.0</td><td>1.5</td><td>-9.6</td><td>5.8E-22</td><td>1.3E-20</td><td>MB231/MB468</td><td>MP</td></tr>
<tr><td>>MI0000542|hsa-mir-320a&WithFlank&8|-|22244960|22245043@48.70.23(+1C)&[MIMAT0000510&hsa-miR-320a-3p&offsets|0|+1(+1C);m-60&8|-|22244975|22244996&offsets|0|+1(+1C)]|AAAAGCTGGGTTGAGAGGGCGAAC</td><td>>MI0000542|hsa-mir-320a&WithFlank&8|-|22244960|22245043</td><td>48</td><td>70</td><td>23(+1C)</td><td>AAAAGCTGGGTTGAGAGGGCGAAC</td><td>5130.4</td><td>11.8</td><td>1.2</td><td>9.5</td><td>1.6E-21</td><td>1.3E-20</td><td>BT20/MB468</td><td>CYTO</td></tr>
<tr><td>>hg38_21_+_8433172_8446622_RNA45SN1_45S_50ntFlanks@12008.12042.35|CCGCCGGTGAAATACCACTACTCTGATCGTTTTTT</td><td>>hg38_21_+_8433172_8446622_RNA45SN1_45S_50ntFlanks</td><td>12008</td><td>12042</td><td>35</td><td>CCGCCGGTGAAATACCACTACTCTGATCGTTTTTT</td><td>2582.9</td><td>-15.1</td><td>1.6</td><td>-9.6</td><td>9.4E-22</td><td>1.3E-20</td><td>MB231/MB468</td><td>MITO</td></tr>
<tr><td>>hg38_21_+_8433172_8446622_RNA45SN1_45S_50ntFlanks@11921.11948.28|TGAGAGGTGTAGAATAAGTGGGAGGCCC</td><td>>hg38_21_+_8433172_8446622_RNA45SN1_45S_50ntFlanks</td><td>11921</td><td>11948</td><td>28</td><td>TGAGAGGTGTAGAATAAGTGGGAGGCCC</td><td>1501.3</td><td>-14.4</td><td>1.5</td><td>-9.6</td><td>5.9E-22</td><td>1.3E-20</td><td>MB231/MB468</td><td>MP</td></tr>
</table>

<table>
<tr><td>>am_trna18_AlaTGC_11_-_50233879_50233950@58.75.18,trna67_AlaAGC_6_+_28678366_28678437@58.75.18,trna87_AlaAGC_1_-_150017382_150017452@57.74.18|TCCCCAGCACCTCCACCA</td><td>>am_trna18_AlaTGC_11_-_50233879_50233950</td><td>58</td><td>75</td><td>18</td><td>TCCCCAGCACCTCCACCA</td><td>439.6</td><td>13.0</td><td>1.4</td><td>9.5</td><td>1.6E-21</td><td>1.3E-20</td><td>BT20/MB468</td><td>CYTO</td></tr>
<tr><td>>hg38_21_+_8433172_8446622_RNA45SN1_45S_50ntFlanks@11767.11792.26|TCTAATTAGTGACGCGCATGAATGGA</td><td>>hg38_21_+_8433172_8446622_RNA45SN1_45S_50ntFlanks</td><td>11767</td><td>11792</td><td>26</td><td>TCTAATTAGTGACGCGCATGAATGGA</td><td>349.7</td><td>-12.9</td><td>1.4</td><td>-9.5</td><td>3.2E-21</td><td>1.3E-20</td><td>BT20/MB468</td><td>NUC</td></tr>
<tr><td>>hg38_21_+_8433172_8446622_RNA45SN1_45S_50ntFlanks@8351.8375.25|AAGTTGAAAAGAACTTTGAAGAGAG</td><td>>hg38_21_+_8433172_8446622_RNA45SN1_45S_50ntFlanks</td><td>8351</td><td>8375</td><td>25</td><td>AAGTTGAAAAGAACTTTGAAGAGAG</td><td>1015.2</td><td>14.4</td><td>1.5</td><td>9.6</td><td>1.2E-21</td><td>1.4E-20</td><td>BT20/MB468</td><td>MITO</td></tr>
<tr><td>>hg38_21_+_8433172_8446622_RNA45SN1_45S_50ntFlanks@7983.8004.22|CAGATCAGACGTGGCGACCCGC</td><td>>hg38_21_+_8433172_8446622_RNA45SN1_45S_50ntFlanks</td><td>7983</td><td>8004</td><td>22</td><td>CAGATCAGACGTGGCGACCCGC</td><td>4323.9</td><td>-7.6</td><td>0.8</td><td>-9.6</td><td>9.6E-22</td><td>1.4E-20</td><td>MB231/MB468</td><td>MITO</td></tr>
<tr><td>>hg38_21_+_8433172_8446622_RNA45SN1_45S_50ntFlanks@8112.8138.27|GGGCGCGGGACATGTGGCGTACGGAAG</td><td>>hg38_21_+_8433172_8446622_RNA45SN1_45S_50ntFlanks</td><td>8112</td><td>8138</td><td>27</td><td>GGGCGCGGGACATGTGGCGTACGGAAG</td><td>2317.9</td><td>-15.0</td><td>1.6</td><td>-9.6</td><td>6.1E-22</td><td>1.4E-20</td><td>MB231/MB468</td><td>MP</td></tr>
<tr><td>>hg38_21_+_8433172_8446622_RNA45SN1_45S_50ntFlanks@4064.4081.18|ATCAACTTTCGATGGTAG</td><td>>hg38_21_+_8433172_8446622_RNA45SN1_45S_50ntFlanks</td><td>4064</td><td>4081</td><td>18</td><td>ATCAACTTTCGATGGTAG</td><td>691.7</td><td>-13.3</td><td>1.4</td><td>-9.6</td><td>6.1E-22</td><td>1.4E-20</td><td>MB231/MB468</td><td>MP</td></tr>
<tr><td>>hg38_21_+_8433172_8446622_RNA45SN1_45S_50ntFlanks@4003.4023.21|ATAACCTCGGGCCGATCGCAC</td><td>>hg38_21_+_8433172_8446622_RNA45SN1_45S_50ntFlanks</td><td>4003</td><td>4023</td><td>21</td><td>ATAACCTCGGGCCGATCGCAC</td><td>4504.8</td><td>-7.8</td><td>0.8</td><td>-9.5</td><td>3.3E-21</td><td>1.4E-20</td><td>BT20/MB468</td><td>NUC</td></tr>
<tr><td>>MI0003628|hsa-mir-615&WithFlank&12|+|54033944|54034051@67.85.19(+1U)&[MIMAT0003283&hsa-miR-615-3p&offsets|0|-3(+1U);m-495&12|+|54034010|54034030&offsets|0|-2(+1U)]|TCCGAGCCTGGGTCTCCCTT</td><td>>MI0003628|hsa-mir-615&WithFlank&12|+|54033944|54034051</td><td>67</td><td>85</td><td>19(+1U)</td><td>TCCGAGCCTGGGTCTCCCTT</td><td>256.6</td><td>8.9</td><td>0.9</td><td>9.5</td><td>1.6E-21</td><td>1.4E-20</td><td>BT20/MB468</td><td>CYTO</td></tr>
<tr><td>>hg38_21_+_8433172_8446622_RNA45SN1_45S_50ntFlanks@6667.6686.20|ATCACTCGGCTCGTGCGTCG</td><td>>hg38_21_+_8433172_8446622_RNA45SN1_45S_50ntFlanks</td><td>6667</td><td>6686</td><td>20</td><td>ATCACTCGGCTCGTGCGTCG</td><td>1164.3</td><td>-14.0</td><td>1.5</td><td>-9.6</td><td>9.7E-22</td><td>1.4E-20</td><td>MB231/MB468</td><td>MITO</td></tr>
<tr><td>>MI0005756|hsa-mir-934&WithFlank&X|+|136550872|136550966@21.42.22&[MIMAT0004977&hsa-miR-934&offsets|0|0;m-943&X|+|136550892|136550913&offsets|0|0]|TGTCTACTACTGGAGACACTGG</td><td>>MI0005756|hsa-mir-934&WithFlank&X|+|136550872|136550966</td><td>21</td><td>42</td><td>22</td><td>TGTCTACTACTGGAGACACTGG</td><td>1298.3</td><td>14.0</td><td>1.4</td><td>9.8</td><td>7.7E-23</td><td>1.4E-20</td><td>BT20/MB231</td><td>MITO</td></tr>
<tr><td>>hg38_21_+_8433172_8446622_RNA45SN1_45S_50ntFlanks@10690.10708.19|CGTCCGGTGAGCTCTCGCT</td><td>>hg38_21_+_8433172_8446622_RNA45SN1_45S_50ntFlanks</td><td>10690</td><td>10708</td><td>19</td><td>CGTCCGGTGAGCTCTCGCT</td><td>7107.1</td><td>-11.9</td><td>1.2</td><td>-9.6</td><td>6.2E-22</td><td>1.4E-20</td><td>MB231/MB468</td><td>MP</td></tr>
<tr><td>>hg38_21_+_8433172_8446622_RNA45SN1_45S_50ntFlanks@11666.11706.41|TCTGCCCAGTGCTCTGAATGTCAAAGTGAAGAAATTCAATG</td><td>>hg38_21_+_8433172_8446622_RNA45SN1_45S_50ntFlanks</td><td>11666</td><td>11706</td><td>41</td><td>TCTGCCCAGTGCTCTGAATGTCAAAGTGAAGAAATTCAATG</td><td>1420.4</td><td>8.1</td><td>0.8</td><td>9.6</td><td>5.0E-22</td><td>1.4E-20</td><td>MB231/MB468</td><td>CYTO</td></tr>
<tr><td>>hg38_21_+_8433172_8446622_RNA45SN1_45S_50ntFlanks@4758.4780.23|GATCAGATACCGTCGTAGTTCCG</td><td>>hg38_21_+_8433172_8446622_RNA45SN1_45S_50ntFlanks</td><td>4758</td><td>4780</td><td>23</td><td>GATCAGATACCGTCGTAGTTCCG</td><td>355.3</td><td>-12.9</td><td>1.4</td><td>-9.5</td><td>3.4E-21</td><td>1.4E-20</td><td>BT20/MB468</td><td>NUC</td></tr>
<tr><td>>hg38_21_+_8433172_8446622_RNA45SN1_45S_50ntFlanks@11233.11254.22|CCCTTCCCGTGGATCGCCCCAG</td><td>>hg38_21_+_8433172_8446622_RNA45SN1_45S_50ntFlanks</td><td>11233</td><td>11254</td><td>22</td><td>CCCTTCCCGTGGATCGCCCCAG</td><td>1836.9</td><td>-13.8</td><td>1.4</td><td>-9.6</td><td>9.1E-22</td><td>1.4E-20</td><td>MB231/MB468</td><td>NUC</td></tr>
<tr><td>>hg38_21_+_8433172_8446622_RNA45SN1_45S_50ntFlanks@9465.9503.39|AGGTGGAGCACGAGCGCACGTGTTAGGACCCGAAAGATG</td><td>>hg38_21_+_8433172_8446622_RNA45SN1_45S_50ntFlanks</td><td>9465</td><td>9503</td><td>39</td><td>AGGTGGAGCACGAGCGCACGTGTTAGGACCCGAAAGATG</td><td>1694.6</td><td>-14.5</td><td>1.5</td><td>-9.6</td><td>9.9E-22</td><td>1.4E-20</td><td>MB231/MB468</td><td>MITO</td></tr>
<tr><td>>hg38_21_+_8433172_8446622_RNA45SN1_45S_50ntFlanks@12539.12568.30|ATGATGTGTTGTTGCCATGGTAATCCTGCT</td><td>>hg38_21_+_8433172_8446622_RNA45SN1_45S_50ntFlanks</td><td>12539</td><td>12568</td><td>30</td><td>ATGATGTGTTGTTGCCATGGTAATCCTGCT</td><td>820.1</td><td>-13.5</td><td>1.4</td><td>-9.6</td><td>6.3E-22</td><td>1.4E-20</td><td>MB231/MB468</td><td>MP</td></tr>
<tr><td>>hg38_21_+_8433172_8446622_RNA45SN1_45S_50ntFlanks@6667.6699.33|ATCACTCGGCTCGTGCGTCGATGAAGAACGCAG</td><td>>hg38_21_+_8433172_8446622_RNA45SN1_45S_50ntFlanks</td><td>6667</td><td>6699</td><td>33</td><td>ATCACTCGGCTCGTGCGTCGATGAAGAACGCAG</td><td>341.3</td><td>-12.8</td><td>1.4</td><td>-9.4</td><td>3.4E-21</td><td>1.4E-20</td><td>BT20/MB468</td><td>NUC</td></tr>
<tr><td>>MI0000255|hsa-mir-30d&WithFlank&8|-|134804870|134804951@12.34.23(+1C)&[MIMAT0000245&hsa-miR-30d-5p&offsets|0|+1(+1C);m-21&8|-|134804919|134804940&offsets|0|+1(+1C)]|TGTAAACATCCCCGACTGGAAGCC</td><td>>MI0000255|hsa-mir-30d&WithFlank&8|-|134804870|134804951</td><td>12</td><td>34</td><td>23(+1C)</td><td>TGTAAACATCCCCGACTGGAAGCC</td><td>2892.1</td><td>3.8</td><td>0.4</td><td>9.4</td><td>3.4E-21</td><td>1.4E-20</td><td>BT20/MB468</td><td>NUC</td></tr>
<tr><td>>hg38_21_+_8433172_8446622_RNA45SN1_45S_50ntFlanks@12892.12935.44|CCCTCGCCCGTCACGCACCGCACGTTCGTGGGGAACCTGGCGCT</td><td>>hg38_21_+_8433172_8446622_RNA45SN1_45S_50ntFlanks</td><td>12892</td><td>12935</td><td>44</td><td>CCCTCGCCCGTCACGCACCGCACGTTCGTGGGGAACCTGGCGCT</td><td>1031.6</td><td>-13.8</td><td>1.4</td><td>-9.6</td><td>1.0E-21</td><td>1.4E-20</td><td>MB231/MB468</td><td>MITO</td></tr>
<tr><td>>hg38_21_+_8433172_8446622_RNA45SN1_45S_50ntFlanks@9545.9566.22|GTGGAGGTCCGTAGCGGTCCTG</td><td>>hg38_21_+_8433172_8446622_RNA45SN1_45S_50ntFlanks</td><td>9545</td><td>9566</td><td>22</td><td>GTGGAGGTCCGTAGCGGTCCTG</td><td>995.6</td><td>-12.9</td><td>1.3</td><td>-9.6</td><td>9.4E-22</td><td>1.4E-20</td><td>MB231/MB468</td><td>NUC</td></tr>
<tr><td>>hg38_21_+_8433172_8446622_RNA45SN1_45S_50ntFlanks@10402.10429.28|AACATGGGTCAGTCGGTCCTGAGAGATG</td><td>>hg38_21_+_8433172_8446622_RNA45SN1_45S_50ntFlanks</td><td>10402</td><td>10429</td><td>28</td><td>AACATGGGTCAGTCGGTCCTGAGAGATG</td><td>6729.7</td><td>-5.9</td><td>0.6</td><td>-9.6</td><td>9.5E-22</td><td>1.5E-20</td><td>MB231/MB468</td><td>NUC</td></tr>
<tr><td>>trnaMT_ValTAC_MT_+_1602_1670@53.71.19|TAACTTGACCGCTCTGACC</td><td>>trnaMT_ValTAC_MT_+_1602_1670</td><td>53</td><td>71</td><td>19</td><td>TAACTTGACCGCTCTGACC</td><td>260.7</td><td>12.0</td><td>1.2</td><td>9.6</td><td>5.2E-22</td><td>1.5E-20</td><td>MB231/MB468</td><td>CYTO</td></tr>
<tr><td>>hg38_21_+_8433172_8446622_RNA45SN1_45S_50ntFlanks@9739.9759.21|AAACGATCTCAACCTATTCTC</td><td>>hg38_21_+_8433172_8446622_RNA45SN1_45S_50ntFlanks</td><td>9739</td><td>9759</td><td>21</td><td>AAACGATCTCAACCTATTCTC</td><td>762.0</td><td>-13.4</td><td>1.4</td><td>-9.6</td><td>6.5E-22</td><td>1.5E-20</td><td>MB231/MB468</td><td>MP</td></tr>
<tr><td>>hg38_21_+_8433172_8446622_RNA45SN1_45S_50ntFlanks@12937.12956.20|AACCATTCGTAGACGACCTG</td><td>>hg38_21_+_8433172_8446622_RNA45SN1_45S_50ntFlanks</td><td>12937</td><td>12956</td><td>20</td><td>AACCATTCGTAGACGACCTG</td><td>3387.0</td><td>-11.3</td><td>1.2</td><td>-9.6</td><td>6.6E-22</td><td>1.5E-20</td><td>MB231/MB468</td><td>MP</td></tr>
<tr><td>>trna119_LysCTT_1_-_145395522_145395594@1.34.34,trna11_LysCTT_5_-_180648979_180649051@1.34.34,trna13_LysCTT_6_+_26556774_26556846@1.34.34,trna7_LysCTT_16_+_3225692_3225764@1.34.34,trna9_LysCTT_5_+_180634755_180634827@1.34.34|GCCCGGCTAGCTCAGTCGGTAGAGCATGAGACTC</td><td>>trna119_LysCTT_1_-_145395522_145395594</td><td>1</td><td>34</td><td>34</td><td>GCCCGGCTAGCTCAGTCGGTAGAGCATGAGACTC</td><td>4118.4</td><td>7.2</td><td>0.8</td><td>9.5</td><td>1.8E-21</td><td>1.5E-20</td><td>BT20/MB468</td><td>CYTO</td></tr>
<tr><td>>MI0000108|hsa-mir-103a-2&WithFlank&20|+|3917488|3917577@54.76.23(+2U);MI0000109|hsa-mir-103a-1&WithFlank&5|-|168560890|168560979@54.76.23(+2U)&[MIMAT0000101&hsa-miR-103a-3p&offsets|0|0(+2U);m-12&20|+|3917541|3917563&offsets|0|0(+2U)];[MIMAT0000101_1&hsa-miR-103a-3p&offsets|0|0(+2U);m-13&5|-|168560904|168560926&offsets|0|0(+2U);m-44220&5|-|168560908|168560926&offsets|0|+4(+2U)]|AGCAGCATTGTACAGGGCTATGATT</td><td>>MI0000108|hsa-mir-103a-2&WithFlank&20|+|3917488|3917577</td><td>54</td><td>76</td><td>23(+2U)</td><td>AGCAGCATTGTACAGGGCTATGATT</td><td>640.7</td><td>6.1</td><td>0.6</td><td>9.5</td><td>1.8E-21</td><td>1.5E-20</td><td>BT20/MB468</td><td>CYTO</td></tr>
<tr><td>>hg38_21_+_8433172_8446622_RNA45SN1_45S_50ntFlanks@11578.11602.25|TAGAACTGGTGCGGACCAGGGGAAT</td><td>>hg38_21_+_8433172_8446622_RNA45SN1_45S_50ntFlanks</td><td>11578</td><td>11602</td><td>25</td><td>TAGAACTGGTGCGGACCAGGGGAAT</td><td>309.7</td><td>-4.4</td><td>0.5</td><td>-9.4</td><td>3.6E-21</td><td>1.5E-20</td><td>BT20/MB468</td><td>NUC</td></tr>
<tr><td>>MI0000086|hsa-mir-28&WithFlank&3|+|188688775|188688872@60.81.22&[MIMAT0004502&hsa-miR-28-3p&offsets|0|0;m-28&3|+|188688834|188688855&offsets|0|0]|CACTAGATTGTGAGCTCCTGGA</td><td>>MI0000086|hsa-mir-28&WithFlank&3|+|188688775|188688872</td><td>60</td><td>81</td><td>22</td><td>CACTAGATTGTGAGCTCCTGGA</td><td>7907.9</td><td>8.6</td><td>0.9</td><td>9.6</td><td>5.4E-22</td><td>1.5E-20</td><td>MB231/MB468</td><td>CYTO</td></tr>
<tr><td>>hg38_21_+_8433172_8446622_RNA45SN1_45S_50ntFlanks@4445.4472.28|CTTGCCTCTCGGCGCCCCCTCGATGCTC</td><td>>hg38_21_+_8433172_8446622_RNA45SN1_45S_50ntFlanks</td><td>4445</td><td>4472</td><td>28</td><td>CTTGCCTCTCGGCGCCCCCTCGATGCTC</td><td>1203.4</td><td>-14.7</td><td>1.6</td><td>-9.4</td><td>3.6E-21</td><td>1.5E-20</td><td>BT20/MB468</td><td>NUC</td></tr>
<tr><td>>hg38_21_+_8433172_8446622_RNA45SN1_45S_50ntFlanks@12414.12444.31|CCTTCGATGTCGGCTCTTCCTATCATTGTGA</td><td>>hg38_21_+_8433172_8446622_RNA45SN1_45S_50ntFlanks</td><td>12414</td><td>12444</td><td>31</td><td>CCTTCGATGTCGGCTCTTCCTATCATTGTGA</td><td>382.3</td><td>5.7</td><td>0.6</td><td>9.5</td><td>1.3E-21</td><td>1.5E-20</td><td>BT20/MB468</td><td>MITO</td></tr>
<tr><td>>hg38_21_+_8433172_8446622_RNA45SN1_45S_50ntFlanks@12419.12463.45|GATGTCGGCTCTTCCTATCATTGTGAAGCAGAATTCACCAAGCGT</td><td>>hg38_21_+_8433172_8446622_RNA45SN1_45S_50ntFlanks</td><td>12419</td><td>12463</td><td>45</td><td>GATGTCGGCTCTTCCTATCATTGTGAAGCAGAATTCACCAAGCGT</td><td>481.4</td><td>12.9</td><td>1.3</td><td>9.6</td><td>5.5E-22</td><td>1.5E-20</td><td>MB231/MB468</td><td>CYTO</td></tr>
<tr><td>>am_MI0000786|hsa-mir-378a&WithFlank&5|+|149732819|149732896@49.69.21(+1A)&[MIMAT0000732&hsa-miR-378a-3p&offsets|0|-1(+1A);m-44&5|+|149732867|149732888&offsets|0|-1(+1A)]|ACTGGACTTGGAGTCAGAAGGA</td><td>>am_MI0000786|hsa-mir-378a&WithFlank&5|+|149732819|149732896</td><td>49</td><td>69</td><td>21(+1A)</td><td>ACTGGACTTGGAGTCAGAAGGA</td><td>1087.9</td><td>5.6</td><td>0.6</td><td>9.5</td><td>1.3E-21</td><td>1.5E-20</td><td>BT20/MB468</td><td>MITO</td></tr>
<tr><td>>hg38_21_+_8433172_8446622_RNA45SN1_45S_50ntFlanks@12289.12318.30|CCGTGAAAGCGGGGCCTCACGATCCTTCTG</td><td>>hg38_21_+_8433172_8446622_RNA45SN1_45S_50ntFlanks</td><td>12289</td><td>12318</td><td>30</td><td>CCGTGAAAGCGGGGCCTCACGATCCTTCTG</td><td>6238.9</td><td>-16.4</td><td>1.7</td><td>-9.6</td><td>1.1E-21</td><td>1.5E-20</td><td>MB231/MB468</td><td>MITO</td></tr>
<tr><td>>MI0000542|hsa-mir-320a&WithFlank&8|-|22244960|22245043@48.70.23(+1U)&[MIMAT0000510&hsa-miR-320a-3p&offsets|0|+1(+1U);m-60&8|-|22244975|22244996&offsets|0|+1(+1U)]|AAAAGCTGGGTTGAGAGGGCGAAT</td><td>>MI0000542|hsa-mir-320a&WithFlank&8|-|22244960|22245043</td><td>48</td><td>70</td><td>23(+1U)</td><td>AAAAGCTGGGTTGAGAGGGCGAAT</td><td>3656.8</td><td>7.7</td><td>0.8</td><td>9.6</td><td>5.6E-22</td><td>1.5E-20</td><td>MB231/MB468</td><td>CYTO</td></tr>
<tr><td>>hg38_21_+_8433172_8446622_RNA45SN1_45S_50ntFlanks@11669.11691.23|GCCCAGTGCTCTGAATGTCAAAG</td><td>>hg38_21_+_8433172_8446622_RNA45SN1_45S_50ntFlanks</td><td>11669</td><td>11691</td><td>23</td><td>GCCCAGTGCTCTGAATGTCAAAG</td><td>245.7</td><td>5.3</td><td>0.6</td><td>9.5</td><td>1.9E-21</td><td>1.6E-20</td><td>BT20/MB468</td><td>CYTO</td></tr>
<tr><td>>trna10_MetCAT_8_-_124169470_124169542@31.51.21,trna22_MetCAT_16_-_87417628_87417700@31.51.21|TCTCATAATCTGAAGGTCGTG</td><td>>trna10_MetCAT_8_-_124169470_124169542</td><td>31</td><td>51</td><td>21</td><td>TCTCATAATCTGAAGGTCGTG</td><td>1056.5</td><td>-13.9</td><td>1.4</td><td>-9.6</td><td>7.0E-22</td><td>1.6E-20</td><td>MB231/MB468</td><td>MP</td></tr>
<tr><td>>hg38_21_+_8433172_8446622_RNA45SN1_45S_50ntFlanks@8170.8196.27|CAAGTCCTTCTGATCGAGGCCCAGCCC</td><td>>hg38_21_+_8433172_8446622_RNA45SN1_45S_50ntFlanks</td><td>8170</td><td>8196</td><td>27</td><td>CAAGTCCTTCTGATCGAGGCCCAGCCC</td><td>587.3</td><td>-13.6</td><td>1.4</td><td>-9.4</td><td>3.8E-21</td><td>1.6E-20</td><td>BT20/MB468</td><td>NUC</td></tr>
<tr><td>>MI0000255|hsa-mir-30d&WithFlank&8|-|134804870|134804951@12.34.23(+1C)&[MIMAT0000245&hsa-miR-30d-5p&offsets|0|+1(+1C);m-21&8|-|134804919|134804940&offsets|0|+1(+1C)]|TGTAAACATCCCCGACTGGAAGCC</td><td>>MI0000255|hsa-mir-30d&WithFlank&8|-|134804870|134804951</td><td>12</td><td>34</td><td>23(+1C)</td><td>TGTAAACATCCCCGACTGGAAGCC</td><td>6391.0</td><td>6.1</td><td>0.6</td><td>9.6</td><td>1.0E-21</td><td>1.6E-20</td><td>MB231/MB468</td><td>NUC</td></tr>
<tr><td>>hg38_21_+_8433172_8446622_RNA45SN1_45S_50ntFlanks@12412.12446.35|ATCCTTCGATGTCGGCTCTTCCTATCATTGTGAAG</td><td>>hg38_21_+_8433172_8446622_RNA45SN1_45S_50ntFlanks</td><td>12412</td><td>12446</td><td>35</td><td>ATCCTTCGATGTCGGCTCTTCCTATCATTGTGAAG</td><td>477.6</td><td>-13.3</td><td>1.4</td><td>-9.4</td><td>3.8E-21</td><td>1.6E-20</td><td>BT20/MB468</td><td>NUC</td></tr>
<tr><td>>hg38_21_+_8433172_8446622_RNA45SN1_45S_50ntFlanks@5492.5514.23|AGAAGACGGTCGAACTTGACTAT</td><td>>hg38_21_+_8433172_8446622_RNA45SN1_45S_50ntFlanks</td><td>5492</td><td>5514</td><td>23</td><td>AGAAGACGGTCGAACTTGACTAT</td><td>11210.8</td><td>-10.7</td><td>1.1</td><td>-9.6</td><td>7.0E-22</td><td>1.6E-20</td><td>MB231/MB468</td><td>MP</td></tr>
<tr><td>>hg38_21_+_8433172_8446622_RNA45SN1_45S_50ntFlanks@11785.11803.19|TGAATGGATGAACGAGATT</td><td>>hg38_21_+_8433172_8446622_RNA45SN1_45S_50ntFlanks</td><td>11785</td><td>11803</td><td>19</td><td>TGAATGGATGAACGAGATT</td><td>13963.5</td><td>-11.4</td><td>1.2</td><td>-9.4</td><td>3.8E-21</td><td>1.6E-20</td><td>BT20/MB468</td><td>NUC</td></tr>
<tr><td>>hg38_21_+_8433172_8446622_RNA45SN1_45S_50ntFlanks@4759.4778.20|ATCAGATACCGTCGTAGTTC</td><td>>hg38_21_+_8433172_8446622_RNA45SN1_45S_50ntFlanks</td><td>4759</td><td>4778</td><td>20</td><td>ATCAGATACCGTCGTAGTTC</td><td>1153.7</td><td>-12.5</td><td>1.3</td><td>-9.5</td><td>1.9E-21</td><td>1.6E-20</td><td>BT20/MB468</td><td>CYTO</td></tr>
<tr><td>>hg38_21_+_8433172_8446622_RNA45SN1_45S_50ntFlanks@10410.10429.20|TCAGTCGGTCCTGAGAGATG</td><td>>hg38_21_+_8433172_8446622_RNA45SN1_45S_50ntFlanks</td><td>10410</td><td>10429</td><td>20</td><td>TCAGTCGGTCCTGAGAGATG</td><td>22300.0</td><td>-11.6</td><td>1.2</td><td>-9.4</td><td>3.9E-21</td><td>1.6E-20</td><td>BT20/MB468</td><td>NUC</td></tr>
<tr><td>>hg38_21_+_8433172_8446622_RNA45SN1_45S_50ntFlanks@12991.13011.21|CTCCCTCGCTGCGATCTATTG</td><td>>hg38_21_+_8433172_8446622_RNA45SN1_45S_50ntFlanks</td><td>12991</td><td>13011</td><td>21</td><td>CTCCCTCGCTGCGATCTATTG</td><td>5577.5</td><td>-8.0</td><td>0.8</td><td>-9.6</td><td>1.1E-21</td><td>1.6E-20</td><td>MB231/MB468</td><td>NUC</td></tr>
<tr><td>>MI0000283|hsa-mir-203a&WithFlank&14|+|104117399|104117520@72.92.21(+1U)&[MIMAT0000264&hsa-miR-203a-3p&offsets|+1|0(+1U);m-79&14|+|104117469|104117490&offsets|+1|0(+1U)]|TGAAATGTTTAGGACCACTAGT</td><td>>MI0000283|hsa-mir-203a&WithFlank&14|+|104117399|104117520</td><td>72</td><td>92</td><td>21(+1U)</td><td>TGAAATGTTTAGGACCACTAGT</td><td>1751.6</td><td>9.5</td><td>1.0</td><td>9.5</td><td>1.9E-21</td><td>1.6E-20</td><td>BT20/MB468</td><td>CYTO</td></tr>
<tr><td>>MI0003188|hsa-mir-503&WithFlank&X|-|134546322|134546404@12.31.20(+1A)&[MIMAT0002874&hsa-miR-503-5p&offsets|0|-3(+1A);m-297&X|-|134546371|134546393&offsets|0|-3(+1A)]|TAGCAGCGGGAACAGTTCTGA</td><td>>MI0003188|hsa-mir-503&WithFlank&X|-|134546322|134546404</td><td>12</td><td>31</td><td>20(+1A)</td><td>TAGCAGCGGGAACAGTTCTGA</td><td>471.0</td><td>13.1</td><td>1.4</td><td>9.5</td><td>1.9E-21</td><td>1.6E-20</td><td>BT20/MB468</td><td>CYTO</td></tr>
<tr><td>>hg38_21_+_8433172_8446622_RNA45SN1_45S_50ntFlanks@6719.6745.27|TGAATTGCAGGACACATTGATCATCGA</td><td>>hg38_21_+_8433172_8446622_RNA45SN1_45S_50ntFlanks</td><td>6719</td><td>6745</td><td>27</td><td>TGAATTGCAGGACACATTGATCATCGA</td><td>1093.5</td><td>-7.4</td><td>0.8</td><td>-9.6</td><td>5.9E-22</td><td>1.6E-20</td><td>MB231/MB468</td><td>CYTO</td></tr>
</table>

| | | | | | | | | | | | | | |
|---|---|---|---|---|---|---|---|---|---|---|---|---|---|
| >MI0000434\|hsa-let-7i&WithFlank&12\|+\|62603680\|62603775@12.33.22(+1U)&[MIMAT0000415&hsa-let-7i-5p&offsets\|0\|0(+1U);m-49&12\|+\|62603691\|62603712&offsets\|0\|0(+1U)]\|TGAGGTAGTAGTTTGTGCTGTTT | >MI0000434\|hsa-let-7i&WithFlank&12\|+\|62603680\|62603775 | 12 | 33 | 22(+1U) | TGAGGTAGTAGTTTGTGCTGTTT | 7657.2 | 6.5 | 0.7 | 9.6 | 1.1E-21 | 1.6E-20 | MB231/MB468 | NUC |
| >MI0000266\|hsa-mir-10a&WithFlank&17\|-\|48579832\|48579953@28.50.23(+1A)&[MIMAT0000253&hsa-miR-10a-5p&offsets\|0\|0(+1A);m-18&17\|-\|48579905\|48579926&offsets\|0\|+1(+1A)]\|TACCCTGTAGATCCGAATTTGTGA | >MI0000266\|hsa-mir-10a&WithFlank&17\|-\|48579832\|48579953 | 28 | 50 | 23(+1A) | TACCCTGTAGATCCGAATTTGTGA | 185.2 | 10.4 | 1.1 | 9.4 | 3.9E-21 | 1.6E-20 | BT20/MB468 | NUC |
| >hg38_21_+_8433172_8446622_RNA45SN1_45S_50ntFlanks@11565.11584.20\|CTAGCAGCCGACTTAGAACT | >hg38_21_+_8433172_8446622_RNA45SN1_45S_50ntFlanks | 11565 | 11584 | 20 | CTAGCAGCCGACTTAGAACT | 384.4 | 5.4 | 0.6 | 9.5 | 1.4E-21 | 1.6E-20 | BT20/MB468 | MITO |
| >hg38_21_+_8433172_8446622_RNA45SN1_45S_50ntFlanks@12419.12461.43\|GATGTCGGCTCTTCCTATCATTGTGAAGCAGAATTCACCAAGC | >hg38_21_+_8433172_8446622_RNA45SN1_45S_50ntFlanks | 12419 | 12461 | 43 | GATGTCGGCTCTTCCTATCATTGTGAAGCAGAATTCACCAAGC | 320.3 | 12.3 | 1.3 | 9.6 | 6.0E-22 | 1.7E-20 | MB231/MB468 | CYTO |
| >hg38_21_+_8433172_8446622_RNA45SN1_45S_50ntFlanks@10866.10891.26\|CTAAGGGCTGGGTCGGTCGGGCTGGG | >hg38_21_+_8433172_8446622_RNA45SN1_45S_50ntFlanks | 10866 | 10891 | 26 | CTAAGGGCTGGGTCGGTCGGGCTGGG | 1193.0 | -14.1 | 1.5 | -9.6 | 7.4E-22 | 1.7E-20 | MB231/MB468 | MP |
| >hg38_21_+_8433172_8446622_RNA45SN1_45S_50ntFlanks@12931.12956.26\|GCGCTAAACCATTCGTAGACGACCTG | >hg38_21_+_8433172_8446622_RNA45SN1_45S_50ntFlanks | 12931 | 12956 | 26 | GCGCTAAACCATTCGTAGACGACCTG | 964.4 | -13.8 | 1.4 | -9.6 | 7.5E-22 | 1.7E-20 | MB231/MB468 | MP |
| >hg38_21_+_8433172_8446622_RNA45SN1_45S_50ntFlanks@4002.4024.23\|GATAACCTCGGGCCGATCGCACG | >hg38_21_+_8433172_8446622_RNA45SN1_45S_50ntFlanks | 4002 | 4024 | 23 | GATAACCTCGGGCCGATCGCACG | 1204.2 | -14.1 | 1.5 | -9.6 | 7.6E-22 | 1.7E-20 | MB231/MB468 | MP |
| >hg38_21_+_8433172_8446622_RNA45SN1_45S_50ntFlanks@4726.4758.33\|TAATCAAGAACGAAAGTCGGAGGTTCGAAGACG | >hg38_21_+_8433172_8446622_RNA45SN1_45S_50ntFlanks | 4726 | 4758 | 33 | TAATCAAGAACGAAAGTCGGAGGTTCGAAGACG | 10128.2 | -10.8 | 1.1 | -9.4 | 4.1E-21 | 1.7E-20 | BT20/MB468 | NUC |
| >hg38_21_+_8433172_8446622_RNA45SN1_45S_50ntFlanks@10392.10435.44\|ACAGCAGTTGAACATGGGTCAGTCGGTCCTGAGAGATGGGCGAG | >hg38_21_+_8433172_8446622_RNA45SN1_45S_50ntFlanks | 10392 | 10435 | 44 | ACAGCAGTTGAACATGGGTCAGTCGGTCCTGAGAGATGGGCGAG | 2938.7 | -15.9 | 1.7 | -9.4 | 4.1E-21 | 1.7E-20 | BT20/MB468 | NUC |
| >hg38_21_+_8433172_8446622_RNA45SN1_45S_50ntFlanks@7985.8006.22\|GATCAGACGTGGCGACCCGCTG | >hg38_21_+_8433172_8446622_RNA45SN1_45S_50ntFlanks | 7985 | 8006 | 22 | GATCAGACGTGGCGACCCGCTG | 8503.2 | -12.1 | 1.3 | -9.6 | 1.2E-21 | 1.7E-20 | MB231/MB468 | MITO |
| >hg38_21_+_8433172_8446622_RNA45SN1_45S_50ntFlanks@6790.6807.18\|CCTGTCTGAGCGTCGCTT | >hg38_21_+_8433172_8446622_RNA45SN1_45S_50ntFlanks | 6790 | 6807 | 18 | CCTGTCTGAGCGTCGCTT | 1312.7 | 4.1 | 0.4 | 9.5 | 2.0E-21 | 1.7E-20 | BT20/MB468 | CYTO |
| >hg38_21_+_8433172_8446622_RNA45SN1_45S_50ntFlanks@7986.8017.32\|ATCAGACGTGGCGACCCGCTGAATTTAAGCAT | >hg38_21_+_8433172_8446622_RNA45SN1_45S_50ntFlanks | 7986 | 8017 | 32 | ATCAGACGTGGCGACCCGCTGAATTTAAGCAT | 995.6 | -13.8 | 1.4 | -9.6 | 7.7E-22 | 1.7E-20 | MB231/MB468 | MP |
| >hg38_21_+_8433172_8446622_RNA45SN1_45S_50ntFlanks@8350.8375.26\|AAAGTTGAAAAGAACTTTGAAGAGAG | >hg38_21_+_8433172_8446622_RNA45SN1_45S_50ntFlanks | 8350 | 8375 | 26 | AAAGTTGAAAAGAACTTTGAAGAGAG | 611.7 | 13.7 | 1.4 | 9.5 | 1.5E-21 | 1.7E-20 | BT20/MB468 | MITO |
| >MI0000103\|hsa-mir-101-1&WithFlank&1\|-\|65058428\|65058514@52.73.22;MI0000739\|hsa-mir-101-2&WithFlank&9\|+\|4850291\|4850381@54.75.22&[MIMAT0000099&hsa-miR-101-3p&offsets\|-1\|0;m-17&1\|-\|65058443\|65058462&offsets\|-1\|+1];[MIMAT0000099_1&hsa-miR-101-3p&offsets\|-1\|0;m-16&9\|+\|4850345\|4850364&offsets\|-1\|+1]\|GTACAGTACTGTGATAACTGAA | >MI0000103\|hsa-mir-101-1&WithFlank&1\|-\|65058428\|65058514 | 52 | 73 | 22 | GTACAGTACTGTGATAACTGAA | 477.1 | 4.7 | 0.5 | 9.5 | 1.5E-21 | 1.7E-20 | BT20/MB468 | MITO |
| >MI0000743\|hsa-mir-34c&WithFlank&11\|+\|111513433\|111513521@19.41.23&[MIMAT0000686&hsa-miR-34c-5p&offsets\|0\|0;m-336&11\|+\|111513451\|111513473&offsets\|0\|0]\|AGGCAGTGTAGTTAGCTGATTGC | >MI0000743\|hsa-mir-34c&WithFlank&11\|+\|111513433\|111513521 | 19 | 41 | 23 | AGGCAGTGTAGTTAGCTGATTGC | 882.4 | 13.0 | 1.4 | 9.6 | 7.8E-22 | 1.7E-20 | MB231/MB468 | MP |
| >trna22_MetCAT_16_-_87417628_87417700@15.42.28,trna75_MetCAT_6_+_28912352_28912424@15.42.28,trna97_MetCAT_6_-_28921042_28921114@15.42.28\|GTAGGCAGCGCGTCAGTCTCATAATCTG | >trna22_MetCAT_16_-_87417628_87417700 | 15 | 42 | 28 | GTAGGCAGCGCGTCAGTCTCATAATCTG | 385.5 | -13.0 | 1.4 | -9.4 | 4.2E-21 | 1.7E-20 | BT20/MB468 | NUC |
| >hg38_21_+_8433172_8446622_RNA45SN1_45S_50ntFlanks@12934.12951.18\|CTAAACCATTCGTAGACG | >hg38_21_+_8433172_8446622_RNA45SN1_45S_50ntFlanks | 12934 | 12951 | 18 | CTAAACCATTCGTAGACG | 1154.8 | -8.4 | 0.9 | -9.6 | 7.9E-22 | 1.7E-20 | MB231/MB468 | MP |
| >hg38_21_+_8433172_8446622_RNA45SN1_45S_50ntFlanks@4045.4063.19\|CCATTCGAACGTCTGCCCT | >hg38_21_+_8433172_8446622_RNA45SN1_45S_50ntFlanks | 4045 | 4063 | 19 | CCATTCGAACGTCTGCCCT | 2028.8 | -5.6 | 0.6 | -9.5 | 2.1E-21 | 1.8E-20 | BT20/MB468 | CYTO |
| >MI0000077\|hsa-mir-21&WithFlank&17\|+\|59841260\|59841343@14.32.19&[MIMAT0000076&hsa-miR-21-5p&offsets\|0\|-3;m-1&17\|+\|59841273\|59841294&offsets\|0\|-3]\|TAGCTTATCAGACTGATGT | >MI0000077\|hsa-mir-21&WithFlank&17\|+\|59841260\|59841343 | 14 | 32 | 19 | TAGCTTATCAGACTGATGT | 2694.0 | 8.5 | 0.9 | 9.5 | 2.1E-21 | 1.8E-20 | BT20/MB468 | CYTO |
| >hg38_21_+_8433172_8446622_RNA45SN1_45S_50ntFlanks@4555.4572.18\|CGCAGCTAGGAATAATGG | >hg38_21_+_8433172_8446622_RNA45SN1_45S_50ntFlanks | 4555 | 4572 | 18 | CGCAGCTAGGAATAATGG | 890.8 | -13.7 | 1.4 | -9.6 | 8.0E-22 | 1.8E-20 | MB231/MB468 | MP |
| >MI0000285\|hsa-mir-205&WithFlank&1\|+\|209432127\|209432248@40.61.22&[MIMAT0000266&hsa-miR-205-5p&offsets\|0\|0;m-351&1\|+\|209432166\|209432187&offsets\|0\|0]\|TCCTTCATTCCACCGGAGTCTG | >MI0000285\|hsa-mir-205&WithFlank&1\|+\|209432127\|209432248 | 40 | 61 | 22 | TCCTTCATTCCACCGGAGTCTG | 8220.3 | -12.4 | 1.3 | -9.6 | 8.0E-22 | 1.8E-20 | MB231/MB468 | MP |
| >hg38_21_+_8433172_8446622_RNA45SN1_45S_50ntFlanks@12300.12318.19\|GGGCCTCACGATCCTTCTG | >hg38_21_+_8433172_8446622_RNA45SN1_45S_50ntFlanks | 12300 | 12318 | 19 | GGGCCTCACGATCCTTCTG | 3279.2 | -15.5 | 1.6 | -9.6 | 1.3E-21 | 1.8E-20 | MB231/MB468 | MITO |
| >MI0000102\|hsa-mir-100&WithFlank&11\|-\|122152223\|122152314@19.38.20&[MIMAT0000098&hsa-miR-100-5p&offsets\|0\|-2;m-27&11\|-\|122152276\|122152296&offsets\|0\|-1]\|AACCCGTAGATCCGAACTTG | >MI0000102\|hsa-mir-100&WithFlank&11\|-\|122152223\|122152314 | 19 | 38 | 20 | AACCCGTAGATCCGAACTTG | 19849.7 | 14.1 | 1.5 | 9.6 | 1.3E-21 | 1.8E-20 | MB231/MB468 | MITO |
| >hg38_21_+_8433172_8446622_RNA45SN1_45S_50ntFlanks@4759.4782.24\|ATCAGATACCGTCGTAGTTCCGAC | >hg38_21_+_8433172_8446622_RNA45SN1_45S_50ntFlanks | 4759 | 4782 | 24 | ATCAGATACCGTCGTAGTTCCGAC | 6560.7 | -16.5 | 1.7 | -9.6 | 1.3E-21 | 1.8E-20 | MB231/MB468 | MITO |
| >MI0000477\|hsa-mir-146a&WithFlank&5\|+\|160485346\|160485456@27.48.22&[MIMAT0000449&hsa-miR-146a-5p&offsets\|0\|0;m-129&5\|+\|160485372\|160485393&offsets\|0\|0]\|TGAGAACTGAATTCCATGGGTT | >MI0000477\|hsa-mir-146a&WithFlank&5\|+\|160485346\|160485456 | 27 | 48 | 22 | TGAGAACTGAATTCCATGGGTT | 96419.8 | 12.4 | 1.3 | 9.6 | 8.1E-22 | 1.8E-20 | MB231/MB468 | MP |
| >hg38_21_+_8433172_8446622_RNA45SN1_45S_50ntFlanks@10544.10567.24\|TCCAGTGCGGTAACGCGACCGATC | >hg38_21_+_8433172_8446622_RNA45SN1_45S_50ntFlanks | 10544 | 10567 | 24 | TCCAGTGCGGTAACGCGACCGATC | 886.2 | -13.7 | 1.4 | -9.6 | 8.1E-22 | 1.8E-20 | MB231/MB468 | MP |
| >hg38_21_+_8433172_8446622_RNA45SN1_45S_50ntFlanks@12072.12095.24\|CCCGAGGGGCTCTCGCTTCTGGCG | >hg38_21_+_8433172_8446622_RNA45SN1_45S_50ntFlanks | 12072 | 12095 | 24 | CCCGAGGGGCTCTCGCTTCTGGCG | 1093.0 | -14.0 | 1.5 | -9.6 | 8.3E-22 | 1.8E-20 | MB231/MB468 | MP |
| >hg38_21_+_8433172_8446622_RNA45SN1_45S_50ntFlanks@485.509.25\|GATGTGGTGACGTCGTGCTCTCCCG | >hg38_21_+_8433172_8446622_RNA45SN1_45S_50ntFlanks | 485 | 509 | 25 | GATGTGGTGACGTCGTGCTCTCCCG | 399.9 | 11.5 | 1.2 | 9.4 | 4.4E-21 | 1.8E-20 | BT20/MB468 | NUC |
| >hg38_21_+_8433172_8446622_RNA45SN1_45S_50ntFlanks@11667.11703.37\|CTGCCCAGTGCTCTGAATGTCAAAGTGAAGAAATTCA | >hg38_21_+_8433172_8446622_RNA45SN1_45S_50ntFlanks | 11667 | 11703 | 37 | CTGCCCAGTGCTCTGAATGTCAAAGTGAAGAAATTCA | 302.7 | 5.8 | 0.6 | 9.5 | 1.6E-21 | 1.8E-20 | BT20/MB468 | MITO |
| >MI0000061\|hsa-let-7a-2&WithFlank&11\|-\|122146516\|122146599@11.33.23;MI0000062\|hsa-let-7a-3&WithFlank&22\|+\|46112743\|46112828@10.32.23;MI0000060\|hsa-let-7a-1&WithFlank&9\|+\|94175951\|94176042@12.34.23&[MIMAT0000062&hsa-let-7a-5p&offsets\|0\|+1;m-6&11\|-\|122146568\|122146589&offsets\|0\|+1];[MIMAT0000062_1&hsa-let-7a-5p&offsets\|0\|+1;m-5&22\|+\|46112752\|46112773&offsets\|0\|+1];[MIMAT0000062_2&hsa-let-7a-5p&offsets\|0\|+1;m-4&9\|+\|94175962\|94175983&offsets\|0\|+1]\|TGAGGTAGTAGGTTGTATAGTTT | >MI0000061\|hsa-let-7a-2&WithFlank&11\|-\|122146516\|122146599 | 11 | 33 | 23 | TGAGGTAGTAGGTTGTATAGTTT | 3308.2 | 5.5 | 0.6 | 9.6 | 1.2E-21 | 1.8E-20 | MB231/MB468 | NUC |
| >hg38_21_+_8433172_8446622_RNA45SN1_45S_50ntFlanks@6747.6771.25\|ACTTCGAACGCACTTGCGGCCCCGG | >hg38_21_+_8433172_8446622_RNA45SN1_45S_50ntFlanks | 6747 | 6771 | 25 | ACTTCGAACGCACTTGCGGCCCCGG | 879.5 | -13.6 | 1.4 | -9.5 | 1.3E-21 | 1.8E-20 | MB231/MB468 | MITO |
| >hg38_21_+_8433172_8446622_RNA45SN1_45S_50ntFlanks@8357.8375.19\|AAAAGAACTTTGAAGAGAG | >hg38_21_+_8433172_8446622_RNA45SN1_45S_50ntFlanks | 8357 | 8375 | 19 | AAAAGAACTTTGAAGAGAG | 3771.1 | -7.6 | 0.8 | -9.5 | 1.3E-21 | 1.8E-20 | MB231/MB468 | MITO |
| >hg38_21_+_8433172_8446622_RNA45SN1_45S_50ntFlanks@9534.9571.38\|AGGAAACTCTGGTGGAGGTCCGTAGCGGTCCTGACGTG | >hg38_21_+_8433172_8446622_RNA45SN1_45S_50ntFlanks | 9534 | 9571 | 38 | AGGAAACTCTGGTGGAGGTCCGTAGCGGTCCTGACGTG | 16867.8 | -8.8 | 0.9 | -9.5 | 1.3E-21 | 1.8E-20 | MB231/MB468 | MITO |
| >hg38_21_+_8433172_8446622_RNA45SN1_45S_50ntFlanks@11575.11603.29\|ACTTAGAACTGGTGCGGACCAGGGGAATC | >hg38_21_+_8433172_8446622_RNA45SN1_45S_50ntFlanks | 11575 | 11603 | 29 | ACTTAGAACTGGTGCGGACCAGGGGAATC | 7365.8 | -5.4 | 0.6 | -9.5 | 2.2E-21 | 1.9E-20 | BT20/MB468 | CYTO |
| >hg38_21_+_8433172_8446622_RNA45SN1_45S_50ntFlanks@10914.10946.33\|CGCCGCGGCTGGACGAGGCGCCGCCGCCCCCCC | >hg38_21_+_8433172_8446622_RNA45SN1_45S_50ntFlanks | 10914 | 10946 | 33 | CGCCGCGGCTGGACGAGGCGCCGCCGCCCCCCC | 816.2 | -14.1 | 1.5 | -9.4 | 4.5E-21 | 1.9E-20 | BT20/MB468 | NUC |
| >MI0000087\|hsa-mir-29a&WithFlank&7\|-\|130876741\|130876816@47.69.23&[MIMAT0000086&hsa-miR-29a-3p&offsets\|-1\|0;m-26&7\|-\|130876749\|130876769&offsets\|-1\|+1]\|CTAGCACCATCTGAAATCGGTTA | >MI0000087\|hsa-mir-29a&WithFlank&7\|-\|130876741\|130876816 | 47 | 69 | 23 | CTAGCACCATCTGAAATCGGTTA | 1176.1 | 9.4 | 1.0 | 9.6 | 1.2E-21 | 1.9E-20 | MB231/MB468 | NUC |
| >hg38_21_+_8433172_8446622_RNA45SN1_45S_50ntFlanks@11874.11892.19\|AGAAGACCCTGTTGAGCTT | >hg38_21_+_8433172_8446622_RNA45SN1_45S_50ntFlanks | 11874 | 11892 | 19 | AGAAGACCCTGTTGAGCTT | 905.1 | -13.7 | 1.4 | -9.6 | 8.6E-22 | 1.9E-20 | MB231/MB468 | MP |
| >am_trna134_GluTTC_1_-_16861774_16861845@1.30.30,trna5_GluTTC_1_+_17199078_17199149@1.30.30,trna84_GluTTC_1_-_161391883_161391954@1.30.30,trna94_GluTTC_1_-_149664355_149664427@1.30.30\|TCCCTGGTGGTCTAGTGGCTAGGATTCGGC | >am_trna134_GluTTC_1_-_16861774_16861845 | 1 | 30 | 30 | TCCCTGGTGGTCTAGTGGCTAGGATTCGGC | 1043.8 | 27.2 | 2.7 | 10.2 | 3.0E-24 | 1.9E-20 | BT20/MB231 | MP |
| >hg38_21_+_8433172_8446622_RNA45SN1_45S_50ntFlanks@4377.4405.29\|GCTCGTAGTTGGATCTTGGGAGCGGGCGG | >hg38_21_+_8433172_8446622_RNA45SN1_45S_50ntFlanks | 4377 | 4405 | 29 | GCTCGTAGTTGGATCTTGGGAGCGGGCGG | 2600.0 | 27.9 | 2.8 | 10.1 | 7.1E-24 | 1.9E-20 | BT20/MB231 | MP |
| >hg38_21_+_8433172_8446622_RNA45SN1_45S_50ntFlanks@5485.5507.23\|AGCGCTGAGAAGACGGTCGAACT | >hg38_21_+_8433172_8446622_RNA45SN1_45S_50ntFlanks | 5485 | 5507 | 23 | AGCGCTGAGAAGACGGTCGAACT | 586.0 | 26.4 | 2.6 | 10.1 | 5.7E-24 | 1.9E-20 | BT20/MB231 | MP |
| >hg38_21_+_8433172_8446622_RNA45SN1_45S_50ntFlanks@11564.11605.42\|CCTAGCAGCCGACTTAGAACTGGTGCGGACCAGGGGAATCCG | >hg38_21_+_8433172_8446622_RNA45SN1_45S_50ntFlanks | 11564 | 11605 | 42 | CCTAGCAGCCGACTTAGAACTGGTGCGGACCAGGGGAATCCG | 1990.9 | -14.7 | 1.5 | -9.5 | 1.4E-21 | 1.9E-20 | MB231/MB468 | MITO |
| >hg38_21_+_8433172_8446622_RNA45SN1_45S_50ntFlanks@10351.10376.26\|CGAGAACTTTGAAGGCCGAAGTGGAG | >hg38_21_+_8433172_8446622_RNA45SN1_45S_50ntFlanks | 10351 | 10376 | 26 | CGAGAACTTTGAAGGCCGAAGTGGAG | 10542.0 | -17.2 | 1.8 | -9.6 | 9.0E-22 | 2.0E-20 | MB231/MB468 | MP |
| >MI0000086\|hsa-mir-28&WithFlank&3\|+\|188688775\|188688872@60.79.20&[MIMAT0004502&hsa-miR-28-3p&offsets\|0\|-2;m-28&3\|+\|188688834\|188688855&offsets\|0\|-2]\|CACTAGATTGTGAGCTCCTG | >MI0000086\|hsa-mir-28&WithFlank&3\|+\|188688775\|188688872 | 60 | 79 | 20 | CACTAGATTGTGAGCTCCTG | 533.1 | 13.3 | 1.4 | 9.5 | 2.4E-21 | 2.0E-20 | BT20/MB468 | CYTO |
| >hg38_21_+_8433172_8446622_RNA45SN1_45S_50ntFlanks@7977.8011.35\|CGACCTCAGATCAGACGTGGCGACCCGCTGAATTT | >hg38_21_+_8433172_8446622_RNA45SN1_45S_50ntFlanks | 7977 | 8011 | 35 | CGACCTCAGATCAGACGTGGCGACCCGCTGAATTT | 3821.9 | 8.5 | 0.9 | 9.6 | 7.3E-22 | 2.0E-20 | MB231/MB468 | CYTO |
| >hg38_21_+_8433172_8446622_RNA45SN1_45S_50ntFlanks@8169.8197.29\|CCAAGTCCTTCTGATCGAGGCCCAGCCCG | >hg38_21_+_8433172_8446622_RNA45SN1_45S_50ntFlanks | 8169 | 8197 | 29 | CCAAGTCCTTCTGATCGAGGCCCAGCCCG | 1203.6 | -14.7 | 1.6 | -9.4 | 4.8E-21 | 2.0E-20 | BT20/MB468 | NUC |
| >MI0000650\|hsa-mir-200c&WithFlank&12\|+\|6963693\|6963772@50.70.21&[MIMAT0000617&hsa-miR-200c-3p&offsets\|0\|-2;m-54&12\|+\|6963742\|6963764&offsets\|0\|-2]\|TAATACTGCCGGGTAATGATG | >MI0000650\|hsa-mir-200c&WithFlank&12\|+\|6963693\|6963772 | 50 | 70 | 21 | TAATACTGCCGGGTAATGATG | 2819.1 | 11.4 | 1.2 | 9.8 | 1.6E-22 | 2.0E-20 | BT20/MB231 | NUC |
| >MI0000440\|hsa-mir-27b&WithFlank&9\|+\|95085439\|95085547@70.87.18&[MIMAT0000419&hsa-miR-27b-3p&offsets\|+3\|0;m-39&9\|+\|95085505\|95085525&offsets\|+3\|0]\|ACAGTGGCTAAGTTCTGC | >MI0000440\|hsa-mir-27b&WithFlank&9\|+\|95085439\|95085547 | 70 | 87 | 18 | ACAGTGGCTAAGTTCTGC | 198.9 | 10.5 | 1.1 | 9.4 | 4.9E-21 | 2.0E-20 | BT20/MB468 | NUC |
| >hg38_21_+_8433172_8446622_RNA45SN1_45S_50ntFlanks@5490.5533.44\|TGAGAAGACGGTCGAACTTGACTATCTAGAGGAAGTAAAAGTCG | >hg38_21_+_8433172_8446622_RNA45SN1_45S_50ntFlanks | 5490 | 5533 | 44 | TGAGAAGACGGTCGAACTTGACTATCTAGAGGAAGTAAAAGTCG | 221.0 | 11.8 | 1.2 | 9.6 | 7.5E-22 | 2.0E-20 | MB231/MB468 | CYTO |

<table>
<tr><th></th><th></th><th></th><th></th><th></th><th></th><th></th><th></th><th></th><th></th><th></th><th></th><th></th><th></th></tr>
<tr><td>>hg38_21_+_8433172_8446622_RNA45SN1_45S_50ntFlanks@12151.12179.29|GTGGGGAGTTTGACTGGGGCGGTACACCT</td><td>>hg38_21_+_8433172_8446622_RNA45SN1_45S_50ntFlanks</td><td>12151</td><td>12179</td><td>29</td><td>GTGGGGAGTTTGACTGGGGCGGTACACCT</td><td>792.7</td><td>-13.5</td><td>1.4</td><td>-9.6</td><td>9.3E-22</td><td>2.0E-20</td><td>MB231/MB468</td><td>MP</td></tr>
<tr><td>>MI0000088|hsa-mir-30a&WithFlank&6|-|71403545|71403627@53.73.21&[MIMAT0000088&hsa-miR-30a-3p&offsets|0|-1;m-38&6|-|71403554|71403575&offsets|0|-1]|CTTTCAGTCGGATGTTTGCAG</td><td>>MI0000088|hsa-mir-30a&WithFlank&6|-|71403545|71403627</td><td>53</td><td>73</td><td>21</td><td>CTTTCAGTCGGATGTTTGCAG</td><td>1306.4</td><td>7.3</td><td>0.8</td><td>9.6</td><td>7.6E-22</td><td>2.0E-20</td><td>MB231/MB468</td><td>CYTO</td></tr>
<tr><td>>trnaMT_GluTTC_MT_-_14674_14742@40.61.22,trnalookalike8_GluTTC_5_-_93905172_93905240@40.61.22|TTGGTCGTGGTTGTAGTCCGTG</td><td>>trnaMT_GluTTC_MT_-_14674_14742</td><td>40</td><td>61</td><td>22</td><td>TTGGTCGTGGTTGTAGTCCGTG</td><td>867.0</td><td>7.4</td><td>0.8</td><td>9.5</td><td>1.8E-21</td><td>2.0E-20</td><td>BT20/MB468</td><td>MITO</td></tr>
<tr><td>>hg38_21_+_8433172_8446622_RNA45SN1_45S_50ntFlanks@10421.10456.36|TGAGAGATGGGCGAGCGCCGTTCCGAAGGGACGGGC</td><td>>hg38_21_+_8433172_8446622_RNA45SN1_45S_50ntFlanks</td><td>10421</td><td>10456</td><td>36</td><td>TGAGAGATGGGCGAGCGCCGTTCCGAAGGGACGGGC</td><td>1874.7</td><td>-6.6</td><td>0.7</td><td>-9.5</td><td>1.4E-21</td><td>2.1E-20</td><td>MB231/MB468</td><td>NUC</td></tr>
<tr><td>>MI0000066|hsa-let-7e&WithFlank&19|+|51692780|51692870@14.35.22(+1U)&[MIMAT0000066&hsa-let-7e-5p&offsets|0|0(+1U);m-50&19|+|51692793|51692814&offsets|0|0(+1U)]|TGAGGTAGGAGGTTGTATAGTTT</td><td>>MI0000066|hsa-let-7e&WithFlank&19|+|51692780|51692870</td><td>14</td><td>35</td><td>22(+1U)</td><td>TGAGGTAGGAGGTTGTATAGTTT</td><td>681.2</td><td>6.5</td><td>0.7</td><td>9.5</td><td>1.8E-21</td><td>2.1E-20</td><td>BT20/MB468</td><td>MITO</td></tr>
<tr><td>>hg38_1_-_228634819_228635039_RNA5S12_5S_50ntFlanks@51.72.22|GTCTACGGCCATACCACCCTGA</td><td>>hg38_1_-_228634819_228635039_RNA5S12_5S_50ntFlanks</td><td>51</td><td>72</td><td>22</td><td>GTCTACGGCCATACCACCCTGA</td><td>478.6</td><td>5.4</td><td>0.6</td><td>9.5</td><td>2.5E-21</td><td>2.1E-20</td><td>BT20/MB468</td><td>CYTO</td></tr>
<tr><td>>hg38_21_+_8433172_8446622_RNA45SN1_45S_50ntFlanks@12539.12563.25|ATGATGTGTTGTTGCCATGGTAATC</td><td>>hg38_21_+_8433172_8446622_RNA45SN1_45S_50ntFlanks</td><td>12539</td><td>12563</td><td>25</td><td>ATGATGTGTTGTTGCCATGGTAATC</td><td>865.3</td><td>-13.6</td><td>1.4</td><td>-9.6</td><td>9.5E-22</td><td>2.1E-20</td><td>MB231/MB468</td><td>MP</td></tr>
<tr><td>>hg38_21_+_8433172_8446622_RNA45SN1_45S_50ntFlanks@12641.12659.19|CCATCTGTGGGATTATGAC</td><td>>hg38_21_+_8433172_8446622_RNA45SN1_45S_50ntFlanks</td><td>12641</td><td>12659</td><td>19</td><td>CCATCTGTGGGATTATGAC</td><td>1988.7</td><td>-14.8</td><td>1.5</td><td>-9.6</td><td>9.5E-22</td><td>2.1E-20</td><td>MB231/MB468</td><td>MP</td></tr>
<tr><td>>hg38_21_+_8433172_8446622_RNA45SN1_45S_50ntFlanks@10735.10755.21|TGTAAATCTCGCGCCGGGCCG</td><td>>hg38_21_+_8433172_8446622_RNA45SN1_45S_50ntFlanks</td><td>10735</td><td>10755</td><td>21</td><td>TGTAAATCTCGCGCCGGGCCG</td><td>7113.6</td><td>-13.0</td><td>1.4</td><td>-9.4</td><td>5.1E-21</td><td>2.1E-20</td><td>BT20/MB468</td><td>NUC</td></tr>
<tr><td>>MI0003591|hsa-mir-584&WithFlank&5|-|149062307|149062415@22.43.22(+1A)&[MIMAT0003249&hsa-miR-584-5p&offsets|0|0(+1A);m-167&5|-|149062374|149062394&offsets|0|+1(+1A)]|TTATGGTTTGCCTGGGACTGAGA</td><td>>MI0003591|hsa-mir-584&WithFlank&5|-|149062307|149062415</td><td>22</td><td>43</td><td>22(+1A)</td><td>TTATGGTTTGCCTGGGACTGAGA</td><td>247.9</td><td>12.2</td><td>1.3</td><td>9.5</td><td>2.5E-21</td><td>2.1E-20</td><td>BT20/MB468</td><td>CYTO</td></tr>
<tr><td>>hg38_21_+_8433172_8446622_RNA45SN1_45S_50ntFlanks@10683.10705.23|CCGGCGGCGTCCGGTGAGCTCTC</td><td>>hg38_21_+_8433172_8446622_RNA45SN1_45S_50ntFlanks</td><td>10683</td><td>10705</td><td>23</td><td>CCGGCGGCGTCCGGTGAGCTCTC</td><td>652.0</td><td>-13.2</td><td>1.4</td><td>-9.6</td><td>9.8E-22</td><td>2.1E-20</td><td>MB231/MB468</td><td>MP</td></tr>
<tr><td>>hg38_21_+_8433172_8446622_RNA45SN1_45S_50ntFlanks@11930.11948.19|TAGAATAAGTGGGAGGCCC</td><td>>hg38_21_+_8433172_8446622_RNA45SN1_45S_50ntFlanks</td><td>11930</td><td>11948</td><td>19</td><td>TAGAATAAGTGGGAGGCCC</td><td>810.3</td><td>-13.5</td><td>1.4</td><td>-9.6</td><td>9.8E-22</td><td>2.1E-20</td><td>MB231/MB468</td><td>MP</td></tr>
<tr><td>>hg38_21_+_8433172_8446622_RNA45SN1_45S_50ntFlanks@10402.10435.34|AACATGGGTCAGTCGGTCCTGAGAGATGGGCGAG</td><td>>hg38_21_+_8433172_8446622_RNA45SN1_45S_50ntFlanks</td><td>10402</td><td>10435</td><td>34</td><td>AACATGGGTCAGTCGGTCCTGAGAGATGGGCGAG</td><td>1188.6</td><td>-14.1</td><td>1.5</td><td>-9.6</td><td>1.0E-21</td><td>2.2E-20</td><td>MB231/MB468</td><td>MP</td></tr>
<tr><td>>hg38_21_+_8433172_8446622_RNA45SN1_45S_50ntFlanks@6738.6772.35|ATCATCGACACTTCGAACGCACTTGCGGCCCCGGG</td><td>>hg38_21_+_8433172_8446622_RNA45SN1_45S_50ntFlanks</td><td>6738</td><td>6772</td><td>35</td><td>ATCATCGACACTTCGAACGCACTTGCGGCCCCGGG</td><td>2004.8</td><td>-13.1</td><td>1.4</td><td>-9.5</td><td>1.9E-21</td><td>2.2E-20</td><td>BT20/MB468</td><td>MITO</td></tr>
<tr><td>>am_MI0015825|hsa-mir-378c&WithFlank&10|-|130962582|130962674@17.35.19;MI0016749|hsa-mir-378d-1&WithFlank&4|-|5923269|5923334@37.55.19;MI0000786|hsa-mir-378a&WithFlank&5|+|149732819|149732896@49.67.19;MI0003840|hsa-mir-378d-2&WithFlank&8|-|93916016|93916125@28.46.19&[MIMAT0016847&hsa-miR-378c&offsets|0|-6;m-130&10|-|130962639|130962658&offsets|0|-1];[MIMAT0018926&hsa-miR-378d&offsets|0|-1;m-228&4|-|5923281|5923298&offsets|0|+1];[MIMAT0000732&hsa-miR-378a-3p&offsets|0|-3;m-44&5|+|149732867|149732888&offsets|0|-3];[MIMAT0018926_1&hsa-miR-378d&offsets|0|-1;m-227&8|-|93916081|93916098&offsets|0|+1]|ACTGGACTTGGAGTCAGAA</td><td>>am_MI0015825|hsa-mir-378c&WithFlank&10|-|130962582|130962674</td><td>17</td><td>35</td><td>19</td><td>ACTGGACTTGGAGTCAGAA</td><td>465.7</td><td>5.9</td><td>0.6</td><td>9.5</td><td>2.7E-21</td><td>2.2E-20</td><td>BT20/MB468</td><td>CYTO</td></tr>
<tr><td>>hg38_21_+_8433172_8446622_RNA45SN1_45S_50ntFlanks@12001.12032.32|CTGCGGGCCGCCGGTGAAATACCACTACTCTG</td><td>>hg38_21_+_8433172_8446622_RNA45SN1_45S_50ntFlanks</td><td>12001</td><td>12032</td><td>32</td><td>CTGCGGGCCGCCGGTGAAATACCACTACTCTG</td><td>9029.5</td><td>-13.1</td><td>1.4</td><td>-9.4</td><td>5.4E-21</td><td>2.2E-20</td><td>BT20/MB468</td><td>NUC</td></tr>
<tr><td>>MI0000108|hsa-mir-103a-2&WithFlank&20|+|3917488|3917577@54.75.22(+1U);MI0000109|hsa-mir-103a-1&WithFlank&5|-|168560890|168560979@54.75.22(+1U)&[MIMAT0000101&hsa-miR-103a-3p&offsets|0|-1(+1U);m-12&20|+|3917541|3917563&offsets|0|-1(+1U)];[MIMAT0000101_1&hsa-miR-103a-3p&offsets|0|-1(+1U);m-13&5|-|168560904|168560926&offsets|0|-1(+1U);m-44220&5|-|168560908|168560926&offsets|0|+3(+1U)]|AGCAGCATTGTACAGGGCTATGT</td><td>>MI0000108|hsa-mir-103a-2&WithFlank&20|+|3917488|3917577</td><td>54</td><td>75</td><td>22(+1U)</td><td>AGCAGCATTGTACAGGGCTATGT</td><td>1111.4</td><td>6.1</td><td>0.6</td><td>9.5</td><td>1.5E-21</td><td>2.2E-20</td><td>MB231/MB468</td><td>NUC</td></tr>
<tr><td>>hg38_21_+_8433172_8446622_RNA45SN1_45S_50ntFlanks@11620.11640.21|ACAAAGCATCGCGAAGGCCCG</td><td>>hg38_21_+_8433172_8446622_RNA45SN1_45S_50ntFlanks</td><td>11620</td><td>11640</td><td>21</td><td>ACAAAGCATCGCGAAGGCCCG</td><td>3592.8</td><td>-11.4</td><td>1.2</td><td>-9.5</td><td>1.6E-21</td><td>2.2E-20</td><td>MB231/MB468</td><td>MITO</td></tr>
<tr><td>>hg38_21_+_8433172_8446622_RNA45SN1_45S_50ntFlanks@12415.12436.22|CTTCGATGTCGGCTCTTCCTAT</td><td>>hg38_21_+_8433172_8446622_RNA45SN1_45S_50ntFlanks</td><td>12415</td><td>12436</td><td>22</td><td>CTTCGATGTCGGCTCTTCCTAT</td><td>682.2</td><td>-13.3</td><td>1.4</td><td>-9.6</td><td>1.0E-21</td><td>2.2E-20</td><td>MB231/MB468</td><td>MP</td></tr>
<tr><td>>MI0000449|hsa-mir-132&WithFlank&17|-|2049902|2050014@65.86.22(+1U)&[MIMAT0000426&hsa-miR-132-3p&offsets|0|0(+1U);m-138&17|-|2049929|2049950&offsets|0|0(+1U)]|TAACAGTCTACAGCCATGGTCGT</td><td>>MI0000449|hsa-mir-132&WithFlank&17|-|2049902|2050014</td><td>65</td><td>86</td><td>22(+1U)</td><td>TAACAGTCTACAGCCATGGTCGT</td><td>865.0</td><td>11.6</td><td>1.2</td><td>9.5</td><td>2.7E-21</td><td>2.2E-20</td><td>BT20/MB468</td><td>CYTO</td></tr>
<tr><td>>MI0000279|hsa-mir-196a-2&WithFlank&12|+|53991732|53991853@31.52.22;MI0000238|hsa-mir-196a-1&WithFlank&17|-|48632484|48632565@13.34.22&[MIMAT0000226&hsa-miR-196a-5p&offsets|0|0;m-261&12|+|53991762|53991783&offsets|0|0];[MIMAT0000226_1&hsa-miR-196a-5p&offsets|0|0;m-268&17|-|48632532|48632553&offsets|0|0]|TAGGTAGTTTCATGTTGTTGGG</td><td>>MI0000279|hsa-mir-196a-2&WithFlank&12|+|53991732|53991853</td><td>31</td><td>52</td><td>22</td><td>TAGGTAGTTTCATGTTGTTGGG</td><td>1102.0</td><td>13.7</td><td>1.4</td><td>9.8</td><td>1.3E-22</td><td>2.2E-20</td><td>BT20/MB231</td><td>MITO</td></tr>
<tr><td>>MI0003560|hsa-mir-92b&WithFlank&1|+|155195171|155195278@67.87.21(+1G)&[MIMAT0003218&hsa-miR-92b-3p&offsets|0|-1(+1G);m-115&1|+|155195237|155195258&offsets|0|-1(+1G)]|TATTGCACTCGTCCCGGCCTCG</td><td>>MI0003560|hsa-mir-92b&WithFlank&1|+|155195171|155195278</td><td>67</td><td>87</td><td>21(+1G)</td><td>TATTGCACTCGTCCCGGCCTCG</td><td>320.8</td><td>12.6</td><td>1.3</td><td>9.5</td><td>2.7E-21</td><td>2.3E-20</td><td>BT20/MB468</td><td>CYTO</td></tr>
<tr><td>>MI0000477|hsa-mir-146a&WithFlank&5|+|160485346|160485456@27.47.21(+1A)&[MIMAT0000449&hsa-miR-146a-5p&offsets|0|-1(+1A);m-129&5|+|160485372|160485393&offsets|0|-1(+1A)]|TGAGAACTGAATTCCATGGGTA</td><td>>MI0000477|hsa-mir-146a&WithFlank&5|+|160485346|160485456</td><td>27</td><td>47</td><td>21(+1A)</td><td>TGAGAACTGAATTCCATGGGTA</td><td>4821.5</td><td>15.5</td><td>1.6</td><td>9.6</td><td>1.0E-21</td><td>2.3E-20</td><td>MB231/MB468</td><td>MP</td></tr>
<tr><td>>MI0000750|hsa-mir-26a-2&WithFlank&12|-|57824603|57824698@20.41.22(+2U);MI0000083|hsa-mir-26a-1&WithFlank&3|+|37969398|37969486@16.37.22(+2U)&[MIMAT0000082&hsa-miR-26a-5p&offsets|0|0(+2U);m-33&12|-|57824658|57824679&offsets|0|0(+2U)];[MIMAT0000082_1&hsa-miR-26a-5p&offsets|0|0(+2U);m-34&3|+|37969413|37969434&offsets|0|0(+2U)]|TTCAAGTAATCCAGGATAGGCTTT</td><td>>MI0000750|hsa-mir-26a-2&WithFlank&12|-|57824603|57824698</td><td>20</td><td>41</td><td>22(+2U)</td><td>TTCAAGTAATCCAGGATAGGCTTT</td><td>2756.7</td><td>6.8</td><td>0.7</td><td>9.5</td><td>1.5E-21</td><td>2.3E-20</td><td>MB231/MB468</td><td>NUC</td></tr>
<tr><td>>hg38_21_+_8433172_8446622_RNA45SN1_45S_50ntFlanks@9304.9336.33|ACCAAGGAGTCTAACACGTGCGCGAGTCGGGGG</td><td>>hg38_21_+_8433172_8446622_RNA45SN1_45S_50ntFlanks</td><td>9304</td><td>9336</td><td>33</td><td>ACCAAGGAGTCTAACACGTGCGCGAGTCGGGGG</td><td>691.9</td><td>-13.3</td><td>1.4</td><td>-9.6</td><td>1.1E-21</td><td>2.3E-20</td><td>MB231/MB468</td><td>MP</td></tr>
<tr><td>>hg38_21_+_8433172_8446622_RNA45SN1_45S_50ntFlanks@6719.6747.29|TGAATTGCAGGACACATTGATCATCGACA</td><td>>hg38_21_+_8433172_8446622_RNA45SN1_45S_50ntFlanks</td><td>6719</td><td>6747</td><td>29</td><td>TGAATTGCAGGACACATTGATCATCGACA</td><td>879.8</td><td>-13.6</td><td>1.4</td><td>-9.6</td><td>1.1E-21</td><td>2.3E-20</td><td>MB231/MB468</td><td>MP</td></tr>
<tr><td>>MI0000767|hsa-mir-365a&WithFlank&16|+|14309279|14309377@62.84.23;MI0000769|hsa-mir-365b&WithFlank&17|+|31575405|31575527@74.96.23&[MIMAT0000710&hsa-miR-365a-3p&offsets|0|+1;m-176&16|+|14309340|14309361&offsets|0|+1];[MIMAT0022834&hsa-miR-365b-3p&offsets|0|+1;m-175&17|+|31575478|31575499&offsets|0|+1]|TAATGCCCCTAAAAATCCTTATT</td><td>>MI0000767|hsa-mir-365a&WithFlank&16|+|14309279|14309377</td><td>62</td><td>84</td><td>23</td><td>TAATGCCCCTAAAAATCCTTATT</td><td>1223.8</td><td>10.4</td><td>1.1</td><td>9.5</td><td>2.8E-21</td><td>2.3E-20</td><td>BT20/MB468</td><td>CYTO</td></tr>
<tr><td>>hg38_21_+_8433172_8446622_RNA45SN1_45S_50ntFlanks@10841.10858.18|CGTAACTTCGGGATAAGG</td><td>>hg38_21_+_8433172_8446622_RNA45SN1_45S_50ntFlanks</td><td>10841</td><td>10858</td><td>18</td><td>CGTAACTTCGGGATAAGG</td><td>635.3</td><td>-13.2</td><td>1.4</td><td>-9.6</td><td>1.1E-21</td><td>2.3E-20</td><td>MB231/MB468</td><td>MP</td></tr>
<tr><td>>hg38_21_+_8433172_8446622_RNA45SN1_45S_50ntFlanks@8106.8143.38|GCGGCGGGGCGCGGGACATGTGGCGTACGGAAGACCCG</td><td>>hg38_21_+_8433172_8446622_RNA45SN1_45S_50ntFlanks</td><td>8106</td><td>8143</td><td>38</td><td>GCGGCGGGGCGCGGGACATGTGGCGTACGGAAGACCCG</td><td>2182.0</td><td>-15.0</td><td>1.6</td><td>-9.6</td><td>1.1E-21</td><td>2.3E-20</td><td>MB231/MB468</td><td>MP</td></tr>
<tr><td>>MI0000816|hsa-mir-335&WithFlank&7|+|130496105|130496210@58.78.21&[MIMAT0004703&hsa-miR-335-3p&offsets|0|-1;m-159&7|+|130496162|130496183&offsets|0|-1]|TTTTTCATTATTGCTCCTGAC</td><td>>MI0000816|hsa-mir-335&WithFlank&7|+|130496105|130496210</td><td>58</td><td>78</td><td>21</td><td>TTTTTCATTATTGCTCCTGAC</td><td>328.3</td><td>5.6</td><td>0.6</td><td>9.5</td><td>1.5E-21</td><td>2.3E-20</td><td>MB231/MB468</td><td>NUC</td></tr>
<tr><td>>hg38_21_+_8433172_8446622_RNA45SN1_45S_50ntFlanks@10362.10383.22|AAGGCCGAAGTGGAGAAGGGTT</td><td>>hg38_21_+_8433172_8446622_RNA45SN1_45S_50ntFlanks</td><td>10362</td><td>10383</td><td>22</td><td>AAGGCCGAAGTGGAGAAGGGTT</td><td>2339.3</td><td>-5.6</td><td>0.6</td><td>-9.6</td><td>8.6E-22</td><td>2.3E-20</td><td>MB231/MB468</td><td>CYTO</td></tr>
<tr><td>>hg38_21_+_8433172_8446622_RNA45SN1_45S_50ntFlanks@4722.4752.31|TCATTAATCAAGAACGAAAGTCGGAGGTTCG</td><td>>hg38_21_+_8433172_8446622_RNA45SN1_45S_50ntFlanks</td><td>4722</td><td>4752</td><td>31</td><td>TCATTAATCAAGAACGAAAGTCGGAGGTTCG</td><td>932.3</td><td>-13.7</td><td>1.4</td><td>-9.6</td><td>1.1E-21</td><td>2.3E-20</td><td>MB231/MB468</td><td>MP</td></tr>
<tr><td>>hg38_21_+_8433172_8446622_RNA45SN1_45S_50ntFlanks@11768.11791.24|CTAATTAGTGACGCGCATGAATGG</td><td>>hg38_21_+_8433172_8446622_RNA45SN1_45S_50ntFlanks</td><td>11768</td><td>11791</td><td>24</td><td>CTAATTAGTGACGCGCATGAATGG</td><td>735.2</td><td>-8.0</td><td>0.8</td><td>-9.6</td><td>8.7E-22</td><td>2.3E-20</td><td>MB231/MB468</td><td>CYTO</td></tr>
<tr><td>>hg38_21_+_8433172_8446622_RNA45SN1_45S_50ntFlanks@9201.9222.22|CTCTCGGGGCCACGCGCGCGTC</td><td>>hg38_21_+_8433172_8446622_RNA45SN1_45S_50ntFlanks</td><td>9201</td><td>9222</td><td>22</td><td>CTCTCGGGGCCACGCGCGCGTC</td><td>1095.5</td><td>-14.0</td><td>1.5</td><td>-9.6</td><td>1.1E-21</td><td>2.3E-20</td><td>MB231/MB468</td><td>MP</td></tr>
<tr><td>>hg38_21_+_8433172_8446622_RNA45SN1_45S_50ntFlanks@11575.11601.27|ACTTAGAACTGGTGCGGACCAGGGGAA</td><td>>hg38_21_+_8433172_8446622_RNA45SN1_45S_50ntFlanks</td><td>11575</td><td>11601</td><td>27</td><td>ACTTAGAACTGGTGCGGACCAGGGGAA</td><td>1599.3</td><td>-8.5</td><td>0.9</td><td>-9.6</td><td>8.8E-22</td><td>2.3E-20</td><td>MB231/MB468</td><td>CYTO</td></tr>
<tr><td>>hg38_21_+_8433172_8446622_RNA45SN1_45S_50ntFlanks@9198.9218.21|GTTCTCTCGGGGCCACGCGCG</td><td>>hg38_21_+_8433172_8446622_RNA45SN1_45S_50ntFlanks</td><td>9198</td><td>9218</td><td>21</td><td>GTTCTCTCGGGGCCACGCGCG</td><td>2140.1</td><td>-14.8</td><td>1.6</td><td>-9.5</td><td>1.7E-21</td><td>2.3E-20</td><td>MB231/MB468</td><td>MITO</td></tr>
<tr><td>>hg38_21_+_8433172_8446622_RNA45SN1_45S_50ntFlanks@12366.12384.19|TAACTGGCTTGTGGCGGCC</td><td>>hg38_21_+_8433172_8446622_RNA45SN1_45S_50ntFlanks</td><td>12366</td><td>12384</td><td>19</td><td>TAACTGGCTTGTGGCGGCC</td><td>893.1</td><td>-13.7</td><td>1.4</td><td>-9.6</td><td>1.1E-21</td><td>2.3E-20</td><td>MB231/MB468</td><td>MP</td></tr>
<tr><td>>hg38_21_+_8433172_8446622_RNA45SN1_45S_50ntFlanks@10794.10823.30|CTGGCATGTTGGAACAATGTAGGTAAGGGA</td><td>>hg38_21_+_8433172_8446622_RNA45SN1_45S_50ntFlanks</td><td>10794</td><td>10823</td><td>30</td><td>CTGGCATGTTGGAACAATGTAGGTAAGGGA</td><td>638.1</td><td>-13.2</td><td>1.4</td><td>-9.6</td><td>1.1E-21</td><td>2.3E-20</td><td>MB231/MB468</td><td>MP</td></tr>
<tr><td>>hg38_21_+_8433172_8446622_RNA45SN1_45S_50ntFlanks@6647.6667.21|CGTACGACTCTTAGCGGTGGA</td><td>>hg38_21_+_8433172_8446622_RNA45SN1_45S_50ntFlanks</td><td>6647</td><td>6667</td><td>21</td><td>CGTACGACTCTTAGCGGTGGA</td><td>9451.1</td><td>6.5</td><td>0.7</td><td>9.5</td><td>2.1E-21</td><td>2.3E-20</td><td>BT20/MB468</td><td>MITO</td></tr>
<tr><td>>hg38_21_+_8433172_8446622_RNA45SN1_45S_50ntFlanks@6731.6772.42|CACATTGATCATCGACACTTCGAACGCACTTGCGGCCCCGGG</td><td>>hg38_21_+_8433172_8446622_RNA45SN1_45S_50ntFlanks</td><td>6731</td><td>6772</td><td>42</td><td>CACATTGATCATCGACACTTCGAACGCACTTGCGGCCCCGGG</td><td>926.0</td><td>-14.3</td><td>1.5</td><td>-9.4</td><td>5.7E-21</td><td>2.4E-20</td><td>BT20/MB468</td><td>NUC</td></tr>
<tr><td>>hg38_21_+_8433172_8446622_RNA45SN1_45S_50ntFlanks@4439.4471.33|CCGCCCCTTGCCTCTCGGCGCCCCCTCGATGCT</td><td>>hg38_21_+_8433172_8446622_RNA45SN1_45S_50ntFlanks</td><td>4439</td><td>4471</td><td>33</td><td>CCGCCCCTTGCCTCTCGGCGCCCCCTCGATGCT</td><td>513.2</td><td>-13.4</td><td>1.4</td><td>-9.4</td><td>5.7E-21</td><td>2.4E-20</td><td>BT20/MB468</td><td>NUC</td></tr>
</table>

<table>
<tr><td>>MI0003581|hsa-mir-574&WithFlank&4|+|38868026|38868133@67.88.22&[MIMAT0003239&hsa-miR-574-3p&offsets|0|0;m-149&4|+|38868092|38868113&offsets|0|0]|CACGCTCATGCACACACCCACA</td><td>>MI0003581|hsa-mir-574&WithFlank&4|+|38868026|38868133</td><td>67</td><td>88</td><td>22</td><td>CACGCTCATGCACACACCCACA</td><td>4256.3</td><td>7.7</td><td>0.8</td><td>9.5</td><td>2.9E-21</td><td>2.4E-20</td><td>BT20/MB468</td><td>CYTO</td></tr>
<tr><td>>MI0000255|hsa-mir-30d&WithFlank&8|-|134804870|134804951@12.35.24&[MIMAT0000245&hsa-miR-30d-5p&offsets|0|+2;m-21&8|-|134804919|134804940&offsets|0|+2]|TGTAAACATCCCCGACTGGAAGCT</td><td>>MI0000255|hsa-mir-30d&WithFlank&8|-|134804870|134804951</td><td>12</td><td>35</td><td>24</td><td>TGTAAACATCCCCGACTGGAAGCT</td><td>47370.7</td><td>6.1</td><td>0.6</td><td>9.5</td><td>1.6E-21</td><td>2.4E-20</td><td>MB231/MB468</td><td>NUC</td></tr>
<tr><td>>hg38_21_+_8433172_8446622_RNA45SN1_45S_50ntFlanks@10410.10430.21|TCAGTCGGTCCTGAGAGATGG</td><td>>hg38_21_+_8433172_8446622_RNA45SN1_45S_50ntFlanks</td><td>10410</td><td>10430</td><td>21</td><td>TCAGTCGGTCCTGAGAGATGG</td><td>17201.8</td><td>-13.7</td><td>1.5</td><td>-9.4</td><td>5.8E-21</td><td>2.4E-20</td><td>BT20/MB468</td><td>NUC</td></tr>
<tr><td>>MI0000081|hsa-mir-24-2&WithFlank&19|-|13836281|13836365@56.74.19(+1U);MI0000080|hsa-mir-24-1&WithFlank&9|+|95086015|95086094@50.68.19(+1U)&[MIMAT0000080&hsa-miR-24-3p&offsets|0|-3(+1U);m-36&19|-|13836288|13836310&offsets|0|-4(+1U)];[MIMAT0000080_1&hsa-miR-24-3p&offsets|0|-3(+1U);m-35&9|+|95086064|95086086&offsets|0|-4(+1U)]|TGGCTCAGTTCAGCAGGAAT</td><td>>MI0000081|hsa-mir-24-2&WithFlank&19|-|13836281|13836365</td><td>56</td><td>74</td><td>19(+1U)</td><td>TGGCTCAGTTCAGCAGGAAT</td><td>1326.5</td><td>6.6</td><td>0.7</td><td>9.5</td><td>2.1E-21</td><td>2.4E-20</td><td>BT20/MB468</td><td>MITO</td></tr>
<tr><td>>hg38_21_+_8433172_8446622_RNA45SN1_45S_50ntFlanks@12386.12414.29|AGCGTTCATAGCGACGTCGCTTTTTGATC</td><td>>hg38_21_+_8433172_8446622_RNA45SN1_45S_50ntFlanks</td><td>12386</td><td>12414</td><td>29</td><td>AGCGTTCATAGCGACGTCGCTTTTTGATC</td><td>617.6</td><td>-13.1</td><td>1.4</td><td>-9.6</td><td>1.1E-21</td><td>2.4E-20</td><td>MB231/MB468</td><td>MP</td></tr>
<tr><td>>hg38_1_-_228634819_228635039_RNA5S12_5S_50ntFlanks@51.89.39|GTCTACGGCCATACCACCCTGAACGCGCCCGATCTCGTC</td><td>>hg38_1_-_228634819_228635039_RNA5S12_5S_50ntFlanks</td><td>51</td><td>89</td><td>39</td><td>GTCTACGGCCATACCACCCTGAACGCGCCCGATCTCGTC</td><td>93298.6</td><td>10.4</td><td>1.1</td><td>9.6</td><td>9.1E-22</td><td>2.4E-20</td><td>MB231/MB468</td><td>CYTO</td></tr>
<tr><td>>hg38_21_+_8433172_8446622_RNA45SN1_45S_50ntFlanks@12624.12659.36|CCAATGGGGCGAAGCTACCATCTGTGGGATTATGAC</td><td>>hg38_21_+_8433172_8446622_RNA45SN1_45S_50ntFlanks</td><td>12624</td><td>12659</td><td>36</td><td>CCAATGGGGCGAAGCTACCATCTGTGGGATTATGAC</td><td>2599.4</td><td>-11.0</td><td>1.2</td><td>-9.5</td><td>1.7E-21</td><td>2.4E-20</td><td>MB231/MB468</td><td>MITO</td></tr>
<tr><td>>hg38_21_+_8433172_8446622_RNA45SN1_45S_50ntFlanks@12008.12039.32|CCGCCGGTGAAATACCACTACTCTGATCGTTT</td><td>>hg38_21_+_8433172_8446622_RNA45SN1_45S_50ntFlanks</td><td>12008</td><td>12039</td><td>32</td><td>CCGCCGGTGAAATACCACTACTCTGATCGTTT</td><td>904.8</td><td>-13.6</td><td>1.4</td><td>-9.5</td><td>1.8E-21</td><td>2.4E-20</td><td>MB231/MB468</td><td>MITO</td></tr>
<tr><td>>hg38_21_+_8433172_8446622_RNA45SN1_45S_50ntFlanks@12937.12956.20|AACCATTCGTAGACGACCTG</td><td>>hg38_21_+_8433172_8446622_RNA45SN1_45S_50ntFlanks</td><td>12937</td><td>12956</td><td>20</td><td>AACCATTCGTAGACGACCTG</td><td>3839.8</td><td>-9.2</td><td>1.0</td><td>-9.5</td><td>1.8E-21</td><td>2.4E-20</td><td>MB231/MB468</td><td>MITO</td></tr>
<tr><td>>hg38_21_+_8433172_8446622_RNA45SN1_45S_50ntFlanks@6645.6669.25|CTCGTACGACTCTTAGCGGTGGATC</td><td>>hg38_21_+_8433172_8446622_RNA45SN1_45S_50ntFlanks</td><td>6645</td><td>6669</td><td>25</td><td>CTCGTACGACTCTTAGCGGTGGATC</td><td>6922.3</td><td>4.8</td><td>0.5</td><td>9.5</td><td>3.0E-21</td><td>2.4E-20</td><td>BT20/MB468</td><td>CYTO</td></tr>
<tr><td>>hg38_21_+_8433172_8446622_RNA45SN1_45S_50ntFlanks@11846.11872.27|GAACGGGCTTGGCGGAATCAGCGGGGA</td><td>>hg38_21_+_8433172_8446622_RNA45SN1_45S_50ntFlanks</td><td>11846</td><td>11872</td><td>27</td><td>GAACGGGCTTGGCGGAATCAGCGGGGA</td><td>593.0</td><td>-13.1</td><td>1.4</td><td>-9.6</td><td>1.2E-21</td><td>2.4E-20</td><td>MB231/MB468</td><td>MP</td></tr>
<tr><td>>hg38_21_+_8433172_8446622_RNA45SN1_45S_50ntFlanks@7984.8004.21|AGATCAGACGTGGCGACCCGC</td><td>>hg38_21_+_8433172_8446622_RNA45SN1_45S_50ntFlanks</td><td>7984</td><td>8004</td><td>21</td><td>AGATCAGACGTGGCGACCCGC</td><td>1182.3</td><td>-14.1</td><td>1.5</td><td>-9.6</td><td>1.2E-21</td><td>2.4E-20</td><td>MB231/MB468</td><td>MP</td></tr>
<tr><td>>hg38_21_+_8433172_8446622_RNA45SN1_45S_50ntFlanks@12405.12431.27|CTTTTTGATCCTTCGATGTCGGCTCTT</td><td>>hg38_21_+_8433172_8446622_RNA45SN1_45S_50ntFlanks</td><td>12405</td><td>12431</td><td>27</td><td>CTTTTTGATCCTTCGATGTCGGCTCTT</td><td>18931.3</td><td>-8.0</td><td>0.8</td><td>-9.5</td><td>3.0E-21</td><td>2.5E-20</td><td>BT20/MB468</td><td>CYTO</td></tr>
<tr><td>>MI0000439|hsa-mir-23b&WithFlank&9|+|95085202|95085310@67.85.19(+1U)&[MIMAT0000418&hsa-miR-23b-3p&offsets|+3|-1(+1U);m-42&9|+|95085265|95085285&offsets|+3|+1(+1U)]|ACATTGCCAGGGATTACCAT</td><td>>MI0000439|hsa-mir-23b&WithFlank&9|+|95085202|95085310</td><td>67</td><td>85</td><td>19(+1U)</td><td>ACATTGCCAGGGATTACCAT</td><td>327.4</td><td>12.8</td><td>1.3</td><td>9.5</td><td>2.2E-21</td><td>2.5E-20</td><td>BT20/MB468</td><td>MITO</td></tr>
<tr><td>>hg38_21_+_8433172_8446622_RNA45SN1_45S_50ntFlanks@9562.9597.36|TCCTGACGTGCAAATCGGTCGTCCGACCTGGGTATA</td><td>>hg38_21_+_8433172_8446622_RNA45SN1_45S_50ntFlanks</td><td>9562</td><td>9597</td><td>36</td><td>TCCTGACGTGCAAATCGGTCGTCCGACCTGGGTATA</td><td>4142.4</td><td>-11.4</td><td>1.2</td><td>-9.5</td><td>1.8E-21</td><td>2.5E-20</td><td>MB231/MB468</td><td>MITO</td></tr>
<tr><td>>hg38_21_+_8433172_8446622_RNA45SN1_45S_50ntFlanks@7974.8009.36|ACGCGACCTCAGATCAGACGTGGCGACCCGCTGAAT</td><td>>hg38_21_+_8433172_8446622_RNA45SN1_45S_50ntFlanks</td><td>7974</td><td>8009</td><td>36</td><td>ACGCGACCTCAGATCAGACGTGGCGACCCGCTGAAT</td><td>88057.9</td><td>5.9</td><td>0.6</td><td>9.5</td><td>2.2E-21</td><td>2.5E-20</td><td>BT20/MB468</td><td>MITO</td></tr>
<tr><td>>hg38_21_+_8433172_8446622_RNA45SN1_45S_50ntFlanks@10922.10946.25|CTGGACGAGGCGCCGCCGCCCCCCC</td><td>>hg38_21_+_8433172_8446622_RNA45SN1_45S_50ntFlanks</td><td>10922</td><td>10946</td><td>25</td><td>CTGGACGAGGCGCCGCCGCCCCCCC</td><td>2157.0</td><td>-14.9</td><td>1.6</td><td>-9.6</td><td>1.2E-21</td><td>2.5E-20</td><td>MB231/MB468</td><td>MP</td></tr>
<tr><td>>MI0000266|hsa-mir-10a&WithFlank&17|-|48579832|48579953@29.49.21(+1A)&[MIMAT0000253&hsa-miR-10a-5p&offsets|+1|-1(+1A);m-18&17|-|48579905|48579926&offsets|+1|0(+1A)]|ACCCTGTAGATCCGAATTTGTA</td><td>>MI0000266|hsa-mir-10a&WithFlank&17|-|48579832|48579953</td><td>29</td><td>49</td><td>21(+1A)</td><td>ACCCTGTAGATCCGAATTTGTA</td><td>316.0</td><td>12.2</td><td>1.3</td><td>9.5</td><td>1.7E-21</td><td>2.5E-20</td><td>MB231/MB468</td><td>NUC</td></tr>
<tr><td>>hg38_21_+_8433172_8446622_RNA45SN1_45S_50ntFlanks@11670.11698.29|CCCAGTGCTCTGAATGTCAAAGTGAAGAA</td><td>>hg38_21_+_8433172_8446622_RNA45SN1_45S_50ntFlanks</td><td>11670</td><td>11698</td><td>29</td><td>CCCAGTGCTCTGAATGTCAAAGTGAAGAA</td><td>650.0</td><td>-5.0</td><td>0.5</td><td>-9.5</td><td>1.7E-21</td><td>2.5E-20</td><td>MB231/MB468</td><td>NUC</td></tr>
<tr><td>>hg38_21_+_8433172_8446622_RNA45SN1_45S_50ntFlanks@9305.9337.33|CCAAGGAGTCTAACACGTGCGCGAGTCGGGGGC</td><td>>hg38_21_+_8433172_8446622_RNA45SN1_45S_50ntFlanks</td><td>9305</td><td>9337</td><td>33</td><td>CCAAGGAGTCTAACACGTGCGCGAGTCGGGGGC</td><td>855.4</td><td>-13.6</td><td>1.4</td><td>-9.6</td><td>1.2E-21</td><td>2.5E-20</td><td>MB231/MB468</td><td>MP</td></tr>
<tr><td>>MI0000471|hsa-mir-126&WithFlank&9|+|136670596|136670692@59.78.20&[MIMAT0000445&hsa-miR-126-3p&offsets|+1|-1;m-40&9|+|136670653|136670674&offsets|+1|-1]|CGTACCGTGAGTAATAATGC</td><td>>MI0000471|hsa-mir-126&WithFlank&9|+|136670596|136670692</td><td>59</td><td>78</td><td>20</td><td>CGTACCGTGAGTAATAATGC</td><td>511.7</td><td>13.5</td><td>1.4</td><td>9.5</td><td>2.3E-21</td><td>2.5E-20</td><td>BT20/MB468</td><td>MITO</td></tr>
<tr><td>>hg38_21_+_8433172_8446622_RNA45SN1_45S_50ntFlanks@9738.9759.22|GAAACGATCTCAACCTATTCTC</td><td>>hg38_21_+_8433172_8446622_RNA45SN1_45S_50ntFlanks</td><td>9738</td><td>9759</td><td>22</td><td>GAAACGATCTCAACCTATTCTC</td><td>850.4</td><td>-13.6</td><td>1.4</td><td>-9.6</td><td>1.2E-21</td><td>2.6E-20</td><td>MB231/MB468</td><td>MP</td></tr>
<tr><td>>hg38_21_+_8433172_8446622_RNA45SN1_45S_50ntFlanks@12000.12032.33|CCTGCGGGCCGCCGGTGAAATACCACTACTCTG</td><td>>hg38_21_+_8433172_8446622_RNA45SN1_45S_50ntFlanks</td><td>12000</td><td>12032</td><td>33</td><td>CCTGCGGGCCGCCGGTGAAATACCACTACTCTG</td><td>948.5</td><td>-13.7</td><td>1.4</td><td>-9.5</td><td>1.9E-21</td><td>2.6E-20</td><td>MB231/MB468</td><td>MITO</td></tr>
<tr><td>>hg38_21_+_8433172_8446622_RNA45SN1_45S_50ntFlanks@8113.8143.31|GGCGCGGGACATGTGGCGTACGGAAGACCCG</td><td>>hg38_21_+_8433172_8446622_RNA45SN1_45S_50ntFlanks</td><td>8113</td><td>8143</td><td>31</td><td>GGCGCGGGACATGTGGCGTACGGAAGACCCG</td><td>1055.9</td><td>-13.8</td><td>1.5</td><td>-9.5</td><td>1.9E-21</td><td>2.6E-20</td><td>MB231/MB468</td><td>MITO</td></tr>
<tr><td>>hg38_21_+_8433172_8446622_RNA45SN1_45S_50ntFlanks@5494.5516.23|AAGACGGTCGAACTTGACTATCT</td><td>>hg38_21_+_8433172_8446622_RNA45SN1_45S_50ntFlanks</td><td>5494</td><td>5516</td><td>23</td><td>AAGACGGTCGAACTTGACTATCT</td><td>5280.8</td><td>6.8</td><td>0.7</td><td>9.6</td><td>9.8E-22</td><td>2.6E-20</td><td>MB231/MB468</td><td>CYTO</td></tr>
<tr><td>>hg38_21_+_8433172_8446622_RNA45SN1_45S_50ntFlanks@11161.11188.28|CACGGTCCCCCGCGAGGGGGGCCCGGGC</td><td>>hg38_21_+_8433172_8446622_RNA45SN1_45S_50ntFlanks</td><td>11161</td><td>11188</td><td>28</td><td>CACGGTCCCCCGCGAGGGGGGCCCGGGC</td><td>1435.2</td><td>-14.3</td><td>1.5</td><td>-9.5</td><td>1.9E-21</td><td>2.6E-20</td><td>MB231/MB468</td><td>MITO</td></tr>
<tr><td>>hg38_21_+_8433172_8446622_RNA45SN1_45S_50ntFlanks@3985.4002.18|CGGCTTTGGTGACTCTAG</td><td>>hg38_21_+_8433172_8446622_RNA45SN1_45S_50ntFlanks</td><td>3985</td><td>4002</td><td>18</td><td>CGGCTTTGGTGACTCTAG</td><td>3621.6</td><td>-7.4</td><td>0.8</td><td>-9.5</td><td>2.3E-21</td><td>2.6E-20</td><td>BT20/MB468</td><td>MITO</td></tr>
<tr><td>>hg38_21_+_8433172_8446622_RNA45SN1_45S_50ntFlanks@10786.10822.37|AACAGCCTCTGGCATGTTGGAACAATGTAGGTAAGGG</td><td>>hg38_21_+_8433172_8446622_RNA45SN1_45S_50ntFlanks</td><td>10786</td><td>10822</td><td>37</td><td>AACAGCCTCTGGCATGTTGGAACAATGTAGGTAAGGG</td><td>2674.0</td><td>-9.2</td><td>1.0</td><td>-9.5</td><td>1.9E-21</td><td>2.6E-20</td><td>MB231/MB468</td><td>MITO</td></tr>
<tr><td>>hg38_21_+_8433172_8446622_RNA45SN1_45S_50ntFlanks@8106.8138.33|GCGGCGGGGCGCGGGACATGTGGCGTACGGAAG</td><td>>hg38_21_+_8433172_8446622_RNA45SN1_45S_50ntFlanks</td><td>8106</td><td>8138</td><td>33</td><td>GCGGCGGGGCGCGGGACATGTGGCGTACGGAAG</td><td>976.0</td><td>-13.8</td><td>1.4</td><td>-9.6</td><td>1.2E-21</td><td>2.6E-20</td><td>MB231/MB468</td><td>MP</td></tr>
<tr><td>>hg38_21_+_8433172_8446622_RNA45SN1_45S_50ntFlanks@12002.12031.30|TGCGGGCCGCCGGTGAAATACCACTACTCT</td><td>>hg38_21_+_8433172_8446622_RNA45SN1_45S_50ntFlanks</td><td>12002</td><td>12031</td><td>30</td><td>TGCGGGCCGCCGGTGAAATACCACTACTCT</td><td>652.5</td><td>-13.2</td><td>1.4</td><td>-9.6</td><td>1.3E-21</td><td>2.6E-20</td><td>MB231/MB468</td><td>MP</td></tr>
<tr><td>>hg38_21_+_8433172_8446622_RNA45SN1_45S_50ntFlanks@6744.6770.27|GACACTTCGAACGCACTTGCGGCCCCG</td><td>>hg38_21_+_8433172_8446622_RNA45SN1_45S_50ntFlanks</td><td>6744</td><td>6770</td><td>27</td><td>GACACTTCGAACGCACTTGCGGCCCCG</td><td>2700.2</td><td>-8.8</td><td>0.9</td><td>-9.5</td><td>1.9E-21</td><td>2.6E-20</td><td>MB231/MB468</td><td>MITO</td></tr>
<tr><td>>hg38_21_+_8433172_8446622_RNA45SN1_45S_50ntFlanks@11161.11179.19|CACGGTCCCCCGCGAGGGG</td><td>>hg38_21_+_8433172_8446622_RNA45SN1_45S_50ntFlanks</td><td>11161</td><td>11179</td><td>19</td><td>CACGGTCCCCCGCGAGGGG</td><td>1015.6</td><td>-13.8</td><td>1.4</td><td>-9.5</td><td>1.9E-21</td><td>2.6E-20</td><td>MB231/MB468</td><td>MITO</td></tr>
<tr><td>>hg38_21_+_8433172_8446622_RNA45SN1_45S_50ntFlanks@12934.12956.23|CTAAACCATTCGTAGACGACCTG</td><td>>hg38_21_+_8433172_8446622_RNA45SN1_45S_50ntFlanks</td><td>12934</td><td>12956</td><td>23</td><td>CTAAACCATTCGTAGACGACCTG</td><td>2767.5</td><td>-4.9</td><td>0.5</td><td>-9.5</td><td>2.4E-21</td><td>2.6E-20</td><td>BT20/MB468</td><td>MITO</td></tr>
<tr><td>>hg38_21_+_8433172_8446622_RNA45SN1_45S_50ntFlanks@11705.11728.24|TGAAGCGCGGGTAAACGGCGGGAG</td><td>>hg38_21_+_8433172_8446622_RNA45SN1_45S_50ntFlanks</td><td>11705</td><td>11728</td><td>24</td><td>TGAAGCGCGGGTAAACGGCGGGAG</td><td>986.0</td><td>-13.8</td><td>1.4</td><td>-9.6</td><td>1.3E-21</td><td>2.6E-20</td><td>MB231/MB468</td><td>MP</td></tr>
<tr><td>>hg38_21_+_8433172_8446622_RNA45SN1_45S_50ntFlanks@11027.11050.24|CTCTCTCTCTCTCTCCCCCGCTCC</td><td>>hg38_21_+_8433172_8446622_RNA45SN1_45S_50ntFlanks</td><td>11027</td><td>11050</td><td>24</td><td>CTCTCTCTCTCTCTCCCCCGCTCC</td><td>1141.8</td><td>-13.9</td><td>1.5</td><td>-9.5</td><td>2.0E-21</td><td>2.6E-20</td><td>MB231/MB468</td><td>MITO</td></tr>
<tr><td>>hg38_21_+_8433172_8446622_RNA45SN1_45S_50ntFlanks@10351.10387.37|CGAGAACTTTGAAGGCCGAAGTGGAGAAGGGTTCCAT</td><td>>hg38_21_+_8433172_8446622_RNA45SN1_45S_50ntFlanks</td><td>10351</td><td>10387</td><td>37</td><td>CGAGAACTTTGAAGGCCGAAGTGGAGAAGGGTTCCAT</td><td>415.7</td><td>-13.1</td><td>1.4</td><td>-9.4</td><td>6.5E-21</td><td>2.7E-20</td><td>BT20/MB468</td><td>NUC</td></tr>
<tr><td>>MI0000087|hsa-mir-29a&WithFlank&7|-|130876741|130876816@47.67.21(+1A)&[MIMAT0000086&hsa-miR-29a-3p&offsets|-1|-2(+1A);m-26&7|-|130876749|130876769&offsets|-1|-1(+1A)]|CTAGCACCATCTGAAATCGGTA</td><td>>MI0000087|hsa-mir-29a&WithFlank&7|-|130876741|130876816</td><td>47</td><td>67</td><td>21(+1A)</td><td>CTAGCACCATCTGAAATCGGTA</td><td>618.5</td><td>13.5</td><td>1.4</td><td>9.5</td><td>3.3E-21</td><td>2.7E-20</td><td>BT20/MB468</td><td>CYTO</td></tr>
<tr><td>>trna5_AspGTC_12_+_98897281_98897352@36.74.39|CACGCGGGAGACCGGGGTTCAATTCCCCGACGGGGAGCC</td><td>>trna5_AspGTC_12_+_98897281_98897352</td><td>36</td><td>74</td><td>39</td><td>CACGCGGGAGACCGGGGTTCAATTCCCCGACGGGGAGCC</td><td>2874.4</td><td>15.5</td><td>1.6</td><td>9.6</td><td>1.0E-21</td><td>2.7E-20</td><td>MB231/MB468</td><td>CYTO</td></tr>
<tr><td>>hg38_21_+_8433172_8446622_RNA45SN1_45S_50ntFlanks@3813.3843.31|TTATGGTTCCTTTGGTCGCTCGCTCCTCTCC</td><td>>hg38_21_+_8433172_8446622_RNA45SN1_45S_50ntFlanks</td><td>3813</td><td>3843</td><td>31</td><td>TTATGGTTCCTTTGGTCGCTCGCTCCTCTCC</td><td>472.7</td><td>-13.3</td><td>1.4</td><td>-9.4</td><td>6.6E-21</td><td>2.7E-20</td><td>BT20/MB468</td><td>NUC</td></tr>
<tr><td>>hg38_21_+_8433172_8446622_RNA45SN1_45S_50ntFlanks@10866.10891.26|CTAAGGGCTGGGTCGGTCGGGCTGGG</td><td>>hg38_21_+_8433172_8446622_RNA45SN1_45S_50ntFlanks</td><td>10866</td><td>10891</td><td>26</td><td>CTAAGGGCTGGGTCGGTCGGGCTGGG</td><td>1105.2</td><td>-13.9</td><td>1.5</td><td>-9.5</td><td>2.0E-21</td><td>2.7E-20</td><td>MB231/MB468</td><td>MITO</td></tr>
<tr><td>>hg38_21_+_8433172_8446622_RNA45SN1_45S_50ntFlanks@12415.12436.22|CTTCGATGTCGGCTCTTCCTAT</td><td>>hg38_21_+_8433172_8446622_RNA45SN1_45S_50ntFlanks</td><td>12415</td><td>12436</td><td>22</td><td>CTTCGATGTCGGCTCTTCCTAT</td><td>14633.2</td><td>-8.1</td><td>0.9</td><td>-9.5</td><td>2.0E-21</td><td>2.7E-20</td><td>MB231/MB468</td><td>MITO</td></tr>
<tr><td>>hg38_21_+_8433172_8446622_RNA45SN1_45S_50ntFlanks@3816.3852.37|TGGTTCCTTTGGTCGCTCGCTCCTCTCCTACTTGGAT</td><td>>hg38_21_+_8433172_8446622_RNA45SN1_45S_50ntFlanks</td><td>3816</td><td>3852</td><td>37</td><td>TGGTTCCTTTGGTCGCTCGCTCCTCTCCTACTTGGAT</td><td>427.3</td><td>-13.2</td><td>1.4</td><td>-9.4</td><td>6.7E-21</td><td>2.7E-20</td><td>BT20/MB468</td><td>NUC</td></tr>
<tr><td>>hg38_21_+_8433172_8446622_RNA45SN1_45S_50ntFlanks@12405.12430.26|CTTTTTGATCCTTCGATGTCGGCTCT</td><td>>hg38_21_+_8433172_8446622_RNA45SN1_45S_50ntFlanks</td><td>12405</td><td>12430</td><td>26</td><td>CTTTTTGATCCTTCGATGTCGGCTCT</td><td>731.8</td><td>-13.4</td><td>1.4</td><td>-9.5</td><td>1.3E-21</td><td>2.7E-20</td><td>MB231/MB468</td><td>MP</td></tr>
<tr><td>>hg38_21_+_8433172_8446622_RNA45SN1_45S_50ntFlanks@4732.4751.20|AGAACGAAAGTCGGAGGTTC</td><td>>hg38_21_+_8433172_8446622_RNA45SN1_45S_50ntFlanks</td><td>4732</td><td>4751</td><td>20</td><td>AGAACGAAAGTCGGAGGTTC</td><td>663.1</td><td>-4.9</td><td>0.5</td><td>-9.4</td><td>6.7E-21</td><td>2.7E-20</td><td>BT20/MB468</td><td>NUC</td></tr>
<tr><td>>MI0000811|hsa-mir-148b&WithFlank&12|+|54337210|54337320@69.90.22&[MIMAT0000759&hsa-miR-148b-3p&offsets|0|0;m-77&12|+|54337278|54337299&offsets|0|0]|TCAGTGCATCACAGAACTTTGT</td><td>>MI0000811|hsa-mir-148b&WithFlank&12|+|54337210|54337320</td><td>69</td><td>90</td><td>22</td><td>TCAGTGCATCACAGAACTTTGT</td><td>2527.2</td><td>6.2</td><td>0.6</td><td>9.5</td><td>2.5E-21</td><td>2.8E-20</td><td>BT20/MB468</td><td>MITO</td></tr>
<tr><td>>MI0000088|hsa-mir-30a&WithFlank&6|-|71403545|71403627@54.75.22&[MIMAT0000088&hsa-miR-30a-3p&offsets|+1|+1;m-38&6|-|71403554|71403575&offsets|+1|+1]|TTTCAGTCGGATGTTTGCAGCT</td><td>>MI0000088|hsa-mir-30a&WithFlank&6|-|71403545|71403627</td><td>54</td><td>75</td><td>22</td><td>TTTCAGTCGGATGTTTGCAGCT</td><td>1027.7</td><td>7.2</td><td>0.8</td><td>9.5</td><td>1.9E-21</td><td>2.8E-20</td><td>MB231/MB468</td><td>NUC</td></tr>
<tr><td>>hg38_21_+_8433172_8446622_RNA45SN1_45S_50ntFlanks@12405.12431.27|CTTTTTGATCCTTCGATGTCGGCTCTT</td><td>>hg38_21_+_8433172_8446622_RNA45SN1_45S_50ntFlanks</td><td>12405</td><td>12431</td><td>27</td><td>CTTTTTGATCCTTCGATGTCGGCTCTT</td><td>12336.2</td><td>-10.2</td><td>1.1</td><td>-9.5</td><td>1.9E-21</td><td>2.8E-20</td><td>MB231/MB468</td><td>NUC</td></tr>
<tr><td>>hg38_21_+_8433172_8446622_RNA45SN1_45S_50ntFlanks@9572.9595.24|CAAATCGGTCGTCCGACCTGGGTA</td><td>>hg38_21_+_8433172_8446622_RNA45SN1_45S_50ntFlanks</td><td>9572</td><td>9595</td><td>24</td><td>CAAATCGGTCGTCCGACCTGGGTA</td><td>7576.4</td><td>-10.8</td><td>1.1</td><td>-9.5</td><td>2.1E-21</td><td>2.8E-20</td><td>MB231/MB468</td><td>MITO</td></tr>
</table>

| | | | | | | | | | | | | | |
|---|---|---|---|---|---|---|---|---|---|---|---|---|---|
| >hg38_21_+_8433172_8446622_RNA45SN1_45S_50ntFlanks@4567.4587.21\|TAATGGAATAGGACCGCGGTT | >hg38_21_+_8433172_8446622_RNA45SN1_45S_50ntFlanks | 4567 | 4587 | 21 | TAATGGAATAGGACCGCGGTT | 501.6 | -4.3 | 0.5 | -9.4 | 3.4E-21 | 2.8E-20 | BT20/MB468 | CYTO |
| >hg38_21_+_8433172_8446622_RNA45SN1_45S_50ntFlanks@6744.6771.28\|GACACTTCGAACGCACTTGCGGCCCCGG | >hg38_21_+_8433172_8446622_RNA45SN1_45S_50ntFlanks | 6744 | 6771 | 28 | GACACTTCGAACGCACTTGCGGCCCCGG | 2040.4 | -14.9 | 1.6 | -9.5 | 1.4E-21 | 2.8E-20 | MB231/MB468 | MP |
| >MI0000301\|hsa-mir-224&WithFlank&X\|-\|151958572\|151958664@14.35.22&[MIMAT0000281&hsa-miR-224-5p&offsets\|+1\|-2;m-238&X\|-\|151958630\|151958651&offsets\|0\|0]\|CAAGTCACTAGTGGTTCCGTTT | >MI0000301\|hsa-mir-224&WithFlank&X\|-\|151958572\|151958664 | 14 | 35 | 22 | CAAGTCACTAGTGGTTCCGTTT | 10336.1 | 13.0 | 1.4 | 9.5 | 2.5E-21 | 2.8E-20 | BT20/MB468 | MITO |
| >hg38_21_+_8433172_8446622_RNA45SN1_45S_50ntFlanks@11316.11340.25\|CACCCCACGTCTCGTCGCGCGCGCG | >hg38_21_+_8433172_8446622_RNA45SN1_45S_50ntFlanks | 11316 | 11340 | 25 | CACCCCACGTCTCGTCGCGCGCGCG | 1112.2 | -13.1 | 1.4 | -9.5 | 1.9E-21 | 2.8E-20 | MB231/MB468 | NUC |
| >hg38_21_+_8433172_8446622_RNA45SN1_45S_50ntFlanks@9563.9596.34\|CCTGACGTGCAAATCGGTCGTCCGACCTGGGTAT | >hg38_21_+_8433172_8446622_RNA45SN1_45S_50ntFlanks | 9563 | 9596 | 34 | CCTGACGTGCAAATCGGTCGTCCGACCTGGGTAT | 1886.3 | -5.8 | 0.6 | -9.5 | 1.9E-21 | 2.9E-20 | MB231/MB468 | NUC |
| >hg38_21_+_8433172_8446622_RNA45SN1_45S_50ntFlanks@12627.12652.26\|ATGGGGCGAAGCTACCATCTGTGGGA | >hg38_21_+_8433172_8446622_RNA45SN1_45S_50ntFlanks | 12627 | 12652 | 26 | ATGGGGCGAAGCTACCATCTGTGGGA | 842.4 | -13.5 | 1.4 | -9.5 | 2.1E-21 | 2.9E-20 | MB231/MB468 | MITO |
| >MI0000298\|hsa-mir-221&WithFlank&X\|-\|45746151\|45746272@71.93.23&[MIMAT0000278&hsa-miR-221-3p&offsets\|0\|0;m-91&X\|-\|45746181\|45746202&offsets\|0\|+1]\|AGCTACATTGTCTGCTGGGTTTC | >MI0000298\|hsa-mir-221&WithFlank&X\|-\|45746151\|45746272 | 71 | 93 | 23 | AGCTACATTGTCTGCTGGGTTTC | 26863.8 | 5.4 | 0.6 | 9.4 | 3.5E-21 | 2.9E-20 | BT20/MB468 | CYTO |
| >hg38_21_+_8433172_8446622_RNA45SN1_45S_50ntFlanks@12617.12654.38\|TGAGGAGCCAATGGGGCGAAGCTACCATCTGTGGGATT | >hg38_21_+_8433172_8446622_RNA45SN1_45S_50ntFlanks | 12617 | 12654 | 38 | TGAGGAGCCAATGGGGCGAAGCTACCATCTGTGGGATT | 1640.9 | -7.1 | 0.8 | -9.5 | 2.0E-21 | 2.9E-20 | MB231/MB468 | NUC |
| >hg38_21_+_8433172_8446622_RNA45SN1_45S_50ntFlanks@12973.12991.19\|TCGTACGTAGCAGAGCAGC | >hg38_21_+_8433172_8446622_RNA45SN1_45S_50ntFlanks | 12973 | 12991 | 19 | TCGTACGTAGCAGAGCAGC | 5218.8 | -6.3 | 0.7 | -9.4 | 3.6E-21 | 2.9E-20 | BT20/MB468 | CYTO |
| >hg38_21_+_8433172_8446622_RNA45SN1_45S_50ntFlanks@10402.10420.19\|AACATGGGTCAGTCGGTCC | >hg38_21_+_8433172_8446622_RNA45SN1_45S_50ntFlanks | 10402 | 10420 | 19 | AACATGGGTCAGTCGGTCC | 1857.0 | -7.5 | 0.8 | -9.6 | 1.1E-21 | 3.0E-20 | MB231/MB468 | CYTO |
| >hg38_21_+_8433172_8446622_RNA45SN1_45S_50ntFlanks@11704.11728.25\|ATGAAGCGCGGGTAAACGGCGGGAG | >hg38_21_+_8433172_8446622_RNA45SN1_45S_50ntFlanks | 11704 | 11728 | 25 | ATGAAGCGCGGGTAAACGGCGGGAG | 1959.7 | -8.8 | 0.9 | -9.5 | 2.1E-21 | 3.0E-20 | MB231/MB468 | NUC |
| >MI0000088\|hsa-mir-30a&WithFlank&6\|-\|71403545\|71403627@12.33.22(+1A)&[MIMAT0000087&hsa-miR-30a-5p&offsets\|0\|0(+1A);m-15&6\|-\|71403595\|71403616&offsets\|0\|0(+1A)]\|TGTAAACATCCTCGACTGGAAGA | >MI0000088\|hsa-mir-30a&WithFlank&6\|-\|71403545\|71403627 | 12 | 33 | 22(+1A) | TGTAAACATCCTCGACTGGAAGA | 697.3 | 6.3 | 0.7 | 9.5 | 2.7E-21 | 3.0E-20 | BT20/MB468 | MITO |
| >hg38_21_+_8433172_8446622_RNA45SN1_45S_50ntFlanks@9567.9594.28\|ACGTGCAAATCGGTCGTCCGACCTGGGT | >hg38_21_+_8433172_8446622_RNA45SN1_45S_50ntFlanks | 9567 | 9594 | 28 | ACGTGCAAATCGGTCGTCCGACCTGGGT | 784.3 | -13.5 | 1.4 | -9.5 | 1.4E-21 | 3.0E-20 | MB231/MB468 | MP |
| >hg38_21_+_8433172_8446622_RNA45SN1_45S_50ntFlanks@12457.12489.33\|CAAGCGTTGGATTGTTCACCCACTAATAGGGAA | >hg38_21_+_8433172_8446622_RNA45SN1_45S_50ntFlanks | 12457 | 12489 | 33 | CAAGCGTTGGATTGTTCACCCACTAATAGGGAA | 367.1 | -12.9 | 1.4 | -9.4 | 7.4E-21 | 3.0E-20 | BT20/MB468 | NUC |
| >hg38_21_+_8433172_8446622_RNA45SN1_45S_50ntFlanks@8115.8136.22\|CGCGGGACATGTGGCGTACGGA | >hg38_21_+_8433172_8446622_RNA45SN1_45S_50ntFlanks | 8115 | 8136 | 22 | CGCGGGACATGTGGCGTACGGA | 1247.1 | -14.1 | 1.5 | -9.5 | 1.5E-21 | 3.0E-20 | MB231/MB468 | MP |
| >MI0000102\|hsa-mir-100&WithFlank&11\|-\|122152223\|122152314@19.40.22&[MIMAT0000098&hsa-miR-100-5p&offsets\|0\|0;m-27&11\|-\|122152276\|122152296&offsets\|0\|+1]\|AACCCGTAGATCCGAACTTGTG | >MI0000102\|hsa-mir-100&WithFlank&11\|-\|122152223\|122152314 | 19 | 40 | 22 | AACCCGTAGATCCGAACTTGTG | 713584.9 | -7.4 | 0.8 | -9.7 | 1.9E-22 | 3.0E-20 | BT20/MB231 | MITO |
| >MI0000285\|hsa-mir-205&WithFlank&1\|+\|209432127\|209432248@40.60.21&[MIMAT0000266&hsa-miR-205-5p&offsets\|0\|-1;m-351&1\|+\|209432166\|209432187&offsets\|0\|-1]\|TCCTTCATTCCACCGGAGTCT | >MI0000285\|hsa-mir-205&WithFlank&1\|+\|209432127\|209432248 | 40 | 60 | 21 | TCCTTCATTCCACCGGAGTCT | 1395.7 | -14.3 | 1.5 | -9.5 | 1.5E-21 | 3.0E-20 | MB231/MB468 | MP |
| >hg38_21_+_8433172_8446622_RNA45SN1_45S_50ntFlanks@9032.9059.28\|CCTCGTGGGGGGGCCGGGCCACCCCTCC | >hg38_21_+_8433172_8446622_RNA45SN1_45S_50ntFlanks | 9032 | 9059 | 28 | CCTCGTGGGGGGGCCGGGCCACCCCTCC | 773.0 | -13.5 | 1.4 | -9.5 | 1.5E-21 | 3.0E-20 | MB231/MB468 | MP |
| >hg38_21_+_8433172_8446622_RNA45SN1_45S_50ntFlanks@11205.11225.21\|GCGGCGGCGACTCTGGACGCG | >hg38_21_+_8433172_8446622_RNA45SN1_45S_50ntFlanks | 11205 | 11225 | 21 | GCGGCGGCGACTCTGGACGCG | 911.1 | -13.7 | 1.4 | -9.5 | 1.5E-21 | 3.0E-20 | MB231/MB468 | MP |
| >MI0000266\|hsa-mir-10a&WithFlank&17\|-\|48579832\|48579953@28.51.24&[MIMAT0000253&hsa-miR-10a-5p&offsets\|0\|+1;m-18&17\|-\|48579905\|48579926&offsets\|0\|+2]\|TACCCTGTAGATCCGAATTTGTGT | >MI0000266\|hsa-mir-10a&WithFlank&17\|-\|48579832\|48579953 | 28 | 51 | 24 | TACCCTGTAGATCCGAATTTGTGT | 773.6 | 13.6 | 1.4 | 9.6 | 1.2E-21 | 3.0E-20 | MB231/MB468 | CYTO |
| >hg38_21_+_8433172_8446622_RNA45SN1_45S_50ntFlanks@4759.4782.24\|ATCAGATACCGTCGTAGTTCCGAC | >hg38_21_+_8433172_8446622_RNA45SN1_45S_50ntFlanks | 4759 | 4782 | 24 | ATCAGATACCGTCGTAGTTCCGAC | 3376.3 | -14.8 | 1.5 | -9.9 | 2.7E-23 | 3.1E-20 | BT20/MB468 | MP |
| >hg38_21_+_8433172_8446622_RNA45SN1_45S_50ntFlanks@10866.10892.27\|CTAAGGGCTGGGTCGGTCGGGCTGGGG | >hg38_21_+_8433172_8446622_RNA45SN1_45S_50ntFlanks | 10866 | 10892 | 27 | CTAAGGGCTGGGTCGGTCGGGCTGGGG | 1230.6 | -14.0 | 1.5 | -9.5 | 2.3E-21 | 3.1E-20 | MB231/MB468 | MITO |
| >hg38_21_+_8433172_8446622_RNA45SN1_45S_50ntFlanks@4759.4778.20\|ATCAGATACCGTCGTAGTTC | >hg38_21_+_8433172_8446622_RNA45SN1_45S_50ntFlanks | 4759 | 4778 | 20 | ATCAGATACCGTCGTAGTTC | 1879.2 | -14.7 | 1.5 | -9.5 | 2.3E-21 | 3.1E-20 | MB231/MB468 | MITO |
| >MI0001448\|hsa-mir-425&WithFlank&3\|-\|49020142\|49020240@20.44.25&[MIMAT0003393&hsa-miR-425-5p&offsets\|0\|+2;m-95&3\|-\|49020200\|49020221&offsets\|0\|+3]\|AATGACACGATCACTCCCGTTGAGT | >MI0001448\|hsa-mir-425&WithFlank&3\|-\|49020142\|49020240 | 20 | 44 | 25 | AATGACACGATCACTCCCGTTGAGT | 5523.2 | 6.2 | 0.6 | 9.5 | 2.1E-21 | 3.1E-20 | MB231/MB468 | NUC |
| >MI0000483\|hsa-mir-186&WithFlank&1\|-\|71067625\|71067722@21.42.22&[MIMAT0000456&hsa-miR-186-5p&offsets\|0\|0;m-84&1\|-\|71067681\|71067702&offsets\|0\|0]\|CAAAGAATTCTCCTTTTGGGCT | >MI0000483\|hsa-mir-186&WithFlank&1\|-\|71067625\|71067722 | 21 | 42 | 22 | CAAAGAATTCTCCTTTTGGGCT | 2511.1 | 7.1 | 0.7 | 9.4 | 3.8E-21 | 3.1E-20 | BT20/MB468 | CYTO |
| >hg38_21_+_8433172_8446622_RNA45SN1_45S_50ntFlanks@11873.11903.31\|AAGAAGACCCTGTTGAGCTTGACTCTAGTCT | >hg38_21_+_8433172_8446622_RNA45SN1_45S_50ntFlanks | 11873 | 11903 | 31 | AAGAAGACCCTGTTGAGCTTGACTCTAGTCT | 647.5 | -13.2 | 1.4 | -9.5 | 1.5E-21 | 3.2E-20 | MB231/MB468 | MP |
| >hg38_21_+_8433172_8446622_RNA45SN1_45S_50ntFlanks@10771.10797.27\|CAGGTCTCCAAGGTGAACAGCCTCTGG | >hg38_21_+_8433172_8446622_RNA45SN1_45S_50ntFlanks | 10771 | 10797 | 27 | CAGGTCTCCAAGGTGAACAGCCTCTGG | 701.7 | -8.5 | 0.9 | -9.6 | 1.2E-21 | 3.2E-20 | MB231/MB468 | CYTO |
| >hg38_21_+_8433172_8446622_RNA45SN1_45S_50ntFlanks@10391.10409.19\|AACAGCAGTTGAACATGGG | >hg38_21_+_8433172_8446622_RNA45SN1_45S_50ntFlanks | 10391 | 10409 | 19 | AACAGCAGTTGAACATGGG | 1129.0 | -14.0 | 1.5 | -9.5 | 1.5E-21 | 3.2E-20 | MB231/MB468 | MP |
| >hg38_1_-_228634819_228635039_RNA5S12_5S_50ntFlanks@146.169.24\|TGGGAATACCGGGTGCTGTAGGCT | >hg38_1_-_228634819_228635039_RNA5S12_5S_50ntFlanks | 146 | 169 | 24 | TGGGAATACCGGGTGCTGTAGGCT | 1148.3 | 6.7 | 0.7 | 9.6 | 1.2E-21 | 3.2E-20 | MB231/MB468 | CYTO |
| >MI0000447\|hsa-mir-128-1&WithFlank&2\|+\|135665391\|135665484@56.78.23&[MIMAT0000424&hsa-miR-128-3p&offsets\|0\|+2;m-108&2\|+\|135665446\|135665466&offsets\|0\|+2]\|TCACAGTGAACCGGTCTCTTTTT | >MI0000447\|hsa-mir-128-1&WithFlank&2\|+\|135665391\|135665484 | 56 | 78 | 23 | TCACAGTGAACCGGTCTCTTTTT | 381.1 | 5.0 | 0.5 | 9.4 | 3.9E-21 | 3.2E-20 | BT20/MB468 | CYTO |
| >hg38_21_+_8433172_8446622_RNA45SN1_45S_50ntFlanks@8170.8201.32\|CAAGTCCTTCTGATCGAGGCCCAGCCCGTGGA | >hg38_21_+_8433172_8446622_RNA45SN1_45S_50ntFlanks | 8170 | 8201 | 32 | CAAGTCCTTCTGATCGAGGCCCAGCCCGTGGA | 972.4 | -14.4 | 1.5 | -9.4 | 8.0E-21 | 3.3E-20 | BT20/MB468 | NUC |
| >MI0000482\|hsa-mir-185&WithFlank&22\|+\|20033133\|20033226@21.43.23&[MIMAT0000455&hsa-miR-185-5p&offsets\|0\|+1;m-145&22\|+\|20033153\|20033174&offsets\|0\|+1]\|TGGAGAGAAAGGCAGTTCCTGAT | >MI0000482\|hsa-mir-185&WithFlank&22\|+\|20033133\|20033226 | 21 | 43 | 23 | TGGAGAGAAAGGCAGTTCCTGAT | 897.0 | 7.3 | 0.8 | 9.4 | 4.1E-21 | 3.3E-20 | BT20/MB468 | CYTO |
| >hg38_21_+_8433172_8446622_RNA45SN1_45S_50ntFlanks@6721.6739.19\|AATTGCAGGACACATTGAT | >hg38_21_+_8433172_8446622_RNA45SN1_45S_50ntFlanks | 6721 | 6739 | 19 | AATTGCAGGACACATTGAT | 1021.3 | -13.9 | 1.5 | -9.5 | 1.6E-21 | 3.3E-20 | MB231/MB468 | MP |
| >hg38_21_+_8433172_8446622_RNA45SN1_45S_50ntFlanks@9567.9595.29\|ACGTGCAAATCGGTCGTCCGACCTGGGTA | >hg38_21_+_8433172_8446622_RNA45SN1_45S_50ntFlanks | 9567 | 9595 | 29 | ACGTGCAAATCGGTCGTCCGACCTGGGTA | 3187.3 | -5.4 | 0.6 | -9.4 | 4.1E-21 | 3.3E-20 | BT20/MB468 | CYTO |
| >hg38_21_+_8433172_8446622_RNA45SN1_45S_50ntFlanks@3900.3919.20\|CTTCGCGGGGGGGATGCGTG | >hg38_21_+_8433172_8446622_RNA45SN1_45S_50ntFlanks | 3900 | 3919 | 20 | CTTCGCGGGGGGGATGCGTG | 1040.5 | -13.8 | 1.5 | -9.5 | 2.5E-21 | 3.3E-20 | MB231/MB468 | MITO |
| >hg38_21_+_8433172_8446622_RNA45SN1_45S_50ntFlanks@11669.11687.19\|GCCCAGTGCTCTGAATGTC | >hg38_21_+_8433172_8446622_RNA45SN1_45S_50ntFlanks | 11669 | 11687 | 19 | GCCCAGTGCTCTGAATGTC | 943.4 | 4.9 | 0.5 | 9.4 | 4.1E-21 | 3.3E-20 | BT20/MB468 | CYTO |
| >hg38_21_+_8433172_8446622_RNA45SN1_45S_50ntFlanks@12456.12478.23\|CCAAGCGTTGGATTGTTCACCCA | >hg38_21_+_8433172_8446622_RNA45SN1_45S_50ntFlanks | 12456 | 12478 | 23 | CCAAGCGTTGGATTGTTCACCCA | 22467.9 | -11.4 | 1.2 | -9.4 | 8.2E-21 | 3.3E-20 | BT20/MB468 | NUC |
| >hg38_21_+_8433172_8446622_RNA45SN1_45S_50ntFlanks@10411.10445.35\|CAGTCGGTCCTGAGAGATGGGCGAGCGCCGTTCCG | >hg38_21_+_8433172_8446622_RNA45SN1_45S_50ntFlanks | 10411 | 10445 | 35 | CAGTCGGTCCTGAGAGATGGGCGAGCGCCGTTCCG | 869.0 | -13.5 | 1.4 | -9.5 | 2.5E-21 | 3.4E-20 | MB231/MB468 | MITO |
| >MI0000746\|hsa-mir-99b&WithFlank&19\|+\|51692606\|51692687@14.32.19(+1U)&[MIMAT0000689&hsa-miR-99b-5p&offsets\|+1\|-2(+1U);m-10&19\|+\|51692618\|51692639&offsets\|+1\|-2(+1U)]\|ACCCGTAGAACCGACCTTGT | >MI0000746\|hsa-mir-99b&WithFlank&19\|+\|51692606\|51692687 | 14 | 32 | 19(+1U) | ACCCGTAGAACCGACCTTGT | 418.5 | 13.0 | 1.4 | 9.4 | 4.2E-21 | 3.4E-20 | BT20/MB468 | CYTO |
| >hg38_21_+_8433172_8446622_RNA45SN1_45S_50ntFlanks@11576.11601.26\|CTTAGAACTGGTGCGGACCAGGGGAA | >hg38_21_+_8433172_8446622_RNA45SN1_45S_50ntFlanks | 11576 | 11601 | 26 | CTTAGAACTGGTGCGGACCAGGGGAA | 1023.8 | -13.9 | 1.5 | -9.5 | 1.7E-21 | 3.4E-20 | MB231/MB468 | MP |
| >hg38_21_+_8433172_8446622_RNA45SN1_45S_50ntFlanks@12003.12043.41\|GCGGGCCGCCGGTGAAATACCACTACTCTGATCGTTTTTTC | >hg38_21_+_8433172_8446622_RNA45SN1_45S_50ntFlanks | 12003 | 12043 | 41 | GCGGGCCGCCGGTGAAATACCACTACTCTGATCGTTTTTTC | 2091.4 | -14.8 | 1.6 | -9.5 | 2.6E-21 | 3.4E-20 | MB231/MB468 | MITO |
| >hg38_21_+_8433172_8446622_RNA45SN1_45S_50ntFlanks@12365.12382.18\|ATAACTGGCTTGTGGCGG | >hg38_21_+_8433172_8446622_RNA45SN1_45S_50ntFlanks | 12365 | 12382 | 18 | ATAACTGGCTTGTGGCGG | 792.6 | -13.5 | 1.4 | -9.5 | 1.7E-21 | 3.5E-20 | MB231/MB468 | MP |
| >MI0000760\|hsa-mir-361&WithFlank&X\|-\|85903630\|85903713@51.72.22&[MIMAT0004682&hsa-miR-361-3p&offsets\|0\|-1;m-124&X\|-\|85903642\|85903663&offsets\|0\|0]\|TCCCCCAGGTGTGATTCTGATT | >MI0000760\|hsa-mir-361&WithFlank&X\|-\|85903630\|85903713 | 51 | 72 | 22 | TCCCCCAGGTGTGATTCTGATT | 376.7 | 9.3 | 1.0 | 9.4 | 4.3E-21 | 3.5E-20 | BT20/MB468 | CYTO |
| >hg38_21_+_8433172_8446622_RNA45SN1_45S_50ntFlanks@4724.4768.45\|ATTAATCAAGAACGAAAGTCGGAGGTTCGAAGACGATCAGATACC | >hg38_21_+_8433172_8446622_RNA45SN1_45S_50ntFlanks | 4724 | 4768 | 45 | ATTAATCAAGAACGAAAGTCGGAGGTTCGAAGACGATCAGATACC | 325.2 | 12.3 | 1.3 | 9.5 | 1.4E-21 | 3.5E-20 | MB231/MB468 | CYTO |
| >hg38_21_+_8433172_8446622_RNA45SN1_45S_50ntFlanks@10798.10817.20\|CATGTTGGAACAATGTAGGT | >hg38_21_+_8433172_8446622_RNA45SN1_45S_50ntFlanks | 10798 | 10817 | 20 | CATGTTGGAACAATGTAGGT | 861.1 | -13.6 | 1.4 | -9.5 | 1.7E-21 | 3.5E-20 | MB231/MB468 | MP |
| >hg38_21_+_8433172_8446622_RNA45SN1_45S_50ntFlanks@6772.6803.32\|GTTCCTCCCGGGGCTACGCCTGTCTGAGCGTC | >hg38_21_+_8433172_8446622_RNA45SN1_45S_50ntFlanks | 6772 | 6803 | 32 | GTTCCTCCCGGGGCTACGCCTGTCTGAGCGTC | 2320.6 | -8.8 | 0.9 | -9.5 | 2.6E-21 | 3.5E-20 | MB231/MB468 | MITO |
| >hg38_1_-_228634819_228635039_RNA5S12_5S_50ntFlanks@84.101.18\|CTCGTCTGATCTCGGAAG | >hg38_1_-_228634819_228635039_RNA5S12_5S_50ntFlanks | 84 | 101 | 18 | CTCGTCTGATCTCGGAAG | 879.9 | -13.6 | 1.4 | -9.5 | 2.6E-21 | 3.5E-20 | MB231/MB468 | MITO |
| >trnaMT_PheGAA_MT_+_577_647@35.59.25\|GAAAATGTTTAGACGGGCTCACATC | >trnaMT_PheGAA_MT_+_577_647 | 35 | 59 | 25 | GAAAATGTTTAGACGGGCTCACATC | 1915.0 | 8.3 | 0.9 | 9.5 | 3.2E-21 | 3.5E-20 | BT20/MB468 | MITO |
| >MI0000077\|hsa-mir-21&WithFlank&17\|+\|59841260\|59841343@14.36.23(+1G)&[MIMAT0000076&hsa-miR-21-5p&offsets\|0\|+1(+1G);m-1&17\|+\|59841273\|59841294&offsets\|0\|+1(+1G)]\|TAGCTTATCAGACTGATGTTGACG | >MI0000077\|hsa-mir-21&WithFlank&17\|+\|59841260\|59841343 | 14 | 36 | 23(+1G) | TAGCTTATCAGACTGATGTTGACG | 9524.4 | 6.4 | 0.7 | 9.5 | 2.4E-21 | 3.5E-20 | MB231/MB468 | NUC |
| >hg38_21_+_8433172_8446622_RNA45SN1_45S_50ntFlanks@10574.10608.35\|AAGCCGGCGGGAGCCCCGGGGAGAGTTCTCTTTTC | >hg38_21_+_8433172_8446622_RNA45SN1_45S_50ntFlanks | 10574 | 10608 | 35 | AAGCCGGCGGGAGCCCCGGGGAGAGTTCTCTTTTC | 1679.1 | -7.9 | 0.8 | -9.5 | 1.4E-21 | 3.5E-20 | MB231/MB468 | CYTO |

<table>
<tr><td>>hg38_21_+_8433172_8446622_RNA45SN1_45S_50ntFlanks@10425.10457.33|AGATGGGCGAGCGCCGTTCCGAAGGGACGGGCG</td><td>>hg38_21_+_8433172_8446622_RNA45SN1_45S_50ntFlanks</td><td>10425</td><td>10457</td><td>33</td><td>AGATGGGCGAGCGCCGTTCCGAAGGGACGGGCG</td><td>495.0</td><td>-13.4</td><td>1.4</td><td>-9.4</td><td>8.7E-21</td><td>3.6E-20</td><td>BT20/MB468</td><td>NUC</td></tr>
<tr><td>>hg38_21_+_8433172_8446622_RNA45SN1_45S_50ntFlanks@5472.5500.29|ACGGCCCTGGCGGAGCGCTGAGAAGACGG</td><td>>hg38_21_+_8433172_8446622_RNA45SN1_45S_50ntFlanks</td><td>5472</td><td>5500</td><td>29</td><td>ACGGCCCTGGCGGAGCGCTGAGAAGACGG</td><td>2828.1</td><td>6.4</td><td>0.7</td><td>9.5</td><td>1.4E-21</td><td>3.6E-20</td><td>MB231/MB468</td><td>CYTO</td></tr>
<tr><td>>hg38_21_+_8433172_8446622_RNA45SN1_45S_50ntFlanks@485.521.37|GATGTGGTGACGTCGTGCTCTCCCGGGCCGGGTCCGA</td><td>>hg38_21_+_8433172_8446622_RNA45SN1_45S_50ntFlanks</td><td>485</td><td>521</td><td>37</td><td>GATGTGGTGACGTCGTGCTCTCCCGGGCCGGGTCCGA</td><td>902.3</td><td>12.7</td><td>1.4</td><td>9.3</td><td>8.8E-21</td><td>3.6E-20</td><td>BT20/MB468</td><td>NUC</td></tr>
<tr><td>>MI0000102|hsa-mir-100&WithFlank&11|-|122152223|122152314@20.40.21&[MIMAT0000098&hsa-miR-100-5p&offsets|+1|0;m-27&11|-|122152276|122152296&offsets|+1|+1]|ACCCGTAGATCCGAACTTGTG</td><td>>MI0000102|hsa-mir-100&WithFlank&11|-|122152223|122152314</td><td>20</td><td>40</td><td>21</td><td>ACCCGTAGATCCGAACTTGTG</td><td>1000.3</td><td>13.2</td><td>1.4</td><td>9.5</td><td>1.8E-21</td><td>3.6E-20</td><td>MB231/MB468</td><td>MP</td></tr>
<tr><td>>hg38_21_+_8433172_8446622_RNA45SN1_45S_50ntFlanks@10727.10763.37|GGAGAGGGTGTAAATCTCGCGCCGGGCCGTACCCATA</td><td>>hg38_21_+_8433172_8446622_RNA45SN1_45S_50ntFlanks</td><td>10727</td><td>10763</td><td>37</td><td>GGAGAGGGTGTAAATCTCGCGCCGGGCCGTACCCATA</td><td>342.3</td><td>-12.8</td><td>1.4</td><td>-9.3</td><td>8.8E-21</td><td>3.6E-20</td><td>BT20/MB468</td><td>NUC</td></tr>
<tr><td>>MI0000255|hsa-mir-30d&WithFlank&8|-|134804870|134804951@12.34.23(+1G)&[MIMAT0000245&hsa-miR-30d-5p&offsets|0|+1(+1G);m-21&8|-|134804919|134804940&offsets|0|+1(+1G)]|TGTAAACATCCCCGACTGGAAGCG</td><td>>MI0000255|hsa-mir-30d&WithFlank&8|-|134804870|134804951</td><td>12</td><td>34</td><td>23(+1G)</td><td>TGTAAACATCCCCGACTGGAAGCG</td><td>3781.9</td><td>7.9</td><td>0.8</td><td>9.4</td><td>4.5E-21</td><td>3.6E-20</td><td>BT20/MB468</td><td>CYTO</td></tr>
<tr><td>>MI0000255|hsa-mir-30d&WithFlank&8|-|134804870|134804951@12.34.23(+1A)&[MIMAT0000245&hsa-miR-30d-5p&offsets|0|+1(+1A);m-21&8|-|134804919|134804940&offsets|0|+1(+1A)]|TGTAAACATCCCCGACTGGAAGCA</td><td>>MI0000255|hsa-mir-30d&WithFlank&8|-|134804870|134804951</td><td>12</td><td>34</td><td>23(+1A)</td><td>TGTAAACATCCCCGACTGGAAGCA</td><td>3084.1</td><td>6.4</td><td>0.7</td><td>9.5</td><td>2.5E-21</td><td>3.7E-20</td><td>MB231/MB468</td><td>NUC</td></tr>
<tr><td>>MI0000266|hsa-mir-10a&WithFlank&17|-|48579832|48579953@28.51.24&[MIMAT0000253&hsa-miR-10a-5p&offsets|0|+1;m-18&17|-|48579905|48579926&offsets|0|+2]|TACCCTGTAGATCCGAATTTGTGT</td><td>>MI0000266|hsa-mir-10a&WithFlank&17|-|48579832|48579953</td><td>28</td><td>51</td><td>24</td><td>TACCCTGTAGATCCGAATTTGTGT</td><td>922.4</td><td>14.3</td><td>1.5</td><td>9.5</td><td>3.3E-21</td><td>3.7E-20</td><td>BT20/MB468</td><td>MITO</td></tr>
<tr><td>>hg38_21_+_8433172_8446622_RNA45SN1_45S_50ntFlanks@11573.11603.31|CGACTTAGAACTGGTGCGGACCAGGGGAATC</td><td>>hg38_21_+_8433172_8446622_RNA45SN1_45S_50ntFlanks</td><td>11573</td><td>11603</td><td>31</td><td>CGACTTAGAACTGGTGCGGACCAGGGGAATC</td><td>1181.2</td><td>-5.0</td><td>0.5</td><td>-9.5</td><td>2.6E-21</td><td>3.7E-20</td><td>MB231/MB468</td><td>NUC</td></tr>
<tr><td>>hg38_21_+_8433172_8446622_RNA45SN1_45S_50ntFlanks@4449.4471.23|CCTCTCGGCGCCCCCTCGATGCT</td><td>>hg38_21_+_8433172_8446622_RNA45SN1_45S_50ntFlanks</td><td>4449</td><td>4471</td><td>23</td><td>CCTCTCGGCGCCCCCTCGATGCT</td><td>701.1</td><td>-13.3</td><td>1.4</td><td>-9.5</td><td>1.8E-21</td><td>3.7E-20</td><td>MB231/MB468</td><td>MP</td></tr>
<tr><td>>am_trna117_GlyTCC_1_-_145397864_145397935@1.30.30,trna45_GlyTCC_1_+_161500903_161500974@1.30.30,trna70_GlyTCC_1_-_161439547_161439618@1.30.30,trna73_GlyTCC_1_-_161432166_161432237@1.30.30,trna76_GlyTCC_1_-_161424756_161424827@1.30.30,trna79_GlyTCC_1_-_161417375_161417446@1.30.30|GCGTTGGTGGTATAGTGGTGAGCATAGCTG</td><td>>am_trna117_GlyTCC_1_-_145397864_145397935</td><td>1</td><td>30</td><td>30</td><td>GCGTTGGTGGTATAGTGGTGAGCATAGCTG</td><td>1559.9</td><td>-14.4</td><td>1.5</td><td>-9.5</td><td>2.8E-21</td><td>3.7E-20</td><td>MB231/MB468</td><td>MITO</td></tr>
<tr><td>>hg38_21_+_8433172_8446622_RNA45SN1_45S_50ntFlanks@9180.9197.18|CCGTCGGGCCCGGGGGAG</td><td>>hg38_21_+_8433172_8446622_RNA45SN1_45S_50ntFlanks</td><td>9180</td><td>9197</td><td>18</td><td>CCGTCGGGCCCGGGGGAG</td><td>2013.8</td><td>-14.8</td><td>1.6</td><td>-9.5</td><td>1.8E-21</td><td>3.8E-20</td><td>MB231/MB468</td><td>MP</td></tr>
<tr><td>>hg38_21_+_8433172_8446622_RNA45SN1_45S_50ntFlanks@6680.6718.39|TGCGTCGATGAAGAACGCAGCTAGCTGCGAGAATTAATG</td><td>>hg38_21_+_8433172_8446622_RNA45SN1_45S_50ntFlanks</td><td>6680</td><td>6718</td><td>39</td><td>TGCGTCGATGAAGAACGCAGCTAGCTGCGAGAATTAATG</td><td>381.5</td><td>-13.0</td><td>1.4</td><td>-9.3</td><td>9.2E-21</td><td>3.8E-20</td><td>BT20/MB468</td><td>NUC</td></tr>
<tr><td>>hg38_21_+_8433172_8446622_RNA45SN1_45S_50ntFlanks@10392.10430.39|ACAGCAGTTGAACATGGGTCAGTCGGTCCTGAGAGATGG</td><td>>hg38_21_+_8433172_8446622_RNA45SN1_45S_50ntFlanks</td><td>10392</td><td>10430</td><td>39</td><td>ACAGCAGTTGAACATGGGTCAGTCGGTCCTGAGAGATGG</td><td>1681.1</td><td>-7.3</td><td>0.8</td><td>-9.5</td><td>2.6E-21</td><td>3.8E-20</td><td>MB231/MB468</td><td>NUC</td></tr>
<tr><td>>MI0000542|hsa-mir-320a&WithFlank&8|-|22244960|22245043@48.72.25&[MIMAT0000510&hsa-miR-320a-3p&offsets|0|+3;m-60&8|-|22244975|22244996&offsets|0|+3]|AAAAGCTGGGTTGAGAGGGCGAAAA</td><td>>MI0000542|hsa-mir-320a&WithFlank&8|-|22244960|22245043</td><td>48</td><td>72</td><td>25</td><td>AAAAGCTGGGTTGAGAGGGCGAAAA</td><td>1253.5</td><td>12.2</td><td>1.3</td><td>9.4</td><td>4.7E-21</td><td>3.8E-20</td><td>BT20/MB468</td><td>CYTO</td></tr>
<tr><td>>hg38_21_+_8433172_8446622_RNA45SN1_45S_50ntFlanks@8170.8197.28|CAAGTCCTTCTGATCGAGGCCCAGCCCG</td><td>>hg38_21_+_8433172_8446622_RNA45SN1_45S_50ntFlanks</td><td>8170</td><td>8197</td><td>28</td><td>CAAGTCCTTCTGATCGAGGCCCAGCCCG</td><td>9724.5</td><td>-12.7</td><td>1.3</td><td>-9.5</td><td>2.9E-21</td><td>3.8E-20</td><td>MB231/MB468</td><td>MITO</td></tr>
<tr><td>>MI0000272|hsa-mir-182&WithFlank&7|-|129770377|129770498@29.52.24(+1U)&[MIMAT0000259&hsa-miR-182-5p&offsets|0|0(+1U);m-22&7|-|129770449|129770470&offsets|0|+2(+1U)]|TTTGGCAATGGTAGAACTCACACTT</td><td>>MI0000272|hsa-mir-182&WithFlank&7|-|129770377|129770498</td><td>29</td><td>52</td><td>24(+1U)</td><td>TTTGGCAATGGTAGAACTCACACTT</td><td>86.7</td><td>3.5</td><td>0.4</td><td>9.3</td><td>9.4E-21</td><td>3.8E-20</td><td>BT20/MB468</td><td>NUC</td></tr>
<tr><td>>hg38_21_+_8433172_8446622_RNA45SN1_45S_50ntFlanks@7986.8017.32|ATCAGACGTGGCGACCCGCTGAATTTAAGCAT</td><td>>hg38_21_+_8433172_8446622_RNA45SN1_45S_50ntFlanks</td><td>7986</td><td>8017</td><td>32</td><td>ATCAGACGTGGCGACCCGCTGAATTTAAGCAT</td><td>509.7</td><td>-13.4</td><td>1.4</td><td>-9.3</td><td>9.4E-21</td><td>3.8E-20</td><td>BT20/MB468</td><td>NUC</td></tr>
<tr><td>>hg38_21_+_8433172_8446622_RNA45SN1_45S_50ntFlanks@10771.10805.35|CAGGTCTCCAAGGTGAACAGCCTCTGGCATGTTGG</td><td>>hg38_21_+_8433172_8446622_RNA45SN1_45S_50ntFlanks</td><td>10771</td><td>10805</td><td>35</td><td>CAGGTCTCCAAGGTGAACAGCCTCTGGCATGTTGG</td><td>853.7</td><td>-13.6</td><td>1.4</td><td>-9.5</td><td>1.9E-21</td><td>3.8E-20</td><td>MB231/MB468</td><td>MP</td></tr>
<tr><td>>hg38_21_+_8433172_8446622_RNA45SN1_45S_50ntFlanks@10682.10706.25|TCCGGCGGCGTCCGGTGAGCTCTCG</td><td>>hg38_21_+_8433172_8446622_RNA45SN1_45S_50ntFlanks</td><td>10682</td><td>10706</td><td>25</td><td>TCCGGCGGCGTCCGGTGAGCTCTCG</td><td>1177.1</td><td>-14.1</td><td>1.5</td><td>-9.5</td><td>1.9E-21</td><td>3.8E-20</td><td>MB231/MB468</td><td>MP</td></tr>
<tr><td>>hg38_21_+_8433172_8446622_RNA45SN1_45S_50ntFlanks@10350.10368.19|ACGAGAACTTTGAAGGCCG</td><td>>hg38_21_+_8433172_8446622_RNA45SN1_45S_50ntFlanks</td><td>10350</td><td>10368</td><td>19</td><td>ACGAGAACTTTGAAGGCCG</td><td>5882.3</td><td>-5.9</td><td>0.6</td><td>-9.5</td><td>2.7E-21</td><td>3.9E-20</td><td>MB231/MB468</td><td>NUC</td></tr>
<tr><td>>trna111_HisGTG_1_-_147774845_147774916@-1G.35.36,trna118_HisGTG_1_-_145396881_145396952@-1G.35.36,trna16_HisGTG_1_+_146544773_146544844@-1G.35.36,trna1_HisGTG_15_+_45493349_45493420@-1G.35.36,trna21_HisGTG_1_+_147753471_147753542@-1G.35.36,trna33_HisGTG_6_+_27125906_27125977@-1G.35.36,trna7_HisGTG_9_-_14433938_14434009@-1G.35.36,trna8_HisGTG_15_-_45492611_45492682@-1G.35.36,trna9_HisGTG_15_-_45490804_45490875@-1G.35.36|GGCCGTGATCGTATAGTGGTTAGTACTCTGCGTTGT</td><td>>trna111_HisGTG_1_-_147774845_147774916</td><td>-1G</td><td>35</td><td>36</td><td>GGCCGTGATCGTATAGTGGTTAGTACTCTGCGTTGT</td><td>6370.6</td><td>14.0</td><td>1.5</td><td>9.5</td><td>1.5E-21</td><td>3.9E-20</td><td>MB231/MB468</td><td>CYTO</td></tr>
<tr><td>>hg38_21_+_8433172_8446622_RNA45SN1_45S_50ntFlanks@9554.9571.18|CGTAGCGGTCCTGACGTG</td><td>>hg38_21_+_8433172_8446622_RNA45SN1_45S_50ntFlanks</td><td>9554</td><td>9571</td><td>18</td><td>CGTAGCGGTCCTGACGTG</td><td>85629.1</td><td>-9.0</td><td>0.9</td><td>-9.5</td><td>1.9E-21</td><td>3.9E-20</td><td>MB231/MB468</td><td>MP</td></tr>
<tr><td>>hg38_21_+_8433172_8446622_RNA45SN1_45S_50ntFlanks@10735.10755.21|TGTAAATCTCGCGCCGGGCCG</td><td>>hg38_21_+_8433172_8446622_RNA45SN1_45S_50ntFlanks</td><td>10735</td><td>10755</td><td>21</td><td>TGTAAATCTCGCGCCGGGCCG</td><td>1333.4</td><td>-14.2</td><td>1.5</td><td>-9.5</td><td>3.0E-21</td><td>3.9E-20</td><td>MB231/MB468</td><td>MITO</td></tr>
<tr><td>>hg38_MT_+_1621_3279_RNR2_16S_50ntFlanks@493.517.25|GCTCTTTGGACACTAGGAAAAAACC</td><td>>hg38_MT_+_1621_3279_RNR2_16S_50ntFlanks</td><td>493</td><td>517</td><td>25</td><td>GCTCTTTGGACACTAGGAAAAAACC</td><td>452.5</td><td>12.8</td><td>1.3</td><td>9.5</td><td>1.5E-21</td><td>3.9E-20</td><td>MB231/MB468</td><td>CYTO</td></tr>
<tr><td>>MI0000734|hsa-mir-106b&WithFlank&7|-|100093987|100094080@58.78.21(+1A)&[MIMAT0004672&hsa-miR-106b-3p&offsets|0|-1(+1A);m-87&7|-|100094003|100094023&offsets|0|0(+1A)]|CCGCACTGTGGGTACTTGCTGA</td><td>>MI0000734|hsa-mir-106b&WithFlank&7|-|100093987|100094080</td><td>58</td><td>78</td><td>21(+1A)</td><td>CCGCACTGTGGGTACTTGCTGA</td><td>1283.2</td><td>8.3</td><td>0.9</td><td>9.4</td><td>4.9E-21</td><td>3.9E-20</td><td>BT20/MB468</td><td>CYTO</td></tr>
<tr><td>>hg38_21_+_8433172_8446622_RNA45SN1_45S_50ntFlanks@12459.12487.29|AGCGTTGGATTGTTCACCCACTAATAGGG</td><td>>hg38_21_+_8433172_8446622_RNA45SN1_45S_50ntFlanks</td><td>12459</td><td>12487</td><td>29</td><td>AGCGTTGGATTGTTCACCCACTAATAGGG</td><td>1277.1</td><td>-13.3</td><td>1.4</td><td>-9.5</td><td>2.8E-21</td><td>4.0E-20</td><td>MB231/MB468</td><td>NUC</td></tr>
<tr><td>>hg38_21_+_8433172_8446622_RNA45SN1_45S_50ntFlanks@11564.11582.19|CCTAGCAGCCGACTTAGAA</td><td>>hg38_21_+_8433172_8446622_RNA45SN1_45S_50ntFlanks</td><td>11564</td><td>11582</td><td>19</td><td>CCTAGCAGCCGACTTAGAA</td><td>3859.5</td><td>-9.3</td><td>1.0</td><td>-9.5</td><td>3.1E-21</td><td>4.0E-20</td><td>MB231/MB468</td><td>MITO</td></tr>
<tr><td>>MI0000750|hsa-mir-26a-2&WithFlank&12|-|57824603|57824698@20.40.21(+1C);MI0000083|hsa-mir-26a-1&WithFlank&3|+|37969398|37969486@16.36.21(+1C)&[MIMAT0000082&hsa-miR-26a-5p&offsets|0|-1(+1C);m-33&12|-|57824658|57824679&offsets|0|-1(+1C)];[MIMAT0000082_1&hsa-miR-26a-5p&offsets|0|-1(+1C);m-34&3|+|37969413|37969434&offsets|0|-1(+1C)]|TTCAAGTAATCCAGGATAGGCC</td><td>>MI0000750|hsa-mir-26a-2&WithFlank&12|-|57824603|57824698</td><td>20</td><td>40</td><td>21(+1C)</td><td>TTCAAGTAATCCAGGATAGGCC</td><td>14345.5</td><td>6.4</td><td>0.7</td><td>9.4</td><td>5.0E-21</td><td>4.0E-20</td><td>BT20/MB468</td><td>CYTO</td></tr>
<tr><td>>hg38_21_+_8433172_8446622_RNA45SN1_45S_50ntFlanks@12445.12484.40|AGCAGAATTCACCAAGCGTTGGATTGTTCACCCACTAATA</td><td>>hg38_21_+_8433172_8446622_RNA45SN1_45S_50ntFlanks</td><td>12445</td><td>12484</td><td>40</td><td>AGCAGAATTCACCAAGCGTTGGATTGTTCACCCACTAATA</td><td>401.9</td><td>-13.1</td><td>1.4</td><td>-9.3</td><td>1.0E-20</td><td>4.1E-20</td><td>BT20/MB468</td><td>NUC</td></tr>
<tr><td>>hg38_21_+_8433172_8446622_RNA45SN1_45S_50ntFlanks@11318.11338.21|CCCCACGTCTCGTCGCGCGCG</td><td>>hg38_21_+_8433172_8446622_RNA45SN1_45S_50ntFlanks</td><td>11318</td><td>11338</td><td>21</td><td>CCCCACGTCTCGTCGCGCGCG</td><td>1629.6</td><td>-14.5</td><td>1.5</td><td>-9.5</td><td>3.2E-21</td><td>4.2E-20</td><td>MB231/MB468</td><td>MITO</td></tr>
<tr><td>>hg38_21_+_8433172_8446622_RNA45SN1_45S_50ntFlanks@8154.8182.29|CCGCTCGTGGGGGGCCCAAGTCCTTCTGA</td><td>>hg38_21_+_8433172_8446622_RNA45SN1_45S_50ntFlanks</td><td>8154</td><td>8182</td><td>29</td><td>CCGCTCGTGGGGGGCCCAAGTCCTTCTGA</td><td>1598.2</td><td>-14.4</td><td>1.5</td><td>-9.5</td><td>3.2E-21</td><td>4.2E-20</td><td>MB231/MB468</td><td>MITO</td></tr>
<tr><td>>hg38_21_+_8433172_8446622_RNA45SN1_45S_50ntFlanks@9166.9197.32|CCGGGCGGGTCGCGCCGTCGGGCCCGGGGGAG</td><td>>hg38_21_+_8433172_8446622_RNA45SN1_45S_50ntFlanks</td><td>9166</td><td>9197</td><td>32</td><td>CCGGGCGGGTCGCGCCGTCGGGCCCGGGGGAG</td><td>2027.8</td><td>-14.8</td><td>1.6</td><td>-9.5</td><td>2.1E-21</td><td>4.2E-20</td><td>MB231/MB468</td><td>MP</td></tr>
<tr><td>>hg38_21_+_8433172_8446622_RNA45SN1_45S_50ntFlanks@12996.13017.22|TCGCTGCGATCTATTGAAAGTC</td><td>>hg38_21_+_8433172_8446622_RNA45SN1_45S_50ntFlanks</td><td>12996</td><td>13017</td><td>22</td><td>TCGCTGCGATCTATTGAAAGTC</td><td>1839.2</td><td>7.0</td><td>0.7</td><td>9.4</td><td>5.2E-21</td><td>4.2E-20</td><td>BT20/MB468</td><td>CYTO</td></tr>
<tr><td>>hg38_21_+_8433172_8446622_RNA45SN1_45S_50ntFlanks@5487.5513.27|CGCTGAGAAGACGGTCGAACTTGACTA</td><td>>hg38_21_+_8433172_8446622_RNA45SN1_45S_50ntFlanks</td><td>5487</td><td>5513</td><td>27</td><td>CGCTGAGAAGACGGTCGAACTTGACTA</td><td>1103.4</td><td>-14.0</td><td>1.5</td><td>-9.5</td><td>2.1E-21</td><td>4.2E-20</td><td>MB231/MB468</td><td>MP</td></tr>
<tr><td>>hg38_21_+_8433172_8446622_RNA45SN1_45S_50ntFlanks@9504.9527.24|GTGAACTATGCCTGGGCAGGGCGA</td><td>>hg38_21_+_8433172_8446622_RNA45SN1_45S_50ntFlanks</td><td>9504</td><td>9527</td><td>24</td><td>GTGAACTATGCCTGGGCAGGGCGA</td><td>2290.9</td><td>-15.0</td><td>1.6</td><td>-9.5</td><td>2.1E-21</td><td>4.2E-20</td><td>MB231/MB468</td><td>MP</td></tr>
<tr><td>>hg38_21_+_8433172_8446622_RNA45SN1_45S_50ntFlanks@11562.11592.31|CGCCTAGCAGCCGACTTAGAACTGGTGCGGA</td><td>>hg38_21_+_8433172_8446622_RNA45SN1_45S_50ntFlanks</td><td>11562</td><td>11592</td><td>31</td><td>CGCCTAGCAGCCGACTTAGAACTGGTGCGGA</td><td>1460.7</td><td>-6.4</td><td>0.7</td><td>-9.5</td><td>3.2E-21</td><td>4.2E-20</td><td>MB231/MB468</td><td>MITO</td></tr>
<tr><td>>hg38_21_+_8433172_8446622_RNA45SN1_45S_50ntFlanks@11572.11600.29|CCGACTTAGAACTGGTGCGGACCAGGGGA</td><td>>hg38_21_+_8433172_8446622_RNA45SN1_45S_50ntFlanks</td><td>11572</td><td>11600</td><td>29</td><td>CCGACTTAGAACTGGTGCGGACCAGGGGA</td><td>6665.3</td><td>-5.8</td><td>0.6</td><td>-9.5</td><td>2.9E-21</td><td>4.2E-20</td><td>MB231/MB468</td><td>NUC</td></tr>
<tr><td>>hg38_21_+_8433172_8446622_RNA45SN1_45S_50ntFlanks@7984.8003.20|AGATCAGACGTGGCGACCCG</td><td>>hg38_21_+_8433172_8446622_RNA45SN1_45S_50ntFlanks</td><td>7984</td><td>8003</td><td>20</td><td>AGATCAGACGTGGCGACCCG</td><td>970.5</td><td>-13.8</td><td>1.5</td><td>-9.5</td><td>2.1E-21</td><td>4.3E-20</td><td>MB231/MB468</td><td>MP</td></tr>
<tr><td>>hg38_21_+_8433172_8446622_RNA45SN1_45S_50ntFlanks@11874.11906.33|AGAAGACCCTGTTGAGCTTGACTCTAGTCTGGC</td><td>>hg38_21_+_8433172_8446622_RNA45SN1_45S_50ntFlanks</td><td>11874</td><td>11906</td><td>33</td><td>AGAAGACCCTGTTGAGCTTGACTCTAGTCTGGC</td><td>1308.5</td><td>-7.5</td><td>0.8</td><td>-9.5</td><td>1.7E-21</td><td>4.3E-20</td><td>MB231/MB468</td><td>CYTO</td></tr>
<tr><td>>MI0000255|hsa-mir-30d&WithFlank&8|-|134804870|134804951@12.33.22(+1A)&[MIMAT0000245&hsa-miR-30d-5p&offsets|0|0(+1A);m-21&8|-|134804919|134804940&offsets|0|0(+1A)]|TGTAAACATCCCCGACTGGAAGA</td><td>>MI0000255|hsa-mir-30d&WithFlank&8|-|134804870|134804951</td><td>12</td><td>33</td><td>22(+1A)</td><td>TGTAAACATCCCCGACTGGAAGA</td><td>5223.4</td><td>10.0</td><td>1.1</td><td>9.4</td><td>5.3E-21</td><td>4.3E-20</td><td>BT20/MB468</td><td>CYTO</td></tr>
<tr><td>>hg38_21_+_8433172_8446622_RNA45SN1_45S_50ntFlanks@12244.12268.25|AAGGGCAAAAGCTCGCTTGATCTTG</td><td>>hg38_21_+_8433172_8446622_RNA45SN1_45S_50ntFlanks</td><td>12244</td><td>12268</td><td>25</td><td>AAGGGCAAAAGCTCGCTTGATCTTG</td><td>899.9</td><td>-13.7</td><td>1.4</td><td>-9.5</td><td>2.1E-21</td><td>4.3E-20</td><td>MB231/MB468</td><td>MP</td></tr>
<tr><td>>MI0000434|hsa-let-7i&WithFlank&12|+|62603680|62603775@12.33.22(+1C)&[MIMAT0000415&hsa-let-7i-5p&offsets|0|0(+1C);m-49&12|+|62603691|62603712&offsets|0|0(+1C)]|TGAGGTAGTAGTTTGTGCTGTTC</td><td>>MI0000434|hsa-let-7i&WithFlank&12|+|62603680|62603775</td><td>12</td><td>33</td><td>22(+1C)</td><td>TGAGGTAGTAGTTTGTGCTGTTC</td><td>794.2</td><td>6.6</td><td>0.7</td><td>9.5</td><td>3.0E-21</td><td>4.3E-20</td><td>MB231/MB468</td><td>NUC</td></tr>
</table>

<table>
<tr><td>>MI0000650|hsa-mir-200c&WithFlank&12|+|6963693|6963772@50.72.23(+1C)&[MIMAT0000617&hsa-miR-200c-3p&offsets|0|0(+1C);m-54&12|+|6963742|6963764&offsets|0|0(+1C)]|TAATACTGCCGGGTAATGATGGAC</td><td>>MI0000650|hsa-mir-200c&WithFlank&12|+|6963693|6963772</td><td>50</td><td>72</td><td>23(+1C)</td><td>TAATACTGCCGGGTAATGATGGAC</td><td>927.1</td><td>13.5</td><td>1.4</td><td>9.7</td><td>2.8E-22</td><td>4.4E-20</td><td>BT20/MB231</td><td>MITO</td></tr>
<tr><td>>hg38_21_+_8433172_8446622_RNA45SN1_45S_50ntFlanks@4722.4751.30|TCATTAATCAAGAACGAAAGTCGGAGGTTC</td><td>>hg38_21_+_8433172_8446622_RNA45SN1_45S_50ntFlanks</td><td>4722</td><td>4751</td><td>30</td><td>TCATTAATCAAGAACGAAAGTCGGAGGTTC</td><td>877.6</td><td>-13.6</td><td>1.4</td><td>-9.5</td><td>3.4E-21</td><td>4.4E-20</td><td>MB231/MB468</td><td>MITO</td></tr>
<tr><td>>hg38_21_+_8433172_8446622_RNA45SN1_45S_50ntFlanks@4724.4752.29|ATTAATCAAGAACGAAAGTCGGAGGTTCG</td><td>>hg38_21_+_8433172_8446622_RNA45SN1_45S_50ntFlanks</td><td>4724</td><td>4752</td><td>29</td><td>ATTAATCAAGAACGAAAGTCGGAGGTTCG</td><td>707.0</td><td>-12.4</td><td>1.3</td><td>-9.5</td><td>3.1E-21</td><td>4.4E-20</td><td>MB231/MB468</td><td>NUC</td></tr>
<tr><td>>hg38_21_+_8433172_8446622_RNA45SN1_45S_50ntFlanks@12433.12455.23|CTATCATTGTGAAGCAGAATTCA</td><td>>hg38_21_+_8433172_8446622_RNA45SN1_45S_50ntFlanks</td><td>12433</td><td>12455</td><td>23</td><td>CTATCATTGTGAAGCAGAATTCA</td><td>312.8</td><td>-12.7</td><td>1.4</td><td>-9.3</td><td>1.1E-20</td><td>4.4E-20</td><td>BT20/MB468</td><td>NUC</td></tr>
<tr><td>>hg38_21_+_8433172_8446622_RNA45SN1_45S_50ntFlanks@9572.9597.26|CAAATCGGTCGTCCGACCTGGGTATA</td><td>>hg38_21_+_8433172_8446622_RNA45SN1_45S_50ntFlanks</td><td>9572</td><td>9597</td><td>26</td><td>CAAATCGGTCGTCCGACCTGGGTATA</td><td>5294.2</td><td>-8.9</td><td>0.9</td><td>-9.5</td><td>1.8E-21</td><td>4.4E-20</td><td>MB231/MB468</td><td>CYTO</td></tr>
<tr><td>>hg38_21_+_8433172_8446622_RNA45SN1_45S_50ntFlanks@9033.9076.44|CTCGTGGGGGGGCCGGGCCACCCCTCCCACGGCGCGACCGCTCT</td><td>>hg38_21_+_8433172_8446622_RNA45SN1_45S_50ntFlanks</td><td>9033</td><td>9076</td><td>44</td><td>CTCGTGGGGGGGCCGGGCCACCCCTCCCACGGCGCGACCGCTCT</td><td>1985.9</td><td>-14.7</td><td>1.6</td><td>-9.4</td><td>3.4E-21</td><td>4.4E-20</td><td>MB231/MB468</td><td>MITO</td></tr>
<tr><td>>hg38_21_+_8433172_8446622_RNA45SN1_45S_50ntFlanks@10680.10709.30|GTTCCGGCGGCGTCCGGTGAGCTCTCGCTG</td><td>>hg38_21_+_8433172_8446622_RNA45SN1_45S_50ntFlanks</td><td>10680</td><td>10709</td><td>30</td><td>GTTCCGGCGGCGTCCGGTGAGCTCTCGCTG</td><td>3700.1</td><td>-8.3</td><td>0.9</td><td>-9.5</td><td>1.8E-21</td><td>4.4E-20</td><td>MB231/MB468</td><td>CYTO</td></tr>
<tr><td>>MI0005756|hsa-mir-934&WithFlank&X|+|136550872|136550966@21.43.23&[MIMAT0004977&hsa-miR-934&offsets|0|+1;m-943&X|+|136550892|136550913&offsets|0|+1]|TGTCTACTACTGGAGACACTGGT</td><td>>MI0005756|hsa-mir-934&WithFlank&X|+|136550872|136550966</td><td>21</td><td>43</td><td>23</td><td>TGTCTACTACTGGAGACACTGGT</td><td>1444.2</td><td>14.1</td><td>1.5</td><td>9.7</td><td>3.0E-22</td><td>4.4E-20</td><td>BT20/MB231</td><td>MITO</td></tr>
<tr><td>>hg38_21_+_8433172_8446622_RNA45SN1_45S_50ntFlanks@7985.8012.28|GATCAGACGTGGCGACCCGCTGAATTTA</td><td>>hg38_21_+_8433172_8446622_RNA45SN1_45S_50ntFlanks</td><td>7985</td><td>8012</td><td>28</td><td>GATCAGACGTGGCGACCCGCTGAATTTA</td><td>638.7</td><td>-13.2</td><td>1.4</td><td>-9.5</td><td>2.2E-21</td><td>4.5E-20</td><td>MB231/MB468</td><td>MP</td></tr>
<tr><td>>MI0000446|hsa-mir-125b-1&WithFlank&11|-|122099751|122099850@21.40.20(+1A);MI0000470|hsa-mir-125b-2&WithFlank&21|+|16590231|16590331@23.42.20(+1A)&[MIMAT0000423&hsa-miR-125b-5p&offsets|0|-2(+1A);m-67&11|-|122099809|122099830&offsets|0|-2(+1A)];[MIMAT0000423_1&hsa-miR-125b-5p&offsets|0|-2(+1A);m-68&21|+|16590253|16590274&offsets|0|-2(+1A)]|TCCCTGAGACCCTAACTTGTA</td><td>>MI0000446|hsa-mir-125b-1&WithFlank&11|-|122099751|122099850</td><td>21</td><td>40</td><td>20(+1A)</td><td>TCCCTGAGACCCTAACTTGTA</td><td>819.5</td><td>7.1</td><td>0.8</td><td>9.5</td><td>3.1E-21</td><td>4.5E-20</td><td>MB231/MB468</td><td>NUC</td></tr>
<tr><td>>hg38_21_+_8433172_8446622_RNA45SN1_45S_50ntFlanks@8170.8212.43|CAAGTCCTTCTGATCGAGGCCCAGCCCGTGGACGGTGTGAGGC</td><td>>hg38_21_+_8433172_8446622_RNA45SN1_45S_50ntFlanks</td><td>8170</td><td>8212</td><td>43</td><td>CAAGTCCTTCTGATCGAGGCCCAGCCCGTGGACGGTGTGAGGC</td><td>10748.5</td><td>-10.2</td><td>1.1</td><td>-9.4</td><td>3.5E-21</td><td>4.5E-20</td><td>MB231/MB468</td><td>MITO</td></tr>
<tr><td>>hg38_21_+_8433172_8446622_RNA45SN1_45S_50ntFlanks@12324.12344.21|TTGGGTTTTAAGCAGGAGGTG</td><td>>hg38_21_+_8433172_8446622_RNA45SN1_45S_50ntFlanks</td><td>12324</td><td>12344</td><td>21</td><td>TTGGGTTTTAAGCAGGAGGTG</td><td>1210.8</td><td>-14.0</td><td>1.5</td><td>-9.4</td><td>3.5E-21</td><td>4.5E-20</td><td>MB231/MB468</td><td>MITO</td></tr>
<tr><td>>hg38_21_+_8433172_8446622_RNA45SN1_45S_50ntFlanks@3899.3920.22|CCTTCGCGGGGGGGATGCGTGC</td><td>>hg38_21_+_8433172_8446622_RNA45SN1_45S_50ntFlanks</td><td>3899</td><td>3920</td><td>22</td><td>CCTTCGCGGGGGGGATGCGTGC</td><td>1638.5</td><td>-14.5</td><td>1.5</td><td>-9.5</td><td>2.2E-21</td><td>4.5E-20</td><td>MB231/MB468</td><td>MP</td></tr>
<tr><td>>hg38_21_+_8433172_8446622_RNA45SN1_45S_50ntFlanks@3854.3898.45|ACTGTGGTAATTCTAGAGCTAATACATGCCGACGGGCGCTGACCC</td><td>>hg38_21_+_8433172_8446622_RNA45SN1_45S_50ntFlanks</td><td>3854</td><td>3898</td><td>45</td><td>ACTGTGGTAATTCTAGAGCTAATACATGCCGACGGGCGCTGACCC</td><td>3189.7</td><td>-16.1</td><td>1.7</td><td>-9.3</td><td>1.1E-20</td><td>4.5E-20</td><td>BT20/MB468</td><td>NUC</td></tr>
<tr><td>>trna101_AlaAGC_6_-_28831462_28831533@58.75.18,trna102_AlaAGC_6_-_28806221_28806292@58.75.18,trna110_AlaTGC_6_-_28757547_28757618@58.75.18,trna113_AlaTGC_6_-_28726141_28726212@58.75.18,trna65_AlaAGC_6_+_28574933_28575004@58.75.18|TCCCCGGCACCTCCACCA</td><td>>trna101_AlaAGC_6_-_28831462_28831533</td><td>58</td><td>75</td><td>18</td><td>TCCCCGGCACCTCCACCA</td><td>1501.0</td><td>6.3</td><td>0.7</td><td>9.4</td><td>4.1E-21</td><td>4.5E-20</td><td>BT20/MB468</td><td>MITO</td></tr>
<tr><td>>hg38_21_+_8433172_8446622_RNA45SN1_45S_50ntFlanks@6645.6670.26|CTCGTACGACTCTTAGCGGTGGATCA</td><td>>hg38_21_+_8433172_8446622_RNA45SN1_45S_50ntFlanks</td><td>6645</td><td>6670</td><td>26</td><td>CTCGTACGACTCTTAGCGGTGGATCA</td><td>904.4</td><td>-3.3</td><td>0.4</td><td>-9.3</td><td>1.1E-20</td><td>4.5E-20</td><td>BT20/MB468</td><td>NUC</td></tr>
<tr><td>>hg38_21_+_8433172_8446622_RNA45SN1_45S_50ntFlanks@3830.3852.23|GCTCGCTCCTCTCCTACTTGGAT</td><td>>hg38_21_+_8433172_8446622_RNA45SN1_45S_50ntFlanks</td><td>3830</td><td>3852</td><td>23</td><td>GCTCGCTCCTCTCCTACTTGGAT</td><td>483.5</td><td>-13.3</td><td>1.4</td><td>-9.3</td><td>1.1E-20</td><td>4.5E-20</td><td>BT20/MB468</td><td>NUC</td></tr>
<tr><td>>MI0000082|hsa-mir-25&WithFlank&7|-|100093554|100093649@58.79.22(+1U)&[MIMAT0000081&hsa-miR-25-3p&offsets|0|0(+1U);m-23&7|-|100093571|100093592&offsets|0|0(+1U)]|CATTGCACTTGTCTCGGTCTGAT</td><td>>MI0000082|hsa-mir-25&WithFlank&7|-|100093554|100093649</td><td>58</td><td>79</td><td>22(+1U)</td><td>CATTGCACTTGTCTCGGTCTGAT</td><td>722.8</td><td>6.5</td><td>0.7</td><td>9.4</td><td>4.1E-21</td><td>4.6E-20</td><td>BT20/MB468</td><td>MITO</td></tr>
<tr><td>>hg38_21_+_8433172_8446622_RNA45SN1_45S_50ntFlanks@12416.12462.47|TTCGATGTCGGCTCTTCCTATCATTGTGAAGCAGAATTCACCAAGCG</td><td>>hg38_21_+_8433172_8446622_RNA45SN1_45S_50ntFlanks</td><td>12416</td><td>12462</td><td>47</td><td>TTCGATGTCGGCTCTTCCTATCATTGTGAAGCAGAATTCACCAAGCG</td><td>594.2</td><td>13.2</td><td>1.4</td><td>9.5</td><td>1.8E-21</td><td>4.6E-20</td><td>MB231/MB468</td><td>CYTO</td></tr>
<tr><td>>hg38_21_+_8433172_8446622_RNA45SN1_45S_50ntFlanks@10866.10884.19|CTAAGGGCTGGGTCGGTCG</td><td>>hg38_21_+_8433172_8446622_RNA45SN1_45S_50ntFlanks</td><td>10866</td><td>10884</td><td>19</td><td>CTAAGGGCTGGGTCGGTCG</td><td>903.8</td><td>-13.7</td><td>1.4</td><td>-9.5</td><td>2.3E-21</td><td>4.6E-20</td><td>MB231/MB468</td><td>MP</td></tr>
<tr><td>>hg38_21_+_8433172_8446622_RNA45SN1_45S_50ntFlanks@9198.9218.21|GTTCTCTCGGGGCCACGCGCG</td><td>>hg38_21_+_8433172_8446622_RNA45SN1_45S_50ntFlanks</td><td>9198</td><td>9218</td><td>21</td><td>GTTCTCTCGGGGCCACGCGCG</td><td>892.9</td><td>-13.7</td><td>1.4</td><td>-9.5</td><td>2.3E-21</td><td>4.6E-20</td><td>MB231/MB468</td><td>MP</td></tr>
<tr><td>>MI0000102|hsa-mir-100&WithFlank&11|-|122152223|122152314@19.40.22(+2A)&[MIMAT0000098&hsa-miR-100-5p&offsets|0|0(+2A);m-27&11|-|122152276|122152296&offsets|0|+1(+2A)]|AACCCGTAGATCCGAACTTGTGAA</td><td>>MI0000102|hsa-mir-100&WithFlank&11|-|122152223|122152314</td><td>19</td><td>40</td><td>22(+2A)</td><td>AACCCGTAGATCCGAACTTGTGAA</td><td>807.8</td><td>12.9</td><td>1.4</td><td>9.5</td><td>2.3E-21</td><td>4.6E-20</td><td>MB231/MB468</td><td>MP</td></tr>
<tr><td>>am_trna106_HisGTG_1_-_149155828_149155899@7.29.23,trna111_HisGTG_1_-_147774845_147774916@7.29.23,trna118_HisGTG_1_-_145396881_145396952@7.29.23,trna16_HisGTG_1_+_146544773_146544844@7.29.23,trna1_HisGTG_15_+_45493349_45493420@7.29.23,trna21_HisGTG_1_+_147753471_147753542@7.29.23,trna33_HisGTG_6_+_27125906_27125977@7.29.23,trna7_HisGTG_9_-_14433938_14434009@7.29.23,trna8_HisGTG_15_-_45492611_45492682@7.29.23,trna9_HisGTG_15_-_45490804_45490875@7.29.23|ATCGTATAGTGGTTAGTACTCTG</td><td>>am_trna106_HisGTG_1_-_149155828_149155899</td><td>7</td><td>29</td><td>23</td><td>ATCGTATAGTGGTTAGTACTCTG</td><td>743.4</td><td>-13.4</td><td>1.4</td><td>-9.5</td><td>2.3E-21</td><td>4.6E-20</td><td>MB231/MB468</td><td>MP</td></tr>
<tr><td>>hg38_21_+_8433172_8446622_RNA45SN1_45S_50ntFlanks@11873.11897.25|AAGAAGACCCTGTTGAGCTTGACTC</td><td>>hg38_21_+_8433172_8446622_RNA45SN1_45S_50ntFlanks</td><td>11873</td><td>11897</td><td>25</td><td>AAGAAGACCCTGTTGAGCTTGACTC</td><td>667.5</td><td>-13.2</td><td>1.4</td><td>-9.5</td><td>2.3E-21</td><td>4.6E-20</td><td>MB231/MB468</td><td>MP</td></tr>
<tr><td>>hg38_21_+_8433172_8446622_RNA45SN1_45S_50ntFlanks@12405.12422.18|CTTTTTGATCCTTCGATG</td><td>>hg38_21_+_8433172_8446622_RNA45SN1_45S_50ntFlanks</td><td>12405</td><td>12422</td><td>18</td><td>CTTTTTGATCCTTCGATG</td><td>783.6</td><td>-13.5</td><td>1.4</td><td>-9.5</td><td>2.3E-21</td><td>4.6E-20</td><td>MB231/MB468</td><td>MP</td></tr>
<tr><td>>hg38_21_+_8433172_8446622_RNA45SN1_45S_50ntFlanks@9504.9524.21|GTGAACTATGCCTGGGCAGGG</td><td>>hg38_21_+_8433172_8446622_RNA45SN1_45S_50ntFlanks</td><td>9504</td><td>9524</td><td>21</td><td>GTGAACTATGCCTGGGCAGGG</td><td>1148.4</td><td>-14.0</td><td>1.5</td><td>-9.5</td><td>2.3E-21</td><td>4.7E-20</td><td>MB231/MB468</td><td>MP</td></tr>
<tr><td>>hg38_21_+_8433172_8446622_RNA45SN1_45S_50ntFlanks@9305.9336.32|CCAAGGAGTCTAACACGTGCGCGAGTCGGGGG</td><td>>hg38_21_+_8433172_8446622_RNA45SN1_45S_50ntFlanks</td><td>9305</td><td>9336</td><td>32</td><td>CCAAGGAGTCTAACACGTGCGCGAGTCGGGGG</td><td>934.1</td><td>-13.7</td><td>1.4</td><td>-9.4</td><td>3.6E-21</td><td>4.7E-20</td><td>MB231/MB468</td><td>MITO</td></tr>
<tr><td>>hg38_21_+_8433172_8446622_RNA45SN1_45S_50ntFlanks@10410.10431.22|TCAGTCGGTCCTGAGAGATGGG</td><td>>hg38_21_+_8433172_8446622_RNA45SN1_45S_50ntFlanks</td><td>10410</td><td>10431</td><td>22</td><td>TCAGTCGGTCCTGAGAGATGGG</td><td>1271.4</td><td>-14.2</td><td>1.5</td><td>-9.5</td><td>2.3E-21</td><td>4.7E-20</td><td>MB231/MB468</td><td>MP</td></tr>
<tr><td>>hg38_21_+_8433172_8446622_RNA45SN1_45S_50ntFlanks@4444.4471.28|CCTTGCCTCTCGGCGCCCCCTCGATGCT</td><td>>hg38_21_+_8433172_8446622_RNA45SN1_45S_50ntFlanks</td><td>4444</td><td>4471</td><td>28</td><td>CCTTGCCTCTCGGCGCCCCCTCGATGCT</td><td>808.9</td><td>-13.4</td><td>1.4</td><td>-9.4</td><td>3.6E-21</td><td>4.7E-20</td><td>MB231/MB468</td><td>MITO</td></tr>
<tr><td>>hg38_21_+_8433172_8446622_RNA45SN1_45S_50ntFlanks@8356.8378.23|GAAAAGAACTTTGAAGAGAGAGT</td><td>>hg38_21_+_8433172_8446622_RNA45SN1_45S_50ntFlanks</td><td>8356</td><td>8378</td><td>23</td><td>GAAAAGAACTTTGAAGAGAGAGT</td><td>10274.9</td><td>-8.5</td><td>0.9</td><td>-9.5</td><td>2.4E-21</td><td>4.7E-20</td><td>MB231/MB468</td><td>MP</td></tr>
<tr><td>>hg38_21_+_8433172_8446622_RNA45SN1_45S_50ntFlanks@6721.6746.26|AATTGCAGGACACATTGATCATCGAC</td><td>>hg38_21_+_8433172_8446622_RNA45SN1_45S_50ntFlanks</td><td>6721</td><td>6746</td><td>26</td><td>AATTGCAGGACACATTGATCATCGAC</td><td>2456.5</td><td>-8.8</td><td>0.9</td><td>-9.4</td><td>3.7E-21</td><td>4.7E-20</td><td>MB231/MB468</td><td>MITO</td></tr>
<tr><td>>hg38_21_+_8433172_8446622_RNA45SN1_45S_50ntFlanks@7975.7997.23|CGCGACCTCAGATCAGACGTGGC</td><td>>hg38_21_+_8433172_8446622_RNA45SN1_45S_50ntFlanks</td><td>7975</td><td>7997</td><td>23</td><td>CGCGACCTCAGATCAGACGTGGC</td><td>255917.2</td><td>7.6</td><td>0.8</td><td>9.4</td><td>4.3E-21</td><td>4.7E-20</td><td>BT20/MB468</td><td>MITO</td></tr>
<tr><td>>hg38_21_+_8433172_8446622_RNA45SN1_45S_50ntFlanks@485.505.21|GATGTGGTGACGTCGTGCTCT</td><td>>hg38_21_+_8433172_8446622_RNA45SN1_45S_50ntFlanks</td><td>485</td><td>505</td><td>21</td><td>GATGTGGTGACGTCGTGCTCT</td><td>5102.0</td><td>16.1</td><td>1.6</td><td>9.9</td><td>4.5E-23</td><td>4.7E-20</td><td>BT20/MB468</td><td>MP</td></tr>
<tr><td>>trnaMT_PheGAA_MT_+_577_647@36.57.22|AAAATGTTTAGACGGGCTCACA</td><td>>trnaMT_PheGAA_MT_+_577_647</td><td>36</td><td>57</td><td>22</td><td>AAAATGTTTAGACGGGCTCACA</td><td>2077.3</td><td>7.7</td><td>0.8</td><td>9.4</td><td>4.3E-21</td><td>4.7E-20</td><td>BT20/MB468</td><td>MITO</td></tr>
<tr><td>>hg38_21_+_8433172_8446622_RNA45SN1_45S_50ntFlanks@9202.9220.19|TCTCGGGGCCACGCGCGCG</td><td>>hg38_21_+_8433172_8446622_RNA45SN1_45S_50ntFlanks</td><td>9202</td><td>9220</td><td>19</td><td>TCTCGGGGCCACGCGCGCG</td><td>1239.6</td><td>-14.1</td><td>1.5</td><td>-9.5</td><td>2.4E-21</td><td>4.8E-20</td><td>MB231/MB468</td><td>MP</td></tr>
<tr><td>>hg38_21_+_8433172_8446622_RNA45SN1_45S_50ntFlanks@7983.8008.26|CAGATCAGACGTGGCGACCCGCTGAA</td><td>>hg38_21_+_8433172_8446622_RNA45SN1_45S_50ntFlanks</td><td>7983</td><td>8008</td><td>26</td><td>CAGATCAGACGTGGCGACCCGCTGAA</td><td>10514.7</td><td>-7.1</td><td>0.8</td><td>-9.4</td><td>3.7E-21</td><td>4.8E-20</td><td>MB231/MB468</td><td>MITO</td></tr>
<tr><td>>hg38_21_+_8433172_8446622_RNA45SN1_45S_50ntFlanks@10463.10497.35|CTCCGTTGCCCTCGGCCGATCGAAAGGGAGTCGGG</td><td>>hg38_21_+_8433172_8446622_RNA45SN1_45S_50ntFlanks</td><td>10463</td><td>10497</td><td>35</td><td>CTCCGTTGCCCTCGGCCGATCGAAAGGGAGTCGGG</td><td>574.5</td><td>-13.0</td><td>1.4</td><td>-9.5</td><td>2.4E-21</td><td>4.8E-20</td><td>MB231/MB468</td><td>MP</td></tr>
<tr><td>>trnaMT_CysGCA_MT_-_5761_5826@47.69.23|CTTCAAACCTGCCGGGGCTTCCA</td><td>>trnaMT_CysGCA_MT_-_5761_5826</td><td>47</td><td>69</td><td>23</td><td>CTTCAAACCTGCCGGGGCTTCCA</td><td>541.0</td><td>-13.5</td><td>1.4</td><td>-9.3</td><td>1.2E-20</td><td>4.8E-20</td><td>BT20/MB468</td><td>NUC</td></tr>
<tr><td>>MI0000446|hsa-mir-125b-1&WithFlank&11|-|122099751|122099850@21.42.22;MI0000470|hsa-mir-125b-2&WithFlank&21|+|16590231|16590331@23.44.22&[MIMAT0000423&hsa-miR-125b-5p&offsets|0|0;m-67&11|-|122099809|122099830&offsets|0|0];[MIMAT0000423_1&hsa-miR-125b-5p&offsets|0|0;m-68&21|+|16590253|16590274&offsets|0|0]|TCCCTGAGACCCTAACTTGTGA</td><td>>MI0000446|hsa-mir-125b-1&WithFlank&11|-|122099751|122099850</td><td>21</td><td>42</td><td>22</td><td>TCCCTGAGACCCTAACTTGTGA</td><td>16266.2</td><td>9.0</td><td>0.9</td><td>9.5</td><td>2.0E-21</td><td>4.9E-20</td><td>MB231/MB468</td><td>CYTO</td></tr>
<tr><td>>MI0000285|hsa-mir-205&WithFlank&1|+|209432127|209432248@40.61.22(+1G)&[MIMAT0000266&hsa-miR-205-5p&offsets|0|0(+1G);m-351&1|+|209432166|209432187&offsets|0|0(+1G)]|TCCTTCATTCCACCGGAGTCTGG</td><td>>MI0000285|hsa-mir-205&WithFlank&1|+|209432127|209432248</td><td>40</td><td>61</td><td>22(+1G)</td><td>TCCTTCATTCCACCGGAGTCTGG</td><td>2073.7</td><td>14.8</td><td>1.5</td><td>9.7</td><td>4.1E-22</td><td>4.9E-20</td><td>BT20/MB231</td><td>NUC</td></tr>
<tr><td>>MI0000067|hsa-let-7f-1&WithFlank&9|+|94176341|94176439@13.30.18;MI0000068|hsa-let-7f-2&WithFlank&X|-|53557186|53557280@14.31.18&[MIMAT0000067&hsa-let-7f-5p&offsets|0|-4;m-20&9|+|94176353|94176374&offsets|0|-4];[MIMAT0000067_1&hsa-let-7f-5p&offsets|0|-4;m-19&X|-|53557246|53557267&offsets|0|-4]|TGAGGTAGTAGATTGTAT</td><td>>MI0000067|hsa-let-7f-1&WithFlank&9|+|94176341|94176439</td><td>13</td><td>30</td><td>18</td><td>TGAGGTAGTAGATTGTAT</td><td>9623.8</td><td>7.7</td><td>0.8</td><td>9.4</td><td>6.1E-21</td><td>4.9E-20</td><td>BT20/MB468</td><td>CYTO</td></tr>
<tr><td>>hg38_21_+_8433172_8446622_RNA45SN1_45S_50ntFlanks@12954.12973.20|CTGCTTCTGGGTCGGGGTTT</td><td>>hg38_21_+_8433172_8446622_RNA45SN1_45S_50ntFlanks</td><td>12954</td><td>12973</td><td>20</td><td>CTGCTTCTGGGTCGGGGTTT</td><td>463.2</td><td>-13.3</td><td>1.4</td><td>-9.3</td><td>1.2E-20</td><td>4.9E-20</td><td>BT20/MB468</td><td>NUC</td></tr>
<tr><td>>hg38_21_+_8433172_8446622_RNA45SN1_45S_50ntFlanks@9712.9737.26|AGCGAATGATTAGAGGTCTTGGGGCC</td><td>>hg38_21_+_8433172_8446622_RNA45SN1_45S_50ntFlanks</td><td>9712</td><td>9737</td><td>26</td><td>AGCGAATGATTAGAGGTCTTGGGGCC</td><td>1836.4</td><td>-14.6</td><td>1.5</td><td>-9.4</td><td>3.8E-21</td><td>4.9E-20</td><td>MB231/MB468</td><td>MITO</td></tr>
</table>

<table>
<tr><td></td><td></td><td></td><td></td><td></td><td></td><td></td><td></td><td></td><td></td><td></td><td></td><td></td><td></td></tr>
<tr><td>>hg38_21_+_8433172_8446622_RNA45SN1_45S_50ntFlanks@9532.9576.45|AGAGGAAACTCTGGTGGAGGTCCGTAGCGGTCCTGACGTGCAAAT</td><td>>hg38_21_+_8433172_8446622_RNA45SN1_45S_50ntFlanks</td><td>9532</td><td>9576</td><td>45</td><td>AGAGGAAACTCTGGTGGAGGTCCGTAGCGGTCCTGACGTGCAAAT</td><td>160.0</td><td>11.3</td><td>1.2</td><td>9.5</td><td>2.0E-21</td><td>4.9E-20</td><td>MB231/MB468</td><td>CYTO</td></tr>
<tr><td>>hg38_1_-_228634819_228635039_RNA5S12_5S_50ntFlanks@149.171.23|GAATACCGGGTGCTGTAGGCTTT</td><td>>hg38_1_-_228634819_228635039_RNA5S12_5S_50ntFlanks</td><td>149</td><td>171</td><td>23</td><td>GAATACCGGGTGCTGTAGGCTTT</td><td>4173.0</td><td>5.7</td><td>0.6</td><td>9.4</td><td>6.2E-21</td><td>4.9E-20</td><td>BT20/MB468</td><td>CYTO</td></tr>
<tr><td>>hg38_21_+_8433172_8446622_RNA45SN1_45S_50ntFlanks@9203.9225.23|CTCGGGGCCACGCGCGCGTCCCC</td><td>>hg38_21_+_8433172_8446622_RNA45SN1_45S_50ntFlanks</td><td>9203</td><td>9225</td><td>23</td><td>CTCGGGGCCACGCGCGCGTCCCC</td><td>878.8</td><td>-13.6</td><td>1.4</td><td>-9.5</td><td>2.5E-21</td><td>5.0E-20</td><td>MB231/MB468</td><td>MP</td></tr>
<tr><td>>hg38_21_+_8433172_8446622_RNA45SN1_45S_50ntFlanks@11786.11803.18|GAATGGATGAACGAGATT</td><td>>hg38_21_+_8433172_8446622_RNA45SN1_45S_50ntFlanks</td><td>11786</td><td>11803</td><td>18</td><td>GAATGGATGAACGAGATT</td><td>1340.3</td><td>-5.8</td><td>0.6</td><td>-9.5</td><td>2.5E-21</td><td>5.0E-20</td><td>MB231/MB468</td><td>MP</td></tr>
<tr><td>>MI0003776|hsa-mir-320b-1&WithFlank&1|+|116671740|116671823@49.68.20;MI0003839|hsa-mir-320b-2&WithFlank&1|-|224257034|224257116@48.67.20&[MIMAT0005792&hsa-miR-320b&offsets|+1|-1;m-183&1|+|116671787|116671806&offsets|+1|+1];[MIMAT0005792_1&hsa-miR-320b&offsets|+1|-1;m-183&1|-|224257051|224257070&offsets|+1|+1]|AAAGCTGGGTTGAGAGGGCA</td><td>>MI0003776|hsa-mir-320b-1&WithFlank&1|+|116671740|116671823</td><td>49</td><td>68</td><td>20</td><td>AAAGCTGGGTTGAGAGGGCA</td><td>403.7</td><td>12.9</td><td>1.4</td><td>9.4</td><td>6.2E-21</td><td>5.0E-20</td><td>BT20/MB468</td><td>CYTO</td></tr>
<tr><td>>hg38_21_+_8433172_8446622_RNA45SN1_45S_50ntFlanks@12539.12563.25|ATGATGTGTTGTTGCCATGGTAATC</td><td>>hg38_21_+_8433172_8446622_RNA45SN1_45S_50ntFlanks</td><td>12539</td><td>12563</td><td>25</td><td>ATGATGTGTTGTTGCCATGGTAATC</td><td>436.5</td><td>-6.8</td><td>0.7</td><td>-9.5</td><td>2.0E-21</td><td>5.0E-20</td><td>MB231/MB468</td><td>CYTO</td></tr>
<tr><td>>hg38_21_+_8433172_8446622_RNA45SN1_45S_50ntFlanks@9574.9595.22|AATCGGTCGTCCGACCTGGGTA</td><td>>hg38_21_+_8433172_8446622_RNA45SN1_45S_50ntFlanks</td><td>9574</td><td>9595</td><td>22</td><td>AATCGGTCGTCCGACCTGGGTA</td><td>11364.2</td><td>-9.0</td><td>1.0</td><td>-9.5</td><td>2.5E-21</td><td>5.0E-20</td><td>MB231/MB468</td><td>MP</td></tr>
<tr><td>>hg38_21_+_8433172_8446622_RNA45SN1_45S_50ntFlanks@5421.5443.23|CGATTGGATGGTTTAGTGAGGCC</td><td>>hg38_21_+_8433172_8446622_RNA45SN1_45S_50ntFlanks</td><td>5421</td><td>5443</td><td>23</td><td>CGATTGGATGGTTTAGTGAGGCC</td><td>2127.0</td><td>-14.9</td><td>1.6</td><td>-9.5</td><td>2.6E-21</td><td>5.0E-20</td><td>MB231/MB468</td><td>MP</td></tr>
<tr><td>>hg38_21_+_8433172_8446622_RNA45SN1_45S_50ntFlanks@11323.11344.22|CGTCTCGTCGCGCGCGCGTCCG</td><td>>hg38_21_+_8433172_8446622_RNA45SN1_45S_50ntFlanks</td><td>11323</td><td>11344</td><td>22</td><td>CGTCTCGTCGCGCGCGCGTCCG</td><td>19318.4</td><td>-10.5</td><td>1.1</td><td>-9.4</td><td>3.6E-21</td><td>5.1E-20</td><td>MB231/MB468</td><td>NUC</td></tr>
<tr><td>>hg38_21_+_8433172_8446622_RNA45SN1_45S_50ntFlanks@12438.12455.18|ATTGTGAAGCAGAATTCA</td><td>>hg38_21_+_8433172_8446622_RNA45SN1_45S_50ntFlanks</td><td>12438</td><td>12455</td><td>18</td><td>ATTGTGAAGCAGAATTCA</td><td>7588.0</td><td>-10.6</td><td>1.1</td><td>-9.5</td><td>2.6E-21</td><td>5.1E-20</td><td>MB231/MB468</td><td>MP</td></tr>
<tr><td>>hg38_21_+_8433172_8446622_RNA45SN1_45S_50ntFlanks@6670.6696.27|ACTCGGCTCGTGCGTCGATGAAGAACG</td><td>>hg38_21_+_8433172_8446622_RNA45SN1_45S_50ntFlanks</td><td>6670</td><td>6696</td><td>27</td><td>ACTCGGCTCGTGCGTCGATGAAGAACG</td><td>868.4</td><td>-5.4</td><td>0.6</td><td>-9.5</td><td>2.6E-21</td><td>5.2E-20</td><td>MB231/MB468</td><td>MP</td></tr>
<tr><td>>am_trnaMT_GluTTC_MT_-_14674_14742@27.48.22,trnalookalike8_GluTTC_5_-_93905172_93905240@27.48.22|GTTTTTCATATCATTGGTCGTG</td><td>>am_trnaMT_GluTTC_MT_-_14674_14742</td><td>27</td><td>48</td><td>22</td><td>GTTTTTCATATCATTGGTCGTG</td><td>338.3</td><td>-12.8</td><td>1.4</td><td>-9.3</td><td>1.3E-20</td><td>5.2E-20</td><td>BT20/MB468</td><td>NUC</td></tr>
<tr><td>>hg38_21_+_8433172_8446622_RNA45SN1_45S_50ntFlanks@4047.4081.35|ATTCGAACGTCTGCCCTATCAACTTTCGATGGTAG</td><td>>hg38_21_+_8433172_8446622_RNA45SN1_45S_50ntFlanks</td><td>4047</td><td>4081</td><td>35</td><td>ATTCGAACGTCTGCCCTATCAACTTTCGATGGTAG</td><td>1294.0</td><td>-14.1</td><td>1.5</td><td>-9.4</td><td>4.1E-21</td><td>5.2E-20</td><td>MB231/MB468</td><td>MITO</td></tr>
<tr><td>>hg38_21_+_8433172_8446622_RNA45SN1_45S_50ntFlanks@11270.11295.26|CGGCCGCCCCCGGGGAGCCCGGCGGG</td><td>>hg38_21_+_8433172_8446622_RNA45SN1_45S_50ntFlanks</td><td>11270</td><td>11295</td><td>26</td><td>CGGCCGCCCCCGGGGAGCCCGGCGGG</td><td>3130.0</td><td>-14.6</td><td>1.5</td><td>-9.4</td><td>3.7E-21</td><td>5.2E-20</td><td>MB231/MB468</td><td>NUC</td></tr>
<tr><td>>hg38_21_+_8433172_8446622_RNA45SN1_45S_50ntFlanks@5470.5514.45|CCACGGCCCTGGCGGAGCGCTGAGAAGACGGTCGAACTTGACTAT</td><td>>hg38_21_+_8433172_8446622_RNA45SN1_45S_50ntFlanks</td><td>5470</td><td>5514</td><td>45</td><td>CCACGGCCCTGGCGGAGCGCTGAGAAGACGGTCGAACTTGACTAT</td><td>1157.1</td><td>-14.6</td><td>1.6</td><td>-9.3</td><td>1.3E-20</td><td>5.2E-20</td><td>BT20/MB468</td><td>NUC</td></tr>
<tr><td>>hg38_21_+_8433172_8446622_RNA45SN1_45S_50ntFlanks@6771.6799.29|GGTTCCTCCCGGGGCTACGCCTGTCTGAG</td><td>>hg38_21_+_8433172_8446622_RNA45SN1_45S_50ntFlanks</td><td>6771</td><td>6799</td><td>29</td><td>GGTTCCTCCCGGGGCTACGCCTGTCTGAG</td><td>990.5</td><td>-13.7</td><td>1.5</td><td>-9.4</td><td>4.1E-21</td><td>5.3E-20</td><td>MB231/MB468</td><td>MITO</td></tr>
<tr><td>>hg38_21_+_8433172_8446622_RNA45SN1_45S_50ntFlanks@10794.10826.33|CTGGCATGTTGGAACAATGTAGGTAAGGGAAGT</td><td>>hg38_21_+_8433172_8446622_RNA45SN1_45S_50ntFlanks</td><td>10794</td><td>10826</td><td>33</td><td>CTGGCATGTTGGAACAATGTAGGTAAGGGAAGT</td><td>759.8</td><td>-13.4</td><td>1.4</td><td>-9.5</td><td>2.7E-21</td><td>5.3E-20</td><td>MB231/MB468</td><td>MP</td></tr>
<tr><td>>hg38_21_+_8433172_8446622_RNA45SN1_45S_50ntFlanks@10389.10426.38|TGAACAGCAGTTGAACATGGGTCAGTCGGTCCTGAGAG</td><td>>hg38_21_+_8433172_8446622_RNA45SN1_45S_50ntFlanks</td><td>10389</td><td>10426</td><td>38</td><td>TGAACAGCAGTTGAACATGGGTCAGTCGGTCCTGAGAG</td><td>584.3</td><td>-13.6</td><td>1.5</td><td>-9.3</td><td>1.3E-20</td><td>5.3E-20</td><td>BT20/MB468</td><td>NUC</td></tr>
<tr><td>>MI0000067|hsa-let-7f-1&WithFlank&9|+|94176341|94176439@13.32.20(+1C);MI0000068|hsa-let-7f-2&WithFlank&X|-|53557186|53557280@14.33.20(+1C)&[MIMAT0000067&hsa-let-7f-5p&offsets|0|-2(+1C);m-20&9|+|94176353|94176374&offsets|0|-2(+1C)];[MIMAT0000067_1&hsa-let-7f-5p&offsets|0|-2(+1C);m-19&X|-|53557246|53557267&offsets|0|-2(+1C)]|TGAGGTAGTAGATTGTATAGC</td><td>>MI0000067|hsa-let-7f-1&WithFlank&9|+|94176341|94176439</td><td>13</td><td>32</td><td>20(+1C)</td><td>TGAGGTAGTAGATTGTATAGC</td><td>1220.6</td><td>4.9</td><td>0.5</td><td>9.4</td><td>3.7E-21</td><td>5.3E-20</td><td>MB231/MB468</td><td>NUC</td></tr>
<tr><td>>hg38_21_+_8433172_8446622_RNA45SN1_45S_50ntFlanks@6745.6770.26|ACACTTCGAACGCACTTGCGGCCCCG</td><td>>hg38_21_+_8433172_8446622_RNA45SN1_45S_50ntFlanks</td><td>6745</td><td>6770</td><td>26</td><td>ACACTTCGAACGCACTTGCGGCCCCG</td><td>11637.9</td><td>-12.3</td><td>1.3</td><td>-9.5</td><td>2.7E-21</td><td>5.3E-20</td><td>MB231/MB468</td><td>MP</td></tr>
<tr><td>>hg38_21_+_8433172_8446622_RNA45SN1_45S_50ntFlanks@4657.4688.32|CTAGAGGTGAAATTCTTGGACCGGCGCAAGAC</td><td>>hg38_21_+_8433172_8446622_RNA45SN1_45S_50ntFlanks</td><td>4657</td><td>4688</td><td>32</td><td>CTAGAGGTGAAATTCTTGGACCGGCGCAAGAC</td><td>2546.2</td><td>-15.1</td><td>1.6</td><td>-9.4</td><td>4.2E-21</td><td>5.3E-20</td><td>MB231/MB468</td><td>MITO</td></tr>
<tr><td>>trna111_HisGTG_1_-_147774845_147774916@1.34.34,trna118_HisGTG_1_-_145396881_145396952@1.34.34,trna16_HisGTG_1_+_146544773_146544844@1.34.34,trna1_HisGTG_15_+_45493349_45493420@1.34.34,trna21_HisGTG_1_+_147753471_147753542@1.34.34,trna33_HisGTG_6_+_27125906_27125977@1.34.34,trna7_HisGTG_9_-_14433938_14434009@1.34.34,trna8_HisGTG_15_-_45492611_45492682@1.34.34,trna9_HisGTG_15_-_45490804_45490875@1.34.34|GCCGTGATCGTATAGTGGTTAGTACTCTGCGTTG</td><td>>trna111_HisGTG_1_-_147774845_147774916</td><td>1</td><td>34</td><td>34</td><td>GCCGTGATCGTATAGTGGTTAGTACTCTGCGTTG</td><td>11017.7</td><td>8.9</td><td>0.9</td><td>9.5</td><td>2.2E-21</td><td>5.4E-20</td><td>MB231/MB468</td><td>CYTO</td></tr>
<tr><td>>MI0000086|hsa-mir-28&WithFlank&3|+|188688775|188688872@60.81.22(+1A)&[MIMAT0004502&hsa-miR-28-3p&offsets|0|0(+1A);m-28&3|+|188688834|188688855&offsets|0|0(+1A)]|CACTAGATTGTGAGCTCCTGGAA</td><td>>MI0000086|hsa-mir-28&WithFlank&3|+|188688775|188688872</td><td>60</td><td>81</td><td>22(+1A)</td><td>CACTAGATTGTGAGCTCCTGGAA</td><td>1182.0</td><td>10.3</td><td>1.1</td><td>9.4</td><td>4.9E-21</td><td>5.4E-20</td><td>BT20/MB468</td><td>MITO</td></tr>
<tr><td>>hg38_21_+_8433172_8446622_RNA45SN1_45S_50ntFlanks@5472.5494.23|ACGGCCCTGGCGGAGCGCTGAGA</td><td>>hg38_21_+_8433172_8446622_RNA45SN1_45S_50ntFlanks</td><td>5472</td><td>5494</td><td>23</td><td>ACGGCCCTGGCGGAGCGCTGAGA</td><td>3047.3</td><td>7.5</td><td>0.8</td><td>9.5</td><td>2.2E-21</td><td>5.4E-20</td><td>MB231/MB468</td><td>CYTO</td></tr>
<tr><td>>MI0003776|hsa-mir-320b-1&WithFlank&1|+|116671740|116671823@48.69.22;MI0003839|hsa-mir-320b-2&WithFlank&1|-|224257034|224257116@47.68.22&[MIMAT0005792&hsa-miR-320b&offsets|0|0;m-183&1|+|116671787|116671806&offsets|0|+2];[MIMAT0005792_1&hsa-miR-320b&offsets|0|0;m-183&1|-|224257051|224257070&offsets|0|+2]|AAAAGCTGGGTTGAGAGGGCAA</td><td>>MI0003776|hsa-mir-320b-1&WithFlank&1|+|116671740|116671823</td><td>48</td><td>69</td><td>22</td><td>AAAAGCTGGGTTGAGAGGGCAA</td><td>277.2</td><td>12.4</td><td>1.3</td><td>9.4</td><td>6.8E-21</td><td>5.4E-20</td><td>BT20/MB468</td><td>CYTO</td></tr>
<tr><td>>hg38_21_+_8433172_8446622_RNA45SN1_45S_50ntFlanks@11729.11765.37|TAACTATGACTCTCTTAAGGTAGCCAAATGCCTCGTC</td><td>>hg38_21_+_8433172_8446622_RNA45SN1_45S_50ntFlanks</td><td>11729</td><td>11765</td><td>37</td><td>TAACTATGACTCTCTTAAGGTAGCCAAATGCCTCGTC</td><td>8403.0</td><td>-7.3</td><td>0.8</td><td>-9.4</td><td>3.9E-21</td><td>5.4E-20</td><td>MB231/MB468</td><td>NUC</td></tr>
<tr><td>>hg38_21_+_8433172_8446622_RNA45SN1_45S_50ntFlanks@8110.8140.31|CGGGGCGCGGGACATGTGGCGTACGGAAGAC</td><td>>hg38_21_+_8433172_8446622_RNA45SN1_45S_50ntFlanks</td><td>8110</td><td>8140</td><td>31</td><td>CGGGGCGCGGGACATGTGGCGTACGGAAGAC</td><td>646.0</td><td>-13.2</td><td>1.4</td><td>-9.5</td><td>2.8E-21</td><td>5.4E-20</td><td>MB231/MB468</td><td>MP</td></tr>
<tr><td>>hg38_21_+_8433172_8446622_RNA45SN1_45S_50ntFlanks@11933.11954.22|AATAAGTGGGAGGCCCCCGGCG</td><td>>hg38_21_+_8433172_8446622_RNA45SN1_45S_50ntFlanks</td><td>11933</td><td>11954</td><td>22</td><td>AATAAGTGGGAGGCCCCCGGCG</td><td>915.9</td><td>-13.7</td><td>1.4</td><td>-9.5</td><td>2.8E-21</td><td>5.5E-20</td><td>MB231/MB468</td><td>MP</td></tr>
<tr><td>>hg38_21_+_8433172_8446622_RNA45SN1_45S_50ntFlanks@12539.12573.35|ATGATGTGTTGTTGCCATGGTAATCCTGCTCAGTA</td><td>>hg38_21_+_8433172_8446622_RNA45SN1_45S_50ntFlanks</td><td>12539</td><td>12573</td><td>35</td><td>ATGATGTGTTGTTGCCATGGTAATCCTGCTCAGTA</td><td>887.9</td><td>-13.7</td><td>1.4</td><td>-9.5</td><td>2.8E-21</td><td>5.5E-20</td><td>MB231/MB468</td><td>MP</td></tr>
<tr><td>>hg38_21_+_8433172_8446622_RNA45SN1_45S_50ntFlanks@4824.4859.36|TGACCCGCCGGGCAGCTTCCGGGAAACCAAAGTCTT</td><td>>hg38_21_+_8433172_8446622_RNA45SN1_45S_50ntFlanks</td><td>4824</td><td>4859</td><td>36</td><td>TGACCCGCCGGGCAGCTTCCGGGAAACCAAAGTCTT</td><td>359.9</td><td>12.7</td><td>1.4</td><td>9.4</td><td>6.9E-21</td><td>5.5E-20</td><td>BT20/MB468</td><td>CYTO</td></tr>
<tr><td>>hg38_21_+_8433172_8446622_RNA45SN1_45S_50ntFlanks@3776.3805.30|CCGGTACAGTGAAACTGCGAATGGCTCATT</td><td>>hg38_21_+_8433172_8446622_RNA45SN1_45S_50ntFlanks</td><td>3776</td><td>3805</td><td>30</td><td>CCGGTACAGTGAAACTGCGAATGGCTCATT</td><td>1376.4</td><td>-6.0</td><td>0.6</td><td>-9.3</td><td>1.4E-20</td><td>5.5E-20</td><td>BT20/MB468</td><td>NUC</td></tr>
<tr><td>>hg38_21_+_8433172_8446622_RNA45SN1_45S_50ntFlanks@11212.11232.21|CGACTCTGGACGCGAGCCGGG</td><td>>hg38_21_+_8433172_8446622_RNA45SN1_45S_50ntFlanks</td><td>11212</td><td>11232</td><td>21</td><td>CGACTCTGGACGCGAGCCGGG</td><td>1029.6</td><td>-13.8</td><td>1.5</td><td>-9.4</td><td>4.4E-21</td><td>5.6E-20</td><td>MB231/MB468</td><td>MITO</td></tr>
<tr><td>>hg38_21_+_8433172_8446622_RNA45SN1_45S_50ntFlanks@4588.4614.27|CTATTTTGTTGGTTTTCGGAACTGAGG</td><td>>hg38_21_+_8433172_8446622_RNA45SN1_45S_50ntFlanks</td><td>4588</td><td>4614</td><td>27</td><td>CTATTTTGTTGGTTTTCGGAACTGAGG</td><td>989.7</td><td>-6.9</td><td>0.7</td><td>-9.5</td><td>2.9E-21</td><td>5.6E-20</td><td>MB231/MB468</td><td>MP</td></tr>
<tr><td>>MI0000082|hsa-mir-25&WithFlank&7|-|100093554|100093649@58.78.21(+1G)&[MIMAT0000081&hsa-miR-25-3p&offsets|0|-1(+1G);m-23&7|-|100093571|100093592&offsets|0|-1(+1G)]|CATTGCACTTGTCTCGGTCTGG</td><td>>MI0000082|hsa-mir-25&WithFlank&7|-|100093554|100093649</td><td>58</td><td>78</td><td>21(+1G)</td><td>CATTGCACTTGTCTCGGTCTGG</td><td>708.2</td><td>6.8</td><td>0.7</td><td>9.4</td><td>5.1E-21</td><td>5.6E-20</td><td>BT20/MB468</td><td>MITO</td></tr>
<tr><td>>MI0001445|hsa-mir-423&WithFlank&17|+|30117073|30117178@58.81.24(+1U)&[MIMAT0001340&hsa-miR-423-3p&offsets|-1|0(+1U);m-100&17|+|30117131|30117153&offsets|-1|0(+1U)]|AAGCTCGGTCTGAGGCCCCTCAGTT</td><td>>MI0001445|hsa-mir-423&WithFlank&17|+|30117073|30117178</td><td>58</td><td>81</td><td>24(+1U)</td><td>AAGCTCGGTCTGAGGCCCCTCAGTT</td><td>659.1</td><td>9.0</td><td>1.0</td><td>9.4</td><td>5.2E-21</td><td>5.7E-20</td><td>BT20/MB468</td><td>MITO</td></tr>
<tr><td>>hg38_21_+_8433172_8446622_RNA45SN1_45S_50ntFlanks@7988.8011.24|CAGACGTGGCGACCCGCTGAATTT</td><td>>hg38_21_+_8433172_8446622_RNA45SN1_45S_50ntFlanks</td><td>7988</td><td>8011</td><td>24</td><td>CAGACGTGGCGACCCGCTGAATTT</td><td>1880.9</td><td>-5.7</td><td>0.6</td><td>-9.5</td><td>2.9E-21</td><td>5.7E-20</td><td>MB231/MB468</td><td>MP</td></tr>
<tr><td>>hg38_21_+_8433172_8446622_RNA45SN1_45S_50ntFlanks@3838.3868.31|CTCTCCTACTTGGATAACTGTGGTAATTCTA</td><td>>hg38_21_+_8433172_8446622_RNA45SN1_45S_50ntFlanks</td><td>3838</td><td>3868</td><td>31</td><td>CTCTCCTACTTGGATAACTGTGGTAATTCTA</td><td>1120.4</td><td>-6.2</td><td>0.7</td><td>-9.3</td><td>1.4E-20</td><td>5.7E-20</td><td>BT20/MB468</td><td>NUC</td></tr>
<tr><td>>hg38_21_+_8433172_8446622_RNA45SN1_45S_50ntFlanks@8339.8375.37|TACCGTAAGGGAAAGTTGAAAAGAACTTTGAAGAGAG</td><td>>hg38_21_+_8433172_8446622_RNA45SN1_45S_50ntFlanks</td><td>8339</td><td>8375</td><td>37</td><td>TACCGTAAGGGAAAGTTGAAAAGAACTTTGAAGAGAG</td><td>540.4</td><td>13.5</td><td>1.4</td><td>9.4</td><td>5.2E-21</td><td>5.7E-20</td><td>BT20/MB468</td><td>MITO</td></tr>
<tr><td>>hg38_21_+_8433172_8446622_RNA45SN1_45S_50ntFlanks@12417.12436.20|TCGATGTCGGCTCTTCCTAT</td><td>>hg38_21_+_8433172_8446622_RNA45SN1_45S_50ntFlanks</td><td>12417</td><td>12436</td><td>20</td><td>TCGATGTCGGCTCTTCCTAT</td><td>4309.9</td><td>-5.5</td><td>0.6</td><td>-9.4</td><td>5.2E-21</td><td>5.7E-20</td><td>BT20/MB468</td><td>MITO</td></tr>
<tr><td>>hg38_21_+_8433172_8446622_RNA45SN1_45S_50ntFlanks@11586.11611.26|GTGCGGACCAGGGGAATCCGACTGTT</td><td>>hg38_21_+_8433172_8446622_RNA45SN1_45S_50ntFlanks</td><td>11586</td><td>11611</td><td>26</td><td>GTGCGGACCAGGGGAATCCGACTGTT</td><td>3340.3</td><td>-5.4</td><td>0.6</td><td>-9.4</td><td>4.5E-21</td><td>5.8E-20</td><td>MB231/MB468</td><td>MITO</td></tr>
<tr><td>>hg38_21_+_8433172_8446622_RNA45SN1_45S_50ntFlanks@12418.12436.19|CGATGTCGGCTCTTCCTAT</td><td>>hg38_21_+_8433172_8446622_RNA45SN1_45S_50ntFlanks</td><td>12418</td><td>12436</td><td>19</td><td>CGATGTCGGCTCTTCCTAT</td><td>2806.5</td><td>-9.2</td><td>1.0</td><td>-9.5</td><td>3.0E-21</td><td>5.8E-20</td><td>MB231/MB468</td><td>MP</td></tr>
<tr><td>>hg38_21_+_8433172_8446622_RNA45SN1_45S_50ntFlanks@11805.11827.23|CCACTGTCCCTACCTACTATCCA</td><td>>hg38_21_+_8433172_8446622_RNA45SN1_45S_50ntFlanks</td><td>11805</td><td>11827</td><td>23</td><td>CCACTGTCCCTACCTACTATCCA</td><td>339.9</td><td>-12.8</td><td>1.4</td><td>-9.3</td><td>1.4E-20</td><td>5.8E-20</td><td>BT20/MB468</td><td>NUC</td></tr>
<tr><td>>hg38_21_+_8433172_8446622_RNA45SN1_45S_50ntFlanks@8380.8399.20|CAAGAGGGCGTGAAACCGTT</td><td>>hg38_21_+_8433172_8446622_RNA45SN1_45S_50ntFlanks</td><td>8380</td><td>8399</td><td>20</td><td>CAAGAGGGCGTGAAACCGTT</td><td>607.9</td><td>-7.4</td><td>0.8</td><td>-9.5</td><td>2.4E-21</td><td>5.8E-20</td><td>MB231/MB468</td><td>CYTO</td></tr>
<tr><td>>MI0000462|hsa-mir-152&WithFlank&17|-|48037155|48037253@60.80.21&[MIMAT0000438&hsa-miR-152-3p&offsets|0|0;m-97&17|-|48037174|48037194&offsets|0|0]|TCAGTGCATGACAGAACTTGG</td><td>>MI0000462|hsa-mir-152&WithFlank&17|-|48037155|48037253</td><td>60</td><td>80</td><td>21</td><td>TCAGTGCATGACAGAACTTGG</td><td>342.4</td><td>12.9</td><td>1.4</td><td>9.4</td><td>5.3E-21</td><td>5.8E-20</td><td>BT20/MB468</td><td>MITO</td></tr>
<tr><td>>hg38_1_-_228634819_228635039_RNA5S12_5S_50ntFlanks@135.170.36|GGGAGACCGCCTGGGAATACCGGGTGCTGTAGGCTT</td><td>>hg38_1_-_228634819_228635039_RNA5S12_5S_50ntFlanks</td><td>135</td><td>170</td><td>36</td><td>GGGAGACCGCCTGGGAATACCGGGTGCTGTAGGCTT</td><td>23728.4</td><td>5.8</td><td>0.6</td><td>9.4</td><td>7.4E-21</td><td>5.9E-20</td><td>BT20/MB468</td><td>CYTO</td></tr>
<tr><td>>hg38_21_+_8433172_8446622_RNA45SN1_45S_50ntFlanks@6745.6772.28|ACACTTCGAACGCACTTGCGGCCCCGGG</td><td>>hg38_21_+_8433172_8446622_RNA45SN1_45S_50ntFlanks</td><td>6745</td><td>6772</td><td>28</td><td>ACACTTCGAACGCACTTGCGGCCCCGGG</td><td>1379.1</td><td>-14.3</td><td>1.5</td><td>-9.5</td><td>3.1E-21</td><td>6.0E-20</td><td>MB231/MB468</td><td>MP</td></tr>
</table>

| | | | | | | | | | | | | | |
|---|---|---|---|---|---|---|---|---|---|---|---|---|---|
| >hg38_21_+_8433172_8446622_RNA45SN1_45S_50ntFlanks@11579.11603.25\|AGAACTGGTGCGGACCAGGGGAATC | >hg38_21_+_8433172_8446622_RNA45SN1_45S_50ntFlanks | 11579 | 11603 | 25 | AGAACTGGTGCGGACCAGGGGAATC | 2446.1 | -6.0 | 0.6 | -9.4 | 4.3E-21 | 6.0E-20 | MB231/MB468 | NUC |
| >MI0000264\|hsa-mir-7-2&WithFlank&15\|+\|88611819\|88611940@38.60.23;MI0000265\|hsa-mir-7-3&WithFlank&19\|+\|4770664\|4770785@37.59.23;MI0000263\|hsa-mir-7-1&WithFlank&9\|-\|83969742\|83969863@30.52.23&[MIMAT0000252&hsa-miR-7-5p&offsets\|0\|-1;m-400&15\|+\|88611856\|88611879&offsets\|0\|-1];[MIMAT0000252_1&hsa-miR-7-5p&offsets\|0\|-1;m-402&19\|+\|4770700\|4770723&offsets\|0\|-1];[MIMAT0000252_2&hsa-miR-7-5p&offsets\|0\|-1;m-401&9\|-\|83969811\|83969834&offsets\|0\|-1]\|TGGAAGACTAGTGATTTTGTTGT | >MI0000264\|hsa-mir-7-2&WithFlank&15\|+\|88611819\|88611940 | 38 | 60 | 23 | TGGAAGACTAGTGATTTTGTTGT | 5505.9 | 5.6 | 0.6 | 9.4 | 4.3E-21 | 6.0E-20 | MB231/MB468 | NUC |
| >hg38_21_+_8433172_8446622_RNA45SN1_45S_50ntFlanks@12627.12659.33\|ATGGGGCGAAGCTACCATCTGTGGGATTATGAC | >hg38_21_+_8433172_8446622_RNA45SN1_45S_50ntFlanks | 12627 | 12659 | 33 | ATGGGGCGAAGCTACCATCTGTGGGATTATGAC | 3718.1 | -15.7 | 1.7 | -9.5 | 3.1E-21 | 6.0E-20 | MB231/MB468 | MP |
| >MI0000272\|hsa-mir-182&WithFlank&7\|-\|129770377\|129770498@29.51.23(+1G)&[MIMAT0000259&hsa-miR-182-5p&offsets\|0\|-1(+1G);m-22&7\|-\|129770449\|129770470&offsets\|0\|+1(+1G)]\|TTTGGCAATGGTAGAACTCACACG | >MI0000272\|hsa-mir-182&WithFlank&7\|-\|129770377\|129770498 | 29 | 51 | 23(+1G) | TTTGGCAATGGTAGAACTCACACG | 5913.8 | 7.6 | 0.8 | 9.4 | 7.6E-21 | 6.0E-20 | BT20/MB468 | CYTO |
| >MI0000264\|hsa-mir-7-2&WithFlank&15\|+\|88611819\|88611940@38.58.21;MI0000265\|hsa-mir-7-3&WithFlank&19\|+\|4770664\|4770785@37.57.21;MI0000263\|hsa-mir-7-1&WithFlank&9\|-\|83969742\|83969863@30.50.21&[MIMAT0000252&hsa-miR-7-5p&offsets\|0\|-3;m-400&15\|+\|88611856\|88611879&offsets\|0\|-3];[MIMAT0000252_1&hsa-miR-7-5p&offsets\|0\|-3;m-402&19\|+\|4770700\|4770723&offsets\|0\|-3];[MIMAT0000252_2&hsa-miR-7-5p&offsets\|0\|-3;m-401&9\|-\|83969811\|83969834&offsets\|0\|-3]\|TGGAAGACTAGTGATTTTGTT | >MI0000264\|hsa-mir-7-2&WithFlank&15\|+\|88611819\|88611940 | 38 | 58 | 21 | TGGAAGACTAGTGATTTTGTT | 1217.5 | 6.2 | 0.7 | 9.4 | 4.3E-21 | 6.1E-20 | MB231/MB468 | NUC |
| >hg38_21_+_8433172_8446622_RNA45SN1_45S_50ntFlanks@12476.12509.34\|CCACTAATAGGGAACGTGAGCTGGGTTTAGACCG | >hg38_21_+_8433172_8446622_RNA45SN1_45S_50ntFlanks | 12476 | 12509 | 34 | CCACTAATAGGGAACGTGAGCTGGGTTTAGACCG | 471.0 | -13.3 | 1.4 | -9.3 | 1.5E-20 | 6.1E-20 | BT20/MB468 | NUC |
| >MI0000477\|hsa-mir-146a&WithFlank&5\|+\|160485346\|160485456@27.47.21&[MIMAT0000449&hsa-miR-146a-5p&offsets\|0\|-1;m-129&5\|+\|160485372\|160485393&offsets\|0\|-1]\|TGAGAACTGAATTCCATGGGT | >MI0000477\|hsa-mir-146a&WithFlank&5\|+\|160485346\|160485456 | 27 | 47 | 21 | TGAGAACTGAATTCCATGGGT | 1683.7 | 14.7 | 1.6 | 9.5 | 2.5E-21 | 6.1E-20 | MB231/MB468 | CYTO |
| >hg38_21_+_8433172_8446622_RNA45SN1_45S_50ntFlanks@11668.11685.18\|TGCCCAGTGCTCTGAATG | >hg38_21_+_8433172_8446622_RNA45SN1_45S_50ntFlanks | 11668 | 11685 | 18 | TGCCCAGTGCTCTGAATG | 991.2 | -5.0 | 0.5 | -9.3 | 1.5E-20 | 6.1E-20 | BT20/MB468 | NUC |
| >hg38_21_+_8433172_8446622_RNA45SN1_45S_50ntFlanks@12324.12355.32\|TTGGGTTTTAAGCAGGAGGTGTCAGAAAAGTT | >hg38_21_+_8433172_8446622_RNA45SN1_45S_50ntFlanks | 12324 | 12355 | 32 | TTGGGTTTTAAGCAGGAGGTGTCAGAAAAGTT | 472.5 | -13.3 | 1.4 | -9.3 | 1.5E-20 | 6.1E-20 | BT20/MB468 | NUC |
| >MI0003643\|hsa-mir-629&WithFlank&15\|-\|70079366\|70079474@28.49.22&[MIMAT0004810&hsa-miR-629-5p&offsets\|0\|+1;m-150&15\|-\|70079427\|70079447&offsets\|0\|+1]\|TGGGTTTACGTTGGGAGAACTT | >MI0003643\|hsa-mir-629&WithFlank&15\|-\|70079366\|70079474 | 28 | 49 | 22 | TGGGTTTACGTTGGGAGAACTT | 805.8 | 13.7 | 1.4 | 9.5 | 2.6E-21 | 6.2E-20 | MB231/MB468 | CYTO |
| >hg38_21_+_8433172_8446622_RNA45SN1_45S_50ntFlanks@4472.4494.23\|CTTAGCTGAGTGTCCCGCGGGGC | >hg38_21_+_8433172_8446622_RNA45SN1_45S_50ntFlanks | 4472 | 4494 | 23 | CTTAGCTGAGTGTCCCGCGGGGC | 1065.8 | -13.9 | 1.5 | -9.5 | 3.2E-21 | 6.2E-20 | MB231/MB468 | MP |
| >hg38_21_+_8433172_8446622_RNA45SN1_45S_50ntFlanks@10841.10863.23\|CGTAACTTCGGGATAAGGATTGG | >hg38_21_+_8433172_8446622_RNA45SN1_45S_50ntFlanks | 10841 | 10863 | 23 | CGTAACTTCGGGATAAGGATTGG | 5007.5 | -6.0 | 0.6 | -9.4 | 7.9E-21 | 6.3E-20 | BT20/MB468 | CYTO |
| >hg38_21_+_8433172_8446622_RNA45SN1_45S_50ntFlanks@10408.10453.46\|GGTCAGTCGGTCCTGAGAGATGGGCGAGCGCCGTTCCGAAGGGACG | >hg38_21_+_8433172_8446622_RNA45SN1_45S_50ntFlanks | 10408 | 10453 | 46 | GGTCAGTCGGTCCTGAGAGATGGGCGAGCGCCGTTCCGAAGGGACG | 758.7 | -14.0 | 1.5 | -9.3 | 1.6E-20 | 6.3E-20 | BT20/MB468 | NUC |
| >hg38_21_+_8433172_8446622_RNA45SN1_45S_50ntFlanks@11620.11640.21\|ACAAAGCATCGCGAAGGCCCG | >hg38_21_+_8433172_8446622_RNA45SN1_45S_50ntFlanks | 11620 | 11640 | 21 | ACAAAGCATCGCGAAGGCCCG | 388.8 | -13.0 | 1.4 | -9.3 | 1.6E-20 | 6.3E-20 | BT20/MB468 | NUC |
| >MI0000809\|hsa-mir-151a&WithFlank&8\|-\|140732558\|140732659@51.72.22(+1A)&[MIMAT0000757&hsa-miR-151a-3p&offsets\|-2\|-1(+1A);m-30&8\|-\|140732587\|140732607&offsets\|-2\|-1(+1A)]\|TACTAGACTGAAGCTCCTTGAGA | >MI0000809\|hsa-mir-151a&WithFlank&8\|-\|140732558\|140732659 | 51 | 72 | 22(+1A) | TACTAGACTGAAGCTCCTTGAGA | 370.5 | 5.0 | 0.5 | 9.4 | 4.5E-21 | 6.3E-20 | MB231/MB468 | NUC |
| >hg38_21_+_8433172_8446622_RNA45SN1_45S_50ntFlanks@6738.6756.19\|ATCATCGACACTTCGAACG | >hg38_21_+_8433172_8446622_RNA45SN1_45S_50ntFlanks | 6738 | 6756 | 19 | ATCATCGACACTTCGAACG | 905.4 | -13.7 | 1.4 | -9.5 | 3.3E-21 | 6.3E-20 | MB231/MB468 | MP |
| >hg38_21_+_8433172_8446622_RNA45SN1_45S_50ntFlanks@12456.12485.30\|CCAAGCGTTGGATTGTTCACCCACTAATAG | >hg38_21_+_8433172_8446622_RNA45SN1_45S_50ntFlanks | 12456 | 12485 | 30 | CCAAGCGTTGGATTGTTCACCCACTAATAG | 1216.5 | -12.3 | 1.3 | -9.4 | 5.8E-21 | 6.3E-20 | BT20/MB468 | MITO |
| >hg38_21_+_8433172_8446622_RNA45SN1_45S_50ntFlanks@10771.10806.36\|CAGGTCTCCAAGGTGAACAGCCTCTGGCATGTTGGA | >hg38_21_+_8433172_8446622_RNA45SN1_45S_50ntFlanks | 10771 | 10806 | 36 | CAGGTCTCCAAGGTGAACAGCCTCTGGCATGTTGGA | 1545.5 | -10.5 | 1.1 | -9.4 | 5.0E-21 | 6.3E-20 | MB231/MB468 | MITO |
| >Antisense_hg38_21_-_8433172_8446622_RNA45SN1_45S_50ntFlanks@7674.7693.20\|CTCTCTCTCTCTCTCCCCCG | >Antisense_hg38_21_-_8433172_8446622_RNA45SN1_45S_50ntFlanks | 7674 | 7693 | 20 | CTCTCTCTCTCTCTCCCCCG | 848.4 | -13.5 | 1.4 | -9.4 | 5.0E-21 | 6.4E-20 | MB231/MB468 | MITO |
| >hg38_21_+_8433172_8446622_RNA45SN1_45S_50ntFlanks@11027.11046.20\|CTCTCTCTCTCTCTCCCCCG | >hg38_21_+_8433172_8446622_RNA45SN1_45S_50ntFlanks | 11027 | 11046 | 20 | CTCTCTCTCTCTCTCCCCCG | 848.4 | -13.5 | 1.4 | -9.4 | 5.0E-21 | 6.4E-20 | MB231/MB468 | MITO |
| >hg38_21_+_8433172_8446622_RNA45SN1_45S_50ntFlanks@3899.3937.39\|CCTTCGCGGGGGGGATGCGTGCATTTATCAGATCAAAAC | >hg38_21_+_8433172_8446622_RNA45SN1_45S_50ntFlanks | 3899 | 3937 | 39 | CCTTCGCGGGGGGGATGCGTGCATTTATCAGATCAAAAC | 1365.9 | -14.2 | 1.5 | -9.4 | 5.0E-21 | 6.4E-20 | MB231/MB468 | MITO |
| >hg38_21_+_8433172_8446622_RNA45SN1_45S_50ntFlanks@7981.8006.26\|CTCAGATCAGACGTGGCGACCCGCTG | >hg38_21_+_8433172_8446622_RNA45SN1_45S_50ntFlanks | 7981 | 8006 | 26 | CTCAGATCAGACGTGGCGACCCGCTG | 6578.3 | -5.1 | 0.5 | -9.4 | 5.8E-21 | 6.4E-20 | BT20/MB468 | MITO |
| >hg38_21_+_8433172_8446622_RNA45SN1_45S_50ntFlanks@12989.13013.25\|AGCTCCCTCGCTGCGATCTATTGAA | >hg38_21_+_8433172_8446622_RNA45SN1_45S_50ntFlanks | 12989 | 13013 | 25 | AGCTCCCTCGCTGCGATCTATTGAA | 427.7 | 5.3 | 0.6 | 9.5 | 2.6E-21 | 6.4E-20 | MB231/MB468 | CYTO |
| >hg38_1_-_228634819_228635039_RNA5S12_5S_50ntFlanks@152.170.19\|TACCGGGTGCTGTAGGCTT | >hg38_1_-_228634819_228635039_RNA5S12_5S_50ntFlanks | 152 | 170 | 19 | TACCGGGTGCTGTAGGCTT | 4124.4 | 6.0 | 0.6 | 9.4 | 8.0E-21 | 6.4E-20 | BT20/MB468 | CYTO |
| >hg38_21_+_8433172_8446622_RNA45SN1_45S_50ntFlanks@11270.11295.26\|CGGCCGCCCCCGGGGAGCCCGGCGGG | >hg38_21_+_8433172_8446622_RNA45SN1_45S_50ntFlanks | 11270 | 11295 | 26 | CGGCCGCCCCCGGGGAGCCCGGCGGG | 1464.7 | -14.4 | 1.5 | -9.5 | 3.3E-21 | 6.4E-20 | MB231/MB468 | MP |
| >hg38_21_+_8433172_8446622_RNA45SN1_45S_50ntFlanks@9568.9595.28\|CGTGCAAATCGGTCGTCCGACCTGGGTA | >hg38_21_+_8433172_8446622_RNA45SN1_45S_50ntFlanks | 9568 | 9595 | 28 | CGTGCAAATCGGTCGTCCGACCTGGGTA | 2908.6 | -5.3 | 0.6 | -9.4 | 8.1E-21 | 6.4E-20 | BT20/MB468 | CYTO |
| >hg38_21_+_8433172_8446622_RNA45SN1_45S_50ntFlanks@11037.11060.24\|CTCTCCCCCGCTCCCCGTCCTCCC | >hg38_21_+_8433172_8446622_RNA45SN1_45S_50ntFlanks | 11037 | 11060 | 24 | CTCTCCCCCGCTCCCCGTCCTCCC | 884.9 | -13.6 | 1.4 | -9.4 | 5.1E-21 | 6.5E-20 | MB231/MB468 | MITO |
| >hg38_21_+_8433172_8446622_RNA45SN1_45S_50ntFlanks@12636.12654.19\|AGCTACCATCTGTGGGATT | >hg38_21_+_8433172_8446622_RNA45SN1_45S_50ntFlanks | 12636 | 12654 | 19 | AGCTACCATCTGTGGGATT | 6273.7 | -12.6 | 1.3 | -9.4 | 5.1E-21 | 6.5E-20 | MB231/MB468 | MITO |
| >hg38_21_+_8433172_8446622_RNA45SN1_45S_50ntFlanks@10107.10124.18\|CGTGTGGTGTGCGTCGGA | >hg38_21_+_8433172_8446622_RNA45SN1_45S_50ntFlanks | 10107 | 10124 | 18 | CGTGTGGTGTGCGTCGGA | 453.6 | 26.1 | 2.6 | 9.9 | 3.1E-23 | 6.5E-20 | BT20/MB231 | MP |
| >MI0000270\|hsa-mir-181b-1&WithFlank&1\|-\|198858867\|198858988@42.63.22(+1A);MI0000683\|hsa-mir-181b-2&WithFlank&9\|+\|124693704\|124693804@22.43.22(+1A)&[MIMAT0000257&hsa-miR-181b-5p&offsets\|0\|-1(+1A);m-73&1\|-\|198858925\|198858947&offsets\|0\|-1(+1A)];[MIMAT0000257_1&hsa-miR-181b-5p&offsets\|0\|-1(+1A);m-72&9\|+\|124693725\|124693747&offsets\|0\|-1(+1A)]\|AACATTCATTGCTGTCGGTGGGA | >MI0000270\|hsa-mir-181b-1&WithFlank&1\|-\|198858867\|198858988 | 42 | 63 | 22(+1A) | AACATTCATTGCTGTCGGTGGGA | 541.2 | 6.1 | 0.6 | 9.4 | 8.2E-21 | 6.5E-20 | BT20/MB468 | CYTO |
| >hg38_21_+_8433172_8446622_RNA45SN1_45S_50ntFlanks@12539.12567.29\|ATGATGTGTTGTTGCCATGGTAATCCTGC | >hg38_21_+_8433172_8446622_RNA45SN1_45S_50ntFlanks | 12539 | 12567 | 29 | ATGATGTGTTGTTGCCATGGTAATCCTGC | 1146.7 | -13.9 | 1.5 | -9.4 | 5.2E-21 | 6.5E-20 | MB231/MB468 | MITO |
| >am_MI0000786\|hsa-mir-378a&WithFlank&5\|+\|149732819\|149732896@49.69.21&[MIMAT0000732&hsa-miR-378a-3p&offsets\|0\|-1;m-44&5\|+\|149732867\|149732888&offsets\|0\|-1]\|ACTGGACTTGGAGTCAGAAGG | >am_MI0000786\|hsa-mir-378a&WithFlank&5\|+\|149732819\|149732896 | 49 | 69 | 21 | ACTGGACTTGGAGTCAGAAGG | 3348.0 | 5.2 | 0.6 | 9.4 | 8.3E-21 | 6.5E-20 | BT20/MB468 | CYTO |
| >hg38_21_+_8433172_8446622_RNA45SN1_45S_50ntFlanks@6722.6739.18\|ATTGCAGGACACATTGAT | >hg38_21_+_8433172_8446622_RNA45SN1_45S_50ntFlanks | 6722 | 6739 | 18 | ATTGCAGGACACATTGAT | 403.3 | -13.1 | 1.4 | -9.3 | 1.6E-20 | 6.5E-20 | BT20/MB468 | NUC |
| >MI0000299\|hsa-mir-222&WithFlank&X\|-\|45747009\|45747130@75.98.24(+1G)&[MIMAT0000279&hsa-miR-222-3p&offsets\|0\|+3(+1G);m-140&X\|-\|45747033\|45747056&offsets\|0\|0(+1G)]\|AGCTACATCTGGCTACTGGGTCTCG | >MI0000299\|hsa-mir-222&WithFlank&X\|-\|45747009\|45747130 | 75 | 98 | 24(+1G) | AGCTACATCTGGCTACTGGGTCTCG | 834.7 | 6.0 | 0.6 | 9.4 | 4.7E-21 | 6.5E-20 | MB231/MB468 | NUC |
| >hg38_21_+_8433172_8446622_RNA45SN1_45S_50ntFlanks@11576.11605.30\|CTTAGAACTGGTGCGGACCAGGGGAATCCG | >hg38_21_+_8433172_8446622_RNA45SN1_45S_50ntFlanks | 11576 | 11605 | 30 | CTTAGAACTGGTGCGGACCAGGGGAATCCG | 969.4 | -13.8 | 1.5 | -9.4 | 3.4E-21 | 6.5E-20 | MB231/MB468 | MP |
| >hg38_21_+_8433172_8446622_RNA45SN1_45S_50ntFlanks@11582.11611.30\|ACTGGTGCGGACCAGGGGAATCCGACTGTT | >hg38_21_+_8433172_8446622_RNA45SN1_45S_50ntFlanks | 11582 | 11611 | 30 | ACTGGTGCGGACCAGGGGAATCCGACTGTT | 1499.8 | -5.6 | 0.6 | -9.4 | 3.4E-21 | 6.5E-20 | MB231/MB468 | MP |
| >hg38_21_+_8433172_8446622_RNA45SN1_45S_50ntFlanks@12411.12435.25\|GATCCTTCGATGTCGGCTCTTCCTA | >hg38_21_+_8433172_8446622_RNA45SN1_45S_50ntFlanks | 12411 | 12435 | 25 | GATCCTTCGATGTCGGCTCTTCCTA | 429.3 | -13.2 | 1.4 | -9.3 | 1.6E-20 | 6.5E-20 | BT20/MB468 | NUC |
| >trnaMT_MetCAT_MT_+_4402_4469@37.71.35\|CCGAAAATGTTGGTTATACCCTTCCCGTACTACCA | >trnaMT_MetCAT_MT_+_4402_4469 | 37 | 71 | 35 | CCGAAAATGTTGGTTATACCCTTCCCGTACTACCA | 1074.0 | 14.1 | 1.5 | 9.5 | 2.7E-21 | 6.6E-20 | MB231/MB468 | CYTO |
| >hg38_21_+_8433172_8446622_RNA45SN1_45S_50ntFlanks@11575.11603.29\|ACTTAGAACTGGTGCGGACCAGGGGAATC | >hg38_21_+_8433172_8446622_RNA45SN1_45S_50ntFlanks | 11575 | 11603 | 29 | ACTTAGAACTGGTGCGGACCAGGGGAATC | 4469.6 | -9.7 | 1.0 | -9.4 | 5.2E-21 | 6.6E-20 | MB231/MB468 | MITO |
| >hg38_21_+_8433172_8446622_RNA45SN1_45S_50ntFlanks@4046.4063.18\|CATTCGAACGTCTGCCCT | >hg38_21_+_8433172_8446622_RNA45SN1_45S_50ntFlanks | 4046 | 4063 | 18 | CATTCGAACGTCTGCCCT | 4047.7 | -9.8 | 1.0 | -9.4 | 5.3E-21 | 6.6E-20 | MB231/MB468 | MITO |
| >hg38_21_+_8433172_8446622_RNA45SN1_45S_50ntFlanks@8115.8142.28\|CGCGGGACATGTGGCGTACGGAAGACCC | >hg38_21_+_8433172_8446622_RNA45SN1_45S_50ntFlanks | 8115 | 8142 | 28 | CGCGGGACATGTGGCGTACGGAAGACCC | 831.3 | -13.5 | 1.4 | -9.4 | 5.3E-21 | 6.6E-20 | MB231/MB468 | MITO |
| >hg38_21_+_8433172_8446622_RNA45SN1_45S_50ntFlanks@9567.9595.29\|ACGTGCAAATCGGTCGTCCGACCTGGGTA | >hg38_21_+_8433172_8446622_RNA45SN1_45S_50ntFlanks | 9567 | 9595 | 29 | ACGTGCAAATCGGTCGTCCGACCTGGGTA | 1738.8 | -6.0 | 0.6 | -9.4 | 4.8E-21 | 6.7E-20 | MB231/MB468 | NUC |
| >hg38_21_+_8433172_8446622_RNA45SN1_45S_50ntFlanks@5475.5516.42\|GCCCTGGCGGAGCGCTGAGAAGACGGTCGAACTTGACTATCT | >hg38_21_+_8433172_8446622_RNA45SN1_45S_50ntFlanks | 5475 | 5516 | 42 | GCCCTGGCGGAGCGCTGAGAAGACGGTCGAACTTGACTATCT | 1274.2 | 11.0 | 1.2 | 9.5 | 2.8E-21 | 6.7E-20 | MB231/MB468 | CYTO |
| >hg38_21_+_8433172_8446622_RNA45SN1_45S_50ntFlanks@11668.11710.43\|TGCCCAGTGCTCTGAATGTCAAAGTGAAGAAATTCAATGAAGC | >hg38_21_+_8433172_8446622_RNA45SN1_45S_50ntFlanks | 11668 | 11710 | 43 | TGCCCAGTGCTCTGAATGTCAAAGTGAAGAAATTCAATGAAGC | 798.3 | 13.6 | 1.4 | 9.5 | 2.8E-21 | 6.7E-20 | MB231/MB468 | CYTO |
| >MI0000081\|hsa-mir-24-2&WithFlank&19\|-\|13836281\|13836365@56.75.20;MI0000080\|hsa-mir-24-1&WithFlank&9\|+\|95086015\|95086094@50.69.20&[MIMAT0000080&hsa-miR-24-3p&offsets\|0\|-2;m-36&19\|-\|13836288\|13836310&offsets\|0\|-3];[MIMAT0000080_1&hsa-miR-24-3p&offsets\|0\|-2;m-35&9\|+\|95086064\|95086086&offsets\|0\|-3]\|TGGCTCAGTTCAGCAGGAAC | >MI0000081\|hsa-mir-24-2&WithFlank&19\|-\|13836281\|13836365 | 56 | 75 | 20 | TGGCTCAGTTCAGCAGGAAC | 14885.8 | 6.3 | 0.7 | 9.4 | 8.6E-21 | 6.8E-20 | BT20/MB468 | CYTO |

<table>
<tr><td>>MI0000108|hsa-mir-103a-2&WithFlank&20|+|3917488|3917577@54.75.22;MI0000109|hsa-mir-103a-1&WithFlank&5|-|168560890|168560979@54.75.22&[MIMAT0000101&hsa-miR-103a-3p&offsets|0|-1;m-12&20|+|3917541|3917563&offsets|0|-1];[MIMAT0000101_1&hsa-miR-103a-3p&offsets|0|-1;m-13&5|-|168560904|168560926&offsets|0|-1;m-44220&5|-|168560908|168560926&offsets|0|+3]|AGCAGCATTGTACAGGGCTATG</td><td>>MI0000108|hsa-mir-103a-2&WithFlank&20|+|3917488|3917577</td><td>54</td><td>75</td><td>22</td><td>AGCAGCATTGTACAGGGCTATG</td><td>5618.9</td><td>5.7</td><td>0.6</td><td>9.4</td><td>4.9E-21</td><td>6.8E-20</td><td>MB231/MB468</td><td>NUC</td></tr>
<tr><td>>hg38_21_+_8433172_8446622_RNA45SN1_45S_50ntFlanks@11999.12038.40|CCCTGCGGGCCGCCGGTGAAATACCACTACTCTGATCGTT</td><td>>hg38_21_+_8433172_8446622_RNA45SN1_45S_50ntFlanks</td><td>11999</td><td>12038</td><td>40</td><td>CCCTGCGGGCCGCCGGTGAAATACCACTACTCTGATCGTT</td><td>525.7</td><td>-13.5</td><td>1.5</td><td>-9.3</td><td>1.7E-20</td><td>6.8E-20</td><td>BT20/MB468</td><td>NUC</td></tr>
<tr><td>>hg38_21_+_8433172_8446622_RNA45SN1_45S_50ntFlanks@12944.12972.29|CGTAGACGACCTGCTTCTGGGTCGGGGTT</td><td>>hg38_21_+_8433172_8446622_RNA45SN1_45S_50ntFlanks</td><td>12944</td><td>12972</td><td>29</td><td>CGTAGACGACCTGCTTCTGGGTCGGGGTT</td><td>784.1</td><td>-13.5</td><td>1.4</td><td>-9.4</td><td>3.6E-21</td><td>6.8E-20</td><td>MB231/MB468</td><td>MP</td></tr>
<tr><td>>hg38_21_+_8433172_8446622_RNA45SN1_45S_50ntFlanks@12627.12653.27|ATGGGGCGAAGCTACCATCTGTGGGAT</td><td>>hg38_21_+_8433172_8446622_RNA45SN1_45S_50ntFlanks</td><td>12627</td><td>12653</td><td>27</td><td>ATGGGGCGAAGCTACCATCTGTGGGAT</td><td>2369.4</td><td>-11.0</td><td>1.2</td><td>-9.4</td><td>4.9E-21</td><td>6.8E-20</td><td>MB231/MB468</td><td>NUC</td></tr>
<tr><td>>hg38_21_+_8433172_8446622_RNA45SN1_45S_50ntFlanks@12636.12661.26|AGCTACCATCTGTGGGATTATGACTG</td><td>>hg38_21_+_8433172_8446622_RNA45SN1_45S_50ntFlanks</td><td>12636</td><td>12661</td><td>26</td><td>AGCTACCATCTGTGGGATTATGACTG</td><td>603.8</td><td>26.4</td><td>2.7</td><td>9.9</td><td>4.2E-23</td><td>6.9E-20</td><td>BT20/MB231</td><td>MP</td></tr>
<tr><td>>hg38_21_+_8433172_8446622_RNA45SN1_45S_50ntFlanks@9739.9765.27|AAACGATCTCAACCTATTCTCAAACTT</td><td>>hg38_21_+_8433172_8446622_RNA45SN1_45S_50ntFlanks</td><td>9739</td><td>9765</td><td>27</td><td>AAACGATCTCAACCTATTCTCAAACTT</td><td>12721.2</td><td>-6.5</td><td>0.7</td><td>-9.4</td><td>6.3E-21</td><td>6.9E-20</td><td>BT20/MB468</td><td>MITO</td></tr>
<tr><td>>hg38_21_+_8433172_8446622_RNA45SN1_45S_50ntFlanks@8112.8143.32|GGGCGCGGGACATGTGGCGTACGGAAGACCCG</td><td>>hg38_21_+_8433172_8446622_RNA45SN1_45S_50ntFlanks</td><td>8112</td><td>8143</td><td>32</td><td>GGGCGCGGGACATGTGGCGTACGGAAGACCCG</td><td>1086.7</td><td>-13.9</td><td>1.5</td><td>-9.4</td><td>3.6E-21</td><td>6.9E-20</td><td>MB231/MB468</td><td>MP</td></tr>
<tr><td>>hg38_21_+_8433172_8446622_RNA45SN1_45S_50ntFlanks@8687.8707.21|CGGCTCCGGGACGGCTGGGAA</td><td>>hg38_21_+_8433172_8446622_RNA45SN1_45S_50ntFlanks</td><td>8687</td><td>8707</td><td>21</td><td>CGGCTCCGGGACGGCTGGGAA</td><td>746.9</td><td>13.5</td><td>1.4</td><td>9.5</td><td>2.9E-21</td><td>6.9E-20</td><td>MB231/MB468</td><td>CYTO</td></tr>
<tr><td>>hg38_21_+_8433172_8446622_RNA45SN1_45S_50ntFlanks@5486.5526.41|GCGCTGAGAAGACGGTCGAACTTGACTATCTAGAGGAAGTA</td><td>>hg38_21_+_8433172_8446622_RNA45SN1_45S_50ntFlanks</td><td>5486</td><td>5526</td><td>41</td><td>GCGCTGAGAAGACGGTCGAACTTGACTATCTAGAGGAAGTA</td><td>898.0</td><td>-13.6</td><td>1.4</td><td>-9.4</td><td>5.6E-21</td><td>6.9E-20</td><td>MB231/MB468</td><td>MITO</td></tr>
<tr><td>>hg38_21_+_8433172_8446622_RNA45SN1_45S_50ntFlanks@11617.11640.24|AAAACAAAGCATCGCGAAGGCCCG</td><td>>hg38_21_+_8433172_8446622_RNA45SN1_45S_50ntFlanks</td><td>11617</td><td>11640</td><td>24</td><td>AAAACAAAGCATCGCGAAGGCCCG</td><td>882.5</td><td>-13.6</td><td>1.4</td><td>-9.4</td><td>5.6E-21</td><td>6.9E-20</td><td>MB231/MB468</td><td>MITO</td></tr>
<tr><td>>hg38_21_+_8433172_8446622_RNA45SN1_45S_50ntFlanks@6719.6737.19|TGAATTGCAGGACACATTG</td><td>>hg38_21_+_8433172_8446622_RNA45SN1_45S_50ntFlanks</td><td>6719</td><td>6737</td><td>19</td><td>TGAATTGCAGGACACATTG</td><td>7731.9</td><td>-13.1</td><td>1.4</td><td>-9.3</td><td>1.7E-20</td><td>7.0E-20</td><td>BT20/MB468</td><td>NUC</td></tr>
<tr><td>>hg38_21_+_8433172_8446622_RNA45SN1_45S_50ntFlanks@4641.4662.22|CATTCGTATTGCGCCGCTAGAG</td><td>>hg38_21_+_8433172_8446622_RNA45SN1_45S_50ntFlanks</td><td>4641</td><td>4662</td><td>22</td><td>CATTCGTATTGCGCCGCTAGAG</td><td>473.8</td><td>-12.8</td><td>1.4</td><td>-9.4</td><td>3.7E-21</td><td>7.0E-20</td><td>MB231/MB468</td><td>MP</td></tr>
<tr><td>>hg38_21_+_8433172_8446622_RNA45SN1_45S_50ntFlanks@12447.12485.39|CAGAATTCACCAAGCGTTGGATTGTTCACCCACTAATAG</td><td>>hg38_21_+_8433172_8446622_RNA45SN1_45S_50ntFlanks</td><td>12447</td><td>12485</td><td>39</td><td>CAGAATTCACCAAGCGTTGGATTGTTCACCCACTAATAG</td><td>451.4</td><td>-13.2</td><td>1.4</td><td>-9.3</td><td>1.8E-20</td><td>7.1E-20</td><td>BT20/MB468</td><td>NUC</td></tr>
<tr><td>>hg38_21_+_8433172_8446622_RNA45SN1_45S_50ntFlanks@6719.6744.26|TGAATTGCAGGACACATTGATCATCG</td><td>>hg38_21_+_8433172_8446622_RNA45SN1_45S_50ntFlanks</td><td>6719</td><td>6744</td><td>26</td><td>TGAATTGCAGGACACATTGATCATCG</td><td>15073.9</td><td>-13.5</td><td>1.5</td><td>-9.3</td><td>1.8E-20</td><td>7.1E-20</td><td>BT20/MB468</td><td>NUC</td></tr>
<tr><td>>hg38_21_+_8433172_8446622_RNA45SN1_45S_50ntFlanks@4728.4758.31|ATCAAGAACGAAAGTCGGAGGTTCGAAGACG</td><td>>hg38_21_+_8433172_8446622_RNA45SN1_45S_50ntFlanks</td><td>4728</td><td>4758</td><td>31</td><td>ATCAAGAACGAAAGTCGGAGGTTCGAAGACG</td><td>1292.8</td><td>-12.4</td><td>1.3</td><td>-9.4</td><td>6.6E-21</td><td>7.1E-20</td><td>BT20/MB468</td><td>MITO</td></tr>
<tr><td>>hg38_21_+_8433172_8446622_RNA45SN1_45S_50ntFlanks@3984.4002.19|GCGGCTTTGGTGACTCTAG</td><td>>hg38_21_+_8433172_8446622_RNA45SN1_45S_50ntFlanks</td><td>3984</td><td>4002</td><td>19</td><td>GCGGCTTTGGTGACTCTAG</td><td>3002.9</td><td>-15.4</td><td>1.6</td><td>-9.4</td><td>3.7E-21</td><td>7.1E-20</td><td>MB231/MB468</td><td>MP</td></tr>
<tr><td>>hg38_21_+_8433172_8446622_RNA45SN1_45S_50ntFlanks@10869.10901.33|AGGGCTGGGTCGGTCGGGCTGGGGCGCGAAGCG</td><td>>hg38_21_+_8433172_8446622_RNA45SN1_45S_50ntFlanks</td><td>10869</td><td>10901</td><td>33</td><td>AGGGCTGGGTCGGTCGGGCTGGGGCGCGAAGCG</td><td>1090.6</td><td>-13.9</td><td>1.5</td><td>-9.4</td><td>5.7E-21</td><td>7.1E-20</td><td>MB231/MB468</td><td>MITO</td></tr>
<tr><td>>hg38_21_+_8433172_8446622_RNA45SN1_45S_50ntFlanks@8117.8138.22|CGGGACATGTGGCGTACGGAAG</td><td>>hg38_21_+_8433172_8446622_RNA45SN1_45S_50ntFlanks</td><td>8117</td><td>8138</td><td>22</td><td>CGGGACATGTGGCGTACGGAAG</td><td>1059.5</td><td>-13.9</td><td>1.5</td><td>-9.4</td><td>3.7E-21</td><td>7.1E-20</td><td>MB231/MB468</td><td>MP</td></tr>
<tr><td>>MI0003560|hsa-mir-92b&WithFlank&1|+|155195171|155195278@67.86.20(+1A)&[MIMAT0003218&hsa-miR-92b-3p&offsets|0|-2(+1A);m-115&1|+|155195237|155195258&offsets|0|-2(+1A)]|TATTGCACTCGTCCCGGCCTA</td><td>>MI0003560|hsa-mir-92b&WithFlank&1|+|155195171|155195278</td><td>67</td><td>86</td><td>20(+1A)</td><td>TATTGCACTCGTCCCGGCCTA</td><td>312.3</td><td>12.5</td><td>1.3</td><td>9.3</td><td>9.1E-21</td><td>7.1E-20</td><td>BT20/MB468</td><td>CYTO</td></tr>
<tr><td>>hg38_21_+_8433172_8446622_RNA45SN1_45S_50ntFlanks@9454.9488.35|CCGCGCCGGGGAGGTGGAGCACGAGCGCACGTGTT</td><td>>hg38_21_+_8433172_8446622_RNA45SN1_45S_50ntFlanks</td><td>9454</td><td>9488</td><td>35</td><td>CCGCGCCGGGGAGGTGGAGCACGAGCGCACGTGTT</td><td>978.6</td><td>-13.7</td><td>1.5</td><td>-9.4</td><td>5.8E-21</td><td>7.1E-20</td><td>MB231/MB468</td><td>MITO</td></tr>
<tr><td>>hg38_21_+_8433172_8446622_RNA45SN1_45S_50ntFlanks@11580.11611.32|GAACTGGTGCGGACCAGGGGAATCCGACTGTT</td><td>>hg38_21_+_8433172_8446622_RNA45SN1_45S_50ntFlanks</td><td>11580</td><td>11611</td><td>32</td><td>GAACTGGTGCGGACCAGGGGAATCCGACTGTT</td><td>1777.2</td><td>-5.2</td><td>0.6</td><td>-9.3</td><td>1.8E-20</td><td>7.2E-20</td><td>BT20/MB468</td><td>NUC</td></tr>
<tr><td>>hg38_21_+_8433172_8446622_RNA45SN1_45S_50ntFlanks@12657.12674.18|GACTGAACGCCTCTAAGT</td><td>>hg38_21_+_8433172_8446622_RNA45SN1_45S_50ntFlanks</td><td>12657</td><td>12674</td><td>18</td><td>GACTGAACGCCTCTAAGT</td><td>1025.4</td><td>-5.0</td><td>0.5</td><td>-9.4</td><td>5.2E-21</td><td>7.2E-20</td><td>MB231/MB468</td><td>NUC</td></tr>
<tr><td>>hg38_21_+_8433172_8446622_RNA45SN1_45S_50ntFlanks@12934.12960.27|CTAAACCATTCGTAGACGACCTGCTTC</td><td>>hg38_21_+_8433172_8446622_RNA45SN1_45S_50ntFlanks</td><td>12934</td><td>12960</td><td>27</td><td>CTAAACCATTCGTAGACGACCTGCTTC</td><td>1076.8</td><td>-4.8</td><td>0.5</td><td>-9.4</td><td>5.2E-21</td><td>7.2E-20</td><td>MB231/MB468</td><td>NUC</td></tr>
<tr><td>>MI0000434|hsa-let-7i&WithFlank&12|+|62603680|62603775@12.32.21(+1A)&[MIMAT0000415&hsa-let-7i-5p&offsets|0|-1(+1A);m-49&12|+|62603691|62603712&offsets|0|-1(+1A)]|TGAGGTAGTAGTTTGTGCTGTA</td><td>>MI0000434|hsa-let-7i&WithFlank&12|+|62603680|62603775</td><td>12</td><td>32</td><td>21(+1A)</td><td>TGAGGTAGTAGTTTGTGCTGTA</td><td>4678.6</td><td>6.2</td><td>0.7</td><td>9.4</td><td>5.3E-21</td><td>7.2E-20</td><td>MB231/MB468</td><td>NUC</td></tr>
<tr><td>>MI0017878|hsa-mir-5010&WithFlank&17|+|42514182|42514313@86.107.22(+1U)&[MIMAT0021044&hsa-miR-5010-3p&offsets|0|0(+1U);m-421&17|+|42514267|42514289&offsets|0|-1(+1U)]|TTTTGTGTCTCCCATTCCCCAGT</td><td>>MI0017878|hsa-mir-5010&WithFlank&17|+|42514182|42514313</td><td>86</td><td>107</td><td>22(+1U)</td><td>TTTTGTGTCTCCCATTCCCCAGT</td><td>333.0</td><td>12.6</td><td>1.4</td><td>9.3</td><td>9.2E-21</td><td>7.3E-20</td><td>BT20/MB468</td><td>CYTO</td></tr>
<tr><td>>hg38_1_-_228634819_228635039_RNA5S12_5S_50ntFlanks@131.170.40|GGATGGGAGACCGCCTGGGAATACCGGGTGCTGTAGGCTT</td><td>>hg38_1_-_228634819_228635039_RNA5S12_5S_50ntFlanks</td><td>131</td><td>170</td><td>40</td><td>GGATGGGAGACCGCCTGGGAATACCGGGTGCTGTAGGCTT</td><td>3128.6</td><td>8.9</td><td>1.0</td><td>9.3</td><td>9.3E-21</td><td>7.3E-20</td><td>BT20/MB468</td><td>CYTO</td></tr>
<tr><td>>MI0000077|hsa-mir-21&WithFlank&17|+|59841260|59841343@14.35.22&[MIMAT0000076&hsa-miR-21-5p&offsets|0|0;m-1&17|+|59841273|59841294&offsets|0|0]|TAGCTTATCAGACTGATGTTGA</td><td>>MI0000077|hsa-mir-21&WithFlank&17|+|59841260|59841343</td><td>14</td><td>35</td><td>22</td><td>TAGCTTATCAGACTGATGTTGA</td><td>215581.7</td><td>8.2</td><td>0.9</td><td>9.3</td><td>9.3E-21</td><td>7.3E-20</td><td>BT20/MB468</td><td>CYTO</td></tr>
<tr><td>>hg38_21_+_8433172_8446622_RNA45SN1_45S_50ntFlanks@11873.11907.35|AAGAAGACCCTGTTGAGCTTGACTCTAGTCTGGCA</td><td>>hg38_21_+_8433172_8446622_RNA45SN1_45S_50ntFlanks</td><td>11873</td><td>11907</td><td>35</td><td>AAGAAGACCCTGTTGAGCTTGACTCTAGTCTGGCA</td><td>897.9</td><td>-13.6</td><td>1.4</td><td>-9.4</td><td>5.9E-21</td><td>7.3E-20</td><td>MB231/MB468</td><td>MITO</td></tr>
<tr><td>>hg38_21_+_8433172_8446622_RNA45SN1_45S_50ntFlanks@10686.10706.21|GCGGCGTCCGGTGAGCTCTCG</td><td>>hg38_21_+_8433172_8446622_RNA45SN1_45S_50ntFlanks</td><td>10686</td><td>10706</td><td>21</td><td>GCGGCGTCCGGTGAGCTCTCG</td><td>1213.0</td><td>-14.1</td><td>1.5</td><td>-9.4</td><td>3.8E-21</td><td>7.3E-20</td><td>MB231/MB468</td><td>MP</td></tr>
<tr><td>>hg38_21_+_8433172_8446622_RNA45SN1_45S_50ntFlanks@9736.9754.19|CCGAAACGATCTCAACCTA</td><td>>hg38_21_+_8433172_8446622_RNA45SN1_45S_50ntFlanks</td><td>9736</td><td>9754</td><td>19</td><td>CCGAAACGATCTCAACCTA</td><td>613.3</td><td>-13.1</td><td>1.4</td><td>-9.4</td><td>3.8E-21</td><td>7.3E-20</td><td>MB231/MB468</td><td>MP</td></tr>
<tr><td>>hg38_21_+_8433172_8446622_RNA45SN1_45S_50ntFlanks@10389.10410.22|TGAACAGCAGTTGAACATGGGT</td><td>>hg38_21_+_8433172_8446622_RNA45SN1_45S_50ntFlanks</td><td>10389</td><td>10410</td><td>22</td><td>TGAACAGCAGTTGAACATGGGT</td><td>2109.7</td><td>-5.9</td><td>0.6</td><td>-9.5</td><td>3.1E-21</td><td>7.3E-20</td><td>MB231/MB468</td><td>CYTO</td></tr>
<tr><td>>hg38_1_-_228634819_228635039_RNA5S12_5S_50ntFlanks@125.169.45|GTACTTGGATGGGAGACCGCCTGGGAATACCGGGTGCTGTAGGCT</td><td>>hg38_1_-_228634819_228635039_RNA5S12_5S_50ntFlanks</td><td>125</td><td>169</td><td>45</td><td>GTACTTGGATGGGAGACCGCCTGGGAATACCGGGTGCTGTAGGCT</td><td>359.7</td><td>13.0</td><td>1.4</td><td>9.4</td><td>6.8E-21</td><td>7.3E-20</td><td>BT20/MB468</td><td>MITO</td></tr>
<tr><td>>hg38_21_+_8433172_8446622_RNA45SN1_45S_50ntFlanks@10869.10901.33|AGGGCTGGGTCGGTCGGGCTGGGGCGCGAAGCG</td><td>>hg38_21_+_8433172_8446622_RNA45SN1_45S_50ntFlanks</td><td>10869</td><td>10901</td><td>33</td><td>AGGGCTGGGTCGGTCGGGCTGGGGCGCGAAGCG</td><td>1177.1</td><td>-14.1</td><td>1.5</td><td>-9.4</td><td>3.9E-21</td><td>7.3E-20</td><td>MB231/MB468</td><td>MP</td></tr>
<tr><td>>trna116_GluCTC_1_-_145399233_145399304@32.70.39,trna71_GluCTC_1_-_161439189_161439260@32.70.39,trna74_GluCTC_1_-_161431809_161431880@32.70.39,trna77_GluCTC_1_-_161424398_161424469@32.70.39,trna77_GluCTC_6_+_28949976_28950047@32.70.39,trna80_GluCTC_1_-_161417018_161417089@32.70.39,trna87_GluCTC_6_-_126101393_126101464@32.70.39|CTCTCACCGCCGCGGCCCGGGTTCGATTCCCGGTCAGGG</td><td>>trna116_GluCTC_1_-_145399233_145399304</td><td>32</td><td>70</td><td>39</td><td>CTCTCACCGCCGCGGCCCGGGTTCGATTCCCGGTCAGGG</td><td>2280.6</td><td>-14.9</td><td>1.6</td><td>-9.4</td><td>6.0E-21</td><td>7.4E-20</td><td>MB231/MB468</td><td>MITO</td></tr>
<tr><td>>hg38_21_+_8433172_8446622_RNA45SN1_45S_50ntFlanks@6699.6718.20|GCTAGCTGCGAGAATTAATG</td><td>>hg38_21_+_8433172_8446622_RNA45SN1_45S_50ntFlanks</td><td>6699</td><td>6718</td><td>20</td><td>GCTAGCTGCGAGAATTAATG</td><td>14030.3</td><td>-6.1</td><td>0.6</td><td>-9.3</td><td>9.4E-21</td><td>7.4E-20</td><td>BT20/MB468</td><td>CYTO</td></tr>
<tr><td>>hg38_21_+_8433172_8446622_RNA45SN1_45S_50ntFlanks@6774.6806.33|TCCTCCCGGGGCTACGCCTGTCTGAGCGTCGCT</td><td>>hg38_21_+_8433172_8446622_RNA45SN1_45S_50ntFlanks</td><td>6774</td><td>6806</td><td>33</td><td>TCCTCCCGGGGCTACGCCTGTCTGAGCGTCGCT</td><td>3223.8</td><td>7.6</td><td>0.8</td><td>9.3</td><td>9.5E-21</td><td>7.4E-20</td><td>BT20/MB468</td><td>CYTO</td></tr>
<tr><td>>hg38_21_+_8433172_8446622_RNA45SN1_45S_50ntFlanks@11641.11667.27|CGGCGGGTGTTGACGCGATGTGATTTC</td><td>>hg38_21_+_8433172_8446622_RNA45SN1_45S_50ntFlanks</td><td>11641</td><td>11667</td><td>27</td><td>CGGCGGGTGTTGACGCGATGTGATTTC</td><td>2638.5</td><td>-15.2</td><td>1.6</td><td>-9.4</td><td>3.9E-21</td><td>7.4E-20</td><td>MB231/MB468</td><td>MP</td></tr>
<tr><td>>hg38_21_+_8433172_8446622_RNA45SN1_45S_50ntFlanks@6645.6677.33|CTCGTACGACTCTTAGCGGTGGATCACTCGGCT</td><td>>hg38_21_+_8433172_8446622_RNA45SN1_45S_50ntFlanks</td><td>6645</td><td>6677</td><td>33</td><td>CTCGTACGACTCTTAGCGGTGGATCACTCGGCT</td><td>606.5</td><td>5.2</td><td>0.6</td><td>9.3</td><td>9.6E-21</td><td>7.5E-20</td><td>BT20/MB468</td><td>CYTO</td></tr>
<tr><td>>hg38_21_+_8433172_8446622_RNA45SN1_45S_50ntFlanks@12456.12489.34|CCAAGCGTTGGATTGTTCACCCACTAATAGGGAA</td><td>>hg38_21_+_8433172_8446622_RNA45SN1_45S_50ntFlanks</td><td>12456</td><td>12489</td><td>34</td><td>CCAAGCGTTGGATTGTTCACCCACTAATAGGGAA</td><td>1549.3</td><td>-14.4</td><td>1.5</td><td>-9.4</td><td>6.1E-21</td><td>7.5E-20</td><td>MB231/MB468</td><td>MITO</td></tr>
<tr><td>>hg38_21_+_8433172_8446622_RNA45SN1_45S_50ntFlanks@11641.11668.28|CGGCGGGTGTTGACGCGATGTGATTTCT</td><td>>hg38_21_+_8433172_8446622_RNA45SN1_45S_50ntFlanks</td><td>11641</td><td>11668</td><td>28</td><td>CGGCGGGTGTTGACGCGATGTGATTTCT</td><td>2100.0</td><td>-14.9</td><td>1.6</td><td>-9.4</td><td>4.0E-21</td><td>7.6E-20</td><td>MB231/MB468</td><td>MP</td></tr>
<tr><td>>hg38_21_+_8433172_8446622_RNA45SN1_45S_50ntFlanks@10572.10612.41|AGAAGCCGGCGGGAGCCCCGGGGAGAGTTCTCTTTTCTTTG</td><td>>hg38_21_+_8433172_8446622_RNA45SN1_45S_50ntFlanks</td><td>10572</td><td>10612</td><td>41</td><td>AGAAGCCGGCGGGAGCCCCGGGGAGAGTTCTCTTTTCTTTG</td><td>393.2</td><td>-13.0</td><td>1.4</td><td>-9.3</td><td>1.9E-20</td><td>7.6E-20</td><td>BT20/MB468</td><td>NUC</td></tr>
<tr><td>>hg38_21_+_8433172_8446622_RNA45SN1_45S_50ntFlanks@4367.4398.32|CAGTTAAAAAGCTCGTAGTTGGATCTTGGGAG</td><td>>hg38_21_+_8433172_8446622_RNA45SN1_45S_50ntFlanks</td><td>4367</td><td>4398</td><td>32</td><td>CAGTTAAAAAGCTCGTAGTTGGATCTTGGGAG</td><td>783.9</td><td>-13.4</td><td>1.4</td><td>-9.4</td><td>6.2E-21</td><td>7.6E-20</td><td>MB231/MB468</td><td>MITO</td></tr>
<tr><td>>trnaMT_PheGAA_MT_+_577_647@39.56.18|ATGTTTAGACGGGCTCAC</td><td>>trnaMT_PheGAA_MT_+_577_647</td><td>39</td><td>56</td><td>18</td><td>ATGTTTAGACGGGCTCAC</td><td>2192.0</td><td>7.2</td><td>0.8</td><td>9.4</td><td>5.6E-21</td><td>7.7E-20</td><td>MB231/MB468</td><td>NUC</td></tr>
<tr><td>>hg38_21_+_8433172_8446622_RNA45SN1_45S_50ntFlanks@11586.11611.26|GTGCGGACCAGGGGAATCCGACTGTT</td><td>>hg38_21_+_8433172_8446622_RNA45SN1_45S_50ntFlanks</td><td>11586</td><td>11611</td><td>26</td><td>GTGCGGACCAGGGGAATCCGACTGTT</td><td>3140.3</td><td>-6.6</td><td>0.7</td><td>-9.4</td><td>4.1E-21</td><td>7.7E-20</td><td>MB231/MB468</td><td>MP</td></tr>
<tr><td>>hg38_21_+_8433172_8446622_RNA45SN1_45S_50ntFlanks@11161.11178.18|CACGGTCCCCCGCGAGGG</td><td>>hg38_21_+_8433172_8446622_RNA45SN1_45S_50ntFlanks</td><td>11161</td><td>11178</td><td>18</td><td>CACGGTCCCCCGCGAGGG</td><td>2692.0</td><td>-10.2</td><td>1.1</td><td>-9.5</td><td>3.3E-21</td><td>7.7E-20</td><td>MB231/MB468</td><td>CYTO</td></tr>
<tr><td>>hg38_21_+_8433172_8446622_RNA45SN1_45S_50ntFlanks@12395.12418.24|AGCGACGTCGCTTTTTGATCCTTC</td><td>>hg38_21_+_8433172_8446622_RNA45SN1_45S_50ntFlanks</td><td>12395</td><td>12418</td><td>24</td><td>AGCGACGTCGCTTTTTGATCCTTC</td><td>442.0</td><td>4.7</td><td>0.5</td><td>9.4</td><td>5.7E-21</td><td>7.7E-20</td><td>MB231/MB468</td><td>NUC</td></tr>
<tr><td>>hg38_21_+_8433172_8446622_RNA45SN1_45S_50ntFlanks@10391.10410.20|AACAGCAGTTGAACATGGGT</td><td>>hg38_21_+_8433172_8446622_RNA45SN1_45S_50ntFlanks</td><td>10391</td><td>10410</td><td>20</td><td>AACAGCAGTTGAACATGGGT</td><td>11699.1</td><td>-10.8</td><td>1.2</td><td>-9.3</td><td>2.0E-20</td><td>7.8E-20</td><td>BT20/MB468</td><td>NUC</td></tr>
</table>

<table>
<tr><th></th><th></th><th></th><th></th><th></th><th></th><th></th><th></th><th></th><th></th><th></th><th></th><th></th><th></th></tr>
<tr><td>>MI0000088|hsa-mir-30a&WithFlank&6|-|71403545|71403627@12.34.23&[MIMAT0000087&hsa-miR-30a-5p&offsets|0|+1;m-15&6|-|71403595|71403616&offsets|0|+1]|TGTAAACATCCTCGACTGGAAGC</td><td>>MI0000088|hsa-mir-30a&WithFlank&6|-|71403545|71403627</td><td>12</td><td>34</td><td>23</td><td>TGTAAACATCCTCGACTGGAAGC</td><td>24585.7</td><td>7.0</td><td>0.7</td><td>9.4</td><td>5.7E-21</td><td>7.8E-20</td><td>MB231/MB468</td><td>NUC</td></tr>
<tr><td>>MI0000067|hsa-let-7f-1&WithFlank&9|+|94176341|94176439@13.33.21(+1C);MI0000068|hsa-let-7f-2&WithFlank&X|-|53557186|53557280@14.34.21(+1C)&[MIMAT0000067&hsa-let-7f-5p&offsets|0|-1(+1C);m-20&9|+|94176353|94176374&offsets|0|-1(+1C)];[MIMAT0000067_1&hsa-let-7f-5p&offsets|0|-1(+1C);m-19&X|-|53557246|53557267&offsets|0|-1(+1C)]|TGAGGTAGTAGATTGTATAGTC</td><td>>MI0000067|hsa-let-7f-1&WithFlank&9|+|94176341|94176439</td><td>13</td><td>33</td><td>21(+1C)</td><td>TGAGGTAGTAGATTGTATAGTC</td><td>26328.1</td><td>6.8</td><td>0.7</td><td>9.3</td><td>1.0E-20</td><td>7.8E-20</td><td>BT20/MB468</td><td>CYTO</td></tr>
<tr><td>>hg38_21_+_8433172_8446622_RNA45SN1_45S_50ntFlanks@10794.10825.32|CTGGCATGTTGGAACAATGTAGGTAAGGGAAG</td><td>>hg38_21_+_8433172_8446622_RNA45SN1_45S_50ntFlanks</td><td>10794</td><td>10825</td><td>32</td><td>CTGGCATGTTGGAACAATGTAGGTAAGGGAAG</td><td>607.3</td><td>-13.1</td><td>1.4</td><td>-9.4</td><td>4.2E-21</td><td>7.9E-20</td><td>MB231/MB468</td><td>MP</td></tr>
<tr><td>>hg38_21_+_8433172_8446622_RNA45SN1_45S_50ntFlanks@11602.11620.19|TCCGACTGTTTAATTAAAA</td><td>>hg38_21_+_8433172_8446622_RNA45SN1_45S_50ntFlanks</td><td>11602</td><td>11620</td><td>19</td><td>TCCGACTGTTTAATTAAAA</td><td>506.9</td><td>-12.8</td><td>1.4</td><td>-9.4</td><td>4.2E-21</td><td>7.9E-20</td><td>MB231/MB468</td><td>MP</td></tr>
<tr><td>>MI0000750|hsa-mir-26a-2&WithFlank&12|-|57824603|57824698@20.39.20;MI0000083|hsa-mir-26a-1&WithFlank&3|+|37969398|37969486@16.35.20&[MIMAT0000082&hsa-miR-26a-5p&offsets|0|-2;m-33&12|-|57824658|57824679&offsets|0|-2];[MIMAT0000082_1&hsa-miR-26a-5p&offsets|0|-2;m-34&3|+|37969413|37969434&offsets|0|-2]|TTCAAGTAATCCAGGATAGG</td><td>>MI0000750|hsa-mir-26a-2&WithFlank&12|-|57824603|57824698</td><td>20</td><td>39</td><td>20</td><td>TTCAAGTAATCCAGGATAGG</td><td>495.6</td><td>6.0</td><td>0.6</td><td>9.4</td><td>5.8E-21</td><td>7.9E-20</td><td>MB231/MB468</td><td>NUC</td></tr>
<tr><td>>hg38_21_+_8433172_8446622_RNA45SN1_45S_50ntFlanks@5079.5120.42|GATAACGAACGAGACTCTGGCATGCTAACTAGTTACGCGACC</td><td>>hg38_21_+_8433172_8446622_RNA45SN1_45S_50ntFlanks</td><td>5079</td><td>5120</td><td>42</td><td>GATAACGAACGAGACTCTGGCATGCTAACTAGTTACGCGACC</td><td>172.1</td><td>11.4</td><td>1.2</td><td>9.5</td><td>3.4E-21</td><td>7.9E-20</td><td>MB231/MB468</td><td>CYTO</td></tr>
<tr><td>>hg38_21_+_8433172_8446622_RNA45SN1_45S_50ntFlanks@12386.12414.29|AGCGTTCATAGCGACGTCGCTTTTTGATC</td><td>>hg38_21_+_8433172_8446622_RNA45SN1_45S_50ntFlanks</td><td>12386</td><td>12414</td><td>29</td><td>AGCGTTCATAGCGACGTCGCTTTTTGATC</td><td>2496.7</td><td>-15.1</td><td>1.6</td><td>-9.4</td><td>6.5E-21</td><td>8.0E-20</td><td>MB231/MB468</td><td>MITO</td></tr>
<tr><td>>hg38_21_+_8433172_8446622_RNA45SN1_45S_50ntFlanks@6730.6771.42|ACACATTGATCATCGACACTTCGAACGCACTTGCGGCCCCGG</td><td>>hg38_21_+_8433172_8446622_RNA45SN1_45S_50ntFlanks</td><td>6730</td><td>6771</td><td>42</td><td>ACACATTGATCATCGACACTTCGAACGCACTTGCGGCCCCGG</td><td>1235.7</td><td>-14.7</td><td>1.6</td><td>-9.3</td><td>2.0E-20</td><td>8.0E-20</td><td>BT20/MB468</td><td>NUC</td></tr>
<tr><td>>hg38_21_+_8433172_8446622_RNA45SN1_45S_50ntFlanks@3842.3881.40|CCTACTTGGATAACTGTGGTAATTCTAGAGCTAATACATG</td><td>>hg38_21_+_8433172_8446622_RNA45SN1_45S_50ntFlanks</td><td>3842</td><td>3881</td><td>40</td><td>CCTACTTGGATAACTGTGGTAATTCTAGAGCTAATACATG</td><td>327.1</td><td>-12.8</td><td>1.4</td><td>-9.3</td><td>2.0E-20</td><td>8.0E-20</td><td>BT20/MB468</td><td>NUC</td></tr>
<tr><td>>MI0000273|hsa-mir-183&WithFlank&7|-|129774899|129775020@34.55.22(+1A)&[MIMAT0000261&hsa-miR-183-5p&offsets|+1|+1(+1A);m-41&7|-|129774966|129774987&offsets|0|0(+1A)]|ATGGCACTGGTAGAATTCACTGA</td><td>>MI0000273|hsa-mir-183&WithFlank&7|-|129774899|129775020</td><td>34</td><td>55</td><td>22(+1A)</td><td>ATGGCACTGGTAGAATTCACTGA</td><td>363.5</td><td>12.8</td><td>1.4</td><td>9.3</td><td>1.0E-20</td><td>8.0E-20</td><td>BT20/MB468</td><td>CYTO</td></tr>
<tr><td>>trnaMT_HisGTG_MT_+_12138_12206@26.61.36|AGATTGTGAATCTGACAACAGAGGCTTACGACCCCT</td><td>>trnaMT_HisGTG_MT_+_12138_12206</td><td>26</td><td>61</td><td>36</td><td>AGATTGTGAATCTGACAACAGAGGCTTACGACCCCT</td><td>582.3</td><td>13.6</td><td>1.5</td><td>9.4</td><td>7.5E-21</td><td>8.1E-20</td><td>BT20/MB468</td><td>MITO</td></tr>
<tr><td>>am_MI0000809|hsa-mir-151a&WithFlank&8|-|140732558|140732659@53.73.21&[MIMAT0000757&hsa-miR-151a-3p&offsets|0|0;m-30&8|-|140732587|140732607&offsets|0|0]|CTAGACTGAAGCTCCTTGAGG</td><td>>am_MI0000809|hsa-mir-151a&WithFlank&8|-|140732558|140732659</td><td>53</td><td>73</td><td>21</td><td>CTAGACTGAAGCTCCTTGAGG</td><td>46425.2</td><td>5.5</td><td>0.6</td><td>9.4</td><td>6.0E-21</td><td>8.1E-20</td><td>MB231/MB468</td><td>NUC</td></tr>
<tr><td>>hg38_21_+_8433172_8446622_RNA45SN1_45S_50ntFlanks@11235.11252.18|CTTCCCGTGGATCGCCCC</td><td>>hg38_21_+_8433172_8446622_RNA45SN1_45S_50ntFlanks</td><td>11235</td><td>11252</td><td>18</td><td>CTTCCCGTGGATCGCCCC</td><td>2376.4</td><td>-8.8</td><td>0.9</td><td>-9.4</td><td>6.7E-21</td><td>8.1E-20</td><td>MB231/MB468</td><td>MITO</td></tr>
<tr><td>>hg38_21_+_8433172_8446622_RNA45SN1_45S_50ntFlanks@12931.12956.26|GCGCTAAACCATTCGTAGACGACCTG</td><td>>hg38_21_+_8433172_8446622_RNA45SN1_45S_50ntFlanks</td><td>12931</td><td>12956</td><td>26</td><td>GCGCTAAACCATTCGTAGACGACCTG</td><td>1152.6</td><td>-5.0</td><td>0.5</td><td>-9.4</td><td>6.0E-21</td><td>8.1E-20</td><td>MB231/MB468</td><td>NUC</td></tr>
<tr><td>>hg38_21_+_8433172_8446622_RNA45SN1_45S_50ntFlanks@12405.12435.31|CTTTTTGATCCTTCGATGTCGGCTCTTCCTA</td><td>>hg38_21_+_8433172_8446622_RNA45SN1_45S_50ntFlanks</td><td>12405</td><td>12435</td><td>31</td><td>CTTTTTGATCCTTCGATGTCGGCTCTTCCTA</td><td>21579.2</td><td>-11.2</td><td>1.2</td><td>-9.4</td><td>6.0E-21</td><td>8.1E-20</td><td>MB231/MB468</td><td>NUC</td></tr>
<tr><td>>hg38_21_+_8433172_8446622_RNA45SN1_45S_50ntFlanks@6740.6770.31|CATCGACACTTCGAACGCACTTGCGGCCCCG</td><td>>hg38_21_+_8433172_8446622_RNA45SN1_45S_50ntFlanks</td><td>6740</td><td>6770</td><td>31</td><td>CATCGACACTTCGAACGCACTTGCGGCCCCG</td><td>4969.7</td><td>-11.2</td><td>1.2</td><td>-9.4</td><td>6.7E-21</td><td>8.2E-20</td><td>MB231/MB468</td><td>MITO</td></tr>
<tr><td>>hg38_21_+_8433172_8446622_RNA45SN1_45S_50ntFlanks@12419.12464.46|GATGTCGGCTCTTCCTATCATTGTGAAGCAGAATTCACCAAGCGTT</td><td>>hg38_21_+_8433172_8446622_RNA45SN1_45S_50ntFlanks</td><td>12419</td><td>12464</td><td>46</td><td>GATGTCGGCTCTTCCTATCATTGTGAAGCAGAATTCACCAAGCGTT</td><td>225.5</td><td>11.8</td><td>1.3</td><td>9.4</td><td>3.6E-21</td><td>8.3E-20</td><td>MB231/MB468</td><td>CYTO</td></tr>
<tr><td>>hg38_21_+_8433172_8446622_RNA45SN1_45S_50ntFlanks@6700.6718.19|CTAGCTGCGAGAATTAATG</td><td>>hg38_21_+_8433172_8446622_RNA45SN1_45S_50ntFlanks</td><td>6700</td><td>6718</td><td>19</td><td>CTAGCTGCGAGAATTAATG</td><td>10460.8</td><td>-6.5</td><td>0.7</td><td>-9.4</td><td>6.2E-21</td><td>8.4E-20</td><td>MB231/MB468</td><td>NUC</td></tr>
<tr><td>>hg38_21_+_8433172_8446622_RNA45SN1_45S_50ntFlanks@8154.8181.28|CCGCTCGTGGGGGGCCCAAGTCCTTCTG</td><td>>hg38_21_+_8433172_8446622_RNA45SN1_45S_50ntFlanks</td><td>8154</td><td>8181</td><td>28</td><td>CCGCTCGTGGGGGGCCCAAGTCCTTCTG</td><td>1519.2</td><td>-14.4</td><td>1.5</td><td>-9.4</td><td>4.5E-21</td><td>8.4E-20</td><td>MB231/MB468</td><td>MP</td></tr>
<tr><td>>hg38_21_+_8433172_8446622_RNA45SN1_45S_50ntFlanks@11209.11254.46|CGGCGACTCTGGACGCGAGCCGGGCCCTTCCCGTGGATCGCCCCAG</td><td>>hg38_21_+_8433172_8446622_RNA45SN1_45S_50ntFlanks</td><td>11209</td><td>11254</td><td>46</td><td>CGGCGACTCTGGACGCGAGCCGGGCCCTTCCCGTGGATCGCCCCAG</td><td>39674.2</td><td>-9.5</td><td>1.0</td><td>-9.4</td><td>6.3E-21</td><td>8.5E-20</td><td>MB231/MB468</td><td>NUC</td></tr>
<tr><td>>hg38_21_+_8433172_8446622_RNA45SN1_45S_50ntFlanks@12324.12344.21|TTGGGTTTTAAGCAGGAGGTG</td><td>>hg38_21_+_8433172_8446622_RNA45SN1_45S_50ntFlanks</td><td>12324</td><td>12344</td><td>21</td><td>TTGGGTTTTAAGCAGGAGGTG</td><td>3399.2</td><td>-8.2</td><td>0.9</td><td>-9.4</td><td>3.7E-21</td><td>8.5E-20</td><td>MB231/MB468</td><td>CYTO</td></tr>
<tr><td>>MI0000067|hsa-let-7f-1&WithFlank&9|+|94176341|94176439@13.34.22(+1C);MI0000068|hsa-let-7f-2&WithFlank&X|-|53557186|53557280@14.35.22(+1C)&[MIMAT0000067&hsa-let-7f-5p&offsets|0|0(+1C);m-20&9|+|94176353|94176374&offsets|0|0(+1C)];[MIMAT0000067_1&hsa-let-7f-5p&offsets|0|0(+1C);m-19&X|-|53557246|53557267&offsets|0|0(+1C)]|TGAGGTAGTAGATTGTATAGTTC</td><td>>MI0000067|hsa-let-7f-1&WithFlank&9|+|94176341|94176439</td><td>13</td><td>34</td><td>22(+1C)</td><td>TGAGGTAGTAGATTGTATAGTTC</td><td>282.1</td><td>4.7</td><td>0.5</td><td>9.4</td><td>6.4E-21</td><td>8.6E-20</td><td>MB231/MB468</td><td>NUC</td></tr>
<tr><td>>hg38_21_+_8433172_8446622_RNA45SN1_45S_50ntFlanks@8380.8402.23|CAAGAGGGCGTGAAACCGTTAAG</td><td>>hg38_21_+_8433172_8446622_RNA45SN1_45S_50ntFlanks</td><td>8380</td><td>8402</td><td>23</td><td>CAAGAGGGCGTGAAACCGTTAAG</td><td>2959.9</td><td>-13.6</td><td>1.5</td><td>-9.4</td><td>8.0E-21</td><td>8.6E-20</td><td>BT20/MB468</td><td>MITO</td></tr>
<tr><td>>MI0000449|hsa-mir-132&WithFlank&17|-|2049902|2050014@65.85.21(+1A)&[MIMAT0000426&hsa-miR-132-3p&offsets|0|-1(+1A);m-138&17|-|2049929|2049950&offsets|0|-1(+1A)]|TAACAGTCTACAGCCATGGTCA</td><td>>MI0000449|hsa-mir-132&WithFlank&17|-|2049902|2050014</td><td>65</td><td>85</td><td>21(+1A)</td><td>TAACAGTCTACAGCCATGGTCA</td><td>350.9</td><td>12.7</td><td>1.4</td><td>9.3</td><td>1.1E-20</td><td>8.6E-20</td><td>BT20/MB468</td><td>CYTO</td></tr>
<tr><td>>hg38_21_+_8433172_8446622_RNA45SN1_45S_50ntFlanks@5473.5516.44|CGGCCCTGGCGGAGCGCTGAGAAGACGGTCGAACTTGACTATCT</td><td>>hg38_21_+_8433172_8446622_RNA45SN1_45S_50ntFlanks</td><td>5473</td><td>5516</td><td>44</td><td>CGGCCCTGGCGGAGCGCTGAGAAGACGGTCGAACTTGACTATCT</td><td>3420.4</td><td>7.3</td><td>0.8</td><td>9.3</td><td>1.1E-20</td><td>8.6E-20</td><td>BT20/MB468</td><td>CYTO</td></tr>
<tr><td>>hg38_21_+_8433172_8446622_RNA45SN1_45S_50ntFlanks@12927.12956.30|CCTGGCGCTAAACCATTCGTAGACGACCTG</td><td>>hg38_21_+_8433172_8446622_RNA45SN1_45S_50ntFlanks</td><td>12927</td><td>12956</td><td>30</td><td>CCTGGCGCTAAACCATTCGTAGACGACCTG</td><td>1294.8</td><td>-10.3</td><td>1.1</td><td>-9.4</td><td>7.1E-21</td><td>8.7E-20</td><td>MB231/MB468</td><td>MITO</td></tr>
<tr><td>>hg38_21_+_8433172_8446622_RNA45SN1_45S_50ntFlanks@10458.10497.40|ATGGCCTCCGTTGCCCTCGGCCGATCGAAAGGGAGTCGGG</td><td>>hg38_21_+_8433172_8446622_RNA45SN1_45S_50ntFlanks</td><td>10458</td><td>10497</td><td>40</td><td>ATGGCCTCCGTTGCCCTCGGCCGATCGAAAGGGAGTCGGG</td><td>1051.8</td><td>-13.9</td><td>1.5</td><td>-9.4</td><td>4.6E-21</td><td>8.7E-20</td><td>MB231/MB468</td><td>MP</td></tr>
<tr><td>>hg38_21_+_8433172_8446622_RNA45SN1_45S_50ntFlanks@11933.11950.18|AATAAGTGGGAGGCCCCC</td><td>>hg38_21_+_8433172_8446622_RNA45SN1_45S_50ntFlanks</td><td>11933</td><td>11950</td><td>18</td><td>AATAAGTGGGAGGCCCCC</td><td>1051.3</td><td>-13.9</td><td>1.5</td><td>-9.4</td><td>4.6E-21</td><td>8.7E-20</td><td>MB231/MB468</td><td>MP</td></tr>
<tr><td>>hg38_21_+_8433172_8446622_RNA45SN1_45S_50ntFlanks@6664.6686.23|TGGATCACTCGGCTCGTGCGTCG</td><td>>hg38_21_+_8433172_8446622_RNA45SN1_45S_50ntFlanks</td><td>6664</td><td>6686</td><td>23</td><td>TGGATCACTCGGCTCGTGCGTCG</td><td>622.1</td><td>-13.1</td><td>1.4</td><td>-9.4</td><td>4.6E-21</td><td>8.7E-20</td><td>MB231/MB468</td><td>MP</td></tr>
<tr><td>>hg38_21_+_8433172_8446622_RNA45SN1_45S_50ntFlanks@8182.8224.43|ATCGAGGCCCAGCCCGTGGACGGTGTGAGGCCGGTAGCGGCCC</td><td>>hg38_21_+_8433172_8446622_RNA45SN1_45S_50ntFlanks</td><td>8182</td><td>8224</td><td>43</td><td>ATCGAGGCCCAGCCCGTGGACGGTGTGAGGCCGGTAGCGGCCC</td><td>450.0</td><td>-13.2</td><td>1.4</td><td>-9.3</td><td>2.2E-20</td><td>8.7E-20</td><td>BT20/MB468</td><td>NUC</td></tr>
<tr><td>>MI0000077|hsa-mir-21&WithFlank&17|+|59841260|59841343@14.33.20(+1A)&[MIMAT0000076&hsa-miR-21-5p&offsets|0|-2(+1A);m-1&17|+|59841273|59841294&offsets|0|-2(+1A)]|TAGCTTATCAGACTGATGTTA</td><td>>MI0000077|hsa-mir-21&WithFlank&17|+|59841260|59841343</td><td>14</td><td>33</td><td>20(+1A)</td><td>TAGCTTATCAGACTGATGTTA</td><td>769.2</td><td>6.8</td><td>0.7</td><td>9.4</td><td>6.5E-21</td><td>8.7E-20</td><td>MB231/MB468</td><td>NUC</td></tr>
<tr><td>>hg38_21_+_8433172_8446622_RNA45SN1_45S_50ntFlanks@12927.12956.30|CCTGGCGCTAAACCATTCGTAGACGACCTG</td><td>>hg38_21_+_8433172_8446622_RNA45SN1_45S_50ntFlanks</td><td>12927</td><td>12956</td><td>30</td><td>CCTGGCGCTAAACCATTCGTAGACGACCTG</td><td>515.8</td><td>-12.9</td><td>1.4</td><td>-9.4</td><td>4.7E-21</td><td>8.7E-20</td><td>MB231/MB468</td><td>MP</td></tr>
<tr><td>>MI0000077|hsa-mir-21&WithFlank&17|+|59841260|59841343@14.34.21&[MIMAT0000076&hsa-miR-21-5p&offsets|0|-1;m-1&17|+|59841273|59841294&offsets|0|-1]|TAGCTTATCAGACTGATGTTG</td><td>>MI0000077|hsa-mir-21&WithFlank&17|+|59841260|59841343</td><td>14</td><td>34</td><td>21</td><td>TAGCTTATCAGACTGATGTTG</td><td>9846.5</td><td>6.8</td><td>0.7</td><td>9.4</td><td>6.6E-21</td><td>8.7E-20</td><td>MB231/MB468</td><td>NUC</td></tr>
<tr><td>>MI0000114|hsa-mir-107&WithFlank&10|-|89592741|89592833@56.76.21;MI0000108|hsa-mir-103a-2&WithFlank&20|+|3917488|3917577@54.74.21;MI0000109|hsa-mir-103a-1&WithFlank&5|-|168560890|168560979@54.74.21&[MIMAT0000104&hsa-miR-107&offsets|0|-2;m-78&10|-|89592758|89592778&offsets|0|0];[MIMAT0000101&hsa-miR-103a-3p&offsets|0|-2;m-12&20|+|3917541|3917563&offsets|0|-2];[MIMAT0000101_1&hsa-miR-103a-3p&offsets|0|-2;m-13&5|-|168560904|168560926&offsets|0|-2;m-44220&5|-|168560908|168560926&offsets|0|+2]|AGCAGCATTGTACAGGGCTAT</td><td>>MI0000114|hsa-mir-107&WithFlank&10|-|89592741|89592833</td><td>56</td><td>76</td><td>21</td><td>AGCAGCATTGTACAGGGCTAT</td><td>742.0</td><td>6.8</td><td>0.7</td><td>9.4</td><td>6.6E-21</td><td>8.7E-20</td><td>MB231/MB468</td><td>NUC</td></tr>
<tr><td>>hg38_21_+_8433172_8446622_RNA45SN1_45S_50ntFlanks@12007.12032.26|GCCGCCGGTGAAATACCACTACTCTG</td><td>>hg38_21_+_8433172_8446622_RNA45SN1_45S_50ntFlanks</td><td>12007</td><td>12032</td><td>26</td><td>GCCGCCGGTGAAATACCACTACTCTG</td><td>10488.0</td><td>-8.3</td><td>0.8</td><td>-9.8</td><td>9.0E-23</td><td>8.8E-20</td><td>BT20/MB468</td><td>MP</td></tr>
<tr><td>>hg38_21_+_8433172_8446622_RNA45SN1_45S_50ntFlanks@12973.13001.29|TCGTACGTAGCAGAGCAGCTCCCTCGCTG</td><td>>hg38_21_+_8433172_8446622_RNA45SN1_45S_50ntFlanks</td><td>12973</td><td>13001</td><td>29</td><td>TCGTACGTAGCAGAGCAGCTCCCTCGCTG</td><td>650.0</td><td>-12.3</td><td>1.3</td><td>-9.4</td><td>6.6E-21</td><td>8.8E-20</td><td>MB231/MB468</td><td>NUC</td></tr>
<tr><td>>hg38_21_+_8433172_8446622_RNA45SN1_45S_50ntFlanks@3841.3865.25|TCCTACTTGGATAACTGTGGTAATT</td><td>>hg38_21_+_8433172_8446622_RNA45SN1_45S_50ntFlanks</td><td>3841</td><td>3865</td><td>25</td><td>TCCTACTTGGATAACTGTGGTAATT</td><td>460.9</td><td>5.7</td><td>0.6</td><td>9.3</td><td>1.1E-20</td><td>8.8E-20</td><td>BT20/MB468</td><td>CYTO</td></tr>
<tr><td>>MI0000087|hsa-mir-29a&WithFlank&7|-|130876741|130876816@47.68.22(+1U)&[MIMAT0000086&hsa-miR-29a-3p&offsets|-1|-1(+1U);m-26&7|-|130876749|130876769&offsets|-1|0(+1U)]|CTAGCACCATCTGAAATCGGTTT</td><td>>MI0000087|hsa-mir-29a&WithFlank&7|-|130876741|130876816</td><td>47</td><td>68</td><td>22(+1U)</td><td>CTAGCACCATCTGAAATCGGTTT</td><td>1292.5</td><td>10.7</td><td>1.1</td><td>9.4</td><td>6.7E-21</td><td>8.9E-20</td><td>MB231/MB468</td><td>NUC</td></tr>
<tr><td>>hg38_21_+_8433172_8446622_RNA45SN1_45S_50ntFlanks@10383.10424.42|TCCATGTGAACAGCAGTTGAACATGGGTCAGTCGGTCCTGAG</td><td>>hg38_21_+_8433172_8446622_RNA45SN1_45S_50ntFlanks</td><td>10383</td><td>10424</td><td>42</td><td>TCCATGTGAACAGCAGTTGAACATGGGTCAGTCGGTCCTGAG</td><td>3489.9</td><td>8.8</td><td>0.9</td><td>9.4</td><td>3.9E-21</td><td>8.9E-20</td><td>MB231/MB468</td><td>CYTO</td></tr>
<tr><td>>hg38_21_+_8433172_8446622_RNA45SN1_45S_50ntFlanks@4722.4748.27|TCATTAATCAAGAACGAAAGTCGGAGG</td><td>>hg38_21_+_8433172_8446622_RNA45SN1_45S_50ntFlanks</td><td>4722</td><td>4748</td><td>27</td><td>TCATTAATCAAGAACGAAAGTCGGAGG</td><td>921.8</td><td>-13.6</td><td>1.5</td><td>-9.4</td><td>7.4E-21</td><td>9.0E-20</td><td>MB231/MB468</td><td>MITO</td></tr>
<tr><td>>MI0000088|hsa-mir-30a&WithFlank&6|-|71403545|71403627@53.73.21&[MIMAT0000088&hsa-miR-30a-3p&offsets|0|-1;m-38&6|-|71403554|71403575&offsets|0|-1]|CTTTCAGTCGGATGTTTGCAG</td><td>>MI0000088|hsa-mir-30a&WithFlank&6|-|71403545|71403627</td><td>53</td><td>73</td><td>21</td><td>CTTTCAGTCGGATGTTTGCAG</td><td>1581.5</td><td>5.7</td><td>0.6</td><td>9.4</td><td>6.8E-21</td><td>9.1E-20</td><td>MB231/MB468</td><td>NUC</td></tr>
<tr><td>>hg38_21_+_8433172_8446622_RNA45SN1_45S_50ntFlanks@11666.11705.40|TCTGCCCAGTGCTCTGAATGTCAAAGTGAAGAAATTCAAT</td><td>>hg38_21_+_8433172_8446622_RNA45SN1_45S_50ntFlanks</td><td>11666</td><td>11705</td><td>40</td><td>TCTGCCCAGTGCTCTGAATGTCAAAGTGAAGAAATTCAAT</td><td>163.4</td><td>11.4</td><td>1.2</td><td>9.4</td><td>3.9E-21</td><td>9.1E-20</td><td>MB231/MB468</td><td>CYTO</td></tr>
<tr><td>>hg38_21_+_8433172_8446622_RNA45SN1_45S_50ntFlanks@11729.11750.22|TAACTATGACTCTCTTAAGGTA</td><td>>hg38_21_+_8433172_8446622_RNA45SN1_45S_50ntFlanks</td><td>11729</td><td>11750</td><td>22</td><td>TAACTATGACTCTCTTAAGGTA</td><td>1607.5</td><td>-4.5</td><td>0.5</td><td>-9.3</td><td>1.2E-20</td><td>9.1E-20</td><td>BT20/MB468</td><td>CYTO</td></tr>
<tr><td>>hg38_21_+_8433172_8446622_RNA45SN1_45S_50ntFlanks@10689.10706.18|GCGTCCGGTGAGCTCTCG</td><td>>hg38_21_+_8433172_8446622_RNA45SN1_45S_50ntFlanks</td><td>10689</td><td>10706</td><td>18</td><td>GCGTCCGGTGAGCTCTCG</td><td>2831.1</td><td>-14.4</td><td>1.5</td><td>-9.4</td><td>7.0E-21</td><td>9.2E-20</td><td>MB231/MB468</td><td>NUC</td></tr>
</table>

<table>
<tr><td></td><td></td><td></td><td></td><td></td><td></td><td></td><td></td><td></td><td></td><td></td><td></td><td></td><td></td></tr>
<tr><td>>MI0000439|hsa-mir-23b&WithFlank&9|+|95085202|95085310@67.86.20(+1A)&[MIMAT0000418&hsa-miR-23b-3p&offsets|+3|0(+1A);m-42&9|+|95085265|95085285&offsets|+3|+2(+1A)]|ACATTGCCAGGGATTACCACA</td><td>>MI0000439|hsa-mir-23b&WithFlank&9|+|95085202|95085310</td><td>67</td><td>86</td><td>20(+1A)</td><td>ACATTGCCAGGGATTACCACA</td><td>133.2</td><td>9.9</td><td>1.1</td><td>9.2</td><td>2.3E-20</td><td>9.3E-20</td><td>BT20/MB468</td><td>NUC</td></tr>
<tr><td>>hg38_21_+_8433172_8446622_RNA45SN1_45S_50ntFlanks@10797.10825.29|GCATGTTGGAACAATGTAGGTAAGGGAAG</td><td>>hg38_21_+_8433172_8446622_RNA45SN1_45S_50ntFlanks</td><td>10797</td><td>10825</td><td>29</td><td>GCATGTTGGAACAATGTAGGTAAGGGAAG</td><td>583.7</td><td>-13.1</td><td>1.4</td><td>-9.4</td><td>5.0E-21</td><td>9.4E-20</td><td>MB231/MB468</td><td>MP</td></tr>
<tr><td>>hg38_21_+_8433172_8446622_RNA45SN1_45S_50ntFlanks@4367.4391.25|CAGTTAAAAAGCTCGTAGTTGGATC</td><td>>hg38_21_+_8433172_8446622_RNA45SN1_45S_50ntFlanks</td><td>4367</td><td>4391</td><td>25</td><td>CAGTTAAAAAGCTCGTAGTTGGATC</td><td>810.1</td><td>-13.4</td><td>1.4</td><td>-9.4</td><td>7.8E-21</td><td>9.5E-20</td><td>MB231/MB468</td><td>MITO</td></tr>
<tr><td>>hg38_21_+_8433172_8446622_RNA45SN1_45S_50ntFlanks@12893.12927.35|CCTCGCCCGTCACGCACCGCACGTTCGTGGGGAAC</td><td>>hg38_21_+_8433172_8446622_RNA45SN1_45S_50ntFlanks</td><td>12893</td><td>12927</td><td>35</td><td>CCTCGCCCGTCACGCACCGCACGTTCGTGGGGAAC</td><td>562.3</td><td>-13.0</td><td>1.4</td><td>-9.4</td><td>5.1E-21</td><td>9.6E-20</td><td>MB231/MB468</td><td>MP</td></tr>
<tr><td>>hg38_21_+_8433172_8446622_RNA45SN1_45S_50ntFlanks@12643.12661.19|ATCTGTGGGATTATGACTG</td><td>>hg38_21_+_8433172_8446622_RNA45SN1_45S_50ntFlanks</td><td>12643</td><td>12661</td><td>19</td><td>ATCTGTGGGATTATGACTG</td><td>1041.3</td><td>-13.9</td><td>1.5</td><td>-9.4</td><td>5.2E-21</td><td>9.6E-20</td><td>MB231/MB468</td><td>MP</td></tr>
<tr><td>>hg38_21_+_8433172_8446622_RNA45SN1_45S_50ntFlanks@9504.9523.20|GTGAACTATGCCTGGGCAGG</td><td>>hg38_21_+_8433172_8446622_RNA45SN1_45S_50ntFlanks</td><td>9504</td><td>9523</td><td>20</td><td>GTGAACTATGCCTGGGCAGG</td><td>5355.3</td><td>-10.3</td><td>1.1</td><td>-9.2</td><td>2.4E-20</td><td>9.6E-20</td><td>BT20/MB468</td><td>NUC</td></tr>
<tr><td>>hg38_21_+_8433172_8446622_RNA45SN1_45S_50ntFlanks@4045.4063.19|CCATTCGAACGTCTGCCCT</td><td>>hg38_21_+_8433172_8446622_RNA45SN1_45S_50ntFlanks</td><td>4045</td><td>4063</td><td>19</td><td>CCATTCGAACGTCTGCCCT</td><td>2542.4</td><td>-11.3</td><td>1.2</td><td>-9.4</td><td>8.0E-21</td><td>9.7E-20</td><td>MB231/MB468</td><td>MITO</td></tr>
<tr><td>>hg38_21_+_8433172_8446622_RNA45SN1_45S_50ntFlanks@10351.10384.34|CGAGAACTTTGAAGGCCGAAGTGGAGAAGGGTTC</td><td>>hg38_21_+_8433172_8446622_RNA45SN1_45S_50ntFlanks</td><td>10351</td><td>10384</td><td>34</td><td>CGAGAACTTTGAAGGCCGAAGTGGAGAAGGGTTC</td><td>2369.7</td><td>-4.7</td><td>0.5</td><td>-9.3</td><td>1.3E-20</td><td>9.8E-20</td><td>BT20/MB468</td><td>CYTO</td></tr>
<tr><td>>hg38_21_+_8433172_8446622_RNA45SN1_45S_50ntFlanks@12347.12365.19|AGAAAAGTTACCACAGGGA</td><td>>hg38_21_+_8433172_8446622_RNA45SN1_45S_50ntFlanks</td><td>12347</td><td>12365</td><td>19</td><td>AGAAAAGTTACCACAGGGA</td><td>782.4</td><td>-9.5</td><td>1.0</td><td>-9.4</td><td>4.3E-21</td><td>9.9E-20</td><td>MB231/MB468</td><td>CYTO</td></tr>
<tr><td>>hg38_21_+_8433172_8446622_RNA45SN1_45S_50ntFlanks@8355.8373.19|TGAAAAGAACTTTGAAGAG</td><td>>hg38_21_+_8433172_8446622_RNA45SN1_45S_50ntFlanks</td><td>8355</td><td>8373</td><td>19</td><td>TGAAAAGAACTTTGAAGAG</td><td>661.2</td><td>13.8</td><td>1.5</td><td>9.3</td><td>9.3E-21</td><td>1.0E-19</td><td>BT20/MB468</td><td>MITO</td></tr>
<tr><td>>hg38_21_+_8433172_8446622_RNA45SN1_45S_50ntFlanks@11768.11792.25|CTAATTAGTGACGCGCATGAATGGA</td><td>>hg38_21_+_8433172_8446622_RNA45SN1_45S_50ntFlanks</td><td>11768</td><td>11792</td><td>25</td><td>CTAATTAGTGACGCGCATGAATGGA</td><td>922.7</td><td>-13.6</td><td>1.5</td><td>-9.4</td><td>8.2E-21</td><td>1.0E-19</td><td>MB231/MB468</td><td>MITO</td></tr>
<tr><td>>hg38_21_+_8433172_8446622_RNA45SN1_45S_50ntFlanks@5472.5507.36|ACGGCCCTGGCGGAGCGCTGAGAAGACGGTCGAACT</td><td>>hg38_21_+_8433172_8446622_RNA45SN1_45S_50ntFlanks</td><td>5472</td><td>5507</td><td>36</td><td>ACGGCCCTGGCGGAGCGCTGAGAAGACGGTCGAACT</td><td>1968.1</td><td>7.0</td><td>0.7</td><td>9.4</td><td>4.4E-21</td><td>1.0E-19</td><td>MB231/MB468</td><td>CYTO</td></tr>
<tr><td>>hg38_21_+_8433172_8446622_RNA45SN1_45S_50ntFlanks@11920.11951.32|ATGAGAGGTGTAGAATAAGTGGGAGGCCCCCG</td><td>>hg38_21_+_8433172_8446622_RNA45SN1_45S_50ntFlanks</td><td>11920</td><td>11951</td><td>32</td><td>ATGAGAGGTGTAGAATAAGTGGGAGGCCCCCG</td><td>840.6</td><td>-13.6</td><td>1.4</td><td>-9.4</td><td>5.4E-21</td><td>1.0E-19</td><td>MB231/MB468</td><td>MP</td></tr>
<tr><td>>hg38_21_+_8433172_8446622_RNA45SN1_45S_50ntFlanks@11315.11338.24|CCACCCCACGTCTCGTCGCGCGCG</td><td>>hg38_21_+_8433172_8446622_RNA45SN1_45S_50ntFlanks</td><td>11315</td><td>11338</td><td>24</td><td>CCACCCCACGTCTCGTCGCGCGCG</td><td>1411.6</td><td>-14.2</td><td>1.5</td><td>-9.4</td><td>8.4E-21</td><td>1.0E-19</td><td>MB231/MB468</td><td>MITO</td></tr>
<tr><td>>hg38_21_+_8433172_8446622_RNA45SN1_45S_50ntFlanks@12991.13010.20|CTCCCTCGCTGCGATCTATT</td><td>>hg38_21_+_8433172_8446622_RNA45SN1_45S_50ntFlanks</td><td>12991</td><td>13010</td><td>20</td><td>CTCCCTCGCTGCGATCTATT</td><td>757.0</td><td>-13.3</td><td>1.4</td><td>-9.4</td><td>8.4E-21</td><td>1.0E-19</td><td>MB231/MB468</td><td>MITO</td></tr>
<tr><td>>hg38_21_+_8433172_8446622_RNA45SN1_45S_50ntFlanks@6751.6770.20|CGAACGCACTTGCGGCCCCG</td><td>>hg38_21_+_8433172_8446622_RNA45SN1_45S_50ntFlanks</td><td>6751</td><td>6770</td><td>20</td><td>CGAACGCACTTGCGGCCCCG</td><td>867.7</td><td>-13.6</td><td>1.4</td><td>-9.4</td><td>5.5E-21</td><td>1.0E-19</td><td>MB231/MB468</td><td>MP</td></tr>
<tr><td>>hg38_21_+_8433172_8446622_RNA45SN1_45S_50ntFlanks@11561.11585.25|GCGCCTAGCAGCCGACTTAGAACTG</td><td>>hg38_21_+_8433172_8446622_RNA45SN1_45S_50ntFlanks</td><td>11561</td><td>11585</td><td>25</td><td>GCGCCTAGCAGCCGACTTAGAACTG</td><td>2360.7</td><td>-10.9</td><td>1.2</td><td>-9.4</td><td>5.5E-21</td><td>1.0E-19</td><td>MB231/MB468</td><td>MP</td></tr>
<tr><td>>MI0000067|hsa-let-7f-1&WithFlank&9|+|94176341|94176439@13.33.21(+1C);MI0000068|hsa-let-7f-2&WithFlank&X|-|53557186|53557280@14.34.21(+1C)&[MIMAT0000067&hsa-let-7f-5p&offsets|0|-1(+1C);m-20&9|+|94176353|94176374&offsets|0|-1(+1C)];[MIMAT0000067_1&hsa-let-7f-5p&offsets|0|-1(+1C);m-19&X|-|53557246|53557267&offsets|0|-1(+1C)]|TGAGGTAGTAGATTGTATAGTC</td><td>>MI0000067|hsa-let-7f-1&WithFlank&9|+|94176341|94176439</td><td>13</td><td>33</td><td>21(+1C)</td><td>TGAGGTAGTAGATTGTATAGTC</td><td>14263.7</td><td>4.5</td><td>0.5</td><td>9.4</td><td>7.8E-21</td><td>1.0E-19</td><td>MB231/MB468</td><td>NUC</td></tr>
<tr><td>>hg38_21_+_8433172_8446622_RNA45SN1_45S_50ntFlanks@12995.13017.23|CTCGCTGCGATCTATTGAAAGTC</td><td>>hg38_21_+_8433172_8446622_RNA45SN1_45S_50ntFlanks</td><td>12995</td><td>13017</td><td>23</td><td>CTCGCTGCGATCTATTGAAAGTC</td><td>577.5</td><td>4.5</td><td>0.5</td><td>9.4</td><td>7.8E-21</td><td>1.0E-19</td><td>MB231/MB468</td><td>NUC</td></tr>
<tr><td>>hg38_21_+_8433172_8446622_RNA45SN1_45S_50ntFlanks@12416.12440.25|TTCGATGTCGGCTCTTCCTATCATT</td><td>>hg38_21_+_8433172_8446622_RNA45SN1_45S_50ntFlanks</td><td>12416</td><td>12440</td><td>25</td><td>TTCGATGTCGGCTCTTCCTATCATT</td><td>3903.8</td><td>6.5</td><td>0.7</td><td>9.4</td><td>4.5E-21</td><td>1.0E-19</td><td>MB231/MB468</td><td>CYTO</td></tr>
<tr><td>>trnaMT_ValTAC_MT_+_1602_1670@48.70.23|CAACTTAACTTGACCGCTCTGAC</td><td>>trnaMT_ValTAC_MT_+_1602_1670</td><td>48</td><td>70</td><td>23</td><td>CAACTTAACTTGACCGCTCTGAC</td><td>897.6</td><td>11.8</td><td>1.3</td><td>9.3</td><td>1.3E-20</td><td>1.0E-19</td><td>BT20/MB468</td><td>CYTO</td></tr>
<tr><td>>hg38_21_+_8433172_8446622_RNA45SN1_45S_50ntFlanks@8380.8403.24|CAAGAGGGCGTGAAACCGTTAAGA</td><td>>hg38_21_+_8433172_8446622_RNA45SN1_45S_50ntFlanks</td><td>8380</td><td>8403</td><td>24</td><td>CAAGAGGGCGTGAAACCGTTAAGA</td><td>3224.9</td><td>-8.4</td><td>0.9</td><td>-9.4</td><td>8.7E-21</td><td>1.0E-19</td><td>MB231/MB468</td><td>MITO</td></tr>
<tr><td>>hg38_21_+_8433172_8446622_RNA45SN1_45S_50ntFlanks@12605.12624.20|TATGTGCTTGGCTGAGGAGC</td><td>>hg38_21_+_8433172_8446622_RNA45SN1_45S_50ntFlanks</td><td>12605</td><td>12624</td><td>20</td><td>TATGTGCTTGGCTGAGGAGC</td><td>6080.0</td><td>-5.3</td><td>0.6</td><td>-9.4</td><td>8.0E-21</td><td>1.1E-19</td><td>MB231/MB468</td><td>NUC</td></tr>
<tr><td>>hg38_21_+_8433172_8446622_RNA45SN1_45S_50ntFlanks@12991.13028.38|CTCCCTCGCTGCGATCTATTGAAAGTCAGCCCTCGACA</td><td>>hg38_21_+_8433172_8446622_RNA45SN1_45S_50ntFlanks</td><td>12991</td><td>13028</td><td>38</td><td>CTCCCTCGCTGCGATCTATTGAAAGTCAGCCCTCGACA</td><td>1297.1</td><td>-14.8</td><td>1.6</td><td>-9.2</td><td>2.7E-20</td><td>1.1E-19</td><td>BT20/MB468</td><td>NUC</td></tr>
<tr><td>>hg38_1_-_228634819_228635039_RNA5S12_5S_50ntFlanks@145.171.27|CTGGGAATACCGGGTGCTGTAGGCTTT</td><td>>hg38_1_-_228634819_228635039_RNA5S12_5S_50ntFlanks</td><td>145</td><td>171</td><td>27</td><td>CTGGGAATACCGGGTGCTGTAGGCTTT</td><td>5088.1</td><td>7.2</td><td>0.8</td><td>9.4</td><td>4.7E-21</td><td>1.1E-19</td><td>MB231/MB468</td><td>CYTO</td></tr>
<tr><td>>MI0000103|hsa-mir-101-1&WithFlank&1|-|65058428|65058514@53.74.22;MI0000739|hsa-mir-101-2&WithFlank&9|+|4850291|4850381@55.76.22&[MIMAT0000099&hsa-miR-101-3p&offsets|0|+1;m-17&1|-|65058443|65058462&offsets|0|+2];[MIMAT0000099_1&hsa-miR-101-3p&offsets|0|+1;m-16&9|+|4850345|4850364&offsets|0|+2]|TACAGTACTGTGATAACTGAAG</td><td>>MI0000103|hsa-mir-101-1&WithFlank&1|-|65058428|65058514</td><td>53</td><td>74</td><td>22</td><td>TACAGTACTGTGATAACTGAAG</td><td>472.1</td><td>5.5</td><td>0.6</td><td>9.3</td><td>9.9E-21</td><td>1.1E-19</td><td>BT20/MB468</td><td>MITO</td></tr>
<tr><td>>hg38_21_+_8433172_8446622_RNA45SN1_45S_50ntFlanks@6647.6672.26|CGTACGACTCTTAGCGGTGGATCACT</td><td>>hg38_21_+_8433172_8446622_RNA45SN1_45S_50ntFlanks</td><td>6647</td><td>6672</td><td>26</td><td>CGTACGACTCTTAGCGGTGGATCACT</td><td>5751.7</td><td>6.9</td><td>0.7</td><td>9.3</td><td>9.9E-21</td><td>1.1E-19</td><td>BT20/MB468</td><td>MITO</td></tr>
<tr><td>>hg38_21_+_8433172_8446622_RNA45SN1_45S_50ntFlanks@10348.10377.30|AAACGAGAACTTTGAAGGCCGAAGTGGAGA</td><td>>hg38_21_+_8433172_8446622_RNA45SN1_45S_50ntFlanks</td><td>10348</td><td>10377</td><td>30</td><td>AAACGAGAACTTTGAAGGCCGAAGTGGAGA</td><td>704.1</td><td>-5.7</td><td>0.6</td><td>-9.2</td><td>2.7E-20</td><td>1.1E-19</td><td>BT20/MB468</td><td>NUC</td></tr>
<tr><td>>hg38_21_+_8433172_8446622_RNA45SN1_45S_50ntFlanks@10771.10815.45|CAGGTCTCCAAGGTGAACAGCCTCTGGCATGTTGGAACAATGTAG</td><td>>hg38_21_+_8433172_8446622_RNA45SN1_45S_50ntFlanks</td><td>10771</td><td>10815</td><td>45</td><td>CAGGTCTCCAAGGTGAACAGCCTCTGGCATGTTGGAACAATGTAG</td><td>713.9</td><td>-13.9</td><td>1.5</td><td>-9.2</td><td>2.7E-20</td><td>1.1E-19</td><td>BT20/MB468</td><td>NUC</td></tr>
<tr><td>>hg38_21_+_8433172_8446622_RNA45SN1_45S_50ntFlanks@12003.12032.30|GCGGGCCGCCGGTGAAATACCACTACTCTG</td><td>>hg38_21_+_8433172_8446622_RNA45SN1_45S_50ntFlanks</td><td>12003</td><td>12032</td><td>30</td><td>GCGGGCCGCCGGTGAAATACCACTACTCTG</td><td>6983.1</td><td>-6.8</td><td>0.7</td><td>-9.3</td><td>9.0E-21</td><td>1.1E-19</td><td>MB231/MB468</td><td>MITO</td></tr>
<tr><td>>hg38_21_+_8433172_8446622_RNA45SN1_45S_50ntFlanks@9178.9197.20|CGCCGTCGGGCCCGGGGGAG</td><td>>hg38_21_+_8433172_8446622_RNA45SN1_45S_50ntFlanks</td><td>9178</td><td>9197</td><td>20</td><td>CGCCGTCGGGCCCGGGGGAG</td><td>2107.2</td><td>-14.8</td><td>1.6</td><td>-9.3</td><td>9.0E-21</td><td>1.1E-19</td><td>MB231/MB468</td><td>MITO</td></tr>
<tr><td>>hg38_21_+_8433172_8446622_RNA45SN1_45S_50ntFlanks@10819.10840.22|AGGGAAGTCGGCAAGCCGGATC</td><td>>hg38_21_+_8433172_8446622_RNA45SN1_45S_50ntFlanks</td><td>10819</td><td>10840</td><td>22</td><td>AGGGAAGTCGGCAAGCCGGATC</td><td>13135.8</td><td>-11.1</td><td>1.2</td><td>-9.2</td><td>2.7E-20</td><td>1.1E-19</td><td>BT20/MB468</td><td>NUC</td></tr>
<tr><td>>hg38_21_+_8433172_8446622_RNA45SN1_45S_50ntFlanks@6646.6672.27|TCGTACGACTCTTAGCGGTGGATCACT</td><td>>hg38_21_+_8433172_8446622_RNA45SN1_45S_50ntFlanks</td><td>6646</td><td>6672</td><td>27</td><td>TCGTACGACTCTTAGCGGTGGATCACT</td><td>45590.4</td><td>7.5</td><td>0.8</td><td>9.3</td><td>1.0E-20</td><td>1.1E-19</td><td>BT20/MB468</td><td>MITO</td></tr>
<tr><td>>hg38_21_+_8433172_8446622_RNA45SN1_45S_50ntFlanks@6653.6676.24|ACTCTTAGCGGTGGATCACTCGGC</td><td>>hg38_21_+_8433172_8446622_RNA45SN1_45S_50ntFlanks</td><td>6653</td><td>6676</td><td>24</td><td>ACTCTTAGCGGTGGATCACTCGGC</td><td>787.6</td><td>6.2</td><td>0.7</td><td>9.3</td><td>1.0E-20</td><td>1.1E-19</td><td>BT20/MB468</td><td>MITO</td></tr>
<tr><td>>hg38_21_+_8433172_8446622_RNA45SN1_45S_50ntFlanks@11577.11603.27|TTAGAACTGGTGCGGACCAGGGGAATC</td><td>>hg38_21_+_8433172_8446622_RNA45SN1_45S_50ntFlanks</td><td>11577</td><td>11603</td><td>27</td><td>TTAGAACTGGTGCGGACCAGGGGAATC</td><td>1397.0</td><td>-14.3</td><td>1.5</td><td>-9.4</td><td>5.9E-21</td><td>1.1E-19</td><td>MB231/MB468</td><td>MP</td></tr>
<tr><td>>MI0016746|hsa-mir-548o-2&WithFlank&20|+|38516557|38516638@51.72.22;MI0006402|hsa-mir-548o&WithFlank&7|-|102405736|102405861@93.114.22&[MIMAT0005919&hsa-miR-548o-3p&offsets|0|0;m-431&20|+|38516607|38516628&offsets|0|0];[MIMAT0005919_1&hsa-miR-548o-3p&offsets|0|0;m-430&7|-|102405748|102405769&offsets|0|0]|CCAAAACTGCAGTTACTTTTGC</td><td>>MI0016746|hsa-mir-548o-2&WithFlank&20|+|38516557|38516638</td><td>51</td><td>72</td><td>22</td><td>CCAAAACTGCAGTTACTTTTGC</td><td>520.8</td><td>7.9</td><td>0.8</td><td>9.3</td><td>1.0E-20</td><td>1.1E-19</td><td>BT20/MB468</td><td>MITO</td></tr>
<tr><td>>trna116_GluCTC_1_-_145399233_145399304@4.33.30,trna59_GluCTC_1_+_249168447_249168518@4.33.30,trna71_GluCTC_1_-_161439189_161439260@4.33.30,trna74_GluCTC_1_-_161431809_161431880@4.33.30,trna77_GluCTC_1_-_161424398_161424469@4.33.30,trna77_GluCTC_6_+_28949976_28950047@4.33.30,trna80_GluCTC_1_-_161417018_161417089@4.33.30,trna87_GluCTC_6_-_126101393_126101464@4.33.30|CTGGTGGTCTAGTGGTTAGGATTCGGCGCT</td><td>>trna116_GluCTC_1_-_145399233_145399304</td><td>4</td><td>33</td><td>30</td><td>CTGGTGGTCTAGTGGTTAGGATTCGGCGCT</td><td>542.5</td><td>6.4</td><td>0.7</td><td>9.3</td><td>1.0E-20</td><td>1.1E-19</td><td>BT20/MB468</td><td>MITO</td></tr>
<tr><td>>hg38_21_+_8433172_8446622_RNA45SN1_45S_50ntFlanks@9166.9183.18|CCGGGCGGGTCGCGCCGT</td><td>>hg38_21_+_8433172_8446622_RNA45SN1_45S_50ntFlanks</td><td>9166</td><td>9183</td><td>18</td><td>CCGGGCGGGTCGCGCCGT</td><td>838.5</td><td>-13.6</td><td>1.4</td><td>-9.4</td><td>5.9E-21</td><td>1.1E-19</td><td>MB231/MB468</td><td>MP</td></tr>
<tr><td>>MI0000446|hsa-mir-125b-1&WithFlank&11|-|122099751|122099850@21.42.22(+1A);MI0000470|hsa-mir-125b-2&WithFlank&21|+|16590231|16590331@23.44.22(+1A)&[MIMAT0000423&hsa-miR-125b-5p&offsets|0|0(+1A);m-67&11|-|122099809|122099830&offsets|0|0(+1A)];[MIMAT0000423_1&hsa-miR-125b-5p&offsets|0|0(+1A);m-68&21|+|16590253|16590274&offsets|0|0(+1A)]|TCCCTGAGACCCTAACTTGTGAA</td><td>>MI0000446|hsa-mir-125b-1&WithFlank&11|-|122099751|122099850</td><td>21</td><td>42</td><td>22(+1A)</td><td>TCCCTGAGACCCTAACTTGTGAA</td><td>6075.7</td><td>9.6</td><td>1.0</td><td>9.4</td><td>4.9E-21</td><td>1.1E-19</td><td>MB231/MB468</td><td>CYTO</td></tr>
<tr><td>>MI0000272|hsa-mir-182&WithFlank&7|-|129770377|129770498@29.53.25&[MIMAT0000259&hsa-miR-182-5p&offsets|0|+1;m-22&7|-|129770449|129770470&offsets|0|+3]|TTTGGCAATGGTAGAACTCACACTG</td><td>>MI0000272|hsa-mir-182&WithFlank&7|-|129770377|129770498</td><td>29</td><td>53</td><td>25</td><td>TTTGGCAATGGTAGAACTCACACTG</td><td>3054.7</td><td>2.6</td><td>0.3</td><td>9.2</td><td>2.8E-20</td><td>1.1E-19</td><td>BT20/MB468</td><td>NUC</td></tr>
<tr><td>>hg38_21_+_8433172_8446622_RNA45SN1_45S_50ntFlanks@485.520.36|GATGTGGTGACGTCGTGCTCTCCCGGGCCGGGTCCG</td><td>>hg38_21_+_8433172_8446622_RNA45SN1_45S_50ntFlanks</td><td>485</td><td>520</td><td>36</td><td>GATGTGGTGACGTCGTGCTCTCCCGGGCCGGGTCCG</td><td>539.7</td><td>11.9</td><td>1.3</td><td>9.2</td><td>2.8E-20</td><td>1.1E-19</td><td>BT20/MB468</td><td>NUC</td></tr>
<tr><td>>hg38_21_+_8433172_8446622_RNA45SN1_45S_50ntFlanks@10410.10431.22|TCAGTCGGTCCTGAGAGATGGG</td><td>>hg38_21_+_8433172_8446622_RNA45SN1_45S_50ntFlanks</td><td>10410</td><td>10431</td><td>22</td><td>TCAGTCGGTCCTGAGAGATGGG</td><td>8788.7</td><td>-13.1</td><td>1.4</td><td>-9.2</td><td>2.8E-20</td><td>1.1E-19</td><td>BT20/MB468</td><td>NUC</td></tr>
<tr><td>>hg38_21_+_8433172_8446622_RNA45SN1_45S_50ntFlanks@9567.9594.28|ACGTGCAAATCGGTCGTCCGACCTGGGT</td><td>>hg38_21_+_8433172_8446622_RNA45SN1_45S_50ntFlanks</td><td>9567</td><td>9594</td><td>28</td><td>ACGTGCAAATCGGTCGTCCGACCTGGGT</td><td>549.7</td><td>-4.1</td><td>0.4</td><td>-9.3</td><td>1.4E-20</td><td>1.1E-19</td><td>BT20/MB468</td><td>CYTO</td></tr>
<tr><td>>hg38_21_+_8433172_8446622_RNA45SN1_45S_50ntFlanks@12419.12443.25|GATGTCGGCTCTTCCTATCATTGTG</td><td>>hg38_21_+_8433172_8446622_RNA45SN1_45S_50ntFlanks</td><td>12419</td><td>12443</td><td>25</td><td>GATGTCGGCTCTTCCTATCATTGTG</td><td>3441.2</td><td>-5.6</td><td>0.6</td><td>-9.2</td><td>2.8E-20</td><td>1.1E-19</td><td>BT20/MB468</td><td>NUC</td></tr>
<tr><td>>MI0000746|hsa-mir-99b&WithFlank&19|+|51692606|51692687@51.70.20(+1U)&[MIMAT0004678&hsa-miR-99b-3p&offsets|0|-2(+1U);m-151&19|+|51692656|51692678&offsets|0|-3(+1U)]|CAAGCTCGTGTCTGTGGGTCT</td><td>>MI0000746|hsa-mir-99b&WithFlank&19|+|51692606|51692687</td><td>51</td><td>70</td><td>20(+1U)</td><td>CAAGCTCGTGTCTGTGGGTCT</td><td>844.9</td><td>10.4</td><td>1.1</td><td>9.3</td><td>1.4E-20</td><td>1.1E-19</td><td>BT20/MB468</td><td>CYTO</td></tr>
</table>

<table>
<tr><td>>hg38_21_+_8433172_8446622_RNA45SN1_45S_50ntFlanks@11568.11601.34|GCAGCCGACTTAGAACTGGTGCGGACCAGGGGAA</td><td>>hg38_21_+_8433172_8446622_RNA45SN1_45S_50ntFlanks</td><td>11568</td><td>11601</td><td>34</td><td>GCAGCCGACTTAGAACTGGTGCGGACCAGGGGAA</td><td>2026.7</td><td>-5.2</td><td>0.6</td><td>-9.4</td><td>8.5E-21</td><td>1.1E-19</td><td>MB231/MB468</td><td>NUC</td></tr>
<tr><td>>hg38_21_+_8433172_8446622_RNA45SN1_45S_50ntFlanks@9806.9838.33|CGTGGAATGCGAGTGCCTAGTGGGCCACTTTTG</td><td>>hg38_21_+_8433172_8446622_RNA45SN1_45S_50ntFlanks</td><td>9806</td><td>9838</td><td>33</td><td>CGTGGAATGCGAGTGCCTAGTGGGCCACTTTTG</td><td>801.0</td><td>-13.5</td><td>1.4</td><td>-9.4</td><td>6.1E-21</td><td>1.1E-19</td><td>MB231/MB468</td><td>MP</td></tr>
<tr><td>>hg38_21_+_8433172_8446622_RNA45SN1_45S_50ntFlanks@4731.4768.38|AAGAACGAAAGTCGGAGGTTCGAAGACGATCAGATACC</td><td>>hg38_21_+_8433172_8446622_RNA45SN1_45S_50ntFlanks</td><td>4731</td><td>4768</td><td>38</td><td>AAGAACGAAAGTCGGAGGTTCGAAGACGATCAGATACC</td><td>3082.2</td><td>10.2</td><td>1.1</td><td>9.4</td><td>5.0E-21</td><td>1.1E-19</td><td>MB231/MB468</td><td>CYTO</td></tr>
<tr><td>>MI0000102|hsa-mir-100&WithFlank&11|-|122152223|122152314@19.37.19(+1A)&[MIMAT0000098&hsa-miR-100-5p&offsets|0|-3(+1A);m-27&11|-|122152276|122152296&offsets|0|-2(+1A)]|AACCCGTAGATCCGAACTTA</td><td>>MI0000102|hsa-mir-100&WithFlank&11|-|122152223|122152314</td><td>19</td><td>37</td><td>19(+1A)</td><td>AACCCGTAGATCCGAACTTA</td><td>3928.1</td><td>15.9</td><td>1.7</td><td>9.4</td><td>5.1E-21</td><td>1.1E-19</td><td>MB231/MB468</td><td>CYTO</td></tr>
<tr><td>>hg38_21_+_8433172_8446622_RNA45SN1_45S_50ntFlanks@11668.11705.38|TGCCCAGTGCTCTGAATGTCAAAGTGAAGAAATTCAAT</td><td>>hg38_21_+_8433172_8446622_RNA45SN1_45S_50ntFlanks</td><td>11668</td><td>11705</td><td>38</td><td>TGCCCAGTGCTCTGAATGTCAAAGTGAAGAAATTCAAT</td><td>491.1</td><td>10.6</td><td>1.1</td><td>9.4</td><td>5.1E-21</td><td>1.1E-19</td><td>MB231/MB468</td><td>CYTO</td></tr>
<tr><td>>hg38_21_+_8433172_8446622_RNA45SN1_45S_50ntFlanks@12456.12490.35|CCAAGCGTTGGATTGTTCACCCACTAATAGGGAAC</td><td>>hg38_21_+_8433172_8446622_RNA45SN1_45S_50ntFlanks</td><td>12456</td><td>12490</td><td>35</td><td>CCAAGCGTTGGATTGTTCACCCACTAATAGGGAAC</td><td>1673.2</td><td>-6.5</td><td>0.7</td><td>-9.3</td><td>9.6E-21</td><td>1.1E-19</td><td>MB231/MB468</td><td>MITO</td></tr>
<tr><td>>MI0000298|hsa-mir-221&WithFlank&X|-|45746151|45746272@71.91.21(+1C)&[MIMAT0000278&hsa-miR-221-3p&offsets|0|-2(+1C);m-91&X|-|45746181|45746202&offsets|0|-1(+1C)]|AGCTACATTGTCTGCTGGGTTC</td><td>>MI0000298|hsa-mir-221&WithFlank&X|-|45746151|45746272</td><td>71</td><td>91</td><td>21(+1C)</td><td>AGCTACATTGTCTGCTGGGTTC</td><td>1512.0</td><td>6.5</td><td>0.7</td><td>9.3</td><td>1.5E-20</td><td>1.1E-19</td><td>BT20/MB468</td><td>CYTO</td></tr>
<tr><td>>hg38_21_+_8433172_8446622_RNA45SN1_45S_50ntFlanks@3984.4001.18|GCGGCTTTGGTGACTCTA</td><td>>hg38_21_+_8433172_8446622_RNA45SN1_45S_50ntFlanks</td><td>3984</td><td>4001</td><td>18</td><td>GCGGCTTTGGTGACTCTA</td><td>2618.9</td><td>-15.2</td><td>1.6</td><td>-9.4</td><td>6.2E-21</td><td>1.1E-19</td><td>MB231/MB468</td><td>MP</td></tr>
<tr><td>>hg38_21_+_8433172_8446622_RNA45SN1_45S_50ntFlanks@4047.4064.18|ATTCGAACGTCTGCCCTA</td><td>>hg38_21_+_8433172_8446622_RNA45SN1_45S_50ntFlanks</td><td>4047</td><td>4064</td><td>18</td><td>ATTCGAACGTCTGCCCTA</td><td>502.1</td><td>-12.8</td><td>1.4</td><td>-9.4</td><td>6.2E-21</td><td>1.1E-19</td><td>MB231/MB468</td><td>MP</td></tr>
<tr><td>>trna111_HisGTG_1_-_147774845_147774916@-1G.30.31,trna118_HisGTG_1_-_145396881_145396952@-1G.30.31,trna16_HisGTG_1_+_146544773_146544844@-1G.30.31,trna1_HisGTG_15_+_45493349_45493420@-1G.30.31,trna21_HisGTG_1_+_147753471_147753542@-1G.30.31,trna33_HisGTG_6_+_27125906_27125977@-1G.30.31,trna7_HisGTG_9_-_14433938_14434009@-1G.30.31,trna8_HisGTG_15_-_45492611_45492682@-1G.30.31,trna9_HisGTG_15_-_45490804_45490875@-1G.30.31|GGCCGTGATCGTATAGTGGTTAGTACTCTGC</td><td>>trna111_HisGTG_1_-_147774845_147774916</td><td>-1G</td><td>30</td><td>31</td><td>GGCCGTGATCGTATAGTGGTTAGTACTCTGC</td><td>3949.1</td><td>-7.0</td><td>0.8</td><td>-9.2</td><td>2.9E-20</td><td>1.2E-19</td><td>BT20/MB468</td><td>NUC</td></tr>
<tr><td>>hg38_21_+_8433172_8446622_RNA45SN1_45S_50ntFlanks@4657.4682.26|CTAGAGGTGAAATTCTTGGACCGGCG</td><td>>hg38_21_+_8433172_8446622_RNA45SN1_45S_50ntFlanks</td><td>4657</td><td>4682</td><td>26</td><td>CTAGAGGTGAAATTCTTGGACCGGCG</td><td>1371.0</td><td>-14.2</td><td>1.5</td><td>-9.3</td><td>9.7E-21</td><td>1.2E-19</td><td>MB231/MB468</td><td>MITO</td></tr>
<tr><td>>hg38_21_+_8433172_8446622_RNA45SN1_45S_50ntFlanks@5527.5549.23|AAAGTCGTAACAAGGTTTCCGTA</td><td>>hg38_21_+_8433172_8446622_RNA45SN1_45S_50ntFlanks</td><td>5527</td><td>5549</td><td>23</td><td>AAAGTCGTAACAAGGTTTCCGTA</td><td>757.7</td><td>-13.4</td><td>1.4</td><td>-9.4</td><td>6.3E-21</td><td>1.2E-19</td><td>MB231/MB468</td><td>MP</td></tr>
<tr><td>>hg38_21_+_8433172_8446622_RNA45SN1_45S_50ntFlanks@10392.10410.19|ACAGCAGTTGAACATGGGT</td><td>>hg38_21_+_8433172_8446622_RNA45SN1_45S_50ntFlanks</td><td>10392</td><td>10410</td><td>19</td><td>ACAGCAGTTGAACATGGGT</td><td>4150.0</td><td>-8.4</td><td>0.9</td><td>-9.4</td><td>5.3E-21</td><td>1.2E-19</td><td>MB231/MB468</td><td>CYTO</td></tr>
<tr><td>>hg38_21_+_8433172_8446622_RNA45SN1_45S_50ntFlanks@10873.10902.30|CTGGGTCGGTCGGGCTGGGGCGCGAAGCGG</td><td>>hg38_21_+_8433172_8446622_RNA45SN1_45S_50ntFlanks</td><td>10873</td><td>10902</td><td>30</td><td>CTGGGTCGGTCGGGCTGGGGCGCGAAGCGG</td><td>758.7</td><td>-7.8</td><td>0.8</td><td>-9.4</td><td>5.3E-21</td><td>1.2E-19</td><td>MB231/MB468</td><td>CYTO</td></tr>
<tr><td>>hg38_21_+_8433172_8446622_RNA45SN1_45S_50ntFlanks@4773.4793.21|TAGTTCCGACCATAAACGATG</td><td>>hg38_21_+_8433172_8446622_RNA45SN1_45S_50ntFlanks</td><td>4773</td><td>4793</td><td>21</td><td>TAGTTCCGACCATAAACGATG</td><td>1794.2</td><td>-4.8</td><td>0.5</td><td>-9.3</td><td>1.5E-20</td><td>1.2E-19</td><td>BT20/MB468</td><td>CYTO</td></tr>
<tr><td>>hg38_21_+_8433172_8446622_RNA45SN1_45S_50ntFlanks@12152.12182.31|TGGGGAGTTTGACTGGGGCGGTACACCTGTC</td><td>>hg38_21_+_8433172_8446622_RNA45SN1_45S_50ntFlanks</td><td>12152</td><td>12182</td><td>31</td><td>TGGGGAGTTTGACTGGGGCGGTACACCTGTC</td><td>545.7</td><td>-13.0</td><td>1.4</td><td>-9.4</td><td>6.4E-21</td><td>1.2E-19</td><td>MB231/MB468</td><td>MP</td></tr>
<tr><td>>hg38_21_+_8433172_8446622_RNA45SN1_45S_50ntFlanks@7984.8007.24|AGATCAGACGTGGCGACCCGCTGA</td><td>>hg38_21_+_8433172_8446622_RNA45SN1_45S_50ntFlanks</td><td>7984</td><td>8007</td><td>24</td><td>AGATCAGACGTGGCGACCCGCTGA</td><td>1008.5</td><td>-13.8</td><td>1.5</td><td>-9.4</td><td>6.4E-21</td><td>1.2E-19</td><td>MB231/MB468</td><td>MP</td></tr>
<tr><td>>hg38_21_+_8433172_8446622_RNA45SN1_45S_50ntFlanks@12934.12972.39|CTAAACCATTCGTAGACGACCTGCTTCTGGGTCGGGGTT</td><td>>hg38_21_+_8433172_8446622_RNA45SN1_45S_50ntFlanks</td><td>12934</td><td>12972</td><td>39</td><td>CTAAACCATTCGTAGACGACCTGCTTCTGGGTCGGGGTT</td><td>2652.2</td><td>-11.0</td><td>1.2</td><td>-9.3</td><td>9.9E-21</td><td>1.2E-19</td><td>MB231/MB468</td><td>MITO</td></tr>
<tr><td>>hg38_21_+_8433172_8446622_RNA45SN1_45S_50ntFlanks@11874.11903.30|AGAAGACCCTGTTGAGCTTGACTCTAGTCT</td><td>>hg38_21_+_8433172_8446622_RNA45SN1_45S_50ntFlanks</td><td>11874</td><td>11903</td><td>30</td><td>AGAAGACCCTGTTGAGCTTGACTCTAGTCT</td><td>653.8</td><td>-13.2</td><td>1.4</td><td>-9.4</td><td>6.5E-21</td><td>1.2E-19</td><td>MB231/MB468</td><td>MP</td></tr>
<tr><td>>MI0000079|hsa-mir-23a&WithFlank&19|-|13836581|13836665@54.74.21&[MIMAT0000078&hsa-miR-23a-3p&offsets|+3|+3;m-31&19|-|13836595|13836615&offsets|+3|+3]|ACATTGCCAGGGATTTCCAAC</td><td>>MI0000079|hsa-mir-23a&WithFlank&19|-|13836581|13836665</td><td>54</td><td>74</td><td>21</td><td>ACATTGCCAGGGATTTCCAAC</td><td>382.9</td><td>12.8</td><td>1.4</td><td>9.3</td><td>1.6E-20</td><td>1.2E-19</td><td>BT20/MB468</td><td>CYTO</td></tr>
<tr><td>>hg38_21_+_8433172_8446622_RNA45SN1_45S_50ntFlanks@12151.12181.31|GTGGGGAGTTTGACTGGGGCGGTACACCTGT</td><td>>hg38_21_+_8433172_8446622_RNA45SN1_45S_50ntFlanks</td><td>12151</td><td>12181</td><td>31</td><td>GTGGGGAGTTTGACTGGGGCGGTACACCTGT</td><td>512.4</td><td>-12.9</td><td>1.4</td><td>-9.4</td><td>6.6E-21</td><td>1.2E-19</td><td>MB231/MB468</td><td>MP</td></tr>
<tr><td>>hg38_21_+_8433172_8446622_RNA45SN1_45S_50ntFlanks@6787.6806.20|ACGCCTGTCTGAGCGTCGCT</td><td>>hg38_21_+_8433172_8446622_RNA45SN1_45S_50ntFlanks</td><td>6787</td><td>6806</td><td>20</td><td>ACGCCTGTCTGAGCGTCGCT</td><td>1570.4</td><td>5.6</td><td>0.6</td><td>9.3</td><td>9.2E-21</td><td>1.2E-19</td><td>MB231/MB468</td><td>NUC</td></tr>
<tr><td>>MI0000478|hsa-mir-149&WithFlank&2|+|240455995|240456095@21.43.23(+1U)&[MIMAT0000450&hsa-miR-149-5p&offsets|0|0(+1U);m-195&2|+|240456015|240456037&offsets|0|0(+1U)]|TCTGGCTCCGTGTCTTCACTCCCT</td><td>>MI0000478|hsa-mir-149&WithFlank&2|+|240455995|240456095</td><td>21</td><td>43</td><td>23(+1U)</td><td>TCTGGCTCCGTGTCTTCACTCCCT</td><td>400.5</td><td>12.9</td><td>1.4</td><td>9.3</td><td>1.6E-20</td><td>1.2E-19</td><td>BT20/MB468</td><td>CYTO</td></tr>
<tr><td>>hg38_21_+_8433172_8446622_RNA45SN1_45S_50ntFlanks@7975.8014.40|CGCGACCTCAGATCAGACGTGGCGACCCGCTGAATTTAAG</td><td>>hg38_21_+_8433172_8446622_RNA45SN1_45S_50ntFlanks</td><td>7975</td><td>8014</td><td>40</td><td>CGCGACCTCAGATCAGACGTGGCGACCCGCTGAATTTAAG</td><td>41109.7</td><td>4.7</td><td>0.5</td><td>9.3</td><td>1.1E-20</td><td>1.2E-19</td><td>BT20/MB468</td><td>MITO</td></tr>
<tr><td>>MI0000542|hsa-mir-320a&WithFlank&8|-|22244960|22245043@49.71.23&[MIMAT0000510&hsa-miR-320a-3p&offsets|+1|+2;m-60&8|-|22244975|22244996&offsets|+1|+2]|AAAGCTGGGTTGAGAGGGCGAAA</td><td>>MI0000542|hsa-mir-320a&WithFlank&8|-|22244960|22245043</td><td>49</td><td>71</td><td>23</td><td>AAAGCTGGGTTGAGAGGGCGAAA</td><td>926.1</td><td>11.5</td><td>1.2</td><td>9.3</td><td>1.6E-20</td><td>1.2E-19</td><td>BT20/MB468</td><td>CYTO</td></tr>
<tr><td>>hg38_21_+_8433172_8446622_RNA45SN1_45S_50ntFlanks@6663.6688.26|GTGGATCACTCGGCTCGTGCGTCGAT</td><td>>hg38_21_+_8433172_8446622_RNA45SN1_45S_50ntFlanks</td><td>6663</td><td>6688</td><td>26</td><td>GTGGATCACTCGGCTCGTGCGTCGAT</td><td>639.8</td><td>-13.2</td><td>1.4</td><td>-9.4</td><td>6.6E-21</td><td>1.2E-19</td><td>MB231/MB468</td><td>MP</td></tr>
<tr><td>>hg38_21_+_8433172_8446622_RNA45SN1_45S_50ntFlanks@11641.11662.22|CGGCGGGTGTTGACGCGATGTG</td><td>>hg38_21_+_8433172_8446622_RNA45SN1_45S_50ntFlanks</td><td>11641</td><td>11662</td><td>22</td><td>CGGCGGGTGTTGACGCGATGTG</td><td>1379.0</td><td>-6.3</td><td>0.7</td><td>-9.3</td><td>9.3E-21</td><td>1.2E-19</td><td>MB231/MB468</td><td>NUC</td></tr>
<tr><td>>am_trna10_LysCTT_16_+_3241501_3241573@31.51.21,trna119_LysCTT_1_-_145395522_145395594@31.51.21,trna11_LysCTT_5_-_180648979_180649051@31.51.21,trna13_LysCTT_16_+_3246153_3246225@31.51.21,trna13_LysCTT_6_+_26556774_26556846@31.51.21,trna30_LysCTT_16_-_3230555_3230627@31.51.21,trna5_LysCTT_16_+_3214939_3215011@31.51.21,trna7_LysCTT_16_+_3225692_3225764@31.51.21,trna8_LysCTT_1_+_39970195_39970267@31.51.21,trna9_LysCTT_5_+_180634755_180634827@31.51.21|ACTCTTAATCTCAGGGTCGTG</td><td>>am_trna10_LysCTT_16_+_3241501_3241573</td><td>31</td><td>51</td><td>21</td><td>ACTCTTAATCTCAGGGTCGTG</td><td>616.3</td><td>-13.7</td><td>1.5</td><td>-9.2</td><td>3.1E-20</td><td>1.2E-19</td><td>BT20/MB468</td><td>NUC</td></tr>
<tr><td>>MI0003667|hsa-mir-652&WithFlank&X|+|110055323|110055432@67.88.22&[MIMAT0003322&hsa-miR-652-3p&offsets|0|+1;m-148&X|+|110055389|110055410&offsets|0|0]|AATGGCGCCACTAGGGTTGTGC</td><td>>MI0003667|hsa-mir-652&WithFlank&X|+|110055323|110055432</td><td>67</td><td>88</td><td>22</td><td>AATGGCGCCACTAGGGTTGTGC</td><td>387.2</td><td>12.9</td><td>1.4</td><td>9.3</td><td>1.6E-20</td><td>1.2E-19</td><td>BT20/MB468</td><td>CYTO</td></tr>
<tr><td>>hg38_21_+_8433172_8446622_RNA45SN1_45S_50ntFlanks@9381.9409.29|CGCTCGCCGGCCGAGGTGGGATCCCGAGG</td><td>>hg38_21_+_8433172_8446622_RNA45SN1_45S_50ntFlanks</td><td>9381</td><td>9409</td><td>29</td><td>CGCTCGCCGGCCGAGGTGGGATCCCGAGG</td><td>5699.6</td><td>-16.3</td><td>1.7</td><td>-9.3</td><td>1.0E-20</td><td>1.2E-19</td><td>MB231/MB468</td><td>MITO</td></tr>
<tr><td>>hg38_21_+_8433172_8446622_RNA45SN1_45S_50ntFlanks@11894.11912.19|ACTCTAGTCTGGCACGGTG</td><td>>hg38_21_+_8433172_8446622_RNA45SN1_45S_50ntFlanks</td><td>11894</td><td>11912</td><td>19</td><td>ACTCTAGTCTGGCACGGTG</td><td>645.9</td><td>-13.2</td><td>1.4</td><td>-9.4</td><td>6.7E-21</td><td>1.2E-19</td><td>MB231/MB468</td><td>MP</td></tr>
<tr><td>>MI0005756|hsa-mir-934&WithFlank&X|+|136550872|136550966@21.42.22(+1A)&[MIMAT0004977&hsa-miR-934&offsets|0|0(+1A);m-943&X|+|136550892|136550913&offsets|0|0(+1A)]|TGTCTACTACTGGAGACACTGGA</td><td>>MI0005756|hsa-mir-934&WithFlank&X|+|136550872|136550966</td><td>21</td><td>42</td><td>22(+1A)</td><td>TGTCTACTACTGGAGACACTGGA</td><td>1349.4</td><td>13.8</td><td>1.4</td><td>9.7</td><td>4.7E-22</td><td>1.2E-19</td><td>BT20/MB231</td><td>CYTO</td></tr>
<tr><td>>MI0000077|hsa-mir-21&WithFlank&17|+|59841260|59841343@14.33.20&[MIMAT0000076&hsa-miR-21-5p&offsets|0|-2;m-1&17|+|59841273|59841294&offsets|0|-2]|TAGCTTATCAGACTGATGTT</td><td>>MI0000077|hsa-mir-21&WithFlank&17|+|59841260|59841343</td><td>14</td><td>33</td><td>20</td><td>TAGCTTATCAGACTGATGTT</td><td>634.8</td><td>6.4</td><td>0.7</td><td>9.3</td><td>1.1E-20</td><td>1.2E-19</td><td>BT20/MB468</td><td>MITO</td></tr>
<tr><td>>MI0000449|hsa-mir-132&WithFlank&17|-|2049902|2050014@29.50.22&[MIMAT0004594&hsa-miR-132-5p&offsets|0|0;m-414&17|-|2049965|2049986&offsets|0|0]|ACCGTGGCTTTCGATTGTTACT</td><td>>MI0000449|hsa-mir-132&WithFlank&17|-|2049902|2050014</td><td>29</td><td>50</td><td>22</td><td>ACCGTGGCTTTCGATTGTTACT</td><td>478.5</td><td>13.4</td><td>1.4</td><td>9.3</td><td>1.2E-20</td><td>1.2E-19</td><td>BT20/MB468</td><td>MITO</td></tr>
<tr><td>>hg38_21_+_8433172_8446622_RNA45SN1_45S_50ntFlanks@12641.12659.19|CCATCTGTGGGATTATGAC</td><td>>hg38_21_+_8433172_8446622_RNA45SN1_45S_50ntFlanks</td><td>12641</td><td>12659</td><td>19</td><td>CCATCTGTGGGATTATGAC</td><td>1077.6</td><td>-14.5</td><td>1.6</td><td>-9.2</td><td>3.1E-20</td><td>1.2E-19</td><td>BT20/MB468</td><td>NUC</td></tr>
<tr><td>>hg38_21_+_8433172_8446622_RNA45SN1_45S_50ntFlanks@10319.10340.22|GCAGATCTTGGTGGTAGTAGCA</td><td>>hg38_21_+_8433172_8446622_RNA45SN1_45S_50ntFlanks</td><td>10319</td><td>10340</td><td>22</td><td>GCAGATCTTGGTGGTAGTAGCA</td><td>435.2</td><td>6.2</td><td>0.7</td><td>9.3</td><td>1.2E-20</td><td>1.2E-19</td><td>BT20/MB468</td><td>MITO</td></tr>
<tr><td>>MI0000764|hsa-mir-363&WithFlank&X|-|134169372|134169458@56.77.22&[MIMAT0000707&hsa-miR-363-3p&offsets|0|0;m-178&X|-|134169381|134169403&offsets|0|-1]|AATTGCACGGTATCCATCTGTA</td><td>>MI0000764|hsa-mir-363&WithFlank&X|-|134169372|134169458</td><td>56</td><td>77</td><td>22</td><td>AATTGCACGGTATCCATCTGTA</td><td>580.6</td><td>-13.0</td><td>1.4</td><td>-9.4</td><td>6.8E-21</td><td>1.2E-19</td><td>MB231/MB468</td><td>MP</td></tr>
<tr><td>>hg38_21_+_8433172_8446622_RNA45SN1_45S_50ntFlanks@6699.6719.21|GCTAGCTGCGAGAATTAATGT</td><td>>hg38_21_+_8433172_8446622_RNA45SN1_45S_50ntFlanks</td><td>6699</td><td>6719</td><td>21</td><td>GCTAGCTGCGAGAATTAATGT</td><td>1502.2</td><td>-5.1</td><td>0.6</td><td>-9.3</td><td>1.6E-20</td><td>1.2E-19</td><td>BT20/MB468</td><td>CYTO</td></tr>
<tr><td>>hg38_21_+_8433172_8446622_RNA45SN1_45S_50ntFlanks@11199.11225.27|CCGGCGGCGGCGGCGACTCTGGACGCG</td><td>>hg38_21_+_8433172_8446622_RNA45SN1_45S_50ntFlanks</td><td>11199</td><td>11225</td><td>27</td><td>CCGGCGGCGGCGGCGACTCTGGACGCG</td><td>1769.5</td><td>-14.7</td><td>1.6</td><td>-9.4</td><td>6.8E-21</td><td>1.2E-19</td><td>MB231/MB468</td><td>MP</td></tr>
<tr><td>>hg38_21_+_8433172_8446622_RNA45SN1_45S_50ntFlanks@6774.6806.33|TCCTCCCGGGGCTACGCCTGTCTGAGCGTCGCT</td><td>>hg38_21_+_8433172_8446622_RNA45SN1_45S_50ntFlanks</td><td>6774</td><td>6806</td><td>33</td><td>TCCTCCCGGGGCTACGCCTGTCTGAGCGTCGCT</td><td>752.4</td><td>3.7</td><td>0.4</td><td>9.2</td><td>3.2E-20</td><td>1.3E-19</td><td>BT20/MB468</td><td>NUC</td></tr>
<tr><td>>hg38_21_+_8433172_8446622_RNA45SN1_45S_50ntFlanks@6690.6720.31|AAGAACGCAGCTAGCTGCGAGAATTAATGTG</td><td>>hg38_21_+_8433172_8446622_RNA45SN1_45S_50ntFlanks</td><td>6690</td><td>6720</td><td>31</td><td>AAGAACGCAGCTAGCTGCGAGAATTAATGTG</td><td>3940.6</td><td>-11.3</td><td>1.2</td><td>-9.3</td><td>1.1E-20</td><td>1.3E-19</td><td>MB231/MB468</td><td>MITO</td></tr>
<tr><td>>hg38_21_+_8433172_8446622_RNA45SN1_45S_50ntFlanks@11570.11601.32|AGCCGACTTAGAACTGGTGCGGACCAGGGGAA</td><td>>hg38_21_+_8433172_8446622_RNA45SN1_45S_50ntFlanks</td><td>11570</td><td>11601</td><td>32</td><td>AGCCGACTTAGAACTGGTGCGGACCAGGGGAA</td><td>3242.3</td><td>-6.0</td><td>0.6</td><td>-9.3</td><td>9.8E-21</td><td>1.3E-19</td><td>MB231/MB468</td><td>NUC</td></tr>
<tr><td>>hg38_21_+_8433172_8446622_RNA45SN1_45S_50ntFlanks@11161.11179.19|CACGGTCCCCCGCGAGGGG</td><td>>hg38_21_+_8433172_8446622_RNA45SN1_45S_50ntFlanks</td><td>11161</td><td>11179</td><td>19</td><td>CACGGTCCCCCGCGAGGGG</td><td>1455.3</td><td>-14.4</td><td>1.5</td><td>-9.4</td><td>7.0E-21</td><td>1.3E-19</td><td>MB231/MB468</td><td>MP</td></tr>
<tr><td>>hg38_21_+_8433172_8446622_RNA45SN1_45S_50ntFlanks@7980.8005.26|CCTCAGATCAGACGTGGCGACCCGCT</td><td>>hg38_21_+_8433172_8446622_RNA45SN1_45S_50ntFlanks</td><td>7980</td><td>8005</td><td>26</td><td>CCTCAGATCAGACGTGGCGACCCGCT</td><td>458.0</td><td>-12.7</td><td>1.4</td><td>-9.4</td><td>7.0E-21</td><td>1.3E-19</td><td>MB231/MB468</td><td>MP</td></tr>
</table>

<table>
<tr><td>>trna111_HisGTG_1_-_147774845_147774916@-1G.33.34,trna118_HisGTG_1_-_145396881_145396952@-1G.33.34,trna16_HisGTG_1_+_146544773_146544844@-1G.33.34,trna1_HisGTG_15_+_45493349_45493420@-1G.33.34,trna21_HisGTG_1_+_147753471_147753542@-1G.33.34,trna33_HisGTG_6_+_27125906_27125977@-1G.33.34,trna7_HisGTG_9_-_14433938_14434009@-1G.33.34,trna8_HisGTG_15_-_45492611_45492682@-1G.33.34,trna9_HisGTG_15_-_45490804_45490875@-1G.33.34|GGCCGTGATCGTATAGTGGTTAGTACTCTGCGTT</td><td>>trna111_HisGTG_1_-_147774845_147774916</td><td>-1G</td><td>33</td><td>34</td><td>GGCCGTGATCGTATAGTGGTTAGTACTCTGCGTT</td><td>43930.2</td><td>10.4</td><td>1.1</td><td>9.3</td><td>1.0E-20</td><td>1.3E-19</td><td>MB231/MB468</td><td>NUC</td></tr>
<tr><td>>hg38_21_+_8433172_8446622_RNA45SN1_45S_50ntFlanks@10687.10710.24|CGGCGTCCGGTGAGCTCTCGCTGG</td><td>>hg38_21_+_8433172_8446622_RNA45SN1_45S_50ntFlanks</td><td>10687</td><td>10710</td><td>24</td><td>CGGCGTCCGGTGAGCTCTCGCTGG</td><td>4970.7</td><td>-11.4</td><td>1.2</td><td>-9.4</td><td>7.2E-21</td><td>1.3E-19</td><td>MB231/MB468</td><td>MP</td></tr>
<tr><td>>hg38_21_+_8433172_8446622_RNA45SN1_45S_50ntFlanks@10807.10825.19|ACAATGTAGGTAAGGGAAG</td><td>>hg38_21_+_8433172_8446622_RNA45SN1_45S_50ntFlanks</td><td>10807</td><td>10825</td><td>19</td><td>ACAATGTAGGTAAGGGAAG</td><td>676.2</td><td>-13.8</td><td>1.5</td><td>-9.2</td><td>3.3E-20</td><td>1.3E-19</td><td>BT20/MB468</td><td>NUC</td></tr>
<tr><td>>hg38_21_+_8433172_8446622_RNA45SN1_45S_50ntFlanks@11564.11588.25|CCTAGCAGCCGACTTAGAACTGGTG</td><td>>hg38_21_+_8433172_8446622_RNA45SN1_45S_50ntFlanks</td><td>11564</td><td>11588</td><td>25</td><td>CCTAGCAGCCGACTTAGAACTGGTG</td><td>933.4</td><td>-4.5</td><td>0.5</td><td>-9.3</td><td>1.7E-20</td><td>1.3E-19</td><td>BT20/MB468</td><td>CYTO</td></tr>
<tr><td>>hg38_21_+_8433172_8446622_RNA45SN1_45S_50ntFlanks@10916.10933.18|CCGCGGCTGGACGAGGCG</td><td>>hg38_21_+_8433172_8446622_RNA45SN1_45S_50ntFlanks</td><td>10916</td><td>10933</td><td>18</td><td>CCGCGGCTGGACGAGGCG</td><td>1016.9</td><td>-13.9</td><td>1.5</td><td>-9.4</td><td>7.3E-21</td><td>1.3E-19</td><td>MB231/MB468</td><td>MP</td></tr>
<tr><td>>hg38_21_+_8433172_8446622_RNA45SN1_45S_50ntFlanks@8170.8187.18|CAAGTCCTTCTGATCGAG</td><td>>hg38_21_+_8433172_8446622_RNA45SN1_45S_50ntFlanks</td><td>8170</td><td>8187</td><td>18</td><td>CAAGTCCTTCTGATCGAG</td><td>676.9</td><td>-13.3</td><td>1.4</td><td>-9.4</td><td>7.3E-21</td><td>1.3E-19</td><td>MB231/MB468</td><td>MP</td></tr>
<tr><td>>hg38_21_+_8433172_8446622_RNA45SN1_45S_50ntFlanks@6733.6772.40|CATTGATCATCGACACTTCGAACGCACTTGCGGCCCCGGG</td><td>>hg38_21_+_8433172_8446622_RNA45SN1_45S_50ntFlanks</td><td>6733</td><td>6772</td><td>40</td><td>CATTGATCATCGACACTTCGAACGCACTTGCGGCCCCGGG</td><td>905.4</td><td>-6.4</td><td>0.7</td><td>-9.2</td><td>3.4E-20</td><td>1.3E-19</td><td>BT20/MB468</td><td>NUC</td></tr>
<tr><td>>hg38_21_+_8433172_8446622_RNA45SN1_45S_50ntFlanks@3996.4025.30|ACTCTAGATAACCTCGGGCCGATCGCACGC</td><td>>hg38_21_+_8433172_8446622_RNA45SN1_45S_50ntFlanks</td><td>3996</td><td>4025</td><td>30</td><td>ACTCTAGATAACCTCGGGCCGATCGCACGC</td><td>561.9</td><td>-13.6</td><td>1.5</td><td>-9.2</td><td>3.4E-20</td><td>1.3E-19</td><td>BT20/MB468</td><td>NUC</td></tr>
<tr><td>>hg38_21_+_8433172_8446622_RNA45SN1_45S_50ntFlanks@9623.9658.36|TAGTAGCTGGTTCCCTCCGAAGTTTCCCTCAGGATA</td><td>>hg38_21_+_8433172_8446622_RNA45SN1_45S_50ntFlanks</td><td>9623</td><td>9658</td><td>36</td><td>TAGTAGCTGGTTCCCTCCGAAGTTTCCCTCAGGATA</td><td>2165.2</td><td>-7.0</td><td>0.7</td><td>-9.3</td><td>1.1E-20</td><td>1.3E-19</td><td>MB231/MB468</td><td>MITO</td></tr>
<tr><td>>hg38_21_+_8433172_8446622_RNA45SN1_45S_50ntFlanks@8552.8577.26|CTCCCCGGAGGGGGCGGGCTCCGGCG</td><td>>hg38_21_+_8433172_8446622_RNA45SN1_45S_50ntFlanks</td><td>8552</td><td>8577</td><td>26</td><td>CTCCCCGGAGGGGGCGGGCTCCGGCG</td><td>784.4</td><td>-13.4</td><td>1.4</td><td>-9.3</td><td>1.1E-20</td><td>1.3E-19</td><td>MB231/MB468</td><td>MITO</td></tr>
<tr><td>>hg38_21_+_8433172_8446622_RNA45SN1_45S_50ntFlanks@11576.11601.26|CTTAGAACTGGTGCGGACCAGGGGAA</td><td>>hg38_21_+_8433172_8446622_RNA45SN1_45S_50ntFlanks</td><td>11576</td><td>11601</td><td>26</td><td>CTTAGAACTGGTGCGGACCAGGGGAA</td><td>3725.8</td><td>-12.1</td><td>1.3</td><td>-9.2</td><td>3.4E-20</td><td>1.3E-19</td><td>BT20/MB468</td><td>NUC</td></tr>
<tr><td>>hg38_21_+_8433172_8446622_RNA45SN1_45S_50ntFlanks@11686.11717.32|TCAAAGTGAAGAAATTCAATGAAGCGCGGGTA</td><td>>hg38_21_+_8433172_8446622_RNA45SN1_45S_50ntFlanks</td><td>11686</td><td>11717</td><td>32</td><td>TCAAAGTGAAGAAATTCAATGAAGCGCGGGTA</td><td>403.6</td><td>-13.1</td><td>1.4</td><td>-9.2</td><td>3.4E-20</td><td>1.3E-19</td><td>BT20/MB468</td><td>NUC</td></tr>
<tr><td>>hg38_21_+_8433172_8446622_RNA45SN1_45S_50ntFlanks@6699.6718.20|GCTAGCTGCGAGAATTAATG</td><td>>hg38_21_+_8433172_8446622_RNA45SN1_45S_50ntFlanks</td><td>6699</td><td>6718</td><td>20</td><td>GCTAGCTGCGAGAATTAATG</td><td>53110.2</td><td>-12.1</td><td>1.3</td><td>-9.2</td><td>3.4E-20</td><td>1.3E-19</td><td>BT20/MB468</td><td>NUC</td></tr>
<tr><td>>hg38_21_+_8433172_8446622_RNA45SN1_45S_50ntFlanks@12615.12653.39|GCTGAGGAGCCAATGGGGCGAAGCTACCATCTGTGGGAT</td><td>>hg38_21_+_8433172_8446622_RNA45SN1_45S_50ntFlanks</td><td>12615</td><td>12653</td><td>39</td><td>GCTGAGGAGCCAATGGGGCGAAGCTACCATCTGTGGGAT</td><td>1465.7</td><td>-14.4</td><td>1.5</td><td>-9.4</td><td>7.5E-21</td><td>1.3E-19</td><td>MB231/MB468</td><td>MP</td></tr>
<tr><td>>trna116_GluCTC_1_-_145399233_145399304@1.34.34,trna59_GluCTC_1_+_249168447_249168518@1.34.34,trna71_GluCTC_1_-_161439189_161439260@1.34.34,trna74_GluCTC_1_-_161431809_161431880@1.34.34,trna77_GluCTC_1_-_161424398_161424469@1.34.34,trna77_GluCTC_6_+_28949976_28950047@1.34.34,trna80_GluCTC_1_-_161417018_161417089@1.34.34,trna87_GluCTC_6_-_126101393_126101464@1.34.34|TCCCTGGTGGTCTAGTGGTTAGGATTCGGCGCTC</td><td>>trna116_GluCTC_1_-_145399233_145399304</td><td>1</td><td>34</td><td>34</td><td>TCCCTGGTGGTCTAGTGGTTAGGATTCGGCGCTC</td><td>20203.2</td><td>8.5</td><td>0.9</td><td>9.3</td><td>1.3E-20</td><td>1.4E-19</td><td>BT20/MB468</td><td>MITO</td></tr>
<tr><td>>MI0000114|hsa-mir-107&WithFlank&10|-|89592741|89592833@56.76.21(+1A);MI0000108|hsa-mir-103a-2&WithFlank&20|+|3917488|3917577@54.74.21(+1A);MI0000109|hsa-mir-103a-1&WithFlank&5|-|168560890|168560979@54.74.21(+1A)&[MIMAT0000104&hsa-miR-107&offsets|0|-2(+1A);m-78&10|-|89592758|89592778&offsets|0|0(+1A)];[MIMAT0000101&hsa-miR-103a-3p&offsets|0|-2(+1A);m-12&20|+|3917541|3917563&offsets|0|-2(+1A)];[MIMAT0000101_1&hsa-miR-103a-3p&offsets|0|-2(+1A);m-13&5|-|168560904|168560926&offsets|0|-2(+1A);m-44220&5|-|168560908|168560926&offsets|0|+2(+1A)]|AGCAGCATTGTACAGGGCTATA</td><td>>MI0000114|hsa-mir-107&WithFlank&10|-|89592741|89592833</td><td>56</td><td>76</td><td>21(+1A)</td><td>AGCAGCATTGTACAGGGCTATA</td><td>376.3</td><td>5.9</td><td>0.6</td><td>9.3</td><td>1.1E-20</td><td>1.4E-19</td><td>MB231/MB468</td><td>NUC</td></tr>
<tr><td>>am_MI0000809|hsa-mir-151a&WithFlank&8|-|140732558|140732659@17.37.21&[MIMAT0004697&hsa-miR-151a-5p&offsets|0|0;m-75&8|-|140732623|140732643&offsets|0|0]|TCGAGGAGCTCACAGTCTAGT</td><td>>am_MI0000809|hsa-mir-151a&WithFlank&8|-|140732558|140732659</td><td>17</td><td>37</td><td>21</td><td>TCGAGGAGCTCACAGTCTAGT</td><td>1189.5</td><td>7.3</td><td>0.8</td><td>9.3</td><td>1.1E-20</td><td>1.4E-19</td><td>MB231/MB468</td><td>NUC</td></tr>
<tr><td>>hg38_21_+_8433172_8446622_RNA45SN1_45S_50ntFlanks@9565.9596.32|TGACGTGCAAATCGGTCGTCCGACCTGGGTAT</td><td>>hg38_21_+_8433172_8446622_RNA45SN1_45S_50ntFlanks</td><td>9565</td><td>9596</td><td>32</td><td>TGACGTGCAAATCGGTCGTCCGACCTGGGTAT</td><td>10739.0</td><td>-6.2</td><td>0.7</td><td>-9.3</td><td>1.1E-20</td><td>1.4E-19</td><td>MB231/MB468</td><td>NUC</td></tr>
<tr><td>>hg38_21_+_8433172_8446622_RNA45SN1_45S_50ntFlanks@6645.6672.28|CTCGTACGACTCTTAGCGGTGGATCACT</td><td>>hg38_21_+_8433172_8446622_RNA45SN1_45S_50ntFlanks</td><td>6645</td><td>6672</td><td>28</td><td>CTCGTACGACTCTTAGCGGTGGATCACT</td><td>6613.3</td><td>6.8</td><td>0.7</td><td>9.3</td><td>1.3E-20</td><td>1.4E-19</td><td>BT20/MB468</td><td>MITO</td></tr>
<tr><td>>MI0003513|hsa-mir-455&WithFlank&9|+|114209428|114209535@60.80.21(+1C)&[MIMAT0004784&hsa-miR-455-3p&offsets|0|0(+1C);m-135&9|+|114209487|114209508&offsets|0|-1(+1C)]|GCAGTCCATGGGCATATACACC</td><td>>MI0003513|hsa-mir-455&WithFlank&9|+|114209428|114209535</td><td>60</td><td>80</td><td>21(+1C)</td><td>GCAGTCCATGGGCATATACACC</td><td>339.3</td><td>12.3</td><td>1.3</td><td>9.3</td><td>1.1E-20</td><td>1.4E-19</td><td>MB231/MB468</td><td>NUC</td></tr>
<tr><td>>hg38_21_+_8433172_8446622_RNA45SN1_45S_50ntFlanks@12076.12095.20|AGGGGCTCTCGCTTCTGGCG</td><td>>hg38_21_+_8433172_8446622_RNA45SN1_45S_50ntFlanks</td><td>12076</td><td>12095</td><td>20</td><td>AGGGGCTCTCGCTTCTGGCG</td><td>2241.0</td><td>-15.0</td><td>1.6</td><td>-9.4</td><td>7.6E-21</td><td>1.4E-19</td><td>MB231/MB468</td><td>MP</td></tr>
<tr><td>>hg38_21_+_8433172_8446622_RNA45SN1_45S_50ntFlanks@12419.12442.24|GATGTCGGCTCTTCCTATCATTGT</td><td>>hg38_21_+_8433172_8446622_RNA45SN1_45S_50ntFlanks</td><td>12419</td><td>12442</td><td>24</td><td>GATGTCGGCTCTTCCTATCATTGT</td><td>3976.4</td><td>6.5</td><td>0.7</td><td>9.4</td><td>6.2E-21</td><td>1.4E-19</td><td>MB231/MB468</td><td>CYTO</td></tr>
<tr><td>>trna152_ValCAC_6_-_27248049_27248121@1.34.34|GCTTCTGTAGTGTAGTGGTTATCACGTTCGCCTC</td><td>>trna152_ValCAC_6_-_27248049_27248121</td><td>1</td><td>34</td><td>34</td><td>GCTTCTGTAGTGTAGTGGTTATCACGTTCGCCTC</td><td>20289.6</td><td>11.1</td><td>1.2</td><td>9.4</td><td>6.3E-21</td><td>1.4E-19</td><td>MB231/MB468</td><td>CYTO</td></tr>
<tr><td>>MI0000466|hsa-mir-9-1&WithFlank&1|-|156420335|156420435@22.44.23;MI0000468|hsa-mir-9-3&WithFlank&15|+|89368011|89368112@22.44.23;MI0000467|hsa-mir-9-2&WithFlank&5|-|88666847|88666945@22.44.23&[MIMAT0000441&hsa-miR-9-5p&offsets|0|0;m-104&1|-|156420392|156420414&offsets|0|0];[MIMAT0000441_1&hsa-miR-9-5p&offsets|0|0;m-105&15|+|89368032|89368054&offsets|0|0];[MIMAT0000441_2&hsa-miR-9-5p&offsets|0|0;m-104&5|-|88666902|88666924&offsets|0|0]|TCTTTGGTTATCTAGCTGTATGA</td><td>>MI0000466|hsa-mir-9-1&WithFlank&1|-|156420335|156420435</td><td>22</td><td>44</td><td>23</td><td>TCTTTGGTTATCTAGCTGTATGA</td><td>1684.7</td><td>9.4</td><td>1.0</td><td>9.3</td><td>1.3E-20</td><td>1.4E-19</td><td>BT20/MB468</td><td>MITO</td></tr>
<tr><td>>hg38_21_+_8433172_8446622_RNA45SN1_45S_50ntFlanks@11321.11338.18|CACGTCTCGTCGCGCGCG</td><td>>hg38_21_+_8433172_8446622_RNA45SN1_45S_50ntFlanks</td><td>11321</td><td>11338</td><td>18</td><td>CACGTCTCGTCGCGCGCG</td><td>1023.1</td><td>-13.8</td><td>1.5</td><td>-9.3</td><td>1.2E-20</td><td>1.4E-19</td><td>MB231/MB468</td><td>MITO</td></tr>
<tr><td>>hg38_21_+_8433172_8446622_RNA45SN1_45S_50ntFlanks@12324.12349.26|TTGGGTTTTAAGCAGGAGGTGTCAGA</td><td>>hg38_21_+_8433172_8446622_RNA45SN1_45S_50ntFlanks</td><td>12324</td><td>12349</td><td>26</td><td>TTGGGTTTTAAGCAGGAGGTGTCAGA</td><td>487.0</td><td>-12.8</td><td>1.4</td><td>-9.4</td><td>7.8E-21</td><td>1.4E-19</td><td>MB231/MB468</td><td>MP</td></tr>
<tr><td>>hg38_21_+_8433172_8446622_RNA45SN1_45S_50ntFlanks@10101.10125.25|CGCGCGCGTGTGGTGTGCGTCGGAG</td><td>>hg38_21_+_8433172_8446622_RNA45SN1_45S_50ntFlanks</td><td>10101</td><td>10125</td><td>25</td><td>CGCGCGCGTGTGGTGTGCGTCGGAG</td><td>1444.8</td><td>-14.3</td><td>1.5</td><td>-9.3</td><td>1.2E-20</td><td>1.4E-19</td><td>MB231/MB468</td><td>MITO</td></tr>
<tr><td>>hg38_21_+_8433172_8446622_RNA45SN1_45S_50ntFlanks@10574.10608.35|AAGCCGGCGGGAGCCCCGGGGAGAGTTCTCTTTTC</td><td>>hg38_21_+_8433172_8446622_RNA45SN1_45S_50ntFlanks</td><td>10574</td><td>10608</td><td>35</td><td>AAGCCGGCGGGAGCCCCGGGGAGAGTTCTCTTTTC</td><td>693.1</td><td>-13.2</td><td>1.4</td><td>-9.3</td><td>1.2E-20</td><td>1.4E-19</td><td>MB231/MB468</td><td>MITO</td></tr>
<tr><td>>hg38_21_+_8433172_8446622_RNA45SN1_45S_50ntFlanks@9570.9596.27|TGCAAATCGGTCGTCCGACCTGGGTAT</td><td>>hg38_21_+_8433172_8446622_RNA45SN1_45S_50ntFlanks</td><td>9570</td><td>9596</td><td>27</td><td>TGCAAATCGGTCGTCCGACCTGGGTAT</td><td>7610.9</td><td>-10.0</td><td>1.1</td><td>-9.3</td><td>1.2E-20</td><td>1.4E-19</td><td>MB231/MB468</td><td>MITO</td></tr>
<tr><td>>MI0000067|hsa-let-7f-1&WithFlank&9|+|94176341|94176439@13.34.22(+1A);MI0000068|hsa-let-7f-2&WithFlank&X|-|53557186|53557280@14.35.22(+1A)&[MIMAT0000067&hsa-let-7f-5p&offsets|0|0(+1A);m-20&9|+|94176353|94176374&offsets|0|0(+1A)];[MIMAT0000067_1&hsa-let-7f-5p&offsets|0|0(+1A);m-19&X|-|53557246|53557267&offsets|0|0(+1A)]|TGAGGTAGTAGATTGTATAGTTA</td><td>>MI0000067|hsa-let-7f-1&WithFlank&9|+|94176341|94176439</td><td>13</td><td>34</td><td>22(+1A)</td><td>TGAGGTAGTAGATTGTATAGTTA</td><td>3732.8</td><td>6.5</td><td>0.7</td><td>9.3</td><td>1.9E-20</td><td>1.4E-19</td><td>BT20/MB468</td><td>CYTO</td></tr>
<tr><td>>hg38_21_+_8433172_8446622_RNA45SN1_45S_50ntFlanks@11881.11906.26|CCTGTTGAGCTTGACTCTAGTCTGGC</td><td>>hg38_21_+_8433172_8446622_RNA45SN1_45S_50ntFlanks</td><td>11881</td><td>11906</td><td>26</td><td>CCTGTTGAGCTTGACTCTAGTCTGGC</td><td>495.0</td><td>-7.4</td><td>0.8</td><td>-9.4</td><td>6.6E-21</td><td>1.4E-19</td><td>MB231/MB468</td><td>CYTO</td></tr>
<tr><td>>hg38_21_+_8433172_8446622_RNA45SN1_45S_50ntFlanks@10362.10391.30|AAGGCCGAAGTGGAGAAGGGTTCCATGTGA</td><td>>hg38_21_+_8433172_8446622_RNA45SN1_45S_50ntFlanks</td><td>10362</td><td>10391</td><td>30</td><td>AAGGCCGAAGTGGAGAAGGGTTCCATGTGA</td><td>531.7</td><td>-13.5</td><td>1.5</td><td>-9.2</td><td>3.7E-20</td><td>1.5E-19</td><td>BT20/MB468</td><td>NUC</td></tr>
<tr><td>>hg38_21_+_8433172_8446622_RNA45SN1_45S_50ntFlanks@10510.10534.25|AATCCGGAGTGGCGGAGATGGGCGC</td><td>>hg38_21_+_8433172_8446622_RNA45SN1_45S_50ntFlanks</td><td>10510</td><td>10534</td><td>25</td><td>AATCCGGAGTGGCGGAGATGGGCGC</td><td>762.2</td><td>-13.4</td><td>1.4</td><td>-9.4</td><td>8.1E-21</td><td>1.5E-19</td><td>MB231/MB468</td><td>MP</td></tr>
<tr><td>>hg38_21_+_8433172_8446622_RNA45SN1_45S_50ntFlanks@475.514.40|CTTCGTGATCGATGTGGTGACGTCGTGCTCTCCCGGGCCG</td><td>>hg38_21_+_8433172_8446622_RNA45SN1_45S_50ntFlanks</td><td>475</td><td>514</td><td>40</td><td>CTTCGTGATCGATGTGGTGACGTCGTGCTCTCCCGGGCCG</td><td>986.0</td><td>9.0</td><td>1.0</td><td>9.3</td><td>1.9E-20</td><td>1.5E-19</td><td>BT20/MB468</td><td>CYTO</td></tr>
<tr><td>>hg38_21_+_8433172_8446622_RNA45SN1_45S_50ntFlanks@12655.12674.20|ATGACTGAACGCCTCTAAGT</td><td>>hg38_21_+_8433172_8446622_RNA45SN1_45S_50ntFlanks</td><td>12655</td><td>12674</td><td>20</td><td>ATGACTGAACGCCTCTAAGT</td><td>2614.6</td><td>-5.4</td><td>0.6</td><td>-9.3</td><td>1.9E-20</td><td>1.5E-19</td><td>BT20/MB468</td><td>CYTO</td></tr>
<tr><td>>hg38_1_-_228634819_228635039_RNA5S12_5S_50ntFlanks@147.170.24|GGGAATACCGGGTGCTGTAGGCTT</td><td>>hg38_1_-_228634819_228635039_RNA5S12_5S_50ntFlanks</td><td>147</td><td>170</td><td>24</td><td>GGGAATACCGGGTGCTGTAGGCTT</td><td>7500.3</td><td>6.0</td><td>0.7</td><td>9.3</td><td>1.9E-20</td><td>1.5E-19</td><td>BT20/MB468</td><td>CYTO</td></tr>
<tr><td>>hg38_21_+_8433172_8446622_RNA45SN1_45S_50ntFlanks@11670.11709.40|CCCAGTGCTCTGAATGTCAAAGTGAAGAAATTCAATGAAG</td><td>>hg38_21_+_8433172_8446622_RNA45SN1_45S_50ntFlanks</td><td>11670</td><td>11709</td><td>40</td><td>CCCAGTGCTCTGAATGTCAAAGTGAAGAAATTCAATGAAG</td><td>449.2</td><td>-13.2</td><td>1.4</td><td>-9.2</td><td>3.7E-20</td><td>1.5E-19</td><td>BT20/MB468</td><td>NUC</td></tr>
<tr><td>>hg38_21_+_8433172_8446622_RNA45SN1_45S_50ntFlanks@10808.10825.18|CAATGTAGGTAAGGGAAG</td><td>>hg38_21_+_8433172_8446622_RNA45SN1_45S_50ntFlanks</td><td>10808</td><td>10825</td><td>18</td><td>CAATGTAGGTAAGGGAAG</td><td>745.2</td><td>-13.4</td><td>1.4</td><td>-9.4</td><td>8.3E-21</td><td>1.5E-19</td><td>MB231/MB468</td><td>MP</td></tr>
<tr><td>>hg38_21_+_8433172_8446622_RNA45SN1_45S_50ntFlanks@10873.10891.19|CTGGGTCGGTCGGGCTGGG</td><td>>hg38_21_+_8433172_8446622_RNA45SN1_45S_50ntFlanks</td><td>10873</td><td>10891</td><td>19</td><td>CTGGGTCGGTCGGGCTGGG</td><td>919.5</td><td>-13.7</td><td>1.5</td><td>-9.4</td><td>8.3E-21</td><td>1.5E-19</td><td>MB231/MB468</td><td>MP</td></tr>
<tr><td>>hg38_21_+_8433172_8446622_RNA45SN1_45S_50ntFlanks@11621.11640.20|CAAAGCATCGCGAAGGCCCG</td><td>>hg38_21_+_8433172_8446622_RNA45SN1_45S_50ntFlanks</td><td>11621</td><td>11640</td><td>20</td><td>CAAAGCATCGCGAAGGCCCG</td><td>366.1</td><td>-12.9</td><td>1.4</td><td>-9.2</td><td>3.8E-20</td><td>1.5E-19</td><td>BT20/MB468</td><td>NUC</td></tr>
</table>

<table>
<tr><td>>MI0000102|hsa-mir-100&WithFlank&11|-|122152223|122152314@19.38.20(+1A)&[MIMAT0000098&hsa-miR-100-5p&offsets|0|-2(+1A);m-27&11|-|122152276|122152296&offsets|0|-1(+1A)]|AACCCGTAGATCCGAACTTGA</td><td>>MI0000102|hsa-mir-100&WithFlank&11|-|122152223|122152314</td><td>19</td><td>38</td><td>20(+1A)</td><td>AACCCGTAGATCCGAACTTGA</td><td>4382.8</td><td>-7.9</td><td>0.8</td><td>-9.6</td><td>1.1E-21</td><td>1.5E-19</td><td>BT20/MB231</td><td>MITO</td></tr>
<tr><td>>MI0000456|hsa-mir-140&WithFlank&16|+|69933075|69933186@69.90.22&[MIMAT0004597&hsa-miR-140-3p&offsets|+1|+2;m-45&16|+|69933143|69933164&offsets|0|0]|ACCACAGGGTAGAACCACGGAC</td><td>>MI0000456|hsa-mir-140&WithFlank&16|+|69933075|69933186</td><td>69</td><td>90</td><td>22</td><td>ACCACAGGGTAGAACCACGGAC</td><td>3351.2</td><td>11.2</td><td>1.2</td><td>9.3</td><td>2.0E-20</td><td>1.5E-19</td><td>BT20/MB468</td><td>CYTO</td></tr>
<tr><td>>hg38_21_+_8433172_8446622_RNA45SN1_45S_50ntFlanks@12936.12953.18|AAACCATTCGTAGACGAC</td><td>>hg38_21_+_8433172_8446622_RNA45SN1_45S_50ntFlanks</td><td>12936</td><td>12953</td><td>18</td><td>AAACCATTCGTAGACGAC</td><td>662.2</td><td>-13.2</td><td>1.4</td><td>-9.4</td><td>8.5E-21</td><td>1.5E-19</td><td>MB231/MB468</td><td>MP</td></tr>
<tr><td>>hg38_21_+_8433172_8446622_RNA45SN1_45S_50ntFlanks@9470.9503.34|GAGCACGAGCGCACGTGTTAGGACCCGAAAGATG</td><td>>hg38_21_+_8433172_8446622_RNA45SN1_45S_50ntFlanks</td><td>9470</td><td>9503</td><td>34</td><td>GAGCACGAGCGCACGTGTTAGGACCCGAAAGATG</td><td>1584.1</td><td>-14.5</td><td>1.5</td><td>-9.4</td><td>8.5E-21</td><td>1.5E-19</td><td>MB231/MB468</td><td>MP</td></tr>
<tr><td>>hg38_21_+_8433172_8446622_RNA45SN1_45S_50ntFlanks@6741.6770.30|ATCGACACTTCGAACGCACTTGCGGCCCCG</td><td>>hg38_21_+_8433172_8446622_RNA45SN1_45S_50ntFlanks</td><td>6741</td><td>6770</td><td>30</td><td>ATCGACACTTCGAACGCACTTGCGGCCCCG</td><td>1724.5</td><td>-7.8</td><td>0.8</td><td>-9.4</td><td>7.0E-21</td><td>1.5E-19</td><td>MB231/MB468</td><td>CYTO</td></tr>
<tr><td>>MI0000088|hsa-mir-30a&WithFlank&6|-|71403545|71403627@12.34.23(+1C)&[MIMAT0000087&hsa-miR-30a-5p&offsets|0|+1(+1C);m-15&6|-|71403595|71403616&offsets|0|+1(+1C)]|TGTAAACATCCTCGACTGGAAGCC</td><td>>MI0000088|hsa-mir-30a&WithFlank&6|-|71403545|71403627</td><td>12</td><td>34</td><td>23(+1C)</td><td>TGTAAACATCCTCGACTGGAAGCC</td><td>11798.7</td><td>7.5</td><td>0.8</td><td>9.3</td><td>1.2E-20</td><td>1.5E-19</td><td>MB231/MB468</td><td>NUC</td></tr>
<tr><td>>MI0000462|hsa-mir-152&WithFlank&17|-|48037155|48037253@60.81.22&[MIMAT0000438&hsa-miR-152-3p&offsets|0|+1;m-97&17|-|48037174|48037194&offsets|0|+1]|TCAGTGCATGACAGAACTTGGG</td><td>>MI0000462|hsa-mir-152&WithFlank&17|-|48037155|48037253</td><td>60</td><td>81</td><td>22</td><td>TCAGTGCATGACAGAACTTGGG</td><td>183.6</td><td>6.9</td><td>0.8</td><td>9.2</td><td>3.9E-20</td><td>1.5E-19</td><td>BT20/MB468</td><td>NUC</td></tr>
<tr><td>>hg38_21_+_8433172_8446622_RNA45SN1_45S_50ntFlanks@10408.10435.28|GGTCAGTCGGTCCTGAGAGATGGGCGAG</td><td>>hg38_21_+_8433172_8446622_RNA45SN1_45S_50ntFlanks</td><td>10408</td><td>10435</td><td>28</td><td>GGTCAGTCGGTCCTGAGAGATGGGCGAG</td><td>2581.3</td><td>-14.3</td><td>1.5</td><td>-9.3</td><td>1.2E-20</td><td>1.5E-19</td><td>MB231/MB468</td><td>NUC</td></tr>
<tr><td>>hg38_21_+_8433172_8446622_RNA45SN1_45S_50ntFlanks@10783.10818.36|GTGAACAGCCTCTGGCATGTTGGAACAATGTAGGTA</td><td>>hg38_21_+_8433172_8446622_RNA45SN1_45S_50ntFlanks</td><td>10783</td><td>10818</td><td>36</td><td>GTGAACAGCCTCTGGCATGTTGGAACAATGTAGGTA</td><td>1325.1</td><td>-14.2</td><td>1.5</td><td>-9.4</td><td>8.6E-21</td><td>1.5E-19</td><td>MB231/MB468</td><td>MP</td></tr>
<tr><td>>hg38_21_+_8433172_8446622_RNA45SN1_45S_50ntFlanks@9033.9055.23|CTCGTGGGGGGGCCGGGCCACCC</td><td>>hg38_21_+_8433172_8446622_RNA45SN1_45S_50ntFlanks</td><td>9033</td><td>9055</td><td>23</td><td>CTCGTGGGGGGGCCGGGCCACCC</td><td>769.0</td><td>-13.5</td><td>1.4</td><td>-9.4</td><td>8.7E-21</td><td>1.5E-19</td><td>MB231/MB468</td><td>MP</td></tr>
<tr><td>>MI0000298|hsa-mir-221&WithFlank&X|-|45746151|45746272@31.54.24&[MIMAT0004568&hsa-miR-221-5p&offsets|0|+2;m-405&X|-|45746220|45746242&offsets|0|+1]|ACCTGGCATACAATGTAGATTTCT</td><td>>MI0000298|hsa-mir-221&WithFlank&X|-|45746151|45746272</td><td>31</td><td>54</td><td>24</td><td>ACCTGGCATACAATGTAGATTTCT</td><td>1743.3</td><td>6.2</td><td>0.7</td><td>9.3</td><td>1.2E-20</td><td>1.6E-19</td><td>MB231/MB468</td><td>NUC</td></tr>
<tr><td>>MI0000065|hsa-let-7d&WithFlank&9|+|94178828|94178926@14.36.23&[MIMAT0000065&hsa-let-7d-5p&offsets|0|+1;m-90&9|+|94178841|94178862&offsets|0|+1]|AGAGGTAGTAGGTTGCATAGTTT</td><td>>MI0000065|hsa-let-7d&WithFlank&9|+|94178828|94178926</td><td>14</td><td>36</td><td>23</td><td>AGAGGTAGTAGGTTGCATAGTTT</td><td>824.2</td><td>7.6</td><td>0.8</td><td>9.3</td><td>2.1E-20</td><td>1.6E-19</td><td>BT20/MB468</td><td>CYTO</td></tr>
<tr><td>>MI0000456|hsa-mir-140&WithFlank&16|+|69933075|69933186@69.91.23&[MIMAT0004597&hsa-miR-140-3p&offsets|+1|+3;m-45&16|+|69933143|69933164&offsets|0|+1]|ACCACAGGGTAGAACCACGGACA</td><td>>MI0000456|hsa-mir-140&WithFlank&16|+|69933075|69933186</td><td>69</td><td>91</td><td>23</td><td>ACCACAGGGTAGAACCACGGACA</td><td>2109.5</td><td>12.2</td><td>1.3</td><td>9.3</td><td>2.1E-20</td><td>1.6E-19</td><td>BT20/MB468</td><td>CYTO</td></tr>
<tr><td>>hg38_21_+_8433172_8446622_RNA45SN1_45S_50ntFlanks@3931.3959.29|TCAAAACCAACCCGGTCAGCCCCTCTCCG</td><td>>hg38_21_+_8433172_8446622_RNA45SN1_45S_50ntFlanks</td><td>3931</td><td>3959</td><td>29</td><td>TCAAAACCAACCCGGTCAGCCCCTCTCCG</td><td>868.6</td><td>-13.5</td><td>1.5</td><td>-9.3</td><td>1.3E-20</td><td>1.6E-19</td><td>MB231/MB468</td><td>MITO</td></tr>
<tr><td>>hg38_21_+_8433172_8446622_RNA45SN1_45S_50ntFlanks@10684.10706.23|CGGCGGCGTCCGGTGAGCTCTCG</td><td>>hg38_21_+_8433172_8446622_RNA45SN1_45S_50ntFlanks</td><td>10684</td><td>10706</td><td>23</td><td>CGGCGGCGTCCGGTGAGCTCTCG</td><td>5951.7</td><td>-7.2</td><td>0.8</td><td>-9.4</td><td>7.2E-21</td><td>1.6E-19</td><td>MB231/MB468</td><td>CYTO</td></tr>
<tr><td>>hg38_21_+_8433172_8446622_RNA45SN1_45S_50ntFlanks@11999.12037.39|CCCTGCGGGCCGCCGGTGAAATACCACTACTCTGATCGT</td><td>>hg38_21_+_8433172_8446622_RNA45SN1_45S_50ntFlanks</td><td>11999</td><td>12037</td><td>39</td><td>CCCTGCGGGCCGCCGGTGAAATACCACTACTCTGATCGT</td><td>370.6</td><td>-13.0</td><td>1.4</td><td>-9.2</td><td>4.0E-20</td><td>1.6E-19</td><td>BT20/MB468</td><td>NUC</td></tr>
<tr><td>>MI0000270|hsa-mir-181b-1&WithFlank&1|-|198858867|198858988@42.64.23;MI0000683|hsa-mir-181b-2&WithFlank&9|+|124693704|124693804@22.44.23&[MIMAT0000257&hsa-miR-181b-5p&offsets|0|0;m-73&1|-|198858925|198858947&offsets|0|0];[MIMAT0000257_1&hsa-miR-181b-5p&offsets|0|0;m-72&9|+|124693725|124693747&offsets|0|0]|AACATTCATTGCTGTCGGTGGGT</td><td>>MI0000270|hsa-mir-181b-1&WithFlank&1|-|198858867|198858988</td><td>42</td><td>64</td><td>23</td><td>AACATTCATTGCTGTCGGTGGGT</td><td>1285.2</td><td>6.1</td><td>0.7</td><td>9.3</td><td>1.5E-20</td><td>1.6E-19</td><td>BT20/MB468</td><td>MITO</td></tr>
<tr><td>>MI0000088|hsa-mir-30a&WithFlank&6|-|71403545|71403627@12.33.22(+1U)&[MIMAT0000087&hsa-miR-30a-5p&offsets|0|0(+1U);m-15&6|-|71403595|71403616&offsets|0|0(+1U)]|TGTAAACATCCTCGACTGGAAGT</td><td>>MI0000088|hsa-mir-30a&WithFlank&6|-|71403545|71403627</td><td>12</td><td>33</td><td>22(+1U)</td><td>TGTAAACATCCTCGACTGGAAGT</td><td>2826.4</td><td>6.5</td><td>0.7</td><td>9.3</td><td>1.5E-20</td><td>1.6E-19</td><td>BT20/MB468</td><td>MITO</td></tr>
<tr><td>>hg38_21_+_8433172_8446622_RNA45SN1_45S_50ntFlanks@10408.10431.24|GGTCAGTCGGTCCTGAGAGATGGG</td><td>>hg38_21_+_8433172_8446622_RNA45SN1_45S_50ntFlanks</td><td>10408</td><td>10431</td><td>24</td><td>GGTCAGTCGGTCCTGAGAGATGGG</td><td>1742.1</td><td>-13.7</td><td>1.5</td><td>-9.3</td><td>1.2E-20</td><td>1.6E-19</td><td>MB231/MB468</td><td>NUC</td></tr>
<tr><td>>hg38_21_+_8433172_8446622_RNA45SN1_45S_50ntFlanks@12932.12972.41|CGCTAAACCATTCGTAGACGACCTGCTTCTGGGTCGGGGTT</td><td>>hg38_21_+_8433172_8446622_RNA45SN1_45S_50ntFlanks</td><td>12932</td><td>12972</td><td>41</td><td>CGCTAAACCATTCGTAGACGACCTGCTTCTGGGTCGGGGTT</td><td>1081.6</td><td>-7.1</td><td>0.8</td><td>-9.2</td><td>4.1E-20</td><td>1.6E-19</td><td>BT20/MB468</td><td>NUC</td></tr>
<tr><td>>hg38_21_+_8433172_8446622_RNA45SN1_45S_50ntFlanks@9032.9058.27|CCTCGTGGGGGGGCCGGGCCACCCCTC</td><td>>hg38_21_+_8433172_8446622_RNA45SN1_45S_50ntFlanks</td><td>9032</td><td>9058</td><td>27</td><td>CCTCGTGGGGGGGCCGGGCCACCCCTC</td><td>678.7</td><td>-13.3</td><td>1.4</td><td>-9.3</td><td>8.9E-21</td><td>1.6E-19</td><td>MB231/MB468</td><td>MP</td></tr>
<tr><td>>hg38_21_+_8433172_8446622_RNA45SN1_45S_50ntFlanks@10683.10710.28|CCGGCGGCGTCCGGTGAGCTCTCGCTGG</td><td>>hg38_21_+_8433172_8446622_RNA45SN1_45S_50ntFlanks</td><td>10683</td><td>10710</td><td>28</td><td>CCGGCGGCGTCCGGTGAGCTCTCGCTGG</td><td>291.2</td><td>-12.6</td><td>1.4</td><td>-9.2</td><td>4.1E-20</td><td>1.6E-19</td><td>BT20/MB468</td><td>NUC</td></tr>
<tr><td>>hg38_21_+_8433172_8446622_RNA45SN1_45S_50ntFlanks@11641.11661.21|CGGCGGGTGTTGACGCGATGT</td><td>>hg38_21_+_8433172_8446622_RNA45SN1_45S_50ntFlanks</td><td>11641</td><td>11661</td><td>21</td><td>CGGCGGGTGTTGACGCGATGT</td><td>1011.2</td><td>-13.8</td><td>1.5</td><td>-9.3</td><td>1.4E-20</td><td>1.6E-19</td><td>MB231/MB468</td><td>MITO</td></tr>
<tr><td>>hg38_21_+_8433172_8446622_RNA45SN1_45S_50ntFlanks@12991.13014.24|CTCCCTCGCTGCGATCTATTGAAA</td><td>>hg38_21_+_8433172_8446622_RNA45SN1_45S_50ntFlanks</td><td>12991</td><td>13014</td><td>24</td><td>CTCCCTCGCTGCGATCTATTGAAA</td><td>3489.1</td><td>-9.3</td><td>1.0</td><td>-9.3</td><td>1.4E-20</td><td>1.6E-19</td><td>MB231/MB468</td><td>MITO</td></tr>
<tr><td>>MI0000077|hsa-mir-21&WithFlank&17|+|59841260|59841343@14.31.18&[MIMAT0000076&hsa-miR-21-5p&offsets|0|-4;m-1&17|+|59841273|59841294&offsets|0|-4]|TAGCTTATCAGACTGATG</td><td>>MI0000077|hsa-mir-21&WithFlank&17|+|59841260|59841343</td><td>14</td><td>31</td><td>18</td><td>TAGCTTATCAGACTGATG</td><td>546.5</td><td>5.9</td><td>0.6</td><td>9.3</td><td>1.3E-20</td><td>1.6E-19</td><td>MB231/MB468</td><td>NUC</td></tr>
<tr><td>>hg38_21_+_8433172_8446622_RNA45SN1_45S_50ntFlanks@10351.10390.40|CGAGAACTTTGAAGGCCGAAGTGGAGAAGGGTTCCATGTG</td><td>>hg38_21_+_8433172_8446622_RNA45SN1_45S_50ntFlanks</td><td>10351</td><td>10390</td><td>40</td><td>CGAGAACTTTGAAGGCCGAAGTGGAGAAGGGTTCCATGTG</td><td>16569.8</td><td>-7.3</td><td>0.8</td><td>-9.3</td><td>1.3E-20</td><td>1.6E-19</td><td>MB231/MB468</td><td>NUC</td></tr>
<tr><td>>hg38_21_+_8433172_8446622_RNA45SN1_45S_50ntFlanks@10727.10755.29|GGAGAGGGTGTAAATCTCGCGCCGGGCCG</td><td>>hg38_21_+_8433172_8446622_RNA45SN1_45S_50ntFlanks</td><td>10727</td><td>10755</td><td>29</td><td>GGAGAGGGTGTAAATCTCGCGCCGGGCCG</td><td>1925.5</td><td>-13.9</td><td>1.5</td><td>-9.3</td><td>1.3E-20</td><td>1.6E-19</td><td>MB231/MB468</td><td>NUC</td></tr>
<tr><td>>hg38_21_+_8433172_8446622_RNA45SN1_45S_50ntFlanks@11321.11342.22|CACGTCTCGTCGCGCGCGCGTC</td><td>>hg38_21_+_8433172_8446622_RNA45SN1_45S_50ntFlanks</td><td>11321</td><td>11342</td><td>22</td><td>CACGTCTCGTCGCGCGCGCGTC</td><td>592.8</td><td>-13.1</td><td>1.4</td><td>-9.3</td><td>9.1E-21</td><td>1.6E-19</td><td>MB231/MB468</td><td>MP</td></tr>
<tr><td>>hg38_21_+_8433172_8446622_RNA45SN1_45S_50ntFlanks@4723.4768.46|CATTAATCAAGAACGAAAGTCGGAGGTTCGAAGACGATCAGATACC</td><td>>hg38_21_+_8433172_8446622_RNA45SN1_45S_50ntFlanks</td><td>4723</td><td>4768</td><td>46</td><td>CATTAATCAAGAACGAAAGTCGGAGGTTCGAAGACGATCAGATACC</td><td>369.9</td><td>12.5</td><td>1.3</td><td>9.4</td><td>7.5E-21</td><td>1.6E-19</td><td>MB231/MB468</td><td>CYTO</td></tr>
<tr><td>>hg38_21_+_8433172_8446622_RNA45SN1_45S_50ntFlanks@4220.4242.23|ACGAAAAATAACAATACAGGACT</td><td>>hg38_21_+_8433172_8446622_RNA45SN1_45S_50ntFlanks</td><td>4220</td><td>4242</td><td>23</td><td>ACGAAAAATAACAATACAGGACT</td><td>730.6</td><td>-5.0</td><td>0.5</td><td>-9.3</td><td>1.6E-20</td><td>1.6E-19</td><td>BT20/MB468</td><td>MITO</td></tr>
<tr><td>>MI0000299|hsa-mir-222&WithFlank&X|-|45747009|45747130@75.96.22&[MIMAT0000279&hsa-miR-222-3p&offsets|0|+1;m-140&X|-|45747033|45747056&offsets|0|-2]|AGCTACATCTGGCTACTGGGTC</td><td>>MI0000299|hsa-mir-222&WithFlank&X|-|45747009|45747130</td><td>75</td><td>96</td><td>22</td><td>AGCTACATCTGGCTACTGGGTC</td><td>1808.8</td><td>7.2</td><td>0.8</td><td>9.4</td><td>7.5E-21</td><td>1.6E-19</td><td>MB231/MB468</td><td>CYTO</td></tr>
<tr><td>>hg38_21_+_8433172_8446622_RNA45SN1_45S_50ntFlanks@10866.10901.36|CTAAGGGCTGGGTCGGTCGGGCTGGGGCGCGAAGCG</td><td>>hg38_21_+_8433172_8446622_RNA45SN1_45S_50ntFlanks</td><td>10866</td><td>10901</td><td>36</td><td>CTAAGGGCTGGGTCGGTCGGGCTGGGGCGCGAAGCG</td><td>678.0</td><td>-13.3</td><td>1.4</td><td>-9.3</td><td>9.2E-21</td><td>1.6E-19</td><td>MB231/MB468</td><td>MP</td></tr>
<tr><td>>hg38_21_+_8433172_8446622_RNA45SN1_45S_50ntFlanks@12081.12110.30|CTCTCGCTTCTGGCGCCAAGCGCCCGGCCG</td><td>>hg38_21_+_8433172_8446622_RNA45SN1_45S_50ntFlanks</td><td>12081</td><td>12110</td><td>30</td><td>CTCTCGCTTCTGGCGCCAAGCGCCCGGCCG</td><td>857.5</td><td>-13.6</td><td>1.5</td><td>-9.3</td><td>9.3E-21</td><td>1.6E-19</td><td>MB231/MB468</td><td>MP</td></tr>
<tr><td>>hg38_21_+_8433172_8446622_RNA45SN1_45S_50ntFlanks@12619.12654.36|AGGAGCCAATGGGGCGAAGCTACCATCTGTGGGATT</td><td>>hg38_21_+_8433172_8446622_RNA45SN1_45S_50ntFlanks</td><td>12619</td><td>12654</td><td>36</td><td>AGGAGCCAATGGGGCGAAGCTACCATCTGTGGGATT</td><td>2111.6</td><td>-6.9</td><td>0.7</td><td>-9.3</td><td>1.3E-20</td><td>1.7E-19</td><td>MB231/MB468</td><td>NUC</td></tr>
<tr><td>>hg38_21_+_8433172_8446622_RNA45SN1_45S_50ntFlanks@9133.9152.20|TCGGGGGGCGGGGCGGACTG</td><td>>hg38_21_+_8433172_8446622_RNA45SN1_45S_50ntFlanks</td><td>9133</td><td>9152</td><td>20</td><td>TCGGGGGGCGGGGCGGACTG</td><td>1820.8</td><td>-14.7</td><td>1.6</td><td>-9.3</td><td>9.4E-21</td><td>1.7E-19</td><td>MB231/MB468</td><td>MP</td></tr>
<tr><td>>MI0000082|hsa-mir-25&WithFlank&7|-|100093554|100093649@58.78.21(+1G)&[MIMAT0000081&hsa-miR-25-3p&offsets|0|-1(+1G);m-23&7|-|100093571|100093592&offsets|0|-1(+1G)]|CATTGCACTTGTCTCGGTCTGG</td><td>>MI0000082|hsa-mir-25&WithFlank&7|-|100093554|100093649</td><td>58</td><td>78</td><td>21(+1G)</td><td>CATTGCACTTGTCTCGGTCTGG</td><td>2213.9</td><td>6.5</td><td>0.7</td><td>9.3</td><td>2.2E-20</td><td>1.7E-19</td><td>BT20/MB468</td><td>CYTO</td></tr>
<tr><td>>hg38_21_+_8433172_8446622_RNA45SN1_45S_50ntFlanks@9575.9595.21|ATCGGTCGTCCGACCTGGGTA</td><td>>hg38_21_+_8433172_8446622_RNA45SN1_45S_50ntFlanks</td><td>9575</td><td>9595</td><td>21</td><td>ATCGGTCGTCCGACCTGGGTA</td><td>899.5</td><td>-12.1</td><td>1.3</td><td>-9.3</td><td>2.2E-20</td><td>1.7E-19</td><td>BT20/MB468</td><td>CYTO</td></tr>
<tr><td>>am_MI0000809|hsa-mir-151a&WithFlank&8|-|140732558|140732659@52.72.21(+1A)&[MIMAT0000757&hsa-miR-151a-3p&offsets|-1|-1(+1A);m-30&8|-|140732587|140732607&offsets|-1|-1(+1A)]|ACTAGACTGAAGCTCCTTGAGA</td><td>>am_MI0000809|hsa-mir-151a&WithFlank&8|-|140732558|140732659</td><td>52</td><td>72</td><td>21(+1A)</td><td>ACTAGACTGAAGCTCCTTGAGA</td><td>697.0</td><td>11.1</td><td>1.2</td><td>9.3</td><td>2.2E-20</td><td>1.7E-19</td><td>BT20/MB468</td><td>CYTO</td></tr>
<tr><td>>hg38_21_+_8433172_8446622_RNA45SN1_45S_50ntFlanks@5492.5516.25|AGAAGACGGTCGAACTTGACTATCT</td><td>>hg38_21_+_8433172_8446622_RNA45SN1_45S_50ntFlanks</td><td>5492</td><td>5516</td><td>25</td><td>AGAAGACGGTCGAACTTGACTATCT</td><td>50466.1</td><td>8.0</td><td>0.9</td><td>9.3</td><td>2.2E-20</td><td>1.7E-19</td><td>BT20/MB468</td><td>CYTO</td></tr>
<tr><td>>MI0000266|hsa-mir-10a&WithFlank&17|-|48579832|48579953@29.51.23&[MIMAT0000253&hsa-miR-10a-5p&offsets|+1|+1;m-18&17|-|48579905|48579926&offsets|+1|+2]|ACCCTGTAGATCCGAATTTGTGT</td><td>>MI0000266|hsa-mir-10a&WithFlank&17|-|48579832|48579953</td><td>29</td><td>51</td><td>23</td><td>ACCCTGTAGATCCGAATTTGTGT</td><td>775.6</td><td>13.6</td><td>1.5</td><td>9.4</td><td>7.7E-21</td><td>1.7E-19</td><td>MB231/MB468</td><td>CYTO</td></tr>
<tr><td>>hg38_21_+_8433172_8446622_RNA45SN1_45S_50ntFlanks@9622.9652.31|CTAGTAGCTGGTTCCCTCCGAAGTTTCCCTC</td><td>>hg38_21_+_8433172_8446622_RNA45SN1_45S_50ntFlanks</td><td>9622</td><td>9652</td><td>31</td><td>CTAGTAGCTGGTTCCCTCCGAAGTTTCCCTC</td><td>720.3</td><td>-13.3</td><td>1.4</td><td>-9.3</td><td>1.4E-20</td><td>1.7E-19</td><td>MB231/MB468</td><td>MITO</td></tr>
<tr><td>>MI0000298|hsa-mir-221&WithFlank&X|-|45746151|45746272@71.92.22(+1A)&[MIMAT0000278&hsa-miR-221-3p&offsets|0|-1(+1A);m-91&X|-|45746181|45746202&offsets|0|0(+1A)]|AGCTACATTGTCTGCTGGGTTTA</td><td>>MI0000298|hsa-mir-221&WithFlank&X|-|45746151|45746272</td><td>71</td><td>92</td><td>22(+1A)</td><td>AGCTACATTGTCTGCTGGGTTTA</td><td>716.6</td><td>6.0</td><td>0.6</td><td>9.3</td><td>1.3E-20</td><td>1.7E-19</td><td>MB231/MB468</td><td>NUC</td></tr>
<tr><td>>hg38_21_+_8433172_8446622_RNA45SN1_45S_50ntFlanks@11104.11123.20|AAGGGTCGGGGCGGCAGGGG</td><td>>hg38_21_+_8433172_8446622_RNA45SN1_45S_50ntFlanks</td><td>11104</td><td>11123</td><td>20</td><td>AAGGGTCGGGGCGGCAGGGG</td><td>1123.1</td><td>-13.1</td><td>1.4</td><td>-9.3</td><td>1.3E-20</td><td>1.7E-19</td><td>MB231/MB468</td><td>NUC</td></tr>
<tr><td>>hg38_21_+_8433172_8446622_RNA45SN1_45S_50ntFlanks@4367.4398.32|CAGTTAAAAAGCTCGTAGTTGGATCTTGGGAG</td><td>>hg38_21_+_8433172_8446622_RNA45SN1_45S_50ntFlanks</td><td>4367</td><td>4398</td><td>32</td><td>CAGTTAAAAAGCTCGTAGTTGGATCTTGGGAG</td><td>1298.9</td><td>-13.3</td><td>1.4</td><td>-9.3</td><td>1.3E-20</td><td>1.7E-19</td><td>MB231/MB468</td><td>NUC</td></tr>
<tr><td>>hg38_21_+_8433172_8446622_RNA45SN1_45S_50ntFlanks@5488.5516.29|GCTGAGAAGACGGTCGAACTTGACTATCT</td><td>>hg38_21_+_8433172_8446622_RNA45SN1_45S_50ntFlanks</td><td>5488</td><td>5516</td><td>29</td><td>GCTGAGAAGACGGTCGAACTTGACTATCT</td><td>1657.4</td><td>5.8</td><td>0.6</td><td>9.4</td><td>7.9E-21</td><td>1.7E-19</td><td>MB231/MB468</td><td>CYTO</td></tr>
<tr><td>>hg38_21_+_8433172_8446622_RNA45SN1_45S_50ntFlanks@10411.10433.23|CAGTCGGTCCTGAGAGATGGGCG</td><td>>hg38_21_+_8433172_8446622_RNA45SN1_45S_50ntFlanks</td><td>10411</td><td>10433</td><td>23</td><td>CAGTCGGTCCTGAGAGATGGGCG</td><td>2234.7</td><td>-14.9</td><td>1.6</td><td>-9.3</td><td>1.5E-20</td><td>1.7E-19</td><td>MB231/MB468</td><td>MITO</td></tr>
<tr><td>>hg38_21_+_8433172_8446622_RNA45SN1_45S_50ntFlanks@12991.13011.21|CTCCCTCGCTGCGATCTATTG</td><td>>hg38_21_+_8433172_8446622_RNA45SN1_45S_50ntFlanks</td><td>12991</td><td>13011</td><td>21</td><td>CTCCCTCGCTGCGATCTATTG</td><td>2107.3</td><td>-10.7</td><td>1.2</td><td>-9.3</td><td>1.5E-20</td><td>1.7E-19</td><td>MB231/MB468</td><td>MITO</td></tr>
</table>

<table>
<tr><td>>hg38_21_+_8433172_8446622_RNA45SN1_45S_50ntFlanks@12419.12459.41|GATGTCGGCTCTTCCTATCATTGTGAAGCAGAATTCACCAA</td><td>>hg38_21_+_8433172_8446622_RNA45SN1_45S_50ntFlanks</td><td>12419</td><td>12459</td><td>41</td><td>GATGTCGGCTCTTCCTATCATTGTGAAGCAGAATTCACCAA</td><td>378.3</td><td>10.2</td><td>1.1</td><td>9.4</td><td>7.9E-21</td><td>1.7E-19</td><td>MB231/MB468</td><td>CYTO</td></tr>
<tr><td>>hg38_21_+_8433172_8446622_RNA45SN1_45S_50ntFlanks@11104.11123.20|AAGGGTCGGGGCGGCAGGGG</td><td>>hg38_21_+_8433172_8446622_RNA45SN1_45S_50ntFlanks</td><td>11104</td><td>11123</td><td>20</td><td>AAGGGTCGGGGCGGCAGGGG</td><td>988.5</td><td>-13.8</td><td>1.5</td><td>-9.3</td><td>9.7E-21</td><td>1.7E-19</td><td>MB231/MB468</td><td>MP</td></tr>
<tr><td>>MI0003776|hsa-mir-320b-1&WithFlank&1|+|116671740|116671823@48.67.20(+1C);MI0003839|hsa-mir-320b-2&WithFlank&1|-|224257034|224257116@47.66.20(+1C);MI0000542|hsa-mir-320a&WithFlank&8|-|22244960|22245043@48.67.20(+1C)&[MIMAT0005792&hsa-miR-320b&offsets|0|-2(+1C);m-183&1|+|116671787|116671806&offsets|0|0(+1C)];[MIMAT0005792_1&hsa-miR-320b&offsets|0|-2(+1C);m-183&1|-|224257051|224257070&offsets|0|0(+1C)];[MIMAT0000510&hsa-miR-320a-3p&offsets|0|-2(+1C);m-60&8|-|22244975|22244996&offsets|0|-2(+1C)]|AAAAGCTGGGTTGAGAGGGCC</td><td>>MI0003776|hsa-mir-320b-1&WithFlank&1|+|116671740|116671823</td><td>48</td><td>67</td><td>20(+1C)</td><td>AAAAGCTGGGTTGAGAGGGCC</td><td>393.7</td><td>12.9</td><td>1.4</td><td>9.2</td><td>2.3E-20</td><td>1.7E-19</td><td>BT20/MB468</td><td>CYTO</td></tr>
<tr><td>>hg38_1_-_228634819_228635039_RNA5S12_5S_50ntFlanks@135.169.35|GGGAGACCGCCTGGGAATACCGGGTGCTGTAGGCT</td><td>>hg38_1_-_228634819_228635039_RNA5S12_5S_50ntFlanks</td><td>135</td><td>169</td><td>35</td><td>GGGAGACCGCCTGGGAATACCGGGTGCTGTAGGCT</td><td>4804.8</td><td>6.7</td><td>0.7</td><td>9.2</td><td>2.3E-20</td><td>1.7E-19</td><td>BT20/MB468</td><td>CYTO</td></tr>
<tr><td>>MI0000086|hsa-mir-28&WithFlank&3|+|188688775|188688872@60.80.21&[MIMAT0004502&hsa-miR-28-3p&offsets|0|-1;m-28&3|+|188688834|188688855&offsets|0|-1]|CACTAGATTGTGAGCTCCTGG</td><td>>MI0000086|hsa-mir-28&WithFlank&3|+|188688775|188688872</td><td>60</td><td>80</td><td>21</td><td>CACTAGATTGTGAGCTCCTGG</td><td>248.3</td><td>3.6</td><td>0.4</td><td>9.2</td><td>4.4E-20</td><td>1.7E-19</td><td>BT20/MB468</td><td>NUC</td></tr>
<tr><td>>hg38_21_+_8433172_8446622_RNA45SN1_45S_50ntFlanks@12073.12095.23|CCGAGGGGCTCTCGCTTCTGGCG</td><td>>hg38_21_+_8433172_8446622_RNA45SN1_45S_50ntFlanks</td><td>12073</td><td>12095</td><td>23</td><td>CCGAGGGGCTCTCGCTTCTGGCG</td><td>5784.5</td><td>-6.6</td><td>0.7</td><td>-9.4</td><td>8.1E-21</td><td>1.7E-19</td><td>MB231/MB468</td><td>CYTO</td></tr>
<tr><td>>hg38_21_+_8433172_8446622_RNA45SN1_45S_50ntFlanks@12158.12180.23|GTTTGACTGGGGCGGTACACCTG</td><td>>hg38_21_+_8433172_8446622_RNA45SN1_45S_50ntFlanks</td><td>12158</td><td>12180</td><td>23</td><td>GTTTGACTGGGGCGGTACACCTG</td><td>610.2</td><td>-13.1</td><td>1.4</td><td>-9.3</td><td>9.8E-21</td><td>1.7E-19</td><td>MB231/MB468</td><td>MP</td></tr>
<tr><td>>hg38_21_+_8433172_8446622_RNA45SN1_45S_50ntFlanks@10234.10262.29|CGGACGCTACGCCGCGACGAGTAGGAGGG</td><td>>hg38_21_+_8433172_8446622_RNA45SN1_45S_50ntFlanks</td><td>10234</td><td>10262</td><td>29</td><td>CGGACGCTACGCCGCGACGAGTAGGAGGG</td><td>518.3</td><td>-12.9</td><td>1.4</td><td>-9.3</td><td>9.8E-21</td><td>1.7E-19</td><td>MB231/MB468</td><td>MP</td></tr>
<tr><td>>hg38_21_+_8433172_8446622_RNA45SN1_45S_50ntFlanks@4759.4783.25|ATCAGATACCGTCGTAGTTCCGACC</td><td>>hg38_21_+_8433172_8446622_RNA45SN1_45S_50ntFlanks</td><td>4759</td><td>4783</td><td>25</td><td>ATCAGATACCGTCGTAGTTCCGACC</td><td>1042.8</td><td>-12.3</td><td>1.3</td><td>-9.2</td><td>2.3E-20</td><td>1.7E-19</td><td>BT20/MB468</td><td>CYTO</td></tr>
<tr><td>>MI0000255|hsa-mir-30d&WithFlank&8|-|134804870|134804951@12.35.24(+2U)&[MIMAT0000245&hsa-miR-30d-5p&offsets|0|+2(+2U);m-21&8|-|134804919|134804940&offsets|0|+2(+2U)]|TGTAAACATCCCCGACTGGAAGCTTT</td><td>>MI0000255|hsa-mir-30d&WithFlank&8|-|134804870|134804951</td><td>12</td><td>35</td><td>24(+2U)</td><td>TGTAAACATCCCCGACTGGAAGCTTT</td><td>678.0</td><td>7.1</td><td>0.8</td><td>9.3</td><td>1.4E-20</td><td>1.7E-19</td><td>MB231/MB468</td><td>NUC</td></tr>
<tr><td>>hg38_21_+_8433172_8446622_RNA45SN1_45S_50ntFlanks@9534.9558.25|AGGAAACTCTGGTGGAGGTCCGTAG</td><td>>hg38_21_+_8433172_8446622_RNA45SN1_45S_50ntFlanks</td><td>9534</td><td>9558</td><td>25</td><td>AGGAAACTCTGGTGGAGGTCCGTAG</td><td>1088.6</td><td>-14.0</td><td>1.5</td><td>-9.3</td><td>9.9E-21</td><td>1.7E-19</td><td>MB231/MB468</td><td>MP</td></tr>
<tr><td>>hg38_21_+_8433172_8446622_RNA45SN1_45S_50ntFlanks@10414.10431.18|TCGGTCCTGAGAGATGGG</td><td>>hg38_21_+_8433172_8446622_RNA45SN1_45S_50ntFlanks</td><td>10414</td><td>10431</td><td>18</td><td>TCGGTCCTGAGAGATGGG</td><td>545.4</td><td>-13.0</td><td>1.4</td><td>-9.3</td><td>1.0E-20</td><td>1.7E-19</td><td>MB231/MB468</td><td>MP</td></tr>
<tr><td>>hg38_21_+_8433172_8446622_RNA45SN1_45S_50ntFlanks@11319.11342.24|CCCACGTCTCGTCGCGCGCGCGTC</td><td>>hg38_21_+_8433172_8446622_RNA45SN1_45S_50ntFlanks</td><td>11319</td><td>11342</td><td>24</td><td>CCCACGTCTCGTCGCGCGCGCGTC</td><td>913.4</td><td>-13.6</td><td>1.5</td><td>-9.3</td><td>1.5E-20</td><td>1.8E-19</td><td>MB231/MB468</td><td>MITO</td></tr>
<tr><td>>hg38_21_+_8433172_8446622_RNA45SN1_45S_50ntFlanks@8380.8404.25|CAAGAGGGCGTGAAACCGTTAAGAG</td><td>>hg38_21_+_8433172_8446622_RNA45SN1_45S_50ntFlanks</td><td>8380</td><td>8404</td><td>25</td><td>CAAGAGGGCGTGAAACCGTTAAGAG</td><td>1647.4</td><td>-14.5</td><td>1.6</td><td>-9.3</td><td>1.5E-20</td><td>1.8E-19</td><td>MB231/MB468</td><td>MITO</td></tr>
<tr><td>>hg38_21_+_8433172_8446622_RNA45SN1_45S_50ntFlanks@9565.9596.32|TGACGTGCAAATCGGTCGTCCGACCTGGGTAT</td><td>>hg38_21_+_8433172_8446622_RNA45SN1_45S_50ntFlanks</td><td>9565</td><td>9596</td><td>32</td><td>TGACGTGCAAATCGGTCGTCCGACCTGGGTAT</td><td>19577.6</td><td>-7.8</td><td>0.8</td><td>-9.3</td><td>1.5E-20</td><td>1.8E-19</td><td>MB231/MB468</td><td>MITO</td></tr>
<tr><td>>trnaMT_SerGCT_MT_+_12207_12265@14.62.49|AGAACTGCTAACTCATGCCCCCATGTCTAACAACATGGCTTTCTCACCA</td><td>>trnaMT_SerGCT_MT_+_12207_12265</td><td>14</td><td>62</td><td>49</td><td>AGAACTGCTAACTCATGCCCCCATGTCTAACAACATGGCTTTCTCACCA</td><td>234.0</td><td>11.9</td><td>1.3</td><td>9.4</td><td>8.3E-21</td><td>1.8E-19</td><td>MB231/MB468</td><td>CYTO</td></tr>
<tr><td>>MI0003591|hsa-mir-584&WithFlank&5|-|149062307|149062415@22.41.20&[MIMAT0003249&hsa-miR-584-5p&offsets|0|-2;m-167&5|-|149062374|149062394&offsets|0|-1]|TTATGGTTTGCCTGGGACTG</td><td>>MI0003591|hsa-mir-584&WithFlank&5|-|149062307|149062415</td><td>22</td><td>41</td><td>20</td><td>TTATGGTTTGCCTGGGACTG</td><td>702.5</td><td>9.8</td><td>1.1</td><td>9.3</td><td>1.7E-20</td><td>1.8E-19</td><td>BT20/MB468</td><td>MITO</td></tr>
<tr><td>>hg38_21_+_8433172_8446622_RNA45SN1_45S_50ntFlanks@11312.11344.33|CCCCCACCCCACGTCTCGTCGCGCGCGCGTCCG</td><td>>hg38_21_+_8433172_8446622_RNA45SN1_45S_50ntFlanks</td><td>11312</td><td>11344</td><td>33</td><td>CCCCCACCCCACGTCTCGTCGCGCGCGCGTCCG</td><td>737.2</td><td>-12.5</td><td>1.3</td><td>-9.3</td><td>1.4E-20</td><td>1.8E-19</td><td>MB231/MB468</td><td>NUC</td></tr>
<tr><td>>hg38_21_+_8433172_8446622_RNA45SN1_45S_50ntFlanks@11576.11609.34|CTTAGAACTGGTGCGGACCAGGGGAATCCGACTG</td><td>>hg38_21_+_8433172_8446622_RNA45SN1_45S_50ntFlanks</td><td>11576</td><td>11609</td><td>34</td><td>CTTAGAACTGGTGCGGACCAGGGGAATCCGACTG</td><td>531.0</td><td>-4.9</td><td>0.5</td><td>-9.3</td><td>1.4E-20</td><td>1.8E-19</td><td>MB231/MB468</td><td>NUC</td></tr>
<tr><td>>hg38_21_+_8433172_8446622_RNA45SN1_45S_50ntFlanks@3705.3739.35|TACCTGGTTGATCCTGCCAGTAGCATATGCTTGTC</td><td>>hg38_21_+_8433172_8446622_RNA45SN1_45S_50ntFlanks</td><td>3705</td><td>3739</td><td>35</td><td>TACCTGGTTGATCCTGCCAGTAGCATATGCTTGTC</td><td>5224.0</td><td>6.2</td><td>0.7</td><td>9.2</td><td>2.4E-20</td><td>1.8E-19</td><td>BT20/MB468</td><td>CYTO</td></tr>
<tr><td>>hg38_21_+_8433172_8446622_RNA45SN1_45S_50ntFlanks@11104.11133.30|AAGGGTCGGGGCGGCAGGGGCCGGCGGCGG</td><td>>hg38_21_+_8433172_8446622_RNA45SN1_45S_50ntFlanks</td><td>11104</td><td>11133</td><td>30</td><td>AAGGGTCGGGGCGGCAGGGGCCGGCGGCGG</td><td>697.2</td><td>-12.4</td><td>1.3</td><td>-9.3</td><td>1.4E-20</td><td>1.8E-19</td><td>MB231/MB468</td><td>NUC</td></tr>
<tr><td>>hg38_21_+_8433172_8446622_RNA45SN1_45S_50ntFlanks@4449.4472.24|CCTCTCGGCGCCCCCTCGATGCTC</td><td>>hg38_21_+_8433172_8446622_RNA45SN1_45S_50ntFlanks</td><td>4449</td><td>4472</td><td>24</td><td>CCTCTCGGCGCCCCCTCGATGCTC</td><td>4263.2</td><td>-16.5</td><td>1.8</td><td>-9.2</td><td>4.6E-20</td><td>1.8E-19</td><td>BT20/MB468</td><td>NUC</td></tr>
<tr><td>>hg38_21_+_8433172_8446622_RNA45SN1_45S_50ntFlanks@12407.12437.31|TTTTGATCCTTCGATGTCGGCTCTTCCTATC</td><td>>hg38_21_+_8433172_8446622_RNA45SN1_45S_50ntFlanks</td><td>12407</td><td>12437</td><td>31</td><td>TTTTGATCCTTCGATGTCGGCTCTTCCTATC</td><td>5200.0</td><td>-7.5</td><td>0.8</td><td>-9.2</td><td>4.6E-20</td><td>1.8E-19</td><td>BT20/MB468</td><td>NUC</td></tr>
<tr><td>>hg38_21_+_8433172_8446622_RNA45SN1_45S_50ntFlanks@6667.6686.20|ATCACTCGGCTCGTGCGTCG</td><td>>hg38_21_+_8433172_8446622_RNA45SN1_45S_50ntFlanks</td><td>6667</td><td>6686</td><td>20</td><td>ATCACTCGGCTCGTGCGTCG</td><td>543.6</td><td>-13.0</td><td>1.4</td><td>-9.3</td><td>1.0E-20</td><td>1.8E-19</td><td>MB231/MB468</td><td>MP</td></tr>
<tr><td>>hg38_21_+_8433172_8446622_RNA45SN1_45S_50ntFlanks@9791.9817.27|TGGCGTGGAGCCGGGCGTGGAATGCGA</td><td>>hg38_21_+_8433172_8446622_RNA45SN1_45S_50ntFlanks</td><td>9791</td><td>9817</td><td>27</td><td>TGGCGTGGAGCCGGGCGTGGAATGCGA</td><td>396.4</td><td>25.9</td><td>2.6</td><td>9.8</td><td>1.3E-22</td><td>1.8E-19</td><td>BT20/MB231</td><td>MP</td></tr>
<tr><td>>hg38_21_+_8433172_8446622_RNA45SN1_45S_50ntFlanks@12973.12991.19|TCGTACGTAGCAGAGCAGC</td><td>>hg38_21_+_8433172_8446622_RNA45SN1_45S_50ntFlanks</td><td>12973</td><td>12991</td><td>19</td><td>TCGTACGTAGCAGAGCAGC</td><td>1474.5</td><td>-5.1</td><td>0.5</td><td>-9.3</td><td>1.7E-20</td><td>1.8E-19</td><td>BT20/MB468</td><td>MITO</td></tr>
<tr><td>>hg38_21_+_8433172_8446622_RNA45SN1_45S_50ntFlanks@11781.11805.25|CGCATGAATGGATGAACGAGATTCC</td><td>>hg38_21_+_8433172_8446622_RNA45SN1_45S_50ntFlanks</td><td>11781</td><td>11805</td><td>25</td><td>CGCATGAATGGATGAACGAGATTCC</td><td>626.6</td><td>-13.2</td><td>1.4</td><td>-9.3</td><td>1.0E-20</td><td>1.8E-19</td><td>MB231/MB468</td><td>MP</td></tr>
<tr><td>>hg38_21_+_8433172_8446622_RNA45SN1_45S_50ntFlanks@12476.12496.21|CCACTAATAGGGAACGTGAGC</td><td>>hg38_21_+_8433172_8446622_RNA45SN1_45S_50ntFlanks</td><td>12476</td><td>12496</td><td>21</td><td>CCACTAATAGGGAACGTGAGC</td><td>2245.2</td><td>-6.2</td><td>0.7</td><td>-9.3</td><td>1.6E-20</td><td>1.8E-19</td><td>MB231/MB468</td><td>MITO</td></tr>
<tr><td>>hg38_21_+_8433172_8446622_RNA45SN1_45S_50ntFlanks@12933.12956.24|GCTAAACCATTCGTAGACGACCTG</td><td>>hg38_21_+_8433172_8446622_RNA45SN1_45S_50ntFlanks</td><td>12933</td><td>12956</td><td>24</td><td>GCTAAACCATTCGTAGACGACCTG</td><td>3271.7</td><td>-6.4</td><td>0.7</td><td>-9.3</td><td>1.5E-20</td><td>1.8E-19</td><td>MB231/MB468</td><td>NUC</td></tr>
<tr><td>>hg38_21_+_8433172_8446622_RNA45SN1_45S_50ntFlanks@10770.10797.28|GCAGGTCTCCAAGGTGAACAGCCTCTGG</td><td>>hg38_21_+_8433172_8446622_RNA45SN1_45S_50ntFlanks</td><td>10770</td><td>10797</td><td>28</td><td>GCAGGTCTCCAAGGTGAACAGCCTCTGG</td><td>246.8</td><td>-12.4</td><td>1.3</td><td>-9.2</td><td>4.7E-20</td><td>1.8E-19</td><td>BT20/MB468</td><td>NUC</td></tr>
<tr><td>>hg38_21_+_8433172_8446622_RNA45SN1_45S_50ntFlanks@9004.9026.23|TCTCCCCCGCGGGGGCGCGCCGG</td><td>>hg38_21_+_8433172_8446622_RNA45SN1_45S_50ntFlanks</td><td>9004</td><td>9026</td><td>23</td><td>TCTCCCCCGCGGGGGCGCGCCGG</td><td>1126.7</td><td>-13.1</td><td>1.4</td><td>-9.3</td><td>1.5E-20</td><td>1.8E-19</td><td>MB231/MB468</td><td>NUC</td></tr>
<tr><td>>MI0003776|hsa-mir-320b-1&WithFlank&1|+|116671740|116671823@48.67.20(+2U);MI0003839|hsa-mir-320b-2&WithFlank&1|-|224257034|224257116@47.66.20(+2U);MI0000542|hsa-mir-320a&WithFlank&8|-|22244960|22245043@48.67.20(+2U)&[MIMAT0005792&hsa-miR-320b&offsets|0|-2(+2U);m-183&1|+|116671787|116671806&offsets|0|0(+2U)];[MIMAT0005792_1&hsa-miR-320b&offsets|0|-2(+2U);m-183&1|-|224257051|224257070&offsets|0|0(+2U)];[MIMAT0000510&hsa-miR-320a-3p&offsets|0|-2(+2U);m-60&8|-|22244975|22244996&offsets|0|-2(+2U)]|AAAAGCTGGGTTGAGAGGGCTT</td><td>>MI0003776|hsa-mir-320b-1&WithFlank&1|+|116671740|116671823</td><td>48</td><td>67</td><td>20(+2U)</td><td>AAAAGCTGGGTTGAGAGGGCTT</td><td>372.3</td><td>12.8</td><td>1.4</td><td>9.2</td><td>2.5E-20</td><td>1.8E-19</td><td>BT20/MB468</td><td>CYTO</td></tr>
<tr><td>>hg38_21_+_8433172_8446622_RNA45SN1_45S_50ntFlanks@7976.8011.36|GCGACCTCAGATCAGACGTGGCGACCCGCTGAATTT</td><td>>hg38_21_+_8433172_8446622_RNA45SN1_45S_50ntFlanks</td><td>7976</td><td>8011</td><td>36</td><td>GCGACCTCAGATCAGACGTGGCGACCCGCTGAATTT</td><td>2015.5</td><td>9.2</td><td>1.0</td><td>9.3</td><td>8.8E-21</td><td>1.9E-19</td><td>MB231/MB468</td><td>CYTO</td></tr>
<tr><td>>hg38_21_+_8433172_8446622_RNA45SN1_45S_50ntFlanks@11583.11605.23|CTGGTGCGGACCAGGGGAATCCG</td><td>>hg38_21_+_8433172_8446622_RNA45SN1_45S_50ntFlanks</td><td>11583</td><td>11605</td><td>23</td><td>CTGGTGCGGACCAGGGGAATCCG</td><td>1902.9</td><td>-10.9</td><td>1.2</td><td>-9.3</td><td>1.6E-20</td><td>1.9E-19</td><td>MB231/MB468</td><td>MITO</td></tr>
<tr><td>>hg38_21_+_8433172_8446622_RNA45SN1_45S_50ntFlanks@12623.12654.32|GCCAATGGGGCGAAGCTACCATCTGTGGGATT</td><td>>hg38_21_+_8433172_8446622_RNA45SN1_45S_50ntFlanks</td><td>12623</td><td>12654</td><td>32</td><td>GCCAATGGGGCGAAGCTACCATCTGTGGGATT</td><td>434.8</td><td>-12.6</td><td>1.4</td><td>-9.3</td><td>1.1E-20</td><td>1.9E-19</td><td>MB231/MB468</td><td>MP</td></tr>
<tr><td>>hg38_21_+_8433172_8446622_RNA45SN1_45S_50ntFlanks@4472.4494.23|CTTAGCTGAGTGTCCCGCGGGGC</td><td>>hg38_21_+_8433172_8446622_RNA45SN1_45S_50ntFlanks</td><td>4472</td><td>4494</td><td>23</td><td>CTTAGCTGAGTGTCCCGCGGGGC</td><td>1205.3</td><td>-8.3</td><td>0.9</td><td>-9.3</td><td>8.9E-21</td><td>1.9E-19</td><td>MB231/MB468</td><td>CYTO</td></tr>
<tr><td>>hg38_21_+_8433172_8446622_RNA45SN1_45S_50ntFlanks@8107.8143.37|CGGCGGGGCGCGGGACATGTGGCGTACGGAAGACCCG</td><td>>hg38_21_+_8433172_8446622_RNA45SN1_45S_50ntFlanks</td><td>8107</td><td>8143</td><td>37</td><td>CGGCGGGGCGCGGGACATGTGGCGTACGGAAGACCCG</td><td>1317.8</td><td>-14.1</td><td>1.5</td><td>-9.3</td><td>1.6E-20</td><td>1.9E-19</td><td>MB231/MB468</td><td>MITO</td></tr>
<tr><td>>hg38_21_+_8433172_8446622_RNA45SN1_45S_50ntFlanks@475.517.43|CTTCGTGATCGATGTGGTGACGTCGTGCTCTCCCGGGCCGGGT</td><td>>hg38_21_+_8433172_8446622_RNA45SN1_45S_50ntFlanks</td><td>475</td><td>517</td><td>43</td><td>CTTCGTGATCGATGTGGTGACGTCGTGCTCTCCCGGGCCGGGT</td><td>1073.6</td><td>14.1</td><td>1.5</td><td>9.3</td><td>9.0E-21</td><td>1.9E-19</td><td>MB231/MB468</td><td>CYTO</td></tr>
<tr><td>>MI0000478|hsa-mir-149&WithFlank&2|+|240455995|240456095@21.41.21(+1A)&[MIMAT0000450&hsa-miR-149-5p&offsets|0|-2(+1A);m-195&2|+|240456015|240456037&offsets|0|-2(+1A)]|TCTGGCTCCGTGTCTTCACTCA</td><td>>MI0000478|hsa-mir-149&WithFlank&2|+|240455995|240456095</td><td>21</td><td>41</td><td>21(+1A)</td><td>TCTGGCTCCGTGTCTTCACTCA</td><td>872.6</td><td>14.0</td><td>1.5</td><td>9.2</td><td>2.5E-20</td><td>1.9E-19</td><td>BT20/MB468</td><td>CYTO</td></tr>
<tr><td>>hg38_21_+_8433172_8446622_RNA45SN1_45S_50ntFlanks@11592.11611.20|ACCAGGGGAATCCGACTGTT</td><td>>hg38_21_+_8433172_8446622_RNA45SN1_45S_50ntFlanks</td><td>11592</td><td>11611</td><td>20</td><td>ACCAGGGGAATCCGACTGTT</td><td>920.4</td><td>-13.6</td><td>1.5</td><td>-9.3</td><td>1.7E-20</td><td>1.9E-19</td><td>MB231/MB468</td><td>MITO</td></tr>
<tr><td>>hg38_21_+_8433172_8446622_RNA45SN1_45S_50ntFlanks@12033.12053.21|ATCGTTTTTTCACTGACCCGG</td><td>>hg38_21_+_8433172_8446622_RNA45SN1_45S_50ntFlanks</td><td>12033</td><td>12053</td><td>21</td><td>ATCGTTTTTTCACTGACCCGG</td><td>720.2</td><td>-13.4</td><td>1.4</td><td>-9.3</td><td>1.1E-20</td><td>1.9E-19</td><td>MB231/MB468</td><td>MP</td></tr>
<tr><td>>MI0000465|hsa-mir-191&WithFlank&3|-|49020612|49020715@22.41.20(+1U)&[MIMAT0000440&hsa-miR-191-5p&offsets|0|-3(+1U);m-48&3|-|49020672|49020694&offsets|0|-3(+1U)]|CAACGGAATCCCAAAAGCAGT</td><td>>MI0000465|hsa-mir-191&WithFlank&3|-|49020612|49020715</td><td>22</td><td>41</td><td>20(+1U)</td><td>CAACGGAATCCCAAAAGCAGT</td><td>361.3</td><td>12.8</td><td>1.4</td><td>9.2</td><td>2.6E-20</td><td>1.9E-19</td><td>BT20/MB468</td><td>CYTO</td></tr>
<tr><td>>hg38_21_+_8433172_8446622_RNA45SN1_45S_50ntFlanks@8168.8215.48|CCCAAGTCCTTCTGATCGAGGCCCAGCCCGTGGACGGTGTGAGGCCGG</td><td>>hg38_21_+_8433172_8446622_RNA45SN1_45S_50ntFlanks</td><td>8168</td><td>8215</td><td>48</td><td>CCCAAGTCCTTCTGATCGAGGCCCAGCCCGTGGACGGTGTGAGGCCGG</td><td>2248.4</td><td>-15.6</td><td>1.7</td><td>-9.2</td><td>4.9E-20</td><td>1.9E-19</td><td>BT20/MB468</td><td>NUC</td></tr>
<tr><td>>hg38_21_+_8433172_8446622_RNA45SN1_45S_50ntFlanks@12476.12493.18|CCACTAATAGGGAACGTG</td><td>>hg38_21_+_8433172_8446622_RNA45SN1_45S_50ntFlanks</td><td>12476</td><td>12493</td><td>18</td><td>CCACTAATAGGGAACGTG</td><td>525.6</td><td>-12.9</td><td>1.4</td><td>-9.3</td><td>1.1E-20</td><td>1.9E-19</td><td>MB231/MB468</td><td>MP</td></tr>
</table>

| | | | | | | | | | | | | | |
|---|---|---|---|---|---|---|---|---|---|---|---|---|---|
| >trna11_GluTTC_15_-_26327381_26327452@1.29.29,trna3_GluTTC_13_-_45492062_45492133@1.29.29\|TCCCACATGGTCTAGCGGTTAGGATTCCT | >trna11_GluTTC_15_-_26327381_26327452 | 1 | 29 | 29 | TCCCACATGGTCTAGCGGTTAGGATTCCT | 3438.9 | -7.6 | 0.8 | -9.2 | 5.0E-20 | 1.9E-19 | BT20/MB468 | NUC |
| >MI0000266\|hsa-mir-10a&WithFlank&17\|-\|48579832\|48579953@28.51.24&[MIMAT0000253&hsa-miR-10a-5p&offsets\|0\|+1;m-18&17\|-\|48579905\|48579926&offsets\|0\|+2]\|TACCCTGTAGATCCGAATTTGTGT | >MI0000266\|hsa-mir-10a&WithFlank&17\|-\|48579832\|48579953 | 28 | 51 | 24 | TACCCTGTAGATCCGAATTTGTGT | 177.3 | 10.3 | 1.1 | 9.2 | 5.0E-20 | 1.9E-19 | BT20/MB468 | NUC |
| >MI0003591\|hsa-mir-584&WithFlank&5\|-\|149062307\|149062415@22.41.20(+1U)&[MIMAT0003249&hsa-miR-584-5p&offsets\|0\|-2(+1U);m-167&5\|-\|149062374\|149062394&offsets\|0\|-1(+1U)]\|TTATGGTTTGCCTGGGACTGT | >MI0003591\|hsa-mir-584&WithFlank&5\|-\|149062307\|149062415 | 22 | 41 | 20(+1U) | TTATGGTTTGCCTGGGACTGT | 592.8 | 7.3 | 0.8 | 9.3 | 1.6E-20 | 1.9E-19 | MB231/MB468 | NUC |
| >hg38_21_+_8433172_8446622_RNA45SN1_45S_50ntFlanks@8687.8706.20\|CGGCTCCGGGACGGCTGGGA | >hg38_21_+_8433172_8446622_RNA45SN1_45S_50ntFlanks | 8687 | 8706 | 20 | CGGCTCCGGGACGGCTGGGA | 1962.1 | 11.8 | 1.3 | 9.3 | 9.3E-21 | 2.0E-19 | MB231/MB468 | CYTO |
| >hg38_21_+_8433172_8446622_RNA45SN1_45S_50ntFlanks@7975.8001.27\|CGCGACCTCAGATCAGACGTGGCGACC | >hg38_21_+_8433172_8446622_RNA45SN1_45S_50ntFlanks | 7975 | 8001 | 27 | CGCGACCTCAGATCAGACGTGGCGACC | 778.3 | 4.5 | 0.5 | 9.2 | 5.0E-20 | 2.0E-19 | BT20/MB468 | NUC |
| >hg38_21_+_8433172_8446622_RNA45SN1_45S_50ntFlanks@9738.9758.21\|GAAACGATCTCAACCTATTCT | >hg38_21_+_8433172_8446622_RNA45SN1_45S_50ntFlanks | 9738 | 9758 | 21 | GAAACGATCTCAACCTATTCT | 2783.2 | -11.1 | 1.2 | -9.3 | 1.1E-20 | 2.0E-19 | MB231/MB468 | MP |
| >hg38_21_+_8433172_8446622_RNA45SN1_45S_50ntFlanks@11576.11611.36\|CTTAGAACTGGTGCGGACCAGGGGAATCCGACTGTT | >hg38_21_+_8433172_8446622_RNA45SN1_45S_50ntFlanks | 11576 | 11611 | 36 | CTTAGAACTGGTGCGGACCAGGGGAATCCGACTGTT | 704.1 | -4.3 | 0.5 | -9.2 | 5.1E-20 | 2.0E-19 | BT20/MB468 | NUC |
| >hg38_21_+_8433172_8446622_RNA45SN1_45S_50ntFlanks@4575.4611.37\|TAGGACCGCGGTTCTATTTTGTTGGTTTTCGGAACTG | >hg38_21_+_8433172_8446622_RNA45SN1_45S_50ntFlanks | 4575 | 4611 | 37 | TAGGACCGCGGTTCTATTTTGTTGGTTTTCGGAACTG | 386.8 | -13.0 | 1.4 | -9.2 | 5.1E-20 | 2.0E-19 | BT20/MB468 | NUC |
| >hg38_21_+_8433172_8446622_RNA45SN1_45S_50ntFlanks@6645.6668.24\|CTCGTACGACTCTTAGCGGTGGAT | >hg38_21_+_8433172_8446622_RNA45SN1_45S_50ntFlanks | 6645 | 6668 | 24 | CTCGTACGACTCTTAGCGGTGGAT | 10450.6 | 6.2 | 0.7 | 9.3 | 1.9E-20 | 2.0E-19 | BT20/MB468 | MITO |
| >hg38_21_+_8433172_8446622_RNA45SN1_45S_50ntFlanks@485.518.34\|GATGTGGTGACGTCGTGCTCTCCCGGGCCGGGTC | >hg38_21_+_8433172_8446622_RNA45SN1_45S_50ntFlanks | 485 | 518 | 34 | GATGTGGTGACGTCGTGCTCTCCCGGGCCGGGTC | 514.9 | 13.5 | 1.5 | 9.3 | 1.9E-20 | 2.0E-19 | BT20/MB468 | MITO |
| >MI0000067\|hsa-let-7f-1&WithFlank&9\|+\|94176341\|94176439@13.31.19;MI0000068\|hsa-let-7f-2&WithFlank&X\|-\|53557186\|53557280@14.32.19&[MIMAT0000067&hsa-let-7f-5p&offsets\|0\|-3;m-20&9\|+\|94176353\|94176374&offsets\|0\|-3];[MIMAT0000067_1&hsa-let-7f-5p&offsets\|0\|-3;m-19&X\|-\|53557246\|53557267&offsets\|0\|-3]\|TGAGGTAGTAGATTGTATA | >MI0000067\|hsa-let-7f-1&WithFlank&9\|+\|94176341\|94176439 | 13 | 31 | 19 | TGAGGTAGTAGATTGTATA | 4692.6 | 7.5 | 0.8 | 9.2 | 2.7E-20 | 2.0E-19 | BT20/MB468 | CYTO |
| >MI0000767\|hsa-mir-365a&WithFlank&16\|+\|14309279\|14309377@22.45.24&[MIMAT0009199&hsa-miR-365a-5p&offsets\|0\|+1;m-595&16\|+\|14309300\|14309322&offsets\|0\|+1]\|AGGGACTTTTGGGGGCAGATGTGT | >MI0000767\|hsa-mir-365a&WithFlank&16\|+\|14309279\|14309377 | 22 | 45 | 24 | AGGGACTTTTGGGGGCAGATGTGT | 273.0 | 12.3 | 1.3 | 9.2 | 2.7E-20 | 2.0E-19 | BT20/MB468 | CYTO |
| >hg38_21_+_8433172_8446622_RNA45SN1_45S_50ntFlanks@4003.4024.22\|ATAACCTCGGGCCGATCGCACG | >hg38_21_+_8433172_8446622_RNA45SN1_45S_50ntFlanks | 4003 | 4024 | 22 | ATAACCTCGGGCCGATCGCACG | 4170.5 | -5.5 | 0.6 | -9.3 | 9.5E-21 | 2.0E-19 | MB231/MB468 | CYTO |
| >hg38_21_+_8433172_8446622_RNA45SN1_45S_50ntFlanks@9570.9594.25\|TGCAAATCGGTCGTCCGACCTGGGT | >hg38_21_+_8433172_8446622_RNA45SN1_45S_50ntFlanks | 9570 | 9594 | 25 | TGCAAATCGGTCGTCCGACCTGGGT | 1188.1 | -14.1 | 1.5 | -9.3 | 1.1E-20 | 2.0E-19 | MB231/MB468 | MP |
| >MI0005756\|hsa-mir-934&WithFlank&X\|+\|136550872\|136550966@21.43.23&[MIMAT0004977&hsa-miR-934&offsets\|0\|+1;m-943&X\|+\|136550892\|136550913&offsets\|0\|+1]\|TGTCTACTACTGGAGACACTGGT | >MI0005756\|hsa-mir-934&WithFlank&X\|+\|136550872\|136550966 | 21 | 43 | 23 | TGTCTACTACTGGAGACACTGGT | 1338.6 | 14.2 | 1.5 | 9.5 | 1.8E-21 | 2.0E-19 | BT20/MB231 | NUC |
| >hg38_21_+_8433172_8446622_RNA45SN1_45S_50ntFlanks@11583.11602.20\|CTGGTGCGGACCAGGGGAAT | >hg38_21_+_8433172_8446622_RNA45SN1_45S_50ntFlanks | 11583 | 11602 | 20 | CTGGTGCGGACCAGGGGAAT | 768.9 | -4.6 | 0.5 | -9.3 | 1.6E-20 | 2.0E-19 | MB231/MB468 | NUC |
| >hg38_21_+_8433172_8446622_RNA45SN1_45S_50ntFlanks@9504.9523.20\|GTGAACTATGCCTGGGCAGG | >hg38_21_+_8433172_8446622_RNA45SN1_45S_50ntFlanks | 9504 | 9523 | 20 | GTGAACTATGCCTGGGCAGG | 2752.2 | -8.1 | 0.9 | -9.3 | 9.7E-21 | 2.0E-19 | MB231/MB468 | CYTO |
| >hg38_21_+_8433172_8446622_RNA45SN1_45S_50ntFlanks@12008.12034.27\|CCGCCGGTGAAATACCACTACTCTGAT | >hg38_21_+_8433172_8446622_RNA45SN1_45S_50ntFlanks | 12008 | 12034 | 27 | CCGCCGGTGAAATACCACTACTCTGAT | 658.1 | -13.1 | 1.4 | -9.3 | 1.8E-20 | 2.0E-19 | MB231/MB468 | MITO |
| >hg38_21_+_8433172_8446622_RNA45SN1_45S_50ntFlanks@8113.8140.28\|GGCGCGGGACATGTGGCGTACGGAAGAC | >hg38_21_+_8433172_8446622_RNA45SN1_45S_50ntFlanks | 8113 | 8140 | 28 | GGCGCGGGACATGTGGCGTACGGAAGAC | 874.5 | -13.6 | 1.5 | -9.3 | 1.8E-20 | 2.0E-19 | MB231/MB468 | MITO |
| >hg38_21_+_8433172_8446622_RNA45SN1_45S_50ntFlanks@10101.10127.27\|CGCGCGCGTGTGGTGTGCGTCGGAGGG | >hg38_21_+_8433172_8446622_RNA45SN1_45S_50ntFlanks | 10101 | 10127 | 27 | CGCGCGCGTGTGGTGTGCGTCGGAGGG | 1029.6 | -13.9 | 1.5 | -9.3 | 1.2E-20 | 2.0E-19 | MB231/MB468 | MP |
| >hg38_21_+_8433172_8446622_RNA45SN1_45S_50ntFlanks@11581.11603.23\|AACTGGTGCGGACCAGGGGAATC | >hg38_21_+_8433172_8446622_RNA45SN1_45S_50ntFlanks | 11581 | 11603 | 23 | AACTGGTGCGGACCAGGGGAATC | 18880.3 | -9.9 | 1.1 | -9.3 | 1.2E-20 | 2.0E-19 | MB231/MB468 | MP |
| >hg38_21_+_8433172_8446622_RNA45SN1_45S_50ntFlanks@8472.8491.20\|CGGCCCGGCGGATCTTTCCC | >hg38_21_+_8433172_8446622_RNA45SN1_45S_50ntFlanks | 8472 | 8491 | 20 | CGGCCCGGCGGATCTTTCCC | 1543.9 | -14.4 | 1.6 | -9.3 | 1.8E-20 | 2.0E-19 | MB231/MB468 | MITO |
| >MI0000079\|hsa-mir-23a&WithFlank&19\|-\|13836581\|13836665@54.73.20(+1A)&[MIMAT0000078&hsa-miR-23a-3p&offsets\|+3\|+2(+1A);m-31&19\|-\|13836595\|13836615&offsets\|+3\|+2(+1A)]\|ACATTGCCAGGGATTTCCAAA | >MI0000079\|hsa-mir-23a&WithFlank&19\|-\|13836581\|13836665 | 54 | 73 | 20(+1A) | ACATTGCCAGGGATTTCCAAA | 283.4 | 12.4 | 1.3 | 9.2 | 2.8E-20 | 2.0E-19 | BT20/MB468 | CYTO |
| >hg38_21_+_8433172_8446622_RNA45SN1_45S_50ntFlanks@11646.11687.42\|GGTGTTGACGCGATGTGATTTCTGCCCAGTGCTCTGAATGTC | >hg38_21_+_8433172_8446622_RNA45SN1_45S_50ntFlanks | 11646 | 11687 | 42 | GGTGTTGACGCGATGTGATTTCTGCCCAGTGCTCTGAATGTC | 174.2 | 11.4 | 1.2 | 9.3 | 9.9E-21 | 2.1E-19 | MB231/MB468 | CYTO |
| >hg38_21_+_8433172_8446622_RNA45SN1_45S_50ntFlanks@3922.3959.38\|TTTATCAGATCAAAACCAACCCGGTCAGCCCCTCTCCG | >hg38_21_+_8433172_8446622_RNA45SN1_45S_50ntFlanks | 3922 | 3959 | 38 | TTTATCAGATCAAAACCAACCCGGTCAGCCCCTCTCCG | 718.2 | -13.9 | 1.5 | -9.2 | 5.3E-20 | 2.1E-19 | BT20/MB468 | NUC |
| >hg38_21_+_8433172_8446622_RNA45SN1_45S_50ntFlanks@11874.11898.25\|AGAAGACCCTGTTGAGCTTGACTCT | >hg38_21_+_8433172_8446622_RNA45SN1_45S_50ntFlanks | 11874 | 11898 | 25 | AGAAGACCCTGTTGAGCTTGACTCT | 2159.7 | -4.5 | 0.5 | -9.3 | 1.0E-20 | 2.1E-19 | MB231/MB468 | CYTO |
| >MI0000102\|hsa-mir-100&WithFlank&11\|-\|122152223\|122152314@19.40.22(+2A)&[MIMAT0000098&hsa-miR-100-5p&offsets\|0\|0(+2A);m-27&11\|-\|122152276\|122152296&offsets\|0\|+1(+2A)]\|AACCCGTAGATCCGAACTTGTGAA | >MI0000102\|hsa-mir-100&WithFlank&11\|-\|122152223\|122152314 | 19 | 40 | 22(+2A) | AACCCGTAGATCCGAACTTGTGAA | 1450.2 | 13.7 | 1.5 | 9.3 | 1.8E-20 | 2.1E-19 | MB231/MB468 | MITO |
| >hg38_21_+_8433172_8446622_RNA45SN1_45S_50ntFlanks@9747.9769.23\|TCAACCTATTCTCAAACTTTAAA | >hg38_21_+_8433172_8446622_RNA45SN1_45S_50ntFlanks | 9747 | 9769 | 23 | TCAACCTATTCTCAAACTTTAAA | 377.2 | 13.0 | 1.4 | 9.3 | 2.0E-20 | 2.1E-19 | BT20/MB468 | MITO |
| >hg38_21_+_8433172_8446622_RNA45SN1_45S_50ntFlanks@11534.11553.20\|GTTCCGCGCGGCGCCTCGCC | >hg38_21_+_8433172_8446622_RNA45SN1_45S_50ntFlanks | 11534 | 11553 | 20 | GTTCCGCGCGGCGCCTCGCC | 859.0 | -13.5 | 1.5 | -9.3 | 1.8E-20 | 2.1E-19 | MB231/MB468 | MITO |
| >MI0000086\|hsa-mir-28&WithFlank&3\|+\|188688775\|188688872@62.82.21(+1U)&[MIMAT0004502&hsa-miR-28-3p&offsets\|+2\|+1(+1U);m-28&3\|+\|188688834\|188688855&offsets\|+2\|+1(+1U)]\|CTAGATTGTGAGCTCCTGGAGT | >MI0000086\|hsa-mir-28&WithFlank&3\|+\|188688775\|188688872 | 62 | 82 | 21(+1U) | CTAGATTGTGAGCTCCTGGAGT | 69.9 | 3.6 | 0.4 | 9.2 | 5.4E-20 | 2.1E-19 | BT20/MB468 | NUC |
| >hg38_21_+_8433172_8446622_RNA45SN1_45S_50ntFlanks@3844.3868.25\|TACTTGGATAACTGTGGTAATTCTA | >hg38_21_+_8433172_8446622_RNA45SN1_45S_50ntFlanks | 3844 | 3868 | 25 | TACTTGGATAACTGTGGTAATTCTA | 416.6 | -4.2 | 0.5 | -9.2 | 5.4E-20 | 2.1E-19 | BT20/MB468 | NUC |
| >hg38_21_+_8433172_8446622_RNA45SN1_45S_50ntFlanks@9504.9529.26\|GTGAACTATGCCTGGGCAGGGCGAAG | >hg38_21_+_8433172_8446622_RNA45SN1_45S_50ntFlanks | 9504 | 9529 | 26 | GTGAACTATGCCTGGGCAGGGCGAAG | 1147.6 | -14.0 | 1.5 | -9.3 | 1.2E-20 | 2.1E-19 | MB231/MB468 | MP |
| >hg38_21_+_8433172_8446622_RNA45SN1_45S_50ntFlanks@12417.12434.18\|TCGATGTCGGCTCTTCCT | >hg38_21_+_8433172_8446622_RNA45SN1_45S_50ntFlanks | 12417 | 12434 | 18 | TCGATGTCGGCTCTTCCT | 1344.9 | -6.1 | 0.7 | -9.3 | 1.2E-20 | 2.1E-19 | MB231/MB468 | MP |
| >hg38_21_+_8433172_8446622_RNA45SN1_45S_50ntFlanks@11886.11904.19\|TGAGCTTGACTCTAGTCTG | >hg38_21_+_8433172_8446622_RNA45SN1_45S_50ntFlanks | 11886 | 11904 | 19 | TGAGCTTGACTCTAGTCTG | 997.2 | -13.7 | 1.5 | -9.3 | 1.9E-20 | 2.1E-19 | MB231/MB468 | MITO |
| >hg38_21_+_8433172_8446622_RNA45SN1_45S_50ntFlanks@12957.12979.23\|CTTCTGGGTCGGGGTTTCGTACG | >hg38_21_+_8433172_8446622_RNA45SN1_45S_50ntFlanks | 12957 | 12979 | 23 | CTTCTGGGTCGGGGTTTCGTACG | 1356.6 | -14.2 | 1.5 | -9.3 | 1.9E-20 | 2.1E-19 | MB231/MB468 | MITO |
| >hg38_21_+_8433172_8446622_RNA45SN1_45S_50ntFlanks@11580.11601.22\|GAACTGGTGCGGACCAGGGGAA | >hg38_21_+_8433172_8446622_RNA45SN1_45S_50ntFlanks | 11580 | 11601 | 22 | GAACTGGTGCGGACCAGGGGAA | 1549.1 | -14.5 | 1.6 | -9.3 | 1.2E-20 | 2.1E-19 | MB231/MB468 | MP |
| >MI0000253\|hsa-mir-148a&WithFlank&7\|-\|25949913\|25949992@50.70.21(+1A)&[MIMAT0000243&hsa-miR-148a-3p&offsets\|0\|-1(+1A);m-7&7\|-\|25949922\|25949943&offsets\|0\|-1(+1A)]\|TCAGTGCACTACAGAACTTTGA | >MI0000253\|hsa-mir-148a&WithFlank&7\|-\|25949913\|25949992 | 50 | 70 | 21(+1A) | TCAGTGCACTACAGAACTTTGA | 3294.7 | 7.3 | 0.8 | 9.3 | 2.1E-20 | 2.2E-19 | BT20/MB468 | MITO |
| >trna10_AspGTC_12_-_125424193_125424264@35.74.40,trna12_AspGTC_12_-_125411891_125411962@35.74.40,trna144_AspGTC_6_-_27551236_27551307@35.74.40,trna38_AspGTC_17_-_8125556_8125627@35.74.40,trna45_AspGTC_6_+_27447453_27447524@35.74.40,trna48_AspGTC_6_+_27471523_27471594@35.74.40,trna4_AspGTC_12_+_96429799_96429870@35.74.40,trna69_AspGTC_1_-_161440205_161440276@35.74.40,trna72_AspGTC_1_-_161432824_161432895@35.74.40,trna75_AspGTC_1_-_161425414_161425485@35.74.40,trna78_AspGTC_1_-_161418033_161418104@35.74.40,trna81_AspGTC_1_-_161410615_161410686@35.74.40\|TCACGCGGGAGACCGGGGTTCGATTCCCCGACGGGGAGCC | >trna10_AspGTC_12_-_125424193_125424264 | 35 | 74 | 40 | TCACGCGGGAGACCGGGGTTCGATTCCCCGACGGGGAGCC | 18961.6 | 12.3 | 1.3 | 9.3 | 1.1E-20 | 2.2E-19 | MB231/MB468 | CYTO |
| >hg38_21_+_8433172_8446622_RNA45SN1_45S_50ntFlanks@10771.10805.35\|CAGGTCTCCAAGGTGAACAGCCTCTGGCATGTTGG | >hg38_21_+_8433172_8446622_RNA45SN1_45S_50ntFlanks | 10771 | 10805 | 35 | CAGGTCTCCAAGGTGAACAGCCTCTGGCATGTTGG | 818.1 | -7.0 | 0.7 | -9.3 | 1.0E-20 | 2.2E-19 | MB231/MB468 | CYTO |
| >hg38_21_+_8433172_8446622_RNA45SN1_45S_50ntFlanks@8380.8401.22\|CAAGAGGGCGTGAAACCGTTAA | >hg38_21_+_8433172_8446622_RNA45SN1_45S_50ntFlanks | 8380 | 8401 | 22 | CAAGAGGGCGTGAAACCGTTAA | 799.1 | -13.5 | 1.5 | -9.3 | 1.3E-20 | 2.2E-19 | MB231/MB468 | MP |
| >hg38_21_+_8433172_8446622_RNA45SN1_45S_50ntFlanks@9306.9338.33\|CAAGGAGTCTAACACGTGCGCGAGTCGGGGGCT | >hg38_21_+_8433172_8446622_RNA45SN1_45S_50ntFlanks | 9306 | 9338 | 33 | CAAGGAGTCTAACACGTGCGCGAGTCGGGGGCT | 2247.9 | -10.8 | 1.2 | -9.3 | 1.3E-20 | 2.2E-19 | MB231/MB468 | MP |
| >hg38_21_+_8433172_8446622_RNA45SN1_45S_50ntFlanks@11729.11767.39\|TAACTATGACTCTCTTAAGGTAGCCAAATGCCTCGTCAT | >hg38_21_+_8433172_8446622_RNA45SN1_45S_50ntFlanks | 11729 | 11767 | 39 | TAACTATGACTCTCTTAAGGTAGCCAAATGCCTCGTCAT | 1565.2 | -14.5 | 1.6 | -9.3 | 1.3E-20 | 2.2E-19 | MB231/MB468 | MP |
| >MI0000089\|hsa-mir-31&WithFlank&9\|-\|21512109\|21512191@14.36.23&[MIMAT0000089&hsa-miR-31-5p&offsets\|0\|+2;m-360&9\|-\|21512157\|21512178&offsets\|0\|+1]\|AGGCAAGATGCTGGCATAGCTGT | >MI0000089\|hsa-mir-31&WithFlank&9\|-\|21512109\|21512191 | 14 | 36 | 23 | AGGCAAGATGCTGGCATAGCTGT | 1520.4 | 14.3 | 1.5 | 9.5 | 2.0E-21 | 2.2E-19 | BT20/MB231 | NUC |
| >hg38_21_+_8433172_8446622_RNA45SN1_45S_50ntFlanks@12712.12747.36\|CGGAGCCTCGGTTGGCCTCGGATAGCCGGTCCCCCG | >hg38_21_+_8433172_8446622_RNA45SN1_45S_50ntFlanks | 12712 | 12747 | 36 | CGGAGCCTCGGTTGGCCTCGGATAGCCGGTCCCCCG | 1096.3 | -13.9 | 1.5 | -9.3 | 2.0E-20 | 2.2E-19 | MB231/MB468 | MITO |
| >hg38_21_+_8433172_8446622_RNA45SN1_45S_50ntFlanks@6670.6699.30\|ACTCGGCTCGTGCGTCGATGAAGAACGCAG | >hg38_21_+_8433172_8446622_RNA45SN1_45S_50ntFlanks | 6670 | 6699 | 30 | ACTCGGCTCGTGCGTCGATGAAGAACGCAG | 485.3 | -5.7 | 0.6 | -9.1 | 5.7E-20 | 2.2E-19 | BT20/MB468 | NUC |

<table>
<tr><th></th><th></th><th></th><th></th><th></th><th></th><th></th><th></th><th></th><th></th><th></th><th></th><th></th><th></th></tr>
<tr><td>>hg38_21_+_8433172_8446622_RNA45SN1_45S_50ntFlanks@11728.11765.38|GTAACTATGACTCTCTTAAGGTAGCCAAATGCCTCGTC</td><td>>hg38_21_+_8433172_8446622_RNA45SN1_45S_50ntFlanks</td><td>11728</td><td>11765</td><td>38</td><td>GTAACTATGACTCTCTTAAGGTAGCCAAATGCCTCGTC</td><td>442.3</td><td>-13.2</td><td>1.4</td><td>-9.1</td><td>5.7E-20</td><td>2.2E-19</td><td>BT20/MB468</td><td>NUC</td></tr>
<tr><td>>hg38_21_+_8433172_8446622_RNA45SN1_45S_50ntFlanks@10410.10453.44|TCAGTCGGTCCTGAGAGATGGGCGAGCGCCGTTCCGAAGGGACG</td><td>>hg38_21_+_8433172_8446622_RNA45SN1_45S_50ntFlanks</td><td>10410</td><td>10453</td><td>44</td><td>TCAGTCGGTCCTGAGAGATGGGCGAGCGCCGTTCCGAAGGGACG</td><td>1580.6</td><td>-15.1</td><td>1.6</td><td>-9.1</td><td>5.8E-20</td><td>2.2E-19</td><td>BT20/MB468</td><td>NUC</td></tr>
<tr><td>>hg38_21_+_8433172_8446622_RNA45SN1_45S_50ntFlanks@6648.6676.29|GTACGACTCTTAGCGGTGGATCACTCGGC</td><td>>hg38_21_+_8433172_8446622_RNA45SN1_45S_50ntFlanks</td><td>6648</td><td>6676</td><td>29</td><td>GTACGACTCTTAGCGGTGGATCACTCGGC</td><td>6948.0</td><td>5.7</td><td>0.6</td><td>9.3</td><td>2.2E-20</td><td>2.2E-19</td><td>BT20/MB468</td><td>MITO</td></tr>
<tr><td>>hg38_21_+_8433172_8446622_RNA45SN1_45S_50ntFlanks@6745.6772.28|ACACTTCGAACGCACTTGCGGCCCCGGG</td><td>>hg38_21_+_8433172_8446622_RNA45SN1_45S_50ntFlanks</td><td>6745</td><td>6772</td><td>28</td><td>ACACTTCGAACGCACTTGCGGCCCCGGG</td><td>1825.6</td><td>-10.5</td><td>1.1</td><td>-9.3</td><td>2.0E-20</td><td>2.3E-19</td><td>MB231/MB468</td><td>MITO</td></tr>
<tr><td>>hg38_21_+_8433172_8446622_RNA45SN1_45S_50ntFlanks@4759.4782.24|ATCAGATACCGTCGTAGTTCCGAC</td><td>>hg38_21_+_8433172_8446622_RNA45SN1_45S_50ntFlanks</td><td>4759</td><td>4782</td><td>24</td><td>ATCAGATACCGTCGTAGTTCCGAC</td><td>1706.0</td><td>-13.0</td><td>1.4</td><td>-9.2</td><td>3.1E-20</td><td>2.3E-19</td><td>BT20/MB468</td><td>CYTO</td></tr>
<tr><td>>hg38_21_+_8433172_8446622_RNA45SN1_45S_50ntFlanks@4735.4767.33|ACGAAAGTCGGAGGTTCGAAGACGATCAGATAC</td><td>>hg38_21_+_8433172_8446622_RNA45SN1_45S_50ntFlanks</td><td>4735</td><td>4767</td><td>33</td><td>ACGAAAGTCGGAGGTTCGAAGACGATCAGATAC</td><td>669.3</td><td>13.4</td><td>1.4</td><td>9.3</td><td>1.1E-20</td><td>2.3E-19</td><td>MB231/MB468</td><td>CYTO</td></tr>
<tr><td>>hg38_21_+_8433172_8446622_RNA45SN1_45S_50ntFlanks@12961.12993.33|TGGGTCGGGGTTTCGTACGTAGCAGAGCAGCTC</td><td>>hg38_21_+_8433172_8446622_RNA45SN1_45S_50ntFlanks</td><td>12961</td><td>12993</td><td>33</td><td>TGGGTCGGGGTTTCGTACGTAGCAGAGCAGCTC</td><td>182.6</td><td>8.2</td><td>0.9</td><td>9.3</td><td>1.1E-20</td><td>2.3E-19</td><td>MB231/MB468</td><td>CYTO</td></tr>
<tr><td>>hg38_21_+_8433172_8446622_RNA45SN1_45S_50ntFlanks@12160.12179.20|TTGACTGGGGCGGTACACCT</td><td>>hg38_21_+_8433172_8446622_RNA45SN1_45S_50ntFlanks</td><td>12160</td><td>12179</td><td>20</td><td>TTGACTGGGGCGGTACACCT</td><td>594.2</td><td>-13.1</td><td>1.4</td><td>-9.3</td><td>1.3E-20</td><td>2.3E-19</td><td>MB231/MB468</td><td>MP</td></tr>
<tr><td>>hg38_21_+_8433172_8446622_RNA45SN1_45S_50ntFlanks@10679.10712.34|GGTTCCGGCGGCGTCCGGTGAGCTCTCGCTGGCC</td><td>>hg38_21_+_8433172_8446622_RNA45SN1_45S_50ntFlanks</td><td>10679</td><td>10712</td><td>34</td><td>GGTTCCGGCGGCGTCCGGTGAGCTCTCGCTGGCC</td><td>1251.4</td><td>-14.1</td><td>1.5</td><td>-9.3</td><td>2.1E-20</td><td>2.3E-19</td><td>MB231/MB468</td><td>MITO</td></tr>
<tr><td>>hg38_21_+_8433172_8446622_RNA45SN1_45S_50ntFlanks@10516.10534.19|GAGTGGCGGAGATGGGCGC</td><td>>hg38_21_+_8433172_8446622_RNA45SN1_45S_50ntFlanks</td><td>10516</td><td>10534</td><td>19</td><td>GAGTGGCGGAGATGGGCGC</td><td>1267.4</td><td>-14.2</td><td>1.5</td><td>-9.3</td><td>1.3E-20</td><td>2.3E-19</td><td>MB231/MB468</td><td>MP</td></tr>
<tr><td>>MI0000102|hsa-mir-100&WithFlank&11|-|122152223|122152314@19.41.23&[MIMAT0000098&hsa-miR-100-5p&offsets|0|+1;m-27&11|-|122152276|122152296&offsets|0|+2]|AACCCGTAGATCCGAACTTGTGG</td><td>>MI0000102|hsa-mir-100&WithFlank&11|-|122152223|122152314</td><td>19</td><td>41</td><td>23</td><td>AACCCGTAGATCCGAACTTGTGG</td><td>2057.8</td><td>14.2</td><td>1.5</td><td>9.3</td><td>2.1E-20</td><td>2.3E-19</td><td>MB231/MB468</td><td>MITO</td></tr>
<tr><td>>hg38_21_+_8433172_8446622_RNA45SN1_45S_50ntFlanks@12420.12464.45|ATGTCGGCTCTTCCTATCATTGTGAAGCAGAATTCACCAAGCGTT</td><td>>hg38_21_+_8433172_8446622_RNA45SN1_45S_50ntFlanks</td><td>12420</td><td>12464</td><td>45</td><td>ATGTCGGCTCTTCCTATCATTGTGAAGCAGAATTCACCAAGCGTT</td><td>223.8</td><td>11.8</td><td>1.3</td><td>9.3</td><td>1.1E-20</td><td>2.3E-19</td><td>MB231/MB468</td><td>CYTO</td></tr>
<tr><td>>hg38_21_+_8433172_8446622_RNA45SN1_45S_50ntFlanks@7977.8010.34|CGACCTCAGATCAGACGTGGCGACCCGCTGAATT</td><td>>hg38_21_+_8433172_8446622_RNA45SN1_45S_50ntFlanks</td><td>7977</td><td>8010</td><td>34</td><td>CGACCTCAGATCAGACGTGGCGACCCGCTGAATT</td><td>866.9</td><td>4.5</td><td>0.5</td><td>9.2</td><td>2.3E-20</td><td>2.3E-19</td><td>BT20/MB468</td><td>MITO</td></tr>
<tr><td>>hg38_21_+_8433172_8446622_RNA45SN1_45S_50ntFlanks@10808.10829.22|CAATGTAGGTAAGGGAAGTCGG</td><td>>hg38_21_+_8433172_8446622_RNA45SN1_45S_50ntFlanks</td><td>10808</td><td>10829</td><td>22</td><td>CAATGTAGGTAAGGGAAGTCGG</td><td>1136.7</td><td>-13.9</td><td>1.5</td><td>-9.3</td><td>2.1E-20</td><td>2.3E-19</td><td>MB231/MB468</td><td>MITO</td></tr>
<tr><td>>hg38_21_+_8433172_8446622_RNA45SN1_45S_50ntFlanks@10808.10830.23|CAATGTAGGTAAGGGAAGTCGGC</td><td>>hg38_21_+_8433172_8446622_RNA45SN1_45S_50ntFlanks</td><td>10808</td><td>10830</td><td>23</td><td>CAATGTAGGTAAGGGAAGTCGGC</td><td>843.1</td><td>-13.5</td><td>1.5</td><td>-9.3</td><td>2.1E-20</td><td>2.3E-19</td><td>MB231/MB468</td><td>MITO</td></tr>
<tr><td>>hg38_21_+_8433172_8446622_RNA45SN1_45S_50ntFlanks@12624.12654.31|CCAATGGGGCGAAGCTACCATCTGTGGGATT</td><td>>hg38_21_+_8433172_8446622_RNA45SN1_45S_50ntFlanks</td><td>12624</td><td>12654</td><td>31</td><td>CCAATGGGGCGAAGCTACCATCTGTGGGATT</td><td>1810.6</td><td>-5.8</td><td>0.6</td><td>-9.2</td><td>3.2E-20</td><td>2.3E-19</td><td>BT20/MB468</td><td>CYTO</td></tr>
<tr><td>>MI0000061|hsa-let-7a-2&WithFlank&11|-|122146516|122146599@11.33.23;MI0000062|hsa-let-7a-3&WithFlank&22|+|46112743|46112828@10.32.23;MI0000060|hsa-let-7a-1&WithFlank&9|+|94175951|94176042@12.34.23&[MIMAT0000062&hsa-let-7a-5p&offsets|0|+1;m-6&11|-|122146568|122146589&offsets|0|+1];[MIMAT0000062_1&hsa-let-7a-5p&offsets|0|+1;m-5&22|+|46112752|46112773&offsets|0|+1];[MIMAT0000062_2&hsa-let-7a-5p&offsets|0|+1;m-4&9|+|94175962|94175983&offsets|0|+1]|TGAGGTAGTAGGTTGTATAGTTT</td><td>>MI0000061|hsa-let-7a-2&WithFlank&11|-|122146516|122146599</td><td>11</td><td>33</td><td>23</td><td>TGAGGTAGTAGGTTGTATAGTTT</td><td>2715.8</td><td>6.8</td><td>0.7</td><td>9.3</td><td>1.2E-20</td><td>2.4E-19</td><td>MB231/MB468</td><td>CYTO</td></tr>
<tr><td>>hg38_21_+_8433172_8446622_RNA45SN1_45S_50ntFlanks@11161.11179.19|CACGGTCCCCCGCGAGGGG</td><td>>hg38_21_+_8433172_8446622_RNA45SN1_45S_50ntFlanks</td><td>11161</td><td>11179</td><td>19</td><td>CACGGTCCCCCGCGAGGGG</td><td>968.4</td><td>-12.9</td><td>1.4</td><td>-9.3</td><td>1.9E-20</td><td>2.4E-19</td><td>MB231/MB468</td><td>NUC</td></tr>
<tr><td>>hg38_21_+_8433172_8446622_RNA45SN1_45S_50ntFlanks@12633.12653.21|CGAAGCTACCATCTGTGGGAT</td><td>>hg38_21_+_8433172_8446622_RNA45SN1_45S_50ntFlanks</td><td>12633</td><td>12653</td><td>21</td><td>CGAAGCTACCATCTGTGGGAT</td><td>1516.2</td><td>-10.4</td><td>1.1</td><td>-9.3</td><td>1.9E-20</td><td>2.4E-19</td><td>MB231/MB468</td><td>NUC</td></tr>
<tr><td>>hg38_21_+_8433172_8446622_RNA45SN1_45S_50ntFlanks@11568.11601.34|GCAGCCGACTTAGAACTGGTGCGGACCAGGGGAA</td><td>>hg38_21_+_8433172_8446622_RNA45SN1_45S_50ntFlanks</td><td>11568</td><td>11601</td><td>34</td><td>GCAGCCGACTTAGAACTGGTGCGGACCAGGGGAA</td><td>1432.6</td><td>-6.1</td><td>0.7</td><td>-9.3</td><td>1.4E-20</td><td>2.4E-19</td><td>MB231/MB468</td><td>MP</td></tr>
<tr><td>>hg38_21_+_8433172_8446622_RNA45SN1_45S_50ntFlanks@9566.9596.31|GACGTGCAAATCGGTCGTCCGACCTGGGTAT</td><td>>hg38_21_+_8433172_8446622_RNA45SN1_45S_50ntFlanks</td><td>9566</td><td>9596</td><td>31</td><td>GACGTGCAAATCGGTCGTCCGACCTGGGTAT</td><td>8232.8</td><td>-7.0</td><td>0.8</td><td>-9.3</td><td>2.1E-20</td><td>2.4E-19</td><td>MB231/MB468</td><td>MITO</td></tr>
<tr><td>>hg38_21_+_8433172_8446622_RNA45SN1_45S_50ntFlanks@11205.11232.28|GCGGCGGCGACTCTGGACGCGAGCCGGG</td><td>>hg38_21_+_8433172_8446622_RNA45SN1_45S_50ntFlanks</td><td>11205</td><td>11232</td><td>28</td><td>GCGGCGGCGACTCTGGACGCGAGCCGGG</td><td>846.1</td><td>-13.6</td><td>1.5</td><td>-9.3</td><td>1.4E-20</td><td>2.4E-19</td><td>MB231/MB468</td><td>MP</td></tr>
<tr><td>>hg38_21_+_8433172_8446622_RNA45SN1_45S_50ntFlanks@11661.11681.21|TGATTTCTGCCCAGTGCTCTG</td><td>>hg38_21_+_8433172_8446622_RNA45SN1_45S_50ntFlanks</td><td>11661</td><td>11681</td><td>21</td><td>TGATTTCTGCCCAGTGCTCTG</td><td>1525.5</td><td>-14.4</td><td>1.6</td><td>-9.3</td><td>1.4E-20</td><td>2.4E-19</td><td>MB231/MB468</td><td>MP</td></tr>
<tr><td>>hg38_21_+_8433172_8446622_RNA45SN1_45S_50ntFlanks@10806.10823.18|AACAATGTAGGTAAGGGA</td><td>>hg38_21_+_8433172_8446622_RNA45SN1_45S_50ntFlanks</td><td>10806</td><td>10823</td><td>18</td><td>AACAATGTAGGTAAGGGA</td><td>653.9</td><td>-13.2</td><td>1.4</td><td>-9.3</td><td>1.4E-20</td><td>2.4E-19</td><td>MB231/MB468</td><td>MP</td></tr>
<tr><td>>hg38_21_+_8433172_8446622_RNA45SN1_45S_50ntFlanks@4002.4024.23|GATAACCTCGGGCCGATCGCACG</td><td>>hg38_21_+_8433172_8446622_RNA45SN1_45S_50ntFlanks</td><td>4002</td><td>4024</td><td>23</td><td>GATAACCTCGGGCCGATCGCACG</td><td>2357.8</td><td>-8.7</td><td>0.9</td><td>-9.3</td><td>2.2E-20</td><td>2.4E-19</td><td>MB231/MB468</td><td>MITO</td></tr>
<tr><td>>hg38_21_+_8433172_8446622_RNA45SN1_45S_50ntFlanks@10726.10755.30|GGGAGAGGGTGTAAATCTCGCGCCGGGCCG</td><td>>hg38_21_+_8433172_8446622_RNA45SN1_45S_50ntFlanks</td><td>10726</td><td>10755</td><td>30</td><td>GGGAGAGGGTGTAAATCTCGCGCCGGGCCG</td><td>747.1</td><td>-14.0</td><td>1.5</td><td>-9.1</td><td>6.3E-20</td><td>2.4E-19</td><td>BT20/MB468</td><td>NUC</td></tr>
<tr><td>>hg38_21_+_8433172_8446622_RNA45SN1_45S_50ntFlanks@10922.10945.24|CTGGACGAGGCGCCGCCGCCCCCC</td><td>>hg38_21_+_8433172_8446622_RNA45SN1_45S_50ntFlanks</td><td>10922</td><td>10945</td><td>24</td><td>CTGGACGAGGCGCCGCCGCCCCCC</td><td>785.1</td><td>-13.4</td><td>1.4</td><td>-9.3</td><td>2.2E-20</td><td>2.5E-19</td><td>MB231/MB468</td><td>MITO</td></tr>
<tr><td>>hg38_21_+_8433172_8446622_RNA45SN1_45S_50ntFlanks@6700.6718.19|CTAGCTGCGAGAATTAATG</td><td>>hg38_21_+_8433172_8446622_RNA45SN1_45S_50ntFlanks</td><td>6700</td><td>6718</td><td>19</td><td>CTAGCTGCGAGAATTAATG</td><td>21815.2</td><td>-13.7</td><td>1.5</td><td>-9.1</td><td>6.5E-20</td><td>2.5E-19</td><td>BT20/MB468</td><td>NUC</td></tr>
<tr><td>>MI0000078|hsa-mir-22&WithFlank&17|-|1713897|1713993@59.77.19&[MIMAT0000077&hsa-miR-22-3p&offsets|0|-3;m-2&17|-|1713914|1713935&offsets|0|-3]|AAGCTGCCAGTTGAAGAAC</td><td>>MI0000078|hsa-mir-22&WithFlank&17|-|1713897|1713993</td><td>59</td><td>77</td><td>19</td><td>AAGCTGCCAGTTGAAGAAC</td><td>520.4</td><td>13.3</td><td>1.4</td><td>9.2</td><td>3.4E-20</td><td>2.5E-19</td><td>BT20/MB468</td><td>CYTO</td></tr>
<tr><td>>MI0000289|hsa-mir-181a-1&WithFlank&1|-|198859038|198859159@70.91.22&[MIMAT0000270&hsa-miR-181a-3p&offsets|0|0;m-125&1|-|198859069|198859090&offsets|0|0]|ACCATCGACCGTTGATTGTACC</td><td>>MI0000289|hsa-mir-181a-1&WithFlank&1|-|198859038|198859159</td><td>70</td><td>91</td><td>22</td><td>ACCATCGACCGTTGATTGTACC</td><td>921.6</td><td>5.9</td><td>0.6</td><td>9.3</td><td>2.0E-20</td><td>2.5E-19</td><td>MB231/MB468</td><td>NUC</td></tr>
<tr><td>>MI0000477|hsa-mir-146a&WithFlank&5|+|160485346|160485456@27.49.23&[MIMAT0000449&hsa-miR-146a-5p&offsets|0|+1;m-129&5|+|160485372|160485393&offsets|0|+1]|TGAGAACTGAATTCCATGGGTTG</td><td>>MI0000477|hsa-mir-146a&WithFlank&5|+|160485346|160485456</td><td>27</td><td>49</td><td>23</td><td>TGAGAACTGAATTCCATGGGTTG</td><td>7860.6</td><td>-16.3</td><td>1.7</td><td>-9.7</td><td>2.1E-22</td><td>2.5E-19</td><td>BT20/MB231</td><td>MP</td></tr>
<tr><td>>MI0003643|hsa-mir-629&WithFlank&15|-|70079366|70079474@28.48.21&[MIMAT0004810&hsa-miR-629-5p&offsets|0|0;m-150&15|-|70079427|70079447&offsets|0|0]|TGGGTTTACGTTGGGAGAACT</td><td>>MI0003643|hsa-mir-629&WithFlank&15|-|70079366|70079474</td><td>28</td><td>48</td><td>21</td><td>TGGGTTTACGTTGGGAGAACT</td><td>982.2</td><td>13.2</td><td>1.4</td><td>9.3</td><td>1.5E-20</td><td>2.5E-19</td><td>MB231/MB468</td><td>MP</td></tr>
<tr><td>>MI0000067|hsa-let-7f-1&WithFlank&9|+|94176341|94176439@13.32.20(+1C);MI0000068|hsa-let-7f-2&WithFlank&X|-|53557186|53557280@14.33.20(+1C)&[MIMAT0000067&hsa-let-7f-5p&offsets|0|-2(+1C);m-20&9|+|94176353|94176374&offsets|0|-2(+1C)];[MIMAT0000067_1&hsa-let-7f-5p&offsets|0|-2(+1C);m-19&X|-|53557246|53557267&offsets|0|-2(+1C)]|TGAGGTAGTAGATTGTATAGC</td><td>>MI0000067|hsa-let-7f-1&WithFlank&9|+|94176341|94176439</td><td>13</td><td>32</td><td>20(+1C)</td><td>TGAGGTAGTAGATTGTATAGC</td><td>1992.2</td><td>7.0</td><td>0.8</td><td>9.2</td><td>3.4E-20</td><td>2.5E-19</td><td>BT20/MB468</td><td>CYTO</td></tr>
<tr><td>>hg38_21_+_8433172_8446622_RNA45SN1_45S_50ntFlanks@12365.12382.18|ATAACTGGCTTGTGGCGG</td><td>>hg38_21_+_8433172_8446622_RNA45SN1_45S_50ntFlanks</td><td>12365</td><td>12382</td><td>18</td><td>ATAACTGGCTTGTGGCGG</td><td>852.5</td><td>-13.5</td><td>1.5</td><td>-9.2</td><td>2.3E-20</td><td>2.6E-19</td><td>MB231/MB468</td><td>MITO</td></tr>
<tr><td>>MI0003643|hsa-mir-629&WithFlank&15|-|70079366|70079474@28.49.22&[MIMAT0004810&hsa-miR-629-5p&offsets|0|+1;m-150&15|-|70079427|70079447&offsets|0|+1]|TGGGTTTACGTTGGGAGAACTT</td><td>>MI0003643|hsa-mir-629&WithFlank&15|-|70079366|70079474</td><td>28</td><td>49</td><td>22</td><td>TGGGTTTACGTTGGGAGAACTT</td><td>319.5</td><td>12.2</td><td>1.3</td><td>9.3</td><td>2.1E-20</td><td>2.6E-19</td><td>MB231/MB468</td><td>NUC</td></tr>
<tr><td>>hg38_21_+_8433172_8446622_RNA45SN1_45S_50ntFlanks@11582.11605.24|ACTGGTGCGGACCAGGGGAATCCG</td><td>>hg38_21_+_8433172_8446622_RNA45SN1_45S_50ntFlanks</td><td>11582</td><td>11605</td><td>24</td><td>ACTGGTGCGGACCAGGGGAATCCG</td><td>453.9</td><td>-3.9</td><td>0.4</td><td>-9.1</td><td>6.6E-20</td><td>2.6E-19</td><td>BT20/MB468</td><td>NUC</td></tr>
<tr><td>>hg38_21_+_8433172_8446622_RNA45SN1_45S_50ntFlanks@485.508.24|GATGTGGTGACGTCGTGCTCTCCC</td><td>>hg38_21_+_8433172_8446622_RNA45SN1_45S_50ntFlanks</td><td>485</td><td>508</td><td>24</td><td>GATGTGGTGACGTCGTGCTCTCCC</td><td>529.7</td><td>13.5</td><td>1.5</td><td>9.2</td><td>2.5E-20</td><td>2.6E-19</td><td>BT20/MB468</td><td>MITO</td></tr>
<tr><td>>hg38_21_+_8433172_8446622_RNA45SN1_45S_50ntFlanks@10458.10477.20|ATGGCCTCCGTTGCCCTCGG</td><td>>hg38_21_+_8433172_8446622_RNA45SN1_45S_50ntFlanks</td><td>10458</td><td>10477</td><td>20</td><td>ATGGCCTCCGTTGCCCTCGG</td><td>896.2</td><td>-11.9</td><td>1.3</td><td>-9.2</td><td>2.5E-20</td><td>2.6E-19</td><td>BT20/MB468</td><td>MITO</td></tr>
<tr><td>>MI0000433|hsa-let-7g&WithFlank&3|-|52268272|52268367@11.33.23&[MIMAT0000414&hsa-let-7g-5p&offsets|0|+1;m-58&3|-|52268336|52268357&offsets|0|+1]|TGAGGTAGTAGTTTGTACAGTTT</td><td>>MI0000433|hsa-let-7g&WithFlank&3|-|52268272|52268367</td><td>11</td><td>33</td><td>23</td><td>TGAGGTAGTAGTTTGTACAGTTT</td><td>2202.2</td><td>5.6</td><td>0.6</td><td>9.3</td><td>2.1E-20</td><td>2.6E-19</td><td>MB231/MB468</td><td>NUC</td></tr>
<tr><td>>hg38_21_+_8433172_8446622_RNA45SN1_45S_50ntFlanks@6702.6720.19|AGCTGCGAGAATTAATGTG</td><td>>hg38_21_+_8433172_8446622_RNA45SN1_45S_50ntFlanks</td><td>6702</td><td>6720</td><td>19</td><td>AGCTGCGAGAATTAATGTG</td><td>649.2</td><td>-5.9</td><td>0.6</td><td>-9.1</td><td>6.7E-20</td><td>2.6E-19</td><td>BT20/MB468</td><td>NUC</td></tr>
<tr><td>>hg38_21_+_8433172_8446622_RNA45SN1_45S_50ntFlanks@10384.10405.22|CCATGTGAACAGCAGTTGAACA</td><td>>hg38_21_+_8433172_8446622_RNA45SN1_45S_50ntFlanks</td><td>10384</td><td>10405</td><td>22</td><td>CCATGTGAACAGCAGTTGAACA</td><td>1112.8</td><td>-14.5</td><td>1.6</td><td>-9.1</td><td>6.7E-20</td><td>2.6E-19</td><td>BT20/MB468</td><td>NUC</td></tr>
<tr><td>>MI0003513|hsa-mir-455&WithFlank&9|+|114209428|114209535@60.81.22&[MIMAT0004784&hsa-miR-455-3p&offsets|0|+1;m-135&9|+|114209487|114209508&offsets|0|0]|GCAGTCCATGGGCATATACACT</td><td>>MI0003513|hsa-mir-455&WithFlank&9|+|114209428|114209535</td><td>60</td><td>81</td><td>22</td><td>GCAGTCCATGGGCATATACACT</td><td>420.3</td><td>12.6</td><td>1.4</td><td>9.3</td><td>2.1E-20</td><td>2.6E-19</td><td>MB231/MB468</td><td>NUC</td></tr>
<tr><td>>hg38_21_+_8433172_8446622_RNA45SN1_45S_50ntFlanks@4555.4573.19|CGCAGCTAGGAATAATGGA</td><td>>hg38_21_+_8433172_8446622_RNA45SN1_45S_50ntFlanks</td><td>4555</td><td>4573</td><td>19</td><td>CGCAGCTAGGAATAATGGA</td><td>622.0</td><td>-13.1</td><td>1.4</td><td>-9.3</td><td>1.5E-20</td><td>2.6E-19</td><td>MB231/MB468</td><td>MP</td></tr>
<tr><td>>hg38_21_+_8433172_8446622_RNA45SN1_45S_50ntFlanks@12488.12521.34|AACGTGAGCTGGGTTTAGACCGTCGTGAGACAGG</td><td>>hg38_21_+_8433172_8446622_RNA45SN1_45S_50ntFlanks</td><td>12488</td><td>12521</td><td>34</td><td>AACGTGAGCTGGGTTTAGACCGTCGTGAGACAGG</td><td>555.9</td><td>-13.0</td><td>1.4</td><td>-9.3</td><td>1.5E-20</td><td>2.6E-19</td><td>MB231/MB468</td><td>MP</td></tr>
<tr><td>>hg38_21_+_8433172_8446622_RNA45SN1_45S_50ntFlanks@7985.8003.19|GATCAGACGTGGCGACCCG</td><td>>hg38_21_+_8433172_8446622_RNA45SN1_45S_50ntFlanks</td><td>7985</td><td>8003</td><td>19</td><td>GATCAGACGTGGCGACCCG</td><td>658.3</td><td>-13.2</td><td>1.4</td><td>-9.3</td><td>1.5E-20</td><td>2.6E-19</td><td>MB231/MB468</td><td>MP</td></tr>
<tr><td>>hg38_21_+_8433172_8446622_RNA45SN1_45S_50ntFlanks@11876.11906.31|AAGACCCTGTTGAGCTTGACTCTAGTCTGGC</td><td>>hg38_21_+_8433172_8446622_RNA45SN1_45S_50ntFlanks</td><td>11876</td><td>11906</td><td>31</td><td>AAGACCCTGTTGAGCTTGACTCTAGTCTGGC</td><td>491.1</td><td>-12.8</td><td>1.4</td><td>-9.3</td><td>1.5E-20</td><td>2.6E-19</td><td>MB231/MB468</td><td>MP</td></tr>
<tr><td>>MI0000266|hsa-mir-10a&WithFlank&17|-|48579832|48579953@28.50.23(+1A)&[MIMAT0000253&hsa-miR-10a-5p&offsets|0|0(+1A);m-18&17|-|48579905|48579926&offsets|0|+1(+1A)]|TACCCTGTAGATCCGAATTTGTGA</td><td>>MI0000266|hsa-mir-10a&WithFlank&17|-|48579832|48579953</td><td>28</td><td>50</td><td>23(+1A)</td><td>TACCCTGTAGATCCGAATTTGTGA</td><td>393.4</td><td>12.6</td><td>1.4</td><td>9.3</td><td>1.3E-20</td><td>2.6E-19</td><td>MB231/MB468</td><td>CYTO</td></tr>
<tr><td>>trnaMT_GluTTC_MT_-_14674_14742@40.71.32,trnalookalike8_GluTTC_5_-_93905172_93905240@40.71.32|TTGGTCGTGGTTGTAGTCCGTGCGAGAATACC</td><td>>trnaMT_GluTTC_MT_-_14674_14742</td><td>40</td><td>71</td><td>32</td><td>TTGGTCGTGGTTGTAGTCCGTGCGAGAATACC</td><td>2325.9</td><td>6.9</td><td>0.7</td><td>9.2</td><td>2.6E-20</td><td>2.6E-19</td><td>BT20/MB468</td><td>MITO</td></tr>
<tr><td>>hg38_21_+_8433172_8446622_RNA45SN1_45S_50ntFlanks@8552.8569.18|CTCCCCGGAGGGGGCGGG</td><td>>hg38_21_+_8433172_8446622_RNA45SN1_45S_50ntFlanks</td><td>8552</td><td>8569</td><td>18</td><td>CTCCCCGGAGGGGGCGGG</td><td>1465.8</td><td>-14.3</td><td>1.5</td><td>-9.2</td><td>2.4E-20</td><td>2.6E-19</td><td>MB231/MB468</td><td>MITO</td></tr>
</table>

<table>
<tr><td>>MI0000434|hsa-let-7i&WithFlank&12|+|62603680|62603775@12.32.21&[MIMAT0000415&hsa-let-7i-5p&offsets|0|-1;m-49&12|+|62603691|62603712&offsets|0|-1]|TGAGGTAGTAGTTTGTGCTGT</td><td>>MI0000434|hsa-let-7i&WithFlank&12|+|62603680|62603775</td><td>12</td><td>32</td><td>21</td><td>TGAGGTAGTAGTTTGTGCTGT</td><td>7883.3</td><td>6.4</td><td>0.7</td><td>9.3</td><td>2.2E-20</td><td>2.6E-19</td><td>MB231/MB468</td><td>NUC</td></tr>
<tr><td>>hg38_21_+_8433172_8446622_RNA45SN1_45S_50ntFlanks@5073.5110.38|AATTCCGATAACGAACGAGACTCTGGCATGCTAACTAG</td><td>>hg38_21_+_8433172_8446622_RNA45SN1_45S_50ntFlanks</td><td>5073</td><td>5110</td><td>38</td><td>AATTCCGATAACGAACGAGACTCTGGCATGCTAACTAG</td><td>1538.3</td><td>-14.4</td><td>1.6</td><td>-9.2</td><td>2.4E-20</td><td>2.6E-19</td><td>MB231/MB468</td><td>MITO</td></tr>
<tr><td>>hg38_21_+_8433172_8446622_RNA45SN1_45S_50ntFlanks@11795.11821.27|AACGAGATTCCCACTGTCCCTACCTAC</td><td>>hg38_21_+_8433172_8446622_RNA45SN1_45S_50ntFlanks</td><td>11795</td><td>11821</td><td>27</td><td>AACGAGATTCCCACTGTCCCTACCTAC</td><td>741.3</td><td>-14.0</td><td>1.5</td><td>-9.1</td><td>6.9E-20</td><td>2.6E-19</td><td>BT20/MB468</td><td>NUC</td></tr>
<tr><td>>trnaMT_GluTTC_MT_-_14674_14742@40.72.33,trnalookalike8_GluTTC_5_-_93905172_93905240@40.72.33|TTGGTCGTGGTTGTAGTCCGTGCGAGAATACCA</td><td>>trnaMT_GluTTC_MT_-_14674_14742</td><td>40</td><td>72</td><td>33</td><td>TTGGTCGTGGTTGTAGTCCGTGCGAGAATACCA</td><td>39718.9</td><td>6.6</td><td>0.7</td><td>9.2</td><td>2.6E-20</td><td>2.6E-19</td><td>BT20/MB468</td><td>MITO</td></tr>
<tr><td>>hg38_21_+_8433172_8446622_RNA45SN1_45S_50ntFlanks@10823.10865.43|AAGTCGGCAAGCCGGATCCGTAACTTCGGGATAAGGATTGGCT</td><td>>hg38_21_+_8433172_8446622_RNA45SN1_45S_50ntFlanks</td><td>10823</td><td>10865</td><td>43</td><td>AAGTCGGCAAGCCGGATCCGTAACTTCGGGATAAGGATTGGCT</td><td>2034.6</td><td>-15.4</td><td>1.7</td><td>-9.1</td><td>7.0E-20</td><td>2.7E-19</td><td>BT20/MB468</td><td>NUC</td></tr>
<tr><td>>hg38_21_+_8433172_8446622_RNA45SN1_45S_50ntFlanks@11619.11640.22|AACAAAGCATCGCGAAGGCCCG</td><td>>hg38_21_+_8433172_8446622_RNA45SN1_45S_50ntFlanks</td><td>11619</td><td>11640</td><td>22</td><td>AACAAAGCATCGCGAAGGCCCG</td><td>1957.8</td><td>-10.8</td><td>1.2</td><td>-9.2</td><td>2.4E-20</td><td>2.7E-19</td><td>MB231/MB468</td><td>MITO</td></tr>
<tr><td>>hg38_21_+_8433172_8446622_RNA45SN1_45S_50ntFlanks@11668.11693.26|TGCCCAGTGCTCTGAATGTCAAAGTG</td><td>>hg38_21_+_8433172_8446622_RNA45SN1_45S_50ntFlanks</td><td>11668</td><td>11693</td><td>26</td><td>TGCCCAGTGCTCTGAATGTCAAAGTG</td><td>721.9</td><td>6.4</td><td>0.7</td><td>9.3</td><td>1.3E-20</td><td>2.7E-19</td><td>MB231/MB468</td><td>CYTO</td></tr>
<tr><td>>hg38_21_+_8433172_8446622_RNA45SN1_45S_50ntFlanks@12676.12703.28|AGAATCCCGCCCAGGCGGAACGATACGG</td><td>>hg38_21_+_8433172_8446622_RNA45SN1_45S_50ntFlanks</td><td>12676</td><td>12703</td><td>28</td><td>AGAATCCCGCCCAGGCGGAACGATACGG</td><td>812.3</td><td>-13.4</td><td>1.5</td><td>-9.2</td><td>2.5E-20</td><td>2.7E-19</td><td>MB231/MB468</td><td>MITO</td></tr>
<tr><td>>MI0003205|hsa-mir-532&WithFlank&X|+|50003142|50003244@26.47.22&[MIMAT0002888&hsa-miR-532-5p&offsets|0|0;m-65&X|+|50003167|50003188&offsets|0|0]|CATGCCTTGAGTGTAGGACCGT</td><td>>MI0003205|hsa-mir-532&WithFlank&X|+|50003142|50003244</td><td>26</td><td>47</td><td>22</td><td>CATGCCTTGAGTGTAGGACCGT</td><td>2835.4</td><td>5.7</td><td>0.6</td><td>9.2</td><td>2.7E-20</td><td>2.7E-19</td><td>BT20/MB468</td><td>MITO</td></tr>
<tr><td>>hg38_21_+_8433172_8446622_RNA45SN1_45S_50ntFlanks@12476.12498.23|CCACTAATAGGGAACGTGAGCTG</td><td>>hg38_21_+_8433172_8446622_RNA45SN1_45S_50ntFlanks</td><td>12476</td><td>12498</td><td>23</td><td>CCACTAATAGGGAACGTGAGCTG</td><td>1512.2</td><td>-10.3</td><td>1.1</td><td>-9.2</td><td>2.5E-20</td><td>2.7E-19</td><td>MB231/MB468</td><td>MITO</td></tr>
<tr><td>>hg38_21_+_8433172_8446622_RNA45SN1_45S_50ntFlanks@12475.12498.24|CCCACTAATAGGGAACGTGAGCTG</td><td>>hg38_21_+_8433172_8446622_RNA45SN1_45S_50ntFlanks</td><td>12475</td><td>12498</td><td>24</td><td>CCCACTAATAGGGAACGTGAGCTG</td><td>718.8</td><td>-5.4</td><td>0.6</td><td>-9.2</td><td>2.3E-20</td><td>2.7E-19</td><td>MB231/MB468</td><td>NUC</td></tr>
<tr><td>>hg38_21_+_8433172_8446622_RNA45SN1_45S_50ntFlanks@10865.10902.38|TCTAAGGGCTGGGTCGGTCGGGCTGGGGCGCGAAGCGG</td><td>>hg38_21_+_8433172_8446622_RNA45SN1_45S_50ntFlanks</td><td>10865</td><td>10902</td><td>38</td><td>TCTAAGGGCTGGGTCGGTCGGGCTGGGGCGCGAAGCGG</td><td>494.1</td><td>-13.4</td><td>1.5</td><td>-9.1</td><td>7.2E-20</td><td>2.7E-19</td><td>BT20/MB468</td><td>NUC</td></tr>
<tr><td>>hg38_21_+_8433172_8446622_RNA45SN1_45S_50ntFlanks@11847.11871.25|AACGGGCTTGGCGGAATCAGCGGGG</td><td>>hg38_21_+_8433172_8446622_RNA45SN1_45S_50ntFlanks</td><td>11847</td><td>11871</td><td>25</td><td>AACGGGCTTGGCGGAATCAGCGGGG</td><td>605.0</td><td>-13.1</td><td>1.4</td><td>-9.3</td><td>1.6E-20</td><td>2.8E-19</td><td>MB231/MB468</td><td>MP</td></tr>
<tr><td>>trnaMT_MetCAT_MT_+_4402_4469@36.71.36|CCCGAAAATGTTGGTTATACCCTTCCCGTACTACCA</td><td>>trnaMT_MetCAT_MT_+_4402_4469</td><td>36</td><td>71</td><td>36</td><td>CCCGAAAATGTTGGTTATACCCTTCCCGTACTACCA</td><td>680.6</td><td>13.4</td><td>1.4</td><td>9.3</td><td>1.4E-20</td><td>2.8E-19</td><td>MB231/MB468</td><td>CYTO</td></tr>
<tr><td>>hg38_21_+_8433172_8446622_RNA45SN1_45S_50ntFlanks@9532.9560.29|AGAGGAAACTCTGGTGGAGGTCCGTAGCG</td><td>>hg38_21_+_8433172_8446622_RNA45SN1_45S_50ntFlanks</td><td>9532</td><td>9560</td><td>29</td><td>AGAGGAAACTCTGGTGGAGGTCCGTAGCG</td><td>174.8</td><td>11.5</td><td>1.2</td><td>9.3</td><td>1.4E-20</td><td>2.8E-19</td><td>MB231/MB468</td><td>CYTO</td></tr>
<tr><td>>MI0001446|hsa-mir-424&WithFlank&X|-|134546608|134546717@54.74.21&[MIMAT0004749&hsa-miR-424-3p&offsets|0|0;m-305&X|-|134546644|134546664&offsets|0|0]|CAAAACGTGAGGCGCTGCTAT</td><td>>MI0001446|hsa-mir-424&WithFlank&X|-|134546608|134546717</td><td>54</td><td>74</td><td>21</td><td>CAAAACGTGAGGCGCTGCTAT</td><td>914.9</td><td>14.3</td><td>1.5</td><td>9.2</td><td>2.7E-20</td><td>2.8E-19</td><td>BT20/MB468</td><td>MITO</td></tr>
<tr><td>>MI0003643|hsa-mir-629&WithFlank&15|-|70079366|70079474@28.49.22&[MIMAT0004810&hsa-miR-629-5p&offsets|0|+1;m-150&15|-|70079427|70079447&offsets|0|+1]|TGGGTTTACGTTGGGAGAACTT</td><td>>MI0003643|hsa-mir-629&WithFlank&15|-|70079366|70079474</td><td>28</td><td>49</td><td>22</td><td>TGGGTTTACGTTGGGAGAACTT</td><td>12860.7</td><td>17.4</td><td>1.8</td><td>9.7</td><td>3.1E-22</td><td>2.8E-19</td><td>BT20/MB468</td><td>MP</td></tr>
<tr><td>>hg38_21_+_8433172_8446622_RNA45SN1_45S_50ntFlanks@11572.11597.26|CCGACTTAGAACTGGTGCGGACCAGG</td><td>>hg38_21_+_8433172_8446622_RNA45SN1_45S_50ntFlanks</td><td>11572</td><td>11597</td><td>26</td><td>CCGACTTAGAACTGGTGCGGACCAGG</td><td>449.8</td><td>-12.7</td><td>1.4</td><td>-9.3</td><td>1.6E-20</td><td>2.8E-19</td><td>MB231/MB468</td><td>MP</td></tr>
<tr><td>>hg38_21_+_8433172_8446622_RNA45SN1_45S_50ntFlanks@8168.8188.21|CCCAAGTCCTTCTGATCGAGG</td><td>>hg38_21_+_8433172_8446622_RNA45SN1_45S_50ntFlanks</td><td>8168</td><td>8188</td><td>21</td><td>CCCAAGTCCTTCTGATCGAGG</td><td>600.4</td><td>-13.1</td><td>1.4</td><td>-9.3</td><td>1.7E-20</td><td>2.8E-19</td><td>MB231/MB468</td><td>MP</td></tr>
<tr><td>>hg38_21_+_8433172_8446622_RNA45SN1_45S_50ntFlanks@6661.6707.47|CGGTGGATCACTCGGCTCGTGCGTCGATGAAGAACGCAGCTAGCTGC</td><td>>hg38_21_+_8433172_8446622_RNA45SN1_45S_50ntFlanks</td><td>6661</td><td>6707</td><td>47</td><td>CGGTGGATCACTCGGCTCGTGCGTCGATGAAGAACGCAGCTAGCTGC</td><td>15510.5</td><td>8.7</td><td>0.9</td><td>9.2</td><td>2.7E-20</td><td>2.8E-19</td><td>BT20/MB468</td><td>MITO</td></tr>
<tr><td>>hg38_21_+_8433172_8446622_RNA45SN1_45S_50ntFlanks@12935.12956.22|TAAACCATTCGTAGACGACCTG</td><td>>hg38_21_+_8433172_8446622_RNA45SN1_45S_50ntFlanks</td><td>12935</td><td>12956</td><td>22</td><td>TAAACCATTCGTAGACGACCTG</td><td>8720.8</td><td>-12.9</td><td>1.4</td><td>-9.1</td><td>7.3E-20</td><td>2.8E-19</td><td>BT20/MB468</td><td>NUC</td></tr>
<tr><td>>MI0000746|hsa-mir-99b&WithFlank&19|+|51692606|51692687@13.32.20(+2U)&[MIMAT0000689&hsa-miR-99b-5p&offsets|0|-2(+2U);m-10&19|+|51692618|51692639&offsets|0|-2(+2U)]|CACCCGTAGAACCGACCTTGTT</td><td>>MI0000746|hsa-mir-99b&WithFlank&19|+|51692606|51692687</td><td>13</td><td>32</td><td>20(+2U)</td><td>CACCCGTAGAACCGACCTTGTT</td><td>2415.7</td><td>11.2</td><td>1.2</td><td>9.6</td><td>1.2E-21</td><td>2.8E-19</td><td>BT20/MB231</td><td>CYTO</td></tr>
<tr><td>>hg38_21_+_8433172_8446622_RNA45SN1_45S_50ntFlanks@12928.12956.29|CTGGCGCTAAACCATTCGTAGACGACCTG</td><td>>hg38_21_+_8433172_8446622_RNA45SN1_45S_50ntFlanks</td><td>12928</td><td>12956</td><td>29</td><td>CTGGCGCTAAACCATTCGTAGACGACCTG</td><td>677.5</td><td>-13.2</td><td>1.4</td><td>-9.2</td><td>2.6E-20</td><td>2.8E-19</td><td>MB231/MB468</td><td>MITO</td></tr>
<tr><td>>hg38_21_+_8433172_8446622_RNA45SN1_45S_50ntFlanks@3994.4023.30|TGACTCTAGATAACCTCGGGCCGATCGCAC</td><td>>hg38_21_+_8433172_8446622_RNA45SN1_45S_50ntFlanks</td><td>3994</td><td>4023</td><td>30</td><td>TGACTCTAGATAACCTCGGGCCGATCGCAC</td><td>338.4</td><td>5.4</td><td>0.6</td><td>9.3</td><td>1.4E-20</td><td>2.8E-19</td><td>MB231/MB468</td><td>CYTO</td></tr>
<tr><td>>am_trnaMT_HisGTG_MT_+_12138_12206@20.51.32|AACATCAGATTGTGAATCTGACAACAGAGGCT</td><td>>am_trnaMT_HisGTG_MT_+_12138_12206</td><td>20</td><td>51</td><td>32</td><td>AACATCAGATTGTGAATCTGACAACAGAGGCT</td><td>1272.6</td><td>8.0</td><td>0.9</td><td>9.2</td><td>2.8E-20</td><td>2.8E-19</td><td>BT20/MB468</td><td>MITO</td></tr>
<tr><td>>hg38_21_+_8433172_8446622_RNA45SN1_45S_50ntFlanks@11923.11948.26|AGAGGTGTAGAATAAGTGGGAGGCCC</td><td>>hg38_21_+_8433172_8446622_RNA45SN1_45S_50ntFlanks</td><td>11923</td><td>11948</td><td>26</td><td>AGAGGTGTAGAATAAGTGGGAGGCCC</td><td>577.6</td><td>-13.0</td><td>1.4</td><td>-9.3</td><td>1.7E-20</td><td>2.9E-19</td><td>MB231/MB468</td><td>MP</td></tr>
<tr><td>>hg38_21_+_8433172_8446622_RNA45SN1_45S_50ntFlanks@7988.8006.19|CAGACGTGGCGACCCGCTG</td><td>>hg38_21_+_8433172_8446622_RNA45SN1_45S_50ntFlanks</td><td>7988</td><td>8006</td><td>19</td><td>CAGACGTGGCGACCCGCTG</td><td>2523.9</td><td>-15.2</td><td>1.6</td><td>-9.3</td><td>1.7E-20</td><td>2.9E-19</td><td>MB231/MB468</td><td>MP</td></tr>
<tr><td>>hg38_21_+_8433172_8446622_RNA45SN1_45S_50ntFlanks@9005.9026.22|CTCCCCCGCGGGGGCGCGCCGG</td><td>>hg38_21_+_8433172_8446622_RNA45SN1_45S_50ntFlanks</td><td>9005</td><td>9026</td><td>22</td><td>CTCCCCCGCGGGGGCGCGCCGG</td><td>1590.8</td><td>-14.4</td><td>1.6</td><td>-9.2</td><td>2.6E-20</td><td>2.9E-19</td><td>MB231/MB468</td><td>MITO</td></tr>
<tr><td>>hg38_21_+_8433172_8446622_RNA45SN1_45S_50ntFlanks@7980.8007.28|CCTCAGATCAGACGTGGCGACCCGCTGA</td><td>>hg38_21_+_8433172_8446622_RNA45SN1_45S_50ntFlanks</td><td>7980</td><td>8007</td><td>28</td><td>CCTCAGATCAGACGTGGCGACCCGCTGA</td><td>416.2</td><td>-12.6</td><td>1.4</td><td>-9.3</td><td>1.7E-20</td><td>2.9E-19</td><td>MB231/MB468</td><td>MP</td></tr>
<tr><td>>hg38_21_+_8433172_8446622_RNA45SN1_45S_50ntFlanks@12949.12989.41|ACGACCTGCTTCTGGGTCGGGGTTTCGTACGTAGCAGAGCA</td><td>>hg38_21_+_8433172_8446622_RNA45SN1_45S_50ntFlanks</td><td>12949</td><td>12989</td><td>41</td><td>ACGACCTGCTTCTGGGTCGGGGTTTCGTACGTAGCAGAGCA</td><td>364.6</td><td>-12.9</td><td>1.4</td><td>-9.1</td><td>7.7E-20</td><td>2.9E-19</td><td>BT20/MB468</td><td>NUC</td></tr>
<tr><td>>hg38_21_+_8433172_8446622_RNA45SN1_45S_50ntFlanks@11729.11748.20|TAACTATGACTCTCTTAAGG</td><td>>hg38_21_+_8433172_8446622_RNA45SN1_45S_50ntFlanks</td><td>11729</td><td>11748</td><td>20</td><td>TAACTATGACTCTCTTAAGG</td><td>2803.2</td><td>-5.8</td><td>0.6</td><td>-9.2</td><td>2.7E-20</td><td>2.9E-19</td><td>MB231/MB468</td><td>MITO</td></tr>
<tr><td>>hg38_21_+_8433172_8446622_RNA45SN1_45S_50ntFlanks@11210.11254.45|GGCGACTCTGGACGCGAGCCGGGCCCTTCCCGTGGATCGCCCCAG</td><td>>hg38_21_+_8433172_8446622_RNA45SN1_45S_50ntFlanks</td><td>11210</td><td>11254</td><td>45</td><td>GGCGACTCTGGACGCGAGCCGGGCCCTTCCCGTGGATCGCCCCAG</td><td>1450.5</td><td>-14.3</td><td>1.5</td><td>-9.2</td><td>2.7E-20</td><td>3.0E-19</td><td>MB231/MB468</td><td>MITO</td></tr>
<tr><td>>am_MI0000786|hsa-mir-378a&WithFlank&5|+|149732819|149732896@49.70.22(+1A)&[MIMAT0000732&hsa-miR-378a-3p&offsets|0|0(+1A);m-44&5|+|149732867|149732888&offsets|0|0(+1A)]|ACTGGACTTGGAGTCAGAAGGCA</td><td>>am_MI0000786|hsa-mir-378a&WithFlank&5|+|149732819|149732896</td><td>49</td><td>70</td><td>22(+1A)</td><td>ACTGGACTTGGAGTCAGAAGGCA</td><td>1365.2</td><td>5.2</td><td>0.6</td><td>9.2</td><td>2.9E-20</td><td>3.0E-19</td><td>BT20/MB468</td><td>MITO</td></tr>
<tr><td>>MI0005756|hsa-mir-934&WithFlank&X|+|136550872|136550966@21.42.22(+2A)&[MIMAT0004977&hsa-miR-934&offsets|0|0(+2A);m-943&X|+|136550892|136550913&offsets|0|0(+2A)]|TGTCTACTACTGGAGACACTGGAA</td><td>>MI0005756|hsa-mir-934&WithFlank&X|+|136550872|136550966</td><td>21</td><td>42</td><td>22(+2A)</td><td>TGTCTACTACTGGAGACACTGGAA</td><td>798.0</td><td>8.5</td><td>0.9</td><td>9.2</td><td>4.0E-20</td><td>3.0E-19</td><td>BT20/MB468</td><td>CYTO</td></tr>
<tr><td>>hg38_21_+_8433172_8446622_RNA45SN1_45S_50ntFlanks@10408.10426.19|GGTCAGTCGGTCCTGAGAG</td><td>>hg38_21_+_8433172_8446622_RNA45SN1_45S_50ntFlanks</td><td>10408</td><td>10426</td><td>19</td><td>GGTCAGTCGGTCCTGAGAG</td><td>1323.6</td><td>-13.3</td><td>1.4</td><td>-9.2</td><td>2.5E-20</td><td>3.0E-19</td><td>MB231/MB468</td><td>NUC</td></tr>
<tr><td>>hg38_21_+_8433172_8446622_RNA45SN1_45S_50ntFlanks@4760.4778.19|TCAGATACCGTCGTAGTTC</td><td>>hg38_21_+_8433172_8446622_RNA45SN1_45S_50ntFlanks</td><td>4760</td><td>4778</td><td>19</td><td>TCAGATACCGTCGTAGTTC</td><td>704.1</td><td>-13.3</td><td>1.4</td><td>-9.3</td><td>1.8E-20</td><td>3.0E-19</td><td>MB231/MB468</td><td>MP</td></tr>
<tr><td>>MI0000650|hsa-mir-200c&WithFlank&12|+|6963693|6963772@50.71.22(+1C)&[MIMAT0000617&hsa-miR-200c-3p&offsets|0|-1(+1C);m-54&12|+|6963742|6963764&offsets|0|-1(+1C)]|TAATACTGCCGGGTAATGATGGC</td><td>>MI0000650|hsa-mir-200c&WithFlank&12|+|6963693|6963772</td><td>50</td><td>71</td><td>22(+1C)</td><td>TAATACTGCCGGGTAATGATGGC</td><td>559.0</td><td>9.9</td><td>1.1</td><td>9.2</td><td>4.1E-20</td><td>3.0E-19</td><td>BT20/MB468</td><td>CYTO</td></tr>
<tr><td>>hg38_21_+_8433172_8446622_RNA45SN1_45S_50ntFlanks@8381.8400.20|AAGAGGGCGTGAAACCGTTA</td><td>>hg38_21_+_8433172_8446622_RNA45SN1_45S_50ntFlanks</td><td>8381</td><td>8400</td><td>20</td><td>AAGAGGGCGTGAAACCGTTA</td><td>1013.3</td><td>-13.8</td><td>1.5</td><td>-9.2</td><td>2.7E-20</td><td>3.0E-19</td><td>MB231/MB468</td><td>MITO</td></tr>
<tr><td>>hg38_21_+_8433172_8446622_RNA45SN1_45S_50ntFlanks@10802.10826.25|TTGGAACAATGTAGGTAAGGGAAGT</td><td>>hg38_21_+_8433172_8446622_RNA45SN1_45S_50ntFlanks</td><td>10802</td><td>10826</td><td>25</td><td>TTGGAACAATGTAGGTAAGGGAAGT</td><td>531.4</td><td>-12.9</td><td>1.4</td><td>-9.3</td><td>1.8E-20</td><td>3.0E-19</td><td>MB231/MB468</td><td>MP</td></tr>
<tr><td>>MI0000456|hsa-mir-140&WithFlank&16|+|69933075|69933186@68.89.22(+1U)&[MIMAT0004597&hsa-miR-140-3p&offsets|0|+1(+1U);m-45&16|+|69933143|69933164&offsets|-1|-1(+1U)]|TACCACAGGGTAGAACCACGGAT</td><td>>MI0000456|hsa-mir-140&WithFlank&16|+|69933075|69933186</td><td>68</td><td>89</td><td>22(+1U)</td><td>TACCACAGGGTAGAACCACGGAT</td><td>339.2</td><td>12.7</td><td>1.4</td><td>9.2</td><td>4.1E-20</td><td>3.0E-19</td><td>BT20/MB468</td><td>CYTO</td></tr>
<tr><td>>hg38_21_+_8433172_8446622_RNA45SN1_45S_50ntFlanks@11583.11601.19|CTGGTGCGGACCAGGGGAA</td><td>>hg38_21_+_8433172_8446622_RNA45SN1_45S_50ntFlanks</td><td>11583</td><td>11601</td><td>19</td><td>CTGGTGCGGACCAGGGGAA</td><td>1278.7</td><td>-5.4</td><td>0.6</td><td>-9.2</td><td>2.5E-20</td><td>3.0E-19</td><td>MB231/MB468</td><td>NUC</td></tr>
<tr><td>>hg38_21_+_8433172_8446622_RNA45SN1_45S_50ntFlanks@10808.10828.21|CAATGTAGGTAAGGGAAGTCG</td><td>>hg38_21_+_8433172_8446622_RNA45SN1_45S_50ntFlanks</td><td>10808</td><td>10828</td><td>21</td><td>CAATGTAGGTAAGGGAAGTCG</td><td>614.7</td><td>-13.1</td><td>1.4</td><td>-9.3</td><td>1.8E-20</td><td>3.0E-19</td><td>MB231/MB468</td><td>MP</td></tr>
<tr><td>>hg38_21_+_8433172_8446622_RNA45SN1_45S_50ntFlanks@10408.10435.28|GGTCAGTCGGTCCTGAGAGATGGGCGAG</td><td>>hg38_21_+_8433172_8446622_RNA45SN1_45S_50ntFlanks</td><td>10408</td><td>10435</td><td>28</td><td>GGTCAGTCGGTCCTGAGAGATGGGCGAG</td><td>2114.0</td><td>-14.8</td><td>1.6</td><td>-9.2</td><td>2.8E-20</td><td>3.0E-19</td><td>MB231/MB468</td><td>MITO</td></tr>
<tr><td>>hg38_21_+_8433172_8446622_RNA45SN1_45S_50ntFlanks@8423.8442.20|CGCAGTCCGCCCGGAGGATT</td><td>>hg38_21_+_8433172_8446622_RNA45SN1_45S_50ntFlanks</td><td>8423</td><td>8442</td><td>20</td><td>CGCAGTCCGCCCGGAGGATT</td><td>10856.2</td><td>-12.6</td><td>1.4</td><td>-9.2</td><td>2.8E-20</td><td>3.0E-19</td><td>MB231/MB468</td><td>MITO</td></tr>
<tr><td>>hg38_21_+_8433172_8446622_RNA45SN1_45S_50ntFlanks@9032.9059.28|CCTCGTGGGGGGGCCGGGCCACCCCTCC</td><td>>hg38_21_+_8433172_8446622_RNA45SN1_45S_50ntFlanks</td><td>9032</td><td>9059</td><td>28</td><td>CCTCGTGGGGGGGCCGGGCCACCCCTCC</td><td>1249.4</td><td>-14.1</td><td>1.5</td><td>-9.2</td><td>2.8E-20</td><td>3.0E-19</td><td>MB231/MB468</td><td>MITO</td></tr>
<tr><td>>hg38_21_+_8433172_8446622_RNA45SN1_45S_50ntFlanks@3985.4005.21|CGGCTTTGGTGACTCTAGATA</td><td>>hg38_21_+_8433172_8446622_RNA45SN1_45S_50ntFlanks</td><td>3985</td><td>4005</td><td>21</td><td>CGGCTTTGGTGACTCTAGATA</td><td>1418.8</td><td>-14.3</td><td>1.5</td><td>-9.3</td><td>1.8E-20</td><td>3.1E-19</td><td>MB231/MB468</td><td>MP</td></tr>
<tr><td>>hg38_21_+_8433172_8446622_RNA45SN1_45S_50ntFlanks@11208.11227.20|GCGGCGACTCTGGACGCGAG</td><td>>hg38_21_+_8433172_8446622_RNA45SN1_45S_50ntFlanks</td><td>11208</td><td>11227</td><td>20</td><td>GCGGCGACTCTGGACGCGAG</td><td>1226.7</td><td>-10.3</td><td>1.1</td><td>-9.2</td><td>2.8E-20</td><td>3.1E-19</td><td>MB231/MB468</td><td>MITO</td></tr>
<tr><td>>hg38_21_+_8433172_8446622_RNA45SN1_45S_50ntFlanks@4449.4472.24|CCTCTCGGCGCCCCCTCGATGCTC</td><td>>hg38_21_+_8433172_8446622_RNA45SN1_45S_50ntFlanks</td><td>4449</td><td>4472</td><td>24</td><td>CCTCTCGGCGCCCCCTCGATGCTC</td><td>547.8</td><td>-13.0</td><td>1.4</td><td>-9.3</td><td>1.8E-20</td><td>3.1E-19</td><td>MB231/MB468</td><td>MP</td></tr>
<tr><td>>trnaMT_ValTAC_MT_+_1602_1670@53.70.18|TAACTTGACCGCTCTGAC</td><td>>trnaMT_ValTAC_MT_+_1602_1670</td><td>53</td><td>70</td><td>18</td><td>TAACTTGACCGCTCTGAC</td><td>319.3</td><td>12.6</td><td>1.4</td><td>9.2</td><td>4.2E-20</td><td>3.1E-19</td><td>BT20/MB468</td><td>CYTO</td></tr>
<tr><td>>MI0000088|hsa-mir-30a&WithFlank&6|-|71403545|71403627@54.73.20(+1U)&[MIMAT0000088&hsa-miR-30a-3p&offsets|+1|-1(+1U);m-38&6|-|71403554|71403575&offsets|+1|-1(+1U)]|TTTCAGTCGGATGTTTGCAGT</td><td>>MI0000088|hsa-mir-30a&WithFlank&6|-|71403545|71403627</td><td>54</td><td>73</td><td>20(+1U)</td><td>TTTCAGTCGGATGTTTGCAGT</td><td>3310.9</td><td>7.5</td><td>0.8</td><td>9.3</td><td>1.5E-20</td><td>3.1E-19</td><td>MB231/MB468</td><td>CYTO</td></tr>
</table>

| | | | | | | | | | | | | | |
|---|---|---|---|---|---|---|---|---|---|---|---|---|---|
| >hg38_21_+_8433172_8446622_RNA45SN1_45S_50ntFlanks@4732.4771.40\|AGAACGAAAGTCGGAGGTTCGAAGACGATCAGATACCGTC | >hg38_21_+_8433172_8446622_RNA45SN1_45S_50ntFlanks | 4732 | 4771 | 40 | AGAACGAAAGTCGGAGGTTCGAAGACGATCAGATACCGTC | 340.4 | 7.4 | 0.8 | 9.3 | 1.5E-20 | 3.1E-19 | MB231/MB468 | CYTO |
| >hg38_21_+_8433172_8446622_RNA45SN1_45S_50ntFlanks@10392.10422.31\|ACAGCAGTTGAACATGGGTCAGTCGGTCCTG | >hg38_21_+_8433172_8446622_RNA45SN1_45S_50ntFlanks | 10392 | 10422 | 31 | ACAGCAGTTGAACATGGGTCAGTCGGTCCTG | 1210.1 | -6.3 | 0.7 | -9.2 | 2.5E-20 | 3.1E-19 | MB231/MB468 | NUC |
| >hg38_21_+_8433172_8446622_RNA45SN1_45S_50ntFlanks@12954.12972.19\|CTGCTTCTGGGTCGGGGTT | >hg38_21_+_8433172_8446622_RNA45SN1_45S_50ntFlanks | 12954 | 12972 | 19 | CTGCTTCTGGGTCGGGGTT | 548.0 | -13.0 | 1.4 | -9.3 | 1.8E-20 | 3.1E-19 | MB231/MB468 | MP |
| >MI0003643\|hsa-mir-629&WithFlank&15\|-\|70079366\|70079474@28.48.21&[MIMAT0004810&hsa-miR-629-5p&offsets\|0\|0;m-150&15\|-\|70079427\|70079447&offsets\|0\|0]\|TGGGTTTACGTTGGGAGAACT | >MI0003643\|hsa-mir-629&WithFlank&15\|-\|70079366\|70079474 | 28 | 48 | 21 | TGGGTTTACGTTGGGAGAACT | 151.5 | 10.1 | 1.1 | 9.1 | 8.1E-20 | 3.1E-19 | BT20/MB468 | NUC |
| >hg38_21_+_8433172_8446622_RNA45SN1_45S_50ntFlanks@8113.8138.26\|GGCGCGGGACATGTGGCGTACGGAAG | >hg38_21_+_8433172_8446622_RNA45SN1_45S_50ntFlanks | 8113 | 8138 | 26 | GGCGCGGGACATGTGGCGTACGGAAG | 1121.9 | -13.9 | 1.5 | -9.2 | 2.8E-20 | 3.1E-19 | MB231/MB468 | MITO |
| >hg38_21_+_8433172_8446622_RNA45SN1_45S_50ntFlanks@11874.11908.35\|AGAAGACCCTGTTGAGCTTGACTCTAGTCTGGCAC | >hg38_21_+_8433172_8446622_RNA45SN1_45S_50ntFlanks | 11874 | 11908 | 35 | AGAAGACCCTGTTGAGCTTGACTCTAGTCTGGCAC | 1714.1 | -8.5 | 0.9 | -9.2 | 2.9E-20 | 3.1E-19 | MB231/MB468 | MITO |
| >am_MI0000786\|hsa-mir-378a&WithFlank&5\|+\|149732819\|149732896@49.69.21&[MIMAT0000732&hsa-miR-378a-3p&offsets\|0\|-1;m-44&5\|+\|149732867\|149732888&offsets\|0\|-1]\|ACTGGACTTGGAGTCAGAAGG | >am_MI0000786\|hsa-mir-378a&WithFlank&5\|+\|149732819\|149732896 | 49 | 69 | 21 | ACTGGACTTGGAGTCAGAAGG | 2124.6 | 5.5 | 0.6 | 9.2 | 3.1E-20 | 3.1E-19 | BT20/MB468 | MITO |
| >hg38_MT_+_1621_3279_RNR2_16S_50ntFlanks@1438.1477.40\|TACGTGATCTGAGTTCAGACCGGAGTAATCCAGGTCGGTT | >hg38_MT_+_1621_3279_RNR2_16S_50ntFlanks | 1438 | 1477 | 40 | TACGTGATCTGAGTTCAGACCGGAGTAATCCAGGTCGGTT | 1065.7 | 6.8 | 0.7 | 9.2 | 3.1E-20 | 3.1E-19 | BT20/MB468 | MITO |
| >hg38_21_+_8433172_8446622_RNA45SN1_45S_50ntFlanks@13004.13027.24\|ATCTATTGAAAGTCAGCCCTCGAC | >hg38_21_+_8433172_8446622_RNA45SN1_45S_50ntFlanks | 13004 | 13027 | 24 | ATCTATTGAAAGTCAGCCCTCGAC | 2557.5 | -13.6 | 1.5 | -9.2 | 4.2E-20 | 3.1E-19 | BT20/MB468 | CYTO |
| >hg38_21_+_8433172_8446622_RNA45SN1_45S_50ntFlanks@10128.10145.18\|CGGCGGCGGCGGCGGCGG | >hg38_21_+_8433172_8446622_RNA45SN1_45S_50ntFlanks | 10128 | 10145 | 18 | CGGCGGCGGCGGCGGCGG | 1317.3 | -14.1 | 1.5 | -9.2 | 2.9E-20 | 3.1E-19 | MB231/MB468 | MITO |
| >hg38_21_+_8433172_8446622_RNA45SN1_45S_50ntFlanks@10131.10148.18\|CGGCGGCGGCGGCGGCGG | >hg38_21_+_8433172_8446622_RNA45SN1_45S_50ntFlanks | 10131 | 10148 | 18 | CGGCGGCGGCGGCGGCGG | 1317.3 | -14.1 | 1.5 | -9.2 | 2.9E-20 | 3.1E-19 | MB231/MB468 | MITO |
| >hg38_21_+_8433172_8446622_RNA45SN1_45S_50ntFlanks@8824.8841.18\|CGGCGGCGGCGGCGGCGG | >hg38_21_+_8433172_8446622_RNA45SN1_45S_50ntFlanks | 8824 | 8841 | 18 | CGGCGGCGGCGGCGGCGG | 1317.3 | -14.1 | 1.5 | -9.2 | 2.9E-20 | 3.1E-19 | MB231/MB468 | MITO |
| >hg38_21_+_8433172_8446622_RNA45SN1_45S_50ntFlanks@8845.8862.18\|CGGCGGCGGCGGCGGCGG | >hg38_21_+_8433172_8446622_RNA45SN1_45S_50ntFlanks | 8845 | 8862 | 18 | CGGCGGCGGCGGCGGCGG | 1317.3 | -14.1 | 1.5 | -9.2 | 2.9E-20 | 3.1E-19 | MB231/MB468 | MITO |
| >hg38_21_+_8433172_8446622_RNA45SN1_45S_50ntFlanks@8848.8865.18\|CGGCGGCGGCGGCGGCGG | >hg38_21_+_8433172_8446622_RNA45SN1_45S_50ntFlanks | 8848 | 8865 | 18 | CGGCGGCGGCGGCGGCGG | 1317.3 | -14.1 | 1.5 | -9.2 | 2.9E-20 | 3.1E-19 | MB231/MB468 | MITO |
| >am_trna11_ArgACG_3_-_45730491_45730563@39.56.18,trna138_ArgACG_6_-_27638344_27638416@39.56.18,trna156_ArgACG_6_-_27181623_27181695@39.56.18,trna36_ArgACG_6_+_27182952_27183024@39.56.18\|TCAGAAGATTCTAGGTTC | >am_trna11_ArgACG_3_-_45730491_45730563 | 39 | 56 | 18 | TCAGAAGATTCTAGGTTC | 284.9 | -12.6 | 1.4 | -9.1 | 8.2E-20 | 3.1E-19 | BT20/MB468 | NUC |
| >hg38_21_+_8433172_8446622_RNA45SN1_45S_50ntFlanks@5478.5516.39\|CTGGCGGAGCGCTGAGAAGACGGTCGAACTTGACTATCT | >hg38_21_+_8433172_8446622_RNA45SN1_45S_50ntFlanks | 5478 | 5516 | 39 | CTGGCGGAGCGCTGAGAAGACGGTCGAACTTGACTATCT | 459.2 | 13.3 | 1.4 | 9.2 | 3.1E-20 | 3.2E-19 | BT20/MB468 | MITO |
| >hg38_21_+_8433172_8446622_RNA45SN1_45S_50ntFlanks@12419.12438.20\|GATGTCGGCTCTTCCTATCA | >hg38_21_+_8433172_8446622_RNA45SN1_45S_50ntFlanks | 12419 | 12438 | 20 | GATGTCGGCTCTTCCTATCA | 3946.1 | 6.3 | 0.7 | 9.3 | 1.6E-20 | 3.2E-19 | MB231/MB468 | CYTO |
| >trnaMT_PheGAA_MT_+_577_647@1.34.34\|GTTTATGTAGCTTACCTCCTCAAAGCAATACACT | >trnaMT_PheGAA_MT_+_577_647 | 1 | 34 | 34 | GTTTATGTAGCTTACCTCCTCAAAGCAATACACT | 178.7 | 6.9 | 0.7 | 9.2 | 2.6E-20 | 3.2E-19 | MB231/MB468 | NUC |
| >hg38_21_+_8433172_8446622_RNA45SN1_45S_50ntFlanks@4445.4472.28\|CTTGCCTCTCGGCGCCCCCTCGATGCTC | >hg38_21_+_8433172_8446622_RNA45SN1_45S_50ntFlanks | 4445 | 4472 | 28 | CTTGCCTCTCGGCGCCCCCTCGATGCTC | 489.7 | -12.8 | 1.4 | -9.3 | 1.9E-20 | 3.2E-19 | MB231/MB468 | MP |
| >hg38_21_+_8433172_8446622_RNA45SN1_45S_50ntFlanks@9306.9334.29\|CAAGGAGTCTAACACGTGCGCGAGTCGGG | >hg38_21_+_8433172_8446622_RNA45SN1_45S_50ntFlanks | 9306 | 9334 | 29 | CAAGGAGTCTAACACGTGCGCGAGTCGGG | 771.5 | -13.5 | 1.5 | -9.3 | 1.9E-20 | 3.2E-19 | MB231/MB468 | MP |
| >hg38_21_+_8433172_8446622_RNA45SN1_45S_50ntFlanks@9339.9370.32\|CGCACGAAAGCCGCCGTGGCGCAATGAAGGTG | >hg38_21_+_8433172_8446622_RNA45SN1_45S_50ntFlanks | 9339 | 9370 | 32 | CGCACGAAAGCCGCCGTGGCGCAATGAAGGTG | 1096.8 | -13.9 | 1.5 | -9.2 | 3.0E-20 | 3.2E-19 | MB231/MB468 | MITO |
| >hg38_21_+_8433172_8446622_RNA45SN1_45S_50ntFlanks@4995.5013.19\|ACAGATTGATAGCTCTTTC | >hg38_21_+_8433172_8446622_RNA45SN1_45S_50ntFlanks | 4995 | 5013 | 19 | ACAGATTGATAGCTCTTTC | 2532.9 | -15.2 | 1.6 | -9.3 | 1.9E-20 | 3.2E-19 | MB231/MB468 | MP |
| >hg38_21_+_8433172_8446622_RNA45SN1_45S_50ntFlanks@12711.12742.32\|GCGGAGCCTCGGTTGGCCTCGGATAGCCGGTC | >hg38_21_+_8433172_8446622_RNA45SN1_45S_50ntFlanks | 12711 | 12742 | 32 | GCGGAGCCTCGGTTGGCCTCGGATAGCCGGTC | 512.7 | -12.9 | 1.4 | -9.3 | 1.9E-20 | 3.2E-19 | MB231/MB468 | MP |
| >hg38_21_+_8433172_8446622_RNA45SN1_45S_50ntFlanks@4122.4145.24\|AGGGTTCGATTCCGGAGAGGGAGC | >hg38_21_+_8433172_8446622_RNA45SN1_45S_50ntFlanks | 4122 | 4145 | 24 | AGGGTTCGATTCCGGAGAGGGAGC | 777.1 | -13.5 | 1.5 | -9.3 | 1.9E-20 | 3.2E-19 | MB231/MB468 | MP |
| >hg38_21_+_8433172_8446622_RNA45SN1_45S_50ntFlanks@10787.10817.31\|ACAGCCTCTGGCATGTTGGAACAATGTAGGT | >hg38_21_+_8433172_8446622_RNA45SN1_45S_50ntFlanks | 10787 | 10817 | 31 | ACAGCCTCTGGCATGTTGGAACAATGTAGGT | 448.8 | -12.7 | 1.4 | -9.3 | 1.9E-20 | 3.2E-19 | MB231/MB468 | MP |
| >MI0000255\|hsa-mir-30d&WithFlank&8\|-\|134804870\|134804951@12.32.21(+1A)&[MIMAT0000245&hsa-miR-30d-5p&offsets\|0\|-1(+1A);m-21&8\|-\|134804919\|134804940&offsets\|0\|-1(+1A)]\|TGTAAACATCCCCGACTGGAAA | >MI0000255\|hsa-mir-30d&WithFlank&8\|-\|134804870\|134804951 | 12 | 32 | 21(+1A) | TGTAAACATCCCCGACTGGAAA | 180.3 | 4.3 | 0.5 | 9.1 | 8.5E-20 | 3.2E-19 | BT20/MB468 | NUC |
| >hg38_21_+_8433172_8446622_RNA45SN1_45S_50ntFlanks@11146.11166.21\|CCCCGGCGGCGGGGGCACGGT | >hg38_21_+_8433172_8446622_RNA45SN1_45S_50ntFlanks | 11146 | 11166 | 21 | CCCCGGCGGCGGGGGCACGGT | 335.8 | -12.8 | 1.4 | -9.1 | 8.5E-20 | 3.2E-19 | BT20/MB468 | NUC |
| >hg38_21_+_8433172_8446622_RNA45SN1_45S_50ntFlanks@5080.5112.33\|ATAACGAACGAGACTCTGGCATGCTAACTAGTT | >hg38_21_+_8433172_8446622_RNA45SN1_45S_50ntFlanks | 5080 | 5112 | 33 | ATAACGAACGAGACTCTGGCATGCTAACTAGTT | 220.1 | 4.3 | 0.5 | 9.3 | 1.7E-20 | 3.3E-19 | MB231/MB468 | CYTO |
| >hg38_21_+_8433172_8446622_RNA45SN1_45S_50ntFlanks@7979.8006.28\|ACCTCAGATCAGACGTGGCGACCCGCTG | >hg38_21_+_8433172_8446622_RNA45SN1_45S_50ntFlanks | 7979 | 8006 | 28 | ACCTCAGATCAGACGTGGCGACCCGCTG | 7342.8 | -11.8 | 1.3 | -9.3 | 2.0E-20 | 3.3E-19 | MB231/MB468 | MP |
| >MI0000650\|hsa-mir-200c&WithFlank&12\|+\|6963693\|6963772@50.72.23(+1C)&[MIMAT0000617&hsa-miR-200c-3p&offsets\|0\|0(+1C);m-54&12\|+\|6963742\|6963764&offsets\|0\|0(+1C)]\|TAATACTGCCGGGTAATGATGGAC | >MI0000650\|hsa-mir-200c&WithFlank&12\|+\|6963693\|6963772 | 50 | 72 | 23(+1C) | TAATACTGCCGGGTAATGATGGAC | 1796.4 | 9.1 | 1.0 | 9.2 | 3.3E-20 | 3.4E-19 | BT20/MB468 | MITO |
| >hg38_21_+_8433172_8446622_RNA45SN1_45S_50ntFlanks@12073.12090.18\|CCGAGGGGCTCTCGCTTC | >hg38_21_+_8433172_8446622_RNA45SN1_45S_50ntFlanks | 12073 | 12090 | 18 | CCGAGGGGCTCTCGCTTC | 701.2 | -13.3 | 1.4 | -9.3 | 2.0E-20 | 3.4E-19 | MB231/MB468 | MP |
| >MI0000077\|hsa-mir-21&WithFlank&17\|+\|59841260\|59841343@13.36.24&[MIMAT0000076&hsa-miR-21-5p&offsets\|-1\|+1;m-1&17\|+\|59841273\|59841294&offsets\|-1\|+1]\|GTAGCTTATCAGACTGATGTTGAC | >MI0000077\|hsa-mir-21&WithFlank&17\|+\|59841260\|59841343 | 13 | 36 | 24 | GTAGCTTATCAGACTGATGTTGAC | 403.0 | 6.7 | 0.7 | 9.2 | 2.8E-20 | 3.4E-19 | MB231/MB468 | NUC |
| >MI0000089\|hsa-mir-31&WithFlank&9\|-\|21512109\|21512191@14.35.22&[MIMAT0000089&hsa-miR-31-5p&offsets\|0\|+1;m-360&9\|-\|21512157\|21512178&offsets\|0\|0]\|AGGCAAGATGCTGGCATAGCTG | >MI0000089\|hsa-mir-31&WithFlank&9\|-\|21512109\|21512191 | 14 | 35 | 22 | AGGCAAGATGCTGGCATAGCTG | 1030.6 | 13.8 | 1.5 | 9.5 | 3.3E-21 | 3.4E-19 | BT20/MB231 | NUC |
| >hg38_21_+_8433172_8446622_RNA45SN1_45S_50ntFlanks@12405.12422.18\|CTTTTTGATCCTTCGATG | >hg38_21_+_8433172_8446622_RNA45SN1_45S_50ntFlanks | 12405 | 12422 | 18 | CTTTTTGATCCTTCGATG | 2052.7 | -8.5 | 0.9 | -9.3 | 1.8E-20 | 3.4E-19 | MB231/MB468 | CYTO |
| >hg38_21_+_8433172_8446622_RNA45SN1_45S_50ntFlanks@10798.10825.28\|CATGTTGGAACAATGTAGGTAAGGGAAG | >hg38_21_+_8433172_8446622_RNA45SN1_45S_50ntFlanks | 10798 | 10825 | 28 | CATGTTGGAACAATGTAGGTAAGGGAAG | 475.0 | -12.8 | 1.4 | -9.3 | 2.1E-20 | 3.5E-19 | MB231/MB468 | MP |
| >hg38_21_+_8433172_8446622_RNA45SN1_45S_50ntFlanks@11641.11660.20\|CGGCGGGTGTTGACGCGATG | >hg38_21_+_8433172_8446622_RNA45SN1_45S_50ntFlanks | 11641 | 11660 | 20 | CGGCGGGTGTTGACGCGATG | 34028.8 | -10.6 | 1.2 | -9.2 | 3.4E-20 | 3.5E-19 | BT20/MB468 | MITO |
| >hg38_21_+_8433172_8446622_RNA45SN1_45S_50ntFlanks@5493.5516.24\|GAAGACGGTCGAACTTGACTATCT | >hg38_21_+_8433172_8446622_RNA45SN1_45S_50ntFlanks | 5493 | 5516 | 24 | GAAGACGGTCGAACTTGACTATCT | 1568.5 | 7.7 | 0.8 | 9.2 | 3.5E-20 | 3.5E-19 | BT20/MB468 | MITO |
| >hg38_21_+_8433172_8446622_RNA45SN1_45S_50ntFlanks@11199.11233.35\|CCGGCGGCGGCGGCGACTCTGGACGCGAGCCGGGC | >hg38_21_+_8433172_8446622_RNA45SN1_45S_50ntFlanks | 11199 | 11233 | 35 | CCGGCGGCGGCGGCGACTCTGGACGCGAGCCGGGC | 400.7 | -13.1 | 1.4 | -9.1 | 9.1E-20 | 3.5E-19 | BT20/MB468 | NUC |
| >hg38_21_+_8433172_8446622_RNA45SN1_45S_50ntFlanks@8192.8212.21\|AGCCCGTGGACGGTGTGAGGC | >hg38_21_+_8433172_8446622_RNA45SN1_45S_50ntFlanks | 8192 | 8212 | 21 | AGCCCGTGGACGGTGTGAGGC | 515.7 | -5.9 | 0.6 | -9.1 | 9.2E-20 | 3.5E-19 | BT20/MB468 | NUC |
| >hg38_21_+_8433172_8446622_RNA45SN1_45S_50ntFlanks@5444.5467.24\|CTCGGATCGGCCCCGCCGGGGTCG | >hg38_21_+_8433172_8446622_RNA45SN1_45S_50ntFlanks | 5444 | 5467 | 24 | CTCGGATCGGCCCCGCCGGGGTCG | 3186.3 | -8.0 | 0.8 | -9.7 | 4.1E-22 | 3.5E-19 | BT20/MB468 | MP |
| >MI0000285\|hsa-mir-205&WithFlank&1\|+\|209432127\|209432248@40.60.21(+1A)&[MIMAT0000266&hsa-miR-205-5p&offsets\|0\|-1(+1A);m-351&1\|+\|209432166\|209432187&offsets\|0\|-1(+1A)]\|TCCTTCATTCCACCGGAGTCTA | >MI0000285\|hsa-mir-205&WithFlank&1\|+\|209432127\|209432248 | 40 | 60 | 21(+1A) | TCCTTCATTCCACCGGAGTCTA | 1030.1 | -13.9 | 1.5 | -9.3 | 2.1E-20 | 3.5E-19 | MB231/MB468 | MP |
| >trna2_GlyGCC_21_-_18827107_18827177@1.33.33,trna35_GlyGCC_1_+_161413094_161413164@1.33.33,trna37_GlyGCC_1_+_161420467_161420537@1.33.33,trna39_GlyGCC_1_+_161427898_161427968@1.33.33,trna41_GlyGCC_1_+_161435258_161435328@1.33.33\|GCATGGGTGGTTCAGTGGTAGAATTCTCGCCTG | >trna2_GlyGCC_21_-_18827107_18827177 | 1 | 33 | 33 | GCATGGGTGGTTCAGTGGTAGAATTCTCGCCTG | 2384.2 | 5.3 | 0.6 | 9.2 | 4.8E-20 | 3.5E-19 | BT20/MB468 | CYTO |
| >hg38_21_+_8433172_8446622_RNA45SN1_45S_50ntFlanks@11323.11343.21\|CGTCTCGTCGCGCGCGCGTCC | >hg38_21_+_8433172_8446622_RNA45SN1_45S_50ntFlanks | 11323 | 11343 | 21 | CGTCTCGTCGCGCGCGCGTCC | 6691.5 | -6.7 | 0.7 | -9.3 | 1.8E-20 | 3.5E-19 | MB231/MB468 | CYTO |
| >MI0000064\|hsa-let-7c&WithFlank&21\|+\|16539822\|16539917@17.38.22&[MIMAT0000064&hsa-let-7c-5p&offsets\|0\|0;m-37&21\|+\|16539838\|16539859&offsets\|0\|0]\|TGAGGTAGTAGGTTGTATGGTT | >MI0000064\|hsa-let-7c&WithFlank&21\|+\|16539822\|16539917 | 17 | 38 | 22 | TGAGGTAGTAGGTTGTATGGTT | 1400.4 | -3.7 | 0.4 | -9.1 | 9.3E-20 | 3.5E-19 | BT20/MB468 | NUC |
| >hg38_21_+_8433172_8446622_RNA45SN1_45S_50ntFlanks@5470.5499.30\|CCACGGCCCTGGCGGAGCGCTGAGAAGACG | >hg38_21_+_8433172_8446622_RNA45SN1_45S_50ntFlanks | 5470 | 5499 | 30 | CCACGGCCCTGGCGGAGCGCTGAGAAGACG | 1173.1 | -14.1 | 1.5 | -9.3 | 2.1E-20 | 3.5E-19 | MB231/MB468 | MP |
| >hg38_21_+_8433172_8446622_RNA45SN1_45S_50ntFlanks@7986.8003.18\|ATCAGACGTGGCGACCCG | >hg38_21_+_8433172_8446622_RNA45SN1_45S_50ntFlanks | 7986 | 8003 | 18 | ATCAGACGTGGCGACCCG | 1718.5 | -13.1 | 1.4 | -9.2 | 4.8E-20 | 3.6E-19 | BT20/MB468 | CYTO |
| >hg38_21_+_8433172_8446622_RNA45SN1_45S_50ntFlanks@10534.10564.31\|CCGCGAGGCGTCCAGTGCGGTAACGCGACCG | >hg38_21_+_8433172_8446622_RNA45SN1_45S_50ntFlanks | 10534 | 10564 | 31 | CCGCGAGGCGTCCAGTGCGGTAACGCGACCG | 601.1 | -13.1 | 1.4 | -9.3 | 2.2E-20 | 3.6E-19 | MB231/MB468 | MP |
| >MI0000077\|hsa-mir-21&WithFlank&17\|+\|59841260\|59841343@13.35.23&[MIMAT0000076&hsa-miR-21-5p&offsets\|-1\|0;m-1&17\|+\|59841273\|59841294&offsets\|-1\|0]\|GTAGCTTATCAGACTGATGTTGA | >MI0000077\|hsa-mir-21&WithFlank&17\|+\|59841260\|59841343 | 13 | 35 | 23 | GTAGCTTATCAGACTGATGTTGA | 510.2 | 6.6 | 0.7 | 9.2 | 3.6E-20 | 3.6E-19 | BT20/MB468 | MITO |
| >hg38_21_+_8433172_8446622_RNA45SN1_45S_50ntFlanks@9545.9563.19\|GTGGAGGTCCGTAGCGGTC | >hg38_21_+_8433172_8446622_RNA45SN1_45S_50ntFlanks | 9545 | 9563 | 19 | GTGGAGGTCCGTAGCGGTC | 19598.4 | -13.9 | 1.5 | -9.1 | 9.4E-20 | 3.6E-19 | BT20/MB468 | NUC |
| >hg38_21_+_8433172_8446622_RNA45SN1_45S_50ntFlanks@6773.6797.25\|TTCCTCCCGGGGCTACGCCTGTCTG | >hg38_21_+_8433172_8446622_RNA45SN1_45S_50ntFlanks | 6773 | 6797 | 25 | TTCCTCCCGGGGCTACGCCTGTCTG | 786.4 | -13.4 | 1.5 | -9.2 | 3.4E-20 | 3.6E-19 | MB231/MB468 | MITO |

| | | | | | | | | | | | | | |
|---|---|---|---|---|---|---|---|---|---|---|---|---|---|
| >hg38_21_+_8433172_8446622_RNA45SN1_45S_50ntFlanks@11581.11601.21\|AACTGGTGCGGACCAGGGGAA | >hg38_21_+_8433172_8446622_RNA45SN1_45S_50ntFlanks | 11581 | 11601 | 21 | AACTGGTGCGGACCAGGGGAA | 1766.2 | -14.7 | 1.6 | -9.3 | 2.2E-20 | 3.6E-19 | MB231/MB468 | MP |
| >hg38_21_+_8433172_8446622_RNA45SN1_45S_50ntFlanks@5420.5445.26\|CCGATTGGATGGTTTAGTGAGGCCCT | >hg38_21_+_8433172_8446622_RNA45SN1_45S_50ntFlanks | 5420 | 5445 | 26 | CCGATTGGATGGTTTAGTGAGGCCCT | 1531.8 | -14.4 | 1.6 | -9.2 | 3.5E-20 | 3.7E-19 | MB231/MB468 | MITO |
| >hg38_21_+_8433172_8446622_RNA45SN1_45S_50ntFlanks@11206.11232.27\|CGGCGGCGACTCTGGACGCGAGCCGGG | >hg38_21_+_8433172_8446622_RNA45SN1_45S_50ntFlanks | 11206 | 11232 | 27 | CGGCGGCGACTCTGGACGCGAGCCGGG | 959.0 | -13.8 | 1.5 | -9.2 | 2.3E-20 | 3.7E-19 | MB231/MB468 | MP |
| >hg38_21_+_8433172_8446622_RNA45SN1_45S_50ntFlanks@9570.9596.27\|TGCAAATCGGTCGTCCGACCTGGGTAT | >hg38_21_+_8433172_8446622_RNA45SN1_45S_50ntFlanks | 9570 | 9596 | 27 | TGCAAATCGGTCGTCCGACCTGGGTAT | 2374.8 | -7.9 | 0.9 | -9.3 | 1.9E-20 | 3.7E-19 | MB231/MB468 | CYTO |
| >MI0000272\|hsa-mir-182&WithFlank&7\|-\|129770377\|129770498@29.53.25(+1A)&[MIMAT0000259&hsa-miR-182-5p&offsets\|0\|+1(+1A);m-22&7\|-\|129770449\|129770470&offsets\|0\|+3(+1A)]\|TTTGGCAATGGTAGAACTCACACTGA | >MI0000272\|hsa-mir-182&WithFlank&7\|-\|129770377\|129770498 | 29 | 53 | 25(+1A) | TTTGGCAATGGTAGAACTCACACTGA | 835.5 | 8.4 | 0.9 | 9.2 | 3.7E-20 | 3.7E-19 | BT20/MB468 | MITO |
| >MI0000734\|hsa-mir-106b&WithFlank&7\|-\|100093987\|100094080@58.77.20(+1A)&[MIMAT0004672&hsa-miR-106b-3p&offsets\|0\|-2(+1A);m-87&7\|-\|100094003\|100094023&offsets\|0\|-1(+1A)]\|CCGCACTGTGGGTACTTGCTA | >MI0000734\|hsa-mir-106b&WithFlank&7\|-\|100093987\|100094080 | 58 | 77 | 20(+1A) | CCGCACTGTGGGTACTTGCTA | 1243.8 | 10.3 | 1.1 | 9.2 | 5.1E-20 | 3.7E-19 | BT20/MB468 | CYTO |
| >MI0000433\|hsa-let-7g&WithFlank&3\|-\|52268272\|52268367@11.30.20(+1C)&[MIMAT0000414&hsa-let-7g-5p&offsets\|0\|-2(+1C);m-58&3\|-\|52268336\|52268357&offsets\|0\|-2(+1C)]\|TGAGGTAGTAGTTTGTACAGC | >MI0000433\|hsa-let-7g&WithFlank&3\|-\|52268272\|52268367 | 11 | 30 | 20(+1C) | TGAGGTAGTAGTTTGTACAGC | 837.4 | 7.4 | 0.8 | 9.2 | 5.1E-20 | 3.7E-19 | BT20/MB468 | CYTO |
| >hg38_21_+_8433172_8446622_RNA45SN1_45S_50ntFlanks@12437.12455.19\|CATTGTGAAGCAGAATTCA | >hg38_21_+_8433172_8446622_RNA45SN1_45S_50ntFlanks | 12437 | 12455 | 19 | CATTGTGAAGCAGAATTCA | 14542.6 | -13.4 | 1.5 | -9.1 | 9.9E-20 | 3.8E-19 | BT20/MB468 | NUC |
| >hg38_21_+_8433172_8446622_RNA45SN1_45S_50ntFlanks@11575.11603.29\|ACTTAGAACTGGTGCGGACCAGGGGAATC | >hg38_21_+_8433172_8446622_RNA45SN1_45S_50ntFlanks | 11575 | 11603 | 29 | ACTTAGAACTGGTGCGGACCAGGGGAATC | 13376.3 | -11.1 | 1.2 | -9.1 | 9.9E-20 | 3.8E-19 | BT20/MB468 | NUC |
| >hg38_21_+_8433172_8446622_RNA45SN1_45S_50ntFlanks@3984.4026.43\|GCGGCTTTGGTGACTCTAGATAACCTCGGGCCGATCGCACGCC | >hg38_21_+_8433172_8446622_RNA45SN1_45S_50ntFlanks | 3984 | 4026 | 43 | GCGGCTTTGGTGACTCTAGATAACCTCGGGCCGATCGCACGCC | 3113.7 | -15.4 | 1.7 | -9.2 | 3.5E-20 | 3.8E-19 | MB231/MB468 | MITO |
| >hg38_21_+_8433172_8446622_RNA45SN1_45S_50ntFlanks@11874.11903.30\|AGAAGACCCTGTTGAGCTTGACTCTAGTCT | >hg38_21_+_8433172_8446622_RNA45SN1_45S_50ntFlanks | 11874 | 11903 | 30 | AGAAGACCCTGTTGAGCTTGACTCTAGTCT | 870.6 | -13.5 | 1.5 | -9.2 | 3.5E-20 | 3.8E-19 | MB231/MB468 | MITO |
| >hg38_21_+_8433172_8446622_RNA45SN1_45S_50ntFlanks@8380.8397.18\|CAAGAGGGCGTGAAACCG | >hg38_21_+_8433172_8446622_RNA45SN1_45S_50ntFlanks | 8380 | 8397 | 18 | CAAGAGGGCGTGAAACCG | 749.3 | -13.3 | 1.4 | -9.2 | 3.6E-20 | 3.8E-19 | MB231/MB468 | MITO |
| >am_trnaMT_PheGAA_MT_+_577_647@41.58.18\|GTTTAGACGGGCTCACAT | >am_trnaMT_PheGAA_MT_+_577_647 | 41 | 58 | 18 | GTTTAGACGGGCTCACAT | 3335.2 | 6.9 | 0.8 | 9.2 | 3.8E-20 | 3.8E-19 | BT20/MB468 | MITO |
| >trna14_LysTTT_11_-_59327808_59327880@1.33.33,trna2_LysTTT_17_+_8022473_8022545@1.33.33,trna54_LysTTT_1_+_204475655_204475727@1.33.33,trna5_LysTTT_11_+_59323902_59323974@1.33.33,trna62_LysTTT_1_-_204476158_204476230@1.33.33,trna76_LysTTT_6_+_28918806_28918878@1.33.33\|GCCCGGATAGCTCAGTCGGTAGAGCATCAGACT | >trna14_LysTTT_11_-_59327808_59327880 | 1 | 33 | 33 | GCCCGGATAGCTCAGTCGGTAGAGCATCAGACT | 2863.7 | 8.5 | 0.9 | 9.2 | 3.2E-20 | 3.8E-19 | MB231/MB468 | NUC |
| >hg38_21_+_8433172_8446622_RNA45SN1_45S_50ntFlanks@12126.12150.25\|CGACCCGCTCCGGGGACAGTGCCAG | >hg38_21_+_8433172_8446622_RNA45SN1_45S_50ntFlanks | 12126 | 12150 | 25 | CGACCCGCTCCGGGGACAGTGCCAG | 824.7 | -13.6 | 1.5 | -9.2 | 2.3E-20 | 3.8E-19 | MB231/MB468 | MP |
| >hg38_21_+_8433172_8446622_RNA45SN1_45S_50ntFlanks@11874.11893.20\|AGAAGACCCTGTTGAGCTTG | >hg38_21_+_8433172_8446622_RNA45SN1_45S_50ntFlanks | 11874 | 11893 | 20 | AGAAGACCCTGTTGAGCTTG | 9001.0 | -10.9 | 1.2 | -9.1 | 1.0E-19 | 3.9E-19 | BT20/MB468 | NUC |
| >hg38_21_+_8433172_8446622_RNA45SN1_45S_50ntFlanks@11881.11904.24\|CCTGTTGAGCTTGACTCTAGTCTG | >hg38_21_+_8433172_8446622_RNA45SN1_45S_50ntFlanks | 11881 | 11904 | 24 | CCTGTTGAGCTTGACTCTAGTCTG | 1616.6 | -7.8 | 0.8 | -9.3 | 2.0E-20 | 3.9E-19 | MB231/MB468 | CYTO |
| >hg38_21_+_8433172_8446622_RNA45SN1_45S_50ntFlanks@11853.11872.20\|CTTGGCGGAATCAGCGGGGA | >hg38_21_+_8433172_8446622_RNA45SN1_45S_50ntFlanks | 11853 | 11872 | 20 | CTTGGCGGAATCAGCGGGGA | 440.5 | -12.6 | 1.4 | -9.2 | 2.4E-20 | 3.9E-19 | MB231/MB468 | MP |
| >hg38_21_+_8433172_8446622_RNA45SN1_45S_50ntFlanks@4443.4471.29\|CCCTTGCCTCTCGGCGCCCCCTCGATGCT | >hg38_21_+_8433172_8446622_RNA45SN1_45S_50ntFlanks | 4443 | 4471 | 29 | CCCTTGCCTCTCGGCGCCCCCTCGATGCT | 744.4 | -13.4 | 1.4 | -9.2 | 2.4E-20 | 3.9E-19 | MB231/MB468 | MP |
| >hg38_21_+_8433172_8446622_RNA45SN1_45S_50ntFlanks@5442.5466.25\|CCCTCGGATCGGCCCCGCCGGGGTC | >hg38_21_+_8433172_8446622_RNA45SN1_45S_50ntFlanks | 5442 | 5466 | 25 | CCCTCGGATCGGCCCCGCCGGGGTC | 703.8 | -13.2 | 1.4 | -9.2 | 3.7E-20 | 3.9E-19 | MB231/MB468 | MITO |
| >hg38_21_+_8433172_8446622_RNA45SN1_45S_50ntFlanks@10841.10865.25\|CGTAACTTCGGGATAAGGATTGGCT | >hg38_21_+_8433172_8446622_RNA45SN1_45S_50ntFlanks | 10841 | 10865 | 25 | CGTAACTTCGGGATAAGGATTGGCT | 7165.5 | -5.6 | 0.6 | -9.2 | 3.9E-20 | 3.9E-19 | BT20/MB468 | MITO |
| >hg38_21_+_8433172_8446622_RNA45SN1_45S_50ntFlanks@12633.12652.20\|CGAAGCTACCATCTGTGGGA | >hg38_21_+_8433172_8446622_RNA45SN1_45S_50ntFlanks | 12633 | 12652 | 20 | CGAAGCTACCATCTGTGGGA | 1281.1 | -10.3 | 1.1 | -9.2 | 3.7E-20 | 3.9E-19 | MB231/MB468 | MITO |
| >hg38_21_+_8433172_8446622_RNA45SN1_45S_50ntFlanks@12997.13037.41\|CGCTGCGATCTATTGAAAGTCAGCCCTCGACACAAGGGTTT | >hg38_21_+_8433172_8446622_RNA45SN1_45S_50ntFlanks | 12997 | 13037 | 41 | CGCTGCGATCTATTGAAAGTCAGCCCTCGACACAAGGGTTT | 1647.7 | 12.0 | 1.3 | 9.3 | 2.0E-20 | 3.9E-19 | MB231/MB468 | CYTO |
| >hg38_21_+_8433172_8446622_RNA45SN1_45S_50ntFlanks@3899.3920.22\|CCTTCGCGGGGGGGATGCGTGC | >hg38_21_+_8433172_8446622_RNA45SN1_45S_50ntFlanks | 3899 | 3920 | 22 | CCTTCGCGGGGGGGATGCGTGC | 1433.5 | -14.3 | 1.6 | -9.2 | 3.7E-20 | 3.9E-19 | MB231/MB468 | MITO |
| >hg38_21_+_8433172_8446622_RNA45SN1_45S_50ntFlanks@6654.6681.28\|CTCTTAGCGGTGGATCACTCGGCTCGTG | >hg38_21_+_8433172_8446622_RNA45SN1_45S_50ntFlanks | 6654 | 6681 | 28 | CTCTTAGCGGTGGATCACTCGGCTCGTG | 1544.8 | -4.5 | 0.5 | -9.1 | 1.0E-19 | 3.9E-19 | BT20/MB468 | NUC |
| >hg38_21_+_8433172_8446622_RNA45SN1_45S_50ntFlanks@476.503.28\|TTCGTGATCGATGTGGTGACGTCGTGCT | >hg38_21_+_8433172_8446622_RNA45SN1_45S_50ntFlanks | 476 | 503 | 28 | TTCGTGATCGATGTGGTGACGTCGTGCT | 403.9 | 12.9 | 1.4 | 9.2 | 5.4E-20 | 3.9E-19 | BT20/MB468 | CYTO |
| >MI0001733\|hsa-mir-452&WithFlank&X\|-\|151959622\|151959718@20.41.22&[MIMAT0001635&hsa-miR-452-5p&offsets\|0\|0;m-188&X\|-\|151959677\|151959699&offsets\|0\|-1]\|AACTGTTTGCAGAGGAAACTGA | >MI0001733\|hsa-mir-452&WithFlank&X\|-\|151959622\|151959718 | 20 | 41 | 22 | AACTGTTTGCAGAGGAAACTGA | 120.7 | 9.8 | 1.1 | 9.1 | 1.0E-19 | 3.9E-19 | BT20/MB468 | NUC |
| >hg38_21_+_8433172_8446622_RNA45SN1_45S_50ntFlanks@6666.6686.21\|GATCACTCGGCTCGTGCGTCG | >hg38_21_+_8433172_8446622_RNA45SN1_45S_50ntFlanks | 6666 | 6686 | 21 | GATCACTCGGCTCGTGCGTCG | 705.3 | -13.2 | 1.4 | -9.2 | 3.7E-20 | 4.0E-19 | MB231/MB468 | MITO |
| >hg38_21_+_8433172_8446622_RNA45SN1_45S_50ntFlanks@9198.9224.27\|GTTCTCTCGGGGCCACGCGCGCGTCCC | >hg38_21_+_8433172_8446622_RNA45SN1_45S_50ntFlanks | 9198 | 9224 | 27 | GTTCTCTCGGGGCCACGCGCGCGTCCC | 774.5 | -13.4 | 1.5 | -9.2 | 3.7E-20 | 4.0E-19 | MB231/MB468 | MITO |
| >hg38_21_+_8433172_8446622_RNA45SN1_45S_50ntFlanks@12152.12180.29\|TGGGGAGTTTGACTGGGGCGGTACACCTG | >hg38_21_+_8433172_8446622_RNA45SN1_45S_50ntFlanks | 12152 | 12180 | 29 | TGGGGAGTTTGACTGGGGCGGTACACCTG | 433.5 | -12.6 | 1.4 | -9.2 | 2.4E-20 | 4.0E-19 | MB231/MB468 | MP |
| >hg38_21_+_8433172_8446622_RNA45SN1_45S_50ntFlanks@11576.11611.36\|CTTAGAACTGGTGCGGACCAGGGGAATCCGACTGTT | >hg38_21_+_8433172_8446622_RNA45SN1_45S_50ntFlanks | 11576 | 11611 | 36 | CTTAGAACTGGTGCGGACCAGGGGAATCCGACTGTT | 1549.3 | -6.2 | 0.7 | -9.2 | 3.8E-20 | 4.0E-19 | MB231/MB468 | MITO |
| >hg38_21_+_8433172_8446622_RNA45SN1_45S_50ntFlanks@9027.9066.40\|CGTCTCCTCGTGGGGGGGCCGGGCCACCCCTCCCACGGCG | >hg38_21_+_8433172_8446622_RNA45SN1_45S_50ntFlanks | 9027 | 9066 | 40 | CGTCTCCTCGTGGGGGGGCCGGGCCACCCCTCCCACGGCG | 1373.7 | -14.2 | 1.5 | -9.2 | 3.8E-20 | 4.0E-19 | MB231/MB468 | MITO |
| >MI0000809\|hsa-mir-151a&WithFlank&8\|-\|140732558\|140732659@53.74.22(+1U)&[MIMAT0000757&hsa-miR-151a-3p&offsets\|0\|+1(+1U);m-30&8\|-\|140732587\|140732607&offsets\|0\|+1(+1U)]\|CTAGACTGAAGCTCCTTGAGGAT | >MI0000809\|hsa-mir-151a&WithFlank&8\|-\|140732558\|140732659 | 53 | 74 | 22(+1U) | CTAGACTGAAGCTCCTTGAGGAT | 2439.2 | 8.8 | 1.0 | 9.2 | 5.5E-20 | 4.0E-19 | BT20/MB468 | CYTO |
| >hg38_21_+_8433172_8446622_RNA45SN1_45S_50ntFlanks@4729.4768.40\|TCAAGAACGAAAGTCGGAGGTTCGAAGACGATCAGATACC | >hg38_21_+_8433172_8446622_RNA45SN1_45S_50ntFlanks | 4729 | 4768 | 40 | TCAAGAACGAAAGTCGGAGGTTCGAAGACGATCAGATACC | 157.2 | 11.3 | 1.2 | 9.3 | 2.1E-20 | 4.0E-19 | MB231/MB468 | CYTO |
| >hg38_21_+_8433172_8446622_RNA45SN1_45S_50ntFlanks@4002.4024.23\|GATAACCTCGGGCCGATCGCACG | >hg38_21_+_8433172_8446622_RNA45SN1_45S_50ntFlanks | 4002 | 4024 | 23 | GATAACCTCGGGCCGATCGCACG | 2893.1 | -4.6 | 0.5 | -9.3 | 2.1E-20 | 4.0E-19 | MB231/MB468 | CYTO |
| >hg38_21_+_8433172_8446622_RNA45SN1_45S_50ntFlanks@7980.8003.24\|CCTCAGATCAGACGTGGCGACCCG | >hg38_21_+_8433172_8446622_RNA45SN1_45S_50ntFlanks | 7980 | 8003 | 24 | CCTCAGATCAGACGTGGCGACCCG | 518.2 | -12.9 | 1.4 | -9.2 | 2.5E-20 | 4.0E-19 | MB231/MB468 | MP |
| >hg38_21_+_8433172_8446622_RNA45SN1_45S_50ntFlanks@10511.10533.23\|ATCCGGAGTGGCGGAGATGGGCG | >hg38_21_+_8433172_8446622_RNA45SN1_45S_50ntFlanks | 10511 | 10533 | 23 | ATCCGGAGTGGCGGAGATGGGCG | 813.4 | -14.1 | 1.6 | -9.1 | 1.1E-19 | 4.1E-19 | BT20/MB468 | NUC |
| >hg38_21_+_8433172_8446622_RNA45SN1_45S_50ntFlanks@4046.4063.18\|CATTCGAACGTCTGCCCT | >hg38_21_+_8433172_8446622_RNA45SN1_45S_50ntFlanks | 4046 | 4063 | 18 | CATTCGAACGTCTGCCCT | 2980.6 | -9.1 | 1.0 | -9.2 | 3.4E-20 | 4.1E-19 | MB231/MB468 | NUC |
| >hg38_21_+_8433172_8446622_RNA45SN1_45S_50ntFlanks@9563.9597.35\|CCTGACGTGCAAATCGGTCGTCCGACCTGGGTATA | >hg38_21_+_8433172_8446622_RNA45SN1_45S_50ntFlanks | 9563 | 9597 | 35 | CCTGACGTGCAAATCGGTCGTCCGACCTGGGTATA | 1571.9 | -4.5 | 0.5 | -9.2 | 4.1E-20 | 4.1E-19 | BT20/MB468 | MITO |
| >hg38_21_+_8433172_8446622_RNA45SN1_45S_50ntFlanks@12044.12072.29\|ACTGACCCGGTGAGGCGGGGGGGCGAGCC | >hg38_21_+_8433172_8446622_RNA45SN1_45S_50ntFlanks | 12044 | 12072 | 29 | ACTGACCCGGTGAGGCGGGGGGGCGAGCC | 872.3 | -13.6 | 1.5 | -9.2 | 2.5E-20 | 4.1E-19 | MB231/MB468 | MP |
| >hg38_21_+_8433172_8446622_RNA45SN1_45S_50ntFlanks@4560.4587.28\|CTAGGAATAATGGAATAGGACCGCGGTT | >hg38_21_+_8433172_8446622_RNA45SN1_45S_50ntFlanks | 4560 | 4587 | 28 | CTAGGAATAATGGAATAGGACCGCGGTT | 734.7 | -13.4 | 1.4 | -9.2 | 2.5E-20 | 4.1E-19 | MB231/MB468 | MP |
| >hg38_21_+_8433172_8446622_RNA45SN1_45S_50ntFlanks@7988.8012.25\|CAGACGTGGCGACCCGCTGAATTTA | >hg38_21_+_8433172_8446622_RNA45SN1_45S_50ntFlanks | 7988 | 8012 | 25 | CAGACGTGGCGACCCGCTGAATTTA | 6456.9 | -9.7 | 1.1 | -9.2 | 2.5E-20 | 4.1E-19 | MB231/MB468 | MP |
| >MI0000804\|hsa-mir-328&WithFlank&16\|-\|67202315\|67202401@54.74.21(+1C)&[MIMAT0000752&hsa-miR-328-3p&offsets\|0\|-1(+1C);m-160&16\|-\|67202327\|67202348&offsets\|0\|-1(+1C)]\|CTGGCCCTCTCTGCCCTTCCGC | >MI0000804\|hsa-mir-328&WithFlank&16\|-\|67202315\|67202401 | 54 | 74 | 21(+1C) | CTGGCCCTCTCTGCCCTTCCGC | 333.0 | 10.3 | 1.1 | 9.2 | 5.7E-20 | 4.1E-19 | BT20/MB468 | CYTO |
| >hg38_21_+_8433172_8446622_RNA45SN1_45S_50ntFlanks@10786.10822.37\|AACAGCCTCTGGCATGTTGGAACAATGTAGGTAAGGG | >hg38_21_+_8433172_8446622_RNA45SN1_45S_50ntFlanks | 10786 | 10822 | 37 | AACAGCCTCTGGCATGTTGGAACAATGTAGGTAAGGG | 1147.8 | -14.0 | 1.5 | -9.2 | 2.6E-20 | 4.1E-19 | MB231/MB468 | MP |
| >hg38_21_+_8433172_8446622_RNA45SN1_45S_50ntFlanks@11270.11293.24\|CGGCCGCCCCCGGGGAGCCCGGCG | >hg38_21_+_8433172_8446622_RNA45SN1_45S_50ntFlanks | 11270 | 11293 | 24 | CGGCCGCCCCCGGGGAGCCCGGCG | 971.5 | -13.7 | 1.5 | -9.2 | 4.0E-20 | 4.2E-19 | MB231/MB468 | MITO |
| >hg38_21_+_8433172_8446622_RNA45SN1_45S_50ntFlanks@11785.11804.20\|TGAATGGATGAACGAGATTC | >hg38_21_+_8433172_8446622_RNA45SN1_45S_50ntFlanks | 11785 | 11804 | 20 | TGAATGGATGAACGAGATTC | 2693.8 | -6.9 | 0.7 | -9.2 | 4.0E-20 | 4.2E-19 | MB231/MB468 | MITO |
| >MI0000102\|hsa-mir-100&WithFlank&11\|-\|122152223\|122152314@19.40.22(+3A)&[MIMAT0000098&hsa-miR-100-5p&offsets\|0\|0(+3A);m-27&11\|-\|122152276\|122152296&offsets\|0\|+1(+3A)]\|AACCCGTAGATCCGAACTTGTGAAA | >MI0000102\|hsa-mir-100&WithFlank&11\|-\|122152223\|122152314 | 19 | 40 | 22(+3A) | AACCCGTAGATCCGAACTTGTGAAA | 355.5 | 12.4 | 1.3 | 9.2 | 3.5E-20 | 4.2E-19 | MB231/MB468 | NUC |
| >hg38_21_+_8433172_8446622_RNA45SN1_45S_50ntFlanks@10402.10433.32\|AACATGGGTCAGTCGGTCCTGAGAGATGGGCG | >hg38_21_+_8433172_8446622_RNA45SN1_45S_50ntFlanks | 10402 | 10433 | 32 | AACATGGGTCAGTCGGTCCTGAGAGATGGGCG | 838.0 | -13.6 | 1.5 | -9.2 | 2.6E-20 | 4.2E-19 | MB231/MB468 | MP |

<table>
<tr><th></th><th></th><th></th><th></th><th></th><th></th><th></th><th></th><th></th><th></th><th></th><th></th><th></th><th></th></tr>
<tr><td>>hg38_21_+_8433172_8446622_RNA45SN1_45S_50ntFlanks@11573.11594.22|CGACTTAGAACTGGTGCGGACC</td><td>>hg38_21_+_8433172_8446622_RNA45SN1_45S_50ntFlanks</td><td>11573</td><td>11594</td><td>22</td><td>CGACTTAGAACTGGTGCGGACC</td><td>617.1</td><td>-5.3</td><td>0.6</td><td>-9.1</td><td>1.1E-19</td><td>4.2E-19</td><td>BT20/MB468</td><td>NUC</td></tr>
<tr><td>>hg38_21_+_8433172_8446622_RNA45SN1_45S_50ntFlanks@12420.12446.27|ATGTCGGCTCTTCCTATCATTGTGAAG</td><td>>hg38_21_+_8433172_8446622_RNA45SN1_45S_50ntFlanks</td><td>12420</td><td>12446</td><td>27</td><td>ATGTCGGCTCTTCCTATCATTGTGAAG</td><td>2600.4</td><td>-6.5</td><td>0.7</td><td>-9.2</td><td>2.6E-20</td><td>4.2E-19</td><td>MB231/MB468</td><td>MP</td></tr>
<tr><td>>MI0005559|hsa-mir-744&WithFlank&17|+|12081893|12082002@17.36.20(+1U)&[MIMAT0004945&hsa-miR-744-5p&offsets|0|-2(+1U);m-172&17|+|12081909|12081929&offsets|0|-1(+1U)]|TGCGGGGCTAGGGCTAACAGT</td><td>>MI0005559|hsa-mir-744&WithFlank&17|+|12081893|12082002</td><td>17</td><td>36</td><td>20(+1U)</td><td>TGCGGGGCTAGGGCTAACAGT</td><td>1142.8</td><td>10.2</td><td>1.1</td><td>9.1</td><td>5.8E-20</td><td>4.2E-19</td><td>BT20/MB468</td><td>CYTO</td></tr>
<tr><td>>hg38_21_+_8433172_8446622_RNA45SN1_45S_50ntFlanks@13002.13029.28|CGATCTATTGAAAGTCAGCCCTCGACAC</td><td>>hg38_21_+_8433172_8446622_RNA45SN1_45S_50ntFlanks</td><td>13002</td><td>13029</td><td>28</td><td>CGATCTATTGAAAGTCAGCCCTCGACAC</td><td>397.5</td><td>-12.5</td><td>1.4</td><td>-9.2</td><td>2.6E-20</td><td>4.3E-19</td><td>MB231/MB468</td><td>MP</td></tr>
<tr><td>>hg38_21_+_8433172_8446622_RNA45SN1_45S_50ntFlanks@12324.12348.25|TTGGGTTTTAAGCAGGAGGTGTCAG</td><td>>hg38_21_+_8433172_8446622_RNA45SN1_45S_50ntFlanks</td><td>12324</td><td>12348</td><td>25</td><td>TTGGGTTTTAAGCAGGAGGTGTCAG</td><td>933.7</td><td>-9.3</td><td>1.0</td><td>-9.3</td><td>2.2E-20</td><td>4.3E-19</td><td>MB231/MB468</td><td>CYTO</td></tr>
<tr><td>>hg38_21_+_8433172_8446622_RNA45SN1_45S_50ntFlanks@10116.10135.20|TGCGTCGGAGGGCGGCGGCG</td><td>>hg38_21_+_8433172_8446622_RNA45SN1_45S_50ntFlanks</td><td>10116</td><td>10135</td><td>20</td><td>TGCGTCGGAGGGCGGCGGCG</td><td>620.0</td><td>-13.1</td><td>1.4</td><td>-9.2</td><td>4.1E-20</td><td>4.3E-19</td><td>MB231/MB468</td><td>MITO</td></tr>
<tr><td>>hg38_21_+_8433172_8446622_RNA45SN1_45S_50ntFlanks@476.505.30|TTCGTGATCGATGTGGTGACGTCGTGCTCT</td><td>>hg38_21_+_8433172_8446622_RNA45SN1_45S_50ntFlanks</td><td>476</td><td>505</td><td>30</td><td>TTCGTGATCGATGTGGTGACGTCGTGCTCT</td><td>2749.7</td><td>7.6</td><td>0.8</td><td>9.2</td><td>4.3E-20</td><td>4.3E-19</td><td>BT20/MB468</td><td>MITO</td></tr>
<tr><td>>MI0000088|hsa-mir-30a&WithFlank&6|-|71403545|71403627@12.31.20&[MIMAT0000087&hsa-miR-30a-5p&offsets|0|-2;m-15&6|-|71403595|71403616&offsets|0|-2]|TGTAAACATCCTCGACTGGA</td><td>>MI0000088|hsa-mir-30a&WithFlank&6|-|71403545|71403627</td><td>12</td><td>31</td><td>20</td><td>TGTAAACATCCTCGACTGGA</td><td>2570.8</td><td>7.5</td><td>0.8</td><td>9.2</td><td>3.6E-20</td><td>4.3E-19</td><td>MB231/MB468</td><td>NUC</td></tr>
<tr><td>>hg38_21_+_8433172_8446622_RNA45SN1_45S_50ntFlanks@12633.12654.22|CGAAGCTACCATCTGTGGGATT</td><td>>hg38_21_+_8433172_8446622_RNA45SN1_45S_50ntFlanks</td><td>12633</td><td>12654</td><td>22</td><td>CGAAGCTACCATCTGTGGGATT</td><td>12454.5</td><td>-6.3</td><td>0.7</td><td>-9.2</td><td>4.3E-20</td><td>4.3E-19</td><td>BT20/MB468</td><td>MITO</td></tr>
<tr><td>>hg38_21_+_8433172_8446622_RNA45SN1_45S_50ntFlanks@11575.11601.27|ACTTAGAACTGGTGCGGACCAGGGGAA</td><td>>hg38_21_+_8433172_8446622_RNA45SN1_45S_50ntFlanks</td><td>11575</td><td>11601</td><td>27</td><td>ACTTAGAACTGGTGCGGACCAGGGGAA</td><td>733.3</td><td>-13.3</td><td>1.4</td><td>-9.2</td><td>4.2E-20</td><td>4.3E-19</td><td>MB231/MB468</td><td>MITO</td></tr>
<tr><td>>hg38_21_+_8433172_8446622_RNA45SN1_45S_50ntFlanks@11999.12030.32|CCCTGCGGGCCGCCGGTGAAATACCACTACTC</td><td>>hg38_21_+_8433172_8446622_RNA45SN1_45S_50ntFlanks</td><td>11999</td><td>12030</td><td>32</td><td>CCCTGCGGGCCGCCGGTGAAATACCACTACTC</td><td>649.4</td><td>-13.1</td><td>1.4</td><td>-9.2</td><td>4.2E-20</td><td>4.3E-19</td><td>MB231/MB468</td><td>MITO</td></tr>
<tr><td>>hg38_21_+_8433172_8446622_RNA45SN1_45S_50ntFlanks@13002.13027.26|CGATCTATTGAAAGTCAGCCCTCGAC</td><td>>hg38_21_+_8433172_8446622_RNA45SN1_45S_50ntFlanks</td><td>13002</td><td>13027</td><td>26</td><td>CGATCTATTGAAAGTCAGCCCTCGAC</td><td>7724.0</td><td>-11.6</td><td>1.3</td><td>-9.2</td><td>4.1E-20</td><td>4.3E-19</td><td>MB231/MB468</td><td>MITO</td></tr>
<tr><td>>hg38_21_+_8433172_8446622_RNA45SN1_45S_50ntFlanks@4728.4758.31|ATCAAGAACGAAAGTCGGAGGTTCGAAGACG</td><td>>hg38_21_+_8433172_8446622_RNA45SN1_45S_50ntFlanks</td><td>4728</td><td>4758</td><td>31</td><td>ATCAAGAACGAAAGTCGGAGGTTCGAAGACG</td><td>1284.3</td><td>-12.6</td><td>1.4</td><td>-9.1</td><td>6.0E-20</td><td>4.3E-19</td><td>BT20/MB468</td><td>CYTO</td></tr>
<tr><td>>hg38_21_+_8433172_8446622_RNA45SN1_45S_50ntFlanks@10690.10710.21|CGTCCGGTGAGCTCTCGCTGG</td><td>>hg38_21_+_8433172_8446622_RNA45SN1_45S_50ntFlanks</td><td>10690</td><td>10710</td><td>21</td><td>CGTCCGGTGAGCTCTCGCTGG</td><td>1355.3</td><td>-13.1</td><td>1.4</td><td>-9.2</td><td>2.3E-20</td><td>4.3E-19</td><td>MB231/MB468</td><td>CYTO</td></tr>
<tr><td>>hg38_21_+_8433172_8446622_RNA45SN1_45S_50ntFlanks@11575.11602.28|ACTTAGAACTGGTGCGGACCAGGGGAAT</td><td>>hg38_21_+_8433172_8446622_RNA45SN1_45S_50ntFlanks</td><td>11575</td><td>11602</td><td>28</td><td>ACTTAGAACTGGTGCGGACCAGGGGAAT</td><td>892.6</td><td>-13.7</td><td>1.5</td><td>-9.2</td><td>2.7E-20</td><td>4.3E-19</td><td>MB231/MB468</td><td>MP</td></tr>
<tr><td>>MI0000102|hsa-mir-100&WithFlank&11|-|122152223|122152314@19.37.19&[MIMAT0000098&hsa-miR-100-5p&offsets|0|-3;m-27&11|-|122152276|122152296&offsets|0|-2]|AACCCGTAGATCCGAACTT</td><td>>MI0000102|hsa-mir-100&WithFlank&11|-|122152223|122152314</td><td>19</td><td>37</td><td>19</td><td>AACCCGTAGATCCGAACTT</td><td>346.3</td><td>12.3</td><td>1.3</td><td>9.2</td><td>3.7E-20</td><td>4.4E-19</td><td>MB231/MB468</td><td>NUC</td></tr>
<tr><td>>MI0000095|hsa-mir-93&WithFlank&7|-|100093762|100093853@17.40.24&[MIMAT0000093&hsa-miR-93-5p&offsets|0|+1;m-29&7|-|100093815|100093837&offsets|0|+1]|CAAAGTGCTGTTCGTGCAGGTAGT</td><td>>MI0000095|hsa-mir-93&WithFlank&7|-|100093762|100093853</td><td>17</td><td>40</td><td>24</td><td>CAAAGTGCTGTTCGTGCAGGTAGT</td><td>2504.5</td><td>5.9</td><td>0.6</td><td>9.1</td><td>6.0E-20</td><td>4.4E-19</td><td>BT20/MB468</td><td>CYTO</td></tr>
<tr><td>>hg38_21_+_8433172_8446622_RNA45SN1_45S_50ntFlanks@12605.12625.21|TATGTGCTTGGCTGAGGAGCC</td><td>>hg38_21_+_8433172_8446622_RNA45SN1_45S_50ntFlanks</td><td>12605</td><td>12625</td><td>21</td><td>TATGTGCTTGGCTGAGGAGCC</td><td>6331.2</td><td>-5.0</td><td>0.6</td><td>-9.1</td><td>6.0E-20</td><td>4.4E-19</td><td>BT20/MB468</td><td>CYTO</td></tr>
<tr><td>>hg38_21_+_8433172_8446622_RNA45SN1_45S_50ntFlanks@11233.11250.18|CCCTTCCCGTGGATCGCC</td><td>>hg38_21_+_8433172_8446622_RNA45SN1_45S_50ntFlanks</td><td>11233</td><td>11250</td><td>18</td><td>CCCTTCCCGTGGATCGCC</td><td>816.2</td><td>-13.5</td><td>1.5</td><td>-9.2</td><td>4.2E-20</td><td>4.4E-19</td><td>MB231/MB468</td><td>MITO</td></tr>
<tr><td>>hg38_21_+_8433172_8446622_RNA45SN1_45S_50ntFlanks@4163.4182.20|CATCCAAGGAAGGCAGCAGG</td><td>>hg38_21_+_8433172_8446622_RNA45SN1_45S_50ntFlanks</td><td>4163</td><td>4182</td><td>20</td><td>CATCCAAGGAAGGCAGCAGG</td><td>1030.9</td><td>-13.9</td><td>1.5</td><td>-9.2</td><td>2.7E-20</td><td>4.4E-19</td><td>MB231/MB468</td><td>MP</td></tr>
<tr><td>>hg38_21_+_8433172_8446622_RNA45SN1_45S_50ntFlanks@12437.12454.18|CATTGTGAAGCAGAATTC</td><td>>hg38_21_+_8433172_8446622_RNA45SN1_45S_50ntFlanks</td><td>12437</td><td>12454</td><td>18</td><td>CATTGTGAAGCAGAATTC</td><td>5520.0</td><td>-10.6</td><td>1.2</td><td>-9.1</td><td>1.2E-19</td><td>4.4E-19</td><td>BT20/MB468</td><td>NUC</td></tr>
<tr><td>>hg38_21_+_8433172_8446622_RNA45SN1_45S_50ntFlanks@12957.12979.23|CTTCTGGGTCGGGGTTTCGTACG</td><td>>hg38_21_+_8433172_8446622_RNA45SN1_45S_50ntFlanks</td><td>12957</td><td>12979</td><td>23</td><td>CTTCTGGGTCGGGGTTTCGTACG</td><td>4968.2</td><td>-12.5</td><td>1.4</td><td>-9.1</td><td>1.2E-19</td><td>4.4E-19</td><td>BT20/MB468</td><td>NUC</td></tr>
<tr><td>>hg38_21_+_8433172_8446622_RNA45SN1_45S_50ntFlanks@10411.10435.25|CAGTCGGTCCTGAGAGATGGGCGAG</td><td>>hg38_21_+_8433172_8446622_RNA45SN1_45S_50ntFlanks</td><td>10411</td><td>10435</td><td>25</td><td>CAGTCGGTCCTGAGAGATGGGCGAG</td><td>1655.6</td><td>-14.5</td><td>1.6</td><td>-9.2</td><td>4.2E-20</td><td>4.4E-19</td><td>MB231/MB468</td><td>MITO</td></tr>
<tr><td>>hg38_21_+_8433172_8446622_RNA45SN1_45S_50ntFlanks@7985.8004.20|GATCAGACGTGGCGACCCGC</td><td>>hg38_21_+_8433172_8446622_RNA45SN1_45S_50ntFlanks</td><td>7985</td><td>8004</td><td>20</td><td>GATCAGACGTGGCGACCCGC</td><td>1037.0</td><td>-13.8</td><td>1.5</td><td>-9.2</td><td>4.2E-20</td><td>4.4E-19</td><td>MB231/MB468</td><td>MITO</td></tr>
<tr><td>>hg38_21_+_8433172_8446622_RNA45SN1_45S_50ntFlanks@7975.8015.41|CGCGACCTCAGATCAGACGTGGCGACCCGCTGAATTTAAGC</td><td>>hg38_21_+_8433172_8446622_RNA45SN1_45S_50ntFlanks</td><td>7975</td><td>8015</td><td>41</td><td>CGCGACCTCAGATCAGACGTGGCGACCCGCTGAATTTAAGC</td><td>9276.4</td><td>7.7</td><td>0.8</td><td>9.2</td><td>2.3E-20</td><td>4.4E-19</td><td>MB231/MB468</td><td>CYTO</td></tr>
<tr><td>>hg38_21_+_8433172_8446622_RNA45SN1_45S_50ntFlanks@3985.4005.21|CGGCTTTGGTGACTCTAGATA</td><td>>hg38_21_+_8433172_8446622_RNA45SN1_45S_50ntFlanks</td><td>3985</td><td>4005</td><td>21</td><td>CGGCTTTGGTGACTCTAGATA</td><td>1359.7</td><td>-14.2</td><td>1.5</td><td>-9.2</td><td>4.3E-20</td><td>4.4E-19</td><td>MB231/MB468</td><td>MITO</td></tr>
<tr><td>>hg38_21_+_8433172_8446622_RNA45SN1_45S_50ntFlanks@8182.8206.25|ATCGAGGCCCAGCCCGTGGACGGTG</td><td>>hg38_21_+_8433172_8446622_RNA45SN1_45S_50ntFlanks</td><td>8182</td><td>8206</td><td>25</td><td>ATCGAGGCCCAGCCCGTGGACGGTG</td><td>804.9</td><td>-13.5</td><td>1.5</td><td>-9.2</td><td>2.8E-20</td><td>4.4E-19</td><td>MB231/MB468</td><td>MP</td></tr>
<tr><td>>hg38_21_+_8433172_8446622_RNA45SN1_45S_50ntFlanks@11564.11581.18|CCTAGCAGCCGACTTAGA</td><td>>hg38_21_+_8433172_8446622_RNA45SN1_45S_50ntFlanks</td><td>11564</td><td>11581</td><td>18</td><td>CCTAGCAGCCGACTTAGA</td><td>451.0</td><td>-12.7</td><td>1.4</td><td>-9.2</td><td>2.8E-20</td><td>4.5E-19</td><td>MB231/MB468</td><td>MP</td></tr>
<tr><td>>hg38_21_+_8433172_8446622_RNA45SN1_45S_50ntFlanks@10351.10389.39|CGAGAACTTTGAAGGCCGAAGTGGAGAAGGGTTCCATGT</td><td>>hg38_21_+_8433172_8446622_RNA45SN1_45S_50ntFlanks</td><td>10351</td><td>10389</td><td>39</td><td>CGAGAACTTTGAAGGCCGAAGTGGAGAAGGGTTCCATGT</td><td>700.9</td><td>-13.9</td><td>1.5</td><td>-9.1</td><td>1.2E-19</td><td>4.6E-19</td><td>BT20/MB468</td><td>NUC</td></tr>
<tr><td>>hg38_21_+_8433172_8446622_RNA45SN1_45S_50ntFlanks@4045.4063.19|CCATTCGAACGTCTGCCCT</td><td>>hg38_21_+_8433172_8446622_RNA45SN1_45S_50ntFlanks</td><td>4045</td><td>4063</td><td>19</td><td>CCATTCGAACGTCTGCCCT</td><td>4367.0</td><td>-12.3</td><td>1.4</td><td>-9.1</td><td>1.2E-19</td><td>4.6E-19</td><td>BT20/MB468</td><td>NUC</td></tr>
<tr><td>>MI0000482|hsa-mir-185&WithFlank&22|+|20033133|20033226@21.42.22&[MIMAT0000455&hsa-miR-185-5p&offsets|0|0;m-145&22|+|20033153|20033174&offsets|0|0]|TGGAGAGAAAGGCAGTTCCTGA</td><td>>MI0000482|hsa-mir-185&WithFlank&22|+|20033133|20033226</td><td>21</td><td>42</td><td>22</td><td>TGGAGAGAAAGGCAGTTCCTGA</td><td>3849.3</td><td>7.0</td><td>0.8</td><td>9.1</td><td>6.3E-20</td><td>4.6E-19</td><td>BT20/MB468</td><td>CYTO</td></tr>
<tr><td>>hg38_21_+_8433172_8446622_RNA45SN1_45S_50ntFlanks@12249.12284.36|CAAAAGCTCGCTTGATCTTGATTTTCAGTACGAATA</td><td>>hg38_21_+_8433172_8446622_RNA45SN1_45S_50ntFlanks</td><td>12249</td><td>12284</td><td>36</td><td>CAAAAGCTCGCTTGATCTTGATTTTCAGTACGAATA</td><td>724.2</td><td>-13.3</td><td>1.4</td><td>-9.2</td><td>4.5E-20</td><td>4.6E-19</td><td>MB231/MB468</td><td>MITO</td></tr>
<tr><td>>hg38_21_+_8433172_8446622_RNA45SN1_45S_50ntFlanks@10572.10591.20|AGAAGCCGGCGGGAGCCCCG</td><td>>hg38_21_+_8433172_8446622_RNA45SN1_45S_50ntFlanks</td><td>10572</td><td>10591</td><td>20</td><td>AGAAGCCGGCGGGAGCCCCG</td><td>800.8</td><td>-13.4</td><td>1.5</td><td>-9.2</td><td>4.5E-20</td><td>4.6E-19</td><td>MB231/MB468</td><td>MITO</td></tr>
<tr><td>>hg38_21_+_8433172_8446622_RNA45SN1_45S_50ntFlanks@12636.12661.26|AGCTACCATCTGTGGGATTATGACTG</td><td>>hg38_21_+_8433172_8446622_RNA45SN1_45S_50ntFlanks</td><td>12636</td><td>12661</td><td>26</td><td>AGCTACCATCTGTGGGATTATGACTG</td><td>8525.4</td><td>-8.4</td><td>0.9</td><td>-9.2</td><td>4.5E-20</td><td>4.6E-19</td><td>MB231/MB468</td><td>MITO</td></tr>
<tr><td>>MI0000542|hsa-mir-320a&WithFlank&8|-|22244960|22245043@50.70.21(+1U)&[MIMAT0000510&hsa-miR-320a-3p&offsets|+2|+1(+1U);m-60&8|-|22244975|22244996&offsets|+2|+1(+1U)]|AAGCTGGGTTGAGAGGGCGAAT</td><td>>MI0000542|hsa-mir-320a&WithFlank&8|-|22244960|22245043</td><td>50</td><td>70</td><td>21(+1U)</td><td>AAGCTGGGTTGAGAGGGCGAAT</td><td>504.8</td><td>10.9</td><td>1.2</td><td>9.1</td><td>6.4E-20</td><td>4.6E-19</td><td>BT20/MB468</td><td>CYTO</td></tr>
<tr><td>>hg38_21_+_8433172_8446622_RNA45SN1_45S_50ntFlanks@11641.11665.25|CGGCGGGTGTTGACGCGATGTGATT</td><td>>hg38_21_+_8433172_8446622_RNA45SN1_45S_50ntFlanks</td><td>11641</td><td>11665</td><td>25</td><td>CGGCGGGTGTTGACGCGATGTGATT</td><td>16514.2</td><td>-12.3</td><td>1.3</td><td>-9.2</td><td>4.5E-20</td><td>4.6E-19</td><td>MB231/MB468</td><td>MITO</td></tr>
<tr><td>>hg38_21_+_8433172_8446622_RNA45SN1_45S_50ntFlanks@8114.8139.26|GCGCGGGACATGTGGCGTACGGAAGA</td><td>>hg38_21_+_8433172_8446622_RNA45SN1_45S_50ntFlanks</td><td>8114</td><td>8139</td><td>26</td><td>GCGCGGGACATGTGGCGTACGGAAGA</td><td>807.1</td><td>-13.4</td><td>1.5</td><td>-9.2</td><td>4.5E-20</td><td>4.6E-19</td><td>MB231/MB468</td><td>MITO</td></tr>
<tr><td>>hg38_21_+_8433172_8446622_RNA45SN1_45S_50ntFlanks@4259.4282.24|ATTGGAATGAGTCCACTTTAAATC</td><td>>hg38_21_+_8433172_8446622_RNA45SN1_45S_50ntFlanks</td><td>4259</td><td>4282</td><td>24</td><td>ATTGGAATGAGTCCACTTTAAATC</td><td>942.8</td><td>-13.7</td><td>1.5</td><td>-9.2</td><td>2.9E-20</td><td>4.7E-19</td><td>MB231/MB468</td><td>MP</td></tr>
<tr><td>>MI0000749|hsa-mir-30e&WithFlank&1|+|40754349|40754452@65.85.21&[MIMAT0000693&hsa-miR-30e-3p&offsets|0|-1;m-25&1|+|40754413|40754434&offsets|0|-1]|CTTTCAGTCGGATGTTTACAG</td><td>>MI0000749|hsa-mir-30e&WithFlank&1|+|40754349|40754452</td><td>65</td><td>85</td><td>21</td><td>CTTTCAGTCGGATGTTTACAG</td><td>570.5</td><td>5.8</td><td>0.6</td><td>9.1</td><td>6.4E-20</td><td>4.7E-19</td><td>BT20/MB468</td><td>CYTO</td></tr>
<tr><td>>hg38_21_+_8433172_8446622_RNA45SN1_45S_50ntFlanks@9540.9571.32|CTCTGGTGGAGGTCCGTAGCGGTCCTGACGTG</td><td>>hg38_21_+_8433172_8446622_RNA45SN1_45S_50ntFlanks</td><td>9540</td><td>9571</td><td>32</td><td>CTCTGGTGGAGGTCCGTAGCGGTCCTGACGTG</td><td>1865.5</td><td>-5.8</td><td>0.6</td><td>-9.2</td><td>3.9E-20</td><td>4.7E-19</td><td>MB231/MB468</td><td>NUC</td></tr>
<tr><td>>MI0006444|hsa-mir-1307&WithFlank&10|-|103394247|103394407@86.107.22(+1U)&[MIMAT0005951&hsa-miR-1307-3p&offsets|0|0(+1U);m-66&10|-|103394300|103394322&offsets|0|-1(+1U)]|ACTCGGCGTGGCGTCGGTCGTGT</td><td>>MI0006444|hsa-mir-1307&WithFlank&10|-|103394247|103394407</td><td>86</td><td>107</td><td>22(+1U)</td><td>ACTCGGCGTGGCGTCGGTCGTGT</td><td>1045.8</td><td>7.4</td><td>0.8</td><td>9.2</td><td>4.7E-20</td><td>4.7E-19</td><td>BT20/MB468</td><td>MITO</td></tr>
<tr><td>>hg38_21_+_8433172_8446622_RNA45SN1_45S_50ntFlanks@12893.12933.41|CCTCGCCCGTCACGCACCGCACGTTCGTGGGGAACCTGGCG</td><td>>hg38_21_+_8433172_8446622_RNA45SN1_45S_50ntFlanks</td><td>12893</td><td>12933</td><td>41</td><td>CCTCGCCCGTCACGCACCGCACGTTCGTGGGGAACCTGGCG</td><td>2078.5</td><td>-14.9</td><td>1.6</td><td>-9.2</td><td>2.9E-20</td><td>4.7E-19</td><td>MB231/MB468</td><td>MP</td></tr>
<tr><td>>hg38_21_+_8433172_8446622_RNA45SN1_45S_50ntFlanks@12416.12459.44|TTCGATGTCGGCTCTTCCTATCATTGTGAAGCAGAATTCACCAA</td><td>>hg38_21_+_8433172_8446622_RNA45SN1_45S_50ntFlanks</td><td>12416</td><td>12459</td><td>44</td><td>TTCGATGTCGGCTCTTCCTATCATTGTGAAGCAGAATTCACCAA</td><td>175.6</td><td>11.5</td><td>1.2</td><td>9.2</td><td>2.5E-20</td><td>4.7E-19</td><td>MB231/MB468</td><td>CYTO</td></tr>
<tr><td>>hg38_21_+_8433172_8446622_RNA45SN1_45S_50ntFlanks@3985.4026.42|CGGCTTTGGTGACTCTAGATAACCTCGGGCCGATCGCACGCC</td><td>>hg38_21_+_8433172_8446622_RNA45SN1_45S_50ntFlanks</td><td>3985</td><td>4026</td><td>42</td><td>CGGCTTTGGTGACTCTAGATAACCTCGGGCCGATCGCACGCC</td><td>2074.8</td><td>-14.9</td><td>1.6</td><td>-9.2</td><td>2.9E-20</td><td>4.7E-19</td><td>MB231/MB468</td><td>MP</td></tr>
<tr><td>>trnaMT_GluTTC_MT_-_14674_14742@38.61.24,trnalookalike8_GluTTC_5_-_93905172_93905240@38.61.24|CATTGGTCGTGGTTGTAGTCCGTG</td><td>>trnaMT_GluTTC_MT_-_14674_14742</td><td>38</td><td>61</td><td>24</td><td>CATTGGTCGTGGTTGTAGTCCGTG</td><td>670.9</td><td>7.2</td><td>0.8</td><td>9.2</td><td>4.7E-20</td><td>4.7E-19</td><td>BT20/MB468</td><td>MITO</td></tr>
<tr><td>>hg38_21_+_8433172_8446622_RNA45SN1_45S_50ntFlanks@11667.11709.43|CTGCCCAGTGCTCTGAATGTCAAAGTGAAGAAATTCAATGAAG</td><td>>hg38_21_+_8433172_8446622_RNA45SN1_45S_50ntFlanks</td><td>11667</td><td>11709</td><td>43</td><td>CTGCCCAGTGCTCTGAATGTCAAAGTGAAGAAATTCAATGAAG</td><td>201.8</td><td>11.7</td><td>1.3</td><td>9.2</td><td>2.5E-20</td><td>4.7E-19</td><td>MB231/MB468</td><td>CYTO</td></tr>
<tr><td>>hg38_21_+_8433172_8446622_RNA45SN1_45S_50ntFlanks@12418.12436.19|CGATGTCGGCTCTTCCTAT</td><td>>hg38_21_+_8433172_8446622_RNA45SN1_45S_50ntFlanks</td><td>12418</td><td>12436</td><td>19</td><td>CGATGTCGGCTCTTCCTAT</td><td>19836.3</td><td>-5.5</td><td>0.6</td><td>-9.2</td><td>2.5E-20</td><td>4.7E-19</td><td>MB231/MB468</td><td>CYTO</td></tr>
<tr><td>>MI0000088|hsa-mir-30a&WithFlank&6|-|71403545|71403627@53.71.19&[MIMAT0000088&hsa-miR-30a-3p&offsets|0|-3;m-38&6|-|71403554|71403575&offsets|0|-3]|CTTTCAGTCGGATGTTTGC</td><td>>MI0000088|hsa-mir-30a&WithFlank&6|-|71403545|71403627</td><td>53</td><td>71</td><td>19</td><td>CTTTCAGTCGGATGTTTGC</td><td>1500.7</td><td>6.7</td><td>0.7</td><td>9.1</td><td>6.6E-20</td><td>4.7E-19</td><td>BT20/MB468</td><td>CYTO</td></tr>
<tr><td>>hg38_1_-_228634819_228635039_RNA5S12_5S_50ntFlanks@139.170.32|GACCGCCTGGGAATACCGGGTGCTGTAGGCTT</td><td>>hg38_1_-_228634819_228635039_RNA5S12_5S_50ntFlanks</td><td>139</td><td>170</td><td>32</td><td>GACCGCCTGGGAATACCGGGTGCTGTAGGCTT</td><td>2261.6</td><td>7.3</td><td>0.8</td><td>9.1</td><td>6.6E-20</td><td>4.7E-19</td><td>BT20/MB468</td><td>CYTO</td></tr>
<tr><td>>hg38_21_+_8433172_8446622_RNA45SN1_45S_50ntFlanks@11234.11254.21|CCTTCCCGTGGATCGCCCCAG</td><td>>hg38_21_+_8433172_8446622_RNA45SN1_45S_50ntFlanks</td><td>11234</td><td>11254</td><td>21</td><td>CCTTCCCGTGGATCGCCCCAG</td><td>691.2</td><td>-13.2</td><td>1.4</td><td>-9.2</td><td>4.6E-20</td><td>4.7E-19</td><td>MB231/MB468</td><td>MITO</td></tr>
<tr><td>>MI0000483|hsa-mir-186&WithFlank&1|-|71067625|71067722@21.43.23&[MIMAT0000456&hsa-miR-186-5p&offsets|0|+1;m-84&1|-|71067681|71067702&offsets|0|+1]|CAAAGAATTCTCCTTTTGGGCTT</td><td>>MI0000483|hsa-mir-186&WithFlank&1|-|71067625|71067722</td><td>21</td><td>43</td><td>23</td><td>CAAAGAATTCTCCTTTTGGGCTT</td><td>844.4</td><td>6.6</td><td>0.7</td><td>9.1</td><td>6.6E-20</td><td>4.8E-19</td><td>BT20/MB468</td><td>CYTO</td></tr>
<tr><td>>hg38_21_+_8433172_8446622_RNA45SN1_45S_50ntFlanks@4759.4779.21|ATCAGATACCGTCGTAGTTCC</td><td>>hg38_21_+_8433172_8446622_RNA45SN1_45S_50ntFlanks</td><td>4759</td><td>4779</td><td>21</td><td>ATCAGATACCGTCGTAGTTCC</td><td>664.1</td><td>-11.7</td><td>1.3</td><td>-9.1</td><td>6.7E-20</td><td>4.8E-19</td><td>BT20/MB468</td><td>CYTO</td></tr>
</table>

| | | | | | | | | | | | | | |
|---|---|---|---|---|---|---|---|---|---|---|---|---|---|
| >hg38_21_+_8433172_8446622_RNA45SN1_45S_50ntFlanks@6741.6771.31\|ATCGACACTTCGAACGCACTTGCGGCCCCGG | >hg38_21_+_8433172_8446622_RNA45SN1_45S_50ntFlanks | 6741 | 6771 | 31 | ATCGACACTTCGAACGCACTTGCGGCCCCGG | 2071.9 | -4.3 | 0.5 | -9.2 | 4.9E-20 | 4.8E-19 | BT20/MB468 | MITO |
| >hg38_21_+_8433172_8446622_RNA45SN1_45S_50ntFlanks@6738.6756.19\|ATCATCGACACTTCGAACG | >hg38_21_+_8433172_8446622_RNA45SN1_45S_50ntFlanks | 6738 | 6756 | 19 | ATCATCGACACTTCGAACG | 278.2 | -12.5 | 1.4 | -9.1 | 1.3E-19 | 4.8E-19 | BT20/MB468 | NUC |
| >hg38_21_+_8433172_8446622_RNA45SN1_45S_50ntFlanks@6777.6806.30\|TCCCGGGGCTACGCCTGTCTGAGCGTCGCT | >hg38_21_+_8433172_8446622_RNA45SN1_45S_50ntFlanks | 6777 | 6806 | 30 | TCCCGGGGCTACGCCTGTCTGAGCGTCGCT | 522.3 | 5.8 | 0.6 | 9.1 | 6.7E-20 | 4.8E-19 | BT20/MB468 | CYTO |
| >hg38_21_+_8433172_8446622_RNA45SN1_45S_50ntFlanks@10679.10709.31\|GGTTCCGGCGGCGTCCGGTGAGCTCTCGCTG | >hg38_21_+_8433172_8446622_RNA45SN1_45S_50ntFlanks | 10679 | 10709 | 31 | GGTTCCGGCGGCGTCCGGTGAGCTCTCGCTG | 5495.7 | -11.6 | 1.3 | -9.2 | 3.1E-20 | 4.9E-19 | MB231/MB468 | MP |
| >MI0001448\|hsa-mir-425&WithFlank&3\|-\|49020142\|49020240@20.43.24(+1C)&[MIMAT0003393&hsa-miR-425-5p&offsets\|0\|+1(+1C);m-95&3\|-\|49020200\|49020221&offsets\|0\|+2(+1C)]\|AATGACACGATCACTCCCGTTGAGC | >MI0001448\|hsa-mir-425&WithFlank&3\|-\|49020142\|49020240 | 20 | 43 | 24(+1C) | AATGACACGATCACTCCCGTTGAGC | 563.0 | 5.9 | 0.6 | 9.2 | 4.2E-20 | 5.0E-19 | MB231/MB468 | NUC |
| >hg38_21_+_8433172_8446622_RNA45SN1_45S_50ntFlanks@12626.12653.28\|AATGGGGCGAAGCTACCATCTGTGGGAT | >hg38_21_+_8433172_8446622_RNA45SN1_45S_50ntFlanks | 12626 | 12653 | 28 | AATGGGGCGAAGCTACCATCTGTGGGAT | 3057.1 | -9.2 | 1.0 | -9.2 | 4.2E-20 | 5.0E-19 | MB231/MB468 | NUC |
| >hg38_21_+_8433172_8446622_RNA45SN1_45S_50ntFlanks@11572.11605.34\|CCGACTTAGAACTGGTGCGGACCAGGGGAATCCG | >hg38_21_+_8433172_8446622_RNA45SN1_45S_50ntFlanks | 11572 | 11605 | 34 | CCGACTTAGAACTGGTGCGGACCAGGGGAATCCG | 10300.7 | -10.0 | 1.1 | -9.2 | 4.9E-20 | 5.0E-19 | MB231/MB468 | MITO |
| >am_trna2_GlyTCC_19_+_4724082_4724153@1.30.30\|GCGTTGGTGGTATAGTGGTTAGCATAGCTG | >am_trna2_GlyTCC_19_+_4724082_4724153 | 1 | 30 | 30 | GCGTTGGTGGTATAGTGGTTAGCATAGCTG | 505.0 | -12.8 | 1.4 | -9.2 | 3.1E-20 | 5.0E-19 | MB231/MB468 | MP |
| >hg38_21_+_8433172_8446622_RNA45SN1_45S_50ntFlanks@11199.11220.22\|CCGGCGGCGGCGGCGACTCTGG | >hg38_21_+_8433172_8446622_RNA45SN1_45S_50ntFlanks | 11199 | 11220 | 22 | CCGGCGGCGGCGGCGACTCTGG | 543.2 | -7.0 | 0.8 | -9.2 | 2.7E-20 | 5.0E-19 | MB231/MB468 | CYTO |
| >hg38_21_+_8433172_8446622_RNA45SN1_45S_50ntFlanks@12419.12439.21\|GATGTCGGCTCTTCCTATCAT | >hg38_21_+_8433172_8446622_RNA45SN1_45S_50ntFlanks | 12419 | 12439 | 21 | GATGTCGGCTCTTCCTATCAT | 537.7 | 4.5 | 0.5 | 9.2 | 5.1E-20 | 5.0E-19 | BT20/MB468 | MITO |
| >hg38_21_+_8433172_8446622_RNA45SN1_45S_50ntFlanks@10350.10368.19\|ACGAGAACTTTGAAGGCCG | >hg38_21_+_8433172_8446622_RNA45SN1_45S_50ntFlanks | 10350 | 10368 | 19 | ACGAGAACTTTGAAGGCCG | 2246.6 | -4.8 | 0.5 | -9.2 | 5.1E-20 | 5.0E-19 | BT20/MB468 | MITO |
| >hg38_21_+_8433172_8446622_RNA45SN1_45S_50ntFlanks@10534.10570.37\|CCGCGAGGCGTCCAGTGCGGTAACGCGACCGATCCCG | >hg38_21_+_8433172_8446622_RNA45SN1_45S_50ntFlanks | 10534 | 10570 | 37 | CCGCGAGGCGTCCAGTGCGGTAACGCGACCGATCCCG | 906.7 | -13.7 | 1.5 | -9.2 | 3.1E-20 | 5.0E-19 | MB231/MB468 | MP |
| >hg38_21_+_8433172_8446622_RNA45SN1_45S_50ntFlanks@11577.11605.29\|TTAGAACTGGTGCGGACCAGGGGAATCCG | >hg38_21_+_8433172_8446622_RNA45SN1_45S_50ntFlanks | 11577 | 11605 | 29 | TTAGAACTGGTGCGGACCAGGGGAATCCG | 654.3 | -12.3 | 1.3 | -9.2 | 4.3E-20 | 5.1E-19 | MB231/MB468 | NUC |
| >hg38_21_+_8433172_8446622_RNA45SN1_45S_50ntFlanks@7986.8003.18\|ATCAGACGTGGCGACCCG | >hg38_21_+_8433172_8446622_RNA45SN1_45S_50ntFlanks | 7986 | 8003 | 18 | ATCAGACGTGGCGACCCG | 2395.2 | -14.3 | 1.5 | -9.6 | 6.3E-22 | 5.1E-19 | BT20/MB468 | MP |
| >hg38_21_+_8433172_8446622_RNA45SN1_45S_50ntFlanks@10727.10755.29\|GGAGAGGGTGTAAATCTCGCGCCGGGCCG | >hg38_21_+_8433172_8446622_RNA45SN1_45S_50ntFlanks | 10727 | 10755 | 29 | GGAGAGGGTGTAAATCTCGCGCCGGGCCG | 875.8 | -13.6 | 1.5 | -9.2 | 5.0E-20 | 5.1E-19 | MB231/MB468 | MITO |
| >hg38_21_+_8433172_8446622_RNA45SN1_45S_50ntFlanks@11233.11254.22\|CCCTTCCCGTGGATCGCCCCAG | >hg38_21_+_8433172_8446622_RNA45SN1_45S_50ntFlanks | 11233 | 11254 | 22 | CCCTTCCCGTGGATCGCCCCAG | 1403.1 | -12.5 | 1.4 | -9.2 | 5.2E-20 | 5.1E-19 | BT20/MB468 | MITO |
| >hg38_21_+_8433172_8446622_RNA45SN1_45S_50ntFlanks@13008.13039.32\|ATTGAAAGTCAGCCCTCGACACAAGGGTTTGT | >hg38_21_+_8433172_8446622_RNA45SN1_45S_50ntFlanks | 13008 | 13039 | 32 | ATTGAAAGTCAGCCCTCGACACAAGGGTTTGT | 15678.5 | 7.2 | 0.8 | 9.2 | 5.2E-20 | 5.1E-19 | BT20/MB468 | MITO |
| >hg38_21_+_8433172_8446622_RNA45SN1_45S_50ntFlanks@12202.12220.19\|CTAAGGCGAGCTCAGGGAG | >hg38_21_+_8433172_8446622_RNA45SN1_45S_50ntFlanks | 12202 | 12220 | 19 | CTAAGGCGAGCTCAGGGAG | 980.2 | -13.8 | 1.5 | -9.2 | 3.2E-20 | 5.1E-19 | MB231/MB468 | MP |
| >hg38_21_+_8433172_8446622_RNA45SN1_45S_50ntFlanks@10680.10709.30\|GTTCCGGCGGCGTCCGGTGAGCTCTCGCTG | >hg38_21_+_8433172_8446622_RNA45SN1_45S_50ntFlanks | 10680 | 10709 | 30 | GTTCCGGCGGCGTCCGGTGAGCTCTCGCTG | 3215.0 | -11.0 | 1.2 | -9.2 | 3.2E-20 | 5.1E-19 | MB231/MB468 | MP |
| >MI0000743\|hsa-mir-34c&WithFlank&11\|+\|111513433\|111513521@19.41.23&[MIMAT0000686&hsa-miR-34c-5p&offsets\|0\|0;m-336&11\|+\|111513451\|111513473&offsets\|0\|0]\|AGGCAGTGTAGTTAGCTGATTGC | >MI0000743\|hsa-mir-34c&WithFlank&11\|+\|111513433\|111513521 | 19 | 41 | 23 | AGGCAGTGTAGTTAGCTGATTGC | 321.2 | 12.2 | 1.3 | 9.2 | 4.4E-20 | 5.1E-19 | MB231/MB468 | NUC |
| >MI0000487\|hsa-mir-193a&WithFlank&17\|+\|31559990\|31560089@27.47.21&[MIMAT0004614&hsa-miR-193a-5p&offsets\|0\|-1;m-123&17\|+\|31560016\|31560037&offsets\|0\|-1]\|TGGGTCTTTGCGGGCGAGATG | >MI0000487\|hsa-mir-193a&WithFlank&17\|+\|31559990\|31560089 | 27 | 47 | 21 | TGGGTCTTTGCGGGCGAGATG | 366.8 | 13.0 | 1.4 | 9.2 | 5.2E-20 | 5.1E-19 | BT20/MB468 | MITO |
| >hg38_21_+_8433172_8446622_RNA45SN1_45S_50ntFlanks@11573.11601.29\|CGACTTAGAACTGGTGCGGACCAGGGGAA | >hg38_21_+_8433172_8446622_RNA45SN1_45S_50ntFlanks | 11573 | 11601 | 29 | CGACTTAGAACTGGTGCGGACCAGGGGAA | 2219.3 | -6.2 | 0.7 | -9.2 | 4.4E-20 | 5.1E-19 | MB231/MB468 | NUC |
| >hg38_21_+_8433172_8446622_RNA45SN1_45S_50ntFlanks@11233.11252.20\|CCCTTCCCGTGGATCGCCCC | >hg38_21_+_8433172_8446622_RNA45SN1_45S_50ntFlanks | 11233 | 11252 | 20 | CCCTTCCCGTGGATCGCCCC | 976.0 | -13.7 | 1.5 | -9.2 | 5.1E-20 | 5.2E-19 | MB231/MB468 | MITO |
| >hg38_21_+_8433172_8446622_RNA45SN1_45S_50ntFlanks@8104.8142.39\|CCGCGGCGGGGCGCGGGACATGTGGCGTACGGAAGACCC | >hg38_21_+_8433172_8446622_RNA45SN1_45S_50ntFlanks | 8104 | 8142 | 39 | CCGCGGCGGGGCGCGGGACATGTGGCGTACGGAAGACCC | 1159.0 | -14.0 | 1.5 | -9.2 | 5.1E-20 | 5.2E-19 | MB231/MB468 | MITO |
| >hg38_21_+_8433172_8446622_RNA45SN1_45S_50ntFlanks@9504.9523.20\|GTGAACTATGCCTGGGCAGG | >hg38_21_+_8433172_8446622_RNA45SN1_45S_50ntFlanks | 9504 | 9523 | 20 | GTGAACTATGCCTGGGCAGG | 4142.1 | -11.7 | 1.3 | -9.2 | 3.3E-20 | 5.2E-19 | MB231/MB468 | MP |
| >MI0000088\|hsa-mir-30a&WithFlank&6\|-\|71403545\|71403627@12.35.24&[MIMAT0000087&hsa-miR-30a-5p&offsets\|0\|+2;m-15&6\|-\|71403595\|71403616&offsets\|0\|+2]\|TGTAAACATCCTCGACTGGAAGCT | >MI0000088\|hsa-mir-30a&WithFlank&6\|-\|71403545\|71403627 | 12 | 35 | 24 | TGTAAACATCCTCGACTGGAAGCT | 89220.0 | 7.5 | 0.8 | 9.2 | 4.5E-20 | 5.2E-19 | MB231/MB468 | NUC |
| >am_MI0003581\|hsa-mir-574&WithFlank&4\|+\|38868026\|38868133@31.55.25&[MIMAT0004795&hsa-miR-574-5p&offsets\|0\|+2;m-213&4\|+\|38868056\|38868078&offsets\|0\|+2]\|TGAGTGTGTGTGTGTGAGTGTGTGT | >am_MI0003581\|hsa-mir-574&WithFlank&4\|+\|38868026\|38868133 | 31 | 55 | 25 | TGAGTGTGTGTGTGTGAGTGTGTGT | 392.3 | 5.5 | 0.6 | 9.2 | 4.5E-20 | 5.2E-19 | MB231/MB468 | NUC |
| >hg38_21_+_8433172_8446622_RNA45SN1_45S_50ntFlanks@10410.10432.23\|TCAGTCGGTCCTGAGAGATGGGC | >hg38_21_+_8433172_8446622_RNA45SN1_45S_50ntFlanks | 10410 | 10432 | 23 | TCAGTCGGTCCTGAGAGATGGGC | 721.8 | -4.7 | 0.5 | -9.2 | 4.5E-20 | 5.2E-19 | MB231/MB468 | NUC |
| >hg38_21_+_8433172_8446622_RNA45SN1_45S_50ntFlanks@5086.5123.38\|AACGAGACTCTGGCATGCTAACTAGTTACGCGACCCCC | >hg38_21_+_8433172_8446622_RNA45SN1_45S_50ntFlanks | 5086 | 5123 | 38 | AACGAGACTCTGGCATGCTAACTAGTTACGCGACCCCC | 228.4 | 11.8 | 1.3 | 9.2 | 2.8E-20 | 5.3E-19 | MB231/MB468 | CYTO |
| >hg38_MT_+_1621_3279_RNR2_16S_50ntFlanks@1349.1388.40\|ACAATAGGGTTTACGACCTCGATGTTGGATCAGGACATCC | >hg38_MT_+_1621_3279_RNR2_16S_50ntFlanks | 1349 | 1388 | 40 | ACAATAGGGTTTACGACCTCGATGTTGGATCAGGACATCC | 255.3 | 11.9 | 1.3 | 9.2 | 4.5E-20 | 5.3E-19 | MB231/MB468 | NUC |
| >MI0000446\|hsa-mir-125b-1&WithFlank&11\|-\|122099751\|122099850@21.42.22(+2A);MI0000470\|hsa-mir-125b-2&WithFlank&21\|+\|16590231\|16590331@23.44.22(+2A)&[MIMAT0000423&hsa-miR-125b-5p&offsets\|0\|0(+2A);m-67&11\|-\|122099809\|122099830&offsets\|0\|0(+2A)];[MIMAT0000423_1&hsa-miR-125b-5p&offsets\|0\|0(+2A);m-68&21\|+\|16590253\|16590274&offsets\|0\|0(+2A)]\|TCCCTGAGACCCTAACTTGTGAAA | >MI0000446\|hsa-mir-125b-1&WithFlank&11\|-\|122099751\|122099850 | 21 | 42 | 22(+2A) | TCCCTGAGACCCTAACTTGTGAAA | 796.1 | 6.5 | 0.7 | 9.2 | 4.6E-20 | 5.3E-19 | MB231/MB468 | NUC |
| >hg38_21_+_8433172_8446622_RNA45SN1_45S_50ntFlanks@11235.11253.19\|CTTCCCGTGGATCGCCCCA | >hg38_21_+_8433172_8446622_RNA45SN1_45S_50ntFlanks | 11235 | 11253 | 19 | CTTCCCGTGGATCGCCCCA | 2070.5 | -10.7 | 1.2 | -9.2 | 5.3E-20 | 5.4E-19 | MB231/MB468 | MITO |
| >hg38_21_+_8433172_8446622_RNA45SN1_45S_50ntFlanks@11876.11893.18\|AAGACCCTGTTGAGCTTG | >hg38_21_+_8433172_8446622_RNA45SN1_45S_50ntFlanks | 11876 | 11893 | 18 | AAGACCCTGTTGAGCTTG | 1644.0 | -7.2 | 0.8 | -9.2 | 5.4E-20 | 5.4E-19 | MB231/MB468 | MITO |
| >hg38_21_+_8433172_8446622_RNA45SN1_45S_50ntFlanks@12991.13012.22\|CTCCCTCGCTGCGATCTATTGA | >hg38_21_+_8433172_8446622_RNA45SN1_45S_50ntFlanks | 12991 | 13012 | 22 | CTCCCTCGCTGCGATCTATTGA | 2644.9 | -8.9 | 1.0 | -9.2 | 5.4E-20 | 5.5E-19 | MB231/MB468 | MITO |
| >hg38_21_+_8433172_8446622_RNA45SN1_45S_50ntFlanks@11233.11251.19\|CCCTTCCCGTGGATCGCCC | >hg38_21_+_8433172_8446622_RNA45SN1_45S_50ntFlanks | 11233 | 11251 | 19 | CCCTTCCCGTGGATCGCCC | 409.8 | -13.1 | 1.4 | -9.0 | 1.5E-19 | 5.5E-19 | BT20/MB468 | NUC |
| >hg38_21_+_8433172_8446622_RNA45SN1_45S_50ntFlanks@9575.9597.23\|ATCGGTCGTCCGACCTGGGTATA | >hg38_21_+_8433172_8446622_RNA45SN1_45S_50ntFlanks | 9575 | 9597 | 23 | ATCGGTCGTCCGACCTGGGTATA | 3075.6 | -13.7 | 1.5 | -9.2 | 5.6E-20 | 5.5E-19 | BT20/MB468 | MITO |
| >hg38_21_+_8433172_8446622_RNA45SN1_45S_50ntFlanks@10866.10884.19\|CTAAGGGCTGGGTCGGTCG | >hg38_21_+_8433172_8446622_RNA45SN1_45S_50ntFlanks | 10866 | 10884 | 19 | CTAAGGGCTGGGTCGGTCG | 598.0 | -13.0 | 1.4 | -9.2 | 5.5E-20 | 5.5E-19 | MB231/MB468 | MITO |
| >hg38_21_+_8433172_8446622_RNA45SN1_45S_50ntFlanks@12436.12455.20\|TCATTGTGAAGCAGAATTCA | >hg38_21_+_8433172_8446622_RNA45SN1_45S_50ntFlanks | 12436 | 12455 | 20 | TCATTGTGAAGCAGAATTCA | 1353.4 | -12.5 | 1.4 | -9.2 | 5.7E-20 | 5.5E-19 | BT20/MB468 | MITO |
| >hg38_21_+_8433172_8446622_RNA45SN1_45S_50ntFlanks@3994.4023.30\|TGACTCTAGATAACCTCGGGCCGATCGCAC | >hg38_21_+_8433172_8446622_RNA45SN1_45S_50ntFlanks | 3994 | 4023 | 30 | TGACTCTAGATAACCTCGGGCCGATCGCAC | 434.7 | 6.0 | 0.7 | 9.1 | 7.7E-20 | 5.6E-19 | BT20/MB468 | CYTO |
| >hg38_21_+_8433172_8446622_RNA45SN1_45S_50ntFlanks@4283.4307.25\|CTTTAACGAGGATCCATTGGAGGGC | >hg38_21_+_8433172_8446622_RNA45SN1_45S_50ntFlanks | 4283 | 4307 | 25 | CTTTAACGAGGATCCATTGGAGGGC | 661.2 | -13.2 | 1.4 | -9.2 | 3.5E-20 | 5.6E-19 | MB231/MB468 | MP |
| >hg38_21_+_8433172_8446622_RNA45SN1_45S_50ntFlanks@8115.8143.29\|CGCGGGACATGTGGCGTACGGAAGACCCG | >hg38_21_+_8433172_8446622_RNA45SN1_45S_50ntFlanks | 8115 | 8143 | 29 | CGCGGGACATGTGGCGTACGGAAGACCCG | 602.9 | -12.2 | 1.3 | -9.2 | 4.8E-20 | 5.6E-19 | MB231/MB468 | NUC |
| >am_trna111_HisGTG_1_-_147774845_147774916@-1G.23.24,trna118_HisGTG_1_-_145396881_145396952@-1G.23.24,trna16_HisGTG_1_+_146544773_146544844@-1G.23.24,trna1_HisGTG_15_+_45493349_45493420@-1G.23.24,trna21_HisGTG_1_+_147753471_147753542@-1G.23.24,trna33_HisGTG_6_+_27125906_27125977@-1G.23.24,trna7_HisGTG_9_-_14433938_14434009@-1G.23.24,trna8_HisGTG_15_-_45492611_45492682@-1G.23.24,trna9_HisGTG_15_-_45490804_45490875@-1G.23.24\|GGCCGTGATCGTATAGTGGTTAGT | >am_trna111_HisGTG_1_-_147774845_147774916 | -1G | 23 | 24 | GGCCGTGATCGTATAGTGGTTAGT | 1811.2 | 8.1 | 0.9 | 9.2 | 4.8E-20 | 5.6E-19 | MB231/MB468 | NUC |
| >hg38_21_+_8433172_8446622_RNA45SN1_45S_50ntFlanks@3835.3860.26\|CTCCTCTCCTACTTGGATAACTGTGG | >hg38_21_+_8433172_8446622_RNA45SN1_45S_50ntFlanks | 3835 | 3860 | 26 | CTCCTCTCCTACTTGGATAACTGTGG | 4891.5 | -8.1 | 0.9 | -9.2 | 5.5E-20 | 5.6E-19 | MB231/MB468 | MITO |
| >hg38_21_+_8433172_8446622_RNA45SN1_45S_50ntFlanks@10689.10708.20\|GCGTCCGGTGAGCTCTCGCT | >hg38_21_+_8433172_8446622_RNA45SN1_45S_50ntFlanks | 10689 | 10708 | 20 | GCGTCCGGTGAGCTCTCGCT | 756.1 | -13.4 | 1.5 | -9.2 | 3.5E-20 | 5.6E-19 | MB231/MB468 | MP |
| >hg38_21_+_8433172_8446622_RNA45SN1_45S_50ntFlanks@10802.10822.21\|TTGGAACAATGTAGGTAAGGG | >hg38_21_+_8433172_8446622_RNA45SN1_45S_50ntFlanks | 10802 | 10822 | 21 | TTGGAACAATGTAGGTAAGGG | 547.8 | -13.0 | 1.4 | -9.2 | 3.5E-20 | 5.6E-19 | MB231/MB468 | MP |
| >hg38_21_+_8433172_8446622_RNA45SN1_45S_50ntFlanks@8110.8138.29\|CGGGGCGCGGGACATGTGGCGTACGGAAG | >hg38_21_+_8433172_8446622_RNA45SN1_45S_50ntFlanks | 8110 | 8138 | 29 | CGGGGCGCGGGACATGTGGCGTACGGAAG | 743.7 | -13.4 | 1.5 | -9.2 | 3.5E-20 | 5.6E-19 | MB231/MB468 | MP |
| >MI0000301\|hsa-mir-224&WithFlank&X\|-\|151958572\|151958664@14.37.24&[MIMAT0000281&hsa-miR-224-5p&offsets\|+1\|0;m-238&X\|-\|151958630\|151958651&offsets\|0\|+2]\|CAAGTCACTAGTGGTTCCGTTTAG | >MI0000301\|hsa-mir-224&WithFlank&X\|-\|151958572\|151958664 | 14 | 37 | 24 | CAAGTCACTAGTGGTTCCGTTTAG | 1206.0 | 11.9 | 1.3 | 9.2 | 3.0E-20 | 5.7E-19 | MB231/MB468 | CYTO |
| >MI0003591\|hsa-mir-584&WithFlank&5\|-\|149062307\|149062415@22.42.21(+1U)&[MIMAT0003249&hsa-miR-584-5p&offsets\|0\|-1(+1U);m-167&5\|-\|149062374\|149062394&offsets\|0\|0(+1U)]\|TTATGGTTTGCCTGGGACTGAT | >MI0003591\|hsa-mir-584&WithFlank&5\|-\|149062307\|149062415 | 22 | 42 | 21(+1U) | TTATGGTTTGCCTGGGACTGAT | 64.9 | 3.3 | 0.4 | 9.0 | 1.5E-19 | 5.7E-19 | BT20/MB468 | NUC |

| | | | | | | | | | | | | | |
|---|---|---|---|---|---|---|---|---|---|---|---|---|---|
| >hg38_21_+_8433172_8446622_RNA45SN1_45S_50ntFlanks@11739.11758.20\|TCTCTTAAGGTAGCCAAATG | >hg38_21_+_8433172_8446622_RNA45SN1_45S_50ntFlanks | 11739 | 11758 | 20 | TCTCTTAAGGTAGCCAAATG | 433.4 | -13.2 | 1.5 | -9.0 | 1.5E-19 | 5.7E-19 | BT20/MB468 | NUC |
| >hg38_21_+_8433172_8446622_RNA45SN1_45S_50ntFlanks@8380.8402.23\|CAAGAGGGCGTGAAACCGTTAAG | >hg38_21_+_8433172_8446622_RNA45SN1_45S_50ntFlanks | 8380 | 8402 | 23 | CAAGAGGGCGTGAAACCGTTAAG | 1530.6 | -7.8 | 0.8 | -9.2 | 3.0E-20 | 5.7E-19 | MB231/MB468 | CYTO |
| >hg38_21_+_8433172_8446622_RNA45SN1_45S_50ntFlanks@9565.9592.28\|TGACGTGCAAATCGGTCGTCCGACCTGG | >hg38_21_+_8433172_8446622_RNA45SN1_45S_50ntFlanks | 9565 | 9592 | 28 | TGACGTGCAAATCGGTCGTCCGACCTGG | 485.0 | -12.8 | 1.4 | -9.2 | 3.6E-20 | 5.7E-19 | MB231/MB468 | MP |
| >hg38_21_+_8433172_8446622_RNA45SN1_45S_50ntFlanks@4367.4389.23\|CAGTTAAAAAGCTCGTAGTTGGA | >hg38_21_+_8433172_8446622_RNA45SN1_45S_50ntFlanks | 4367 | 4389 | 23 | CAGTTAAAAAGCTCGTAGTTGGA | 802.1 | -13.4 | 1.5 | -9.1 | 5.8E-20 | 5.9E-19 | MB231/MB468 | MITO |
| >hg38_21_+_8433172_8446622_RNA45SN1_45S_50ntFlanks@10841.10864.24\|CGTAACTTCGGGATAAGGATTGGC | >hg38_21_+_8433172_8446622_RNA45SN1_45S_50ntFlanks | 10841 | 10864 | 24 | CGTAACTTCGGGATAAGGATTGGC | 6575.9 | -5.5 | 0.6 | -9.2 | 3.2E-20 | 5.9E-19 | MB231/MB468 | CYTO |
| >hg38_21_+_8433172_8446622_RNA45SN1_45S_50ntFlanks@3899.3937.39\|CCTTCGCGGGGGGGATGCGTGCATTTATCAGATCAAAAC | >hg38_21_+_8433172_8446622_RNA45SN1_45S_50ntFlanks | 3899 | 3937 | 39 | CCTTCGCGGGGGGGATGCGTGCATTTATCAGATCAAAAC | 938.8 | -14.3 | 1.6 | -9.0 | 1.6E-19 | 5.9E-19 | BT20/MB468 | NUC |
| >MI0005559\|hsa-mir-744&WithFlank&17\|+\|12081893\|12082002@17.36.20(+1U)&[MIMAT0004945&hsa-miR-744-5p&offsets\|0\|-2(+1U);m-172&17\|+\|12081909\|12081929&offsets\|0\|-1(+1U)]\|TGCGGGGCTAGGGCTAACAGT | >MI0005559\|hsa-mir-744&WithFlank&17\|+\|12081893\|12082002 | 17 | 36 | 20(+1U) | TGCGGGGCTAGGGCTAACAGT | 1041.4 | 10.3 | 1.1 | 9.1 | 6.1E-20 | 5.9E-19 | BT20/MB468 | MITO |
| >hg38_21_+_8433172_8446622_RNA45SN1_45S_50ntFlanks@5490.5516.27\|TGAGAAGACGGTCGAACTTGACTATCT | >hg38_21_+_8433172_8446622_RNA45SN1_45S_50ntFlanks | 5490 | 5516 | 27 | TGAGAAGACGGTCGAACTTGACTATCT | 4898.5 | 10.9 | 1.2 | 9.1 | 6.1E-20 | 5.9E-19 | BT20/MB468 | MITO |
| >hg38_21_+_8433172_8446622_RNA45SN1_45S_50ntFlanks@6719.6752.34\|TGAATTGCAGGACACATTGATCATCGACACTTCG | >hg38_21_+_8433172_8446622_RNA45SN1_45S_50ntFlanks | 6719 | 6752 | 34 | TGAATTGCAGGACACATTGATCATCGACACTTCG | 693.9 | -13.9 | 1.5 | -9.0 | 1.6E-19 | 5.9E-19 | BT20/MB468 | NUC |
| >hg38_21_+_8433172_8446622_RNA45SN1_45S_50ntFlanks@11199.11227.29\|CCGGCGGCGGCGGCGACTCTGGACGCGAG | >hg38_21_+_8433172_8446622_RNA45SN1_45S_50ntFlanks | 11199 | 11227 | 29 | CCGGCGGCGGCGGCGACTCTGGACGCGAG | 659.2 | -13.1 | 1.4 | -9.1 | 5.9E-20 | 5.9E-19 | MB231/MB468 | MITO |
| >hg38_21_+_8433172_8446622_RNA45SN1_45S_50ntFlanks@12874.12891.18\|GCTGGAAAGGCGGCCGCC | >hg38_21_+_8433172_8446622_RNA45SN1_45S_50ntFlanks | 12874 | 12891 | 18 | GCTGGAAAGGCGGCCGCC | 356.8 | -12.9 | 1.4 | -9.0 | 1.6E-19 | 5.9E-19 | BT20/MB468 | NUC |
| >MI0000086\|hsa-mir-28&WithFlank&3\|+\|188688775\|188688872@61.82.22(+1U)&[MIMAT0004502&hsa-miR-28-3p&offsets\|+1\|+1(+1U);m-28&3\|+\|188688834\|188688855&offsets\|+1\|+1(+1U)]\|ACTAGATTGTGAGCTCCTGGAGT | >MI0000086\|hsa-mir-28&WithFlank&3\|+\|188688775\|188688872 | 61 | 82 | 22(+1U) | ACTAGATTGTGAGCTCCTGGAGT | 930.0 | 10.1 | 1.1 | 9.1 | 8.4E-20 | 6.0E-19 | BT20/MB468 | CYTO |
| >MI0005756\|hsa-mir-934&WithFlank&X\|+\|136550872\|136550966@21.43.23&[MIMAT0004977&hsa-miR-934&offsets\|0\|+1;m-943&X\|+\|136550892\|136550913&offsets\|0\|+1]\|TGTCTACTACTGGAGACACTGGT | >MI0005756\|hsa-mir-934&WithFlank&X\|+\|136550872\|136550966 | 21 | 43 | 23 | TGTCTACTACTGGAGACACTGGT | 2927.6 | 6.9 | 0.8 | 9.1 | 8.4E-20 | 6.0E-19 | BT20/MB468 | CYTO |
| >hg38_21_+_8433172_8446622_RNA45SN1_45S_50ntFlanks@7975.8005.31\|CGCGACCTCAGATCAGACGTGGCGACCCGCT | >hg38_21_+_8433172_8446622_RNA45SN1_45S_50ntFlanks | 7975 | 8005 | 31 | CGCGACCTCAGATCAGACGTGGCGACCCGCT | 44976.6 | 5.7 | 0.6 | 9.1 | 8.4E-20 | 6.0E-19 | BT20/MB468 | CYTO |
| >hg38_21_+_8433172_8446622_RNA45SN1_45S_50ntFlanks@12717.12745.29\|CCTCGGTTGGCCTCGGATAGCCGGTCCCC | >hg38_21_+_8433172_8446622_RNA45SN1_45S_50ntFlanks | 12717 | 12745 | 29 | CCTCGGTTGGCCTCGGATAGCCGGTCCCC | 482.3 | -12.8 | 1.4 | -9.2 | 3.8E-20 | 6.0E-19 | MB231/MB468 | MP |
| >hg38_1_-_228634819_228635039_RNA5S12_5S_50ntFlanks@139.169.31\|GACCGCCTGGGAATACCGGGTGCTGTAGGCT | >hg38_1_-_228634819_228635039_RNA5S12_5S_50ntFlanks | 139 | 169 | 31 | GACCGCCTGGGAATACCGGGTGCTGTAGGCT | 795.1 | 4.7 | 0.5 | 9.1 | 6.2E-20 | 6.0E-19 | BT20/MB468 | MITO |
| >MI0000088\|hsa-mir-30a&WithFlank&6\|-\|71403545\|71403627@12.34.23(+1A)&[MIMAT0000087&hsa-miR-30a-5p&offsets\|0\|+1(+1A);m-15&6\|-\|71403595\|71403616&offsets\|0\|+1(+1A)]\|TGTAAACATCCTCGACTGGAAGCA | >MI0000088\|hsa-mir-30a&WithFlank&6\|-\|71403545\|71403627 | 12 | 34 | 23(+1A) | TGTAAACATCCTCGACTGGAAGCA | 2498.7 | 5.9 | 0.6 | 9.1 | 6.2E-20 | 6.0E-19 | BT20/MB468 | MITO |
| >MI0000283\|hsa-mir-203a&WithFlank&14\|+\|104117399\|104117520@72.92.21(+1U)&[MIMAT0000264&hsa-miR-203a-3p&offsets\|+1\|0(+1U);m-79&14\|+\|104117469\|104117490&offsets\|+1\|0(+1U)]\|TGAAATGTTTAGGACCACTAGT | >MI0000283\|hsa-mir-203a&WithFlank&14\|+\|104117399\|104117520 | 72 | 92 | 21(+1U) | TGAAATGTTTAGGACCACTAGT | 1007.7 | 10.1 | 1.1 | 9.5 | 2.7E-21 | 6.1E-19 | BT20/MB231 | CYTO |
| >hg38_21_+_8433172_8446622_RNA45SN1_45S_50ntFlanks@9198.9220.23\|GTTCTCTCGGGGCCACGCGCGCG | >hg38_21_+_8433172_8446622_RNA45SN1_45S_50ntFlanks | 9198 | 9220 | 23 | GTTCTCTCGGGGCCACGCGCGCG | 2122.4 | -14.8 | 1.6 | -9.1 | 6.1E-20 | 6.1E-19 | MB231/MB468 | MITO |
| >MI0000433\|hsa-let-7g&WithFlank&3\|-\|52268272\|52268367@11.28.18&[MIMAT0000414&hsa-let-7g-5p&offsets\|0\|-4;m-58&3\|-\|52268336\|52268357&offsets\|0\|-4]\|TGAGGTAGTAGTTTGTAC | >MI0000433\|hsa-let-7g&WithFlank&3\|-\|52268272\|52268367 | 11 | 28 | 18 | TGAGGTAGTAGTTTGTAC | 4609.5 | 7.9 | 0.9 | 9.1 | 8.6E-20 | 6.1E-19 | BT20/MB468 | CYTO |
| >MI0000079\|hsa-mir-23a&WithFlank&19\|-\|13836581\|13836665@54.73.20(+1U)&[MIMAT0000078&hsa-miR-23a-3p&offsets\|+3\|+2(+1U);m-31&19\|-\|13836595\|13836615&offsets\|+3\|+2(+1U)]\|ACATTGCCAGGGATTTCCAAT | >MI0000079\|hsa-mir-23a&WithFlank&19\|-\|13836581\|13836665 | 54 | 73 | 20(+1U) | ACATTGCCAGGGATTTCCAAT | 712.1 | 13.9 | 1.5 | 9.1 | 6.3E-20 | 6.1E-19 | BT20/MB468 | MITO |
| >hg38_21_+_8433172_8446622_RNA45SN1_45S_50ntFlanks@8845.8867.23\|CGGCGGCGGCGGCGGCGGCGGGA | >hg38_21_+_8433172_8446622_RNA45SN1_45S_50ntFlanks | 8845 | 8867 | 23 | CGGCGGCGGCGGCGGCGGCGGGA | 946.8 | -13.8 | 1.5 | -9.2 | 3.9E-20 | 6.2E-19 | MB231/MB468 | MP |
| >hg38_21_+_8433172_8446622_RNA45SN1_45S_50ntFlanks@4722.4752.31\|TCATTAATCAAGAACGAAAGTCGGAGGTTCG | >hg38_21_+_8433172_8446622_RNA45SN1_45S_50ntFlanks | 4722 | 4752 | 31 | TCATTAATCAAGAACGAAAGTCGGAGGTTCG | 813.4 | -11.8 | 1.3 | -9.1 | 6.4E-20 | 6.2E-19 | BT20/MB468 | MITO |
| >trna130_GlnTTG_6_-_27763640_27763711@1.29.29,trna173_GlnTTG_6_-_26311975_26312046@1.29.29,trna174_GlnTTG_6_-_26311424_26311495@1.29.29\|GGCCCCATGGTGTAATGGTTAGCACTCTG | >trna130_GlnTTG_6_-_27763640_27763711 | 1 | 29 | 29 | GGCCCCATGGTGTAATGGTTAGCACTCTG | 883.0 | -12.7 | 1.4 | -9.2 | 5.4E-20 | 6.2E-19 | MB231/MB468 | NUC |
| >hg38_21_+_8433172_8446622_RNA45SN1_45S_50ntFlanks@12456.12490.35\|CCAAGCGTTGGATTGTTCACCCACTAATAGGGAAC | >hg38_21_+_8433172_8446622_RNA45SN1_45S_50ntFlanks | 12456 | 12490 | 35 | CCAAGCGTTGGATTGTTCACCCACTAATAGGGAAC | 924.6 | -4.1 | 0.4 | -9.1 | 8.8E-20 | 6.3E-19 | BT20/MB468 | CYTO |
| >hg38_21_+_8433172_8446622_RNA45SN1_45S_50ntFlanks@10392.10409.18\|ACAGCAGTTGAACATGGG | >hg38_21_+_8433172_8446622_RNA45SN1_45S_50ntFlanks | 10392 | 10409 | 18 | ACAGCAGTTGAACATGGG | 1045.9 | -13.9 | 1.5 | -9.2 | 4.0E-20 | 6.3E-19 | MB231/MB468 | MP |
| >trnaMT_ValTAC_MT_+_1602_1670@42.71.30\|AGATTTCAACTTAACTTGACCGCTCTGACC | >trnaMT_ValTAC_MT_+_1602_1670 | 42 | 71 | 30 | AGATTTCAACTTAACTTGACCGCTCTGACC | 1325.7 | 5.8 | 0.6 | 9.2 | 5.4E-20 | 6.3E-19 | MB231/MB468 | NUC |
| >hg38_21_+_8433172_8446622_RNA45SN1_45S_50ntFlanks@12626.12652.27\|AATGGGGCGAAGCTACCATCTGTGGGA | >hg38_21_+_8433172_8446622_RNA45SN1_45S_50ntFlanks | 12626 | 12652 | 27 | AATGGGGCGAAGCTACCATCTGTGGGA | 1777.0 | -14.7 | 1.6 | -9.2 | 4.0E-20 | 6.3E-19 | MB231/MB468 | MP |
| >hg38_21_+_8433172_8446622_RNA45SN1_45S_50ntFlanks@11581.11603.23\|AACTGGTGCGGACCAGGGGAATC | >hg38_21_+_8433172_8446622_RNA45SN1_45S_50ntFlanks | 11581 | 11603 | 23 | AACTGGTGCGGACCAGGGGAATC | 8863.2 | -5.3 | 0.6 | -9.2 | 3.4E-20 | 6.4E-19 | MB231/MB468 | CYTO |
| >hg38_21_+_8433172_8446622_RNA45SN1_45S_50ntFlanks@6738.6770.33\|ATCATCGACACTTCGAACGCACTTGCGGCCCCG | >hg38_21_+_8433172_8446622_RNA45SN1_45S_50ntFlanks | 6738 | 6770 | 33 | ATCATCGACACTTCGAACGCACTTGCGGCCCCG | 11372.9 | -11.9 | 1.3 | -9.1 | 6.3E-20 | 6.4E-19 | MB231/MB468 | MITO |
| >MI0000465\|hsa-mir-191&WithFlank&3\|-\|49020612\|49020715@22.42.21(+1A)&[MIMAT0000440&hsa-miR-191-5p&offsets\|0\|-2(+1A);m-48&3\|-\|49020672\|49020694&offsets\|0\|-2(+1A)]\|CAACGGAATCCCAAAAGCAGCA | >MI0000465\|hsa-mir-191&WithFlank&3\|-\|49020612\|49020715 | 22 | 42 | 21(+1A) | CAACGGAATCCCAAAAGCAGCA | 820.5 | 7.3 | 0.8 | 9.1 | 6.6E-20 | 6.4E-19 | BT20/MB468 | MITO |
| >hg38_21_+_8433172_8446622_RNA45SN1_45S_50ntFlanks@6733.6774.42\|CATTGATCATCGACACTTCGAACGCACTTGCGGCCCCGGGTT | >hg38_21_+_8433172_8446622_RNA45SN1_45S_50ntFlanks | 6733 | 6774 | 42 | CATTGATCATCGACACTTCGAACGCACTTGCGGCCCCGGGTT | 1029.3 | 6.8 | 0.7 | 9.1 | 6.6E-20 | 6.4E-19 | BT20/MB468 | MITO |
| >hg38_21_+_8433172_8446622_RNA45SN1_45S_50ntFlanks@9713.9737.25\|GCGAATGATTAGAGGTCTTGGGGCC | >hg38_21_+_8433172_8446622_RNA45SN1_45S_50ntFlanks | 9713 | 9737 | 25 | GCGAATGATTAGAGGTCTTGGGGCC | 1704.1 | -14.5 | 1.6 | -9.1 | 6.4E-20 | 6.4E-19 | MB231/MB468 | MITO |
| >trnaMT_ValTAC_MT_+_1602_1670@46.71.26\|TTCAACTTAACTTGACCGCTCTGACC | >trnaMT_ValTAC_MT_+_1602_1670 | 46 | 71 | 26 | TTCAACTTAACTTGACCGCTCTGACC | 401.0 | 12.6 | 1.4 | 9.2 | 3.5E-20 | 6.5E-19 | MB231/MB468 | CYTO |
| >hg38_21_+_8433172_8446622_RNA45SN1_45S_50ntFlanks@475.504.30\|CTTCGTGATCGATGTGGTGACGTCGTGCTC | >hg38_21_+_8433172_8446622_RNA45SN1_45S_50ntFlanks | 475 | 504 | 30 | CTTCGTGATCGATGTGGTGACGTCGTGCTC | 8322.4 | 8.6 | 0.9 | 9.1 | 6.8E-20 | 6.5E-19 | BT20/MB468 | MITO |
| >hg38_21_+_8433172_8446622_RNA45SN1_45S_50ntFlanks@8114.8142.29\|GCGCGGGACATGTGGCGTACGGAAGACCC | >hg38_21_+_8433172_8446622_RNA45SN1_45S_50ntFlanks | 8114 | 8142 | 29 | GCGCGGGACATGTGGCGTACGGAAGACCC | 694.0 | -13.2 | 1.4 | -9.1 | 6.5E-20 | 6.5E-19 | MB231/MB468 | MITO |
| >hg38_21_+_8433172_8446622_RNA45SN1_45S_50ntFlanks@9202.9220.19\|TCTCGGGGCCACGCGCGCG | >hg38_21_+_8433172_8446622_RNA45SN1_45S_50ntFlanks | 9202 | 9220 | 19 | TCTCGGGGCCACGCGCGCG | 848.0 | -13.5 | 1.5 | -9.1 | 6.5E-20 | 6.5E-19 | MB231/MB468 | MITO |
| >hg38_21_+_8433172_8446622_RNA45SN1_45S_50ntFlanks@3930.3959.30\|ATCAAAACCAACCCGGTCAGCCCCTCTCCG | >hg38_21_+_8433172_8446622_RNA45SN1_45S_50ntFlanks | 3930 | 3959 | 30 | ATCAAAACCAACCCGGTCAGCCCCTCTCCG | 834.5 | -12.7 | 1.4 | -9.2 | 5.7E-20 | 6.5E-19 | MB231/MB468 | NUC |
| >hg38_21_+_8433172_8446622_RNA45SN1_45S_50ntFlanks@9738.9755.18\|GAAACGATCTCAACCTAT | >hg38_21_+_8433172_8446622_RNA45SN1_45S_50ntFlanks | 9738 | 9755 | 18 | GAAACGATCTCAACCTAT | 982.6 | -13.7 | 1.5 | -9.1 | 6.6E-20 | 6.6E-19 | MB231/MB468 | MITO |
| >hg38_21_+_8433172_8446622_RNA45SN1_45S_50ntFlanks@11581.11605.25\|AACTGGTGCGGACCAGGGGAATCCG | >hg38_21_+_8433172_8446622_RNA45SN1_45S_50ntFlanks | 11581 | 11605 | 25 | AACTGGTGCGGACCAGGGGAATCCG | 1364.3 | -3.9 | 0.4 | -9.1 | 6.8E-20 | 6.6E-19 | BT20/MB468 | MITO |
| >hg38_21_+_8433172_8446622_RNA45SN1_45S_50ntFlanks@11564.11603.40\|CCTAGCAGCCGACTTAGAACTGGTGCGGACCAGGGGAATC | >hg38_21_+_8433172_8446622_RNA45SN1_45S_50ntFlanks | 11564 | 11603 | 40 | CCTAGCAGCCGACTTAGAACTGGTGCGGACCAGGGGAATC | 6540.7 | -10.7 | 1.2 | -9.0 | 1.7E-19 | 6.6E-19 | BT20/MB468 | NUC |
| >hg38_21_+_8433172_8446622_RNA45SN1_45S_50ntFlanks@11595.11616.22\|AGGGGAATCCGACTGTTTAATT | >hg38_21_+_8433172_8446622_RNA45SN1_45S_50ntFlanks | 11595 | 11616 | 22 | AGGGGAATCCGACTGTTTAATT | 221.1 | 4.5 | 0.5 | 9.2 | 3.6E-20 | 6.6E-19 | MB231/MB468 | CYTO |
| >hg38_1_-_228634819_228635039_RNA5S12_5S_50ntFlanks@146.170.25\|TGGGAATACCGGGTGCTGTAGGCTT | >hg38_1_-_228634819_228635039_RNA5S12_5S_50ntFlanks | 146 | 170 | 25 | TGGGAATACCGGGTGCTGTAGGCTT | 2705.2 | 5.3 | 0.6 | 9.1 | 6.9E-20 | 6.6E-19 | BT20/MB468 | MITO |
| >MI0000299\|hsa-mir-222&WithFlank&X\|-\|45747009\|45747130@75.94.20(+1C)&[MIMAT0000279&hsa-miR-222-3p&offsets\|0\|-1(+1C);m-140&X\|-\|45747033\|45747056&offsets\|0\|-4(+1C)]\|AGCTACATCTGGCTACTGGGC | >MI0000299\|hsa-mir-222&WithFlank&X\|-\|45747009\|45747130 | 75 | 94 | 20(+1C) | AGCTACATCTGGCTACTGGGC | 238.1 | 5.5 | 0.6 | 9.1 | 5.8E-20 | 6.7E-19 | MB231/MB468 | NUC |
| >hg38_21_+_8433172_8446622_RNA45SN1_45S_50ntFlanks@12616.12654.39\|CTGAGGAGCCAATGGGGCGAAGCTACCATCTGTGGGATT | >hg38_21_+_8433172_8446622_RNA45SN1_45S_50ntFlanks | 12616 | 12654 | 39 | CTGAGGAGCCAATGGGGCGAAGCTACCATCTGTGGGATT | 4736.5 | -9.7 | 1.1 | -9.1 | 6.7E-20 | 6.7E-19 | MB231/MB468 | MITO |
| >MI0000816\|hsa-mir-335&WithFlank&7\|+\|130496105\|130496210@58.78.21&[MIMAT0004703&hsa-miR-335-3p&offsets\|0\|-1;m-159&7\|+\|130496162\|130496183&offsets\|0\|-1]\|TTTTTCATTATTGCTCCTGAC | >MI0000816\|hsa-mir-335&WithFlank&7\|+\|130496105\|130496210 | 58 | 78 | 21 | TTTTTCATTATTGCTCCTGAC | 557.0 | 6.9 | 0.8 | 9.1 | 9.4E-20 | 6.7E-19 | BT20/MB468 | CYTO |
| >MI0000440\|hsa-mir-27b&WithFlank&9\|+\|95085439\|95085547@66.86.21(+1U)&[MIMAT0000419&hsa-miR-27b-3p&offsets\|-1\|-1(+1U);m-39&9\|+\|95085505\|95085525&offsets\|-1\|-1(+1U)]\|GTTCACAGTGGCTAAGTTCTGT | >MI0000440\|hsa-mir-27b&WithFlank&9\|+\|95085439\|95085547 | 66 | 86 | 21(+1U) | GTTCACAGTGGCTAAGTTCTGT | 340.4 | 12.7 | 1.4 | 9.1 | 9.5E-20 | 6.7E-19 | BT20/MB468 | CYTO |
| >MI0000077\|hsa-mir-21&WithFlank&17\|+\|59841260\|59841343@14.34.21&[MIMAT0000076&hsa-miR-21-5p&offsets\|0\|-1;m-1&17\|+\|59841273\|59841294&offsets\|0\|-1]\|TAGCTTATCAGACTGATGTTG | >MI0000077\|hsa-mir-21&WithFlank&17\|+\|59841260\|59841343 | 14 | 34 | 21 | TAGCTTATCAGACTGATGTTG | 28928.4 | 8.0 | 0.9 | 9.1 | 9.6E-20 | 6.8E-19 | BT20/MB468 | CYTO |

<table>
<tr><th></th><th></th><th></th><th></th><th></th><th></th><th></th><th></th><th></th><th></th><th></th><th></th><th></th><th></th></tr>
<tr><td>>MI0000465|hsa-mir-191&WithFlank&3|-|49020612|49020715@22.43.22(+1U)&[MIMAT0000440&hsa-miR-191-5p&offsets|0|-1(+1U);m-48&3|-|49020672|49020694&offsets|0|-1(+1U)]|CAACGGAATCCCAAAAGCAGCTT</td><td>>MI0000465|hsa-mir-191&WithFlank&3|-|49020612|49020715</td><td>22</td><td>43</td><td>22(+1U)</td><td>CAACGGAATCCCAAAAGCAGCTT</td><td>819.3</td><td>8.4</td><td>0.9</td><td>9.1</td><td>9.6E-20</td><td>6.8E-19</td><td>BT20/MB468</td><td>CYTO</td></tr>
<tr><td>>hg38_21_+_8433172_8446622_RNA45SN1_45S_50ntFlanks@10392.10435.44|ACAGCAGTTGAACATGGGTCAGTCGGTCCTGAGAGATGGGCGAG</td><td>>hg38_21_+_8433172_8446622_RNA45SN1_45S_50ntFlanks</td><td>10392</td><td>10435</td><td>44</td><td>ACAGCAGTTGAACATGGGTCAGTCGGTCCTGAGAGATGGGCGAG</td><td>1546.8</td><td>-14.4</td><td>1.6</td><td>-9.1</td><td>6.8E-20</td><td>6.8E-19</td><td>MB231/MB468</td><td>MITO</td></tr>
<tr><td>>hg38_MT_+_598_1651_RNR1_12S_50ntFlanks@908.946.39|ATACTTCAAAGGACATTTAACTAAAACCCCTACGCATTT</td><td>>hg38_MT_+_598_1651_RNR1_12S_50ntFlanks</td><td>908</td><td>946</td><td>39</td><td>ATACTTCAAAGGACATTTAACTAAAACCCCTACGCATTT</td><td>330.1</td><td>12.8</td><td>1.4</td><td>9.1</td><td>7.1E-20</td><td>6.8E-19</td><td>BT20/MB468</td><td>MITO</td></tr>
<tr><td>>hg38_21_+_8433172_8446622_RNA45SN1_45S_50ntFlanks@10866.10902.37|CTAAGGGCTGGGTCGGTCGGGCTGGGGCGCGAAGCGG</td><td>>hg38_21_+_8433172_8446622_RNA45SN1_45S_50ntFlanks</td><td>10866</td><td>10902</td><td>37</td><td>CTAAGGGCTGGGTCGGTCGGGCTGGGGCGCGAAGCGG</td><td>3199.9</td><td>-11.4</td><td>1.2</td><td>-9.1</td><td>5.9E-20</td><td>6.8E-19</td><td>MB231/MB468</td><td>NUC</td></tr>
<tr><td>>hg38_21_+_8433172_8446622_RNA45SN1_45S_50ntFlanks@12405.12425.21|CTTTTTGATCCTTCGATGTCG</td><td>>hg38_21_+_8433172_8446622_RNA45SN1_45S_50ntFlanks</td><td>12405</td><td>12425</td><td>21</td><td>CTTTTTGATCCTTCGATGTCG</td><td>846.5</td><td>-13.5</td><td>1.5</td><td>-9.1</td><td>6.9E-20</td><td>6.8E-19</td><td>MB231/MB468</td><td>MITO</td></tr>
<tr><td>>hg38_21_+_8433172_8446622_RNA45SN1_45S_50ntFlanks@12202.12221.20|CTAAGGCGAGCTCAGGGAGG</td><td>>hg38_21_+_8433172_8446622_RNA45SN1_45S_50ntFlanks</td><td>12202</td><td>12221</td><td>20</td><td>CTAAGGCGAGCTCAGGGAGG</td><td>810.6</td><td>-13.5</td><td>1.5</td><td>-9.2</td><td>4.4E-20</td><td>6.9E-19</td><td>MB231/MB468</td><td>MP</td></tr>
<tr><td>>hg38_21_+_8433172_8446622_RNA45SN1_45S_50ntFlanks@5446.5465.20|CGGATCGGCCCCGCCGGGGT</td><td>>hg38_21_+_8433172_8446622_RNA45SN1_45S_50ntFlanks</td><td>5446</td><td>5465</td><td>20</td><td>CGGATCGGCCCCGCCGGGGT</td><td>533.8</td><td>-12.9</td><td>1.4</td><td>-9.2</td><td>4.4E-20</td><td>6.9E-19</td><td>MB231/MB468</td><td>MP</td></tr>
<tr><td>>hg38_21_+_8433172_8446622_RNA45SN1_45S_50ntFlanks@10783.10801.19|GTGAACAGCCTCTGGCATG</td><td>>hg38_21_+_8433172_8446622_RNA45SN1_45S_50ntFlanks</td><td>10783</td><td>10801</td><td>19</td><td>GTGAACAGCCTCTGGCATG</td><td>760.8</td><td>-13.4</td><td>1.5</td><td>-9.2</td><td>4.4E-20</td><td>6.9E-19</td><td>MB231/MB468</td><td>MP</td></tr>
<tr><td>>hg38_21_+_8433172_8446622_RNA45SN1_45S_50ntFlanks@11703.11728.26|AATGAAGCGCGGGTAAACGGCGGGAG</td><td>>hg38_21_+_8433172_8446622_RNA45SN1_45S_50ntFlanks</td><td>11703</td><td>11728</td><td>26</td><td>AATGAAGCGCGGGTAAACGGCGGGAG</td><td>3783.8</td><td>-11.6</td><td>1.3</td><td>-9.2</td><td>4.4E-20</td><td>6.9E-19</td><td>MB231/MB468</td><td>MP</td></tr>
<tr><td>>hg38_21_+_8433172_8446622_RNA45SN1_45S_50ntFlanks@9345.9370.26|AAAGCCGCCGTGGCGCAATGAAGGTG</td><td>>hg38_21_+_8433172_8446622_RNA45SN1_45S_50ntFlanks</td><td>9345</td><td>9370</td><td>26</td><td>AAAGCCGCCGTGGCGCAATGAAGGTG</td><td>681.2</td><td>-13.2</td><td>1.4</td><td>-9.1</td><td>7.0E-20</td><td>6.9E-19</td><td>MB231/MB468</td><td>MITO</td></tr>
<tr><td>>hg38_21_+_8433172_8446622_RNA45SN1_45S_50ntFlanks@10411.10432.22|CAGTCGGTCCTGAGAGATGGGC</td><td>>hg38_21_+_8433172_8446622_RNA45SN1_45S_50ntFlanks</td><td>10411</td><td>10432</td><td>22</td><td>CAGTCGGTCCTGAGAGATGGGC</td><td>1814.9</td><td>-6.4</td><td>0.7</td><td>-9.2</td><td>3.8E-20</td><td>7.0E-19</td><td>MB231/MB468</td><td>CYTO</td></tr>
<tr><td>>hg38_21_+_8433172_8446622_RNA45SN1_45S_50ntFlanks@10678.10711.34|CGGTTCCGGCGGCGTCCGGTGAGCTCTCGCTGGC</td><td>>hg38_21_+_8433172_8446622_RNA45SN1_45S_50ntFlanks</td><td>10678</td><td>10711</td><td>34</td><td>CGGTTCCGGCGGCGTCCGGTGAGCTCTCGCTGGC</td><td>37472.3</td><td>-8.0</td><td>0.9</td><td>-9.2</td><td>3.8E-20</td><td>7.0E-19</td><td>MB231/MB468</td><td>CYTO</td></tr>
<tr><td>>hg38_21_+_8433172_8446622_RNA45SN1_45S_50ntFlanks@12476.12499.24|CCACTAATAGGGAACGTGAGCTGG</td><td>>hg38_21_+_8433172_8446622_RNA45SN1_45S_50ntFlanks</td><td>12476</td><td>12499</td><td>24</td><td>CCACTAATAGGGAACGTGAGCTGG</td><td>418.4</td><td>-6.6</td><td>0.7</td><td>-9.2</td><td>3.8E-20</td><td>7.0E-19</td><td>MB231/MB468</td><td>CYTO</td></tr>
<tr><td>>hg38_21_+_8433172_8446622_RNA45SN1_45S_50ntFlanks@12625.12653.29|CAATGGGGCGAAGCTACCATCTGTGGGAT</td><td>>hg38_21_+_8433172_8446622_RNA45SN1_45S_50ntFlanks</td><td>12625</td><td>12653</td><td>29</td><td>CAATGGGGCGAAGCTACCATCTGTGGGAT</td><td>4654.4</td><td>-11.1</td><td>1.2</td><td>-9.1</td><td>7.0E-20</td><td>7.0E-19</td><td>MB231/MB468</td><td>MITO</td></tr>
<tr><td>>MI0002470|hsa-mir-486-1&WithFlank&8|-|41660435|41660514@10.31.22(+1U);MI0023622|hsa-mir-486-2&WithFlank&8|+|41660438|41660513@7.28.22(+1U)&[MIMAT0002177&hsa-miR-486-5p&offsets|0|0(+1U);m-107&8|-|41660484|41660505&offsets|0|0(+1U)];[MIMAT0002177_1&hsa-miR-486-5p&offsets|0|0(+1U);TJU_CMC_MD2.ID03138.5p-miR&offsets|0|-2(+1U);m-106&8|+|41660444|41660465&offsets|0|0(+1U)]|TCCTGTACTGAGCTGCCCCGAGT</td><td>>MI0002470|hsa-mir-486-1&WithFlank&8|-|41660435|41660514</td><td>10</td><td>31</td><td>22(+1U)</td><td>TCCTGTACTGAGCTGCCCCGAGT</td><td>645.8</td><td>13.3</td><td>1.5</td><td>9.2</td><td>3.8E-20</td><td>7.0E-19</td><td>MB231/MB468</td><td>CYTO</td></tr>
<tr><td>>hg38_21_+_8433172_8446622_RNA45SN1_45S_50ntFlanks@11784.11803.20|ATGAATGGATGAACGAGATT</td><td>>hg38_21_+_8433172_8446622_RNA45SN1_45S_50ntFlanks</td><td>11784</td><td>11803</td><td>20</td><td>ATGAATGGATGAACGAGATT</td><td>10266.2</td><td>-10.9</td><td>1.2</td><td>-9.0</td><td>1.9E-19</td><td>7.0E-19</td><td>BT20/MB468</td><td>NUC</td></tr>
<tr><td>>hg38_21_+_8433172_8446622_RNA45SN1_45S_50ntFlanks@10410.10428.19|TCAGTCGGTCCTGAGAGAT</td><td>>hg38_21_+_8433172_8446622_RNA45SN1_45S_50ntFlanks</td><td>10410</td><td>10428</td><td>19</td><td>TCAGTCGGTCCTGAGAGAT</td><td>1441.2</td><td>-8.6</td><td>0.9</td><td>-9.1</td><td>6.2E-20</td><td>7.1E-19</td><td>MB231/MB468</td><td>NUC</td></tr>
<tr><td>>hg38_21_+_8433172_8446622_RNA45SN1_45S_50ntFlanks@12008.12030.23|CCGCCGGTGAAATACCACTACTC</td><td>>hg38_21_+_8433172_8446622_RNA45SN1_45S_50ntFlanks</td><td>12008</td><td>12030</td><td>23</td><td>CCGCCGGTGAAATACCACTACTC</td><td>1072.5</td><td>-10.0</td><td>1.1</td><td>-9.2</td><td>3.9E-20</td><td>7.1E-19</td><td>MB231/MB468</td><td>CYTO</td></tr>
<tr><td>>hg38_21_+_8433172_8446622_RNA45SN1_45S_50ntFlanks@9201.9222.22|CTCTCGGGGCCACGCGCGCGTC</td><td>>hg38_21_+_8433172_8446622_RNA45SN1_45S_50ntFlanks</td><td>9201</td><td>9222</td><td>22</td><td>CTCTCGGGGCCACGCGCGCGTC</td><td>1104.3</td><td>-13.9</td><td>1.5</td><td>-9.1</td><td>7.2E-20</td><td>7.1E-19</td><td>MB231/MB468</td><td>MITO</td></tr>
<tr><td>>trna2_GlyGCC_21_-_18827107_18827177@1.32.32,trna35_GlyGCC_1_+_161413094_161413164@1.32.32,trna37_GlyGCC_1_+_161420467_161420537@1.32.32,trna39_GlyGCC_1_+_161427898_161427968@1.32.32,trna41_GlyGCC_1_+_161435258_161435328@1.32.32|GCATGGGTGGTTCAGTGGTAGAATTCTCGCCT</td><td>>trna2_GlyGCC_21_-_18827107_18827177</td><td>1</td><td>32</td><td>32</td><td>GCATGGGTGGTTCAGTGGTAGAATTCTCGCCT</td><td>15074.5</td><td>7.2</td><td>0.8</td><td>9.1</td><td>1.0E-19</td><td>7.1E-19</td><td>BT20/MB468</td><td>CYTO</td></tr>
<tr><td>>hg38_21_+_8433172_8446622_RNA45SN1_45S_50ntFlanks@8423.8471.49|CGCAGTCCGCCCGGAGGATTCAACCCGGCGGCGGGTCCGGCCGTGTCGG</td><td>>hg38_21_+_8433172_8446622_RNA45SN1_45S_50ntFlanks</td><td>8423</td><td>8471</td><td>49</td><td>CGCAGTCCGCCCGGAGGATTCAACCCGGCGGCGGGTCCGGCCGTGTCGG</td><td>1494.0</td><td>-14.3</td><td>1.6</td><td>-9.1</td><td>7.4E-20</td><td>7.3E-19</td><td>MB231/MB468</td><td>MITO</td></tr>
<tr><td>>MI0000102|hsa-mir-100&WithFlank&11|-|122152223|122152314@19.39.21&[MIMAT0000098&hsa-miR-100-5p&offsets|0|-1;m-27&11|-|122152276|122152296&offsets|0|0]|AACCCGTAGATCCGAACTTGT</td><td>>MI0000102|hsa-mir-100&WithFlank&11|-|122152223|122152314</td><td>19</td><td>39</td><td>21</td><td>AACCCGTAGATCCGAACTTGT</td><td>154031.6</td><td>-7.9</td><td>0.8</td><td>-9.4</td><td>5.6E-21</td><td>7.3E-19</td><td>BT20/MB231</td><td>MITO</td></tr>
<tr><td>>MI0000102|hsa-mir-100&WithFlank&11|-|122152223|122152314@20.40.21&[MIMAT0000098&hsa-miR-100-5p&offsets|+1|0;m-27&11|-|122152276|122152296&offsets|+1|+1]|ACCCGTAGATCCGAACTTGTG</td><td>>MI0000102|hsa-mir-100&WithFlank&11|-|122152223|122152314</td><td>20</td><td>40</td><td>21</td><td>ACCCGTAGATCCGAACTTGTG</td><td>1156.3</td><td>-13.4</td><td>1.4</td><td>-9.4</td><td>5.7E-21</td><td>7.3E-19</td><td>BT20/MB231</td><td>MITO</td></tr>
<tr><td>>hg38_21_+_8433172_8446622_RNA45SN1_45S_50ntFlanks@4853.4883.31|AAGTCTTTGGGTTCCGGGGGGAGTATGGTTG</td><td>>hg38_21_+_8433172_8446622_RNA45SN1_45S_50ntFlanks</td><td>4853</td><td>4883</td><td>31</td><td>AAGTCTTTGGGTTCCGGGGGGAGTATGGTTG</td><td>633.4</td><td>-12.3</td><td>1.3</td><td>-9.1</td><td>6.4E-20</td><td>7.3E-19</td><td>MB231/MB468</td><td>NUC</td></tr>
<tr><td>>MI0005756|hsa-mir-934&WithFlank&X|+|136550872|136550966@21.42.22(+1A)&[MIMAT0004977&hsa-miR-934&offsets|0|0(+1A);m-943&X|+|136550892|136550913&offsets|0|0(+1A)]|TGTCTACTACTGGAGACACTGGA</td><td>>MI0005756|hsa-mir-934&WithFlank&X|+|136550872|136550966</td><td>21</td><td>42</td><td>22(+1A)</td><td>TGTCTACTACTGGAGACACTGGA</td><td>804.8</td><td>13.3</td><td>1.4</td><td>9.4</td><td>6.0E-21</td><td>7.4E-19</td><td>BT20/MB231</td><td>MITO</td></tr>
<tr><td>>hg38_21_+_8433172_8446622_RNA45SN1_45S_50ntFlanks@12044.12071.28|ACTGACCCGGTGAGGCGGGGGGGCGAGC</td><td>>hg38_21_+_8433172_8446622_RNA45SN1_45S_50ntFlanks</td><td>12044</td><td>12071</td><td>28</td><td>ACTGACCCGGTGAGGCGGGGGGGCGAGC</td><td>1015.6</td><td>-13.9</td><td>1.5</td><td>-9.2</td><td>4.7E-20</td><td>7.4E-19</td><td>MB231/MB468</td><td>MP</td></tr>
<tr><td>>MI0000477|hsa-mir-146a&WithFlank&5|+|160485346|160485456@27.49.23&[MIMAT0000449&hsa-miR-146a-5p&offsets|0|+1;m-129&5|+|160485372|160485393&offsets|0|+1]|TGAGAACTGAATTCCATGGGTTG</td><td>>MI0000477|hsa-mir-146a&WithFlank&5|+|160485346|160485456</td><td>27</td><td>49</td><td>23</td><td>TGAGAACTGAATTCCATGGGTTG</td><td>1882.0</td><td>-14.1</td><td>1.5</td><td>-9.4</td><td>6.2E-21</td><td>7.4E-19</td><td>BT20/MB231</td><td>MITO</td></tr>
<tr><td>>hg38_21_+_8433172_8446622_RNA45SN1_45S_50ntFlanks@11641.11658.18|CGGCGGGTGTTGACGCGA</td><td>>hg38_21_+_8433172_8446622_RNA45SN1_45S_50ntFlanks</td><td>11641</td><td>11658</td><td>18</td><td>CGGCGGGTGTTGACGCGA</td><td>1577.1</td><td>-14.5</td><td>1.6</td><td>-9.2</td><td>4.7E-20</td><td>7.4E-19</td><td>MB231/MB468</td><td>MP</td></tr>
<tr><td>>MI0000477|hsa-mir-146a&WithFlank&5|+|160485346|160485456@27.48.22(+1A)&[MIMAT0000449&hsa-miR-146a-5p&offsets|0|0(+1A);m-129&5|+|160485372|160485393&offsets|0|0(+1A)]|TGAGAACTGAATTCCATGGGTTA</td><td>>MI0000477|hsa-mir-146a&WithFlank&5|+|160485346|160485456</td><td>27</td><td>48</td><td>22(+1A)</td><td>TGAGAACTGAATTCCATGGGTTA</td><td>1815.0</td><td>14.0</td><td>1.5</td><td>9.2</td><td>4.7E-20</td><td>7.4E-19</td><td>MB231/MB468</td><td>MP</td></tr>
<tr><td>>hg38_21_+_8433172_8446622_RNA45SN1_45S_50ntFlanks@11576.11602.27|CTTAGAACTGGTGCGGACCAGGGGAAT</td><td>>hg38_21_+_8433172_8446622_RNA45SN1_45S_50ntFlanks</td><td>11576</td><td>11602</td><td>27</td><td>CTTAGAACTGGTGCGGACCAGGGGAAT</td><td>610.2</td><td>-13.1</td><td>1.4</td><td>-9.2</td><td>4.8E-20</td><td>7.4E-19</td><td>MB231/MB468</td><td>MP</td></tr>
<tr><td>>hg38_21_+_8433172_8446622_RNA45SN1_45S_50ntFlanks@10362.10390.29|AAGGCCGAAGTGGAGAAGGGTTCCATGTG</td><td>>hg38_21_+_8433172_8446622_RNA45SN1_45S_50ntFlanks</td><td>10362</td><td>10390</td><td>29</td><td>AAGGCCGAAGTGGAGAAGGGTTCCATGTG</td><td>1878.4</td><td>-10.5</td><td>1.1</td><td>-9.1</td><td>7.5E-20</td><td>7.4E-19</td><td>MB231/MB468</td><td>MITO</td></tr>
<tr><td>>hg38_21_+_8433172_8446622_RNA45SN1_45S_50ntFlanks@11235.11253.19|CTTCCCGTGGATCGCCCCA</td><td>>hg38_21_+_8433172_8446622_RNA45SN1_45S_50ntFlanks</td><td>11235</td><td>11253</td><td>19</td><td>CTTCCCGTGGATCGCCCCA</td><td>585.6</td><td>-13.1</td><td>1.4</td><td>-9.2</td><td>4.8E-20</td><td>7.5E-19</td><td>MB231/MB468</td><td>MP</td></tr>
<tr><td>>MI0000279|hsa-mir-196a-2&WithFlank&12|+|53991732|53991853@31.51.21;MI0000238|hsa-mir-196a-1&WithFlank&17|-|48632484|48632565@13.33.21&[MIMAT0000226&hsa-miR-196a-5p&offsets|0|-1;m-261&12|+|53991762|53991783&offsets|0|-1];[MIMAT0000226_1&hsa-miR-196a-5p&offsets|0|-1;m-268&17|-|48632532|48632553&offsets|0|-1]|TAGGTAGTTTCATGTTGTTGG</td><td>>MI0000279|hsa-mir-196a-2&WithFlank&12|+|53991732|53991853</td><td>31</td><td>51</td><td>21</td><td>TAGGTAGTTTCATGTTGTTGG</td><td>2871.4</td><td>7.2</td><td>0.8</td><td>9.1</td><td>1.1E-19</td><td>7.5E-19</td><td>BT20/MB468</td><td>CYTO</td></tr>
<tr><td>>hg38_21_+_8433172_8446622_RNA45SN1_45S_50ntFlanks@8317.8356.40|CACGAGACCGATAGTCAACAAGTACCGTAAGGGAAAGTTG</td><td>>hg38_21_+_8433172_8446622_RNA45SN1_45S_50ntFlanks</td><td>8317</td><td>8356</td><td>40</td><td>CACGAGACCGATAGTCAACAAGTACCGTAAGGGAAAGTTG</td><td>1307.8</td><td>-6.5</td><td>0.7</td><td>-9.1</td><td>7.6E-20</td><td>7.5E-19</td><td>MB231/MB468</td><td>MITO</td></tr>
<tr><td>>hg38_21_+_8433172_8446622_RNA45SN1_45S_50ntFlanks@11561.11594.34|GCGCCTAGCAGCCGACTTAGAACTGGTGCGGACC</td><td>>hg38_21_+_8433172_8446622_RNA45SN1_45S_50ntFlanks</td><td>11561</td><td>11594</td><td>34</td><td>GCGCCTAGCAGCCGACTTAGAACTGGTGCGGACC</td><td>749.0</td><td>-5.9</td><td>0.6</td><td>-9.1</td><td>7.6E-20</td><td>7.5E-19</td><td>MB231/MB468</td><td>MITO</td></tr>
<tr><td>>hg38_21_+_8433172_8446622_RNA45SN1_45S_50ntFlanks@10690.10710.21|CGTCCGGTGAGCTCTCGCTGG</td><td>>hg38_21_+_8433172_8446622_RNA45SN1_45S_50ntFlanks</td><td>10690</td><td>10710</td><td>21</td><td>CGTCCGGTGAGCTCTCGCTGG</td><td>847.3</td><td>-13.5</td><td>1.5</td><td>-9.1</td><td>7.7E-20</td><td>7.6E-19</td><td>MB231/MB468</td><td>MITO</td></tr>
<tr><td>>MI0000471|hsa-mir-126&WithFlank&9|+|136670596|136670692@57.79.23&[MIMAT0000445&hsa-miR-126-3p&offsets|-1|0;m-40&9|+|136670653|136670674&offsets|-1|0]|CTCGTACCGTGAGTAATAATGCG</td><td>>MI0000471|hsa-mir-126&WithFlank&9|+|136670596|136670692</td><td>57</td><td>79</td><td>23</td><td>CTCGTACCGTGAGTAATAATGCG</td><td>478.4</td><td>10.5</td><td>1.1</td><td>9.1</td><td>6.7E-20</td><td>7.6E-19</td><td>MB231/MB468</td><td>NUC</td></tr>
<tr><td>>MI0000809|hsa-mir-151a&WithFlank&8|-|140732558|140732659@51.73.23&[MIMAT0000757&hsa-miR-151a-3p&offsets|-2|0;m-30&8|-|140732587|140732607&offsets|-2|0]|TACTAGACTGAAGCTCCTTGAGG</td><td>>MI0000809|hsa-mir-151a&WithFlank&8|-|140732558|140732659</td><td>51</td><td>73</td><td>23</td><td>TACTAGACTGAAGCTCCTTGAGG</td><td>3654.9</td><td>5.3</td><td>0.6</td><td>9.1</td><td>6.6E-20</td><td>7.6E-19</td><td>MB231/MB468</td><td>NUC</td></tr>
<tr><td>>hg38_21_+_8433172_8446622_RNA45SN1_45S_50ntFlanks@4759.4793.35|ATCAGATACCGTCGTAGTTCCGACCATAAACGATG</td><td>>hg38_21_+_8433172_8446622_RNA45SN1_45S_50ntFlanks</td><td>4759</td><td>4793</td><td>35</td><td>ATCAGATACCGTCGTAGTTCCGACCATAAACGATG</td><td>577.3</td><td>-12.1</td><td>1.3</td><td>-9.1</td><td>6.7E-20</td><td>7.6E-19</td><td>MB231/MB468</td><td>NUC</td></tr>
<tr><td>>hg38_21_+_8433172_8446622_RNA45SN1_45S_50ntFlanks@8378.8402.25|TTCAAGAGGGCGTGAAACCGTTAAG</td><td>>hg38_21_+_8433172_8446622_RNA45SN1_45S_50ntFlanks</td><td>8378</td><td>8402</td><td>25</td><td>TTCAAGAGGGCGTGAAACCGTTAAG</td><td>638.4</td><td>-13.2</td><td>1.4</td><td>-9.2</td><td>4.9E-20</td><td>7.6E-19</td><td>MB231/MB468</td><td>MP</td></tr>
<tr><td>>trnaMT_GluTTC_MT_-_14674_14742@35.72.38,trnalookalike8_GluTTC_5_-_93905172_93905240@35.72.38|TATCATTGGTCGTGGTTGTAGTCCGTGCGAGAATACCA</td><td>>trnaMT_GluTTC_MT_-_14674_14742</td><td>35</td><td>72</td><td>38</td><td>TATCATTGGTCGTGGTTGTAGTCCGTGCGAGAATACCA</td><td>12984.0</td><td>10.9</td><td>1.2</td><td>9.2</td><td>4.2E-20</td><td>7.6E-19</td><td>MB231/MB468</td><td>CYTO</td></tr>
<tr><td>>hg38_21_+_8433172_8446622_RNA45SN1_45S_50ntFlanks@10683.10712.30|CCGGCGGCGTCCGGTGAGCTCTCGCTGGCC</td><td>>hg38_21_+_8433172_8446622_RNA45SN1_45S_50ntFlanks</td><td>10683</td><td>10712</td><td>30</td><td>CCGGCGGCGTCCGGTGAGCTCTCGCTGGCC</td><td>4035.5</td><td>-5.4</td><td>0.6</td><td>-9.1</td><td>7.8E-20</td><td>7.6E-19</td><td>MB231/MB468</td><td>MITO</td></tr>
<tr><td>>hg38_21_+_8433172_8446622_RNA45SN1_45S_50ntFlanks@11876.11910.35|AAGACCCTGTTGAGCTTGACTCTAGTCTGGCACGG</td><td>>hg38_21_+_8433172_8446622_RNA45SN1_45S_50ntFlanks</td><td>11876</td><td>11910</td><td>35</td><td>AAGACCCTGTTGAGCTTGACTCTAGTCTGGCACGG</td><td>625.4</td><td>-13.1</td><td>1.4</td><td>-9.1</td><td>7.8E-20</td><td>7.6E-19</td><td>MB231/MB468</td><td>MITO</td></tr>
<tr><td>>hg38_21_+_8433172_8446622_RNA45SN1_45S_50ntFlanks@10788.10818.31|CAGCCTCTGGCATGTTGGAACAATGTAGGTA</td><td>>hg38_21_+_8433172_8446622_RNA45SN1_45S_50ntFlanks</td><td>10788</td><td>10818</td><td>31</td><td>CAGCCTCTGGCATGTTGGAACAATGTAGGTA</td><td>4469.4</td><td>-7.5</td><td>0.8</td><td>-9.1</td><td>7.8E-20</td><td>7.6E-19</td><td>MB231/MB468</td><td>MITO</td></tr>
<tr><td>>hg38_21_+_8433172_8446622_RNA45SN1_45S_50ntFlanks@9198.9223.26|GTTCTCTCGGGGCCACGCGCGCGTCC</td><td>>hg38_21_+_8433172_8446622_RNA45SN1_45S_50ntFlanks</td><td>9198</td><td>9223</td><td>26</td><td>GTTCTCTCGGGGCCACGCGCGCGTCC</td><td>937.6</td><td>-13.7</td><td>1.5</td><td>-9.1</td><td>7.8E-20</td><td>7.6E-19</td><td>MB231/MB468</td><td>MITO</td></tr>
</table>

| | | | | | | | | | | | | | |
|---|---|---|---|---|---|---|---|---|---|---|---|---|---|
| >hg38_21_+_8433172_8446622_RNA45SN1_45S_50ntFlanks@12939.12956.18\|CCATTCGTAGACGACCTG | >hg38_21_+_8433172_8446622_RNA45SN1_45S_50ntFlanks | 12939 | 12956 | 18 | CCATTCGTAGACGACCTG | 1029.6 | -5.0 | 0.5 | -9.1 | 6.7E-20 | 7.7E-19 | MB231/MB468 | NUC |
| >MI0000816\|hsa-mir-335&WithFlank&7\|+\|130496105\|130496210@22.43.22&[MIMAT0000765&hsa-miR-335-5p&offsets\|0\|-1;m-249&7\|+\|130496126\|130496147&offsets\|0\|0]\|TCAAGAGCAATAACGAAAAATG | >MI0000816\|hsa-mir-335&WithFlank&7\|+\|130496105\|130496210 | 22 | 43 | 22 | TCAAGAGCAATAACGAAAAATG | 262.3 | 5.8 | 0.6 | 9.1 | 6.8E-20 | 7.7E-19 | MB231/MB468 | NUC |
| >am_trna10_LysCTT_16_+_3241501_3241573@31.51.21,trna119_LysCTT_1_-_145395522_145395594@31.51.21,trna11_LysCTT_5_-_180648979_180649051@31.51.21,trna13_LysCTT_16_+_3246153_3246225@31.51.21,trna13_LysCTT_6_+_26556774_26556846@31.51.21,trna30_LysCTT_16_-_3230555_3230627@31.51.21,trna5_LysCTT_16_+_3214939_3215011@31.51.21,trna7_LysCTT_16_+_3225692_3225764@31.51.21,trna8_LysCTT_1_+_39970195_39970267@31.51.21,trna9_LysCTT_5_+_180634755_180634827@31.51.21\|ACTCTTAATCTCAGGGTCGTG | >am_trna10_LysCTT_16_+_3241501_3241573 | 31 | 51 | 21 | ACTCTTAATCTCAGGGTCGTG | 455.3 | -12.7 | 1.4 | -9.2 | 5.0E-20 | 7.8E-19 | MB231/MB468 | MP |
| >MI0000255\|hsa-mir-30d&WithFlank&8\|-\|134804870\|134804951@12.35.24&[MIMAT0000245&hsa-miR-30d-5p&offsets\|0\|+2;m-21&8\|-\|134804919\|134804940&offsets\|0\|+2]\|TGTAAACATCCCCGACTGGAAGCT | >MI0000255\|hsa-mir-30d&WithFlank&8\|-\|134804870\|134804951 | 12 | 35 | 24 | TGTAAACATCCCCGACTGGAAGCT | 17254.7 | 3.4 | 0.4 | 9.0 | 2.1E-19 | 7.8E-19 | BT20/MB468 | NUC |
| >hg38_21_+_8433172_8446622_RNA45SN1_45S_50ntFlanks@5420.5444.25\|CCGATTGGATGGTTTAGTGAGGCCC | >hg38_21_+_8433172_8446622_RNA45SN1_45S_50ntFlanks | 5420 | 5444 | 25 | CCGATTGGATGGTTTAGTGAGGCCC | 1110.9 | -14.5 | 1.6 | -9.0 | 2.1E-19 | 7.9E-19 | BT20/MB468 | NUC |
| >hg38_21_+_8433172_8446622_RNA45SN1_45S_50ntFlanks@5478.5516.39\|CTGGCGGAGCGCTGAGAAGACGGTCGAACTTGACTATCT | >hg38_21_+_8433172_8446622_RNA45SN1_45S_50ntFlanks | 5478 | 5516 | 39 | CTGGCGGAGCGCTGAGAAGACGGTCGAACTTGACTATCT | 487.4 | 12.9 | 1.4 | 9.2 | 4.4E-20 | 7.9E-19 | MB231/MB468 | CYTO |
| >hg38_21_+_8433172_8446622_RNA45SN1_45S_50ntFlanks@8324.8356.33\|CCGATAGTCAACAAGTACCGTAAGGGAAAGTTG | >hg38_21_+_8433172_8446622_RNA45SN1_45S_50ntFlanks | 8324 | 8356 | 33 | CCGATAGTCAACAAGTACCGTAAGGGAAAGTTG | 410.9 | -12.5 | 1.4 | -9.2 | 5.1E-20 | 7.9E-19 | MB231/MB468 | MP |
| >hg38_21_+_8433172_8446622_RNA45SN1_45S_50ntFlanks@7987.8006.20\|TCAGACGTGGCGACCCGCTG | >hg38_21_+_8433172_8446622_RNA45SN1_45S_50ntFlanks | 7987 | 8006 | 20 | TCAGACGTGGCGACCCGCTG | 1146.4 | -14.0 | 1.5 | -9.2 | 5.1E-20 | 7.9E-19 | MB231/MB468 | MP |
| >hg38_21_+_8433172_8446622_RNA45SN1_45S_50ntFlanks@8357.8387.31\|AAAAGAACTTTGAAGAGAGAGTTCAAGAGGG | >hg38_21_+_8433172_8446622_RNA45SN1_45S_50ntFlanks | 8357 | 8387 | 31 | AAAAGAACTTTGAAGAGAGAGTTCAAGAGGG | 890.1 | -13.7 | 1.5 | -9.2 | 5.1E-20 | 7.9E-19 | MB231/MB468 | MP |
| >hg38_21_+_8433172_8446622_RNA45SN1_45S_50ntFlanks@12654.12675.22\|TATGACTGAACGCCTCTAAGTC | >hg38_21_+_8433172_8446622_RNA45SN1_45S_50ntFlanks | 12654 | 12675 | 22 | TATGACTGAACGCCTCTAAGTC | 10867.1 | -13.0 | 1.4 | -9.0 | 2.1E-19 | 8.0E-19 | BT20/MB468 | NUC |
| >MI0000734\|hsa-mir-106b&WithFlank&7\|-\|100093987\|100094080@56.77.22&[MIMAT0004672&hsa-miR-106b-3p&offsets\|-2\|-2;m-87&7\|-\|100094003\|100094023&offsets\|-2\|-1]\|TACCGCACTGTGGGTACTTGCT | >MI0000734\|hsa-mir-106b&WithFlank&7\|-\|100093987\|100094080 | 56 | 77 | 22 | TACCGCACTGTGGGTACTTGCT | 855.9 | 7.3 | 0.8 | 9.1 | 7.1E-20 | 8.0E-19 | MB231/MB468 | NUC |
| >MI0000283\|hsa-mir-203a&WithFlank&14\|+\|104117399\|104117520@72.93.22&[MIMAT0000264&hsa-miR-203a-3p&offsets\|+1\|+1;m-79&14\|+\|104117469\|104117490&offsets\|+1\|+1]\|TGAAATGTTTAGGACCACTAGA | >MI0000283\|hsa-mir-203a&WithFlank&14\|+\|104117399\|104117520 | 72 | 93 | 22 | TGAAATGTTTAGGACCACTAGA | 583.2 | 13.0 | 1.4 | 9.4 | 8.0E-21 | 8.0E-19 | BT20/MB231 | NUC |
| >hg38_21_+_8433172_8446622_RNA45SN1_45S_50ntFlanks@12158.12181.24\|GTTTGACTGGGGCGGTACACCTGT | >hg38_21_+_8433172_8446622_RNA45SN1_45S_50ntFlanks | 12158 | 12181 | 24 | GTTTGACTGGGGCGGTACACCTGT | 470.5 | -12.7 | 1.4 | -9.2 | 5.2E-20 | 8.0E-19 | MB231/MB468 | MP |
| >hg38_21_+_8433172_8446622_RNA45SN1_45S_50ntFlanks@12157.12183.27\|AGTTTGACTGGGGCGGTACACCTGTCA | >hg38_21_+_8433172_8446622_RNA45SN1_45S_50ntFlanks | 12157 | 12183 | 27 | AGTTTGACTGGGGCGGTACACCTGTCA | 801.3 | -13.5 | 1.5 | -9.2 | 5.2E-20 | 8.1E-19 | MB231/MB468 | MP |
| >am_MI0008190\|hsa-mir-320d-1&WithFlank&13\|-\|40727810\|40727893@48.66.19;MI0008192\|hsa-mir-320d-2&WithFlank&X\|-\|140926154\|140926237@48.66.19&[MIMAT0006764&hsa-miR-320d&offsets\|0\|0;m-246&13\|-\|40727829\|40727846&offsets\|0\|+1];[MIMAT0006764_1&hsa-miR-320d&offsets\|0\|0;m-253&X\|-\|140926173\|140926190&offsets\|0\|+1]\|AAAAGCTGGGTTGAGAGGA | >am_MI0008190\|hsa-mir-320d-1&WithFlank&13\|-\|40727810\|40727893 | 48 | 66 | 19 | AAAAGCTGGGTTGAGAGGA | 305.6 | 12.5 | 1.4 | 9.1 | 1.1E-19 | 8.1E-19 | BT20/MB468 | CYTO |
| >hg38_21_+_8433172_8446622_RNA45SN1_45S_50ntFlanks@10107.10127.21\|CGTGTGGTGTGCGTCGGAGGG | >hg38_21_+_8433172_8446622_RNA45SN1_45S_50ntFlanks | 10107 | 10127 | 21 | CGTGTGGTGTGCGTCGGAGGG | 720.1 | -13.3 | 1.5 | -9.1 | 8.3E-20 | 8.1E-19 | MB231/MB468 | MITO |
| >hg38_21_+_8433172_8446622_RNA45SN1_45S_50ntFlanks@12957.12990.34\|CTTCTGGGTCGGGGTTTCGTACGTAGCAGAGCAG | >hg38_21_+_8433172_8446622_RNA45SN1_45S_50ntFlanks | 12957 | 12990 | 34 | CTTCTGGGTCGGGGTTTCGTACGTAGCAGAGCAG | 696.8 | -13.2 | 1.5 | -9.1 | 8.4E-20 | 8.1E-19 | MB231/MB468 | MITO |
| >hg38_21_+_8433172_8446622_RNA45SN1_45S_50ntFlanks@12627.12652.26\|ATGGGGCGAAGCTACCATCTGTGGGA | >hg38_21_+_8433172_8446622_RNA45SN1_45S_50ntFlanks | 12627 | 12652 | 26 | ATGGGGCGAAGCTACCATCTGTGGGA | 1518.4 | -14.4 | 1.6 | -9.2 | 5.3E-20 | 8.2E-19 | MB231/MB468 | MP |
| >hg38_21_+_8433172_8446622_RNA45SN1_45S_50ntFlanks@10727.10757.31\|GGAGAGGGTGTAAATCTCGCGCCGGGCCGTA | >hg38_21_+_8433172_8446622_RNA45SN1_45S_50ntFlanks | 10727 | 10757 | 31 | GGAGAGGGTGTAAATCTCGCGCCGGGCCGTA | 903.3 | -12.8 | 1.4 | -9.1 | 7.2E-20 | 8.2E-19 | MB231/MB468 | NUC |
| >hg38_21_+_8433172_8446622_RNA45SN1_45S_50ntFlanks@11821.11844.24\|CTATCCAGCGAAACCACAGCCAAG | >hg38_21_+_8433172_8446622_RNA45SN1_45S_50ntFlanks | 11821 | 11844 | 24 | CTATCCAGCGAAACCACAGCCAAG | 630.2 | -13.1 | 1.4 | -9.1 | 8.5E-20 | 8.2E-19 | MB231/MB468 | MITO |
| >hg38_21_+_8433172_8446622_RNA45SN1_45S_50ntFlanks@11918.11950.33\|ACATGAGAGGTGTAGAATAAGTGGGAGGCCCCC | >hg38_21_+_8433172_8446622_RNA45SN1_45S_50ntFlanks | 11918 | 11950 | 33 | ACATGAGAGGTGTAGAATAAGTGGGAGGCCCCC | 1900.8 | -6.9 | 0.8 | -9.2 | 4.6E-20 | 8.2E-19 | MB231/MB468 | CYTO |
| >MI0000809\|hsa-mir-151a&WithFlank&8\|-\|140732558\|140732659@51.72.22(+1U)&[MIMAT0000757&hsa-miR-151a-3p&offsets\|-2\|-1(+1U);m-30&8\|-\|140732587\|140732607&offsets\|-2\|-1(+1U)]\|TACTAGACTGAAGCTCCTTGAGT | >MI0000809\|hsa-mir-151a&WithFlank&8\|-\|140732558\|140732659 | 51 | 72 | 22(+1U) | TACTAGACTGAAGCTCCTTGAGT | 553.0 | 6.4 | 0.7 | 9.1 | 7.3E-20 | 8.3E-19 | MB231/MB468 | NUC |
| >hg38_21_+_8433172_8446622_RNA45SN1_45S_50ntFlanks@9572.9598.27\|CAAATCGGTCGTCCGACCTGGGTATAG | >hg38_21_+_8433172_8446622_RNA45SN1_45S_50ntFlanks | 9572 | 9598 | 27 | CAAATCGGTCGTCCGACCTGGGTATAG | 606.7 | -13.0 | 1.4 | -9.1 | 8.5E-20 | 8.3E-19 | MB231/MB468 | MITO |
| >hg38_21_+_8433172_8446622_RNA45SN1_45S_50ntFlanks@11641.11677.37\|CGGCGGGTGTTGACGCGATGTGATTTCTGCCCAGTGC | >hg38_21_+_8433172_8446622_RNA45SN1_45S_50ntFlanks | 11641 | 11677 | 37 | CGGCGGGTGTTGACGCGATGTGATTTCTGCCCAGTGC | 836.3 | -13.5 | 1.5 | -9.1 | 8.6E-20 | 8.3E-19 | MB231/MB468 | MITO |
| >hg38_21_+_8433172_8446622_RNA45SN1_45S_50ntFlanks@6663.6689.27\|GTGGATCACTCGGCTCGTGCGTCGATG | >hg38_21_+_8433172_8446622_RNA45SN1_45S_50ntFlanks | 6663 | 6689 | 27 | GTGGATCACTCGGCTCGTGCGTCGATG | 7649.5 | -9.8 | 1.1 | -9.2 | 5.4E-20 | 8.3E-19 | MB231/MB468 | MP |
| >hg38_21_+_8433172_8446622_RNA45SN1_45S_50ntFlanks@12622.12654.33\|AGCCAATGGGGCGAAGCTACCATCTGTGGGATT | >hg38_21_+_8433172_8446622_RNA45SN1_45S_50ntFlanks | 12622 | 12654 | 33 | AGCCAATGGGGCGAAGCTACCATCTGTGGGATT | 428.2 | -12.6 | 1.4 | -9.2 | 5.4E-20 | 8.3E-19 | MB231/MB468 | MP |
| >hg38_21_+_8433172_8446622_RNA45SN1_45S_50ntFlanks@9621.9658.38\|TCTAGTAGCTGGTTCCCTCCGAAGTTTCCCTCAGGATA | >hg38_21_+_8433172_8446622_RNA45SN1_45S_50ntFlanks | 9621 | 9658 | 38 | TCTAGTAGCTGGTTCCCTCCGAAGTTTCCCTCAGGATA | 412.5 | -13.1 | 1.5 | -9.0 | 2.2E-19 | 8.3E-19 | BT20/MB468 | NUC |
| >hg38_21_+_8433172_8446622_RNA45SN1_45S_50ntFlanks@10678.10706.29\|CGGTTCCGGCGGCGTCCGGTGAGCTCTCG | >hg38_21_+_8433172_8446622_RNA45SN1_45S_50ntFlanks | 10678 | 10706 | 29 | CGGTTCCGGCGGCGTCCGGTGAGCTCTCG | 22217.8 | -8.5 | 0.9 | -9.2 | 4.7E-20 | 8.4E-19 | MB231/MB468 | CYTO |
| >hg38_21_+_8433172_8446622_RNA45SN1_45S_50ntFlanks@11315.11336.22\|CCACCCCACGTCTCGTCGCGCG | >hg38_21_+_8433172_8446622_RNA45SN1_45S_50ntFlanks | 11315 | 11336 | 22 | CCACCCCACGTCTCGTCGCGCG | 1308.8 | -14.1 | 1.6 | -9.1 | 8.7E-20 | 8.4E-19 | MB231/MB468 | MITO |
| >hg38_21_+_8433172_8446622_RNA45SN1_45S_50ntFlanks@9929.9960.32\|GATATAGACAGCAGGACGGTGGCCATGGAAGT | >hg38_21_+_8433172_8446622_RNA45SN1_45S_50ntFlanks | 9929 | 9960 | 32 | GATATAGACAGCAGGACGGTGGCCATGGAAGT | 923.6 | -6.1 | 0.7 | -9.2 | 5.5E-20 | 8.4E-19 | MB231/MB468 | MP |
| >hg38_21_+_8433172_8446622_RNA45SN1_45S_50ntFlanks@10794.10818.25\|CTGGCATGTTGGAACAATGTAGGTA | >hg38_21_+_8433172_8446622_RNA45SN1_45S_50ntFlanks | 10794 | 10818 | 25 | CTGGCATGTTGGAACAATGTAGGTA | 1502.9 | -5.2 | 0.6 | -9.1 | 8.9E-20 | 8.5E-19 | BT20/MB468 | MITO |
| >MI0003188\|hsa-mir-503&WithFlank&X\|-\|134546322\|134546404@12.31.20(+3A)&[MIMAT0002874&hsa-miR-503-5p&offsets\|0\|-3(+3A);m-297&X\|-\|134546371\|134546393&offsets\|0\|-3(+3A)]\|TAGCAGCGGGAACAGTTCTGAAA | >MI0003188\|hsa-mir-503&WithFlank&X\|-\|134546322\|134546404 | 12 | 31 | 20(+3A) | TAGCAGCGGGAACAGTTCTGAAA | 360.1 | 12.7 | 1.4 | 9.1 | 1.2E-19 | 8.5E-19 | BT20/MB468 | CYTO |
| >hg38_21_+_8433172_8446622_RNA45SN1_45S_50ntFlanks@12274.12293.20\|CAGTACGAATACAGACCGTG | >hg38_21_+_8433172_8446622_RNA45SN1_45S_50ntFlanks | 12274 | 12293 | 20 | CAGTACGAATACAGACCGTG | 1151.7 | -14.0 | 1.5 | -9.2 | 5.5E-20 | 8.5E-19 | MB231/MB468 | MP |
| >hg38_21_+_8433172_8446622_RNA45SN1_45S_50ntFlanks@4679.4720.42\|GGCGCAAGACGGACCAGAGCGAAAGCATTTGCCAAGAATGTT | >hg38_21_+_8433172_8446622_RNA45SN1_45S_50ntFlanks | 4679 | 4720 | 42 | GGCGCAAGACGGACCAGAGCGAAAGCATTTGCCAAGAATGTT | 852.0 | 13.7 | 1.5 | 9.2 | 4.8E-20 | 8.5E-19 | MB231/MB468 | CYTO |
| >hg38_21_+_8433172_8446622_RNA45SN1_45S_50ntFlanks@6789.6806.18\|GCCTGTCTGAGCGTCGCT | >hg38_21_+_8433172_8446622_RNA45SN1_45S_50ntFlanks | 6789 | 6806 | 18 | GCCTGTCTGAGCGTCGCT | 3544.3 | 5.9 | 0.6 | 9.2 | 4.8E-20 | 8.5E-19 | MB231/MB468 | CYTO |
| >hg38_21_+_8433172_8446622_RNA45SN1_45S_50ntFlanks@11270.11293.24\|CGGCCGCCCCCGGGGAGCCCGGCG | >hg38_21_+_8433172_8446622_RNA45SN1_45S_50ntFlanks | 11270 | 11293 | 24 | CGGCCGCCCCCGGGGAGCCCGGCG | 2164.1 | -14.9 | 1.6 | -9.2 | 5.6E-20 | 8.6E-19 | MB231/MB468 | MP |
| >hg38_21_+_8433172_8446622_RNA45SN1_45S_50ntFlanks@12416.12457.42\|TTCGATGTCGGCTCTTCCTATCATTGTGAAGCAGAATTCACC | >hg38_21_+_8433172_8446622_RNA45SN1_45S_50ntFlanks | 12416 | 12457 | 42 | TTCGATGTCGGCTCTTCCTATCATTGTGAAGCAGAATTCACC | 2384.0 | 9.1 | 1.0 | 9.2 | 4.9E-20 | 8.6E-19 | MB231/MB468 | CYTO |
| >hg38_21_+_8433172_8446622_RNA45SN1_45S_50ntFlanks@9575.9603.29\|ATCGGTCGTCCGACCTGGGTATAGGGGCG | >hg38_21_+_8433172_8446622_RNA45SN1_45S_50ntFlanks | 9575 | 9603 | 29 | ATCGGTCGTCCGACCTGGGTATAGGGGCG | 1049.1 | -12.1 | 1.3 | -9.1 | 9.0E-20 | 8.6E-19 | BT20/MB468 | MITO |
| >hg38_21_+_8433172_8446622_RNA45SN1_45S_50ntFlanks@10101.10125.25\|CGCGCGCGTGTGGTGTGCGTCGGAG | >hg38_21_+_8433172_8446622_RNA45SN1_45S_50ntFlanks | 10101 | 10125 | 25 | CGCGCGCGTGTGGTGTGCGTCGGAG | 977.3 | -13.8 | 1.5 | -9.2 | 5.6E-20 | 8.6E-19 | MB231/MB468 | MP |
| >hg38_1_-_228634819_228635039_RNA5S12_5S_50ntFlanks@129.170.42\|TTGGATGGGAGACCGCCTGGGAATACCGGGTGCTGTAGGCTT | >hg38_1_-_228634819_228635039_RNA5S12_5S_50ntFlanks | 129 | 170 | 42 | TTGGATGGGAGACCGCCTGGGAATACCGGGTGCTGTAGGCTT | 6496.9 | 10.3 | 1.1 | 9.2 | 4.9E-20 | 8.7E-19 | MB231/MB468 | CYTO |
| >hg38_21_+_8433172_8446622_RNA45SN1_45S_50ntFlanks@11876.11904.29\|AAGACCCTGTTGAGCTTGACTCTAGTCTG | >hg38_21_+_8433172_8446622_RNA45SN1_45S_50ntFlanks | 11876 | 11904 | 29 | AAGACCCTGTTGAGCTTGACTCTAGTCTG | 969.1 | -13.8 | 1.5 | -9.1 | 5.7E-20 | 8.8E-19 | MB231/MB468 | MP |
| >hg38_21_+_8433172_8446622_RNA45SN1_45S_50ntFlanks@10794.10823.30\|CTGGCATGTTGGAACAATGTAGGTAAGGGA | >hg38_21_+_8433172_8446622_RNA45SN1_45S_50ntFlanks | 10794 | 10823 | 30 | CTGGCATGTTGGAACAATGTAGGTAAGGGA | 582.5 | -13.0 | 1.4 | -9.1 | 9.1E-20 | 8.8E-19 | MB231/MB468 | MITO |
| >trna152_ValCAC_6_-_27248049_27248121@1.36.36\|GCTTCTGTAGTGTAGTGGTTATCACGTTCGCCTCAC | >trna152_ValCAC_6_-_27248049_27248121 | 1 | 36 | 36 | GCTTCTGTAGTGTAGTGGTTATCACGTTCGCCTCAC | 2748.4 | 11.6 | 1.3 | 9.2 | 5.0E-20 | 8.8E-19 | MB231/MB468 | CYTO |
| >hg38_21_+_8433172_8446622_RNA45SN1_45S_50ntFlanks@12628.12653.26\|TGGGGCGAAGCTACCATCTGTGGGAT | >hg38_21_+_8433172_8446622_RNA45SN1_45S_50ntFlanks | 12628 | 12653 | 26 | TGGGGCGAAGCTACCATCTGTGGGAT | 794.2 | -12.6 | 1.4 | -9.1 | 7.8E-20 | 8.8E-19 | MB231/MB468 | NUC |
| >hg38_21_+_8433172_8446622_RNA45SN1_45S_50ntFlanks@12444.12486.43\|AAGCAGAATTCACCAAGCGTTGGATTGTTCACCCACTAATAGG | >hg38_21_+_8433172_8446622_RNA45SN1_45S_50ntFlanks | 12444 | 12486 | 43 | AAGCAGAATTCACCAAGCGTTGGATTGTTCACCCACTAATAGG | 1853.2 | -15.3 | 1.7 | -9.0 | 2.4E-19 | 8.9E-19 | BT20/MB468 | NUC |
| >hg38_21_+_8433172_8446622_RNA45SN1_45S_50ntFlanks@11768.11786.19\|CTAATTAGTGACGCGCATG | >hg38_21_+_8433172_8446622_RNA45SN1_45S_50ntFlanks | 11768 | 11786 | 19 | CTAATTAGTGACGCGCATG | 688.6 | -13.2 | 1.5 | -9.1 | 9.2E-20 | 8.9E-19 | MB231/MB468 | MITO |
| >hg38_21_+_8433172_8446622_RNA45SN1_45S_50ntFlanks@11885.11903.19\|TTGAGCTTGACTCTAGTCT | >hg38_21_+_8433172_8446622_RNA45SN1_45S_50ntFlanks | 11885 | 11903 | 19 | TTGAGCTTGACTCTAGTCT | 499.4 | -12.8 | 1.4 | -9.1 | 5.8E-20 | 8.9E-19 | MB231/MB468 | MP |
| >hg38_21_+_8433172_8446622_RNA45SN1_45S_50ntFlanks@9171.9197.27\|CGGGTCGCGCCGTCGGGCCCGGGGGAG | >hg38_21_+_8433172_8446622_RNA45SN1_45S_50ntFlanks | 9171 | 9197 | 27 | CGGGTCGCGCCGTCGGGCCCGGGGGAG | 1604.5 | -14.4 | 1.6 | -9.1 | 9.3E-20 | 9.0E-19 | MB231/MB468 | MITO |

<table>
<tr><th></th><th></th><th></th><th></th><th></th><th></th><th></th><th></th><th></th><th></th><th></th><th></th><th></th><th></th></tr>
<tr><td>>hg38_21_+_8433172_8446622_RNA45SN1_45S_50ntFlanks@12154.12179.26|GGGAGTTTGACTGGGGCGGTACACCT</td><td>>hg38_21_+_8433172_8446622_RNA45SN1_45S_50ntFlanks</td><td>12154</td><td>12179</td><td>26</td><td>GGGAGTTTGACTGGGGCGGTACACCT</td><td>588.7</td><td>-13.1</td><td>1.4</td><td>-9.1</td><td>5.9E-20</td><td>9.1E-19</td><td>MB231/MB468</td><td>MP</td></tr>
<tr><td>>hg38_21_+_8433172_8446622_RNA45SN1_45S_50ntFlanks@11270.11293.24|CGGCCGCCCCCGGGGAGCCCGGCG</td><td>>hg38_21_+_8433172_8446622_RNA45SN1_45S_50ntFlanks</td><td>11270</td><td>11293</td><td>24</td><td>CGGCCGCCCCCGGGGAGCCCGGCG</td><td>686.5</td><td>-12.1</td><td>1.3</td><td>-9.2</td><td>5.2E-20</td><td>9.1E-19</td><td>MB231/MB468</td><td>CYTO</td></tr>
<tr><td>>hg38_21_+_8433172_8446622_RNA45SN1_45S_50ntFlanks@9738.9766.29|GAAACGATCTCAACCTATTCTCAAACTTT</td><td>>hg38_21_+_8433172_8446622_RNA45SN1_45S_50ntFlanks</td><td>9738</td><td>9766</td><td>29</td><td>GAAACGATCTCAACCTATTCTCAAACTTT</td><td>2911.0</td><td>-7.5</td><td>0.8</td><td>-9.1</td><td>9.5E-20</td><td>9.1E-19</td><td>MB231/MB468</td><td>MITO</td></tr>
<tr><td>>hg38_21_+_8433172_8446622_RNA45SN1_45S_50ntFlanks@12007.12030.24|GCCGCCGGTGAAATACCACTACTC</td><td>>hg38_21_+_8433172_8446622_RNA45SN1_45S_50ntFlanks</td><td>12007</td><td>12030</td><td>24</td><td>GCCGCCGGTGAAATACCACTACTC</td><td>652.3</td><td>-13.2</td><td>1.4</td><td>-9.1</td><td>6.0E-20</td><td>9.1E-19</td><td>MB231/MB468</td><td>MP</td></tr>
<tr><td>>MI0000650|hsa-mir-200c&WithFlank&12|+|6963693|6963772@50.72.23(+1C)&[MIMAT0000617&hsa-miR-200c-3p&offsets|0|0(+1C);m-54&12|+|6963742|6963764&offsets|0|0(+1C)]|TAATACTGCCGGGTAATGATGGAC</td><td>>MI0000650|hsa-mir-200c&WithFlank&12|+|6963693|6963772</td><td>50</td><td>72</td><td>23(+1C)</td><td>TAATACTGCCGGGTAATGATGGAC</td><td>1583.8</td><td>10.0</td><td>1.1</td><td>9.4</td><td>4.3E-21</td><td>9.2E-19</td><td>BT20/MB231</td><td>CYTO</td></tr>
<tr><td>>MI0000650|hsa-mir-200c&WithFlank&12|+|6963693|6963772@50.72.23(+1U)&[MIMAT0000617&hsa-miR-200c-3p&offsets|0|0(+1U);m-54&12|+|6963742|6963764&offsets|0|0(+1U)]|TAATACTGCCGGGTAATGATGGAT</td><td>>MI0000650|hsa-mir-200c&WithFlank&12|+|6963693|6963772</td><td>50</td><td>72</td><td>23(+1U)</td><td>TAATACTGCCGGGTAATGATGGAT</td><td>7392.8</td><td>10.1</td><td>1.1</td><td>9.3</td><td>9.5E-21</td><td>9.2E-19</td><td>BT20/MB231</td><td>NUC</td></tr>
<tr><td>>hg38_21_+_8433172_8446622_RNA45SN1_45S_50ntFlanks@7980.8006.27|CCTCAGATCAGACGTGGCGACCCGCTG</td><td>>hg38_21_+_8433172_8446622_RNA45SN1_45S_50ntFlanks</td><td>7980</td><td>8006</td><td>27</td><td>CCTCAGATCAGACGTGGCGACCCGCTG</td><td>8426.5</td><td>-5.4</td><td>0.6</td><td>-9.1</td><td>9.7E-20</td><td>9.2E-19</td><td>BT20/MB468</td><td>MITO</td></tr>
<tr><td>>hg38_21_+_8433172_8446622_RNA45SN1_45S_50ntFlanks@5486.5518.33|GCGCTGAGAAGACGGTCGAACTTGACTATCTAG</td><td>>hg38_21_+_8433172_8446622_RNA45SN1_45S_50ntFlanks</td><td>5486</td><td>5518</td><td>33</td><td>GCGCTGAGAAGACGGTCGAACTTGACTATCTAG</td><td>523.5</td><td>-12.9</td><td>1.4</td><td>-9.1</td><td>6.1E-20</td><td>9.3E-19</td><td>MB231/MB468</td><td>MP</td></tr>
<tr><td>>hg38_21_+_8433172_8446622_RNA45SN1_45S_50ntFlanks@12615.12654.40|GCTGAGGAGCCAATGGGGCGAAGCTACCATCTGTGGGATT</td><td>>hg38_21_+_8433172_8446622_RNA45SN1_45S_50ntFlanks</td><td>12615</td><td>12654</td><td>40</td><td>GCTGAGGAGCCAATGGGGCGAAGCTACCATCTGTGGGATT</td><td>2647.3</td><td>-9.7</td><td>1.1</td><td>-9.1</td><td>6.1E-20</td><td>9.3E-19</td><td>MB231/MB468</td><td>MP</td></tr>
<tr><td>>hg38_21_+_8433172_8446622_RNA45SN1_45S_50ntFlanks@10574.10591.18|AAGCCGGCGGGAGCCCCG</td><td>>hg38_21_+_8433172_8446622_RNA45SN1_45S_50ntFlanks</td><td>10574</td><td>10591</td><td>18</td><td>AAGCCGGCGGGAGCCCCG</td><td>530.3</td><td>-12.9</td><td>1.4</td><td>-9.1</td><td>6.1E-20</td><td>9.3E-19</td><td>MB231/MB468</td><td>MP</td></tr>
<tr><td>>hg38_21_+_8433172_8446622_RNA45SN1_45S_50ntFlanks@11919.11948.30|CATGAGAGGTGTAGAATAAGTGGGAGGCCC</td><td>>hg38_21_+_8433172_8446622_RNA45SN1_45S_50ntFlanks</td><td>11919</td><td>11948</td><td>30</td><td>CATGAGAGGTGTAGAATAAGTGGGAGGCCC</td><td>1087.4</td><td>-14.0</td><td>1.5</td><td>-9.1</td><td>6.1E-20</td><td>9.3E-19</td><td>MB231/MB468</td><td>MP</td></tr>
<tr><td>>hg38_21_+_8433172_8446622_RNA45SN1_45S_50ntFlanks@5087.5123.37|ACGAGACTCTGGCATGCTAACTAGTTACGCGACCCCC</td><td>>hg38_21_+_8433172_8446622_RNA45SN1_45S_50ntFlanks</td><td>5087</td><td>5123</td><td>37</td><td>ACGAGACTCTGGCATGCTAACTAGTTACGCGACCCCC</td><td>268.7</td><td>12.1</td><td>1.3</td><td>9.2</td><td>5.3E-20</td><td>9.3E-19</td><td>MB231/MB468</td><td>CYTO</td></tr>
<tr><td>>MI0000469|hsa-mir-125a&WithFlank&19|+|51693248|51693345@21.43.23(+1U)&[MIMAT0000443&hsa-miR-125a-5p&offsets|0|-1(+1U);m-63&19|+|51693268|51693289&offsets|0|+1(+1U)]|TCCCTGAGACCCTTTAACCTGTGT</td><td>>MI0000469|hsa-mir-125a&WithFlank&19|+|51693248|51693345</td><td>21</td><td>43</td><td>23(+1U)</td><td>TCCCTGAGACCCTTTAACCTGTGT</td><td>2134.4</td><td>7.2</td><td>0.8</td><td>9.1</td><td>9.8E-20</td><td>9.4E-19</td><td>BT20/MB468</td><td>MITO</td></tr>
<tr><td>>MI0000266|hsa-mir-10a&WithFlank&17|-|48579832|48579953@29.50.22(+1A)&[MIMAT0000253&hsa-miR-10a-5p&offsets|+1|0(+1A);m-18&17|-|48579905|48579926&offsets|+1|+1(+1A)]|ACCCTGTAGATCCGAATTTGTGA</td><td>>MI0000266|hsa-mir-10a&WithFlank&17|-|48579832|48579953</td><td>29</td><td>50</td><td>22(+1A)</td><td>ACCCTGTAGATCCGAATTTGTGA</td><td>363.9</td><td>12.5</td><td>1.4</td><td>9.2</td><td>5.4E-20</td><td>9.4E-19</td><td>MB231/MB468</td><td>CYTO</td></tr>
<tr><td>>hg38_21_+_8433172_8446622_RNA45SN1_45S_50ntFlanks@5482.5514.33|CGGAGCGCTGAGAAGACGGTCGAACTTGACTAT</td><td>>hg38_21_+_8433172_8446622_RNA45SN1_45S_50ntFlanks</td><td>5482</td><td>5514</td><td>33</td><td>CGGAGCGCTGAGAAGACGGTCGAACTTGACTAT</td><td>1067.7</td><td>-13.9</td><td>1.5</td><td>-9.1</td><td>6.2E-20</td><td>9.5E-19</td><td>MB231/MB468</td><td>MP</td></tr>
<tr><td>>hg38_21_+_8433172_8446622_RNA45SN1_45S_50ntFlanks@5467.5484.18|GGCCCACGGCCCTGGCGG</td><td>>hg38_21_+_8433172_8446622_RNA45SN1_45S_50ntFlanks</td><td>5467</td><td>5484</td><td>18</td><td>GGCCCACGGCCCTGGCGG</td><td>750.5</td><td>-13.3</td><td>1.5</td><td>-9.1</td><td>1.0E-19</td><td>9.6E-19</td><td>MB231/MB468</td><td>MITO</td></tr>
<tr><td>>hg38_21_+_8433172_8446622_RNA45SN1_45S_50ntFlanks@6775.6806.32|CCTCCCGGGGCTACGCCTGTCTGAGCGTCGCT</td><td>>hg38_21_+_8433172_8446622_RNA45SN1_45S_50ntFlanks</td><td>6775</td><td>6806</td><td>32</td><td>CCTCCCGGGGCTACGCCTGTCTGAGCGTCGCT</td><td>592.3</td><td>5.8</td><td>0.6</td><td>9.1</td><td>1.4E-19</td><td>9.6E-19</td><td>BT20/MB468</td><td>CYTO</td></tr>
<tr><td>>am_trna117_GlyTCC_1_-_145397864_145397935@1.30.30,trna45_GlyTCC_1_+_161500903_161500974@1.30.30,trna70_GlyTCC_1_-_161439547_161439618@1.30.30,trna73_GlyTCC_1_-_161432166_161432237@1.30.30,trna76_GlyTCC_1_-_161424756_161424827@1.30.30,trna79_GlyTCC_1_-_161417375_161417446@1.30.30|GCGTTGGTGGTATAGTGGTGAGCATAGCTG</td><td>>am_trna117_GlyTCC_1_-_145397864_145397935</td><td>1</td><td>30</td><td>30</td><td>GCGTTGGTGGTATAGTGGTGAGCATAGCTG</td><td>2951.7</td><td>-11.0</td><td>1.2</td><td>-9.1</td><td>6.3E-20</td><td>9.6E-19</td><td>MB231/MB468</td><td>MP</td></tr>
<tr><td>>hg38_21_+_8433172_8446622_RNA45SN1_45S_50ntFlanks@11843.11860.18|AGGGAACGGGCTTGGCGG</td><td>>hg38_21_+_8433172_8446622_RNA45SN1_45S_50ntFlanks</td><td>11843</td><td>11860</td><td>18</td><td>AGGGAACGGGCTTGGCGG</td><td>1038.1</td><td>-13.9</td><td>1.5</td><td>-9.1</td><td>6.3E-20</td><td>9.6E-19</td><td>MB231/MB468</td><td>MP</td></tr>
<tr><td>>MI0000301|hsa-mir-224&WithFlank&X|-|151958572|151958664@14.34.21&[MIMAT0000281&hsa-miR-224-5p&offsets|+1|-3;m-238&X|-|151958630|151958651&offsets|0|-1]|CAAGTCACTAGTGGTTCCGTT</td><td>>MI0000301|hsa-mir-224&WithFlank&X|-|151958572|151958664</td><td>14</td><td>34</td><td>21</td><td>CAAGTCACTAGTGGTTCCGTT</td><td>325.7</td><td>8.0</td><td>0.9</td><td>9.0</td><td>2.6E-19</td><td>9.6E-19</td><td>BT20/MB468</td><td>NUC</td></tr>
<tr><td>>hg38_21_+_8433172_8446622_RNA45SN1_45S_50ntFlanks@4439.4471.33|CCGCCCCTTGCCTCTCGGCGCCCCCTCGATGCT</td><td>>hg38_21_+_8433172_8446622_RNA45SN1_45S_50ntFlanks</td><td>4439</td><td>4471</td><td>33</td><td>CCGCCCCTTGCCTCTCGGCGCCCCCTCGATGCT</td><td>576.3</td><td>-13.0</td><td>1.4</td><td>-9.1</td><td>1.0E-19</td><td>9.6E-19</td><td>MB231/MB468</td><td>MITO</td></tr>
<tr><td>>MI0000285|hsa-mir-205&WithFlank&1|+|209432127|209432248@40.60.21(+1U)&[MIMAT0000266&hsa-miR-205-5p&offsets|0|-1(+1U);m-351&1|+|209432166|209432187&offsets|0|-1(+1U)]|TCCTTCATTCCACCGGAGTCTT</td><td>>MI0000285|hsa-mir-205&WithFlank&1|+|209432127|209432248</td><td>40</td><td>60</td><td>21(+1U)</td><td>TCCTTCATTCCACCGGAGTCTT</td><td>1387.0</td><td>13.8</td><td>1.5</td><td>9.4</td><td>4.7E-21</td><td>9.6E-19</td><td>BT20/MB231</td><td>CYTO</td></tr>
<tr><td>>hg38_21_+_8433172_8446622_RNA45SN1_45S_50ntFlanks@6650.6676.27|ACGACTCTTAGCGGTGGATCACTCGGC</td><td>>hg38_21_+_8433172_8446622_RNA45SN1_45S_50ntFlanks</td><td>6650</td><td>6676</td><td>27</td><td>ACGACTCTTAGCGGTGGATCACTCGGC</td><td>6343.0</td><td>6.2</td><td>0.7</td><td>9.1</td><td>1.0E-19</td><td>9.6E-19</td><td>BT20/MB468</td><td>MITO</td></tr>
<tr><td>>hg38_21_+_8433172_8446622_RNA45SN1_45S_50ntFlanks@4283.4317.35|CTTTAACGAGGATCCATTGGAGGGCAAGTCTGGTG</td><td>>hg38_21_+_8433172_8446622_RNA45SN1_45S_50ntFlanks</td><td>4283</td><td>4317</td><td>35</td><td>CTTTAACGAGGATCCATTGGAGGGCAAGTCTGGTG</td><td>2004.0</td><td>-10.8</td><td>1.2</td><td>-9.1</td><td>1.0E-19</td><td>9.7E-19</td><td>MB231/MB468</td><td>MITO</td></tr>
<tr><td>>hg38_21_+_8433172_8446622_RNA45SN1_45S_50ntFlanks@6747.6770.24|ACTTCGAACGCACTTGCGGCCCCG</td><td>>hg38_21_+_8433172_8446622_RNA45SN1_45S_50ntFlanks</td><td>6747</td><td>6770</td><td>24</td><td>ACTTCGAACGCACTTGCGGCCCCG</td><td>1370.1</td><td>-10.3</td><td>1.1</td><td>-9.1</td><td>1.0E-19</td><td>9.7E-19</td><td>MB231/MB468</td><td>MITO</td></tr>
<tr><td>>hg38_21_+_8433172_8446622_RNA45SN1_45S_50ntFlanks@7985.8004.20|GATCAGACGTGGCGACCCGC</td><td>>hg38_21_+_8433172_8446622_RNA45SN1_45S_50ntFlanks</td><td>7985</td><td>8004</td><td>20</td><td>GATCAGACGTGGCGACCCGC</td><td>889.3</td><td>-13.7</td><td>1.5</td><td>-9.1</td><td>6.4E-20</td><td>9.7E-19</td><td>MB231/MB468</td><td>MP</td></tr>
<tr><td>>hg38_21_+_8433172_8446622_RNA45SN1_45S_50ntFlanks@12001.12043.43|CTGCGGGCCGCCGGTGAAATACCACTACTCTGATCGTTTTTTC</td><td>>hg38_21_+_8433172_8446622_RNA45SN1_45S_50ntFlanks</td><td>12001</td><td>12043</td><td>43</td><td>CTGCGGGCCGCCGGTGAAATACCACTACTCTGATCGTTTTTTC</td><td>1128.1</td><td>-14.6</td><td>1.6</td><td>-9.0</td><td>2.6E-19</td><td>9.8E-19</td><td>BT20/MB468</td><td>NUC</td></tr>
<tr><td>>hg38_21_+_8433172_8446622_RNA45SN1_45S_50ntFlanks@10869.10886.18|AGGGCTGGGTCGGTCGGG</td><td>>hg38_21_+_8433172_8446622_RNA45SN1_45S_50ntFlanks</td><td>10869</td><td>10886</td><td>18</td><td>AGGGCTGGGTCGGTCGGG</td><td>902.3</td><td>-13.7</td><td>1.5</td><td>-9.1</td><td>6.5E-20</td><td>9.8E-19</td><td>MB231/MB468</td><td>MP</td></tr>
<tr><td>>hg38_21_+_8433172_8446622_RNA45SN1_45S_50ntFlanks@8154.8175.22|CCGCTCGTGGGGGGCCCAAGTC</td><td>>hg38_21_+_8433172_8446622_RNA45SN1_45S_50ntFlanks</td><td>8154</td><td>8175</td><td>22</td><td>CCGCTCGTGGGGGGCCCAAGTC</td><td>868.8</td><td>-13.5</td><td>1.5</td><td>-9.1</td><td>1.0E-19</td><td>9.8E-19</td><td>MB231/MB468</td><td>MITO</td></tr>
<tr><td>>hg38_21_+_8433172_8446622_RNA45SN1_45S_50ntFlanks@475.513.39|CTTCGTGATCGATGTGGTGACGTCGTGCTCTCCCGGGCC</td><td>>hg38_21_+_8433172_8446622_RNA45SN1_45S_50ntFlanks</td><td>475</td><td>513</td><td>39</td><td>CTTCGTGATCGATGTGGTGACGTCGTGCTCTCCCGGGCC</td><td>338.1</td><td>12.7</td><td>1.4</td><td>9.1</td><td>1.4E-19</td><td>9.8E-19</td><td>BT20/MB468</td><td>CYTO</td></tr>
<tr><td>>MI0000088|hsa-mir-30a&WithFlank&6|-|71403545|71403627@53.74.22&[MIMAT0000088&hsa-miR-30a-3p&offsets|0|0;m-38&6|-|71403554|71403575&offsets|0|0]|CTTTCAGTCGGATGTTTGCAGC</td><td>>MI0000088|hsa-mir-30a&WithFlank&6|-|71403545|71403627</td><td>53</td><td>74</td><td>22</td><td>CTTTCAGTCGGATGTTTGCAGC</td><td>16303.2</td><td>5.4</td><td>0.6</td><td>9.1</td><td>8.8E-20</td><td>9.9E-19</td><td>MB231/MB468</td><td>NUC</td></tr>
<tr><td>>am_MI0000809|hsa-mir-151a&WithFlank&8|-|140732558|140732659@53.71.19&[MIMAT0000757&hsa-miR-151a-3p&offsets|0|-2;m-30&8|-|140732587|140732607&offsets|0|-2]|CTAGACTGAAGCTCCTTGA</td><td>>am_MI0000809|hsa-mir-151a&WithFlank&8|-|140732558|140732659</td><td>53</td><td>71</td><td>19</td><td>CTAGACTGAAGCTCCTTGA</td><td>501.0</td><td>6.2</td><td>0.7</td><td>9.1</td><td>8.9E-20</td><td>1.0E-18</td><td>MB231/MB468</td><td>NUC</td></tr>
<tr><td>>hg38_21_+_8433172_8446622_RNA45SN1_45S_50ntFlanks@10411.10431.21|CAGTCGGTCCTGAGAGATGGG</td><td>>hg38_21_+_8433172_8446622_RNA45SN1_45S_50ntFlanks</td><td>10411</td><td>10431</td><td>21</td><td>CAGTCGGTCCTGAGAGATGGG</td><td>2837.4</td><td>-11.0</td><td>1.2</td><td>-9.1</td><td>1.1E-19</td><td>1.0E-18</td><td>MB231/MB468</td><td>MITO</td></tr>
<tr><td>>hg38_21_+_8433172_8446622_RNA45SN1_45S_50ntFlanks@9171.9197.27|CGGGTCGCGCCGTCGGGCCCGGGGGAG</td><td>>hg38_21_+_8433172_8446622_RNA45SN1_45S_50ntFlanks</td><td>9171</td><td>9197</td><td>27</td><td>CGGGTCGCGCCGTCGGGCCCGGGGGAG</td><td>1311.0</td><td>-14.2</td><td>1.6</td><td>-9.1</td><td>6.7E-20</td><td>1.0E-18</td><td>MB231/MB468</td><td>MP</td></tr>
<tr><td>>MI0000809|hsa-mir-151a&WithFlank&8|-|140732558|140732659@53.74.22(+1A)&[MIMAT0000757&hsa-miR-151a-3p&offsets|0|+1(+1A);m-30&8|-|140732587|140732607&offsets|0|+1(+1A)]|CTAGACTGAAGCTCCTTGAGGAA</td><td>>MI0000809|hsa-mir-151a&WithFlank&8|-|140732558|140732659</td><td>53</td><td>74</td><td>22(+1A)</td><td>CTAGACTGAAGCTCCTTGAGGAA</td><td>2233.2</td><td>4.6</td><td>0.5</td><td>9.1</td><td>9.0E-20</td><td>1.0E-18</td><td>MB231/MB468</td><td>NUC</td></tr>
<tr><td>>hg38_21_+_8433172_8446622_RNA45SN1_45S_50ntFlanks@12628.12661.34|TGGGGCGAAGCTACCATCTGTGGGATTATGACTG</td><td>>hg38_21_+_8433172_8446622_RNA45SN1_45S_50ntFlanks</td><td>12628</td><td>12661</td><td>34</td><td>TGGGGCGAAGCTACCATCTGTGGGATTATGACTG</td><td>2474.9</td><td>-15.1</td><td>1.7</td><td>-9.1</td><td>6.7E-20</td><td>1.0E-18</td><td>MB231/MB468</td><td>MP</td></tr>
<tr><td>>hg38_21_+_8433172_8446622_RNA45SN1_45S_50ntFlanks@12959.12997.39|TCTGGGTCGGGGTTTCGTACGTAGCAGAGCAGCTCCCTC</td><td>>hg38_21_+_8433172_8446622_RNA45SN1_45S_50ntFlanks</td><td>12959</td><td>12997</td><td>39</td><td>TCTGGGTCGGGGTTTCGTACGTAGCAGAGCAGCTCCCTC</td><td>263.7</td><td>12.0</td><td>1.3</td><td>9.1</td><td>5.8E-20</td><td>1.0E-18</td><td>MB231/MB468</td><td>CYTO</td></tr>
<tr><td>>am_trna2_GlyGCC_21_-_18827107_18827177@1.24.24,trna35_GlyGCC_1_+_161413094_161413164@1.24.24,trna37_GlyGCC_1_+_161420467_161420537@1.24.24,trna39_GlyGCC_1_+_161427898_161427968@1.24.24,trna41_GlyGCC_1_+_161435258_161435328@1.24.24|GCATGGGTGGTTCAGTGGTAGAAT</td><td>>am_trna2_GlyGCC_21_-_18827107_18827177</td><td>1</td><td>24</td><td>24</td><td>GCATGGGTGGTTCAGTGGTAGAAT</td><td>423.3</td><td>5.5</td><td>0.6</td><td>9.0</td><td>1.4E-19</td><td>1.0E-18</td><td>BT20/MB468</td><td>CYTO</td></tr>
<tr><td>>hg38_21_+_8433172_8446622_RNA45SN1_45S_50ntFlanks@11315.11332.18|CCACCCCACGTCTCGTCG</td><td>>hg38_21_+_8433172_8446622_RNA45SN1_45S_50ntFlanks</td><td>11315</td><td>11332</td><td>18</td><td>CCACCCCACGTCTCGTCG</td><td>590.6</td><td>-13.0</td><td>1.4</td><td>-9.1</td><td>1.1E-19</td><td>1.0E-18</td><td>MB231/MB468</td><td>MITO</td></tr>
<tr><td>>hg38_21_+_8433172_8446622_RNA45SN1_45S_50ntFlanks@11594.11611.18|CAGGGGAATCCGACTGTT</td><td>>hg38_21_+_8433172_8446622_RNA45SN1_45S_50ntFlanks</td><td>11594</td><td>11611</td><td>18</td><td>CAGGGGAATCCGACTGTT</td><td>4039.0</td><td>-6.9</td><td>0.8</td><td>-9.1</td><td>1.1E-19</td><td>1.0E-18</td><td>MB231/MB468</td><td>MITO</td></tr>
<tr><td>>hg38_21_+_8433172_8446622_RNA45SN1_45S_50ntFlanks@12007.12029.23|GCCGCCGGTGAAATACCACTACT</td><td>>hg38_21_+_8433172_8446622_RNA45SN1_45S_50ntFlanks</td><td>12007</td><td>12029</td><td>23</td><td>GCCGCCGGTGAAATACCACTACT</td><td>736.8</td><td>-8.0</td><td>0.9</td><td>-9.1</td><td>6.7E-20</td><td>1.0E-18</td><td>MB231/MB468</td><td>MP</td></tr>
<tr><td>>MI0000294|hsa-mir-218-1&WithFlank&4|+|20528269|20528390@31.51.21;MI0000295|hsa-mir-218-2&WithFlank&5|-|168768140|168768261@31.51.21&[MIMAT0000275&hsa-miR-218-5p&offsets|0|0;m-185&4|+|20528299|20528319&offsets|0|0];[MIMAT0000275_1&hsa-miR-218-5p&offsets|0|0;m-187&5|-|168768211|168768231&offsets|0|0]|TTGTGCTTGATCTAACCATGT</td><td>>MI0000294|hsa-mir-218-1&WithFlank&4|+|20528269|20528390</td><td>31</td><td>51</td><td>21</td><td>TTGTGCTTGATCTAACCATGT</td><td>451.5</td><td>12.7</td><td>1.4</td><td>9.1</td><td>9.1E-20</td><td>1.0E-18</td><td>MB231/MB468</td><td>NUC</td></tr>
<tr><td>>MI0000102|hsa-mir-100&WithFlank&11|-|122152223|122152314@19.40.22(+1U)&[MIMAT0000098&hsa-miR-100-5p&offsets|0|0(+1U);m-27&11|-|122152276|122152296&offsets|0|+1(+1U)]|AACCCGTAGATCCGAACTTGTGT</td><td>>MI0000102|hsa-mir-100&WithFlank&11|-|122152223|122152314</td><td>19</td><td>40</td><td>22(+1U)</td><td>AACCCGTAGATCCGAACTTGTGT</td><td>35812.3</td><td>11.3</td><td>1.2</td><td>9.1</td><td>6.8E-20</td><td>1.0E-18</td><td>MB231/MB468</td><td>MP</td></tr>
<tr><td>>hg38_21_+_8433172_8446622_RNA45SN1_45S_50ntFlanks@12436.12454.19|TCATTGTGAAGCAGAATTC</td><td>>hg38_21_+_8433172_8446622_RNA45SN1_45S_50ntFlanks</td><td>12436</td><td>12454</td><td>19</td><td>TCATTGTGAAGCAGAATTC</td><td>1116.1</td><td>-4.9</td><td>0.5</td><td>-9.1</td><td>9.2E-20</td><td>1.0E-18</td><td>MB231/MB468</td><td>NUC</td></tr>
<tr><td>>hg38_21_+_8433172_8446622_RNA45SN1_45S_50ntFlanks@6772.6807.36|GTTCCTCCCGGGGCTACGCCTGTCTGAGCGTCGCTT</td><td>>hg38_21_+_8433172_8446622_RNA45SN1_45S_50ntFlanks</td><td>6772</td><td>6807</td><td>36</td><td>GTTCCTCCCGGGGCTACGCCTGTCTGAGCGTCGCTT</td><td>4569.6</td><td>-5.8</td><td>0.6</td><td>-9.1</td><td>1.1E-19</td><td>1.0E-18</td><td>MB231/MB468</td><td>MITO</td></tr>
</table>

| | | | | | | | | | | | | | |
|---|---|---|---|---|---|---|---|---|---|---|---|---|---|
| >am_MI0005763\|hsa-mir-941-1&WithFlank&20\|+\|63919443\|63919526@53.73.21(+1A);MI0005764\|hsa-mir-941-2&WithFlank&20\|+\|63919499\|63919582@53.73.21(+1A);MI0005765\|hsa-mir-941-3&WithFlank&20\|+\|63919555\|63919638@53.73.21(+1A);MI0005766\|hsa-mir-941-4&WithFlank&20\|+\|63919750\|63919833@53.73.21(+1A);MI0031520\|hsa-mir-941-5&WithFlank&20\|+\|63919862\|63919945@53.73.21(+1A)&[MIMAT0004984&hsa-miR-941&offsets\|0\|-2(+1A);m-263&20\|+\|63919495\|63919517&offsets\|0\|-2(+1A)];[MIMAT0004984_1&hsa-miR-941&offsets\|0\|-2(+1A);m-263&20\|+\|63919551\|63919573&offsets\|0\|-2(+1A)];[MIMAT0004984_2&hsa-miR-941&offsets\|0\|-2(+1A);m-263&20\|+\|63919607\|63919629&offsets\|0\|-2(+1A)];[MIMAT0004984_3&hsa-miR-941&offsets\|0\|-2(+1A);m-263&20\|+\|63919802\|63919824&offsets\|0\|-2(+1A)];[MIMAT0004984_4&hsa-miR-941&offsets\|0\|-2(+1A);m-263&20\|+\|63919914\|63919936&offsets\|0\|-2(+1A)]\|CACCCGGCTGTGTGCACATGTA | >am_MI0005763\|hsa-mir-941-1&WithFlank&20\|+\|63919443\|63919526 | 53 | 73 | 21(+1A) | CACCCGGCTGTGTGCACATGTA | 370.4 | 9.5 | 1.0 | 9.0 | 1.5E-19 | 1.0E-18 | BT20/MB468 | CYTO |
| >hg38_21_+_8433172_8446622_RNA45SN1_45S_50ntFlanks@10389.10409.21\|TGAACAGCAGTTGAACATGGG | >hg38_21_+_8433172_8446622_RNA45SN1_45S_50ntFlanks | 10389 | 10409 | 21 | TGAACAGCAGTTGAACATGGG | 998.4 | -7.9 | 0.9 | -9.1 | 5.9E-20 | 1.0E-18 | MB231/MB468 | CYTO |
| >hg38_21_+_8433172_8446622_RNA45SN1_45S_50ntFlanks@12164.12182.19\|CTGGGGCGGTACACCTGTC | >hg38_21_+_8433172_8446622_RNA45SN1_45S_50ntFlanks | 12164 | 12182 | 19 | CTGGGGCGGTACACCTGTC | 560.5 | -13.0 | 1.4 | -9.1 | 6.8E-20 | 1.0E-18 | MB231/MB468 | MP |
| >MI0000809\|hsa-mir-151a&WithFlank&8\|-\|140732558\|140732659@53.74.22(+1G)&[MIMAT0000757&hsa-miR-151a-3p&offsets\|0\|+1(+1G);m-30&8\|-\|140732587\|140732607&offsets\|0\|+1(+1G)]\|CTAGACTGAAGCTCCTTGAGGAG | >MI0000809\|hsa-mir-151a&WithFlank&8\|-\|140732558\|140732659 | 53 | 74 | 22(+1G) | CTAGACTGAAGCTCCTTGAGGAG | 307.8 | 8.9 | 1.0 | 9.0 | 1.5E-19 | 1.0E-18 | BT20/MB468 | CYTO |
| >MI0000683\|hsa-mir-181b-2&WithFlank&9\|+\|124693704\|124693804@22.46.25&[MIMAT0000257_1&hsa-miR-181b-5p&offsets\|0\|+2;m-72&9\|+\|124693725\|124693747&offsets\|0\|+2]\|AACATTCATTGCTGTCGGTGGGTTT | >MI0000683\|hsa-mir-181b-2&WithFlank&9\|+\|124693704\|124693804 | 22 | 46 | 25 | AACATTCATTGCTGTCGGTGGGTTT | 630.0 | 7.5 | 0.8 | 9.1 | 9.2E-20 | 1.0E-18 | MB231/MB468 | NUC |
| >hg38_21_+_8433172_8446622_RNA45SN1_45S_50ntFlanks@10411.10431.21\|CAGTCGGTCCTGAGAGATGGG | >hg38_21_+_8433172_8446622_RNA45SN1_45S_50ntFlanks | 10411 | 10431 | 21 | CAGTCGGTCCTGAGAGATGGG | 5737.5 | -11.3 | 1.2 | -9.1 | 9.3E-20 | 1.0E-18 | MB231/MB468 | NUC |
| >hg38_21_+_8433172_8446622_RNA45SN1_45S_50ntFlanks@10479.10497.19\|CGATCGAAAGGGAGTCGGG | >hg38_21_+_8433172_8446622_RNA45SN1_45S_50ntFlanks | 10479 | 10497 | 19 | CGATCGAAAGGGAGTCGGG | 692.7 | -11.7 | 1.3 | -9.0 | 1.5E-19 | 1.0E-18 | BT20/MB468 | CYTO |
| >MI0000253\|hsa-mir-148a&WithFlank&7\|-\|25949913\|25949992@50.70.21(+1A)&[MIMAT0000243&hsa-miR-148a-3p&offsets\|0\|-1(+1A);m-7&7\|-\|25949922\|25949943&offsets\|0\|-1(+1A)]\|TCAGTGCACTACAGAACTTTGA | >MI0000253\|hsa-mir-148a&WithFlank&7\|-\|25949913\|25949992 | 50 | 70 | 21(+1A) | TCAGTGCACTACAGAACTTTGA | 3736.1 | 4.9 | 0.5 | 9.3 | 1.1E-20 | 1.0E-18 | BT20/MB231 | NUC |
| >hg38_21_+_8433172_8446622_RNA45SN1_45S_50ntFlanks@11643.11660.18\|GCGGGTGTTGACGCGATG | >hg38_21_+_8433172_8446622_RNA45SN1_45S_50ntFlanks | 11643 | 11660 | 18 | GCGGGTGTTGACGCGATG | 6163.3 | -8.6 | 0.9 | -9.1 | 6.9E-20 | 1.0E-18 | MB231/MB468 | MP |
| >hg38_21_+_8433172_8446622_RNA45SN1_45S_50ntFlanks@12712.12742.31\|CGGAGCCTCGGTTGGCCTCGGATAGCCGGTC | >hg38_21_+_8433172_8446622_RNA45SN1_45S_50ntFlanks | 12712 | 12742 | 31 | CGGAGCCTCGGTTGGCCTCGGATAGCCGGTC | 810.0 | -13.4 | 1.5 | -9.1 | 1.1E-19 | 1.0E-18 | MB231/MB468 | MITO |
| >hg38_21_+_8433172_8446622_RNA45SN1_45S_50ntFlanks@12004.12033.30\|CGGGCCGCCGGTGAAATACCACTACTCTGA | >hg38_21_+_8433172_8446622_RNA45SN1_45S_50ntFlanks | 12004 | 12033 | 30 | CGGGCCGCCGGTGAAATACCACTACTCTGA | 2585.9 | 7.6 | 0.8 | 9.1 | 1.1E-19 | 1.0E-18 | BT20/MB468 | MITO |
| >hg38_1_-_228634819_228635039_RNA5S12_5S_50ntFlanks@51.84.34\|GTCTACGGCCATACCACCCTGAACGCGCCCGATC | >hg38_1_-_228634819_228635039_RNA5S12_5S_50ntFlanks | 51 | 84 | 34 | GTCTACGGCCATACCACCCTGAACGCGCCCGATC | 40255.2 | 7.4 | 0.8 | 9.0 | 1.5E-19 | 1.0E-18 | BT20/MB468 | CYTO |
| >hg38_21_+_8433172_8446622_RNA45SN1_45S_50ntFlanks@10729.10755.27\|AGAGGGTGTAAATCTCGCGCCGGGCCG | >hg38_21_+_8433172_8446622_RNA45SN1_45S_50ntFlanks | 10729 | 10755 | 27 | AGAGGGTGTAAATCTCGCGCCGGGCCG | 442.6 | -12.7 | 1.4 | -9.1 | 7.0E-20 | 1.0E-18 | MB231/MB468 | MP |
| >hg38_21_+_8433172_8446622_RNA45SN1_45S_50ntFlanks@4003.4026.24\|ATAACCTCGGGCCGATCGCACGCC | >hg38_21_+_8433172_8446622_RNA45SN1_45S_50ntFlanks | 4003 | 4026 | 24 | ATAACCTCGGGCCGATCGCACGCC | 572.3 | -13.0 | 1.4 | -9.1 | 7.0E-20 | 1.0E-18 | MB231/MB468 | MP |
| >trna116_GluCTC_1_-_145399233_145399304@1.33.33,trna59_GluCTC_1_+_249168447_249168518@1.33.33,trna71_GluCTC_1_-_161439189_161439260@1.33.33,trna74_GluCTC_1_-_161431809_161431880@1.33.33,trna77_GluCTC_1_-_161424398_161424469@1.33.33,trna77_GluCTC_6_+_28949976_28950047@1.33.33,trna80_GluCTC_1_-_161417018_161417089@1.33.33,trna87_GluCTC_6_-_126101393_126101464@1.33.33\|TCCCTGGTGGTCTAGTGGTTAGGATTCGGCGCT | >trna116_GluCTC_1_-_145399233_145399304 | 1 | 33 | 33 | TCCCTGGTGGTCTAGTGGTTAGGATTCGGCGCT | 189374.8 | 8.2 | 0.9 | 9.1 | 6.0E-20 | 1.0E-18 | MB231/MB468 | CYTO |
| >hg38_21_+_8433172_8446622_RNA45SN1_45S_50ntFlanks@10101.10126.26\|CGCGCGCGTGTGGTGTGCGTCGGAGG | >hg38_21_+_8433172_8446622_RNA45SN1_45S_50ntFlanks | 10101 | 10126 | 26 | CGCGCGCGTGTGGTGTGCGTCGGAGG | 1427.0 | -14.3 | 1.6 | -9.1 | 1.1E-19 | 1.1E-18 | MB231/MB468 | MITO |
| >hg38_21_+_8433172_8446622_RNA45SN1_45S_50ntFlanks@12159.12183.25\|TTTGACTGGGGCGGTACACCTGTCA | >hg38_21_+_8433172_8446622_RNA45SN1_45S_50ntFlanks | 12159 | 12183 | 25 | TTTGACTGGGGCGGTACACCTGTCA | 892.2 | -13.7 | 1.5 | -9.1 | 7.1E-20 | 1.1E-18 | MB231/MB468 | MP |
| >hg38_21_+_8433172_8446622_RNA45SN1_45S_50ntFlanks@8154.8178.25\|CCGCTCGTGGGGGGCCCAAGTCCTT | >hg38_21_+_8433172_8446622_RNA45SN1_45S_50ntFlanks | 8154 | 8178 | 25 | CCGCTCGTGGGGGGCCCAAGTCCTT | 806.9 | -13.4 | 1.5 | -9.1 | 1.1E-19 | 1.1E-18 | MB231/MB468 | MITO |
| >hg38_21_+_8433172_8446622_RNA45SN1_45S_50ntFlanks@6778.6807.30\|CCCGGGGCTACGCCTGTCTGAGCGTCGCTT | >hg38_21_+_8433172_8446622_RNA45SN1_45S_50ntFlanks | 6778 | 6807 | 30 | CCCGGGGCTACGCCTGTCTGAGCGTCGCTT | 151.7 | 4.8 | 0.5 | 9.1 | 6.2E-20 | 1.1E-18 | MB231/MB468 | CYTO |
| >hg38_21_+_8433172_8446622_RNA45SN1_45S_50ntFlanks@8179.8211.33\|CTGATCGAGGCCCAGCCCGTGGACGGTGTGAGG | >hg38_21_+_8433172_8446622_RNA45SN1_45S_50ntFlanks | 8179 | 8211 | 33 | CTGATCGAGGCCCAGCCCGTGGACGGTGTGAGG | 1274.4 | -14.2 | 1.6 | -9.1 | 7.2E-20 | 1.1E-18 | MB231/MB468 | MP |
| >trna10_AspGTC_12_-_125424193_125424264@35.75.41,trna12_AspGTC_12_-_125411891_125411962@35.75.41,trna144_AspGTC_6_-_27551236_27551307@35.75.41,trna38_AspGTC_17_-_8125556_8125627@35.75.41,trna45_AspGTC_6_+_27447453_27447524@35.75.41,trna48_AspGTC_6_+_27471523_27471594@35.75.41,trna4_AspGTC_12_+_96429799_96429870@35.75.41,trna69_AspGTC_1_-_161440205_161440276@35.75.41,trna72_AspGTC_1_-_161432824_161432895@35.75.41,trna75_AspGTC_1_-_161425414_161425485@35.75.41,trna78_AspGTC_1_-_161418033_161418104@35.75.41,trna81_AspGTC_1_-_161410615_161410686@35.75.41\|TCACGCGGGAGACCGGGGTTCGATTCCCCGACGGGGAGCCA | >trna10_AspGTC_12_-_125424193_125424264 | 35 | 75 | 41 | TCACGCGGGAGACCGGGGTTCGATTCCCCGACGGGGAGCCA | 61859.1 | 11.3 | 1.2 | 9.1 | 6.2E-20 | 1.1E-18 | MB231/MB468 | CYTO |
| >MI0000264\|hsa-mir-7-2&WithFlank&15\|+\|88611819\|88611940@38.60.23(+1G);MI0000265\|hsa-mir-7-3&WithFlank&19\|+\|4770664\|4770785@37.59.23(+1G);MI0000263\|hsa-mir-7-1&WithFlank&9\|-\|83969742\|83969863@30.52.23(+1G)&[MIMAT0000252&hsa-miR-7-5p&offsets\|0\|-1(+1G);m-400&15\|+\|88611856\|88611879&offsets\|0\|-1(+1G)];[MIMAT0000252_1&hsa-miR-7-5p&offsets\|0\|-1(+1G);m-402&19\|+\|4770700\|4770723&offsets\|0\|-1(+1G)];[MIMAT0000252_2&hsa-miR-7-5p&offsets\|0\|-1(+1G);m-401&9\|-\|83969811\|83969834&offsets\|0\|-1(+1G)]\|TGGAAGACTAGTGATTTTGTTGTG | >MI0000264\|hsa-mir-7-2&WithFlank&15\|+\|88611819\|88611940 | 38 | 60 | 23(+1G) | TGGAAGACTAGTGATTTTGTTGTG | 348.8 | 3.7 | 0.4 | 9.0 | 2.9E-19 | 1.1E-18 | BT20/MB468 | NUC |
| >hg38_21_+_8433172_8446622_RNA45SN1_45S_50ntFlanks@4367.4391.25\|CAGTTAAAAAGCTCGTAGTTGGATC | >hg38_21_+_8433172_8446622_RNA45SN1_45S_50ntFlanks | 4367 | 4391 | 25 | CAGTTAAAAAGCTCGTAGTTGGATC | 1268.0 | -8.5 | 0.9 | -9.1 | 7.2E-20 | 1.1E-18 | MB231/MB468 | MP |
| >hg38_21_+_8433172_8446622_RNA45SN1_45S_50ntFlanks@6645.6664.20\|CTCGTACGACTCTTAGCGGT | >hg38_21_+_8433172_8446622_RNA45SN1_45S_50ntFlanks | 6645 | 6664 | 20 | CTCGTACGACTCTTAGCGGT | 13877.6 | 5.7 | 0.6 | 9.1 | 1.2E-19 | 1.1E-18 | BT20/MB468 | MITO |
| >hg38_21_+_8433172_8446622_RNA45SN1_45S_50ntFlanks@4760.4780.21\|TCAGATACCGTCGTAGTTCCG | >hg38_21_+_8433172_8446622_RNA45SN1_45S_50ntFlanks | 4760 | 4780 | 21 | TCAGATACCGTCGTAGTTCCG | 1992.6 | -13.1 | 1.4 | -9.1 | 1.2E-19 | 1.1E-18 | BT20/MB468 | MITO |
| >MI0000446\|hsa-mir-125b-1&WithFlank&11\|-\|122099751\|122099850@21.43.23&[MIMAT0000423&hsa-miR-125b-5p&offsets\|0\|+1;m-67&11\|-\|122099809\|122099830&offsets\|0\|+1]\|TCCCTGAGACCCTAACTTGTGAT | >MI0000446\|hsa-mir-125b-1&WithFlank&11\|-\|122099751\|122099850 | 21 | 43 | 23 | TCCCTGAGACCCTAACTTGTGAT | 2442.4 | 12.9 | 1.4 | 9.1 | 6.5E-20 | 1.1E-18 | MB231/MB468 | CYTO |
| >hg38_21_+_8433172_8446622_RNA45SN1_45S_50ntFlanks@11857.11884.28\|GCGGAATCAGCGGGGAAAGAAGACCCTG | >hg38_21_+_8433172_8446622_RNA45SN1_45S_50ntFlanks | 11857 | 11884 | 28 | GCGGAATCAGCGGGGAAAGAAGACCCTG | 369.1 | -12.4 | 1.4 | -9.1 | 7.5E-20 | 1.1E-18 | MB231/MB468 | MP |
| >MI0000462\|hsa-mir-152&WithFlank&17\|-\|48037155\|48037253@60.80.21(+1U)&[MIMAT0000438&hsa-miR-152-3p&offsets\|0\|0(+1U);m-97&17\|-\|48037174\|48037194&offsets\|0\|0(+1U)]\|TCAGTGCATGACAGAACTTGGT | >MI0000462\|hsa-mir-152&WithFlank&17\|-\|48037155\|48037253 | 60 | 80 | 21(+1U) | TCAGTGCATGACAGAACTTGGT | 595.5 | 13.5 | 1.5 | 9.0 | 1.6E-19 | 1.1E-18 | BT20/MB468 | CYTO |
| >MI0001445\|hsa-mir-423&WithFlank&17\|+\|30117073\|30117178@23.42.20(+1C)&[MIMAT0004748&hsa-miR-423-5p&offsets\|0\|-3(+1C);m-141&17\|+\|30117095\|30117117&offsets\|0\|-3(+1C)]\|TGAGGGGCAGAGAGCGAGACC | >MI0001445\|hsa-mir-423&WithFlank&17\|+\|30117073\|30117178 | 23 | 42 | 20(+1C) | TGAGGGGCAGAGAGCGAGACC | 578.8 | 11.1 | 1.2 | 9.0 | 1.6E-19 | 1.1E-18 | BT20/MB468 | CYTO |
| >hg38_21_+_8433172_8446622_RNA45SN1_45S_50ntFlanks@4122.4144.23\|AGGGTTCGATTCCGGAGAGGGAG | >hg38_21_+_8433172_8446622_RNA45SN1_45S_50ntFlanks | 4122 | 4144 | 23 | AGGGTTCGATTCCGGAGAGGGAG | 396.2 | -12.5 | 1.4 | -9.1 | 7.7E-20 | 1.1E-18 | MB231/MB468 | MP |
| >hg38_21_+_8433172_8446622_RNA45SN1_45S_50ntFlanks@11561.11586.26\|GCGCCTAGCAGCCGACTTAGAACTGG | >hg38_21_+_8433172_8446622_RNA45SN1_45S_50ntFlanks | 11561 | 11586 | 26 | GCGCCTAGCAGCCGACTTAGAACTGG | 1472.4 | -8.4 | 0.9 | -9.1 | 7.7E-20 | 1.1E-18 | MB231/MB468 | MP |
| >hg38_21_+_8433172_8446622_RNA45SN1_45S_50ntFlanks@11564.11603.40\|CCTAGCAGCCGACTTAGAACTGGTGCGGACCAGGGGAATC | >hg38_21_+_8433172_8446622_RNA45SN1_45S_50ntFlanks | 11564 | 11603 | 40 | CCTAGCAGCCGACTTAGAACTGGTGCGGACCAGGGGAATC | 2385.3 | -15.1 | 1.7 | -9.1 | 7.7E-20 | 1.1E-18 | MB231/MB468 | MP |
| >MI0000466\|hsa-mir-9-1&WithFlank&1\|-\|156420335\|156420435@22.43.22;MI0000468\|hsa-mir-9-3&WithFlank&15\|+\|89368011\|89368112@22.43.22;MI0000467\|hsa-mir-9-2&WithFlank&5\|-\|88666847\|88666945@22.43.22&[MIMAT0000441&hsa-miR-9-5p&offsets\|0\|-1;m-104&1\|-\|156420392\|156420414&offsets\|0\|-1];[MIMAT0000441_1&hsa-miR-9-5p&offsets\|0\|-1;m-105&15\|+\|89368032\|89368054&offsets\|0\|-1];[MIMAT0000441_2&hsa-miR-9-5p&offsets\|0\|-1;m-104&5\|-\|88666902\|88666924&offsets\|0\|-1]\|TCTTTGGTTATCTAGCTGTATG | >MI0000466\|hsa-mir-9-1&WithFlank&1\|-\|156420335\|156420435 | 22 | 43 | 22 | TCTTTGGTTATCTAGCTGTATG | 2057.2 | 11.1 | 1.2 | 9.0 | 1.6E-19 | 1.1E-18 | BT20/MB468 | CYTO |

<table>
<tr><th></th><th></th><th></th><th></th><th></th><th></th><th></th><th></th><th></th><th></th><th></th><th></th><th></th><th></th></tr>
<tr><td>>hg38_21_+_8433172_8446622_RNA45SN1_45S_50ntFlanks@6650.6673.24|ACGACTCTTAGCGGTGGATCACTC</td><td>>hg38_21_+_8433172_8446622_RNA45SN1_45S_50ntFlanks</td><td>6650</td><td>6673</td><td>24</td><td>ACGACTCTTAGCGGTGGATCACTC</td><td>1882.6</td><td>6.2</td><td>0.7</td><td>9.1</td><td>1.2E-19</td><td>1.1E-18</td><td>BT20/MB468</td><td>MITO</td></tr>
<tr><td>>hg38_21_+_8433172_8446622_RNA45SN1_45S_50ntFlanks@6650.6672.23|ACGACTCTTAGCGGTGGATCACT</td><td>>hg38_21_+_8433172_8446622_RNA45SN1_45S_50ntFlanks</td><td>6650</td><td>6672</td><td>23</td><td>ACGACTCTTAGCGGTGGATCACT</td><td>5215.5</td><td>7.2</td><td>0.8</td><td>9.1</td><td>1.2E-19</td><td>1.2E-18</td><td>BT20/MB468</td><td>MITO</td></tr>
<tr><td>>hg38_21_+_8433172_8446622_RNA45SN1_45S_50ntFlanks@4443.4470.28|CCCTTGCCTCTCGGCGCCCCCTCGATGC</td><td>>hg38_21_+_8433172_8446622_RNA45SN1_45S_50ntFlanks</td><td>4443</td><td>4470</td><td>28</td><td>CCCTTGCCTCTCGGCGCCCCCTCGATGC</td><td>635.3</td><td>-13.1</td><td>1.4</td><td>-9.1</td><td>1.2E-19</td><td>1.2E-18</td><td>MB231/MB468</td><td>MITO</td></tr>
<tr><td>>hg38_21_+_8433172_8446622_RNA45SN1_45S_50ntFlanks@12997.13016.20|CGCTGCGATCTATTGAAAGT</td><td>>hg38_21_+_8433172_8446622_RNA45SN1_45S_50ntFlanks</td><td>12997</td><td>13016</td><td>20</td><td>CGCTGCGATCTATTGAAAGT</td><td>332.3</td><td>5.3</td><td>0.6</td><td>9.0</td><td>1.7E-19</td><td>1.2E-18</td><td>BT20/MB468</td><td>CYTO</td></tr>
<tr><td>>hg38_21_+_8433172_8446622_RNA45SN1_45S_50ntFlanks@10679.10712.34|GGTTCCGGCGGCGTCCGGTGAGCTCTCGCTGGCC</td><td>>hg38_21_+_8433172_8446622_RNA45SN1_45S_50ntFlanks</td><td>10679</td><td>10712</td><td>34</td><td>GGTTCCGGCGGCGTCCGGTGAGCTCTCGCTGGCC</td><td>3013.5</td><td>-9.6</td><td>1.0</td><td>-9.1</td><td>7.9E-20</td><td>1.2E-18</td><td>MB231/MB468</td><td>MP</td></tr>
<tr><td>>hg38_21_+_8433172_8446622_RNA45SN1_45S_50ntFlanks@11613.11640.28|AATTAAAACAAAGCATCGCGAAGGCCCG</td><td>>hg38_21_+_8433172_8446622_RNA45SN1_45S_50ntFlanks</td><td>11613</td><td>11640</td><td>28</td><td>AATTAAAACAAAGCATCGCGAAGGCCCG</td><td>1938.9</td><td>-13.0</td><td>1.4</td><td>-9.1</td><td>1.2E-19</td><td>1.2E-18</td><td>BT20/MB468</td><td>MITO</td></tr>
<tr><td>>hg38_21_+_8433172_8446622_RNA45SN1_45S_50ntFlanks@10679.10708.30|GGTTCCGGCGGCGTCCGGTGAGCTCTCGCT</td><td>>hg38_21_+_8433172_8446622_RNA45SN1_45S_50ntFlanks</td><td>10679</td><td>10708</td><td>30</td><td>GGTTCCGGCGGCGTCCGGTGAGCTCTCGCT</td><td>3268.4</td><td>-8.0</td><td>0.9</td><td>-9.1</td><td>8.0E-20</td><td>1.2E-18</td><td>MB231/MB468</td><td>MP</td></tr>
<tr><td>>hg38_21_+_8433172_8446622_RNA45SN1_45S_50ntFlanks@4979.4999.21|GACACGGACAGGATTGACAGA</td><td>>hg38_21_+_8433172_8446622_RNA45SN1_45S_50ntFlanks</td><td>4979</td><td>4999</td><td>21</td><td>GACACGGACAGGATTGACAGA</td><td>805.8</td><td>-13.5</td><td>1.5</td><td>-9.1</td><td>8.0E-20</td><td>1.2E-18</td><td>MB231/MB468</td><td>MP</td></tr>
<tr><td>>hg38_21_+_8433172_8446622_RNA45SN1_45S_50ntFlanks@11785.11803.19|TGAATGGATGAACGAGATT</td><td>>hg38_21_+_8433172_8446622_RNA45SN1_45S_50ntFlanks</td><td>11785</td><td>11803</td><td>19</td><td>TGAATGGATGAACGAGATT</td><td>6959.7</td><td>-7.0</td><td>0.8</td><td>-9.1</td><td>1.1E-19</td><td>1.2E-18</td><td>MB231/MB468</td><td>NUC</td></tr>
<tr><td>>MI0000102|hsa-mir-100&WithFlank&11|-|122152223|122152314@19.40.22(+3A)&[MIMAT0000098&hsa-miR-100-5p&offsets|0|0(+3A);m-27&11|-|122152276|122152296&offsets|0|+1(+3A)]|AACCCGTAGATCCGAACTTGTGAAA</td><td>>MI0000102|hsa-mir-100&WithFlank&11|-|122152223|122152314</td><td>19</td><td>40</td><td>22(+3A)</td><td>AACCCGTAGATCCGAACTTGTGAAA</td><td>269.2</td><td>12.1</td><td>1.3</td><td>9.1</td><td>6.9E-20</td><td>1.2E-18</td><td>MB231/MB468</td><td>CYTO</td></tr>
<tr><td>>hg38_21_+_8433172_8446622_RNA45SN1_45S_50ntFlanks@9533.9576.44|GAGGAAACTCTGGTGGAGGTCCGTAGCGGTCCTGACGTGCAAAT</td><td>>hg38_21_+_8433172_8446622_RNA45SN1_45S_50ntFlanks</td><td>9533</td><td>9576</td><td>44</td><td>GAGGAAACTCTGGTGGAGGTCCGTAGCGGTCCTGACGTGCAAAT</td><td>297.7</td><td>12.2</td><td>1.3</td><td>9.1</td><td>6.9E-20</td><td>1.2E-18</td><td>MB231/MB468</td><td>CYTO</td></tr>
<tr><td>>hg38_21_+_8433172_8446622_RNA45SN1_45S_50ntFlanks@9576.9596.21|TCGGTCGTCCGACCTGGGTAT</td><td>>hg38_21_+_8433172_8446622_RNA45SN1_45S_50ntFlanks</td><td>9576</td><td>9596</td><td>21</td><td>TCGGTCGTCCGACCTGGGTAT</td><td>780.3</td><td>-13.5</td><td>1.5</td><td>-9.1</td><td>8.0E-20</td><td>1.2E-18</td><td>MB231/MB468</td><td>MP</td></tr>
<tr><td>>hg38_21_+_8433172_8446622_RNA45SN1_45S_50ntFlanks@10781.10818.38|AGGTGAACAGCCTCTGGCATGTTGGAACAATGTAGGTA</td><td>>hg38_21_+_8433172_8446622_RNA45SN1_45S_50ntFlanks</td><td>10781</td><td>10818</td><td>38</td><td>AGGTGAACAGCCTCTGGCATGTTGGAACAATGTAGGTA</td><td>2927.8</td><td>-8.8</td><td>1.0</td><td>-9.1</td><td>1.3E-19</td><td>1.2E-18</td><td>MB231/MB468</td><td>MITO</td></tr>
<tr><td>>hg38_21_+_8433172_8446622_RNA45SN1_45S_50ntFlanks@10684.10710.27|CGGCGGCGTCCGGTGAGCTCTCGCTGG</td><td>>hg38_21_+_8433172_8446622_RNA45SN1_45S_50ntFlanks</td><td>10684</td><td>10710</td><td>27</td><td>CGGCGGCGTCCGGTGAGCTCTCGCTGG</td><td>12425.3</td><td>-6.4</td><td>0.7</td><td>-9.1</td><td>1.1E-19</td><td>1.2E-18</td><td>MB231/MB468</td><td>NUC</td></tr>
<tr><td>>hg38_21_+_8433172_8446622_RNA45SN1_45S_50ntFlanks@11027.11050.24|CTCTCTCTCTCTCTCCCCCGCTCC</td><td>>hg38_21_+_8433172_8446622_RNA45SN1_45S_50ntFlanks</td><td>11027</td><td>11050</td><td>24</td><td>CTCTCTCTCTCTCTCCCCCGCTCC</td><td>1917.8</td><td>-13.9</td><td>1.5</td><td>-9.1</td><td>1.1E-19</td><td>1.2E-18</td><td>MB231/MB468</td><td>NUC</td></tr>
<tr><td>>hg38_21_+_8433172_8446622_RNA45SN1_45S_50ntFlanks@11891.11912.22|TTGACTCTAGTCTGGCACGGTG</td><td>>hg38_21_+_8433172_8446622_RNA45SN1_45S_50ntFlanks</td><td>11891</td><td>11912</td><td>22</td><td>TTGACTCTAGTCTGGCACGGTG</td><td>2266.4</td><td>-5.8</td><td>0.6</td><td>-9.1</td><td>7.0E-20</td><td>1.2E-18</td><td>MB231/MB468</td><td>CYTO</td></tr>
<tr><td>>MI0000434|hsa-let-7i&WithFlank&12|+|62603680|62603775@12.32.21(+1C)&[MIMAT0000415&hsa-let-7i-5p&offsets|0|-1(+1C);m-49&12|+|62603691|62603712&offsets|0|-1(+1C)]|TGAGGTAGTAGTTTGTGCTGTC</td><td>>MI0000434|hsa-let-7i&WithFlank&12|+|62603680|62603775</td><td>12</td><td>32</td><td>21(+1C)</td><td>TGAGGTAGTAGTTTGTGCTGTC</td><td>11711.4</td><td>7.4</td><td>0.8</td><td>9.1</td><td>1.3E-19</td><td>1.2E-18</td><td>BT20/MB468</td><td>MITO</td></tr>
<tr><td>>hg38_21_+_8433172_8446622_RNA45SN1_45S_50ntFlanks@12940.12972.33|CATTCGTAGACGACCTGCTTCTGGGTCGGGGTT</td><td>>hg38_21_+_8433172_8446622_RNA45SN1_45S_50ntFlanks</td><td>12940</td><td>12972</td><td>33</td><td>CATTCGTAGACGACCTGCTTCTGGGTCGGGGTT</td><td>583.1</td><td>-13.1</td><td>1.4</td><td>-9.1</td><td>8.2E-20</td><td>1.2E-18</td><td>MB231/MB468</td><td>MP</td></tr>
<tr><td>>hg38_21_+_8433172_8446622_RNA45SN1_45S_50ntFlanks@9565.9597.33|TGACGTGCAAATCGGTCGTCCGACCTGGGTATA</td><td>>hg38_21_+_8433172_8446622_RNA45SN1_45S_50ntFlanks</td><td>9565</td><td>9597</td><td>33</td><td>TGACGTGCAAATCGGTCGTCCGACCTGGGTATA</td><td>2287.6</td><td>-5.3</td><td>0.6</td><td>-9.0</td><td>1.7E-19</td><td>1.2E-18</td><td>BT20/MB468</td><td>CYTO</td></tr>
<tr><td>>MI0000087|hsa-mir-29a&WithFlank&7|-|130876741|130876816@48.69.22&[MIMAT0000086&hsa-miR-29a-3p&offsets|0|0;m-26&7|-|130876749|130876769&offsets|0|+1]|TAGCACCATCTGAAATCGGTTA</td><td>>MI0000087|hsa-mir-29a&WithFlank&7|-|130876741|130876816</td><td>48</td><td>69</td><td>22</td><td>TAGCACCATCTGAAATCGGTTA</td><td>6430.2</td><td>8.8</td><td>1.0</td><td>9.1</td><td>1.3E-19</td><td>1.2E-18</td><td>BT20/MB468</td><td>MITO</td></tr>
<tr><td>>hg38_21_+_8433172_8446622_RNA45SN1_45S_50ntFlanks@9504.9523.20|GTGAACTATGCCTGGGCAGG</td><td>>hg38_21_+_8433172_8446622_RNA45SN1_45S_50ntFlanks</td><td>9504</td><td>9523</td><td>20</td><td>GTGAACTATGCCTGGGCAGG</td><td>764.0</td><td>-3.6</td><td>0.4</td><td>-9.1</td><td>1.3E-19</td><td>1.2E-18</td><td>BT20/MB468</td><td>MITO</td></tr>
<tr><td>>MI0000255|hsa-mir-30d&WithFlank&8|-|134804870|134804951@12.34.23(+1G)&[MIMAT0000245&hsa-miR-30d-5p&offsets|0|+1(+1G);m-21&8|-|134804919|134804940&offsets|0|+1(+1G)]|TGTAAACATCCCCGACTGGAAGCG</td><td>>MI0000255|hsa-mir-30d&WithFlank&8|-|134804870|134804951</td><td>12</td><td>34</td><td>23(+1G)</td><td>TGTAAACATCCCCGACTGGAAGCG</td><td>1170.2</td><td>7.1</td><td>0.8</td><td>9.1</td><td>1.3E-19</td><td>1.2E-18</td><td>BT20/MB468</td><td>MITO</td></tr>
<tr><td>>hg38_21_+_8433172_8446622_RNA45SN1_45S_50ntFlanks@12126.12148.23|CGACCCGCTCCGGGGACAGTGCC</td><td>>hg38_21_+_8433172_8446622_RNA45SN1_45S_50ntFlanks</td><td>12126</td><td>12148</td><td>23</td><td>CGACCCGCTCCGGGGACAGTGCC</td><td>1066.4</td><td>-13.8</td><td>1.5</td><td>-9.1</td><td>1.3E-19</td><td>1.2E-18</td><td>MB231/MB468</td><td>MITO</td></tr>
<tr><td>>hg38_21_+_8433172_8446622_RNA45SN1_45S_50ntFlanks@11593.11612.20|CCAGGGGAATCCGACTGTTT</td><td>>hg38_21_+_8433172_8446622_RNA45SN1_45S_50ntFlanks</td><td>11593</td><td>11612</td><td>20</td><td>CCAGGGGAATCCGACTGTTT</td><td>2042.2</td><td>-6.7</td><td>0.7</td><td>-9.0</td><td>3.3E-19</td><td>1.2E-18</td><td>BT20/MB468</td><td>NUC</td></tr>
<tr><td>>hg38_21_+_8433172_8446622_RNA45SN1_45S_50ntFlanks@12676.12708.33|AGAATCCCGCCCAGGCGGAACGATACGGCAGCG</td><td>>hg38_21_+_8433172_8446622_RNA45SN1_45S_50ntFlanks</td><td>12676</td><td>12708</td><td>33</td><td>AGAATCCCGCCCAGGCGGAACGATACGGCAGCG</td><td>586.3</td><td>-13.0</td><td>1.4</td><td>-9.1</td><td>1.3E-19</td><td>1.2E-18</td><td>MB231/MB468</td><td>MITO</td></tr>
<tr><td>>hg38_21_+_8433172_8446622_RNA45SN1_45S_50ntFlanks@10402.10435.34|AACATGGGTCAGTCGGTCCTGAGAGATGGGCGAG</td><td>>hg38_21_+_8433172_8446622_RNA45SN1_45S_50ntFlanks</td><td>10402</td><td>10435</td><td>34</td><td>AACATGGGTCAGTCGGTCCTGAGAGATGGGCGAG</td><td>1337.5</td><td>-10.3</td><td>1.1</td><td>-9.1</td><td>1.3E-19</td><td>1.2E-18</td><td>MB231/MB468</td><td>MITO</td></tr>
<tr><td>>hg38_21_+_8433172_8446622_RNA45SN1_45S_50ntFlanks@5490.5511.22|TGAGAAGACGGTCGAACTTGAC</td><td>>hg38_21_+_8433172_8446622_RNA45SN1_45S_50ntFlanks</td><td>5490</td><td>5511</td><td>22</td><td>TGAGAAGACGGTCGAACTTGAC</td><td>200.2</td><td>11.7</td><td>1.3</td><td>9.1</td><td>7.3E-20</td><td>1.2E-18</td><td>MB231/MB468</td><td>CYTO</td></tr>
<tr><td>>hg38_21_+_8433172_8446622_RNA45SN1_45S_50ntFlanks@11564.11601.38|CCTAGCAGCCGACTTAGAACTGGTGCGGACCAGGGGAA</td><td>>hg38_21_+_8433172_8446622_RNA45SN1_45S_50ntFlanks</td><td>11564</td><td>11601</td><td>38</td><td>CCTAGCAGCCGACTTAGAACTGGTGCGGACCAGGGGAA</td><td>9761.7</td><td>-10.9</td><td>1.2</td><td>-9.0</td><td>3.4E-19</td><td>1.3E-18</td><td>BT20/MB468</td><td>NUC</td></tr>
<tr><td>>hg38_21_+_8433172_8446622_RNA45SN1_45S_50ntFlanks@11345.11362.18|CTGGGGGCGGGGAGCGGT</td><td>>hg38_21_+_8433172_8446622_RNA45SN1_45S_50ntFlanks</td><td>11345</td><td>11362</td><td>18</td><td>CTGGGGGCGGGGAGCGGT</td><td>1493.2</td><td>-14.4</td><td>1.6</td><td>-9.1</td><td>8.5E-20</td><td>1.3E-18</td><td>MB231/MB468</td><td>MP</td></tr>
<tr><td>>hg38_21_+_8433172_8446622_RNA45SN1_45S_50ntFlanks@4734.4768.35|AACGAAAGTCGGAGGTTCGAAGACGATCAGATACC</td><td>>hg38_21_+_8433172_8446622_RNA45SN1_45S_50ntFlanks</td><td>4734</td><td>4768</td><td>35</td><td>AACGAAAGTCGGAGGTTCGAAGACGATCAGATACC</td><td>339.6</td><td>12.4</td><td>1.4</td><td>9.1</td><td>7.4E-20</td><td>1.3E-18</td><td>MB231/MB468</td><td>CYTO</td></tr>
<tr><td>>hg38_21_+_8433172_8446622_RNA45SN1_45S_50ntFlanks@10408.10456.49|GGTCAGTCGGTCCTGAGAGATGGGCGAGCGCCGTTCCGAAGGGACGGGC</td><td>>hg38_21_+_8433172_8446622_RNA45SN1_45S_50ntFlanks</td><td>10408</td><td>10456</td><td>49</td><td>GGTCAGTCGGTCCTGAGAGATGGGCGAGCGCCGTTCCGAAGGGACGGGC</td><td>893.7</td><td>-14.2</td><td>1.6</td><td>-9.0</td><td>3.4E-19</td><td>1.3E-18</td><td>BT20/MB468</td><td>NUC</td></tr>
<tr><td>>hg38_21_+_8433172_8446622_RNA45SN1_45S_50ntFlanks@9554.9597.44|CGTAGCGGTCCTGACGTGCAAATCGGTCGTCCGACCTGGGTATA</td><td>>hg38_21_+_8433172_8446622_RNA45SN1_45S_50ntFlanks</td><td>9554</td><td>9597</td><td>44</td><td>CGTAGCGGTCCTGACGTGCAAATCGGTCGTCCGACCTGGGTATA</td><td>5846.2</td><td>-16.9</td><td>1.9</td><td>-9.0</td><td>3.4E-19</td><td>1.3E-18</td><td>BT20/MB468</td><td>NUC</td></tr>
<tr><td>>hg38_21_+_8433172_8446622_RNA45SN1_45S_50ntFlanks@10786.10818.33|AACAGCCTCTGGCATGTTGGAACAATGTAGGTA</td><td>>hg38_21_+_8433172_8446622_RNA45SN1_45S_50ntFlanks</td><td>10786</td><td>10818</td><td>33</td><td>AACAGCCTCTGGCATGTTGGAACAATGTAGGTA</td><td>8998.8</td><td>-9.6</td><td>1.1</td><td>-9.1</td><td>1.4E-19</td><td>1.3E-18</td><td>MB231/MB468</td><td>MITO</td></tr>
<tr><td>>hg38_21_+_8433172_8446622_RNA45SN1_45S_50ntFlanks@12407.12431.25|TTTTGATCCTTCGATGTCGGCTCTT</td><td>>hg38_21_+_8433172_8446622_RNA45SN1_45S_50ntFlanks</td><td>12407</td><td>12431</td><td>25</td><td>TTTTGATCCTTCGATGTCGGCTCTT</td><td>1402.5</td><td>-6.6</td><td>0.7</td><td>-9.0</td><td>3.4E-19</td><td>1.3E-18</td><td>BT20/MB468</td><td>NUC</td></tr>
<tr><td>>hg38_21_+_8433172_8446622_RNA45SN1_45S_50ntFlanks@5489.5515.27|CTGAGAAGACGGTCGAACTTGACTATC</td><td>>hg38_21_+_8433172_8446622_RNA45SN1_45S_50ntFlanks</td><td>5489</td><td>5515</td><td>27</td><td>CTGAGAAGACGGTCGAACTTGACTATC</td><td>899.2</td><td>-5.3</td><td>0.6</td><td>-9.1</td><td>8.7E-20</td><td>1.3E-18</td><td>MB231/MB468</td><td>MP</td></tr>
<tr><td>>MI0000266|hsa-mir-10a&WithFlank&17|-|48579832|48579953@29.48.20(+1A)&[MIMAT0000253&hsa-miR-10a-5p&offsets|+1|-2(+1A);m-18&17|-|48579905|48579926&offsets|+1|-1(+1A)]|ACCCTGTAGATCCGAATTTGA</td><td>>MI0000266|hsa-mir-10a&WithFlank&17|-|48579832|48579953</td><td>29</td><td>48</td><td>20(+1A)</td><td>ACCCTGTAGATCCGAATTTGA</td><td>349.4</td><td>12.7</td><td>1.4</td><td>9.0</td><td>1.8E-19</td><td>1.3E-18</td><td>BT20/MB468</td><td>CYTO</td></tr>
<tr><td>>hg38_21_+_8433172_8446622_RNA45SN1_45S_50ntFlanks@12324.12348.25|TTGGGTTTTAAGCAGGAGGTGTCAG</td><td>>hg38_21_+_8433172_8446622_RNA45SN1_45S_50ntFlanks</td><td>12324</td><td>12348</td><td>25</td><td>TTGGGTTTTAAGCAGGAGGTGTCAG</td><td>627.2</td><td>-12.3</td><td>1.4</td><td>-9.1</td><td>1.2E-19</td><td>1.3E-18</td><td>MB231/MB468</td><td>NUC</td></tr>
<tr><td>>MI0000279|hsa-mir-196a-2&WithFlank&12|+|53991732|53991853@31.51.21(+1A);MI0000238|hsa-mir-196a-1&WithFlank&17|-|48632484|48632565@13.33.21(+1A)&[MIMAT0000226&hsa-miR-196a-5p&offsets|0|-1(+1A);m-261&12|+|53991762|53991783&offsets|0|-1(+1A)];[MIMAT0000226_1&hsa-miR-196a-5p&offsets|0|-1(+1A);m-268&17|-|48632532|48632553&offsets|0|-1(+1A)]|TAGGTAGTTTCATGTTGTTGGA</td><td>>MI0000279|hsa-mir-196a-2&WithFlank&12|+|53991732|53991853</td><td>31</td><td>51</td><td>21(+1A)</td><td>TAGGTAGTTTCATGTTGTTGGA</td><td>614.9</td><td>13.0</td><td>1.4</td><td>9.3</td><td>1.4E-20</td><td>1.3E-18</td><td>BT20/MB231</td><td>NUC</td></tr>
<tr><td>>hg38_21_+_8433172_8446622_RNA45SN1_45S_50ntFlanks@9198.9223.26|GTTCTCTCGGGGCCACGCGCGCGTCC</td><td>>hg38_21_+_8433172_8446622_RNA45SN1_45S_50ntFlanks</td><td>9198</td><td>9223</td><td>26</td><td>GTTCTCTCGGGGCCACGCGCGCGTCC</td><td>812.0</td><td>-13.5</td><td>1.5</td><td>-9.1</td><td>8.8E-20</td><td>1.3E-18</td><td>MB231/MB468</td><td>MP</td></tr>
<tr><td>>hg38_21_+_8433172_8446622_RNA45SN1_45S_50ntFlanks@11027.11061.35|CTCTCTCTCTCTCTCCCCCGCTCCCCGTCCTCCCC</td><td>>hg38_21_+_8433172_8446622_RNA45SN1_45S_50ntFlanks</td><td>11027</td><td>11061</td><td>35</td><td>CTCTCTCTCTCTCTCCCCCGCTCCCCGTCCTCCCC</td><td>621.8</td><td>-13.1</td><td>1.4</td><td>-9.1</td><td>1.4E-19</td><td>1.3E-18</td><td>MB231/MB468</td><td>MITO</td></tr>
<tr><td>>hg38_21_+_8433172_8446622_RNA45SN1_45S_50ntFlanks@11580.11601.22|GAACTGGTGCGGACCAGGGGAA</td><td>>hg38_21_+_8433172_8446622_RNA45SN1_45S_50ntFlanks</td><td>11580</td><td>11601</td><td>22</td><td>GAACTGGTGCGGACCAGGGGAA</td><td>640.9</td><td>-13.1</td><td>1.4</td><td>-9.1</td><td>1.4E-19</td><td>1.3E-18</td><td>MB231/MB468</td><td>MITO</td></tr>
<tr><td>>hg38_21_+_8433172_8446622_RNA45SN1_45S_50ntFlanks@5472.5490.19|ACGGCCCTGGCGGAGCGCT</td><td>>hg38_21_+_8433172_8446622_RNA45SN1_45S_50ntFlanks</td><td>5472</td><td>5490</td><td>19</td><td>ACGGCCCTGGCGGAGCGCT</td><td>457.8</td><td>7.6</td><td>0.8</td><td>9.1</td><td>1.4E-19</td><td>1.3E-18</td><td>BT20/MB468</td><td>MITO</td></tr>
<tr><td>>hg38_21_+_8433172_8446622_RNA45SN1_45S_50ntFlanks@10841.10865.25|CGTAACTTCGGGATAAGGATTGGCT</td><td>>hg38_21_+_8433172_8446622_RNA45SN1_45S_50ntFlanks</td><td>10841</td><td>10865</td><td>25</td><td>CGTAACTTCGGGATAAGGATTGGCT</td><td>108850.6</td><td>-12.1</td><td>1.4</td><td>-9.0</td><td>3.5E-19</td><td>1.3E-18</td><td>BT20/MB468</td><td>NUC</td></tr>
<tr><td>>hg38_21_+_8433172_8446622_RNA45SN1_45S_50ntFlanks@10845.10868.24|ACTTCGGGATAAGGATTGGCTCTA</td><td>>hg38_21_+_8433172_8446622_RNA45SN1_45S_50ntFlanks</td><td>10845</td><td>10868</td><td>24</td><td>ACTTCGGGATAAGGATTGGCTCTA</td><td>1756.7</td><td>-11.1</td><td>1.2</td><td>-9.1</td><td>1.2E-19</td><td>1.3E-18</td><td>MB231/MB468</td><td>NUC</td></tr>
<tr><td>>MI0000542|hsa-mir-320a&WithFlank&8|-|22244960|22245043@48.69.22(+2U)&[MIMAT0000510&hsa-miR-320a-3p&offsets|0|0(+2U);m-60&8|-|22244975|22244996&offsets|0|0(+2U)]|AAAAGCTGGGTTGAGAGGGCGATT</td><td>>MI0000542|hsa-mir-320a&WithFlank&8|-|22244960|22245043</td><td>48</td><td>69</td><td>22(+2U)</td><td>AAAAGCTGGGTTGAGAGGGCGATT</td><td>1691.0</td><td>7.8</td><td>0.9</td><td>9.1</td><td>7.9E-20</td><td>1.3E-18</td><td>MB231/MB468</td><td>CYTO</td></tr>
<tr><td>>hg38_21_+_8433172_8446622_RNA45SN1_45S_50ntFlanks@12635.12657.23|AAGCTACCATCTGTGGGATTATG</td><td>>hg38_21_+_8433172_8446622_RNA45SN1_45S_50ntFlanks</td><td>12635</td><td>12657</td><td>23</td><td>AAGCTACCATCTGTGGGATTATG</td><td>496.8</td><td>-13.4</td><td>1.5</td><td>-8.9</td><td>3.6E-19</td><td>1.3E-18</td><td>BT20/MB468</td><td>NUC</td></tr>
<tr><td>>hg38_21_+_8433172_8446622_RNA45SN1_45S_50ntFlanks@10436.10453.18|CGCCGTTCCGAAGGGACG</td><td>>hg38_21_+_8433172_8446622_RNA45SN1_45S_50ntFlanks</td><td>10436</td><td>10453</td><td>18</td><td>CGCCGTTCCGAAGGGACG</td><td>554.5</td><td>-13.0</td><td>1.4</td><td>-9.1</td><td>9.1E-20</td><td>1.3E-18</td><td>MB231/MB468</td><td>MP</td></tr>
<tr><td>>hg38_21_+_8433172_8446622_RNA45SN1_45S_50ntFlanks@6659.6681.23|AGCGGTGGATCACTCGGCTCGTG</td><td>>hg38_21_+_8433172_8446622_RNA45SN1_45S_50ntFlanks</td><td>6659</td><td>6681</td><td>23</td><td>AGCGGTGGATCACTCGGCTCGTG</td><td>2895.7</td><td>-9.6</td><td>1.1</td><td>-9.1</td><td>9.1E-20</td><td>1.3E-18</td><td>MB231/MB468</td><td>MP</td></tr>
</table>

| | | | | | | | | | | | | | |
|---|---|---|---|---|---|---|---|---|---|---|---|---|---|
| >hg38_21_+_8433172_8446622_RNA45SN1_45S_50ntFlanks@7986.8014.29\|ATCAGACGTGGCGACCCGCTGAATTTAAG | >hg38_21_+_8433172_8446622_RNA45SN1_45S_50ntFlanks | 7986 | 8014 | 29 | ATCAGACGTGGCGACCCGCTGAATTTAAG | 854.9 | -13.5 | 1.5 | -9.0 | 1.4E-19 | 1.3E-18 | MB231/MB468 | MITO |
| >MI0000446\|hsa-mir-125b-1&WithFlank&11\|-\|122099751\|122099850@21.41.21(+1U);MI0000470\|hsa-mir-125b-2&WithFlank&21\|+\|16590231\|16590331@23.43.21(+1U)&[MIMAT0000423&hsa-miR-125b-5p&offsets\|0\|-1(+1U);m-67&11\|-\|122099809\|122099830&offsets\|0\|-1(+1U)];[MIMAT0000423_1&hsa-miR-125b-5p&offsets\|0\|-1(+1U);m-68&21\|+\|16590253\|16590274&offsets\|0\|-1(+1U)]\|TCCCTGAGACCCTAACTTGTGT | >MI0000446\|hsa-mir-125b-1&WithFlank&11\|-\|122099751\|122099850 | 21 | 41 | 21(+1U) | TCCCTGAGACCCTAACTTGTGT | 1008.0 | 14.0 | 1.5 | 9.1 | 8.0E-20 | 1.3E-18 | MB231/MB468 | CYTO |
| >hg38_21_+_8433172_8446622_RNA45SN1_45S_50ntFlanks@12008.12038.31\|CCGCCGGTGAAATACCACTACTCTGATCGTT | >hg38_21_+_8433172_8446622_RNA45SN1_45S_50ntFlanks | 12008 | 12038 | 31 | CCGCCGGTGAAATACCACTACTCTGATCGTT | 745.0 | -13.3 | 1.5 | -9.0 | 1.4E-19 | 1.4E-18 | MB231/MB468 | MITO |
| >hg38_21_+_8433172_8446622_RNA45SN1_45S_50ntFlanks@12934.12961.28\|CTAAACCATTCGTAGACGACCTGCTTCT | >hg38_21_+_8433172_8446622_RNA45SN1_45S_50ntFlanks | 12934 | 12961 | 28 | CTAAACCATTCGTAGACGACCTGCTTCT | 3486.4 | -12.0 | 1.3 | -8.9 | 3.7E-19 | 1.4E-18 | BT20/MB468 | NUC |
| >hg38_21_+_8433172_8446622_RNA45SN1_45S_50ntFlanks@6647.6668.22\|CGTACGACTCTTAGCGGTGGAT | >hg38_21_+_8433172_8446622_RNA45SN1_45S_50ntFlanks | 6647 | 6668 | 22 | CGTACGACTCTTAGCGGTGGAT | 10446.0 | 6.3 | 0.7 | 9.0 | 1.5E-19 | 1.4E-18 | BT20/MB468 | MITO |
| >hg38_21_+_8433172_8446622_RNA45SN1_45S_50ntFlanks@10781.10818.38\|AGGTGAACAGCCTCTGGCATGTTGGAACAATGTAGGTA | >hg38_21_+_8433172_8446622_RNA45SN1_45S_50ntFlanks | 10781 | 10818 | 38 | AGGTGAACAGCCTCTGGCATGTTGGAACAATGTAGGTA | 1305.4 | -14.2 | 1.6 | -9.1 | 9.3E-20 | 1.4E-18 | MB231/MB468 | MP |
| >hg38_21_+_8433172_8446622_RNA45SN1_45S_50ntFlanks@7988.8006.19\|CAGACGTGGCGACCCGCTG | >hg38_21_+_8433172_8446622_RNA45SN1_45S_50ntFlanks | 7988 | 8006 | 19 | CAGACGTGGCGACCCGCTG | 875.8 | -13.6 | 1.5 | -9.0 | 1.5E-19 | 1.4E-18 | MB231/MB468 | MITO |
| >hg38_21_+_8433172_8446622_RNA45SN1_45S_50ntFlanks@8552.8569.18\|CTCCCCGGAGGGGGCGGG | >hg38_21_+_8433172_8446622_RNA45SN1_45S_50ntFlanks | 8552 | 8569 | 18 | CTCCCCGGAGGGGGCGGG | 748.4 | -12.3 | 1.3 | -9.1 | 8.1E-20 | 1.4E-18 | MB231/MB468 | CYTO |
| >hg38_21_+_8433172_8446622_RNA45SN1_45S_50ntFlanks@4051.4070.20\|GAACGTCTGCCCTATCAACT | >hg38_21_+_8433172_8446622_RNA45SN1_45S_50ntFlanks | 4051 | 4070 | 20 | GAACGTCTGCCCTATCAACT | 273.8 | 12.1 | 1.3 | 9.1 | 8.2E-20 | 1.4E-18 | MB231/MB468 | CYTO |
| >hg38_21_+_8433172_8446622_RNA45SN1_45S_50ntFlanks@4588.4616.29\|CTATTTTGTTGGTTTTCGGAACTGAGGCC | >hg38_21_+_8433172_8446622_RNA45SN1_45S_50ntFlanks | 4588 | 4616 | 29 | CTATTTTGTTGGTTTTCGGAACTGAGGCC | 2141.6 | -5.0 | 0.6 | -9.0 | 2.0E-19 | 1.4E-18 | BT20/MB468 | CYTO |
| >hg38_21_+_8433172_8446622_RNA45SN1_45S_50ntFlanks@10479.10512.34\|CGATCGAAAGGGAGTCGGGTTCAGATCCCCGAAT | >hg38_21_+_8433172_8446622_RNA45SN1_45S_50ntFlanks | 10479 | 10512 | 34 | CGATCGAAAGGGAGTCGGGTTCAGATCCCCGAAT | 686.6 | -11.7 | 1.3 | -9.0 | 2.0E-19 | 1.4E-18 | BT20/MB468 | CYTO |
| >trna143_LysTTT_6_-_27559593_27559665@14.31.18,trna149_LysTTT_6_-_27302769_27302841@14.31.18,trna14_LysTTT_11_-_59327808_59327880@14.31.18,trna2_LysTTT_17_+_8022473_8022545@14.31.18,trna54_LysTTT_1_+_204475655_204475727@14.31.18,trna5_LysTTT_11_+_59323902_59323974@14.31.18,trna62_LysTTT_1_-_204476158_204476230@14.31.18,trna76_LysTTT_6_+_28918806_28918878@14.31.18\|AGTCGGTAGAGCATCAGA | >trna143_LysTTT_6_-_27559593_27559665 | 14 | 31 | 18 | AGTCGGTAGAGCATCAGA | 387.1 | -13.0 | 1.5 | -8.9 | 3.7E-19 | 1.4E-18 | BT20/MB468 | NUC |
| >trna20_GluTTC_2_-_131094701_131094772@1.33.33,trna5_GluTTC_13_-_41634874_41634945@1.33.33\|TCCCATATGGTCTAGCGGTTAGGATTCCTGGTT | >trna20_GluTTC_2_-_131094701_131094772 | 1 | 33 | 33 | TCCCATATGGTCTAGCGGTTAGGATTCCTGGTT | 1185.1 | 10.3 | 1.1 | 9.0 | 2.0E-19 | 1.4E-18 | BT20/MB468 | CYTO |
| >hg38_21_+_8433172_8446622_RNA45SN1_45S_50ntFlanks@4163.4184.22\|CATCCAAGGAAGGCAGCAGGCG | >hg38_21_+_8433172_8446622_RNA45SN1_45S_50ntFlanks | 4163 | 4184 | 22 | CATCCAAGGAAGGCAGCAGGCG | 648.6 | -13.1 | 1.5 | -9.0 | 1.5E-19 | 1.4E-18 | MB231/MB468 | MITO |
| >MI0000764\|hsa-mir-363&WithFlank&X\|-\|134169372\|134169458@56.77.22&[MIMAT0000707&hsa-miR-363-3p&offsets\|0\|0;m-178&X\|-\|134169381\|134169403&offsets\|0\|-1]\|AATTGCACGGTATCCATCTGTA | >MI0000764\|hsa-mir-363&WithFlank&X\|-\|134169372\|134169458 | 56 | 77 | 22 | AATTGCACGGTATCCATCTGTA | 1154.1 | 13.9 | 1.5 | 9.3 | 1.6E-20 | 1.4E-18 | BT20/MB231 | NUC |
| >hg38_21_+_8433172_8446622_RNA45SN1_45S_50ntFlanks@11580.11605.26\|GAACTGGTGCGGACCAGGGGAATCCG | >hg38_21_+_8433172_8446622_RNA45SN1_45S_50ntFlanks | 11580 | 11605 | 26 | GAACTGGTGCGGACCAGGGGAATCCG | 1777.3 | -5.4 | 0.6 | -9.0 | 2.0E-19 | 1.4E-18 | BT20/MB468 | CYTO |
| >hg38_21_+_8433172_8446622_RNA45SN1_45S_50ntFlanks@12073.12110.38\|CCGAGGGGCTCTCGCTTCTGGCGCCAAGCGCCCGGCCG | >hg38_21_+_8433172_8446622_RNA45SN1_45S_50ntFlanks | 12073 | 12110 | 38 | CCGAGGGGCTCTCGCTTCTGGCGCCAAGCGCCCGGCCG | 1748.5 | -10.4 | 1.1 | -9.0 | 1.5E-19 | 1.4E-18 | MB231/MB468 | MITO |
| >trnaMT_GluTTC_MT_-_14674_14742@38.71.34,trnalookalike8_GluTTC_5_-_93905172_93905240@38.71.34\|CATTGGTCGTGGTTGTAGTCCGTGCGAGAATACC | >trnaMT_GluTTC_MT_-_14674_14742 | 38 | 71 | 34 | CATTGGTCGTGGTTGTAGTCCGTGCGAGAATACC | 437.8 | 12.8 | 1.4 | 9.1 | 8.4E-20 | 1.4E-18 | MB231/MB468 | CYTO |
| >hg38_21_+_8433172_8446622_RNA45SN1_45S_50ntFlanks@12133.12150.18\|CTCCGGGGACAGTGCCAG | >hg38_21_+_8433172_8446622_RNA45SN1_45S_50ntFlanks | 12133 | 12150 | 18 | CTCCGGGGACAGTGCCAG | 1274.7 | -14.2 | 1.6 | -9.1 | 9.6E-20 | 1.4E-18 | MB231/MB468 | MP |
| >hg38_21_+_8433172_8446622_RNA45SN1_45S_50ntFlanks@9575.9596.22\|ATCGGTCGTCCGACCTGGGTAT | >hg38_21_+_8433172_8446622_RNA45SN1_45S_50ntFlanks | 9575 | 9596 | 22 | ATCGGTCGTCCGACCTGGGTAT | 7642.7 | -16.0 | 1.7 | -9.5 | 1.9E-21 | 1.4E-18 | BT20/MB468 | MP |
| >trna119_LysCTT_1_-_145395522_145395594@4.35.32,trna11_LysCTT_5_-_180648979_180649051@4.35.32,trna13_LysCTT_6_+_26556774_26556846@4.35.32,trna7_LysCTT_16_+_3225692_3225764@4.35.32,trna9_LysCTT_5_+_180634755_180634827@4.35.32\|CGGCTAGCTCAGTCGGTAGAGCATGAGACTCT | >trna119_LysCTT_1_-_145395522_145395594 | 4 | 35 | 32 | CGGCTAGCTCAGTCGGTAGAGCATGAGACTCT | 1286.5 | 7.7 | 0.8 | 9.1 | 1.3E-19 | 1.4E-18 | MB231/MB468 | NUC |
| >hg38_21_+_8433172_8446622_RNA45SN1_45S_50ntFlanks@11785.11804.20\|TGAATGGATGAACGAGATTC | >hg38_21_+_8433172_8446622_RNA45SN1_45S_50ntFlanks | 11785 | 11804 | 20 | TGAATGGATGAACGAGATTC | 14897.0 | -7.3 | 0.8 | -9.1 | 1.3E-19 | 1.4E-18 | MB231/MB468 | NUC |
| >hg38_21_+_8433172_8446622_RNA45SN1_45S_50ntFlanks@6733.6756.24\|CATTGATCATCGACACTTCGAACG | >hg38_21_+_8433172_8446622_RNA45SN1_45S_50ntFlanks | 6733 | 6756 | 24 | CATTGATCATCGACACTTCGAACG | 2156.5 | -6.3 | 0.7 | -9.1 | 9.7E-20 | 1.4E-18 | MB231/MB468 | MP |
| >hg38_21_+_8433172_8446622_RNA45SN1_45S_50ntFlanks@9567.9584.18\|ACGTGCAAATCGGTCGTC | >hg38_21_+_8433172_8446622_RNA45SN1_45S_50ntFlanks | 9567 | 9584 | 18 | ACGTGCAAATCGGTCGTC | 778.3 | -13.5 | 1.5 | -9.1 | 9.7E-20 | 1.4E-18 | MB231/MB468 | MP |
| >hg38_21_+_8433172_8446622_RNA45SN1_45S_50ntFlanks@12991.13015.25\|CTCCCTCGCTGCGATCTATTGAAAG | >hg38_21_+_8433172_8446622_RNA45SN1_45S_50ntFlanks | 12991 | 13015 | 25 | CTCCCTCGCTGCGATCTATTGAAAG | 9153.5 | -10.0 | 1.1 | -9.0 | 1.5E-19 | 1.4E-18 | MB231/MB468 | MITO |
| >hg38_21_+_8433172_8446622_RNA45SN1_45S_50ntFlanks@11621.11660.40\|CAAAGCATCGCGAAGGCCCGCGGCGGGTGTTGACGCGATG | >hg38_21_+_8433172_8446622_RNA45SN1_45S_50ntFlanks | 11621 | 11660 | 40 | CAAAGCATCGCGAAGGCCCGCGGCGGGTGTTGACGCGATG | 1141.6 | -13.9 | 1.5 | -9.0 | 1.5E-19 | 1.4E-18 | MB231/MB468 | MITO |
| >hg38_21_+_8433172_8446622_RNA45SN1_45S_50ntFlanks@10786.10818.33\|AACAGCCTCTGGCATGTTGGAACAATGTAGGTA | >hg38_21_+_8433172_8446622_RNA45SN1_45S_50ntFlanks | 10786 | 10818 | 33 | AACAGCCTCTGGCATGTTGGAACAATGTAGGTA | 4342.4 | -5.4 | 0.6 | -9.0 | 2.1E-19 | 1.4E-18 | BT20/MB468 | CYTO |
| >MI0000301\|hsa-mir-224&WithFlank&X\|-\|151958572\|151958664@14.38.25&[MIMAT0000281&hsa-miR-224-5p&offsets\|+1\|+1;m-238&X\|-\|151958630\|151958651&offsets\|0\|+3]\|CAAGTCACTAGTGGTTCCGTTTAGT | >MI0000301\|hsa-mir-224&WithFlank&X\|-\|151958572\|151958664 | 14 | 38 | 25 | CAAGTCACTAGTGGTTCCGTTTAGT | 719.5 | 14.0 | 1.5 | 9.0 | 1.5E-19 | 1.4E-18 | BT20/MB468 | MITO |
| >hg38_21_+_8433172_8446622_RNA45SN1_45S_50ntFlanks@4442.4471.30\|CCCCTTGCCTCTCGGCGCCCCCTCGATGCT | >hg38_21_+_8433172_8446622_RNA45SN1_45S_50ntFlanks | 4442 | 4471 | 30 | CCCCTTGCCTCTCGGCGCCCCCTCGATGCT | 563.2 | -12.9 | 1.4 | -9.0 | 1.6E-19 | 1.4E-18 | MB231/MB468 | MITO |
| >MI0000433\|hsa-let-7g&WithFlank&3\|-\|52268272\|52268367@11.29.19&[MIMAT0000414&hsa-let-7g-5p&offsets\|0\|-3;m-58&3\|-\|52268336\|52268357&offsets\|0\|-3]\|TGAGGTAGTAGTTTGTACA | >MI0000433\|hsa-let-7g&WithFlank&3\|-\|52268272\|52268367 | 11 | 29 | 19 | TGAGGTAGTAGTTTGTACA | 716.1 | 8.2 | 0.9 | 9.0 | 2.1E-19 | 1.4E-18 | BT20/MB468 | CYTO |
| >hg38_21_+_8433172_8446622_RNA45SN1_45S_50ntFlanks@3930.3959.30\|ATCAAAACCAACCCGGTCAGCCCCTCTCCG | >hg38_21_+_8433172_8446622_RNA45SN1_45S_50ntFlanks | 3930 | 3959 | 30 | ATCAAAACCAACCCGGTCAGCCCCTCTCCG | 624.8 | -11.6 | 1.3 | -9.0 | 2.1E-19 | 1.4E-18 | BT20/MB468 | CYTO |
| >MI0000108\|hsa-mir-103a-2&WithFlank&20\|+\|3917488\|3917577@54.76.23;MI0000109\|hsa-mir-103a-1&WithFlank&5\|-\|168560890\|168560979@54.76.23&[MIMAT0000101&hsa-miR-103a-3p&offsets\|0\|0;m-12&20\|+\|3917541\|3917563&offsets\|0\|0];[MIMAT0000101_1&hsa-miR-103a-3p&offsets\|0\|0;m-13&5\|-\|168560904\|168560926&offsets\|0\|0;m-44220&5\|-\|168560908\|168560926&offsets\|0\|+4]\|AGCAGCATTGTACAGGGCTATGA | >MI0000108\|hsa-mir-103a-2&WithFlank&20\|+\|3917488\|3917577 | 54 | 76 | 23 | AGCAGCATTGTACAGGGCTATGA | 22246.8 | 6.6 | 0.7 | 9.0 | 1.6E-19 | 1.5E-18 | BT20/MB468 | MITO |
| >hg38_21_+_8433172_8446622_RNA45SN1_45S_50ntFlanks@12628.12659.32\|TGGGGCGAAGCTACCATCTGTGGGATTATGAC | >hg38_21_+_8433172_8446622_RNA45SN1_45S_50ntFlanks | 12628 | 12659 | 32 | TGGGGCGAAGCTACCATCTGTGGGATTATGAC | 3507.1 | -15.6 | 1.7 | -9.0 | 1.6E-19 | 1.5E-18 | MB231/MB468 | MITO |
| >hg38_21_+_8433172_8446622_RNA45SN1_45S_50ntFlanks@6658.6681.24\|TAGCGGTGGATCACTCGGCTCGTG | >hg38_21_+_8433172_8446622_RNA45SN1_45S_50ntFlanks | 6658 | 6681 | 24 | TAGCGGTGGATCACTCGGCTCGTG | 464.5 | -12.7 | 1.4 | -9.1 | 1.0E-19 | 1.5E-18 | MB231/MB468 | MP |
| >hg38_21_+_8433172_8446622_RNA45SN1_45S_50ntFlanks@10782.10818.37\|GGTGAACAGCCTCTGGCATGTTGGAACAATGTAGGTA | >hg38_21_+_8433172_8446622_RNA45SN1_45S_50ntFlanks | 10782 | 10818 | 37 | GGTGAACAGCCTCTGGCATGTTGGAACAATGTAGGTA | 627.5 | -13.1 | 1.4 | -9.0 | 1.6E-19 | 1.5E-18 | MB231/MB468 | MITO |
| >hg38_21_+_8433172_8446622_RNA45SN1_45S_50ntFlanks@10807.10825.19\|ACAATGTAGGTAAGGGAAG | >hg38_21_+_8433172_8446622_RNA45SN1_45S_50ntFlanks | 10807 | 10825 | 19 | ACAATGTAGGTAAGGGAAG | 671.4 | -13.2 | 1.5 | -9.0 | 1.6E-19 | 1.5E-18 | MB231/MB468 | MITO |
| >hg38_21_+_8433172_8446622_RNA45SN1_45S_50ntFlanks@11564.11601.38\|CCTAGCAGCCGACTTAGAACTGGTGCGGACCAGGGGAA | >hg38_21_+_8433172_8446622_RNA45SN1_45S_50ntFlanks | 11564 | 11601 | 38 | CCTAGCAGCCGACTTAGAACTGGTGCGGACCAGGGGAA | 4878.0 | -7.2 | 0.8 | -9.1 | 1.3E-19 | 1.5E-18 | MB231/MB468 | NUC |
| >hg38_21_+_8433172_8446622_RNA45SN1_45S_50ntFlanks@12447.12486.40\|CAGAATTCACCAAGCGTTGGATTGTTCACCCACTAATAGG | >hg38_21_+_8433172_8446622_RNA45SN1_45S_50ntFlanks | 12447 | 12486 | 40 | CAGAATTCACCAAGCGTTGGATTGTTCACCCACTAATAGG | 751.5 | -14.0 | 1.6 | -8.9 | 4.0E-19 | 1.5E-18 | BT20/MB468 | NUC |
| >am_MI0000786\|hsa-mir-378a&WithFlank&5\|+\|149732819\|149732896@49.70.22(+1G)&[MIMAT0000732&hsa-miR-378a-3p&offsets\|0\|0(+1G);m-44&5\|+\|149732867\|149732888&offsets\|0\|0(+1G)]\|ACTGGACTTGGAGTCAGAAGGCG | >am_MI0000786\|hsa-mir-378a&WithFlank&5\|+\|149732819\|149732896 | 49 | 70 | 22(+1G) | ACTGGACTTGGAGTCAGAAGGCG | 755.9 | 5.1 | 0.6 | 9.0 | 2.2E-19 | 1.5E-18 | BT20/MB468 | CYTO |
| >hg38_21_+_8433172_8446622_RNA45SN1_45S_50ntFlanks@6721.6743.23\|AATTGCAGGACACATTGATCATC | >hg38_21_+_8433172_8446622_RNA45SN1_45S_50ntFlanks | 6721 | 6743 | 23 | AATTGCAGGACACATTGATCATC | 9533.0 | -6.6 | 0.7 | -9.1 | 1.4E-19 | 1.5E-18 | MB231/MB468 | NUC |
| >hg38_21_+_8433172_8446622_RNA45SN1_45S_50ntFlanks@4443.4472.30\|CCCTTGCCTCTCGGCGCCCCCTCGATGCTC | >hg38_21_+_8433172_8446622_RNA45SN1_45S_50ntFlanks | 4443 | 4472 | 30 | CCCTTGCCTCTCGGCGCCCCCTCGATGCTC | 490.7 | -12.8 | 1.4 | -9.1 | 1.0E-19 | 1.5E-18 | MB231/MB468 | MP |
| >hg38_21_+_8433172_8446622_RNA45SN1_45S_50ntFlanks@12008.12032.25\|CCGCCGGTGAAATACCACTACTCTG | >hg38_21_+_8433172_8446622_RNA45SN1_45S_50ntFlanks | 12008 | 12032 | 25 | CCGCCGGTGAAATACCACTACTCTG | 17538.8 | -10.0 | 1.1 | -9.1 | 8.9E-20 | 1.5E-18 | MB231/MB468 | CYTO |
| >MI0000809\|hsa-mir-151a&WithFlank&8\|-\|140732558\|140732659@51.73.23(+1U)&[MIMAT0000757&hsa-miR-151a-3p&offsets\|-2\|0(+1U);m-30&8\|-\|140732587\|140732607&offsets\|-2\|0(+1U)]\|TACTAGACTGAAGCTCCTTGAGGT | >MI0000809\|hsa-mir-151a&WithFlank&8\|-\|140732558\|140732659 | 51 | 73 | 23(+1U) | TACTAGACTGAAGCTCCTTGAGGT | 388.9 | 5.4 | 0.6 | 9.1 | 1.4E-19 | 1.5E-18 | MB231/MB468 | NUC |
| >hg38_21_+_8433172_8446622_RNA45SN1_45S_50ntFlanks@10922.10939.18\|CTGGACGAGGCGCCGCCG | >hg38_21_+_8433172_8446622_RNA45SN1_45S_50ntFlanks | 10922 | 10939 | 18 | CTGGACGAGGCGCCGCCG | 3168.1 | -11.3 | 1.3 | -9.0 | 1.6E-19 | 1.5E-18 | MB231/MB468 | MITO |
| >hg38_21_+_8433172_8446622_RNA45SN1_45S_50ntFlanks@475.504.30\|CTTCGTGATCGATGTGGTGACGTCGTGCTC | >hg38_21_+_8433172_8446622_RNA45SN1_45S_50ntFlanks | 475 | 504 | 30 | CTTCGTGATCGATGTGGTGACGTCGTGCTC | 5896.9 | 9.6 | 1.1 | 9.0 | 2.2E-19 | 1.5E-18 | BT20/MB468 | CYTO |
| >hg38_21_+_8433172_8446622_RNA45SN1_45S_50ntFlanks@12405.12429.25\|CTTTTTGATCCTTCGATGTCGGCTC | >hg38_21_+_8433172_8446622_RNA45SN1_45S_50ntFlanks | 12405 | 12429 | 25 | CTTTTTGATCCTTCGATGTCGGCTC | 5752.5 | -6.2 | 0.7 | -9.0 | 2.2E-19 | 1.5E-18 | BT20/MB468 | CYTO |

<table>
<tr><th></th><th></th><th></th><th></th><th></th><th></th><th></th><th></th><th></th><th></th><th></th><th></th><th></th><th></th></tr>
<tr><td>>hg38_21_+_8433172_8446622_RNA45SN1_45S_50ntFlanks@11592.11611.20|ACCAGGGGAATCCGACTGTT</td><td>>hg38_21_+_8433172_8446622_RNA45SN1_45S_50ntFlanks</td><td>11592</td><td>11611</td><td>20</td><td>ACCAGGGGAATCCGACTGTT</td><td>1381.4</td><td>-14.3</td><td>1.6</td><td>-9.1</td><td>1.0E-19</td><td>1.5E-18</td><td>MB231/MB468</td><td>MP</td></tr>
<tr><td>>hg38_21_+_8433172_8446622_RNA45SN1_45S_50ntFlanks@10684.10709.26|CGGCGGCGTCCGGTGAGCTCTCGCTG</td><td>>hg38_21_+_8433172_8446622_RNA45SN1_45S_50ntFlanks</td><td>10684</td><td>10709</td><td>26</td><td>CGGCGGCGTCCGGTGAGCTCTCGCTG</td><td>1159.3</td><td>-5.1</td><td>0.6</td><td>-9.1</td><td>1.4E-19</td><td>1.5E-18</td><td>MB231/MB468</td><td>NUC</td></tr>
<tr><td>>hg38_21_+_8433172_8446622_RNA45SN1_45S_50ntFlanks@4977.4999.23|CGGACACGGACAGGATTGACAGA</td><td>>hg38_21_+_8433172_8446622_RNA45SN1_45S_50ntFlanks</td><td>4977</td><td>4999</td><td>23</td><td>CGGACACGGACAGGATTGACAGA</td><td>515.2</td><td>-12.9</td><td>1.4</td><td>-9.1</td><td>1.0E-19</td><td>1.5E-18</td><td>MB231/MB468</td><td>MP</td></tr>
<tr><td>>MI0000650|hsa-mir-200c&WithFlank&12|+|6963693|6963772@49.71.23&[MIMAT0000617&hsa-miR-200c-3p&offsets|-1|-1;m-54&12|+|6963742|6963764&offsets|-1|-1]|CTAATACTGCCGGGTAATGATGG</td><td>>MI0000650|hsa-mir-200c&WithFlank&12|+|6963693|6963772</td><td>49</td><td>71</td><td>23</td><td>CTAATACTGCCGGGTAATGATGG</td><td>1001.5</td><td>10.0</td><td>1.1</td><td>9.0</td><td>2.2E-19</td><td>1.5E-18</td><td>BT20/MB468</td><td>CYTO</td></tr>
<tr><td>>hg38_21_+_8433172_8446622_RNA45SN1_45S_50ntFlanks@8169.8215.47|CCAAGTCCTTCTGATCGAGGCCCAGCCCGTGGACGGTGTGAGGCCGG</td><td>>hg38_21_+_8433172_8446622_RNA45SN1_45S_50ntFlanks</td><td>8169</td><td>8215</td><td>47</td><td>CCAAGTCCTTCTGATCGAGGCCCAGCCCGTGGACGGTGTGAGGCCGG</td><td>821.0</td><td>-14.1</td><td>1.6</td><td>-8.9</td><td>4.1E-19</td><td>1.5E-18</td><td>BT20/MB468</td><td>NUC</td></tr>
<tr><td>>hg38_21_+_8433172_8446622_RNA45SN1_45S_50ntFlanks@4443.4472.30|CCCTTGCCTCTCGGCGCCCCCTCGATGCTC</td><td>>hg38_21_+_8433172_8446622_RNA45SN1_45S_50ntFlanks</td><td>4443</td><td>4472</td><td>30</td><td>CCCTTGCCTCTCGGCGCCCCCTCGATGCTC</td><td>1718.4</td><td>-15.2</td><td>1.7</td><td>-8.9</td><td>4.1E-19</td><td>1.5E-18</td><td>BT20/MB468</td><td>NUC</td></tr>
<tr><td>>MI0000255|hsa-mir-30d&WithFlank&8|-|134804870|134804951@12.33.22&[MIMAT0000245&hsa-miR-30d-5p&offsets|0|0;m-21&8|-|134804919|134804940&offsets|0|0]|TGTAAACATCCCCGACTGGAAG</td><td>>MI0000255|hsa-mir-30d&WithFlank&8|-|134804870|134804951</td><td>12</td><td>33</td><td>22</td><td>TGTAAACATCCCCGACTGGAAG</td><td>1956.4</td><td>5.8</td><td>0.6</td><td>9.1</td><td>1.4E-19</td><td>1.5E-18</td><td>MB231/MB468</td><td>NUC</td></tr>
<tr><td>>MI0000255|hsa-mir-30d&WithFlank&8|-|134804870|134804951@12.33.22(+1G)&[MIMAT0000245&hsa-miR-30d-5p&offsets|0|0(+1G);m-21&8|-|134804919|134804940&offsets|0|0(+1G)]|TGTAAACATCCCCGACTGGAAGG</td><td>>MI0000255|hsa-mir-30d&WithFlank&8|-|134804870|134804951</td><td>12</td><td>33</td><td>22(+1G)</td><td>TGTAAACATCCCCGACTGGAAGG</td><td>316.7</td><td>12.6</td><td>1.4</td><td>9.0</td><td>2.2E-19</td><td>1.5E-18</td><td>BT20/MB468</td><td>CYTO</td></tr>
<tr><td>>hg38_21_+_8433172_8446622_RNA45SN1_45S_50ntFlanks@11233.11253.21|CCCTTCCCGTGGATCGCCCCA</td><td>>hg38_21_+_8433172_8446622_RNA45SN1_45S_50ntFlanks</td><td>11233</td><td>11253</td><td>21</td><td>CCCTTCCCGTGGATCGCCCCA</td><td>859.4</td><td>-13.5</td><td>1.5</td><td>-9.0</td><td>1.7E-19</td><td>1.5E-18</td><td>MB231/MB468</td><td>MITO</td></tr>
<tr><td>>MI0000456|hsa-mir-140&WithFlank&16|+|69933075|69933186@69.91.23&[MIMAT0004597&hsa-miR-140-3p&offsets|+1|+3;m-45&16|+|69933143|69933164&offsets|0|+1]|ACCACAGGGTAGAACCACGGACA</td><td>>MI0000456|hsa-mir-140&WithFlank&16|+|69933075|69933186</td><td>69</td><td>91</td><td>23</td><td>ACCACAGGGTAGAACCACGGACA</td><td>296.0</td><td>7.1</td><td>0.8</td><td>9.1</td><td>1.4E-19</td><td>1.5E-18</td><td>MB231/MB468</td><td>NUC</td></tr>
<tr><td>>hg38_21_+_8433172_8446622_RNA45SN1_45S_50ntFlanks@10866.10891.26|CTAAGGGCTGGGTCGGTCGGGCTGGG</td><td>>hg38_21_+_8433172_8446622_RNA45SN1_45S_50ntFlanks</td><td>10866</td><td>10891</td><td>26</td><td>CTAAGGGCTGGGTCGGTCGGGCTGGG</td><td>578.2</td><td>-12.1</td><td>1.3</td><td>-9.1</td><td>1.4E-19</td><td>1.5E-18</td><td>MB231/MB468</td><td>NUC</td></tr>
<tr><td>>hg38_21_+_8433172_8446622_RNA45SN1_45S_50ntFlanks@10903.10935.33|GGCTGGGCGCGCGCCGCGGCTGGACGAGGCGCC</td><td>>hg38_21_+_8433172_8446622_RNA45SN1_45S_50ntFlanks</td><td>10903</td><td>10935</td><td>33</td><td>GGCTGGGCGCGCGCCGCGGCTGGACGAGGCGCC</td><td>592.5</td><td>-13.6</td><td>1.5</td><td>-8.9</td><td>4.1E-19</td><td>1.5E-18</td><td>BT20/MB468</td><td>NUC</td></tr>
<tr><td>>hg38_21_+_8433172_8446622_RNA45SN1_45S_50ntFlanks@11578.11603.26|TAGAACTGGTGCGGACCAGGGGAATC</td><td>>hg38_21_+_8433172_8446622_RNA45SN1_45S_50ntFlanks</td><td>11578</td><td>11603</td><td>26</td><td>TAGAACTGGTGCGGACCAGGGGAATC</td><td>764.3</td><td>-5.3</td><td>0.6</td><td>-9.1</td><td>1.4E-19</td><td>1.5E-18</td><td>MB231/MB468</td><td>NUC</td></tr>
<tr><td>>hg38_21_+_8433172_8446622_RNA45SN1_45S_50ntFlanks@12957.12982.26|CTTCTGGGTCGGGGTTTCGTACGTAG</td><td>>hg38_21_+_8433172_8446622_RNA45SN1_45S_50ntFlanks</td><td>12957</td><td>12982</td><td>26</td><td>CTTCTGGGTCGGGGTTTCGTACGTAG</td><td>12412.7</td><td>-13.2</td><td>1.5</td><td>-8.9</td><td>4.1E-19</td><td>1.5E-18</td><td>BT20/MB468</td><td>NUC</td></tr>
<tr><td>>hg38_21_+_8433172_8446622_RNA45SN1_45S_50ntFlanks@11873.11909.37|AAGAAGACCCTGTTGAGCTTGACTCTAGTCTGGCACG</td><td>>hg38_21_+_8433172_8446622_RNA45SN1_45S_50ntFlanks</td><td>11873</td><td>11909</td><td>37</td><td>AAGAAGACCCTGTTGAGCTTGACTCTAGTCTGGCACG</td><td>766.6</td><td>-13.4</td><td>1.5</td><td>-9.0</td><td>1.7E-19</td><td>1.5E-18</td><td>MB231/MB468</td><td>MITO</td></tr>
<tr><td>>hg38_21_+_8433172_8446622_RNA45SN1_45S_50ntFlanks@11163.11188.26|CGGTCCCCCGCGAGGGGGGCCCGGGC</td><td>>hg38_21_+_8433172_8446622_RNA45SN1_45S_50ntFlanks</td><td>11163</td><td>11188</td><td>26</td><td>CGGTCCCCCGCGAGGGGGGCCCGGGC</td><td>382.6</td><td>-13.0</td><td>1.5</td><td>-8.9</td><td>4.2E-19</td><td>1.5E-18</td><td>BT20/MB468</td><td>NUC</td></tr>
<tr><td>>trnaMT_MetCAT_MT_+_4402_4469@35.71.37|CCCCGAAAATGTTGGTTATACCCTTCCCGTACTACCA</td><td>>trnaMT_MetCAT_MT_+_4402_4469</td><td>35</td><td>71</td><td>37</td><td>CCCCGAAAATGTTGGTTATACCCTTCCCGTACTACCA</td><td>452.6</td><td>12.8</td><td>1.4</td><td>9.1</td><td>9.3E-20</td><td>1.5E-18</td><td>MB231/MB468</td><td>CYTO</td></tr>
<tr><td>>hg38_21_+_8433172_8446622_RNA45SN1_45S_50ntFlanks@8170.8197.28|CAAGTCCTTCTGATCGAGGCCCAGCCCG</td><td>>hg38_21_+_8433172_8446622_RNA45SN1_45S_50ntFlanks</td><td>8170</td><td>8197</td><td>28</td><td>CAAGTCCTTCTGATCGAGGCCCAGCCCG</td><td>4065.1</td><td>-8.1</td><td>0.9</td><td>-9.0</td><td>1.4E-19</td><td>1.5E-18</td><td>MB231/MB468</td><td>NUC</td></tr>
<tr><td>>hg38_21_+_8433172_8446622_RNA45SN1_45S_50ntFlanks@11918.11951.34|ACATGAGAGGTGTAGAATAAGTGGGAGGCCCCCG</td><td>>hg38_21_+_8433172_8446622_RNA45SN1_45S_50ntFlanks</td><td>11918</td><td>11951</td><td>34</td><td>ACATGAGAGGTGTAGAATAAGTGGGAGGCCCCCG</td><td>1026.7</td><td>-13.9</td><td>1.5</td><td>-9.1</td><td>1.1E-19</td><td>1.5E-18</td><td>MB231/MB468</td><td>MP</td></tr>
<tr><td>>MI0001733|hsa-mir-452&WithFlank&X|-|151959622|151959718@23.44.22&[MIMAT0001635&hsa-miR-452-5p&offsets|+3|+3;m-188&X|-|151959677|151959699&offsets|+3|+2]|TGTTTGCAGAGGAAACTGAGAC</td><td>>MI0001733|hsa-mir-452&WithFlank&X|-|151959622|151959718</td><td>23</td><td>44</td><td>22</td><td>TGTTTGCAGAGGAAACTGAGAC</td><td>555.1</td><td>13.4</td><td>1.5</td><td>9.0</td><td>2.3E-19</td><td>1.5E-18</td><td>BT20/MB468</td><td>CYTO</td></tr>
<tr><td>>hg38_21_+_8433172_8446622_RNA45SN1_45S_50ntFlanks@12405.12431.27|CTTTTTGATCCTTCGATGTCGGCTCTT</td><td>>hg38_21_+_8433172_8446622_RNA45SN1_45S_50ntFlanks</td><td>12405</td><td>12431</td><td>27</td><td>CTTTTTGATCCTTCGATGTCGGCTCTT</td><td>25575.9</td><td>-11.3</td><td>1.3</td><td>-8.9</td><td>4.2E-19</td><td>1.6E-18</td><td>BT20/MB468</td><td>NUC</td></tr>
<tr><td>>MI0000067|hsa-let-7f-1&WithFlank&9|+|94176341|94176439@13.34.22;MI0000068|hsa-let-7f-2&WithFlank&X|-|53557186|53557280@14.35.22&[MIMAT0000067&hsa-let-7f-5p&offsets|0|0;m-20&9|+|94176353|94176374&offsets|0|0];[MIMAT0000067_1&hsa-let-7f-5p&offsets|0|0;m-19&X|-|53557246|53557267&offsets|0|0]|TGAGGTAGTAGATTGTATAGTT</td><td>>MI0000067|hsa-let-7f-1&WithFlank&9|+|94176341|94176439</td><td>13</td><td>34</td><td>22</td><td>TGAGGTAGTAGATTGTATAGTT</td><td>141033.3</td><td>4.4</td><td>0.5</td><td>9.0</td><td>1.5E-19</td><td>1.6E-18</td><td>MB231/MB468</td><td>NUC</td></tr>
<tr><td>>MI0000650|hsa-mir-200c&WithFlank&12|+|6963693|6963772@50.70.21&[MIMAT0000617&hsa-miR-200c-3p&offsets|0|-2;m-54&12|+|6963742|6963764&offsets|0|-2]|TAATACTGCCGGGTAATGATG</td><td>>MI0000650|hsa-mir-200c&WithFlank&12|+|6963693|6963772</td><td>50</td><td>70</td><td>21</td><td>TAATACTGCCGGGTAATGATG</td><td>3507.3</td><td>15.1</td><td>1.6</td><td>9.5</td><td>1.5E-21</td><td>1.6E-18</td><td>BT20/MB231</td><td>MP</td></tr>
<tr><td>>trna111_HisGTG_1_-_147774845_147774916@-1T.29.30,trna118_HisGTG_1_-_145396881_145396952@-1T.29.30,trna16_HisGTG_1_+_146544773_146544844@-1T.29.30,trna1_HisGTG_15_+_45493349_45493420@-1T.29.30,trna21_HisGTG_1_+_147753471_147753542@-1T.29.30,trna33_HisGTG_6_+_27125906_27125977@-1T.29.30,trna7_HisGTG_9_-_14433938_14434009@-1T.29.30,trna8_HisGTG_15_-_45492611_45492682@-1T.29.30,trna9_HisGTG_15_-_45490804_45490875@-1T.29.30|TGCCGTGATCGTATAGTGGTTAGTACTCTG</td><td>>trna111_HisGTG_1_-_147774845_147774916</td><td>-1T</td><td>29</td><td>30</td><td>TGCCGTGATCGTATAGTGGTTAGTACTCTG</td><td>3409.6</td><td>-7.6</td><td>0.8</td><td>-9.1</td><td>1.1E-19</td><td>1.6E-18</td><td>MB231/MB468</td><td>MP</td></tr>
<tr><td>>hg38_21_+_8433172_8446622_RNA45SN1_45S_50ntFlanks@4759.4782.24|ATCAGATACCGTCGTAGTTCCGAC</td><td>>hg38_21_+_8433172_8446622_RNA45SN1_45S_50ntFlanks</td><td>4759</td><td>4782</td><td>24</td><td>ATCAGATACCGTCGTAGTTCCGAC</td><td>535.7</td><td>-12.0</td><td>1.3</td><td>-9.0</td><td>1.5E-19</td><td>1.6E-18</td><td>MB231/MB468</td><td>NUC</td></tr>
<tr><td>>hg38_21_+_8433172_8446622_RNA45SN1_45S_50ntFlanks@4726.4751.26|TAATCAAGAACGAAAGTCGGAGGTTC</td><td>>hg38_21_+_8433172_8446622_RNA45SN1_45S_50ntFlanks</td><td>4726</td><td>4751</td><td>26</td><td>TAATCAAGAACGAAAGTCGGAGGTTC</td><td>585.1</td><td>-13.1</td><td>1.4</td><td>-9.1</td><td>1.1E-19</td><td>1.6E-18</td><td>MB231/MB468</td><td>MP</td></tr>
<tr><td>>hg38_21_+_8433172_8446622_RNA45SN1_45S_50ntFlanks@485.521.37|GATGTGGTGACGTCGTGCTCTCCCGGGCCGGGTCCGA</td><td>>hg38_21_+_8433172_8446622_RNA45SN1_45S_50ntFlanks</td><td>485</td><td>521</td><td>37</td><td>GATGTGGTGACGTCGTGCTCTCCCGGGCCGGGTCCGA</td><td>369.6</td><td>12.8</td><td>1.4</td><td>9.0</td><td>2.3E-19</td><td>1.6E-18</td><td>BT20/MB468</td><td>CYTO</td></tr>
<tr><td>>hg38_21_+_8433172_8446622_RNA45SN1_45S_50ntFlanks@11579.11603.25|AGAACTGGTGCGGACCAGGGGAATC</td><td>>hg38_21_+_8433172_8446622_RNA45SN1_45S_50ntFlanks</td><td>11579</td><td>11603</td><td>25</td><td>AGAACTGGTGCGGACCAGGGGAATC</td><td>5731.4</td><td>-5.4</td><td>0.6</td><td>-9.1</td><td>9.5E-20</td><td>1.6E-18</td><td>MB231/MB468</td><td>CYTO</td></tr>
<tr><td>>trna2_GlyGCC_21_-_18827107_18827177@1.31.31,trna35_GlyGCC_1_+_161413094_161413164@1.31.31,trna37_GlyGCC_1_+_161420467_161420537@1.31.31,trna39_GlyGCC_1_+_161427898_161427968@1.31.31,trna41_GlyGCC_1_+_161435258_161435328@1.31.31|GCATGGGTGGTTCAGTGGTAGAATTCTCGCC</td><td>>trna2_GlyGCC_21_-_18827107_18827177</td><td>1</td><td>31</td><td>31</td><td>GCATGGGTGGTTCAGTGGTAGAATTCTCGCC</td><td>78635.4</td><td>10.3</td><td>1.1</td><td>9.0</td><td>1.7E-19</td><td>1.6E-18</td><td>BT20/MB468</td><td>MITO</td></tr>
<tr><td>>MI0000433|hsa-let-7g&WithFlank&3|-|52268272|52268367@11.31.21(+1G)&[MIMAT0000414&hsa-let-7g-5p&offsets|0|-1(+1G);m-58&3|-|52268336|52268357&offsets|0|-1(+1G)]|TGAGGTAGTAGTTTGTACAGTG</td><td>>MI0000433|hsa-let-7g&WithFlank&3|-|52268272|52268367</td><td>11</td><td>31</td><td>21(+1G)</td><td>TGAGGTAGTAGTTTGTACAGTG</td><td>490.9</td><td>6.0</td><td>0.7</td><td>9.0</td><td>1.7E-19</td><td>1.6E-18</td><td>BT20/MB468</td><td>MITO</td></tr>
<tr><td>>hg38_21_+_8433172_8446622_RNA45SN1_45S_50ntFlanks@9575.9596.22|ATCGGTCGTCCGACCTGGGTAT</td><td>>hg38_21_+_8433172_8446622_RNA45SN1_45S_50ntFlanks</td><td>9575</td><td>9596</td><td>22</td><td>ATCGGTCGTCCGACCTGGGTAT</td><td>2247.9</td><td>-13.2</td><td>1.5</td><td>-9.0</td><td>1.7E-19</td><td>1.6E-18</td><td>BT20/MB468</td><td>MITO</td></tr>
<tr><td>>MI0000077|hsa-mir-21&WithFlank&17|+|59841260|59841343@14.39.26(+1U)&[MIMAT0000076&hsa-miR-21-5p&offsets|0|+4(+1U);m-1&17|+|59841273|59841294&offsets|0|+4(+1U)]|TAGCTTATCAGACTGATGTTGACTGTT</td><td>>MI0000077|hsa-mir-21&WithFlank&17|+|59841260|59841343</td><td>14</td><td>39</td><td>26(+1U)</td><td>TAGCTTATCAGACTGATGTTGACTGTT</td><td>393.5</td><td>6.5</td><td>0.7</td><td>9.0</td><td>1.7E-19</td><td>1.6E-18</td><td>BT20/MB468</td><td>MITO</td></tr>
<tr><td>>hg38_21_+_8433172_8446622_RNA45SN1_45S_50ntFlanks@12479.12496.18|CTAATAGGGAACGTGAGC</td><td>>hg38_21_+_8433172_8446622_RNA45SN1_45S_50ntFlanks</td><td>12479</td><td>12496</td><td>18</td><td>CTAATAGGGAACGTGAGC</td><td>1414.3</td><td>-5.3</td><td>0.6</td><td>-8.9</td><td>4.4E-19</td><td>1.6E-18</td><td>BT20/MB468</td><td>NUC</td></tr>
<tr><td>>am_trnaMT_ValTAC_MT_+_1602_1670@28.52.25|AACTTACACTTAGGAGATTTCAACT</td><td>>am_trnaMT_ValTAC_MT_+_1602_1670</td><td>28</td><td>52</td><td>25</td><td>AACTTACACTTAGGAGATTTCAACT</td><td>466.2</td><td>10.8</td><td>1.2</td><td>9.0</td><td>2.4E-19</td><td>1.6E-18</td><td>BT20/MB468</td><td>CYTO</td></tr>
<tr><td>>MI0000264|hsa-mir-7-2&WithFlank&15|+|88611819|88611940@38.58.21;MI0000265|hsa-mir-7-3&WithFlank&19|+|4770664|4770785@37.57.21;MI0000263|hsa-mir-7-1&WithFlank&9|-|83969742|83969863@30.50.21&[MIMAT0000252&hsa-miR-7-5p&offsets|0|-3;m-400&15|+|88611856|88611879&offsets|0|-3];[MIMAT0000252_1&hsa-miR-7-5p&offsets|0|-3;m-402&19|+|4770700|4770723&offsets|0|-3];[MIMAT0000252_2&hsa-miR-7-5p&offsets|0|-3;m-401&9|-|83969811|83969834&offsets|0|-3]|TGGAAGACTAGTGATTTTGTT</td><td>>MI0000264|hsa-mir-7-2&WithFlank&15|+|88611819|88611940</td><td>38</td><td>58</td><td>21</td><td>TGGAAGACTAGTGATTTTGTT</td><td>1615.0</td><td>7.5</td><td>0.8</td><td>9.0</td><td>2.4E-19</td><td>1.6E-18</td><td>BT20/MB468</td><td>CYTO</td></tr>
<tr><td>>hg38_21_+_8433172_8446622_RNA45SN1_45S_50ntFlanks@11767.11803.37|TCTAATTAGTGACGCGCATGAATGGATGAACGAGATT</td><td>>hg38_21_+_8433172_8446622_RNA45SN1_45S_50ntFlanks</td><td>11767</td><td>11803</td><td>37</td><td>TCTAATTAGTGACGCGCATGAATGGATGAACGAGATT</td><td>757.6</td><td>-4.5</td><td>0.5</td><td>-9.0</td><td>1.5E-19</td><td>1.6E-18</td><td>MB231/MB468</td><td>NUC</td></tr>
<tr><td>>hg38_21_+_8433172_8446622_RNA45SN1_45S_50ntFlanks@12464.12496.33|TGGATTGTTCACCCACTAATAGGGAACGTGAGC</td><td>>hg38_21_+_8433172_8446622_RNA45SN1_45S_50ntFlanks</td><td>12464</td><td>12496</td><td>33</td><td>TGGATTGTTCACCCACTAATAGGGAACGTGAGC</td><td>201.0</td><td>4.2</td><td>0.5</td><td>9.1</td><td>9.9E-20</td><td>1.6E-18</td><td>MB231/MB468</td><td>CYTO</td></tr>
<tr><td>>hg38_21_+_8433172_8446622_RNA45SN1_45S_50ntFlanks@3899.3927.29|CCTTCGCGGGGGGGATGCGTGCATTTATC</td><td>>hg38_21_+_8433172_8446622_RNA45SN1_45S_50ntFlanks</td><td>3899</td><td>3927</td><td>29</td><td>CCTTCGCGGGGGGGATGCGTGCATTTATC</td><td>657.9</td><td>-13.2</td><td>1.5</td><td>-9.1</td><td>1.1E-19</td><td>1.6E-18</td><td>MB231/MB468</td><td>MP</td></tr>
<tr><td>>hg38_21_+_8433172_8446622_RNA45SN1_45S_50ntFlanks@12942.12960.19|TTCGTAGACGACCTGCTTC</td><td>>hg38_21_+_8433172_8446622_RNA45SN1_45S_50ntFlanks</td><td>12942</td><td>12960</td><td>19</td><td>TTCGTAGACGACCTGCTTC</td><td>549.3</td><td>-4.8</td><td>0.5</td><td>-8.9</td><td>4.5E-19</td><td>1.7E-18</td><td>BT20/MB468</td><td>NUC</td></tr>
<tr><td>>hg38_21_+_8433172_8446622_RNA45SN1_45S_50ntFlanks@11027.11051.25|CTCTCTCTCTCTCTCCCCCGCTCCC</td><td>>hg38_21_+_8433172_8446622_RNA45SN1_45S_50ntFlanks</td><td>11027</td><td>11051</td><td>25</td><td>CTCTCTCTCTCTCTCCCCCGCTCCC</td><td>1685.5</td><td>-13.7</td><td>1.5</td><td>-9.0</td><td>1.5E-19</td><td>1.7E-18</td><td>MB231/MB468</td><td>NUC</td></tr>
<tr><td>>MI0000266|hsa-mir-10a&WithFlank&17|-|48579832|48579953@28.49.22(+1A)&[MIMAT0000253&hsa-miR-10a-5p&offsets|0|-1(+1A);m-18&17|-|48579905|48579926&offsets|0|0(+1A)]|TACCCTGTAGATCCGAATTTGTA</td><td>>MI0000266|hsa-mir-10a&WithFlank&17|-|48579832|48579953</td><td>28</td><td>49</td><td>22(+1A)</td><td>TACCCTGTAGATCCGAATTTGTA</td><td>514.2</td><td>13.0</td><td>1.4</td><td>9.1</td><td>1.0E-19</td><td>1.7E-18</td><td>MB231/MB468</td><td>CYTO</td></tr>
<tr><td>>hg38_21_+_8433172_8446622_RNA45SN1_45S_50ntFlanks@12420.12457.38|ATGTCGGCTCTTCCTATCATTGTGAAGCAGAATTCACC</td><td>>hg38_21_+_8433172_8446622_RNA45SN1_45S_50ntFlanks</td><td>12420</td><td>12457</td><td>38</td><td>ATGTCGGCTCTTCCTATCATTGTGAAGCAGAATTCACC</td><td>2781.3</td><td>5.2</td><td>0.6</td><td>9.1</td><td>1.0E-19</td><td>1.7E-18</td><td>MB231/MB468</td><td>CYTO</td></tr>
<tr><td>>trnaMT_SerGCT_MT_+_12207_12265@6.38.33|AGCTCACAAGAACTGCTAACTCATGCCCCCATG</td><td>>trnaMT_SerGCT_MT_+_12207_12265</td><td>6</td><td>38</td><td>33</td><td>AGCTCACAAGAACTGCTAACTCATGCCCCCATG</td><td>710.0</td><td>-13.9</td><td>1.6</td><td>-8.9</td><td>4.5E-19</td><td>1.7E-18</td><td>BT20/MB468</td><td>NUC</td></tr>
<tr><td>>hg38_21_+_8433172_8446622_RNA45SN1_45S_50ntFlanks@12615.12661.47|GCTGAGGAGCCAATGGGGCGAAGCTACCATCTGTGGGATTATGACTG</td><td>>hg38_21_+_8433172_8446622_RNA45SN1_45S_50ntFlanks</td><td>12615</td><td>12661</td><td>47</td><td>GCTGAGGAGCCAATGGGGCGAAGCTACCATCTGTGGGATTATGACTG</td><td>1909.8</td><td>-15.3</td><td>1.7</td><td>-8.9</td><td>4.5E-19</td><td>1.7E-18</td><td>BT20/MB468</td><td>NUC</td></tr>
</table>

<table>
<tr><td>>MI0000079|hsa-mir-23a&WithFlank&19|-|13836581|13836665@54.71.18(+1U)&[MIMAT0000078&hsa-miR-23a-3p&offsets|+3|0(+1U);m-31&19|-|13836595|13836615&offsets|+3|0(+1U)]|ACATTGCCAGGGATTTCCT</td><td>>MI0000079|hsa-mir-23a&WithFlank&19|-|13836581|13836665</td><td>54</td><td>71</td><td>18(+1U)</td><td>ACATTGCCAGGGATTTCCT</td><td>347.1</td><td>12.7</td><td>1.4</td><td>9.0</td><td>2.4E-19</td><td>1.7E-18</td><td>BT20/MB468</td><td>CYTO</td></tr>
<tr><td>>hg38_21_+_8433172_8446622_RNA45SN1_45S_50ntFlanks@10403.10424.22|ACATGGGTCAGTCGGTCCTGAG</td><td>>hg38_21_+_8433172_8446622_RNA45SN1_45S_50ntFlanks</td><td>10403</td><td>10424</td><td>22</td><td>ACATGGGTCAGTCGGTCCTGAG</td><td>1295.5</td><td>4.1</td><td>0.5</td><td>9.0</td><td>2.5E-19</td><td>1.7E-18</td><td>BT20/MB468</td><td>CYTO</td></tr>
<tr><td>>hg38_21_+_8433172_8446622_RNA45SN1_45S_50ntFlanks@10393.10410.18|CAGCAGTTGAACATGGGT</td><td>>hg38_21_+_8433172_8446622_RNA45SN1_45S_50ntFlanks</td><td>10393</td><td>10410</td><td>18</td><td>CAGCAGTTGAACATGGGT</td><td>2068.1</td><td>-8.8</td><td>1.0</td><td>-9.1</td><td>1.2E-19</td><td>1.7E-18</td><td>MB231/MB468</td><td>MP</td></tr>
<tr><td>>MI0003190|hsa-mir-505&WithFlank&X|-|139924142|139924237@57.80.24&[MIMAT0002876&hsa-miR-505-3p&offsets|0|+2;m-196&X|-|139924160|139924181&offsets|0|+2]|CGTCAACACTTGCTGGTTTCCTCT</td><td>>MI0003190|hsa-mir-505&WithFlank&X|-|139924142|139924237</td><td>57</td><td>80</td><td>24</td><td>CGTCAACACTTGCTGGTTTCCTCT</td><td>282.7</td><td>8.8</td><td>1.0</td><td>9.0</td><td>2.5E-19</td><td>1.7E-18</td><td>BT20/MB468</td><td>CYTO</td></tr>
<tr><td>>trnaMT_PheGAA_MT_+_577_647@35.57.23|GAAAATGTTTAGACGGGCTCACA</td><td>>trnaMT_PheGAA_MT_+_577_647</td><td>35</td><td>57</td><td>23</td><td>GAAAATGTTTAGACGGGCTCACA</td><td>281.5</td><td>7.0</td><td>0.8</td><td>9.0</td><td>1.6E-19</td><td>1.7E-18</td><td>MB231/MB468</td><td>NUC</td></tr>
<tr><td>>hg38_21_+_8433172_8446622_RNA45SN1_45S_50ntFlanks@9027.9057.31|CGTCTCCTCGTGGGGGGGCCGGGCCACCCCT</td><td>>hg38_21_+_8433172_8446622_RNA45SN1_45S_50ntFlanks</td><td>9027</td><td>9057</td><td>31</td><td>CGTCTCCTCGTGGGGGGGCCGGGCCACCCCT</td><td>595.1</td><td>-13.0</td><td>1.4</td><td>-9.0</td><td>1.8E-19</td><td>1.7E-18</td><td>MB231/MB468</td><td>MITO</td></tr>
<tr><td>>am_trna134_GluTTC_1_-_16861774_16861845@1.28.28,trna5_GluTTC_1_+_17199078_17199149@1.28.28,trna84_GluTTC_1_-_161391883_161391954@1.28.28,trna94_GluTTC_1_-_149664355_149664427@1.28.28|TCCCTGGTGGTCTAGTGGCTAGGATTCG</td><td>>am_trna134_GluTTC_1_-_16861774_16861845</td><td>1</td><td>28</td><td>28</td><td>TCCCTGGTGGTCTAGTGGCTAGGATTCG</td><td>1151.4</td><td>-7.2</td><td>0.8</td><td>-9.1</td><td>1.0E-19</td><td>1.7E-18</td><td>MB231/MB468</td><td>CYTO</td></tr>
<tr><td>>MI0000477|hsa-mir-146a&WithFlank&5|+|160485346|160485456@27.50.24&[MIMAT0000449&hsa-miR-146a-5p&offsets|0|+2;m-129&5|+|160485372|160485393&offsets|0|+2]|TGAGAACTGAATTCCATGGGTTGT</td><td>>MI0000477|hsa-mir-146a&WithFlank&5|+|160485346|160485456</td><td>27</td><td>50</td><td>24</td><td>TGAGAACTGAATTCCATGGGTTGT</td><td>1186.2</td><td>13.4</td><td>1.5</td><td>9.1</td><td>1.2E-19</td><td>1.7E-18</td><td>MB231/MB468</td><td>MP</td></tr>
<tr><td>>hg38_21_+_8433172_8446622_RNA45SN1_45S_50ntFlanks@9929.9959.31|GATATAGACAGCAGGACGGTGGCCATGGAAG</td><td>>hg38_21_+_8433172_8446622_RNA45SN1_45S_50ntFlanks</td><td>9929</td><td>9959</td><td>31</td><td>GATATAGACAGCAGGACGGTGGCCATGGAAG</td><td>1483.8</td><td>-14.3</td><td>1.6</td><td>-9.0</td><td>1.9E-19</td><td>1.7E-18</td><td>MB231/MB468</td><td>MITO</td></tr>
<tr><td>>trna16_GlnTTG_17_+_47269890_47269961@1.29.29,trna64_GlnTTG_6_+_28557156_28557227@1.29.29,trna84_GlnTTG_6_+_145503859_145503930@1.29.29|GGTCCCATGGTGTAATGGTTAGCACTCTG</td><td>>trna16_GlnTTG_17_+_47269890_47269961</td><td>1</td><td>29</td><td>29</td><td>GGTCCCATGGTGTAATGGTTAGCACTCTG</td><td>5092.7</td><td>-9.1</td><td>1.0</td><td>-9.1</td><td>1.0E-19</td><td>1.7E-18</td><td>MB231/MB468</td><td>CYTO</td></tr>
<tr><td>>hg38_21_+_8433172_8446622_RNA45SN1_45S_50ntFlanks@12459.12486.28|AGCGTTGGATTGTTCACCCACTAATAGG</td><td>>hg38_21_+_8433172_8446622_RNA45SN1_45S_50ntFlanks</td><td>12459</td><td>12486</td><td>28</td><td>AGCGTTGGATTGTTCACCCACTAATAGG</td><td>270.0</td><td>-12.5</td><td>1.4</td><td>-8.9</td><td>4.6E-19</td><td>1.7E-18</td><td>BT20/MB468</td><td>NUC</td></tr>
<tr><td>>MI0000285|hsa-mir-205&WithFlank&1|+|209432127|209432248@40.59.20(+1C)&[MIMAT0000266&hsa-miR-205-5p&offsets|0|-2(+1C);m-351&1|+|209432166|209432187&offsets|0|-2(+1C)]|TCCTTCATTCCACCGGAGTCC</td><td>>MI0000285|hsa-mir-205&WithFlank&1|+|209432127|209432248</td><td>40</td><td>59</td><td>20(+1C)</td><td>TCCTTCATTCCACCGGAGTCC</td><td>905.6</td><td>10.5</td><td>1.1</td><td>9.3</td><td>8.9E-21</td><td>1.7E-18</td><td>BT20/MB231</td><td>CYTO</td></tr>
<tr><td>>hg38_21_+_8433172_8446622_RNA45SN1_45S_50ntFlanks@12126.12151.26|CGACCCGCTCCGGGGACAGTGCCAGG</td><td>>hg38_21_+_8433172_8446622_RNA45SN1_45S_50ntFlanks</td><td>12126</td><td>12151</td><td>26</td><td>CGACCCGCTCCGGGGACAGTGCCAGG</td><td>1272.6</td><td>-14.1</td><td>1.6</td><td>-9.0</td><td>1.9E-19</td><td>1.7E-18</td><td>MB231/MB468</td><td>MITO</td></tr>
<tr><td>>MI0006444|hsa-mir-1307&WithFlank&10|-|103394247|103394407@86.107.22&[MIMAT0005951&hsa-miR-1307-3p&offsets|0|0;m-66&10|-|103394300|103394322&offsets|0|-1]|ACTCGGCGTGGCGTCGGTCGTG</td><td>>MI0006444|hsa-mir-1307&WithFlank&10|-|103394247|103394407</td><td>86</td><td>107</td><td>22</td><td>ACTCGGCGTGGCGTCGGTCGTG</td><td>812.9</td><td>6.3</td><td>0.7</td><td>9.0</td><td>1.9E-19</td><td>1.7E-18</td><td>BT20/MB468</td><td>MITO</td></tr>
<tr><td>>MI0006444|hsa-mir-1307&WithFlank&10|-|103394247|103394407@86.109.24&[MIMAT0005951&hsa-miR-1307-3p&offsets|0|+2;m-66&10|-|103394300|103394322&offsets|0|+1]|ACTCGGCGTGGCGTCGGTCGTGGT</td><td>>MI0006444|hsa-mir-1307&WithFlank&10|-|103394247|103394407</td><td>86</td><td>109</td><td>24</td><td>ACTCGGCGTGGCGTCGGTCGTGGT</td><td>654.1</td><td>7.5</td><td>0.8</td><td>9.0</td><td>1.9E-19</td><td>1.7E-18</td><td>BT20/MB468</td><td>MITO</td></tr>
<tr><td>>hg38_21_+_8433172_8446622_RNA45SN1_45S_50ntFlanks@12678.12708.31|AATCCCGCCCAGGCGGAACGATACGGCAGCG</td><td>>hg38_21_+_8433172_8446622_RNA45SN1_45S_50ntFlanks</td><td>12678</td><td>12708</td><td>31</td><td>AATCCCGCCCAGGCGGAACGATACGGCAGCG</td><td>482.2</td><td>-12.8</td><td>1.4</td><td>-9.1</td><td>1.2E-19</td><td>1.7E-18</td><td>MB231/MB468</td><td>MP</td></tr>
<tr><td>>hg38_21_+_8433172_8446622_RNA45SN1_45S_50ntFlanks@10682.10706.25|TCCGGCGGCGTCCGGTGAGCTCTCG</td><td>>hg38_21_+_8433172_8446622_RNA45SN1_45S_50ntFlanks</td><td>10682</td><td>10706</td><td>25</td><td>TCCGGCGGCGTCCGGTGAGCTCTCG</td><td>2950.2</td><td>-6.4</td><td>0.7</td><td>-9.0</td><td>1.6E-19</td><td>1.7E-18</td><td>MB231/MB468</td><td>NUC</td></tr>
<tr><td>>hg38_21_+_8433172_8446622_RNA45SN1_45S_50ntFlanks@11323.11342.20|CGTCTCGTCGCGCGCGCGTC</td><td>>hg38_21_+_8433172_8446622_RNA45SN1_45S_50ntFlanks</td><td>11323</td><td>11342</td><td>20</td><td>CGTCTCGTCGCGCGCGCGTC</td><td>11761.6</td><td>-10.9</td><td>1.2</td><td>-9.0</td><td>1.9E-19</td><td>1.7E-18</td><td>MB231/MB468</td><td>MITO</td></tr>
<tr><td>>hg38_21_+_8433172_8446622_RNA45SN1_45S_50ntFlanks@9545.9572.28|GTGGAGGTCCGTAGCGGTCCTGACGTGC</td><td>>hg38_21_+_8433172_8446622_RNA45SN1_45S_50ntFlanks</td><td>9545</td><td>9572</td><td>28</td><td>GTGGAGGTCCGTAGCGGTCCTGACGTGC</td><td>1176.7</td><td>-5.6</td><td>0.6</td><td>-9.0</td><td>1.9E-19</td><td>1.7E-18</td><td>MB231/MB468</td><td>MITO</td></tr>
<tr><td>>hg38_21_+_8433172_8446622_RNA45SN1_45S_50ntFlanks@11541.11583.43|GCGGCGCCTCGCCTCGGCCGGCGCCTAGCAGCCGACTTAGAAC</td><td>>hg38_21_+_8433172_8446622_RNA45SN1_45S_50ntFlanks</td><td>11541</td><td>11583</td><td>43</td><td>GCGGCGCCTCGCCTCGGCCGGCGCCTAGCAGCCGACTTAGAAC</td><td>203.8</td><td>9.3</td><td>1.0</td><td>9.1</td><td>1.1E-19</td><td>1.7E-18</td><td>MB231/MB468</td><td>CYTO</td></tr>
<tr><td>>trnaMT_GluTTC_MT_-_14674_14742@40.71.32,trnalookalike8_GluTTC_5_-_93905172_93905240@40.71.32|TTGGTCGTGGTTGTAGTCCGTGCGAGAATACC</td><td>>trnaMT_GluTTC_MT_-_14674_14742</td><td>40</td><td>71</td><td>32</td><td>TTGGTCGTGGTTGTAGTCCGTGCGAGAATACC</td><td>1826.9</td><td>8.9</td><td>1.0</td><td>9.1</td><td>1.1E-19</td><td>1.7E-18</td><td>MB231/MB468</td><td>CYTO</td></tr>
<tr><td>>hg38_21_+_8433172_8446622_RNA45SN1_45S_50ntFlanks@5379.5399.21|ATTAAGTCCCTGCCCTTTGTA</td><td>>hg38_21_+_8433172_8446622_RNA45SN1_45S_50ntFlanks</td><td>5379</td><td>5399</td><td>21</td><td>ATTAAGTCCCTGCCCTTTGTA</td><td>3533.5</td><td>-15.6</td><td>1.7</td><td>-9.0</td><td>1.9E-19</td><td>1.7E-18</td><td>MB231/MB468</td><td>MITO</td></tr>
<tr><td>>hg38_21_+_8433172_8446622_RNA45SN1_45S_50ntFlanks@11027.11060.34|CTCTCTCTCTCTCTCCCCCGCTCCCCGTCCTCCC</td><td>>hg38_21_+_8433172_8446622_RNA45SN1_45S_50ntFlanks</td><td>11027</td><td>11060</td><td>34</td><td>CTCTCTCTCTCTCTCCCCCGCTCCCCGTCCTCCC</td><td>3363.9</td><td>-14.7</td><td>1.6</td><td>-9.0</td><td>1.6E-19</td><td>1.7E-18</td><td>MB231/MB468</td><td>NUC</td></tr>
<tr><td>>MI0000816|hsa-mir-335&WithFlank&7|+|130496105|130496210@58.79.22&[MIMAT0004703&hsa-miR-335-3p&offsets|0|0;m-159&7|+|130496162|130496183&offsets|0|0]|TTTTTCATTATTGCTCCTGACC</td><td>>MI0000816|hsa-mir-335&WithFlank&7|+|130496105|130496210</td><td>58</td><td>79</td><td>22</td><td>TTTTTCATTATTGCTCCTGACC</td><td>4358.0</td><td>7.3</td><td>0.8</td><td>9.1</td><td>1.1E-19</td><td>1.7E-18</td><td>MB231/MB468</td><td>CYTO</td></tr>
<tr><td>>hg38_21_+_8433172_8446622_RNA45SN1_45S_50ntFlanks@10684.10709.26|CGGCGGCGTCCGGTGAGCTCTCGCTG</td><td>>hg38_21_+_8433172_8446622_RNA45SN1_45S_50ntFlanks</td><td>10684</td><td>10709</td><td>26</td><td>CGGCGGCGTCCGGTGAGCTCTCGCTG</td><td>1667.9</td><td>-7.0</td><td>0.8</td><td>-9.1</td><td>1.1E-19</td><td>1.7E-18</td><td>MB231/MB468</td><td>CYTO</td></tr>
<tr><td>>hg38_21_+_8433172_8446622_RNA45SN1_45S_50ntFlanks@7985.8007.23|GATCAGACGTGGCGACCCGCTGA</td><td>>hg38_21_+_8433172_8446622_RNA45SN1_45S_50ntFlanks</td><td>7985</td><td>8007</td><td>23</td><td>GATCAGACGTGGCGACCCGCTGA</td><td>589.1</td><td>-13.1</td><td>1.4</td><td>-9.1</td><td>1.2E-19</td><td>1.8E-18</td><td>MB231/MB468</td><td>MP</td></tr>
<tr><td>>hg38_21_+_8433172_8446622_RNA45SN1_45S_50ntFlanks@12936.12956.21|AAACCATTCGTAGACGACCTG</td><td>>hg38_21_+_8433172_8446622_RNA45SN1_45S_50ntFlanks</td><td>12936</td><td>12956</td><td>21</td><td>AAACCATTCGTAGACGACCTG</td><td>6065.5</td><td>-4.9</td><td>0.5</td><td>-9.1</td><td>1.1E-19</td><td>1.8E-18</td><td>MB231/MB468</td><td>CYTO</td></tr>
<tr><td>>hg38_21_+_8433172_8446622_RNA45SN1_45S_50ntFlanks@11320.11344.25|CCACGTCTCGTCGCGCGCGCGTCCG</td><td>>hg38_21_+_8433172_8446622_RNA45SN1_45S_50ntFlanks</td><td>11320</td><td>11344</td><td>25</td><td>CCACGTCTCGTCGCGCGCGCGTCCG</td><td>585.5</td><td>-13.1</td><td>1.4</td><td>-9.1</td><td>1.2E-19</td><td>1.8E-18</td><td>MB231/MB468</td><td>MP</td></tr>
<tr><td>>hg38_21_+_8433172_8446622_RNA45SN1_45S_50ntFlanks@11873.11905.33|AAGAAGACCCTGTTGAGCTTGACTCTAGTCTGG</td><td>>hg38_21_+_8433172_8446622_RNA45SN1_45S_50ntFlanks</td><td>11873</td><td>11905</td><td>33</td><td>AAGAAGACCCTGTTGAGCTTGACTCTAGTCTGG</td><td>311.2</td><td>-6.6</td><td>0.7</td><td>-9.1</td><td>1.1E-19</td><td>1.8E-18</td><td>MB231/MB468</td><td>CYTO</td></tr>
<tr><td>>hg38_21_+_8433172_8446622_RNA45SN1_45S_50ntFlanks@3838.3860.23|CTCTCCTACTTGGATAACTGTGG</td><td>>hg38_21_+_8433172_8446622_RNA45SN1_45S_50ntFlanks</td><td>3838</td><td>3860</td><td>23</td><td>CTCTCCTACTTGGATAACTGTGG</td><td>1748.0</td><td>-4.9</td><td>0.5</td><td>-9.0</td><td>1.7E-19</td><td>1.8E-18</td><td>MB231/MB468</td><td>NUC</td></tr>
<tr><td>>MI0000066|hsa-let-7e&WithFlank&19|+|51692780|51692870@14.36.23&[MIMAT0000066&hsa-let-7e-5p&offsets|0|+1;m-50&19|+|51692793|51692814&offsets|0|+1]|TGAGGTAGGAGGTTGTATAGTTG</td><td>>MI0000066|hsa-let-7e&WithFlank&19|+|51692780|51692870</td><td>14</td><td>36</td><td>23</td><td>TGAGGTAGGAGGTTGTATAGTTG</td><td>1676.6</td><td>8.8</td><td>1.0</td><td>9.0</td><td>2.6E-19</td><td>1.8E-18</td><td>BT20/MB468</td><td>CYTO</td></tr>
<tr><td>>MI0000088|hsa-mir-30a&WithFlank&6|-|71403545|71403627@54.74.21&[MIMAT0000088&hsa-miR-30a-3p&offsets|+1|0;m-38&6|-|71403554|71403575&offsets|+1|0]|TTTCAGTCGGATGTTTGCAGC</td><td>>MI0000088|hsa-mir-30a&WithFlank&6|-|71403545|71403627</td><td>54</td><td>74</td><td>21</td><td>TTTCAGTCGGATGTTTGCAGC</td><td>4941.6</td><td>7.2</td><td>0.8</td><td>9.1</td><td>1.1E-19</td><td>1.8E-18</td><td>MB231/MB468</td><td>CYTO</td></tr>
<tr><td>>hg38_21_+_8433172_8446622_RNA45SN1_45S_50ntFlanks@7974.8015.42|ACGCGACCTCAGATCAGACGTGGCGACCCGCTGAATTTAAGC</td><td>>hg38_21_+_8433172_8446622_RNA45SN1_45S_50ntFlanks</td><td>7974</td><td>8015</td><td>42</td><td>ACGCGACCTCAGATCAGACGTGGCGACCCGCTGAATTTAAGC</td><td>1517.3</td><td>7.9</td><td>0.9</td><td>9.1</td><td>1.1E-19</td><td>1.8E-18</td><td>MB231/MB468</td><td>CYTO</td></tr>
<tr><td>>hg38_21_+_8433172_8446622_RNA45SN1_45S_50ntFlanks@11847.11873.27|AACGGGCTTGGCGGAATCAGCGGGGAA</td><td>>hg38_21_+_8433172_8446622_RNA45SN1_45S_50ntFlanks</td><td>11847</td><td>11873</td><td>27</td><td>AACGGGCTTGGCGGAATCAGCGGGGAA</td><td>616.3</td><td>-13.1</td><td>1.4</td><td>-9.1</td><td>1.2E-19</td><td>1.8E-18</td><td>MB231/MB468</td><td>MP</td></tr>
<tr><td>>hg38_21_+_8433172_8446622_RNA45SN1_45S_50ntFlanks@8168.8185.18|CCCAAGTCCTTCTGATCG</td><td>>hg38_21_+_8433172_8446622_RNA45SN1_45S_50ntFlanks</td><td>8168</td><td>8185</td><td>18</td><td>CCCAAGTCCTTCTGATCG</td><td>853.2</td><td>-13.6</td><td>1.5</td><td>-9.1</td><td>1.2E-19</td><td>1.8E-18</td><td>MB231/MB468</td><td>MP</td></tr>
<tr><td>>hg38_21_+_8433172_8446622_RNA45SN1_45S_50ntFlanks@6666.6699.34|GATCACTCGGCTCGTGCGTCGATGAAGAACGCAG</td><td>>hg38_21_+_8433172_8446622_RNA45SN1_45S_50ntFlanks</td><td>6666</td><td>6699</td><td>34</td><td>GATCACTCGGCTCGTGCGTCGATGAAGAACGCAG</td><td>578.6</td><td>-13.0</td><td>1.4</td><td>-9.0</td><td>2.0E-19</td><td>1.8E-18</td><td>MB231/MB468</td><td>MITO</td></tr>
<tr><td>>MI0000264|hsa-mir-7-2&WithFlank&15|+|88611819|88611940@38.61.24(+1A);MI0000265|hsa-mir-7-3&WithFlank&19|+|4770664|4770785@37.60.24(+1A);MI0000263|hsa-mir-7-1&WithFlank&9|-|83969742|83969863@30.53.24(+1A)&[MIMAT0000252&hsa-miR-7-5p&offsets|0|0(+1A);m-400&15|+|88611856|88611879&offsets|0|0(+1A)];[MIMAT0000252_1&hsa-miR-7-5p&offsets|0|0(+1A);m-402&19|+|4770700|4770723&offsets|0|0(+1A)];[MIMAT0000252_2&hsa-miR-7-5p&offsets|0|0(+1A);m-401&9|-|83969811|83969834&offsets|0|0(+1A)]|TGGAAGACTAGTGATTTTGTTGTTA</td><td>>MI0000264|hsa-mir-7-2&WithFlank&15|+|88611819|88611940</td><td>38</td><td>61</td><td>24(+1A)</td><td>TGGAAGACTAGTGATTTTGTTGTTA</td><td>1560.6</td><td>7.7</td><td>0.9</td><td>9.0</td><td>2.6E-19</td><td>1.8E-18</td><td>BT20/MB468</td><td>CYTO</td></tr>
<tr><td>>hg38_21_+_8433172_8446622_RNA45SN1_45S_50ntFlanks@7975.8002.28|CGCGACCTCAGATCAGACGTGGCGACCC</td><td>>hg38_21_+_8433172_8446622_RNA45SN1_45S_50ntFlanks</td><td>7975</td><td>8002</td><td>28</td><td>CGCGACCTCAGATCAGACGTGGCGACCC</td><td>2424.6</td><td>4.2</td><td>0.5</td><td>8.9</td><td>4.9E-19</td><td>1.8E-18</td><td>BT20/MB468</td><td>NUC</td></tr>
<tr><td>>MI0000650|hsa-mir-200c&WithFlank&12|+|6963693|6963772@50.72.23&[MIMAT0000617&hsa-miR-200c-3p&offsets|0|0;m-54&12|+|6963742|6963764&offsets|0|0]|TAATACTGCCGGGTAATGATGGA</td><td>>MI0000650|hsa-mir-200c&WithFlank&12|+|6963693|6963772</td><td>50</td><td>72</td><td>23</td><td>TAATACTGCCGGGTAATGATGGA</td><td>70081.8</td><td>8.6</td><td>0.9</td><td>9.5</td><td>2.0E-21</td><td>1.8E-18</td><td>BT20/MB231</td><td>MP</td></tr>
<tr><td>>MI0001445|hsa-mir-423&WithFlank&17|+|30117073|30117178@23.43.21(+1C)&[MIMAT0004748&hsa-miR-423-5p&offsets|0|-2(+1C);m-141&17|+|30117095|30117117&offsets|0|-2(+1C)]|TGAGGGGCAGAGAGCGAGACTC</td><td>>MI0001445|hsa-mir-423&WithFlank&17|+|30117073|30117178</td><td>23</td><td>43</td><td>21(+1C)</td><td>TGAGGGGCAGAGAGCGAGACTC</td><td>361.5</td><td>10.4</td><td>1.2</td><td>9.0</td><td>2.7E-19</td><td>1.8E-18</td><td>BT20/MB468</td><td>CYTO</td></tr>
<tr><td>>MI0000809|hsa-mir-151a&WithFlank&8|-|140732558|140732659@51.72.22&[MIMAT0000757&hsa-miR-151a-3p&offsets|-2|-1;m-30&8|-|140732587|140732607&offsets|-2|-1]|TACTAGACTGAAGCTCCTTGAG</td><td>>MI0000809|hsa-mir-151a&WithFlank&8|-|140732558|140732659</td><td>51</td><td>72</td><td>22</td><td>TACTAGACTGAAGCTCCTTGAG</td><td>1432.4</td><td>8.3</td><td>0.9</td><td>9.0</td><td>2.7E-19</td><td>1.8E-18</td><td>BT20/MB468</td><td>CYTO</td></tr>
<tr><td>>hg38_21_+_8433172_8446622_RNA45SN1_45S_50ntFlanks@7983.8008.26|CAGATCAGACGTGGCGACCCGCTGAA</td><td>>hg38_21_+_8433172_8446622_RNA45SN1_45S_50ntFlanks</td><td>7983</td><td>8008</td><td>26</td><td>CAGATCAGACGTGGCGACCCGCTGAA</td><td>1083.9</td><td>-5.4</td><td>0.6</td><td>-9.0</td><td>1.7E-19</td><td>1.8E-18</td><td>MB231/MB468</td><td>NUC</td></tr>
<tr><td>>hg38_21_+_8433172_8446622_RNA45SN1_45S_50ntFlanks@10841.10873.33|CGTAACTTCGGGATAAGGATTGGCTCTAAGGGC</td><td>>hg38_21_+_8433172_8446622_RNA45SN1_45S_50ntFlanks</td><td>10841</td><td>10873</td><td>33</td><td>CGTAACTTCGGGATAAGGATTGGCTCTAAGGGC</td><td>1150.5</td><td>-5.3</td><td>0.6</td><td>-9.0</td><td>1.7E-19</td><td>1.9E-18</td><td>MB231/MB468</td><td>NUC</td></tr>
<tr><td>>trnaMT_ValTAC_MT_+_1602_1670@48.72.25|CAACTTAACTTGACCGCTCTGACCA</td><td>>trnaMT_ValTAC_MT_+_1602_1670</td><td>48</td><td>72</td><td>25</td><td>CAACTTAACTTGACCGCTCTGACCA</td><td>3301.4</td><td>10.2</td><td>1.1</td><td>9.0</td><td>2.7E-19</td><td>1.9E-18</td><td>BT20/MB468</td><td>CYTO</td></tr>
<tr><td>>MI0000089|hsa-mir-31&WithFlank&9|-|21512109|21512191@14.36.23&[MIMAT0000089&hsa-miR-31-5p&offsets|0|+2;m-360&9|-|21512157|21512178&offsets|0|+1]|AGGCAAGATGCTGGCATAGCTGT</td><td>>MI0000089|hsa-mir-31&WithFlank&9|-|21512109|21512191</td><td>14</td><td>36</td><td>23</td><td>AGGCAAGATGCTGGCATAGCTGT</td><td>1364.4</td><td>10.8</td><td>1.2</td><td>9.3</td><td>1.1E-20</td><td>1.9E-18</td><td>BT20/MB231</td><td>CYTO</td></tr>
<tr><td>>MI0005756|hsa-mir-934&WithFlank&X|+|136550872|136550966@21.42.22&[MIMAT0004977&hsa-miR-934&offsets|0|0;m-943&X|+|136550892|136550913&offsets|0|0]|TGTCTACTACTGGAGACACTGG</td><td>>MI0005756|hsa-mir-934&WithFlank&X|+|136550872|136550966</td><td>21</td><td>42</td><td>22</td><td>TGTCTACTACTGGAGACACTGG</td><td>994.0</td><td>13.3</td><td>1.4</td><td>9.3</td><td>1.0E-20</td><td>1.9E-18</td><td>BT20/MB231</td><td>CYTO</td></tr>
</table>

<table>
<tr><td>>hg38_21_+_8433172_8446622_RNA45SN1_45S_50ntFlanks@11873.11907.35|AAGAAGACCCTGTTGAGCTTGACTCTAGTCTGGCA</td><td>>hg38_21_+_8433172_8446622_RNA45SN1_45S_50ntFlanks</td><td>11873</td><td>11907</td><td>35</td><td>AAGAAGACCCTGTTGAGCTTGACTCTAGTCTGGCA</td><td>328.5</td><td>-6.7</td><td>0.7</td><td>-9.1</td><td>1.2E-19</td><td>1.9E-18</td><td>MB231/MB468</td><td>CYTO</td></tr>
<tr><td>>hg38_21_+_8433172_8446622_RNA45SN1_45S_50ntFlanks@9201.9220.20|CTCTCGGGGCCACGCGCGCG</td><td>>hg38_21_+_8433172_8446622_RNA45SN1_45S_50ntFlanks</td><td>9201</td><td>9220</td><td>20</td><td>CTCTCGGGGCCACGCGCGCG</td><td>1092.9</td><td>-13.9</td><td>1.5</td><td>-9.0</td><td>2.1E-19</td><td>1.9E-18</td><td>MB231/MB468</td><td>MITO</td></tr>
<tr><td>>trnaMT_GluTTC_MT_-_14674_14742@46.72.27,trnalookalike8_GluTTC_5_-_93905172_93905240@46.72.27|GTGGTTGTAGTCCGTGCGAGAATACCA</td><td>>trnaMT_GluTTC_MT_-_14674_14742</td><td>46</td><td>72</td><td>27</td><td>GTGGTTGTAGTCCGTGCGAGAATACCA</td><td>599.3</td><td>6.2</td><td>0.7</td><td>9.0</td><td>1.8E-19</td><td>1.9E-18</td><td>MB231/MB468</td><td>NUC</td></tr>
<tr><td>>trna10_AspGTC_12_-_125424193_125424264@32.71.40,trna12_AspGTC_12_-_125411891_125411962@32.71.40,trna144_AspGTC_6_-_27551236_27551307@32.71.40,trna38_AspGTC_17_-_8125556_8125627@32.71.40,trna45_AspGTC_6_+_27447453_27447524@32.71.40,trna48_AspGTC_6_+_27471523_27471594@32.71.40,trna4_AspGTC_12_+_96429799_96429870@32.71.40,trna69_AspGTC_1_-_161440205_161440276@32.71.40,trna72_AspGTC_1_-_161432824_161432895@32.71.40,trna75_AspGTC_1_-_161425414_161425485@32.71.40,trna78_AspGTC_1_-_161418033_161418104@32.71.40,trna81_AspGTC_1_-_161410615_161410686@32.71.40|CTGTCACGCGGGAGACCGGGGTTCGATTCCCCGACGGGGA</td><td>>trna10_AspGTC_12_-_125424193_125424264</td><td>32</td><td>71</td><td>40</td><td>CTGTCACGCGGGAGACCGGGGTTCGATTCCCCGACGGGGA</td><td>477.9</td><td>-13.3</td><td>1.5</td><td>-8.9</td><td>5.1E-19</td><td>1.9E-18</td><td>BT20/MB468</td><td>NUC</td></tr>
<tr><td>>hg38_21_+_8433172_8446622_RNA45SN1_45S_50ntFlanks@12991.13028.38|CTCCCTCGCTGCGATCTATTGAAAGTCAGCCCTCGACA</td><td>>hg38_21_+_8433172_8446622_RNA45SN1_45S_50ntFlanks</td><td>12991</td><td>13028</td><td>38</td><td>CTCCCTCGCTGCGATCTATTGAAAGTCAGCCCTCGACA</td><td>1153.1</td><td>-14.0</td><td>1.5</td><td>-9.0</td><td>2.1E-19</td><td>1.9E-18</td><td>MB231/MB468</td><td>MITO</td></tr>
<tr><td>>am_trna128_GlyGCC_6_-_27870686_27870756@1.28.28,trna133_GlyCCC_1_-_16872434_16872504@1.28.28,trna18_GlyGCC_16_+_70822597_70822667@1.28.28,trna19_GlyGCC_16_+_70823410_70823480@1.28.28,trna19_GlyGCC_2_-_157257659_157257729@1.28.28,trna24_GlyGCC_16_-_70812942_70813012@1.28.28,trna25_GlyGCC_16_-_70812114_70812184@1.28.28,trna4_GlyCCC_1_+_17188416_17188486@1.28.28,trna5_GlyGCC_17_+_8029064_8029134@1.28.28,trna68_GlyGCC_1_-_161493637_161493707@1.28.28|GCATTGGTGGTTCAGTGGTAGAATTCTC</td><td>>am_trna128_GlyGCC_6_-_27870686_27870756</td><td>1</td><td>28</td><td>28</td><td>GCATTGGTGGTTCAGTGGTAGAATTCTC</td><td>104063.4</td><td>-9.7</td><td>1.1</td><td>-8.9</td><td>5.1E-19</td><td>1.9E-18</td><td>BT20/MB468</td><td>NUC</td></tr>
<tr><td>>hg38_21_+_8433172_8446622_RNA45SN1_45S_50ntFlanks@4726.4758.33|TAATCAAGAACGAAAGTCGGAGGTTCGAAGACG</td><td>>hg38_21_+_8433172_8446622_RNA45SN1_45S_50ntFlanks</td><td>4726</td><td>4758</td><td>33</td><td>TAATCAAGAACGAAAGTCGGAGGTTCGAAGACG</td><td>2429.6</td><td>-5.7</td><td>0.6</td><td>-9.0</td><td>2.0E-19</td><td>1.9E-18</td><td>BT20/MB468</td><td>MITO</td></tr>
<tr><td>>hg38_21_+_8433172_8446622_RNA45SN1_45S_50ntFlanks@6739.6770.32|TCATCGACACTTCGAACGCACTTGCGGCCCCG</td><td>>hg38_21_+_8433172_8446622_RNA45SN1_45S_50ntFlanks</td><td>6739</td><td>6770</td><td>32</td><td>TCATCGACACTTCGAACGCACTTGCGGCCCCG</td><td>644.8</td><td>-13.2</td><td>1.5</td><td>-9.1</td><td>1.3E-19</td><td>1.9E-18</td><td>MB231/MB468</td><td>MP</td></tr>
<tr><td>>hg38_21_+_8433172_8446622_RNA45SN1_45S_50ntFlanks@11575.11592.18|ACTTAGAACTGGTGCGGA</td><td>>hg38_21_+_8433172_8446622_RNA45SN1_45S_50ntFlanks</td><td>11575</td><td>11592</td><td>18</td><td>ACTTAGAACTGGTGCGGA</td><td>5562.3</td><td>-12.4</td><td>1.4</td><td>-8.9</td><td>5.1E-19</td><td>1.9E-18</td><td>BT20/MB468</td><td>NUC</td></tr>
<tr><td>>hg38_21_+_8433172_8446622_RNA45SN1_45S_50ntFlanks@10408.10433.26|GGTCAGTCGGTCCTGAGAGATGGGCG</td><td>>hg38_21_+_8433172_8446622_RNA45SN1_45S_50ntFlanks</td><td>10408</td><td>10433</td><td>26</td><td>GGTCAGTCGGTCCTGAGAGATGGGCG</td><td>1366.6</td><td>-14.3</td><td>1.6</td><td>-9.1</td><td>1.3E-19</td><td>1.9E-18</td><td>MB231/MB468</td><td>MP</td></tr>
<tr><td>>hg38_21_+_8433172_8446622_RNA45SN1_45S_50ntFlanks@11641.11662.22|CGGCGGGTGTTGACGCGATGTG</td><td>>hg38_21_+_8433172_8446622_RNA45SN1_45S_50ntFlanks</td><td>11641</td><td>11662</td><td>22</td><td>CGGCGGGTGTTGACGCGATGTG</td><td>12865.4</td><td>-12.2</td><td>1.3</td><td>-9.1</td><td>1.3E-19</td><td>1.9E-18</td><td>MB231/MB468</td><td>MP</td></tr>
<tr><td>>hg38_21_+_8433172_8446622_RNA45SN1_45S_50ntFlanks@9204.9223.20|TCGGGGCCACGCGCGCGTCC</td><td>>hg38_21_+_8433172_8446622_RNA45SN1_45S_50ntFlanks</td><td>9204</td><td>9223</td><td>20</td><td>TCGGGGCCACGCGCGCGTCC</td><td>714.3</td><td>-13.3</td><td>1.5</td><td>-9.1</td><td>1.3E-19</td><td>1.9E-18</td><td>MB231/MB468</td><td>MP</td></tr>
<tr><td>>hg38_21_+_8433172_8446622_RNA45SN1_45S_50ntFlanks@11615.11640.26|TTAAAACAAAGCATCGCGAAGGCCCG</td><td>>hg38_21_+_8433172_8446622_RNA45SN1_45S_50ntFlanks</td><td>11615</td><td>11640</td><td>26</td><td>TTAAAACAAAGCATCGCGAAGGCCCG</td><td>371.5</td><td>-12.4</td><td>1.4</td><td>-9.1</td><td>1.3E-19</td><td>1.9E-18</td><td>MB231/MB468</td><td>MP</td></tr>
<tr><td>>hg38_21_+_8433172_8446622_RNA45SN1_45S_50ntFlanks@11613.11640.28|AATTAAAACAAAGCATCGCGAAGGCCCG</td><td>>hg38_21_+_8433172_8446622_RNA45SN1_45S_50ntFlanks</td><td>11613</td><td>11640</td><td>28</td><td>AATTAAAACAAAGCATCGCGAAGGCCCG</td><td>2358.1</td><td>-14.3</td><td>1.5</td><td>-9.5</td><td>2.6E-21</td><td>1.9E-18</td><td>BT20/MB468</td><td>MP</td></tr>
<tr><td>>hg38_21_+_8433172_8446622_RNA45SN1_45S_50ntFlanks@6771.6799.29|GGTTCCTCCCGGGGCTACGCCTGTCTGAG</td><td>>hg38_21_+_8433172_8446622_RNA45SN1_45S_50ntFlanks</td><td>6771</td><td>6799</td><td>29</td><td>GGTTCCTCCCGGGGCTACGCCTGTCTGAG</td><td>699.0</td><td>-13.3</td><td>1.5</td><td>-9.1</td><td>1.3E-19</td><td>1.9E-18</td><td>MB231/MB468</td><td>MP</td></tr>
<tr><td>>hg38_21_+_8433172_8446622_RNA45SN1_45S_50ntFlanks@12008.12034.27|CCGCCGGTGAAATACCACTACTCTGAT</td><td>>hg38_21_+_8433172_8446622_RNA45SN1_45S_50ntFlanks</td><td>12008</td><td>12034</td><td>27</td><td>CCGCCGGTGAAATACCACTACTCTGAT</td><td>438.5</td><td>-12.6</td><td>1.4</td><td>-9.1</td><td>1.3E-19</td><td>1.9E-18</td><td>MB231/MB468</td><td>MP</td></tr>
<tr><td>>hg38_21_+_8433172_8446622_RNA45SN1_45S_50ntFlanks@10346.10374.29|TCAAACGAGAACTTTGAAGGCCGAAGTGG</td><td>>hg38_21_+_8433172_8446622_RNA45SN1_45S_50ntFlanks</td><td>10346</td><td>10374</td><td>29</td><td>TCAAACGAGAACTTTGAAGGCCGAAGTGG</td><td>2865.3</td><td>-8.5</td><td>0.9</td><td>-9.0</td><td>2.1E-19</td><td>1.9E-18</td><td>MB231/MB468</td><td>MITO</td></tr>
<tr><td>>MI0000650|hsa-mir-200c&WithFlank&12|+|6963693|6963772@50.72.23(+2A)&[MIMAT0000617&hsa-miR-200c-3p&offsets|0|0(+2A);m-54&12|+|6963742|6963764&offsets|0|0(+2A)]|TAATACTGCCGGGTAATGATGGAAA</td><td>>MI0000650|hsa-mir-200c&WithFlank&12|+|6963693|6963772</td><td>50</td><td>72</td><td>23(+2A)</td><td>TAATACTGCCGGGTAATGATGGAAA</td><td>685.0</td><td>13.0</td><td>1.4</td><td>9.3</td><td>1.7E-20</td><td>1.9E-18</td><td>BT20/MB231</td><td>MITO</td></tr>
<tr><td>>MI0005756|hsa-mir-934&WithFlank&X|+|136550872|136550966@21.43.23&[MIMAT0004977&hsa-miR-934&offsets|0|+1;m-943&X|+|136550892|136550913&offsets|0|+1]|TGTCTACTACTGGAGACACTGGT</td><td>>MI0005756|hsa-mir-934&WithFlank&X|+|136550872|136550966</td><td>21</td><td>43</td><td>23</td><td>TGTCTACTACTGGAGACACTGGT</td><td>1628.9</td><td>10.8</td><td>1.2</td><td>9.3</td><td>1.2E-20</td><td>1.9E-18</td><td>BT20/MB231</td><td>CYTO</td></tr>
<tr><td>>hg38_21_+_8433172_8446622_RNA45SN1_45S_50ntFlanks@9567.9594.28|ACGTGCAAATCGGTCGTCCGACCTGGGT</td><td>>hg38_21_+_8433172_8446622_RNA45SN1_45S_50ntFlanks</td><td>9567</td><td>9594</td><td>28</td><td>ACGTGCAAATCGGTCGTCCGACCTGGGT</td><td>1185.4</td><td>-7.0</td><td>0.8</td><td>-9.0</td><td>2.1E-19</td><td>1.9E-18</td><td>MB231/MB468</td><td>MITO</td></tr>
<tr><td>>hg38_21_+_8433172_8446622_RNA45SN1_45S_50ntFlanks@6741.6772.32|ATCGACACTTCGAACGCACTTGCGGCCCCGGG</td><td>>hg38_21_+_8433172_8446622_RNA45SN1_45S_50ntFlanks</td><td>6741</td><td>6772</td><td>32</td><td>ATCGACACTTCGAACGCACTTGCGGCCCCGGG</td><td>2307.1</td><td>-8.9</td><td>1.0</td><td>-9.0</td><td>2.1E-19</td><td>1.9E-18</td><td>MB231/MB468</td><td>MITO</td></tr>
<tr><td>>am_trna106_HisGTG_1_-_149155828_149155899@7.29.23,trna111_HisGTG_1_-_147774845_147774916@7.29.23,trna118_HisGTG_1_-_145396881_145396952@7.29.23,trna16_HisGTG_1_+_146544773_146544844@7.29.23,trna1_HisGTG_15_+_45493349_45493420@7.29.23,trna21_HisGTG_1_+_147753471_147753542@7.29.23,trna33_HisGTG_6_+_27125906_27125977@7.29.23,trna7_HisGTG_9_-_14433938_14434009@7.29.23,trna8_HisGTG_15_-_45492611_45492682@7.29.23,trna9_HisGTG_15_-_45490804_45490875@7.29.23|ATCGTATAGTGGTTAGTACTCTG</td><td>>am_trna106_HisGTG_1_-_149155828_149155899</td><td>7</td><td>29</td><td>23</td><td>ATCGTATAGTGGTTAGTACTCTG</td><td>572.5</td><td>-12.9</td><td>1.4</td><td>-9.0</td><td>2.1E-19</td><td>1.9E-18</td><td>MB231/MB468</td><td>MITO</td></tr>
<tr><td>>hg38_21_+_8433172_8446622_RNA45SN1_45S_50ntFlanks@6664.6688.25|TGGATCACTCGGCTCGTGCGTCGAT</td><td>>hg38_21_+_8433172_8446622_RNA45SN1_45S_50ntFlanks</td><td>6664</td><td>6688</td><td>25</td><td>TGGATCACTCGGCTCGTGCGTCGAT</td><td>457.0</td><td>-12.7</td><td>1.4</td><td>-9.1</td><td>1.4E-19</td><td>1.9E-18</td><td>MB231/MB468</td><td>MP</td></tr>
<tr><td>>MI0000286|hsa-mir-210&WithFlank&11|-|568083|568204@72.93.22(+1A)&[MIMAT0000267&hsa-miR-210-3p&offsets|0|0(+1A);m-102&11|-|568112|568133&offsets|0|0(+1A)]|CTGTGCGTGTGACAGCGGCTGAA</td><td>>MI0000286|hsa-mir-210&WithFlank&11|-|568083|568204</td><td>72</td><td>93</td><td>22(+1A)</td><td>CTGTGCGTGTGACAGCGGCTGAA</td><td>1265.5</td><td>14.6</td><td>1.6</td><td>9.0</td><td>2.9E-19</td><td>1.9E-18</td><td>BT20/MB468</td><td>CYTO</td></tr>
<tr><td>>hg38_21_+_8433172_8446622_RNA45SN1_45S_50ntFlanks@11849.11873.25|CGGGCTTGGCGGAATCAGCGGGGAA</td><td>>hg38_21_+_8433172_8446622_RNA45SN1_45S_50ntFlanks</td><td>11849</td><td>11873</td><td>25</td><td>CGGGCTTGGCGGAATCAGCGGGGAA</td><td>596.4</td><td>-13.1</td><td>1.4</td><td>-9.1</td><td>1.4E-19</td><td>1.9E-18</td><td>MB231/MB468</td><td>MP</td></tr>
<tr><td>>MI0000750|hsa-mir-26a-2&WithFlank&12|-|57824603|57824698@23.41.19;MI0000083|hsa-mir-26a-1&WithFlank&3|+|37969398|37969486@19.37.19&[MIMAT0000082&hsa-miR-26a-5p&offsets|+3|0;m-33&12|-|57824658|57824679&offsets|+3|0];[MIMAT0000082_1&hsa-miR-26a-5p&offsets|+3|0;m-34&3|+|37969413|37969434&offsets|+3|0]|AAGTAATCCAGGATAGGCT</td><td>>MI0000750|hsa-mir-26a-2&WithFlank&12|-|57824603|57824698</td><td>23</td><td>41</td><td>19</td><td>AAGTAATCCAGGATAGGCT</td><td>195.6</td><td>10.4</td><td>1.2</td><td>8.9</td><td>5.3E-19</td><td>1.9E-18</td><td>BT20/MB468</td><td>NUC</td></tr>
<tr><td>>hg38_21_+_8433172_8446622_RNA45SN1_45S_50ntFlanks@10683.10710.28|CCGGCGGCGTCCGGTGAGCTCTCGCTGG</td><td>>hg38_21_+_8433172_8446622_RNA45SN1_45S_50ntFlanks</td><td>10683</td><td>10710</td><td>28</td><td>CCGGCGGCGTCCGGTGAGCTCTCGCTGG</td><td>461.2</td><td>-12.7</td><td>1.4</td><td>-9.1</td><td>1.4E-19</td><td>1.9E-18</td><td>MB231/MB468</td><td>MP</td></tr>
<tr><td>>hg38_21_+_8433172_8446622_RNA45SN1_45S_50ntFlanks@10351.10388.38|CGAGAACTTTGAAGGCCGAAGTGGAGAAGGGTTCCATG</td><td>>hg38_21_+_8433172_8446622_RNA45SN1_45S_50ntFlanks</td><td>10351</td><td>10388</td><td>38</td><td>CGAGAACTTTGAAGGCCGAAGTGGAGAAGGGTTCCATG</td><td>8326.6</td><td>-7.7</td><td>0.9</td><td>-9.0</td><td>1.9E-19</td><td>2.0E-18</td><td>MB231/MB468</td><td>NUC</td></tr>
<tr><td>>hg38_21_+_8433172_8446622_RNA45SN1_45S_50ntFlanks@12457.12478.22|CAAGCGTTGGATTGTTCACCCA</td><td>>hg38_21_+_8433172_8446622_RNA45SN1_45S_50ntFlanks</td><td>12457</td><td>12478</td><td>22</td><td>CAAGCGTTGGATTGTTCACCCA</td><td>3308.3</td><td>-6.2</td><td>0.7</td><td>-9.0</td><td>1.9E-19</td><td>2.0E-18</td><td>MB231/MB468</td><td>NUC</td></tr>
<tr><td>>hg38_21_+_8433172_8446622_RNA45SN1_45S_50ntFlanks@12624.12659.36|CCAATGGGGCGAAGCTACCATCTGTGGGATTATGAC</td><td>>hg38_21_+_8433172_8446622_RNA45SN1_45S_50ntFlanks</td><td>12624</td><td>12659</td><td>36</td><td>CCAATGGGGCGAAGCTACCATCTGTGGGATTATGAC</td><td>1087.6</td><td>-12.2</td><td>1.4</td><td>-9.0</td><td>2.1E-19</td><td>2.0E-18</td><td>BT20/MB468</td><td>MITO</td></tr>
<tr><td>>hg38_21_+_8433172_8446622_RNA45SN1_45S_50ntFlanks@12678.12703.26|AATCCCGCCCAGGCGGAACGATACGG</td><td>>hg38_21_+_8433172_8446622_RNA45SN1_45S_50ntFlanks</td><td>12678</td><td>12703</td><td>26</td><td>AATCCCGCCCAGGCGGAACGATACGG</td><td>680.8</td><td>-13.2</td><td>1.5</td><td>-9.0</td><td>2.2E-19</td><td>2.0E-18</td><td>MB231/MB468</td><td>MITO</td></tr>
<tr><td>>hg38_21_+_8433172_8446622_RNA45SN1_45S_50ntFlanks@10348.10365.18|AAACGAGAACTTTGAAGG</td><td>>hg38_21_+_8433172_8446622_RNA45SN1_45S_50ntFlanks</td><td>10348</td><td>10365</td><td>18</td><td>AAACGAGAACTTTGAAGG</td><td>588.1</td><td>-6.5</td><td>0.7</td><td>-8.9</td><td>5.4E-19</td><td>2.0E-18</td><td>BT20/MB468</td><td>NUC</td></tr>
<tr><td>>hg38_21_+_8433172_8446622_RNA45SN1_45S_50ntFlanks@7978.8017.40|GACCTCAGATCAGACGTGGCGACCCGCTGAATTTAAGCAT</td><td>>hg38_21_+_8433172_8446622_RNA45SN1_45S_50ntFlanks</td><td>7978</td><td>8017</td><td>40</td><td>GACCTCAGATCAGACGTGGCGACCCGCTGAATTTAAGCAT</td><td>430.9</td><td>12.8</td><td>1.4</td><td>9.1</td><td>1.2E-19</td><td>2.0E-18</td><td>MB231/MB468</td><td>CYTO</td></tr>
<tr><td>>MI0002468|hsa-mir-484&WithFlank&16|+|15643288|15643378@14.33.20(+2U)&[MIMAT0002174&hsa-miR-484&offsets|0|-2(+2U);m-127&16|+|15643301|15643321&offsets|0|-1(+2U)]|TCAGGCTCAGTCCCCTCCCGTT</td><td>>MI0002468|hsa-mir-484&WithFlank&16|+|15643288|15643378</td><td>14</td><td>33</td><td>20(+2U)</td><td>TCAGGCTCAGTCCCCTCCCGTT</td><td>347.6</td><td>9.2</td><td>1.0</td><td>9.0</td><td>2.9E-19</td><td>2.0E-18</td><td>BT20/MB468</td><td>CYTO</td></tr>
<tr><td>>hg38_21_+_8433172_8446622_RNA45SN1_45S_50ntFlanks@3802.3842.41|CATTAAATCAGTTATGGTTCCTTTGGTCGCTCGCTCCTCTC</td><td>>hg38_21_+_8433172_8446622_RNA45SN1_45S_50ntFlanks</td><td>3802</td><td>3842</td><td>41</td><td>CATTAAATCAGTTATGGTTCCTTTGGTCGCTCGCTCCTCTC</td><td>608.9</td><td>-13.7</td><td>1.5</td><td>-8.9</td><td>5.5E-19</td><td>2.0E-18</td><td>BT20/MB468</td><td>NUC</td></tr>
<tr><td>>hg38_21_+_8433172_8446622_RNA45SN1_45S_50ntFlanks@4724.4758.35|ATTAATCAAGAACGAAAGTCGGAGGTTCGAAGACG</td><td>>hg38_21_+_8433172_8446622_RNA45SN1_45S_50ntFlanks</td><td>4724</td><td>4758</td><td>35</td><td>ATTAATCAAGAACGAAAGTCGGAGGTTCGAAGACG</td><td>31801.9</td><td>-11.3</td><td>1.3</td><td>-8.9</td><td>5.5E-19</td><td>2.0E-18</td><td>BT20/MB468</td><td>NUC</td></tr>
<tr><td>>hg38_21_+_8433172_8446622_RNA45SN1_45S_50ntFlanks@10808.10830.23|CAATGTAGGTAAGGGAAGTCGGC</td><td>>hg38_21_+_8433172_8446622_RNA45SN1_45S_50ntFlanks</td><td>10808</td><td>10830</td><td>23</td><td>CAATGTAGGTAAGGGAAGTCGGC</td><td>511.3</td><td>-12.9</td><td>1.4</td><td>-9.1</td><td>1.4E-19</td><td>2.0E-18</td><td>MB231/MB468</td><td>MP</td></tr>
<tr><td>>hg38_21_+_8433172_8446622_RNA45SN1_45S_50ntFlanks@4725.4758.34|TTAATCAAGAACGAAAGTCGGAGGTTCGAAGACG</td><td>>hg38_21_+_8433172_8446622_RNA45SN1_45S_50ntFlanks</td><td>4725</td><td>4758</td><td>34</td><td>TTAATCAAGAACGAAAGTCGGAGGTTCGAAGACG</td><td>1893.3</td><td>-4.6</td><td>0.5</td><td>-9.0</td><td>1.9E-19</td><td>2.0E-18</td><td>MB231/MB468</td><td>NUC</td></tr>
<tr><td>>hg38_21_+_8433172_8446622_RNA45SN1_45S_50ntFlanks@5489.5518.30|CTGAGAAGACGGTCGAACTTGACTATCTAG</td><td>>hg38_21_+_8433172_8446622_RNA45SN1_45S_50ntFlanks</td><td>5489</td><td>5518</td><td>30</td><td>CTGAGAAGACGGTCGAACTTGACTATCTAG</td><td>777.2</td><td>-13.5</td><td>1.5</td><td>-9.1</td><td>1.4E-19</td><td>2.0E-18</td><td>MB231/MB468</td><td>MP</td></tr>
<tr><td>>hg38_21_+_8433172_8446622_RNA45SN1_45S_50ntFlanks@7977.8019.43|CGACCTCAGATCAGACGTGGCGACCCGCTGAATTTAAGCATAT</td><td>>hg38_21_+_8433172_8446622_RNA45SN1_45S_50ntFlanks</td><td>7977</td><td>8019</td><td>43</td><td>CGACCTCAGATCAGACGTGGCGACCCGCTGAATTTAAGCATAT</td><td>724.5</td><td>13.5</td><td>1.5</td><td>9.1</td><td>1.3E-19</td><td>2.0E-18</td><td>MB231/MB468</td><td>CYTO</td></tr>
<tr><td>>hg38_21_+_8433172_8446622_RNA45SN1_45S_50ntFlanks@11876.11905.30|AAGACCCTGTTGAGCTTGACTCTAGTCTGG</td><td>>hg38_21_+_8433172_8446622_RNA45SN1_45S_50ntFlanks</td><td>11876</td><td>11905</td><td>30</td><td>AAGACCCTGTTGAGCTTGACTCTAGTCTGG</td><td>1014.5</td><td>-13.8</td><td>1.5</td><td>-9.0</td><td>2.3E-19</td><td>2.0E-18</td><td>MB231/MB468</td><td>MITO</td></tr>
</table>

| | | | | | | | | | | | | | |
|---|---|---|---|---|---|---|---|---|---|---|---|---|---|
| >MI0000077\|hsa-mir-21&WithFlank&17\|+\|59841260\|59841343@15.36.22&[MIMAT0000076&hsa-miR-21-5p&offsets\|+1\|+1;m-1&17\|+\|59841273\|59841294&offsets\|+1\|+1]\|AGCTTATCAGACTGATGTTGAC | >MI0000077\|hsa-mir-21&WithFlank&17\|+\|59841260\|59841343 | 15 | 36 | 22 | AGCTTATCAGACTGATGTTGAC | 962.1 | 6.4 | 0.7 | 9.0 | 1.9E-19 | 2.0E-18 | MB231/MB468 | NUC |
| >MI0003667\|hsa-mir-652&WithFlank&X\|+\|110055323\|110055432@67.87.21(+1U)&[MIMAT0003322&hsa-miR-652-3p&offsets\|0\|0(+1U);m-148&X\|+\|110055389\|110055410&offsets\|0\|-1(+1U)]\|AATGGCGCCACTAGGGTTGTGT | >MI0003667\|hsa-mir-652&WithFlank&X\|+\|110055323\|110055432 | 67 | 87 | 21(+1U) | AATGGCGCCACTAGGGTTGTGT | 2050.8 | 8.9 | 1.0 | 9.0 | 3.0E-19 | 2.0E-18 | BT20/MB468 | CYTO |
| >hg38_21_+_8433172_8446622_RNA45SN1_45S_50ntFlanks@10845.10868.24\|ACTTCGGGATAAGGATTGGCTCTA | >hg38_21_+_8433172_8446622_RNA45SN1_45S_50ntFlanks | 10845 | 10868 | 24 | ACTTCGGGATAAGGATTGGCTCTA | 860.9 | -13.6 | 1.5 | -9.0 | 1.4E-19 | 2.0E-18 | MB231/MB468 | MP |
| >hg38_21_+_8433172_8446622_RNA45SN1_45S_50ntFlanks@4297.4316.20\|CATTGGAGGGCAAGTCTGGT | >hg38_21_+_8433172_8446622_RNA45SN1_45S_50ntFlanks | 4297 | 4316 | 20 | CATTGGAGGGCAAGTCTGGT | 596.2 | -5.5 | 0.6 | -8.9 | 5.6E-19 | 2.1E-18 | BT20/MB468 | NUC |
| >hg38_21_+_8433172_8446622_RNA45SN1_45S_50ntFlanks@11579.11605.27\|AGAACTGGTGCGGACCAGGGGAATCCG | >hg38_21_+_8433172_8446622_RNA45SN1_45S_50ntFlanks | 11579 | 11605 | 27 | AGAACTGGTGCGGACCAGGGGAATCCG | 1677.2 | -4.9 | 0.5 | -9.0 | 3.1E-19 | 2.1E-18 | BT20/MB468 | CYTO |
| >hg38_21_+_8433172_8446622_RNA45SN1_45S_50ntFlanks@3882.3899.18\|CCGACGGGCGCTGACCCC | >hg38_21_+_8433172_8446622_RNA45SN1_45S_50ntFlanks | 3882 | 3899 | 18 | CCGACGGGCGCTGACCCC | 4987.3 | -11.4 | 1.3 | -9.0 | 1.5E-19 | 2.1E-18 | MB231/MB468 | MP |
| >hg38_21_+_8433172_8446622_RNA45SN1_45S_50ntFlanks@11849.11871.23\|CGGGCTTGGCGGAATCAGCGGGG | >hg38_21_+_8433172_8446622_RNA45SN1_45S_50ntFlanks | 11849 | 11871 | 23 | CGGGCTTGGCGGAATCAGCGGGG | 570.7 | -13.0 | 1.4 | -9.0 | 1.5E-19 | 2.1E-18 | MB231/MB468 | MP |
| >hg38_21_+_8433172_8446622_RNA45SN1_45S_50ntFlanks@10351.10383.33\|CGAGAACTTTGAAGGCCGAAGTGGAGAAGGGTT | >hg38_21_+_8433172_8446622_RNA45SN1_45S_50ntFlanks | 10351 | 10383 | 33 | CGAGAACTTTGAAGGCCGAAGTGGAGAAGGGTT | 15277.6 | -5.1 | 0.6 | -9.0 | 3.1E-19 | 2.1E-18 | BT20/MB468 | CYTO |
| >hg38_21_+_8433172_8446622_RNA45SN1_45S_50ntFlanks@9621.9652.32\|TCTAGTAGCTGGTTCCCTCCGAAGTTTCCCTC | >hg38_21_+_8433172_8446622_RNA45SN1_45S_50ntFlanks | 9621 | 9652 | 32 | TCTAGTAGCTGGTTCCCTCCGAAGTTTCCCTC | 657.0 | -13.1 | 1.5 | -9.0 | 2.3E-19 | 2.1E-18 | MB231/MB468 | MITO |
| >MI0003591\|hsa-mir-584&WithFlank&5\|-\|149062307\|149062415@22.41.20&[MIMAT0003249&hsa-miR-584-5p&offsets\|0\|-2;m-167&5\|-\|149062374\|149062394&offsets\|0\|-1]\|TTATGGTTTGCCTGGGACTG | >MI0003591\|hsa-mir-584&WithFlank&5\|-\|149062307\|149062415 | 22 | 41 | 20 | TTATGGTTTGCCTGGGACTG | 328.0 | 5.9 | 0.7 | 9.0 | 2.0E-19 | 2.1E-18 | MB231/MB468 | NUC |
| >hg38_21_+_8433172_8446622_RNA45SN1_45S_50ntFlanks@4722.4748.27\|TCATTAATCAAGAACGAAAGTCGGAGG | >hg38_21_+_8433172_8446622_RNA45SN1_45S_50ntFlanks | 4722 | 4748 | 27 | TCATTAATCAAGAACGAAAGTCGGAGG | 573.9 | -13.0 | 1.4 | -9.0 | 1.5E-19 | 2.1E-18 | MB231/MB468 | MP |
| >hg38_21_+_8433172_8446622_RNA45SN1_45S_50ntFlanks@10351.10379.29\|CGAGAACTTTGAAGGCCGAAGTGGAGAAG | >hg38_21_+_8433172_8446622_RNA45SN1_45S_50ntFlanks | 10351 | 10379 | 29 | CGAGAACTTTGAAGGCCGAAGTGGAGAAG | 1218.0 | -4.9 | 0.5 | -9.0 | 2.0E-19 | 2.1E-18 | MB231/MB468 | NUC |
| >hg38_21_+_8433172_8446622_RNA45SN1_45S_50ntFlanks@12658.12675.18\|ACTGAACGCCTCTAAGTC | >hg38_21_+_8433172_8446622_RNA45SN1_45S_50ntFlanks | 12658 | 12675 | 18 | ACTGAACGCCTCTAAGTC | 634.4 | -13.2 | 1.5 | -9.0 | 1.5E-19 | 2.1E-18 | MB231/MB468 | MP |
| >hg38_21_+_8433172_8446622_RNA45SN1_45S_50ntFlanks@9570.9595.26\|TGCAAATCGGTCGTCCGACCTGGGTA | >hg38_21_+_8433172_8446622_RNA45SN1_45S_50ntFlanks | 9570 | 9595 | 26 | TGCAAATCGGTCGTCCGACCTGGGTA | 3361.9 | -9.9 | 1.1 | -9.0 | 1.5E-19 | 2.1E-18 | MB231/MB468 | MP |
| >hg38_21_+_8433172_8446622_RNA45SN1_45S_50ntFlanks@9459.9503.45\|CCGGGGAGGTGGAGCACGAGCGCACGTGTTAGGACCCGAAAGATG | >hg38_21_+_8433172_8446622_RNA45SN1_45S_50ntFlanks | 9459 | 9503 | 45 | CCGGGGAGGTGGAGCACGAGCGCACGTGTTAGGACCCGAAAGATG | 581.1 | -13.6 | 1.5 | -8.9 | 5.8E-19 | 2.1E-18 | BT20/MB468 | NUC |
| >hg38_21_+_8433172_8446622_RNA45SN1_45S_50ntFlanks@7985.8006.22\|GATCAGACGTGGCGACCCGCTG | >hg38_21_+_8433172_8446622_RNA45SN1_45S_50ntFlanks | 7985 | 8006 | 22 | GATCAGACGTGGCGACCCGCTG | 824.9 | -12.0 | 1.3 | -9.0 | 3.2E-19 | 2.1E-18 | BT20/MB468 | CYTO |
| >hg38_21_+_8433172_8446622_RNA45SN1_45S_50ntFlanks@12306.12323.18\|CACGATCCTTCTGACCTT | >hg38_21_+_8433172_8446622_RNA45SN1_45S_50ntFlanks | 12306 | 12323 | 18 | CACGATCCTTCTGACCTT | 9296.8 | -12.8 | 1.4 | -9.0 | 2.4E-19 | 2.1E-18 | MB231/MB468 | MITO |
| >MI0000272\|hsa-mir-182&WithFlank&7\|-\|129770377\|129770498@29.49.21&[MIMAT0000259&hsa-miR-182-5p&offsets\|0\|-3;m-22&7\|-\|129770449\|129770470&offsets\|0\|-1]\|TTTGGCAATGGTAGAACTCAC | >MI0000272\|hsa-mir-182&WithFlank&7\|-\|129770377\|129770498 | 29 | 49 | 21 | TTTGGCAATGGTAGAACTCAC | 165.3 | 2.7 | 0.3 | 8.9 | 5.9E-19 | 2.1E-18 | BT20/MB468 | NUC |
| >MI0010635\|hsa-mir-2116&WithFlank&15\|-\|59171177\|59171268@56.76.21&[MIMAT0011161&hsa-miR-2116-3p&offsets\|-1\|-1;m-656&15\|-\|59171193\|59171213&offsets\|0\|0]\|TCCTCCCATGCCAAGAACTCC | >MI0010635\|hsa-mir-2116&WithFlank&15\|-\|59171177\|59171268 | 56 | 76 | 21 | TCCTCCCATGCCAAGAACTCC | 310.5 | 12.5 | 1.4 | 9.0 | 3.2E-19 | 2.2E-18 | BT20/MB468 | CYTO |
| >hg38_21_+_8433172_8446622_RNA45SN1_45S_50ntFlanks@7974.7991.18\|ACGCGACCTCAGATCAGA | >hg38_21_+_8433172_8446622_RNA45SN1_45S_50ntFlanks | 7974 | 7991 | 18 | ACGCGACCTCAGATCAGA | 22264.4 | 6.7 | 0.7 | 9.0 | 2.4E-19 | 2.2E-18 | BT20/MB468 | MITO |
| >hg38_21_+_8433172_8446622_RNA45SN1_45S_50ntFlanks@12632.12653.22\|GCGAAGCTACCATCTGTGGGAT | >hg38_21_+_8433172_8446622_RNA45SN1_45S_50ntFlanks | 12632 | 12653 | 22 | GCGAAGCTACCATCTGTGGGAT | 2579.7 | -10.9 | 1.2 | -9.0 | 2.4E-19 | 2.2E-18 | MB231/MB468 | MITO |
| >MI0000086\|hsa-mir-28&WithFlank&3\|+\|188688775\|188688872@62.82.21(+1U)&[MIMAT0004502&hsa-miR-28-3p&offsets\|+2\|+1(+1U);m-28&3\|+\|188688834\|188688855&offsets\|+2\|+1(+1U)]\|CTAGATTGTGAGCTCCTGGAGT | >MI0000086\|hsa-mir-28&WithFlank&3\|+\|188688775\|188688872 | 62 | 82 | 21(+1U) | CTAGATTGTGAGCTCCTGGAGT | 778.9 | 11.2 | 1.2 | 9.0 | 3.2E-19 | 2.2E-18 | BT20/MB468 | CYTO |
| >MI0000062\|hsa-let-7a-3&WithFlank&22\|+\|46112743\|46112828@58.78.21(+1U);MI0000060\|hsa-let-7a-1&WithFlank&9\|+\|94175951\|94176042@63.83.21(+1U)&[MIMAT0004481&hsa-let-7a-3p&offsets\|0\|0(+1U);m-180&22\|+\|46112800\|46112821&offsets\|0\|-1(+1U)];[MIMAT0004481_1&hsa-let-7a-3p&offsets\|0\|0(+1U);m-181&9\|+\|94176013\|94176034&offsets\|0\|-1(+1U)]\|CTATACAATCTACTGTCTTTCT | >MI0000062\|hsa-let-7a-3&WithFlank&22\|+\|46112743\|46112828 | 58 | 78 | 21(+1U) | CTATACAATCTACTGTCTTTCT | 1561.0 | 5.3 | 0.6 | 9.0 | 2.1E-19 | 2.2E-18 | MB231/MB468 | NUC |
| >hg38_21_+_8433172_8446622_RNA45SN1_45S_50ntFlanks@11452.11478.27\|CTCCTCGGCGCGCGGCGGCGGCGG | >hg38_21_+_8433172_8446622_RNA45SN1_45S_50ntFlanks | 11452 | 11478 | 27 | CTCCTCGGCGCGCGGCGGCGGCGG | 597.6 | -13.0 | 1.4 | -9.0 | 2.5E-19 | 2.2E-18 | MB231/MB468 | MITO |
| >MI0000487\|hsa-mir-193a&WithFlank&17\|+\|31559990\|31560089@27.48.22&[MIMAT0004614&hsa-miR-193a-5p&offsets\|0\|0;m-123&17\|+\|31560016\|31560037&offsets\|0\|0]\|TGGGTCTTTGCGGGCGAGATGA | >MI0000487\|hsa-mir-193a&WithFlank&17\|+\|31559990\|31560089 | 27 | 48 | 22 | TGGGTCTTTGCGGGCGAGATGA | 807.1 | 8.8 | 1.0 | 9.0 | 3.3E-19 | 2.2E-18 | BT20/MB468 | CYTO |
| >hg38_21_+_8433172_8446622_RNA45SN1_45S_50ntFlanks@12006.12032.27\|GGCCGCCGGTGAAATACCACTACTCTG | >hg38_21_+_8433172_8446622_RNA45SN1_45S_50ntFlanks | 12006 | 12032 | 27 | GGCCGCCGGTGAAATACCACTACTCTG | 876.6 | -13.6 | 1.5 | -9.0 | 1.6E-19 | 2.2E-18 | MB231/MB468 | MP |
| >MI0000102\|hsa-mir-100&WithFlank&11\|-\|122152223\|122152314@19.37.19(+1A)&[MIMAT0000098&hsa-miR-100-5p&offsets\|0\|-3(+1A);m-27&11\|-\|122152276\|122152296&offsets\|0\|-2(+1A)]\|AACCCGTAGATCCGAACTTA | >MI0000102\|hsa-mir-100&WithFlank&11\|-\|122152223\|122152314 | 19 | 37 | 19(+1A) | AACCCGTAGATCCGAACTTA | 2322.4 | 14.4 | 1.6 | 9.0 | 2.5E-19 | 2.2E-18 | MB231/MB468 | MITO |
| >hg38_21_+_8433172_8446622_RNA45SN1_45S_50ntFlanks@12893.12926.34\|CCTCGCCCGTCACGCACCGCACGTTCGTGGGGAA | >hg38_21_+_8433172_8446622_RNA45SN1_45S_50ntFlanks | 12893 | 12926 | 34 | CCTCGCCCGTCACGCACCGCACGTTCGTGGGGAA | 529.6 | -12.9 | 1.4 | -9.0 | 1.6E-19 | 2.2E-18 | MB231/MB468 | MP |
| >hg38_21_+_8433172_8446622_RNA45SN1_45S_50ntFlanks@4742.4767.26\|TCGGAGGTTCGAAGACGATCAGATAC | >hg38_21_+_8433172_8446622_RNA45SN1_45S_50ntFlanks | 4742 | 4767 | 26 | TCGGAGGTTCGAAGACGATCAGATAC | 190.1 | 11.6 | 1.3 | 9.1 | 1.4E-19 | 2.2E-18 | MB231/MB468 | CYTO |
| >hg38_21_+_8433172_8446622_RNA45SN1_45S_50ntFlanks@12405.12425.21\|CTTTTTGATCCTTCGATGTCG | >hg38_21_+_8433172_8446622_RNA45SN1_45S_50ntFlanks | 12405 | 12425 | 21 | CTTTTTGATCCTTCGATGTCG | 1654.9 | -7.4 | 0.8 | -9.1 | 1.4E-19 | 2.2E-18 | MB231/MB468 | CYTO |
| >hg38_21_+_8433172_8446622_RNA45SN1_45S_50ntFlanks@8109.8143.35\|GCGGGGCGCGGGACATGTGGCGTACGGAAGACCCG | >hg38_21_+_8433172_8446622_RNA45SN1_45S_50ntFlanks | 8109 | 8143 | 35 | GCGGGGCGCGGGACATGTGGCGTACGGAAGACCCG | 438.7 | -12.6 | 1.4 | -9.0 | 1.6E-19 | 2.3E-18 | MB231/MB468 | MP |
| >hg38_21_+_8433172_8446622_RNA45SN1_45S_50ntFlanks@8168.8190.23\|CCCAAGTCCTTCTGATCGAGGCC | >hg38_21_+_8433172_8446622_RNA45SN1_45S_50ntFlanks | 8168 | 8190 | 23 | CCCAAGTCCTTCTGATCGAGGCC | 400.3 | -12.5 | 1.4 | -9.0 | 1.6E-19 | 2.3E-18 | MB231/MB468 | MP |
| >hg38_21_+_8433172_8446622_RNA45SN1_45S_50ntFlanks@12456.12488.33\|CCAAGCGTTGGATTGTTCACCCACTAATAGGGA | >hg38_21_+_8433172_8446622_RNA45SN1_45S_50ntFlanks | 12456 | 12488 | 33 | CCAAGCGTTGGATTGTTCACCCACTAATAGGGA | 2450.1 | -6.0 | 0.7 | -9.0 | 2.5E-19 | 2.3E-18 | BT20/MB468 | MITO |
| >hg38_21_+_8433172_8446622_RNA45SN1_45S_50ntFlanks@10346.10368.23\|TCAAACGAGAACTTTGAAGGCCG | >hg38_21_+_8433172_8446622_RNA45SN1_45S_50ntFlanks | 10346 | 10368 | 23 | TCAAACGAGAACTTTGAAGGCCG | 2562.6 | -3.5 | 0.4 | -9.0 | 3.4E-19 | 2.3E-18 | BT20/MB468 | CYTO |
| >trna116_GluCTC_1_-_145399233_145399304@32.71.40,trna71_GluCTC_1_-_161439189_161439260@32.71.40,trna74_GluCTC_1_-_161431809_161431880@32.71.40,trna77_GluCTC_1_-_161424398_161424469@32.71.40,trna77_GluCTC_6_+_28949976_28950047@32.71.40,trna80_GluCTC_1_-_161417018_161417089@32.71.40,trna87_GluCTC_6_-_126101393_126101464@32.71.40\|CTCTCACCGCCGCGGCCCGGGTTCGATTCCCGGTCAGGGA | >trna116_GluCTC_1_-_145399233_145399304 | 32 | 71 | 40 | CTCTCACCGCCGCGGCCCGGGTTCGATTCCCGGTCAGGGA | 895.6 | -12.8 | 1.4 | -9.0 | 2.2E-19 | 2.3E-18 | MB231/MB468 | NUC |
| >hg38_21_+_8433172_8446622_RNA45SN1_45S_50ntFlanks@5421.5443.23\|CGATTGGATGGTTTAGTGAGGCC | >hg38_21_+_8433172_8446622_RNA45SN1_45S_50ntFlanks | 5421 | 5443 | 23 | CGATTGGATGGTTTAGTGAGGCC | 883.7 | -13.6 | 1.5 | -9.0 | 2.6E-19 | 2.3E-18 | MB231/MB468 | MITO |
| >hg38_21_+_8433172_8446622_RNA45SN1_45S_50ntFlanks@12419.12439.21\|GATGTCGGCTCTTCCTATCAT | >hg38_21_+_8433172_8446622_RNA45SN1_45S_50ntFlanks | 12419 | 12439 | 21 | GATGTCGGCTCTTCCTATCAT | 1327.5 | 4.1 | 0.5 | 9.0 | 3.5E-19 | 2.3E-18 | BT20/MB468 | CYTO |
| >hg38_21_+_8433172_8446622_RNA45SN1_45S_50ntFlanks@11317.11340.24\|ACCCCACGTCTCGTCGCGCGCGCG | >hg38_21_+_8433172_8446622_RNA45SN1_45S_50ntFlanks | 11317 | 11340 | 24 | ACCCCACGTCTCGTCGCGCGCGCG | 1426.3 | -14.3 | 1.6 | -9.0 | 2.6E-19 | 2.3E-18 | MB231/MB468 | MITO |
| >am_MI0000809\|hsa-mir-151a&WithFlank&8\|-\|140732558\|140732659@17.38.22&[MIMAT0004697&hsa-miR-151a-5p&offsets\|0\|+1;m-75&8\|-\|140732623\|140732643&offsets\|0\|+1]\|TCGAGGAGCTCACAGTCTAGTA | >am_MI0000809\|hsa-mir-151a&WithFlank&8\|-\|140732558\|140732659 | 17 | 38 | 22 | TCGAGGAGCTCACAGTCTAGTA | 344.2 | 4.8 | 0.5 | 8.9 | 6.4E-19 | 2.3E-18 | BT20/MB468 | NUC |
| >MI0000077\|hsa-mir-21&WithFlank&17\|+\|59841260\|59841343@14.35.22&[MIMAT0000076&hsa-miR-21-5p&offsets\|0\|0;m-1&17\|+\|59841273\|59841294&offsets\|0\|0]\|TAGCTTATCAGACTGATGTTGA | >MI0000077\|hsa-mir-21&WithFlank&17\|+\|59841260\|59841343 | 14 | 35 | 22 | TAGCTTATCAGACTGATGTTGA | 54203.8 | 6.5 | 0.7 | 9.0 | 2.6E-19 | 2.3E-18 | BT20/MB468 | MITO |
| >trna10_GlyTCC_17_+_8124866_8124937@1.30.30\|GCGTTGGTGGTATAGTGGTAAGCATAGCTG | >trna10_GlyTCC_17_+_8124866_8124937 | 1 | 30 | 30 | GCGTTGGTGGTATAGTGGTAAGCATAGCTG | 475.9 | -12.8 | 1.4 | -9.0 | 1.7E-19 | 2.3E-18 | MB231/MB468 | MP |
| >MI0000264\|hsa-mir-7-2&WithFlank&15\|+\|88611819\|88611940@38.61.24;MI0000265\|hsa-mir-7-3&WithFlank&19\|+\|4770664\|4770785@37.60.24;MI0000263\|hsa-mir-7-1&WithFlank&9\|-\|83969742\|83969863@30.53.24&[MIMAT0000252&hsa-miR-7-5p&offsets\|0\|0;m-400&15\|+\|88611856\|88611879&offsets\|0\|0];[MIMAT0000252_1&hsa-miR-7-5p&offsets\|0\|0;m-402&19\|+\|4770700\|4770723&offsets\|0\|0];[MIMAT0000252_2&hsa-miR-7-5p&offsets\|0\|0;m-401&9\|-\|83969811\|83969834&offsets\|0\|0]\|TGGAAGACTAGTGATTTTGTTGTT | >MI0000264\|hsa-mir-7-2&WithFlank&15\|+\|88611819\|88611940 | 38 | 61 | 24 | TGGAAGACTAGTGATTTTGTTGTT | 167792.9 | 7.5 | 0.8 | 9.0 | 3.5E-19 | 2.3E-18 | BT20/MB468 | CYTO |
| >hg38_21_+_8433172_8446622_RNA45SN1_45S_50ntFlanks@9306.9323.18\|CAAGGAGTCTAACACGTG | >hg38_21_+_8433172_8446622_RNA45SN1_45S_50ntFlanks | 9306 | 9323 | 18 | CAAGGAGTCTAACACGTG | 606.4 | -13.0 | 1.5 | -9.0 | 2.6E-19 | 2.3E-18 | MB231/MB468 | MITO |
| >hg38_21_+_8433172_8446622_RNA45SN1_45S_50ntFlanks@3861.3898.38\|TAATTCTAGAGCTAATACATGCCGACGGGCGCTGACCC | >hg38_21_+_8433172_8446622_RNA45SN1_45S_50ntFlanks | 3861 | 3898 | 38 | TAATTCTAGAGCTAATACATGCCGACGGGCGCTGACCC | 569.0 | -12.9 | 1.4 | -9.0 | 2.7E-19 | 2.4E-18 | MB231/MB468 | MITO |
| >hg38_21_+_8433172_8446622_RNA45SN1_45S_50ntFlanks@9201.9220.20\|CTCTCGGGGCCACGCGCGCG | >hg38_21_+_8433172_8446622_RNA45SN1_45S_50ntFlanks | 9201 | 9220 | 20 | CTCTCGGGGCCACGCGCGCG | 680.8 | -13.3 | 1.5 | -9.0 | 1.7E-19 | 2.4E-18 | MB231/MB468 | MP |
| >hg38_21_+_8433172_8446622_RNA45SN1_45S_50ntFlanks@3802.3838.37\|CATTAAATCAGTTATGGTTCCTTTGGTCGCTCGCTCC | >hg38_21_+_8433172_8446622_RNA45SN1_45S_50ntFlanks | 3802 | 3838 | 37 | CATTAAATCAGTTATGGTTCCTTTGGTCGCTCGCTCC | 400.2 | -13.1 | 1.5 | -8.9 | 6.5E-19 | 2.4E-18 | BT20/MB468 | NUC |
| >hg38_21_+_8433172_8446622_RNA45SN1_45S_50ntFlanks@5446.5467.22\|CGGATCGGCCCCGCCGGGGTCG | >hg38_21_+_8433172_8446622_RNA45SN1_45S_50ntFlanks | 5446 | 5467 | 22 | CGGATCGGCCCCGCCGGGGTCG | 779.9 | -13.4 | 1.5 | -9.0 | 2.7E-19 | 2.4E-18 | MB231/MB468 | MITO |

| | | | | | | | | | | | | | |
|---|---|---|---|---|---|---|---|---|---|---|---|---|---|
| >MI0000289\|hsa-mir-181a-1&WithFlank&1\|-\|198859038\|198859159@70.91.22&[MIMAT0000270&hsa-miR-181a-3p&offsets\|0\|0;m-125&1\|-\|198859069\|198859090&offsets\|0\|0]\|ACCATCGACCGTTGATTGTACC | >MI0000289\|hsa-mir-181a-1&WithFlank&1\|-\|198859038\|198859159 | 70 | 91 | 22 | ACCATCGACCGTTGATTGTACC | 1114.3 | -13.4 | 1.4 | -9.3 | 2.2E-20 | 2.4E-18 | BT20/MB231 | MITO |
| >MI0000285\|hsa-mir-205&WithFlank&1\|+\|209432127\|209432248@40.59.20(+1A)&[MIMAT0000266&hsa-miR-205-5p&offsets\|0\|-2(+1A);m-351&1\|+\|209432166\|209432187&offsets\|0\|-2(+1A)]\|TCCTTCATTCCACCGGAGTCA | >MI0000285\|hsa-mir-205&WithFlank&1\|+\|209432127\|209432248 | 40 | 59 | 20(+1A) | TCCTTCATTCCACCGGAGTCA | 885.8 | 13.2 | 1.4 | 9.3 | 1.5E-20 | 2.4E-18 | BT20/MB231 | CYTO |
| >hg38_21_+_8433172_8446622_RNA45SN1_45S_50ntFlanks@9203.9223.21\|CTCGGGGCCACGCGCGCGTCC | >hg38_21_+_8433172_8446622_RNA45SN1_45S_50ntFlanks | 9203 | 9223 | 21 | CTCGGGGCCACGCGCGCGTCC | 714.7 | -13.3 | 1.5 | -9.0 | 1.7E-19 | 2.4E-18 | MB231/MB468 | MP |
| >hg38_21_+_8433172_8446622_RNA45SN1_45S_50ntFlanks@4724.4751.28\|ATTAATCAAGAACGAAAGTCGGAGGTTC | >hg38_21_+_8433172_8446622_RNA45SN1_45S_50ntFlanks | 4724 | 4751 | 28 | ATTAATCAAGAACGAAAGTCGGAGGTTC | 1102.3 | -13.9 | 1.5 | -9.0 | 2.7E-19 | 2.4E-18 | MB231/MB468 | MITO |
| >hg38_21_+_8433172_8446622_RNA45SN1_45S_50ntFlanks@12626.12659.34\|AATGGGGCGAAGCTACCATCTGTGGGATTATGAC | >hg38_21_+_8433172_8446622_RNA45SN1_45S_50ntFlanks | 12626 | 12659 | 34 | AATGGGGCGAAGCTACCATCTGTGGGATTATGAC | 1960.0 | -14.8 | 1.6 | -9.0 | 1.7E-19 | 2.4E-18 | MB231/MB468 | MP |
| >trna119_LysCTT_1_-_145395522_145395594@1.33.33,trna11_LysCTT_5_-_180648979_180649051@1.33.33,trna13_LysCTT_6_+_26556774_26556846@1.33.33,trna7_LysCTT_16_+_3225692_3225764@1.33.33,trna9_LysCTT_5_+_180634755_180634827@1.33.33\|GCCCGGCTAGCTCAGTCGGTAGAGCATGAGACT | >trna119_LysCTT_1_-_145395522_145395594 | 1 | 33 | 33 | GCCCGGCTAGCTCAGTCGGTAGAGCATGAGACT | 6050.9 | 5.5 | 0.6 | 8.9 | 3.6E-19 | 2.4E-18 | BT20/MB468 | CYTO |
| >hg38_21_+_8433172_8446622_RNA45SN1_45S_50ntFlanks@6690.6720.31\|AAGAACGCAGCTAGCTGCGAGAATTAATGTG | >hg38_21_+_8433172_8446622_RNA45SN1_45S_50ntFlanks | 6690 | 6720 | 31 | AAGAACGCAGCTAGCTGCGAGAATTAATGTG | 2901.5 | -7.8 | 0.9 | -9.0 | 1.5E-19 | 2.4E-18 | MB231/MB468 | CYTO |
| >am_MI0000809\|hsa-mir-151a&WithFlank&8\|-\|140732558\|140732659@53.72.20(+1U)&[MIMAT0000757&hsa-miR-151a-3p&offsets\|0\|-1(+1U);m-30&8\|-\|140732587\|140732607&offsets\|0\|-1(+1U)]\|CTAGACTGAAGCTCCTTGAGT | >am_MI0000809\|hsa-mir-151a&WithFlank&8\|-\|140732558\|140732659 | 53 | 72 | 20(+1U) | CTAGACTGAAGCTCCTTGAGT | 1936.1 | 7.1 | 0.8 | 9.0 | 1.5E-19 | 2.4E-18 | MB231/MB468 | CYTO |
| >hg38_21_+_8433172_8446622_RNA45SN1_45S_50ntFlanks@11669.11707.39\|GCCCAGTGCTCTGAATGTCAAAGTGAAGAAATTCAATGA | >hg38_21_+_8433172_8446622_RNA45SN1_45S_50ntFlanks | 11669 | 11707 | 39 | GCCCAGTGCTCTGAATGTCAAAGTGAAGAAATTCAATGA | 1703.6 | 10.1 | 1.1 | 9.0 | 1.5E-19 | 2.4E-18 | MB231/MB468 | CYTO |
| >hg38_21_+_8433172_8446622_RNA45SN1_45S_50ntFlanks@5420.5445.26\|CCGATTGGATGGTTTAGTGAGGCCCT | >hg38_21_+_8433172_8446622_RNA45SN1_45S_50ntFlanks | 5420 | 5445 | 26 | CCGATTGGATGGTTTAGTGAGGCCCT | 1108.7 | -14.0 | 1.5 | -9.0 | 1.7E-19 | 2.4E-18 | MB231/MB468 | MP |
| >hg38_21_+_8433172_8446622_RNA45SN1_45S_50ntFlanks@7973.7992.20\|GACGCGACCTCAGATCAGAC | >hg38_21_+_8433172_8446622_RNA45SN1_45S_50ntFlanks | 7973 | 7992 | 20 | GACGCGACCTCAGATCAGAC | 4927.0 | 5.9 | 0.7 | 9.0 | 2.7E-19 | 2.4E-18 | BT20/MB468 | MITO |
| >MI0000446\|hsa-mir-125b-1&WithFlank&11\|-\|122099751\|122099850@21.40.20;MI0000470\|hsa-mir-125b-2&WithFlank&21\|+\|16590231\|16590331@23.42.20&[MIMAT0000423&hsa-miR-125b-5p&offsets\|0\|-2;m-67&11\|-\|122099809\|122099830&offsets\|0\|-2];[MIMAT0000423_1&hsa-miR-125b-5p&offsets\|0\|-2;m-68&21\|+\|16590253\|16590274&offsets\|0\|-2]\|TCCCTGAGACCCTAACTTGT | >MI0000446\|hsa-mir-125b-1&WithFlank&11\|-\|122099751\|122099850 | 21 | 40 | 20 | TCCCTGAGACCCTAACTTGT | 3137.9 | 9.9 | 1.1 | 9.0 | 1.6E-19 | 2.4E-18 | MB231/MB468 | CYTO |
| >hg38_21_+_8433172_8446622_RNA45SN1_45S_50ntFlanks@12937.12956.20\|AACCATTCGTAGACGACCTG | >hg38_21_+_8433172_8446622_RNA45SN1_45S_50ntFlanks | 12937 | 12956 | 20 | AACCATTCGTAGACGACCTG | 20593.8 | -13.5 | 1.5 | -8.9 | 6.7E-19 | 2.4E-18 | BT20/MB468 | NUC |
| >hg38_21_+_8433172_8446622_RNA45SN1_45S_50ntFlanks@8814.8835.22\|CGGGTCGGGGCGGCGGCGGCGG | >hg38_21_+_8433172_8446622_RNA45SN1_45S_50ntFlanks | 8814 | 8835 | 22 | CGGGTCGGGGCGGCGGCGGCGG | 1292.1 | -14.2 | 1.6 | -9.0 | 1.8E-19 | 2.4E-18 | MB231/MB468 | MP |
| >hg38_21_+_8433172_8446622_RNA45SN1_45S_50ntFlanks@11707.11728.22\|AAGCGCGGGTAAACGGCGGGAG | >hg38_21_+_8433172_8446622_RNA45SN1_45S_50ntFlanks | 11707 | 11728 | 22 | AAGCGCGGGTAAACGGCGGGAG | 530.5 | -12.9 | 1.4 | -9.0 | 1.8E-19 | 2.4E-18 | MB231/MB468 | MP |
| >hg38_21_+_8433172_8446622_RNA45SN1_45S_50ntFlanks@4472.4497.26\|CTTAGCTGAGTGTCCCGCGGGGCCCG | >hg38_21_+_8433172_8446622_RNA45SN1_45S_50ntFlanks | 4472 | 4497 | 26 | CTTAGCTGAGTGTCCCGCGGGGCCCG | 772.6 | -13.4 | 1.5 | -9.0 | 2.8E-19 | 2.5E-18 | MB231/MB468 | MITO |
| >hg38_21_+_8433172_8446622_RNA45SN1_45S_50ntFlanks@4367.4406.40\|CAGTTAAAAAGCTCGTAGTTGGATCTTGGGAGCGGGCGGG | >hg38_21_+_8433172_8446622_RNA45SN1_45S_50ntFlanks | 4367 | 4406 | 40 | CAGTTAAAAAGCTCGTAGTTGGATCTTGGGAGCGGGCGGG | 509.8 | -13.4 | 1.5 | -8.9 | 6.7E-19 | 2.5E-18 | BT20/MB468 | NUC |
| >MI0001445\|hsa-mir-423&WithFlank&17\|+\|30117073\|30117178@23.45.23&[MIMAT0004748&hsa-miR-423-5p&offsets\|0\|0;m-141&17\|+\|30117095\|30117117&offsets\|0\|0]\|TGAGGGGCAGAGAGCGAGACTTT | >MI0001445\|hsa-mir-423&WithFlank&17\|+\|30117073\|30117178 | 23 | 45 | 23 | TGAGGGGCAGAGAGCGAGACTTT | 6970.1 | 7.9 | 0.9 | 9.0 | 1.6E-19 | 2.5E-18 | MB231/MB468 | CYTO |
| >MI0000438\|hsa-mir-15b&WithFlank&3\|+\|160404582\|160404691@64.84.21&[MIMAT0004586&hsa-miR-15b-3p&offsets\|0\|-1;m-204&3\|+\|160404645\|160404665&offsets\|0\|0]\|CGAATCATTATTTGCTGCTCT | >MI0000438\|hsa-mir-15b&WithFlank&3\|+\|160404582\|160404691 | 64 | 84 | 21 | CGAATCATTATTTGCTGCTCT | 566.9 | 6.1 | 0.7 | 8.9 | 3.7E-19 | 2.5E-18 | BT20/MB468 | CYTO |
| >hg38_21_+_8433172_8446622_RNA45SN1_45S_50ntFlanks@9790.9820.31\|CTGGCGTGGAGCCGGGCGTGGAATGCGAGTG | >hg38_21_+_8433172_8446622_RNA45SN1_45S_50ntFlanks | 9790 | 9820 | 31 | CTGGCGTGGAGCCGGGCGTGGAATGCGAGTG | 488.5 | -12.8 | 1.4 | -9.0 | 1.8E-19 | 2.5E-18 | MB231/MB468 | MP |
| >MI0003560\|hsa-mir-92b&WithFlank&1\|+\|155195171\|155195278@67.86.20(+1U)&[MIMAT0003218&hsa-miR-92b-3p&offsets\|0\|-2(+1U);m-115&1\|+\|155195237\|155195258&offsets\|0\|-2(+1U)]\|TATTGCACTCGTCCCGGCCTT | >MI0003560\|hsa-mir-92b&WithFlank&1\|+\|155195171\|155195278 | 67 | 86 | 20(+1U) | TATTGCACTCGTCCCGGCCTT | 470.1 | 12.9 | 1.4 | 9.0 | 1.6E-19 | 2.5E-18 | MB231/MB468 | CYTO |
| >hg38_21_+_8433172_8446622_RNA45SN1_45S_50ntFlanks@8552.8579.28\|CTCCCCGGAGGGGGCGGGCTCCGGCGGG | >hg38_21_+_8433172_8446622_RNA45SN1_45S_50ntFlanks | 8552 | 8579 | 28 | CTCCCCGGAGGGGGCGGGCTCCGGCGGG | 395.1 | -13.1 | 1.5 | -8.9 | 6.9E-19 | 2.5E-18 | BT20/MB468 | NUC |
| >hg38_21_+_8433172_8446622_RNA45SN1_45S_50ntFlanks@4472.4496.25\|CTTAGCTGAGTGTCCCGCGGGGCCC | >hg38_21_+_8433172_8446622_RNA45SN1_45S_50ntFlanks | 4472 | 4496 | 25 | CTTAGCTGAGTGTCCCGCGGGGCCC | 548.6 | -12.9 | 1.4 | -9.0 | 2.9E-19 | 2.5E-18 | MB231/MB468 | MITO |
| >hg38_21_+_8433172_8446622_RNA45SN1_45S_50ntFlanks@11575.11600.26\|ACTTAGAACTGGTGCGGACCAGGGGA | >hg38_21_+_8433172_8446622_RNA45SN1_45S_50ntFlanks | 11575 | 11600 | 26 | ACTTAGAACTGGTGCGGACCAGGGGA | 898.7 | -13.7 | 1.5 | -9.0 | 1.8E-19 | 2.5E-18 | MB231/MB468 | MP |
| >hg38_21_+_8433172_8446622_RNA45SN1_45S_50ntFlanks@10866.10892.27\|CTAAGGGCTGGGTCGGTCGGGCTGGGG | >hg38_21_+_8433172_8446622_RNA45SN1_45S_50ntFlanks | 10866 | 10892 | 27 | CTAAGGGCTGGGTCGGTCGGGCTGGGG | 567.5 | -12.1 | 1.3 | -9.0 | 2.4E-19 | 2.5E-18 | MB231/MB468 | NUC |
| >hg38_21_+_8433172_8446622_RNA45SN1_45S_50ntFlanks@4679.4720.42\|GGCGCAAGACGGACCAGAGCGAAAGCATTTGCCAAGAATGTT | >hg38_21_+_8433172_8446622_RNA45SN1_45S_50ntFlanks | 4679 | 4720 | 42 | GGCGCAAGACGGACCAGAGCGAAAGCATTTGCCAAGAATGTT | 331.2 | 12.6 | 1.4 | 8.9 | 3.8E-19 | 2.5E-18 | BT20/MB468 | CYTO |
| >hg38_21_+_8433172_8446622_RNA45SN1_45S_50ntFlanks@3831.3872.42\|CTCGCTCCTCTCCTACTTGGATAACTGTGGTAATTCTAGAGC | >hg38_21_+_8433172_8446622_RNA45SN1_45S_50ntFlanks | 3831 | 3872 | 42 | CTCGCTCCTCTCCTACTTGGATAACTGTGGTAATTCTAGAGC | 506.0 | -13.4 | 1.5 | -8.9 | 6.9E-19 | 2.5E-18 | BT20/MB468 | NUC |
| >hg38_21_+_8433172_8446622_RNA45SN1_45S_50ntFlanks@12151.12180.30\|GTGGGGAGTTTGACTGGGGCGGTACACCTG | >hg38_21_+_8433172_8446622_RNA45SN1_45S_50ntFlanks | 12151 | 12180 | 30 | GTGGGGAGTTTGACTGGGGCGGTACACCTG | 642.8 | -13.1 | 1.5 | -9.0 | 2.9E-19 | 2.5E-18 | MB231/MB468 | MITO |
| >hg38_21_+_8433172_8446622_RNA45SN1_45S_50ntFlanks@8115.8141.27\|CGCGGGACATGTGGCGTACGGAAGACC | >hg38_21_+_8433172_8446622_RNA45SN1_45S_50ntFlanks | 8115 | 8141 | 27 | CGCGGGACATGTGGCGTACGGAAGACC | 4211.2 | -11.4 | 1.3 | -9.0 | 1.8E-19 | 2.5E-18 | MB231/MB468 | MP |
| >MI0000102\|hsa-mir-100&WithFlank&11\|-\|122152223\|122152314@19.39.21(+1C)&[MIMAT0000098&hsa-miR-100-5p&offsets\|0\|-1(+1C);m-27&11\|-\|122152276\|122152296&offsets\|0\|0(+1C)]\|AACCCGTAGATCCGAACTTGTC | >MI0000102\|hsa-mir-100&WithFlank&11\|-\|122152223\|122152314 | 19 | 39 | 21(+1C) | AACCCGTAGATCCGAACTTGTC | 2595.5 | 14.6 | 1.6 | 9.0 | 2.9E-19 | 2.5E-18 | MB231/MB468 | MITO |
| >MI0015866\|hsa-mir-4326&WithFlank&20\|+\|63286802\|63286872@17.39.23&[MIMAT0016888&hsa-miR-4326&offsets\|0\|+3;m-339&20\|+\|63286818\|63286841&offsets\|0\|-1]\|TGTTCCTCTGTCTCCCAGACTCT | >MI0015866\|hsa-mir-4326&WithFlank&20\|+\|63286802\|63286872 | 17 | 39 | 23 | TGTTCCTCTGTCTCCCAGACTCT | 619.9 | 10.9 | 1.2 | 8.9 | 3.9E-19 | 2.6E-18 | BT20/MB468 | CYTO |
| >hg38_21_+_8433172_8446622_RNA45SN1_45S_50ntFlanks@10873.10902.30\|CTGGGTCGGTCGGGCTGGGGCGCGAAGCGG | >hg38_21_+_8433172_8446622_RNA45SN1_45S_50ntFlanks | 10873 | 10902 | 30 | CTGGGTCGGTCGGGCTGGGGCGCGAAGCGG | 734.3 | -13.3 | 1.5 | -9.0 | 2.9E-19 | 2.6E-18 | MB231/MB468 | MITO |
| >hg38_21_+_8433172_8446622_RNA45SN1_45S_50ntFlanks@10687.10712.26\|CGGCGTCCGGTGAGCTCTCGCTGGCC | >hg38_21_+_8433172_8446622_RNA45SN1_45S_50ntFlanks | 10687 | 10712 | 26 | CGGCGTCCGGTGAGCTCTCGCTGGCC | 1945.1 | -6.0 | 0.7 | -9.0 | 1.7E-19 | 2.6E-18 | MB231/MB468 | CYTO |
| >hg38_21_+_8433172_8446622_RNA45SN1_45S_50ntFlanks@12636.12653.18\|AGCTACCATCTGTGGGAT | >hg38_21_+_8433172_8446622_RNA45SN1_45S_50ntFlanks | 12636 | 12653 | 18 | AGCTACCATCTGTGGGAT | 1603.5 | -14.4 | 1.6 | -9.0 | 3.0E-19 | 2.6E-18 | MB231/MB468 | MITO |
| >hg38_21_+_8433172_8446622_RNA45SN1_45S_50ntFlanks@9554.9572.19\|CGTAGCGGTCCTGACGTGC | >hg38_21_+_8433172_8446622_RNA45SN1_45S_50ntFlanks | 9554 | 9572 | 19 | CGTAGCGGTCCTGACGTGC | 16358.0 | -3.7 | 0.4 | -8.9 | 3.9E-19 | 2.6E-18 | BT20/MB468 | CYTO |
| >hg38_21_+_8433172_8446622_RNA45SN1_45S_50ntFlanks@3876.3898.23\|TACATGCCGACGGGCGCTGACCC | >hg38_21_+_8433172_8446622_RNA45SN1_45S_50ntFlanks | 3876 | 3898 | 23 | TACATGCCGACGGGCGCTGACCC | 2069.9 | -8.4 | 0.9 | -9.0 | 3.0E-19 | 2.6E-18 | MB231/MB468 | MITO |
| >trna108_AlaAGC_6_-_28763741_28763812@58.75.18\|TCCCCAGTACCTCCACCA | >trna108_AlaAGC_6_-_28763741_28763812 | 58 | 75 | 18 | TCCCCAGTACCTCCACCA | 4335.9 | 7.9 | 0.9 | 8.9 | 4.0E-19 | 2.6E-18 | BT20/MB468 | CYTO |
| >hg38_21_+_8433172_8446622_RNA45SN1_45S_50ntFlanks@10350.10374.25\|ACGAGAACTTTGAAGGCCGAAGTGG | >hg38_21_+_8433172_8446622_RNA45SN1_45S_50ntFlanks | 10350 | 10374 | 25 | ACGAGAACTTTGAAGGCCGAAGTGG | 1489.3 | -4.6 | 0.5 | -9.0 | 2.9E-19 | 2.6E-18 | BT20/MB468 | MITO |
| >hg38_21_+_8433172_8446622_RNA45SN1_45S_50ntFlanks@12405.12422.18\|CTTTTTGATCCTTCGATG | >hg38_21_+_8433172_8446622_RNA45SN1_45S_50ntFlanks | 12405 | 12422 | 18 | CTTTTTGATCCTTCGATG | 642.4 | -13.1 | 1.5 | -9.0 | 3.0E-19 | 2.6E-18 | MB231/MB468 | MITO |
| >hg38_21_+_8433172_8446622_RNA45SN1_45S_50ntFlanks@12456.12485.30\|CCAAGCGTTGGATTGTTCACCCACTAATAG | >hg38_21_+_8433172_8446622_RNA45SN1_45S_50ntFlanks | 12456 | 12485 | 30 | CCAAGCGTTGGATTGTTCACCCACTAATAG | 2830.2 | -10.6 | 1.2 | -9.0 | 3.0E-19 | 2.6E-18 | MB231/MB468 | MITO |
| >hg38_21_+_8433172_8446622_RNA45SN1_45S_50ntFlanks@10131.10149.19\|CGGCGGCGGCGGCGGCGGG | >hg38_21_+_8433172_8446622_RNA45SN1_45S_50ntFlanks | 10131 | 10149 | 19 | CGGCGGCGGCGGCGGCGGG | 1276.2 | -14.1 | 1.6 | -9.0 | 3.1E-19 | 2.7E-18 | MB231/MB468 | MITO |
| >hg38_21_+_8433172_8446622_RNA45SN1_45S_50ntFlanks@5527.5550.24\|AAAGTCGTAACAAGGTTTCCGTAG | >hg38_21_+_8433172_8446622_RNA45SN1_45S_50ntFlanks | 5527 | 5550 | 24 | AAAGTCGTAACAAGGTTTCCGTAG | 720.4 | -13.3 | 1.5 | -9.0 | 3.1E-19 | 2.7E-18 | MB231/MB468 | MITO |
| >hg38_21_+_8433172_8446622_RNA45SN1_45S_50ntFlanks@8848.8866.19\|CGGCGGCGGCGGCGGCGGG | >hg38_21_+_8433172_8446622_RNA45SN1_45S_50ntFlanks | 8848 | 8866 | 19 | CGGCGGCGGCGGCGGCGGG | 1276.2 | -14.1 | 1.6 | -9.0 | 3.1E-19 | 2.7E-18 | MB231/MB468 | MITO |
| >hg38_21_+_8433172_8446622_RNA45SN1_45S_50ntFlanks@12934.12970.37\|CTAAACCATTCGTAGACGACCTGCTTCTGGGTCGGGG | >hg38_21_+_8433172_8446622_RNA45SN1_45S_50ntFlanks | 12934 | 12970 | 37 | CTAAACCATTCGTAGACGACCTGCTTCTGGGTCGGGG | 685.5 | -8.2 | 0.9 | -9.0 | 1.7E-19 | 2.7E-18 | MB231/MB468 | CYTO |
| >hg38_21_+_8433172_8446622_RNA45SN1_45S_50ntFlanks@11561.11582.22\|GCGCCTAGCAGCCGACTTAGAA | >hg38_21_+_8433172_8446622_RNA45SN1_45S_50ntFlanks | 11561 | 11582 | 22 | GCGCCTAGCAGCCGACTTAGAA | 1003.6 | -13.8 | 1.5 | -9.0 | 3.1E-19 | 2.7E-18 | MB231/MB468 | MITO |
| >MI0000095\|hsa-mir-93&WithFlank&7\|-\|100093762\|100093853@17.38.22&[MIMAT0000093&hsa-miR-93-5p&offsets\|0\|-1;m-29&7\|-\|100093815\|100093837&offsets\|0\|-1]\|CAAAGTGCTGTTCGTGCAGGTA | >MI0000095\|hsa-mir-93&WithFlank&7\|-\|100093762\|100093853 | 17 | 38 | 22 | CAAAGTGCTGTTCGTGCAGGTA | 890.9 | 6.5 | 0.7 | 8.9 | 4.0E-19 | 2.7E-18 | BT20/MB468 | CYTO |
| >hg38_21_+_8433172_8446622_RNA45SN1_45S_50ntFlanks@12627.12657.31\|ATGGGGCGAAGCTACCATCTGTGGGATTATG | >hg38_21_+_8433172_8446622_RNA45SN1_45S_50ntFlanks | 12627 | 12657 | 31 | ATGGGGCGAAGCTACCATCTGTGGGATTATG | 4894.1 | -11.6 | 1.3 | -9.0 | 3.1E-19 | 2.7E-18 | MB231/MB468 | MITO |
| >hg38_21_+_8433172_8446622_RNA45SN1_45S_50ntFlanks@5485.5510.26\|AGCGCTGAGAAGACGGTCGAACTTGA | >hg38_21_+_8433172_8446622_RNA45SN1_45S_50ntFlanks | 5485 | 5510 | 26 | AGCGCTGAGAAGACGGTCGAACTTGA | 614.5 | -13.0 | 1.5 | -9.0 | 3.1E-19 | 2.7E-18 | MB231/MB468 | MITO |
| >hg38_1_-_228634819_228635039_RNA5S12_5S_50ntFlanks@152.171.20\|TACCGGGTGCTGTAGGCTTT | >hg38_1_-_228634819_228635039_RNA5S12_5S_50ntFlanks | 152 | 171 | 20 | TACCGGGTGCTGTAGGCTTT | 7313.8 | 6.5 | 0.7 | 9.0 | 1.7E-19 | 2.7E-18 | MB231/MB468 | CYTO |

| | | | | | | | | | | | | | |
|---|---|---|---|---|---|---|---|---|---|---|---|---|---|
| >hg38_21_+_8433172_8446622_RNA45SN1_45S_50ntFlanks@11886.11904.19\|TGAGCTTGACTCTAGTCTG | >hg38_21_+_8433172_8446622_RNA45SN1_45S_50ntFlanks | 11886 | 11904 | 19 | TGAGCTTGACTCTAGTCTG | 1518.1 | -13.3 | 1.5 | -9.0 | 1.7E-19 | 2.7E-18 | MB231/MB468 | CYTO |
| >hg38_21_+_8433172_8446622_RNA45SN1_45S_50ntFlanks@10350.10374.25\|ACGAGAACTTTGAAGGCCGAAGTGG | >hg38_21_+_8433172_8446622_RNA45SN1_45S_50ntFlanks | 10350 | 10374 | 25 | ACGAGAACTTTGAAGGCCGAAGTGG | 5475.7 | -12.4 | 1.4 | -8.9 | 7.4E-19 | 2.7E-18 | BT20/MB468 | NUC |
| >hg38_21_+_8433172_8446622_RNA45SN1_45S_50ntFlanks@11586.11612.27\|GTGCGGACCAGGGGAATCCGACTGTTT | >hg38_21_+_8433172_8446622_RNA45SN1_45S_50ntFlanks | 11586 | 11612 | 27 | GTGCGGACCAGGGGAATCCGACTGTTT | 1917.9 | -5.3 | 0.6 | -9.0 | 3.1E-19 | 2.7E-18 | MB231/MB468 | MITO |
| >MI0000261\|hsa-mir-139&WithFlank&11\|-\|72615057\|72615136@13.35.23&[MIMAT0000250&hsa-miR-139-5p&offsets\|0\|0;m-186&11\|-\|72615102\|72615124&offsets\|0\|0]\|TCTACAGTGCACGTGTCTCCAGT | >MI0000261\|hsa-mir-139&WithFlank&11\|-\|72615057\|72615136 | 13 | 35 | 23 | TCTACAGTGCACGTGTCTCCAGT | 1811.0 | 14.0 | 1.6 | 9.0 | 3.2E-19 | 2.7E-18 | MB231/MB468 | MITO |
| >hg38_21_+_8433172_8446622_RNA45SN1_45S_50ntFlanks@12073.12090.18\|CCGAGGGGCTCTCGCTTC | >hg38_21_+_8433172_8446622_RNA45SN1_45S_50ntFlanks | 12073 | 12090 | 18 | CCGAGGGGCTCTCGCTTC | 636.5 | -13.1 | 1.5 | -9.0 | 3.2E-19 | 2.7E-18 | MB231/MB468 | MITO |
| >hg38_21_+_8433172_8446622_RNA45SN1_45S_50ntFlanks@8439.8471.33\|GATTCAACCCGGCGGCGGGTCCGGCCGTGTCGG | >hg38_21_+_8433172_8446622_RNA45SN1_45S_50ntFlanks | 8439 | 8471 | 33 | GATTCAACCCGGCGGCGGGTCCGGCCGTGTCGG | 382.4 | -12.4 | 1.4 | -9.0 | 2.0E-19 | 2.7E-18 | MB231/MB468 | MP |
| >hg38_21_+_8433172_8446622_RNA45SN1_45S_50ntFlanks@6789.6806.18\|GCCTGTCTGAGCGTCGCT | >hg38_21_+_8433172_8446622_RNA45SN1_45S_50ntFlanks | 6789 | 6806 | 18 | GCCTGTCTGAGCGTCGCT | 1938.8 | 4.7 | 0.5 | 9.0 | 2.7E-19 | 2.8E-18 | MB231/MB468 | NUC |
| >hg38_21_+_8433172_8446622_RNA45SN1_45S_50ntFlanks@12357.12376.20\|CCACAGGGATAACTGGCTTG | >hg38_21_+_8433172_8446622_RNA45SN1_45S_50ntFlanks | 12357 | 12376 | 20 | CCACAGGGATAACTGGCTTG | 1029.0 | -13.8 | 1.5 | -9.0 | 3.2E-19 | 2.8E-18 | MB231/MB468 | MITO |
| >MI0000102\|hsa-mir-100&WithFlank&11\|-\|122152223\|122152314@20.39.20&[MIMAT0000098&hsa-miR-100-5p&offsets\|+1\|-1;m-27&11\|-\|122152276\|122152296&offsets\|+1\|0]\|ACCCGTAGATCCGAACTTGT | >MI0000102\|hsa-mir-100&WithFlank&11\|-\|122152223\|122152314 | 20 | 39 | 20 | ACCCGTAGATCCGAACTTGT | 425.2 | 12.6 | 1.4 | 9.0 | 2.7E-19 | 2.8E-18 | MB231/MB468 | NUC |
| >hg38_21_+_8433172_8446622_RNA45SN1_45S_50ntFlanks@10869.10901.33\|AGGGCTGGGTCGGTCGGGCTGGGGCGCGAAGCG | >hg38_21_+_8433172_8446622_RNA45SN1_45S_50ntFlanks | 10869 | 10901 | 33 | AGGGCTGGGTCGGTCGGGCTGGGGCGCGAAGCG | 563.9 | -7.4 | 0.8 | -9.0 | 1.8E-19 | 2.8E-18 | MB231/MB468 | CYTO |
| >hg38_21_+_8433172_8446622_RNA45SN1_45S_50ntFlanks@6719.6743.25\|TGAATTGCAGGACACATTGATCATC | >hg38_21_+_8433172_8446622_RNA45SN1_45S_50ntFlanks | 6719 | 6743 | 25 | TGAATTGCAGGACACATTGATCATC | 5400.0 | -9.5 | 1.1 | -9.0 | 2.0E-19 | 2.8E-18 | MB231/MB468 | MP |
| >MI0000439\|hsa-mir-23b&WithFlank&9\|+\|95085202\|95085310@67.85.19(+1A)&[MIMAT0000418&hsa-miR-23b-3p&offsets\|+3\|-1(+1A);m-42&9\|+\|95085265\|95085285&offsets\|+3\|+1(+1A)]\|ACATTGCCAGGGATTACCAA | >MI0000439\|hsa-mir-23b&WithFlank&9\|+\|95085202\|95085310 | 67 | 85 | 19(+1A) | ACATTGCCAGGGATTACCAA | 515.9 | 13.3 | 1.5 | 8.9 | 4.3E-19 | 2.8E-18 | BT20/MB468 | CYTO |
| >hg38_21_+_8433172_8446622_RNA45SN1_45S_50ntFlanks@475.509.35\|CTTCGTGATCGATGTGGTGACGTCGTGCTCTCCCG | >hg38_21_+_8433172_8446622_RNA45SN1_45S_50ntFlanks | 475 | 509 | 35 | CTTCGTGATCGATGTGGTGACGTCGTGCTCTCCCG | 1503.4 | 8.7 | 1.0 | 8.9 | 4.3E-19 | 2.8E-18 | BT20/MB468 | CYTO |
| >hg38_21_+_8433172_8446622_RNA45SN1_45S_50ntFlanks@12948.12989.42\|GACGACCTGCTTCTGGGTCGGGGTTTCGTACGTAGCAGAGCA | >hg38_21_+_8433172_8446622_RNA45SN1_45S_50ntFlanks | 12948 | 12989 | 42 | GACGACCTGCTTCTGGGTCGGGGTTTCGTACGTAGCAGAGCA | 431.3 | -13.2 | 1.5 | -8.9 | 7.8E-19 | 2.8E-18 | BT20/MB468 | NUC |
| >am_MI0000809\|hsa-mir-151a&WithFlank&8\|-\|140732558\|140732659@53.73.21(+3U)&[MIMAT0000757&hsa-miR-151a-3p&offsets\|0\|0(+3U);m-30&8\|-\|140732587\|140732607&offsets\|0\|0(+3U)]\|CTAGACTGAAGCTCCTTGAGGTTT | >am_MI0000809\|hsa-mir-151a&WithFlank&8\|-\|140732558\|140732659 | 53 | 73 | 21(+3U) | CTAGACTGAAGCTCCTTGAGGTTT | 259.6 | 12.0 | 1.3 | 9.0 | 1.9E-19 | 2.9E-18 | MB231/MB468 | CYTO |
| >hg38_21_+_8433172_8446622_RNA45SN1_45S_50ntFlanks@12418.12435.18\|CGATGTCGGCTCTTCCTA | >hg38_21_+_8433172_8446622_RNA45SN1_45S_50ntFlanks | 12418 | 12435 | 18 | CGATGTCGGCTCTTCCTA | 5266.4 | -5.7 | 0.6 | -9.0 | 3.1E-19 | 2.9E-18 | BT20/MB468 | MITO |
| >hg38_21_+_8433172_8446622_RNA45SN1_45S_50ntFlanks@6646.6685.40\|TCGTACGACTCTTAGCGGTGGATCACTCGGCTCGTGCGTC | >hg38_21_+_8433172_8446622_RNA45SN1_45S_50ntFlanks | 6646 | 6685 | 40 | TCGTACGACTCTTAGCGGTGGATCACTCGGCTCGTGCGTC | 12398.2 | 8.0 | 0.9 | 8.9 | 4.3E-19 | 2.9E-18 | BT20/MB468 | CYTO |
| >MI0000266\|hsa-mir-10a&WithFlank&17\|-\|48579832\|48579953@29.50.22&[MIMAT0000253&hsa-miR-10a-5p&offsets\|+1\|0;m-18&17\|-\|48579905\|48579926&offsets\|+1\|+1]\|ACCCTGTAGATCCGAATTTGTG | >MI0000266\|hsa-mir-10a&WithFlank&17\|-\|48579832\|48579953 | 29 | 50 | 22 | ACCCTGTAGATCCGAATTTGTG | 3680.5 | 12.3 | 1.4 | 9.0 | 2.8E-19 | 2.9E-18 | MB231/MB468 | NUC |
| >hg38_21_+_8433172_8446622_RNA45SN1_45S_50ntFlanks@9504.9533.30\|GTGAACTATGCCTGGGCAGGGCGAAGCCAG | >hg38_21_+_8433172_8446622_RNA45SN1_45S_50ntFlanks | 9504 | 9533 | 30 | GTGAACTATGCCTGGGCAGGGCGAAGCCAG | 999.1 | -12.1 | 1.3 | -9.0 | 3.2E-19 | 2.9E-18 | BT20/MB468 | MITO |
| >hg38_21_+_8433172_8446622_RNA45SN1_45S_50ntFlanks@8170.8197.28\|CAAGTCCTTCTGATCGAGGCCCAGCCCG | >hg38_21_+_8433172_8446622_RNA45SN1_45S_50ntFlanks | 8170 | 8197 | 28 | CAAGTCCTTCTGATCGAGGCCCAGCCCG | 844.9 | -13.6 | 1.5 | -9.0 | 2.1E-19 | 2.9E-18 | MB231/MB468 | MP |
| >trnaMT_ValTAC_MT_+_1602_1670@46.72.27\|TTCAACTTAACTTGACCGCTCTGACCA | >trnaMT_ValTAC_MT_+_1602_1670 | 46 | 72 | 27 | TTCAACTTAACTTGACCGCTCTGACCA | 350.4 | 12.5 | 1.4 | 9.0 | 1.9E-19 | 2.9E-18 | MB231/MB468 | CYTO |
| >hg38_21_+_8433172_8446622_RNA45SN1_45S_50ntFlanks@9306.9338.33\|CAAGGAGTCTAACACGTGCGCGAGTCGGGGGCT | >hg38_21_+_8433172_8446622_RNA45SN1_45S_50ntFlanks | 9306 | 9338 | 33 | CAAGGAGTCTAACACGTGCGCGAGTCGGGGGCT | 1797.0 | -10.5 | 1.2 | -9.0 | 3.4E-19 | 2.9E-18 | MB231/MB468 | MITO |
| >hg38_21_+_8433172_8446622_RNA45SN1_45S_50ntFlanks@10461.10497.37\|GCCTCCGTTGCCCTCGGCCGATCGAAAGGGAGTCGGG | >hg38_21_+_8433172_8446622_RNA45SN1_45S_50ntFlanks | 10461 | 10497 | 37 | GCCTCCGTTGCCCTCGGCCGATCGAAAGGGAGTCGGG | 534.3 | -12.8 | 1.4 | -9.0 | 3.4E-19 | 2.9E-18 | MB231/MB468 | MITO |
| >hg38_21_+_8433172_8446622_RNA45SN1_45S_50ntFlanks@12073.12096.24\|CCGAGGGGCTCTCGCTTCTGGCGC | >hg38_21_+_8433172_8446622_RNA45SN1_45S_50ntFlanks | 12073 | 12096 | 24 | CCGAGGGGCTCTCGCTTCTGGCGC | 804.2 | -13.5 | 1.5 | -9.0 | 2.1E-19 | 2.9E-18 | MB231/MB468 | MP |
| >hg38_21_+_8433172_8446622_RNA45SN1_45S_50ntFlanks@12008.12043.36\|CCGCCGGTGAAATACCACTACTCTGATCGTTTTTTC | >hg38_21_+_8433172_8446622_RNA45SN1_45S_50ntFlanks | 12008 | 12043 | 36 | CCGCCGGTGAAATACCACTACTCTGATCGTTTTTTC | 5178.3 | -16.2 | 1.8 | -9.0 | 2.1E-19 | 2.9E-18 | MB231/MB468 | MP |
| >MI0000809\|hsa-mir-151a&WithFlank&8\|-\|140732558\|140732659@51.72.22&[MIMAT0000757&hsa-miR-151a-3p&offsets\|-2\|-1;m-30&8\|-\|140732587\|140732607&offsets\|-2\|-1]\|TACTAGACTGAAGCTCCTTGAG | >MI0000809\|hsa-mir-151a&WithFlank&8\|-\|140732558\|140732659 | 51 | 72 | 22 | TACTAGACTGAAGCTCCTTGAG | 541.6 | 5.7 | 0.6 | 9.0 | 2.8E-19 | 2.9E-18 | MB231/MB468 | NUC |
| >MI0000102\|hsa-mir-100&WithFlank&11\|-\|122152223\|122152314@19.38.20(+2G)&[MIMAT0000098&hsa-miR-100-5p&offsets\|0\|-2(+2G);m-27&11\|-\|122152276\|122152296&offsets\|0\|-1(+2G)]\|AACCCGTAGATCCGAACTTGGG | >MI0000102\|hsa-mir-100&WithFlank&11\|-\|122152223\|122152314 | 19 | 38 | 20(+2G) | AACCCGTAGATCCGAACTTGGG | 1123.0 | 14.1 | 1.6 | 9.0 | 1.9E-19 | 3.0E-18 | MB231/MB468 | CYTO |
| >hg38_21_+_8433172_8446622_RNA45SN1_45S_50ntFlanks@5473.5494.22\|CGGCCCTGGCGGAGCGCTGAGA | >hg38_21_+_8433172_8446622_RNA45SN1_45S_50ntFlanks | 5473 | 5494 | 22 | CGGCCCTGGCGGAGCGCTGAGA | 296.2 | 5.2 | 0.6 | 9.0 | 1.9E-19 | 3.0E-18 | MB231/MB468 | CYTO |
| >hg38_21_+_8433172_8446622_RNA45SN1_45S_50ntFlanks@12008.12031.24\|CCGCCGGTGAAATACCACTACTCT | >hg38_21_+_8433172_8446622_RNA45SN1_45S_50ntFlanks | 12008 | 12031 | 24 | CCGCCGGTGAAATACCACTACTCT | 2520.0 | -6.8 | 0.8 | -9.0 | 1.9E-19 | 3.0E-18 | MB231/MB468 | CYTO |
| >hg38_21_+_8433172_8446622_RNA45SN1_45S_50ntFlanks@9564.9594.31\|CTGACGTGCAAATCGGTCGTCCGACCTGGGT | >hg38_21_+_8433172_8446622_RNA45SN1_45S_50ntFlanks | 9564 | 9594 | 31 | CTGACGTGCAAATCGGTCGTCCGACCTGGGT | 397.2 | -12.5 | 1.4 | -9.0 | 2.1E-19 | 3.0E-18 | MB231/MB468 | MP |
| >hg38_21_+_8433172_8446622_RNA45SN1_45S_50ntFlanks@11561.11590.30\|GCGCCTAGCAGCCGACTTAGAACTGGTGCG | >hg38_21_+_8433172_8446622_RNA45SN1_45S_50ntFlanks | 11561 | 11590 | 30 | GCGCCTAGCAGCCGACTTAGAACTGGTGCG | 445.3 | -12.7 | 1.4 | -9.0 | 2.2E-19 | 3.0E-18 | MB231/MB468 | MP |
| >hg38_21_+_8433172_8446622_RNA45SN1_45S_50ntFlanks@5472.5507.36\|ACGGCCCTGGCGGAGCGCTGAGAAGACGGTCGAACT | >hg38_21_+_8433172_8446622_RNA45SN1_45S_50ntFlanks | 5472 | 5507 | 36 | ACGGCCCTGGCGGAGCGCTGAGAAGACGGTCGAACT | 4435.4 | 8.5 | 1.0 | 8.9 | 4.5E-19 | 3.0E-18 | BT20/MB468 | CYTO |
| >hg38_21_+_8433172_8446622_RNA45SN1_45S_50ntFlanks@8472.8494.23\|CGGCCCGGCGGATCTTTCCCGCC | >hg38_21_+_8433172_8446622_RNA45SN1_45S_50ntFlanks | 8472 | 8494 | 23 | CGGCCCGGCGGATCTTTCCCGCC | 2977.3 | -15.3 | 1.7 | -9.0 | 3.5E-19 | 3.0E-18 | MB231/MB468 | MITO |
| >hg38_21_+_8433172_8446622_RNA45SN1_45S_50ntFlanks@12073.12092.20\|CCGAGGGGCTCTCGCTTCTG | >hg38_21_+_8433172_8446622_RNA45SN1_45S_50ntFlanks | 12073 | 12092 | 20 | CCGAGGGGCTCTCGCTTCTG | 414.8 | -12.6 | 1.4 | -9.0 | 2.2E-19 | 3.0E-18 | MB231/MB468 | MP |
| >hg38_21_+_8433172_8446622_RNA45SN1_45S_50ntFlanks@11768.11791.24\|CTAATTAGTGACGCGCATGAATGG | >hg38_21_+_8433172_8446622_RNA45SN1_45S_50ntFlanks | 11768 | 11791 | 24 | CTAATTAGTGACGCGCATGAATGG | 550.0 | -12.9 | 1.4 | -9.0 | 3.5E-19 | 3.0E-18 | MB231/MB468 | MITO |
| >hg38_21_+_8433172_8446622_RNA45SN1_45S_50ntFlanks@6721.6745.25\|AATTGCAGGACACATTGATCATCGA | >hg38_21_+_8433172_8446622_RNA45SN1_45S_50ntFlanks | 6721 | 6745 | 25 | AATTGCAGGACACATTGATCATCGA | 736.3 | -13.4 | 1.5 | -9.0 | 2.2E-19 | 3.0E-18 | MB231/MB468 | MP |
| >hg38_21_+_8433172_8446622_RNA45SN1_45S_50ntFlanks@11580.11603.24\|GAACTGGTGCGGACCAGGGGAATC | >hg38_21_+_8433172_8446622_RNA45SN1_45S_50ntFlanks | 11580 | 11603 | 24 | GAACTGGTGCGGACCAGGGGAATC | 7866.4 | -4.6 | 0.5 | -9.0 | 2.0E-19 | 3.0E-18 | MB231/MB468 | CYTO |
| >hg38_21_+_8433172_8446622_RNA45SN1_45S_50ntFlanks@12357.12375.19\|CCACAGGGATAACTGGCTT | >hg38_21_+_8433172_8446622_RNA45SN1_45S_50ntFlanks | 12357 | 12375 | 19 | CCACAGGGATAACTGGCTT | 479.0 | -12.8 | 1.4 | -9.0 | 2.2E-19 | 3.0E-18 | MB231/MB468 | MP |
| >hg38_21_+_8433172_8446622_RNA45SN1_45S_50ntFlanks@10351.10382.32\|CGAGAACTTTGAAGGCCGAAGTGGAGAAGGGT | >hg38_21_+_8433172_8446622_RNA45SN1_45S_50ntFlanks | 10351 | 10382 | 32 | CGAGAACTTTGAAGGCCGAAGTGGAGAAGGGT | 4285.3 | -6.9 | 0.8 | -9.0 | 2.2E-19 | 3.0E-18 | MB231/MB468 | MP |
| >hg38_21_+_8433172_8446622_RNA45SN1_45S_50ntFlanks@10805.10826.22\|GAACAATGTAGGTAAGGGAAGT | >hg38_21_+_8433172_8446622_RNA45SN1_45S_50ntFlanks | 10805 | 10826 | 22 | GAACAATGTAGGTAAGGGAAGT | 531.0 | -12.8 | 1.4 | -9.0 | 3.5E-19 | 3.0E-18 | MB231/MB468 | MITO |
| >hg38_21_+_8433172_8446622_RNA45SN1_45S_50ntFlanks@12151.12182.32\|GTGGGGAGTTTGACTGGGGCGGTACACCTGTC | >hg38_21_+_8433172_8446622_RNA45SN1_45S_50ntFlanks | 12151 | 12182 | 32 | GTGGGGAGTTTGACTGGGGCGGTACACCTGTC | 700.0 | -13.2 | 1.5 | -8.9 | 3.6E-19 | 3.0E-18 | MB231/MB468 | MITO |
| >hg38_21_+_8433172_8446622_RNA45SN1_45S_50ntFlanks@11452.11478.27\|CTCCTCGGCGCGCGGCGGCGGCGG | >hg38_21_+_8433172_8446622_RNA45SN1_45S_50ntFlanks | 11452 | 11478 | 27 | CTCCTCGGCGCGCGGCGGCGGCGG | 983.8 | -12.9 | 1.4 | -9.0 | 3.0E-19 | 3.0E-18 | MB231/MB468 | NUC |
| >hg38_21_+_8433172_8446622_RNA45SN1_45S_50ntFlanks@12160.12181.22\|TTGACTGGGGCGGTACACCTGT | >hg38_21_+_8433172_8446622_RNA45SN1_45S_50ntFlanks | 12160 | 12181 | 22 | TTGACTGGGGCGGTACACCTGT | 338.6 | -12.3 | 1.4 | -9.0 | 2.2E-19 | 3.1E-18 | MB231/MB468 | MP |
| >MI0001445\|hsa-mir-423&WithFlank&17\|+\|30117073\|30117178@58.80.23(+1C)&[MIMAT0001340&hsa-miR-423-3p&offsets\|-1\|-1(+1C);m-100&17\|+\|30117131\|30117153&offsets\|-1\|-1(+1C)]\|AAGCTCGGTCTGAGGCCCCTCAGC | >MI0001445\|hsa-mir-423&WithFlank&17\|+\|30117073\|30117178 | 58 | 80 | 23(+1C) | AAGCTCGGTCTGAGGCCCCTCAGC | 704.8 | 9.6 | 1.1 | 8.9 | 4.7E-19 | 3.1E-18 | BT20/MB468 | CYTO |
| >hg38_21_+_8433172_8446622_RNA45SN1_45S_50ntFlanks@10408.10433.26\|GGTCAGTCGGTCCTGAGAGATGGGCG | >hg38_21_+_8433172_8446622_RNA45SN1_45S_50ntFlanks | 10408 | 10433 | 26 | GGTCAGTCGGTCCTGAGAGATGGGCG | 1804.2 | -7.4 | 0.8 | -8.9 | 4.7E-19 | 3.1E-18 | BT20/MB468 | CYTO |
| >hg38_21_+_8433172_8446622_RNA45SN1_45S_50ntFlanks@8213.8235.23\|CGGTAGCGGCCCCCGGCGCGCCG | >hg38_21_+_8433172_8446622_RNA45SN1_45S_50ntFlanks | 8213 | 8235 | 23 | CGGTAGCGGCCCCCGGCGCGCCG | 362.4 | -12.4 | 1.4 | -9.0 | 2.3E-19 | 3.1E-18 | MB231/MB468 | MP |
| >MI0000285\|hsa-mir-205&WithFlank&1\|+\|209432127\|209432248@40.61.22&[MIMAT0000266&hsa-miR-205-5p&offsets\|0\|0;m-351&1\|+\|209432166\|209432187&offsets\|0\|0]\|TCCTTCATTCCACCGGAGTCTG | >MI0000285\|hsa-mir-205&WithFlank&1\|+\|209432127\|209432248 | 40 | 61 | 22 | TCCTTCATTCCACCGGAGTCTG | 949.0 | -10.2 | 1.1 | -9.0 | 3.0E-19 | 3.1E-18 | MB231/MB468 | NUC |
| >trna116_GluCTC_1_-_145399233_145399304@1.35.35,trna59_GluCTC_1_+_249168447_249168518@1.35.35,trna71_GluCTC_1_-_161439189_161439260@1.35.35,trna74_GluCTC_1_-_161431809_161431880@1.35.35,trna77_GluCTC_1_-_161424398_161424469@1.35.35,trna77_GluCTC_6_+_28949976_28950047@1.35.35,trna80_GluCTC_1_-_161417018_161417089@1.35.35,trna87_GluCTC_6_-_126101393_126101464@1.35.35\|TCCCTGGTGGTCTAGTGGTTAGGATTCGGCGCTCT | >trna116_GluCTC_1_-_145399233_145399304 | 1 | 35 | 35 | TCCCTGGTGGTCTAGTGGTTAGGATTCGGCGCTCT | 16318.4 | 7.8 | 0.9 | 9.0 | 2.1E-19 | 3.1E-18 | MB231/MB468 | CYTO |

| | | | | | | | | | | | | | |
|---|---|---|---|---|---|---|---|---|---|---|---|---|---|
| >hg38_21_+_8433172_8446622_RNA45SN1_45S_50ntFlanks@11561.11602.42\|GCGCCTAGCAGCCGACTTAGAACTGGTGCGGACCAGGGGAAT | >hg38_21_+_8433172_8446622_RNA45SN1_45S_50ntFlanks | 11561 | 11602 | 42 | GCGCCTAGCAGCCGACTTAGAACTGGTGCGGACCAGGGGAAT | 4179.5 | -9.5 | 1.1 | -8.9 | 3.6E-19 | 3.1E-18 | MB231/MB468 | MITO |
| >hg38_21_+_8433172_8446622_RNA45SN1_45S_50ntFlanks@11783.11820.38\|CATGAATGGATGAACGAGATTCCCACTGTCCCTACCTA | >hg38_21_+_8433172_8446622_RNA45SN1_45S_50ntFlanks | 11783 | 11820 | 38 | CATGAATGGATGAACGAGATTCCCACTGTCCCTACCTA | 1080.6 | -13.9 | 1.5 | -8.9 | 3.6E-19 | 3.1E-18 | MB231/MB468 | MITO |
| >hg38_21_+_8433172_8446622_RNA45SN1_45S_50ntFlanks@11787.11804.18\|AATGGATGAACGAGATTC | >hg38_21_+_8433172_8446622_RNA45SN1_45S_50ntFlanks | 11787 | 11804 | 18 | AATGGATGAACGAGATTC | 2599.3 | -6.3 | 0.7 | -8.9 | 3.7E-19 | 3.1E-18 | MB231/MB468 | MITO |
| >hg38_21_+_8433172_8446622_RNA45SN1_45S_50ntFlanks@4588.4611.24\|CTATTTTGTTGGTTTTCGGAACTG | >hg38_21_+_8433172_8446622_RNA45SN1_45S_50ntFlanks | 4588 | 4611 | 24 | CTATTTTGTTGGTTTTCGGAACTG | 607.3 | -13.1 | 1.5 | -9.0 | 2.3E-19 | 3.1E-18 | MB231/MB468 | MP |
| >hg38_21_+_8433172_8446622_RNA45SN1_45S_50ntFlanks@9532.9571.40\|AGAGGAAACTCTGGTGGAGGTCCGTAGCGGTCCTGACGTG | >hg38_21_+_8433172_8446622_RNA45SN1_45S_50ntFlanks | 9532 | 9571 | 40 | AGAGGAAACTCTGGTGGAGGTCCGTAGCGGTCCTGACGTG | 1446.8 | 8.3 | 0.9 | 9.0 | 2.1E-19 | 3.1E-18 | MB231/MB468 | CYTO |
| >hg38_21_+_8433172_8446622_RNA45SN1_45S_50ntFlanks@9467.9503.37\|GTGGAGCACGAGCGCACGTGTTAGGACCCGAAAGATG | >hg38_21_+_8433172_8446622_RNA45SN1_45S_50ntFlanks | 9467 | 9503 | 37 | GTGGAGCACGAGCGCACGTGTTAGGACCCGAAAGATG | 1814.6 | -10.8 | 1.2 | -8.9 | 3.7E-19 | 3.2E-18 | MB231/MB468 | MITO |
| >hg38_21_+_8433172_8446622_RNA45SN1_45S_50ntFlanks@8380.8401.22\|CAAGAGGGCGTGAAACCGTTAA | >hg38_21_+_8433172_8446622_RNA45SN1_45S_50ntFlanks | 8380 | 8401 | 22 | CAAGAGGGCGTGAAACCGTTAA | 800.2 | -13.4 | 1.5 | -8.9 | 3.7E-19 | 3.2E-18 | MB231/MB468 | MITO |
| >hg38_21_+_8433172_8446622_RNA45SN1_45S_50ntFlanks@12836.12869.34\|CGGTGCGGAGTGCCCTTCGTCCTGGGAAACGGGG | >hg38_21_+_8433172_8446622_RNA45SN1_45S_50ntFlanks | 12836 | 12869 | 34 | CGGTGCGGAGTGCCCTTCGTCCTGGGAAACGGGG | 386.6 | -12.5 | 1.4 | -9.0 | 2.3E-19 | 3.2E-18 | MB231/MB468 | MP |
| >hg38_21_+_8433172_8446622_RNA45SN1_45S_50ntFlanks@12639.12661.23\|TACCATCTGTGGGATTATGACTG | >hg38_21_+_8433172_8446622_RNA45SN1_45S_50ntFlanks | 12639 | 12661 | 23 | TACCATCTGTGGGATTATGACTG | 972.3 | -13.8 | 1.5 | -9.0 | 2.3E-19 | 3.2E-18 | MB231/MB468 | MP |
| >hg38_21_+_8433172_8446622_RNA45SN1_45S_50ntFlanks@9572.9592.21\|CAAATCGGTCGTCCGACCTGG | >hg38_21_+_8433172_8446622_RNA45SN1_45S_50ntFlanks | 9572 | 9592 | 21 | CAAATCGGTCGTCCGACCTGG | 1634.9 | -14.5 | 1.6 | -9.0 | 2.3E-19 | 3.2E-18 | MB231/MB468 | MP |
| >MI0000446\|hsa-mir-125b-1&WithFlank&11\|-\|122099751\|122099850@21.41.21;MI0000470\|hsa-mir-125b-2&WithFlank&21\|+\|16590231\|16590331@23.43.21&[MIMAT0000423&hsa-miR-125b-5p&offsets\|0\|-1;m-67&11\|-\|122099809\|122099830&offsets\|0\|-1];[MIMAT0000423_1&hsa-miR-125b-5p&offsets\|0\|-1;m-68&21\|+\|16590253\|16590274&offsets\|0\|-1]\|TCCCTGAGACCCTAACTTGTG | >MI0000446\|hsa-mir-125b-1&WithFlank&11\|-\|122099751\|122099850 | 21 | 41 | 21 | TCCCTGAGACCCTAACTTGTG | 14554.4 | 8.3 | 0.9 | 9.0 | 2.1E-19 | 3.2E-18 | MB231/MB468 | CYTO |
| >hg38_21_+_8433172_8446622_RNA45SN1_45S_50ntFlanks@11315.11344.30\|CCACCCCACGTCTCGTCGCGCGCGCGTCCG | >hg38_21_+_8433172_8446622_RNA45SN1_45S_50ntFlanks | 11315 | 11344 | 30 | CCACCCCACGTCTCGTCGCGCGCGCGTCCG | 407.2 | -12.5 | 1.4 | -9.0 | 2.3E-19 | 3.2E-18 | MB231/MB468 | MP |
| >hg38_21_+_8433172_8446622_RNA45SN1_45S_50ntFlanks@11115.11133.19\|CGGCAGGGGCCGGCGGCGG | >hg38_21_+_8433172_8446622_RNA45SN1_45S_50ntFlanks | 11115 | 11133 | 19 | CGGCAGGGGCCGGCGGCGG | 899.8 | -12.8 | 1.4 | -9.0 | 3.1E-19 | 3.2E-18 | MB231/MB468 | NUC |
| >hg38_21_+_8433172_8446622_RNA45SN1_45S_50ntFlanks@4367.4407.41\|CAGTTAAAAAGCTCGTAGTTGGATCTTGGGAGCGGGCGGGC | >hg38_21_+_8433172_8446622_RNA45SN1_45S_50ntFlanks | 4367 | 4407 | 41 | CAGTTAAAAAGCTCGTAGTTGGATCTTGGGAGCGGGCGGGC | 402.1 | -13.1 | 1.5 | -8.9 | 8.7E-19 | 3.2E-18 | BT20/MB468 | NUC |
| >hg38_21_+_8433172_8446622_RNA45SN1_45S_50ntFlanks@9552.9571.20\|TCCGTAGCGGTCCTGACGTG | >hg38_21_+_8433172_8446622_RNA45SN1_45S_50ntFlanks | 9552 | 9571 | 20 | TCCGTAGCGGTCCTGACGTG | 758.7 | -4.3 | 0.5 | -8.9 | 4.8E-19 | 3.2E-18 | BT20/MB468 | CYTO |
| >hg38_21_+_8433172_8446622_RNA45SN1_45S_50ntFlanks@9562.9597.36\|TCCTGACGTGCAAATCGGTCGTCCGACCTGGGTATA | >hg38_21_+_8433172_8446622_RNA45SN1_45S_50ntFlanks | 9562 | 9597 | 36 | TCCTGACGTGCAAATCGGTCGTCCGACCTGGGTATA | 1804.7 | -5.8 | 0.6 | -9.0 | 3.5E-19 | 3.2E-18 | BT20/MB468 | MITO |
| >MI0000078\|hsa-mir-22&WithFlank&17\|-\|1713897\|1713993@21.42.22&[MIMAT0004495&hsa-miR-22-5p&offsets\|0\|0;m-217&17\|-\|1713953\|1713973&offsets\|0\|+1]\|AGTTCTTCAGTGGCAAGCTTTA | >MI0000078\|hsa-mir-22&WithFlank&17\|-\|1713897\|1713993 | 21 | 42 | 22 | AGTTCTTCAGTGGCAAGCTTTA | 299.6 | 12.5 | 1.4 | 8.9 | 4.9E-19 | 3.2E-18 | BT20/MB468 | CYTO |
| >hg38_21_+_8433172_8446622_RNA45SN1_45S_50ntFlanks@4731.4748.18\|AAGAACGAAAGTCGGAGG | >hg38_21_+_8433172_8446622_RNA45SN1_45S_50ntFlanks | 4731 | 4748 | 18 | AAGAACGAAAGTCGGAGG | 378.7 | -4.7 | 0.5 | -8.8 | 8.9E-19 | 3.2E-18 | BT20/MB468 | NUC |
| >hg38_21_+_8433172_8446622_RNA45SN1_45S_50ntFlanks@11705.11728.24\|TGAAGCGCGGGTAAACGGCGGGAG | >hg38_21_+_8433172_8446622_RNA45SN1_45S_50ntFlanks | 11705 | 11728 | 24 | TGAAGCGCGGGTAAACGGCGGGAG | 3040.7 | -11.8 | 1.3 | -8.8 | 8.9E-19 | 3.2E-18 | BT20/MB468 | NUC |
| >hg38_21_+_8433172_8446622_RNA45SN1_45S_50ntFlanks@10805.10823.19\|GAACAATGTAGGTAAGGGA | >hg38_21_+_8433172_8446622_RNA45SN1_45S_50ntFlanks | 10805 | 10823 | 19 | GAACAATGTAGGTAAGGGA | 524.1 | -12.8 | 1.4 | -8.9 | 3.8E-19 | 3.2E-18 | MB231/MB468 | MITO |
| >hg38_21_+_8433172_8446622_RNA45SN1_45S_50ntFlanks@11729.11752.24\|TAACTATGACTCTCTTAAGGTAGC | >hg38_21_+_8433172_8446622_RNA45SN1_45S_50ntFlanks | 11729 | 11752 | 24 | TAACTATGACTCTCTTAAGGTAGC | 571.5 | -13.0 | 1.4 | -9.0 | 2.4E-19 | 3.2E-18 | MB231/MB468 | MP |
| >hg38_21_+_8433172_8446622_RNA45SN1_45S_50ntFlanks@6646.6664.19\|TCGTACGACTCTTAGCGGT | >hg38_21_+_8433172_8446622_RNA45SN1_45S_50ntFlanks | 6646 | 6664 | 19 | TCGTACGACTCTTAGCGGT | 37250.6 | 6.0 | 0.7 | 8.9 | 4.9E-19 | 3.2E-18 | BT20/MB468 | CYTO |
| >hg38_21_+_8433172_8446622_RNA45SN1_45S_50ntFlanks@11235.11254.20\|CTTCCCGTGGATCGCCCCAG | >hg38_21_+_8433172_8446622_RNA45SN1_45S_50ntFlanks | 11235 | 11254 | 20 | CTTCCCGTGGATCGCCCCAG | 3732.1 | -10.5 | 1.2 | -9.0 | 2.2E-19 | 3.2E-18 | MB231/MB468 | CYTO |
| >hg38_21_+_8433172_8446622_RNA45SN1_45S_50ntFlanks@6739.6770.32\|TCATCGACACTTCGAACGCACTTGCGGCCCCG | >hg38_21_+_8433172_8446622_RNA45SN1_45S_50ntFlanks | 6739 | 6770 | 32 | TCATCGACACTTCGAACGCACTTGCGGCCCCG | 1499.5 | -10.2 | 1.1 | -8.9 | 3.8E-19 | 3.2E-18 | MB231/MB468 | MITO |
| >hg38_21_+_8433172_8446622_RNA45SN1_45S_50ntFlanks@11894.11912.19\|ACTCTAGTCTGGCACGGTG | >hg38_21_+_8433172_8446622_RNA45SN1_45S_50ntFlanks | 11894 | 11912 | 19 | ACTCTAGTCTGGCACGGTG | 576.5 | -13.0 | 1.4 | -8.9 | 3.8E-19 | 3.2E-18 | MB231/MB468 | MITO |
| >hg38_21_+_8433172_8446622_RNA45SN1_45S_50ntFlanks@10866.10889.24\|CTAAGGGCTGGGTCGGTCGGGCTG | >hg38_21_+_8433172_8446622_RNA45SN1_45S_50ntFlanks | 10866 | 10889 | 24 | CTAAGGGCTGGGTCGGTCGGGCTG | 508.4 | -9.2 | 1.0 | -9.0 | 2.2E-19 | 3.2E-18 | MB231/MB468 | CYTO |
| >MI0000285\|hsa-mir-205&WithFlank&1\|+\|209432127\|209432248@40.62.23(+1A)&[MIMAT0000266&hsa-miR-205-5p&offsets\|0\|+1(+1A);m-351&1\|+\|209432166\|209432187&offsets\|0\|+1(+1A)]\|TCCTTCATTCCACCGGAGTCTGTA | >MI0000285\|hsa-mir-205&WithFlank&1\|+\|209432127\|209432248 | 40 | 62 | 23(+1A) | TCCTTCATTCCACCGGAGTCTGTA | 1075.4 | 13.7 | 1.5 | 9.2 | 3.1E-20 | 3.3E-18 | BT20/MB231 | MITO |
| >hg38_21_+_8433172_8446622_RNA45SN1_45S_50ntFlanks@9203.9220.18\|CTCGGGGCCACGCGCGCG | >hg38_21_+_8433172_8446622_RNA45SN1_45S_50ntFlanks | 9203 | 9220 | 18 | CTCGGGGCCACGCGCGCG | 675.5 | -13.2 | 1.5 | -8.9 | 3.9E-19 | 3.3E-18 | MB231/MB468 | MITO |
| >MI0000434\|hsa-let-7i&WithFlank&12\|+\|62603680\|62603775@12.32.21(+1A)&[MIMAT0000415&hsa-let-7i-5p&offsets\|0\|-1(+1A);m-49&12\|+\|62603691\|62603712&offsets\|0\|-1(+1A)]\|TGAGGTAGTAGTTTGTGCTGTA | >MI0000434\|hsa-let-7i&WithFlank&12\|+\|62603680\|62603775 | 12 | 32 | 21(+1A) | TGAGGTAGTAGTTTGTGCTGTA | 6651.8 | 7.2 | 0.8 | 8.9 | 5.0E-19 | 3.3E-18 | BT20/MB468 | CYTO |
| >hg38_21_+_8433172_8446622_RNA45SN1_45S_50ntFlanks@12628.12657.30\|TGGGGCGAAGCTACCATCTGTGGGATTATG | >hg38_21_+_8433172_8446622_RNA45SN1_45S_50ntFlanks | 12628 | 12657 | 30 | TGGGGCGAAGCTACCATCTGTGGGATTATG | 1372.6 | -14.2 | 1.6 | -8.9 | 3.9E-19 | 3.3E-18 | MB231/MB468 | MITO |
| >hg38_21_+_8433172_8446622_RNA45SN1_45S_50ntFlanks@12748.12774.27\|CCTGTCCCCGCCGGCGGGCCGCCCCCC | >hg38_21_+_8433172_8446622_RNA45SN1_45S_50ntFlanks | 12748 | 12774 | 27 | CCTGTCCCCGCCGGCGGGCCGCCCCCC | 566.1 | -12.9 | 1.4 | -8.9 | 3.9E-19 | 3.3E-18 | MB231/MB468 | MITO |
| >hg38_21_+_8433172_8446622_RNA45SN1_45S_50ntFlanks@3782.3801.20\|CAGTGAAACTGCGAATGGCT | >hg38_21_+_8433172_8446622_RNA45SN1_45S_50ntFlanks | 3782 | 3801 | 20 | CAGTGAAACTGCGAATGGCT | 5324.1 | -8.2 | 0.9 | -8.9 | 4.0E-19 | 3.3E-18 | MB231/MB468 | MITO |
| >MI0001445\|hsa-mir-423&WithFlank&17\|+\|30117073\|30117178@23.45.23(+1A)&[MIMAT0004748&hsa-miR-423-5p&offsets\|0\|0(+1A);m-141&17\|+\|30117095\|30117117&offsets\|0\|0(+1A)]\|TGAGGGGCAGAGAGCGAGACTTTA | >MI0001445\|hsa-mir-423&WithFlank&17\|+\|30117073\|30117178 | 23 | 45 | 23(+1A) | TGAGGGGCAGAGAGCGAGACTTTA | 298.8 | 10.2 | 1.1 | 8.9 | 5.1E-19 | 3.3E-18 | BT20/MB468 | CYTO |
| >hg38_21_+_8433172_8446622_RNA45SN1_45S_50ntFlanks@6748.6770.23\|CTTCGAACGCACTTGCGGCCCCG | >hg38_21_+_8433172_8446622_RNA45SN1_45S_50ntFlanks | 6748 | 6770 | 23 | CTTCGAACGCACTTGCGGCCCCG | 1023.7 | -8.2 | 0.9 | -8.9 | 4.0E-19 | 3.4E-18 | MB231/MB468 | MITO |
| >hg38_21_+_8433172_8446622_RNA45SN1_45S_50ntFlanks@8170.8214.45\|CAAGTCCTTCTGATCGAGGCCCAGCCCGTGGACGGTGTGAGGCCG | >hg38_21_+_8433172_8446622_RNA45SN1_45S_50ntFlanks | 8170 | 8214 | 45 | CAAGTCCTTCTGATCGAGGCCCAGCCCGTGGACGGTGTGAGGCCG | 8656.7 | -9.0 | 1.0 | -8.9 | 4.0E-19 | 3.4E-18 | MB231/MB468 | MITO |
| >hg38_21_+_8433172_8446622_RNA45SN1_45S_50ntFlanks@4207.4242.36\|CGGGGAGGTAGTGACGAAAAATAACAATACAGGACT | >hg38_21_+_8433172_8446622_RNA45SN1_45S_50ntFlanks | 4207 | 4242 | 36 | CGGGGAGGTAGTGACGAAAAATAACAATACAGGACT | 246.6 | 12.0 | 1.3 | 9.0 | 2.2E-19 | 3.4E-18 | MB231/MB468 | CYTO |
| >hg38_21_+_8433172_8446622_RNA45SN1_45S_50ntFlanks@9504.9523.20\|GTGAACTATGCCTGGGCAGG | >hg38_21_+_8433172_8446622_RNA45SN1_45S_50ntFlanks | 9504 | 9523 | 20 | GTGAACTATGCCTGGGCAGG | 2601.4 | -8.8 | 1.0 | -9.0 | 3.3E-19 | 3.4E-18 | MB231/MB468 | NUC |
| >hg38_21_+_8433172_8446622_RNA45SN1_45S_50ntFlanks@10726.10755.30\|GGGAGAGGGTGTAAATCTCGCGCCGGGCCG | >hg38_21_+_8433172_8446622_RNA45SN1_45S_50ntFlanks | 10726 | 10755 | 30 | GGGAGAGGGTGTAAATCTCGCGCCGGGCCG | 432.1 | -12.6 | 1.4 | -9.0 | 2.5E-19 | 3.4E-18 | MB231/MB468 | MP |
| >trnaMT_SerGCT_MT_+_12207_12265@8.38.31\|CTCACAAGAACTGCTAACTCATGCCCCCATG | >trnaMT_SerGCT_MT_+_12207_12265 | 8 | 38 | 31 | CTCACAAGAACTGCTAACTCATGCCCCCATG | 1845.2 | -13.8 | 1.5 | -9.0 | 3.3E-19 | 3.4E-18 | MB231/MB468 | NUC |
| >MI0000283\|hsa-mir-203a&WithFlank&14\|+\|104117399\|104117520@72.90.19&[MIMAT0000264&hsa-miR-203a-3p&offsets\|+1\|-2;m-79&14\|+\|104117469\|104117490&offsets\|+1\|-2]\|TGAAATGTTTAGGACCACT | >MI0000283\|hsa-mir-203a&WithFlank&14\|+\|104117399\|104117520 | 72 | 90 | 19 | TGAAATGTTTAGGACCACT | 1261.0 | 9.1 | 1.0 | 8.9 | 5.2E-19 | 3.4E-18 | BT20/MB468 | CYTO |
| >hg38_1_-_228634819_228635039_RNA5S12_5S_50ntFlanks@131.171.41\|GGATGGGAGACCGCCTGGGAATACCGGGTGCTGTAGGCTTT | >hg38_1_-_228634819_228635039_RNA5S12_5S_50ntFlanks | 131 | 171 | 41 | GGATGGGAGACCGCCTGGGAATACCGGGTGCTGTAGGCTTT | 3092.6 | 9.3 | 1.0 | 9.0 | 2.3E-19 | 3.4E-18 | MB231/MB468 | CYTO |
| >hg38_21_+_8433172_8446622_RNA45SN1_45S_50ntFlanks@12159.12181.23\|TTTGACTGGGGCGGTACACCTGT | >hg38_21_+_8433172_8446622_RNA45SN1_45S_50ntFlanks | 12159 | 12181 | 23 | TTTGACTGGGGCGGTACACCTGT | 1285.8 | -5.7 | 0.6 | -8.9 | 3.7E-19 | 3.4E-18 | BT20/MB468 | MITO |
| >hg38_21_+_8433172_8446622_RNA45SN1_45S_50ntFlanks@11207.11232.26\|GGCGGCGACTCTGGACGCGAGCCGGG | >hg38_21_+_8433172_8446622_RNA45SN1_45S_50ntFlanks | 11207 | 11232 | 26 | GGCGGCGACTCTGGACGCGAGCCGGG | 696.2 | -13.3 | 1.5 | -9.0 | 2.5E-19 | 3.4E-18 | MB231/MB468 | MP |
| >hg38_21_+_8433172_8446622_RNA45SN1_45S_50ntFlanks@13002.13023.22\|CGATCTATTGAAAGTCAGCCCT | >hg38_21_+_8433172_8446622_RNA45SN1_45S_50ntFlanks | 13002 | 13023 | 22 | CGATCTATTGAAAGTCAGCCCT | 834.3 | -12.7 | 1.4 | -9.0 | 3.3E-19 | 3.4E-18 | MB231/MB468 | NUC |
| >hg38_21_+_8433172_8446622_RNA45SN1_45S_50ntFlanks@9575.9596.22\|ATCGGTCGTCCGACCTGGGTAT | >hg38_21_+_8433172_8446622_RNA45SN1_45S_50ntFlanks | 9575 | 9596 | 22 | ATCGGTCGTCCGACCTGGGTAT | 10860.6 | -12.5 | 1.4 | -9.0 | 2.5E-19 | 3.4E-18 | MB231/MB468 | MP |
| >hg38_21_+_8433172_8446622_RNA45SN1_45S_50ntFlanks@9545.9566.22\|GTGGAGGTCCGTAGCGGTCCTG | >hg38_21_+_8433172_8446622_RNA45SN1_45S_50ntFlanks | 9545 | 9566 | 22 | GTGGAGGTCCGTAGCGGTCCTG | 720.8 | -13.3 | 1.5 | -8.9 | 4.1E-19 | 3.4E-18 | MB231/MB468 | MITO |
| >hg38_21_+_8433172_8446622_RNA45SN1_45S_50ntFlanks@3771.3792.22\|CACGGCCGGTACAGTGAAACTG | >hg38_21_+_8433172_8446622_RNA45SN1_45S_50ntFlanks | 3771 | 3792 | 22 | CACGGCCGGTACAGTGAAACTG | 1745.0 | -14.6 | 1.6 | -8.9 | 4.1E-19 | 3.5E-18 | MB231/MB468 | MITO |
| >hg38_21_+_8433172_8446622_RNA45SN1_45S_50ntFlanks@11881.11904.24\|CCTGTTGAGCTTGACTCTAGTCTG | >hg38_21_+_8433172_8446622_RNA45SN1_45S_50ntFlanks | 11881 | 11904 | 24 | CCTGTTGAGCTTGACTCTAGTCTG | 1776.5 | -10.5 | 1.2 | -8.9 | 4.1E-19 | 3.5E-18 | MB231/MB468 | MITO |
| >hg38_21_+_8433172_8446622_RNA45SN1_45S_50ntFlanks@5487.5518.32\|CGCTGAGAAGACGGTCGAACTTGACTATCTAG | >hg38_21_+_8433172_8446622_RNA45SN1_45S_50ntFlanks | 5487 | 5518 | 32 | CGCTGAGAAGACGGTCGAACTTGACTATCTAG | 1013.8 | -13.8 | 1.5 | -9.0 | 2.6E-19 | 3.5E-18 | MB231/MB468 | MP |
| >MI0002470\|hsa-mir-486-1&WithFlank&8\|-\|41660435\|41660514@10.31.22;MI0023622\|hsa-mir-486-2&WithFlank&8\|+\|41660438\|41660513@7.28.22&[MIMAT0002177&hsa-miR-486-5p&offsets\|0\|0;m-107&8\|-\|41660484\|41660505&offsets\|0\|0];[MIMAT0002177_1&hsa-miR-486-5p&offsets\|0\|0;TJU_CMC_MD2.ID03138.5p-miR&offsets\|0\|-2;m-106&8\|+\|41660444\|41660465&offsets\|0\|0]\|TCCTGTACTGAGCTGCCCCGAG | >MI0002470\|hsa-mir-486-1&WithFlank&8\|-\|41660435\|41660514 | 10 | 31 | 22 | TCCTGTACTGAGCTGCCCCGAG | 383.7 | 12.6 | 1.4 | 9.0 | 2.3E-19 | 3.5E-18 | MB231/MB468 | CYTO |

<table>
<thead><tr><th></th><th></th><th></th><th></th><th></th><th></th><th></th><th></th><th></th><th></th><th></th><th></th><th></th><th></th></tr></thead>
<tbody>
<tr><td>>hg38_21_+_8433172_8446622_RNA45SN1_45S_50ntFlanks@11576.11601.26|CTTAGAACTGGTGCGGACCAGGGGAA</td><td>>hg38_21_+_8433172_8446622_RNA45SN1_45S_50ntFlanks</td><td>11576</td><td>11601</td><td>26</td><td>CTTAGAACTGGTGCGGACCAGGGGAA</td><td>536.9</td><td>-12.9</td><td>1.4</td><td>-8.9</td><td>4.2E-19</td><td>3.5E-18</td><td>MB231/MB468</td><td>MITO</td></tr>
<tr><td>>trna100_LeuCAA_6_-_28864000_28864105@65.86.22,trna140_LeuCAA_6_-_27573417_27573524@66.87.22,trna141_LeuCAA_6_-_27570348_27570454@66.87.22,trna17_LeuCAG_16_+_57333863_57333945@65.86.22,trna26_LeuCAG_16_-_57334392_57334474@65.86.22,trna58_LeuCAA_1_+_249168054_249168159@66.87.22,trna74_LeuCAA_6_+_28908830_28908934@65.86.22|TCGAATCCCACTTCTGACACCA</td><td>>trna100_LeuCAA_6_-_28864000_28864105</td><td>65</td><td>86</td><td>22</td><td>TCGAATCCCACTTCTGACACCA</td><td>284.5</td><td>12.4</td><td>1.4</td><td>8.9</td><td>5.4E-19</td><td>3.5E-18</td><td>BT20/MB468</td><td>CYTO</td></tr>
<tr><td>>hg38_21_+_8433172_8446622_RNA45SN1_45S_50ntFlanks@10411.10429.19|CAGTCGGTCCTGAGAGATG</td><td>>hg38_21_+_8433172_8446622_RNA45SN1_45S_50ntFlanks</td><td>10411</td><td>10429</td><td>19</td><td>CAGTCGGTCCTGAGAGATG</td><td>8835.6</td><td>-11.9</td><td>1.3</td><td>-9.0</td><td>2.6E-19</td><td>3.5E-18</td><td>MB231/MB468</td><td>MP</td></tr>
<tr><td>>trna116_GluCTC_1_-_145399233_145399304@32.69.38,trna59_GluCTC_1_+_249168447_249168518@32.69.38,trna71_GluCTC_1_-_161439189_161439260@32.69.38,trna74_GluCTC_1_-_161431809_161431880@32.69.38,trna77_GluCTC_1_-_161424398_161424469@32.69.38,trna77_GluCTC_6_+_28949976_28950047@32.69.38,trna80_GluCTC_1_-_161417018_161417089@32.69.38,trna87_GluCTC_6_-_126101393_126101464@32.69.38|CTCTCACCGCCGCGGCCCGGGTTCGATTCCCGGTCAGG</td><td>>trna116_GluCTC_1_-_145399233_145399304</td><td>32</td><td>69</td><td>38</td><td>CTCTCACCGCCGCGGCCCGGGTTCGATTCCCGGTCAGG</td><td>793.2</td><td>-12.6</td><td>1.4</td><td>-9.0</td><td>3.4E-19</td><td>3.5E-18</td><td>MB231/MB468</td><td>NUC</td></tr>
<tr><td>>MI0003513|hsa-mir-455&WithFlank&9|+|114209428|114209535@22.44.23&[MIMAT0003150&hsa-miR-455-5p&offsets|0|+1;m-232&9|+|114209449|114209470&offsets|0|+1]|TATGTGCCTTTGGACTACATCGT</td><td>>MI0003513|hsa-mir-455&WithFlank&9|+|114209428|114209535</td><td>22</td><td>44</td><td>23</td><td>TATGTGCCTTTGGACTACATCGT</td><td>307.2</td><td>12.3</td><td>1.4</td><td>9.0</td><td>2.4E-19</td><td>3.5E-18</td><td>MB231/MB468</td><td>CYTO</td></tr>
<tr><td>>hg38_21_+_8433172_8446622_RNA45SN1_45S_50ntFlanks@6738.6770.33|ATCATCGACACTTCGAACGCACTTGCGGCCCCG</td><td>>hg38_21_+_8433172_8446622_RNA45SN1_45S_50ntFlanks</td><td>6738</td><td>6770</td><td>33</td><td>ATCATCGACACTTCGAACGCACTTGCGGCCCCG</td><td>3979.0</td><td>-15.0</td><td>1.6</td><td>-9.4</td><td>5.2E-21</td><td>3.6E-18</td><td>BT20/MB468</td><td>MP</td></tr>
<tr><td>>MI0000764|hsa-mir-363&WithFlank&X|-|134169372|134169458@56.77.22&[MIMAT0000707&hsa-miR-363-3p&offsets|0|0;m-178&X|-|134169381|134169403&offsets|0|-1]|AATTGCACGGTATCCATCTGTA</td><td>>MI0000764|hsa-mir-363&WithFlank&X|-|134169372|134169458</td><td>56</td><td>77</td><td>22</td><td>AATTGCACGGTATCCATCTGTA</td><td>2267.1</td><td>5.6</td><td>0.6</td><td>8.9</td><td>3.9E-19</td><td>3.6E-18</td><td>BT20/MB468</td><td>MITO</td></tr>
<tr><td>>hg38_21_+_8433172_8446622_RNA45SN1_45S_50ntFlanks@475.505.31|CTTCGTGATCGATGTGGTGACGTCGTGCTCT</td><td>>hg38_21_+_8433172_8446622_RNA45SN1_45S_50ntFlanks</td><td>475</td><td>505</td><td>31</td><td>CTTCGTGATCGATGTGGTGACGTCGTGCTCT</td><td>10118.3</td><td>8.0</td><td>0.9</td><td>8.9</td><td>4.0E-19</td><td>3.6E-18</td><td>BT20/MB468</td><td>MITO</td></tr>
<tr><td>>hg38_21_+_8433172_8446622_RNA45SN1_45S_50ntFlanks@10572.10608.37|AGAAGCCGGCGGGAGCCCCGGGGAGAGTTCTCTTTTC</td><td>>hg38_21_+_8433172_8446622_RNA45SN1_45S_50ntFlanks</td><td>10572</td><td>10608</td><td>37</td><td>AGAAGCCGGCGGGAGCCCCGGGGAGAGTTCTCTTTTC</td><td>656.3</td><td>-13.1</td><td>1.5</td><td>-8.9</td><td>4.3E-19</td><td>3.6E-18</td><td>MB231/MB468</td><td>MITO</td></tr>
<tr><td>>hg38_21_+_8433172_8446622_RNA45SN1_45S_50ntFlanks@9305.9323.19|CCAAGGAGTCTAACACGTG</td><td>>hg38_21_+_8433172_8446622_RNA45SN1_45S_50ntFlanks</td><td>9305</td><td>9323</td><td>19</td><td>CCAAGGAGTCTAACACGTG</td><td>842.0</td><td>-13.5</td><td>1.5</td><td>-8.9</td><td>4.3E-19</td><td>3.6E-18</td><td>MB231/MB468</td><td>MITO</td></tr>
<tr><td>>hg38_21_+_8433172_8446622_RNA45SN1_45S_50ntFlanks@12008.12041.34|CCGCCGGTGAAATACCACTACTCTGATCGTTTTT</td><td>>hg38_21_+_8433172_8446622_RNA45SN1_45S_50ntFlanks</td><td>12008</td><td>12041</td><td>34</td><td>CCGCCGGTGAAATACCACTACTCTGATCGTTTTT</td><td>1249.5</td><td>-14.1</td><td>1.6</td><td>-8.9</td><td>4.3E-19</td><td>3.6E-18</td><td>MB231/MB468</td><td>MITO</td></tr>
<tr><td>>hg38_21_+_8433172_8446622_RNA45SN1_45S_50ntFlanks@8799.8823.25|CCTCGGGAGGGCGCGCGGGTCGGGG</td><td>>hg38_21_+_8433172_8446622_RNA45SN1_45S_50ntFlanks</td><td>8799</td><td>8823</td><td>25</td><td>CCTCGGGAGGGCGCGCGGGTCGGGG</td><td>615.2</td><td>-13.1</td><td>1.5</td><td>-9.0</td><td>2.7E-19</td><td>3.6E-18</td><td>MB231/MB468</td><td>MP</td></tr>
<tr><td>>am_trna117_GlyTCC_1_-_145397864_145397935@1.29.29,trna45_GlyTCC_1_+_161500903_161500974@1.29.29,trna70_GlyTCC_1_-_161439547_161439618@1.29.29,trna73_GlyTCC_1_-_161432166_161432237@1.29.29,trna76_GlyTCC_1_-_161424756_161424827@1.29.29,trna79_GlyTCC_1_-_161417375_161417446@1.29.29|GCGTTGGTGGTATAGTGGTGAGCATAGCT</td><td>>am_trna117_GlyTCC_1_-_145397864_145397935</td><td>1</td><td>29</td><td>29</td><td>GCGTTGGTGGTATAGTGGTGAGCATAGCT</td><td>1611.9</td><td>-8.3</td><td>0.9</td><td>-9.0</td><td>2.7E-19</td><td>3.6E-18</td><td>MB231/MB468</td><td>MP</td></tr>
<tr><td>>hg38_21_+_8433172_8446622_RNA45SN1_45S_50ntFlanks@11594.11612.19|CAGGGGAATCCGACTGTTT</td><td>>hg38_21_+_8433172_8446622_RNA45SN1_45S_50ntFlanks</td><td>11594</td><td>11612</td><td>19</td><td>CAGGGGAATCCGACTGTTT</td><td>1589.7</td><td>-6.0</td><td>0.7</td><td>-8.8</td><td>1.0E-18</td><td>3.6E-18</td><td>BT20/MB468</td><td>NUC</td></tr>
<tr><td>>hg38_21_+_8433172_8446622_RNA45SN1_45S_50ntFlanks@12417.12435.19|TCGATGTCGGCTCTTCCTA</td><td>>hg38_21_+_8433172_8446622_RNA45SN1_45S_50ntFlanks</td><td>12417</td><td>12435</td><td>19</td><td>TCGATGTCGGCTCTTCCTA</td><td>1604.3</td><td>-4.2</td><td>0.5</td><td>-8.9</td><td>4.0E-19</td><td>3.6E-18</td><td>BT20/MB468</td><td>MITO</td></tr>
<tr><td>>hg38_21_+_8433172_8446622_RNA45SN1_45S_50ntFlanks@10427.10457.31|ATGGGCGAGCGCCGTTCCGAAGGGACGGGCG</td><td>>hg38_21_+_8433172_8446622_RNA45SN1_45S_50ntFlanks</td><td>10427</td><td>10457</td><td>31</td><td>ATGGGCGAGCGCCGTTCCGAAGGGACGGGCG</td><td>576.8</td><td>-13.6</td><td>1.5</td><td>-8.8</td><td>1.0E-18</td><td>3.7E-18</td><td>BT20/MB468</td><td>NUC</td></tr>
<tr><td>>hg38_21_+_8433172_8446622_RNA45SN1_45S_50ntFlanks@12412.12443.32|ATCCTTCGATGTCGGCTCTTCCTATCATTGTG</td><td>>hg38_21_+_8433172_8446622_RNA45SN1_45S_50ntFlanks</td><td>12412</td><td>12443</td><td>32</td><td>ATCCTTCGATGTCGGCTCTTCCTATCATTGTG</td><td>756.8</td><td>-11.9</td><td>1.3</td><td>-8.9</td><td>5.7E-19</td><td>3.7E-18</td><td>BT20/MB468</td><td>CYTO</td></tr>
<tr><td>>MI0000102|hsa-mir-100&WithFlank&11|-|122152223|122152314@19.39.21&[MIMAT0000098&hsa-miR-100-5p&offsets|0|-1;m-27&11|-|122152276|122152296&offsets|0|0]|AACCCGTAGATCCGAACTTGT</td><td>>MI0000102|hsa-mir-100&WithFlank&11|-|122152223|122152314</td><td>19</td><td>39</td><td>21</td><td>AACCCGTAGATCCGAACTTGT</td><td>18112.2</td><td>12.9</td><td>1.4</td><td>8.9</td><td>5.7E-19</td><td>3.7E-18</td><td>BT20/MB468</td><td>CYTO</td></tr>
<tr><td>>hg38_21_+_8433172_8446622_RNA45SN1_45S_50ntFlanks@11580.11602.23|GAACTGGTGCGGACCAGGGGAAT</td><td>>hg38_21_+_8433172_8446622_RNA45SN1_45S_50ntFlanks</td><td>11580</td><td>11602</td><td>23</td><td>GAACTGGTGCGGACCAGGGGAAT</td><td>1176.4</td><td>-14.1</td><td>1.6</td><td>-9.0</td><td>2.8E-19</td><td>3.7E-18</td><td>MB231/MB468</td><td>MP</td></tr>
<tr><td>>MI0000746|hsa-mir-99b&WithFlank&19|+|51692606|51692687@13.32.20(+1U)&[MIMAT0000689&hsa-miR-99b-5p&offsets|0|-2(+1U);m-10&19|+|51692618|51692639&offsets|0|-2(+1U)]|CACCCGTAGAACCGACCTTGT</td><td>>MI0000746|hsa-mir-99b&WithFlank&19|+|51692606|51692687</td><td>13</td><td>32</td><td>20(+1U)</td><td>CACCCGTAGAACCGACCTTGT</td><td>2662.6</td><td>4.0</td><td>0.5</td><td>8.8</td><td>1.0E-18</td><td>3.7E-18</td><td>BT20/MB468</td><td>NUC</td></tr>
<tr><td>>hg38_21_+_8433172_8446622_RNA45SN1_45S_50ntFlanks@5443.5466.24|CCTCGGATCGGCCCCGCCGGGGTC</td><td>>hg38_21_+_8433172_8446622_RNA45SN1_45S_50ntFlanks</td><td>5443</td><td>5466</td><td>24</td><td>CCTCGGATCGGCCCCGCCGGGGTC</td><td>1639.4</td><td>-8.3</td><td>0.9</td><td>-8.9</td><td>4.5E-19</td><td>3.7E-18</td><td>MB231/MB468</td><td>MITO</td></tr>
<tr><td>>hg38_21_+_8433172_8446622_RNA45SN1_45S_50ntFlanks@10186.10207.22|TCCTCCCCTCCTCCCGCCCACG</td><td>>hg38_21_+_8433172_8446622_RNA45SN1_45S_50ntFlanks</td><td>10186</td><td>10207</td><td>22</td><td>TCCTCCCCTCCTCCCGCCCACG</td><td>551.3</td><td>-12.9</td><td>1.4</td><td>-8.9</td><td>4.5E-19</td><td>3.8E-18</td><td>MB231/MB468</td><td>MITO</td></tr>
<tr><td>>hg38_21_+_8433172_8446622_RNA45SN1_45S_50ntFlanks@9167.9197.31|CGGGCGGGTCGCGCCGTCGGGCCCGGGGGAG</td><td>>hg38_21_+_8433172_8446622_RNA45SN1_45S_50ntFlanks</td><td>9167</td><td>9197</td><td>31</td><td>CGGGCGGGTCGCGCCGTCGGGCCCGGGGGAG</td><td>1498.1</td><td>-14.4</td><td>1.6</td><td>-9.0</td><td>2.8E-19</td><td>3.8E-18</td><td>MB231/MB468</td><td>MP</td></tr>
<tr><td>>hg38_21_+_8433172_8446622_RNA45SN1_45S_50ntFlanks@12415.12435.21|CTTCGATGTCGGCTCTTCCTA</td><td>>hg38_21_+_8433172_8446622_RNA45SN1_45S_50ntFlanks</td><td>12415</td><td>12435</td><td>21</td><td>CTTCGATGTCGGCTCTTCCTA</td><td>4924.7</td><td>-7.6</td><td>0.8</td><td>-8.9</td><td>4.6E-19</td><td>3.8E-18</td><td>MB231/MB468</td><td>MITO</td></tr>
<tr><td>>hg38_21_+_8433172_8446622_RNA45SN1_45S_50ntFlanks@4128.4162.35|CGATTCCGGAGAGGGAGCCTGAGAAACGGCTACCA</td><td>>hg38_21_+_8433172_8446622_RNA45SN1_45S_50ntFlanks</td><td>4128</td><td>4162</td><td>35</td><td>CGATTCCGGAGAGGGAGCCTGAGAAACGGCTACCA</td><td>815.0</td><td>-13.5</td><td>1.5</td><td>-8.9</td><td>4.6E-19</td><td>3.8E-18</td><td>MB231/MB468</td><td>MITO</td></tr>
<tr><td>>hg38_21_+_8433172_8446622_RNA45SN1_45S_50ntFlanks@12044.12072.29|ACTGACCCGGTGAGGCGGGGGGGCGAGCC</td><td>>hg38_21_+_8433172_8446622_RNA45SN1_45S_50ntFlanks</td><td>12044</td><td>12072</td><td>29</td><td>ACTGACCCGGTGAGGCGGGGGGGCGAGCC</td><td>911.3</td><td>-12.8</td><td>1.4</td><td>-8.9</td><td>3.8E-19</td><td>3.8E-18</td><td>MB231/MB468</td><td>NUC</td></tr>
<tr><td>>MI0000433|hsa-let-7g&WithFlank&3|-|52268272|52268367@11.32.22&[MIMAT0000414&hsa-let-7g-5p&offsets|0|0;m-58&3|-|52268336|52268357&offsets|0|0]|TGAGGTAGTAGTTTGTACAGTT</td><td>>MI0000433|hsa-let-7g&WithFlank&3|-|52268272|52268367</td><td>11</td><td>32</td><td>22</td><td>TGAGGTAGTAGTTTGTACAGTT</td><td>113950.3</td><td>7.0</td><td>0.8</td><td>8.9</td><td>5.9E-19</td><td>3.8E-18</td><td>BT20/MB468</td><td>CYTO</td></tr>
<tr><td>>MI0000102|hsa-mir-100&WithFlank&11|-|122152223|122152314@19.40.22(+2U)&[MIMAT0000098&hsa-miR-100-5p&offsets|0|0(+2U);m-27&11|-|122152276|122152296&offsets|0|+1(+2U)]|AACCCGTAGATCCGAACTTGTGTT</td><td>>MI0000102|hsa-mir-100&WithFlank&11|-|122152223|122152314</td><td>19</td><td>40</td><td>22(+2U)</td><td>AACCCGTAGATCCGAACTTGTGTT</td><td>620.1</td><td>13.3</td><td>1.5</td><td>9.0</td><td>2.6E-19</td><td>3.9E-18</td><td>MB231/MB468</td><td>CYTO</td></tr>
<tr><td>>MI0000098|hsa-mir-96&WithFlank&7|-|129774686|129774775@15.36.22(+1C)&[MIMAT0000095&hsa-miR-96-5p&offsets|0|-1(+1C);m-248&7|-|129774739|129774761&offsets|0|-1(+1C)]|TTTGGCACTAGCACATTTTTGCC</td><td>>MI0000098|hsa-mir-96&WithFlank&7|-|129774686|129774775</td><td>15</td><td>36</td><td>22(+1C)</td><td>TTTGGCACTAGCACATTTTTGCC</td><td>729.4</td><td>6.9</td><td>0.8</td><td>8.9</td><td>5.9E-19</td><td>3.9E-18</td><td>BT20/MB468</td><td>CYTO</td></tr>
<tr><td>>hg38_21_+_8433172_8446622_RNA45SN1_45S_50ntFlanks@12044.12070.27|ACTGACCCGGTGAGGCGGGGGGGCGAG</td><td>>hg38_21_+_8433172_8446622_RNA45SN1_45S_50ntFlanks</td><td>12044</td><td>12070</td><td>27</td><td>ACTGACCCGGTGAGGCGGGGGGGCGAG</td><td>1469.7</td><td>-10.8</td><td>1.2</td><td>-8.9</td><td>3.8E-19</td><td>3.9E-18</td><td>MB231/MB468</td><td>NUC</td></tr>
<tr><td>>hg38_21_+_8433172_8446622_RNA45SN1_45S_50ntFlanks@10391.10409.19|AACAGCAGTTGAACATGGG</td><td>>hg38_21_+_8433172_8446622_RNA45SN1_45S_50ntFlanks</td><td>10391</td><td>10409</td><td>19</td><td>AACAGCAGTTGAACATGGG</td><td>2273.3</td><td>-10.6</td><td>1.2</td><td>-8.9</td><td>3.8E-19</td><td>3.9E-18</td><td>MB231/MB468</td><td>NUC</td></tr>
<tr><td>>hg38_21_+_8433172_8446622_RNA45SN1_45S_50ntFlanks@9454.9503.50|CCGCGCCGGGGAGGTGGAGCACGAGCGCACGTGTTAGGACCCGAAAGATG</td><td>>hg38_21_+_8433172_8446622_RNA45SN1_45S_50ntFlanks</td><td>9454</td><td>9503</td><td>50</td><td>CCGCGCCGGGGAGGTGGAGCACGAGCGCACGTGTTAGGACCCGAAAGATG</td><td>3122.4</td><td>-14.6</td><td>1.6</td><td>-8.9</td><td>3.8E-19</td><td>3.9E-18</td><td>MB231/MB468</td><td>NUC</td></tr>
<tr><td>>MI0000283|hsa-mir-203a&WithFlank&14|+|104117399|104117520@72.92.21(+1U)&[MIMAT0000264&hsa-miR-203a-3p&offsets|+1|0(+1U);m-79&14|+|104117469|104117490&offsets|+1|0(+1U)]|TGAAATGTTTAGGACCACTAGT</td><td>>MI0000283|hsa-mir-203a&WithFlank&14|+|104117399|104117520</td><td>72</td><td>92</td><td>21(+1U)</td><td>TGAAATGTTTAGGACCACTAGT</td><td>247.2</td><td>4.2</td><td>0.5</td><td>8.8</td><td>1.1E-18</td><td>3.9E-18</td><td>BT20/MB468</td><td>NUC</td></tr>
<tr><td>>hg38_21_+_8433172_8446622_RNA45SN1_45S_50ntFlanks@6786.6806.21|TACGCCTGTCTGAGCGTCGCT</td><td>>hg38_21_+_8433172_8446622_RNA45SN1_45S_50ntFlanks</td><td>6786</td><td>6806</td><td>21</td><td>TACGCCTGTCTGAGCGTCGCT</td><td>2105.5</td><td>6.2</td><td>0.7</td><td>9.0</td><td>2.7E-19</td><td>3.9E-18</td><td>MB231/MB468</td><td>CYTO</td></tr>
<tr><td>>hg38_21_+_8433172_8446622_RNA45SN1_45S_50ntFlanks@4443.4470.28|CCCTTGCCTCTCGGCGCCCCCTCGATGC</td><td>>hg38_21_+_8433172_8446622_RNA45SN1_45S_50ntFlanks</td><td>4443</td><td>4470</td><td>28</td><td>CCCTTGCCTCTCGGCGCCCCCTCGATGC</td><td>974.2</td><td>-14.4</td><td>1.6</td><td>-8.8</td><td>1.1E-18</td><td>3.9E-18</td><td>BT20/MB468</td><td>NUC</td></tr>
<tr><td>>hg38_21_+_8433172_8446622_RNA45SN1_45S_50ntFlanks@11317.11336.20|ACCCCACGTCTCGTCGCGCG</td><td>>hg38_21_+_8433172_8446622_RNA45SN1_45S_50ntFlanks</td><td>11317</td><td>11336</td><td>20</td><td>ACCCCACGTCTCGTCGCGCG</td><td>1019.2</td><td>-12.7</td><td>1.4</td><td>-9.0</td><td>2.7E-19</td><td>3.9E-18</td><td>MB231/MB468</td><td>CYTO</td></tr>
<tr><td>>hg38_21_+_8433172_8446622_RNA45SN1_45S_50ntFlanks@476.516.41|TTCGTGATCGATGTGGTGACGTCGTGCTCTCCCGGGCCGGG</td><td>>hg38_21_+_8433172_8446622_RNA45SN1_45S_50ntFlanks</td><td>476</td><td>516</td><td>41</td><td>TTCGTGATCGATGTGGTGACGTCGTGCTCTCCCGGGCCGGG</td><td>833.1</td><td>11.0</td><td>1.2</td><td>9.0</td><td>2.7E-19</td><td>4.0E-18</td><td>MB231/MB468</td><td>CYTO</td></tr>
<tr><td>>hg38_21_+_8433172_8446622_RNA45SN1_45S_50ntFlanks@10458.10488.31|ATGGCCTCCGTTGCCCTCGGCCGATCGAAAG</td><td>>hg38_21_+_8433172_8446622_RNA45SN1_45S_50ntFlanks</td><td>10458</td><td>10488</td><td>31</td><td>ATGGCCTCCGTTGCCCTCGGCCGATCGAAAG</td><td>353.4</td><td>-12.3</td><td>1.4</td><td>-9.0</td><td>3.0E-19</td><td>4.0E-18</td><td>MB231/MB468</td><td>MP</td></tr>
<tr><td>>hg38_21_+_8433172_8446622_RNA45SN1_45S_50ntFlanks@4760.4782.23|TCAGATACCGTCGTAGTTCCGAC</td><td>>hg38_21_+_8433172_8446622_RNA45SN1_45S_50ntFlanks</td><td>4760</td><td>4782</td><td>23</td><td>TCAGATACCGTCGTAGTTCCGAC</td><td>3022.8</td><td>-15.3</td><td>1.7</td><td>-8.9</td><td>4.9E-19</td><td>4.0E-18</td><td>MB231/MB468</td><td>MITO</td></tr>
<tr><td>>MI0000650|hsa-mir-200c&WithFlank&12|+|6963693|6963772@50.69.20&[MIMAT0000617&hsa-miR-200c-3p&offsets|0|-3;m-54&12|+|6963742|6963764&offsets|0|-3]|TAATACTGCCGGGTAATGAT</td><td>>MI0000650|hsa-mir-200c&WithFlank&12|+|6963693|6963772</td><td>50</td><td>69</td><td>20</td><td>TAATACTGCCGGGTAATGAT</td><td>1747.8</td><td>9.2</td><td>1.0</td><td>9.2</td><td>2.7E-20</td><td>4.0E-18</td><td>BT20/MB231</td><td>CYTO</td></tr>
<tr><td>>MI0003628|hsa-mir-615&WithFlank&12|+|54033944|54034051@67.88.22&[MIMAT0003283&hsa-miR-615-3p&offsets|0|0;m-495&12|+|54034010|54034030&offsets|0|+1]|TCCGAGCCTGGGTCTCCCTCTT</td><td>>MI0003628|hsa-mir-615&WithFlank&12|+|54033944|54034051</td><td>67</td><td>88</td><td>22</td><td>TCCGAGCCTGGGTCTCCCTCTT</td><td>1585.3</td><td>6.7</td><td>0.7</td><td>9.2</td><td>2.8E-20</td><td>4.0E-18</td><td>BT20/MB231</td><td>CYTO</td></tr>
<tr><td>>hg38_21_+_8433172_8446622_RNA45SN1_45S_50ntFlanks@3993.4026.34|GTGACTCTAGATAACCTCGGGCCGATCGCACGCC</td><td>>hg38_21_+_8433172_8446622_RNA45SN1_45S_50ntFlanks</td><td>3993</td><td>4026</td><td>34</td><td>GTGACTCTAGATAACCTCGGGCCGATCGCACGCC</td><td>516.0</td><td>-12.9</td><td>1.4</td><td>-9.0</td><td>3.0E-19</td><td>4.0E-18</td><td>MB231/MB468</td><td>MP</td></tr>
<tr><td>>hg38_21_+_8433172_8446622_RNA45SN1_45S_50ntFlanks@12126.12146.21|CGACCCGCTCCGGGGACAGTG</td><td>>hg38_21_+_8433172_8446622_RNA45SN1_45S_50ntFlanks</td><td>12126</td><td>12146</td><td>21</td><td>CGACCCGCTCCGGGGACAGTG</td><td>1793.2</td><td>-14.6</td><td>1.6</td><td>-8.9</td><td>4.9E-19</td><td>4.1E-18</td><td>MB231/MB468</td><td>MITO</td></tr>
</tbody>
</table>

<table>
<tr><th></th><th></th><th></th><th></th><th></th><th></th><th></th><th></th><th></th><th></th><th></th><th></th><th></th><th></th></tr>
<tr><td>>hg38_21_+_8433172_8446622_RNA45SN1_45S_50ntFlanks@12624.12652.29|CCAATGGGGCGAAGCTACCATCTGTGGGA</td><td>>hg38_21_+_8433172_8446622_RNA45SN1_45S_50ntFlanks</td><td>12624</td><td>12652</td><td>29</td><td>CCAATGGGGCGAAGCTACCATCTGTGGGA</td><td>502.8</td><td>-12.8</td><td>1.4</td><td>-8.9</td><td>4.9E-19</td><td>4.1E-18</td><td>MB231/MB468</td><td>MITO</td></tr>
<tr><td>>hg38_21_+_8433172_8446622_RNA45SN1_45S_50ntFlanks@5087.5124.38|ACGAGACTCTGGCATGCTAACTAGTTACGCGACCCCCG</td><td>>hg38_21_+_8433172_8446622_RNA45SN1_45S_50ntFlanks</td><td>5087</td><td>5124</td><td>38</td><td>ACGAGACTCTGGCATGCTAACTAGTTACGCGACCCCCG</td><td>189.8</td><td>11.6</td><td>1.3</td><td>9.0</td><td>2.8E-19</td><td>4.1E-18</td><td>MB231/MB468</td><td>CYTO</td></tr>
<tr><td>>hg38_21_+_8433172_8446622_RNA45SN1_45S_50ntFlanks@12162.12180.19|GACTGGGGCGGTACACCTG</td><td>>hg38_21_+_8433172_8446622_RNA45SN1_45S_50ntFlanks</td><td>12162</td><td>12180</td><td>19</td><td>GACTGGGGCGGTACACCTG</td><td>1115.9</td><td>-13.9</td><td>1.6</td><td>-8.9</td><td>5.0E-19</td><td>4.1E-18</td><td>MB231/MB468</td><td>MITO</td></tr>
<tr><td>>hg38_21_+_8433172_8446622_RNA45SN1_45S_50ntFlanks@9738.9759.22|GAAACGATCTCAACCTATTCTC</td><td>>hg38_21_+_8433172_8446622_RNA45SN1_45S_50ntFlanks</td><td>9738</td><td>9759</td><td>22</td><td>GAAACGATCTCAACCTATTCTC</td><td>6755.4</td><td>-10.3</td><td>1.2</td><td>-8.9</td><td>5.0E-19</td><td>4.1E-18</td><td>MB231/MB468</td><td>MITO</td></tr>
<tr><td>>hg38_21_+_8433172_8446622_RNA45SN1_45S_50ntFlanks@11564.11592.29|CCTAGCAGCCGACTTAGAACTGGTGCGGA</td><td>>hg38_21_+_8433172_8446622_RNA45SN1_45S_50ntFlanks</td><td>11564</td><td>11592</td><td>29</td><td>CCTAGCAGCCGACTTAGAACTGGTGCGGA</td><td>2589.8</td><td>-15.2</td><td>1.7</td><td>-9.0</td><td>3.0E-19</td><td>4.1E-18</td><td>MB231/MB468</td><td>MP</td></tr>
<tr><td>>MI0000077|hsa-mir-21&WithFlank&17|+|59841260|59841343@14.33.20&[MIMAT0000076&hsa-miR-21-5p&offsets|0|-2;m-1&17|+|59841273|59841294&offsets|0|-2]|TAGCTTATCAGACTGATGTT</td><td>>MI0000077|hsa-mir-21&WithFlank&17|+|59841260|59841343</td><td>14</td><td>33</td><td>20</td><td>TAGCTTATCAGACTGATGTT</td><td>2701.7</td><td>8.1</td><td>0.9</td><td>8.9</td><td>6.3E-19</td><td>4.1E-18</td><td>BT20/MB468</td><td>CYTO</td></tr>
<tr><td>>hg38_21_+_8433172_8446622_RNA45SN1_45S_50ntFlanks@12366.12384.19|TAACTGGCTTGTGGCGGCC</td><td>>hg38_21_+_8433172_8446622_RNA45SN1_45S_50ntFlanks</td><td>12366</td><td>12384</td><td>19</td><td>TAACTGGCTTGTGGCGGCC</td><td>1000.7</td><td>-13.7</td><td>1.5</td><td>-8.9</td><td>5.0E-19</td><td>4.1E-18</td><td>MB231/MB468</td><td>MITO</td></tr>
<tr><td>>hg38_21_+_8433172_8446622_RNA45SN1_45S_50ntFlanks@10392.10410.19|ACAGCAGTTGAACATGGGT</td><td>>hg38_21_+_8433172_8446622_RNA45SN1_45S_50ntFlanks</td><td>10392</td><td>10410</td><td>19</td><td>ACAGCAGTTGAACATGGGT</td><td>12221.2</td><td>-13.0</td><td>1.5</td><td>-8.8</td><td>1.1E-18</td><td>4.1E-18</td><td>BT20/MB468</td><td>NUC</td></tr>
<tr><td>>hg38_21_+_8433172_8446622_RNA45SN1_45S_50ntFlanks@8848.8867.20|CGGCGGCGGCGGCGGCGGGA</td><td>>hg38_21_+_8433172_8446622_RNA45SN1_45S_50ntFlanks</td><td>8848</td><td>8867</td><td>20</td><td>CGGCGGCGGCGGCGGCGGGA</td><td>978.5</td><td>-13.7</td><td>1.5</td><td>-8.9</td><td>5.0E-19</td><td>4.1E-18</td><td>MB231/MB468</td><td>MITO</td></tr>
<tr><td>>trnaMT_GluTTC_MT_-_14674_14742@39.72.34,trnalookalike8_GluTTC_5_-_93905172_93905240@39.72.34|ATTGGTCGTGGTTGTAGTCCGTGCGAGAATACCA</td><td>>trnaMT_GluTTC_MT_-_14674_14742</td><td>39</td><td>72</td><td>34</td><td>ATTGGTCGTGGTTGTAGTCCGTGCGAGAATACCA</td><td>44964.8</td><td>7.1</td><td>0.8</td><td>8.9</td><td>4.6E-19</td><td>4.1E-18</td><td>BT20/MB468</td><td>MITO</td></tr>
<tr><td>>hg38_21_+_8433172_8446622_RNA45SN1_45S_50ntFlanks@11729.11766.38|TAACTATGACTCTCTTAAGGTAGCCAAATGCCTCGTCA</td><td>>hg38_21_+_8433172_8446622_RNA45SN1_45S_50ntFlanks</td><td>11729</td><td>11766</td><td>38</td><td>TAACTATGACTCTCTTAAGGTAGCCAAATGCCTCGTCA</td><td>855.7</td><td>-14.2</td><td>1.6</td><td>-8.8</td><td>1.1E-18</td><td>4.1E-18</td><td>BT20/MB468</td><td>NUC</td></tr>
<tr><td>>hg38_21_+_8433172_8446622_RNA45SN1_45S_50ntFlanks@12631.12654.24|GGCGAAGCTACCATCTGTGGGATT</td><td>>hg38_21_+_8433172_8446622_RNA45SN1_45S_50ntFlanks</td><td>12631</td><td>12654</td><td>24</td><td>GGCGAAGCTACCATCTGTGGGATT</td><td>646.3</td><td>-13.2</td><td>1.5</td><td>-9.0</td><td>3.1E-19</td><td>4.2E-18</td><td>MB231/MB468</td><td>MP</td></tr>
<tr><td>>hg38_21_+_8433172_8446622_RNA45SN1_45S_50ntFlanks@9306.9335.30|CAAGGAGTCTAACACGTGCGCGAGTCGGGG</td><td>>hg38_21_+_8433172_8446622_RNA45SN1_45S_50ntFlanks</td><td>9306</td><td>9335</td><td>30</td><td>CAAGGAGTCTAACACGTGCGCGAGTCGGGG</td><td>780.7</td><td>-13.4</td><td>1.5</td><td>-8.9</td><td>5.1E-19</td><td>4.2E-18</td><td>MB231/MB468</td><td>MITO</td></tr>
<tr><td>>hg38_21_+_8433172_8446622_RNA45SN1_45S_50ntFlanks@12786.12805.20|CCCGCGCGCGCGGGAGGGCG</td><td>>hg38_21_+_8433172_8446622_RNA45SN1_45S_50ntFlanks</td><td>12786</td><td>12805</td><td>20</td><td>CCCGCGCGCGCGGGAGGGCG</td><td>1128.1</td><td>-13.1</td><td>1.5</td><td>-8.9</td><td>4.2E-19</td><td>4.2E-18</td><td>MB231/MB468</td><td>NUC</td></tr>
<tr><td>>MI0000088|hsa-mir-30a&WithFlank&6|-|71403545|71403627@53.71.19&[MIMAT0000088&hsa-miR-30a-3p&offsets|0|-3;m-38&6|-|71403554|71403575&offsets|0|-3]|CTTTCAGTCGGATGTTTGC</td><td>>MI0000088|hsa-mir-30a&WithFlank&6|-|71403545|71403627</td><td>53</td><td>71</td><td>19</td><td>CTTTCAGTCGGATGTTTGC</td><td>2886.5</td><td>7.4</td><td>0.8</td><td>9.0</td><td>2.9E-19</td><td>4.2E-18</td><td>MB231/MB468</td><td>CYTO</td></tr>
<tr><td>>hg38_21_+_8433172_8446622_RNA45SN1_45S_50ntFlanks@11873.11906.34|AAGAAGACCCTGTTGAGCTTGACTCTAGTCTGGC</td><td>>hg38_21_+_8433172_8446622_RNA45SN1_45S_50ntFlanks</td><td>11873</td><td>11906</td><td>34</td><td>AAGAAGACCCTGTTGAGCTTGACTCTAGTCTGGC</td><td>1212.3</td><td>-7.8</td><td>0.9</td><td>-9.0</td><td>2.9E-19</td><td>4.2E-18</td><td>MB231/MB468</td><td>CYTO</td></tr>
<tr><td>>hg38_21_+_8433172_8446622_RNA45SN1_45S_50ntFlanks@12633.12653.21|CGAAGCTACCATCTGTGGGAT</td><td>>hg38_21_+_8433172_8446622_RNA45SN1_45S_50ntFlanks</td><td>12633</td><td>12653</td><td>21</td><td>CGAAGCTACCATCTGTGGGAT</td><td>6808.7</td><td>-7.0</td><td>0.8</td><td>-8.9</td><td>4.7E-19</td><td>4.2E-18</td><td>BT20/MB468</td><td>MITO</td></tr>
<tr><td>>hg38_21_+_8433172_8446622_RNA45SN1_45S_50ntFlanks@11207.11225.19|GGCGGCGACTCTGGACGCG</td><td>>hg38_21_+_8433172_8446622_RNA45SN1_45S_50ntFlanks</td><td>11207</td><td>11225</td><td>19</td><td>GGCGGCGACTCTGGACGCG</td><td>683.3</td><td>-9.6</td><td>1.1</td><td>-9.0</td><td>2.9E-19</td><td>4.3E-18</td><td>MB231/MB468</td><td>CYTO</td></tr>
<tr><td>>hg38_21_+_8433172_8446622_RNA45SN1_45S_50ntFlanks@3843.3861.19|CTACTTGGATAACTGTGGT</td><td>>hg38_21_+_8433172_8446622_RNA45SN1_45S_50ntFlanks</td><td>3843</td><td>3861</td><td>19</td><td>CTACTTGGATAACTGTGGT</td><td>1166.7</td><td>-5.3</td><td>0.6</td><td>-8.8</td><td>1.2E-18</td><td>4.3E-18</td><td>BT20/MB468</td><td>NUC</td></tr>
<tr><td>>MI0005531|hsa-mir-450b&WithFlank&X|-|134540179|134540268@17.38.22&[MIMAT0004909&hsa-miR-450b-5p&offsets|0|0;m-266&X|-|134540232|134540252&offsets|0|+1]|TTTTGCAATATGTTCCTGAATA</td><td>>MI0005531|hsa-mir-450b&WithFlank&X|-|134540179|134540268</td><td>17</td><td>38</td><td>22</td><td>TTTTGCAATATGTTCCTGAATA</td><td>477.2</td><td>13.2</td><td>1.5</td><td>8.9</td><td>6.6E-19</td><td>4.3E-18</td><td>BT20/MB468</td><td>CYTO</td></tr>
<tr><td>>trna11_GluTTC_15_-_26327381_26327452@1.32.32,trna3_GluTTC_13_-_45492062_45492133@1.32.32|TCCCACATGGTCTAGCGGTTAGGATTCCTGGT</td><td>>trna11_GluTTC_15_-_26327381_26327452</td><td>1</td><td>32</td><td>32</td><td>TCCCACATGGTCTAGCGGTTAGGATTCCTGGT</td><td>501.9</td><td>8.1</td><td>0.9</td><td>8.9</td><td>6.6E-19</td><td>4.3E-18</td><td>BT20/MB468</td><td>CYTO</td></tr>
<tr><td>>hg38_21_+_8433172_8446622_RNA45SN1_45S_50ntFlanks@12786.12805.20|CCCGCGCGCGCGGGAGGGCG</td><td>>hg38_21_+_8433172_8446622_RNA45SN1_45S_50ntFlanks</td><td>12786</td><td>12805</td><td>20</td><td>CCCGCGCGCGCGGGAGGGCG</td><td>589.3</td><td>-13.0</td><td>1.5</td><td>-8.9</td><td>5.2E-19</td><td>4.3E-18</td><td>MB231/MB468</td><td>MITO</td></tr>
<tr><td>>MI0000434|hsa-let-7i&WithFlank&12|+|62603680|62603775@12.32.21(+1G)&[MIMAT0000415&hsa-let-7i-5p&offsets|0|-1(+1G);m-49&12|+|62603691|62603712&offsets|0|-1(+1G)]|TGAGGTAGTAGTTTGTGCTGTG</td><td>>MI0000434|hsa-let-7i&WithFlank&12|+|62603680|62603775</td><td>12</td><td>32</td><td>21(+1G)</td><td>TGAGGTAGTAGTTTGTGCTGTG</td><td>2709.3</td><td>6.2</td><td>0.7</td><td>8.9</td><td>4.3E-19</td><td>4.3E-18</td><td>MB231/MB468</td><td>NUC</td></tr>
<tr><td>>hg38_21_+_8433172_8446622_RNA45SN1_45S_50ntFlanks@485.504.20|GATGTGGTGACGTCGTGCTC</td><td>>hg38_21_+_8433172_8446622_RNA45SN1_45S_50ntFlanks</td><td>485</td><td>504</td><td>20</td><td>GATGTGGTGACGTCGTGCTC</td><td>553.1</td><td>13.6</td><td>1.5</td><td>8.9</td><td>4.8E-19</td><td>4.3E-18</td><td>BT20/MB468</td><td>MITO</td></tr>
<tr><td>>hg38_21_+_8433172_8446622_RNA45SN1_45S_50ntFlanks@12418.12462.45|CGATGTCGGCTCTTCCTATCATTGTGAAGCAGAATTCACCAAGCG</td><td>>hg38_21_+_8433172_8446622_RNA45SN1_45S_50ntFlanks</td><td>12418</td><td>12462</td><td>45</td><td>CGATGTCGGCTCTTCCTATCATTGTGAAGCAGAATTCACCAAGCG</td><td>235.2</td><td>11.9</td><td>1.3</td><td>9.0</td><td>3.0E-19</td><td>4.3E-18</td><td>MB231/MB468</td><td>CYTO</td></tr>
<tr><td>>hg38_1_-_228634819_228635039_RNA5S12_5S_50ntFlanks@128.171.44|CTTGGATGGGAGACCGCCTGGGAATACCGGGTGCTGTAGGCTTT</td><td>>hg38_1_-_228634819_228635039_RNA5S12_5S_50ntFlanks</td><td>128</td><td>171</td><td>44</td><td>CTTGGATGGGAGACCGCCTGGGAATACCGGGTGCTGTAGGCTTT</td><td>3198.2</td><td>8.0</td><td>0.9</td><td>9.0</td><td>3.0E-19</td><td>4.3E-18</td><td>MB231/MB468</td><td>CYTO</td></tr>
<tr><td>>hg38_21_+_8433172_8446622_RNA45SN1_45S_50ntFlanks@11921.11951.31|TGAGAGGTGTAGAATAAGTGGGAGGCCCCCG</td><td>>hg38_21_+_8433172_8446622_RNA45SN1_45S_50ntFlanks</td><td>11921</td><td>11951</td><td>31</td><td>TGAGAGGTGTAGAATAAGTGGGAGGCCCCCG</td><td>591.2</td><td>-13.1</td><td>1.5</td><td>-9.0</td><td>3.2E-19</td><td>4.3E-18</td><td>MB231/MB468</td><td>MP</td></tr>
<tr><td>>hg38_1_-_228634819_228635039_RNA5S12_5S_50ntFlanks@152.171.20|TACCGGGTGCTGTAGGCTTT</td><td>>hg38_1_-_228634819_228635039_RNA5S12_5S_50ntFlanks</td><td>152</td><td>171</td><td>20</td><td>TACCGGGTGCTGTAGGCTTT</td><td>3344.8</td><td>5.6</td><td>0.6</td><td>8.9</td><td>6.7E-19</td><td>4.3E-18</td><td>BT20/MB468</td><td>CYTO</td></tr>
<tr><td>>hg38_21_+_8433172_8446622_RNA45SN1_45S_50ntFlanks@12567.12587.21|CTCAGTACGAGAGGAACCGCA</td><td>>hg38_21_+_8433172_8446622_RNA45SN1_45S_50ntFlanks</td><td>12567</td><td>12587</td><td>21</td><td>CTCAGTACGAGAGGAACCGCA</td><td>535.9</td><td>-12.9</td><td>1.4</td><td>-9.0</td><td>3.2E-19</td><td>4.3E-18</td><td>MB231/MB468</td><td>MP</td></tr>
<tr><td>>MI0000434|hsa-let-7i&WithFlank&12|+|62603680|62603775@12.31.20(+1C)&[MIMAT0000415&hsa-let-7i-5p&offsets|0|-2(+1C);m-49&12|+|62603691|62603712&offsets|0|-2(+1C)]|TGAGGTAGTAGTTTGTGCTGC</td><td>>MI0000434|hsa-let-7i&WithFlank&12|+|62603680|62603775</td><td>12</td><td>31</td><td>20(+1C)</td><td>TGAGGTAGTAGTTTGTGCTGC</td><td>861.2</td><td>6.4</td><td>0.7</td><td>8.9</td><td>4.3E-19</td><td>4.4E-18</td><td>MB231/MB468</td><td>NUC</td></tr>
<tr><td>>hg38_21_+_8433172_8446622_RNA45SN1_45S_50ntFlanks@12133.12150.18|CTCCGGGGACAGTGCCAG</td><td>>hg38_21_+_8433172_8446622_RNA45SN1_45S_50ntFlanks</td><td>12133</td><td>12150</td><td>18</td><td>CTCCGGGGACAGTGCCAG</td><td>5186.9</td><td>-11.7</td><td>1.3</td><td>-8.9</td><td>5.3E-19</td><td>4.4E-18</td><td>MB231/MB468</td><td>MITO</td></tr>
<tr><td>>hg38_21_+_8433172_8446622_RNA45SN1_45S_50ntFlanks@10823.10865.43|AAGTCGGCAAGCCGGATCCGTAACTTCGGGATAAGGATTGGCT</td><td>>hg38_21_+_8433172_8446622_RNA45SN1_45S_50ntFlanks</td><td>10823</td><td>10865</td><td>43</td><td>AAGTCGGCAAGCCGGATCCGTAACTTCGGGATAAGGATTGGCT</td><td>567.6</td><td>-12.9</td><td>1.5</td><td>-8.9</td><td>5.4E-19</td><td>4.4E-18</td><td>MB231/MB468</td><td>MITO</td></tr>
<tr><td>>hg38_21_+_8433172_8446622_RNA45SN1_45S_50ntFlanks@11564.11582.19|CCTAGCAGCCGACTTAGAA</td><td>>hg38_21_+_8433172_8446622_RNA45SN1_45S_50ntFlanks</td><td>11564</td><td>11582</td><td>19</td><td>CCTAGCAGCCGACTTAGAA</td><td>5992.7</td><td>-12.5</td><td>1.4</td><td>-8.8</td><td>1.2E-18</td><td>4.4E-18</td><td>BT20/MB468</td><td>NUC</td></tr>
<tr><td>>hg38_21_+_8433172_8446622_RNA45SN1_45S_50ntFlanks@9305.9335.31|CCAAGGAGTCTAACACGTGCGCGAGTCGGGG</td><td>>hg38_21_+_8433172_8446622_RNA45SN1_45S_50ntFlanks</td><td>9305</td><td>9335</td><td>31</td><td>CCAAGGAGTCTAACACGTGCGCGAGTCGGGG</td><td>700.7</td><td>-13.2</td><td>1.5</td><td>-8.9</td><td>5.4E-19</td><td>4.4E-18</td><td>MB231/MB468</td><td>MITO</td></tr>
<tr><td>>hg38_21_+_8433172_8446622_RNA45SN1_45S_50ntFlanks@12405.12437.33|CTTTTTGATCCTTCGATGTCGGCTCTTCCTATC</td><td>>hg38_21_+_8433172_8446622_RNA45SN1_45S_50ntFlanks</td><td>12405</td><td>12437</td><td>33</td><td>CTTTTTGATCCTTCGATGTCGGCTCTTCCTATC</td><td>36657.8</td><td>-12.0</td><td>1.4</td><td>-8.8</td><td>1.2E-18</td><td>4.4E-18</td><td>BT20/MB468</td><td>NUC</td></tr>
<tr><td>>hg38_21_+_8433172_8446622_RNA45SN1_45S_50ntFlanks@4045.4063.19|CCATTCGAACGTCTGCCCT</td><td>>hg38_21_+_8433172_8446622_RNA45SN1_45S_50ntFlanks</td><td>4045</td><td>4063</td><td>19</td><td>CCATTCGAACGTCTGCCCT</td><td>2149.7</td><td>-8.7</td><td>1.0</td><td>-8.9</td><td>4.4E-19</td><td>4.4E-18</td><td>MB231/MB468</td><td>NUC</td></tr>
<tr><td>>MI0000273|hsa-mir-183&WithFlank&7|-|129774899|129775020@34.54.21(+1A)&[MIMAT0000261&hsa-miR-183-5p&offsets|+1|0(+1A);m-41&7|-|129774966|129774987&offsets|0|-1(+1A)]|ATGGCACTGGTAGAATTCACTA</td><td>>MI0000273|hsa-mir-183&WithFlank&7|-|129774899|129775020</td><td>34</td><td>54</td><td>21(+1A)</td><td>ATGGCACTGGTAGAATTCACTA</td><td>1238.6</td><td>8.7</td><td>1.0</td><td>8.9</td><td>6.8E-19</td><td>4.4E-18</td><td>BT20/MB468</td><td>CYTO</td></tr>
<tr><td>>hg38_21_+_8433172_8446622_RNA45SN1_45S_50ntFlanks@7973.8005.33|GACGCGACCTCAGATCAGACGTGGCGACCCGCT</td><td>>hg38_21_+_8433172_8446622_RNA45SN1_45S_50ntFlanks</td><td>7973</td><td>8005</td><td>33</td><td>GACGCGACCTCAGATCAGACGTGGCGACCCGCT</td><td>9230.3</td><td>6.9</td><td>0.8</td><td>8.9</td><td>4.9E-19</td><td>4.4E-18</td><td>BT20/MB468</td><td>MITO</td></tr>
<tr><td>>hg38_21_+_8433172_8446622_RNA45SN1_45S_50ntFlanks@10819.10840.22|AGGGAAGTCGGCAAGCCGGATC</td><td>>hg38_21_+_8433172_8446622_RNA45SN1_45S_50ntFlanks</td><td>10819</td><td>10840</td><td>22</td><td>AGGGAAGTCGGCAAGCCGGATC</td><td>1813.2</td><td>-4.3</td><td>0.5</td><td>-8.9</td><td>4.9E-19</td><td>4.4E-18</td><td>BT20/MB468</td><td>MITO</td></tr>
<tr><td>>MI0000285|hsa-mir-205&WithFlank&1|+|209432127|209432248@40.62.23(+1A)&[MIMAT0000266&hsa-miR-205-5p&offsets|0|+1(+1A);m-351&1|+|209432166|209432187&offsets|0|+1(+1A)]|TCCTTCATTCCACCGGAGTCTGTA</td><td>>MI0000285|hsa-mir-205&WithFlank&1|+|209432127|209432248</td><td>40</td><td>62</td><td>23(+1A)</td><td>TCCTTCATTCCACCGGAGTCTGTA</td><td>1337.9</td><td>14.2</td><td>1.5</td><td>9.2</td><td>5.3E-20</td><td>4.4E-18</td><td>BT20/MB231</td><td>NUC</td></tr>
<tr><td>>MI0000478|hsa-mir-149&WithFlank&2|+|240455995|240456095@21.43.23(+1A)&[MIMAT0000450&hsa-miR-149-5p&offsets|0|0(+1A);m-195&2|+|240456015|240456037&offsets|0|0(+1A)]|TCTGGCTCCGTGTCTTCACTCCCA</td><td>>MI0000478|hsa-mir-149&WithFlank&2|+|240455995|240456095</td><td>21</td><td>43</td><td>23(+1A)</td><td>TCTGGCTCCGTGTCTTCACTCCCA</td><td>480.8</td><td>13.2</td><td>1.5</td><td>8.9</td><td>6.9E-19</td><td>4.5E-18</td><td>BT20/MB468</td><td>CYTO</td></tr>
<tr><td>>hg38_21_+_8433172_8446622_RNA45SN1_45S_50ntFlanks@4557.4587.31|CAGCTAGGAATAATGGAATAGGACCGCGGTT</td><td>>hg38_21_+_8433172_8446622_RNA45SN1_45S_50ntFlanks</td><td>4557</td><td>4587</td><td>31</td><td>CAGCTAGGAATAATGGAATAGGACCGCGGTT</td><td>543.3</td><td>-12.0</td><td>1.3</td><td>-8.9</td><td>4.4E-19</td><td>4.5E-18</td><td>MB231/MB468</td><td>NUC</td></tr>
<tr><td>>hg38_21_+_8433172_8446622_RNA45SN1_45S_50ntFlanks@6738.6772.35|ATCATCGACACTTCGAACGCACTTGCGGCCCCGGG</td><td>>hg38_21_+_8433172_8446622_RNA45SN1_45S_50ntFlanks</td><td>6738</td><td>6772</td><td>35</td><td>ATCATCGACACTTCGAACGCACTTGCGGCCCCGGG</td><td>864.7</td><td>-12.7</td><td>1.4</td><td>-8.9</td><td>4.5E-19</td><td>4.5E-18</td><td>MB231/MB468</td><td>NUC</td></tr>
<tr><td>>hg38_21_+_8433172_8446622_RNA45SN1_45S_50ntFlanks@6744.6771.28|GACACTTCGAACGCACTTGCGGCCCCGG</td><td>>hg38_21_+_8433172_8446622_RNA45SN1_45S_50ntFlanks</td><td>6744</td><td>6771</td><td>28</td><td>GACACTTCGAACGCACTTGCGGCCCCGG</td><td>2126.3</td><td>-8.3</td><td>0.9</td><td>-8.9</td><td>5.5E-19</td><td>4.5E-18</td><td>MB231/MB468</td><td>MITO</td></tr>
<tr><td>>hg38_21_+_8433172_8446622_RNA45SN1_45S_50ntFlanks@5515.5533.19|CTAGAGGAAGTAAAAGTCG</td><td>>hg38_21_+_8433172_8446622_RNA45SN1_45S_50ntFlanks</td><td>5515</td><td>5533</td><td>19</td><td>CTAGAGGAAGTAAAAGTCG</td><td>443.2</td><td>-12.7</td><td>1.4</td><td>-9.0</td><td>3.3E-19</td><td>4.5E-18</td><td>MB231/MB468</td><td>MP</td></tr>
<tr><td>>hg38_21_+_8433172_8446622_RNA45SN1_45S_50ntFlanks@12973.12990.18|TCGTACGTAGCAGAGCAG</td><td>>hg38_21_+_8433172_8446622_RNA45SN1_45S_50ntFlanks</td><td>12973</td><td>12990</td><td>18</td><td>TCGTACGTAGCAGAGCAG</td><td>1906.5</td><td>-9.3</td><td>1.0</td><td>-8.9</td><td>5.0E-19</td><td>4.5E-18</td><td>BT20/MB468</td><td>MITO</td></tr>
<tr><td>>MI0000088|hsa-mir-30a&WithFlank&6|-|71403545|71403627@12.31.20&[MIMAT0000087&hsa-miR-30a-5p&offsets|0|-2;m-15&6|-|71403595|71403616&offsets|0|-2]|TGTAAACATCCTCGACTGGA</td><td>>MI0000088|hsa-mir-30a&WithFlank&6|-|71403545|71403627</td><td>12</td><td>31</td><td>20</td><td>TGTAAACATCCTCGACTGGA</td><td>924.4</td><td>5.9</td><td>0.7</td><td>8.9</td><td>5.0E-19</td><td>4.5E-18</td><td>BT20/MB468</td><td>MITO</td></tr>
<tr><td>>MI0000447|hsa-mir-128-1&WithFlank&2|+|135665391|135665484@56.75.20(+1C);MI0000727|hsa-mir-128-2&WithFlank&3|+|35744470|35744565@58.77.20(+1C)&[MIMAT0000424&hsa-miR-128-3p&offsets|0|-1(+1C);m-108&2|+|135665446|135665466&offsets|0|-1(+1C)];[MIMAT0000424_1&hsa-miR-128-3p&offsets|0|-1(+1C);m-117&3|+|35744527|35744547&offsets|0|-1(+1C)]|TCACAGTGAACCGGTCTCTTC</td><td>>MI0000447|hsa-mir-128-1&WithFlank&2|+|135665391|135665484</td><td>56</td><td>75</td><td>20(+1C)</td><td>TCACAGTGAACCGGTCTCTTC</td><td>1391.3</td><td>5.2</td><td>0.6</td><td>8.9</td><td>5.0E-19</td><td>4.5E-18</td><td>BT20/MB468</td><td>MITO</td></tr>
</table>

| | | | | | | | | | | | | | |
|---|---|---|---|---|---|---|---|---|---|---|---|---|---|
| >am_MI0000809\|hsa-mir-151a&WithFlank&8\|-\|140732558\|140732659@53.71.19&[MIMAT0000757&hsa-miR-151a-3p&offsets\|0\|-2;m-30&8\|-\|140732587\|140732607&offsets\|0\|-2]\|CTAGACTGAAGCTCCTTGA | >am_MI0000809\|hsa-mir-151a&WithFlank&8\|-\|140732558\|140732659 | 53 | 71 | 19 | CTAGACTGAAGCTCCTTGA | 1853.7 | 10.6 | 1.2 | 8.9 | 5.0E-19 | 4.5E-18 | BT20/MB468 | MITO |
| >hg38_21_+_8433172_8446622_RNA45SN1_45S_50ntFlanks@476.518.43\|TTCGTGATCGATGTGGTGACGTCGTGCTCTCCCGGGCCGGGTC | >hg38_21_+_8433172_8446622_RNA45SN1_45S_50ntFlanks | 476 | 518 | 43 | TTCGTGATCGATGTGGTGACGTCGTGCTCTCCCGGGCCGGGTC | 458.3 | 12.8 | 1.4 | 9.0 | 3.1E-19 | 4.5E-18 | MB231/MB468 | CYTO |
| >hg38_21_+_8433172_8446622_RNA45SN1_45S_50ntFlanks@11233.11254.22\|CCCTTCCCGTGGATCGCCCCAG | >hg38_21_+_8433172_8446622_RNA45SN1_45S_50ntFlanks | 11233 | 11254 | 22 | CCCTTCCCGTGGATCGCCCCAG | 986.0 | -13.8 | 1.5 | -9.0 | 3.4E-19 | 4.5E-18 | MB231/MB468 | MP |
| >hg38_21_+_8433172_8446622_RNA45SN1_45S_50ntFlanks@7984.8012.29\|AGATCAGACGTGGCGACCCGCTGAATTTA | >hg38_21_+_8433172_8446622_RNA45SN1_45S_50ntFlanks | 7984 | 8012 | 29 | AGATCAGACGTGGCGACCCGCTGAATTTA | 506.2 | -12.8 | 1.4 | -9.0 | 3.4E-19 | 4.5E-18 | MB231/MB468 | MP |
| >hg38_21_+_8433172_8446622_RNA45SN1_45S_50ntFlanks@475.495.21\|CTTCGTGATCGATGTGGTGAC | >hg38_21_+_8433172_8446622_RNA45SN1_45S_50ntFlanks | 475 | 495 | 21 | CTTCGTGATCGATGTGGTGAC | 451.1 | 12.8 | 1.4 | 9.0 | 3.1E-19 | 4.5E-18 | MB231/MB468 | CYTO |
| >hg38_21_+_8433172_8446622_RNA45SN1_45S_50ntFlanks@11874.11912.39\|AGAAGACCCTGTTGAGCTTGACTCTAGTCTGGCACGGTG | >hg38_21_+_8433172_8446622_RNA45SN1_45S_50ntFlanks | 11874 | 11912 | 39 | AGAAGACCCTGTTGAGCTTGACTCTAGTCTGGCACGGTG | 813.0 | -13.5 | 1.5 | -8.9 | 5.6E-19 | 4.6E-18 | MB231/MB468 | MITO |
| >hg38_21_+_8433172_8446622_RNA45SN1_45S_50ntFlanks@10513.10533.21\|CCGGAGTGGCGGAGATGGGCG | >hg38_21_+_8433172_8446622_RNA45SN1_45S_50ntFlanks | 10513 | 10533 | 21 | CCGGAGTGGCGGAGATGGGCG | 3246.3 | -11.9 | 1.3 | -8.9 | 4.6E-19 | 4.6E-18 | MB231/MB468 | NUC |
| >hg38_21_+_8433172_8446622_RNA45SN1_45S_50ntFlanks@9566.9596.31\|GACGTGCAAATCGGTCGTCCGACCTGGGTAT | >hg38_21_+_8433172_8446622_RNA45SN1_45S_50ntFlanks | 9566 | 9596 | 31 | GACGTGCAAATCGGTCGTCCGACCTGGGTAT | 3533.8 | -4.9 | 0.5 | -8.9 | 4.6E-19 | 4.6E-18 | MB231/MB468 | NUC |
| >hg38_21_+_8433172_8446622_RNA45SN1_45S_50ntFlanks@9534.9575.42\|AGGAAACTCTGGTGGAGGTCCGTAGCGGTCCTGACGTGCAAA | >hg38_21_+_8433172_8446622_RNA45SN1_45S_50ntFlanks | 9534 | 9575 | 42 | AGGAAACTCTGGTGGAGGTCCGTAGCGGTCCTGACGTGCAAA | 887.1 | -14.2 | 1.6 | -8.8 | 1.3E-18 | 4.6E-18 | BT20/MB468 | NUC |
| >hg38_21_+_8433172_8446622_RNA45SN1_45S_50ntFlanks@10534.10567.34\|CCGCGAGGCGTCCAGTGCGGTAACGCGACCGATC | >hg38_21_+_8433172_8446622_RNA45SN1_45S_50ntFlanks | 10534 | 10567 | 34 | CCGCGAGGCGTCCAGTGCGGTAACGCGACCGATC | 671.1 | -13.2 | 1.5 | -8.9 | 5.7E-19 | 4.6E-18 | MB231/MB468 | MITO |
| >hg38_21_+_8433172_8446622_RNA45SN1_45S_50ntFlanks@11783.11805.23\|CATGAATGGATGAACGAGATTCC | >hg38_21_+_8433172_8446622_RNA45SN1_45S_50ntFlanks | 11783 | 11805 | 23 | CATGAATGGATGAACGAGATTCC | 1491.6 | -6.0 | 0.7 | -8.9 | 5.7E-19 | 4.6E-18 | MB231/MB468 | MITO |
| >MI0000061\|hsa-let-7a-2&WithFlank&11\|-\|122146516\|122146599@11.32.22;MI0000062\|hsa-let-7a-3&WithFlank&22\|+\|46112743\|46112828@10.31.22;MI0000060\|hsa-let-7a-1&WithFlank&9\|+\|94175951\|94176042@12.33.22&[MIMAT0000062&hsa-let-7a-5p&offsets\|0\|0;m-6&11\|-\|122146568\|122146589&offsets\|0\|0];[MIMAT0000062_1&hsa-let-7a-5p&offsets\|0\|0;m-5&22\|+\|46112752\|46112773&offsets\|0\|0];[MIMAT0000062_2&hsa-let-7a-5p&offsets\|0\|0;m-4&9\|+\|94175962\|94175983&offsets\|0\|0]\|TGAGGTAGTAGGTTGTATAGTT | >MI0000061\|hsa-let-7a-2&WithFlank&11\|-\|122146516\|122146599 | 11 | 32 | 22 | TGAGGTAGTAGGTTGTATAGTT | 118482.7 | 5.2 | 0.6 | 8.9 | 4.6E-19 | 4.6E-18 | MB231/MB468 | NUC |
| >hg38_21_+_8433172_8446622_RNA45SN1_45S_50ntFlanks@12570.12587.18\|AGTACGAGAGGAACCGCA | >hg38_21_+_8433172_8446622_RNA45SN1_45S_50ntFlanks | 12570 | 12587 | 18 | AGTACGAGAGGAACCGCA | 598.4 | -12.2 | 1.4 | -8.9 | 4.6E-19 | 4.6E-18 | MB231/MB468 | NUC |
| >hg38_21_+_8433172_8446622_RNA45SN1_45S_50ntFlanks@13004.13025.22\|ATCTATTGAAAGTCAGCCCTCG | >hg38_21_+_8433172_8446622_RNA45SN1_45S_50ntFlanks | 13004 | 13025 | 22 | ATCTATTGAAAGTCAGCCCTCG | 1220.9 | -12.6 | 1.4 | -8.9 | 7.2E-19 | 4.6E-18 | BT20/MB468 | CYTO |
| >hg38_21_+_8433172_8446622_RNA45SN1_45S_50ntFlanks@12406.12436.31\|TTTTTGATCCTTCGATGTCGGCTCTTCCTAT | >hg38_21_+_8433172_8446622_RNA45SN1_45S_50ntFlanks | 12406 | 12436 | 31 | TTTTTGATCCTTCGATGTCGGCTCTTCCTAT | 2019.4 | -6.5 | 0.7 | -8.9 | 4.7E-19 | 4.6E-18 | MB231/MB468 | NUC |
| >hg38_21_+_8433172_8446622_RNA45SN1_45S_50ntFlanks@12419.12439.21\|GATGTCGGCTCTTCCTATCAT | >hg38_21_+_8433172_8446622_RNA45SN1_45S_50ntFlanks | 12419 | 12439 | 21 | GATGTCGGCTCTTCCTATCAT | 3339.4 | 5.2 | 0.6 | 9.0 | 3.2E-19 | 4.6E-18 | MB231/MB468 | CYTO |
| >hg38_21_+_8433172_8446622_RNA45SN1_45S_50ntFlanks@12539.12566.28\|ATGATGTGTTGTTGCCATGGTAATCCTG | >hg38_21_+_8433172_8446622_RNA45SN1_45S_50ntFlanks | 12539 | 12566 | 28 | ATGATGTGTTGTTGCCATGGTAATCCTG | 3045.8 | -11.8 | 1.3 | -8.8 | 1.3E-18 | 4.7E-18 | BT20/MB468 | NUC |
| >hg38_21_+_8433172_8446622_RNA45SN1_45S_50ntFlanks@13002.13039.38\|CGATCTATTGAAAGTCAGCCCTCGACACAAGGGTTTGT | >hg38_21_+_8433172_8446622_RNA45SN1_45S_50ntFlanks | 13002 | 13039 | 38 | CGATCTATTGAAAGTCAGCCCTCGACACAAGGGTTTGT | 3746.5 | -14.0 | 1.6 | -8.9 | 5.3E-19 | 4.7E-18 | BT20/MB468 | MITO |
| >hg38_21_+_8433172_8446622_RNA45SN1_45S_50ntFlanks@9454.9493.40\|CCGCGCCGGGGAGGTGGAGCACGAGCGCACGTGTTAGGAC | >hg38_21_+_8433172_8446622_RNA45SN1_45S_50ntFlanks | 9454 | 9493 | 40 | CCGCGCCGGGGAGGTGGAGCACGAGCGCACGTGTTAGGAC | 671.0 | -13.2 | 1.5 | -8.9 | 5.8E-19 | 4.7E-18 | MB231/MB468 | MITO |
| >hg38_21_+_8433172_8446622_RNA45SN1_45S_50ntFlanks@3984.4026.43\|GCGGCTTTGGTGACTCTAGATAACCTCGGGCCGATCGCACGCC | >hg38_21_+_8433172_8446622_RNA45SN1_45S_50ntFlanks | 3984 | 4026 | 43 | GCGGCTTTGGTGACTCTAGATAACCTCGGGCCGATCGCACGCC | 1292.6 | -14.2 | 1.6 | -9.0 | 3.5E-19 | 4.7E-18 | MB231/MB468 | MP |
| >hg38_21_+_8433172_8446622_RNA45SN1_45S_50ntFlanks@10841.10859.19\|CGTAACTTCGGGATAAGGA | >hg38_21_+_8433172_8446622_RNA45SN1_45S_50ntFlanks | 10841 | 10859 | 19 | CGTAACTTCGGGATAAGGA | 413.8 | -12.6 | 1.4 | -9.0 | 3.5E-19 | 4.7E-18 | MB231/MB468 | MP |
| >hg38_21_+_8433172_8446622_RNA45SN1_45S_50ntFlanks@12715.12751.37\|AGCCTCGGTTGGCCTCGGATAGCCGGTCCCCCGCCTG | >hg38_21_+_8433172_8446622_RNA45SN1_45S_50ntFlanks | 12715 | 12751 | 37 | AGCCTCGGTTGGCCTCGGATAGCCGGTCCCCCGCCTG | 640.4 | -13.1 | 1.5 | -8.9 | 5.8E-19 | 4.7E-18 | MB231/MB468 | MITO |
| >hg38_21_+_8433172_8446622_RNA45SN1_45S_50ntFlanks@5494.5513.20\|AAGACGGTCGAACTTGACTA | >hg38_21_+_8433172_8446622_RNA45SN1_45S_50ntFlanks | 5494 | 5513 | 20 | AAGACGGTCGAACTTGACTA | 493.1 | -12.8 | 1.4 | -8.9 | 3.6E-19 | 4.8E-18 | MB231/MB468 | MP |
| >hg38_21_+_8433172_8446622_RNA45SN1_45S_50ntFlanks@8437.8469.33\|AGGATTCAACCCGGCGGCGGGTCCGGCCGTGTC | >hg38_21_+_8433172_8446622_RNA45SN1_45S_50ntFlanks | 8437 | 8469 | 33 | AGGATTCAACCCGGCGGCGGGTCCGGCCGTGTC | 380.8 | -12.4 | 1.4 | -8.9 | 3.6E-19 | 4.8E-18 | MB231/MB468 | MP |
| >hg38_21_+_8433172_8446622_RNA45SN1_45S_50ntFlanks@10389.10430.42\|TGAACAGCAGTTGAACATGGGTCAGTCGGTCCTGAGAGATGG | >hg38_21_+_8433172_8446622_RNA45SN1_45S_50ntFlanks | 10389 | 10430 | 42 | TGAACAGCAGTTGAACATGGGTCAGTCGGTCCTGAGAGATGG | 1067.8 | -14.5 | 1.6 | -8.8 | 1.3E-18 | 4.8E-18 | BT20/MB468 | NUC |
| >hg38_21_+_8433172_8446622_RNA45SN1_45S_50ntFlanks@11620.11640.21\|ACAAAGCATCGCGAAGGCCCG | >hg38_21_+_8433172_8446622_RNA45SN1_45S_50ntFlanks | 11620 | 11640 | 21 | ACAAAGCATCGCGAAGGCCCG | 2951.3 | -14.6 | 1.6 | -9.4 | 7.4E-21 | 4.8E-18 | BT20/MB468 | MP |
| >am_trna117_GlyTCC_1_-_145397864_145397935@1.30.30,trna45_GlyTCC_1_+_161500903_161500974@1.30.30,trna70_GlyTCC_1_-_161439547_161439618@1.30.30,trna73_GlyTCC_1_-_161432166_161432237@1.30.30,trna76_GlyTCC_1_-_161424756_161424827@1.30.30,trna79_GlyTCC_1_-_161417375_161417446@1.30.30\|GCGTTGGTGGTATAGTGGTGAGCATAGCTG | >am_trna117_GlyTCC_1_-_145397864_145397935 | 1 | 30 | 30 | GCGTTGGTGGTATAGTGGTGAGCATAGCTG | 5235.7 | -5.4 | 0.6 | -8.9 | 7.4E-19 | 4.8E-18 | BT20/MB468 | CYTO |
| >hg38_21_+_8433172_8446622_RNA45SN1_45S_50ntFlanks@8114.8134.21\|GCGCGGGACATGTGGCGTACG | >hg38_21_+_8433172_8446622_RNA45SN1_45S_50ntFlanks | 8114 | 8134 | 21 | GCGCGGGACATGTGGCGTACG | 432.9 | -12.6 | 1.4 | -8.9 | 3.6E-19 | 4.8E-18 | MB231/MB468 | MP |
| >hg38_21_+_8433172_8446622_RNA45SN1_45S_50ntFlanks@8190.8213.24\|CCAGCCCGTGGACGGTGTGAGGCC | >hg38_21_+_8433172_8446622_RNA45SN1_45S_50ntFlanks | 8190 | 8213 | 24 | CCAGCCCGTGGACGGTGTGAGGCC | 816.7 | -6.0 | 0.7 | -8.8 | 1.3E-18 | 4.8E-18 | BT20/MB468 | NUC |
| >hg38_21_+_8433172_8446622_RNA45SN1_45S_50ntFlanks@10408.10430.23\|GGTCAGTCGGTCCTGAGAGATGG | >hg38_21_+_8433172_8446622_RNA45SN1_45S_50ntFlanks | 10408 | 10430 | 23 | GGTCAGTCGGTCCTGAGAGATGG | 1151.1 | -14.0 | 1.6 | -8.9 | 3.7E-19 | 4.9E-18 | MB231/MB468 | MP |
| >MI0000456\|hsa-mir-140&WithFlank&16\|+\|69933075\|69933186@69.89.21&[MIMAT0004597&hsa-miR-140-3p&offsets\|+1\|+1;m-45&16\|+\|69933143\|69933164&offsets\|0\|-1]\|ACCACAGGGTAGAACCACGGA | >MI0000456\|hsa-mir-140&WithFlank&16\|+\|69933075\|69933186 | 69 | 89 | 21 | ACCACAGGGTAGAACCACGGA | 1547.1 | 14.6 | 1.6 | 9.0 | 3.4E-19 | 4.9E-18 | MB231/MB468 | CYTO |
| >hg38_21_+_8433172_8446622_RNA45SN1_45S_50ntFlanks@6663.6699.37\|GTGGATCACTCGGCTCGTGCGTCGATGAAGAACGCAG | >hg38_21_+_8433172_8446622_RNA45SN1_45S_50ntFlanks | 6663 | 6699 | 37 | GTGGATCACTCGGCTCGTGCGTCGATGAAGAACGCAG | 433.8 | -5.2 | 0.6 | -8.8 | 1.4E-18 | 4.9E-18 | BT20/MB468 | NUC |
| >hg38_21_+_8433172_8446622_RNA45SN1_45S_50ntFlanks@6782.6803.22\|GGGCTACGCCTGTCTGAGCGTC | >hg38_21_+_8433172_8446622_RNA45SN1_45S_50ntFlanks | 6782 | 6803 | 22 | GGGCTACGCCTGTCTGAGCGTC | 657.3 | -13.2 | 1.5 | -8.9 | 3.7E-19 | 4.9E-18 | MB231/MB468 | MP |
| >hg38_21_+_8433172_8446622_RNA45SN1_45S_50ntFlanks@9454.9494.41\|CCGCGCCGGGGAGGTGGAGCACGAGCGCACGTGTTAGGACC | >hg38_21_+_8433172_8446622_RNA45SN1_45S_50ntFlanks | 9454 | 9494 | 41 | CCGCGCCGGGGAGGTGGAGCACGAGCGCACGTGTTAGGACC | 791.9 | -13.4 | 1.5 | -8.9 | 6.1E-19 | 5.0E-18 | MB231/MB468 | MITO |
| >hg38_21_+_8433172_8446622_RNA45SN1_45S_50ntFlanks@10466.10497.32\|CGTTGCCCTCGGCCGATCGAAAGGGAGTCGGG | >hg38_21_+_8433172_8446622_RNA45SN1_45S_50ntFlanks | 10466 | 10497 | 32 | CGTTGCCCTCGGCCGATCGAAAGGGAGTCGGG | 720.1 | -11.6 | 1.3 | -8.9 | 5.6E-19 | 5.0E-18 | BT20/MB468 | MITO |
| >hg38_21_+_8433172_8446622_RNA45SN1_45S_50ntFlanks@9534.9563.30\|AGGAAACTCTGGTGGAGGTCCGTAGCGGTC | >hg38_21_+_8433172_8446622_RNA45SN1_45S_50ntFlanks | 9534 | 9563 | 30 | AGGAAACTCTGGTGGAGGTCCGTAGCGGTC | 3524.8 | -5.1 | 0.6 | -8.9 | 5.6E-19 | 5.0E-18 | BT20/MB468 | MITO |
| >MI0000077\|hsa-mir-21&WithFlank&17\|+\|59841260\|59841343@13.35.23&[MIMAT0000076&hsa-miR-21-5p&offsets\|-1\|0;m-1&17\|+\|59841273\|59841294&offsets\|-1\|0]\|GTAGCTTATCAGACTGATGTTGA | >MI0000077\|hsa-mir-21&WithFlank&17\|+\|59841260\|59841343 | 13 | 35 | 23 | GTAGCTTATCAGACTGATGTTGA | 1277.9 | 7.5 | 0.8 | 8.9 | 7.7E-19 | 5.0E-18 | BT20/MB468 | CYTO |
| >MI0000469\|hsa-mir-125a&WithFlank&19\|+\|51693248\|51693345@59.78.20(+1U)&[MIMAT0004602&hsa-miR-125a-3p&offsets\|0\|-2(+1U);m-333&19\|+\|51693306\|51693326&offsets\|0\|-1(+1U)]\|ACAGGTGAGGTTCTTGGGAGT | >MI0000469\|hsa-mir-125a&WithFlank&19\|+\|51693248\|51693345 | 59 | 78 | 20(+1U) | ACAGGTGAGGTTCTTGGGAGT | 390.3 | 10.5 | 1.2 | 8.9 | 7.7E-19 | 5.0E-18 | BT20/MB468 | CYTO |
| >MI0016887\|hsa-mir-4521&WithFlank&17\|+\|8186939\|8187010@6.31.26(+3U)&[MIMAT0019058&hsa-miR-4521&offsets\|-4\|0(+3U);m-765&17\|+\|8186948\|8186969&offsets\|-4\|0(+3U)]\|GTCGGCTAAGGAAGTCCTGTGCTCAGTTT | >MI0016887\|hsa-mir-4521&WithFlank&17\|+\|8186939\|8187010 | 6 | 31 | 26(+3U) | GTCGGCTAAGGAAGTCCTGTGCTCAGTTT | 323.8 | 12.6 | 1.4 | 8.9 | 7.7E-19 | 5.0E-18 | BT20/MB468 | CYTO |
| >hg38_21_+_8433172_8446622_RNA45SN1_45S_50ntFlanks@9575.9596.22\|ATCGGTCGTCCGACCTGGGTAT | >hg38_21_+_8433172_8446622_RNA45SN1_45S_50ntFlanks | 9575 | 9596 | 22 | ATCGGTCGTCCGACCTGGGTAT | 1263.4 | -12.6 | 1.4 | -8.9 | 7.7E-19 | 5.0E-18 | BT20/MB468 | CYTO |
| >hg38_21_+_8433172_8446622_RNA45SN1_45S_50ntFlanks@6651.6681.31\|CGACTCTTAGCGGTGGATCACTCGGCTCGTG | >hg38_21_+_8433172_8446622_RNA45SN1_45S_50ntFlanks | 6651 | 6681 | 31 | CGACTCTTAGCGGTGGATCACTCGGCTCGTG | 70399.2 | -6.7 | 0.7 | -8.9 | 3.8E-19 | 5.0E-18 | MB231/MB468 | MP |
| >MI0000270\|hsa-mir-181b-1&WithFlank&1\|-\|198858867\|198858988@42.65.24;MI0000683\|hsa-mir-181b-2&WithFlank&9\|+\|124693704\|124693804@22.45.24&[MIMAT0000257&hsa-miR-181b-5p&offsets\|0\|+1;m-73&1\|-\|198858925\|198858947&offsets\|0\|+1];[MIMAT0000257_1&hsa-miR-181b-5p&offsets\|0\|+1;m-72&9\|+\|124693725\|124693747&offsets\|0\|+1]\|AACATTCATTGCTGTCGGTGGGTT | >MI0000270\|hsa-mir-181b-1&WithFlank&1\|-\|198858867\|198858988 | 42 | 65 | 24 | AACATTCATTGCTGTCGGTGGGTT | 974.7 | 5.4 | 0.6 | 8.9 | 7.8E-19 | 5.0E-18 | BT20/MB468 | CYTO |
| >hg38_21_+_8433172_8446622_RNA45SN1_45S_50ntFlanks@8437.8467.31\|AGGATTCAACCCGGCGGCGGGTCCGGCCGTG | >hg38_21_+_8433172_8446622_RNA45SN1_45S_50ntFlanks | 8437 | 8467 | 31 | AGGATTCAACCCGGCGGCGGGTCCGGCCGTG | 491.5 | -12.8 | 1.4 | -8.9 | 3.8E-19 | 5.0E-18 | MB231/MB468 | MP |
| >hg38_21_+_8433172_8446622_RNA45SN1_45S_50ntFlanks@12992.13013.22\|TCCCTCGCTGCGATCTATTGAA | >hg38_21_+_8433172_8446622_RNA45SN1_45S_50ntFlanks | 12992 | 13013 | 22 | TCCCTCGCTGCGATCTATTGAA | 9065.3 | -12.9 | 1.5 | -8.8 | 1.4E-18 | 5.0E-18 | BT20/MB468 | NUC |
| >hg38_21_+_8433172_8446622_RNA45SN1_45S_50ntFlanks@10866.10884.19\|CTAAGGGCTGGGTCGGTCG | >hg38_21_+_8433172_8446622_RNA45SN1_45S_50ntFlanks | 10866 | 10884 | 19 | CTAAGGGCTGGGTCGGTCG | 926.2 | -12.8 | 1.4 | -8.9 | 5.0E-19 | 5.0E-18 | MB231/MB468 | NUC |

| | | | | | | | | | | | | | |
|---|---|---|---|---|---|---|---|---|---|---|---|---|---|
| >hg38_21_+_8433172_8446622_RNA45SN1_45S_50ntFlanks@11845.11872.28\|GGAACGGGCTTGGCGGAATCAGCGGGGA | >hg38_21_+_8433172_8446622_RNA45SN1_45S_50ntFlanks | 11845 | 11872 | 28 | GGAACGGGCTTGGCGGAATCAGCGGGGA | 1143.9 | -10.2 | 1.1 | -8.9 | 6.2E-19 | 5.0E-18 | MB231/MB468 | MITO |
| >hg38_21_+_8433172_8446622_RNA45SN1_45S_50ntFlanks@5484.5513.30\|GAGCGCTGAGAAGACGGTCGAACTTGACTA | >hg38_21_+_8433172_8446622_RNA45SN1_45S_50ntFlanks | 5484 | 5513 | 30 | GAGCGCTGAGAAGACGGTCGAACTTGACTA | 464.5 | -12.7 | 1.4 | -8.9 | 3.8E-19 | 5.0E-18 | MB231/MB468 | MP |
| >MI0000466\|hsa-mir-9-1&WithFlank&1\|-\|156420335\|156420435@22.42.21(+1A);MI0000468\|hsa-mir-9-3&WithFlank&15\|+\|89368011\|89368112@22.42.21(+1A);MI0000467\|hsa-mir-9-2&WithFlank&5\|-\|88666847\|88666945@22.42.21(+1A)&[MIMAT0000441&hsa-miR-9-5p&offsets\|0\|-2(+1A);m-104&1\|-\|156420392\|156420414&offsets\|0\|-2(+1A)];[MIMAT0000441_1&hsa-miR-9-5p&offsets\|0\|-2(+1A);m-105&15\|+\|89368032\|89368054&offsets\|0\|-2(+1A)];[MIMAT0000441_2&hsa-miR-9-5p&offsets\|0\|-2(+1A);m-104&5\|-\|88666902\|88666924&offsets\|0\|-2(+1A)]\|TCTTTGGTTATCTAGCTGTATA | >MI0000466\|hsa-mir-9-1&WithFlank&1\|-\|156420335\|156420435 | 22 | 42 | 21(+1A) | TCTTTGGTTATCTAGCTGTATA | 272.2 | 12.3 | 1.4 | 8.9 | 7.9E-19 | 5.0E-18 | BT20/MB468 | CYTO |
| >hg38_21_+_8433172_8446622_RNA45SN1_45S_50ntFlanks@10830.10875.46\|CAAGCCGGATCCGTAACTTCGGGATAAGGATTGGCTCTAAGGGCTG | >hg38_21_+_8433172_8446622_RNA45SN1_45S_50ntFlanks | 10830 | 10875 | 46 | CAAGCCGGATCCGTAACTTCGGGATAAGGATTGGCTCTAAGGGCTG | 4171.9 | -16.5 | 1.9 | -8.8 | 1.4E-18 | 5.1E-18 | BT20/MB468 | NUC |
| >trna111_HisGTG_1_-_147774845_147774916@-1T.33.34,trna118_HisGTG_1_-_145396881_145396952@-1T.33.34,trna16_HisGTG_1_+_146544773_146544844@-1T.33.34,trna1_HisGTG_15_+_45493349_45493420@-1T.33.34,trna21_HisGTG_1_+_147753471_147753542@-1T.33.34,trna33_HisGTG_6_+_27125906_27125977@-1T.33.34,trna7_HisGTG_9_-_14433938_14434009@-1T.33.34,trna8_HisGTG_15_-_45492611_45492682@-1T.33.34,trna9_HisGTG_15_-_45490804_45490875@-1T.33.34\|TGCCGTGATCGTATAGTGGTTAGTACTCTGCGTT | >trna111_HisGTG_1_-_147774845_147774916 | -1T | 33 | 34 | TGCCGTGATCGTATAGTGGTTAGTACTCTGCGTT | 6175.7 | 8.8 | 1.0 | 8.9 | 3.6E-19 | 5.1E-18 | MB231/MB468 | CYTO |
| >MI0000746\|hsa-mir-99b&WithFlank&19\|+\|51692606\|51692687@13.32.20(+2U)&[MIMAT0000689&hsa-miR-99b-5p&offsets\|0\|-2(+2U);m-10&19\|+\|51692618\|51692639&offsets\|0\|-2(+2U)]\|CACCCGTAGAACCGACCTTGTT | >MI0000746\|hsa-mir-99b&WithFlank&19\|+\|51692606\|51692687 | 13 | 32 | 20(+2U) | CACCCGTAGAACCGACCTTGTT | 209.0 | 10.5 | 1.2 | 8.8 | 1.4E-18 | 5.1E-18 | BT20/MB468 | NUC |
| >hg38_21_+_8433172_8446622_RNA45SN1_45S_50ntFlanks@9156.9187.32\|CCAGTGCGCCCCGGGCGGGTCGCGCCGTCGGG | >hg38_21_+_8433172_8446622_RNA45SN1_45S_50ntFlanks | 9156 | 9187 | 32 | CCAGTGCGCCCCGGGCGGGTCGCGCCGTCGGG | 495.3 | -12.7 | 1.4 | -8.9 | 6.4E-19 | 5.1E-18 | MB231/MB468 | MITO |
| >hg38_21_+_8433172_8446622_RNA45SN1_45S_50ntFlanks@7974.8013.40\|ACGCGACCTCAGATCAGACGTGGCGACCCGCTGAATTTAA | >hg38_21_+_8433172_8446622_RNA45SN1_45S_50ntFlanks | 7974 | 8013 | 40 | ACGCGACCTCAGATCAGACGTGGCGACCCGCTGAATTTAA | 2902.5 | 5.0 | 0.6 | 8.9 | 5.8E-19 | 5.2E-18 | BT20/MB468 | MITO |
| >hg38_21_+_8433172_8446622_RNA45SN1_45S_50ntFlanks@9567.9595.29\|ACGTGCAAATCGGTCGTCCGACCTGGGTA | >hg38_21_+_8433172_8446622_RNA45SN1_45S_50ntFlanks | 9567 | 9595 | 29 | ACGTGCAAATCGGTCGTCCGACCTGGGTA | 3807.8 | -6.0 | 0.7 | -8.9 | 3.6E-19 | 5.2E-18 | MB231/MB468 | CYTO |
| >MI0000074\|hsa-mir-19b-1&WithFlank&13\|+\|91351186\|91351284@60.81.22;MI0000075\|hsa-mir-19b-2&WithFlank&X\|-\|134169665\|134169772@68.89.22&[MIMAT0000074&hsa-miR-19b-3p&offsets\|0\|-1;m-122&13\|+\|91351245\|91351267&offsets\|0\|-1];[MIMAT0000074_1&hsa-miR-19b-3p&offsets\|0\|-1;m-121&X\|-\|134169683\|134169705&offsets\|0\|-1]\|TGTGCAAATCCATGCAAAACTG | >MI0000074\|hsa-mir-19b-1&WithFlank&13\|+\|91351186\|91351284 | 60 | 81 | 22 | TGTGCAAATCCATGCAAAACTG | 88.4 | 3.0 | 0.3 | 8.8 | 1.5E-18 | 5.2E-18 | BT20/MB468 | NUC |
| >hg38_21_+_8433172_8446622_RNA45SN1_45S_50ntFlanks@9545.9571.27\|GTGGAGGTCCGTAGCGGTCCTGACGTG | >hg38_21_+_8433172_8446622_RNA45SN1_45S_50ntFlanks | 9545 | 9571 | 27 | GTGGAGGTCCGTAGCGGTCCTGACGTG | 15056.5 | -6.1 | 0.7 | -8.9 | 5.3E-19 | 5.2E-18 | MB231/MB468 | NUC |
| >hg38_21_+_8433172_8446622_RNA45SN1_45S_50ntFlanks@12456.12480.25\|CCAAGCGTTGGATTGTTCACCCACT | >hg38_21_+_8433172_8446622_RNA45SN1_45S_50ntFlanks | 12456 | 12480 | 25 | CCAAGCGTTGGATTGTTCACCCACT | 1688.5 | -6.5 | 0.7 | -8.9 | 6.6E-19 | 5.3E-18 | MB231/MB468 | MITO |
| >trna10_AspGTC_12_-_125424193_125424264@36.74.39,trna12_AspGTC_12_-_125411891_125411962@36.74.39,trna144_AspGTC_6_-_27551236_27551307@36.74.39,trna38_AspGTC_17_-_8125556_8125627@36.74.39,trna45_AspGTC_6_+_27447453_27447524@36.74.39,trna48_AspGTC_6_+_27471523_27471594@36.74.39,trna4_AspGTC_12_+_96429799_96429870@36.74.39,trna69_AspGTC_1_-_161440205_161440276@36.74.39,trna72_AspGTC_1_-_161432824_161432895@36.74.39,trna75_AspGTC_1_-_161425414_161425485@36.74.39,trna78_AspGTC_1_-_161418033_161418104@36.74.39,trna81_AspGTC_1_-_161410615_161410686@36.74.39\|CACGCGGGAGACCGGGGTTCGATTCCCCGACGGGGAGCC | >trna10_AspGTC_12_-_125424193_125424264 | 36 | 74 | 39 | CACGCGGGAGACCGGGGTTCGATTCCCCGACGGGGAGCC | 17466.4 | 11.7 | 1.3 | 8.9 | 3.7E-19 | 5.3E-18 | MB231/MB468 | CYTO |
| >hg38_21_+_8433172_8446622_RNA45SN1_45S_50ntFlanks@11577.11605.29\|TTAGAACTGGTGCGGACCAGGGGAATCCG | >hg38_21_+_8433172_8446622_RNA45SN1_45S_50ntFlanks | 11577 | 11605 | 29 | TTAGAACTGGTGCGGACCAGGGGAATCCG | 806.9 | -6.7 | 0.8 | -8.9 | 3.7E-19 | 5.3E-18 | MB231/MB468 | CYTO |
| >trna116_GluCTC_1_-_145399233_145399304@32.73.42,trna71_GluCTC_1_-_161439189_161439260@32.73.42,trna74_GluCTC_1_-_161431809_161431880@32.73.42,trna77_GluCTC_1_-_161424398_161424469@32.73.42,trna77_GluCTC_6_+_28949976_28950047@32.73.42,trna80_GluCTC_1_-_161417018_161417089@32.73.42,trna87_GluCTC_6_-_126101393_126101464@32.73.42\|CTCTCACCGCCGCGGCCCGGGTTCGATTCCCGGTCAGGGAAC | >trna116_GluCTC_1_-_145399233_145399304 | 32 | 73 | 42 | CTCTCACCGCCGCGGCCCGGGTTCGATTCCCGGTCAGGGAAC | 811.1 | -12.6 | 1.4 | -8.9 | 5.4E-19 | 5.3E-18 | MB231/MB468 | NUC |
| >MI0003513\|hsa-mir-455&WithFlank&9\|+\|114209428\|114209535@60.80.21(+1C)&[MIMAT0004784&hsa-miR-455-3p&offsets\|0\|0(+1C);m-135&9\|+\|114209487\|114209508&offsets\|0\|-1(+1C)]\|GCAGTCCATGGGCATATACACC | >MI0003513\|hsa-mir-455&WithFlank&9\|+\|114209428\|114209535 | 60 | 80 | 21(+1C) | GCAGTCCATGGGCATATACACC | 316.9 | 12.3 | 1.4 | 8.9 | 3.8E-19 | 5.3E-18 | MB231/MB468 | CYTO |
| >MI0000750\|hsa-mir-26a-2&WithFlank&12\|-\|57824603\|57824698@20.41.22(+1C);MI0000083\|hsa-mir-26a-1&WithFlank&3\|+\|37969398\|37969486@16.37.22(+1C)&[MIMAT0000082&hsa-miR-26a-5p&offsets\|0\|0(+1C);m-33&12\|-\|57824658\|57824679&offsets\|0\|0(+1C)];[MIMAT0000082_1&hsa-miR-26a-5p&offsets\|0\|0(+1C);m-34&3\|+\|37969413\|37969434&offsets\|0\|0(+1C)]\|TTCAAGTAATCCAGGATAGGCTC | >MI0000750\|hsa-mir-26a-2&WithFlank&12\|-\|57824603\|57824698 | 20 | 41 | 22(+1C) | TTCAAGTAATCCAGGATAGGCTC | 780.2 | 7.2 | 0.8 | 8.9 | 8.3E-19 | 5.3E-18 | BT20/MB468 | CYTO |
| >hg38_21_+_8433172_8446622_RNA45SN1_45S_50ntFlanks@12994.13039.46\|CCTCGCTGCGATCTATTGAAAGTCAGCCCTCGACACAAGGGTTTGT | >hg38_21_+_8433172_8446622_RNA45SN1_45S_50ntFlanks | 12994 | 13039 | 46 | CCTCGCTGCGATCTATTGAAAGTCAGCCCTCGACACAAGGGTTTGT | 41118.4 | 7.4 | 0.8 | 8.9 | 3.8E-19 | 5.3E-18 | MB231/MB468 | CYTO |
| >hg38_21_+_8433172_8446622_RNA45SN1_45S_50ntFlanks@10351.10374.24\|CGAGAACTTTGAAGGCCGAAGTGG | >hg38_21_+_8433172_8446622_RNA45SN1_45S_50ntFlanks | 10351 | 10374 | 24 | CGAGAACTTTGAAGGCCGAAGTGG | 2364.7 | -5.3 | 0.6 | -8.9 | 3.8E-19 | 5.3E-18 | MB231/MB468 | CYTO |
| >hg38_21_+_8433172_8446622_RNA45SN1_45S_50ntFlanks@10866.10890.25\|CTAAGGGCTGGGTCGGTCGGGCTGG | >hg38_21_+_8433172_8446622_RNA45SN1_45S_50ntFlanks | 10866 | 10890 | 25 | CTAAGGGCTGGGTCGGTCGGGCTGG | 872.9 | -7.0 | 0.8 | -8.9 | 3.8E-19 | 5.3E-18 | MB231/MB468 | CYTO |
| >hg38_21_+_8433172_8446622_RNA45SN1_45S_50ntFlanks@11318.11336.19\|CCCCACGTCTCGTCGCGCG | >hg38_21_+_8433172_8446622_RNA45SN1_45S_50ntFlanks | 11318 | 11336 | 19 | CCCCACGTCTCGTCGCGCG | 1611.0 | -14.4 | 1.6 | -8.9 | 6.7E-19 | 5.4E-18 | MB231/MB468 | MITO |
| >MI0000542\|hsa-mir-320a&WithFlank&8\|-\|22244960\|22245043@48.70.23&[MIMAT0000510&hsa-miR-320a-3p&offsets\|0\|+1;m-60&8\|-\|22244975\|22244996&offsets\|0\|+1]\|AAAAGCTGGGTTGAGAGGGCGAA | >MI0000542\|hsa-mir-320a&WithFlank&8\|-\|22244960\|22245043 | 48 | 70 | 23 | AAAAGCTGGGTTGAGAGGGCGAA | 2020.6 | 3.0 | 0.3 | 8.8 | 1.5E-18 | 5.4E-18 | BT20/MB468 | NUC |
| >hg38_21_+_8433172_8446622_RNA45SN1_45S_50ntFlanks@4729.4767.39\|TCAAGAACGAAAGTCGGAGGTTCGAAGACGATCAGATAC | >hg38_21_+_8433172_8446622_RNA45SN1_45S_50ntFlanks | 4729 | 4767 | 39 | TCAAGAACGAAAGTCGGAGGTTCGAAGACGATCAGATAC | 615.4 | 10.9 | 1.2 | 8.9 | 3.9E-19 | 5.4E-18 | MB231/MB468 | CYTO |
| >hg38_21_+_8433172_8446622_RNA45SN1_45S_50ntFlanks@6731.6771.41\|CACATTGATCATCGACACTTCGAACGCACTTGCGGCCCCGG | >hg38_21_+_8433172_8446622_RNA45SN1_45S_50ntFlanks | 6731 | 6771 | 41 | CACATTGATCATCGACACTTCGAACGCACTTGCGGCCCCGG | 1873.1 | -5.1 | 0.6 | -8.9 | 6.1E-19 | 5.4E-18 | BT20/MB468 | MITO |
| >hg38_21_+_8433172_8446622_RNA45SN1_45S_50ntFlanks@8178.8203.26\|TCTGATCGAGGCCCAGCCCGTGGACG | >hg38_21_+_8433172_8446622_RNA45SN1_45S_50ntFlanks | 8178 | 8203 | 26 | TCTGATCGAGGCCCAGCCCGTGGACG | 816.5 | -5.2 | 0.6 | -8.8 | 1.5E-18 | 5.4E-18 | BT20/MB468 | NUC |
| >hg38_21_+_8433172_8446622_RNA45SN1_45S_50ntFlanks@9203.9224.22\|CTCGGGGCCACGCGCGCGTCCC | >hg38_21_+_8433172_8446622_RNA45SN1_45S_50ntFlanks | 9203 | 9224 | 22 | CTCGGGGCCACGCGCGCGTCCC | 603.3 | -13.1 | 1.5 | -8.9 | 4.1E-19 | 5.4E-18 | MB231/MB468 | MP |
| >hg38_21_+_8433172_8446622_RNA45SN1_45S_50ntFlanks@10808.10829.22\|CAATGTAGGTAAGGGAAGTCGG | >hg38_21_+_8433172_8446622_RNA45SN1_45S_50ntFlanks | 10808 | 10829 | 22 | CAATGTAGGTAAGGGAAGTCGG | 472.2 | -8.3 | 0.9 | -8.9 | 3.9E-19 | 5.5E-18 | MB231/MB468 | CYTO |
| >hg38_21_+_8433172_8446622_RNA45SN1_45S_50ntFlanks@10768.10797.30\|CAGCAGGTCTCCAAGGTGAACAGCCTCTGG | >hg38_21_+_8433172_8446622_RNA45SN1_45S_50ntFlanks | 10768 | 10797 | 30 | CAGCAGGTCTCCAAGGTGAACAGCCTCTGG | 201.1 | -5.8 | 0.7 | -8.9 | 3.9E-19 | 5.5E-18 | MB231/MB468 | CYTO |
| >hg38_21_+_8433172_8446622_RNA45SN1_45S_50ntFlanks@12456.12477.22\|CCAAGCGTTGGATTGTTCACCC | >hg38_21_+_8433172_8446622_RNA45SN1_45S_50ntFlanks | 12456 | 12477 | 22 | CCAAGCGTTGGATTGTTCACCC | 2675.0 | -4.9 | 0.5 | -8.9 | 6.2E-19 | 5.5E-18 | BT20/MB468 | MITO |
| >MI0000266\|hsa-mir-10a&WithFlank&17\|-\|48579832\|48579953@29.49.21(+1A)&[MIMAT0000253&hsa-miR-10a-5p&offsets\|+1\|-1(+1A);m-18&17\|-\|48579905\|48579926&offsets\|+1\|0(+1A)]\|ACCCTGTAGATCCGAATTTGTA | >MI0000266\|hsa-mir-10a&WithFlank&17\|-\|48579832\|48579953 | 29 | 49 | 21(+1A) | ACCCTGTAGATCCGAATTTGTA | 442.9 | 12.8 | 1.4 | 8.9 | 3.9E-19 | 5.5E-18 | MB231/MB468 | CYTO |
| >hg38_MT_+_1621_3279_RNR2_16S_50ntFlanks@51.87.37\|GCTAAACCTAGCCCCAAACCCACTCCACCTTACTACC | >hg38_MT_+_1621_3279_RNR2_16S_50ntFlanks | 51 | 87 | 37 | GCTAAACCTAGCCCCAAACCCACTCCACCTTACTACC | 456.7 | 12.8 | 1.4 | 8.9 | 3.9E-19 | 5.5E-18 | MB231/MB468 | CYTO |
| >MI0003591\|hsa-mir-584&WithFlank&5\|-\|149062307\|149062415@22.42.21(+1U)&[MIMAT0003249&hsa-miR-584-5p&offsets\|0\|-1(+1U);m-167&5\|-\|149062374\|149062394&offsets\|0\|0(+1U)]\|TTATGGTTTGCCTGGGACTGAT | >MI0003591\|hsa-mir-584&WithFlank&5\|-\|149062307\|149062415 | 22 | 42 | 21(+1U) | TTATGGTTTGCCTGGGACTGAT | 311.4 | 6.8 | 0.8 | 8.9 | 5.6E-19 | 5.5E-18 | MB231/MB468 | NUC |
| >hg38_21_+_8433172_8446622_RNA45SN1_45S_50ntFlanks@7986.8018.33\|ATCAGACGTGGCGACCCGCTGAATTTAAGCATA | >hg38_21_+_8433172_8446622_RNA45SN1_45S_50ntFlanks | 7986 | 8018 | 33 | ATCAGACGTGGCGACCCGCTGAATTTAAGCATA | 342.4 | -12.8 | 1.5 | -8.8 | 1.5E-18 | 5.5E-18 | BT20/MB468 | NUC |
| >hg38_21_+_8433172_8446622_RNA45SN1_45S_50ntFlanks@3705.3741.37\|TACCTGGTTGATCCTGCCAGTAGCATATGCTTGTCTC | >hg38_21_+_8433172_8446622_RNA45SN1_45S_50ntFlanks | 3705 | 3741 | 37 | TACCTGGTTGATCCTGCCAGTAGCATATGCTTGTCTC | 9230.7 | 6.6 | 0.8 | 8.9 | 8.7E-19 | 5.5E-18 | BT20/MB468 | CYTO |
| >MI0000266\|hsa-mir-10a&WithFlank&17\|-\|48579832\|48579953@29.49.21&[MIMAT0000253&hsa-miR-10a-5p&offsets\|+1\|-1;m-18&17\|-\|48579905\|48579926&offsets\|+1\|0]\|ACCCTGTAGATCCGAATTTGT | >MI0000266\|hsa-mir-10a&WithFlank&17\|-\|48579832\|48579953 | 29 | 49 | 21 | ACCCTGTAGATCCGAATTTGT | 963.4 | 13.1 | 1.5 | 8.9 | 6.9E-19 | 5.5E-18 | MB231/MB468 | MITO |
| >hg38_1_-_228634819_228635039_RNA5S12_5S_50ntFlanks@145.170.26\|CTGGGAATACCGGGTGCTGTAGGCTT | >hg38_1_-_228634819_228635039_RNA5S12_5S_50ntFlanks | 145 | 170 | 26 | CTGGGAATACCGGGTGCTGTAGGCTT | 4177.6 | 7.1 | 0.8 | 8.9 | 4.0E-19 | 5.6E-18 | MB231/MB468 | CYTO |

<table>
<tr><td>>MI0000289|hsa-mir-181a-1&WithFlank&1|-|198859038|198859159@30.53.24;MI0000269|hsa-mir-181a-2&WithFlank&9|+|124692436|124692557@45.68.24&[MIMAT0000256&hsa-miR-181a-5p&offsets|0|+1;m-32&1|-|198859108|198859130&offsets|0|+1];[MIMAT0000256_1&hsa-miR-181a-5p&offsets|0|+1;m-32&9|+|124692480|124692502&offsets|0|+1]|AACATTCAACGCTGTCGGTGAGTT</td><td>>MI0000289|hsa-mir-181a-1&WithFlank&1|-|198859038|198859159</td><td>30</td><td>53</td><td>24</td><td>AACATTCAACGCTGTCGGTGAGTT</td><td>1056.7</td><td>7.8</td><td>0.9</td><td>8.9</td><td>5.7E-19</td><td>5.6E-18</td><td>MB231/MB468</td><td>NUC</td></tr>
<tr><td>>MI0002468|hsa-mir-484&WithFlank&16|+|15643288|15643378@14.36.23&[MIMAT0002174&hsa-miR-484&offsets|0|+1;m-127&16|+|15643301|15643321&offsets|0|+2]|TCAGGCTCAGTCCCCTCCCGATA</td><td>>MI0002468|hsa-mir-484&WithFlank&16|+|15643288|15643378</td><td>14</td><td>36</td><td>23</td><td>TCAGGCTCAGTCCCCTCCCGATA</td><td>347.1</td><td>7.2</td><td>0.8</td><td>8.9</td><td>8.7E-19</td><td>5.6E-18</td><td>BT20/MB468</td><td>CYTO</td></tr>
<tr><td>>hg38_21_+_8433172_8446622_RNA45SN1_45S_50ntFlanks@10782.10801.20|GGTGAACAGCCTCTGGCATG</td><td>>hg38_21_+_8433172_8446622_RNA45SN1_45S_50ntFlanks</td><td>10782</td><td>10801</td><td>20</td><td>GGTGAACAGCCTCTGGCATG</td><td>597.3</td><td>-13.1</td><td>1.5</td><td>-8.9</td><td>4.2E-19</td><td>5.6E-18</td><td>MB231/MB468</td><td>MP</td></tr>
<tr><td>>hg38_21_+_8433172_8446622_RNA45SN1_45S_50ntFlanks@6645.6674.30|CTCGTACGACTCTTAGCGGTGGATCACTCG</td><td>>hg38_21_+_8433172_8446622_RNA45SN1_45S_50ntFlanks</td><td>6645</td><td>6674</td><td>30</td><td>CTCGTACGACTCTTAGCGGTGGATCACTCG</td><td>5361.0</td><td>4.7</td><td>0.5</td><td>8.9</td><td>6.3E-19</td><td>5.6E-18</td><td>BT20/MB468</td><td>MITO</td></tr>
<tr><td>>trnaMT_PheGAA_MT_+_577_647@35.55.21|GAAAATGTTTAGACGGGCTCA</td><td>>trnaMT_PheGAA_MT_+_577_647</td><td>35</td><td>55</td><td>21</td><td>GAAAATGTTTAGACGGGCTCA</td><td>365.6</td><td>12.8</td><td>1.4</td><td>8.8</td><td>8.8E-19</td><td>5.6E-18</td><td>BT20/MB468</td><td>CYTO</td></tr>
<tr><td>>MI0000440|hsa-mir-27b&WithFlank&9|+|95085439|95085547@67.86.20&[MIMAT0000419&hsa-miR-27b-3p&offsets|0|-1;m-39&9|+|95085505|95085525&offsets|0|-1]|TTCACAGTGGCTAAGTTCTG</td><td>>MI0000440|hsa-mir-27b&WithFlank&9|+|95085439|95085547</td><td>67</td><td>86</td><td>20</td><td>TTCACAGTGGCTAAGTTCTG</td><td>1933.3</td><td>3.5</td><td>0.4</td><td>8.8</td><td>1.6E-18</td><td>5.6E-18</td><td>BT20/MB468</td><td>NUC</td></tr>
<tr><td>>trnaMT_ValTAC_MT_+_1602_1670@46.72.27|TTCAACTTAACTTGACCGCTCTGACCA</td><td>>trnaMT_ValTAC_MT_+_1602_1670</td><td>46</td><td>72</td><td>27</td><td>TTCAACTTAACTTGACCGCTCTGACCA</td><td>405.5</td><td>12.9</td><td>1.5</td><td>8.8</td><td>8.8E-19</td><td>5.6E-18</td><td>BT20/MB468</td><td>CYTO</td></tr>
<tr><td>>hg38_21_+_8433172_8446622_RNA45SN1_45S_50ntFlanks@12996.13016.21|TCGCTGCGATCTATTGAAAGT</td><td>>hg38_21_+_8433172_8446622_RNA45SN1_45S_50ntFlanks</td><td>12996</td><td>13016</td><td>21</td><td>TCGCTGCGATCTATTGAAAGT</td><td>546.6</td><td>5.4</td><td>0.6</td><td>8.9</td><td>4.1E-19</td><td>5.6E-18</td><td>MB231/MB468</td><td>CYTO</td></tr>
<tr><td>>hg38_21_+_8433172_8446622_RNA45SN1_45S_50ntFlanks@11660.11684.25|GTGATTTCTGCCCAGTGCTCTGAAT</td><td>>hg38_21_+_8433172_8446622_RNA45SN1_45S_50ntFlanks</td><td>11660</td><td>11684</td><td>25</td><td>GTGATTTCTGCCCAGTGCTCTGAAT</td><td>312.2</td><td>24.9</td><td>2.7</td><td>9.4</td><td>6.9E-21</td><td>5.6E-18</td><td>BT20/MB231</td><td>MP</td></tr>
<tr><td>>hg38_21_+_8433172_8446622_RNA45SN1_45S_50ntFlanks@10689.10708.20|GCGTCCGGTGAGCTCTCGCT</td><td>>hg38_21_+_8433172_8446622_RNA45SN1_45S_50ntFlanks</td><td>10689</td><td>10708</td><td>20</td><td>GCGTCCGGTGAGCTCTCGCT</td><td>1225.9</td><td>-6.3</td><td>0.7</td><td>-8.9</td><td>7.1E-19</td><td>5.7E-18</td><td>MB231/MB468</td><td>MITO</td></tr>
<tr><td>>hg38_21_+_8433172_8446622_RNA45SN1_45S_50ntFlanks@6666.6689.24|GATCACTCGGCTCGTGCGTCGATG</td><td>>hg38_21_+_8433172_8446622_RNA45SN1_45S_50ntFlanks</td><td>6666</td><td>6689</td><td>24</td><td>GATCACTCGGCTCGTGCGTCGATG</td><td>546.3</td><td>-12.9</td><td>1.5</td><td>-8.9</td><td>7.1E-19</td><td>5.7E-18</td><td>MB231/MB468</td><td>MITO</td></tr>
<tr><td>>hg38_21_+_8433172_8446622_RNA45SN1_45S_50ntFlanks@12617.12653.37|TGAGGAGCCAATGGGGCGAAGCTACCATCTGTGGGAT</td><td>>hg38_21_+_8433172_8446622_RNA45SN1_45S_50ntFlanks</td><td>12617</td><td>12653</td><td>37</td><td>TGAGGAGCCAATGGGGCGAAGCTACCATCTGTGGGAT</td><td>640.7</td><td>-13.2</td><td>1.5</td><td>-8.9</td><td>4.3E-19</td><td>5.7E-18</td><td>MB231/MB468</td><td>MP</td></tr>
<tr><td>>hg38_21_+_8433172_8446622_RNA45SN1_45S_50ntFlanks@11233.11269.37|CCCTTCCCGTGGATCGCCCCAGCTGCGGCGGGCGTCG</td><td>>hg38_21_+_8433172_8446622_RNA45SN1_45S_50ntFlanks</td><td>11233</td><td>11269</td><td>37</td><td>CCCTTCCCGTGGATCGCCCCAGCTGCGGCGGGCGTCG</td><td>470.0</td><td>-13.3</td><td>1.5</td><td>-8.8</td><td>1.6E-18</td><td>5.7E-18</td><td>BT20/MB468</td><td>NUC</td></tr>
<tr><td>>hg38_21_+_8433172_8446622_RNA45SN1_45S_50ntFlanks@12405.12444.40|CTTTTTGATCCTTCGATGTCGGCTCTTCCTATCATTGTGA</td><td>>hg38_21_+_8433172_8446622_RNA45SN1_45S_50ntFlanks</td><td>12405</td><td>12444</td><td>40</td><td>CTTTTTGATCCTTCGATGTCGGCTCTTCCTATCATTGTGA</td><td>14568.3</td><td>-11.8</td><td>1.3</td><td>-8.8</td><td>1.6E-18</td><td>5.7E-18</td><td>BT20/MB468</td><td>NUC</td></tr>
<tr><td>>hg38_21_+_8433172_8446622_RNA45SN1_45S_50ntFlanks@11921.11967.47|TGAGAGGTGTAGAATAAGTGGGAGGCCCCCGGCGCCCCCCCGGTGTC</td><td>>hg38_21_+_8433172_8446622_RNA45SN1_45S_50ntFlanks</td><td>11921</td><td>11967</td><td>47</td><td>TGAGAGGTGTAGAATAAGTGGGAGGCCCCCGGCGCCCCCCCGGTGTC</td><td>1406.1</td><td>-14.9</td><td>1.7</td><td>-8.8</td><td>1.6E-18</td><td>5.7E-18</td><td>BT20/MB468</td><td>NUC</td></tr>
<tr><td>>hg38_21_+_8433172_8446622_RNA45SN1_45S_50ntFlanks@10097.10117.21|CGCGCGCGCGCGTGTGGTGTG</td><td>>hg38_21_+_8433172_8446622_RNA45SN1_45S_50ntFlanks</td><td>10097</td><td>10117</td><td>21</td><td>CGCGCGCGCGCGTGTGGTGTG</td><td>424.1</td><td>-12.6</td><td>1.4</td><td>-8.9</td><td>4.3E-19</td><td>5.7E-18</td><td>MB231/MB468</td><td>MP</td></tr>
<tr><td>>hg38_21_+_8433172_8446622_RNA45SN1_45S_50ntFlanks@10107.10125.19|CGTGTGGTGTGCGTCGGAG</td><td>>hg38_21_+_8433172_8446622_RNA45SN1_45S_50ntFlanks</td><td>10107</td><td>10125</td><td>19</td><td>CGTGTGGTGTGCGTCGGAG</td><td>549.0</td><td>-12.9</td><td>1.5</td><td>-8.9</td><td>7.2E-19</td><td>5.7E-18</td><td>MB231/MB468</td><td>MITO</td></tr>
<tr><td>>hg38_21_+_8433172_8446622_RNA45SN1_45S_50ntFlanks@10866.10889.24|CTAAGGGCTGGGTCGGTCGGGCTG</td><td>>hg38_21_+_8433172_8446622_RNA45SN1_45S_50ntFlanks</td><td>10866</td><td>10889</td><td>24</td><td>CTAAGGGCTGGGTCGGTCGGGCTG</td><td>714.9</td><td>-13.3</td><td>1.5</td><td>-8.9</td><td>7.2E-19</td><td>5.8E-18</td><td>MB231/MB468</td><td>MITO</td></tr>
<tr><td>>hg38_21_+_8433172_8446622_RNA45SN1_45S_50ntFlanks@5484.5518.35|GAGCGCTGAGAAGACGGTCGAACTTGACTATCTAG</td><td>>hg38_21_+_8433172_8446622_RNA45SN1_45S_50ntFlanks</td><td>5484</td><td>5518</td><td>35</td><td>GAGCGCTGAGAAGACGGTCGAACTTGACTATCTAG</td><td>536.3</td><td>-12.9</td><td>1.4</td><td>-8.9</td><td>4.4E-19</td><td>5.8E-18</td><td>MB231/MB468</td><td>MP</td></tr>
<tr><td>>hg38_21_+_8433172_8446622_RNA45SN1_45S_50ntFlanks@10843.10868.26|TAACTTCGGGATAAGGATTGGCTCTA</td><td>>hg38_21_+_8433172_8446622_RNA45SN1_45S_50ntFlanks</td><td>10843</td><td>10868</td><td>26</td><td>TAACTTCGGGATAAGGATTGGCTCTA</td><td>1233.7</td><td>-8.5</td><td>1.0</td><td>-8.9</td><td>5.9E-19</td><td>5.8E-18</td><td>MB231/MB468</td><td>NUC</td></tr>
<tr><td>>hg38_21_+_8433172_8446622_RNA45SN1_45S_50ntFlanks@3854.3898.45|ACTGTGGTAATTCTAGAGCTAATACATGCCGACGGGCGCTGACCC</td><td>>hg38_21_+_8433172_8446622_RNA45SN1_45S_50ntFlanks</td><td>3854</td><td>3898</td><td>45</td><td>ACTGTGGTAATTCTAGAGCTAATACATGCCGACGGGCGCTGACCC</td><td>1227.1</td><td>-14.0</td><td>1.6</td><td>-8.9</td><td>7.3E-19</td><td>5.8E-18</td><td>MB231/MB468</td><td>MITO</td></tr>
<tr><td>>hg38_21_+_8433172_8446622_RNA45SN1_45S_50ntFlanks@10806.10826.21|AACAATGTAGGTAAGGGAAGT</td><td>>hg38_21_+_8433172_8446622_RNA45SN1_45S_50ntFlanks</td><td>10806</td><td>10826</td><td>21</td><td>AACAATGTAGGTAAGGGAAGT</td><td>405.2</td><td>-12.5</td><td>1.4</td><td>-8.9</td><td>4.4E-19</td><td>5.8E-18</td><td>MB231/MB468</td><td>MP</td></tr>
<tr><td>>hg38_21_+_8433172_8446622_RNA45SN1_45S_50ntFlanks@11885.11904.20|TTGAGCTTGACTCTAGTCTG</td><td>>hg38_21_+_8433172_8446622_RNA45SN1_45S_50ntFlanks</td><td>11885</td><td>11904</td><td>20</td><td>TTGAGCTTGACTCTAGTCTG</td><td>1829.1</td><td>-10.6</td><td>1.2</td><td>-8.9</td><td>7.4E-19</td><td>5.9E-18</td><td>MB231/MB468</td><td>MITO</td></tr>
<tr><td>>hg38_21_+_8433172_8446622_RNA45SN1_45S_50ntFlanks@3866.3898.33|CTAGAGCTAATACATGCCGACGGGCGCTGACCC</td><td>>hg38_21_+_8433172_8446622_RNA45SN1_45S_50ntFlanks</td><td>3866</td><td>3898</td><td>33</td><td>CTAGAGCTAATACATGCCGACGGGCGCTGACCC</td><td>599.0</td><td>-13.7</td><td>1.6</td><td>-8.8</td><td>1.7E-18</td><td>5.9E-18</td><td>BT20/MB468</td><td>NUC</td></tr>
<tr><td>>hg38_21_+_8433172_8446622_RNA45SN1_45S_50ntFlanks@12624.12654.31|CCAATGGGGCGAAGCTACCATCTGTGGGATT</td><td>>hg38_21_+_8433172_8446622_RNA45SN1_45S_50ntFlanks</td><td>12624</td><td>12654</td><td>31</td><td>CCAATGGGGCGAAGCTACCATCTGTGGGATT</td><td>2139.2</td><td>-7.7</td><td>0.9</td><td>-8.9</td><td>4.3E-19</td><td>5.9E-18</td><td>MB231/MB468</td><td>CYTO</td></tr>
<tr><td>>hg38_21_+_8433172_8446622_RNA45SN1_45S_50ntFlanks@11890.11912.23|CTTGACTCTAGTCTGGCACGGTG</td><td>>hg38_21_+_8433172_8446622_RNA45SN1_45S_50ntFlanks</td><td>11890</td><td>11912</td><td>23</td><td>CTTGACTCTAGTCTGGCACGGTG</td><td>1664.2</td><td>-6.4</td><td>0.7</td><td>-8.9</td><td>4.3E-19</td><td>6.0E-18</td><td>MB231/MB468</td><td>CYTO</td></tr>
<tr><td>>hg38_1_-_228634819_228635039_RNA5S12_5S_50ntFlanks@139.171.33|GACCGCCTGGGAATACCGGGTGCTGTAGGCTTT</td><td>>hg38_1_-_228634819_228635039_RNA5S12_5S_50ntFlanks</td><td>139</td><td>171</td><td>33</td><td>GACCGCCTGGGAATACCGGGTGCTGTAGGCTTT</td><td>2050.1</td><td>7.1</td><td>0.8</td><td>8.8</td><td>9.5E-19</td><td>6.0E-18</td><td>BT20/MB468</td><td>CYTO</td></tr>
<tr><td>>hg38_21_+_8433172_8446622_RNA45SN1_45S_50ntFlanks@12616.12653.38|CTGAGGAGCCAATGGGGCGAAGCTACCATCTGTGGGAT</td><td>>hg38_21_+_8433172_8446622_RNA45SN1_45S_50ntFlanks</td><td>12616</td><td>12653</td><td>38</td><td>CTGAGGAGCCAATGGGGCGAAGCTACCATCTGTGGGAT</td><td>1748.0</td><td>-10.8</td><td>1.2</td><td>-8.9</td><td>7.6E-19</td><td>6.1E-18</td><td>MB231/MB468</td><td>MITO</td></tr>
<tr><td>>hg38_21_+_8433172_8446622_RNA45SN1_45S_50ntFlanks@10414.10431.18|TCGGTCCTGAGAGATGGG</td><td>>hg38_21_+_8433172_8446622_RNA45SN1_45S_50ntFlanks</td><td>10414</td><td>10431</td><td>18</td><td>TCGGTCCTGAGAGATGGG</td><td>983.4</td><td>-8.1</td><td>0.9</td><td>-8.9</td><td>4.4E-19</td><td>6.1E-18</td><td>MB231/MB468</td><td>CYTO</td></tr>
<tr><td>>MI0000266|hsa-mir-10a&WithFlank&17|-|48579832|48579953@28.50.23&[MIMAT0000253&hsa-miR-10a-5p&offsets|0|0;m-18&17|-|48579905|48579926&offsets|0|+1]|TACCCTGTAGATCCGAATTTGTG</td><td>>MI0000266|hsa-mir-10a&WithFlank&17|-|48579832|48579953</td><td>28</td><td>50</td><td>23</td><td>TACCCTGTAGATCCGAATTTGTG</td><td>2031.8</td><td>11.0</td><td>1.3</td><td>8.8</td><td>1.7E-18</td><td>6.1E-18</td><td>BT20/MB468</td><td>NUC</td></tr>
<tr><td>>hg38_21_+_8433172_8446622_RNA45SN1_45S_50ntFlanks@10792.10818.27|CTCTGGCATGTTGGAACAATGTAGGTA</td><td>>hg38_21_+_8433172_8446622_RNA45SN1_45S_50ntFlanks</td><td>10792</td><td>10818</td><td>27</td><td>CTCTGGCATGTTGGAACAATGTAGGTA</td><td>477.3</td><td>-12.8</td><td>1.4</td><td>-8.9</td><td>4.6E-19</td><td>6.1E-18</td><td>MB231/MB468</td><td>MP</td></tr>
<tr><td>>hg38_21_+_8433172_8446622_RNA45SN1_45S_50ntFlanks@9623.9650.28|TAGTAGCTGGTTCCCTCCGAAGTTTCCC</td><td>>hg38_21_+_8433172_8446622_RNA45SN1_45S_50ntFlanks</td><td>9623</td><td>9650</td><td>28</td><td>TAGTAGCTGGTTCCCTCCGAAGTTTCCC</td><td>329.2</td><td>-12.2</td><td>1.4</td><td>-8.9</td><td>4.7E-19</td><td>6.2E-18</td><td>MB231/MB468</td><td>MP</td></tr>
<tr><td>>hg38_21_+_8433172_8446622_RNA45SN1_45S_50ntFlanks@10389.10409.21|TGAACAGCAGTTGAACATGGG</td><td>>hg38_21_+_8433172_8446622_RNA45SN1_45S_50ntFlanks</td><td>10389</td><td>10409</td><td>21</td><td>TGAACAGCAGTTGAACATGGG</td><td>617.8</td><td>-13.1</td><td>1.5</td><td>-8.9</td><td>4.7E-19</td><td>6.2E-18</td><td>MB231/MB468</td><td>MP</td></tr>
<tr><td>>hg38_21_+_8433172_8446622_RNA45SN1_45S_50ntFlanks@9313.9338.26|TCTAACACGTGCGCGAGTCGGGGGCT</td><td>>hg38_21_+_8433172_8446622_RNA45SN1_45S_50ntFlanks</td><td>9313</td><td>9338</td><td>26</td><td>TCTAACACGTGCGCGAGTCGGGGGCT</td><td>959.3</td><td>-13.8</td><td>1.5</td><td>-8.9</td><td>4.7E-19</td><td>6.2E-18</td><td>MB231/MB468</td><td>MP</td></tr>
<tr><td>>trnaMT_MetCAT_MT_+_4402_4469@34.71.38|ACCCCGAAAATGTTGGTTATACCCTTCCCGTACTACCA</td><td>>trnaMT_MetCAT_MT_+_4402_4469</td><td>34</td><td>71</td><td>38</td><td>ACCCCGAAAATGTTGGTTATACCCTTCCCGTACTACCA</td><td>975.1</td><td>13.9</td><td>1.6</td><td>8.9</td><td>4.5E-19</td><td>6.2E-18</td><td>MB231/MB468</td><td>CYTO</td></tr>
<tr><td>>hg38_21_+_8433172_8446622_RNA45SN1_45S_50ntFlanks@10866.10891.26|CTAAGGGCTGGGTCGGTCGGGCTGGG</td><td>>hg38_21_+_8433172_8446622_RNA45SN1_45S_50ntFlanks</td><td>10866</td><td>10891</td><td>26</td><td>CTAAGGGCTGGGTCGGTCGGGCTGGG</td><td>664.4</td><td>-8.8</td><td>1.0</td><td>-8.9</td><td>4.5E-19</td><td>6.2E-18</td><td>MB231/MB468</td><td>CYTO</td></tr>
<tr><td>>MI0000815|hsa-mir-339&WithFlank&7|-|1022927|1023032@56.76.21&[MIMAT0004702&hsa-miR-339-3p&offsets|0|-2;m-235&7|-|1022957|1022977&offsets|0|0]|TGAGCGCCTCGACGACAGAGC</td><td>>MI0000815|hsa-mir-339&WithFlank&7|-|1022927|1023032</td><td>56</td><td>76</td><td>21</td><td>TGAGCGCCTCGACGACAGAGC</td><td>471.8</td><td>8.2</td><td>0.9</td><td>8.9</td><td>7.1E-19</td><td>6.2E-18</td><td>BT20/MB468</td><td>MITO</td></tr>
<tr><td>>hg38_21_+_8433172_8446622_RNA45SN1_45S_50ntFlanks@4730.4767.38|CAAGAACGAAAGTCGGAGGTTCGAAGACGATCAGATAC</td><td>>hg38_21_+_8433172_8446622_RNA45SN1_45S_50ntFlanks</td><td>4730</td><td>4767</td><td>38</td><td>CAAGAACGAAAGTCGGAGGTTCGAAGACGATCAGATAC</td><td>1693.1</td><td>10.2</td><td>1.1</td><td>8.9</td><td>4.6E-19</td><td>6.2E-18</td><td>MB231/MB468</td><td>CYTO</td></tr>
<tr><td>>hg38_21_+_8433172_8446622_RNA45SN1_45S_50ntFlanks@9574.9595.22|AATCGGTCGTCCGACCTGGGTA</td><td>>hg38_21_+_8433172_8446622_RNA45SN1_45S_50ntFlanks</td><td>9574</td><td>9595</td><td>22</td><td>AATCGGTCGTCCGACCTGGGTA</td><td>3320.6</td><td>-5.0</td><td>0.6</td><td>-8.9</td><td>4.6E-19</td><td>6.2E-18</td><td>MB231/MB468</td><td>CYTO</td></tr>
<tr><td>>MI0000285|hsa-mir-205&WithFlank&1|+|209432127|209432248@40.61.22(+2A)&[MIMAT0000266&hsa-miR-205-5p&offsets|0|0(+2A);m-351&1|+|209432166|209432187&offsets|0|0(+2A)]|TCCTTCATTCCACCGGAGTCTGAA</td><td>>MI0000285|hsa-mir-205&WithFlank&1|+|209432127|209432248</td><td>40</td><td>61</td><td>22(+2A)</td><td>TCCTTCATTCCACCGGAGTCTGAA</td><td>1023.7</td><td>13.8</td><td>1.5</td><td>9.1</td><td>7.7E-20</td><td>6.3E-18</td><td>BT20/MB231</td><td>NUC</td></tr>
<tr><td>>hg38_21_+_8433172_8446622_RNA45SN1_45S_50ntFlanks@12941.12972.32|ATTCGTAGACGACCTGCTTCTGGGTCGGGGTT</td><td>>hg38_21_+_8433172_8446622_RNA45SN1_45S_50ntFlanks</td><td>12941</td><td>12972</td><td>32</td><td>ATTCGTAGACGACCTGCTTCTGGGTCGGGGTT</td><td>561.0</td><td>-13.0</td><td>1.5</td><td>-8.9</td><td>4.8E-19</td><td>6.3E-18</td><td>MB231/MB468</td><td>MP</td></tr>
<tr><td>>hg38_21_+_8433172_8446622_RNA45SN1_45S_50ntFlanks@7988.8007.20|CAGACGTGGCGACCCGCTGA</td><td>>hg38_21_+_8433172_8446622_RNA45SN1_45S_50ntFlanks</td><td>7988</td><td>8007</td><td>20</td><td>CAGACGTGGCGACCCGCTGA</td><td>397.1</td><td>-5.5</td><td>0.6</td><td>-8.8</td><td>1.8E-18</td><td>6.3E-18</td><td>BT20/MB468</td><td>NUC</td></tr>
<tr><td>>am_MI0005763|hsa-mir-941-1&WithFlank&20|+|63919443|63919526@53.74.22(+1U);MI0005764|hsa-mir-941-2&WithFlank&20|+|63919499|63919582@53.74.22(+1U);MI0005765|hsa-mir-941-3&WithFlank&20|+|63919555|63919638@53.74.22(+1U);MI0005766|hsa-mir-941-4&WithFlank&20|+|63919750|63919833@53.74.22(+1U);MI0031520|hsa-mir-941-5&WithFlank&20|+|63919862|63919945@53.74.22(+1U)&[MIMAT0004984&hsa-miR-941&offsets|0|-1(+1U);m-263&20|+|63919495|63919517&offsets|0|-1(+1U)];[MIMAT0004984_1&hsa-miR-941&offsets|0|-1(+1U);m-263&20|+|63919551|63919573&offsets|0|-1(+1U)];[MIMAT0004984_2&hsa-miR-941&offsets|0|-1(+1U);m-263&20|+|63919607|63919629&offsets|0|-1(+1U)];[MIMAT0004984_3&hsa-miR-941&offsets|0|-1(+1U);m-263&20|+|63919802|63919824&offsets|0|-1(+1U)];[MIMAT0004984_4&hsa-miR-941&offsets|0|-1(+1U);m-263&20|+|63919914|63919936&offsets|0|-1(+1U)]|CACCCGGCTGTGTGCACATGTGT</td><td>>am_MI0005763|hsa-mir-941-1&WithFlank&20|+|63919443|63919526</td><td>53</td><td>74</td><td>22(+1U)</td><td>CACCCGGCTGTGTGCACATGTGT</td><td>2445.0</td><td>5.4</td><td>0.6</td><td>8.9</td><td>6.4E-19</td><td>6.3E-18</td><td>MB231/MB468</td><td>NUC</td></tr>
<tr><td>>hg38_21_+_8433172_8446622_RNA45SN1_45S_50ntFlanks@4445.4472.28|CTTGCCTCTCGGCGCCCCCTCGATGCTC</td><td>>hg38_21_+_8433172_8446622_RNA45SN1_45S_50ntFlanks</td><td>4445</td><td>4472</td><td>28</td><td>CTTGCCTCTCGGCGCCCCCTCGATGCTC</td><td>644.9</td><td>-13.1</td><td>1.5</td><td>-8.9</td><td>7.9E-19</td><td>6.3E-18</td><td>MB231/MB468</td><td>MITO</td></tr>
<tr><td>>hg38_21_+_8433172_8446622_RNA45SN1_45S_50ntFlanks@10351.10371.21|CGAGAACTTTGAAGGCCGAAG</td><td>>hg38_21_+_8433172_8446622_RNA45SN1_45S_50ntFlanks</td><td>10351</td><td>10371</td><td>21</td><td>CGAGAACTTTGAAGGCCGAAG</td><td>2685.7</td><td>-6.6</td><td>0.7</td><td>-8.9</td><td>4.8E-19</td><td>6.3E-18</td><td>MB231/MB468</td><td>MP</td></tr>
<tr><td>>hg38_21_+_8433172_8446622_RNA45SN1_45S_50ntFlanks@9305.9334.30|CCAAGGAGTCTAACACGTGCGCGAGTCGGG</td><td>>hg38_21_+_8433172_8446622_RNA45SN1_45S_50ntFlanks</td><td>9305</td><td>9334</td><td>30</td><td>CCAAGGAGTCTAACACGTGCGCGAGTCGGG</td><td>573.8</td><td>-13.0</td><td>1.5</td><td>-8.9</td><td>4.9E-19</td><td>6.3E-18</td><td>MB231/MB468</td><td>MP</td></tr>
</table>

<table>
<tr><td>>am_MI0005763|hsa-mir-941-1&WithFlank&20|+|63919443|63919526@53.74.22(+1A);MI0005764|hsa-mir-941-2&WithFlank&20|+|63919499|63919582@53.74.22(+1A);MI0005765|hsa-mir-941-3&WithFlank&20|+|63919555|63919638@53.74.22(+1A);MI0005766|hsa-mir-941-4&WithFlank&20|+|63919750|63919833@53.74.22(+1A);MI0031520|hsa-mir-941-5&WithFlank&20|+|63919862|63919945@53.74.22(+1A)&[MIMAT0004984&hsa-miR-941&offsets|0|-1(+1A);m-263&20|+|63919495|63919517&offsets|0|-1(+1A)];[MIMAT0004984_1&hsa-miR-941&offsets|0|-1(+1A);m-263&20|+|63919551|63919573&offsets|0|-1(+1A)];[MIMAT0004984_2&hsa-miR-941&offsets|0|-1(+1A);m-263&20|+|63919607|63919629&offsets|0|-1(+1A)];[MIMAT0004984_3&hsa-miR-941&offsets|0|-1(+1A);m-263&20|+|63919802|63919824&offsets|0|-1(+1A)];[MIMAT0004984_4&hsa-miR-941&offsets|0|-1(+1A);m-263&20|+|63919914|63919936&offsets|0|-1(+1A)]|CACCCGGCTGTGTGCACATGTGA</td><td>>am_MI0005763|hsa-mir-941-1&WithFlank&20|+|63919443|63919526</td><td>53</td><td>74</td><td>22(+1A)</td><td>CACCCGGCTGTGTGCACATGTGA</td><td>994.9</td><td>8.8</td><td>1.0</td><td>8.9</td><td>7.2E-19</td><td>6.4E-18</td><td>BT20/MB468</td><td>MITO</td></tr>
<tr><td>>hg38_21_+_8433172_8446622_RNA45SN1_45S_50ntFlanks@11233.11269.37|CCCTTCCCGTGGATCGCCCCAGCTGCGGCGGGCGTCG</td><td>>hg38_21_+_8433172_8446622_RNA45SN1_45S_50ntFlanks</td><td>11233</td><td>11269</td><td>37</td><td>CCCTTCCCGTGGATCGCCCCAGCTGCGGCGGGCGTCG</td><td>501.3</td><td>-12.8</td><td>1.4</td><td>-8.9</td><td>8.0E-19</td><td>6.4E-18</td><td>MB231/MB468</td><td>MITO</td></tr>
<tr><td>>hg38_21_+_8433172_8446622_RNA45SN1_45S_50ntFlanks@3920.3959.40|CATTTATCAGATCAAAACCAACCCGGTCAGCCCCTCTCCG</td><td>>hg38_21_+_8433172_8446622_RNA45SN1_45S_50ntFlanks</td><td>3920</td><td>3959</td><td>40</td><td>CATTTATCAGATCAAAACCAACCCGGTCAGCCCCTCTCCG</td><td>880.8</td><td>-13.6</td><td>1.5</td><td>-8.9</td><td>8.0E-19</td><td>6.4E-18</td><td>MB231/MB468</td><td>MITO</td></tr>
<tr><td>>hg38_21_+_8433172_8446622_RNA45SN1_45S_50ntFlanks@7986.8012.27|ATCAGACGTGGCGACCCGCTGAATTTA</td><td>>hg38_21_+_8433172_8446622_RNA45SN1_45S_50ntFlanks</td><td>7986</td><td>8012</td><td>27</td><td>ATCAGACGTGGCGACCCGCTGAATTTA</td><td>1639.1</td><td>-12.8</td><td>1.4</td><td>-8.9</td><td>7.3E-19</td><td>6.4E-18</td><td>BT20/MB468</td><td>MITO</td></tr>
<tr><td>>hg38_21_+_8433172_8446622_RNA45SN1_45S_50ntFlanks@6699.6720.22|GCTAGCTGCGAGAATTAATGTG</td><td>>hg38_21_+_8433172_8446622_RNA45SN1_45S_50ntFlanks</td><td>6699</td><td>6720</td><td>22</td><td>GCTAGCTGCGAGAATTAATGTG</td><td>31993.5</td><td>-5.0</td><td>0.6</td><td>-8.8</td><td>1.0E-18</td><td>6.4E-18</td><td>BT20/MB468</td><td>CYTO</td></tr>
<tr><td>>hg38_21_+_8433172_8446622_RNA45SN1_45S_50ntFlanks@11235.11268.34|CTTCCCGTGGATCGCCCCAGCTGCGGCGGGCGTC</td><td>>hg38_21_+_8433172_8446622_RNA45SN1_45S_50ntFlanks</td><td>11235</td><td>11268</td><td>34</td><td>CTTCCCGTGGATCGCCCCAGCTGCGGCGGGCGTC</td><td>611.7</td><td>-13.0</td><td>1.5</td><td>-8.9</td><td>8.1E-19</td><td>6.4E-18</td><td>MB231/MB468</td><td>MITO</td></tr>
<tr><td>>MI0000446|hsa-mir-125b-1&WithFlank&11|-|122099751|122099850@61.80.20(+1U)&[MIMAT0004592&hsa-miR-125b-1-3p&offsets|0|-2(+1U);m-485&11|-|122099769|122099790&offsets|0|-2(+1U)]|ACGGGTTAGGCTCTTGGGAGT</td><td>>MI0000446|hsa-mir-125b-1&WithFlank&11|-|122099751|122099850</td><td>61</td><td>80</td><td>20(+1U)</td><td>ACGGGTTAGGCTCTTGGGAGT</td><td>676.1</td><td>12.6</td><td>1.4</td><td>8.9</td><td>4.9E-19</td><td>6.4E-18</td><td>MB231/MB468</td><td>MP</td></tr>
<tr><td>>MI0000465|hsa-mir-191&WithFlank&3|-|49020612|49020715@22.44.23(+1C)&[MIMAT0000440&hsa-miR-191-5p&offsets|0|0(+1C);m-48&3|-|49020672|49020694&offsets|0|0(+1C)]|CAACGGAATCCCAAAAGCAGCTGC</td><td>>MI0000465|hsa-mir-191&WithFlank&3|-|49020612|49020715</td><td>22</td><td>44</td><td>23(+1C)</td><td>CAACGGAATCCCAAAAGCAGCTGC</td><td>526.4</td><td>7.1</td><td>0.8</td><td>8.8</td><td>1.0E-18</td><td>6.4E-18</td><td>BT20/MB468</td><td>CYTO</td></tr>
<tr><td>>hg38_21_+_8433172_8446622_RNA45SN1_45S_50ntFlanks@3931.3959.29|TCAAAACCAACCCGGTCAGCCCCTCTCCG</td><td>>hg38_21_+_8433172_8446622_RNA45SN1_45S_50ntFlanks</td><td>3931</td><td>3959</td><td>29</td><td>TCAAAACCAACCCGGTCAGCCCCTCTCCG</td><td>521.6</td><td>-12.0</td><td>1.3</td><td>-8.9</td><td>6.5E-19</td><td>6.4E-18</td><td>MB231/MB468</td><td>NUC</td></tr>
<tr><td>>hg38_21_+_8433172_8446622_RNA45SN1_45S_50ntFlanks@12405.12432.28|CTTTTTGATCCTTCGATGTCGGCTCTTC</td><td>>hg38_21_+_8433172_8446622_RNA45SN1_45S_50ntFlanks</td><td>12405</td><td>12432</td><td>28</td><td>CTTTTTGATCCTTCGATGTCGGCTCTTC</td><td>25589.2</td><td>-11.2</td><td>1.3</td><td>-8.8</td><td>1.8E-18</td><td>6.4E-18</td><td>BT20/MB468</td><td>NUC</td></tr>
<tr><td>>hg38_21_+_8433172_8446622_RNA45SN1_45S_50ntFlanks@11933.11951.19|AATAAGTGGGAGGCCCCCG</td><td>>hg38_21_+_8433172_8446622_RNA45SN1_45S_50ntFlanks</td><td>11933</td><td>11951</td><td>19</td><td>AATAAGTGGGAGGCCCCCG</td><td>893.7</td><td>-13.7</td><td>1.5</td><td>-8.9</td><td>5.0E-19</td><td>6.4E-18</td><td>MB231/MB468</td><td>MP</td></tr>
<tr><td>>hg38_21_+_8433172_8446622_RNA45SN1_45S_50ntFlanks@5472.5512.41|ACGGCCCTGGCGGAGCGCTGAGAAGACGGTCGAACTTGACT</td><td>>hg38_21_+_8433172_8446622_RNA45SN1_45S_50ntFlanks</td><td>5472</td><td>5512</td><td>41</td><td>ACGGCCCTGGCGGAGCGCTGAGAAGACGGTCGAACTTGACT</td><td>5017.2</td><td>7.1</td><td>0.8</td><td>8.8</td><td>1.0E-18</td><td>6.5E-18</td><td>BT20/MB468</td><td>CYTO</td></tr>
<tr><td>>hg38_21_+_8433172_8446622_RNA45SN1_45S_50ntFlanks@10866.10886.21|CTAAGGGCTGGGTCGGTCGGG</td><td>>hg38_21_+_8433172_8446622_RNA45SN1_45S_50ntFlanks</td><td>10866</td><td>10886</td><td>21</td><td>CTAAGGGCTGGGTCGGTCGGG</td><td>550.3</td><td>-13.0</td><td>1.5</td><td>-8.9</td><td>5.0E-19</td><td>6.5E-18</td><td>MB231/MB468</td><td>MP</td></tr>
<tr><td>>hg38_21_+_8433172_8446622_RNA45SN1_45S_50ntFlanks@12433.12455.23|CTATCATTGTGAAGCAGAATTCA</td><td>>hg38_21_+_8433172_8446622_RNA45SN1_45S_50ntFlanks</td><td>12433</td><td>12455</td><td>23</td><td>CTATCATTGTGAAGCAGAATTCA</td><td>683.9</td><td>-13.3</td><td>1.5</td><td>-8.9</td><td>5.0E-19</td><td>6.5E-18</td><td>MB231/MB468</td><td>MP</td></tr>
<tr><td>>hg38_21_+_8433172_8446622_RNA45SN1_45S_50ntFlanks@12934.12972.39|CTAAACCATTCGTAGACGACCTGCTTCTGGGTCGGGGTT</td><td>>hg38_21_+_8433172_8446622_RNA45SN1_45S_50ntFlanks</td><td>12934</td><td>12972</td><td>39</td><td>CTAAACCATTCGTAGACGACCTGCTTCTGGGTCGGGGTT</td><td>1584.5</td><td>-14.5</td><td>1.6</td><td>-8.9</td><td>5.0E-19</td><td>6.5E-18</td><td>MB231/MB468</td><td>MP</td></tr>
<tr><td>>MI0000081|hsa-mir-24-2&WithFlank&19|-|13836281|13836365@56.77.22(+1U);MI0000080|hsa-mir-24-1&WithFlank&9|+|95086015|95086094@50.71.22(+1U)&[MIMAT0000080&hsa-miR-24-3p&offsets|0|0(+1U);m-36&19|-|13836288|13836310&offsets|0|-1(+1U)];[MIMAT0000080_1&hsa-miR-24-3p&offsets|0|0(+1U);m-35&9|+|95086064|95086086&offsets|0|-1(+1U)]|TGGCTCAGTTCAGCAGGAACAGT</td><td>>MI0000081|hsa-mir-24-2&WithFlank&19|-|13836281|13836365</td><td>56</td><td>77</td><td>22(+1U)</td><td>TGGCTCAGTTCAGCAGGAACAGT</td><td>4562.3</td><td>5.8</td><td>0.7</td><td>8.9</td><td>7.4E-19</td><td>6.5E-18</td><td>BT20/MB468</td><td>MITO</td></tr>
<tr><td>>hg38_21_+_8433172_8446622_RNA45SN1_45S_50ntFlanks@9202.9222.21|TCTCGGGGCCACGCGCGCGTC</td><td>>hg38_21_+_8433172_8446622_RNA45SN1_45S_50ntFlanks</td><td>9202</td><td>9222</td><td>21</td><td>TCTCGGGGCCACGCGCGCGTC</td><td>669.2</td><td>-13.2</td><td>1.5</td><td>-8.9</td><td>8.2E-19</td><td>6.5E-18</td><td>MB231/MB468</td><td>MITO</td></tr>
<tr><td>>hg38_21_+_8433172_8446622_RNA45SN1_45S_50ntFlanks@12456.12489.34|CCAAGCGTTGGATTGTTCACCCACTAATAGGGAA</td><td>>hg38_21_+_8433172_8446622_RNA45SN1_45S_50ntFlanks</td><td>12456</td><td>12489</td><td>34</td><td>CCAAGCGTTGGATTGTTCACCCACTAATAGGGAA</td><td>1708.2</td><td>-14.6</td><td>1.6</td><td>-8.9</td><td>5.0E-19</td><td>6.5E-18</td><td>MB231/MB468</td><td>MP</td></tr>
<tr><td>>hg38_21_+_8433172_8446622_RNA45SN1_45S_50ntFlanks@10234.10260.27|CGGACGCTACGCCGCGACGAGTAGGAG</td><td>>hg38_21_+_8433172_8446622_RNA45SN1_45S_50ntFlanks</td><td>10234</td><td>10260</td><td>27</td><td>CGGACGCTACGCCGCGACGAGTAGGAG</td><td>454.8</td><td>-12.7</td><td>1.4</td><td>-8.9</td><td>5.0E-19</td><td>6.5E-18</td><td>MB231/MB468</td><td>MP</td></tr>
<tr><td>>MI0000284|hsa-mir-204&WithFlank&9|-|70809969|70810090@39.60.22&[MIMAT0000265&hsa-miR-204-5p&offsets|0|0;m-296&9|-|70810031|70810052&offsets|0|0]|TTCCCTTTGTCATCCTATGCCT</td><td>>MI0000284|hsa-mir-204&WithFlank&9|-|70809969|70810090</td><td>39</td><td>60</td><td>22</td><td>TTCCCTTTGTCATCCTATGCCT</td><td>220.8</td><td>11.7</td><td>1.3</td><td>8.9</td><td>6.7E-19</td><td>6.6E-18</td><td>MB231/MB468</td><td>NUC</td></tr>
<tr><td>>hg38_21_+_8433172_8446622_RNA45SN1_45S_50ntFlanks@4002.4025.24|GATAACCTCGGGCCGATCGCACGC</td><td>>hg38_21_+_8433172_8446622_RNA45SN1_45S_50ntFlanks</td><td>4002</td><td>4025</td><td>24</td><td>GATAACCTCGGGCCGATCGCACGC</td><td>751.7</td><td>-13.3</td><td>1.5</td><td>-8.9</td><td>8.3E-19</td><td>6.6E-18</td><td>MB231/MB468</td><td>MITO</td></tr>
<tr><td>>hg38_21_+_8433172_8446622_RNA45SN1_45S_50ntFlanks@11848.11872.25|ACGGGCTTGGCGGAATCAGCGGGGA</td><td>>hg38_21_+_8433172_8446622_RNA45SN1_45S_50ntFlanks</td><td>11848</td><td>11872</td><td>25</td><td>ACGGGCTTGGCGGAATCAGCGGGGA</td><td>624.1</td><td>-13.1</td><td>1.5</td><td>-8.9</td><td>8.4E-19</td><td>6.6E-18</td><td>MB231/MB468</td><td>MITO</td></tr>
<tr><td>>hg38_21_+_8433172_8446622_RNA45SN1_45S_50ntFlanks@12991.13011.21|CTCCCTCGCTGCGATCTATTG</td><td>>hg38_21_+_8433172_8446622_RNA45SN1_45S_50ntFlanks</td><td>12991</td><td>13011</td><td>21</td><td>CTCCCTCGCTGCGATCTATTG</td><td>880.0</td><td>-11.9</td><td>1.3</td><td>-8.9</td><td>7.6E-19</td><td>6.7E-18</td><td>BT20/MB468</td><td>MITO</td></tr>
<tr><td>>MI0000299|hsa-mir-222&WithFlank&X|-|45747009|45747130@75.99.25&[MIMAT0000279&hsa-miR-222-3p&offsets|0|+4;m-140&X|-|45747033|45747056&offsets|0|+1]|AGCTACATCTGGCTACTGGGTCTCT</td><td>>MI0000299|hsa-mir-222&WithFlank&X|-|45747009|45747130</td><td>75</td><td>99</td><td>25</td><td>AGCTACATCTGGCTACTGGGTCTCT</td><td>14629.3</td><td>6.8</td><td>0.8</td><td>8.9</td><td>7.7E-19</td><td>6.7E-18</td><td>BT20/MB468</td><td>MITO</td></tr>
<tr><td>>hg38_21_+_8433172_8446622_RNA45SN1_45S_50ntFlanks@13022.13039.18|CTCGACACAAGGGTTTGT</td><td>>hg38_21_+_8433172_8446622_RNA45SN1_45S_50ntFlanks</td><td>13022</td><td>13039</td><td>18</td><td>CTCGACACAAGGGTTTGT</td><td>655.0</td><td>4.0</td><td>0.4</td><td>8.9</td><td>6.8E-19</td><td>6.7E-18</td><td>MB231/MB468</td><td>NUC</td></tr>
<tr><td>>hg38_21_+_8433172_8446622_RNA45SN1_45S_50ntFlanks@8191.8211.21|CAGCCCGTGGACGGTGTGAGG</td><td>>hg38_21_+_8433172_8446622_RNA45SN1_45S_50ntFlanks</td><td>8191</td><td>8211</td><td>21</td><td>CAGCCCGTGGACGGTGTGAGG</td><td>1272.0</td><td>-14.2</td><td>1.6</td><td>-8.9</td><td>5.2E-19</td><td>6.7E-18</td><td>MB231/MB468</td><td>MP</td></tr>
<tr><td>>hg38_21_+_8433172_8446622_RNA45SN1_45S_50ntFlanks@11729.11749.21|TAACTATGACTCTCTTAAGGT</td><td>>hg38_21_+_8433172_8446622_RNA45SN1_45S_50ntFlanks</td><td>11729</td><td>11749</td><td>21</td><td>TAACTATGACTCTCTTAAGGT</td><td>15043.4</td><td>-10.7</td><td>1.2</td><td>-8.8</td><td>1.9E-18</td><td>6.7E-18</td><td>BT20/MB468</td><td>NUC</td></tr>
<tr><td>>hg38_21_+_8433172_8446622_RNA45SN1_45S_50ntFlanks@12717.12755.39|CCTCGGTTGGCCTCGGATAGCCGGTCCCCCGCCTGTCCC</td><td>>hg38_21_+_8433172_8446622_RNA45SN1_45S_50ntFlanks</td><td>12717</td><td>12755</td><td>39</td><td>CCTCGGTTGGCCTCGGATAGCCGGTCCCCCGCCTGTCCC</td><td>628.2</td><td>-13.1</td><td>1.5</td><td>-8.9</td><td>8.6E-19</td><td>6.8E-18</td><td>MB231/MB468</td><td>MITO</td></tr>
<tr><td>>trna10_AspGTC_12_-_125424193_125424264@36.75.40,trna12_AspGTC_12_-_125411891_125411962@36.75.40,trna144_AspGTC_6_-_27551236_27551307@36.75.40,trna38_AspGTC_17_-_8125556_8125627@36.75.40,trna45_AspGTC_6_+_27447453_27447524@36.75.40,trna48_AspGTC_6_+_27471523_27471594@36.75.40,trna4_AspGTC_12_+_96429799_96429870@36.75.40,trna69_AspGTC_1_-_161440205_161440276@36.75.40,trna72_AspGTC_1_-_161432824_161432895@36.75.40,trna75_AspGTC_1_-_161425414_161425485@36.75.40,trna78_AspGTC_1_-_161418033_161418104@36.75.40,trna81_AspGTC_1_-_161410615_161410686@36.75.40|CACGCGGGAGACCGGGGTTCGATTCCCCGACGGGGAGCCA</td><td>>trna10_AspGTC_12_-_125424193_125424264</td><td>36</td><td>75</td><td>40</td><td>CACGCGGGAGACCGGGGTTCGATTCCCCGACGGGGAGCCA</td><td>35693.6</td><td>10.5</td><td>1.2</td><td>8.9</td><td>5.0E-19</td><td>6.8E-18</td><td>MB231/MB468</td><td>CYTO</td></tr>
<tr><td>>hg38_21_+_8433172_8446622_RNA45SN1_45S_50ntFlanks@11874.11908.35|AGAAGACCCTGTTGAGCTTGACTCTAGTCTGGCAC</td><td>>hg38_21_+_8433172_8446622_RNA45SN1_45S_50ntFlanks</td><td>11874</td><td>11908</td><td>35</td><td>AGAAGACCCTGTTGAGCTTGACTCTAGTCTGGCAC</td><td>446.4</td><td>-6.4</td><td>0.7</td><td>-8.9</td><td>5.0E-19</td><td>6.8E-18</td><td>MB231/MB468</td><td>CYTO</td></tr>
<tr><td>>hg38_21_+_8433172_8446622_RNA45SN1_45S_50ntFlanks@9454.9486.33|CCGCGCCGGGGAGGTGGAGCACGAGCGCACGTG</td><td>>hg38_21_+_8433172_8446622_RNA45SN1_45S_50ntFlanks</td><td>9454</td><td>9486</td><td>33</td><td>CCGCGCCGGGGAGGTGGAGCACGAGCGCACGTG</td><td>396.9</td><td>-12.5</td><td>1.4</td><td>-8.9</td><td>5.2E-19</td><td>6.8E-18</td><td>MB231/MB468</td><td>MP</td></tr>
<tr><td>>hg38_21_+_8433172_8446622_RNA45SN1_45S_50ntFlanks@12162.12179.18|GACTGGGGCGGTACACCT</td><td>>hg38_21_+_8433172_8446622_RNA45SN1_45S_50ntFlanks</td><td>12162</td><td>12179</td><td>18</td><td>GACTGGGGCGGTACACCT</td><td>917.4</td><td>-13.7</td><td>1.5</td><td>-8.9</td><td>5.2E-19</td><td>6.8E-18</td><td>MB231/MB468</td><td>MP</td></tr>
<tr><td>>hg38_21_+_8433172_8446622_RNA45SN1_45S_50ntFlanks@12423.12462.40|TCGGCTCTTCCTATCATTGTGAAGCAGAATTCACCAAGCG</td><td>>hg38_21_+_8433172_8446622_RNA45SN1_45S_50ntFlanks</td><td>12423</td><td>12462</td><td>40</td><td>TCGGCTCTTCCTATCATTGTGAAGCAGAATTCACCAAGCG</td><td>153.5</td><td>11.3</td><td>1.3</td><td>8.9</td><td>5.0E-19</td><td>6.8E-18</td><td>MB231/MB468</td><td>CYTO</td></tr>
<tr><td>>hg38_21_+_8433172_8446622_RNA45SN1_45S_50ntFlanks@10683.10706.24|CCGGCGGCGTCCGGTGAGCTCTCG</td><td>>hg38_21_+_8433172_8446622_RNA45SN1_45S_50ntFlanks</td><td>10683</td><td>10706</td><td>24</td><td>CCGGCGGCGTCCGGTGAGCTCTCG</td><td>7165.4</td><td>-5.6</td><td>0.6</td><td>-8.9</td><td>7.8E-19</td><td>6.8E-18</td><td>BT20/MB468</td><td>MITO</td></tr>
<tr><td>>MI0003599|hsa-mir-589&WithFlank&7|-|5495813|5495923@30.51.22(+1U)&[MIMAT0004799&hsa-miR-589-5p&offsets|0|0(+1U);m-162&7|-|5495874|5495894&offsets|0|+1(+1U)]|TGAGAACCACGTCTGCTCTGAGT</td><td>>MI0003599|hsa-mir-589&WithFlank&7|-|5495813|5495923</td><td>30</td><td>51</td><td>22(+1U)</td><td>TGAGAACCACGTCTGCTCTGAGT</td><td>541.8</td><td>10.7</td><td>1.2</td><td>8.8</td><td>1.1E-18</td><td>6.8E-18</td><td>BT20/MB468</td><td>CYTO</td></tr>
<tr><td>>hg38_21_+_8433172_8446622_RNA45SN1_45S_50ntFlanks@12941.12966.26|ATTCGTAGACGACCTGCTTCTGGGTC</td><td>>hg38_21_+_8433172_8446622_RNA45SN1_45S_50ntFlanks</td><td>12941</td><td>12966</td><td>26</td><td>ATTCGTAGACGACCTGCTTCTGGGTC</td><td>816.8</td><td>-13.5</td><td>1.5</td><td>-8.9</td><td>5.3E-19</td><td>6.8E-18</td><td>MB231/MB468</td><td>MP</td></tr>
<tr><td>>hg38_21_+_8433172_8446622_RNA45SN1_45S_50ntFlanks@12283.12318.36|TACAGACCGTGAAAGCGGGGCCTCACGATCCTTCTG</td><td>>hg38_21_+_8433172_8446622_RNA45SN1_45S_50ntFlanks</td><td>12283</td><td>12318</td><td>36</td><td>TACAGACCGTGAAAGCGGGGCCTCACGATCCTTCTG</td><td>473.1</td><td>-12.7</td><td>1.4</td><td>-8.9</td><td>8.7E-19</td><td>6.8E-18</td><td>MB231/MB468</td><td>MITO</td></tr>
<tr><td>>hg38_21_+_8433172_8446622_RNA45SN1_45S_50ntFlanks@9573.9596.24|AAATCGGTCGTCCGACCTGGGTAT</td><td>>hg38_21_+_8433172_8446622_RNA45SN1_45S_50ntFlanks</td><td>9573</td><td>9596</td><td>24</td><td>AAATCGGTCGTCCGACCTGGGTAT</td><td>8296.3</td><td>-8.6</td><td>1.0</td><td>-8.9</td><td>8.7E-19</td><td>6.8E-18</td><td>MB231/MB468</td><td>MITO</td></tr>
<tr><td>>hg38_21_+_8433172_8446622_RNA45SN1_45S_50ntFlanks@11199.11223.25|CCGGCGGCGGCGGCGACTCTGGACG</td><td>>hg38_21_+_8433172_8446622_RNA45SN1_45S_50ntFlanks</td><td>11199</td><td>11223</td><td>25</td><td>CCGGCGGCGGCGGCGACTCTGGACG</td><td>620.3</td><td>-13.1</td><td>1.5</td><td>-8.9</td><td>8.7E-19</td><td>6.9E-18</td><td>MB231/MB468</td><td>MITO</td></tr>
<tr><td>>trnaMT_GluTTC_MT_-_14674_14742@34.71.38,trnalookalike8_GluTTC_5_-_93905172_93905240@34.71.38|ATATCATTGGTCGTGGTTGTAGTCCGTGCGAGAATACC</td><td>>trnaMT_GluTTC_MT_-_14674_14742</td><td>34</td><td>71</td><td>38</td><td>ATATCATTGGTCGTGGTTGTAGTCCGTGCGAGAATACC</td><td>996.8</td><td>14.0</td><td>1.6</td><td>8.9</td><td>5.1E-19</td><td>6.9E-18</td><td>MB231/MB468</td><td>CYTO</td></tr>
</table>

| | | | | | | | | | | | | | |
|---|---|---|---|---|---|---|---|---|---|---|---|---|---|
| >hg38_21_+_8433172_8446622_RNA45SN1_45S_50ntFlanks@6651.6672.22\|CGACTCTTAGCGGTGGATCACT | >hg38_21_+_8433172_8446622_RNA45SN1_45S_50ntFlanks | 6651 | 6672 | 22 | CGACTCTTAGCGGTGGATCACT | 5045.2 | 6.0 | 0.7 | 8.8 | 1.1E-18 | 6.9E-18 | BT20/MB468 | CYTO |
| >hg38_21_+_8433172_8446622_RNA45SN1_45S_50ntFlanks@12574.12604.31\|CGAGAGGAACCGCAGGTTCAGACATTTGGTG | >hg38_21_+_8433172_8446622_RNA45SN1_45S_50ntFlanks | 12574 | 12604 | 31 | CGAGAGGAACCGCAGGTTCAGACATTTGGTG | 452.7 | -12.6 | 1.4 | -8.8 | 8.9E-19 | 7.0E-18 | MB231/MB468 | MITO |
| >hg38_21_+_8433172_8446622_RNA45SN1_45S_50ntFlanks@4724.4748.25\|ATTAATCAAGAACGAAAGTCGGAGG | >hg38_21_+_8433172_8446622_RNA45SN1_45S_50ntFlanks | 4724 | 4748 | 25 | ATTAATCAAGAACGAAAGTCGGAGG | 1062.6 | -13.8 | 1.6 | -8.8 | 8.9E-19 | 7.0E-18 | MB231/MB468 | MITO |
| >hg38_21_+_8433172_8446622_RNA45SN1_45S_50ntFlanks@8437.8471.35\|AGGATTCAACCCGGCGGCGGGTCCGGCCGTGTCGG | >hg38_21_+_8433172_8446622_RNA45SN1_45S_50ntFlanks | 8437 | 8471 | 35 | AGGATTCAACCCGGCGGCGGGTCCGGCCGTGTCGG | 461.9 | -12.7 | 1.4 | -8.9 | 5.4E-19 | 7.0E-18 | MB231/MB468 | MP |
| >hg38_21_+_8433172_8446622_RNA45SN1_45S_50ntFlanks@12625.12653.29\|CAATGGGGCGAAGCTACCATCTGTGGGAT | >hg38_21_+_8433172_8446622_RNA45SN1_45S_50ntFlanks | 12625 | 12653 | 29 | CAATGGGGCGAAGCTACCATCTGTGGGAT | 2482.6 | -10.6 | 1.2 | -8.9 | 5.4E-19 | 7.0E-18 | MB231/MB468 | MP |
| >hg38_21_+_8433172_8446622_RNA45SN1_45S_50ntFlanks@12420.12446.27\|ATGTCGGCTCTTCCTATCATTGTGAAG | >hg38_21_+_8433172_8446622_RNA45SN1_45S_50ntFlanks | 12420 | 12446 | 27 | ATGTCGGCTCTTCCTATCATTGTGAAG | 11536.9 | -5.5 | 0.6 | -8.9 | 7.2E-19 | 7.0E-18 | MB231/MB468 | NUC |
| >hg38_21_+_8433172_8446622_RNA45SN1_45S_50ntFlanks@9305.9337.33\|CCAAGGAGTCTAACACGTGCGCGAGTCGGGGGC | >hg38_21_+_8433172_8446622_RNA45SN1_45S_50ntFlanks | 9305 | 9337 | 33 | CCAAGGAGTCTAACACGTGCGCGAGTCGGGGGC | 697.7 | -13.2 | 1.5 | -8.8 | 9.0E-19 | 7.0E-18 | MB231/MB468 | MITO |
| >hg38_21_+_8433172_8446622_RNA45SN1_45S_50ntFlanks@9306.9337.32\|CAAGGAGTCTAACACGTGCGCGAGTCGGGGGC | >hg38_21_+_8433172_8446622_RNA45SN1_45S_50ntFlanks | 9306 | 9337 | 32 | CAAGGAGTCTAACACGTGCGCGAGTCGGGGGC | 509.8 | -12.9 | 1.4 | -8.9 | 5.5E-19 | 7.0E-18 | MB231/MB468 | MP |
| >hg38_21_+_8433172_8446622_RNA45SN1_45S_50ntFlanks@9534.9563.30\|AGGAAACTCTGGTGGAGGTCCGTAGCGGTC | >hg38_21_+_8433172_8446622_RNA45SN1_45S_50ntFlanks | 9534 | 9563 | 30 | AGGAAACTCTGGTGGAGGTCCGTAGCGGTC | 17320.2 | -6.0 | 0.7 | -8.8 | 1.1E-18 | 7.0E-18 | BT20/MB468 | CYTO |
| >hg38_21_+_8433172_8446622_RNA45SN1_45S_50ntFlanks@9564.9597.34\|CTGACGTGCAAATCGGTCGTCCGACCTGGGTATA | >hg38_21_+_8433172_8446622_RNA45SN1_45S_50ntFlanks | 9564 | 9597 | 34 | CTGACGTGCAAATCGGTCGTCCGACCTGGGTATA | 3326.5 | -9.6 | 1.1 | -8.9 | 5.5E-19 | 7.1E-18 | MB231/MB468 | MP |
| >MI0000067\|hsa-let-7f-1&WithFlank&9\|+\|94176341\|94176439@13.33.21;MI0000068\|hsa-let-7f-2&WithFlank&X\|-\|53557186\|53557280@14.34.21&[MIMAT0000067&hsa-let-7f-5p&offsets\|0\|-1;m-20&9\|+\|94176353\|94176374&offsets\|0\|-1];[MIMAT0000067_1&hsa-let-7f-5p&offsets\|0\|-1;m-19&X\|-\|53557246\|53557267&offsets\|0\|-1]\|TGAGGTAGTAGATTGTATAGT | >MI0000067\|hsa-let-7f-1&WithFlank&9\|+\|94176341\|94176439 | 13 | 33 | 21 | TGAGGTAGTAGATTGTATAGT | 12732.6 | 4.9 | 0.5 | 8.9 | 7.3E-19 | 7.1E-18 | MB231/MB468 | NUC |
| >hg38_21_+_8433172_8446622_RNA45SN1_45S_50ntFlanks@10512.10533.22\|TCCGGAGTGGCGGAGATGGGCG | >hg38_21_+_8433172_8446622_RNA45SN1_45S_50ntFlanks | 10512 | 10533 | 22 | TCCGGAGTGGCGGAGATGGGCG | 3072.8 | -14.5 | 1.6 | -8.9 | 7.3E-19 | 7.1E-18 | MB231/MB468 | NUC |
| >hg38_21_+_8433172_8446622_RNA45SN1_45S_50ntFlanks@12161.12180.20\|TGACTGGGGCGGTACACCTG | >hg38_21_+_8433172_8446622_RNA45SN1_45S_50ntFlanks | 12161 | 12180 | 20 | TGACTGGGGCGGTACACCTG | 890.7 | -13.6 | 1.5 | -8.8 | 9.1E-19 | 7.1E-18 | MB231/MB468 | MITO |
| >hg38_21_+_8433172_8446622_RNA45SN1_45S_50ntFlanks@12008.12030.23\|CCGCCGGTGAAATACCACTACTC | >hg38_21_+_8433172_8446622_RNA45SN1_45S_50ntFlanks | 12008 | 12030 | 23 | CCGCCGGTGAAATACCACTACTC | 3499.3 | -8.3 | 0.9 | -8.9 | 8.2E-19 | 7.1E-18 | BT20/MB468 | MITO |
| >hg38_21_+_8433172_8446622_RNA45SN1_45S_50ntFlanks@3802.3841.40\|CATTAAATCAGTTATGGTTCCTTTGGTCGCTCGCTCCTCT | >hg38_21_+_8433172_8446622_RNA45SN1_45S_50ntFlanks | 3802 | 3841 | 40 | CATTAAATCAGTTATGGTTCCTTTGGTCGCTCGCTCCTCT | 384.9 | -13.0 | 1.5 | -8.8 | 2.0E-18 | 7.1E-18 | BT20/MB468 | NUC |
| >MI0000102\|hsa-mir-100&WithFlank&11\|-\|122152223\|122152314@19.36.18&[MIMAT0000098&hsa-miR-100-5p&offsets\|0\|-4;m-27&11\|-\|122152276\|122152296&offsets\|0\|-3]\|AACCCGTAGATCCGAACT | >MI0000102\|hsa-mir-100&WithFlank&11\|-\|122152223\|122152314 | 19 | 36 | 18 | AACCCGTAGATCCGAACT | 362.2 | 12.5 | 1.4 | 8.9 | 5.3E-19 | 7.2E-18 | MB231/MB468 | CYTO |
| >hg38_21_+_8433172_8446622_RNA45SN1_45S_50ntFlanks@6700.6720.21\|CTAGCTGCGAGAATTAATGTG | >hg38_21_+_8433172_8446622_RNA45SN1_45S_50ntFlanks | 6700 | 6720 | 21 | CTAGCTGCGAGAATTAATGTG | 4336.2 | -4.6 | 0.5 | -8.9 | 8.3E-19 | 7.2E-18 | BT20/MB468 | MITO |
| >hg38_21_+_8433172_8446622_RNA45SN1_45S_50ntFlanks@10846.10868.23\|CTTCGGGATAAGGATTGGCTCTA | >hg38_21_+_8433172_8446622_RNA45SN1_45S_50ntFlanks | 10846 | 10868 | 23 | CTTCGGGATAAGGATTGGCTCTA | 576.6 | -13.0 | 1.5 | -8.9 | 5.6E-19 | 7.2E-18 | MB231/MB468 | MP |
| >MI0000285\|hsa-mir-205&WithFlank&1\|+\|209432127\|209432248@40.61.22(+2A)&[MIMAT0000266&hsa-miR-205-5p&offsets\|0\|0(+2A);m-351&1\|+\|209432166\|209432187&offsets\|0\|0(+2A)]\|TCCTTCATTCCACCGGAGTCTGAA | >MI0000285\|hsa-mir-205&WithFlank&1\|+\|209432127\|209432248 | 40 | 61 | 22(+2A) | TCCTTCATTCCACCGGAGTCTGAA | 746.7 | 13.2 | 1.4 | 9.1 | 7.1E-20 | 7.3E-18 | BT20/MB231 | MITO |
| >MI0016887\|hsa-mir-4521&WithFlank&17\|+\|8186939\|8187010@12.34.23&[MIMAT0019058&hsa-miR-4521&offsets\|+2\|+3;m-765&17\|+\|8186948\|8186969&offsets\|+2\|+3]\|TAAGGAAGTCCTGTGCTCAGTTT | >MI0016887\|hsa-mir-4521&WithFlank&17\|+\|8186939\|8187010 | 12 | 34 | 23 | TAAGGAAGTCCTGTGCTCAGTTT | 490.5 | 13.2 | 1.5 | 8.8 | 1.2E-18 | 7.3E-18 | BT20/MB468 | CYTO |
| >hg38_21_+_8433172_8446622_RNA45SN1_45S_50ntFlanks@11318.11338.21\|CCCCACGTCTCGTCGCGCGCG | >hg38_21_+_8433172_8446622_RNA45SN1_45S_50ntFlanks | 11318 | 11338 | 21 | CCCCACGTCTCGTCGCGCGCG | 890.5 | -12.8 | 1.4 | -8.9 | 7.5E-19 | 7.3E-18 | MB231/MB468 | NUC |
| >hg38_21_+_8433172_8446622_RNA45SN1_45S_50ntFlanks@10402.10430.29\|AACATGGGTCAGTCGGTCCTGAGAGATGG | >hg38_21_+_8433172_8446622_RNA45SN1_45S_50ntFlanks | 10402 | 10430 | 29 | AACATGGGTCAGTCGGTCCTGAGAGATGG | 2083.3 | -5.2 | 0.6 | -8.9 | 5.4E-19 | 7.3E-18 | MB231/MB468 | CYTO |
| >hg38_MT_+_1621_3279_RNR2_16S_50ntFlanks@51.92.42\|GCTAAACCTAGCCCCAAACCCACTCCACCTTACTACCAGACA | >hg38_MT_+_1621_3279_RNR2_16S_50ntFlanks | 51 | 92 | 42 | GCTAAACCTAGCCCCAAACCCACTCCACCTTACTACCAGACA | 821.8 | 13.7 | 1.5 | 8.9 | 5.5E-19 | 7.4E-18 | MB231/MB468 | CYTO |
| >hg38_21_+_8433172_8446622_RNA45SN1_45S_50ntFlanks@12154.12181.28\|GGGAGTTTGACTGGGGCGGTACACCTGT | >hg38_21_+_8433172_8446622_RNA45SN1_45S_50ntFlanks | 12154 | 12181 | 28 | GGGAGTTTGACTGGGGCGGTACACCTGT | 378.4 | -12.4 | 1.4 | -8.9 | 5.8E-19 | 7.4E-18 | MB231/MB468 | MP |
| >hg38_21_+_8433172_8446622_RNA45SN1_45S_50ntFlanks@7975.8010.36\|CGCGACCTCAGATCAGACGTGGCGACCCGCTGAATT | >hg38_21_+_8433172_8446622_RNA45SN1_45S_50ntFlanks | 7975 | 8010 | 36 | CGCGACCTCAGATCAGACGTGGCGACCCGCTGAATT | 128456.8 | 4.2 | 0.5 | 8.8 | 2.1E-18 | 7.4E-18 | BT20/MB468 | NUC |
| >hg38_21_+_8433172_8446622_RNA45SN1_45S_50ntFlanks@11621.11640.20\|CAAAGCATCGCGAAGGCCCG | >hg38_21_+_8433172_8446622_RNA45SN1_45S_50ntFlanks | 11621 | 11640 | 20 | CAAAGCATCGCGAAGGCCCG | 1720.5 | -12.8 | 1.5 | -8.9 | 8.5E-19 | 7.4E-18 | BT20/MB468 | MITO |
| >trnaMT_GluTTC_MT_-_14674_14742@39.65.27,trnalookalike8_GluTTC_5_-_93905172_93905240@39.65.27\|ATTGGTCGTGGTTGTAGTCCGTGCGAG | >trnaMT_GluTTC_MT_-_14674_14742 | 39 | 65 | 27 | ATTGGTCGTGGTTGTAGTCCGTGCGAG | 1478.7 | 8.3 | 0.9 | 8.9 | 8.6E-19 | 7.4E-18 | BT20/MB468 | MITO |
| >hg38_21_+_8433172_8446622_RNA45SN1_45S_50ntFlanks@6719.6740.22\|TGAATTGCAGGACACATTGATC | >hg38_21_+_8433172_8446622_RNA45SN1_45S_50ntFlanks | 6719 | 6740 | 22 | TGAATTGCAGGACACATTGATC | 2922.0 | -5.6 | 0.6 | -8.8 | 1.2E-18 | 7.4E-18 | BT20/MB468 | CYTO |
| >hg38_21_+_8433172_8446622_RNA45SN1_45S_50ntFlanks@12033.12055.23\|ATCGTTTTTTCACTGACCCGGTG | >hg38_21_+_8433172_8446622_RNA45SN1_45S_50ntFlanks | 12033 | 12055 | 23 | ATCGTTTTTTCACTGACCCGGTG | 505.3 | -12.8 | 1.4 | -8.9 | 5.8E-19 | 7.4E-18 | MB231/MB468 | MP |
| >hg38_21_+_8433172_8446622_RNA45SN1_45S_50ntFlanks@475.516.42\|CTTCGTGATCGATGTGGTGACGTCGTGCTCTCCCGGGCCGGG | >hg38_21_+_8433172_8446622_RNA45SN1_45S_50ntFlanks | 475 | 516 | 42 | CTTCGTGATCGATGTGGTGACGTCGTGCTCTCCCGGGCCGGG | 586.2 | 8.2 | 0.9 | 8.8 | 1.2E-18 | 7.5E-18 | BT20/MB468 | CYTO |
| >hg38_21_+_8433172_8446622_RNA45SN1_45S_50ntFlanks@11210.11227.18\|GGCGACTCTGGACGCGAG | >hg38_21_+_8433172_8446622_RNA45SN1_45S_50ntFlanks | 11210 | 11227 | 18 | GGCGACTCTGGACGCGAG | 431.0 | -12.6 | 1.4 | -8.9 | 5.8E-19 | 7.5E-18 | MB231/MB468 | MP |
| >hg38_21_+_8433172_8446622_RNA45SN1_45S_50ntFlanks@6773.6799.27\|TTCCTCCCGGGGCTACGCCTGTCTGAG | >hg38_21_+_8433172_8446622_RNA45SN1_45S_50ntFlanks | 6773 | 6799 | 27 | TTCCTCCCGGGGCTACGCCTGTCTGAG | 386.7 | -12.5 | 1.4 | -8.9 | 5.8E-19 | 7.5E-18 | MB231/MB468 | MP |
| >hg38_21_+_8433172_8446622_RNA45SN1_45S_50ntFlanks@9505.9526.22\|TGAACTATGCCTGGGCAGGGCG | >hg38_21_+_8433172_8446622_RNA45SN1_45S_50ntFlanks | 9505 | 9526 | 22 | TGAACTATGCCTGGGCAGGGCG | 1695.7 | -14.6 | 1.6 | -8.9 | 5.9E-19 | 7.6E-18 | MB231/MB468 | MP |
| >hg38_21_+_8433172_8446622_RNA45SN1_45S_50ntFlanks@9602.9622.21\|CGAAAGACTAATCGAACCATC | >hg38_21_+_8433172_8446622_RNA45SN1_45S_50ntFlanks | 9602 | 9622 | 21 | CGAAAGACTAATCGAACCATC | 1005.6 | -13.8 | 1.6 | -8.9 | 5.9E-19 | 7.6E-18 | MB231/MB468 | MP |
| >hg38_21_+_8433172_8446622_RNA45SN1_45S_50ntFlanks@9198.9220.23\|GTTCTCTCGGGGCCACGCGCGCG | >hg38_21_+_8433172_8446622_RNA45SN1_45S_50ntFlanks | 9198 | 9220 | 23 | GTTCTCTCGGGGCCACGCGCGCG | 580.1 | -13.0 | 1.5 | -8.9 | 5.9E-19 | 7.6E-18 | MB231/MB468 | MP |
| >hg38_21_+_8433172_8446622_RNA45SN1_45S_50ntFlanks@11576.11603.28\|CTTAGAACTGGTGCGGACCAGGGGAATC | >hg38_21_+_8433172_8446622_RNA45SN1_45S_50ntFlanks | 11576 | 11603 | 28 | CTTAGAACTGGTGCGGACCAGGGGAATC | 4244.0 | -9.0 | 1.0 | -8.9 | 7.8E-19 | 7.6E-18 | MB231/MB468 | NUC |
| >hg38_21_+_8433172_8446622_RNA45SN1_45S_50ntFlanks@12712.12742.31\|CGGAGCCTCGGTTGGCCTCGGATAGCCGGTC | >hg38_21_+_8433172_8446622_RNA45SN1_45S_50ntFlanks | 12712 | 12742 | 31 | CGGAGCCTCGGTTGGCCTCGGATAGCCGGTC | 3707.3 | -14.9 | 1.6 | -9.3 | 1.2E-20 | 7.6E-18 | BT20/MB468 | MP |
| >hg38_21_+_8433172_8446622_RNA45SN1_45S_50ntFlanks@12156.12180.25\|GAGTTTGACTGGGGCGGTACACCTG | >hg38_21_+_8433172_8446622_RNA45SN1_45S_50ntFlanks | 12156 | 12180 | 25 | GAGTTTGACTGGGGCGGTACACCTG | 460.4 | -12.7 | 1.4 | -8.9 | 6.0E-19 | 7.6E-18 | MB231/MB468 | MP |
| >hg38_21_+_8433172_8446622_RNA45SN1_45S_50ntFlanks@9835.9875.41\|TTTGGTAAGCAGAACTGGCGCTGCGGGATGAACCGAACGCC | >hg38_21_+_8433172_8446622_RNA45SN1_45S_50ntFlanks | 9835 | 9875 | 41 | TTTGGTAAGCAGAACTGGCGCTGCGGGATGAACCGAACGCC | 382.1 | 13.0 | 1.5 | 8.8 | 8.8E-19 | 7.6E-18 | BT20/MB468 | MITO |
| >hg38_21_+_8433172_8446622_RNA45SN1_45S_50ntFlanks@10845.10868.24\|ACTTCGGGATAAGGATTGGCTCTA | >hg38_21_+_8433172_8446622_RNA45SN1_45S_50ntFlanks | 10845 | 10868 | 24 | ACTTCGGGATAAGGATTGGCTCTA | 462.5 | -12.6 | 1.4 | -8.8 | 9.8E-19 | 7.6E-18 | MB231/MB468 | MITO |
| >hg38_21_+_8433172_8446622_RNA45SN1_45S_50ntFlanks@10771.10806.36\|CAGGTCTCCAAGGTGAACAGCCTCTGGCATGTTGGA | >hg38_21_+_8433172_8446622_RNA45SN1_45S_50ntFlanks | 10771 | 10806 | 36 | CAGGTCTCCAAGGTGAACAGCCTCTGGCATGTTGGA | 775.4 | -13.5 | 1.5 | -8.9 | 6.0E-19 | 7.7E-18 | MB231/MB468 | MP |
| >hg38_21_+_8433172_8446622_RNA45SN1_45S_50ntFlanks@11316.11343.28\|CACCCCACGTCTCGTCGCGCGCGCGTCC | >hg38_21_+_8433172_8446622_RNA45SN1_45S_50ntFlanks | 11316 | 11343 | 28 | CACCCCACGTCTCGTCGCGCGCGCGTCC | 712.8 | -9.2 | 1.0 | -8.9 | 5.7E-19 | 7.7E-18 | MB231/MB468 | CYTO |
| >hg38_21_+_8433172_8446622_RNA45SN1_45S_50ntFlanks@5086.5124.39\|AACGAGACTCTGGCATGCTAACTAGTTACGCGACCCCCG | >hg38_21_+_8433172_8446622_RNA45SN1_45S_50ntFlanks | 5086 | 5124 | 39 | AACGAGACTCTGGCATGCTAACTAGTTACGCGACCCCCG | 258.8 | 8.5 | 1.0 | 8.9 | 5.8E-19 | 7.7E-18 | MB231/MB468 | CYTO |
| >hg38_21_+_8433172_8446622_RNA45SN1_45S_50ntFlanks@8195.8213.19\|CCGTGGACGGTGTGAGGCC | >hg38_21_+_8433172_8446622_RNA45SN1_45S_50ntFlanks | 8195 | 8213 | 19 | CCGTGGACGGTGTGAGGCC | 2114.3 | 5.0 | 0.6 | 8.9 | 5.8E-19 | 7.7E-18 | MB231/MB468 | CYTO |
| >hg38_21_+_8433172_8446622_RNA45SN1_45S_50ntFlanks@11580.11603.24\|GAACTGGTGCGGACCAGGGGAATC | >hg38_21_+_8433172_8446622_RNA45SN1_45S_50ntFlanks | 11580 | 11603 | 24 | GAACTGGTGCGGACCAGGGGAATC | 3682.2 | -8.8 | 1.0 | -8.8 | 9.9E-19 | 7.7E-18 | MB231/MB468 | MITO |
| >hg38_21_+_8433172_8446622_RNA45SN1_45S_50ntFlanks@9545.9570.26\|GTGGAGGTCCGTAGCGGTCCTGACGT | >hg38_21_+_8433172_8446622_RNA45SN1_45S_50ntFlanks | 9545 | 9570 | 26 | GTGGAGGTCCGTAGCGGTCCTGACGT | 1972.4 | -4.2 | 0.5 | -8.8 | 1.2E-18 | 7.7E-18 | BT20/MB468 | CYTO |
| >hg38_21_+_8433172_8446622_RNA45SN1_45S_50ntFlanks@11845.11872.28\|GGAACGGGCTTGGCGGAATCAGCGGGGA | >hg38_21_+_8433172_8446622_RNA45SN1_45S_50ntFlanks | 11845 | 11872 | 28 | GGAACGGGCTTGGCGGAATCAGCGGGGA | 449.9 | -11.8 | 1.3 | -8.9 | 8.0E-19 | 7.8E-18 | MB231/MB468 | NUC |
| >hg38_21_+_8433172_8446622_RNA45SN1_45S_50ntFlanks@8117.8140.24\|CGGGACATGTGGCGTACGGAAGAC | >hg38_21_+_8433172_8446622_RNA45SN1_45S_50ntFlanks | 8117 | 8140 | 24 | CGGGACATGTGGCGTACGGAAGAC | 648.0 | -13.2 | 1.5 | -8.9 | 6.1E-19 | 7.8E-18 | MB231/MB468 | MP |
| >hg38_21_+_8433172_8446622_RNA45SN1_45S_50ntFlanks@9534.9569.36\|AGGAAACTCTGGTGGAGGTCCGTAGCGGTCCTGACG | >hg38_21_+_8433172_8446622_RNA45SN1_45S_50ntFlanks | 9534 | 9569 | 36 | AGGAAACTCTGGTGGAGGTCCGTAGCGGTCCTGACG | 9292.4 | -7.2 | 0.8 | -8.9 | 8.1E-19 | 7.8E-18 | MB231/MB468 | NUC |
| >hg38_21_+_8433172_8446622_RNA45SN1_45S_50ntFlanks@10910.10932.23\|CGCGCGCCGCGGCTGGACGAGGC | >hg38_21_+_8433172_8446622_RNA45SN1_45S_50ntFlanks | 10910 | 10932 | 23 | CGCGCGCCGCGGCTGGACGAGGC | 493.4 | -12.8 | 1.4 | -8.9 | 6.1E-19 | 7.8E-18 | MB231/MB468 | MP |

<table>
<tr><td></td><td></td><td></td><td></td><td></td><td></td><td></td><td></td><td></td><td></td><td></td><td></td><td></td><td></td></tr>
<tr><td>>MI0000746|hsa-mir-99b&WithFlank&19|+|51692606|51692687@51.72.22(+1A)&[MIMAT0004678&hsa-miR-99b-3p&offsets|0|0(+1A);m-151&19|+|51692656|51692678&offsets|0|-1(+1A)]|CAAGCTCGTGTCTGTGGGTCCGA</td><td>>MI0000746|hsa-mir-99b&WithFlank&19|+|51692606|51692687</td><td>51</td><td>72</td><td>22(+1A)</td><td>CAAGCTCGTGTCTGTGGGTCCGA</td><td>571.6</td><td>7.6</td><td>0.9</td><td>8.8</td><td>9.0E-19</td><td>7.8E-18</td><td>BT20/MB468</td><td>MITO</td></tr>
<tr><td>>hg38_21_+_8433172_8446622_RNA45SN1_45S_50ntFlanks@10914.10946.33|CGCCGCGGCTGGACGAGGCGCCGCCGCCCCCCC</td><td>>hg38_21_+_8433172_8446622_RNA45SN1_45S_50ntFlanks</td><td>10914</td><td>10946</td><td>33</td><td>CGCCGCGGCTGGACGAGGCGCCGCCGCCCCCCC</td><td>1016.4</td><td>-13.8</td><td>1.6</td><td>-8.8</td><td>1.0E-18</td><td>7.9E-18</td><td>MB231/MB468</td><td>MITO</td></tr>
<tr><td>>hg38_21_+_8433172_8446622_RNA45SN1_45S_50ntFlanks@12324.12346.23|TTGGGTTTTAAGCAGGAGGTGTC</td><td>>hg38_21_+_8433172_8446622_RNA45SN1_45S_50ntFlanks</td><td>12324</td><td>12346</td><td>23</td><td>TTGGGTTTTAAGCAGGAGGTGTC</td><td>2640.3</td><td>-7.4</td><td>0.8</td><td>-8.9</td><td>5.9E-19</td><td>7.9E-18</td><td>MB231/MB468</td><td>CYTO</td></tr>
<tr><td>>hg38_21_+_8433172_8446622_RNA45SN1_45S_50ntFlanks@12617.12653.37|TGAGGAGCCAATGGGGCGAAGCTACCATCTGTGGGAT</td><td>>hg38_21_+_8433172_8446622_RNA45SN1_45S_50ntFlanks</td><td>12617</td><td>12653</td><td>37</td><td>TGAGGAGCCAATGGGGCGAAGCTACCATCTGTGGGAT</td><td>581.5</td><td>-12.1</td><td>1.4</td><td>-8.9</td><td>8.2E-19</td><td>7.9E-18</td><td>MB231/MB468</td><td>NUC</td></tr>
<tr><td>>MI0000239|hsa-mir-197&WithFlank&1|+|109598887|109598973@54.74.21&[MIMAT0000227&hsa-miR-197-3p&offsets|0|-1;m-99&1|+|109598940|109598961&offsets|0|-1]|TTCACCACCTTCTCCACCCAG</td><td>>MI0000239|hsa-mir-197&WithFlank&1|+|109598887|109598973</td><td>54</td><td>74</td><td>21</td><td>TTCACCACCTTCTCCACCCAG</td><td>1159.4</td><td>10.4</td><td>1.2</td><td>8.8</td><td>1.3E-18</td><td>7.9E-18</td><td>BT20/MB468</td><td>CYTO</td></tr>
<tr><td>>hg38_21_+_8433172_8446622_RNA45SN1_45S_50ntFlanks@12997.13038.42|CGCTGCGATCTATTGAAAGTCAGCCCTCGACACAAGGGTTTG</td><td>>hg38_21_+_8433172_8446622_RNA45SN1_45S_50ntFlanks</td><td>12997</td><td>13038</td><td>42</td><td>CGCTGCGATCTATTGAAAGTCAGCCCTCGACACAAGGGTTTG</td><td>5589.8</td><td>10.0</td><td>1.1</td><td>8.9</td><td>6.0E-19</td><td>7.9E-18</td><td>MB231/MB468</td><td>CYTO</td></tr>
<tr><td>>hg38_21_+_8433172_8446622_RNA45SN1_45S_50ntFlanks@8919.8936.18|TCCCGGGGCCGAGGGAGC</td><td>>hg38_21_+_8433172_8446622_RNA45SN1_45S_50ntFlanks</td><td>8919</td><td>8936</td><td>18</td><td>TCCCGGGGCCGAGGGAGC</td><td>447.1</td><td>12.8</td><td>1.4</td><td>8.9</td><td>6.0E-19</td><td>7.9E-18</td><td>MB231/MB468</td><td>CYTO</td></tr>
<tr><td>>hg38_21_+_8433172_8446622_RNA45SN1_45S_50ntFlanks@9741.9765.25|ACGATCTCAACCTATTCTCAAACTT</td><td>>hg38_21_+_8433172_8446622_RNA45SN1_45S_50ntFlanks</td><td>9741</td><td>9765</td><td>25</td><td>ACGATCTCAACCTATTCTCAAACTT</td><td>1241.4</td><td>-14.1</td><td>1.6</td><td>-8.8</td><td>1.0E-18</td><td>8.0E-18</td><td>MB231/MB468</td><td>MITO</td></tr>
<tr><td>>hg38_21_+_8433172_8446622_RNA45SN1_45S_50ntFlanks@10819.10840.22|AGGGAAGTCGGCAAGCCGGATC</td><td>>hg38_21_+_8433172_8446622_RNA45SN1_45S_50ntFlanks</td><td>10819</td><td>10840</td><td>22</td><td>AGGGAAGTCGGCAAGCCGGATC</td><td>4205.7</td><td>-8.8</td><td>1.0</td><td>-8.8</td><td>1.0E-18</td><td>8.0E-18</td><td>MB231/MB468</td><td>MITO</td></tr>
<tr><td>>hg38_21_+_8433172_8446622_RNA45SN1_45S_50ntFlanks@9003.9026.24|GTCTCCCCCGCGGGGGCGCGCCGG</td><td>>hg38_21_+_8433172_8446622_RNA45SN1_45S_50ntFlanks</td><td>9003</td><td>9026</td><td>24</td><td>GTCTCCCCCGCGGGGGCGCGCCGG</td><td>702.3</td><td>-13.2</td><td>1.5</td><td>-8.8</td><td>1.0E-18</td><td>8.0E-18</td><td>MB231/MB468</td><td>MITO</td></tr>
<tr><td>>hg38_21_+_8433172_8446622_RNA45SN1_45S_50ntFlanks@5484.5510.27|GAGCGCTGAGAAGACGGTCGAACTTGA</td><td>>hg38_21_+_8433172_8446622_RNA45SN1_45S_50ntFlanks</td><td>5484</td><td>5510</td><td>27</td><td>GAGCGCTGAGAAGACGGTCGAACTTGA</td><td>517.8</td><td>-12.9</td><td>1.4</td><td>-8.9</td><td>6.3E-19</td><td>8.0E-18</td><td>MB231/MB468</td><td>MP</td></tr>
<tr><td>>hg38_21_+_8433172_8446622_RNA45SN1_45S_50ntFlanks@10905.10936.32|CTGGGCGCGCGCCGCGGCTGGACGAGGCGCCG</td><td>>hg38_21_+_8433172_8446622_RNA45SN1_45S_50ntFlanks</td><td>10905</td><td>10936</td><td>32</td><td>CTGGGCGCGCGCCGCGGCTGGACGAGGCGCCG</td><td>401.4</td><td>-12.5</td><td>1.4</td><td>-8.9</td><td>6.3E-19</td><td>8.0E-18</td><td>MB231/MB468</td><td>MP</td></tr>
<tr><td>>hg38_21_+_8433172_8446622_RNA45SN1_45S_50ntFlanks@11571.11594.24|GCCGACTTAGAACTGGTGCGGACC</td><td>>hg38_21_+_8433172_8446622_RNA45SN1_45S_50ntFlanks</td><td>11571</td><td>11594</td><td>24</td><td>GCCGACTTAGAACTGGTGCGGACC</td><td>384.7</td><td>4.1</td><td>0.5</td><td>8.8</td><td>9.3E-19</td><td>8.1E-18</td><td>BT20/MB468</td><td>MITO</td></tr>
<tr><td>>trna116_GluCTC_1_-_145399233_145399304@31.70.40,trna71_GluCTC_1_-_161439189_161439260@31.70.40,trna74_GluCTC_1_-_161431809_161431880@31.70.40,trna77_GluCTC_1_-_161424398_161424469@31.70.40,trna77_GluCTC_6_+_28949976_28950047@31.70.40,trna80_GluCTC_1_-_161417018_161417089@31.70.40,trna87_GluCTC_6_-_126101393_126101464@31.70.40|GCTCTCACCGCCGCGGCCCGGGTTCGATTCCCGGTCAGGG</td><td>>trna116_GluCTC_1_-_145399233_145399304</td><td>31</td><td>70</td><td>40</td><td>GCTCTCACCGCCGCGGCCCGGGTTCGATTCCCGGTCAGGG</td><td>431.4</td><td>-11.7</td><td>1.3</td><td>-8.9</td><td>8.4E-19</td><td>8.1E-18</td><td>MB231/MB468</td><td>NUC</td></tr>
<tr><td>>hg38_21_+_8433172_8446622_RNA45SN1_45S_50ntFlanks@10808.10827.20|CAATGTAGGTAAGGGAAGTC</td><td>>hg38_21_+_8433172_8446622_RNA45SN1_45S_50ntFlanks</td><td>10808</td><td>10827</td><td>20</td><td>CAATGTAGGTAAGGGAAGTC</td><td>453.8</td><td>-12.7</td><td>1.4</td><td>-8.9</td><td>6.4E-19</td><td>8.1E-18</td><td>MB231/MB468</td><td>MP</td></tr>
<tr><td>>MI0000650|hsa-mir-200c&WithFlank&12|+|6963693|6963772@50.72.23(+2A)&[MIMAT0000617&hsa-miR-200c-3p&offsets|0|0(+2A);m-54&12|+|6963742|6963764&offsets|0|0(+2A)]|TAATACTGCCGGGTAATGATGGAAA</td><td>>MI0000650|hsa-mir-200c&WithFlank&12|+|6963693|6963772</td><td>50</td><td>72</td><td>23(+2A)</td><td>TAATACTGCCGGGTAATGATGGAAA</td><td>2226.5</td><td>9.0</td><td>1.0</td><td>8.8</td><td>1.3E-18</td><td>8.1E-18</td><td>BT20/MB468</td><td>CYTO</td></tr>
<tr><td>>hg38_21_+_8433172_8446622_RNA45SN1_45S_50ntFlanks@10391.10431.41|AACAGCAGTTGAACATGGGTCAGTCGGTCCTGAGAGATGGG</td><td>>hg38_21_+_8433172_8446622_RNA45SN1_45S_50ntFlanks</td><td>10391</td><td>10431</td><td>41</td><td>AACAGCAGTTGAACATGGGTCAGTCGGTCCTGAGAGATGGG</td><td>618.1</td><td>-13.7</td><td>1.6</td><td>-8.7</td><td>2.3E-18</td><td>8.1E-18</td><td>BT20/MB468</td><td>NUC</td></tr>
<tr><td>>hg38_21_+_8433172_8446622_RNA45SN1_45S_50ntFlanks@10873.10892.20|CTGGGTCGGTCGGGCTGGGG</td><td>>hg38_21_+_8433172_8446622_RNA45SN1_45S_50ntFlanks</td><td>10873</td><td>10892</td><td>20</td><td>CTGGGTCGGTCGGGCTGGGG</td><td>504.1</td><td>-12.8</td><td>1.4</td><td>-8.8</td><td>1.1E-18</td><td>8.2E-18</td><td>MB231/MB468</td><td>MITO</td></tr>
<tr><td>>MI0000102|hsa-mir-100&WithFlank&11|-|122152223|122152314@19.39.21(+1A)&[MIMAT0000098&hsa-miR-100-5p&offsets|0|-1(+1A);m-27&11|-|122152276|122152296&offsets|0|0(+1A)]|AACCCGTAGATCCGAACTTGTA</td><td>>MI0000102|hsa-mir-100&WithFlank&11|-|122152223|122152314</td><td>19</td><td>39</td><td>21(+1A)</td><td>AACCCGTAGATCCGAACTTGTA</td><td>94311.4</td><td>-7.8</td><td>0.9</td><td>-9.1</td><td>8.3E-20</td><td>8.2E-18</td><td>BT20/MB231</td><td>MITO</td></tr>
<tr><td>>MI0000542|hsa-mir-320a&WithFlank&8|-|22244960|22245043@49.68.20(+1U)&[MIMAT0000510&hsa-miR-320a-3p&offsets|+1|-1(+1U);m-60&8|-|22244975|22244996&offsets|+1|-1(+1U)]|AAAGCTGGGTTGAGAGGGCGT</td><td>>MI0000542|hsa-mir-320a&WithFlank&8|-|22244960|22245043</td><td>49</td><td>68</td><td>20(+1U)</td><td>AAAGCTGGGTTGAGAGGGCGT</td><td>1159.1</td><td>11.4</td><td>1.3</td><td>8.8</td><td>1.3E-18</td><td>8.2E-18</td><td>BT20/MB468</td><td>CYTO</td></tr>
<tr><td>>MI0000811|hsa-mir-148b&WithFlank&12|+|54337210|54337320@69.90.22(+1U)&[MIMAT0000759&hsa-miR-148b-3p&offsets|0|0(+1U);m-77&12|+|54337278|54337299&offsets|0|0(+1U)]|TCAGTGCATCACAGAACTTTGTT</td><td>>MI0000811|hsa-mir-148b&WithFlank&12|+|54337210|54337320</td><td>69</td><td>90</td><td>22(+1U)</td><td>TCAGTGCATCACAGAACTTTGTT</td><td>1873.0</td><td>6.0</td><td>0.7</td><td>8.8</td><td>9.6E-19</td><td>8.3E-18</td><td>BT20/MB468</td><td>MITO</td></tr>
<tr><td>>trnaMT_PheGAA_MT_+_577_647@39.58.20|ATGTTTAGACGGGCTCACAT</td><td>>trnaMT_PheGAA_MT_+_577_647</td><td>39</td><td>58</td><td>20</td><td>ATGTTTAGACGGGCTCACAT</td><td>965.7</td><td>7.1</td><td>0.8</td><td>8.8</td><td>9.7E-19</td><td>8.3E-18</td><td>BT20/MB468</td><td>MITO</td></tr>
<tr><td>>hg38_21_+_8433172_8446622_RNA45SN1_45S_50ntFlanks@5084.5105.22|CGAACGAGACTCTGGCATGCTA</td><td>>hg38_21_+_8433172_8446622_RNA45SN1_45S_50ntFlanks</td><td>5084</td><td>5105</td><td>22</td><td>CGAACGAGACTCTGGCATGCTA</td><td>1301.9</td><td>-14.8</td><td>1.7</td><td>-8.7</td><td>2.4E-18</td><td>8.4E-18</td><td>BT20/MB468</td><td>NUC</td></tr>
<tr><td>>hg38_21_+_8433172_8446622_RNA45SN1_45S_50ntFlanks@7991.8008.18|ACGTGGCGACCCGCTGAA</td><td>>hg38_21_+_8433172_8446622_RNA45SN1_45S_50ntFlanks</td><td>7991</td><td>8008</td><td>18</td><td>ACGTGGCGACCCGCTGAA</td><td>47296.5</td><td>-7.8</td><td>0.9</td><td>-8.8</td><td>1.1E-18</td><td>8.4E-18</td><td>MB231/MB468</td><td>MITO</td></tr>
<tr><td>>hg38_21_+_8433172_8446622_RNA45SN1_45S_50ntFlanks@5492.5526.35|AGAAGACGGTCGAACTTGACTATCTAGAGGAAGTA</td><td>>hg38_21_+_8433172_8446622_RNA45SN1_45S_50ntFlanks</td><td>5492</td><td>5526</td><td>35</td><td>AGAAGACGGTCGAACTTGACTATCTAGAGGAAGTA</td><td>1632.4</td><td>-6.4</td><td>0.7</td><td>-8.9</td><td>6.6E-19</td><td>8.4E-18</td><td>MB231/MB468</td><td>MP</td></tr>
<tr><td>>hg38_21_+_8433172_8446622_RNA45SN1_45S_50ntFlanks@11317.11344.28|ACCCCACGTCTCGTCGCGCGCGCGTCCG</td><td>>hg38_21_+_8433172_8446622_RNA45SN1_45S_50ntFlanks</td><td>11317</td><td>11344</td><td>28</td><td>ACCCCACGTCTCGTCGCGCGCGCGTCCG</td><td>464.3</td><td>-12.7</td><td>1.4</td><td>-8.9</td><td>6.6E-19</td><td>8.4E-18</td><td>MB231/MB468</td><td>MP</td></tr>
<tr><td>>hg38_21_+_8433172_8446622_RNA45SN1_45S_50ntFlanks@11312.11344.33|CCCCCACCCCACGTCTCGTCGCGCGCGCGTCCG</td><td>>hg38_21_+_8433172_8446622_RNA45SN1_45S_50ntFlanks</td><td>11312</td><td>11344</td><td>33</td><td>CCCCCACCCCACGTCTCGTCGCGCGCGCGTCCG</td><td>621.9</td><td>-13.1</td><td>1.5</td><td>-8.8</td><td>1.1E-18</td><td>8.5E-18</td><td>MB231/MB468</td><td>MITO</td></tr>
<tr><td>>MI0003129|hsa-mir-146b&WithFlank&10|+|102436506|102436590@15.38.24&[MIMAT0002809&hsa-miR-146b-5p&offsets|0|+1;m-70&10|+|102436520|102436543&offsets|0|0]|TGAGAACTGAATTCCATAGGCTGT</td><td>>MI0003129|hsa-mir-146b&WithFlank&10|+|102436506|102436590</td><td>15</td><td>38</td><td>24</td><td>TGAGAACTGAATTCCATAGGCTGT</td><td>582.2</td><td>6.0</td><td>0.7</td><td>8.8</td><td>1.4E-18</td><td>8.5E-18</td><td>BT20/MB468</td><td>CYTO</td></tr>
<tr><td>>hg38_21_+_8433172_8446622_RNA45SN1_45S_50ntFlanks@5490.5525.36|TGAGAAGACGGTCGAACTTGACTATCTAGAGGAAGT</td><td>>hg38_21_+_8433172_8446622_RNA45SN1_45S_50ntFlanks</td><td>5490</td><td>5525</td><td>36</td><td>TGAGAAGACGGTCGAACTTGACTATCTAGAGGAAGT</td><td>302.5</td><td>9.6</td><td>1.1</td><td>8.9</td><td>6.5E-19</td><td>8.6E-18</td><td>MB231/MB468</td><td>CYTO</td></tr>
<tr><td>>hg38_21_+_8433172_8446622_RNA45SN1_45S_50ntFlanks@12126.12150.25|CGACCCGCTCCGGGGACAGTGCCAG</td><td>>hg38_21_+_8433172_8446622_RNA45SN1_45S_50ntFlanks</td><td>12126</td><td>12150</td><td>25</td><td>CGACCCGCTCCGGGGACAGTGCCAG</td><td>1286.1</td><td>-12.4</td><td>1.4</td><td>-8.8</td><td>1.0E-18</td><td>8.6E-18</td><td>BT20/MB468</td><td>MITO</td></tr>
<tr><td>>hg38_1_-_228634819_228635039_RNA5S12_5S_50ntFlanks@130.169.40|TGGATGGGAGACCGCCTGGGAATACCGGGTGCTGTAGGCT</td><td>>hg38_1_-_228634819_228635039_RNA5S12_5S_50ntFlanks</td><td>130</td><td>169</td><td>40</td><td>TGGATGGGAGACCGCCTGGGAATACCGGGTGCTGTAGGCT</td><td>1081.4</td><td>8.4</td><td>0.9</td><td>8.9</td><td>6.5E-19</td><td>8.6E-18</td><td>MB231/MB468</td><td>CYTO</td></tr>
<tr><td>>MI0001445|hsa-mir-423&WithFlank&17|+|30117073|30117178@59.77.19(+1U)&[MIMAT0001340&hsa-miR-423-3p&offsets|0|-4(+1U);m-100&17|+|30117131|30117153&offsets|0|-4(+1U)]|AGCTCGGTCTGAGGCCCCTT</td><td>>MI0001445|hsa-mir-423&WithFlank&17|+|30117073|30117178</td><td>59</td><td>77</td><td>19(+1U)</td><td>AGCTCGGTCTGAGGCCCCTT</td><td>901.6</td><td>9.8</td><td>1.1</td><td>8.8</td><td>1.4E-18</td><td>8.6E-18</td><td>BT20/MB468</td><td>CYTO</td></tr>
<tr><td>>hg38_21_+_8433172_8446622_RNA45SN1_45S_50ntFlanks@5495.5516.22|AGACGGTCGAACTTGACTATCT</td><td>>hg38_21_+_8433172_8446622_RNA45SN1_45S_50ntFlanks</td><td>5495</td><td>5516</td><td>22</td><td>AGACGGTCGAACTTGACTATCT</td><td>2518.0</td><td>9.4</td><td>1.1</td><td>8.8</td><td>1.4E-18</td><td>8.6E-18</td><td>BT20/MB468</td><td>CYTO</td></tr>
<tr><td>>hg38_21_+_8433172_8446622_RNA45SN1_45S_50ntFlanks@12655.12673.19|ATGACTGAACGCCTCTAAG</td><td>>hg38_21_+_8433172_8446622_RNA45SN1_45S_50ntFlanks</td><td>12655</td><td>12673</td><td>19</td><td>ATGACTGAACGCCTCTAAG</td><td>502.9</td><td>-12.8</td><td>1.4</td><td>-8.9</td><td>6.8E-19</td><td>8.6E-18</td><td>MB231/MB468</td><td>MP</td></tr>
<tr><td>>hg38_21_+_8433172_8446622_RNA45SN1_45S_50ntFlanks@10781.10818.38|AGGTGAACAGCCTCTGGCATGTTGGAACAATGTAGGTA</td><td>>hg38_21_+_8433172_8446622_RNA45SN1_45S_50ntFlanks</td><td>10781</td><td>10818</td><td>38</td><td>AGGTGAACAGCCTCTGGCATGTTGGAACAATGTAGGTA</td><td>2512.1</td><td>-6.1</td><td>0.7</td><td>-8.9</td><td>6.6E-19</td><td>8.7E-18</td><td>MB231/MB468</td><td>CYTO</td></tr>
<tr><td>>hg38_21_+_8433172_8446622_RNA45SN1_45S_50ntFlanks@11212.11231.20|CGACTCTGGACGCGAGCCGG</td><td>>hg38_21_+_8433172_8446622_RNA45SN1_45S_50ntFlanks</td><td>11212</td><td>11231</td><td>20</td><td>CGACTCTGGACGCGAGCCGG</td><td>382.6</td><td>-12.4</td><td>1.4</td><td>-8.9</td><td>6.9E-19</td><td>8.7E-18</td><td>MB231/MB468</td><td>MP</td></tr>
<tr><td>>trna116_GluCTC_1_-_145399233_145399304@32.72.41,trna71_GluCTC_1_-_161439189_161439260@32.72.41,trna74_GluCTC_1_-_161431809_161431880@32.72.41,trna77_GluCTC_1_-_161424398_161424469@32.72.41,trna77_GluCTC_6_+_28949976_28950047@32.72.41,trna80_GluCTC_1_-_161417018_161417089@32.72.41,trna87_GluCTC_6_-_126101393_126101464@32.72.41|CTCTCACCGCCGCGGCCCGGGTTCGATTCCCGGTCAGGGAA</td><td>>trna116_GluCTC_1_-_145399233_145399304</td><td>32</td><td>72</td><td>41</td><td>CTCTCACCGCCGCGGCCCGGGTTCGATTCCCGGTCAGGGAA</td><td>772.1</td><td>-12.3</td><td>1.4</td><td>-8.9</td><td>6.7E-19</td><td>8.7E-18</td><td>MB231/MB468</td><td>CYTO</td></tr>
<tr><td>>hg38_21_+_8433172_8446622_RNA45SN1_45S_50ntFlanks@11199.11220.22|CCGGCGGCGGCGGCGACTCTGG</td><td>>hg38_21_+_8433172_8446622_RNA45SN1_45S_50ntFlanks</td><td>11199</td><td>11220</td><td>22</td><td>CCGGCGGCGGCGGCGACTCTGG</td><td>582.3</td><td>-13.0</td><td>1.5</td><td>-8.8</td><td>1.1E-18</td><td>8.8E-18</td><td>MB231/MB468</td><td>MITO</td></tr>
<tr><td>>hg38_21_+_8433172_8446622_RNA45SN1_45S_50ntFlanks@9504.9524.21|GTGAACTATGCCTGGGCAGGG</td><td>>hg38_21_+_8433172_8446622_RNA45SN1_45S_50ntFlanks</td><td>9504</td><td>9524</td><td>21</td><td>GTGAACTATGCCTGGGCAGGG</td><td>537.7</td><td>-12.0</td><td>1.4</td><td>-8.8</td><td>9.1E-19</td><td>8.8E-18</td><td>MB231/MB468</td><td>NUC</td></tr>
<tr><td>>hg38_21_+_8433172_8446622_RNA45SN1_45S_50ntFlanks@8403.8422.20|AGGTAAACGGGTGGGGTCCG</td><td>>hg38_21_+_8433172_8446622_RNA45SN1_45S_50ntFlanks</td><td>8403</td><td>8422</td><td>20</td><td>AGGTAAACGGGTGGGGTCCG</td><td>540.0</td><td>-12.9</td><td>1.5</td><td>-8.8</td><td>1.1E-18</td><td>8.8E-18</td><td>MB231/MB468</td><td>MITO</td></tr>
<tr><td>>hg38_21_+_8433172_8446622_RNA45SN1_45S_50ntFlanks@11576.11603.28|CTTAGAACTGGTGCGGACCAGGGGAATC</td><td>>hg38_21_+_8433172_8446622_RNA45SN1_45S_50ntFlanks</td><td>11576</td><td>11603</td><td>28</td><td>CTTAGAACTGGTGCGGACCAGGGGAATC</td><td>6280.0</td><td>-6.3</td><td>0.7</td><td>-8.9</td><td>6.7E-19</td><td>8.8E-18</td><td>MB231/MB468</td><td>CYTO</td></tr>
<tr><td>>MI0000446|hsa-mir-125b-1&WithFlank&11|-|122099751|122099850@21.43.23&[MIMAT0000423&hsa-miR-125b-5p&offsets|0|+1;m-67&11|-|122099809|122099830&offsets|0|+1]|TCCCTGAGACCCTAACTTGTGAT</td><td>>MI0000446|hsa-mir-125b-1&WithFlank&11|-|122099751|122099850</td><td>21</td><td>43</td><td>23</td><td>TCCCTGAGACCCTAACTTGTGAT</td><td>2743.2</td><td>14.6</td><td>1.7</td><td>8.8</td><td>1.1E-18</td><td>8.8E-18</td><td>MB231/MB468</td><td>MITO</td></tr>
<tr><td>>hg38_21_+_8433172_8446622_RNA45SN1_45S_50ntFlanks@4760.4778.19|TCAGATACCGTCGTAGTTC</td><td>>hg38_21_+_8433172_8446622_RNA45SN1_45S_50ntFlanks</td><td>4760</td><td>4778</td><td>19</td><td>TCAGATACCGTCGTAGTTC</td><td>710.9</td><td>-13.3</td><td>1.5</td><td>-8.8</td><td>1.1E-18</td><td>8.8E-18</td><td>MB231/MB468</td><td>MITO</td></tr>
<tr><td>>hg38_21_+_8433172_8446622_RNA45SN1_45S_50ntFlanks@9739.9757.19|AAACGATCTCAACCTATTC</td><td>>hg38_21_+_8433172_8446622_RNA45SN1_45S_50ntFlanks</td><td>9739</td><td>9757</td><td>19</td><td>AAACGATCTCAACCTATTC</td><td>723.9</td><td>-4.4</td><td>0.5</td><td>-8.9</td><td>6.8E-19</td><td>8.9E-18</td><td>MB231/MB468</td><td>CYTO</td></tr>
<tr><td>>hg38_21_+_8433172_8446622_RNA45SN1_45S_50ntFlanks@10351.10377.27|CGAGAACTTTGAAGGCCGAAGTGGAGA</td><td>>hg38_21_+_8433172_8446622_RNA45SN1_45S_50ntFlanks</td><td>10351</td><td>10377</td><td>27</td><td>CGAGAACTTTGAAGGCCGAAGTGGAGA</td><td>18513.0</td><td>-11.0</td><td>1.3</td><td>-8.7</td><td>2.5E-18</td><td>8.9E-18</td><td>BT20/MB468</td><td>NUC</td></tr>
</table>

| | | | | | | | | | | | | | |
|---|---|---|---|---|---|---|---|---|---|---|---|---|---|
| >MI0000446\|hsa-mir-125b-1&WithFlank&11\|-\|122099751\|122099850@22.42.21;MI0000470\|hsa-mir-125b-2&WithFlank&21\|+\|16590231\|16590331@24.44.21&[MIMAT0000423&hsa-miR-125b-5p&offsets\|+1\|0;m-67&11\|-\|122099809\|122099830&offsets\|+1\|0];[MIMAT0000423_1&hsa-miR-125b-5p&offsets\|+1\|0;m-68&21\|+\|16590253\|16590274&offsets\|+1\|0]\|CCCTGAGACCCTAACTTGTGA | >MI0000446\|hsa-mir-125b-1&WithFlank&11\|-\|122099751\|122099850 | 22 | 42 | 21 | CCCTGAGACCCTAACTTGTGA | 351.9 | 12.4 | 1.4 | 8.8 | 9.3E-19 | 8.9E-18 | MB231/MB468 | NUC |
| >hg38_21_+_8433172_8446622_RNA45SN1_45S_50ntFlanks@10391.10433.43\|AACAGCAGTTGAACATGGGTCAGTCGGTCCTGAGAGATGGGCG | >hg38_21_+_8433172_8446622_RNA45SN1_45S_50ntFlanks | 10391 | 10433 | 43 | AACAGCAGTTGAACATGGGTCAGTCGGTCCTGAGAGATGGGCG | 1266.3 | -14.7 | 1.7 | -8.7 | 2.5E-18 | 8.9E-18 | BT20/MB468 | NUC |
| >hg38_21_+_8433172_8446622_RNA45SN1_45S_50ntFlanks@6778.6802.25\|CCCGGGGCTACGCCTGTCTGAGCGT | >hg38_21_+_8433172_8446622_RNA45SN1_45S_50ntFlanks | 6778 | 6802 | 25 | CCCGGGGCTACGCCTGTCTGAGCGT | 3112.7 | 6.3 | 0.7 | 8.8 | 1.0E-18 | 8.9E-18 | BT20/MB468 | MITO |
| >hg38_21_+_8433172_8446622_RNA45SN1_45S_50ntFlanks@12033.12053.21\|ATCGTTTTTTCACTGACCCGG | >hg38_21_+_8433172_8446622_RNA45SN1_45S_50ntFlanks | 12033 | 12053 | 21 | ATCGTTTTTTCACTGACCCGG | 791.5 | -11.7 | 1.3 | -8.8 | 1.0E-18 | 8.9E-18 | BT20/MB468 | MITO |
| >hg38_21_+_8433172_8446622_RNA45SN1_45S_50ntFlanks@9623.9652.30\|TAGTAGCTGGTTCCCTCCGAAGTTTCCCTC | >hg38_21_+_8433172_8446622_RNA45SN1_45S_50ntFlanks | 9623 | 9652 | 30 | TAGTAGCTGGTTCCCTCCGAAGTTTCCCTC | 344.8 | -12.3 | 1.4 | -8.9 | 7.1E-19 | 9.0E-18 | MB231/MB468 | MP |
| >hg38_21_+_8433172_8446622_RNA45SN1_45S_50ntFlanks@10101.10126.26\|CGCGCGCGTGTGGTGTGCGTCGGAGG | >hg38_21_+_8433172_8446622_RNA45SN1_45S_50ntFlanks | 10101 | 10126 | 26 | CGCGCGCGTGTGGTGTGCGTCGGAGG | 681.0 | -13.3 | 1.5 | -8.9 | 7.1E-19 | 9.0E-18 | MB231/MB468 | MP |
| >hg38_21_+_8433172_8446622_RNA45SN1_45S_50ntFlanks@5472.5490.19\|ACGGCCCTGGCGGAGCGCT | >hg38_21_+_8433172_8446622_RNA45SN1_45S_50ntFlanks | 5472 | 5490 | 19 | ACGGCCCTGGCGGAGCGCT | 3449.9 | 8.7 | 1.0 | 8.8 | 1.4E-18 | 9.0E-18 | BT20/MB468 | CYTO |
| >hg38_21_+_8433172_8446622_RNA45SN1_45S_50ntFlanks@5487.5506.20\|CGCTGAGAAGACGGTCGAAC | >hg38_21_+_8433172_8446622_RNA45SN1_45S_50ntFlanks | 5487 | 5506 | 20 | CGCTGAGAAGACGGTCGAAC | 334.6 | -12.3 | 1.4 | -8.9 | 7.2E-19 | 9.1E-18 | MB231/MB468 | MP |
| >hg38_21_+_8433172_8446622_RNA45SN1_45S_50ntFlanks@6742.6771.30\|TCGACACTTCGAACGCACTTGCGGCCCCGG | >hg38_21_+_8433172_8446622_RNA45SN1_45S_50ntFlanks | 6742 | 6771 | 30 | TCGACACTTCGAACGCACTTGCGGCCCCGG | 420.7 | -12.6 | 1.4 | -8.9 | 7.3E-19 | 9.1E-18 | MB231/MB468 | MP |
| >hg38_21_+_8433172_8446622_RNA45SN1_45S_50ntFlanks@12959.12988.30\|TCTGGGTCGGGGTTTCGTACGTAGCAGAGC | >hg38_21_+_8433172_8446622_RNA45SN1_45S_50ntFlanks | 12959 | 12988 | 30 | TCTGGGTCGGGGTTTCGTACGTAGCAGAGC | 223.6 | 4.9 | 0.6 | 8.9 | 7.0E-19 | 9.1E-18 | MB231/MB468 | CYTO |
| >hg38_21_+_8433172_8446622_RNA45SN1_45S_50ntFlanks@10351.10368.18\|CGAGAACTTTGAAGGCCG | >hg38_21_+_8433172_8446622_RNA45SN1_45S_50ntFlanks | 10351 | 10368 | 18 | CGAGAACTTTGAAGGCCG | 17773.5 | -9.9 | 1.1 | -8.8 | 1.2E-18 | 9.2E-18 | MB231/MB468 | MITO |
| >hg38_21_+_8433172_8446622_RNA45SN1_45S_50ntFlanks@4449.4466.18\|CCTCTCGGCGCCCCCTCG | >hg38_21_+_8433172_8446622_RNA45SN1_45S_50ntFlanks | 4449 | 4466 | 18 | CCTCTCGGCGCCCCCTCG | 758.1 | -13.3 | 1.5 | -8.8 | 1.2E-18 | 9.2E-18 | MB231/MB468 | MITO |
| >hg38_21_+_8433172_8446622_RNA45SN1_45S_50ntFlanks@9203.9222.20\|CTCGGGGCCACGCGCGCGTC | >hg38_21_+_8433172_8446622_RNA45SN1_45S_50ntFlanks | 9203 | 9222 | 20 | CTCGGGGCCACGCGCGCGTC | 612.6 | -13.0 | 1.5 | -8.8 | 1.2E-18 | 9.2E-18 | MB231/MB468 | MITO |
| >hg38_21_+_8433172_8446622_RNA45SN1_45S_50ntFlanks@4003.4025.23\|ATAACCTCGGGCCGATCGCACGC | >hg38_21_+_8433172_8446622_RNA45SN1_45S_50ntFlanks | 4003 | 4025 | 23 | ATAACCTCGGGCCGATCGCACGC | 1904.4 | -10.8 | 1.2 | -8.8 | 1.2E-18 | 9.3E-18 | MB231/MB468 | MITO |
| >hg38_21_+_8433172_8446622_RNA45SN1_45S_50ntFlanks@5489.5513.25\|CTGAGAAGACGGTCGAACTTGACTA | >hg38_21_+_8433172_8446622_RNA45SN1_45S_50ntFlanks | 5489 | 5513 | 25 | CTGAGAAGACGGTCGAACTTGACTA | 815.7 | -8.0 | 0.9 | -8.9 | 7.4E-19 | 9.3E-18 | MB231/MB468 | MP |
| >hg38_21_+_8433172_8446622_RNA45SN1_45S_50ntFlanks@10781.10804.24\|AGGTGAACAGCCTCTGGCATGTTG | >hg38_21_+_8433172_8446622_RNA45SN1_45S_50ntFlanks | 10781 | 10804 | 24 | AGGTGAACAGCCTCTGGCATGTTG | 521.1 | -12.9 | 1.5 | -8.9 | 7.4E-19 | 9.3E-18 | MB231/MB468 | MP |
| >hg38_21_+_8433172_8446622_RNA45SN1_45S_50ntFlanks@12008.12038.31\|CCGCCGGTGAAATACCACTACTCTGATCGTT | >hg38_21_+_8433172_8446622_RNA45SN1_45S_50ntFlanks | 12008 | 12038 | 31 | CCGCCGGTGAAATACCACTACTCTGATCGTT | 610.3 | -13.1 | 1.5 | -8.9 | 7.4E-19 | 9.3E-18 | MB231/MB468 | MP |
| >hg38_21_+_8433172_8446622_RNA45SN1_45S_50ntFlanks@12479.12509.31\|CTAATAGGGAACGTGAGCTGGGTTTAGACCG | >hg38_21_+_8433172_8446622_RNA45SN1_45S_50ntFlanks | 12479 | 12509 | 31 | CTAATAGGGAACGTGAGCTGGGTTTAGACCG | 578.7 | -13.6 | 1.6 | -8.7 | 2.6E-18 | 9.3E-18 | BT20/MB468 | NUC |
| >hg38_21_+_8433172_8446622_RNA45SN1_45S_50ntFlanks@4726.4752.27\|TAATCAAGAACGAAAGTCGGAGGTTCG | >hg38_21_+_8433172_8446622_RNA45SN1_45S_50ntFlanks | 4726 | 4752 | 27 | TAATCAAGAACGAAAGTCGGAGGTTCG | 422.7 | -12.6 | 1.4 | -8.9 | 7.5E-19 | 9.3E-18 | MB231/MB468 | MP |
| >hg38_21_+_8433172_8446622_RNA45SN1_45S_50ntFlanks@6698.6719.22\|AGCTAGCTGCGAGAATTAATGT | >hg38_21_+_8433172_8446622_RNA45SN1_45S_50ntFlanks | 6698 | 6719 | 22 | AGCTAGCTGCGAGAATTAATGT | 691.4 | -4.1 | 0.5 | -8.8 | 1.5E-18 | 9.4E-18 | BT20/MB468 | CYTO |
| >hg38_21_+_8433172_8446622_RNA45SN1_45S_50ntFlanks@13004.13025.22\|ATCTATTGAAAGTCAGCCCTCG | >hg38_21_+_8433172_8446622_RNA45SN1_45S_50ntFlanks | 13004 | 13025 | 22 | ATCTATTGAAAGTCAGCCCTCG | 477.6 | -12.7 | 1.4 | -8.8 | 1.2E-18 | 9.4E-18 | MB231/MB468 | MITO |
| >hg38_21_+_8433172_8446622_RNA45SN1_45S_50ntFlanks@5467.5491.25\|GGCCCACGGCCCTGGCGGAGCGCTG | >hg38_21_+_8433172_8446622_RNA45SN1_45S_50ntFlanks | 5467 | 5491 | 25 | GGCCCACGGCCCTGGCGGAGCGCTG | 457.2 | -12.6 | 1.4 | -8.8 | 1.2E-18 | 9.4E-18 | MB231/MB468 | MITO |
| >trna116_GluCTC_1_-_145399233_145399304@32.71.40,trna71_GluCTC_1_-_161439189_161439260@32.71.40,trna74_GluCTC_1_-_161431809_161431880@32.71.40,trna77_GluCTC_1_-_161424398_161424469@32.71.40,trna77_GluCTC_6_+_28949976_28950047@32.71.40,trna80_GluCTC_1_-_161417018_161417089@32.71.40,trna87_GluCTC_6_-_126101393_126101464@32.71.40\|CTCTCACCGCCGCGGCCCGGGTTCGATTCCCGGTCAGGGA | >trna116_GluCTC_1_-_145399233_145399304 | 32 | 71 | 40 | CTCTCACCGCCGCGGCCCGGGTTCGATTCCCGGTCAGGGA | 1944.5 | -9.4 | 1.1 | -8.9 | 7.3E-19 | 9.5E-18 | MB231/MB468 | CYTO |
| >MI0005756\|hsa-mir-934&WithFlank&X\|+\|136550872\|136550966@21.41.21&[MIMAT0004977&hsa-miR-934&offsets\|0\|-1;m-943&X\|+\|136550892\|136550913&offsets\|0\|-1]\|TGTCTACTACTGGAGACACTG | >MI0005756\|hsa-mir-934&WithFlank&X\|+\|136550872\|136550966 | 21 | 41 | 21 | TGTCTACTACTGGAGACACTG | 843.3 | 13.1 | 1.4 | 9.1 | 6.8E-20 | 9.5E-18 | BT20/MB231 | CYTO |
| >hg38_21_+_8433172_8446622_RNA45SN1_45S_50ntFlanks@10866.10892.27\|CTAAGGGCTGGGTCGGTCGGGCTGGGG | >hg38_21_+_8433172_8446622_RNA45SN1_45S_50ntFlanks | 10866 | 10892 | 27 | CTAAGGGCTGGGTCGGTCGGGCTGGGG | 554.4 | -11.8 | 1.3 | -8.9 | 7.3E-19 | 9.5E-18 | MB231/MB468 | CYTO |
| >hg38_21_+_8433172_8446622_RNA45SN1_45S_50ntFlanks@11208.11239.32\|GCGGCGACTCTGGACGCGAGCCGGGCCCTTCC | >hg38_21_+_8433172_8446622_RNA45SN1_45S_50ntFlanks | 11208 | 11239 | 32 | GCGGCGACTCTGGACGCGAGCCGGGCCCTTCC | 662.1 | -7.5 | 0.8 | -8.9 | 7.3E-19 | 9.5E-18 | MB231/MB468 | CYTO |
| >hg38_21_+_8433172_8446622_RNA45SN1_45S_50ntFlanks@9575.9597.23\|ATCGGTCGTCCGACCTGGGTATA | >hg38_21_+_8433172_8446622_RNA45SN1_45S_50ntFlanks | 9575 | 9597 | 23 | ATCGGTCGTCCGACCTGGGTATA | 4436.6 | -15.2 | 1.6 | -9.3 | 1.6E-20 | 9.5E-18 | BT20/MB468 | MP |
| >hg38_21_+_8433172_8446622_RNA45SN1_45S_50ntFlanks@7976.8005.30\|GCGACCTCAGATCAGACGTGGCGACCCGCT | >hg38_21_+_8433172_8446622_RNA45SN1_45S_50ntFlanks | 7976 | 8005 | 30 | GCGACCTCAGATCAGACGTGGCGACCCGCT | 1494.6 | 7.0 | 0.8 | 8.8 | 1.1E-18 | 9.6E-18 | BT20/MB468 | MITO |
| >am_trnaMT_LysTTT_MT_+_8295_8364@31.52.22,trnalookalike7_LysTTT_1_+_568844_568913@31.52.22\|TAAGTTAAAGATTAAGAGAACC | >am_trnaMT_LysTTT_MT_+_8295_8364 | 31 | 52 | 22 | TAAGTTAAAGATTAAGAGAACC | 422.8 | 13.0 | 1.5 | 8.8 | 1.5E-18 | 9.6E-18 | BT20/MB468 | CYTO |
| >hg38_21_+_8433172_8446622_RNA45SN1_45S_50ntFlanks@10420.10456.37\|CTGAGAGATGGGCGAGCGCCGTTCCGAAGGGACGGGC | >hg38_21_+_8433172_8446622_RNA45SN1_45S_50ntFlanks | 10420 | 10456 | 37 | CTGAGAGATGGGCGAGCGCCGTTCCGAAGGGACGGGC | 4500.9 | -7.0 | 0.8 | -8.8 | 1.0E-18 | 9.7E-18 | MB231/MB468 | NUC |
| >hg38_21_+_8433172_8446622_RNA45SN1_45S_50ntFlanks@4759.4779.21\|ATCAGATACCGTCGTAGTTCC | >hg38_21_+_8433172_8446622_RNA45SN1_45S_50ntFlanks | 4759 | 4779 | 21 | ATCAGATACCGTCGTAGTTCC | 954.2 | -13.7 | 1.6 | -8.8 | 1.3E-18 | 9.7E-18 | MB231/MB468 | MITO |
| >hg38_21_+_8433172_8446622_RNA45SN1_45S_50ntFlanks@3813.3842.30\|TTATGGTTCCTTTGGTCGCTCGCTCCTCTC | >hg38_21_+_8433172_8446622_RNA45SN1_45S_50ntFlanks | 3813 | 3842 | 30 | TTATGGTTCCTTTGGTCGCTCGCTCCTCTC | 616.1 | -13.0 | 1.5 | -8.8 | 1.3E-18 | 9.7E-18 | MB231/MB468 | MITO |
| >hg38_21_+_8433172_8446622_RNA45SN1_45S_50ntFlanks@10466.10497.32\|CGTTGCCCTCGGCCGATCGAAAGGGAGTCGGG | >hg38_21_+_8433172_8446622_RNA45SN1_45S_50ntFlanks | 10466 | 10497 | 32 | CGTTGCCCTCGGCCGATCGAAAGGGAGTCGGG | 1713.5 | -10.2 | 1.2 | -8.8 | 1.3E-18 | 9.8E-18 | MB231/MB468 | MITO |
| >hg38_21_+_8433172_8446622_RNA45SN1_45S_50ntFlanks@3996.4026.31\|ACTCTAGATAACCTCGGGCCGATCGCACGCC | >hg38_21_+_8433172_8446622_RNA45SN1_45S_50ntFlanks | 3996 | 4026 | 31 | ACTCTAGATAACCTCGGGCCGATCGCACGCC | 615.3 | -13.7 | 1.6 | -8.7 | 2.8E-18 | 9.8E-18 | BT20/MB468 | NUC |
| >hg38_21_+_8433172_8446622_RNA45SN1_45S_50ntFlanks@5485.5522.38\|AGCGCTGAGAAGACGGTCGAACTTGACTATCTAGAGGA | >hg38_21_+_8433172_8446622_RNA45SN1_45S_50ntFlanks | 5485 | 5522 | 38 | AGCGCTGAGAAGACGGTCGAACTTGACTATCTAGAGGA | 483.4 | -12.7 | 1.4 | -8.8 | 1.3E-18 | 9.8E-18 | MB231/MB468 | MITO |
| >hg38_21_+_8433172_8446622_RNA45SN1_45S_50ntFlanks@9903.9928.26\|ATCAGACCCCAGAAAAGGTGTTGGTT | >hg38_21_+_8433172_8446622_RNA45SN1_45S_50ntFlanks | 9903 | 9928 | 26 | ATCAGACCCCAGAAAAGGTGTTGGTT | 456.8 | -13.3 | 1.5 | -8.7 | 2.8E-18 | 9.8E-18 | BT20/MB468 | NUC |
| >MI0006444\|hsa-mir-1307&WithFlank&10\|-\|103394247\|103394407@87.107.21(+1U)&[MIMAT0005951&hsa-miR-1307-3p&offsets\|+1\|0(+1U);m-66&10\|-\|103394300\|103394322&offsets\|+1\|-1(+1U)]\|CTCGGCGTGGCGTCGGTCGTGT | >MI0006444\|hsa-mir-1307&WithFlank&10\|-\|103394247\|103394407 | 87 | 107 | 21(+1U) | CTCGGCGTGGCGTCGGTCGTGT | 329.9 | 6.0 | 0.7 | 8.9 | 7.6E-19 | 9.8E-18 | MB231/MB468 | CYTO |
| >hg38_21_+_8433172_8446622_RNA45SN1_45S_50ntFlanks@11729.11753.25\|TAACTATGACTCTCTTAAGGTAGCC | >hg38_21_+_8433172_8446622_RNA45SN1_45S_50ntFlanks | 11729 | 11753 | 25 | TAACTATGACTCTCTTAAGGTAGCC | 2303.7 | -8.0 | 0.9 | -8.9 | 7.7E-19 | 9.9E-18 | MB231/MB468 | CYTO |
| >hg38_21_+_8433172_8446622_RNA45SN1_45S_50ntFlanks@4002.4020.19\|GATAACCTCGGGCCGATCG | >hg38_21_+_8433172_8446622_RNA45SN1_45S_50ntFlanks | 4002 | 4020 | 19 | GATAACCTCGGGCCGATCG | 773.9 | -13.4 | 1.5 | -8.8 | 1.3E-18 | 9.9E-18 | MB231/MB468 | MITO |
| >hg38_21_+_8433172_8446622_RNA45SN1_45S_50ntFlanks@6645.6676.32\|CTCGTACGACTCTTAGCGGTGGATCACTCGGC | >hg38_21_+_8433172_8446622_RNA45SN1_45S_50ntFlanks | 6645 | 6676 | 32 | CTCGTACGACTCTTAGCGGTGGATCACTCGGC | 4007.5 | 5.7 | 0.6 | 8.8 | 1.2E-18 | 9.9E-18 | BT20/MB468 | MITO |
| >hg38_21_+_8433172_8446622_RNA45SN1_45S_50ntFlanks@12934.12960.27\|CTAAACCATTCGTAGACGACCTGCTTC | >hg38_21_+_8433172_8446622_RNA45SN1_45S_50ntFlanks | 12934 | 12960 | 27 | CTAAACCATTCGTAGACGACCTGCTTC | 343.2 | -12.3 | 1.4 | -8.9 | 7.9E-19 | 9.9E-18 | MB231/MB468 | MP |
| >hg38_21_+_8433172_8446622_RNA45SN1_45S_50ntFlanks@4472.4497.26\|CTTAGCTGAGTGTCCCGCGGGGCCCG | >hg38_21_+_8433172_8446622_RNA45SN1_45S_50ntFlanks | 4472 | 4497 | 26 | CTTAGCTGAGTGTCCCGCGGGGCCCG | 731.7 | -13.4 | 1.5 | -8.9 | 8.0E-19 | 1.0E-17 | MB231/MB468 | MP |
| >hg38_21_+_8433172_8446622_RNA45SN1_45S_50ntFlanks@4367.4408.42\|CAGTTAAAAAGCTCGTAGTTGGATCTTGGGAGCGGGCGGGCG | >hg38_21_+_8433172_8446622_RNA45SN1_45S_50ntFlanks | 4367 | 4408 | 42 | CAGTTAAAAAGCTCGTAGTTGGATCTTGGGAGCGGGCGGGCG | 518.0 | -13.4 | 1.5 | -8.7 | 2.8E-18 | 1.0E-17 | BT20/MB468 | NUC |
| >hg38_21_+_8433172_8446622_RNA45SN1_45S_50ntFlanks@485.519.35\|GATGTGGTGACGTCGTGCTCTCCCGGGCCGGGTCC | >hg38_21_+_8433172_8446622_RNA45SN1_45S_50ntFlanks | 485 | 519 | 35 | GATGTGGTGACGTCGTGCTCTCCCGGGCCGGGTCC | 204.5 | 10.5 | 1.2 | 8.7 | 2.8E-18 | 1.0E-17 | BT20/MB468 | NUC |
| >MI0000269\|hsa-mir-181a-2&WithFlank&9\|+\|124692436\|124692557@83.104.22&[MIMAT0004558&hsa-miR-181a-2-3p&offsets\|0\|0;m-71&9\|+\|124692518\|124692539&offsets\|0\|0]\|ACCACTGACCGTTGACTGTACC | >MI0000269\|hsa-mir-181a-2&WithFlank&9\|+\|124692436\|124692557 | 83 | 104 | 22 | ACCACTGACCGTTGACTGTACC | 439.1 | 6.6 | 0.7 | 8.8 | 1.2E-18 | 1.0E-17 | BT20/MB468 | MITO |
| >hg38_21_+_8433172_8446622_RNA45SN1_45S_50ntFlanks@10839.10865.27\|TCCGTAACTTCGGGATAAGGATTGGCT | >hg38_21_+_8433172_8446622_RNA45SN1_45S_50ntFlanks | 10839 | 10865 | 27 | TCCGTAACTTCGGGATAAGGATTGGCT | 522.4 | -12.9 | 1.5 | -8.9 | 8.1E-19 | 1.0E-17 | MB231/MB468 | MP |
| >hg38_21_+_8433172_8446622_RNA45SN1_45S_50ntFlanks@12934.12959.26\|CTAAACCATTCGTAGACGACCTGCTT | >hg38_21_+_8433172_8446622_RNA45SN1_45S_50ntFlanks | 12934 | 12959 | 26 | CTAAACCATTCGTAGACGACCTGCTT | 3962.1 | -11.9 | 1.4 | -8.7 | 2.9E-18 | 1.0E-17 | BT20/MB468 | NUC |
| >hg38_21_+_8433172_8446622_RNA45SN1_45S_50ntFlanks@10362.10383.22\|AAGGCCGAAGTGGAGAAGGGTT | >hg38_21_+_8433172_8446622_RNA45SN1_45S_50ntFlanks | 10362 | 10383 | 22 | AAGGCCGAAGTGGAGAAGGGTT | 2368.1 | -8.5 | 1.0 | -8.8 | 1.3E-18 | 1.0E-17 | MB231/MB468 | MITO |
| >MI0000255\|hsa-mir-30d&WithFlank&8\|-\|134804870\|134804951@12.33.22(+1A)&[MIMAT0000245&hsa-miR-30d-5p&offsets\|0\|0(+1A);m-21&8\|-\|134804919\|134804940&offsets\|0\|0(+1A)]\|TGTAAACATCCCCGACTGGAAGA | >MI0000255\|hsa-mir-30d&WithFlank&8\|-\|134804870\|134804951 | 12 | 33 | 22(+1A) | TGTAAACATCCCCGACTGGAAGA | 683.0 | 5.2 | 0.6 | 8.8 | 1.1E-18 | 1.0E-17 | MB231/MB468 | NUC |

| | | | | | | | | | | | | | |
|---|---|---|---|---|---|---|---|---|---|---|---|---|---|
| >hg38_21_+_8433172_8446622_RNA45SN1_45S_50ntFlanks@11930.11950.21\|TAGAATAAGTGGGAGGCCCCC | >hg38_21_+_8433172_8446622_RNA45SN1_45S_50ntFlanks | 11930 | 11950 | 21 | TAGAATAAGTGGGAGGCCCCC | 554.6 | -13.0 | 1.5 | -8.9 | 8.2E-19 | 1.0E-17 | MB231/MB468 | MP |
| >hg38_21_+_8433172_8446622_RNA45SN1_45S_50ntFlanks@11561.11605.45\|GCGCCTAGCAGCCGACTTAGAACTGGTGCGGACCAGGGGAATCCG | >hg38_21_+_8433172_8446622_RNA45SN1_45S_50ntFlanks | 11561 | 11605 | 45 | GCGCCTAGCAGCCGACTTAGAACTGGTGCGGACCAGGGGAATCCG | 976.9 | -13.7 | 1.6 | -8.8 | 1.4E-18 | 1.0E-17 | MB231/MB468 | MITO |
| >hg38_21_+_8433172_8446622_RNA45SN1_45S_50ntFlanks@12991.13019.29\|CTCCCTCGCTGCGATCTATTGAAAGTCAG | >hg38_21_+_8433172_8446622_RNA45SN1_45S_50ntFlanks | 12991 | 13019 | 29 | CTCCCTCGCTGCGATCTATTGAAAGTCAG | 560.4 | -12.9 | 1.5 | -8.8 | 1.4E-18 | 1.0E-17 | MB231/MB468 | MITO |
| >hg38_21_+_8433172_8446622_RNA45SN1_45S_50ntFlanks@4002.4042.41\|GATAACCTCGGGCCGATCGCACGCCCCCCGTGGCGGCGACG | >hg38_21_+_8433172_8446622_RNA45SN1_45S_50ntFlanks | 4002 | 4042 | 41 | GATAACCTCGGGCCGATCGCACGCCCCCCGTGGCGGCGACG | 512.6 | -12.8 | 1.5 | -8.8 | 1.4E-18 | 1.0E-17 | MB231/MB468 | MITO |
| >hg38_21_+_8433172_8446622_RNA45SN1_45S_50ntFlanks@9033.9057.25\|CTCGTGGGGGGGCCGGGCCACCCCT | >hg38_21_+_8433172_8446622_RNA45SN1_45S_50ntFlanks | 9033 | 9057 | 25 | CTCGTGGGGGGGCCGGGCCACCCCT | 704.4 | -13.3 | 1.5 | -8.9 | 8.3E-19 | 1.0E-17 | MB231/MB468 | MP |
| >hg38_21_+_8433172_8446622_RNA45SN1_45S_50ntFlanks@3813.3836.24\|TTATGGTTCCTTTGGTCGCTCGCT | >hg38_21_+_8433172_8446622_RNA45SN1_45S_50ntFlanks | 3813 | 3836 | 24 | TTATGGTTCCTTTGGTCGCTCGCT | 893.6 | -5.2 | 0.6 | -8.7 | 2.9E-18 | 1.0E-17 | BT20/MB468 | NUC |
| >hg38_21_+_8433172_8446622_RNA45SN1_45S_50ntFlanks@12992.13015.24\|TCCCTCGCTGCGATCTATTGAAAG | >hg38_21_+_8433172_8446622_RNA45SN1_45S_50ntFlanks | 12992 | 13015 | 24 | TCCCTCGCTGCGATCTATTGAAAG | 1919.5 | -6.0 | 0.7 | -8.8 | 1.1E-18 | 1.0E-17 | MB231/MB468 | NUC |
| >hg38_21_+_8433172_8446622_RNA45SN1_45S_50ntFlanks@9554.9584.31\|CGTAGCGGTCCTGACGTGCAAATCGGTCGTC | >hg38_21_+_8433172_8446622_RNA45SN1_45S_50ntFlanks | 9554 | 9584 | 31 | CGTAGCGGTCCTGACGTGCAAATCGGTCGTC | 4538.0 | -5.6 | 0.6 | -8.8 | 1.1E-18 | 1.0E-17 | MB231/MB468 | NUC |
| >hg38_21_+_8433172_8446622_RNA45SN1_45S_50ntFlanks@10916.10936.21\|CCGCGGCTGGACGAGGCGCCG | >hg38_21_+_8433172_8446622_RNA45SN1_45S_50ntFlanks | 10916 | 10936 | 21 | CCGCGGCTGGACGAGGCGCCG | 432.2 | -12.6 | 1.4 | -8.9 | 8.4E-19 | 1.0E-17 | MB231/MB468 | MP |
| >hg38_21_+_8433172_8446622_RNA45SN1_45S_50ntFlanks@12164.12181.18\|CTGGGGCGGTACACCTGT | >hg38_21_+_8433172_8446622_RNA45SN1_45S_50ntFlanks | 12164 | 12181 | 18 | CTGGGGCGGTACACCTGT | 370.1 | -12.4 | 1.4 | -8.9 | 8.4E-19 | 1.0E-17 | MB231/MB468 | MP |
| >hg38_21_+_8433172_8446622_RNA45SN1_45S_50ntFlanks@9252.9274.23\|CACGGGGTCGGCGGCGACGTCGG | >hg38_21_+_8433172_8446622_RNA45SN1_45S_50ntFlanks | 9252 | 9274 | 23 | CACGGGGTCGGCGGCGACGTCGG | 559.9 | -13.0 | 1.5 | -8.9 | 8.4E-19 | 1.0E-17 | MB231/MB468 | MP |
| >hg38_21_+_8433172_8446622_RNA45SN1_45S_50ntFlanks@10794.10815.22\|CTGGCATGTTGGAACAATGTAG | >hg38_21_+_8433172_8446622_RNA45SN1_45S_50ntFlanks | 10794 | 10815 | 22 | CTGGCATGTTGGAACAATGTAG | 315.2 | -12.2 | 1.4 | -8.9 | 8.4E-19 | 1.0E-17 | MB231/MB468 | MP |
| >hg38_21_+_8433172_8446622_RNA45SN1_45S_50ntFlanks@11781.11804.24\|CGCATGAATGGATGAACGAGATTC | >hg38_21_+_8433172_8446622_RNA45SN1_45S_50ntFlanks | 11781 | 11804 | 24 | CGCATGAATGGATGAACGAGATTC | 14249.4 | -12.8 | 1.5 | -8.7 | 3.0E-18 | 1.0E-17 | BT20/MB468 | NUC |
| >hg38_21_+_8433172_8446622_RNA45SN1_45S_50ntFlanks@12973.13001.29\|TCGTACGTAGCAGAGCAGCTCCCTCGCTG | >hg38_21_+_8433172_8446622_RNA45SN1_45S_50ntFlanks | 12973 | 13001 | 29 | TCGTACGTAGCAGAGCAGCTCCCTCGCTG | 462.3 | -12.6 | 1.4 | -8.8 | 1.4E-18 | 1.1E-17 | MB231/MB468 | MITO |
| >hg38_21_+_8433172_8446622_RNA45SN1_45S_50ntFlanks@9552.9571.20\|TCCGTAGCGGTCCTGACGTG | >hg38_21_+_8433172_8446622_RNA45SN1_45S_50ntFlanks | 9552 | 9571 | 20 | TCCGTAGCGGTCCTGACGTG | 579.1 | -3.7 | 0.4 | -8.8 | 1.1E-18 | 1.1E-17 | MB231/MB468 | NUC |
| >hg38_21_+_8433172_8446622_RNA45SN1_45S_50ntFlanks@11161.11189.29\|CACGGTCCCCCGCGAGGGGGGCCCGGGCA | >hg38_21_+_8433172_8446622_RNA45SN1_45S_50ntFlanks | 11161 | 11189 | 29 | CACGGTCCCCCGCGAGGGGGGCCCGGGCA | 642.1 | -13.8 | 1.6 | -8.7 | 3.0E-18 | 1.1E-17 | BT20/MB468 | NUC |
| >hg38_21_+_8433172_8446622_RNA45SN1_45S_50ntFlanks@10350.10384.35\|ACGAGAACTTTGAAGGCCGAAGTGGAGAAGGGTTC | >hg38_21_+_8433172_8446622_RNA45SN1_45S_50ntFlanks | 10350 | 10384 | 35 | ACGAGAACTTTGAAGGCCGAAGTGGAGAAGGGTTC | 9496.4 | -10.6 | 1.2 | -8.7 | 3.0E-18 | 1.1E-17 | BT20/MB468 | NUC |
| >hg38_21_+_8433172_8446622_RNA45SN1_45S_50ntFlanks@12933.12956.24\|GCTAAACCATTCGTAGACGACCTG | >hg38_21_+_8433172_8446622_RNA45SN1_45S_50ntFlanks | 12933 | 12956 | 24 | GCTAAACCATTCGTAGACGACCTG | 504.5 | -12.8 | 1.5 | -8.9 | 8.7E-19 | 1.1E-17 | MB231/MB468 | MP |
| >hg38_21_+_8433172_8446622_RNA45SN1_45S_50ntFlanks@6667.6689.23\|ATCACTCGGCTCGTGCGTCGATG | >hg38_21_+_8433172_8446622_RNA45SN1_45S_50ntFlanks | 6667 | 6689 | 23 | ATCACTCGGCTCGTGCGTCGATG | 845.2 | -13.6 | 1.5 | -8.9 | 8.7E-19 | 1.1E-17 | MB231/MB468 | MP |
| >hg38_21_+_8433172_8446622_RNA45SN1_45S_50ntFlanks@4486.4511.26\|CCGCGGGGCCCGAAGCGTTTACTTTG | >hg38_21_+_8433172_8446622_RNA45SN1_45S_50ntFlanks | 4486 | 4511 | 26 | CCGCGGGGCCCGAAGCGTTTACTTTG | 1076.1 | -14.5 | 1.7 | -8.7 | 3.0E-18 | 1.1E-17 | BT20/MB468 | NUC |
| >trnaMT_GluTTC_MT_-_14674_14742@38.72.35,trnalookalike8_GluTTC_5_-_93905172_93905240@38.72.35\|CATTGGTCGTGGTTGTAGTCCGTGCGAGAATACCA | >trnaMT_GluTTC_MT_-_14674_14742 | 38 | 72 | 35 | CATTGGTCGTGGTTGTAGTCCGTGCGAGAATACCA | 1117.0 | 8.5 | 1.0 | 8.8 | 1.7E-18 | 1.1E-17 | BT20/MB468 | CYTO |
| >hg38_21_+_8433172_8446622_RNA45SN1_45S_50ntFlanks@11318.11338.21\|CCCCACGTCTCGTCGCGCGCG | >hg38_21_+_8433172_8446622_RNA45SN1_45S_50ntFlanks | 11318 | 11338 | 21 | CCCCACGTCTCGTCGCGCGCG | 859.4 | -9.5 | 1.1 | -8.9 | 8.4E-19 | 1.1E-17 | MB231/MB468 | CYTO |
| >hg38_21_+_8433172_8446622_RNA45SN1_45S_50ntFlanks@10404.10430.27\|CATGGGTCAGTCGGTCCTGAGAGATGG | >hg38_21_+_8433172_8446622_RNA45SN1_45S_50ntFlanks | 10404 | 10430 | 27 | CATGGGTCAGTCGGTCCTGAGAGATGG | 467.9 | -12.7 | 1.4 | -8.8 | 8.8E-19 | 1.1E-17 | MB231/MB468 | MP |
| >hg38_21_+_8433172_8446622_RNA45SN1_45S_50ntFlanks@11669.11703.35\|GCCCAGTGCTCTGAATGTCAAAGTGAAGAAATTCA | >hg38_21_+_8433172_8446622_RNA45SN1_45S_50ntFlanks | 11669 | 11703 | 35 | GCCCAGTGCTCTGAATGTCAAAGTGAAGAAATTCA | 1474.9 | 6.3 | 0.7 | 8.9 | 8.5E-19 | 1.1E-17 | MB231/MB468 | CYTO |
| >hg38_21_+_8433172_8446622_RNA45SN1_45S_50ntFlanks@11620.11640.21\|ACAAAGCATCGCGAAGGCCCG | >hg38_21_+_8433172_8446622_RNA45SN1_45S_50ntFlanks | 11620 | 11640 | 21 | ACAAAGCATCGCGAAGGCCCG | 1584.2 | -12.7 | 1.4 | -8.8 | 1.3E-18 | 1.1E-17 | BT20/MB468 | MITO |
| >hg38_21_+_8433172_8446622_RNA45SN1_45S_50ntFlanks@11768.11791.24\|CTAATTAGTGACGCGCATGAATGG | >hg38_21_+_8433172_8446622_RNA45SN1_45S_50ntFlanks | 11768 | 11791 | 24 | CTAATTAGTGACGCGCATGAATGG | 376.2 | -12.4 | 1.4 | -8.8 | 8.8E-19 | 1.1E-17 | MB231/MB468 | MP |
| >hg38_21_+_8433172_8446622_RNA45SN1_45S_50ntFlanks@11233.11254.22\|CCCTTCCCGTGGATCGCCCCAG | >hg38_21_+_8433172_8446622_RNA45SN1_45S_50ntFlanks | 11233 | 11254 | 22 | CCCTTCCCGTGGATCGCCCCAG | 1615.9 | -9.9 | 1.1 | -8.9 | 8.5E-19 | 1.1E-17 | MB231/MB468 | CYTO |
| >hg38_21_+_8433172_8446622_RNA45SN1_45S_50ntFlanks@6647.6669.23\|CGTACGACTCTTAGCGGTGGATC | >hg38_21_+_8433172_8446622_RNA45SN1_45S_50ntFlanks | 6647 | 6669 | 23 | CGTACGACTCTTAGCGGTGGATC | 28425.1 | 4.4 | 0.5 | 8.8 | 1.3E-18 | 1.1E-17 | BT20/MB468 | MITO |
| >hg38_21_+_8433172_8446622_RNA45SN1_45S_50ntFlanks@9338.9370.33\|TCGCACGAAAGCCGCCGTGGCGCAATGAAGGTG | >hg38_21_+_8433172_8446622_RNA45SN1_45S_50ntFlanks | 9338 | 9370 | 33 | TCGCACGAAAGCCGCCGTGGCGCAATGAAGGTG | 716.3 | -13.3 | 1.5 | -8.8 | 8.8E-19 | 1.1E-17 | MB231/MB468 | MP |
| >hg38_21_+_8433172_8446622_RNA45SN1_45S_50ntFlanks@10410.10430.21\|TCAGTCGGTCCTGAGAGATGG | >hg38_21_+_8433172_8446622_RNA45SN1_45S_50ntFlanks | 10410 | 10430 | 21 | TCAGTCGGTCCTGAGAGATGG | 2836.3 | -10.6 | 1.2 | -8.8 | 1.4E-18 | 1.1E-17 | MB231/MB468 | MITO |
| >hg38_21_+_8433172_8446622_RNA45SN1_45S_50ntFlanks@3921.3959.39\|ATTTATCAGATCAAAACCAACCCGGTCAGCCCCTCTCCG | >hg38_21_+_8433172_8446622_RNA45SN1_45S_50ntFlanks | 3921 | 3959 | 39 | ATTTATCAGATCAAAACCAACCCGGTCAGCCCCTCTCCG | 733.7 | -13.9 | 1.6 | -8.7 | 3.1E-18 | 1.1E-17 | BT20/MB468 | NUC |
| >MI0000542\|hsa-mir-320a&WithFlank&8\|-\|22244960\|22245043@48.70.23(+1G)&[MIMAT0000510&hsa-miR-320a-3p&offsets\|0\|+1(+1G);m-60&8\|-\|22244975\|22244996&offsets\|0\|+1(+1G)]\|AAAAGCTGGGTTGAGAGGGCGAAG | >MI0000542\|hsa-mir-320a&WithFlank&8\|-\|22244960\|22245043 | 48 | 70 | 23(+1G) | AAAAGCTGGGTTGAGAGGGCGAAG | 135.4 | 3.6 | 0.4 | 8.7 | 3.1E-18 | 1.1E-17 | BT20/MB468 | NUC |
| >hg38_21_+_8433172_8446622_RNA45SN1_45S_50ntFlanks@12957.12990.34\|CTTCTGGGTCGGGGTTTCGTACGTAGCAGAGCAG | >hg38_21_+_8433172_8446622_RNA45SN1_45S_50ntFlanks | 12957 | 12990 | 34 | CTTCTGGGTCGGGGTTTCGTACGTAGCAGAGCAG | 4523.4 | -12.1 | 1.4 | -8.7 | 3.2E-18 | 1.1E-17 | BT20/MB468 | NUC |
| >MI0000477\|hsa-mir-146a&WithFlank&5\|+\|160485346\|160485456@27.47.21(+1C)&[MIMAT0000449&hsa-miR-146a-5p&offsets\|0\|-1(+1C);m-129&5\|+\|160485372\|160485393&offsets\|0\|-1(+1C)]\|TGAGAACTGAATTCCATGGGTC | >MI0000477\|hsa-mir-146a&WithFlank&5\|+\|160485346\|160485456 | 27 | 47 | 21(+1C) | TGAGAACTGAATTCCATGGGTC | 1488.7 | 14.5 | 1.6 | 8.9 | 8.6E-19 | 1.1E-17 | MB231/MB468 | CYTO |
| >MI0000088\|hsa-mir-30a&WithFlank&6\|-\|71403545\|71403627@53.73.21(+1U)&[MIMAT0000088&hsa-miR-30a-3p&offsets\|0\|-1(+1U);m-38&6\|-\|71403554\|71403575&offsets\|0\|-1(+1U)]\|CTTTCAGTCGGATGTTTGCAGT | >MI0000088\|hsa-mir-30a&WithFlank&6\|-\|71403545\|71403627 | 53 | 73 | 21(+1U) | CTTTCAGTCGGATGTTTGCAGT | 9337.5 | 7.8 | 0.9 | 8.9 | 8.7E-19 | 1.1E-17 | MB231/MB468 | CYTO |
| >hg38_21_+_8433172_8446622_RNA45SN1_45S_50ntFlanks@11784.11804.21\|ATGAATGGATGAACGAGATTC | >hg38_21_+_8433172_8446622_RNA45SN1_45S_50ntFlanks | 11784 | 11804 | 21 | ATGAATGGATGAACGAGATTC | 21661.0 | -11.0 | 1.3 | -8.7 | 3.2E-18 | 1.1E-17 | BT20/MB468 | NUC |
| >hg38_21_+_8433172_8446622_RNA45SN1_45S_50ntFlanks@12605.12640.36\|TATGTGCTTGGCTGAGGAGCCAATGGGGCGAAGCTA | >hg38_21_+_8433172_8446622_RNA45SN1_45S_50ntFlanks | 12605 | 12640 | 36 | TATGTGCTTGGCTGAGGAGCCAATGGGGCGAAGCTA | 319.0 | -12.7 | 1.5 | -8.7 | 3.2E-18 | 1.1E-17 | BT20/MB468 | NUC |
| >hg38_21_+_8433172_8446622_RNA45SN1_45S_50ntFlanks@10463.10497.35\|CTCCGTTGCCCTCGGCCGATCGAAAGGGAGTCGGG | >hg38_21_+_8433172_8446622_RNA45SN1_45S_50ntFlanks | 10463 | 10497 | 35 | CTCCGTTGCCCTCGGCCGATCGAAAGGGAGTCGGG | 500.6 | -12.8 | 1.5 | -8.8 | 1.5E-18 | 1.1E-17 | MB231/MB468 | MITO |
| >hg38_21_+_8433172_8446622_RNA45SN1_45S_50ntFlanks@6699.6720.22\|GCTAGCTGCGAGAATTAATGTG | >hg38_21_+_8433172_8446622_RNA45SN1_45S_50ntFlanks | 6699 | 6720 | 22 | GCTAGCTGCGAGAATTAATGTG | 35859.4 | -8.3 | 0.9 | -8.8 | 1.5E-18 | 1.1E-17 | MB231/MB468 | MITO |
| >hg38_21_+_8433172_8446622_RNA45SN1_45S_50ntFlanks@9167.9197.31\|CGGGCGGGTCGCGCCGTCGGGCCCGGGGGAG | >hg38_21_+_8433172_8446622_RNA45SN1_45S_50ntFlanks | 9167 | 9197 | 31 | CGGGCGGGTCGCGCCGTCGGGCCCGGGGGAG | 2034.4 | -14.8 | 1.7 | -8.8 | 1.5E-18 | 1.1E-17 | MB231/MB468 | MITO |
| >hg38_21_+_8433172_8446622_RNA45SN1_45S_50ntFlanks@7986.8007.22\|ATCAGACGTGGCGACCCGCTGA | >hg38_21_+_8433172_8446622_RNA45SN1_45S_50ntFlanks | 7986 | 8007 | 22 | ATCAGACGTGGCGACCCGCTGA | 972.8 | -13.8 | 1.6 | -8.8 | 9.1E-19 | 1.1E-17 | MB231/MB468 | MP |
| >MI0000746\|hsa-mir-99b&WithFlank&19\|+\|51692606\|51692687@14.34.21&[MIMAT0000689&hsa-miR-99b-5p&offsets\|+1\|0;m-10&19\|+\|51692618\|51692639&offsets\|+1\|0]\|ACCCGTAGAACCGACCTTGCG | >MI0000746\|hsa-mir-99b&WithFlank&19\|+\|51692606\|51692687 | 14 | 34 | 21 | ACCCGTAGAACCGACCTTGCG | 956.6 | 8.7 | 1.0 | 8.8 | 1.8E-18 | 1.1E-17 | BT20/MB468 | CYTO |
| >hg38_21_+_8433172_8446622_RNA45SN1_45S_50ntFlanks@6646.6668.23\|TCGTACGACTCTTAGCGGTGGAT | >hg38_21_+_8433172_8446622_RNA45SN1_45S_50ntFlanks | 6646 | 6668 | 23 | TCGTACGACTCTTAGCGGTGGAT | 60008.9 | 6.5 | 0.7 | 8.8 | 1.3E-18 | 1.1E-17 | BT20/MB468 | MITO |
| >hg38_21_+_8433172_8446622_RNA45SN1_45S_50ntFlanks@10391.10408.18\|AACAGCAGTTGAACATGG | >hg38_21_+_8433172_8446622_RNA45SN1_45S_50ntFlanks | 10391 | 10408 | 18 | AACAGCAGTTGAACATGG | 592.5 | -13.1 | 1.5 | -8.8 | 9.3E-19 | 1.1E-17 | MB231/MB468 | MP |
| >MI0003137\|hsa-mir-193b&WithFlank&16\|+\|14303961\|14304055@57.79.23&[MIMAT0002819&hsa-miR-193b-3p&offsets\|0\|+1;m-166&16\|+\|14304017\|14304038&offsets\|0\|+1]\|AACTGGCCCTCAAAGTCCCGCTT | >MI0003137\|hsa-mir-193b&WithFlank&16\|+\|14303961\|14304055 | 57 | 79 | 23 | AACTGGCCCTCAAAGTCCCGCTT | 315.0 | 12.6 | 1.4 | 8.8 | 1.8E-18 | 1.1E-17 | BT20/MB468 | CYTO |
| >am_MI0000786\|hsa-mir-378a&WithFlank&5\|+\|149732819\|149732896@49.70.22&[MIMAT0000732&hsa-miR-378a-3p&offsets\|0\|0;m-44&5\|+\|149732867\|149732888&offsets\|0\|0]\|ACTGGACTTGGAGTCAGAAGGC | >am_MI0000786\|hsa-mir-378a&WithFlank&5\|+\|149732819\|149732896 | 49 | 70 | 22 | ACTGGACTTGGAGTCAGAAGGC | 13822.7 | 5.5 | 0.6 | 8.8 | 1.4E-18 | 1.1E-17 | BT20/MB468 | MITO |
| >hg38_21_+_8433172_8446622_RNA45SN1_45S_50ntFlanks@10679.10705.27\|GGTTCCGGCGGCGTCCGGTGAGCTCTC | >hg38_21_+_8433172_8446622_RNA45SN1_45S_50ntFlanks | 10679 | 10705 | 27 | GGTTCCGGCGGCGTCCGGTGAGCTCTC | 552.8 | -7.5 | 0.8 | -8.8 | 9.0E-19 | 1.1E-17 | MB231/MB468 | CYTO |
| >MI0000279\|hsa-mir-196a-2&WithFlank&12\|+\|53991732\|53991853@31.51.21(+1A);MI0000238\|hsa-mir-196a-1&WithFlank&17\|-\|48632484\|48632565@13.33.21(+1A)&[MIMAT0000226&hsa-miR-196a-5p&offsets\|0\|-1(+1A);m-261&12\|+\|53991762\|53991783&offsets\|0\|-1(+1A)];[MIMAT0000226_1&hsa-miR-196a-5p&offsets\|0\|-1(+1A);m-268&17\|-\|48632532\|48632553&offsets\|0\|-1(+1A)]\|TAGGTAGTTTCATGTTGTTGGA | >MI0000279\|hsa-mir-196a-2&WithFlank&12\|+\|53991732\|53991853 | 31 | 51 | 21(+1A) | TAGGTAGTTTCATGTTGTTGGA | 1051.2 | 10.8 | 1.2 | 9.1 | 8.8E-20 | 1.1E-17 | BT20/MB231 | CYTO |

| | | | | | | | | | | | | | |
|---|---|---|---|---|---|---|---|---|---|---|---|---|---|
| >MI0000283\|hsa-mir-203a&WithFlank&14\|+\|104117399\|104117520@72.90.19&[MIMAT0000264&hsa-miR-203a-3p&offsets\|+1\|-2;m-79&14\|+\|104117469\|104117490&offsets\|+1\|-2]\|TGAAATGTTTAGGACCACT | >MI0000283\|hsa-mir-203a&WithFlank&14\|+\|104117399\|104117520 | 72 | 90 | 19 | TGAAATGTTTAGGACCACT | 720.6 | 9.9 | 1.1 | 9.1 | 8.7E-20 | 1.1E-17 | BT20/MB231 | CYTO |
| >hg38_21_+_8433172_8446622_RNA45SN1_45S_50ntFlanks@11316.11338.23\|CACCCCACGTCTCGTCGCGCGCG | >hg38_21_+_8433172_8446622_RNA45SN1_45S_50ntFlanks | 11316 | 11338 | 23 | CACCCCACGTCTCGTCGCGCGCG | 491.8 | -12.8 | 1.4 | -8.8 | 9.3E-19 | 1.2E-17 | MB231/MB468 | MP |
| >hg38_21_+_8433172_8446622_RNA45SN1_45S_50ntFlanks@4099.4121.23\|GTGACCACGGGTGACGGGGAATC | >hg38_21_+_8433172_8446622_RNA45SN1_45S_50ntFlanks | 4099 | 4121 | 23 | GTGACCACGGGTGACGGGGAATC | 517.3 | -12.9 | 1.5 | -8.8 | 9.4E-19 | 1.2E-17 | MB231/MB468 | MP |
| >hg38_21_+_8433172_8446622_RNA45SN1_45S_50ntFlanks@11566.11584.19\|TAGCAGCCGACTTAGAACT | >hg38_21_+_8433172_8446622_RNA45SN1_45S_50ntFlanks | 11566 | 11584 | 19 | TAGCAGCCGACTTAGAACT | 1065.0 | 6.3 | 0.7 | 8.8 | 1.9E-18 | 1.2E-17 | BT20/MB468 | CYTO |
| >MI0000102\|hsa-mir-100&WithFlank&11\|-\|122152223\|122152314@21.40.20&[MIMAT0000098&hsa-miR-100-5p&offsets\|+2\|0;m-27&11\|-\|122152276\|122152296&offsets\|+2\|+1]\|CCCGTAGATCCGAACTTGTG | >MI0000102\|hsa-mir-100&WithFlank&11\|-\|122152223\|122152314 | 21 | 40 | 20 | CCCGTAGATCCGAACTTGTG | 316.7 | 12.3 | 1.4 | 8.8 | 9.2E-19 | 1.2E-17 | MB231/MB468 | CYTO |
| >MI0000650\|hsa-mir-200c&WithFlank&12\|+\|6963693\|6963772@50.72.23(+1A)&[MIMAT0000617&hsa-miR-200c-3p&offsets\|0\|0(+1A);m-54&12\|+\|6963742\|6963764&offsets\|0\|0(+1A)]\|TAATACTGCCGGGTAATGATGGAA | >MI0000650\|hsa-mir-200c&WithFlank&12\|+\|6963693\|6963772 | 50 | 72 | 23(+1A) | TAATACTGCCGGGTAATGATGGAA | 3285.2 | 15.0 | 1.6 | 9.3 | 1.6E-20 | 1.2E-17 | BT20/MB231 | MP |
| >hg38_21_+_8433172_8446622_RNA45SN1_45S_50ntFlanks@5490.5516.27\|TGAGAAGACGGTCGAACTTGACTATCT | >hg38_21_+_8433172_8446622_RNA45SN1_45S_50ntFlanks | 5490 | 5516 | 27 | TGAGAAGACGGTCGAACTTGACTATCT | 6632.2 | 9.2 | 1.0 | 8.8 | 9.2E-19 | 1.2E-17 | MB231/MB468 | CYTO |
| >hg38_21_+_8433172_8446622_RNA45SN1_45S_50ntFlanks@8179.8223.45\|CTGATCGAGGCCCAGCCCGTGGACGGTGTGAGGCCGGTAGCGGCC | >hg38_21_+_8433172_8446622_RNA45SN1_45S_50ntFlanks | 8179 | 8223 | 45 | CTGATCGAGGCCCAGCCCGTGGACGGTGTGAGGCCGGTAGCGGCC | 421.9 | -13.1 | 1.5 | -8.7 | 3.3E-18 | 1.2E-17 | BT20/MB468 | NUC |
| >hg38_21_+_8433172_8446622_RNA45SN1_45S_50ntFlanks@11666.11709.44\|TCTGCCCAGTGCTCTGAATGTCAAAGTGAAGAAATTCAATGAAG | >hg38_21_+_8433172_8446622_RNA45SN1_45S_50ntFlanks | 11666 | 11709 | 44 | TCTGCCCAGTGCTCTGAATGTCAAAGTGAAGAAATTCAATGAAG | 310.1 | 12.3 | 1.4 | 8.8 | 9.3E-19 | 1.2E-17 | MB231/MB468 | CYTO |
| >MI0000286\|hsa-mir-210&WithFlank&11\|-\|568083\|568204@72.93.22(+3A)&[MIMAT0000267&hsa-miR-210-3p&offsets\|0\|0(+3A);m-102&11\|-\|568112\|568133&offsets\|0\|0(+3A)]\|CTGTGCGTGTGACAGCGGCTGAAAA | >MI0000286\|hsa-mir-210&WithFlank&11\|-\|568083\|568204 | 72 | 93 | 22(+3A) | CTGTGCGTGTGACAGCGGCTGAAAA | 519.4 | 13.5 | 1.5 | 8.8 | 1.4E-18 | 1.2E-17 | BT20/MB468 | MITO |
| >hg38_21_+_8433172_8446622_RNA45SN1_45S_50ntFlanks@7986.8016.31\|ATCAGACGTGGCGACCCGCTGAATTTAAGCA | >hg38_21_+_8433172_8446622_RNA45SN1_45S_50ntFlanks | 7986 | 8016 | 31 | ATCAGACGTGGCGACCCGCTGAATTTAAGCA | 437.2 | -13.2 | 1.5 | -8.7 | 3.4E-18 | 1.2E-17 | BT20/MB468 | NUC |
| >hg38_21_+_8433172_8446622_RNA45SN1_45S_50ntFlanks@7977.8007.31\|CGACCTCAGATCAGACGTGGCGACCCGCTGA | >hg38_21_+_8433172_8446622_RNA45SN1_45S_50ntFlanks | 7977 | 8007 | 31 | CGACCTCAGATCAGACGTGGCGACCCGCTGA | 315.3 | -3.9 | 0.5 | -8.7 | 3.4E-18 | 1.2E-17 | BT20/MB468 | NUC |
| >MI0000802\|hsa-mir-340&WithFlank&5\|-\|180015297\|180015403@22.43.22&[MIMAT0004692&hsa-miR-340-5p&offsets\|0\|0;m-220&5\|-\|180015361\|180015382&offsets\|0\|0]\|TTATAAAGCAATGAGACTGATT | >MI0000802\|hsa-mir-340&WithFlank&5\|-\|180015297\|180015403 | 22 | 43 | 22 | TTATAAAGCAATGAGACTGATT | 838.5 | 4.8 | 0.5 | 8.8 | 1.4E-18 | 1.2E-17 | BT20/MB468 | MITO |
| >hg38_21_+_8433172_8446622_RNA45SN1_45S_50ntFlanks@10680.10711.32\|GTTCCGGCGGCGTCCGGTGAGCTCTCGCTGGC | >hg38_21_+_8433172_8446622_RNA45SN1_45S_50ntFlanks | 10680 | 10711 | 32 | GTTCCGGCGGCGTCCGGTGAGCTCTCGCTGGC | 3673.7 | -11.1 | 1.3 | -8.8 | 9.7E-19 | 1.2E-17 | MB231/MB468 | MP |
| >MI0000471\|hsa-mir-126&WithFlank&9\|+\|136670596\|136670692@21.41.21&[MIMAT0000444&hsa-miR-126-5p&offsets\|0\|0;m-114&9\|+\|136670616\|136670636&offsets\|0\|0]\|CATTATTACTTTTGGTACGCG | >MI0000471\|hsa-mir-126&WithFlank&9\|+\|136670596\|136670692 | 21 | 41 | 21 | CATTATTACTTTTGGTACGCG | 406.4 | 12.9 | 1.5 | 8.8 | 1.9E-18 | 1.2E-17 | BT20/MB468 | CYTO |
| >hg38_21_+_8433172_8446622_RNA45SN1_45S_50ntFlanks@12539.12565.27\|ATGATGTGTTGTTGCCATGGTAATCCT | >hg38_21_+_8433172_8446622_RNA45SN1_45S_50ntFlanks | 12539 | 12565 | 27 | ATGATGTGTTGTTGCCATGGTAATCCT | 452.3 | -12.7 | 1.4 | -8.8 | 9.7E-19 | 1.2E-17 | MB231/MB468 | MP |
| >hg38_21_+_8433172_8446622_RNA45SN1_45S_50ntFlanks@12324.12348.25\|TTGGGTTTTAAGCAGGAGGTGTCAG | >hg38_21_+_8433172_8446622_RNA45SN1_45S_50ntFlanks | 12324 | 12348 | 25 | TTGGGTTTTAAGCAGGAGGTGTCAG | 524.2 | -12.8 | 1.5 | -8.8 | 1.6E-18 | 1.2E-17 | MB231/MB468 | MITO |
| >MI0000477\|hsa-mir-146a&WithFlank&5\|+\|160485346\|160485456@27.48.22(+1U)&[MIMAT0000449&hsa-miR-146a-5p&offsets\|0\|0(+1U);m-129&5\|+\|160485372\|160485393&offsets\|0\|0(+1U)]\|TGAGAACTGAATTCCATGGGTTT | >MI0000477\|hsa-mir-146a&WithFlank&5\|+\|160485346\|160485456 | 27 | 48 | 22(+1U) | TGAGAACTGAATTCCATGGGTTT | 1249.7 | 14.2 | 1.6 | 8.8 | 1.3E-18 | 1.2E-17 | MB231/MB468 | NUC |
| >hg38_21_+_8433172_8446622_RNA45SN1_45S_50ntFlanks@12615.12654.40\|GCTGAGGAGCCAATGGGGCGAAGCTACCATCTGTGGGATT | >hg38_21_+_8433172_8446622_RNA45SN1_45S_50ntFlanks | 12615 | 12654 | 40 | GCTGAGGAGCCAATGGGGCGAAGCTACCATCTGTGGGATT | 10453.2 | -8.3 | 0.9 | -8.8 | 1.3E-18 | 1.2E-17 | MB231/MB468 | NUC |
| >hg38_21_+_8433172_8446622_RNA45SN1_45S_50ntFlanks@10905.10933.29\|CTGGGCGCGCGCCGCGGCTGGACGAGGCG | >hg38_21_+_8433172_8446622_RNA45SN1_45S_50ntFlanks | 10905 | 10933 | 29 | CTGGGCGCGCGCCGCGGCTGGACGAGGCG | 535.2 | -12.9 | 1.5 | -8.8 | 9.8E-19 | 1.2E-17 | MB231/MB468 | MP |
| >hg38_21_+_8433172_8446622_RNA45SN1_45S_50ntFlanks@10727.10757.31\|GGAGAGGGTGTAAATCTCGCGCCGGGCCGTA | >hg38_21_+_8433172_8446622_RNA45SN1_45S_50ntFlanks | 10727 | 10757 | 31 | GGAGAGGGTGTAAATCTCGCGCCGGGCCGTA | 347.5 | -12.3 | 1.4 | -8.8 | 9.9E-19 | 1.2E-17 | MB231/MB468 | MP |
| >hg38_21_+_8433172_8446622_RNA45SN1_45S_50ntFlanks@6719.6739.21\|TGAATTGCAGGACACATTGAT | >hg38_21_+_8433172_8446622_RNA45SN1_45S_50ntFlanks | 6719 | 6739 | 21 | TGAATTGCAGGACACATTGAT | 460.9 | -12.7 | 1.4 | -8.8 | 9.9E-19 | 1.2E-17 | MB231/MB468 | MP |
| >MI0000434\|hsa-let-7i&WithFlank&12\|+\|62603680\|62603775@12.34.23&[MIMAT0000415&hsa-let-7i-5p&offsets\|0\|+1;m-49&12\|+\|62603691\|62603712&offsets\|0\|+1]\|TGAGGTAGTAGTTTGTGCTGTTG | >MI0000434\|hsa-let-7i&WithFlank&12\|+\|62603680\|62603775 | 12 | 34 | 23 | TGAGGTAGTAGTTTGTGCTGTTG | 348.2 | 6.2 | 0.7 | 8.8 | 2.0E-18 | 1.2E-17 | BT20/MB468 | CYTO |
| >am_trnaMT_ValTAC_MT_+_1602_1670@50.68.19\|ACTTAACTTGACCGCTCTG | >am_trnaMT_ValTAC_MT_+_1602_1670 | 50 | 68 | 19 | ACTTAACTTGACCGCTCTG | 22744.4 | -8.6 | 1.0 | -8.8 | 1.3E-18 | 1.2E-17 | MB231/MB468 | NUC |
| >hg38_21_+_8433172_8446622_RNA45SN1_45S_50ntFlanks@11579.11605.27\|AGAACTGGTGCGGACCAGGGGAATCCG | >hg38_21_+_8433172_8446622_RNA45SN1_45S_50ntFlanks | 11579 | 11605 | 27 | AGAACTGGTGCGGACCAGGGGAATCCG | 1978.8 | -5.5 | 0.6 | -8.8 | 9.6E-19 | 1.2E-17 | MB231/MB468 | CYTO |
| >MI0016597\|hsa-mir-3940&WithFlank&19\|-\|6416404\|6416517@63.83.21&[MIMAT0018356&hsa-miR-3940-3p&offsets\|0\|-1;m-605&19\|-\|6416435\|6416455&offsets\|0\|0]\|CAGCCCGGATCCCAGCCCACT | >MI0016597\|hsa-mir-3940&WithFlank&19\|-\|6416404\|6416517 | 63 | 83 | 21 | CAGCCCGGATCCCAGCCCACT | 431.5 | 13.0 | 1.5 | 8.8 | 2.0E-18 | 1.2E-17 | BT20/MB468 | CYTO |
| >hg38_21_+_8433172_8446622_RNA45SN1_45S_50ntFlanks@12588.12610.23\|GGTTCAGACATTTGGTGTATGTG | >hg38_21_+_8433172_8446622_RNA45SN1_45S_50ntFlanks | 12588 | 12610 | 23 | GGTTCAGACATTTGGTGTATGTG | 603.2 | -13.1 | 1.5 | -8.8 | 1.0E-18 | 1.2E-17 | MB231/MB468 | MP |
| >MI0000085\|hsa-mir-27a&WithFlank&19\|-\|13836434\|13836523@57.75.19&[MIMAT0000084&hsa-miR-27a-3p&offsets\|0\|-2;m-43&19\|-\|13836447\|13836467&offsets\|0\|-2]\|TTCACAGTGGCTAAGTTCC | >MI0000085\|hsa-mir-27a&WithFlank&19\|-\|13836434\|13836523 | 57 | 75 | 19 | TTCACAGTGGCTAAGTTCC | 699.8 | 6.6 | 0.8 | 8.8 | 2.0E-18 | 1.2E-17 | BT20/MB468 | CYTO |
| >hg38_21_+_8433172_8446622_RNA45SN1_45S_50ntFlanks@6659.6707.49\|AGCGGTGGATCACTCGGCTCGTGCGTCGATGAAGAACGCAGCTAGCTGC | >hg38_21_+_8433172_8446622_RNA45SN1_45S_50ntFlanks | 6659 | 6707 | 49 | AGCGGTGGATCACTCGGCTCGTGCGTCGATGAAGAACGCAGCTAGCTGC | 505.7 | 13.4 | 1.5 | 8.8 | 1.5E-18 | 1.2E-17 | BT20/MB468 | MITO |
| >hg38_21_+_8433172_8446622_RNA45SN1_45S_50ntFlanks@12625.12654.30\|CAATGGGGCGAAGCTACCATCTGTGGGATT | >hg38_21_+_8433172_8446622_RNA45SN1_45S_50ntFlanks | 12625 | 12654 | 30 | CAATGGGGCGAAGCTACCATCTGTGGGATT | 2585.5 | -4.0 | 0.5 | -8.8 | 2.0E-18 | 1.2E-17 | BT20/MB468 | CYTO |
| >hg38_21_+_8433172_8446622_RNA45SN1_45S_50ntFlanks@12715.12742.28\|AGCCTCGGTTGGCCTCGGATAGCCGGTC | >hg38_21_+_8433172_8446622_RNA45SN1_45S_50ntFlanks | 12715 | 12742 | 28 | AGCCTCGGTTGGCCTCGGATAGCCGGTC | 398.2 | -12.5 | 1.4 | -8.8 | 1.0E-18 | 1.2E-17 | MB231/MB468 | MP |
| >hg38_21_+_8433172_8446622_RNA45SN1_45S_50ntFlanks@4919.4940.22\|CCACCAGGAGTGGAGCCTGCGG | >hg38_21_+_8433172_8446622_RNA45SN1_45S_50ntFlanks | 4919 | 4940 | 22 | CCACCAGGAGTGGAGCCTGCGG | 371.1 | -12.4 | 1.4 | -8.8 | 1.0E-18 | 1.2E-17 | MB231/MB468 | MP |
| >MI0000255\|hsa-mir-30d&WithFlank&8\|-\|134804870\|134804951@12.33.22(+1U)&[MIMAT0000245&hsa-miR-30d-5p&offsets\|0\|0(+1U);m-21&8\|-\|134804919\|134804940&offsets\|0\|0(+1U)]\|TGTAAACATCCCCGACTGGAAGT | >MI0000255\|hsa-mir-30d&WithFlank&8\|-\|134804870\|134804951 | 12 | 33 | 22(+1U) | TGTAAACATCCCCGACTGGAAGT | 3254.1 | 5.4 | 0.6 | 8.8 | 1.3E-18 | 1.2E-17 | MB231/MB468 | NUC |
| >hg38_21_+_8433172_8446622_RNA45SN1_45S_50ntFlanks@8115.8134.20\|CGCGGGACATGTGGCGTACG | >hg38_21_+_8433172_8446622_RNA45SN1_45S_50ntFlanks | 8115 | 8134 | 20 | CGCGGGACATGTGGCGTACG | 475.8 | -12.8 | 1.4 | -8.8 | 1.0E-18 | 1.2E-17 | MB231/MB468 | MP |
| >hg38_21_+_8433172_8446622_RNA45SN1_45S_50ntFlanks@4759.4780.22\|ATCAGATACCGTCGTAGTTCCG | >hg38_21_+_8433172_8446622_RNA45SN1_45S_50ntFlanks | 4759 | 4780 | 22 | ATCAGATACCGTCGTAGTTCCG | 5098.5 | -15.4 | 1.7 | -9.3 | 2.2E-20 | 1.2E-17 | BT20/MB468 | MP |
| >hg38_21_+_8433172_8446622_RNA45SN1_45S_50ntFlanks@11076.11103.28\|CGCCGCGTGGGGGCGGCGGCGGGGGGAG | >hg38_21_+_8433172_8446622_RNA45SN1_45S_50ntFlanks | 11076 | 11103 | 28 | CGCCGCGTGGGGGCGGCGGCGGGGGGAG | 970.5 | -12.9 | 1.5 | -8.8 | 1.3E-18 | 1.3E-17 | MB231/MB468 | NUC |
| >hg38_21_+_8433172_8446622_RNA45SN1_45S_50ntFlanks@5486.5510.25\|GCGCTGAGAAGACGGTCGAACTTGA | >hg38_21_+_8433172_8446622_RNA45SN1_45S_50ntFlanks | 5486 | 5510 | 25 | GCGCTGAGAAGACGGTCGAACTTGA | 372.2 | -12.4 | 1.4 | -8.8 | 1.0E-18 | 1.3E-17 | MB231/MB468 | MP |
| >hg38_21_+_8433172_8446622_RNA45SN1_45S_50ntFlanks@7982.8008.27\|TCAGATCAGACGTGGCGACCCGCTGAA | >hg38_21_+_8433172_8446622_RNA45SN1_45S_50ntFlanks | 7982 | 8008 | 27 | TCAGATCAGACGTGGCGACCCGCTGAA | 1466.0 | -8.3 | 0.9 | -8.8 | 1.0E-18 | 1.3E-17 | MB231/MB468 | MP |
| >hg38_21_+_8433172_8446622_RNA45SN1_45S_50ntFlanks@5490.5516.27\|TGAGAAGACGGTCGAACTTGACTATCT | >hg38_21_+_8433172_8446622_RNA45SN1_45S_50ntFlanks | 5490 | 5516 | 27 | TGAGAAGACGGTCGAACTTGACTATCT | 16451.8 | 10.7 | 1.2 | 8.8 | 2.0E-18 | 1.3E-17 | BT20/MB468 | CYTO |
| >hg38_21_+_8433172_8446622_RNA45SN1_45S_50ntFlanks@10444.10462.19\|CGAAGGGACGGGCGATGGC | >hg38_21_+_8433172_8446622_RNA45SN1_45S_50ntFlanks | 10444 | 10462 | 19 | CGAAGGGACGGGCGATGGC | 511.8 | -12.0 | 1.4 | -8.8 | 1.3E-18 | 1.3E-17 | MB231/MB468 | NUC |
| >trna10_AspGTC_12_-_125424193_125424264@37.74.38,trna12_AspGTC_12_-_125411891_125411962@37.74.38,trna144_AspGTC_6_-_27551236_27551307@37.74.38,trna38_AspGTC_17_-_8125556_8125627@37.74.38,trna45_AspGTC_6_+_27447453_27447524@37.74.38,trna48_AspGTC_6_+_27471523_27471594@37.74.38,trna4_AspGTC_12_+_96429799_96429870@37.74.38,trna69_AspGTC_1_-_161440205_161440276@37.74.38,trna72_AspGTC_1_-_161432824_161432895@37.74.38,trna75_AspGTC_1_-_161425414_161425485@37.74.38,trna78_AspGTC_1_-_161418033_161418104@37.74.38,trna81_AspGTC_1_-_161410615_161410686@37.74.38\|ACGCGGGAGACCGGGGTTCGATTCCCCGACGGGGAGCC | >trna10_AspGTC_12_-_125424193_125424264 | 37 | 74 | 38 | ACGCGGGAGACCGGGGTTCGATTCCCCGACGGGGAGCC | 40869.9 | 11.3 | 1.3 | 8.8 | 1.0E-18 | 1.3E-17 | MB231/MB468 | CYTO |
| >MI0000462\|hsa-mir-152&WithFlank&17\|-\|48037155\|48037253@60.80.21&[MIMAT0000438&hsa-miR-152-3p&offsets\|0\|0;m-97&17\|-\|48037174\|48037194&offsets\|0\|0]\|TCAGTGCATGACAGAACTTGG | >MI0000462\|hsa-mir-152&WithFlank&17\|-\|48037155\|48037253 | 60 | 80 | 21 | TCAGTGCATGACAGAACTTGG | 683.9 | 11.0 | 1.3 | 8.8 | 1.4E-18 | 1.3E-17 | MB231/MB468 | NUC |
| >hg38_21_+_8433172_8446622_RNA45SN1_45S_50ntFlanks@8168.8197.30\|CCCAAGTCCTTCTGATCGAGGCCCAGCCCG | >hg38_21_+_8433172_8446622_RNA45SN1_45S_50ntFlanks | 8168 | 8197 | 30 | CCCAAGTCCTTCTGATCGAGGCCCAGCCCG | 1430.8 | -14.3 | 1.6 | -8.8 | 1.7E-18 | 1.3E-17 | MB231/MB468 | MITO |
| >MI0000734\|hsa-mir-106b&WithFlank&7\|-\|100093987\|100094080@58.78.21&[MIMAT0004672&hsa-miR-106b-3p&offsets\|0\|-1;m-87&7\|-\|100094003\|100094023&offsets\|0\|0]\|CCGCACTGTGGGTACTTGCTG | >MI0000734\|hsa-mir-106b&WithFlank&7\|-\|100093987\|100094080 | 58 | 78 | 21 | CCGCACTGTGGGTACTTGCTG | 3348.5 | 7.0 | 0.8 | 8.8 | 1.0E-18 | 1.3E-17 | MB231/MB468 | CYTO |

| | | | | | | | | | | | | | |
|---|---|---|---|---|---|---|---|---|---|---|---|---|---|
| >hg38_21_+_8433172_8446622_RNA45SN1_45S_50ntFlanks@9622.9652.31\|CTAGTAGCTGGTTCCCTCCGAAGTTTCCCTC | >hg38_21_+_8433172_8446622_RNA45SN1_45S_50ntFlanks | 9622 | 9652 | 31 | CTAGTAGCTGGTTCCCTCCGAAGTTTCCCTC | 331.4 | -12.2 | 1.4 | -8.8 | 1.1E-18 | 1.3E-17 | MB231/MB468 | MP |
| >MI0000433\|hsa-let-7g&WithFlank&3\|-\|52268272\|52268367@11.28.18&[MIMAT0000414&hsa-let-7g-5p&offsets\|0\|-4;m-58&3\|-\|52268336\|52268357&offsets\|0\|-4]\|TGAGGTAGTAGTTTGTAC | >MI0000433\|hsa-let-7g&WithFlank&3\|-\|52268272\|52268367 | 11 | 28 | 18 | TGAGGTAGTAGTTTGTAC | 2592.5 | 7.3 | 0.8 | 8.8 | 1.5E-18 | 1.3E-17 | BT20/MB468 | MITO |
| >hg38_21_+_8433172_8446622_RNA45SN1_45S_50ntFlanks@9504.9533.30\|GTGAACTATGCCTGGGCAGGGCGAAGCCAG | >hg38_21_+_8433172_8446622_RNA45SN1_45S_50ntFlanks | 9504 | 9533 | 30 | GTGAACTATGCCTGGGCAGGGCGAAGCCAG | 889.8 | -12.8 | 1.5 | -8.8 | 1.4E-18 | 1.3E-17 | MB231/MB468 | NUC |
| >trna116_GluCTC_1_-_145399233_145399304@1.33.33,trna59_GluCTC_1_+_249168447_249168518@1.33.33,trna71_GluCTC_1_-_161439189_161439260@1.33.33,trna74_GluCTC_1_-_161431809_161431880@1.33.33,trna77_GluCTC_1_-_161424398_161424469@1.33.33,trna77_GluCTC_6_+_28949976_28950047@1.33.33,trna80_GluCTC_1_-_161417018_161417089@1.33.33,trna87_GluCTC_6_-_126101393_126101464@1.33.33\|TCCCTGGTGGTCTAGTGGTTAGGATTCGGCGCT | >trna116_GluCTC_1_-_145399233_145399304 | 1 | 33 | 33 | TCCCTGGTGGTCTAGTGGTTAGGATTCGGCGCT | 69614.2 | 8.7 | 1.0 | 8.8 | 1.6E-18 | 1.3E-17 | BT20/MB468 | MITO |
| >hg38_21_+_8433172_8446622_RNA45SN1_45S_50ntFlanks@8357.8391.35\|AAAAGAACTTTGAAGAGAGAGTTCAAGAGGGCGTG | >hg38_21_+_8433172_8446622_RNA45SN1_45S_50ntFlanks | 8357 | 8391 | 35 | AAAAGAACTTTGAAGAGAGAGTTCAAGAGGGCGTG | 1275.6 | -4.8 | 0.5 | -8.8 | 1.4E-18 | 1.3E-17 | MB231/MB468 | NUC |
| >hg38_21_+_8433172_8446622_RNA45SN1_45S_50ntFlanks@4823.4858.36\|ATGACCCGCCGGGCAGCTTCCGGGAAACCAAAGTCT | >hg38_21_+_8433172_8446622_RNA45SN1_45S_50ntFlanks | 4823 | 4858 | 36 | ATGACCCGCCGGGCAGCTTCCGGGAAACCAAAGTCT | 367.5 | 12.8 | 1.5 | 8.8 | 2.1E-18 | 1.3E-17 | BT20/MB468 | CYTO |
| >hg38_21_+_8433172_8446622_RNA45SN1_45S_50ntFlanks@11323.11342.20\|CGTCTCGTCGCGCGCGCGTC | >hg38_21_+_8433172_8446622_RNA45SN1_45S_50ntFlanks | 11323 | 11342 | 20 | CGTCTCGTCGCGCGCGCGTC | 9113.9 | -9.7 | 1.1 | -8.8 | 1.4E-18 | 1.3E-17 | MB231/MB468 | NUC |
| >MI0000087\|hsa-mir-29a&WithFlank&7\|-\|130876741\|130876816@48.67.20&[MIMAT0000086&hsa-miR-29a-3p&offsets\|0\|-2;m-26&7\|-\|130876749\|130876769&offsets\|0\|-1]\|TAGCACCATCTGAAATCGGT | >MI0000087\|hsa-mir-29a&WithFlank&7\|-\|130876741\|130876816 | 48 | 67 | 20 | TAGCACCATCTGAAATCGGT | 514.6 | 13.3 | 1.5 | 8.7 | 2.1E-18 | 1.3E-17 | BT20/MB468 | CYTO |
| >hg38_21_+_8433172_8446622_RNA45SN1_45S_50ntFlanks@10873.10894.22\|CTGGGTCGGTCGGGCTGGGGCG | >hg38_21_+_8433172_8446622_RNA45SN1_45S_50ntFlanks | 10873 | 10894 | 22 | CTGGGTCGGTCGGGCTGGGGCG | 448.2 | -12.6 | 1.4 | -8.8 | 1.8E-18 | 1.3E-17 | MB231/MB468 | MITO |
| >MI0000746\|hsa-mir-99b&WithFlank&19\|+\|51692606\|51692687@51.72.22(+1A)&[MIMAT0004678&hsa-miR-99b-3p&offsets\|0\|0(+1A);m-151&19\|+\|51692656\|51692678&offsets\|0\|-1(+1A)]\|CAAGCTCGTGTCTGTGGGTCCGA | >MI0000746\|hsa-mir-99b&WithFlank&19\|+\|51692606\|51692687 | 51 | 72 | 22(+1A) | CAAGCTCGTGTCTGTGGGTCCGA | 149.1 | 3.3 | 0.4 | 8.7 | 3.8E-18 | 1.3E-17 | BT20/MB468 | NUC |
| >hg38_21_+_8433172_8446622_RNA45SN1_45S_50ntFlanks@6785.6803.19\|CTACGCCTGTCTGAGCGTC | >hg38_21_+_8433172_8446622_RNA45SN1_45S_50ntFlanks | 6785 | 6803 | 19 | CTACGCCTGTCTGAGCGTC | 865.1 | -13.6 | 1.5 | -8.8 | 1.1E-18 | 1.3E-17 | MB231/MB468 | MP |
| >hg38_1_-_228634819_228635039_RNA5S12_5S_50ntFlanks@84.103.20\|CTCGTCTGATCTCGGAAGCT | >hg38_1_-_228634819_228635039_RNA5S12_5S_50ntFlanks | 84 | 103 | 20 | CTCGTCTGATCTCGGAAGCT | 492.4 | -12.7 | 1.5 | -8.8 | 1.8E-18 | 1.3E-17 | MB231/MB468 | MITO |
| >am_trna10_ValCAC_5_-_180649395_180649467@1.32.32,trna12_ValAAC_5_-_180645270_180645342@1.32.32,trna132_ValAAC_6_-_27721179_27721251@1.32.32,trna136_ValAAC_6_-_27648885_27648957@1.32.32,trna139_ValAAC_6_-_27618707_27618779@1.32.32,trna18_ValCAC_5_-_180529253_180529325@1.32.32,trna2_ValAAC_3_+_169490018_169490090@1.32.32,trna2_ValCAC_5_+_180524070_180524142@1.32.32,trna4_ValAAC_5_+_180591154_180591226@1.32.32,trna5_ValAAC_5_+_180596610_180596682@1.32.32,trna6_ValCAC_5_+_180600650_180600722@1.32.32,trna85_ValCAC_1_-_161369490_161369562@1.32.32,trna90_ValCAC_1_-_149684088_149684161@1.32.32,trna98_ValCAC_1_-_149298555_149298627@1.32.32,trna9_ValCAC_6_+_26538282_26538354@1.32.32\|GTTTCCGTAGTGTAGTGGTTATCACGTTCGCC | >am_trna10_ValCAC_5_-_180649395_180649467 | 1 | 32 | 32 | GTTTCCGTAGTGTAGTGGTTATCACGTTCGCC | 5384.2 | 7.0 | 0.8 | 8.7 | 2.2E-18 | 1.3E-17 | BT20/MB468 | CYTO |
| >hg38_21_+_8433172_8446622_RNA45SN1_45S_50ntFlanks@12626.12654.29\|AATGGGGCGAAGCTACCATCTGTGGGATT | >hg38_21_+_8433172_8446622_RNA45SN1_45S_50ntFlanks | 12626 | 12654 | 29 | AATGGGGCGAAGCTACCATCTGTGGGATT | 6331.9 | -4.5 | 0.5 | -8.8 | 1.1E-18 | 1.3E-17 | MB231/MB468 | CYTO |
| >hg38_21_+_8433172_8446622_RNA45SN1_45S_50ntFlanks@12712.12743.32\|CGGAGCCTCGGTTGGCCTCGGATAGCCGGTCC | >hg38_21_+_8433172_8446622_RNA45SN1_45S_50ntFlanks | 12712 | 12743 | 32 | CGGAGCCTCGGTTGGCCTCGGATAGCCGGTCC | 442.2 | -12.7 | 1.4 | -8.8 | 1.1E-18 | 1.3E-17 | MB231/MB468 | MP |
| >hg38_21_+_8433172_8446622_RNA45SN1_45S_50ntFlanks@10346.10369.24\|TCAAACGAGAACTTTGAAGGCCGA | >hg38_21_+_8433172_8446622_RNA45SN1_45S_50ntFlanks | 10346 | 10369 | 24 | TCAAACGAGAACTTTGAAGGCCGA | 1759.1 | -6.1 | 0.7 | -8.8 | 1.1E-18 | 1.3E-17 | MB231/MB468 | MP |
| >hg38_21_+_8433172_8446622_RNA45SN1_45S_50ntFlanks@10409.10429.21\|GTCAGTCGGTCCTGAGAGATG | >hg38_21_+_8433172_8446622_RNA45SN1_45S_50ntFlanks | 10409 | 10429 | 21 | GTCAGTCGGTCCTGAGAGATG | 2110.7 | -4.5 | 0.5 | -8.8 | 1.4E-18 | 1.4E-17 | MB231/MB468 | NUC |
| >MI0000089\|hsa-mir-31&WithFlank&9\|-\|21512109\|21512191@14.35.22&[MIMAT0000089&hsa-miR-31-5p&offsets\|0\|+1;m-360&9\|-\|21512157\|21512178&offsets\|0\|0]\|AGGCAAGATGCTGGCATAGCTG | >MI0000089\|hsa-mir-31&WithFlank&9\|-\|21512109\|21512191 | 14 | 35 | 22 | AGGCAAGATGCTGGCATAGCTG | 1147.7 | 8.6 | 1.0 | 8.7 | 2.2E-18 | 1.4E-17 | BT20/MB468 | CYTO |
| >MI0000077\|hsa-mir-21&WithFlank&17\|+\|59841260\|59841343@14.36.23(+1G)&[MIMAT0000076&hsa-miR-21-5p&offsets\|0\|+1(+1G);m-1&17\|+\|59841273\|59841294&offsets\|0\|+1(+1G)]\|TAGCTTATCAGACTGATGTTGACG | >MI0000077\|hsa-mir-21&WithFlank&17\|+\|59841260\|59841343 | 14 | 36 | 23(+1G) | TAGCTTATCAGACTGATGTTGACG | 4338.6 | 6.2 | 0.7 | 8.8 | 1.6E-18 | 1.4E-17 | BT20/MB468 | MITO |
| >trnaMT_LeuTAA_MT_+_3230_3304@16.44.29\|CCCGGTAATCGCATAAAACTTAAAACTTT | >trnaMT_LeuTAA_MT_+_3230_3304 | 16 | 44 | 29 | CCCGGTAATCGCATAAAACTTAAAACTTT | 5322.8 | -12.5 | 1.4 | -8.7 | 3.9E-18 | 1.4E-17 | BT20/MB468 | NUC |
| >hg38_21_+_8433172_8446622_RNA45SN1_45S_50ntFlanks@3940.3959.20\|ACCCGGTCAGCCCCTCTCCG | >hg38_21_+_8433172_8446622_RNA45SN1_45S_50ntFlanks | 3940 | 3959 | 20 | ACCCGGTCAGCCCCTCTCCG | 498.4 | -12.7 | 1.5 | -8.8 | 1.8E-18 | 1.4E-17 | MB231/MB468 | MITO |
| >hg38_21_+_8433172_8446622_RNA45SN1_45S_50ntFlanks@4257.4275.19\|TAATTGGAATGAGTCCACT | >hg38_21_+_8433172_8446622_RNA45SN1_45S_50ntFlanks | 4257 | 4275 | 19 | TAATTGGAATGAGTCCACT | 322.2 | -12.2 | 1.4 | -8.8 | 1.1E-18 | 1.4E-17 | MB231/MB468 | MP |
| >hg38_21_+_8433172_8446622_RNA45SN1_45S_50ntFlanks@6748.6771.24\|CTTCGAACGCACTTGCGGCCCCGG | >hg38_21_+_8433172_8446622_RNA45SN1_45S_50ntFlanks | 6748 | 6771 | 24 | CTTCGAACGCACTTGCGGCCCCGG | 454.6 | -12.6 | 1.4 | -8.8 | 1.8E-18 | 1.4E-17 | MB231/MB468 | MITO |
| >trna10_MetCAT_8_-_124169470_124169542@31.51.21,trna22_MetCAT_16_-_87417628_87417700@31.51.21\|TCTCATAATCTGAAGGTCGTG | >trna10_MetCAT_8_-_124169470_124169542 | 31 | 51 | 21 | TCTCATAATCTGAAGGTCGTG | 479.1 | -12.7 | 1.4 | -8.8 | 1.8E-18 | 1.4E-17 | MB231/MB468 | MITO |
| >hg38_21_+_8433172_8446622_RNA45SN1_45S_50ntFlanks@3782.3801.20\|CAGTGAAACTGCGAATGGCT | >hg38_21_+_8433172_8446622_RNA45SN1_45S_50ntFlanks | 3782 | 3801 | 20 | CAGTGAAACTGCGAATGGCT | 3955.2 | -10.2 | 1.2 | -8.7 | 3.9E-18 | 1.4E-17 | BT20/MB468 | NUC |
| >hg38_21_+_8433172_8446622_RNA45SN1_45S_50ntFlanks@3870.3898.29\|AGCTAATACATGCCGACGGGCGCTGACCC | >hg38_21_+_8433172_8446622_RNA45SN1_45S_50ntFlanks | 3870 | 3898 | 29 | AGCTAATACATGCCGACGGGCGCTGACCC | 738.5 | -14.0 | 1.6 | -8.7 | 3.9E-18 | 1.4E-17 | BT20/MB468 | NUC |
| >hg38_21_+_8433172_8446622_RNA45SN1_45S_50ntFlanks@6731.6752.22\|CACATTGATCATCGACACTTCG | >hg38_21_+_8433172_8446622_RNA45SN1_45S_50ntFlanks | 6731 | 6752 | 22 | CACATTGATCATCGACACTTCG | 354.2 | -12.3 | 1.4 | -8.8 | 1.1E-18 | 1.4E-17 | MB231/MB468 | MP |
| >MI0000477\|hsa-mir-146a&WithFlank&5\|+\|160485346\|160485456@27.47.21(+1C)&[MIMAT0000449&hsa-miR-146a-5p&offsets\|0\|-1(+1C);m-129&5\|+\|160485372\|160485393&offsets\|0\|-1(+1C)]\|TGAGAACTGAATTCCATGGGTC | >MI0000477\|hsa-mir-146a&WithFlank&5\|+\|160485346\|160485456 | 27 | 47 | 21(+1C) | TGAGAACTGAATTCCATGGGTC | 3062.8 | 14.8 | 1.7 | 8.8 | 1.9E-18 | 1.4E-17 | MB231/MB468 | MITO |
| >hg38_21_+_8433172_8446622_RNA45SN1_45S_50ntFlanks@9562.9596.35\|TCCTGACGTGCAAATCGGTCGTCCGACCTGGGTAT | >hg38_21_+_8433172_8446622_RNA45SN1_45S_50ntFlanks | 9562 | 9596 | 35 | TCCTGACGTGCAAATCGGTCGTCCGACCTGGGTAT | 1100.1 | -7.8 | 0.9 | -8.8 | 1.1E-18 | 1.4E-17 | MB231/MB468 | CYTO |
| >hg38_21_+_8433172_8446622_RNA45SN1_45S_50ntFlanks@11575.11605.31\|ACTTAGAACTGGTGCGGACCAGGGGAATCCG | >hg38_21_+_8433172_8446622_RNA45SN1_45S_50ntFlanks | 11575 | 11605 | 31 | ACTTAGAACTGGTGCGGACCAGGGGAATCCG | 2210.6 | -5.6 | 0.6 | -8.7 | 2.3E-18 | 1.4E-17 | BT20/MB468 | CYTO |
| >hg38_21_+_8433172_8446622_RNA45SN1_45S_50ntFlanks@12712.12745.34\|CGGAGCCTCGGTTGGCCTCGGATAGCCGGTCCCC | >hg38_21_+_8433172_8446622_RNA45SN1_45S_50ntFlanks | 12712 | 12745 | 34 | CGGAGCCTCGGTTGGCCTCGGATAGCCGGTCCCC | 743.3 | -13.3 | 1.5 | -8.8 | 1.9E-18 | 1.4E-17 | MB231/MB468 | MITO |
| >am_MI0015825\|hsa-mir-378c&WithFlank&10\|-\|130962582\|130962674@17.37.21&[MIMAT0016847&hsa-miR-378c&offsets\|0\|-4;m-130&10\|-\|130962639\|130962658&offsets\|0\|+1]\|ACTGGACTTGGAGTCAGAAGA | >am_MI0015825\|hsa-mir-378c&WithFlank&10\|-\|130962582\|130962674 | 17 | 37 | 21 | ACTGGACTTGGAGTCAGAAGA | 592.8 | 6.0 | 0.7 | 8.7 | 2.3E-18 | 1.4E-17 | BT20/MB468 | CYTO |
| >MI0000471\|hsa-mir-126&WithFlank&9\|+\|136670596\|136670692@59.78.20&[MIMAT0000445&hsa-miR-126-3p&offsets\|+1\|-1;m-40&9\|+\|136670653\|136670674&offsets\|+1\|-1]\|CGTACCGTGAGTAATAATGC | >MI0000471\|hsa-mir-126&WithFlank&9\|+\|136670596\|136670692 | 59 | 78 | 20 | CGTACCGTGAGTAATAATGC | 455.6 | 10.4 | 1.2 | 8.8 | 1.5E-18 | 1.4E-17 | MB231/MB468 | NUC |
| >hg38_21_+_8433172_8446622_RNA45SN1_45S_50ntFlanks@8552.8577.26\|CTCCCCGGAGGGGGCGGGCTCCGGCG | >hg38_21_+_8433172_8446622_RNA45SN1_45S_50ntFlanks | 8552 | 8577 | 26 | CTCCCCGGAGGGGGCGGGCTCCGGCG | 556.5 | -11.8 | 1.3 | -8.8 | 1.1E-18 | 1.4E-17 | MB231/MB468 | CYTO |
| >hg38_21_+_8433172_8446622_RNA45SN1_45S_50ntFlanks@7986.8007.22\|ATCAGACGTGGCGACCCGCTGA | >hg38_21_+_8433172_8446622_RNA45SN1_45S_50ntFlanks | 7986 | 8007 | 22 | ATCAGACGTGGCGACCCGCTGA | 683.0 | -13.2 | 1.5 | -8.8 | 1.9E-18 | 1.4E-17 | MB231/MB468 | MITO |
| >hg38_21_+_8433172_8446622_RNA45SN1_45S_50ntFlanks@11027.11049.23\|CTCTCTCTCTCTCTCCCCCGCTC | >hg38_21_+_8433172_8446622_RNA45SN1_45S_50ntFlanks | 11027 | 11049 | 23 | CTCTCTCTCTCTCTCCCCCGCTC | 730.9 | -13.3 | 1.5 | -8.8 | 1.9E-18 | 1.4E-17 | MB231/MB468 | MITO |
| >MI0000477\|hsa-mir-146a&WithFlank&5\|+\|160485346\|160485456@27.48.22(+2U)&[MIMAT0000449&hsa-miR-146a-5p&offsets\|0\|0(+2U);m-129&5\|+\|160485372\|160485393&offsets\|0\|0(+2U)]\|TGAGAACTGAATTCCATGGGTTTT | >MI0000477\|hsa-mir-146a&WithFlank&5\|+\|160485346\|160485456 | 27 | 48 | 22(+2U) | TGAGAACTGAATTCCATGGGTTTT | 1001.7 | 13.2 | 1.5 | 8.8 | 1.2E-18 | 1.4E-17 | MB231/MB468 | MP |
| >MI0003643\|hsa-mir-629&WithFlank&15\|-\|70079366\|70079474@28.48.21&[MIMAT0004810&hsa-miR-629-5p&offsets\|0\|0;m-150&15\|-\|70079427\|70079447&offsets\|0\|0]\|TGGGTTTACGTTGGGAGAACT | >MI0003643\|hsa-mir-629&WithFlank&15\|-\|70079366\|70079474 | 28 | 48 | 21 | TGGGTTTACGTTGGGAGAACT | 5425.3 | 16.2 | 1.7 | 9.2 | 2.6E-20 | 1.4E-17 | BT20/MB468 | MP |
| >hg38_21_+_8433172_8446622_RNA45SN1_45S_50ntFlanks@6789.6806.18\|GCCTGTCTGAGCGTCGCT | >hg38_21_+_8433172_8446622_RNA45SN1_45S_50ntFlanks | 6789 | 6806 | 18 | GCCTGTCTGAGCGTCGCT | 5643.6 | 5.5 | 0.6 | 8.8 | 1.7E-18 | 1.4E-17 | BT20/MB468 | MITO |
| >hg38_21_+_8433172_8446622_RNA45SN1_45S_50ntFlanks@12126.12146.21\|CGACCCGCTCCGGGGACAGTG | >hg38_21_+_8433172_8446622_RNA45SN1_45S_50ntFlanks | 12126 | 12146 | 21 | CGACCCGCTCCGGGGACAGTG | 2345.8 | -14.3 | 1.5 | -9.2 | 2.7E-20 | 1.4E-17 | BT20/MB468 | MP |
| >hg38_21_+_8433172_8446622_RNA45SN1_45S_50ntFlanks@12405.12428.24\|CTTTTTGATCCTTCGATGTCGGCT | >hg38_21_+_8433172_8446622_RNA45SN1_45S_50ntFlanks | 12405 | 12428 | 24 | CTTTTTGATCCTTCGATGTCGGCT | 300.6 | -6.0 | 0.7 | -8.8 | 1.1E-18 | 1.4E-17 | MB231/MB468 | CYTO |
| >hg38_21_+_8433172_8446622_RNA45SN1_45S_50ntFlanks@11572.11605.34\|CCGACTTAGAACTGGTGCGGACCAGGGGAATCCG | >hg38_21_+_8433172_8446622_RNA45SN1_45S_50ntFlanks | 11572 | 11605 | 34 | CCGACTTAGAACTGGTGCGGACCAGGGGAATCCG | 3504.4 | -11.4 | 1.3 | -8.8 | 1.2E-18 | 1.4E-17 | MB231/MB468 | MP |
| >hg38_21_+_8433172_8446622_RNA45SN1_45S_50ntFlanks@12457.12475.19\|CAAGCGTTGGATTGTTCAC | >hg38_21_+_8433172_8446622_RNA45SN1_45S_50ntFlanks | 12457 | 12475 | 19 | CAAGCGTTGGATTGTTCAC | 457.9 | -12.7 | 1.4 | -8.8 | 1.2E-18 | 1.4E-17 | MB231/MB468 | MP |
| >MI0000273\|hsa-mir-183&WithFlank&7\|-\|129774899\|129775020@33.54.22(+1A)&[MIMAT0000261&hsa-miR-183-5p&offsets\|0\|0(+1A);m-41&7\|-\|129774966\|129774987&offsets\|-1\|-1(+1A)]\|TATGGCACTGGTAGAATTCACTA | >MI0000273\|hsa-mir-183&WithFlank&7\|-\|129774899\|129775020 | 33 | 54 | 22(+1A) | TATGGCACTGGTAGAATTCACTA | 1326.4 | 8.6 | 1.0 | 8.7 | 2.3E-18 | 1.4E-17 | BT20/MB468 | CYTO |

<table>
<tr><td>>MI0000078|hsa-mir-22&WithFlank&17|-|1713897|1713993@59.79.21(+1A)&[MIMAT0000077&hsa-miR-22-3p&offsets|0|-1(+1A);m-2&17|-|1713914|1713935&offsets|0|-1(+1A)]|AAGCTGCCAGTTGAAGAACTGA</td><td>>MI0000078|hsa-mir-22&WithFlank&17|-|1713897|1713993</td><td>59</td><td>79</td><td>21(+1A)</td><td>AAGCTGCCAGTTGAAGAACTGA</td><td>1147.3</td><td>14.4</td><td>1.6</td><td>8.7</td><td>2.3E-18</td><td>1.4E-17</td><td>BT20/MB468</td><td>CYTO</td></tr>
<tr><td>>hg38_21_+_8433172_8446622_RNA45SN1_45S_50ntFlanks@12892.12932.41|CCCTCGCCCGTCACGCACCGCACGTTCGTGGGGAACCTGGC</td><td>>hg38_21_+_8433172_8446622_RNA45SN1_45S_50ntFlanks</td><td>12892</td><td>12932</td><td>41</td><td>CCCTCGCCCGTCACGCACCGCACGTTCGTGGGGAACCTGGC</td><td>8539.4</td><td>-9.5</td><td>1.1</td><td>-8.8</td><td>1.9E-18</td><td>1.4E-17</td><td>MB231/MB468</td><td>MITO</td></tr>
<tr><td>>hg38_21_+_8433172_8446622_RNA45SN1_45S_50ntFlanks@12942.12973.32|TTCGTAGACGACCTGCTTCTGGGTCGGGGTTT</td><td>>hg38_21_+_8433172_8446622_RNA45SN1_45S_50ntFlanks</td><td>12942</td><td>12973</td><td>32</td><td>TTCGTAGACGACCTGCTTCTGGGTCGGGGTTT</td><td>377.1</td><td>-12.4</td><td>1.4</td><td>-8.8</td><td>1.2E-18</td><td>1.4E-17</td><td>MB231/MB468</td><td>MP</td></tr>
<tr><td>>hg38_21_+_8433172_8446622_RNA45SN1_45S_50ntFlanks@12438.12473.36|ATTGTGAAGCAGAATTCACCAAGCGTTGGATTGTTC</td><td>>hg38_21_+_8433172_8446622_RNA45SN1_45S_50ntFlanks</td><td>12438</td><td>12473</td><td>36</td><td>ATTGTGAAGCAGAATTCACCAAGCGTTGGATTGTTC</td><td>907.5</td><td>-5.5</td><td>0.6</td><td>-8.7</td><td>4.1E-18</td><td>1.4E-17</td><td>BT20/MB468</td><td>NUC</td></tr>
<tr><td>>hg38_21_+_8433172_8446622_RNA45SN1_45S_50ntFlanks@12456.12489.34|CCAAGCGTTGGATTGTTCACCCACTAATAGGGAA</td><td>>hg38_21_+_8433172_8446622_RNA45SN1_45S_50ntFlanks</td><td>12456</td><td>12489</td><td>34</td><td>CCAAGCGTTGGATTGTTCACCCACTAATAGGGAA</td><td>2339.4</td><td>-9.2</td><td>1.0</td><td>-8.8</td><td>1.5E-18</td><td>1.4E-17</td><td>MB231/MB468</td><td>NUC</td></tr>
<tr><td>>MI0000285|hsa-mir-205&WithFlank&1|+|209432127|209432248@40.59.20(+1C)&[MIMAT0000266&hsa-miR-205-5p&offsets|0|-2(+1C);m-351&1|+|209432166|209432187&offsets|0|-2(+1C)]|TCCTTCATTCCACCGGAGTCC</td><td>>MI0000285|hsa-mir-205&WithFlank&1|+|209432127|209432248</td><td>40</td><td>59</td><td>20(+1C)</td><td>TCCTTCATTCCACCGGAGTCC</td><td>1031.4</td><td>13.8</td><td>1.5</td><td>9.0</td><td>1.9E-19</td><td>1.4E-17</td><td>BT20/MB231</td><td>NUC</td></tr>
<tr><td>>MI0000102|hsa-mir-100&WithFlank&11|-|122152223|122152314@19.38.20(+2G)&[MIMAT0000098&hsa-miR-100-5p&offsets|0|-2(+2G);m-27&11|-|122152276|122152296&offsets|0|-1(+2G)]|AACCCGTAGATCCGAACTTGGG</td><td>>MI0000102|hsa-mir-100&WithFlank&11|-|122152223|122152314</td><td>19</td><td>38</td><td>20(+2G)</td><td>AACCCGTAGATCCGAACTTGGG</td><td>882.3</td><td>-12.9</td><td>1.4</td><td>-9.0</td><td>1.9E-19</td><td>1.4E-17</td><td>BT20/MB231</td><td>NUC</td></tr>
<tr><td>>hg38_21_+_8433172_8446622_RNA45SN1_45S_50ntFlanks@12071.12095.25|CCCCGAGGGGCTCTCGCTTCTGGCG</td><td>>hg38_21_+_8433172_8446622_RNA45SN1_45S_50ntFlanks</td><td>12071</td><td>12095</td><td>25</td><td>CCCCGAGGGGCTCTCGCTTCTGGCG</td><td>2155.9</td><td>-10.6</td><td>1.2</td><td>-8.8</td><td>2.0E-18</td><td>1.4E-17</td><td>MB231/MB468</td><td>MITO</td></tr>
<tr><td>>hg38_21_+_8433172_8446622_RNA45SN1_45S_50ntFlanks@10830.10858.29|CAAGCCGGATCCGTAACTTCGGGATAAGG</td><td>>hg38_21_+_8433172_8446622_RNA45SN1_45S_50ntFlanks</td><td>10830</td><td>10858</td><td>29</td><td>CAAGCCGGATCCGTAACTTCGGGATAAGG</td><td>1205.6</td><td>-14.1</td><td>1.6</td><td>-8.8</td><td>1.2E-18</td><td>1.5E-17</td><td>MB231/MB468</td><td>MP</td></tr>
<tr><td>>hg38_21_+_8433172_8446622_RNA45SN1_45S_50ntFlanks@10786.10829.44|AACAGCCTCTGGCATGTTGGAACAATGTAGGTAAGGGAAGTCGG</td><td>>hg38_21_+_8433172_8446622_RNA45SN1_45S_50ntFlanks</td><td>10786</td><td>10829</td><td>44</td><td>AACAGCCTCTGGCATGTTGGAACAATGTAGGTAAGGGAAGTCGG</td><td>1087.2</td><td>-13.9</td><td>1.6</td><td>-8.8</td><td>2.0E-18</td><td>1.5E-17</td><td>MB231/MB468</td><td>MITO</td></tr>
<tr><td>>MI0000760|hsa-mir-361&WithFlank&X|-|85903630|85903713@51.73.23&[MIMAT0004682&hsa-miR-361-3p&offsets|0|0;m-124&X|-|85903642|85903663&offsets|0|+1]|TCCCCCAGGTGTGATTCTGATTT</td><td>>MI0000760|hsa-mir-361&WithFlank&X|-|85903630|85903713</td><td>51</td><td>73</td><td>23</td><td>TCCCCCAGGTGTGATTCTGATTT</td><td>1198.0</td><td>8.1</td><td>0.9</td><td>8.7</td><td>2.4E-18</td><td>1.5E-17</td><td>BT20/MB468</td><td>CYTO</td></tr>
<tr><td>>MI0000433|hsa-let-7g&WithFlank&3|-|52268272|52268367@11.31.21&[MIMAT0000414&hsa-let-7g-5p&offsets|0|-1;m-58&3|-|52268336|52268357&offsets|0|-1]|TGAGGTAGTAGTTTGTACAGT</td><td>>MI0000433|hsa-let-7g&WithFlank&3|-|52268272|52268367</td><td>11</td><td>31</td><td>21</td><td>TGAGGTAGTAGTTTGTACAGT</td><td>6179.1</td><td>5.4</td><td>0.6</td><td>8.8</td><td>1.6E-18</td><td>1.5E-17</td><td>MB231/MB468</td><td>NUC</td></tr>
<tr><td>>MI0000264|hsa-mir-7-2&WithFlank&15|+|88611819|88611940@38.60.23;MI0000265|hsa-mir-7-3&WithFlank&19|+|4770664|4770785@37.59.23;MI0000263|hsa-mir-7-1&WithFlank&9|-|83969742|83969863@30.52.23&[MIMAT0000252&hsa-miR-7-5p&offsets|0|-1;m-400&15|+|88611856|88611879&offsets|0|-1];[MIMAT0000252_1&hsa-miR-7-5p&offsets|0|-1;m-402&19|+|4770700|4770723&offsets|0|-1];[MIMAT0000252_2&hsa-miR-7-5p&offsets|0|-1;m-401&9|-|83969811|83969834&offsets|0|-1]|TGGAAGACTAGTGATTTTGTTGT</td><td>>MI0000264|hsa-mir-7-2&WithFlank&15|+|88611819|88611940</td><td>38</td><td>60</td><td>23</td><td>TGGAAGACTAGTGATTTTGTTGT</td><td>18285.7</td><td>7.9</td><td>0.9</td><td>8.7</td><td>2.4E-18</td><td>1.5E-17</td><td>BT20/MB468</td><td>CYTO</td></tr>
<tr><td>>hg38_21_+_8433172_8446622_RNA45SN1_45S_50ntFlanks@3872.3899.28|CTAATACATGCCGACGGGCGCTGACCCC</td><td>>hg38_21_+_8433172_8446622_RNA45SN1_45S_50ntFlanks</td><td>3872</td><td>3899</td><td>28</td><td>CTAATACATGCCGACGGGCGCTGACCCC</td><td>505.8</td><td>-12.8</td><td>1.5</td><td>-8.8</td><td>2.0E-18</td><td>1.5E-17</td><td>MB231/MB468</td><td>MITO</td></tr>
<tr><td>>hg38_21_+_8433172_8446622_RNA45SN1_45S_50ntFlanks@12686.12711.26|CCAGGCGGAACGATACGGCAGCGCCG</td><td>>hg38_21_+_8433172_8446622_RNA45SN1_45S_50ntFlanks</td><td>12686</td><td>12711</td><td>26</td><td>CCAGGCGGAACGATACGGCAGCGCCG</td><td>300.8</td><td>-12.1</td><td>1.4</td><td>-8.8</td><td>1.2E-18</td><td>1.5E-17</td><td>MB231/MB468</td><td>MP</td></tr>
<tr><td>>hg38_21_+_8433172_8446622_RNA45SN1_45S_50ntFlanks@8324.8356.33|CCGATAGTCAACAAGTACCGTAAGGGAAAGTTG</td><td>>hg38_21_+_8433172_8446622_RNA45SN1_45S_50ntFlanks</td><td>8324</td><td>8356</td><td>33</td><td>CCGATAGTCAACAAGTACCGTAAGGGAAAGTTG</td><td>681.7</td><td>-11.7</td><td>1.3</td><td>-8.7</td><td>2.4E-18</td><td>1.5E-17</td><td>BT20/MB468</td><td>CYTO</td></tr>
<tr><td>>hg38_21_+_8433172_8446622_RNA45SN1_45S_50ntFlanks@10866.10889.24|CTAAGGGCTGGGTCGGTCGGGCTG</td><td>>hg38_21_+_8433172_8446622_RNA45SN1_45S_50ntFlanks</td><td>10866</td><td>10889</td><td>24</td><td>CTAAGGGCTGGGTCGGTCGGGCTG</td><td>611.3</td><td>-13.1</td><td>1.5</td><td>-8.8</td><td>1.2E-18</td><td>1.5E-17</td><td>MB231/MB468</td><td>MP</td></tr>
<tr><td>>hg38_21_+_8433172_8446622_RNA45SN1_45S_50ntFlanks@10398.10424.27|GTTGAACATGGGTCAGTCGGTCCTGAG</td><td>>hg38_21_+_8433172_8446622_RNA45SN1_45S_50ntFlanks</td><td>10398</td><td>10424</td><td>27</td><td>GTTGAACATGGGTCAGTCGGTCCTGAG</td><td>1456.5</td><td>7.1</td><td>0.8</td><td>8.8</td><td>1.2E-18</td><td>1.5E-17</td><td>MB231/MB468</td><td>CYTO</td></tr>
<tr><td>>hg38_21_+_8433172_8446622_RNA45SN1_45S_50ntFlanks@6698.6718.21|AGCTAGCTGCGAGAATTAATG</td><td>>hg38_21_+_8433172_8446622_RNA45SN1_45S_50ntFlanks</td><td>6698</td><td>6718</td><td>21</td><td>AGCTAGCTGCGAGAATTAATG</td><td>5631.4</td><td>-4.8</td><td>0.6</td><td>-8.7</td><td>2.4E-18</td><td>1.5E-17</td><td>BT20/MB468</td><td>CYTO</td></tr>
<tr><td>>hg38_21_+_8433172_8446622_RNA45SN1_45S_50ntFlanks@6671.6696.26|CTCGGCTCGTGCGTCGATGAAGAACG</td><td>>hg38_21_+_8433172_8446622_RNA45SN1_45S_50ntFlanks</td><td>6671</td><td>6696</td><td>26</td><td>CTCGGCTCGTGCGTCGATGAAGAACG</td><td>1243.1</td><td>-5.8</td><td>0.7</td><td>-8.7</td><td>4.3E-18</td><td>1.5E-17</td><td>BT20/MB468</td><td>NUC</td></tr>
<tr><td>>MI0003643|hsa-mir-629&WithFlank&15|-|70079366|70079474@28.48.21&[MIMAT0004810&hsa-miR-629-5p&offsets|0|0;m-150&15|-|70079427|70079447&offsets|0|0]|TGGGTTTACGTTGGGAGAACT</td><td>>MI0003643|hsa-mir-629&WithFlank&15|-|70079366|70079474</td><td>28</td><td>48</td><td>21</td><td>TGGGTTTACGTTGGGAGAACT</td><td>262.2</td><td>12.0</td><td>1.4</td><td>8.8</td><td>1.2E-18</td><td>1.5E-17</td><td>MB231/MB468</td><td>CYTO</td></tr>
<tr><td>>hg38_21_+_8433172_8446622_RNA45SN1_45S_50ntFlanks@12974.12991.18|CGTACGTAGCAGAGCAGC</td><td>>hg38_21_+_8433172_8446622_RNA45SN1_45S_50ntFlanks</td><td>12974</td><td>12991</td><td>18</td><td>CGTACGTAGCAGAGCAGC</td><td>2946.5</td><td>-5.0</td><td>0.6</td><td>-8.7</td><td>2.5E-18</td><td>1.5E-17</td><td>BT20/MB468</td><td>CYTO</td></tr>
<tr><td>>hg38_21_+_8433172_8446622_RNA45SN1_45S_50ntFlanks@7974.8014.41|ACGCGACCTCAGATCAGACGTGGCGACCCGCTGAATTTAAG</td><td>>hg38_21_+_8433172_8446622_RNA45SN1_45S_50ntFlanks</td><td>7974</td><td>8014</td><td>41</td><td>ACGCGACCTCAGATCAGACGTGGCGACCCGCTGAATTTAAG</td><td>930.4</td><td>5.4</td><td>0.6</td><td>8.8</td><td>1.2E-18</td><td>1.5E-17</td><td>MB231/MB468</td><td>CYTO</td></tr>
<tr><td>>MI0000081|hsa-mir-24-2&WithFlank&19|-|13836281|13836365@56.77.22(+1U);MI0000080|hsa-mir-24-1&WithFlank&9|+|95086015|95086094@50.71.22(+1U)&[MIMAT0000080&hsa-miR-24-3p&offsets|0|0(+1U);m-36&19|-|13836288|13836310&offsets|0|-1(+1U)];[MIMAT0000080_1&hsa-miR-24-3p&offsets|0|0(+1U);m-35&9|+|95086064|95086086&offsets|0|-1(+1U)]|TGGCTCAGTTCAGCAGGAACAGT</td><td>>MI0000081|hsa-mir-24-2&WithFlank&19|-|13836281|13836365</td><td>56</td><td>77</td><td>22(+1U)</td><td>TGGCTCAGTTCAGCAGGAACAGT</td><td>5082.9</td><td>6.2</td><td>0.7</td><td>8.7</td><td>2.5E-18</td><td>1.5E-17</td><td>BT20/MB468</td><td>CYTO</td></tr>
<tr><td>>MI0000283|hsa-mir-203a&WithFlank&14|+|104117399|104117520@71.90.20&[MIMAT0000264&hsa-miR-203a-3p&offsets|0|-2;m-79&14|+|104117469|104117490&offsets|0|-2]|GTGAAATGTTTAGGACCACT</td><td>>MI0000283|hsa-mir-203a&WithFlank&14|+|104117399|104117520</td><td>71</td><td>90</td><td>20</td><td>GTGAAATGTTTAGGACCACT</td><td>150.4</td><td>4.0</td><td>0.5</td><td>8.7</td><td>4.3E-18</td><td>1.5E-17</td><td>BT20/MB468</td><td>NUC</td></tr>
<tr><td>>hg38_21_+_8433172_8446622_RNA45SN1_45S_50ntFlanks@9572.9601.30|CAAATCGGTCGTCCGACCTGGGTATAGGGG</td><td>>hg38_21_+_8433172_8446622_RNA45SN1_45S_50ntFlanks</td><td>9572</td><td>9601</td><td>30</td><td>CAAATCGGTCGTCCGACCTGGGTATAGGGG</td><td>1369.5</td><td>-5.8</td><td>0.7</td><td>-8.8</td><td>2.1E-18</td><td>1.5E-17</td><td>MB231/MB468</td><td>MITO</td></tr>
<tr><td>>hg38_21_+_8433172_8446622_RNA45SN1_45S_50ntFlanks@485.522.38|GATGTGGTGACGTCGTGCTCTCCCGGGCCGGGTCCGAG</td><td>>hg38_21_+_8433172_8446622_RNA45SN1_45S_50ntFlanks</td><td>485</td><td>522</td><td>38</td><td>GATGTGGTGACGTCGTGCTCTCCCGGGCCGGGTCCGAG</td><td>949.9</td><td>12.7</td><td>1.5</td><td>8.7</td><td>4.4E-18</td><td>1.5E-17</td><td>BT20/MB468</td><td>NUC</td></tr>
<tr><td>>hg38_21_+_8433172_8446622_RNA45SN1_45S_50ntFlanks@10572.10607.36|AGAAGCCGGCGGGAGCCCCGGGGAGAGTTCTCTTTT</td><td>>hg38_21_+_8433172_8446622_RNA45SN1_45S_50ntFlanks</td><td>10572</td><td>10607</td><td>36</td><td>AGAAGCCGGCGGGAGCCCCGGGGAGAGTTCTCTTTT</td><td>776.2</td><td>-12.6</td><td>1.4</td><td>-8.8</td><td>1.7E-18</td><td>1.5E-17</td><td>MB231/MB468</td><td>NUC</td></tr>
<tr><td>>hg38_21_+_8433172_8446622_RNA45SN1_45S_50ntFlanks@10116.10139.24|TGCGTCGGAGGGCGGCGGCGGCGG</td><td>>hg38_21_+_8433172_8446622_RNA45SN1_45S_50ntFlanks</td><td>10116</td><td>10139</td><td>24</td><td>TGCGTCGGAGGGCGGCGGCGGCGG</td><td>497.8</td><td>-12.7</td><td>1.5</td><td>-8.8</td><td>2.1E-18</td><td>1.5E-17</td><td>MB231/MB468</td><td>MITO</td></tr>
<tr><td>>hg38_21_+_8433172_8446622_RNA45SN1_45S_50ntFlanks@12161.12179.19|TGACTGGGGCGGTACACCT</td><td>>hg38_21_+_8433172_8446622_RNA45SN1_45S_50ntFlanks</td><td>12161</td><td>12179</td><td>19</td><td>TGACTGGGGCGGTACACCT</td><td>1077.1</td><td>-13.9</td><td>1.6</td><td>-8.8</td><td>2.1E-18</td><td>1.6E-17</td><td>MB231/MB468</td><td>MITO</td></tr>
<tr><td>>MI0000434|hsa-let-7i&WithFlank&12|+|62603680|62603775@12.33.22(+1C)&[MIMAT0000415&hsa-let-7i-5p&offsets|0|0(+1C);m-49&12|+|62603691|62603712&offsets|0|0(+1C)]|TGAGGTAGTAGTTTGTGCTGTTC</td><td>>MI0000434|hsa-let-7i&WithFlank&12|+|62603680|62603775</td><td>12</td><td>33</td><td>22(+1C)</td><td>TGAGGTAGTAGTTTGTGCTGTTC</td><td>479.6</td><td>6.5</td><td>0.7</td><td>8.7</td><td>2.6E-18</td><td>1.6E-17</td><td>BT20/MB468</td><td>CYTO</td></tr>
<tr><td>>MI0000462|hsa-mir-152&WithFlank&17|-|48037155|48037253@60.81.22(+1U)&[MIMAT0000438&hsa-miR-152-3p&offsets|0|+1(+1U);m-97&17|-|48037174|48037194&offsets|0|+1(+1U)]|TCAGTGCATGACAGAACTTGGGT</td><td>>MI0000462|hsa-mir-152&WithFlank&17|-|48037155|48037253</td><td>60</td><td>81</td><td>22(+1U)</td><td>TCAGTGCATGACAGAACTTGGGT</td><td>293.7</td><td>12.1</td><td>1.4</td><td>8.8</td><td>1.7E-18</td><td>1.6E-17</td><td>MB231/MB468</td><td>NUC</td></tr>
<tr><td>>hg38_21_+_8433172_8446622_RNA45SN1_45S_50ntFlanks@10807.10840.34|ACAATGTAGGTAAGGGAAGTCGGCAAGCCGGATC</td><td>>hg38_21_+_8433172_8446622_RNA45SN1_45S_50ntFlanks</td><td>10807</td><td>10840</td><td>34</td><td>ACAATGTAGGTAAGGGAAGTCGGCAAGCCGGATC</td><td>497.7</td><td>-12.8</td><td>1.5</td><td>-8.8</td><td>1.3E-18</td><td>1.6E-17</td><td>MB231/MB468</td><td>MP</td></tr>
<tr><td>>trnaMT_ValTAC_MT_+_1602_1670@47.71.25|TCAACTTAACTTGACCGCTCTGACC</td><td>>trnaMT_ValTAC_MT_+_1602_1670</td><td>47</td><td>71</td><td>25</td><td>TCAACTTAACTTGACCGCTCTGACC</td><td>312.9</td><td>12.3</td><td>1.4</td><td>8.8</td><td>1.3E-18</td><td>1.6E-17</td><td>MB231/MB468</td><td>CYTO</td></tr>
<tr><td>>hg38_21_+_8433172_8446622_RNA45SN1_45S_50ntFlanks@12008.12036.29|CCGCCGGTGAAATACCACTACTCTGATCG</td><td>>hg38_21_+_8433172_8446622_RNA45SN1_45S_50ntFlanks</td><td>12008</td><td>12036</td><td>29</td><td>CCGCCGGTGAAATACCACTACTCTGATCG</td><td>496.0</td><td>-12.8</td><td>1.5</td><td>-8.8</td><td>1.3E-18</td><td>1.6E-17</td><td>MB231/MB468</td><td>MP</td></tr>
<tr><td>>hg38_21_+_8433172_8446622_RNA45SN1_45S_50ntFlanks@6732.6756.25|ACATTGATCATCGACACTTCGAACG</td><td>>hg38_21_+_8433172_8446622_RNA45SN1_45S_50ntFlanks</td><td>6732</td><td>6756</td><td>25</td><td>ACATTGATCATCGACACTTCGAACG</td><td>958.1</td><td>-5.1</td><td>0.6</td><td>-8.7</td><td>4.5E-18</td><td>1.6E-17</td><td>BT20/MB468</td><td>NUC</td></tr>
<tr><td>>hg38_21_+_8433172_8446622_RNA45SN1_45S_50ntFlanks@9533.9563.31|GAGGAAACTCTGGTGGAGGTCCGTAGCGGTC</td><td>>hg38_21_+_8433172_8446622_RNA45SN1_45S_50ntFlanks</td><td>9533</td><td>9563</td><td>31</td><td>GAGGAAACTCTGGTGGAGGTCCGTAGCGGTC</td><td>4923.0</td><td>-6.8</td><td>0.8</td><td>-8.8</td><td>1.7E-18</td><td>1.6E-17</td><td>MB231/MB468</td><td>NUC</td></tr>
<tr><td>>MI0000289|hsa-mir-181a-1&WithFlank&1|-|198859038|198859159@30.54.25;MI0000269|hsa-mir-181a-2&WithFlank&9|+|124692436|124692557@45.69.25&[MIMAT0000256&hsa-miR-181a-5p&offsets|0|+2;m-32&1|-|198859108|198859130&offsets|0|+2];[MIMAT0000256_1&hsa-miR-181a-5p&offsets|0|+2;m-32&9|+|124692480|124692502&offsets|0|+2]|AACATTCAACGCTGTCGGTGAGTTT</td><td>>MI0000289|hsa-mir-181a-1&WithFlank&1|-|198859038|198859159</td><td>30</td><td>54</td><td>25</td><td>AACATTCAACGCTGTCGGTGAGTTT</td><td>1277.5</td><td>7.6</td><td>0.9</td><td>8.8</td><td>1.7E-18</td><td>1.6E-17</td><td>MB231/MB468</td><td>NUC</td></tr>
<tr><td>>MI0000078|hsa-mir-22&WithFlank&17|-|1713897|1713993@59.80.22(+1A)&[MIMAT0000077&hsa-miR-22-3p&offsets|0|0(+1A);m-2&17|-|1713914|1713935&offsets|0|0(+1A)]|AAGCTGCCAGTTGAAGAACTGTA</td><td>>MI0000078|hsa-mir-22&WithFlank&17|-|1713897|1713993</td><td>59</td><td>80</td><td>22(+1A)</td><td>AAGCTGCCAGTTGAAGAACTGTA</td><td>207.1</td><td>6.4</td><td>0.7</td><td>8.8</td><td>1.7E-18</td><td>1.6E-17</td><td>MB231/MB468</td><td>NUC</td></tr>
<tr><td>>hg38_21_+_8433172_8446622_RNA45SN1_45S_50ntFlanks@6721.6747.27|AATTGCAGGACACATTGATCATCGACA</td><td>>hg38_21_+_8433172_8446622_RNA45SN1_45S_50ntFlanks</td><td>6721</td><td>6747</td><td>27</td><td>AATTGCAGGACACATTGATCATCGACA</td><td>1735.0</td><td>-5.0</td><td>0.6</td><td>-8.8</td><td>1.7E-18</td><td>1.6E-17</td><td>MB231/MB468</td><td>NUC</td></tr>
<tr><td>>hg38_21_+_8433172_8446622_RNA45SN1_45S_50ntFlanks@11933.11967.35|AATAAGTGGGAGGCCCCCGGCGCCCCCCCGGTGTC</td><td>>hg38_21_+_8433172_8446622_RNA45SN1_45S_50ntFlanks</td><td>11933</td><td>11967</td><td>35</td><td>AATAAGTGGGAGGCCCCCGGCGCCCCCCCGGTGTC</td><td>530.2</td><td>-12.9</td><td>1.5</td><td>-8.8</td><td>1.3E-18</td><td>1.6E-17</td><td>MB231/MB468</td><td>MP</td></tr>
<tr><td>>MI0000285|hsa-mir-205&WithFlank&1|+|209432127|209432248@40.60.21&[MIMAT0000266&hsa-miR-205-5p&offsets|0|-1;m-351&1|+|209432166|209432187&offsets|0|-1]|TCCTTCATTCCACCGGAGTCT</td><td>>MI0000285|hsa-mir-205&WithFlank&1|+|209432127|209432248</td><td>40</td><td>60</td><td>21</td><td>TCCTTCATTCCACCGGAGTCT</td><td>6308.6</td><td>9.4</td><td>1.0</td><td>9.1</td><td>1.3E-19</td><td>1.6E-17</td><td>BT20/MB231</td><td>CYTO</td></tr>
<tr><td>>hg38_21_+_8433172_8446622_RNA45SN1_45S_50ntFlanks@11661.11681.21|TGATTTCTGCCCAGTGCTCTG</td><td>>hg38_21_+_8433172_8446622_RNA45SN1_45S_50ntFlanks</td><td>11661</td><td>11681</td><td>21</td><td>TGATTTCTGCCCAGTGCTCTG</td><td>5061.4</td><td>-6.3</td><td>0.7</td><td>-8.7</td><td>2.7E-18</td><td>1.6E-17</td><td>BT20/MB468</td><td>CYTO</td></tr>
<tr><td>>hg38_21_+_8433172_8446622_RNA45SN1_45S_50ntFlanks@12539.12557.19|ATGATGTGTTGTTGCCATG</td><td>>hg38_21_+_8433172_8446622_RNA45SN1_45S_50ntFlanks</td><td>12539</td><td>12557</td><td>19</td><td>ATGATGTGTTGTTGCCATG</td><td>704.2</td><td>-13.2</td><td>1.5</td><td>-8.7</td><td>2.2E-18</td><td>1.6E-17</td><td>MB231/MB468</td><td>MITO</td></tr>
</table>

| | | | | | | | | | | | | | |
|---|---|---|---|---|---|---|---|---|---|---|---|---|---|
| >MI0005756\|hsa-mir-934&WithFlank&X\|+\|136550872\|136550966@21.41.21&[MIMAT0004977&hsa-miR-934&offsets\|0\|-1;m-943&X\|+\|136550892\|136550913&offsets\|0\|-1]\|TGTCTACTACTGGAGACACTG | >MI0005756\|hsa-mir-934&WithFlank&X\|+\|136550872\|136550966 | 21 | 41 | 21 | TGTCTACTACTGGAGACACTG | 1261.4 | -14.2 | 1.6 | -8.8 | 1.3E-18 | 1.6E-17 | MB231/MB468 | MP |
| >am_MI0000809\|hsa-mir-151a&WithFlank&8\|-\|140732558\|140732659@52.73.22&[MIMAT0000757&hsa-miR-151a-3p&offsets\|-1\|0;m-30&8\|-\|140732587\|140732607&offsets\|-1\|0]\|ACTAGACTGAAGCTCCTTGAGG | >am_MI0000809\|hsa-mir-151a&WithFlank&8\|-\|140732558\|140732659 | 52 | 73 | 22 | ACTAGACTGAAGCTCCTTGAGG | 1241.4 | 5.1 | 0.6 | 8.8 | 1.8E-18 | 1.6E-17 | MB231/MB468 | NUC |
| >hg38_21_+_8433172_8446622_RNA45SN1_45S_50ntFlanks@5525.5550.26\|TAAAAGTCGTAACAAGGTTTCCGTAG | >hg38_21_+_8433172_8446622_RNA45SN1_45S_50ntFlanks | 5525 | 5550 | 26 | TAAAAGTCGTAACAAGGTTTCCGTAG | 684.2 | -12.4 | 1.4 | -8.8 | 1.8E-18 | 1.6E-17 | MB231/MB468 | NUC |
| >hg38_21_+_8433172_8446622_RNA45SN1_45S_50ntFlanks@5485.5524.40\|AGCGCTGAGAAGACGGTCGAACTTGACTATCTAGAGGAAG | >hg38_21_+_8433172_8446622_RNA45SN1_45S_50ntFlanks | 5485 | 5524 | 40 | AGCGCTGAGAAGACGGTCGAACTTGACTATCTAGAGGAAG | 486.4 | -12.7 | 1.5 | -8.7 | 2.2E-18 | 1.6E-17 | MB231/MB468 | MITO |
| >hg38_21_+_8433172_8446622_RNA45SN1_45S_50ntFlanks@485.514.30\|GATGTGGTGACGTCGTGCTCTCCCGGGCCG | >hg38_21_+_8433172_8446622_RNA45SN1_45S_50ntFlanks | 485 | 514 | 30 | GATGTGGTGACGTCGTGCTCTCCCGGGCCG | 491.4 | 8.1 | 0.9 | 8.7 | 4.7E-18 | 1.6E-17 | BT20/MB468 | NUC |
| >hg38_21_+_8433172_8446622_RNA45SN1_45S_50ntFlanks@11874.11900.27\|AGAAGACCCTGTTGAGCTTGACTCTAG | >hg38_21_+_8433172_8446622_RNA45SN1_45S_50ntFlanks | 11874 | 11900 | 27 | AGAAGACCCTGTTGAGCTTGACTCTAG | 1065.8 | -7.4 | 0.8 | -8.8 | 1.3E-18 | 1.6E-17 | MB231/MB468 | CYTO |
| >hg38_21_+_8433172_8446622_RNA45SN1_45S_50ntFlanks@12624.12653.30\|CCAATGGGGCGAAGCTACCATCTGTGGGAT | >hg38_21_+_8433172_8446622_RNA45SN1_45S_50ntFlanks | 12624 | 12653 | 30 | CCAATGGGGCGAAGCTACCATCTGTGGGAT | 919.1 | -11.9 | 1.4 | -8.8 | 2.0E-18 | 1.6E-17 | BT20/MB468 | MITO |
| >hg38_21_+_8433172_8446622_RNA45SN1_45S_50ntFlanks@7986.8005.20\|ATCAGACGTGGCGACCCGCT | >hg38_21_+_8433172_8446622_RNA45SN1_45S_50ntFlanks | 7986 | 8005 | 20 | ATCAGACGTGGCGACCCGCT | 2207.0 | -14.2 | 1.5 | -9.2 | 3.3E-20 | 1.6E-17 | BT20/MB468 | MP |
| >hg38_21_+_8433172_8446622_RNA45SN1_45S_50ntFlanks@11321.11338.18\|CACGTCTCGTCGCGCGCG | >hg38_21_+_8433172_8446622_RNA45SN1_45S_50ntFlanks | 11321 | 11338 | 18 | CACGTCTCGTCGCGCGCG | 310.0 | -12.1 | 1.4 | -8.8 | 1.4E-18 | 1.6E-17 | MB231/MB468 | MP |
| >hg38_21_+_8433172_8446622_RNA45SN1_45S_50ntFlanks@9203.9225.23\|CTCGGGGCCACGCGCGCGTCCCC | >hg38_21_+_8433172_8446622_RNA45SN1_45S_50ntFlanks | 9203 | 9225 | 23 | CTCGGGGCCACGCGCGCGTCCCC | 593.3 | -13.0 | 1.5 | -8.7 | 2.2E-18 | 1.6E-17 | MB231/MB468 | MITO |
| >MI0000440\|hsa-mir-27b&WithFlank&9\|+\|95085439\|95085547@67.88.22&[MIMAT0000419&hsa-miR-27b-3p&offsets\|0\|+1;m-39&9\|+\|95085505\|95085525&offsets\|0\|+1]\|TTCACAGTGGCTAAGTTCTGCA | >MI0000440\|hsa-mir-27b&WithFlank&9\|+\|95085439\|95085547 | 67 | 88 | 22 | TTCACAGTGGCTAAGTTCTGCA | 5936.7 | 5.8 | 0.7 | 8.8 | 1.8E-18 | 1.7E-17 | MB231/MB468 | NUC |
| >hg38_21_+_8433172_8446622_RNA45SN1_45S_50ntFlanks@10350.10384.35\|ACGAGAACTTTGAAGGCCGAAGTGGAGAAGGGTTC | >hg38_21_+_8433172_8446622_RNA45SN1_45S_50ntFlanks | 10350 | 10384 | 35 | ACGAGAACTTTGAAGGCCGAAGTGGAGAAGGGTTC | 4534.6 | -6.7 | 0.8 | -8.8 | 1.8E-18 | 1.7E-17 | MB231/MB468 | NUC |
| >hg38_21_+_8433172_8446622_RNA45SN1_45S_50ntFlanks@6726.6743.18\|CAGGACACATTGATCATC | >hg38_21_+_8433172_8446622_RNA45SN1_45S_50ntFlanks | 6726 | 6743 | 18 | CAGGACACATTGATCATC | 313.1 | -12.7 | 1.5 | -8.7 | 4.8E-18 | 1.7E-17 | BT20/MB468 | NUC |
| >hg38_21_+_8433172_8446622_RNA45SN1_45S_50ntFlanks@4241.4258.18\|CTCTTTCGAGGCCCTGTA | >hg38_21_+_8433172_8446622_RNA45SN1_45S_50ntFlanks | 4241 | 4258 | 18 | CTCTTTCGAGGCCCTGTA | 617.4 | -13.1 | 1.5 | -8.7 | 2.3E-18 | 1.7E-17 | MB231/MB468 | MITO |
| >hg38_21_+_8433172_8446622_RNA45SN1_45S_50ntFlanks@12478.12498.21\|ACTAATAGGGAACGTGAGCTG | >hg38_21_+_8433172_8446622_RNA45SN1_45S_50ntFlanks | 12478 | 12498 | 21 | ACTAATAGGGAACGTGAGCTG | 863.5 | -5.2 | 0.6 | -8.8 | 1.8E-18 | 1.7E-17 | MB231/MB468 | NUC |
| >trna10_AspGTC_12_-_125424193_125424264@32.70.39,trna12_AspGTC_12_-_125411891_125411962@32.70.39,trna144_AspGTC_6_-_27551236_27551307@32.70.39,trna38_AspGTC_17_-_8125556_8125627@32.70.39,trna45_AspGTC_6_+_27447453_27447524@32.70.39,trna48_AspGTC_6_+_27471523_27471594@32.70.39,trna4_AspGTC_12_+_96429799_96429870@32.70.39,trna69_AspGTC_1_-_161440205_161440276@32.70.39,trna72_AspGTC_1_-_161432824_161432895@32.70.39,trna75_AspGTC_1_-_161425414_161425485@32.70.39,trna78_AspGTC_1_-_161418033_161418104@32.70.39,trna81_AspGTC_1_-_161410615_161410686@32.70.39\|CTGTCACGCGGGAGACCGGGGTTCGATTCCCCGACGGGG | >trna10_AspGTC_12_-_125424193_125424264 | 32 | 70 | 39 | CTGTCACGCGGGAGACCGGGGTTCGATTCCCCGACGGGG | 561.9 | -11.9 | 1.3 | -8.8 | 1.3E-18 | 1.7E-17 | MB231/MB468 | CYTO |
| >hg38_21_+_8433172_8446622_RNA45SN1_45S_50ntFlanks@12456.12476.21\|CCAAGCGTTGGATTGTTCACC | >hg38_21_+_8433172_8446622_RNA45SN1_45S_50ntFlanks | 12456 | 12476 | 21 | CCAAGCGTTGGATTGTTCACC | 16821.6 | -6.9 | 0.8 | -8.8 | 1.8E-18 | 1.7E-17 | MB231/MB468 | NUC |
| >hg38_21_+_8433172_8446622_RNA45SN1_45S_50ntFlanks@11885.11912.28\|TTGAGCTTGACTCTAGTCTGGCACGGTG | >hg38_21_+_8433172_8446622_RNA45SN1_45S_50ntFlanks | 11885 | 11912 | 28 | TTGAGCTTGACTCTAGTCTGGCACGGTG | 677.3 | -13.3 | 1.5 | -8.8 | 1.4E-18 | 1.7E-17 | MB231/MB468 | MP |
| >hg38_MT_+_1621_3279_RNR2_16S_50ntFlanks@51.72.22\|GCTAAACCTAGCCCCAAACCCA | >hg38_MT_+_1621_3279_RNR2_16S_50ntFlanks | 51 | 72 | 22 | GCTAAACCTAGCCCCAAACCCA | 321.9 | 5.6 | 0.6 | 8.8 | 1.8E-18 | 1.7E-17 | MB231/MB468 | NUC |
| >hg38_21_+_8433172_8446622_RNA45SN1_45S_50ntFlanks@11321.11340.20\|CACGTCTCGTCGCGCGCGCG | >hg38_21_+_8433172_8446622_RNA45SN1_45S_50ntFlanks | 11321 | 11340 | 20 | CACGTCTCGTCGCGCGCGCG | 749.4 | -13.3 | 1.5 | -8.7 | 2.3E-18 | 1.7E-17 | MB231/MB468 | MITO |
| >hg38_21_+_8433172_8446622_RNA45SN1_45S_50ntFlanks@12324.12349.26\|TTGGGTTTTAAGCAGGAGGTGTCAGA | >hg38_21_+_8433172_8446622_RNA45SN1_45S_50ntFlanks | 12324 | 12349 | 26 | TTGGGTTTTAAGCAGGAGGTGTCAGA | 212.1 | -6.5 | 0.7 | -8.8 | 1.4E-18 | 1.7E-17 | MB231/MB468 | CYTO |
| >hg38_21_+_8433172_8446622_RNA45SN1_45S_50ntFlanks@9568.9596.29\|CGTGCAAATCGGTCGTCCGACCTGGGTAT | >hg38_21_+_8433172_8446622_RNA45SN1_45S_50ntFlanks | 9568 | 9596 | 29 | CGTGCAAATCGGTCGTCCGACCTGGGTAT | 6069.1 | -12.4 | 1.4 | -8.7 | 4.8E-18 | 1.7E-17 | BT20/MB468 | NUC |
| >hg38_21_+_8433172_8446622_RNA45SN1_45S_50ntFlanks@8342.8380.39\|CGTAAGGGAAAGTTGAAAAGAACTTTGAAGAGAGAGTTC | >hg38_21_+_8433172_8446622_RNA45SN1_45S_50ntFlanks | 8342 | 8380 | 39 | CGTAAGGGAAAGTTGAAAAGAACTTTGAAGAGAGAGTTC | 267.8 | 12.1 | 1.4 | 8.8 | 1.4E-18 | 1.7E-17 | MB231/MB468 | CYTO |
| >hg38_21_+_8433172_8446622_RNA45SN1_45S_50ntFlanks@12934.12972.39\|CTAAACCATTCGTAGACGACCTGCTTCTGGGTCGGGGTT | >hg38_21_+_8433172_8446622_RNA45SN1_45S_50ntFlanks | 12934 | 12972 | 39 | CTAAACCATTCGTAGACGACCTGCTTCTGGGTCGGGGTT | 7894.8 | -9.2 | 1.0 | -8.8 | 1.4E-18 | 1.7E-17 | MB231/MB468 | CYTO |
| >hg38_21_+_8433172_8446622_RNA45SN1_45S_50ntFlanks@5421.5445.25\|CGATTGGATGGTTTAGTGAGGCCCT | >hg38_21_+_8433172_8446622_RNA45SN1_45S_50ntFlanks | 5421 | 5445 | 25 | CGATTGGATGGTTTAGTGAGGCCCT | 890.2 | -13.7 | 1.6 | -8.8 | 1.4E-18 | 1.7E-17 | MB231/MB468 | MP |
| >hg38_21_+_8433172_8446622_RNA45SN1_45S_50ntFlanks@476.509.34\|TTCGTGATCGATGTGGTGACGTCGTGCTCTCCCG | >hg38_21_+_8433172_8446622_RNA45SN1_45S_50ntFlanks | 476 | 509 | 34 | TTCGTGATCGATGTGGTGACGTCGTGCTCTCCCG | 526.7 | 8.0 | 0.9 | 8.7 | 2.8E-18 | 1.7E-17 | BT20/MB468 | CYTO |
| >hg38_21_+_8433172_8446622_RNA45SN1_45S_50ntFlanks@11890.11912.23\|CTTGACTCTAGTCTGGCACGGTG | >hg38_21_+_8433172_8446622_RNA45SN1_45S_50ntFlanks | 11890 | 11912 | 23 | CTTGACTCTAGTCTGGCACGGTG | 1050.8 | -10.0 | 1.1 | -8.7 | 2.3E-18 | 1.7E-17 | MB231/MB468 | MITO |
| >hg38_21_+_8433172_8446622_RNA45SN1_45S_50ntFlanks@9504.9523.20\|GTGAACTATGCCTGGGCAGG | >hg38_21_+_8433172_8446622_RNA45SN1_45S_50ntFlanks | 9504 | 9523 | 20 | GTGAACTATGCCTGGGCAGG | 1699.2 | -10.1 | 1.2 | -8.7 | 2.3E-18 | 1.7E-17 | MB231/MB468 | MITO |
| >hg38_21_+_8433172_8446622_RNA45SN1_45S_50ntFlanks@4735.4758.24\|ACGAAAGTCGGAGGTTCGAAGACG | >hg38_21_+_8433172_8446622_RNA45SN1_45S_50ntFlanks | 4735 | 4758 | 24 | ACGAAAGTCGGAGGTTCGAAGACG | 2973.0 | -6.8 | 0.8 | -8.7 | 4.9E-18 | 1.7E-17 | BT20/MB468 | NUC |
| >hg38_21_+_8433172_8446622_RNA45SN1_45S_50ntFlanks@11208.11225.18\|GCGGCGACTCTGGACGCG | >hg38_21_+_8433172_8446622_RNA45SN1_45S_50ntFlanks | 11208 | 11225 | 18 | GCGGCGACTCTGGACGCG | 6274.0 | -6.7 | 0.8 | -8.7 | 2.8E-18 | 1.7E-17 | BT20/MB468 | CYTO |
| >hg38_21_+_8433172_8446622_RNA45SN1_45S_50ntFlanks@10681.10711.31\|TTCCGGCGGCGTCCGGTGAGCTCTCGCTGGC | >hg38_21_+_8433172_8446622_RNA45SN1_45S_50ntFlanks | 10681 | 10711 | 31 | TTCCGGCGGCGTCCGGTGAGCTCTCGCTGGC | 316.7 | -12.2 | 1.4 | -8.8 | 1.4E-18 | 1.7E-17 | MB231/MB468 | MP |
| >hg38_21_+_8433172_8446622_RNA45SN1_45S_50ntFlanks@12935.12959.25\|TAAACCATTCGTAGACGACCTGCTT | >hg38_21_+_8433172_8446622_RNA45SN1_45S_50ntFlanks | 12935 | 12959 | 25 | TAAACCATTCGTAGACGACCTGCTT | 363.1 | -4.1 | 0.5 | -8.8 | 1.9E-18 | 1.7E-17 | MB231/MB468 | NUC |
| >hg38_21_+_8433172_8446622_RNA45SN1_45S_50ntFlanks@13004.13025.22\|ATCTATTGAAAGTCAGCCCTCG | >hg38_21_+_8433172_8446622_RNA45SN1_45S_50ntFlanks | 13004 | 13025 | 22 | ATCTATTGAAAGTCAGCCCTCG | 1152.1 | -13.1 | 1.5 | -8.8 | 1.9E-18 | 1.7E-17 | MB231/MB468 | NUC |
| >hg38_21_+_8433172_8446622_RNA45SN1_45S_50ntFlanks@11821.11844.24\|CTATCCAGCGAAACCACAGCCAAG | >hg38_21_+_8433172_8446622_RNA45SN1_45S_50ntFlanks | 11821 | 11844 | 24 | CTATCCAGCGAAACCACAGCCAAG | 889.8 | -13.7 | 1.6 | -8.8 | 1.4E-18 | 1.7E-17 | MB231/MB468 | MP |
| >hg38_21_+_8433172_8446622_RNA45SN1_45S_50ntFlanks@12539.12568.30\|ATGATGTGTTGTTGCCATGGTAATCCTGCT | >hg38_21_+_8433172_8446622_RNA45SN1_45S_50ntFlanks | 12539 | 12568 | 30 | ATGATGTGTTGTTGCCATGGTAATCCTGCT | 440.9 | -12.6 | 1.4 | -8.7 | 2.4E-18 | 1.7E-17 | MB231/MB468 | MITO |
| >hg38_21_+_8433172_8446622_RNA45SN1_45S_50ntFlanks@10351.10377.27\|CGAGAACTTTGAAGGCCGAAGTGGAGA | >hg38_21_+_8433172_8446622_RNA45SN1_45S_50ntFlanks | 10351 | 10377 | 27 | CGAGAACTTTGAAGGCCGAAGTGGAGA | 32715.0 | -11.3 | 1.3 | -8.8 | 1.4E-18 | 1.7E-17 | MB231/MB468 | MP |
| >hg38_21_+_8433172_8446622_RNA45SN1_45S_50ntFlanks@3853.3898.46\|AACTGTGGTAATTCTAGAGCTAATACATGCCGACGGGCGCTGACCC | >hg38_21_+_8433172_8446622_RNA45SN1_45S_50ntFlanks | 3853 | 3898 | 46 | AACTGTGGTAATTCTAGAGCTAATACATGCCGACGGGCGCTGACCC | 1514.4 | -15.0 | 1.7 | -8.7 | 5.0E-18 | 1.7E-17 | BT20/MB468 | NUC |
| >hg38_21_+_8433172_8446622_RNA45SN1_45S_50ntFlanks@12511.12538.28\|CGTGAGACAGGTTAGTTTTACCCTACTG | >hg38_21_+_8433172_8446622_RNA45SN1_45S_50ntFlanks | 12511 | 12538 | 28 | CGTGAGACAGGTTAGTTTTACCCTACTG | 325.6 | -12.2 | 1.4 | -8.8 | 1.5E-18 | 1.7E-17 | MB231/MB468 | MP |
| >MI0000061\|hsa-let-7a-2&WithFlank&11\|-\|122146516\|122146599@11.31.21;MI0000062\|hsa-let-7a-3&WithFlank&22\|+\|46112743\|46112828@10.30.21;MI0000060\|hsa-let-7a-1&WithFlank&9\|+\|94175951\|94176042@12.32.21&[MIMAT0000062&hsa-let-7a-5p&offsets\|0\|-1;m-6&11\|-\|122146568\|122146589&offsets\|0\|-1];[MIMAT0000062_1&hsa-let-7a-5p&offsets\|0\|-1;m-5&22\|+\|46112752\|46112773&offsets\|0\|-1];[MIMAT0000062_2&hsa-let-7a-5p&offsets\|0\|-1;m-4&9\|+\|94175962\|94175983&offsets\|0\|-1]\|TGAGGTAGTAGGTTGTATAGT | >MI0000061\|hsa-let-7a-2&WithFlank&11\|-\|122146516\|122146599 | 11 | 31 | 21 | TGAGGTAGTAGGTTGTATAGT | 8183.0 | 6.6 | 0.7 | 8.8 | 1.4E-18 | 1.7E-17 | MB231/MB468 | CYTO |
| >hg38_21_+_8433172_8446622_RNA45SN1_45S_50ntFlanks@11884.11903.20\|GTTGAGCTTGACTCTAGTCT | >hg38_21_+_8433172_8446622_RNA45SN1_45S_50ntFlanks | 11884 | 11903 | 20 | GTTGAGCTTGACTCTAGTCT | 1368.3 | 6.7 | 0.8 | 8.8 | 2.1E-18 | 1.7E-17 | BT20/MB468 | MITO |
| >hg38_21_+_8433172_8446622_RNA45SN1_45S_50ntFlanks@5449.5466.18\|ATCGGCCCCGCCGGGGTC | >hg38_21_+_8433172_8446622_RNA45SN1_45S_50ntFlanks | 5449 | 5466 | 18 | ATCGGCCCCGCCGGGGTC | 491.7 | -12.7 | 1.5 | -8.7 | 2.4E-18 | 1.8E-17 | MB231/MB468 | MITO |
| >hg38_21_+_8433172_8446622_RNA45SN1_45S_50ntFlanks@11576.11600.25\|CTTAGAACTGGTGCGGACCAGGGGA | >hg38_21_+_8433172_8446622_RNA45SN1_45S_50ntFlanks | 11576 | 11600 | 25 | CTTAGAACTGGTGCGGACCAGGGGA | 596.3 | -13.1 | 1.5 | -8.8 | 1.5E-18 | 1.8E-17 | MB231/MB468 | MP |
| >trna130_GlnTTG_6_-_27763640_27763711@1.29.29,trna173_GlnTTG_6_-_26311975_26312046@1.29.29,trna174_GlnTTG_6_-_26311424_26311495@1.29.29\|GGCCCCATGGTGTAATGGTTAGCACTCTG | >trna130_GlnTTG_6_-_27763640_27763711 | 1 | 29 | 29 | GGCCCCATGGTGTAATGGTTAGCACTCTG | 1864.9 | -8.1 | 0.9 | -8.8 | 1.4E-18 | 1.8E-17 | MB231/MB468 | CYTO |
| >MI0000301\|hsa-mir-224&WithFlank&X\|-\|151958572\|151958664@14.37.24&[MIMAT0000281&hsa-miR-224-5p&offsets\|+1\|0;m-238&X\|-\|151958630\|151958651&offsets\|0\|+2]\|CAAGTCACTAGTGGTTCCGTTTAG | >MI0000301\|hsa-mir-224&WithFlank&X\|-\|151958572\|151958664 | 14 | 37 | 24 | CAAGTCACTAGTGGTTCCGTTTAG | 4784.7 | 13.0 | 1.5 | 8.8 | 2.1E-18 | 1.8E-17 | BT20/MB468 | MITO |
| >MI0000283\|hsa-mir-203a&WithFlank&14\|+\|104117399\|104117520@72.93.22&[MIMAT0000264&hsa-miR-203a-3p&offsets\|+1\|+1;m-79&14\|+\|104117469\|104117490&offsets\|+1\|+1]\|TGAAATGTTTAGGACCACTAGA | >MI0000283\|hsa-mir-203a&WithFlank&14\|+\|104117399\|104117520 | 72 | 93 | 22 | TGAAATGTTTAGGACCACTAGA | 721.0 | 12.9 | 1.4 | 9.0 | 1.5E-19 | 1.8E-17 | BT20/MB231 | CYTO |
| >hg38_21_+_8433172_8446622_RNA45SN1_45S_50ntFlanks@5484.5526.43\|GAGCGCTGAGAAGACGGTCGAACTTGACTATCTAGAGGAAGTA | >hg38_21_+_8433172_8446622_RNA45SN1_45S_50ntFlanks | 5484 | 5526 | 43 | GAGCGCTGAGAAGACGGTCGAACTTGACTATCTAGAGGAAGTA | 534.0 | -12.8 | 1.5 | -8.7 | 2.4E-18 | 1.8E-17 | MB231/MB468 | MITO |

<table>
<tr><td>>hg38_21_+_8433172_8446622_RNA45SN1_45S_50ntFlanks@5467.5499.33|GGCCCACGGCCCTGGCGGAGCGCTGAGAAGACG</td><td>>hg38_21_+_8433172_8446622_RNA45SN1_45S_50ntFlanks</td><td>5467</td><td>5499</td><td>33</td><td>GGCCCACGGCCCTGGCGGAGCGCTGAGAAGACG</td><td>3154.2</td><td>-15.5</td><td>1.8</td><td>-8.8</td><td>1.5E-18</td><td>1.8E-17</td><td>MB231/MB468</td><td>MP</td></tr>
<tr><td>>hg38_21_+_8433172_8446622_RNA45SN1_45S_50ntFlanks@10349.10366.18|AACGAGAACTTTGAAGGC</td><td>>hg38_21_+_8433172_8446622_RNA45SN1_45S_50ntFlanks</td><td>10349</td><td>10366</td><td>18</td><td>AACGAGAACTTTGAAGGC</td><td>2102.8</td><td>-6.1</td><td>0.7</td><td>-8.6</td><td>5.2E-18</td><td>1.8E-17</td><td>BT20/MB468</td><td>NUC</td></tr>
<tr><td>>hg38_21_+_8433172_8446622_RNA45SN1_45S_50ntFlanks@11572.11609.38|CCGACTTAGAACTGGTGCGGACCAGGGGAATCCGACTG</td><td>>hg38_21_+_8433172_8446622_RNA45SN1_45S_50ntFlanks</td><td>11572</td><td>11609</td><td>38</td><td>CCGACTTAGAACTGGTGCGGACCAGGGGAATCCGACTG</td><td>1044.2</td><td>-5.8</td><td>0.7</td><td>-8.6</td><td>5.2E-18</td><td>1.8E-17</td><td>BT20/MB468</td><td>NUC</td></tr>
<tr><td>>hg38_21_+_8433172_8446622_RNA45SN1_45S_50ntFlanks@11592.11612.21|ACCAGGGGAATCCGACTGTTT</td><td>>hg38_21_+_8433172_8446622_RNA45SN1_45S_50ntFlanks</td><td>11592</td><td>11612</td><td>21</td><td>ACCAGGGGAATCCGACTGTTT</td><td>816.2</td><td>-13.5</td><td>1.5</td><td>-8.8</td><td>1.5E-18</td><td>1.8E-17</td><td>MB231/MB468</td><td>MP</td></tr>
<tr><td>>MI0000272|hsa-mir-182&WithFlank&7|-|129770377|129770498@29.48.20&[MIMAT0000259&hsa-miR-182-5p&offsets|0|-4;m-22&7|-|129770449|129770470&offsets|0|-2]|TTTGGCAATGGTAGAACTCA</td><td>>MI0000272|hsa-mir-182&WithFlank&7|-|129770377|129770498</td><td>29</td><td>48</td><td>20</td><td>TTTGGCAATGGTAGAACTCA</td><td>388.0</td><td>12.9</td><td>1.5</td><td>8.7</td><td>3.0E-18</td><td>1.8E-17</td><td>BT20/MB468</td><td>CYTO</td></tr>
<tr><td>>MI0016887|hsa-mir-4521&WithFlank&17|+|8186939|8187010@10.35.26&[MIMAT0019058&hsa-miR-4521&offsets|0|+4;m-765&17|+|8186948|8186969&offsets|0|+4]|GCTAAGGAAGTCCTGTGCTCAGTTTT</td><td>>MI0016887|hsa-mir-4521&WithFlank&17|+|8186939|8187010</td><td>10</td><td>35</td><td>26</td><td>GCTAAGGAAGTCCTGTGCTCAGTTTT</td><td>338.8</td><td>5.6</td><td>0.6</td><td>8.8</td><td>2.0E-18</td><td>1.8E-17</td><td>MB231/MB468</td><td>NUC</td></tr>
<tr><td>>MI0000477|hsa-mir-146a&WithFlank&5|+|160485346|160485456@27.47.21&[MIMAT0000449&hsa-miR-146a-5p&offsets|0|-1;m-129&5|+|160485372|160485393&offsets|0|-1]|TGAGAACTGAATTCCATGGGT</td><td>>MI0000477|hsa-mir-146a&WithFlank&5|+|160485346|160485456</td><td>27</td><td>47</td><td>21</td><td>TGAGAACTGAATTCCATGGGT</td><td>1988.9</td><td>14.2</td><td>1.6</td><td>8.7</td><td>2.5E-18</td><td>1.8E-17</td><td>MB231/MB468</td><td>MITO</td></tr>
<tr><td>>am_trna128_GlyGCC_6_-_27870686_27870756@1.35.35,trna18_GlyGCC_16_+_70822597_70822667@1.35.35,trna19_GlyGCC_16_+_70823410_70823480@1.35.35,trna19_GlyGCC_2_-_157257659_157257729@1.35.35,trna24_GlyGCC_16_-_70812942_70813012@1.35.35,trna25_GlyGCC_16_-_70812114_70812184@1.35.35,trna5_GlyGCC_17_+_8029064_8029134@1.35.35,trna68_GlyGCC_1_-_161493637_161493707@1.35.35|GCATTGGTGGTTCAGTGGTAGAATTCTCGCCTGCC</td><td>>am_trna128_GlyGCC_6_-_27870686_27870756</td><td>1</td><td>35</td><td>35</td><td>GCATTGGTGGTTCAGTGGTAGAATTCTCGCCTGCC</td><td>4118.4</td><td>9.0</td><td>1.0</td><td>8.8</td><td>2.0E-18</td><td>1.8E-17</td><td>MB231/MB468</td><td>NUC</td></tr>
<tr><td>>MI0001733|hsa-mir-452&WithFlank&X|-|151959622|151959718@23.44.22&[MIMAT0001635&hsa-miR-452-5p&offsets|+3|+3;m-188&X|-|151959677|151959699&offsets|+3|+2]|TGTTTGCAGAGGAAACTGAGAC</td><td>>MI0001733|hsa-mir-452&WithFlank&X|-|151959622|151959718</td><td>23</td><td>44</td><td>22</td><td>TGTTTGCAGAGGAAACTGAGAC</td><td>110.4</td><td>9.6</td><td>1.1</td><td>8.6</td><td>5.2E-18</td><td>1.8E-17</td><td>BT20/MB468</td><td>NUC</td></tr>
<tr><td>>MI0000077|hsa-mir-21&WithFlank&17|+|59841260|59841343@14.35.22(+1A)&[MIMAT0000076&hsa-miR-21-5p&offsets|0|0(+1A);m-1&17|+|59841273|59841294&offsets|0|0(+1A)]|TAGCTTATCAGACTGATGTTGAA</td><td>>MI0000077|hsa-mir-21&WithFlank&17|+|59841260|59841343</td><td>14</td><td>35</td><td>22(+1A)</td><td>TAGCTTATCAGACTGATGTTGAA</td><td>7113.6</td><td>6.3</td><td>0.7</td><td>8.7</td><td>2.2E-18</td><td>1.8E-17</td><td>BT20/MB468</td><td>MITO</td></tr>
<tr><td>>hg38_21_+_8433172_8446622_RNA45SN1_45S_50ntFlanks@3876.3898.23|TACATGCCGACGGGCGCTGACCC</td><td>>hg38_21_+_8433172_8446622_RNA45SN1_45S_50ntFlanks</td><td>3876</td><td>3898</td><td>23</td><td>TACATGCCGACGGGCGCTGACCC</td><td>633.2</td><td>-13.2</td><td>1.5</td><td>-8.8</td><td>1.5E-18</td><td>1.8E-17</td><td>MB231/MB468</td><td>MP</td></tr>
<tr><td>>MI0000102|hsa-mir-100&WithFlank&11|-|122152223|122152314@19.40.22(+2U)&[MIMAT0000098&hsa-miR-100-5p&offsets|0|0(+2U);m-27&11|-|122152276|122152296&offsets|0|+1(+2U)]|AACCCGTAGATCCGAACTTGTGTT</td><td>>MI0000102|hsa-mir-100&WithFlank&11|-|122152223|122152314</td><td>19</td><td>40</td><td>22(+2U)</td><td>AACCCGTAGATCCGAACTTGTGTT</td><td>881.2</td><td>-13.0</td><td>1.4</td><td>-9.0</td><td>1.9E-19</td><td>1.8E-17</td><td>BT20/MB231</td><td>MITO</td></tr>
<tr><td>>hg38_21_+_8433172_8446622_RNA45SN1_45S_50ntFlanks@7988.8014.27|CAGACGTGGCGACCCGCTGAATTTAAG</td><td>>hg38_21_+_8433172_8446622_RNA45SN1_45S_50ntFlanks</td><td>7988</td><td>8014</td><td>27</td><td>CAGACGTGGCGACCCGCTGAATTTAAG</td><td>1615.1</td><td>-10.2</td><td>1.2</td><td>-8.8</td><td>1.6E-18</td><td>1.9E-17</td><td>MB231/MB468</td><td>MP</td></tr>
<tr><td>>hg38_21_+_8433172_8446622_RNA45SN1_45S_50ntFlanks@4731.4757.27|AAGAACGAAAGTCGGAGGTTCGAAGAC</td><td>>hg38_21_+_8433172_8446622_RNA45SN1_45S_50ntFlanks</td><td>4731</td><td>4757</td><td>27</td><td>AAGAACGAAAGTCGGAGGTTCGAAGAC</td><td>351.0</td><td>-3.5</td><td>0.4</td><td>-8.6</td><td>5.4E-18</td><td>1.9E-17</td><td>BT20/MB468</td><td>NUC</td></tr>
<tr><td>>MI0000108|hsa-mir-103a-2&WithFlank&20|+|3917488|3917577@54.76.23(+1U);MI0000109|hsa-mir-103a-1&WithFlank&5|-|168560890|168560979@54.76.23(+1U)&[MIMAT0000101&hsa-miR-103a-3p&offsets|0|0(+1U);m-12&20|+|3917541|3917563&offsets|0|0(+1U)];[MIMAT0000101_1&hsa-miR-103a-3p&offsets|0|0(+1U);m-13&5|-|168560904|168560926&offsets|0|0(+1U);m-44220&5|-|168560908|168560926&offsets|0|+4(+1U)]|AGCAGCATTGTACAGGGCTATGAT</td><td>>MI0000108|hsa-mir-103a-2&WithFlank&20|+|3917488|3917577</td><td>54</td><td>76</td><td>23(+1U)</td><td>AGCAGCATTGTACAGGGCTATGAT</td><td>2903.2</td><td>5.7</td><td>0.7</td><td>8.8</td><td>2.1E-18</td><td>1.9E-17</td><td>MB231/MB468</td><td>NUC</td></tr>
<tr><td>>hg38_21_+_8433172_8446622_RNA45SN1_45S_50ntFlanks@5472.5516.45|ACGGCCCTGGCGGAGCGCTGAGAAGACGGTCGAACTTGACTATCT</td><td>>hg38_21_+_8433172_8446622_RNA45SN1_45S_50ntFlanks</td><td>5472</td><td>5516</td><td>45</td><td>ACGGCCCTGGCGGAGCGCTGAGAAGACGGTCGAACTTGACTATCT</td><td>5891.1</td><td>7.2</td><td>0.8</td><td>8.7</td><td>3.1E-18</td><td>1.9E-17</td><td>BT20/MB468</td><td>CYTO</td></tr>
<tr><td>>hg38_21_+_8433172_8446622_RNA45SN1_45S_50ntFlanks@9306.9336.31|CAAGGAGTCTAACACGTGCGCGAGTCGGGGG</td><td>>hg38_21_+_8433172_8446622_RNA45SN1_45S_50ntFlanks</td><td>9306</td><td>9336</td><td>31</td><td>CAAGGAGTCTAACACGTGCGCGAGTCGGGGG</td><td>518.1</td><td>-12.8</td><td>1.5</td><td>-8.7</td><td>2.6E-18</td><td>1.9E-17</td><td>MB231/MB468</td><td>MITO</td></tr>
<tr><td>>hg38_21_+_8433172_8446622_RNA45SN1_45S_50ntFlanks@10508.10533.26|CGAATCCGGAGTGGCGGAGATGGGCG</td><td>>hg38_21_+_8433172_8446622_RNA45SN1_45S_50ntFlanks</td><td>10508</td><td>10533</td><td>26</td><td>CGAATCCGGAGTGGCGGAGATGGGCG</td><td>653.5</td><td>-13.1</td><td>1.5</td><td>-8.7</td><td>2.6E-18</td><td>1.9E-17</td><td>MB231/MB468</td><td>MITO</td></tr>
<tr><td>>hg38_21_+_8433172_8446622_RNA45SN1_45S_50ntFlanks@4729.4752.24|TCAAGAACGAAAGTCGGAGGTTCG</td><td>>hg38_21_+_8433172_8446622_RNA45SN1_45S_50ntFlanks</td><td>4729</td><td>4752</td><td>24</td><td>TCAAGAACGAAAGTCGGAGGTTCG</td><td>316.2</td><td>-12.2</td><td>1.4</td><td>-8.8</td><td>1.6E-18</td><td>1.9E-17</td><td>MB231/MB468</td><td>MP</td></tr>
<tr><td>>hg38_21_+_8433172_8446622_RNA45SN1_45S_50ntFlanks@10432.10453.22|CGAGCGCCGTTCCGAAGGGACG</td><td>>hg38_21_+_8433172_8446622_RNA45SN1_45S_50ntFlanks</td><td>10432</td><td>10453</td><td>22</td><td>CGAGCGCCGTTCCGAAGGGACG</td><td>532.7</td><td>-12.9</td><td>1.5</td><td>-8.8</td><td>1.6E-18</td><td>1.9E-17</td><td>MB231/MB468</td><td>MP</td></tr>
<tr><td>>hg38_21_+_8433172_8446622_RNA45SN1_45S_50ntFlanks@10116.10138.23|TGCGTCGGAGGGCGGCGGCGGCG</td><td>>hg38_21_+_8433172_8446622_RNA45SN1_45S_50ntFlanks</td><td>10116</td><td>10138</td><td>23</td><td>TGCGTCGGAGGGCGGCGGCGGCG</td><td>439.6</td><td>-12.6</td><td>1.4</td><td>-8.7</td><td>2.7E-18</td><td>1.9E-17</td><td>MB231/MB468</td><td>MITO</td></tr>
<tr><td>>hg38_21_+_8433172_8446622_RNA45SN1_45S_50ntFlanks@6667.6689.23|ATCACTCGGCTCGTGCGTCGATG</td><td>>hg38_21_+_8433172_8446622_RNA45SN1_45S_50ntFlanks</td><td>6667</td><td>6689</td><td>23</td><td>ATCACTCGGCTCGTGCGTCGATG</td><td>485.9</td><td>-12.7</td><td>1.5</td><td>-8.7</td><td>2.7E-18</td><td>1.9E-17</td><td>MB231/MB468</td><td>MITO</td></tr>
<tr><td>>hg38_21_+_8433172_8446622_RNA45SN1_45S_50ntFlanks@4472.4496.25|CTTAGCTGAGTGTCCCGCGGGGCCC</td><td>>hg38_21_+_8433172_8446622_RNA45SN1_45S_50ntFlanks</td><td>4472</td><td>4496</td><td>25</td><td>CTTAGCTGAGTGTCCCGCGGGGCCC</td><td>366.6</td><td>-6.6</td><td>0.8</td><td>-8.8</td><td>1.6E-18</td><td>1.9E-17</td><td>MB231/MB468</td><td>CYTO</td></tr>
<tr><td>>MI0000084|hsa-mir-26b&WithFlank&2|+|218402640|218402728@18.39.22&[MIMAT0000083&hsa-miR-26b-5p&offsets|0|+1;m-53&2|+|218402657|218402678&offsets|0|0]|TTCAAGTAATTCAGGATAGGTT</td><td>>MI0000084|hsa-mir-26b&WithFlank&2|+|218402640|218402728</td><td>18</td><td>39</td><td>22</td><td>TTCAAGTAATTCAGGATAGGTT</td><td>2572.9</td><td>4.7</td><td>0.5</td><td>8.7</td><td>2.3E-18</td><td>1.9E-17</td><td>BT20/MB468</td><td>MITO</td></tr>
<tr><td>>hg38_21_+_8433172_8446622_RNA45SN1_45S_50ntFlanks@9714.9737.24|CGAATGATTAGAGGTCTTGGGGCC</td><td>>hg38_21_+_8433172_8446622_RNA45SN1_45S_50ntFlanks</td><td>9714</td><td>9737</td><td>24</td><td>CGAATGATTAGAGGTCTTGGGGCC</td><td>885.4</td><td>-10.1</td><td>1.2</td><td>-8.7</td><td>2.2E-18</td><td>2.0E-17</td><td>MB231/MB468</td><td>NUC</td></tr>
<tr><td>>am_MI0000809|hsa-mir-151a&WithFlank&8|-|140732558|140732659@53.71.19&[MIMAT0000757&hsa-miR-151a-3p&offsets|0|-2;m-30&8|-|140732587|140732607&offsets|0|-2]|CTAGACTGAAGCTCCTTGA</td><td>>am_MI0000809|hsa-mir-151a&WithFlank&8|-|140732558|140732659</td><td>53</td><td>71</td><td>19</td><td>CTAGACTGAAGCTCCTTGA</td><td>1722.2</td><td>10.5</td><td>1.2</td><td>8.7</td><td>3.2E-18</td><td>2.0E-17</td><td>BT20/MB468</td><td>CYTO</td></tr>
<tr><td>>hg38_21_+_8433172_8446622_RNA45SN1_45S_50ntFlanks@6652.6684.33|GACTCTTAGCGGTGGATCACTCGGCTCGTGCGT</td><td>>hg38_21_+_8433172_8446622_RNA45SN1_45S_50ntFlanks</td><td>6652</td><td>6684</td><td>33</td><td>GACTCTTAGCGGTGGATCACTCGGCTCGTGCGT</td><td>734.3</td><td>5.7</td><td>0.7</td><td>8.7</td><td>3.3E-18</td><td>2.0E-17</td><td>BT20/MB468</td><td>CYTO</td></tr>
<tr><td>>MI0000746|hsa-mir-99b&WithFlank&19|+|51692606|51692687@13.32.20(+1A)&[MIMAT0000689&hsa-miR-99b-5p&offsets|0|-2(+1A);m-10&19|+|51692618|51692639&offsets|0|-2(+1A)]|CACCCGTAGAACCGACCTTGA</td><td>>MI0000746|hsa-mir-99b&WithFlank&19|+|51692606|51692687</td><td>13</td><td>32</td><td>20(+1A)</td><td>CACCCGTAGAACCGACCTTGA</td><td>4720.7</td><td>10.0</td><td>1.1</td><td>8.7</td><td>3.3E-18</td><td>2.0E-17</td><td>BT20/MB468</td><td>CYTO</td></tr>
<tr><td>>hg38_21_+_8433172_8446622_RNA45SN1_45S_50ntFlanks@10808.10828.21|CAATGTAGGTAAGGGAAGTCG</td><td>>hg38_21_+_8433172_8446622_RNA45SN1_45S_50ntFlanks</td><td>10808</td><td>10828</td><td>21</td><td>CAATGTAGGTAAGGGAAGTCG</td><td>2834.8</td><td>-10.9</td><td>1.2</td><td>-8.7</td><td>2.7E-18</td><td>2.0E-17</td><td>MB231/MB468</td><td>MITO</td></tr>
<tr><td>>hg38_21_+_8433172_8446622_RNA45SN1_45S_50ntFlanks@9714.9738.25|CGAATGATTAGAGGTCTTGGGGCCG</td><td>>hg38_21_+_8433172_8446622_RNA45SN1_45S_50ntFlanks</td><td>9714</td><td>9738</td><td>25</td><td>CGAATGATTAGAGGTCTTGGGGCCG</td><td>1240.5</td><td>-14.1</td><td>1.6</td><td>-8.7</td><td>2.7E-18</td><td>2.0E-17</td><td>MB231/MB468</td><td>MITO</td></tr>
<tr><td>>hg38_21_+_8433172_8446622_RNA45SN1_45S_50ntFlanks@11212.11254.43|CGACTCTGGACGCGAGCCGGGCCCTTCCCGTGGATCGCCCCAG</td><td>>hg38_21_+_8433172_8446622_RNA45SN1_45S_50ntFlanks</td><td>11212</td><td>11254</td><td>43</td><td>CGACTCTGGACGCGAGCCGGGCCCTTCCCGTGGATCGCCCCAG</td><td>1549.4</td><td>-15.0</td><td>1.7</td><td>-8.6</td><td>5.7E-18</td><td>2.0E-17</td><td>BT20/MB468</td><td>NUC</td></tr>
<tr><td>>hg38_21_+_8433172_8446622_RNA45SN1_45S_50ntFlanks@6772.6793.22|GTTCCTCCCGGGGCTACGCCTG</td><td>>hg38_21_+_8433172_8446622_RNA45SN1_45S_50ntFlanks</td><td>6772</td><td>6793</td><td>22</td><td>GTTCCTCCCGGGGCTACGCCTG</td><td>734.1</td><td>-13.3</td><td>1.5</td><td>-8.7</td><td>2.8E-18</td><td>2.0E-17</td><td>MB231/MB468</td><td>MITO</td></tr>
<tr><td>>hg38_21_+_8433172_8446622_RNA45SN1_45S_50ntFlanks@4444.4472.29|CCTTGCCTCTCGGCGCCCCCTCGATGCTC</td><td>>hg38_21_+_8433172_8446622_RNA45SN1_45S_50ntFlanks</td><td>4444</td><td>4472</td><td>29</td><td>CCTTGCCTCTCGGCGCCCCCTCGATGCTC</td><td>593.2</td><td>-13.6</td><td>1.6</td><td>-8.6</td><td>5.7E-18</td><td>2.0E-17</td><td>BT20/MB468</td><td>NUC</td></tr>
<tr><td>>MI0000077|hsa-mir-21&WithFlank&17|+|59841260|59841343@14.35.22(+2A)&[MIMAT0000076&hsa-miR-21-5p&offsets|0|0(+2A);m-1&17|+|59841273|59841294&offsets|0|0(+2A)]|TAGCTTATCAGACTGATGTTGAAA</td><td>>MI0000077|hsa-mir-21&WithFlank&17|+|59841260|59841343</td><td>14</td><td>35</td><td>22(+2A)</td><td>TAGCTTATCAGACTGATGTTGAAA</td><td>2342.2</td><td>7.5</td><td>0.9</td><td>8.7</td><td>3.3E-18</td><td>2.0E-17</td><td>BT20/MB468</td><td>CYTO</td></tr>
<tr><td>>hg38_21_+_8433172_8446622_RNA45SN1_45S_50ntFlanks@3853.3871.19|AACTGTGGTAATTCTAGAG</td><td>>hg38_21_+_8433172_8446622_RNA45SN1_45S_50ntFlanks</td><td>3853</td><td>3871</td><td>19</td><td>AACTGTGGTAATTCTAGAG</td><td>530.0</td><td>-12.9</td><td>1.5</td><td>-8.8</td><td>1.7E-18</td><td>2.0E-17</td><td>MB231/MB468</td><td>MP</td></tr>
<tr><td>>hg38_21_+_8433172_8446622_RNA45SN1_45S_50ntFlanks@10513.10534.22|CCGGAGTGGCGGAGATGGGCGC</td><td>>hg38_21_+_8433172_8446622_RNA45SN1_45S_50ntFlanks</td><td>10513</td><td>10534</td><td>22</td><td>CCGGAGTGGCGGAGATGGGCGC</td><td>859.6</td><td>-13.5</td><td>1.6</td><td>-8.7</td><td>2.8E-18</td><td>2.0E-17</td><td>MB231/MB468</td><td>MITO</td></tr>
<tr><td>>hg38_MT_+_598_1651_RNR1_12S_50ntFlanks@428.468.41|AAAGTGGCTTTAACATATCTGAACACACAATAGCTAAGACC</td><td>>hg38_MT_+_598_1651_RNR1_12S_50ntFlanks</td><td>428</td><td>468</td><td>41</td><td>AAAGTGGCTTTAACATATCTGAACACACAATAGCTAAGACC</td><td>305.5</td><td>12.3</td><td>1.4</td><td>8.8</td><td>1.7E-18</td><td>2.0E-17</td><td>MB231/MB468</td><td>CYTO</td></tr>
<tr><td>>hg38_21_+_8433172_8446622_RNA45SN1_45S_50ntFlanks@12971.12989.19|TTTCGTACGTAGCAGAGCA</td><td>>hg38_21_+_8433172_8446622_RNA45SN1_45S_50ntFlanks</td><td>12971</td><td>12989</td><td>19</td><td>TTTCGTACGTAGCAGAGCA</td><td>537.5</td><td>-4.8</td><td>0.5</td><td>-8.7</td><td>2.2E-18</td><td>2.0E-17</td><td>MB231/MB468</td><td>NUC</td></tr>
<tr><td>>hg38_21_+_8433172_8446622_RNA45SN1_45S_50ntFlanks@11918.11950.33|ACATGAGAGGTGTAGAATAAGTGGGAGGCCCCC</td><td>>hg38_21_+_8433172_8446622_RNA45SN1_45S_50ntFlanks</td><td>11918</td><td>11950</td><td>33</td><td>ACATGAGAGGTGTAGAATAAGTGGGAGGCCCCC</td><td>999.8</td><td>-13.8</td><td>1.6</td><td>-8.8</td><td>1.7E-18</td><td>2.0E-17</td><td>MB231/MB468</td><td>MP</td></tr>
<tr><td>>hg38_21_+_8433172_8446622_RNA45SN1_45S_50ntFlanks@8113.8136.24|GGCGCGGGACATGTGGCGTACGGA</td><td>>hg38_21_+_8433172_8446622_RNA45SN1_45S_50ntFlanks</td><td>8113</td><td>8136</td><td>24</td><td>GGCGCGGGACATGTGGCGTACGGA</td><td>1687.6</td><td>-9.2</td><td>1.0</td><td>-8.8</td><td>1.7E-18</td><td>2.0E-17</td><td>MB231/MB468</td><td>MP</td></tr>
<tr><td>>MI0000750|hsa-mir-26a-2&WithFlank&12|-|57824603|57824698@20.40.21(+1A);MI0000083|hsa-mir-26a-1&WithFlank&3|+|37969398|37969486@16.36.21(+1A)&[MIMAT0000082&hsa-miR-26a-5p&offsets|0|-1(+1A);m-33&12|-|57824658|57824679&offsets|0|-1(+1A)];[MIMAT0000082_1&hsa-miR-26a-5p&offsets|0|-1(+1A);m-34&3|+|37969413|37969434&offsets|0|-1(+1A)]|TTCAAGTAATCCAGGATAGGCA</td><td>>MI0000750|hsa-mir-26a-2&WithFlank&12|-|57824603|57824698</td><td>20</td><td>40</td><td>21(+1A)</td><td>TTCAAGTAATCCAGGATAGGCA</td><td>5958.5</td><td>5.9</td><td>0.7</td><td>8.7</td><td>2.4E-18</td><td>2.0E-17</td><td>BT20/MB468</td><td>MITO</td></tr>
<tr><td>>hg38_21_+_8433172_8446622_RNA45SN1_45S_50ntFlanks@4048.4070.23|TTCGAACGTCTGCCCTATCAACT</td><td>>hg38_21_+_8433172_8446622_RNA45SN1_45S_50ntFlanks</td><td>4048</td><td>4070</td><td>23</td><td>TTCGAACGTCTGCCCTATCAACT</td><td>148.8</td><td>8.9</td><td>1.0</td><td>8.8</td><td>1.7E-18</td><td>2.0E-17</td><td>MB231/MB468</td><td>CYTO</td></tr>
<tr><td>>hg38_21_+_8433172_8446622_RNA45SN1_45S_50ntFlanks@11718.11751.34|AACGGCGGGAGTAACTATGACTCTCTTAAGGTAG</td><td>>hg38_21_+_8433172_8446622_RNA45SN1_45S_50ntFlanks</td><td>11718</td><td>11751</td><td>34</td><td>AACGGCGGGAGTAACTATGACTCTCTTAAGGTAG</td><td>429.9</td><td>-12.5</td><td>1.4</td><td>-8.7</td><td>2.9E-18</td><td>2.1E-17</td><td>MB231/MB468</td><td>MITO</td></tr>
</table>

| | | | | | | | | | | | | | |
|---|---|---|---|---|---|---|---|---|---|---|---|---|---|
| >hg38_21_+_8433172_8446622_RNA45SN1_45S_50ntFlanks@13004.13028.25\|ATCTATTGAAAGTCAGCCCTCGACA | >hg38_21_+_8433172_8446622_RNA45SN1_45S_50ntFlanks | 13004 | 13028 | 25 | ATCTATTGAAAGTCAGCCCTCGACA | 598.5 | -12.2 | 1.4 | -8.7 | 2.3E-18 | 2.1E-17 | MB231/MB468 | NUC |
| >MI0000299\|hsa-mir-222&WithFlank&X\|-\|45747009\|45747130@75.98.24&[MIMAT0000279&hsa-miR-222-3p&offsets\|0\|+3;m-140&X\|-\|45747033\|45747056&offsets\|0\|0]\|AGCTACATCTGGCTACTGGGTCTC | >MI0000299\|hsa-mir-222&WithFlank&X\|-\|45747009\|45747130 | 75 | 98 | 24 | AGCTACATCTGGCTACTGGGTCTC | 11329.5 | 6.9 | 0.8 | 8.7 | 2.5E-18 | 2.1E-17 | BT20/MB468 | MITO |
| >hg38_21_+_8433172_8446622_RNA45SN1_45S_50ntFlanks@9141.9164.24\|CGGGGCGGACTGTCCCCAGTGCGC | >hg38_21_+_8433172_8446622_RNA45SN1_45S_50ntFlanks | 9141 | 9164 | 24 | CGGGGCGGACTGTCCCCAGTGCGC | 320.2 | -12.2 | 1.4 | -8.8 | 1.7E-18 | 2.1E-17 | MB231/MB468 | MP |
| >hg38_21_+_8433172_8446622_RNA45SN1_45S_50ntFlanks@10768.10797.30\|CAGCAGGTCTCCAAGGTGAACAGCCTCTGG | >hg38_21_+_8433172_8446622_RNA45SN1_45S_50ntFlanks | 10768 | 10797 | 30 | CAGCAGGTCTCCAAGGTGAACAGCCTCTGG | 361.1 | -12.4 | 1.4 | -8.8 | 1.7E-18 | 2.1E-17 | MB231/MB468 | MP |
| >MI0000803\|hsa-mir-330&WithFlank&19\|-\|45638988\|45639093@63.86.24(+1U)&[MIMAT0000751&hsa-miR-330-3p&offsets\|0\|+1(+1U);m-347&19\|-\|45639009\|45639030&offsets\|-1\|+1(+1U)]\|GCAAAGCACACGGCCTGCAGAGAGT | >MI0000803\|hsa-mir-330&WithFlank&19\|-\|45638988\|45639093 | 63 | 86 | 24(+1U) | GCAAAGCACACGGCCTGCAGAGAGT | 462.1 | 6.5 | 0.7 | 8.7 | 2.5E-18 | 2.1E-17 | BT20/MB468 | MITO |
| >hg38_21_+_8433172_8446622_RNA45SN1_45S_50ntFlanks@8198.8225.28\|TGGACGGTGTGAGGCCGGTAGCGGCCCC | >hg38_21_+_8433172_8446622_RNA45SN1_45S_50ntFlanks | 8198 | 8225 | 28 | TGGACGGTGTGAGGCCGGTAGCGGCCCC | 401.5 | -12.5 | 1.4 | -8.8 | 1.8E-18 | 2.1E-17 | MB231/MB468 | MP |
| >hg38_21_+_8433172_8446622_RNA45SN1_45S_50ntFlanks@4760.4786.27\|TCAGATACCGTCGTAGTTCCGACCATA | >hg38_21_+_8433172_8446622_RNA45SN1_45S_50ntFlanks | 4760 | 4786 | 27 | TCAGATACCGTCGTAGTTCCGACCATA | 958.3 | -13.7 | 1.6 | -8.7 | 2.9E-18 | 2.1E-17 | MB231/MB468 | MITO |
| >hg38_21_+_8433172_8446622_RNA45SN1_45S_50ntFlanks@12617.12654.38\|TGAGGAGCCAATGGGGCGAAGCTACCATCTGTGGGATT | >hg38_21_+_8433172_8446622_RNA45SN1_45S_50ntFlanks | 12617 | 12654 | 38 | TGAGGAGCCAATGGGGCGAAGCTACCATCTGTGGGATT | 3598.2 | -9.4 | 1.1 | -8.7 | 2.9E-18 | 2.1E-17 | MB231/MB468 | MITO |
| >MI0000088\|hsa-mir-30a&WithFlank&6\|-\|71403545\|71403627@53.74.22&[MIMAT0000088&hsa-miR-30a-3p&offsets\|0\|0;m-38&6\|-\|71403554\|71403575&offsets\|0\|0]\|CTTTCAGTCGGATGTTTGCAGC | >MI0000088\|hsa-mir-30a&WithFlank&6\|-\|71403545\|71403627 | 53 | 74 | 22 | CTTTCAGTCGGATGTTTGCAGC | 19093.3 | 7.3 | 0.8 | 8.8 | 1.7E-18 | 2.1E-17 | MB231/MB468 | CYTO |
| >MI0000433\|hsa-let-7g&WithFlank&3\|-\|52268272\|52268367@11.31.21(+1C)&[MIMAT0000414&hsa-let-7g-5p&offsets\|0\|-1(+1C);m-58&3\|-\|52268336\|52268357&offsets\|0\|-1(+1C)]\|TGAGGTAGTAGTTTGTACAGTC | >MI0000433\|hsa-let-7g&WithFlank&3\|-\|52268272\|52268367 | 11 | 31 | 21(+1C) | TGAGGTAGTAGTTTGTACAGTC | 14774.6 | 7.1 | 0.8 | 8.7 | 3.5E-18 | 2.1E-17 | BT20/MB468 | CYTO |
| >hg38_21_+_8433172_8446622_RNA45SN1_45S_50ntFlanks@10408.10430.23\|GGTCAGTCGGTCCTGAGAGATGG | >hg38_21_+_8433172_8446622_RNA45SN1_45S_50ntFlanks | 10408 | 10430 | 23 | GGTCAGTCGGTCCTGAGAGATGG | 632.9 | -13.1 | 1.5 | -8.7 | 2.9E-18 | 2.1E-17 | MB231/MB468 | MITO |
| >hg38_21_+_8433172_8446622_RNA45SN1_45S_50ntFlanks@10806.10829.24\|AACAATGTAGGTAAGGGAAGTCGG | >hg38_21_+_8433172_8446622_RNA45SN1_45S_50ntFlanks | 10806 | 10829 | 24 | AACAATGTAGGTAAGGGAAGTCGG | 450.5 | -12.6 | 1.4 | -8.7 | 2.9E-18 | 2.1E-17 | MB231/MB468 | MITO |
| >MI0000078\|hsa-mir-22&WithFlank&17\|-\|1713897\|1713993@59.78.20&[MIMAT0000077&hsa-miR-22-3p&offsets\|0\|-2;m-2&17\|-\|1713914\|1713935&offsets\|0\|-2]\|AAGCTGCCAGTTGAAGAACT | >MI0000078\|hsa-mir-22&WithFlank&17\|-\|1713897\|1713993 | 59 | 78 | 20 | AAGCTGCCAGTTGAAGAACT | 528.2 | 6.5 | 0.7 | 8.7 | 2.3E-18 | 2.1E-17 | MB231/MB468 | NUC |
| >MI0000084\|hsa-mir-26b&WithFlank&2\|+\|218402640\|218402728@18.38.21(+1A)&[MIMAT0000083&hsa-miR-26b-5p&offsets\|0\|0(+1A);m-53&2\|+\|218402657\|218402678&offsets\|0\|-1(+1A)]\|TTCAAGTAATTCAGGATAGGTA | >MI0000084\|hsa-mir-26b&WithFlank&2\|+\|218402640\|218402728 | 18 | 38 | 21(+1A) | TTCAAGTAATTCAGGATAGGTA | 83.4 | 3.3 | 0.4 | 8.6 | 6.1E-18 | 2.1E-17 | BT20/MB468 | NUC |
| >MI0000086\|hsa-mir-28&WithFlank&3\|+\|188688775\|188688872@60.81.22(+1U)&[MIMAT0004502&hsa-miR-28-3p&offsets\|0\|0(+1U);m-28&3\|+\|188688834\|188688855&offsets\|0\|0(+1U)]\|CACTAGATTGTGAGCTCCTGGAT | >MI0000086\|hsa-mir-28&WithFlank&3\|+\|188688775\|188688872 | 60 | 81 | 22(+1U) | CACTAGATTGTGAGCTCCTGGAT | 460.2 | 6.7 | 0.8 | 8.7 | 2.4E-18 | 2.1E-17 | MB231/MB468 | NUC |
| >MI0000088\|hsa-mir-30a&WithFlank&6\|-\|71403545\|71403627@13.35.23(+1U)&[MIMAT0000087&hsa-miR-30a-5p&offsets\|+1\|+2(+1U);m-15&6\|-\|71403595\|71403616&offsets\|+1\|+2(+1U)]\|GTAAACATCCTCGACTGGAAGCTT | >MI0000088\|hsa-mir-30a&WithFlank&6\|-\|71403545\|71403627 | 13 | 35 | 23(+1U) | GTAAACATCCTCGACTGGAAGCTT | 247.9 | 11.8 | 1.4 | 8.7 | 2.4E-18 | 2.1E-17 | MB231/MB468 | NUC |
| >trnaMT_GluTTC_MT_-_14674_14742@40.58.19,trnalookalike8_GluTTC_5_-_93905172_93905240@40.58.19\|TTGGTCGTGGTTGTAGTCC | >trnaMT_GluTTC_MT_-_14674_14742 | 40 | 58 | 19 | TTGGTCGTGGTTGTAGTCC | 350.5 | 6.0 | 0.7 | 8.7 | 2.4E-18 | 2.1E-17 | MB231/MB468 | NUC |
| >hg38_21_+_8433172_8446622_RNA45SN1_45S_50ntFlanks@5471.5499.29\|CACGGCCCTGGCGGAGCGCTGAGAAGACG | >hg38_21_+_8433172_8446622_RNA45SN1_45S_50ntFlanks | 5471 | 5499 | 29 | CACGGCCCTGGCGGAGCGCTGAGAAGACG | 4345.4 | -9.9 | 1.1 | -8.8 | 1.8E-18 | 2.1E-17 | MB231/MB468 | MP |
| >hg38_21_+_8433172_8446622_RNA45SN1_45S_50ntFlanks@9532.9570.39\|AGAGGAAACTCTGGTGGAGGTCCGTAGCGGTCCTGACGT | >hg38_21_+_8433172_8446622_RNA45SN1_45S_50ntFlanks | 9532 | 9570 | 39 | AGAGGAAACTCTGGTGGAGGTCCGTAGCGGTCCTGACGT | 280.3 | 9.5 | 1.1 | 8.8 | 1.8E-18 | 2.1E-17 | MB231/MB468 | CYTO |
| >hg38_21_+_8433172_8446622_RNA45SN1_45S_50ntFlanks@5421.5445.25\|CGATTGGATGGTTTAGTGAGGCCCT | >hg38_21_+_8433172_8446622_RNA45SN1_45S_50ntFlanks | 5421 | 5445 | 25 | CGATTGGATGGTTTAGTGAGGCCCT | 1249.6 | -14.1 | 1.6 | -8.7 | 3.0E-18 | 2.1E-17 | MB231/MB468 | MITO |
| >hg38_21_+_8433172_8446622_RNA45SN1_45S_50ntFlanks@12285.12318.34\|CAGACCGTGAAAGCGGGGCCTCACGATCCTTCTG | >hg38_21_+_8433172_8446622_RNA45SN1_45S_50ntFlanks | 12285 | 12318 | 34 | CAGACCGTGAAAGCGGGGCCTCACGATCCTTCTG | 901.0 | -13.6 | 1.6 | -8.7 | 3.0E-18 | 2.2E-17 | MB231/MB468 | MITO |
| >MI0000298\|hsa-mir-221&WithFlank&X\|-\|45746151\|45746272@71.91.21&[MIMAT0000278&hsa-miR-221-3p&offsets\|0\|-2;m-91&X\|-\|45746181\|45746202&offsets\|0\|-1]\|AGCTACATTGTCTGCTGGGTT | >MI0000298\|hsa-mir-221&WithFlank&X\|-\|45746151\|45746272 | 71 | 91 | 21 | AGCTACATTGTCTGCTGGGTT | 1842.0 | 6.7 | 0.8 | 8.7 | 2.4E-18 | 2.2E-17 | MB231/MB468 | NUC |
| >hg38_21_+_8433172_8446622_RNA45SN1_45S_50ntFlanks@10458.10477.20\|ATGGCCTCCGTTGCCCTCGG | >hg38_21_+_8433172_8446622_RNA45SN1_45S_50ntFlanks | 10458 | 10477 | 20 | ATGGCCTCCGTTGCCCTCGG | 1581.1 | -10.4 | 1.2 | -8.7 | 2.4E-18 | 2.2E-17 | MB231/MB468 | NUC |
| >hg38_21_+_8433172_8446622_RNA45SN1_45S_50ntFlanks@3593.3610.18\|CCCGTCCGTCCGTCCGTC | >hg38_21_+_8433172_8446622_RNA45SN1_45S_50ntFlanks | 3593 | 3610 | 18 | CCCGTCCGTCCGTCCGTC | 481.2 | -12.7 | 1.5 | -8.7 | 3.0E-18 | 2.2E-17 | MB231/MB468 | MITO |
| >hg38_21_+_8433172_8446622_RNA45SN1_45S_50ntFlanks@8737.8754.18\|CCCGTCCGTCCGTCCGTC | >hg38_21_+_8433172_8446622_RNA45SN1_45S_50ntFlanks | 8737 | 8754 | 18 | CCCGTCCGTCCGTCCGTC | 481.2 | -12.7 | 1.5 | -8.7 | 3.0E-18 | 2.2E-17 | MB231/MB468 | MITO |
| >hg38_21_+_8433172_8446622_RNA45SN1_45S_50ntFlanks@10729.10752.24\|AGAGGGTGTAAATCTCGCGCCGGG | >hg38_21_+_8433172_8446622_RNA45SN1_45S_50ntFlanks | 10729 | 10752 | 24 | AGAGGGTGTAAATCTCGCGCCGGG | 308.4 | -12.1 | 1.4 | -8.8 | 1.8E-18 | 2.2E-17 | MB231/MB468 | MP |
| >hg38_21_+_8433172_8446622_RNA45SN1_45S_50ntFlanks@12259.12282.24\|CTTGATCTTGATTTTCAGTACGAA | >hg38_21_+_8433172_8446622_RNA45SN1_45S_50ntFlanks | 12259 | 12282 | 24 | CTTGATCTTGATTTTCAGTACGAA | 981.1 | -13.8 | 1.6 | -8.8 | 1.8E-18 | 2.2E-17 | MB231/MB468 | MP |
| >MI0000087\|hsa-mir-29a&WithFlank&7\|-\|130876741\|130876816@48.67.20(+1C)&[MIMAT0000086&hsa-miR-29a-3p&offsets\|0\|-2(+1C);m-26&7\|-\|130876749\|130876769&offsets\|0\|-1(+1C)]\|TAGCACCATCTGAAATCGGTC | >MI0000087\|hsa-mir-29a&WithFlank&7\|-\|130876741\|130876816 | 48 | 67 | 20(+1C) | TAGCACCATCTGAAATCGGTC | 1160.3 | 12.1 | 1.4 | 8.7 | 3.6E-18 | 2.2E-17 | BT20/MB468 | CYTO |
| >hg38_21_+_8433172_8446622_RNA45SN1_45S_50ntFlanks@12418.12442.25\|CGATGTCGGCTCTTCCTATCATTGT | >hg38_21_+_8433172_8446622_RNA45SN1_45S_50ntFlanks | 12418 | 12442 | 25 | CGATGTCGGCTCTTCCTATCATTGT | 860.1 | -5.4 | 0.6 | -8.6 | 6.3E-18 | 2.2E-17 | BT20/MB468 | NUC |
| >hg38_21_+_8433172_8446622_RNA45SN1_45S_50ntFlanks@11579.11601.23\|AGAACTGGTGCGGACCAGGGGAA | >hg38_21_+_8433172_8446622_RNA45SN1_45S_50ntFlanks | 11579 | 11601 | 23 | AGAACTGGTGCGGACCAGGGGAA | 1010.4 | -13.8 | 1.6 | -8.8 | 1.9E-18 | 2.2E-17 | MB231/MB468 | MP |
| >hg38_21_+_8433172_8446622_RNA45SN1_45S_50ntFlanks@12366.12384.19\|TAACTGGCTTGTGGCGGCC | >hg38_21_+_8433172_8446622_RNA45SN1_45S_50ntFlanks | 12366 | 12384 | 19 | TAACTGGCTTGTGGCGGCC | 908.4 | -12.5 | 1.4 | -8.8 | 1.8E-18 | 2.2E-17 | MB231/MB468 | CYTO |
| >MI0000086\|hsa-mir-28&WithFlank&3\|+\|188688775\|188688872@20.40.21&[MIMAT0000085&hsa-miR-28-5p&offsets\|0\|-1;m-133&3\|+\|188688794\|188688815&offsets\|0\|-1]\|AAGGAGCTCACAGTCTATTGA | >MI0000086\|hsa-mir-28&WithFlank&3\|+\|188688775\|188688872 | 20 | 40 | 21 | AAGGAGCTCACAGTCTATTGA | 542.0 | 11.0 | 1.3 | 8.7 | 3.7E-18 | 2.2E-17 | BT20/MB468 | CYTO |
| >hg38_21_+_8433172_8446622_RNA45SN1_45S_50ntFlanks@4208.4242.35\|GGGGAGGTAGTGACGAAAAATAACAATACAGGACT | >hg38_21_+_8433172_8446622_RNA45SN1_45S_50ntFlanks | 4208 | 4242 | 35 | GGGGAGGTAGTGACGAAAAATAACAATACAGGACT | 266.2 | 12.1 | 1.4 | 8.8 | 1.9E-18 | 2.2E-17 | MB231/MB468 | CYTO |
| >hg38_21_+_8433172_8446622_RNA45SN1_45S_50ntFlanks@11209.11233.25\|CGGCGACTCTGGACGCGAGCCGGGC | >hg38_21_+_8433172_8446622_RNA45SN1_45S_50ntFlanks | 11209 | 11233 | 25 | CGGCGACTCTGGACGCGAGCCGGGC | 2852.4 | -6.5 | 0.7 | -8.8 | 1.9E-18 | 2.2E-17 | MB231/MB468 | CYTO |
| >hg38_21_+_8433172_8446622_RNA45SN1_45S_50ntFlanks@12405.12431.27\|CTTTTTGATCCTTCGATGTCGGCTCTT | >hg38_21_+_8433172_8446622_RNA45SN1_45S_50ntFlanks | 12405 | 12431 | 27 | CTTTTTGATCCTTCGATGTCGGCTCTT | 22276.9 | -7.4 | 0.8 | -8.8 | 1.9E-18 | 2.2E-17 | MB231/MB468 | CYTO |
| >hg38_21_+_8433172_8446622_RNA45SN1_45S_50ntFlanks@11572.11605.34\|CCGACTTAGAACTGGTGCGGACCAGGGGAATCCG | >hg38_21_+_8433172_8446622_RNA45SN1_45S_50ntFlanks | 11572 | 11605 | 34 | CCGACTTAGAACTGGTGCGGACCAGGGGAATCCG | 2335.7 | -4.6 | 0.5 | -8.7 | 3.8E-18 | 2.3E-17 | BT20/MB468 | CYTO |
| >hg38_21_+_8433172_8446622_RNA45SN1_45S_50ntFlanks@9736.9757.22\|CCGAAACGATCTCAACCTATTC | >hg38_21_+_8433172_8446622_RNA45SN1_45S_50ntFlanks | 9736 | 9757 | 22 | CCGAAACGATCTCAACCTATTC | 1400.8 | -14.2 | 1.6 | -8.7 | 3.2E-18 | 2.3E-17 | MB231/MB468 | MITO |
| >hg38_21_+_8433172_8446622_RNA45SN1_45S_50ntFlanks@10410.10428.19\|TCAGTCGGTCCTGAGAGAT | >hg38_21_+_8433172_8446622_RNA45SN1_45S_50ntFlanks | 10410 | 10428 | 19 | TCAGTCGGTCCTGAGAGAT | 2992.3 | -11.7 | 1.4 | -8.6 | 6.5E-18 | 2.3E-17 | BT20/MB468 | NUC |
| >MI0003560\|hsa-mir-92b&WithFlank&1\|+\|155195171\|155195278@67.88.22(+3U)&[MIMAT0003218&hsa-miR-92b-3p&offsets\|0\|0(+3U);m-115&1\|+\|155195237\|155195258&offsets\|0\|0(+3U)]\|TATTGCACTCGTCCCGGCCTCCTTT | >MI0003560\|hsa-mir-92b&WithFlank&1\|+\|155195171\|155195278 | 67 | 88 | 22(+3U) | TATTGCACTCGTCCCGGCCTCCTTT | 415.1 | 12.7 | 1.4 | 8.8 | 1.9E-18 | 2.3E-17 | MB231/MB468 | CYTO |
| >hg38_21_+_8433172_8446622_RNA45SN1_45S_50ntFlanks@9240.9274.35\|GCGGAGCGAGCGCACGGGGTCGGCGGCGACGTCGG | >hg38_21_+_8433172_8446622_RNA45SN1_45S_50ntFlanks | 9240 | 9274 | 35 | GCGGAGCGAGCGCACGGGGTCGGCGGCGACGTCGG | 454.0 | -12.7 | 1.4 | -8.8 | 1.9E-18 | 2.3E-17 | MB231/MB468 | MP |
| >hg38_21_+_8433172_8446622_RNA45SN1_45S_50ntFlanks@12457.12487.31\|CAAGCGTTGGATTGTTCACCCACTAATAGGG | >hg38_21_+_8433172_8446622_RNA45SN1_45S_50ntFlanks | 12457 | 12487 | 31 | CAAGCGTTGGATTGTTCACCCACTAATAGGG | 839.8 | -13.6 | 1.5 | -8.8 | 2.0E-18 | 2.3E-17 | MB231/MB468 | MP |
| >hg38_21_+_8433172_8446622_RNA45SN1_45S_50ntFlanks@9459.9488.30\|CCGGGGAGGTGGAGCACGAGCGCACGTGTT | >hg38_21_+_8433172_8446622_RNA45SN1_45S_50ntFlanks | 9459 | 9488 | 30 | CCGGGGAGGTGGAGCACGAGCGCACGTGTT | 496.6 | -12.8 | 1.5 | -8.8 | 2.0E-18 | 2.3E-17 | MB231/MB468 | MP |
| >hg38_21_+_8433172_8446622_RNA45SN1_45S_50ntFlanks@6722.6744.23\|ATTGCAGGACACATTGATCATCG | >hg38_21_+_8433172_8446622_RNA45SN1_45S_50ntFlanks | 6722 | 6744 | 23 | ATTGCAGGACACATTGATCATCG | 1546.0 | -6.0 | 0.7 | -8.7 | 2.6E-18 | 2.3E-17 | MB231/MB468 | NUC |
| >MI0000446\|hsa-mir-125b-1&WithFlank&11\|-\|122099751\|122099850@61.81.21&[MIMAT0004592&hsa-miR-125b-1-3p&offsets\|0\|-1;m-485&11\|-\|122099769\|122099790&offsets\|0\|-1]\|ACGGGTTAGGCTCTTGGGAGC | >MI0000446\|hsa-mir-125b-1&WithFlank&11\|-\|122099751\|122099850 | 61 | 81 | 21 | ACGGGTTAGGCTCTTGGGAGC | 211.4 | 11.6 | 1.3 | 8.7 | 2.6E-18 | 2.3E-17 | MB231/MB468 | NUC |
| >hg38_21_+_8433172_8446622_RNA45SN1_45S_50ntFlanks@8357.8378.22\|AAAAGAACTTTGAAGAGAGAGT | >hg38_21_+_8433172_8446622_RNA45SN1_45S_50ntFlanks | 8357 | 8378 | 22 | AAAAGAACTTTGAAGAGAGAGT | 14227.3 | -6.9 | 0.8 | -8.7 | 2.6E-18 | 2.3E-17 | MB231/MB468 | NUC |
| >hg38_21_+_8433172_8446622_RNA45SN1_45S_50ntFlanks@4547.4565.19\|CTGGATACCGCAGCTAGGA | >hg38_21_+_8433172_8446622_RNA45SN1_45S_50ntFlanks | 4547 | 4565 | 19 | CTGGATACCGCAGCTAGGA | 833.5 | -13.6 | 1.5 | -8.8 | 2.0E-18 | 2.3E-17 | MB231/MB468 | MP |
| >hg38_21_+_8433172_8446622_RNA45SN1_45S_50ntFlanks@11207.11227.21\|GGCGGCGACTCTGGACGCGAG | >hg38_21_+_8433172_8446622_RNA45SN1_45S_50ntFlanks | 11207 | 11227 | 21 | GGCGGCGACTCTGGACGCGAG | 396.1 | -12.5 | 1.4 | -8.8 | 2.0E-18 | 2.3E-17 | MB231/MB468 | MP |

<table>
<tr><td>>hg38_21_+_8433172_8446622_RNA45SN1_45S_50ntFlanks@12606.12626.21|ATGTGCTTGGCTGAGGAGCCA</td><td>>hg38_21_+_8433172_8446622_RNA45SN1_45S_50ntFlanks</td><td>12606</td><td>12626</td><td>21</td><td>ATGTGCTTGGCTGAGGAGCCA</td><td>414.7</td><td>-12.6</td><td>1.4</td><td>-8.8</td><td>2.0E-18</td><td>2.3E-17</td><td>MB231/MB468</td><td>MP</td></tr>
<tr><td>>hg38_21_+_8433172_8446622_RNA45SN1_45S_50ntFlanks@12347.12365.19|AGAAAAGTTACCACAGGGA</td><td>>hg38_21_+_8433172_8446622_RNA45SN1_45S_50ntFlanks</td><td>12347</td><td>12365</td><td>19</td><td>AGAAAAGTTACCACAGGGA</td><td>660.2</td><td>-9.4</td><td>1.1</td><td>-8.7</td><td>3.9E-18</td><td>2.3E-17</td><td>BT20/MB468</td><td>CYTO</td></tr>
<tr><td>>hg38_21_+_8433172_8446622_RNA45SN1_45S_50ntFlanks@7973.8005.33|GACGCGACCTCAGATCAGACGTGGCGACCCGCT</td><td>>hg38_21_+_8433172_8446622_RNA45SN1_45S_50ntFlanks</td><td>7973</td><td>8005</td><td>33</td><td>GACGCGACCTCAGATCAGACGTGGCGACCCGCT</td><td>433.1</td><td>4.8</td><td>0.5</td><td>8.7</td><td>3.9E-18</td><td>2.4E-17</td><td>BT20/MB468</td><td>CYTO</td></tr>
<tr><td>>hg38_21_+_8433172_8446622_RNA45SN1_45S_50ntFlanks@3846.3868.23|CTTGGATAACTGTGGTAATTCTA</td><td>>hg38_21_+_8433172_8446622_RNA45SN1_45S_50ntFlanks</td><td>3846</td><td>3868</td><td>23</td><td>CTTGGATAACTGTGGTAATTCTA</td><td>928.8</td><td>-5.6</td><td>0.6</td><td>-8.6</td><td>6.8E-18</td><td>2.4E-17</td><td>BT20/MB468</td><td>NUC</td></tr>
<tr><td>>hg38_21_+_8433172_8446622_RNA45SN1_45S_50ntFlanks@11316.11342.27|CACCCCACGTCTCGTCGCGCGCGCGTC</td><td>>hg38_21_+_8433172_8446622_RNA45SN1_45S_50ntFlanks</td><td>11316</td><td>11342</td><td>27</td><td>CACCCCACGTCTCGTCGCGCGCGCGTC</td><td>1440.6</td><td>-14.4</td><td>1.6</td><td>-8.8</td><td>2.0E-18</td><td>2.4E-17</td><td>MB231/MB468</td><td>MP</td></tr>
<tr><td>>hg38_21_+_8433172_8446622_RNA45SN1_45S_50ntFlanks@11561.11585.25|GCGCCTAGCAGCCGACTTAGAACTG</td><td>>hg38_21_+_8433172_8446622_RNA45SN1_45S_50ntFlanks</td><td>11561</td><td>11585</td><td>25</td><td>GCGCCTAGCAGCCGACTTAGAACTG</td><td>4679.3</td><td>-9.0</td><td>1.0</td><td>-8.7</td><td>3.3E-18</td><td>2.4E-17</td><td>MB231/MB468</td><td>MITO</td></tr>
<tr><td>>trnaMT_PheGAA_MT_+_577_647@35.56.22|GAAAATGTTTAGACGGGCTCAC</td><td>>trnaMT_PheGAA_MT_+_577_647</td><td>35</td><td>56</td><td>22</td><td>GAAAATGTTTAGACGGGCTCAC</td><td>319.8</td><td>12.3</td><td>1.4</td><td>8.8</td><td>2.0E-18</td><td>2.4E-17</td><td>MB231/MB468</td><td>CYTO</td></tr>
<tr><td>>hg38_21_+_8433172_8446622_RNA45SN1_45S_50ntFlanks@10351.10383.33|CGAGAACTTTGAAGGCCGAAGTGGAGAAGGGTT</td><td>>hg38_21_+_8433172_8446622_RNA45SN1_45S_50ntFlanks</td><td>10351</td><td>10383</td><td>33</td><td>CGAGAACTTTGAAGGCCGAAGTGGAGAAGGGTT</td><td>11462.5</td><td>-5.6</td><td>0.6</td><td>-8.7</td><td>2.9E-18</td><td>2.4E-17</td><td>BT20/MB468</td><td>MITO</td></tr>
<tr><td>>hg38_21_+_8433172_8446622_RNA45SN1_45S_50ntFlanks@9306.9323.18|CAAGGAGTCTAACACGTG</td><td>>hg38_21_+_8433172_8446622_RNA45SN1_45S_50ntFlanks</td><td>9306</td><td>9323</td><td>18</td><td>CAAGGAGTCTAACACGTG</td><td>1347.5</td><td>-14.3</td><td>1.6</td><td>-8.8</td><td>2.0E-18</td><td>2.4E-17</td><td>MB231/MB468</td><td>MP</td></tr>
<tr><td>>hg38_21_+_8433172_8446622_RNA45SN1_45S_50ntFlanks@12539.12565.27|ATGATGTGTTGTTGCCATGGTAATCCT</td><td>>hg38_21_+_8433172_8446622_RNA45SN1_45S_50ntFlanks</td><td>12539</td><td>12565</td><td>27</td><td>ATGATGTGTTGTTGCCATGGTAATCCT</td><td>517.1</td><td>-12.8</td><td>1.5</td><td>-8.7</td><td>3.4E-18</td><td>2.4E-17</td><td>MB231/MB468</td><td>MITO</td></tr>
<tr><td>>hg38_21_+_8433172_8446622_RNA45SN1_45S_50ntFlanks@12432.12455.24|CCTATCATTGTGAAGCAGAATTCA</td><td>>hg38_21_+_8433172_8446622_RNA45SN1_45S_50ntFlanks</td><td>12432</td><td>12455</td><td>24</td><td>CCTATCATTGTGAAGCAGAATTCA</td><td>706.3</td><td>-13.3</td><td>1.5</td><td>-8.8</td><td>2.1E-18</td><td>2.4E-17</td><td>MB231/MB468</td><td>MP</td></tr>
<tr><td>>hg38_21_+_8433172_8446622_RNA45SN1_45S_50ntFlanks@12940.12966.27|CATTCGTAGACGACCTGCTTCTGGGTC</td><td>>hg38_21_+_8433172_8446622_RNA45SN1_45S_50ntFlanks</td><td>12940</td><td>12966</td><td>27</td><td>CATTCGTAGACGACCTGCTTCTGGGTC</td><td>664.2</td><td>-13.2</td><td>1.5</td><td>-8.8</td><td>2.1E-18</td><td>2.4E-17</td><td>MB231/MB468</td><td>MP</td></tr>
<tr><td>>hg38_21_+_8433172_8446622_RNA45SN1_45S_50ntFlanks@12003.12032.30|GCGGGCCGCCGGTGAAATACCACTACTCTG</td><td>>hg38_21_+_8433172_8446622_RNA45SN1_45S_50ntFlanks</td><td>12003</td><td>12032</td><td>30</td><td>GCGGGCCGCCGGTGAAATACCACTACTCTG</td><td>3059.8</td><td>-3.8</td><td>0.4</td><td>-8.7</td><td>2.9E-18</td><td>2.4E-17</td><td>BT20/MB468</td><td>MITO</td></tr>
<tr><td>>hg38_21_+_8433172_8446622_RNA45SN1_45S_50ntFlanks@9504.9532.29|GTGAACTATGCCTGGGCAGGGCGAAGCCA</td><td>>hg38_21_+_8433172_8446622_RNA45SN1_45S_50ntFlanks</td><td>9504</td><td>9532</td><td>29</td><td>GTGAACTATGCCTGGGCAGGGCGAAGCCA</td><td>479.7</td><td>-12.7</td><td>1.5</td><td>-8.7</td><td>3.4E-18</td><td>2.4E-17</td><td>MB231/MB468</td><td>MITO</td></tr>
<tr><td>>hg38_21_+_8433172_8446622_RNA45SN1_45S_50ntFlanks@12955.12988.34|TGCTTCTGGGTCGGGGTTTCGTACGTAGCAGAGC</td><td>>hg38_21_+_8433172_8446622_RNA45SN1_45S_50ntFlanks</td><td>12955</td><td>12988</td><td>34</td><td>TGCTTCTGGGTCGGGGTTTCGTACGTAGCAGAGC</td><td>246.1</td><td>4.8</td><td>0.5</td><td>8.8</td><td>2.0E-18</td><td>2.4E-17</td><td>MB231/MB468</td><td>CYTO</td></tr>
<tr><td>>hg38_21_+_8433172_8446622_RNA45SN1_45S_50ntFlanks@5473.5512.40|CGGCCCTGGCGGAGCGCTGAGAAGACGGTCGAACTTGACT</td><td>>hg38_21_+_8433172_8446622_RNA45SN1_45S_50ntFlanks</td><td>5473</td><td>5512</td><td>40</td><td>CGGCCCTGGCGGAGCGCTGAGAAGACGGTCGAACTTGACT</td><td>1744.9</td><td>6.7</td><td>0.8</td><td>8.8</td><td>2.0E-18</td><td>2.4E-17</td><td>MB231/MB468</td><td>CYTO</td></tr>
<tr><td>>hg38_21_+_8433172_8446622_RNA45SN1_45S_50ntFlanks@8475.8497.23|CCCGGCGGATCTTTCCCGCCCCC</td><td>>hg38_21_+_8433172_8446622_RNA45SN1_45S_50ntFlanks</td><td>8475</td><td>8497</td><td>23</td><td>CCCGGCGGATCTTTCCCGCCCCC</td><td>691.5</td><td>-13.3</td><td>1.5</td><td>-8.8</td><td>2.1E-18</td><td>2.4E-17</td><td>MB231/MB468</td><td>MP</td></tr>
<tr><td>>hg38_21_+_8433172_8446622_RNA45SN1_45S_50ntFlanks@12405.12426.22|CTTTTTGATCCTTCGATGTCGG</td><td>>hg38_21_+_8433172_8446622_RNA45SN1_45S_50ntFlanks</td><td>12405</td><td>12426</td><td>22</td><td>CTTTTTGATCCTTCGATGTCGG</td><td>2692.1</td><td>-10.5</td><td>1.2</td><td>-8.7</td><td>3.4E-18</td><td>2.4E-17</td><td>MB231/MB468</td><td>MITO</td></tr>
<tr><td>>MI0000087|hsa-mir-29a&WithFlank&7|-|130876741|130876816@48.67.20(+1A)&[MIMAT0000086&hsa-miR-29a-3p&offsets|0|-2(+1A);m-26&7|-|130876749|130876769&offsets|0|-1(+1A)]|TAGCACCATCTGAAATCGGTA</td><td>>MI0000087|hsa-mir-29a&WithFlank&7|-|130876741|130876816</td><td>48</td><td>67</td><td>20(+1A)</td><td>TAGCACCATCTGAAATCGGTA</td><td>441.0</td><td>13.0</td><td>1.5</td><td>8.7</td><td>4.1E-18</td><td>2.4E-17</td><td>BT20/MB468</td><td>CYTO</td></tr>
<tr><td>>hg38_21_+_8433172_8446622_RNA45SN1_45S_50ntFlanks@10391.10410.20|AACAGCAGTTGAACATGGGT</td><td>>hg38_21_+_8433172_8446622_RNA45SN1_45S_50ntFlanks</td><td>10391</td><td>10410</td><td>20</td><td>AACAGCAGTTGAACATGGGT</td><td>3868.7</td><td>-5.6</td><td>0.6</td><td>-8.7</td><td>4.1E-18</td><td>2.4E-17</td><td>BT20/MB468</td><td>CYTO</td></tr>
<tr><td>>hg38_21_+_8433172_8446622_RNA45SN1_45S_50ntFlanks@4163.4182.20|CATCCAAGGAAGGCAGCAGG</td><td>>hg38_21_+_8433172_8446622_RNA45SN1_45S_50ntFlanks</td><td>4163</td><td>4182</td><td>20</td><td>CATCCAAGGAAGGCAGCAGG</td><td>496.1</td><td>-12.7</td><td>1.5</td><td>-8.7</td><td>3.4E-18</td><td>2.4E-17</td><td>MB231/MB468</td><td>MITO</td></tr>
<tr><td>>MI0000301|hsa-mir-224&WithFlank&X|-|151958572|151958664@14.38.25&[MIMAT0000281&hsa-miR-224-5p&offsets|+1|+1;m-238&X|-|151958630|151958651&offsets|0|+3]|CAAGTCACTAGTGGTTCCGTTTAGT</td><td>>MI0000301|hsa-mir-224&WithFlank&X|-|151958572|151958664</td><td>14</td><td>38</td><td>25</td><td>CAAGTCACTAGTGGTTCCGTTTAGT</td><td>1420.3</td><td>12.4</td><td>1.4</td><td>8.7</td><td>4.1E-18</td><td>2.4E-17</td><td>BT20/MB468</td><td>CYTO</td></tr>
<tr><td>>hg38_21_+_8433172_8446622_RNA45SN1_45S_50ntFlanks@10468.10497.30|TTGCCCTCGGCCGATCGAAAGGGAGTCGGG</td><td>>hg38_21_+_8433172_8446622_RNA45SN1_45S_50ntFlanks</td><td>10468</td><td>10497</td><td>30</td><td>TTGCCCTCGGCCGATCGAAAGGGAGTCGGG</td><td>358.6</td><td>-12.4</td><td>1.4</td><td>-8.8</td><td>2.1E-18</td><td>2.5E-17</td><td>MB231/MB468</td><td>MP</td></tr>
<tr><td>>hg38_21_+_8433172_8446622_RNA45SN1_45S_50ntFlanks@6738.6769.32|ATCATCGACACTTCGAACGCACTTGCGGCCCC</td><td>>hg38_21_+_8433172_8446622_RNA45SN1_45S_50ntFlanks</td><td>6738</td><td>6769</td><td>32</td><td>ATCATCGACACTTCGAACGCACTTGCGGCCCC</td><td>409.8</td><td>-12.5</td><td>1.4</td><td>-8.7</td><td>3.5E-18</td><td>2.5E-17</td><td>MB231/MB468</td><td>MITO</td></tr>
<tr><td>>hg38_21_+_8433172_8446622_RNA45SN1_45S_50ntFlanks@6644.6665.22|CCTCGTACGACTCTTAGCGGTG</td><td>>hg38_21_+_8433172_8446622_RNA45SN1_45S_50ntFlanks</td><td>6644</td><td>6665</td><td>22</td><td>CCTCGTACGACTCTTAGCGGTG</td><td>2048.4</td><td>6.0</td><td>0.7</td><td>8.7</td><td>3.0E-18</td><td>2.5E-17</td><td>BT20/MB468</td><td>MITO</td></tr>
<tr><td>>hg38_21_+_8433172_8446622_RNA45SN1_45S_50ntFlanks@12628.12654.27|TGGGGCGAAGCTACCATCTGTGGGATT</td><td>>hg38_21_+_8433172_8446622_RNA45SN1_45S_50ntFlanks</td><td>12628</td><td>12654</td><td>27</td><td>TGGGGCGAAGCTACCATCTGTGGGATT</td><td>7018.6</td><td>-12.4</td><td>1.4</td><td>-8.6</td><td>7.1E-18</td><td>2.5E-17</td><td>BT20/MB468</td><td>NUC</td></tr>
<tr><td>>hg38_21_+_8433172_8446622_RNA45SN1_45S_50ntFlanks@9570.9595.26|TGCAAATCGGTCGTCCGACCTGGGTA</td><td>>hg38_21_+_8433172_8446622_RNA45SN1_45S_50ntFlanks</td><td>9570</td><td>9595</td><td>26</td><td>TGCAAATCGGTCGTCCGACCTGGGTA</td><td>891.0</td><td>-5.3</td><td>0.6</td><td>-8.7</td><td>2.8E-18</td><td>2.5E-17</td><td>MB231/MB468</td><td>NUC</td></tr>
<tr><td>>MI0000283|hsa-mir-203a&WithFlank&14|+|104117399|104117520@72.90.19&[MIMAT0000264&hsa-miR-203a-3p&offsets|+1|-2;m-79&14|+|104117469|104117490&offsets|+1|-2]|TGAAATGTTTAGGACCACT</td><td>>MI0000283|hsa-mir-203a&WithFlank&14|+|104117399|104117520</td><td>72</td><td>90</td><td>19</td><td>TGAAATGTTTAGGACCACT</td><td>429.6</td><td>12.5</td><td>1.4</td><td>9.0</td><td>3.3E-19</td><td>2.5E-17</td><td>BT20/MB231</td><td>NUC</td></tr>
<tr><td>>hg38_21_+_8433172_8446622_RNA45SN1_45S_50ntFlanks@3775.3799.25|GCCGGTACAGTGAAACTGCGAATGG</td><td>>hg38_21_+_8433172_8446622_RNA45SN1_45S_50ntFlanks</td><td>3775</td><td>3799</td><td>25</td><td>GCCGGTACAGTGAAACTGCGAATGG</td><td>381.0</td><td>-12.4</td><td>1.4</td><td>-8.7</td><td>2.1E-18</td><td>2.5E-17</td><td>MB231/MB468</td><td>MP</td></tr>
<tr><td>>MI0000264|hsa-mir-7-2&WithFlank&15|+|88611819|88611940@38.61.24;MI0000265|hsa-mir-7-3&WithFlank&19|+|4770664|4770785@37.60.24;MI0000263|hsa-mir-7-1&WithFlank&9|-|83969742|83969863@30.53.24&[MIMAT0000252&hsa-miR-7-5p&offsets|0|0;m-400&15|+|88611856|88611879&offsets|0|0];[MIMAT0000252_1&hsa-miR-7-5p&offsets|0|0;m-402&19|+|4770700|4770723&offsets|0|0];[MIMAT0000252_2&hsa-miR-7-5p&offsets|0|0;m-401&9|-|83969811|83969834&offsets|0|0]|TGGAAGACTAGTGATTTTGTTGTT</td><td>>MI0000264|hsa-mir-7-2&WithFlank&15|+|88611819|88611940</td><td>38</td><td>61</td><td>24</td><td>TGGAAGACTAGTGATTTTGTTGTT</td><td>43806.6</td><td>5.2</td><td>0.6</td><td>8.7</td><td>2.8E-18</td><td>2.5E-17</td><td>MB231/MB468</td><td>NUC</td></tr>
<tr><td>>am_trnaMT_SerGCT_MT_+_12207_12265@18.38.21|CTGCTAACTCATGCCCCCATG</td><td>>am_trnaMT_SerGCT_MT_+_12207_12265</td><td>18</td><td>38</td><td>21</td><td>CTGCTAACTCATGCCCCCATG</td><td>531.0</td><td>-13.5</td><td>1.6</td><td>-8.6</td><td>7.2E-18</td><td>2.5E-17</td><td>BT20/MB468</td><td>NUC</td></tr>
<tr><td>>hg38_21_+_8433172_8446622_RNA45SN1_45S_50ntFlanks@9533.9553.21|GAGGAAACTCTGGTGGAGGTC</td><td>>hg38_21_+_8433172_8446622_RNA45SN1_45S_50ntFlanks</td><td>9533</td><td>9553</td><td>21</td><td>GAGGAAACTCTGGTGGAGGTC</td><td>5795.2</td><td>-6.9</td><td>0.8</td><td>-8.7</td><td>2.2E-18</td><td>2.5E-17</td><td>MB231/MB468</td><td>MP</td></tr>
<tr><td>>am_MI0000809|hsa-mir-151a&WithFlank&8|-|140732558|140732659@17.38.22&[MIMAT0004697&hsa-miR-151a-5p&offsets|0|+1;m-75&8|-|140732623|140732643&offsets|0|+1]|TCGAGGAGCTCACAGTCTAGTA</td><td>>am_MI0000809|hsa-mir-151a&WithFlank&8|-|140732558|140732659</td><td>17</td><td>38</td><td>22</td><td>TCGAGGAGCTCACAGTCTAGTA</td><td>466.0</td><td>6.4</td><td>0.7</td><td>8.7</td><td>2.8E-18</td><td>2.5E-17</td><td>MB231/MB468</td><td>NUC</td></tr>
<tr><td>>hg38_21_+_8433172_8446622_RNA45SN1_45S_50ntFlanks@9033.9058.26|CTCGTGGGGGGGCCGGGCCACCCCTC</td><td>>hg38_21_+_8433172_8446622_RNA45SN1_45S_50ntFlanks</td><td>9033</td><td>9058</td><td>26</td><td>CTCGTGGGGGGGCCGGGCCACCCCTC</td><td>2167.7</td><td>-14.2</td><td>1.5</td><td>-9.2</td><td>5.1E-20</td><td>2.5E-17</td><td>BT20/MB468</td><td>MP</td></tr>
<tr><td>>hg38_21_+_8433172_8446622_RNA45SN1_45S_50ntFlanks@9739.9766.28|AAACGATCTCAACCTATTCTCAAACTTT</td><td>>hg38_21_+_8433172_8446622_RNA45SN1_45S_50ntFlanks</td><td>9739</td><td>9766</td><td>28</td><td>AAACGATCTCAACCTATTCTCAAACTTT</td><td>2163.6</td><td>-7.2</td><td>0.8</td><td>-8.7</td><td>3.5E-18</td><td>2.5E-17</td><td>MB231/MB468</td><td>MITO</td></tr>
<tr><td>>hg38_21_+_8433172_8446622_RNA45SN1_45S_50ntFlanks@10731.10755.25|AGGGTGTAAATCTCGCGCCGGGCCG</td><td>>hg38_21_+_8433172_8446622_RNA45SN1_45S_50ntFlanks</td><td>10731</td><td>10755</td><td>25</td><td>AGGGTGTAAATCTCGCGCCGGGCCG</td><td>554.0</td><td>-12.9</td><td>1.5</td><td>-8.7</td><td>3.6E-18</td><td>2.5E-17</td><td>MB231/MB468</td><td>MITO</td></tr>
<tr><td>>hg38_21_+_8433172_8446622_RNA45SN1_45S_50ntFlanks@7975.7994.20|CGCGACCTCAGATCAGACGT</td><td>>hg38_21_+_8433172_8446622_RNA45SN1_45S_50ntFlanks</td><td>7975</td><td>7994</td><td>20</td><td>CGCGACCTCAGATCAGACGT</td><td>759686.3</td><td>5.8</td><td>0.7</td><td>8.7</td><td>3.0E-18</td><td>2.5E-17</td><td>BT20/MB468</td><td>MITO</td></tr>
<tr><td>>hg38_21_+_8433172_8446622_RNA45SN1_45S_50ntFlanks@12033.12068.36|ATCGTTTTTTTCACTGACCCGGTGAGGCGGGGGGGCG</td><td>>hg38_21_+_8433172_8446622_RNA45SN1_45S_50ntFlanks</td><td>12033</td><td>12068</td><td>36</td><td>ATCGTTTTTTTCACTGACCCGGTGAGGCGGGGGGGCG</td><td>589.2</td><td>-13.0</td><td>1.5</td><td>-8.7</td><td>3.6E-18</td><td>2.5E-17</td><td>MB231/MB468</td><td>MITO</td></tr>
<tr><td>>hg38_21_+_8433172_8446622_RNA45SN1_45S_50ntFlanks@11203.11225.23|CGGCGGCGGCGACTCTGGACGCG</td><td>>hg38_21_+_8433172_8446622_RNA45SN1_45S_50ntFlanks</td><td>11203</td><td>11225</td><td>23</td><td>CGGCGGCGGCGACTCTGGACGCG</td><td>365.1</td><td>-12.4</td><td>1.4</td><td>-8.7</td><td>2.2E-18</td><td>2.5E-17</td><td>MB231/MB468</td><td>MP</td></tr>
<tr><td>>hg38_21_+_8433172_8446622_RNA45SN1_45S_50ntFlanks@12678.12703.26|AATCCCGCCCAGGCGGAACGATACGG</td><td>>hg38_21_+_8433172_8446622_RNA45SN1_45S_50ntFlanks</td><td>12678</td><td>12703</td><td>26</td><td>AATCCCGCCCAGGCGGAACGATACGG</td><td>2612.4</td><td>-11.1</td><td>1.3</td><td>-8.7</td><td>2.2E-18</td><td>2.5E-17</td><td>MB231/MB468</td><td>MP</td></tr>
<tr><td>>hg38_21_+_8433172_8446622_RNA45SN1_45S_50ntFlanks@5082.5104.23|AACGAACGAGACTCTGGCATGCT</td><td>>hg38_21_+_8433172_8446622_RNA45SN1_45S_50ntFlanks</td><td>5082</td><td>5104</td><td>23</td><td>AACGAACGAGACTCTGGCATGCT</td><td>509.5</td><td>5.7</td><td>0.7</td><td>8.7</td><td>4.3E-18</td><td>2.5E-17</td><td>BT20/MB468</td><td>CYTO</td></tr>
<tr><td>>hg38_21_+_8433172_8446622_RNA45SN1_45S_50ntFlanks@7981.8007.27|CTCAGATCAGACGTGGCGACCCGCTGA</td><td>>hg38_21_+_8433172_8446622_RNA45SN1_45S_50ntFlanks</td><td>7981</td><td>8007</td><td>27</td><td>CTCAGATCAGACGTGGCGACCCGCTGA</td><td>294.6</td><td>-12.1</td><td>1.4</td><td>-8.7</td><td>2.2E-18</td><td>2.5E-17</td><td>MB231/MB468</td><td>MP</td></tr>
<tr><td>>hg38_21_+_8433172_8446622_RNA45SN1_45S_50ntFlanks@9342.9370.29|ACGAAAGCCGCCGTGGCGCAATGAAGGTG</td><td>>hg38_21_+_8433172_8446622_RNA45SN1_45S_50ntFlanks</td><td>9342</td><td>9370</td><td>29</td><td>ACGAAAGCCGCCGTGGCGCAATGAAGGTG</td><td>532.7</td><td>-12.9</td><td>1.5</td><td>-8.7</td><td>2.2E-18</td><td>2.5E-17</td><td>MB231/MB468</td><td>MP</td></tr>
<tr><td>>MI0000279|hsa-mir-196a-2&WithFlank&12|+|53991732|53991853@31.51.21;MI0000238|hsa-mir-196a-1&WithFlank&17|-|48632484|48632565@13.33.21&[MIMAT0000226&hsa-miR-196a-5p&offsets|0|-1;m-261&12|+|53991762|53991783&offsets|0|-1];[MIMAT0000226_1&hsa-miR-196a-5p&offsets|0|-1;m-268&17|-|48632532|48632553&offsets|0|-1]|TAGGTAGTTTCATGTTGTTGG</td><td>>MI0000279|hsa-mir-196a-2&WithFlank&12|+|53991732|53991853</td><td>31</td><td>51</td><td>21</td><td>TAGGTAGTTTCATGTTGTTGG</td><td>1550.3</td><td>10.4</td><td>1.2</td><td>9.0</td><td>2.2E-19</td><td>2.5E-17</td><td>BT20/MB231</td><td>CYTO</td></tr>
<tr><td>>hg38_21_+_8433172_8446622_RNA45SN1_45S_50ntFlanks@12641.12659.19|CCATCTGTGGGATTATGAC</td><td>>hg38_21_+_8433172_8446622_RNA45SN1_45S_50ntFlanks</td><td>12641</td><td>12659</td><td>19</td><td>CCATCTGTGGGATTATGAC</td><td>8964.9</td><td>-10.3</td><td>1.2</td><td>-8.7</td><td>3.6E-18</td><td>2.6E-17</td><td>MB231/MB468</td><td>MITO</td></tr>
<tr><td>>hg38_21_+_8433172_8446622_RNA45SN1_45S_50ntFlanks@7987.8006.20|TCAGACGTGGCGACCCGCTG</td><td>>hg38_21_+_8433172_8446622_RNA45SN1_45S_50ntFlanks</td><td>7987</td><td>8006</td><td>20</td><td>TCAGACGTGGCGACCCGCTG</td><td>681.8</td><td>-13.2</td><td>1.5</td><td>-8.7</td><td>3.6E-18</td><td>2.6E-17</td><td>MB231/MB468</td><td>MITO</td></tr>
<tr><td>>hg38_1_-_228634819_228635039_RNA5S12_5S_50ntFlanks@126.170.45|TACTTGGATGGGAGACCGCCTGGGAATACCGGGTGCTGTAGGCTT</td><td>>hg38_1_-_228634819_228635039_RNA5S12_5S_50ntFlanks</td><td>126</td><td>170</td><td>45</td><td>TACTTGGATGGGAGACCGCCTGGGAATACCGGGTGCTGTAGGCTT</td><td>4672.6</td><td>8.4</td><td>1.0</td><td>8.7</td><td>2.2E-18</td><td>2.6E-17</td><td>MB231/MB468</td><td>CYTO</td></tr>
<tr><td>>hg38_21_+_8433172_8446622_RNA45SN1_45S_50ntFlanks@4380.4427.48|CGTAGTTGGATCTTGGGAGCGGGCGGGCGGTCCGCCGCGAGGCGAGCC</td><td>>hg38_21_+_8433172_8446622_RNA45SN1_45S_50ntFlanks</td><td>4380</td><td>4427</td><td>48</td><td>CGTAGTTGGATCTTGGGAGCGGGCGGGCGGTCCGCCGCGAGGCGAGCC</td><td>174.8</td><td>11.5</td><td>1.3</td><td>8.7</td><td>2.2E-18</td><td>2.6E-17</td><td>MB231/MB468</td><td>CYTO</td></tr>
<tr><td>>hg38_21_+_8433172_8446622_RNA45SN1_45S_50ntFlanks@10458.10478.21|ATGGCCTCCGTTGCCCTCGGC</td><td>>hg38_21_+_8433172_8446622_RNA45SN1_45S_50ntFlanks</td><td>10458</td><td>10478</td><td>21</td><td>ATGGCCTCCGTTGCCCTCGGC</td><td>416.2</td><td>-12.6</td><td>1.4</td><td>-8.7</td><td>2.2E-18</td><td>2.6E-17</td><td>MB231/MB468</td><td>MP</td></tr>
</table>

| | | | | | | | | | | | | | |
|---|---|---|---|---|---|---|---|---|---|---|---|---|---|
| >hg38_21_+_8433172_8446622_RNA45SN1_45S_50ntFlanks@10821.10865.45\|GGAAGTCGGCAAGCCGGATCCGTAACTTCGGGATAAGGATTGGCT | >hg38_21_+_8433172_8446622_RNA45SN1_45S_50ntFlanks | 10821 | 10865 | 45 | GGAAGTCGGCAAGCCGGATCCGTAACTTCGGGATAAGGATTGGCT | 2345.5 | -15.6 | 1.8 | -8.6 | 7.5E-18 | 2.6E-17 | BT20/MB468 | NUC |
| >hg38_21_+_8433172_8446622_RNA45SN1_45S_50ntFlanks@12074.12095.22\|CGAGGGGCTCTCGCTTCTGGCG | >hg38_21_+_8433172_8446622_RNA45SN1_45S_50ntFlanks | 12074 | 12095 | 22 | CGAGGGGCTCTCGCTTCTGGCG | 604.0 | -4.7 | 0.5 | -8.7 | 2.9E-18 | 2.6E-17 | MB231/MB468 | NUC |
| >hg38_21_+_8433172_8446622_RNA45SN1_45S_50ntFlanks@8114.8136.23\|GCGCGGGACATGTGGCGTACGGA | >hg38_21_+_8433172_8446622_RNA45SN1_45S_50ntFlanks | 8114 | 8136 | 23 | GCGCGGGACATGTGGCGTACGGA | 3680.8 | -11.5 | 1.3 | -8.7 | 2.3E-18 | 2.6E-17 | MB231/MB468 | MP |
| >MI0000542\|hsa-mir-320a&WithFlank&8\|-\|22244960\|22245043@48.69.22(+1U)&[MIMAT0000510&hsa-miR-320a-3p&offsets\|0\|0(+1U);m-60&8\|-\|22244975\|22244996&offsets\|0\|0(+1U)]\|AAAAGCTGGGTTGAGAGGGCGAT | >MI0000542\|hsa-mir-320a&WithFlank&8\|-\|22244960\|22245043 | 48 | 69 | 22(+1U) | AAAAGCTGGGTTGAGAGGGCGAT | 2908.3 | 7.7 | 0.9 | 8.7 | 2.2E-18 | 2.6E-17 | MB231/MB468 | CYTO |
| >trna59_GluCTC_1_+_249168447_249168518@32.70.39\|CTCTCACCGCCGCGGCCCGGGTTCGATTCCCGGTCAGGA | >trna59_GluCTC_1_+_249168447_249168518 | 32 | 70 | 39 | CTCTCACCGCCGCGGCCCGGGTTCGATTCCCGGTCAGGA | 580.2 | -11.9 | 1.4 | -8.7 | 2.2E-18 | 2.6E-17 | MB231/MB468 | CYTO |
| >trna116_GluCTC_1_-_145399233_145399304@1.32.32,trna59_GluCTC_1_+_249168447_249168518@1.32.32,trna71_GluCTC_1_-_161439189_161439260@1.32.32,trna74_GluCTC_1_-_161431809_161431880@1.32.32,trna77_GluCTC_1_-_161424398_161424469@1.32.32,trna77_GluCTC_6_+_28949976_28950047@1.32.32,trna80_GluCTC_1_-_161417018_161417089@1.32.32,trna87_GluCTC_6_-_126101393_126101464@1.32.32\|TCCCTGGTGGTCTAGTGGTTAGGATTCGGCGC | >trna116_GluCTC_1_-_145399233_145399304 | 1 | 32 | 32 | TCCCTGGTGGTCTAGTGGTTAGGATTCGGCGC | 8966.7 | 9.4 | 1.1 | 8.7 | 3.2E-18 | 2.6E-17 | BT20/MB468 | MITO |
| >hg38_21_+_8433172_8446622_RNA45SN1_45S_50ntFlanks@4003.4024.22\|ATAACCTCGGGCCGATCGCACG | >hg38_21_+_8433172_8446622_RNA45SN1_45S_50ntFlanks | 4003 | 4024 | 22 | ATAACCTCGGGCCGATCGCACG | 1903.5 | -10.5 | 1.2 | -8.7 | 2.3E-18 | 2.6E-17 | MB231/MB468 | MP |
| >hg38_21_+_8433172_8446622_RNA45SN1_45S_50ntFlanks@5237.5282.46\|ACTGACTGGCTCAGCGTGTGCCTACCCTACGCCGGCAGGCGCGGGT | >hg38_21_+_8433172_8446622_RNA45SN1_45S_50ntFlanks | 5237 | 5282 | 46 | ACTGACTGGCTCAGCGTGTGCCTACCCTACGCCGGCAGGCGCGGGT | 353.7 | 6.0 | 0.7 | 8.7 | 3.2E-18 | 2.6E-17 | BT20/MB468 | MITO |
| >hg38_21_+_8433172_8446622_RNA45SN1_45S_50ntFlanks@12712.12733.22\|CGGAGCCTCGGTTGGCCTCGGA | >hg38_21_+_8433172_8446622_RNA45SN1_45S_50ntFlanks | 12712 | 12733 | 22 | CGGAGCCTCGGTTGGCCTCGGA | 798.6 | -13.5 | 1.5 | -8.7 | 2.3E-18 | 2.7E-17 | MB231/MB468 | MP |
| >MI0000650\|hsa-mir-200c&WithFlank&12\|+\|6963693\|6963772@50.71.22(+1U)&[MIMAT0000617&hsa-miR-200c-3p&offsets\|0\|-1(+1U);m-54&12\|+\|6963742\|6963764&offsets\|0\|-1(+1U)]\|TAATACTGCCGGGTAATGATGGT | >MI0000650\|hsa-mir-200c&WithFlank&12\|+\|6963693\|6963772 | 50 | 71 | 22(+1U) | TAATACTGCCGGGTAATGATGGT | 1093.8 | 4.5 | 0.5 | 8.6 | 7.8E-18 | 2.7E-17 | BT20/MB468 | NUC |
| >hg38_21_+_8433172_8446622_RNA45SN1_45S_50ntFlanks@11672.11706.35\|CAGTGCTCTGAATGTCAAAGTGAAGAAATTCAATG | >hg38_21_+_8433172_8446622_RNA45SN1_45S_50ntFlanks | 11672 | 11706 | 35 | CAGTGCTCTGAATGTCAAAGTGAAGAAATTCAATG | 1537.1 | -5.9 | 0.7 | -8.6 | 7.8E-18 | 2.7E-17 | BT20/MB468 | NUC |
| >hg38_21_+_8433172_8446622_RNA45SN1_45S_50ntFlanks@11641.11662.22\|CGGCGGGTGTTGACGCGATGTG | >hg38_21_+_8433172_8446622_RNA45SN1_45S_50ntFlanks | 11641 | 11662 | 22 | CGGCGGGTGTTGACGCGATGTG | 3989.8 | -6.0 | 0.7 | -8.7 | 3.3E-18 | 2.7E-17 | BT20/MB468 | MITO |
| >hg38_21_+_8433172_8446622_RNA45SN1_45S_50ntFlanks@7974.7997.24\|ACGCGACCTCAGATCAGACGTGGC | >hg38_21_+_8433172_8446622_RNA45SN1_45S_50ntFlanks | 7974 | 7997 | 24 | ACGCGACCTCAGATCAGACGTGGC | 24382.7 | 7.1 | 0.8 | 8.7 | 3.3E-18 | 2.7E-17 | BT20/MB468 | MITO |
| >MI0000433\|hsa-let-7g&WithFlank&3\|-\|52268272\|52268367@11.30.20(+1C)&[MIMAT0000414&hsa-let-7g-5p&offsets\|0\|-2(+1C);m-58&3\|-\|52268336\|52268357&offsets\|0\|-2(+1C)]\|TGAGGTAGTAGTTTGTACAGC | >MI0000433\|hsa-let-7g&WithFlank&3\|-\|52268272\|52268367 | 11 | 30 | 20(+1C) | TGAGGTAGTAGTTTGTACAGC | 612.5 | 5.6 | 0.6 | 8.7 | 3.1E-18 | 2.7E-17 | MB231/MB468 | NUC |
| >hg38_21_+_8433172_8446622_RNA45SN1_45S_50ntFlanks@12045.12070.26\|CTGACCCGGTGAGGCGGGGGGGCGAG | >hg38_21_+_8433172_8446622_RNA45SN1_45S_50ntFlanks | 12045 | 12070 | 26 | CTGACCCGGTGAGGCGGGGGGGCGAG | 1097.2 | -13.9 | 1.6 | -8.7 | 3.8E-18 | 2.7E-17 | MB231/MB468 | MITO |
| >hg38_21_+_8433172_8446622_RNA45SN1_45S_50ntFlanks@8194.8213.20\|CCCGTGGACGGTGTGAGGCC | >hg38_21_+_8433172_8446622_RNA45SN1_45S_50ntFlanks | 8194 | 8213 | 20 | CCCGTGGACGGTGTGAGGCC | 2118.9 | -6.5 | 0.7 | -8.7 | 2.4E-18 | 2.7E-17 | MB231/MB468 | MP |
| >hg38_21_+_8433172_8446622_RNA45SN1_45S_50ntFlanks@10798.10816.19\|CATGTTGGAACAATGTAGG | >hg38_21_+_8433172_8446622_RNA45SN1_45S_50ntFlanks | 10798 | 10816 | 19 | CATGTTGGAACAATGTAGG | 338.8 | -12.3 | 1.4 | -8.7 | 2.4E-18 | 2.7E-17 | MB231/MB468 | MP |
| >hg38_21_+_8433172_8446622_RNA45SN1_45S_50ntFlanks@9504.9533.30\|GTGAACTATGCCTGGGCAGGGCGAAGCCAG | >hg38_21_+_8433172_8446622_RNA45SN1_45S_50ntFlanks | 9504 | 9533 | 30 | GTGAACTATGCCTGGGCAGGGCGAAGCCAG | 1249.5 | -10.0 | 1.1 | -8.7 | 2.4E-18 | 2.8E-17 | MB231/MB468 | CYTO |
| >hg38_MT_+_1621_3279_RNR2_16S_50ntFlanks@51.95.45\|GCTAAACCTAGCCCCAAACCCACTCCACCTTACTACCAGACAACC | >hg38_MT_+_1621_3279_RNR2_16S_50ntFlanks | 51 | 95 | 45 | GCTAAACCTAGCCCCAAACCCACTCCACCTTACTACCAGACAACC | 393.0 | 12.6 | 1.4 | 8.7 | 2.4E-18 | 2.8E-17 | MB231/MB468 | CYTO |
| >hg38_21_+_8433172_8446622_RNA45SN1_45S_50ntFlanks@11623.11640.18\|AAGCATCGCGAAGGCCCG | >hg38_21_+_8433172_8446622_RNA45SN1_45S_50ntFlanks | 11623 | 11640 | 18 | AAGCATCGCGAAGGCCCG | 466.8 | -12.7 | 1.5 | -8.7 | 2.4E-18 | 2.8E-17 | MB231/MB468 | MP |
| >hg38_21_+_8433172_8446622_RNA45SN1_45S_50ntFlanks@8552.8581.30\|CTCCCCGGAGGGGGCGGGCTCCGGCGGGTG | >hg38_21_+_8433172_8446622_RNA45SN1_45S_50ntFlanks | 8552 | 8581 | 30 | CTCCCCGGAGGGGGCGGGCTCCGGCGGGTG | 526.1 | -12.8 | 1.5 | -8.7 | 4.0E-18 | 2.8E-17 | MB231/MB468 | MITO |
| >hg38_21_+_8433172_8446622_RNA45SN1_45S_50ntFlanks@12004.12037.34\|CGGGCCGCCGGTGAAATACCACTACTCTGATCGT | >hg38_21_+_8433172_8446622_RNA45SN1_45S_50ntFlanks | 12004 | 12037 | 34 | CGGGCCGCCGGTGAAATACCACTACTCTGATCGT | 434.3 | 6.4 | 0.7 | 8.7 | 3.4E-18 | 2.8E-17 | BT20/MB468 | MITO |
| >hg38_21_+_8433172_8446622_RNA45SN1_45S_50ntFlanks@5472.5489.18\|ACGGCCCTGGCGGAGCGC | >hg38_21_+_8433172_8446622_RNA45SN1_45S_50ntFlanks | 5472 | 5489 | 18 | ACGGCCCTGGCGGAGCGC | 369.3 | 6.8 | 0.8 | 8.7 | 3.4E-18 | 2.8E-17 | BT20/MB468 | MITO |
| >hg38_21_+_8433172_8446622_RNA45SN1_45S_50ntFlanks@6737.6771.35\|GATCATCGACACTTCGAACGCACTTGCGGCCCCGG | >hg38_21_+_8433172_8446622_RNA45SN1_45S_50ntFlanks | 6737 | 6771 | 35 | GATCATCGACACTTCGAACGCACTTGCGGCCCCGG | 831.7 | -11.8 | 1.4 | -8.7 | 3.4E-18 | 2.8E-17 | BT20/MB468 | MITO |
| >hg38_21_+_8433172_8446622_RNA45SN1_45S_50ntFlanks@10679.10708.30\|GGTTCCGGCGGCGTCCGGTGAGCTCTCGCT | >hg38_21_+_8433172_8446622_RNA45SN1_45S_50ntFlanks | 10679 | 10708 | 30 | GGTTCCGGCGGCGTCCGGTGAGCTCTCGCT | 4070.6 | -6.6 | 0.8 | -8.7 | 2.4E-18 | 2.8E-17 | MB231/MB468 | CYTO |
| >MI0005559\|hsa-mir-744&WithFlank&17\|+\|12081893\|12082002@17.37.21&[MIMAT0004945&hsa-miR-744-5p&offsets\|0\|-1;m-172&17\|+\|12081909\|12081929&offsets\|0\|0]\|TGCGGGGCTAGGGCTAACAGC | >MI0005559\|hsa-mir-744&WithFlank&17\|+\|12081893\|12082002 | 17 | 37 | 21 | TGCGGGGCTAGGGCTAACAGC | 1600.6 | 6.6 | 0.8 | 8.7 | 2.4E-18 | 2.8E-17 | MB231/MB468 | CYTO |
| >MI0000095\|hsa-mir-93&WithFlank&7\|-\|100093762\|100093853@17.38.22(+1A)&[MIMAT0000093&hsa-miR-93-5p&offsets\|0\|-1(+1A);m-29&7\|-\|100093815\|100093837&offsets\|0\|-1(+1A)]\|CAAAGTGCTGTTCGTGCAGGTAA | >MI0000095\|hsa-mir-93&WithFlank&7\|-\|100093762\|100093853 | 17 | 38 | 22(+1A) | CAAAGTGCTGTTCGTGCAGGTAA | 2433.9 | 7.2 | 0.8 | 8.7 | 4.8E-18 | 2.9E-17 | BT20/MB468 | CYTO |
| >hg38_21_+_8433172_8446622_RNA45SN1_45S_50ntFlanks@12539.12557.19\|ATGATGTGTTGTTGCCATG | >hg38_21_+_8433172_8446622_RNA45SN1_45S_50ntFlanks | 12539 | 12557 | 19 | ATGATGTGTTGTTGCCATG | 781.8 | -13.5 | 1.5 | -8.7 | 2.5E-18 | 2.9E-17 | MB231/MB468 | MP |
| >hg38_21_+_8433172_8446622_RNA45SN1_45S_50ntFlanks@11886.11905.20\|TGAGCTTGACTCTAGTCTGG | >hg38_21_+_8433172_8446622_RNA45SN1_45S_50ntFlanks | 11886 | 11905 | 20 | TGAGCTTGACTCTAGTCTGG | 440.2 | -12.6 | 1.4 | -8.7 | 2.5E-18 | 2.9E-17 | MB231/MB468 | MP |
| >hg38_21_+_8433172_8446622_RNA45SN1_45S_50ntFlanks@6719.6737.19\|TGAATTGCAGGACACATTG | >hg38_21_+_8433172_8446622_RNA45SN1_45S_50ntFlanks | 6719 | 6737 | 19 | TGAATTGCAGGACACATTG | 1423.7 | -10.3 | 1.2 | -8.7 | 4.1E-18 | 2.9E-17 | MB231/MB468 | MITO |
| >hg38_21_+_8433172_8446622_RNA45SN1_45S_50ntFlanks@10869.10904.36\|AGGGCTGGGTCGGTCGGGCTGGGGCGCGAAGCGGGG | >hg38_21_+_8433172_8446622_RNA45SN1_45S_50ntFlanks | 10869 | 10904 | 36 | AGGGCTGGGTCGGTCGGGCTGGGGCGCGAAGCGGGG | 619.2 | -13.1 | 1.5 | -8.7 | 2.5E-18 | 2.9E-17 | MB231/MB468 | MP |
| >hg38_21_+_8433172_8446622_RNA45SN1_45S_50ntFlanks@5420.5443.24\|CCGATTGGATGGTTTAGTGAGGCC | >hg38_21_+_8433172_8446622_RNA45SN1_45S_50ntFlanks | 5420 | 5443 | 24 | CCGATTGGATGGTTTAGTGAGGCC | 1683.0 | -10.5 | 1.2 | -8.7 | 3.3E-18 | 2.9E-17 | MB231/MB468 | NUC |
| >MI0000433\|hsa-let-7g&WithFlank&3\|-\|52268272\|52268367@11.31.21(+1C)&[MIMAT0000414&hsa-let-7g-5p&offsets\|0\|-1(+1C);m-58&3\|-\|52268336\|52268357&offsets\|0\|-1(+1C)]\|TGAGGTAGTAGTTTGTACAGTC | >MI0000433\|hsa-let-7g&WithFlank&3\|-\|52268272\|52268367 | 11 | 31 | 21(+1C) | TGAGGTAGTAGTTTGTACAGTC | 11973.1 | 5.1 | 0.6 | 8.7 | 3.3E-18 | 2.9E-17 | MB231/MB468 | NUC |
| >hg38_21_+_8433172_8446622_RNA45SN1_45S_50ntFlanks@10392.10420.29\|ACAGCAGTTGAACATGGGTCAGTCGGTCC | >hg38_21_+_8433172_8446622_RNA45SN1_45S_50ntFlanks | 10392 | 10420 | 29 | ACAGCAGTTGAACATGGGTCAGTCGGTCC | 561.0 | -13.0 | 1.5 | -8.7 | 2.5E-18 | 2.9E-17 | MB231/MB468 | MP |
| >hg38_21_+_8433172_8446622_RNA45SN1_45S_50ntFlanks@3813.3853.41\|TTATGGTTCCTTTGGTCGCTCGCTCCTCTCCTACTTGGATA | >hg38_21_+_8433172_8446622_RNA45SN1_45S_50ntFlanks | 3813 | 3853 | 41 | TTATGGTTCCTTTGGTCGCTCGCTCCTCTCCTACTTGGATA | 448.1 | -13.2 | 1.5 | -8.6 | 8.4E-18 | 2.9E-17 | BT20/MB468 | NUC |
| >hg38_21_+_8433172_8446622_RNA45SN1_45S_50ntFlanks@6699.6718.20\|GCTAGCTGCGAGAATTAATG | >hg38_21_+_8433172_8446622_RNA45SN1_45S_50ntFlanks | 6699 | 6718 | 20 | GCTAGCTGCGAGAATTAATG | 20333.4 | -9.8 | 1.1 | -8.7 | 4.1E-18 | 2.9E-17 | MB231/MB468 | MITO |
| >hg38_21_+_8433172_8446622_RNA45SN1_45S_50ntFlanks@4678.4720.43\|CGGCGCAAGACGGACCAGAGCGAAAGCATTTGCCAAGAATGTT | >hg38_21_+_8433172_8446622_RNA45SN1_45S_50ntFlanks | 4678 | 4720 | 43 | CGGCGCAAGACGGACCAGAGCGAAAGCATTTGCCAAGAATGTT | 617.9 | 13.3 | 1.5 | 8.7 | 2.5E-18 | 2.9E-17 | MB231/MB468 | CYTO |
| >MI0000750\|hsa-mir-26a-2&WithFlank&12\|-\|57824603\|57824698@20.41.22;MI0000083\|hsa-mir-26a-1&WithFlank&3\|+\|37969398\|37969486@16.37.22&[MIMAT0000082&hsa-miR-26a-5p&offsets\|0\|0;m-33&12\|-\|57824658\|57824679&offsets\|0\|0];[MIMAT0000082_1&hsa-miR-26a-5p&offsets\|0\|0;m-34&3\|+\|37969413\|37969434&offsets\|0\|0]\|TTCAAGTAATCCAGGATAGGCT | >MI0000750\|hsa-mir-26a-2&WithFlank&12\|-\|57824603\|57824698 | 20 | 41 | 22 | TTCAAGTAATCCAGGATAGGCT | 71410.1 | 5.8 | 0.7 | 8.7 | 3.6E-18 | 2.9E-17 | BT20/MB468 | MITO |
| >hg38_21_+_8433172_8446622_RNA45SN1_45S_50ntFlanks@11318.11343.26\|CCCCACGTCTCGTCGCGCGCGCGTCC | >hg38_21_+_8433172_8446622_RNA45SN1_45S_50ntFlanks | 11318 | 11343 | 26 | CCCCACGTCTCGTCGCGCGCGCGTCC | 512.6 | -11.7 | 1.3 | -8.7 | 2.5E-18 | 2.9E-17 | MB231/MB468 | CYTO |
| >hg38_21_+_8433172_8446622_RNA45SN1_45S_50ntFlanks@6646.6677.32\|TCGTACGACTCTTAGCGGTGGATCACTCGGCT | >hg38_21_+_8433172_8446622_RNA45SN1_45S_50ntFlanks | 6646 | 6677 | 32 | TCGTACGACTCTTAGCGGTGGATCACTCGGCT | 3707.1 | 5.3 | 0.6 | 8.7 | 4.9E-18 | 2.9E-17 | BT20/MB468 | CYTO |
| >trna10_AspGTC_12_-_125424193_125424264@32.72.41,trna12_AspGTC_12_-_125411891_125411962@32.72.41,trna144_AspGTC_6_-_27551236_27551307@32.72.41,trna38_AspGTC_17_-_8125556_8125627@32.72.41,trna45_AspGTC_6_+_27447453_27447524@32.72.41,trna48_AspGTC_6_+_27471523_27471594@32.72.41,trna4_AspGTC_12_+_96429799_96429870@32.72.41,trna69_AspGTC_1_-_161440205_161440276@32.72.41,trna72_AspGTC_1_-_161432824_161432895@32.72.41,trna75_AspGTC_1_-_161425414_161425485@32.72.41,trna78_AspGTC_1_-_161418033_161418104@32.72.41,trna81_AspGTC_1_-_161410615_161410686@32.72.41\|CTGTCACGCGGGAGACCGGGGTTCGATTCCCCGACGGGGAG | >trna10_AspGTC_12_-_125424193_125424264 | 32 | 72 | 41 | CTGTCACGCGGGAGACCGGGGTTCGATTCCCCGACGGGGAG | 1601.7 | -13.6 | 1.6 | -8.7 | 3.3E-18 | 2.9E-17 | MB231/MB468 | NUC |
| >hg38_21_+_8433172_8446622_RNA45SN1_45S_50ntFlanks@5420.5438.19\|CCGATTGGATGGTTTAGTG | >hg38_21_+_8433172_8446622_RNA45SN1_45S_50ntFlanks | 5420 | 5438 | 19 | CCGATTGGATGGTTTAGTG | 615.6 | -13.1 | 1.5 | -8.7 | 2.6E-18 | 2.9E-17 | MB231/MB468 | MP |
| >hg38_21_+_8433172_8446622_RNA45SN1_45S_50ntFlanks@12589.12614.26\|GTTCAGACATTTGGTGTATGTGCTTG | >hg38_21_+_8433172_8446622_RNA45SN1_45S_50ntFlanks | 12589 | 12614 | 26 | GTTCAGACATTTGGTGTATGTGCTTG | 519.2 | -12.8 | 1.5 | -8.7 | 4.2E-18 | 2.9E-17 | MB231/MB468 | MITO |
| >am_MI0000809\|hsa-mir-151a&WithFlank&8\|-\|140732558\|140732659@17.37.21&[MIMAT0004697&hsa-miR-151a-5p&offsets\|0\|0;m-75&8\|-\|140732623\|140732643&offsets\|0\|0]\|TCGAGGAGCTCACAGTCTAGT | >am_MI0000809\|hsa-mir-151a&WithFlank&8\|-\|140732558\|140732659 | 17 | 37 | 21 | TCGAGGAGCTCACAGTCTAGT | 1047.8 | 8.0 | 0.9 | 8.7 | 3.6E-18 | 2.9E-17 | BT20/MB468 | MITO |

| | | | | | | | | | | | | | |
|---|---|---|---|---|---|---|---|---|---|---|---|---|---|
| >hg38_21_+_8433172_8446622_RNA45SN1_45S_50ntFlanks@8079.8105.27\|AAGAGCCCAGCGCCGAATCCCCGCCCC | >hg38_21_+_8433172_8446622_RNA45SN1_45S_50ntFlanks | 8079 | 8105 | 27 | AAGAGCCCAGCGCCGAATCCCCGCCCC | 540.2 | -12.9 | 1.5 | -8.7 | 4.2E-18 | 3.0E-17 | MB231/MB468 | MITO |
| >trna152_ValCAC_6_-_27248049_27248121@1.34.34\|GCTTCTGTAGTGTAGTGGTTATCACGTTCGCCTC | >trna152_ValCAC_6_-_27248049_27248121 | 1 | 34 | 34 | GCTTCTGTAGTGTAGTGGTTATCACGTTCGCCTC | 1305.4 | 7.3 | 0.8 | 8.7 | 5.0E-18 | 3.0E-17 | BT20/MB468 | CYTO |
| >hg38_21_+_8433172_8446622_RNA45SN1_45S_50ntFlanks@12419.12448.30\|GATGTCGGCTCTTCCTATCATTGTGAAGCA | >hg38_21_+_8433172_8446622_RNA45SN1_45S_50ntFlanks | 12419 | 12448 | 30 | GATGTCGGCTCTTCCTATCATTGTGAAGCA | 7574.8 | 7.3 | 0.8 | 8.7 | 2.6E-18 | 3.0E-17 | MB231/MB468 | CYTO |
| >hg38_21_+_8433172_8446622_RNA45SN1_45S_50ntFlanks@9572.9598.27\|CAAATCGGTCGTCCGACCTGGGTATAG | >hg38_21_+_8433172_8446622_RNA45SN1_45S_50ntFlanks | 9572 | 9598 | 27 | CAAATCGGTCGTCCGACCTGGGTATAG | 330.8 | -12.2 | 1.4 | -8.7 | 2.6E-18 | 3.0E-17 | MB231/MB468 | MP |
| >hg38_21_+_8433172_8446622_RNA45SN1_45S_50ntFlanks@5515.5534.20\|CTAGAGGAAGTAAAAGTCGT | >hg38_21_+_8433172_8446622_RNA45SN1_45S_50ntFlanks | 5515 | 5534 | 20 | CTAGAGGAAGTAAAAGTCGT | 400.4 | -12.5 | 1.4 | -8.7 | 2.6E-18 | 3.0E-17 | MB231/MB468 | MP |
| >hg38_21_+_8433172_8446622_RNA45SN1_45S_50ntFlanks@6645.6667.23\|CTCGTACGACTCTTAGCGGTGGA | >hg38_21_+_8433172_8446622_RNA45SN1_45S_50ntFlanks | 6645 | 6667 | 23 | CTCGTACGACTCTTAGCGGTGGA | 4479.9 | 6.5 | 0.8 | 8.7 | 5.0E-18 | 3.0E-17 | BT20/MB468 | CYTO |
| >hg38_21_+_8433172_8446622_RNA45SN1_45S_50ntFlanks@13006.13025.20\|CTATTGAAAGTCAGCCCTCG | >hg38_21_+_8433172_8446622_RNA45SN1_45S_50ntFlanks | 13006 | 13025 | 20 | CTATTGAAAGTCAGCCCTCG | 876.6 | -12.7 | 1.5 | -8.7 | 3.4E-18 | 3.0E-17 | MB231/MB468 | NUC |
| >hg38_21_+_8433172_8446622_RNA45SN1_45S_50ntFlanks@10535.10567.33\|CGCGAGGCGTCCAGTGCGGTAACGCGACCGATC | >hg38_21_+_8433172_8446622_RNA45SN1_45S_50ntFlanks | 10535 | 10567 | 33 | CGCGAGGCGTCCAGTGCGGTAACGCGACCGATC | 333.2 | -12.2 | 1.4 | -8.7 | 2.6E-18 | 3.0E-17 | MB231/MB468 | MP |
| >hg38_21_+_8433172_8446622_RNA45SN1_45S_50ntFlanks@11577.11603.27\|TTAGAACTGGTGCGGACCAGGGGAATC | >hg38_21_+_8433172_8446622_RNA45SN1_45S_50ntFlanks | 11577 | 11603 | 27 | TTAGAACTGGTGCGGACCAGGGGAATC | 1810.4 | -4.6 | 0.5 | -8.7 | 5.1E-18 | 3.0E-17 | BT20/MB468 | CYTO |
| >MI0000063\|hsa-let-7b&WithFlank&22\|+\|46113680\|46113774@12.31.20&[MIMAT0000063&hsa-let-7b-5p&offsets\|0\|-2;m-14&22\|+\|46113691\|46113712&offsets\|0\|-2]\|TGAGGTAGTAGGTTGTGTGG | >MI0000063\|hsa-let-7b&WithFlank&22\|+\|46113680\|46113774 | 12 | 31 | 20 | TGAGGTAGTAGGTTGTGTGG | 449.0 | 8.1 | 0.9 | 8.7 | 3.7E-18 | 3.0E-17 | BT20/MB468 | MITO |
| >hg38_21_+_8433172_8446622_RNA45SN1_45S_50ntFlanks@4725.4765.41\|TTAATCAAGAACGAAAGTCGGAGGTTCGAAGACGATCAGAT | >hg38_21_+_8433172_8446622_RNA45SN1_45S_50ntFlanks | 4725 | 4765 | 41 | TTAATCAAGAACGAAAGTCGGAGGTTCGAAGACGATCAGAT | 551.4 | 7.5 | 0.9 | 8.7 | 2.6E-18 | 3.0E-17 | MB231/MB468 | CYTO |
| >hg38_21_+_8433172_8446622_RNA45SN1_45S_50ntFlanks@10846.10863.18\|CTTCGGGATAAGGATTGG | >hg38_21_+_8433172_8446622_RNA45SN1_45S_50ntFlanks | 10846 | 10863 | 18 | CTTCGGGATAAGGATTGG | 426.9 | -12.6 | 1.4 | -8.7 | 2.7E-18 | 3.0E-17 | MB231/MB468 | MP |
| >MI0000086\|hsa-mir-28&WithFlank&3\|+\|188688775\|188688872@20.41.22&[MIMAT0000085&hsa-miR-28-5p&offsets\|0\|0;m-133&3\|+\|188688794\|188688815&offsets\|0\|0]\|AAGGAGCTCACAGTCTATTGAG | >MI0000086\|hsa-mir-28&WithFlank&3\|+\|188688775\|188688872 | 20 | 41 | 22 | AAGGAGCTCACAGTCTATTGAG | 969.7 | 10.4 | 1.2 | 8.7 | 5.1E-18 | 3.1E-17 | BT20/MB468 | CYTO |
| >hg38_21_+_8433172_8446622_RNA45SN1_45S_50ntFlanks@4759.4793.35\|ATCAGATACCGTCGTAGTTCCGACCATAAACGATG | >hg38_21_+_8433172_8446622_RNA45SN1_45S_50ntFlanks | 4759 | 4793 | 35 | ATCAGATACCGTCGTAGTTCCGACCATAAACGATG | 793.6 | -11.9 | 1.4 | -8.7 | 5.1E-18 | 3.1E-17 | BT20/MB468 | CYTO |
| >hg38_21_+_8433172_8446622_RNA45SN1_45S_50ntFlanks@11561.11605.45\|GCGCCTAGCAGCCGACTTAGAACTGGTGCGGACCAGGGGAATCCG | >hg38_21_+_8433172_8446622_RNA45SN1_45S_50ntFlanks | 11561 | 11605 | 45 | GCGCCTAGCAGCCGACTTAGAACTGGTGCGGACCAGGGGAATCCG | 603.1 | -13.7 | 1.6 | -8.6 | 8.9E-18 | 3.1E-17 | BT20/MB468 | NUC |
| >hg38_21_+_8433172_8446622_RNA45SN1_45S_50ntFlanks@12996.13039.44\|TCGCTGCGATCTATTGAAAGTCAGCCCTCGACACAAGGGTTTGT | >hg38_21_+_8433172_8446622_RNA45SN1_45S_50ntFlanks | 12996 | 13039 | 44 | TCGCTGCGATCTATTGAAAGTCAGCCCTCGACACAAGGGTTTGT | 395010.5 | 8.2 | 0.9 | 8.7 | 2.6E-18 | 3.1E-17 | MB231/MB468 | CYTO |
| >hg38_21_+_8433172_8446622_RNA45SN1_45S_50ntFlanks@12942.12972.31\|TTCGTAGACGACCTGCTTCTGGGTCGGGGTT | >hg38_21_+_8433172_8446622_RNA45SN1_45S_50ntFlanks | 12942 | 12972 | 31 | TTCGTAGACGACCTGCTTCTGGGTCGGGGTT | 1074.9 | -9.8 | 1.1 | -8.7 | 2.6E-18 | 3.1E-17 | MB231/MB468 | CYTO |
| >hg38_21_+_8433172_8446622_RNA45SN1_45S_50ntFlanks@11568.11603.36\|GCAGCCGACTTAGAACTGGTGCGGACCAGGGGAATC | >hg38_21_+_8433172_8446622_RNA45SN1_45S_50ntFlanks | 11568 | 11603 | 36 | GCAGCCGACTTAGAACTGGTGCGGACCAGGGGAATC | 3026.6 | -6.7 | 0.8 | -8.7 | 4.4E-18 | 3.1E-17 | MB231/MB468 | MITO |
| >MI0016005\|hsa-mir-3615&WithFlank&17\|+\|74748607\|74748705@57.78.22(+1U)&[MIMAT0017994&hsa-miR-3615&offsets\|0\|+1(+1U);m-271&17\|+\|74748663\|74748684&offsets\|0\|0(+1U)]\|TCTCTCGGCTCCTCGCGGCTCGT | >MI0016005\|hsa-mir-3615&WithFlank&17\|+\|74748607\|74748705 | 57 | 78 | 22(+1U) | TCTCTCGGCTCCTCGCGGCTCGT | 368.3 | 7.7 | 0.9 | 8.6 | 5.2E-18 | 3.1E-17 | BT20/MB468 | CYTO |
| >hg38_21_+_8433172_8446622_RNA45SN1_45S_50ntFlanks@4241.4258.18\|CTCTTTCGAGGCCCTGTA | >hg38_21_+_8433172_8446622_RNA45SN1_45S_50ntFlanks | 4241 | 4258 | 18 | CTCTTTCGAGGCCCTGTA | 1370.8 | -5.8 | 0.7 | -8.6 | 5.2E-18 | 3.1E-17 | BT20/MB468 | CYTO |
| >hg38_21_+_8433172_8446622_RNA45SN1_45S_50ntFlanks@4661.4689.29\|AGGTGAAATTCTTGGACCGGCGCAAGACG | >hg38_21_+_8433172_8446622_RNA45SN1_45S_50ntFlanks | 4661 | 4689 | 29 | AGGTGAAATTCTTGGACCGGCGCAAGACG | 496.2 | -12.7 | 1.5 | -8.7 | 4.5E-18 | 3.1E-17 | MB231/MB468 | MITO |
| >hg38_21_+_8433172_8446622_RNA45SN1_45S_50ntFlanks@11579.11603.25\|AGAACTGGTGCGGACCAGGGGAATC | >hg38_21_+_8433172_8446622_RNA45SN1_45S_50ntFlanks | 11579 | 11603 | 25 | AGAACTGGTGCGGACCAGGGGAATC | 1746.0 | -4.9 | 0.6 | -8.7 | 3.8E-18 | 3.1E-17 | BT20/MB468 | MITO |
| >MI0000650\|hsa-mir-200c&WithFlank&12\|+\|6963693\|6963772@50.72.23(+2A)&[MIMAT0000617&hsa-miR-200c-3p&offsets\|0\|0(+2A);m-54&12\|+\|6963742\|6963764&offsets\|0\|0(+2A)]\|TAATACTGCCGGGTAATGATGGAAA | >MI0000650\|hsa-mir-200c&WithFlank&12\|+\|6963693\|6963772 | 50 | 72 | 23(+2A) | TAATACTGCCGGGTAATGATGGAAA | 758.1 | 10.2 | 1.1 | 8.9 | 4.3E-19 | 3.1E-17 | BT20/MB231 | NUC |
| >hg38_21_+_8433172_8446622_RNA45SN1_45S_50ntFlanks@12936.12955.20\|AAACCATTCGTAGACGACCT | >hg38_21_+_8433172_8446622_RNA45SN1_45S_50ntFlanks | 12936 | 12955 | 20 | AAACCATTCGTAGACGACCT | 736.5 | -5.2 | 0.6 | -8.7 | 3.6E-18 | 3.1E-17 | MB231/MB468 | NUC |
| >hg38_1_-_228634819_228635039_RNA5S12_5S_50ntFlanks@153.171.19\|ACCGGGTGCTGTAGGCTTT | >hg38_1_-_228634819_228635039_RNA5S12_5S_50ntFlanks | 153 | 171 | 19 | ACCGGGTGCTGTAGGCTTT | 33082.6 | 7.4 | 0.8 | 8.7 | 2.7E-18 | 3.1E-17 | MB231/MB468 | CYTO |
| >hg38_21_+_8433172_8446622_RNA45SN1_45S_50ntFlanks@9573.9597.25\|AAATCGGTCGTCCGACCTGGGTATA | >hg38_21_+_8433172_8446622_RNA45SN1_45S_50ntFlanks | 9573 | 9597 | 25 | AAATCGGTCGTCCGACCTGGGTATA | 1979.8 | -8.1 | 0.9 | -8.7 | 2.7E-18 | 3.1E-17 | MB231/MB468 | CYTO |
| >hg38_21_+_8433172_8446622_RNA45SN1_45S_50ntFlanks@11575.11611.37\|ACTTAGAACTGGTGCGGACCAGGGGAATCCGACTGTT | >hg38_21_+_8433172_8446622_RNA45SN1_45S_50ntFlanks | 11575 | 11611 | 37 | ACTTAGAACTGGTGCGGACCAGGGGAATCCGACTGTT | 975.4 | -4.5 | 0.5 | -8.6 | 9.2E-18 | 3.1E-17 | BT20/MB468 | NUC |
| >MI0000272\|hsa-mir-182&WithFlank&7\|-\|129770377\|129770498@29.54.26(+1A)&[MIMAT0000259&hsa-miR-182-5p&offsets\|0\|+2(+1A);m-22&7\|-\|129770449\|129770470&offsets\|0\|+4(+1A)]\|TTTGGCAATGGTAGAACTCACACTGGA | >MI0000272\|hsa-mir-182&WithFlank&7\|-\|129770377\|129770498 | 29 | 54 | 26(+1A) | TTTGGCAATGGTAGAACTCACACTGGA | 427.2 | 9.2 | 1.1 | 8.6 | 5.3E-18 | 3.1E-17 | BT20/MB468 | CYTO |
| >hg38_21_+_8433172_8446622_RNA45SN1_45S_50ntFlanks@12638.12659.22\|CTACCATCTGTGGGATTATGAC | >hg38_21_+_8433172_8446622_RNA45SN1_45S_50ntFlanks | 12638 | 12659 | 22 | CTACCATCTGTGGGATTATGAC | 7412.6 | -10.4 | 1.2 | -8.7 | 4.5E-18 | 3.2E-17 | MB231/MB468 | MITO |
| >hg38_21_+_8433172_8446622_RNA45SN1_45S_50ntFlanks@10806.10840.35\|AACAATGTAGGTAAGGGAAGTCGGCAAGCCGGATC | >hg38_21_+_8433172_8446622_RNA45SN1_45S_50ntFlanks | 10806 | 10840 | 35 | AACAATGTAGGTAAGGGAAGTCGGCAAGCCGGATC | 405.0 | -3.7 | 0.4 | -8.6 | 5.3E-18 | 3.2E-17 | BT20/MB468 | CYTO |
| >MI0000298\|hsa-mir-221&WithFlank&X\|-\|45746151\|45746272@71.94.24&[MIMAT0000278&hsa-miR-221-3p&offsets\|0\|+1;m-91&X\|-\|45746181\|45746202&offsets\|0\|+2]\|AGCTACATTGTCTGCTGGGTTTCA | >MI0000298\|hsa-mir-221&WithFlank&X\|-\|45746151\|45746272 | 71 | 94 | 24 | AGCTACATTGTCTGCTGGGTTTCA | 2404.8 | 5.9 | 0.7 | 8.6 | 5.4E-18 | 3.2E-17 | BT20/MB468 | CYTO |
| >hg38_21_+_8433172_8446622_RNA45SN1_45S_50ntFlanks@11768.11794.27\|CTAATTAGTGACGCGCATGAATGGATG | >hg38_21_+_8433172_8446622_RNA45SN1_45S_50ntFlanks | 11768 | 11794 | 27 | CTAATTAGTGACGCGCATGAATGGATG | 1383.1 | -7.9 | 0.9 | -8.7 | 4.5E-18 | 3.2E-17 | MB231/MB468 | MITO |
| >hg38_21_+_8433172_8446622_RNA45SN1_45S_50ntFlanks@11843.11872.30\|AGGGAACGGGCTTGGCGGAATCAGCGGGGA | >hg38_21_+_8433172_8446622_RNA45SN1_45S_50ntFlanks | 11843 | 11872 | 30 | AGGGAACGGGCTTGGCGGAATCAGCGGGGA | 532.5 | -12.8 | 1.5 | -8.7 | 4.6E-18 | 3.2E-17 | MB231/MB468 | MITO |
| >hg38_21_+_8433172_8446622_RNA45SN1_45S_50ntFlanks@11920.11967.48\|ATGAGAGGTGTAGAATAAGTGGGAGGCCCCCGGCGCCCCCCCGGTGTC | >hg38_21_+_8433172_8446622_RNA45SN1_45S_50ntFlanks | 11920 | 11967 | 48 | ATGAGAGGTGTAGAATAAGTGGGAGGCCCCCGGCGCCCCCCCGGTGTC | 1541.1 | -15.0 | 1.7 | -8.6 | 9.3E-18 | 3.2E-17 | BT20/MB468 | NUC |
| >hg38_21_+_8433172_8446622_RNA45SN1_45S_50ntFlanks@9714.9735.22\|CGAATGATTAGAGGTCTTGGGG | >hg38_21_+_8433172_8446622_RNA45SN1_45S_50ntFlanks | 9714 | 9735 | 22 | CGAATGATTAGAGGTCTTGGGG | 544.2 | -13.0 | 1.5 | -8.7 | 2.8E-18 | 3.2E-17 | MB231/MB468 | MP |
| >hg38_21_+_8433172_8446622_RNA45SN1_45S_50ntFlanks@4919.4937.19\|CCACCAGGAGTGGAGCCTG | >hg38_21_+_8433172_8446622_RNA45SN1_45S_50ntFlanks | 4919 | 4937 | 19 | CCACCAGGAGTGGAGCCTG | 611.5 | -13.1 | 1.5 | -8.7 | 2.8E-18 | 3.2E-17 | MB231/MB468 | MP |
| >hg38_21_+_8433172_8446622_RNA45SN1_45S_50ntFlanks@4730.4767.38\|CAAGAACGAAAGTCGGAGGTTCGAAGACGATCAGATAC | >hg38_21_+_8433172_8446622_RNA45SN1_45S_50ntFlanks | 4730 | 4767 | 38 | CAAGAACGAAAGTCGGAGGTTCGAAGACGATCAGATAC | 2018.3 | 10.8 | 1.2 | 8.6 | 5.5E-18 | 3.2E-17 | BT20/MB468 | CYTO |
| >hg38_21_+_8433172_8446622_RNA45SN1_45S_50ntFlanks@3872.3898.27\|CTAATACATGCCGACGGGCGCTGACCC | >hg38_21_+_8433172_8446622_RNA45SN1_45S_50ntFlanks | 3872 | 3898 | 27 | CTAATACATGCCGACGGGCGCTGACCC | 503.9 | -12.8 | 1.5 | -8.7 | 2.9E-18 | 3.2E-17 | MB231/MB468 | MP |
| >hg38_21_+_8433172_8446622_RNA45SN1_45S_50ntFlanks@10403.10420.18\|ACATGGGTCAGTCGGTCC | >hg38_21_+_8433172_8446622_RNA45SN1_45S_50ntFlanks | 10403 | 10420 | 18 | ACATGGGTCAGTCGGTCC | 202.4 | -6.1 | 0.7 | -8.7 | 2.8E-18 | 3.2E-17 | MB231/MB468 | CYTO |
| >hg38_21_+_8433172_8446622_RNA45SN1_45S_50ntFlanks@11578.11605.28\|TAGAACTGGTGCGGACCAGGGGAATCCG | >hg38_21_+_8433172_8446622_RNA45SN1_45S_50ntFlanks | 11578 | 11605 | 28 | TAGAACTGGTGCGGACCAGGGGAATCCG | 446.2 | -12.6 | 1.5 | -8.7 | 4.7E-18 | 3.3E-17 | MB231/MB468 | MITO |
| >hg38_21_+_8433172_8446622_RNA45SN1_45S_50ntFlanks@12412.12432.21\|ATCCTTCGATGTCGGCTCTTC | >hg38_21_+_8433172_8446622_RNA45SN1_45S_50ntFlanks | 12412 | 12432 | 21 | ATCCTTCGATGTCGGCTCTTC | 1195.6 | -12.5 | 1.5 | -8.6 | 5.5E-18 | 3.3E-17 | BT20/MB468 | CYTO |
| >MI0000102\|hsa-mir-100&WithFlank&11\|-\|122152223\|122152314@19.38.20(+2G)&[MIMAT0000098&hsa-miR-100-5p&offsets\|0\|-2(+2G);m-27&11\|-\|122152276\|122152296&offsets\|0\|-1(+2G)]\|AACCCGTAGATCCGAACTTGGG | >MI0000102\|hsa-mir-100&WithFlank&11\|-\|122152223\|122152314 | 19 | 38 | 20(+2G) | AACCCGTAGATCCGAACTTGGG | 1064.2 | -13.3 | 1.5 | -9.0 | 3.5E-19 | 3.3E-17 | BT20/MB231 | MITO |
| >hg38_21_+_8433172_8446622_RNA45SN1_45S_50ntFlanks@5487.5526.40\|CGCTGAGAAGACGGTCGAACTTGACTATCTAGAGGAAGTA | >hg38_21_+_8433172_8446622_RNA45SN1_45S_50ntFlanks | 5487 | 5526 | 40 | CGCTGAGAAGACGGTCGAACTTGACTATCTAGAGGAAGTA | 950.1 | -13.8 | 1.6 | -8.7 | 2.9E-18 | 3.3E-17 | MB231/MB468 | MP |
| >hg38_21_+_8433172_8446622_RNA45SN1_45S_50ntFlanks@11581.11611.31\|AACTGGTGCGGACCAGGGGAATCCGACTGTT | >hg38_21_+_8433172_8446622_RNA45SN1_45S_50ntFlanks | 11581 | 11611 | 31 | AACTGGTGCGGACCAGGGGAATCCGACTGTT | 1259.2 | -5.3 | 0.6 | -8.6 | 9.5E-18 | 3.3E-17 | BT20/MB468 | NUC |
| >hg38_21_+_8433172_8446622_RNA45SN1_45S_50ntFlanks@13004.13028.25\|ATCTATTGAAAGTCAGCCCTCGACA | >hg38_21_+_8433172_8446622_RNA45SN1_45S_50ntFlanks | 13004 | 13028 | 25 | ATCTATTGAAAGTCAGCCCTCGACA | 555.8 | -12.9 | 1.5 | -8.7 | 4.7E-18 | 3.3E-17 | MB231/MB468 | MITO |
| >MI0005566\|hsa-mir-374b&WithFlank&X\|-\|74218541\|74218624@17.38.22&[MIMAT0004955&hsa-miR-374b-5p&offsets\|0\|0;m-153&X\|-\|74218587\|74218608&offsets\|0\|0]\|ATATAATACAACCTGCTAAGTG | >MI0005566\|hsa-mir-374b&WithFlank&X\|-\|74218541\|74218624 | 17 | 38 | 22 | ATATAATACAACCTGCTAAGTG | 810.4 | 6.5 | 0.8 | 8.6 | 5.6E-18 | 3.3E-17 | BT20/MB468 | CYTO |
| >trna2_GlyGCC_21_-_18827107_18827177@1.28.28,trna35_GlyGCC_1_+_161413094_161413164@1.28.28,trna37_GlyGCC_1_+_161420467_161420537@1.28.28,trna39_GlyGCC_1_+_161427898_161427968@1.28.28,trna41_GlyGCC_1_+_161435258_161435328@1.28.28\|GCATGGGTGGTTCAGTGGTAGAATTCTC | >trna2_GlyGCC_21_-_18827107_18827177 | 1 | 28 | 28 | GCATGGGTGGTTCAGTGGTAGAATTCTC | 24494.9 | -12.7 | 1.5 | -8.7 | 3.8E-18 | 3.3E-17 | MB231/MB468 | NUC |
| >hg38_21_+_8433172_8446622_RNA45SN1_45S_50ntFlanks@482.525.44\|ATCGATGTGGTGACGTCGTGCTCTCCCGGGCCGGGTCCGAGCCG | >hg38_21_+_8433172_8446622_RNA45SN1_45S_50ntFlanks | 482 | 525 | 44 | ATCGATGTGGTGACGTCGTGCTCTCCCGGGCCGGGTCCGAGCCG | 852.8 | -14.2 | 1.7 | -8.6 | 9.6E-18 | 3.3E-17 | BT20/MB468 | NUC |
| >hg38_21_+_8433172_8446622_RNA45SN1_45S_50ntFlanks@10346.10374.29\|TCAAACGAGAACTTTGAAGGCCGAAGTGG | >hg38_21_+_8433172_8446622_RNA45SN1_45S_50ntFlanks | 10346 | 10374 | 29 | TCAAACGAGAACTTTGAAGGCCGAAGTGG | 2382.0 | -11.4 | 1.3 | -8.6 | 9.7E-18 | 3.3E-17 | BT20/MB468 | NUC |
| >hg38_21_+_8433172_8446622_RNA45SN1_45S_50ntFlanks@8342.8385.44\|CGTAAGGGAAAGTTGAAAAGAACTTTGAAGAGAGAGTTCAAGAG | >hg38_21_+_8433172_8446622_RNA45SN1_45S_50ntFlanks | 8342 | 8385 | 44 | CGTAAGGGAAAGTTGAAAAGAACTTTGAAGAGAGAGTTCAAGAG | 145.9 | 11.2 | 1.3 | 8.7 | 2.9E-18 | 3.3E-17 | MB231/MB468 | CYTO |

<table>
<tr><td>>hg38_21_+_8433172_8446622_RNA45SN1_45S_50ntFlanks@11051.11075.25|CCGTCCTCCCCCCTCCCCGGGGGAG</td><td>>hg38_21_+_8433172_8446622_RNA45SN1_45S_50ntFlanks</td><td>11051</td><td>11075</td><td>25</td><td>CCGTCCTCCCCCCTCCCCGGGGGAG</td><td>485.6</td><td>-11.6</td><td>1.3</td><td>-8.7</td><td>2.9E-18</td><td>3.3E-17</td><td>MB231/MB468</td><td>CYTO</td></tr>
<tr><td>>hg38_21_+_8433172_8446622_RNA45SN1_45S_50ntFlanks@10926.10946.21|ACGAGGCGCCGCCGCCCCCCC</td><td>>hg38_21_+_8433172_8446622_RNA45SN1_45S_50ntFlanks</td><td>10926</td><td>10946</td><td>21</td><td>ACGAGGCGCCGCCGCCCCCCC</td><td>321.2</td><td>-12.8</td><td>1.5</td><td>-8.6</td><td>9.8E-18</td><td>3.3E-17</td><td>BT20/MB468</td><td>NUC</td></tr>
<tr><td>>hg38_21_+_8433172_8446622_RNA45SN1_45S_50ntFlanks@7988.8005.18|CAGACGTGGCGACCCGCT</td><td>>hg38_21_+_8433172_8446622_RNA45SN1_45S_50ntFlanks</td><td>7988</td><td>8005</td><td>18</td><td>CAGACGTGGCGACCCGCT</td><td>528.3</td><td>-12.9</td><td>1.5</td><td>-8.7</td><td>3.0E-18</td><td>3.4E-17</td><td>MB231/MB468</td><td>MP</td></tr>
<tr><td>>hg38_21_+_8433172_8446622_RNA45SN1_45S_50ntFlanks@12405.12443.39|CTTTTTGATCCTTCGATGTCGGCTCTTCCTATCATTGTG</td><td>>hg38_21_+_8433172_8446622_RNA45SN1_45S_50ntFlanks</td><td>12405</td><td>12443</td><td>39</td><td>CTTTTTGATCCTTCGATGTCGGCTCTTCCTATCATTGTG</td><td>5245.9</td><td>-5.4</td><td>0.6</td><td>-8.7</td><td>4.2E-18</td><td>3.4E-17</td><td>BT20/MB468</td><td>MITO</td></tr>
<tr><td>>hg38_21_+_8433172_8446622_RNA45SN1_45S_50ntFlanks@12934.12955.22|CTAAACCATTCGTAGACGACCT</td><td>>hg38_21_+_8433172_8446622_RNA45SN1_45S_50ntFlanks</td><td>12934</td><td>12955</td><td>22</td><td>CTAAACCATTCGTAGACGACCT</td><td>424.2</td><td>-12.5</td><td>1.4</td><td>-8.7</td><td>4.9E-18</td><td>3.4E-17</td><td>MB231/MB468</td><td>MITO</td></tr>
<tr><td>>MI0003628|hsa-mir-615&WithFlank&12|+|54033944|54034051@67.88.22&[MIMAT0003283&hsa-miR-615-3p&offsets|0|0;m-495&12|+|54034010|54034030&offsets|0|+1]|TCCGAGCCTGGGTCTCCCTCTT</td><td>>MI0003628|hsa-mir-615&WithFlank&12|+|54033944|54034051</td><td>67</td><td>88</td><td>22</td><td>TCCGAGCCTGGGTCTCCCTCTT</td><td>2105.8</td><td>5.8</td><td>0.6</td><td>8.9</td><td>3.8E-19</td><td>3.4E-17</td><td>BT20/MB231</td><td>MITO</td></tr>
<tr><td>>hg38_21_+_8433172_8446622_RNA45SN1_45S_50ntFlanks@5491.5512.22|GAGAAGACGGTCGAACTTGACT</td><td>>hg38_21_+_8433172_8446622_RNA45SN1_45S_50ntFlanks</td><td>5491</td><td>5512</td><td>22</td><td>GAGAAGACGGTCGAACTTGACT</td><td>5575.0</td><td>10.0</td><td>1.2</td><td>8.6</td><td>5.8E-18</td><td>3.4E-17</td><td>BT20/MB468</td><td>CYTO</td></tr>
<tr><td>>hg38_21_+_8433172_8446622_RNA45SN1_45S_50ntFlanks@6777.6806.30|TCCCGGGGCTACGCCTGTCTGAGCGTCGCT</td><td>>hg38_21_+_8433172_8446622_RNA45SN1_45S_50ntFlanks</td><td>6777</td><td>6806</td><td>30</td><td>TCCCGGGGCTACGCCTGTCTGAGCGTCGCT</td><td>381.2</td><td>4.2</td><td>0.5</td><td>8.7</td><td>3.9E-18</td><td>3.4E-17</td><td>MB231/MB468</td><td>NUC</td></tr>
<tr><td>>hg38_21_+_8433172_8446622_RNA45SN1_45S_50ntFlanks@10572.10591.20|AGAAGCCGGCGGGAGCCCCG</td><td>>hg38_21_+_8433172_8446622_RNA45SN1_45S_50ntFlanks</td><td>10572</td><td>10591</td><td>20</td><td>AGAAGCCGGCGGGAGCCCCG</td><td>507.3</td><td>-12.9</td><td>1.5</td><td>-8.7</td><td>3.0E-18</td><td>3.4E-17</td><td>MB231/MB468</td><td>MP</td></tr>
<tr><td>>MI0000272|hsa-mir-182&WithFlank&7|-|129770377|129770498@29.47.19&[MIMAT0000259&hsa-miR-182-5p&offsets|0|-5;m-22&7|-|129770449|129770470&offsets|0|-3]|TTTGGCAATGGTAGAACTC</td><td>>MI0000272|hsa-mir-182&WithFlank&7|-|129770377|129770498</td><td>29</td><td>47</td><td>19</td><td>TTTGGCAATGGTAGAACTC</td><td>293.8</td><td>10.1</td><td>1.2</td><td>8.6</td><td>5.9E-18</td><td>3.4E-17</td><td>BT20/MB468</td><td>CYTO</td></tr>
<tr><td>>hg38_21_+_8433172_8446622_RNA45SN1_45S_50ntFlanks@10535.10570.36|CGCGAGGCGTCCAGTGCGGTAACGCGACCGATCCCG</td><td>>hg38_21_+_8433172_8446622_RNA45SN1_45S_50ntFlanks</td><td>10535</td><td>10570</td><td>36</td><td>CGCGAGGCGTCCAGTGCGGTAACGCGACCGATCCCG</td><td>372.1</td><td>-12.4</td><td>1.4</td><td>-8.7</td><td>3.1E-18</td><td>3.5E-17</td><td>MB231/MB468</td><td>MP</td></tr>
<tr><td>>trna116_GluCTC_1_-_145399233_145399304@4.35.32,trna59_GluCTC_1_+_249168447_249168518@4.35.32,trna71_GluCTC_1_-_161439189_161439260@4.35.32,trna74_GluCTC_1_-_161431809_161431880@4.35.32,trna77_GluCTC_1_-_161424398_161424469@4.35.32,trna77_GluCTC_6_+_28949976_28950047@4.35.32,trna80_GluCTC_1_-_161417018_161417089@4.35.32,trna87_GluCTC_6_-_126101393_126101464@4.35.32|CTGGTGGTCTAGTGGTTAGGATTCGGCGCTCT</td><td>>trna116_GluCTC_1_-_145399233_145399304</td><td>4</td><td>35</td><td>32</td><td>CTGGTGGTCTAGTGGTTAGGATTCGGCGCTCT</td><td>342.2</td><td>6.9</td><td>0.8</td><td>8.7</td><td>4.3E-18</td><td>3.5E-17</td><td>BT20/MB468</td><td>MITO</td></tr>
<tr><td>>MI0000650|hsa-mir-200c&WithFlank&12|+|6963693|6963772@49.71.23&[MIMAT0000617&hsa-miR-200c-3p&offsets|-1|-1;m-54&12|+|6963742|6963764&offsets|-1|-1]|CTAATACTGCCGGGTAATGATGG</td><td>>MI0000650|hsa-mir-200c&WithFlank&12|+|6963693|6963772</td><td>49</td><td>71</td><td>23</td><td>CTAATACTGCCGGGTAATGATGG</td><td>468.1</td><td>12.6</td><td>1.4</td><td>8.9</td><td>4.9E-19</td><td>3.5E-17</td><td>BT20/MB231</td><td>NUC</td></tr>
<tr><td>>hg38_21_+_8433172_8446622_RNA45SN1_45S_50ntFlanks@4283.4307.25|CTTTAACGAGGATCCATTGGAGGGC</td><td>>hg38_21_+_8433172_8446622_RNA45SN1_45S_50ntFlanks</td><td>4283</td><td>4307</td><td>25</td><td>CTTTAACGAGGATCCATTGGAGGGC</td><td>460.6</td><td>-11.8</td><td>1.4</td><td>-8.7</td><td>4.0E-18</td><td>3.5E-17</td><td>MB231/MB468</td><td>NUC</td></tr>
<tr><td>>MI0003560|hsa-mir-92b&WithFlank&1|+|155195171|155195278@67.88.22(+1U)&[MIMAT0003218&hsa-miR-92b-3p&offsets|0|0(+1U);m-115&1|+|155195237|155195258&offsets|0|0(+1U)]|TATTGCACTCGTCCCGGCCTCCT</td><td>>MI0003560|hsa-mir-92b&WithFlank&1|+|155195171|155195278</td><td>67</td><td>88</td><td>22(+1U)</td><td>TATTGCACTCGTCCCGGCCTCCT</td><td>1333.5</td><td>10.3</td><td>1.2</td><td>8.6</td><td>6.0E-18</td><td>3.5E-17</td><td>BT20/MB468</td><td>CYTO</td></tr>
<tr><td>>MI0000477|hsa-mir-146a&WithFlank&5|+|160485346|160485456@27.49.23&[MIMAT0000449&hsa-miR-146a-5p&offsets|0|+1;m-129&5|+|160485372|160485393&offsets|0|+1]|TGAGAACTGAATTCCATGGGTTG</td><td>>MI0000477|hsa-mir-146a&WithFlank&5|+|160485346|160485456</td><td>27</td><td>49</td><td>23</td><td>TGAGAACTGAATTCCATGGGTTG</td><td>770.7</td><td>13.6</td><td>1.6</td><td>8.7</td><td>3.1E-18</td><td>3.5E-17</td><td>MB231/MB468</td><td>CYTO</td></tr>
<tr><td>>hg38_21_+_8433172_8446622_RNA45SN1_45S_50ntFlanks@5467.5514.48|GGCCCACGGCCCTGGCGGAGCGCTGAGAAGACGGTCGAACTTGACTAT</td><td>>hg38_21_+_8433172_8446622_RNA45SN1_45S_50ntFlanks</td><td>5467</td><td>5514</td><td>48</td><td>GGCCCACGGCCCTGGCGGAGCGCTGAGAAGACGGTCGAACTTGACTAT</td><td>198.3</td><td>-24.7</td><td>2.8</td><td>-8.7</td><td>3.1E-18</td><td>3.5E-17</td><td>MB231/MB468</td><td>MP</td></tr>
<tr><td>>hg38_21_+_8433172_8446622_RNA45SN1_45S_50ntFlanks@12488.12518.31|AACGTGAGCTGGGTTTAGACCGTCGTGAGAC</td><td>>hg38_21_+_8433172_8446622_RNA45SN1_45S_50ntFlanks</td><td>12488</td><td>12518</td><td>31</td><td>AACGTGAGCTGGGTTTAGACCGTCGTGAGAC</td><td>432.9</td><td>-12.6</td><td>1.4</td><td>-8.7</td><td>3.1E-18</td><td>3.5E-17</td><td>MB231/MB468</td><td>MP</td></tr>
<tr><td>>hg38_21_+_8433172_8446622_RNA45SN1_45S_50ntFlanks@8403.8442.40|AGGTAAACGGGTGGGGTCCGCGCAGTCCGCCCGGAGGATT</td><td>>hg38_21_+_8433172_8446622_RNA45SN1_45S_50ntFlanks</td><td>8403</td><td>8442</td><td>40</td><td>AGGTAAACGGGTGGGGTCCGCGCAGTCCGCCCGGAGGATT</td><td>427.2</td><td>-12.5</td><td>1.4</td><td>-8.7</td><td>5.1E-18</td><td>3.5E-17</td><td>MB231/MB468</td><td>MITO</td></tr>
<tr><td>>hg38_21_+_8433172_8446622_RNA45SN1_45S_50ntFlanks@9131.9152.22|GGTCGGGGGGCGGGGCGGACTG</td><td>>hg38_21_+_8433172_8446622_RNA45SN1_45S_50ntFlanks</td><td>9131</td><td>9152</td><td>22</td><td>GGTCGGGGGGCGGGGCGGACTG</td><td>627.5</td><td>-13.1</td><td>1.5</td><td>-8.7</td><td>5.1E-18</td><td>3.5E-17</td><td>MB231/MB468</td><td>MITO</td></tr>
<tr><td>>hg38_21_+_8433172_8446622_RNA45SN1_45S_50ntFlanks@11787.11804.18|AATGGATGAACGAGATTC</td><td>>hg38_21_+_8433172_8446622_RNA45SN1_45S_50ntFlanks</td><td>11787</td><td>11804</td><td>18</td><td>AATGGATGAACGAGATTC</td><td>8339.8</td><td>-6.4</td><td>0.7</td><td>-8.7</td><td>4.1E-18</td><td>3.5E-17</td><td>MB231/MB468</td><td>NUC</td></tr>
<tr><td>>hg38_21_+_8433172_8446622_RNA45SN1_45S_50ntFlanks@9554.9576.23|CGTAGCGGTCCTGACGTGCAAAT</td><td>>hg38_21_+_8433172_8446622_RNA45SN1_45S_50ntFlanks</td><td>9554</td><td>9576</td><td>23</td><td>CGTAGCGGTCCTGACGTGCAAAT</td><td>1463.2</td><td>2.8</td><td>0.3</td><td>8.6</td><td>6.1E-18</td><td>3.5E-17</td><td>BT20/MB468</td><td>CYTO</td></tr>
<tr><td>>trna10_LysCTT_16_+_3241501_3241573@1.28.28,trna119_LysCTT_1_-_145395522_145395594@1.28.28,trna11_LysCTT_5_-_180648979_180649051@1.28.28,trna13_LysCTT_14_-_58706613_58706685@1.28.28,trna13_LysCTT_6_+_26556774_26556846@1.28.28,trna2_LysCTT_15_+_79152904_79152976@1.28.28,trna32_LysCTT_16_-_3207406_3207478@1.28.28,trna7_LysCTT_16_+_3225692_3225764@1.28.28,trna9_LysCTT_5_+_180634755_180634827@1.28.28|GCCCGGCTAGCTCAGTCGGTAGAGCATG</td><td>>trna10_LysCTT_16_+_3241501_3241573</td><td>1</td><td>28</td><td>28</td><td>GCCCGGCTAGCTCAGTCGGTAGAGCATG</td><td>3991.1</td><td>-10.0</td><td>1.2</td><td>-8.6</td><td>1.0E-17</td><td>3.6E-17</td><td>BT20/MB468</td><td>NUC</td></tr>
<tr><td>>hg38_21_+_8433172_8446622_RNA45SN1_45S_50ntFlanks@9158.9187.30|AGTGCGCCCCGGGCGGGTCGCGCCGTCGGG</td><td>>hg38_21_+_8433172_8446622_RNA45SN1_45S_50ntFlanks</td><td>9158</td><td>9187</td><td>30</td><td>AGTGCGCCCCGGGCGGGTCGCGCCGTCGGG</td><td>581.9</td><td>-13.0</td><td>1.5</td><td>-8.6</td><td>5.2E-18</td><td>3.6E-17</td><td>MB231/MB468</td><td>MITO</td></tr>
<tr><td>>hg38_21_+_8433172_8446622_RNA45SN1_45S_50ntFlanks@4050.4070.21|CGAACGTCTGCCCTATCAACT</td><td>>hg38_21_+_8433172_8446622_RNA45SN1_45S_50ntFlanks</td><td>4050</td><td>4070</td><td>21</td><td>CGAACGTCTGCCCTATCAACT</td><td>263.4</td><td>9.4</td><td>1.1</td><td>8.7</td><td>3.1E-18</td><td>3.6E-17</td><td>MB231/MB468</td><td>CYTO</td></tr>
<tr><td>>MI0000746|hsa-mir-99b&WithFlank&19|+|51692606|51692687@13.35.23&[MIMAT0000689&hsa-miR-99b-5p&offsets|0|+1;m-10&19|+|51692618|51692639&offsets|0|+1]|CACCCGTAGAACCGACCTTGCGG</td><td>>MI0000746|hsa-mir-99b&WithFlank&19|+|51692606|51692687</td><td>13</td><td>35</td><td>23</td><td>CACCCGTAGAACCGACCTTGCGG</td><td>343.3</td><td>6.3</td><td>0.7</td><td>8.6</td><td>6.1E-18</td><td>3.6E-17</td><td>BT20/MB468</td><td>CYTO</td></tr>
<tr><td>>hg38_21_+_8433172_8446622_RNA45SN1_45S_50ntFlanks@9720.9737.18|ATTAGAGGTCTTGGGGCC</td><td>>hg38_21_+_8433172_8446622_RNA45SN1_45S_50ntFlanks</td><td>9720</td><td>9737</td><td>18</td><td>ATTAGAGGTCTTGGGGCC</td><td>1192.3</td><td>-14.0</td><td>1.6</td><td>-8.6</td><td>5.2E-18</td><td>3.6E-17</td><td>MB231/MB468</td><td>MITO</td></tr>
<tr><td>>hg38_21_+_8433172_8446622_RNA45SN1_45S_50ntFlanks@10384.10404.21|CCATGTGAACAGCAGTTGAAC</td><td>>hg38_21_+_8433172_8446622_RNA45SN1_45S_50ntFlanks</td><td>10384</td><td>10404</td><td>21</td><td>CCATGTGAACAGCAGTTGAAC</td><td>315.7</td><td>-12.7</td><td>1.5</td><td>-8.6</td><td>1.1E-17</td><td>3.6E-17</td><td>BT20/MB468</td><td>NUC</td></tr>
<tr><td>>hg38_21_+_8433172_8446622_RNA45SN1_45S_50ntFlanks@11934.11954.21|ATAAGTGGGAGGCCCCCGGCG</td><td>>hg38_21_+_8433172_8446622_RNA45SN1_45S_50ntFlanks</td><td>11934</td><td>11954</td><td>21</td><td>ATAAGTGGGAGGCCCCCGGCG</td><td>349.5</td><td>-12.3</td><td>1.4</td><td>-8.7</td><td>3.2E-18</td><td>3.6E-17</td><td>MB231/MB468</td><td>MP</td></tr>
<tr><td>>hg38_21_+_8433172_8446622_RNA45SN1_45S_50ntFlanks@12367.12384.18|AACTGGCTTGTGGCGGCC</td><td>>hg38_21_+_8433172_8446622_RNA45SN1_45S_50ntFlanks</td><td>12367</td><td>12384</td><td>18</td><td>AACTGGCTTGTGGCGGCC</td><td>846.7</td><td>-13.6</td><td>1.6</td><td>-8.7</td><td>3.2E-18</td><td>3.6E-17</td><td>MB231/MB468</td><td>MP</td></tr>
<tr><td>>trna111_HisGTG_1_-_147774845_147774916@-1T.34.35,trna118_HisGTG_1_-_145396881_145396952@-1T.34.35,trna16_HisGTG_1_+_146544773_146544844@-1T.34.35,trna1_HisGTG_15_+_45493349_45493420@-1T.34.35,trna21_HisGTG_1_+_147753471_147753542@-1T.34.35,trna33_HisGTG_6_+_27125906_27125977@-1T.34.35,trna7_HisGTG_9_-_14433938_14434009@-1T.34.35,trna8_HisGTG_15_-_45492611_45492682@-1T.34.35,trna9_HisGTG_15_-_45490804_45490875@-1T.34.35|TGCCGTGATCGTATAGTGGTTAGTACTCTGCGTTG</td><td>>trna111_HisGTG_1_-_147774845_147774916</td><td>-1T</td><td>34</td><td>35</td><td>TGCCGTGATCGTATAGTGGTTAGTACTCTGCGTTG</td><td>1398.7</td><td>7.3</td><td>0.8</td><td>8.6</td><td>6.2E-18</td><td>3.6E-17</td><td>BT20/MB468</td><td>CYTO</td></tr>
<tr><td>>hg38_21_+_8433172_8446622_RNA45SN1_45S_50ntFlanks@3993.4023.31|GTGACTCTAGATAACCTCGGGCCGATCGCAC</td><td>>hg38_21_+_8433172_8446622_RNA45SN1_45S_50ntFlanks</td><td>3993</td><td>4023</td><td>31</td><td>GTGACTCTAGATAACCTCGGGCCGATCGCAC</td><td>1245.2</td><td>5.2</td><td>0.6</td><td>8.6</td><td>6.2E-18</td><td>3.6E-17</td><td>BT20/MB468</td><td>CYTO</td></tr>
<tr><td>>hg38_21_+_8433172_8446622_RNA45SN1_45S_50ntFlanks@3802.3834.33|CATTAAATCAGTTATGGTTCCTTTGGTCGCTCG</td><td>>hg38_21_+_8433172_8446622_RNA45SN1_45S_50ntFlanks</td><td>3802</td><td>3834</td><td>33</td><td>CATTAAATCAGTTATGGTTCCTTTGGTCGCTCG</td><td>892.6</td><td>-13.6</td><td>1.6</td><td>-8.6</td><td>5.3E-18</td><td>3.6E-17</td><td>MB231/MB468</td><td>MITO</td></tr>
<tr><td>>hg38_21_+_8433172_8446622_RNA45SN1_45S_50ntFlanks@12488.12521.34|AACGTGAGCTGGGTTTAGACCGTCGTGAGACAGG</td><td>>hg38_21_+_8433172_8446622_RNA45SN1_45S_50ntFlanks</td><td>12488</td><td>12521</td><td>34</td><td>AACGTGAGCTGGGTTTAGACCGTCGTGAGACAGG</td><td>541.4</td><td>-12.0</td><td>1.4</td><td>-8.7</td><td>4.2E-18</td><td>3.7E-17</td><td>MB231/MB468</td><td>NUC</td></tr>
<tr><td>>hg38_21_+_8433172_8446622_RNA45SN1_45S_50ntFlanks@12632.12657.26|GCGAAGCTACCATCTGTGGGATTATG</td><td>>hg38_21_+_8433172_8446622_RNA45SN1_45S_50ntFlanks</td><td>12632</td><td>12657</td><td>26</td><td>GCGAAGCTACCATCTGTGGGATTATG</td><td>560.9</td><td>-13.0</td><td>1.5</td><td>-8.7</td><td>3.2E-18</td><td>3.7E-17</td><td>MB231/MB468</td><td>MP</td></tr>
<tr><td>>hg38_21_+_8433172_8446622_RNA45SN1_45S_50ntFlanks@6697.6718.22|CAGCTAGCTGCGAGAATTAATG</td><td>>hg38_21_+_8433172_8446622_RNA45SN1_45S_50ntFlanks</td><td>6697</td><td>6718</td><td>22</td><td>CAGCTAGCTGCGAGAATTAATG</td><td>4703.5</td><td>-8.6</td><td>1.0</td><td>-8.6</td><td>5.3E-18</td><td>3.7E-17</td><td>MB231/MB468</td><td>MITO</td></tr>
<tr><td>>hg38_21_+_8433172_8446622_RNA45SN1_45S_50ntFlanks@8336.8375.40|AAGTACCGTAAGGGAAAGTTGAAAAGAACTTTGAAGAGAG</td><td>>hg38_21_+_8433172_8446622_RNA45SN1_45S_50ntFlanks</td><td>8336</td><td>8375</td><td>40</td><td>AAGTACCGTAAGGGAAAGTTGAAAAGAACTTTGAAGAGAG</td><td>437.0</td><td>13.2</td><td>1.5</td><td>8.7</td><td>4.5E-18</td><td>3.7E-17</td><td>BT20/MB468</td><td>MITO</td></tr>
<tr><td>>hg38_21_+_8433172_8446622_RNA45SN1_45S_50ntFlanks@6741.6771.31|ATCGACACTTCGAACGCACTTGCGGCCCCGG</td><td>>hg38_21_+_8433172_8446622_RNA45SN1_45S_50ntFlanks</td><td>6741</td><td>6771</td><td>31</td><td>ATCGACACTTCGAACGCACTTGCGGCCCCGG</td><td>4703.5</td><td>-10.7</td><td>1.2</td><td>-8.6</td><td>5.3E-18</td><td>3.7E-17</td><td>MB231/MB468</td><td>MITO</td></tr>
<tr><td>>hg38_21_+_8433172_8446622_RNA45SN1_45S_50ntFlanks@9575.9597.23|ATCGGTCGTCCGACCTGGGTATA</td><td>>hg38_21_+_8433172_8446622_RNA45SN1_45S_50ntFlanks</td><td>9575</td><td>9597</td><td>23</td><td>ATCGGTCGTCCGACCTGGGTATA</td><td>6222.6</td><td>-11.7</td><td>1.3</td><td>-8.7</td><td>3.3E-18</td><td>3.7E-17</td><td>MB231/MB468</td><td>MP</td></tr>
<tr><td>>hg38_21_+_8433172_8446622_RNA45SN1_45S_50ntFlanks@6782.6799.18|GGGCTACGCCTGTCTGAG</td><td>>hg38_21_+_8433172_8446622_RNA45SN1_45S_50ntFlanks</td><td>6782</td><td>6799</td><td>18</td><td>GGGCTACGCCTGTCTGAG</td><td>468.1</td><td>-12.7</td><td>1.5</td><td>-8.7</td><td>3.3E-18</td><td>3.7E-17</td><td>MB231/MB468</td><td>MP</td></tr>
<tr><td>>MI0000283|hsa-mir-203a&WithFlank&14|+|104117399|104117520@72.90.19&[MIMAT0000264&hsa-miR-203a-3p&offsets|+1|-2;m-79&14|+|104117469|104117490&offsets|+1|-2]|TGAAATGTTTAGGACCACT</td><td>>MI0000283|hsa-mir-203a&WithFlank&14|+|104117399|104117520</td><td>72</td><td>90</td><td>19</td><td>TGAAATGTTTAGGACCACT</td><td>139.6</td><td>3.8</td><td>0.4</td><td>8.6</td><td>1.1E-17</td><td>3.7E-17</td><td>BT20/MB468</td><td>NUC</td></tr>
<tr><td>>hg38_21_+_8433172_8446622_RNA45SN1_45S_50ntFlanks@10678.10705.28|CGGTTCCGGCGGCGTCCGGTGAGCTCTC</td><td>>hg38_21_+_8433172_8446622_RNA45SN1_45S_50ntFlanks</td><td>10678</td><td>10705</td><td>28</td><td>CGGTTCCGGCGGCGTCCGGTGAGCTCTC</td><td>1509.9</td><td>-10.3</td><td>1.2</td><td>-8.7</td><td>3.3E-18</td><td>3.7E-17</td><td>MB231/MB468</td><td>MP</td></tr>
<tr><td>>hg38_21_+_8433172_8446622_RNA45SN1_45S_50ntFlanks@8155.8178.24|CGCTCGTGGGGGGCCCAAGTCCTT</td><td>>hg38_21_+_8433172_8446622_RNA45SN1_45S_50ntFlanks</td><td>8155</td><td>8178</td><td>24</td><td>CGCTCGTGGGGGGCCCAAGTCCTT</td><td>659.3</td><td>-13.1</td><td>1.5</td><td>-8.6</td><td>5.4E-18</td><td>3.7E-17</td><td>MB231/MB468</td><td>MITO</td></tr>
<tr><td>>hg38_21_+_8433172_8446622_RNA45SN1_45S_50ntFlanks@8359.8379.21|AAGAACTTTGAAGAGAGAGTT</td><td>>hg38_21_+_8433172_8446622_RNA45SN1_45S_50ntFlanks</td><td>8359</td><td>8379</td><td>21</td><td>AAGAACTTTGAAGAGAGAGTT</td><td>1101.4</td><td>-7.1</td><td>0.8</td><td>-8.6</td><td>1.1E-17</td><td>3.8E-17</td><td>BT20/MB468</td><td>NUC</td></tr>
</table>

<table>
<tr><td>>hg38_21_+_8433172_8446622_RNA45SN1_45S_50ntFlanks@3899.3935.37|CCTTCGCGGGGGGGATGCGTGCATTTATCAGATCAAA</td><td>>hg38_21_+_8433172_8446622_RNA45SN1_45S_50ntFlanks</td><td>3899</td><td>3935</td><td>37</td><td>CCTTCGCGGGGGGGATGCGTGCATTTATCAGATCAAA</td><td>513.1</td><td>-12.8</td><td>1.5</td><td>-8.6</td><td>5.5E-18</td><td>3.8E-17</td><td>MB231/MB468</td><td>MITO</td></tr>
<tr><td>>hg38_21_+_8433172_8446622_RNA45SN1_45S_50ntFlanks@12415.12447.33|CTTCGATGTCGGCTCTTCCTATCATTGTGAAGC</td><td>>hg38_21_+_8433172_8446622_RNA45SN1_45S_50ntFlanks</td><td>12415</td><td>12447</td><td>33</td><td>CTTCGATGTCGGCTCTTCCTATCATTGTGAAGC</td><td>388.5</td><td>3.5</td><td>0.4</td><td>8.6</td><td>6.5E-18</td><td>3.8E-17</td><td>BT20/MB468</td><td>CYTO</td></tr>
<tr><td>>hg38_21_+_8433172_8446622_RNA45SN1_45S_50ntFlanks@11792.11820.29|ATGAACGAGATTCCCACTGTCCCTACCTA</td><td>>hg38_21_+_8433172_8446622_RNA45SN1_45S_50ntFlanks</td><td>11792</td><td>11820</td><td>29</td><td>ATGAACGAGATTCCCACTGTCCCTACCTA</td><td>442.3</td><td>-12.6</td><td>1.5</td><td>-8.6</td><td>5.5E-18</td><td>3.8E-17</td><td>MB231/MB468</td><td>MITO</td></tr>
<tr><td>>hg38_21_+_8433172_8446622_RNA45SN1_45S_50ntFlanks@5524.5550.27|GTAAAAGTCGTAACAAGGTTTCCGTAG</td><td>>hg38_21_+_8433172_8446622_RNA45SN1_45S_50ntFlanks</td><td>5524</td><td>5550</td><td>27</td><td>GTAAAAGTCGTAACAAGGTTTCCGTAG</td><td>605.2</td><td>-12.2</td><td>1.4</td><td>-8.7</td><td>4.4E-18</td><td>3.8E-17</td><td>MB231/MB468</td><td>NUC</td></tr>
<tr><td>>MI0000079|hsa-mir-23a&WithFlank&19|-|13836581|13836665@54.71.18(+1C)&[MIMAT0000078&hsa-miR-23a-3p&offsets|+3|0(+1C);m-31&19|-|13836595|13836615&offsets|+3|0(+1C)]|ACATTGCCAGGGATTTCCC</td><td>>MI0000079|hsa-mir-23a&WithFlank&19|-|13836581|13836665</td><td>54</td><td>71</td><td>18(+1C)</td><td>ACATTGCCAGGGATTTCCC</td><td>307.7</td><td>12.5</td><td>1.5</td><td>8.6</td><td>6.5E-18</td><td>3.8E-17</td><td>BT20/MB468</td><td>CYTO</td></tr>
<tr><td>>hg38_21_+_8433172_8446622_RNA45SN1_45S_50ntFlanks@10737.10755.19|TAAATCTCGCGCCGGGCCG</td><td>>hg38_21_+_8433172_8446622_RNA45SN1_45S_50ntFlanks</td><td>10737</td><td>10755</td><td>19</td><td>TAAATCTCGCGCCGGGCCG</td><td>1784.7</td><td>-10.5</td><td>1.2</td><td>-8.6</td><td>5.6E-18</td><td>3.8E-17</td><td>MB231/MB468</td><td>MITO</td></tr>
<tr><td>>hg38_21_+_8433172_8446622_RNA45SN1_45S_50ntFlanks@10425.10457.33|AGATGGGCGAGCGCCGTTCCGAAGGGACGGGCG</td><td>>hg38_21_+_8433172_8446622_RNA45SN1_45S_50ntFlanks</td><td>10425</td><td>10457</td><td>33</td><td>AGATGGGCGAGCGCCGTTCCGAAGGGACGGGCG</td><td>573.0</td><td>-6.7</td><td>0.8</td><td>-8.7</td><td>3.3E-18</td><td>3.8E-17</td><td>MB231/MB468</td><td>CYTO</td></tr>
<tr><td>>MI0000114|hsa-mir-107&WithFlank&10|-|89592741|89592833@56.76.21(+1A);MI0000108|hsa-mir-103a-2&WithFlank&20|+|3917488|3917577@54.74.21(+1A);MI0000109|hsa-mir-103a-1&WithFlank&5|-|168560890|168560979@54.74.21(+1A)&[MIMAT0000104&hsa-miR-107&offsets|0|-2(+1A);m-78&10|-|89592758|89592778&offsets|0|0(+1A)];[MIMAT0000101&hsa-miR-103a-3p&offsets|0|-2(+1A);m-12&20|+|3917541|3917563&offsets|0|-2(+1A)];[MIMAT0000101_1&hsa-miR-103a-3p&offsets|0|-2(+1A);m-13&5|-|168560904|168560926&offsets|0|-2(+1A);m-44220&5|-|168560908|168560926&offsets|0|+2(+1A)]|AGCAGCATTGTACAGGGCTATA</td><td>>MI0000114|hsa-mir-107&WithFlank&10|-|89592741|89592833</td><td>56</td><td>76</td><td>21(+1A)</td><td>AGCAGCATTGTACAGGGCTATA</td><td>1080.8</td><td>7.6</td><td>0.9</td><td>8.6</td><td>6.6E-18</td><td>3.8E-17</td><td>BT20/MB468</td><td>CYTO</td></tr>
<tr><td>>hg38_21_+_8433172_8446622_RNA45SN1_45S_50ntFlanks@12073.12095.23|CCGAGGGGCTCTCGCTTCTGGCG</td><td>>hg38_21_+_8433172_8446622_RNA45SN1_45S_50ntFlanks</td><td>12073</td><td>12095</td><td>23</td><td>CCGAGGGGCTCTCGCTTCTGGCG</td><td>3789.7</td><td>-11.1</td><td>1.3</td><td>-8.7</td><td>3.4E-18</td><td>3.8E-17</td><td>MB231/MB468</td><td>MP</td></tr>
<tr><td>>hg38_21_+_8433172_8446622_RNA45SN1_45S_50ntFlanks@4260.4282.23|TTGGAATGAGTCCACTTTAAATC</td><td>>hg38_21_+_8433172_8446622_RNA45SN1_45S_50ntFlanks</td><td>4260</td><td>4282</td><td>23</td><td>TTGGAATGAGTCCACTTTAAATC</td><td>310.0</td><td>-12.1</td><td>1.4</td><td>-8.7</td><td>3.4E-18</td><td>3.8E-17</td><td>MB231/MB468</td><td>MP</td></tr>
<tr><td>>hg38_21_+_8433172_8446622_RNA45SN1_45S_50ntFlanks@6663.6699.37|GTGGATCACTCGGCTCGTGCGTCGATGAAGAACGCAG</td><td>>hg38_21_+_8433172_8446622_RNA45SN1_45S_50ntFlanks</td><td>6663</td><td>6699</td><td>37</td><td>GTGGATCACTCGGCTCGTGCGTCGATGAAGAACGCAG</td><td>658.6</td><td>-13.2</td><td>1.5</td><td>-8.7</td><td>3.4E-18</td><td>3.8E-17</td><td>MB231/MB468</td><td>MP</td></tr>
<tr><td>>hg38_21_+_8433172_8446622_RNA45SN1_45S_50ntFlanks@10680.10710.31|GTTCCGGCGGCGTCCGGTGAGCTCTCGCTGG</td><td>>hg38_21_+_8433172_8446622_RNA45SN1_45S_50ntFlanks</td><td>10680</td><td>10710</td><td>31</td><td>GTTCCGGCGGCGTCCGGTGAGCTCTCGCTGG</td><td>22193.1</td><td>-9.5</td><td>1.1</td><td>-8.7</td><td>3.4E-18</td><td>3.8E-17</td><td>MB231/MB468</td><td>CYTO</td></tr>
<tr><td>>MI0000542|hsa-mir-320a&WithFlank&8|-|22244960|22245043@48.68.21(+1U)&[MIMAT0000510&hsa-miR-320a-3p&offsets|0|-1(+1U);m-60&8|-|22244975|22244996&offsets|0|-1(+1U)]|AAAAGCTGGGTTGAGAGGGCGT</td><td>>MI0000542|hsa-mir-320a&WithFlank&8|-|22244960|22245043</td><td>48</td><td>68</td><td>21(+1U)</td><td>AAAAGCTGGGTTGAGAGGGCGT</td><td>2285.5</td><td>7.3</td><td>0.8</td><td>8.7</td><td>3.4E-18</td><td>3.8E-17</td><td>MB231/MB468</td><td>CYTO</td></tr>
<tr><td>>hg38_21_+_8433172_8446622_RNA45SN1_45S_50ntFlanks@9573.9597.25|AAATCGGTCGTCCGACCTGGGTATA</td><td>>hg38_21_+_8433172_8446622_RNA45SN1_45S_50ntFlanks</td><td>9573</td><td>9597</td><td>25</td><td>AAATCGGTCGTCCGACCTGGGTATA</td><td>6799.5</td><td>-11.7</td><td>1.3</td><td>-8.7</td><td>3.4E-18</td><td>3.9E-17</td><td>MB231/MB468</td><td>MP</td></tr>
<tr><td>>hg38_21_+_8433172_8446622_RNA45SN1_45S_50ntFlanks@8196.8214.19|CGTGGACGGTGTGAGGCCG</td><td>>hg38_21_+_8433172_8446622_RNA45SN1_45S_50ntFlanks</td><td>8196</td><td>8214</td><td>19</td><td>CGTGGACGGTGTGAGGCCG</td><td>4545.5</td><td>-6.4</td><td>0.7</td><td>-8.7</td><td>3.4E-18</td><td>3.9E-17</td><td>MB231/MB468</td><td>MP</td></tr>
<tr><td>>hg38_21_+_8433172_8446622_RNA45SN1_45S_50ntFlanks@10841.10864.24|CGTAACTTCGGGATAAGGATTGGC</td><td>>hg38_21_+_8433172_8446622_RNA45SN1_45S_50ntFlanks</td><td>10841</td><td>10864</td><td>24</td><td>CGTAACTTCGGGATAAGGATTGGC</td><td>9905.4</td><td>-10.4</td><td>1.2</td><td>-8.6</td><td>1.1E-17</td><td>3.9E-17</td><td>BT20/MB468</td><td>NUC</td></tr>
<tr><td>>hg38_21_+_8433172_8446622_RNA45SN1_45S_50ntFlanks@12995.13012.18|CTCGCTGCGATCTATTGA</td><td>>hg38_21_+_8433172_8446622_RNA45SN1_45S_50ntFlanks</td><td>12995</td><td>13012</td><td>18</td><td>CTCGCTGCGATCTATTGA</td><td>641.7</td><td>-5.1</td><td>0.6</td><td>-8.6</td><td>1.1E-17</td><td>3.9E-17</td><td>BT20/MB468</td><td>NUC</td></tr>
<tr><td>>hg38_21_+_8433172_8446622_RNA45SN1_45S_50ntFlanks@10771.10818.48|CAGGTCTCCAAGGTGAACAGCCTCTGGCATGTTGGAACAATGTAGGTA</td><td>>hg38_21_+_8433172_8446622_RNA45SN1_45S_50ntFlanks</td><td>10771</td><td>10818</td><td>48</td><td>CAGGTCTCCAAGGTGAACAGCCTCTGGCATGTTGGAACAATGTAGGTA</td><td>1576.3</td><td>-15.0</td><td>1.8</td><td>-8.6</td><td>1.1E-17</td><td>3.9E-17</td><td>BT20/MB468</td><td>NUC</td></tr>
<tr><td>>MI0000650|hsa-mir-200c&WithFlank&12|+|6963693|6963772@50.73.24&[MIMAT0000617&hsa-miR-200c-3p&offsets|0|+1;m-54&12|+|6963742|6963764&offsets|0|+1]|TAATACTGCCGGGTAATGATGGAG</td><td>>MI0000650|hsa-mir-200c&WithFlank&12|+|6963693|6963772</td><td>50</td><td>73</td><td>24</td><td>TAATACTGCCGGGTAATGATGGAG</td><td>641.1</td><td>12.7</td><td>1.4</td><td>9.0</td><td>3.4E-19</td><td>3.9E-17</td><td>BT20/MB231</td><td>CYTO</td></tr>
<tr><td>>MI0000809|hsa-mir-151a&WithFlank&8|-|140732558|140732659@53.74.22(+2U)&[MIMAT0000757&hsa-miR-151a-3p&offsets|0|+1(+2U);m-30&8|-|140732587|140732607&offsets|0|+1(+2U)]|CTAGACTGAAGCTCCTTGAGGATT</td><td>>MI0000809|hsa-mir-151a&WithFlank&8|-|140732558|140732659</td><td>53</td><td>74</td><td>22(+2U)</td><td>CTAGACTGAAGCTCCTTGAGGATT</td><td>98.1</td><td>2.2</td><td>0.3</td><td>8.6</td><td>1.1E-17</td><td>3.9E-17</td><td>BT20/MB468</td><td>NUC</td></tr>
<tr><td>>hg38_21_+_8433172_8446622_RNA45SN1_45S_50ntFlanks@9178.9197.20|CGCCGTCGGGCCCGGGGGAG</td><td>>hg38_21_+_8433172_8446622_RNA45SN1_45S_50ntFlanks</td><td>9178</td><td>9197</td><td>20</td><td>CGCCGTCGGGCCCGGGGGAG</td><td>786.4</td><td>-13.5</td><td>1.6</td><td>-8.7</td><td>3.5E-18</td><td>3.9E-17</td><td>MB231/MB468</td><td>MP</td></tr>
<tr><td>>hg38_21_+_8433172_8446622_RNA45SN1_45S_50ntFlanks@9352.9371.20|CCGTGGCGCAATGAAGGTGA</td><td>>hg38_21_+_8433172_8446622_RNA45SN1_45S_50ntFlanks</td><td>9352</td><td>9371</td><td>20</td><td>CCGTGGCGCAATGAAGGTGA</td><td>532.7</td><td>-12.8</td><td>1.5</td><td>-8.6</td><td>5.7E-18</td><td>3.9E-17</td><td>MB231/MB468</td><td>MITO</td></tr>
<tr><td>>hg38_21_+_8433172_8446622_RNA45SN1_45S_50ntFlanks@8182.8211.30|ATCGAGGCCCAGCCCGTGGACGGTGTGAGG</td><td>>hg38_21_+_8433172_8446622_RNA45SN1_45S_50ntFlanks</td><td>8182</td><td>8211</td><td>30</td><td>ATCGAGGCCCAGCCCGTGGACGGTGTGAGG</td><td>574.5</td><td>-13.0</td><td>1.5</td><td>-8.7</td><td>3.5E-18</td><td>3.9E-17</td><td>MB231/MB468</td><td>MP</td></tr>
<tr><td>>hg38_21_+_8433172_8446622_RNA45SN1_45S_50ntFlanks@4735.4759.25|ACGAAAGTCGGAGGTTCGAAGACGA</td><td>>hg38_21_+_8433172_8446622_RNA45SN1_45S_50ntFlanks</td><td>4735</td><td>4759</td><td>25</td><td>ACGAAAGTCGGAGGTTCGAAGACGA</td><td>1090.3</td><td>-6.3</td><td>0.7</td><td>-8.6</td><td>1.1E-17</td><td>3.9E-17</td><td>BT20/MB468</td><td>NUC</td></tr>
<tr><td>>hg38_21_+_8433172_8446622_RNA45SN1_45S_50ntFlanks@12406.12446.41|TTTTTGATCCTTCGATGTCGGCTCTTCCTATCATTGTGAAG</td><td>>hg38_21_+_8433172_8446622_RNA45SN1_45S_50ntFlanks</td><td>12406</td><td>12446</td><td>41</td><td>TTTTTGATCCTTCGATGTCGGCTCTTCCTATCATTGTGAAG</td><td>649.2</td><td>-13.8</td><td>1.6</td><td>-8.6</td><td>1.2E-17</td><td>3.9E-17</td><td>BT20/MB468</td><td>NUC</td></tr>
<tr><td>>hg38_21_+_8433172_8446622_RNA45SN1_45S_50ntFlanks@11998.12032.35|GCCCTGCGGGCCGCCGGTGAAATACCACTACTCTG</td><td>>hg38_21_+_8433172_8446622_RNA45SN1_45S_50ntFlanks</td><td>11998</td><td>12032</td><td>35</td><td>GCCCTGCGGGCCGCCGGTGAAATACCACTACTCTG</td><td>595.6</td><td>-13.0</td><td>1.5</td><td>-8.6</td><td>5.7E-18</td><td>3.9E-17</td><td>MB231/MB468</td><td>MITO</td></tr>
<tr><td>>hg38_21_+_8433172_8446622_RNA45SN1_45S_50ntFlanks@8380.8406.27|CAAGAGGGCGTGAAACCGTTAAGAGGT</td><td>>hg38_21_+_8433172_8446622_RNA45SN1_45S_50ntFlanks</td><td>8380</td><td>8406</td><td>27</td><td>CAAGAGGGCGTGAAACCGTTAAGAGGT</td><td>2704.0</td><td>-7.3</td><td>0.8</td><td>-8.6</td><td>5.7E-18</td><td>3.9E-17</td><td>MB231/MB468</td><td>MITO</td></tr>
<tr><td>>hg38_21_+_8433172_8446622_RNA45SN1_45S_50ntFlanks@12324.12345.22|TTGGGTTTTAAGCAGGAGGTGT</td><td>>hg38_21_+_8433172_8446622_RNA45SN1_45S_50ntFlanks</td><td>12324</td><td>12345</td><td>22</td><td>TTGGGTTTTAAGCAGGAGGTGT</td><td>350.5</td><td>-12.3</td><td>1.4</td><td>-8.7</td><td>3.5E-18</td><td>3.9E-17</td><td>MB231/MB468</td><td>MP</td></tr>
<tr><td>>hg38_21_+_8433172_8446622_RNA45SN1_45S_50ntFlanks@4486.4511.26|CCGCGGGGCCCGAAGCGTTTACTTTG</td><td>>hg38_21_+_8433172_8446622_RNA45SN1_45S_50ntFlanks</td><td>4486</td><td>4511</td><td>26</td><td>CCGCGGGGCCCGAAGCGTTTACTTTG</td><td>1046.9</td><td>-13.8</td><td>1.6</td><td>-8.6</td><td>5.8E-18</td><td>3.9E-17</td><td>MB231/MB468</td><td>MITO</td></tr>
<tr><td>>hg38_21_+_8433172_8446622_RNA45SN1_45S_50ntFlanks@12157.12178.22|AGTTTGACTGGGGCGGTACACC</td><td>>hg38_21_+_8433172_8446622_RNA45SN1_45S_50ntFlanks</td><td>12157</td><td>12178</td><td>22</td><td>AGTTTGACTGGGGCGGTACACC</td><td>503.5</td><td>-12.8</td><td>1.5</td><td>-8.7</td><td>3.5E-18</td><td>3.9E-17</td><td>MB231/MB468</td><td>MP</td></tr>
<tr><td>>am_trna162_MetCAT_6_-_26758550_26758622@34.54.21,trna164_MetCAT_6_-_26735574_26735646@34.54.21,trna20_MetCAT_16_+_71460396_71460468@34.54.21,trna21_MetCAT_6_+_26701712_26701784@34.54.21,trna27_MetCAT_6_+_26766444_26766516@34.54.21,trna75_MetCAT_6_+_28912352_28912424@34.54.21,trna97_MetCAT_6_-_28921042_28921114@34.54.21|CATAATCTGAAGGTCCTGAGT</td><td>>am_trna162_MetCAT_6_-_26758550_26758622</td><td>34</td><td>54</td><td>21</td><td>CATAATCTGAAGGTCCTGAGT</td><td>969.5</td><td>4.7</td><td>0.5</td><td>8.6</td><td>6.8E-18</td><td>4.0E-17</td><td>BT20/MB468</td><td>CYTO</td></tr>
<tr><td>>trna10_AspGTC_12_-_125424193_125424264@1.29.29,trna12_AspGTC_12_-_125411891_125411962@1.29.29,trna38_AspGTC_17_-_8125556_8125627@1.29.29,trna45_AspGTC_6_+_27447453_27447524@1.29.29,trna48_AspGTC_6_+_27471523_27471594@1.29.29,trna4_AspGTC_12_+_96429799_96429870@1.29.29,trna69_AspGTC_1_-_161440205_161440276@1.29.29,trna72_AspGTC_1_-_161432824_161432895@1.29.29,trna75_AspGTC_1_-_161425414_161425485@1.29.29,trna78_AspGTC_1_-_161418033_161418104@1.29.29,trna81_AspGTC_1_-_161410615_161410686@1.29.29|TCCTCGTTAGTATAGTGGTGAGTATCCCC</td><td>>trna10_AspGTC_12_-_125424193_125424264</td><td>1</td><td>29</td><td>29</td><td>TCCTCGTTAGTATAGTGGTGAGTATCCCC</td><td>316.0</td><td>-12.2</td><td>1.4</td><td>-8.7</td><td>3.6E-18</td><td>4.0E-17</td><td>MB231/MB468</td><td>MP</td></tr>
<tr><td>>hg38_21_+_8433172_8446622_RNA45SN1_45S_50ntFlanks@9504.9521.18|GTGAACTATGCCTGGGCA</td><td>>hg38_21_+_8433172_8446622_RNA45SN1_45S_50ntFlanks</td><td>9504</td><td>9521</td><td>18</td><td>GTGAACTATGCCTGGGCA</td><td>3013.5</td><td>-6.2</td><td>0.7</td><td>-8.7</td><td>3.5E-18</td><td>4.0E-17</td><td>MB231/MB468</td><td>CYTO</td></tr>
<tr><td>>hg38_21_+_8433172_8446622_RNA45SN1_45S_50ntFlanks@10806.10825.20|AACAATGTAGGTAAGGGAAG</td><td>>hg38_21_+_8433172_8446622_RNA45SN1_45S_50ntFlanks</td><td>10806</td><td>10825</td><td>20</td><td>AACAATGTAGGTAAGGGAAG</td><td>361.8</td><td>-12.4</td><td>1.4</td><td>-8.7</td><td>3.6E-18</td><td>4.0E-17</td><td>MB231/MB468</td><td>MP</td></tr>
<tr><td>>hg38_21_+_8433172_8446622_RNA45SN1_45S_50ntFlanks@5428.5450.23|ATGGTTTAGTGAGGCCCTCGGAT</td><td>>hg38_21_+_8433172_8446622_RNA45SN1_45S_50ntFlanks</td><td>5428</td><td>5450</td><td>23</td><td>ATGGTTTAGTGAGGCCCTCGGAT</td><td>131.7</td><td>11.0</td><td>1.3</td><td>8.7</td><td>3.6E-18</td><td>4.0E-17</td><td>MB231/MB468</td><td>CYTO</td></tr>
<tr><td>>hg38_21_+_8433172_8446622_RNA45SN1_45S_50ntFlanks@12405.12426.22|CTTTTTGATCCTTCGATGTCGG</td><td>>hg38_21_+_8433172_8446622_RNA45SN1_45S_50ntFlanks</td><td>12405</td><td>12426</td><td>22</td><td>CTTTTTGATCCTTCGATGTCGG</td><td>3000.2</td><td>-8.7</td><td>1.0</td><td>-8.7</td><td>4.7E-18</td><td>4.0E-17</td><td>MB231/MB468</td><td>NUC</td></tr>
<tr><td>>trna10_AspGTC_12_-_125424193_125424264@31.72.42,trna12_AspGTC_12_-_125411891_125411962@31.72.42,trna38_AspGTC_17_-_8125556_8125627@31.72.42,trna45_AspGTC_6_+_27447453_27447524@31.72.42,trna48_AspGTC_6_+_27471523_27471594@31.72.42,trna4_AspGTC_12_+_96429799_96429870@31.72.42,trna69_AspGTC_1_-_161440205_161440276@31.72.42,trna72_AspGTC_1_-_161432824_161432895@31.72.42,trna75_AspGTC_1_-_161425414_161425485@31.72.42,trna78_AspGTC_1_-_161418033_161418104@31.72.42,trna81_AspGTC_1_-_161410615_161410686@31.72.42|CCTGTCACGCGGGAGACCGGGGTTCGATTCCCCGACGGGGAG</td><td>>trna10_AspGTC_12_-_125424193_125424264</td><td>31</td><td>72</td><td>42</td><td>CCTGTCACGCGGGAGACCGGGGTTCGATTCCCCGACGGGGAG</td><td>1722.1</td><td>-9.7</td><td>1.1</td><td>-8.7</td><td>3.6E-18</td><td>4.0E-17</td><td>MB231/MB468</td><td>CYTO</td></tr>
<tr><td>>hg38_21_+_8433172_8446622_RNA45SN1_45S_50ntFlanks@9623.9650.28|TAGTAGCTGGTTCCCTCCGAAGTTTCCC</td><td>>hg38_21_+_8433172_8446622_RNA45SN1_45S_50ntFlanks</td><td>9623</td><td>9650</td><td>28</td><td>TAGTAGCTGGTTCCCTCCGAAGTTTCCC</td><td>1925.0</td><td>-10.1</td><td>1.2</td><td>-8.6</td><td>5.9E-18</td><td>4.0E-17</td><td>MB231/MB468</td><td>MITO</td></tr>
<tr><td>>hg38_21_+_8433172_8446622_RNA45SN1_45S_50ntFlanks@11324.11347.24|GTCTCGTCGCGCGCGCGTCCGCTG</td><td>>hg38_21_+_8433172_8446622_RNA45SN1_45S_50ntFlanks</td><td>11324</td><td>11347</td><td>24</td><td>GTCTCGTCGCGCGCGCGTCCGCTG</td><td>3664.0</td><td>16.3</td><td>1.9</td><td>8.7</td><td>5.0E-18</td><td>4.0E-17</td><td>BT20/MB468</td><td>MITO</td></tr>
<tr><td>>hg38_21_+_8433172_8446622_RNA45SN1_45S_50ntFlanks@12420.12462.43|ATGTCGGCTCTTCCTATCATTGTGAAGCAGAATTCACCAAGCG</td><td>>hg38_21_+_8433172_8446622_RNA45SN1_45S_50ntFlanks</td><td>12420</td><td>12462</td><td>43</td><td>ATGTCGGCTCTTCCTATCATTGTGAAGCAGAATTCACCAAGCG</td><td>1263.6</td><td>8.8</td><td>1.0</td><td>8.7</td><td>3.6E-18</td><td>4.1E-17</td><td>MB231/MB468</td><td>CYTO</td></tr>
<tr><td>>hg38_21_+_8433172_8446622_RNA45SN1_45S_50ntFlanks@3835.3871.37|CTCCTCTCCTACTTGGATAACTGTGGTAATTCTAGAG</td><td>>hg38_21_+_8433172_8446622_RNA45SN1_45S_50ntFlanks</td><td>3835</td><td>3871</td><td>37</td><td>CTCCTCTCCTACTTGGATAACTGTGGTAATTCTAGAG</td><td>829.4</td><td>-7.9</td><td>0.9</td><td>-8.6</td><td>6.0E-18</td><td>4.1E-17</td><td>MB231/MB468</td><td>MITO</td></tr>
</table>

<table>
<tr><td></td><td></td><td></td><td></td><td></td><td></td><td></td><td></td><td></td><td></td><td></td><td></td><td></td><td></td></tr>
<tr><td>>hg38_21_+_8433172_8446622_RNA45SN1_45S_50ntFlanks@12615.12657.43|GCTGAGGAGCCAATGGGGCGAAGCTACCATCTGTGGGATTATG</td><td>>hg38_21_+_8433172_8446622_RNA45SN1_45S_50ntFlanks</td><td>12615</td><td>12657</td><td>43</td><td>GCTGAGGAGCCAATGGGGCGAAGCTACCATCTGTGGGATTATG</td><td>900.2</td><td>-14.2</td><td>1.7</td><td>-8.6</td><td>1.2E-17</td><td>4.1E-17</td><td>BT20/MB468</td><td>NUC</td></tr>
<tr><td>>hg38_21_+_8433172_8446622_RNA45SN1_45S_50ntFlanks@9504.9525.22|GTGAACTATGCCTGGGCAGGGC</td><td>>hg38_21_+_8433172_8446622_RNA45SN1_45S_50ntFlanks</td><td>9504</td><td>9525</td><td>22</td><td>GTGAACTATGCCTGGGCAGGGC</td><td>3889.4</td><td>-9.8</td><td>1.1</td><td>-8.7</td><td>3.7E-18</td><td>4.1E-17</td><td>MB231/MB468</td><td>MP</td></tr>
<tr><td>>hg38_21_+_8433172_8446622_RNA45SN1_45S_50ntFlanks@10680.10711.32|GTTCCGGCGGCGTCCGGTGAGCTCTCGCTGGC</td><td>>hg38_21_+_8433172_8446622_RNA45SN1_45S_50ntFlanks</td><td>10680</td><td>10711</td><td>32</td><td>GTTCCGGCGGCGTCCGGTGAGCTCTCGCTGGC</td><td>8415.2</td><td>-5.7</td><td>0.7</td><td>-8.7</td><td>5.1E-18</td><td>4.1E-17</td><td>BT20/MB468</td><td>MITO</td></tr>
<tr><td>>hg38_21_+_8433172_8446622_RNA45SN1_45S_50ntFlanks@7991.8016.26|ACGTGGCGACCCGCTGAATTTAAGCA</td><td>>hg38_21_+_8433172_8446622_RNA45SN1_45S_50ntFlanks</td><td>7991</td><td>8016</td><td>26</td><td>ACGTGGCGACCCGCTGAATTTAAGCA</td><td>796.0</td><td>-5.5</td><td>0.6</td><td>-8.7</td><td>3.7E-18</td><td>4.1E-17</td><td>MB231/MB468</td><td>MP</td></tr>
<tr><td>>trna119_LysCTT_1_-_145395522_145395594@1.30.30,trna11_LysCTT_5_-_180648979_180649051@1.30.30,trna13_LysCTT_6_+_26556774_26556846@1.30.30,trna32_LysCTT_16_-_3207406_3207478@1.30.30,trna7_LysCTT_16_+_3225692_3225764@1.30.30,trna9_LysCTT_5_+_180634755_180634827@1.30.30|GCCCGGCTAGCTCAGTCGGTAGAGCATGAG</td><td>>trna119_LysCTT_1_-_145395522_145395594</td><td>1</td><td>30</td><td>30</td><td>GCCCGGCTAGCTCAGTCGGTAGAGCATGAG</td><td>10270.0</td><td>-6.3</td><td>0.7</td><td>-8.7</td><td>4.8E-18</td><td>4.1E-17</td><td>MB231/MB468</td><td>NUC</td></tr>
<tr><td>>hg38_21_+_8433172_8446622_RNA45SN1_45S_50ntFlanks@8170.8212.43|CAAGTCCTTCTGATCGAGGCCCAGCCCGTGGACGGTGTGAGGC</td><td>>hg38_21_+_8433172_8446622_RNA45SN1_45S_50ntFlanks</td><td>8170</td><td>8212</td><td>43</td><td>CAAGTCCTTCTGATCGAGGCCCAGCCCGTGGACGGTGTGAGGC</td><td>6954.6</td><td>-7.3</td><td>0.8</td><td>-8.7</td><td>4.8E-18</td><td>4.1E-17</td><td>MB231/MB468</td><td>NUC</td></tr>
<tr><td>>MI0000081|hsa-mir-24-2&WithFlank&19|-|13836281|13836365@56.77.22(+1C);MI0000080|hsa-mir-24-1&WithFlank&9|+|95086015|95086094@50.71.22(+1C)&[MIMAT0000080&hsa-miR-24-3p&offsets|0|0(+1C);m-36&19|-|13836288|13836310&offsets|0|-1(+1C)];[MIMAT0000080_1&hsa-miR-24-3p&offsets|0|0(+1C);m-35&9|+|95086064|95086086&offsets|0|-1(+1C)]|TGGCTCAGTTCAGCAGGAACAGC</td><td>>MI0000081|hsa-mir-24-2&WithFlank&19|-|13836281|13836365</td><td>56</td><td>77</td><td>22(+1C)</td><td>TGGCTCAGTTCAGCAGGAACAGC</td><td>627.0</td><td>6.1</td><td>0.7</td><td>8.7</td><td>5.1E-18</td><td>4.1E-17</td><td>BT20/MB468</td><td>MITO</td></tr>
<tr><td>>hg38_21_+_8433172_8446622_RNA45SN1_45S_50ntFlanks@9127.9152.26|CGCGGGTCGGGGGGCGGGGCGGACTG</td><td>>hg38_21_+_8433172_8446622_RNA45SN1_45S_50ntFlanks</td><td>9127</td><td>9152</td><td>26</td><td>CGCGGGTCGGGGGGCGGGGCGGACTG</td><td>636.4</td><td>-13.1</td><td>1.5</td><td>-8.6</td><td>6.1E-18</td><td>4.2E-17</td><td>MB231/MB468</td><td>MITO</td></tr>
<tr><td>>am_MI0003581|hsa-mir-574&WithFlank&4|+|38868026|38868133@31.53.23&[MIMAT0004795&hsa-miR-574-5p&offsets|0|0;m-213&4|+|38868056|38868078&offsets|0|0]|TGAGTGTGTGTGTGTGAGTGTGT</td><td>>am_MI0003581|hsa-mir-574&WithFlank&4|+|38868026|38868133</td><td>31</td><td>53</td><td>23</td><td>TGAGTGTGTGTGTGTGAGTGTGT</td><td>542.5</td><td>5.1</td><td>0.6</td><td>8.7</td><td>4.8E-18</td><td>4.2E-17</td><td>MB231/MB468</td><td>NUC</td></tr>
<tr><td>>hg38_21_+_8433172_8446622_RNA45SN1_45S_50ntFlanks@10516.10533.18|GAGTGGCGGAGATGGGCG</td><td>>hg38_21_+_8433172_8446622_RNA45SN1_45S_50ntFlanks</td><td>10516</td><td>10533</td><td>18</td><td>GAGTGGCGGAGATGGGCG</td><td>5985.7</td><td>-11.9</td><td>1.4</td><td>-8.7</td><td>3.7E-18</td><td>4.2E-17</td><td>MB231/MB468</td><td>MP</td></tr>
<tr><td>>hg38_21_+_8433172_8446622_RNA45SN1_45S_50ntFlanks@11718.11751.34|AACGGCGGGAGTAACTATGACTCTCTTAAGGTAG</td><td>>hg38_21_+_8433172_8446622_RNA45SN1_45S_50ntFlanks</td><td>11718</td><td>11751</td><td>34</td><td>AACGGCGGGAGTAACTATGACTCTCTTAAGGTAG</td><td>479.4</td><td>-12.8</td><td>1.5</td><td>-8.7</td><td>3.8E-18</td><td>4.2E-17</td><td>MB231/MB468</td><td>MP</td></tr>
<tr><td>>hg38_21_+_8433172_8446622_RNA45SN1_45S_50ntFlanks@12202.12222.21|CTAAGGCGAGCTCAGGGAGGA</td><td>>hg38_21_+_8433172_8446622_RNA45SN1_45S_50ntFlanks</td><td>12202</td><td>12222</td><td>21</td><td>CTAAGGCGAGCTCAGGGAGGA</td><td>455.3</td><td>-12.7</td><td>1.5</td><td>-8.7</td><td>3.8E-18</td><td>4.2E-17</td><td>MB231/MB468</td><td>MP</td></tr>
<tr><td>>hg38_21_+_8433172_8446622_RNA45SN1_45S_50ntFlanks@11621.11640.20|CAAAGCATCGCGAAGGCCCG</td><td>>hg38_21_+_8433172_8446622_RNA45SN1_45S_50ntFlanks</td><td>11621</td><td>11640</td><td>20</td><td>CAAAGCATCGCGAAGGCCCG</td><td>2531.9</td><td>-14.4</td><td>1.6</td><td>-9.1</td><td>9.2E-20</td><td>4.2E-17</td><td>BT20/MB468</td><td>MP</td></tr>
<tr><td>>hg38_21_+_8433172_8446622_RNA45SN1_45S_50ntFlanks@13002.13027.26|CGATCTATTGAAAGTCAGCCCTCGAC</td><td>>hg38_21_+_8433172_8446622_RNA45SN1_45S_50ntFlanks</td><td>13002</td><td>13027</td><td>26</td><td>CGATCTATTGAAAGTCAGCCCTCGAC</td><td>1388.5</td><td>-13.5</td><td>1.5</td><td>-9.1</td><td>9.0E-20</td><td>4.2E-17</td><td>BT20/MB468</td><td>MP</td></tr>
<tr><td>>hg38_21_+_8433172_8446622_RNA45SN1_45S_50ntFlanks@11847.11872.26|AACGGGCTTGGCGGAATCAGCGGGGA</td><td>>hg38_21_+_8433172_8446622_RNA45SN1_45S_50ntFlanks</td><td>11847</td><td>11872</td><td>26</td><td>AACGGGCTTGGCGGAATCAGCGGGGA</td><td>1050.0</td><td>-9.9</td><td>1.2</td><td>-8.6</td><td>6.1E-18</td><td>4.2E-17</td><td>MB231/MB468</td><td>MITO</td></tr>
<tr><td>>hg38_21_+_8433172_8446622_RNA45SN1_45S_50ntFlanks@10786.10823.38|AACAGCCTCTGGCATGTTGGAACAATGTAGGTAAGGGA</td><td>>hg38_21_+_8433172_8446622_RNA45SN1_45S_50ntFlanks</td><td>10786</td><td>10823</td><td>38</td><td>AACAGCCTCTGGCATGTTGGAACAATGTAGGTAAGGGA</td><td>898.8</td><td>-13.7</td><td>1.6</td><td>-8.7</td><td>3.8E-18</td><td>4.2E-17</td><td>MB231/MB468</td><td>MP</td></tr>
<tr><td>>MI0003513|hsa-mir-455&WithFlank&9|+|114209428|114209535@22.45.24&[MIMAT0003150&hsa-miR-455-5p&offsets|0|+2;m-232&9|+|114209449|114209470&offsets|0|+2]|TATGTGCCTTTGGACTACATCGTG</td><td>>MI0003513|hsa-mir-455&WithFlank&9|+|114209428|114209535</td><td>22</td><td>45</td><td>24</td><td>TATGTGCCTTTGGACTACATCGTG</td><td>209.5</td><td>11.6</td><td>1.3</td><td>8.7</td><td>4.9E-18</td><td>4.2E-17</td><td>MB231/MB468</td><td>NUC</td></tr>
<tr><td>>hg38_MT_+_1621_3279_RNR2_16S_50ntFlanks@51.88.38|GCTAAACCTAGCCCCAAACCCACTCCACCTTACTACCA</td><td>>hg38_MT_+_1621_3279_RNR2_16S_50ntFlanks</td><td>51</td><td>88</td><td>38</td><td>GCTAAACCTAGCCCCAAACCCACTCCACCTTACTACCA</td><td>442.2</td><td>12.8</td><td>1.5</td><td>8.7</td><td>3.7E-18</td><td>4.2E-17</td><td>MB231/MB468</td><td>CYTO</td></tr>
<tr><td>>hg38_21_+_8433172_8446622_RNA45SN1_45S_50ntFlanks@6652.6669.18|GACTCTTAGCGGTGGATC</td><td>>hg38_21_+_8433172_8446622_RNA45SN1_45S_50ntFlanks</td><td>6652</td><td>6669</td><td>18</td><td>GACTCTTAGCGGTGGATC</td><td>677.5</td><td>4.7</td><td>0.5</td><td>8.6</td><td>7.3E-18</td><td>4.2E-17</td><td>BT20/MB468</td><td>CYTO</td></tr>
<tr><td>>hg38_21_+_8433172_8446622_RNA45SN1_45S_50ntFlanks@6737.6770.34|GATCATCGACACTTCGAACGCACTTGCGGCCCCG</td><td>>hg38_21_+_8433172_8446622_RNA45SN1_45S_50ntFlanks</td><td>6737</td><td>6770</td><td>34</td><td>GATCATCGACACTTCGAACGCACTTGCGGCCCCG</td><td>766.3</td><td>-11.7</td><td>1.3</td><td>-8.6</td><td>5.2E-18</td><td>4.2E-17</td><td>BT20/MB468</td><td>MITO</td></tr>
<tr><td>>MI0000108|hsa-mir-103a-2&WithFlank&20|+|3917488|3917577@54.76.23(+1C);MI0000109|hsa-mir-103a-1&WithFlank&5|-|168560890|168560979@54.76.23(+1C)&[MIMAT0000101&hsa-miR-103a-3p&offsets|0|0(+1C);m-12&20|+|3917541|3917563&offsets|0|0(+1C)];[MIMAT0000101_1&hsa-miR-103a-3p&offsets|0|0(+1C);m-13&5|-|168560904|168560926&offsets|0|0(+1C);m-44220&5|-|168560908|168560926&offsets|0|+4(+1C)]|AGCAGCATTGTACAGGGCTATGAC</td><td>>MI0000108|hsa-mir-103a-2&WithFlank&20|+|3917488|3917577</td><td>54</td><td>76</td><td>23(+1C)</td><td>AGCAGCATTGTACAGGGCTATGAC</td><td>259.8</td><td>5.8</td><td>0.7</td><td>8.7</td><td>4.9E-18</td><td>4.2E-17</td><td>MB231/MB468</td><td>NUC</td></tr>
<tr><td>>hg38_21_+_8433172_8446622_RNA45SN1_45S_50ntFlanks@12456.12491.36|CCAAGCGTTGGATTGTTCACCCACTAATAGGGAACG</td><td>>hg38_21_+_8433172_8446622_RNA45SN1_45S_50ntFlanks</td><td>12456</td><td>12491</td><td>36</td><td>CCAAGCGTTGGATTGTTCACCCACTAATAGGGAACG</td><td>405.4</td><td>-12.4</td><td>1.4</td><td>-8.6</td><td>6.2E-18</td><td>4.2E-17</td><td>MB231/MB468</td><td>MITO</td></tr>
<tr><td>>hg38_21_+_8433172_8446622_RNA45SN1_45S_50ntFlanks@10802.10823.22|TTGGAACAATGTAGGTAAGGGA</td><td>>hg38_21_+_8433172_8446622_RNA45SN1_45S_50ntFlanks</td><td>10802</td><td>10823</td><td>22</td><td>TTGGAACAATGTAGGTAAGGGA</td><td>403.7</td><td>-12.5</td><td>1.4</td><td>-8.7</td><td>3.8E-18</td><td>4.2E-17</td><td>MB231/MB468</td><td>MP</td></tr>
<tr><td>>MI0000255|hsa-mir-30d&WithFlank&8|-|134804870|134804951@12.32.21(+1A)&[MIMAT0000245&hsa-miR-30d-5p&offsets|0|-1(+1A);m-21&8|-|134804919|134804940&offsets|0|-1(+1A)]|TGTAAACATCCCCGACTGGAAA</td><td>>MI0000255|hsa-mir-30d&WithFlank&8|-|134804870|134804951</td><td>12</td><td>32</td><td>21(+1A)</td><td>TGTAAACATCCCCGACTGGAAA</td><td>1515.7</td><td>10.4</td><td>1.2</td><td>8.6</td><td>7.3E-18</td><td>4.3E-17</td><td>BT20/MB468</td><td>CYTO</td></tr>
<tr><td>>hg38_21_+_8433172_8446622_RNA45SN1_45S_50ntFlanks@9576.9597.22|TCGGTCGTCCGACCTGGGTATA</td><td>>hg38_21_+_8433172_8446622_RNA45SN1_45S_50ntFlanks</td><td>9576</td><td>9597</td><td>22</td><td>TCGGTCGTCCGACCTGGGTATA</td><td>607.8</td><td>-13.0</td><td>1.5</td><td>-8.6</td><td>6.3E-18</td><td>4.3E-17</td><td>MB231/MB468</td><td>MITO</td></tr>
<tr><td>>hg38_21_+_8433172_8446622_RNA45SN1_45S_50ntFlanks@4002.4026.25|GATAACCTCGGGCCGATCGCACGCC</td><td>>hg38_21_+_8433172_8446622_RNA45SN1_45S_50ntFlanks</td><td>4002</td><td>4026</td><td>25</td><td>GATAACCTCGGGCCGATCGCACGCC</td><td>562.1</td><td>-12.9</td><td>1.5</td><td>-8.6</td><td>6.3E-18</td><td>4.3E-17</td><td>MB231/MB468</td><td>MITO</td></tr>
<tr><td>>hg38_21_+_8433172_8446622_RNA45SN1_45S_50ntFlanks@12991.13009.19|CTCCCTCGCTGCGATCTAT</td><td>>hg38_21_+_8433172_8446622_RNA45SN1_45S_50ntFlanks</td><td>12991</td><td>13009</td><td>19</td><td>CTCCCTCGCTGCGATCTAT</td><td>514.9</td><td>-12.8</td><td>1.5</td><td>-8.6</td><td>6.3E-18</td><td>4.3E-17</td><td>MB231/MB468</td><td>MITO</td></tr>
<tr><td>>hg38_21_+_8433172_8446622_RNA45SN1_45S_50ntFlanks@11569.11597.29|CAGCCGACTTAGAACTGGTGCGGACCAGG</td><td>>hg38_21_+_8433172_8446622_RNA45SN1_45S_50ntFlanks</td><td>11569</td><td>11597</td><td>29</td><td>CAGCCGACTTAGAACTGGTGCGGACCAGG</td><td>318.7</td><td>-12.2</td><td>1.4</td><td>-8.7</td><td>3.9E-18</td><td>4.3E-17</td><td>MB231/MB468</td><td>MP</td></tr>
<tr><td>>hg38_21_+_8433172_8446622_RNA45SN1_45S_50ntFlanks@10787.10818.32|ACAGCCTCTGGCATGTTGGAACAATGTAGGTA</td><td>>hg38_21_+_8433172_8446622_RNA45SN1_45S_50ntFlanks</td><td>10787</td><td>10818</td><td>32</td><td>ACAGCCTCTGGCATGTTGGAACAATGTAGGTA</td><td>2108.6</td><td>-10.4</td><td>1.2</td><td>-8.7</td><td>3.9E-18</td><td>4.3E-17</td><td>MB231/MB468</td><td>MP</td></tr>
<tr><td>>hg38_21_+_8433172_8446622_RNA45SN1_45S_50ntFlanks@11567.11585.19|AGCAGCCGACTTAGAACTG</td><td>>hg38_21_+_8433172_8446622_RNA45SN1_45S_50ntFlanks</td><td>11567</td><td>11585</td><td>19</td><td>AGCAGCCGACTTAGAACTG</td><td>1685.4</td><td>-4.2</td><td>0.5</td><td>-8.5</td><td>1.3E-17</td><td>4.3E-17</td><td>BT20/MB468</td><td>NUC</td></tr>
<tr><td>>hg38_21_+_8433172_8446622_RNA45SN1_45S_50ntFlanks@4858.4890.33|TTTGGGTTCCGGGGGGAGTATGGTTGCAAAGCT</td><td>>hg38_21_+_8433172_8446622_RNA45SN1_45S_50ntFlanks</td><td>4858</td><td>4890</td><td>33</td><td>TTTGGGTTCCGGGGGGAGTATGGTTGCAAAGCT</td><td>132.9</td><td>11.1</td><td>1.3</td><td>8.7</td><td>3.8E-18</td><td>4.3E-17</td><td>MB231/MB468</td><td>CYTO</td></tr>
<tr><td>>hg38_21_+_8433172_8446622_RNA45SN1_45S_50ntFlanks@8971.8990.20|CCCACCCCCGCGGGGAATCC</td><td>>hg38_21_+_8433172_8446622_RNA45SN1_45S_50ntFlanks</td><td>8971</td><td>8990</td><td>20</td><td>CCCACCCCCGCGGGGAATCC</td><td>838.9</td><td>-13.5</td><td>1.6</td><td>-8.6</td><td>6.4E-18</td><td>4.3E-17</td><td>MB231/MB468</td><td>MITO</td></tr>
<tr><td>>hg38_21_+_8433172_8446622_RNA45SN1_45S_50ntFlanks@12008.12032.25|CCGCCGGTGAAATACCACTACTCTG</td><td>>hg38_21_+_8433172_8446622_RNA45SN1_45S_50ntFlanks</td><td>12008</td><td>12032</td><td>25</td><td>CCGCCGGTGAAATACCACTACTCTG</td><td>24685.0</td><td>-9.4</td><td>1.1</td><td>-8.6</td><td>5.4E-18</td><td>4.3E-17</td><td>BT20/MB468</td><td>MITO</td></tr>
<tr><td>>hg38_21_+_8433172_8446622_RNA45SN1_45S_50ntFlanks@10411.10440.30|CAGTCGGTCCTGAGAGATGGGCGAGCGCCG</td><td>>hg38_21_+_8433172_8446622_RNA45SN1_45S_50ntFlanks</td><td>10411</td><td>10440</td><td>30</td><td>CAGTCGGTCCTGAGAGATGGGCGAGCGCCG</td><td>469.4</td><td>-12.7</td><td>1.5</td><td>-8.6</td><td>6.4E-18</td><td>4.3E-17</td><td>MB231/MB468</td><td>MITO</td></tr>
<tr><td>>hg38_21_+_8433172_8446622_RNA45SN1_45S_50ntFlanks@3939.3957.19|AACCCGGTCAGCCCCTCTC</td><td>>hg38_21_+_8433172_8446622_RNA45SN1_45S_50ntFlanks</td><td>3939</td><td>3957</td><td>19</td><td>AACCCGGTCAGCCCCTCTC</td><td>411.0</td><td>-12.5</td><td>1.4</td><td>-8.6</td><td>6.4E-18</td><td>4.3E-17</td><td>MB231/MB468</td><td>MITO</td></tr>
<tr><td>>am_trna116_GluCTC_1_-_145399233_145399304@1.27.27,trna59_GluCTC_1_+_249168447_249168518@1.27.27,trna71_GluCTC_1_-_161439189_161439260@1.27.27,trna74_GluCTC_1_-_161431809_161431880@1.27.27,trna77_GluCTC_1_-_161424398_161424469@1.27.27,trna77_GluCTC_6_+_28949976_28950047@1.27.27,trna80_GluCTC_1_-_161417018_161417089@1.27.27,trna87_GluCTC_6_-_126101393_126101464@1.27.27|TCCCTGGTGGTCTAGTGGTTAGGATTC</td><td>>am_trna116_GluCTC_1_-_145399233_145399304</td><td>1</td><td>27</td><td>27</td><td>TCCCTGGTGGTCTAGTGGTTAGGATTC</td><td>6169.6</td><td>-6.4</td><td>0.8</td><td>-8.5</td><td>1.3E-17</td><td>4.3E-17</td><td>BT20/MB468</td><td>NUC</td></tr>
<tr><td>>MI0000456|hsa-mir-140&WithFlank&16|+|69933075|69933186@69.89.21&[MIMAT0004597&hsa-miR-140-3p&offsets|+1|+1;m-45&16|+|69933143|69933164&offsets|0|-1]|ACCACAGGGTAGAACCACGGA</td><td>>MI0000456|hsa-mir-140&WithFlank&16|+|69933075|69933186</td><td>69</td><td>89</td><td>21</td><td>ACCACAGGGTAGAACCACGGA</td><td>700.6</td><td>9.7</td><td>1.1</td><td>8.7</td><td>5.1E-18</td><td>4.4E-17</td><td>MB231/MB468</td><td>NUC</td></tr>
<tr><td>>hg38_21_+_8433172_8446622_RNA45SN1_45S_50ntFlanks@6651.6684.34|CGACTCTTAGCGGTGGATCACTCGGCTCGTGCGT</td><td>>hg38_21_+_8433172_8446622_RNA45SN1_45S_50ntFlanks</td><td>6651</td><td>6684</td><td>34</td><td>CGACTCTTAGCGGTGGATCACTCGGCTCGTGCGT</td><td>3070.5</td><td>5.5</td><td>0.6</td><td>8.6</td><td>7.5E-18</td><td>4.4E-17</td><td>BT20/MB468</td><td>CYTO</td></tr>
<tr><td>>hg38_21_+_8433172_8446622_RNA45SN1_45S_50ntFlanks@12202.12220.19|CTAAGGCGAGCTCAGGGAG</td><td>>hg38_21_+_8433172_8446622_RNA45SN1_45S_50ntFlanks</td><td>12202</td><td>12220</td><td>19</td><td>CTAAGGCGAGCTCAGGGAG</td><td>521.1</td><td>-12.0</td><td>1.4</td><td>-8.7</td><td>5.1E-18</td><td>4.4E-17</td><td>MB231/MB468</td><td>NUC</td></tr>
<tr><td>>MI0000477|hsa-mir-146a&WithFlank&5|+|160485346|160485456@27.49.23&[MIMAT0000449&hsa-miR-146a-5p&offsets|0|+1;m-129&5|+|160485372|160485393&offsets|0|+1]|TGAGAACTGAATTCCATGGGTTG</td><td>>MI0000477|hsa-mir-146a&WithFlank&5|+|160485346|160485456</td><td>27</td><td>49</td><td>23</td><td>TGAGAACTGAATTCCATGGGTTG</td><td>1329.9</td><td>13.6</td><td>1.6</td><td>8.6</td><td>6.5E-18</td><td>4.4E-17</td><td>MB231/MB468</td><td>MITO</td></tr>
<tr><td>>hg38_21_+_8433172_8446622_RNA45SN1_45S_50ntFlanks@10905.10936.32|CTGGGCGCGCGCCGCGGCTGGACGAGGCGCCG</td><td>>hg38_21_+_8433172_8446622_RNA45SN1_45S_50ntFlanks</td><td>10905</td><td>10936</td><td>32</td><td>CTGGGCGCGCGCCGCGGCTGGACGAGGCGCCG</td><td>498.7</td><td>-12.7</td><td>1.5</td><td>-8.6</td><td>6.5E-18</td><td>4.4E-17</td><td>MB231/MB468</td><td>MITO</td></tr>
<tr><td>>hg38_21_+_8433172_8446622_RNA45SN1_45S_50ntFlanks@9033.9070.38|CTCGTGGGGGGGCCGGGCCACCCCTCCCACGGCGCGAC</td><td>>hg38_21_+_8433172_8446622_RNA45SN1_45S_50ntFlanks</td><td>9033</td><td>9070</td><td>38</td><td>CTCGTGGGGGGGCCGGGCCACCCCTCCCACGGCGCGAC</td><td>464.9</td><td>-12.6</td><td>1.5</td><td>-8.6</td><td>6.6E-18</td><td>4.4E-17</td><td>MB231/MB468</td><td>MITO</td></tr>
<tr><td>>hg38_21_+_8433172_8446622_RNA45SN1_45S_50ntFlanks@476.501.26|TTCGTGATCGATGTGGTGACGTCGTG</td><td>>hg38_21_+_8433172_8446622_RNA45SN1_45S_50ntFlanks</td><td>476</td><td>501</td><td>26</td><td>TTCGTGATCGATGTGGTGACGTCGTG</td><td>853.7</td><td>6.9</td><td>0.8</td><td>8.6</td><td>5.5E-18</td><td>4.4E-17</td><td>BT20/MB468</td><td>MITO</td></tr>
<tr><td>>hg38_21_+_8433172_8446622_RNA45SN1_45S_50ntFlanks@12001.12043.43|CTGCGGGCCGCCGGTGAAATACCACTACTCTGATCGTTTTTTC</td><td>>hg38_21_+_8433172_8446622_RNA45SN1_45S_50ntFlanks</td><td>12001</td><td>12043</td><td>43</td><td>CTGCGGGCCGCCGGTGAAATACCACTACTCTGATCGTTTTTTC</td><td>942.1</td><td>-13.7</td><td>1.6</td><td>-8.6</td><td>6.6E-18</td><td>4.4E-17</td><td>MB231/MB468</td><td>MITO</td></tr>
</table>

<table>
<tr><td>>hg38_21_+_8433172_8446622_RNA45SN1_45S_50ntFlanks@10678.10705.28|CGGTTCCGGCGGCGTCCGGTGAGCTCTC</td><td>>hg38_21_+_8433172_8446622_RNA45SN1_45S_50ntFlanks</td><td>10678</td><td>10705</td><td>28</td><td>CGGTTCCGGCGGCGTCCGGTGAGCTCTC</td><td>1136.0</td><td>-7.3</td><td>0.8</td><td>-8.7</td><td>3.9E-18</td><td>4.4E-17</td><td>MB231/MB468</td><td>CYTO</td></tr>
<tr><td>>MI0000483|hsa-mir-186&WithFlank&1|-|71067625|71067722@21.41.21&[MIMAT0000456&hsa-miR-186-5p&offsets|0|-1;m-84&1|-|71067681|71067702&offsets|0|-1]|CAAAGAATTCTCCTTTTGGGC</td><td>>MI0000483|hsa-mir-186&WithFlank&1|-|71067625|71067722</td><td>21</td><td>41</td><td>21</td><td>CAAAGAATTCTCCTTTTGGGC</td><td>244.6</td><td>5.2</td><td>0.6</td><td>8.6</td><td>5.2E-18</td><td>4.5E-17</td><td>MB231/MB468</td><td>NUC</td></tr>
<tr><td>>trna10_AspGTC_12_-_125424193_125424264@1.30.30,trna12_AspGTC_12_-_125411891_125411962@1.30.30,trna38_AspGTC_17_-_8125556_8125627@1.30.30,trna45_AspGTC_6_+_27447453_27447524@1.30.30,trna48_AspGTC_6_+_27471523_27471594@1.30.30,trna4_AspGTC_12_+_96429799_96429870@1.30.30,trna69_AspGTC_1_-_161440205_161440276@1.30.30,trna72_AspGTC_1_-_161432824_161432895@1.30.30,trna75_AspGTC_1_-_161425414_161425485@1.30.30,trna78_AspGTC_1_-_161418033_161418104@1.30.30,trna81_AspGTC_1_-_161410615_161410686@1.30.30|TCCTCGTTAGTATAGTGGTGAGTATCCCCG</td><td>>trna10_AspGTC_12_-_125424193_125424264</td><td>1</td><td>30</td><td>30</td><td>TCCTCGTTAGTATAGTGGTGAGTATCCCCG</td><td>314.6</td><td>-12.2</td><td>1.4</td><td>-8.7</td><td>4.1E-18</td><td>4.5E-17</td><td>MB231/MB468</td><td>MP</td></tr>
<tr><td>>MI0003513|hsa-mir-455&WithFlank&9|+|114209428|114209535@60.81.22&[MIMAT0004784&hsa-miR-455-3p&offsets|0|+1;m-135&9|+|114209487|114209508&offsets|0|0]|GCAGTCCATGGGCATATACACT</td><td>>MI0003513|hsa-mir-455&WithFlank&9|+|114209428|114209535</td><td>60</td><td>81</td><td>22</td><td>GCAGTCCATGGGCATATACACT</td><td>419.5</td><td>12.7</td><td>1.5</td><td>8.7</td><td>4.0E-18</td><td>4.5E-17</td><td>MB231/MB468</td><td>CYTO</td></tr>
<tr><td>>am_MI0005763|hsa-mir-941-1&WithFlank&20|+|63919443|63919526@53.74.22(+1A);MI0005764|hsa-mir-941-2&WithFlank&20|+|63919499|63919582@53.74.22(+1A);MI0005765|hsa-mir-941-3&WithFlank&20|+|63919555|63919638@53.74.22(+1A);MI0005766|hsa-mir-941-4&WithFlank&20|+|63919750|63919833@53.74.22(+1A);MI0031520|hsa-mir-941-5&WithFlank&20|+|63919862|63919945@53.74.22(+1A)&[MIMAT0004984&hsa-miR-941&offsets|0|-1(+1A);m-263&20|+|63919495|63919517&offsets|0|-1(+1A)];[MIMAT0004984_1&hsa-miR-941&offsets|0|-1(+1A);m-263&20|+|63919551|63919573&offsets|0|-1(+1A)];[MIMAT0004984_2&hsa-miR-941&offsets|0|-1(+1A);m-263&20|+|63919607|63919629&offsets|0|-1(+1A)];[MIMAT0004984_3&hsa-miR-941&offsets|0|-1(+1A);m-263&20|+|63919802|63919824&offsets|0|-1(+1A)];[MIMAT0004984_4&hsa-miR-941&offsets|0|-1(+1A);m-263&20|+|63919914|63919936&offsets|0|-1(+1A)]|CACCCGGCTGTGTGCACATGTGA</td><td>>am_MI0005763|hsa-mir-941-1&WithFlank&20|+|63919443|63919526</td><td>53</td><td>74</td><td>22(+1A)</td><td>CACCCGGCTGTGTGCACATGTGA</td><td>1009.5</td><td>7.0</td><td>0.8</td><td>8.7</td><td>4.0E-18</td><td>4.5E-17</td><td>MB231/MB468</td><td>CYTO</td></tr>
<tr><td>>MI0000447|hsa-mir-128-1&WithFlank&2|+|135665391|135665484@56.76.21(+1A);MI0000727|hsa-mir-128-2&WithFlank&3|+|35744470|35744565@58.78.21(+1A)&[MIMAT0000424&hsa-miR-128-3p&offsets|0|0(+1A);m-108&2|+|135665446|135665466&offsets|0|0(+1A)];[MIMAT0000424_1&hsa-miR-128-3p&offsets|0|0(+1A);m-117&3|+|35744527|35744547&offsets|0|0(+1A)]|TCACAGTGAACCGGTCTCTTTA</td><td>>MI0000447|hsa-mir-128-1&WithFlank&2|+|135665391|135665484</td><td>56</td><td>76</td><td>21(+1A)</td><td>TCACAGTGAACCGGTCTCTTTA</td><td>1757.1</td><td>5.0</td><td>0.6</td><td>8.6</td><td>5.6E-18</td><td>4.5E-17</td><td>BT20/MB468</td><td>MITO</td></tr>
<tr><td>>hg38_21_+_8433172_8446622_RNA45SN1_45S_50ntFlanks@9573.9597.25|AAATCGGTCGTCCGACCTGGGTATA</td><td>>hg38_21_+_8433172_8446622_RNA45SN1_45S_50ntFlanks</td><td>9573</td><td>9597</td><td>25</td><td>AAATCGGTCGTCCGACCTGGGTATA</td><td>10342.7</td><td>-9.8</td><td>1.1</td><td>-8.6</td><td>6.8E-18</td><td>4.5E-17</td><td>MB231/MB468</td><td>MITO</td></tr>
<tr><td>>hg38_21_+_8433172_8446622_RNA45SN1_45S_50ntFlanks@4728.4752.25|ATCAAGAACGAAAGTCGGAGGTTCG</td><td>>hg38_21_+_8433172_8446622_RNA45SN1_45S_50ntFlanks</td><td>4728</td><td>4752</td><td>25</td><td>ATCAAGAACGAAAGTCGGAGGTTCG</td><td>392.6</td><td>-12.4</td><td>1.4</td><td>-8.6</td><td>6.8E-18</td><td>4.5E-17</td><td>MB231/MB468</td><td>MITO</td></tr>
<tr><td>>MI0000289|hsa-mir-181a-1&WithFlank&1|-|198859038|198859159@30.52.23;MI0000269|hsa-mir-181a-2&WithFlank&9|+|124692436|124692557@45.67.23&[MIMAT0000256&hsa-miR-181a-5p&offsets|0|0;m-32&1|-|198859108|198859130&offsets|0|0];[MIMAT0000256_1&hsa-miR-181a-5p&offsets|0|0;m-32&9|+|124692480|124692502&offsets|0|0]|AACATTCAACGCTGTCGGTGAGT</td><td>>MI0000289|hsa-mir-181a-1&WithFlank&1|-|198859038|198859159</td><td>30</td><td>52</td><td>23</td><td>AACATTCAACGCTGTCGGTGAGT</td><td>2100.8</td><td>4.8</td><td>0.6</td><td>8.6</td><td>7.8E-18</td><td>4.5E-17</td><td>BT20/MB468</td><td>CYTO</td></tr>
<tr><td>>hg38_21_+_8433172_8446622_RNA45SN1_45S_50ntFlanks@9337.9370.34|CTCGCACGAAAGCCGCCGTGGCGCAATGAAGGTG</td><td>>hg38_21_+_8433172_8446622_RNA45SN1_45S_50ntFlanks</td><td>9337</td><td>9370</td><td>34</td><td>CTCGCACGAAAGCCGCCGTGGCGCAATGAAGGTG</td><td>473.0</td><td>-12.7</td><td>1.5</td><td>-8.7</td><td>4.1E-18</td><td>4.6E-17</td><td>MB231/MB468</td><td>MP</td></tr>
<tr><td>>hg38_21_+_8433172_8446622_RNA45SN1_45S_50ntFlanks@10808.10829.22|CAATGTAGGTAAGGGAAGTCGG</td><td>>hg38_21_+_8433172_8446622_RNA45SN1_45S_50ntFlanks</td><td>10808</td><td>10829</td><td>22</td><td>CAATGTAGGTAAGGGAAGTCGG</td><td>435.5</td><td>-12.6</td><td>1.5</td><td>-8.7</td><td>4.1E-18</td><td>4.6E-17</td><td>MB231/MB468</td><td>MP</td></tr>
<tr><td>>hg38_21_+_8433172_8446622_RNA45SN1_45S_50ntFlanks@11594.11612.19|CAGGGGAATCCGACTGTTT</td><td>>hg38_21_+_8433172_8446622_RNA45SN1_45S_50ntFlanks</td><td>11594</td><td>11612</td><td>19</td><td>CAGGGGAATCCGACTGTTT</td><td>1825.4</td><td>-6.5</td><td>0.8</td><td>-8.6</td><td>6.8E-18</td><td>4.6E-17</td><td>MB231/MB468</td><td>MITO</td></tr>
<tr><td>>hg38_21_+_8433172_8446622_RNA45SN1_45S_50ntFlanks@12950.12990.41|CGACCTGCTTCTGGGTCGGGGTTTCGTACGTAGCAGAGCAG</td><td>>hg38_21_+_8433172_8446622_RNA45SN1_45S_50ntFlanks</td><td>12950</td><td>12990</td><td>41</td><td>CGACCTGCTTCTGGGTCGGGGTTTCGTACGTAGCAGAGCAG</td><td>342.1</td><td>-12.8</td><td>1.5</td><td>-8.5</td><td>1.4E-17</td><td>4.6E-17</td><td>BT20/MB468</td><td>NUC</td></tr>
<tr><td>>hg38_21_+_8433172_8446622_RNA45SN1_45S_50ntFlanks@7986.8018.33|ATCAGACGTGGCGACCCGCTGAATTTAAGCATA</td><td>>hg38_21_+_8433172_8446622_RNA45SN1_45S_50ntFlanks</td><td>7986</td><td>8018</td><td>33</td><td>ATCAGACGTGGCGACCCGCTGAATTTAAGCATA</td><td>588.0</td><td>-13.1</td><td>1.5</td><td>-8.7</td><td>4.2E-18</td><td>4.6E-17</td><td>MB231/MB468</td><td>MP</td></tr>
<tr><td>>trna116_GluCTC_1_-_145399233_145399304@4.34.31,trna59_GluCTC_1_+_249168447_249168518@4.34.31,trna71_GluCTC_1_-_161439189_161439260@4.34.31,trna74_GluCTC_1_-_161431809_161431880@4.34.31,trna77_GluCTC_1_-_161424398_161424469@4.34.31,trna77_GluCTC_6_+_28949976_28950047@4.34.31,trna80_GluCTC_1_-_161417018_161417089@4.34.31,trna87_GluCTC_6_-_126101393_126101464@4.34.31|CTGGTGGTCTAGTGGTTAGGATTCGGCGCTC</td><td>>trna116_GluCTC_1_-_145399233_145399304</td><td>4</td><td>34</td><td>31</td><td>CTGGTGGTCTAGTGGTTAGGATTCGGCGCTC</td><td>371.7</td><td>6.7</td><td>0.8</td><td>8.6</td><td>5.7E-18</td><td>4.6E-17</td><td>BT20/MB468</td><td>MITO</td></tr>
<tr><td>>hg38_21_+_8433172_8446622_RNA45SN1_45S_50ntFlanks@11321.11342.22|CACGTCTCGTCGCGCGCGCGTC</td><td>>hg38_21_+_8433172_8446622_RNA45SN1_45S_50ntFlanks</td><td>11321</td><td>11342</td><td>22</td><td>CACGTCTCGTCGCGCGCGCGTC</td><td>2366.2</td><td>-7.9</td><td>0.9</td><td>-8.7</td><td>4.1E-18</td><td>4.6E-17</td><td>MB231/MB468</td><td>CYTO</td></tr>
<tr><td>>hg38_21_+_8433172_8446622_RNA45SN1_45S_50ntFlanks@11890.11910.21|CTTGACTCTAGTCTGGCACGG</td><td>>hg38_21_+_8433172_8446622_RNA45SN1_45S_50ntFlanks</td><td>11890</td><td>11910</td><td>21</td><td>CTTGACTCTAGTCTGGCACGG</td><td>184.9</td><td>-5.8</td><td>0.7</td><td>-8.7</td><td>4.1E-18</td><td>4.6E-17</td><td>MB231/MB468</td><td>CYTO</td></tr>
<tr><td>>hg38_21_+_8433172_8446622_RNA45SN1_45S_50ntFlanks@9575.9597.23|ATCGGTCGTCCGACCTGGGTATA</td><td>>hg38_21_+_8433172_8446622_RNA45SN1_45S_50ntFlanks</td><td>9575</td><td>9597</td><td>23</td><td>ATCGGTCGTCCGACCTGGGTATA</td><td>1610.0</td><td>-9.9</td><td>1.1</td><td>-8.7</td><td>4.2E-18</td><td>4.6E-17</td><td>MB231/MB468</td><td>CYTO</td></tr>
<tr><td>>hg38_21_+_8433172_8446622_RNA45SN1_45S_50ntFlanks@8170.8213.44|CAAGTCCTTCTGATCGAGGCCCAGCCCGTGGACGGTGTGAGGCC</td><td>>hg38_21_+_8433172_8446622_RNA45SN1_45S_50ntFlanks</td><td>8170</td><td>8213</td><td>44</td><td>CAAGTCCTTCTGATCGAGGCCCAGCCCGTGGACGGTGTGAGGCC</td><td>1695.6</td><td>-6.8</td><td>0.8</td><td>-8.6</td><td>6.9E-18</td><td>4.6E-17</td><td>MB231/MB468</td><td>MITO</td></tr>
<tr><td>>hg38_21_+_8433172_8446622_RNA45SN1_45S_50ntFlanks@10845.10865.21|ACTTCGGGATAAGGATTGGCT</td><td>>hg38_21_+_8433172_8446622_RNA45SN1_45S_50ntFlanks</td><td>10845</td><td>10865</td><td>21</td><td>ACTTCGGGATAAGGATTGGCT</td><td>962.8</td><td>-5.1</td><td>0.6</td><td>-8.6</td><td>5.8E-18</td><td>4.6E-17</td><td>BT20/MB468</td><td>MITO</td></tr>
<tr><td>>hg38_21_+_8433172_8446622_RNA45SN1_45S_50ntFlanks@4657.4686.30|CTAGAGGTGAAATTCTTGGACCGGCGCAAG</td><td>>hg38_21_+_8433172_8446622_RNA45SN1_45S_50ntFlanks</td><td>4657</td><td>4686</td><td>30</td><td>CTAGAGGTGAAATTCTTGGACCGGCGCAAG</td><td>878.1</td><td>-13.6</td><td>1.6</td><td>-8.6</td><td>6.9E-18</td><td>4.6E-17</td><td>MB231/MB468</td><td>MITO</td></tr>
<tr><td>>MI0003188|hsa-mir-503&WithFlank&X|-|134546322|134546404@12.31.20&[MIMAT0002874&hsa-miR-503-5p&offsets|0|-3;m-297&X|-|134546371|134546393&offsets|0|-3]|TAGCAGCGGGAACAGTTCTG</td><td>>MI0003188|hsa-mir-503&WithFlank&X|-|134546322|134546404</td><td>12</td><td>31</td><td>20</td><td>TAGCAGCGGGAACAGTTCTG</td><td>85.4</td><td>6.8</td><td>0.8</td><td>8.5</td><td>1.4E-17</td><td>4.7E-17</td><td>BT20/MB468</td><td>NUC</td></tr>
<tr><td>>hg38_21_+_8433172_8446622_RNA45SN1_45S_50ntFlanks@12008.12030.23|CCGCCGGTGAAATACCACTACTC</td><td>>hg38_21_+_8433172_8446622_RNA45SN1_45S_50ntFlanks</td><td>12008</td><td>12030</td><td>23</td><td>CCGCCGGTGAAATACCACTACTC</td><td>564.4</td><td>-12.1</td><td>1.4</td><td>-8.6</td><td>5.5E-18</td><td>4.7E-17</td><td>MB231/MB468</td><td>NUC</td></tr>
<tr><td>>hg38_21_+_8433172_8446622_RNA45SN1_45S_50ntFlanks@4445.4471.27|CTTGCCTCTCGGCGCCCCCTCGATGCT</td><td>>hg38_21_+_8433172_8446622_RNA45SN1_45S_50ntFlanks</td><td>4445</td><td>4471</td><td>27</td><td>CTTGCCTCTCGGCGCCCCCTCGATGCT</td><td>724.2</td><td>-12.5</td><td>1.4</td><td>-8.6</td><td>5.5E-18</td><td>4.7E-17</td><td>MB231/MB468</td><td>NUC</td></tr>
<tr><td>>hg38_21_+_8433172_8446622_RNA45SN1_45S_50ntFlanks@11668.11687.20|TGCCCAGTGCTCTGAATGTC</td><td>>hg38_21_+_8433172_8446622_RNA45SN1_45S_50ntFlanks</td><td>11668</td><td>11687</td><td>20</td><td>TGCCCAGTGCTCTGAATGTC</td><td>2427.6</td><td>6.4</td><td>0.7</td><td>8.7</td><td>4.2E-18</td><td>4.7E-17</td><td>MB231/MB468</td><td>CYTO</td></tr>
<tr><td>>hg38_21_+_8433172_8446622_RNA45SN1_45S_50ntFlanks@3900.3920.21|CTTCGCGGGGGGGATGCGTGC</td><td>>hg38_21_+_8433172_8446622_RNA45SN1_45S_50ntFlanks</td><td>3900</td><td>3920</td><td>21</td><td>CTTCGCGGGGGGGATGCGTGC</td><td>566.4</td><td>-13.0</td><td>1.5</td><td>-8.7</td><td>4.3E-18</td><td>4.7E-17</td><td>MB231/MB468</td><td>MP</td></tr>
<tr><td>>hg38_21_+_8433172_8446622_RNA45SN1_45S_50ntFlanks@9574.9598.25|AATCGGTCGTCCGACCTGGGTATAG</td><td>>hg38_21_+_8433172_8446622_RNA45SN1_45S_50ntFlanks</td><td>9574</td><td>9598</td><td>25</td><td>AATCGGTCGTCCGACCTGGGTATAG</td><td>337.1</td><td>-12.3</td><td>1.4</td><td>-8.7</td><td>4.3E-18</td><td>4.7E-17</td><td>MB231/MB468</td><td>MP</td></tr>
<tr><td>>hg38_21_+_8433172_8446622_RNA45SN1_45S_50ntFlanks@10726.10752.27|GGGAGAGGGTGTAAATCTCGCGCCGGG</td><td>>hg38_21_+_8433172_8446622_RNA45SN1_45S_50ntFlanks</td><td>10726</td><td>10752</td><td>27</td><td>GGGAGAGGGTGTAAATCTCGCGCCGGG</td><td>387.3</td><td>-12.5</td><td>1.4</td><td>-8.7</td><td>4.3E-18</td><td>4.7E-17</td><td>MB231/MB468</td><td>MP</td></tr>
<tr><td>>MI0000767|hsa-mir-365a&WithFlank&16|+|14309279|14309377@62.82.21;MI0000769|hsa-mir-365b&WithFlank&17|+|31575405|31575527@74.94.21&[MIMAT0000710&hsa-miR-365a-3p&offsets|0|-1;m-176&16|+|14309340|14309361&offsets|0|-1];[MIMAT0022834&hsa-miR-365b-3p&offsets|0|-1;m-175&17|+|31575478|31575499&offsets|0|-1]|TAATGCCCCTAAAAATCCTTA</td><td>>MI0000767|hsa-mir-365a&WithFlank&16|+|14309279|14309377</td><td>62</td><td>82</td><td>21</td><td>TAATGCCCCTAAAAATCCTTA</td><td>941.7</td><td>11.5</td><td>1.3</td><td>8.6</td><td>8.2E-18</td><td>4.7E-17</td><td>BT20/MB468</td><td>CYTO</td></tr>
<tr><td>>MI0000102|hsa-mir-100&WithFlank&11|-|122152223|122152314@19.38.20(+1G)&[MIMAT0000098&hsa-miR-100-5p&offsets|0|-2(+1G);m-27&11|-|122152276|122152296&offsets|0|-1(+1G)]|AACCCGTAGATCCGAACTTGG</td><td>>MI0000102|hsa-mir-100&WithFlank&11|-|122152223|122152314</td><td>19</td><td>38</td><td>20(+1G)</td><td>AACCCGTAGATCCGAACTTGG</td><td>1691.3</td><td>13.9</td><td>1.6</td><td>8.6</td><td>7.1E-18</td><td>4.7E-17</td><td>MB231/MB468</td><td>MITO</td></tr>
<tr><td>>hg38_MT_+_598_1651_RNR1_12S_50ntFlanks@916.950.35|AAGGACATTTAACTAAAACCCCTACGCATTTATAT</td><td>>hg38_MT_+_598_1651_RNR1_12S_50ntFlanks</td><td>916</td><td>950</td><td>35</td><td>AAGGACATTTAACTAAAACCCCTACGCATTTATAT</td><td>412.3</td><td>13.1</td><td>1.5</td><td>8.6</td><td>6.0E-18</td><td>4.8E-17</td><td>BT20/MB468</td><td>MITO</td></tr>
<tr><td>>MI0000462|hsa-mir-152&WithFlank&17|-|48037155|48037253@60.81.22&[MIMAT0000438&hsa-miR-152-3p&offsets|0|+1;m-97&17|-|48037174|48037194&offsets|0|+1]|TCAGTGCATGACAGAACTTGGG</td><td>>MI0000462|hsa-mir-152&WithFlank&17|-|48037155|48037253</td><td>60</td><td>81</td><td>22</td><td>TCAGTGCATGACAGAACTTGGG</td><td>421.7</td><td>12.7</td><td>1.5</td><td>8.7</td><td>4.3E-18</td><td>4.8E-17</td><td>MB231/MB468</td><td>CYTO</td></tr>
<tr><td>>hg38_21_+_8433172_8446622_RNA45SN1_45S_50ntFlanks@12973.13001.29|TCGTACGTAGCAGAGCAGCTCCCTCGCTG</td><td>>hg38_21_+_8433172_8446622_RNA45SN1_45S_50ntFlanks</td><td>12973</td><td>13001</td><td>29</td><td>TCGTACGTAGCAGAGCAGCTCCCTCGCTG</td><td>281.3</td><td>-12.0</td><td>1.4</td><td>-8.7</td><td>4.4E-18</td><td>4.8E-17</td><td>MB231/MB468</td><td>MP</td></tr>
<tr><td>>hg38_21_+_8433172_8446622_RNA45SN1_45S_50ntFlanks@5379.5419.41|ATTAAGTCCCTGCCCTTTGTACACACCGCCCGTCGCTACTA</td><td>>hg38_21_+_8433172_8446622_RNA45SN1_45S_50ntFlanks</td><td>5379</td><td>5419</td><td>41</td><td>ATTAAGTCCCTGCCCTTTGTACACACCGCCCGTCGCTACTA</td><td>745.3</td><td>-13.3</td><td>1.5</td><td>-8.6</td><td>7.2E-18</td><td>4.8E-17</td><td>MB231/MB468</td><td>MITO</td></tr>
<tr><td>>hg38_21_+_8433172_8446622_RNA45SN1_45S_50ntFlanks@9714.9735.22|CGAATGATTAGAGGTCTTGGGG</td><td>>hg38_21_+_8433172_8446622_RNA45SN1_45S_50ntFlanks</td><td>9714</td><td>9735</td><td>22</td><td>CGAATGATTAGAGGTCTTGGGG</td><td>1045.0</td><td>-13.8</td><td>1.6</td><td>-8.6</td><td>7.3E-18</td><td>4.8E-17</td><td>MB231/MB468</td><td>MITO</td></tr>
<tr><td>>MI0000299|hsa-mir-222&WithFlank&X|-|45747009|45747130@75.97.23&[MIMAT0000279&hsa-miR-222-3p&offsets|0|+2;m-140&X|-|45747033|45747056&offsets|0|-1]|AGCTACATCTGGCTACTGGGTCT</td><td>>MI0000299|hsa-mir-222&WithFlank&X|-|45747009|45747130</td><td>75</td><td>97</td><td>23</td><td>AGCTACATCTGGCTACTGGGTCT</td><td>5214.6</td><td>7.2</td><td>0.8</td><td>8.7</td><td>4.4E-18</td><td>4.8E-17</td><td>MB231/MB468</td><td>CYTO</td></tr>
</table>

<table>
<tr><td>>hg38_21_+_8433172_8446622_RNA45SN1_45S_50ntFlanks@10351.10382.32|CGAGAACTTTGAAGGCCGAAGTGGAGAAGGGT</td><td>>hg38_21_+_8433172_8446622_RNA45SN1_45S_50ntFlanks</td><td>10351</td><td>10382</td><td>32</td><td>CGAGAACTTTGAAGGCCGAAGTGGAGAAGGGT</td><td>2471.9</td><td>-5.1</td><td>0.6</td><td>-8.6</td><td>5.7E-18</td><td>4.8E-17</td><td>MB231/MB468</td><td>NUC</td></tr>
<tr><td>>MI0005566|hsa-mir-374b&WithFlank&X|-|74218541|74218624@17.37.21&[MIMAT0004955&hsa-miR-374b-5p&offsets|0|-1;m-153&X|-|74218587|74218608&offsets|0|-1]|ATATAATACAACCTGCTAAGT</td><td>>MI0005566|hsa-mir-374b&WithFlank&X|-|74218541|74218624</td><td>17</td><td>37</td><td>21</td><td>ATATAATACAACCTGCTAAGT</td><td>1055.7</td><td>7.2</td><td>0.8</td><td>8.6</td><td>8.4E-18</td><td>4.9E-17</td><td>BT20/MB468</td><td>CYTO</td></tr>
<tr><td>>hg38_21_+_8433172_8446622_RNA45SN1_45S_50ntFlanks@8154.8183.30|CCGCTCGTGGGGGGCCCAAGTCCTTCTGAT</td><td>>hg38_21_+_8433172_8446622_RNA45SN1_45S_50ntFlanks</td><td>8154</td><td>8183</td><td>30</td><td>CCGCTCGTGGGGGGCCCAAGTCCTTCTGAT</td><td>520.9</td><td>-12.8</td><td>1.5</td><td>-8.6</td><td>7.3E-18</td><td>4.9E-17</td><td>MB231/MB468</td><td>MITO</td></tr>
<tr><td>>hg38_21_+_8433172_8446622_RNA45SN1_45S_50ntFlanks@10073.10092.20|GACGGGAGCGGCGGGGGCGG</td><td>>hg38_21_+_8433172_8446622_RNA45SN1_45S_50ntFlanks</td><td>10073</td><td>10092</td><td>20</td><td>GACGGGAGCGGCGGGGGCGG</td><td>587.1</td><td>-13.1</td><td>1.5</td><td>-8.7</td><td>4.4E-18</td><td>4.9E-17</td><td>MB231/MB468</td><td>MP</td></tr>
<tr><td>>hg38_21_+_8433172_8446622_RNA45SN1_45S_50ntFlanks@4050.4081.32|CGAACGTCTGCCCTATCAACTTTCGATGGTAG</td><td>>hg38_21_+_8433172_8446622_RNA45SN1_45S_50ntFlanks</td><td>4050</td><td>4081</td><td>32</td><td>CGAACGTCTGCCCTATCAACTTTCGATGGTAG</td><td>526.0</td><td>-12.8</td><td>1.5</td><td>-8.6</td><td>7.4E-18</td><td>4.9E-17</td><td>MB231/MB468</td><td>MITO</td></tr>
<tr><td>>hg38_21_+_8433172_8446622_RNA45SN1_45S_50ntFlanks@6700.6720.21|CTAGCTGCGAGAATTAATGTG</td><td>>hg38_21_+_8433172_8446622_RNA45SN1_45S_50ntFlanks</td><td>6700</td><td>6720</td><td>21</td><td>CTAGCTGCGAGAATTAATGTG</td><td>21231.3</td><td>-7.3</td><td>0.8</td><td>-8.7</td><td>4.5E-18</td><td>4.9E-17</td><td>MB231/MB468</td><td>MP</td></tr>
<tr><td>>MI0000650|hsa-mir-200c&WithFlank&12|+|6963693|6963772@50.70.21(+1U)&[MIMAT0000617&hsa-miR-200c-3p&offsets|0|-2(+1U);m-54&12|+|6963742|6963764&offsets|0|-2(+1U)]|TAATACTGCCGGGTAATGATGT</td><td>>MI0000650|hsa-mir-200c&WithFlank&12|+|6963693|6963772</td><td>50</td><td>70</td><td>21(+1U)</td><td>TAATACTGCCGGGTAATGATGT</td><td>545.9</td><td>12.9</td><td>1.4</td><td>8.9</td><td>7.2E-19</td><td>4.9E-17</td><td>BT20/MB231</td><td>NUC</td></tr>
<tr><td>>MI0000068|hsa-let-7f-2&WithFlank&X|-|53557186|53557280@14.36.23&[MIMAT0000067_1&hsa-let-7f-5p&offsets|0|+1;m-19&X|-|53557246|53557267&offsets|0|+1]|TGAGGTAGTAGATTGTATAGTTT</td><td>>MI0000068|hsa-let-7f-2&WithFlank&X|-|53557186|53557280</td><td>14</td><td>36</td><td>23</td><td>TGAGGTAGTAGATTGTATAGTTT</td><td>1800.7</td><td>4.4</td><td>0.5</td><td>8.7</td><td>4.5E-18</td><td>4.9E-17</td><td>MB231/MB468</td><td>CYTO</td></tr>
<tr><td>>hg38_21_+_8433172_8446622_RNA45SN1_45S_50ntFlanks@12437.12455.19|CATTGTGAAGCAGAATTCA</td><td>>hg38_21_+_8433172_8446622_RNA45SN1_45S_50ntFlanks</td><td>12437</td><td>12455</td><td>19</td><td>CATTGTGAAGCAGAATTCA</td><td>26753.2</td><td>-12.6</td><td>1.5</td><td>-8.7</td><td>4.5E-18</td><td>5.0E-17</td><td>MB231/MB468</td><td>MP</td></tr>
<tr><td>>hg38_21_+_8433172_8446622_RNA45SN1_45S_50ntFlanks@5072.5112.41|TAATTCCGATAACGAACGAGACTCTGGCATGCTAACTAGTT</td><td>>hg38_21_+_8433172_8446622_RNA45SN1_45S_50ntFlanks</td><td>5072</td><td>5112</td><td>41</td><td>TAATTCCGATAACGAACGAGACTCTGGCATGCTAACTAGTT</td><td>270.2</td><td>12.1</td><td>1.4</td><td>8.7</td><td>4.5E-18</td><td>5.0E-17</td><td>MB231/MB468</td><td>CYTO</td></tr>
<tr><td>>hg38_21_+_8433172_8446622_RNA45SN1_45S_50ntFlanks@5484.5518.35|GAGCGCTGAGAAGACGGTCGAACTTGACTATCTAG</td><td>>hg38_21_+_8433172_8446622_RNA45SN1_45S_50ntFlanks</td><td>5484</td><td>5518</td><td>35</td><td>GAGCGCTGAGAAGACGGTCGAACTTGACTATCTAG</td><td>456.0</td><td>-12.6</td><td>1.5</td><td>-8.6</td><td>7.5E-18</td><td>5.0E-17</td><td>MB231/MB468</td><td>MITO</td></tr>
<tr><td>>hg38_21_+_8433172_8446622_RNA45SN1_45S_50ntFlanks@11768.11786.19|CTAATTAGTGACGCGCATG</td><td>>hg38_21_+_8433172_8446622_RNA45SN1_45S_50ntFlanks</td><td>11768</td><td>11786</td><td>19</td><td>CTAATTAGTGACGCGCATG</td><td>841.4</td><td>-12.7</td><td>1.5</td><td>-8.6</td><td>5.8E-18</td><td>5.0E-17</td><td>MB231/MB468</td><td>NUC</td></tr>
<tr><td>>hg38_21_+_8433172_8446622_RNA45SN1_45S_50ntFlanks@3705.3742.38|TACCTGGTTGATCCTGCCAGTAGCATATGCTTGTCTCA</td><td>>hg38_21_+_8433172_8446622_RNA45SN1_45S_50ntFlanks</td><td>3705</td><td>3742</td><td>38</td><td>TACCTGGTTGATCCTGCCAGTAGCATATGCTTGTCTCA</td><td>1577.6</td><td>5.3</td><td>0.6</td><td>8.7</td><td>4.5E-18</td><td>5.0E-17</td><td>MB231/MB468</td><td>CYTO</td></tr>
<tr><td>>hg38_21_+_8433172_8446622_RNA45SN1_45S_50ntFlanks@11768.11790.23|CTAATTAGTGACGCGCATGAATG</td><td>>hg38_21_+_8433172_8446622_RNA45SN1_45S_50ntFlanks</td><td>11768</td><td>11790</td><td>23</td><td>CTAATTAGTGACGCGCATGAATG</td><td>2128.2</td><td>-7.1</td><td>0.8</td><td>-8.7</td><td>4.6E-18</td><td>5.0E-17</td><td>MB231/MB468</td><td>CYTO</td></tr>
<tr><td>>hg38_21_+_8433172_8446622_RNA45SN1_45S_50ntFlanks@476.506.31|TTCGTGATCGATGTGGTGACGTCGTGCTCTC</td><td>>hg38_21_+_8433172_8446622_RNA45SN1_45S_50ntFlanks</td><td>476</td><td>506</td><td>31</td><td>TTCGTGATCGATGTGGTGACGTCGTGCTCTC</td><td>1451.8</td><td>8.5</td><td>1.0</td><td>8.6</td><td>8.7E-18</td><td>5.0E-17</td><td>BT20/MB468</td><td>CYTO</td></tr>
<tr><td>>hg38_21_+_8433172_8446622_RNA45SN1_45S_50ntFlanks@3830.3872.43|GCTCGCTCCTCTCCTACTTGGATAACTGTGGTAATTCTAGAGC</td><td>>hg38_21_+_8433172_8446622_RNA45SN1_45S_50ntFlanks</td><td>3830</td><td>3872</td><td>43</td><td>GCTCGCTCCTCTCCTACTTGGATAACTGTGGTAATTCTAGAGC</td><td>580.5</td><td>-13.6</td><td>1.6</td><td>-8.5</td><td>1.5E-17</td><td>5.0E-17</td><td>BT20/MB468</td><td>NUC</td></tr>
<tr><td>>hg38_21_+_8433172_8446622_RNA45SN1_45S_50ntFlanks@3927.3959.33|CAGATCAAAACCAACCCGGTCAGCCCCTCTCCG</td><td>>hg38_21_+_8433172_8446622_RNA45SN1_45S_50ntFlanks</td><td>3927</td><td>3959</td><td>33</td><td>CAGATCAAAACCAACCCGGTCAGCCCCTCTCCG</td><td>440.3</td><td>-12.6</td><td>1.5</td><td>-8.6</td><td>7.6E-18</td><td>5.1E-17</td><td>MB231/MB468</td><td>MITO</td></tr>
<tr><td>>hg38_21_+_8433172_8446622_RNA45SN1_45S_50ntFlanks@7986.8009.24|ATCAGACGTGGCGACCCGCTGAAT</td><td>>hg38_21_+_8433172_8446622_RNA45SN1_45S_50ntFlanks</td><td>7986</td><td>8009</td><td>24</td><td>ATCAGACGTGGCGACCCGCTGAAT</td><td>506.7</td><td>-12.8</td><td>1.5</td><td>-8.6</td><td>7.6E-18</td><td>5.1E-17</td><td>MB231/MB468</td><td>MITO</td></tr>
<tr><td>>hg38_21_+_8433172_8446622_RNA45SN1_45S_50ntFlanks@8357.8391.35|AAAAGAACTTTGAAGAGAGAGTTCAAGAGGGCGTG</td><td>>hg38_21_+_8433172_8446622_RNA45SN1_45S_50ntFlanks</td><td>8357</td><td>8391</td><td>35</td><td>AAAAGAACTTTGAAGAGAGAGTTCAAGAGGGCGTG</td><td>979.9</td><td>-6.5</td><td>0.8</td><td>-8.7</td><td>4.6E-18</td><td>5.1E-17</td><td>MB231/MB468</td><td>MP</td></tr>
<tr><td>>hg38_21_+_8433172_8446622_RNA45SN1_45S_50ntFlanks@7986.8005.20|ATCAGACGTGGCGACCCGCT</td><td>>hg38_21_+_8433172_8446622_RNA45SN1_45S_50ntFlanks</td><td>7986</td><td>8005</td><td>20</td><td>ATCAGACGTGGCGACCCGCT</td><td>1228.9</td><td>-12.4</td><td>1.4</td><td>-8.6</td><td>6.4E-18</td><td>5.1E-17</td><td>BT20/MB468</td><td>MITO</td></tr>
<tr><td>>hg38_21_+_8433172_8446622_RNA45SN1_45S_50ntFlanks@10349.10376.28|AACGAGAACTTTGAAGGCCGAAGTGGAG</td><td>>hg38_21_+_8433172_8446622_RNA45SN1_45S_50ntFlanks</td><td>10349</td><td>10376</td><td>28</td><td>AACGAGAACTTTGAAGGCCGAAGTGGAG</td><td>1151.2</td><td>-4.8</td><td>0.6</td><td>-8.6</td><td>6.0E-18</td><td>5.1E-17</td><td>MB231/MB468</td><td>NUC</td></tr>
<tr><td>>hg38_21_+_8433172_8446622_RNA45SN1_45S_50ntFlanks@10571.10591.21|GAGAAGCCGGCGGGAGCCCCG</td><td>>hg38_21_+_8433172_8446622_RNA45SN1_45S_50ntFlanks</td><td>10571</td><td>10591</td><td>21</td><td>GAGAAGCCGGCGGGAGCCCCG</td><td>598.3</td><td>-13.0</td><td>1.5</td><td>-8.6</td><td>7.7E-18</td><td>5.1E-17</td><td>MB231/MB468</td><td>MITO</td></tr>
<tr><td>>trnaMT_ValTAC_MT_+_1602_1670@45.72.28|TTTCAACTTAACTTGACCGCTCTGACCA</td><td>>trnaMT_ValTAC_MT_+_1602_1670</td><td>45</td><td>72</td><td>28</td><td>TTTCAACTTAACTTGACCGCTCTGACCA</td><td>424.3</td><td>12.7</td><td>1.5</td><td>8.7</td><td>4.6E-18</td><td>5.1E-17</td><td>MB231/MB468</td><td>CYTO</td></tr>
<tr><td>>hg38_21_+_8433172_8446622_RNA45SN1_45S_50ntFlanks@9564.9596.33|CTGACGTGCAAATCGGTCGTCCGACCTGGGTAT</td><td>>hg38_21_+_8433172_8446622_RNA45SN1_45S_50ntFlanks</td><td>9564</td><td>9596</td><td>33</td><td>CTGACGTGCAAATCGGTCGTCCGACCTGGGTAT</td><td>14627.3</td><td>-6.8</td><td>0.8</td><td>-8.6</td><td>8.8E-18</td><td>5.1E-17</td><td>BT20/MB468</td><td>CYTO</td></tr>
<tr><td>>MI0000477|hsa-mir-146a&WithFlank&5|+|160485346|160485456@27.47.21&[MIMAT0000449&hsa-miR-146a-5p&offsets|0|-1;m-129&5|+|160485372|160485393&offsets|0|-1]|TGAGAACTGAATTCCATGGGT</td><td>>MI0000477|hsa-mir-146a&WithFlank&5|+|160485346|160485456</td><td>27</td><td>47</td><td>21</td><td>TGAGAACTGAATTCCATGGGT</td><td>1991.4</td><td>-14.0</td><td>1.6</td><td>-8.9</td><td>7.6E-19</td><td>5.1E-17</td><td>BT20/MB231</td><td>NUC</td></tr>
<tr><td>>MI0000086|hsa-mir-28&WithFlank&3|+|188688775|188688872@61.81.21(+1A)&[MIMAT0004502&hsa-miR-28-3p&offsets|+1|0(+1A);m-28&3|+|188688834|188688855&offsets|+1|0(+1A)]|ACTAGATTGTGAGCTCCTGGAA</td><td>>MI0000086|hsa-mir-28&WithFlank&3|+|188688775|188688872</td><td>61</td><td>81</td><td>21(+1A)</td><td>ACTAGATTGTGAGCTCCTGGAA</td><td>417.2</td><td>13.0</td><td>1.5</td><td>8.6</td><td>8.9E-18</td><td>5.1E-17</td><td>BT20/MB468</td><td>CYTO</td></tr>
<tr><td>>hg38_1_-_228634819_228635039_RNA5S12_5S_50ntFlanks@130.169.40|TGGATGGGAGACCGCCTGGGAATACCGGGTGCTGTAGGCT</td><td>>hg38_1_-_228634819_228635039_RNA5S12_5S_50ntFlanks</td><td>130</td><td>169</td><td>40</td><td>TGGATGGGAGACCGCCTGGGAATACCGGGTGCTGTAGGCT</td><td>2973.2</td><td>10.2</td><td>1.2</td><td>8.6</td><td>8.9E-18</td><td>5.1E-17</td><td>BT20/MB468</td><td>CYTO</td></tr>
<tr><td>>trnaMT_ValTAC_MT_+_1602_1670@45.71.27|TTTCAACTTAACTTGACCGCTCTGACC</td><td>>trnaMT_ValTAC_MT_+_1602_1670</td><td>45</td><td>71</td><td>27</td><td>TTTCAACTTAACTTGACCGCTCTGACC</td><td>418.0</td><td>12.7</td><td>1.5</td><td>8.7</td><td>4.7E-18</td><td>5.2E-17</td><td>MB231/MB468</td><td>CYTO</td></tr>
<tr><td>>hg38_21_+_8433172_8446622_RNA45SN1_45S_50ntFlanks@11668.11704.37|TGCCCAGTGCTCTGAATGTCAAAGTGAAGAAATTCAA</td><td>>hg38_21_+_8433172_8446622_RNA45SN1_45S_50ntFlanks</td><td>11668</td><td>11704</td><td>37</td><td>TGCCCAGTGCTCTGAATGTCAAAGTGAAGAAATTCAA</td><td>625.3</td><td>5.0</td><td>0.6</td><td>8.7</td><td>4.7E-18</td><td>5.2E-17</td><td>MB231/MB468</td><td>CYTO</td></tr>
<tr><td>>trnaMT_ProTGG_MT_-_15956_16023@44.70.27|GTGGAGTTAAAGACTTTTTCTCTGACC</td><td>>trnaMT_ProTGG_MT_-_15956_16023</td><td>44</td><td>70</td><td>27</td><td>GTGGAGTTAAAGACTTTTTCTCTGACC</td><td>177.1</td><td>11.4</td><td>1.3</td><td>8.6</td><td>6.1E-18</td><td>5.2E-17</td><td>MB231/MB468</td><td>NUC</td></tr>
<tr><td>>hg38_21_+_8433172_8446622_RNA45SN1_45S_50ntFlanks@7980.8005.26|CCTCAGATCAGACGTGGCGACCCGCT</td><td>>hg38_21_+_8433172_8446622_RNA45SN1_45S_50ntFlanks</td><td>7980</td><td>8005</td><td>26</td><td>CCTCAGATCAGACGTGGCGACCCGCT</td><td>797.9</td><td>-4.8</td><td>0.6</td><td>-8.5</td><td>1.5E-17</td><td>5.2E-17</td><td>BT20/MB468</td><td>NUC</td></tr>
<tr><td>>trnaMT_PheGAA_MT_+_577_647@36.55.20|AAAATGTTTAGACGGGCTCA</td><td>>trnaMT_PheGAA_MT_+_577_647</td><td>36</td><td>55</td><td>20</td><td>AAAATGTTTAGACGGGCTCA</td><td>1204.1</td><td>9.0</td><td>1.0</td><td>8.6</td><td>6.5E-18</td><td>5.2E-17</td><td>BT20/MB468</td><td>MITO</td></tr>
<tr><td>>hg38_21_+_8433172_8446622_RNA45SN1_45S_50ntFlanks@4995.5022.28|ACAGATTGATAGCTCTTTCTCGATTCCG</td><td>>hg38_21_+_8433172_8446622_RNA45SN1_45S_50ntFlanks</td><td>4995</td><td>5022</td><td>28</td><td>ACAGATTGATAGCTCTTTCTCGATTCCG</td><td>1884.1</td><td>-15.3</td><td>1.8</td><td>-8.5</td><td>1.5E-17</td><td>5.2E-17</td><td>BT20/MB468</td><td>NUC</td></tr>
<tr><td>>hg38_21_+_8433172_8446622_RNA45SN1_45S_50ntFlanks@8105.8141.37|CGCGGCGGGGCGCGGGACATGTGGCGTACGGAAGACC</td><td>>hg38_21_+_8433172_8446622_RNA45SN1_45S_50ntFlanks</td><td>8105</td><td>8141</td><td>37</td><td>CGCGGCGGGGCGCGGGACATGTGGCGTACGGAAGACC</td><td>2223.4</td><td>-7.5</td><td>0.9</td><td>-8.6</td><td>7.9E-18</td><td>5.2E-17</td><td>MB231/MB468</td><td>MITO</td></tr>
<tr><td>>MI0001445|hsa-mir-423&WithFlank&17|+|30117073|30117178@59.77.19(+1A)&[MIMAT0001340&hsa-miR-423-3p&offsets|0|-4(+1A);m-100&17|+|30117131|30117153&offsets|0|-4(+1A)]|AGCTCGGTCTGAGGCCCCTA</td><td>>MI0001445|hsa-mir-423&WithFlank&17|+|30117073|30117178</td><td>59</td><td>77</td><td>19(+1A)</td><td>AGCTCGGTCTGAGGCCCCTA</td><td>426.2</td><td>10.4</td><td>1.2</td><td>8.6</td><td>9.1E-18</td><td>5.2E-17</td><td>BT20/MB468</td><td>CYTO</td></tr>
<tr><td>>hg38_21_+_8433172_8446622_RNA45SN1_45S_50ntFlanks@12630.12654.25|GGGCGAAGCTACCATCTGTGGGATT</td><td>>hg38_21_+_8433172_8446622_RNA45SN1_45S_50ntFlanks</td><td>12630</td><td>12654</td><td>25</td><td>GGGCGAAGCTACCATCTGTGGGATT</td><td>741.2</td><td>-13.3</td><td>1.5</td><td>-8.6</td><td>7.9E-18</td><td>5.2E-17</td><td>MB231/MB468</td><td>MITO</td></tr>
<tr><td>>hg38_21_+_8433172_8446622_RNA45SN1_45S_50ntFlanks@11672.11701.30|CAGTGCTCTGAATGTCAAAGTGAAGAAATT</td><td>>hg38_21_+_8433172_8446622_RNA45SN1_45S_50ntFlanks</td><td>11672</td><td>11701</td><td>30</td><td>CAGTGCTCTGAATGTCAAAGTGAAGAAATT</td><td>1225.8</td><td>-5.5</td><td>0.6</td><td>-8.5</td><td>1.6E-17</td><td>5.3E-17</td><td>BT20/MB468</td><td>NUC</td></tr>
<tr><td>>hg38_21_+_8433172_8446622_RNA45SN1_45S_50ntFlanks@5490.5531.42|TGAGAAGACGGTCGAACTTGACTATCTAGAGGAAGTAAAAGT</td><td>>hg38_21_+_8433172_8446622_RNA45SN1_45S_50ntFlanks</td><td>5490</td><td>5531</td><td>42</td><td>TGAGAAGACGGTCGAACTTGACTATCTAGAGGAAGTAAAAGT</td><td>730.3</td><td>13.5</td><td>1.6</td><td>8.7</td><td>4.8E-18</td><td>5.3E-17</td><td>MB231/MB468</td><td>CYTO</td></tr>
<tr><td>>hg38_21_+_8433172_8446622_RNA45SN1_45S_50ntFlanks@12437.12454.18|CATTGTGAAGCAGAATTC</td><td>>hg38_21_+_8433172_8446622_RNA45SN1_45S_50ntFlanks</td><td>12437</td><td>12454</td><td>18</td><td>CATTGTGAAGCAGAATTC</td><td>2566.2</td><td>-6.0</td><td>0.7</td><td>-8.6</td><td>6.2E-18</td><td>5.3E-17</td><td>MB231/MB468</td><td>NUC</td></tr>
<tr><td>>MI0000285|hsa-mir-205&WithFlank&1|+|209432127|209432248@40.61.22(+1G)&[MIMAT0000266&hsa-miR-205-5p&offsets|0|0(+1G);m-351&1|+|209432166|209432187&offsets|0|0(+1G)]|TCCTTCATTCCACCGGAGTCTGG</td><td>>MI0000285|hsa-mir-205&WithFlank&1|+|209432127|209432248</td><td>40</td><td>61</td><td>22(+1G)</td><td>TCCTTCATTCCACCGGAGTCTGG</td><td>700.4</td><td>13.1</td><td>1.5</td><td>8.9</td><td>6.0E-19</td><td>5.3E-17</td><td>BT20/MB231</td><td>MITO</td></tr>
<tr><td>>hg38_21_+_8433172_8446622_RNA45SN1_45S_50ntFlanks@12126.12148.23|CGACCCGCTCCGGGGACAGTGCC</td><td>>hg38_21_+_8433172_8446622_RNA45SN1_45S_50ntFlanks</td><td>12126</td><td>12148</td><td>23</td><td>CGACCCGCTCCGGGGACAGTGCC</td><td>589.4</td><td>-13.1</td><td>1.5</td><td>-8.7</td><td>4.8E-18</td><td>5.3E-17</td><td>MB231/MB468</td><td>MP</td></tr>
<tr><td>>MI0000767|hsa-mir-365a&WithFlank&16|+|14309279|14309377@62.84.23;MI0000769|hsa-mir-365b&WithFlank&17|+|31575405|31575527@74.96.23&[MIMAT0000710&hsa-miR-365a-3p&offsets|0|+1;m-176&16|+|14309340|14309361&offsets|0|+1];[MIMAT0022834&hsa-miR-365b-3p&offsets|0|+1;m-175&17|+|31575478|31575499&offsets|0|+1]|TAATGCCCCTAAAAATCCTTATT</td><td>>MI0000767|hsa-mir-365a&WithFlank&16|+|14309279|14309377</td><td>62</td><td>84</td><td>23</td><td>TAATGCCCCTAAAAATCCTTATT</td><td>696.0</td><td>8.0</td><td>0.9</td><td>8.6</td><td>6.6E-18</td><td>5.3E-17</td><td>BT20/MB468</td><td>MITO</td></tr>
<tr><td>>hg38_21_+_8433172_8446622_RNA45SN1_45S_50ntFlanks@10687.10712.26|CGGCGTCCGGTGAGCTCTCGCTGGCC</td><td>>hg38_21_+_8433172_8446622_RNA45SN1_45S_50ntFlanks</td><td>10687</td><td>10712</td><td>26</td><td>CGGCGTCCGGTGAGCTCTCGCTGGCC</td><td>1349.5</td><td>-6.6</td><td>0.8</td><td>-8.6</td><td>8.0E-18</td><td>5.3E-17</td><td>MB231/MB468</td><td>MITO</td></tr>
<tr><td>>hg38_21_+_8433172_8446622_RNA45SN1_45S_50ntFlanks@9204.9222.19|TCGGGGCCACGCGCGCGTC</td><td>>hg38_21_+_8433172_8446622_RNA45SN1_45S_50ntFlanks</td><td>9204</td><td>9222</td><td>19</td><td>TCGGGGCCACGCGCGCGTC</td><td>460.1</td><td>-12.6</td><td>1.5</td><td>-8.6</td><td>8.0E-18</td><td>5.3E-17</td><td>MB231/MB468</td><td>MITO</td></tr>
<tr><td>>hg38_21_+_8433172_8446622_RNA45SN1_45S_50ntFlanks@10679.10711.33|GGTTCCGGCGGCGTCCGGTGAGCTCTCGCTGGC</td><td>>hg38_21_+_8433172_8446622_RNA45SN1_45S_50ntFlanks</td><td>10679</td><td>10711</td><td>33</td><td>GGTTCCGGCGGCGTCCGGTGAGCTCTCGCTGGC</td><td>9929.4</td><td>-6.6</td><td>0.8</td><td>-8.6</td><td>6.7E-18</td><td>5.3E-17</td><td>BT20/MB468</td><td>MITO</td></tr>
<tr><td>>MI0000088|hsa-mir-30a&WithFlank&6|-|71403545|71403627@12.32.21&[MIMAT0000087&hsa-miR-30a-5p&offsets|0|-1;m-15&6|-|71403595|71403616&offsets|0|-1]|TGTAAACATCCTCGACTGGAA</td><td>>MI0000088|hsa-mir-30a&WithFlank&6|-|71403545|71403627</td><td>12</td><td>32</td><td>21</td><td>TGTAAACATCCTCGACTGGAA</td><td>327.1</td><td>5.3</td><td>0.6</td><td>8.6</td><td>6.7E-18</td><td>5.3E-17</td><td>BT20/MB468</td><td>MITO</td></tr>
<tr><td>>hg38_21_+_8433172_8446622_RNA45SN1_45S_50ntFlanks@12633.12654.22|CGAAGCTACCATCTGTGGGATT</td><td>>hg38_21_+_8433172_8446622_RNA45SN1_45S_50ntFlanks</td><td>12633</td><td>12654</td><td>22</td><td>CGAAGCTACCATCTGTGGGATT</td><td>5823.9</td><td>-10.0</td><td>1.2</td><td>-8.5</td><td>1.6E-17</td><td>5.3E-17</td><td>BT20/MB468</td><td>NUC</td></tr>
<tr><td>>MI0000078|hsa-mir-22&WithFlank&17|-|1713897|1713993@59.79.21(+1C)&[MIMAT0000077&hsa-miR-22-3p&offsets|0|-1(+1C);m-2&17|-|1713914|1713935&offsets|0|-1(+1C)]|AAGCTGCCAGTTGAAGAACTGC</td><td>>MI0000078|hsa-mir-22&WithFlank&17|-|1713897|1713993</td><td>59</td><td>79</td><td>21(+1C)</td><td>AAGCTGCCAGTTGAAGAACTGC</td><td>478.1</td><td>5.3</td><td>0.6</td><td>8.5</td><td>1.6E-17</td><td>5.3E-17</td><td>BT20/MB468</td><td>NUC</td></tr>
<tr><td>>hg38_21_+_8433172_8446622_RNA45SN1_45S_50ntFlanks@10791.10818.28|CCTCTGGCATGTTGGAACAATGTAGGTA</td><td>>hg38_21_+_8433172_8446622_RNA45SN1_45S_50ntFlanks</td><td>10791</td><td>10818</td><td>28</td><td>CCTCTGGCATGTTGGAACAATGTAGGTA</td><td>292.4</td><td>-12.1</td><td>1.4</td><td>-8.7</td><td>4.9E-18</td><td>5.4E-17</td><td>MB231/MB468</td><td>MP</td></tr>
<tr><td>>hg38_21_+_8433172_8446622_RNA45SN1_45S_50ntFlanks@5484.5514.31|GAGCGCTGAGAAGACGGTCGAACTTGACTAT</td><td>>hg38_21_+_8433172_8446622_RNA45SN1_45S_50ntFlanks</td><td>5484</td><td>5514</td><td>31</td><td>GAGCGCTGAGAAGACGGTCGAACTTGACTAT</td><td>12472.4</td><td>-12.1</td><td>1.4</td><td>-8.7</td><td>4.9E-18</td><td>5.4E-17</td><td>MB231/MB468</td><td>MP</td></tr>
</table>

<table>
<tr><td>>hg38_21_+_8433172_8446622_RNA45SN1_45S_50ntFlanks@11641.11662.22|CGGCGGGTGTTGACGCGATGTG</td><td>>hg38_21_+_8433172_8446622_RNA45SN1_45S_50ntFlanks</td><td>11641</td><td>11662</td><td>22</td><td>CGGCGGGTGTTGACGCGATGTG</td><td>9356.1</td><td>-11.5</td><td>1.3</td><td>-8.6</td><td>8.1E-18</td><td>5.4E-17</td><td>MB231/MB468</td><td>MITO</td></tr>
<tr><td>>hg38_21_+_8433172_8446622_RNA45SN1_45S_50ntFlanks@3831.3871.41|CTCGCTCCTCTCCTACTTGGATAACTGTGGTAATTCTAGAG</td><td>>hg38_21_+_8433172_8446622_RNA45SN1_45S_50ntFlanks</td><td>3831</td><td>3871</td><td>41</td><td>CTCGCTCCTCTCCTACTTGGATAACTGTGGTAATTCTAGAG</td><td>599.5</td><td>-13.7</td><td>1.6</td><td>-8.5</td><td>1.6E-17</td><td>5.4E-17</td><td>BT20/MB468</td><td>NUC</td></tr>
<tr><td>>hg38_21_+_8433172_8446622_RNA45SN1_45S_50ntFlanks@9576.9596.21|TCGGTCGTCCGACCTGGGTAT</td><td>>hg38_21_+_8433172_8446622_RNA45SN1_45S_50ntFlanks</td><td>9576</td><td>9596</td><td>21</td><td>TCGGTCGTCCGACCTGGGTAT</td><td>524.4</td><td>-12.8</td><td>1.5</td><td>-8.6</td><td>8.2E-18</td><td>5.4E-17</td><td>MB231/MB468</td><td>MITO</td></tr>
<tr><td>>hg38_21_+_8433172_8446622_RNA45SN1_45S_50ntFlanks@10411.10442.32|CAGTCGGTCCTGAGAGATGGGCGAGCGCCGTT</td><td>>hg38_21_+_8433172_8446622_RNA45SN1_45S_50ntFlanks</td><td>10411</td><td>10442</td><td>32</td><td>CAGTCGGTCCTGAGAGATGGGCGAGCGCCGTT</td><td>748.8</td><td>-5.1</td><td>0.6</td><td>-8.7</td><td>5.0E-18</td><td>5.4E-17</td><td>MB231/MB468</td><td>CYTO</td></tr>
<tr><td>>hg38_21_+_8433172_8446622_RNA45SN1_45S_50ntFlanks@11569.11605.37|CAGCCGACTTAGAACTGGTGCGGACCAGGGGAATCCG</td><td>>hg38_21_+_8433172_8446622_RNA45SN1_45S_50ntFlanks</td><td>11569</td><td>11605</td><td>37</td><td>CAGCCGACTTAGAACTGGTGCGGACCAGGGGAATCCG</td><td>1314.7</td><td>-14.2</td><td>1.6</td><td>-8.7</td><td>5.0E-18</td><td>5.4E-17</td><td>MB231/MB468</td><td>MP</td></tr>
<tr><td>>hg38_21_+_8433172_8446622_RNA45SN1_45S_50ntFlanks@12939.12956.18|CCATTCGTAGACGACCTG</td><td>>hg38_21_+_8433172_8446622_RNA45SN1_45S_50ntFlanks</td><td>12939</td><td>12956</td><td>18</td><td>CCATTCGTAGACGACCTG</td><td>1032.4</td><td>-7.8</td><td>0.9</td><td>-8.6</td><td>8.2E-18</td><td>5.4E-17</td><td>MB231/MB468</td><td>MITO</td></tr>
<tr><td>>hg38_21_+_8433172_8446622_RNA45SN1_45S_50ntFlanks@5527.5548.22|AAAGTCGTAACAAGGTTTCCGT</td><td>>hg38_21_+_8433172_8446622_RNA45SN1_45S_50ntFlanks</td><td>5527</td><td>5548</td><td>22</td><td>AAAGTCGTAACAAGGTTTCCGT</td><td>408.3</td><td>-12.5</td><td>1.4</td><td>-8.7</td><td>5.0E-18</td><td>5.4E-17</td><td>MB231/MB468</td><td>MP</td></tr>
<tr><td>>hg38_21_+_8433172_8446622_RNA45SN1_45S_50ntFlanks@12073.12095.23|CCGAGGGGCTCTCGCTTCTGGCG</td><td>>hg38_21_+_8433172_8446622_RNA45SN1_45S_50ntFlanks</td><td>12073</td><td>12095</td><td>23</td><td>CCGAGGGGCTCTCGCTTCTGGCG</td><td>1120.6</td><td>-9.9</td><td>1.2</td><td>-8.6</td><td>6.4E-18</td><td>5.5E-17</td><td>MB231/MB468</td><td>NUC</td></tr>
<tr><td>>hg38_21_+_8433172_8446622_RNA45SN1_45S_50ntFlanks@11215.11234.20|CTCTGGACGCGAGCCGGGCC</td><td>>hg38_21_+_8433172_8446622_RNA45SN1_45S_50ntFlanks</td><td>11215</td><td>11234</td><td>20</td><td>CTCTGGACGCGAGCCGGGCC</td><td>451.5</td><td>-8.3</td><td>1.0</td><td>-8.7</td><td>5.0E-18</td><td>5.5E-17</td><td>MB231/MB468</td><td>CYTO</td></tr>
<tr><td>>MI0000081|hsa-mir-24-2&WithFlank&19|-|13836281|13836365@56.77.22(+1U);MI0000080|hsa-mir-24-1&WithFlank&9|+|95086015|95086094@50.71.22(+1U)&[MIMAT0000080&hsa-miR-24-3p&offsets|0|0(+1U);m-36&19|-|13836288|13836310&offsets|0|-1(+1U)];[MIMAT0000080_1&hsa-miR-24-3p&offsets|0|0(+1U);m-35&9|+|95086064|95086086&offsets|0|-1(+1U)]|TGGCTCAGTTCAGCAGGAACAGT</td><td>>MI0000081|hsa-mir-24-2&WithFlank&19|-|13836281|13836365</td><td>56</td><td>77</td><td>22(+1U)</td><td>TGGCTCAGTTCAGCAGGAACAGT</td><td>6527.4</td><td>5.5</td><td>0.6</td><td>8.6</td><td>6.5E-18</td><td>5.5E-17</td><td>MB231/MB468</td><td>NUC</td></tr>
<tr><td>>MI0000447|hsa-mir-128-1&WithFlank&2|+|135665391|135665484@56.77.22&[MIMAT0000424&hsa-miR-128-3p&offsets|0|+1;m-108&2|+|135665446|135665466&offsets|0|+1]|TCACAGTGAACCGGTCTCTTTT</td><td>>MI0000447|hsa-mir-128-1&WithFlank&2|+|135665391|135665484</td><td>56</td><td>77</td><td>22</td><td>TCACAGTGAACCGGTCTCTTTT</td><td>3082.1</td><td>5.1</td><td>0.6</td><td>8.6</td><td>6.9E-18</td><td>5.5E-17</td><td>BT20/MB468</td><td>MITO</td></tr>
<tr><td>>hg38_21_+_8433172_8446622_RNA45SN1_45S_50ntFlanks@4297.4321.25|CATTGGAGGGCAAGTCTGGTGCCAG</td><td>>hg38_21_+_8433172_8446622_RNA45SN1_45S_50ntFlanks</td><td>4297</td><td>4321</td><td>25</td><td>CATTGGAGGGCAAGTCTGGTGCCAG</td><td>334.8</td><td>-11.3</td><td>1.3</td><td>-8.6</td><td>6.5E-18</td><td>5.5E-17</td><td>MB231/MB468</td><td>NUC</td></tr>
<tr><td>>hg38_21_+_8433172_8446622_RNA45SN1_45S_50ntFlanks@10805.10826.22|GAACAATGTAGGTAAGGGAAGT</td><td>>hg38_21_+_8433172_8446622_RNA45SN1_45S_50ntFlanks</td><td>10805</td><td>10826</td><td>22</td><td>GAACAATGTAGGTAAGGGAAGT</td><td>195.6</td><td>-5.8</td><td>0.7</td><td>-8.7</td><td>5.1E-18</td><td>5.5E-17</td><td>MB231/MB468</td><td>CYTO</td></tr>
<tr><td>>MI0000087|hsa-mir-29a&WithFlank&7|-|130876741|130876816@47.68.22&[MIMAT0000086&hsa-miR-29a-3p&offsets|-1|-1;m-26&7|-|130876749|130876769&offsets|-1|0]|CTAGCACCATCTGAAATCGGTT</td><td>>MI0000087|hsa-mir-29a&WithFlank&7|-|130876741|130876816</td><td>47</td><td>68</td><td>22</td><td>CTAGCACCATCTGAAATCGGTT</td><td>3053.4</td><td>5.0</td><td>0.6</td><td>8.5</td><td>1.6E-17</td><td>5.5E-17</td><td>BT20/MB468</td><td>NUC</td></tr>
<tr><td>>hg38_21_+_8433172_8446622_RNA45SN1_45S_50ntFlanks@12934.12951.18|CTAAACCATTCGTAGACG</td><td>>hg38_21_+_8433172_8446622_RNA45SN1_45S_50ntFlanks</td><td>12934</td><td>12951</td><td>18</td><td>CTAAACCATTCGTAGACG</td><td>1600.1</td><td>-5.4</td><td>0.6</td><td>-8.7</td><td>5.1E-18</td><td>5.5E-17</td><td>MB231/MB468</td><td>CYTO</td></tr>
<tr><td>>MI0000433|hsa-let-7g&WithFlank&3|-|52268272|52268367@10.32.23&[MIMAT0000414&hsa-let-7g-5p&offsets|-1|0;m-58&3|-|52268336|52268357&offsets|-1|0]|CTGAGGTAGTAGTTTGTACAGTT</td><td>>MI0000433|hsa-let-7g&WithFlank&3|-|52268272|52268367</td><td>10</td><td>32</td><td>23</td><td>CTGAGGTAGTAGTTTGTACAGTT</td><td>519.9</td><td>6.4</td><td>0.7</td><td>8.6</td><td>9.6E-18</td><td>5.5E-17</td><td>BT20/MB468</td><td>CYTO</td></tr>
<tr><td>>hg38_21_+_8433172_8446622_RNA45SN1_45S_50ntFlanks@8737.8758.22|CCCGTCCGTCCGTCCGTCCGTC</td><td>>hg38_21_+_8433172_8446622_RNA45SN1_45S_50ntFlanks</td><td>8737</td><td>8758</td><td>22</td><td>CCCGTCCGTCCGTCCGTCCGTC</td><td>443.2</td><td>-12.6</td><td>1.5</td><td>-8.6</td><td>8.4E-18</td><td>5.5E-17</td><td>MB231/MB468</td><td>MITO</td></tr>
<tr><td>>hg38_21_+_8433172_8446622_RNA45SN1_45S_50ntFlanks@12456.12499.44|CCAAGCGTTGGATTGTTCACCCACTAATAGGGAACGTGAGCTGG</td><td>>hg38_21_+_8433172_8446622_RNA45SN1_45S_50ntFlanks</td><td>12456</td><td>12499</td><td>44</td><td>CCAAGCGTTGGATTGTTCACCCACTAATAGGGAACGTGAGCTGG</td><td>877.3</td><td>-14.2</td><td>1.7</td><td>-8.5</td><td>1.6E-17</td><td>5.6E-17</td><td>BT20/MB468</td><td>NUC</td></tr>
<tr><td>>am_trna134_GluTTC_1_-_16861774_16861845@1.29.29,trna5_GluTTC_1_+_17199078_17199149@1.29.29,trna84_GluTTC_1_-_161391883_161391954@1.29.29,trna94_GluTTC_1_-_149664355_149664427@1.29.29|TCCCTGGTGGTCTAGTGGCTAGGATTCGG</td><td>>am_trna134_GluTTC_1_-_16861774_16861845</td><td>1</td><td>29</td><td>29</td><td>TCCCTGGTGGTCTAGTGGCTAGGATTCGG</td><td>675.6</td><td>-13.3</td><td>1.5</td><td>-8.7</td><td>5.1E-18</td><td>5.6E-17</td><td>MB231/MB468</td><td>MP</td></tr>
<tr><td>>hg38_21_+_8433172_8446622_RNA45SN1_45S_50ntFlanks@11316.11336.21|CACCCCACGTCTCGTCGCGCG</td><td>>hg38_21_+_8433172_8446622_RNA45SN1_45S_50ntFlanks</td><td>11316</td><td>11336</td><td>21</td><td>CACCCCACGTCTCGTCGCGCG</td><td>430.9</td><td>-12.6</td><td>1.5</td><td>-8.7</td><td>5.1E-18</td><td>5.6E-17</td><td>MB231/MB468</td><td>MP</td></tr>
<tr><td>>hg38_21_+_8433172_8446622_RNA45SN1_45S_50ntFlanks@12633.12659.27|CGAAGCTACCATCTGTGGGATTATGAC</td><td>>hg38_21_+_8433172_8446622_RNA45SN1_45S_50ntFlanks</td><td>12633</td><td>12659</td><td>27</td><td>CGAAGCTACCATCTGTGGGATTATGAC</td><td>9618.9</td><td>-10.0</td><td>1.2</td><td>-8.6</td><td>8.5E-18</td><td>5.6E-17</td><td>MB231/MB468</td><td>MITO</td></tr>
<tr><td>>MI0000102|hsa-mir-100&WithFlank&11|-|122152223|122152314@19.40.22(+2U)&[MIMAT0000098&hsa-miR-100-5p&offsets|0|0(+2U);m-27&11|-|122152276|122152296&offsets|0|+1(+2U)]|AACCCGTAGATCCGAACTTGTGTT</td><td>>MI0000102|hsa-mir-100&WithFlank&11|-|122152223|122152314</td><td>19</td><td>40</td><td>22(+2U)</td><td>AACCCGTAGATCCGAACTTGTGTT</td><td>585.3</td><td>-12.3</td><td>1.4</td><td>-8.9</td><td>8.7E-19</td><td>5.7E-17</td><td>BT20/MB231</td><td>NUC</td></tr>
<tr><td>>hg38_21_+_8433172_8446622_RNA45SN1_45S_50ntFlanks@7973.8010.38|GACGCGACCTCAGATCAGACGTGGCGACCCGCTGAATT</td><td>>hg38_21_+_8433172_8446622_RNA45SN1_45S_50ntFlanks</td><td>7973</td><td>8010</td><td>38</td><td>GACGCGACCTCAGATCAGACGTGGCGACCCGCTGAATT</td><td>1747.4</td><td>8.1</td><td>0.9</td><td>8.6</td><td>5.3E-18</td><td>5.7E-17</td><td>MB231/MB468</td><td>CYTO</td></tr>
<tr><td>>hg38_21_+_8433172_8446622_RNA45SN1_45S_50ntFlanks@12419.12444.26|GATGTCGGCTCTTCCTATCATTGTGA</td><td>>hg38_21_+_8433172_8446622_RNA45SN1_45S_50ntFlanks</td><td>12419</td><td>12444</td><td>26</td><td>GATGTCGGCTCTTCCTATCATTGTGA</td><td>1153.9</td><td>-4.0</td><td>0.5</td><td>-8.5</td><td>1.7E-17</td><td>5.7E-17</td><td>BT20/MB468</td><td>NUC</td></tr>
<tr><td>>hg38_21_+_8433172_8446622_RNA45SN1_45S_50ntFlanks@11575.11603.29|ACTTAGAACTGGTGCGGACCAGGGGAATC</td><td>>hg38_21_+_8433172_8446622_RNA45SN1_45S_50ntFlanks</td><td>11575</td><td>11603</td><td>29</td><td>ACTTAGAACTGGTGCGGACCAGGGGAATC</td><td>8755.5</td><td>-5.8</td><td>0.7</td><td>-8.6</td><td>5.3E-18</td><td>5.7E-17</td><td>MB231/MB468</td><td>CYTO</td></tr>
<tr><td>>hg38_21_+_8433172_8446622_RNA45SN1_45S_50ntFlanks@11561.11593.33|GCGCCTAGCAGCCGACTTAGAACTGGTGCGGAC</td><td>>hg38_21_+_8433172_8446622_RNA45SN1_45S_50ntFlanks</td><td>11561</td><td>11593</td><td>33</td><td>GCGCCTAGCAGCCGACTTAGAACTGGTGCGGAC</td><td>552.1</td><td>-13.0</td><td>1.5</td><td>-8.6</td><td>5.3E-18</td><td>5.7E-17</td><td>MB231/MB468</td><td>MP</td></tr>
<tr><td>>hg38_21_+_8433172_8446622_RNA45SN1_45S_50ntFlanks@10802.10825.24|TTGGAACAATGTAGGTAAGGGAAG</td><td>>hg38_21_+_8433172_8446622_RNA45SN1_45S_50ntFlanks</td><td>10802</td><td>10825</td><td>24</td><td>TTGGAACAATGTAGGTAAGGGAAG</td><td>326.8</td><td>-12.2</td><td>1.4</td><td>-8.6</td><td>5.3E-18</td><td>5.7E-17</td><td>MB231/MB468</td><td>MP</td></tr>
<tr><td>>MI0003591|hsa-mir-584&WithFlank&5|-|149062307|149062415@22.41.20(+1U)&[MIMAT0003249&hsa-miR-584-5p&offsets|0|-2(+1U);m-167&5|-|149062374|149062394&offsets|0|-1(+1U)]|TTATGGTTTGCCTGGGACTGT</td><td>>MI0003591|hsa-mir-584&WithFlank&5|-|149062307|149062415</td><td>22</td><td>41</td><td>20(+1U)</td><td>TTATGGTTTGCCTGGGACTGT</td><td>879.5</td><td>9.8</td><td>1.1</td><td>8.6</td><td>1.0E-17</td><td>5.7E-17</td><td>BT20/MB468</td><td>CYTO</td></tr>
<tr><td>>hg38_21_+_8433172_8446622_RNA45SN1_45S_50ntFlanks@12709.12747.39|CCGCGGAGCCTCGGTTGGCCTCGGATAGCCGGTCCCCCG</td><td>>hg38_21_+_8433172_8446622_RNA45SN1_45S_50ntFlanks</td><td>12709</td><td>12747</td><td>39</td><td>CCGCGGAGCCTCGGTTGGCCTCGGATAGCCGGTCCCCCG</td><td>557.7</td><td>-12.9</td><td>1.5</td><td>-8.6</td><td>8.8E-18</td><td>5.8E-17</td><td>MB231/MB468</td><td>MITO</td></tr>
<tr><td>>MI0000289|hsa-mir-181a-1&WithFlank&1|-|198859038|198859159@70.91.22&[MIMAT0000270&hsa-miR-181a-3p&offsets|0|0;m-125&1|-|198859069|198859090&offsets|0|0]|ACCATCGACCGTTGATTGTACC</td><td>>MI0000289|hsa-mir-181a-1&WithFlank&1|-|198859038|198859159</td><td>70</td><td>91</td><td>22</td><td>ACCATCGACCGTTGATTGTACC</td><td>622.6</td><td>-12.4</td><td>1.4</td><td>-8.8</td><td>9.1E-19</td><td>5.8E-17</td><td>BT20/MB231</td><td>NUC</td></tr>
<tr><td>>hg38_21_+_8433172_8446622_RNA45SN1_45S_50ntFlanks@11729.11753.25|TAACTATGACTCTCTTAAGGTAGCC</td><td>>hg38_21_+_8433172_8446622_RNA45SN1_45S_50ntFlanks</td><td>11729</td><td>11753</td><td>25</td><td>TAACTATGACTCTCTTAAGGTAGCC</td><td>2194.0</td><td>-10.8</td><td>1.3</td><td>-8.6</td><td>5.4E-18</td><td>5.8E-17</td><td>MB231/MB468</td><td>MP</td></tr>
<tr><td>>hg38_21_+_8433172_8446622_RNA45SN1_45S_50ntFlanks@12626.12651.26|AATGGGGCGAAGCTACCATCTGTGGG</td><td>>hg38_21_+_8433172_8446622_RNA45SN1_45S_50ntFlanks</td><td>12626</td><td>12651</td><td>26</td><td>AATGGGGCGAAGCTACCATCTGTGGG</td><td>518.9</td><td>-12.9</td><td>1.5</td><td>-8.6</td><td>5.4E-18</td><td>5.9E-17</td><td>MB231/MB468</td><td>MP</td></tr>
<tr><td>>MI0000269|hsa-mir-181a-2&WithFlank&9|+|124692436|124692557@83.103.21(+1A)&[MIMAT0004558&hsa-miR-181a-2-3p&offsets|0|-1(+1A);m-71&9|+|124692518|124692539&offsets|0|-1(+1A)]|ACCACTGACCGTTGACTGTACA</td><td>>MI0000269|hsa-mir-181a-2&WithFlank&9|+|124692436|124692557</td><td>83</td><td>103</td><td>21(+1A)</td><td>ACCACTGACCGTTGACTGTACA</td><td>349.6</td><td>12.5</td><td>1.4</td><td>8.6</td><td>5.5E-18</td><td>5.9E-17</td><td>MB231/MB468</td><td>CYTO</td></tr>
<tr><td>>hg38_21_+_8433172_8446622_RNA45SN1_45S_50ntFlanks@11586.11612.27|GTGCGGACCAGGGGAATCCGACTGTTT</td><td>>hg38_21_+_8433172_8446622_RNA45SN1_45S_50ntFlanks</td><td>11586</td><td>11612</td><td>27</td><td>GTGCGGACCAGGGGAATCCGACTGTTT</td><td>2213.4</td><td>-5.2</td><td>0.6</td><td>-8.5</td><td>1.8E-17</td><td>5.9E-17</td><td>BT20/MB468</td><td>NUC</td></tr>
<tr><td>>hg38_21_+_8433172_8446622_RNA45SN1_45S_50ntFlanks@7982.8007.26|TCAGATCAGACGTGGCGACCCGCTGA</td><td>>hg38_21_+_8433172_8446622_RNA45SN1_45S_50ntFlanks</td><td>7982</td><td>8007</td><td>26</td><td>TCAGATCAGACGTGGCGACCCGCTGA</td><td>282.7</td><td>-12.0</td><td>1.4</td><td>-8.6</td><td>5.5E-18</td><td>5.9E-17</td><td>MB231/MB468</td><td>MP</td></tr>
<tr><td>>hg38_21_+_8433172_8446622_RNA45SN1_45S_50ntFlanks@10683.10711.29|CCGGCGGCGTCCGGTGAGCTCTCGCTGGC</td><td>>hg38_21_+_8433172_8446622_RNA45SN1_45S_50ntFlanks</td><td>10683</td><td>10711</td><td>29</td><td>CCGGCGGCGTCCGGTGAGCTCTCGCTGGC</td><td>1281.0</td><td>-9.9</td><td>1.1</td><td>-8.6</td><td>5.5E-18</td><td>6.0E-17</td><td>MB231/MB468</td><td>MP</td></tr>
<tr><td>>hg38_21_+_8433172_8446622_RNA45SN1_45S_50ntFlanks@11641.11676.36|CGGCGGGTGTTGACGCGATGTGATTTCTGCCCAGTG</td><td>>hg38_21_+_8433172_8446622_RNA45SN1_45S_50ntFlanks</td><td>11641</td><td>11676</td><td>36</td><td>CGGCGGGTGTTGACGCGATGTGATTTCTGCCCAGTG</td><td>402.3</td><td>-12.5</td><td>1.4</td><td>-8.6</td><td>5.5E-18</td><td>6.0E-17</td><td>MB231/MB468</td><td>MP</td></tr>
<tr><td>>hg38_21_+_8433172_8446622_RNA45SN1_45S_50ntFlanks@8403.8422.20|AGGTAAACGGGTGGGGTCCG</td><td>>hg38_21_+_8433172_8446622_RNA45SN1_45S_50ntFlanks</td><td>8403</td><td>8422</td><td>20</td><td>AGGTAAACGGGTGGGGTCCG</td><td>457.5</td><td>-12.7</td><td>1.5</td><td>-8.6</td><td>5.5E-18</td><td>6.0E-17</td><td>MB231/MB468</td><td>MP</td></tr>
<tr><td>>hg38_21_+_8433172_8446622_RNA45SN1_45S_50ntFlanks@3873.3898.26|TAATACATGCCGACGGGCGCTGACCC</td><td>>hg38_21_+_8433172_8446622_RNA45SN1_45S_50ntFlanks</td><td>3873</td><td>3898</td><td>26</td><td>TAATACATGCCGACGGGCGCTGACCC</td><td>2280.1</td><td>-5.4</td><td>0.6</td><td>-8.6</td><td>1.1E-17</td><td>6.0E-17</td><td>BT20/MB468</td><td>CYTO</td></tr>
<tr><td>>MI0000272|hsa-mir-182&WithFlank&7|-|129770377|129770498@29.53.25(+1A)&[MIMAT0000259&hsa-miR-182-5p&offsets|0|+1(+1A);m-22&7|-|129770449|129770470&offsets|0|+3(+1A)]|TTTGGCAATGGTAGAACTCACACTGA</td><td>>MI0000272|hsa-mir-182&WithFlank&7|-|129770377|129770498</td><td>29</td><td>53</td><td>25(+1A)</td><td>TTTGGCAATGGTAGAACTCACACTGA</td><td>298.1</td><td>3.0</td><td>0.3</td><td>8.5</td><td>1.8E-17</td><td>6.1E-17</td><td>BT20/MB468</td><td>NUC</td></tr>
<tr><td>>MI0000650|hsa-mir-200c&WithFlank&12|+|6963693|6963772@50.69.20(+1A)&[MIMAT0000617&hsa-miR-200c-3p&offsets|0|-3(+1A);m-54&12|+|6963742|6963764&offsets|0|-3(+1A)]|TAATACTGCCGGGTAATGATA</td><td>>MI0000650|hsa-mir-200c&WithFlank&12|+|6963693|6963772</td><td>50</td><td>69</td><td>20(+1A)</td><td>TAATACTGCCGGGTAATGATA</td><td>663.4</td><td>10.1</td><td>1.1</td><td>8.9</td><td>5.5E-19</td><td>6.1E-17</td><td>BT20/MB231</td><td>CYTO</td></tr>
<tr><td>>hg38_21_+_8433172_8446622_RNA45SN1_45S_50ntFlanks@7981.8005.25|CTCAGATCAGACGTGGCGACCCGCT</td><td>>hg38_21_+_8433172_8446622_RNA45SN1_45S_50ntFlanks</td><td>7981</td><td>8005</td><td>25</td><td>CTCAGATCAGACGTGGCGACCCGCT</td><td>351.0</td><td>-12.3</td><td>1.4</td><td>-8.6</td><td>5.7E-18</td><td>6.1E-17</td><td>MB231/MB468</td><td>MP</td></tr>
<tr><td>>MI0000283|hsa-mir-203a&WithFlank&14|+|104117399|104117520@72.92.21&[MIMAT0000264&hsa-miR-203a-3p&offsets|+1|0;m-79&14|+|104117469|104117490&offsets|+1|0]|TGAAATGTTTAGGACCACTAG</td><td>>MI0000283|hsa-mir-203a&WithFlank&14|+|104117399|104117520</td><td>72</td><td>92</td><td>21</td><td>TGAAATGTTTAGGACCACTAG</td><td>456.2</td><td>12.5</td><td>1.4</td><td>8.9</td><td>7.2E-19</td><td>6.1E-17</td><td>BT20/MB231</td><td>MITO</td></tr>
<tr><td>>MI0003628|hsa-mir-615&WithFlank&12|+|54033944|54034051@67.87.21&[MIMAT0003283&hsa-miR-615-3p&offsets|0|-1;m-495&12|+|54034010|54034030&offsets|0|0]|TCCGAGCCTGGGTCTCCCTCT</td><td>>MI0003628|hsa-mir-615&WithFlank&12|+|54033944|54034051</td><td>67</td><td>87</td><td>21</td><td>TCCGAGCCTGGGTCTCCCTCT</td><td>3584.3</td><td>5.8</td><td>0.7</td><td>8.9</td><td>7.4E-19</td><td>6.1E-17</td><td>BT20/MB231</td><td>MITO</td></tr>
<tr><td>>hg38_21_+_8433172_8446622_RNA45SN1_45S_50ntFlanks@476.505.30|TTCGTGATCGATGTGGTGACGTCGTGCTCT</td><td>>hg38_21_+_8433172_8446622_RNA45SN1_45S_50ntFlanks</td><td>476</td><td>505</td><td>30</td><td>TTCGTGATCGATGTGGTGACGTCGTGCTCT</td><td>1926.2</td><td>9.0</td><td>1.1</td><td>8.6</td><td>1.1E-17</td><td>6.1E-17</td><td>BT20/MB468</td><td>CYTO</td></tr>
</table>

| | | | | | | | | | | | | | |
|---|---|---|---|---|---|---|---|---|---|---|---|---|---|
| >hg38_21_+_8433172_8446622_RNA45SN1_45S_50ntFlanks@11843.11871.29\|AGGGAACGGGCTTGGCGGAATCAGCGGGG | >hg38_21_+_8433172_8446622_RNA45SN1_45S_50ntFlanks | 11843 | 11871 | 29 | AGGGAACGGGCTTGGCGGAATCAGCGGGG | 413.1 | -12.6 | 1.5 | -8.6 | 5.7E-18 | 6.1E-17 | MB231/MB468 | MP |
| >hg38_21_+_8433172_8446622_RNA45SN1_45S_50ntFlanks@12625.12654.30\|CAATGGGGCGAAGCTACCATCTGTGGGATT | >hg38_21_+_8433172_8446622_RNA45SN1_45S_50ntFlanks | 12625 | 12654 | 30 | CAATGGGGCGAAGCTACCATCTGTGGGATT | 5438.6 | -9.5 | 1.1 | -8.6 | 5.7E-18 | 6.1E-17 | MB231/MB468 | MP |
| >hg38_21_+_8433172_8446622_RNA45SN1_45S_50ntFlanks@485.513.29\|GATGTGGTGACGTCGTGCTCTCCCGGGCC | >hg38_21_+_8433172_8446622_RNA45SN1_45S_50ntFlanks | 485 | 513 | 29 | GATGTGGTGACGTCGTGCTCTCCCGGGCC | 120.2 | 9.7 | 1.1 | 8.5 | 1.8E-17 | 6.1E-17 | BT20/MB468 | NUC |
| >hg38_21_+_8433172_8446622_RNA45SN1_45S_50ntFlanks@10348.10374.27\|AAACGAGAACTTTGAAGGCCGAAGTGG | >hg38_21_+_8433172_8446622_RNA45SN1_45S_50ntFlanks | 10348 | 10374 | 27 | AAACGAGAACTTTGAAGGCCGAAGTGG | 2256.2 | -9.5 | 1.1 | -8.5 | 1.8E-17 | 6.1E-17 | BT20/MB468 | NUC |
| >hg38_21_+_8433172_8446622_RNA45SN1_45S_50ntFlanks@11999.12033.35\|CCCTGCGGGCCGCCGGTGAAATACCACTACTCTGA | >hg38_21_+_8433172_8446622_RNA45SN1_45S_50ntFlanks | 11999 | 12033 | 35 | CCCTGCGGGCCGCCGGTGAAATACCACTACTCTGA | 782.5 | -9.9 | 1.2 | -8.6 | 7.3E-18 | 6.2E-17 | MB231/MB468 | NUC |
| >trna112_GlnCTG_1_-_147737382_147737453@30.50.21,trna131_GlnCTG_6_-_27759135_27759206@30.50.21,trna146_GlnCTG_6_-_27515531_27515602@30.50.21,trna15_GlnCTG_1_+_145963304_145963375@30.50.21,trna19_GlnCTG_1_+_147505038_147505109@30.50.21,trna1_GlnCTG_6_+_18836402_18836473@30.50.21,trna28_GlnCTG_1_+_149186125_149186196@30.50.21,trna3_GlnCTG_17_+_8023070_8023141@30.50.21,trna49_GlnCTG_6_+_27487308_27487379@30.50.21,trna7_GlnCTG_15_-_66161400_66161471@30.50.21,trna99_GlnCTG_6_-_28909378_28909449@30.50.21\|GACTCTGAATCCAGCGATCCG | >trna112_GlnCTG_1_-_147737382_147737453 | 30 | 50 | 21 | GACTCTGAATCCAGCGATCCG | 680.2 | -13.2 | 1.5 | -8.6 | 9.4E-18 | 6.2E-17 | MB231/MB468 | MITO |
| >trna20_GluTTC_2_-_131094701_131094772@1.36.36,trna5_GluTTC_13_-_41634874_41634945@1.36.36\|TCCCATATGGTCTAGCGGTTAGGATTCCTGGTTTTC | >trna20_GluTTC_2_-_131094701_131094772 | 1 | 36 | 36 | TCCCATATGGTCTAGCGGTTAGGATTCCTGGTTTTC | 1437.6 | 14.5 | 1.7 | 8.6 | 5.8E-18 | 6.2E-17 | MB231/MB468 | CYTO |
| >hg38_21_+_8433172_8446622_RNA45SN1_45S_50ntFlanks@4045.4062.18\|CCATTCGAACGTCTGCCC | >hg38_21_+_8433172_8446622_RNA45SN1_45S_50ntFlanks | 4045 | 4062 | 18 | CCATTCGAACGTCTGCCC | 279.0 | -12.0 | 1.4 | -8.6 | 5.8E-18 | 6.2E-17 | MB231/MB468 | MP |
| >hg38_21_+_8433172_8446622_RNA45SN1_45S_50ntFlanks@11930.11951.22\|TAGAATAAGTGGGAGGCCCCCG | >hg38_21_+_8433172_8446622_RNA45SN1_45S_50ntFlanks | 11930 | 11951 | 22 | TAGAATAAGTGGGAGGCCCCCG | 716.9 | -13.3 | 1.5 | -8.6 | 5.8E-18 | 6.3E-17 | MB231/MB468 | MP |
| >hg38_21_+_8433172_8446622_RNA45SN1_45S_50ntFlanks@12632.12654.23\|GCGAAGCTACCATCTGTGGGATT | >hg38_21_+_8433172_8446622_RNA45SN1_45S_50ntFlanks | 12632 | 12654 | 23 | GCGAAGCTACCATCTGTGGGATT | 761.9 | -4.3 | 0.5 | -8.6 | 1.1E-17 | 6.3E-17 | BT20/MB468 | CYTO |
| >hg38_21_+_8433172_8446622_RNA45SN1_45S_50ntFlanks@11592.11612.21\|ACCAGGGGAATCCGACTGTTT | >hg38_21_+_8433172_8446622_RNA45SN1_45S_50ntFlanks | 11592 | 11612 | 21 | ACCAGGGGAATCCGACTGTTT | 834.7 | -8.2 | 1.0 | -8.6 | 9.5E-18 | 6.3E-17 | MB231/MB468 | MITO |
| >hg38_21_+_8433172_8446622_RNA45SN1_45S_50ntFlanks@12407.12435.29\|TTTTGATCCTTCGATGTCGGCTCTTCCTA | >hg38_21_+_8433172_8446622_RNA45SN1_45S_50ntFlanks | 12407 | 12435 | 29 | TTTTGATCCTTCGATGTCGGCTCTTCCTA | 1788.2 | -6.4 | 0.7 | -8.6 | 7.5E-18 | 6.3E-17 | MB231/MB468 | NUC |
| >hg38_21_+_8433172_8446622_RNA45SN1_45S_50ntFlanks@10391.10435.45\|AACAGCAGTTGAACATGGGTCAGTCGGTCCTGAGAGATGGGCGAG | >hg38_21_+_8433172_8446622_RNA45SN1_45S_50ntFlanks | 10391 | 10435 | 45 | AACAGCAGTTGAACATGGGTCAGTCGGTCCTGAGAGATGGGCGAG | 1768.0 | -15.2 | 1.8 | -8.5 | 1.9E-17 | 6.3E-17 | BT20/MB468 | NUC |
| >hg38_21_+_8433172_8446622_RNA45SN1_45S_50ntFlanks@11874.11902.29\|AGAAGACCCTGTTGAGCTTGACTCTAGTC | >hg38_21_+_8433172_8446622_RNA45SN1_45S_50ntFlanks | 11874 | 11902 | 29 | AGAAGACCCTGTTGAGCTTGACTCTAGTC | 295.6 | -12.1 | 1.4 | -8.6 | 5.9E-18 | 6.3E-17 | MB231/MB468 | MP |
| >hg38_21_+_8433172_8446622_RNA45SN1_45S_50ntFlanks@6648.6665.18\|GTACGACTCTTAGCGGTG | >hg38_21_+_8433172_8446622_RNA45SN1_45S_50ntFlanks | 6648 | 6665 | 18 | GTACGACTCTTAGCGGTG | 4760.3 | 6.1 | 0.7 | 8.6 | 8.0E-18 | 6.3E-17 | BT20/MB468 | MITO |
| >am_trna117_GlyTCC_1_-_145397864_145397935@1.29.29,trna45_GlyTCC_1_+_161500903_161500974@1.29.29,trna70_GlyTCC_1_-_161439547_161439618@1.29.29,trna73_GlyTCC_1_-_161432166_161432237@1.29.29,trna76_GlyTCC_1_-_161424756_161424827@1.29.29,trna79_GlyTCC_1_-_161417375_161417446@1.29.29\|GCGTTGGTGGTATAGTGGTGAGCATAGCT | >am_trna117_GlyTCC_1_-_145397864_145397935 | 1 | 29 | 29 | GCGTTGGTGGTATAGTGGTGAGCATAGCT | 3325.6 | -5.4 | 0.6 | -8.6 | 1.1E-17 | 6.3E-17 | BT20/MB468 | CYTO |
| >hg38_21_+_8433172_8446622_RNA45SN1_45S_50ntFlanks@5487.5513.27\|CGCTGAGAAGACGGTCGAACTTGACTA | >hg38_21_+_8433172_8446622_RNA45SN1_45S_50ntFlanks | 5487 | 5513 | 27 | CGCTGAGAAGACGGTCGAACTTGACTA | 379.8 | -12.4 | 1.4 | -8.6 | 9.7E-18 | 6.3E-17 | MB231/MB468 | MITO |
| >trna111_HisGTG_1_-_147774845_147774916@-1A.34.35,trna118_HisGTG_1_-_145396881_145396952@-1A.34.35,trna16_HisGTG_1_+_146544773_146544844@-1A.34.35,trna1_HisGTG_15_+_45493349_45493420@-1A.34.35,trna21_HisGTG_1_+_147753471_147753542@-1A.34.35,trna33_HisGTG_6_+_27125906_27125977@-1A.34.35,trna7_HisGTG_9_-_14433938_14434009@-1A.34.35,trna8_HisGTG_15_-_45492611_45492682@-1A.34.35,trna9_HisGTG_15_-_45490804_45490875@-1A.34.35\|AGCCGTGATCGTATAGTGGTTAGTACTCTGCGTTG | >trna111_HisGTG_1_-_147774845_147774916 | -1A | 34 | 35 | AGCCGTGATCGTATAGTGGTTAGTACTCTGCGTTG | 1420.9 | 10.9 | 1.3 | 8.6 | 5.9E-18 | 6.3E-17 | MB231/MB468 | CYTO |
| >hg38_21_+_8433172_8446622_RNA45SN1_45S_50ntFlanks@4000.4024.25\|TAGATAACCTCGGGCCGATCGCACG | >hg38_21_+_8433172_8446622_RNA45SN1_45S_50ntFlanks | 4000 | 4024 | 25 | TAGATAACCTCGGGCCGATCGCACG | 729.5 | -4.7 | 0.5 | -8.6 | 7.6E-18 | 6.4E-17 | MB231/MB468 | NUC |
| >MI0000264\|hsa-mir-7-2&WithFlank&15\|+\|88611819\|88611940@38.60.23(+1C);MI0000265\|hsa-mir-7-3&WithFlank&19\|+\|4770664\|4770785@37.59.23(+1C);MI0000263\|hsa-mir-7-1&WithFlank&9\|-\|83969742\|83969863@30.52.23(+1C)&[MIMAT0000252&hsa-miR-7-5p&offsets\|0\|-1(+1C);m-400&15\|+\|88611856\|88611879&offsets\|0\|-1(+1C)];[MIMAT0000252_1&hsa-miR-7-5p&offsets\|0\|-1(+1C);m-402&19\|+\|4770700\|4770723&offsets\|0\|-1(+1C)];[MIMAT0000252_2&hsa-miR-7-5p&offsets\|0\|-1(+1C);m-401&9\|-\|83969811\|83969834&offsets\|0\|-1(+1C)]\|TGGAAGACTAGTGATTTTGTTGTC | >MI0000264\|hsa-mir-7-2&WithFlank&15\|+\|88611819\|88611940 | 38 | 60 | 23(+1C) | TGGAAGACTAGTGATTTTGTTGTC | 24492.2 | 7.9 | 0.9 | 8.6 | 1.1E-17 | 6.4E-17 | BT20/MB468 | CYTO |
| >hg38_21_+_8433172_8446622_RNA45SN1_45S_50ntFlanks@9571.9596.26\|GCAAATCGGTCGTCCGACCTGGGTAT | >hg38_21_+_8433172_8446622_RNA45SN1_45S_50ntFlanks | 9571 | 9596 | 26 | GCAAATCGGTCGTCCGACCTGGGTAT | 534.3 | -12.9 | 1.5 | -8.6 | 6.0E-18 | 6.4E-17 | MB231/MB468 | MP |
| >hg38_21_+_8433172_8446622_RNA45SN1_45S_50ntFlanks@9534.9567.34\|AGGAAACTCTGGTGGAGGTCCGTAGCGGTCCTGA | >hg38_21_+_8433172_8446622_RNA45SN1_45S_50ntFlanks | 9534 | 9567 | 34 | AGGAAACTCTGGTGGAGGTCCGTAGCGGTCCTGA | 621.7 | -13.1 | 1.5 | -8.6 | 6.0E-18 | 6.4E-17 | MB231/MB468 | MP |
| >hg38_21_+_8433172_8446622_RNA45SN1_45S_50ntFlanks@10402.10431.30\|AACATGGGTCAGTCGGTCCTGAGAGATGGG | >hg38_21_+_8433172_8446622_RNA45SN1_45S_50ntFlanks | 10402 | 10431 | 30 | AACATGGGTCAGTCGGTCCTGAGAGATGGG | 1085.1 | -8.6 | 1.0 | -8.6 | 6.0E-18 | 6.4E-17 | MB231/MB468 | MP |
| >hg38_21_+_8433172_8446622_RNA45SN1_45S_50ntFlanks@10805.10823.19\|GAACAATGTAGGTAAGGGA | >hg38_21_+_8433172_8446622_RNA45SN1_45S_50ntFlanks | 10805 | 10823 | 19 | GAACAATGTAGGTAAGGGA | 321.2 | -12.2 | 1.4 | -8.6 | 6.0E-18 | 6.4E-17 | MB231/MB468 | MP |
| >hg38_21_+_8433172_8446622_RNA45SN1_45S_50ntFlanks@12456.12489.34\|CCAAGCGTTGGATTGTTCACCCACTAATAGGGAA | >hg38_21_+_8433172_8446622_RNA45SN1_45S_50ntFlanks | 12456 | 12489 | 34 | CCAAGCGTTGGATTGTTCACCCACTAATAGGGAA | 1174.9 | -7.2 | 0.8 | -8.6 | 6.0E-18 | 6.4E-17 | MB231/MB468 | CYTO |
| >hg38_21_+_8433172_8446622_RNA45SN1_45S_50ntFlanks@12155.12180.26\|GGAGTTTGACTGGGGCGGTACACCTG | >hg38_21_+_8433172_8446622_RNA45SN1_45S_50ntFlanks | 12155 | 12180 | 26 | GGAGTTTGACTGGGGCGGTACACCTG | 381.7 | -12.4 | 1.4 | -8.6 | 6.0E-18 | 6.5E-17 | MB231/MB468 | MP |
| >hg38_21_+_8433172_8446622_RNA45SN1_45S_50ntFlanks@10781.10807.27\|AGGTGAACAGCCTCTGGCATGTTGGAA | >hg38_21_+_8433172_8446622_RNA45SN1_45S_50ntFlanks | 10781 | 10807 | 27 | AGGTGAACAGCCTCTGGCATGTTGGAA | 405.5 | -11.6 | 1.4 | -8.6 | 7.7E-18 | 6.5E-17 | MB231/MB468 | NUC |
| >hg38_21_+_8433172_8446622_RNA45SN1_45S_50ntFlanks@13006.13025.20\|CTATTGAAAGTCAGCCCTCG | >hg38_21_+_8433172_8446622_RNA45SN1_45S_50ntFlanks | 13006 | 13025 | 20 | CTATTGAAAGTCAGCCCTCG | 387.6 | -12.4 | 1.4 | -8.6 | 9.9E-18 | 6.5E-17 | MB231/MB468 | MITO |
| >hg38_21_+_8433172_8446622_RNA45SN1_45S_50ntFlanks@10735.10755.21\|TGTAAATCTCGCGCCGGGCCG | >hg38_21_+_8433172_8446622_RNA45SN1_45S_50ntFlanks | 10735 | 10755 | 21 | TGTAAATCTCGCGCCGGGCCG | 322.5 | -12.2 | 1.4 | -8.6 | 6.1E-18 | 6.5E-17 | MB231/MB468 | MP |
| >hg38_21_+_8433172_8446622_RNA45SN1_45S_50ntFlanks@3875.3898.24\|ATACATGCCGACGGGCGCTGACCC | >hg38_21_+_8433172_8446622_RNA45SN1_45S_50ntFlanks | 3875 | 3898 | 24 | ATACATGCCGACGGGCGCTGACCC | 3547.6 | -5.2 | 0.6 | -8.6 | 1.1E-17 | 6.5E-17 | BT20/MB468 | CYTO |
| >hg38_21_+_8433172_8446622_RNA45SN1_45S_50ntFlanks@10186.10207.22\|TCCTCCCCTCCTCCCGCCCACG | >hg38_21_+_8433172_8446622_RNA45SN1_45S_50ntFlanks | 10186 | 10207 | 22 | TCCTCCCCTCCTCCCGCCCACG | 1191.0 | -13.2 | 1.5 | -8.6 | 7.8E-18 | 6.5E-17 | MB231/MB468 | NUC |
| >hg38_21_+_8433172_8446622_RNA45SN1_45S_50ntFlanks@4099.4117.19\|GTGACCACGGGTGACGGGG | >hg38_21_+_8433172_8446622_RNA45SN1_45S_50ntFlanks | 4099 | 4117 | 19 | GTGACCACGGGTGACGGGG | 545.5 | -13.0 | 1.5 | -8.6 | 6.1E-18 | 6.5E-17 | MB231/MB468 | MP |
| >hg38_21_+_8433172_8446622_RNA45SN1_45S_50ntFlanks@4823.4844.22\|ATGACCCGCCGGGCAGCTTCCG | >hg38_21_+_8433172_8446622_RNA45SN1_45S_50ntFlanks | 4823 | 4844 | 22 | ATGACCCGCCGGGCAGCTTCCG | 1183.5 | 5.9 | 0.7 | 8.6 | 6.1E-18 | 6.5E-17 | MB231/MB468 | CYTO |
| >hg38_21_+_8433172_8446622_RNA45SN1_45S_50ntFlanks@9572.9596.25\|CAAATCGGTCGTCCGACCTGGGTAT | >hg38_21_+_8433172_8446622_RNA45SN1_45S_50ntFlanks | 9572 | 9596 | 25 | CAAATCGGTCGTCCGACCTGGGTAT | 10459.8 | -11.4 | 1.3 | -8.6 | 8.3E-18 | 6.6E-17 | BT20/MB468 | MITO |
| >MI0000301\|hsa-mir-224&WithFlank&X\|-\|151958572\|151958664@14.34.21&[MIMAT0000281&hsa-miR-224-5p&offsets\|+1\|-3;m-238&X\|-\|151958630\|151958651&offsets\|0\|-1]\|CAAGTCACTAGTGGTTCCGTT | >MI0000301\|hsa-mir-224&WithFlank&X\|-\|151958572\|151958664 | 14 | 34 | 21 | CAAGTCACTAGTGGTTCCGTT | 341.9 | 9.1 | 1.1 | 8.6 | 7.9E-18 | 6.6E-17 | MB231/MB468 | NUC |
| >MI0000272\|hsa-mir-182&WithFlank&7\|-\|129770377\|129770498@29.50.22(+1A)&[MIMAT0000259&hsa-miR-182-5p&offsets\|0\|-2(+1A);m-22&7\|-\|129770449\|129770470&offsets\|0\|0(+1A)]\|TTTGGCAATGGTAGAACTCACAA | >MI0000272\|hsa-mir-182&WithFlank&7\|-\|129770377\|129770498 | 29 | 50 | 22(+1A) | TTTGGCAATGGTAGAACTCACAA | 392.3 | 12.9 | 1.5 | 8.6 | 1.2E-17 | 6.6E-17 | BT20/MB468 | CYTO |
| >hg38_21_+_8433172_8446622_RNA45SN1_45S_50ntFlanks@6744.6770.27\|GACACTTCGAACGCACTTGCGGCCCCG | >hg38_21_+_8433172_8446622_RNA45SN1_45S_50ntFlanks | 6744 | 6770 | 27 | GACACTTCGAACGCACTTGCGGCCCCG | 1199.4 | -3.8 | 0.4 | -8.6 | 8.4E-18 | 6.6E-17 | BT20/MB468 | MITO |
| >hg38_21_+_8433172_8446622_RNA45SN1_45S_50ntFlanks@10914.10939.26\|CGCCGCGGCTGGACGAGGCGCCGCCG | >hg38_21_+_8433172_8446622_RNA45SN1_45S_50ntFlanks | 10914 | 10939 | 26 | CGCCGCGGCTGGACGAGGCGCCGCCG | 463.4 | -12.7 | 1.5 | -8.6 | 6.2E-18 | 6.6E-17 | MB231/MB468 | MP |
| >hg38_21_+_8433172_8446622_RNA45SN1_45S_50ntFlanks@11200.11232.33\|CGGCGGCGGCGGCGACTCTGGACGCGAGCCGGG | >hg38_21_+_8433172_8446622_RNA45SN1_45S_50ntFlanks | 11200 | 11232 | 33 | CGGCGGCGGCGGCGACTCTGGACGCGAGCCGGG | 4037.6 | -4.9 | 0.6 | -8.6 | 1.2E-17 | 6.6E-17 | BT20/MB468 | CYTO |
| >hg38_21_+_8433172_8446622_RNA45SN1_45S_50ntFlanks@7988.8008.21\|CAGACGTGGCGACCCGCTGAA | >hg38_21_+_8433172_8446622_RNA45SN1_45S_50ntFlanks | 7988 | 8008 | 21 | CAGACGTGGCGACCCGCTGAA | 7896.4 | -5.1 | 0.6 | -8.6 | 8.5E-18 | 6.7E-17 | BT20/MB468 | MITO |

| | | | | | | | | | | | | | |
|---|---|---|---|---|---|---|---|---|---|---|---|---|---|
| >trna119_LysCTT_1_-_145395522_145395594@1.35.35,trna11_LysCTT_5_-_180648979_180649051@1.35.35,trna13_LysCTT_6_+_26556774_26556846@1.35.35,trna7_LysCTT_16_+_3225692_3225764@1.35.35,trna9_LysCTT_5_+_180634755_180634827@1.35.35\|GCCCGGCTAGCTCAGTCGGTAGAGCATGAGACTCT | >trna119_LysCTT_1_-_145395522_145395594 | 1 | 35 | 35 | GCCCGGCTAGCTCAGTCGGTAGAGCATGAGACTCT | 7117.2 | 7.7 | 0.9 | 8.6 | 6.3E-18 | 6.7E-17 | MB231/MB468 | CYTO |
| >hg38_21_+_8433172_8446622_RNA45SN1_45S_50ntFlanks@6772.6803.32\|GTTCCTCCCGGGGCTACGCCTGTCTGAGCGTC | >hg38_21_+_8433172_8446622_RNA45SN1_45S_50ntFlanks | 6772 | 6803 | 32 | GTTCCTCCCGGGGCTACGCCTGTCTGAGCGTC | 1415.9 | -10.1 | 1.2 | -8.6 | 6.3E-18 | 6.7E-17 | MB231/MB468 | MP |
| >MI0000734\|hsa-mir-106b&WithFlank&7\|-\|100093987\|100094080@58.78.21(+1U)&[MIMAT0004672&hsa-miR-106b-3p&offsets\|0\|-1(+1U);m-87&7\|-\|100094003\|100094023&offsets\|0\|0(+1U)]\|CCGCACTGTGGGTACTTGCTGT | >MI0000734\|hsa-mir-106b&WithFlank&7\|-\|100093987\|100094080 | 58 | 78 | 21(+1U) | CCGCACTGTGGGTACTTGCTGT | 1817.1 | 6.4 | 0.7 | 8.6 | 6.3E-18 | 6.8E-17 | MB231/MB468 | CYTO |
| >hg38_21_+_8433172_8446622_RNA45SN1_45S_50ntFlanks@12954.12972.19\|CTGCTTCTGGGTCGGGGTT | >hg38_21_+_8433172_8446622_RNA45SN1_45S_50ntFlanks | 12954 | 12972 | 19 | CTGCTTCTGGGTCGGGGTT | 499.3 | -12.7 | 1.5 | -8.6 | 1.0E-17 | 6.8E-17 | MB231/MB468 | MITO |
| >hg38_21_+_8433172_8446622_RNA45SN1_45S_50ntFlanks@10914.10932.19\|CGCCGCGGCTGGACGAGGC | >hg38_21_+_8433172_8446622_RNA45SN1_45S_50ntFlanks | 10914 | 10932 | 19 | CGCCGCGGCTGGACGAGGC | 633.0 | -13.1 | 1.5 | -8.6 | 1.0E-17 | 6.8E-17 | MB231/MB468 | MITO |
| >hg38_21_+_8433172_8446622_RNA45SN1_45S_50ntFlanks@6721.6758.38\|AATTGCAGGACACATTGATCATCGACACTTCGAACGCA | >hg38_21_+_8433172_8446622_RNA45SN1_45S_50ntFlanks | 6721 | 6758 | 38 | AATTGCAGGACACATTGATCATCGACACTTCGAACGCA | 4087.1 | -6.7 | 0.8 | -8.6 | 8.2E-18 | 6.8E-17 | MB231/MB468 | NUC |
| >hg38_21_+_8433172_8446622_RNA45SN1_45S_50ntFlanks@11564.11581.18\|CCTAGCAGCCGACTTAGA | >hg38_21_+_8433172_8446622_RNA45SN1_45S_50ntFlanks | 11564 | 11581 | 18 | CCTAGCAGCCGACTTAGA | 2645.6 | -8.4 | 1.0 | -8.6 | 1.1E-17 | 6.9E-17 | MB231/MB468 | MITO |
| >trnaMT_ValTAC_MT_+_1602_1670@47.72.26\|TCAACTTAACTTGACCGCTCTGACCA | >trnaMT_ValTAC_MT_+_1602_1670 | 47 | 72 | 26 | TCAACTTAACTTGACCGCTCTGACCA | 384.6 | 12.8 | 1.5 | 8.6 | 1.2E-17 | 6.9E-17 | BT20/MB468 | CYTO |
| >hg38_21_+_8433172_8446622_RNA45SN1_45S_50ntFlanks@4927.4944.18\|AGTGGAGCCTGCGGCTTA | >hg38_21_+_8433172_8446622_RNA45SN1_45S_50ntFlanks | 4927 | 4944 | 18 | AGTGGAGCCTGCGGCTTA | 348.0 | 5.2 | 0.6 | 8.6 | 1.2E-17 | 6.9E-17 | BT20/MB468 | CYTO |
| >hg38_21_+_8433172_8446622_RNA45SN1_45S_50ntFlanks@12456.12474.19\|CCAAGCGTTGGATTGTTCA | >hg38_21_+_8433172_8446622_RNA45SN1_45S_50ntFlanks | 12456 | 12474 | 19 | CCAAGCGTTGGATTGTTCA | 1885.6 | -10.2 | 1.2 | -8.6 | 1.1E-17 | 6.9E-17 | MB231/MB468 | MITO |
| >trnaMT_MetCAT_MT_+_4402_4469@32.71.40\|ATACCCCGAAAATGTTGGTTATACCCTTCCCGTACTACCA | >trnaMT_MetCAT_MT_+_4402_4469 | 32 | 71 | 40 | ATACCCCGAAAATGTTGGTTATACCCTTCCCGTACTACCA | 635.4 | 13.3 | 1.5 | 8.6 | 6.5E-18 | 6.9E-17 | MB231/MB468 | CYTO |
| >hg38_21_+_8433172_8446622_RNA45SN1_45S_50ntFlanks@11999.12036.38\|CCCTGCGGGCCGCCGGTGAAATACCACTACTCTGATCG | >hg38_21_+_8433172_8446622_RNA45SN1_45S_50ntFlanks | 11999 | 12036 | 38 | CCCTGCGGGCCGCCGGTGAAATACCACTACTCTGATCG | 413.4 | -12.5 | 1.5 | -8.6 | 1.1E-17 | 6.9E-17 | MB231/MB468 | MITO |
| >MI0000289\|hsa-mir-181a-1&WithFlank&1\|-\|198859038\|198859159@30.51.22(+1A);MI0000269\|hsa-mir-181a-2&WithFlank&9\|+\|124692436\|124692557@45.66.22(+1A)&[MIMAT0000256&hsa-miR-181a-5p&offsets\|0\|-1(+1A);m-32&1\|-\|198859108\|198859130&offsets\|0\|-1(+1A)];[MIMAT0000256_1&hsa-miR-181a-5p&offsets\|0\|-1(+1A);m-32&9\|+\|124692480\|124692502&offsets\|0\|-1(+1A)]\|AACATTCAACGCTGTCGGTGAGA | >MI0000289\|hsa-mir-181a-1&WithFlank&1\|-\|198859038\|198859159 | 30 | 51 | 22(+1A) | AACATTCAACGCTGTCGGTGAGA | 305.5 | 5.2 | 0.6 | 8.6 | 1.2E-17 | 7.0E-17 | BT20/MB468 | CYTO |
| >hg38_21_+_8433172_8446622_RNA45SN1_45S_50ntFlanks@11857.11887.31\|GCGGAATCAGCGGGGAAAGAAGACCCTGTTG | >hg38_21_+_8433172_8446622_RNA45SN1_45S_50ntFlanks | 11857 | 11887 | 31 | GCGGAATCAGCGGGGAAAGAAGACCCTGTTG | 314.5 | -12.2 | 1.4 | -8.6 | 6.5E-18 | 7.0E-17 | MB231/MB468 | MP |
| >hg38_21_+_8433172_8446622_RNA45SN1_45S_50ntFlanks@11873.11894.22\|AAGAAGACCCTGTTGAGCTTGA | >hg38_21_+_8433172_8446622_RNA45SN1_45S_50ntFlanks | 11873 | 11894 | 22 | AAGAAGACCCTGTTGAGCTTGA | 299.3 | -12.1 | 1.4 | -8.6 | 6.6E-18 | 7.0E-17 | MB231/MB468 | MP |
| >hg38_21_+_8433172_8446622_RNA45SN1_45S_50ntFlanks@5472.5515.44\|ACGGCCCTGGCGGAGCGCTGAGAAGACGGTCGAACTTGACTATC | >hg38_21_+_8433172_8446622_RNA45SN1_45S_50ntFlanks | 5472 | 5515 | 44 | ACGGCCCTGGCGGAGCGCTGAGAAGACGGTCGAACTTGACTATC | 2916.7 | 7.3 | 0.8 | 8.6 | 6.6E-18 | 7.0E-17 | MB231/MB468 | CYTO |
| >hg38_21_+_8433172_8446622_RNA45SN1_45S_50ntFlanks@12356.12376.21\|ACCACAGGGATAACTGGCTTG | >hg38_21_+_8433172_8446622_RNA45SN1_45S_50ntFlanks | 12356 | 12376 | 21 | ACCACAGGGATAACTGGCTTG | 327.3 | -12.2 | 1.4 | -8.6 | 6.6E-18 | 7.0E-17 | MB231/MB468 | MP |
| >hg38_21_+_8433172_8446622_RNA45SN1_45S_50ntFlanks@11323.11342.20\|CGTCTCGTCGCGCGCGCGTC | >hg38_21_+_8433172_8446622_RNA45SN1_45S_50ntFlanks | 11323 | 11342 | 20 | CGTCTCGTCGCGCGCGCGTC | 11022.5 | -8.1 | 0.9 | -8.6 | 6.6E-18 | 7.0E-17 | MB231/MB468 | CYTO |
| >hg38_21_+_8433172_8446622_RNA45SN1_45S_50ntFlanks@11668.11688.21\|TGCCCAGTGCTCTGAATGTCA | >hg38_21_+_8433172_8446622_RNA45SN1_45S_50ntFlanks | 11668 | 11688 | 21 | TGCCCAGTGCTCTGAATGTCA | 764.1 | 6.3 | 0.7 | 8.6 | 6.7E-18 | 7.0E-17 | MB231/MB468 | CYTO |
| >hg38_21_+_8433172_8446622_RNA45SN1_45S_50ntFlanks@5072.5115.44\|TAATTCCGATAACGAACGAGACTCTGGCATGCTAACTAGTTACG | >hg38_21_+_8433172_8446622_RNA45SN1_45S_50ntFlanks | 5072 | 5115 | 44 | TAATTCCGATAACGAACGAGACTCTGGCATGCTAACTAGTTACG | 342.7 | 12.4 | 1.4 | 8.6 | 6.7E-18 | 7.0E-17 | MB231/MB468 | CYTO |
| >MI0000078\|hsa-mir-22&WithFlank&17\|-\|1713897\|1713993@21.41.21&[MIMAT0004495&hsa-miR-22-5p&offsets\|0\|-1;m-217&17\|-\|1713953\|1713973&offsets\|0\|0]\|AGTTCTTCAGTGGCAAGCTTT | >MI0000078\|hsa-mir-22&WithFlank&17\|-\|1713897\|1713993 | 21 | 41 | 21 | AGTTCTTCAGTGGCAAGCTTT | 693.5 | 6.8 | 0.8 | 8.6 | 8.5E-18 | 7.1E-17 | MB231/MB468 | NUC |
| >MI0000298\|hsa-mir-221&WithFlank&X\|-\|45746151\|45746272@71.93.23(+1G)&[MIMAT0000278&hsa-miR-221-3p&offsets\|0\|0(+1G);m-91&X\|-\|45746181\|45746202&offsets\|0\|+1(+1G)]\|AGCTACATTGTCTGCTGGGTTTCG | >MI0000298\|hsa-mir-221&WithFlank&X\|-\|45746151\|45746272 | 71 | 93 | 23(+1G) | AGCTACATTGTCTGCTGGGTTTCG | 422.6 | 5.1 | 0.6 | 8.6 | 8.5E-18 | 7.1E-17 | MB231/MB468 | NUC |
| >hg38_21_+_8433172_8446622_RNA45SN1_45S_50ntFlanks@9339.9369.31\|CGCACGAAAGCCGCCGTGGCGCAATGAAGGT | >hg38_21_+_8433172_8446622_RNA45SN1_45S_50ntFlanks | 9339 | 9369 | 31 | CGCACGAAAGCCGCCGTGGCGCAATGAAGGT | 670.2 | -13.3 | 1.5 | -8.6 | 6.6E-18 | 7.1E-17 | MB231/MB468 | MP |
| >hg38_21_+_8433172_8446622_RNA45SN1_45S_50ntFlanks@3984.4005.22\|GCGGCTTTGGTGACTCTAGATA | >hg38_21_+_8433172_8446622_RNA45SN1_45S_50ntFlanks | 3984 | 4005 | 22 | GCGGCTTTGGTGACTCTAGATA | 650.5 | -13.2 | 1.5 | -8.6 | 6.7E-18 | 7.1E-17 | MB231/MB468 | MP |
| >hg38_21_+_8433172_8446622_RNA45SN1_45S_50ntFlanks@10840.10865.26\|CCGTAACTTCGGGATAAGGATTGGCT | >hg38_21_+_8433172_8446622_RNA45SN1_45S_50ntFlanks | 10840 | 10865 | 26 | CCGTAACTTCGGGATAAGGATTGGCT | 4636.0 | -9.7 | 1.1 | -8.5 | 2.1E-17 | 7.1E-17 | BT20/MB468 | NUC |
| >hg38_21_+_8433172_8446622_RNA45SN1_45S_50ntFlanks@8189.8208.20\|CCCAGCCCGTGGACGGTGTG | >hg38_21_+_8433172_8446622_RNA45SN1_45S_50ntFlanks | 8189 | 8208 | 20 | CCCAGCCCGTGGACGGTGTG | 447.3 | -12.7 | 1.5 | -8.6 | 6.7E-18 | 7.1E-17 | MB231/MB468 | MP |
| >hg38_21_+_8433172_8446622_RNA45SN1_45S_50ntFlanks@11663.11706.44\|ATTTCTGCCCAGTGCTCTGAATGTCAAAGTGAAGAAATTCAATG | >hg38_21_+_8433172_8446622_RNA45SN1_45S_50ntFlanks | 11663 | 11706 | 44 | ATTTCTGCCCAGTGCTCTGAATGTCAAAGTGAAGAAATTCAATG | 194.5 | 8.9 | 1.0 | 8.6 | 6.7E-18 | 7.1E-17 | MB231/MB468 | CYTO |
| >hg38_21_+_8433172_8446622_RNA45SN1_45S_50ntFlanks@10391.10420.30\|AACAGCAGTTGAACATGGGTCAGTCGGTCC | >hg38_21_+_8433172_8446622_RNA45SN1_45S_50ntFlanks | 10391 | 10420 | 30 | AACAGCAGTTGAACATGGGTCAGTCGGTCC | 442.6 | -12.7 | 1.5 | -8.6 | 6.7E-18 | 7.1E-17 | MB231/MB468 | MP |
| >hg38_21_+_8433172_8446622_RNA45SN1_45S_50ntFlanks@11315.11343.29\|CCACCCCACGTCTCGTCGCGCGCGCGTCC | >hg38_21_+_8433172_8446622_RNA45SN1_45S_50ntFlanks | 11315 | 11343 | 29 | CCACCCCACGTCTCGTCGCGCGCGCGTCC | 465.0 | -12.6 | 1.5 | -8.6 | 1.1E-17 | 7.1E-17 | MB231/MB468 | MITO |
| >hg38_21_+_8433172_8446622_RNA45SN1_45S_50ntFlanks@5078.5102.25\|CGATAACGAACGAGACTCTGGCATG | >hg38_21_+_8433172_8446622_RNA45SN1_45S_50ntFlanks | 5078 | 5102 | 25 | CGATAACGAACGAGACTCTGGCATG | 1560.1 | -14.5 | 1.7 | -8.6 | 6.7E-18 | 7.1E-17 | MB231/MB468 | MP |
| >hg38_21_+_8433172_8446622_RNA45SN1_45S_50ntFlanks@11999.12035.37\|CCCTGCGGGCCGCCGGTGAAATACCACTACTCTGATC | >hg38_21_+_8433172_8446622_RNA45SN1_45S_50ntFlanks | 11999 | 12035 | 37 | CCCTGCGGGCCGCCGGTGAAATACCACTACTCTGATC | 210.1 | -6.2 | 0.7 | -8.6 | 6.8E-18 | 7.2E-17 | MB231/MB468 | CYTO |
| >hg38_21_+_8433172_8446622_RNA45SN1_45S_50ntFlanks@4884.4906.23\|CAAAGCTGAAACTTAAAGGAATT | >hg38_21_+_8433172_8446622_RNA45SN1_45S_50ntFlanks | 4884 | 4906 | 23 | CAAAGCTGAAACTTAAAGGAATT | 292.3 | -12.1 | 1.4 | -8.6 | 6.8E-18 | 7.2E-17 | MB231/MB468 | MP |
| >trnaMT_GluTTC_MT_-_14674_14742@46.71.26,trnalookalike8_GluTTC_5_-_93905172_93905240@46.71.26\|GTGGTTGTAGTCCGTGCGAGAATACC | >trnaMT_GluTTC_MT_-_14674_14742 | 46 | 71 | 26 | GTGGTTGTAGTCCGTGCGAGAATACC | 299.0 | 12.2 | 1.4 | 8.6 | 6.8E-18 | 7.2E-17 | MB231/MB468 | CYTO |
| >hg38_21_+_8433172_8446622_RNA45SN1_45S_50ntFlanks@10664.10710.47\|CTTGGAAAGCGTCGCGGTTCCGGCGGCGTCCGGTGAGCTCTCGCTGG | >hg38_21_+_8433172_8446622_RNA45SN1_45S_50ntFlanks | 10664 | 10710 | 47 | CTTGGAAAGCGTCGCGGTTCCGGCGGCGTCCGGTGAGCTCTCGCTGG | 951.4 | -13.7 | 1.6 | -8.6 | 1.1E-17 | 7.2E-17 | MB231/MB468 | MITO |
| >hg38_21_+_8433172_8446622_RNA45SN1_45S_50ntFlanks@10349.10369.21\|AACGAGAACTTTGAAGGCCGA | >hg38_21_+_8433172_8446622_RNA45SN1_45S_50ntFlanks | 10349 | 10369 | 21 | AACGAGAACTTTGAAGGCCGA | 1034.6 | 4.0 | 0.5 | 8.6 | 9.2E-18 | 7.2E-17 | BT20/MB468 | MITO |
| >hg38_21_+_8433172_8446622_RNA45SN1_45S_50ntFlanks@12001.12031.31\|CTGCGGGCCGCCGGTGAAATACCACTACTCT | >hg38_21_+_8433172_8446622_RNA45SN1_45S_50ntFlanks | 12001 | 12031 | 31 | CTGCGGGCCGCCGGTGAAATACCACTACTCT | 310.4 | -12.1 | 1.4 | -8.6 | 6.8E-18 | 7.2E-17 | MB231/MB468 | MP |
| >hg38_21_+_8433172_8446622_RNA45SN1_45S_50ntFlanks@7978.8019.42\|GACCTCAGATCAGACGTGGCGACCCGCTGAATTTAAGCATAT | >hg38_21_+_8433172_8446622_RNA45SN1_45S_50ntFlanks | 7978 | 8019 | 42 | GACCTCAGATCAGACGTGGCGACCCGCTGAATTTAAGCATAT | 477.2 | 12.9 | 1.5 | 8.6 | 6.9E-18 | 7.3E-17 | MB231/MB468 | CYTO |
| >hg38_21_+_8433172_8446622_RNA45SN1_45S_50ntFlanks@11235.11253.19\|CTTCCCGTGGATCGCCCCA | >hg38_21_+_8433172_8446622_RNA45SN1_45S_50ntFlanks | 11235 | 11253 | 19 | CTTCCCGTGGATCGCCCCA | 1634.0 | -8.3 | 1.0 | -8.6 | 7.0E-18 | 7.3E-17 | MB231/MB468 | CYTO |
| >MI0000088\|hsa-mir-30a&WithFlank&6\|-\|71403545\|71403627@53.72.20&[MIMAT0000088&hsa-miR-30a-3p&offsets\|0\|-2;m-38&6\|-\|71403554\|71403575&offsets\|0\|-2]\|CTTTCAGTCGGATGTTTGCA | >MI0000088\|hsa-mir-30a&WithFlank&6\|-\|71403545\|71403627 | 53 | 72 | 20 | CTTTCAGTCGGATGTTTGCA | 266.3 | 5.9 | 0.7 | 8.6 | 8.8E-18 | 7.3E-17 | MB231/MB468 | NUC |
| >hg38_21_+_8433172_8446622_RNA45SN1_45S_50ntFlanks@12412.12431.20\|ATCCTTCGATGTCGGCTCTT | >hg38_21_+_8433172_8446622_RNA45SN1_45S_50ntFlanks | 12412 | 12431 | 20 | ATCCTTCGATGTCGGCTCTT | 371.3 | -11.5 | 1.3 | -8.6 | 8.8E-18 | 7.3E-17 | MB231/MB468 | NUC |
| >hg38_21_+_8433172_8446622_RNA45SN1_45S_50ntFlanks@11613.11640.28\|AATTAAAACAAAGCATCGCGAAGGCCCG | >hg38_21_+_8433172_8446622_RNA45SN1_45S_50ntFlanks | 11613 | 11640 | 28 | AATTAAAACAAAGCATCGCGAAGGCCCG | 394.1 | -11.6 | 1.3 | -8.6 | 8.8E-18 | 7.3E-17 | MB231/MB468 | NUC |
| >hg38_21_+_8433172_8446622_RNA45SN1_45S_50ntFlanks@12997.13016.20\|CGCTGCGATCTATTGAAAGT | >hg38_21_+_8433172_8446622_RNA45SN1_45S_50ntFlanks | 12997 | 13016 | 20 | CGCTGCGATCTATTGAAAGT | 452.4 | 5.5 | 0.6 | 8.6 | 7.0E-18 | 7.4E-17 | MB231/MB468 | CYTO |
| >hg38_21_+_8433172_8446622_RNA45SN1_45S_50ntFlanks@12992.13015.24\|TCCCTCGCTGCGATCTATTGAAAG | >hg38_21_+_8433172_8446622_RNA45SN1_45S_50ntFlanks | 12992 | 13015 | 24 | TCCCTCGCTGCGATCTATTGAAAG | 3848.2 | -11.9 | 1.4 | -8.5 | 2.2E-17 | 7.4E-17 | BT20/MB468 | NUC |
| >MI0000081\|hsa-mir-24-2&WithFlank&19\|-\|13836281\|13836365@56.76.21(+1A);MI0000080\|hsa-mir-24-1&WithFlank&9\|+\|95086015\|95086094@50.70.21(+1A)&[MIMAT0000080&hsa-miR-24-3p&offsets\|0\|-1(+1A);m-36&19\|-\|13836288\|13836310&offsets\|0\|-2(+1A)];[MIMAT0000080_1&hsa-miR-24-3p&offsets\|0\|-1(+1A);m-35&9\|+\|95086064\|95086086&offsets\|0\|-2(+1A)]\|TGGCTCAGTTCAGCAGGAACAA | >MI0000081\|hsa-mir-24-2&WithFlank&19\|-\|13836281\|13836365 | 56 | 76 | 21(+1A) | TGGCTCAGTTCAGCAGGAACAA | 1832.1 | 6.3 | 0.7 | 8.5 | 1.3E-17 | 7.4E-17 | BT20/MB468 | CYTO |
| >hg38_21_+_8433172_8446622_RNA45SN1_45S_50ntFlanks@12942.12972.31\|TTCGTAGACGACCTGCTTCTGGGTCGGGGTT | >hg38_21_+_8433172_8446622_RNA45SN1_45S_50ntFlanks | 12942 | 12972 | 31 | TTCGTAGACGACCTGCTTCTGGGTCGGGGTT | 424.5 | -12.5 | 1.5 | -8.6 | 1.1E-17 | 7.5E-17 | MB231/MB468 | MITO |
| >hg38_21_+_8433172_8446622_RNA45SN1_45S_50ntFlanks@11318.11344.27\|CCCCACGTCTCGTCGCGCGCGCGTCCG | >hg38_21_+_8433172_8446622_RNA45SN1_45S_50ntFlanks | 11318 | 11344 | 27 | CCCCACGTCTCGTCGCGCGCGCGTCCG | 16446.6 | -11.9 | 1.4 | -8.6 | 9.0E-18 | 7.5E-17 | MB231/MB468 | NUC |
| >hg38_1_-_228634819_228635039_RNA5S12_5S_50ntFlanks@127.171.45\|ACTTGGATGGGAGACCGCCTGGGAATACCGGGTGCTGTAGGCTTT | >hg38_1_-_228634819_228635039_RNA5S12_5S_50ntFlanks | 127 | 171 | 45 | ACTTGGATGGGAGACCGCCTGGGAATACCGGGTGCTGTAGGCTTT | 3337.3 | 10.9 | 1.3 | 8.6 | 7.2E-18 | 7.5E-17 | MB231/MB468 | CYTO |
| >trnaMT_GluTTC_MT_-_14674_14742@39.66.28,trnalookalike8_GluTTC_5_-_93905172_93905240@39.66.28\|ATTGGTCGTGGTTGTAGTCCGTGCGAGA | >trnaMT_GluTTC_MT_-_14674_14742 | 39 | 66 | 28 | ATTGGTCGTGGTTGTAGTCCGTGCGAGA | 1741.4 | 9.3 | 1.1 | 8.6 | 9.5E-18 | 7.5E-17 | BT20/MB468 | MITO |

| | | | | | | | | | | | | | |
|---|---|---|---|---|---|---|---|---|---|---|---|---|---|
| >hg38_21_+_8433172_8446622_RNA45SN1_45S_50ntFlanks@4160.4182.23\|CCACATCCAAGGAAGGCAGCAGG | >hg38_21_+_8433172_8446622_RNA45SN1_45S_50ntFlanks | 4160 | 4182 | 23 | CCACATCCAAGGAAGGCAGCAGG | 390.9 | -11.6 | 1.3 | -8.6 | 9.2E-18 | 7.6E-17 | MB231/MB468 | NUC |
| >hg38_21_+_8433172_8446622_RNA45SN1_45S_50ntFlanks@12991.13014.24\|CTCCCTCGCTGCGATCTATTGAAA | >hg38_21_+_8433172_8446622_RNA45SN1_45S_50ntFlanks | 12991 | 13014 | 24 | CTCCCTCGCTGCGATCTATTGAAA | 386.1 | -12.5 | 1.4 | -8.6 | 7.2E-18 | 7.6E-17 | MB231/MB468 | MP |
| >hg38_21_+_8433172_8446622_RNA45SN1_45S_50ntFlanks@10512.10534.23\|TCCGGAGTGGCGGAGATGGGCGC | >hg38_21_+_8433172_8446622_RNA45SN1_45S_50ntFlanks | 10512 | 10534 | 23 | TCCGGAGTGGCGGAGATGGGCGC | 585.7 | -13.0 | 1.5 | -8.6 | 1.2E-17 | 7.6E-17 | MB231/MB468 | MITO |
| >hg38_21_+_8433172_8446622_RNA45SN1_45S_50ntFlanks@9175.9197.23\|TCGCGCCGTCGGGCCCGGGGGAG | >hg38_21_+_8433172_8446622_RNA45SN1_45S_50ntFlanks | 9175 | 9197 | 23 | TCGCGCCGTCGGGCCCGGGGGAG | 894.6 | -13.6 | 1.6 | -8.6 | 1.2E-17 | 7.6E-17 | MB231/MB468 | MITO |
| >trnaMT_GluTTC_MT_-_14674_14742@38.72.35,trnalookalike8_GluTTC_5_-_93905172_93905240@38.72.35\|CATTGGTCGTGGTTGTAGTCCGTGCGAGAATACCA | >trnaMT_GluTTC_MT_-_14674_14742 | 38 | 72 | 35 | CATTGGTCGTGGTTGTAGTCCGTGCGAGAATACCA | 5414.3 | 5.9 | 0.7 | 8.6 | 9.3E-18 | 7.6E-17 | MB231/MB468 | NUC |
| >hg38_21_+_8433172_8446622_RNA45SN1_45S_50ntFlanks@12992.13013.22\|TCCCTCGCTGCGATCTATTGAA | >hg38_21_+_8433172_8446622_RNA45SN1_45S_50ntFlanks | 12992 | 13013 | 22 | TCCCTCGCTGCGATCTATTGAA | 4584.1 | -6.2 | 0.7 | -8.6 | 9.2E-18 | 7.6E-17 | MB231/MB468 | NUC |
| >hg38_21_+_8433172_8446622_RNA45SN1_45S_50ntFlanks@6737.6770.34\|GATCATCGACACTTCGAACGCACTTGCGGCCCCG | >hg38_21_+_8433172_8446622_RNA45SN1_45S_50ntFlanks | 6737 | 6770 | 34 | GATCATCGACACTTCGAACGCACTTGCGGCCCCG | 629.2 | -13.2 | 1.5 | -8.6 | 7.2E-18 | 7.7E-17 | MB231/MB468 | MP |
| >hg38_21_+_8433172_8446622_RNA45SN1_45S_50ntFlanks@11027.11075.49\|CTCTCTCTCTCTCTCCCCCGCTCCCCGTCCTCCCCCCTCCCCGGGGGAG | >hg38_21_+_8433172_8446622_RNA45SN1_45S_50ntFlanks | 11027 | 11075 | 49 | CTCTCTCTCTCTCTCCCCCGCTCCCCGTCCTCCCCCCTCCCCGGGGGAG | 2553.8 | -15.7 | 1.9 | -8.5 | 2.3E-17 | 7.7E-17 | BT20/MB468 | NUC |
| >hg38_21_+_8433172_8446622_RNA45SN1_45S_50ntFlanks@12004.12033.30\|CGGGCCGCCGGTGAAATACCACTACTCTGA | >hg38_21_+_8433172_8446622_RNA45SN1_45S_50ntFlanks | 12004 | 12033 | 30 | CGGGCCGCCGGTGAAATACCACTACTCTGA | 1145.7 | 7.5 | 0.9 | 8.5 | 1.4E-17 | 7.7E-17 | BT20/MB468 | CYTO |
| >trnaMT_GluTTC_MT_-_14674_14742@38.66.29,trnalookalike8_GluTTC_5_-_93905172_93905240@38.66.29\|CATTGGTCGTGGTTGTAGTCCGTGCGAGA | >trnaMT_GluTTC_MT_-_14674_14742 | 38 | 66 | 29 | CATTGGTCGTGGTTGTAGTCCGTGCGAGA | 1216.3 | 7.4 | 0.9 | 8.6 | 9.8E-18 | 7.7E-17 | BT20/MB468 | MITO |
| >hg38_21_+_8433172_8446622_RNA45SN1_45S_50ntFlanks@6646.6672.27\|TCGTACGACTCTTAGCGGTGGATCACT | >hg38_21_+_8433172_8446622_RNA45SN1_45S_50ntFlanks | 6646 | 6672 | 27 | TCGTACGACTCTTAGCGGTGGATCACT | 11074.0 | 7.0 | 0.8 | 8.5 | 1.4E-17 | 7.7E-17 | BT20/MB468 | CYTO |
| >MI0000434\|hsa-let-7i&WithFlank&12\|+\|62603680\|62603775@12.31.20(+1C)&[MIMAT0000415&hsa-let-7i-5p&offsets\|0\|-2(+1C);m-49&12\|+\|62603691\|62603712&offsets\|0\|-2(+1C)]\|TGAGGTAGTAGTTTGTGCTGC | >MI0000434\|hsa-let-7i&WithFlank&12\|+\|62603680\|62603775 | 12 | 31 | 20(+1C) | TGAGGTAGTAGTTTGTGCTGC | 149.5 | 2.6 | 0.3 | 8.5 | 2.3E-17 | 7.7E-17 | BT20/MB468 | NUC |
| >hg38_21_+_8433172_8446622_RNA45SN1_45S_50ntFlanks@11561.11589.29\|GCGCCTAGCAGCCGACTTAGAACTGGTGC | >hg38_21_+_8433172_8446622_RNA45SN1_45S_50ntFlanks | 11561 | 11589 | 29 | GCGCCTAGCAGCCGACTTAGAACTGGTGC | 302.0 | -12.1 | 1.4 | -8.6 | 7.3E-18 | 7.7E-17 | MB231/MB468 | MP |
| >hg38_21_+_8433172_8446622_RNA45SN1_45S_50ntFlanks@8380.8401.22\|CAAGAGGGCGTGAAACCGTTAA | >hg38_21_+_8433172_8446622_RNA45SN1_45S_50ntFlanks | 8380 | 8401 | 22 | CAAGAGGGCGTGAAACCGTTAA | 543.2 | -7.1 | 0.8 | -8.6 | 7.4E-18 | 7.7E-17 | MB231/MB468 | CYTO |
| >hg38_21_+_8433172_8446622_RNA45SN1_45S_50ntFlanks@9554.9572.19\|CGTAGCGGTCCTGACGTGC | >hg38_21_+_8433172_8446622_RNA45SN1_45S_50ntFlanks | 9554 | 9572 | 19 | CGTAGCGGTCCTGACGTGC | 7790.0 | -5.7 | 0.7 | -8.6 | 1.2E-17 | 7.7E-17 | MB231/MB468 | MITO |
| >hg38_1_-_228634819_228635039_RNA5S12_5S_50ntFlanks@139.171.33\|GACCGCCTGGGAATACCGGGTGCTGTAGGCTTT | >hg38_1_-_228634819_228635039_RNA5S12_5S_50ntFlanks | 139 | 171 | 33 | GACCGCCTGGGAATACCGGGTGCTGTAGGCTTT | 1849.3 | 6.7 | 0.8 | 8.6 | 7.5E-18 | 7.8E-17 | MB231/MB468 | CYTO |
| >hg38_21_+_8433172_8446622_RNA45SN1_45S_50ntFlanks@10375.10407.33\|AGAAGGGTTCCATGTGAACAGCAGTTGAACATG | >hg38_21_+_8433172_8446622_RNA45SN1_45S_50ntFlanks | 10375 | 10407 | 33 | AGAAGGGTTCCATGTGAACAGCAGTTGAACATG | 574.5 | -7.0 | 0.8 | -8.6 | 7.5E-18 | 7.8E-17 | MB231/MB468 | CYTO |
| >hg38_21_+_8433172_8446622_RNA45SN1_45S_50ntFlanks@10680.10711.32\|GTTCCGGCGGCGTCCGGTGAGCTCTCGCTGGC | >hg38_21_+_8433172_8446622_RNA45SN1_45S_50ntFlanks | 10680 | 10711 | 32 | GTTCCGGCGGCGTCCGGTGAGCTCTCGCTGGC | 12229.3 | -9.2 | 1.1 | -8.6 | 7.5E-18 | 7.8E-17 | MB231/MB468 | CYTO |
| >hg38_21_+_8433172_8446622_RNA45SN1_45S_50ntFlanks@12625.12653.29\|CAATGGGGCGAAGCTACCATCTGTGGGAT | >hg38_21_+_8433172_8446622_RNA45SN1_45S_50ntFlanks | 12625 | 12653 | 29 | CAATGGGGCGAAGCTACCATCTGTGGGAT | 1048.6 | -7.9 | 0.9 | -8.6 | 9.5E-18 | 7.8E-17 | MB231/MB468 | NUC |
| >hg38_21_+_8433172_8446622_RNA45SN1_45S_50ntFlanks@7981.8005.25\|CTCAGATCAGACGTGGCGACCCGCT | >hg38_21_+_8433172_8446622_RNA45SN1_45S_50ntFlanks | 7981 | 8005 | 25 | CTCAGATCAGACGTGGCGACCCGCT | 695.9 | -4.4 | 0.5 | -8.5 | 2.3E-17 | 7.8E-17 | BT20/MB468 | NUC |
| >MI0000750\|hsa-mir-26a-2&WithFlank&12\|-\|57824603\|57824698@20.41.22(+1U);MI0000083\|hsa-mir-26a-1&WithFlank&3\|+\|37969398\|37969486@16.37.22(+1U)&[MIMAT0000082&hsa-miR-26a-5p&offsets\|0\|0(+1U);m-33&12\|-\|57824658\|57824679&offsets\|0\|0(+1U)];[MIMAT0000082_1&hsa-miR-26a-5p&offsets\|0\|0(+1U);m-34&3\|+\|37969413\|37969434&offsets\|0\|0(+1U)]\|TTCAAGTAATCCAGGATAGGCTT | >MI0000750\|hsa-mir-26a-2&WithFlank&12\|-\|57824603\|57824698 | 20 | 41 | 22(+1U) | TTCAAGTAATCCAGGATAGGCTT | 1926.1 | 6.3 | 0.7 | 8.6 | 1.0E-17 | 7.8E-17 | BT20/MB468 | MITO |
| >hg38_21_+_8433172_8446622_RNA45SN1_45S_50ntFlanks@9252.9273.22\|CACGGGGTCGGCGGCGACGTCG | >hg38_21_+_8433172_8446622_RNA45SN1_45S_50ntFlanks | 9252 | 9273 | 22 | CACGGGGTCGGCGGCGACGTCG | 565.6 | -13.0 | 1.5 | -8.6 | 7.4E-18 | 7.8E-17 | MB231/MB468 | MP |
| >hg38_21_+_8433172_8446622_RNA45SN1_45S_50ntFlanks@9534.9567.34\|AGGAAACTCTGGTGGAGGTCCGTAGCGGTCCTGA | >hg38_21_+_8433172_8446622_RNA45SN1_45S_50ntFlanks | 9534 | 9567 | 34 | AGGAAACTCTGGTGGAGGTCCGTAGCGGTCCTGA | 467.7 | -12.7 | 1.5 | -8.5 | 1.2E-17 | 8.0E-17 | MB231/MB468 | MITO |
| >MI0000650\|hsa-mir-200c&WithFlank&12\|+\|6963693\|6963772@50.70.21(+1U)&[MIMAT0000617&hsa-miR-200c-3p&offsets\|0\|-2(+1U);m-54&12\|+\|6963742\|6963764&offsets\|0\|-2(+1U)]\|TAATACTGCCGGGTAATGATGT | >MI0000650\|hsa-mir-200c&WithFlank&12\|+\|6963693\|6963772 | 50 | 70 | 21(+1U) | TAATACTGCCGGGTAATGATGT | 1132.9 | 10.1 | 1.2 | 8.5 | 1.4E-17 | 8.0E-17 | BT20/MB468 | CYTO |
| >hg38_21_+_8433172_8446622_RNA45SN1_45S_50ntFlanks@6690.6720.31\|AAGAACGCAGCTAGCTGCGAGAATTAATGTG | >hg38_21_+_8433172_8446622_RNA45SN1_45S_50ntFlanks | 6690 | 6720 | 31 | AAGAACGCAGCTAGCTGCGAGAATTAATGTG | 1887.6 | -11.1 | 1.3 | -8.5 | 2.4E-17 | 8.0E-17 | BT20/MB468 | NUC |
| >hg38_21_+_8433172_8446622_RNA45SN1_45S_50ntFlanks@12957.12987.31\|CTTCTGGGTCGGGGTTTCGTACGTAGCAGAG | >hg38_21_+_8433172_8446622_RNA45SN1_45S_50ntFlanks | 12957 | 12987 | 31 | CTTCTGGGTCGGGGTTTCGTACGTAGCAGAG | 385.8 | -12.4 | 1.4 | -8.5 | 1.2E-17 | 8.0E-17 | MB231/MB468 | MITO |
| >hg38_21_+_8433172_8446622_RNA45SN1_45S_50ntFlanks@10393.10410.18\|CAGCAGTTGAACATGGGT | >hg38_21_+_8433172_8446622_RNA45SN1_45S_50ntFlanks | 10393 | 10410 | 18 | CAGCAGTTGAACATGGGT | 6752.3 | -10.2 | 1.2 | -8.5 | 2.4E-17 | 8.1E-17 | BT20/MB468 | NUC |
| >hg38_21_+_8433172_8446622_RNA45SN1_45S_50ntFlanks@11540.11583.44\|CGCGGCGCCTCGCCTCGGCCGGCGCCTAGCAGCCGACTTAGAAC | >hg38_21_+_8433172_8446622_RNA45SN1_45S_50ntFlanks | 11540 | 11583 | 44 | CGCGGCGCCTCGCCTCGGCCGGCGCCTAGCAGCCGACTTAGAAC | 1832.8 | 8.3 | 1.0 | 8.6 | 7.9E-18 | 8.2E-17 | MB231/MB468 | CYTO |
| >trna152_ValCAC_6_-_27248049_27248121@1.32.32\|GCTTCTGTAGTGTAGTGGTTATCACGTTCGCC | >trna152_ValCAC_6_-_27248049_27248121 | 1 | 32 | 32 | GCTTCTGTAGTGTAGTGGTTATCACGTTCGCC | 11914.5 | 9.6 | 1.1 | 8.6 | 7.9E-18 | 8.2E-17 | MB231/MB468 | CYTO |
| >hg38_21_+_8433172_8446622_RNA45SN1_45S_50ntFlanks@9198.9225.28\|GTTCTCTCGGGGCCACGCGCGCGTCCCC | >hg38_21_+_8433172_8446622_RNA45SN1_45S_50ntFlanks | 9198 | 9225 | 28 | GTTCTCTCGGGGCCACGCGCGCGTCCCC | 499.1 | -12.8 | 1.5 | -8.6 | 7.8E-18 | 8.2E-17 | MB231/MB468 | MP |
| >hg38_21_+_8433172_8446622_RNA45SN1_45S_50ntFlanks@10571.10591.21\|GAGAAGCCGGCGGGAGCCCCG | >hg38_21_+_8433172_8446622_RNA45SN1_45S_50ntFlanks | 10571 | 10591 | 21 | GAGAAGCCGGCGGGAGCCCCG | 509.2 | -12.9 | 1.5 | -8.6 | 7.8E-18 | 8.2E-17 | MB231/MB468 | MP |
| >MI0000089\|hsa-mir-31&WithFlank&9\|-\|21512109\|21512191@14.36.23&[MIMAT0000089&hsa-miR-31-5p&offsets\|0\|+2;m-360&9\|-\|21512157\|21512178&offsets\|0\|+1]\|AGGCAAGATGCTGGCATAGCTGT | >MI0000089\|hsa-mir-31&WithFlank&9\|-\|21512109\|21512191 | 14 | 36 | 23 | AGGCAAGATGCTGGCATAGCTGT | 680.8 | 13.0 | 1.5 | 8.8 | 1.0E-18 | 8.2E-17 | BT20/MB231 | MITO |
| >hg38_21_+_8433172_8446622_RNA45SN1_45S_50ntFlanks@11795.11820.26\|AACGAGATTCCCACTGTCCCTACCTA | >hg38_21_+_8433172_8446622_RNA45SN1_45S_50ntFlanks | 11795 | 11820 | 26 | AACGAGATTCCCACTGTCCCTACCTA | 428.7 | -12.5 | 1.5 | -8.5 | 1.3E-17 | 8.3E-17 | MB231/MB468 | MITO |
| >hg38_21_+_8433172_8446622_RNA45SN1_45S_50ntFlanks@10786.10815.30\|AACAGCCTCTGGCATGTTGGAACAATGTAG | >hg38_21_+_8433172_8446622_RNA45SN1_45S_50ntFlanks | 10786 | 10815 | 30 | AACAGCCTCTGGCATGTTGGAACAATGTAG | 363.2 | -12.4 | 1.4 | -8.6 | 7.9E-18 | 8.3E-17 | MB231/MB468 | MP |
| >hg38_21_+_8433172_8446622_RNA45SN1_45S_50ntFlanks@12415.12447.33\|CTTCGATGTCGGCTCTTCCTATCATTGTGAAGC | >hg38_21_+_8433172_8446622_RNA45SN1_45S_50ntFlanks | 12415 | 12447 | 33 | CTTCGATGTCGGCTCTTCCTATCATTGTGAAGC | 1008.3 | 4.7 | 0.6 | 8.6 | 8.1E-18 | 8.3E-17 | MB231/MB468 | CYTO |
| >hg38_21_+_8433172_8446622_RNA45SN1_45S_50ntFlanks@4733.4766.34\|GAACGAAAGTCGGAGGTTCGAAGACGATCAGATA | >hg38_21_+_8433172_8446622_RNA45SN1_45S_50ntFlanks | 4733 | 4766 | 34 | GAACGAAAGTCGGAGGTTCGAAGACGATCAGATA | 588.5 | 7.2 | 0.8 | 8.6 | 8.1E-18 | 8.4E-17 | MB231/MB468 | CYTO |
| >hg38_21_+_8433172_8446622_RNA45SN1_45S_50ntFlanks@11999.12039.41\|CCCTGCGGGCCGCCGGTGAAATACCACTACTCTGATCGTTT | >hg38_21_+_8433172_8446622_RNA45SN1_45S_50ntFlanks | 11999 | 12039 | 41 | CCCTGCGGGCCGCCGGTGAAATACCACTACTCTGATCGTTT | 583.9 | -13.6 | 1.6 | -8.5 | 2.5E-17 | 8.4E-17 | BT20/MB468 | NUC |
| >trnaMT_GluTTC_MT_-_14674_14742@21.58.38,trnalookalike8_GluTTC_5_-_93905172_93905240@21.58.38\|ACGATGGTTTTTCATATCATTGGTCGTGGTTGTAGTCC | >trnaMT_GluTTC_MT_-_14674_14742 | 21 | 58 | 38 | ACGATGGTTTTTCATATCATTGGTCGTGGTTGTAGTCC | 1078.7 | 8.6 | 1.0 | 8.6 | 1.1E-17 | 8.4E-17 | BT20/MB468 | MITO |
| >am_trna2_GlyTCC_19_+_4724082_4724153@1.30.30\|GCGTTGGTGGTATAGTGGTTAGCATAGCTG | >am_trna2_GlyTCC_19_+_4724082_4724153 | 1 | 30 | 30 | GCGTTGGTGGTATAGTGGTTAGCATAGCTG | 1082.4 | -4.8 | 0.6 | -8.5 | 1.5E-17 | 8.4E-17 | BT20/MB468 | CYTO |
| >hg38_21_+_8433172_8446622_RNA45SN1_45S_50ntFlanks@10408.10426.19\|GGTCAGTCGGTCCTGAGAG | >hg38_21_+_8433172_8446622_RNA45SN1_45S_50ntFlanks | 10408 | 10426 | 19 | GGTCAGTCGGTCCTGAGAG | 547.7 | -13.0 | 1.5 | -8.6 | 8.0E-18 | 8.5E-17 | MB231/MB468 | MP |
| >hg38_21_+_8433172_8446622_RNA45SN1_45S_50ntFlanks@12360.12382.23\|CAGGGATAACTGGCTTGTGGCGG | >hg38_21_+_8433172_8446622_RNA45SN1_45S_50ntFlanks | 12360 | 12382 | 23 | CAGGGATAACTGGCTTGTGGCGG | 337.8 | -12.3 | 1.4 | -8.6 | 8.1E-18 | 8.5E-17 | MB231/MB468 | MP |
| >MI0000088\|hsa-mir-30a&WithFlank&6\|-\|71403545\|71403627@12.33.22&[MIMAT0000087&hsa-miR-30a-5p&offsets\|0\|0;m-15&6\|-\|71403595\|71403616&offsets\|0\|0]\|TGTAAACATCCTCGACTGGAAG | >MI0000088\|hsa-mir-30a&WithFlank&6\|-\|71403545\|71403627 | 12 | 33 | 22 | TGTAAACATCCTCGACTGGAAG | 3319.8 | 6.7 | 0.8 | 8.6 | 1.0E-17 | 8.5E-17 | MB231/MB468 | NUC |
| >MI0001652\|hsa-mir-450a-1&WithFlank&X\|-\|134540335\|134540437@24.43.20;MI0003187\|hsa-mir-450a-2&WithFlank&X\|-\|134540502\|134540613@29.48.20&[MIMAT0001545&hsa-miR-450a-5p&offsets\|0\|-2;m-288&X\|-\|134540393\|134540414&offsets\|0\|-2];[MIMAT0001545_1&hsa-miR-450a-5p&offsets\|0\|-2;m-287&X\|-\|134540564\|134540585&offsets\|0\|-2]\|TTTTGCGATGTGTTCCTAAT | >MI0001652\|hsa-mir-450a-1&WithFlank&X\|-\|134540335\|134540437 | 24 | 43 | 20 | TTTTGCGATGTGTTCCTAAT | 352.7 | 12.7 | 1.5 | 8.5 | 1.5E-17 | 8.5E-17 | BT20/MB468 | CYTO |
| >hg38_1_-_228634819_228635039_RNA5S12_5S_50ntFlanks@152.170.19\|TACCGGGTGCTGTAGGCTT | >hg38_1_-_228634819_228635039_RNA5S12_5S_50ntFlanks | 152 | 170 | 19 | TACCGGGTGCTGTAGGCTT | 6462.0 | 6.4 | 0.7 | 8.6 | 8.3E-18 | 8.5E-17 | MB231/MB468 | CYTO |
| >hg38_21_+_8433172_8446622_RNA45SN1_45S_50ntFlanks@4189.4207.19\|AATTACCCACTCCCGACCC | >hg38_21_+_8433172_8446622_RNA45SN1_45S_50ntFlanks | 4189 | 4207 | 19 | AATTACCCACTCCCGACCC | 212.8 | 11.7 | 1.4 | 8.6 | 8.3E-18 | 8.5E-17 | MB231/MB468 | CYTO |
| >hg38_21_+_8433172_8446622_RNA45SN1_45S_50ntFlanks@5436.5466.31\|GTGAGGCCCTCGGATCGGCCCCGCCGGGGTC | >hg38_21_+_8433172_8446622_RNA45SN1_45S_50ntFlanks | 5436 | 5466 | 31 | GTGAGGCCCTCGGATCGGCCCCGCCGGGGTC | 328.9 | -6.2 | 0.7 | -8.6 | 8.3E-18 | 8.5E-17 | MB231/MB468 | CYTO |
| >hg38_21_+_8433172_8446622_RNA45SN1_45S_50ntFlanks@12329.12346.18\|TTTTAAGCAGGAGGTGTC | >hg38_21_+_8433172_8446622_RNA45SN1_45S_50ntFlanks | 12329 | 12346 | 18 | TTTTAAGCAGGAGGTGTC | 453.3 | -11.8 | 1.4 | -8.6 | 1.1E-17 | 8.6E-17 | MB231/MB468 | NUC |
| >am_MI0003772\|hsa-mir-151b&WithFlank&14\|-\|100109413\|100109520@66.85.20(+1C);MI0000809\|hsa-mir-151a&WithFlank&8\|-\|140732558\|140732659@17.36.20(+1C)&[MIMAT0010214&hsa-miR-151b&offsets\|0\|+2(+1C);m-179&14\|-\|100109436\|100109455&offsets\|0\|0(+1C)];[MIMAT0004697&hsa-miR-151a-5p&offsets\|0\|-1(+1C);m-75&8\|-\|140732623\|140732643&offsets\|0\|-1(+1C)]\|TCGAGGAGCTCACAGTCTAGC | >am_MI0003772\|hsa-mir-151b&WithFlank&14\|-\|100109413\|100109520 | 66 | 85 | 20(+1C) | TCGAGGAGCTCACAGTCTAGC | 588.4 | 11.1 | 1.3 | 8.5 | 1.5E-17 | 8.7E-17 | BT20/MB468 | CYTO |

| | | | | | | | | | | | | | |
|---|---|---|---|---|---|---|---|---|---|---|---|---|---|
| >hg38_21_+_8433172_8446622_RNA45SN1_45S_50ntFlanks@8079.8105.27\|AAGAGCCCAGCGCCGAATCCCCGCCCC | >hg38_21_+_8433172_8446622_RNA45SN1_45S_50ntFlanks | 8079 | 8105 | 27 | AAGAGCCCAGCGCCGAATCCCCGCCCC | 914.4 | -13.7 | 1.6 | -8.6 | 8.3E-18 | 8.7E-17 | MB231/MB468 | MP |
| >hg38_21_+_8433172_8446622_RNA45SN1_45S_50ntFlanks@12973.12992.20\|TCGTACGTAGCAGAGCAGCT | >hg38_21_+_8433172_8446622_RNA45SN1_45S_50ntFlanks | 12973 | 12992 | 20 | TCGTACGTAGCAGAGCAGCT | 1361.9 | -3.7 | 0.4 | -8.6 | 8.5E-18 | 8.7E-17 | MB231/MB468 | CYTO |
| >hg38_21_+_8433172_8446622_RNA45SN1_45S_50ntFlanks@11843.11873.31\|AGGGAACGGGCTTGGCGGAATCAGCGGGGAA | >hg38_21_+_8433172_8446622_RNA45SN1_45S_50ntFlanks | 11843 | 11873 | 31 | AGGGAACGGGCTTGGCGGAATCAGCGGGGAA | 449.2 | -12.7 | 1.5 | -8.6 | 8.3E-18 | 8.7E-17 | MB231/MB468 | MP |
| >am_MI0000809\|hsa-mir-151a&WithFlank&8\|-\|140732558\|140732659@53.70.18&[MIMAT0000757&hsa-miR-151a-3p&offsets\|0\|-3;m-30&8\|-\|140732587\|140732607&offsets\|0\|-3]\|CTAGACTGAAGCTCCTTG | >am_MI0000809\|hsa-mir-151a&WithFlank&8\|-\|140732558\|140732659 | 53 | 70 | 18 | CTAGACTGAAGCTCCTTG | 358.0 | 3.7 | 0.4 | 8.5 | 2.6E-17 | 8.7E-17 | BT20/MB468 | NUC |
| >hg38_21_+_8433172_8446622_RNA45SN1_45S_50ntFlanks@4734.4767.34\|AACGAAAGTCGGAGGTTCGAAGACGATCAGATAC | >hg38_21_+_8433172_8446622_RNA45SN1_45S_50ntFlanks | 4734 | 4767 | 34 | AACGAAAGTCGGAGGTTCGAAGACGATCAGATAC | 532.3 | 10.7 | 1.2 | 8.6 | 8.5E-18 | 8.7E-17 | MB231/MB468 | CYTO |
| >hg38_21_+_8433172_8446622_RNA45SN1_45S_50ntFlanks@10351.10381.31\|CGAGAACTTTGAAGGCCGAAGTGGAGAAGGG | >hg38_21_+_8433172_8446622_RNA45SN1_45S_50ntFlanks | 10351 | 10381 | 31 | CGAGAACTTTGAAGGCCGAAGTGGAGAAGGG | 2932.3 | -9.3 | 1.1 | -8.5 | 2.6E-17 | 8.7E-17 | BT20/MB468 | NUC |
| >hg38_21_+_8433172_8446622_RNA45SN1_45S_50ntFlanks@9352.9377.26\|CCGTGGCGCAATGAAGGTGAAGGCCG | >hg38_21_+_8433172_8446622_RNA45SN1_45S_50ntFlanks | 9352 | 9377 | 26 | CCGTGGCGCAATGAAGGTGAAGGCCG | 582.4 | -13.0 | 1.5 | -8.5 | 1.4E-17 | 8.7E-17 | MB231/MB468 | MITO |
| >MI0000440\|hsa-mir-27b&WithFlank&9\|+\|95085439\|95085547@67.87.21(+1U)&[MIMAT0000419&hsa-miR-27b-3p&offsets\|0\|0(+1U);m-39&9\|+\|95085505\|95085525&offsets\|0\|0(+1U)]\|TTCACAGTGGCTAAGTTCTGCT | >MI0000440\|hsa-mir-27b&WithFlank&9\|+\|95085439\|95085547 | 67 | 87 | 21(+1U) | TTCACAGTGGCTAAGTTCTGCT | 907.4 | 6.5 | 0.8 | 8.6 | 1.1E-17 | 8.7E-17 | BT20/MB468 | MITO |
| >MI0000650\|hsa-mir-200c&WithFlank&12\|+\|6963693\|6963772@50.73.24&[MIMAT0000617&hsa-miR-200c-3p&offsets\|0\|+1;m-54&12\|+\|6963742\|6963764&offsets\|0\|+1]\|TAATACTGCCGGGTAATGATGGAG | >MI0000650\|hsa-mir-200c&WithFlank&12\|+\|6963693\|6963772 | 50 | 73 | 24 | TAATACTGCCGGGTAATGATGGAG | 492.3 | 12.7 | 1.4 | 8.8 | 1.4E-18 | 8.8E-17 | BT20/MB231 | NUC |
| >hg38_21_+_8433172_8446622_RNA45SN1_45S_50ntFlanks@9623.9654.32\|TAGTAGCTGGTTCCCTCCGAAGTTTCCCTCAG | >hg38_21_+_8433172_8446622_RNA45SN1_45S_50ntFlanks | 9623 | 9654 | 32 | TAGTAGCTGGTTCCCTCCGAAGTTTCCCTCAG | 413.5 | -12.5 | 1.5 | -8.5 | 1.4E-17 | 8.8E-17 | MB231/MB468 | MITO |
| >hg38_21_+_8433172_8446622_RNA45SN1_45S_50ntFlanks@3783.3801.19\|AGTGAAACTGCGAATGGCT | >hg38_21_+_8433172_8446622_RNA45SN1_45S_50ntFlanks | 3783 | 3801 | 19 | AGTGAAACTGCGAATGGCT | 858.2 | -6.8 | 0.8 | -8.5 | 2.6E-17 | 8.8E-17 | BT20/MB468 | NUC |
| >MI0005756\|hsa-mir-934&WithFlank&X\|+\|136550872\|136550966@21.40.20(+1A)&[MIMAT0004977&hsa-miR-934&offsets\|0\|-2(+1A);m-943&X\|+\|136550892\|136550913&offsets\|0\|-2(+1A)]\|TGTCTACTACTGGAGACACTA | >MI0005756\|hsa-mir-934&WithFlank&X\|+\|136550872\|136550966 | 21 | 40 | 20(+1A) | TGTCTACTACTGGAGACACTA | 307.7 | 12.5 | 1.5 | 8.5 | 1.6E-17 | 8.8E-17 | BT20/MB468 | CYTO |
| >hg38_21_+_8433172_8446622_RNA45SN1_45S_50ntFlanks@9454.9500.47\|CCGCGCCGGGGAGGTGGAGCACGAGCGCACGTGTTAGGACCCGAAAG | >hg38_21_+_8433172_8446622_RNA45SN1_45S_50ntFlanks | 9454 | 9500 | 47 | CCGCGCCGGGGAGGTGGAGCACGAGCGCACGTGTTAGGACCCGAAAG | 545.7 | -12.9 | 1.5 | -8.5 | 1.4E-17 | 8.9E-17 | MB231/MB468 | MITO |
| >hg38_21_+_8433172_8446622_RNA45SN1_45S_50ntFlanks@9714.9737.24\|CGAATGATTAGAGGTCTTGGGGCC | >hg38_21_+_8433172_8446622_RNA45SN1_45S_50ntFlanks | 9714 | 9737 | 24 | CGAATGATTAGAGGTCTTGGGGCC | 2756.5 | -6.9 | 0.8 | -8.6 | 1.1E-17 | 8.9E-17 | BT20/MB468 | MITO |
| >MI0000736\|hsa-mir-30c-1&WithFlank&1\|+\|40757278\|40757378@23.46.24(+2U);MI0000254\|hsa-mir-30c-2&WithFlank&6\|-\|71376954\|71377037@13.36.24(+2U)&[MIMAT0000244&hsa-miR-30c-5p&offsets\|0\|+1(+2U);m-57&1\|+\|40757300\|40757323&offsets\|0\|0(+2U)];[MIMAT0000244_1&hsa-miR-30c-5p&offsets\|0\|+1(+2U);m-56&6\|-\|71377002\|71377025&offsets\|0\|0(+2U)]\|TGTAAACATCCTACACTCTCAGCTTT | >MI0000736\|hsa-mir-30c-1&WithFlank&1\|+\|40757278\|40757378 | 23 | 46 | 24(+2U) | TGTAAACATCCTACACTCTCAGCTTT | 403.4 | 6.0 | 0.7 | 8.6 | 1.1E-17 | 8.9E-17 | MB231/MB468 | NUC |
| >am_trna13_LysCTT_14_-_58706613_58706685@31.51.21,trna2_LysCTT_15_+_79152904_79152976@31.51.21,trna4_LysCTT_18_-_43669271_43669343@31.51.21\|ACTCTTAATCCCAGGGTCGTG | >am_trna13_LysCTT_14_-_58706613_58706685 | 31 | 51 | 21 | ACTCTTAATCCCAGGGTCGTG | 391.1 | -12.5 | 1.5 | -8.6 | 8.5E-18 | 8.9E-17 | MB231/MB468 | MP |
| >hg38_21_+_8433172_8446622_RNA45SN1_45S_50ntFlanks@10351.10388.38\|CGAGAACTTTGAAGGCCGAAGTGGAGAAGGGTTCCATG | >hg38_21_+_8433172_8446622_RNA45SN1_45S_50ntFlanks | 10351 | 10388 | 38 | CGAGAACTTTGAAGGCCGAAGTGGAGAAGGGTTCCATG | 16422.9 | -11.2 | 1.3 | -8.5 | 2.7E-17 | 9.0E-17 | BT20/MB468 | NUC |
| >hg38_21_+_8433172_8446622_RNA45SN1_45S_50ntFlanks@10689.10706.18\|GCGTCCGGTGAGCTCTCG | >hg38_21_+_8433172_8446622_RNA45SN1_45S_50ntFlanks | 10689 | 10706 | 18 | GCGTCCGGTGAGCTCTCG | 5541.2 | -12.6 | 1.5 | -8.5 | 2.7E-17 | 9.0E-17 | BT20/MB468 | NUC |
| >hg38_21_+_8433172_8446622_RNA45SN1_45S_50ntFlanks@11577.11605.29\|TTAGAACTGGTGCGGACCAGGGGAATCCG | >hg38_21_+_8433172_8446622_RNA45SN1_45S_50ntFlanks | 11577 | 11605 | 29 | TTAGAACTGGTGCGGACCAGGGGAATCCG | 492.0 | -12.7 | 1.5 | -8.5 | 1.4E-17 | 9.0E-17 | MB231/MB468 | MITO |
| >MI0000285\|hsa-mir-205&WithFlank&1\|+\|209432127\|209432248@40.62.23(+1U)&[MIMAT0000266&hsa-miR-205-5p&offsets\|0\|+1(+1U);m-351&1\|+\|209432166\|209432187&offsets\|0\|+1(+1U)]\|TCCTTCATTCCACCGGAGTCTGTT | >MI0000285\|hsa-mir-205&WithFlank&1\|+\|209432127\|209432248 | 40 | 62 | 23(+1U) | TCCTTCATTCCACCGGAGTCTGTT | 1079.7 | 13.8 | 1.6 | 8.8 | 1.5E-18 | 9.0E-17 | BT20/MB231 | NUC |
| >hg38_21_+_8433172_8446622_RNA45SN1_45S_50ntFlanks@10128.10145.18\|CGGCGGCGGCGGCGGCGG | >hg38_21_+_8433172_8446622_RNA45SN1_45S_50ntFlanks | 10128 | 10145 | 18 | CGGCGGCGGCGGCGGCGG | 458.7 | -12.7 | 1.5 | -8.6 | 8.6E-18 | 9.0E-17 | MB231/MB468 | MP |
| >hg38_21_+_8433172_8446622_RNA45SN1_45S_50ntFlanks@10131.10148.18\|CGGCGGCGGCGGCGGCGG | >hg38_21_+_8433172_8446622_RNA45SN1_45S_50ntFlanks | 10131 | 10148 | 18 | CGGCGGCGGCGGCGGCGG | 458.7 | -12.7 | 1.5 | -8.6 | 8.6E-18 | 9.0E-17 | MB231/MB468 | MP |
| >hg38_21_+_8433172_8446622_RNA45SN1_45S_50ntFlanks@8824.8841.18\|CGGCGGCGGCGGCGGCGG | >hg38_21_+_8433172_8446622_RNA45SN1_45S_50ntFlanks | 8824 | 8841 | 18 | CGGCGGCGGCGGCGGCGG | 458.7 | -12.7 | 1.5 | -8.6 | 8.6E-18 | 9.0E-17 | MB231/MB468 | MP |
| >hg38_21_+_8433172_8446622_RNA45SN1_45S_50ntFlanks@8845.8862.18\|CGGCGGCGGCGGCGGCGG | >hg38_21_+_8433172_8446622_RNA45SN1_45S_50ntFlanks | 8845 | 8862 | 18 | CGGCGGCGGCGGCGGCGG | 458.7 | -12.7 | 1.5 | -8.6 | 8.6E-18 | 9.0E-17 | MB231/MB468 | MP |
| >hg38_21_+_8433172_8446622_RNA45SN1_45S_50ntFlanks@8848.8865.18\|CGGCGGCGGCGGCGGCGG | >hg38_21_+_8433172_8446622_RNA45SN1_45S_50ntFlanks | 8848 | 8865 | 18 | CGGCGGCGGCGGCGGCGG | 458.7 | -12.7 | 1.5 | -8.6 | 8.6E-18 | 9.0E-17 | MB231/MB468 | MP |
| >hg38_21_+_8433172_8446622_RNA45SN1_45S_50ntFlanks@10376.10424.49\|GAAGGGTTCCATGTGAACAGCAGTTGAACATGGGTCAGTCGGTCCTGAG | >hg38_21_+_8433172_8446622_RNA45SN1_45S_50ntFlanks | 10376 | 10424 | 49 | GAAGGGTTCCATGTGAACAGCAGTTGAACATGGGTCAGTCGGTCCTGAG | 1392.2 | 14.9 | 1.7 | 8.6 | 1.2E-17 | 9.0E-17 | BT20/MB468 | MITO |
| >hg38_21_+_8433172_8446622_RNA45SN1_45S_50ntFlanks@6645.6665.21\|CTCGTACGACTCTTAGCGGTG | >hg38_21_+_8433172_8446622_RNA45SN1_45S_50ntFlanks | 6645 | 6665 | 21 | CTCGTACGACTCTTAGCGGTG | 14832.0 | 5.7 | 0.7 | 8.5 | 1.6E-17 | 9.1E-17 | BT20/MB468 | CYTO |
| >hg38_21_+_8433172_8446622_RNA45SN1_45S_50ntFlanks@6719.6743.25\|TGAATTGCAGGACACATTGATCATC | >hg38_21_+_8433172_8446622_RNA45SN1_45S_50ntFlanks | 6719 | 6743 | 25 | TGAATTGCAGGACACATTGATCATC | 4524.2 | -6.0 | 0.7 | -8.5 | 1.6E-17 | 9.1E-17 | BT20/MB468 | CYTO |
| >hg38_21_+_8433172_8446622_RNA45SN1_45S_50ntFlanks@8112.8141.30\|GGGCGCGGGACATGTGGCGTACGGAAGACC | >hg38_21_+_8433172_8446622_RNA45SN1_45S_50ntFlanks | 8112 | 8141 | 30 | GGGCGCGGGACATGTGGCGTACGGAAGACC | 722.3 | -13.4 | 1.6 | -8.6 | 8.7E-18 | 9.1E-17 | MB231/MB468 | MP |
| >hg38_21_+_8433172_8446622_RNA45SN1_45S_50ntFlanks@8179.8206.28\|CTGATCGAGGCCCAGCCCGTGGACGGTG | >hg38_21_+_8433172_8446622_RNA45SN1_45S_50ntFlanks | 8179 | 8206 | 28 | CTGATCGAGGCCCAGCCCGTGGACGGTG | 362.5 | -12.4 | 1.4 | -8.6 | 8.8E-18 | 9.1E-17 | MB231/MB468 | MP |
| >hg38_21_+_8433172_8446622_RNA45SN1_45S_50ntFlanks@11212.11230.19\|CGACTCTGGACGCGAGCCG | >hg38_21_+_8433172_8446622_RNA45SN1_45S_50ntFlanks | 11212 | 11230 | 19 | CGACTCTGGACGCGAGCCG | 326.9 | -12.2 | 1.4 | -8.6 | 8.8E-18 | 9.1E-17 | MB231/MB468 | MP |
| >hg38_21_+_8433172_8446622_RNA45SN1_45S_50ntFlanks@11930.11954.25\|TAGAATAAGTGGGAGGCCCCCGGCG | >hg38_21_+_8433172_8446622_RNA45SN1_45S_50ntFlanks | 11930 | 11954 | 25 | TAGAATAAGTGGGAGGCCCCCGGCG | 347.5 | -12.3 | 1.4 | -8.6 | 8.8E-18 | 9.1E-17 | MB231/MB468 | MP |
| >hg38_21_+_8433172_8446622_RNA45SN1_45S_50ntFlanks@12157.12180.24\|AGTTTGACTGGGGCGGTACACCTG | >hg38_21_+_8433172_8446622_RNA45SN1_45S_50ntFlanks | 12157 | 12180 | 24 | AGTTTGACTGGGGCGGTACACCTG | 562.9 | -12.1 | 1.4 | -8.6 | 1.1E-17 | 9.2E-17 | MB231/MB468 | NUC |
| >MI0000478\|hsa-mir-149&WithFlank&2\|+\|240455995\|240456095@21.42.22&[MIMAT0000450&hsa-miR-149-5p&offsets\|0\|-1;m-195&2\|+\|240456015\|240456037&offsets\|0\|-1]\|TCTGGCTCCGTGTCTTCACTCC | >MI0000478\|hsa-mir-149&WithFlank&2\|+\|240455995\|240456095 | 21 | 42 | 22 | TCTGGCTCCGTGTCTTCACTCC | 368.9 | 5.3 | 0.6 | 8.5 | 2.8E-17 | 9.2E-17 | BT20/MB468 | NUC |
| >hg38_21_+_8433172_8446622_RNA45SN1_45S_50ntFlanks@9563.9596.34\|CCTGACGTGCAAATCGGTCGTCCGACCTGGGTAT | >hg38_21_+_8433172_8446622_RNA45SN1_45S_50ntFlanks | 9563 | 9596 | 34 | CCTGACGTGCAAATCGGTCGTCCGACCTGGGTAT | 3790.3 | -11.9 | 1.4 | -8.5 | 2.8E-17 | 9.2E-17 | BT20/MB468 | NUC |
| >hg38_21_+_8433172_8446622_RNA45SN1_45S_50ntFlanks@9574.9596.23\|AATCGGTCGTCCGACCTGGGTAT | >hg38_21_+_8433172_8446622_RNA45SN1_45S_50ntFlanks | 9574 | 9596 | 23 | AATCGGTCGTCCGACCTGGGTAT | 24226.0 | -10.7 | 1.2 | -8.6 | 8.8E-18 | 9.2E-17 | MB231/MB468 | MP |
| >hg38_21_+_8433172_8446622_RNA45SN1_45S_50ntFlanks@12002.12031.30\|TGCGGGCCGCCGGTGAAATACCACTACTCT | >hg38_21_+_8433172_8446622_RNA45SN1_45S_50ntFlanks | 12002 | 12031 | 30 | TGCGGGCCGCCGGTGAAATACCACTACTCT | 1410.0 | -5.9 | 0.7 | -8.5 | 1.4E-17 | 9.3E-17 | MB231/MB468 | MITO |
| >hg38_21_+_8433172_8446622_RNA45SN1_45S_50ntFlanks@10808.10828.21\|CAATGTAGGTAAGGGAAGTCG | >hg38_21_+_8433172_8446622_RNA45SN1_45S_50ntFlanks | 10808 | 10828 | 21 | CAATGTAGGTAAGGGAAGTCG | 1030.0 | -13.0 | 1.5 | -8.6 | 1.1E-17 | 9.3E-17 | MB231/MB468 | NUC |
| >hg38_21_+_8433172_8446622_RNA45SN1_45S_50ntFlanks@12489.12509.21\|ACGTGAGCTGGGTTTAGACCG | >hg38_21_+_8433172_8446622_RNA45SN1_45S_50ntFlanks | 12489 | 12509 | 21 | ACGTGAGCTGGGTTTAGACCG | 567.6 | -13.0 | 1.5 | -8.6 | 9.0E-18 | 9.3E-17 | MB231/MB468 | MP |
| >hg38_21_+_8433172_8446622_RNA45SN1_45S_50ntFlanks@11581.11612.32\|AACTGGTGCGGACCAGGGGAATCCGACTGTTT | >hg38_21_+_8433172_8446622_RNA45SN1_45S_50ntFlanks | 11581 | 11612 | 32 | AACTGGTGCGGACCAGGGGAATCCGACTGTTT | 2591.9 | -6.3 | 0.7 | -8.6 | 9.0E-18 | 9.4E-17 | MB231/MB468 | MP |
| >hg38_21_+_8433172_8446622_RNA45SN1_45S_50ntFlanks@9035.9059.25\|CGTGGGGGGGCCGGGCCACCCCTCC | >hg38_21_+_8433172_8446622_RNA45SN1_45S_50ntFlanks | 9035 | 9059 | 25 | CGTGGGGGGGCCGGGCCACCCCTCC | 536.4 | -12.9 | 1.5 | -8.6 | 9.0E-18 | 9.4E-17 | MB231/MB468 | MP |
| >MI0000746\|hsa-mir-99b&WithFlank&19\|+\|51692606\|51692687@13.32.20(+2U)&[MIMAT0000689&hsa-miR-99b-5p&offsets\|0\|-2(+2U);m-10&19\|+\|51692618\|51692639&offsets\|0\|-2(+2U)]\|CACCCGTAGAACCGACCTTGTT | >MI0000746\|hsa-mir-99b&WithFlank&19\|+\|51692606\|51692687 | 13 | 32 | 20(+2U) | CACCCGTAGAACCGACCTTGTT | 731.5 | 10.8 | 1.2 | 8.8 | 1.6E-18 | 9.4E-17 | BT20/MB231 | NUC |
| >hg38_21_+_8433172_8446622_RNA45SN1_45S_50ntFlanks@12710.12745.36\|CGCGGAGCCTCGGTTGGCCTCGGATAGCCGGTCCCC | >hg38_21_+_8433172_8446622_RNA45SN1_45S_50ntFlanks | 12710 | 12745 | 36 | CGCGGAGCCTCGGTTGGCCTCGGATAGCCGGTCCCC | 429.3 | -12.6 | 1.5 | -8.6 | 9.1E-18 | 9.4E-17 | MB231/MB468 | MP |
| >MI0000434\|hsa-let-7i&WithFlank&12\|+\|62603680\|62603775@12.32.21(+1C)&[MIMAT0000415&hsa-let-7i-5p&offsets\|0\|-1(+1C);m-49&12\|+\|62603691\|62603712&offsets\|0\|-1(+1C)]\|TGAGGTAGTAGTTTGTGCTGTC | >MI0000434\|hsa-let-7i&WithFlank&12\|+\|62603680\|62603775 | 12 | 32 | 21(+1C) | TGAGGTAGTAGTTTGTGCTGTC | 21540.7 | 6.0 | 0.7 | 8.6 | 1.2E-17 | 9.4E-17 | MB231/MB468 | NUC |
| >hg38_21_+_8433172_8446622_RNA45SN1_45S_50ntFlanks@4444.4472.29\|CCTTGCCTCTCGGCGCCCCCTCGATGCTC | >hg38_21_+_8433172_8446622_RNA45SN1_45S_50ntFlanks | 4444 | 4472 | 29 | CCTTGCCTCTCGGCGCCCCCTCGATGCTC | 317.8 | -12.2 | 1.4 | -8.6 | 9.1E-18 | 9.5E-17 | MB231/MB468 | MP |
| >hg38_21_+_8433172_8446622_RNA45SN1_45S_50ntFlanks@10830.10852.23\|CAAGCCGGATCCGTAACTTCGGG | >hg38_21_+_8433172_8446622_RNA45SN1_45S_50ntFlanks | 10830 | 10852 | 23 | CAAGCCGGATCCGTAACTTCGGG | 305.6 | -12.1 | 1.4 | -8.6 | 9.2E-18 | 9.5E-17 | MB231/MB468 | MP |
| >MI0000088\|hsa-mir-30a&WithFlank&6\|-\|71403545\|71403627@54.74.21&[MIMAT0000088&hsa-miR-30a-3p&offsets\|+1\|0;m-38&6\|-\|71403554\|71403575&offsets\|+1\|0]\|TTTCAGTCGGATGTTTGCAGC | >MI0000088\|hsa-mir-30a&WithFlank&6\|-\|71403545\|71403627 | 54 | 74 | 21 | TTTCAGTCGGATGTTTGCAGC | 4977.5 | 6.5 | 0.8 | 8.6 | 1.2E-17 | 9.5E-17 | MB231/MB468 | NUC |
| >hg38_21_+_8433172_8446622_RNA45SN1_45S_50ntFlanks@11452.11472.21\|CTCCTCGGCGCGCGGCGGCGG | >hg38_21_+_8433172_8446622_RNA45SN1_45S_50ntFlanks | 11452 | 11472 | 21 | CTCCTCGGCGCGCGGCGGCGG | 516.5 | -12.8 | 1.5 | -8.5 | 1.5E-17 | 9.5E-17 | MB231/MB468 | MITO |

<table>
<tr><td>>MI0000239|hsa-mir-197&WithFlank&1|+|109598887|109598973@54.76.23&[MIMAT0000227&hsa-miR-197-3p&offsets|0|+1;m-99&1|+|109598940|109598961&offsets|0|+1]|TTCACCACCTTCTCCACCCAGCA</td><td>>MI0000239|hsa-mir-197&WithFlank&1|+|109598887|109598973</td><td>54</td><td>76</td><td>23</td><td>TTCACCACCTTCTCCACCCAGCA</td><td>670.6</td><td>11.0</td><td>1.3</td><td>8.5</td><td>1.7E-17</td><td>9.6E-17</td><td>BT20/MB468</td><td>CYTO</td></tr>
<tr><td>>hg38_21_+_8433172_8446622_RNA45SN1_45S_50ntFlanks@13004.13028.25|ATCTATTGAAAGTCAGCCCTCGACA</td><td>>hg38_21_+_8433172_8446622_RNA45SN1_45S_50ntFlanks</td><td>13004</td><td>13028</td><td>25</td><td>ATCTATTGAAAGTCAGCCCTCGACA</td><td>1365.0</td><td>-12.7</td><td>1.5</td><td>-8.5</td><td>1.7E-17</td><td>9.6E-17</td><td>BT20/MB468</td><td>CYTO</td></tr>
<tr><td>>hg38_21_+_8433172_8446622_RNA45SN1_45S_50ntFlanks@11885.11905.21|TTGAGCTTGACTCTAGTCTGG</td><td>>hg38_21_+_8433172_8446622_RNA45SN1_45S_50ntFlanks</td><td>11885</td><td>11905</td><td>21</td><td>TTGAGCTTGACTCTAGTCTGG</td><td>733.8</td><td>-13.3</td><td>1.6</td><td>-8.5</td><td>1.5E-17</td><td>9.6E-17</td><td>MB231/MB468</td><td>MITO</td></tr>
<tr><td>>hg38_21_+_8433172_8446622_RNA45SN1_45S_50ntFlanks@10574.10592.19|AAGCCGGCGGGAGCCCCGG</td><td>>hg38_21_+_8433172_8446622_RNA45SN1_45S_50ntFlanks</td><td>10574</td><td>10592</td><td>19</td><td>AAGCCGGCGGGAGCCCCGG</td><td>274.8</td><td>-12.0</td><td>1.4</td><td>-8.6</td><td>9.3E-18</td><td>9.6E-17</td><td>MB231/MB468</td><td>MP</td></tr>
<tr><td>>hg38_21_+_8433172_8446622_RNA45SN1_45S_50ntFlanks@12324.12350.27|TTGGGTTTTAAGCAGGAGGTGTCAGAA</td><td>>hg38_21_+_8433172_8446622_RNA45SN1_45S_50ntFlanks</td><td>12324</td><td>12350</td><td>27</td><td>TTGGGTTTTAAGCAGGAGGTGTCAGAA</td><td>281.8</td><td>-12.0</td><td>1.4</td><td>-8.6</td><td>9.3E-18</td><td>9.6E-17</td><td>MB231/MB468</td><td>MP</td></tr>
<tr><td>>am_trnaMT_PheGAA_MT_+_577_647@40.62.23|TGTTTAGACGGGCTCACATCACC</td><td>>am_trnaMT_PheGAA_MT_+_577_647</td><td>40</td><td>62</td><td>23</td><td>TGTTTAGACGGGCTCACATCACC</td><td>938.7</td><td>8.4</td><td>1.0</td><td>8.5</td><td>1.2E-17</td><td>9.6E-17</td><td>BT20/MB468</td><td>MITO</td></tr>
<tr><td>>hg38_21_+_8433172_8446622_RNA45SN1_45S_50ntFlanks@12892.12931.40|CCCTCGCCCGTCACGCACCGCACGTTCGTGGGGAACCTGG</td><td>>hg38_21_+_8433172_8446622_RNA45SN1_45S_50ntFlanks</td><td>12892</td><td>12931</td><td>40</td><td>CCCTCGCCCGTCACGCACCGCACGTTCGTGGGGAACCTGG</td><td>441.7</td><td>-12.6</td><td>1.5</td><td>-8.5</td><td>1.5E-17</td><td>9.7E-17</td><td>MB231/MB468</td><td>MITO</td></tr>
<tr><td>>hg38_21_+_8433172_8446622_RNA45SN1_45S_50ntFlanks@12126.12150.25|CGACCCGCTCCGGGGACAGTGCCAG</td><td>>hg38_21_+_8433172_8446622_RNA45SN1_45S_50ntFlanks</td><td>12126</td><td>12150</td><td>25</td><td>CGACCCGCTCCGGGGACAGTGCCAG</td><td>521.5</td><td>-12.0</td><td>1.4</td><td>-8.6</td><td>1.2E-17</td><td>9.7E-17</td><td>MB231/MB468</td><td>NUC</td></tr>
<tr><td>>hg38_21_+_8433172_8446622_RNA45SN1_45S_50ntFlanks@6719.6737.19|TGAATTGCAGGACACATTG</td><td>>hg38_21_+_8433172_8446622_RNA45SN1_45S_50ntFlanks</td><td>6719</td><td>6737</td><td>19</td><td>TGAATTGCAGGACACATTG</td><td>3592.9</td><td>-9.0</td><td>1.1</td><td>-8.6</td><td>1.2E-17</td><td>9.7E-17</td><td>MB231/MB468</td><td>NUC</td></tr>
<tr><td>>MI0001448|hsa-mir-425&WithFlank&3|-|49020142|49020240@20.43.24&[MIMAT0003393&hsa-miR-425-5p&offsets|0|+1;m-95&3|-|49020200|49020221&offsets|0|+2]|AATGACACGATCACTCCCGTTGAG</td><td>>MI0001448|hsa-mir-425&WithFlank&3|-|49020142|49020240</td><td>20</td><td>43</td><td>24</td><td>AATGACACGATCACTCCCGTTGAG</td><td>1762.9</td><td>5.6</td><td>0.7</td><td>8.6</td><td>1.2E-17</td><td>9.7E-17</td><td>MB231/MB468</td><td>NUC</td></tr>
<tr><td>>hg38_21_+_8433172_8446622_RNA45SN1_45S_50ntFlanks@6666.6686.21|GATCACTCGGCTCGTGCGTCG</td><td>>hg38_21_+_8433172_8446622_RNA45SN1_45S_50ntFlanks</td><td>6666</td><td>6686</td><td>21</td><td>GATCACTCGGCTCGTGCGTCG</td><td>354.1</td><td>-12.3</td><td>1.4</td><td>-8.6</td><td>9.4E-18</td><td>9.7E-17</td><td>MB231/MB468</td><td>MP</td></tr>
<tr><td>>hg38_21_+_8433172_8446622_RNA45SN1_45S_50ntFlanks@12456.12486.31|CCAAGCGTTGGATTGTTCACCCACTAATAGG</td><td>>hg38_21_+_8433172_8446622_RNA45SN1_45S_50ntFlanks</td><td>12456</td><td>12486</td><td>31</td><td>CCAAGCGTTGGATTGTTCACCCACTAATAGG</td><td>30781.4</td><td>-13.6</td><td>1.6</td><td>-8.4</td><td>2.9E-17</td><td>9.7E-17</td><td>BT20/MB468</td><td>NUC</td></tr>
<tr><td>>hg38_21_+_8433172_8446622_RNA45SN1_45S_50ntFlanks@12476.12497.22|CCACTAATAGGGAACGTGAGCT</td><td>>hg38_21_+_8433172_8446622_RNA45SN1_45S_50ntFlanks</td><td>12476</td><td>12497</td><td>22</td><td>CCACTAATAGGGAACGTGAGCT</td><td>1704.2</td><td>-4.0</td><td>0.5</td><td>-8.6</td><td>1.2E-17</td><td>9.8E-17</td><td>MB231/MB468</td><td>NUC</td></tr>
<tr><td>>hg38_21_+_8433172_8446622_RNA45SN1_45S_50ntFlanks@11317.11334.18|ACCCCACGTCTCGTCGCG</td><td>>hg38_21_+_8433172_8446622_RNA45SN1_45S_50ntFlanks</td><td>11317</td><td>11334</td><td>18</td><td>ACCCCACGTCTCGTCGCG</td><td>902.9</td><td>-13.6</td><td>1.6</td><td>-8.5</td><td>1.5E-17</td><td>9.8E-17</td><td>MB231/MB468</td><td>MITO</td></tr>
<tr><td>>hg38_21_+_8433172_8446622_RNA45SN1_45S_50ntFlanks@8194.8213.20|CCCGTGGACGGTGTGAGGCC</td><td>>hg38_21_+_8433172_8446622_RNA45SN1_45S_50ntFlanks</td><td>8194</td><td>8213</td><td>20</td><td>CCCGTGGACGGTGTGAGGCC</td><td>1323.3</td><td>-5.6</td><td>0.7</td><td>-8.5</td><td>1.5E-17</td><td>9.8E-17</td><td>MB231/MB468</td><td>MITO</td></tr>
<tr><td>>hg38_21_+_8433172_8446622_RNA45SN1_45S_50ntFlanks@6719.6744.26|TGAATTGCAGGACACATTGATCATCG</td><td>>hg38_21_+_8433172_8446622_RNA45SN1_45S_50ntFlanks</td><td>6719</td><td>6744</td><td>26</td><td>TGAATTGCAGGACACATTGATCATCG</td><td>15979.2</td><td>-7.6</td><td>0.9</td><td>-8.5</td><td>1.8E-17</td><td>9.8E-17</td><td>BT20/MB468</td><td>CYTO</td></tr>
<tr><td>>hg38_21_+_8433172_8446622_RNA45SN1_45S_50ntFlanks@4728.4751.24|ATCAAGAACGAAAGTCGGAGGTTC</td><td>>hg38_21_+_8433172_8446622_RNA45SN1_45S_50ntFlanks</td><td>4728</td><td>4751</td><td>24</td><td>ATCAAGAACGAAAGTCGGAGGTTC</td><td>475.0</td><td>-11.2</td><td>1.3</td><td>-8.5</td><td>1.8E-17</td><td>9.9E-17</td><td>BT20/MB468</td><td>CYTO</td></tr>
<tr><td>>hg38_21_+_8433172_8446622_RNA45SN1_45S_50ntFlanks@5417.5443.27|CTACCGATTGGATGGTTTAGTGAGGCC</td><td>>hg38_21_+_8433172_8446622_RNA45SN1_45S_50ntFlanks</td><td>5417</td><td>5443</td><td>27</td><td>CTACCGATTGGATGGTTTAGTGAGGCC</td><td>351.5</td><td>-12.3</td><td>1.4</td><td>-8.6</td><td>9.6E-18</td><td>9.9E-17</td><td>MB231/MB468</td><td>MP</td></tr>
<tr><td>>MI0000542|hsa-mir-320a&WithFlank&8|-|22244960|22245043@49.68.20&[MIMAT0000510&hsa-miR-320a-3p&offsets|+1|-1;m-60&8|-|22244975|22244996&offsets|+1|-1]|AAAGCTGGGTTGAGAGGGCG</td><td>>MI0000542|hsa-mir-320a&WithFlank&8|-|22244960|22245043</td><td>49</td><td>68</td><td>20</td><td>AAAGCTGGGTTGAGAGGGCG</td><td>1216.9</td><td>10.0</td><td>1.2</td><td>8.5</td><td>1.8E-17</td><td>9.9E-17</td><td>BT20/MB468</td><td>CYTO</td></tr>
<tr><td>>hg38_21_+_8433172_8446622_RNA45SN1_45S_50ntFlanks@6721.6740.20|AATTGCAGGACACATTGATC</td><td>>hg38_21_+_8433172_8446622_RNA45SN1_45S_50ntFlanks</td><td>6721</td><td>6740</td><td>20</td><td>AATTGCAGGACACATTGATC</td><td>20583.4</td><td>-11.2</td><td>1.3</td><td>-8.4</td><td>3.0E-17</td><td>9.9E-17</td><td>BT20/MB468</td><td>NUC</td></tr>
<tr><td>>MI0000088|hsa-mir-30a&WithFlank&6|-|71403545|71403627@53.73.21(+1A)&[MIMAT0000088&hsa-miR-30a-3p&offsets|0|-1(+1A);m-38&6|-|71403554|71403575&offsets|0|-1(+1A)]|CTTTCAGTCGGATGTTTGCAGA</td><td>>MI0000088|hsa-mir-30a&WithFlank&6|-|71403545|71403627</td><td>53</td><td>73</td><td>21(+1A)</td><td>CTTTCAGTCGGATGTTTGCAGA</td><td>655.2</td><td>5.6</td><td>0.7</td><td>8.6</td><td>1.2E-17</td><td>9.9E-17</td><td>MB231/MB468</td><td>NUC</td></tr>
<tr><td>>MI0000483|hsa-mir-186&WithFlank&1|-|71067625|71067722@21.44.24&[MIMAT0000456&hsa-miR-186-5p&offsets|0|+2;m-84&1|-|71067681|71067702&offsets|0|+2]|CAAAGAATTCTCCTTTTGGGCTTT</td><td>>MI0000483|hsa-mir-186&WithFlank&1|-|71067625|71067722</td><td>21</td><td>44</td><td>24</td><td>CAAAGAATTCTCCTTTTGGGCTTT</td><td>293.3</td><td>6.5</td><td>0.8</td><td>8.5</td><td>1.2E-17</td><td>1.0E-16</td><td>MB231/MB468</td><td>NUC</td></tr>
<tr><td>>hg38_21_+_8433172_8446622_RNA45SN1_45S_50ntFlanks@12202.12220.19|CTAAGGCGAGCTCAGGGAG</td><td>>hg38_21_+_8433172_8446622_RNA45SN1_45S_50ntFlanks</td><td>12202</td><td>12220</td><td>19</td><td>CTAAGGCGAGCTCAGGGAG</td><td>1509.8</td><td>-14.3</td><td>1.7</td><td>-8.5</td><td>1.6E-17</td><td>1.0E-16</td><td>MB231/MB468</td><td>MITO</td></tr>
<tr><td>>hg38_21_+_8433172_8446622_RNA45SN1_45S_50ntFlanks@12539.12560.22|ATGATGTGTTGTTGCCATGGTA</td><td>>hg38_21_+_8433172_8446622_RNA45SN1_45S_50ntFlanks</td><td>12539</td><td>12560</td><td>22</td><td>ATGATGTGTTGTTGCCATGGTA</td><td>464.9</td><td>-12.6</td><td>1.5</td><td>-8.5</td><td>1.6E-17</td><td>1.0E-16</td><td>MB231/MB468</td><td>MITO</td></tr>
<tr><td>>hg38_21_+_8433172_8446622_RNA45SN1_45S_50ntFlanks@5484.5509.26|GAGCGCTGAGAAGACGGTCGAACTTG</td><td>>hg38_21_+_8433172_8446622_RNA45SN1_45S_50ntFlanks</td><td>5484</td><td>5509</td><td>26</td><td>GAGCGCTGAGAAGACGGTCGAACTTG</td><td>364.7</td><td>-12.3</td><td>1.4</td><td>-8.5</td><td>1.6E-17</td><td>1.0E-16</td><td>MB231/MB468</td><td>MITO</td></tr>
<tr><td>>hg38_21_+_8433172_8446622_RNA45SN1_45S_50ntFlanks@9738.9757.20|GAAACGATCTCAACCTATTC</td><td>>hg38_21_+_8433172_8446622_RNA45SN1_45S_50ntFlanks</td><td>9738</td><td>9757</td><td>20</td><td>GAAACGATCTCAACCTATTC</td><td>852.5</td><td>-13.6</td><td>1.6</td><td>-8.6</td><td>9.7E-18</td><td>1.0E-16</td><td>MB231/MB468</td><td>MP</td></tr>
<tr><td>>hg38_21_+_8433172_8446622_RNA45SN1_45S_50ntFlanks@12045.12068.24|CTGACCCGGTGAGGCGGGGGGGCG</td><td>>hg38_21_+_8433172_8446622_RNA45SN1_45S_50ntFlanks</td><td>12045</td><td>12068</td><td>24</td><td>CTGACCCGGTGAGGCGGGGGGGCG</td><td>411.6</td><td>-12.5</td><td>1.5</td><td>-8.6</td><td>9.8E-18</td><td>1.0E-16</td><td>MB231/MB468</td><td>MP</td></tr>
<tr><td>>hg38_21_+_8433172_8446622_RNA45SN1_45S_50ntFlanks@5436.5466.31|GTGAGGCCCTCGGATCGGCCCCGCCGGGGTC</td><td>>hg38_21_+_8433172_8446622_RNA45SN1_45S_50ntFlanks</td><td>5436</td><td>5466</td><td>31</td><td>GTGAGGCCCTCGGATCGGCCCCGCCGGGGTC</td><td>450.0</td><td>-12.6</td><td>1.5</td><td>-8.5</td><td>1.6E-17</td><td>1.0E-16</td><td>MB231/MB468</td><td>MITO</td></tr>
<tr><td>>hg38_21_+_8433172_8446622_RNA45SN1_45S_50ntFlanks@4008.4026.19|CTCGGGCCGATCGCACGCC</td><td>>hg38_21_+_8433172_8446622_RNA45SN1_45S_50ntFlanks</td><td>4008</td><td>4026</td><td>19</td><td>CTCGGGCCGATCGCACGCC</td><td>382.1</td><td>-12.4</td><td>1.5</td><td>-8.6</td><td>9.9E-18</td><td>1.0E-16</td><td>MB231/MB468</td><td>MP</td></tr>
<tr><td>>hg38_21_+_8433172_8446622_RNA45SN1_45S_50ntFlanks@11766.11791.26|ATCTAATTAGTGACGCGCATGAATGG</td><td>>hg38_21_+_8433172_8446622_RNA45SN1_45S_50ntFlanks</td><td>11766</td><td>11791</td><td>26</td><td>ATCTAATTAGTGACGCGCATGAATGG</td><td>616.0</td><td>-12.2</td><td>1.4</td><td>-8.5</td><td>1.3E-17</td><td>1.0E-16</td><td>MB231/MB468</td><td>NUC</td></tr>
<tr><td>>MI0000283|hsa-mir-203a&WithFlank&14|+|104117399|104117520@71.92.22&[MIMAT0000264&hsa-miR-203a-3p&offsets|0|0;m-79&14|+|104117469|104117490&offsets|0|0]|GTGAAATGTTTAGGACCACTAG</td><td>>MI0000283|hsa-mir-203a&WithFlank&14|+|104117399|104117520</td><td>71</td><td>92</td><td>22</td><td>GTGAAATGTTTAGGACCACTAG</td><td>417.4</td><td>12.3</td><td>1.4</td><td>8.8</td><td>1.3E-18</td><td>1.0E-16</td><td>BT20/MB231</td><td>MITO</td></tr>
<tr><td>>trnaMT_GluTTC_MT_-_14674_14742@30.72.43,trnalookalike8_GluTTC_5_-_93905172_93905240@30.72.43|TTTCATATCATTGGTCGTGGTTGTAGTCCGTGCGAGAATACCA</td><td>>trnaMT_GluTTC_MT_-_14674_14742</td><td>30</td><td>72</td><td>43</td><td>TTTCATATCATTGGTCGTGGTTGTAGTCCGTGCGAGAATACCA</td><td>320.3</td><td>12.3</td><td>1.4</td><td>8.6</td><td>1.0E-17</td><td>1.0E-16</td><td>MB231/MB468</td><td>CYTO</td></tr>
<tr><td>>hg38_21_+_8433172_8446622_RNA45SN1_45S_50ntFlanks@10781.10805.25|AGGTGAACAGCCTCTGGCATGTTGG</td><td>>hg38_21_+_8433172_8446622_RNA45SN1_45S_50ntFlanks</td><td>10781</td><td>10805</td><td>25</td><td>AGGTGAACAGCCTCTGGCATGTTGG</td><td>390.4</td><td>-12.5</td><td>1.5</td><td>-8.6</td><td>1.0E-17</td><td>1.0E-16</td><td>MB231/MB468</td><td>MP</td></tr>
<tr><td>>hg38_21_+_8433172_8446622_RNA45SN1_45S_50ntFlanks@5445.5465.21|TCGGATCGGCCCCGCCGGGGT</td><td>>hg38_21_+_8433172_8446622_RNA45SN1_45S_50ntFlanks</td><td>5445</td><td>5465</td><td>21</td><td>TCGGATCGGCCCCGCCGGGGT</td><td>314.9</td><td>-12.2</td><td>1.4</td><td>-8.6</td><td>1.0E-17</td><td>1.0E-16</td><td>MB231/MB468</td><td>MP</td></tr>
<tr><td>>MI0000750|hsa-mir-26a-2&WithFlank&12|-|57824603|57824698@20.40.21(+1C);MI0000083|hsa-mir-26a-1&WithFlank&3|+|37969398|37969486@16.36.21(+1C)&[MIMAT0000082&hsa-miR-26a-5p&offsets|0|-1(+1C);m-33&12|-|57824658|57824679&offsets|0|-1(+1C)];[MIMAT0000082_1&hsa-miR-26a-5p&offsets|0|-1(+1C);m-34&3|+|37969413|37969434&offsets|0|-1(+1C)]|TTCAAGTAATCCAGGATAGGCC</td><td>>MI0000750|hsa-mir-26a-2&WithFlank&12|-|57824603|57824698</td><td>20</td><td>40</td><td>21(+1C)</td><td>TTCAAGTAATCCAGGATAGGCC</td><td>10778.9</td><td>5.9</td><td>0.7</td><td>8.5</td><td>1.3E-17</td><td>1.0E-16</td><td>BT20/MB468</td><td>MITO</td></tr>
<tr><td>>hg38_21_+_8433172_8446622_RNA45SN1_45S_50ntFlanks@476.506.31|TTCGTGATCGATGTGGTGACGTCGTGCTCTC</td><td>>hg38_21_+_8433172_8446622_RNA45SN1_45S_50ntFlanks</td><td>476</td><td>506</td><td>31</td><td>TTCGTGATCGATGTGGTGACGTCGTGCTCTC</td><td>1707.5</td><td>7.1</td><td>0.8</td><td>8.5</td><td>1.3E-17</td><td>1.0E-16</td><td>BT20/MB468</td><td>MITO</td></tr>
<tr><td>>hg38_21_+_8433172_8446622_RNA45SN1_45S_50ntFlanks@4224.4243.20|AAAATAACAATACAGGACTC</td><td>>hg38_21_+_8433172_8446622_RNA45SN1_45S_50ntFlanks</td><td>4224</td><td>4243</td><td>20</td><td>AAAATAACAATACAGGACTC</td><td>388.1</td><td>-12.5</td><td>1.5</td><td>-8.6</td><td>1.0E-17</td><td>1.0E-16</td><td>MB231/MB468</td><td>MP</td></tr>
<tr><td>>MI0000294|hsa-mir-218-1&WithFlank&4|+|20528269|20528390@31.52.22;MI0000295|hsa-mir-218-2&WithFlank&5|-|168768140|168768261@31.52.22&[MIMAT0000275&hsa-miR-218-5p&offsets|0|+1;m-185&4|+|20528299|20528319&offsets|0|+1];[MIMAT0000275_1&hsa-miR-218-5p&offsets|0|+1;m-187&5|-|168768211|168768231&offsets|0|+1]|TTGTGCTTGATCTAACCATGTG</td><td>>MI0000294|hsa-mir-218-1&WithFlank&4|+|20528269|20528390</td><td>31</td><td>52</td><td>22</td><td>TTGTGCTTGATCTAACCATGTG</td><td>243.1</td><td>11.8</td><td>1.4</td><td>8.5</td><td>1.3E-17</td><td>1.0E-16</td><td>MB231/MB468</td><td>NUC</td></tr>
<tr><td>>hg38_MT_+_598_1651_RNR1_12S_50ntFlanks@428.468.41|AAAGTGGCTTTAACATATCTGAACACACAATAGCTAAGACC</td><td>>hg38_MT_+_598_1651_RNR1_12S_50ntFlanks</td><td>428</td><td>468</td><td>41</td><td>AAAGTGGCTTTAACATATCTGAACACACAATAGCTAAGACC</td><td>206.8</td><td>11.6</td><td>1.4</td><td>8.5</td><td>1.3E-17</td><td>1.0E-16</td><td>MB231/MB468</td><td>NUC</td></tr>
<tr><td>>hg38_21_+_8433172_8446622_RNA45SN1_45S_50ntFlanks@9202.9225.24|TCTCGGGGCCACGCGCGCGTCCCC</td><td>>hg38_21_+_8433172_8446622_RNA45SN1_45S_50ntFlanks</td><td>9202</td><td>9225</td><td>24</td><td>TCTCGGGGCCACGCGCGCGTCCCC</td><td>578.4</td><td>-13.0</td><td>1.5</td><td>-8.6</td><td>1.0E-17</td><td>1.0E-16</td><td>MB231/MB468</td><td>MP</td></tr>
<tr><td>>MI0003205|hsa-mir-532&WithFlank&X|+|50003142|50003244@26.46.21(+1C)&[MIMAT0002888&hsa-miR-532-5p&offsets|0|-1(+1C);m-65&X|+|50003167|50003188&offsets|0|-1(+1C)]|CATGCCTTGAGTGTAGGACCGC</td><td>>MI0003205|hsa-mir-532&WithFlank&X|+|50003142|50003244</td><td>26</td><td>46</td><td>21(+1C)</td><td>CATGCCTTGAGTGTAGGACCGC</td><td>371.5</td><td>5.7</td><td>0.7</td><td>8.5</td><td>1.3E-17</td><td>1.0E-16</td><td>BT20/MB468</td><td>MITO</td></tr>
<tr><td>>hg38_21_+_8433172_8446622_RNA45SN1_45S_50ntFlanks@10513.10533.21|CCGGAGTGGCGGAGATGGGCG</td><td>>hg38_21_+_8433172_8446622_RNA45SN1_45S_50ntFlanks</td><td>10513</td><td>10533</td><td>21</td><td>CCGGAGTGGCGGAGATGGGCG</td><td>20985.7</td><td>-12.7</td><td>1.5</td><td>-8.6</td><td>1.0E-17</td><td>1.0E-16</td><td>MB231/MB468</td><td>MP</td></tr>
<tr><td>>hg38_21_+_8433172_8446622_RNA45SN1_45S_50ntFlanks@12044.12068.25|ACTGACCCGGTGAGGCGGGGGGGCG</td><td>>hg38_21_+_8433172_8446622_RNA45SN1_45S_50ntFlanks</td><td>12044</td><td>12068</td><td>25</td><td>ACTGACCCGGTGAGGCGGGGGGGCG</td><td>809.1</td><td>-12.6</td><td>1.5</td><td>-8.5</td><td>1.3E-17</td><td>1.1E-16</td><td>MB231/MB468</td><td>NUC</td></tr>
<tr><td>>hg38_21_+_8433172_8446622_RNA45SN1_45S_50ntFlanks@11641.11666.26|CGGCGGGTGTTGACGCGATGTGATTT</td><td>>hg38_21_+_8433172_8446622_RNA45SN1_45S_50ntFlanks</td><td>11641</td><td>11666</td><td>26</td><td>CGGCGGGTGTTGACGCGATGTGATTT</td><td>400.9</td><td>-12.5</td><td>1.5</td><td>-8.6</td><td>1.0E-17</td><td>1.1E-16</td><td>MB231/MB468</td><td>MP</td></tr>
<tr><td>>hg38_21_+_8433172_8446622_RNA45SN1_45S_50ntFlanks@12453.12477.25|TCACCAAGCGTTGGATTGTTCACCC</td><td>>hg38_21_+_8433172_8446622_RNA45SN1_45S_50ntFlanks</td><td>12453</td><td>12477</td><td>25</td><td>TCACCAAGCGTTGGATTGTTCACCC</td><td>429.9</td><td>6.2</td><td>0.7</td><td>8.6</td><td>1.0E-17</td><td>1.1E-16</td><td>MB231/MB468</td><td>CYTO</td></tr>
<tr><td>>hg38_21_+_8433172_8446622_RNA45SN1_45S_50ntFlanks@10914.10946.33|CGCCGCGGCTGGACGAGGCGCCGCCGCCCCCCC</td><td>>hg38_21_+_8433172_8446622_RNA45SN1_45S_50ntFlanks</td><td>10914</td><td>10946</td><td>33</td><td>CGCCGCGGCTGGACGAGGCGCCGCCGCCCCCCC</td><td>453.7</td><td>-12.7</td><td>1.5</td><td>-8.6</td><td>1.0E-17</td><td>1.1E-16</td><td>MB231/MB468</td><td>MP</td></tr>
<tr><td>>hg38_21_+_8433172_8446622_RNA45SN1_45S_50ntFlanks@6737.6770.34|GATCATCGACACTTCGAACGCACTTGCGGCCCCG</td><td>>hg38_21_+_8433172_8446622_RNA45SN1_45S_50ntFlanks</td><td>6737</td><td>6770</td><td>34</td><td>GATCATCGACACTTCGAACGCACTTGCGGCCCCG</td><td>674.6</td><td>-12.4</td><td>1.4</td><td>-8.5</td><td>1.3E-17</td><td>1.1E-16</td><td>MB231/MB468</td><td>NUC</td></tr>
<tr><td>>hg38_21_+_8433172_8446622_RNA45SN1_45S_50ntFlanks@4122.4146.25|AGGGTTCGATTCCGGAGAGGGAGCC</td><td>>hg38_21_+_8433172_8446622_RNA45SN1_45S_50ntFlanks</td><td>4122</td><td>4146</td><td>25</td><td>AGGGTTCGATTCCGGAGAGGGAGCC</td><td>292.4</td><td>-12.1</td><td>1.4</td><td>-8.6</td><td>1.0E-17</td><td>1.1E-16</td><td>MB231/MB468</td><td>MP</td></tr>
<tr><td>>hg38_21_+_8433172_8446622_RNA45SN1_45S_50ntFlanks@5496.5516.21|GACGGTCGAACTTGACTATCT</td><td>>hg38_21_+_8433172_8446622_RNA45SN1_45S_50ntFlanks</td><td>5496</td><td>5516</td><td>21</td><td>GACGGTCGAACTTGACTATCT</td><td>972.5</td><td>10.3</td><td>1.2</td><td>8.5</td><td>1.9E-17</td><td>1.1E-16</td><td>BT20/MB468</td><td>CYTO</td></tr>
<tr><td>>hg38_21_+_8433172_8446622_RNA45SN1_45S_50ntFlanks@12941.12966.26|ATTCGTAGACGACCTGCTTCTGGGTC</td><td>>hg38_21_+_8433172_8446622_RNA45SN1_45S_50ntFlanks</td><td>12941</td><td>12966</td><td>26</td><td>ATTCGTAGACGACCTGCTTCTGGGTC</td><td>1466.6</td><td>-7.7</td><td>0.9</td><td>-8.6</td><td>1.1E-17</td><td>1.1E-16</td><td>MB231/MB468</td><td>CYTO</td></tr>
<tr><td>>hg38_21_+_8433172_8446622_RNA45SN1_45S_50ntFlanks@12624.12659.36|CCAATGGGGCGAAGCTACCATCTGTGGGATTATGAC</td><td>>hg38_21_+_8433172_8446622_RNA45SN1_45S_50ntFlanks</td><td>12624</td><td>12659</td><td>36</td><td>CCAATGGGGCGAAGCTACCATCTGTGGGATTATGAC</td><td>532.4</td><td>-12.9</td><td>1.5</td><td>-8.6</td><td>1.1E-17</td><td>1.1E-16</td><td>MB231/MB468</td><td>MP</td></tr>
</table>

<table>
<tr><th></th><th></th><th></th><th></th><th></th><th></th><th></th><th></th><th></th><th></th><th></th><th></th><th></th><th></th></tr>
<tr><td>>hg38_21_+_8433172_8446622_RNA45SN1_45S_50ntFlanks@9567.9586.20|ACGTGCAAATCGGTCGTCCG</td><td>>hg38_21_+_8433172_8446622_RNA45SN1_45S_50ntFlanks</td><td>9567</td><td>9586</td><td>20</td><td>ACGTGCAAATCGGTCGTCCG</td><td>310.1</td><td>-12.1</td><td>1.4</td><td>-8.6</td><td>1.1E-17</td><td>1.1E-16</td><td>MB231/MB468</td><td>MP</td></tr>
<tr><td>>hg38_21_+_8433172_8446622_RNA45SN1_45S_50ntFlanks@12360.12382.23|CAGGGATAACTGGCTTGTGGCGG</td><td>>hg38_21_+_8433172_8446622_RNA45SN1_45S_50ntFlanks</td><td>12360</td><td>12382</td><td>23</td><td>CAGGGATAACTGGCTTGTGGCGG</td><td>517.9</td><td>-12.0</td><td>1.4</td><td>-8.5</td><td>1.4E-17</td><td>1.1E-16</td><td>MB231/MB468</td><td>NUC</td></tr>
<tr><td>>hg38_21_+_8433172_8446622_RNA45SN1_45S_50ntFlanks@12006.12043.38|GGCCGCCGGTGAAATACCACTACTCTGATCGTTTTTTC</td><td>>hg38_21_+_8433172_8446622_RNA45SN1_45S_50ntFlanks</td><td>12006</td><td>12043</td><td>38</td><td>GGCCGCCGGTGAAATACCACTACTCTGATCGTTTTTTC</td><td>665.8</td><td>-13.2</td><td>1.5</td><td>-8.5</td><td>1.7E-17</td><td>1.1E-16</td><td>MB231/MB468</td><td>MITO</td></tr>
<tr><td>>hg38_21_+_8433172_8446622_RNA45SN1_45S_50ntFlanks@12991.13016.26|CTCCCTCGCTGCGATCTATTGAAAGT</td><td>>hg38_21_+_8433172_8446622_RNA45SN1_45S_50ntFlanks</td><td>12991</td><td>13016</td><td>26</td><td>CTCCCTCGCTGCGATCTATTGAAAGT</td><td>5101.8</td><td>-9.8</td><td>1.2</td><td>-8.4</td><td>3.3E-17</td><td>1.1E-16</td><td>BT20/MB468</td><td>NUC</td></tr>
<tr><td>>hg38_21_+_8433172_8446622_RNA45SN1_45S_50ntFlanks@3873.3898.26|TAATACATGCCGACGGGCGCTGACCC</td><td>>hg38_21_+_8433172_8446622_RNA45SN1_45S_50ntFlanks</td><td>3873</td><td>3898</td><td>26</td><td>TAATACATGCCGACGGGCGCTGACCC</td><td>586.9</td><td>-13.1</td><td>1.5</td><td>-8.6</td><td>1.1E-17</td><td>1.1E-16</td><td>MB231/MB468</td><td>MP</td></tr>
<tr><td>>hg38_21_+_8433172_8446622_RNA45SN1_45S_50ntFlanks@9576.9597.22|TCGGTCGTCCGACCTGGGTATA</td><td>>hg38_21_+_8433172_8446622_RNA45SN1_45S_50ntFlanks</td><td>9576</td><td>9597</td><td>22</td><td>TCGGTCGTCCGACCTGGGTATA</td><td>430.9</td><td>-12.6</td><td>1.5</td><td>-8.6</td><td>1.1E-17</td><td>1.1E-16</td><td>MB231/MB468</td><td>MP</td></tr>
<tr><td>>MI0000087|hsa-mir-29a&WithFlank&7|-|130876741|130876816@47.67.21(+1C)&[MIMAT0000086&hsa-miR-29a-3p&offsets|-1|-2(+1C);m-26&7|-|130876749|130876769&offsets|-1|-1(+1C)]|CTAGCACCATCTGAAATCGGTC</td><td>>MI0000087|hsa-mir-29a&WithFlank&7|-|130876741|130876816</td><td>47</td><td>67</td><td>21(+1C)</td><td>CTAGCACCATCTGAAATCGGTC</td><td>1758.1</td><td>9.6</td><td>1.1</td><td>8.5</td><td>2.0E-17</td><td>1.1E-16</td><td>BT20/MB468</td><td>CYTO</td></tr>
<tr><td>>hg38_21_+_8433172_8446622_RNA45SN1_45S_50ntFlanks@10905.10932.28|CTGGGCGCGCGCCGCGGCTGGACGAGGC</td><td>>hg38_21_+_8433172_8446622_RNA45SN1_45S_50ntFlanks</td><td>10905</td><td>10932</td><td>28</td><td>CTGGGCGCGCGCCGCGGCTGGACGAGGC</td><td>356.7</td><td>-12.3</td><td>1.4</td><td>-8.6</td><td>1.1E-17</td><td>1.1E-16</td><td>MB231/MB468</td><td>MP</td></tr>
<tr><td>>MI0000264|hsa-mir-7-2&WithFlank&15|+|88611819|88611940@38.60.23(+1A);MI0000265|hsa-mir-7-3&WithFlank&19|+|4770664|4770785@37.59.23(+1A);MI0000263|hsa-mir-7-1&WithFlank&9|-|83969742|83969863@30.52.23(+1A)&[MIMAT0000252&hsa-miR-7-5p&offsets|0|-1(+1A);m-400&15|+|88611856|88611879&offsets|0|-1(+1A)];[MIMAT0000252_1&hsa-miR-7-5p&offsets|0|-1(+1A);m-402&19|+|4770700|4770723&offsets|0|-1(+1A)];[MIMAT0000252_2&hsa-miR-7-5p&offsets|0|-1(+1A);m-401&9|-|83969811|83969834&offsets|0|-1(+1A)]|TGGAAGACTAGTGATTTTGTTGTA</td><td>>MI0000264|hsa-mir-7-2&WithFlank&15|+|88611819|88611940</td><td>38</td><td>60</td><td>23(+1A)</td><td>TGGAAGACTAGTGATTTTGTTGTA</td><td>10822.6</td><td>8.9</td><td>1.0</td><td>8.5</td><td>2.0E-17</td><td>1.1E-16</td><td>BT20/MB468</td><td>CYTO</td></tr>
<tr><td>>trna11_GluTTC_15_-_26327381_26327452@1.33.33,trna3_GluTTC_13_-_45492062_45492133@1.33.33|TCCCACATGGTCTAGCGGTTAGGATTCCTGGTT</td><td>>trna11_GluTTC_15_-_26327381_26327452</td><td>1</td><td>33</td><td>33</td><td>TCCCACATGGTCTAGCGGTTAGGATTCCTGGTT</td><td>5814.3</td><td>10.2</td><td>1.2</td><td>8.5</td><td>1.4E-17</td><td>1.1E-16</td><td>BT20/MB468</td><td>MITO</td></tr>
<tr><td>>hg38_21_+_8433172_8446622_RNA45SN1_45S_50ntFlanks@9471.9503.33|AGCACGAGCGCACGTGTTAGGACCCGAAAGATG</td><td>>hg38_21_+_8433172_8446622_RNA45SN1_45S_50ntFlanks</td><td>9471</td><td>9503</td><td>33</td><td>AGCACGAGCGCACGTGTTAGGACCCGAAAGATG</td><td>1905.7</td><td>-10.2</td><td>1.2</td><td>-8.5</td><td>1.8E-17</td><td>1.1E-16</td><td>MB231/MB468</td><td>MITO</td></tr>
<tr><td>>hg38_21_+_8433172_8446622_RNA45SN1_45S_50ntFlanks@11641.11663.23|CGGCGGGTGTTGACGCGATGTGA</td><td>>hg38_21_+_8433172_8446622_RNA45SN1_45S_50ntFlanks</td><td>11641</td><td>11663</td><td>23</td><td>CGGCGGGTGTTGACGCGATGTGA</td><td>1761.8</td><td>-10.6</td><td>1.2</td><td>-8.5</td><td>1.8E-17</td><td>1.1E-16</td><td>MB231/MB468</td><td>MITO</td></tr>
<tr><td>>hg38_21_+_8433172_8446622_RNA45SN1_45S_50ntFlanks@9622.9650.29|CTAGTAGCTGGTTCCCTCCGAAGTTTCCC</td><td>>hg38_21_+_8433172_8446622_RNA45SN1_45S_50ntFlanks</td><td>9622</td><td>9650</td><td>29</td><td>CTAGTAGCTGGTTCCCTCCGAAGTTTCCC</td><td>332.4</td><td>-12.2</td><td>1.4</td><td>-8.6</td><td>1.1E-17</td><td>1.1E-16</td><td>MB231/MB468</td><td>MP</td></tr>
<tr><td>>MI0001445|hsa-mir-423&WithFlank&17|+|30117073|30117178@59.80.22(+1G)&[MIMAT0001340&hsa-miR-423-3p&offsets|0|-1(+1G);m-100&17|+|30117131|30117153&offsets|0|-1(+1G)]|AGCTCGGTCTGAGGCCCCTCAGG</td><td>>MI0001445|hsa-mir-423&WithFlank&17|+|30117073|30117178</td><td>59</td><td>80</td><td>22(+1G)</td><td>AGCTCGGTCTGAGGCCCCTCAGG</td><td>125.4</td><td>3.5</td><td>0.4</td><td>8.4</td><td>3.4E-17</td><td>1.1E-16</td><td>BT20/MB468</td><td>NUC</td></tr>
<tr><td>>am_trna2_GlyGCC_21_-_18827107_18827177@57.74.18,trna35_GlyGCC_1_+_161413094_161413164@57.74.18,trna37_GlyGCC_1_+_161420467_161420537@57.74.18,trna39_GlyGCC_1_+_161427898_161427968@57.74.18,trna41_GlyGCC_1_+_161435258_161435328@57.74.18|TTCCCGGCCCATGCACCA</td><td>>am_trna2_GlyGCC_21_-_18827107_18827177</td><td>57</td><td>74</td><td>18</td><td>TTCCCGGCCCATGCACCA</td><td>1189.0</td><td>8.2</td><td>0.9</td><td>8.8</td><td>1.0E-18</td><td>1.1E-16</td><td>BT20/MB231</td><td>CYTO</td></tr>
<tr><td>>am_MI0000809|hsa-mir-151a&WithFlank&8|-|140732558|140732659@53.73.21(+2U)&[MIMAT0000757&hsa-miR-151a-3p&offsets|0|0(+2U);m-30&8|-|140732587|140732607&offsets|0|0(+2U)]|CTAGACTGAAGCTCCTTGAGGTT</td><td>>am_MI0000809|hsa-mir-151a&WithFlank&8|-|140732558|140732659</td><td>53</td><td>73</td><td>21(+2U)</td><td>CTAGACTGAAGCTCCTTGAGGTT</td><td>1652.8</td><td>10.2</td><td>1.2</td><td>8.5</td><td>2.0E-17</td><td>1.1E-16</td><td>BT20/MB468</td><td>CYTO</td></tr>
<tr><td>>hg38_21_+_8433172_8446622_RNA45SN1_45S_50ntFlanks@6787.6806.20|ACGCCTGTCTGAGCGTCGCT</td><td>>hg38_21_+_8433172_8446622_RNA45SN1_45S_50ntFlanks</td><td>6787</td><td>6806</td><td>20</td><td>ACGCCTGTCTGAGCGTCGCT</td><td>2098.6</td><td>5.4</td><td>0.6</td><td>8.5</td><td>2.0E-17</td><td>1.1E-16</td><td>BT20/MB468</td><td>CYTO</td></tr>
<tr><td>>hg38_21_+_8433172_8446622_RNA45SN1_45S_50ntFlanks@11212.11254.43|CGACTCTGGACGCGAGCCGGGCCCTTCCCGTGGATCGCCCCAG</td><td>>hg38_21_+_8433172_8446622_RNA45SN1_45S_50ntFlanks</td><td>11212</td><td>11254</td><td>43</td><td>CGACTCTGGACGCGAGCCGGGCCCTTCCCGTGGATCGCCCCAG</td><td>1380.9</td><td>-14.2</td><td>1.7</td><td>-8.5</td><td>1.8E-17</td><td>1.1E-16</td><td>MB231/MB468</td><td>MITO</td></tr>
<tr><td>>am_MI0003639|hsa-mir-625&WithFlank&14|+|65471096|65471192@58.79.22&[MIMAT0004808&hsa-miR-625-3p&offsets|0|0;m-137&14|+|65471153|65471174&offsets|0|0]|GACTATAGAACTTTCCCCCTCA</td><td>>am_MI0003639|hsa-mir-625&WithFlank&14|+|65471096|65471192</td><td>58</td><td>79</td><td>22</td><td>GACTATAGAACTTTCCCCCTCA</td><td>249.8</td><td>12.0</td><td>1.4</td><td>8.6</td><td>1.1E-17</td><td>1.1E-16</td><td>MB231/MB468</td><td>CYTO</td></tr>
<tr><td>>hg38_21_+_8433172_8446622_RNA45SN1_45S_50ntFlanks@7980.8007.28|CCTCAGATCAGACGTGGCGACCCGCTGA</td><td>>hg38_21_+_8433172_8446622_RNA45SN1_45S_50ntFlanks</td><td>7980</td><td>8007</td><td>28</td><td>CCTCAGATCAGACGTGGCGACCCGCTGA</td><td>789.2</td><td>-8.0</td><td>0.9</td><td>-8.5</td><td>1.8E-17</td><td>1.1E-16</td><td>MB231/MB468</td><td>MITO</td></tr>
<tr><td>>hg38_21_+_8433172_8446622_RNA45SN1_45S_50ntFlanks@12151.12181.31|GTGGGGAGTTTGACTGGGGCGGTACACCTGT</td><td>>hg38_21_+_8433172_8446622_RNA45SN1_45S_50ntFlanks</td><td>12151</td><td>12181</td><td>31</td><td>GTGGGGAGTTTGACTGGGGCGGTACACCTGT</td><td>393.1</td><td>-12.4</td><td>1.5</td><td>-8.5</td><td>1.8E-17</td><td>1.1E-16</td><td>MB231/MB468</td><td>MITO</td></tr>
<tr><td>>hg38_21_+_8433172_8446622_RNA45SN1_45S_50ntFlanks@10131.10149.19|CGGCGGCGGCGGCGGCGGG</td><td>>hg38_21_+_8433172_8446622_RNA45SN1_45S_50ntFlanks</td><td>10131</td><td>10149</td><td>19</td><td>CGGCGGCGGCGGCGGCGGG</td><td>581.1</td><td>-13.0</td><td>1.5</td><td>-8.6</td><td>1.1E-17</td><td>1.1E-16</td><td>MB231/MB468</td><td>MP</td></tr>
<tr><td>>hg38_21_+_8433172_8446622_RNA45SN1_45S_50ntFlanks@10826.10858.33|TCGGCAAGCCGGATCCGTAACTTCGGGATAAGG</td><td>>hg38_21_+_8433172_8446622_RNA45SN1_45S_50ntFlanks</td><td>10826</td><td>10858</td><td>33</td><td>TCGGCAAGCCGGATCCGTAACTTCGGGATAAGG</td><td>551.1</td><td>-13.0</td><td>1.5</td><td>-8.6</td><td>1.1E-17</td><td>1.1E-16</td><td>MB231/MB468</td><td>MP</td></tr>
<tr><td>>hg38_21_+_8433172_8446622_RNA45SN1_45S_50ntFlanks@8848.8866.19|CGGCGGCGGCGGCGGCGGG</td><td>>hg38_21_+_8433172_8446622_RNA45SN1_45S_50ntFlanks</td><td>8848</td><td>8866</td><td>19</td><td>CGGCGGCGGCGGCGGCGGG</td><td>581.1</td><td>-13.0</td><td>1.5</td><td>-8.6</td><td>1.1E-17</td><td>1.1E-16</td><td>MB231/MB468</td><td>MP</td></tr>
<tr><td>>hg38_21_+_8433172_8446622_RNA45SN1_45S_50ntFlanks@10797.10818.22|GCATGTTGGAACAATGTAGGTA</td><td>>hg38_21_+_8433172_8446622_RNA45SN1_45S_50ntFlanks</td><td>10797</td><td>10818</td><td>22</td><td>GCATGTTGGAACAATGTAGGTA</td><td>3637.4</td><td>-8.9</td><td>1.0</td><td>-8.6</td><td>1.1E-17</td><td>1.1E-16</td><td>MB231/MB468</td><td>MP</td></tr>
<tr><td>>hg38_21_+_8433172_8446622_RNA45SN1_45S_50ntFlanks@12045.12072.28|CTGACCCGGTGAGGCGGGGGGGCGAGCC</td><td>>hg38_21_+_8433172_8446622_RNA45SN1_45S_50ntFlanks</td><td>12045</td><td>12072</td><td>28</td><td>CTGACCCGGTGAGGCGGGGGGGCGAGCC</td><td>800.5</td><td>-13.4</td><td>1.6</td><td>-8.5</td><td>1.8E-17</td><td>1.1E-16</td><td>MB231/MB468</td><td>MITO</td></tr>
<tr><td>>hg38_21_+_8433172_8446622_RNA45SN1_45S_50ntFlanks@12083.12110.28|CTCGCTTCTGGCGCCAAGCGCCCGGCCG</td><td>>hg38_21_+_8433172_8446622_RNA45SN1_45S_50ntFlanks</td><td>12083</td><td>12110</td><td>28</td><td>CTCGCTTCTGGCGCCAAGCGCCCGGCCG</td><td>388.7</td><td>-12.4</td><td>1.5</td><td>-8.5</td><td>1.8E-17</td><td>1.2E-16</td><td>MB231/MB468</td><td>MITO</td></tr>
<tr><td>>hg38_21_+_8433172_8446622_RNA45SN1_45S_50ntFlanks@11190.11207.18|CCCGGGGGGCCGGCGGCG</td><td>>hg38_21_+_8433172_8446622_RNA45SN1_45S_50ntFlanks</td><td>11190</td><td>11207</td><td>18</td><td>CCCGGGGGGCCGGCGGCG</td><td>779.9</td><td>-13.5</td><td>1.6</td><td>-8.6</td><td>1.1E-17</td><td>1.2E-16</td><td>MB231/MB468</td><td>MP</td></tr>
<tr><td>>hg38_21_+_8433172_8446622_RNA45SN1_45S_50ntFlanks@12003.12031.29|GCGGGCCGCCGGTGAAATACCACTACTCT</td><td>>hg38_21_+_8433172_8446622_RNA45SN1_45S_50ntFlanks</td><td>12003</td><td>12031</td><td>29</td><td>GCGGGCCGCCGGTGAAATACCACTACTCT</td><td>1059.1</td><td>-5.4</td><td>0.6</td><td>-8.4</td><td>3.5E-17</td><td>1.2E-16</td><td>BT20/MB468</td><td>NUC</td></tr>
<tr><td>>hg38_21_+_8433172_8446622_RNA45SN1_45S_50ntFlanks@3926.3959.34|TCAGATCAAAACCAACCCGGTCAGCCCCTCTCCG</td><td>>hg38_21_+_8433172_8446622_RNA45SN1_45S_50ntFlanks</td><td>3926</td><td>3959</td><td>34</td><td>TCAGATCAAAACCAACCCGGTCAGCCCCTCTCCG</td><td>397.8</td><td>-12.4</td><td>1.5</td><td>-8.5</td><td>1.9E-17</td><td>1.2E-16</td><td>MB231/MB468</td><td>MITO</td></tr>
<tr><td>>hg38_21_+_8433172_8446622_RNA45SN1_45S_50ntFlanks@10798.10818.21|CATGTTGGAACAATGTAGGTA</td><td>>hg38_21_+_8433172_8446622_RNA45SN1_45S_50ntFlanks</td><td>10798</td><td>10818</td><td>21</td><td>CATGTTGGAACAATGTAGGTA</td><td>1960.9</td><td>-8.6</td><td>1.0</td><td>-8.5</td><td>1.9E-17</td><td>1.2E-16</td><td>MB231/MB468</td><td>MITO</td></tr>
<tr><td>>trnaMT_ValTAC_MT_+_1602_1670@28.72.45|AACTTACACTTAGGAGATTTCAACTTAACTTGACCGCTCTGACCA</td><td>>trnaMT_ValTAC_MT_+_1602_1670</td><td>28</td><td>72</td><td>45</td><td>AACTTACACTTAGGAGATTTCAACTTAACTTGACCGCTCTGACCA</td><td>253.5</td><td>12.0</td><td>1.4</td><td>8.6</td><td>1.2E-17</td><td>1.2E-16</td><td>MB231/MB468</td><td>CYTO</td></tr>
<tr><td>>MI0000294|hsa-mir-218-1&WithFlank&4|+|20528269|20528390@31.52.22(+1U);MI0000295|hsa-mir-218-2&WithFlank&5|-|168768140|168768261@31.52.22(+1U)&[MIMAT0000275&hsa-miR-218-5p&offsets|0|+1(+1U);m-185&4|+|20528299|20528319&offsets|0|+1(+1U)];[MIMAT0000275_1&hsa-miR-218-5p&offsets|0|+1(+1U);m-187&5|-|168768211|168768231&offsets|0|+1(+1U)]|TTGTGCTTGATCTAACCATGTGT</td><td>>MI0000294|hsa-mir-218-1&WithFlank&4|+|20528269|20528390</td><td>31</td><td>52</td><td>22(+1U)</td><td>TTGTGCTTGATCTAACCATGTGT</td><td>222.8</td><td>11.7</td><td>1.4</td><td>8.5</td><td>1.5E-17</td><td>1.2E-16</td><td>MB231/MB468</td><td>NUC</td></tr>
<tr><td>>hg38_21_+_8433172_8446622_RNA45SN1_45S_50ntFlanks@13002.13025.24|CGATCTATTGAAAGTCAGCCCTCG</td><td>>hg38_21_+_8433172_8446622_RNA45SN1_45S_50ntFlanks</td><td>13002</td><td>13025</td><td>24</td><td>CGATCTATTGAAAGTCAGCCCTCG</td><td>298.7</td><td>-11.2</td><td>1.3</td><td>-8.5</td><td>1.5E-17</td><td>1.2E-16</td><td>MB231/MB468</td><td>NUC</td></tr>
<tr><td>>hg38_21_+_8433172_8446622_RNA45SN1_45S_50ntFlanks@11873.11900.28|AAGAAGACCCTGTTGAGCTTGACTCTAG</td><td>>hg38_21_+_8433172_8446622_RNA45SN1_45S_50ntFlanks</td><td>11873</td><td>11900</td><td>28</td><td>AAGAAGACCCTGTTGAGCTTGACTCTAG</td><td>915.3</td><td>-9.6</td><td>1.1</td><td>-8.5</td><td>1.9E-17</td><td>1.2E-16</td><td>MB231/MB468</td><td>MITO</td></tr>
<tr><td>>hg38_21_+_8433172_8446622_RNA45SN1_45S_50ntFlanks@10409.10435.27|GTCAGTCGGTCCTGAGAGATGGGCGAG</td><td>>hg38_21_+_8433172_8446622_RNA45SN1_45S_50ntFlanks</td><td>10409</td><td>10435</td><td>27</td><td>GTCAGTCGGTCCTGAGAGATGGGCGAG</td><td>346.7</td><td>-12.3</td><td>1.4</td><td>-8.6</td><td>1.2E-17</td><td>1.2E-16</td><td>MB231/MB468</td><td>MP</td></tr>
<tr><td>>hg38_21_+_8433172_8446622_RNA45SN1_45S_50ntFlanks@11562.11592.31|CGCCTAGCAGCCGACTTAGAACTGGTGCGGA</td><td>>hg38_21_+_8433172_8446622_RNA45SN1_45S_50ntFlanks</td><td>11562</td><td>11592</td><td>31</td><td>CGCCTAGCAGCCGACTTAGAACTGGTGCGGA</td><td>748.4</td><td>-4.2</td><td>0.5</td><td>-8.5</td><td>2.1E-17</td><td>1.2E-16</td><td>BT20/MB468</td><td>CYTO</td></tr>
<tr><td>>MI0000088|hsa-mir-30a&WithFlank&6|-|71403545|71403627@12.33.22(+1A)&[MIMAT0000087&hsa-miR-30a-5p&offsets|0|0(+1A);m-15&6|-|71403595|71403616&offsets|0|0(+1A)]|TGTAAACATCCTCGACTGGAAGA</td><td>>MI0000088|hsa-mir-30a&WithFlank&6|-|71403545|71403627</td><td>12</td><td>33</td><td>22(+1A)</td><td>TGTAAACATCCTCGACTGGAAGA</td><td>1025.8</td><td>6.7</td><td>0.8</td><td>8.5</td><td>1.5E-17</td><td>1.2E-16</td><td>MB231/MB468</td><td>NUC</td></tr>
<tr><td>>MI0000298|hsa-mir-221&WithFlank&X|-|45746151|45746272@71.93.23(+1U)&[MIMAT0000278&hsa-miR-221-3p&offsets|0|0(+1U);m-91&X|-|45746181|45746202&offsets|0|+1(+1U)]|AGCTACATTGTCTGCTGGGTTTCT</td><td>>MI0000298|hsa-mir-221&WithFlank&X|-|45746151|45746272</td><td>71</td><td>93</td><td>23(+1U)</td><td>AGCTACATTGTCTGCTGGGTTTCT</td><td>3832.9</td><td>5.8</td><td>0.7</td><td>8.5</td><td>1.5E-17</td><td>1.2E-16</td><td>BT20/MB468</td><td>MITO</td></tr>
<tr><td>>hg38_21_+_8433172_8446622_RNA45SN1_45S_50ntFlanks@11667.11703.37|CTGCCCAGTGCTCTGAATGTCAAAGTGAAGAAATTCA</td><td>>hg38_21_+_8433172_8446622_RNA45SN1_45S_50ntFlanks</td><td>11667</td><td>11703</td><td>37</td><td>CTGCCCAGTGCTCTGAATGTCAAAGTGAAGAAATTCA</td><td>533.8</td><td>6.1</td><td>0.7</td><td>8.6</td><td>1.2E-17</td><td>1.2E-16</td><td>MB231/MB468</td><td>CYTO</td></tr>
<tr><td>>hg38_21_+_8433172_8446622_RNA45SN1_45S_50ntFlanks@10116.10135.20|TGCGTCGGAGGGCGGCGGCG</td><td>>hg38_21_+_8433172_8446622_RNA45SN1_45S_50ntFlanks</td><td>10116</td><td>10135</td><td>20</td><td>TGCGTCGGAGGGCGGCGGCG</td><td>401.5</td><td>-11.4</td><td>1.3</td><td>-8.6</td><td>1.2E-17</td><td>1.2E-16</td><td>MB231/MB468</td><td>CYTO</td></tr>
<tr><td>>hg38_21_+_8433172_8446622_RNA45SN1_45S_50ntFlanks@10680.10706.27|GTTCCGGCGGCGTCCGGTGAGCTCTCG</td><td>>hg38_21_+_8433172_8446622_RNA45SN1_45S_50ntFlanks</td><td>10680</td><td>10706</td><td>27</td><td>GTTCCGGCGGCGTCCGGTGAGCTCTCG</td><td>11332.9</td><td>-9.8</td><td>1.2</td><td>-8.5</td><td>1.9E-17</td><td>1.2E-16</td><td>MB231/MB468</td><td>MITO</td></tr>
<tr><td>>MI0000433|hsa-let-7g&WithFlank&3|-|52268272|52268367@11.31.21(+1A)&[MIMAT0000414&hsa-let-7g-5p&offsets|0|-1(+1A);m-58&3|-|52268336|52268357&offsets|0|-1(+1A)]|TGAGGTAGTAGTTTGTACAGTA</td><td>>MI0000433|hsa-let-7g&WithFlank&3|-|52268272|52268367</td><td>11</td><td>31</td><td>21(+1A)</td><td>TGAGGTAGTAGTTTGTACAGTA</td><td>4826.8</td><td>7.5</td><td>0.9</td><td>8.5</td><td>2.1E-17</td><td>1.2E-16</td><td>BT20/MB468</td><td>CYTO</td></tr>
</table>

<table>
<tr><td>>trna111_HisGTG_1_-_147774845_147774916@-1G.33.34,trna118_HisGTG_1_-_145396881_145396952@-1G.33.34,trna16_HisGTG_1_+_146544773_146544844@-1G.33.34,trna1_HisGTG_15_+_45493349_45493420@-1G.33.34,trna21_HisGTG_1_+_147753471_147753542@-1G.33.34,trna33_HisGTG_6_+_27125906_27125977@-1G.33.34,trna7_HisGTG_9_-_14433938_14434009@-1G.33.34,trna8_HisGTG_15_-_45492611_45492682@-1G.33.34,trna9_HisGTG_15_-_45490804_45490875@-1G.33.34|GGCCGTGATCGTATAGTGGTTAGTACTCTGCGTT</td><td>>trna111_HisGTG_1_-_147774845_147774916</td><td>-1G</td><td>33</td><td>34</td><td>GGCCGTGATCGTATAGTGGTTAGTACTCTGCGTT</td><td>7839.7</td><td>8.5</td><td>1.0</td><td>8.5</td><td>2.1E-17</td><td>1.2E-16</td><td>BT20/MB468</td><td>CYTO</td></tr>
<tr><td>>hg38_21_+_8433172_8446622_RNA45SN1_45S_50ntFlanks@12002.12030.29|TGCGGGCCGCCGGTGAAATACCACTACTC</td><td>>hg38_21_+_8433172_8446622_RNA45SN1_45S_50ntFlanks</td><td>12002</td><td>12030</td><td>29</td><td>TGCGGGCCGCCGGTGAAATACCACTACTC</td><td>359.3</td><td>-12.4</td><td>1.4</td><td>-8.6</td><td>1.2E-17</td><td>1.2E-16</td><td>MB231/MB468</td><td>MP</td></tr>
<tr><td>>hg38_21_+_8433172_8446622_RNA45SN1_45S_50ntFlanks@6660.6678.19|GCGGTGGATCACTCGGCTC</td><td>>hg38_21_+_8433172_8446622_RNA45SN1_45S_50ntFlanks</td><td>6660</td><td>6678</td><td>19</td><td>GCGGTGGATCACTCGGCTC</td><td>464.8</td><td>6.0</td><td>0.7</td><td>8.5</td><td>1.5E-17</td><td>1.2E-16</td><td>BT20/MB468</td><td>MITO</td></tr>
<tr><td>>hg38_21_+_8433172_8446622_RNA45SN1_45S_50ntFlanks@11881.11904.24|CCTGTTGAGCTTGACTCTAGTCTG</td><td>>hg38_21_+_8433172_8446622_RNA45SN1_45S_50ntFlanks</td><td>11881</td><td>11904</td><td>24</td><td>CCTGTTGAGCTTGACTCTAGTCTG</td><td>2210.0</td><td>-11.3</td><td>1.3</td><td>-8.4</td><td>3.6E-17</td><td>1.2E-16</td><td>BT20/MB468</td><td>NUC</td></tr>
<tr><td>>hg38_21_+_8433172_8446622_RNA45SN1_45S_50ntFlanks@11729.11753.25|TAACTATGACTCTCTTAAGGTAGCC</td><td>>hg38_21_+_8433172_8446622_RNA45SN1_45S_50ntFlanks</td><td>11729</td><td>11753</td><td>25</td><td>TAACTATGACTCTCTTAAGGTAGCC</td><td>661.3</td><td>-13.2</td><td>1.5</td><td>-8.5</td><td>1.9E-17</td><td>1.2E-16</td><td>MB231/MB468</td><td>MITO</td></tr>
<tr><td>>MI0000298|hsa-mir-221&WithFlank&X|-|45746151|45746272@71.93.23&[MIMAT0000278&hsa-miR-221-3p&offsets|0|0;m-91&X|-|45746181|45746202&offsets|0|+1]|AGCTACATTGTCTGCTGGGTTTC</td><td>>MI0000298|hsa-mir-221&WithFlank&X|-|45746151|45746272</td><td>71</td><td>93</td><td>23</td><td>AGCTACATTGTCTGCTGGGTTTC</td><td>15415.9</td><td>5.8</td><td>0.7</td><td>8.5</td><td>1.6E-17</td><td>1.2E-16</td><td>BT20/MB468</td><td>MITO</td></tr>
<tr><td>>trna10_AspGTC_12_-_125424193_125424264@31.75.45,trna12_AspGTC_12_-_125411891_125411962@31.75.45,trna38_AspGTC_17_-_8125556_8125627@31.75.45,trna45_AspGTC_6_+_27447453_27447524@31.75.45,trna48_AspGTC_6_+_27471523_27471594@31.75.45,trna4_AspGTC_12_+_96429799_96429870@31.75.45,trna69_AspGTC_1_-_161440205_161440276@31.75.45,trna72_AspGTC_1_-_161432824_161432895@31.75.45,trna75_AspGTC_1_-_161425414_161425485@31.75.45,trna78_AspGTC_1_-_161418033_161418104@31.75.45,trna81_AspGTC_1_-_161410615_161410686@31.75.45|CCTGTCACGCGGGAGACCGGGGTTCGATTCCCCGACGGGGAGCCA</td><td>>trna10_AspGTC_12_-_125424193_125424264</td><td>31</td><td>75</td><td>45</td><td>CCTGTCACGCGGGAGACCGGGGTTCGATTCCCCGACGGGGAGCCA</td><td>964.7</td><td>-14.3</td><td>1.7</td><td>-8.4</td><td>3.6E-17</td><td>1.2E-16</td><td>BT20/MB468</td><td>NUC</td></tr>
<tr><td>>hg38_21_+_8433172_8446622_RNA45SN1_45S_50ntFlanks@3815.3836.22|ATGGTTCCTTTGGTCGCTCGCT</td><td>>hg38_21_+_8433172_8446622_RNA45SN1_45S_50ntFlanks</td><td>3815</td><td>3836</td><td>22</td><td>ATGGTTCCTTTGGTCGCTCGCT</td><td>365.8</td><td>-4.6</td><td>0.5</td><td>-8.4</td><td>3.6E-17</td><td>1.2E-16</td><td>BT20/MB468</td><td>NUC</td></tr>
<tr><td>>MI0005756|hsa-mir-934&WithFlank&X|+|136550872|136550966@21.42.22(+2A)&[MIMAT0004977&hsa-miR-934&offsets|0|0(+2A);m-943&X|+|136550892|136550913&offsets|0|0(+2A)]|TGTCTACTACTGGAGACACTGGAA</td><td>>MI0005756|hsa-mir-934&WithFlank&X|+|136550872|136550966</td><td>21</td><td>42</td><td>22(+2A)</td><td>TGTCTACTACTGGAGACACTGGAA</td><td>584.7</td><td>13.0</td><td>1.5</td><td>8.8</td><td>2.1E-18</td><td>1.2E-16</td><td>BT20/MB231</td><td>NUC</td></tr>
<tr><td>>hg38_21_+_8433172_8446622_RNA45SN1_45S_50ntFlanks@12360.12382.23|CAGGGATAACTGGCTTGTGGCGG</td><td>>hg38_21_+_8433172_8446622_RNA45SN1_45S_50ntFlanks</td><td>12360</td><td>12382</td><td>23</td><td>CAGGGATAACTGGCTTGTGGCGG</td><td>392.0</td><td>-12.4</td><td>1.5</td><td>-8.5</td><td>1.9E-17</td><td>1.2E-16</td><td>MB231/MB468</td><td>MITO</td></tr>
<tr><td>>hg38_21_+_8433172_8446622_RNA45SN1_45S_50ntFlanks@5469.5499.31|CCCACGGCCCTGGCGGAGCGCTGAGAAGACG</td><td>>hg38_21_+_8433172_8446622_RNA45SN1_45S_50ntFlanks</td><td>5469</td><td>5499</td><td>31</td><td>CCCACGGCCCTGGCGGAGCGCTGAGAAGACG</td><td>416.8</td><td>-12.5</td><td>1.5</td><td>-8.5</td><td>1.9E-17</td><td>1.2E-16</td><td>MB231/MB468</td><td>MITO</td></tr>
<tr><td>>MI0000816|hsa-mir-335&WithFlank&7|+|130496105|130496210@58.79.22&[MIMAT0004703&hsa-miR-335-3p&offsets|0|0;m-159&7|+|130496162|130496183&offsets|0|0]|TTTTTCATTATTGCTCCTGACC</td><td>>MI0000816|hsa-mir-335&WithFlank&7|+|130496105|130496210</td><td>58</td><td>79</td><td>22</td><td>TTTTTCATTATTGCTCCTGACC</td><td>2245.1</td><td>5.1</td><td>0.6</td><td>8.5</td><td>1.5E-17</td><td>1.2E-16</td><td>MB231/MB468</td><td>NUC</td></tr>
<tr><td>>hg38_21_+_8433172_8446622_RNA45SN1_45S_50ntFlanks@13002.13020.19|CGATCTATTGAAAGTCAGC</td><td>>hg38_21_+_8433172_8446622_RNA45SN1_45S_50ntFlanks</td><td>13002</td><td>13020</td><td>19</td><td>CGATCTATTGAAAGTCAGC</td><td>352.5</td><td>-12.3</td><td>1.4</td><td>-8.6</td><td>1.2E-17</td><td>1.2E-16</td><td>MB231/MB468</td><td>MP</td></tr>
<tr><td>>trnaMT_LeuTAA_MT_+_3230_3304@16.44.29|CCCGGTAATCGCATAAAACTTAAAACTTT</td><td>>trnaMT_LeuTAA_MT_+_3230_3304</td><td>16</td><td>44</td><td>29</td><td>CCCGGTAATCGCATAAAACTTAAAACTTT</td><td>2109.4</td><td>-8.8</td><td>1.0</td><td>-8.5</td><td>1.9E-17</td><td>1.2E-16</td><td>MB231/MB468</td><td>MITO</td></tr>
<tr><td>>hg38_21_+_8433172_8446622_RNA45SN1_45S_50ntFlanks@11641.11658.18|CGGCGGGTGTTGACGCGA</td><td>>hg38_21_+_8433172_8446622_RNA45SN1_45S_50ntFlanks</td><td>11641</td><td>11658</td><td>18</td><td>CGGCGGGTGTTGACGCGA</td><td>740.1</td><td>-13.3</td><td>1.6</td><td>-8.5</td><td>2.0E-17</td><td>1.2E-16</td><td>MB231/MB468</td><td>MITO</td></tr>
<tr><td>>trnaMT_SerGCT_MT_+_12207_12265@9.38.30|TCACAAGAACTGCTAACTCATGCCCCCATG</td><td>>trnaMT_SerGCT_MT_+_12207_12265</td><td>9</td><td>38</td><td>30</td><td>TCACAAGAACTGCTAACTCATGCCCCCATG</td><td>446.3</td><td>-13.2</td><td>1.6</td><td>-8.4</td><td>3.7E-17</td><td>1.2E-16</td><td>BT20/MB468</td><td>NUC</td></tr>
<tr><td>>hg38_21_+_8433172_8446622_RNA45SN1_45S_50ntFlanks@11572.11601.30|CCGACTTAGAACTGGTGCGGACCAGGGGAA</td><td>>hg38_21_+_8433172_8446622_RNA45SN1_45S_50ntFlanks</td><td>11572</td><td>11601</td><td>30</td><td>CCGACTTAGAACTGGTGCGGACCAGGGGAA</td><td>41053.2</td><td>-11.5</td><td>1.4</td><td>-8.4</td><td>3.7E-17</td><td>1.2E-16</td><td>BT20/MB468</td><td>NUC</td></tr>
<tr><td>>hg38_21_+_8433172_8446622_RNA45SN1_45S_50ntFlanks@6722.6744.23|ATTGCAGGACACATTGATCATCG</td><td>>hg38_21_+_8433172_8446622_RNA45SN1_45S_50ntFlanks</td><td>6722</td><td>6744</td><td>23</td><td>ATTGCAGGACACATTGATCATCG</td><td>3180.3</td><td>-11.6</td><td>1.4</td><td>-8.4</td><td>3.7E-17</td><td>1.2E-16</td><td>BT20/MB468</td><td>NUC</td></tr>
<tr><td>>MI0000650|hsa-mir-200c&WithFlank&12|+|6963693|6963772@49.72.24&[MIMAT0000617&hsa-miR-200c-3p&offsets|-1|0;m-54&12|+|6963742|6963764&offsets|-1|0]|CTAATACTGCCGGGTAATGATGGA</td><td>>MI0000650|hsa-mir-200c&WithFlank&12|+|6963693|6963772</td><td>49</td><td>72</td><td>24</td><td>CTAATACTGCCGGGTAATGATGGA</td><td>409.5</td><td>12.4</td><td>1.4</td><td>8.7</td><td>2.2E-18</td><td>1.2E-16</td><td>BT20/MB231</td><td>NUC</td></tr>
<tr><td>>hg38_21_+_8433172_8446622_RNA45SN1_45S_50ntFlanks@9570.9597.28|TGCAAATCGGTCGTCCGACCTGGGTATA</td><td>>hg38_21_+_8433172_8446622_RNA45SN1_45S_50ntFlanks</td><td>9570</td><td>9597</td><td>28</td><td>TGCAAATCGGTCGTCCGACCTGGGTATA</td><td>3411.0</td><td>-11.1</td><td>1.3</td><td>-8.6</td><td>1.2E-17</td><td>1.2E-16</td><td>MB231/MB468</td><td>MP</td></tr>
<tr><td>>hg38_1_-_228634819_228635039_RNA5S12_5S_50ntFlanks@138.170.33|AGACCGCCTGGGAATACCGGGTGCTGTAGGCTT</td><td>>hg38_1_-_228634819_228635039_RNA5S12_5S_50ntFlanks</td><td>138</td><td>170</td><td>33</td><td>AGACCGCCTGGGAATACCGGGTGCTGTAGGCTT</td><td>1624.8</td><td>6.7</td><td>0.8</td><td>8.6</td><td>1.2E-17</td><td>1.2E-16</td><td>MB231/MB468</td><td>CYTO</td></tr>
<tr><td>>hg38_21_+_8433172_8446622_RNA45SN1_45S_50ntFlanks@12615.12659.45|GCTGAGGAGCCAATGGGGCGAAGCTACCATCTGTGGGATTATGAC</td><td>>hg38_21_+_8433172_8446622_RNA45SN1_45S_50ntFlanks</td><td>12615</td><td>12659</td><td>45</td><td>GCTGAGGAGCCAATGGGGCGAAGCTACCATCTGTGGGATTATGAC</td><td>629.5</td><td>-13.7</td><td>1.6</td><td>-8.4</td><td>3.7E-17</td><td>1.2E-16</td><td>BT20/MB468</td><td>NUC</td></tr>
<tr><td>>trnaMT_GluTTC_MT_-_14674_14742@39.68.30,trnalookalike8_GluTTC_5_-_93905172_93905240@39.68.30|ATTGGTCGTGGTTGTAGTCCGTGCGAGAAT</td><td>>trnaMT_GluTTC_MT_-_14674_14742</td><td>39</td><td>68</td><td>30</td><td>ATTGGTCGTGGTTGTAGTCCGTGCGAGAAT</td><td>1044.7</td><td>7.7</td><td>0.9</td><td>8.5</td><td>1.6E-17</td><td>1.2E-16</td><td>BT20/MB468</td><td>MITO</td></tr>
<tr><td>>hg38_21_+_8433172_8446622_RNA45SN1_45S_50ntFlanks@12405.12433.29|CTTTTTGATCCTTCGATGTCGGCTCTTCC</td><td>>hg38_21_+_8433172_8446622_RNA45SN1_45S_50ntFlanks</td><td>12405</td><td>12433</td><td>29</td><td>CTTTTTGATCCTTCGATGTCGGCTCTTCC</td><td>1322.1</td><td>-8.3</td><td>1.0</td><td>-8.6</td><td>1.2E-17</td><td>1.2E-16</td><td>MB231/MB468</td><td>MP</td></tr>
<tr><td>>hg38_21_+_8433172_8446622_RNA45SN1_45S_50ntFlanks@12634.12667.34|GAAGCTACCATCTGTGGGATTATGACTGAACGCC</td><td>>hg38_21_+_8433172_8446622_RNA45SN1_45S_50ntFlanks</td><td>12634</td><td>12667</td><td>34</td><td>GAAGCTACCATCTGTGGGATTATGACTGAACGCC</td><td>153.1</td><td>11.3</td><td>1.3</td><td>8.5</td><td>1.2E-17</td><td>1.2E-16</td><td>MB231/MB468</td><td>CYTO</td></tr>
<tr><td>>hg38_21_+_8433172_8446622_RNA45SN1_45S_50ntFlanks@10116.10136.21|TGCGTCGGAGGGCGGCGGCGG</td><td>>hg38_21_+_8433172_8446622_RNA45SN1_45S_50ntFlanks</td><td>10116</td><td>10136</td><td>21</td><td>TGCGTCGGAGGGCGGCGGCGG</td><td>466.8</td><td>-11.6</td><td>1.4</td><td>-8.5</td><td>1.2E-17</td><td>1.2E-16</td><td>MB231/MB468</td><td>CYTO</td></tr>
<tr><td>>hg38_21_+_8433172_8446622_RNA45SN1_45S_50ntFlanks@11581.11602.22|AACTGGTGCGGACCAGGGGAAT</td><td>>hg38_21_+_8433172_8446622_RNA45SN1_45S_50ntFlanks</td><td>11581</td><td>11602</td><td>22</td><td>AACTGGTGCGGACCAGGGGAAT</td><td>909.0</td><td>-5.7</td><td>0.7</td><td>-8.4</td><td>3.8E-17</td><td>1.2E-16</td><td>BT20/MB468</td><td>NUC</td></tr>
<tr><td>>hg38_21_+_8433172_8446622_RNA45SN1_45S_50ntFlanks@5487.5509.23|CGCTGAGAAGACGGTCGAACTTG</td><td>>hg38_21_+_8433172_8446622_RNA45SN1_45S_50ntFlanks</td><td>5487</td><td>5509</td><td>23</td><td>CGCTGAGAAGACGGTCGAACTTG</td><td>377.1</td><td>-12.3</td><td>1.5</td><td>-8.5</td><td>2.0E-17</td><td>1.2E-16</td><td>MB231/MB468</td><td>MITO</td></tr>
<tr><td>>hg38_21_+_8433172_8446622_RNA45SN1_45S_50ntFlanks@9381.9410.30|CGCTCGCCGGCCGAGGTGGGATCCCGAGGC</td><td>>hg38_21_+_8433172_8446622_RNA45SN1_45S_50ntFlanks</td><td>9381</td><td>9410</td><td>30</td><td>CGCTCGCCGGCCGAGGTGGGATCCCGAGGC</td><td>1843.3</td><td>-14.6</td><td>1.7</td><td>-8.5</td><td>2.0E-17</td><td>1.2E-16</td><td>MB231/MB468</td><td>MITO</td></tr>
<tr><td>>hg38_21_+_8433172_8446622_RNA45SN1_45S_50ntFlanks@11873.11890.18|AAGAAGACCCTGTTGAGC</td><td>>hg38_21_+_8433172_8446622_RNA45SN1_45S_50ntFlanks</td><td>11873</td><td>11890</td><td>18</td><td>AAGAAGACCCTGTTGAGC</td><td>855.2</td><td>-4.8</td><td>0.6</td><td>-8.5</td><td>1.2E-17</td><td>1.2E-16</td><td>MB231/MB468</td><td>CYTO</td></tr>
<tr><td>>hg38_21_+_8433172_8446622_RNA45SN1_45S_50ntFlanks@8157.8178.22|CTCGTGGGGGGCCCAAGTCCTT</td><td>>hg38_21_+_8433172_8446622_RNA45SN1_45S_50ntFlanks</td><td>8157</td><td>8178</td><td>22</td><td>CTCGTGGGGGGCCCAAGTCCTT</td><td>2207.0</td><td>-11.2</td><td>1.3</td><td>-8.5</td><td>2.0E-17</td><td>1.3E-16</td><td>MB231/MB468</td><td>MITO</td></tr>
<tr><td>>hg38_21_+_8433172_8446622_RNA45SN1_45S_50ntFlanks@3793.3814.22|CGAATGGCTCATTAAATCAGTT</td><td>>hg38_21_+_8433172_8446622_RNA45SN1_45S_50ntFlanks</td><td>3793</td><td>3814</td><td>22</td><td>CGAATGGCTCATTAAATCAGTT</td><td>272.5</td><td>-12.0</td><td>1.4</td><td>-8.5</td><td>1.2E-17</td><td>1.3E-16</td><td>MB231/MB468</td><td>MP</td></tr>
<tr><td>>hg38_21_+_8433172_8446622_RNA45SN1_45S_50ntFlanks@6649.6681.33|TACGACTCTTAGCGGTGGATCACTCGGCTCGTG</td><td>>hg38_21_+_8433172_8446622_RNA45SN1_45S_50ntFlanks</td><td>6649</td><td>6681</td><td>33</td><td>TACGACTCTTAGCGGTGGATCACTCGGCTCGTG</td><td>2603.8</td><td>-6.7</td><td>0.8</td><td>-8.5</td><td>1.2E-17</td><td>1.3E-16</td><td>MB231/MB468</td><td>MP</td></tr>
<tr><td>>hg38_21_+_8433172_8446622_RNA45SN1_45S_50ntFlanks@12419.12440.22|GATGTCGGCTCTTCCTATCATT</td><td>>hg38_21_+_8433172_8446622_RNA45SN1_45S_50ntFlanks</td><td>12419</td><td>12440</td><td>22</td><td>GATGTCGGCTCTTCCTATCATT</td><td>333.9</td><td>4.7</td><td>0.6</td><td>8.5</td><td>1.6E-17</td><td>1.3E-16</td><td>BT20/MB468</td><td>MITO</td></tr>
<tr><td>>trna11_GluTTC_15_-_26327381_26327452@1.33.33,trna3_GluTTC_13_-_45492062_45492133@1.33.33|TCCCACATGGTCTAGCGGTTAGGATTCCTGGTT</td><td>>trna11_GluTTC_15_-_26327381_26327452</td><td>1</td><td>33</td><td>33</td><td>TCCCACATGGTCTAGCGGTTAGGATTCCTGGTT</td><td>2383.0</td><td>8.2</td><td>1.0</td><td>8.5</td><td>1.6E-17</td><td>1.3E-16</td><td>MB231/MB468</td><td>NUC</td></tr>
<tr><td>>hg38_21_+_8433172_8446622_RNA45SN1_45S_50ntFlanks@11235.11269.35|CTTCCCGTGGATCGCCCCAGCTGCGGCGGGCGTCG</td><td>>hg38_21_+_8433172_8446622_RNA45SN1_45S_50ntFlanks</td><td>11235</td><td>11269</td><td>35</td><td>CTTCCCGTGGATCGCCCCAGCTGCGGCGGGCGTCG</td><td>1484.3</td><td>-13.5</td><td>1.6</td><td>-8.5</td><td>1.6E-17</td><td>1.3E-16</td><td>MB231/MB468</td><td>NUC</td></tr>
<tr><td>>hg38_21_+_8433172_8446622_RNA45SN1_45S_50ntFlanks@10681.10710.30|TTCCGGCGGCGTCCGGTGAGCTCTCGCTGG</td><td>>hg38_21_+_8433172_8446622_RNA45SN1_45S_50ntFlanks</td><td>10681</td><td>10710</td><td>30</td><td>TTCCGGCGGCGTCCGGTGAGCTCTCGCTGG</td><td>2320.1</td><td>-8.9</td><td>1.0</td><td>-8.5</td><td>1.3E-17</td><td>1.3E-16</td><td>MB231/MB468</td><td>CYTO</td></tr>
<tr><td>>hg38_21_+_8433172_8446622_RNA45SN1_45S_50ntFlanks@10682.10706.25|TCCGGCGGCGTCCGGTGAGCTCTCG</td><td>>hg38_21_+_8433172_8446622_RNA45SN1_45S_50ntFlanks</td><td>10682</td><td>10706</td><td>25</td><td>TCCGGCGGCGTCCGGTGAGCTCTCG</td><td>2327.4</td><td>-6.7</td><td>0.8</td><td>-8.5</td><td>2.0E-17</td><td>1.3E-16</td><td>MB231/MB468</td><td>MITO</td></tr>
<tr><td>>hg38_21_+_8433172_8446622_RNA45SN1_45S_50ntFlanks@11874.11907.34|AGAAGACCCTGTTGAGCTTGACTCTAGTCTGGCA</td><td>>hg38_21_+_8433172_8446622_RNA45SN1_45S_50ntFlanks</td><td>11874</td><td>11907</td><td>34</td><td>AGAAGACCCTGTTGAGCTTGACTCTAGTCTGGCA</td><td>554.6</td><td>-4.3</td><td>0.5</td><td>-8.5</td><td>1.7E-17</td><td>1.3E-16</td><td>BT20/MB468</td><td>MITO</td></tr>
<tr><td>>hg38_21_+_8433172_8446622_RNA45SN1_45S_50ntFlanks@12002.12043.42|TGCGGGCCGCCGGTGAAATACCACTACTCTGATCGTTTTTTC</td><td>>hg38_21_+_8433172_8446622_RNA45SN1_45S_50ntFlanks</td><td>12002</td><td>12043</td><td>42</td><td>TGCGGGCCGCCGGTGAAATACCACTACTCTGATCGTTTTTTC</td><td>2258.1</td><td>-14.1</td><td>1.7</td><td>-8.5</td><td>1.6E-17</td><td>1.3E-16</td><td>MB231/MB468</td><td>NUC</td></tr>
<tr><td>>hg38_21_+_8433172_8446622_RNA45SN1_45S_50ntFlanks@9805.9838.34|GCGTGGAATGCGAGTGCCTAGTGGGCCACTTTTG</td><td>>hg38_21_+_8433172_8446622_RNA45SN1_45S_50ntFlanks</td><td>9805</td><td>9838</td><td>34</td><td>GCGTGGAATGCGAGTGCCTAGTGGGCCACTTTTG</td><td>313.2</td><td>-12.2</td><td>1.4</td><td>-8.5</td><td>1.3E-17</td><td>1.3E-16</td><td>MB231/MB468</td><td>MP</td></tr>
<tr><td>>hg38_21_+_8433172_8446622_RNA45SN1_45S_50ntFlanks@10879.10902.24|CGGTCGGGCTGGGGCGCGAAGCGG</td><td>>hg38_21_+_8433172_8446622_RNA45SN1_45S_50ntFlanks</td><td>10879</td><td>10902</td><td>24</td><td>CGGTCGGGCTGGGGCGCGAAGCGG</td><td>384.2</td><td>-12.4</td><td>1.5</td><td>-8.5</td><td>1.3E-17</td><td>1.3E-16</td><td>MB231/MB468</td><td>MP</td></tr>
<tr><td>>hg38_21_+_8433172_8446622_RNA45SN1_45S_50ntFlanks@4657.4686.30|CTAGAGGTGAAATTCTTGGACCGGCGCAAG</td><td>>hg38_21_+_8433172_8446622_RNA45SN1_45S_50ntFlanks</td><td>4657</td><td>4686</td><td>30</td><td>CTAGAGGTGAAATTCTTGGACCGGCGCAAG</td><td>470.7</td><td>-12.7</td><td>1.5</td><td>-8.5</td><td>1.3E-17</td><td>1.3E-16</td><td>MB231/MB468</td><td>MP</td></tr>
<tr><td>>hg38_21_+_8433172_8446622_RNA45SN1_45S_50ntFlanks@5436.5466.31|GTGAGGCCCTCGGATCGGCCCCGCCGGGGTC</td><td>>hg38_21_+_8433172_8446622_RNA45SN1_45S_50ntFlanks</td><td>5436</td><td>5466</td><td>31</td><td>GTGAGGCCCTCGGATCGGCCCCGCCGGGGTC</td><td>317.1</td><td>-12.2</td><td>1.4</td><td>-8.5</td><td>1.3E-17</td><td>1.3E-16</td><td>MB231/MB468</td><td>MP</td></tr>
<tr><td>>hg38_21_+_8433172_8446622_RNA45SN1_45S_50ntFlanks@12893.12933.41|CCTCGCCCGTCACGCACCGCACGTTCGTGGGGAACCTGGCG</td><td>>hg38_21_+_8433172_8446622_RNA45SN1_45S_50ntFlanks</td><td>12893</td><td>12933</td><td>41</td><td>CCTCGCCCGTCACGCACCGCACGTTCGTGGGGAACCTGGCG</td><td>535.0</td><td>-12.0</td><td>1.4</td><td>-8.5</td><td>1.6E-17</td><td>1.3E-16</td><td>MB231/MB468</td><td>NUC</td></tr>
</table>

<table>
<tr><td>>hg38_21_+_8433172_8446622_RNA45SN1_45S_50ntFlanks@6733.6770.38|CATTGATCATCGACACTTCGAACGCACTTGCGGCCCCG</td><td>>hg38_21_+_8433172_8446622_RNA45SN1_45S_50ntFlanks</td><td>6733</td><td>6770</td><td>38</td><td>CATTGATCATCGACACTTCGAACGCACTTGCGGCCCCG</td><td>2666.1</td><td>-4.3</td><td>0.5</td><td>-8.5</td><td>1.7E-17</td><td>1.3E-16</td><td>BT20/MB468</td><td>MITO</td></tr>
<tr><td>>MI0003137|hsa-mir-193b&WithFlank&16|+|14303961|14304055@57.77.21(+1A)&[MIMAT0002819&hsa-miR-193b-3p&offsets|0|-1(+1A);m-166&16|+|14304017|14304038&offsets|0|-1(+1A)]|AACTGGCCCTCAAAGTCCCGCA</td><td>>MI0003137|hsa-mir-193b&WithFlank&16|+|14303961|14304055</td><td>57</td><td>77</td><td>21(+1A)</td><td>AACTGGCCCTCAAAGTCCCGCA</td><td>605.1</td><td>11.2</td><td>1.3</td><td>8.5</td><td>2.3E-17</td><td>1.3E-16</td><td>BT20/MB468</td><td>CYTO</td></tr>
<tr><td>>hg38_21_+_8433172_8446622_RNA45SN1_45S_50ntFlanks@10692.10709.18|TCCGGTGAGCTCTCGCTG</td><td>>hg38_21_+_8433172_8446622_RNA45SN1_45S_50ntFlanks</td><td>10692</td><td>10709</td><td>18</td><td>TCCGGTGAGCTCTCGCTG</td><td>384.6</td><td>-12.5</td><td>1.5</td><td>-8.5</td><td>1.3E-17</td><td>1.3E-16</td><td>MB231/MB468</td><td>MP</td></tr>
<tr><td>>hg38_21_+_8433172_8446622_RNA45SN1_45S_50ntFlanks@6646.6669.24|TCGTACGACTCTTAGCGGTGGATC</td><td>>hg38_21_+_8433172_8446622_RNA45SN1_45S_50ntFlanks</td><td>6646</td><td>6669</td><td>24</td><td>TCGTACGACTCTTAGCGGTGGATC</td><td>201858.1</td><td>4.9</td><td>0.6</td><td>8.5</td><td>1.7E-17</td><td>1.3E-16</td><td>BT20/MB468</td><td>MITO</td></tr>
<tr><td>>MI0000095|hsa-mir-93&WithFlank&7|-|100093762|100093853@17.37.21&[MIMAT0000093&hsa-miR-93-5p&offsets|0|-2;m-29&7|-|100093815|100093837&offsets|0|-2]|CAAAGTGCTGTTCGTGCAGGT</td><td>>MI0000095|hsa-mir-93&WithFlank&7|-|100093762|100093853</td><td>17</td><td>37</td><td>21</td><td>CAAAGTGCTGTTCGTGCAGGT</td><td>1062.5</td><td>6.1</td><td>0.7</td><td>8.5</td><td>2.3E-17</td><td>1.3E-16</td><td>BT20/MB468</td><td>CYTO</td></tr>
<tr><td>>hg38_21_+_8433172_8446622_RNA45SN1_45S_50ntFlanks@7982.8005.24|TCAGATCAGACGTGGCGACCCGCT</td><td>>hg38_21_+_8433172_8446622_RNA45SN1_45S_50ntFlanks</td><td>7982</td><td>8005</td><td>24</td><td>TCAGATCAGACGTGGCGACCCGCT</td><td>434.6</td><td>-3.9</td><td>0.5</td><td>-8.4</td><td>3.9E-17</td><td>1.3E-16</td><td>BT20/MB468</td><td>NUC</td></tr>
<tr><td>>hg38_21_+_8433172_8446622_RNA45SN1_45S_50ntFlanks@12072.12095.24|CCCGAGGGGCTCTCGCTTCTGGCG</td><td>>hg38_21_+_8433172_8446622_RNA45SN1_45S_50ntFlanks</td><td>12072</td><td>12095</td><td>24</td><td>CCCGAGGGGCTCTCGCTTCTGGCG</td><td>1378.9</td><td>-10.0</td><td>1.2</td><td>-8.5</td><td>2.1E-17</td><td>1.3E-16</td><td>MB231/MB468</td><td>MITO</td></tr>
<tr><td>>MI0000299|hsa-mir-222&WithFlank&X|-|45747009|45747130@75.99.25(+4U)&[MIMAT0000279&hsa-miR-222-3p&offsets|0|+4(+4U);m-140&X|-|45747033|45747056&offsets|0|+1(+4U)]|AGCTACATCTGGCTACTGGGTCTCTTTTT</td><td>>MI0000299|hsa-mir-222&WithFlank&X|-|45747009|45747130</td><td>75</td><td>99</td><td>25(+4U)</td><td>AGCTACATCTGGCTACTGGGTCTCTTTTT</td><td>353.3</td><td>6.8</td><td>0.8</td><td>8.5</td><td>1.6E-17</td><td>1.3E-16</td><td>MB231/MB468</td><td>NUC</td></tr>
<tr><td>>hg38_21_+_8433172_8446622_RNA45SN1_45S_50ntFlanks@11707.11728.22|AAGCGCGGGTAAACGGCGGGAG</td><td>>hg38_21_+_8433172_8446622_RNA45SN1_45S_50ntFlanks</td><td>11707</td><td>11728</td><td>22</td><td>AAGCGCGGGTAAACGGCGGGAG</td><td>426.7</td><td>-12.5</td><td>1.5</td><td>-8.5</td><td>2.1E-17</td><td>1.3E-16</td><td>MB231/MB468</td><td>MITO</td></tr>
<tr><td>>hg38_21_+_8433172_8446622_RNA45SN1_45S_50ntFlanks@12624.12652.29|CCAATGGGGCGAAGCTACCATCTGTGGGA</td><td>>hg38_21_+_8433172_8446622_RNA45SN1_45S_50ntFlanks</td><td>12624</td><td>12652</td><td>29</td><td>CCAATGGGGCGAAGCTACCATCTGTGGGA</td><td>751.5</td><td>-13.4</td><td>1.6</td><td>-8.5</td><td>1.3E-17</td><td>1.3E-16</td><td>MB231/MB468</td><td>MP</td></tr>
<tr><td>>MI0000469|hsa-mir-125a&WithFlank&19|+|51693248|51693345@21.42.22(+1U)&[MIMAT0000443&hsa-miR-125a-5p&offsets|0|-2(+1U);m-63&19|+|51693268|51693289&offsets|0|0(+1U)]|TCCCTGAGACCCTTTAACCTGTT</td><td>>MI0000469|hsa-mir-125a&WithFlank&19|+|51693248|51693345</td><td>21</td><td>42</td><td>22(+1U)</td><td>TCCCTGAGACCCTTTAACCTGTT</td><td>702.6</td><td>9.8</td><td>1.2</td><td>8.5</td><td>2.4E-17</td><td>1.3E-16</td><td>BT20/MB468</td><td>CYTO</td></tr>
<tr><td>>hg38_21_+_8433172_8446622_RNA45SN1_45S_50ntFlanks@485.518.34|GATGTGGTGACGTCGTGCTCTCCCGGGCCGGGTC</td><td>>hg38_21_+_8433172_8446622_RNA45SN1_45S_50ntFlanks</td><td>485</td><td>518</td><td>34</td><td>GATGTGGTGACGTCGTGCTCTCCCGGGCCGGGTC</td><td>396.5</td><td>11.5</td><td>1.4</td><td>8.4</td><td>4.0E-17</td><td>1.3E-16</td><td>BT20/MB468</td><td>NUC</td></tr>
<tr><td>>MI0000108|hsa-mir-103a-2&WithFlank&20|+|3917488|3917577@54.76.23;MI0000109|hsa-mir-103a-1&WithFlank&5|-|168560890|168560979@54.76.23&[MIMAT0000101&hsa-miR-103a-3p&offsets|0|0;m-12&20|+|3917541|3917563&offsets|0|0];[MIMAT0000101_1&hsa-miR-103a-3p&offsets|0|0;m-13&5|-|168560904|168560926&offsets|0|0;m-44220&5|-|168560908|168560926&offsets|0|+4]|AGCAGCATTGTACAGGGCTATGA</td><td>>MI0000108|hsa-mir-103a-2&WithFlank&20|+|3917488|3917577</td><td>54</td><td>76</td><td>23</td><td>AGCAGCATTGTACAGGGCTATGA</td><td>24973.4</td><td>5.4</td><td>0.6</td><td>8.5</td><td>1.7E-17</td><td>1.3E-16</td><td>MB231/MB468</td><td>NUC</td></tr>
<tr><td>>hg38_21_+_8433172_8446622_RNA45SN1_45S_50ntFlanks@9457.9503.47|CGCCGGGGAGGTGGAGCACGAGCGCACGTGTTAGGACCCGAAAGATG</td><td>>hg38_21_+_8433172_8446622_RNA45SN1_45S_50ntFlanks</td><td>9457</td><td>9503</td><td>47</td><td>CGCCGGGGAGGTGGAGCACGAGCGCACGTGTTAGGACCCGAAAGATG</td><td>704.5</td><td>-13.9</td><td>1.7</td><td>-8.4</td><td>4.0E-17</td><td>1.3E-16</td><td>BT20/MB468</td><td>NUC</td></tr>
<tr><td>>hg38_21_+_8433172_8446622_RNA45SN1_45S_50ntFlanks@475.512.38|CTTCGTGATCGATGTGGTGACGTCGTGCTCTCCCGGGC</td><td>>hg38_21_+_8433172_8446622_RNA45SN1_45S_50ntFlanks</td><td>475</td><td>512</td><td>38</td><td>CTTCGTGATCGATGTGGTGACGTCGTGCTCTCCCGGGC</td><td>318.2</td><td>10.3</td><td>1.2</td><td>8.5</td><td>2.4E-17</td><td>1.3E-16</td><td>BT20/MB468</td><td>CYTO</td></tr>
<tr><td>>hg38_21_+_8433172_8446622_RNA45SN1_45S_50ntFlanks@12488.12520.33|AACGTGAGCTGGGTTTAGACCGTCGTGAGACAG</td><td>>hg38_21_+_8433172_8446622_RNA45SN1_45S_50ntFlanks</td><td>12488</td><td>12520</td><td>33</td><td>AACGTGAGCTGGGTTTAGACCGTCGTGAGACAG</td><td>280.7</td><td>-12.0</td><td>1.4</td><td>-8.5</td><td>1.3E-17</td><td>1.3E-16</td><td>MB231/MB468</td><td>MP</td></tr>
<tr><td>>MI0000477|hsa-mir-146a&WithFlank&5|+|160485346|160485456@27.50.24&[MIMAT0000449&hsa-miR-146a-5p&offsets|0|+2;m-129&5|+|160485372|160485393&offsets|0|+2]|TGAGAACTGAATTCCATGGGTTGT</td><td>>MI0000477|hsa-mir-146a&WithFlank&5|+|160485346|160485456</td><td>27</td><td>50</td><td>24</td><td>TGAGAACTGAATTCCATGGGTTGT</td><td>511.9</td><td>12.9</td><td>1.5</td><td>8.5</td><td>1.7E-17</td><td>1.3E-16</td><td>MB231/MB468</td><td>NUC</td></tr>
<tr><td>>hg38_21_+_8433172_8446622_RNA45SN1_45S_50ntFlanks@10534.10569.36|CCGCGAGGCGTCCAGTGCGGTAACGCGACCGATCCC</td><td>>hg38_21_+_8433172_8446622_RNA45SN1_45S_50ntFlanks</td><td>10534</td><td>10569</td><td>36</td><td>CCGCGAGGCGTCCAGTGCGGTAACGCGACCGATCCC</td><td>447.5</td><td>-12.6</td><td>1.5</td><td>-8.5</td><td>2.1E-17</td><td>1.3E-16</td><td>MB231/MB468</td><td>MITO</td></tr>
<tr><td>>trna116_GluCTC_1_-_145399233_145399304@1.36.36,trna59_GluCTC_1_+_249168447_249168518@1.36.36,trna71_GluCTC_1_-_161439189_161439260@1.36.36,trna74_GluCTC_1_-_161431809_161431880@1.36.36,trna77_GluCTC_1_-_161424398_161424469@1.36.36,trna77_GluCTC_6_+_28949976_28950047@1.36.36,trna80_GluCTC_1_-_161417018_161417089@1.36.36,trna87_GluCTC_6_-_126101393_126101464@1.36.36|TCCCTGGTGGTCTAGTGGTTAGGATTCGGCGCTCTC</td><td>>trna116_GluCTC_1_-_145399233_145399304</td><td>1</td><td>36</td><td>36</td><td>TCCCTGGTGGTCTAGTGGTTAGGATTCGGCGCTCTC</td><td>4978.2</td><td>7.5</td><td>0.9</td><td>8.5</td><td>1.7E-17</td><td>1.3E-16</td><td>BT20/MB468</td><td>MITO</td></tr>
<tr><td>>hg38_21_+_8433172_8446622_RNA45SN1_45S_50ntFlanks@10681.10709.29|TTCCGGCGGCGTCCGGTGAGCTCTCGCTG</td><td>>hg38_21_+_8433172_8446622_RNA45SN1_45S_50ntFlanks</td><td>10681</td><td>10709</td><td>29</td><td>TTCCGGCGGCGTCCGGTGAGCTCTCGCTG</td><td>254.9</td><td>-11.9</td><td>1.4</td><td>-8.5</td><td>1.3E-17</td><td>1.3E-16</td><td>MB231/MB468</td><td>MP</td></tr>
<tr><td>>hg38_21_+_8433172_8446622_RNA45SN1_45S_50ntFlanks@4759.4780.22|ATCAGATACCGTCGTAGTTCCG</td><td>>hg38_21_+_8433172_8446622_RNA45SN1_45S_50ntFlanks</td><td>4759</td><td>4780</td><td>22</td><td>ATCAGATACCGTCGTAGTTCCG</td><td>8204.4</td><td>-12.2</td><td>1.4</td><td>-8.5</td><td>1.3E-17</td><td>1.3E-16</td><td>MB231/MB468</td><td>MP</td></tr>
<tr><td>>hg38_21_+_8433172_8446622_RNA45SN1_45S_50ntFlanks@10841.10862.22|CGTAACTTCGGGATAAGGATTG</td><td>>hg38_21_+_8433172_8446622_RNA45SN1_45S_50ntFlanks</td><td>10841</td><td>10862</td><td>22</td><td>CGTAACTTCGGGATAAGGATTG</td><td>4752.5</td><td>-9.7</td><td>1.2</td><td>-8.4</td><td>4.0E-17</td><td>1.3E-16</td><td>BT20/MB468</td><td>NUC</td></tr>
<tr><td>>hg38_21_+_8433172_8446622_RNA45SN1_45S_50ntFlanks@11452.11472.21|CTCCTCGGCGCGCGGCGGCGG</td><td>>hg38_21_+_8433172_8446622_RNA45SN1_45S_50ntFlanks</td><td>11452</td><td>11472</td><td>21</td><td>CTCCTCGGCGCGCGGCGGCGG</td><td>657.7</td><td>-8.6</td><td>1.0</td><td>-8.5</td><td>1.3E-17</td><td>1.3E-16</td><td>MB231/MB468</td><td>CYTO</td></tr>
<tr><td>>hg38_21_+_8433172_8446622_RNA45SN1_45S_50ntFlanks@12003.12030.28|GCGGGCCGCCGGTGAAATACCACTACTC</td><td>>hg38_21_+_8433172_8446622_RNA45SN1_45S_50ntFlanks</td><td>12003</td><td>12030</td><td>28</td><td>GCGGGCCGCCGGTGAAATACCACTACTC</td><td>297.0</td><td>-12.1</td><td>1.4</td><td>-8.5</td><td>1.3E-17</td><td>1.3E-16</td><td>MB231/MB468</td><td>MP</td></tr>
<tr><td>>trna116_GluCTC_1_-_145399233_145399304@32.69.38,trna59_GluCTC_1_+_249168447_249168518@32.69.38,trna71_GluCTC_1_-_161439189_161439260@32.69.38,trna74_GluCTC_1_-_161431809_161431880@32.69.38,trna77_GluCTC_1_-_161424398_161424469@32.69.38,trna77_GluCTC_6_+_28949976_28950047@32.69.38,trna80_GluCTC_1_-_161417018_161417089@32.69.38,trna87_GluCTC_6_-_126101393_126101464@32.69.38|CTCTCACCGCCGCGGCCCGGGTTCGATTCCCGGTCAGG</td><td>>trna116_GluCTC_1_-_145399233_145399304</td><td>32</td><td>69</td><td>38</td><td>CTCTCACCGCCGCGGCCCGGGTTCGATTCCCGGTCAGG</td><td>570.8</td><td>-11.9</td><td>1.4</td><td>-8.5</td><td>1.3E-17</td><td>1.3E-16</td><td>MB231/MB468</td><td>CYTO</td></tr>
<tr><td>>hg38_21_+_8433172_8446622_RNA45SN1_45S_50ntFlanks@10681.10711.31|TTCCGGCGGCGTCCGGTGAGCTCTCGCTGGC</td><td>>hg38_21_+_8433172_8446622_RNA45SN1_45S_50ntFlanks</td><td>10681</td><td>10711</td><td>31</td><td>TTCCGGCGGCGTCCGGTGAGCTCTCGCTGGC</td><td>1576.7</td><td>-8.7</td><td>1.0</td><td>-8.5</td><td>1.3E-17</td><td>1.3E-16</td><td>MB231/MB468</td><td>CYTO</td></tr>
<tr><td>>hg38_21_+_8433172_8446622_RNA45SN1_45S_50ntFlanks@6719.6740.22|TGAATTGCAGGACACATTGATC</td><td>>hg38_21_+_8433172_8446622_RNA45SN1_45S_50ntFlanks</td><td>6719</td><td>6740</td><td>22</td><td>TGAATTGCAGGACACATTGATC</td><td>3199.9</td><td>-8.8</td><td>1.0</td><td>-8.5</td><td>1.3E-17</td><td>1.3E-16</td><td>MB231/MB468</td><td>MP</td></tr>
<tr><td>>MI0000088|hsa-mir-30a&WithFlank&6|-|71403545|71403627@53.74.22&[MIMAT0000088&hsa-miR-30a-3p&offsets|0|0;m-38&6|-|71403554|71403575&offsets|0|0]|CTTTCAGTCGGATGTTTGCAGC</td><td>>MI0000088|hsa-mir-30a&WithFlank&6|-|71403545|71403627</td><td>53</td><td>74</td><td>22</td><td>CTTTCAGTCGGATGTTTGCAGC</td><td>3054.6</td><td>5.8</td><td>0.7</td><td>8.5</td><td>1.8E-17</td><td>1.3E-16</td><td>BT20/MB468</td><td>MITO</td></tr>
<tr><td>>hg38_21_+_8433172_8446622_RNA45SN1_45S_50ntFlanks@11768.11786.19|CTAATTAGTGACGCGCATG</td><td>>hg38_21_+_8433172_8446622_RNA45SN1_45S_50ntFlanks</td><td>11768</td><td>11786</td><td>19</td><td>CTAATTAGTGACGCGCATG</td><td>354.3</td><td>-12.3</td><td>1.4</td><td>-8.5</td><td>1.4E-17</td><td>1.4E-16</td><td>MB231/MB468</td><td>MP</td></tr>
<tr><td>>hg38_21_+_8433172_8446622_RNA45SN1_45S_50ntFlanks@11612.11640.29|TAATTAAAACAAAGCATCGCGAAGGCCCG</td><td>>hg38_21_+_8433172_8446622_RNA45SN1_45S_50ntFlanks</td><td>11612</td><td>11640</td><td>29</td><td>TAATTAAAACAAAGCATCGCGAAGGCCCG</td><td>1478.3</td><td>-12.6</td><td>1.5</td><td>-8.5</td><td>1.8E-17</td><td>1.4E-16</td><td>BT20/MB468</td><td>MITO</td></tr>
<tr><td>>MI0000102|hsa-mir-100&WithFlank&11|-|122152223|122152314@20.40.21&[MIMAT0000098&hsa-miR-100-5p&offsets|+1|0;m-27&11|-|122152276|122152296&offsets|+1|+1]|ACCCGTAGATCCGAACTTGTG</td><td>>MI0000102|hsa-mir-100&WithFlank&11|-|122152223|122152314</td><td>20</td><td>40</td><td>21</td><td>ACCCGTAGATCCGAACTTGTG</td><td>1036.9</td><td>13.2</td><td>1.6</td><td>8.5</td><td>2.2E-17</td><td>1.4E-16</td><td>MB231/MB468</td><td>MITO</td></tr>
<tr><td>>hg38_21_+_8433172_8446622_RNA45SN1_45S_50ntFlanks@9166.9183.18|CCGGGCGGGTCGCGCCGT</td><td>>hg38_21_+_8433172_8446622_RNA45SN1_45S_50ntFlanks</td><td>9166</td><td>9183</td><td>18</td><td>CCGGGCGGGTCGCGCCGT</td><td>595.2</td><td>-13.0</td><td>1.5</td><td>-8.5</td><td>2.2E-17</td><td>1.4E-16</td><td>MB231/MB468</td><td>MITO</td></tr>
<tr><td>>hg38_21_+_8433172_8446622_RNA45SN1_45S_50ntFlanks@11562.11585.24|CGCCTAGCAGCCGACTTAGAACTG</td><td>>hg38_21_+_8433172_8446622_RNA45SN1_45S_50ntFlanks</td><td>11562</td><td>11585</td><td>24</td><td>CGCCTAGCAGCCGACTTAGAACTG</td><td>291.7</td><td>-12.1</td><td>1.4</td><td>-8.5</td><td>1.4E-17</td><td>1.4E-16</td><td>MB231/MB468</td><td>MP</td></tr>
<tr><td>>trna15_ValAAC_5_-_180615416_180615488@1.29.29|GTTTCCGTAGTGTAGTGGTCATCACGTTC</td><td>>trna15_ValAAC_5_-_180615416_180615488</td><td>1</td><td>29</td><td>29</td><td>GTTTCCGTAGTGTAGTGGTCATCACGTTC</td><td>5539.5</td><td>-7.0</td><td>0.8</td><td>-8.5</td><td>1.4E-17</td><td>1.4E-16</td><td>MB231/MB468</td><td>MP</td></tr>
<tr><td>>hg38_21_+_8433172_8446622_RNA45SN1_45S_50ntFlanks@11534.11552.19|GTTCCGCGCGGCGCCTCGC</td><td>>hg38_21_+_8433172_8446622_RNA45SN1_45S_50ntFlanks</td><td>11534</td><td>11552</td><td>19</td><td>GTTCCGCGCGGCGCCTCGC</td><td>1480.0</td><td>-8.8</td><td>1.0</td><td>-8.5</td><td>2.2E-17</td><td>1.4E-16</td><td>MB231/MB468</td><td>MITO</td></tr>
<tr><td>>MI0000283|hsa-mir-203a&WithFlank&14|+|104117399|104117520@71.90.20&[MIMAT0000264&hsa-miR-203a-3p&offsets|0|-2;m-79&14|+|104117469|104117490&offsets|0|-2]|GTGAAATGTTTAGGACCACT</td><td>>MI0000283|hsa-mir-203a&WithFlank&14|+|104117399|104117520</td><td>71</td><td>90</td><td>20</td><td>GTGAAATGTTTAGGACCACT</td><td>624.4</td><td>12.7</td><td>1.4</td><td>8.8</td><td>1.3E-18</td><td>1.4E-16</td><td>BT20/MB231</td><td>CYTO</td></tr>
<tr><td>>hg38_21_+_8433172_8446622_RNA45SN1_45S_50ntFlanks@11208.11233.26|GCGGCGACTCTGGACGCGAGCCGGGC</td><td>>hg38_21_+_8433172_8446622_RNA45SN1_45S_50ntFlanks</td><td>11208</td><td>11233</td><td>26</td><td>GCGGCGACTCTGGACGCGAGCCGGGC</td><td>6902.0</td><td>-7.1</td><td>0.8</td><td>-8.5</td><td>1.4E-17</td><td>1.4E-16</td><td>MB231/MB468</td><td>CYTO</td></tr>
<tr><td>>hg38_21_+_8433172_8446622_RNA45SN1_45S_50ntFlanks@10869.10894.26|AGGGCTGGGTCGGTCGGGCTGGGGCG</td><td>>hg38_21_+_8433172_8446622_RNA45SN1_45S_50ntFlanks</td><td>10869</td><td>10894</td><td>26</td><td>AGGGCTGGGTCGGTCGGGCTGGGGCG</td><td>1600.1</td><td>-6.2</td><td>0.7</td><td>-8.5</td><td>1.4E-17</td><td>1.4E-16</td><td>MB231/MB468</td><td>MP</td></tr>
<tr><td>>hg38_21_+_8433172_8446622_RNA45SN1_45S_50ntFlanks@11781.11806.26|CGCATGAATGGATGAACGAGATTCCC</td><td>>hg38_21_+_8433172_8446622_RNA45SN1_45S_50ntFlanks</td><td>11781</td><td>11806</td><td>26</td><td>CGCATGAATGGATGAACGAGATTCCC</td><td>1116.7</td><td>-9.7</td><td>1.1</td><td>-8.5</td><td>2.2E-17</td><td>1.4E-16</td><td>MB231/MB468</td><td>MITO</td></tr>
<tr><td>>hg38_21_+_8433172_8446622_RNA45SN1_45S_50ntFlanks@12996.13038.43|TCGCTGCGATCTATTGAAAGTCAGCCCTCGACACAAGGGTTTG</td><td>>hg38_21_+_8433172_8446622_RNA45SN1_45S_50ntFlanks</td><td>12996</td><td>13038</td><td>43</td><td>TCGCTGCGATCTATTGAAAGTCAGCCCTCGACACAAGGGTTTG</td><td>4444.8</td><td>9.6</td><td>1.1</td><td>8.5</td><td>1.4E-17</td><td>1.4E-16</td><td>MB231/MB468</td><td>CYTO</td></tr>
<tr><td>>hg38_21_+_8433172_8446622_RNA45SN1_45S_50ntFlanks@10841.10865.25|CGTAACTTCGGGATAAGGATTGGCT</td><td>>hg38_21_+_8433172_8446622_RNA45SN1_45S_50ntFlanks</td><td>10841</td><td>10865</td><td>25</td><td>CGTAACTTCGGGATAAGGATTGGCT</td><td>88107.1</td><td>-6.5</td><td>0.8</td><td>-8.5</td><td>1.4E-17</td><td>1.4E-16</td><td>MB231/MB468</td><td>CYTO</td></tr>
<tr><td>>hg38_21_+_8433172_8446622_RNA45SN1_45S_50ntFlanks@11729.11765.37|TAACTATGACTCTCTTAAGGTAGCCAAATGCCTCGTC</td><td>>hg38_21_+_8433172_8446622_RNA45SN1_45S_50ntFlanks</td><td>11729</td><td>11765</td><td>37</td><td>TAACTATGACTCTCTTAAGGTAGCCAAATGCCTCGTC</td><td>3291.4</td><td>-8.8</td><td>1.0</td><td>-8.5</td><td>2.2E-17</td><td>1.4E-16</td><td>MB231/MB468</td><td>MITO</td></tr>
<tr><td>>hg38_21_+_8433172_8446622_RNA45SN1_45S_50ntFlanks@12636.12653.18|AGCTACCATCTGTGGGAT</td><td>>hg38_21_+_8433172_8446622_RNA45SN1_45S_50ntFlanks</td><td>12636</td><td>12653</td><td>18</td><td>AGCTACCATCTGTGGGAT</td><td>1820.1</td><td>-13.9</td><td>1.6</td><td>-9.0</td><td>3.1E-19</td><td>1.4E-16</td><td>BT20/MB468</td><td>MP</td></tr>
</table>

| | | | | | | | | | | | | | |
|---|---|---|---|---|---|---|---|---|---|---|---|---|---|
| >hg38_21_+_8433172_8446622_RNA45SN1_45S_50ntFlanks@11564.11603.40\|CCTAGCAGCCGACTTAGAACTGGTGCGGACCAGGGGAATC | >hg38_21_+_8433172_8446622_RNA45SN1_45S_50ntFlanks | 11564 | 11603 | 40 | CCTAGCAGCCGACTTAGAACTGGTGCGGACCAGGGGAATC | 3184.6 | -6.7 | 0.8 | -8.5 | 1.8E-17 | 1.4E-16 | MB231/MB468 | NUC |
| >hg38_21_+_8433172_8446622_RNA45SN1_45S_50ntFlanks@11576.11605.30\|CTTAGAACTGGTGCGGACCAGGGGAATCCG | >hg38_21_+_8433172_8446622_RNA45SN1_45S_50ntFlanks | 11576 | 11605 | 30 | CTTAGAACTGGTGCGGACCAGGGGAATCCG | 1521.8 | -5.6 | 0.7 | -8.5 | 2.5E-17 | 1.4E-16 | BT20/MB468 | CYTO |
| >hg38_21_+_8433172_8446622_RNA45SN1_45S_50ntFlanks@13002.13025.24\|CGATCTATTGAAAGTCAGCCCTCG | >hg38_21_+_8433172_8446622_RNA45SN1_45S_50ntFlanks | 13002 | 13025 | 24 | CGATCTATTGAAAGTCAGCCCTCG | 553.2 | -11.4 | 1.3 | -8.5 | 2.5E-17 | 1.4E-16 | BT20/MB468 | CYTO |
| >am_trna134_GluTTC_1_-_16861774_16861845@1.30.30,trna5_GluTTC_1_+_17199078_17199149@1.30.30,trna84_GluTTC_1_-_161391883_161391954@1.30.30,trna94_GluTTC_1_-_149664355_149664427@1.30.30\|TCCCTGGTGGTCTAGTGGCTAGGATTCGGC | >am_trna134_GluTTC_1_-_16861774_16861845 | 1 | 30 | 30 | TCCCTGGTGGTCTAGTGGCTAGGATTCGGC | 2880.3 | -7.1 | 0.8 | -8.4 | 4.2E-17 | 1.4E-16 | BT20/MB468 | NUC |
| >hg38_21_+_8433172_8446622_RNA45SN1_45S_50ntFlanks@12457.12476.20\|CAAGCGTTGGATTGTTCACC | >hg38_21_+_8433172_8446622_RNA45SN1_45S_50ntFlanks | 12457 | 12476 | 20 | CAAGCGTTGGATTGTTCACC | 3883.9 | -6.8 | 0.8 | -8.5 | 1.4E-17 | 1.4E-16 | MB231/MB468 | MP |
| >hg38_21_+_8433172_8446622_RNA45SN1_45S_50ntFlanks@10822.10865.44\|GAAGTCGGCAAGCCGGATCCGTAACTTCGGGATAAGGATTGGCT | >hg38_21_+_8433172_8446622_RNA45SN1_45S_50ntFlanks | 10822 | 10865 | 44 | GAAGTCGGCAAGCCGGATCCGTAACTTCGGGATAAGGATTGGCT | 1578.5 | -15.1 | 1.8 | -8.4 | 4.2E-17 | 1.4E-16 | BT20/MB468 | NUC |
| >hg38_21_+_8433172_8446622_RNA45SN1_45S_50ntFlanks@12289.12317.29\|CCGTGAAAGCGGGGCCTCACGATCCTTCT | >hg38_21_+_8433172_8446622_RNA45SN1_45S_50ntFlanks | 12289 | 12317 | 29 | CCGTGAAAGCGGGGCCTCACGATCCTTCT | 1487.3 | -14.3 | 1.7 | -8.5 | 2.3E-17 | 1.4E-16 | MB231/MB468 | MITO |
| >hg38_21_+_8433172_8446622_RNA45SN1_45S_50ntFlanks@7986.8008.23\|ATCAGACGTGGCGACCCGCTGAA | >hg38_21_+_8433172_8446622_RNA45SN1_45S_50ntFlanks | 7986 | 8008 | 23 | ATCAGACGTGGCGACCCGCTGAA | 1246.0 | -12.4 | 1.5 | -8.5 | 1.8E-17 | 1.4E-16 | BT20/MB468 | MITO |
| >hg38_21_+_8433172_8446622_RNA45SN1_45S_50ntFlanks@6721.6770.50\|AATTGCAGGACACATTGATCATCGACACTTCGAACGCACTTGCGGCCCCG | >hg38_21_+_8433172_8446622_RNA45SN1_45S_50ntFlanks | 6721 | 6770 | 50 | AATTGCAGGACACATTGATCATCGACACTTCGAACGCACTTGCGGCCCCG | 1967.2 | -15.4 | 1.8 | -8.4 | 4.3E-17 | 1.4E-16 | BT20/MB468 | NUC |
| >hg38_21_+_8433172_8446622_RNA45SN1_45S_50ntFlanks@11844.11873.30\|GGGAACGGGCTTGGCGGAATCAGCGGGGAA | >hg38_21_+_8433172_8446622_RNA45SN1_45S_50ntFlanks | 11844 | 11873 | 30 | GGGAACGGGCTTGGCGGAATCAGCGGGGAA | 343.0 | -12.3 | 1.4 | -8.5 | 1.4E-17 | 1.4E-16 | MB231/MB468 | MP |
| >hg38_21_+_8433172_8446622_RNA45SN1_45S_50ntFlanks@10914.10946.33\|CGCCGCGGCTGGACGAGGCGCCGCCGCCCCCCC | >hg38_21_+_8433172_8446622_RNA45SN1_45S_50ntFlanks | 10914 | 10946 | 33 | CGCCGCGGCTGGACGAGGCGCCGCCGCCCCCCC | 353.5 | -11.2 | 1.3 | -8.5 | 1.4E-17 | 1.4E-16 | MB231/MB468 | CYTO |
| >hg38_21_+_8433172_8446622_RNA45SN1_45S_50ntFlanks@11235.11269.35\|CTTCCCGTGGATCGCCCCAGCTGCGGCGGGCGTCG | >hg38_21_+_8433172_8446622_RNA45SN1_45S_50ntFlanks | 11235 | 11269 | 35 | CTTCCCGTGGATCGCCCCAGCTGCGGCGGGCGTCG | 585.6 | -11.9 | 1.4 | -8.5 | 1.4E-17 | 1.4E-16 | MB231/MB468 | CYTO |
| >MI0003834\|hsa-mir-769&WithFlank&19\|+\|46018926\|46019055@36.57.22&[MIMAT0003886&hsa-miR-769-5p&offsets\|0\|0;m-216&19\|+\|46018961\|46018982&offsets\|0\|0]\|TGAGACCTCTGGGTTCTGAGCT | >MI0003834\|hsa-mir-769&WithFlank&19\|+\|46018926\|46019055 | 36 | 57 | 22 | TGAGACCTCTGGGTTCTGAGCT | 321.5 | 2.5 | 0.3 | 8.4 | 4.3E-17 | 1.4E-16 | BT20/MB468 | NUC |
| >MI0000086\|hsa-mir-28&WithFlank&3\|+\|188688775\|188688872@60.81.22(+1U)&[MIMAT0004502&hsa-miR-28-3p&offsets\|0\|0(+1U);m-28&3\|+\|188688834\|188688855&offsets\|0\|0(+1U)]\|CACTAGATTGTGAGCTCCTGGAT | >MI0000086\|hsa-mir-28&WithFlank&3\|+\|188688775\|188688872 | 60 | 81 | 22(+1U) | CACTAGATTGTGAGCTCCTGGAT | 1497.2 | 10.5 | 1.2 | 8.5 | 2.6E-17 | 1.4E-16 | BT20/MB468 | CYTO |
| >MI0000266\|hsa-mir-10a&WithFlank&17\|-\|48579832\|48579953@28.48.21(+1A)&[MIMAT0000253&hsa-miR-10a-5p&offsets\|0\|-2(+1A);m-18&17\|-\|48579905\|48579926&offsets\|0\|-1(+1A)]\|TACCCTGTAGATCCGAATTTGA | >MI0000266\|hsa-mir-10a&WithFlank&17\|-\|48579832\|48579953 | 28 | 48 | 21(+1A) | TACCCTGTAGATCCGAATTTGA | 78.4 | 9.1 | 1.1 | 8.4 | 4.3E-17 | 1.4E-16 | BT20/MB468 | NUC |
| >MI0000446\|hsa-mir-125b-1&WithFlank&11\|-\|122099751\|122099850@21.43.23(+1U)&[MIMAT0000423&hsa-miR-125b-5p&offsets\|0\|+1(+1U);m-67&11\|-\|122099809\|122099830&offsets\|0\|+1(+1U)]\|TCCCTGAGACCCTAACTTGTGATT | >MI0000446\|hsa-mir-125b-1&WithFlank&11\|-\|122099751\|122099850 | 21 | 43 | 23(+1U) | TCCCTGAGACCCTAACTTGTGATT | 310.7 | 12.2 | 1.4 | 8.5 | 1.8E-17 | 1.4E-16 | MB231/MB468 | NUC |
| >hg38_21_+_8433172_8446622_RNA45SN1_45S_50ntFlanks@6772.6807.36\|GTTCCTCCCGGGGCTACGCCTGTCTGAGCGTCGCTT | >hg38_21_+_8433172_8446622_RNA45SN1_45S_50ntFlanks | 6772 | 6807 | 36 | GTTCCTCCCGGGGCTACGCCTGTCTGAGCGTCGCTT | 1255.9 | -5.3 | 0.6 | -8.4 | 4.4E-17 | 1.4E-16 | BT20/MB468 | NUC |
| >hg38_21_+_8433172_8446622_RNA45SN1_45S_50ntFlanks@12616.12653.38\|CTGAGGAGCCAATGGGGCGAAGCTACCATCTGTGGGAT | >hg38_21_+_8433172_8446622_RNA45SN1_45S_50ntFlanks | 12616 | 12653 | 38 | CTGAGGAGCCAATGGGGCGAAGCTACCATCTGTGGGAT | 614.4 | -13.1 | 1.5 | -8.5 | 1.4E-17 | 1.4E-16 | MB231/MB468 | MP |
| >hg38_21_+_8433172_8446622_RNA45SN1_45S_50ntFlanks@10727.10757.31\|GGAGAGGGTGTAAATCTCGCGCCGGGCCGTA | >hg38_21_+_8433172_8446622_RNA45SN1_45S_50ntFlanks | 10727 | 10757 | 31 | GGAGAGGGTGTAAATCTCGCGCCGGGCCGTA | 443.4 | -12.6 | 1.5 | -8.5 | 2.3E-17 | 1.4E-16 | MB231/MB468 | MITO |
| >hg38_21_+_8433172_8446622_RNA45SN1_45S_50ntFlanks@13006.13027.22\|CTATTGAAAGTCAGCCCTCGAC | >hg38_21_+_8433172_8446622_RNA45SN1_45S_50ntFlanks | 13006 | 13027 | 22 | CTATTGAAAGTCAGCCCTCGAC | 471.1 | -11.8 | 1.4 | -8.5 | 1.8E-17 | 1.4E-16 | MB231/MB468 | NUC |
| >hg38_21_+_8433172_8446622_RNA45SN1_45S_50ntFlanks@12147.12169.23\|CCAGGTGGGGAGTTTGACTGGGG | >hg38_21_+_8433172_8446622_RNA45SN1_45S_50ntFlanks | 12147 | 12169 | 23 | CCAGGTGGGGAGTTTGACTGGGG | 351.0 | -12.3 | 1.4 | -8.5 | 1.4E-17 | 1.4E-16 | MB231/MB468 | MP |
| >hg38_21_+_8433172_8446622_RNA45SN1_45S_50ntFlanks@6647.6664.18\|CGTACGACTCTTAGCGGT | >hg38_21_+_8433172_8446622_RNA45SN1_45S_50ntFlanks | 6647 | 6664 | 18 | CGTACGACTCTTAGCGGT | 3792.3 | 5.6 | 0.7 | 8.5 | 2.6E-17 | 1.4E-16 | BT20/MB468 | CYTO |
| >MI0000462\|hsa-mir-152&WithFlank&17\|-\|48037155\|48037253@60.81.22(+1U)&[MIMAT0000438&hsa-miR-152-3p&offsets\|0\|+1(+1U);m-97&17\|-\|48037174\|48037194&offsets\|0\|+1(+1U)]\|TCAGTGCATGACAGAACTTGGGT | >MI0000462\|hsa-mir-152&WithFlank&17\|-\|48037155\|48037253 | 60 | 81 | 22(+1U) | TCAGTGCATGACAGAACTTGGGT | 438.5 | 12.8 | 1.5 | 8.5 | 1.5E-17 | 1.4E-16 | MB231/MB468 | CYTO |
| >hg38_21_+_8433172_8446622_RNA45SN1_45S_50ntFlanks@12934.12959.26\|CTAAACCATTCGTAGACGACCTGCTT | >hg38_21_+_8433172_8446622_RNA45SN1_45S_50ntFlanks | 12934 | 12959 | 26 | CTAAACCATTCGTAGACGACCTGCTT | 249.3 | -11.8 | 1.4 | -8.5 | 1.5E-17 | 1.4E-16 | MB231/MB468 | MP |
| >hg38_21_+_8433172_8446622_RNA45SN1_45S_50ntFlanks@12045.12068.24\|CTGACCCGGTGAGGCGGGGGGGCG | >hg38_21_+_8433172_8446622_RNA45SN1_45S_50ntFlanks | 12045 | 12068 | 24 | CTGACCCGGTGAGGCGGGGGGGCG | 787.3 | -13.4 | 1.6 | -8.5 | 2.3E-17 | 1.4E-16 | MB231/MB468 | MITO |
| >hg38_21_+_8433172_8446622_RNA45SN1_45S_50ntFlanks@10681.10712.32\|TTCCGGCGGCGTCCGGTGAGCTCTCGCTGGCC | >hg38_21_+_8433172_8446622_RNA45SN1_45S_50ntFlanks | 10681 | 10712 | 32 | TTCCGGCGGCGTCCGGTGAGCTCTCGCTGGCC | 2901.2 | -5.9 | 0.7 | -8.5 | 2.3E-17 | 1.4E-16 | MB231/MB468 | MITO |
| >MI0000279\|hsa-mir-196a-2&WithFlank&12\|+\|53991732\|53991853@31.52.22(+1U);MI0000238\|hsa-mir-196a-1&WithFlank&17\|-\|48632484\|48632565@13.34.22(+1U)&[MIMAT0000226&hsa-miR-196a-5p&offsets\|0\|0(+1U);m-261&12\|+\|53991762\|53991783&offsets\|0\|0(+1U)];[MIMAT0000226_1&hsa-miR-196a-5p&offsets\|0\|0(+1U);m-268&17\|-\|48632532\|48632553&offsets\|0\|0(+1U)]\|TAGGTAGTTTCATGTTGTTGGGT | >MI0000279\|hsa-mir-196a-2&WithFlank&12\|+\|53991732\|53991853 | 31 | 52 | 22(+1U) | TAGGTAGTTTCATGTTGTTGGGT | 484.0 | 12.7 | 1.5 | 8.7 | 2.6E-18 | 1.5E-16 | BT20/MB231 | NUC |
| >hg38_21_+_8433172_8446622_RNA45SN1_45S_50ntFlanks@12928.12956.29\|CTGGCGCTAAACCATTCGTAGACGACCTG | >hg38_21_+_8433172_8446622_RNA45SN1_45S_50ntFlanks | 12928 | 12956 | 29 | CTGGCGCTAAACCATTCGTAGACGACCTG | 271.8 | -12.0 | 1.4 | -8.5 | 1.5E-17 | 1.5E-16 | MB231/MB468 | MP |
| >hg38_21_+_8433172_8446622_RNA45SN1_45S_50ntFlanks@11876.11900.25\|AAGACCCTGTTGAGCTTGACTCTAG | >hg38_21_+_8433172_8446622_RNA45SN1_45S_50ntFlanks | 11876 | 11900 | 25 | AAGACCCTGTTGAGCTTGACTCTAG | 338.8 | -12.3 | 1.4 | -8.5 | 1.5E-17 | 1.5E-16 | MB231/MB468 | MP |
| >hg38_21_+_8433172_8446622_RNA45SN1_45S_50ntFlanks@12717.12739.23\|CCTCGGTTGGCCTCGGATAGCCG | >hg38_21_+_8433172_8446622_RNA45SN1_45S_50ntFlanks | 12717 | 12739 | 23 | CCTCGGTTGGCCTCGGATAGCCG | 311.0 | -12.1 | 1.4 | -8.5 | 1.5E-17 | 1.5E-16 | MB231/MB468 | MP |
| >hg38_21_+_8433172_8446622_RNA45SN1_45S_50ntFlanks@9305.9323.19\|CCAAGGAGTCTAACACGTG | >hg38_21_+_8433172_8446622_RNA45SN1_45S_50ntFlanks | 9305 | 9323 | 19 | CCAAGGAGTCTAACACGTG | 728.8 | -13.4 | 1.6 | -8.5 | 1.5E-17 | 1.5E-16 | MB231/MB468 | MP |
| >hg38_21_+_8433172_8446622_RNA45SN1_45S_50ntFlanks@10350.10370.21\|ACGAGAACTTTGAAGGCCGAA | >hg38_21_+_8433172_8446622_RNA45SN1_45S_50ntFlanks | 10350 | 10370 | 21 | ACGAGAACTTTGAAGGCCGAA | 355.9 | -5.2 | 0.6 | -8.4 | 4.5E-17 | 1.5E-16 | BT20/MB468 | NUC |
| >hg38_21_+_8433172_8446622_RNA45SN1_45S_50ntFlanks@11027.11051.25\|CTCTCTCTCTCTCTCCCCCGCTCCC | >hg38_21_+_8433172_8446622_RNA45SN1_45S_50ntFlanks | 11027 | 11051 | 25 | CTCTCTCTCTCTCTCCCCCGCTCCC | 1852.8 | -9.2 | 1.1 | -8.5 | 1.5E-17 | 1.5E-16 | MB231/MB468 | CYTO |
| >hg38_21_+_8433172_8446622_RNA45SN1_45S_50ntFlanks@11572.11611.40\|CCGACTTAGAACTGGTGCGGACCAGGGGAATCCGACTGTT | >hg38_21_+_8433172_8446622_RNA45SN1_45S_50ntFlanks | 11572 | 11611 | 40 | CCGACTTAGAACTGGTGCGGACCAGGGGAATCCGACTGTT | 6273.0 | -6.8 | 0.8 | -8.5 | 2.4E-17 | 1.5E-16 | MB231/MB468 | MITO |
| >hg38_21_+_8433172_8446622_RNA45SN1_45S_50ntFlanks@11199.11223.25\|CCGGCGGCGGCGGCGACTCTGGACG | >hg38_21_+_8433172_8446622_RNA45SN1_45S_50ntFlanks | 11199 | 11223 | 25 | CCGGCGGCGGCGGCGACTCTGGACG | 546.1 | -13.0 | 1.5 | -8.5 | 1.5E-17 | 1.5E-16 | MB231/MB468 | MP |
| >hg38_21_+_8433172_8446622_RNA45SN1_45S_50ntFlanks@10679.10711.33\|GGTTCCGGCGGCGTCCGGTGAGCTCTCGCTGGC | >hg38_21_+_8433172_8446622_RNA45SN1_45S_50ntFlanks | 10679 | 10711 | 33 | GGTTCCGGCGGCGTCCGGTGAGCTCTCGCTGGC | 5414.9 | -8.7 | 1.0 | -8.5 | 1.5E-17 | 1.5E-16 | MB231/MB468 | CYTO |
| >hg38_21_+_8433172_8446622_RNA45SN1_45S_50ntFlanks@12405.12436.32\|CTTTTTGATCCTTCGATGTCGGCTCTTCCTAT | >hg38_21_+_8433172_8446622_RNA45SN1_45S_50ntFlanks | 12405 | 12436 | 32 | CTTTTTGATCCTTCGATGTCGGCTCTTCCTAT | 10299.6 | -9.0 | 1.1 | -8.5 | 1.9E-17 | 1.5E-16 | BT20/MB468 | MITO |
| >hg38_21_+_8433172_8446622_RNA45SN1_45S_50ntFlanks@10869.10904.36\|AGGGCTGGGTCGGTCGGGCTGGGGCGCGAAGCGGGG | >hg38_21_+_8433172_8446622_RNA45SN1_45S_50ntFlanks | 10869 | 10904 | 36 | AGGGCTGGGTCGGTCGGGCTGGGGCGCGAAGCGGGG | 381.5 | -8.5 | 1.0 | -8.5 | 1.5E-17 | 1.5E-16 | MB231/MB468 | CYTO |
| >hg38_21_+_8433172_8446622_RNA45SN1_45S_50ntFlanks@4763.4780.18\|GATACCGTCGTAGTTCCG | >hg38_21_+_8433172_8446622_RNA45SN1_45S_50ntFlanks | 4763 | 4780 | 18 | GATACCGTCGTAGTTCCG | 813.5 | -9.7 | 1.1 | -8.5 | 2.4E-17 | 1.5E-16 | MB231/MB468 | MITO |
| >hg38_21_+_8433172_8446622_RNA45SN1_45S_50ntFlanks@12083.12110.28\|CTCGCTTCTGGCGCCAAGCGCCCGGCCG | >hg38_21_+_8433172_8446622_RNA45SN1_45S_50ntFlanks | 12083 | 12110 | 28 | CTCGCTTCTGGCGCCAAGCGCCCGGCCG | 275.7 | -12.0 | 1.4 | -8.5 | 1.5E-17 | 1.5E-16 | MB231/MB468 | MP |
| >hg38_21_+_8433172_8446622_RNA45SN1_45S_50ntFlanks@11873.11902.30\|AAGAAGACCCTGTTGAGCTTGACTCTAGTC | >hg38_21_+_8433172_8446622_RNA45SN1_45S_50ntFlanks | 11873 | 11902 | 30 | AAGAAGACCCTGTTGAGCTTGACTCTAGTC | 265.4 | -11.9 | 1.4 | -8.5 | 1.5E-17 | 1.5E-16 | MB231/MB468 | MP |
| >hg38_21_+_8433172_8446622_RNA45SN1_45S_50ntFlanks@11534.11560.27\|GTTCCGCGCGGCGCCTCGCCTCGGCCG | >hg38_21_+_8433172_8446622_RNA45SN1_45S_50ntFlanks | 11534 | 11560 | 27 | GTTCCGCGCGGCGCCTCGCCTCGGCCG | 711.6 | -12.2 | 1.4 | -8.5 | 1.5E-17 | 1.5E-16 | MB231/MB468 | CYTO |
| >hg38_21_+_8433172_8446622_RNA45SN1_45S_50ntFlanks@10410.10435.26\|TCAGTCGGTCCTGAGAGATGGGCGAG | >hg38_21_+_8433172_8446622_RNA45SN1_45S_50ntFlanks | 10410 | 10435 | 26 | TCAGTCGGTCCTGAGAGATGGGCGAG | 608.4 | -13.0 | 1.5 | -8.5 | 2.5E-17 | 1.5E-16 | MB231/MB468 | MITO |
| >hg38_21_+_8433172_8446622_RNA45SN1_45S_50ntFlanks@10878.10902.25\|TCGGTCGGGCTGGGGCGCGAAGCGG | >hg38_21_+_8433172_8446622_RNA45SN1_45S_50ntFlanks | 10878 | 10902 | 25 | TCGGTCGGGCTGGGGCGCGAAGCGG | 424.8 | -12.6 | 1.5 | -8.5 | 1.5E-17 | 1.5E-16 | MB231/MB468 | MP |
| >MI0000108\|hsa-mir-103a-2&WithFlank&20\|+\|3917488\|3917577@54.77.24;MI0000109\|hsa-mir-103a-1&WithFlank&5\|-\|168560890\|168560979@54.77.24&[MIMAT0000101&hsa-miR-103a-3p&offsets\|0\|+1;m-12&20\|+\|3917541\|3917563&offsets\|0\|+1];[MIMAT0000101_1&hsa-miR-103a-3p&offsets\|0\|+1;m-13&5\|-\|168560904\|168560926&offsets\|0\|+1;m-44220&5\|-\|168560908\|168560926&offsets\|0\|+5]\|AGCAGCATTGTACAGGGCTATGAA | >MI0000108\|hsa-mir-103a-2&WithFlank&20\|+\|3917488\|3917577 | 54 | 77 | 24 | AGCAGCATTGTACAGGGCTATGAA | 500.4 | 6.4 | 0.8 | 8.5 | 2.0E-17 | 1.5E-16 | BT20/MB468 | MITO |
| >hg38_1_-_228634819_228635039_RNA5S12_5S_50ntFlanks@151.170.20\|ATACCGGGTGCTGTAGGCTT | >hg38_1_-_228634819_228635039_RNA5S12_5S_50ntFlanks | 151 | 170 | 20 | ATACCGGGTGCTGTAGGCTT | 4157.0 | 5.7 | 0.7 | 8.5 | 2.7E-17 | 1.5E-16 | BT20/MB468 | CYTO |
| >hg38_21_+_8433172_8446622_RNA45SN1_45S_50ntFlanks@12804.12821.18\|CGCGTGCCCCGCCGCGCG | >hg38_21_+_8433172_8446622_RNA45SN1_45S_50ntFlanks | 12804 | 12821 | 18 | CGCGTGCCCCGCCGCGCG | 449.2 | -12.6 | 1.5 | -8.5 | 2.5E-17 | 1.5E-16 | MB231/MB468 | MITO |

<table>
<tr><td>>MI0000255|hsa-mir-30d&WithFlank&8|-|134804870|134804951@12.35.24(+3U)&[MIMAT0000245&hsa-miR-30d-5p&offsets|0|+2(+3U);m-21&8|-|134804919|134804940&offsets|0|+2(+3U)]|TGTAAACATCCCCGACTGGAAGCTTTT</td><td>>MI0000255|hsa-mir-30d&WithFlank&8|-|134804870|134804951</td><td>12</td><td>35</td><td>24(+3U)</td><td>TGTAAACATCCCCGACTGGAAGCTTTT</td><td>445.3</td><td>6.5</td><td>0.8</td><td>8.5</td><td>1.9E-17</td><td>1.5E-16</td><td>MB231/MB468</td><td>NUC</td></tr>
<tr><td>>MI0002470|hsa-mir-486-1&WithFlank&8|-|41660435|41660514@10.31.22(+1U);MI0023622|hsa-mir-486-2&WithFlank&8|+|41660438|41660513@7.28.22(+1U)&[MIMAT0002177&hsa-miR-486-5p&offsets|0|0(+1U);m-107&8|-|41660484|41660505&offsets|0|0(+1U)];[MIMAT0002177_1&hsa-miR-486-5p&offsets|0|0(+1U);TJU_CMC_MD2.ID03138.5p-miR&offsets|0|-2(+1U);m-106&8|+|41660444|41660465&offsets|0|0(+1U)]|TCCTGTACTGAGCTGCCCCGAGT</td><td>>MI0002470|hsa-mir-486-1&WithFlank&8|-|41660435|41660514</td><td>10</td><td>31</td><td>22(+1U)</td><td>TCCTGTACTGAGCTGCCCCGAGT</td><td>283.0</td><td>12.0</td><td>1.4</td><td>8.5</td><td>1.9E-17</td><td>1.5E-16</td><td>MB231/MB468</td><td>NUC</td></tr>
<tr><td>>hg38_21_+_8433172_8446622_RNA45SN1_45S_50ntFlanks@11209.11234.26|CGGCGACTCTGGACGCGAGCCGGGCC</td><td>>hg38_21_+_8433172_8446622_RNA45SN1_45S_50ntFlanks</td><td>11209</td><td>11234</td><td>26</td><td>CGGCGACTCTGGACGCGAGCCGGGCC</td><td>2197.6</td><td>-8.7</td><td>1.0</td><td>-8.5</td><td>2.5E-17</td><td>1.5E-16</td><td>MB231/MB468</td><td>MITO</td></tr>
<tr><td>>hg38_21_+_8433172_8446622_RNA45SN1_45S_50ntFlanks@12658.12675.18|ACTGAACGCCTCTAAGTC</td><td>>hg38_21_+_8433172_8446622_RNA45SN1_45S_50ntFlanks</td><td>12658</td><td>12675</td><td>18</td><td>ACTGAACGCCTCTAAGTC</td><td>1314.4</td><td>-8.2</td><td>1.0</td><td>-8.5</td><td>2.5E-17</td><td>1.5E-16</td><td>MB231/MB468</td><td>MITO</td></tr>
<tr><td>>hg38_21_+_8433172_8446622_RNA45SN1_45S_50ntFlanks@5445.5466.22|TCGGATCGGCCCCGCCGGGGTC</td><td>>hg38_21_+_8433172_8446622_RNA45SN1_45S_50ntFlanks</td><td>5445</td><td>5466</td><td>22</td><td>TCGGATCGGCCCCGCCGGGGTC</td><td>7436.7</td><td>-9.7</td><td>1.1</td><td>-8.5</td><td>2.5E-17</td><td>1.5E-16</td><td>MB231/MB468</td><td>MITO</td></tr>
<tr><td>>hg38_21_+_8433172_8446622_RNA45SN1_45S_50ntFlanks@11619.11640.22|AACAAAGCATCGCGAAGGCCCG</td><td>>hg38_21_+_8433172_8446622_RNA45SN1_45S_50ntFlanks</td><td>11619</td><td>11640</td><td>22</td><td>AACAAAGCATCGCGAAGGCCCG</td><td>1821.0</td><td>-13.9</td><td>1.6</td><td>-8.9</td><td>3.6E-19</td><td>1.5E-16</td><td>BT20/MB468</td><td>MP</td></tr>
<tr><td>>hg38_21_+_8433172_8446622_RNA45SN1_45S_50ntFlanks@4367.4394.28|CAGTTAAAAAGCTCGTAGTTGGATCTTG</td><td>>hg38_21_+_8433172_8446622_RNA45SN1_45S_50ntFlanks</td><td>4367</td><td>4394</td><td>28</td><td>CAGTTAAAAAGCTCGTAGTTGGATCTTG</td><td>295.9</td><td>-12.1</td><td>1.4</td><td>-8.5</td><td>1.6E-17</td><td>1.5E-16</td><td>MB231/MB468</td><td>MP</td></tr>
<tr><td>>hg38_21_+_8433172_8446622_RNA45SN1_45S_50ntFlanks@7979.8006.28|ACCTCAGATCAGACGTGGCGACCCGCTG</td><td>>hg38_21_+_8433172_8446622_RNA45SN1_45S_50ntFlanks</td><td>7979</td><td>8006</td><td>28</td><td>ACCTCAGATCAGACGTGGCGACCCGCTG</td><td>9031.6</td><td>-5.0</td><td>0.6</td><td>-8.5</td><td>2.0E-17</td><td>1.5E-16</td><td>BT20/MB468</td><td>MITO</td></tr>
<tr><td>>MI0000254|hsa-mir-30c-2&WithFlank&6|-|71376954|71377037@53.74.22&[MIMAT0004550&hsa-miR-30c-2-3p&offsets|0|0;m-206&6|-|71376964|71376985&offsets|0|0]|CTGGGAGAAGGCTGTTTACTCT</td><td>>MI0000254|hsa-mir-30c-2&WithFlank&6|-|71376954|71377037</td><td>53</td><td>74</td><td>22</td><td>CTGGGAGAAGGCTGTTTACTCT</td><td>1289.0</td><td>5.5</td><td>0.6</td><td>8.5</td><td>2.0E-17</td><td>1.5E-16</td><td>MB231/MB468</td><td>NUC</td></tr>
<tr><td>>hg38_21_+_8433172_8446622_RNA45SN1_45S_50ntFlanks@12625.12657.33|CAATGGGGCGAAGCTACCATCTGTGGGATTATG</td><td>>hg38_21_+_8433172_8446622_RNA45SN1_45S_50ntFlanks</td><td>12625</td><td>12657</td><td>33</td><td>CAATGGGGCGAAGCTACCATCTGTGGGATTATG</td><td>438.0</td><td>-12.6</td><td>1.5</td><td>-8.5</td><td>1.6E-17</td><td>1.5E-16</td><td>MB231/MB468</td><td>MP</td></tr>
<tr><td>>hg38_21_+_8433172_8446622_RNA45SN1_45S_50ntFlanks@11579.11607.29|AGAACTGGTGCGGACCAGGGGAATCCGAC</td><td>>hg38_21_+_8433172_8446622_RNA45SN1_45S_50ntFlanks</td><td>11579</td><td>11607</td><td>29</td><td>AGAACTGGTGCGGACCAGGGGAATCCGAC</td><td>675.9</td><td>-4.3</td><td>0.5</td><td>-8.4</td><td>4.7E-17</td><td>1.6E-16</td><td>BT20/MB468</td><td>NUC</td></tr>
<tr><td>>hg38_21_+_8433172_8446622_RNA45SN1_45S_50ntFlanks@3854.3881.28|ACTGTGGTAATTCTAGAGCTAATACATG</td><td>>hg38_21_+_8433172_8446622_RNA45SN1_45S_50ntFlanks</td><td>3854</td><td>3881</td><td>28</td><td>ACTGTGGTAATTCTAGAGCTAATACATG</td><td>1972.6</td><td>-10.8</td><td>1.3</td><td>-8.5</td><td>2.5E-17</td><td>1.6E-16</td><td>MB231/MB468</td><td>MITO</td></tr>
<tr><td>>hg38_21_+_8433172_8446622_RNA45SN1_45S_50ntFlanks@10839.10865.27|TCCGTAACTTCGGGATAAGGATTGGCT</td><td>>hg38_21_+_8433172_8446622_RNA45SN1_45S_50ntFlanks</td><td>10839</td><td>10865</td><td>27</td><td>TCCGTAACTTCGGGATAAGGATTGGCT</td><td>384.6</td><td>-12.4</td><td>1.5</td><td>-8.5</td><td>2.6E-17</td><td>1.6E-16</td><td>MB231/MB468</td><td>MITO</td></tr>
<tr><td>>hg38_21_+_8433172_8446622_RNA45SN1_45S_50ntFlanks@3782.3812.31|CAGTGAAACTGCGAATGGCTCATTAAATCAG</td><td>>hg38_21_+_8433172_8446622_RNA45SN1_45S_50ntFlanks</td><td>3782</td><td>3812</td><td>31</td><td>CAGTGAAACTGCGAATGGCTCATTAAATCAG</td><td>964.7</td><td>-13.7</td><td>1.6</td><td>-8.5</td><td>2.6E-17</td><td>1.6E-16</td><td>MB231/MB468</td><td>MITO</td></tr>
<tr><td>>hg38_21_+_8433172_8446622_RNA45SN1_45S_50ntFlanks@6748.6770.23|CTTCGAACGCACTTGCGGCCCCG</td><td>>hg38_21_+_8433172_8446622_RNA45SN1_45S_50ntFlanks</td><td>6748</td><td>6770</td><td>23</td><td>CTTCGAACGCACTTGCGGCCCCG</td><td>384.7</td><td>-6.4</td><td>0.7</td><td>-8.5</td><td>1.6E-17</td><td>1.6E-16</td><td>MB231/MB468</td><td>CYTO</td></tr>
<tr><td>>hg38_21_+_8433172_8446622_RNA45SN1_45S_50ntFlanks@7977.8009.33|CGACCTCAGATCAGACGTGGCGACCCGCTGAAT</td><td>>hg38_21_+_8433172_8446622_RNA45SN1_45S_50ntFlanks</td><td>7977</td><td>8009</td><td>33</td><td>CGACCTCAGATCAGACGTGGCGACCCGCTGAAT</td><td>387.6</td><td>5.2</td><td>0.6</td><td>8.5</td><td>2.9E-17</td><td>1.6E-16</td><td>BT20/MB468</td><td>CYTO</td></tr>
<tr><td>>hg38_21_+_8433172_8446622_RNA45SN1_45S_50ntFlanks@10786.10828.43|AACAGCCTCTGGCATGTTGGAACAATGTAGGTAAGGGAAGTCG</td><td>>hg38_21_+_8433172_8446622_RNA45SN1_45S_50ntFlanks</td><td>10786</td><td>10828</td><td>43</td><td>AACAGCCTCTGGCATGTTGGAACAATGTAGGTAAGGGAAGTCG</td><td>536.8</td><td>-13.5</td><td>1.6</td><td>-8.4</td><td>4.8E-17</td><td>1.6E-16</td><td>BT20/MB468</td><td>NUC</td></tr>
<tr><td>>hg38_21_+_8433172_8446622_RNA45SN1_45S_50ntFlanks@12578.12604.27|AGGAACCGCAGGTTCAGACATTTGGTG</td><td>>hg38_21_+_8433172_8446622_RNA45SN1_45S_50ntFlanks</td><td>12578</td><td>12604</td><td>27</td><td>AGGAACCGCAGGTTCAGACATTTGGTG</td><td>393.0</td><td>-12.4</td><td>1.5</td><td>-8.5</td><td>2.6E-17</td><td>1.6E-16</td><td>MB231/MB468</td><td>MITO</td></tr>
<tr><td>>hg38_21_+_8433172_8446622_RNA45SN1_45S_50ntFlanks@9419.9453.35|TCCGCCGAGGGCGCACCACCGGCCCGTCTCGCCCG</td><td>>hg38_21_+_8433172_8446622_RNA45SN1_45S_50ntFlanks</td><td>9419</td><td>9453</td><td>35</td><td>TCCGCCGAGGGCGCACCACCGGCCCGTCTCGCCCG</td><td>455.1</td><td>-12.6</td><td>1.5</td><td>-8.5</td><td>2.6E-17</td><td>1.6E-16</td><td>MB231/MB468</td><td>MITO</td></tr>
<tr><td>>hg38_21_+_8433172_8446622_RNA45SN1_45S_50ntFlanks@5420.5445.26|CCGATTGGATGGTTTAGTGAGGCCCT</td><td>>hg38_21_+_8433172_8446622_RNA45SN1_45S_50ntFlanks</td><td>5420</td><td>5445</td><td>26</td><td>CCGATTGGATGGTTTAGTGAGGCCCT</td><td>2207.8</td><td>-8.0</td><td>0.9</td><td>-8.5</td><td>1.6E-17</td><td>1.6E-16</td><td>MB231/MB468</td><td>CYTO</td></tr>
<tr><td>>hg38_21_+_8433172_8446622_RNA45SN1_45S_50ntFlanks@10128.10148.21|CGGCGGCGGCGGCGGCGGCGG</td><td>>hg38_21_+_8433172_8446622_RNA45SN1_45S_50ntFlanks</td><td>10128</td><td>10148</td><td>21</td><td>CGGCGGCGGCGGCGGCGGCGG</td><td>816.7</td><td>-13.5</td><td>1.6</td><td>-8.5</td><td>2.6E-17</td><td>1.6E-16</td><td>MB231/MB468</td><td>MITO</td></tr>
<tr><td>>hg38_21_+_8433172_8446622_RNA45SN1_45S_50ntFlanks@8845.8865.21|CGGCGGCGGCGGCGGCGGCGG</td><td>>hg38_21_+_8433172_8446622_RNA45SN1_45S_50ntFlanks</td><td>8845</td><td>8865</td><td>21</td><td>CGGCGGCGGCGGCGGCGGCGG</td><td>816.7</td><td>-13.5</td><td>1.6</td><td>-8.5</td><td>2.6E-17</td><td>1.6E-16</td><td>MB231/MB468</td><td>MITO</td></tr>
<tr><td>>hg38_21_+_8433172_8446622_RNA45SN1_45S_50ntFlanks@4443.4471.29|CCCTTGCCTCTCGGCGCCCCCTCGATGCT</td><td>>hg38_21_+_8433172_8446622_RNA45SN1_45S_50ntFlanks</td><td>4443</td><td>4471</td><td>29</td><td>CCCTTGCCTCTCGGCGCCCCCTCGATGCT</td><td>2126.0</td><td>-14.0</td><td>1.7</td><td>-8.5</td><td>2.0E-17</td><td>1.6E-16</td><td>MB231/MB468</td><td>NUC</td></tr>
<tr><td>>hg38_21_+_8433172_8446622_RNA45SN1_45S_50ntFlanks@9545.9567.23|GTGGAGGTCCGTAGCGGTCCTGA</td><td>>hg38_21_+_8433172_8446622_RNA45SN1_45S_50ntFlanks</td><td>9545</td><td>9567</td><td>23</td><td>GTGGAGGTCCGTAGCGGTCCTGA</td><td>509.6</td><td>-7.3</td><td>0.9</td><td>-8.5</td><td>1.6E-17</td><td>1.6E-16</td><td>MB231/MB468</td><td>CYTO</td></tr>
<tr><td>>hg38_21_+_8433172_8446622_RNA45SN1_45S_50ntFlanks@10678.10712.35|CGGTTCCGGCGGCGTCCGGTGAGCTCTCGCTGGCC</td><td>>hg38_21_+_8433172_8446622_RNA45SN1_45S_50ntFlanks</td><td>10678</td><td>10712</td><td>35</td><td>CGGTTCCGGCGGCGTCCGGTGAGCTCTCGCTGGCC</td><td>41398.7</td><td>-9.5</td><td>1.1</td><td>-8.4</td><td>4.8E-17</td><td>1.6E-16</td><td>BT20/MB468</td><td>NUC</td></tr>
<tr><td>>hg38_21_+_8433172_8446622_RNA45SN1_45S_50ntFlanks@4731.4758.28|AAGAACGAAAGTCGGAGGTTCGAAGACG</td><td>>hg38_21_+_8433172_8446622_RNA45SN1_45S_50ntFlanks</td><td>4731</td><td>4758</td><td>28</td><td>AAGAACGAAAGTCGGAGGTTCGAAGACG</td><td>19125.9</td><td>-5.6</td><td>0.7</td><td>-8.5</td><td>1.6E-17</td><td>1.6E-16</td><td>MB231/MB468</td><td>MP</td></tr>
<tr><td>>hg38_21_+_8433172_8446622_RNA45SN1_45S_50ntFlanks@9505.9526.22|TGAACTATGCCTGGGCAGGGCG</td><td>>hg38_21_+_8433172_8446622_RNA45SN1_45S_50ntFlanks</td><td>9505</td><td>9526</td><td>22</td><td>TGAACTATGCCTGGGCAGGGCG</td><td>503.0</td><td>-12.8</td><td>1.5</td><td>-8.5</td><td>2.6E-17</td><td>1.6E-16</td><td>MB231/MB468</td><td>MITO</td></tr>
<tr><td>>hg38_21_+_8433172_8446622_RNA45SN1_45S_50ntFlanks@12628.12649.22|TGGGGCGAAGCTACCATCTGTG</td><td>>hg38_21_+_8433172_8446622_RNA45SN1_45S_50ntFlanks</td><td>12628</td><td>12649</td><td>22</td><td>TGGGGCGAAGCTACCATCTGTG</td><td>328.4</td><td>-12.2</td><td>1.4</td><td>-8.5</td><td>1.6E-17</td><td>1.6E-16</td><td>MB231/MB468</td><td>MP</td></tr>
<tr><td>>hg38_21_+_8433172_8446622_RNA45SN1_45S_50ntFlanks@11668.11693.26|TGCCCAGTGCTCTGAATGTCAAAGTG</td><td>>hg38_21_+_8433172_8446622_RNA45SN1_45S_50ntFlanks</td><td>11668</td><td>11693</td><td>26</td><td>TGCCCAGTGCTCTGAATGTCAAAGTG</td><td>366.8</td><td>5.6</td><td>0.7</td><td>8.5</td><td>2.9E-17</td><td>1.6E-16</td><td>BT20/MB468</td><td>CYTO</td></tr>
<tr><td>>hg38_21_+_8433172_8446622_RNA45SN1_45S_50ntFlanks@4104.4121.18|CACGGGTGACGGGGAATC</td><td>>hg38_21_+_8433172_8446622_RNA45SN1_45S_50ntFlanks</td><td>4104</td><td>4121</td><td>18</td><td>CACGGGTGACGGGGAATC</td><td>487.1</td><td>-12.8</td><td>1.5</td><td>-8.5</td><td>1.6E-17</td><td>1.6E-16</td><td>MB231/MB468</td><td>MP</td></tr>
<tr><td>>MI0003820|hsa-mir-454&WithFlank&17|-|59137752|59137878@70.93.24&[MIMAT0003885&hsa-miR-454-3p&offsets|0|+1;m-272&17|-|59137787|59137809&offsets|0|+1]|TAGTGCAATATTGCTTATAGGGTT</td><td>>MI0003820|hsa-mir-454&WithFlank&17|-|59137752|59137878</td><td>70</td><td>93</td><td>24</td><td>TAGTGCAATATTGCTTATAGGGTT</td><td>510.1</td><td>5.4</td><td>0.6</td><td>8.5</td><td>2.0E-17</td><td>1.6E-16</td><td>MB231/MB468</td><td>NUC</td></tr>
<tr><td>>MI0000750|hsa-mir-26a-2&WithFlank&12|-|57824603|57824698@20.39.20;MI0000083|hsa-mir-26a-1&WithFlank&3|+|37969398|37969486@16.35.20&[MIMAT0000082&hsa-miR-26a-5p&offsets|0|-2;m-33&12|-|57824658|57824679&offsets|0|-2];[MIMAT0000082_1&hsa-miR-26a-5p&offsets|0|-2;m-34&3|+|37969413|37969434&offsets|0|-2]|TTCAAGTAATCCAGGATAGG</td><td>>MI0000750|hsa-mir-26a-2&WithFlank&12|-|57824603|57824698</td><td>20</td><td>39</td><td>20</td><td>TTCAAGTAATCCAGGATAGG</td><td>101.5</td><td>2.3</td><td>0.3</td><td>8.4</td><td>4.9E-17</td><td>1.6E-16</td><td>BT20/MB468</td><td>NUC</td></tr>
<tr><td>>hg38_21_+_8433172_8446622_RNA45SN1_45S_50ntFlanks@9201.9225.25|CTCTCGGGGCCACGCGCGCGTCCCC</td><td>>hg38_21_+_8433172_8446622_RNA45SN1_45S_50ntFlanks</td><td>9201</td><td>9225</td><td>25</td><td>CTCTCGGGGCCACGCGCGCGTCCCC</td><td>431.3</td><td>-12.6</td><td>1.5</td><td>-8.5</td><td>1.6E-17</td><td>1.6E-16</td><td>MB231/MB468</td><td>MP</td></tr>
<tr><td>>hg38_21_+_8433172_8446622_RNA45SN1_45S_50ntFlanks@9567.9595.29|ACGTGCAAATCGGTCGTCCGACCTGGGTA</td><td>>hg38_21_+_8433172_8446622_RNA45SN1_45S_50ntFlanks</td><td>9567</td><td>9595</td><td>29</td><td>ACGTGCAAATCGGTCGTCCGACCTGGGTA</td><td>5183.6</td><td>-9.0</td><td>1.1</td><td>-8.5</td><td>2.7E-17</td><td>1.6E-16</td><td>MB231/MB468</td><td>MITO</td></tr>
<tr><td>>hg38_21_+_8433172_8446622_RNA45SN1_45S_50ntFlanks@7986.8008.23|ATCAGACGTGGCGACCCGCTGAA</td><td>>hg38_21_+_8433172_8446622_RNA45SN1_45S_50ntFlanks</td><td>7986</td><td>8008</td><td>23</td><td>ATCAGACGTGGCGACCCGCTGAA</td><td>3289.4</td><td>-14.8</td><td>1.7</td><td>-8.9</td><td>3.9E-19</td><td>1.6E-16</td><td>BT20/MB468</td><td>MP</td></tr>
<tr><td>>hg38_21_+_8433172_8446622_RNA45SN1_45S_50ntFlanks@8388.8405.18|CGTGAAACCGTTAAGAGG</td><td>>hg38_21_+_8433172_8446622_RNA45SN1_45S_50ntFlanks</td><td>8388</td><td>8405</td><td>18</td><td>CGTGAAACCGTTAAGAGG</td><td>297.2</td><td>-12.1</td><td>1.4</td><td>-8.5</td><td>1.6E-17</td><td>1.6E-16</td><td>MB231/MB468</td><td>MP</td></tr>
<tr><td>>MI0000299|hsa-mir-222&WithFlank&X|-|45747009|45747130@75.97.23(+1A)&[MIMAT0000279&hsa-miR-222-3p&offsets|0|+2(+1A);m-140&X|-|45747033|45747056&offsets|0|-1(+1A)]|AGCTACATCTGGCTACTGGGTCTA</td><td>>MI0000299|hsa-mir-222&WithFlank&X|-|45747009|45747130</td><td>75</td><td>97</td><td>23(+1A)</td><td>AGCTACATCTGGCTACTGGGTCTA</td><td>617.4</td><td>6.8</td><td>0.8</td><td>8.5</td><td>2.1E-17</td><td>1.6E-16</td><td>BT20/MB468</td><td>MITO</td></tr>
<tr><td>>hg38_21_+_8433172_8446622_RNA45SN1_45S_50ntFlanks@11316.11338.23|CACCCCACGTCTCGTCGCGCGCG</td><td>>hg38_21_+_8433172_8446622_RNA45SN1_45S_50ntFlanks</td><td>11316</td><td>11338</td><td>23</td><td>CACCCCACGTCTCGTCGCGCGCG</td><td>4766.7</td><td>-11.1</td><td>1.3</td><td>-8.5</td><td>2.1E-17</td><td>1.6E-16</td><td>MB231/MB468</td><td>NUC</td></tr>
<tr><td>>hg38_21_+_8433172_8446622_RNA45SN1_45S_50ntFlanks@4380.4408.29|CGTAGTTGGATCTTGGGAGCGGGCGGGCG</td><td>>hg38_21_+_8433172_8446622_RNA45SN1_45S_50ntFlanks</td><td>4380</td><td>4408</td><td>29</td><td>CGTAGTTGGATCTTGGGAGCGGGCGGGCG</td><td>302.4</td><td>-11.2</td><td>1.3</td><td>-8.5</td><td>2.1E-17</td><td>1.6E-16</td><td>MB231/MB468</td><td>NUC</td></tr>
<tr><td>>hg38_1_-_228634819_228635039_RNA5S12_5S_50ntFlanks@134.171.38|TGGGAGACCGCCTGGGAATACCGGGTGCTGTAGGCTTT</td><td>>hg38_1_-_228634819_228635039_RNA5S12_5S_50ntFlanks</td><td>134</td><td>171</td><td>38</td><td>TGGGAGACCGCCTGGGAATACCGGGTGCTGTAGGCTTT</td><td>60585.7</td><td>8.5</td><td>1.0</td><td>8.5</td><td>1.7E-17</td><td>1.6E-16</td><td>MB231/MB468</td><td>CYTO</td></tr>
<tr><td>>hg38_21_+_8433172_8446622_RNA45SN1_45S_50ntFlanks@11586.11605.20|GTGCGGACCAGGGGAATCCG</td><td>>hg38_21_+_8433172_8446622_RNA45SN1_45S_50ntFlanks</td><td>11586</td><td>11605</td><td>20</td><td>GTGCGGACCAGGGGAATCCG</td><td>878.2</td><td>-13.6</td><td>1.6</td><td>-8.5</td><td>1.7E-17</td><td>1.6E-16</td><td>MB231/MB468</td><td>MP</td></tr>
<tr><td>>MI0000462|hsa-mir-152&WithFlank&17|-|48037155|48037253@60.80.21(+1A)&[MIMAT0000438&hsa-miR-152-3p&offsets|0|0(+1A);m-97&17|-|48037174|48037194&offsets|0|0(+1A)]|TCAGTGCATGACAGAACTTGGA</td><td>>MI0000462|hsa-mir-152&WithFlank&17|-|48037155|48037253</td><td>60</td><td>80</td><td>21(+1A)</td><td>TCAGTGCATGACAGAACTTGGA</td><td>378.4</td><td>12.8</td><td>1.5</td><td>8.4</td><td>3.0E-17</td><td>1.6E-16</td><td>BT20/MB468</td><td>CYTO</td></tr>
<tr><td>>hg38_21_+_8433172_8446622_RNA45SN1_45S_50ntFlanks@9570.9597.28|TGCAAATCGGTCGTCCGACCTGGGTATA</td><td>>hg38_21_+_8433172_8446622_RNA45SN1_45S_50ntFlanks</td><td>9570</td><td>9597</td><td>28</td><td>TGCAAATCGGTCGTCCGACCTGGGTATA</td><td>8388.6</td><td>-9.5</td><td>1.1</td><td>-8.5</td><td>2.7E-17</td><td>1.6E-16</td><td>MB231/MB468</td><td>MITO</td></tr>
<tr><td>>hg38_21_+_8433172_8446622_RNA45SN1_45S_50ntFlanks@9564.9595.32|CTGACGTGCAAATCGGTCGTCCGACCTGGGTA</td><td>>hg38_21_+_8433172_8446622_RNA45SN1_45S_50ntFlanks</td><td>9564</td><td>9595</td><td>32</td><td>CTGACGTGCAAATCGGTCGTCCGACCTGGGTA</td><td>1784.6</td><td>-4.8</td><td>0.6</td><td>-8.4</td><td>3.0E-17</td><td>1.6E-16</td><td>BT20/MB468</td><td>CYTO</td></tr>
<tr><td>>MI0000650|hsa-mir-200c&WithFlank&12|+|6963693|6963772@49.70.22&[MIMAT0000617&hsa-miR-200c-3p&offsets|-1|-2;m-54&12|+|6963742|6963764&offsets|-1|-2]|CTAATACTGCCGGGTAATGATG</td><td>>MI0000650|hsa-mir-200c&WithFlank&12|+|6963693|6963772</td><td>49</td><td>70</td><td>22</td><td>CTAATACTGCCGGGTAATGATG</td><td>358.9</td><td>12.3</td><td>1.4</td><td>8.7</td><td>3.0E-18</td><td>1.6E-16</td><td>BT20/MB231</td><td>NUC</td></tr>
<tr><td>>hg38_21_+_8433172_8446622_RNA45SN1_45S_50ntFlanks@12459.12487.29|AGCGTTGGATTGTTCACCCACTAATAGGG</td><td>>hg38_21_+_8433172_8446622_RNA45SN1_45S_50ntFlanks</td><td>12459</td><td>12487</td><td>29</td><td>AGCGTTGGATTGTTCACCCACTAATAGGG</td><td>520.1</td><td>-8.5</td><td>1.0</td><td>-8.5</td><td>1.7E-17</td><td>1.7E-16</td><td>MB231/MB468</td><td>CYTO</td></tr>
<tr><td>>hg38_21_+_8433172_8446622_RNA45SN1_45S_50ntFlanks@9534.9551.18|AGGAAACTCTGGTGGAGG</td><td>>hg38_21_+_8433172_8446622_RNA45SN1_45S_50ntFlanks</td><td>9534</td><td>9551</td><td>18</td><td>AGGAAACTCTGGTGGAGG</td><td>4658.5</td><td>-9.0</td><td>1.1</td><td>-8.5</td><td>2.1E-17</td><td>1.7E-16</td><td>MB231/MB468</td><td>NUC</td></tr>
</table>

<table>
<tr><td>>MI0000736|hsa-mir-30c-1&WithFlank&1|+|40757278|40757378@23.45.23(+1A);MI0000254|hsa-mir-30c-2&WithFlank&6|-|71376954|71377037@13.35.23(+1A)&[MIMAT0000244&hsa-miR-30c-5p&offsets|0|0(+1A);m-57&1|+|40757300|40757323&offsets|0|-1(+1A)];[MIMAT0000244_1&hsa-miR-30c-5p&offsets|0|0(+1A);m-56&6|-|71377002|71377025&offsets|0|-1(+1A)]|TGTAAACATCCTACACTCTCAGCA</td><td>>MI0000736|hsa-mir-30c-1&WithFlank&1|+|40757278|40757378</td><td>23</td><td>45</td><td>23(+1A)</td><td>TGTAAACATCCTACACTCTCAGCA</td><td>2199.7</td><td>6.1</td><td>0.7</td><td>8.4</td><td>3.0E-17</td><td>1.7E-16</td><td>BT20/MB468</td><td>CYTO</td></tr>
<tr><td>>hg38_1_-_228634819_228635039_RNA5S12_5S_50ntFlanks@84.103.20|CTCGTCTGATCTCGGAAGCT</td><td>>hg38_1_-_228634819_228635039_RNA5S12_5S_50ntFlanks</td><td>84</td><td>103</td><td>20</td><td>CTCGTCTGATCTCGGAAGCT</td><td>273.7</td><td>-12.0</td><td>1.4</td><td>-8.5</td><td>1.7E-17</td><td>1.7E-16</td><td>MB231/MB468</td><td>MP</td></tr>
<tr><td>>hg38_21_+_8433172_8446622_RNA45SN1_45S_50ntFlanks@7977.8006.30|CGACCTCAGATCAGACGTGGCGACCCGCTG</td><td>>hg38_21_+_8433172_8446622_RNA45SN1_45S_50ntFlanks</td><td>7977</td><td>8006</td><td>30</td><td>CGACCTCAGATCAGACGTGGCGACCCGCTG</td><td>1998.6</td><td>-5.0</td><td>0.6</td><td>-8.5</td><td>2.2E-17</td><td>1.7E-16</td><td>MB231/MB468</td><td>NUC</td></tr>
<tr><td>>hg38_21_+_8433172_8446622_RNA45SN1_45S_50ntFlanks@9252.9272.21|CACGGGGTCGGCGGCGACGTC</td><td>>hg38_21_+_8433172_8446622_RNA45SN1_45S_50ntFlanks</td><td>9252</td><td>9272</td><td>21</td><td>CACGGGGTCGGCGGCGACGTC</td><td>291.1</td><td>-12.0</td><td>1.4</td><td>-8.5</td><td>1.7E-17</td><td>1.7E-16</td><td>MB231/MB468</td><td>MP</td></tr>
<tr><td>>hg38_21_+_8433172_8446622_RNA45SN1_45S_50ntFlanks@5485.5521.37|AGCGCTGAGAAGACGGTCGAACTTGACTATCTAGAGG</td><td>>hg38_21_+_8433172_8446622_RNA45SN1_45S_50ntFlanks</td><td>5485</td><td>5521</td><td>37</td><td>AGCGCTGAGAAGACGGTCGAACTTGACTATCTAGAGG</td><td>419.0</td><td>-12.6</td><td>1.5</td><td>-8.5</td><td>1.7E-17</td><td>1.7E-16</td><td>MB231/MB468</td><td>MP</td></tr>
<tr><td>>hg38_21_+_8433172_8446622_RNA45SN1_45S_50ntFlanks@10728.10755.28|GAGAGGGTGTAAATCTCGCGCCGGGCCG</td><td>>hg38_21_+_8433172_8446622_RNA45SN1_45S_50ntFlanks</td><td>10728</td><td>10755</td><td>28</td><td>GAGAGGGTGTAAATCTCGCGCCGGGCCG</td><td>691.8</td><td>-12.4</td><td>1.5</td><td>-8.5</td><td>2.2E-17</td><td>1.7E-16</td><td>MB231/MB468</td><td>NUC</td></tr>
<tr><td>>hg38_21_+_8433172_8446622_RNA45SN1_45S_50ntFlanks@11561.11600.40|GCGCCTAGCAGCCGACTTAGAACTGGTGCGGACCAGGGGA</td><td>>hg38_21_+_8433172_8446622_RNA45SN1_45S_50ntFlanks</td><td>11561</td><td>11600</td><td>40</td><td>GCGCCTAGCAGCCGACTTAGAACTGGTGCGGACCAGGGGA</td><td>1724.5</td><td>-8.6</td><td>1.0</td><td>-8.5</td><td>2.8E-17</td><td>1.7E-16</td><td>MB231/MB468</td><td>MITO</td></tr>
<tr><td>>hg38_21_+_8433172_8446622_RNA45SN1_45S_50ntFlanks@11534.11553.20|GTTCCGCGCGGCGCCTCGCC</td><td>>hg38_21_+_8433172_8446622_RNA45SN1_45S_50ntFlanks</td><td>11534</td><td>11553</td><td>20</td><td>GTTCCGCGCGGCGCCTCGCC</td><td>970.3</td><td>-12.9</td><td>1.5</td><td>-8.5</td><td>2.2E-17</td><td>1.7E-16</td><td>MB231/MB468</td><td>NUC</td></tr>
<tr><td>>hg38_21_+_8433172_8446622_RNA45SN1_45S_50ntFlanks@9027.9059.33|CGTCTCCTCGTGGGGGGGCCGGGCCACCCCTCC</td><td>>hg38_21_+_8433172_8446622_RNA45SN1_45S_50ntFlanks</td><td>9027</td><td>9059</td><td>33</td><td>CGTCTCCTCGTGGGGGGGCCGGGCCACCCCTCC</td><td>334.9</td><td>-12.3</td><td>1.4</td><td>-8.5</td><td>1.7E-17</td><td>1.7E-16</td><td>MB231/MB468</td><td>MP</td></tr>
<tr><td>>hg38_21_+_8433172_8446622_RNA45SN1_45S_50ntFlanks@3847.3866.20|TTGGATAACTGTGGTAATTC</td><td>>hg38_21_+_8433172_8446622_RNA45SN1_45S_50ntFlanks</td><td>3847</td><td>3866</td><td>20</td><td>TTGGATAACTGTGGTAATTC</td><td>224.0</td><td>5.0</td><td>0.6</td><td>8.5</td><td>1.8E-17</td><td>1.7E-16</td><td>MB231/MB468</td><td>CYTO</td></tr>
<tr><td>>hg38_21_+_8433172_8446622_RNA45SN1_45S_50ntFlanks@7978.8011.34|GACCTCAGATCAGACGTGGCGACCCGCTGAATTT</td><td>>hg38_21_+_8433172_8446622_RNA45SN1_45S_50ntFlanks</td><td>7978</td><td>8011</td><td>34</td><td>GACCTCAGATCAGACGTGGCGACCCGCTGAATTT</td><td>1934.0</td><td>10.1</td><td>1.2</td><td>8.5</td><td>1.8E-17</td><td>1.7E-16</td><td>MB231/MB468</td><td>CYTO</td></tr>
<tr><td>>trnaMT_ProTGG_MT_-_15956_16023@50.70.21|TTAAAGACTTTTTCTCTGACC</td><td>>trnaMT_ProTGG_MT_-_15956_16023</td><td>50</td><td>70</td><td>21</td><td>TTAAAGACTTTTTCTCTGACC</td><td>265.2</td><td>11.9</td><td>1.4</td><td>8.5</td><td>2.2E-17</td><td>1.7E-16</td><td>MB231/MB468</td><td>NUC</td></tr>
<tr><td>>hg38_21_+_8433172_8446622_RNA45SN1_45S_50ntFlanks@10869.10886.18|AGGGCTGGGTCGGTCGGG</td><td>>hg38_21_+_8433172_8446622_RNA45SN1_45S_50ntFlanks</td><td>10869</td><td>10886</td><td>18</td><td>AGGGCTGGGTCGGTCGGG</td><td>739.4</td><td>-8.6</td><td>1.0</td><td>-8.5</td><td>1.8E-17</td><td>1.7E-16</td><td>MB231/MB468</td><td>CYTO</td></tr>
<tr><td>>hg38_21_+_8433172_8446622_RNA45SN1_45S_50ntFlanks@9349.9370.22|CCGCCGTGGCGCAATGAAGGTG</td><td>>hg38_21_+_8433172_8446622_RNA45SN1_45S_50ntFlanks</td><td>9349</td><td>9370</td><td>22</td><td>CCGCCGTGGCGCAATGAAGGTG</td><td>414.3</td><td>-12.6</td><td>1.5</td><td>-8.5</td><td>1.8E-17</td><td>1.7E-16</td><td>MB231/MB468</td><td>MP</td></tr>
<tr><td>>hg38_21_+_8433172_8446622_RNA45SN1_45S_50ntFlanks@12934.12965.32|CTAAACCATTCGTAGACGACCTGCTTCTGGGT</td><td>>hg38_21_+_8433172_8446622_RNA45SN1_45S_50ntFlanks</td><td>12934</td><td>12965</td><td>32</td><td>CTAAACCATTCGTAGACGACCTGCTTCTGGGT</td><td>670.2</td><td>-13.3</td><td>1.6</td><td>-8.5</td><td>1.8E-17</td><td>1.7E-16</td><td>MB231/MB468</td><td>MP</td></tr>
<tr><td>>hg38_21_+_8433172_8446622_RNA45SN1_45S_50ntFlanks@12894.12930.37|CTCGCCCGTCACGCACCGCACGTTCGTGGGGAACCTG</td><td>>hg38_21_+_8433172_8446622_RNA45SN1_45S_50ntFlanks</td><td>12894</td><td>12930</td><td>37</td><td>CTCGCCCGTCACGCACCGCACGTTCGTGGGGAACCTG</td><td>543.0</td><td>-12.9</td><td>1.5</td><td>-8.5</td><td>2.8E-17</td><td>1.7E-16</td><td>MB231/MB468</td><td>MITO</td></tr>
<tr><td>>hg38_21_+_8433172_8446622_RNA45SN1_45S_50ntFlanks@12081.12108.28|CTCTCGCTTCTGGCGCCAAGCGCCCGGC</td><td>>hg38_21_+_8433172_8446622_RNA45SN1_45S_50ntFlanks</td><td>12081</td><td>12108</td><td>28</td><td>CTCTCGCTTCTGGCGCCAAGCGCCCGGC</td><td>337.9</td><td>-12.3</td><td>1.4</td><td>-8.5</td><td>1.8E-17</td><td>1.7E-16</td><td>MB231/MB468</td><td>MP</td></tr>
<tr><td>>trna111_HisGTG_1_-_147774845_147774916@1.33.33,trna118_HisGTG_1_-_145396881_145396952@1.33.33,trna16_HisGTG_1_+_146544773_146544844@1.33.33,trna1_HisGTG_15_+_45493349_45493420@1.33.33,trna21_HisGTG_1_+_147753471_147753542@1.33.33,trna33_HisGTG_6_+_27125906_27125977@1.33.33,trna7_HisGTG_9_-_14433938_14434009@1.33.33,trna8_HisGTG_15_-_45492611_45492682@1.33.33,trna9_HisGTG_15_-_45490804_45490875@1.33.33|GCCGTGATCGTATAGTGGTTAGTACTCTGCGTT</td><td>>trna111_HisGTG_1_-_147774845_147774916</td><td>1</td><td>33</td><td>33</td><td>GCCGTGATCGTATAGTGGTTAGTACTCTGCGTT</td><td>2360.4</td><td>7.6</td><td>0.9</td><td>8.5</td><td>2.3E-17</td><td>1.7E-16</td><td>BT20/MB468</td><td>MITO</td></tr>
<tr><td>>hg38_21_+_8433172_8446622_RNA45SN1_45S_50ntFlanks@8570.8593.24|CTCCGGCGGGTGCGGGGGTGGGCG</td><td>>hg38_21_+_8433172_8446622_RNA45SN1_45S_50ntFlanks</td><td>8570</td><td>8593</td><td>24</td><td>CTCCGGCGGGTGCGGGGGTGGGCG</td><td>369.2</td><td>-12.4</td><td>1.5</td><td>-8.5</td><td>1.8E-17</td><td>1.7E-16</td><td>MB231/MB468</td><td>MP</td></tr>
<tr><td>>hg38_21_+_8433172_8446622_RNA45SN1_45S_50ntFlanks@6733.6756.24|CATTGATCATCGACACTTCGAACG</td><td>>hg38_21_+_8433172_8446622_RNA45SN1_45S_50ntFlanks</td><td>6733</td><td>6756</td><td>24</td><td>CATTGATCATCGACACTTCGAACG</td><td>1280.9</td><td>-5.9</td><td>0.7</td><td>-8.4</td><td>5.3E-17</td><td>1.7E-16</td><td>BT20/MB468</td><td>NUC</td></tr>
<tr><td>>hg38_21_+_8433172_8446622_RNA45SN1_45S_50ntFlanks@12405.12446.42|CTTTTTGATCCTTCGATGTCGGCTCTTCCTATCATTGTGAAG</td><td>>hg38_21_+_8433172_8446622_RNA45SN1_45S_50ntFlanks</td><td>12405</td><td>12446</td><td>42</td><td>CTTTTTGATCCTTCGATGTCGGCTCTTCCTATCATTGTGAAG</td><td>16777.9</td><td>-14.0</td><td>1.7</td><td>-8.4</td><td>5.3E-17</td><td>1.7E-16</td><td>BT20/MB468</td><td>NUC</td></tr>
<tr><td>>hg38_21_+_8433172_8446622_RNA45SN1_45S_50ntFlanks@12935.12956.22|TAAACCATTCGTAGACGACCTG</td><td>>hg38_21_+_8433172_8446622_RNA45SN1_45S_50ntFlanks</td><td>12935</td><td>12956</td><td>22</td><td>TAAACCATTCGTAGACGACCTG</td><td>1455.1</td><td>-7.9</td><td>0.9</td><td>-8.5</td><td>2.9E-17</td><td>1.7E-16</td><td>MB231/MB468</td><td>MITO</td></tr>
<tr><td>>MI0000086|hsa-mir-28&WithFlank&3|+|188688775|188688872@61.82.22(+1A)&[MIMAT0004502&hsa-miR-28-3p&offsets|+1|+1(+1A);m-28&3|+|188688834|188688855&offsets|+1|+1(+1A)]|ACTAGATTGTGAGCTCCTGGAGA</td><td>>MI0000086|hsa-mir-28&WithFlank&3|+|188688775|188688872</td><td>61</td><td>82</td><td>22(+1A)</td><td>ACTAGATTGTGAGCTCCTGGAGA</td><td>465.9</td><td>10.8</td><td>1.3</td><td>8.4</td><td>3.2E-17</td><td>1.8E-16</td><td>BT20/MB468</td><td>CYTO</td></tr>
<tr><td>>hg38_21_+_8433172_8446622_RNA45SN1_45S_50ntFlanks@4051.4069.19|GAACGTCTGCCCTATCAAC</td><td>>hg38_21_+_8433172_8446622_RNA45SN1_45S_50ntFlanks</td><td>4051</td><td>4069</td><td>19</td><td>GAACGTCTGCCCTATCAAC</td><td>144.8</td><td>11.2</td><td>1.3</td><td>8.5</td><td>1.8E-17</td><td>1.8E-16</td><td>MB231/MB468</td><td>CYTO</td></tr>
<tr><td>>MI0000102|hsa-mir-100&WithFlank&11|-|122152223|122152314@19.38.20(+1C)&[MIMAT0000098&hsa-miR-100-5p&offsets|0|-2(+1C);m-27&11|-|122152276|122152296&offsets|0|-1(+1C)]|AACCCGTAGATCCGAACTTGC</td><td>>MI0000102|hsa-mir-100&WithFlank&11|-|122152223|122152314</td><td>19</td><td>38</td><td>20(+1C)</td><td>AACCCGTAGATCCGAACTTGC</td><td>22288.5</td><td>-8.0</td><td>0.9</td><td>-8.7</td><td>2.3E-18</td><td>1.8E-16</td><td>BT20/MB231</td><td>MITO</td></tr>
<tr><td>>MI0000077|hsa-mir-21&WithFlank&17|+|59841260|59841343@14.32.19&[MIMAT0000076&hsa-miR-21-5p&offsets|0|-3;m-1&17|+|59841273|59841294&offsets|0|-3]|TAGCTTATCAGACTGATGT</td><td>>MI0000077|hsa-mir-21&WithFlank&17|+|59841260|59841343</td><td>14</td><td>32</td><td>19</td><td>TAGCTTATCAGACTGATGT</td><td>593.4</td><td>6.3</td><td>0.7</td><td>8.5</td><td>2.3E-17</td><td>1.8E-16</td><td>MB231/MB468</td><td>NUC</td></tr>
<tr><td>>hg38_21_+_8433172_8446622_RNA45SN1_45S_50ntFlanks@6643.6683.41|ACCTCGTACGACTCTTAGCGGTGGATCACTCGGCTCGTGCG</td><td>>hg38_21_+_8433172_8446622_RNA45SN1_45S_50ntFlanks</td><td>6643</td><td>6683</td><td>41</td><td>ACCTCGTACGACTCTTAGCGGTGGATCACTCGGCTCGTGCG</td><td>875.1</td><td>9.0</td><td>1.1</td><td>8.5</td><td>2.3E-17</td><td>1.8E-16</td><td>MB231/MB468</td><td>NUC</td></tr>
<tr><td>>hg38_21_+_8433172_8446622_RNA45SN1_45S_50ntFlanks@9821.9838.18|CCTAGTGGGCCACTTTTG</td><td>>hg38_21_+_8433172_8446622_RNA45SN1_45S_50ntFlanks</td><td>9821</td><td>9838</td><td>18</td><td>CCTAGTGGGCCACTTTTG</td><td>642.3</td><td>-13.1</td><td>1.6</td><td>-8.4</td><td>2.9E-17</td><td>1.8E-16</td><td>MB231/MB468</td><td>MITO</td></tr>
<tr><td>>hg38_21_+_8433172_8446622_RNA45SN1_45S_50ntFlanks@10408.10431.24|GGTCAGTCGGTCCTGAGAGATGGG</td><td>>hg38_21_+_8433172_8446622_RNA45SN1_45S_50ntFlanks</td><td>10408</td><td>10431</td><td>24</td><td>GGTCAGTCGGTCCTGAGAGATGGG</td><td>804.1</td><td>-8.7</td><td>1.0</td><td>-8.4</td><td>3.2E-17</td><td>1.8E-16</td><td>BT20/MB468</td><td>CYTO</td></tr>
<tr><td>>hg38_21_+_8433172_8446622_RNA45SN1_45S_50ntFlanks@4047.4064.18|ATTCGAACGTCTGCCCTA</td><td>>hg38_21_+_8433172_8446622_RNA45SN1_45S_50ntFlanks</td><td>4047</td><td>4064</td><td>18</td><td>ATTCGAACGTCTGCCCTA</td><td>1018.0</td><td>-13.8</td><td>1.6</td><td>-8.4</td><td>2.9E-17</td><td>1.8E-16</td><td>MB231/MB468</td><td>MITO</td></tr>
<tr><td>>hg38_21_+_8433172_8446622_RNA45SN1_45S_50ntFlanks@9027.9066.40|CGTCTCCTCGTGGGGGGGCCGGGCCACCCCTCCCACGGCG</td><td>>hg38_21_+_8433172_8446622_RNA45SN1_45S_50ntFlanks</td><td>9027</td><td>9066</td><td>40</td><td>CGTCTCCTCGTGGGGGGGCCGGGCCACCCCTCCCACGGCG</td><td>327.0</td><td>-12.2</td><td>1.4</td><td>-8.5</td><td>1.8E-17</td><td>1.8E-16</td><td>MB231/MB468</td><td>MP</td></tr>
<tr><td>>MI0000477|hsa-mir-146a&WithFlank&5|+|160485346|160485456@27.46.20(+1C)&[MIMAT0000449&hsa-miR-146a-5p&offsets|0|-2(+1C);m-129&5|+|160485372|160485393&offsets|0|-2(+1C)]|TGAGAACTGAATTCCATGGGC</td><td>>MI0000477|hsa-mir-146a&WithFlank&5|+|160485346|160485456</td><td>27</td><td>46</td><td>20(+1C)</td><td>TGAGAACTGAATTCCATGGGC</td><td>1432.7</td><td>13.7</td><td>1.6</td><td>8.5</td><td>1.8E-17</td><td>1.8E-16</td><td>MB231/MB468</td><td>MP</td></tr>
<tr><td>>hg38_21_+_8433172_8446622_RNA45SN1_45S_50ntFlanks@12416.12442.27|TTCGATGTCGGCTCTTCCTATCATTGT</td><td>>hg38_21_+_8433172_8446622_RNA45SN1_45S_50ntFlanks</td><td>12416</td><td>12442</td><td>27</td><td>TTCGATGTCGGCTCTTCCTATCATTGT</td><td>986.4</td><td>5.1</td><td>0.6</td><td>8.4</td><td>3.3E-17</td><td>1.8E-16</td><td>BT20/MB468</td><td>CYTO</td></tr>
<tr><td>>MI0000093|hsa-mir-92a-1&WithFlank&13|+|91351308|91351397@54.76.23&[MIMAT0000092&hsa-miR-92a-3p&offsets|0|+1;m-9&13|+|91351361|91351382&offsets|0|+1]|TATTGCACTTGTCCCGGCCTGTT</td><td>>MI0000093|hsa-mir-92a-1&WithFlank&13|+|91351308|91351397</td><td>54</td><td>76</td><td>23</td><td>TATTGCACTTGTCCCGGCCTGTT</td><td>7762.1</td><td>6.6</td><td>0.8</td><td>8.5</td><td>1.9E-17</td><td>1.8E-16</td><td>MB231/MB468</td><td>CYTO</td></tr>
<tr><td>>hg38_21_+_8433172_8446622_RNA45SN1_45S_50ntFlanks@9035.9056.22|CGTGGGGGGGCCGGGCCACCCC</td><td>>hg38_21_+_8433172_8446622_RNA45SN1_45S_50ntFlanks</td><td>9035</td><td>9056</td><td>22</td><td>CGTGGGGGGGCCGGGCCACCCC</td><td>597.7</td><td>-13.1</td><td>1.5</td><td>-8.5</td><td>1.8E-17</td><td>1.8E-16</td><td>MB231/MB468</td><td>MP</td></tr>
<tr><td>>hg38_21_+_8433172_8446622_RNA45SN1_45S_50ntFlanks@9413.9453.41|CTCCAGTCCGCCGAGGGCGCACCACCGGCCCGTCTCGCCCG</td><td>>hg38_21_+_8433172_8446622_RNA45SN1_45S_50ntFlanks</td><td>9413</td><td>9453</td><td>41</td><td>CTCCAGTCCGCCGAGGGCGCACCACCGGCCCGTCTCGCCCG</td><td>376.3</td><td>-12.3</td><td>1.5</td><td>-8.4</td><td>3.0E-17</td><td>1.8E-16</td><td>MB231/MB468</td><td>MITO</td></tr>
<tr><td>>hg38_21_+_8433172_8446622_RNA45SN1_45S_50ntFlanks@4884.4918.35|CAAAGCTGAAACTTAAAGGAATTGACGGAAGGGCA</td><td>>hg38_21_+_8433172_8446622_RNA45SN1_45S_50ntFlanks</td><td>4884</td><td>4918</td><td>35</td><td>CAAAGCTGAAACTTAAAGGAATTGACGGAAGGGCA</td><td>403.5</td><td>-12.4</td><td>1.5</td><td>-8.4</td><td>3.0E-17</td><td>1.8E-16</td><td>MB231/MB468</td><td>MITO</td></tr>
<tr><td>>hg38_21_+_8433172_8446622_RNA45SN1_45S_50ntFlanks@10351.10388.38|CGAGAACTTTGAAGGCCGAAGTGGAGAAGGGTTCCATG</td><td>>hg38_21_+_8433172_8446622_RNA45SN1_45S_50ntFlanks</td><td>10351</td><td>10388</td><td>38</td><td>CGAGAACTTTGAAGGCCGAAGTGGAGAAGGGTTCCATG</td><td>8250.7</td><td>-11.8</td><td>1.4</td><td>-8.5</td><td>1.9E-17</td><td>1.8E-16</td><td>MB231/MB468</td><td>MP</td></tr>
<tr><td>>am_trna128_GlyGCC_6_-_27870686_27870756@1.32.32,trna133_GlyCCC_1_-_16872434_16872504@1.32.32,trna18_GlyGCC_16_+_70822597_70822667@1.32.32,trna19_GlyGCC_16_+_70823410_70823480@1.32.32,trna19_GlyGCC_2_-_157257659_157257729@1.32.32,trna24_GlyGCC_16_-_70812942_70813012@1.32.32,trna25_GlyGCC_16_-_70812114_70812184@1.32.32,trna4_GlyCCC_1_+_17188416_17188486@1.32.32,trna5_GlyGCC_17_+_8029064_8029134@1.32.32,trna68_GlyGCC_1_-_161493637_161493707@1.32.32|GCATTGGTGGTTCAGTGGTAGAATTCTCGCCT</td><td>>am_trna128_GlyGCC_6_-_27870686_27870756</td><td>1</td><td>32</td><td>32</td><td>GCATTGGTGGTTCAGTGGTAGAATTCTCGCCT</td><td>8887.2</td><td>8.8</td><td>1.0</td><td>8.5</td><td>2.4E-17</td><td>1.8E-16</td><td>MB231/MB468</td><td>NUC</td></tr>
<tr><td>>MI0000114|hsa-mir-107&WithFlank&10|-|89592741|89592833@56.77.22&[MIMAT0000104&hsa-miR-107&offsets|0|-1;m-78&10|-|89592758|89592778&offsets|0|+1]|AGCAGCATTGTACAGGGCTATC</td><td>>MI0000114|hsa-mir-107&WithFlank&10|-|89592741|89592833</td><td>56</td><td>77</td><td>22</td><td>AGCAGCATTGTACAGGGCTATC</td><td>589.9</td><td>5.6</td><td>0.7</td><td>8.5</td><td>2.4E-17</td><td>1.8E-16</td><td>MB231/MB468</td><td>NUC</td></tr>
<tr><td>>trna10_MetCAT_8_-_124169470_124169542@31.51.21,trna22_MetCAT_16_-_87417628_87417700@31.51.21|TCTCATAATCTGAAGGTCGTG</td><td>>trna10_MetCAT_8_-_124169470_124169542</td><td>31</td><td>51</td><td>21</td><td>TCTCATAATCTGAAGGTCGTG</td><td>1010.0</td><td>-10.0</td><td>1.2</td><td>-8.4</td><td>3.4E-17</td><td>1.8E-16</td><td>BT20/MB468</td><td>CYTO</td></tr>
<tr><td>>hg38_21_+_8433172_8446622_RNA45SN1_45S_50ntFlanks@8472.8491.20|CGGCCCGGCGGATCTTTCCC</td><td>>hg38_21_+_8433172_8446622_RNA45SN1_45S_50ntFlanks</td><td>8472</td><td>8491</td><td>20</td><td>CGGCCCGGCGGATCTTTCCC</td><td>772.2</td><td>-12.6</td><td>1.5</td><td>-8.5</td><td>2.4E-17</td><td>1.8E-16</td><td>MB231/MB468</td><td>NUC</td></tr>
<tr><td>>MI0000272|hsa-mir-182&WithFlank&7|-|129770377|129770498@29.51.23(+1A)&[MIMAT0000259&hsa-miR-182-5p&offsets|0|-1(+1A);m-22&7|-|129770449|129770470&offsets|0|+1(+1A)]|TTTGGCAATGGTAGAACTCACACA</td><td>>MI0000272|hsa-mir-182&WithFlank&7|-|129770377|129770498</td><td>29</td><td>51</td><td>23(+1A)</td><td>TTTGGCAATGGTAGAACTCACACA</td><td>3340.1</td><td>4.1</td><td>0.5</td><td>8.7</td><td>3.5E-18</td><td>1.8E-16</td><td>BT20/MB231</td><td>NUC</td></tr>
<tr><td>>hg38_21_+_8433172_8446622_RNA45SN1_45S_50ntFlanks@12991.13008.18|CTCCCTCGCTGCGATCTA</td><td>>hg38_21_+_8433172_8446622_RNA45SN1_45S_50ntFlanks</td><td>12991</td><td>13008</td><td>18</td><td>CTCCCTCGCTGCGATCTA</td><td>404.7</td><td>-12.4</td><td>1.5</td><td>-8.4</td><td>3.1E-17</td><td>1.9E-16</td><td>MB231/MB468</td><td>MITO</td></tr>
<tr><td>>hg38_21_+_8433172_8446622_RNA45SN1_45S_50ntFlanks@11873.11907.35|AAGAAGACCCTGTTGAGCTTGACTCTAGTCTGGCA</td><td>>hg38_21_+_8433172_8446622_RNA45SN1_45S_50ntFlanks</td><td>11873</td><td>11907</td><td>35</td><td>AAGAAGACCCTGTTGAGCTTGACTCTAGTCTGGCA</td><td>347.5</td><td>-12.3</td><td>1.4</td><td>-8.5</td><td>1.9E-17</td><td>1.9E-16</td><td>MB231/MB468</td><td>MP</td></tr>
</table>

<table>
<tr><td></td><td></td><td></td><td></td><td></td><td></td><td></td><td></td><td></td><td></td><td></td><td></td><td></td><td></td></tr>
<tr><td>>hg38_21_+_8433172_8446622_RNA45SN1_45S_50ntFlanks@6661.6695.35|CGGTGGATCACTCGGCTCGTGCGTCGATGAAGAAC</td><td>>hg38_21_+_8433172_8446622_RNA45SN1_45S_50ntFlanks</td><td>6661</td><td>6695</td><td>35</td><td>CGGTGGATCACTCGGCTCGTGCGTCGATGAAGAAC</td><td>967.0</td><td>8.0</td><td>0.9</td><td>8.4</td><td>3.4E-17</td><td>1.9E-16</td><td>BT20/MB468</td><td>CYTO</td></tr>
<tr><td>>hg38_21_+_8433172_8446622_RNA45SN1_45S_50ntFlanks@11517.11539.23|CCGCCCCCGGGGCCGCGGTTCCG</td><td>>hg38_21_+_8433172_8446622_RNA45SN1_45S_50ntFlanks</td><td>11517</td><td>11539</td><td>23</td><td>CCGCCCCCGGGGCCGCGGTTCCG</td><td>493.8</td><td>-12.7</td><td>1.5</td><td>-8.4</td><td>3.1E-17</td><td>1.9E-16</td><td>MB231/MB468</td><td>MITO</td></tr>
<tr><td>>hg38_21_+_8433172_8446622_RNA45SN1_45S_50ntFlanks@9035.9059.25|CGTGGGGGGGCCGGGCCACCCCTCC</td><td>>hg38_21_+_8433172_8446622_RNA45SN1_45S_50ntFlanks</td><td>9035</td><td>9059</td><td>25</td><td>CGTGGGGGGGCCGGGCCACCCCTCC</td><td>814.4</td><td>-13.5</td><td>1.6</td><td>-8.4</td><td>3.1E-17</td><td>1.9E-16</td><td>MB231/MB468</td><td>MITO</td></tr>
<tr><td>>hg38_21_+_8433172_8446622_RNA45SN1_45S_50ntFlanks@9738.9760.23|GAAACGATCTCAACCTATTCTCA</td><td>>hg38_21_+_8433172_8446622_RNA45SN1_45S_50ntFlanks</td><td>9738</td><td>9760</td><td>23</td><td>GAAACGATCTCAACCTATTCTCA</td><td>892.5</td><td>-13.6</td><td>1.6</td><td>-8.4</td><td>3.1E-17</td><td>1.9E-16</td><td>MB231/MB468</td><td>MITO</td></tr>
<tr><td>>hg38_21_+_8433172_8446622_RNA45SN1_45S_50ntFlanks@10845.10865.21|ACTTCGGGATAAGGATTGGCT</td><td>>hg38_21_+_8433172_8446622_RNA45SN1_45S_50ntFlanks</td><td>10845</td><td>10865</td><td>21</td><td>ACTTCGGGATAAGGATTGGCT</td><td>3613.1</td><td>-11.1</td><td>1.3</td><td>-8.5</td><td>1.9E-17</td><td>1.9E-16</td><td>MB231/MB468</td><td>MP</td></tr>
<tr><td>>hg38_21_+_8433172_8446622_RNA45SN1_45S_50ntFlanks@5493.5516.24|GAAGACGGTCGAACTTGACTATCT</td><td>>hg38_21_+_8433172_8446622_RNA45SN1_45S_50ntFlanks</td><td>5493</td><td>5516</td><td>24</td><td>GAAGACGGTCGAACTTGACTATCT</td><td>222.1</td><td>4.3</td><td>0.5</td><td>8.5</td><td>2.5E-17</td><td>1.9E-16</td><td>MB231/MB468</td><td>NUC</td></tr>
<tr><td>>hg38_21_+_8433172_8446622_RNA45SN1_45S_50ntFlanks@9740.9758.19|AACGATCTCAACCTATTCT</td><td>>hg38_21_+_8433172_8446622_RNA45SN1_45S_50ntFlanks</td><td>9740</td><td>9758</td><td>19</td><td>AACGATCTCAACCTATTCT</td><td>250.0</td><td>-11.8</td><td>1.4</td><td>-8.5</td><td>2.0E-17</td><td>1.9E-16</td><td>MB231/MB468</td><td>MP</td></tr>
<tr><td>>hg38_21_+_8433172_8446622_RNA45SN1_45S_50ntFlanks@12934.12966.33|CTAAACCATTCGTAGACGACCTGCTTCTGGGTC</td><td>>hg38_21_+_8433172_8446622_RNA45SN1_45S_50ntFlanks</td><td>12934</td><td>12966</td><td>33</td><td>CTAAACCATTCGTAGACGACCTGCTTCTGGGTC</td><td>1312.2</td><td>-7.3</td><td>0.9</td><td>-8.5</td><td>2.0E-17</td><td>1.9E-16</td><td>MB231/MB468</td><td>CYTO</td></tr>
<tr><td>>hg38_21_+_8433172_8446622_RNA45SN1_45S_50ntFlanks@12437.12473.37|CATTGTGAAGCAGAATTCACCAAGCGTTGGATTGTTC</td><td>>hg38_21_+_8433172_8446622_RNA45SN1_45S_50ntFlanks</td><td>12437</td><td>12473</td><td>37</td><td>CATTGTGAAGCAGAATTCACCAAGCGTTGGATTGTTC</td><td>4362.7</td><td>-10.0</td><td>1.2</td><td>-8.4</td><td>5.8E-17</td><td>1.9E-16</td><td>BT20/MB468</td><td>NUC</td></tr>
<tr><td>>hg38_21_+_8433172_8446622_RNA45SN1_45S_50ntFlanks@9545.9570.26|GTGGAGGTCCGTAGCGGTCCTGACGT</td><td>>hg38_21_+_8433172_8446622_RNA45SN1_45S_50ntFlanks</td><td>9545</td><td>9570</td><td>26</td><td>GTGGAGGTCCGTAGCGGTCCTGACGT</td><td>2004.9</td><td>-7.4</td><td>0.9</td><td>-8.4</td><td>5.9E-17</td><td>1.9E-16</td><td>BT20/MB468</td><td>NUC</td></tr>
<tr><td>>am_trna111_HisGTG_1_-_147774845_147774916@1.29.29,trna118_HisGTG_1_-_145396881_145396952@1.29.29,trna16_HisGTG_1_+_146544773_146544844@1.29.29,trna1_HisGTG_15_+_45493349_45493420@1.29.29,trna21_HisGTG_1_+_147753471_147753542@1.29.29,trna33_HisGTG_6_+_27125906_27125977@1.29.29,trna7_HisGTG_9_-_14433938_14434009@1.29.29,trna8_HisGTG_15_-_45492611_45492682@1.29.29,trna9_HisGTG_15_-_45490804_45490875@1.29.29|GCCGTGATCGTATAGTGGTTAGTACTCTG</td><td>>am_trna111_HisGTG_1_-_147774845_147774916</td><td>1</td><td>29</td><td>29</td><td>GCCGTGATCGTATAGTGGTTAGTACTCTG</td><td>18270.4</td><td>-5.9</td><td>0.7</td><td>-8.4</td><td>3.5E-17</td><td>1.9E-16</td><td>BT20/MB468</td><td>CYTO</td></tr>
<tr><td>>hg38_21_+_8433172_8446622_RNA45SN1_45S_50ntFlanks@6646.6674.29|TCGTACGACTCTTAGCGGTGGATCACTCG</td><td>>hg38_21_+_8433172_8446622_RNA45SN1_45S_50ntFlanks</td><td>6646</td><td>6674</td><td>29</td><td>TCGTACGACTCTTAGCGGTGGATCACTCG</td><td>14230.7</td><td>5.3</td><td>0.6</td><td>8.5</td><td>2.5E-17</td><td>1.9E-16</td><td>BT20/MB468</td><td>MITO</td></tr>
<tr><td>>hg38_21_+_8433172_8446622_RNA45SN1_45S_50ntFlanks@5420.5445.26|CCGATTGGATGGTTTAGTGAGGCCCT</td><td>>hg38_21_+_8433172_8446622_RNA45SN1_45S_50ntFlanks</td><td>5420</td><td>5445</td><td>26</td><td>CCGATTGGATGGTTTAGTGAGGCCCT</td><td>1405.6</td><td>-13.4</td><td>1.6</td><td>-8.5</td><td>2.5E-17</td><td>1.9E-16</td><td>MB231/MB468</td><td>NUC</td></tr>
<tr><td>>hg38_21_+_8433172_8446622_RNA45SN1_45S_50ntFlanks@5449.5466.18|ATCGGCCCCGCCGGGGTC</td><td>>hg38_21_+_8433172_8446622_RNA45SN1_45S_50ntFlanks</td><td>5449</td><td>5466</td><td>18</td><td>ATCGGCCCCGCCGGGGTC</td><td>2545.0</td><td>-14.4</td><td>1.6</td><td>-8.9</td><td>4.8E-19</td><td>1.9E-16</td><td>BT20/MB468</td><td>MP</td></tr>
<tr><td>>hg38_21_+_8433172_8446622_RNA45SN1_45S_50ntFlanks@12357.12375.19|CCACAGGGATAACTGGCTT</td><td>>hg38_21_+_8433172_8446622_RNA45SN1_45S_50ntFlanks</td><td>12357</td><td>12375</td><td>19</td><td>CCACAGGGATAACTGGCTT</td><td>511.7</td><td>-12.8</td><td>1.5</td><td>-8.4</td><td>3.2E-17</td><td>1.9E-16</td><td>MB231/MB468</td><td>MITO</td></tr>
<tr><td>>am_MI0005763|hsa-mir-941-1&WithFlank&20|+|63919443|63919526@53.74.22;MI0005764|hsa-mir-941-2&WithFlank&20|+|63919499|63919582@53.74.22;MI0005765|hsa-mir-941-3&WithFlank&20|+|63919555|63919638@53.74.22;MI0005766|hsa-mir-941-4&WithFlank&20|+|63919750|63919833@53.74.22;MI0031520|hsa-mir-941-5&WithFlank&20|+|63919862|63919945@53.74.22&[MIMAT0004984&hsa-miR-941&offsets|0|-1;m-263&20|+|63919495|63919517&offsets|0|-1];[MIMAT0004984_1&hsa-miR-941&offsets|0|-1;m-263&20|+|63919551|63919573&offsets|0|-1];[MIMAT0004984_2&hsa-miR-941&offsets|0|-1;m-263&20|+|63919607|63919629&offsets|0|-1];[MIMAT0004984_3&hsa-miR-941&offsets|0|-1;m-263&20|+|63919802|63919824&offsets|0|-1];[MIMAT0004984_4&hsa-miR-941&offsets|0|-1;m-263&20|+|63919914|63919936&offsets|0|-1]|CACCCGGCTGTGTGCACATGTG</td><td>>am_MI0005763|hsa-mir-941-1&WithFlank&20|+|63919443|63919526</td><td>53</td><td>74</td><td>22</td><td>CACCCGGCTGTGTGCACATGTG</td><td>508.7</td><td>5.9</td><td>0.7</td><td>8.5</td><td>2.5E-17</td><td>1.9E-16</td><td>MB231/MB468</td><td>NUC</td></tr>
<tr><td>>hg38_21_+_8433172_8446622_RNA45SN1_45S_50ntFlanks@11270.11293.24|CGGCCGCCCCCGGGGAGCCCGGCG</td><td>>hg38_21_+_8433172_8446622_RNA45SN1_45S_50ntFlanks</td><td>11270</td><td>11293</td><td>24</td><td>CGGCCGCCCCCGGGGAGCCCGGCG</td><td>624.5</td><td>-12.2</td><td>1.4</td><td>-8.5</td><td>2.5E-17</td><td>1.9E-16</td><td>MB231/MB468</td><td>NUC</td></tr>
<tr><td>>hg38_21_+_8433172_8446622_RNA45SN1_45S_50ntFlanks@11564.11585.22|CCTAGCAGCCGACTTAGAACTG</td><td>>hg38_21_+_8433172_8446622_RNA45SN1_45S_50ntFlanks</td><td>11564</td><td>11585</td><td>22</td><td>CCTAGCAGCCGACTTAGAACTG</td><td>1724.6</td><td>-5.2</td><td>0.6</td><td>-8.4</td><td>3.6E-17</td><td>1.9E-16</td><td>BT20/MB468</td><td>CYTO</td></tr>
<tr><td>>hg38_21_+_8433172_8446622_RNA45SN1_45S_50ntFlanks@10882.10901.20|TCGGGCTGGGGCGCGAAGCG</td><td>>hg38_21_+_8433172_8446622_RNA45SN1_45S_50ntFlanks</td><td>10882</td><td>10901</td><td>20</td><td>TCGGGCTGGGGCGCGAAGCG</td><td>141.4</td><td>-5.4</td><td>0.6</td><td>-8.5</td><td>2.0E-17</td><td>1.9E-16</td><td>MB231/MB468</td><td>CYTO</td></tr>
<tr><td>>hg38_21_+_8433172_8446622_RNA45SN1_45S_50ntFlanks@12836.12868.33|CGGTGCGGAGTGCCCTTCGTCCTGGGAAACGGG</td><td>>hg38_21_+_8433172_8446622_RNA45SN1_45S_50ntFlanks</td><td>12836</td><td>12868</td><td>33</td><td>CGGTGCGGAGTGCCCTTCGTCCTGGGAAACGGG</td><td>276.8</td><td>-12.0</td><td>1.4</td><td>-8.5</td><td>2.0E-17</td><td>2.0E-16</td><td>MB231/MB468</td><td>MP</td></tr>
<tr><td>>hg38_21_+_8433172_8446622_RNA45SN1_45S_50ntFlanks@10392.10409.18|ACAGCAGTTGAACATGGG</td><td>>hg38_21_+_8433172_8446622_RNA45SN1_45S_50ntFlanks</td><td>10392</td><td>10409</td><td>18</td><td>ACAGCAGTTGAACATGGG</td><td>2418.5</td><td>-10.6</td><td>1.3</td><td>-8.5</td><td>2.6E-17</td><td>2.0E-16</td><td>MB231/MB468</td><td>NUC</td></tr>
<tr><td>>hg38_21_+_8433172_8446622_RNA45SN1_45S_50ntFlanks@6661.6694.34|CGGTGGATCACTCGGCTCGTGCGTCGATGAAGAA</td><td>>hg38_21_+_8433172_8446622_RNA45SN1_45S_50ntFlanks</td><td>6661</td><td>6694</td><td>34</td><td>CGGTGGATCACTCGGCTCGTGCGTCGATGAAGAA</td><td>899.5</td><td>4.7</td><td>0.6</td><td>8.5</td><td>2.6E-17</td><td>2.0E-16</td><td>BT20/MB468</td><td>MITO</td></tr>
<tr><td>>hg38_21_+_8433172_8446622_RNA45SN1_45S_50ntFlanks@11581.11605.25|AACTGGTGCGGACCAGGGGAATCCG</td><td>>hg38_21_+_8433172_8446622_RNA45SN1_45S_50ntFlanks</td><td>11581</td><td>11605</td><td>25</td><td>AACTGGTGCGGACCAGGGGAATCCG</td><td>2492.2</td><td>-5.1</td><td>0.6</td><td>-8.4</td><td>3.6E-17</td><td>2.0E-16</td><td>BT20/MB468</td><td>CYTO</td></tr>
<tr><td>>hg38_21_+_8433172_8446622_RNA45SN1_45S_50ntFlanks@11581.11600.20|AACTGGTGCGGACCAGGGGA</td><td>>hg38_21_+_8433172_8446622_RNA45SN1_45S_50ntFlanks</td><td>11581</td><td>11600</td><td>20</td><td>AACTGGTGCGGACCAGGGGA</td><td>880.8</td><td>-13.6</td><td>1.6</td><td>-8.5</td><td>2.0E-17</td><td>2.0E-16</td><td>MB231/MB468</td><td>MP</td></tr>
<tr><td>>hg38_21_+_8433172_8446622_RNA45SN1_45S_50ntFlanks@9304.9335.32|ACCAAGGAGTCTAACACGTGCGCGAGTCGGGG</td><td>>hg38_21_+_8433172_8446622_RNA45SN1_45S_50ntFlanks</td><td>9304</td><td>9335</td><td>32</td><td>ACCAAGGAGTCTAACACGTGCGCGAGTCGGGG</td><td>309.9</td><td>-12.1</td><td>1.4</td><td>-8.5</td><td>2.0E-17</td><td>2.0E-16</td><td>MB231/MB468</td><td>MP</td></tr>
<tr><td>>hg38_21_+_8433172_8446622_RNA45SN1_45S_50ntFlanks@6652.6691.40|GACTCTTAGCGGTGGATCACTCGGCTCGTGCGTCGATGAA</td><td>>hg38_21_+_8433172_8446622_RNA45SN1_45S_50ntFlanks</td><td>6652</td><td>6691</td><td>40</td><td>GACTCTTAGCGGTGGATCACTCGGCTCGTGCGTCGATGAA</td><td>604.6</td><td>4.1</td><td>0.5</td><td>8.5</td><td>2.6E-17</td><td>2.0E-16</td><td>BT20/MB468</td><td>MITO</td></tr>
<tr><td>>trna129_MetCAT_6_-_27870271_27870342@31.51.21,trna142_MetCAT_6_-_27560600_27560671@31.51.21,trna150_MetCAT_6_-_27300764_27300835@31.51.21,trna169_MetCAT_6_-_26330529_26330600@31.51.21,trna171_MetCAT_6_-_26313352_26313423@31.51.21,trna20_MetCAT_17_-_80452597_80452668@31.51.21,trna2_MetCAT_6_+_26286754_26286825@31.51.21,trna32_MetCAT_1_+_153643726_153643797@31.51.21,trna61_MetCAT_6_+_27745664_27745735@31.51.21|CCCATAACCCAGAGGTCGATG</td><td>>trna129_MetCAT_6_-_27870271_27870342</td><td>31</td><td>51</td><td>21</td><td>CCCATAACCCAGAGGTCGATG</td><td>581.5</td><td>-12.1</td><td>1.4</td><td>-8.5</td><td>2.6E-17</td><td>2.0E-16</td><td>MB231/MB468</td><td>NUC</td></tr>
<tr><td>>hg38_21_+_8433172_8446622_RNA45SN1_45S_50ntFlanks@13002.13023.22|CGATCTATTGAAAGTCAGCCCT</td><td>>hg38_21_+_8433172_8446622_RNA45SN1_45S_50ntFlanks</td><td>13002</td><td>13023</td><td>22</td><td>CGATCTATTGAAAGTCAGCCCT</td><td>716.2</td><td>-11.8</td><td>1.4</td><td>-8.4</td><td>3.6E-17</td><td>2.0E-16</td><td>BT20/MB468</td><td>CYTO</td></tr>
<tr><td>>MI0000093|hsa-mir-92a-1&WithFlank&13|+|91351308|91351397@54.74.21(+1A);MI0000094|hsa-mir-92a-2&WithFlank&X|-|134169532|134169618@54.74.21(+1A)&[MIMAT0000092&hsa-miR-92a-3p&offsets|0|-1(+1A);m-9&13|+|91351361|91351382&offsets|0|-1(+1A)];[MIMAT0000092_1&hsa-miR-92a-3p&offsets|0|-1(+1A);m-11&X|-|134169544|134169565&offsets|0|-1(+1A)]|TATTGCACTTGTCCCGGCCTGA</td><td>>MI0000093|hsa-mir-92a-1&WithFlank&13|+|91351308|91351397</td><td>54</td><td>74</td><td>21(+1A)</td><td>TATTGCACTTGTCCCGGCCTGA</td><td>1275.9</td><td>5.6</td><td>0.7</td><td>8.5</td><td>2.6E-17</td><td>2.0E-16</td><td>BT20/MB468</td><td>MITO</td></tr>
<tr><td>>hg38_21_+_8433172_8446622_RNA45SN1_45S_50ntFlanks@12621.12654.34|GAGCCAATGGGGCGAAGCTACCATCTGTGGGATT</td><td>>hg38_21_+_8433172_8446622_RNA45SN1_45S_50ntFlanks</td><td>12621</td><td>12654</td><td>34</td><td>GAGCCAATGGGGCGAAGCTACCATCTGTGGGATT</td><td>260.3</td><td>-11.9</td><td>1.4</td><td>-8.5</td><td>2.1E-17</td><td>2.0E-16</td><td>MB231/MB468</td><td>MP</td></tr>
<tr><td>>hg38_21_+_8433172_8446622_RNA45SN1_45S_50ntFlanks@11581.11604.24|AACTGGTGCGGACCAGGGGAATCC</td><td>>hg38_21_+_8433172_8446622_RNA45SN1_45S_50ntFlanks</td><td>11581</td><td>11604</td><td>24</td><td>AACTGGTGCGGACCAGGGGAATCC</td><td>470.5</td><td>-5.7</td><td>0.7</td><td>-8.4</td><td>6.2E-17</td><td>2.0E-16</td><td>BT20/MB468</td><td>NUC</td></tr>
<tr><td>>MI0000084|hsa-mir-26b&WithFlank&2|+|218402640|218402728@18.39.22&[MIMAT0000083&hsa-miR-26b-5p&offsets|0|+1;m-53&2|+|218402657|218402678&offsets|0|0]|TTCAAGTAATTCAGGATAGGTT</td><td>>MI0000084|hsa-mir-26b&WithFlank&2|+|218402640|218402728</td><td>18</td><td>39</td><td>22</td><td>TTCAAGTAATTCAGGATAGGTT</td><td>12226.5</td><td>6.4</td><td>0.8</td><td>8.4</td><td>3.7E-17</td><td>2.0E-16</td><td>BT20/MB468</td><td>CYTO</td></tr>
<tr><td>>hg38_21_+_8433172_8446622_RNA45SN1_45S_50ntFlanks@10845.10863.19|ACTTCGGGATAAGGATTGG</td><td>>hg38_21_+_8433172_8446622_RNA45SN1_45S_50ntFlanks</td><td>10845</td><td>10863</td><td>19</td><td>ACTTCGGGATAAGGATTGG</td><td>474.5</td><td>-12.8</td><td>1.5</td><td>-8.5</td><td>2.1E-17</td><td>2.0E-16</td><td>MB231/MB468</td><td>MP</td></tr>
<tr><td>>hg38_21_+_8433172_8446622_RNA45SN1_45S_50ntFlanks@9575.9597.23|ATCGGTCGTCCGACCTGGGTATA</td><td>>hg38_21_+_8433172_8446622_RNA45SN1_45S_50ntFlanks</td><td>9575</td><td>9597</td><td>23</td><td>ATCGGTCGTCCGACCTGGGTATA</td><td>1372.9</td><td>-12.7</td><td>1.5</td><td>-8.4</td><td>3.7E-17</td><td>2.0E-16</td><td>BT20/MB468</td><td>CYTO</td></tr>
<tr><td>>hg38_21_+_8433172_8446622_RNA45SN1_45S_50ntFlanks@12931.12956.26|GCGCTAAACCATTCGTAGACGACCTG</td><td>>hg38_21_+_8433172_8446622_RNA45SN1_45S_50ntFlanks</td><td>12931</td><td>12956</td><td>26</td><td>GCGCTAAACCATTCGTAGACGACCTG</td><td>1734.6</td><td>-7.9</td><td>0.9</td><td>-8.4</td><td>3.4E-17</td><td>2.0E-16</td><td>MB231/MB468</td><td>MITO</td></tr>
<tr><td>>MI0000447|hsa-mir-128-1&WithFlank&2|+|135665391|135665484@56.76.21;MI0000727|hsa-mir-128-2&WithFlank&3|+|35744470|35744565@58.78.21&[MIMAT0000424&hsa-miR-128-3p&offsets|0|0;m-108&2|+|135665446|135665466&offsets|0|0];[MIMAT0000424_1&hsa-miR-128-3p&offsets|0|0;m-117&3|+|35744527|35744547&offsets|0|0]|TCACAGTGAACCGGTCTCTTT</td><td>>MI0000447|hsa-mir-128-1&WithFlank&2|+|135665391|135665484</td><td>56</td><td>76</td><td>21</td><td>TCACAGTGAACCGGTCTCTTT</td><td>12321.6</td><td>5.2</td><td>0.6</td><td>8.5</td><td>2.7E-17</td><td>2.0E-16</td><td>BT20/MB468</td><td>MITO</td></tr>
<tr><td>>MI0000102|hsa-mir-100&WithFlank&11|-|122152223|122152314@19.37.19&[MIMAT0000098&hsa-miR-100-5p&offsets|0|-3;m-27&11|-|122152276|122152296&offsets|0|-2]|AACCCGTAGATCCGAACTT</td><td>>MI0000102|hsa-mir-100&WithFlank&11|-|122152223|122152314</td><td>19</td><td>37</td><td>19</td><td>AACCCGTAGATCCGAACTT</td><td>510.4</td><td>12.2</td><td>1.4</td><td>8.5</td><td>2.1E-17</td><td>2.0E-16</td><td>MB231/MB468</td><td>MP</td></tr>
<tr><td>>hg38_21_+_8433172_8446622_RNA45SN1_45S_50ntFlanks@11855.11872.18|TGGCGGAATCAGCGGGGA</td><td>>hg38_21_+_8433172_8446622_RNA45SN1_45S_50ntFlanks</td><td>11855</td><td>11872</td><td>18</td><td>TGGCGGAATCAGCGGGGA</td><td>287.9</td><td>-12.0</td><td>1.4</td><td>-8.5</td><td>2.1E-17</td><td>2.0E-16</td><td>MB231/MB468</td><td>MP</td></tr>
</table>

| | | | | | | | | | | | | | |
|---|---|---|---|---|---|---|---|---|---|---|---|---|---|
| >MI0000061\|hsa-let-7a-2&WithFlank&11\|-\|122146516\|122146599@11.28.18;MI0000064\|hsa-let-7c&WithFlank&21\|+\|16539822\|16539917@17.34.18;MI0000062\|hsa-let-7a-3&WithFlank&22\|+\|46112743\|46112828@10.27.18;MI0000060\|hsa-let-7a-1&WithFlank&9\|+\|94175951\|94176042@12.29.18&[MIMAT0000062&hsa-let-7a-5p&offsets\|0\|-4;m-6&11\|-\|122146568\|122146589&offsets\|0\|-4];[MIMAT0000064&hsa-let-7c-5p&offsets\|0\|-4;m-37&21\|+\|16539838\|16539859&offsets\|0\|-4];[MIMAT0000062_1&hsa-let-7a-5p&offsets\|0\|-4;m-5&22\|+\|46112752\|46112773&offsets\|0\|-4];[MIMAT0000062_2&hsa-let-7a-5p&offsets\|0\|-4;m-4&9\|+\|94175962\|94175983&offsets\|0\|-4]\|TGAGGTAGTAGGTTGTAT | >MI0000061\|hsa-let-7a-2&WithFlank&11\|-\|122146516\|122146599 | 11 | 28 | 18 | TGAGGTAGTAGGTTGTAT | 3246.0 | 6.8 | 0.8 | 8.5 | 2.1E-17 | 2.0E-16 | MB231/MB468 | CYTO |
| >hg38_21_+_8433172_8446622_RNA45SN1_45S_50ntFlanks@8342.8377.36\|CGTAAGGGAAAGTTGAAAAGAACTTTGAAGAGAGAG | >hg38_21_+_8433172_8446622_RNA45SN1_45S_50ntFlanks | 8342 | 8377 | 36 | CGTAAGGGAAAGTTGAAAAGAACTTTGAAGAGAGAG | 524.6 | 10.4 | 1.2 | 8.5 | 2.1E-17 | 2.0E-16 | MB231/MB468 | CYTO |
| >MI0000440\|hsa-mir-27b&WithFlank&9\|+\|95085439\|95085547@67.88.22(+1U)&[MIMAT0000419&hsa-miR-27b-3p&offsets\|0\|+1(+1U);m-39&9\|+\|95085505\|95085525&offsets\|0\|+1(+1U)]\|TTCACAGTGGCTAAGTTCTGCAT | >MI0000440\|hsa-mir-27b&WithFlank&9\|+\|95085439\|95085547 | 67 | 88 | 22(+1U) | TTCACAGTGGCTAAGTTCTGCAT | 263.7 | 5.9 | 0.7 | 8.5 | 2.7E-17 | 2.0E-16 | MB231/MB468 | NUC |
| >hg38_21_+_8433172_8446622_RNA45SN1_45S_50ntFlanks@4995.5022.28\|ACAGATTGATAGCTCTTTCTCGATTCCG | >hg38_21_+_8433172_8446622_RNA45SN1_45S_50ntFlanks | 4995 | 5022 | 28 | ACAGATTGATAGCTCTTTCTCGATTCCG | 1704.7 | -14.5 | 1.7 | -8.4 | 3.4E-17 | 2.1E-16 | MB231/MB468 | MITO |
| >hg38_21_+_8433172_8446622_RNA45SN1_45S_50ntFlanks@12463.12496.34\|TTGGATTGTTCACCCACTAATAGGGAACGTGAGC | >hg38_21_+_8433172_8446622_RNA45SN1_45S_50ntFlanks | 12463 | 12496 | 34 | TTGGATTGTTCACCCACTAATAGGGAACGTGAGC | 599.9 | 5.2 | 0.6 | 8.5 | 2.1E-17 | 2.1E-16 | MB231/MB468 | CYTO |
| >hg38_21_+_8433172_8446622_RNA45SN1_45S_50ntFlanks@10914.10935.22\|CGCCGCGGCTGGACGAGGCGCC | >hg38_21_+_8433172_8446622_RNA45SN1_45S_50ntFlanks | 10914 | 10935 | 22 | CGCCGCGGCTGGACGAGGCGCC | 355.7 | -12.3 | 1.5 | -8.5 | 2.1E-17 | 2.1E-16 | MB231/MB468 | MP |
| >hg38_21_+_8433172_8446622_RNA45SN1_45S_50ntFlanks@9533.9553.21\|GAGGAAACTCTGGTGGAGGTC | >hg38_21_+_8433172_8446622_RNA45SN1_45S_50ntFlanks | 9533 | 9553 | 21 | GAGGAAACTCTGGTGGAGGTC | 1697.2 | -5.9 | 0.7 | -8.4 | 3.4E-17 | 2.1E-16 | MB231/MB468 | MITO |
| >hg38_21_+_8433172_8446622_RNA45SN1_45S_50ntFlanks@12081.12108.28\|CTCTCGCTTCTGGCGCCAAGCGCCCGGC | >hg38_21_+_8433172_8446622_RNA45SN1_45S_50ntFlanks | 12081 | 12108 | 28 | CTCTCGCTTCTGGCGCCAAGCGCCCGGC | 642.2 | -7.8 | 0.9 | -8.5 | 2.2E-17 | 2.1E-16 | MB231/MB468 | CYTO |
| >hg38_21_+_8433172_8446622_RNA45SN1_45S_50ntFlanks@11641.11677.37\|CGGCGGGTGTTGACGCGATGTGATTTCTGCCCAGTGC | >hg38_21_+_8433172_8446622_RNA45SN1_45S_50ntFlanks | 11641 | 11677 | 37 | CGGCGGGTGTTGACGCGATGTGATTTCTGCCCAGTGC | 416.1 | -12.6 | 1.5 | -8.5 | 2.1E-17 | 2.1E-16 | MB231/MB468 | MP |
| >MI0000446\|hsa-mir-125b-1&WithFlank&11\|-\|122099751\|122099850@21.40.20(+1A);MI0000470\|hsa-mir-125b-2&WithFlank&21\|+\|16590231\|16590331@23.42.20(+1A)&[MIMAT0000423&hsa-miR-125b-5p&offsets\|0\|-2(+1A);m-67&11\|-\|122099809\|122099830&offsets\|0\|-2(+1A)];[MIMAT0000423_1&hsa-miR-125b-5p&offsets\|0\|-2(+1A);m-68&21\|+\|16590253\|16590274&offsets\|0\|-2(+1A)]\|TCCCTGAGACCCTAACTTGTA | >MI0000446\|hsa-mir-125b-1&WithFlank&11\|-\|122099751\|122099850 | 21 | 40 | 20(+1A) | TCCCTGAGACCCTAACTTGTA | 1852.6 | 8.4 | 1.0 | 8.5 | 2.2E-17 | 2.1E-16 | MB231/MB468 | CYTO |
| >hg38_21_+_8433172_8446622_RNA45SN1_45S_50ntFlanks@12875.12892.18\|CTGGAAAGGCGGCCGCCC | >hg38_21_+_8433172_8446622_RNA45SN1_45S_50ntFlanks | 12875 | 12892 | 18 | CTGGAAAGGCGGCCGCCC | 481.5 | -12.8 | 1.5 | -8.5 | 2.2E-17 | 2.1E-16 | MB231/MB468 | MP |
| >MI0000477\|hsa-mir-146a&WithFlank&5\|+\|160485346\|160485456@27.47.21(+1A)&[MIMAT0000449&hsa-miR-146a-5p&offsets\|0\|-1(+1A);m-129&5\|+\|160485372\|160485393&offsets\|0\|-1(+1A)]\|TGAGAACTGAATTCCATGGGTA | >MI0000477\|hsa-mir-146a&WithFlank&5\|+\|160485346\|160485456 | 27 | 47 | 21(+1A) | TGAGAACTGAATTCCATGGGTA | 691.5 | 13.3 | 1.6 | 8.5 | 2.7E-17 | 2.1E-16 | MB231/MB468 | NUC |
| >hg38_21_+_8433172_8446622_RNA45SN1_45S_50ntFlanks@12435.12455.21\|ATCATTGTGAAGCAGAATTCA | >hg38_21_+_8433172_8446622_RNA45SN1_45S_50ntFlanks | 12435 | 12455 | 21 | ATCATTGTGAAGCAGAATTCA | 2029.4 | -14.1 | 1.6 | -8.9 | 5.4E-19 | 2.1E-16 | BT20/MB468 | MP |
| >hg38_21_+_8433172_8446622_RNA45SN1_45S_50ntFlanks@11615.11640.26\|TTAAAACAAAGCATCGCGAAGGCCCG | >hg38_21_+_8433172_8446622_RNA45SN1_45S_50ntFlanks | 11615 | 11640 | 26 | TTAAAACAAAGCATCGCGAAGGCCCG | 440.4 | -12.6 | 1.5 | -8.4 | 3.5E-17 | 2.1E-16 | MB231/MB468 | MITO |
| >hg38_21_+_8433172_8446622_RNA45SN1_45S_50ntFlanks@8170.8199.30\|CAAGTCCTTCTGATCGAGGCCCAGCCCGTG | >hg38_21_+_8433172_8446622_RNA45SN1_45S_50ntFlanks | 8170 | 8199 | 30 | CAAGTCCTTCTGATCGAGGCCCAGCCCGTG | 329.6 | -12.2 | 1.4 | -8.5 | 2.2E-17 | 2.1E-16 | MB231/MB468 | MP |
| >MI0000077\|hsa-mir-21&WithFlank&17\|+\|59841260\|59841343@14.35.22(+1G)&[MIMAT0000076&hsa-miR-21-5p&offsets\|0\|0(+1G);m-1&17\|+\|59841273\|59841294&offsets\|0\|0(+1G)]\|TAGCTTATCAGACTGATGTTGAG | >MI0000077\|hsa-mir-21&WithFlank&17\|+\|59841260\|59841343 | 14 | 35 | 22(+1G) | TAGCTTATCAGACTGATGTTGAG | 1837.6 | 7.4 | 0.9 | 8.5 | 2.8E-17 | 2.1E-16 | MB231/MB468 | NUC |
| >hg38_21_+_8433172_8446622_RNA45SN1_45S_50ntFlanks@12416.12442.27\|TTCGATGTCGGCTCTTCCTATCATTGT | >hg38_21_+_8433172_8446622_RNA45SN1_45S_50ntFlanks | 12416 | 12442 | 27 | TTCGATGTCGGCTCTTCCTATCATTGT | 2839.8 | 6.4 | 0.8 | 8.5 | 2.2E-17 | 2.1E-16 | MB231/MB468 | CYTO |
| >hg38_21_+_8433172_8446622_RNA45SN1_45S_50ntFlanks@11564.11585.22\|CCTAGCAGCCGACTTAGAACTG | >hg38_21_+_8433172_8446622_RNA45SN1_45S_50ntFlanks | 11564 | 11585 | 22 | CCTAGCAGCCGACTTAGAACTG | 2550.3 | -8.6 | 1.0 | -8.5 | 2.2E-17 | 2.1E-16 | MB231/MB468 | MP |
| >hg38_21_+_8433172_8446622_RNA45SN1_45S_50ntFlanks@9339.9367.29\|CGCACGAAAGCCGCCGTGGCGCAATGAAG | >hg38_21_+_8433172_8446622_RNA45SN1_45S_50ntFlanks | 9339 | 9367 | 29 | CGCACGAAAGCCGCCGTGGCGCAATGAAG | 716.6 | -13.3 | 1.6 | -8.5 | 2.2E-17 | 2.1E-16 | MB231/MB468 | MP |
| >hg38_21_+_8433172_8446622_RNA45SN1_45S_50ntFlanks@9622.9655.34\|CTAGTAGCTGGTTCCCTCCGAAGTTTCCCTCAGG | >hg38_21_+_8433172_8446622_RNA45SN1_45S_50ntFlanks | 9622 | 9655 | 34 | CTAGTAGCTGGTTCCCTCCGAAGTTTCCCTCAGG | 376.6 | -12.3 | 1.5 | -8.4 | 3.6E-17 | 2.1E-16 | MB231/MB468 | MITO |
| >trnaMT_GluTTC_MT_-_14674_14742@31.71.41,trnalookalike8_GluTTC_5_-_93905172_93905240@31.71.41\|TTCATATCATTGGTCGTGGTTGTAGTCCGTGCGAGAATACC | >trnaMT_GluTTC_MT_-_14674_14742 | 31 | 71 | 41 | TTCATATCATTGGTCGTGGTTGTAGTCCGTGCGAGAATACC | 413.2 | 12.7 | 1.5 | 8.5 | 2.2E-17 | 2.1E-16 | MB231/MB468 | CYTO |
| >hg38_21_+_8433172_8446622_RNA45SN1_45S_50ntFlanks@12940.12966.27\|CATTCGTAGACGACCTGCTTCTGGGTC | >hg38_21_+_8433172_8446622_RNA45SN1_45S_50ntFlanks | 12940 | 12966 | 27 | CATTCGTAGACGACCTGCTTCTGGGTC | 1572.5 | -7.4 | 0.9 | -8.5 | 2.2E-17 | 2.1E-16 | MB231/MB468 | CYTO |
| >hg38_21_+_8433172_8446622_RNA45SN1_45S_50ntFlanks@11848.11873.26\|ACGGGCTTGGCGGAATCAGCGGGGAA | >hg38_21_+_8433172_8446622_RNA45SN1_45S_50ntFlanks | 11848 | 11873 | 26 | ACGGGCTTGGCGGAATCAGCGGGGAA | 358.9 | -12.4 | 1.5 | -8.5 | 2.2E-17 | 2.1E-16 | MB231/MB468 | MP |
| >hg38_21_+_8433172_8446622_RNA45SN1_45S_50ntFlanks@5078.5098.21\|CGATAACGAACGAGACTCTGG | >hg38_21_+_8433172_8446622_RNA45SN1_45S_50ntFlanks | 5078 | 5098 | 21 | CGATAACGAACGAGACTCTGG | 1866.7 | -14.7 | 1.7 | -8.5 | 2.2E-17 | 2.1E-16 | MB231/MB468 | MP |
| >am_trna134_GluTTC_1_-_16861774_16861845@1.29.29,trna5_GluTTC_1_+_17199078_17199149@1.29.29,trna84_GluTTC_1_-_161391883_161391954@1.29.29,trna94_GluTTC_1_-_149664355_149664427@1.29.29\|TCCCTGGTGGTCTAGTGGCTAGGATTCGG | >am_trna134_GluTTC_1_-_16861774_16861845 | 1 | 29 | 29 | TCCCTGGTGGTCTAGTGGCTAGGATTCGG | 503.0 | -12.8 | 1.5 | -8.4 | 3.6E-17 | 2.1E-16 | MB231/MB468 | MITO |
| >hg38_21_+_8433172_8446622_RNA45SN1_45S_50ntFlanks@10841.10864.24\|CGTAACTTCGGGATAAGGATTGGC | >hg38_21_+_8433172_8446622_RNA45SN1_45S_50ntFlanks | 10841 | 10864 | 24 | CGTAACTTCGGGATAAGGATTGGC | 5533.6 | -5.4 | 0.6 | -8.4 | 3.9E-17 | 2.1E-16 | BT20/MB468 | CYTO |
| >hg38_21_+_8433172_8446622_RNA45SN1_45S_50ntFlanks@4005.4026.22\|AACCTCGGGCCGATCGCACGCC | >hg38_21_+_8433172_8446622_RNA45SN1_45S_50ntFlanks | 4005 | 4026 | 22 | AACCTCGGGCCGATCGCACGCC | 708.8 | -13.3 | 1.6 | -8.4 | 3.6E-17 | 2.1E-16 | MB231/MB468 | MITO |
| >MI0000102\|hsa-mir-100&WithFlank&11\|-\|122152223\|122152314@19.38.20(+2G)&[MIMAT0000098&hsa-miR-100-5p&offsets\|0\|-2(+2G);m-27&11\|-\|122152276\|122152296&offsets\|0\|-1(+2G)]\|AACCCGTAGATCCGAACTTGGG | >MI0000102\|hsa-mir-100&WithFlank&11\|-\|122152223\|122152314 | 19 | 38 | 20(+2G) | AACCCGTAGATCCGAACTTGGG | 447.4 | 12.0 | 1.4 | 8.5 | 2.2E-17 | 2.1E-16 | MB231/MB468 | MP |
| >hg38_21_+_8433172_8446622_RNA45SN1_45S_50ntFlanks@5467.5499.33\|GGCCCACGGCCCTGGCGGAGCGCTGAGAAGACG | >hg38_21_+_8433172_8446622_RNA45SN1_45S_50ntFlanks | 5467 | 5499 | 33 | GGCCCACGGCCCTGGCGGAGCGCTGAGAAGACG | 4152.9 | -11.3 | 1.3 | -8.4 | 3.6E-17 | 2.1E-16 | MB231/MB468 | MITO |
| >hg38_21_+_8433172_8446622_RNA45SN1_45S_50ntFlanks@3921.3959.39\|ATTTATCAGATCAAAACCAACCCGGTCAGCCCCTCTCCG | >hg38_21_+_8433172_8446622_RNA45SN1_45S_50ntFlanks | 3921 | 3959 | 39 | ATTTATCAGATCAAAACCAACCCGGTCAGCCCCTCTCCG | 473.9 | -12.7 | 1.5 | -8.4 | 3.6E-17 | 2.2E-16 | MB231/MB468 | MITO |
| >hg38_21_+_8433172_8446622_RNA45SN1_45S_50ntFlanks@6656.6681.26\|CTTAGCGGTGGATCACTCGGCTCGTG | >hg38_21_+_8433172_8446622_RNA45SN1_45S_50ntFlanks | 6656 | 6681 | 26 | CTTAGCGGTGGATCACTCGGCTCGTG | 1775.2 | -4.6 | 0.5 | -8.4 | 4.0E-17 | 2.2E-16 | BT20/MB468 | CYTO |
| >hg38_21_+_8433172_8446622_RNA45SN1_45S_50ntFlanks@12156.12181.26\|GAGTTTGACTGGGGCGGTACACCTGT | >hg38_21_+_8433172_8446622_RNA45SN1_45S_50ntFlanks | 12156 | 12181 | 26 | GAGTTTGACTGGGGCGGTACACCTGT | 301.2 | -12.1 | 1.4 | -8.5 | 2.3E-17 | 2.2E-16 | MB231/MB468 | MP |
| >hg38_21_+_8433172_8446622_RNA45SN1_45S_50ntFlanks@4724.4751.28\|ATTAATCAAGAACGAAAGTCGGAGGTTC | >hg38_21_+_8433172_8446622_RNA45SN1_45S_50ntFlanks | 4724 | 4751 | 28 | ATTAATCAAGAACGAAAGTCGGAGGTTC | 1882.1 | -10.3 | 1.2 | -8.5 | 2.3E-17 | 2.2E-16 | MB231/MB468 | MP |
| >hg38_21_+_8433172_8446622_RNA45SN1_45S_50ntFlanks@10402.10429.28\|AACATGGGTCAGTCGGTCCTGAGAGATG | >hg38_21_+_8433172_8446622_RNA45SN1_45S_50ntFlanks | 10402 | 10429 | 28 | AACATGGGTCAGTCGGTCCTGAGAGATG | 4414.7 | -6.2 | 0.7 | -8.4 | 3.6E-17 | 2.2E-16 | MB231/MB468 | MITO |
| >hg38_21_+_8433172_8446622_RNA45SN1_45S_50ntFlanks@12511.12538.28\|CGTGAGACAGGTTAGTTTTACCCTACTG | >hg38_21_+_8433172_8446622_RNA45SN1_45S_50ntFlanks | 12511 | 12538 | 28 | CGTGAGACAGGTTAGTTTTACCCTACTG | 464.7 | -12.6 | 1.5 | -8.4 | 3.7E-17 | 2.2E-16 | MB231/MB468 | MITO |
| >hg38_21_+_8433172_8446622_RNA45SN1_45S_50ntFlanks@10392.10426.35\|ACAGCAGTTGAACATGGGTCAGTCGGTCCTGAGAG | >hg38_21_+_8433172_8446622_RNA45SN1_45S_50ntFlanks | 10392 | 10426 | 35 | ACAGCAGTTGAACATGGGTCAGTCGGTCCTGAGAG | 594.3 | -13.1 | 1.5 | -8.5 | 2.3E-17 | 2.2E-16 | MB231/MB468 | MP |
| >hg38_21_+_8433172_8446622_RNA45SN1_45S_50ntFlanks@11930.11949.20\|TAGAATAAGTGGGAGGCCCC | >hg38_21_+_8433172_8446622_RNA45SN1_45S_50ntFlanks | 11930 | 11949 | 20 | TAGAATAAGTGGGAGGCCCC | 350.6 | -12.3 | 1.5 | -8.5 | 2.3E-17 | 2.2E-16 | MB231/MB468 | MP |
| >hg38_21_+_8433172_8446622_RNA45SN1_45S_50ntFlanks@9568.9596.29\|CGTGCAAATCGGTCGTCCGACCTGGGTAT | >hg38_21_+_8433172_8446622_RNA45SN1_45S_50ntFlanks | 9568 | 9596 | 29 | CGTGCAAATCGGTCGTCCGACCTGGGTAT | 4452.4 | -8.0 | 0.9 | -8.5 | 2.3E-17 | 2.2E-16 | MB231/MB468 | CYTO |
| >hg38_21_+_8433172_8446622_RNA45SN1_45S_50ntFlanks@4163.4182.20\|CATCCAAGGAAGGCAGCAGG | >hg38_21_+_8433172_8446622_RNA45SN1_45S_50ntFlanks | 4163 | 4182 | 20 | CATCCAAGGAAGGCAGCAGG | 3130.0 | -11.6 | 1.4 | -8.4 | 6.7E-17 | 2.2E-16 | BT20/MB468 | NUC |
| >MI0003513\|hsa-mir-455&WithFlank&9\|+\|114209428\|114209535@22.44.23&[MIMAT0003150&hsa-miR-455-5p&offsets\|0\|+1;m-232&9\|+\|114209449\|114209470&offsets\|0\|+1]\|TATGTGCCTTTGGACTACATCGT | >MI0003513\|hsa-mir-455&WithFlank&9\|+\|114209428\|114209535 | 22 | 44 | 23 | TATGTGCCTTTGGACTACATCGT | 1144.0 | 13.4 | 1.6 | 8.4 | 3.7E-17 | 2.2E-16 | MB231/MB468 | MITO |
| >hg38_21_+_8433172_8446622_RNA45SN1_45S_50ntFlanks@12934.12952.19\|CTAAACCATTCGTAGACGA | >hg38_21_+_8433172_8446622_RNA45SN1_45S_50ntFlanks | 12934 | 12952 | 19 | CTAAACCATTCGTAGACGA | 432.2 | -12.5 | 1.5 | -8.4 | 3.7E-17 | 2.2E-16 | MB231/MB468 | MITO |
| >MI0000279\|hsa-mir-196a-2&WithFlank&12\|+\|53991732\|53991853@31.53.23&[MIMAT0000226&hsa-miR-196a-5p&offsets\|0\|+1;m-261&12\|+\|53991762\|53991783&offsets\|0\|+1]\|TAGGTAGTTTCATGTTGTTGGGA | >MI0000279\|hsa-mir-196a-2&WithFlank&12\|+\|53991732\|53991853 | 31 | 53 | 23 | TAGGTAGTTTCATGTTGTTGGGA | 365.3 | 12.1 | 1.4 | 8.7 | 2.9E-18 | 2.2E-16 | BT20/MB231 | MITO |
| >hg38_21_+_8433172_8446622_RNA45SN1_45S_50ntFlanks@12008.12037.30\|CCGCCGGTGAAATACCACTACTCTGATCGT | >hg38_21_+_8433172_8446622_RNA45SN1_45S_50ntFlanks | 12008 | 12037 | 30 | CCGCCGGTGAAATACCACTACTCTGATCGT | 394.9 | -12.5 | 1.5 | -8.5 | 2.3E-17 | 2.2E-16 | MB231/MB468 | MP |
| >hg38_21_+_8433172_8446622_RNA45SN1_45S_50ntFlanks@12627.12659.33\|ATGGGGCGAAGCTACCATCTGTGGGATTATGAC | >hg38_21_+_8433172_8446622_RNA45SN1_45S_50ntFlanks | 12627 | 12659 | 33 | ATGGGGCGAAGCTACCATCTGTGGGATTATGAC | 15573.0 | -12.6 | 1.5 | -8.4 | 3.7E-17 | 2.2E-16 | MB231/MB468 | MITO |

| | | | | | | | | | | | | | |
|---|---|---|---|---|---|---|---|---|---|---|---|---|---|
| >hg38_21_+_8433172_8446622_RNA45SN1_45S_50ntFlanks@11845.11873.29\|GGAACGGGCTTGGCGGAATCAGCGGGGAA | >hg38_21_+_8433172_8446622_RNA45SN1_45S_50ntFlanks | 11845 | 11873 | 29 | GGAACGGGCTTGGCGGAATCAGCGGGGAA | 345.2 | -12.2 | 1.5 | -8.4 | 3.8E-17 | 2.2E-16 | MB231/MB468 | MITO |
| >hg38_21_+_8433172_8446622_RNA45SN1_45S_50ntFlanks@5493.5514.22\|GAAGACGGTCGAACTTGACTAT | >hg38_21_+_8433172_8446622_RNA45SN1_45S_50ntFlanks | 5493 | 5514 | 22 | GAAGACGGTCGAACTTGACTAT | 571.2 | -13.0 | 1.5 | -8.5 | 2.3E-17 | 2.2E-16 | MB231/MB468 | MP |
| >hg38_21_+_8433172_8446622_RNA45SN1_45S_50ntFlanks@5071.5110.40\|TTAATTCCGATAACGAACGAGACTCTGGCATGCTAACTAG | >hg38_21_+_8433172_8446622_RNA45SN1_45S_50ntFlanks | 5071 | 5110 | 40 | TTAATTCCGATAACGAACGAGACTCTGGCATGCTAACTAG | 517.2 | -12.8 | 1.5 | -8.4 | 3.8E-17 | 2.2E-16 | MB231/MB468 | MITO |
| >MI0000650\|hsa-mir-200c&WithFlank&12\|+\|6963693\|6963772@49.72.24&[MIMAT0000617&hsa-miR-200c-3p&offsets\|-1\|0;m-54&12\|+\|6963742\|6963764&offsets\|-1\|0]\|CTAATACTGCCGGGTAATGATGGA | >MI0000650\|hsa-mir-200c&WithFlank&12\|+\|6963693\|6963772 | 49 | 72 | 24 | CTAATACTGCCGGGTAATGATGGA | 379.3 | 12.2 | 1.4 | 8.7 | 3.0E-18 | 2.2E-16 | BT20/MB231 | MITO |
| >hg38_21_+_8433172_8446622_RNA45SN1_45S_50ntFlanks@9621.9648.28\|TCTAGTAGCTGGTTCCCTCCGAAGTTTC | >hg38_21_+_8433172_8446622_RNA45SN1_45S_50ntFlanks | 9621 | 9648 | 28 | TCTAGTAGCTGGTTCCCTCCGAAGTTTC | 1231.5 | -9.8 | 1.2 | -8.4 | 3.8E-17 | 2.2E-16 | MB231/MB468 | MITO |
| >hg38_21_+_8433172_8446622_RNA45SN1_45S_50ntFlanks@9033.9056.24\|CTCGTGGGGGGGCCGGGCCACCCC | >hg38_21_+_8433172_8446622_RNA45SN1_45S_50ntFlanks | 9033 | 9056 | 24 | CTCGTGGGGGGGCCGGGCCACCCC | 944.8 | -13.7 | 1.6 | -8.4 | 3.8E-17 | 2.2E-16 | MB231/MB468 | MITO |
| >hg38_21_+_8433172_8446622_RNA45SN1_45S_50ntFlanks@4721.4767.47\|TTCATTAATCAAGAACGAAAGTCGGAGGTTCGAAGACGATCAGATAC | >hg38_21_+_8433172_8446622_RNA45SN1_45S_50ntFlanks | 4721 | 4767 | 47 | TTCATTAATCAAGAACGAAAGTCGGAGGTTCGAAGACGATCAGATAC | 263.5 | 12.0 | 1.4 | 8.5 | 2.4E-17 | 2.3E-16 | MB231/MB468 | CYTO |
| >hg38_21_+_8433172_8446622_RNA45SN1_45S_50ntFlanks@12974.12991.18\|CGTACGTAGCAGAGCAGC | >hg38_21_+_8433172_8446622_RNA45SN1_45S_50ntFlanks | 12974 | 12991 | 18 | CGTACGTAGCAGAGCAGC | 2854.0 | -8.4 | 1.0 | -8.4 | 3.8E-17 | 2.3E-16 | MB231/MB468 | MITO |
| >hg38_21_+_8433172_8446622_RNA45SN1_45S_50ntFlanks@3988.4024.37\|CTTTGGTGACTCTAGATAACCTCGGGCCGATCGCACG | >hg38_21_+_8433172_8446622_RNA45SN1_45S_50ntFlanks | 3988 | 4024 | 37 | CTTTGGTGACTCTAGATAACCTCGGGCCGATCGCACG | 1113.2 | -14.0 | 1.7 | -8.5 | 2.4E-17 | 2.3E-16 | MB231/MB468 | MP |
| >hg38_21_+_8433172_8446622_RNA45SN1_45S_50ntFlanks@8472.8497.26\|CGGCCCGGCGGATCTTTCCCGCCCCC | >hg38_21_+_8433172_8446622_RNA45SN1_45S_50ntFlanks | 8472 | 8497 | 26 | CGGCCCGGCGGATCTTTCCCGCCCCC | 708.1 | -11.6 | 1.4 | -8.4 | 3.0E-17 | 2.3E-16 | BT20/MB468 | MITO |
| >hg38_21_+_8433172_8446622_RNA45SN1_45S_50ntFlanks@8465.8497.33\|GTGTCGGCGGCCCGGCGGATCTTTCCCGCCCCC | >hg38_21_+_8433172_8446622_RNA45SN1_45S_50ntFlanks | 8465 | 8497 | 33 | GTGTCGGCGGCCCGGCGGATCTTTCCCGCCCCC | 1194.4 | -14.0 | 1.7 | -8.4 | 3.8E-17 | 2.3E-16 | MB231/MB468 | MITO |
| >hg38_21_+_8433172_8446622_RNA45SN1_45S_50ntFlanks@11710.11728.19\|CGCGGGTAAACGGCGGGAG | >hg38_21_+_8433172_8446622_RNA45SN1_45S_50ntFlanks | 11710 | 11728 | 19 | CGCGGGTAAACGGCGGGAG | 485.7 | -12.7 | 1.5 | -8.4 | 3.8E-17 | 2.3E-16 | MB231/MB468 | MITO |
| >hg38_21_+_8433172_8446622_RNA45SN1_45S_50ntFlanks@12785.12817.33\|CCCCGCGCGCGCGGGAGGGCGCGTGCCCCGCCG | >hg38_21_+_8433172_8446622_RNA45SN1_45S_50ntFlanks | 12785 | 12817 | 33 | CCCCGCGCGCGCGGGAGGGCGCGTGCCCCGCCG | 476.4 | -12.7 | 1.5 | -8.4 | 3.9E-17 | 2.3E-16 | MB231/MB468 | MITO |
| >hg38_21_+_8433172_8446622_RNA45SN1_45S_50ntFlanks@11534.11551.18\|GTTCCGCGCGGCGCCTCG | >hg38_21_+_8433172_8446622_RNA45SN1_45S_50ntFlanks | 11534 | 11551 | 18 | GTTCCGCGCGGCGCCTCG | 4663.7 | -8.2 | 1.0 | -8.4 | 3.0E-17 | 2.3E-16 | MB231/MB468 | NUC |
| >MI0000298\|hsa-mir-221&WithFlank&X\|-\|45746151\|45746272@71.93.23&[MIMAT0000278&hsa-miR-221-3p&offsets\|0\|0;m-91&X\|-\|45746181\|45746202&offsets\|0\|+1]\|AGCTACATTGTCTGCTGGGTTTC | >MI0000298\|hsa-mir-221&WithFlank&X\|-\|45746151\|45746272 | 71 | 93 | 23 | AGCTACATTGTCTGCTGGGTTTC | 36164.5 | 5.8 | 0.7 | 8.4 | 3.0E-17 | 2.3E-16 | MB231/MB468 | NUC |
| >hg38_21_+_8433172_8446622_RNA45SN1_45S_50ntFlanks@4445.4471.27\|CTTGCCTCTCGGCGCCCCCTCGATGCT | >hg38_21_+_8433172_8446622_RNA45SN1_45S_50ntFlanks | 4445 | 4471 | 27 | CTTGCCTCTCGGCGCCCCCTCGATGCT | 862.2 | -13.6 | 1.6 | -8.5 | 2.4E-17 | 2.3E-16 | MB231/MB468 | MP |
| >hg38_21_+_8433172_8446622_RNA45SN1_45S_50ntFlanks@5482.5514.33\|CGGAGCGCTGAGAAGACGGTCGAACTTGACTAT | >hg38_21_+_8433172_8446622_RNA45SN1_45S_50ntFlanks | 5482 | 5514 | 33 | CGGAGCGCTGAGAAGACGGTCGAACTTGACTAT | 633.4 | -13.1 | 1.6 | -8.4 | 3.9E-17 | 2.3E-16 | MB231/MB468 | MITO |
| >MI0000266\|hsa-mir-10a&WithFlank&17\|-\|48579832\|48579953@28.48.21(+1C)&[MIMAT0000253&hsa-miR-10a-5p&offsets\|0\|-2(+1C);m-18&17\|-\|48579905\|48579926&offsets\|0\|-1(+1C)]\|TACCCTGTAGATCCGAATTTGC | >MI0000266\|hsa-mir-10a&WithFlank&17\|-\|48579832\|48579953 | 28 | 48 | 21(+1C) | TACCCTGTAGATCCGAATTTGC | 948.0 | 13.1 | 1.6 | 8.4 | 3.9E-17 | 2.3E-16 | MB231/MB468 | MITO |
| >hg38_21_+_8433172_8446622_RNA45SN1_45S_50ntFlanks@9029.9059.31\|TCTCCTCGTGGGGGGGCCGGGCCACCCCTCC | >hg38_21_+_8433172_8446622_RNA45SN1_45S_50ntFlanks | 9029 | 9059 | 31 | TCTCCTCGTGGGGGGGCCGGGCCACCCCTCC | 630.6 | -13.1 | 1.6 | -8.4 | 3.9E-17 | 2.3E-16 | MB231/MB468 | MITO |
| >hg38_21_+_8433172_8446622_RNA45SN1_45S_50ntFlanks@12457.12487.31\|CAAGCGTTGGATTGTTCACCCACTAATAGGG | >hg38_21_+_8433172_8446622_RNA45SN1_45S_50ntFlanks | 12457 | 12487 | 31 | CAAGCGTTGGATTGTTCACCCACTAATAGGG | 3053.3 | -8.7 | 1.0 | -8.4 | 3.0E-17 | 2.3E-16 | MB231/MB468 | NUC |
| >hg38_21_+_8433172_8446622_RNA45SN1_45S_50ntFlanks@9739.9758.20\|AAACGATCTCAACCTATTCT | >hg38_21_+_8433172_8446622_RNA45SN1_45S_50ntFlanks | 9739 | 9758 | 20 | AAACGATCTCAACCTATTCT | 2157.7 | -3.6 | 0.4 | -8.4 | 3.0E-17 | 2.3E-16 | MB231/MB468 | NUC |
| >hg38_21_+_8433172_8446622_RNA45SN1_45S_50ntFlanks@6737.6771.35\|GATCATCGACACTTCGAACGCACTTGCGGCCCCGG | >hg38_21_+_8433172_8446622_RNA45SN1_45S_50ntFlanks | 6737 | 6771 | 35 | GATCATCGACACTTCGAACGCACTTGCGGCCCCGG | 545.0 | -12.1 | 1.4 | -8.4 | 3.0E-17 | 2.3E-16 | MB231/MB468 | NUC |
| >MI0000650\|hsa-mir-200c&WithFlank&12\|+\|6963693\|6963772@50.71.22&[MIMAT0000617&hsa-miR-200c-3p&offsets\|0\|-1;m-54&12\|+\|6963742\|6963764&offsets\|0\|-1]\|TAATACTGCCGGGTAATGATGG | >MI0000650\|hsa-mir-200c&WithFlank&12\|+\|6963693\|6963772 | 50 | 71 | 22 | TAATACTGCCGGGTAATGATGG | 17070.4 | 8.6 | 1.0 | 9.0 | 3.4E-19 | 2.3E-16 | BT20/MB231 | MP |
| >hg38_21_+_8433172_8446622_RNA45SN1_45S_50ntFlanks@11661.11685.25\|TGATTTCTGCCCAGTGCTCTGAATG | >hg38_21_+_8433172_8446622_RNA45SN1_45S_50ntFlanks | 11661 | 11685 | 25 | TGATTTCTGCCCAGTGCTCTGAATG | 15083.6 | -10.6 | 1.3 | -8.3 | 7.1E-17 | 2.3E-16 | BT20/MB468 | NUC |
| >hg38_21_+_8433172_8446622_RNA45SN1_45S_50ntFlanks@10866.10904.39\|CTAAGGGCTGGGTCGGTCGGGCTGGGGCGCGAAGCGGGG | >hg38_21_+_8433172_8446622_RNA45SN1_45S_50ntFlanks | 10866 | 10904 | 39 | CTAAGGGCTGGGTCGGTCGGGCTGGGGCGCGAAGCGGGG | 469.6 | -12.7 | 1.5 | -8.4 | 4.0E-17 | 2.3E-16 | MB231/MB468 | MITO |
| >hg38_21_+_8433172_8446622_RNA45SN1_45S_50ntFlanks@4734.4751.18\|AACGAAAGTCGGAGGTTC | >hg38_21_+_8433172_8446622_RNA45SN1_45S_50ntFlanks | 4734 | 4751 | 18 | AACGAAAGTCGGAGGTTC | 1777.2 | -6.0 | 0.7 | -8.3 | 7.1E-17 | 2.3E-16 | BT20/MB468 | NUC |
| >MI0000079\|hsa-mir-23a&WithFlank&19\|-\|13836581\|13836665@54.73.20(+1U)&[MIMAT0000078&hsa-miR-23a-3p&offsets\|+3\|+2(+1U);m-31&19\|-\|13836595\|13836615&offsets\|+3\|+2(+1U)]\|ACATTGCCAGGGATTTCCAAT | >MI0000079\|hsa-mir-23a&WithFlank&19\|-\|13836581\|13836665 | 54 | 73 | 20(+1U) | ACATTGCCAGGGATTTCCAAT | 250.8 | 10.8 | 1.3 | 8.3 | 7.1E-17 | 2.3E-16 | BT20/MB468 | NUC |
| >hg38_21_+_8433172_8446622_RNA45SN1_45S_50ntFlanks@10806.10826.21\|AACAATGTAGGTAAGGGAAGT | >hg38_21_+_8433172_8446622_RNA45SN1_45S_50ntFlanks | 10806 | 10826 | 21 | AACAATGTAGGTAAGGGAAGT | 374.2 | -6.4 | 0.8 | -8.5 | 2.5E-17 | 2.3E-16 | MB231/MB468 | CYTO |
| >hg38_21_+_8433172_8446622_RNA45SN1_45S_50ntFlanks@7980.7998.19\|CCTCAGATCAGACGTGGCG | >hg38_21_+_8433172_8446622_RNA45SN1_45S_50ntFlanks | 7980 | 7998 | 19 | CCTCAGATCAGACGTGGCG | 249.3 | -11.8 | 1.4 | -8.5 | 2.5E-17 | 2.3E-16 | MB231/MB468 | MP |
| >hg38_21_+_8433172_8446622_RNA45SN1_45S_50ntFlanks@8379.8405.27\|TCAAGAGGGCGTGAAACCGTTAAGAGG | >hg38_21_+_8433172_8446622_RNA45SN1_45S_50ntFlanks | 8379 | 8405 | 27 | TCAAGAGGGCGTGAAACCGTTAAGAGG | 328.5 | -12.2 | 1.4 | -8.5 | 2.5E-17 | 2.3E-16 | MB231/MB468 | MP |
| >hg38_21_+_8433172_8446622_RNA45SN1_45S_50ntFlanks@6649.6681.33\|TACGACTCTTAGCGGTGGATCACTCGGCTCGTG | >hg38_21_+_8433172_8446622_RNA45SN1_45S_50ntFlanks | 6649 | 6681 | 33 | TACGACTCTTAGCGGTGGATCACTCGGCTCGTG | 692.9 | -3.5 | 0.4 | -8.3 | 7.2E-17 | 2.3E-16 | BT20/MB468 | NUC |
| >hg38_21_+_8433172_8446622_RNA45SN1_45S_50ntFlanks@11583.11605.23\|CTGGTGCGGACCAGGGGAATCCG | >hg38_21_+_8433172_8446622_RNA45SN1_45S_50ntFlanks | 11583 | 11605 | 23 | CTGGTGCGGACCAGGGGAATCCG | 1069.8 | -5.2 | 0.6 | -8.5 | 2.5E-17 | 2.4E-16 | MB231/MB468 | CYTO |
| >hg38_1_-_228634819_228635039_RNA5S12_5S_50ntFlanks@138.171.34\|AGACCGCCTGGGAATACCGGGTGCTGTAGGCTTT | >hg38_1_-_228634819_228635039_RNA5S12_5S_50ntFlanks | 138 | 171 | 34 | AGACCGCCTGGGAATACCGGGTGCTGTAGGCTTT | 2200.1 | 6.9 | 0.8 | 8.5 | 2.5E-17 | 2.4E-16 | MB231/MB468 | CYTO |
| >MI0000066\|hsa-let-7e&WithFlank&19\|+\|51692780\|51692870@14.35.22(+1A)&[MIMAT0000066&hsa-let-7e-5p&offsets\|0\|0(+1A);m-50&19\|+\|51692793\|51692814&offsets\|0\|0(+1A)]\|TGAGGTAGGAGGTTGTATAGTTA | >MI0000066\|hsa-let-7e&WithFlank&19\|+\|51692780\|51692870 | 14 | 35 | 22(+1A) | TGAGGTAGGAGGTTGTATAGTTA | 1449.9 | 8.5 | 1.0 | 8.4 | 4.4E-17 | 2.4E-16 | BT20/MB468 | CYTO |
| >hg38_21_+_8433172_8446622_RNA45SN1_45S_50ntFlanks@12126.12146.21\|CGACCCGCTCCGGGGACAGTG | >hg38_21_+_8433172_8446622_RNA45SN1_45S_50ntFlanks | 12126 | 12146 | 21 | CGACCCGCTCCGGGGACAGTG | 562.3 | -11.9 | 1.4 | -8.5 | 2.5E-17 | 2.4E-16 | MB231/MB468 | CYTO |
| >hg38_21_+_8433172_8446622_RNA45SN1_45S_50ntFlanks@12151.12183.33\|GTGGGGAGTTTGACTGGGGCGGTACACCTGTCA | >hg38_21_+_8433172_8446622_RNA45SN1_45S_50ntFlanks | 12151 | 12183 | 33 | GTGGGGAGTTTGACTGGGGCGGTACACCTGTCA | 419.7 | -12.6 | 1.5 | -8.5 | 2.5E-17 | 2.4E-16 | MB231/MB468 | MP |
| >hg38_21_+_8433172_8446622_RNA45SN1_45S_50ntFlanks@11919.11951.33\|CATGAGAGGTGTAGAATAAGTGGGAGGCCCCCG | >hg38_21_+_8433172_8446622_RNA45SN1_45S_50ntFlanks | 11919 | 11951 | 33 | CATGAGAGGTGTAGAATAAGTGGGAGGCCCCCG | 414.3 | -12.6 | 1.5 | -8.5 | 2.5E-17 | 2.4E-16 | MB231/MB468 | MP |
| >hg38_21_+_8433172_8446622_RNA45SN1_45S_50ntFlanks@12710.12742.33\|CGCGGAGCCTCGGTTGGCCTCGGATAGCCGGTC | >hg38_21_+_8433172_8446622_RNA45SN1_45S_50ntFlanks | 12710 | 12742 | 33 | CGCGGAGCCTCGGTTGGCCTCGGATAGCCGGTC | 389.9 | -12.4 | 1.5 | -8.4 | 4.1E-17 | 2.4E-16 | MB231/MB468 | MITO |
| >MI0000743\|hsa-mir-34c&WithFlank&11\|+\|111513433\|111513521@19.41.23&[MIMAT0000686&hsa-miR-34c-5p&offsets\|0\|0;m-336&11\|+\|111513451\|111513473&offsets\|0\|0]\|AGGCAGTGTAGTTAGCTGATTGC | >MI0000743\|hsa-mir-34c&WithFlank&11\|+\|111513433\|111513521 | 19 | 41 | 23 | AGGCAGTGTAGTTAGCTGATTGC | 468.4 | 12.9 | 1.5 | 8.5 | 2.5E-17 | 2.4E-16 | MB231/MB468 | CYTO |
| >hg38_21_+_8433172_8446622_RNA45SN1_45S_50ntFlanks@10794.10825.32\|CTGGCATGTTGGAACAATGTAGGTAAGGGAAG | >hg38_21_+_8433172_8446622_RNA45SN1_45S_50ntFlanks | 10794 | 10825 | 32 | CTGGCATGTTGGAACAATGTAGGTAAGGGAAG | 878.3 | -9.8 | 1.2 | -8.4 | 4.1E-17 | 2.4E-16 | MB231/MB468 | MITO |
| >MI0000446\|hsa-mir-125b-1&WithFlank&11\|-\|122099751\|122099850@21.39.19;MI0000470\|hsa-mir-125b-2&WithFlank&21\|+\|16590231\|16590331@23.41.19&[MIMAT0000423&hsa-miR-125b-5p&offsets\|0\|-3;m-67&11\|-\|122099809\|122099830&offsets\|0\|-3];[MIMAT0000423_1&hsa-miR-125b-5p&offsets\|0\|-3;m-68&21\|+\|16590253\|16590274&offsets\|0\|-3]\|TCCCTGAGACCCTAACTTG | >MI0000446\|hsa-mir-125b-1&WithFlank&11\|-\|122099751\|122099850 | 21 | 39 | 19 | TCCCTGAGACCCTAACTTG | 1269.8 | 13.5 | 1.6 | 8.4 | 4.1E-17 | 2.4E-16 | MB231/MB468 | MITO |
| >hg38_21_+_8433172_8446622_RNA45SN1_45S_50ntFlanks@4005.4024.20\|AACCTCGGGCCGATCGCACG | >hg38_21_+_8433172_8446622_RNA45SN1_45S_50ntFlanks | 4005 | 4024 | 20 | AACCTCGGGCCGATCGCACG | 1110.4 | -3.8 | 0.4 | -8.4 | 4.4E-17 | 2.4E-16 | BT20/MB468 | CYTO |
| >MI0000299\|hsa-mir-222&WithFlank&X\|-\|45747009\|45747130@75.97.23(+1U)&[MIMAT0000279&hsa-miR-222-3p&offsets\|0\|+2(+1U);m-140&X\|-\|45747033\|45747056&offsets\|0\|-1(+1U)]\|AGCTACATCTGGCTACTGGGTCTT | >MI0000299\|hsa-mir-222&WithFlank&X\|-\|45747009\|45747130 | 75 | 97 | 23(+1U) | AGCTACATCTGGCTACTGGGTCTT | 2650.7 | 6.0 | 0.7 | 8.4 | 3.2E-17 | 2.4E-16 | MB231/MB468 | NUC |
| >hg38_21_+_8433172_8446622_RNA45SN1_45S_50ntFlanks@6784.6803.20\|GCTACGCCTGTCTGAGCGTC | >hg38_21_+_8433172_8446622_RNA45SN1_45S_50ntFlanks | 6784 | 6803 | 20 | GCTACGCCTGTCTGAGCGTC | 2270.4 | -14.2 | 1.6 | -8.9 | 6.4E-19 | 2.4E-16 | BT20/MB468 | MP |
| >hg38_21_+_8433172_8446622_RNA45SN1_45S_50ntFlanks@8357.8377.21\|AAAAGAACTTTGAAGAGAGAG | >hg38_21_+_8433172_8446622_RNA45SN1_45S_50ntFlanks | 8357 | 8377 | 21 | AAAAGAACTTTGAAGAGAGAG | 4767.2 | -6.3 | 0.7 | -8.4 | 3.2E-17 | 2.4E-16 | MB231/MB468 | NUC |
| >hg38_21_+_8433172_8446622_RNA45SN1_45S_50ntFlanks@11874.11909.36\|AGAAGACCCTGTTGAGCTTGACTCTAGTCTGGCACG | >hg38_21_+_8433172_8446622_RNA45SN1_45S_50ntFlanks | 11874 | 11909 | 36 | AGAAGACCCTGTTGAGCTTGACTCTAGTCTGGCACG | 1242.6 | -9.8 | 1.2 | -8.4 | 4.2E-17 | 2.4E-16 | MB231/MB468 | MITO |
| >hg38_21_+_8433172_8446622_RNA45SN1_45S_50ntFlanks@8158.8188.31\|TCGTGGGGGGCCCAAGTCCTTCTGATCGAGG | >hg38_21_+_8433172_8446622_RNA45SN1_45S_50ntFlanks | 8158 | 8188 | 31 | TCGTGGGGGGCCCAAGTCCTTCTGATCGAGG | 382.4 | -12.4 | 1.5 | -8.5 | 2.6E-17 | 2.4E-16 | MB231/MB468 | MP |
| >hg38_21_+_8433172_8446622_RNA45SN1_45S_50ntFlanks@10805.10825.21\|GAACAATGTAGGTAAGGGAAG | >hg38_21_+_8433172_8446622_RNA45SN1_45S_50ntFlanks | 10805 | 10825 | 21 | GAACAATGTAGGTAAGGGAAG | 401.5 | -12.4 | 1.5 | -8.4 | 4.2E-17 | 2.5E-16 | MB231/MB468 | MITO |
| >hg38_21_+_8433172_8446622_RNA45SN1_45S_50ntFlanks@10041.10072.32\|CCATACCCGGCCGTCGCCGGCAGTCGAGAGTG | >hg38_21_+_8433172_8446622_RNA45SN1_45S_50ntFlanks | 10041 | 10072 | 32 | CCATACCCGGCCGTCGCCGGCAGTCGAGAGTG | 391.5 | -12.4 | 1.5 | -8.4 | 4.2E-17 | 2.5E-16 | MB231/MB468 | MITO |
| >hg38_21_+_8433172_8446622_RNA45SN1_45S_50ntFlanks@12973.12991.19\|TCGTACGTAGCAGAGCAGC | >hg38_21_+_8433172_8446622_RNA45SN1_45S_50ntFlanks | 12973 | 12991 | 19 | TCGTACGTAGCAGAGCAGC | 2242.1 | -8.3 | 1.0 | -8.5 | 2.6E-17 | 2.5E-16 | MB231/MB468 | MP |

<table>
<tr><td>>hg38_21_+_8433172_8446622_RNA45SN1_45S_50ntFlanks@7980.8012.33|CCTCAGATCAGACGTGGCGACCCGCTGAATTTA</td><td>>hg38_21_+_8433172_8446622_RNA45SN1_45S_50ntFlanks</td><td>7980</td><td>8012</td><td>33</td><td>CCTCAGATCAGACGTGGCGACCCGCTGAATTTA</td><td>484.5</td><td>-12.8</td><td>1.5</td><td>-8.5</td><td>2.6E-17</td><td>2.5E-16</td><td>MB231/MB468</td><td>MP</td></tr>
<tr><td>>hg38_21_+_8433172_8446622_RNA45SN1_45S_50ntFlanks@10781.10803.23|AGGTGAACAGCCTCTGGCATGTT</td><td>>hg38_21_+_8433172_8446622_RNA45SN1_45S_50ntFlanks</td><td>10781</td><td>10803</td><td>23</td><td>AGGTGAACAGCCTCTGGCATGTT</td><td>263.9</td><td>-11.9</td><td>1.4</td><td>-8.5</td><td>2.6E-17</td><td>2.5E-16</td><td>MB231/MB468</td><td>MP</td></tr>
<tr><td>>hg38_21_+_8433172_8446622_RNA45SN1_45S_50ntFlanks@4736.4759.24|CGAAAGTCGGAGGTTCGAAGACGA</td><td>>hg38_21_+_8433172_8446622_RNA45SN1_45S_50ntFlanks</td><td>4736</td><td>4759</td><td>24</td><td>CGAAAGTCGGAGGTTCGAAGACGA</td><td>302.5</td><td>-12.1</td><td>1.4</td><td>-8.5</td><td>2.6E-17</td><td>2.5E-16</td><td>MB231/MB468</td><td>MP</td></tr>
<tr><td>>hg38_21_+_8433172_8446622_RNA45SN1_45S_50ntFlanks@8110.8143.34|CGGGGCGCGGGACATGTGGCGTACGGAAGACCCG</td><td>>hg38_21_+_8433172_8446622_RNA45SN1_45S_50ntFlanks</td><td>8110</td><td>8143</td><td>34</td><td>CGGGGCGCGGGACATGTGGCGTACGGAAGACCCG</td><td>313.8</td><td>-12.2</td><td>1.4</td><td>-8.5</td><td>2.6E-17</td><td>2.5E-16</td><td>MB231/MB468</td><td>MP</td></tr>
<tr><td>>hg38_21_+_8433172_8446622_RNA45SN1_45S_50ntFlanks@9806.9831.26|CGTGGAATGCGAGTGCCTAGTGGGCC</td><td>>hg38_21_+_8433172_8446622_RNA45SN1_45S_50ntFlanks</td><td>9806</td><td>9831</td><td>26</td><td>CGTGGAATGCGAGTGCCTAGTGGGCC</td><td>535.9</td><td>-12.9</td><td>1.5</td><td>-8.5</td><td>2.6E-17</td><td>2.5E-16</td><td>MB231/MB468</td><td>MP</td></tr>
<tr><td>>hg38_21_+_8433172_8446622_RNA45SN1_45S_50ntFlanks@12073.12095.23|CCGAGGGGCTCTCGCTTCTGGCG</td><td>>hg38_21_+_8433172_8446622_RNA45SN1_45S_50ntFlanks</td><td>12073</td><td>12095</td><td>23</td><td>CCGAGGGGCTCTCGCTTCTGGCG</td><td>2786.0</td><td>-10.5</td><td>1.2</td><td>-8.4</td><td>4.3E-17</td><td>2.5E-16</td><td>MB231/MB468</td><td>MITO</td></tr>
<tr><td>>hg38_21_+_8433172_8446622_RNA45SN1_45S_50ntFlanks@5487.5525.39|CGCTGAGAAGACGGTCGAACTTGACTATCTAGAGGAAGT</td><td>>hg38_21_+_8433172_8446622_RNA45SN1_45S_50ntFlanks</td><td>5487</td><td>5525</td><td>39</td><td>CGCTGAGAAGACGGTCGAACTTGACTATCTAGAGGAAGT</td><td>358.9</td><td>-12.3</td><td>1.5</td><td>-8.4</td><td>4.3E-17</td><td>2.5E-16</td><td>MB231/MB468</td><td>MITO</td></tr>
<tr><td>>MI0000255|hsa-mir-30d&WithFlank&8|-|134804870|134804951@12.35.24(+1U)&[MIMAT0000245&hsa-miR-30d-5p&offsets|0|+2(+1U);m-21&8|-|134804919|134804940&offsets|0|+2(+1U)]|TGTAAACATCCCCGACTGGAAGCTT</td><td>>MI0000255|hsa-mir-30d&WithFlank&8|-|134804870|134804951</td><td>12</td><td>35</td><td>24(+1U)</td><td>TGTAAACATCCCCGACTGGAAGCTT</td><td>1426.1</td><td>7.0</td><td>0.8</td><td>8.5</td><td>2.7E-17</td><td>2.5E-16</td><td>MB231/MB468</td><td>CYTO</td></tr>
<tr><td>>MI0000107|hsa-mir-29b-2&WithFlank&1|-|207802437|207802529@58.80.23;MI0000105|hsa-mir-29b-1&WithFlank&7|-|130877453|130877545@57.79.23&[MIMAT0000100&hsa-miR-29b-3p&offsets|0|0;m-81&1|-|207802450|207802472&offsets|0|0];[MIMAT0000100_1&hsa-miR-29b-3p&offsets|0|0;m-82&7|-|130877467|130877489&offsets|0|0]|TAGCACCATTTGAAATCAGTGTT</td><td>>MI0000107|hsa-mir-29b-2&WithFlank&1|-|207802437|207802529</td><td>58</td><td>80</td><td>23</td><td>TAGCACCATTTGAAATCAGTGTT</td><td>772.3</td><td>9.9</td><td>1.2</td><td>8.4</td><td>3.3E-17</td><td>2.5E-16</td><td>MB231/MB468</td><td>NUC</td></tr>
<tr><td>>hg38_21_+_8433172_8446622_RNA45SN1_45S_50ntFlanks@11161.11188.28|CACGGTCCCCCGCGAGGGGGGCCCGGGC</td><td>>hg38_21_+_8433172_8446622_RNA45SN1_45S_50ntFlanks</td><td>11161</td><td>11188</td><td>28</td><td>CACGGTCCCCCGCGAGGGGGGCCCGGGC</td><td>907.7</td><td>-13.7</td><td>1.6</td><td>-8.5</td><td>2.7E-17</td><td>2.5E-16</td><td>MB231/MB468</td><td>MP</td></tr>
<tr><td>>hg38_21_+_8433172_8446622_RNA45SN1_45S_50ntFlanks@11781.11804.24|CGCATGAATGGATGAACGAGATTC</td><td>>hg38_21_+_8433172_8446622_RNA45SN1_45S_50ntFlanks</td><td>11781</td><td>11804</td><td>24</td><td>CGCATGAATGGATGAACGAGATTC</td><td>7029.3</td><td>-9.3</td><td>1.1</td><td>-8.4</td><td>3.4E-17</td><td>2.5E-16</td><td>MB231/MB468</td><td>NUC</td></tr>
<tr><td>>MI0006444|hsa-mir-1307&WithFlank&10|-|103394247|103394407@87.108.22&[MIMAT0005951&hsa-miR-1307-3p&offsets|+1|+1;m-66&10|-|103394300|103394322&offsets|+1|0]|CTCGGCGTGGCGTCGGTCGTGG</td><td>>MI0006444|hsa-mir-1307&WithFlank&10|-|103394247|103394407</td><td>87</td><td>108</td><td>22</td><td>CTCGGCGTGGCGTCGGTCGTGG</td><td>1461.1</td><td>5.7</td><td>0.7</td><td>8.5</td><td>2.7E-17</td><td>2.5E-16</td><td>MB231/MB468</td><td>CYTO</td></tr>
<tr><td>>hg38_21_+_8433172_8446622_RNA45SN1_45S_50ntFlanks@11170.11188.19|CCGCGAGGGGGGCCCGGGC</td><td>>hg38_21_+_8433172_8446622_RNA45SN1_45S_50ntFlanks</td><td>11170</td><td>11188</td><td>19</td><td>CCGCGAGGGGGGCCCGGGC</td><td>1074.7</td><td>9.8</td><td>1.2</td><td>8.5</td><td>2.7E-17</td><td>2.5E-16</td><td>MB231/MB468</td><td>CYTO</td></tr>
<tr><td>>hg38_21_+_8433172_8446622_RNA45SN1_45S_50ntFlanks@4687.4723.37|ACGGACCAGAGCGAAAGCATTTGCCAAGAATGTTTTC</td><td>>hg38_21_+_8433172_8446622_RNA45SN1_45S_50ntFlanks</td><td>4687</td><td>4723</td><td>37</td><td>ACGGACCAGAGCGAAAGCATTTGCCAAGAATGTTTTC</td><td>1319.1</td><td>-14.1</td><td>1.7</td><td>-8.4</td><td>4.4E-17</td><td>2.5E-16</td><td>MB231/MB468</td><td>MITO</td></tr>
<tr><td>>hg38_21_+_8433172_8446622_RNA45SN1_45S_50ntFlanks@6699.6718.20|GCTAGCTGCGAGAATTAATG</td><td>>hg38_21_+_8433172_8446622_RNA45SN1_45S_50ntFlanks</td><td>6699</td><td>6718</td><td>20</td><td>GCTAGCTGCGAGAATTAATG</td><td>16576.9</td><td>-5.7</td><td>0.7</td><td>-8.5</td><td>2.7E-17</td><td>2.5E-16</td><td>MB231/MB468</td><td>CYTO</td></tr>
<tr><td>>hg38_21_+_8433172_8446622_RNA45SN1_45S_50ntFlanks@6700.6718.19|CTAGCTGCGAGAATTAATG</td><td>>hg38_21_+_8433172_8446622_RNA45SN1_45S_50ntFlanks</td><td>6700</td><td>6718</td><td>19</td><td>CTAGCTGCGAGAATTAATG</td><td>12595.3</td><td>-9.5</td><td>1.1</td><td>-8.4</td><td>4.4E-17</td><td>2.5E-16</td><td>MB231/MB468</td><td>MITO</td></tr>
<tr><td>>hg38_21_+_8433172_8446622_RNA45SN1_45S_50ntFlanks@9202.9223.22|TCTCGGGGCCACGCGCGCGTCC</td><td>>hg38_21_+_8433172_8446622_RNA45SN1_45S_50ntFlanks</td><td>9202</td><td>9223</td><td>22</td><td>TCTCGGGGCCACGCGCGCGTCC</td><td>470.9</td><td>-12.7</td><td>1.5</td><td>-8.5</td><td>2.7E-17</td><td>2.5E-16</td><td>MB231/MB468</td><td>MP</td></tr>
<tr><td>>hg38_21_+_8433172_8446622_RNA45SN1_45S_50ntFlanks@11847.11872.26|AACGGGCTTGGCGGAATCAGCGGGGA</td><td>>hg38_21_+_8433172_8446622_RNA45SN1_45S_50ntFlanks</td><td>11847</td><td>11872</td><td>26</td><td>AACGGGCTTGGCGGAATCAGCGGGGA</td><td>241.7</td><td>-5.9</td><td>0.7</td><td>-8.5</td><td>2.7E-17</td><td>2.5E-16</td><td>MB231/MB468</td><td>CYTO</td></tr>
<tr><td>>MI0000459|hsa-mir-143&WithFlank&5|+|149428912|149429029@67.87.21&[MIMAT0000435&hsa-miR-143-3p&offsets|0|0;m-3&5|+|149428978|149428997&offsets|0|+1]|TGAGATGAAGCACTGTAGCTC</td><td>>MI0000459|hsa-mir-143&WithFlank&5|+|149428912|149429029</td><td>67</td><td>87</td><td>21</td><td>TGAGATGAAGCACTGTAGCTC</td><td>723.4</td><td>-8.0</td><td>0.9</td><td>-8.5</td><td>2.7E-17</td><td>2.5E-16</td><td>MB231/MB468</td><td>MP</td></tr>
<tr><td>>hg38_21_+_8433172_8446622_RNA45SN1_45S_50ntFlanks@9454.9491.38|CCGCGCCGGGGAGGTGGAGCACGAGCGCACGTGTTAGG</td><td>>hg38_21_+_8433172_8446622_RNA45SN1_45S_50ntFlanks</td><td>9454</td><td>9491</td><td>38</td><td>CCGCGCCGGGGAGGTGGAGCACGAGCGCACGTGTTAGG</td><td>279.9</td><td>-12.0</td><td>1.4</td><td>-8.5</td><td>2.7E-17</td><td>2.5E-16</td><td>MB231/MB468</td><td>MP</td></tr>
<tr><td>>hg38_21_+_8433172_8446622_RNA45SN1_45S_50ntFlanks@12636.12659.24|AGCTACCATCTGTGGGATTATGAC</td><td>>hg38_21_+_8433172_8446622_RNA45SN1_45S_50ntFlanks</td><td>12636</td><td>12659</td><td>24</td><td>AGCTACCATCTGTGGGATTATGAC</td><td>7184.4</td><td>-10.2</td><td>1.2</td><td>-8.4</td><td>4.4E-17</td><td>2.6E-16</td><td>MB231/MB468</td><td>MITO</td></tr>
<tr><td>>hg38_21_+_8433172_8446622_RNA45SN1_45S_50ntFlanks@10574.10591.18|AAGCCGGCGGGAGCCCCG</td><td>>hg38_21_+_8433172_8446622_RNA45SN1_45S_50ntFlanks</td><td>10574</td><td>10591</td><td>18</td><td>AAGCCGGCGGGAGCCCCG</td><td>442.5</td><td>-12.6</td><td>1.5</td><td>-8.4</td><td>4.4E-17</td><td>2.6E-16</td><td>MB231/MB468</td><td>MITO</td></tr>
<tr><td>>hg38_21_+_8433172_8446622_RNA45SN1_45S_50ntFlanks@5492.5516.25|AGAAGACGGTCGAACTTGACTATCT</td><td>>hg38_21_+_8433172_8446622_RNA45SN1_45S_50ntFlanks</td><td>5492</td><td>5516</td><td>25</td><td>AGAAGACGGTCGAACTTGACTATCT</td><td>11230.5</td><td>5.5</td><td>0.7</td><td>8.5</td><td>2.7E-17</td><td>2.6E-16</td><td>MB231/MB468</td><td>CYTO</td></tr>
<tr><td>>hg38_21_+_8433172_8446622_RNA45SN1_45S_50ntFlanks@485.524.40|GATGTGGTGACGTCGTGCTCTCCCGGGCCGGGTCCGAGCC</td><td>>hg38_21_+_8433172_8446622_RNA45SN1_45S_50ntFlanks</td><td>485</td><td>524</td><td>40</td><td>GATGTGGTGACGTCGTGCTCTCCCGGGCCGGGTCCGAGCC</td><td>388.9</td><td>11.4</td><td>1.4</td><td>8.3</td><td>7.9E-17</td><td>2.6E-16</td><td>BT20/MB468</td><td>NUC</td></tr>
<tr><td>>MI0000086|hsa-mir-28&WithFlank&3|+|188688775|188688872@60.80.21(+1U)&[MIMAT0004502&hsa-miR-28-3p&offsets|0|-1(+1U);m-28&3|+|188688834|188688855&offsets|0|-1(+1U)]|CACTAGATTGTGAGCTCCTGGT</td><td>>MI0000086|hsa-mir-28&WithFlank&3|+|188688775|188688872</td><td>60</td><td>80</td><td>21(+1U)</td><td>CACTAGATTGTGAGCTCCTGGT</td><td>382.5</td><td>12.6</td><td>1.5</td><td>8.5</td><td>2.8E-17</td><td>2.6E-16</td><td>MB231/MB468</td><td>CYTO</td></tr>
<tr><td>>MI0000086|hsa-mir-28&WithFlank&3|+|188688775|188688872@60.80.21(+1G)&[MIMAT0004502&hsa-miR-28-3p&offsets|0|-1(+1G);m-28&3|+|188688834|188688855&offsets|0|-1(+1G)]|CACTAGATTGTGAGCTCCTGGG</td><td>>MI0000086|hsa-mir-28&WithFlank&3|+|188688775|188688872</td><td>60</td><td>80</td><td>21(+1G)</td><td>CACTAGATTGTGAGCTCCTGGG</td><td>1324.9</td><td>12.0</td><td>1.4</td><td>8.4</td><td>4.8E-17</td><td>2.6E-16</td><td>BT20/MB468</td><td>CYTO</td></tr>
<tr><td>>hg38_21_+_8433172_8446622_RNA45SN1_45S_50ntFlanks@10820.10840.21|GGGAAGTCGGCAAGCCGGATC</td><td>>hg38_21_+_8433172_8446622_RNA45SN1_45S_50ntFlanks</td><td>10820</td><td>10840</td><td>21</td><td>GGGAAGTCGGCAAGCCGGATC</td><td>1691.4</td><td>-5.7</td><td>0.7</td><td>-8.4</td><td>3.4E-17</td><td>2.6E-16</td><td>MB231/MB468</td><td>NUC</td></tr>
<tr><td>>MI0000746|hsa-mir-99b&WithFlank&19|+|51692606|51692687@51.71.21&[MIMAT0004678&hsa-miR-99b-3p&offsets|0|-1;m-151&19|+|51692656|51692678&offsets|0|-2]|CAAGCTCGTGTCTGTGGGTCC</td><td>>MI0000746|hsa-mir-99b&WithFlank&19|+|51692606|51692687</td><td>51</td><td>71</td><td>21</td><td>CAAGCTCGTGTCTGTGGGTCC</td><td>1166.5</td><td>8.2</td><td>1.0</td><td>8.4</td><td>4.8E-17</td><td>2.6E-16</td><td>BT20/MB468</td><td>CYTO</td></tr>
<tr><td>>hg38_21_+_8433172_8446622_RNA45SN1_45S_50ntFlanks@10805.10822.18|GAACAATGTAGGTAAGGG</td><td>>hg38_21_+_8433172_8446622_RNA45SN1_45S_50ntFlanks</td><td>10805</td><td>10822</td><td>18</td><td>GAACAATGTAGGTAAGGG</td><td>390.3</td><td>-12.4</td><td>1.5</td><td>-8.4</td><td>4.5E-17</td><td>2.6E-16</td><td>MB231/MB468</td><td>MITO</td></tr>
<tr><td>>hg38_21_+_8433172_8446622_RNA45SN1_45S_50ntFlanks@11207.11232.26|GGCGGCGACTCTGGACGCGAGCCGGG</td><td>>hg38_21_+_8433172_8446622_RNA45SN1_45S_50ntFlanks</td><td>11207</td><td>11232</td><td>26</td><td>GGCGGCGACTCTGGACGCGAGCCGGG</td><td>359.1</td><td>-12.3</td><td>1.5</td><td>-8.4</td><td>4.5E-17</td><td>2.6E-16</td><td>MB231/MB468</td><td>MITO</td></tr>
<tr><td>>hg38_21_+_8433172_8446622_RNA45SN1_45S_50ntFlanks@3853.3898.46|AACTGTGGTAATTCTAGAGCTAATACATGCCGACGGGCGCTGACCC</td><td>>hg38_21_+_8433172_8446622_RNA45SN1_45S_50ntFlanks</td><td>3853</td><td>3898</td><td>46</td><td>AACTGTGGTAATTCTAGAGCTAATACATGCCGACGGGCGCTGACCC</td><td>796.8</td><td>-13.4</td><td>1.6</td><td>-8.4</td><td>4.5E-17</td><td>2.6E-16</td><td>MB231/MB468</td><td>MITO</td></tr>
<tr><td>>hg38_21_+_8433172_8446622_RNA45SN1_45S_50ntFlanks@10664.10711.48|CTTGGAAAGCGTCGCGGTTCCGGCGGCGTCCGGTGAGCTCTCGCTGGC</td><td>>hg38_21_+_8433172_8446622_RNA45SN1_45S_50ntFlanks</td><td>10664</td><td>10711</td><td>48</td><td>CTTGGAAAGCGTCGCGGTTCCGGCGGCGTCCGGTGAGCTCTCGCTGGC</td><td>830.9</td><td>-13.5</td><td>1.6</td><td>-8.4</td><td>4.6E-17</td><td>2.6E-16</td><td>MB231/MB468</td><td>MITO</td></tr>
<tr><td>>hg38_21_+_8433172_8446622_RNA45SN1_45S_50ntFlanks@8114.8136.23|GCGCGGGACATGTGGCGTACGGA</td><td>>hg38_21_+_8433172_8446622_RNA45SN1_45S_50ntFlanks</td><td>8114</td><td>8136</td><td>23</td><td>GCGCGGGACATGTGGCGTACGGA</td><td>670.0</td><td>-13.2</td><td>1.6</td><td>-8.4</td><td>4.6E-17</td><td>2.6E-16</td><td>MB231/MB468</td><td>MITO</td></tr>
<tr><td>>hg38_21_+_8433172_8446622_RNA45SN1_45S_50ntFlanks@11874.11905.32|AGAAGACCCTGTTGAGCTTGACTCTAGTCTGG</td><td>>hg38_21_+_8433172_8446622_RNA45SN1_45S_50ntFlanks</td><td>11874</td><td>11905</td><td>32</td><td>AGAAGACCCTGTTGAGCTTGACTCTAGTCTGG</td><td>250.0</td><td>-6.7</td><td>0.8</td><td>-8.5</td><td>2.8E-17</td><td>2.6E-16</td><td>MB231/MB468</td><td>CYTO</td></tr>
<tr><td>>MI0000093|hsa-mir-92a-1&WithFlank&13|+|91351308|91351397@54.75.22;MI0000094|hsa-mir-92a-2&WithFlank&X|-|134169532|134169618@54.75.22&[MIMAT0000092&hsa-miR-92a-3p&offsets|0|0;m-9&13|+|91351361|91351382&offsets|0|0];[MIMAT0000092_1&hsa-miR-92a-3p&offsets|0|0;m-11&X|-|134169544|134169565&offsets|0|0]|TATTGCACTTGTCCCGGCCTGT</td><td>>MI0000093|hsa-mir-92a-1&WithFlank&13|+|91351308|91351397</td><td>54</td><td>75</td><td>22</td><td>TATTGCACTTGTCCCGGCCTGT</td><td>40107.1</td><td>5.7</td><td>0.7</td><td>8.4</td><td>3.5E-17</td><td>2.7E-16</td><td>BT20/MB468</td><td>MITO</td></tr>
<tr><td>>am_trna128_GlyGCC_6_-_27870686_27870756@1.33.33,trna18_GlyGCC_16_+_70822597_70822667@1.33.33,trna19_GlyGCC_16_+_70823410_70823480@1.33.33,trna19_GlyGCC_2_-_157257659_157257729@1.33.33,trna24_GlyGCC_16_-_70812942_70813012@1.33.33,trna25_GlyGCC_16_-_70812114_70812184@1.33.33,trna5_GlyGCC_17_+_8029064_8029134@1.33.33,trna68_GlyGCC_1_-_161493637_161493707@1.33.33|GCATTGGTGGTTCAGTGGTAGAATTCTCGCCTG</td><td>>am_trna128_GlyGCC_6_-_27870686_27870756</td><td>1</td><td>33</td><td>33</td><td>GCATTGGTGGTTCAGTGGTAGAATTCTCGCCTG</td><td>3783.1</td><td>5.4</td><td>0.6</td><td>8.4</td><td>4.9E-17</td><td>2.7E-16</td><td>BT20/MB468</td><td>CYTO</td></tr>
<tr><td>>hg38_21_+_8433172_8446622_RNA45SN1_45S_50ntFlanks@12617.12653.37|TGAGGAGCCAATGGGGCGAAGCTACCATCTGTGGGAT</td><td>>hg38_21_+_8433172_8446622_RNA45SN1_45S_50ntFlanks</td><td>12617</td><td>12653</td><td>37</td><td>TGAGGAGCCAATGGGGCGAAGCTACCATCTGTGGGAT</td><td>1315.2</td><td>-10.4</td><td>1.2</td><td>-8.4</td><td>4.6E-17</td><td>2.7E-16</td><td>MB231/MB468</td><td>MITO</td></tr>
<tr><td>>hg38_1_-_228634819_228635039_RNA5S12_5S_50ntFlanks@139.170.32|GACCGCCTGGGAATACCGGGTGCTGTAGGCTT</td><td>>hg38_1_-_228634819_228635039_RNA5S12_5S_50ntFlanks</td><td>139</td><td>170</td><td>32</td><td>GACCGCCTGGGAATACCGGGTGCTGTAGGCTT</td><td>1570.4</td><td>6.5</td><td>0.8</td><td>8.5</td><td>2.9E-17</td><td>2.7E-16</td><td>MB231/MB468</td><td>CYTO</td></tr>
<tr><td>>MI0000650|hsa-mir-200c&WithFlank&12|+|6963693|6963772@50.69.20&[MIMAT0000617&hsa-miR-200c-3p&offsets|0|-3;m-54&12|+|6963742|6963764&offsets|0|-3]|TAATACTGCCGGGTAATGAT</td><td>>MI0000650|hsa-mir-200c&WithFlank&12|+|6963693|6963772</td><td>50</td><td>69</td><td>20</td><td>TAATACTGCCGGGTAATGAT</td><td>3008.5</td><td>9.9</td><td>1.2</td><td>8.4</td><td>4.9E-17</td><td>2.7E-16</td><td>BT20/MB468</td><td>CYTO</td></tr>
<tr><td>>hg38_21_+_8433172_8446622_RNA45SN1_45S_50ntFlanks@3849.3866.18|GGATAACTGTGGTAATTC</td><td>>hg38_21_+_8433172_8446622_RNA45SN1_45S_50ntFlanks</td><td>3849</td><td>3866</td><td>18</td><td>GGATAACTGTGGTAATTC</td><td>335.4</td><td>4.6</td><td>0.5</td><td>8.4</td><td>4.9E-17</td><td>2.7E-16</td><td>BT20/MB468</td><td>CYTO</td></tr>
<tr><td>>MI0003560|hsa-mir-92b&WithFlank&1|+|155195171|155195278@67.87.21(+1A)&[MIMAT0003218&hsa-miR-92b-3p&offsets|0|-1(+1A);m-115&1|+|155195237|155195258&offsets|0|-1(+1A)]|TATTGCACTCGTCCCGGCCTCA</td><td>>MI0003560|hsa-mir-92b&WithFlank&1|+|155195171|155195278</td><td>67</td><td>87</td><td>21(+1A)</td><td>TATTGCACTCGTCCCGGCCTCA</td><td>1427.0</td><td>9.5</td><td>1.1</td><td>8.4</td><td>3.5E-17</td><td>2.7E-16</td><td>BT20/MB468</td><td>MITO</td></tr>
<tr><td>>hg38_21_+_8433172_8446622_RNA45SN1_45S_50ntFlanks@10511.10533.23|ATCCGGAGTGGCGGAGATGGGCG</td><td>>hg38_21_+_8433172_8446622_RNA45SN1_45S_50ntFlanks</td><td>10511</td><td>10533</td><td>23</td><td>ATCCGGAGTGGCGGAGATGGGCG</td><td>674.0</td><td>-13.2</td><td>1.6</td><td>-8.4</td><td>4.7E-17</td><td>2.7E-16</td><td>MB231/MB468</td><td>MITO</td></tr>
<tr><td>>hg38_21_+_8433172_8446622_RNA45SN1_45S_50ntFlanks@12072.12096.25|CCCGAGGGGCTCTCGCTTCTGGCGC</td><td>>hg38_21_+_8433172_8446622_RNA45SN1_45S_50ntFlanks</td><td>12072</td><td>12096</td><td>25</td><td>CCCGAGGGGCTCTCGCTTCTGGCGC</td><td>374.8</td><td>-12.3</td><td>1.5</td><td>-8.4</td><td>4.7E-17</td><td>2.7E-16</td><td>MB231/MB468</td><td>MITO</td></tr>
<tr><td>>hg38_21_+_8433172_8446622_RNA45SN1_45S_50ntFlanks@6772.6797.26|GTTCCTCCCGGGGCTACGCCTGTCTG</td><td>>hg38_21_+_8433172_8446622_RNA45SN1_45S_50ntFlanks</td><td>6772</td><td>6797</td><td>26</td><td>GTTCCTCCCGGGGCTACGCCTGTCTG</td><td>924.9</td><td>-9.8</td><td>1.2</td><td>-8.4</td><td>4.7E-17</td><td>2.7E-16</td><td>MB231/MB468</td><td>MITO</td></tr>
<tr><td>>hg38_21_+_8433172_8446622_RNA45SN1_45S_50ntFlanks@3939.3959.21|AACCCGGTCAGCCCCTCTCCG</td><td>>hg38_21_+_8433172_8446622_RNA45SN1_45S_50ntFlanks</td><td>3939</td><td>3959</td><td>21</td><td>AACCCGGTCAGCCCCTCTCCG</td><td>1164.4</td><td>-8.6</td><td>1.0</td><td>-8.5</td><td>2.9E-17</td><td>2.7E-16</td><td>MB231/MB468</td><td>CYTO</td></tr>
<tr><td>>hg38_21_+_8433172_8446622_RNA45SN1_45S_50ntFlanks@12274.12306.33|CAGTACGAATACAGACCGTGAAAGCGGGGCCTC</td><td>>hg38_21_+_8433172_8446622_RNA45SN1_45S_50ntFlanks</td><td>12274</td><td>12306</td><td>33</td><td>CAGTACGAATACAGACCGTGAAAGCGGGGCCTC</td><td>379.9</td><td>-12.4</td><td>1.5</td><td>-8.5</td><td>2.9E-17</td><td>2.7E-16</td><td>MB231/MB468</td><td>MP</td></tr>
</table>

| | | | | | | | | | | | | | |
|---|---|---|---|---|---|---|---|---|---|---|---|---|---|
| >hg38_21_+_8433172_8446622_RNA45SN1_45S_50ntFlanks@10727.10756.30\|GGAGAGGGTGTAAATCTCGCGCCGGGCCGT | >hg38_21_+_8433172_8446622_RNA45SN1_45S_50ntFlanks | 10727 | 10756 | 30 | GGAGAGGGTGTAAATCTCGCGCCGGGCCGT | 234.9 | -11.7 | 1.4 | -8.5 | 2.9E-17 | 2.7E-16 | MB231/MB468 | MP |
| >hg38_21_+_8433172_8446622_RNA45SN1_45S_50ntFlanks@6744.6771.28\|GACACTTCGAACGCACTTGCGGCCCCGG | >hg38_21_+_8433172_8446622_RNA45SN1_45S_50ntFlanks | 6744 | 6771 | 28 | GACACTTCGAACGCACTTGCGGCCCCGG | 964.7 | -3.3 | 0.4 | -8.4 | 3.6E-17 | 2.7E-16 | BT20/MB468 | MITO |
| >hg38_1_-_228634819_228635039_RNA5S12_5S_50ntFlanks@147.171.25\|GGGAATACCGGGTGCTGTAGGCTTT | >hg38_1_-_228634819_228635039_RNA5S12_5S_50ntFlanks | 147 | 171 | 25 | GGGAATACCGGGTGCTGTAGGCTTT | 4048.4 | 5.6 | 0.7 | 8.4 | 5.0E-17 | 2.7E-16 | BT20/MB468 | CYTO |
| >hg38_21_+_8433172_8446622_RNA45SN1_45S_50ntFlanks@3838.3873.36\|CTCTCCTACTTGGATAACTGTGGTAATTCTAGAGCT | >hg38_21_+_8433172_8446622_RNA45SN1_45S_50ntFlanks | 3838 | 3873 | 36 | CTCTCCTACTTGGATAACTGTGGTAATTCTAGAGCT | 419.5 | -4.0 | 0.5 | -8.4 | 3.6E-17 | 2.7E-16 | MB231/MB468 | NUC |
| >hg38_21_+_8433172_8446622_RNA45SN1_45S_50ntFlanks@4160.4184.25\|CCACATCCAAGGAAGGCAGCAGGCG | >hg38_21_+_8433172_8446622_RNA45SN1_45S_50ntFlanks | 4160 | 4184 | 25 | CCACATCCAAGGAAGGCAGCAGGCG | 376.7 | -11.5 | 1.4 | -8.4 | 3.6E-17 | 2.7E-16 | MB231/MB468 | NUC |
| >MI0000102\|hsa-mir-100&WithFlank&11\|-\|122152223\|122152314@20.39.20&[MIMAT0000098&hsa-miR-100-5p&offsets\|+1\|-1;m-27&11\|-\|122152276\|122152296&offsets\|+1\|0]\|ACCCGTAGATCCGAACTTGT | >MI0000102\|hsa-mir-100&WithFlank&11\|-\|122152223\|122152314 | 20 | 39 | 20 | ACCCGTAGATCCGAACTTGT | 483.9 | 12.9 | 1.5 | 8.4 | 3.0E-17 | 2.7E-16 | MB231/MB468 | CYTO |
| >hg38_21_+_8433172_8446622_RNA45SN1_45S_50ntFlanks@12081.12110.30\|CTCTCGCTTCTGGCGCCAAGCGCCCGGCCG | >hg38_21_+_8433172_8446622_RNA45SN1_45S_50ntFlanks | 12081 | 12110 | 30 | CTCTCGCTTCTGGCGCCAAGCGCCCGGCCG | 381.0 | -11.5 | 1.4 | -8.4 | 3.7E-17 | 2.7E-16 | MB231/MB468 | NUC |
| >hg38_21_+_8433172_8446622_RNA45SN1_45S_50ntFlanks@10391.10435.45\|AACAGCAGTTGAACATGGGTCAGTCGGTCCTGAGAGATGGGCGAG | >hg38_21_+_8433172_8446622_RNA45SN1_45S_50ntFlanks | 10391 | 10435 | 45 | AACAGCAGTTGAACATGGGTCAGTCGGTCCTGAGAGATGGGCGAG | 1193.0 | -14.0 | 1.7 | -8.4 | 4.8E-17 | 2.7E-16 | MB231/MB468 | MITO |
| >hg38_21_+_8433172_8446622_RNA45SN1_45S_50ntFlanks@12405.12434.30\|CTTTTTGATCCTTCGATGTCGGCTCTTCCT | >hg38_21_+_8433172_8446622_RNA45SN1_45S_50ntFlanks | 12405 | 12434 | 30 | CTTTTTGATCCTTCGATGTCGGCTCTTCCT | 2737.6 | -10.6 | 1.3 | -8.5 | 2.9E-17 | 2.7E-16 | MB231/MB468 | MP |
| >hg38_1_-_228634819_228635039_RNA5S12_5S_50ntFlanks@153.170.18\|ACCGGGTGCTGTAGGCTT | >hg38_1_-_228634819_228635039_RNA5S12_5S_50ntFlanks | 153 | 170 | 18 | ACCGGGTGCTGTAGGCTT | 31113.5 | 7.2 | 0.9 | 8.4 | 5.1E-17 | 2.8E-16 | BT20/MB468 | CYTO |
| >hg38_21_+_8433172_8446622_RNA45SN1_45S_50ntFlanks@8357.8377.21\|AAAAGAACTTTGAAGAGAGAG | >hg38_21_+_8433172_8446622_RNA45SN1_45S_50ntFlanks | 8357 | 8377 | 21 | AAAAGAACTTTGAAGAGAGAG | 12907.7 | -10.7 | 1.3 | -8.4 | 2.9E-17 | 2.8E-16 | MB231/MB468 | MP |
| >hg38_21_+_8433172_8446622_RNA45SN1_45S_50ntFlanks@11918.11949.32\|ACATGAGAGGTGTAGAATAAGTGGGAGGCCCC | >hg38_21_+_8433172_8446622_RNA45SN1_45S_50ntFlanks | 11918 | 11949 | 32 | ACATGAGAGGTGTAGAATAAGTGGGAGGCCCC | 568.2 | -13.0 | 1.5 | -8.4 | 2.9E-17 | 2.8E-16 | MB231/MB468 | MP |
| >hg38_21_+_8433172_8446622_RNA45SN1_45S_50ntFlanks@10403.10420.18\|ACATGGGTCAGTCGGTCC | >hg38_21_+_8433172_8446622_RNA45SN1_45S_50ntFlanks | 10403 | 10420 | 18 | ACATGGGTCAGTCGGTCC | 265.9 | -11.9 | 1.4 | -8.4 | 2.9E-17 | 2.8E-16 | MB231/MB468 | MP |
| >hg38_21_+_8433172_8446622_RNA45SN1_45S_50ntFlanks@11584.11611.28\|TGGTGCGGACCAGGGGAATCCGACTGTT | >hg38_21_+_8433172_8446622_RNA45SN1_45S_50ntFlanks | 11584 | 11611 | 28 | TGGTGCGGACCAGGGGAATCCGACTGTT | 1552.0 | 5.4 | 0.6 | 8.4 | 3.7E-17 | 2.8E-16 | BT20/MB468 | MITO |
| >hg38_21_+_8433172_8446622_RNA45SN1_45S_50ntFlanks@12405.12443.39\|CTTTTTGATCCTTCGATGTCGGCTCTTCCTATCATTGTG | >hg38_21_+_8433172_8446622_RNA45SN1_45S_50ntFlanks | 12405 | 12443 | 39 | CTTTTTGATCCTTCGATGTCGGCTCTTCCTATCATTGTG | 1323.5 | -14.2 | 1.7 | -8.4 | 3.0E-17 | 2.8E-16 | MB231/MB468 | MP |
| >hg38_21_+_8433172_8446622_RNA45SN1_45S_50ntFlanks@11804.11821.18\|CCCACTGTCCCTACCTAC | >hg38_21_+_8433172_8446622_RNA45SN1_45S_50ntFlanks | 11804 | 11821 | 18 | CCCACTGTCCCTACCTAC | 1116.3 | -13.9 | 1.7 | -8.4 | 4.9E-17 | 2.8E-16 | MB231/MB468 | MITO |
| >MI0000439\|hsa-mir-23b&WithFlank&9\|+\|95085202\|95085310@65.86.22(+1U)&[MIMAT0000418&hsa-miR-23b-3p&offsets\|+1\|0(+1U);m-42&9\|+\|95085265\|95085285&offsets\|+1\|+2(+1U)]\|TCACATTGCCAGGGATTACCACT | >MI0000439\|hsa-mir-23b&WithFlank&9\|+\|95085202\|95085310 | 65 | 86 | 22(+1U) | TCACATTGCCAGGGATTACCACT | 445.7 | 7.0 | 0.8 | 8.4 | 5.2E-17 | 2.8E-16 | BT20/MB468 | CYTO |
| >hg38_21_+_8433172_8446622_RNA45SN1_45S_50ntFlanks@6780.6802.23\|CGGGGCTACGCCTGTCTGAGCGT | >hg38_21_+_8433172_8446622_RNA45SN1_45S_50ntFlanks | 6780 | 6802 | 23 | CGGGGCTACGCCTGTCTGAGCGT | 636.9 | 5.2 | 0.6 | 8.4 | 3.8E-17 | 2.8E-16 | BT20/MB468 | MITO |
| >hg38_21_+_8433172_8446622_RNA45SN1_45S_50ntFlanks@10808.10840.33\|CAATGTAGGTAAGGGAAGTCGGCAAGCCGGATC | >hg38_21_+_8433172_8446622_RNA45SN1_45S_50ntFlanks | 10808 | 10840 | 33 | CAATGTAGGTAAGGGAAGTCGGCAAGCCGGATC | 5234.9 | -10.9 | 1.3 | -8.4 | 4.9E-17 | 2.8E-16 | MB231/MB468 | MITO |
| >hg38_21_+_8433172_8446622_RNA45SN1_45S_50ntFlanks@9031.9059.29\|TCCTCGTGGGGGGGCCGGGCCACCCCTCC | >hg38_21_+_8433172_8446622_RNA45SN1_45S_50ntFlanks | 9031 | 9059 | 29 | TCCTCGTGGGGGGGCCGGGCCACCCCTCC | 547.3 | -12.9 | 1.5 | -8.4 | 4.9E-17 | 2.8E-16 | MB231/MB468 | MITO |
| >hg38_21_+_8433172_8446622_RNA45SN1_45S_50ntFlanks@8176.8205.30\|CTTCTGATCGAGGCCCAGCCCGTGGACGGT | >hg38_21_+_8433172_8446622_RNA45SN1_45S_50ntFlanks | 8176 | 8205 | 30 | CTTCTGATCGAGGCCCAGCCCGTGGACGGT | 747.5 | -5.0 | 0.6 | -8.3 | 8.8E-17 | 2.8E-16 | BT20/MB468 | NUC |
| >hg38_21_+_8433172_8446622_RNA45SN1_45S_50ntFlanks@5473.5516.44\|CGGCCCTGGCGGAGCGCTGAGAAGACGGTCGAACTTGACTATCT | >hg38_21_+_8433172_8446622_RNA45SN1_45S_50ntFlanks | 5473 | 5516 | 44 | CGGCCCTGGCGGAGCGCTGAGAAGACGGTCGAACTTGACTATCT | 18048.5 | 10.6 | 1.3 | 8.4 | 3.8E-17 | 2.8E-16 | BT20/MB468 | MITO |
| >hg38_21_+_8433172_8446622_RNA45SN1_45S_50ntFlanks@10352.10369.18\|GAGAACTTTGAAGGCCGA | >hg38_21_+_8433172_8446622_RNA45SN1_45S_50ntFlanks | 10352 | 10369 | 18 | GAGAACTTTGAAGGCCGA | 514.2 | 5.5 | 0.7 | 8.4 | 3.8E-17 | 2.9E-16 | MB231/MB468 | NUC |
| >hg38_21_+_8433172_8446622_RNA45SN1_45S_50ntFlanks@12616.12654.39\|CTGAGGAGCCAATGGGGCGAAGCTACCATCTGTGGGATT | >hg38_21_+_8433172_8446622_RNA45SN1_45S_50ntFlanks | 12616 | 12654 | 39 | CTGAGGAGCCAATGGGGCGAAGCTACCATCTGTGGGATT | 1678.2 | -6.6 | 0.8 | -8.4 | 3.8E-17 | 2.9E-16 | MB231/MB468 | NUC |
| >MI0000746\|hsa-mir-99b&WithFlank&19\|+\|51692606\|51692687@51.72.22(+1A)&[MIMAT0004678&hsa-miR-99b-3p&offsets\|0\|0(+1A);m-151&19\|+\|51692656\|51692678&offsets\|0\|-1(+1A)]\|CAAGCTCGTGTCTGTGGGTCCGA | >MI0000746\|hsa-mir-99b&WithFlank&19\|+\|51692606\|51692687 | 51 | 72 | 22(+1A) | CAAGCTCGTGTCTGTGGGTCCGA | 2590.3 | 8.7 | 1.0 | 8.4 | 5.3E-17 | 2.9E-16 | BT20/MB468 | CYTO |
| >hg38_21_+_8433172_8446622_RNA45SN1_45S_50ntFlanks@11570.11600.31\|AGCCGACTTAGAACTGGTGCGGACCAGGGGA | >hg38_21_+_8433172_8446622_RNA45SN1_45S_50ntFlanks | 11570 | 11600 | 31 | AGCCGACTTAGAACTGGTGCGGACCAGGGGA | 733.0 | -5.3 | 0.6 | -8.4 | 3.0E-17 | 2.9E-16 | MB231/MB468 | MP |
| >hg38_21_+_8433172_8446622_RNA45SN1_45S_50ntFlanks@5467.5514.48\|GGCCCACGGCCCTGGCGGAGCGCTGAGAAGACGGTCGAACTTGACTAT | >hg38_21_+_8433172_8446622_RNA45SN1_45S_50ntFlanks | 5467 | 5514 | 48 | GGCCCACGGCCCTGGCGGAGCGCTGAGAAGACGGTCGAACTTGACTAT | 895.4 | -13.6 | 1.6 | -8.4 | 5.0E-17 | 2.9E-16 | MB231/MB468 | MITO |
| >hg38_21_+_8433172_8446622_RNA45SN1_45S_50ntFlanks@8238.8266.29\|CCCGGGTCTTCCCGGAGTCGGGTTGCTTG | >hg38_21_+_8433172_8446622_RNA45SN1_45S_50ntFlanks | 8238 | 8266 | 29 | CCCGGGTCTTCCCGGAGTCGGGTTGCTTG | 375.5 | -12.3 | 1.5 | -8.4 | 5.0E-17 | 2.9E-16 | MB231/MB468 | MITO |
| >hg38_21_+_8433172_8446622_RNA45SN1_45S_50ntFlanks@8498.8523.26\|CGTTCCTCCCGACCCCTCCACCCGCC | >hg38_21_+_8433172_8446622_RNA45SN1_45S_50ntFlanks | 8498 | 8523 | 26 | CGTTCCTCCCGACCCCTCCACCCGCC | 356.9 | -12.3 | 1.5 | -8.4 | 5.0E-17 | 2.9E-16 | MB231/MB468 | MITO |
| >hg38_21_+_8433172_8446622_RNA45SN1_45S_50ntFlanks@11874.11907.34\|AGAAGACCCTGTTGAGCTTGACTCTAGTCTGGCA | >hg38_21_+_8433172_8446622_RNA45SN1_45S_50ntFlanks | 11874 | 11907 | 34 | AGAAGACCCTGTTGAGCTTGACTCTAGTCTGGCA | 366.1 | -12.4 | 1.5 | -8.4 | 3.1E-17 | 2.9E-16 | MB231/MB468 | MP |
| >hg38_21_+_8433172_8446622_RNA45SN1_45S_50ntFlanks@3846.3869.24\|CTTGGATAACTGTGGTAATTCTAG | >hg38_21_+_8433172_8446622_RNA45SN1_45S_50ntFlanks | 3846 | 3869 | 24 | CTTGGATAACTGTGGTAATTCTAG | 778.9 | -8.2 | 1.0 | -8.4 | 3.1E-17 | 2.9E-16 | MB231/MB468 | MP |
| >trna10_AspGTC_12_-_125424193_125424264@32.72.41,trna12_AspGTC_12_-_125411891_125411962@32.72.41,trna144_AspGTC_6_-_27551236_27551307@32.72.41,trna38_AspGTC_17_-_8125556_8125627@32.72.41,trna45_AspGTC_6_+_27447453_27447524@32.72.41,trna48_AspGTC_6_+_27471523_27471594@32.72.41,trna4_AspGTC_12_+_96429799_96429870@32.72.41,trna69_AspGTC_1_-_161440205_161440276@32.72.41,trna72_AspGTC_1_-_161432824_161432895@32.72.41,trna75_AspGTC_1_-_161425414_161425485@32.72.41,trna78_AspGTC_1_-_161418033_161418104@32.72.41,trna81_AspGTC_1_-_161410615_161410686@32.72.41\|CTGTCACGCGGGAGACCGGGGTTCGATTCCCCGACGGGGAG | >trna10_AspGTC_12_-_125424193_125424264 | 32 | 72 | 41 | CTGTCACGCGGGAGACCGGGGTTCGATTCCCCGACGGGGAG | 1968.8 | -9.2 | 1.1 | -8.4 | 3.1E-17 | 2.9E-16 | MB231/MB468 | CYTO |
| >hg38_21_+_8433172_8446622_RNA45SN1_45S_50ntFlanks@12108.12125.18\|CCGCGCGCCGGCCGGGCG | >hg38_21_+_8433172_8446622_RNA45SN1_45S_50ntFlanks | 12108 | 12125 | 18 | CCGCGCGCCGGCCGGGCG | 359.9 | -6.5 | 0.8 | -8.4 | 3.1E-17 | 2.9E-16 | MB231/MB468 | CYTO |
| >hg38_21_+_8433172_8446622_RNA45SN1_45S_50ntFlanks@10808.10838.31\|CAATGTAGGTAAGGGAAGTCGGCAAGCCGGA | >hg38_21_+_8433172_8446622_RNA45SN1_45S_50ntFlanks | 10808 | 10838 | 31 | CAATGTAGGTAAGGGAAGTCGGCAAGCCGGA | 431.2 | -12.5 | 1.5 | -8.4 | 5.0E-17 | 2.9E-16 | MB231/MB468 | MITO |
| >hg38_21_+_8433172_8446622_RNA45SN1_45S_50ntFlanks@12417.12443.27\|TCGATGTCGGCTCTTCCTATCATTGTG | >hg38_21_+_8433172_8446622_RNA45SN1_45S_50ntFlanks | 12417 | 12443 | 27 | TCGATGTCGGCTCTTCCTATCATTGTG | 6918.8 | -3.9 | 0.5 | -8.4 | 5.3E-17 | 2.9E-16 | BT20/MB468 | CYTO |
| >hg38_21_+_8433172_8446622_RNA45SN1_45S_50ntFlanks@10361.10388.28\|GAAGGCCGAAGTGGAGAAGGGTTCCATG | >hg38_21_+_8433172_8446622_RNA45SN1_45S_50ntFlanks | 10361 | 10388 | 28 | GAAGGCCGAAGTGGAGAAGGGTTCCATG | 927.0 | -13.7 | 1.6 | -8.4 | 3.1E-17 | 2.9E-16 | MB231/MB468 | MP |
| >MI0000063\|hsa-let-7b&WithFlank&22\|+\|46113680\|46113774@12.30.19(+1A)&[MIMAT0000063&hsa-let-7b-5p&offsets\|0\|-3(+1A);m-14&22\|+\|46113691\|46113712&offsets\|0\|-3(+1A)]\|TGAGGTAGTAGGTTGTGTGA | >MI0000063\|hsa-let-7b&WithFlank&22\|+\|46113680\|46113774 | 12 | 30 | 19(+1A) | TGAGGTAGTAGGTTGTGTGA | 434.4 | 10.7 | 1.3 | 8.4 | 5.4E-17 | 2.9E-16 | BT20/MB468 | CYTO |
| >am_trnaMT_PheGAA_MT_+_577_647@41.59.19\|GTTTAGACGGGCTCACATC | >am_trnaMT_PheGAA_MT_+_577_647 | 41 | 59 | 19 | GTTTAGACGGGCTCACATC | 1564.8 | 7.9 | 0.9 | 8.4 | 3.9E-17 | 2.9E-16 | BT20/MB468 | MITO |
| >MI0000103\|hsa-mir-101-1&WithFlank&1\|-\|65058428\|65058514@53.72.20;MI0000739\|hsa-mir-101-2&WithFlank&9\|+\|4850291\|4850381@55.74.20&[MIMAT0000099&hsa-miR-101-3p&offsets\|0\|-1;m-17&1\|-\|65058443\|65058462&offsets\|0\|0];[MIMAT0000099_1&hsa-miR-101-3p&offsets\|0\|-1;m-16&9\|+\|4850345\|4850364&offsets\|0\|0]\|TACAGTACTGTGATAACTGA | >MI0000103\|hsa-mir-101-1&WithFlank&1\|-\|65058428\|65058514 | 53 | 72 | 20 | TACAGTACTGTGATAACTGA | 552.0 | 5.7 | 0.7 | 8.4 | 3.9E-17 | 2.9E-16 | BT20/MB468 | MITO |
| >hg38_21_+_8433172_8446622_RNA45SN1_45S_50ntFlanks@10391.10419.29\|AACAGCAGTTGAACATGGGTCAGTCGGTC | >hg38_21_+_8433172_8446622_RNA45SN1_45S_50ntFlanks | 10391 | 10419 | 29 | AACAGCAGTTGAACATGGGTCAGTCGGTC | 424.3 | -12.6 | 1.5 | -8.4 | 3.1E-17 | 2.9E-16 | MB231/MB468 | MP |
| >MI0000477\|hsa-mir-146a&WithFlank&5\|+\|160485346\|160485456@27.47.21(+1C)&[MIMAT0000449&hsa-miR-146a-5p&offsets\|0\|-1(+1C);m-129&5\|+\|160485372\|160485393&offsets\|0\|-1(+1C)]\|TGAGAACTGAATTCCATGGGTC | >MI0000477\|hsa-mir-146a&WithFlank&5\|+\|160485346\|160485456 | 27 | 47 | 21(+1C) | TGAGAACTGAATTCCATGGGTC | 2201.7 | -14.2 | 1.6 | -8.6 | 5.5E-18 | 2.9E-16 | BT20/MB231 | NUC |
| >hg38_21_+_8433172_8446622_RNA45SN1_45S_50ntFlanks@10576.10595.20\|GCCGGCGGGAGCCCCGGGGA | >hg38_21_+_8433172_8446622_RNA45SN1_45S_50ntFlanks | 10576 | 10595 | 20 | GCCGGCGGGAGCCCCGGGGA | 703.4 | 13.5 | 1.6 | 8.4 | 3.2E-17 | 2.9E-16 | MB231/MB468 | CYTO |
| >hg38_21_+_8433172_8446622_RNA45SN1_45S_50ntFlanks@11918.11952.35\|ACATGAGAGGTGTAGAATAAGTGGGAGGCCCCCGG | >hg38_21_+_8433172_8446622_RNA45SN1_45S_50ntFlanks | 11918 | 11952 | 35 | ACATGAGAGGTGTAGAATAAGTGGGAGGCCCCCGG | 426.6 | -7.2 | 0.9 | -8.4 | 3.2E-17 | 2.9E-16 | MB231/MB468 | CYTO |
| >hg38_21_+_8433172_8446622_RNA45SN1_45S_50ntFlanks@4884.4907.24\|CAAAGCTGAAACTTAAAGGAATTG | >hg38_21_+_8433172_8446622_RNA45SN1_45S_50ntFlanks | 4884 | 4907 | 24 | CAAAGCTGAAACTTAAAGGAATTG | 246.4 | -11.8 | 1.4 | -8.4 | 3.1E-17 | 2.9E-16 | MB231/MB468 | MP |
| >hg38_21_+_8433172_8446622_RNA45SN1_45S_50ntFlanks@8168.8187.20\|CCCAAGTCCTTCTGATCGAG | >hg38_21_+_8433172_8446622_RNA45SN1_45S_50ntFlanks | 8168 | 8187 | 20 | CCCAAGTCCTTCTGATCGAG | 258.5 | -11.9 | 1.4 | -8.4 | 3.1E-17 | 2.9E-16 | MB231/MB468 | MP |
| >hg38_21_+_8433172_8446622_RNA45SN1_45S_50ntFlanks@12927.12973.47\|CCTGGCGCTAAACCATTCGTAGACGACCTGCTTCTGGGTCGGGGTTT | >hg38_21_+_8433172_8446622_RNA45SN1_45S_50ntFlanks | 12927 | 12973 | 47 | CCTGGCGCTAAACCATTCGTAGACGACCTGCTTCTGGGTCGGGGTTT | 725.0 | -13.3 | 1.6 | -8.4 | 5.1E-17 | 2.9E-16 | MB231/MB468 | MITO |

<table>
<tr><th></th><th></th><th></th><th></th><th></th><th></th><th></th><th></th><th></th><th></th><th></th><th></th><th></th><th></th></tr>
<tr><td>>MI0000285|hsa-mir-205&WithFlank&1|+|209432127|209432248@40.60.21(+1U)&[MIMAT0000266&hsa-miR-205-5p&offsets|0|-1(+1U);m-351&1|+|209432166|209432187&offsets|0|-1(+1U)]|TCCTTCATTCCACCGGAGTCTT</td><td>>MI0000285|hsa-mir-205&WithFlank&1|+|209432127|209432248</td><td>40</td><td>60</td><td>21(+1U)</td><td>TCCTTCATTCCACCGGAGTCTT</td><td>912.9</td><td>13.6</td><td>1.6</td><td>8.6</td><td>5.7E-18</td><td>2.9E-16</td><td>BT20/MB231</td><td>NUC</td></tr>
<tr><td>>hg38_21_+_8433172_8446622_RNA45SN1_45S_50ntFlanks@11876.11893.18|AAGACCCTGTTGAGCTTG</td><td>>hg38_21_+_8433172_8446622_RNA45SN1_45S_50ntFlanks</td><td>11876</td><td>11893</td><td>18</td><td>AAGACCCTGTTGAGCTTG</td><td>4169.5</td><td>-5.7</td><td>0.7</td><td>-8.4</td><td>3.2E-17</td><td>2.9E-16</td><td>MB231/MB468</td><td>CYTO</td></tr>
<tr><td>>hg38_21_+_8433172_8446622_RNA45SN1_45S_50ntFlanks@9352.9380.29|CCGTGGCGCAATGAAGGTGAAGGCCGGCG</td><td>>hg38_21_+_8433172_8446622_RNA45SN1_45S_50ntFlanks</td><td>9352</td><td>9380</td><td>29</td><td>CCGTGGCGCAATGAAGGTGAAGGCCGGCG</td><td>410.9</td><td>-12.5</td><td>1.5</td><td>-8.4</td><td>5.1E-17</td><td>2.9E-16</td><td>MB231/MB468</td><td>MITO</td></tr>
<tr><td>>hg38_21_+_8433172_8446622_RNA45SN1_45S_50ntFlanks@8176.8215.40|CTTCTGATCGAGGCCCAGCCCGTGGACGGTGTGAGGCCGG</td><td>>hg38_21_+_8433172_8446622_RNA45SN1_45S_50ntFlanks</td><td>8176</td><td>8215</td><td>40</td><td>CTTCTGATCGAGGCCCAGCCCGTGGACGGTGTGAGGCCGG</td><td>1157.5</td><td>-14.0</td><td>1.7</td><td>-8.4</td><td>3.1E-17</td><td>2.9E-16</td><td>MB231/MB468</td><td>MP</td></tr>
<tr><td>>hg38_21_+_8433172_8446622_RNA45SN1_45S_50ntFlanks@10845.10862.18|ACTTCGGGATAAGGATTG</td><td>>hg38_21_+_8433172_8446622_RNA45SN1_45S_50ntFlanks</td><td>10845</td><td>10862</td><td>18</td><td>ACTTCGGGATAAGGATTG</td><td>344.5</td><td>-12.3</td><td>1.5</td><td>-8.4</td><td>3.1E-17</td><td>2.9E-16</td><td>MB231/MB468</td><td>MP</td></tr>
<tr><td>>hg38_21_+_8433172_8446622_RNA45SN1_45S_50ntFlanks@12957.12981.25|CTTCTGGGTCGGGGTTTCGTACGTA</td><td>>hg38_21_+_8433172_8446622_RNA45SN1_45S_50ntFlanks</td><td>12957</td><td>12981</td><td>25</td><td>CTTCTGGGTCGGGGTTTCGTACGTA</td><td>382.7</td><td>-12.4</td><td>1.5</td><td>-8.4</td><td>5.2E-17</td><td>3.0E-16</td><td>MB231/MB468</td><td>MITO</td></tr>
<tr><td>>hg38_21_+_8433172_8446622_RNA45SN1_45S_50ntFlanks@10663.10710.48|CCTTGGAAAGCGTCGCGGTTCCGGCGGCGTCCGGTGAGCTCTCGCTGG</td><td>>hg38_21_+_8433172_8446622_RNA45SN1_45S_50ntFlanks</td><td>10663</td><td>10710</td><td>48</td><td>CCTTGGAAAGCGTCGCGGTTCCGGCGGCGTCCGGTGAGCTCTCGCTGG</td><td>902.4</td><td>-13.6</td><td>1.6</td><td>-8.4</td><td>5.2E-17</td><td>3.0E-16</td><td>MB231/MB468</td><td>MITO</td></tr>
<tr><td>>hg38_21_+_8433172_8446622_RNA45SN1_45S_50ntFlanks@5482.5499.18|CGGAGCGCTGAGAAGACG</td><td>>hg38_21_+_8433172_8446622_RNA45SN1_45S_50ntFlanks</td><td>5482</td><td>5499</td><td>18</td><td>CGGAGCGCTGAGAAGACG</td><td>3133.6</td><td>-9.0</td><td>1.1</td><td>-8.4</td><td>3.2E-17</td><td>3.0E-16</td><td>MB231/MB468</td><td>MP</td></tr>
<tr><td>>hg38_21_+_8433172_8446622_RNA45SN1_45S_50ntFlanks@9027.9064.38|CGTCTCCTCGTGGGGGGGCCGGGCCACCCCTCCCACGG</td><td>>hg38_21_+_8433172_8446622_RNA45SN1_45S_50ntFlanks</td><td>9027</td><td>9064</td><td>38</td><td>CGTCTCCTCGTGGGGGGGCCGGGCCACCCCTCCCACGG</td><td>389.6</td><td>-12.5</td><td>1.5</td><td>-8.4</td><td>3.2E-17</td><td>3.0E-16</td><td>MB231/MB468</td><td>MP</td></tr>
<tr><td>>hg38_21_+_8433172_8446622_RNA45SN1_45S_50ntFlanks@6663.6680.18|GTGGATCACTCGGCTCGT</td><td>>hg38_21_+_8433172_8446622_RNA45SN1_45S_50ntFlanks</td><td>6663</td><td>6680</td><td>18</td><td>GTGGATCACTCGGCTCGT</td><td>395.1</td><td>-12.5</td><td>1.5</td><td>-8.4</td><td>3.2E-17</td><td>3.0E-16</td><td>MB231/MB468</td><td>MP</td></tr>
<tr><td>>MI0000279|hsa-mir-196a-2&WithFlank&12|+|53991732|53991853@31.52.22;MI0000238|hsa-mir-196a-1&WithFlank&17|-|48632484|48632565@13.34.22&[MIMAT0000226&hsa-miR-196a-5p&offsets|0|0;m-261&12|+|53991762|53991783&offsets|0|0];[MIMAT0000226_1&hsa-miR-196a-5p&offsets|0|0;m-268&17|-|48632532|48632553&offsets|0|0]|TAGGTAGTTTCATGTTGTTGGG</td><td>>MI0000279|hsa-mir-196a-2&WithFlank&12|+|53991732|53991853</td><td>31</td><td>52</td><td>22</td><td>TAGGTAGTTTCATGTTGTTGGG</td><td>2981.1</td><td>14.9</td><td>1.7</td><td>8.9</td><td>4.9E-19</td><td>3.0E-16</td><td>BT20/MB231</td><td>MP</td></tr>
<tr><td>>MI0000285|hsa-mir-205&WithFlank&1|+|209432127|209432248@40.61.22(+1A)&[MIMAT0000266&hsa-miR-205-5p&offsets|0|0(+1A);m-351&1|+|209432166|209432187&offsets|0|0(+1A)]|TCCTTCATTCCACCGGAGTCTGA</td><td>>MI0000285|hsa-mir-205&WithFlank&1|+|209432127|209432248</td><td>40</td><td>61</td><td>22(+1A)</td><td>TCCTTCATTCCACCGGAGTCTGA</td><td>3109.5</td><td>14.9</td><td>1.7</td><td>8.9</td><td>5.1E-19</td><td>3.0E-16</td><td>BT20/MB231</td><td>MP</td></tr>
<tr><td>>hg38_21_+_8433172_8446622_RNA45SN1_45S_50ntFlanks@11564.11585.22|CCTAGCAGCCGACTTAGAACTG</td><td>>hg38_21_+_8433172_8446622_RNA45SN1_45S_50ntFlanks</td><td>11564</td><td>11585</td><td>22</td><td>CCTAGCAGCCGACTTAGAACTG</td><td>2061.8</td><td>-4.8</td><td>0.6</td><td>-8.4</td><td>3.3E-17</td><td>3.0E-16</td><td>MB231/MB468</td><td>CYTO</td></tr>
<tr><td>>hg38_21_+_8433172_8446622_RNA45SN1_45S_50ntFlanks@12007.12032.26|GCCGCCGGTGAAATACCACTACTCTG</td><td>>hg38_21_+_8433172_8446622_RNA45SN1_45S_50ntFlanks</td><td>12007</td><td>12032</td><td>26</td><td>GCCGCCGGTGAAATACCACTACTCTG</td><td>15750.3</td><td>-12.0</td><td>1.4</td><td>-8.4</td><td>3.2E-17</td><td>3.0E-16</td><td>MB231/MB468</td><td>MP</td></tr>
<tr><td>>hg38_21_+_8433172_8446622_RNA45SN1_45S_50ntFlanks@12045.12070.26|CTGACCCGGTGAGGCGGGGGGGCGAG</td><td>>hg38_21_+_8433172_8446622_RNA45SN1_45S_50ntFlanks</td><td>12045</td><td>12070</td><td>26</td><td>CTGACCCGGTGAGGCGGGGGGGCGAG</td><td>423.0</td><td>-12.6</td><td>1.5</td><td>-8.4</td><td>3.2E-17</td><td>3.0E-16</td><td>MB231/MB468</td><td>MP</td></tr>
<tr><td>>hg38_21_+_8433172_8446622_RNA45SN1_45S_50ntFlanks@6745.6769.25|ACACTTCGAACGCACTTGCGGCCCC</td><td>>hg38_21_+_8433172_8446622_RNA45SN1_45S_50ntFlanks</td><td>6745</td><td>6769</td><td>25</td><td>ACACTTCGAACGCACTTGCGGCCCC</td><td>366.0</td><td>-12.4</td><td>1.5</td><td>-8.4</td><td>3.2E-17</td><td>3.0E-16</td><td>MB231/MB468</td><td>MP</td></tr>
<tr><td>>hg38_21_+_8433172_8446622_RNA45SN1_45S_50ntFlanks@12627.12651.25|ATGGGGCGAAGCTACCATCTGTGGG</td><td>>hg38_21_+_8433172_8446622_RNA45SN1_45S_50ntFlanks</td><td>12627</td><td>12651</td><td>25</td><td>ATGGGGCGAAGCTACCATCTGTGGG</td><td>422.2</td><td>-12.6</td><td>1.5</td><td>-8.4</td><td>3.2E-17</td><td>3.0E-16</td><td>MB231/MB468</td><td>MP</td></tr>
<tr><td>>hg38_21_+_8433172_8446622_RNA45SN1_45S_50ntFlanks@10873.10901.29|CTGGGTCGGTCGGGCTGGGGCGCGAAGCG</td><td>>hg38_21_+_8433172_8446622_RNA45SN1_45S_50ntFlanks</td><td>10873</td><td>10901</td><td>29</td><td>CTGGGTCGGTCGGGCTGGGGCGCGAAGCG</td><td>451.5</td><td>-12.7</td><td>1.5</td><td>-8.4</td><td>3.2E-17</td><td>3.0E-16</td><td>MB231/MB468</td><td>MP</td></tr>
<tr><td>>am_MI0000809|hsa-mir-151a&WithFlank&8|-|140732558|140732659@53.72.20(+1A)&[MIMAT0000757&hsa-miR-151a-3p&offsets|0|-1(+1A);m-30&8|-|140732587|140732607&offsets|0|-1(+1A)]|CTAGACTGAAGCTCCTTGAGA</td><td>>am_MI0000809|hsa-mir-151a&WithFlank&8|-|140732558|140732659</td><td>53</td><td>72</td><td>20(+1A)</td><td>CTAGACTGAAGCTCCTTGAGA</td><td>4870.2</td><td>7.1</td><td>0.8</td><td>8.4</td><td>3.3E-17</td><td>3.0E-16</td><td>MB231/MB468</td><td>CYTO</td></tr>
<tr><td>>hg38_21_+_8433172_8446622_RNA45SN1_45S_50ntFlanks@10348.10366.19|AAACGAGAACTTTGAAGGC</td><td>>hg38_21_+_8433172_8446622_RNA45SN1_45S_50ntFlanks</td><td>10348</td><td>10366</td><td>19</td><td>AAACGAGAACTTTGAAGGC</td><td>1371.4</td><td>-5.5</td><td>0.7</td><td>-8.3</td><td>9.3E-17</td><td>3.0E-16</td><td>BT20/MB468</td><td>NUC</td></tr>
<tr><td>>MI0000462|hsa-mir-152&WithFlank&17|-|48037155|48037253@60.81.22(+1U)&[MIMAT0000438&hsa-miR-152-3p&offsets|0|+1(+1U);m-97&17|-|48037174|48037194&offsets|0|+1(+1U)]|TCAGTGCATGACAGAACTTGGGT</td><td>>MI0000462|hsa-mir-152&WithFlank&17|-|48037155|48037253</td><td>60</td><td>81</td><td>22(+1U)</td><td>TCAGTGCATGACAGAACTTGGGT</td><td>72.5</td><td>9.0</td><td>1.1</td><td>8.3</td><td>9.3E-17</td><td>3.0E-16</td><td>BT20/MB468</td><td>NUC</td></tr>
<tr><td>>hg38_21_+_8433172_8446622_RNA45SN1_45S_50ntFlanks@11534.11560.27|GTTCCGCGCGGCGCCTCGCCTCGGCCG</td><td>>hg38_21_+_8433172_8446622_RNA45SN1_45S_50ntFlanks</td><td>11534</td><td>11560</td><td>27</td><td>GTTCCGCGCGGCGCCTCGCCTCGGCCG</td><td>564.4</td><td>-12.1</td><td>1.4</td><td>-8.4</td><td>4.1E-17</td><td>3.0E-16</td><td>MB231/MB468</td><td>NUC</td></tr>
<tr><td>>hg38_21_+_8433172_8446622_RNA45SN1_45S_50ntFlanks@3846.3869.24|CTTGGATAACTGTGGTAATTCTAG</td><td>>hg38_21_+_8433172_8446622_RNA45SN1_45S_50ntFlanks</td><td>3846</td><td>3869</td><td>24</td><td>CTTGGATAACTGTGGTAATTCTAG</td><td>351.2</td><td>-3.9</td><td>0.5</td><td>-8.4</td><td>4.1E-17</td><td>3.0E-16</td><td>MB231/MB468</td><td>NUC</td></tr>
<tr><td>>hg38_21_+_8433172_8446622_RNA45SN1_45S_50ntFlanks@10869.10891.23|AGGGCTGGGTCGGTCGGGCTGGG</td><td>>hg38_21_+_8433172_8446622_RNA45SN1_45S_50ntFlanks</td><td>10869</td><td>10891</td><td>23</td><td>AGGGCTGGGTCGGTCGGGCTGGG</td><td>751.5</td><td>-13.3</td><td>1.6</td><td>-8.4</td><td>5.3E-17</td><td>3.0E-16</td><td>MB231/MB468</td><td>MITO</td></tr>
<tr><td>>hg38_21_+_8433172_8446622_RNA45SN1_45S_50ntFlanks@4006.4026.21|ACCTCGGGCCGATCGCACGCC</td><td>>hg38_21_+_8433172_8446622_RNA45SN1_45S_50ntFlanks</td><td>4006</td><td>4026</td><td>21</td><td>ACCTCGGGCCGATCGCACGCC</td><td>571.4</td><td>-12.9</td><td>1.5</td><td>-8.4</td><td>5.4E-17</td><td>3.0E-16</td><td>MB231/MB468</td><td>MITO</td></tr>
<tr><td>>hg38_21_+_8433172_8446622_RNA45SN1_45S_50ntFlanks@7978.8010.33|GACCTCAGATCAGACGTGGCGACCCGCTGAATT</td><td>>hg38_21_+_8433172_8446622_RNA45SN1_45S_50ntFlanks</td><td>7978</td><td>8010</td><td>33</td><td>GACCTCAGATCAGACGTGGCGACCCGCTGAATT</td><td>693.5</td><td>9.3</td><td>1.1</td><td>8.4</td><td>3.4E-17</td><td>3.1E-16</td><td>MB231/MB468</td><td>CYTO</td></tr>
<tr><td>>hg38_21_+_8433172_8446622_RNA45SN1_45S_50ntFlanks@6774.6807.34|TCCTCCCGGGGCTACGCCTGTCTGAGCGTCGCTT</td><td>>hg38_21_+_8433172_8446622_RNA45SN1_45S_50ntFlanks</td><td>6774</td><td>6807</td><td>34</td><td>TCCTCCCGGGGCTACGCCTGTCTGAGCGTCGCTT</td><td>1388.0</td><td>6.8</td><td>0.8</td><td>8.4</td><td>4.1E-17</td><td>3.1E-16</td><td>BT20/MB468</td><td>MITO</td></tr>
<tr><td>>hg38_21_+_8433172_8446622_RNA45SN1_45S_50ntFlanks@11575.11592.18|ACTTAGAACTGGTGCGGA</td><td>>hg38_21_+_8433172_8446622_RNA45SN1_45S_50ntFlanks</td><td>11575</td><td>11592</td><td>18</td><td>ACTTAGAACTGGTGCGGA</td><td>4196.3</td><td>-8.3</td><td>1.0</td><td>-8.4</td><td>3.4E-17</td><td>3.1E-16</td><td>MB231/MB468</td><td>CYTO</td></tr>
<tr><td>>hg38_21_+_8433172_8446622_RNA45SN1_45S_50ntFlanks@3985.4002.18|CGGCTTTGGTGACTCTAG</td><td>>hg38_21_+_8433172_8446622_RNA45SN1_45S_50ntFlanks</td><td>3985</td><td>4002</td><td>18</td><td>CGGCTTTGGTGACTCTAG</td><td>6518.6</td><td>-11.1</td><td>1.3</td><td>-8.4</td><td>4.2E-17</td><td>3.1E-16</td><td>MB231/MB468</td><td>NUC</td></tr>
<tr><td>>trnaMT_LeuTAA_MT_+_3230_3304@18.42.25|CGGTAATCGCATAAAACTTAAAACT</td><td>>trnaMT_LeuTAA_MT_+_3230_3304</td><td>18</td><td>42</td><td>25</td><td>CGGTAATCGCATAAAACTTAAAACT</td><td>704.7</td><td>11.1</td><td>1.3</td><td>8.4</td><td>5.8E-17</td><td>3.1E-16</td><td>BT20/MB468</td><td>CYTO</td></tr>
<tr><td>>hg38_21_+_8433172_8446622_RNA45SN1_45S_50ntFlanks@11769.11794.26|TAATTAGTGACGCGCATGAATGGATG</td><td>>hg38_21_+_8433172_8446622_RNA45SN1_45S_50ntFlanks</td><td>11769</td><td>11794</td><td>26</td><td>TAATTAGTGACGCGCATGAATGGATG</td><td>2819.3</td><td>-5.3</td><td>0.6</td><td>-8.4</td><td>4.2E-17</td><td>3.1E-16</td><td>MB231/MB468</td><td>NUC</td></tr>
<tr><td>>MI0000650|hsa-mir-200c&WithFlank&12|+|6963693|6963772@50.72.23(+1A)&[MIMAT0000617&hsa-miR-200c-3p&offsets|0|0(+1A);m-54&12|+|6963742|6963764&offsets|0|0(+1A)]|TAATACTGCCGGGTAATGATGGAA</td><td>>MI0000650|hsa-mir-200c&WithFlank&12|+|6963693|6963772</td><td>50</td><td>72</td><td>23(+1A)</td><td>TAATACTGCCGGGTAATGATGGAA</td><td>4096.9</td><td>3.3</td><td>0.4</td><td>8.3</td><td>9.6E-17</td><td>3.1E-16</td><td>BT20/MB468</td><td>NUC</td></tr>
<tr><td>>hg38_21_+_8433172_8446622_RNA45SN1_45S_50ntFlanks@8114.8140.27|GCGCGGGACATGTGGCGTACGGAAGAC</td><td>>hg38_21_+_8433172_8446622_RNA45SN1_45S_50ntFlanks</td><td>8114</td><td>8140</td><td>27</td><td>GCGCGGGACATGTGGCGTACGGAAGAC</td><td>411.0</td><td>-11.6</td><td>1.4</td><td>-8.4</td><td>4.2E-17</td><td>3.1E-16</td><td>MB231/MB468</td><td>NUC</td></tr>
<tr><td>>hg38_21_+_8433172_8446622_RNA45SN1_45S_50ntFlanks@11207.11232.26|GGCGGCGACTCTGGACGCGAGCCGGG</td><td>>hg38_21_+_8433172_8446622_RNA45SN1_45S_50ntFlanks</td><td>11207</td><td>11232</td><td>26</td><td>GGCGGCGACTCTGGACGCGAGCCGGG</td><td>986.0</td><td>-7.8</td><td>0.9</td><td>-8.4</td><td>3.4E-17</td><td>3.1E-16</td><td>MB231/MB468</td><td>CYTO</td></tr>
<tr><td>>hg38_21_+_8433172_8446622_RNA45SN1_45S_50ntFlanks@11881.11905.25|CCTGTTGAGCTTGACTCTAGTCTGG</td><td>>hg38_21_+_8433172_8446622_RNA45SN1_45S_50ntFlanks</td><td>11881</td><td>11905</td><td>25</td><td>CCTGTTGAGCTTGACTCTAGTCTGG</td><td>308.5</td><td>-12.1</td><td>1.4</td><td>-8.4</td><td>3.4E-17</td><td>3.1E-16</td><td>MB231/MB468</td><td>MP</td></tr>
<tr><td>>MI0001733|hsa-mir-452&WithFlank&X|-|151959622|151959718@20.42.23(+1U)&[MIMAT0001635&hsa-miR-452-5p&offsets|0|+1(+1U);m-188&X|-|151959677|151959699&offsets|0|0(+1U)]|AACTGTTTGCAGAGGAAACTGAGT</td><td>>MI0001733|hsa-mir-452&WithFlank&X|-|151959622|151959718</td><td>20</td><td>42</td><td>23(+1U)</td><td>AACTGTTTGCAGAGGAAACTGAGT</td><td>66.5</td><td>8.9</td><td>1.1</td><td>8.3</td><td>9.7E-17</td><td>3.1E-16</td><td>BT20/MB468</td><td>NUC</td></tr>
<tr><td>>hg38_21_+_8433172_8446622_RNA45SN1_45S_50ntFlanks@6663.6686.24|GTGGATCACTCGGCTCGTGCGTCG</td><td>>hg38_21_+_8433172_8446622_RNA45SN1_45S_50ntFlanks</td><td>6663</td><td>6686</td><td>24</td><td>GTGGATCACTCGGCTCGTGCGTCG</td><td>1118.0</td><td>-5.2</td><td>0.6</td><td>-8.3</td><td>9.7E-17</td><td>3.1E-16</td><td>BT20/MB468</td><td>NUC</td></tr>
<tr><td>>am_MI0000809|hsa-mir-151a&WithFlank&8|-|140732558|140732659@53.70.18&[MIMAT0000757&hsa-miR-151a-3p&offsets|0|-3;m-30&8|-|140732587|140732607&offsets|0|-3]|CTAGACTGAAGCTCCTTG</td><td>>am_MI0000809|hsa-mir-151a&WithFlank&8|-|140732558|140732659</td><td>53</td><td>70</td><td>18</td><td>CTAGACTGAAGCTCCTTG</td><td>2638.2</td><td>10.7</td><td>1.3</td><td>8.4</td><td>5.9E-17</td><td>3.1E-16</td><td>BT20/MB468</td><td>CYTO</td></tr>
<tr><td>>hg38_21_+_8433172_8446622_RNA45SN1_45S_50ntFlanks@12283.12323.41|TACAGACCGTGAAAGCGGGGCCTCACGATCCTTCTGACCTT</td><td>>hg38_21_+_8433172_8446622_RNA45SN1_45S_50ntFlanks</td><td>12283</td><td>12323</td><td>41</td><td>TACAGACCGTGAAAGCGGGGCCTCACGATCCTTCTGACCTT</td><td>389.8</td><td>-12.4</td><td>1.5</td><td>-8.4</td><td>5.5E-17</td><td>3.1E-16</td><td>MB231/MB468</td><td>MITO</td></tr>
<tr><td>>am_MI0000786|hsa-mir-378a&WithFlank&5|+|149732819|149732896@50.69.20(+1A)&[MIMAT0000732&hsa-miR-378a-3p&offsets|+1|-1(+1A);m-44&5|+|149732867|149732888&offsets|+1|-1(+1A)]|CTGGACTTGGAGTCAGAAGGA</td><td>>am_MI0000786|hsa-mir-378a&WithFlank&5|+|149732819|149732896</td><td>50</td><td>69</td><td>20(+1A)</td><td>CTGGACTTGGAGTCAGAAGGA</td><td>438.1</td><td>7.8</td><td>0.9</td><td>8.4</td><td>5.9E-17</td><td>3.2E-16</td><td>BT20/MB468</td><td>CYTO</td></tr>
<tr><td>>hg38_21_+_8433172_8446622_RNA45SN1_45S_50ntFlanks@6649.6679.31|TACGACTCTTAGCGGTGGATCACTCGGCTCG</td><td>>hg38_21_+_8433172_8446622_RNA45SN1_45S_50ntFlanks</td><td>6649</td><td>6679</td><td>31</td><td>TACGACTCTTAGCGGTGGATCACTCGGCTCG</td><td>1520.4</td><td>-3.8</td><td>0.5</td><td>-8.3</td><td>9.7E-17</td><td>3.2E-16</td><td>BT20/MB468</td><td>NUC</td></tr>
<tr><td>>hg38_21_+_8433172_8446622_RNA45SN1_45S_50ntFlanks@4452.4472.21|CTCGGCGCCCCCTCGATGCTC</td><td>>hg38_21_+_8433172_8446622_RNA45SN1_45S_50ntFlanks</td><td>4452</td><td>4472</td><td>21</td><td>CTCGGCGCCCCCTCGATGCTC</td><td>286.3</td><td>-12.0</td><td>1.4</td><td>-8.4</td><td>3.4E-17</td><td>3.2E-16</td><td>MB231/MB468</td><td>MP</td></tr>
<tr><td>>hg38_21_+_8433172_8446622_RNA45SN1_45S_50ntFlanks@9624.9650.27|AGTAGCTGGTTCCCTCCGAAGTTTCCC</td><td>>hg38_21_+_8433172_8446622_RNA45SN1_45S_50ntFlanks</td><td>9624</td><td>9650</td><td>27</td><td>AGTAGCTGGTTCCCTCCGAAGTTTCCC</td><td>389.8</td><td>-12.4</td><td>1.5</td><td>-8.4</td><td>5.6E-17</td><td>3.2E-16</td><td>MB231/MB468</td><td>MITO</td></tr>
<tr><td>>MI0000285|hsa-mir-205&WithFlank&1|+|209432127|209432248@40.59.20(+1A)&[MIMAT0000266&hsa-miR-205-5p&offsets|0|-2(+1A);m-351&1|+|209432166|209432187&offsets|0|-2(+1A)]|TCCTTCATTCCACCGGAGTCA</td><td>>MI0000285|hsa-mir-205&WithFlank&1|+|209432127|209432248</td><td>40</td><td>59</td><td>20(+1A)</td><td>TCCTTCATTCCACCGGAGTCA</td><td>679.9</td><td>13.2</td><td>1.5</td><td>8.6</td><td>6.3E-18</td><td>3.2E-16</td><td>BT20/MB231</td><td>NUC</td></tr>
<tr><td>>hg38_21_+_8433172_8446622_RNA45SN1_45S_50ntFlanks@11051.11075.25|CCGTCCTCCCCCCTCCCCGGGGGAG</td><td>>hg38_21_+_8433172_8446622_RNA45SN1_45S_50ntFlanks</td><td>11051</td><td>11075</td><td>25</td><td>CCGTCCTCCCCCCTCCCCGGGGGAG</td><td>405.4</td><td>-8.7</td><td>1.0</td><td>-8.4</td><td>5.9E-17</td><td>3.2E-16</td><td>BT20/MB468</td><td>CYTO</td></tr>
<tr><td>>hg38_21_+_8433172_8446622_RNA45SN1_45S_50ntFlanks@11208.11233.26|GCGGCGACTCTGGACGCGAGCCGGGC</td><td>>hg38_21_+_8433172_8446622_RNA45SN1_45S_50ntFlanks</td><td>11208</td><td>11233</td><td>26</td><td>GCGGCGACTCTGGACGCGAGCCGGGC</td><td>2902.8</td><td>-10.6</td><td>1.3</td><td>-8.4</td><td>3.4E-17</td><td>3.2E-16</td><td>MB231/MB468</td><td>MP</td></tr>
<tr><td>>hg38_21_+_8433172_8446622_RNA45SN1_45S_50ntFlanks@9571.9597.27|GCAAATCGGTCGTCCGACCTGGGTATA</td><td>>hg38_21_+_8433172_8446622_RNA45SN1_45S_50ntFlanks</td><td>9571</td><td>9597</td><td>27</td><td>GCAAATCGGTCGTCCGACCTGGGTATA</td><td>396.3</td><td>-12.5</td><td>1.5</td><td>-8.4</td><td>3.5E-17</td><td>3.2E-16</td><td>MB231/MB468</td><td>MP</td></tr>
</table>

<table>
<tr><th></th><th></th><th></th><th></th><th></th><th></th><th></th><th></th><th></th><th></th><th></th><th></th><th></th><th></th></tr>
<tr><td>>MI0000469|hsa-mir-125a&WithFlank&19|+|51693248|51693345@21.41.21(+1A)&[MIMAT0000443&hsa-miR-125a-5p&offsets|0|-3(+1A);m-63&19|+|51693268|51693289&offsets|0|-1(+1A)]|TCCCTGAGACCCTTTAACCTGA</td><td>>MI0000469|hsa-mir-125a&WithFlank&19|+|51693248|51693345</td><td>21</td><td>41</td><td>21(+1A)</td><td>TCCCTGAGACCCTTTAACCTGA</td><td>472.0</td><td>10.8</td><td>1.3</td><td>8.4</td><td>6.0E-17</td><td>3.2E-16</td><td>BT20/MB468</td><td>CYTO</td></tr>
<tr><td>>MI0000087|hsa-mir-29a&WithFlank&7|-|130876741|130876816@48.68.21(+1U)&[MIMAT0000086&hsa-miR-29a-3p&offsets|0|-1(+1U);m-26&7|-|130876749|130876769&offsets|0|0(+1U)]|TAGCACCATCTGAAATCGGTTT</td><td>>MI0000087|hsa-mir-29a&WithFlank&7|-|130876741|130876816</td><td>48</td><td>68</td><td>21(+1U)</td><td>TAGCACCATCTGAAATCGGTTT</td><td>283.1</td><td>12.0</td><td>1.4</td><td>8.4</td><td>4.3E-17</td><td>3.2E-16</td><td>MB231/MB468</td><td>NUC</td></tr>
<tr><td>>hg38_21_+_8433172_8446622_RNA45SN1_45S_50ntFlanks@12934.12958.25|CTAAACCATTCGTAGACGACCTGCT</td><td>>hg38_21_+_8433172_8446622_RNA45SN1_45S_50ntFlanks</td><td>12934</td><td>12958</td><td>25</td><td>CTAAACCATTCGTAGACGACCTGCT</td><td>715.8</td><td>-4.6</td><td>0.5</td><td>-8.4</td><td>4.3E-17</td><td>3.2E-16</td><td>MB231/MB468</td><td>NUC</td></tr>
<tr><td>>hg38_21_+_8433172_8446622_RNA45SN1_45S_50ntFlanks@3866.3899.34|CTAGAGCTAATACATGCCGACGGGCGCTGACCCC</td><td>>hg38_21_+_8433172_8446622_RNA45SN1_45S_50ntFlanks</td><td>3866</td><td>3899</td><td>34</td><td>CTAGAGCTAATACATGCCGACGGGCGCTGACCCC</td><td>337.3</td><td>-12.2</td><td>1.5</td><td>-8.4</td><td>5.7E-17</td><td>3.2E-16</td><td>MB231/MB468</td><td>MITO</td></tr>
<tr><td>>MI0016887|hsa-mir-4521&WithFlank&17|+|8186939|8187010@12.35.24&[MIMAT0019058&hsa-miR-4521&offsets|+2|+4;m-765&17|+|8186948|8186969&offsets|+2|+4]|TAAGGAAGTCCTGTGCTCAGTTTT</td><td>>MI0016887|hsa-mir-4521&WithFlank&17|+|8186939|8187010</td><td>12</td><td>35</td><td>24</td><td>TAAGGAAGTCCTGTGCTCAGTTTT</td><td>436.4</td><td>13.0</td><td>1.6</td><td>8.4</td><td>6.0E-17</td><td>3.2E-16</td><td>BT20/MB468</td><td>CYTO</td></tr>
<tr><td>>hg38_21_+_8433172_8446622_RNA45SN1_45S_50ntFlanks@13002.13039.38|CGATCTATTGAAAGTCAGCCCTCGACACAAGGGTTTGT</td><td>>hg38_21_+_8433172_8446622_RNA45SN1_45S_50ntFlanks</td><td>13002</td><td>13039</td><td>38</td><td>CGATCTATTGAAAGTCAGCCCTCGACACAAGGGTTTGT</td><td>2420.3</td><td>-13.5</td><td>1.6</td><td>-8.4</td><td>6.0E-17</td><td>3.2E-16</td><td>BT20/MB468</td><td>CYTO</td></tr>
<tr><td>>MI0000072|hsa-mir-18a&WithFlank&13|+|91350745|91350827@12.34.23&[MIMAT0000072&hsa-miR-18a-5p&offsets|0|0;m-262&13|+|91350756|91350778&offsets|0|0]|TAAGGTGCATCTAGTGCAGATAG</td><td>>MI0000072|hsa-mir-18a&WithFlank&13|+|91350745|91350827</td><td>12</td><td>34</td><td>23</td><td>TAAGGTGCATCTAGTGCAGATAG</td><td>455.5</td><td>6.4</td><td>0.8</td><td>8.4</td><td>4.4E-17</td><td>3.2E-16</td><td>MB231/MB468</td><td>NUC</td></tr>
<tr><td>>hg38_21_+_8433172_8446622_RNA45SN1_45S_50ntFlanks@11669.11692.24|GCCCAGTGCTCTGAATGTCAAAGT</td><td>>hg38_21_+_8433172_8446622_RNA45SN1_45S_50ntFlanks</td><td>11669</td><td>11692</td><td>24</td><td>GCCCAGTGCTCTGAATGTCAAAGT</td><td>264.5</td><td>4.0</td><td>0.5</td><td>8.4</td><td>4.4E-17</td><td>3.2E-16</td><td>MB231/MB468</td><td>NUC</td></tr>
<tr><td>>hg38_21_+_8433172_8446622_RNA45SN1_45S_50ntFlanks@4859.4890.32|TTGGGTTCCGGGGGGAGTATGGTTGCAAAGCT</td><td>>hg38_21_+_8433172_8446622_RNA45SN1_45S_50ntFlanks</td><td>4859</td><td>4890</td><td>32</td><td>TTGGGTTCCGGGGGGAGTATGGTTGCAAAGCT</td><td>135.5</td><td>11.1</td><td>1.3</td><td>8.4</td><td>3.6E-17</td><td>3.3E-16</td><td>MB231/MB468</td><td>CYTO</td></tr>
<tr><td>>hg38_21_+_8433172_8446622_RNA45SN1_45S_50ntFlanks@10099.10127.29|CGCGCGCGCGTGTGGTGTGCGTCGGAGGG</td><td>>hg38_21_+_8433172_8446622_RNA45SN1_45S_50ntFlanks</td><td>10099</td><td>10127</td><td>29</td><td>CGCGCGCGCGTGTGGTGTGCGTCGGAGGG</td><td>625.8</td><td>-13.1</td><td>1.6</td><td>-8.4</td><td>5.8E-17</td><td>3.3E-16</td><td>MB231/MB468</td><td>MITO</td></tr>
<tr><td>>MI0000285|hsa-mir-205&WithFlank&1|+|209432127|209432248@40.62.23(+1U)&[MIMAT0000266&hsa-miR-205-5p&offsets|0|+1(+1U);m-351&1|+|209432166|209432187&offsets|0|+1(+1U)]|TCCTTCATTCCACCGGAGTCTGTT</td><td>>MI0000285|hsa-mir-205&WithFlank&1|+|209432127|209432248</td><td>40</td><td>62</td><td>23(+1U)</td><td>TCCTTCATTCCACCGGAGTCTGTT</td><td>730.2</td><td>13.1</td><td>1.5</td><td>8.7</td><td>4.6E-18</td><td>3.3E-16</td><td>BT20/MB231</td><td>MITO</td></tr>
<tr><td>>hg38_21_+_8433172_8446622_RNA45SN1_45S_50ntFlanks@10806.10826.21|AACAATGTAGGTAAGGGAAGT</td><td>>hg38_21_+_8433172_8446622_RNA45SN1_45S_50ntFlanks</td><td>10806</td><td>10826</td><td>21</td><td>AACAATGTAGGTAAGGGAAGT</td><td>1108.3</td><td>-9.7</td><td>1.2</td><td>-8.4</td><td>5.8E-17</td><td>3.3E-16</td><td>MB231/MB468</td><td>MITO</td></tr>
<tr><td>>hg38_21_+_8433172_8446622_RNA45SN1_45S_50ntFlanks@11669.11709.41|GCCCAGTGCTCTGAATGTCAAAGTGAAGAAATTCAATGAAG</td><td>>hg38_21_+_8433172_8446622_RNA45SN1_45S_50ntFlanks</td><td>11669</td><td>11709</td><td>41</td><td>GCCCAGTGCTCTGAATGTCAAAGTGAAGAAATTCAATGAAG</td><td>1076.3</td><td>10.8</td><td>1.3</td><td>8.4</td><td>3.6E-17</td><td>3.3E-16</td><td>MB231/MB468</td><td>CYTO</td></tr>
<tr><td>>hg38_21_+_8433172_8446622_RNA45SN1_45S_50ntFlanks@11578.11603.26|TAGAACTGGTGCGGACCAGGGGAATC</td><td>>hg38_21_+_8433172_8446622_RNA45SN1_45S_50ntFlanks</td><td>11578</td><td>11603</td><td>26</td><td>TAGAACTGGTGCGGACCAGGGGAATC</td><td>1534.9</td><td>-4.3</td><td>0.5</td><td>-8.4</td><td>3.6E-17</td><td>3.3E-16</td><td>MB231/MB468</td><td>CYTO</td></tr>
<tr><td>>trnaMT_LeuTAA_MT_+_3230_3304@19.42.24|GGTAATCGCATAAAACTTAAAACT</td><td>>trnaMT_LeuTAA_MT_+_3230_3304</td><td>19</td><td>42</td><td>24</td><td>GGTAATCGCATAAAACTTAAAACT</td><td>510.8</td><td>11.0</td><td>1.3</td><td>8.4</td><td>6.2E-17</td><td>3.3E-16</td><td>BT20/MB468</td><td>CYTO</td></tr>
<tr><td>>hg38_21_+_8433172_8446622_RNA45SN1_45S_50ntFlanks@10866.10904.39|CTAAGGGCTGGGTCGGTCGGGCTGGGGCGCGAAGCGGGG</td><td>>hg38_21_+_8433172_8446622_RNA45SN1_45S_50ntFlanks</td><td>10866</td><td>10904</td><td>39</td><td>CTAAGGGCTGGGTCGGTCGGGCTGGGGCGCGAAGCGGGG</td><td>844.0</td><td>-12.7</td><td>1.5</td><td>-8.4</td><td>4.5E-17</td><td>3.3E-16</td><td>MB231/MB468</td><td>NUC</td></tr>
<tr><td>>MI0000077|hsa-mir-21&WithFlank&17|+|59841260|59841343@14.35.22(+2A)&[MIMAT0000076&hsa-miR-21-5p&offsets|0|0(+2A);m-1&17|+|59841273|59841294&offsets|0|0(+2A)]|TAGCTTATCAGACTGATGTTGAAA</td><td>>MI0000077|hsa-mir-21&WithFlank&17|+|59841260|59841343</td><td>14</td><td>35</td><td>22(+2A)</td><td>TAGCTTATCAGACTGATGTTGAAA</td><td>632.6</td><td>5.1</td><td>0.6</td><td>8.4</td><td>4.5E-17</td><td>3.3E-16</td><td>MB231/MB468</td><td>NUC</td></tr>
<tr><td>>hg38_21_+_8433172_8446622_RNA45SN1_45S_50ntFlanks@11718.11742.25|AACGGCGGGAGTAACTATGACTCTC</td><td>>hg38_21_+_8433172_8446622_RNA45SN1_45S_50ntFlanks</td><td>11718</td><td>11742</td><td>25</td><td>AACGGCGGGAGTAACTATGACTCTC</td><td>709.3</td><td>-6.2</td><td>0.7</td><td>-8.4</td><td>5.9E-17</td><td>3.3E-16</td><td>MB231/MB468</td><td>MITO</td></tr>
<tr><td>>hg38_21_+_8433172_8446622_RNA45SN1_45S_50ntFlanks@10808.10825.18|CAATGTAGGTAAGGGAAG</td><td>>hg38_21_+_8433172_8446622_RNA45SN1_45S_50ntFlanks</td><td>10808</td><td>10825</td><td>18</td><td>CAATGTAGGTAAGGGAAG</td><td>1612.4</td><td>-13.6</td><td>1.6</td><td>-8.4</td><td>4.6E-17</td><td>3.4E-16</td><td>MB231/MB468</td><td>NUC</td></tr>
<tr><td>>hg38_21_+_8433172_8446622_RNA45SN1_45S_50ntFlanks@11874.11912.39|AGAAGACCCTGTTGAGCTTGACTCTAGTCTGGCACGGTG</td><td>>hg38_21_+_8433172_8446622_RNA45SN1_45S_50ntFlanks</td><td>11874</td><td>11912</td><td>39</td><td>AGAAGACCCTGTTGAGCTTGACTCTAGTCTGGCACGGTG</td><td>572.8</td><td>-13.0</td><td>1.5</td><td>-8.4</td><td>3.6E-17</td><td>3.4E-16</td><td>MB231/MB468</td><td>MP</td></tr>
<tr><td>>MI0000811|hsa-mir-148b&WithFlank&12|+|54337210|54337320@69.90.22(+2U)&[MIMAT0000759&hsa-miR-148b-3p&offsets|0|0(+2U);m-77&12|+|54337278|54337299&offsets|0|0(+2U)]|TCAGTGCATCACAGAACTTTGTTT</td><td>>MI0000811|hsa-mir-148b&WithFlank&12|+|54337210|54337320</td><td>69</td><td>90</td><td>22(+2U)</td><td>TCAGTGCATCACAGAACTTTGTTT</td><td>325.8</td><td>6.6</td><td>0.8</td><td>8.4</td><td>4.6E-17</td><td>3.4E-16</td><td>MB231/MB468</td><td>NUC</td></tr>
<tr><td>>trna2_GlyGCC_21_-_18827107_18827177@1.29.29,trna35_GlyGCC_1_+_161413094_161413164@1.29.29,trna37_GlyGCC_1_+_161420467_161420537@1.29.29,trna39_GlyGCC_1_+_161427898_161427968@1.29.29,trna41_GlyGCC_1_+_161435258_161435328@1.29.29|GCATGGGTGGTTCAGTGGTAGAATTCTCG</td><td>>trna2_GlyGCC_21_-_18827107_18827177</td><td>1</td><td>29</td><td>29</td><td>GCATGGGTGGTTCAGTGGTAGAATTCTCG</td><td>16196.2</td><td>-11.9</td><td>1.4</td><td>-8.4</td><td>4.6E-17</td><td>3.4E-16</td><td>MB231/MB468</td><td>NUC</td></tr>
<tr><td>>trna2_GlyGCC_21_-_18827107_18827177@1.29.29,trna35_GlyGCC_1_+_161413094_161413164@1.29.29,trna37_GlyGCC_1_+_161420467_161420537@1.29.29,trna39_GlyGCC_1_+_161427898_161427968@1.29.29,trna41_GlyGCC_1_+_161435258_161435328@1.29.29|GCATGGGTGGTTCAGTGGTAGAATTCTCG</td><td>>trna2_GlyGCC_21_-_18827107_18827177</td><td>1</td><td>29</td><td>29</td><td>GCATGGGTGGTTCAGTGGTAGAATTCTCG</td><td>4570.8</td><td>-11.1</td><td>1.3</td><td>-8.4</td><td>6.0E-17</td><td>3.4E-16</td><td>MB231/MB468</td><td>MITO</td></tr>
<tr><td>>hg38_21_+_8433172_8446622_RNA45SN1_45S_50ntFlanks@4884.4917.34|CAAAGCTGAAACTTAAAGGAATTGACGGAAGGGC</td><td>>hg38_21_+_8433172_8446622_RNA45SN1_45S_50ntFlanks</td><td>4884</td><td>4917</td><td>34</td><td>CAAAGCTGAAACTTAAAGGAATTGACGGAAGGGC</td><td>405.9</td><td>-12.4</td><td>1.5</td><td>-8.4</td><td>6.0E-17</td><td>3.4E-16</td><td>MB231/MB468</td><td>MITO</td></tr>
<tr><td>>MI0000093|hsa-mir-92a-1&WithFlank&13|+|91351308|91351397@54.74.21(+1C);MI0000094|hsa-mir-92a-2&WithFlank&X|-|134169532|134169618@54.74.21(+1C)&[MIMAT0000092&hsa-miR-92a-3p&offsets|0|-1(+1C);m-9&13|+|91351361|91351382&offsets|0|-1(+1C)];[MIMAT0000092_1&hsa-miR-92a-3p&offsets|0|-1(+1C);m-11&X|-|134169544|134169565&offsets|0|-1(+1C)]|TATTGCACTTGTCCCGGCCTGC</td><td>>MI0000093|hsa-mir-92a-1&WithFlank&13|+|91351308|91351397</td><td>54</td><td>74</td><td>21(+1C)</td><td>TATTGCACTTGTCCCGGCCTGC</td><td>5885.4</td><td>5.9</td><td>0.7</td><td>8.4</td><td>4.5E-17</td><td>3.4E-16</td><td>BT20/MB468</td><td>MITO</td></tr>
<tr><td>>hg38_21_+_8433172_8446622_RNA45SN1_45S_50ntFlanks@8189.8206.18|CCCAGCCCGTGGACGGTG</td><td>>hg38_21_+_8433172_8446622_RNA45SN1_45S_50ntFlanks</td><td>8189</td><td>8206</td><td>18</td><td>CCCAGCCCGTGGACGGTG</td><td>275.9</td><td>-12.0</td><td>1.4</td><td>-8.4</td><td>3.7E-17</td><td>3.4E-16</td><td>MB231/MB468</td><td>MP</td></tr>
<tr><td>>hg38_21_+_8433172_8446622_RNA45SN1_45S_50ntFlanks@10687.10705.19|CGGCGTCCGGTGAGCTCTC</td><td>>hg38_21_+_8433172_8446622_RNA45SN1_45S_50ntFlanks</td><td>10687</td><td>10705</td><td>19</td><td>CGGCGTCCGGTGAGCTCTC</td><td>345.3</td><td>-12.2</td><td>1.5</td><td>-8.4</td><td>6.0E-17</td><td>3.4E-16</td><td>MB231/MB468</td><td>MITO</td></tr>
<tr><td>>hg38_21_+_8433172_8446622_RNA45SN1_45S_50ntFlanks@4738.4759.22|AAAGTCGGAGGTTCGAAGACGA</td><td>>hg38_21_+_8433172_8446622_RNA45SN1_45S_50ntFlanks</td><td>4738</td><td>4759</td><td>22</td><td>AAAGTCGGAGGTTCGAAGACGA</td><td>2703.8</td><td>-5.6</td><td>0.7</td><td>-8.4</td><td>3.7E-17</td><td>3.4E-16</td><td>MB231/MB468</td><td>MP</td></tr>
<tr><td>>hg38_21_+_8433172_8446622_RNA45SN1_45S_50ntFlanks@5056.5094.39|GAGCGATTTGTCTGGTTAATTCCGATAACGAACGAGACT</td><td>>hg38_21_+_8433172_8446622_RNA45SN1_45S_50ntFlanks</td><td>5056</td><td>5094</td><td>39</td><td>GAGCGATTTGTCTGGTTAATTCCGATAACGAACGAGACT</td><td>136.0</td><td>-24.1</td><td>2.9</td><td>-8.4</td><td>3.7E-17</td><td>3.4E-16</td><td>MB231/MB468</td><td>MP</td></tr>
<tr><td>>hg38_21_+_8433172_8446622_RNA45SN1_45S_50ntFlanks@6699.6720.22|GCTAGCTGCGAGAATTAATGTG</td><td>>hg38_21_+_8433172_8446622_RNA45SN1_45S_50ntFlanks</td><td>6699</td><td>6720</td><td>22</td><td>GCTAGCTGCGAGAATTAATGTG</td><td>71196.0</td><td>-7.9</td><td>0.9</td><td>-8.4</td><td>3.7E-17</td><td>3.4E-16</td><td>MB231/MB468</td><td>MP</td></tr>
<tr><td>>hg38_21_+_8433172_8446622_RNA45SN1_45S_50ntFlanks@10916.10933.18|CCGCGGCTGGACGAGGCG</td><td>>hg38_21_+_8433172_8446622_RNA45SN1_45S_50ntFlanks</td><td>10916</td><td>10933</td><td>18</td><td>CCGCGGCTGGACGAGGCG</td><td>564.3</td><td>-7.2</td><td>0.9</td><td>-8.4</td><td>3.8E-17</td><td>3.4E-16</td><td>MB231/MB468</td><td>CYTO</td></tr>
<tr><td>>hg38_21_+_8433172_8446622_RNA45SN1_45S_50ntFlanks@8115.8139.25|CGCGGGACATGTGGCGTACGGAAGA</td><td>>hg38_21_+_8433172_8446622_RNA45SN1_45S_50ntFlanks</td><td>8115</td><td>8139</td><td>25</td><td>CGCGGGACATGTGGCGTACGGAAGA</td><td>397.6</td><td>-12.4</td><td>1.5</td><td>-8.4</td><td>6.1E-17</td><td>3.4E-16</td><td>MB231/MB468</td><td>MITO</td></tr>
<tr><td>>MI0000089|hsa-mir-31&WithFlank&9|-|21512109|21512191@14.35.22&[MIMAT0000089&hsa-miR-31-5p&offsets|0|+1;m-360&9|-|21512157|21512178&offsets|0|0]|AGGCAAGATGCTGGCATAGCTG</td><td>>MI0000089|hsa-mir-31&WithFlank&9|-|21512109|21512191</td><td>14</td><td>35</td><td>22</td><td>AGGCAAGATGCTGGCATAGCTG</td><td>663.4</td><td>12.8</td><td>1.5</td><td>8.7</td><td>3.4E-18</td><td>3.4E-16</td><td>BT20/MB231</td><td>CYTO</td></tr>
<tr><td>>hg38_21_+_8433172_8446622_RNA45SN1_45S_50ntFlanks@3939.3959.21|AACCCGGTCAGCCCCTCTCCG</td><td>>hg38_21_+_8433172_8446622_RNA45SN1_45S_50ntFlanks</td><td>3939</td><td>3959</td><td>21</td><td>AACCCGGTCAGCCCCTCTCCG</td><td>400.8</td><td>-12.5</td><td>1.5</td><td>-8.4</td><td>3.7E-17</td><td>3.4E-16</td><td>MB231/MB468</td><td>MP</td></tr>
<tr><td>>hg38_21_+_8433172_8446622_RNA45SN1_45S_50ntFlanks@11768.11803.36|CTAATTAGTGACGCGCATGAATGGATGAACGAGATT</td><td>>hg38_21_+_8433172_8446622_RNA45SN1_45S_50ntFlanks</td><td>11768</td><td>11803</td><td>36</td><td>CTAATTAGTGACGCGCATGAATGGATGAACGAGATT</td><td>1174.9</td><td>-4.9</td><td>0.6</td><td>-8.4</td><td>6.5E-17</td><td>3.4E-16</td><td>BT20/MB468</td><td>CYTO</td></tr>
<tr><td>>MI0006444|hsa-mir-1307&WithFlank&10|-|103394247|103394407@86.107.22(+1A)&[MIMAT0005951&hsa-miR-1307-3p&offsets|0|0(+1A);m-66&10|-|103394300|103394322&offsets|0|-1(+1A)]|ACTCGGCGTGGCGTCGGTCGTGA</td><td>>MI0006444|hsa-mir-1307&WithFlank&10|-|103394247|103394407</td><td>86</td><td>107</td><td>22(+1A)</td><td>ACTCGGCGTGGCGTCGGTCGTGA</td><td>696.3</td><td>6.4</td><td>0.8</td><td>8.4</td><td>3.8E-17</td><td>3.5E-16</td><td>MB231/MB468</td><td>CYTO</td></tr>
<tr><td>>hg38_21_+_8433172_8446622_RNA45SN1_45S_50ntFlanks@12627.12652.26|ATGGGGCGAAGCTACCATCTGTGGGA</td><td>>hg38_21_+_8433172_8446622_RNA45SN1_45S_50ntFlanks</td><td>12627</td><td>12652</td><td>26</td><td>ATGGGGCGAAGCTACCATCTGTGGGA</td><td>349.5</td><td>-6.1</td><td>0.7</td><td>-8.4</td><td>3.8E-17</td><td>3.5E-16</td><td>MB231/MB468</td><td>CYTO</td></tr>
<tr><td>>trna111_HisGTG_1_-_147774845_147774916@-1A.29.30,trna118_HisGTG_1_-_145396881_145396952@-1A.29.30,trna16_HisGTG_1_+_146544773_146544844@-1A.29.30,trna1_HisGTG_15_+_45493349_45493420@-1A.29.30,trna21_HisGTG_1_+_147753471_147753542@-1A.29.30,trna33_HisGTG_6_+_27125906_27125977@-1A.29.30,trna7_HisGTG_9_-_14433938_14434009@-1A.29.30,trna8_HisGTG_15_-_45492611_45492682@-1A.29.30,trna9_HisGTG_15_-_45490804_45490875@-1A.29.30|AGCCGTGATCGTATAGTGGTTAGTACTCTG</td><td>>trna111_HisGTG_1_-_147774845_147774916</td><td>-1A</td><td>29</td><td>30</td><td>AGCCGTGATCGTATAGTGGTTAGTACTCTG</td><td>371.0</td><td>-12.4</td><td>1.5</td><td>-8.4</td><td>3.8E-17</td><td>3.5E-16</td><td>MB231/MB468</td><td>MP</td></tr>
<tr><td>>hg38_21_+_8433172_8446622_RNA45SN1_45S_50ntFlanks@11618.11640.23|AAACAAAGCATCGCGAAGGCCCG</td><td>>hg38_21_+_8433172_8446622_RNA45SN1_45S_50ntFlanks</td><td>11618</td><td>11640</td><td>23</td><td>AAACAAAGCATCGCGAAGGCCCG</td><td>1536.6</td><td>-10.0</td><td>1.2</td><td>-8.4</td><td>6.1E-17</td><td>3.5E-16</td><td>MB231/MB468</td><td>MITO</td></tr>
<tr><td>>hg38_21_+_8433172_8446622_RNA45SN1_45S_50ntFlanks@4722.4755.34|TCATTAATCAAGAACGAAAGTCGGAGGTTCGAAG</td><td>>hg38_21_+_8433172_8446622_RNA45SN1_45S_50ntFlanks</td><td>4722</td><td>4755</td><td>34</td><td>TCATTAATCAAGAACGAAAGTCGGAGGTTCGAAG</td><td>444.3</td><td>-12.6</td><td>1.5</td><td>-8.4</td><td>6.2E-17</td><td>3.5E-16</td><td>MB231/MB468</td><td>MITO</td></tr>
<tr><td>>hg38_21_+_8433172_8446622_RNA45SN1_45S_50ntFlanks@11667.11702.36|CTGCCCAGTGCTCTGAATGTCAAAGTGAAGAAATTC</td><td>>hg38_21_+_8433172_8446622_RNA45SN1_45S_50ntFlanks</td><td>11667</td><td>11702</td><td>36</td><td>CTGCCCAGTGCTCTGAATGTCAAAGTGAAGAAATTC</td><td>2152.8</td><td>6.4</td><td>0.8</td><td>8.4</td><td>3.8E-17</td><td>3.5E-16</td><td>MB231/MB468</td><td>CYTO</td></tr>
<tr><td>>hg38_21_+_8433172_8446622_RNA45SN1_45S_50ntFlanks@10684.10710.27|CGGCGGCGTCCGGTGAGCTCTCGCTGG</td><td>>hg38_21_+_8433172_8446622_RNA45SN1_45S_50ntFlanks</td><td>10684</td><td>10710</td><td>27</td><td>CGGCGGCGTCCGGTGAGCTCTCGCTGG</td><td>18328.5</td><td>-7.5</td><td>0.9</td><td>-8.4</td><td>6.2E-17</td><td>3.5E-16</td><td>MB231/MB468</td><td>MITO</td></tr>
<tr><td>>hg38_21_+_8433172_8446622_RNA45SN1_45S_50ntFlanks@4064.4101.38|ATCAACTTTCGATGGTAGTCGCCGTGCCTACCATGGTG</td><td>>hg38_21_+_8433172_8446622_RNA45SN1_45S_50ntFlanks</td><td>4064</td><td>4101</td><td>38</td><td>ATCAACTTTCGATGGTAGTCGCCGTGCCTACCATGGTG</td><td>262.3</td><td>-11.0</td><td>1.3</td><td>-8.4</td><td>4.8E-17</td><td>3.5E-16</td><td>MB231/MB468</td><td>NUC</td></tr>
</table>

| | | | | | | | | | | | | | |
|---|---|---|---|---|---|---|---|---|---|---|---|---|---|
| >hg38_21_+_8433172_8446622_RNA45SN1_45S_50ntFlanks@6731.6753.23\|CACATTGATCATCGACACTTCGA | >hg38_21_+_8433172_8446622_RNA45SN1_45S_50ntFlanks | 6731 | 6753 | 23 | CACATTGATCATCGACACTTCGA | 272.5 | -12.0 | 1.4 | -8.4 | 3.8E-17 | 3.5E-16 | MB231/MB468 | MP |
| >hg38_21_+_8433172_8446622_RNA45SN1_45S_50ntFlanks@6721.6744.24\|AATTGCAGGACACATTGATCATCG | >hg38_21_+_8433172_8446622_RNA45SN1_45S_50ntFlanks | 6721 | 6744 | 24 | AATTGCAGGACACATTGATCATCG | 32793.3 | -7.0 | 0.8 | -8.4 | 3.9E-17 | 3.5E-16 | MB231/MB468 | CYTO |
| >hg38_21_+_8433172_8446622_RNA45SN1_45S_50ntFlanks@8191.8215.25\|CAGCCCGTGGACGGTGTGAGGCCGG | >hg38_21_+_8433172_8446622_RNA45SN1_45S_50ntFlanks | 8191 | 8215 | 25 | CAGCCCGTGGACGGTGTGAGGCCGG | 1150.4 | -7.2 | 0.9 | -8.4 | 3.9E-17 | 3.5E-16 | MB231/MB468 | CYTO |
| >hg38_21_+_8433172_8446622_RNA45SN1_45S_50ntFlanks@12000.12032.33\|CCTGCGGGCCGCCGGTGAAATACCACTACTCTG | >hg38_21_+_8433172_8446622_RNA45SN1_45S_50ntFlanks | 12000 | 12032 | 33 | CCTGCGGGCCGCCGGTGAAATACCACTACTCTG | 272.0 | -12.0 | 1.4 | -8.4 | 3.8E-17 | 3.5E-16 | MB231/MB468 | MP |
| >hg38_21_+_8433172_8446622_RNA45SN1_45S_50ntFlanks@12418.12446.29\|CGATGTCGGCTCTTCCTATCATTGTGAAG | >hg38_21_+_8433172_8446622_RNA45SN1_45S_50ntFlanks | 12418 | 12446 | 29 | CGATGTCGGCTCTTCCTATCATTGTGAAG | 1368.2 | -4.1 | 0.5 | -8.4 | 4.7E-17 | 3.5E-16 | BT20/MB468 | MITO |
| >hg38_21_+_8433172_8446622_RNA45SN1_45S_50ntFlanks@5073.5112.40\|AATTCCGATAACGAACGAGACTCTGGCATGCTAACTAGTT | >hg38_21_+_8433172_8446622_RNA45SN1_45S_50ntFlanks | 5073 | 5112 | 40 | AATTCCGATAACGAACGAGACTCTGGCATGCTAACTAGTT | 128.1 | -24.1 | 2.9 | -8.4 | 3.8E-17 | 3.5E-16 | MB231/MB468 | MP |
| >hg38_21_+_8433172_8446622_RNA45SN1_45S_50ntFlanks@12635.12653.19\|AAGCTACCATCTGTGGGAT | >hg38_21_+_8433172_8446622_RNA45SN1_45S_50ntFlanks | 12635 | 12653 | 19 | AAGCTACCATCTGTGGGAT | 1973.3 | -13.0 | 1.6 | -8.4 | 4.7E-17 | 3.5E-16 | BT20/MB468 | MITO |
| >am_MI0000809\|hsa-mir-151a&WithFlank&8\|-\|140732558\|140732659@53.72.20(+1A)&[MIMAT0000757&hsa-miR-151a-3p&offsets\|0\|-1(+1A);m-30&8\|-\|140732587\|140732607&offsets\|0\|-1(+1A)]\|CTAGACTGAAGCTCCTTGAGA | >am_MI0000809\|hsa-mir-151a&WithFlank&8\|-\|140732558\|140732659 | 53 | 72 | 20(+1A) | CTAGACTGAAGCTCCTTGAGA | 2747.5 | 3.8 | 0.5 | 8.3 | 1.1E-16 | 3.5E-16 | BT20/MB468 | NUC |
| >hg38_21_+_8433172_8446622_RNA45SN1_45S_50ntFlanks@12081.12107.27\|CTCTCGCTTCTGGCGCCAAGCGCCCGG | >hg38_21_+_8433172_8446622_RNA45SN1_45S_50ntFlanks | 12081 | 12107 | 27 | CTCTCGCTTCTGGCGCCAAGCGCCCGG | 291.3 | -12.0 | 1.4 | -8.4 | 3.8E-17 | 3.5E-16 | MB231/MB468 | MP |
| >MI0000253\|hsa-mir-148a&WithFlank&7\|-\|25949913\|25949992@50.70.21(+1C)&[MIMAT0000243&hsa-miR-148a-3p&offsets\|0\|-1(+1C);m-7&7\|-\|25949922\|25949943&offsets\|0\|-1(+1C)]\|TCAGTGCACTACAGAACTTTGC | >MI0000253\|hsa-mir-148a&WithFlank&7\|-\|25949913\|25949992 | 50 | 70 | 21(+1C) | TCAGTGCACTACAGAACTTTGC | 21500.1 | 4.7 | 0.5 | 8.6 | 7.2E-18 | 3.5E-16 | BT20/MB231 | NUC |
| >hg38_21_+_8433172_8446622_RNA45SN1_45S_50ntFlanks@4657.4684.28\|CTAGAGGTGAAATTCTTGGACCGGCGCA | >hg38_21_+_8433172_8446622_RNA45SN1_45S_50ntFlanks | 4657 | 4684 | 28 | CTAGAGGTGAAATTCTTGGACCGGCGCA | 295.1 | -11.2 | 1.3 | -8.4 | 4.8E-17 | 3.5E-16 | MB231/MB468 | NUC |
| >hg38_21_+_8433172_8446622_RNA45SN1_45S_50ntFlanks@10916.10936.21\|CCGCGGCTGGACGAGGCGCCG | >hg38_21_+_8433172_8446622_RNA45SN1_45S_50ntFlanks | 10916 | 10936 | 21 | CCGCGGCTGGACGAGGCGCCG | 560.0 | -12.9 | 1.5 | -8.4 | 6.3E-17 | 3.5E-16 | MB231/MB468 | MITO |
| >am_trnaMT_ValTAC_MT_+_1602_1670@50.68.19\|ACTTAACTTGACCGCTCTG | >am_trnaMT_ValTAC_MT_+_1602_1670 | 50 | 68 | 19 | ACTTAACTTGACCGCTCTG | 4358.2 | -6.1 | 0.7 | -8.4 | 6.4E-17 | 3.6E-16 | MB231/MB468 | MITO |
| >hg38_21_+_8433172_8446622_RNA45SN1_45S_50ntFlanks@11583.11612.30\|CTGGTGCGGACCAGGGGAATCCGACTGTTT | >hg38_21_+_8433172_8446622_RNA45SN1_45S_50ntFlanks | 11583 | 11612 | 30 | CTGGTGCGGACCAGGGGAATCCGACTGTTT | 2899.6 | -5.3 | 0.6 | -8.4 | 6.4E-17 | 3.6E-16 | MB231/MB468 | MITO |
| >hg38_21_+_8433172_8446622_RNA45SN1_45S_50ntFlanks@10841.10863.23\|CGTAACTTCGGGATAAGGATTGG | >hg38_21_+_8433172_8446622_RNA45SN1_45S_50ntFlanks | 10841 | 10863 | 23 | CGTAACTTCGGGATAAGGATTGG | 5051.6 | -9.8 | 1.2 | -8.3 | 1.1E-16 | 3.6E-16 | BT20/MB468 | NUC |
| >MI0000477\|hsa-mir-146a&WithFlank&5\|+\|160485346\|160485456@27.48.22(+1C)&[MIMAT0000449&hsa-miR-146a-5p&offsets\|0\|0(+1C);m-129&5\|+\|160485372\|160485393&offsets\|0\|0(+1C)]\|TGAGAACTGAATTCCATGGGTTC | >MI0000477\|hsa-mir-146a&WithFlank&5\|+\|160485346\|160485456 | 27 | 48 | 22(+1C) | TGAGAACTGAATTCCATGGGTTC | 1245.3 | 13.5 | 1.6 | 8.4 | 3.9E-17 | 3.6E-16 | MB231/MB468 | MP |
| >hg38_21_+_8433172_8446622_RNA45SN1_45S_50ntFlanks@11581.11609.29\|AACTGGTGCGGACCAGGGGAATCCGACTG | >hg38_21_+_8433172_8446622_RNA45SN1_45S_50ntFlanks | 11581 | 11609 | 29 | AACTGGTGCGGACCAGGGGAATCCGACTG | 2653.1 | -5.5 | 0.7 | -8.4 | 6.5E-17 | 3.6E-16 | MB231/MB468 | MITO |
| >trna10_LysCTT_16_+_3241501_3241573@1.30.30,trna13_LysCTT_14_-_58706613_58706685@1.30.30,trna2_LysCTT_15_+_79152904_79152976@1.30.30\|GCCCGGCTAGCTCAGTCGGTAGAGCATGGG | >trna10_LysCTT_16_+_3241501_3241573 | 1 | 30 | 30 | GCCCGGCTAGCTCAGTCGGTAGAGCATGGG | 616.8 | -12.0 | 1.4 | -8.4 | 4.0E-17 | 3.6E-16 | MB231/MB468 | CYTO |
| >hg38_21_+_8433172_8446622_RNA45SN1_45S_50ntFlanks@12158.12182.25\|GTTTGACTGGGGCGGTACACCTGTC | >hg38_21_+_8433172_8446622_RNA45SN1_45S_50ntFlanks | 12158 | 12182 | 25 | GTTTGACTGGGGCGGTACACCTGTC | 536.6 | -12.9 | 1.5 | -8.4 | 4.0E-17 | 3.6E-16 | MB231/MB468 | MP |
| >hg38_21_+_8433172_8446622_RNA45SN1_45S_50ntFlanks@13006.13027.22\|CTATTGAAAGTCAGCCCTCGAC | >hg38_21_+_8433172_8446622_RNA45SN1_45S_50ntFlanks | 13006 | 13027 | 22 | CTATTGAAAGTCAGCCCTCGAC | 361.3 | -12.3 | 1.5 | -8.4 | 6.5E-17 | 3.6E-16 | MB231/MB468 | MITO |
| >hg38_21_+_8433172_8446622_RNA45SN1_45S_50ntFlanks@12957.12979.23\|CTTCTGGGTCGGGGTTTCGTACG | >hg38_21_+_8433172_8446622_RNA45SN1_45S_50ntFlanks | 12957 | 12979 | 23 | CTTCTGGGTCGGGGTTTCGTACG | 463.1 | -12.7 | 1.5 | -8.4 | 4.0E-17 | 3.6E-16 | MB231/MB468 | MP |
| >MI0000743\|hsa-mir-34c&WithFlank&11\|+\|111513433\|111513521@19.41.23&[MIMAT0000686&hsa-miR-34c-5p&offsets\|0\|0;m-336&11\|+\|111513451\|111513473&offsets\|0\|0]\|AGGCAGTGTAGTTAGCTGATTGC | >MI0000743\|hsa-mir-34c&WithFlank&11\|+\|111513433\|111513521 | 19 | 41 | 23 | AGGCAGTGTAGTTAGCTGATTGC | 850.7 | -13.0 | 1.5 | -8.6 | 5.2E-18 | 3.6E-16 | BT20/MB231 | MITO |
| >MI0000283\|hsa-mir-203a&WithFlank&14\|+\|104117399\|104117520@72.92.21(+1U)&[MIMAT0000264&hsa-miR-203a-3p&offsets\|+1\|0(+1U);m-79&14\|+\|104117469\|104117490&offsets\|+1\|0(+1U)]\|TGAAATGTTTAGGACCACTAGT | >MI0000283\|hsa-mir-203a&WithFlank&14\|+\|104117399\|104117520 | 72 | 92 | 21(+1U) | TGAAATGTTTAGGACCACTAGT | 346.2 | 12.1 | 1.4 | 8.6 | 5.3E-18 | 3.6E-16 | BT20/MB231 | MITO |
| >hg38_21_+_8433172_8446622_RNA45SN1_45S_50ntFlanks@10681.10712.32\|TTCCGGCGGCGTCCGGTGAGCTCTCGCTGGCC | >hg38_21_+_8433172_8446622_RNA45SN1_45S_50ntFlanks | 10681 | 10712 | 32 | TTCCGGCGGCGTCCGGTGAGCTCTCGCTGGCC | 3221.8 | -5.8 | 0.7 | -8.4 | 5.0E-17 | 3.7E-16 | MB231/MB468 | NUC |
| >hg38_21_+_8433172_8446622_RNA45SN1_45S_50ntFlanks@4453.4473.21\|TCGGCGCCCCCTCGATGCTCT | >hg38_21_+_8433172_8446622_RNA45SN1_45S_50ntFlanks | 4453 | 4473 | 21 | TCGGCGCCCCCTCGATGCTCT | 1026.7 | 11.4 | 1.4 | 8.4 | 4.1E-17 | 3.7E-16 | MB231/MB468 | CYTO |
| >hg38_21_+_8433172_8446622_RNA45SN1_45S_50ntFlanks@6651.6676.26\|CGACTCTTAGCGGTGGATCACTCGGC | >hg38_21_+_8433172_8446622_RNA45SN1_45S_50ntFlanks | 6651 | 6676 | 26 | CGACTCTTAGCGGTGGATCACTCGGC | 75872.1 | 6.0 | 0.7 | 8.4 | 4.9E-17 | 3.7E-16 | BT20/MB468 | MITO |
| >hg38_21_+_8433172_8446622_RNA45SN1_45S_50ntFlanks@3777.3799.23\|CGGTACAGTGAAACTGCGAATGG | >hg38_21_+_8433172_8446622_RNA45SN1_45S_50ntFlanks | 3777 | 3799 | 23 | CGGTACAGTGAAACTGCGAATGG | 299.3 | -12.1 | 1.4 | -8.4 | 4.0E-17 | 3.7E-16 | MB231/MB468 | MP |
| >hg38_21_+_8433172_8446622_RNA45SN1_45S_50ntFlanks@6731.6771.41\|CACATTGATCATCGACACTTCGAACGCACTTGCGGCCCCGG | >hg38_21_+_8433172_8446622_RNA45SN1_45S_50ntFlanks | 6731 | 6771 | 41 | CACATTGATCATCGACACTTCGAACGCACTTGCGGCCCCGG | 4364.1 | -8.7 | 1.0 | -8.4 | 6.6E-17 | 3.7E-16 | MB231/MB468 | MITO |
| >hg38_21_+_8433172_8446622_RNA45SN1_45S_50ntFlanks@7986.8005.20\|ATCAGACGTGGCGACCCGCT | >hg38_21_+_8433172_8446622_RNA45SN1_45S_50ntFlanks | 7986 | 8005 | 20 | ATCAGACGTGGCGACCCGCT | 284.5 | -11.1 | 1.3 | -8.4 | 5.1E-17 | 3.7E-16 | MB231/MB468 | NUC |
| >hg38_21_+_8433172_8446622_RNA45SN1_45S_50ntFlanks@10806.10828.23\|AACAATGTAGGTAAGGGAAGTCG | >hg38_21_+_8433172_8446622_RNA45SN1_45S_50ntFlanks | 10806 | 10828 | 23 | AACAATGTAGGTAAGGGAAGTCG | 295.7 | -12.1 | 1.4 | -8.4 | 4.1E-17 | 3.7E-16 | MB231/MB468 | MP |
| >hg38_21_+_8433172_8446622_RNA45SN1_45S_50ntFlanks@11918.11967.50\|ACATGAGAGGTGTAGAATAAGTGGGAGGCCCCCGGCGCCCCCCCGGTGTC | >hg38_21_+_8433172_8446622_RNA45SN1_45S_50ntFlanks | 11918 | 11967 | 50 | ACATGAGAGGTGTAGAATAAGTGGGAGGCCCCCGGCGCCCCCCCGGTGTC | 2019.6 | -15.4 | 1.9 | -8.3 | 1.1E-16 | 3.7E-16 | BT20/MB468 | NUC |
| >hg38_21_+_8433172_8446622_RNA45SN1_45S_50ntFlanks@9503.9526.24\|GGTGAACTATGCCTGGGCAGGGCG | >hg38_21_+_8433172_8446622_RNA45SN1_45S_50ntFlanks | 9503 | 9526 | 24 | GGTGAACTATGCCTGGGCAGGGCG | 993.1 | -13.8 | 1.6 | -8.4 | 4.1E-17 | 3.7E-16 | MB231/MB468 | MP |
| >MI0000273\|hsa-mir-183&WithFlank&7\|-\|129774899\|129775020@33.54.22(+1A)&[MIMAT0000261&hsa-miR-183-5p&offsets\|0\|0(+1A);m-41&7\|-\|129774966\|129774987&offsets\|-1\|-1(+1A)]\|TATGGCACTGGTAGAATTCACTA | >MI0000273\|hsa-mir-183&WithFlank&7\|-\|129774899\|129775020 | 33 | 54 | 22(+1A) | TATGGCACTGGTAGAATTCACTA | 166.8 | 2.7 | 0.3 | 8.3 | 1.2E-16 | 3.7E-16 | BT20/MB468 | NUC |
| >hg38_21_+_8433172_8446622_RNA45SN1_45S_50ntFlanks@9805.9838.34\|GCGTGGAATGCGAGTGCCTAGTGGGCCACTTTTG | >hg38_21_+_8433172_8446622_RNA45SN1_45S_50ntFlanks | 9805 | 9838 | 34 | GCGTGGAATGCGAGTGCCTAGTGGGCCACTTTTG | 397.1 | -12.4 | 1.5 | -8.4 | 6.7E-17 | 3.7E-16 | MB231/MB468 | MITO |
| >MI0001445\|hsa-mir-423&WithFlank&17\|+\|30117073\|30117178@59.80.22(+1A)&[MIMAT0001340&hsa-miR-423-3p&offsets\|0\|-1(+1A);m-100&17\|+\|30117131\|30117153&offsets\|0\|-1(+1A)]\|AGCTCGGTCTGAGGCCCCTCAGA | >MI0001445\|hsa-mir-423&WithFlank&17\|+\|30117073\|30117178 | 59 | 80 | 22(+1A) | AGCTCGGTCTGAGGCCCCTCAGA | 2032.7 | 8.4 | 1.0 | 8.3 | 7.1E-17 | 3.8E-16 | BT20/MB468 | CYTO |
| >hg38_21_+_8433172_8446622_RNA45SN1_45S_50ntFlanks@10107.10126.20\|CGTGTGGTGTGCGTCGGAGG | >hg38_21_+_8433172_8446622_RNA45SN1_45S_50ntFlanks | 10107 | 10126 | 20 | CGTGTGGTGTGCGTCGGAGG | 359.0 | -12.4 | 1.5 | -8.4 | 4.1E-17 | 3.8E-16 | MB231/MB468 | MP |
| >trnaMT_ValTAC_MT_+_1602_1670@42.72.31\|AGATTTCAACTTAACTTGACCGCTCTGACCA | >trnaMT_ValTAC_MT_+_1602_1670 | 42 | 72 | 31 | AGATTTCAACTTAACTTGACCGCTCTGACCA | 5119.3 | 5.9 | 0.7 | 8.4 | 5.2E-17 | 3.8E-16 | MB231/MB468 | NUC |
| >hg38_21_+_8433172_8446622_RNA45SN1_45S_50ntFlanks@6647.6665.19\|CGTACGACTCTTAGCGGTG | >hg38_21_+_8433172_8446622_RNA45SN1_45S_50ntFlanks | 6647 | 6665 | 19 | CGTACGACTCTTAGCGGTG | 7826.7 | 5.6 | 0.7 | 8.3 | 7.1E-17 | 3.8E-16 | BT20/MB468 | CYTO |
| >MI0000734\|hsa-mir-106b&WithFlank&7\|-\|100093987\|100094080@18.38.21&[MIMAT0000680&hsa-miR-106b-5p&offsets\|0\|0;m-103&7\|-\|100094043\|100094063&offsets\|0\|0]\|TAAAGTGCTGACAGTGCAGAT | >MI0000734\|hsa-mir-106b&WithFlank&7\|-\|100093987\|100094080 | 18 | 38 | 21 | TAAAGTGCTGACAGTGCAGAT | 387.4 | 6.2 | 0.7 | 8.3 | 7.1E-17 | 3.8E-16 | BT20/MB468 | CYTO |
| >hg38_21_+_8433172_8446622_RNA45SN1_45S_50ntFlanks@11604.11623.20\|CGACTGTTTAATTAAAACAA | >hg38_21_+_8433172_8446622_RNA45SN1_45S_50ntFlanks | 11604 | 11623 | 20 | CGACTGTTTAATTAAAACAA | 1321.7 | 5.8 | 0.7 | 8.3 | 7.1E-17 | 3.8E-16 | BT20/MB468 | CYTO |
| >hg38_21_+_8433172_8446622_RNA45SN1_45S_50ntFlanks@8552.8579.28\|CTCCCCGGAGGGGGCGGGCTCCGGCGGG | >hg38_21_+_8433172_8446622_RNA45SN1_45S_50ntFlanks | 8552 | 8579 | 28 | CTCCCCGGAGGGGGCGGGCTCCGGCGGG | 345.0 | -12.2 | 1.5 | -8.4 | 6.8E-17 | 3.8E-16 | MB231/MB468 | MITO |
| >hg38_21_+_8433172_8446622_RNA45SN1_45S_50ntFlanks@10498.10516.19\|TTCAGATCCCCGAATCCGG | >hg38_21_+_8433172_8446622_RNA45SN1_45S_50ntFlanks | 10498 | 10516 | 19 | TTCAGATCCCCGAATCCGG | 491.1 | -11.7 | 1.4 | -8.4 | 4.3E-17 | 3.8E-16 | MB231/MB468 | CYTO |
| >hg38_21_+_8433172_8446622_RNA45SN1_45S_50ntFlanks@13004.13027.24\|ATCTATTGAAAGTCAGCCCTCGAC | >hg38_21_+_8433172_8446622_RNA45SN1_45S_50ntFlanks | 13004 | 13027 | 24 | ATCTATTGAAAGTCAGCCCTCGAC | 585.2 | -11.3 | 1.3 | -8.4 | 5.1E-17 | 3.8E-16 | BT20/MB468 | MITO |
| >hg38_21_+_8433172_8446622_RNA45SN1_45S_50ntFlanks@11671.11691.21\|CCAGTGCTCTGAATGTCAAAG | >hg38_21_+_8433172_8446622_RNA45SN1_45S_50ntFlanks | 11671 | 11691 | 21 | CCAGTGCTCTGAATGTCAAAG | 1506.4 | -6.2 | 0.8 | -8.3 | 1.2E-16 | 3.8E-16 | BT20/MB468 | NUC |
| >hg38_21_+_8433172_8446622_RNA45SN1_45S_50ntFlanks@10107.10127.21\|CGTGTGGTGTGCGTCGGAGGG | >hg38_21_+_8433172_8446622_RNA45SN1_45S_50ntFlanks | 10107 | 10127 | 21 | CGTGTGGTGTGCGTCGGAGGG | 504.4 | -11.9 | 1.4 | -8.4 | 5.3E-17 | 3.8E-16 | MB231/MB468 | NUC |
| >hg38_21_+_8433172_8446622_RNA45SN1_45S_50ntFlanks@8176.8204.29\|CTTCTGATCGAGGCCCAGCCCGTGGACGG | >hg38_21_+_8433172_8446622_RNA45SN1_45S_50ntFlanks | 8176 | 8204 | 29 | CTTCTGATCGAGGCCCAGCCCGTGGACGG | 1658.0 | -5.0 | 0.6 | -8.4 | 5.3E-17 | 3.9E-16 | MB231/MB468 | NUC |
| >MI0000071\|hsa-mir-17&WithFlank&13\|+\|91350599\|91350694@20.43.24&[MIMAT0000070&hsa-miR-17-5p&offsets\|0\|+1;m-88&13\|+\|91350618\|91350640&offsets\|0\|+1]\|CAAAGTGCTTACAGTGCAGGTAGT | >MI0000071\|hsa-mir-17&WithFlank&13\|+\|91350599\|91350694 | 20 | 43 | 24 | CAAAGTGCTTACAGTGCAGGTAGT | 2213.6 | 5.5 | 0.7 | 8.4 | 5.4E-17 | 3.9E-16 | MB231/MB468 | NUC |
| >MI0000077\|hsa-mir-21&WithFlank&17\|+\|59841260\|59841343@14.34.21(+1G)&[MIMAT0000076&hsa-miR-21-5p&offsets\|0\|-1(+1G);m-1&17\|+\|59841273\|59841294&offsets\|0\|-1(+1G)]\|TAGCTTATCAGACTGATGTTGG | >MI0000077\|hsa-mir-21&WithFlank&17\|+\|59841260\|59841343 | 14 | 34 | 21(+1G) | TAGCTTATCAGACTGATGTTGG | 1174.6 | 6.4 | 0.8 | 8.4 | 5.4E-17 | 3.9E-16 | MB231/MB468 | NUC |

| | | | | | | | | | | | | | |
|---|---|---|---|---|---|---|---|---|---|---|---|---|---|
| >MI0000433\|hsa-let-7g&WithFlank&3\|-\|52268272\|52268367@11.32.22&[MIMAT0000414&hsa-let-7g-5p&offsets\|0\|0;m-58&3\|-\|52268336\|52268357&offsets\|0\|0]\|TGAGGTAGTAGTTTGTACAGTT | >MI0000433\|hsa-let-7g&WithFlank&3\|-\|52268272\|52268367 | 11 | 32 | 22 | TGAGGTAGTAGTTTGTACAGTT | 112903.8 | 5.0 | 0.6 | 8.4 | 5.4E-17 | 3.9E-16 | MB231/MB468 | NUC |
| >hg38_21_+_8433172_8446622_RNA45SN1_45S_50ntFlanks@9736.9757.22\|CCGAAACGATCTCAACCTATTC | >hg38_21_+_8433172_8446622_RNA45SN1_45S_50ntFlanks | 9736 | 9757 | 22 | CCGAAACGATCTCAACCTATTC | 532.3 | -12.9 | 1.5 | -8.4 | 4.3E-17 | 3.9E-16 | MB231/MB468 | MP |
| >hg38_21_+_8433172_8446622_RNA45SN1_45S_50ntFlanks@7980.7999.20\|CCTCAGATCAGACGTGGCGA | >hg38_21_+_8433172_8446622_RNA45SN1_45S_50ntFlanks | 7980 | 7999 | 20 | CCTCAGATCAGACGTGGCGA | 228.0 | -11.7 | 1.4 | -8.4 | 4.3E-17 | 3.9E-16 | MB231/MB468 | MP |
| >hg38_21_+_8433172_8446622_RNA45SN1_45S_50ntFlanks@13008.13025.18\|ATTGAAAGTCAGCCCTCG | >hg38_21_+_8433172_8446622_RNA45SN1_45S_50ntFlanks | 13008 | 13025 | 18 | ATTGAAAGTCAGCCCTCG | 432.2 | -12.6 | 1.5 | -8.4 | 4.3E-17 | 3.9E-16 | MB231/MB468 | MP |
| >hg38_21_+_8433172_8446622_RNA45SN1_45S_50ntFlanks@9167.9187.21\|CGGGCGGGTCGCGCCGTCGGG | >hg38_21_+_8433172_8446622_RNA45SN1_45S_50ntFlanks | 9167 | 9187 | 21 | CGGGCGGGTCGCGCCGTCGGG | 495.7 | -12.8 | 1.5 | -8.4 | 4.3E-17 | 4.0E-16 | MB231/MB468 | MP |
| >hg38_21_+_8433172_8446622_RNA45SN1_45S_50ntFlanks@10737.10755.19\|TAAATCTCGCGCCGGGCCG | >hg38_21_+_8433172_8446622_RNA45SN1_45S_50ntFlanks | 10737 | 10755 | 19 | TAAATCTCGCGCCGGGCCG | 8601.1 | -12.5 | 1.5 | -8.3 | 1.2E-16 | 4.0E-16 | BT20/MB468 | NUC |
| >hg38_21_+_8433172_8446622_RNA45SN1_45S_50ntFlanks@11575.11593.19\|ACTTAGAACTGGTGCGGAC | >hg38_21_+_8433172_8446622_RNA45SN1_45S_50ntFlanks | 11575 | 11593 | 19 | ACTTAGAACTGGTGCGGAC | 2155.5 | -5.0 | 0.6 | -8.3 | 7.5E-17 | 4.0E-16 | BT20/MB468 | CYTO |
| >hg38_21_+_8433172_8446622_RNA45SN1_45S_50ntFlanks@13002.13023.22\|CGATCTATTGAAAGTCAGCCCT | >hg38_21_+_8433172_8446622_RNA45SN1_45S_50ntFlanks | 13002 | 13023 | 22 | CGATCTATTGAAAGTCAGCCCT | 527.5 | -12.8 | 1.5 | -8.3 | 7.1E-17 | 4.0E-16 | MB231/MB468 | MITO |
| >hg38_21_+_8433172_8446622_RNA45SN1_45S_50ntFlanks@9032.9076.45\|CCTCGTGGGGGGGCCGGGCCACCCCTCCCACGGCGCGACCGCTCT | >hg38_21_+_8433172_8446622_RNA45SN1_45S_50ntFlanks | 9032 | 9076 | 45 | CCTCGTGGGGGGGCCGGGCCACCCCTCCCACGGCGCGACCGCTCT | 824.3 | -13.5 | 1.6 | -8.3 | 7.1E-17 | 4.0E-16 | MB231/MB468 | MITO |
| >hg38_21_+_8433172_8446622_RNA45SN1_45S_50ntFlanks@11569.11591.23\|CAGCCGACTTAGAACTGGTGCGG | >hg38_21_+_8433172_8446622_RNA45SN1_45S_50ntFlanks | 11569 | 11591 | 23 | CAGCCGACTTAGAACTGGTGCGG | 1287.3 | -14.2 | 1.7 | -8.4 | 4.4E-17 | 4.0E-16 | MB231/MB468 | MP |
| >hg38_21_+_8433172_8446622_RNA45SN1_45S_50ntFlanks@11208.11254.47\|GCGGCGACTCTGGACGCGAGCCGGGCCCTTCCCGTGGATCGCCCCAG | >hg38_21_+_8433172_8446622_RNA45SN1_45S_50ntFlanks | 11208 | 11254 | 47 | GCGGCGACTCTGGACGCGAGCCGGGCCCTTCCCGTGGATCGCCCCAG | 66318.4 | -12.4 | 1.5 | -8.3 | 1.2E-16 | 4.0E-16 | BT20/MB468 | NUC |
| >hg38_21_+_8433172_8446622_RNA45SN1_45S_50ntFlanks@10830.10874.45\|CAAGCCGGATCCGTAACTTCGGGATAAGGATTGGCTCTAAGGGCT | >hg38_21_+_8433172_8446622_RNA45SN1_45S_50ntFlanks | 10830 | 10874 | 45 | CAAGCCGGATCCGTAACTTCGGGATAAGGATTGGCTCTAAGGGCT | 2574.0 | -15.8 | 1.9 | -8.3 | 1.2E-16 | 4.0E-16 | BT20/MB468 | NUC |
| >hg38_21_+_8433172_8446622_RNA45SN1_45S_50ntFlanks@6721.6740.20\|AATTGCAGGACACATTGATC | >hg38_21_+_8433172_8446622_RNA45SN1_45S_50ntFlanks | 6721 | 6740 | 20 | AATTGCAGGACACATTGATC | 7037.7 | -6.3 | 0.8 | -8.3 | 7.5E-17 | 4.0E-16 | BT20/MB468 | CYTO |
| >trnaMT_PheGAA_MT_+_577_647@35.55.21\|GAAAATGTTTAGACGGGCTCA | >trnaMT_PheGAA_MT_+_577_647 | 35 | 55 | 21 | GAAAATGTTTAGACGGGCTCA | 2993.7 | 9.5 | 1.1 | 8.4 | 5.4E-17 | 4.0E-16 | BT20/MB468 | MITO |
| >hg38_21_+_8433172_8446622_RNA45SN1_45S_50ntFlanks@11665.11687.23\|TTCTGCCCAGTGCTCTGAATGTC | >hg38_21_+_8433172_8446622_RNA45SN1_45S_50ntFlanks | 11665 | 11687 | 23 | TTCTGCCCAGTGCTCTGAATGTC | 758.9 | 5.5 | 0.7 | 8.4 | 4.5E-17 | 4.0E-16 | MB231/MB468 | CYTO |
| >hg38_21_+_8433172_8446622_RNA45SN1_45S_50ntFlanks@8380.8404.25\|CAAGAGGGCGTGAAACCGTTAAGAG | >hg38_21_+_8433172_8446622_RNA45SN1_45S_50ntFlanks | 8380 | 8404 | 25 | CAAGAGGGCGTGAAACCGTTAAGAG | 578.8 | -6.9 | 0.8 | -8.4 | 4.5E-17 | 4.0E-16 | MB231/MB468 | CYTO |
| >MI0000102\|hsa-mir-100&WithFlank&11\|-\|122152223\|122152314@19.36.18&[MIMAT0000098&hsa-miR-100-5p&offsets\|0\|-4;m-27&11\|-\|122152276\|122152296&offsets\|0\|-3]\|AACCCGTAGATCCGAACT | >MI0000102\|hsa-mir-100&WithFlank&11\|-\|122152223\|122152314 | 19 | 36 | 18 | AACCCGTAGATCCGAACT | 399.9 | 11.9 | 1.4 | 8.4 | 4.4E-17 | 4.0E-16 | MB231/MB468 | MP |
| >MI0000811\|hsa-mir-148b&WithFlank&12\|+\|54337210\|54337320@69.89.21(+1C)&[MIMAT0000759&hsa-miR-148b-3p&offsets\|0\|-1(+1C);m-77&12\|+\|54337278\|54337299&offsets\|0\|-1(+1C)]\|TCAGTGCATCACAGAACTTTGC | >MI0000811\|hsa-mir-148b&WithFlank&12\|+\|54337210\|54337320 | 69 | 89 | 21(+1C) | TCAGTGCATCACAGAACTTTGC | 438.8 | 5.7 | 0.7 | 8.4 | 5.6E-17 | 4.0E-16 | MB231/MB468 | NUC |
| >hg38_21_+_8433172_8446622_RNA45SN1_45S_50ntFlanks@10572.10607.36\|AGAAGCCGGCGGGAGCCCCGGGGAGAGTTCTCTTTT | >hg38_21_+_8433172_8446622_RNA45SN1_45S_50ntFlanks | 10572 | 10607 | 36 | AGAAGCCGGCGGGAGCCCCGGGGAGAGTTCTCTTTT | 384.9 | -12.4 | 1.5 | -8.3 | 7.2E-17 | 4.0E-16 | MB231/MB468 | MITO |
| >hg38_21_+_8433172_8446622_RNA45SN1_45S_50ntFlanks@9353.9370.18\|CGTGGCGCAATGAAGGTG | >hg38_21_+_8433172_8446622_RNA45SN1_45S_50ntFlanks | 9353 | 9370 | 18 | CGTGGCGCAATGAAGGTG | 378.0 | -12.3 | 1.5 | -8.3 | 7.2E-17 | 4.0E-16 | MB231/MB468 | MITO |
| >MI0000088\|hsa-mir-30a&WithFlank&6\|-\|71403545\|71403627@12.33.22(+1U)&[MIMAT0000087&hsa-miR-30a-5p&offsets\|0\|0(+1U);m-15&6\|-\|71403595\|71403616&offsets\|0\|0(+1U)]\|TGTAAACATCCTCGACTGGAAGT | >MI0000088\|hsa-mir-30a&WithFlank&6\|-\|71403545\|71403627 | 12 | 33 | 22(+1U) | TGTAAACATCCTCGACTGGAAGT | 4247.4 | 7.0 | 0.8 | 8.4 | 5.6E-17 | 4.0E-16 | MB231/MB468 | NUC |
| >hg38_21_+_8433172_8446622_RNA45SN1_45S_50ntFlanks@12161.12181.21\|TGACTGGGGCGGTACACCTGT | >hg38_21_+_8433172_8446622_RNA45SN1_45S_50ntFlanks | 12161 | 12181 | 21 | TGACTGGGGCGGTACACCTGT | 502.5 | -12.8 | 1.5 | -8.3 | 7.3E-17 | 4.0E-16 | MB231/MB468 | MITO |
| >hg38_21_+_8433172_8446622_RNA45SN1_45S_50ntFlanks@11805.11823.19\|CCACTGTCCCTACCTACTA | >hg38_21_+_8433172_8446622_RNA45SN1_45S_50ntFlanks | 11805 | 11823 | 19 | CCACTGTCCCTACCTACTA | 399.5 | -12.4 | 1.5 | -8.3 | 7.3E-17 | 4.0E-16 | MB231/MB468 | MITO |
| >hg38_21_+_8433172_8446622_RNA45SN1_45S_50ntFlanks@11768.11792.25\|CTAATTAGTGACGCGCATGAATGGA | >hg38_21_+_8433172_8446622_RNA45SN1_45S_50ntFlanks | 11768 | 11792 | 25 | CTAATTAGTGACGCGCATGAATGGA | 2435.5 | -6.0 | 0.7 | -8.4 | 4.5E-17 | 4.0E-16 | MB231/MB468 | CYTO |
| >trna112_GlnCTG_1_-_147737382_147737453@30.50.21,trna131_GlnCTG_6_-_27759135_27759206@30.50.21,trna146_GlnCTG_6_-_27515531_27515602@30.50.21,trna15_GlnCTG_1_+_145963304_145963375@30.50.21,trna19_GlnCTG_1_+_147505038_147505109@30.50.21,trna1_GlnCTG_6_+_18836402_18836473@30.50.21,trna28_GlnCTG_1_+_149186125_149186196@30.50.21,trna3_GlnCTG_17_+_8023070_8023141@30.50.21,trna49_GlnCTG_6_+_27487308_27487379@30.50.21,trna7_GlnCTG_15_-_66161400_66161471@30.50.21,trna99_GlnCTG_6_-_28909378_28909449@30.50.21\|GACTCTGAATCCAGCGATCCG | >trna112_GlnCTG_1_-_147737382_147737453 | 30 | 50 | 21 | GACTCTGAATCCAGCGATCCG | 318.1 | -12.2 | 1.4 | -8.4 | 4.5E-17 | 4.1E-16 | MB231/MB468 | MP |
| >hg38_21_+_8433172_8446622_RNA45SN1_45S_50ntFlanks@8472.8492.21\|CGGCCCGGCGGATCTTTCCCG | >hg38_21_+_8433172_8446622_RNA45SN1_45S_50ntFlanks | 8472 | 8492 | 21 | CGGCCCGGCGGATCTTTCCCG | 1373.4 | -12.5 | 1.5 | -8.4 | 5.5E-17 | 4.1E-16 | BT20/MB468 | MITO |
| >MI0000088\|hsa-mir-30a&WithFlank&6\|-\|71403545\|71403627@54.74.21(+1A)&[MIMAT0000088&hsa-miR-30a-3p&offsets\|+1\|0(+1A);m-38&6\|-\|71403554\|71403575&offsets\|+1\|0(+1A)]\|TTTCAGTCGGATGTTTGCAGCA | >MI0000088\|hsa-mir-30a&WithFlank&6\|-\|71403545\|71403627 | 54 | 74 | 21(+1A) | TTTCAGTCGGATGTTTGCAGCA | 331.8 | 9.0 | 1.1 | 8.3 | 7.7E-17 | 4.1E-16 | BT20/MB468 | CYTO |
| >hg38_21_+_8433172_8446622_RNA45SN1_45S_50ntFlanks@7991.8008.18\|ACGTGGCGACCCGCTGAA | >hg38_21_+_8433172_8446622_RNA45SN1_45S_50ntFlanks | 7991 | 8008 | 18 | ACGTGGCGACCCGCTGAA | 4311.2 | -6.0 | 0.7 | -8.3 | 1.3E-16 | 4.1E-16 | BT20/MB468 | NUC |
| >hg38_21_+_8433172_8446622_RNA45SN1_45S_50ntFlanks@5035.5072.38\|CATGGCCGTTCTTAGTTGGTGGAGCGATTTGTCTGGTT | >hg38_21_+_8433172_8446622_RNA45SN1_45S_50ntFlanks | 5035 | 5072 | 38 | CATGGCCGTTCTTAGTTGGTGGAGCGATTTGTCTGGTT | 1044.8 | -13.8 | 1.7 | -8.3 | 7.4E-17 | 4.1E-16 | MB231/MB468 | MITO |
| >hg38_21_+_8433172_8446622_RNA45SN1_45S_50ntFlanks@8155.8181.27\|CGCTCGTGGGGGGCCCAAGTCCTTCTG | >hg38_21_+_8433172_8446622_RNA45SN1_45S_50ntFlanks | 8155 | 8181 | 27 | CGCTCGTGGGGGGCCCAAGTCCTTCTG | 828.7 | -13.6 | 1.6 | -8.4 | 4.5E-17 | 4.1E-16 | MB231/MB468 | MP |
| >hg38_21_+_8433172_8446622_RNA45SN1_45S_50ntFlanks@11586.11603.18\|GTGCGGACCAGGGGAATC | >hg38_21_+_8433172_8446622_RNA45SN1_45S_50ntFlanks | 11586 | 11603 | 18 | GTGCGGACCAGGGGAATC | 1433.6 | -5.1 | 0.6 | -8.3 | 7.8E-17 | 4.1E-16 | BT20/MB468 | CYTO |
| >hg38_21_+_8433172_8446622_RNA45SN1_45S_50ntFlanks@6666.6688.23\|GATCACTCGGCTCGTGCGTCGAT | >hg38_21_+_8433172_8446622_RNA45SN1_45S_50ntFlanks | 6666 | 6688 | 23 | GATCACTCGGCTCGTGCGTCGAT | 261.0 | -11.9 | 1.4 | -8.4 | 4.5E-17 | 4.1E-16 | MB231/MB468 | MP |
| >hg38_21_+_8433172_8446622_RNA45SN1_45S_50ntFlanks@9457.9488.32\|CGCCGGGGAGGTGGAGCACGAGCGCACGTGTT | >hg38_21_+_8433172_8446622_RNA45SN1_45S_50ntFlanks | 9457 | 9488 | 32 | CGCCGGGGAGGTGGAGCACGAGCGCACGTGTT | 356.8 | -12.3 | 1.5 | -8.3 | 7.4E-17 | 4.1E-16 | MB231/MB468 | MITO |
| >hg38_21_+_8433172_8446622_RNA45SN1_45S_50ntFlanks@11876.11907.32\|AAGACCCTGTTGAGCTTGACTCTAGTCTGGCA | >hg38_21_+_8433172_8446622_RNA45SN1_45S_50ntFlanks | 11876 | 11907 | 32 | AAGACCCTGTTGAGCTTGACTCTAGTCTGGCA | 341.4 | -12.2 | 1.5 | -8.3 | 7.5E-17 | 4.1E-16 | MB231/MB468 | MITO |
| >hg38_21_+_8433172_8446622_RNA45SN1_45S_50ntFlanks@9504.9527.24\|GTGAACTATGCCTGGGCAGGGCGA | >hg38_21_+_8433172_8446622_RNA45SN1_45S_50ntFlanks | 9504 | 9527 | 24 | GTGAACTATGCCTGGGCAGGGCGA | 437.6 | -11.5 | 1.4 | -8.4 | 4.7E-17 | 4.1E-16 | MB231/MB468 | CYTO |
| >hg38_21_+_8433172_8446622_RNA45SN1_45S_50ntFlanks@10849.10868.20\|CGGGATAAGGATTGGCTCTA | >hg38_21_+_8433172_8446622_RNA45SN1_45S_50ntFlanks | 10849 | 10868 | 20 | CGGGATAAGGATTGGCTCTA | 396.8 | -12.5 | 1.5 | -8.4 | 4.6E-17 | 4.1E-16 | MB231/MB468 | MP |
| >MI0000264\|hsa-mir-7-2&WithFlank&15\|+\|88611819\|88611940@38.59.22(+1C);MI0000265\|hsa-mir-7-3&WithFlank&19\|+\|4770664\|4770785@37.58.22(+1C);MI0000263\|hsa-mir-7-1&WithFlank&9\|-\|83969742\|83969863@30.51.22(+1C)&[MIMAT0000252&hsa-miR-7-5p&offsets\|0\|-2(+1C);m-400&15\|+\|88611856\|88611879&offsets\|0\|-2(+1C)];[MIMAT0000252_1&hsa-miR-7-5p&offsets\|0\|-2(+1C);m-402&19\|+\|4770700\|4770723&offsets\|0\|-2(+1C)];[MIMAT0000252_2&hsa-miR-7-5p&offsets\|0\|-2(+1C);m-401&9\|-\|83969811\|83969834&offsets\|0\|-2(+1C)]\|TGGAAGACTAGTGATTTTGTTGC | >MI0000264\|hsa-mir-7-2&WithFlank&15\|+\|88611819\|88611940 | 38 | 59 | 22(+1C) | TGGAAGACTAGTGATTTTGTTGC | 2596.8 | 8.0 | 1.0 | 8.3 | 7.9E-17 | 4.2E-16 | BT20/MB468 | CYTO |
| >hg38_21_+_8433172_8446622_RNA45SN1_45S_50ntFlanks@10841.10863.23\|CGTAACTTCGGGATAAGGATTGG | >hg38_21_+_8433172_8446622_RNA45SN1_45S_50ntFlanks | 10841 | 10863 | 23 | CGTAACTTCGGGATAAGGATTGG | 5912.5 | -5.8 | 0.7 | -8.4 | 4.7E-17 | 4.2E-16 | MB231/MB468 | CYTO |
| >MI0000061\|hsa-let-7a-2&WithFlank&11\|-\|122146516\|122146599@11.31.21(+1A);MI0000062\|hsa-let-7a-3&WithFlank&22\|+\|46112743\|46112828@10.30.21(+1A);MI0000060\|hsa-let-7a-1&WithFlank&9\|+\|94175951\|94176042@12.32.21(+1A)&[MIMAT0000062&hsa-let-7a-5p&offsets\|0\|-1(+1A);m-6&11\|-\|122146568\|122146589&offsets\|0\|-1(+1A)];[MIMAT0000062_1&hsa-let-7a-5p&offsets\|0\|-1(+1A);m-5&22\|+\|46112752\|46112773&offsets\|0\|-1(+1A)];[MIMAT0000062_2&hsa-let-7a-5p&offsets\|0\|-1(+1A);m-4&9\|+\|94175962\|94175983&offsets\|0\|-1(+1A)]\|TGAGGTAGTAGGTTGTATAGTA | >MI0000061\|hsa-let-7a-2&WithFlank&11\|-\|122146516\|122146599 | 11 | 31 | 21(+1A) | TGAGGTAGTAGGTTGTATAGTA | 1692.7 | 5.2 | 0.6 | 8.4 | 5.8E-17 | 4.2E-16 | MB231/MB468 | NUC |
| >hg38_21_+_8433172_8446622_RNA45SN1_45S_50ntFlanks@9821.9850.30\|CCTAGTGGGCCACTTTTGGTAAGCAGAACT | >hg38_21_+_8433172_8446622_RNA45SN1_45S_50ntFlanks | 9821 | 9850 | 30 | CCTAGTGGGCCACTTTTGGTAAGCAGAACT | 456.6 | -12.7 | 1.5 | -8.4 | 4.6E-17 | 4.2E-16 | MB231/MB468 | MP |
| >MI0000086\|hsa-mir-28&WithFlank&3\|+\|188688775\|188688872@60.82.23&[MIMAT0004502&hsa-miR-28-3p&offsets\|0\|+1;m-28&3\|+\|188688834\|188688855&offsets\|0\|+1]\|CACTAGATTGTGAGCTCCTGGAG | >MI0000086\|hsa-mir-28&WithFlank&3\|+\|188688775\|188688872 | 60 | 82 | 23 | CACTAGATTGTGAGCTCCTGGAG | 209.5 | 6.1 | 0.7 | 8.4 | 5.8E-17 | 4.2E-16 | MB231/MB468 | NUC |
| >hg38_21_+_8433172_8446622_RNA45SN1_45S_50ntFlanks@10781.10817.37\|AGGTGAACAGCCTCTGGCATGTTGGAACAATGTAGGT | >hg38_21_+_8433172_8446622_RNA45SN1_45S_50ntFlanks | 10781 | 10817 | 37 | AGGTGAACAGCCTCTGGCATGTTGGAACAATGTAGGT | 421.6 | -12.6 | 1.5 | -8.4 | 4.6E-17 | 4.2E-16 | MB231/MB468 | MP |
| >hg38_21_+_8433172_8446622_RNA45SN1_45S_50ntFlanks@5487.5514.28\|CGCTGAGAAGACGGTCGAACTTGACTAT | >hg38_21_+_8433172_8446622_RNA45SN1_45S_50ntFlanks | 5487 | 5514 | 28 | CGCTGAGAAGACGGTCGAACTTGACTAT | 3628.3 | -4.1 | 0.5 | -8.3 | 7.9E-17 | 4.2E-16 | BT20/MB468 | CYTO |
| >hg38_21_+_8433172_8446622_RNA45SN1_45S_50ntFlanks@11363.11383.21\|CGGGCGGCGGCGGTCGGCGGG | >hg38_21_+_8433172_8446622_RNA45SN1_45S_50ntFlanks | 11363 | 11383 | 21 | CGGGCGGCGGCGGTCGGCGGG | 702.7 | -13.3 | 1.6 | -8.4 | 4.6E-17 | 4.2E-16 | MB231/MB468 | MP |
| >hg38_21_+_8433172_8446622_RNA45SN1_45S_50ntFlanks@475.503.29\|CTTCGTGATCGATGTGGTGACGTCGTGCT | >hg38_21_+_8433172_8446622_RNA45SN1_45S_50ntFlanks | 475 | 503 | 29 | CTTCGTGATCGATGTGGTGACGTCGTGCT | 3693.4 | 7.8 | 0.9 | 8.4 | 5.7E-17 | 4.2E-16 | BT20/MB468 | MITO |

| | | | | | | | | | | | | | |
|---|---|---|---|---|---|---|---|---|---|---|---|---|---|
| >hg38_21_+_8433172_8446622_RNA45SN1_45S_50ntFlanks@12973.13003.31\|TCGTACGTAGCAGAGCAGCTCCCTCGCTGCG | >hg38_21_+_8433172_8446622_RNA45SN1_45S_50ntFlanks | 12973 | 13003 | 31 | TCGTACGTAGCAGAGCAGCTCCCTCGCTGCG | 251.2 | -6.8 | 0.8 | -8.4 | 4.8E-17 | 4.2E-16 | MB231/MB468 | CYTO |
| >hg38_21_+_8433172_8446622_RNA45SN1_45S_50ntFlanks@10375.10407.33\|AGAAGGGTTCCATGTGAACAGCAGTTGAACATG | >hg38_21_+_8433172_8446622_RNA45SN1_45S_50ntFlanks | 10375 | 10407 | 33 | AGAAGGGTTCCATGTGAACAGCAGTTGAACATG | 346.0 | -12.2 | 1.5 | -8.3 | 7.7E-17 | 4.2E-16 | MB231/MB468 | MITO |
| >hg38_21_+_8433172_8446622_RNA45SN1_45S_50ntFlanks@12934.12959.26\|CTAAACCATTCGTAGACGACCTGCTT | >hg38_21_+_8433172_8446622_RNA45SN1_45S_50ntFlanks | 12934 | 12959 | 26 | CTAAACCATTCGTAGACGACCTGCTT | 820.8 | -7.5 | 0.9 | -8.3 | 7.7E-17 | 4.2E-16 | MB231/MB468 | MITO |
| >hg38_21_+_8433172_8446622_RNA45SN1_45S_50ntFlanks@8114.8137.24\|GCGCGGGACATGTGGCGTACGGAA | >hg38_21_+_8433172_8446622_RNA45SN1_45S_50ntFlanks | 8114 | 8137 | 24 | GCGCGGGACATGTGGCGTACGGAA | 600.1 | -13.1 | 1.6 | -8.4 | 4.7E-17 | 4.2E-16 | MB231/MB468 | MP |
| >hg38_21_+_8433172_8446622_RNA45SN1_45S_50ntFlanks@12605.12626.22\|TATGTGCTTGGCTGAGGAGCCA | >hg38_21_+_8433172_8446622_RNA45SN1_45S_50ntFlanks | 12605 | 12626 | 22 | TATGTGCTTGGCTGAGGAGCCA | 1975.0 | -10.5 | 1.3 | -8.3 | 7.7E-17 | 4.2E-16 | MB231/MB468 | MITO |
| >hg38_21_+_8433172_8446622_RNA45SN1_45S_50ntFlanks@9507.9526.20\|AACTATGCCTGGGCAGGGCG | >hg38_21_+_8433172_8446622_RNA45SN1_45S_50ntFlanks | 9507 | 9526 | 20 | AACTATGCCTGGGCAGGGCG | 1503.4 | -14.4 | 1.7 | -8.4 | 4.7E-17 | 4.2E-16 | MB231/MB468 | MP |
| >hg38_21_+_8433172_8446622_RNA45SN1_45S_50ntFlanks@10383.10424.42\|TCCATGTGAACAGCAGTTGAACATGGGTCAGTCGGTCCTGAG | >hg38_21_+_8433172_8446622_RNA45SN1_45S_50ntFlanks | 10383 | 10424 | 42 | TCCATGTGAACAGCAGTTGAACATGGGTCAGTCGGTCCTGAG | 8217.5 | 8.5 | 1.0 | 8.4 | 5.8E-17 | 4.2E-16 | BT20/MB468 | MITO |
| >hg38_21_+_8433172_8446622_RNA45SN1_45S_50ntFlanks@5087.5122.36\|ACGAGACTCTGGCATGCTAACTAGTTACGCGACCCC | >hg38_21_+_8433172_8446622_RNA45SN1_45S_50ntFlanks | 5087 | 5122 | 36 | ACGAGACTCTGGCATGCTAACTAGTTACGCGACCCC | 418.7 | 10.4 | 1.2 | 8.4 | 4.8E-17 | 4.2E-16 | MB231/MB468 | CYTO |
| >hg38_21_+_8433172_8446622_RNA45SN1_45S_50ntFlanks@11679.11702.24\|CTGAATGTCAAAGTGAAGAAATTC | >hg38_21_+_8433172_8446622_RNA45SN1_45S_50ntFlanks | 11679 | 11702 | 24 | CTGAATGTCAAAGTGAAGAAATTC | 265.9 | -11.9 | 1.4 | -8.4 | 4.7E-17 | 4.3E-16 | MB231/MB468 | MP |
| >hg38_21_+_8433172_8446622_RNA45SN1_45S_50ntFlanks@9308.9336.29\|AGGAGTCTAACACGTGCGCGAGTCGGGGG | >hg38_21_+_8433172_8446622_RNA45SN1_45S_50ntFlanks | 9308 | 9336 | 29 | AGGAGTCTAACACGTGCGCGAGTCGGGGG | 433.7 | -12.5 | 1.5 | -8.3 | 7.7E-17 | 4.3E-16 | MB231/MB468 | MITO |
| >MI0003836\|hsa-mir-766&WithFlank&X\|-\|119646732\|119646854@71.92.22&[MIMAT0003888&hsa-miR-766-3p&offsets\|0\|0;m-286&X\|-\|119646763\|119646784&offsets\|0\|0]\|ACTCCAGCCCCACAGCCTCAGC | >MI0003836\|hsa-mir-766&WithFlank&X\|-\|119646732\|119646854 | 71 | 92 | 22 | ACTCCAGCCCCACAGCCTCAGC | 227.1 | 9.8 | 1.2 | 8.3 | 8.1E-17 | 4.3E-16 | BT20/MB468 | CYTO |
| >hg38_21_+_8433172_8446622_RNA45SN1_45S_50ntFlanks@8357.8402.46\|AAAAGAACTTTGAAGAGAGAGTTCAAGAGGGCGTGAAACCGTTAAG | >hg38_21_+_8433172_8446622_RNA45SN1_45S_50ntFlanks | 8357 | 8402 | 46 | AAAAGAACTTTGAAGAGAGAGTTCAAGAGGGCGTGAAACCGTTAAG | 642.2 | -13.1 | 1.6 | -8.3 | 7.7E-17 | 4.3E-16 | MB231/MB468 | MITO |
| >hg38_21_+_8433172_8446622_RNA45SN1_45S_50ntFlanks@12974.12991.18\|CGTACGTAGCAGAGCAGC | >hg38_21_+_8433172_8446622_RNA45SN1_45S_50ntFlanks | 12974 | 12991 | 18 | CGTACGTAGCAGAGCAGC | 1227.8 | -4.6 | 0.6 | -8.4 | 5.8E-17 | 4.3E-16 | BT20/MB468 | MITO |
| >hg38_1_-_228634819_228635039_RNA5S12_5S_50ntFlanks@147.171.25\|GGGAATACCGGGTGCTGTAGGCTTT | >hg38_1_-_228634819_228635039_RNA5S12_5S_50ntFlanks | 147 | 171 | 25 | GGGAATACCGGGTGCTGTAGGCTTT | 5695.4 | 5.9 | 0.7 | 8.4 | 4.8E-17 | 4.3E-16 | MB231/MB468 | CYTO |
| >hg38_21_+_8433172_8446622_RNA45SN1_45S_50ntFlanks@4727.4766.40\|AATCAAGAACGAAAGTCGGAGGTTCGAAGACGATCAGATA | >hg38_21_+_8433172_8446622_RNA45SN1_45S_50ntFlanks | 4727 | 4766 | 40 | AATCAAGAACGAAAGTCGGAGGTTCGAAGACGATCAGATA | 807.4 | 8.1 | 1.0 | 8.4 | 4.9E-17 | 4.3E-16 | MB231/MB468 | CYTO |
| >hg38_21_+_8433172_8446622_RNA45SN1_45S_50ntFlanks@11589.11611.23\|CGGACCAGGGGAATCCGACTGTT | >hg38_21_+_8433172_8446622_RNA45SN1_45S_50ntFlanks | 11589 | 11611 | 23 | CGGACCAGGGGAATCCGACTGTT | 1199.8 | -5.9 | 0.7 | -8.4 | 4.7E-17 | 4.3E-16 | MB231/MB468 | MP |
| >hg38_21_+_8433172_8446622_RNA45SN1_45S_50ntFlanks@11621.11660.40\|CAAAGCATCGCGAAGGCCCGCGGCGGGTGTTGACGCGATG | >hg38_21_+_8433172_8446622_RNA45SN1_45S_50ntFlanks | 11621 | 11660 | 40 | CAAAGCATCGCGAAGGCCCGCGGCGGGTGTTGACGCGATG | 328.0 | -12.2 | 1.5 | -8.4 | 4.7E-17 | 4.3E-16 | MB231/MB468 | MP |
| >hg38_21_+_8433172_8446622_RNA45SN1_45S_50ntFlanks@12996.13017.22\|TCGCTGCGATCTATTGAAAGTC | >hg38_21_+_8433172_8446622_RNA45SN1_45S_50ntFlanks | 12996 | 13017 | 22 | TCGCTGCGATCTATTGAAAGTC | 1070.4 | 5.9 | 0.7 | 8.4 | 5.9E-17 | 4.3E-16 | MB231/MB468 | NUC |
| >hg38_21_+_8433172_8446622_RNA45SN1_45S_50ntFlanks@10408.10433.26\|GGTCAGTCGGTCCTGAGAGATGGGCG | >hg38_21_+_8433172_8446622_RNA45SN1_45S_50ntFlanks | 10408 | 10433 | 26 | GGTCAGTCGGTCCTGAGAGATGGGCG | 1036.2 | -13.8 | 1.7 | -8.3 | 7.8E-17 | 4.3E-16 | MB231/MB468 | MITO |
| >MI0000255\|hsa-mir-30d&WithFlank&8\|-\|134804870\|134804951@12.34.23&[MIMAT0000245&hsa-miR-30d-5p&offsets\|0\|+1;m-21&8\|-\|134804919\|134804940&offsets\|0\|+1]\|TGTAAACATCCCCGACTGGAAGC | >MI0000255\|hsa-mir-30d&WithFlank&8\|-\|134804870\|134804951 | 12 | 34 | 23 | TGTAAACATCCCCGACTGGAAGC | 19274.9 | 5.3 | 0.6 | 8.4 | 6.0E-17 | 4.3E-16 | MB231/MB468 | NUC |
| >hg38_21_+_8433172_8446622_RNA45SN1_45S_50ntFlanks@11874.11899.26\|AGAAGACCCTGTTGAGCTTGACTCTA | >hg38_21_+_8433172_8446622_RNA45SN1_45S_50ntFlanks | 11874 | 11899 | 26 | AGAAGACCCTGTTGAGCTTGACTCTA | 269.4 | -11.9 | 1.4 | -8.4 | 4.8E-17 | 4.3E-16 | MB231/MB468 | MP |
| >hg38_21_+_8433172_8446622_RNA45SN1_45S_50ntFlanks@9574.9596.23\|AATCGGTCGTCCGACCTGGGTAT | >hg38_21_+_8433172_8446622_RNA45SN1_45S_50ntFlanks | 9574 | 9596 | 23 | AATCGGTCGTCCGACCTGGGTAT | 6036.1 | -6.6 | 0.8 | -8.4 | 5.9E-17 | 4.3E-16 | BT20/MB468 | MITO |
| >hg38_21_+_8433172_8446622_RNA45SN1_45S_50ntFlanks@9621.9650.30\|TCTAGTAGCTGGTTCCCTCCGAAGTTTCCC | >hg38_21_+_8433172_8446622_RNA45SN1_45S_50ntFlanks | 9621 | 9650 | 30 | TCTAGTAGCTGGTTCCCTCCGAAGTTTCCC | 216.1 | -11.6 | 1.4 | -8.4 | 4.8E-17 | 4.3E-16 | MB231/MB468 | MP |
| >hg38_21_+_8433172_8446622_RNA45SN1_45S_50ntFlanks@11190.11207.18\|CCCGGGGGGCCGGCGGCG | >hg38_21_+_8433172_8446622_RNA45SN1_45S_50ntFlanks | 11190 | 11207 | 18 | CCCGGGGGGCCGGCGGCG | 675.2 | -13.2 | 1.6 | -8.3 | 7.9E-17 | 4.4E-16 | MB231/MB468 | MITO |
| >hg38_21_+_8433172_8446622_RNA45SN1_45S_50ntFlanks@9133.9152.20\|TCGGGGGGCGGGGCGGACTG | >hg38_21_+_8433172_8446622_RNA45SN1_45S_50ntFlanks | 9133 | 9152 | 20 | TCGGGGGGCGGGGCGGACTG | 487.0 | -12.7 | 1.5 | -8.3 | 8.0E-17 | 4.4E-16 | MB231/MB468 | MITO |
| >hg38_21_+_8433172_8446622_RNA45SN1_45S_50ntFlanks@11575.11592.18\|ACTTAGAACTGGTGCGGA | >hg38_21_+_8433172_8446622_RNA45SN1_45S_50ntFlanks | 11575 | 11592 | 18 | ACTTAGAACTGGTGCGGA | 1793.7 | -9.9 | 1.2 | -8.3 | 8.0E-17 | 4.4E-16 | MB231/MB468 | MITO |
| >hg38_21_+_8433172_8446622_RNA45SN1_45S_50ntFlanks@12961.12994.34\|TGGGTCGGGGTTTCGTACGTAGCAGAGCAGCTCC | >hg38_21_+_8433172_8446622_RNA45SN1_45S_50ntFlanks | 12961 | 12994 | 34 | TGGGTCGGGGTTTCGTACGTAGCAGAGCAGCTCC | 143.5 | 8.5 | 1.0 | 8.4 | 5.0E-17 | 4.4E-16 | MB231/MB468 | CYTO |
| >hg38_21_+_8433172_8446622_RNA45SN1_45S_50ntFlanks@485.504.20\|GATGTGGTGACGTCGTGCTC | >hg38_21_+_8433172_8446622_RNA45SN1_45S_50ntFlanks | 485 | 504 | 20 | GATGTGGTGACGTCGTGCTC | 298.2 | 11.1 | 1.3 | 8.3 | 1.4E-16 | 4.4E-16 | BT20/MB468 | NUC |
| >hg38_21_+_8433172_8446622_RNA45SN1_45S_50ntFlanks@11845.11871.27\|GGAACGGGCTTGGCGGAATCAGCGGGG | >hg38_21_+_8433172_8446622_RNA45SN1_45S_50ntFlanks | 11845 | 11871 | 27 | GGAACGGGCTTGGCGGAATCAGCGGGG | 849.5 | -6.3 | 0.8 | -8.4 | 4.9E-17 | 4.4E-16 | MB231/MB468 | MP |
| >hg38_21_+_8433172_8446622_RNA45SN1_45S_50ntFlanks@6721.6738.18\|AATTGCAGGACACATTGA | >hg38_21_+_8433172_8446622_RNA45SN1_45S_50ntFlanks | 6721 | 6738 | 18 | AATTGCAGGACACATTGA | 356.6 | -12.3 | 1.5 | -8.4 | 4.9E-17 | 4.4E-16 | MB231/MB468 | MP |
| >hg38_21_+_8433172_8446622_RNA45SN1_45S_50ntFlanks@9621.9652.32\|TCTAGTAGCTGGTTCCCTCCGAAGTTTCCCTC | >hg38_21_+_8433172_8446622_RNA45SN1_45S_50ntFlanks | 9621 | 9652 | 32 | TCTAGTAGCTGGTTCCCTCCGAAGTTTCCCTC | 233.6 | -11.7 | 1.4 | -8.4 | 4.9E-17 | 4.4E-16 | MB231/MB468 | MP |
| >hg38_21_+_8433172_8446622_RNA45SN1_45S_50ntFlanks@10239.10259.21\|GCTACGCCGCGACGAGTAGGA | >hg38_21_+_8433172_8446622_RNA45SN1_45S_50ntFlanks | 10239 | 10259 | 21 | GCTACGCCGCGACGAGTAGGA | 513.1 | 6.6 | 0.8 | 8.4 | 6.0E-17 | 4.4E-16 | BT20/MB468 | MITO |
| >hg38_21_+_8433172_8446622_RNA45SN1_45S_50ntFlanks@11646.11671.26\|GGTGTTGACGCGATGTGATTTCTGCC | >hg38_21_+_8433172_8446622_RNA45SN1_45S_50ntFlanks | 11646 | 11671 | 26 | GGTGTTGACGCGATGTGATTTCTGCC | 323.6 | -12.2 | 1.5 | -8.4 | 4.9E-17 | 4.4E-16 | MB231/MB468 | MP |
| >hg38_21_+_8433172_8446622_RNA45SN1_45S_50ntFlanks@11668.11687.20\|TGCCCAGTGCTCTGAATGTC | >hg38_21_+_8433172_8446622_RNA45SN1_45S_50ntFlanks | 11668 | 11687 | 20 | TGCCCAGTGCTCTGAATGTC | 315.1 | 4.8 | 0.6 | 8.4 | 6.1E-17 | 4.4E-16 | BT20/MB468 | MITO |
| >hg38_21_+_8433172_8446622_RNA45SN1_45S_50ntFlanks@10679.10705.27\|GGTTCCGGCGGCGTCCGGTGAGCTCTC | >hg38_21_+_8433172_8446622_RNA45SN1_45S_50ntFlanks | 10679 | 10705 | 27 | GGTTCCGGCGGCGTCCGGTGAGCTCTC | 504.9 | -12.8 | 1.5 | -8.3 | 8.1E-17 | 4.4E-16 | MB231/MB468 | MITO |
| >hg38_21_+_8433172_8446622_RNA45SN1_45S_50ntFlanks@9339.9368.30\|CGCACGAAAGCCGCCGTGGCGCAATGAAGG | >hg38_21_+_8433172_8446622_RNA45SN1_45S_50ntFlanks | 9339 | 9368 | 30 | CGCACGAAAGCCGCCGTGGCGCAATGAAGG | 738.2 | -13.4 | 1.6 | -8.4 | 5.0E-17 | 4.4E-16 | MB231/MB468 | MP |
| >hg38_21_+_8433172_8446622_RNA45SN1_45S_50ntFlanks@13004.13027.24\|ATCTATTGAAAGTCAGCCCTCGAC | >hg38_21_+_8433172_8446622_RNA45SN1_45S_50ntFlanks | 13004 | 13027 | 24 | ATCTATTGAAAGTCAGCCCTCGAC | 243.1 | -11.8 | 1.4 | -8.4 | 5.0E-17 | 4.5E-16 | MB231/MB468 | MP |
| >hg38_21_+_8433172_8446622_RNA45SN1_45S_50ntFlanks@7985.8008.24\|GATCAGACGTGGCGACCCGCTGAA | >hg38_21_+_8433172_8446622_RNA45SN1_45S_50ntFlanks | 7985 | 8008 | 24 | GATCAGACGTGGCGACCCGCTGAA | 1690.1 | -13.8 | 1.6 | -8.8 | 1.2E-18 | 4.5E-16 | BT20/MB468 | MP |
| >hg38_21_+_8433172_8446622_RNA45SN1_45S_50ntFlanks@12324.12342.19\|TTGGGTTTTAAGCAGGAGG | >hg38_21_+_8433172_8446622_RNA45SN1_45S_50ntFlanks | 12324 | 12342 | 19 | TTGGGTTTTAAGCAGGAGG | 343.5 | -12.3 | 1.5 | -8.4 | 5.0E-17 | 4.5E-16 | MB231/MB468 | MP |
| >hg38_1_-_228634819_228635039_RNA5S12_5S_50ntFlanks@148.169.22\|GGAATACCGGGTGCTGTAGGCT | >hg38_1_-_228634819_228635039_RNA5S12_5S_50ntFlanks | 148 | 169 | 22 | GGAATACCGGGTGCTGTAGGCT | 5967.9 | -6.1 | 0.7 | -8.4 | 5.0E-17 | 4.5E-16 | MB231/MB468 | MP |
| >hg38_21_+_8433172_8446622_RNA45SN1_45S_50ntFlanks@7992.8012.21\|CGTGGCGACCCGCTGAATTTA | >hg38_21_+_8433172_8446622_RNA45SN1_45S_50ntFlanks | 7992 | 8012 | 21 | CGTGGCGACCCGCTGAATTTA | 3264.5 | -6.3 | 0.8 | -8.3 | 8.1E-17 | 4.5E-16 | MB231/MB468 | MITO |
| >hg38_21_+_8433172_8446622_RNA45SN1_45S_50ntFlanks@12045.12071.27\|CTGACCCGGTGAGGCGGGGGGGCGAGC | >hg38_21_+_8433172_8446622_RNA45SN1_45S_50ntFlanks | 12045 | 12071 | 27 | CTGACCCGGTGAGGCGGGGGGGCGAGC | 632.4 | -13.1 | 1.6 | -8.3 | 8.2E-17 | 4.5E-16 | MB231/MB468 | MITO |
| >hg38_21_+_8433172_8446622_RNA45SN1_45S_50ntFlanks@10411.10443.33\|CAGTCGGTCCTGAGAGATGGGCGAGCGCCGTTC | >hg38_21_+_8433172_8446622_RNA45SN1_45S_50ntFlanks | 10411 | 10443 | 33 | CAGTCGGTCCTGAGAGATGGGCGAGCGCCGTTC | 5848.2 | -9.8 | 1.2 | -8.4 | 5.0E-17 | 4.5E-16 | MB231/MB468 | MP |
| >hg38_21_+_8433172_8446622_RNA45SN1_45S_50ntFlanks@4472.4492.21\|CTTAGCTGAGTGTCCCGCGGG | >hg38_21_+_8433172_8446622_RNA45SN1_45S_50ntFlanks | 4472 | 4492 | 21 | CTTAGCTGAGTGTCCCGCGGG | 515.5 | -12.9 | 1.5 | -8.4 | 5.0E-17 | 4.5E-16 | MB231/MB468 | MP |
| >hg38_21_+_8433172_8446622_RNA45SN1_45S_50ntFlanks@11577.11605.29\|TTAGAACTGGTGCGGACCAGGGGAATCCG | >hg38_21_+_8433172_8446622_RNA45SN1_45S_50ntFlanks | 11577 | 11605 | 29 | TTAGAACTGGTGCGGACCAGGGGAATCCG | 347.4 | -12.3 | 1.5 | -8.4 | 5.0E-17 | 4.5E-16 | MB231/MB468 | MP |
| >hg38_21_+_8433172_8446622_RNA45SN1_45S_50ntFlanks@10866.10901.36\|CTAAGGGCTGGGTCGGTCGGGCTGGGGCGCGAAGCG | >hg38_21_+_8433172_8446622_RNA45SN1_45S_50ntFlanks | 10866 | 10901 | 36 | CTAAGGGCTGGGTCGGTCGGGCTGGGGCGCGAAGCG | 1918.0 | -10.7 | 1.3 | -8.4 | 6.3E-17 | 4.5E-16 | MB231/MB468 | NUC |
| >hg38_21_+_8433172_8446622_RNA45SN1_45S_50ntFlanks@12939.12956.18\|CCATTCGTAGACGACCTG | >hg38_21_+_8433172_8446622_RNA45SN1_45S_50ntFlanks | 12939 | 12956 | 18 | CCATTCGTAGACGACCTG | 1741.6 | -4.2 | 0.5 | -8.4 | 5.1E-17 | 4.5E-16 | MB231/MB468 | CYTO |
| >hg38_21_+_8433172_8446622_RNA45SN1_45S_50ntFlanks@3985.4026.42\|CGGCTTTGGTGACTCTAGATAACCTCGGGCCGATCGCACGCC | >hg38_21_+_8433172_8446622_RNA45SN1_45S_50ntFlanks | 3985 | 4026 | 42 | CGGCTTTGGTGACTCTAGATAACCTCGGGCCGATCGCACGCC | 2329.6 | -13.3 | 1.6 | -8.4 | 6.2E-17 | 4.5E-16 | BT20/MB468 | MITO |
| >MI0000071\|hsa-mir-17&WithFlank&13\|+\|91350599\|91350694@20.42.23&[MIMAT0000070&hsa-miR-17-5p&offsets\|0\|0;m-88&13\|+\|91350618\|91350640&offsets\|0\|0]\|CAAAGTGCTTACAGTGCAGGTAG | >MI0000071\|hsa-mir-17&WithFlank&13\|+\|91350599\|91350694 | 20 | 42 | 23 | CAAAGTGCTTACAGTGCAGGTAG | 8785.2 | 4.9 | 0.6 | 8.4 | 6.3E-17 | 4.5E-16 | MB231/MB468 | NUC |
| >hg38_21_+_8433172_8446622_RNA45SN1_45S_50ntFlanks@12997.13036.40\|CGCTGCGATCTATTGAAAGTCAGCCCTCGACACAAGGGTT | >hg38_21_+_8433172_8446622_RNA45SN1_45S_50ntFlanks | 12997 | 13036 | 40 | CGCTGCGATCTATTGAAAGTCAGCCCTCGACACAAGGGTT | 5038.5 | 8.3 | 1.0 | 8.3 | 8.6E-17 | 4.5E-16 | BT20/MB468 | CYTO |
| >hg38_21_+_8433172_8446622_RNA45SN1_45S_50ntFlanks@4002.4026.25\|GATAACCTCGGGCCGATCGCACGCC | >hg38_21_+_8433172_8446622_RNA45SN1_45S_50ntFlanks | 4002 | 4026 | 25 | GATAACCTCGGGCCGATCGCACGCC | 348.2 | -12.3 | 1.5 | -8.4 | 5.1E-17 | 4.6E-16 | MB231/MB468 | MP |
| >MI0000650\|hsa-mir-200c&WithFlank&12\|+\|6963693\|6963772@50.73.24&[MIMAT0000617&hsa-miR-200c-3p&offsets\|0\|+1;m-54&12\|+\|6963742\|6963764&offsets\|0\|+1]\|TAATACTGCCGGGTAATGATGGAG | >MI0000650\|hsa-mir-200c&WithFlank&12\|+\|6963693\|6963772 | 50 | 73 | 24 | TAATACTGCCGGGTAATGATGGAG | 1126.1 | 8.4 | 1.0 | 8.3 | 8.7E-17 | 4.6E-16 | BT20/MB468 | CYTO |

<table>
<tr><th></th><th></th><th></th><th></th><th></th><th></th><th></th><th></th><th></th><th></th><th></th><th></th><th></th><th></th><th></th></tr>
<tr><td>>MI0000816|hsa-mir-335&WithFlank&7|+|130496105|130496210@58.78.21(+1U)&[MIMAT0004703&hsa-miR-335-3p&offsets|0|-1(+1U);m-159&7|+|130496162|130496183&offsets|0|-1(+1U)]|TTTTTCATTATTGCTCCTGACT</td><td>>MI0000816|hsa-mir-335&WithFlank&7|+|130496105|130496210</td><td>58</td><td>78</td><td>21(+1U)</td><td>TTTTTCATTATTGCTCCTGACT</td><td>1329.3</td><td>7.2</td><td>0.9</td><td>8.4</td><td>5.2E-17</td><td>4.6E-16</td><td>MB231/MB468</td><td>CYTO</td><td></td></tr>
<tr><td>>hg38_21_+_8433172_8446622_RNA45SN1_45S_50ntFlanks@12388.12437.50|CGTTCATAGCGACGTCGCTTTTTGATCCTTCGATGTCGGCTCTTCCTATC</td><td>>hg38_21_+_8433172_8446622_RNA45SN1_45S_50ntFlanks</td><td>12388</td><td>12437</td><td>50</td><td>CGTTCATAGCGACGTCGCTTTTTGATCCTTCGATGTCGGCTCTTCCTATC</td><td>204.2</td><td>8.7</td><td>1.0</td><td>8.4</td><td>5.2E-17</td><td>4.6E-16</td><td>MB231/MB468</td><td>CYTO</td><td></td></tr>
<tr><td>>hg38_21_+_8433172_8446622_RNA45SN1_45S_50ntFlanks@13006.13028.23|CTATTGAAAGTCAGCCCTCGACA</td><td>>hg38_21_+_8433172_8446622_RNA45SN1_45S_50ntFlanks</td><td>13006</td><td>13028</td><td>23</td><td>CTATTGAAAGTCAGCCCTCGACA</td><td>1334.7</td><td>-8.1</td><td>1.0</td><td>-8.4</td><td>5.2E-17</td><td>4.6E-16</td><td>MB231/MB468</td><td>CYTO</td><td></td></tr>
<tr><td>>hg38_21_+_8433172_8446622_RNA45SN1_45S_50ntFlanks@13002.13025.24|CGATCTATTGAAAGTCAGCCCTCG</td><td>>hg38_21_+_8433172_8446622_RNA45SN1_45S_50ntFlanks</td><td>13002</td><td>13025</td><td>24</td><td>CGATCTATTGAAAGTCAGCCCTCG</td><td>361.8</td><td>-12.4</td><td>1.5</td><td>-8.4</td><td>5.1E-17</td><td>4.6E-16</td><td>MB231/MB468</td><td>MP</td><td></td></tr>
<tr><td>>hg38_21_+_8433172_8446622_RNA45SN1_45S_50ntFlanks@10388.10410.23|GTGAACAGCAGTTGAACATGGGT</td><td>>hg38_21_+_8433172_8446622_RNA45SN1_45S_50ntFlanks</td><td>10388</td><td>10410</td><td>23</td><td>GTGAACAGCAGTTGAACATGGGT</td><td>470.1</td><td>-12.7</td><td>1.5</td><td>-8.4</td><td>5.2E-17</td><td>4.6E-16</td><td>MB231/MB468</td><td>MP</td><td></td></tr>
<tr><td>>hg38_21_+_8433172_8446622_RNA45SN1_45S_50ntFlanks@9572.9596.25|CAAATCGGTCGTCCGACCTGGGTAT</td><td>>hg38_21_+_8433172_8446622_RNA45SN1_45S_50ntFlanks</td><td>9572</td><td>9596</td><td>25</td><td>CAAATCGGTCGTCCGACCTGGGTAT</td><td>5836.6</td><td>-8.5</td><td>1.0</td><td>-8.4</td><td>5.3E-17</td><td>4.6E-16</td><td>MB231/MB468</td><td>CYTO</td><td></td></tr>
<tr><td>>hg38_21_+_8433172_8446622_RNA45SN1_45S_50ntFlanks@5417.5445.29|CTACCGATTGGATGGTTTAGTGAGGCCCT</td><td>>hg38_21_+_8433172_8446622_RNA45SN1_45S_50ntFlanks</td><td>5417</td><td>5445</td><td>29</td><td>CTACCGATTGGATGGTTTAGTGAGGCCCT</td><td>643.2</td><td>-13.1</td><td>1.6</td><td>-8.3</td><td>8.4E-17</td><td>4.6E-16</td><td>MB231/MB468</td><td>MITO</td><td></td></tr>
<tr><td>>hg38_21_+_8433172_8446622_RNA45SN1_45S_50ntFlanks@12405.12429.25|CTTTTTGATCCTTCGATGTCGGCTC</td><td>>hg38_21_+_8433172_8446622_RNA45SN1_45S_50ntFlanks</td><td>12405</td><td>12429</td><td>25</td><td>CTTTTTGATCCTTCGATGTCGGCTC</td><td>6861.7</td><td>-5.3</td><td>0.6</td><td>-8.4</td><td>5.3E-17</td><td>4.6E-16</td><td>MB231/MB468</td><td>CYTO</td><td></td></tr>
<tr><td>>hg38_21_+_8433172_8446622_RNA45SN1_45S_50ntFlanks@12893.12928.36|CCTCGCCCGTCACGCACCGCACGTTCGTGGGGAACC</td><td>>hg38_21_+_8433172_8446622_RNA45SN1_45S_50ntFlanks</td><td>12893</td><td>12928</td><td>36</td><td>CCTCGCCCGTCACGCACCGCACGTTCGTGGGGAACC</td><td>244.5</td><td>-11.8</td><td>1.4</td><td>-8.4</td><td>5.2E-17</td><td>4.6E-16</td><td>MB231/MB468</td><td>MP</td><td></td></tr>
<tr><td>>hg38_21_+_8433172_8446622_RNA45SN1_45S_50ntFlanks@8168.8186.19|CCCAAGTCCTTCTGATCGA</td><td>>hg38_21_+_8433172_8446622_RNA45SN1_45S_50ntFlanks</td><td>8168</td><td>8186</td><td>19</td><td>CCCAAGTCCTTCTGATCGA</td><td>1169.2</td><td>-14.1</td><td>1.7</td><td>-8.4</td><td>5.2E-17</td><td>4.7E-16</td><td>MB231/MB468</td><td>MP</td><td></td></tr>
<tr><td>>hg38_21_+_8433172_8446622_RNA45SN1_45S_50ntFlanks@9198.9224.27|GTTCTCTCGGGGCCACGCGCGCGTCCC</td><td>>hg38_21_+_8433172_8446622_RNA45SN1_45S_50ntFlanks</td><td>9198</td><td>9224</td><td>27</td><td>GTTCTCTCGGGGCCACGCGCGCGTCCC</td><td>447.6</td><td>-12.7</td><td>1.5</td><td>-8.4</td><td>5.2E-17</td><td>4.7E-16</td><td>MB231/MB468</td><td>MP</td><td></td></tr>
<tr><td>>hg38_1_-_228634819_228635039_RNA5S12_5S_50ntFlanks@144.170.27|CCTGGGAATACCGGGTGCTGTAGGCTT</td><td>>hg38_1_-_228634819_228635039_RNA5S12_5S_50ntFlanks</td><td>144</td><td>170</td><td>27</td><td>CCTGGGAATACCGGGTGCTGTAGGCTT</td><td>2948.2</td><td>5.8</td><td>0.7</td><td>8.3</td><td>8.9E-17</td><td>4.7E-16</td><td>BT20/MB468</td><td>CYTO</td><td></td></tr>
<tr><td>>trna59_GluCTC_1_+_249168447_249168518@32.70.39|CTCTCACCGCCGCGGCCCGGGTTCGATTCCCGGTCAGGA</td><td>>trna59_GluCTC_1_+_249168447_249168518</td><td>32</td><td>70</td><td>39</td><td>CTCTCACCGCCGCGGCCCGGGTTCGATTCCCGGTCAGGA</td><td>417.7</td><td>-11.7</td><td>1.4</td><td>-8.4</td><td>6.5E-17</td><td>4.7E-16</td><td>MB231/MB468</td><td>NUC</td><td></td></tr>
<tr><td>>hg38_21_+_8433172_8446622_RNA45SN1_45S_50ntFlanks@7985.8008.24|GATCAGACGTGGCGACCCGCTGAA</td><td>>hg38_21_+_8433172_8446622_RNA45SN1_45S_50ntFlanks</td><td>7985</td><td>8008</td><td>24</td><td>GATCAGACGTGGCGACCCGCTGAA</td><td>951.1</td><td>-12.0</td><td>1.4</td><td>-8.4</td><td>6.4E-17</td><td>4.7E-16</td><td>BT20/MB468</td><td>MITO</td><td></td></tr>
<tr><td>>MI0000734|hsa-mir-106b&WithFlank&7|-|100093987|100094080@18.38.21&[MIMAT0000680&hsa-miR-106b-5p&offsets|0|0;m-103&7|-|100094043|100094063&offsets|0|0]|TAAAGTGCTGACAGTGCAGAT</td><td>>MI0000734|hsa-mir-106b&WithFlank&7|-|100093987|100094080</td><td>18</td><td>38</td><td>21</td><td>TAAAGTGCTGACAGTGCAGAT</td><td>989.7</td><td>6.2</td><td>0.7</td><td>8.4</td><td>6.6E-17</td><td>4.7E-16</td><td>MB231/MB468</td><td>NUC</td><td></td></tr>
<tr><td>>MI0000254|hsa-mir-30c-2&WithFlank&6|-|71376954|71377037@53.75.23&[MIMAT0004550&hsa-miR-30c-2-3p&offsets|0|+1;m-206&6|-|71376964|71376985&offsets|0|+1]|CTGGGAGAAGGCTGTTTACTCTT</td><td>>MI0000254|hsa-mir-30c-2&WithFlank&6|-|71376954|71377037</td><td>53</td><td>75</td><td>23</td><td>CTGGGAGAAGGCTGTTTACTCTT</td><td>357.5</td><td>5.9</td><td>0.7</td><td>8.4</td><td>6.6E-17</td><td>4.7E-16</td><td>MB231/MB468</td><td>NUC</td><td></td></tr>
<tr><td>>MI0000253|hsa-mir-148a&WithFlank&7|-|25949913|25949992@50.70.21(+1G)&[MIMAT0000243&hsa-miR-148a-3p&offsets|0|-1(+1G);m-7&7|-|25949922|25949943&offsets|0|-1(+1G)]|TCAGTGCACTACAGAACTTTGG</td><td>>MI0000253|hsa-mir-148a&WithFlank&7|-|25949913|25949992</td><td>50</td><td>70</td><td>21(+1G)</td><td>TCAGTGCACTACAGAACTTTGG</td><td>3819.3</td><td>5.0</td><td>0.6</td><td>8.6</td><td>9.7E-18</td><td>4.7E-16</td><td>BT20/MB231</td><td>NUC</td><td></td></tr>
<tr><td>>MI0003628|hsa-mir-615&WithFlank&12|+|54033944|54034051@67.88.22&[MIMAT0003283&hsa-miR-615-3p&offsets|0|0;m-495&12|+|54034010|54034030&offsets|0|+1]|TCCGAGCCTGGGTCTCCCTCTT</td><td>>MI0003628|hsa-mir-615&WithFlank&12|+|54033944|54034051</td><td>67</td><td>88</td><td>22</td><td>TCCGAGCCTGGGTCTCCCTCTT</td><td>1123.6</td><td>5.2</td><td>0.6</td><td>8.6</td><td>9.9E-18</td><td>4.7E-16</td><td>BT20/MB231</td><td>NUC</td><td></td></tr>
<tr><td>>hg38_21_+_8433172_8446622_RNA45SN1_45S_50ntFlanks@12163.12183.21|ACTGGGGCGGTACACCTGTCA</td><td>>hg38_21_+_8433172_8446622_RNA45SN1_45S_50ntFlanks</td><td>12163</td><td>12183</td><td>21</td><td>ACTGGGGCGGTACACCTGTCA</td><td>1299.1</td><td>-10.1</td><td>1.2</td><td>-8.3</td><td>8.7E-17</td><td>4.8E-16</td><td>MB231/MB468</td><td>MITO</td><td></td></tr>
<tr><td>>hg38_21_+_8433172_8446622_RNA45SN1_45S_50ntFlanks@4731.4765.35|AAGAACGAAAGTCGGAGGTTCGAAGACGATCAGAT</td><td>>hg38_21_+_8433172_8446622_RNA45SN1_45S_50ntFlanks</td><td>4731</td><td>4765</td><td>35</td><td>AAGAACGAAAGTCGGAGGTTCGAAGACGATCAGAT</td><td>4676.5</td><td>7.2</td><td>0.9</td><td>8.3</td><td>9.1E-17</td><td>4.8E-16</td><td>BT20/MB468</td><td>CYTO</td><td></td></tr>
<tr><td>>MI0000809|hsa-mir-151a&WithFlank&8|-|140732558|140732659@53.74.22&[MIMAT0000757&hsa-miR-151a-3p&offsets|0|+1;m-30&8|-|140732587|140732607&offsets|0|+1]|CTAGACTGAAGCTCCTTGAGGA</td><td>>MI0000809|hsa-mir-151a&WithFlank&8|-|140732558|140732659</td><td>53</td><td>74</td><td>22</td><td>CTAGACTGAAGCTCCTTGAGGA</td><td>6601.3</td><td>6.1</td><td>0.7</td><td>8.4</td><td>5.5E-17</td><td>4.8E-16</td><td>MB231/MB468</td><td>CYTO</td><td></td></tr>
<tr><td>>hg38_21_+_8433172_8446622_RNA45SN1_45S_50ntFlanks@11752.11780.29|CCAAATGCCTCGTCATCTAATTAGTGACG</td><td>>hg38_21_+_8433172_8446622_RNA45SN1_45S_50ntFlanks</td><td>11752</td><td>11780</td><td>29</td><td>CCAAATGCCTCGTCATCTAATTAGTGACG</td><td>462.9</td><td>-11.6</td><td>1.4</td><td>-8.4</td><td>5.5E-17</td><td>4.8E-16</td><td>MB231/MB468</td><td>CYTO</td><td></td></tr>
<tr><td>>MI0000477|hsa-mir-146a&WithFlank&5|+|160485346|160485456@27.47.21&[MIMAT0000449&hsa-miR-146a-5p&offsets|0|-1;m-129&5|+|160485372|160485393&offsets|0|-1]|TGAGAACTGAATTCCATGGGT</td><td>>MI0000477|hsa-mir-146a&WithFlank&5|+|160485346|160485456</td><td>27</td><td>47</td><td>21</td><td>TGAGAACTGAATTCCATGGGT</td><td>8642.2</td><td>11.8</td><td>1.4</td><td>8.4</td><td>5.4E-17</td><td>4.8E-16</td><td>MB231/MB468</td><td>MP</td><td></td></tr>
<tr><td>>MI0001445|hsa-mir-423&WithFlank&17|+|30117073|30117178@59.78.20&[MIMAT0001340&hsa-miR-423-3p&offsets|0|-3;m-100&17|+|30117131|30117153&offsets|0|-3]|AGCTCGGTCTGAGGCCCCTC</td><td>>MI0001445|hsa-mir-423&WithFlank&17|+|30117073|30117178</td><td>59</td><td>78</td><td>20</td><td>AGCTCGGTCTGAGGCCCCTC</td><td>2502.7</td><td>6.7</td><td>0.8</td><td>8.4</td><td>5.5E-17</td><td>4.8E-16</td><td>MB231/MB468</td><td>CYTO</td><td></td></tr>
<tr><td>>hg38_21_+_8433172_8446622_RNA45SN1_45S_50ntFlanks@9572.9601.30|CAAATCGGTCGTCCGACCTGGGTATAGGGG</td><td>>hg38_21_+_8433172_8446622_RNA45SN1_45S_50ntFlanks</td><td>9572</td><td>9601</td><td>30</td><td>CAAATCGGTCGTCCGACCTGGGTATAGGGG</td><td>1064.2</td><td>-6.3</td><td>0.7</td><td>-8.4</td><td>5.4E-17</td><td>4.8E-16</td><td>MB231/MB468</td><td>MP</td><td></td></tr>
<tr><td>>MI0000102|hsa-mir-100&WithFlank&11|-|122152223|122152314@19.40.22(+1U)&[MIMAT0000098&hsa-miR-100-5p&offsets|0|0(+1U);m-27&11|-|122152276|122152296&offsets|0|+1(+1U)]|AACCCGTAGATCCGAACTTGTGT</td><td>>MI0000102|hsa-mir-100&WithFlank&11|-|122152223|122152314</td><td>19</td><td>40</td><td>22(+1U)</td><td>AACCCGTAGATCCGAACTTGTGT</td><td>6614.9</td><td>16.9</td><td>2.0</td><td>8.3</td><td>9.2E-17</td><td>4.8E-16</td><td>BT20/MB468</td><td>CYTO</td><td></td></tr>
<tr><td>>hg38_21_+_8433172_8446622_RNA45SN1_45S_50ntFlanks@10805.10822.18|GAACAATGTAGGTAAGGG</td><td>>hg38_21_+_8433172_8446622_RNA45SN1_45S_50ntFlanks</td><td>10805</td><td>10822</td><td>18</td><td>GAACAATGTAGGTAAGGG</td><td>237.5</td><td>-11.8</td><td>1.4</td><td>-8.4</td><td>5.4E-17</td><td>4.8E-16</td><td>MB231/MB468</td><td>MP</td><td></td></tr>
<tr><td>>MI0000087|hsa-mir-29a&WithFlank&7|-|130876741|130876816@47.68.22&[MIMAT0000086&hsa-miR-29a-3p&offsets|-1|-1;m-26&7|-|130876749|130876769&offsets|-1|0]|CTAGCACCATCTGAAATCGGTT</td><td>>MI0000087|hsa-mir-29a&WithFlank&7|-|130876741|130876816</td><td>47</td><td>68</td><td>22</td><td>CTAGCACCATCTGAAATCGGTT</td><td>13689.5</td><td>7.5</td><td>0.9</td><td>8.3</td><td>8.9E-17</td><td>4.8E-16</td><td>MB231/MB468</td><td>MITO</td><td></td></tr>
<tr><td>>hg38_21_+_8433172_8446622_RNA45SN1_45S_50ntFlanks@12974.13001.28|CGTACGTAGCAGAGCAGCTCCCTCGCTG</td><td>>hg38_21_+_8433172_8446622_RNA45SN1_45S_50ntFlanks</td><td>12974</td><td>13001</td><td>28</td><td>CGTACGTAGCAGAGCAGCTCCCTCGCTG</td><td>328.5</td><td>-12.1</td><td>1.5</td><td>-8.3</td><td>8.9E-17</td><td>4.8E-16</td><td>MB231/MB468</td><td>MITO</td><td></td></tr>
<tr><td>>hg38_21_+_8433172_8446622_RNA45SN1_45S_50ntFlanks@4560.4579.20|CTAGGAATAATGGAATAGGA</td><td>>hg38_21_+_8433172_8446622_RNA45SN1_45S_50ntFlanks</td><td>4560</td><td>4579</td><td>20</td><td>CTAGGAATAATGGAATAGGA</td><td>345.9</td><td>-12.3</td><td>1.5</td><td>-8.4</td><td>5.5E-17</td><td>4.8E-16</td><td>MB231/MB468</td><td>MP</td><td></td></tr>
<tr><td>>hg38_21_+_8433172_8446622_RNA45SN1_45S_50ntFlanks@11612.11640.29|TAATTAAAACAAAGCATCGCGAAGGCCCG</td><td>>hg38_21_+_8433172_8446622_RNA45SN1_45S_50ntFlanks</td><td>11612</td><td>11640</td><td>29</td><td>TAATTAAAACAAAGCATCGCGAAGGCCCG</td><td>334.8</td><td>-11.4</td><td>1.4</td><td>-8.4</td><td>6.8E-17</td><td>4.8E-16</td><td>MB231/MB468</td><td>NUC</td><td></td></tr>
<tr><td>>hg38_21_+_8433172_8446622_RNA45SN1_45S_50ntFlanks@5472.5494.23|ACGGCCCTGGCGGAGCGCTGAGA</td><td>>hg38_21_+_8433172_8446622_RNA45SN1_45S_50ntFlanks</td><td>5472</td><td>5494</td><td>23</td><td>ACGGCCCTGGCGGAGCGCTGAGA</td><td>8192.2</td><td>9.2</td><td>1.1</td><td>8.3</td><td>9.3E-17</td><td>4.9E-16</td><td>BT20/MB468</td><td>CYTO</td><td></td></tr>
<tr><td>>hg38_21_+_8433172_8446622_RNA45SN1_45S_50ntFlanks@9736.9758.23|CCGAAACGATCTCAACCTATTCT</td><td>>hg38_21_+_8433172_8446622_RNA45SN1_45S_50ntFlanks</td><td>9736</td><td>9758</td><td>23</td><td>CCGAAACGATCTCAACCTATTCT</td><td>937.4</td><td>-13.7</td><td>1.6</td><td>-8.3</td><td>8.9E-17</td><td>4.9E-16</td><td>MB231/MB468</td><td>MITO</td><td></td></tr>
<tr><td>>hg38_21_+_8433172_8446622_RNA45SN1_45S_50ntFlanks@9201.9218.18|CTCTCGGGGCCACGCGCG</td><td>>hg38_21_+_8433172_8446622_RNA45SN1_45S_50ntFlanks</td><td>9201</td><td>9218</td><td>18</td><td>CTCTCGGGGCCACGCGCG</td><td>1284.6</td><td>-10.4</td><td>1.2</td><td>-8.3</td><td>9.0E-17</td><td>4.9E-16</td><td>MB231/MB468</td><td>MITO</td><td></td></tr>
<tr><td>>hg38_21_+_8433172_8446622_RNA45SN1_45S_50ntFlanks@8107.8139.33|CGGCGGGGCGCGGGACATGTGGCGTACGGAAGA</td><td>>hg38_21_+_8433172_8446622_RNA45SN1_45S_50ntFlanks</td><td>8107</td><td>8139</td><td>33</td><td>CGGCGGGGCGCGGGACATGTGGCGTACGGAAGA</td><td>253.7</td><td>-11.9</td><td>1.4</td><td>-8.4</td><td>5.5E-17</td><td>4.9E-16</td><td>MB231/MB468</td><td>MP</td><td></td></tr>
<tr><td>>hg38_21_+_8433172_8446622_RNA45SN1_45S_50ntFlanks@6772.6798.27|GTTCCTCCCGGGGCTACGCCTGTCTGA</td><td>>hg38_21_+_8433172_8446622_RNA45SN1_45S_50ntFlanks</td><td>6772</td><td>6798</td><td>27</td><td>GTTCCTCCCGGGGCTACGCCTGTCTGA</td><td>391.6</td><td>-12.4</td><td>1.5</td><td>-8.3</td><td>9.0E-17</td><td>4.9E-16</td><td>MB231/MB468</td><td>MITO</td><td></td></tr>
<tr><td>>hg38_21_+_8433172_8446622_RNA45SN1_45S_50ntFlanks@8779.8796.18|CCCCCCGGCCCCGCGTCC</td><td>>hg38_21_+_8433172_8446622_RNA45SN1_45S_50ntFlanks</td><td>8779</td><td>8796</td><td>18</td><td>CCCCCCGGCCCCGCGTCC</td><td>390.0</td><td>-12.4</td><td>1.5</td><td>-8.3</td><td>9.0E-17</td><td>4.9E-16</td><td>MB231/MB468</td><td>MITO</td><td></td></tr>
<tr><td>>hg38_21_+_8433172_8446622_RNA45SN1_45S_50ntFlanks@6661.6693.33|CGGTGGATCACTCGGCTCGTGCGTCGATGAAGA</td><td>>hg38_21_+_8433172_8446622_RNA45SN1_45S_50ntFlanks</td><td>6661</td><td>6693</td><td>33</td><td>CGGTGGATCACTCGGCTCGTGCGTCGATGAAGA</td><td>1867.8</td><td>5.4</td><td>0.6</td><td>8.4</td><td>6.7E-17</td><td>4.9E-16</td><td>BT20/MB468</td><td>MITO</td><td></td></tr>
<tr><td>>hg38_21_+_8433172_8446622_RNA45SN1_45S_50ntFlanks@10806.10823.18|AACAATGTAGGTAAGGGA</td><td>>hg38_21_+_8433172_8446622_RNA45SN1_45S_50ntFlanks</td><td>10806</td><td>10823</td><td>18</td><td>AACAATGTAGGTAAGGGA</td><td>572.3</td><td>-7.3</td><td>0.9</td><td>-8.4</td><td>5.7E-17</td><td>4.9E-16</td><td>MB231/MB468</td><td>CYTO</td><td></td></tr>
<tr><td>>MI0000086|hsa-mir-28&WithFlank&3|+|188688775|188688872@61.82.22&[MIMAT0004502&hsa-miR-28-3p&offsets|+1|+1;m-28&3|+|188688834|188688855&offsets|+1|+1]|ACTAGATTGTGAGCTCCTGGAG</td><td>>MI0000086|hsa-mir-28&WithFlank&3|+|188688775|188688872</td><td>61</td><td>82</td><td>22</td><td>ACTAGATTGTGAGCTCCTGGAG</td><td>175.2</td><td>5.5</td><td>0.7</td><td>8.3</td><td>6.9E-17</td><td>4.9E-16</td><td>MB231/MB468</td><td>NUC</td><td></td></tr>
<tr><td>>MI0000076|hsa-mir-20a&WithFlank&13|+|91351059|91351141@14.36.23&[MIMAT0000075&hsa-miR-20a-5p&offsets|0|0;m-96&13|+|91351072|91351094&offsets|0|0]|TAAAGTGCTTATAGTGCAGGTAG</td><td>>MI0000076|hsa-mir-20a&WithFlank&13|+|91351059|91351141</td><td>14</td><td>36</td><td>23</td><td>TAAAGTGCTTATAGTGCAGGTAG</td><td>16572.3</td><td>5.7</td><td>0.7</td><td>8.3</td><td>6.9E-17</td><td>4.9E-16</td><td>MB231/MB468</td><td>NUC</td><td></td></tr>
<tr><td>>hg38_21_+_8433172_8446622_RNA45SN1_45S_50ntFlanks@12709.12743.35|CCGCGGAGCCTCGGTTGGCCTCGGATAGCCGGTCC</td><td>>hg38_21_+_8433172_8446622_RNA45SN1_45S_50ntFlanks</td><td>12709</td><td>12743</td><td>35</td><td>CCGCGGAGCCTCGGTTGGCCTCGGATAGCCGGTCC</td><td>349.2</td><td>-12.2</td><td>1.5</td><td>-8.3</td><td>9.1E-17</td><td>4.9E-16</td><td>MB231/MB468</td><td>MITO</td><td></td></tr>
<tr><td>>hg38_21_+_8433172_8446622_RNA45SN1_45S_50ntFlanks@5448.5466.19|GATCGGCCCCGCCGGGGTC</td><td>>hg38_21_+_8433172_8446622_RNA45SN1_45S_50ntFlanks</td><td>5448</td><td>5466</td><td>19</td><td>GATCGGCCCCGCCGGGGTC</td><td>805.0</td><td>-13.5</td><td>1.6</td><td>-8.4</td><td>5.6E-17</td><td>4.9E-16</td><td>MB231/MB468</td><td>MP</td><td></td></tr>
<tr><td>>hg38_21_+_8433172_8446622_RNA45SN1_45S_50ntFlanks@11199.11233.35|CCGGCGGCGGCGGCGACTCTGGACGCGAGCCGGGC</td><td>>hg38_21_+_8433172_8446622_RNA45SN1_45S_50ntFlanks</td><td>11199</td><td>11233</td><td>35</td><td>CCGGCGGCGGCGGCGACTCTGGACGCGAGCCGGGC</td><td>312.6</td><td>-8.0</td><td>1.0</td><td>-8.4</td><td>5.7E-17</td><td>5.0E-16</td><td>MB231/MB468</td><td>CYTO</td><td></td></tr>
<tr><td>>hg38_21_+_8433172_8446622_RNA45SN1_45S_50ntFlanks@12249.12273.25|CAAAAGCTCGCTTGATCTTGATTTT</td><td>>hg38_21_+_8433172_8446622_RNA45SN1_45S_50ntFlanks</td><td>12249</td><td>12273</td><td>25</td><td>CAAAAGCTCGCTTGATCTTGATTTT</td><td>1160.8</td><td>-14.0</td><td>1.7</td><td>-8.3</td><td>9.1E-17</td><td>5.0E-16</td><td>MB231/MB468</td><td>MITO</td><td></td></tr>
<tr><td>>hg38_21_+_8433172_8446622_RNA45SN1_45S_50ntFlanks@11885.11904.20|TTGAGCTTGACTCTAGTCTG</td><td>>hg38_21_+_8433172_8446622_RNA45SN1_45S_50ntFlanks</td><td>11885</td><td>11904</td><td>20</td><td>TTGAGCTTGACTCTAGTCTG</td><td>2835.8</td><td>-8.7</td><td>1.0</td><td>-8.4</td><td>5.7E-17</td><td>5.0E-16</td><td>MB231/MB468</td><td>CYTO</td><td></td></tr>
<tr><td>>hg38_21_+_8433172_8446622_RNA45SN1_45S_50ntFlanks@11568.11603.36|GCAGCCGACTTAGAACTGGTGCGGACCAGGGGAATC</td><td>>hg38_21_+_8433172_8446622_RNA45SN1_45S_50ntFlanks</td><td>11568</td><td>11603</td><td>36</td><td>GCAGCCGACTTAGAACTGGTGCGGACCAGGGGAATC</td><td>1824.6</td><td>-5.5</td><td>0.7</td><td>-8.3</td><td>7.0E-17</td><td>5.0E-16</td><td>MB231/MB468</td><td>NUC</td><td></td></tr>
<tr><td>>hg38_21_+_8433172_8446622_RNA45SN1_45S_50ntFlanks@12633.12661.29|CGAAGCTACCATCTGTGGGATTATGACTG</td><td>>hg38_21_+_8433172_8446622_RNA45SN1_45S_50ntFlanks</td><td>12633</td><td>12661</td><td>29</td><td>CGAAGCTACCATCTGTGGGATTATGACTG</td><td>1447.4</td><td>-4.8</td><td>0.6</td><td>-8.3</td><td>7.0E-17</td><td>5.0E-16</td><td>MB231/MB468</td><td>NUC</td><td></td></tr>
<tr><td>>hg38_21_+_8433172_8446622_RNA45SN1_45S_50ntFlanks@12006.12043.38|GGCCGCCGGTGAAATACCACTACTCTGATCGTTTTTTC</td><td>>hg38_21_+_8433172_8446622_RNA45SN1_45S_50ntFlanks</td><td>12006</td><td>12043</td><td>38</td><td>GGCCGCCGGTGAAATACCACTACTCTGATCGTTTTTTC</td><td>118.1</td><td>-24.0</td><td>2.9</td><td>-8.4</td><td>5.6E-17</td><td>5.0E-16</td><td>MB231/MB468</td><td>MP</td><td></td></tr>
<tr><td>>hg38_21_+_8433172_8446622_RNA45SN1_45S_50ntFlanks@10866.10901.36|CTAAGGGCTGGGTCGGTCGGGCTGGGGCGCGAAGCG</td><td>>hg38_21_+_8433172_8446622_RNA45SN1_45S_50ntFlanks</td><td>10866</td><td>10901</td><td>36</td><td>CTAAGGGCTGGGTCGGTCGGGCTGGGGCGCGAAGCG</td><td>1626.7</td><td>-10.0</td><td>1.2</td><td>-8.3</td><td>9.2E-17</td><td>5.0E-16</td><td>MB231/MB468</td><td>MITO</td><td></td></tr>
</table>

<table>
<tr><th></th><th></th><th></th><th></th><th></th><th></th><th></th><th></th><th></th><th></th><th></th><th></th><th></th><th></th></tr>
<tr><td>>trna101_AlaAGC_6_-_28831462_28831533@58.75.18,trna102_AlaAGC_6_-_28806221_28806292@58.75.18,trna110_AlaTGC_6_-_28757547_28757618@58.75.18,trna113_AlaTGC_6_-_28726141_28726212@58.75.18,trna65_AlaAGC_6_+_28574933_28575004@58.75.18|TCCCCGGCACCTCCACCA</td><td>>trna101_AlaAGC_6_-_28831462_28831533</td><td>58</td><td>75</td><td>18</td><td>TCCCCGGCACCTCCACCA</td><td>14007.6</td><td>7.4</td><td>0.9</td><td>8.4</td><td>5.8E-17</td><td>5.0E-16</td><td>MB231/MB468</td><td>CYTO</td></tr>
<tr><td>>am_trna13_LysCTT_14_-_58706613_58706685@31.51.21,trna2_LysCTT_15_+_79152904_79152976@31.51.21,trna4_LysCTT_18_-_43669271_43669343@31.51.21|ACTCTTAATCCCAGGGTCGTG</td><td>>am_trna13_LysCTT_14_-_58706613_58706685</td><td>31</td><td>51</td><td>21</td><td>ACTCTTAATCCCAGGGTCGTG</td><td>657.4</td><td>-12.1</td><td>1.4</td><td>-8.4</td><td>5.8E-17</td><td>5.0E-16</td><td>MB231/MB468</td><td>CYTO</td></tr>
<tr><td>>MI0000286|hsa-mir-210&WithFlank&11|-|568083|568204@72.93.22&[MIMAT0000267&hsa-miR-210-3p&offsets|0|0;m-102&11|-|568112|568133&offsets|0|0]|CTGTGCGTGTGACAGCGGCTGA</td><td>>MI0000286|hsa-mir-210&WithFlank&11|-|568083|568204</td><td>72</td><td>93</td><td>22</td><td>CTGTGCGTGTGACAGCGGCTGA</td><td>1338.9</td><td>12.4</td><td>1.5</td><td>8.3</td><td>9.5E-17</td><td>5.0E-16</td><td>BT20/MB468</td><td>CYTO</td></tr>
<tr><td>>hg38_21_+_8433172_8446622_RNA45SN1_45S_50ntFlanks@9131.9152.22|GGTCGGGGGGCGGGGCGGACTG</td><td>>hg38_21_+_8433172_8446622_RNA45SN1_45S_50ntFlanks</td><td>9131</td><td>9152</td><td>22</td><td>GGTCGGGGGGCGGGGCGGACTG</td><td>1090.4</td><td>-14.0</td><td>1.7</td><td>-8.4</td><td>5.7E-17</td><td>5.0E-16</td><td>MB231/MB468</td><td>MP</td></tr>
<tr><td>>hg38_21_+_8433172_8446622_RNA45SN1_45S_50ntFlanks@4723.4759.37|CATTAATCAAGAACGAAAGTCGGAGGTTCGAAGACGA</td><td>>hg38_21_+_8433172_8446622_RNA45SN1_45S_50ntFlanks</td><td>4723</td><td>4759</td><td>37</td><td>CATTAATCAAGAACGAAAGTCGGAGGTTCGAAGACGA</td><td>2286.4</td><td>-6.4</td><td>0.8</td><td>-8.3</td><td>7.1E-17</td><td>5.0E-16</td><td>MB231/MB468</td><td>NUC</td></tr>
<tr><td>>hg38_21_+_8433172_8446622_RNA45SN1_45S_50ntFlanks@12717.12743.27|CCTCGGTTGGCCTCGGATAGCCGGTCC</td><td>>hg38_21_+_8433172_8446622_RNA45SN1_45S_50ntFlanks</td><td>12717</td><td>12743</td><td>27</td><td>CCTCGGTTGGCCTCGGATAGCCGGTCC</td><td>248.2</td><td>-11.8</td><td>1.4</td><td>-8.4</td><td>5.7E-17</td><td>5.0E-16</td><td>MB231/MB468</td><td>MP</td></tr>
<tr><td>>trna2_GlyGCC_21_-_18827107_18827177@53.74.22,trna35_GlyGCC_1_+_161413094_161413164@53.74.22,trna37_GlyGCC_1_+_161420467_161420537@53.74.22,trna39_GlyGCC_1_+_161427898_161427968@53.74.22,trna41_GlyGCC_1_+_161435258_161435328@53.74.22|TCGATTCCCGGCCCATGCACCA</td><td>>trna2_GlyGCC_21_-_18827107_18827177</td><td>53</td><td>74</td><td>22</td><td>TCGATTCCCGGCCCATGCACCA</td><td>558.6</td><td>11.1</td><td>1.3</td><td>8.3</td><td>9.6E-17</td><td>5.0E-16</td><td>BT20/MB468</td><td>CYTO</td></tr>
<tr><td>>hg38_21_+_8433172_8446622_RNA45SN1_45S_50ntFlanks@11768.11804.37|CTAATTAGTGACGCGCATGAATGGATGAACGAGATTC</td><td>>hg38_21_+_8433172_8446622_RNA45SN1_45S_50ntFlanks</td><td>11768</td><td>11804</td><td>37</td><td>CTAATTAGTGACGCGCATGAATGGATGAACGAGATTC</td><td>642.2</td><td>-13.2</td><td>1.6</td><td>-8.4</td><td>5.7E-17</td><td>5.0E-16</td><td>MB231/MB468</td><td>MP</td></tr>
<tr><td>>hg38_21_+_8433172_8446622_RNA45SN1_45S_50ntFlanks@4297.4321.25|CATTGGAGGGCAAGTCTGGTGCCAG</td><td>>hg38_21_+_8433172_8446622_RNA45SN1_45S_50ntFlanks</td><td>4297</td><td>4321</td><td>25</td><td>CATTGGAGGGCAAGTCTGGTGCCAG</td><td>336.1</td><td>-12.3</td><td>1.5</td><td>-8.4</td><td>5.7E-17</td><td>5.0E-16</td><td>MB231/MB468</td><td>MP</td></tr>
<tr><td>>hg38_21_+_8433172_8446622_RNA45SN1_45S_50ntFlanks@9471.9503.33|AGCACGAGCGCACGTGTTAGGACCCGAAAGATG</td><td>>hg38_21_+_8433172_8446622_RNA45SN1_45S_50ntFlanks</td><td>9471</td><td>9503</td><td>33</td><td>AGCACGAGCGCACGTGTTAGGACCCGAAAGATG</td><td>1576.2</td><td>-10.3</td><td>1.2</td><td>-8.4</td><td>5.7E-17</td><td>5.0E-16</td><td>MB231/MB468</td><td>MP</td></tr>
<tr><td>>hg38_21_+_8433172_8446622_RNA45SN1_45S_50ntFlanks@10392.10424.33|ACAGCAGTTGAACATGGGTCAGTCGGTCCTGAG</td><td>>hg38_21_+_8433172_8446622_RNA45SN1_45S_50ntFlanks</td><td>10392</td><td>10424</td><td>33</td><td>ACAGCAGTTGAACATGGGTCAGTCGGTCCTGAG</td><td>6986.9</td><td>-5.0</td><td>0.6</td><td>-8.3</td><td>1.6E-16</td><td>5.0E-16</td><td>BT20/MB468</td><td>NUC</td></tr>
<tr><td>>hg38_21_+_8433172_8446622_RNA45SN1_45S_50ntFlanks@10789.10818.30|AGCCTCTGGCATGTTGGAACAATGTAGGTA</td><td>>hg38_21_+_8433172_8446622_RNA45SN1_45S_50ntFlanks</td><td>10789</td><td>10818</td><td>30</td><td>AGCCTCTGGCATGTTGGAACAATGTAGGTA</td><td>760.8</td><td>-7.7</td><td>0.9</td><td>-8.4</td><td>5.7E-17</td><td>5.0E-16</td><td>MB231/MB468</td><td>MP</td></tr>
<tr><td>>MI0000266|hsa-mir-10a&WithFlank&17|-|48579832|48579953@28.49.22(+1A)&[MIMAT0000253&hsa-miR-10a-5p&offsets|0|-1(+1A);m-18&17|-|48579905|48579926&offsets|0|0(+1A)]|TACCCTGTAGATCCGAATTTGTA</td><td>>MI0000266|hsa-mir-10a&WithFlank&17|-|48579832|48579953</td><td>28</td><td>49</td><td>22(+1A)</td><td>TACCCTGTAGATCCGAATTTGTA</td><td>395.8</td><td>11.8</td><td>1.4</td><td>8.4</td><td>5.7E-17</td><td>5.1E-16</td><td>MB231/MB468</td><td>MP</td></tr>
<tr><td>>MI0000286|hsa-mir-210&WithFlank&11|-|568083|568204@72.94.23&[MIMAT0000267&hsa-miR-210-3p&offsets|0|+1;m-102&11|-|568112|568133&offsets|0|+1]|CTGTGCGTGTGACAGCGGCTGAT</td><td>>MI0000286|hsa-mir-210&WithFlank&11|-|568083|568204</td><td>72</td><td>94</td><td>23</td><td>CTGTGCGTGTGACAGCGGCTGAT</td><td>585.8</td><td>13.4</td><td>1.6</td><td>8.3</td><td>9.7E-17</td><td>5.1E-16</td><td>BT20/MB468</td><td>CYTO</td></tr>
<tr><td>>MI0000456|hsa-mir-140&WithFlank&16|+|69933075|69933186@69.90.22(+1U)&[MIMAT0004597&hsa-miR-140-3p&offsets|+1|+2(+1U);m-45&16|+|69933143|69933164&offsets|0|0(+1U)]|ACCACAGGGTAGAACCACGGACT</td><td>>MI0000456|hsa-mir-140&WithFlank&16|+|69933075|69933186</td><td>69</td><td>90</td><td>22(+1U)</td><td>ACCACAGGGTAGAACCACGGACT</td><td>1290.1</td><td>11.3</td><td>1.4</td><td>8.3</td><td>9.7E-17</td><td>5.1E-16</td><td>BT20/MB468</td><td>CYTO</td></tr>
<tr><td>>hg38_21_+_8433172_8446622_RNA45SN1_45S_50ntFlanks@4773.4793.21|TAGTTCCGACCATAAACGATG</td><td>>hg38_21_+_8433172_8446622_RNA45SN1_45S_50ntFlanks</td><td>4773</td><td>4793</td><td>21</td><td>TAGTTCCGACCATAAACGATG</td><td>1258.8</td><td>-7.9</td><td>0.9</td><td>-8.3</td><td>9.3E-17</td><td>5.1E-16</td><td>MB231/MB468</td><td>MITO</td></tr>
<tr><td>>hg38_21_+_8433172_8446622_RNA45SN1_45S_50ntFlanks@6646.6667.22|TCGTACGACTCTTAGCGGTGGA</td><td>>hg38_21_+_8433172_8446622_RNA45SN1_45S_50ntFlanks</td><td>6646</td><td>6667</td><td>22</td><td>TCGTACGACTCTTAGCGGTGGA</td><td>69173.1</td><td>6.9</td><td>0.8</td><td>8.3</td><td>6.9E-17</td><td>5.1E-16</td><td>BT20/MB468</td><td>MITO</td></tr>
<tr><td>>hg38_21_+_8433172_8446622_RNA45SN1_45S_50ntFlanks@10912.10931.20|CGCGCCGCGGCTGGACGAGG</td><td>>hg38_21_+_8433172_8446622_RNA45SN1_45S_50ntFlanks</td><td>10912</td><td>10931</td><td>20</td><td>CGCGCCGCGGCTGGACGAGG</td><td>407.8</td><td>-12.5</td><td>1.5</td><td>-8.4</td><td>5.8E-17</td><td>5.1E-16</td><td>MB231/MB468</td><td>MP</td></tr>
<tr><td>>hg38_21_+_8433172_8446622_RNA45SN1_45S_50ntFlanks@10787.10816.30|ACAGCCTCTGGCATGTTGGAACAATGTAGG</td><td>>hg38_21_+_8433172_8446622_RNA45SN1_45S_50ntFlanks</td><td>10787</td><td>10816</td><td>30</td><td>ACAGCCTCTGGCATGTTGGAACAATGTAGG</td><td>358.0</td><td>-12.3</td><td>1.5</td><td>-8.4</td><td>5.8E-17</td><td>5.1E-16</td><td>MB231/MB468</td><td>MP</td></tr>
<tr><td>>MI0000082|hsa-mir-25&WithFlank&7|-|100093554|100093649@58.78.21(+1U)&[MIMAT0000081&hsa-miR-25-3p&offsets|0|-1(+1U);m-23&7|-|100093571|100093592&offsets|0|-1(+1U)]|CATTGCACTTGTCTCGGTCTGT</td><td>>MI0000082|hsa-mir-25&WithFlank&7|-|100093554|100093649</td><td>58</td><td>78</td><td>21(+1U)</td><td>CATTGCACTTGTCTCGGTCTGT</td><td>1821.9</td><td>5.1</td><td>0.6</td><td>8.3</td><td>7.2E-17</td><td>5.1E-16</td><td>MB231/MB468</td><td>NUC</td></tr>
<tr><td>>hg38_21_+_8433172_8446622_RNA45SN1_45S_50ntFlanks@9304.9336.33|ACCAAGGAGTCTAACACGTGCGCGAGTCGGGGG</td><td>>hg38_21_+_8433172_8446622_RNA45SN1_45S_50ntFlanks</td><td>9304</td><td>9336</td><td>33</td><td>ACCAAGGAGTCTAACACGTGCGCGAGTCGGGGG</td><td>351.5</td><td>-12.2</td><td>1.5</td><td>-8.3</td><td>9.4E-17</td><td>5.1E-16</td><td>MB231/MB468</td><td>MITO</td></tr>
<tr><td>>hg38_21_+_8433172_8446622_RNA45SN1_45S_50ntFlanks@11641.11676.36|CGGCGGGTGTTGACGCGATGTGATTTCTGCCCAGTG</td><td>>hg38_21_+_8433172_8446622_RNA45SN1_45S_50ntFlanks</td><td>11641</td><td>11676</td><td>36</td><td>CGGCGGGTGTTGACGCGATGTGATTTCTGCCCAGTG</td><td>477.2</td><td>-12.7</td><td>1.5</td><td>-8.3</td><td>9.5E-17</td><td>5.1E-16</td><td>MB231/MB468</td><td>MITO</td></tr>
<tr><td>>hg38_21_+_8433172_8446622_RNA45SN1_45S_50ntFlanks@10350.10383.34|ACGAGAACTTTGAAGGCCGAAGTGGAGAAGGGTT</td><td>>hg38_21_+_8433172_8446622_RNA45SN1_45S_50ntFlanks</td><td>10350</td><td>10383</td><td>34</td><td>ACGAGAACTTTGAAGGCCGAAGTGGAGAAGGGTT</td><td>7247.3</td><td>-4.0</td><td>0.5</td><td>-8.4</td><td>6.0E-17</td><td>5.1E-16</td><td>MB231/MB468</td><td>CYTO</td></tr>
<tr><td>>hg38_21_+_8433172_8446622_RNA45SN1_45S_50ntFlanks@3873.3898.26|TAATACATGCCGACGGGCGCTGACCC</td><td>>hg38_21_+_8433172_8446622_RNA45SN1_45S_50ntFlanks</td><td>3873</td><td>3898</td><td>26</td><td>TAATACATGCCGACGGGCGCTGACCC</td><td>2688.5</td><td>-5.4</td><td>0.6</td><td>-8.4</td><td>6.0E-17</td><td>5.1E-16</td><td>MB231/MB468</td><td>CYTO</td></tr>
<tr><td>>hg38_21_+_8433172_8446622_RNA45SN1_45S_50ntFlanks@9545.9567.23|GTGGAGGTCCGTAGCGGTCCTGA</td><td>>hg38_21_+_8433172_8446622_RNA45SN1_45S_50ntFlanks</td><td>9545</td><td>9567</td><td>23</td><td>GTGGAGGTCCGTAGCGGTCCTGA</td><td>267.2</td><td>-11.0</td><td>1.3</td><td>-8.3</td><td>7.3E-17</td><td>5.1E-16</td><td>MB231/MB468</td><td>NUC</td></tr>
<tr><td>>hg38_21_+_8433172_8446622_RNA45SN1_45S_50ntFlanks@4130.4162.33|ATTCCGGAGAGGGAGCCTGAGAAACGGCTACCA</td><td>>hg38_21_+_8433172_8446622_RNA45SN1_45S_50ntFlanks</td><td>4130</td><td>4162</td><td>33</td><td>ATTCCGGAGAGGGAGCCTGAGAAACGGCTACCA</td><td>465.5</td><td>-12.7</td><td>1.5</td><td>-8.4</td><td>5.9E-17</td><td>5.2E-16</td><td>MB231/MB468</td><td>MP</td></tr>
<tr><td>>MI0000471|hsa-mir-126&WithFlank&9|+|136670596|136670692@58.78.21(+1A)&[MIMAT0000445&hsa-miR-126-3p&offsets|0|-1(+1A);m-40&9|+|136670653|136670674&offsets|0|-1(+1A)]|TCGTACCGTGAGTAATAATGCA</td><td>>MI0000471|hsa-mir-126&WithFlank&9|+|136670596|136670692</td><td>58</td><td>78</td><td>21(+1A)</td><td>TCGTACCGTGAGTAATAATGCA</td><td>404.3</td><td>9.9</td><td>1.2</td><td>8.3</td><td>7.3E-17</td><td>5.2E-16</td><td>MB231/MB468</td><td>NUC</td></tr>
<tr><td>>hg38_21_+_8433172_8446622_RNA45SN1_45S_50ntFlanks@8104.8141.38|CCGCGGCGGGGCGCGGGACATGTGGCGTACGGAAGACC</td><td>>hg38_21_+_8433172_8446622_RNA45SN1_45S_50ntFlanks</td><td>8104</td><td>8141</td><td>38</td><td>CCGCGGCGGGGCGCGGGACATGTGGCGTACGGAAGACC</td><td>2124.6</td><td>-7.5</td><td>0.9</td><td>-8.3</td><td>9.6E-17</td><td>5.2E-16</td><td>MB231/MB468</td><td>MITO</td></tr>
<tr><td>>hg38_21_+_8433172_8446622_RNA45SN1_45S_50ntFlanks@12626.12649.24|AATGGGGCGAAGCTACCATCTGTG</td><td>>hg38_21_+_8433172_8446622_RNA45SN1_45S_50ntFlanks</td><td>12626</td><td>12649</td><td>24</td><td>AATGGGGCGAAGCTACCATCTGTG</td><td>369.8</td><td>-12.4</td><td>1.5</td><td>-8.4</td><td>5.9E-17</td><td>5.2E-16</td><td>MB231/MB468</td><td>MP</td></tr>
<tr><td>>MI0001445|hsa-mir-423&WithFlank&17|+|30117073|30117178@59.82.24&[MIMAT0001340&hsa-miR-423-3p&offsets|0|+1;m-100&17|+|30117131|30117153&offsets|0|+1]|AGCTCGGTCTGAGGCCCCTCAGTC</td><td>>MI0001445|hsa-mir-423&WithFlank&17|+|30117073|30117178</td><td>59</td><td>82</td><td>24</td><td>AGCTCGGTCTGAGGCCCCTCAGTC</td><td>581.1</td><td>8.4</td><td>1.0</td><td>8.3</td><td>7.1E-17</td><td>5.2E-16</td><td>BT20/MB468</td><td>MITO</td></tr>
<tr><td>>MI0000434|hsa-let-7i&WithFlank&12|+|62603680|62603775@12.31.20&[MIMAT0000415&hsa-let-7i-5p&offsets|0|-2;m-49&12|+|62603691|62603712&offsets|0|-2]|TGAGGTAGTAGTTTGTGCTG</td><td>>MI0000434|hsa-let-7i&WithFlank&12|+|62603680|62603775</td><td>12</td><td>31</td><td>20</td><td>TGAGGTAGTAGTTTGTGCTG</td><td>521.3</td><td>6.0</td><td>0.7</td><td>8.3</td><td>7.4E-17</td><td>5.2E-16</td><td>MB231/MB468</td><td>NUC</td></tr>
<tr><td>>hg38_21_+_8433172_8446622_RNA45SN1_45S_50ntFlanks@11718.11740.23|AACGGCGGGAGTAACTATGACTC</td><td>>hg38_21_+_8433172_8446622_RNA45SN1_45S_50ntFlanks</td><td>11718</td><td>11740</td><td>23</td><td>AACGGCGGGAGTAACTATGACTC</td><td>7068.8</td><td>-9.7</td><td>1.2</td><td>-8.3</td><td>7.4E-17</td><td>5.2E-16</td><td>MB231/MB468</td><td>NUC</td></tr>
<tr><td>>hg38_21_+_8433172_8446622_RNA45SN1_45S_50ntFlanks@12677.12703.27|GAATCCCGCCCAGGCGGAACGATACGG</td><td>>hg38_21_+_8433172_8446622_RNA45SN1_45S_50ntFlanks</td><td>12677</td><td>12703</td><td>27</td><td>GAATCCCGCCCAGGCGGAACGATACGG</td><td>483.8</td><td>-12.8</td><td>1.5</td><td>-8.4</td><td>5.9E-17</td><td>5.2E-16</td><td>MB231/MB468</td><td>MP</td></tr>
<tr><td>>hg38_21_+_8433172_8446622_RNA45SN1_45S_50ntFlanks@10411.10440.30|CAGTCGGTCCTGAGAGATGGGCGAGCGCCG</td><td>>hg38_21_+_8433172_8446622_RNA45SN1_45S_50ntFlanks</td><td>10411</td><td>10440</td><td>30</td><td>CAGTCGGTCCTGAGAGATGGGCGAGCGCCG</td><td>499.9</td><td>-12.8</td><td>1.5</td><td>-8.4</td><td>6.0E-17</td><td>5.2E-16</td><td>MB231/MB468</td><td>MP</td></tr>
<tr><td>>hg38_21_+_8433172_8446622_RNA45SN1_45S_50ntFlanks@12405.12432.28|CTTTTTGATCCTTCGATGTCGGCTCTTC</td><td>>hg38_21_+_8433172_8446622_RNA45SN1_45S_50ntFlanks</td><td>12405</td><td>12432</td><td>28</td><td>CTTTTTGATCCTTCGATGTCGGCTCTTC</td><td>2465.9</td><td>-5.0</td><td>0.6</td><td>-8.3</td><td>7.2E-17</td><td>5.3E-16</td><td>BT20/MB468</td><td>MITO</td></tr>
<tr><td>>hg38_21_+_8433172_8446622_RNA45SN1_45S_50ntFlanks@10350.10390.41|ACGAGAACTTTGAAGGCCGAAGTGGAGAAGGGTTCCATGTG</td><td>>hg38_21_+_8433172_8446622_RNA45SN1_45S_50ntFlanks</td><td>10350</td><td>10390</td><td>41</td><td>ACGAGAACTTTGAAGGCCGAAGTGGAGAAGGGTTCCATGTG</td><td>2396.4</td><td>-6.7</td><td>0.8</td><td>-8.4</td><td>6.0E-17</td><td>5.3E-16</td><td>MB231/MB468</td><td>MP</td></tr>
<tr><td>>hg38_21_+_8433172_8446622_RNA45SN1_45S_50ntFlanks@10680.10708.29|GTTCCGGCGGCGTCCGGTGAGCTCTCGCT</td><td>>hg38_21_+_8433172_8446622_RNA45SN1_45S_50ntFlanks</td><td>10680</td><td>10708</td><td>29</td><td>GTTCCGGCGGCGTCCGGTGAGCTCTCGCT</td><td>46384.4</td><td>-9.0</td><td>1.1</td><td>-8.2</td><td>1.6E-16</td><td>5.3E-16</td><td>BT20/MB468</td><td>NUC</td></tr>
<tr><td>>hg38_21_+_8433172_8446622_RNA45SN1_45S_50ntFlanks@11641.11678.38|CGGCGGGTGTTGACGCGATGTGATTTCTGCCCAGTGCT</td><td>>hg38_21_+_8433172_8446622_RNA45SN1_45S_50ntFlanks</td><td>11641</td><td>11678</td><td>38</td><td>CGGCGGGTGTTGACGCGATGTGATTTCTGCCCAGTGCT</td><td>609.2</td><td>-13.0</td><td>1.6</td><td>-8.3</td><td>9.8E-17</td><td>5.3E-16</td><td>MB231/MB468</td><td>MITO</td></tr>
<tr><td>>hg38_21_+_8433172_8446622_RNA45SN1_45S_50ntFlanks@4283.4314.32|CTTTAACGAGGATCCATTGGAGGGCAAGTCTG</td><td>>hg38_21_+_8433172_8446622_RNA45SN1_45S_50ntFlanks</td><td>4283</td><td>4314</td><td>32</td><td>CTTTAACGAGGATCCATTGGAGGGCAAGTCTG</td><td>792.5</td><td>-13.5</td><td>1.6</td><td>-8.4</td><td>6.0E-17</td><td>5.3E-16</td><td>MB231/MB468</td><td>MP</td></tr>
<tr><td>>hg38_21_+_8433172_8446622_RNA45SN1_45S_50ntFlanks@10597.10617.21|AGTTCTCTTTTCTTTGTGAAG</td><td>>hg38_21_+_8433172_8446622_RNA45SN1_45S_50ntFlanks</td><td>10597</td><td>10617</td><td>21</td><td>AGTTCTCTTTTCTTTGTGAAG</td><td>245.2</td><td>-12.4</td><td>1.5</td><td>-8.2</td><td>1.7E-16</td><td>5.3E-16</td><td>BT20/MB468</td><td>NUC</td></tr>
<tr><td>>MI0000071|hsa-mir-17&WithFlank&13|+|91350599|91350694@20.43.24&[MIMAT0000070&hsa-miR-17-5p&offsets|0|+1;m-88&13|+|91350618|91350640&offsets|0|+1]|CAAAGTGCTTACAGTGCAGGTAGT</td><td>>MI0000071|hsa-mir-17&WithFlank&13|+|91350599|91350694</td><td>20</td><td>43</td><td>24</td><td>CAAAGTGCTTACAGTGCAGGTAGT</td><td>857.6</td><td>4.8</td><td>0.6</td><td>8.3</td><td>1.0E-16</td><td>5.3E-16</td><td>BT20/MB468</td><td>CYTO</td></tr>
<tr><td>>hg38_21_+_8433172_8446622_RNA45SN1_45S_50ntFlanks@3920.3959.40|CATTTATCAGATCAAAACCAACCCGGTCAGCCCCTCTCCG</td><td>>hg38_21_+_8433172_8446622_RNA45SN1_45S_50ntFlanks</td><td>3920</td><td>3959</td><td>40</td><td>CATTTATCAGATCAAAACCAACCCGGTCAGCCCCTCTCCG</td><td>994.4</td><td>-14.4</td><td>1.7</td><td>-8.2</td><td>1.7E-16</td><td>5.3E-16</td><td>BT20/MB468</td><td>NUC</td></tr>
<tr><td>>hg38_21_+_8433172_8446622_RNA45SN1_45S_50ntFlanks@11209.11233.25|CGGCGACTCTGGACGCGAGCCGGGC</td><td>>hg38_21_+_8433172_8446622_RNA45SN1_45S_50ntFlanks</td><td>11209</td><td>11233</td><td>25</td><td>CGGCGACTCTGGACGCGAGCCGGGC</td><td>1476.6</td><td>-10.1</td><td>1.2</td><td>-8.4</td><td>6.1E-17</td><td>5.3E-16</td><td>MB231/MB468</td><td>MP</td></tr>
<tr><td>>hg38_21_+_8433172_8446622_RNA45SN1_45S_50ntFlanks@10420.10457.38|CTGAGAGATGGGCGAGCGCCGTTCCGAAGGGACGGGCG</td><td>>hg38_21_+_8433172_8446622_RNA45SN1_45S_50ntFlanks</td><td>10420</td><td>10457</td><td>38</td><td>CTGAGAGATGGGCGAGCGCCGTTCCGAAGGGACGGGCG</td><td>799.4</td><td>-12.6</td><td>1.5</td><td>-8.3</td><td>7.6E-17</td><td>5.4E-16</td><td>MB231/MB468</td><td>NUC</td></tr>
<tr><td>>hg38_21_+_8433172_8446622_RNA45SN1_45S_50ntFlanks@10348.10371.24|AAACGAGAACTTTGAAGGCCGAAG</td><td>>hg38_21_+_8433172_8446622_RNA45SN1_45S_50ntFlanks</td><td>10348</td><td>10371</td><td>24</td><td>AAACGAGAACTTTGAAGGCCGAAG</td><td>779.6</td><td>-3.5</td><td>0.4</td><td>-8.2</td><td>1.7E-16</td><td>5.4E-16</td><td>BT20/MB468</td><td>NUC</td></tr>
</table>

| | | | | | | | | | | | | | |
|---|---|---|---|---|---|---|---|---|---|---|---|---|---|
| >hg38_21_+_8433172_8446622_RNA45SN1_45S_50ntFlanks@4995.5024.30\|ACAGATTGATAGCTCTTTCTCGATTCCGTG | >hg38_21_+_8433172_8446622_RNA45SN1_45S_50ntFlanks | 4995 | 5024 | 30 | ACAGATTGATAGCTCTTTCTCGATTCCGTG | 2424.9 | -15.1 | 1.8 | -8.4 | 6.2E-17 | 5.4E-16 | MB231/MB468 | MP |
| >hg38_21_+_8433172_8446622_RNA45SN1_45S_50ntFlanks@11208.11253.46\|GCGGCGACTCTGGACGCGAGCCGGGCCCTTCCCGTGGATCGCCCCA | >hg38_21_+_8433172_8446622_RNA45SN1_45S_50ntFlanks | 11208 | 11253 | 46 | GCGGCGACTCTGGACGCGAGCCGGGCCCTTCCCGTGGATCGCCCCA | 820.8 | -14.1 | 1.7 | -8.2 | 1.7E-16 | 5.4E-16 | BT20/MB468 | NUC |
| >hg38_21_+_8433172_8446622_RNA45SN1_45S_50ntFlanks@10914.10936.23\|CGCCGCGGCTGGACGAGGCGCCG | >hg38_21_+_8433172_8446622_RNA45SN1_45S_50ntFlanks | 10914 | 10936 | 23 | CGCCGCGGCTGGACGAGGCGCCG | 630.2 | -12.0 | 1.4 | -8.4 | 6.3E-17 | 5.4E-16 | MB231/MB468 | CYTO |
| >hg38_21_+_8433172_8446622_RNA45SN1_45S_50ntFlanks@8168.8186.19\|CCCAAGTCCTTCTGATCGA | >hg38_21_+_8433172_8446622_RNA45SN1_45S_50ntFlanks | 8168 | 8186 | 19 | CCCAAGTCCTTCTGATCGA | 1552.1 | -6.0 | 0.7 | -8.3 | 7.7E-17 | 5.4E-16 | MB231/MB468 | NUC |
| >hg38_21_+_8433172_8446622_RNA45SN1_45S_50ntFlanks@11759.11777.19\|CCTCGTCATCTAATTAGTG | >hg38_21_+_8433172_8446622_RNA45SN1_45S_50ntFlanks | 11759 | 11777 | 19 | CCTCGTCATCTAATTAGTG | 369.7 | -12.3 | 1.5 | -8.3 | 1.0E-16 | 5.4E-16 | MB231/MB468 | MITO |
| >MI0000065\|hsa-let-7d&WithFlank&9\|+\|94178828\|94178926@68.89.22&[MIMAT0004484&hsa-let-7d-3p&offsets\|0\|0;m-98&9\|+\|94178895\|94178916&offsets\|0\|0]\|CTATACGACCTGCTGCCTTTCT | >MI0000065\|hsa-let-7d&WithFlank&9\|+\|94178828\|94178926 | 68 | 89 | 22 | CTATACGACCTGCTGCCTTTCT | 1996.1 | 6.3 | 0.8 | 8.4 | 6.3E-17 | 5.4E-16 | MB231/MB468 | CYTO |
| >trna10_AspGTC_12_-_125424193_125424264@32.70.39,trna12_AspGTC_12_-_125411891_125411962@32.70.39,trna144_AspGTC_6_-_27551236_27551307@32.70.39,trna38_AspGTC_17_-_8125556_8125627@32.70.39,trna45_AspGTC_6_+_27447453_27447524@32.70.39,trna48_AspGTC_6_+_27471523_27471594@32.70.39,trna4_AspGTC_12_+_96429799_96429870@32.70.39,trna69_AspGTC_1_-_161440205_161440276@32.70.39,trna72_AspGTC_1_-_161432824_161432895@32.70.39,trna75_AspGTC_1_-_161425414_161425485@32.70.39,trna78_AspGTC_1_-_161418033_161418104@32.70.39,trna81_AspGTC_1_-_161410615_161410686@32.70.39\|CTGTCACGCGGGAGACCGGGGTTCGATTCCCCGACGGGG | >trna10_AspGTC_12_-_125424193_125424264 | 32 | 70 | 39 | CTGTCACGCGGGAGACCGGGGTTCGATTCCCCGACGGGG | 416.0 | -12.6 | 1.5 | -8.4 | 6.2E-17 | 5.5E-16 | MB231/MB468 | MP |
| >hg38_21_+_8433172_8446622_RNA45SN1_45S_50ntFlanks@8104.8136.33\|CCGCGGCGGGGCGCGGGACATGTGGCGTACGGA | >hg38_21_+_8433172_8446622_RNA45SN1_45S_50ntFlanks | 8104 | 8136 | 33 | CCGCGGCGGGGCGCGGGACATGTGGCGTACGGA | 1920.1 | -14.8 | 1.8 | -8.4 | 6.3E-17 | 5.5E-16 | MB231/MB468 | MP |
| >hg38_21_+_8433172_8446622_RNA45SN1_45S_50ntFlanks@11161.11189.29\|CACGGTCCCCCGCGAGGGGGGCCCGGGCA | >hg38_21_+_8433172_8446622_RNA45SN1_45S_50ntFlanks | 11161 | 11189 | 29 | CACGGTCCCCCGCGAGGGGGGCCCGGGCA | 605.2 | -13.0 | 1.6 | -8.3 | 1.0E-16 | 5.5E-16 | MB231/MB468 | MITO |
| >hg38_21_+_8433172_8446622_RNA45SN1_45S_50ntFlanks@10843.10868.26\|TAACTTCGGGATAAGGATTGGCTCTA | >hg38_21_+_8433172_8446622_RNA45SN1_45S_50ntFlanks | 10843 | 10868 | 26 | TAACTTCGGGATAAGGATTGGCTCTA | 405.5 | -12.5 | 1.5 | -8.4 | 6.3E-17 | 5.5E-16 | MB231/MB468 | MP |
| >hg38_21_+_8433172_8446622_RNA45SN1_45S_50ntFlanks@9573.9596.24\|AAATCGGTCGTCCGACCTGGGTAT | >hg38_21_+_8433172_8446622_RNA45SN1_45S_50ntFlanks | 9573 | 9596 | 24 | AAATCGGTCGTCCGACCTGGGTAT | 1695.9 | -6.0 | 0.7 | -8.3 | 1.0E-16 | 5.5E-16 | BT20/MB468 | CYTO |
| >hg38_21_+_8433172_8446622_RNA45SN1_45S_50ntFlanks@10341.10361.21\|AATATTCAAACGAGAACTTTG | >hg38_21_+_8433172_8446622_RNA45SN1_45S_50ntFlanks | 10341 | 10361 | 21 | AATATTCAAACGAGAACTTTG | 249.0 | -11.8 | 1.4 | -8.4 | 6.3E-17 | 5.5E-16 | MB231/MB468 | MP |
| >MI0000085\|hsa-mir-27a&WithFlank&19\|-\|13836434\|13836523@57.74.18(+1A);MI0000440\|hsa-mir-27b&WithFlank&9\|+\|95085439\|95085547@67.84.18(+1A)&[MIMAT0000084&hsa-miR-27a-3p&offsets\|0\|-3(+1A);m-43&19\|-\|13836447\|13836467&offsets\|0\|-3(+1A)];[MIMAT0000419&hsa-miR-27b-3p&offsets\|0\|-3(+1A);m-39&9\|+\|95085505\|95085525&offsets\|0\|-3(+1A)]\|TTCACAGTGGCTAAGTTCA | >MI0000085\|hsa-mir-27a&WithFlank&19\|-\|13836434\|13836523 | 57 | 74 | 18(+1A) | TTCACAGTGGCTAAGTTCA | 395.7 | 12.9 | 1.6 | 8.3 | 1.1E-16 | 5.5E-16 | BT20/MB468 | CYTO |
| >trna116_GluCTC_1_-_145399233_145399304@32.75.44,trna71_GluCTC_1_-_161439189_161439260@32.75.44,trna74_GluCTC_1_-_161431809_161431880@32.75.44,trna77_GluCTC_1_-_161424398_161424469@32.75.44,trna77_GluCTC_6_+_28949976_28950047@32.75.44,trna80_GluCTC_1_-_161417018_161417089@32.75.44,trna87_GluCTC_6_-_126101393_126101464@32.75.44\|CTCTCACCGCCGCGGCCCGGGTTCGATTCCCGGTCAGGGAACCA | >trna116_GluCTC_1_-_145399233_145399304 | 32 | 75 | 44 | CTCTCACCGCCGCGGCCCGGGTTCGATTCCCGGTCAGGGAACCA | 1417.5 | -13.4 | 1.6 | -8.3 | 7.8E-17 | 5.5E-16 | MB231/MB468 | NUC |
| >hg38_21_+_8433172_8446622_RNA45SN1_45S_50ntFlanks@8116.8143.28\|GCGGGACATGTGGCGTACGGAAGACCCG | >hg38_21_+_8433172_8446622_RNA45SN1_45S_50ntFlanks | 8116 | 8143 | 28 | GCGGGACATGTGGCGTACGGAAGACCCG | 449.2 | -12.7 | 1.5 | -8.4 | 6.3E-17 | 5.5E-16 | MB231/MB468 | MP |
| >hg38_21_+_8433172_8446622_RNA45SN1_45S_50ntFlanks@475.497.23\|CTTCGTGATCGATGTGGTGACGT | >hg38_21_+_8433172_8446622_RNA45SN1_45S_50ntFlanks | 475 | 497 | 23 | CTTCGTGATCGATGTGGTGACGT | 55731.0 | 6.6 | 0.8 | 8.3 | 7.8E-17 | 5.5E-16 | MB231/MB468 | NUC |
| >hg38_21_+_8433172_8446622_RNA45SN1_45S_50ntFlanks@4444.4472.29\|CCTTGCCTCTCGGCGCCCCCTCGATGCTC | >hg38_21_+_8433172_8446622_RNA45SN1_45S_50ntFlanks | 4444 | 4472 | 29 | CCTTGCCTCTCGGCGCCCCCTCGATGCTC | 388.6 | -12.4 | 1.5 | -8.3 | 1.0E-16 | 5.5E-16 | MB231/MB468 | MITO |
| >MI0000077\|hsa-mir-21&WithFlank&17\|+\|59841260\|59841343@14.33.20(+1A)&[MIMAT0000076&hsa-miR-21-5p&offsets\|0\|-2(+1A);m-1&17\|+\|59841273\|59841294&offsets\|0\|-2(+1A)]\|TAGCTTATCAGACTGATGTTA | >MI0000077\|hsa-mir-21&WithFlank&17\|+\|59841260\|59841343 | 14 | 33 | 20(+1A) | TAGCTTATCAGACTGATGTTA | 730.8 | 6.3 | 0.8 | 8.3 | 7.6E-17 | 5.5E-16 | BT20/MB468 | MITO |
| >hg38_21_+_8433172_8446622_RNA45SN1_45S_50ntFlanks@5485.5519.35\|AGCGCTGAGAAGACGGTCGAACTTGACTATCTAGA | >hg38_21_+_8433172_8446622_RNA45SN1_45S_50ntFlanks | 5485 | 5519 | 35 | AGCGCTGAGAAGACGGTCGAACTTGACTATCTAGA | 396.6 | -12.5 | 1.5 | -8.4 | 6.4E-17 | 5.5E-16 | MB231/MB468 | MP |
| >trnaMT_GluTTC_MT_-_14674_14742@39.58.20,trnalookalike8_GluTTC_5_-_93905172_93905240@39.58.20\|ATTGGTCGTGGTTGTAGTCC | >trnaMT_GluTTC_MT_-_14674_14742 | 39 | 58 | 20 | ATTGGTCGTGGTTGTAGTCC | 279.7 | 5.6 | 0.7 | 8.3 | 7.9E-17 | 5.5E-16 | MB231/MB468 | NUC |
| >hg38_21_+_8433172_8446622_RNA45SN1_45S_50ntFlanks@11844.11871.28\|GGGAACGGGCTTGGCGGAATCAGCGGGG | >hg38_21_+_8433172_8446622_RNA45SN1_45S_50ntFlanks | 11844 | 11871 | 28 | GGGAACGGGCTTGGCGGAATCAGCGGGG | 331.3 | -12.2 | 1.5 | -8.4 | 6.4E-17 | 5.6E-16 | MB231/MB468 | MP |
| >hg38_21_+_8433172_8446622_RNA45SN1_45S_50ntFlanks@11646.11669.24\|GGTGTTGACGCGATGTGATTTCTG | >hg38_21_+_8433172_8446622_RNA45SN1_45S_50ntFlanks | 11646 | 11669 | 24 | GGTGTTGACGCGATGTGATTTCTG | 455.3 | -12.7 | 1.5 | -8.4 | 6.4E-17 | 5.6E-16 | MB231/MB468 | MP |
| >hg38_21_+_8433172_8446622_RNA45SN1_45S_50ntFlanks@4763.4782.20\|GATACCGTCGTAGTTCCGAC | >hg38_21_+_8433172_8446622_RNA45SN1_45S_50ntFlanks | 4763 | 4782 | 20 | GATACCGTCGTAGTTCCGAC | 271.6 | -11.9 | 1.4 | -8.4 | 6.4E-17 | 5.6E-16 | MB231/MB468 | MP |
| >hg38_21_+_8433172_8446622_RNA45SN1_45S_50ntFlanks@10866.10883.18\|CTAAGGGCTGGGTCGGTC | >hg38_21_+_8433172_8446622_RNA45SN1_45S_50ntFlanks | 10866 | 10883 | 18 | CTAAGGGCTGGGTCGGTC | 296.2 | -12.1 | 1.4 | -8.4 | 6.4E-17 | 5.6E-16 | MB231/MB468 | MP |
| >hg38_21_+_8433172_8446622_RNA45SN1_45S_50ntFlanks@13002.13030.29\|CGATCTATTGAAAGTCAGCCCTCGACACA | >hg38_21_+_8433172_8446622_RNA45SN1_45S_50ntFlanks | 13002 | 13030 | 29 | CGATCTATTGAAAGTCAGCCCTCGACACA | 359.1 | -12.3 | 1.5 | -8.3 | 1.0E-16 | 5.6E-16 | MB231/MB468 | MITO |
| >hg38_21_+_8433172_8446622_RNA45SN1_45S_50ntFlanks@5491.5532.42\|GAGAAGACGGTCGAACTTGACTATCTAGAGGAAGTAAAAGTC | >hg38_21_+_8433172_8446622_RNA45SN1_45S_50ntFlanks | 5491 | 5532 | 42 | GAGAAGACGGTCGAACTTGACTATCTAGAGGAAGTAAAAGTC | 897.5 | 10.7 | 1.3 | 8.4 | 6.5E-17 | 5.6E-16 | MB231/MB468 | CYTO |
| >hg38_21_+_8433172_8446622_RNA45SN1_45S_50ntFlanks@4160.4184.25\|CCACATCCAAGGAAGGCAGCAGGCG | >hg38_21_+_8433172_8446622_RNA45SN1_45S_50ntFlanks | 4160 | 4184 | 25 | CCACATCCAAGGAAGGCAGCAGGCG | 582.1 | -13.0 | 1.6 | -8.4 | 6.4E-17 | 5.6E-16 | MB231/MB468 | MP |
| >hg38_21_+_8433172_8446622_RNA45SN1_45S_50ntFlanks@10683.10709.27\|CCGGCGGCGTCCGGTGAGCTCTCGCTG | >hg38_21_+_8433172_8446622_RNA45SN1_45S_50ntFlanks | 10683 | 10709 | 27 | CCGGCGGCGTCCGGTGAGCTCTCGCTG | 633.0 | -6.9 | 0.8 | -8.4 | 6.5E-17 | 5.6E-16 | MB231/MB468 | CYTO |
| >MI0000253\|hsa-mir-148a&WithFlank&7\|-\|25949913\|25949992@50.70.21(+1G)&[MIMAT0000243&hsa-miR-148a-3p&offsets\|0\|-1(+1G);m-7&7\|-\|25949922\|25949943&offsets\|0\|-1(+1G)]\|TCAGTGCACTACAGAACTTTGG | >MI0000253\|hsa-mir-148a&WithFlank&7\|-\|25949913\|25949992 | 50 | 70 | 21(+1G) | TCAGTGCACTACAGAACTTTGG | 5959.8 | 6.4 | 0.8 | 8.3 | 1.1E-16 | 5.6E-16 | BT20/MB468 | CYTO |
| >hg38_21_+_8433172_8446622_RNA45SN1_45S_50ntFlanks@10684.10711.28\|CGGCGGCGTCCGGTGAGCTCTCGCTGGC | >hg38_21_+_8433172_8446622_RNA45SN1_45S_50ntFlanks | 10684 | 10711 | 28 | CGGCGGCGTCCGGTGAGCTCTCGCTGGC | 6420.8 | -5.8 | 0.7 | -8.3 | 8.0E-17 | 5.6E-16 | MB231/MB468 | NUC |
| >hg38_21_+_8433172_8446622_RNA45SN1_45S_50ntFlanks@9570.9597.28\|TGCAAATCGGTCGTCCGACCTGGGTATA | >hg38_21_+_8433172_8446622_RNA45SN1_45S_50ntFlanks | 9570 | 9597 | 28 | TGCAAATCGGTCGTCCGACCTGGGTATA | 1886.6 | -7.6 | 0.9 | -8.4 | 6.6E-17 | 5.6E-16 | MB231/MB468 | CYTO |
| >hg38_21_+_8433172_8446622_RNA45SN1_45S_50ntFlanks@5443.5466.24\|CCTCGGATCGGCCCCGCCGGGGTC | >hg38_21_+_8433172_8446622_RNA45SN1_45S_50ntFlanks | 5443 | 5466 | 24 | CCTCGGATCGGCCCCGCCGGGGTC | 368.2 | -12.4 | 1.5 | -8.4 | 6.5E-17 | 5.6E-16 | MB231/MB468 | MP |
| >trnaMT_PheGAA_MT_+_577_647@35.57.23\|GAAAATGTTTAGACGGGCTCACA | >trnaMT_PheGAA_MT_+_577_647 | 35 | 57 | 23 | GAAAATGTTTAGACGGGCTCACA | 841.9 | 11.7 | 1.4 | 8.3 | 1.1E-16 | 5.6E-16 | BT20/MB468 | CYTO |
| >hg38_21_+_8433172_8446622_RNA45SN1_45S_50ntFlanks@5469.5514.46\|CCCACGGCCCTGGCGGAGCGCTGAGAAGACGGTCGAACTTGACTAT | >hg38_21_+_8433172_8446622_RNA45SN1_45S_50ntFlanks | 5469 | 5514 | 46 | CCCACGGCCCTGGCGGAGCGCTGAGAAGACGGTCGAACTTGACTAT | 463.4 | -13.3 | 1.6 | -8.2 | 1.8E-16 | 5.7E-16 | BT20/MB468 | NUC |
| >hg38_21_+_8433172_8446622_RNA45SN1_45S_50ntFlanks@11208.11238.31\|GCGGCGACTCTGGACGCGAGCCGGGCCCTTC | >hg38_21_+_8433172_8446622_RNA45SN1_45S_50ntFlanks | 11208 | 11238 | 31 | GCGGCGACTCTGGACGCGAGCCGGGCCCTTC | 424.5 | -6.9 | 0.8 | -8.4 | 6.6E-17 | 5.7E-16 | MB231/MB468 | CYTO |
| >hg38_21_+_8433172_8446622_RNA45SN1_45S_50ntFlanks@12926.12956.31\|ACCTGGCGCTAAACCATTCGTAGACGACCTG | >hg38_21_+_8433172_8446622_RNA45SN1_45S_50ntFlanks | 12926 | 12956 | 31 | ACCTGGCGCTAAACCATTCGTAGACGACCTG | 227.7 | -11.7 | 1.4 | -8.4 | 6.5E-17 | 5.7E-16 | MB231/MB468 | MP |
| >am_trna128_GlyGCC_6_-_27870686_27870756@1.32.32,trna133_GlyCCC_1_-_16872434_16872504@1.32.32,trna18_GlyGCC_16_+_70822597_70822667@1.32.32,trna19_GlyGCC_16_+_70823410_70823480@1.32.32,trna19_GlyGCC_2_-_157257659_157257729@1.32.32,trna24_GlyGCC_16_-_70812942_70813012@1.32.32,trna25_GlyGCC_16_-_70812114_70812184@1.32.32,trna4_GlyCCC_1_+_17188416_17188486@1.32.32,trna5_GlyGCC_17_+_8029064_8029134@1.32.32,trna68_GlyGCC_1_-_161493637_161493707@1.32.32\|GCATTGGTGGTTCAGTGGTAGAATTCTCGCCT | >am_trna128_GlyGCC_6_-_27870686_27870756 | 1 | 32 | 32 | GCATTGGTGGTTCAGTGGTAGAATTCTCGCCT | 44378.4 | 8.4 | 1.0 | 8.4 | 6.7E-17 | 5.7E-16 | MB231/MB468 | CYTO |
| >hg38_21_+_8433172_8446622_RNA45SN1_45S_50ntFlanks@7988.8008.21\|CAGACGTGGCGACCCGCTGAA | >hg38_21_+_8433172_8446622_RNA45SN1_45S_50ntFlanks | 7988 | 8008 | 21 | CAGACGTGGCGACCCGCTGAA | 17781.3 | -6.9 | 0.8 | -8.3 | 1.1E-16 | 5.7E-16 | MB231/MB468 | MITO |
| >MI0000542\|hsa-mir-320a&WithFlank&8\|-\|22244960\|22245043@48.70.23&[MIMAT0000510&hsa-miR-320a-3p&offsets\|0\|+1;m-60&8\|-\|22244975\|22244996&offsets\|0\|+1]\|AAAAGCTGGGTTGAGAGGGCGAA | >MI0000542\|hsa-mir-320a&WithFlank&8\|-\|22244960\|22245043 | 48 | 70 | 23 | AAAAGCTGGGTTGAGAGGGCGAA | 2255.6 | 7.0 | 0.8 | 8.4 | 6.7E-17 | 5.7E-16 | MB231/MB468 | CYTO |
| >hg38_21_+_8433172_8446622_RNA45SN1_45S_50ntFlanks@11913.11951.39\|AAGAGACATGAGAGGTGTAGAATAAGTGGGAGGCCCCCG | >hg38_21_+_8433172_8446622_RNA45SN1_45S_50ntFlanks | 11913 | 11951 | 39 | AAGAGACATGAGAGGTGTAGAATAAGTGGGAGGCCCCCG | 207.6 | -6.4 | 0.8 | -8.4 | 6.8E-17 | 5.7E-16 | MB231/MB468 | CYTO |
| >hg38_21_+_8433172_8446622_RNA45SN1_45S_50ntFlanks@9736.9755.20\|CCGAAACGATCTCAACCTAT | >hg38_21_+_8433172_8446622_RNA45SN1_45S_50ntFlanks | 9736 | 9755 | 20 | CCGAAACGATCTCAACCTAT | 608.6 | -13.0 | 1.6 | -8.3 | 1.1E-16 | 5.8E-16 | MB231/MB468 | MITO |
| >hg38_21_+_8433172_8446622_RNA45SN1_45S_50ntFlanks@6647.6673.27\|CGTACGACTCTTAGCGGTGGATCACTC | >hg38_21_+_8433172_8446622_RNA45SN1_45S_50ntFlanks | 6647 | 6673 | 27 | CGTACGACTCTTAGCGGTGGATCACTC | 1180.8 | 5.4 | 0.7 | 8.3 | 7.9E-17 | 5.8E-16 | BT20/MB468 | MITO |

<table>
<tr><th></th><th></th><th></th><th></th><th></th><th></th><th></th><th></th><th></th><th></th><th></th><th></th><th></th><th></th></tr>
<tr><td>>hg38_21_+_8433172_8446622_RNA45SN1_45S_50ntFlanks@8106.8140.35|GCGGCGGGGCGCGGGACATGTGGCGTACGGAAGAC</td><td>>hg38_21_+_8433172_8446622_RNA45SN1_45S_50ntFlanks</td><td>8106</td><td>8140</td><td>35</td><td>GCGGCGGGGCGCGGGACATGTGGCGTACGGAAGAC</td><td>478.3</td><td>-12.7</td><td>1.5</td><td>-8.3</td><td>1.1E-16</td><td>5.8E-16</td><td>MB231/MB468</td><td>MITO</td></tr>
<tr><td>>hg38_21_+_8433172_8446622_RNA45SN1_45S_50ntFlanks@12588.12614.27|GGTTCAGACATTTGGTGTATGTGCTTG</td><td>>hg38_21_+_8433172_8446622_RNA45SN1_45S_50ntFlanks</td><td>12588</td><td>12614</td><td>27</td><td>GGTTCAGACATTTGGTGTATGTGCTTG</td><td>547.6</td><td>-13.0</td><td>1.6</td><td>-8.4</td><td>6.7E-17</td><td>5.8E-16</td><td>MB231/MB468</td><td>MP</td></tr>
<tr><td>>hg38_21_+_8433172_8446622_RNA45SN1_45S_50ntFlanks@11918.11949.32|ACATGAGAGGTGTAGAATAAGTGGGAGGCCCC</td><td>>hg38_21_+_8433172_8446622_RNA45SN1_45S_50ntFlanks</td><td>11918</td><td>11949</td><td>32</td><td>ACATGAGAGGTGTAGAATAAGTGGGAGGCCCC</td><td>582.7</td><td>-7.8</td><td>0.9</td><td>-8.4</td><td>6.8E-17</td><td>5.8E-16</td><td>MB231/MB468</td><td>CYTO</td></tr>
<tr><td>>hg38_21_+_8433172_8446622_RNA45SN1_45S_50ntFlanks@11921.11969.49|TGAGAGGTGTAGAATAAGTGGGAGGCCCCCGGCGCCCCCCCGGTGTCCC</td><td>>hg38_21_+_8433172_8446622_RNA45SN1_45S_50ntFlanks</td><td>11921</td><td>11969</td><td>49</td><td>TGAGAGGTGTAGAATAAGTGGGAGGCCCCCGGCGCCCCCCCGGTGTCCC</td><td>499.1</td><td>-13.4</td><td>1.6</td><td>-8.2</td><td>1.8E-16</td><td>5.8E-16</td><td>BT20/MB468</td><td>NUC</td></tr>
<tr><td>>hg38_21_+_8433172_8446622_RNA45SN1_45S_50ntFlanks@12942.12966.25|TTCGTAGACGACCTGCTTCTGGGTC</td><td>>hg38_21_+_8433172_8446622_RNA45SN1_45S_50ntFlanks</td><td>12942</td><td>12966</td><td>25</td><td>TTCGTAGACGACCTGCTTCTGGGTC</td><td>582.1</td><td>-13.0</td><td>1.6</td><td>-8.4</td><td>6.7E-17</td><td>5.8E-16</td><td>MB231/MB468</td><td>MP</td></tr>
<tr><td>>MI0000473|hsa-mir-129-2&WithFlank&11|+|43581388|43581489@21.41.21;MI0000252|hsa-mir-129-1&WithFlank&7|+|128207866|128207949@11.31.21&[MIMAT0000242&hsa-miR-129-5p&offsets|0|0;m-434&11|+|43581408|43581428&offsets|0|0];[MIMAT0000242_1&hsa-miR-129-5p&offsets|0|0;m-434&7|+|128207876|128207896&offsets|0|0]|CTTTTTGCGGTCTGGGCTTGC</td><td>>MI0000473|hsa-mir-129-2&WithFlank&11|+|43581388|43581489</td><td>21</td><td>41</td><td>21</td><td>CTTTTTGCGGTCTGGGCTTGC</td><td>273.6</td><td>8.6</td><td>1.0</td><td>8.3</td><td>1.1E-16</td><td>5.9E-16</td><td>BT20/MB468</td><td>CYTO</td></tr>
<tr><td>>hg38_21_+_8433172_8446622_RNA45SN1_45S_50ntFlanks@9805.9830.26|GCGTGGAATGCGAGTGCCTAGTGGGC</td><td>>hg38_21_+_8433172_8446622_RNA45SN1_45S_50ntFlanks</td><td>9805</td><td>9830</td><td>26</td><td>GCGTGGAATGCGAGTGCCTAGTGGGC</td><td>401.7</td><td>-12.5</td><td>1.5</td><td>-8.4</td><td>6.8E-17</td><td>5.9E-16</td><td>MB231/MB468</td><td>MP</td></tr>
<tr><td>>hg38_21_+_8433172_8446622_RNA45SN1_45S_50ntFlanks@12152.12179.28|TGGGGAGTTTGACTGGGGCGGTACACCT</td><td>>hg38_21_+_8433172_8446622_RNA45SN1_45S_50ntFlanks</td><td>12152</td><td>12179</td><td>28</td><td>TGGGGAGTTTGACTGGGGCGGTACACCT</td><td>239.1</td><td>-11.8</td><td>1.4</td><td>-8.4</td><td>6.8E-17</td><td>5.9E-16</td><td>MB231/MB468</td><td>MP</td></tr>
<tr><td>>MI0000811|hsa-mir-148b&WithFlank&12|+|54337210|54337320@69.90.22&[MIMAT0000759&hsa-miR-148b-3p&offsets|0|0;m-77&12|+|54337278|54337299&offsets|0|0]|TCAGTGCATCACAGAACTTTGT</td><td>>MI0000811|hsa-mir-148b&WithFlank&12|+|54337210|54337320</td><td>69</td><td>90</td><td>22</td><td>TCAGTGCATCACAGAACTTTGT</td><td>4357.0</td><td>5.6</td><td>0.7</td><td>8.3</td><td>8.4E-17</td><td>5.9E-16</td><td>MB231/MB468</td><td>NUC</td></tr>
<tr><td>>hg38_21_+_8433172_8446622_RNA45SN1_45S_50ntFlanks@11768.11794.27|CTAATTAGTGACGCGCATGAATGGATG</td><td>>hg38_21_+_8433172_8446622_RNA45SN1_45S_50ntFlanks</td><td>11768</td><td>11794</td><td>27</td><td>CTAATTAGTGACGCGCATGAATGGATG</td><td>2499.3</td><td>-3.9</td><td>0.5</td><td>-8.3</td><td>1.1E-16</td><td>5.9E-16</td><td>BT20/MB468</td><td>CYTO</td></tr>
<tr><td>>hg38_21_+_8433172_8446622_RNA45SN1_45S_50ntFlanks@12290.12323.34|CGTGAAAGCGGGGCCTCACGATCCTTCTGACCTT</td><td>>hg38_21_+_8433172_8446622_RNA45SN1_45S_50ntFlanks</td><td>12290</td><td>12323</td><td>34</td><td>CGTGAAAGCGGGGCCTCACGATCCTTCTGACCTT</td><td>699.5</td><td>-13.2</td><td>1.6</td><td>-8.3</td><td>1.1E-16</td><td>5.9E-16</td><td>MB231/MB468</td><td>MITO</td></tr>
<tr><td>>hg38_21_+_8433172_8446622_RNA45SN1_45S_50ntFlanks@11576.11603.28|CTTAGAACTGGTGCGGACCAGGGGAATC</td><td>>hg38_21_+_8433172_8446622_RNA45SN1_45S_50ntFlanks</td><td>11576</td><td>11603</td><td>28</td><td>CTTAGAACTGGTGCGGACCAGGGGAATC</td><td>5366.8</td><td>-5.4</td><td>0.7</td><td>-8.3</td><td>1.1E-16</td><td>5.9E-16</td><td>BT20/MB468</td><td>CYTO</td></tr>
<tr><td>>hg38_21_+_8433172_8446622_RNA45SN1_45S_50ntFlanks@5472.5489.18|ACGGCCCTGGCGGAGCGC</td><td>>hg38_21_+_8433172_8446622_RNA45SN1_45S_50ntFlanks</td><td>5472</td><td>5489</td><td>18</td><td>ACGGCCCTGGCGGAGCGC</td><td>2884.0</td><td>10.5</td><td>1.3</td><td>8.3</td><td>7.0E-17</td><td>5.9E-16</td><td>MB231/MB468</td><td>CYTO</td></tr>
<tr><td>>hg38_21_+_8433172_8446622_RNA45SN1_45S_50ntFlanks@12672.12703.32|AGTCAGAATCCCGCCCAGGCGGAACGATACGG</td><td>>hg38_21_+_8433172_8446622_RNA45SN1_45S_50ntFlanks</td><td>12672</td><td>12703</td><td>32</td><td>AGTCAGAATCCCGCCCAGGCGGAACGATACGG</td><td>299.7</td><td>-12.1</td><td>1.4</td><td>-8.3</td><td>6.9E-17</td><td>5.9E-16</td><td>MB231/MB468</td><td>MP</td></tr>
<tr><td>>hg38_21_+_8433172_8446622_RNA45SN1_45S_50ntFlanks@12893.12926.34|CCTCGCCCGTCACGCACCGCACGTTCGTGGGGAA</td><td>>hg38_21_+_8433172_8446622_RNA45SN1_45S_50ntFlanks</td><td>12893</td><td>12926</td><td>34</td><td>CCTCGCCCGTCACGCACCGCACGTTCGTGGGGAA</td><td>366.1</td><td>-12.3</td><td>1.5</td><td>-8.3</td><td>1.1E-16</td><td>5.9E-16</td><td>MB231/MB468</td><td>MITO</td></tr>
<tr><td>>hg38_21_+_8433172_8446622_RNA45SN1_45S_50ntFlanks@11618.11640.23|AAACAAAGCATCGCGAAGGCCCG</td><td>>hg38_21_+_8433172_8446622_RNA45SN1_45S_50ntFlanks</td><td>11618</td><td>11640</td><td>23</td><td>AAACAAAGCATCGCGAAGGCCCG</td><td>1479.8</td><td>-13.6</td><td>1.6</td><td>-8.8</td><td>1.7E-18</td><td>6.0E-16</td><td>BT20/MB468</td><td>MP</td></tr>
<tr><td>>MI0005757|hsa-mir-935&WithFlank&19|+|53982301|53982403@63.84.22&[MIMAT0004978&hsa-miR-935&offsets|+1|0;m-549&19|+|53982363|53982384&offsets|0|0]|CAGTTACCGCTTCCGCTACCGC</td><td>>MI0005757|hsa-mir-935&WithFlank&19|+|53982301|53982403</td><td>63</td><td>84</td><td>22</td><td>CAGTTACCGCTTCCGCTACCGC</td><td>54.2</td><td>8.6</td><td>1.0</td><td>8.2</td><td>1.9E-16</td><td>6.0E-16</td><td>BT20/MB468</td><td>NUC</td></tr>
<tr><td>>hg38_21_+_8433172_8446622_RNA45SN1_45S_50ntFlanks@5073.5113.41|AATTCCGATAACGAACGAGACTCTGGCATGCTAACTAGTTA</td><td>>hg38_21_+_8433172_8446622_RNA45SN1_45S_50ntFlanks</td><td>5073</td><td>5113</td><td>41</td><td>AATTCCGATAACGAACGAGACTCTGGCATGCTAACTAGTTA</td><td>489.7</td><td>-12.7</td><td>1.5</td><td>-8.3</td><td>1.1E-16</td><td>6.0E-16</td><td>MB231/MB468</td><td>MITO</td></tr>
<tr><td>>hg38_21_+_8433172_8446622_RNA45SN1_45S_50ntFlanks@8117.8143.27|CGGGACATGTGGCGTACGGAAGACCCG</td><td>>hg38_21_+_8433172_8446622_RNA45SN1_45S_50ntFlanks</td><td>8117</td><td>8143</td><td>27</td><td>CGGGACATGTGGCGTACGGAAGACCCG</td><td>617.2</td><td>-13.1</td><td>1.6</td><td>-8.3</td><td>6.9E-17</td><td>6.0E-16</td><td>MB231/MB468</td><td>MP</td></tr>
<tr><td>>hg38_21_+_8433172_8446622_RNA45SN1_45S_50ntFlanks@11873.11892.20|AAGAAGACCCTGTTGAGCTT</td><td>>hg38_21_+_8433172_8446622_RNA45SN1_45S_50ntFlanks</td><td>11873</td><td>11892</td><td>20</td><td>AAGAAGACCCTGTTGAGCTT</td><td>476.8</td><td>-3.5</td><td>0.4</td><td>-8.3</td><td>1.2E-16</td><td>6.0E-16</td><td>BT20/MB468</td><td>CYTO</td></tr>
<tr><td>>hg38_21_+_8433172_8446622_RNA45SN1_45S_50ntFlanks@8325.8356.32|CGATAGTCAACAAGTACCGTAAGGGAAAGTTG</td><td>>hg38_21_+_8433172_8446622_RNA45SN1_45S_50ntFlanks</td><td>8325</td><td>8356</td><td>32</td><td>CGATAGTCAACAAGTACCGTAAGGGAAAGTTG</td><td>246.4</td><td>-6.7</td><td>0.8</td><td>-8.3</td><td>7.1E-17</td><td>6.0E-16</td><td>MB231/MB468</td><td>CYTO</td></tr>
<tr><td>>hg38_21_+_8433172_8446622_RNA45SN1_45S_50ntFlanks@9501.9526.26|ATGGTGAACTATGCCTGGGCAGGGCG</td><td>>hg38_21_+_8433172_8446622_RNA45SN1_45S_50ntFlanks</td><td>9501</td><td>9526</td><td>26</td><td>ATGGTGAACTATGCCTGGGCAGGGCG</td><td>632.2</td><td>-13.2</td><td>1.6</td><td>-8.3</td><td>7.0E-17</td><td>6.0E-16</td><td>MB231/MB468</td><td>MP</td></tr>
<tr><td>>hg38_21_+_8433172_8446622_RNA45SN1_45S_50ntFlanks@12420.12461.42|ATGTCGGCTCTTCCTATCATTGTGAAGCAGAATTCACCAAGC</td><td>>hg38_21_+_8433172_8446622_RNA45SN1_45S_50ntFlanks</td><td>12420</td><td>12461</td><td>42</td><td>ATGTCGGCTCTTCCTATCATTGTGAAGCAGAATTCACCAAGC</td><td>253.5</td><td>9.6</td><td>1.2</td><td>8.3</td><td>7.1E-17</td><td>6.0E-16</td><td>MB231/MB468</td><td>CYTO</td></tr>
<tr><td>>MI0000082|hsa-mir-25&WithFlank&7|-|100093554|100093649@58.78.21&[MIMAT0000081&hsa-miR-25-3p&offsets|0|-1;m-23&7|-|100093571|100093592&offsets|0|-1]|CATTGCACTTGTCTCGGTCTG</td><td>>MI0000082|hsa-mir-25&WithFlank&7|-|100093554|100093649</td><td>58</td><td>78</td><td>21</td><td>CATTGCACTTGTCTCGGTCTG</td><td>5926.4</td><td>4.4</td><td>0.5</td><td>8.3</td><td>8.6E-17</td><td>6.0E-16</td><td>MB231/MB468</td><td>NUC</td></tr>
<tr><td>>MI0000298|hsa-mir-221&WithFlank&X|-|45746151|45746272@31.55.25&[MIMAT0004568&hsa-miR-221-5p&offsets|0|+3;m-405&X|-|45746220|45746242&offsets|0|+2]|ACCTGGCATACAATGTAGATTTCTG</td><td>>MI0000298|hsa-mir-221&WithFlank&X|-|45746151|45746272</td><td>31</td><td>55</td><td>25</td><td>ACCTGGCATACAATGTAGATTTCTG</td><td>300.1</td><td>5.6</td><td>0.7</td><td>8.3</td><td>8.6E-17</td><td>6.0E-16</td><td>MB231/MB468</td><td>NUC</td></tr>
<tr><td>>MI0000089|hsa-mir-31&WithFlank&9|-|21512109|21512191@14.35.22&[MIMAT0000089&hsa-miR-31-5p&offsets|0|+1;m-360&9|-|21512157|21512178&offsets|0|0]|AGGCAAGATGCTGGCATAGCTG</td><td>>MI0000089|hsa-mir-31&WithFlank&9|-|21512109|21512191</td><td>14</td><td>35</td><td>22</td><td>AGGCAAGATGCTGGCATAGCTG</td><td>365.8</td><td>12.1</td><td>1.4</td><td>8.6</td><td>9.0E-18</td><td>6.0E-16</td><td>BT20/MB231</td><td>MITO</td></tr>
<tr><td>>hg38_21_+_8433172_8446622_RNA45SN1_45S_50ntFlanks@7977.8009.33|CGACCTCAGATCAGACGTGGCGACCCGCTGAAT</td><td>>hg38_21_+_8433172_8446622_RNA45SN1_45S_50ntFlanks</td><td>7977</td><td>8009</td><td>33</td><td>CGACCTCAGATCAGACGTGGCGACCCGCTGAAT</td><td>742.3</td><td>5.9</td><td>0.7</td><td>8.3</td><td>7.2E-17</td><td>6.0E-16</td><td>MB231/MB468</td><td>CYTO</td></tr>
<tr><td>>hg38_21_+_8433172_8446622_RNA45SN1_45S_50ntFlanks@12583.12604.22|CCGCAGGTTCAGACATTTGGTG</td><td>>hg38_21_+_8433172_8446622_RNA45SN1_45S_50ntFlanks</td><td>12583</td><td>12604</td><td>22</td><td>CCGCAGGTTCAGACATTTGGTG</td><td>227.2</td><td>-11.7</td><td>1.4</td><td>-8.3</td><td>7.0E-17</td><td>6.1E-16</td><td>MB231/MB468</td><td>MP</td></tr>
<tr><td>>hg38_21_+_8433172_8446622_RNA45SN1_45S_50ntFlanks@10806.10823.18|AACAATGTAGGTAAGGGA</td><td>>hg38_21_+_8433172_8446622_RNA45SN1_45S_50ntFlanks</td><td>10806</td><td>10823</td><td>18</td><td>AACAATGTAGGTAAGGGA</td><td>573.1</td><td>-12.1</td><td>1.5</td><td>-8.3</td><td>8.7E-17</td><td>6.1E-16</td><td>MB231/MB468</td><td>NUC</td></tr>
<tr><td>>hg38_21_+_8433172_8446622_RNA45SN1_45S_50ntFlanks@11918.11950.33|ACATGAGAGGTGTAGAATAAGTGGGAGGCCCCC</td><td>>hg38_21_+_8433172_8446622_RNA45SN1_45S_50ntFlanks</td><td>11918</td><td>11950</td><td>33</td><td>ACATGAGAGGTGTAGAATAAGTGGGAGGCCCCC</td><td>535.3</td><td>-12.8</td><td>1.5</td><td>-8.3</td><td>1.1E-16</td><td>6.1E-16</td><td>MB231/MB468</td><td>MITO</td></tr>
<tr><td>>trna111_HisGTG_1_-_147774845_147774916@-1T.35.36,trna118_HisGTG_1_-_145396881_145396952@-1T.35.36,trna16_HisGTG_1_+_146544773_146544844@-1T.35.36,trna1_HisGTG_15_+_45493349_45493420@-1T.35.36,trna21_HisGTG_1_+_147753471_147753542@-1T.35.36,trna33_HisGTG_6_+_27125906_27125977@-1T.35.36,trna7_HisGTG_9_-_14433938_14434009@-1T.35.36,trna8_HisGTG_15_-_45492611_45492682@-1T.35.36,trna9_HisGTG_15_-_45490804_45490875@-1T.35.36|TGCCGTGATCGTATAGTGGTTAGTACTCTGCGTTGT</td><td>>trna111_HisGTG_1_-_147774845_147774916</td><td>-1T</td><td>35</td><td>36</td><td>TGCCGTGATCGTATAGTGGTTAGTACTCTGCGTTGT</td><td>917.5</td><td>13.8</td><td>1.7</td><td>8.3</td><td>7.2E-17</td><td>6.1E-16</td><td>MB231/MB468</td><td>CYTO</td></tr>
<tr><td>>MI0000088|hsa-mir-30a&WithFlank&6|-|71403545|71403627@12.31.20&[MIMAT0000087&hsa-miR-30a-5p&offsets|0|-2;m-15&6|-|71403595|71403616&offsets|0|-2]|TGTAAACATCCTCGACTGGA</td><td>>MI0000088|hsa-mir-30a&WithFlank&6|-|71403545|71403627</td><td>12</td><td>31</td><td>20</td><td>TGTAAACATCCTCGACTGGA</td><td>969.6</td><td>6.8</td><td>0.8</td><td>8.3</td><td>1.2E-16</td><td>6.1E-16</td><td>BT20/MB468</td><td>CYTO</td></tr>
<tr><td>>hg38_21_+_8433172_8446622_RNA45SN1_45S_50ntFlanks@11537.11560.24|CCGCGCGGCGCCTCGCCTCGGCCG</td><td>>hg38_21_+_8433172_8446622_RNA45SN1_45S_50ntFlanks</td><td>11537</td><td>11560</td><td>24</td><td>CCGCGCGGCGCCTCGCCTCGGCCG</td><td>487.8</td><td>-12.7</td><td>1.5</td><td>-8.3</td><td>1.1E-16</td><td>6.2E-16</td><td>MB231/MB468</td><td>MITO</td></tr>
<tr><td>>MI0000067|hsa-let-7f-1&WithFlank&9|+|94176341|94176439@13.33.21(+1A);MI0000068|hsa-let-7f-2&WithFlank&X|-|53557186|53557280@14.34.21(+1A)&[MIMAT0000067&hsa-let-7f-5p&offsets|0|-1(+1A);m-20&9|+|94176353|94176374&offsets|0|-1(+1A)];[MIMAT0000067_1&hsa-let-7f-5p&offsets|0|-1(+1A);m-19&X|-|53557246|53557267&offsets|0|-1(+1A)]|TGAGGTAGTAGATTGTATAGTA</td><td>>MI0000067|hsa-let-7f-1&WithFlank&9|+|94176341|94176439</td><td>13</td><td>33</td><td>21(+1A)</td><td>TGAGGTAGTAGATTGTATAGTA</td><td>7784.0</td><td>7.3</td><td>0.9</td><td>8.3</td><td>1.2E-16</td><td>6.2E-16</td><td>BT20/MB468</td><td>CYTO</td></tr>
<tr><td>>hg38_21_+_8433172_8446622_RNA45SN1_45S_50ntFlanks@9313.9336.24|TCTAACACGTGCGCGAGTCGGGGG</td><td>>hg38_21_+_8433172_8446622_RNA45SN1_45S_50ntFlanks</td><td>9313</td><td>9336</td><td>24</td><td>TCTAACACGTGCGCGAGTCGGGGG</td><td>529.6</td><td>-12.9</td><td>1.5</td><td>-8.3</td><td>7.1E-17</td><td>6.2E-16</td><td>MB231/MB468</td><td>MP</td></tr>
<tr><td>>hg38_21_+_8433172_8446622_RNA45SN1_45S_50ntFlanks@10410.10443.34|TCAGTCGGTCCTGAGAGATGGGCGAGCGCCGTTC</td><td>>hg38_21_+_8433172_8446622_RNA45SN1_45S_50ntFlanks</td><td>10410</td><td>10443</td><td>34</td><td>TCAGTCGGTCCTGAGAGATGGGCGAGCGCCGTTC</td><td>1211.8</td><td>-8.6</td><td>1.0</td><td>-8.3</td><td>7.2E-17</td><td>6.2E-16</td><td>MB231/MB468</td><td>MP</td></tr>
<tr><td>>MI0000439|hsa-mir-23b&WithFlank&9|+|95085202|95085310@67.86.20(+1U)&[MIMAT0000418&hsa-miR-23b-3p&offsets|+3|0(+1U);m-42&9|+|95085265|95085285&offsets|+3|+2(+1U)]|ACATTGCCAGGGATTACCACT</td><td>>MI0000439|hsa-mir-23b&WithFlank&9|+|95085202|95085310</td><td>67</td><td>86</td><td>20(+1U)</td><td>ACATTGCCAGGGATTACCACT</td><td>1321.5</td><td>14.1</td><td>1.6</td><td>8.8</td><td>1.8E-18</td><td>6.2E-16</td><td>BT20/MB468</td><td>MP</td></tr>
<tr><td>>MI0000081|hsa-mir-24-2&WithFlank&19|-|13836281|13836365@56.77.22;MI0000080|hsa-mir-24-1&WithFlank&9|+|95086015|95086094@50.71.22&[MIMAT0000080&hsa-miR-24-3p&offsets|0|0;m-36&19|-|13836288|13836310&offsets|0|-1];[MIMAT0000080_1&hsa-miR-24-3p&offsets|0|0;m-35&9|+|95086064|95086086&offsets|0|-1]|TGGCTCAGTTCAGCAGGAACAG</td><td>>MI0000081|hsa-mir-24-2&WithFlank&19|-|13836281|13836365</td><td>56</td><td>77</td><td>22</td><td>TGGCTCAGTTCAGCAGGAACAG</td><td>10900.6</td><td>5.3</td><td>0.6</td><td>8.3</td><td>1.2E-16</td><td>6.2E-16</td><td>BT20/MB468</td><td>CYTO</td></tr>
<tr><td>>hg38_21_+_8433172_8446622_RNA45SN1_45S_50ntFlanks@12632.12653.22|GCGAAGCTACCATCTGTGGGAT</td><td>>hg38_21_+_8433172_8446622_RNA45SN1_45S_50ntFlanks</td><td>12632</td><td>12653</td><td>22</td><td>GCGAAGCTACCATCTGTGGGAT</td><td>1085.8</td><td>-5.4</td><td>0.7</td><td>-8.3</td><td>8.6E-17</td><td>6.2E-16</td><td>BT20/MB468</td><td>MITO</td></tr>
<tr><td>>hg38_21_+_8433172_8446622_RNA45SN1_45S_50ntFlanks@10408.10429.22|GGTCAGTCGGTCCTGAGAGATG</td><td>>hg38_21_+_8433172_8446622_RNA45SN1_45S_50ntFlanks</td><td>10408</td><td>10429</td><td>22</td><td>GGTCAGTCGGTCCTGAGAGATG</td><td>3347.9</td><td>-5.7</td><td>0.7</td><td>-8.3</td><td>7.4E-17</td><td>6.2E-16</td><td>MB231/MB468</td><td>CYTO</td></tr>
<tr><td>>hg38_21_+_8433172_8446622_RNA45SN1_45S_50ntFlanks@11576.11601.26|CTTAGAACTGGTGCGGACCAGGGGAA</td><td>>hg38_21_+_8433172_8446622_RNA45SN1_45S_50ntFlanks</td><td>11576</td><td>11601</td><td>26</td><td>CTTAGAACTGGTGCGGACCAGGGGAA</td><td>1761.2</td><td>-8.3</td><td>1.0</td><td>-8.3</td><td>7.4E-17</td><td>6.2E-16</td><td>MB231/MB468</td><td>CYTO</td></tr>
<tr><td>>hg38_21_+_8433172_8446622_RNA45SN1_45S_50ntFlanks@8079.8104.26|AAGAGCCCAGCGCCGAATCCCCGCCC</td><td>>hg38_21_+_8433172_8446622_RNA45SN1_45S_50ntFlanks</td><td>8079</td><td>8104</td><td>26</td><td>AAGAGCCCAGCGCCGAATCCCCGCCC</td><td>4216.8</td><td>-11.8</td><td>1.4</td><td>-8.3</td><td>7.2E-17</td><td>6.2E-16</td><td>MB231/MB468</td><td>MP</td></tr>
<tr><td>>hg38_21_+_8433172_8446622_RNA45SN1_45S_50ntFlanks@12456.12491.36|CCAAGCGTTGGATTGTTCACCCACTAATAGGGAACG</td><td>>hg38_21_+_8433172_8446622_RNA45SN1_45S_50ntFlanks</td><td>12456</td><td>12491</td><td>36</td><td>CCAAGCGTTGGATTGTTCACCCACTAATAGGGAACG</td><td>587.1</td><td>-13.6</td><td>1.7</td><td>-8.2</td><td>1.9E-16</td><td>6.2E-16</td><td>BT20/MB468</td><td>NUC</td></tr>
<tr><td>>MI0000803|hsa-mir-330&WithFlank&19|-|45638988|45639093@64.87.24&[MIMAT0000751&hsa-miR-330-3p&offsets|+1|+2;m-347&19|-|45639009|45639030&offsets|0|+2]|CAAAGCACACGGCCTGCAGAGAGG</td><td>>MI0000803|hsa-mir-330&WithFlank&19|-|45638988|45639093</td><td>64</td><td>87</td><td>24</td><td>CAAAGCACACGGCCTGCAGAGAGG</td><td>313.4</td><td>8.6</td><td>1.0</td><td>8.3</td><td>1.2E-16</td><td>6.2E-16</td><td>BT20/MB468</td><td>CYTO</td></tr>
</table>

| | | | | | | | | | | | | | |
|---|---|---|---|---|---|---|---|---|---|---|---|---|---|
| >MI0000078\|hsa-mir-22&WithFlank&17\|-\|1713897\|1713993@59.78.20&[MIMAT0000077&hsa-miR-22-3p&offsets\|0\|-2;m-2&17\|-\|1713914\|1713935&offsets\|0\|-2]\|AAGCTGCCAGTTGAAGAACT | >MI0000078\|hsa-mir-22&WithFlank&17\|-\|1713897\|1713993 | 59 | 78 | 20 | AAGCTGCCAGTTGAAGAACT | 1502.5 | 10.5 | 1.3 | 8.3 | 1.2E-16 | 6.3E-16 | BT20/MB468 | CYTO |
| >hg38_21_+_8433172_8446622_RNA45SN1_45S_50ntFlanks@12157.12181.25\|AGTTTGACTGGGGCGGTACACCTGT | >hg38_21_+_8433172_8446622_RNA45SN1_45S_50ntFlanks | 12157 | 12181 | 25 | AGTTTGACTGGGGCGGTACACCTGT | 826.2 | -5.2 | 0.6 | -8.3 | 8.7E-17 | 6.3E-16 | BT20/MB468 | MITO |
| >hg38_21_+_8433172_8446622_RNA45SN1_45S_50ntFlanks@11729.11765.37\|TAACTATGACTCTCTTAAGGTAGCCAAATGCCTCGTC | >hg38_21_+_8433172_8446622_RNA45SN1_45S_50ntFlanks | 11729 | 11765 | 37 | TAACTATGACTCTCTTAAGGTAGCCAAATGCCTCGTC | 10226.0 | -6.5 | 0.8 | -8.3 | 7.5E-17 | 6.3E-16 | MB231/MB468 | CYTO |
| >MI0000102\|hsa-mir-100&WithFlank&11\|-\|122152223\|122152314@19.37.19&[MIMAT0000098&hsa-miR-100-5p&offsets\|0\|-3;m-27&11\|-\|122152276\|122152296&offsets\|0\|-2]\|AACCCGTAGATCCGAACTT | >MI0000102\|hsa-mir-100&WithFlank&11\|-\|122152223\|122152314 | 19 | 37 | 19 | AACCCGTAGATCCGAACTT | 346.1 | 12.4 | 1.5 | 8.3 | 7.5E-17 | 6.3E-16 | MB231/MB468 | CYTO |
| >hg38_21_+_8433172_8446622_RNA45SN1_45S_50ntFlanks@3984.4005.22\|GCGGCTTTGGTGACTCTAGATA | >hg38_21_+_8433172_8446622_RNA45SN1_45S_50ntFlanks | 3984 | 4005 | 22 | GCGGCTTTGGTGACTCTAGATA | 755.8 | -13.3 | 1.6 | -8.3 | 1.2E-16 | 6.3E-16 | MB231/MB468 | MITO |
| >hg38_21_+_8433172_8446622_RNA45SN1_45S_50ntFlanks@4047.4067.21\|ATTCGAACGTCTGCCCTATCA | >hg38_21_+_8433172_8446622_RNA45SN1_45S_50ntFlanks | 4047 | 4067 | 21 | ATTCGAACGTCTGCCCTATCA | 211.9 | -11.6 | 1.4 | -8.3 | 7.4E-17 | 6.4E-16 | MB231/MB468 | MP |
| >hg38_21_+_8433172_8446622_RNA45SN1_45S_50ntFlanks@8155.8175.21\|CGCTCGTGGGGGGCCCAAGTC | >hg38_21_+_8433172_8446622_RNA45SN1_45S_50ntFlanks | 8155 | 8175 | 21 | CGCTCGTGGGGGGCCCAAGTC | 649.6 | -13.1 | 1.6 | -8.3 | 1.2E-16 | 6.4E-16 | MB231/MB468 | MITO |
| >hg38_21_+_8433172_8446622_RNA45SN1_45S_50ntFlanks@9203.9224.22\|CTCGGGGCCACGCGCGCGTCCC | >hg38_21_+_8433172_8446622_RNA45SN1_45S_50ntFlanks | 9203 | 9224 | 22 | CTCGGGGCCACGCGCGCGTCCC | 441.8 | -12.6 | 1.5 | -8.3 | 1.2E-16 | 6.4E-16 | MB231/MB468 | MITO |
| >hg38_21_+_8433172_8446622_RNA45SN1_45S_50ntFlanks@6650.6668.19\|ACGACTCTTAGCGGTGGAT | >hg38_21_+_8433172_8446622_RNA45SN1_45S_50ntFlanks | 6650 | 6668 | 19 | ACGACTCTTAGCGGTGGAT | 1204.7 | 4.5 | 0.5 | 8.3 | 1.2E-16 | 6.4E-16 | BT20/MB468 | CYTO |
| >MI0000750\|hsa-mir-26a-2&WithFlank&12\|-\|57824603\|57824698@20.40.21;MI0000083\|hsa-mir-26a-1&WithFlank&3\|+\|37969398\|37969486@16.36.21&[MIMAT0000082&hsa-miR-26a-5p&offsets\|0\|-1;m-33&12\|-\|57824658\|57824679&offsets\|0\|-1];[MIMAT0000082_1&hsa-miR-26a-5p&offsets\|0\|-1;m-34&3\|+\|37969413\|37969434&offsets\|0\|-1]\|TTCAAGTAATCCAGGATAGGC | >MI0000750\|hsa-mir-26a-2&WithFlank&12\|-\|57824603\|57824698 | 20 | 40 | 21 | TTCAAGTAATCCAGGATAGGC | 1533.5 | 5.4 | 0.6 | 8.3 | 9.2E-17 | 6.4E-16 | MB231/MB468 | NUC |
| >MI0000811\|hsa-mir-148b&WithFlank&12\|+\|54337210\|54337320@69.91.23&[MIMAT0000759&hsa-miR-148b-3p&offsets\|0\|+1;m-77&12\|+\|54337278\|54337299&offsets\|0\|+1]\|TCAGTGCATCACAGAACTTTGTC | >MI0000811\|hsa-mir-148b&WithFlank&12\|+\|54337210\|54337320 | 69 | 91 | 23 | TCAGTGCATCACAGAACTTTGTC | 356.4 | 5.6 | 0.7 | 8.3 | 9.2E-17 | 6.4E-16 | MB231/MB468 | NUC |
| >hg38_21_+_8433172_8446622_RNA45SN1_45S_50ntFlanks@11515.11539.25\|GTCCGCCCCCGGGGCCGCGGTTCCG | >hg38_21_+_8433172_8446622_RNA45SN1_45S_50ntFlanks | 11515 | 11539 | 25 | GTCCGCCCCCGGGGCCGCGGTTCCG | 431.4 | -11.7 | 1.4 | -8.3 | 9.2E-17 | 6.4E-16 | MB231/MB468 | NUC |
| >hg38_21_+_8433172_8446622_RNA45SN1_45S_50ntFlanks@11580.11605.26\|GAACTGGTGCGGACCAGGGGAATCCG | >hg38_21_+_8433172_8446622_RNA45SN1_45S_50ntFlanks | 11580 | 11605 | 26 | GAACTGGTGCGGACCAGGGGAATCCG | 2302.1 | -10.8 | 1.3 | -8.3 | 7.5E-17 | 6.4E-16 | MB231/MB468 | MP |
| >hg38_21_+_8433172_8446622_RNA45SN1_45S_50ntFlanks@11873.11890.18\|AAGAAGACCCTGTTGAGC | >hg38_21_+_8433172_8446622_RNA45SN1_45S_50ntFlanks | 11873 | 11890 | 18 | AAGAAGACCCTGTTGAGC | 3263.7 | -9.5 | 1.2 | -8.2 | 2.0E-16 | 6.4E-16 | BT20/MB468 | NUC |
| >hg38_21_+_8433172_8446622_RNA45SN1_45S_50ntFlanks@11583.11603.21\|CTGGTGCGGACCAGGGGAATC | >hg38_21_+_8433172_8446622_RNA45SN1_45S_50ntFlanks | 11583 | 11603 | 21 | CTGGTGCGGACCAGGGGAATC | 2377.6 | -10.2 | 1.2 | -8.3 | 1.2E-16 | 6.5E-16 | MB231/MB468 | MITO |
| >hg38_21_+_8433172_8446622_RNA45SN1_45S_50ntFlanks@11881.11903.23\|CCTGTTGAGCTTGACTCTAGTCT | >hg38_21_+_8433172_8446622_RNA45SN1_45S_50ntFlanks | 11881 | 11903 | 23 | CCTGTTGAGCTTGACTCTAGTCT | 389.2 | -12.5 | 1.5 | -8.3 | 7.5E-17 | 6.5E-16 | MB231/MB468 | MP |
| >MI0000434\|hsa-let-7i&WithFlank&12\|+\|62603680\|62603775@12.33.22&[MIMAT0000415&hsa-let-7i-5p&offsets\|0\|0;m-49&12\|+\|62603691\|62603712&offsets\|0\|0]\|TGAGGTAGTAGTTTGTGCTGTT | >MI0000434\|hsa-let-7i&WithFlank&12\|+\|62603680\|62603775 | 12 | 33 | 22 | TGAGGTAGTAGTTTGTGCTGTT | 92950.8 | 7.1 | 0.9 | 8.3 | 8.9E-17 | 6.5E-16 | BT20/MB468 | MITO |
| >hg38_21_+_8433172_8446622_RNA45SN1_45S_50ntFlanks@7986.8010.25\|ATCAGACGTGGCGACCCGCTGAATT | >hg38_21_+_8433172_8446622_RNA45SN1_45S_50ntFlanks | 7986 | 8010 | 25 | ATCAGACGTGGCGACCCGCTGAATT | 507.4 | -12.9 | 1.5 | -8.3 | 7.5E-17 | 6.5E-16 | MB231/MB468 | MP |
| >hg38_21_+_8433172_8446622_RNA45SN1_45S_50ntFlanks@7986.8012.27\|ATCAGACGTGGCGACCCGCTGAATTTA | >hg38_21_+_8433172_8446622_RNA45SN1_45S_50ntFlanks | 7986 | 8012 | 27 | ATCAGACGTGGCGACCCGCTGAATTTA | 1737.7 | -13.8 | 1.6 | -8.8 | 1.9E-18 | 6.5E-16 | BT20/MB468 | MP |
| >MI0000088\|hsa-mir-30a&WithFlank&6\|-\|71403545\|71403627@12.35.24(+1C)&[MIMAT0000087&hsa-miR-30a-5p&offsets\|0\|+2(+1C);m-15&6\|-\|71403595\|71403616&offsets\|0\|+2(+1C)]\|TGTAAACATCCTCGACTGGAAGCTC | >MI0000088\|hsa-mir-30a&WithFlank&6\|-\|71403545\|71403627 | 12 | 35 | 24(+1C) | TGTAAACATCCTCGACTGGAAGCTC | 486.3 | 8.2 | 1.0 | 8.3 | 9.4E-17 | 6.5E-16 | MB231/MB468 | NUC |
| >MI0000102\|hsa-mir-100&WithFlank&11\|-\|122152223\|122152314@19.40.22(+3U)&[MIMAT0000098&hsa-miR-100-5p&offsets\|0\|0(+3U);m-27&11\|-\|122152276\|122152296&offsets\|0\|+1(+3U)]\|AACCCGTAGATCCGAACTTGTGTTT | >MI0000102\|hsa-mir-100&WithFlank&11\|-\|122152223\|122152314 | 19 | 40 | 22(+3U) | AACCCGTAGATCCGAACTTGTGTTT | 202.9 | 11.6 | 1.4 | 8.3 | 9.4E-17 | 6.5E-16 | MB231/MB468 | NUC |
| >trna116_GluCTC_1_-_145399233_145399304@30.70.41,trna71_GluCTC_1_-_161439189_161439260@30.70.41,trna74_GluCTC_1_-_161431809_161431880@30.70.41,trna77_GluCTC_1_-_161424398_161424469@30.70.41,trna77_GluCTC_6_+_28949976_28950047@30.70.41,trna80_GluCTC_1_-_161417018_161417089@30.70.41,trna87_GluCTC_6_-_126101393_126101464@30.70.41\|CGCTCTCACCGCCGCGGCCCGGGTTCGATTCCCGGTCAGGG | >trna116_GluCTC_1_-_145399233_145399304 | 30 | 70 | 41 | CGCTCTCACCGCCGCGGCCCGGGTTCGATTCCCGGTCAGGG | 336.6 | -11.4 | 1.4 | -8.3 | 9.4E-17 | 6.5E-16 | MB231/MB468 | NUC |
| >hg38_21_+_8433172_8446622_RNA45SN1_45S_50ntFlanks@11670.11702.33\|CCCAGTGCTCTGAATGTCAAAGTGAAGAAATTC | >hg38_21_+_8433172_8446622_RNA45SN1_45S_50ntFlanks | 11670 | 11702 | 33 | CCCAGTGCTCTGAATGTCAAAGTGAAGAAATTC | 7163.2 | -5.5 | 0.7 | -8.3 | 9.4E-17 | 6.5E-16 | MB231/MB468 | NUC |
| >hg38_21_+_8433172_8446622_RNA45SN1_45S_50ntFlanks@8423.8467.45\|CGCAGTCCGCCCGGAGGATTCAACCCGGCGGCGGGTCCGGCCGTG | >hg38_21_+_8433172_8446622_RNA45SN1_45S_50ntFlanks | 8423 | 8467 | 45 | CGCAGTCCGCCCGGAGGATTCAACCCGGCGGCGGGTCCGGCCGTG | 658.7 | -13.1 | 1.6 | -8.3 | 1.2E-16 | 6.5E-16 | MB231/MB468 | MITO |
| >hg38_21_+_8433172_8446622_RNA45SN1_45S_50ntFlanks@8105.8142.38\|CGCGGCGGGGCGCGGGACATGTGGCGTACGGAAGACCC | >hg38_21_+_8433172_8446622_RNA45SN1_45S_50ntFlanks | 8105 | 8142 | 38 | CGCGGCGGGGCGCGGGACATGTGGCGTACGGAAGACCC | 1590.0 | -10.4 | 1.2 | -8.3 | 1.2E-16 | 6.5E-16 | MB231/MB468 | MITO |
| >am_trnaMT_SerGCT_MT_+_12207_12265@16.50.35\|AACTGCTAACTCATGCCCCCATGTCTAACAACATG | >am_trnaMT_SerGCT_MT_+_12207_12265 | 16 | 50 | 35 | AACTGCTAACTCATGCCCCCATGTCTAACAACATG | 422.7 | -13.1 | 1.6 | -8.2 | 2.0E-16 | 6.5E-16 | BT20/MB468 | NUC |
| >MI0000088\|hsa-mir-30a&WithFlank&6\|-\|71403545\|71403627@54.73.20(+1U)&[MIMAT0000088&hsa-miR-30a-3p&offsets\|+1\|-1(+1U);m-38&6\|-\|71403554\|71403575&offsets\|+1\|-1(+1U)]\|TTTCAGTCGGATGTTTGCAGT | >MI0000088\|hsa-mir-30a&WithFlank&6\|-\|71403545\|71403627 | 54 | 73 | 20(+1U) | TTTCAGTCGGATGTTTGCAGT | 3264.1 | 7.6 | 0.9 | 8.3 | 1.2E-16 | 6.6E-16 | MB231/MB468 | MITO |
| >hg38_21_+_8433172_8446622_RNA45SN1_45S_50ntFlanks@12405.12427.23\|CTTTTTGATCCTTCGATGTCGGC | >hg38_21_+_8433172_8446622_RNA45SN1_45S_50ntFlanks | 12405 | 12427 | 23 | CTTTTTGATCCTTCGATGTCGGC | 267.8 | -8.3 | 1.0 | -8.3 | 7.9E-17 | 6.6E-16 | MB231/MB468 | CYTO |
| >hg38_21_+_8433172_8446622_RNA45SN1_45S_50ntFlanks@8105.8136.32\|CGCGGCGGGGCGCGGGACATGTGGCGTACGGA | >hg38_21_+_8433172_8446622_RNA45SN1_45S_50ntFlanks | 8105 | 8136 | 32 | CGCGGCGGGGCGCGGGACATGTGGCGTACGGA | 901.0 | -13.6 | 1.6 | -8.3 | 1.2E-16 | 6.6E-16 | MB231/MB468 | MITO |
| >hg38_21_+_8433172_8446622_RNA45SN1_45S_50ntFlanks@12273.12293.21\|TCAGTACGAATACAGACCGTG | >hg38_21_+_8433172_8446622_RNA45SN1_45S_50ntFlanks | 12273 | 12293 | 21 | TCAGTACGAATACAGACCGTG | 405.3 | -12.5 | 1.5 | -8.3 | 7.7E-17 | 6.6E-16 | MB231/MB468 | MP |
| >hg38_21_+_8433172_8446622_RNA45SN1_45S_50ntFlanks@10819.10840.22\|AGGGAAGTCGGCAAGCCGGATC | >hg38_21_+_8433172_8446622_RNA45SN1_45S_50ntFlanks | 10819 | 10840 | 22 | AGGGAAGTCGGCAAGCCGGATC | 5107.2 | -5.1 | 0.6 | -8.3 | 7.9E-17 | 6.6E-16 | MB231/MB468 | CYTO |
| >trnaMT_PheGAA_MT_+_577_647@31.53.23\|CACTGAAAATGTTTAGACGGGCT | >trnaMT_PheGAA_MT_+_577_647 | 31 | 53 | 23 | CACTGAAAATGTTTAGACGGGCT | 596.4 | -13.6 | 1.7 | -8.2 | 2.1E-16 | 6.6E-16 | BT20/MB468 | NUC |
| >hg38_21_+_8433172_8446622_RNA45SN1_45S_50ntFlanks@3900.3938.39\|CTTCGCGGGGGGGATGCGTGCATTTATCAGATCAAAACC | >hg38_21_+_8433172_8446622_RNA45SN1_45S_50ntFlanks | 3900 | 3938 | 39 | CTTCGCGGGGGGGATGCGTGCATTTATCAGATCAAAACC | 613.3 | -13.0 | 1.6 | -8.3 | 1.3E-16 | 6.6E-16 | MB231/MB468 | MITO |
| >hg38_21_+_8433172_8446622_RNA45SN1_45S_50ntFlanks@10805.10822.18\|GAACAATGTAGGTAAGGG | >hg38_21_+_8433172_8446622_RNA45SN1_45S_50ntFlanks | 10805 | 10822 | 18 | GAACAATGTAGGTAAGGG | 336.4 | -11.1 | 1.3 | -8.3 | 8.0E-17 | 6.6E-16 | MB231/MB468 | CYTO |
| >hg38_21_+_8433172_8446622_RNA45SN1_45S_50ntFlanks@11575.11602.28\|ACTTAGAACTGGTGCGGACCAGGGGAAT | >hg38_21_+_8433172_8446622_RNA45SN1_45S_50ntFlanks | 11575 | 11602 | 28 | ACTTAGAACTGGTGCGGACCAGGGGAAT | 730.4 | -6.5 | 0.8 | -8.3 | 1.3E-16 | 6.7E-16 | MB231/MB468 | MITO |
| >MI0000449\|hsa-mir-132&WithFlank&17\|-\|2049902\|2050014@65.86.22&[MIMAT0000426&hsa-miR-132-3p&offsets\|0\|0;m-138&17\|-\|2049929\|2049950&offsets\|0\|0]\|TAACAGTCTACAGCCATGGTCG | >MI0000449\|hsa-mir-132&WithFlank&17\|-\|2049902\|2050014 | 65 | 86 | 22 | TAACAGTCTACAGCCATGGTCG | 307.0 | 7.7 | 0.9 | 8.3 | 9.7E-17 | 6.7E-16 | MB231/MB468 | NUC |
| >hg38_21_+_8433172_8446622_RNA45SN1_45S_50ntFlanks@11575.11605.31\|ACTTAGAACTGGTGCGGACCAGGGGAATCCG | >hg38_21_+_8433172_8446622_RNA45SN1_45S_50ntFlanks | 11575 | 11605 | 31 | ACTTAGAACTGGTGCGGACCAGGGGAATCCG | 1841.7 | -10.3 | 1.2 | -8.3 | 1.3E-16 | 6.7E-16 | MB231/MB468 | MITO |
| >Antisense_hg38_21_-_8433172_8446622_RNA45SN1_45S_50ntFlanks@7123.7140.18\|GCGGCGGCGGCGGCGGCG | >Antisense_hg38_21_-_8433172_8446622_RNA45SN1_45S_50ntFlanks | 7123 | 7140 | 18 | GCGGCGGCGGCGGCGGCG | 563.6 | -12.9 | 1.6 | -8.3 | 1.3E-16 | 6.7E-16 | MB231/MB468 | MITO |
| >hg38_21_+_8433172_8446622_RNA45SN1_45S_50ntFlanks@10127.10144.18\|GCGGCGGCGGCGGCGGCG | >hg38_21_+_8433172_8446622_RNA45SN1_45S_50ntFlanks | 10127 | 10144 | 18 | GCGGCGGCGGCGGCGGCG | 563.6 | -12.9 | 1.6 | -8.3 | 1.3E-16 | 6.7E-16 | MB231/MB468 | MITO |
| >hg38_21_+_8433172_8446622_RNA45SN1_45S_50ntFlanks@10130.10147.18\|GCGGCGGCGGCGGCGGCG | >hg38_21_+_8433172_8446622_RNA45SN1_45S_50ntFlanks | 10130 | 10147 | 18 | GCGGCGGCGGCGGCGGCG | 563.6 | -12.9 | 1.6 | -8.3 | 1.3E-16 | 6.7E-16 | MB231/MB468 | MITO |
| >hg38_21_+_8433172_8446622_RNA45SN1_45S_50ntFlanks@8823.8840.18\|GCGGCGGCGGCGGCGGCG | >hg38_21_+_8433172_8446622_RNA45SN1_45S_50ntFlanks | 8823 | 8840 | 18 | GCGGCGGCGGCGGCGGCG | 563.6 | -12.9 | 1.6 | -8.3 | 1.3E-16 | 6.7E-16 | MB231/MB468 | MITO |
| >hg38_21_+_8433172_8446622_RNA45SN1_45S_50ntFlanks@8844.8861.18\|GCGGCGGCGGCGGCGGCG | >hg38_21_+_8433172_8446622_RNA45SN1_45S_50ntFlanks | 8844 | 8861 | 18 | GCGGCGGCGGCGGCGGCG | 563.6 | -12.9 | 1.6 | -8.3 | 1.3E-16 | 6.7E-16 | MB231/MB468 | MITO |
| >hg38_21_+_8433172_8446622_RNA45SN1_45S_50ntFlanks@8847.8864.18\|GCGGCGGCGGCGGCGGCG | >hg38_21_+_8433172_8446622_RNA45SN1_45S_50ntFlanks | 8847 | 8864 | 18 | GCGGCGGCGGCGGCGGCG | 563.6 | -12.9 | 1.6 | -8.3 | 1.3E-16 | 6.7E-16 | MB231/MB468 | MITO |
| >MI0000077\|hsa-mir-21&WithFlank&17\|+\|59841260\|59841343@14.33.20&[MIMAT0000076&hsa-miR-21-5p&offsets\|0\|-2;m-1&17\|+\|59841273\|59841294&offsets\|0\|-2]\|TAGCTTATCAGACTGATGTT | >MI0000077\|hsa-mir-21&WithFlank&17\|+\|59841260\|59841343 | 14 | 33 | 20 | TAGCTTATCAGACTGATGTT | 622.1 | 6.4 | 0.8 | 8.3 | 9.7E-17 | 6.7E-16 | MB231/MB468 | NUC |
| >hg38_21_+_8433172_8446622_RNA45SN1_45S_50ntFlanks@4918.4940.23\|ACCACCAGGAGTGGAGCCTGCGG | >hg38_21_+_8433172_8446622_RNA45SN1_45S_50ntFlanks | 4918 | 4940 | 23 | ACCACCAGGAGTGGAGCCTGCGG | 363.6 | -12.4 | 1.5 | -8.3 | 7.8E-17 | 6.7E-16 | MB231/MB468 | MP |
| >MI0000078\|hsa-mir-22&WithFlank&17\|-\|1713897\|1713993@59.78.20&[MIMAT0000077&hsa-miR-22-3p&offsets\|0\|-2;m-2&17\|-\|1713914\|1713935&offsets\|0\|-2]\|AAGCTGCCAGTTGAAGAACT | >MI0000078\|hsa-mir-22&WithFlank&17\|-\|1713897\|1713993 | 59 | 78 | 20 | AAGCTGCCAGTTGAAGAACT | 777.8 | 9.7 | 1.2 | 8.3 | 9.3E-17 | 6.7E-16 | BT20/MB468 | MITO |
| >hg38_21_+_8433172_8446622_RNA45SN1_45S_50ntFlanks@11561.11594.34\|GCGCCTAGCAGCCGACTTAGAACTGGTGCGGACC | >hg38_21_+_8433172_8446622_RNA45SN1_45S_50ntFlanks | 11561 | 11594 | 34 | GCGCCTAGCAGCCGACTTAGAACTGGTGCGGACC | 295.9 | -12.1 | 1.4 | -8.3 | 7.8E-17 | 6.7E-16 | MB231/MB468 | MP |
| >hg38_21_+_8433172_8446622_RNA45SN1_45S_50ntFlanks@9534.9554.21\|AGGAAACTCTGGTGGAGGTCC | >hg38_21_+_8433172_8446622_RNA45SN1_45S_50ntFlanks | 9534 | 9554 | 21 | AGGAAACTCTGGTGGAGGTCC | 711.2 | -13.3 | 1.6 | -8.3 | 7.8E-17 | 6.7E-16 | MB231/MB468 | MP |
| >hg38_21_+_8433172_8446622_RNA45SN1_45S_50ntFlanks@10841.10862.22\|CGTAACTTCGGGATAAGGATTG | >hg38_21_+_8433172_8446622_RNA45SN1_45S_50ntFlanks | 10841 | 10862 | 22 | CGTAACTTCGGGATAAGGATTG | 6602.2 | -7.0 | 0.8 | -8.3 | 8.1E-17 | 6.8E-16 | MB231/MB468 | CYTO |
| >hg38_21_+_8433172_8446622_RNA45SN1_45S_50ntFlanks@4588.4613.26\|CTATTTTGTTGGTTTTCGGAACTGAG | >hg38_21_+_8433172_8446622_RNA45SN1_45S_50ntFlanks | 4588 | 4613 | 26 | CTATTTTGTTGGTTTTCGGAACTGAG | 300.0 | -12.0 | 1.5 | -8.3 | 1.3E-16 | 6.8E-16 | MB231/MB468 | MITO |

<table>
<tr><td>>hg38_21_+_8433172_8446622_RNA45SN1_45S_50ntFlanks@8192.8213.22|AGCCCGTGGACGGTGTGAGGCC</td><td>>hg38_21_+_8433172_8446622_RNA45SN1_45S_50ntFlanks</td><td>8192</td><td>8213</td><td>22</td><td>AGCCCGTGGACGGTGTGAGGCC</td><td>550.8</td><td>-4.7</td><td>0.6</td><td>-8.2</td><td>2.1E-16</td><td>6.8E-16</td><td>BT20/MB468</td><td>NUC</td></tr>
<tr><td>>MI0000079|hsa-mir-23a&WithFlank&19|-|13836581|13836665@54.71.18&[MIMAT0000078&hsa-miR-23a-3p&offsets|+3|0;m-31&19|-|13836595|13836615&offsets|+3|0]|ACATTGCCAGGGATTTCC</td><td>>MI0000079|hsa-mir-23a&WithFlank&19|-|13836581|13836665</td><td>54</td><td>71</td><td>18</td><td>ACATTGCCAGGGATTTCC</td><td>5791.6</td><td>16.2</td><td>1.9</td><td>8.8</td><td>2.0E-18</td><td>6.8E-16</td><td>BT20/MB468</td><td>MP</td></tr>
<tr><td>>hg38_21_+_8433172_8446622_RNA45SN1_45S_50ntFlanks@8339.8375.37|TACCGTAAGGGAAAGTTGAAAAGAACTTTGAAGAGAG</td><td>>hg38_21_+_8433172_8446622_RNA45SN1_45S_50ntFlanks</td><td>8339</td><td>8375</td><td>37</td><td>TACCGTAAGGGAAAGTTGAAAAGAACTTTGAAGAGAG</td><td>1477.7</td><td>14.3</td><td>1.6</td><td>8.8</td><td>2.1E-18</td><td>6.8E-16</td><td>BT20/MB468</td><td>MP</td></tr>
<tr><td>>hg38_21_+_8433172_8446622_RNA45SN1_45S_50ntFlanks@12633.12654.22|CGAAGCTACCATCTGTGGGATT</td><td>>hg38_21_+_8433172_8446622_RNA45SN1_45S_50ntFlanks</td><td>12633</td><td>12654</td><td>22</td><td>CGAAGCTACCATCTGTGGGATT</td><td>18504.0</td><td>-7.6</td><td>0.9</td><td>-8.7</td><td>2.1E-18</td><td>6.8E-16</td><td>BT20/MB468</td><td>MP</td></tr>
<tr><td>>hg38_21_+_8433172_8446622_RNA45SN1_45S_50ntFlanks@4657.4684.28|CTAGAGGTGAAATTCTTGGACCGGCGCA</td><td>>hg38_21_+_8433172_8446622_RNA45SN1_45S_50ntFlanks</td><td>4657</td><td>4684</td><td>28</td><td>CTAGAGGTGAAATTCTTGGACCGGCGCA</td><td>1438.3</td><td>-14.3</td><td>1.7</td><td>-8.3</td><td>1.3E-16</td><td>6.8E-16</td><td>MB231/MB468</td><td>MITO</td></tr>
<tr><td>>MI0000063|hsa-let-7b&WithFlank&22|+|46113680|46113774@12.31.20(+1A)&[MIMAT0000063&hsa-let-7b-5p&offsets|0|-2(+1A);m-14&22|+|46113691|46113712&offsets|0|-2(+1A)]|TGAGGTAGTAGGTTGTGTGGA</td><td>>MI0000063|hsa-let-7b&WithFlank&22|+|46113680|46113774</td><td>12</td><td>31</td><td>20(+1A)</td><td>TGAGGTAGTAGGTTGTGTGGA</td><td>868.1</td><td>11.4</td><td>1.4</td><td>8.3</td><td>1.3E-16</td><td>6.8E-16</td><td>BT20/MB468</td><td>CYTO</td></tr>
<tr><td>>MI0000301|hsa-mir-224&WithFlank&X|-|151958572|151958664@14.35.22&[MIMAT0000281&hsa-miR-224-5p&offsets|+1|-2;m-238&X|-|151958630|151958651&offsets|0|0]|CAAGTCACTAGTGGTTCCGTTT</td><td>>MI0000301|hsa-mir-224&WithFlank&X|-|151958572|151958664</td><td>14</td><td>35</td><td>22</td><td>CAAGTCACTAGTGGTTCCGTTT</td><td>1630.9</td><td>10.2</td><td>1.2</td><td>8.3</td><td>8.2E-17</td><td>6.8E-16</td><td>MB231/MB468</td><td>CYTO</td></tr>
<tr><td>>hg38_21_+_8433172_8446622_RNA45SN1_45S_50ntFlanks@11585.11616.32|GGTGCGGACCAGGGGAATCCGACTGTTTAATT</td><td>>hg38_21_+_8433172_8446622_RNA45SN1_45S_50ntFlanks</td><td>11585</td><td>11616</td><td>32</td><td>GGTGCGGACCAGGGGAATCCGACTGTTTAATT</td><td>276.1</td><td>8.1</td><td>1.0</td><td>8.3</td><td>8.2E-17</td><td>6.8E-16</td><td>MB231/MB468</td><td>CYTO</td></tr>
<tr><td>>hg38_21_+_8433172_8446622_RNA45SN1_45S_50ntFlanks@8020.8046.27|TAGTCAGCGGAGGAGAAGAAACTAACC</td><td>>hg38_21_+_8433172_8446622_RNA45SN1_45S_50ntFlanks</td><td>8020</td><td>8046</td><td>27</td><td>TAGTCAGCGGAGGAGAAGAAACTAACC</td><td>580.0</td><td>10.5</td><td>1.3</td><td>8.3</td><td>8.2E-17</td><td>6.8E-16</td><td>MB231/MB468</td><td>CYTO</td></tr>
<tr><td>>hg38_21_+_8433172_8446622_RNA45SN1_45S_50ntFlanks@9573.9596.24|AAATCGGTCGTCCGACCTGGGTAT</td><td>>hg38_21_+_8433172_8446622_RNA45SN1_45S_50ntFlanks</td><td>9573</td><td>9596</td><td>24</td><td>AAATCGGTCGTCCGACCTGGGTAT</td><td>2002.9</td><td>-6.1</td><td>0.7</td><td>-8.3</td><td>8.2E-17</td><td>6.8E-16</td><td>MB231/MB468</td><td>CYTO</td></tr>
<tr><td>>hg38_21_+_8433172_8446622_RNA45SN1_45S_50ntFlanks@6737.6772.36|GATCATCGACACTTCGAACGCACTTGCGGCCCCGGG</td><td>>hg38_21_+_8433172_8446622_RNA45SN1_45S_50ntFlanks</td><td>6737</td><td>6772</td><td>36</td><td>GATCATCGACACTTCGAACGCACTTGCGGCCCCGGG</td><td>554.3</td><td>-12.9</td><td>1.6</td><td>-8.3</td><td>1.3E-16</td><td>6.8E-16</td><td>MB231/MB468</td><td>MITO</td></tr>
<tr><td>>am_MI0000786|hsa-mir-378a&WithFlank&5|+|149732819|149732896@50.70.21&[MIMAT0000732&hsa-miR-378a-3p&offsets|+1|0;m-44&5|+|149732867|149732888&offsets|+1|0]|CTGGACTTGGAGTCAGAAGGC</td><td>>am_MI0000786|hsa-mir-378a&WithFlank&5|+|149732819|149732896</td><td>50</td><td>70</td><td>21</td><td>CTGGACTTGGAGTCAGAAGGC</td><td>1517.3</td><td>5.3</td><td>0.6</td><td>8.3</td><td>9.4E-17</td><td>6.8E-16</td><td>BT20/MB468</td><td>MITO</td></tr>
<tr><td>>hg38_21_+_8433172_8446622_RNA45SN1_45S_50ntFlanks@12676.12704.29|AGAATCCCGCCCAGGCGGAACGATACGGC</td><td>>hg38_21_+_8433172_8446622_RNA45SN1_45S_50ntFlanks</td><td>12676</td><td>12704</td><td>29</td><td>AGAATCCCGCCCAGGCGGAACGATACGGC</td><td>608.9</td><td>-13.1</td><td>1.6</td><td>-8.3</td><td>8.0E-17</td><td>6.8E-16</td><td>MB231/MB468</td><td>MP</td></tr>
<tr><td>>hg38_21_+_8433172_8446622_RNA45SN1_45S_50ntFlanks@12347.12365.19|AGAAAAGTTACCACAGGGA</td><td>>hg38_21_+_8433172_8446622_RNA45SN1_45S_50ntFlanks</td><td>12347</td><td>12365</td><td>19</td><td>AGAAAAGTTACCACAGGGA</td><td>274.5</td><td>-11.1</td><td>1.3</td><td>-8.3</td><td>9.9E-17</td><td>6.8E-16</td><td>MB231/MB468</td><td>NUC</td></tr>
<tr><td>>hg38_21_+_8433172_8446622_RNA45SN1_45S_50ntFlanks@11873.11903.31|AAGAAGACCCTGTTGAGCTTGACTCTAGTCT</td><td>>hg38_21_+_8433172_8446622_RNA45SN1_45S_50ntFlanks</td><td>11873</td><td>11903</td><td>31</td><td>AAGAAGACCCTGTTGAGCTTGACTCTAGTCT</td><td>771.4</td><td>-9.6</td><td>1.2</td><td>-8.3</td><td>1.3E-16</td><td>6.8E-16</td><td>MB231/MB468</td><td>MITO</td></tr>
<tr><td>>hg38_21_+_8433172_8446622_RNA45SN1_45S_50ntFlanks@11209.11233.25|CGGCGACTCTGGACGCGAGCCGGGC</td><td>>hg38_21_+_8433172_8446622_RNA45SN1_45S_50ntFlanks</td><td>11209</td><td>11233</td><td>25</td><td>CGGCGACTCTGGACGCGAGCCGGGC</td><td>1269.6</td><td>-7.8</td><td>0.9</td><td>-8.3</td><td>1.3E-16</td><td>6.8E-16</td><td>MB231/MB468</td><td>MITO</td></tr>
<tr><td>>hg38_21_+_8433172_8446622_RNA45SN1_45S_50ntFlanks@9504.9529.26|GTGAACTATGCCTGGGCAGGGCGAAG</td><td>>hg38_21_+_8433172_8446622_RNA45SN1_45S_50ntFlanks</td><td>9504</td><td>9529</td><td>26</td><td>GTGAACTATGCCTGGGCAGGGCGAAG</td><td>473.9</td><td>-12.7</td><td>1.5</td><td>-8.3</td><td>1.3E-16</td><td>6.8E-16</td><td>MB231/MB468</td><td>MITO</td></tr>
<tr><td>>hg38_21_+_8433172_8446622_RNA45SN1_45S_50ntFlanks@3988.4023.36|CTTTGGTGACTCTAGATAACCTCGGGCCGATCGCAC</td><td>>hg38_21_+_8433172_8446622_RNA45SN1_45S_50ntFlanks</td><td>3988</td><td>4023</td><td>36</td><td>CTTTGGTGACTCTAGATAACCTCGGGCCGATCGCAC</td><td>1043.1</td><td>5.1</td><td>0.6</td><td>8.3</td><td>9.5E-17</td><td>6.8E-16</td><td>BT20/MB468</td><td>MITO</td></tr>
<tr><td>>trna112_GlnCTG_1_-_147737382_147737453@31.50.20,trna131_GlnCTG_6_-_27759135_27759206@31.50.20,trna146_GlnCTG_6_-_27515531_27515602@31.50.20,trna15_GlnCTG_1_+_145963304_145963375@31.50.20,trna19_GlnCTG_1_+_147505038_147505109@31.50.20,trna1_GlnCTG_6_+_18836402_18836473@31.50.20,trna28_GlnCTG_1_+_149186125_149186196@31.50.20,trna3_GlnCTG_17_+_8023070_8023141@31.50.20,trna49_GlnCTG_6_+_27487308_27487379@31.50.20,trna7_GlnCTG_15_-_66161400_66161471@31.50.20,trna99_GlnCTG_6_-_28909378_28909449@31.50.20|ACTCTGAATCCAGCGATCCG</td><td>>trna112_GlnCTG_1_-_147737382_147737453</td><td>31</td><td>50</td><td>20</td><td>ACTCTGAATCCAGCGATCCG</td><td>155.7</td><td>-6.0</td><td>0.7</td><td>-8.3</td><td>8.3E-17</td><td>6.9E-16</td><td>MB231/MB468</td><td>CYTO</td></tr>
<tr><td>>hg38_1_-_228634819_228635039_RNA5S12_5S_50ntFlanks@147.170.24|GGGAATACCGGGTGCTGTAGGCTT</td><td>>hg38_1_-_228634819_228635039_RNA5S12_5S_50ntFlanks</td><td>147</td><td>170</td><td>24</td><td>GGGAATACCGGGTGCTGTAGGCTT</td><td>6720.2</td><td>5.6</td><td>0.7</td><td>8.3</td><td>8.3E-17</td><td>6.9E-16</td><td>MB231/MB468</td><td>CYTO</td></tr>
<tr><td>>MI0006273|hsa-mir-1180&WithFlank&17|-|19344500|19344580@47.66.20&[MIMAT0005825&hsa-miR-1180-3p&offsets|0|-2;m-214&17|-|19344513|19344534&offsets|0|-2]|TTTCCGGCTCGCGTGGGTGT</td><td>>MI0006273|hsa-mir-1180&WithFlank&17|-|19344500|19344580</td><td>47</td><td>66</td><td>20</td><td>TTTCCGGCTCGCGTGGGTGT</td><td>1441.2</td><td>10.1</td><td>1.2</td><td>8.3</td><td>1.3E-16</td><td>6.9E-16</td><td>BT20/MB468</td><td>CYTO</td></tr>
<tr><td>>hg38_21_+_8433172_8446622_RNA45SN1_45S_50ntFlanks@10393.10420.28|CAGCAGTTGAACATGGGTCAGTCGGTCC</td><td>>hg38_21_+_8433172_8446622_RNA45SN1_45S_50ntFlanks</td><td>10393</td><td>10420</td><td>28</td><td>CAGCAGTTGAACATGGGTCAGTCGGTCC</td><td>350.7</td><td>-12.3</td><td>1.5</td><td>-8.3</td><td>8.0E-17</td><td>6.9E-16</td><td>MB231/MB468</td><td>MP</td></tr>
<tr><td>>hg38_21_+_8433172_8446622_RNA45SN1_45S_50ntFlanks@10681.10710.30|TTCCGGCGGCGTCCGGTGAGCTCTCGCTGG</td><td>>hg38_21_+_8433172_8446622_RNA45SN1_45S_50ntFlanks</td><td>10681</td><td>10710</td><td>30</td><td>TTCCGGCGGCGTCCGGTGAGCTCTCGCTGG</td><td>3118.1</td><td>-10.2</td><td>1.2</td><td>-8.3</td><td>1.3E-16</td><td>6.9E-16</td><td>MB231/MB468</td><td>MITO</td></tr>
<tr><td>>MI0000298|hsa-mir-221&WithFlank&X|-|45746151|45746272@71.91.21&[MIMAT0000278&hsa-miR-221-3p&offsets|0|-2;m-91&X|-|45746181|45746202&offsets|0|-1]|AGCTACATTGTCTGCTGGGTT</td><td>>MI0000298|hsa-mir-221&WithFlank&X|-|45746151|45746272</td><td>71</td><td>91</td><td>21</td><td>AGCTACATTGTCTGCTGGGTT</td><td>920.6</td><td>6.3</td><td>0.8</td><td>8.3</td><td>9.6E-17</td><td>6.9E-16</td><td>BT20/MB468</td><td>MITO</td></tr>
<tr><td>>MI0000809|hsa-mir-151a&WithFlank&8|-|140732558|140732659@51.73.23&[MIMAT0000757&hsa-miR-151a-3p&offsets|-2|0;m-30&8|-|140732587|140732607&offsets|-2|0]|TACTAGACTGAAGCTCCTTGAGG</td><td>>MI0000809|hsa-mir-151a&WithFlank&8|-|140732558|140732659</td><td>51</td><td>73</td><td>23</td><td>TACTAGACTGAAGCTCCTTGAGG</td><td>3210.8</td><td>6.3</td><td>0.8</td><td>8.3</td><td>8.3E-17</td><td>6.9E-16</td><td>MB231/MB468</td><td>CYTO</td></tr>
<tr><td>>hg38_21_+_8433172_8446622_RNA45SN1_45S_50ntFlanks@12081.12108.28|CTCTCGCTTCTGGCGCCAAGCGCCCGGC</td><td>>hg38_21_+_8433172_8446622_RNA45SN1_45S_50ntFlanks</td><td>12081</td><td>12108</td><td>28</td><td>CTCTCGCTTCTGGCGCCAAGCGCCCGGC</td><td>1337.1</td><td>-7.8</td><td>0.9</td><td>-8.3</td><td>1.3E-16</td><td>6.9E-16</td><td>MB231/MB468</td><td>MITO</td></tr>
<tr><td>>hg38_21_+_8433172_8446622_RNA45SN1_45S_50ntFlanks@5056.5098.43|GAGCGATTTGTCTGGTTAATTCCGATAACGAACGAGACTCTGG</td><td>>hg38_21_+_8433172_8446622_RNA45SN1_45S_50ntFlanks</td><td>5056</td><td>5098</td><td>43</td><td>GAGCGATTTGTCTGGTTAATTCCGATAACGAACGAGACTCTGG</td><td>111.8</td><td>-23.9</td><td>2.9</td><td>-8.3</td><td>8.1E-17</td><td>6.9E-16</td><td>MB231/MB468</td><td>MP</td></tr>
<tr><td>>hg38_21_+_8433172_8446622_RNA45SN1_45S_50ntFlanks@8170.8214.45|CAAGTCCTTCTGATCGAGGCCCAGCCCGTGGACGGTGTGAGGCCG</td><td>>hg38_21_+_8433172_8446622_RNA45SN1_45S_50ntFlanks</td><td>8170</td><td>8214</td><td>45</td><td>CAAGTCCTTCTGATCGAGGCCCAGCCCGTGGACGGTGTGAGGCCG</td><td>972.9</td><td>-13.8</td><td>1.7</td><td>-8.3</td><td>8.1E-17</td><td>6.9E-16</td><td>MB231/MB468</td><td>MP</td></tr>
<tr><td>>hg38_21_+_8433172_8446622_RNA45SN1_45S_50ntFlanks@10783.10818.36|GTGAACAGCCTCTGGCATGTTGGAACAATGTAGGTA</td><td>>hg38_21_+_8433172_8446622_RNA45SN1_45S_50ntFlanks</td><td>10783</td><td>10818</td><td>36</td><td>GTGAACAGCCTCTGGCATGTTGGAACAATGTAGGTA</td><td>1639.0</td><td>-8.2</td><td>1.0</td><td>-8.3</td><td>1.3E-16</td><td>7.0E-16</td><td>MB231/MB468</td><td>MITO</td></tr>
<tr><td>>hg38_21_+_8433172_8446622_RNA45SN1_45S_50ntFlanks@11205.11225.21|GCGGCGGCGACTCTGGACGCG</td><td>>hg38_21_+_8433172_8446622_RNA45SN1_45S_50ntFlanks</td><td>11205</td><td>11225</td><td>21</td><td>GCGGCGGCGACTCTGGACGCG</td><td>619.4</td><td>-9.2</td><td>1.1</td><td>-8.3</td><td>8.4E-17</td><td>7.0E-16</td><td>MB231/MB468</td><td>CYTO</td></tr>
<tr><td>>hg38_21_+_8433172_8446622_RNA45SN1_45S_50ntFlanks@4723.4752.30|CATTAATCAAGAACGAAAGTCGGAGGTTCG</td><td>>hg38_21_+_8433172_8446622_RNA45SN1_45S_50ntFlanks</td><td>4723</td><td>4752</td><td>30</td><td>CATTAATCAAGAACGAAAGTCGGAGGTTCG</td><td>1011.0</td><td>-12.1</td><td>1.5</td><td>-8.3</td><td>9.7E-17</td><td>7.0E-16</td><td>BT20/MB468</td><td>MITO</td></tr>
<tr><td>>hg38_21_+_8433172_8446622_RNA45SN1_45S_50ntFlanks@10410.10431.22|TCAGTCGGTCCTGAGAGATGGG</td><td>>hg38_21_+_8433172_8446622_RNA45SN1_45S_50ntFlanks</td><td>10410</td><td>10431</td><td>22</td><td>TCAGTCGGTCCTGAGAGATGGG</td><td>1210.8</td><td>-10.2</td><td>1.2</td><td>-8.3</td><td>1.3E-16</td><td>7.0E-16</td><td>MB231/MB468</td><td>MITO</td></tr>
<tr><td>>trna10_LysCTT_16_+_3241501_3241573@4.33.30,trna13_LysCTT_14_-_58706613_58706685@4.33.30,trna2_LysCTT_15_+_79152904_79152976@4.33.30|CGGCTAGCTCAGTCGGTAGAGCATGGGACT</td><td>>trna10_LysCTT_16_+_3241501_3241573</td><td>4</td><td>33</td><td>30</td><td>CGGCTAGCTCAGTCGGTAGAGCATGGGACT</td><td>422.8</td><td>5.5</td><td>0.7</td><td>8.3</td><td>9.8E-17</td><td>7.0E-16</td><td>BT20/MB468</td><td>MITO</td></tr>
<tr><td>>hg38_21_+_8433172_8446622_RNA45SN1_45S_50ntFlanks@10882.10902.21|TCGGGCTGGGGCGCGAAGCGG</td><td>>hg38_21_+_8433172_8446622_RNA45SN1_45S_50ntFlanks</td><td>10882</td><td>10902</td><td>21</td><td>TCGGGCTGGGGCGCGAAGCGG</td><td>371.5</td><td>-12.4</td><td>1.5</td><td>-8.3</td><td>8.2E-17</td><td>7.0E-16</td><td>MB231/MB468</td><td>MP</td></tr>
<tr><td>>hg38_21_+_8433172_8446622_RNA45SN1_45S_50ntFlanks@12249.12272.24|CAAAAGCTCGCTTGATCTTGATTT</td><td>>hg38_21_+_8433172_8446622_RNA45SN1_45S_50ntFlanks</td><td>12249</td><td>12272</td><td>24</td><td>CAAAAGCTCGCTTGATCTTGATTT</td><td>417.4</td><td>-12.6</td><td>1.5</td><td>-8.3</td><td>8.2E-17</td><td>7.0E-16</td><td>MB231/MB468</td><td>MP</td></tr>
<tr><td>>hg38_21_+_8433172_8446622_RNA45SN1_45S_50ntFlanks@10794.10826.33|CTGGCATGTTGGAACAATGTAGGTAAGGGAAGT</td><td>>hg38_21_+_8433172_8446622_RNA45SN1_45S_50ntFlanks</td><td>10794</td><td>10826</td><td>33</td><td>CTGGCATGTTGGAACAATGTAGGTAAGGGAAGT</td><td>828.8</td><td>-9.5</td><td>1.1</td><td>-8.3</td><td>1.3E-16</td><td>7.0E-16</td><td>MB231/MB468</td><td>MITO</td></tr>
<tr><td>>hg38_21_+_8433172_8446622_RNA45SN1_45S_50ntFlanks@10841.10862.22|CGTAACTTCGGGATAAGGATTG</td><td>>hg38_21_+_8433172_8446622_RNA45SN1_45S_50ntFlanks</td><td>10841</td><td>10862</td><td>22</td><td>CGTAACTTCGGGATAAGGATTG</td><td>5636.7</td><td>-6.8</td><td>0.8</td><td>-8.3</td><td>1.4E-16</td><td>7.0E-16</td><td>BT20/MB468</td><td>CYTO</td></tr>
<tr><td>>hg38_21_+_8433172_8446622_RNA45SN1_45S_50ntFlanks@12418.12441.24|CGATGTCGGCTCTTCCTATCATTG</td><td>>hg38_21_+_8433172_8446622_RNA45SN1_45S_50ntFlanks</td><td>12418</td><td>12441</td><td>24</td><td>CGATGTCGGCTCTTCCTATCATTG</td><td>1488.0</td><td>-4.8</td><td>0.6</td><td>-8.3</td><td>1.4E-16</td><td>7.0E-16</td><td>BT20/MB468</td><td>CYTO</td></tr>
<tr><td>>hg38_21_+_8433172_8446622_RNA45SN1_45S_50ntFlanks@8425.8469.45|CAGTCCGCCCGGAGGATTCAACCCGGCGGCGGGTCCGGCCGTGTC</td><td>>hg38_21_+_8433172_8446622_RNA45SN1_45S_50ntFlanks</td><td>8425</td><td>8469</td><td>45</td><td>CAGTCCGCCCGGAGGATTCAACCCGGCGGCGGGTCCGGCCGTGTC</td><td>494.4</td><td>-12.7</td><td>1.5</td><td>-8.3</td><td>1.3E-16</td><td>7.0E-16</td><td>MB231/MB468</td><td>MITO</td></tr>
<tr><td>>hg38_21_+_8433172_8446622_RNA45SN1_45S_50ntFlanks@11874.11894.21|AGAAGACCCTGTTGAGCTTGA</td><td>>hg38_21_+_8433172_8446622_RNA45SN1_45S_50ntFlanks</td><td>11874</td><td>11894</td><td>21</td><td>AGAAGACCCTGTTGAGCTTGA</td><td>270.1</td><td>-11.9</td><td>1.4</td><td>-8.3</td><td>8.3E-17</td><td>7.1E-16</td><td>MB231/MB468</td><td>MP</td></tr>
<tr><td>>hg38_21_+_8433172_8446622_RNA45SN1_45S_50ntFlanks@4975.5001.27|CCCGGACACGGACAGGATTGACAGATT</td><td>>hg38_21_+_8433172_8446622_RNA45SN1_45S_50ntFlanks</td><td>4975</td><td>5001</td><td>27</td><td>CCCGGACACGGACAGGATTGACAGATT</td><td>289.8</td><td>6.9</td><td>0.8</td><td>8.3</td><td>1.4E-16</td><td>7.1E-16</td><td>BT20/MB468</td><td>CYTO</td></tr>
<tr><td>>MI0000650|hsa-mir-200c&WithFlank&12|+|6963693|6963772@49.71.23&[MIMAT0000617&hsa-miR-200c-3p&offsets|-1|-1;m-54&12|+|6963742|6963764&offsets|-1|-1]|CTAATACTGCCGGGTAATGATGG</td><td>>MI0000650|hsa-mir-200c&WithFlank&12|+|6963693|6963772</td><td>49</td><td>71</td><td>23</td><td>CTAATACTGCCGGGTAATGATGG</td><td>574.6</td><td>9.1</td><td>1.1</td><td>8.6</td><td>7.2E-18</td><td>7.1E-16</td><td>BT20/MB231</td><td>CYTO</td></tr>
<tr><td>>hg38_21_+_8433172_8446622_RNA45SN1_45S_50ntFlanks@10866.10902.37|CTAAGGGCTGGGTCGGTCGGGCTGGGGCGCGAAGCGG</td><td>>hg38_21_+_8433172_8446622_RNA45SN1_45S_50ntFlanks</td><td>10866</td><td>10902</td><td>37</td><td>CTAAGGGCTGGGTCGGTCGGGCTGGGGCGCGAAGCGG</td><td>1005.4</td><td>-9.0</td><td>1.1</td><td>-8.3</td><td>8.6E-17</td><td>7.1E-16</td><td>MB231/MB468</td><td>CYTO</td></tr>
<tr><td>>hg38_21_+_8433172_8446622_RNA45SN1_45S_50ntFlanks@12407.12436.30|TTTTGATCCTTCGATGTCGGCTCTTCCTAT</td><td>>hg38_21_+_8433172_8446622_RNA45SN1_45S_50ntFlanks</td><td>12407</td><td>12436</td><td>30</td><td>TTTTGATCCTTCGATGTCGGCTCTTCCTAT</td><td>1660.1</td><td>-5.5</td><td>0.7</td><td>-8.3</td><td>9.9E-17</td><td>7.1E-16</td><td>BT20/MB468</td><td>MITO</td></tr>
<tr><td>>hg38_21_+_8433172_8446622_RNA45SN1_45S_50ntFlanks@12419.12437.19|GATGTCGGCTCTTCCTATC</td><td>>hg38_21_+_8433172_8446622_RNA45SN1_45S_50ntFlanks</td><td>12419</td><td>12437</td><td>19</td><td>GATGTCGGCTCTTCCTATC</td><td>3445.0</td><td>-3.8</td><td>0.5</td><td>-8.2</td><td>2.2E-16</td><td>7.1E-16</td><td>BT20/MB468</td><td>NUC</td></tr>
<tr><td>>hg38_21_+_8433172_8446622_RNA45SN1_45S_50ntFlanks@8176.8195.20|CTTCTGATCGAGGCCCAGCC</td><td>>hg38_21_+_8433172_8446622_RNA45SN1_45S_50ntFlanks</td><td>8176</td><td>8195</td><td>20</td><td>CTTCTGATCGAGGCCCAGCC</td><td>5277.9</td><td>-5.6</td><td>0.7</td><td>-8.2</td><td>2.2E-16</td><td>7.1E-16</td><td>BT20/MB468</td><td>NUC</td></tr>
</table>

| | | | | | | | | | | | | | |
|---|---|---|---|---|---|---|---|---|---|---|---|---|---|
| >hg38_21_+_8433172_8446622_RNA45SN1_45S_50ntFlanks@10729.10755.27\|AGAGGGTGTAAATCTCGCGCCGGGCCG | >hg38_21_+_8433172_8446622_RNA45SN1_45S_50ntFlanks | 10729 | 10755 | 27 | AGAGGGTGTAAATCTCGCGCCGGGCCG | 347.1 | -12.2 | 1.5 | -8.3 | 1.4E-16 | 7.1E-16 | MB231/MB468 | MITO |
| >hg38_21_+_8433172_8446622_RNA45SN1_45S_50ntFlanks@6682.6718.37\|CGTCGATGAAGAACGCAGCTAGCTGCGAGAATTAATG | >hg38_21_+_8433172_8446622_RNA45SN1_45S_50ntFlanks | 6682 | 6718 | 37 | CGTCGATGAAGAACGCAGCTAGCTGCGAGAATTAATG | 824.6 | -13.5 | 1.6 | -8.3 | 1.4E-16 | 7.2E-16 | MB231/MB468 | MITO |
| >hg38_21_+_8433172_8446622_RNA45SN1_45S_50ntFlanks@11313.11344.32\|CCCCACCCCACGTCTCGTCGCGCGCGCGTCCG | >hg38_21_+_8433172_8446622_RNA45SN1_45S_50ntFlanks | 11313 | 11344 | 32 | CCCCACCCCACGTCTCGTCGCGCGCGCGTCCG | 419.4 | -12.5 | 1.5 | -8.3 | 1.4E-16 | 7.2E-16 | MB231/MB468 | MITO |
| >hg38_21_+_8433172_8446622_RNA45SN1_45S_50ntFlanks@8110.8141.32\|CGGGGCGCGGGACATGTGGCGTACGGAAGACC | >hg38_21_+_8433172_8446622_RNA45SN1_45S_50ntFlanks | 8110 | 8141 | 32 | CGGGGCGCGGGACATGTGGCGTACGGAAGACC | 291.8 | -12.1 | 1.4 | -8.3 | 8.4E-17 | 7.2E-16 | MB231/MB468 | MP |
| >hg38_21_+_8433172_8446622_RNA45SN1_45S_50ntFlanks@10684.10705.22\|CGGCGGCGTCCGGTGAGCTCTC | >hg38_21_+_8433172_8446622_RNA45SN1_45S_50ntFlanks | 10684 | 10705 | 22 | CGGCGGCGTCCGGTGAGCTCTC | 273.2 | -12.0 | 1.4 | -8.3 | 8.5E-17 | 7.2E-16 | MB231/MB468 | MP |
| >hg38_21_+_8433172_8446622_RNA45SN1_45S_50ntFlanks@9574.9594.21\|AATCGGTCGTCCGACCTGGGT | >hg38_21_+_8433172_8446622_RNA45SN1_45S_50ntFlanks | 9574 | 9594 | 21 | AATCGGTCGTCCGACCTGGGT | 5622.8 | -7.1 | 0.9 | -8.3 | 8.5E-17 | 7.2E-16 | MB231/MB468 | MP |
| >hg38_21_+_8433172_8446622_RNA45SN1_45S_50ntFlanks@12974.13001.28\|CGTACGTAGCAGAGCAGCTCCCTCGCTG | >hg38_21_+_8433172_8446622_RNA45SN1_45S_50ntFlanks | 12974 | 13001 | 28 | CGTACGTAGCAGAGCAGCTCCCTCGCTG | 259.0 | -11.9 | 1.4 | -8.3 | 8.5E-17 | 7.2E-16 | MB231/MB468 | MP |
| >hg38_21_+_8433172_8446622_RNA45SN1_45S_50ntFlanks@9489.9506.18\|AGGACCCGAAAGATGGTG | >hg38_21_+_8433172_8446622_RNA45SN1_45S_50ntFlanks | 9489 | 9506 | 18 | AGGACCCGAAAGATGGTG | 513.9 | -12.9 | 1.5 | -8.3 | 8.5E-17 | 7.2E-16 | MB231/MB468 | MP |
| >hg38_21_+_8433172_8446622_RNA45SN1_45S_50ntFlanks@9862.9893.32\|ATGAACCGAACGCCGGGTTAAGGCGCCCGATG | >hg38_21_+_8433172_8446622_RNA45SN1_45S_50ntFlanks | 9862 | 9893 | 32 | ATGAACCGAACGCCGGGTTAAGGCGCCCGATG | 1007.1 | -13.8 | 1.7 | -8.3 | 8.5E-17 | 7.2E-16 | MB231/MB468 | MP |
| >hg38_21_+_8433172_8446622_RNA45SN1_45S_50ntFlanks@9027.9058.32\|CGTCTCCTCGTGGGGGGGCCGGGCCACCCCTC | >hg38_21_+_8433172_8446622_RNA45SN1_45S_50ntFlanks | 9027 | 9058 | 32 | CGTCTCCTCGTGGGGGGGCCGGGCCACCCCTC | 617.7 | -11.4 | 1.4 | -8.3 | 1.0E-16 | 7.3E-16 | BT20/MB468 | MITO |
| >hg38_21_+_8433172_8446622_RNA45SN1_45S_50ntFlanks@9566.9596.31\|GACGTGCAAATCGGTCGTCCGACCTGGGTAT | >hg38_21_+_8433172_8446622_RNA45SN1_45S_50ntFlanks | 9566 | 9596 | 31 | GACGTGCAAATCGGTCGTCCGACCTGGGTAT | 3591.6 | -4.0 | 0.5 | -8.3 | 1.0E-16 | 7.3E-16 | BT20/MB468 | MITO |
| >hg38_21_+_8433172_8446622_RNA45SN1_45S_50ntFlanks@4684.4726.43\|AAGACGGACCAGAGCGAAAGCATTTGCCAAGAATGTTTTCATT | >hg38_21_+_8433172_8446622_RNA45SN1_45S_50ntFlanks | 4684 | 4726 | 43 | AAGACGGACCAGAGCGAAAGCATTTGCCAAGAATGTTTTCATT | 454.5 | 9.2 | 1.1 | 8.3 | 8.8E-17 | 7.3E-16 | MB231/MB468 | CYTO |
| >hg38_21_+_8433172_8446622_RNA45SN1_45S_50ntFlanks@9032.9058.27\|CCTCGTGGGGGGGCCGGGCCACCCCTC | >hg38_21_+_8433172_8446622_RNA45SN1_45S_50ntFlanks | 9032 | 9058 | 27 | CCTCGTGGGGGGGCCGGGCCACCCCTC | 426.1 | -12.5 | 1.5 | -8.3 | 1.4E-16 | 7.3E-16 | MB231/MB468 | MITO |
| >MI0000434\|hsa-let-7i&WithFlank&12\|+\|62603680\|62603775@12.32.21&[MIMAT0000415&hsa-let-7i-5p&offsets\|0\|-1;m-49&12\|+\|62603691\|62603712&offsets\|0\|-1]\|TGAGGTAGTAGTTTGTGCTGT | >MI0000434\|hsa-let-7i&WithFlank&12\|+\|62603680\|62603775 | 12 | 32 | 21 | TGAGGTAGTAGTTTGTGCTGT | 3990.2 | 6.3 | 0.8 | 8.3 | 8.9E-17 | 7.3E-16 | MB231/MB468 | CYTO |
| >hg38_21_+_8433172_8446622_RNA45SN1_45S_50ntFlanks@11874.11907.34\|AGAAGACCCTGTTGAGCTTGACTCTAGTCTGGCA | >hg38_21_+_8433172_8446622_RNA45SN1_45S_50ntFlanks | 11874 | 11907 | 34 | AGAAGACCCTGTTGAGCTTGACTCTAGTCTGGCA | 331.1 | -11.1 | 1.3 | -8.3 | 8.9E-17 | 7.3E-16 | MB231/MB468 | CYTO |
| >hg38_21_+_8433172_8446622_RNA45SN1_45S_50ntFlanks@10510.10534.25\|AATCCGGAGTGGCGGAGATGGGCGC | >hg38_21_+_8433172_8446622_RNA45SN1_45S_50ntFlanks | 10510 | 10534 | 25 | AATCCGGAGTGGCGGAGATGGGCGC | 404.4 | -12.4 | 1.5 | -8.3 | 1.4E-16 | 7.3E-16 | MB231/MB468 | MITO |
| >hg38_21_+_8433172_8446622_RNA45SN1_45S_50ntFlanks@9930.9950.21\|ATATAGACAGCAGGACGGTGG | >hg38_21_+_8433172_8446622_RNA45SN1_45S_50ntFlanks | 9930 | 9950 | 21 | ATATAGACAGCAGGACGGTGG | 283.3 | -12.0 | 1.4 | -8.3 | 8.7E-17 | 7.4E-16 | MB231/MB468 | MP |
| >hg38_21_+_8433172_8446622_RNA45SN1_45S_50ntFlanks@476.511.36\|TTCGTGATCGATGTGGTGACGTCGTGCTCTCCCGGG | >hg38_21_+_8433172_8446622_RNA45SN1_45S_50ntFlanks | 476 | 511 | 36 | TTCGTGATCGATGTGGTGACGTCGTGCTCTCCCGGG | 446.1 | 10.2 | 1.2 | 8.3 | 9.0E-17 | 7.4E-16 | MB231/MB468 | CYTO |
| >hg38_21_+_8433172_8446622_RNA45SN1_45S_50ntFlanks@10410.10428.19\|TCAGTCGGTCCTGAGAGAT | >hg38_21_+_8433172_8446622_RNA45SN1_45S_50ntFlanks | 10410 | 10428 | 19 | TCAGTCGGTCCTGAGAGAT | 473.6 | -12.8 | 1.5 | -8.3 | 8.7E-17 | 7.4E-16 | MB231/MB468 | MP |
| >hg38_21_+_8433172_8446622_RNA45SN1_45S_50ntFlanks@9352.9371.20\|CCGTGGCGCAATGAAGGTGA | >hg38_21_+_8433172_8446622_RNA45SN1_45S_50ntFlanks | 9352 | 9371 | 20 | CCGTGGCGCAATGAAGGTGA | 392.3 | -12.5 | 1.5 | -8.3 | 8.7E-17 | 7.4E-16 | MB231/MB468 | MP |
| >hg38_21_+_8433172_8446622_RNA45SN1_45S_50ntFlanks@12366.12404.39\|TAACTGGCTTGTGGCGGCCAAGCGTTCATAGCGACGTCG | >hg38_21_+_8433172_8446622_RNA45SN1_45S_50ntFlanks | 12366 | 12404 | 39 | TAACTGGCTTGTGGCGGCCAAGCGTTCATAGCGACGTCG | 341.4 | -12.2 | 1.5 | -8.3 | 1.4E-16 | 7.4E-16 | MB231/MB468 | MITO |
| >hg38_21_+_8433172_8446622_RNA45SN1_45S_50ntFlanks@10517.10534.18\|AGTGGCGGAGATGGGCGC | >hg38_21_+_8433172_8446622_RNA45SN1_45S_50ntFlanks | 10517 | 10534 | 18 | AGTGGCGGAGATGGGCGC | 330.5 | -12.2 | 1.5 | -8.3 | 1.4E-16 | 7.4E-16 | MB231/MB468 | MITO |
| >hg38_21_+_8433172_8446622_RNA45SN1_45S_50ntFlanks@13002.13026.25\|CGATCTATTGAAAGTCAGCCCTCGA | >hg38_21_+_8433172_8446622_RNA45SN1_45S_50ntFlanks | 13002 | 13026 | 25 | CGATCTATTGAAAGTCAGCCCTCGA | 370.7 | -11.3 | 1.4 | -8.3 | 9.1E-17 | 7.4E-16 | MB231/MB468 | CYTO |
| >hg38_21_+_8433172_8446622_RNA45SN1_45S_50ntFlanks@3835.3862.28\|CTCCTCTCCTACTTGGATAACTGTGGTA | >hg38_21_+_8433172_8446622_RNA45SN1_45S_50ntFlanks | 3835 | 3862 | 28 | CTCCTCTCCTACTTGGATAACTGTGGTA | 1509.4 | -14.3 | 1.7 | -8.3 | 1.4E-16 | 7.4E-16 | MB231/MB468 | MITO |
| >hg38_21_+_8433172_8446622_RNA45SN1_45S_50ntFlanks@9505.9526.22\|TGAACTATGCCTGGGCAGGGCG | >hg38_21_+_8433172_8446622_RNA45SN1_45S_50ntFlanks | 9505 | 9526 | 22 | TGAACTATGCCTGGGCAGGGCG | 281.2 | -10.9 | 1.3 | -8.3 | 9.1E-17 | 7.4E-16 | MB231/MB468 | CYTO |
| >hg38_21_+_8433172_8446622_RNA45SN1_45S_50ntFlanks@4283.4321.39\|CTTTAACGAGGATCCATTGGAGGGCAAGTCTGGTGCCAG | >hg38_21_+_8433172_8446622_RNA45SN1_45S_50ntFlanks | 4283 | 4321 | 39 | CTTTAACGAGGATCCATTGGAGGGCAAGTCTGGTGCCAG | 540.4 | -12.0 | 1.5 | -8.3 | 1.1E-16 | 7.5E-16 | MB231/MB468 | NUC |
| >hg38_21_+_8433172_8446622_RNA45SN1_45S_50ntFlanks@12686.12703.18\|CCAGGCGGAACGATACGG | >hg38_21_+_8433172_8446622_RNA45SN1_45S_50ntFlanks | 12686 | 12703 | 18 | CCAGGCGGAACGATACGG | 256.7 | -11.9 | 1.4 | -8.3 | 8.8E-17 | 7.5E-16 | MB231/MB468 | MP |
| >hg38_21_+_8433172_8446622_RNA45SN1_45S_50ntFlanks@3874.3899.26\|AATACATGCCGACGGGCGCTGACCCC | >hg38_21_+_8433172_8446622_RNA45SN1_45S_50ntFlanks | 3874 | 3899 | 26 | AATACATGCCGACGGGCGCTGACCCC | 993.1 | -7.6 | 0.9 | -8.3 | 1.4E-16 | 7.5E-16 | MB231/MB468 | MITO |
| >MI0000746\|hsa-mir-99b&WithFlank&19\|+\|51692606\|51692687@13.32.20(+2U)&[MIMAT0000689&hsa-miR-99b-5p&offsets\|0\|-2(+2U);m-10&19\|+\|51692618\|51692639&offsets\|0\|-2(+2U)]\|CACCCGTAGAACCGACCTTGTT | >MI0000746\|hsa-mir-99b&WithFlank&19\|+\|51692606\|51692687 | 13 | 32 | 20(+2U) | CACCCGTAGAACCGACCTTGTT | 355.5 | 12.1 | 1.4 | 8.6 | 1.1E-17 | 7.5E-16 | BT20/MB231 | MITO |
| >hg38_21_+_8433172_8446622_RNA45SN1_45S_50ntFlanks@4367.4391.25\|CAGTTAAAAAGCTCGTAGTTGGATC | >hg38_21_+_8433172_8446622_RNA45SN1_45S_50ntFlanks | 4367 | 4391 | 25 | CAGTTAAAAAGCTCGTAGTTGGATC | 1695.5 | -4.2 | 0.5 | -8.3 | 9.2E-17 | 7.5E-16 | MB231/MB468 | CYTO |
| >MI0000299\|hsa-mir-222&WithFlank&X\|-\|45747009\|45747130@75.98.24(+1A)&[MIMAT0000279&hsa-miR-222-3p&offsets\|0\|+3(+1A);m-140&X\|-\|45747033\|45747056&offsets\|0\|0(+1A)]\|AGCTACATCTGGCTACTGGGTCTCA | >MI0000299\|hsa-mir-222&WithFlank&X\|-\|45747009\|45747130 | 75 | 98 | 24(+1A) | AGCTACATCTGGCTACTGGGTCTCA | 3179.8 | 5.7 | 0.7 | 8.3 | 1.1E-16 | 7.5E-16 | MB231/MB468 | NUC |
| >hg38_21_+_8433172_8446622_RNA45SN1_45S_50ntFlanks@12324.12343.20\|TTGGGTTTTAAGCAGGAGGT | >hg38_21_+_8433172_8446622_RNA45SN1_45S_50ntFlanks | 12324 | 12343 | 20 | TTGGGTTTTAAGCAGGAGGT | 289.9 | -10.9 | 1.3 | -8.3 | 9.2E-17 | 7.5E-16 | MB231/MB468 | CYTO |
| >hg38_21_+_8433172_8446622_RNA45SN1_45S_50ntFlanks@4687.4707.21\|ACGGACCAGAGCGAAAGCATT | >hg38_21_+_8433172_8446622_RNA45SN1_45S_50ntFlanks | 4687 | 4707 | 21 | ACGGACCAGAGCGAAAGCATT | 312.9 | -12.2 | 1.5 | -8.3 | 8.9E-17 | 7.5E-16 | MB231/MB468 | MP |
| >hg38_21_+_8433172_8446622_RNA45SN1_45S_50ntFlanks@12717.12747.31\|CCTCGGTTGGCCTCGGATAGCCGGTCCCCCG | >hg38_21_+_8433172_8446622_RNA45SN1_45S_50ntFlanks | 12717 | 12747 | 31 | CCTCGGTTGGCCTCGGATAGCCGGTCCCCCG | 396.4 | -12.4 | 1.5 | -8.3 | 1.5E-16 | 7.6E-16 | MB231/MB468 | MITO |
| >trna129_MetCAT_6_-_27870271_27870342@32.51.20,trna142_MetCAT_6_-_27560600_27560671@32.51.20,trna150_MetCAT_6_-_27300764_27300835@32.51.20,trna169_MetCAT_6_-_26330529_26330600@32.51.20,trna171_MetCAT_6_-_26313352_26313423@32.51.20,trna20_MetCAT_17_-_80452597_80452668@32.51.20,trna2_MetCAT_6_+_26286754_26286825@32.51.20,trna32_MetCAT_1_+_153643726_153643797@32.51.20,trna61_MetCAT_6_+_27745664_27745735@32.51.20\|CCATAACCCAGAGGTCGATG | >trna129_MetCAT_6_-_27870271_27870342 | 32 | 51 | 20 | CCATAACCCAGAGGTCGATG | 644.4 | -13.2 | 1.6 | -8.3 | 9.0E-17 | 7.6E-16 | MB231/MB468 | MP |
| >MI0000440\|hsa-mir-27b&WithFlank&9\|+\|95085439\|95085547@66.87.22&[MIMAT0000419&hsa-miR-27b-3p&offsets\|-1\|0;m-39&9\|+\|95085505\|95085525&offsets\|-1\|0]\|GTTCACAGTGGCTAAGTTCTGC | >MI0000440\|hsa-mir-27b&WithFlank&9\|+\|95085439\|95085547 | 66 | 87 | 22 | GTTCACAGTGGCTAAGTTCTGC | 138.0 | 3.3 | 0.4 | 8.2 | 2.4E-16 | 7.6E-16 | BT20/MB468 | NUC |
| >hg38_21_+_8433172_8446622_RNA45SN1_45S_50ntFlanks@11849.11873.25\|CGGGCTTGGCGGAATCAGCGGGGAA | >hg38_21_+_8433172_8446622_RNA45SN1_45S_50ntFlanks | 11849 | 11873 | 25 | CGGGCTTGGCGGAATCAGCGGGGAA | 356.1 | -12.3 | 1.5 | -8.3 | 1.5E-16 | 7.6E-16 | MB231/MB468 | MITO |
| >hg38_21_+_8433172_8446622_RNA45SN1_45S_50ntFlanks@7979.8007.29\|ACCTCAGATCAGACGTGGCGACCCGCTGA | >hg38_21_+_8433172_8446622_RNA45SN1_45S_50ntFlanks | 7979 | 8007 | 29 | ACCTCAGATCAGACGTGGCGACCCGCTGA | 959.4 | -5.5 | 0.7 | -8.3 | 1.5E-16 | 7.6E-16 | MB231/MB468 | MITO |
| >hg38_21_+_8433172_8446622_RNA45SN1_45S_50ntFlanks@9033.9075.43\|CTCGTGGGGGGGCCGGGCCACCCCTCCCACGGCGCGACCGCTC | >hg38_21_+_8433172_8446622_RNA45SN1_45S_50ntFlanks | 9033 | 9075 | 43 | CTCGTGGGGGGGCCGGGCCACCCCTCCCACGGCGCGACCGCTC | 681.0 | -13.2 | 1.6 | -8.3 | 1.5E-16 | 7.6E-16 | MB231/MB468 | MITO |
| >hg38_21_+_8433172_8446622_RNA45SN1_45S_50ntFlanks@5476.5516.41\|CCCTGGCGGAGCGCTGAGAAGACGGTCGAACTTGACTATCT | >hg38_21_+_8433172_8446622_RNA45SN1_45S_50ntFlanks | 5476 | 5516 | 41 | CCCTGGCGGAGCGCTGAGAAGACGGTCGAACTTGACTATCT | 1947.1 | 11.0 | 1.3 | 8.3 | 1.5E-16 | 7.6E-16 | BT20/MB468 | CYTO |
| >hg38_21_+_8433172_8446622_RNA45SN1_45S_50ntFlanks@10127.10147.21\|GCGGCGGCGGCGGCGGCGGCG | >hg38_21_+_8433172_8446622_RNA45SN1_45S_50ntFlanks | 10127 | 10147 | 21 | GCGGCGGCGGCGGCGGCGGCG | 507.6 | -12.8 | 1.5 | -8.3 | 1.5E-16 | 7.6E-16 | MB231/MB468 | MITO |
| >hg38_21_+_8433172_8446622_RNA45SN1_45S_50ntFlanks@8844.8864.21\|GCGGCGGCGGCGGCGGCGGCG | >hg38_21_+_8433172_8446622_RNA45SN1_45S_50ntFlanks | 8844 | 8864 | 21 | GCGGCGGCGGCGGCGGCGGCG | 507.6 | -12.8 | 1.5 | -8.3 | 1.5E-16 | 7.6E-16 | MB231/MB468 | MITO |
| >hg38_21_+_8433172_8446622_RNA45SN1_45S_50ntFlanks@11918.11951.34\|ACATGAGAGGTGTAGAATAAGTGGGAGGCCCCCG | >hg38_21_+_8433172_8446622_RNA45SN1_45S_50ntFlanks | 11918 | 11951 | 34 | ACATGAGAGGTGTAGAATAAGTGGGAGGCCCCCG | 465.4 | -12.6 | 1.5 | -8.3 | 1.5E-16 | 7.7E-16 | MB231/MB468 | MITO |
| >trna129_MetCAT_6_-_27870271_27870342@31.51.21,trna142_MetCAT_6_-_27560600_27560671@31.51.21,trna150_MetCAT_6_-_27300764_27300835@31.51.21,trna169_MetCAT_6_-_26330529_26330600@31.51.21,trna171_MetCAT_6_-_26313352_26313423@31.51.21,trna20_MetCAT_17_-_80452597_80452668@31.51.21,trna2_MetCAT_6_+_26286754_26286825@31.51.21,trna32_MetCAT_1_+_153643726_153643797@31.51.21,trna61_MetCAT_6_+_27745664_27745735@31.51.21\|CCCATAACCCAGAGGTCGATG | >trna129_MetCAT_6_-_27870271_27870342 | 31 | 51 | 21 | CCCATAACCCAGAGGTCGATG | 580.9 | -13.0 | 1.6 | -8.3 | 1.5E-16 | 7.7E-16 | MB231/MB468 | MITO |
| >hg38_21_+_8433172_8446622_RNA45SN1_45S_50ntFlanks@11885.11912.28\|TTGAGCTTGACTCTAGTCTGGCACGGTG | >hg38_21_+_8433172_8446622_RNA45SN1_45S_50ntFlanks | 11885 | 11912 | 28 | TTGAGCTTGACTCTAGTCTGGCACGGTG | 350.6 | -12.2 | 1.5 | -8.3 | 1.5E-16 | 7.7E-16 | MB231/MB468 | MITO |

<table>
<tr><td></td><td></td><td></td><td></td><td></td><td></td><td></td><td></td><td></td><td></td><td></td><td></td><td></td><td></td></tr>
<tr><td>>hg38_21_+_8433172_8446622_RNA45SN1_45S_50ntFlanks@4588.4615.28|CTATTTTGTTGGTTTTCGGAACTGAGGC</td><td>>hg38_21_+_8433172_8446622_RNA45SN1_45S_50ntFlanks</td><td>4588</td><td>4615</td><td>28</td><td>CTATTTTGTTGGTTTTCGGAACTGAGGC</td><td>323.1</td><td>-12.2</td><td>1.5</td><td>-8.3</td><td>9.1E-17</td><td>7.7E-16</td><td>MB231/MB468</td><td>MP</td></tr>
<tr><td>>MI0006444|hsa-mir-1307&WithFlank&10|-|103394247|103394407@86.106.21&[MIMAT0005951&hsa-miR-1307-3p&offsets|0|-1;m-66&10|-|103394300|103394322&offsets|0|-2]|ACTCGGCGTGGCGTCGGTCGT</td><td>>MI0006444|hsa-mir-1307&WithFlank&10|-|103394247|103394407</td><td>86</td><td>106</td><td>21</td><td>ACTCGGCGTGGCGTCGGTCGT</td><td>486.1</td><td>9.6</td><td>1.2</td><td>8.3</td><td>1.5E-16</td><td>7.7E-16</td><td>BT20/MB468</td><td>CYTO</td></tr>
<tr><td>>hg38_21_+_8433172_8446622_RNA45SN1_45S_50ntFlanks@10687.10709.23|CGGCGTCCGGTGAGCTCTCGCTG</td><td>>hg38_21_+_8433172_8446622_RNA45SN1_45S_50ntFlanks</td><td>10687</td><td>10709</td><td>23</td><td>CGGCGTCCGGTGAGCTCTCGCTG</td><td>3376.3</td><td>-7.8</td><td>0.9</td><td>-8.3</td><td>9.5E-17</td><td>7.7E-16</td><td>MB231/MB468</td><td>CYTO</td></tr>
<tr><td>>hg38_21_+_8433172_8446622_RNA45SN1_45S_50ntFlanks@12619.12653.35|AGGAGCCAATGGGGCGAAGCTACCATCTGTGGGAT</td><td>>hg38_21_+_8433172_8446622_RNA45SN1_45S_50ntFlanks</td><td>12619</td><td>12653</td><td>35</td><td>AGGAGCCAATGGGGCGAAGCTACCATCTGTGGGAT</td><td>1630.8</td><td>-10.9</td><td>1.3</td><td>-8.2</td><td>2.4E-16</td><td>7.8E-16</td><td>BT20/MB468</td><td>NUC</td></tr>
<tr><td>>hg38_1_-_228634819_228635039_RNA5S12_5S_50ntFlanks@124.147.24|AGTACTTGGATGGGAGACCGCCTG</td><td>>hg38_1_-_228634819_228635039_RNA5S12_5S_50ntFlanks</td><td>124</td><td>147</td><td>24</td><td>AGTACTTGGATGGGAGACCGCCTG</td><td>310.7</td><td>-12.1</td><td>1.5</td><td>-8.3</td><td>1.5E-16</td><td>7.8E-16</td><td>MB231/MB468</td><td>MITO</td></tr>
<tr><td>>hg38_21_+_8433172_8446622_RNA45SN1_45S_50ntFlanks@11873.11893.21|AAGAAGACCCTGTTGAGCTTG</td><td>>hg38_21_+_8433172_8446622_RNA45SN1_45S_50ntFlanks</td><td>11873</td><td>11893</td><td>21</td><td>AAGAAGACCCTGTTGAGCTTG</td><td>4609.6</td><td>-6.7</td><td>0.8</td><td>-8.3</td><td>9.6E-17</td><td>7.8E-16</td><td>MB231/MB468</td><td>CYTO</td></tr>
<tr><td>>hg38_21_+_8433172_8446622_RNA45SN1_45S_50ntFlanks@11663.11685.23|ATTTCTGCCCAGTGCTCTGAATG</td><td>>hg38_21_+_8433172_8446622_RNA45SN1_45S_50ntFlanks</td><td>11663</td><td>11685</td><td>23</td><td>ATTTCTGCCCAGTGCTCTGAATG</td><td>584.3</td><td>-3.6</td><td>0.4</td><td>-8.3</td><td>1.5E-16</td><td>7.8E-16</td><td>BT20/MB468</td><td>CYTO</td></tr>
<tr><td>>hg38_21_+_8433172_8446622_RNA45SN1_45S_50ntFlanks@12456.12479.24|CCAAGCGTTGGATTGTTCACCCAC</td><td>>hg38_21_+_8433172_8446622_RNA45SN1_45S_50ntFlanks</td><td>12456</td><td>12479</td><td>24</td><td>CCAAGCGTTGGATTGTTCACCCAC</td><td>875.4</td><td>-7.6</td><td>0.9</td><td>-8.3</td><td>1.5E-16</td><td>7.8E-16</td><td>MB231/MB468</td><td>MITO</td></tr>
<tr><td>>hg38_21_+_8433172_8446622_RNA45SN1_45S_50ntFlanks@10689.10709.21|GCGTCCGGTGAGCTCTCGCTG</td><td>>hg38_21_+_8433172_8446622_RNA45SN1_45S_50ntFlanks</td><td>10689</td><td>10709</td><td>21</td><td>GCGTCCGGTGAGCTCTCGCTG</td><td>267.5</td><td>-11.9</td><td>1.4</td><td>-8.3</td><td>9.3E-17</td><td>7.9E-16</td><td>MB231/MB468</td><td>MP</td></tr>
<tr><td>>hg38_21_+_8433172_8446622_RNA45SN1_45S_50ntFlanks@5084.5105.22|CGAACGAGACTCTGGCATGCTA</td><td>>hg38_21_+_8433172_8446622_RNA45SN1_45S_50ntFlanks</td><td>5084</td><td>5105</td><td>22</td><td>CGAACGAGACTCTGGCATGCTA</td><td>1115.5</td><td>-14.0</td><td>1.7</td><td>-8.3</td><td>9.3E-17</td><td>7.9E-16</td><td>MB231/MB468</td><td>MP</td></tr>
<tr><td>>MI0000809|hsa-mir-151a&WithFlank&8|-|140732558|140732659@53.74.22(+2A)&[MIMAT0000757&hsa-miR-151a-3p&offsets|0|+1(+2A);m-30&8|-|140732587|140732607&offsets|0|+1(+2A)]|CTAGACTGAAGCTCCTTGAGGAAA</td><td>>MI0000809|hsa-mir-151a&WithFlank&8|-|140732558|140732659</td><td>53</td><td>74</td><td>22(+2A)</td><td>CTAGACTGAAGCTCCTTGAGGAAA</td><td>2056.7</td><td>8.4</td><td>1.0</td><td>8.3</td><td>1.5E-16</td><td>7.9E-16</td><td>BT20/MB468</td><td>CYTO</td></tr>
<tr><td>>hg38_21_+_8433172_8446622_RNA45SN1_45S_50ntFlanks@6734.6753.20|ATTGATCATCGACACTTCGA</td><td>>hg38_21_+_8433172_8446622_RNA45SN1_45S_50ntFlanks</td><td>6734</td><td>6753</td><td>20</td><td>ATTGATCATCGACACTTCGA</td><td>1048.5</td><td>4.8</td><td>0.6</td><td>8.3</td><td>1.1E-16</td><td>7.9E-16</td><td>BT20/MB468</td><td>MITO</td></tr>
<tr><td>>hg38_21_+_8433172_8446622_RNA45SN1_45S_50ntFlanks@12158.12179.22|GTTTGACTGGGGCGGTACACCT</td><td>>hg38_21_+_8433172_8446622_RNA45SN1_45S_50ntFlanks</td><td>12158</td><td>12179</td><td>22</td><td>GTTTGACTGGGGCGGTACACCT</td><td>449.5</td><td>-12.7</td><td>1.5</td><td>-8.3</td><td>9.3E-17</td><td>7.9E-16</td><td>MB231/MB468</td><td>MP</td></tr>
<tr><td>>hg38_21_+_8433172_8446622_RNA45SN1_45S_50ntFlanks@4220.4241.22|ACGAAAAATAACAATACAGGAC</td><td>>hg38_21_+_8433172_8446622_RNA45SN1_45S_50ntFlanks</td><td>4220</td><td>4241</td><td>22</td><td>ACGAAAAATAACAATACAGGAC</td><td>215.9</td><td>-11.6</td><td>1.4</td><td>-8.3</td><td>9.4E-17</td><td>7.9E-16</td><td>MB231/MB468</td><td>MP</td></tr>
<tr><td>>hg38_21_+_8433172_8446622_RNA45SN1_45S_50ntFlanks@10687.10711.25|CGGCGTCCGGTGAGCTCTCGCTGGC</td><td>>hg38_21_+_8433172_8446622_RNA45SN1_45S_50ntFlanks</td><td>10687</td><td>10711</td><td>25</td><td>CGGCGTCCGGTGAGCTCTCGCTGGC</td><td>5293.8</td><td>-5.3</td><td>0.6</td><td>-8.3</td><td>1.5E-16</td><td>7.9E-16</td><td>BT20/MB468</td><td>CYTO</td></tr>
<tr><td>>MI0000283|hsa-mir-203a&WithFlank&14|+|104117399|104117520@71.93.23&[MIMAT0000264&hsa-miR-203a-3p&offsets|0|+1;m-79&14|+|104117469|104117490&offsets|0|+1]|GTGAAATGTTTAGGACCACTAGA</td><td>>MI0000283|hsa-mir-203a&WithFlank&14|+|104117399|104117520</td><td>71</td><td>93</td><td>23</td><td>GTGAAATGTTTAGGACCACTAGA</td><td>427.6</td><td>12.1</td><td>1.4</td><td>8.6</td><td>8.2E-18</td><td>7.9E-16</td><td>BT20/MB231</td><td>CYTO</td></tr>
<tr><td>>hg38_21_+_8433172_8446622_RNA45SN1_45S_50ntFlanks@10574.10591.18|AAGCCGGCGGGAGCCCCG</td><td>>hg38_21_+_8433172_8446622_RNA45SN1_45S_50ntFlanks</td><td>10574</td><td>10591</td><td>18</td><td>AAGCCGGCGGGAGCCCCG</td><td>485.1</td><td>-11.6</td><td>1.4</td><td>-8.3</td><td>9.8E-17</td><td>7.9E-16</td><td>MB231/MB468</td><td>CYTO</td></tr>
<tr><td>>am_trna11_ProAGG_16_+_3241989_3242060@1.28.28,trna12_ProAGG_6_+_26555498_26555569@1.28.28,trna12_ProTGG_11_-_75946869_75946940@1.28.28,trna14_ProTGG_5_-_180615854_180615925@1.28.28,trna22_ProAGG_14_-_21081560_21081631@1.28.28,trna23_ProAGG_14_-_21077495_21077566@1.28.28,trna28_ProTGG_16_-_3234133_3234204@1.28.28,trna29_ProAGG_16_-_3232635_3232706@1.28.28,trna2_ProAGG_7_+_128423504_128423575@1.28.28,trna30_ProCGG_6_+_27059521_27059592@1.28.28,trna37_ProCGG_17_-_8126151_8126222@1.28.28,trna3_ProTGG_16_+_3208923_3208994@1.28.28,trna52_ProCGG_1_+_167683962_167684033@1.28.28,trna65_ProAGG_1_-_167684725_167684796@1.28.28,trna6_ProCGG_16_+_3222049_3222120@1.28.28,trna6_ProTGG_14_+_21152175_21152246@1.28.28,trna8_ProTGG_16_+_3238094_3238165@1.28.28,trna9_ProAGG_11_+_75946557_75946628@1.28.28,trna9_ProAGG_16_+_3239634_3239705@1.28.28|GGCTCGTTGGTCTAGGGGTATGATTCTC</td><td>>am_trna11_ProAGG_16_+_3241989_3242060</td><td>1</td><td>28</td><td>28</td><td>GGCTCGTTGGTCTAGGGGTATGATTCTC</td><td>517.7</td><td>-12.8</td><td>1.6</td><td>-8.3</td><td>1.5E-16</td><td>8.0E-16</td><td>MB231/MB468</td><td>MITO</td></tr>
<tr><td>>hg38_21_+_8433172_8446622_RNA45SN1_45S_50ntFlanks@11885.11903.19|TTGAGCTTGACTCTAGTCT</td><td>>hg38_21_+_8433172_8446622_RNA45SN1_45S_50ntFlanks</td><td>11885</td><td>11903</td><td>19</td><td>TTGAGCTTGACTCTAGTCT</td><td>486.1</td><td>-12.7</td><td>1.5</td><td>-8.3</td><td>1.6E-16</td><td>8.0E-16</td><td>MB231/MB468</td><td>MITO</td></tr>
<tr><td>>trnaMT_PheGAA_MT_+_577_647@35.56.22|GAAAATGTTTAGACGGGCTCAC</td><td>>trnaMT_PheGAA_MT_+_577_647</td><td>35</td><td>56</td><td>22</td><td>GAAAATGTTTAGACGGGCTCAC</td><td>3019.8</td><td>8.3</td><td>1.0</td><td>8.3</td><td>1.1E-16</td><td>8.0E-16</td><td>BT20/MB468</td><td>MITO</td></tr>
<tr><td>>hg38_21_+_8433172_8446622_RNA45SN1_45S_50ntFlanks@5073.5098.26|AATTCCGATAACGAACGAGACTCTGG</td><td>>hg38_21_+_8433172_8446622_RNA45SN1_45S_50ntFlanks</td><td>5073</td><td>5098</td><td>26</td><td>AATTCCGATAACGAACGAGACTCTGG</td><td>5258.3</td><td>-16.2</td><td>2.0</td><td>-8.3</td><td>9.5E-17</td><td>8.0E-16</td><td>MB231/MB468</td><td>MP</td></tr>
<tr><td>>hg38_21_+_8433172_8446622_RNA45SN1_45S_50ntFlanks@12997.13017.21|CGCTGCGATCTATTGAAAGTC</td><td>>hg38_21_+_8433172_8446622_RNA45SN1_45S_50ntFlanks</td><td>12997</td><td>13017</td><td>21</td><td>CGCTGCGATCTATTGAAAGTC</td><td>2489.4</td><td>6.9</td><td>0.8</td><td>8.3</td><td>1.6E-16</td><td>8.0E-16</td><td>BT20/MB468</td><td>CYTO</td></tr>
<tr><td>>hg38_21_+_8433172_8446622_RNA45SN1_45S_50ntFlanks@3922.3959.38|TTTATCAGATCAAAACCAACCCGGTCAGCCCCTCTCCG</td><td>>hg38_21_+_8433172_8446622_RNA45SN1_45S_50ntFlanks</td><td>3922</td><td>3959</td><td>38</td><td>TTTATCAGATCAAAACCAACCCGGTCAGCCCCTCTCCG</td><td>391.5</td><td>-12.4</td><td>1.5</td><td>-8.3</td><td>1.6E-16</td><td>8.1E-16</td><td>MB231/MB468</td><td>MITO</td></tr>
<tr><td>>hg38_21_+_8433172_8446622_RNA45SN1_45S_50ntFlanks@12710.12745.36|CGCGGAGCCTCGGTTGGCCTCGGATAGCCGGTCCCC</td><td>>hg38_21_+_8433172_8446622_RNA45SN1_45S_50ntFlanks</td><td>12710</td><td>12745</td><td>36</td><td>CGCGGAGCCTCGGTTGGCCTCGGATAGCCGGTCCCC</td><td>397.2</td><td>-12.4</td><td>1.5</td><td>-8.3</td><td>1.6E-16</td><td>8.1E-16</td><td>MB231/MB468</td><td>MITO</td></tr>
<tr><td>>hg38_21_+_8433172_8446622_RNA45SN1_45S_50ntFlanks@12630.12654.25|GGGCGAAGCTACCATCTGTGGGATT</td><td>>hg38_21_+_8433172_8446622_RNA45SN1_45S_50ntFlanks</td><td>12630</td><td>12654</td><td>25</td><td>GGGCGAAGCTACCATCTGTGGGATT</td><td>400.5</td><td>-12.5</td><td>1.5</td><td>-8.3</td><td>9.7E-17</td><td>8.1E-16</td><td>MB231/MB468</td><td>MP</td></tr>
<tr><td>>hg38_21_+_8433172_8446622_RNA45SN1_45S_50ntFlanks@4367.4393.27|CAGTTAAAAAGCTCGTAGTTGGATCTT</td><td>>hg38_21_+_8433172_8446622_RNA45SN1_45S_50ntFlanks</td><td>4367</td><td>4393</td><td>27</td><td>CAGTTAAAAAGCTCGTAGTTGGATCTT</td><td>246.8</td><td>-12.4</td><td>1.5</td><td>-8.2</td><td>2.6E-16</td><td>8.1E-16</td><td>BT20/MB468</td><td>NUC</td></tr>
<tr><td>>MI0000266|hsa-mir-10a&WithFlank&17|-|48579832|48579953@69.89.21&[MIMAT0004555&hsa-miR-10a-3p&offsets|0|-1;m-468&17|-|48579865|48579885&offsets|0|0]|CAAATTCGTATCTAGGGGAAT</td><td>>MI0000266|hsa-mir-10a&WithFlank&17|-|48579832|48579953</td><td>69</td><td>89</td><td>21</td><td>CAAATTCGTATCTAGGGGAAT</td><td>172.4</td><td>11.3</td><td>1.4</td><td>8.3</td><td>1.2E-16</td><td>8.2E-16</td><td>MB231/MB468</td><td>NUC</td></tr>
<tr><td>>MI0000433|hsa-let-7g&WithFlank&3|-|52268272|52268367@11.32.22(+1A)&[MIMAT0000414&hsa-let-7g-5p&offsets|0|0(+1A);m-58&3|-|52268336|52268357&offsets|0|0(+1A)]|TGAGGTAGTAGTTTGTACAGTTA</td><td>>MI0000433|hsa-let-7g&WithFlank&3|-|52268272|52268367</td><td>11</td><td>32</td><td>22(+1A)</td><td>TGAGGTAGTAGTTTGTACAGTTA</td><td>2772.0</td><td>6.9</td><td>0.8</td><td>8.2</td><td>1.6E-16</td><td>8.2E-16</td><td>BT20/MB468</td><td>CYTO</td></tr>
<tr><td>>hg38_21_+_8433172_8446622_RNA45SN1_45S_50ntFlanks@5227.5244.18|CGCGCGCTACACTGACTG</td><td>>hg38_21_+_8433172_8446622_RNA45SN1_45S_50ntFlanks</td><td>5227</td><td>5244</td><td>18</td><td>CGCGCGCTACACTGACTG</td><td>3778.8</td><td>-15.7</td><td>1.9</td><td>-8.2</td><td>1.6E-16</td><td>8.2E-16</td><td>MB231/MB468</td><td>MITO</td></tr>
<tr><td>>hg38_21_+_8433172_8446622_RNA45SN1_45S_50ntFlanks@9604.9622.19|AAAGACTAATCGAACCATC</td><td>>hg38_21_+_8433172_8446622_RNA45SN1_45S_50ntFlanks</td><td>9604</td><td>9622</td><td>19</td><td>AAAGACTAATCGAACCATC</td><td>1105.1</td><td>-5.9</td><td>0.7</td><td>-8.2</td><td>1.6E-16</td><td>8.2E-16</td><td>MB231/MB468</td><td>MITO</td></tr>
<tr><td>>hg38_21_+_8433172_8446622_RNA45SN1_45S_50ntFlanks@12928.12953.26|CTGGCGCTAAACCATTCGTAGACGAC</td><td>>hg38_21_+_8433172_8446622_RNA45SN1_45S_50ntFlanks</td><td>12928</td><td>12953</td><td>26</td><td>CTGGCGCTAAACCATTCGTAGACGAC</td><td>338.4</td><td>-12.2</td><td>1.5</td><td>-8.2</td><td>1.6E-16</td><td>8.2E-16</td><td>MB231/MB468</td><td>MITO</td></tr>
<tr><td>>hg38_21_+_8433172_8446622_RNA45SN1_45S_50ntFlanks@6753.6770.18|AACGCACTTGCGGCCCCG</td><td>>hg38_21_+_8433172_8446622_RNA45SN1_45S_50ntFlanks</td><td>6753</td><td>6770</td><td>18</td><td>AACGCACTTGCGGCCCCG</td><td>377.9</td><td>-12.4</td><td>1.5</td><td>-8.3</td><td>9.8E-17</td><td>8.3E-16</td><td>MB231/MB468</td><td>MP</td></tr>
<tr><td>>hg38_21_+_8433172_8446622_RNA45SN1_45S_50ntFlanks@10678.10705.28|CGGTTCCGGCGGCGTCCGGTGAGCTCTC</td><td>>hg38_21_+_8433172_8446622_RNA45SN1_45S_50ntFlanks</td><td>10678</td><td>10705</td><td>28</td><td>CGGTTCCGGCGGCGTCCGGTGAGCTCTC</td><td>1105.2</td><td>-9.8</td><td>1.2</td><td>-8.2</td><td>1.6E-16</td><td>8.3E-16</td><td>MB231/MB468</td><td>MITO</td></tr>
<tr><td>>hg38_21_+_8433172_8446622_RNA45SN1_45S_50ntFlanks@11200.11223.24|CGGCGGCGGCGGCGACTCTGGACG</td><td>>hg38_21_+_8433172_8446622_RNA45SN1_45S_50ntFlanks</td><td>11200</td><td>11223</td><td>24</td><td>CGGCGGCGGCGGCGACTCTGGACG</td><td>409.8</td><td>-12.5</td><td>1.5</td><td>-8.3</td><td>9.9E-17</td><td>8.3E-16</td><td>MB231/MB468</td><td>MP</td></tr>
<tr><td>>hg38_21_+_8433172_8446622_RNA45SN1_45S_50ntFlanks@11209.11254.46|CGGCGACTCTGGACGCGAGCCGGGCCCTTCCCGTGGATCGCCCCAG</td><td>>hg38_21_+_8433172_8446622_RNA45SN1_45S_50ntFlanks</td><td>11209</td><td>11254</td><td>46</td><td>CGGCGACTCTGGACGCGAGCCGGGCCCTTCCCGTGGATCGCCCCAG</td><td>1582.6</td><td>-14.5</td><td>1.7</td><td>-8.3</td><td>9.9E-17</td><td>8.3E-16</td><td>MB231/MB468</td><td>MP</td></tr>
<tr><td>>hg38_21_+_8433172_8446622_RNA45SN1_45S_50ntFlanks@6734.6770.37|ATTGATCATCGACACTTCGAACGCACTTGCGGCCCCG</td><td>>hg38_21_+_8433172_8446622_RNA45SN1_45S_50ntFlanks</td><td>6734</td><td>6770</td><td>37</td><td>ATTGATCATCGACACTTCGAACGCACTTGCGGCCCCG</td><td>2316.4</td><td>-6.2</td><td>0.8</td><td>-8.2</td><td>2.6E-16</td><td>8.3E-16</td><td>BT20/MB468</td><td>NUC</td></tr>
<tr><td>>hg38_21_+_8433172_8446622_RNA45SN1_45S_50ntFlanks@12991.13017.27|CTCCCTCGCTGCGATCTATTGAAAGTC</td><td>>hg38_21_+_8433172_8446622_RNA45SN1_45S_50ntFlanks</td><td>12991</td><td>13017</td><td>27</td><td>CTCCCTCGCTGCGATCTATTGAAAGTC</td><td>3274.8</td><td>-11.5</td><td>1.4</td><td>-8.2</td><td>2.6E-16</td><td>8.4E-16</td><td>BT20/MB468</td><td>NUC</td></tr>
<tr><td>>hg38_21_+_8433172_8446622_RNA45SN1_45S_50ntFlanks@11778.11798.21|ACGCGCATGAATGGATGAACG</td><td>>hg38_21_+_8433172_8446622_RNA45SN1_45S_50ntFlanks</td><td>11778</td><td>11798</td><td>21</td><td>ACGCGCATGAATGGATGAACG</td><td>336.7</td><td>-12.3</td><td>1.5</td><td>-8.3</td><td>9.9E-17</td><td>8.4E-16</td><td>MB231/MB468</td><td>MP</td></tr>
<tr><td>>hg38_21_+_8433172_8446622_RNA45SN1_45S_50ntFlanks@11619.11640.22|AACAAAGCATCGCGAAGGCCCG</td><td>>hg38_21_+_8433172_8446622_RNA45SN1_45S_50ntFlanks</td><td>11619</td><td>11640</td><td>22</td><td>AACAAAGCATCGCGAAGGCCCG</td><td>860.5</td><td>-11.8</td><td>1.4</td><td>-8.3</td><td>1.2E-16</td><td>8.4E-16</td><td>BT20/MB468</td><td>MITO</td></tr>
<tr><td>>hg38_21_+_8433172_8446622_RNA45SN1_45S_50ntFlanks@3785.3812.28|TGAAACTGCGAATGGCTCATTAAATCAG</td><td>>hg38_21_+_8433172_8446622_RNA45SN1_45S_50ntFlanks</td><td>3785</td><td>3812</td><td>28</td><td>TGAAACTGCGAATGGCTCATTAAATCAG</td><td>981.4</td><td>-13.7</td><td>1.7</td><td>-8.2</td><td>1.7E-16</td><td>8.4E-16</td><td>MB231/MB468</td><td>MITO</td></tr>
<tr><td>>hg38_21_+_8433172_8446622_RNA45SN1_45S_50ntFlanks@11792.11810.19|ATGAACGAGATTCCCACTG</td><td>>hg38_21_+_8433172_8446622_RNA45SN1_45S_50ntFlanks</td><td>11792</td><td>11810</td><td>19</td><td>ATGAACGAGATTCCCACTG</td><td>593.2</td><td>-13.1</td><td>1.6</td><td>-8.3</td><td>1.0E-16</td><td>8.5E-16</td><td>MB231/MB468</td><td>MP</td></tr>
<tr><td>>MI0000087|hsa-mir-29a&WithFlank&7|-|130876741|130876816@48.68.21&[MIMAT0000086&hsa-miR-29a-3p&offsets|0|-1;m-26&7|-|130876749|130876769&offsets|0|0]|TAGCACCATCTGAAATCGGTT</td><td>>MI0000087|hsa-mir-29a&WithFlank&7|-|130876741|130876816</td><td>48</td><td>68</td><td>21</td><td>TAGCACCATCTGAAATCGGTT</td><td>14090.3</td><td>8.6</td><td>1.0</td><td>8.2</td><td>1.7E-16</td><td>8.5E-16</td><td>MB231/MB468</td><td>MITO</td></tr>
<tr><td>>hg38_21_+_8433172_8446622_RNA45SN1_45S_50ntFlanks@4443.4472.30|CCCTTGCCTCTCGGCGCCCCCTCGATGCTC</td><td>>hg38_21_+_8433172_8446622_RNA45SN1_45S_50ntFlanks</td><td>4443</td><td>4472</td><td>30</td><td>CCCTTGCCTCTCGGCGCCCCCTCGATGCTC</td><td>542.8</td><td>-11.2</td><td>1.3</td><td>-8.3</td><td>1.2E-16</td><td>8.5E-16</td><td>BT20/MB468</td><td>MITO</td></tr>
<tr><td>>hg38_21_+_8433172_8446622_RNA45SN1_45S_50ntFlanks@12081.12107.27|CTCTCGCTTCTGGCGCCAAGCGCCCGG</td><td>>hg38_21_+_8433172_8446622_RNA45SN1_45S_50ntFlanks</td><td>12081</td><td>12107</td><td>27</td><td>CTCTCGCTTCTGGCGCCAAGCGCCCGG</td><td>1129.8</td><td>-7.7</td><td>0.9</td><td>-8.2</td><td>1.7E-16</td><td>8.5E-16</td><td>MB231/MB468</td><td>MITO</td></tr>
<tr><td>>hg38_21_+_8433172_8446622_RNA45SN1_45S_50ntFlanks@4995.5013.19|ACAGATTGATAGCTCTTTC</td><td>>hg38_21_+_8433172_8446622_RNA45SN1_45S_50ntFlanks</td><td>4995</td><td>5013</td><td>19</td><td>ACAGATTGATAGCTCTTTC</td><td>1389.2</td><td>-14.2</td><td>1.7</td><td>-8.2</td><td>1.7E-16</td><td>8.5E-16</td><td>MB231/MB468</td><td>MITO</td></tr>
<tr><td>>hg38_21_+_8433172_8446622_RNA45SN1_45S_50ntFlanks@6656.6681.26|CTTAGCGGTGGATCACTCGGCTCGTG</td><td>>hg38_21_+_8433172_8446622_RNA45SN1_45S_50ntFlanks</td><td>6656</td><td>6681</td><td>26</td><td>CTTAGCGGTGGATCACTCGGCTCGTG</td><td>5900.5</td><td>-7.0</td><td>0.9</td><td>-8.2</td><td>1.7E-16</td><td>8.6E-16</td><td>MB231/MB468</td><td>MITO</td></tr>
<tr><td>>hg38_21_+_8433172_8446622_RNA45SN1_45S_50ntFlanks@4512.4537.26|AAAAAATTAGAGTGTTCAAAGCAGGC</td><td>>hg38_21_+_8433172_8446622_RNA45SN1_45S_50ntFlanks</td><td>4512</td><td>4537</td><td>26</td><td>AAAAAATTAGAGTGTTCAAAGCAGGC</td><td>239.1</td><td>-10.9</td><td>1.3</td><td>-8.3</td><td>1.3E-16</td><td>8.6E-16</td><td>MB231/MB468</td><td>NUC</td></tr>
<tr><td>>hg38_21_+_8433172_8446622_RNA45SN1_45S_50ntFlanks@11886.11903.18|TGAGCTTGACTCTAGTCT</td><td>>hg38_21_+_8433172_8446622_RNA45SN1_45S_50ntFlanks</td><td>11886</td><td>11903</td><td>18</td><td>TGAGCTTGACTCTAGTCT</td><td>240.1</td><td>-11.8</td><td>1.4</td><td>-8.3</td><td>1.0E-16</td><td>8.6E-16</td><td>MB231/MB468</td><td>MP</td></tr>
</table>

| | | | | | | | | | | | | | |
|---|---|---|---|---|---|---|---|---|---|---|---|---|---|
| >hg38_21_+_8433172_8446622_RNA45SN1_45S_50ntFlanks@8109.8143.35\|GCGGGGCGCGGGACATGTGGCGTACGGAAGACCCG | >hg38_21_+_8433172_8446622_RNA45SN1_45S_50ntFlanks | 8109 | 8143 | 35 | GCGGGGCGCGGGACATGTGGCGTACGGAAGACCCG | 380.9 | -12.4 | 1.5 | -8.2 | 1.7E-16 | 8.6E-16 | MB231/MB468 | MITO |
| >hg38_21_+_8433172_8446622_RNA45SN1_45S_50ntFlanks@9027.9063.37\|CGTCTCCTCGTGGGGGGGCCGGGCCACCCCTCCCACG | >hg38_21_+_8433172_8446622_RNA45SN1_45S_50ntFlanks | 9027 | 9063 | 37 | CGTCTCCTCGTGGGGGGGCCGGGCCACCCCTCCCACG | 676.0 | -13.2 | 1.6 | -8.2 | 1.7E-16 | 8.6E-16 | MB231/MB468 | MITO |
| >MI0000089\|hsa-mir-31&WithFlank&9\|-\|21512109\|21512191@14.34.21&[MIMAT0000089&hsa-miR-31-5p&offsets\|0\|0;m-360&9\|-\|21512157\|21512178&offsets\|0\|-1]\|AGGCAAGATGCTGGCATAGCT | >MI0000089\|hsa-mir-31&WithFlank&9\|-\|21512109\|21512191 | 14 | 34 | 21 | AGGCAAGATGCTGGCATAGCT | 522.0 | 12.8 | 1.5 | 8.5 | 1.9E-17 | 8.7E-16 | BT20/MB231 | NUC |
| >hg38_21_+_8433172_8446622_RNA45SN1_45S_50ntFlanks@5485.5521.37\|AGCGCTGAGAAGACGGTCGAACTTGACTATCTAGAGG | >hg38_21_+_8433172_8446622_RNA45SN1_45S_50ntFlanks | 5485 | 5521 | 37 | AGCGCTGAGAAGACGGTCGAACTTGACTATCTAGAGG | 337.4 | -12.2 | 1.5 | -8.2 | 1.7E-16 | 8.7E-16 | MB231/MB468 | MITO |
| >hg38_21_+_8433172_8446622_RNA45SN1_45S_50ntFlanks@12159.12179.21\|TTTGACTGGGGCGGTACACCT | >hg38_21_+_8433172_8446622_RNA45SN1_45S_50ntFlanks | 12159 | 12179 | 21 | TTTGACTGGGGCGGTACACCT | 1948.8 | -5.5 | 0.7 | -8.3 | 1.2E-16 | 8.7E-16 | BT20/MB468 | MITO |
| >MI0000100\|hsa-mir-98&WithFlank&X\|-\|53556217\|53556347@28.49.22&[MIMAT0000096&hsa-miR-98-5p&offsets\|0\|0;m-143&X\|-\|53556299\|53556320&offsets\|0\|0]\|TGAGGTAGTAAGTTGTATTGTT | >MI0000100\|hsa-mir-98&WithFlank&X\|-\|53556217\|53556347 | 28 | 49 | 22 | TGAGGTAGTAAGTTGTATTGTT | 2266.7 | 5.0 | 0.6 | 8.3 | 1.2E-16 | 8.7E-16 | BT20/MB468 | MITO |
| >MI0000650\|hsa-mir-200c&WithFlank&12\|+\|6963693\|6963772@49.70.22&[MIMAT0000617&hsa-miR-200c-3p&offsets\|-1\|-2;m-54&12\|+\|6963742\|6963764&offsets\|-1\|-2]\|CTAATACTGCCGGGTAATGATG | >MI0000650\|hsa-mir-200c&WithFlank&12\|+\|6963693\|6963772 | 49 | 70 | 22 | CTAATACTGCCGGGTAATGATG | 439.9 | 9.5 | 1.1 | 8.6 | 9.3E-18 | 8.7E-16 | BT20/MB231 | CYTO |
| >MI0000077\|hsa-mir-21&WithFlank&17\|+\|59841260\|59841343@14.35.22(+1A)&[MIMAT0000076&hsa-miR-21-5p&offsets\|0\|0(+1A);m-1&17\|+\|59841273\|59841294&offsets\|0\|0(+1A)]\|TAGCTTATCAGACTGATGTTGAA | >MI0000077\|hsa-mir-21&WithFlank&17\|+\|59841260\|59841343 | 14 | 35 | 22(+1A) | TAGCTTATCAGACTGATGTTGAA | 65400.7 | 8.8 | 1.1 | 8.2 | 1.7E-16 | 8.7E-16 | BT20/MB468 | CYTO |
| >hg38_21_+_8433172_8446622_RNA45SN1_45S_50ntFlanks@12615.12661.47\|GCTGAGGAGCCAATGGGGCGAAGCTACCATCTGTGGGATTATGACTG | >hg38_21_+_8433172_8446622_RNA45SN1_45S_50ntFlanks | 12615 | 12661 | 47 | GCTGAGGAGCCAATGGGGCGAAGCTACCATCTGTGGGATTATGACTG | 2878.0 | -9.8 | 1.2 | -8.2 | 1.7E-16 | 8.7E-16 | MB231/MB468 | MITO |
| >hg38_21_+_8433172_8446622_RNA45SN1_45S_50ntFlanks@12328.12346.19\|GTTTTAAGCAGGAGGTGTC | >hg38_21_+_8433172_8446622_RNA45SN1_45S_50ntFlanks | 12328 | 12346 | 19 | GTTTTAAGCAGGAGGTGTC | 268.9 | -11.0 | 1.3 | -8.3 | 1.3E-16 | 8.7E-16 | MB231/MB468 | NUC |
| >MI0000749\|hsa-mir-30e&WithFlank&1\|+\|40754349\|40754452@65.85.21(+2U)&[MIMAT0000693&hsa-miR-30e-3p&offsets\|0\|-1(+2U);m-25&1\|+\|40754413\|40754434&offsets\|0\|-1(+2U)]\|CTTTCAGTCGGATGTTTACAGTT | >MI0000749\|hsa-mir-30e&WithFlank&1\|+\|40754349\|40754452 | 65 | 85 | 21(+2U) | CTTTCAGTCGGATGTTTACAGTT | 903.3 | 5.9 | 0.7 | 8.2 | 1.7E-16 | 8.8E-16 | BT20/MB468 | CYTO |
| >hg38_21_+_8433172_8446622_RNA45SN1_45S_50ntFlanks@5485.5509.25\|AGCGCTGAGAAGACGGTCGAACTTG | >hg38_21_+_8433172_8446622_RNA45SN1_45S_50ntFlanks | 5485 | 5509 | 25 | AGCGCTGAGAAGACGGTCGAACTTG | 903.6 | -9.5 | 1.2 | -8.2 | 1.7E-16 | 8.8E-16 | MB231/MB468 | MITO |
| >MI0000088\|hsa-mir-30a&WithFlank&6\|-\|71403545\|71403627@13.34.22(+1C)&[MIMAT0000087&hsa-miR-30a-5p&offsets\|+1\|+1(+1C);m-15&6\|-\|71403595\|71403616&offsets\|+1\|+1(+1C)]\|GTAAACATCCTCGACTGGAAGCC | >MI0000088\|hsa-mir-30a&WithFlank&6\|-\|71403545\|71403627 | 13 | 34 | 22(+1C) | GTAAACATCCTCGACTGGAAGCC | 179.2 | 11.4 | 1.4 | 8.3 | 1.3E-16 | 8.9E-16 | MB231/MB468 | NUC |
| >hg38_21_+_8433172_8446622_RNA45SN1_45S_50ntFlanks@3705.3728.24\|TACCTGGTTGATCCTGCCAGTAGC | >hg38_21_+_8433172_8446622_RNA45SN1_45S_50ntFlanks | 3705 | 3728 | 24 | TACCTGGTTGATCCTGCCAGTAGC | 1345.8 | 5.0 | 0.6 | 8.3 | 1.1E-16 | 8.9E-16 | MB231/MB468 | CYTO |
| >hg38_21_+_8433172_8446622_RNA45SN1_45S_50ntFlanks@10391.10409.19\|AACAGCAGTTGAACATGGG | >hg38_21_+_8433172_8446622_RNA45SN1_45S_50ntFlanks | 10391 | 10409 | 19 | AACAGCAGTTGAACATGGG | 3411.1 | -8.1 | 1.0 | -8.3 | 1.1E-16 | 8.9E-16 | MB231/MB468 | CYTO |
| >hg38_21_+_8433172_8446622_RNA45SN1_45S_50ntFlanks@485.525.41\|GATGTGGTGACGTCGTGCTCTCCCGGGCCGGGTCCGAGCCG | >hg38_21_+_8433172_8446622_RNA45SN1_45S_50ntFlanks | 485 | 525 | 41 | GATGTGGTGACGTCGTGCTCTCCCGGGCCGGGTCCGAGCCG | 645.1 | 12.2 | 1.5 | 8.2 | 2.8E-16 | 8.9E-16 | BT20/MB468 | NUC |
| >hg38_21_+_8433172_8446622_RNA45SN1_45S_50ntFlanks@10807.10829.23\|ACAATGTAGGTAAGGGAAGTCGG | >hg38_21_+_8433172_8446622_RNA45SN1_45S_50ntFlanks | 10807 | 10829 | 23 | ACAATGTAGGTAAGGGAAGTCGG | 335.7 | -12.2 | 1.5 | -8.2 | 1.8E-16 | 9.0E-16 | MB231/MB468 | MITO |
| >hg38_21_+_8433172_8446622_RNA45SN1_45S_50ntFlanks@11661.11682.22\|TGATTTCTGCCCAGTGCTCTGA | >hg38_21_+_8433172_8446622_RNA45SN1_45S_50ntFlanks | 11661 | 11682 | 22 | TGATTTCTGCCCAGTGCTCTGA | 313.7 | -12.2 | 1.5 | -8.3 | 1.1E-16 | 9.0E-16 | MB231/MB468 | MP |
| >hg38_21_+_8433172_8446622_RNA45SN1_45S_50ntFlanks@10391.10422.32\|AACAGCAGTTGAACATGGGTCAGTCGGTCCTG | >hg38_21_+_8433172_8446622_RNA45SN1_45S_50ntFlanks | 10391 | 10422 | 32 | AACAGCAGTTGAACATGGGTCAGTCGGTCCTG | 508.7 | -4.7 | 0.6 | -8.3 | 1.3E-16 | 9.0E-16 | MB231/MB468 | NUC |
| >hg38_21_+_8433172_8446622_RNA45SN1_45S_50ntFlanks@12071.12095.25\|CCCCGAGGGGCTCTCGCTTCTGGCG | >hg38_21_+_8433172_8446622_RNA45SN1_45S_50ntFlanks | 12071 | 12095 | 25 | CCCCGAGGGGCTCTCGCTTCTGGCG | 1204.5 | -14.1 | 1.7 | -8.3 | 1.1E-16 | 9.0E-16 | MB231/MB468 | MP |
| >hg38_21_+_8433172_8446622_RNA45SN1_45S_50ntFlanks@11576.11593.18\|CTTAGAACTGGTGCGGAC | >hg38_21_+_8433172_8446622_RNA45SN1_45S_50ntFlanks | 11576 | 11593 | 18 | CTTAGAACTGGTGCGGAC | 1963.5 | -4.9 | 0.6 | -8.3 | 1.1E-16 | 9.0E-16 | MB231/MB468 | CYTO |
| >hg38_21_+_8433172_8446622_RNA45SN1_45S_50ntFlanks@10535.10564.30\|CGCGAGGCGTCCAGTGCGGTAACGCGACCG | >hg38_21_+_8433172_8446622_RNA45SN1_45S_50ntFlanks | 10535 | 10564 | 30 | CGCGAGGCGTCCAGTGCGGTAACGCGACCG | 240.2 | -11.8 | 1.4 | -8.3 | 1.1E-16 | 9.1E-16 | MB231/MB468 | MP |
| >hg38_21_+_8433172_8446622_RNA45SN1_45S_50ntFlanks@11891.11912.22\|TTGACTCTAGTCTGGCACGGTG | >hg38_21_+_8433172_8446622_RNA45SN1_45S_50ntFlanks | 11891 | 11912 | 22 | TTGACTCTAGTCTGGCACGGTG | 2675.0 | -11.4 | 1.4 | -8.2 | 2.9E-16 | 9.1E-16 | BT20/MB468 | NUC |
| >hg38_21_+_8433172_8446622_RNA45SN1_45S_50ntFlanks@11998.12032.35\|GCCCTGCGGGCCGCCGGTGAAATACCACTACTCTG | >hg38_21_+_8433172_8446622_RNA45SN1_45S_50ntFlanks | 11998 | 12032 | 35 | GCCCTGCGGGCCGCCGGTGAAATACCACTACTCTG | 443.0 | -11.8 | 1.4 | -8.3 | 1.3E-16 | 9.1E-16 | MB231/MB468 | NUC |
| >hg38_21_+_8433172_8446622_RNA45SN1_45S_50ntFlanks@5485.5507.23\|AGCGCTGAGAAGACGGTCGAACT | >hg38_21_+_8433172_8446622_RNA45SN1_45S_50ntFlanks | 5485 | 5507 | 23 | AGCGCTGAGAAGACGGTCGAACT | 208.3 | -11.6 | 1.4 | -8.3 | 1.1E-16 | 9.2E-16 | MB231/MB468 | MP |
| >hg38_21_+_8433172_8446622_RNA45SN1_45S_50ntFlanks@12631.12653.23\|GGCGAAGCTACCATCTGTGGGAT | >hg38_21_+_8433172_8446622_RNA45SN1_45S_50ntFlanks | 12631 | 12653 | 23 | GGCGAAGCTACCATCTGTGGGAT | 233.8 | -11.7 | 1.4 | -8.3 | 1.1E-16 | 9.2E-16 | MB231/MB468 | MP |
| >hg38_21_+_8433172_8446622_RNA45SN1_45S_50ntFlanks@11235.11253.19\|CTTCCCGTGGATCGCCCCA | >hg38_21_+_8433172_8446622_RNA45SN1_45S_50ntFlanks | 11235 | 11253 | 19 | CTTCCCGTGGATCGCCCCA | 995.7 | -10.2 | 1.2 | -8.3 | 1.4E-16 | 9.2E-16 | MB231/MB468 | NUC |
| >hg38_21_+_8433172_8446622_RNA45SN1_45S_50ntFlanks@12971.12988.18\|TTTCGTACGTAGCAGAGC | >hg38_21_+_8433172_8446622_RNA45SN1_45S_50ntFlanks | 12971 | 12988 | 18 | TTTCGTACGTAGCAGAGC | 379.8 | -4.5 | 0.5 | -8.3 | 1.4E-16 | 9.2E-16 | MB231/MB468 | NUC |
| >MI0000477\|hsa-mir-146a&WithFlank&5\|+\|160485346\|160485456@27.47.21(+1C)&[MIMAT0000449&hsa-miR-146a-5p&offsets\|0\|-1(+1C);m-129&5\|+\|160485372\|160485393&offsets\|0\|-1(+1C)]\|TGAGAACTGAATTCCATGGGTC | >MI0000477\|hsa-mir-146a&WithFlank&5\|+\|160485346\|160485456 | 27 | 47 | 21(+1C) | TGAGAACTGAATTCCATGGGTC | 16941.0 | 13.7 | 1.7 | 8.3 | 1.1E-16 | 9.2E-16 | MB231/MB468 | MP |
| >hg38_21_+_8433172_8446622_RNA45SN1_45S_50ntFlanks@8357.8391.35\|AAAAGAACTTTGAAGAGAGAGTTCAAGAGGGCGTG | >hg38_21_+_8433172_8446622_RNA45SN1_45S_50ntFlanks | 8357 | 8391 | 35 | AAAAGAACTTTGAAGAGAGAGTTCAAGAGGGCGTG | 1092.7 | -5.6 | 0.7 | -8.2 | 1.8E-16 | 9.3E-16 | MB231/MB468 | MITO |
| >hg38_21_+_8433172_8446622_RNA45SN1_45S_50ntFlanks@9567.9588.22\|ACGTGCAAATCGGTCGTCCGAC | >hg38_21_+_8433172_8446622_RNA45SN1_45S_50ntFlanks | 9567 | 9588 | 22 | ACGTGCAAATCGGTCGTCCGAC | 267.4 | -11.9 | 1.4 | -8.3 | 1.1E-16 | 9.3E-16 | MB231/MB468 | MP |
| >hg38_21_+_8433172_8446622_RNA45SN1_45S_50ntFlanks@9031.9058.28\|TCCTCGTGGGGGGGCCGGGCCACCCCTC | >hg38_21_+_8433172_8446622_RNA45SN1_45S_50ntFlanks | 9031 | 9058 | 28 | TCCTCGTGGGGGGGCCGGGCCACCCCTC | 272.0 | -12.0 | 1.4 | -8.3 | 1.1E-16 | 9.3E-16 | MB231/MB468 | MP |
| >hg38_21_+_8433172_8446622_RNA45SN1_45S_50ntFlanks@8380.8399.20\|CAAGAGGGCGTGAAACCGTT | >hg38_21_+_8433172_8446622_RNA45SN1_45S_50ntFlanks | 8380 | 8399 | 20 | CAAGAGGGCGTGAAACCGTT | 2111.7 | -10.8 | 1.3 | -8.3 | 1.1E-16 | 9.3E-16 | MB231/MB468 | MP |
| >MI0000743\|hsa-mir-34c&WithFlank&11\|+\|111513433\|111513521@19.40.22(+1U)&[MIMAT0000686&hsa-miR-34c-5p&offsets\|0\|-1(+1U);m-336&11\|+\|111513451\|111513473&offsets\|0\|-1(+1U)]\|AGGCAGTGTAGTTAGCTGATTGT | >MI0000743\|hsa-mir-34c&WithFlank&11\|+\|111513433\|111513521 | 19 | 40 | 22(+1U) | AGGCAGTGTAGTTAGCTGATTGT | 273.9 | 11.3 | 1.4 | 8.3 | 1.1E-16 | 9.4E-16 | MB231/MB468 | MP |
| >MI0000087\|hsa-mir-29a&WithFlank&7\|-\|130876741\|130876816@47.69.23&[MIMAT0000086&hsa-miR-29a-3p&offsets\|-1\|0;m-26&7\|-\|130876749\|130876769&offsets\|-1\|+1]\|CTAGCACCATCTGAAATCGGTTA | >MI0000087\|hsa-mir-29a&WithFlank&7\|-\|130876741\|130876816 | 47 | 69 | 23 | CTAGCACCATCTGAAATCGGTTA | 981.3 | 11.9 | 1.4 | 8.2 | 1.8E-16 | 9.4E-16 | BT20/MB468 | CYTO |
| >MI0003599\|hsa-mir-589&WithFlank&7\|-\|5495813\|5495923@30.50.21&[MIMAT0004799&hsa-miR-589-5p&offsets\|0\|-1;m-162&7\|-\|5495874\|5495894&offsets\|0\|0]\|TGAGAACCACGTCTGCTCTGA | >MI0003599\|hsa-mir-589&WithFlank&7\|-\|5495813\|5495923 | 30 | 50 | 21 | TGAGAACCACGTCTGCTCTGA | 444.2 | 8.9 | 1.1 | 8.2 | 1.8E-16 | 9.4E-16 | BT20/MB468 | CYTO |
| >hg38_21_+_8433172_8446622_RNA45SN1_45S_50ntFlanks@9563.9596.34\|CCTGACGTGCAAATCGGTCGTCCGACCTGGGTAT | >hg38_21_+_8433172_8446622_RNA45SN1_45S_50ntFlanks | 9563 | 9596 | 34 | CCTGACGTGCAAATCGGTCGTCCGACCTGGGTAT | 618.4 | -13.1 | 1.6 | -8.3 | 1.1E-16 | 9.4E-16 | MB231/MB468 | MP |
| >MI0000811\|hsa-mir-148b&WithFlank&12\|+\|54337210\|54337320@69.90.22(+1U)&[MIMAT0000759&hsa-miR-148b-3p&offsets\|0\|0(+1U);m-77&12\|+\|54337278\|54337299&offsets\|0\|0(+1U)]\|TCAGTGCATCACAGAACTTTGTT | >MI0000811\|hsa-mir-148b&WithFlank&12\|+\|54337210\|54337320 | 69 | 90 | 22(+1U) | TCAGTGCATCACAGAACTTTGTT | 3502.5 | 5.8 | 0.7 | 8.3 | 1.4E-16 | 9.4E-16 | MB231/MB468 | NUC |
| >hg38_21_+_8433172_8446622_RNA45SN1_45S_50ntFlanks@11104.11133.30\|AAGGGTCGGGGCGGCAGGGGCCGGCGGCGG | >hg38_21_+_8433172_8446622_RNA45SN1_45S_50ntFlanks | 11104 | 11133 | 30 | AAGGGTCGGGGCGGCAGGGGCCGGCGGCGG | 595.8 | -13.1 | 1.6 | -8.3 | 1.1E-16 | 9.4E-16 | MB231/MB468 | MP |
| >hg38_21_+_8433172_8446622_RNA45SN1_45S_50ntFlanks@10571.10591.21\|GAGAAGCCGGCGGGAGCCCCG | >hg38_21_+_8433172_8446622_RNA45SN1_45S_50ntFlanks | 10571 | 10591 | 21 | GAGAAGCCGGCGGGAGCCCCG | 884.6 | -12.8 | 1.5 | -8.3 | 1.4E-16 | 9.4E-16 | MB231/MB468 | NUC |
| >hg38_21_+_8433172_8446622_RNA45SN1_45S_50ntFlanks@10663.10706.44\|CCTTGGAAAGCGTCGCGGTTCCGGCGGCGTCCGGTGAGCTCTCG | >hg38_21_+_8433172_8446622_RNA45SN1_45S_50ntFlanks | 10663 | 10706 | 44 | CCTTGGAAAGCGTCGCGGTTCCGGCGGCGTCCGGTGAGCTCTCG | 593.2 | -13.0 | 1.6 | -8.2 | 1.9E-16 | 9.5E-16 | MB231/MB468 | MITO |
| >hg38_21_+_8433172_8446622_RNA45SN1_45S_50ntFlanks@5496.5516.21\|GACGGTCGAACTTGACTATCT | >hg38_21_+_8433172_8446622_RNA45SN1_45S_50ntFlanks | 5496 | 5516 | 21 | GACGGTCGAACTTGACTATCT | 212.5 | 7.8 | 0.9 | 8.3 | 1.2E-16 | 9.5E-16 | MB231/MB468 | CYTO |
| >MI0000440\|hsa-mir-27b&WithFlank&9\|+\|95085439\|95085547@66.87.22&[MIMAT0000419&hsa-miR-27b-3p&offsets\|-1\|0;m-39&9\|+\|95085505\|95085525&offsets\|-1\|0]\|GTTCACAGTGGCTAAGTTCTGC | >MI0000440\|hsa-mir-27b&WithFlank&9\|+\|95085439\|95085547 | 66 | 87 | 22 | GTTCACAGTGGCTAAGTTCTGC | 1257.9 | 8.6 | 1.0 | 8.2 | 1.9E-16 | 9.5E-16 | BT20/MB468 | CYTO |
| >MI0000079\|hsa-mir-23a&WithFlank&19\|-\|13836581\|13836665@54.72.19&[MIMAT0000078&hsa-miR-23a-3p&offsets\|+3\|+1;m-31&19\|-\|13836595\|13836615&offsets\|+3\|+1]\|ACATTGCCAGGGATTTCCA | >MI0000079\|hsa-mir-23a&WithFlank&19\|-\|13836581\|13836665 | 54 | 72 | 19 | ACATTGCCAGGGATTTCCA | 4671.6 | 30.0 | 3.6 | 8.3 | 1.4E-16 | 9.6E-16 | MB231/MB468 | NUC |
| >hg38_21_+_8433172_8446622_RNA45SN1_45S_50ntFlanks@485.507.23\|GATGTGGTGACGTCGTGCTCTCC | >hg38_21_+_8433172_8446622_RNA45SN1_45S_50ntFlanks | 485 | 507 | 23 | GATGTGGTGACGTCGTGCTCTCC | 11749.1 | 30.0 | 3.6 | 8.3 | 1.4E-16 | 9.6E-16 | MB231/MB468 | NUC |
| >hg38_21_+_8433172_8446622_RNA45SN1_45S_50ntFlanks@485.517.33\|GATGTGGTGACGTCGTGCTCTCCCGGGCCGGGT | >hg38_21_+_8433172_8446622_RNA45SN1_45S_50ntFlanks | 485 | 517 | 33 | GATGTGGTGACGTCGTGCTCTCCCGGGCCGGGT | 5541.7 | 30.0 | 3.6 | 8.3 | 1.4E-16 | 9.6E-16 | MB231/MB468 | NUC |
| >hg38_21_+_8433172_8446622_RNA45SN1_45S_50ntFlanks@485.520.36\|GATGTGGTGACGTCGTGCTCTCCCGGGCCGGGTCCG | >hg38_21_+_8433172_8446622_RNA45SN1_45S_50ntFlanks | 485 | 520 | 36 | GATGTGGTGACGTCGTGCTCTCCCGGGCCGGGTCCG | 8717.5 | 30.0 | 3.6 | 8.3 | 1.4E-16 | 9.6E-16 | MB231/MB468 | NUC |
| >hg38_21_+_8433172_8446622_RNA45SN1_45S_50ntFlanks@485.521.37\|GATGTGGTGACGTCGTGCTCTCCCGGGCCGGGTCCGA | >hg38_21_+_8433172_8446622_RNA45SN1_45S_50ntFlanks | 485 | 521 | 37 | GATGTGGTGACGTCGTGCTCTCCCGGGCCGGGTCCGA | 7560.7 | 30.0 | 3.6 | 8.3 | 1.4E-16 | 9.6E-16 | MB231/MB468 | NUC |

| | | | | | | | | | | | | | |
|---|---|---|---|---|---|---|---|---|---|---|---|---|---|
| >hg38_21_+_8433172_8446622_RNA45SN1_45S_50ntFlanks@485.522.38\|GATGTGGTGACGTCGTGCTCTCCCGGGCCGGGTCCGAG | >hg38_21_+_8433172_8446622_RNA45SN1_45S_50ntFlanks | 485 | 522 | 38 | GATGTGGTGACGTCGTGCTCTCCCGGGCCGGGTCCGAG | 6663.3 | 30.0 | 3.6 | 8.3 | 1.4E-16 | 9.6E-16 | MB231/MB468 | NUC |
| >hg38_21_+_8433172_8446622_RNA45SN1_45S_50ntFlanks@485.525.41\|GATGTGGTGACGTCGTGCTCTCCCGGGCCGGGTCCGAGCCG | >hg38_21_+_8433172_8446622_RNA45SN1_45S_50ntFlanks | 485 | 525 | 41 | GATGTGGTGACGTCGTGCTCTCCCGGGCCGGGTCCGAGCCG | 7068.9 | 30.0 | 3.6 | 8.3 | 1.4E-16 | 9.6E-16 | MB231/MB468 | NUC |
| >hg38_21_+_8433172_8446622_RNA45SN1_45S_50ntFlanks@10420.10443.24\|CTGAGAGATGGGCGAGCGCCGTTC | >hg38_21_+_8433172_8446622_RNA45SN1_45S_50ntFlanks | 10420 | 10443 | 24 | CTGAGAGATGGGCGAGCGCCGTTC | 524.5 | -9.4 | 1.1 | -8.3 | 1.4E-16 | 9.6E-16 | MB231/MB468 | NUC |
| >hg38_21_+_8433172_8446622_RNA45SN1_45S_50ntFlanks@12539.12587.49\|ATGATGTGTTGTTGCCATGGTAATCCTGCTCAGTACGAGAGGAACCGCA | >hg38_21_+_8433172_8446622_RNA45SN1_45S_50ntFlanks | 12539 | 12587 | 49 | ATGATGTGTTGTTGCCATGGTAATCCTGCTCAGTACGAGAGGAACCGCA | 1548.6 | -15.0 | 1.8 | -8.2 | 3.0E-16 | 9.6E-16 | BT20/MB468 | NUC |
| >hg38_21_+_8433172_8446622_RNA45SN1_45S_50ntFlanks@9033.9076.44\|CTCGTGGGGGGGCCGGGCCACCCCTCCCACGGCGCGACCGCTCT | >hg38_21_+_8433172_8446622_RNA45SN1_45S_50ntFlanks | 9033 | 9076 | 44 | CTCGTGGGGGGGCCGGGCCACCCCTCCCACGGCGCGACCGCTCT | 107.4 | -23.9 | 2.9 | -8.3 | 1.2E-16 | 9.6E-16 | MB231/MB468 | MP |
| >hg38_21_+_8433172_8446622_RNA45SN1_45S_50ntFlanks@5186.5226.41\|CAATAACAGGTCTGTGATGCCCTTAGATGTCCGGGGCTGCA | >hg38_21_+_8433172_8446622_RNA45SN1_45S_50ntFlanks | 5186 | 5226 | 41 | CAATAACAGGTCTGTGATGCCCTTAGATGTCCGGGGCTGCA | 791.5 | -13.4 | 1.6 | -8.2 | 1.9E-16 | 9.6E-16 | MB231/MB468 | MITO |
| >hg38_21_+_8433172_8446622_RNA45SN1_45S_50ntFlanks@12539.12560.22\|ATGATGTGTTGTTGCCATGGTA | >hg38_21_+_8433172_8446622_RNA45SN1_45S_50ntFlanks | 12539 | 12560 | 22 | ATGATGTGTTGTTGCCATGGTA | 265.0 | -11.0 | 1.3 | -8.3 | 1.4E-16 | 9.6E-16 | MB231/MB468 | NUC |
| >hg38_21_+_8433172_8446622_RNA45SN1_45S_50ntFlanks@8107.8143.37\|CGGCGGGGCGCGGGACATGTGGCGTACGGAAGACCCG | >hg38_21_+_8433172_8446622_RNA45SN1_45S_50ntFlanks | 8107 | 8143 | 37 | CGGCGGGGCGCGGGACATGTGGCGTACGGAAGACCCG | 230.3 | -10.8 | 1.3 | -8.3 | 1.4E-16 | 9.6E-16 | MB231/MB468 | NUC |
| >hg38_21_+_8433172_8446622_RNA45SN1_45S_50ntFlanks@5078.5102.25\|CGATAACGAACGAGACTCTGGCATG | >hg38_21_+_8433172_8446622_RNA45SN1_45S_50ntFlanks | 5078 | 5102 | 25 | CGATAACGAACGAGACTCTGGCATG | 1116.0 | -13.9 | 1.7 | -8.2 | 1.9E-16 | 9.6E-16 | MB231/MB468 | MITO |
| >hg38_21_+_8433172_8446622_RNA45SN1_45S_50ntFlanks@8176.8206.31\|CTTCTGATCGAGGCCCAGCCCGTGGACGGTG | >hg38_21_+_8433172_8446622_RNA45SN1_45S_50ntFlanks | 8176 | 8206 | 31 | CTTCTGATCGAGGCCCAGCCCGTGGACGGTG | 1426.6 | -4.5 | 0.5 | -8.3 | 1.4E-16 | 9.6E-16 | MB231/MB468 | NUC |
| >am_trna21_ThrTGT_14_-_21081949_21082021@59.76.18\|TTCTCGCTGGGGCCTCCA | >am_trna21_ThrTGT_14_-_21081949_21082021 | 59 | 76 | 18 | TTCTCGCTGGGGCCTCCA | 253.8 | 8.5 | 1.0 | 8.3 | 1.2E-16 | 9.7E-16 | MB231/MB468 | CYTO |
| >MI0000734\|hsa-mir-106b&WithFlank&7\|-\|100093987\|100094080@58.77.20&[MIMAT0004672&hsa-miR-106b-3p&offsets\|0\|-2;m-87&7\|-\|100094003\|100094023&offsets\|0\|-1]\|CCGCACTGTGGGTACTTGCT | >MI0000734\|hsa-mir-106b&WithFlank&7\|-\|100093987\|100094080 | 58 | 77 | 20 | CCGCACTGTGGGTACTTGCT | 1347.6 | 6.9 | 0.8 | 8.3 | 1.2E-16 | 9.7E-16 | MB231/MB468 | CYTO |
| >hg38_21_+_8433172_8446622_RNA45SN1_45S_50ntFlanks@10571.10608.38\|GAGAAGCCGGCGGGAGCCCCGGGGAGAGTTCTCTTTTC | >hg38_21_+_8433172_8446622_RNA45SN1_45S_50ntFlanks | 10571 | 10608 | 38 | GAGAAGCCGGCGGGAGCCCCGGGGAGAGTTCTCTTTTC | 385.2 | -11.3 | 1.4 | -8.3 | 1.2E-16 | 9.7E-16 | MB231/MB468 | CYTO |
| >hg38_21_+_8433172_8446622_RNA45SN1_45S_50ntFlanks@5414.5445.32\|CTACTACCGATTGGATGGTTTAGTGAGGCCCT | >hg38_21_+_8433172_8446622_RNA45SN1_45S_50ntFlanks | 5414 | 5445 | 32 | CTACTACCGATTGGATGGTTTAGTGAGGCCCT | 165.2 | -5.9 | 0.7 | -8.3 | 1.2E-16 | 9.7E-16 | MB231/MB468 | CYTO |
| >hg38_21_+_8433172_8446622_RNA45SN1_45S_50ntFlanks@4367.4396.30\|CAGTTAAAAAGCTCGTAGTTGGATCTTGGG | >hg38_21_+_8433172_8446622_RNA45SN1_45S_50ntFlanks | 4367 | 4396 | 30 | CAGTTAAAAAGCTCGTAGTTGGATCTTGGG | 416.5 | -9.0 | 1.1 | -8.3 | 1.4E-16 | 9.7E-16 | MB231/MB468 | NUC |
| >hg38_21_+_8433172_8446622_RNA45SN1_45S_50ntFlanks@12893.12925.33\|CCTCGCCCGTCACGCACCGCACGTTCGTGGGGA | >hg38_21_+_8433172_8446622_RNA45SN1_45S_50ntFlanks | 12893 | 12925 | 33 | CCTCGCCCGTCACGCACCGCACGTTCGTGGGGA | 469.0 | -12.7 | 1.5 | -8.3 | 1.2E-16 | 9.7E-16 | MB231/MB468 | MP |
| >hg38_21_+_8433172_8446622_RNA45SN1_45S_50ntFlanks@11569.11601.33\|CAGCCGACTTAGAACTGGTGCGGACCAGGGGAA | >hg38_21_+_8433172_8446622_RNA45SN1_45S_50ntFlanks | 11569 | 11601 | 33 | CAGCCGACTTAGAACTGGTGCGGACCAGGGGAA | 32574.8 | -11.2 | 1.4 | -8.2 | 3.1E-16 | 9.7E-16 | BT20/MB468 | NUC |
| >hg38_21_+_8433172_8446622_RNA45SN1_45S_50ntFlanks@9504.9524.21\|GTGAACTATGCCTGGGCAGGG | >hg38_21_+_8433172_8446622_RNA45SN1_45S_50ntFlanks | 9504 | 9524 | 21 | GTGAACTATGCCTGGGCAGGG | 350.3 | -12.2 | 1.5 | -8.2 | 1.9E-16 | 9.7E-16 | MB231/MB468 | MITO |
| >hg38_21_+_8433172_8446622_RNA45SN1_45S_50ntFlanks@10409.10431.23\|GTCAGTCGGTCCTGAGAGATGGG | >hg38_21_+_8433172_8446622_RNA45SN1_45S_50ntFlanks | 10409 | 10431 | 23 | GTCAGTCGGTCCTGAGAGATGGG | 621.3 | -5.7 | 0.7 | -8.3 | 1.5E-16 | 9.8E-16 | MB231/MB468 | NUC |
| >hg38_21_+_8433172_8446622_RNA45SN1_45S_50ntFlanks@5467.5514.48\|GGCCCACGGCCCTGGCGGAGCGCTGAGAAGACGGTCGAACTTGACTAT | >hg38_21_+_8433172_8446622_RNA45SN1_45S_50ntFlanks | 5467 | 5514 | 48 | GGCCCACGGCCCTGGCGGAGCGCTGAGAAGACGGTCGAACTTGACTAT | 2126.8 | -14.0 | 1.7 | -8.3 | 1.5E-16 | 9.8E-16 | MB231/MB468 | NUC |
| >hg38_21_+_8433172_8446622_RNA45SN1_45S_50ntFlanks@7978.8011.34\|GACCTCAGATCAGACGTGGCGACCCGCTGAATTT | >hg38_21_+_8433172_8446622_RNA45SN1_45S_50ntFlanks | 7978 | 8011 | 34 | GACCTCAGATCAGACGTGGCGACCCGCTGAATTT | 1279.0 | 5.8 | 0.7 | 8.3 | 1.4E-16 | 9.9E-16 | BT20/MB468 | MITO |
| >hg38_21_+_8433172_8446622_RNA45SN1_45S_50ntFlanks@12151.12180.30\|GTGGGGAGTTTGACTGGGGCGGTACACCTG | >hg38_21_+_8433172_8446622_RNA45SN1_45S_50ntFlanks | 12151 | 12180 | 30 | GTGGGGAGTTTGACTGGGGCGGTACACCTG | 437.0 | -11.7 | 1.4 | -8.3 | 1.5E-16 | 9.9E-16 | MB231/MB468 | NUC |
| >hg38_21_+_8433172_8446622_RNA45SN1_45S_50ntFlanks@12641.12661.21\|CCATCTGTGGGATTATGACTG | >hg38_21_+_8433172_8446622_RNA45SN1_45S_50ntFlanks | 12641 | 12661 | 21 | CCATCTGTGGGATTATGACTG | 5039.8 | -9.2 | 1.1 | -8.2 | 2.0E-16 | 9.9E-16 | MB231/MB468 | MITO |
| >MI0006444\|hsa-mir-1307&WithFlank&10\|-\|103394247\|103394407@86.108.23&[MIMAT0005951&hsa-miR-1307-3p&offsets\|0\|+1;m-66&10\|-\|103394300\|103394322&offsets\|0\|0]\|ACTCGGCGTGGCGTCGGTCGTGG | >MI0006444\|hsa-mir-1307&WithFlank&10\|-\|103394247\|103394407 | 86 | 108 | 23 | ACTCGGCGTGGCGTCGGTCGTGG | 3962.6 | 6.1 | 0.7 | 8.3 | 1.2E-16 | 9.9E-16 | MB231/MB468 | CYTO |
| >MI0033423\|hsa-mir-10399&WithFlank&7\|-\|139044094\|139044163@43.63.21(+1U)&[MIMAT0041630&hsa-miR-10399-3p&offsets\|-1\|-1(+1U);TJU_CMC_MD2.ID03061.3p-miR&offsets\|+3\|+2(+1U);m-2657&7\|-\|139044103\|139044124&offsets\|+3\|+2(+1U)]\|TCTCTCGGACAAGCTGTAGGTT | >MI0033423\|hsa-mir-10399&WithFlank&7\|-\|139044094\|139044163 | 43 | 63 | 21(+1U) | TCTCTCGGACAAGCTGTAGGTT | 302.2 | 7.8 | 0.9 | 8.3 | 1.5E-16 | 1.0E-15 | MB231/MB468 | NUC |
| >MI0001445\|hsa-mir-423&WithFlank&17\|+\|30117073\|30117178@58.80.23&[MIMAT0001340&hsa-miR-423-3p&offsets\|-1\|-1;m-100&17\|+\|30117131\|30117153&offsets\|-1\|-1]\|AAGCTCGGTCTGAGGCCCCTCAG | >MI0001445\|hsa-mir-423&WithFlank&17\|+\|30117073\|30117178 | 58 | 80 | 23 | AAGCTCGGTCTGAGGCCCCTCAG | 643.8 | 8.3 | 1.0 | 8.3 | 1.4E-16 | 1.0E-15 | BT20/MB468 | MITO |
| >hg38_21_+_8433172_8446622_RNA45SN1_45S_50ntFlanks@12625.12652.28\|CAATGGGGCGAAGCTACCATCTGTGGGA | >hg38_21_+_8433172_8446622_RNA45SN1_45S_50ntFlanks | 12625 | 12652 | 28 | CAATGGGGCGAAGCTACCATCTGTGGGA | 430.4 | -12.6 | 1.5 | -8.3 | 1.2E-16 | 1.0E-15 | MB231/MB468 | MP |
| >hg38_21_+_8433172_8446622_RNA45SN1_45S_50ntFlanks@12892.12925.34\|CCCTCGCCCGTCACGCACCGCACGTTCGTGGGGA | >hg38_21_+_8433172_8446622_RNA45SN1_45S_50ntFlanks | 12892 | 12925 | 34 | CCCTCGCCCGTCACGCACCGCACGTTCGTGGGGA | 236.3 | -11.7 | 1.4 | -8.3 | 1.2E-16 | 1.0E-15 | MB231/MB468 | MP |
| >MI0000446\|hsa-mir-125b-1&WithFlank&11\|-\|122099751\|122099850@21.42.22(+1A);MI0000470\|hsa-mir-125b-2&WithFlank&21\|+\|16590231\|16590331@23.44.22(+1A)&[MIMAT0000423&hsa-miR-125b-5p&offsets\|0\|0(+1A);m-67&11\|-\|122099809\|122099830&offsets\|0\|0(+1A)];[MIMAT0000423_1&hsa-miR-125b-5p&offsets\|0\|0(+1A);m-68&21\|+\|16590253\|16590274&offsets\|0\|0(+1A)]\|TCCCTGAGACCCTAACTTGTGAA | >MI0000446\|hsa-mir-125b-1&WithFlank&11\|-\|122099751\|122099850 | 21 | 42 | 22(+1A) | TCCCTGAGACCCTAACTTGTGAA | 5650.1 | 8.7 | 1.1 | 8.2 | 2.0E-16 | 1.0E-15 | MB231/MB468 | MITO |
| >hg38_21_+_8433172_8446622_RNA45SN1_45S_50ntFlanks@12633.12661.29\|CGAAGCTACCATCTGTGGGATTATGACTG | >hg38_21_+_8433172_8446622_RNA45SN1_45S_50ntFlanks | 12633 | 12661 | 29 | CGAAGCTACCATCTGTGGGATTATGACTG | 13756.3 | -9.5 | 1.2 | -8.2 | 2.0E-16 | 1.0E-15 | MB231/MB468 | MITO |
| >hg38_21_+_8433172_8446622_RNA45SN1_45S_50ntFlanks@10817.10840.24\|TAAGGGAAGTCGGCAAGCCGGATC | >hg38_21_+_8433172_8446622_RNA45SN1_45S_50ntFlanks | 10817 | 10840 | 24 | TAAGGGAAGTCGGCAAGCCGGATC | 298.2 | -12.1 | 1.5 | -8.3 | 1.2E-16 | 1.0E-15 | MB231/MB468 | MP |
| >hg38_21_+_8433172_8446622_RNA45SN1_45S_50ntFlanks@4555.4572.18\|CGCAGCTAGGAATAATGG | >hg38_21_+_8433172_8446622_RNA45SN1_45S_50ntFlanks | 4555 | 4572 | 18 | CGCAGCTAGGAATAATGG | 499.1 | -12.7 | 1.6 | -8.2 | 2.0E-16 | 1.0E-15 | MB231/MB468 | MITO |
| >hg38_21_+_8433172_8446622_RNA45SN1_45S_50ntFlanks@12456.12475.20\|CCAAGCGTTGGATTGTTCAC | >hg38_21_+_8433172_8446622_RNA45SN1_45S_50ntFlanks | 12456 | 12475 | 20 | CCAAGCGTTGGATTGTTCAC | 17781.0 | -13.2 | 1.6 | -8.2 | 3.2E-16 | 1.0E-15 | BT20/MB468 | NUC |
| >hg38_21_+_8433172_8446622_RNA45SN1_45S_50ntFlanks@11821.11856.36\|CTATCCAGCGAAACCACAGCCAAGGGAACGGGCTTG | >hg38_21_+_8433172_8446622_RNA45SN1_45S_50ntFlanks | 11821 | 11856 | 36 | CTATCCAGCGAAACCACAGCCAAGGGAACGGGCTTG | 303.4 | -12.0 | 1.5 | -8.2 | 2.0E-16 | 1.0E-15 | MB231/MB468 | MITO |
| >hg38_21_+_8433172_8446622_RNA45SN1_45S_50ntFlanks@12456.12478.23\|CCAAGCGTTGGATTGTTCACCCA | >hg38_21_+_8433172_8446622_RNA45SN1_45S_50ntFlanks | 12456 | 12478 | 23 | CCAAGCGTTGGATTGTTCACCCA | 11840.0 | -6.1 | 0.7 | -8.3 | 1.3E-16 | 1.0E-15 | MB231/MB468 | CYTO |
| >hg38_21_+_8433172_8446622_RNA45SN1_45S_50ntFlanks@10679.10712.34\|GGTTCCGGCGGCGTCCGGTGAGCTCTCGCTGGCC | >hg38_21_+_8433172_8446622_RNA45SN1_45S_50ntFlanks | 10679 | 10712 | 34 | GGTTCCGGCGGCGTCCGGTGAGCTCTCGCTGGCC | 345.7 | -7.9 | 1.0 | -8.3 | 1.3E-16 | 1.0E-15 | MB231/MB468 | CYTO |
| >hg38_21_+_8433172_8446622_RNA45SN1_45S_50ntFlanks@10663.10711.49\|CCTTGGAAAGCGTCGCGGTTCCGGCGGCGTCCGGTGAGCTCTCGCTGGC | >hg38_21_+_8433172_8446622_RNA45SN1_45S_50ntFlanks | 10663 | 10711 | 49 | CCTTGGAAAGCGTCGCGGTTCCGGCGGCGTCCGGTGAGCTCTCGCTGGC | 887.5 | -13.6 | 1.7 | -8.2 | 2.0E-16 | 1.0E-15 | MB231/MB468 | MITO |
| >hg38_21_+_8433172_8446622_RNA45SN1_45S_50ntFlanks@6651.6673.23\|CGACTCTTAGCGGTGGATCACTC | >hg38_21_+_8433172_8446622_RNA45SN1_45S_50ntFlanks | 6651 | 6673 | 23 | CGACTCTTAGCGGTGGATCACTC | 16811.3 | 5.6 | 0.7 | 8.3 | 1.4E-16 | 1.0E-15 | BT20/MB468 | MITO |
| >hg38_21_+_8433172_8446622_RNA45SN1_45S_50ntFlanks@4224.4242.19\|AAAATAACAATACAGGACT | >hg38_21_+_8433172_8446622_RNA45SN1_45S_50ntFlanks | 4224 | 4242 | 19 | AAAATAACAATACAGGACT | 1004.6 | -7.8 | 0.9 | -8.2 | 2.0E-16 | 1.0E-15 | MB231/MB468 | MITO |
| >hg38_21_+_8433172_8446622_RNA45SN1_45S_50ntFlanks@3885.3915.31\|ACGGGCGCTGACCCCCTTCGCGGGGGGGATG | >hg38_21_+_8433172_8446622_RNA45SN1_45S_50ntFlanks | 3885 | 3915 | 31 | ACGGGCGCTGACCCCCTTCGCGGGGGGGATG | 231.4 | -11.7 | 1.4 | -8.3 | 1.2E-16 | 1.0E-15 | MB231/MB468 | MP |
| >hg38_21_+_8433172_8446622_RNA45SN1_45S_50ntFlanks@10384.10424.41\|CCATGTGAACAGCAGTTGAACATGGGTCAGTCGGTCCTGAG | >hg38_21_+_8433172_8446622_RNA45SN1_45S_50ntFlanks | 10384 | 10424 | 41 | CCATGTGAACAGCAGTTGAACATGGGTCAGTCGGTCCTGAG | 2339.4 | -5.3 | 0.7 | -8.2 | 3.2E-16 | 1.0E-15 | BT20/MB468 | NUC |
| >hg38_21_+_8433172_8446622_RNA45SN1_45S_50ntFlanks@4728.4751.24\|ATCAAGAACGAAAGTCGGAGGTTC | >hg38_21_+_8433172_8446622_RNA45SN1_45S_50ntFlanks | 4728 | 4751 | 24 | ATCAAGAACGAAAGTCGGAGGTTC | 1246.9 | -13.4 | 1.5 | -8.7 | 3.3E-18 | 1.0E-15 | BT20/MB468 | MP |
| >hg38_21_+_8433172_8446622_RNA45SN1_45S_50ntFlanks@11665.11702.38\|TTCTGCCCAGTGCTCTGAATGTCAAAGTGAAGAAATTC | >hg38_21_+_8433172_8446622_RNA45SN1_45S_50ntFlanks | 11665 | 11702 | 38 | TTCTGCCCAGTGCTCTGAATGTCAAAGTGAAGAAATTC | 1597.3 | 6.2 | 0.8 | 8.3 | 1.3E-16 | 1.0E-15 | MB231/MB468 | CYTO |
| >hg38_21_+_8433172_8446622_RNA45SN1_45S_50ntFlanks@9339.9367.29\|CGCACGAAAGCCGCCGTGGCGCAATGAAG | >hg38_21_+_8433172_8446622_RNA45SN1_45S_50ntFlanks | 9339 | 9367 | 29 | CGCACGAAAGCCGCCGTGGCGCAATGAAG | 574.6 | -12.9 | 1.6 | -8.2 | 2.0E-16 | 1.0E-15 | MB231/MB468 | MITO |
| >hg38_MT_+_1621_3279_RNR2_16S_50ntFlanks@51.90.40\|GCTAAACCTAGCCCCAAACCCACTCCACCTTACTACCAGA | >hg38_MT_+_1621_3279_RNR2_16S_50ntFlanks | 51 | 90 | 40 | GCTAAACCTAGCCCCAAACCCACTCCACCTTACTACCAGA | 294.0 | 12.2 | 1.5 | 8.3 | 1.3E-16 | 1.0E-15 | MB231/MB468 | CYTO |
| >hg38_21_+_8433172_8446622_RNA45SN1_45S_50ntFlanks@10404.10435.32\|CATGGGTCAGTCGGTCCTGAGAGATGGGCGAG | >hg38_21_+_8433172_8446622_RNA45SN1_45S_50ntFlanks | 10404 | 10435 | 32 | CATGGGTCAGTCGGTCCTGAGAGATGGGCGAG | 335.4 | -12.2 | 1.5 | -8.2 | 2.0E-16 | 1.0E-15 | MB231/MB468 | MITO |
| >hg38_21_+_8433172_8446622_RNA45SN1_45S_50ntFlanks@5084.5110.27\|CGAACGAGACTCTGGCATGCTAACTAG | >hg38_21_+_8433172_8446622_RNA45SN1_45S_50ntFlanks | 5084 | 5110 | 27 | CGAACGAGACTCTGGCATGCTAACTAG | 5779.3 | -9.3 | 1.1 | -8.2 | 2.1E-16 | 1.0E-15 | MB231/MB468 | MITO |
| >MI0001445\|hsa-mir-423&WithFlank&17\|+\|30117073\|30117178@58.81.24(+1U)&[MIMAT0001340&hsa-miR-423-3p&offsets\|-1\|0(+1U);m-100&17\|+\|30117131\|30117153&offsets\|-1\|0(+1U)]\|AAGCTCGGTCTGAGGCCCCTCAGTT | >MI0001445\|hsa-mir-423&WithFlank&17\|+\|30117073\|30117178 | 58 | 81 | 24(+1U) | AAGCTCGGTCTGAGGCCCCTCAGTT | 371.0 | 10.2 | 1.2 | 8.2 | 2.0E-16 | 1.0E-15 | BT20/MB468 | CYTO |
| >am_trnaMT_ProTGG_MT_-_15956_16023@52.70.19\|AAAGACTTTTTCTCTGACC | >am_trnaMT_ProTGG_MT_-_15956_16023 | 52 | 70 | 19 | AAAGACTTTTTCTCTGACC | 374.8 | 10.2 | 1.2 | 8.2 | 2.0E-16 | 1.0E-15 | BT20/MB468 | CYTO |

| | | | | | | | | | | | | | |
|---|---|---|---|---|---|---|---|---|---|---|---|---|---|
| >hg38_21_+_8433172_8446622_RNA45SN1_45S_50ntFlanks@6666.6696.31\|GATCACTCGGCTCGTGCGTCGATGAAGAACG | >hg38_21_+_8433172_8446622_RNA45SN1_45S_50ntFlanks | 6666 | 6696 | 31 | GATCACTCGGCTCGTGCGTCGATGAAGAACG | 207.6 | -11.6 | 1.4 | -8.3 | 1.2E-16 | 1.0E-15 | MB231/MB468 | MP |
| >hg38_21_+_8433172_8446622_RNA45SN1_45S_50ntFlanks@11999.12043.45\|CCCTGCGGGCCGCCGGTGAAATACCACTACTCTGATCGTTTTTTC | >hg38_21_+_8433172_8446622_RNA45SN1_45S_50ntFlanks | 11999 | 12043 | 45 | CCCTGCGGGCCGCCGGTGAAATACCACTACTCTGATCGTTTTTTC | 2469.8 | -14.2 | 1.7 | -8.3 | 1.5E-16 | 1.0E-15 | MB231/MB468 | NUC |
| >hg38_21_+_8433172_8446622_RNA45SN1_45S_50ntFlanks@10479.10497.19\|CGATCGAAAGGGAGTCGGG | >hg38_21_+_8433172_8446622_RNA45SN1_45S_50ntFlanks | 10479 | 10497 | 19 | CGATCGAAAGGGAGTCGGG | 1251.5 | -13.4 | 1.5 | -8.7 | 3.4E-18 | 1.0E-15 | BT20/MB468 | MP |
| >hg38_21_+_8433172_8446622_RNA45SN1_45S_50ntFlanks@11570.11591.22\|AGCCGACTTAGAACTGGTGCGG | >hg38_21_+_8433172_8446622_RNA45SN1_45S_50ntFlanks | 11570 | 11591 | 22 | AGCCGACTTAGAACTGGTGCGG | 1315.2 | -8.3 | 1.0 | -8.3 | 1.2E-16 | 1.0E-15 | MB231/MB468 | MP |
| >hg38_21_+_8433172_8446622_RNA45SN1_45S_50ntFlanks@11561.11588.28\|GCGCCTAGCAGCCGACTTAGAACTGGTG | >hg38_21_+_8433172_8446622_RNA45SN1_45S_50ntFlanks | 11561 | 11588 | 28 | GCGCCTAGCAGCCGACTTAGAACTGGTG | 1066.5 | -7.7 | 0.9 | -8.2 | 2.1E-16 | 1.0E-15 | MB231/MB468 | MITO |
| >hg38_21_+_8433172_8446622_RNA45SN1_45S_50ntFlanks@11579.11605.27\|AGAACTGGTGCGGACCAGGGGAATCCG | >hg38_21_+_8433172_8446622_RNA45SN1_45S_50ntFlanks | 11579 | 11605 | 27 | AGAACTGGTGCGGACCAGGGGAATCCG | 2227.4 | -10.0 | 1.2 | -8.2 | 2.1E-16 | 1.1E-15 | MB231/MB468 | MITO |
| >hg38_21_+_8433172_8446622_RNA45SN1_45S_50ntFlanks@8552.8577.26\|CTCCCCGGAGGGGGCGGGCTCCGGCG | >hg38_21_+_8433172_8446622_RNA45SN1_45S_50ntFlanks | 8552 | 8577 | 26 | CTCCCCGGAGGGGGCGGGCTCCGGCG | 444.8 | -11.8 | 1.4 | -8.3 | 1.6E-16 | 1.1E-15 | MB231/MB468 | NUC |
| >hg38_21_+_8433172_8446622_RNA45SN1_45S_50ntFlanks@11873.11900.28\|AAGAAGACCCTGTTGAGCTTGACTCTAG | >hg38_21_+_8433172_8446622_RNA45SN1_45S_50ntFlanks | 11873 | 11900 | 28 | AAGAAGACCCTGTTGAGCTTGACTCTAG | 2437.0 | -11.2 | 1.4 | -8.2 | 3.3E-16 | 1.1E-15 | BT20/MB468 | NUC |
| >hg38_21_+_8433172_8446622_RNA45SN1_45S_50ntFlanks@10406.10424.19\|TGGGTCAGTCGGTCCTGAG | >hg38_21_+_8433172_8446622_RNA45SN1_45S_50ntFlanks | 10406 | 10424 | 19 | TGGGTCAGTCGGTCCTGAG | 418.0 | 4.8 | 0.6 | 8.2 | 2.1E-16 | 1.1E-15 | BT20/MB468 | CYTO |
| >hg38_21_+_8433172_8446622_RNA45SN1_45S_50ntFlanks@10423.10457.35\|AGAGATGGGCGAGCGCCGTTCCGAAGGGACGGGCG | >hg38_21_+_8433172_8446622_RNA45SN1_45S_50ntFlanks | 10423 | 10457 | 35 | AGAGATGGGCGAGCGCCGTTCCGAAGGGACGGGCG | 717.5 | -12.4 | 1.5 | -8.2 | 1.6E-16 | 1.1E-15 | MB231/MB468 | NUC |
| >hg38_21_+_8433172_8446622_RNA45SN1_45S_50ntFlanks@3837.3861.25\|CCTCTCCTACTTGGATAACTGTGGT | >hg38_21_+_8433172_8446622_RNA45SN1_45S_50ntFlanks | 3837 | 3861 | 25 | CCTCTCCTACTTGGATAACTGTGGT | 773.1 | -7.5 | 0.9 | -8.2 | 3.3E-16 | 1.1E-15 | BT20/MB468 | NUC |
| >hg38_21_+_8433172_8446622_RNA45SN1_45S_50ntFlanks@12655.12674.20\|ATGACTGAACGCCTCTAAGT | >hg38_21_+_8433172_8446622_RNA45SN1_45S_50ntFlanks | 12655 | 12674 | 20 | ATGACTGAACGCCTCTAAGT | 2177.5 | -6.4 | 0.8 | -8.2 | 2.1E-16 | 1.1E-15 | MB231/MB468 | MITO |
| >am_trna2_ArgCCT_16_+_3202901_3202973@14.31.18,trna3_ArgCCT_7_+_139025446_139025518@14.31.18\|AATGGATAAGGCATTGGC | >am_trna2_ArgCCT_16_+_3202901_3202973 | 14 | 31 | 18 | AATGGATAAGGCATTGGC | 613.0 | -12.2 | 1.5 | -8.2 | 1.6E-16 | 1.1E-15 | MB231/MB468 | NUC |
| >hg38_21_+_8433172_8446622_RNA45SN1_45S_50ntFlanks@9575.9603.29\|ATCGGTCGTCCGACCTGGGTATAGGGGCG | >hg38_21_+_8433172_8446622_RNA45SN1_45S_50ntFlanks | 9575 | 9603 | 29 | ATCGGTCGTCCGACCTGGGTATAGGGGCG | 293.3 | -10.5 | 1.3 | -8.2 | 2.1E-16 | 1.1E-15 | BT20/MB468 | CYTO |
| >hg38_21_+_8433172_8446622_RNA45SN1_45S_50ntFlanks@8406.8442.37\|TAAACGGGTGGGGTCCGCGCAGTCCGCCCGGAGGATT | >hg38_21_+_8433172_8446622_RNA45SN1_45S_50ntFlanks | 8406 | 8442 | 37 | TAAACGGGTGGGGTCCGCGCAGTCCGCCCGGAGGATT | 296.0 | -12.0 | 1.5 | -8.2 | 2.1E-16 | 1.1E-15 | MB231/MB468 | MITO |
| >hg38_21_+_8433172_8446622_RNA45SN1_45S_50ntFlanks@10412.10430.19\|AGTCGGTCCTGAGAGATGG | >hg38_21_+_8433172_8446622_RNA45SN1_45S_50ntFlanks | 10412 | 10430 | 19 | AGTCGGTCCTGAGAGATGG | 396.9 | -12.5 | 1.5 | -8.3 | 1.3E-16 | 1.1E-15 | MB231/MB468 | MP |
| >hg38_21_+_8433172_8446622_RNA45SN1_45S_50ntFlanks@11210.11233.24\|GGCGACTCTGGACGCGAGCCGGGC | >hg38_21_+_8433172_8446622_RNA45SN1_45S_50ntFlanks | 11210 | 11233 | 24 | GGCGACTCTGGACGCGAGCCGGGC | 283.2 | -12.0 | 1.5 | -8.3 | 1.3E-16 | 1.1E-15 | MB231/MB468 | MP |
| >hg38_21_+_8433172_8446622_RNA45SN1_45S_50ntFlanks@8113.8141.29\|GGCGCGGGACATGTGGCGTACGGAAGACC | >hg38_21_+_8433172_8446622_RNA45SN1_45S_50ntFlanks | 8113 | 8141 | 29 | GGCGCGGGACATGTGGCGTACGGAAGACC | 368.6 | -12.3 | 1.5 | -8.2 | 2.2E-16 | 1.1E-15 | MB231/MB468 | MITO |
| >am_trna10_ValCAC_5_-_180649395_180649467@1.32.32,trna12_ValAAC_5_-_180645270_180645342@1.32.32,trna132_ValAAC_6_-_27721179_27721251@1.32.32,trna136_ValAAC_6_-_27648885_27648957@1.32.32,trna139_ValAAC_6_-_27618707_27618779@1.32.32,trna18_ValCAC_5_-_180529253_180529325@1.32.32,trna2_ValAAC_3_+_169490018_169490090@1.32.32,trna2_ValCAC_5_+_180524070_180524142@1.32.32,trna4_ValAAC_5_+_180591154_180591226@1.32.32,trna5_ValAAC_5_+_180596610_180596682@1.32.32,trna6_ValCAC_5_+_180600650_180600722@1.32.32,trna85_ValCAC_1_-_161369490_161369562@1.32.32,trna90_ValCAC_1_-_149684088_149684161@1.32.32,trna98_ValCAC_1_-_149298555_149298627@1.32.32,trna9_ValCAC_6_+_26538282_26538354@1.32.32\|GTTTCCGTAGTGTAGTGGTTATCACGTTCGCC | >am_trna10_ValCAC_5_-_180649395_180649467 | 1 | 32 | 32 | GTTTCCGTAGTGTAGTGGTTATCACGTTCGCC | 30701.8 | 9.3 | 1.1 | 8.3 | 1.4E-16 | 1.1E-15 | MB231/MB468 | CYTO |
| >MI0000478\|hsa-mir-149&WithFlank&2\|+\|240455995\|240456095@21.43.23&[MIMAT0000450&hsa-miR-149-5p&offsets\|0\|0;m-195&2\|+\|240456015\|240456037&offsets\|0\|0]\|TCTGGCTCCGTGTCTTCACTCCC | >MI0000478\|hsa-mir-149&WithFlank&2\|+\|240455995\|240456095 | 21 | 43 | 23 | TCTGGCTCCGTGTCTTCACTCCC | 600.5 | 5.7 | 0.7 | 8.2 | 3.4E-16 | 1.1E-15 | BT20/MB468 | NUC |
| >hg38_21_+_8433172_8446622_RNA45SN1_45S_50ntFlanks@11768.11804.37\|CTAATTAGTGACGCGCATGAATGGATGAACGAGATTC | >hg38_21_+_8433172_8446622_RNA45SN1_45S_50ntFlanks | 11768 | 11804 | 37 | CTAATTAGTGACGCGCATGAATGGATGAACGAGATTC | 462.9 | -6.3 | 0.8 | -8.3 | 1.4E-16 | 1.1E-15 | MB231/MB468 | CYTO |
| >hg38_21_+_8433172_8446622_RNA45SN1_45S_50ntFlanks@12072.12089.18\|CCCGAGGGGCTCTCGCTT | >hg38_21_+_8433172_8446622_RNA45SN1_45S_50ntFlanks | 12072 | 12089 | 18 | CCCGAGGGGCTCTCGCTT | 157.6 | -5.5 | 0.7 | -8.3 | 1.4E-16 | 1.1E-15 | MB231/MB468 | CYTO |
| >am_trna13_LysCTT_14_-_58706613_58706685@32.51.20,trna2_LysCTT_15_+_79152904_79152976@32.51.20,trna4_LysCTT_18_-_43669271_43669343@32.51.20\|CTCTTAATCCCAGGGTCGTG | >am_trna13_LysCTT_14_-_58706613_58706685 | 32 | 51 | 20 | CTCTTAATCCCAGGGTCGTG | 341.4 | -12.3 | 1.5 | -8.3 | 1.3E-16 | 1.1E-15 | MB231/MB468 | MP |
| >hg38_21_+_8433172_8446622_RNA45SN1_45S_50ntFlanks@12959.12993.35\|TCTGGGTCGGGGTTTCGTACGTAGCAGAGCAGCTC | >hg38_21_+_8433172_8446622_RNA45SN1_45S_50ntFlanks | 12959 | 12993 | 35 | TCTGGGTCGGGGTTTCGTACGTAGCAGAGCAGCTC | 725.8 | 6.9 | 0.8 | 8.2 | 2.2E-16 | 1.1E-15 | BT20/MB468 | CYTO |
| >hg38_21_+_8433172_8446622_RNA45SN1_45S_50ntFlanks@6666.6688.23\|GATCACTCGGCTCGTGCGTCGAT | >hg38_21_+_8433172_8446622_RNA45SN1_45S_50ntFlanks | 6666 | 6688 | 23 | GATCACTCGGCTCGTGCGTCGAT | 325.9 | -12.1 | 1.5 | -8.2 | 2.2E-16 | 1.1E-15 | MB231/MB468 | MITO |
| >hg38_21_+_8433172_8446622_RNA45SN1_45S_50ntFlanks@10458.10490.33\|ATGGCCTCCGTTGCCCTCGGCCGATCGAAAGGG | >hg38_21_+_8433172_8446622_RNA45SN1_45S_50ntFlanks | 10458 | 10490 | 33 | ATGGCCTCCGTTGCCCTCGGCCGATCGAAAGGG | 235.7 | -11.7 | 1.4 | -8.3 | 1.3E-16 | 1.1E-15 | MB231/MB468 | MP |
| >hg38_21_+_8433172_8446622_RNA45SN1_45S_50ntFlanks@12241.12269.29\|CAGAAGGGCAAAAGCTCGCTTGATCTTGA | >hg38_21_+_8433172_8446622_RNA45SN1_45S_50ntFlanks | 12241 | 12269 | 29 | CAGAAGGGCAAAAGCTCGCTTGATCTTGA | 320.3 | -12.2 | 1.5 | -8.3 | 1.3E-16 | 1.1E-15 | MB231/MB468 | MP |
| >hg38_21_+_8433172_8446622_RNA45SN1_45S_50ntFlanks@8113.8142.30\|GGCGCGGGACATGTGGCGTACGGAAGACCC | >hg38_21_+_8433172_8446622_RNA45SN1_45S_50ntFlanks | 8113 | 8142 | 30 | GGCGCGGGACATGTGGCGTACGGAAGACCC | 330.7 | -12.2 | 1.5 | -8.3 | 1.3E-16 | 1.1E-15 | MB231/MB468 | MP |
| >hg38_21_+_8433172_8446622_RNA45SN1_45S_50ntFlanks@9787.9820.34\|TCGCTGGCGTGGAGCCGGGCGTGGAATGCGAGTG | >hg38_21_+_8433172_8446622_RNA45SN1_45S_50ntFlanks | 9787 | 9820 | 34 | TCGCTGGCGTGGAGCCGGGCGTGGAATGCGAGTG | 270.0 | -11.9 | 1.4 | -8.3 | 1.3E-16 | 1.1E-15 | MB231/MB468 | MP |
| >hg38_21_+_8433172_8446622_RNA45SN1_45S_50ntFlanks@10869.10889.21\|AGGGCTGGGTCGGTCGGGCTG | >hg38_21_+_8433172_8446622_RNA45SN1_45S_50ntFlanks | 10869 | 10889 | 21 | AGGGCTGGGTCGGTCGGGCTG | 601.9 | -13.0 | 1.6 | -8.2 | 2.2E-16 | 1.1E-15 | MB231/MB468 | MITO |
| >hg38_21_+_8433172_8446622_RNA45SN1_45S_50ntFlanks@7985.8003.19\|GATCAGACGTGGCGACCCG | >hg38_21_+_8433172_8446622_RNA45SN1_45S_50ntFlanks | 7985 | 8003 | 19 | GATCAGACGTGGCGACCCG | 334.8 | -12.2 | 1.5 | -8.2 | 2.2E-16 | 1.1E-15 | MB231/MB468 | MITO |
| >hg38_21_+_8433172_8446622_RNA45SN1_45S_50ntFlanks@6646.6673.28\|TCGTACGACTCTTAGCGGTGGATCACTC | >hg38_21_+_8433172_8446622_RNA45SN1_45S_50ntFlanks | 6646 | 6673 | 28 | TCGTACGACTCTTAGCGGTGGATCACTC | 8784.8 | 5.9 | 0.7 | 8.3 | 1.6E-16 | 1.1E-15 | BT20/MB468 | MITO |
| >hg38_21_+_8433172_8446622_RNA45SN1_45S_50ntFlanks@12542.12566.25\|ATGTGTTGTTGCCATGGTAATCCTG | >hg38_21_+_8433172_8446622_RNA45SN1_45S_50ntFlanks | 12542 | 12566 | 25 | ATGTGTTGTTGCCATGGTAATCCTG | 242.8 | -11.8 | 1.4 | -8.3 | 1.3E-16 | 1.1E-15 | MB231/MB468 | MP |
| >hg38_21_+_8433172_8446622_RNA45SN1_45S_50ntFlanks@7975.8014.40\|CGCGACCTCAGATCAGACGTGGCGACCCGCTGAATTTAAG | >hg38_21_+_8433172_8446622_RNA45SN1_45S_50ntFlanks | 7975 | 8014 | 40 | CGCGACCTCAGATCAGACGTGGCGACCCGCTGAATTTAAG | 7931.9 | 5.6 | 0.7 | 8.3 | 1.4E-16 | 1.1E-15 | MB231/MB468 | CYTO |
| >am_MI0005763\|hsa-mir-941-1&WithFlank&20\|+\|63919443\|63919526@53.75.23;MI0005764\|hsa-mir-941-2&WithFlank&20\|+\|63919499\|63919582@53.75.23;MI0005765\|hsa-mir-941-3&WithFlank&20\|+\|63919555\|63919638@53.75.23;MI0005766\|hsa-mir-941-4&WithFlank&20\|+\|63919750\|63919833@53.75.23;MI0031520\|hsa-mir-941-5&WithFlank&20\|+\|63919862\|63919945@53.75.23&[MIMAT0004984&hsa-miR-941&offsets\|0\|0;m-263&20\|+\|63919495\|63919517&offsets\|0\|0];[MIMAT0004984_1&hsa-miR-941&offsets\|0\|0;m-263&20\|+\|63919551\|63919573&offsets\|0\|0];[MIMAT0004984_2&hsa-miR-941&offsets\|0\|0;m-263&20\|+\|63919607\|63919629&offsets\|0\|0];[MIMAT0004984_3&hsa-miR-941&offsets\|0\|0;m-263&20\|+\|63919802\|63919824&offsets\|0\|0];[MIMAT0004984_4&hsa-miR-941&offsets\|0\|0;m-263&20\|+\|63919914\|63919936&offsets\|0\|0]\|CACCCGGCTGTGTGCACATGTGC | >am_MI0005763\|hsa-mir-941-1&WithFlank&20\|+\|63919443\|63919526 | 53 | 75 | 23 | CACCCGGCTGTGTGCACATGTGC | 16154.7 | 6.6 | 0.8 | 8.3 | 1.4E-16 | 1.1E-15 | MB231/MB468 | CYTO |
| >hg38_21_+_8433172_8446622_RNA45SN1_45S_50ntFlanks@12365.12383.19\|ATAACTGGCTTGTGGCGGC | >hg38_21_+_8433172_8446622_RNA45SN1_45S_50ntFlanks | 12365 | 12383 | 19 | ATAACTGGCTTGTGGCGGC | 348.6 | -7.9 | 1.0 | -8.3 | 1.4E-16 | 1.1E-15 | MB231/MB468 | CYTO |
| >hg38_21_+_8433172_8446622_RNA45SN1_45S_50ntFlanks@4067.4095.29\|AACTTTCGATGGTAGTCGCCGTGCCTACC | >hg38_21_+_8433172_8446622_RNA45SN1_45S_50ntFlanks | 4067 | 4095 | 29 | AACTTTCGATGGTAGTCGCCGTGCCTACC | 334.3 | -12.2 | 1.5 | -8.3 | 1.3E-16 | 1.1E-15 | MB231/MB468 | MP |
| >MI0000079\|hsa-mir-23a&WithFlank&19\|-\|13836581\|13836665@54.71.18&[MIMAT0000078&hsa-miR-23a-3p&offsets\|+3\|0;m-31&19\|-\|13836595\|13836615&offsets\|+3\|0]\|ACATTGCCAGGGATTTCC | >MI0000079\|hsa-mir-23a&WithFlank&19\|-\|13836581\|13836665 | 54 | 71 | 18 | ACATTGCCAGGGATTTCC | 20922.5 | 30.0 | 3.6 | 8.3 | 1.4E-16 | 1.1E-15 | MB231/MB468 | CYTO |
| >MI0000079\|hsa-mir-23a&WithFlank&19\|-\|13836581\|13836665@54.72.19&[MIMAT0000078&hsa-miR-23a-3p&offsets\|+3\|+1;m-31&19\|-\|13836595\|13836615&offsets\|+3\|+1]\|ACATTGCCAGGGATTTCCA | >MI0000079\|hsa-mir-23a&WithFlank&19\|-\|13836581\|13836665 | 54 | 72 | 19 | ACATTGCCAGGGATTTCCA | 19306.1 | 30.0 | 3.6 | 8.3 | 1.4E-16 | 1.1E-15 | MB231/MB468 | CYTO |
| >MI0000079\|hsa-mir-23a&WithFlank&19\|-\|13836581\|13836665@54.73.20&[MIMAT0000078&hsa-miR-23a-3p&offsets\|+3\|+2;m-31&19\|-\|13836595\|13836615&offsets\|+3\|+2]\|ACATTGCCAGGGATTTCCAA | >MI0000079\|hsa-mir-23a&WithFlank&19\|-\|13836581\|13836665 | 54 | 73 | 20 | ACATTGCCAGGGATTTCCAA | 6999.9 | 30.0 | 3.6 | 8.3 | 1.4E-16 | 1.1E-15 | MB231/MB468 | CYTO |
| >MI0000079\|hsa-mir-23a&WithFlank&19\|-\|13836581\|13836665@54.73.20(+1U)&[MIMAT0000078&hsa-miR-23a-3p&offsets\|+3\|+2(+1U);m-31&19\|-\|13836595\|13836615&offsets\|+3\|+2(+1U)]\|ACATTGCCAGGGATTTCCAAT | >MI0000079\|hsa-mir-23a&WithFlank&19\|-\|13836581\|13836665 | 54 | 73 | 20(+1U) | ACATTGCCAGGGATTTCCAAT | 9257.0 | 30.0 | 3.6 | 8.3 | 1.4E-16 | 1.1E-15 | MB231/MB468 | CYTO |
| >MI0000439\|hsa-mir-23b&WithFlank&9\|+\|95085202\|95085310@67.86.20&[MIMAT0000418&hsa-miR-23b-3p&offsets\|+3\|0;m-42&9\|+\|95085265\|95085285&offsets\|+3\|+2]\|ACATTGCCAGGGATTACCAC | >MI0000439\|hsa-mir-23b&WithFlank&9\|+\|95085202\|95085310 | 67 | 86 | 20 | ACATTGCCAGGGATTACCAC | 5668.2 | 30.0 | 3.6 | 8.3 | 1.4E-16 | 1.1E-15 | MB231/MB468 | CYTO |

| | | | | | | | | | | | | | |
|---|---|---|---|---|---|---|---|---|---|---|---|---|---|
| >MI0000439\|hsa-mir-23b&WithFlank&9\|+\|95085202\|95085310@67.86.20(+1U)&[MIMAT0000418&hsa-miR-23b-3p&offsets\|+3\|0(+1U);m-42&9\|+\|95085265\|95085285&offsets\|+3\|+2(+1U)]\|ACATTGCCAGGGATTACCACT | >MI0000439\|hsa-mir-23b&WithFlank&9\|+\|95085202\|95085310 | 67 | 86 | 20(+1U) | ACATTGCCAGGGATTACCACT | 9725.6 | 30.0 | 3.6 | 8.3 | 1.4E-16 | 1.1E-15 | MB231/MB468 | CYTO |
| >trnaMT_PheGAA_MT_+_577_647@35.54.20\|GAAAATGTTTAGACGGGCTC | >trnaMT_PheGAA_MT_+_577_647 | 35 | 54 | 20 | GAAAATGTTTAGACGGGCTC | 271.6 | 12.1 | 1.5 | 8.3 | 1.4E-16 | 1.1E-15 | MB231/MB468 | CYTO |
| >hg38_21_+_8433172_8446622_RNA45SN1_45S_50ntFlanks@9532.9563.32\|AGAGGAAACTCTGGTGGAGGTCCGTAGCGGTC | >hg38_21_+_8433172_8446622_RNA45SN1_45S_50ntFlanks | 9532 | 9563 | 32 | AGAGGAAACTCTGGTGGAGGTCCGTAGCGGTC | 276.5 | 5.8 | 0.7 | 8.3 | 1.4E-16 | 1.1E-15 | MB231/MB468 | CYTO |
| >hg38_21_+_8433172_8446622_RNA45SN1_45S_50ntFlanks@6719.6744.26\|TGAATTGCAGGACACATTGATCATCG | >hg38_21_+_8433172_8446622_RNA45SN1_45S_50ntFlanks | 6719 | 6744 | 26 | TGAATTGCAGGACACATTGATCATCG | 19061.7 | -6.7 | 0.8 | -8.3 | 1.4E-16 | 1.1E-15 | MB231/MB468 | CYTO |
| >hg38_21_+_8433172_8446622_RNA45SN1_45S_50ntFlanks@12510.12527.18\|TCGTGAGACAGGTTAGTT | >hg38_21_+_8433172_8446622_RNA45SN1_45S_50ntFlanks | 12510 | 12527 | 18 | TCGTGAGACAGGTTAGTT | 320.2 | -12.1 | 1.5 | -8.2 | 2.3E-16 | 1.1E-15 | MB231/MB468 | MITO |
| >hg38_21_+_8433172_8446622_RNA45SN1_45S_50ntFlanks@9129.9151.23\|CGGGTCGGGGGGCGGGGCGGACT | >hg38_21_+_8433172_8446622_RNA45SN1_45S_50ntFlanks | 9129 | 9151 | 23 | CGGGTCGGGGGGCGGGGCGGACT | 453.7 | -12.6 | 1.5 | -8.2 | 2.3E-16 | 1.1E-15 | MB231/MB468 | MITO |
| >hg38_21_+_8433172_8446622_RNA45SN1_45S_50ntFlanks@9465.9503.39\|AGGTGGAGCACGAGCGCACGTGTTAGGACCCGAAAGATG | >hg38_21_+_8433172_8446622_RNA45SN1_45S_50ntFlanks | 9465 | 9503 | 39 | AGGTGGAGCACGAGCGCACGTGTTAGGACCCGAAAGATG | 624.8 | -13.2 | 1.6 | -8.3 | 1.4E-16 | 1.1E-15 | MB231/MB468 | MP |
| >hg38_21_+_8433172_8446622_RNA45SN1_45S_50ntFlanks@8155.8181.27\|CGCTCGTGGGGGGCCCAAGTCCTTCTG | >hg38_21_+_8433172_8446622_RNA45SN1_45S_50ntFlanks | 8155 | 8181 | 27 | CGCTCGTGGGGGGCCCAAGTCCTTCTG | 7243.7 | -9.4 | 1.1 | -8.2 | 2.3E-16 | 1.1E-15 | MB231/MB468 | MITO |
| >hg38_21_+_8433172_8446622_RNA45SN1_45S_50ntFlanks@4724.4755.32\|ATTAATCAAGAACGAAAGTCGGAGGTTCGAAG | >hg38_21_+_8433172_8446622_RNA45SN1_45S_50ntFlanks | 4724 | 4755 | 32 | ATTAATCAAGAACGAAAGTCGGAGGTTCGAAG | 382.2 | -12.4 | 1.5 | -8.2 | 2.3E-16 | 1.1E-15 | MB231/MB468 | MITO |
| >hg38_21_+_8433172_8446622_RNA45SN1_45S_50ntFlanks@12634.12659.26\|GAAGCTACCATCTGTGGGATTATGAC | >hg38_21_+_8433172_8446622_RNA45SN1_45S_50ntFlanks | 12634 | 12659 | 26 | GAAGCTACCATCTGTGGGATTATGAC | 534.9 | -12.9 | 1.6 | -8.3 | 1.4E-16 | 1.1E-15 | MB231/MB468 | MP |
| >hg38_21_+_8433172_8446622_RNA45SN1_45S_50ntFlanks@8380.8397.18\|CAAGAGGGCGTGAAACCG | >hg38_21_+_8433172_8446622_RNA45SN1_45S_50ntFlanks | 8380 | 8397 | 18 | CAAGAGGGCGTGAAACCG | 2238.8 | -10.3 | 1.3 | -8.3 | 1.4E-16 | 1.1E-15 | MB231/MB468 | MP |
| >hg38_21_+_8433172_8446622_RNA45SN1_45S_50ntFlanks@10421.10457.37\|TGAGAGATGGGCGAGCGCCGTTCCGAAGGGACGGGCG | >hg38_21_+_8433172_8446622_RNA45SN1_45S_50ntFlanks | 10421 | 10457 | 37 | TGAGAGATGGGCGAGCGCCGTTCCGAAGGGACGGGCG | 535.3 | -12.0 | 1.5 | -8.2 | 1.7E-16 | 1.1E-15 | MB231/MB468 | NUC |
| >MI0000650\|hsa-mir-200c&WithFlank&12\|+\|6963693\|6963772@50.71.22(+1C)&[MIMAT0000617&hsa-miR-200c-3p&offsets\|0\|-1(+1C);m-54&12\|+\|6963742\|6963764&offsets\|0\|-1(+1C)]\|TAATACTGCCGGGTAATGATGGC | >MI0000650\|hsa-mir-200c&WithFlank&12\|+\|6963693\|6963772 | 50 | 71 | 22(+1C) | TAATACTGCCGGGTAATGATGGC | 449.9 | 12.6 | 1.5 | 8.5 | 2.5E-17 | 1.1E-15 | BT20/MB231 | NUC |
| >hg38_21_+_8433172_8446622_RNA45SN1_45S_50ntFlanks@11999.12031.33\|CCCTGCGGGCCGCCGGTGAAATACCACTACTCT | >hg38_21_+_8433172_8446622_RNA45SN1_45S_50ntFlanks | 11999 | 12031 | 33 | CCCTGCGGGCCGCCGGTGAAATACCACTACTCT | 250.6 | -11.8 | 1.4 | -8.3 | 1.4E-16 | 1.1E-15 | MB231/MB468 | MP |
| >hg38_21_+_8433172_8446622_RNA45SN1_45S_50ntFlanks@7983.8008.26\|CAGATCAGACGTGGCGACCCGCTGAA | >hg38_21_+_8433172_8446622_RNA45SN1_45S_50ntFlanks | 7983 | 8008 | 26 | CAGATCAGACGTGGCGACCCGCTGAA | 2322.0 | -6.5 | 0.8 | -8.2 | 3.6E-16 | 1.1E-15 | BT20/MB468 | NUC |
| >MI0000650\|hsa-mir-200c&WithFlank&12\|+\|6963693\|6963772@50.72.23(+2A)&[MIMAT0000617&hsa-miR-200c-3p&offsets\|0\|0(+2A);m-54&12\|+\|6963742\|6963764&offsets\|0\|0(+2A)]\|TAATACTGCCGGGTAATGATGGAAA | >MI0000650\|hsa-mir-200c&WithFlank&12\|+\|6963693\|6963772 | 50 | 72 | 23(+2A) | TAATACTGCCGGGTAATGATGGAAA | 1259.9 | 9.3 | 1.1 | 8.5 | 1.2E-17 | 1.1E-15 | BT20/MB231 | CYTO |
| >hg38_21_+_8433172_8446622_RNA45SN1_45S_50ntFlanks@10690.10710.21\|CGTCCGGTGAGCTCTCGCTGG | >hg38_21_+_8433172_8446622_RNA45SN1_45S_50ntFlanks | 10690 | 10710 | 21 | CGTCCGGTGAGCTCTCGCTGG | 382.7 | -11.5 | 1.4 | -8.2 | 1.7E-16 | 1.1E-15 | MB231/MB468 | NUC |
| >hg38_21_+_8433172_8446622_RNA45SN1_45S_50ntFlanks@12157.12181.25\|AGTTTGACTGGGGCGGTACACCTGT | >hg38_21_+_8433172_8446622_RNA45SN1_45S_50ntFlanks | 12157 | 12181 | 25 | AGTTTGACTGGGGCGGTACACCTGT | 352.4 | -11.4 | 1.4 | -8.2 | 1.7E-16 | 1.1E-15 | MB231/MB468 | NUC |
| >trna116_GluCTC_1_-_145399233_145399304@2.33.32,trna59_GluCTC_1_+_249168447_249168518@2.33.32,trna71_GluCTC_1_-_161439189_161439260@2.33.32,trna74_GluCTC_1_-_161431809_161431880@2.33.32,trna77_GluCTC_1_-_161424398_161424469@2.33.32,trna77_GluCTC_6_+_28949976_28950047@2.33.32,trna80_GluCTC_1_-_161417018_161417089@2.33.32,trna87_GluCTC_6_-_126101393_126101464@2.33.32\|CCCTGGTGGTCTAGTGGTTAGGATTCGGCGCT | >trna116_GluCTC_1_-_145399233_145399304 | 2 | 33 | 32 | CCCTGGTGGTCTAGTGGTTAGGATTCGGCGCT | 870.8 | 14.2 | 1.7 | 8.2 | 1.6E-16 | 1.1E-15 | BT20/MB468 | MITO |
| >hg38_21_+_8433172_8446622_RNA45SN1_45S_50ntFlanks@4512.4529.18\|AAAAAATTAGAGTGTTCA | >hg38_21_+_8433172_8446622_RNA45SN1_45S_50ntFlanks | 4512 | 4529 | 18 | AAAAAATTAGAGTGTTCA | 272.9 | -12.0 | 1.4 | -8.3 | 1.4E-16 | 1.1E-15 | MB231/MB468 | MP |
| >hg38_21_+_8433172_8446622_RNA45SN1_45S_50ntFlanks@11567.11601.35\|AGCAGCCGACTTAGAACTGGTGCGGACCAGGGGAA | >hg38_21_+_8433172_8446622_RNA45SN1_45S_50ntFlanks | 11567 | 11601 | 35 | AGCAGCCGACTTAGAACTGGTGCGGACCAGGGGAA | 1387.3 | -4.2 | 0.5 | -8.2 | 1.7E-16 | 1.1E-15 | MB231/MB468 | NUC |
| >hg38_21_+_8433172_8446622_RNA45SN1_45S_50ntFlanks@6743.6772.30\|CGACACTTCGAACGCACTTGCGGCCCCGGG | >hg38_21_+_8433172_8446622_RNA45SN1_45S_50ntFlanks | 6743 | 6772 | 30 | CGACACTTCGAACGCACTTGCGGCCCCGGG | 1774.6 | -10.4 | 1.3 | -8.2 | 2.3E-16 | 1.2E-15 | MB231/MB468 | MITO |
| >hg38_21_+_8433172_8446622_RNA45SN1_45S_50ntFlanks@476.511.36\|TTCGTGATCGATGTGGTGACGTCGTGCTCTCCCGGG | >hg38_21_+_8433172_8446622_RNA45SN1_45S_50ntFlanks | 476 | 511 | 36 | TTCGTGATCGATGTGGTGACGTCGTGCTCTCCCGGG | 524.6 | 10.7 | 1.3 | 8.2 | 2.3E-16 | 1.2E-15 | BT20/MB468 | CYTO |
| >MI0001445\|hsa-mir-423&WithFlank&17\|+\|30117073\|30117178@23.44.22(+1C)&[MIMAT0004748&hsa-miR-423-5p&offsets\|0\|-1(+1C);m-141&17\|+\|30117095\|30117117&offsets\|0\|-1(+1C)]\|TGAGGGGCAGAGAGCGAGACTTC | >MI0001445\|hsa-mir-423&WithFlank&17\|+\|30117073\|30117178 | 23 | 44 | 22(+1C) | TGAGGGGCAGAGAGCGAGACTTC | 1854.0 | 10.1 | 1.2 | 8.2 | 2.3E-16 | 1.2E-15 | BT20/MB468 | CYTO |
| >hg38_21_+_8433172_8446622_RNA45SN1_45S_50ntFlanks@5472.5489.18\|ACGGCCCTGGCGGAGCGC | >hg38_21_+_8433172_8446622_RNA45SN1_45S_50ntFlanks | 5472 | 5489 | 18 | ACGGCCCTGGCGGAGCGC | 716.4 | 8.3 | 1.0 | 8.2 | 1.8E-16 | 1.2E-15 | MB231/MB468 | NUC |
| >hg38_21_+_8433172_8446622_RNA45SN1_45S_50ntFlanks@4241.4258.18\|CTCTTTCGAGGCCCTGTA | >hg38_21_+_8433172_8446622_RNA45SN1_45S_50ntFlanks | 4241 | 4258 | 18 | CTCTTTCGAGGCCCTGTA | 272.3 | -12.0 | 1.4 | -8.3 | 1.4E-16 | 1.2E-15 | MB231/MB468 | MP |
| >hg38_21_+_8433172_8446622_RNA45SN1_45S_50ntFlanks@4977.4994.18\|CGGACACGGACAGGATTG | >hg38_21_+_8433172_8446622_RNA45SN1_45S_50ntFlanks | 4977 | 4994 | 18 | CGGACACGGACAGGATTG | 427.9 | -12.6 | 1.5 | -8.3 | 1.4E-16 | 1.2E-15 | MB231/MB468 | MP |
| >hg38_21_+_8433172_8446622_RNA45SN1_45S_50ntFlanks@9568.9597.30\|CGTGCAAATCGGTCGTCCGACCTGGGTATA | >hg38_21_+_8433172_8446622_RNA45SN1_45S_50ntFlanks | 9568 | 9597 | 30 | CGTGCAAATCGGTCGTCCGACCTGGGTATA | 3672.5 | -7.9 | 1.0 | -8.3 | 1.5E-16 | 1.2E-15 | MB231/MB468 | CYTO |
| >MI0000299\|hsa-mir-222&WithFlank&X\|-\|45747009\|45747130@75.99.25(+1C)&[MIMAT0000279&hsa-miR-222-3p&offsets\|0\|+4(+1C);m-140&X\|-\|45747033\|45747056&offsets\|0\|+1(+1C)]\|AGCTACATCTGGCTACTGGGTCTCTC | >MI0000299\|hsa-mir-222&WithFlank&X\|-\|45747009\|45747130 | 75 | 99 | 25(+1C) | AGCTACATCTGGCTACTGGGTCTCTC | 715.5 | 9.5 | 1.1 | 8.3 | 1.5E-16 | 1.2E-15 | MB231/MB468 | CYTO |
| >hg38_21_+_8433172_8446622_RNA45SN1_45S_50ntFlanks@11104.11121.18\|AAGGGTCGGGGCGGCAGG | >hg38_21_+_8433172_8446622_RNA45SN1_45S_50ntFlanks | 11104 | 11121 | 18 | AAGGGTCGGGGCGGCAGG | 369.4 | -11.5 | 1.4 | -8.2 | 1.8E-16 | 1.2E-15 | MB231/MB468 | NUC |
| >hg38_21_+_8433172_8446622_RNA45SN1_45S_50ntFlanks@475.506.32\|CTTCGTGATCGATGTGGTGACGTCGTGCTCTC | >hg38_21_+_8433172_8446622_RNA45SN1_45S_50ntFlanks | 475 | 506 | 32 | CTTCGTGATCGATGTGGTGACGTCGTGCTCTC | 6456.9 | 7.8 | 0.9 | 8.2 | 1.7E-16 | 1.2E-15 | BT20/MB468 | MITO |
| >am_MI0000815\|hsa-mir-339&WithFlank&7\|-\|1022927\|1023032@21.41.21&[MIMAT0000764&hsa-miR-339-5p&offsets\|0\|-2;m-163&7\|-\|1022990\|1023012&offsets\|0\|-2]\|TCCCTGTCCTCCAGGAGCTCA | >am_MI0000815\|hsa-mir-339&WithFlank&7\|-\|1022927\|1023032 | 21 | 41 | 21 | TCCCTGTCCTCCAGGAGCTCA | 416.5 | 7.9 | 1.0 | 8.2 | 2.3E-16 | 1.2E-15 | BT20/MB468 | CYTO |
| >hg38_21_+_8433172_8446622_RNA45SN1_45S_50ntFlanks@10116.10136.21\|TGCGTCGGAGGGCGGCGGCGG | >hg38_21_+_8433172_8446622_RNA45SN1_45S_50ntFlanks | 10116 | 10136 | 21 | TGCGTCGGAGGGCGGCGGCGG | 654.3 | -13.2 | 1.6 | -8.3 | 1.4E-16 | 1.2E-15 | MB231/MB468 | MP |
| >hg38_21_+_8433172_8446622_RNA45SN1_45S_50ntFlanks@6751.6771.21\|CGAACGCACTTGCGGCCCCGG | >hg38_21_+_8433172_8446622_RNA45SN1_45S_50ntFlanks | 6751 | 6771 | 21 | CGAACGCACTTGCGGCCCCGG | 333.9 | -12.2 | 1.5 | -8.3 | 1.4E-16 | 1.2E-15 | MB231/MB468 | MP |
| >hg38_21_+_8433172_8446622_RNA45SN1_45S_50ntFlanks@9027.9065.39\|CGTCTCCTCGTGGGGGGGCCGGGCCACCCCTCCCACGGC | >hg38_21_+_8433172_8446622_RNA45SN1_45S_50ntFlanks | 9027 | 9065 | 39 | CGTCTCCTCGTGGGGGGGCCGGGCCACCCCTCCCACGGC | 355.3 | -12.3 | 1.5 | -8.3 | 1.5E-16 | 1.2E-15 | MB231/MB468 | MP |
| >hg38_21_+_8433172_8446622_RNA45SN1_45S_50ntFlanks@8114.8143.30\|GCGCGGGACATGTGGCGTACGGAAGACCCG | >hg38_21_+_8433172_8446622_RNA45SN1_45S_50ntFlanks | 8114 | 8143 | 30 | GCGCGGGACATGTGGCGTACGGAAGACCCG | 482.2 | -9.2 | 1.1 | -8.2 | 1.8E-16 | 1.2E-15 | MB231/MB468 | NUC |
| >MI0000102\|hsa-mir-100&WithFlank&11\|-\|122152223\|122152314@19.40.22(+1A)&[MIMAT0000098&hsa-miR-100-5p&offsets\|0\|0(+1A);m-27&11\|-\|122152276\|122152296&offsets\|0\|+1(+1A)]\|AACCCGTAGATCCGAACTTGTGA | >MI0000102\|hsa-mir-100&WithFlank&11\|-\|122152223\|122152314 | 19 | 40 | 22(+1A) | AACCCGTAGATCCGAACTTGTGA | 24603.3 | 10.5 | 1.3 | 8.3 | 1.5E-16 | 1.2E-15 | MB231/MB468 | MP |
| >hg38_21_+_8433172_8446622_RNA45SN1_45S_50ntFlanks@10389.10407.19\|TGAACAGCAGTTGAACATG | >hg38_21_+_8433172_8446622_RNA45SN1_45S_50ntFlanks | 10389 | 10407 | 19 | TGAACAGCAGTTGAACATG | 1919.5 | -10.2 | 1.2 | -8.3 | 1.5E-16 | 1.2E-15 | MB231/MB468 | MP |
| >hg38_21_+_8433172_8446622_RNA45SN1_45S_50ntFlanks@6654.6689.36\|CTCTTAGCGGTGGATCACTCGGCTCGTGCGTCGATG | >hg38_21_+_8433172_8446622_RNA45SN1_45S_50ntFlanks | 6654 | 6689 | 36 | CTCTTAGCGGTGGATCACTCGGCTCGTGCGTCGATG | 2625.6 | -5.9 | 0.7 | -8.2 | 2.4E-16 | 1.2E-15 | MB231/MB468 | MITO |
| >hg38_21_+_8433172_8446622_RNA45SN1_45S_50ntFlanks@9180.9210.31\|CCGTCGGGCCCGGGGGAGGTTCTCTCGGGGC | >hg38_21_+_8433172_8446622_RNA45SN1_45S_50ntFlanks | 9180 | 9210 | 31 | CCGTCGGGCCCGGGGGAGGTTCTCTCGGGGC | 421.6 | -12.6 | 1.5 | -8.3 | 1.5E-16 | 1.2E-15 | MB231/MB468 | MP |
| >hg38_21_+_8433172_8446622_RNA45SN1_45S_50ntFlanks@4555.4587.33\|CGCAGCTAGGAATAATGGAATAGGACCGCGGTT | >hg38_21_+_8433172_8446622_RNA45SN1_45S_50ntFlanks | 4555 | 4587 | 33 | CGCAGCTAGGAATAATGGAATAGGACCGCGGTT | 294.4 | -12.0 | 1.5 | -8.2 | 2.4E-16 | 1.2E-15 | MB231/MB468 | MITO |
| >MI0000299\|hsa-mir-222&WithFlank&X\|-\|45747009\|45747130@75.97.23(+1U)&[MIMAT0000279&hsa-miR-222-3p&offsets\|0\|+2(+1U);m-140&X\|-\|45747033\|45747056&offsets\|0\|-1(+1U)]\|AGCTACATCTGGCTACTGGGTCTT | >MI0000299\|hsa-mir-222&WithFlank&X\|-\|45747009\|45747130 | 75 | 97 | 23(+1U) | AGCTACATCTGGCTACTGGGTCTT | 2173.2 | 6.7 | 0.8 | 8.2 | 1.7E-16 | 1.2E-15 | BT20/MB468 | MITO |
| >hg38_21_+_8433172_8446622_RNA45SN1_45S_50ntFlanks@11768.11803.36\|CTAATTAGTGACGCGCATGAATGGATGAACGAGATT | >hg38_21_+_8433172_8446622_RNA45SN1_45S_50ntFlanks | 11768 | 11803 | 36 | CTAATTAGTGACGCGCATGAATGGATGAACGAGATT | 4498.3 | -11.8 | 1.4 | -8.1 | 3.8E-16 | 1.2E-15 | BT20/MB468 | NUC |
| >hg38_21_+_8433172_8446622_RNA45SN1_45S_50ntFlanks@10362.10383.22\|AAGGCCGAAGTGGAGAAGGGTT | >hg38_21_+_8433172_8446622_RNA45SN1_45S_50ntFlanks | 10362 | 10383 | 22 | AAGGCCGAAGTGGAGAAGGGTT | 1038.9 | -4.8 | 0.6 | -8.2 | 1.7E-16 | 1.2E-15 | BT20/MB468 | MITO |
| >hg38_21_+_8433172_8446622_RNA45SN1_45S_50ntFlanks@12959.12995.37\|TCTGGGTCGGGGTTTCGTACGTAGCAGAGCAGCTCCC | >hg38_21_+_8433172_8446622_RNA45SN1_45S_50ntFlanks | 12959 | 12995 | 37 | TCTGGGTCGGGGTTTCGTACGTAGCAGAGCAGCTCCC | 314.4 | 8.3 | 1.0 | 8.3 | 1.6E-16 | 1.2E-15 | MB231/MB468 | CYTO |
| >hg38_21_+_8433172_8446622_RNA45SN1_45S_50ntFlanks@10806.10830.25\|AACAATGTAGGTAAGGGAAGTCGGC | >hg38_21_+_8433172_8446622_RNA45SN1_45S_50ntFlanks | 10806 | 10830 | 25 | AACAATGTAGGTAAGGGAAGTCGGC | 294.4 | -12.0 | 1.5 | -8.2 | 2.5E-16 | 1.2E-15 | MB231/MB468 | MITO |
| >hg38_21_+_8433172_8446622_RNA45SN1_45S_50ntFlanks@12712.12751.40\|CGGAGCCTCGGTTGGCCTCGGATAGCCGGTCCCCCGCCTG | >hg38_21_+_8433172_8446622_RNA45SN1_45S_50ntFlanks | 12712 | 12751 | 40 | CGGAGCCTCGGTTGGCCTCGGATAGCCGGTCCCCCGCCTG | 1044.4 | -12.1 | 1.5 | -8.2 | 1.7E-16 | 1.2E-15 | BT20/MB468 | MITO |
| >hg38_21_+_8433172_8446622_RNA45SN1_45S_50ntFlanks@3866.3898.33\|CTAGAGCTAATACATGCCGACGGGCGCTGACCC | >hg38_21_+_8433172_8446622_RNA45SN1_45S_50ntFlanks | 3866 | 3898 | 33 | CTAGAGCTAATACATGCCGACGGGCGCTGACCC | 505.6 | -11.1 | 1.3 | -8.2 | 1.7E-16 | 1.2E-15 | BT20/MB468 | MITO |
| >hg38_21_+_8433172_8446622_RNA45SN1_45S_50ntFlanks@3738.3756.19\|TCTCAAAGATTAAGCCATG | >hg38_21_+_8433172_8446622_RNA45SN1_45S_50ntFlanks | 3738 | 3756 | 19 | TCTCAAAGATTAAGCCATG | 471.4 | -12.7 | 1.5 | -8.2 | 2.5E-16 | 1.2E-15 | MB231/MB468 | MITO |

<table>
<tr><td></td><td></td><td></td><td></td><td></td><td></td><td></td><td></td><td></td><td></td><td></td><td></td><td></td><td></td></tr>
<tr><td>>MI0000477|hsa-mir-146a&WithFlank&5|+|160485346|160485456@27.48.22(+1A)&[MIMAT0000449&hsa-miR-146a-5p&offsets|0|0(+1A);m-129&5|+|160485372|160485393&offsets|0|0(+1A)]|TGAGAACTGAATTCCATGGGTTA</td><td>>MI0000477|hsa-mir-146a&WithFlank&5|+|160485346|160485456</td><td>27</td><td>48</td><td>22(+1A)</td><td>TGAGAACTGAATTCCATGGGTTA</td><td>646.1</td><td>13.2</td><td>1.6</td><td>8.2</td><td>1.9E-16</td><td>1.2E-15</td><td>MB231/MB468</td><td>NUC</td></tr>
<tr><td>>hg38_21_+_8433172_8446622_RNA45SN1_45S_50ntFlanks@11729.11765.37|TAACTATGACTCTCTTAAGGTAGCCAAATGCCTCGTC</td><td>>hg38_21_+_8433172_8446622_RNA45SN1_45S_50ntFlanks</td><td>11729</td><td>11765</td><td>37</td><td>TAACTATGACTCTCTTAAGGTAGCCAAATGCCTCGTC</td><td>17247.6</td><td>-13.3</td><td>1.6</td><td>-8.1</td><td>3.9E-16</td><td>1.2E-15</td><td>BT20/MB468</td><td>NUC</td></tr>
<tr><td>>hg38_21_+_8433172_8446622_RNA45SN1_45S_50ntFlanks@11452.11475.24|CTCCTCGGCGCGCGGCGGCGGCGG</td><td>>hg38_21_+_8433172_8446622_RNA45SN1_45S_50ntFlanks</td><td>11452</td><td>11475</td><td>24</td><td>CTCCTCGGCGCGCGGCGGCGGCGG</td><td>245.9</td><td>-11.8</td><td>1.4</td><td>-8.3</td><td>1.5E-16</td><td>1.2E-15</td><td>MB231/MB468</td><td>MP</td></tr>
<tr><td>>hg38_21_+_8433172_8446622_RNA45SN1_45S_50ntFlanks@9564.9592.29|CTGACGTGCAAATCGGTCGTCCGACCTGG</td><td>>hg38_21_+_8433172_8446622_RNA45SN1_45S_50ntFlanks</td><td>9564</td><td>9592</td><td>29</td><td>CTGACGTGCAAATCGGTCGTCCGACCTGG</td><td>1370.6</td><td>-4.9</td><td>0.6</td><td>-8.2</td><td>1.9E-16</td><td>1.2E-15</td><td>MB231/MB468</td><td>NUC</td></tr>
<tr><td>>am_MI0000809|hsa-mir-151a&WithFlank&8|-|140732558|140732659@53.72.20&[MIMAT0000757&hsa-miR-151a-3p&offsets|0|-1;m-30&8|-|140732587|140732607&offsets|0|-1]|CTAGACTGAAGCTCCTTGAG</td><td>>am_MI0000809|hsa-mir-151a&WithFlank&8|-|140732558|140732659</td><td>53</td><td>72</td><td>20</td><td>CTAGACTGAAGCTCCTTGAG</td><td>974.7</td><td>6.4</td><td>0.8</td><td>8.2</td><td>1.6E-16</td><td>1.2E-15</td><td>MB231/MB468</td><td>CYTO</td></tr>
<tr><td>>MI0000079|hsa-mir-23a&WithFlank&19|-|13836581|13836665@54.71.18&[MIMAT0000078&hsa-miR-23a-3p&offsets|+3|0;m-31&19|-|13836595|13836615&offsets|+3|0]|ACATTGCCAGGGATTTCC</td><td>>MI0000079|hsa-mir-23a&WithFlank&19|-|13836581|13836665</td><td>54</td><td>71</td><td>18</td><td>ACATTGCCAGGGATTTCC</td><td>3829.9</td><td>29.9</td><td>3.6</td><td>8.2</td><td>1.9E-16</td><td>1.2E-15</td><td>MB231/MB468</td><td>NUC</td></tr>
<tr><td>>hg38_21_+_8433172_8446622_RNA45SN1_45S_50ntFlanks@6721.6750.30|AATTGCAGGACACATTGATCATCGACACTT</td><td>>hg38_21_+_8433172_8446622_RNA45SN1_45S_50ntFlanks</td><td>6721</td><td>6750</td><td>30</td><td>AATTGCAGGACACATTGATCATCGACACTT</td><td>502.3</td><td>-4.5</td><td>0.5</td><td>-8.2</td><td>1.9E-16</td><td>1.2E-15</td><td>MB231/MB468</td><td>NUC</td></tr>
<tr><td>>hg38_21_+_8433172_8446622_RNA45SN1_45S_50ntFlanks@5495.5516.22|AGACGGTCGAACTTGACTATCT</td><td>>hg38_21_+_8433172_8446622_RNA45SN1_45S_50ntFlanks</td><td>5495</td><td>5516</td><td>22</td><td>AGACGGTCGAACTTGACTATCT</td><td>1113.5</td><td>7.9</td><td>1.0</td><td>8.2</td><td>1.6E-16</td><td>1.3E-15</td><td>MB231/MB468</td><td>CYTO</td></tr>
<tr><td>>hg38_21_+_8433172_8446622_RNA45SN1_45S_50ntFlanks@3985.4004.20|CGGCTTTGGTGACTCTAGAT</td><td>>hg38_21_+_8433172_8446622_RNA45SN1_45S_50ntFlanks</td><td>3985</td><td>4004</td><td>20</td><td>CGGCTTTGGTGACTCTAGAT</td><td>656.3</td><td>-13.1</td><td>1.6</td><td>-8.2</td><td>2.5E-16</td><td>1.3E-15</td><td>MB231/MB468</td><td>MITO</td></tr>
<tr><td>>MI0000477|hsa-mir-146a&WithFlank&5|+|160485346|160485456@27.49.23&[MIMAT0000449&hsa-miR-146a-5p&offsets|0|+1;m-129&5|+|160485372|160485393&offsets|0|+1]|TGAGAACTGAATTCCATGGGTTG</td><td>>MI0000477|hsa-mir-146a&WithFlank&5|+|160485346|160485456</td><td>27</td><td>49</td><td>23</td><td>TGAGAACTGAATTCCATGGGTTG</td><td>1944.3</td><td>12.2</td><td>1.5</td><td>8.2</td><td>1.9E-16</td><td>1.3E-15</td><td>MB231/MB468</td><td>NUC</td></tr>
<tr><td>>MI0000253|hsa-mir-148a&WithFlank&7|-|25949913|25949992@50.70.21(+1G)&[MIMAT0000243&hsa-miR-148a-3p&offsets|0|-1(+1G);m-7&7|-|25949922|25949943&offsets|0|-1(+1G)]|TCAGTGCACTACAGAACTTTGG</td><td>>MI0000253|hsa-mir-148a&WithFlank&7|-|25949913|25949992</td><td>50</td><td>70</td><td>21(+1G)</td><td>TCAGTGCACTACAGAACTTTGG</td><td>1541.6</td><td>6.2</td><td>0.8</td><td>8.2</td><td>1.8E-16</td><td>1.3E-15</td><td>BT20/MB468</td><td>MITO</td></tr>
<tr><td>>MI0000439|hsa-mir-23b&WithFlank&9|+|95085202|95085310@67.86.20(+1U)&[MIMAT0000418&hsa-miR-23b-3p&offsets|+3|0(+1U);m-42&9|+|95085265|95085285&offsets|+3|+2(+1U)]|ACATTGCCAGGGATTACCACT</td><td>>MI0000439|hsa-mir-23b&WithFlank&9|+|95085202|95085310</td><td>67</td><td>86</td><td>20(+1U)</td><td>ACATTGCCAGGGATTACCACT</td><td>5499.7</td><td>29.7</td><td>3.6</td><td>8.2</td><td>2.6E-16</td><td>1.3E-15</td><td>MB231/MB468</td><td>MITO</td></tr>
<tr><td>>hg38_21_+_8433172_8446622_RNA45SN1_45S_50ntFlanks@11578.11605.28|TAGAACTGGTGCGGACCAGGGGAATCCG</td><td>>hg38_21_+_8433172_8446622_RNA45SN1_45S_50ntFlanks</td><td>11578</td><td>11605</td><td>28</td><td>TAGAACTGGTGCGGACCAGGGGAATCCG</td><td>303.1</td><td>-12.1</td><td>1.5</td><td>-8.3</td><td>1.5E-16</td><td>1.3E-15</td><td>MB231/MB468</td><td>MP</td></tr>
<tr><td>>hg38_21_+_8433172_8446622_RNA45SN1_45S_50ntFlanks@11641.11674.34|CGGCGGGTGTTGACGCGATGTGATTTCTGCCCAG</td><td>>hg38_21_+_8433172_8446622_RNA45SN1_45S_50ntFlanks</td><td>11641</td><td>11674</td><td>34</td><td>CGGCGGGTGTTGACGCGATGTGATTTCTGCCCAG</td><td>371.3</td><td>-12.4</td><td>1.5</td><td>-8.3</td><td>1.5E-16</td><td>1.3E-15</td><td>MB231/MB468</td><td>MP</td></tr>
<tr><td>>hg38_21_+_8433172_8446622_RNA45SN1_45S_50ntFlanks@4735.4768.34|ACGAAAGTCGGAGGTTCGAAGACGATCAGATACC</td><td>>hg38_21_+_8433172_8446622_RNA45SN1_45S_50ntFlanks</td><td>4735</td><td>4768</td><td>34</td><td>ACGAAAGTCGGAGGTTCGAAGACGATCAGATACC</td><td>638.4</td><td>10.7</td><td>1.3</td><td>8.2</td><td>1.6E-16</td><td>1.3E-15</td><td>MB231/MB468</td><td>CYTO</td></tr>
<tr><td>>hg38_21_+_8433172_8446622_RNA45SN1_45S_50ntFlanks@8357.8387.31|AAAAGAACTTTGAAGAGAGAGTTCAAGAGGG</td><td>>hg38_21_+_8433172_8446622_RNA45SN1_45S_50ntFlanks</td><td>8357</td><td>8387</td><td>31</td><td>AAAAGAACTTTGAAGAGAGAGTTCAAGAGGG</td><td>342.5</td><td>-11.1</td><td>1.4</td><td>-8.2</td><td>1.6E-16</td><td>1.3E-15</td><td>MB231/MB468</td><td>CYTO</td></tr>
<tr><td>>hg38_21_+_8433172_8446622_RNA45SN1_45S_50ntFlanks@12151.12179.29|GTGGGGAGTTTGACTGGGGCGGTACACCT</td><td>>hg38_21_+_8433172_8446622_RNA45SN1_45S_50ntFlanks</td><td>12151</td><td>12179</td><td>29</td><td>GTGGGGAGTTTGACTGGGGCGGTACACCT</td><td>368.8</td><td>-12.3</td><td>1.5</td><td>-8.2</td><td>2.6E-16</td><td>1.3E-15</td><td>MB231/MB468</td><td>MITO</td></tr>
<tr><td>>hg38_21_+_8433172_8446622_RNA45SN1_45S_50ntFlanks@4853.4883.31|AAGTCTTTGGGTTCCGGGGGGAGTATGGTTG</td><td>>hg38_21_+_8433172_8446622_RNA45SN1_45S_50ntFlanks</td><td>4853</td><td>4883</td><td>31</td><td>AAGTCTTTGGGTTCCGGGGGGAGTATGGTTG</td><td>416.5</td><td>-12.5</td><td>1.5</td><td>-8.2</td><td>2.6E-16</td><td>1.3E-15</td><td>MB231/MB468</td><td>MITO</td></tr>
<tr><td>>hg38_21_+_8433172_8446622_RNA45SN1_45S_50ntFlanks@12243.12275.33|GAAGGGCAAAAGCTCGCTTGATCTTGATTTTCA</td><td>>hg38_21_+_8433172_8446622_RNA45SN1_45S_50ntFlanks</td><td>12243</td><td>12275</td><td>33</td><td>GAAGGGCAAAAGCTCGCTTGATCTTGATTTTCA</td><td>1797.7</td><td>7.4</td><td>0.9</td><td>8.2</td><td>1.8E-16</td><td>1.3E-15</td><td>BT20/MB468</td><td>MITO</td></tr>
<tr><td>>hg38_21_+_8433172_8446622_RNA45SN1_45S_50ntFlanks@3899.3936.38|CCTTCGCGGGGGGGATGCGTGCATTTATCAGATCAAAA</td><td>>hg38_21_+_8433172_8446622_RNA45SN1_45S_50ntFlanks</td><td>3899</td><td>3936</td><td>38</td><td>CCTTCGCGGGGGGGATGCGTGCATTTATCAGATCAAAA</td><td>392.7</td><td>-12.4</td><td>1.5</td><td>-8.2</td><td>2.6E-16</td><td>1.3E-15</td><td>MB231/MB468</td><td>MITO</td></tr>
<tr><td>>hg38_21_+_8433172_8446622_RNA45SN1_45S_50ntFlanks@10680.10706.27|GTTCCGGCGGCGTCCGGTGAGCTCTCG</td><td>>hg38_21_+_8433172_8446622_RNA45SN1_45S_50ntFlanks</td><td>10680</td><td>10706</td><td>27</td><td>GTTCCGGCGGCGTCCGGTGAGCTCTCG</td><td>13413.1</td><td>-7.0</td><td>0.9</td><td>-8.2</td><td>1.6E-16</td><td>1.3E-15</td><td>MB231/MB468</td><td>CYTO</td></tr>
<tr><td>>hg38_21_+_8433172_8446622_RNA45SN1_45S_50ntFlanks@9122.9140.19|CCGCGCGCGGGTCGGGGGG</td><td>>hg38_21_+_8433172_8446622_RNA45SN1_45S_50ntFlanks</td><td>9122</td><td>9140</td><td>19</td><td>CCGCGCGCGGGTCGGGGGG</td><td>452.2</td><td>-12.6</td><td>1.5</td><td>-8.2</td><td>2.6E-16</td><td>1.3E-15</td><td>MB231/MB468</td><td>MITO</td></tr>
<tr><td>>MI0005559|hsa-mir-744&WithFlank&17|+|12081893|12082002@17.34.18&[MIMAT0004945&hsa-miR-744-5p&offsets|0|-4;m-172&17|+|12081909|12081929&offsets|0|-3]|TGCGGGGCTAGGGCTAAC</td><td>>MI0005559|hsa-mir-744&WithFlank&17|+|12081893|12082002</td><td>17</td><td>34</td><td>18</td><td>TGCGGGGCTAGGGCTAAC</td><td>319.7</td><td>10.0</td><td>1.2</td><td>8.2</td><td>2.5E-16</td><td>1.3E-15</td><td>BT20/MB468</td><td>CYTO</td></tr>
<tr><td>>hg38_21_+_8433172_8446622_RNA45SN1_45S_50ntFlanks@4297.4320.24|CATTGGAGGGCAAGTCTGGTGCCA</td><td>>hg38_21_+_8433172_8446622_RNA45SN1_45S_50ntFlanks</td><td>4297</td><td>4320</td><td>24</td><td>CATTGGAGGGCAAGTCTGGTGCCA</td><td>231.0</td><td>-10.8</td><td>1.3</td><td>-8.2</td><td>2.0E-16</td><td>1.3E-15</td><td>MB231/MB468</td><td>NUC</td></tr>
<tr><td>>MI0000439|hsa-mir-23b&WithFlank&9|+|95085202|95085310@67.86.20(+1U)&[MIMAT0000418&hsa-miR-23b-3p&offsets|+3|0(+1U);m-42&9|+|95085265|95085285&offsets|+3|+2(+1U)]|ACATTGCCAGGGATTACCACT</td><td>>MI0000439|hsa-mir-23b&WithFlank&9|+|95085202|95085310</td><td>67</td><td>86</td><td>20(+1U)</td><td>ACATTGCCAGGGATTACCACT</td><td>3778.2</td><td>29.9</td><td>3.6</td><td>8.2</td><td>2.0E-16</td><td>1.3E-15</td><td>MB231/MB468</td><td>NUC</td></tr>
<tr><td>>hg38_21_+_8433172_8446622_RNA45SN1_45S_50ntFlanks@12159.12176.18|TTTGACTGGGGCGGTACA</td><td>>hg38_21_+_8433172_8446622_RNA45SN1_45S_50ntFlanks</td><td>12159</td><td>12176</td><td>18</td><td>TTTGACTGGGGCGGTACA</td><td>231.8</td><td>-11.7</td><td>1.4</td><td>-8.3</td><td>1.6E-16</td><td>1.3E-15</td><td>MB231/MB468</td><td>MP</td></tr>
<tr><td>>hg38_21_+_8433172_8446622_RNA45SN1_45S_50ntFlanks@10781.10806.26|AGGTGAACAGCCTCTGGCATGTTGGA</td><td>>hg38_21_+_8433172_8446622_RNA45SN1_45S_50ntFlanks</td><td>10781</td><td>10806</td><td>26</td><td>AGGTGAACAGCCTCTGGCATGTTGGA</td><td>336.5</td><td>-12.3</td><td>1.5</td><td>-8.3</td><td>1.6E-16</td><td>1.3E-15</td><td>MB231/MB468</td><td>MP</td></tr>
<tr><td>>hg38_21_+_8433172_8446622_RNA45SN1_45S_50ntFlanks@10408.10431.24|GGTCAGTCGGTCCTGAGAGATGGG</td><td>>hg38_21_+_8433172_8446622_RNA45SN1_45S_50ntFlanks</td><td>10408</td><td>10431</td><td>24</td><td>GGTCAGTCGGTCCTGAGAGATGGG</td><td>513.4</td><td>-12.9</td><td>1.6</td><td>-8.2</td><td>1.6E-16</td><td>1.3E-15</td><td>MB231/MB468</td><td>MP</td></tr>
<tr><td>>hg38_21_+_8433172_8446622_RNA45SN1_45S_50ntFlanks@10914.10938.25|CGCCGCGGCTGGACGAGGCGCCGCC</td><td>>hg38_21_+_8433172_8446622_RNA45SN1_45S_50ntFlanks</td><td>10914</td><td>10938</td><td>25</td><td>CGCCGCGGCTGGACGAGGCGCCGCC</td><td>233.5</td><td>-11.7</td><td>1.4</td><td>-8.2</td><td>1.6E-16</td><td>1.3E-15</td><td>MB231/MB468</td><td>MP</td></tr>
<tr><td>>hg38_21_+_8433172_8446622_RNA45SN1_45S_50ntFlanks@12347.12365.19|AGAAAAGTTACCACAGGGA</td><td>>hg38_21_+_8433172_8446622_RNA45SN1_45S_50ntFlanks</td><td>12347</td><td>12365</td><td>19</td><td>AGAAAAGTTACCACAGGGA</td><td>513.1</td><td>-12.9</td><td>1.6</td><td>-8.2</td><td>1.6E-16</td><td>1.3E-15</td><td>MB231/MB468</td><td>MP</td></tr>
<tr><td>>hg38_21_+_8433172_8446622_RNA45SN1_45S_50ntFlanks@9129.9149.21|CGGGTCGGGGGGCGGGGCGGA</td><td>>hg38_21_+_8433172_8446622_RNA45SN1_45S_50ntFlanks</td><td>9129</td><td>9149</td><td>21</td><td>CGGGTCGGGGGGCGGGGCGGA</td><td>677.1</td><td>-13.3</td><td>1.6</td><td>-8.2</td><td>1.6E-16</td><td>1.3E-15</td><td>MB231/MB468</td><td>MP</td></tr>
<tr><td>>hg38_21_+_8433172_8446622_RNA45SN1_45S_50ntFlanks@12073.12106.34|CCGAGGGGCTCTCGCTTCTGGCGCCAAGCGCCCG</td><td>>hg38_21_+_8433172_8446622_RNA45SN1_45S_50ntFlanks</td><td>12073</td><td>12106</td><td>34</td><td>CCGAGGGGCTCTCGCTTCTGGCGCCAAGCGCCCG</td><td>1096.0</td><td>-7.7</td><td>0.9</td><td>-8.2</td><td>2.6E-16</td><td>1.3E-15</td><td>MB231/MB468</td><td>MITO</td></tr>
<tr><td>>hg38_21_+_8433172_8446622_RNA45SN1_45S_50ntFlanks@6721.6741.21|AATTGCAGGACACATTGATCA</td><td>>hg38_21_+_8433172_8446622_RNA45SN1_45S_50ntFlanks</td><td>6721</td><td>6741</td><td>21</td><td>AATTGCAGGACACATTGATCA</td><td>287.5</td><td>-12.0</td><td>1.5</td><td>-8.2</td><td>1.6E-16</td><td>1.3E-15</td><td>MB231/MB468</td><td>MP</td></tr>
<tr><td>>hg38_21_+_8433172_8446622_RNA45SN1_45S_50ntFlanks@6741.6770.30|ATCGACACTTCGAACGCACTTGCGGCCCCG</td><td>>hg38_21_+_8433172_8446622_RNA45SN1_45S_50ntFlanks</td><td>6741</td><td>6770</td><td>30</td><td>ATCGACACTTCGAACGCACTTGCGGCCCCG</td><td>1585.8</td><td>-4.5</td><td>0.5</td><td>-8.2</td><td>1.9E-16</td><td>1.3E-15</td><td>BT20/MB468</td><td>MITO</td></tr>
<tr><td>>MI0000102|hsa-mir-100&WithFlank&11|-|122152223|122152314@19.39.21(+1U)&[MIMAT0000098&hsa-miR-100-5p&offsets|0|-1(+1U);m-27&11|-|122152276|122152296&offsets|0|0(+1U)]|AACCCGTAGATCCGAACTTGTT</td><td>>MI0000102|hsa-mir-100&WithFlank&11|-|122152223|122152314</td><td>19</td><td>39</td><td>21(+1U)</td><td>AACCCGTAGATCCGAACTTGTT</td><td>7288.1</td><td>-7.8</td><td>0.9</td><td>-8.5</td><td>2.0E-17</td><td>1.3E-15</td><td>BT20/MB231</td><td>MITO</td></tr>
<tr><td>>hg38_21_+_8433172_8446622_RNA45SN1_45S_50ntFlanks@6746.6772.27|CACTTCGAACGCACTTGCGGCCCCGGG</td><td>>hg38_21_+_8433172_8446622_RNA45SN1_45S_50ntFlanks</td><td>6746</td><td>6772</td><td>27</td><td>CACTTCGAACGCACTTGCGGCCCCGGG</td><td>318.2</td><td>-12.1</td><td>1.5</td><td>-8.2</td><td>2.6E-16</td><td>1.3E-15</td><td>MB231/MB468</td><td>MITO</td></tr>
<tr><td>>hg38_21_+_8433172_8446622_RNA45SN1_45S_50ntFlanks@9205.9222.18|CGGGGCCACGCGCGCGTC</td><td>>hg38_21_+_8433172_8446622_RNA45SN1_45S_50ntFlanks</td><td>9205</td><td>9222</td><td>18</td><td>CGGGGCCACGCGCGCGTC</td><td>365.9</td><td>-12.4</td><td>1.5</td><td>-8.2</td><td>1.6E-16</td><td>1.3E-15</td><td>MB231/MB468</td><td>MP</td></tr>
<tr><td>>MI0000264|hsa-mir-7-2&WithFlank&15|+|88611819|88611940@38.59.22(+1C);MI0000265|hsa-mir-7-3&WithFlank&19|+|4770664|4770785@37.58.22(+1C);MI0000263|hsa-mir-7-1&WithFlank&9|-|83969742|83969863@30.51.22(+1C)&[MIMAT0000252&hsa-miR-7-5p&offsets|0|-2(+1C);m-400&15|+|88611856|88611879&offsets|0|-2(+1C)];[MIMAT0000252_1&hsa-miR-7-5p&offsets|0|-2(+1C);m-402&19|+|4770700|4770723&offsets|0|-2(+1C)];[MIMAT0000252_2&hsa-miR-7-5p&offsets|0|-2(+1C);m-401&9|-|83969811|83969834&offsets|0|-2(+1C)]|TGGAAGACTAGTGATTTTGTTGC</td><td>>MI0000264|hsa-mir-7-2&WithFlank&15|+|88611819|88611940</td><td>38</td><td>59</td><td>22(+1C)</td><td>TGGAAGACTAGTGATTTTGTTGC</td><td>378.8</td><td>3.9</td><td>0.5</td><td>8.1</td><td>4.1E-16</td><td>1.3E-15</td><td>BT20/MB468</td><td>NUC</td></tr>
<tr><td>>MI0000077|hsa-mir-21&WithFlank&17|+|59841260|59841343@14.35.22(+1U)&[MIMAT0000076&hsa-miR-21-5p&offsets|0|0(+1U);m-1&17|+|59841273|59841294&offsets|0|0(+1U)]|TAGCTTATCAGACTGATGTTGAT</td><td>>MI0000077|hsa-mir-21&WithFlank&17|+|59841260|59841343</td><td>14</td><td>35</td><td>22(+1U)</td><td>TAGCTTATCAGACTGATGTTGAT</td><td>282336.9</td><td>8.7</td><td>1.1</td><td>8.2</td><td>2.6E-16</td><td>1.3E-15</td><td>BT20/MB468</td><td>CYTO</td></tr>
<tr><td>>hg38_21_+_8433172_8446622_RNA45SN1_45S_50ntFlanks@11885.11904.20|TTGAGCTTGACTCTAGTCTG</td><td>>hg38_21_+_8433172_8446622_RNA45SN1_45S_50ntFlanks</td><td>11885</td><td>11904</td><td>20</td><td>TTGAGCTTGACTCTAGTCTG</td><td>2410.0</td><td>-10.6</td><td>1.3</td><td>-8.2</td><td>2.6E-16</td><td>1.3E-15</td><td>BT20/MB468</td><td>CYTO</td></tr>
<tr><td>>hg38_21_+_8433172_8446622_RNA45SN1_45S_50ntFlanks@10787.10825.39|ACAGCCTCTGGCATGTTGGAACAATGTAGGTAAGGGAAG</td><td>>hg38_21_+_8433172_8446622_RNA45SN1_45S_50ntFlanks</td><td>10787</td><td>10825</td><td>39</td><td>ACAGCCTCTGGCATGTTGGAACAATGTAGGTAAGGGAAG</td><td>384.5</td><td>-12.4</td><td>1.5</td><td>-8.2</td><td>2.7E-16</td><td>1.3E-15</td><td>MB231/MB468</td><td>MITO</td></tr>
<tr><td>>hg38_21_+_8433172_8446622_RNA45SN1_45S_50ntFlanks@4560.4577.18|CTAGGAATAATGGAATAG</td><td>>hg38_21_+_8433172_8446622_RNA45SN1_45S_50ntFlanks</td><td>4560</td><td>4577</td><td>18</td><td>CTAGGAATAATGGAATAG</td><td>277.8</td><td>-12.0</td><td>1.5</td><td>-8.2</td><td>1.6E-16</td><td>1.3E-15</td><td>MB231/MB468</td><td>MP</td></tr>
<tr><td>>hg38_21_+_8433172_8446622_RNA45SN1_45S_50ntFlanks@12995.13036.42|CTCGCTGCGATCTATTGAAAGTCAGCCCTCGACACAAGGGTT</td><td>>hg38_21_+_8433172_8446622_RNA45SN1_45S_50ntFlanks</td><td>12995</td><td>13036</td><td>42</td><td>CTCGCTGCGATCTATTGAAAGTCAGCCCTCGACACAAGGGTT</td><td>8093.3</td><td>9.4</td><td>1.1</td><td>8.2</td><td>2.6E-16</td><td>1.3E-15</td><td>BT20/MB468</td><td>CYTO</td></tr>
<tr><td>>hg38_21_+_8433172_8446622_RNA45SN1_45S_50ntFlanks@12073.12092.20|CCGAGGGGCTCTCGCTTCTG</td><td>>hg38_21_+_8433172_8446622_RNA45SN1_45S_50ntFlanks</td><td>12073</td><td>12092</td><td>20</td><td>CCGAGGGGCTCTCGCTTCTG</td><td>353.1</td><td>-12.2</td><td>1.5</td><td>-8.2</td><td>2.7E-16</td><td>1.3E-15</td><td>MB231/MB468</td><td>MITO</td></tr>
<tr><td>>hg38_21_+_8433172_8446622_RNA45SN1_45S_50ntFlanks@8799.8823.25|CCTCGGGAGGGCGCGCGGGTCGGGG</td><td>>hg38_21_+_8433172_8446622_RNA45SN1_45S_50ntFlanks</td><td>8799</td><td>8823</td><td>25</td><td>CCTCGGGAGGGCGCGCGGGTCGGGG</td><td>423.0</td><td>-12.5</td><td>1.5</td><td>-8.2</td><td>2.7E-16</td><td>1.3E-15</td><td>MB231/MB468</td><td>MITO</td></tr>
<tr><td>>hg38_21_+_8433172_8446622_RNA45SN1_45S_50ntFlanks@10362.10383.22|AAGGCCGAAGTGGAGAAGGGTT</td><td>>hg38_21_+_8433172_8446622_RNA45SN1_45S_50ntFlanks</td><td>10362</td><td>10383</td><td>22</td><td>AAGGCCGAAGTGGAGAAGGGTT</td><td>9075.4</td><td>-12.4</td><td>1.5</td><td>-8.1</td><td>4.2E-16</td><td>1.3E-15</td><td>BT20/MB468</td><td>NUC</td></tr>
</table>

| | | | | | | | | | | | | | |
|---|---|---|---|---|---|---|---|---|---|---|---|---|---|
| >hg38_21_+_8433172_8446622_RNA45SN1_45S_50ntFlanks@12974.13001.28\|CGTACGTAGCAGAGCAGCTCCCTCGCTG | >hg38_21_+_8433172_8446622_RNA45SN1_45S_50ntFlanks | 12974 | 13001 | 28 | CGTACGTAGCAGAGCAGCTCCCTCGCTG | 277.6 | -11.1 | 1.3 | -8.2 | 2.0E-16 | 1.3E-15 | MB231/MB468 | NUC |
| >hg38_21_+_8433172_8446622_RNA45SN1_45S_50ntFlanks@5082.5112.31\|AACGAACGAGACTCTGGCATGCTAACTAGTT | >hg38_21_+_8433172_8446622_RNA45SN1_45S_50ntFlanks | 5082 | 5112 | 31 | AACGAACGAGACTCTGGCATGCTAACTAGTT | 437.9 | 5.1 | 0.6 | 8.2 | 1.7E-16 | 1.3E-15 | MB231/MB468 | CYTO |
| >hg38_21_+_8433172_8446622_RNA45SN1_45S_50ntFlanks@10411.10435.25\|CAGTCGGTCCTGAGAGATGGGCGAG | >hg38_21_+_8433172_8446622_RNA45SN1_45S_50ntFlanks | 10411 | 10435 | 25 | CAGTCGGTCCTGAGAGATGGGCGAG | 395.1 | -11.3 | 1.4 | -8.2 | 1.7E-16 | 1.3E-15 | MB231/MB468 | CYTO |
| >MI0003560\|hsa-mir-92b&WithFlank&1\|+\|155195171\|155195278@67.87.21(+1U)&[MIMAT0003218&hsa-miR-92b-3p&offsets\|0\|-1(+1U);m-115&1\|+\|155195237\|155195258&offsets\|0\|-1(+1U)]\|TATTGCACTCGTCCCGGCCTCT | >MI0003560\|hsa-mir-92b&WithFlank&1\|+\|155195171\|155195278 | 67 | 87 | 21(+1U) | TATTGCACTCGTCCCGGCCTCT | 1353.0 | 5.6 | 0.7 | 8.2 | 2.0E-16 | 1.3E-15 | MB231/MB468 | NUC |
| >hg38_21_+_8433172_8446622_RNA45SN1_45S_50ntFlanks@4917.4939.23\|CACCACCAGGAGTGGAGCCTGCG | >hg38_21_+_8433172_8446622_RNA45SN1_45S_50ntFlanks | 4917 | 4939 | 23 | CACCACCAGGAGTGGAGCCTGCG | 206.9 | -11.6 | 1.4 | -8.2 | 1.6E-16 | 1.3E-15 | MB231/MB468 | MP |
| >hg38_21_+_8433172_8446622_RNA45SN1_45S_50ntFlanks@6649.6672.24\|TACGACTCTTAGCGGTGGATCACT | >hg38_21_+_8433172_8446622_RNA45SN1_45S_50ntFlanks | 6649 | 6672 | 24 | TACGACTCTTAGCGGTGGATCACT | 2396.9 | 6.6 | 0.8 | 8.2 | 1.9E-16 | 1.3E-15 | BT20/MB468 | MITO |
| >hg38_21_+_8433172_8446622_RNA45SN1_45S_50ntFlanks@10687.10711.25\|CGGCGTCCGGTGAGCTCTCGCTGGC | >hg38_21_+_8433172_8446622_RNA45SN1_45S_50ntFlanks | 10687 | 10711 | 25 | CGGCGTCCGGTGAGCTCTCGCTGGC | 6223.6 | -6.2 | 0.8 | -8.2 | 1.7E-16 | 1.3E-15 | MB231/MB468 | CYTO |
| >hg38_21_+_8433172_8446622_RNA45SN1_45S_50ntFlanks@9759.9789.31\|CAAACTTTAAATGGGTAAGAAGCCCGGCTCG | >hg38_21_+_8433172_8446622_RNA45SN1_45S_50ntFlanks | 9759 | 9789 | 31 | CAAACTTTAAATGGGTAAGAAGCCCGGCTCG | 281.4 | -11.9 | 1.5 | -8.2 | 2.7E-16 | 1.3E-15 | MB231/MB468 | MITO |
| >hg38_21_+_8433172_8446622_RNA45SN1_45S_50ntFlanks@9564.9595.32\|CTGACGTGCAAATCGGTCGTCCGACCTGGGTA | >hg38_21_+_8433172_8446622_RNA45SN1_45S_50ntFlanks | 9564 | 9595 | 32 | CTGACGTGCAAATCGGTCGTCCGACCTGGGTA | 3215.4 | -5.6 | 0.7 | -8.2 | 2.1E-16 | 1.3E-15 | MB231/MB468 | NUC |
| >hg38_21_+_8433172_8446622_RNA45SN1_45S_50ntFlanks@10408.10426.19\|GGTCAGTCGGTCCTGAGAG | >hg38_21_+_8433172_8446622_RNA45SN1_45S_50ntFlanks | 10408 | 10426 | 19 | GGTCAGTCGGTCCTGAGAG | 439.2 | -12.6 | 1.5 | -8.2 | 2.7E-16 | 1.3E-15 | MB231/MB468 | MITO |
| >hg38_21_+_8433172_8446622_RNA45SN1_45S_50ntFlanks@10572.10589.18\|AGAAGCCGGCGGGAGCCC | >hg38_21_+_8433172_8446622_RNA45SN1_45S_50ntFlanks | 10572 | 10589 | 18 | AGAAGCCGGCGGGAGCCC | 303.4 | -11.0 | 1.3 | -8.2 | 1.7E-16 | 1.3E-15 | MB231/MB468 | CYTO |
| >hg38_21_+_8433172_8446622_RNA45SN1_45S_50ntFlanks@11318.11342.25\|CCCCACGTCTCGTCGCGCGCGCGTC | >hg38_21_+_8433172_8446622_RNA45SN1_45S_50ntFlanks | 11318 | 11342 | 25 | CCCCACGTCTCGTCGCGCGCGCGTC | 1457.7 | -8.9 | 1.1 | -8.2 | 1.7E-16 | 1.3E-15 | MB231/MB468 | CYTO |
| >hg38_21_+_8433172_8446622_RNA45SN1_45S_50ntFlanks@6772.6793.22\|GTTCCTCCCGGGGCTACGCCTG | >hg38_21_+_8433172_8446622_RNA45SN1_45S_50ntFlanks | 6772 | 6793 | 22 | GTTCCTCCCGGGGCTACGCCTG | 181.7 | -5.8 | 0.7 | -8.2 | 1.8E-16 | 1.3E-15 | MB231/MB468 | CYTO |
| >MI0000087\|hsa-mir-29a&WithFlank&7\|-\|130876741\|130876816@47.67.21(+1C)&[MIMAT0000086&hsa-miR-29a-3p&offsets\|-1\|-2(+1C);m-26&7\|-\|130876749\|130876769&offsets\|-1\|-1(+1C)]\|CTAGCACCATCTGAAATCGGTC | >MI0000087\|hsa-mir-29a&WithFlank&7\|-\|130876741\|130876816 | 47 | 67 | 21(+1C) | CTAGCACCATCTGAAATCGGTC | 1910.0 | 8.1 | 1.0 | 8.2 | 2.8E-16 | 1.3E-15 | MB231/MB468 | MITO |
| >hg38_21_+_8433172_8446622_RNA45SN1_45S_50ntFlanks@10351.10378.28\|CGAGAACTTTGAAGGCCGAAGTGGAGAA | >hg38_21_+_8433172_8446622_RNA45SN1_45S_50ntFlanks | 10351 | 10378 | 28 | CGAGAACTTTGAAGGCCGAAGTGGAGAA | 645.4 | -4.9 | 0.6 | -8.2 | 1.8E-16 | 1.3E-15 | MB231/MB468 | CYTO |
| >trna5_AspGTC_12_+_98897281_98897352@36.75.40\|CACGCGGGAGACCGGGGTTCAATTCCCCGACGGGGAGCCA | >trna5_AspGTC_12_+_98897281_98897352 | 36 | 75 | 40 | CACGCGGGAGACCGGGGTTCAATTCCCCGACGGGGAGCCA | 5463.7 | 10.9 | 1.3 | 8.2 | 1.8E-16 | 1.4E-15 | MB231/MB468 | CYTO |
| >hg38_1_-_228634819_228635039_RNA5S12_5S_50ntFlanks@51.88.38\|GTCTACGGCCATACCACCCTGAACGCGCCCGATCTCGT | >hg38_1_-_228634819_228635039_RNA5S12_5S_50ntFlanks | 51 | 88 | 38 | GTCTACGGCCATACCACCCTGAACGCGCCCGATCTCGT | 2636.7 | 6.3 | 0.8 | 8.2 | 1.9E-16 | 1.4E-15 | BT20/MB468 | MITO |
| >hg38_21_+_8433172_8446622_RNA45SN1_45S_50ntFlanks@3985.4004.20\|CGGCTTTGGTGACTCTAGAT | >hg38_21_+_8433172_8446622_RNA45SN1_45S_50ntFlanks | 3985 | 4004 | 20 | CGGCTTTGGTGACTCTAGAT | 396.3 | -12.5 | 1.5 | -8.2 | 1.7E-16 | 1.4E-15 | MB231/MB468 | MP |
| >hg38_21_+_8433172_8446622_RNA45SN1_45S_50ntFlanks@8836.8864.29\|CGGCGGTGGCGGCGGCGGCGGCGGCGGCG | >hg38_21_+_8433172_8446622_RNA45SN1_45S_50ntFlanks | 8836 | 8864 | 29 | CGGCGGTGGCGGCGGCGGCGGCGGCGGCG | 635.8 | -13.1 | 1.6 | -8.2 | 2.8E-16 | 1.4E-15 | MB231/MB468 | MITO |
| >MI0003763\|hsa-mir-767&WithFlank&X\|-\|152393415\|152393535@33.56.24&[MIMAT0003882&hsa-miR-767-5p&offsets\|0\|+1;m-788&X\|-\|152393481\|152393503&offsets\|0\|+1]\|TGCACCATGGTTGTCTGAGCATGC | >MI0003763\|hsa-mir-767&WithFlank&X\|-\|152393415\|152393535 | 33 | 56 | 24 | TGCACCATGGTTGTCTGAGCATGC | 675.7 | 9.6 | 1.1 | 8.5 | 1.5E-17 | 1.4E-15 | BT20/MB231 | CYTO |
| >hg38_21_+_8433172_8446622_RNA45SN1_45S_50ntFlanks@4047.4066.20\|ATTCGAACGTCTGCCCTATC | >hg38_21_+_8433172_8446622_RNA45SN1_45S_50ntFlanks | 4047 | 4066 | 20 | ATTCGAACGTCTGCCCTATC | 9070.4 | -7.9 | 1.0 | -8.2 | 2.8E-16 | 1.4E-15 | MB231/MB468 | MITO |
| >hg38_21_+_8433172_8446622_RNA45SN1_45S_50ntFlanks@6719.6741.23\|TGAATTGCAGGACACATTGATCA | >hg38_21_+_8433172_8446622_RNA45SN1_45S_50ntFlanks | 6719 | 6741 | 23 | TGAATTGCAGGACACATTGATCA | 271.9 | -12.0 | 1.4 | -8.2 | 1.7E-16 | 1.4E-15 | MB231/MB468 | MP |
| >trna116_GluCTC_1_-_145399233_145399304@1.32.32,trna59_GluCTC_1_+_249168447_249168518@1.32.32,trna71_GluCTC_1_-_161439189_161439260@1.32.32,trna74_GluCTC_1_-_161431809_161431880@1.32.32,trna77_GluCTC_1_-_161424398_161424469@1.32.32,trna77_GluCTC_6_+_28949976_28950047@1.32.32,trna80_GluCTC_1_-_161417018_161417089@1.32.32,trna87_GluCTC_6_-_126101393_126101464@1.32.32\|TCCCTGGTGGTCTAGTGGTTAGGATTCGGCGC | >trna116_GluCTC_1_-_145399233_145399304 | 1 | 32 | 32 | TCCCTGGTGGTCTAGTGGTTAGGATTCGGCGC | 9425.3 | 6.5 | 0.8 | 8.2 | 1.8E-16 | 1.4E-15 | MB231/MB468 | CYTO |
| >trnaMT_LeuTAA_MT_+_3230_3304@16.45.30\|CCCGGTAATCGCATAAAACTTAAAACTTTA | >trnaMT_LeuTAA_MT_+_3230_3304 | 16 | 45 | 30 | CCCGGTAATCGCATAAAACTTAAAACTTTA | 468.0 | -12.7 | 1.5 | -8.2 | 2.8E-16 | 1.4E-15 | MB231/MB468 | MITO |
| >hg38_21_+_8433172_8446622_RNA45SN1_45S_50ntFlanks@6741.6770.30\|ATCGACACTTCGAACGCACTTGCGGCCCCG | >hg38_21_+_8433172_8446622_RNA45SN1_45S_50ntFlanks | 6741 | 6770 | 30 | ATCGACACTTCGAACGCACTTGCGGCCCCG | 3584.6 | -10.3 | 1.3 | -8.2 | 2.8E-16 | 1.4E-15 | MB231/MB468 | MITO |
| >hg38_21_+_8433172_8446622_RNA45SN1_45S_50ntFlanks@11562.11592.31\|CGCCTAGCAGCCGACTTAGAACTGGTGCGGA | >hg38_21_+_8433172_8446622_RNA45SN1_45S_50ntFlanks | 11562 | 11592 | 31 | CGCCTAGCAGCCGACTTAGAACTGGTGCGGA | 3847.3 | -11.4 | 1.4 | -8.2 | 1.7E-16 | 1.4E-15 | MB231/MB468 | MP |
| >hg38_21_+_8433172_8446622_RNA45SN1_45S_50ntFlanks@11792.11821.30\|ATGAACGAGATTCCCACTGTCCCTACCTAC | >hg38_21_+_8433172_8446622_RNA45SN1_45S_50ntFlanks | 11792 | 11821 | 30 | ATGAACGAGATTCCCACTGTCCCTACCTAC | 452.7 | -12.6 | 1.5 | -8.2 | 2.8E-16 | 1.4E-15 | MB231/MB468 | MITO |
| >hg38_21_+_8433172_8446622_RNA45SN1_45S_50ntFlanks@485.509.25\|GATGTGGTGACGTCGTGCTCTCCCG | >hg38_21_+_8433172_8446622_RNA45SN1_45S_50ntFlanks | 485 | 509 | 25 | GATGTGGTGACGTCGTGCTCTCCCG | 3563.5 | 29.8 | 3.6 | 8.2 | 2.1E-16 | 1.4E-15 | MB231/MB468 | NUC |
| >MI0000750\|hsa-mir-26a-2&WithFlank&12\|-\|57824603\|57824698@20.42.23;MI0000083\|hsa-mir-26a-1&WithFlank&3\|+\|37969398\|37969486@16.38.23&[MIMAT0000082&hsa-miR-26a-5p&offsets\|0\|+1;m-33&12\|-\|57824658\|57824679&offsets\|0\|+1];[MIMAT0000082_1&hsa-miR-26a-5p&offsets\|0\|+1;m-34&3\|+\|37969413\|37969434&offsets\|0\|+1]\|TTCAAGTAATCCAGGATAGGCTG | >MI0000750\|hsa-mir-26a-2&WithFlank&12\|-\|57824603\|57824698 | 20 | 42 | 23 | TTCAAGTAATCCAGGATAGGCTG | 430.3 | 6.5 | 0.8 | 8.2 | 2.7E-16 | 1.4E-15 | BT20/MB468 | CYTO |
| >hg38_21_+_8433172_8446622_RNA45SN1_45S_50ntFlanks@12004.12029.26\|CGGGCCGCCGGTGAAATACCACTACT | >hg38_21_+_8433172_8446622_RNA45SN1_45S_50ntFlanks | 12004 | 12029 | 26 | CGGGCCGCCGGTGAAATACCACTACT | 345.6 | 5.1 | 0.6 | 8.2 | 2.7E-16 | 1.4E-15 | BT20/MB468 | CYTO |
| >hg38_21_+_8433172_8446622_RNA45SN1_45S_50ntFlanks@10678.10696.19\|CGGTTCCGGCGGCGTCCGG | >hg38_21_+_8433172_8446622_RNA45SN1_45S_50ntFlanks | 10678 | 10696 | 19 | CGGTTCCGGCGGCGTCCGG | 321.9 | -12.2 | 1.5 | -8.2 | 1.7E-16 | 1.4E-15 | MB231/MB468 | MP |
| >hg38_21_+_8433172_8446622_RNA45SN1_45S_50ntFlanks@5479.5512.34\|TGGCGGAGCGCTGAGAAGACGGTCGAACTTGACT | >hg38_21_+_8433172_8446622_RNA45SN1_45S_50ntFlanks | 5479 | 5512 | 34 | TGGCGGAGCGCTGAGAAGACGGTCGAACTTGACT | 207.4 | 8.8 | 1.1 | 8.2 | 1.8E-16 | 1.4E-15 | MB231/MB468 | CYTO |
| >hg38_21_+_8433172_8446622_RNA45SN1_45S_50ntFlanks@10829.10875.47\|GCAAGCCGGATCCGTAACTTCGGGATAAGGATTGGCTCTAAGGGCTG | >hg38_21_+_8433172_8446622_RNA45SN1_45S_50ntFlanks | 10829 | 10875 | 47 | GCAAGCCGGATCCGTAACTTCGGGATAAGGATTGGCTCTAAGGGCTG | 2450.6 | -15.7 | 1.9 | -8.1 | 4.4E-16 | 1.4E-15 | BT20/MB468 | NUC |
| >hg38_1_-_228634819_228635039_RNA5S12_5S_50ntFlanks@149.171.23\|GAATACCGGGTGCTGTAGGCTTT | >hg38_1_-_228634819_228635039_RNA5S12_5S_50ntFlanks | 149 | 171 | 23 | GAATACCGGGTGCTGTAGGCTTT | 6770.1 | 6.2 | 0.7 | 8.2 | 1.8E-16 | 1.4E-15 | MB231/MB468 | CYTO |
| >MI0000279\|hsa-mir-196a-2&WithFlank&12\|+\|53991732\|53991853@31.51.21;MI0000238\|hsa-mir-196a-1&WithFlank&17\|-\|48632484\|48632565@13.33.21&[MIMAT0000226&hsa-miR-196a-5p&offsets\|0\|-1;m-261&12\|+\|53991762\|53991783&offsets\|0\|-1];[MIMAT0000226_1&hsa-miR-196a-5p&offsets\|0\|-1;m-268&17\|-\|48632532\|48632553&offsets\|0\|-1]\|TAGGTAGTTTCATGTTGTTGG | >MI0000279\|hsa-mir-196a-2&WithFlank&12\|+\|53991732\|53991853 | 31 | 51 | 21 | TAGGTAGTTTCATGTTGTTGG | 455.8 | 7.3 | 0.9 | 8.2 | 2.0E-16 | 1.4E-15 | BT20/MB468 | MITO |
| >hg38_21_+_8433172_8446622_RNA45SN1_45S_50ntFlanks@12934.12952.19\|CTAAACCATTCGTAGACGA | >hg38_21_+_8433172_8446622_RNA45SN1_45S_50ntFlanks | 12934 | 12952 | 19 | CTAAACCATTCGTAGACGA | 924.9 | -6.8 | 0.8 | -8.2 | 1.8E-16 | 1.4E-15 | MB231/MB468 | CYTO |
| >hg38_21_+_8433172_8446622_RNA45SN1_45S_50ntFlanks@4472.4497.26\|CTTAGCTGAGTGTCCCGCGGGGCCCG | >hg38_21_+_8433172_8446622_RNA45SN1_45S_50ntFlanks | 4472 | 4497 | 26 | CTTAGCTGAGTGTCCCGCGGGGCCCG | 444.6 | -11.5 | 1.4 | -8.2 | 1.8E-16 | 1.4E-15 | MB231/MB468 | CYTO |
| >hg38_21_+_8433172_8446622_RNA45SN1_45S_50ntFlanks@8475.8492.18\|CCCGGCGGATCTTTCCCG | >hg38_21_+_8433172_8446622_RNA45SN1_45S_50ntFlanks | 8475 | 8492 | 18 | CCCGGCGGATCTTTCCCG | 563.6 | -12.9 | 1.6 | -8.2 | 2.8E-16 | 1.4E-15 | MB231/MB468 | MITO |
| >hg38_21_+_8433172_8446622_RNA45SN1_45S_50ntFlanks@11919.11951.33\|CATGAGAGGTGTAGAATAAGTGGGAGGCCCCCG | >hg38_21_+_8433172_8446622_RNA45SN1_45S_50ntFlanks | 11919 | 11951 | 33 | CATGAGAGGTGTAGAATAAGTGGGAGGCCCCCG | 320.1 | -11.0 | 1.3 | -8.2 | 1.8E-16 | 1.4E-15 | MB231/MB468 | CYTO |
| >hg38_21_+_8433172_8446622_RNA45SN1_45S_50ntFlanks@11873.11912.40\|AAGAAGACCCTGTTGAGCTTGACTCTAGTCTGGCACGGTG | >hg38_21_+_8433172_8446622_RNA45SN1_45S_50ntFlanks | 11873 | 11912 | 40 | AAGAAGACCCTGTTGAGCTTGACTCTAGTCTGGCACGGTG | 471.2 | -12.7 | 1.5 | -8.2 | 1.7E-16 | 1.4E-15 | MB231/MB468 | MP |
| >hg38_21_+_8433172_8446622_RNA45SN1_45S_50ntFlanks@11321.11338.18\|CACGTCTCGTCGCGCGCG | >hg38_21_+_8433172_8446622_RNA45SN1_45S_50ntFlanks | 11321 | 11338 | 18 | CACGTCTCGTCGCGCGCG | 1268.2 | -7.6 | 0.9 | -8.2 | 1.8E-16 | 1.4E-15 | MB231/MB468 | CYTO |
| >hg38_21_+_8433172_8446622_RNA45SN1_45S_50ntFlanks@4983.5015.33\|CGGACAGGATTGACAGATTGATAGCTCTTTCTC | >hg38_21_+_8433172_8446622_RNA45SN1_45S_50ntFlanks | 4983 | 5015 | 33 | CGGACAGGATTGACAGATTGATAGCTCTTTCTC | 1777.8 | -14.6 | 1.8 | -8.2 | 2.9E-16 | 1.4E-15 | MB231/MB468 | MITO |
| >trna10_MetCAT_8_-_124169470_124169542@31.51.21,trna22_MetCAT_16_-_87417628_87417700@31.51.21\|TCTCATAATCTGAAGGTCGTG | >trna10_MetCAT_8_-_124169470_124169542 | 31 | 51 | 21 | TCTCATAATCTGAAGGTCGTG | 394.3 | -11.6 | 1.4 | -8.2 | 2.1E-16 | 1.4E-15 | MB231/MB468 | NUC |
| >hg38_21_+_8433172_8446622_RNA45SN1_45S_50ntFlanks@9714.9735.22\|CGAATGATTAGAGGTCTTGGGG | >hg38_21_+_8433172_8446622_RNA45SN1_45S_50ntFlanks | 9714 | 9735 | 22 | CGAATGATTAGAGGTCTTGGGG | 266.2 | -11.0 | 1.3 | -8.2 | 2.1E-16 | 1.4E-15 | MB231/MB468 | NUC |
| >hg38_21_+_8433172_8446622_RNA45SN1_45S_50ntFlanks@12476.12499.24\|CCACTAATAGGGAACGTGAGCTGG | >hg38_21_+_8433172_8446622_RNA45SN1_45S_50ntFlanks | 12476 | 12499 | 24 | CCACTAATAGGGAACGTGAGCTGG | 296.5 | -12.1 | 1.5 | -8.2 | 1.7E-16 | 1.4E-15 | MB231/MB468 | MP |
| >hg38_21_+_8433172_8446622_RNA45SN1_45S_50ntFlanks@9534.9552.19\|AGGAAACTCTGGTGGAGGT | >hg38_21_+_8433172_8446622_RNA45SN1_45S_50ntFlanks | 9534 | 9552 | 19 | AGGAAACTCTGGTGGAGGT | 3003.0 | -7.2 | 0.9 | -8.2 | 1.7E-16 | 1.4E-15 | MB231/MB468 | MP |
| >MI0001445\|hsa-mir-423&WithFlank&17\|+\|30117073\|30117178@58.80.23(+1A)&[MIMAT0001340&hsa-miR-423-3p&offsets\|-1\|-1(+1A);m-100&17\|+\|30117131\|30117153&offsets\|-1\|-1(+1A)]\|AAGCTCGGTCTGAGGCCCCTCAGA | >MI0001445\|hsa-mir-423&WithFlank&17\|+\|30117073\|30117178 | 58 | 80 | 23(+1A) | AAGCTCGGTCTGAGGCCCCTCAGA | 386.3 | 10.2 | 1.2 | 8.2 | 2.8E-16 | 1.4E-15 | BT20/MB468 | CYTO |
| >hg38_21_+_8433172_8446622_RNA45SN1_45S_50ntFlanks@3850.3869.20\|GATAACTGTGGTAATTCTAG | >hg38_21_+_8433172_8446622_RNA45SN1_45S_50ntFlanks | 3850 | 3869 | 20 | GATAACTGTGGTAATTCTAG | 230.5 | -11.7 | 1.4 | -8.2 | 1.7E-16 | 1.4E-15 | MB231/MB468 | MP |

<table>
<tr><td>>hg38_21_+_8433172_8446622_RNA45SN1_45S_50ntFlanks@5443.5467.25|CCTCGGATCGGCCCCGCCGGGGTCG</td><td>>hg38_21_+_8433172_8446622_RNA45SN1_45S_50ntFlanks</td><td>5443</td><td>5467</td><td>25</td><td>CCTCGGATCGGCCCCGCCGGGGTCG</td><td>288.0</td><td>-12.0</td><td>1.5</td><td>-8.2</td><td>1.7E-16</td><td>1.4E-15</td><td>MB231/MB468</td><td>MP</td></tr>
<tr><td>>hg38_21_+_8433172_8446622_RNA45SN1_45S_50ntFlanks@5467.5484.18|GGCCCACGGCCCTGGCGG</td><td>>hg38_21_+_8433172_8446622_RNA45SN1_45S_50ntFlanks</td><td>5467</td><td>5484</td><td>18</td><td>GGCCCACGGCCCTGGCGG</td><td>280.0</td><td>-12.0</td><td>1.5</td><td>-8.2</td><td>1.7E-16</td><td>1.4E-15</td><td>MB231/MB468</td><td>MP</td></tr>
<tr><td>>hg38_21_+_8433172_8446622_RNA45SN1_45S_50ntFlanks@5492.5509.18|AGAAGACGGTCGAACTTG</td><td>>hg38_21_+_8433172_8446622_RNA45SN1_45S_50ntFlanks</td><td>5492</td><td>5509</td><td>18</td><td>AGAAGACGGTCGAACTTG</td><td>1628.6</td><td>-10.3</td><td>1.2</td><td>-8.2</td><td>1.7E-16</td><td>1.4E-15</td><td>MB231/MB468</td><td>MP</td></tr>
<tr><td>>hg38_21_+_8433172_8446622_RNA45SN1_45S_50ntFlanks@12399.12436.38|ACGTCGCTTTTTGATCCTTCGATGTCGGCTCTTCCTAT</td><td>>hg38_21_+_8433172_8446622_RNA45SN1_45S_50ntFlanks</td><td>12399</td><td>12436</td><td>38</td><td>ACGTCGCTTTTTGATCCTTCGATGTCGGCTCTTCCTAT</td><td>803.3</td><td>-14.1</td><td>1.7</td><td>-8.1</td><td>4.4E-16</td><td>1.4E-15</td><td>BT20/MB468</td><td>NUC</td></tr>
<tr><td>>hg38_21_+_8433172_8446622_RNA45SN1_45S_50ntFlanks@10921.10939.19|GCTGGACGAGGCGCCGCCG</td><td>>hg38_21_+_8433172_8446622_RNA45SN1_45S_50ntFlanks</td><td>10921</td><td>10939</td><td>19</td><td>GCTGGACGAGGCGCCGCCG</td><td>501.9</td><td>-12.8</td><td>1.6</td><td>-8.2</td><td>2.9E-16</td><td>1.4E-15</td><td>MB231/MB468</td><td>MITO</td></tr>
<tr><td>>MI0000088|hsa-mir-30a&WithFlank&6|-|71403545|71403627@53.75.23&[MIMAT0000088&hsa-miR-30a-3p&offsets|0|+1;m-38&6|-|71403554|71403575&offsets|0|+1]|CTTTCAGTCGGATGTTTGCAGCT</td><td>>MI0000088|hsa-mir-30a&WithFlank&6|-|71403545|71403627</td><td>53</td><td>75</td><td>23</td><td>CTTTCAGTCGGATGTTTGCAGCT</td><td>396.0</td><td>5.7</td><td>0.7</td><td>8.2</td><td>2.2E-16</td><td>1.4E-15</td><td>MB231/MB468</td><td>NUC</td></tr>
<tr><td>>hg38_21_+_8433172_8446622_RNA45SN1_45S_50ntFlanks@11569.11593.25|CAGCCGACTTAGAACTGGTGCGGAC</td><td>>hg38_21_+_8433172_8446622_RNA45SN1_45S_50ntFlanks</td><td>11569</td><td>11593</td><td>25</td><td>CAGCCGACTTAGAACTGGTGCGGAC</td><td>1015.0</td><td>-13.9</td><td>1.7</td><td>-8.2</td><td>1.8E-16</td><td>1.4E-15</td><td>MB231/MB468</td><td>MP</td></tr>
<tr><td>>hg38_21_+_8433172_8446622_RNA45SN1_45S_50ntFlanks@7983.8007.25|CAGATCAGACGTGGCGACCCGCTGA</td><td>>hg38_21_+_8433172_8446622_RNA45SN1_45S_50ntFlanks</td><td>7983</td><td>8007</td><td>25</td><td>CAGATCAGACGTGGCGACCCGCTGA</td><td>2049.6</td><td>-10.2</td><td>1.2</td><td>-8.2</td><td>2.9E-16</td><td>1.4E-15</td><td>MB231/MB468</td><td>MITO</td></tr>
<tr><td>>hg38_21_+_8433172_8446622_RNA45SN1_45S_50ntFlanks@11999.12034.36|CCCTGCGGGCCGCCGGTGAAATACCACTACTCTGAT</td><td>>hg38_21_+_8433172_8446622_RNA45SN1_45S_50ntFlanks</td><td>11999</td><td>12034</td><td>36</td><td>CCCTGCGGGCCGCCGGTGAAATACCACTACTCTGAT</td><td>404.6</td><td>-12.4</td><td>1.5</td><td>-8.2</td><td>2.9E-16</td><td>1.4E-15</td><td>MB231/MB468</td><td>MITO</td></tr>
<tr><td>>hg38_21_+_8433172_8446622_RNA45SN1_45S_50ntFlanks@12162.12181.20|GACTGGGGCGGTACACCTGT</td><td>>hg38_21_+_8433172_8446622_RNA45SN1_45S_50ntFlanks</td><td>12162</td><td>12181</td><td>20</td><td>GACTGGGGCGGTACACCTGT</td><td>509.4</td><td>-12.8</td><td>1.6</td><td>-8.2</td><td>2.9E-16</td><td>1.4E-15</td><td>MB231/MB468</td><td>MITO</td></tr>
<tr><td>>MI0000734|hsa-mir-106b&WithFlank&7|-|100093987|100094080@58.79.22(+1A)&[MIMAT0004672&hsa-miR-106b-3p&offsets|0|0(+1A);m-87&7|-|100094003|100094023&offsets|0|+1(+1A)]|CCGCACTGTGGGTACTTGCTGCA</td><td>>MI0000734|hsa-mir-106b&WithFlank&7|-|100093987|100094080</td><td>58</td><td>79</td><td>22(+1A)</td><td>CCGCACTGTGGGTACTTGCTGCA</td><td>541.2</td><td>7.2</td><td>0.9</td><td>8.2</td><td>2.8E-16</td><td>1.4E-15</td><td>BT20/MB468</td><td>CYTO</td></tr>
<tr><td>>hg38_21_+_8433172_8446622_RNA45SN1_45S_50ntFlanks@4860.4890.31|TGGGTTCCGGGGGGAGTATGGTTGCAAAGCT</td><td>>hg38_21_+_8433172_8446622_RNA45SN1_45S_50ntFlanks</td><td>4860</td><td>4890</td><td>31</td><td>TGGGTTCCGGGGGGAGTATGGTTGCAAAGCT</td><td>131.6</td><td>11.0</td><td>1.3</td><td>8.2</td><td>1.9E-16</td><td>1.4E-15</td><td>MB231/MB468</td><td>CYTO</td></tr>
<tr><td>>MI0000434|hsa-let-7i&WithFlank&12|+|62603680|62603775@12.33.22(+1U)&[MIMAT0000415&hsa-let-7i-5p&offsets|0|0(+1U);m-49&12|+|62603691|62603712&offsets|0|0(+1U)]|TGAGGTAGTAGTTTGTGCTGTTT</td><td>>MI0000434|hsa-let-7i&WithFlank&12|+|62603680|62603775</td><td>12</td><td>33</td><td>22(+1U)</td><td>TGAGGTAGTAGTTTGTGCTGTTT</td><td>958.3</td><td>2.1</td><td>0.3</td><td>8.1</td><td>4.5E-16</td><td>1.4E-15</td><td>BT20/MB468</td><td>NUC</td></tr>
<tr><td>>hg38_21_+_8433172_8446622_RNA45SN1_45S_50ntFlanks@4726.4752.27|TAATCAAGAACGAAAGTCGGAGGTTCG</td><td>>hg38_21_+_8433172_8446622_RNA45SN1_45S_50ntFlanks</td><td>4726</td><td>4752</td><td>27</td><td>TAATCAAGAACGAAAGTCGGAGGTTCG</td><td>475.6</td><td>-12.7</td><td>1.6</td><td>-8.2</td><td>3.0E-16</td><td>1.4E-15</td><td>MB231/MB468</td><td>MITO</td></tr>
<tr><td>>hg38_21_+_8433172_8446622_RNA45SN1_45S_50ntFlanks@11576.11603.28|CTTAGAACTGGTGCGGACCAGGGGAATC</td><td>>hg38_21_+_8433172_8446622_RNA45SN1_45S_50ntFlanks</td><td>11576</td><td>11603</td><td>28</td><td>CTTAGAACTGGTGCGGACCAGGGGAATC</td><td>3729.5</td><td>-10.9</td><td>1.3</td><td>-8.2</td><td>1.8E-16</td><td>1.4E-15</td><td>MB231/MB468</td><td>MP</td></tr>
<tr><td>>am_trna117_GlyTCC_1_-_145397864_145397935@1.30.30,trna45_GlyTCC_1_+_161500903_161500974@1.30.30,trna70_GlyTCC_1_-_161439547_161439618@1.30.30,trna73_GlyTCC_1_-_161432166_161432237@1.30.30,trna76_GlyTCC_1_-_161424756_161424827@1.30.30,trna79_GlyTCC_1_-_161417375_161417446@1.30.30|GCGTTGGTGGTATAGTGGTGAGCATAGCTG</td><td>>am_trna117_GlyTCC_1_-_145397864_145397935</td><td>1</td><td>30</td><td>30</td><td>GCGTTGGTGGTATAGTGGTGAGCATAGCTG</td><td>3884.7</td><td>-11.9</td><td>1.5</td><td>-8.1</td><td>4.6E-16</td><td>1.4E-15</td><td>BT20/MB468</td><td>NUC</td></tr>
<tr><td>>MI0000070|hsa-mir-16-1&WithFlank&13|-|50048967|50049067@20.39.20(+1A);MI0000115|hsa-mir-16-2&WithFlank&3|+|160404739|160404831@16.35.20(+1A)&[MIMAT0000069&hsa-miR-16-5p&offsets|0|-2(+1A);m-62&13|-|50049027|50049048&offsets|0|-2(+1A)];[MIMAT0000069_1&hsa-miR-16-5p&offsets|0|-2(+1A);m-61&3|+|160404754|160404775&offsets|0|-2(+1A)]|TAGCAGCACGTAAATATTGGA</td><td>>MI0000070|hsa-mir-16-1&WithFlank&13|-|50048967|50049067</td><td>20</td><td>39</td><td>20(+1A)</td><td>TAGCAGCACGTAAATATTGGA</td><td>348.5</td><td>12.7</td><td>1.6</td><td>8.2</td><td>2.9E-16</td><td>1.4E-15</td><td>BT20/MB468</td><td>CYTO</td></tr>
<tr><td>>hg38_21_+_8433172_8446622_RNA45SN1_45S_50ntFlanks@6721.6745.25|AATTGCAGGACACATTGATCATCGA</td><td>>hg38_21_+_8433172_8446622_RNA45SN1_45S_50ntFlanks</td><td>6721</td><td>6745</td><td>25</td><td>AATTGCAGGACACATTGATCATCGA</td><td>2629.3</td><td>-7.5</td><td>0.9</td><td>-8.2</td><td>1.9E-16</td><td>1.4E-15</td><td>MB231/MB468</td><td>CYTO</td></tr>
<tr><td>>hg38_21_+_8433172_8446622_RNA45SN1_45S_50ntFlanks@11921.11968.48|TGAGAGGTGTAGAATAAGTGGGAGGCCCCCGGCGCCCCCCCGGTGTCC</td><td>>hg38_21_+_8433172_8446622_RNA45SN1_45S_50ntFlanks</td><td>11921</td><td>11968</td><td>48</td><td>TGAGAGGTGTAGAATAAGTGGGAGGCCCCCGGCGCCCCCCCGGTGTCC</td><td>622.0</td><td>-13.7</td><td>1.7</td><td>-8.1</td><td>4.6E-16</td><td>1.4E-15</td><td>BT20/MB468</td><td>NUC</td></tr>
<tr><td>>hg38_21_+_8433172_8446622_RNA45SN1_45S_50ntFlanks@6733.6752.20|CATTGATCATCGACACTTCG</td><td>>hg38_21_+_8433172_8446622_RNA45SN1_45S_50ntFlanks</td><td>6733</td><td>6752</td><td>20</td><td>CATTGATCATCGACACTTCG</td><td>662.7</td><td>-5.2</td><td>0.6</td><td>-8.1</td><td>4.6E-16</td><td>1.4E-15</td><td>BT20/MB468</td><td>NUC</td></tr>
<tr><td>>hg38_21_+_8433172_8446622_RNA45SN1_45S_50ntFlanks@7982.8008.27|TCAGATCAGACGTGGCGACCCGCTGAA</td><td>>hg38_21_+_8433172_8446622_RNA45SN1_45S_50ntFlanks</td><td>7982</td><td>8008</td><td>27</td><td>TCAGATCAGACGTGGCGACCCGCTGAA</td><td>717.2</td><td>-5.2</td><td>0.6</td><td>-8.1</td><td>4.6E-16</td><td>1.4E-15</td><td>BT20/MB468</td><td>NUC</td></tr>
<tr><td>>MI0000087|hsa-mir-29a&WithFlank&7|-|130876741|130876816@47.68.22&[MIMAT0000086&hsa-miR-29a-3p&offsets|-1|-1;m-26&7|-|130876749|130876769&offsets|-1|0]|CTAGCACCATCTGAAATCGGTT</td><td>>MI0000087|hsa-mir-29a&WithFlank&7|-|130876741|130876816</td><td>47</td><td>68</td><td>22</td><td>CTAGCACCATCTGAAATCGGTT</td><td>4787.3</td><td>7.4</td><td>0.9</td><td>8.2</td><td>1.9E-16</td><td>1.4E-15</td><td>MB231/MB468</td><td>CYTO</td></tr>
<tr><td>>hg38_21_+_8433172_8446622_RNA45SN1_45S_50ntFlanks@12874.12891.18|GCTGGAAAGGCGGCCGCC</td><td>>hg38_21_+_8433172_8446622_RNA45SN1_45S_50ntFlanks</td><td>12874</td><td>12891</td><td>18</td><td>GCTGGAAAGGCGGCCGCC</td><td>269.4</td><td>-11.9</td><td>1.4</td><td>-8.2</td><td>1.8E-16</td><td>1.4E-15</td><td>MB231/MB468</td><td>MP</td></tr>
<tr><td>>hg38_21_+_8433172_8446622_RNA45SN1_45S_50ntFlanks@9574.9597.24|AATCGGTCGTCCGACCTGGGTATA</td><td>>hg38_21_+_8433172_8446622_RNA45SN1_45S_50ntFlanks</td><td>9574</td><td>9597</td><td>24</td><td>AATCGGTCGTCCGACCTGGGTATA</td><td>15028.5</td><td>-9.9</td><td>1.2</td><td>-8.2</td><td>1.8E-16</td><td>1.4E-15</td><td>MB231/MB468</td><td>MP</td></tr>
<tr><td>>MI0000471|hsa-mir-126&WithFlank&9|+|136670596|136670692@58.78.21&[MIMAT0000445&hsa-miR-126-3p&offsets|0|-1;m-40&9|+|136670653|136670674&offsets|0|-1]|TCGTACCGTGAGTAATAATGC</td><td>>MI0000471|hsa-mir-126&WithFlank&9|+|136670596|136670692</td><td>58</td><td>78</td><td>21</td><td>TCGTACCGTGAGTAATAATGC</td><td>1882.5</td><td>9.0</td><td>1.1</td><td>8.2</td><td>3.0E-16</td><td>1.5E-15</td><td>MB231/MB468</td><td>MITO</td></tr>
<tr><td>>hg38_21_+_8433172_8446622_RNA45SN1_45S_50ntFlanks@6733.6770.38|CATTGATCATCGACACTTCGAACGCACTTGCGGCCCCG</td><td>>hg38_21_+_8433172_8446622_RNA45SN1_45S_50ntFlanks</td><td>6733</td><td>6770</td><td>38</td><td>CATTGATCATCGACACTTCGAACGCACTTGCGGCCCCG</td><td>2895.1</td><td>-5.8</td><td>0.7</td><td>-8.2</td><td>2.2E-16</td><td>1.5E-15</td><td>MB231/MB468</td><td>NUC</td></tr>
<tr><td>>hg38_1_-_228634819_228635039_RNA5S12_5S_50ntFlanks@134.170.37|TGGGAGACCGCCTGGGAATACCGGGTGCTGTAGGCTT</td><td>>hg38_1_-_228634819_228635039_RNA5S12_5S_50ntFlanks</td><td>134</td><td>170</td><td>37</td><td>TGGGAGACCGCCTGGGAATACCGGGTGCTGTAGGCTT</td><td>143951.5</td><td>7.9</td><td>1.0</td><td>8.2</td><td>1.9E-16</td><td>1.5E-15</td><td>MB231/MB468</td><td>CYTO</td></tr>
<tr><td>>am_trnaMT_GluTTC_MT_-_14674_14742@41.58.18,trnalookalike8_GluTTC_5_-_93905172_93905240@41.58.18|TGGTCGTGGTTGTAGTCC</td><td>>am_trnaMT_GluTTC_MT_-_14674_14742</td><td>41</td><td>58</td><td>18</td><td>TGGTCGTGGTTGTAGTCC</td><td>292.8</td><td>9.3</td><td>1.1</td><td>8.2</td><td>1.9E-16</td><td>1.5E-15</td><td>MB231/MB468</td><td>CYTO</td></tr>
<tr><td>>hg38_21_+_8433172_8446622_RNA45SN1_45S_50ntFlanks@11729.11767.39|TAACTATGACTCTCTTAAGGTAGCCAAATGCCTCGTCAT</td><td>>hg38_21_+_8433172_8446622_RNA45SN1_45S_50ntFlanks</td><td>11729</td><td>11767</td><td>39</td><td>TAACTATGACTCTCTTAAGGTAGCCAAATGCCTCGTCAT</td><td>835.3</td><td>-11.8</td><td>1.4</td><td>-8.2</td><td>2.1E-16</td><td>1.5E-15</td><td>BT20/MB468</td><td>MITO</td></tr>
<tr><td>>hg38_21_+_8433172_8446622_RNA45SN1_45S_50ntFlanks@8171.8188.18|AAGTCCTTCTGATCGAGG</td><td>>hg38_21_+_8433172_8446622_RNA45SN1_45S_50ntFlanks</td><td>8171</td><td>8188</td><td>18</td><td>AAGTCCTTCTGATCGAGG</td><td>313.1</td><td>-12.2</td><td>1.5</td><td>-8.2</td><td>1.8E-16</td><td>1.5E-15</td><td>MB231/MB468</td><td>MP</td></tr>
<tr><td>>MI0000767|hsa-mir-365a&WithFlank&16|+|14309279|14309377@62.82.21(+1C);MI0000769|hsa-mir-365b&WithFlank&17|+|31575405|31575527@74.94.21(+1C)&[MIMAT0000710&hsa-miR-365a-3p&offsets|0|-1(+1C);m-176&16|+|14309340|14309361&offsets|0|-1(+1C)];[MIMAT0022834&hsa-miR-365b-3p&offsets|0|-1(+1C);m-175&17|+|31575478|31575499&offsets|0|-1(+1C)]|TAATGCCCCTAAAAATCCTTAC</td><td>>MI0000767|hsa-mir-365a&WithFlank&16|+|14309279|14309377</td><td>62</td><td>82</td><td>21(+1C)</td><td>TAATGCCCCTAAAAATCCTTAC</td><td>63.0</td><td>4.3</td><td>0.5</td><td>8.1</td><td>4.7E-16</td><td>1.5E-15</td><td>BT20/MB468</td><td>NUC</td></tr>
<tr><td>>hg38_21_+_8433172_8446622_RNA45SN1_45S_50ntFlanks@4723.4752.30|CATTAATCAAGAACGAAAGTCGGAGGTTCG</td><td>>hg38_21_+_8433172_8446622_RNA45SN1_45S_50ntFlanks</td><td>4723</td><td>4752</td><td>30</td><td>CATTAATCAAGAACGAAAGTCGGAGGTTCG</td><td>611.4</td><td>-9.1</td><td>1.1</td><td>-8.2</td><td>2.3E-16</td><td>1.5E-15</td><td>MB231/MB468</td><td>NUC</td></tr>
<tr><td>>MI0000102|hsa-mir-100&WithFlank&11|-|122152223|122152314@22.40.19&[MIMAT0000098&hsa-miR-100-5p&offsets|+3|0;m-27&11|-|122152276|122152296&offsets|+3|+1]|CCGTAGATCCGAACTTGTG</td><td>>MI0000102|hsa-mir-100&WithFlank&11|-|122152223|122152314</td><td>22</td><td>40</td><td>19</td><td>CCGTAGATCCGAACTTGTG</td><td>906.5</td><td>13.8</td><td>1.7</td><td>8.2</td><td>1.9E-16</td><td>1.5E-15</td><td>MB231/MB468</td><td>CYTO</td></tr>
<tr><td>>MI0000087|hsa-mir-29a&WithFlank&7|-|130876741|130876816@47.68.22(+1U)&[MIMAT0000086&hsa-miR-29a-3p&offsets|-1|-1(+1U);m-26&7|-|130876749|130876769&offsets|-1|0(+1U)]|CTAGCACCATCTGAAATCGGTTT</td><td>>MI0000087|hsa-mir-29a&WithFlank&7|-|130876741|130876816</td><td>47</td><td>68</td><td>22(+1U)</td><td>CTAGCACCATCTGAAATCGGTTT</td><td>3340.8</td><td>6.6</td><td>0.8</td><td>8.2</td><td>1.8E-16</td><td>1.5E-15</td><td>MB231/MB468</td><td>MP</td></tr>
<tr><td>>hg38_21_+_8433172_8446622_RNA45SN1_45S_50ntFlanks@4759.4778.20|ATCAGATACCGTCGTAGTTC</td><td>>hg38_21_+_8433172_8446622_RNA45SN1_45S_50ntFlanks</td><td>4759</td><td>4778</td><td>20</td><td>ATCAGATACCGTCGTAGTTC</td><td>1195.1</td><td>-13.3</td><td>1.5</td><td>-8.7</td><td>5.1E-18</td><td>1.5E-15</td><td>BT20/MB468</td><td>MP</td></tr>
<tr><td>>hg38_21_+_8433172_8446622_RNA45SN1_45S_50ntFlanks@9572.9595.24|CAAATCGGTCGTCCGACCTGGGTA</td><td>>hg38_21_+_8433172_8446622_RNA45SN1_45S_50ntFlanks</td><td>9572</td><td>9595</td><td>24</td><td>CAAATCGGTCGTCCGACCTGGGTA</td><td>8065.6</td><td>-8.8</td><td>1.0</td><td>-8.7</td><td>5.0E-18</td><td>1.5E-15</td><td>BT20/MB468</td><td>MP</td></tr>
<tr><td>>hg38_21_+_8433172_8446622_RNA45SN1_45S_50ntFlanks@11318.11336.19|CCCCACGTCTCGTCGCGCG</td><td>>hg38_21_+_8433172_8446622_RNA45SN1_45S_50ntFlanks</td><td>11318</td><td>11336</td><td>19</td><td>CCCCACGTCTCGTCGCGCG</td><td>630.3</td><td>-12.3</td><td>1.5</td><td>-8.2</td><td>2.3E-16</td><td>1.5E-15</td><td>MB231/MB468</td><td>NUC</td></tr>
<tr><td>>hg38_21_+_8433172_8446622_RNA45SN1_45S_50ntFlanks@6700.6720.21|CTAGCTGCGAGAATTAATGTG</td><td>>hg38_21_+_8433172_8446622_RNA45SN1_45S_50ntFlanks</td><td>6700</td><td>6720</td><td>21</td><td>CTAGCTGCGAGAATTAATGTG</td><td>10186.1</td><td>-7.0</td><td>0.9</td><td>-8.2</td><td>3.1E-16</td><td>1.5E-15</td><td>MB231/MB468</td><td>MITO</td></tr>
<tr><td>>hg38_21_+_8433172_8446622_RNA45SN1_45S_50ntFlanks@12937.12966.30|AACCATTCGTAGACGACCTGCTTCTGGGTC</td><td>>hg38_21_+_8433172_8446622_RNA45SN1_45S_50ntFlanks</td><td>12937</td><td>12966</td><td>30</td><td>AACCATTCGTAGACGACCTGCTTCTGGGTC</td><td>573.4</td><td>-6.8</td><td>0.8</td><td>-8.2</td><td>2.0E-16</td><td>1.5E-15</td><td>MB231/MB468</td><td>CYTO</td></tr>
<tr><td>>hg38_21_+_8433172_8446622_RNA45SN1_45S_50ntFlanks@3593.3610.18|CCCGTCCGTCCGTCCGTC</td><td>>hg38_21_+_8433172_8446622_RNA45SN1_45S_50ntFlanks</td><td>3593</td><td>3610</td><td>18</td><td>CCCGTCCGTCCGTCCGTC</td><td>431.8</td><td>-8.5</td><td>1.0</td><td>-8.2</td><td>2.0E-16</td><td>1.5E-15</td><td>MB231/MB468</td><td>CYTO</td></tr>
<tr><td>>hg38_21_+_8433172_8446622_RNA45SN1_45S_50ntFlanks@8737.8754.18|CCCGTCCGTCCGTCCGTC</td><td>>hg38_21_+_8433172_8446622_RNA45SN1_45S_50ntFlanks</td><td>8737</td><td>8754</td><td>18</td><td>CCCGTCCGTCCGTCCGTC</td><td>431.8</td><td>-8.5</td><td>1.0</td><td>-8.2</td><td>2.0E-16</td><td>1.5E-15</td><td>MB231/MB468</td><td>CYTO</td></tr>
<tr><td>>hg38_21_+_8433172_8446622_RNA45SN1_45S_50ntFlanks@11325.11344.20|TCTCGTCGCGCGCGCGTCCG</td><td>>hg38_21_+_8433172_8446622_RNA45SN1_45S_50ntFlanks</td><td>11325</td><td>11344</td><td>20</td><td>TCTCGTCGCGCGCGCGTCCG</td><td>483.4</td><td>-12.8</td><td>1.6</td><td>-8.2</td><td>1.9E-16</td><td>1.5E-15</td><td>MB231/MB468</td><td>MP</td></tr>
<tr><td>>hg38_21_+_8433172_8446622_RNA45SN1_45S_50ntFlanks@6775.6803.29|CCTCCCGGGGCTACGCCTGTCTGAGCGTC</td><td>>hg38_21_+_8433172_8446622_RNA45SN1_45S_50ntFlanks</td><td>6775</td><td>6803</td><td>29</td><td>CCTCCCGGGGCTACGCCTGTCTGAGCGTC</td><td>237.6</td><td>-11.8</td><td>1.4</td><td>-8.2</td><td>1.9E-16</td><td>1.5E-15</td><td>MB231/MB468</td><td>MP</td></tr>
<tr><td>>trna2_GlyGCC_21_-_18827107_18827177@1.32.32,trna35_GlyGCC_1_+_161413094_161413164@1.32.32,trna37_GlyGCC_1_+_161420467_161420537@1.32.32,trna39_GlyGCC_1_+_161427898_161427968@1.32.32,trna41_GlyGCC_1_+_161435258_161435328@1.32.32|GCATGGGTGGTTCAGTGGTAGAATTCTCGCCT</td><td>>trna2_GlyGCC_21_-_18827107_18827177</td><td>1</td><td>32</td><td>32</td><td>GCATGGGTGGTTCAGTGGTAGAATTCTCGCCT</td><td>498.7</td><td>4.0</td><td>0.5</td><td>8.1</td><td>4.8E-16</td><td>1.5E-15</td><td>BT20/MB468</td><td>NUC</td></tr>
<tr><td>>hg38_21_+_8433172_8446622_RNA45SN1_45S_50ntFlanks@5422.5450.29|GATTGGATGGTTTAGTGAGGCCCTCGGAT</td><td>>hg38_21_+_8433172_8446622_RNA45SN1_45S_50ntFlanks</td><td>5422</td><td>5450</td><td>29</td><td>GATTGGATGGTTTAGTGAGGCCCTCGGAT</td><td>332.7</td><td>7.2</td><td>0.9</td><td>8.2</td><td>3.0E-16</td><td>1.5E-15</td><td>BT20/MB468</td><td>CYTO</td></tr>
<tr><td>>hg38_21_+_8433172_8446622_RNA45SN1_45S_50ntFlanks@10819.10840.22|AGGGAAGTCGGCAAGCCGGATC</td><td>>hg38_21_+_8433172_8446622_RNA45SN1_45S_50ntFlanks</td><td>10819</td><td>10840</td><td>22</td><td>AGGGAAGTCGGCAAGCCGGATC</td><td>10442.1</td><td>-9.5</td><td>1.2</td><td>-8.2</td><td>1.9E-16</td><td>1.5E-15</td><td>MB231/MB468</td><td>MP</td></tr>
<tr><td>>hg38_21_+_8433172_8446622_RNA45SN1_45S_50ntFlanks@12244.12275.32|AAGGGCAAAAGCTCGCTTGATCTTGATTTTCA</td><td>>hg38_21_+_8433172_8446622_RNA45SN1_45S_50ntFlanks</td><td>12244</td><td>12275</td><td>32</td><td>AAGGGCAAAAGCTCGCTTGATCTTGATTTTCA</td><td>322.5</td><td>4.8</td><td>0.6</td><td>8.2</td><td>2.1E-16</td><td>1.5E-15</td><td>BT20/MB468</td><td>MITO</td></tr>
</table>

| | | | | | | | | | | | | | |
|---|---|---|---|---|---|---|---|---|---|---|---|---|---|
| >hg38_21_+_8433172_8446622_RNA45SN1_45S_50ntFlanks@485.524.40\|GATGTGGTGACGTCGTGCTCTCCCGGGCCGGGTCCGAGCC | >hg38_21_+_8433172_8446622_RNA45SN1_45S_50ntFlanks | 485 | 524 | 40 | GATGTGGTGACGTCGTGCTCTCCCGGGCCGGGTCCGAGCC | 6523.4 | 29.8 | 3.6 | 8.2 | 2.3E-16 | 1.5E-15 | MB231/MB468 | NUC |
| >MI0000061\|hsa-let-7a-2&WithFlank&11\|-\|122146516\|122146599@11.32.22;MI0000062\|hsa-let-7a-3&WithFlank&22\|+\|46112743\|46112828@10.31.22;MI0000060\|hsa-let-7a-1&WithFlank&9\|+\|94175951\|94176042@12.33.22&[MIMAT0000062&hsa-let-7a-5p&offsets\|0\|0;m-6&11\|-\|122146568\|122146589&offsets\|0\|0];[MIMAT0000062_1&hsa-let-7a-5p&offsets\|0\|0;m-5&22\|+\|46112752\|46112773&offsets\|0\|0];[MIMAT0000062_2&hsa-let-7a-5p&offsets\|0\|0;m-4&9\|+\|94175962\|94175983&offsets\|0\|0]\|TGAGGTAGTAGGTTGTATAGTT | >MI0000061\|hsa-let-7a-2&WithFlank&11\|-\|122146516\|122146599 | 11 | 32 | 22 | TGAGGTAGTAGGTTGTATAGTT | 80657.3 | 6.2 | 0.7 | 8.2 | 2.0E-16 | 1.5E-15 | MB231/MB468 | CYTO |
| >hg38_21_+_8433172_8446622_RNA45SN1_45S_50ntFlanks@10678.10695.18\|CGGTTCCGGCGGCGTCCG | >hg38_21_+_8433172_8446622_RNA45SN1_45S_50ntFlanks | 10678 | 10695 | 18 | CGGTTCCGGCGGCGTCCG | 236.1 | -11.7 | 1.4 | -8.2 | 1.9E-16 | 1.5E-15 | MB231/MB468 | MP |
| >hg38_21_+_8433172_8446622_RNA45SN1_45S_50ntFlanks@6784.6803.20\|GCTACGCCTGTCTGAGCGTC | >hg38_21_+_8433172_8446622_RNA45SN1_45S_50ntFlanks | 6784 | 6803 | 20 | GCTACGCCTGTCTGAGCGTC | 882.0 | -13.6 | 1.7 | -8.2 | 3.1E-16 | 1.5E-15 | MB231/MB468 | MITO |
| >MI0000542\|hsa-mir-320a&WithFlank&8\|-\|22244960\|22245043@48.71.24&[MIMAT0000510&hsa-miR-320a-3p&offsets\|0\|+2;m-60&8\|-\|22244975\|22244996&offsets\|0\|+2]\|AAAAGCTGGGTTGAGAGGGCGAAA | >MI0000542\|hsa-mir-320a&WithFlank&8\|-\|22244960\|22245043 | 48 | 71 | 24 | AAAAGCTGGGTTGAGAGGGCGAAA | 1470.6 | 10.3 | 1.3 | 8.2 | 2.2E-16 | 1.5E-15 | BT20/MB468 | MITO |
| >hg38_21_+_8433172_8446622_RNA45SN1_45S_50ntFlanks@11104.11123.20\|AAGGGTCGGGGCGGCAGGGG | >hg38_21_+_8433172_8446622_RNA45SN1_45S_50ntFlanks | 11104 | 11123 | 20 | AAGGGTCGGGGCGGCAGGGG | 359.1 | -12.3 | 1.5 | -8.2 | 3.1E-16 | 1.5E-15 | MB231/MB468 | MITO |
| >hg38_21_+_8433172_8446622_RNA45SN1_45S_50ntFlanks@4763.4780.18\|GATACCGTCGTAGTTCCG | >hg38_21_+_8433172_8446622_RNA45SN1_45S_50ntFlanks | 4763 | 4780 | 18 | GATACCGTCGTAGTTCCG | 402.3 | -12.5 | 1.5 | -8.2 | 1.9E-16 | 1.5E-15 | MB231/MB468 | MP |
| >hg38_21_+_8433172_8446622_RNA45SN1_45S_50ntFlanks@11207.11254.48\|GGCGGCGACTCTGGACGCGAGCCGGGCCCTTCCCGTGGATCGCCCCAG | >hg38_21_+_8433172_8446622_RNA45SN1_45S_50ntFlanks | 11207 | 11254 | 48 | GGCGGCGACTCTGGACGCGAGCCGGGCCCTTCCCGTGGATCGCCCCAG | 632.9 | -13.1 | 1.6 | -8.2 | 3.1E-16 | 1.5E-15 | MB231/MB468 | MITO |
| >hg38_21_+_8433172_8446622_RNA45SN1_45S_50ntFlanks@3991.4026.36\|TGGTGACTCTAGATAACCTCGGGCCGATCGCACGCC | >hg38_21_+_8433172_8446622_RNA45SN1_45S_50ntFlanks | 3991 | 4026 | 36 | TGGTGACTCTAGATAACCTCGGGCCGATCGCACGCC | 710.1 | -7.5 | 0.9 | -8.2 | 3.1E-16 | 1.5E-15 | MB231/MB468 | MITO |
| >hg38_21_+_8433172_8446622_RNA45SN1_45S_50ntFlanks@12274.12299.26\|CAGTACGAATACAGACCGTGAAAGCG | >hg38_21_+_8433172_8446622_RNA45SN1_45S_50ntFlanks | 12274 | 12299 | 26 | CAGTACGAATACAGACCGTGAAAGCG | 271.0 | -11.9 | 1.5 | -8.2 | 1.9E-16 | 1.5E-15 | MB231/MB468 | MP |
| >hg38_21_+_8433172_8446622_RNA45SN1_45S_50ntFlanks@9569.9609.41\|GTGCAAATCGGTCGTCCGACCTGGGTATAGGGGCGAAAGAC | >hg38_21_+_8433172_8446622_RNA45SN1_45S_50ntFlanks | 9569 | 9609 | 41 | GTGCAAATCGGTCGTCCGACCTGGGTATAGGGGCGAAAGAC | 159.0 | 11.3 | 1.4 | 8.2 | 2.0E-16 | 1.5E-15 | MB231/MB468 | CYTO |
| >hg38_21_+_8433172_8446622_RNA45SN1_45S_50ntFlanks@10771.10817.47\|CAGGTCTCCAAGGTGAACAGCCTCTGGCATGTTGGAACAATGTAGGT | >hg38_21_+_8433172_8446622_RNA45SN1_45S_50ntFlanks | 10771 | 10817 | 47 | CAGGTCTCCAAGGTGAACAGCCTCTGGCATGTTGGAACAATGTAGGT | 561.5 | -13.6 | 1.7 | -8.1 | 4.9E-16 | 1.5E-15 | BT20/MB468 | NUC |
| >am_trnaMT_SerGCT_MT_+_12207_12265@14.34.21\|AGAACTGCTAACTCATGCCCC | >am_trnaMT_SerGCT_MT_+_12207_12265 | 14 | 34 | 21 | AGAACTGCTAACTCATGCCCC | 257.5 | 10.0 | 1.2 | 8.2 | 3.1E-16 | 1.5E-15 | BT20/MB468 | CYTO |
| >MI0005531\|hsa-mir-450b&WithFlank&X\|-\|134540179\|134540268@17.37.21&[MIMAT0004909&hsa-miR-450b-5p&offsets\|0\|-1;m-266&X\|-\|134540232\|134540252&offsets\|0\|0]\|TTTTGCAATATGTTCCTGAAT | >MI0005531\|hsa-mir-450b&WithFlank&X\|-\|134540179\|134540268 | 17 | 37 | 21 | TTTTGCAATATGTTCCTGAAT | 1013.0 | 12.0 | 1.5 | 8.2 | 3.1E-16 | 1.5E-15 | BT20/MB468 | CYTO |
| >hg38_21_+_8433172_8446622_RNA45SN1_45S_50ntFlanks@8439.8471.33\|GATTCAACCCGGCGGCGGGTCCGGCCGTGTCGG | >hg38_21_+_8433172_8446622_RNA45SN1_45S_50ntFlanks | 8439 | 8471 | 33 | GATTCAACCCGGCGGCGGGTCCGGCCGTGTCGG | 357.6 | -12.3 | 1.5 | -8.2 | 3.2E-16 | 1.5E-15 | MB231/MB468 | MITO |
| >hg38_21_+_8433172_8446622_RNA45SN1_45S_50ntFlanks@12152.12181.30\|TGGGGAGTTTGACTGGGGCGGTACACCTGT | >hg38_21_+_8433172_8446622_RNA45SN1_45S_50ntFlanks | 12152 | 12181 | 30 | TGGGGAGTTTGACTGGGGCGGTACACCTGT | 215.7 | -11.6 | 1.4 | -8.2 | 1.9E-16 | 1.5E-15 | MB231/MB468 | MP |
| >MI0000299\|hsa-mir-222&WithFlank&X\|-\|45747009\|45747130@75.97.23(+1A)&[MIMAT0000279&hsa-miR-222-3p&offsets\|0\|+2(+1A);m-140&X\|-\|45747033\|45747056&offsets\|0\|-1(+1A)]\|AGCTACATCTGGCTACTGGGTCTA | >MI0000299\|hsa-mir-222&WithFlank&X\|-\|45747009\|45747130 | 75 | 97 | 23(+1A) | AGCTACATCTGGCTACTGGGTCTA | 809.3 | 6.0 | 0.7 | 8.2 | 2.4E-16 | 1.5E-15 | MB231/MB468 | NUC |
| >hg38_21_+_8433172_8446622_RNA45SN1_45S_50ntFlanks@12976.12997.22\|TACGTAGCAGAGCAGCTCCCTC | >hg38_21_+_8433172_8446622_RNA45SN1_45S_50ntFlanks | 12976 | 12997 | 22 | TACGTAGCAGAGCAGCTCCCTC | 331.6 | 7.2 | 0.9 | 8.2 | 2.0E-16 | 1.5E-15 | MB231/MB468 | CYTO |
| >hg38_21_+_8433172_8446622_RNA45SN1_45S_50ntFlanks@8527.8551.25\|CCTTCCCCCGCCGCCCCTCCTCCTC | >hg38_21_+_8433172_8446622_RNA45SN1_45S_50ntFlanks | 8527 | 8551 | 25 | CCTTCCCCCGCCGCCCCTCCTCCTC | 423.5 | -12.5 | 1.5 | -8.2 | 3.2E-16 | 1.5E-15 | MB231/MB468 | MITO |
| >hg38_21_+_8433172_8446622_RNA45SN1_45S_50ntFlanks@5152.5185.34\|AGGGACAAGTGGCGTTCAGCCACCCGAGATTGAG | >hg38_21_+_8433172_8446622_RNA45SN1_45S_50ntFlanks | 5152 | 5185 | 34 | AGGGACAAGTGGCGTTCAGCCACCCGAGATTGAG | 2169.5 | -14.9 | 1.8 | -8.2 | 3.2E-16 | 1.5E-15 | MB231/MB468 | MITO |
| >hg38_21_+_8433172_8446622_RNA45SN1_45S_50ntFlanks@5472.5506.35\|ACGGCCCTGGCGGAGCGCTGAGAAGACGGTCGAAC | >hg38_21_+_8433172_8446622_RNA45SN1_45S_50ntFlanks | 5472 | 5506 | 35 | ACGGCCCTGGCGGAGCGCTGAGAAGACGGTCGAAC | 3030.9 | 7.1 | 0.9 | 8.2 | 2.1E-16 | 1.5E-15 | MB231/MB468 | CYTO |
| >MI0000285\|hsa-mir-205&WithFlank&1\|+\|209432127\|209432248@40.62.23&[MIMAT0000266&hsa-miR-205-5p&offsets\|0\|+1;m-351&1\|+\|209432166\|209432187&offsets\|0\|+1]\|TCCTTCATTCCACCGGAGTCTGT | >MI0000285\|hsa-mir-205&WithFlank&1\|+\|209432127\|209432248 | 40 | 62 | 23 | TCCTTCATTCCACCGGAGTCTGT | 1133.5 | -9.6 | 1.2 | -8.2 | 2.4E-16 | 1.5E-15 | MB231/MB468 | NUC |
| >hg38_21_+_8433172_8446622_RNA45SN1_45S_50ntFlanks@12893.12934.42\|CCTCGCCCGTCACGCACCGCACGTTCGTGGGGAACCTGGCGC | >hg38_21_+_8433172_8446622_RNA45SN1_45S_50ntFlanks | 12893 | 12934 | 42 | CCTCGCCCGTCACGCACCGCACGTTCGTGGGGAACCTGGCGC | 411.5 | -12.5 | 1.5 | -8.2 | 3.2E-16 | 1.5E-15 | MB231/MB468 | MITO |
| >hg38_21_+_8433172_8446622_RNA45SN1_45S_50ntFlanks@10843.10865.23\|TAACTTCGGGATAAGGATTGGCT | >hg38_21_+_8433172_8446622_RNA45SN1_45S_50ntFlanks | 10843 | 10865 | 23 | TAACTTCGGGATAAGGATTGGCT | 297.0 | -12.0 | 1.5 | -8.2 | 3.2E-16 | 1.6E-15 | MB231/MB468 | MITO |
| >hg38_21_+_8433172_8446622_RNA45SN1_45S_50ntFlanks@12300.12323.24\|GGGCCTCACGATCCTTCTGACCTT | >hg38_21_+_8433172_8446622_RNA45SN1_45S_50ntFlanks | 12300 | 12323 | 24 | GGGCCTCACGATCCTTCTGACCTT | 2104.6 | -14.8 | 1.8 | -8.2 | 3.2E-16 | 1.6E-15 | MB231/MB468 | MITO |
| >hg38_21_+_8433172_8446622_RNA45SN1_45S_50ntFlanks@10869.10892.24\|AGGGCTGGGTCGGTCGGGCTGGGG | >hg38_21_+_8433172_8446622_RNA45SN1_45S_50ntFlanks | 10869 | 10892 | 24 | AGGGCTGGGTCGGTCGGGCTGGGG | 820.3 | -13.5 | 1.6 | -8.2 | 3.2E-16 | 1.6E-15 | MB231/MB468 | MITO |
| >hg38_21_+_8433172_8446622_RNA45SN1_45S_50ntFlanks@9198.9226.29\|GTTCTCTCGGGGCCACGCGCGCGTCCCCC | >hg38_21_+_8433172_8446622_RNA45SN1_45S_50ntFlanks | 9198 | 9226 | 29 | GTTCTCTCGGGGCCACGCGCGCGTCCCCC | 259.1 | -11.9 | 1.4 | -8.2 | 2.0E-16 | 1.6E-15 | MB231/MB468 | MP |
| >hg38_21_+_8433172_8446622_RNA45SN1_45S_50ntFlanks@10350.10374.25\|ACGAGAACTTTGAAGGCCGAAGTGG | >hg38_21_+_8433172_8446622_RNA45SN1_45S_50ntFlanks | 10350 | 10374 | 25 | ACGAGAACTTTGAAGGCCGAAGTGG | 3417.5 | -8.2 | 1.0 | -8.2 | 3.2E-16 | 1.6E-15 | MB231/MB468 | MITO |
| >hg38_21_+_8433172_8446622_RNA45SN1_45S_50ntFlanks@9033.9068.36\|CTCGTGGGGGGGCCGGGCCACCCCTCCCACGGCGCG | >hg38_21_+_8433172_8446622_RNA45SN1_45S_50ntFlanks | 9033 | 9068 | 36 | CTCGTGGGGGGGCCGGGCCACCCCTCCCACGGCGCG | 385.9 | -12.4 | 1.5 | -8.2 | 3.2E-16 | 1.6E-15 | MB231/MB468 | MITO |
| >hg38_21_+_8433172_8446622_RNA45SN1_45S_50ntFlanks@10788.10822.35\|CAGCCTCTGGCATGTTGGAACAATGTAGGTAAGGG | >hg38_21_+_8433172_8446622_RNA45SN1_45S_50ntFlanks | 10788 | 10822 | 35 | CAGCCTCTGGCATGTTGGAACAATGTAGGTAAGGG | 278.4 | -12.0 | 1.5 | -8.2 | 2.0E-16 | 1.6E-15 | MB231/MB468 | MP |
| >hg38_21_+_8433172_8446622_RNA45SN1_45S_50ntFlanks@12415.12455.41\|CTTCGATGTCGGCTCTTCCTATCATTGTGAAGCAGAATTCA | >hg38_21_+_8433172_8446622_RNA45SN1_45S_50ntFlanks | 12415 | 12455 | 41 | CTTCGATGTCGGCTCTTCCTATCATTGTGAAGCAGAATTCA | 733.6 | -13.9 | 1.7 | -8.1 | 5.0E-16 | 1.6E-15 | BT20/MB468 | NUC |
| >MI0000650\|hsa-mir-200c&WithFlank&12\|+\|6963693\|6963772@50.69.20(+1A)&[MIMAT0000617&hsa-miR-200c-3p&offsets\|0\|-3(+1A);m-54&12\|+\|6963742\|6963764&offsets\|0\|-3(+1A)]\|TAATACTGCCGGGTAATGATA | >MI0000650\|hsa-mir-200c&WithFlank&12\|+\|6963693\|6963772 | 50 | 69 | 20(+1A) | TAATACTGCCGGGTAATGATA | 280.3 | 11.7 | 1.4 | 8.5 | 2.5E-17 | 1.6E-15 | BT20/MB231 | MITO |
| >hg38_21_+_8433172_8446622_RNA45SN1_45S_50ntFlanks@12407.12426.20\|TTTTGATCCTTCGATGTCGG | >hg38_21_+_8433172_8446622_RNA45SN1_45S_50ntFlanks | 12407 | 12426 | 20 | TTTTGATCCTTCGATGTCGG | 526.9 | -12.9 | 1.6 | -8.2 | 2.0E-16 | 1.6E-15 | MB231/MB468 | MP |
| >hg38_21_+_8433172_8446622_RNA45SN1_45S_50ntFlanks@10687.10705.19\|CGGCGTCCGGTGAGCTCTC | >hg38_21_+_8433172_8446622_RNA45SN1_45S_50ntFlanks | 10687 | 10705 | 19 | CGGCGTCCGGTGAGCTCTC | 275.3 | -6.0 | 0.7 | -8.2 | 2.1E-16 | 1.6E-15 | MB231/MB468 | CYTO |
| >hg38_21_+_8433172_8446622_RNA45SN1_45S_50ntFlanks@10873.10891.19\|CTGGGTCGGTCGGGCTGGG | >hg38_21_+_8433172_8446622_RNA45SN1_45S_50ntFlanks | 10873 | 10891 | 19 | CTGGGTCGGTCGGGCTGGG | 354.2 | -12.3 | 1.5 | -8.2 | 3.3E-16 | 1.6E-15 | MB231/MB468 | MITO |
| >hg38_21_+_8433172_8446622_RNA45SN1_45S_50ntFlanks@10807.10826.20\|ACAATGTAGGTAAGGGAAGT | >hg38_21_+_8433172_8446622_RNA45SN1_45S_50ntFlanks | 10807 | 10826 | 20 | ACAATGTAGGTAAGGGAAGT | 247.3 | -11.8 | 1.4 | -8.2 | 2.0E-16 | 1.6E-15 | MB231/MB468 | MP |
| >MI0000269\|hsa-mir-181a-2&WithFlank&9\|+\|124692436\|124692557@83.103.21(+1U)&[MIMAT0004558&hsa-miR-181a-2-3p&offsets\|0\|-1(+1U);m-71&9\|+\|124692518\|124692539&offsets\|0\|-1(+1U)]\|ACCACTGACCGTTGACTGTACT | >MI0000269\|hsa-mir-181a-2&WithFlank&9\|+\|124692436\|124692557 | 83 | 103 | 21(+1U) | ACCACTGACCGTTGACTGTACT | 1193.3 | 10.1 | 1.2 | 8.2 | 3.1E-16 | 1.6E-15 | BT20/MB468 | CYTO |
| >hg38_21_+_8433172_8446622_RNA45SN1_45S_50ntFlanks@12451.12476.26\|ATTCACCAAGCGTTGGATTGTTCACC | >hg38_21_+_8433172_8446622_RNA45SN1_45S_50ntFlanks | 12451 | 12476 | 26 | ATTCACCAAGCGTTGGATTGTTCACC | 369.7 | 4.8 | 0.6 | 8.2 | 2.1E-16 | 1.6E-15 | MB231/MB468 | CYTO |
| >hg38_21_+_8433172_8446622_RNA45SN1_45S_50ntFlanks@11873.11912.40\|AAGAAGACCCTGTTGAGCTTGACTCTAGTCTGGCACGGTG | >hg38_21_+_8433172_8446622_RNA45SN1_45S_50ntFlanks | 11873 | 11912 | 40 | AAGAAGACCCTGTTGAGCTTGACTCTAGTCTGGCACGGTG | 605.1 | -13.0 | 1.6 | -8.2 | 3.3E-16 | 1.6E-15 | MB231/MB468 | MITO |
| >am_trna129_MetCAT_6_-_27870271_27870342@34.51.18,trna142_MetCAT_6_-_27560600_27560671@34.51.18,trna150_MetCAT_6_-_27300764_27300835@34.51.18,trna169_MetCAT_6_-_26330529_26330600@34.51.18,trna171_MetCAT_6_-_26313352_26313423@34.51.18,trna20_MetCAT_17_-_80452597_80452668@34.51.18,trna2_MetCAT_6_+_26286754_26286825@34.51.18,trna32_MetCAT_1_+_153643726_153643797@34.51.18,trna61_MetCAT_6_+_27745664_27745735@34.51.18\|ATAACCCAGAGGTCGATG | >am_trna129_MetCAT_6_-_27870271_27870342 | 34 | 51 | 18 | ATAACCCAGAGGTCGATG | 598.0 | -13.1 | 1.6 | -8.2 | 2.0E-16 | 1.6E-15 | MB231/MB468 | MP |
| >hg38_21_+_8433172_8446622_RNA45SN1_45S_50ntFlanks@8179.8210.32\|CTGATCGAGGCCCAGCCCGTGGACGGTGTGAG | >hg38_21_+_8433172_8446622_RNA45SN1_45S_50ntFlanks | 8179 | 8210 | 32 | CTGATCGAGGCCCAGCCCGTGGACGGTGTGAG | 315.8 | -12.2 | 1.5 | -8.2 | 2.0E-16 | 1.6E-15 | MB231/MB468 | MP |
| >hg38_21_+_8433172_8446622_RNA45SN1_45S_50ntFlanks@6786.6806.21\|TACGCCTGTCTGAGCGTCGCT | >hg38_21_+_8433172_8446622_RNA45SN1_45S_50ntFlanks | 6786 | 6806 | 21 | TACGCCTGTCTGAGCGTCGCT | 1239.8 | 5.2 | 0.6 | 8.2 | 2.5E-16 | 1.6E-15 | MB231/MB468 | NUC |
| >MI0016746\|hsa-mir-548o-2&WithFlank&20\|+\|38516557\|38516638@51.72.22;MI0006402\|hsa-mir-548o&WithFlank&7\|-\|102405736\|102405861@93.114.22&[MIMAT0005919&hsa-miR-548o-3p&offsets\|0\|0;m-431&20\|+\|38516607\|38516628&offsets\|0\|0];[MIMAT0005919_1&hsa-miR-548o-3p&offsets\|0\|0;m-430&7\|-\|102405748\|102405769&offsets\|0\|0]\|CCAAAACTGCAGTTACTTTTGC | >MI0016746\|hsa-mir-548o-2&WithFlank&20\|+\|38516557\|38516638 | 51 | 72 | 22 | CCAAAACTGCAGTTACTTTTGC | 2002.8 | 8.5 | 1.0 | 8.2 | 3.2E-16 | 1.6E-15 | BT20/MB468 | CYTO |
| >hg38_21_+_8433172_8446622_RNA45SN1_45S_50ntFlanks@11235.11269.35\|CTTCCCGTGGATCGCCCCAGCTGCGGCGGGCGTCG | >hg38_21_+_8433172_8446622_RNA45SN1_45S_50ntFlanks | 11235 | 11269 | 35 | CTTCCCGTGGATCGCCCCAGCTGCGGCGGGCGTCG | 299.1 | -12.1 | 1.5 | -8.2 | 2.0E-16 | 1.6E-15 | MB231/MB468 | MP |
| >hg38_21_+_8433172_8446622_RNA45SN1_45S_50ntFlanks@12155.12179.25\|GGAGTTTGACTGGGGCGGTACACCT | >hg38_21_+_8433172_8446622_RNA45SN1_45S_50ntFlanks | 12155 | 12179 | 25 | GGAGTTTGACTGGGGCGGTACACCT | 274.9 | -12.0 | 1.5 | -8.2 | 2.0E-16 | 1.6E-15 | MB231/MB468 | MP |

| | | | | | | | | | | | | | |
|---|---|---|---|---|---|---|---|---|---|---|---|---|---|
| >MI0000650\|hsa-mir-200c&WithFlank&12\|+\|6963693\|6963772@50.69.20(+1A)&[MIMAT0000617&hsa-miR-200c-3p&offsets\|0\|-3(+1A);m-54&12\|+\|6963742\|6963764&offsets\|0\|-3(+1A)]\|TAATACTGCCGGGTAATGATA | >MI0000650\|hsa-mir-200c&WithFlank&12\|+\|6963693\|6963772 | 50 | 69 | 20(+1A) | TAATACTGCCGGGTAATGATA | 365.5 | 12.3 | 1.5 | 8.4 | 3.5E-17 | 1.6E-15 | BT20/MB231 | NUC |
| >hg38_21_+_8433172_8446622_RNA45SN1_45S_50ntFlanks@7986.8010.25\|ATCAGACGTGGCGACCCGCTGAATT | >hg38_21_+_8433172_8446622_RNA45SN1_45S_50ntFlanks | 7986 | 8010 | 25 | ATCAGACGTGGCGACCCGCTGAATT | 414.0 | -12.5 | 1.5 | -8.2 | 3.3E-16 | 1.6E-15 | MB231/MB468 | MITO |
| >MI0000746\|hsa-mir-99b&WithFlank&19\|+\|51692606\|51692687@13.32.20(+1G)&[MIMAT0000689&hsa-miR-99b-5p&offsets\|0\|-2(+1G);m-10&19\|+\|51692618\|51692639&offsets\|0\|-2(+1G)]\|CACCCGTAGAACCGACCTTGG | >MI0000746\|hsa-mir-99b&WithFlank&19\|+\|51692606\|51692687 | 13 | 32 | 20(+1G) | CACCCGTAGAACCGACCTTGG | 1439.2 | 11.9 | 1.5 | 8.2 | 3.2E-16 | 1.6E-15 | BT20/MB468 | CYTO |
| >hg38_MT_+_598_1651_RNR1_12S_50ntFlanks@971.1004.34\|TGGTAAGTGTACTGGAAAGTGCACTTGGACGAAC | >hg38_MT_+_598_1651_RNR1_12S_50ntFlanks | 971 | 1004 | 34 | TGGTAAGTGTACTGGAAAGTGCACTTGGACGAAC | 227.2 | 11.7 | 1.4 | 8.2 | 2.5E-16 | 1.6E-15 | MB231/MB468 | NUC |
| >hg38_21_+_8433172_8446622_RNA45SN1_45S_50ntFlanks@12281.12321.41\|AATACAGACCGTGAAAGCGGGGCCTCACGATCCTTCTGACC | >hg38_21_+_8433172_8446622_RNA45SN1_45S_50ntFlanks | 12281 | 12321 | 41 | AATACAGACCGTGAAAGCGGGGCCTCACGATCCTTCTGACC | 192.1 | 11.6 | 1.4 | 8.2 | 2.1E-16 | 1.6E-15 | MB231/MB468 | CYTO |
| >hg38_21_+_8433172_8446622_RNA45SN1_45S_50ntFlanks@12324.12343.20\|TTGGGTTTTAAGCAGGAGGT | >hg38_21_+_8433172_8446622_RNA45SN1_45S_50ntFlanks | 12324 | 12343 | 20 | TTGGGTTTTAAGCAGGAGGT | 227.8 | -11.7 | 1.4 | -8.2 | 2.0E-16 | 1.6E-15 | MB231/MB468 | MP |
| >hg38_21_+_8433172_8446622_RNA45SN1_45S_50ntFlanks@12083.12106.24\|CTCGCTTCTGGCGCCAAGCGCCCG | >hg38_21_+_8433172_8446622_RNA45SN1_45S_50ntFlanks | 12083 | 12106 | 24 | CTCGCTTCTGGCGCCAAGCGCCCG | 262.9 | -11.9 | 1.4 | -8.2 | 2.0E-16 | 1.6E-15 | MB231/MB468 | MP |
| >hg38_21_+_8433172_8446622_RNA45SN1_45S_50ntFlanks@10845.10864.20\|ACTTCGGGATAAGGATTGGC | >hg38_21_+_8433172_8446622_RNA45SN1_45S_50ntFlanks | 10845 | 10864 | 20 | ACTTCGGGATAAGGATTGGC | 495.1 | -12.8 | 1.6 | -8.2 | 2.0E-16 | 1.6E-15 | MB231/MB468 | MP |
| >MI0000067\|hsa-let-7f-1&WithFlank&9\|+\|94176341\|94176439@13.33.21(+1A);MI0000068\|hsa-let-7f-2&WithFlank&X\|-\|53557186\|53557280@14.34.21(+1A)&[MIMAT0000067&hsa-let-7f-5p&offsets\|0\|-1(+1A);m-20&9\|+\|94176353\|94176374&offsets\|0\|-1(+1A)];[MIMAT0000067_1&hsa-let-7f-5p&offsets\|0\|-1(+1A);m-19&X\|-\|53557246\|53557267&offsets\|0\|-1(+1A)]\|TGAGGTAGTAGATTGTATAGTA | >MI0000067\|hsa-let-7f-1&WithFlank&9\|+\|94176341\|94176439 | 13 | 33 | 21(+1A) | TGAGGTAGTAGATTGTATAGTA | 1620.0 | 6.9 | 0.8 | 8.2 | 2.3E-16 | 1.6E-15 | BT20/MB468 | MITO |
| >MI0000255\|hsa-mir-30d&WithFlank&8\|-\|134804870\|134804951@12.32.21(+1A)&[MIMAT0000245&hsa-miR-30d-5p&offsets\|0\|-1(+1A);m-21&8\|-\|134804919\|134804940&offsets\|0\|-1(+1A)]\|TGTAAACATCCCCGACTGGAAA | >MI0000255\|hsa-mir-30d&WithFlank&8\|-\|134804870\|134804951 | 12 | 32 | 21(+1A) | TGTAAACATCCCCGACTGGAAA | 718.7 | 9.4 | 1.1 | 8.2 | 2.3E-16 | 1.6E-15 | BT20/MB468 | MITO |
| >MI0000070\|hsa-mir-16-1&WithFlank&13\|-\|50048967\|50049067@20.40.21;MI0000115\|hsa-mir-16-2&WithFlank&3\|+\|160404739\|160404831@16.36.21&[MIMAT0000069&hsa-miR-16-5p&offsets\|0\|-1;m-62&13\|-\|50049027\|50049048&offsets\|0\|-1];[MIMAT0000069_1&hsa-miR-16-5p&offsets\|0\|-1;m-61&3\|+\|160404754\|160404775&offsets\|0\|-1]\|TAGCAGCACGTAAATATTGGC | >MI0000070\|hsa-mir-16-1&WithFlank&13\|-\|50048967\|50049067 | 20 | 40 | 21 | TAGCAGCACGTAAATATTGGC | 988.6 | 4.6 | 0.6 | 8.2 | 2.5E-16 | 1.6E-15 | MB231/MB468 | NUC |
| >hg38_21_+_8433172_8446622_RNA45SN1_45S_50ntFlanks@485.515.31\|GATGTGGTGACGTCGTGCTCTCCCGGGCCGG | >hg38_21_+_8433172_8446622_RNA45SN1_45S_50ntFlanks | 485 | 515 | 31 | GATGTGGTGACGTCGTGCTCTCCCGGGCCGG | 5271.0 | 29.8 | 3.6 | 8.2 | 2.5E-16 | 1.6E-15 | MB231/MB468 | NUC |
| >hg38_21_+_8433172_8446622_RNA45SN1_45S_50ntFlanks@11874.11893.20\|AGAAGACCCTGTTGAGCTTG | >hg38_21_+_8433172_8446622_RNA45SN1_45S_50ntFlanks | 11874 | 11893 | 20 | AGAAGACCCTGTTGAGCTTG | 3627.9 | -9.3 | 1.1 | -8.2 | 3.3E-16 | 1.6E-15 | MB231/MB468 | MITO |
| >hg38_21_+_8433172_8446622_RNA45SN1_45S_50ntFlanks@11891.11909.19\|TTGACTCTAGTCTGGCACG | >hg38_21_+_8433172_8446622_RNA45SN1_45S_50ntFlanks | 11891 | 11909 | 19 | TTGACTCTAGTCTGGCACG | 231.8 | -11.7 | 1.4 | -8.2 | 2.0E-16 | 1.6E-15 | MB231/MB468 | MP |
| >hg38_21_+_8433172_8446622_RNA45SN1_45S_50ntFlanks@12033.12055.23\|ATCGTTTTTTCACTGACCCGGTG | >hg38_21_+_8433172_8446622_RNA45SN1_45S_50ntFlanks | 12033 | 12055 | 23 | ATCGTTTTTTCACTGACCCGGTG | 313.1 | -12.1 | 1.5 | -8.2 | 3.4E-16 | 1.6E-15 | MB231/MB468 | MITO |
| >hg38_21_+_8433172_8446622_RNA45SN1_45S_50ntFlanks@4027.4045.19\|CCCCGTGGCGGCGACGACC | >hg38_21_+_8433172_8446622_RNA45SN1_45S_50ntFlanks | 4027 | 4045 | 19 | CCCCGTGGCGGCGACGACC | 663.2 | -13.2 | 1.6 | -8.2 | 3.4E-16 | 1.6E-15 | MB231/MB468 | MITO |
| >hg38_21_+_8433172_8446622_RNA45SN1_45S_50ntFlanks@4297.4321.25\|CATTGGAGGGCAAGTCTGGTGCCAG | >hg38_21_+_8433172_8446622_RNA45SN1_45S_50ntFlanks | 4297 | 4321 | 25 | CATTGGAGGGCAAGTCTGGTGCCAG | 507.9 | -12.8 | 1.6 | -8.2 | 3.4E-16 | 1.6E-15 | MB231/MB468 | MITO |
| >hg38_21_+_8433172_8446622_RNA45SN1_45S_50ntFlanks@3930.3959.30\|ATCAAAACCAACCCGGTCAGCCCCTCTCCG | >hg38_21_+_8433172_8446622_RNA45SN1_45S_50ntFlanks | 3930 | 3959 | 30 | ATCAAAACCAACCCGGTCAGCCCCTCTCCG | 465.5 | -11.0 | 1.3 | -8.2 | 2.3E-16 | 1.6E-15 | BT20/MB468 | MITO |
| >hg38_21_+_8433172_8446622_RNA45SN1_45S_50ntFlanks@11873.11906.34\|AAGAAGACCCTGTTGAGCTTGACTCTAGTCTGGC | >hg38_21_+_8433172_8446622_RNA45SN1_45S_50ntFlanks | 11873 | 11906 | 34 | AAGAAGACCCTGTTGAGCTTGACTCTAGTCTGGC | 1046.7 | -5.3 | 0.6 | -8.2 | 3.3E-16 | 1.6E-15 | BT20/MB468 | CYTO |
| >hg38_21_+_8433172_8446622_RNA45SN1_45S_50ntFlanks@6690.6718.29\|AAGAACGCAGCTAGCTGCGAGAATTAATG | >hg38_21_+_8433172_8446622_RNA45SN1_45S_50ntFlanks | 6690 | 6718 | 29 | AAGAACGCAGCTAGCTGCGAGAATTAATG | 402.4 | -12.4 | 1.5 | -8.2 | 3.4E-16 | 1.6E-15 | MB231/MB468 | MITO |
| >hg38_21_+_8433172_8446622_RNA45SN1_45S_50ntFlanks@6789.6806.18\|GCCTGTCTGAGCGTCGCT | >hg38_21_+_8433172_8446622_RNA45SN1_45S_50ntFlanks | 6789 | 6806 | 18 | GCCTGTCTGAGCGTCGCT | 1843.9 | 5.2 | 0.6 | 8.2 | 3.3E-16 | 1.6E-15 | BT20/MB468 | CYTO |
| >hg38_21_+_8433172_8446622_RNA45SN1_45S_50ntFlanks@9565.9592.28\|TGACGTGCAAATCGGTCGTCCGACCTGG | >hg38_21_+_8433172_8446622_RNA45SN1_45S_50ntFlanks | 9565 | 9592 | 28 | TGACGTGCAAATCGGTCGTCCGACCTGG | 1214.3 | -4.6 | 0.6 | -8.2 | 3.3E-16 | 1.6E-15 | BT20/MB468 | CYTO |
| >hg38_21_+_8433172_8446622_RNA45SN1_45S_50ntFlanks@12912.12933.22\|CACGTTCGTGGGGAACCTGGCG | >hg38_21_+_8433172_8446622_RNA45SN1_45S_50ntFlanks | 12912 | 12933 | 22 | CACGTTCGTGGGGAACCTGGCG | 1311.8 | -8.0 | 1.0 | -8.2 | 3.4E-16 | 1.6E-15 | MB231/MB468 | MITO |
| >hg38_21_+_8433172_8446622_RNA45SN1_45S_50ntFlanks@9622.9650.29\|CTAGTAGCTGGTTCCCTCCGAAGTTTCCC | >hg38_21_+_8433172_8446622_RNA45SN1_45S_50ntFlanks | 9622 | 9650 | 29 | CTAGTAGCTGGTTCCCTCCGAAGTTTCCC | 433.0 | -10.8 | 1.3 | -8.2 | 2.4E-16 | 1.6E-15 | BT20/MB468 | MITO |
| >hg38_21_+_8433172_8446622_RNA45SN1_45S_50ntFlanks@12993.13017.25\|CCCTCGCTGCGATCTATTGAAAGTC | >hg38_21_+_8433172_8446622_RNA45SN1_45S_50ntFlanks | 12993 | 13017 | 25 | CCCTCGCTGCGATCTATTGAAAGTC | 274.2 | 4.8 | 0.6 | 8.2 | 2.2E-16 | 1.6E-15 | MB231/MB468 | CYTO |
| >hg38_21_+_8433172_8446622_RNA45SN1_45S_50ntFlanks@12973.12991.19\|TCGTACGTAGCAGAGCAGC | >hg38_21_+_8433172_8446622_RNA45SN1_45S_50ntFlanks | 12973 | 12991 | 19 | TCGTACGTAGCAGAGCAGC | 3406.3 | -10.2 | 1.3 | -8.2 | 3.5E-16 | 1.6E-15 | MB231/MB468 | MITO |
| >hg38_21_+_8433172_8446622_RNA45SN1_45S_50ntFlanks@10458.10478.21\|ATGGCCTCCGTTGCCCTCGGC | >hg38_21_+_8433172_8446622_RNA45SN1_45S_50ntFlanks | 10458 | 10478 | 21 | ATGGCCTCCGTTGCCCTCGGC | 3561.7 | -5.9 | 0.7 | -8.2 | 2.2E-16 | 1.6E-15 | MB231/MB468 | CYTO |
| >hg38_21_+_8433172_8446622_RNA45SN1_45S_50ntFlanks@8689.8706.18\|GCTCCGGGACGGCTGGGA | >hg38_21_+_8433172_8446622_RNA45SN1_45S_50ntFlanks | 8689 | 8706 | 18 | GCTCCGGGACGGCTGGGA | 428.8 | 10.1 | 1.2 | 8.2 | 2.2E-16 | 1.7E-15 | MB231/MB468 | CYTO |
| >MI0000285\|hsa-mir-205&WithFlank&1\|+\|209432127\|209432248@40.60.21(+1A)&[MIMAT0000266&hsa-miR-205-5p&offsets\|0\|-1(+1A);m-351&1\|+\|209432166\|209432187&offsets\|0\|-1(+1A)]\|TCCTTCATTCCACCGGAGTCTA | >MI0000285\|hsa-mir-205&WithFlank&1\|+\|209432127\|209432248 | 40 | 60 | 21(+1A) | TCCTTCATTCCACCGGAGTCTA | 3030.9 | 14.9 | 1.7 | 8.7 | 3.4E-18 | 1.7E-15 | BT20/MB231 | MP |
| >MI0000476\|hsa-mir-138-1&WithFlank&3\|+\|44114206\|44114316@29.52.24&[MIMAT0000430_1&hsa-miR-138-5p&offsets\|0\|+1;m-497&3\|+\|44114234\|44114256&offsets\|0\|+1]\|AGCTGGTGTTGTGAATCAGGCCGT | >MI0000476\|hsa-mir-138-1&WithFlank&3\|+\|44114206\|44114316 | 29 | 52 | 24 | AGCTGGTGTTGTGAATCAGGCCGT | 1616.1 | -14.1 | 1.6 | -8.7 | 3.2E-18 | 1.7E-15 | BT20/MB231 | MP |
| >MI0000477\|hsa-mir-146a&WithFlank&5\|+\|160485346\|160485456@27.48.22(+1A)&[MIMAT0000449&hsa-miR-146a-5p&offsets\|0\|0(+1A);m-129&5\|+\|160485372\|160485393&offsets\|0\|0(+1A)]\|TGAGAACTGAATTCCATGGGTTA | >MI0000477\|hsa-mir-146a&WithFlank&5\|+\|160485346\|160485456 | 27 | 48 | 22(+1A) | TGAGAACTGAATTCCATGGGTTA | 3167.1 | -15.0 | 1.7 | -8.7 | 3.3E-18 | 1.7E-15 | BT20/MB231 | MP |
| >MI0000439\|hsa-mir-23b&WithFlank&9\|+\|95085202\|95085310@67.85.19(+2U)&[MIMAT0000418&hsa-miR-23b-3p&offsets\|+3\|-1(+2U);m-42&9\|+\|95085265\|95085285&offsets\|+3\|+1(+2U)]\|ACATTGCCAGGGATTACCATT | >MI0000439\|hsa-mir-23b&WithFlank&9\|+\|95085202\|95085310 | 67 | 85 | 19(+2U) | ACATTGCCAGGGATTACCATT | 98.4 | 9.5 | 1.2 | 8.1 | 5.3E-16 | 1.7E-15 | BT20/MB468 | NUC |
| >hg38_21_+_8433172_8446622_RNA45SN1_45S_50ntFlanks@7988.8014.27\|CAGACGTGGCGACCCGCTGAATTTAAG | >hg38_21_+_8433172_8446622_RNA45SN1_45S_50ntFlanks | 7988 | 8014 | 27 | CAGACGTGGCGACCCGCTGAATTTAAG | 2186.1 | -6.0 | 0.7 | -8.2 | 3.5E-16 | 1.7E-15 | MB231/MB468 | MITO |
| >hg38_21_+_8433172_8446622_RNA45SN1_45S_50ntFlanks@4853.4883.31\|AAGTCTTTGGGTTCCGGGGGGAGTATGGTTG | >hg38_21_+_8433172_8446622_RNA45SN1_45S_50ntFlanks | 4853 | 4883 | 31 | AAGTCTTTGGGTTCCGGGGGGAGTATGGTTG | 270.7 | -11.9 | 1.5 | -8.2 | 2.1E-16 | 1.7E-15 | MB231/MB468 | MP |
| >hg38_21_+_8433172_8446622_RNA45SN1_45S_50ntFlanks@11667.11692.26\|CTGCCCAGTGCTCTGAATGTCAAAGT | >hg38_21_+_8433172_8446622_RNA45SN1_45S_50ntFlanks | 11667 | 11692 | 26 | CTGCCCAGTGCTCTGAATGTCAAAGT | 671.6 | 9.7 | 1.2 | 8.2 | 2.4E-16 | 1.7E-15 | BT20/MB468 | MITO |
| >hg38_21_+_8433172_8446622_RNA45SN1_45S_50ntFlanks@6667.6699.33\|ATCACTCGGCTCGTGCGTCGATGAAGAACGCAG | >hg38_21_+_8433172_8446622_RNA45SN1_45S_50ntFlanks | 6667 | 6699 | 33 | ATCACTCGGCTCGTGCGTCGATGAAGAACGCAG | 426.0 | -10.8 | 1.3 | -8.2 | 2.4E-16 | 1.7E-15 | BT20/MB468 | MITO |
| >hg38_21_+_8433172_8446622_RNA45SN1_45S_50ntFlanks@10878.10901.24\|TCGGTCGGGCTGGGGCGCGAAGCG | >hg38_21_+_8433172_8446622_RNA45SN1_45S_50ntFlanks | 10878 | 10901 | 24 | TCGGTCGGGCTGGGGCGCGAAGCG | 404.1 | -6.1 | 0.7 | -8.2 | 2.3E-16 | 1.7E-15 | MB231/MB468 | CYTO |
| >hg38_21_+_8433172_8446622_RNA45SN1_45S_50ntFlanks@12417.12435.19\|TCGATGTCGGCTCTTCCTA | >hg38_21_+_8433172_8446622_RNA45SN1_45S_50ntFlanks | 12417 | 12435 | 19 | TCGATGTCGGCTCTTCCTA | 6369.4 | -4.4 | 0.5 | -8.2 | 2.3E-16 | 1.7E-15 | MB231/MB468 | CYTO |
| >hg38_21_+_8433172_8446622_RNA45SN1_45S_50ntFlanks@11999.12037.39\|CCCTGCGGGCCGCCGGTGAAATACCACTACTCTGATCGT | >hg38_21_+_8433172_8446622_RNA45SN1_45S_50ntFlanks | 11999 | 12037 | 39 | CCCTGCGGGCCGCCGGTGAAATACCACTACTCTGATCGT | 294.4 | -12.0 | 1.5 | -8.2 | 3.6E-16 | 1.7E-15 | MB231/MB468 | MITO |
| >hg38_21_+_8433172_8446622_RNA45SN1_45S_50ntFlanks@12004.12043.40\|CGGGCCGCCGGTGAAATACCACTACTCTGATCGTTTTTTC | >hg38_21_+_8433172_8446622_RNA45SN1_45S_50ntFlanks | 12004 | 12043 | 40 | CGGGCCGCCGGTGAAATACCACTACTCTGATCGTTTTTTC | 640.1 | -13.1 | 1.6 | -8.2 | 3.6E-16 | 1.7E-15 | MB231/MB468 | MITO |
| >hg38_21_+_8433172_8446622_RNA45SN1_45S_50ntFlanks@8475.8492.18\|CCCGGCGGATCTTTCCCG | >hg38_21_+_8433172_8446622_RNA45SN1_45S_50ntFlanks | 8475 | 8492 | 18 | CCCGGCGGATCTTTCCCG | 227.9 | -11.7 | 1.4 | -8.2 | 2.1E-16 | 1.7E-15 | MB231/MB468 | MP |
| >hg38_21_+_8433172_8446622_RNA45SN1_45S_50ntFlanks@9313.9335.23\|TCTAACACGTGCGCGAGTCGGGG | >hg38_21_+_8433172_8446622_RNA45SN1_45S_50ntFlanks | 9313 | 9335 | 23 | TCTAACACGTGCGCGAGTCGGGG | 500.8 | -12.8 | 1.6 | -8.2 | 2.1E-16 | 1.7E-15 | MB231/MB468 | MP |
| >hg38_21_+_8433172_8446622_RNA45SN1_45S_50ntFlanks@10380.10424.45\|GGTTCCATGTGAACAGCAGTTGAACATGGGTCAGTCGGTCCTGAG | >hg38_21_+_8433172_8446622_RNA45SN1_45S_50ntFlanks | 10380 | 10424 | 45 | GGTTCCATGTGAACAGCAGTTGAACATGGGTCAGTCGGTCCTGAG | 708.8 | 7.5 | 0.9 | 8.2 | 2.3E-16 | 1.7E-15 | MB231/MB468 | CYTO |
| >hg38_21_+_8433172_8446622_RNA45SN1_45S_50ntFlanks@6735.6770.36\|TTGATCATCGACACTTCGAACGCACTTGCGGCCCCG | >hg38_21_+_8433172_8446622_RNA45SN1_45S_50ntFlanks | 6735 | 6770 | 36 | TTGATCATCGACACTTCGAACGCACTTGCGGCCCCG | 400.4 | -12.5 | 1.5 | -8.2 | 2.2E-16 | 1.7E-15 | MB231/MB468 | MP |
| >MI0000108\|hsa-mir-103a-2&WithFlank&20\|+\|3917488\|3917577@54.76.23(+2U);MI0000109\|hsa-mir-103a-1&WithFlank&5\|-\|168560890\|168560979@54.76.23(+2U)&[MIMAT0000101&hsa-miR-103a-3p&offsets\|0\|0(+2U);m-12&20\|+\|3917541\|3917563&offsets\|0\|0(+2U)];[MIMAT0000101_1&hsa-miR-103a-3p&offsets\|0\|0(+2U);m-13&5\|-\|168560904\|168560926&offsets\|0\|0(+2U);m-44220&5\|-\|168560908\|168560926&offsets\|0\|+4(+2U)]\|AGCAGCATTGTACAGGGCTATGATT | >MI0000108\|hsa-mir-103a-2&WithFlank&20\|+\|3917488\|3917577 | 54 | 76 | 23(+2U) | AGCAGCATTGTACAGGGCTATGATT | 833.7 | 5.8 | 0.7 | 8.2 | 2.7E-16 | 1.7E-15 | MB231/MB468 | NUC |

| | | | | | | | | | | | | | |
|---|---|---|---|---|---|---|---|---|---|---|---|---|---|
| >am_MI0000809\|hsa-mir-151a&WithFlank&8\|-\|140732558\|140732659@53.72.20(+1C)&[MIMAT0000757&hsa-miR-151a-3p&offsets\|0\|-1(+1C);m-30&8\|-\|140732587\|140732607&offsets\|0\|-1(+1C)]\|CTAGACTGAAGCTCCTTGAGC | >am_MI0000809\|hsa-mir-151a&WithFlank&8\|-\|140732558\|140732659 | 53 | 72 | 20(+1C) | CTAGACTGAAGCTCCTTGAGC | 308.6 | 8.6 | 1.1 | 8.2 | 2.3E-16 | 1.7E-15 | MB231/MB468 | CYTO |
| >hg38_21_+_8433172_8446622_RNA45SN1_45S_50ntFlanks@8552.8581.30\|CTCCCCGGAGGGGGCGGGCTCCGGCGGGTG | >hg38_21_+_8433172_8446622_RNA45SN1_45S_50ntFlanks | 8552 | 8581 | 30 | CTCCCCGGAGGGGGCGGGCTCCGGCGGGTG | 236.6 | -10.6 | 1.3 | -8.2 | 2.3E-16 | 1.7E-15 | MB231/MB468 | CYTO |
| >MI0000279\|hsa-mir-196a-2&WithFlank&12\|+\|53991732\|53991853@31.52.22(+1U);MI0000238\|hsa-mir-196a-1&WithFlank&17\|-\|48632484\|48632565@13.34.22(+1U)&[MIMAT0000226&hsa-miR-196a-5p&offsets\|0\|0(+1U);m-261&12\|+\|53991762\|53991783&offsets\|0\|0(+1U)];[MIMAT0000226_1&hsa-miR-196a-5p&offsets\|0\|0(+1U);m-268&17\|-\|48632532\|48632553&offsets\|0\|0(+1U)]\|TAGGTAGTTTCATGTTGTTGGGT | >MI0000279\|hsa-mir-196a-2&WithFlank&12\|+\|53991732\|53991853 | 31 | 52 | 22(+1U) | TAGGTAGTTTCATGTTGTTGGGT | 483.5 | 12.3 | 1.4 | 8.5 | 2.0E-17 | 1.7E-15 | BT20/MB231 | CYTO |
| >hg38_21_+_8433172_8446622_RNA45SN1_45S_50ntFlanks@9534.9574.41\|AGGAAACTCTGGTGGAGGTCCGTAGCGGTCCTGACGTGCAA | >hg38_21_+_8433172_8446622_RNA45SN1_45S_50ntFlanks | 9534 | 9574 | 41 | AGGAAACTCTGGTGGAGGTCCGTAGCGGTCCTGACGTGCAA | 2226.8 | -6.6 | 0.8 | -8.2 | 2.7E-16 | 1.7E-15 | MB231/MB468 | NUC |
| >hg38_21_+_8433172_8446622_RNA45SN1_45S_50ntFlanks@12073.12106.34\|CCGAGGGGCTCTCGCTTCTGGCGCCAAGCGCCCG | >hg38_21_+_8433172_8446622_RNA45SN1_45S_50ntFlanks | 12073 | 12106 | 34 | CCGAGGGGCTCTCGCTTCTGGCGCCAAGCGCCCG | 272.1 | -12.0 | 1.5 | -8.2 | 2.2E-16 | 1.7E-15 | MB231/MB468 | MP |
| >MI0000076\|hsa-mir-20a&WithFlank&13\|+\|91351059\|91351141@14.35.22(+1A)&[MIMAT0000075&hsa-miR-20a-5p&offsets\|0\|-1(+1A);m-96&13\|+\|91351072\|91351094&offsets\|0\|-1(+1A)]\|TAAAGTGCTTATAGTGCAGGTAA | >MI0000076\|hsa-mir-20a&WithFlank&13\|+\|91351059\|91351141 | 14 | 35 | 22(+1A) | TAAAGTGCTTATAGTGCAGGTAA | 1079.8 | 5.9 | 0.7 | 8.2 | 2.7E-16 | 1.7E-15 | MB231/MB468 | NUC |
| >hg38_21_+_8433172_8446622_RNA45SN1_45S_50ntFlanks@11583.11603.21\|CTGGTGCGGACCAGGGGAATC | >hg38_21_+_8433172_8446622_RNA45SN1_45S_50ntFlanks | 11583 | 11603 | 21 | CTGGTGCGGACCAGGGGAATC | 6620.2 | -9.2 | 1.1 | -8.2 | 2.2E-16 | 1.7E-15 | MB231/MB468 | MP |
| >hg38_21_+_8433172_8446622_RNA45SN1_45S_50ntFlanks@9903.9928.26\|ATCAGACCCCAGAAAAGGTGTTGGTT | >hg38_21_+_8433172_8446622_RNA45SN1_45S_50ntFlanks | 9903 | 9928 | 26 | ATCAGACCCCAGAAAAGGTGTTGGTT | 416.8 | -12.5 | 1.5 | -8.1 | 3.7E-16 | 1.7E-15 | MB231/MB468 | MITO |
| >MI0000266\|hsa-mir-10a&WithFlank&17\|-\|48579832\|48579953@28.48.21&[MIMAT0000253&hsa-miR-10a-5p&offsets\|0\|-2;m-18&17\|-\|48579905\|48579926&offsets\|0\|-1]\|TACCCTGTAGATCCGAATTTG | >MI0000266\|hsa-mir-10a&WithFlank&17\|-\|48579832\|48579953 | 28 | 48 | 21 | TACCCTGTAGATCCGAATTTG | 207.6 | 11.6 | 1.4 | 8.2 | 2.7E-16 | 1.7E-15 | MB231/MB468 | NUC |
| >hg38_21_+_8433172_8446622_RNA45SN1_45S_50ntFlanks@12676.12708.33\|AGAATCCCGCCCAGGCGGAACGATACGGCAGCG | >hg38_21_+_8433172_8446622_RNA45SN1_45S_50ntFlanks | 12676 | 12708 | 33 | AGAATCCCGCCCAGGCGGAACGATACGGCAGCG | 253.7 | -10.9 | 1.3 | -8.2 | 2.7E-16 | 1.7E-15 | MB231/MB468 | NUC |
| >hg38_21_+_8433172_8446622_RNA45SN1_45S_50ntFlanks@6746.6770.25\|CACTTCGAACGCACTTGCGGCCCCG | >hg38_21_+_8433172_8446622_RNA45SN1_45S_50ntFlanks | 6746 | 6770 | 25 | CACTTCGAACGCACTTGCGGCCCCG | 683.6 | -9.7 | 1.2 | -8.2 | 2.7E-16 | 1.7E-15 | MB231/MB468 | NUC |
| >hg38_21_+_8433172_8446622_RNA45SN1_45S_50ntFlanks@11593.11613.21\|CCAGGGGAATCCGACTGTTTA | >hg38_21_+_8433172_8446622_RNA45SN1_45S_50ntFlanks | 11593 | 11613 | 21 | CCAGGGGAATCCGACTGTTTA | 1210.8 | -8.0 | 1.0 | -8.1 | 3.7E-16 | 1.7E-15 | MB231/MB468 | MITO |
| >trnaMT_ValTAC_MT_+_1602_1670@48.70.23\|CAACTTAACTTGACCGCTCTGAC | >trnaMT_ValTAC_MT_+_1602_1670 | 48 | 70 | 23 | CAACTTAACTTGACCGCTCTGAC | 286.8 | 9.8 | 1.2 | 8.2 | 2.4E-16 | 1.8E-15 | MB231/MB468 | CYTO |
| >hg38_21_+_8433172_8446622_RNA45SN1_45S_50ntFlanks@5469.5499.31\|CCCACGGCCCTGGCGGAGCGCTGAGAAGACG | >hg38_21_+_8433172_8446622_RNA45SN1_45S_50ntFlanks | 5469 | 5499 | 31 | CCCACGGCCCTGGCGGAGCGCTGAGAAGACG | 425.3 | -12.6 | 1.5 | -8.2 | 2.2E-16 | 1.8E-15 | MB231/MB468 | MP |
| >MI0000077\|hsa-mir-21&WithFlank&17\|+\|59841260\|59841343@14.35.22(+1U)&[MIMAT0000076&hsa-miR-21-5p&offsets\|0\|0(+1U);m-1&17\|+\|59841273\|59841294&offsets\|0\|0(+1U)]\|TAGCTTATCAGACTGATGTTGAT | >MI0000077\|hsa-mir-21&WithFlank&17\|+\|59841260\|59841343 | 14 | 35 | 22(+1U) | TAGCTTATCAGACTGATGTTGAT | 40168.9 | 6.5 | 0.8 | 8.2 | 2.6E-16 | 1.8E-15 | BT20/MB468 | MITO |
| >hg38_21_+_8433172_8446622_RNA45SN1_45S_50ntFlanks@3862.3898.37\|AATTCTAGAGCTAATACATGCCGACGGGCGCTGACCC | >hg38_21_+_8433172_8446622_RNA45SN1_45S_50ntFlanks | 3862 | 3898 | 37 | AATTCTAGAGCTAATACATGCCGACGGGCGCTGACCC | 298.8 | -12.0 | 1.5 | -8.1 | 3.7E-16 | 1.8E-15 | MB231/MB468 | MITO |
| >hg38_21_+_8433172_8446622_RNA45SN1_45S_50ntFlanks@12946.12990.45\|TAGACGACCTGCTTCTGGGTCGGGGTTTCGTACGTAGCAGAGCAG | >hg38_21_+_8433172_8446622_RNA45SN1_45S_50ntFlanks | 12946 | 12990 | 45 | TAGACGACCTGCTTCTGGGTCGGGGTTTCGTACGTAGCAGAGCAG | 1216.7 | -14.7 | 1.8 | -8.1 | 5.6E-16 | 1.8E-15 | BT20/MB468 | NUC |
| >hg38_21_+_8433172_8446622_RNA45SN1_45S_50ntFlanks@10830.10853.24\|CAAGCCGGATCCGTAACTTCGGGA | >hg38_21_+_8433172_8446622_RNA45SN1_45S_50ntFlanks | 10830 | 10853 | 24 | CAAGCCGGATCCGTAACTTCGGGA | 362.3 | -12.4 | 1.5 | -8.2 | 2.2E-16 | 1.8E-15 | MB231/MB468 | MP |
| >hg38_21_+_8433172_8446622_RNA45SN1_45S_50ntFlanks@12637.12654.18\|GCTACCATCTGTGGGATT | >hg38_21_+_8433172_8446622_RNA45SN1_45S_50ntFlanks | 12637 | 12654 | 18 | GCTACCATCTGTGGGATT | 2265.1 | -10.7 | 1.3 | -8.1 | 3.8E-16 | 1.8E-15 | MB231/MB468 | MITO |
| >hg38_21_+_8433172_8446622_RNA45SN1_45S_50ntFlanks@12635.12652.18\|AAGCTACCATCTGTGGGA | >hg38_21_+_8433172_8446622_RNA45SN1_45S_50ntFlanks | 12635 | 12652 | 18 | AAGCTACCATCTGTGGGA | 395.1 | -12.4 | 1.5 | -8.1 | 3.8E-16 | 1.8E-15 | MB231/MB468 | MITO |
| >hg38_21_+_8433172_8446622_RNA45SN1_45S_50ntFlanks@10361.10390.30\|GAAGGCCGAAGTGGAGAAGGGTTCCATGTG | >hg38_21_+_8433172_8446622_RNA45SN1_45S_50ntFlanks | 10361 | 10390 | 30 | GAAGGCCGAAGTGGAGAAGGGTTCCATGTG | 319.2 | -12.1 | 1.5 | -8.1 | 3.8E-16 | 1.8E-15 | MB231/MB468 | MITO |
| >hg38_21_+_8433172_8446622_RNA45SN1_45S_50ntFlanks@6719.6745.27\|TGAATTGCAGGACACATTGATCATCGA | >hg38_21_+_8433172_8446622_RNA45SN1_45S_50ntFlanks | 6719 | 6745 | 27 | TGAATTGCAGGACACATTGATCATCGA | 326.4 | -12.1 | 1.5 | -8.1 | 3.8E-16 | 1.8E-15 | MB231/MB468 | MITO |
| >MI0000477\|hsa-mir-146a&WithFlank&5\|+\|160485346\|160485456@27.47.21(+1A)&[MIMAT0000449&hsa-miR-146a-5p&offsets\|0\|-1(+1A);m-129&5\|+\|160485372\|160485393&offsets\|0\|-1(+1A)]\|TGAGAACTGAATTCCATGGGTA | >MI0000477\|hsa-mir-146a&WithFlank&5\|+\|160485346\|160485456 | 27 | 47 | 21(+1A) | TGAGAACTGAATTCCATGGGTA | 845.4 | -13.0 | 1.5 | -8.5 | 2.9E-17 | 1.8E-15 | BT20/MB231 | MITO |
| >hg38_21_+_8433172_8446622_RNA45SN1_45S_50ntFlanks@9574.9596.23\|AATCGGTCGTCCGACCTGGGTAT | >hg38_21_+_8433172_8446622_RNA45SN1_45S_50ntFlanks | 9574 | 9596 | 23 | AATCGGTCGTCCGACCTGGGTAT | 17096.3 | -8.4 | 1.0 | -8.6 | 6.3E-18 | 1.8E-15 | BT20/MB468 | MP |
| >MI0000081\|hsa-mir-24-2&WithFlank&19\|-\|13836281\|13836365@56.76.21(+1A);MI0000080\|hsa-mir-24-1&WithFlank&9\|+\|95086015\|95086094@50.70.21(+1A)&[MIMAT0000080&hsa-miR-24-3p&offsets\|0\|-1(+1A);m-36&19\|-\|13836288\|13836310&offsets\|0\|-2(+1A)];[MIMAT0000080_1&hsa-miR-24-3p&offsets\|0\|-1(+1A);m-35&9\|+\|95086064\|95086086&offsets\|0\|-2(+1A)]\|TGGCTCAGTTCAGCAGGAACAA | >MI0000081\|hsa-mir-24-2&WithFlank&19\|-\|13836281\|13836365 | 56 | 76 | 21(+1A) | TGGCTCAGTTCAGCAGGAACAA | 875.9 | 4.5 | 0.6 | 8.2 | 2.8E-16 | 1.8E-15 | MB231/MB468 | NUC |
| >MI0000736\|hsa-mir-30c-1&WithFlank&1\|+\|40757278\|40757378@23.45.23;MI0000254\|hsa-mir-30c-2&WithFlank&6\|-\|71376954\|71377037@13.35.23&[MIMAT0000244&hsa-miR-30c-5p&offsets\|0\|0;m-57&1\|+\|40757300\|40757323&offsets\|0\|-1];[MIMAT0000244_1&hsa-miR-30c-5p&offsets\|0\|0;m-56&6\|-\|71377002\|71377025&offsets\|0\|-1]\|TGTAAACATCCTACACTCTCAGC | >MI0000736\|hsa-mir-30c-1&WithFlank&1\|+\|40757278\|40757378 | 23 | 45 | 23 | TGTAAACATCCTACACTCTCAGC | 37061.2 | 4.9 | 0.6 | 8.2 | 2.8E-16 | 1.8E-15 | MB231/MB468 | NUC |
| >hg38_21_+_8433172_8446622_RNA45SN1_45S_50ntFlanks@9740.9757.18\|AACGATCTCAACCTATTC | >hg38_21_+_8433172_8446622_RNA45SN1_45S_50ntFlanks | 9740 | 9757 | 18 | AACGATCTCAACCTATTC | 279.3 | -12.0 | 1.5 | -8.2 | 2.3E-16 | 1.8E-15 | MB231/MB468 | MP |
| >hg38_21_+_8433172_8446622_RNA45SN1_45S_50ntFlanks@8737.8756.20\|CCCGTCCGTCCGTCCGTCCG | >hg38_21_+_8433172_8446622_RNA45SN1_45S_50ntFlanks | 8737 | 8756 | 20 | CCCGTCCGTCCGTCCGTCCG | 368.4 | -12.3 | 1.5 | -8.1 | 3.8E-16 | 1.8E-15 | MB231/MB468 | MITO |
| >MI0000266\|hsa-mir-10a&WithFlank&17\|-\|48579832\|48579953@28.48.21&[MIMAT0000253&hsa-miR-10a-5p&offsets\|0\|-2;m-18&17\|-\|48579905\|48579926&offsets\|0\|-1]\|TACCCTGTAGATCCGAATTTG | >MI0000266\|hsa-mir-10a&WithFlank&17\|-\|48579832\|48579953 | 28 | 48 | 21 | TACCCTGTAGATCCGAATTTG | 276.8 | 12.1 | 1.5 | 8.2 | 2.5E-16 | 1.8E-15 | MB231/MB468 | CYTO |
| >hg38_21_+_8433172_8446622_RNA45SN1_45S_50ntFlanks@11669.11688.20\|GCCCAGTGCTCTGAATGTCA | >hg38_21_+_8433172_8446622_RNA45SN1_45S_50ntFlanks | 11669 | 11688 | 20 | GCCCAGTGCTCTGAATGTCA | 688.4 | 6.1 | 0.7 | 8.2 | 2.5E-16 | 1.8E-15 | MB231/MB468 | CYTO |
| >MI0000266\|hsa-mir-10a&WithFlank&17\|-\|48579832\|48579953@28.51.24&[MIMAT0000253&hsa-miR-10a-5p&offsets\|0\|+1;m-18&17\|-\|48579905\|48579926&offsets\|0\|+2]\|TACCCTGTAGATCCGAATTTGTGT | >MI0000266\|hsa-mir-10a&WithFlank&17\|-\|48579832\|48579953 | 28 | 51 | 24 | TACCCTGTAGATCCGAATTTGTGT | 303.0 | 11.5 | 1.4 | 8.2 | 2.3E-16 | 1.8E-15 | MB231/MB468 | MP |
| >am_trna128_GlyGCC_6_-_27870686_27870756@1.32.32,trna133_GlyCCC_1_-_16872434_16872504@1.32.32,trna18_GlyGCC_16_+_70822597_70822667@1.32.32,trna19_GlyGCC_16_+_70823410_70823480@1.32.32,trna19_GlyGCC_2_-_157257659_157257729@1.32.32,trna24_GlyGCC_16_-_70812942_70813012@1.32.32,trna25_GlyGCC_16_-_70812114_70812184@1.32.32,trna4_GlyCCC_1_+_17188416_17188486@1.32.32,trna5_GlyGCC_17_+_8029064_8029134@1.32.32,trna68_GlyGCC_1_-_161493637_161493707@1.32.32\|GCATTGGTGGTTCAGTGGTAGAATTCTCGCCT | >am_trna128_GlyGCC_6_-_27870686_27870756 | 1 | 32 | 32 | GCATTGGTGGTTCAGTGGTAGAATTCTCGCCT | 14968.2 | 7.4 | 0.9 | 8.1 | 3.8E-16 | 1.8E-15 | MB231/MB468 | MITO |
| >hg38_21_+_8433172_8446622_RNA45SN1_45S_50ntFlanks@11653.11687.35\|ACGCGATGTGATTTCTGCCCAGTGCTCTGAATGTC | >hg38_21_+_8433172_8446622_RNA45SN1_45S_50ntFlanks | 11653 | 11687 | 35 | ACGCGATGTGATTTCTGCCCAGTGCTCTGAATGTC | 177.0 | 7.3 | 0.9 | 8.2 | 2.5E-16 | 1.8E-15 | MB231/MB468 | CYTO |
| >hg38_21_+_8433172_8446622_RNA45SN1_45S_50ntFlanks@7981.8003.23\|CTCAGATCAGACGTGGCGACCCG | >hg38_21_+_8433172_8446622_RNA45SN1_45S_50ntFlanks | 7981 | 8003 | 23 | CTCAGATCAGACGTGGCGACCCG | 195.2 | -11.5 | 1.4 | -8.2 | 2.3E-16 | 1.8E-15 | MB231/MB468 | MP |
| >hg38_21_+_8433172_8446622_RNA45SN1_45S_50ntFlanks@10411.10442.32\|CAGTCGGTCCTGAGAGATGGGCGAGCGCCGTT | >hg38_21_+_8433172_8446622_RNA45SN1_45S_50ntFlanks | 10411 | 10442 | 32 | CAGTCGGTCCTGAGAGATGGGCGAGCGCCGTT | 1554.7 | -7.3 | 0.9 | -8.2 | 2.3E-16 | 1.8E-15 | MB231/MB468 | MP |
| >hg38_21_+_8433172_8446622_RNA45SN1_45S_50ntFlanks@11661.11679.19\|TGATTTCTGCCCAGTGCTC | >hg38_21_+_8433172_8446622_RNA45SN1_45S_50ntFlanks | 11661 | 11679 | 19 | TGATTTCTGCCCAGTGCTC | 1000.0 | -9.6 | 1.2 | -8.1 | 3.9E-16 | 1.8E-15 | MB231/MB468 | MITO |
| >hg38_21_+_8433172_8446622_RNA45SN1_45S_50ntFlanks@5485.5526.42\|AGCGCTGAGAAGACGGTCGAACTTGACTATCTAGAGGAAGTA | >hg38_21_+_8433172_8446622_RNA45SN1_45S_50ntFlanks | 5485 | 5526 | 42 | AGCGCTGAGAAGACGGTCGAACTTGACTATCTAGAGGAAGTA | 684.5 | -13.3 | 1.6 | -8.2 | 2.3E-16 | 1.8E-15 | MB231/MB468 | MP |
| >hg38_21_+_8433172_8446622_RNA45SN1_45S_50ntFlanks@10791.10816.26\|CCTCTGGCATGTTGGAACAATGTAGG | >hg38_21_+_8433172_8446622_RNA45SN1_45S_50ntFlanks | 10791 | 10816 | 26 | CCTCTGGCATGTTGGAACAATGTAGG | 250.1 | -11.8 | 1.4 | -8.2 | 2.3E-16 | 1.8E-15 | MB231/MB468 | MP |
| >hg38_21_+_8433172_8446622_RNA45SN1_45S_50ntFlanks@11318.11344.27\|CCCCACGTCTCGTCGCGCGCGCGTCCG | >hg38_21_+_8433172_8446622_RNA45SN1_45S_50ntFlanks | 11318 | 11344 | 27 | CCCCACGTCTCGTCGCGCGCGCGTCCG | 300.9 | -12.1 | 1.5 | -8.2 | 2.3E-16 | 1.8E-15 | MB231/MB468 | MP |
| >hg38_21_+_8433172_8446622_RNA45SN1_45S_50ntFlanks@9504.9521.18\|GTGAACTATGCCTGGGCA | >hg38_21_+_8433172_8446622_RNA45SN1_45S_50ntFlanks | 9504 | 9521 | 18 | GTGAACTATGCCTGGGCA | 2578.7 | -6.0 | 0.7 | -8.1 | 3.7E-16 | 1.8E-15 | BT20/MB468 | CYTO |
| >hg38_21_+_8433172_8446622_RNA45SN1_45S_50ntFlanks@8182.8212.31\|ATCGAGGCCCAGCCCGTGGACGGTGTGAGGC | >hg38_21_+_8433172_8446622_RNA45SN1_45S_50ntFlanks | 8182 | 8212 | 31 | ATCGAGGCCCAGCCCGTGGACGGTGTGAGGC | 539.9 | -12.9 | 1.6 | -8.2 | 2.3E-16 | 1.8E-15 | MB231/MB468 | MP |
| >hg38_21_+_8433172_8446622_RNA45SN1_45S_50ntFlanks@12990.13013.24\|GCTCCCTCGCTGCGATCTATTGAA | >hg38_21_+_8433172_8446622_RNA45SN1_45S_50ntFlanks | 12990 | 13013 | 24 | GCTCCCTCGCTGCGATCTATTGAA | 14708.3 | -10.8 | 1.3 | -8.1 | 5.9E-16 | 1.8E-15 | BT20/MB468 | NUC |
| >hg38_MT_+_598_1651_RNR1_12S_50ntFlanks@608.633.26\|TGTTCTGTAATCGATAAACCCCGATC | >hg38_MT_+_598_1651_RNR1_12S_50ntFlanks | 608 | 633 | 26 | TGTTCTGTAATCGATAAACCCCGATC | 202.5 | 8.5 | 1.0 | 8.2 | 2.5E-16 | 1.8E-15 | MB231/MB468 | CYTO |
| >MI0005756\|hsa-mir-934&WithFlank&X\|+\|136550872\|136550966@21.42.22(+2A)&[MIMAT0004977&hsa-miR-934&offsets\|0\|0(+2A);m-943&X\|+\|136550892\|136550913&offsets\|0\|0(+2A)]\|TGTCTACTACTGGAGACACTGGAA | >MI0005756\|hsa-mir-934&WithFlank&X\|+\|136550872\|136550966 | 21 | 42 | 22(+2A) | TGTCTACTACTGGAGACACTGGAA | 435.6 | 12.1 | 1.4 | 8.5 | 2.2E-17 | 1.8E-15 | BT20/MB231 | CYTO |

<table>
<tr><td></td><td></td><td></td><td></td><td></td><td></td><td></td><td></td><td></td><td></td><td></td><td></td><td></td><td></td></tr>
<tr><td>>hg38_21_+_8433172_8446622_RNA45SN1_45S_50ntFlanks@6775.6806.32|CCTCCCGGGGCTACGCCTGTCTGAGCGTCGCT</td><td>>hg38_21_+_8433172_8446622_RNA45SN1_45S_50ntFlanks</td><td>6775</td><td>6806</td><td>32</td><td>CCTCCCGGGGCTACGCCTGTCTGAGCGTCGCT</td><td>225.6</td><td>3.9</td><td>0.5</td><td>8.2</td><td>2.9E-16</td><td>1.8E-15</td><td>MB231/MB468</td><td>NUC</td></tr>
<tr><td>>hg38_21_+_8433172_8446622_RNA45SN1_45S_50ntFlanks@12407.12436.30|TTTTGATCCTTCGATGTCGGCTCTTCCTAT</td><td>>hg38_21_+_8433172_8446622_RNA45SN1_45S_50ntFlanks</td><td>12407</td><td>12436</td><td>30</td><td>TTTTGATCCTTCGATGTCGGCTCTTCCTAT</td><td>235.3</td><td>-11.7</td><td>1.4</td><td>-8.2</td><td>2.4E-16</td><td>1.8E-15</td><td>MB231/MB468</td><td>MP</td></tr>
<tr><td>>am_trna162_MetCAT_6_-_26758550_26758622@31.51.21,trna164_MetCAT_6_-_26735574_26735646@31.51.21,trna20_MetCAT_16_+_71460396_71460468@31.51.21,trna21_MetCAT_6_+_26701712_26701784@31.51.21,trna27_MetCAT_6_+_26766444_26766516@31.51.21,trna75_MetCAT_6_+_28912352_28912424@31.51.21,trna97_MetCAT_6_-_28921042_28921114@31.51.21|TCTCATAATCTGAAGGTCCTG</td><td>>am_trna162_MetCAT_6_-_26758550_26758622</td><td>31</td><td>51</td><td>21</td><td>TCTCATAATCTGAAGGTCCTG</td><td>250.2</td><td>-11.8</td><td>1.4</td><td>-8.2</td><td>2.4E-16</td><td>1.8E-15</td><td>MB231/MB468</td><td>MP</td></tr>
<tr><td>>hg38_21_+_8433172_8446622_RNA45SN1_45S_50ntFlanks@7986.8012.27|ATCAGACGTGGCGACCCGCTGAATTTA</td><td>>hg38_21_+_8433172_8446622_RNA45SN1_45S_50ntFlanks</td><td>7986</td><td>8012</td><td>27</td><td>ATCAGACGTGGCGACCCGCTGAATTTA</td><td>3746.2</td><td>-10.6</td><td>1.3</td><td>-8.1</td><td>3.9E-16</td><td>1.9E-15</td><td>MB231/MB468</td><td>MITO</td></tr>
<tr><td>>hg38_21_+_8433172_8446622_RNA45SN1_45S_50ntFlanks@10735.10752.18|TGTAAATCTCGCGCCGGG</td><td>>hg38_21_+_8433172_8446622_RNA45SN1_45S_50ntFlanks</td><td>10735</td><td>10752</td><td>18</td><td>TGTAAATCTCGCGCCGGG</td><td>194.1</td><td>-11.5</td><td>1.4</td><td>-8.2</td><td>2.4E-16</td><td>1.9E-15</td><td>MB231/MB468</td><td>MP</td></tr>
<tr><td>>hg38_21_+_8433172_8446622_RNA45SN1_45S_50ntFlanks@10845.10863.19|ACTTCGGGATAAGGATTGG</td><td>>hg38_21_+_8433172_8446622_RNA45SN1_45S_50ntFlanks</td><td>10845</td><td>10863</td><td>19</td><td>ACTTCGGGATAAGGATTGG</td><td>1366.7</td><td>-5.2</td><td>0.6</td><td>-8.2</td><td>2.5E-16</td><td>1.9E-15</td><td>MB231/MB468</td><td>CYTO</td></tr>
<tr><td>>MI0000446|hsa-mir-125b-1&WithFlank&11|-|122099751|122099850@21.39.19(+1C);MI0000470|hsa-mir-125b-2&WithFlank&21|+|16590231|16590331@23.41.19(+1C)&[MIMAT0000423&hsa-miR-125b-5p&offsets|0|-3(+1C);m-67&11|-|122099809|122099830&offsets|0|-3(+1C)];[MIMAT0000423_1&hsa-miR-125b-5p&offsets|0|-3(+1C);m-68&21|+|16590253|16590274&offsets|0|-3(+1C)]|TCCCTGAGACCCTAACTTGC</td><td>>MI0000446|hsa-mir-125b-1&WithFlank&11|-|122099751|122099850</td><td>21</td><td>39</td><td>19(+1C)</td><td>TCCCTGAGACCCTAACTTGC</td><td>349.3</td><td>12.4</td><td>1.5</td><td>8.2</td><td>2.5E-16</td><td>1.9E-15</td><td>MB231/MB468</td><td>CYTO</td></tr>
<tr><td>>hg38_21_+_8433172_8446622_RNA45SN1_45S_50ntFlanks@6655.6678.24|TCTTAGCGGTGGATCACTCGGCTC</td><td>>hg38_21_+_8433172_8446622_RNA45SN1_45S_50ntFlanks</td><td>6655</td><td>6678</td><td>24</td><td>TCTTAGCGGTGGATCACTCGGCTC</td><td>581.5</td><td>5.6</td><td>0.7</td><td>8.2</td><td>2.7E-16</td><td>1.9E-15</td><td>BT20/MB468</td><td>MITO</td></tr>
<tr><td>>hg38_21_+_8433172_8446622_RNA45SN1_45S_50ntFlanks@10391.10432.42|AACAGCAGTTGAACATGGGTCAGTCGGTCCTGAGAGATGGGC</td><td>>hg38_21_+_8433172_8446622_RNA45SN1_45S_50ntFlanks</td><td>10391</td><td>10432</td><td>42</td><td>AACAGCAGTTGAACATGGGTCAGTCGGTCCTGAGAGATGGGC</td><td>466.1</td><td>-13.3</td><td>1.6</td><td>-8.1</td><td>6.0E-16</td><td>1.9E-15</td><td>BT20/MB468</td><td>NUC</td></tr>
<tr><td>>hg38_21_+_8433172_8446622_RNA45SN1_45S_50ntFlanks@10795.10817.23|TGGCATGTTGGAACAATGTAGGT</td><td>>hg38_21_+_8433172_8446622_RNA45SN1_45S_50ntFlanks</td><td>10795</td><td>10817</td><td>23</td><td>TGGCATGTTGGAACAATGTAGGT</td><td>263.7</td><td>-11.9</td><td>1.5</td><td>-8.2</td><td>2.4E-16</td><td>1.9E-15</td><td>MB231/MB468</td><td>MP</td></tr>
<tr><td>>hg38_21_+_8433172_8446622_RNA45SN1_45S_50ntFlanks@11602.11620.19|TCCGACTGTTTAATTAAAA</td><td>>hg38_21_+_8433172_8446622_RNA45SN1_45S_50ntFlanks</td><td>11602</td><td>11620</td><td>19</td><td>TCCGACTGTTTAATTAAAA</td><td>735.9</td><td>-5.3</td><td>0.6</td><td>-8.1</td><td>4.0E-16</td><td>1.9E-15</td><td>MB231/MB468</td><td>MITO</td></tr>
<tr><td>>hg38_21_+_8433172_8446622_RNA45SN1_45S_50ntFlanks@12635.12659.25|AAGCTACCATCTGTGGGATTATGAC</td><td>>hg38_21_+_8433172_8446622_RNA45SN1_45S_50ntFlanks</td><td>12635</td><td>12659</td><td>25</td><td>AAGCTACCATCTGTGGGATTATGAC</td><td>13802.2</td><td>-10.1</td><td>1.2</td><td>-8.1</td><td>4.0E-16</td><td>1.9E-15</td><td>MB231/MB468</td><td>MITO</td></tr>
<tr><td>>hg38_21_+_8433172_8446622_RNA45SN1_45S_50ntFlanks@9034.9076.43|TCGTGGGGGGGCCGGGCCACCCCTCCCACGGCGCGACCGCTCT</td><td>>hg38_21_+_8433172_8446622_RNA45SN1_45S_50ntFlanks</td><td>9034</td><td>9076</td><td>43</td><td>TCGTGGGGGGGCCGGGCCACCCCTCCCACGGCGCGACCGCTCT</td><td>545.2</td><td>-12.9</td><td>1.6</td><td>-8.1</td><td>4.0E-16</td><td>1.9E-15</td><td>MB231/MB468</td><td>MITO</td></tr>
<tr><td>>hg38_21_+_8433172_8446622_RNA45SN1_45S_50ntFlanks@9454.9489.36|CCGCGCCGGGGAGGTGGAGCACGAGCGCACGTGTTA</td><td>>hg38_21_+_8433172_8446622_RNA45SN1_45S_50ntFlanks</td><td>9454</td><td>9489</td><td>36</td><td>CCGCGCCGGGGAGGTGGAGCACGAGCGCACGTGTTA</td><td>272.8</td><td>-11.9</td><td>1.5</td><td>-8.1</td><td>4.0E-16</td><td>1.9E-15</td><td>MB231/MB468</td><td>MITO</td></tr>
<tr><td>>hg38_21_+_8433172_8446622_RNA45SN1_45S_50ntFlanks@10535.10571.37|CGCGAGGCGTCCAGTGCGGTAACGCGACCGATCCCGG</td><td>>hg38_21_+_8433172_8446622_RNA45SN1_45S_50ntFlanks</td><td>10535</td><td>10571</td><td>37</td><td>CGCGAGGCGTCCAGTGCGGTAACGCGACCGATCCCGG</td><td>296.4</td><td>-12.1</td><td>1.5</td><td>-8.2</td><td>2.4E-16</td><td>1.9E-15</td><td>MB231/MB468</td><td>MP</td></tr>
<tr><td>>hg38_21_+_8433172_8446622_RNA45SN1_45S_50ntFlanks@4283.4308.26|CTTTAACGAGGATCCATTGGAGGGCA</td><td>>hg38_21_+_8433172_8446622_RNA45SN1_45S_50ntFlanks</td><td>4283</td><td>4308</td><td>26</td><td>CTTTAACGAGGATCCATTGGAGGGCA</td><td>355.2</td><td>-12.3</td><td>1.5</td><td>-8.2</td><td>2.4E-16</td><td>1.9E-15</td><td>MB231/MB468</td><td>MP</td></tr>
<tr><td>>hg38_21_+_8433172_8446622_RNA45SN1_45S_50ntFlanks@4451.4479.29|TCTCGGCGCCCCCTCGATGCTCTTAGCTG</td><td>>hg38_21_+_8433172_8446622_RNA45SN1_45S_50ntFlanks</td><td>4451</td><td>4479</td><td>29</td><td>TCTCGGCGCCCCCTCGATGCTCTTAGCTG</td><td>139.5</td><td>11.1</td><td>1.4</td><td>8.2</td><td>2.6E-16</td><td>1.9E-15</td><td>MB231/MB468</td><td>CYTO</td></tr>
<tr><td>>hg38_21_+_8433172_8446622_RNA45SN1_45S_50ntFlanks@6723.6744.22|TTGCAGGACACATTGATCATCG</td><td>>hg38_21_+_8433172_8446622_RNA45SN1_45S_50ntFlanks</td><td>6723</td><td>6744</td><td>22</td><td>TTGCAGGACACATTGATCATCG</td><td>294.7</td><td>-12.1</td><td>1.5</td><td>-8.2</td><td>2.4E-16</td><td>1.9E-15</td><td>MB231/MB468</td><td>MP</td></tr>
<tr><td>>hg38_21_+_8433172_8446622_RNA45SN1_45S_50ntFlanks@12141.12179.39|ACAGTGCCAGGTGGGGAGTTTGACTGGGGCGGTACACCT</td><td>>hg38_21_+_8433172_8446622_RNA45SN1_45S_50ntFlanks</td><td>12141</td><td>12179</td><td>39</td><td>ACAGTGCCAGGTGGGGAGTTTGACTGGGGCGGTACACCT</td><td>397.0</td><td>-12.4</td><td>1.5</td><td>-8.1</td><td>4.0E-16</td><td>1.9E-15</td><td>MB231/MB468</td><td>MITO</td></tr>
<tr><td>>hg38_21_+_8433172_8446622_RNA45SN1_45S_50ntFlanks@11104.11133.30|AAGGGTCGGGGCGGCAGGGGCCGGCGGCGG</td><td>>hg38_21_+_8433172_8446622_RNA45SN1_45S_50ntFlanks</td><td>11104</td><td>11133</td><td>30</td><td>AAGGGTCGGGGCGGCAGGGGCCGGCGGCGG</td><td>284.1</td><td>-10.9</td><td>1.3</td><td>-8.2</td><td>2.6E-16</td><td>1.9E-15</td><td>MB231/MB468</td><td>CYTO</td></tr>
<tr><td>>hg38_21_+_8433172_8446622_RNA45SN1_45S_50ntFlanks@5485.5503.19|AGCGCTGAGAAGACGGTCG</td><td>>hg38_21_+_8433172_8446622_RNA45SN1_45S_50ntFlanks</td><td>5485</td><td>5503</td><td>19</td><td>AGCGCTGAGAAGACGGTCG</td><td>350.3</td><td>-12.3</td><td>1.5</td><td>-8.2</td><td>2.4E-16</td><td>1.9E-15</td><td>MB231/MB468</td><td>MP</td></tr>
<tr><td>>hg38_21_+_8433172_8446622_RNA45SN1_45S_50ntFlanks@6666.6699.34|GATCACTCGGCTCGTGCGTCGATGAAGAACGCAG</td><td>>hg38_21_+_8433172_8446622_RNA45SN1_45S_50ntFlanks</td><td>6666</td><td>6699</td><td>34</td><td>GATCACTCGGCTCGTGCGTCGATGAAGAACGCAG</td><td>345.0</td><td>-12.3</td><td>1.5</td><td>-8.2</td><td>2.4E-16</td><td>1.9E-15</td><td>MB231/MB468</td><td>MP</td></tr>
<tr><td>>hg38_21_+_8433172_8446622_RNA45SN1_45S_50ntFlanks@8182.8211.30|ATCGAGGCCCAGCCCGTGGACGGTGTGAGG</td><td>>hg38_21_+_8433172_8446622_RNA45SN1_45S_50ntFlanks</td><td>8182</td><td>8211</td><td>30</td><td>ATCGAGGCCCAGCCCGTGGACGGTGTGAGG</td><td>580.1</td><td>-12.1</td><td>1.5</td><td>-8.2</td><td>3.0E-16</td><td>1.9E-15</td><td>MB231/MB468</td><td>NUC</td></tr>
<tr><td>>trnaMT_PheGAA_MT_+_577_647@27.57.31|AATACACTGAAAATGTTTAGACGGGCTCACA</td><td>>trnaMT_PheGAA_MT_+_577_647</td><td>27</td><td>57</td><td>31</td><td>AATACACTGAAAATGTTTAGACGGGCTCACA</td><td>335.8</td><td>12.9</td><td>1.6</td><td>8.2</td><td>2.8E-16</td><td>1.9E-15</td><td>BT20/MB468</td><td>MITO</td></tr>
<tr><td>>trnaMT_ThrTGT_MT_+_15888_15953@31.52.22|TTGTAAACCGGAGATGAAAACC</td><td>>trnaMT_ThrTGT_MT_+_15888_15953</td><td>31</td><td>52</td><td>22</td><td>TTGTAAACCGGAGATGAAAACC</td><td>428.4</td><td>10.7</td><td>1.3</td><td>8.1</td><td>3.8E-16</td><td>1.9E-15</td><td>BT20/MB468</td><td>CYTO</td></tr>
<tr><td>>hg38_21_+_8433172_8446622_RNA45SN1_45S_50ntFlanks@5485.5522.38|AGCGCTGAGAAGACGGTCGAACTTGACTATCTAGAGGA</td><td>>hg38_21_+_8433172_8446622_RNA45SN1_45S_50ntFlanks</td><td>5485</td><td>5522</td><td>38</td><td>AGCGCTGAGAAGACGGTCGAACTTGACTATCTAGAGGA</td><td>388.4</td><td>-12.5</td><td>1.5</td><td>-8.2</td><td>2.4E-16</td><td>1.9E-15</td><td>MB231/MB468</td><td>MP</td></tr>
<tr><td>>MI0000446|hsa-mir-125b-1&WithFlank&11|-|122099751|122099850@21.42.22;MI0000470|hsa-mir-125b-2&WithFlank&21|+|16590231|16590331@23.44.22&[MIMAT0000423&hsa-miR-125b-5p&offsets|0|0;m-67&11|-|122099809|122099830&offsets|0|0];[MIMAT0000423_1&hsa-miR-125b-5p&offsets|0|0;m-68&21|+|16590253|16590274&offsets|0|0]|TCCCTGAGACCCTAACTTGTGA</td><td>>MI0000446|hsa-mir-125b-1&WithFlank&11|-|122099751|122099850</td><td>21</td><td>42</td><td>22</td><td>TCCCTGAGACCCTAACTTGTGA</td><td>22781.1</td><td>7.9</td><td>1.0</td><td>8.1</td><td>4.1E-16</td><td>1.9E-15</td><td>MB231/MB468</td><td>MITO</td></tr>
<tr><td>>hg38_21_+_8433172_8446622_RNA45SN1_45S_50ntFlanks@11199.11225.27|CCGGCGGCGGCGGCGACTCTGGACGCG</td><td>>hg38_21_+_8433172_8446622_RNA45SN1_45S_50ntFlanks</td><td>11199</td><td>11225</td><td>27</td><td>CCGGCGGCGGCGGCGACTCTGGACGCG</td><td>3437.2</td><td>-9.4</td><td>1.1</td><td>-8.2</td><td>2.6E-16</td><td>1.9E-15</td><td>MB231/MB468</td><td>CYTO</td></tr>
<tr><td>>hg38_21_+_8433172_8446622_RNA45SN1_45S_50ntFlanks@11535.11552.18|TTCCGCGCGGCGCCTCGC</td><td>>hg38_21_+_8433172_8446622_RNA45SN1_45S_50ntFlanks</td><td>11535</td><td>11552</td><td>18</td><td>TTCCGCGCGGCGCCTCGC</td><td>326.5</td><td>-12.1</td><td>1.5</td><td>-8.1</td><td>4.1E-16</td><td>1.9E-15</td><td>MB231/MB468</td><td>MITO</td></tr>
<tr><td>>hg38_21_+_8433172_8446622_RNA45SN1_45S_50ntFlanks@6670.6698.29|ACTCGGCTCGTGCGTCGATGAAGAACGCA</td><td>>hg38_21_+_8433172_8446622_RNA45SN1_45S_50ntFlanks</td><td>6670</td><td>6698</td><td>29</td><td>ACTCGGCTCGTGCGTCGATGAAGAACGCA</td><td>347.0</td><td>-12.3</td><td>1.5</td><td>-8.2</td><td>2.5E-16</td><td>1.9E-15</td><td>MB231/MB468</td><td>MP</td></tr>
<tr><td>>hg38_21_+_8433172_8446622_RNA45SN1_45S_50ntFlanks@11052.11075.24|CGTCCTCCCCCCTCCCCGGGGGAG</td><td>>hg38_21_+_8433172_8446622_RNA45SN1_45S_50ntFlanks</td><td>11052</td><td>11075</td><td>24</td><td>CGTCCTCCCCCCTCCCCGGGGGAG</td><td>254.0</td><td>-10.7</td><td>1.3</td><td>-8.2</td><td>2.6E-16</td><td>1.9E-15</td><td>MB231/MB468</td><td>CYTO</td></tr>
<tr><td>>am_trna134_GluTTC_1_-_16861774_16861845@1.29.29,trna5_GluTTC_1_+_17199078_17199149@1.29.29,trna84_GluTTC_1_-_161391883_161391954@1.29.29,trna94_GluTTC_1_-_149664355_149664427@1.29.29|TCCCTGGTGGTCTAGTGGCTAGGATTCGG</td><td>>am_trna134_GluTTC_1_-_16861774_16861845</td><td>1</td><td>29</td><td>29</td><td>TCCCTGGTGGTCTAGTGGCTAGGATTCGG</td><td>2915.9</td><td>-10.9</td><td>1.3</td><td>-8.2</td><td>3.0E-16</td><td>1.9E-15</td><td>MB231/MB468</td><td>NUC</td></tr>
<tr><td>>hg38_21_+_8433172_8446622_RNA45SN1_45S_50ntFlanks@4590.4614.25|ATTTTGTTGGTTTTCGGAACTGAGG</td><td>>hg38_21_+_8433172_8446622_RNA45SN1_45S_50ntFlanks</td><td>4590</td><td>4614</td><td>25</td><td>ATTTTGTTGGTTTTCGGAACTGAGG</td><td>279.7</td><td>-12.0</td><td>1.5</td><td>-8.2</td><td>2.5E-16</td><td>1.9E-15</td><td>MB231/MB468</td><td>MP</td></tr>
<tr><td>>hg38_21_+_8433172_8446622_RNA45SN1_45S_50ntFlanks@12463.12496.34|TTGGATTGTTCACCCACTAATAGGGAACGTGAGC</td><td>>hg38_21_+_8433172_8446622_RNA45SN1_45S_50ntFlanks</td><td>12463</td><td>12496</td><td>34</td><td>TTGGATTGTTCACCCACTAATAGGGAACGTGAGC</td><td>237.3</td><td>5.0</td><td>0.6</td><td>8.2</td><td>3.1E-16</td><td>1.9E-15</td><td>MB231/MB468</td><td>NUC</td></tr>
<tr><td>>hg38_21_+_8433172_8446622_RNA45SN1_45S_50ntFlanks@11718.11740.23|AACGGCGGGAGTAACTATGACTC</td><td>>hg38_21_+_8433172_8446622_RNA45SN1_45S_50ntFlanks</td><td>11718</td><td>11740</td><td>23</td><td>AACGGCGGGAGTAACTATGACTC</td><td>558.3</td><td>-13.0</td><td>1.6</td><td>-8.2</td><td>2.5E-16</td><td>1.9E-15</td><td>MB231/MB468</td><td>MP</td></tr>
<tr><td>>trnaMT_SerGCT_MT_+_12207_12265@8.31.24|CTCACAAGAACTGCTAACTCATGC</td><td>>trnaMT_SerGCT_MT_+_12207_12265</td><td>8</td><td>31</td><td>24</td><td>CTCACAAGAACTGCTAACTCATGC</td><td>969.4</td><td>-14.3</td><td>1.8</td><td>-8.1</td><td>6.2E-16</td><td>1.9E-15</td><td>BT20/MB468</td><td>NUC</td></tr>
<tr><td>>hg38_21_+_8433172_8446622_RNA45SN1_45S_50ntFlanks@6719.6745.27|TGAATTGCAGGACACATTGATCATCGA</td><td>>hg38_21_+_8433172_8446622_RNA45SN1_45S_50ntFlanks</td><td>6719</td><td>6745</td><td>27</td><td>TGAATTGCAGGACACATTGATCATCGA</td><td>374.7</td><td>-12.4</td><td>1.5</td><td>-8.2</td><td>2.5E-16</td><td>1.9E-15</td><td>MB231/MB468</td><td>MP</td></tr>
<tr><td>>hg38_21_+_8433172_8446622_RNA45SN1_45S_50ntFlanks@12324.12356.33|TTGGGTTTTAAGCAGGAGGTGTCAGAAAAGTTA</td><td>>hg38_21_+_8433172_8446622_RNA45SN1_45S_50ntFlanks</td><td>12324</td><td>12356</td><td>33</td><td>TTGGGTTTTAAGCAGGAGGTGTCAGAAAAGTTA</td><td>433.8</td><td>-12.6</td><td>1.5</td><td>-8.2</td><td>2.5E-16</td><td>1.9E-15</td><td>MB231/MB468</td><td>MP</td></tr>
<tr><td>>hg38_21_+_8433172_8446622_RNA45SN1_45S_50ntFlanks@9822.9850.29|CTAGTGGGCCACTTTTGGTAAGCAGAACT</td><td>>hg38_21_+_8433172_8446622_RNA45SN1_45S_50ntFlanks</td><td>9822</td><td>9850</td><td>29</td><td>CTAGTGGGCCACTTTTGGTAAGCAGAACT</td><td>244.8</td><td>-11.8</td><td>1.4</td><td>-8.2</td><td>2.5E-16</td><td>1.9E-15</td><td>MB231/MB468</td><td>MP</td></tr>
<tr><td>>hg38_21_+_8433172_8446622_RNA45SN1_45S_50ntFlanks@12892.12934.43|CCCTCGCCCGTCACGCACCGCACGTTCGTGGGGAACCTGGCGC</td><td>>hg38_21_+_8433172_8446622_RNA45SN1_45S_50ntFlanks</td><td>12892</td><td>12934</td><td>43</td><td>CCCTCGCCCGTCACGCACCGCACGTTCGTGGGGAACCTGGCGC</td><td>321.5</td><td>-12.1</td><td>1.5</td><td>-8.1</td><td>4.2E-16</td><td>1.9E-15</td><td>MB231/MB468</td><td>MITO</td></tr>
<tr><td>>hg38_21_+_8433172_8446622_RNA45SN1_45S_50ntFlanks@11534.11553.20|GTTCCGCGCGGCGCCTCGCC</td><td>>hg38_21_+_8433172_8446622_RNA45SN1_45S_50ntFlanks</td><td>11534</td><td>11553</td><td>20</td><td>GTTCCGCGCGGCGCCTCGCC</td><td>326.1</td><td>-12.2</td><td>1.5</td><td>-8.2</td><td>2.5E-16</td><td>1.9E-15</td><td>MB231/MB468</td><td>MP</td></tr>
<tr><td>>hg38_21_+_8433172_8446622_RNA45SN1_45S_50ntFlanks@12622.12654.33|AGCCAATGGGGCGAAGCTACCATCTGTGGGATT</td><td>>hg38_21_+_8433172_8446622_RNA45SN1_45S_50ntFlanks</td><td>12622</td><td>12654</td><td>33</td><td>AGCCAATGGGGCGAAGCTACCATCTGTGGGATT</td><td>1152.6</td><td>-9.6</td><td>1.2</td><td>-8.1</td><td>4.2E-16</td><td>2.0E-15</td><td>MB231/MB468</td><td>MITO</td></tr>
<tr><td>>hg38_21_+_8433172_8446622_RNA45SN1_45S_50ntFlanks@7977.8009.33|CGACCTCAGATCAGACGTGGCGACCCGCTGAAT</td><td>>hg38_21_+_8433172_8446622_RNA45SN1_45S_50ntFlanks</td><td>7977</td><td>8009</td><td>33</td><td>CGACCTCAGATCAGACGTGGCGACCCGCTGAAT</td><td>745.1</td><td>3.1</td><td>0.4</td><td>8.2</td><td>2.8E-16</td><td>2.0E-15</td><td>BT20/MB468</td><td>MITO</td></tr>
<tr><td>>hg38_21_+_8433172_8446622_RNA45SN1_45S_50ntFlanks@3899.3932.34|CCTTCGCGGGGGGGATGCGTGCATTTATCAGATC</td><td>>hg38_21_+_8433172_8446622_RNA45SN1_45S_50ntFlanks</td><td>3899</td><td>3932</td><td>34</td><td>CCTTCGCGGGGGGGATGCGTGCATTTATCAGATC</td><td>322.6</td><td>-12.1</td><td>1.5</td><td>-8.1</td><td>4.2E-16</td><td>2.0E-15</td><td>MB231/MB468</td><td>MITO</td></tr>
<tr><td>>hg38_21_+_8433172_8446622_RNA45SN1_45S_50ntFlanks@9454.9491.38|CCGCGCCGGGGAGGTGGAGCACGAGCGCACGTGTTAGG</td><td>>hg38_21_+_8433172_8446622_RNA45SN1_45S_50ntFlanks</td><td>9454</td><td>9491</td><td>38</td><td>CCGCGCCGGGGAGGTGGAGCACGAGCGCACGTGTTAGG</td><td>345.1</td><td>-12.2</td><td>1.5</td><td>-8.1</td><td>4.2E-16</td><td>2.0E-15</td><td>MB231/MB468</td><td>MITO</td></tr>
<tr><td>>hg38_21_+_8433172_8446622_RNA45SN1_45S_50ntFlanks@12991.13018.28|CTCCCTCGCTGCGATCTATTGAAAGTCA</td><td>>hg38_21_+_8433172_8446622_RNA45SN1_45S_50ntFlanks</td><td>12991</td><td>13018</td><td>28</td><td>CTCCCTCGCTGCGATCTATTGAAAGTCA</td><td>347.0</td><td>-11.4</td><td>1.4</td><td>-8.2</td><td>3.1E-16</td><td>2.0E-15</td><td>MB231/MB468</td><td>NUC</td></tr>
<tr><td>>hg38_21_+_8433172_8446622_RNA45SN1_45S_50ntFlanks@12159.12183.25|TTTGACTGGGGCGGTACACCTGTCA</td><td>>hg38_21_+_8433172_8446622_RNA45SN1_45S_50ntFlanks</td><td>12159</td><td>12183</td><td>25</td><td>TTTGACTGGGGCGGTACACCTGTCA</td><td>460.5</td><td>-12.6</td><td>1.6</td><td>-8.1</td><td>4.2E-16</td><td>2.0E-15</td><td>MB231/MB468</td><td>MITO</td></tr>
<tr><td>>hg38_21_+_8433172_8446622_RNA45SN1_45S_50ntFlanks@11190.11225.36|CCCGGGGGGCCGGCGGCGGCGGCGACTCTGGACGCG</td><td>>hg38_21_+_8433172_8446622_RNA45SN1_45S_50ntFlanks</td><td>11190</td><td>11225</td><td>36</td><td>CCCGGGGGGCCGGCGGCGGCGGCGACTCTGGACGCG</td><td>1250.6</td><td>-13.3</td><td>1.6</td><td>-8.2</td><td>3.1E-16</td><td>2.0E-15</td><td>MB231/MB468</td><td>NUC</td></tr>
<tr><td>>hg38_21_+_8433172_8446622_RNA45SN1_45S_50ntFlanks@12957.12980.24|CTTCTGGGTCGGGGTTTCGTACGT</td><td>>hg38_21_+_8433172_8446622_RNA45SN1_45S_50ntFlanks</td><td>12957</td><td>12980</td><td>24</td><td>CTTCTGGGTCGGGGTTTCGTACGT</td><td>649.6</td><td>-9.6</td><td>1.2</td><td>-8.2</td><td>3.1E-16</td><td>2.0E-15</td><td>MB231/MB468</td><td>NUC</td></tr>
</table>

<table>
<tr><td>>hg38_21_+_8433172_8446622_RNA45SN1_45S_50ntFlanks@6650.6674.25|ACGACTCTTAGCGGTGGATCACTCG</td><td>>hg38_21_+_8433172_8446622_RNA45SN1_45S_50ntFlanks</td><td>6650</td><td>6674</td><td>25</td><td>ACGACTCTTAGCGGTGGATCACTCG</td><td>4865.2</td><td>5.9</td><td>0.7</td><td>8.2</td><td>2.9E-16</td><td>2.0E-15</td><td>BT20/MB468</td><td>MITO</td></tr>
<tr><td>>hg38_21_+_8433172_8446622_RNA45SN1_45S_50ntFlanks@5489.5509.21|CTGAGAAGACGGTCGAACTTG</td><td>>hg38_21_+_8433172_8446622_RNA45SN1_45S_50ntFlanks</td><td>5489</td><td>5509</td><td>21</td><td>CTGAGAAGACGGTCGAACTTG</td><td>284.3</td><td>-11.9</td><td>1.5</td><td>-8.1</td><td>4.2E-16</td><td>2.0E-15</td><td>MB231/MB468</td><td>MITO</td></tr>
<tr><td>>hg38_21_+_8433172_8446622_RNA45SN1_45S_50ntFlanks@11564.11593.30|CCTAGCAGCCGACTTAGAACTGGTGCGGAC</td><td>>hg38_21_+_8433172_8446622_RNA45SN1_45S_50ntFlanks</td><td>11564</td><td>11593</td><td>30</td><td>CCTAGCAGCCGACTTAGAACTGGTGCGGAC</td><td>279.0</td><td>-11.9</td><td>1.5</td><td>-8.1</td><td>4.3E-16</td><td>2.0E-15</td><td>MB231/MB468</td><td>MITO</td></tr>
<tr><td>>hg38_21_+_8433172_8446622_RNA45SN1_45S_50ntFlanks@12033.12070.38|ATCGTTTTTTCACTGACCCGGTGAGGCGGGGGGGCGAG</td><td>>hg38_21_+_8433172_8446622_RNA45SN1_45S_50ntFlanks</td><td>12033</td><td>12070</td><td>38</td><td>ATCGTTTTTTCACTGACCCGGTGAGGCGGGGGGGCGAG</td><td>421.2</td><td>-12.5</td><td>1.5</td><td>-8.1</td><td>4.3E-16</td><td>2.0E-15</td><td>MB231/MB468</td><td>MITO</td></tr>
<tr><td>>hg38_21_+_8433172_8446622_RNA45SN1_45S_50ntFlanks@6646.6683.38|TCGTACGACTCTTAGCGGTGGATCACTCGGCTCGTGCG</td><td>>hg38_21_+_8433172_8446622_RNA45SN1_45S_50ntFlanks</td><td>6646</td><td>6683</td><td>38</td><td>TCGTACGACTCTTAGCGGTGGATCACTCGGCTCGTGCG</td><td>2174.9</td><td>7.5</td><td>0.9</td><td>8.2</td><td>3.2E-16</td><td>2.0E-15</td><td>MB231/MB468</td><td>NUC</td></tr>
<tr><td>>MI0000298|hsa-mir-221&WithFlank&X|-|45746151|45746272@31.56.26&[MIMAT0004568&hsa-miR-221-5p&offsets|0|+4;m-405&X|-|45746220|45746242&offsets|0|+3]|ACCTGGCATACAATGTAGATTTCTGT</td><td>>MI0000298|hsa-mir-221&WithFlank&X|-|45746151|45746272</td><td>31</td><td>56</td><td>26</td><td>ACCTGGCATACAATGTAGATTTCTGT</td><td>1454.3</td><td>5.8</td><td>0.7</td><td>8.1</td><td>4.0E-16</td><td>2.0E-15</td><td>BT20/MB468</td><td>CYTO</td></tr>
<tr><td>>hg38_21_+_8433172_8446622_RNA45SN1_45S_50ntFlanks@12157.12183.27|AGTTTGACTGGGGCGGTACACCTGTCA</td><td>>hg38_21_+_8433172_8446622_RNA45SN1_45S_50ntFlanks</td><td>12157</td><td>12183</td><td>27</td><td>AGTTTGACTGGGGCGGTACACCTGTCA</td><td>340.4</td><td>-12.2</td><td>1.5</td><td>-8.1</td><td>4.3E-16</td><td>2.0E-15</td><td>MB231/MB468</td><td>MITO</td></tr>
<tr><td>>hg38_21_+_8433172_8446622_RNA45SN1_45S_50ntFlanks@9454.9493.40|CCGCGCCGGGGAGGTGGAGCACGAGCGCACGTGTTAGGAC</td><td>>hg38_21_+_8433172_8446622_RNA45SN1_45S_50ntFlanks</td><td>9454</td><td>9493</td><td>40</td><td>CCGCGCCGGGGAGGTGGAGCACGAGCGCACGTGTTAGGAC</td><td>491.6</td><td>-12.8</td><td>1.6</td><td>-8.2</td><td>2.6E-16</td><td>2.0E-15</td><td>MB231/MB468</td><td>MP</td></tr>
<tr><td>>hg38_21_+_8433172_8446622_RNA45SN1_45S_50ntFlanks@12044.12070.27|ACTGACCCGGTGAGGCGGGGGGGCGAG</td><td>>hg38_21_+_8433172_8446622_RNA45SN1_45S_50ntFlanks</td><td>12044</td><td>12070</td><td>27</td><td>ACTGACCCGGTGAGGCGGGGGGGCGAG</td><td>315.3</td><td>-11.0</td><td>1.3</td><td>-8.2</td><td>2.8E-16</td><td>2.0E-15</td><td>MB231/MB468</td><td>CYTO</td></tr>
<tr><td>>hg38_21_+_8433172_8446622_RNA45SN1_45S_50ntFlanks@12363.12382.20|GGATAACTGGCTTGTGGCGG</td><td>>hg38_21_+_8433172_8446622_RNA45SN1_45S_50ntFlanks</td><td>12363</td><td>12382</td><td>20</td><td>GGATAACTGGCTTGTGGCGG</td><td>539.6</td><td>-12.0</td><td>1.5</td><td>-8.2</td><td>3.2E-16</td><td>2.0E-15</td><td>MB231/MB468</td><td>NUC</td></tr>
<tr><td>>hg38_21_+_8433172_8446622_RNA45SN1_45S_50ntFlanks@9162.9187.26|CGCCCCGGGCGGGTCGCGCCGTCGGG</td><td>>hg38_21_+_8433172_8446622_RNA45SN1_45S_50ntFlanks</td><td>9162</td><td>9187</td><td>26</td><td>CGCCCCGGGCGGGTCGCGCCGTCGGG</td><td>351.5</td><td>-12.2</td><td>1.5</td><td>-8.1</td><td>4.3E-16</td><td>2.0E-15</td><td>MB231/MB468</td><td>MITO</td></tr>
<tr><td>>hg38_21_+_8433172_8446622_RNA45SN1_45S_50ntFlanks@3984.4001.18|GCGGCTTTGGTGACTCTA</td><td>>hg38_21_+_8433172_8446622_RNA45SN1_45S_50ntFlanks</td><td>3984</td><td>4001</td><td>18</td><td>GCGGCTTTGGTGACTCTA</td><td>4380.8</td><td>-9.2</td><td>1.1</td><td>-8.2</td><td>3.2E-16</td><td>2.0E-15</td><td>MB231/MB468</td><td>NUC</td></tr>
<tr><td>>hg38_21_+_8433172_8446622_RNA45SN1_45S_50ntFlanks@8198.8215.18|TGGACGGTGTGAGGCCGG</td><td>>hg38_21_+_8433172_8446622_RNA45SN1_45S_50ntFlanks</td><td>8198</td><td>8215</td><td>18</td><td>TGGACGGTGTGAGGCCGG</td><td>3469.7</td><td>-8.7</td><td>1.1</td><td>-8.2</td><td>2.6E-16</td><td>2.0E-15</td><td>MB231/MB468</td><td>MP</td></tr>
<tr><td>>hg38_21_+_8433172_8446622_RNA45SN1_45S_50ntFlanks@8154.8179.26|CCGCTCGTGGGGGGCCCAAGTCCTTC</td><td>>hg38_21_+_8433172_8446622_RNA45SN1_45S_50ntFlanks</td><td>8154</td><td>8179</td><td>26</td><td>CCGCTCGTGGGGGGCCCAAGTCCTTC</td><td>426.7</td><td>-12.5</td><td>1.5</td><td>-8.1</td><td>4.4E-16</td><td>2.0E-15</td><td>MB231/MB468</td><td>MITO</td></tr>
<tr><td>>hg38_21_+_8433172_8446622_RNA45SN1_45S_50ntFlanks@10764.10785.22|TCCGCAGCAGGTCTCCAAGGTG</td><td>>hg38_21_+_8433172_8446622_RNA45SN1_45S_50ntFlanks</td><td>10764</td><td>10785</td><td>22</td><td>TCCGCAGCAGGTCTCCAAGGTG</td><td>283.3</td><td>-11.9</td><td>1.5</td><td>-8.1</td><td>4.4E-16</td><td>2.0E-15</td><td>MB231/MB468</td><td>MITO</td></tr>
<tr><td>>MI0000269|hsa-mir-181a-2&WithFlank&9|+|124692436|124692557@83.103.21&[MIMAT0004558&hsa-miR-181a-2-3p&offsets|0|-1;m-71&9|+|124692518|124692539&offsets|0|-1]|ACCACTGACCGTTGACTGTAC</td><td>>MI0000269|hsa-mir-181a-2&WithFlank&9|+|124692436|124692557</td><td>83</td><td>103</td><td>21</td><td>ACCACTGACCGTTGACTGTAC</td><td>376.2</td><td>10.2</td><td>1.3</td><td>8.1</td><td>4.1E-16</td><td>2.0E-15</td><td>BT20/MB468</td><td>CYTO</td></tr>
<tr><td>>hg38_21_+_8433172_8446622_RNA45SN1_45S_50ntFlanks@11873.11890.18|AAGAAGACCCTGTTGAGC</td><td>>hg38_21_+_8433172_8446622_RNA45SN1_45S_50ntFlanks</td><td>11873</td><td>11890</td><td>18</td><td>AAGAAGACCCTGTTGAGC</td><td>727.4</td><td>-3.9</td><td>0.5</td><td>-8.1</td><td>4.1E-16</td><td>2.0E-15</td><td>BT20/MB468</td><td>CYTO</td></tr>
<tr><td>>MI0000299|hsa-mir-222&WithFlank&X|-|45747009|45747130@75.98.24(+1C)&[MIMAT0000279&hsa-miR-222-3p&offsets|0|+3(+1C);m-140&X|-|45747033|45747056&offsets|0|0(+1C)]|AGCTACATCTGGCTACTGGGTCTCC</td><td>>MI0000299|hsa-mir-222&WithFlank&X|-|45747009|45747130</td><td>75</td><td>98</td><td>24(+1C)</td><td>AGCTACATCTGGCTACTGGGTCTCC</td><td>3643.2</td><td>6.6</td><td>0.8</td><td>8.2</td><td>3.0E-16</td><td>2.0E-15</td><td>BT20/MB468</td><td>MITO</td></tr>
<tr><td>>hg38_21_+_8433172_8446622_RNA45SN1_45S_50ntFlanks@4069.4089.21|CTTTCGATGGTAGTCGCCGTG</td><td>>hg38_21_+_8433172_8446622_RNA45SN1_45S_50ntFlanks</td><td>4069</td><td>4089</td><td>21</td><td>CTTTCGATGGTAGTCGCCGTG</td><td>279.2</td><td>-6.5</td><td>0.8</td><td>-8.2</td><td>2.8E-16</td><td>2.0E-15</td><td>MB231/MB468</td><td>CYTO</td></tr>
<tr><td>>hg38_21_+_8433172_8446622_RNA45SN1_45S_50ntFlanks@4443.4466.24|CCCTTGCCTCTCGGCGCCCCCTCG</td><td>>hg38_21_+_8433172_8446622_RNA45SN1_45S_50ntFlanks</td><td>4443</td><td>4466</td><td>24</td><td>CCCTTGCCTCTCGGCGCCCCCTCG</td><td>303.9</td><td>-12.0</td><td>1.5</td><td>-8.1</td><td>4.4E-16</td><td>2.0E-15</td><td>MB231/MB468</td><td>MITO</td></tr>
<tr><td>>hg38_21_+_8433172_8446622_RNA45SN1_45S_50ntFlanks@12567.12587.21|CTCAGTACGAGAGGAACCGCA</td><td>>hg38_21_+_8433172_8446622_RNA45SN1_45S_50ntFlanks</td><td>12567</td><td>12587</td><td>21</td><td>CTCAGTACGAGAGGAACCGCA</td><td>607.8</td><td>-4.3</td><td>0.5</td><td>-8.2</td><td>3.0E-16</td><td>2.1E-15</td><td>BT20/MB468</td><td>MITO</td></tr>
<tr><td>>MI0000446|hsa-mir-125b-1&WithFlank&11|-|122099751|122099850@21.42.22(+1C);MI0000470|hsa-mir-125b-2&WithFlank&21|+|16590231|16590331@23.44.22(+1C)&[MIMAT0000423&hsa-miR-125b-5p&offsets|0|0(+1C);m-67&11|-|122099809|122099830&offsets|0|0(+1C)];[MIMAT0000423_1&hsa-miR-125b-5p&offsets|0|0(+1C);m-68&21|+|16590253|16590274&offsets|0|0(+1C)]|TCCCTGAGACCCTAACTTGTGAC</td><td>>MI0000446|hsa-mir-125b-1&WithFlank&11|-|122099751|122099850</td><td>21</td><td>42</td><td>22(+1C)</td><td>TCCCTGAGACCCTAACTTGTGAC</td><td>405.6</td><td>12.7</td><td>1.5</td><td>8.2</td><td>2.9E-16</td><td>2.1E-15</td><td>MB231/MB468</td><td>CYTO</td></tr>
<tr><td>>hg38_21_+_8433172_8446622_RNA45SN1_45S_50ntFlanks@10349.10374.26|AACGAGAACTTTGAAGGCCGAAGTGG</td><td>>hg38_21_+_8433172_8446622_RNA45SN1_45S_50ntFlanks</td><td>10349</td><td>10374</td><td>26</td><td>AACGAGAACTTTGAAGGCCGAAGTGG</td><td>2513.1</td><td>-9.1</td><td>1.1</td><td>-8.1</td><td>6.6E-16</td><td>2.1E-15</td><td>BT20/MB468</td><td>NUC</td></tr>
<tr><td>>hg38_21_+_8433172_8446622_RNA45SN1_45S_50ntFlanks@10786.10818.33|AACAGCCTCTGGCATGTTGGAACAATGTAGGTA</td><td>>hg38_21_+_8433172_8446622_RNA45SN1_45S_50ntFlanks</td><td>10786</td><td>10818</td><td>33</td><td>AACAGCCTCTGGCATGTTGGAACAATGTAGGTA</td><td>5218.9</td><td>-7.0</td><td>0.8</td><td>-8.6</td><td>7.5E-18</td><td>2.1E-15</td><td>BT20/MB468</td><td>MP</td></tr>
<tr><td>>hg38_21_+_8433172_8446622_RNA45SN1_45S_50ntFlanks@12108.12125.18|CCGCGCGCCGGCCGGGCG</td><td>>hg38_21_+_8433172_8446622_RNA45SN1_45S_50ntFlanks</td><td>12108</td><td>12125</td><td>18</td><td>CCGCGCGCCGGCCGGGCG</td><td>325.8</td><td>-12.2</td><td>1.5</td><td>-8.2</td><td>2.7E-16</td><td>2.1E-15</td><td>MB231/MB468</td><td>MP</td></tr>
<tr><td>>MI0001445|hsa-mir-423&WithFlank&17|+|30117073|30117178@23.46.24&[MIMAT0004748&hsa-miR-423-5p&offsets|0|+1;m-141&17|+|30117095|30117117&offsets|0|+1]|TGAGGGGCAGAGAGCGAGACTTTT</td><td>>MI0001445|hsa-mir-423&WithFlank&17|+|30117073|30117178</td><td>23</td><td>46</td><td>24</td><td>TGAGGGGCAGAGAGCGAGACTTTT</td><td>848.4</td><td>9.7</td><td>1.2</td><td>8.2</td><td>3.0E-16</td><td>2.1E-15</td><td>BT20/MB468</td><td>MITO</td></tr>
<tr><td>>hg38_21_+_8433172_8446622_RNA45SN1_45S_50ntFlanks@10479.10497.19|CGATCGAAAGGGAGTCGGG</td><td>>hg38_21_+_8433172_8446622_RNA45SN1_45S_50ntFlanks</td><td>10479</td><td>10497</td><td>19</td><td>CGATCGAAAGGGAGTCGGG</td><td>822.2</td><td>-8.4</td><td>1.0</td><td>-8.2</td><td>2.9E-16</td><td>2.1E-15</td><td>MB231/MB468</td><td>CYTO</td></tr>
<tr><td>>hg38_21_+_8433172_8446622_RNA45SN1_45S_50ntFlanks@12627.12647.21|ATGGGGCGAAGCTACCATCTG</td><td>>hg38_21_+_8433172_8446622_RNA45SN1_45S_50ntFlanks</td><td>12627</td><td>12647</td><td>21</td><td>ATGGGGCGAAGCTACCATCTG</td><td>346.3</td><td>-12.3</td><td>1.5</td><td>-8.2</td><td>2.7E-16</td><td>2.1E-15</td><td>MB231/MB468</td><td>MP</td></tr>
<tr><td>>hg38_21_+_8433172_8446622_RNA45SN1_45S_50ntFlanks@10392.10429.38|ACAGCAGTTGAACATGGGTCAGTCGGTCCTGAGAGATG</td><td>>hg38_21_+_8433172_8446622_RNA45SN1_45S_50ntFlanks</td><td>10392</td><td>10429</td><td>38</td><td>ACAGCAGTTGAACATGGGTCAGTCGGTCCTGAGAGATG</td><td>4077.9</td><td>-6.2</td><td>0.8</td><td>-8.2</td><td>3.3E-16</td><td>2.1E-15</td><td>MB231/MB468</td><td>NUC</td></tr>
<tr><td>>hg38_21_+_8433172_8446622_RNA45SN1_45S_50ntFlanks@11671.11700.30|CCAGTGCTCTGAATGTCAAAGTGAAGAAAT</td><td>>hg38_21_+_8433172_8446622_RNA45SN1_45S_50ntFlanks</td><td>11671</td><td>11700</td><td>30</td><td>CCAGTGCTCTGAATGTCAAAGTGAAGAAAT</td><td>549.3</td><td>-4.2</td><td>0.5</td><td>-8.2</td><td>3.3E-16</td><td>2.1E-15</td><td>MB231/MB468</td><td>NUC</td></tr>
<tr><td>>hg38_21_+_8433172_8446622_RNA45SN1_45S_50ntFlanks@12954.12982.29|CTGCTTCTGGGTCGGGGTTTCGTACGTAG</td><td>>hg38_21_+_8433172_8446622_RNA45SN1_45S_50ntFlanks</td><td>12954</td><td>12982</td><td>29</td><td>CTGCTTCTGGGTCGGGGTTTCGTACGTAG</td><td>476.1</td><td>-11.9</td><td>1.5</td><td>-8.2</td><td>3.3E-16</td><td>2.1E-15</td><td>MB231/MB468</td><td>NUC</td></tr>
<tr><td>>hg38_21_+_8433172_8446622_RNA45SN1_45S_50ntFlanks@11600.11623.24|AATCCGACTGTTTAATTAAAACAA</td><td>>hg38_21_+_8433172_8446622_RNA45SN1_45S_50ntFlanks</td><td>11600</td><td>11623</td><td>24</td><td>AATCCGACTGTTTAATTAAAACAA</td><td>2621.2</td><td>10.7</td><td>1.3</td><td>8.2</td><td>3.1E-16</td><td>2.1E-15</td><td>BT20/MB468</td><td>MITO</td></tr>
<tr><td>>hg38_21_+_8433172_8446622_RNA45SN1_45S_50ntFlanks@12991.13040.50|CTCCCTCGCTGCGATCTATTGAAAGTCAGCCCTCGACACAAGGGTTTGTC</td><td>>hg38_21_+_8433172_8446622_RNA45SN1_45S_50ntFlanks</td><td>12991</td><td>13040</td><td>50</td><td>CTCCCTCGCTGCGATCTATTGAAAGTCAGCCCTCGACACAAGGGTTTGTC</td><td>532.6</td><td>-13.5</td><td>1.7</td><td>-8.1</td><td>6.7E-16</td><td>2.1E-15</td><td>BT20/MB468</td><td>NUC</td></tr>
<tr><td>>hg38_21_+_8433172_8446622_RNA45SN1_45S_50ntFlanks@9352.9369.18|CCGTGGCGCAATGAAGGT</td><td>>hg38_21_+_8433172_8446622_RNA45SN1_45S_50ntFlanks</td><td>9352</td><td>9369</td><td>18</td><td>CCGTGGCGCAATGAAGGT</td><td>324.1</td><td>-12.1</td><td>1.5</td><td>-8.1</td><td>4.5E-16</td><td>2.1E-15</td><td>MB231/MB468</td><td>MITO</td></tr>
<tr><td>>hg38_21_+_8433172_8446622_RNA45SN1_45S_50ntFlanks@8472.8490.19|CGGCCCGGCGGATCTTTCC</td><td>>hg38_21_+_8433172_8446622_RNA45SN1_45S_50ntFlanks</td><td>8472</td><td>8490</td><td>19</td><td>CGGCCCGGCGGATCTTTCC</td><td>585.7</td><td>-13.0</td><td>1.6</td><td>-8.1</td><td>4.5E-16</td><td>2.1E-15</td><td>MB231/MB468</td><td>MITO</td></tr>
<tr><td>>hg38_21_+_8433172_8446622_RNA45SN1_45S_50ntFlanks@12938.12956.19|ACCATTCGTAGACGACCTG</td><td>>hg38_21_+_8433172_8446622_RNA45SN1_45S_50ntFlanks</td><td>12938</td><td>12956</td><td>19</td><td>ACCATTCGTAGACGACCTG</td><td>3064.9</td><td>-5.3</td><td>0.7</td><td>-8.2</td><td>3.3E-16</td><td>2.1E-15</td><td>MB231/MB468</td><td>NUC</td></tr>
<tr><td>>hg38_21_+_8433172_8446622_RNA45SN1_45S_50ntFlanks@9573.9605.33|AAATCGGTCGTCCGACCTGGGTATAGGGGCGAA</td><td>>hg38_21_+_8433172_8446622_RNA45SN1_45S_50ntFlanks</td><td>9573</td><td>9605</td><td>33</td><td>AAATCGGTCGTCCGACCTGGGTATAGGGGCGAA</td><td>333.8</td><td>-4.2</td><td>0.5</td><td>-8.1</td><td>6.7E-16</td><td>2.1E-15</td><td>BT20/MB468</td><td>NUC</td></tr>
<tr><td>>hg38_21_+_8433172_8446622_RNA45SN1_45S_50ntFlanks@12457.12478.22|CAAGCGTTGGATTGTTCACCCA</td><td>>hg38_21_+_8433172_8446622_RNA45SN1_45S_50ntFlanks</td><td>12457</td><td>12478</td><td>22</td><td>CAAGCGTTGGATTGTTCACCCA</td><td>6674.2</td><td>-9.7</td><td>1.2</td><td>-8.1</td><td>6.7E-16</td><td>2.1E-15</td><td>BT20/MB468</td><td>NUC</td></tr>
<tr><td>>hg38_21_+_8433172_8446622_RNA45SN1_45S_50ntFlanks@12607.12626.20|TGTGCTTGGCTGAGGAGCCA</td><td>>hg38_21_+_8433172_8446622_RNA45SN1_45S_50ntFlanks</td><td>12607</td><td>12626</td><td>20</td><td>TGTGCTTGGCTGAGGAGCCA</td><td>250.8</td><td>-11.8</td><td>1.4</td><td>-8.2</td><td>2.7E-16</td><td>2.1E-15</td><td>MB231/MB468</td><td>MP</td></tr>
<tr><td>>hg38_21_+_8433172_8446622_RNA45SN1_45S_50ntFlanks@9033.9072.40|CTCGTGGGGGGGCCGGGCCACCCCTCCCACGGCGCGACCG</td><td>>hg38_21_+_8433172_8446622_RNA45SN1_45S_50ntFlanks</td><td>9033</td><td>9072</td><td>40</td><td>CTCGTGGGGGGGCCGGGCCACCCCTCCCACGGCGCGACCG</td><td>417.5</td><td>-12.5</td><td>1.5</td><td>-8.1</td><td>4.6E-16</td><td>2.1E-15</td><td>MB231/MB468</td><td>MITO</td></tr>
<tr><td>>hg38_21_+_8433172_8446622_RNA45SN1_45S_50ntFlanks@6721.6739.19|AATTGCAGGACACATTGAT</td><td>>hg38_21_+_8433172_8446622_RNA45SN1_45S_50ntFlanks</td><td>6721</td><td>6739</td><td>19</td><td>AATTGCAGGACACATTGAT</td><td>1713.0</td><td>-7.6</td><td>0.9</td><td>-8.2</td><td>2.9E-16</td><td>2.1E-15</td><td>MB231/MB468</td><td>CYTO</td></tr>
<tr><td>>hg38_21_+_8433172_8446622_RNA45SN1_45S_50ntFlanks@10351.10378.28|CGAGAACTTTGAAGGCCGAAGTGGAGAA</td><td>>hg38_21_+_8433172_8446622_RNA45SN1_45S_50ntFlanks</td><td>10351</td><td>10378</td><td>28</td><td>CGAGAACTTTGAAGGCCGAAGTGGAGAA</td><td>3434.2</td><td>-9.7</td><td>1.2</td><td>-8.2</td><td>2.7E-16</td><td>2.1E-15</td><td>MB231/MB468</td><td>MP</td></tr>
<tr><td>>hg38_21_+_8433172_8446622_RNA45SN1_45S_50ntFlanks@9568.9595.28|CGTGCAAATCGGTCGTCCGACCTGGGTA</td><td>>hg38_21_+_8433172_8446622_RNA45SN1_45S_50ntFlanks</td><td>9568</td><td>9595</td><td>28</td><td>CGTGCAAATCGGTCGTCCGACCTGGGTA</td><td>3167.4</td><td>-9.5</td><td>1.2</td><td>-8.1</td><td>6.8E-16</td><td>2.1E-15</td><td>BT20/MB468</td><td>NUC</td></tr>
<tr><td>>hg38_21_+_8433172_8446622_RNA45SN1_45S_50ntFlanks@8194.8215.22|CCCGTGGACGGTGTGAGGCCGG</td><td>>hg38_21_+_8433172_8446622_RNA45SN1_45S_50ntFlanks</td><td>8194</td><td>8215</td><td>22</td><td>CCCGTGGACGGTGTGAGGCCGG</td><td>6003.6</td><td>-12.2</td><td>1.5</td><td>-8.1</td><td>6.8E-16</td><td>2.1E-15</td><td>BT20/MB468</td><td>NUC</td></tr>
<tr><td>>hg38_21_+_8433172_8446622_RNA45SN1_45S_50ntFlanks@9621.9648.28|TCTAGTAGCTGGTTCCCTCCGAAGTTTC</td><td>>hg38_21_+_8433172_8446622_RNA45SN1_45S_50ntFlanks</td><td>9621</td><td>9648</td><td>28</td><td>TCTAGTAGCTGGTTCCCTCCGAAGTTTC</td><td>331.5</td><td>-12.2</td><td>1.5</td><td>-8.2</td><td>2.8E-16</td><td>2.1E-15</td><td>MB231/MB468</td><td>MP</td></tr>
<tr><td>>hg38_21_+_8433172_8446622_RNA45SN1_45S_50ntFlanks@8737.8758.22|CCCGTCCGTCCGTCCGTCCGTC</td><td>>hg38_21_+_8433172_8446622_RNA45SN1_45S_50ntFlanks</td><td>8737</td><td>8758</td><td>22</td><td>CCCGTCCGTCCGTCCGTCCGTC</td><td>341.8</td><td>-11.1</td><td>1.4</td><td>-8.2</td><td>3.0E-16</td><td>2.1E-15</td><td>MB231/MB468</td><td>CYTO</td></tr>
<tr><td>>hg38_21_+_8433172_8446622_RNA45SN1_45S_50ntFlanks@12324.12355.32|TTGGGTTTTAAGCAGGAGGTGTCAGAAAAGTT</td><td>>hg38_21_+_8433172_8446622_RNA45SN1_45S_50ntFlanks</td><td>12324</td><td>12355</td><td>32</td><td>TTGGGTTTTAAGCAGGAGGTGTCAGAAAAGTT</td><td>348.8</td><td>-12.3</td><td>1.5</td><td>-8.2</td><td>2.8E-16</td><td>2.1E-15</td><td>MB231/MB468</td><td>MP</td></tr>
<tr><td>>hg38_21_+_8433172_8446622_RNA45SN1_45S_50ntFlanks@11873.11899.27|AAGAAGACCCTGTTGAGCTTGACTCTA</td><td>>hg38_21_+_8433172_8446622_RNA45SN1_45S_50ntFlanks</td><td>11873</td><td>11899</td><td>27</td><td>AAGAAGACCCTGTTGAGCTTGACTCTA</td><td>241.4</td><td>-11.8</td><td>1.4</td><td>-8.2</td><td>2.8E-16</td><td>2.1E-15</td><td>MB231/MB468</td><td>MP</td></tr>
<tr><td>>MI0000342|hsa-mir-200b&WithFlank&1|+|1167098|1167204@63.85.23(+1A)&[MIMAT0000318&hsa-miR-200b-3p&offsets|0|+1(+1A);m-93&1|+|1167160|1167182&offsets|0|0(+1A)]|TAATACTGCCTGGTAATGATGACA</td><td>>MI0000342|hsa-mir-200b&WithFlank&1|+|1167098|1167204</td><td>63</td><td>85</td><td>23(+1A)</td><td>TAATACTGCCTGGTAATGATGACA</td><td>719.9</td><td>-5.6</td><td>0.7</td><td>-8.2</td><td>2.8E-16</td><td>2.1E-15</td><td>MB231/MB468</td><td>MP</td></tr>
<tr><td>>hg38_21_+_8433172_8446622_RNA45SN1_45S_50ntFlanks@12628.12659.32|TGGGGCGAAGCTACCATCTGTGGGATTATGAC</td><td>>hg38_21_+_8433172_8446622_RNA45SN1_45S_50ntFlanks</td><td>12628</td><td>12659</td><td>32</td><td>TGGGGCGAAGCTACCATCTGTGGGATTATGAC</td><td>948.9</td><td>-13.8</td><td>1.7</td><td>-8.2</td><td>2.8E-16</td><td>2.1E-15</td><td>MB231/MB468</td><td>MP</td></tr>
<tr><td>>hg38_21_+_8433172_8446622_RNA45SN1_45S_50ntFlanks@12874.12891.18|GCTGGAAAGGCGGCCGCC</td><td>>hg38_21_+_8433172_8446622_RNA45SN1_45S_50ntFlanks</td><td>12874</td><td>12891</td><td>18</td><td>GCTGGAAAGGCGGCCGCC</td><td>390.2</td><td>-11.3</td><td>1.4</td><td>-8.2</td><td>3.0E-16</td><td>2.1E-15</td><td>MB231/MB468</td><td>CYTO</td></tr>
</table>

| | | | | | | | | | | | | | |
|---|---|---|---|---|---|---|---|---|---|---|---|---|---|
| >hg38_21_+_8433172_8446622_RNA45SN1_45S_50ntFlanks@6652.6693.42\|GACTCTTAGCGGTGGATCACTCGGCTCGTGCGTCGATGAAGA | >hg38_21_+_8433172_8446622_RNA45SN1_45S_50ntFlanks | 6652 | 6693 | 42 | GACTCTTAGCGGTGGATCACTCGGCTCGTGCGTCGATGAAGA | 383.6 | 5.2 | 0.6 | 8.2 | 3.0E-16 | 2.1E-15 | MB231/MB468 | CYTO |
| >trna10_AspGTC_12_-_125424193_125424264@30.72.43,trna12_AspGTC_12_-_125411891_125411962@30.72.43,trna38_AspGTC_17_-_8125556_8125627@30.72.43,trna45_AspGTC_6_+_27447453_27447524@30.72.43,trna48_AspGTC_6_+_27471523_27471594@30.72.43,trna4_AspGTC_12_+_96429799_96429870@30.72.43,trna69_AspGTC_1_-_161440205_161440276@30.72.43,trna72_AspGTC_1_-_161432824_161432895@30.72.43,trna75_AspGTC_1_-_161425414_161425485@30.72.43,trna78_AspGTC_1_-_161418033_161418104@30.72.43,trna81_AspGTC_1_-_161410615_161410686@30.72.43\|GCCTGTCACGCGGGAGACCGGGGTTCGATTCCCCGACGGGGAG | >trna10_AspGTC_12_-_125424193_125424264 | 30 | 72 | 43 | GCCTGTCACGCGGGAGACCGGGGTTCGATTCCCCGACGGGGAG | 402.6 | -11.4 | 1.4 | -8.2 | 3.0E-16 | 2.1E-15 | MB231/MB468 | CYTO |
| >hg38_21_+_8433172_8446622_RNA45SN1_45S_50ntFlanks@8169.8188.20\|CCAAGTCCTTCTGATCGAGG | >hg38_21_+_8433172_8446622_RNA45SN1_45S_50ntFlanks | 8169 | 8188 | 20 | CCAAGTCCTTCTGATCGAGG | 267.0 | -11.9 | 1.5 | -8.2 | 2.8E-16 | 2.1E-15 | MB231/MB468 | MP |
| >hg38_MT_+_598_1651_RNR1_12S_50ntFlanks@428.469.42\|AAAGTGGCTTTAACATATCTGAACACACAATAGCTAAGACCC | >hg38_MT_+_598_1651_RNR1_12S_50ntFlanks | 428 | 469 | 42 | AAAGTGGCTTTAACATATCTGAACACACAATAGCTAAGACCC | 248.9 | 12.0 | 1.5 | 8.2 | 3.0E-16 | 2.1E-15 | MB231/MB468 | CYTO |
| >hg38_21_+_8433172_8446622_RNA45SN1_45S_50ntFlanks@9573.9596.24\|AAATCGGTCGTCCGACCTGGGTAT | >hg38_21_+_8433172_8446622_RNA45SN1_45S_50ntFlanks | 9573 | 9596 | 24 | AAATCGGTCGTCCGACCTGGGTAT | 3675.2 | -7.2 | 0.9 | -8.2 | 3.2E-16 | 2.2E-15 | BT20/MB468 | MITO |
| >hg38_21_+_8433172_8446622_RNA45SN1_45S_50ntFlanks@10410.10428.19\|TCAGTCGGTCCTGAGAGAT | >hg38_21_+_8433172_8446622_RNA45SN1_45S_50ntFlanks | 10410 | 10428 | 19 | TCAGTCGGTCCTGAGAGAT | 731.4 | -4.9 | 0.6 | -8.1 | 4.4E-16 | 2.2E-15 | BT20/MB468 | CYTO |
| >hg38_21_+_8433172_8446622_RNA45SN1_45S_50ntFlanks@10436.10453.18\|CGCCGTTCCGAAGGGACG | >hg38_21_+_8433172_8446622_RNA45SN1_45S_50ntFlanks | 10436 | 10453 | 18 | CGCCGTTCCGAAGGGACG | 636.4 | -12.0 | 1.5 | -8.2 | 3.0E-16 | 2.2E-15 | MB231/MB468 | CYTO |
| >hg38_21_+_8433172_8446622_RNA45SN1_45S_50ntFlanks@5472.5519.48\|ACGGCCCTGGCGGAGCGCTGAGAAGACGGTCGAACTTGACTATCTAGA | >hg38_21_+_8433172_8446622_RNA45SN1_45S_50ntFlanks | 5472 | 5519 | 48 | ACGGCCCTGGCGGAGCGCTGAGAAGACGGTCGAACTTGACTATCTAGA | 796.9 | 10.7 | 1.3 | 8.2 | 3.0E-16 | 2.2E-15 | MB231/MB468 | CYTO |
| >hg38_21_+_8433172_8446622_RNA45SN1_45S_50ntFlanks@11572.11590.19\|CCGACTTAGAACTGGTGCG | >hg38_21_+_8433172_8446622_RNA45SN1_45S_50ntFlanks | 11572 | 11590 | 19 | CCGACTTAGAACTGGTGCG | 2111.3 | -4.2 | 0.5 | -8.1 | 4.4E-16 | 2.2E-15 | BT20/MB468 | CYTO |
| >hg38_21_+_8433172_8446622_RNA45SN1_45S_50ntFlanks@12991.13015.25\|CTCCCTCGCTGCGATCTATTGAAAG | >hg38_21_+_8433172_8446622_RNA45SN1_45S_50ntFlanks | 12991 | 13015 | 25 | CTCCCTCGCTGCGATCTATTGAAAG | 264.6 | -11.9 | 1.5 | -8.2 | 2.8E-16 | 2.2E-15 | MB231/MB468 | MP |
| >am_MI0000809\|hsa-mir-151a&WithFlank&8\|-\|140732558\|140732659@52.73.22(+1U)&[MIMAT0000757&hsa-miR-151a-3p&offsets\|-1\|0(+1U);m-30&8\|-\|140732587\|140732607&offsets\|-1\|0(+1U)]\|ACTAGACTGAAGCTCCTTGAGGT | >am_MI0000809\|hsa-mir-151a&WithFlank&8\|-\|140732558\|140732659 | 52 | 73 | 22(+1U) | ACTAGACTGAAGCTCCTTGAGGT | 287.4 | 8.5 | 1.1 | 8.1 | 4.4E-16 | 2.2E-15 | BT20/MB468 | CYTO |
| >hg38_21_+_8433172_8446622_RNA45SN1_45S_50ntFlanks@10186.10208.23\|TCCTCCCCTCCTCCCGCCCACGC | >hg38_21_+_8433172_8446622_RNA45SN1_45S_50ntFlanks | 10186 | 10208 | 23 | TCCTCCCCTCCTCCCGCCCACGC | 514.7 | -12.0 | 1.5 | -8.2 | 3.5E-16 | 2.2E-15 | MB231/MB468 | NUC |
| >hg38_21_+_8433172_8446622_RNA45SN1_45S_50ntFlanks@8839.8859.21\|CGGTGGCGGCGGCGGCGGCGG | >hg38_21_+_8433172_8446622_RNA45SN1_45S_50ntFlanks | 8839 | 8859 | 21 | CGGTGGCGGCGGCGGCGGCGG | 390.9 | -12.4 | 1.5 | -8.1 | 4.7E-16 | 2.2E-15 | MB231/MB468 | MITO |
| >hg38_21_+_8433172_8446622_RNA45SN1_45S_50ntFlanks@485.526.42\|GATGTGGTGACGTCGTGCTCTCCCGGGCCGGGTCCGAGCCGC | >hg38_21_+_8433172_8446622_RNA45SN1_45S_50ntFlanks | 485 | 526 | 42 | GATGTGGTGACGTCGTGCTCTCCCGGGCCGGGTCCGAGCCGC | 3144.2 | 29.6 | 3.6 | 8.2 | 3.5E-16 | 2.2E-15 | MB231/MB468 | NUC |
| >hg38_21_+_8433172_8446622_RNA45SN1_45S_50ntFlanks@11641.11685.45\|CGGCGGGTGTTGACGCGATGTGATTTCTGCCCAGTGCTCTGAATG | >hg38_21_+_8433172_8446622_RNA45SN1_45S_50ntFlanks | 11641 | 11685 | 45 | CGGCGGGTGTTGACGCGATGTGATTTCTGCCCAGTGCTCTGAATG | 11476.0 | -9.9 | 1.2 | -8.1 | 4.7E-16 | 2.2E-15 | MB231/MB468 | MITO |
| >hg38_21_+_8433172_8446622_RNA45SN1_45S_50ntFlanks@11584.11616.33\|TGGTGCGGACCAGGGGAATCCGACTGTTTAATT | >hg38_21_+_8433172_8446622_RNA45SN1_45S_50ntFlanks | 11584 | 11616 | 33 | TGGTGCGGACCAGGGGAATCCGACTGTTTAATT | 543.3 | 6.6 | 0.8 | 8.2 | 3.2E-16 | 2.2E-15 | BT20/MB468 | MITO |
| >hg38_21_+_8433172_8446622_RNA45SN1_45S_50ntFlanks@9201.9223.23\|CTCTCGGGGCCACGCGCGCGTCC | >hg38_21_+_8433172_8446622_RNA45SN1_45S_50ntFlanks | 9201 | 9223 | 23 | CTCTCGGGGCCACGCGCGCGTCC | 344.6 | -12.3 | 1.5 | -8.2 | 2.8E-16 | 2.2E-15 | MB231/MB468 | MP |
| >hg38_21_+_8433172_8446622_RNA45SN1_45S_50ntFlanks@6773.6803.31\|TTCCTCCCGGGGCTACGCCTGTCTGAGCGTC | >hg38_21_+_8433172_8446622_RNA45SN1_45S_50ntFlanks | 6773 | 6803 | 31 | TTCCTCCCGGGGCTACGCCTGTCTGAGCGTC | 6822.7 | -8.7 | 1.1 | -8.1 | 4.7E-16 | 2.2E-15 | MB231/MB468 | MITO |
| >hg38_21_+_8433172_8446622_RNA45SN1_45S_50ntFlanks@8472.8492.21\|CGGCCCGGCGGATCTTTCCCG | >hg38_21_+_8433172_8446622_RNA45SN1_45S_50ntFlanks | 8472 | 8492 | 21 | CGGCCCGGCGGATCTTTCCCG | 579.2 | -12.1 | 1.5 | -8.2 | 3.5E-16 | 2.2E-15 | MB231/MB468 | NUC |
| >MI0000446\|hsa-mir-125b-1&WithFlank&11\|-\|122099751\|122099850@21.43.23(+1U)&[MIMAT0000423&hsa-miR-125b-5p&offsets\|0\|+1(+1U);m-67&11\|-\|122099809\|122099830&offsets\|0\|+1(+1U)]\|TCCCTGAGACCCTAACTTGTGATT | >MI0000446\|hsa-mir-125b-1&WithFlank&11\|-\|122099751\|122099850 | 21 | 43 | 23(+1U) | TCCCTGAGACCCTAACTTGTGATT | 347.7 | 12.4 | 1.5 | 8.2 | 3.1E-16 | 2.2E-15 | MB231/MB468 | CYTO |
| >hg38_21_+_8433172_8446622_RNA45SN1_45S_50ntFlanks@10351.10388.38\|CGAGAACTTTGAAGGCCGAAGTGGAGAAGGGTTCCATG | >hg38_21_+_8433172_8446622_RNA45SN1_45S_50ntFlanks | 10351 | 10388 | 38 | CGAGAACTTTGAAGGCCGAAGTGGAGAAGGGTTCCATG | 5633.2 | -5.0 | 0.6 | -8.2 | 3.3E-16 | 2.2E-15 | BT20/MB468 | MITO |
| >hg38_21_+_8433172_8446622_RNA45SN1_45S_50ntFlanks@4995.5020.26\|ACAGATTGATAGCTCTTTCTCGATTC | >hg38_21_+_8433172_8446622_RNA45SN1_45S_50ntFlanks | 4995 | 5020 | 26 | ACAGATTGATAGCTCTTTCTCGATTC | 234.6 | -11.7 | 1.4 | -8.2 | 2.9E-16 | 2.2E-15 | MB231/MB468 | MP |
| >hg38_21_+_8433172_8446622_RNA45SN1_45S_50ntFlanks@11115.11132.18\|CGGCAGGGGCCGGCGGCG | >hg38_21_+_8433172_8446622_RNA45SN1_45S_50ntFlanks | 11115 | 11132 | 18 | CGGCAGGGGCCGGCGGCG | 277.4 | -10.8 | 1.3 | -8.2 | 3.1E-16 | 2.2E-15 | MB231/MB468 | CYTO |
| >hg38_21_+_8433172_8446622_RNA45SN1_45S_50ntFlanks@12991.13012.22\|CTCCCTCGCTGCGATCTATTGA | >hg38_21_+_8433172_8446622_RNA45SN1_45S_50ntFlanks | 12991 | 13012 | 22 | CTCCCTCGCTGCGATCTATTGA | 229.1 | -11.7 | 1.4 | -8.2 | 2.9E-16 | 2.2E-15 | MB231/MB468 | MP |
| >hg38_21_+_8433172_8446622_RNA45SN1_45S_50ntFlanks@12934.12962.29\|CTAAACCATTCGTAGACGACCTGCTTCTG | >hg38_21_+_8433172_8446622_RNA45SN1_45S_50ntFlanks | 12934 | 12962 | 29 | CTAAACCATTCGTAGACGACCTGCTTCTG | 222.2 | -11.7 | 1.4 | -8.2 | 2.9E-16 | 2.2E-15 | MB231/MB468 | MP |
| >hg38_21_+_8433172_8446622_RNA45SN1_45S_50ntFlanks@11821.11856.36\|CTATCCAGCGAAACCACAGCCAAGGGAACGGGCTTG | >hg38_21_+_8433172_8446622_RNA45SN1_45S_50ntFlanks | 11821 | 11856 | 36 | CTATCCAGCGAAACCACAGCCAAGGGAACGGGCTTG | 303.9 | -12.1 | 1.5 | -8.2 | 2.9E-16 | 2.2E-15 | MB231/MB468 | MP |
| >hg38_21_+_8433172_8446622_RNA45SN1_45S_50ntFlanks@3860.3898.39\|GTAATTCTAGAGCTAATACATGCCGACGGGCGCTGACCC | >hg38_21_+_8433172_8446622_RNA45SN1_45S_50ntFlanks | 3860 | 3898 | 39 | GTAATTCTAGAGCTAATACATGCCGACGGGCGCTGACCC | 315.5 | -12.1 | 1.5 | -8.1 | 4.9E-16 | 2.2E-15 | MB231/MB468 | MITO |
| >hg38_21_+_8433172_8446622_RNA45SN1_45S_50ntFlanks@12412.12446.35\|ATCCTTCGATGTCGGCTCTTCCTATCATTGTGAAG | >hg38_21_+_8433172_8446622_RNA45SN1_45S_50ntFlanks | 12412 | 12446 | 35 | ATCCTTCGATGTCGGCTCTTCCTATCATTGTGAAG | 333.8 | -10.7 | 1.3 | -8.1 | 4.5E-16 | 2.2E-15 | BT20/MB468 | CYTO |
| >MI0000067\|hsa-let-7f-1&WithFlank&9\|+\|94176341\|94176439@13.32.20;MI0000068\|hsa-let-7f-2&WithFlank&X\|-\|53557186\|53557280@14.33.20&[MIMAT0000067&hsa-let-7f-5p&offsets\|0\|-2;m-20&9\|+\|94176353\|94176374&offsets\|0\|-2];[MIMAT0000067_1&hsa-let-7f-5p&offsets\|0\|-2;m-19&X\|-\|53557246\|53557267&offsets\|0\|-2]\|TGAGGTAGTAGATTGTATAG | >MI0000067\|hsa-let-7f-1&WithFlank&9\|+\|94176341\|94176439 | 13 | 32 | 20 | TGAGGTAGTAGATTGTATAG | 239.4 | 5.3 | 0.7 | 8.2 | 3.6E-16 | 2.2E-15 | MB231/MB468 | NUC |
| >hg38_21_+_8433172_8446622_RNA45SN1_45S_50ntFlanks@11234.11254.21\|CCTTCCCGTGGATCGCCCCAG | >hg38_21_+_8433172_8446622_RNA45SN1_45S_50ntFlanks | 11234 | 11254 | 21 | CCTTCCCGTGGATCGCCCCAG | 271.7 | -11.0 | 1.4 | -8.2 | 3.6E-16 | 2.2E-15 | MB231/MB468 | NUC |
| >hg38_21_+_8433172_8446622_RNA45SN1_45S_50ntFlanks@10846.10864.19\|CTTCGGGATAAGGATTGGC | >hg38_21_+_8433172_8446622_RNA45SN1_45S_50ntFlanks | 10846 | 10864 | 19 | CTTCGGGATAAGGATTGGC | 419.0 | -12.6 | 1.5 | -8.2 | 2.9E-16 | 2.2E-15 | MB231/MB468 | MP |
| >MI0000088\|hsa-mir-30a&WithFlank&6\|-\|71403545\|71403627@54.75.22&[MIMAT0000088&hsa-miR-30a-3p&offsets\|+1\|+1;m-38&6\|-\|71403554\|71403575&offsets\|+1\|+1]\|TTTCAGTCGGATGTTTGCAGCT | >MI0000088\|hsa-mir-30a&WithFlank&6\|-\|71403545\|71403627 | 54 | 75 | 22 | TTTCAGTCGGATGTTTGCAGCT | 740.2 | 7.7 | 0.9 | 8.1 | 4.6E-16 | 2.3E-15 | BT20/MB468 | CYTO |
| >hg38_21_+_8433172_8446622_RNA45SN1_45S_50ntFlanks@11821.11842.22\|CTATCCAGCGAAACCACAGCCA | >hg38_21_+_8433172_8446622_RNA45SN1_45S_50ntFlanks | 11821 | 11842 | 22 | CTATCCAGCGAAACCACAGCCA | 426.2 | -12.6 | 1.5 | -8.2 | 2.9E-16 | 2.3E-15 | MB231/MB468 | MP |
| >hg38_21_+_8433172_8446622_RNA45SN1_45S_50ntFlanks@12933.12956.24\|GCTAAACCATTCGTAGACGACCTG | >hg38_21_+_8433172_8446622_RNA45SN1_45S_50ntFlanks | 12933 | 12956 | 24 | GCTAAACCATTCGTAGACGACCTG | 1000.6 | -4.7 | 0.6 | -8.1 | 4.6E-16 | 2.3E-15 | BT20/MB468 | CYTO |
| >MI0000234\|hsa-mir-192&WithFlank&11\|-\|64891131\|64891252@30.49.20(+1U)&[MIMAT0000222&hsa-miR-192-5p&offsets\|0\|-1(+1U);m-59&11\|-\|64891203\|64891223&offsets\|0\|-1(+1U)]\|CTGACCTATGAATTGACAGCT | >MI0000234\|hsa-mir-192&WithFlank&11\|-\|64891131\|64891252 | 30 | 49 | 20(+1U) | CTGACCTATGAATTGACAGCT | 551.4 | 6.8 | 0.8 | 8.1 | 4.6E-16 | 2.3E-15 | BT20/MB468 | CYTO |
| >MI0000102\|hsa-mir-100&WithFlank&11\|-\|122152223\|122152314@19.40.22(+2U)&[MIMAT0000098&hsa-miR-100-5p&offsets\|0\|0(+2U);m-27&11\|-\|122152276\|122152296&offsets\|0\|+1(+2U)]\|AACCCGTAGATCCGAACTTGTGTT | >MI0000102\|hsa-mir-100&WithFlank&11\|-\|122152223\|122152314 | 19 | 40 | 22(+2U) | AACCCGTAGATCCGAACTTGTGTT | 802.7 | 12.9 | 1.6 | 8.1 | 4.9E-16 | 2.3E-15 | MB231/MB468 | MITO |
| >hg38_21_+_8433172_8446622_RNA45SN1_45S_50ntFlanks@3855.3898.44\|CTGTGGTAATTCTAGAGCTAATACATGCCGACGGGCGCTGACCC | >hg38_21_+_8433172_8446622_RNA45SN1_45S_50ntFlanks | 3855 | 3898 | 44 | CTGTGGTAATTCTAGAGCTAATACATGCCGACGGGCGCTGACCC | 551.3 | -13.5 | 1.7 | -8.1 | 7.3E-16 | 2.3E-15 | BT20/MB468 | NUC |
| >hg38_21_+_8433172_8446622_RNA45SN1_45S_50ntFlanks@11821.11842.22\|CTATCCAGCGAAACCACAGCCA | >hg38_21_+_8433172_8446622_RNA45SN1_45S_50ntFlanks | 11821 | 11842 | 22 | CTATCCAGCGAAACCACAGCCA | 238.4 | -10.9 | 1.3 | -8.2 | 3.6E-16 | 2.3E-15 | MB231/MB468 | NUC |
| >hg38_21_+_8433172_8446622_RNA45SN1_45S_50ntFlanks@4659.4678.20\|AGAGGTGAAATTCTTGGACC | >hg38_21_+_8433172_8446622_RNA45SN1_45S_50ntFlanks | 4659 | 4678 | 20 | AGAGGTGAAATTCTTGGACC | 306.6 | 10.2 | 1.3 | 8.1 | 4.6E-16 | 2.3E-15 | BT20/MB468 | CYTO |
| >hg38_21_+_8433172_8446622_RNA45SN1_45S_50ntFlanks@10510.10533.24\|AATCCGGAGTGGCGGAGATGGGCG | >hg38_21_+_8433172_8446622_RNA45SN1_45S_50ntFlanks | 10510 | 10533 | 24 | AATCCGGAGTGGCGGAGATGGGCG | 2060.9 | -11.2 | 1.4 | -8.1 | 3.7E-16 | 2.3E-15 | MB231/MB468 | NUC |
| >hg38_21_+_8433172_8446622_RNA45SN1_45S_50ntFlanks@9177.9197.21\|GCGCCGTCGGGCCCGGGGGAG | >hg38_21_+_8433172_8446622_RNA45SN1_45S_50ntFlanks | 9177 | 9197 | 21 | GCGCCGTCGGGCCCGGGGGAG | 809.1 | -13.4 | 1.7 | -8.1 | 4.9E-16 | 2.3E-15 | MB231/MB468 | MITO |
| >trna119_LysCTT_1_-_145395522_145395594@1.31.31,trna11_LysCTT_5_-_180648979_180649051@1.31.31,trna13_LysCTT_6_+_26556774_26556846@1.31.31,trna32_LysCTT_16_-_3207406_3207478@1.31.31,trna7_LysCTT_16_+_3225692_3225764@1.31.31,trna9_LysCTT_5_+_180634755_180634827@1.31.31\|GCCCGGCTAGCTCAGTCGGTAGAGCATGAGA | >trna119_LysCTT_1_-_145395522_145395594 | 1 | 31 | 31 | GCCCGGCTAGCTCAGTCGGTAGAGCATGAGA | 4881.1 | -9.9 | 1.2 | -8.1 | 7.3E-16 | 2.3E-15 | BT20/MB468 | NUC |
| >hg38_21_+_8433172_8446622_RNA45SN1_45S_50ntFlanks@10878.10901.24\|TCGGTCGGGCTGGGGCGCGAAGCG | >hg38_21_+_8433172_8446622_RNA45SN1_45S_50ntFlanks | 10878 | 10901 | 24 | TCGGTCGGGCTGGGGCGCGAAGCG | 340.8 | -12.3 | 1.5 | -8.2 | 3.0E-16 | 2.3E-15 | MB231/MB468 | MP |
| >hg38_21_+_8433172_8446622_RNA45SN1_45S_50ntFlanks@10350.10384.35\|ACGAGAACTTTGAAGGCCGAAGTGGAGAAGGGTTC | >hg38_21_+_8433172_8446622_RNA45SN1_45S_50ntFlanks | 10350 | 10384 | 35 | ACGAGAACTTTGAAGGCCGAAGTGGAGAAGGGTTC | 1237.7 | -3.7 | 0.5 | -8.1 | 4.7E-16 | 2.3E-15 | BT20/MB468 | CYTO |
| >hg38_21_+_8433172_8446622_RNA45SN1_45S_50ntFlanks@11894.11912.19\|ACTCTAGTCTGGCACGGTG | >hg38_21_+_8433172_8446622_RNA45SN1_45S_50ntFlanks | 11894 | 11912 | 19 | ACTCTAGTCTGGCACGGTG | 1501.6 | -5.6 | 0.7 | -8.1 | 4.6E-16 | 2.3E-15 | BT20/MB468 | CYTO |

<table>
<tr><th></th><th></th><th></th><th></th><th></th><th></th><th></th><th></th><th></th><th></th><th></th><th></th><th></th><th></th></tr>
<tr><td>>trna116_GluCTC_1_-_145399233_145399304@3.35.33,trna59_GluCTC_1_+_249168447_249168518@3.35.33,trna71_GluCTC_1_-_161439189_161439260@3.35.33,trna74_GluCTC_1_-_161431809_161431880@3.35.33,trna77_GluCTC_1_-_161424398_161424469@3.35.33,trna77_GluCTC_6_+_28949976_28950047@3.35.33,trna80_GluCTC_1_-_161417018_161417089@3.35.33,trna87_GluCTC_6_-_126101393_126101464@3.35.33|CCTGGTGGTCTAGTGGTTAGGATTCGGCGCTCT</td><td>>trna116_GluCTC_1_-_145399233_145399304</td><td>3</td><td>35</td><td>33</td><td>CCTGGTGGTCTAGTGGTTAGGATTCGGCGCTCT</td><td>260.4</td><td>7.8</td><td>1.0</td><td>8.2</td><td>3.2E-16</td><td>2.3E-15</td><td>MB231/MB468</td><td>CYTO</td></tr>
<tr><td>>hg38_21_+_8433172_8446622_RNA45SN1_45S_50ntFlanks@11849.11872.24|CGGGCTTGGCGGAATCAGCGGGGA</td><td>>hg38_21_+_8433172_8446622_RNA45SN1_45S_50ntFlanks</td><td>11849</td><td>11872</td><td>24</td><td>CGGGCTTGGCGGAATCAGCGGGGA</td><td>287.1</td><td>-7.5</td><td>0.9</td><td>-8.2</td><td>3.2E-16</td><td>2.3E-15</td><td>MB231/MB468</td><td>CYTO</td></tr>
<tr><td>>hg38_21_+_8433172_8446622_RNA45SN1_45S_50ntFlanks@11885.11903.19|TTGAGCTTGACTCTAGTCT</td><td>>hg38_21_+_8433172_8446622_RNA45SN1_45S_50ntFlanks</td><td>11885</td><td>11903</td><td>19</td><td>TTGAGCTTGACTCTAGTCT</td><td>727.8</td><td>-12.2</td><td>1.5</td><td>-8.2</td><td>3.2E-16</td><td>2.3E-15</td><td>MB231/MB468</td><td>CYTO</td></tr>
<tr><td>>hg38_21_+_8433172_8446622_RNA45SN1_45S_50ntFlanks@6745.6770.26|ACACTTCGAACGCACTTGCGGCCCCG</td><td>>hg38_21_+_8433172_8446622_RNA45SN1_45S_50ntFlanks</td><td>6745</td><td>6770</td><td>26</td><td>ACACTTCGAACGCACTTGCGGCCCCG</td><td>6029.8</td><td>-7.7</td><td>0.9</td><td>-8.2</td><td>3.2E-16</td><td>2.3E-15</td><td>MB231/MB468</td><td>CYTO</td></tr>
<tr><td>>hg38_21_+_8433172_8446622_RNA45SN1_45S_50ntFlanks@4283.4314.32|CTTTAACGAGGATCCATTGGAGGGCAAGTCTG</td><td>>hg38_21_+_8433172_8446622_RNA45SN1_45S_50ntFlanks</td><td>4283</td><td>4314</td><td>32</td><td>CTTTAACGAGGATCCATTGGAGGGCAAGTCTG</td><td>426.4</td><td>-12.5</td><td>1.5</td><td>-8.1</td><td>5.0E-16</td><td>2.3E-15</td><td>MB231/MB468</td><td>MITO</td></tr>
<tr><td>>hg38_21_+_8433172_8446622_RNA45SN1_45S_50ntFlanks@12436.12455.20|TCATTGTGAAGCAGAATTCA</td><td>>hg38_21_+_8433172_8446622_RNA45SN1_45S_50ntFlanks</td><td>12436</td><td>12455</td><td>20</td><td>TCATTGTGAAGCAGAATTCA</td><td>3922.4</td><td>-10.6</td><td>1.3</td><td>-8.2</td><td>3.2E-16</td><td>2.3E-15</td><td>MB231/MB468</td><td>CYTO</td></tr>
<tr><td>>hg38_21_+_8433172_8446622_RNA45SN1_45S_50ntFlanks@12927.12972.46|CCTGGCGCTAAACCATTCGTAGACGACCTGCTTCTGGGTCGGGGTT</td><td>>hg38_21_+_8433172_8446622_RNA45SN1_45S_50ntFlanks</td><td>12927</td><td>12972</td><td>46</td><td>CCTGGCGCTAAACCATTCGTAGACGACCTGCTTCTGGGTCGGGGTT</td><td>544.2</td><td>-12.9</td><td>1.6</td><td>-8.1</td><td>5.0E-16</td><td>2.3E-15</td><td>MB231/MB468</td><td>MITO</td></tr>
<tr><td>>hg38_21_+_8433172_8446622_RNA45SN1_45S_50ntFlanks@11161.11188.28|CACGGTCCCCCGCGAGGGGGGCCCGGGC</td><td>>hg38_21_+_8433172_8446622_RNA45SN1_45S_50ntFlanks</td><td>11161</td><td>11188</td><td>28</td><td>CACGGTCCCCCGCGAGGGGGGCCCGGGC</td><td>638.3</td><td>-12.0</td><td>1.5</td><td>-8.2</td><td>3.3E-16</td><td>2.3E-15</td><td>MB231/MB468</td><td>CYTO</td></tr>
<tr><td>>hg38_21_+_8433172_8446622_RNA45SN1_45S_50ntFlanks@11038.11060.23|TCTCCCCCGCTCCCCGTCCTCCC</td><td>>hg38_21_+_8433172_8446622_RNA45SN1_45S_50ntFlanks</td><td>11038</td><td>11060</td><td>23</td><td>TCTCCCCCGCTCCCCGTCCTCCC</td><td>291.5</td><td>-12.0</td><td>1.5</td><td>-8.1</td><td>5.0E-16</td><td>2.3E-15</td><td>MB231/MB468</td><td>MITO</td></tr>
<tr><td>>hg38_21_+_8433172_8446622_RNA45SN1_45S_50ntFlanks@5422.5450.29|GATTGGATGGTTTAGTGAGGCCCTCGGAT</td><td>>hg38_21_+_8433172_8446622_RNA45SN1_45S_50ntFlanks</td><td>5422</td><td>5450</td><td>29</td><td>GATTGGATGGTTTAGTGAGGCCCTCGGAT</td><td>447.9</td><td>7.4</td><td>0.9</td><td>8.2</td><td>3.3E-16</td><td>2.3E-15</td><td>MB231/MB468</td><td>CYTO</td></tr>
<tr><td>>hg38_21_+_8433172_8446622_RNA45SN1_45S_50ntFlanks@3854.3899.46|ACTGTGGTAATTCTAGAGCTAATACATGCCGACGGGCGCTGACCCC</td><td>>hg38_21_+_8433172_8446622_RNA45SN1_45S_50ntFlanks</td><td>3854</td><td>3899</td><td>46</td><td>ACTGTGGTAATTCTAGAGCTAATACATGCCGACGGGCGCTGACCCC</td><td>449.7</td><td>-13.2</td><td>1.6</td><td>-8.1</td><td>7.5E-16</td><td>2.3E-15</td><td>BT20/MB468</td><td>NUC</td></tr>
<tr><td>>MI0000272|hsa-mir-182&WithFlank&7|-|129770377|129770498@29.48.20(+1U)&[MIMAT0000259&hsa-miR-182-5p&offsets|0|-4(+1U);m-22&7|-|129770449|129770470&offsets|0|-2(+1U)]|TTTGGCAATGGTAGAACTCAT</td><td>>MI0000272|hsa-mir-182&WithFlank&7|-|129770377|129770498</td><td>29</td><td>48</td><td>20(+1U)</td><td>TTTGGCAATGGTAGAACTCAT</td><td>360.2</td><td>12.7</td><td>1.6</td><td>8.1</td><td>4.7E-16</td><td>2.3E-15</td><td>BT20/MB468</td><td>CYTO</td></tr>
<tr><td>>MI0005561|hsa-mir-877&WithFlank&6|+|30584326|30584423@7.26.20&[MIMAT0004949&hsa-miR-877-5p&offsets|0|0;m-382&6|+|30584332|30584354&offsets|0|-3]|GTAGAGGAGATGGCGCAGGG</td><td>>MI0005561|hsa-mir-877&WithFlank&6|+|30584326|30584423</td><td>7</td><td>26</td><td>20</td><td>GTAGAGGAGATGGCGCAGGG</td><td>390.9</td><td>10.5</td><td>1.3</td><td>8.1</td><td>4.8E-16</td><td>2.3E-15</td><td>BT20/MB468</td><td>CYTO</td></tr>
<tr><td>>trna10_AspGTC_12_-_125424193_125424264@32.70.39,trna12_AspGTC_12_-_125411891_125411962@32.70.39,trna144_AspGTC_6_-_27551236_27551307@32.70.39,trna38_AspGTC_17_-_8125556_8125627@32.70.39,trna45_AspGTC_6_+_27447453_27447524@32.70.39,trna48_AspGTC_6_+_27471523_27471594@32.70.39,trna4_AspGTC_12_+_96429799_96429870@32.70.39,trna69_AspGTC_1_-_161440205_161440276@32.70.39,trna72_AspGTC_1_-_161432824_161432895@32.70.39,trna75_AspGTC_1_-_161425414_161425485@32.70.39,trna78_AspGTC_1_-_161418033_161418104@32.70.39,trna81_AspGTC_1_-_161410615_161410686@32.70.39|CTGTCACGCGGGAGACCGGGGTTCGATTCCCCGACGGGG</td><td>>trna10_AspGTC_12_-_125424193_125424264</td><td>32</td><td>70</td><td>39</td><td>CTGTCACGCGGGAGACCGGGGTTCGATTCCCCGACGGGG</td><td>536.5</td><td>-12.0</td><td>1.5</td><td>-8.1</td><td>3.8E-16</td><td>2.3E-15</td><td>MB231/MB468</td><td>NUC</td></tr>
<tr><td>>trnaMT_PheGAA_MT_+_577_647@39.59.21|ATGTTTAGACGGGCTCACATC</td><td>>trnaMT_PheGAA_MT_+_577_647</td><td>39</td><td>59</td><td>21</td><td>ATGTTTAGACGGGCTCACATC</td><td>1463.0</td><td>7.3</td><td>0.9</td><td>8.2</td><td>3.4E-16</td><td>2.3E-15</td><td>BT20/MB468</td><td>MITO</td></tr>
<tr><td>>hg38_21_+_8433172_8446622_RNA45SN1_45S_50ntFlanks@10427.10457.31|ATGGGCGAGCGCCGTTCCGAAGGGACGGGCG</td><td>>hg38_21_+_8433172_8446622_RNA45SN1_45S_50ntFlanks</td><td>10427</td><td>10457</td><td>31</td><td>ATGGGCGAGCGCCGTTCCGAAGGGACGGGCG</td><td>462.7</td><td>-6.6</td><td>0.8</td><td>-8.2</td><td>3.3E-16</td><td>2.3E-15</td><td>MB231/MB468</td><td>CYTO</td></tr>
<tr><td>>hg38_21_+_8433172_8446622_RNA45SN1_45S_50ntFlanks@4064.4101.38|ATCAACTTTCGATGGTAGTCGCCGTGCCTACCATGGTG</td><td>>hg38_21_+_8433172_8446622_RNA45SN1_45S_50ntFlanks</td><td>4064</td><td>4101</td><td>38</td><td>ATCAACTTTCGATGGTAGTCGCCGTGCCTACCATGGTG</td><td>553.8</td><td>-11.4</td><td>1.4</td><td>-8.1</td><td>4.8E-16</td><td>2.3E-15</td><td>BT20/MB468</td><td>CYTO</td></tr>
<tr><td>>hg38_1_-_228634819_228635039_RNA5S12_5S_50ntFlanks@129.171.43|TTGGATGGGAGACCGCCTGGGAATACCGGGTGCTGTAGGCTTT</td><td>>hg38_1_-_228634819_228635039_RNA5S12_5S_50ntFlanks</td><td>129</td><td>171</td><td>43</td><td>TTGGATGGGAGACCGCCTGGGAATACCGGGTGCTGTAGGCTTT</td><td>5033.4</td><td>10.7</td><td>1.3</td><td>8.2</td><td>3.3E-16</td><td>2.3E-15</td><td>MB231/MB468</td><td>CYTO</td></tr>
<tr><td>>hg38_21_+_8433172_8446622_RNA45SN1_45S_50ntFlanks@12459.12488.30|AGCGTTGGATTGTTCACCCACTAATAGGGA</td><td>>hg38_21_+_8433172_8446622_RNA45SN1_45S_50ntFlanks</td><td>12459</td><td>12488</td><td>30</td><td>AGCGTTGGATTGTTCACCCACTAATAGGGA</td><td>268.6</td><td>-11.0</td><td>1.4</td><td>-8.1</td><td>3.8E-16</td><td>2.3E-15</td><td>MB231/MB468</td><td>NUC</td></tr>
<tr><td>>hg38_21_+_8433172_8446622_RNA45SN1_45S_50ntFlanks@11877.11906.30|AGACCCTGTTGAGCTTGACTCTAGTCTGGC</td><td>>hg38_21_+_8433172_8446622_RNA45SN1_45S_50ntFlanks</td><td>11877</td><td>11906</td><td>30</td><td>AGACCCTGTTGAGCTTGACTCTAGTCTGGC</td><td>305.6</td><td>-12.0</td><td>1.5</td><td>-8.1</td><td>5.1E-16</td><td>2.4E-15</td><td>MB231/MB468</td><td>MITO</td></tr>
<tr><td>>MI0000285|hsa-mir-205&WithFlank&1|+|209432127|209432248@40.59.20(+1C)&[MIMAT0000266&hsa-miR-205-5p&offsets|0|-2(+1C);m-351&1|+|209432166|209432187&offsets|0|-2(+1C)]|TCCTTCATTCCACCGGAGTCC</td><td>>MI0000285|hsa-mir-205&WithFlank&1|+|209432127|209432248</td><td>40</td><td>59</td><td>20(+1C)</td><td>TCCTTCATTCCACCGGAGTCC</td><td>424.6</td><td>12.3</td><td>1.5</td><td>8.4</td><td>3.9E-17</td><td>2.4E-15</td><td>BT20/MB231</td><td>MITO</td></tr>
<tr><td>>MI0006444|hsa-mir-1307&WithFlank&10|-|103394247|103394407@86.107.22(+1U)&[MIMAT0005951&hsa-miR-1307-3p&offsets|0|0(+1U);m-66&10|-|103394300|103394322&offsets|0|-1(+1U)]|ACTCGGCGTGGCGTCGGTCGTGT</td><td>>MI0006444|hsa-mir-1307&WithFlank&10|-|103394247|103394407</td><td>86</td><td>107</td><td>22(+1U)</td><td>ACTCGGCGTGGCGTCGGTCGTGT</td><td>854.3</td><td>6.1</td><td>0.7</td><td>8.1</td><td>3.8E-16</td><td>2.4E-15</td><td>MB231/MB468</td><td>NUC</td></tr>
<tr><td>>hg38_21_+_8433172_8446622_RNA45SN1_45S_50ntFlanks@9158.9187.30|AGTGCGCCCCGGGCGGGTCGCGCCGTCGGG</td><td>>hg38_21_+_8433172_8446622_RNA45SN1_45S_50ntFlanks</td><td>9158</td><td>9187</td><td>30</td><td>AGTGCGCCCCGGGCGGGTCGCGCCGTCGGG</td><td>295.8</td><td>-12.1</td><td>1.5</td><td>-8.2</td><td>3.1E-16</td><td>2.4E-15</td><td>MB231/MB468</td><td>MP</td></tr>
<tr><td>>hg38_21_+_8433172_8446622_RNA45SN1_45S_50ntFlanks@9339.9377.39|CGCACGAAAGCCGCCGTGGCGCAATGAAGGTGAAGGCCG</td><td>>hg38_21_+_8433172_8446622_RNA45SN1_45S_50ntFlanks</td><td>9339</td><td>9377</td><td>39</td><td>CGCACGAAAGCCGCCGTGGCGCAATGAAGGTGAAGGCCG</td><td>746.4</td><td>-13.3</td><td>1.6</td><td>-8.1</td><td>5.1E-16</td><td>2.4E-15</td><td>MB231/MB468</td><td>MITO</td></tr>
<tr><td>>hg38_21_+_8433172_8446622_RNA45SN1_45S_50ntFlanks@10401.10424.24|GAACATGGGTCAGTCGGTCCTGAG</td><td>>hg38_21_+_8433172_8446622_RNA45SN1_45S_50ntFlanks</td><td>10401</td><td>10424</td><td>24</td><td>GAACATGGGTCAGTCGGTCCTGAG</td><td>1911.3</td><td>4.5</td><td>0.6</td><td>8.1</td><td>4.8E-16</td><td>2.4E-15</td><td>BT20/MB468</td><td>CYTO</td></tr>
<tr><td>>hg38_21_+_8433172_8446622_RNA45SN1_45S_50ntFlanks@11322.11340.19|ACGTCTCGTCGCGCGCGCG</td><td>>hg38_21_+_8433172_8446622_RNA45SN1_45S_50ntFlanks</td><td>11322</td><td>11340</td><td>19</td><td>ACGTCTCGTCGCGCGCGCG</td><td>206.6</td><td>-11.6</td><td>1.4</td><td>-8.2</td><td>3.1E-16</td><td>2.4E-15</td><td>MB231/MB468</td><td>MP</td></tr>
<tr><td>>hg38_21_+_8433172_8446622_RNA45SN1_45S_50ntFlanks@10786.10825.40|AACAGCCTCTGGCATGTTGGAACAATGTAGGTAAGGGAAG</td><td>>hg38_21_+_8433172_8446622_RNA45SN1_45S_50ntFlanks</td><td>10786</td><td>10825</td><td>40</td><td>AACAGCCTCTGGCATGTTGGAACAATGTAGGTAAGGGAAG</td><td>2648.9</td><td>-10.3</td><td>1.3</td><td>-8.1</td><td>5.2E-16</td><td>2.4E-15</td><td>MB231/MB468</td><td>MITO</td></tr>
<tr><td>>hg38_21_+_8433172_8446622_RNA45SN1_45S_50ntFlanks@5084.5105.22|CGAACGAGACTCTGGCATGCTA</td><td>>hg38_21_+_8433172_8446622_RNA45SN1_45S_50ntFlanks</td><td>5084</td><td>5105</td><td>22</td><td>CGAACGAGACTCTGGCATGCTA</td><td>1520.3</td><td>-14.4</td><td>1.8</td><td>-8.1</td><td>5.2E-16</td><td>2.4E-15</td><td>MB231/MB468</td><td>MITO</td></tr>
<tr><td>>hg38_21_+_8433172_8446622_RNA45SN1_45S_50ntFlanks@8157.8181.25|CTCGTGGGGGGCCCAAGTCCTTCTG</td><td>>hg38_21_+_8433172_8446622_RNA45SN1_45S_50ntFlanks</td><td>8157</td><td>8181</td><td>25</td><td>CTCGTGGGGGGCCCAAGTCCTTCTG</td><td>4743.3</td><td>-7.8</td><td>1.0</td><td>-8.1</td><td>5.2E-16</td><td>2.4E-15</td><td>MB231/MB468</td><td>MITO</td></tr>
<tr><td>>hg38_21_+_8433172_8446622_RNA45SN1_45S_50ntFlanks@11766.11790.25|ATCTAATTAGTGACGCGCATGAATG</td><td>>hg38_21_+_8433172_8446622_RNA45SN1_45S_50ntFlanks</td><td>11766</td><td>11790</td><td>25</td><td>ATCTAATTAGTGACGCGCATGAATG</td><td>342.1</td><td>-10.7</td><td>1.3</td><td>-8.1</td><td>4.9E-16</td><td>2.4E-15</td><td>BT20/MB468</td><td>CYTO</td></tr>
<tr><td>>hg38_21_+_8433172_8446622_RNA45SN1_45S_50ntFlanks@11317.11338.22|ACCCCACGTCTCGTCGCGCGCG</td><td>>hg38_21_+_8433172_8446622_RNA45SN1_45S_50ntFlanks</td><td>11317</td><td>11338</td><td>22</td><td>ACCCCACGTCTCGTCGCGCGCG</td><td>2325.3</td><td>-10.3</td><td>1.3</td><td>-8.1</td><td>3.9E-16</td><td>2.4E-15</td><td>MB231/MB468</td><td>NUC</td></tr>
<tr><td>>hg38_21_+_8433172_8446622_RNA45SN1_45S_50ntFlanks@10794.10827.34|CTGGCATGTTGGAACAATGTAGGTAAGGGAAGTC</td><td>>hg38_21_+_8433172_8446622_RNA45SN1_45S_50ntFlanks</td><td>10794</td><td>10827</td><td>34</td><td>CTGGCATGTTGGAACAATGTAGGTAAGGGAAGTC</td><td>235.7</td><td>-11.7</td><td>1.4</td><td>-8.2</td><td>3.1E-16</td><td>2.4E-15</td><td>MB231/MB468</td><td>MP</td></tr>
<tr><td>>trnaMT_GlnTTG_MT_-_4329_4400@1.33.33,trnalookalike2_GlnTTG_1_-_564879_564950@1.33.33|TAGGATGGGGTGTGATAGGTGGCACGGAGAATT</td><td>>trnaMT_GlnTTG_MT_-_4329_4400</td><td>1</td><td>33</td><td>33</td><td>TAGGATGGGGTGTGATAGGTGGCACGGAGAATT</td><td>1658.0</td><td>15.0</td><td>1.8</td><td>8.1</td><td>4.9E-16</td><td>2.4E-15</td><td>BT20/MB468</td><td>CYTO</td></tr>
<tr><td>>hg38_21_+_8433172_8446622_RNA45SN1_45S_50ntFlanks@3938.3959.22|CAACCCGGTCAGCCCCTCTCCG</td><td>>hg38_21_+_8433172_8446622_RNA45SN1_45S_50ntFlanks</td><td>3938</td><td>3959</td><td>22</td><td>CAACCCGGTCAGCCCCTCTCCG</td><td>355.8</td><td>-12.3</td><td>1.5</td><td>-8.1</td><td>5.3E-16</td><td>2.4E-15</td><td>MB231/MB468</td><td>MITO</td></tr>
<tr><td>>hg38_21_+_8433172_8446622_RNA45SN1_45S_50ntFlanks@4560.4587.28|CTAGGAATAATGGAATAGGACCGCGGTT</td><td>>hg38_21_+_8433172_8446622_RNA45SN1_45S_50ntFlanks</td><td>4560</td><td>4587</td><td>28</td><td>CTAGGAATAATGGAATAGGACCGCGGTT</td><td>363.9</td><td>-12.3</td><td>1.5</td><td>-8.1</td><td>5.3E-16</td><td>2.4E-15</td><td>MB231/MB468</td><td>MITO</td></tr>
<tr><td>>hg38_21_+_8433172_8446622_RNA45SN1_45S_50ntFlanks@12126.12151.26|CGACCCGCTCCGGGGACAGTGCCAGG</td><td>>hg38_21_+_8433172_8446622_RNA45SN1_45S_50ntFlanks</td><td>12126</td><td>12151</td><td>26</td><td>CGACCCGCTCCGGGGACAGTGCCAGG</td><td>259.4</td><td>-11.9</td><td>1.5</td><td>-8.2</td><td>3.2E-16</td><td>2.4E-15</td><td>MB231/MB468</td><td>MP</td></tr>
<tr><td>>hg38_21_+_8433172_8446622_RNA45SN1_45S_50ntFlanks@12357.12376.20|CCACAGGGATAACTGGCTTG</td><td>>hg38_21_+_8433172_8446622_RNA45SN1_45S_50ntFlanks</td><td>12357</td><td>12376</td><td>20</td><td>CCACAGGGATAACTGGCTTG</td><td>605.8</td><td>-8.7</td><td>1.1</td><td>-8.2</td><td>3.4E-16</td><td>2.4E-15</td><td>MB231/MB468</td><td>CYTO</td></tr>
<tr><td>>trnaMT_PheGAA_MT_+_577_647@55.74.20|ACATCACCCCATAAACACCA</td><td>>trnaMT_PheGAA_MT_+_577_647</td><td>55</td><td>74</td><td>20</td><td>ACATCACCCCATAAACACCA</td><td>587.9</td><td>13.2</td><td>1.6</td><td>8.2</td><td>3.4E-16</td><td>2.4E-15</td><td>MB231/MB468</td><td>CYTO</td></tr>
<tr><td>>MI0000086|hsa-mir-28&WithFlank&3|+|188688775|188688872@60.82.23&[MIMAT0004502&hsa-miR-28-3p&offsets|0|+1;m-28&3|+|188688834|188688855&offsets|0|+1]|CACTAGATTGTGAGCTCCTGGAG</td><td>>MI0000086|hsa-mir-28&WithFlank&3|+|188688775|188688872</td><td>60</td><td>82</td><td>23</td><td>CACTAGATTGTGAGCTCCTGGAG</td><td>2745.9</td><td>12.3</td><td>1.5</td><td>8.1</td><td>5.0E-16</td><td>2.4E-15</td><td>BT20/MB468</td><td>CYTO</td></tr>
<tr><td>>hg38_21_+_8433172_8446622_RNA45SN1_45S_50ntFlanks@3993.4024.32|GTGACTCTAGATAACCTCGGGCCGATCGCACG</td><td>>hg38_21_+_8433172_8446622_RNA45SN1_45S_50ntFlanks</td><td>3993</td><td>4024</td><td>32</td><td>GTGACTCTAGATAACCTCGGGCCGATCGCACG</td><td>1829.5</td><td>-6.3</td><td>0.8</td><td>-8.1</td><td>7.8E-16</td><td>2.4E-15</td><td>BT20/MB468</td><td>NUC</td></tr>
<tr><td>>hg38_21_+_8433172_8446622_RNA45SN1_45S_50ntFlanks@12405.12427.23|CTTTTTGATCCTTCGATGTCGGC</td><td>>hg38_21_+_8433172_8446622_RNA45SN1_45S_50ntFlanks</td><td>12405</td><td>12427</td><td>23</td><td>CTTTTTGATCCTTCGATGTCGGC</td><td>236.8</td><td>-10.9</td><td>1.3</td><td>-8.1</td><td>3.9E-16</td><td>2.4E-15</td><td>MB231/MB468</td><td>NUC</td></tr>
<tr><td>>hg38_21_+_8433172_8446622_RNA45SN1_45S_50ntFlanks@11641.11659.19|CGGCGGGTGTTGACGCGAT</td><td>>hg38_21_+_8433172_8446622_RNA45SN1_45S_50ntFlanks</td><td>11641</td><td>11659</td><td>19</td><td>CGGCGGGTGTTGACGCGAT</td><td>966.1</td><td>-12.0</td><td>1.5</td><td>-8.2</td><td>3.6E-16</td><td>2.4E-15</td><td>BT20/MB468</td><td>MITO</td></tr>
<tr><td>>hg38_21_+_8433172_8446622_RNA45SN1_45S_50ntFlanks@9570.9597.28|TGCAAATCGGTCGTCCGACCTGGGTATA</td><td>>hg38_21_+_8433172_8446622_RNA45SN1_45S_50ntFlanks</td><td>9570</td><td>9597</td><td>28</td><td>TGCAAATCGGTCGTCCGACCTGGGTATA</td><td>3719.9</td><td>-7.1</td><td>0.9</td><td>-8.2</td><td>3.6E-16</td><td>2.4E-15</td><td>BT20/MB468</td><td>MITO</td></tr>
</table>

<table>
<tr><td>>trna10_AlaCGC_6_+_26553731_26553802@58.75.18,trna119_AlaCGC_6_-_28641613_28641684@58.75.18,trna13_AlaCGC_2_+_157257281_157257352@58.75.18,trna13_AlaTGC_12_-_125406301_125406372@58.75.18,trna66_AlaTGC_6_+_28611222_28611293@58.75.18,trna8_AlaTGC_12_+_125424512_125424583@58.75.18,trna8_AlaTGC_5_+_180633868_180633939@58.75.18|TCCCCGGCATCTCCACCA</td><td>>trna10_AlaCGC_6_+_26553731_26553802</td><td>58</td><td>75</td><td>18</td><td>TCCCCGGCATCTCCACCA</td><td>853.3</td><td>5.9</td><td>0.7</td><td>8.2</td><td>3.6E-16</td><td>2.4E-15</td><td>BT20/MB468</td><td>MITO</td></tr>
<tr><td>>MI0000102|hsa-mir-100&WithFlank&11|-|122152223|122152314@20.39.20(+1A)&[MIMAT0000098&hsa-miR-100-5p&offsets|+1|-1(+1A);m-27&11|-|122152276|122152296&offsets|+1|0(+1A)]|ACCCGTAGATCCGAACTTGTA</td><td>>MI0000102|hsa-mir-100&WithFlank&11|-|122152223|122152314</td><td>20</td><td>39</td><td>20(+1A)</td><td>ACCCGTAGATCCGAACTTGTA</td><td>278.3</td><td>12.1</td><td>1.5</td><td>8.2</td><td>3.5E-16</td><td>2.4E-15</td><td>MB231/MB468</td><td>CYTO</td></tr>
<tr><td>>hg38_1_-_228634819_228635039_RNA5S12_5S_50ntFlanks@61.89.29|ATACCACCCTGAACGCGCCCGATCTCGTC</td><td>>hg38_1_-_228634819_228635039_RNA5S12_5S_50ntFlanks</td><td>61</td><td>89</td><td>29</td><td>ATACCACCCTGAACGCGCCCGATCTCGTC</td><td>1698.6</td><td>11.8</td><td>1.5</td><td>8.2</td><td>3.5E-16</td><td>2.4E-15</td><td>MB231/MB468</td><td>CYTO</td></tr>
<tr><td>>hg38_21_+_8433172_8446622_RNA45SN1_45S_50ntFlanks@11537.11560.24|CCGCGCGGCGCCTCGCCTCGGCCG</td><td>>hg38_21_+_8433172_8446622_RNA45SN1_45S_50ntFlanks</td><td>11537</td><td>11560</td><td>24</td><td>CCGCGCGGCGCCTCGCCTCGGCCG</td><td>254.0</td><td>-10.7</td><td>1.3</td><td>-8.2</td><td>3.5E-16</td><td>2.4E-15</td><td>MB231/MB468</td><td>CYTO</td></tr>
<tr><td>>hg38_21_+_8433172_8446622_RNA45SN1_45S_50ntFlanks@11821.11842.22|CTATCCAGCGAAACCACAGCCA</td><td>>hg38_21_+_8433172_8446622_RNA45SN1_45S_50ntFlanks</td><td>11821</td><td>11842</td><td>22</td><td>CTATCCAGCGAAACCACAGCCA</td><td>406.6</td><td>-12.5</td><td>1.5</td><td>-8.1</td><td>5.3E-16</td><td>2.4E-15</td><td>MB231/MB468</td><td>MITO</td></tr>
<tr><td>>hg38_21_+_8433172_8446622_RNA45SN1_45S_50ntFlanks@5473.5494.22|CGGCCCTGGCGGAGCGCTGAGA</td><td>>hg38_21_+_8433172_8446622_RNA45SN1_45S_50ntFlanks</td><td>5473</td><td>5494</td><td>22</td><td>CGGCCCTGGCGGAGCGCTGAGA</td><td>3175.0</td><td>9.0</td><td>1.1</td><td>8.1</td><td>5.0E-16</td><td>2.5E-15</td><td>BT20/MB468</td><td>CYTO</td></tr>
<tr><td>>MI0000087|hsa-mir-29a&WithFlank&7|-|130876741|130876816@47.70.24&[MIMAT0000086&hsa-miR-29a-3p&offsets|-1|+1;m-26&7|-|130876749|130876769&offsets|-1|+2]|CTAGCACCATCTGAAATCGGTTAT</td><td>>MI0000087|hsa-mir-29a&WithFlank&7|-|130876741|130876816</td><td>47</td><td>70</td><td>24</td><td>CTAGCACCATCTGAAATCGGTTAT</td><td>239.0</td><td>11.8</td><td>1.4</td><td>8.1</td><td>4.0E-16</td><td>2.5E-15</td><td>MB231/MB468</td><td>NUC</td></tr>
<tr><td>>hg38_21_+_8433172_8446622_RNA45SN1_45S_50ntFlanks@10731.10752.22|AGGGTGTAAATCTCGCGCCGGG</td><td>>hg38_21_+_8433172_8446622_RNA45SN1_45S_50ntFlanks</td><td>10731</td><td>10752</td><td>22</td><td>AGGGTGTAAATCTCGCGCCGGG</td><td>355.3</td><td>-7.9</td><td>1.0</td><td>-8.2</td><td>3.5E-16</td><td>2.5E-15</td><td>MB231/MB468</td><td>CYTO</td></tr>
<tr><td>>MI0000255|hsa-mir-30d&WithFlank&8|-|134804870|134804951@12.34.23(+1A)&[MIMAT0000245&hsa-miR-30d-5p&offsets|0|+1(+1A);m-21&8|-|134804919|134804940&offsets|0|+1(+1A)]|TGTAAACATCCCCGACTGGAAGCA</td><td>>MI0000255|hsa-mir-30d&WithFlank&8|-|134804870|134804951</td><td>12</td><td>34</td><td>23(+1A)</td><td>TGTAAACATCCCCGACTGGAAGCA</td><td>1680.2</td><td>4.4</td><td>0.5</td><td>8.1</td><td>8.0E-16</td><td>2.5E-15</td><td>BT20/MB468</td><td>NUC</td></tr>
<tr><td>>am_trna116_GluCTC_1_-_145399233_145399304@1.28.28,trna59_GluCTC_1_+_249168447_249168518@1.28.28,trna71_GluCTC_1_-_161439189_161439260@1.28.28,trna74_GluCTC_1_-_161431809_161431880@1.28.28,trna77_GluCTC_1_-_161424398_161424469@1.28.28,trna77_GluCTC_6_+_28949976_28950047@1.28.28,trna80_GluCTC_1_-_161417018_161417089@1.28.28,trna87_GluCTC_6_-_126101393_126101464@1.28.28|TCCCTGGTGGTCTAGTGGTTAGGATTCG</td><td>>am_trna116_GluCTC_1_-_145399233_145399304</td><td>1</td><td>28</td><td>28</td><td>TCCCTGGTGGTCTAGTGGTTAGGATTCG</td><td>37063.0</td><td>-8.0</td><td>1.0</td><td>-8.1</td><td>8.0E-16</td><td>2.5E-15</td><td>BT20/MB468</td><td>NUC</td></tr>
<tr><td>>hg38_21_+_8433172_8446622_RNA45SN1_45S_50ntFlanks@6671.6688.18|CTCGGCTCGTGCGTCGAT</td><td>>hg38_21_+_8433172_8446622_RNA45SN1_45S_50ntFlanks</td><td>6671</td><td>6688</td><td>18</td><td>CTCGGCTCGTGCGTCGAT</td><td>310.5</td><td>-12.1</td><td>1.5</td><td>-8.2</td><td>3.2E-16</td><td>2.5E-15</td><td>MB231/MB468</td><td>MP</td></tr>
<tr><td>>hg38_21_+_8433172_8446622_RNA45SN1_45S_50ntFlanks@8357.8376.20|AAAAGAACTTTGAAGAGAGA</td><td>>hg38_21_+_8433172_8446622_RNA45SN1_45S_50ntFlanks</td><td>8357</td><td>8376</td><td>20</td><td>AAAAGAACTTTGAAGAGAGA</td><td>1148.7</td><td>-6.5</td><td>0.8</td><td>-8.1</td><td>8.0E-16</td><td>2.5E-15</td><td>BT20/MB468</td><td>NUC</td></tr>
<tr><td>>hg38_21_+_8433172_8446622_RNA45SN1_45S_50ntFlanks@12457.12488.32|CAAGCGTTGGATTGTTCACCCACTAATAGGGA</td><td>>hg38_21_+_8433172_8446622_RNA45SN1_45S_50ntFlanks</td><td>12457</td><td>12488</td><td>32</td><td>CAAGCGTTGGATTGTTCACCCACTAATAGGGA</td><td>343.4</td><td>-12.3</td><td>1.5</td><td>-8.2</td><td>3.3E-16</td><td>2.5E-15</td><td>MB231/MB468</td><td>MP</td></tr>
<tr><td>>hg38_21_+_8433172_8446622_RNA45SN1_45S_50ntFlanks@8350.8373.24|AAAGTTGAAAAGAACTTTGAAGAG</td><td>>hg38_21_+_8433172_8446622_RNA45SN1_45S_50ntFlanks</td><td>8350</td><td>8373</td><td>24</td><td>AAAGTTGAAAAGAACTTTGAAGAG</td><td>965.5</td><td>13.7</td><td>1.6</td><td>8.6</td><td>9.2E-18</td><td>2.5E-15</td><td>BT20/MB468</td><td>MP</td></tr>
<tr><td>>hg38_21_+_8433172_8446622_RNA45SN1_45S_50ntFlanks@12678.12703.26|AATCCCGCCCAGGCGGAACGATACGG</td><td>>hg38_21_+_8433172_8446622_RNA45SN1_45S_50ntFlanks</td><td>12678</td><td>12703</td><td>26</td><td>AATCCCGCCCAGGCGGAACGATACGG</td><td>1846.9</td><td>-13.9</td><td>1.6</td><td>-8.6</td><td>9.3E-18</td><td>2.5E-15</td><td>BT20/MB468</td><td>MP</td></tr>
<tr><td>>hg38_21_+_8433172_8446622_RNA45SN1_45S_50ntFlanks@485.519.35|GATGTGGTGACGTCGTGCTCTCCCGGGCCGGGTCC</td><td>>hg38_21_+_8433172_8446622_RNA45SN1_45S_50ntFlanks</td><td>485</td><td>519</td><td>35</td><td>GATGTGGTGACGTCGTGCTCTCCCGGGCCGGGTCC</td><td>3338.7</td><td>29.5</td><td>3.6</td><td>8.1</td><td>4.0E-16</td><td>2.5E-15</td><td>MB231/MB468</td><td>NUC</td></tr>
<tr><td>>hg38_21_+_8433172_8446622_RNA45SN1_45S_50ntFlanks@5445.5467.23|TCGGATCGGCCCCGCCGGGGTCG</td><td>>hg38_21_+_8433172_8446622_RNA45SN1_45S_50ntFlanks</td><td>5445</td><td>5467</td><td>23</td><td>TCGGATCGGCCCCGCCGGGGTCG</td><td>351.1</td><td>-12.2</td><td>1.5</td><td>-8.1</td><td>5.5E-16</td><td>2.5E-15</td><td>MB231/MB468</td><td>MITO</td></tr>
<tr><td>>MI0000441|hsa-mir-30b&WithFlank&8|-|134800514|134800613@23.43.21(+1C)&[MIMAT0000420&hsa-miR-30b-5p&offsets|0|-1(+1C);m-76&8|-|134800570|134800591&offsets|0|-1(+1C)]|TGTAAACATCCTACACTCAGCC</td><td>>MI0000441|hsa-mir-30b&WithFlank&8|-|134800514|134800613</td><td>23</td><td>43</td><td>21(+1C)</td><td>TGTAAACATCCTACACTCAGCC</td><td>410.6</td><td>6.2</td><td>0.8</td><td>8.1</td><td>4.0E-16</td><td>2.5E-15</td><td>MB231/MB468</td><td>NUC</td></tr>
<tr><td>>hg38_21_+_8433172_8446622_RNA45SN1_45S_50ntFlanks@11316.11334.19|CACCCCACGTCTCGTCGCG</td><td>>hg38_21_+_8433172_8446622_RNA45SN1_45S_50ntFlanks</td><td>11316</td><td>11334</td><td>19</td><td>CACCCCACGTCTCGTCGCG</td><td>256.4</td><td>-11.9</td><td>1.5</td><td>-8.2</td><td>3.3E-16</td><td>2.5E-15</td><td>MB231/MB468</td><td>MP</td></tr>
<tr><td>>hg38_21_+_8433172_8446622_RNA45SN1_45S_50ntFlanks@12943.12972.30|TCGTAGACGACCTGCTTCTGGGTCGGGGTT</td><td>>hg38_21_+_8433172_8446622_RNA45SN1_45S_50ntFlanks</td><td>12943</td><td>12972</td><td>30</td><td>TCGTAGACGACCTGCTTCTGGGTCGGGGTT</td><td>237.9</td><td>-11.8</td><td>1.4</td><td>-8.2</td><td>3.3E-16</td><td>2.5E-15</td><td>MB231/MB468</td><td>MP</td></tr>
<tr><td>>hg38_21_+_8433172_8446622_RNA45SN1_45S_50ntFlanks@12141.12162.22|ACAGTGCCAGGTGGGGAGTTTG</td><td>>hg38_21_+_8433172_8446622_RNA45SN1_45S_50ntFlanks</td><td>12141</td><td>12162</td><td>22</td><td>ACAGTGCCAGGTGGGGAGTTTG</td><td>246.8</td><td>-11.8</td><td>1.4</td><td>-8.2</td><td>3.3E-16</td><td>2.5E-15</td><td>MB231/MB468</td><td>MP</td></tr>
<tr><td>>trna111_HisGTG_1_-_147774845_147774916@-1A.23.24,trna118_HisGTG_1_-_145396881_145396952@-1A.23.24,trna16_HisGTG_1_+_146544773_146544844@-1A.23.24,trna1_HisGTG_15_+_45493349_45493420@-1A.23.24,trna21_HisGTG_1_+_147753471_147753542@-1A.23.24,trna33_HisGTG_6_+_27125906_27125977@-1A.23.24,trna7_HisGTG_9_-_14433938_14434009@-1A.23.24,trna8_HisGTG_15_-_45492611_45492682@-1A.23.24,trna9_HisGTG_15_-_45490804_45490875@-1A.23.24|AGCCGTGATCGTATAGTGGTTAGT</td><td>>trna111_HisGTG_1_-_147774845_147774916</td><td>-1A</td><td>23</td><td>24</td><td>AGCCGTGATCGTATAGTGGTTAGT</td><td>713.8</td><td>11.1</td><td>1.4</td><td>8.1</td><td>4.1E-16</td><td>2.5E-15</td><td>MB231/MB468</td><td>NUC</td></tr>
<tr><td>>hg38_21_+_8433172_8446622_RNA45SN1_45S_50ntFlanks@12957.12979.23|CTTCTGGGTCGGGGTTTCGTACG</td><td>>hg38_21_+_8433172_8446622_RNA45SN1_45S_50ntFlanks</td><td>12957</td><td>12979</td><td>23</td><td>CTTCTGGGTCGGGGTTTCGTACG</td><td>2478.5</td><td>-10.4</td><td>1.3</td><td>-8.1</td><td>4.1E-16</td><td>2.5E-15</td><td>MB231/MB468</td><td>NUC</td></tr>
<tr><td>>MI0000082|hsa-mir-25&WithFlank&7|-|100093554|100093649@58.78.21(+1U)&[MIMAT0000081&hsa-miR-25-3p&offsets|0|-1(+1U);m-23&7|-|100093571|100093592&offsets|0|-1(+1U)]|CATTGCACTTGTCTCGGTCTGT</td><td>>MI0000082|hsa-mir-25&WithFlank&7|-|100093554|100093649</td><td>58</td><td>78</td><td>21(+1U)</td><td>CATTGCACTTGTCTCGGTCTGT</td><td>1875.9</td><td>6.6</td><td>0.8</td><td>8.2</td><td>3.6E-16</td><td>2.5E-15</td><td>MB231/MB468</td><td>CYTO</td></tr>
<tr><td>>hg38_21_+_8433172_8446622_RNA45SN1_45S_50ntFlanks@6731.6770.40|CACATTGATCATCGACACTTCGAACGCACTTGCGGCCCCG</td><td>>hg38_21_+_8433172_8446622_RNA45SN1_45S_50ntFlanks</td><td>6731</td><td>6770</td><td>40</td><td>CACATTGATCATCGACACTTCGAACGCACTTGCGGCCCCG</td><td>1815.9</td><td>-5.2</td><td>0.6</td><td>-8.1</td><td>3.7E-16</td><td>2.5E-15</td><td>BT20/MB468</td><td>MITO</td></tr>
<tr><td>>hg38_21_+_8433172_8446622_RNA45SN1_45S_50ntFlanks@8472.8489.18|CGGCCCGGCGGATCTTTC</td><td>>hg38_21_+_8433172_8446622_RNA45SN1_45S_50ntFlanks</td><td>8472</td><td>8489</td><td>18</td><td>CGGCCCGGCGGATCTTTC</td><td>305.5</td><td>-11.2</td><td>1.4</td><td>-8.1</td><td>4.1E-16</td><td>2.5E-15</td><td>MB231/MB468</td><td>NUC</td></tr>
<tr><td>>hg38_21_+_8433172_8446622_RNA45SN1_45S_50ntFlanks@11576.11593.18|CTTAGAACTGGTGCGGAC</td><td>>hg38_21_+_8433172_8446622_RNA45SN1_45S_50ntFlanks</td><td>11576</td><td>11593</td><td>18</td><td>CTTAGAACTGGTGCGGAC</td><td>1663.1</td><td>-4.9</td><td>0.6</td><td>-8.1</td><td>5.2E-16</td><td>2.5E-15</td><td>BT20/MB468</td><td>CYTO</td></tr>
<tr><td>>hg38_21_+_8433172_8446622_RNA45SN1_45S_50ntFlanks@9533.9571.39|GAGGAAACTCTGGTGGAGGTCCGTAGCGGTCCTGACGTG</td><td>>hg38_21_+_8433172_8446622_RNA45SN1_45S_50ntFlanks</td><td>9533</td><td>9571</td><td>39</td><td>GAGGAAACTCTGGTGGAGGTCCGTAGCGGTCCTGACGTG</td><td>22601.4</td><td>-8.7</td><td>1.1</td><td>-8.1</td><td>8.2E-16</td><td>2.5E-15</td><td>BT20/MB468</td><td>NUC</td></tr>
<tr><td>>hg38_21_+_8433172_8446622_RNA45SN1_45S_50ntFlanks@4995.5034.40|ACAGATTGATAGCTCTTTCTCGATTCCGTGGGTGGTGGTG</td><td>>hg38_21_+_8433172_8446622_RNA45SN1_45S_50ntFlanks</td><td>4995</td><td>5034</td><td>40</td><td>ACAGATTGATAGCTCTTTCTCGATTCCGTGGGTGGTGGTG</td><td>430.5</td><td>-12.5</td><td>1.5</td><td>-8.1</td><td>5.6E-16</td><td>2.5E-15</td><td>MB231/MB468</td><td>MITO</td></tr>
<tr><td>>trnaMT_PheGAA_MT_+_577_647@39.57.19|ATGTTTAGACGGGCTCACA</td><td>>trnaMT_PheGAA_MT_+_577_647</td><td>39</td><td>57</td><td>19</td><td>ATGTTTAGACGGGCTCACA</td><td>2549.1</td><td>5.8</td><td>0.7</td><td>8.1</td><td>3.8E-16</td><td>2.5E-15</td><td>BT20/MB468</td><td>MITO</td></tr>
<tr><td>>hg38_21_+_8433172_8446622_RNA45SN1_45S_50ntFlanks@11874.11904.31|AGAAGACCCTGTTGAGCTTGACTCTAGTCTG</td><td>>hg38_21_+_8433172_8446622_RNA45SN1_45S_50ntFlanks</td><td>11874</td><td>11904</td><td>31</td><td>AGAAGACCCTGTTGAGCTTGACTCTAGTCTG</td><td>573.7</td><td>-11.3</td><td>1.4</td><td>-8.1</td><td>3.8E-16</td><td>2.5E-15</td><td>BT20/MB468</td><td>MITO</td></tr>
<tr><td>>MI0000111|hsa-mir-105-1&WithFlank&X|-|152392213|152392305@19.40.22;MI0000112|hsa-mir-105-2&WithFlank&X|-|152394406|152394498@19.40.22&[MIMAT0000102&hsa-miR-105-5p&offsets|0|-1;m-735&X|-|152392266|152392287&offsets|0|0];[MIMAT0000102_1&hsa-miR-105-5p&offsets|0|-1;m-735&X|-|152394459|152394480&offsets|0|0]|TCAAATGCTCAGACTCCTGTGG</td><td>>MI0000111|hsa-mir-105-1&WithFlank&X|-|152392213|152392305</td><td>19</td><td>40</td><td>22</td><td>TCAAATGCTCAGACTCCTGTGG</td><td>264.8</td><td>11.7</td><td>1.4</td><td>8.4</td><td>4.3E-17</td><td>2.6E-15</td><td>BT20/MB231</td><td>MITO</td></tr>
<tr><td>>hg38_21_+_8433172_8446622_RNA45SN1_45S_50ntFlanks@11768.11792.25|CTAATTAGTGACGCGCATGAATGGA</td><td>>hg38_21_+_8433172_8446622_RNA45SN1_45S_50ntFlanks</td><td>11768</td><td>11792</td><td>25</td><td>CTAATTAGTGACGCGCATGAATGGA</td><td>3705.5</td><td>-10.5</td><td>1.3</td><td>-8.1</td><td>4.2E-16</td><td>2.6E-15</td><td>MB231/MB468</td><td>NUC</td></tr>
<tr><td>>MI0000253|hsa-mir-148a&WithFlank&7|-|25949913|25949992@50.73.24&[MIMAT0000243&hsa-miR-148a-3p&offsets|0|+2;m-7&7|-|25949922|25949943&offsets|0|+2]|TCAGTGCACTACAGAACTTTGTCT</td><td>>MI0000253|hsa-mir-148a&WithFlank&7|-|25949913|25949992</td><td>50</td><td>73</td><td>24</td><td>TCAGTGCACTACAGAACTTTGTCT</td><td>486.7</td><td>7.8</td><td>1.0</td><td>8.1</td><td>5.2E-16</td><td>2.6E-15</td><td>BT20/MB468</td><td>CYTO</td></tr>
<tr><td>>hg38_21_+_8433172_8446622_RNA45SN1_45S_50ntFlanks@6732.6777.46|ACATTGATCATCGACACTTCGAACGCACTTGCGGCCCCGGGTTCCT</td><td>>hg38_21_+_8433172_8446622_RNA45SN1_45S_50ntFlanks</td><td>6732</td><td>6777</td><td>46</td><td>ACATTGATCATCGACACTTCGAACGCACTTGCGGCCCCGGGTTCCT</td><td>164.4</td><td>11.4</td><td>1.4</td><td>8.1</td><td>3.6E-16</td><td>2.6E-15</td><td>MB231/MB468</td><td>CYTO</td></tr>
<tr><td>>hg38_21_+_8433172_8446622_RNA45SN1_45S_50ntFlanks@9736.9758.23|CCGAAACGATCTCAACCTATTCT</td><td>>hg38_21_+_8433172_8446622_RNA45SN1_45S_50ntFlanks</td><td>9736</td><td>9758</td><td>23</td><td>CCGAAACGATCTCAACCTATTCT</td><td>239.8</td><td>-11.8</td><td>1.4</td><td>-8.2</td><td>3.4E-16</td><td>2.6E-15</td><td>MB231/MB468</td><td>MP</td></tr>
<tr><td>>hg38_21_+_8433172_8446622_RNA45SN1_45S_50ntFlanks@12539.12587.49|ATGATGTGTTGTTGCCATGGTAATCCTGCTCAGTACGAGAGGAACCGCA</td><td>>hg38_21_+_8433172_8446622_RNA45SN1_45S_50ntFlanks</td><td>12539</td><td>12587</td><td>49</td><td>ATGATGTGTTGTTGCCATGGTAATCCTGCTCAGTACGAGAGGAACCGCA</td><td>1240.8</td><td>-14.1</td><td>1.7</td><td>-8.1</td><td>5.6E-16</td><td>2.6E-15</td><td>MB231/MB468</td><td>MITO</td></tr>
<tr><td>>hg38_21_+_8433172_8446622_RNA45SN1_45S_50ntFlanks@6740.6770.31|CATCGACACTTCGAACGCACTTGCGGCCCCG</td><td>>hg38_21_+_8433172_8446622_RNA45SN1_45S_50ntFlanks</td><td>6740</td><td>6770</td><td>31</td><td>CATCGACACTTCGAACGCACTTGCGGCCCCG</td><td>1009.1</td><td>-6.5</td><td>0.8</td><td>-8.1</td><td>3.7E-16</td><td>2.6E-15</td><td>MB231/MB468</td><td>CYTO</td></tr>
<tr><td>>hg38_21_+_8433172_8446622_RNA45SN1_45S_50ntFlanks@10097.10115.19|CGCGCGCGCGCGTGTGGTG</td><td>>hg38_21_+_8433172_8446622_RNA45SN1_45S_50ntFlanks</td><td>10097</td><td>10115</td><td>19</td><td>CGCGCGCGCGCGTGTGGTG</td><td>249.3</td><td>-11.8</td><td>1.4</td><td>-8.2</td><td>3.4E-16</td><td>2.6E-15</td><td>MB231/MB468</td><td>MP</td></tr>
<tr><td>>hg38_21_+_8433172_8446622_RNA45SN1_45S_50ntFlanks@9545.9567.23|GTGGAGGTCCGTAGCGGTCCTGA</td><td>>hg38_21_+_8433172_8446622_RNA45SN1_45S_50ntFlanks</td><td>9545</td><td>9567</td><td>23</td><td>GTGGAGGTCCGTAGCGGTCCTGA</td><td>236.4</td><td>-11.7</td><td>1.4</td><td>-8.2</td><td>3.4E-16</td><td>2.6E-15</td><td>MB231/MB468</td><td>MP</td></tr>
<tr><td>>MI0000542|hsa-mir-320a&WithFlank&8|-|22244960|22245043@48.68.21(+1U)&[MIMAT0000510&hsa-miR-320a-3p&offsets|0|-1(+1U);m-60&8|-|22244975|22244996&offsets|0|-1(+1U)]|AAAAGCTGGGTTGAGAGGGCGT</td><td>>MI0000542|hsa-mir-320a&WithFlank&8|-|22244960|22245043</td><td>48</td><td>68</td><td>21(+1U)</td><td>AAAAGCTGGGTTGAGAGGGCGT</td><td>1098.2</td><td>3.6</td><td>0.4</td><td>8.0</td><td>8.3E-16</td><td>2.6E-15</td><td>BT20/MB468</td><td>NUC</td></tr>
<tr><td>>trna32_LysCTT_16_-_3207406_3207478@1.33.33|GCCCGGCTAGCTCAGTCGGTAGAGCATGAGACC</td><td>>trna32_LysCTT_16_-_3207406_3207478</td><td>1</td><td>33</td><td>33</td><td>GCCCGGCTAGCTCAGTCGGTAGAGCATGAGACC</td><td>7373.1</td><td>8.5</td><td>1.0</td><td>8.1</td><td>3.7E-16</td><td>2.6E-15</td><td>MB231/MB468</td><td>CYTO</td></tr>
<tr><td>>hg38_21_+_8433172_8446622_RNA45SN1_45S_50ntFlanks@11320.11342.23|CCACGTCTCGTCGCGCGCGCGTC</td><td>>hg38_21_+_8433172_8446622_RNA45SN1_45S_50ntFlanks</td><td>11320</td><td>11342</td><td>23</td><td>CCACGTCTCGTCGCGCGCGCGTC</td><td>432.0</td><td>-12.6</td><td>1.5</td><td>-8.2</td><td>3.4E-16</td><td>2.6E-15</td><td>MB231/MB468</td><td>MP</td></tr>
</table>

<table>
<tr><td>>hg38_21_+_8433172_8446622_RNA45SN1_45S_50ntFlanks@10771.10805.35|CAGGTCTCCAAGGTGAACAGCCTCTGGCATGTTGG</td><td>>hg38_21_+_8433172_8446622_RNA45SN1_45S_50ntFlanks</td><td>10771</td><td>10805</td><td>35</td><td>CAGGTCTCCAAGGTGAACAGCCTCTGGCATGTTGG</td><td>1615.9</td><td>-9.7</td><td>1.2</td><td>-8.1</td><td>5.7E-16</td><td>2.6E-15</td><td>MB231/MB468</td><td>MITO</td></tr>
<tr><td>>hg38_21_+_8433172_8446622_RNA45SN1_45S_50ntFlanks@12324.12347.24|TTGGGTTTTAAGCAGGAGGTGTCA</td><td>>hg38_21_+_8433172_8446622_RNA45SN1_45S_50ntFlanks</td><td>12324</td><td>12347</td><td>24</td><td>TTGGGTTTTAAGCAGGAGGTGTCA</td><td>219.2</td><td>-11.6</td><td>1.4</td><td>-8.2</td><td>3.4E-16</td><td>2.6E-15</td><td>MB231/MB468</td><td>MP</td></tr>
<tr><td>>hg38_21_+_8433172_8446622_RNA45SN1_45S_50ntFlanks@10805.10823.19|GAACAATGTAGGTAAGGGA</td><td>>hg38_21_+_8433172_8446622_RNA45SN1_45S_50ntFlanks</td><td>10805</td><td>10823</td><td>19</td><td>GAACAATGTAGGTAAGGGA</td><td>283.3</td><td>-10.9</td><td>1.3</td><td>-8.1</td><td>3.7E-16</td><td>2.6E-15</td><td>MB231/MB468</td><td>CYTO</td></tr>
<tr><td>>hg38_21_+_8433172_8446622_RNA45SN1_45S_50ntFlanks@12676.12708.33|AGAATCCCGCCCAGGCGGAACGATACGGCAGCG</td><td>>hg38_21_+_8433172_8446622_RNA45SN1_45S_50ntFlanks</td><td>12676</td><td>12708</td><td>33</td><td>AGAATCCCGCCCAGGCGGAACGATACGGCAGCG</td><td>272.7</td><td>-10.8</td><td>1.3</td><td>-8.1</td><td>3.7E-16</td><td>2.6E-15</td><td>MB231/MB468</td><td>CYTO</td></tr>
<tr><td>>hg38_21_+_8433172_8446622_RNA45SN1_45S_50ntFlanks@9120.9138.19|TGCCGCGCGCGGGTCGGGG</td><td>>hg38_21_+_8433172_8446622_RNA45SN1_45S_50ntFlanks</td><td>9120</td><td>9138</td><td>19</td><td>TGCCGCGCGCGGGTCGGGG</td><td>310.8</td><td>-12.1</td><td>1.5</td><td>-8.1</td><td>5.8E-16</td><td>2.6E-15</td><td>MB231/MB468</td><td>MITO</td></tr>
<tr><td>>MI0000650|hsa-mir-200c&WithFlank&12|+|6963693|6963772@50.72.23(+2U)&[MIMAT0000617&hsa-miR-200c-3p&offsets|0|0(+2U);m-54&12|+|6963742|6963764&offsets|0|0(+2U)]|TAATACTGCCGGGTAATGATGGATT</td><td>>MI0000650|hsa-mir-200c&WithFlank&12|+|6963693|6963772</td><td>50</td><td>72</td><td>23(+2U)</td><td>TAATACTGCCGGGTAATGATGGATT</td><td>282.8</td><td>11.8</td><td>1.4</td><td>8.4</td><td>4.6E-17</td><td>2.6E-15</td><td>BT20/MB231</td><td>MITO</td></tr>
<tr><td>>hg38_21_+_8433172_8446622_RNA45SN1_45S_50ntFlanks@10797.10816.20|GCATGTTGGAACAATGTAGG</td><td>>hg38_21_+_8433172_8446622_RNA45SN1_45S_50ntFlanks</td><td>10797</td><td>10816</td><td>20</td><td>GCATGTTGGAACAATGTAGG</td><td>217.6</td><td>-11.6</td><td>1.4</td><td>-8.2</td><td>3.5E-16</td><td>2.7E-15</td><td>MB231/MB468</td><td>MP</td></tr>
<tr><td>>hg38_21_+_8433172_8446622_RNA45SN1_45S_50ntFlanks@485.508.24|GATGTGGTGACGTCGTGCTCTCCC</td><td>>hg38_21_+_8433172_8446622_RNA45SN1_45S_50ntFlanks</td><td>485</td><td>508</td><td>24</td><td>GATGTGGTGACGTCGTGCTCTCCC</td><td>2990.2</td><td>29.5</td><td>3.6</td><td>8.1</td><td>4.3E-16</td><td>2.7E-15</td><td>MB231/MB468</td><td>NUC</td></tr>
<tr><td>>hg38_21_+_8433172_8446622_RNA45SN1_45S_50ntFlanks@3771.3792.22|CACGGCCGGTACAGTGAAACTG</td><td>>hg38_21_+_8433172_8446622_RNA45SN1_45S_50ntFlanks</td><td>3771</td><td>3792</td><td>22</td><td>CACGGCCGGTACAGTGAAACTG</td><td>1072.9</td><td>-7.9</td><td>1.0</td><td>-8.1</td><td>4.3E-16</td><td>2.7E-15</td><td>MB231/MB468</td><td>NUC</td></tr>
<tr><td>>hg38_21_+_8433172_8446622_RNA45SN1_45S_50ntFlanks@8107.8138.32|CGGCGGGGCGCGGGACATGTGGCGTACGGAAG</td><td>>hg38_21_+_8433172_8446622_RNA45SN1_45S_50ntFlanks</td><td>8107</td><td>8138</td><td>32</td><td>CGGCGGGGCGCGGGACATGTGGCGTACGGAAG</td><td>548.0</td><td>-12.9</td><td>1.6</td><td>-8.1</td><td>5.8E-16</td><td>2.7E-15</td><td>MB231/MB468</td><td>MITO</td></tr>
<tr><td>>MI0000286|hsa-mir-210&WithFlank&11|-|568083|568204@72.92.21(+1U)&[MIMAT0000267&hsa-miR-210-3p&offsets|0|-1(+1U);m-102&11|-|568112|568133&offsets|0|-1(+1U)]|CTGTGCGTGTGACAGCGGCTGT</td><td>>MI0000286|hsa-mir-210&WithFlank&11|-|568083|568204</td><td>72</td><td>92</td><td>21(+1U)</td><td>CTGTGCGTGTGACAGCGGCTGT</td><td>364.2</td><td>12.8</td><td>1.6</td><td>8.1</td><td>5.5E-16</td><td>2.7E-15</td><td>BT20/MB468</td><td>CYTO</td></tr>
<tr><td>>hg38_21_+_8433172_8446622_RNA45SN1_45S_50ntFlanks@10109.10127.19|TGTGGTGTGCGTCGGAGGG</td><td>>hg38_21_+_8433172_8446622_RNA45SN1_45S_50ntFlanks</td><td>10109</td><td>10127</td><td>19</td><td>TGTGGTGTGCGTCGGAGGG</td><td>273.7</td><td>-12.0</td><td>1.5</td><td>-8.2</td><td>3.5E-16</td><td>2.7E-15</td><td>MB231/MB468</td><td>MP</td></tr>
<tr><td>>hg38_21_+_8433172_8446622_RNA45SN1_45S_50ntFlanks@9738.9757.20|GAAACGATCTCAACCTATTC</td><td>>hg38_21_+_8433172_8446622_RNA45SN1_45S_50ntFlanks</td><td>9738</td><td>9757</td><td>20</td><td>GAAACGATCTCAACCTATTC</td><td>548.8</td><td>-12.9</td><td>1.6</td><td>-8.1</td><td>5.9E-16</td><td>2.7E-15</td><td>MB231/MB468</td><td>MITO</td></tr>
<tr><td>>MI0001445|hsa-mir-423&WithFlank&17|+|30117073|30117178@58.81.24&[MIMAT0001340&hsa-miR-423-3p&offsets|-1|0;m-100&17|+|30117131|30117153&offsets|-1|0]|AAGCTCGGTCTGAGGCCCCTCAGT</td><td>>MI0001445|hsa-mir-423&WithFlank&17|+|30117073|30117178</td><td>58</td><td>81</td><td>24</td><td>AAGCTCGGTCTGAGGCCCCTCAGT</td><td>778.3</td><td>3.3</td><td>0.4</td><td>8.0</td><td>8.6E-16</td><td>2.7E-15</td><td>BT20/MB468</td><td>NUC</td></tr>
<tr><td>>trna11_GluTTC_15_-_26327381_26327452@1.35.35,trna3_GluTTC_13_-_45492062_45492133@1.35.35|TCCCACATGGTCTAGCGGTTAGGATTCCTGGTTTT</td><td>>trna11_GluTTC_15_-_26327381_26327452</td><td>1</td><td>35</td><td>35</td><td>TCCCACATGGTCTAGCGGTTAGGATTCCTGGTTTT</td><td>443.7</td><td>10.4</td><td>1.3</td><td>8.1</td><td>5.5E-16</td><td>2.7E-15</td><td>BT20/MB468</td><td>CYTO</td></tr>
<tr><td>>hg38_21_+_8433172_8446622_RNA45SN1_45S_50ntFlanks@8379.8400.22|TCAAGAGGGCGTGAAACCGTTA</td><td>>hg38_21_+_8433172_8446622_RNA45SN1_45S_50ntFlanks</td><td>8379</td><td>8400</td><td>22</td><td>TCAAGAGGGCGTGAAACCGTTA</td><td>402.7</td><td>-12.5</td><td>1.5</td><td>-8.2</td><td>3.5E-16</td><td>2.7E-15</td><td>MB231/MB468</td><td>MP</td></tr>
<tr><td>>MI0000102|hsa-mir-100&WithFlank&11|-|122152223|122152314@19.40.22(+1U)&[MIMAT0000098&hsa-miR-100-5p&offsets|0|0(+1U);m-27&11|-|122152276|122152296&offsets|0|+1(+1U)]|AACCCGTAGATCCGAACTTGTGT</td><td>>MI0000102|hsa-mir-100&WithFlank&11|-|122152223|122152314</td><td>19</td><td>40</td><td>22(+1U)</td><td>AACCCGTAGATCCGAACTTGTGT</td><td>57343.1</td><td>-8.2</td><td>1.0</td><td>-8.4</td><td>4.7E-17</td><td>2.7E-15</td><td>BT20/MB231</td><td>MITO</td></tr>
<tr><td>>hg38_21_+_8433172_8446622_RNA45SN1_45S_50ntFlanks@6783.6803.21|GGCTACGCCTGTCTGAGCGTC</td><td>>hg38_21_+_8433172_8446622_RNA45SN1_45S_50ntFlanks</td><td>6783</td><td>6803</td><td>21</td><td>GGCTACGCCTGTCTGAGCGTC</td><td>251.2</td><td>-11.8</td><td>1.5</td><td>-8.2</td><td>3.5E-16</td><td>2.7E-15</td><td>MB231/MB468</td><td>MP</td></tr>
<tr><td>>hg38_21_+_8433172_8446622_RNA45SN1_45S_50ntFlanks@10511.10533.23|ATCCGGAGTGGCGGAGATGGGCG</td><td>>hg38_21_+_8433172_8446622_RNA45SN1_45S_50ntFlanks</td><td>10511</td><td>10533</td><td>23</td><td>ATCCGGAGTGGCGGAGATGGGCG</td><td>1691.4</td><td>-13.8</td><td>1.6</td><td>-8.6</td><td>1.0E-17</td><td>2.7E-15</td><td>BT20/MB468</td><td>MP</td></tr>
<tr><td>>hg38_21_+_8433172_8446622_RNA45SN1_45S_50ntFlanks@13002.13028.27|CGATCTATTGAAAGTCAGCCCTCGACA</td><td>>hg38_21_+_8433172_8446622_RNA45SN1_45S_50ntFlanks</td><td>13002</td><td>13028</td><td>27</td><td>CGATCTATTGAAAGTCAGCCCTCGACA</td><td>1040.6</td><td>-13.1</td><td>1.5</td><td>-8.6</td><td>1.1E-17</td><td>2.7E-15</td><td>BT20/MB468</td><td>MP</td></tr>
<tr><td>>hg38_21_+_8433172_8446622_RNA45SN1_45S_50ntFlanks@9575.9603.29|ATCGGTCGTCCGACCTGGGTATAGGGGCG</td><td>>hg38_21_+_8433172_8446622_RNA45SN1_45S_50ntFlanks</td><td>9575</td><td>9603</td><td>29</td><td>ATCGGTCGTCCGACCTGGGTATAGGGGCG</td><td>2078.9</td><td>-14.1</td><td>1.6</td><td>-8.6</td><td>1.1E-17</td><td>2.7E-15</td><td>BT20/MB468</td><td>MP</td></tr>
<tr><td>>MI0000087|hsa-mir-29a&WithFlank&7|-|130876741|130876816@47.67.21(+1G)&[MIMAT0000086&hsa-miR-29a-3p&offsets|-1|-2(+1G);m-26&7|-|130876749|130876769&offsets|-1|-1(+1G)]|CTAGCACCATCTGAAATCGGTG</td><td>>MI0000087|hsa-mir-29a&WithFlank&7|-|130876741|130876816</td><td>47</td><td>67</td><td>21(+1G)</td><td>CTAGCACCATCTGAAATCGGTG</td><td>173.3</td><td>11.3</td><td>1.4</td><td>8.1</td><td>4.4E-16</td><td>2.7E-15</td><td>MB231/MB468</td><td>NUC</td></tr>
<tr><td>>hg38_21_+_8433172_8446622_RNA45SN1_45S_50ntFlanks@12626.12661.36|AATGGGGCGAAGCTACCATCTGTGGGATTATGACTG</td><td>>hg38_21_+_8433172_8446622_RNA45SN1_45S_50ntFlanks</td><td>12626</td><td>12661</td><td>36</td><td>AATGGGGCGAAGCTACCATCTGTGGGATTATGACTG</td><td>1415.2</td><td>-4.5</td><td>0.6</td><td>-8.1</td><td>4.4E-16</td><td>2.7E-15</td><td>MB231/MB468</td><td>NUC</td></tr>
<tr><td>>hg38_21_+_8433172_8446622_RNA45SN1_45S_50ntFlanks@9534.9566.33|AGGAAACTCTGGTGGAGGTCCGTAGCGGTCCTG</td><td>>hg38_21_+_8433172_8446622_RNA45SN1_45S_50ntFlanks</td><td>9534</td><td>9566</td><td>33</td><td>AGGAAACTCTGGTGGAGGTCCGTAGCGGTCCTG</td><td>9171.4</td><td>-12.4</td><td>1.5</td><td>-8.0</td><td>8.8E-16</td><td>2.7E-15</td><td>BT20/MB468</td><td>NUC</td></tr>
<tr><td>>MI0000102|hsa-mir-100&WithFlank&11|-|122152223|122152314@19.39.21(+1U)&[MIMAT0000098&hsa-miR-100-5p&offsets|0|-1(+1U);m-27&11|-|122152276|122152296&offsets|0|0(+1U)]|AACCCGTAGATCCGAACTTGTT</td><td>>MI0000102|hsa-mir-100&WithFlank&11|-|122152223|122152314</td><td>19</td><td>39</td><td>21(+1U)</td><td>AACCCGTAGATCCGAACTTGTT</td><td>6818.0</td><td>12.4</td><td>1.5</td><td>8.1</td><td>6.0E-16</td><td>2.7E-15</td><td>MB231/MB468</td><td>MITO</td></tr>
<tr><td>>hg38_21_+_8433172_8446622_RNA45SN1_45S_50ntFlanks@10664.10712.49|CTTGGAAAGCGTCGCGGTTCCGGCGGCGTCCGGTGAGCTCTCGCTGGCC</td><td>>hg38_21_+_8433172_8446622_RNA45SN1_45S_50ntFlanks</td><td>10664</td><td>10712</td><td>49</td><td>CTTGGAAAGCGTCGCGGTTCCGGCGGCGTCCGGTGAGCTCTCGCTGGCC</td><td>454.3</td><td>-12.6</td><td>1.6</td><td>-8.1</td><td>6.0E-16</td><td>2.7E-15</td><td>MB231/MB468</td><td>MITO</td></tr>
<tr><td>>hg38_21_+_8433172_8446622_RNA45SN1_45S_50ntFlanks@8570.8589.20|CTCCGGCGGGTGCGGGGGTG</td><td>>hg38_21_+_8433172_8446622_RNA45SN1_45S_50ntFlanks</td><td>8570</td><td>8589</td><td>20</td><td>CTCCGGCGGGTGCGGGGGTG</td><td>253.1</td><td>-11.8</td><td>1.5</td><td>-8.2</td><td>3.6E-16</td><td>2.7E-15</td><td>MB231/MB468</td><td>MP</td></tr>
<tr><td>>hg38_21_+_8433172_8446622_RNA45SN1_45S_50ntFlanks@12893.12932.40|CCTCGCCCGTCACGCACCGCACGTTCGTGGGGAACCTGGC</td><td>>hg38_21_+_8433172_8446622_RNA45SN1_45S_50ntFlanks</td><td>12893</td><td>12932</td><td>40</td><td>CCTCGCCCGTCACGCACCGCACGTTCGTGGGGAACCTGGC</td><td>1445.5</td><td>-14.4</td><td>1.8</td><td>-8.2</td><td>3.6E-16</td><td>2.7E-15</td><td>MB231/MB468</td><td>MP</td></tr>
<tr><td>>hg38_21_+_8433172_8446622_RNA45SN1_45S_50ntFlanks@8381.8399.19|AAGAGGGCGTGAAACCGTT</td><td>>hg38_21_+_8433172_8446622_RNA45SN1_45S_50ntFlanks</td><td>8381</td><td>8399</td><td>19</td><td>AAGAGGGCGTGAAACCGTT</td><td>195.5</td><td>-5.8</td><td>0.7</td><td>-8.1</td><td>3.9E-16</td><td>2.7E-15</td><td>MB231/MB468</td><td>CYTO</td></tr>
<tr><td>>MI0000471|hsa-mir-126&WithFlank&9|+|136670596|136670692@58.78.21(+1A)&[MIMAT0000445&hsa-miR-126-3p&offsets|0|-1(+1A);m-40&9|+|136670653|136670674&offsets|0|-1(+1A)]|TCGTACCGTGAGTAATAATGCA</td><td>>MI0000471|hsa-mir-126&WithFlank&9|+|136670596|136670692</td><td>58</td><td>78</td><td>21(+1A)</td><td>TCGTACCGTGAGTAATAATGCA</td><td>1716.6</td><td>12.4</td><td>1.5</td><td>8.1</td><td>5.6E-16</td><td>2.7E-15</td><td>BT20/MB468</td><td>CYTO</td></tr>
<tr><td>>MI0000815|hsa-mir-339&WithFlank&7|-|1022927|1023032@21.43.23&[MIMAT0000764&hsa-miR-339-5p&offsets|0|0;m-163&7|-|1022990|1023012&offsets|0|0]|TCCCTGTCCTCCAGGAGCTCACG</td><td>>MI0000815|hsa-mir-339&WithFlank&7|-|1022927|1023032</td><td>21</td><td>43</td><td>23</td><td>TCCCTGTCCTCCAGGAGCTCACG</td><td>766.0</td><td>6.9</td><td>0.9</td><td>8.1</td><td>4.0E-16</td><td>2.7E-15</td><td>BT20/MB468</td><td>MITO</td></tr>
<tr><td>>hg38_21_+_8433172_8446622_RNA45SN1_45S_50ntFlanks@12621.12654.34|GAGCCAATGGGGCGAAGCTACCATCTGTGGGATT</td><td>>hg38_21_+_8433172_8446622_RNA45SN1_45S_50ntFlanks</td><td>12621</td><td>12654</td><td>34</td><td>GAGCCAATGGGGCGAAGCTACCATCTGTGGGATT</td><td>609.1</td><td>-9.3</td><td>1.1</td><td>-8.1</td><td>6.0E-16</td><td>2.7E-15</td><td>MB231/MB468</td><td>MITO</td></tr>
<tr><td>>hg38_21_+_8433172_8446622_RNA45SN1_45S_50ntFlanks@10116.10134.19|TGCGTCGGAGGGCGGCGGC</td><td>>hg38_21_+_8433172_8446622_RNA45SN1_45S_50ntFlanks</td><td>10116</td><td>10134</td><td>19</td><td>TGCGTCGGAGGGCGGCGGC</td><td>309.1</td><td>-12.1</td><td>1.5</td><td>-8.1</td><td>6.0E-16</td><td>2.7E-15</td><td>MB231/MB468</td><td>MITO</td></tr>
<tr><td>>hg38_21_+_8433172_8446622_RNA45SN1_45S_50ntFlanks@12841.12862.22|CGGAGTGCCCTTCGTCCTGGGA</td><td>>hg38_21_+_8433172_8446622_RNA45SN1_45S_50ntFlanks</td><td>12841</td><td>12862</td><td>22</td><td>CGGAGTGCCCTTCGTCCTGGGA</td><td>307.7</td><td>-12.0</td><td>1.5</td><td>-8.1</td><td>6.0E-16</td><td>2.7E-15</td><td>MB231/MB468</td><td>MITO</td></tr>
<tr><td>>MI0000542|hsa-mir-320a&WithFlank&8|-|22244960|22245043@48.69.22&[MIMAT0000510&hsa-miR-320a-3p&offsets|0|0;m-60&8|-|22244975|22244996&offsets|0|0]|AAAAGCTGGGTTGAGAGGGCGA</td><td>>MI0000542|hsa-mir-320a&WithFlank&8|-|22244960|22245043</td><td>48</td><td>69</td><td>22</td><td>AAAAGCTGGGTTGAGAGGGCGA</td><td>3545.3</td><td>3.8</td><td>0.5</td><td>8.0</td><td>8.9E-16</td><td>2.7E-15</td><td>BT20/MB468</td><td>NUC</td></tr>
<tr><td>>hg38_21_+_8433172_8446622_RNA45SN1_45S_50ntFlanks@12717.12742.26|CCTCGGTTGGCCTCGGATAGCCGGTC</td><td>>hg38_21_+_8433172_8446622_RNA45SN1_45S_50ntFlanks</td><td>12717</td><td>12742</td><td>26</td><td>CCTCGGTTGGCCTCGGATAGCCGGTC</td><td>675.9</td><td>-13.2</td><td>1.6</td><td>-8.1</td><td>6.1E-16</td><td>2.7E-15</td><td>MB231/MB468</td><td>MITO</td></tr>
<tr><td>>hg38_1_-_228634819_228635039_RNA5S12_5S_50ntFlanks@138.171.34|AGACCGCCTGGGAATACCGGGTGCTGTAGGCTTT</td><td>>hg38_1_-_228634819_228635039_RNA5S12_5S_50ntFlanks</td><td>138</td><td>171</td><td>34</td><td>AGACCGCCTGGGAATACCGGGTGCTGTAGGCTTT</td><td>1663.9</td><td>6.8</td><td>0.8</td><td>8.1</td><td>5.7E-16</td><td>2.7E-15</td><td>BT20/MB468</td><td>CYTO</td></tr>
<tr><td>>hg38_21_+_8433172_8446622_RNA45SN1_45S_50ntFlanks@4854.4883.30|AGTCTTTGGGTTCCGGGGGGAGTATGGTTG</td><td>>hg38_21_+_8433172_8446622_RNA45SN1_45S_50ntFlanks</td><td>4854</td><td>4883</td><td>30</td><td>AGTCTTTGGGTTCCGGGGGGAGTATGGTTG</td><td>274.7</td><td>-11.1</td><td>1.4</td><td>-8.1</td><td>4.5E-16</td><td>2.8E-15</td><td>MB231/MB468</td><td>NUC</td></tr>
<tr><td>>hg38_21_+_8433172_8446622_RNA45SN1_45S_50ntFlanks@3985.4005.21|CGGCTTTGGTGACTCTAGATA</td><td>>hg38_21_+_8433172_8446622_RNA45SN1_45S_50ntFlanks</td><td>3985</td><td>4005</td><td>21</td><td>CGGCTTTGGTGACTCTAGATA</td><td>1238.6</td><td>-9.7</td><td>1.2</td><td>-8.1</td><td>4.5E-16</td><td>2.8E-15</td><td>MB231/MB468</td><td>NUC</td></tr>
<tr><td>>hg38_21_+_8433172_8446622_RNA45SN1_45S_50ntFlanks@12412.12432.21|ATCCTTCGATGTCGGCTCTTC</td><td>>hg38_21_+_8433172_8446622_RNA45SN1_45S_50ntFlanks</td><td>12412</td><td>12432</td><td>21</td><td>ATCCTTCGATGTCGGCTCTTC</td><td>294.1</td><td>-12.1</td><td>1.5</td><td>-8.2</td><td>3.6E-16</td><td>2.8E-15</td><td>MB231/MB468</td><td>MP</td></tr>
<tr><td>>hg38_21_+_8433172_8446622_RNA45SN1_45S_50ntFlanks@11616.11640.25|TAAAACAAAGCATCGCGAAGGCCCG</td><td>>hg38_21_+_8433172_8446622_RNA45SN1_45S_50ntFlanks</td><td>11616</td><td>11640</td><td>25</td><td>TAAAACAAAGCATCGCGAAGGCCCG</td><td>187.9</td><td>-11.4</td><td>1.4</td><td>-8.2</td><td>3.6E-16</td><td>2.8E-15</td><td>MB231/MB468</td><td>MP</td></tr>
<tr><td>>hg38_21_+_8433172_8446622_RNA45SN1_45S_50ntFlanks@12998.13017.20|GCTGCGATCTATTGAAAGTC</td><td>>hg38_21_+_8433172_8446622_RNA45SN1_45S_50ntFlanks</td><td>12998</td><td>13017</td><td>20</td><td>GCTGCGATCTATTGAAAGTC</td><td>248.5</td><td>7.8</td><td>1.0</td><td>8.1</td><td>4.0E-16</td><td>2.8E-15</td><td>MB231/MB468</td><td>CYTO</td></tr>
<tr><td>>MI0000477|hsa-mir-146a&WithFlank&5|+|160485346|160485456@27.47.21(+1G)&[MIMAT0000449&hsa-miR-146a-5p&offsets|0|-1(+1G);m-129&5|+|160485372|160485393&offsets|0|-1(+1G)]|TGAGAACTGAATTCCATGGGTG</td><td>>MI0000477|hsa-mir-146a&WithFlank&5|+|160485346|160485456</td><td>27</td><td>47</td><td>21(+1G)</td><td>TGAGAACTGAATTCCATGGGTG</td><td>1311.5</td><td>13.6</td><td>1.7</td><td>8.1</td><td>3.6E-16</td><td>2.8E-15</td><td>MB231/MB468</td><td>MP</td></tr>
<tr><td>>hg38_21_+_8433172_8446622_RNA45SN1_45S_50ntFlanks@9602.9622.21|CGAAAGACTAATCGAACCATC</td><td>>hg38_21_+_8433172_8446622_RNA45SN1_45S_50ntFlanks</td><td>9602</td><td>9622</td><td>21</td><td>CGAAAGACTAATCGAACCATC</td><td>267.5</td><td>-11.8</td><td>1.5</td><td>-8.1</td><td>6.1E-16</td><td>2.8E-15</td><td>MB231/MB468</td><td>MITO</td></tr>
<tr><td>>MI0000446|hsa-mir-125b-1&WithFlank&11|-|122099751|122099850@21.39.19;MI0000470|hsa-mir-125b-2&WithFlank&21|+|16590231|16590331@23.41.19&[MIMAT0000423&hsa-miR-125b-5p&offsets|0|-3;m-67&11|-|122099809|122099830&offsets|0|-3];[MIMAT0000423_1&hsa-miR-125b-5p&offsets|0|-3;m-68&21|+|16590253|16590274&offsets|0|-3]|TCCCTGAGACCCTAACTTG</td><td>>MI0000446|hsa-mir-125b-1&WithFlank&11|-|122099751|122099850</td><td>21</td><td>39</td><td>19</td><td>TCCCTGAGACCCTAACTTG</td><td>2280.2</td><td>12.3</td><td>1.5</td><td>8.1</td><td>4.0E-16</td><td>2.8E-15</td><td>MB231/MB468</td><td>CYTO</td></tr>
<tr><td>>hg38_21_+_8433172_8446622_RNA45SN1_45S_50ntFlanks@9419.9453.35|TCCGCCGAGGGCGCACCACCGGCCCGTCTCGCCCG</td><td>>hg38_21_+_8433172_8446622_RNA45SN1_45S_50ntFlanks</td><td>9419</td><td>9453</td><td>35</td><td>TCCGCCGAGGGCGCACCACCGGCCCGTCTCGCCCG</td><td>198.7</td><td>-11.5</td><td>1.4</td><td>-8.1</td><td>3.7E-16</td><td>2.8E-15</td><td>MB231/MB468</td><td>MP</td></tr>
</table>

| | | | | | | | | | | | | | |
|---|---|---|---|---|---|---|---|---|---|---|---|---|---|
| >MI0005756\|hsa-mir-934&WithFlank&X\|+\|136550872\|136550966@21.42.22(+1A)&[MIMAT0004977&hsa-miR-934&offsets\|0\|0(+1A);m-943&X\|+\|136550892\|136550913&offsets\|0\|0(+1A)]\|TGTCTACTACTGGAGACACTGGA | >MI0005756\|hsa-mir-934&WithFlank&X\|+\|136550872\|136550966 | 21 | 42 | 22(+1A) | TGTCTACTACTGGAGACACTGGA | 1742.4 | 7.7 | 1.0 | 8.1 | 4.1E-16 | 2.8E-15 | BT20/MB468 | MITO |
| >hg38_21_+_8433172_8446622_RNA45SN1_45S_50ntFlanks@11843.11872.30\|AGGGAACGGGCTTGGCGGAATCAGCGGGGA | >hg38_21_+_8433172_8446622_RNA45SN1_45S_50ntFlanks | 11843 | 11872 | 30 | AGGGAACGGGCTTGGCGGAATCAGCGGGGA | 279.5 | -11.1 | 1.4 | -8.1 | 4.5E-16 | 2.8E-15 | MB231/MB468 | NUC |
| >MI0000061\|hsa-let-7a-2&WithFlank&11\|-\|122146516\|122146599@11.30.20(+1A);MI0000062\|hsa-let-7a-3&WithFlank&22\|+\|46112743\|46112828@10.29.20(+1A);MI0000060\|hsa-let-7a-1&WithFlank&9\|+\|94175951\|94176042@12.31.20(+1A)&[MIMAT0000062&hsa-let-7a-5p&offsets\|0\|-2(+1A);m-6&11\|-\|122146568\|122146589&offsets\|0\|-2(+1A)];[MIMAT0000062_1&hsa-let-7a-5p&offsets\|0\|-2(+1A);m-5&22\|+\|46112752\|46112773&offsets\|0\|-2(+1A)];[MIMAT0000062_2&hsa-let-7a-5p&offsets\|0\|-2(+1A);m-4&9\|+\|94175962\|94175983&offsets\|0\|-2(+1A)]\|TGAGGTAGTAGGTTGTATAGA | >MI0000061\|hsa-let-7a-2&WithFlank&11\|-\|122146516\|122146599 | 11 | 30 | 20(+1A) | TGAGGTAGTAGGTTGTATAGA | 510.6 | 9.6 | 1.2 | 8.1 | 5.7E-16 | 2.8E-15 | BT20/MB468 | CYTO |
| >MI0000088\|hsa-mir-30a&WithFlank&6\|-\|71403545\|71403627@12.35.24&[MIMAT0000087&hsa-miR-30a-5p&offsets\|0\|+2;m-15&6\|-\|71403595\|71403616&offsets\|0\|+2]\|TGTAAACATCCTCGACTGGAAGCT | >MI0000088\|hsa-mir-30a&WithFlank&6\|-\|71403545\|71403627 | 12 | 35 | 24 | TGTAAACATCCTCGACTGGAAGCT | 27886.2 | 6.5 | 0.8 | 8.1 | 5.7E-16 | 2.8E-15 | BT20/MB468 | CYTO |
| >MI0000079\|hsa-mir-23a&WithFlank&19\|-\|13836581\|13836665@54.72.19(+1U)&[MIMAT0000078&hsa-miR-23a-3p&offsets\|+3\|+1(+1U);m-31&19\|-\|13836595\|13836615&offsets\|+3\|+1(+1U)]\|ACATTGCCAGGGATTTCCAT | >MI0000079\|hsa-mir-23a&WithFlank&19\|-\|13836581\|13836665 | 54 | 72 | 19(+1U) | ACATTGCCAGGGATTTCCAT | 146.8 | 10.0 | 1.2 | 8.0 | 9.0E-16 | 2.8E-15 | BT20/MB468 | NUC |
| >hg38_21_+_8433172_8446622_RNA45SN1_45S_50ntFlanks@6649.6676.28\|TACGACTCTTAGCGGTGGATCACTCGGC | >hg38_21_+_8433172_8446622_RNA45SN1_45S_50ntFlanks | 6649 | 6676 | 28 | TACGACTCTTAGCGGTGGATCACTCGGC | 1783.2 | 5.5 | 0.7 | 8.1 | 4.1E-16 | 2.8E-15 | BT20/MB468 | MITO |
| >hg38_21_+_8433172_8446622_RNA45SN1_45S_50ntFlanks@9544.9571.28\|GGTGGAGGTCCGTAGCGGTCCTGACGTG | >hg38_21_+_8433172_8446622_RNA45SN1_45S_50ntFlanks | 9544 | 9571 | 28 | GGTGGAGGTCCGTAGCGGTCCTGACGTG | 12529.7 | -8.2 | 1.0 | -8.0 | 9.0E-16 | 2.8E-15 | BT20/MB468 | NUC |
| >MI0000650\|hsa-mir-200c&WithFlank&12\|+\|6963693\|6963772@50.70.21(+1U)&[MIMAT0000617&hsa-miR-200c-3p&offsets\|0\|-2(+1U);m-54&12\|+\|6963742\|6963764&offsets\|0\|-2(+1U)]\|TAATACTGCCGGGTAATGATGT | >MI0000650\|hsa-mir-200c&WithFlank&12\|+\|6963693\|6963772 | 50 | 70 | 21(+1U) | TAATACTGCCGGGTAATGATGT | 635.1 | 9.1 | 1.1 | 8.4 | 3.4E-17 | 2.8E-15 | BT20/MB231 | CYTO |
| >hg38_21_+_8433172_8446622_RNA45SN1_45S_50ntFlanks@4682.4720.39\|GCAAGACGGACCAGAGCGAAAGCATTTGCCAAGAATGTT | >hg38_21_+_8433172_8446622_RNA45SN1_45S_50ntFlanks | 4682 | 4720 | 39 | GCAAGACGGACCAGAGCGAAAGCATTTGCCAAGAATGTT | 451.0 | 9.3 | 1.1 | 8.1 | 4.1E-16 | 2.8E-15 | MB231/MB468 | CYTO |
| >hg38_21_+_8433172_8446622_RNA45SN1_45S_50ntFlanks@10463.10497.35\|CTCCGTTGCCCTCGGCCGATCGAAAGGGAGTCGGG | >hg38_21_+_8433172_8446622_RNA45SN1_45S_50ntFlanks | 10463 | 10497 | 35 | CTCCGTTGCCCTCGGCCGATCGAAAGGGAGTCGGG | 293.4 | -10.9 | 1.3 | -8.1 | 4.1E-16 | 2.8E-15 | MB231/MB468 | CYTO |
| >hg38_21_+_8433172_8446622_RNA45SN1_45S_50ntFlanks@8844.8867.24\|GCGGCGGCGGCGGCGGCGGCGGGA | >hg38_21_+_8433172_8446622_RNA45SN1_45S_50ntFlanks | 8844 | 8867 | 24 | GCGGCGGCGGCGGCGGCGGCGGGA | 507.1 | -12.8 | 1.6 | -8.1 | 6.2E-16 | 2.8E-15 | MB231/MB468 | MITO |
| >hg38_1_-_228634819_228635039_RNA5S12_5S_50ntFlanks@137.170.34\|GAGACCGCCTGGGAATACCGGGTGCTGTAGGCTT | >hg38_1_-_228634819_228635039_RNA5S12_5S_50ntFlanks | 137 | 170 | 34 | GAGACCGCCTGGGAATACCGGGTGCTGTAGGCTT | 999.9 | 8.2 | 1.0 | 8.1 | 5.8E-16 | 2.8E-15 | BT20/MB468 | CYTO |
| >hg38_1_-_228634819_228635039_RNA5S12_5S_50ntFlanks@84.103.20\|CTCGTCTGATCTCGGAAGCT | >hg38_1_-_228634819_228635039_RNA5S12_5S_50ntFlanks | 84 | 103 | 20 | CTCGTCTGATCTCGGAAGCT | 956.7 | -5.0 | 0.6 | -8.1 | 5.8E-16 | 2.8E-15 | BT20/MB468 | CYTO |
| >MI0000089\|hsa-mir-31&WithFlank&9\|-\|21512109\|21512191@14.36.23&[MIMAT0000089&hsa-miR-31-5p&offsets\|0\|+2;m-360&9\|-\|21512157\|21512178&offsets\|0\|+1]\|AGGCAAGATGCTGGCATAGCTGT | >MI0000089\|hsa-mir-31&WithFlank&9\|-\|21512109\|21512191 | 14 | 36 | 23 | AGGCAAGATGCTGGCATAGCTGT | 1699.8 | 7.9 | 1.0 | 8.1 | 4.2E-16 | 2.8E-15 | BT20/MB468 | MITO |
| >hg38_21_+_8433172_8446622_RNA45SN1_45S_50ntFlanks@4122.4143.22\|AGGGTTCGATTCCGGAGAGGGA | >hg38_21_+_8433172_8446622_RNA45SN1_45S_50ntFlanks | 4122 | 4143 | 22 | AGGGTTCGATTCCGGAGAGGGA | 223.4 | -11.7 | 1.4 | -8.1 | 3.7E-16 | 2.8E-15 | MB231/MB468 | MP |
| >hg38_21_+_8433172_8446622_RNA45SN1_45S_50ntFlanks@8325.8356.32\|CGATAGTCAACAAGTACCGTAAGGGAAAGTTG | >hg38_21_+_8433172_8446622_RNA45SN1_45S_50ntFlanks | 8325 | 8356 | 32 | CGATAGTCAACAAGTACCGTAAGGGAAAGTTG | 195.7 | -11.5 | 1.4 | -8.1 | 3.7E-16 | 2.8E-15 | MB231/MB468 | MP |
| >hg38_21_+_8433172_8446622_RNA45SN1_45S_50ntFlanks@9027.9058.32\|CGTCTCCTCGTGGGGGGGCCGGGCCACCCCTC | >hg38_21_+_8433172_8446622_RNA45SN1_45S_50ntFlanks | 9027 | 9058 | 32 | CGTCTCCTCGTGGGGGGGCCGGGCCACCCCTC | 313.3 | -12.2 | 1.5 | -8.1 | 3.7E-16 | 2.8E-15 | MB231/MB468 | MP |
| >hg38_21_+_8433172_8446622_RNA45SN1_45S_50ntFlanks@5472.5506.35\|ACGGCCCTGGCGGAGCGCTGAGAAGACGGTCGAAC | >hg38_21_+_8433172_8446622_RNA45SN1_45S_50ntFlanks | 5472 | 5506 | 35 | ACGGCCCTGGCGGAGCGCTGAGAAGACGGTCGAAC | 2845.4 | 7.2 | 0.9 | 8.1 | 5.9E-16 | 2.8E-15 | BT20/MB468 | CYTO |
| >hg38_21_+_8433172_8446622_RNA45SN1_45S_50ntFlanks@9352.9380.29\|CCGTGGCGCAATGAAGGTGAAGGCCGGCG | >hg38_21_+_8433172_8446622_RNA45SN1_45S_50ntFlanks | 9352 | 9380 | 29 | CCGTGGCGCAATGAAGGTGAAGGCCGGCG | 647.0 | -13.2 | 1.6 | -8.1 | 3.8E-16 | 2.8E-15 | MB231/MB468 | MP |
| >trna10_ValCAC_5_-_180649395_180649467@59.76.18,trna12_ValAAC_5_-_180645270_180645342@59.76.18,trna132_ValAAC_6_-_27721179_27721251@59.76.18,trna136_ValAAC_6_-_27648885_27648957@59.76.18,trna15_ValAAC_5_-_180615416_180615488@59.76.18,trna18_ValCAC_5_-_180529253_180529325@59.76.18,trna2_ValAAC_3_+_169490018_169490090@59.76.18,trna2_ValCAC_5_+_180524070_180524142@59.76.18,trna4_ValAAC_5_+_180591154_180591226@59.76.18,trna5_ValAAC_5_+_180596610_180596682@59.76.18,trna6_ValCAC_5_+_180600650_180600722@59.76.18,trna85_ValCAC_1_-_161369490_161369562@59.76.18,trna90_ValCAC_1_-_149684088_149684161@60.77.18,trna9_ValCAC_6_+_26538282_26538354@59.76.18\|AACCGGGCGGAAACACCA | >trna10_ValCAC_5_-_180649395_180649467 | 59 | 76 | 18 | AACCGGGCGGAAACACCA | 113.3 | 2.9 | 0.4 | 8.0 | 9.2E-16 | 2.9E-15 | BT20/MB468 | NUC |
| >hg38_21_+_8433172_8446622_RNA45SN1_45S_50ntFlanks@10351.10390.40\|CGAGAACTTTGAAGGCCGAAGTGGAGAAGGGTTCCATGTG | >hg38_21_+_8433172_8446622_RNA45SN1_45S_50ntFlanks | 10351 | 10390 | 40 | CGAGAACTTTGAAGGCCGAAGTGGAGAAGGGTTCCATGTG | 9803.8 | -5.2 | 0.6 | -8.1 | 4.1E-16 | 2.9E-15 | MB231/MB468 | CYTO |
| >hg38_21_+_8433172_8446622_RNA45SN1_45S_50ntFlanks@12712.12741.30\|CGGAGCCTCGGTTGGCCTCGGATAGCCGGT | >hg38_21_+_8433172_8446622_RNA45SN1_45S_50ntFlanks | 12712 | 12741 | 30 | CGGAGCCTCGGTTGGCCTCGGATAGCCGGT | 218.1 | -11.6 | 1.4 | -8.1 | 3.8E-16 | 2.9E-15 | MB231/MB468 | MP |
| >hg38_21_+_8433172_8446622_RNA45SN1_45S_50ntFlanks@4884.4911.28\|CAAAGCTGAAACTTAAAGGAATTGACGG | >hg38_21_+_8433172_8446622_RNA45SN1_45S_50ntFlanks | 4884 | 4911 | 28 | CAAAGCTGAAACTTAAAGGAATTGACGG | 278.9 | -12.0 | 1.5 | -8.1 | 3.8E-16 | 2.9E-15 | MB231/MB468 | MP |
| >MI0000455\|hsa-mir-138-2&WithFlank&16\|+\|56858512\|56858607@16.38.23;MI0000476\|hsa-mir-138-1&WithFlank&3\|+\|44114206\|44114316@29.51.23&[MIMAT0000430&hsa-miR-138-5p&offsets\|0\|0;m-509&16\|+\|56858527\|56858549&offsets\|0\|0];[MIMAT0000430_1&hsa-miR-138-5p&offsets\|0\|0;m-497&3\|+\|44114234\|44114256&offsets\|0\|0]\|AGCTGGTGTTGTGAATCAGGCCG | >MI0000455\|hsa-mir-138-2&WithFlank&16\|+\|56858512\|56858607 | 16 | 38 | 23 | AGCTGGTGTTGTGAATCAGGCCG | 271.8 | 11.3 | 1.4 | 8.1 | 3.8E-16 | 2.9E-15 | MB231/MB468 | MP |
| >trnaMT_PheGAA_MT_+_577_647@39.57.19\|ATGTTTAGACGGGCTCACA | >trnaMT_PheGAA_MT_+_577_647 | 39 | 57 | 19 | ATGTTTAGACGGGCTCACA | 340.9 | 5.9 | 0.7 | 8.1 | 4.7E-16 | 2.9E-15 | MB231/MB468 | NUC |
| >hg38_21_+_8433172_8446622_RNA45SN1_45S_50ntFlanks@11569.11588.20\|CAGCCGACTTAGAACTGGTG | >hg38_21_+_8433172_8446622_RNA45SN1_45S_50ntFlanks | 11569 | 11588 | 20 | CAGCCGACTTAGAACTGGTG | 558.5 | -13.0 | 1.6 | -8.1 | 3.8E-16 | 2.9E-15 | MB231/MB468 | MP |
| >MI0000301\|hsa-mir-224&WithFlank&X\|-\|151958572\|151958664@14.36.23&[MIMAT0000281&hsa-miR-224-5p&offsets\|+1\|-1;m-238&X\|-\|151958630\|151958651&offsets\|0\|+1]\|CAAGTCACTAGTGGTTCCGTTTA | >MI0000301\|hsa-mir-224&WithFlank&X\|-\|151958572\|151958664 | 14 | 36 | 23 | CAAGTCACTAGTGGTTCCGTTTA | 352.8 | 12.5 | 1.5 | 8.1 | 4.1E-16 | 2.9E-15 | MB231/MB468 | CYTO |
| >hg38_21_+_8433172_8446622_RNA45SN1_45S_50ntFlanks@11161.11188.28\|CACGGTCCCCCGCGAGGGGGGCCCGGGC | >hg38_21_+_8433172_8446622_RNA45SN1_45S_50ntFlanks | 11161 | 11188 | 28 | CACGGTCCCCCGCGAGGGGGGCCCGGGC | 1053.8 | -13.0 | 1.6 | -8.1 | 4.7E-16 | 2.9E-15 | MB231/MB468 | NUC |
| >MI0000440\|hsa-mir-27b&WithFlank&9\|+\|95085439\|95085547@67.86.20(+1U)&[MIMAT0000419&hsa-miR-27b-3p&offsets\|0\|-1(+1U);m-39&9\|+\|95085505\|95085525&offsets\|0\|-1(+1U)]\|TTCACAGTGGCTAAGTTCTGT | >MI0000440\|hsa-mir-27b&WithFlank&9\|+\|95085439\|95085547 | 67 | 86 | 20(+1U) | TTCACAGTGGCTAAGTTCTGT | 2951.3 | 4.5 | 0.6 | 8.0 | 9.3E-16 | 2.9E-15 | BT20/MB468 | NUC |
| >MI0000085\|hsa-mir-27a&WithFlank&19\|-\|13836434\|13836523@57.77.21&[MIMAT0000084&hsa-miR-27a-3p&offsets\|0\|0;m-43&19\|-\|13836447\|13836467&offsets\|0\|0]\|TTCACAGTGGCTAAGTTCCGC | >MI0000085\|hsa-mir-27a&WithFlank&19\|-\|13836434\|13836523 | 57 | 77 | 21 | TTCACAGTGGCTAAGTTCCGC | 1615.8 | 5.9 | 0.7 | 8.1 | 4.8E-16 | 2.9E-15 | MB231/MB468 | NUC |
| >hg38_21_+_8433172_8446622_RNA45SN1_45S_50ntFlanks@4122.4145.24\|AGGGTTCGATTCCGGAGAGGGAGC | >hg38_21_+_8433172_8446622_RNA45SN1_45S_50ntFlanks | 4122 | 4145 | 24 | AGGGTTCGATTCCGGAGAGGGAGC | 323.6 | -11.3 | 1.4 | -8.1 | 4.8E-16 | 2.9E-15 | MB231/MB468 | NUC |
| >hg38_21_+_8433172_8446622_RNA45SN1_45S_50ntFlanks@10511.10534.24\|ATCCGGAGTGGCGGAGATGGGCGC | >hg38_21_+_8433172_8446622_RNA45SN1_45S_50ntFlanks | 10511 | 10534 | 24 | ATCCGGAGTGGCGGAGATGGGCGC | 302.2 | -12.1 | 1.5 | -8.1 | 3.8E-16 | 2.9E-15 | MB231/MB468 | MP |
| >hg38_21_+_8433172_8446622_RNA45SN1_45S_50ntFlanks@4003.4020.18\|ATAACCTCGGGCCGATCG | >hg38_21_+_8433172_8446622_RNA45SN1_45S_50ntFlanks | 4003 | 4020 | 18 | ATAACCTCGGGCCGATCG | 243.1 | -11.8 | 1.4 | -8.1 | 3.8E-16 | 2.9E-15 | MB231/MB468 | MP |
| >MI0000462\|hsa-mir-152&WithFlank&17\|-\|48037155\|48037253@60.80.21&[MIMAT0000438&hsa-miR-152-3p&offsets\|0\|0;m-97&17\|-\|48037174\|48037194&offsets\|0\|0]\|TCAGTGCATGACAGAACTTGG | >MI0000462\|hsa-mir-152&WithFlank&17\|-\|48037155\|48037253 | 60 | 80 | 21 | TCAGTGCATGACAGAACTTGG | 273.1 | 8.6 | 1.1 | 8.0 | 9.4E-16 | 2.9E-15 | BT20/MB468 | NUC |
| >hg38_21_+_8433172_8446622_RNA45SN1_45S_50ntFlanks@4602.4619.18\|TTCGGAACTGAGGCCATG | >hg38_21_+_8433172_8446622_RNA45SN1_45S_50ntFlanks | 4602 | 4619 | 18 | TTCGGAACTGAGGCCATG | 289.6 | -12.0 | 1.5 | -8.1 | 3.9E-16 | 2.9E-15 | MB231/MB468 | MP |
| >MI0000081\|hsa-mir-24-2&WithFlank&19\|-\|13836281\|13836365@56.77.22(+1C);MI0000080\|hsa-mir-24-1&WithFlank&9\|+\|95086015\|95086094@50.71.22(+1C)&[MIMAT0000080&hsa-miR-24-3p&offsets\|0\|0(+1C);m-36&19\|-\|13836288\|13836310&offsets\|0\|-1(+1C)];[MIMAT0000080_1&hsa-miR-24-3p&offsets\|0\|0(+1C);m-35&9\|+\|95086064\|95086086&offsets\|0\|-1(+1C)]\|TGGCTCAGTTCAGCAGGAACAGC | >MI0000081\|hsa-mir-24-2&WithFlank&19\|-\|13836281\|13836365 | 56 | 77 | 22(+1C) | TGGCTCAGTTCAGCAGGAACAGC | 689.2 | 5.5 | 0.7 | 8.1 | 4.8E-16 | 2.9E-15 | MB231/MB468 | NUC |
| >hg38_21_+_8433172_8446622_RNA45SN1_45S_50ntFlanks@9806.9831.26\|CGTGGAATGCGAGTGCCTAGTGGGCC | >hg38_21_+_8433172_8446622_RNA45SN1_45S_50ntFlanks | 9806 | 9831 | 26 | CGTGGAATGCGAGTGCCTAGTGGGCC | 297.5 | -12.0 | 1.5 | -8.1 | 6.4E-16 | 2.9E-15 | MB231/MB468 | MITO |
| >hg38_21_+_8433172_8446622_RNA45SN1_45S_50ntFlanks@10786.10826.41\|AACAGCCTCTGGCATGTTGGAACAATGTAGGTAAGGGAAGT | >hg38_21_+_8433172_8446622_RNA45SN1_45S_50ntFlanks | 10786 | 10826 | 41 | AACAGCCTCTGGCATGTTGGAACAATGTAGGTAAGGGAAGT | 744.2 | -13.4 | 1.6 | -8.1 | 3.9E-16 | 2.9E-15 | MB231/MB468 | MP |
| >hg38_21_+_8433172_8446622_RNA45SN1_45S_50ntFlanks@12006.12043.38\|GGCCGCCGGTGAAATACCACTACTCTGATCGTTTTTTC | >hg38_21_+_8433172_8446622_RNA45SN1_45S_50ntFlanks | 12006 | 12043 | 38 | GGCCGCCGGTGAAATACCACTACTCTGATCGTTTTTTC | 554.8 | -13.5 | 1.7 | -8.0 | 9.4E-16 | 2.9E-15 | BT20/MB468 | NUC |
| >hg38_21_+_8433172_8446622_RNA45SN1_45S_50ntFlanks@12636.12653.18\|AGCTACCATCTGTGGGAT | >hg38_21_+_8433172_8446622_RNA45SN1_45S_50ntFlanks | 12636 | 12653 | 18 | AGCTACCATCTGTGGGAT | 242.6 | -7.6 | 0.9 | -8.1 | 4.2E-16 | 2.9E-15 | MB231/MB468 | CYTO |

| | | | | | | | | | | | | | |
|---|---|---|---|---|---|---|---|---|---|---|---|---|---|
| >trna129_MetCAT_6_-_27870271_27870342@31.51.21,trna142_MetCAT_6_-_27560600_27560671@31.51.21,trna150_MetCAT_6_-_27300764_27300835@31.51.21,trna169_MetCAT_6_-_26330529_26330600@31.51.21,trna171_MetCAT_6_-_26313352_26313423@31.51.21,trna20_MetCAT_17_-_80452597_80452668@31.51.21,trna2_MetCAT_6_+_26286754_26286825@31.51.21,trna32_MetCAT_1_+_153643726_153643797@31.51.21,trna61_MetCAT_6_+_27745664_27745735@31.51.21\|CCCATAACCCAGAGGTCGATG | >trna129_MetCAT_6_-_27870271_27870342 | 31 | 51 | 21 | CCCATAACCCAGAGGTCGATG | 740.3 | -9.3 | 1.1 | -8.1 | 4.2E-16 | 2.9E-15 | MB231/MB468 | CYTO |
| >hg38_21_+_8433172_8446622_RNA45SN1_45S_50ntFlanks@6741.6771.31\|ATCGACACTTCGAACGCACTTGCGGCCCCGG | >hg38_21_+_8433172_8446622_RNA45SN1_45S_50ntFlanks | 6741 | 6771 | 31 | ATCGACACTTCGAACGCACTTGCGGCCCCGG | 641.2 | -6.5 | 0.8 | -8.1 | 4.2E-16 | 2.9E-15 | MB231/MB468 | CYTO |
| >MI0000764\|hsa-mir-363&WithFlank&X\|-\|134169372\|134169458@56.77.22&[MIMAT0000707&hsa-miR-363-3p&offsets\|0\|0;m-178&X\|-\|134169381\|134169403&offsets\|0\|-1]\|AATTGCACGGTATCCATCTGTA | >MI0000764\|hsa-mir-363&WithFlank&X\|-\|134169372\|134169458 | 56 | 77 | 22 | AATTGCACGGTATCCATCTGTA | 853.3 | 9.7 | 1.2 | 8.4 | 3.6E-17 | 2.9E-15 | BT20/MB231 | CYTO |
| >MI0000088\|hsa-mir-30a&WithFlank&6\|-\|71403545\|71403627@53.70.18(+1U)&[MIMAT0000088&hsa-miR-30a-3p&offsets\|0\|-4(+1U);m-38&6\|-\|71403554\|71403575&offsets\|0\|-4(+1U)]\|CTTTCAGTCGGATGTTTGT | >MI0000088\|hsa-mir-30a&WithFlank&6\|-\|71403545\|71403627 | 53 | 70 | 18(+1U) | CTTTCAGTCGGATGTTTGT | 453.2 | 5.0 | 0.6 | 8.1 | 4.8E-16 | 2.9E-15 | MB231/MB468 | NUC |
| >hg38_21_+_8433172_8446622_RNA45SN1_45S_50ntFlanks@6731.6770.40\|CACATTGATCATCGACACTTCGAACGCACTTGCGGCCCCG | >hg38_21_+_8433172_8446622_RNA45SN1_45S_50ntFlanks | 6731 | 6770 | 40 | CACATTGATCATCGACACTTCGAACGCACTTGCGGCCCCG | 961.0 | -13.8 | 1.7 | -8.1 | 3.9E-16 | 2.9E-15 | MB231/MB468 | MP |
| >MI0000466\|hsa-mir-9-1&WithFlank&1\|-\|156420335\|156420435@22.43.22;MI0000468\|hsa-mir-9-3&WithFlank&15\|+\|89368011\|89368112@22.43.22;MI0000467\|hsa-mir-9-2&WithFlank&5\|-\|88666847\|88666945@22.43.22&[MIMAT0000441&hsa-miR-9-5p&offsets\|0\|-1;m-104&1\|-\|156420392\|156420414&offsets\|0\|-1];[MIMAT0000441_1&hsa-miR-9-5p&offsets\|0\|-1;m-105&15\|+\|89368032\|89368054&offsets\|0\|-1];[MIMAT0000441_2&hsa-miR-9-5p&offsets\|0\|-1;m-104&5\|-\|88666902\|88666924&offsets\|0\|-1]\|TCTTTGGTTATCTAGCTGTATG | >MI0000466\|hsa-mir-9-1&WithFlank&1\|-\|156420335\|156420435 | 22 | 43 | 22 | TCTTTGGTTATCTAGCTGTATG | 279.2 | 6.1 | 0.8 | 8.1 | 4.8E-16 | 2.9E-15 | MB231/MB468 | NUC |
| >hg38_21_+_8433172_8446622_RNA45SN1_45S_50ntFlanks@9471.9500.30\|AGCACGAGCGCACGTGTTAGGACCCGAAAG | >hg38_21_+_8433172_8446622_RNA45SN1_45S_50ntFlanks | 9471 | 9500 | 30 | AGCACGAGCGCACGTGTTAGGACCCGAAAG | 264.0 | -11.8 | 1.5 | -8.1 | 6.5E-16 | 2.9E-15 | MB231/MB468 | MITO |
| >hg38_21_+_8433172_8446622_RNA45SN1_45S_50ntFlanks@4367.4398.32\|CAGTTAAAAAGCTCGTAGTTGGATCTTGGGAG | >hg38_21_+_8433172_8446622_RNA45SN1_45S_50ntFlanks | 4367 | 4398 | 32 | CAGTTAAAAAGCTCGTAGTTGGATCTTGGGAG | 493.4 | -12.8 | 1.6 | -8.1 | 3.9E-16 | 2.9E-15 | MB231/MB468 | MP |
| >hg38_21_+_8433172_8446622_RNA45SN1_45S_50ntFlanks@5059.5094.36\|CGATTTGTCTGGTTAATTCCGATAACGAACGAGACT | >hg38_21_+_8433172_8446622_RNA45SN1_45S_50ntFlanks | 5059 | 5094 | 36 | CGATTTGTCTGGTTAATTCCGATAACGAACGAGACT | 88.6 | -23.6 | 2.9 | -8.1 | 3.9E-16 | 2.9E-15 | MB231/MB468 | MP |
| >hg38_21_+_8433172_8446622_RNA45SN1_45S_50ntFlanks@11208.11226.19\|GCGGCGACTCTGGACGCGA | >hg38_21_+_8433172_8446622_RNA45SN1_45S_50ntFlanks | 11208 | 11226 | 19 | GCGGCGACTCTGGACGCGA | 274.3 | -12.0 | 1.5 | -8.1 | 3.9E-16 | 2.9E-15 | MB231/MB468 | MP |
| >hg38_21_+_8433172_8446622_RNA45SN1_45S_50ntFlanks@7982.8008.27\|TCAGATCAGACGTGGCGACCCGCTGAA | >hg38_21_+_8433172_8446622_RNA45SN1_45S_50ntFlanks | 7982 | 8008 | 27 | TCAGATCAGACGTGGCGACCCGCTGAA | 2712.6 | -5.8 | 0.7 | -8.1 | 6.5E-16 | 3.0E-15 | MB231/MB468 | MITO |
| >hg38_21_+_8433172_8446622_RNA45SN1_45S_50ntFlanks@11920.11949.30\|ATGAGAGGTGTAGAATAAGTGGGAGGCCCC | >hg38_21_+_8433172_8446622_RNA45SN1_45S_50ntFlanks | 11920 | 11949 | 30 | ATGAGAGGTGTAGAATAAGTGGGAGGCCCC | 465.8 | -12.7 | 1.6 | -8.1 | 4.0E-16 | 3.0E-15 | MB231/MB468 | MP |
| >MI0000254\|hsa-mir-30c-2&WithFlank&6\|-\|71376954\|71377037@53.73.21(+1A)&[MIMAT0004550&hsa-miR-30c-2-3p&offsets\|0\|-1(+1A);m-206&6\|-\|71376964\|71376985&offsets\|0\|-1(+1A)]\|CTGGGAGAAGGCTGTTTACTCA | >MI0000254\|hsa-mir-30c-2&WithFlank&6\|-\|71376954\|71377037 | 53 | 73 | 21(+1A) | CTGGGAGAAGGCTGTTTACTCA | 418.1 | 9.5 | 1.2 | 8.1 | 6.1E-16 | 3.0E-15 | BT20/MB468 | CYTO |
| >trna111_HisGTG_1_-_147774845_147774916@16.33.18,trna118_HisGTG_1_-_145396881_145396952@16.33.18,trna16_HisGTG_1_+_146544773_146544844@16.33.18,trna1_HisGTG_15_+_45493349_45493420@16.33.18,trna21_HisGTG_1_+_147753471_147753542@16.33.18,trna33_HisGTG_6_+_27125906_27125977@16.33.18,trna7_HisGTG_9_-_14433938_14434009@16.33.18,trna8_HisGTG_15_-_45492611_45492682@16.33.18,trna9_HisGTG_15_-_45490804_45490875@16.33.18\|TGGTTAGTACTCTGCGTT | >trna111_HisGTG_1_-_147774845_147774916 | 16 | 33 | 18 | TGGTTAGTACTCTGCGTT | 846.9 | 6.0 | 0.7 | 8.1 | 6.1E-16 | 3.0E-15 | BT20/MB468 | CYTO |
| >hg38_21_+_8433172_8446622_RNA45SN1_45S_50ntFlanks@11564.11581.18\|CCTAGCAGCCGACTTAGA | >hg38_21_+_8433172_8446622_RNA45SN1_45S_50ntFlanks | 11564 | 11581 | 18 | CCTAGCAGCCGACTTAGA | 2629.0 | -11.3 | 1.4 | -8.0 | 9.7E-16 | 3.0E-15 | BT20/MB468 | NUC |
| >MI0000809\|hsa-mir-151a&WithFlank&8\|-\|140732558\|140732659@51.72.22&[MIMAT0000757&hsa-miR-151a-3p&offsets\|-2\|-1;m-30&8\|-\|140732587\|140732607&offsets\|-2\|-1]\|TACTAGACTGAAGCTCCTTGAG | >MI0000809\|hsa-mir-151a&WithFlank&8\|-\|140732558\|140732659 | 51 | 72 | 22 | TACTAGACTGAAGCTCCTTGAG | 127.8 | 2.2 | 0.3 | 8.0 | 9.7E-16 | 3.0E-15 | BT20/MB468 | NUC |
| >hg38_21_+_8433172_8446622_RNA45SN1_45S_50ntFlanks@7978.8013.36\|GACCTCAGATCAGACGTGGCGACCCGCTGAATTTAA | >hg38_21_+_8433172_8446622_RNA45SN1_45S_50ntFlanks | 7978 | 8013 | 36 | GACCTCAGATCAGACGTGGCGACCCGCTGAATTTAA | 271.7 | 12.1 | 1.5 | 8.1 | 4.4E-16 | 3.0E-15 | MB231/MB468 | CYTO |
| >hg38_21_+_8433172_8446622_RNA45SN1_45S_50ntFlanks@13006.13025.20\|CTATTGAAAGTCAGCCCTCG | >hg38_21_+_8433172_8446622_RNA45SN1_45S_50ntFlanks | 13006 | 13025 | 20 | CTATTGAAAGTCAGCCCTCG | 223.2 | -11.7 | 1.4 | -8.1 | 4.0E-16 | 3.0E-15 | MB231/MB468 | MP |
| >hg38_21_+_8433172_8446622_RNA45SN1_45S_50ntFlanks@11115.11132.18\|CGGCAGGGGCCGGCGGCG | >hg38_21_+_8433172_8446622_RNA45SN1_45S_50ntFlanks | 11115 | 11132 | 18 | CGGCAGGGGCCGGCGGCG | 356.7 | -12.3 | 1.5 | -8.1 | 4.0E-16 | 3.0E-15 | MB231/MB468 | MP |
| >hg38_21_+_8433172_8446622_RNA45SN1_45S_50ntFlanks@5414.5445.32\|CTACTACCGATTGGATGGTTTAGTGAGGCCCT | >hg38_21_+_8433172_8446622_RNA45SN1_45S_50ntFlanks | 5414 | 5445 | 32 | CTACTACCGATTGGATGGTTTAGTGAGGCCCT | 486.4 | -12.7 | 1.6 | -8.1 | 6.7E-16 | 3.0E-15 | MB231/MB468 | MITO |
| >hg38_21_+_8433172_8446622_RNA45SN1_45S_50ntFlanks@6671.6697.27\|CTCGGCTCGTGCGTCGATGAAGAACGC | >hg38_21_+_8433172_8446622_RNA45SN1_45S_50ntFlanks | 6671 | 6697 | 27 | CTCGGCTCGTGCGTCGATGAAGAACGC | 1276.6 | -10.0 | 1.2 | -8.1 | 4.1E-16 | 3.0E-15 | MB231/MB468 | MP |
| >hg38_21_+_8433172_8446622_RNA45SN1_45S_50ntFlanks@12927.12973.47\|CCTGGCGCTAAACCATTCGTAGACGACCTGCTTCTGGGTCGGGGTTT | >hg38_21_+_8433172_8446622_RNA45SN1_45S_50ntFlanks | 12927 | 12973 | 47 | CCTGGCGCTAAACCATTCGTAGACGACCTGCTTCTGGGTCGGGGTTT | 566.1 | -13.6 | 1.7 | -8.0 | 9.9E-16 | 3.0E-15 | BT20/MB468 | NUC |
| >trnaMT_GluTTC_MT_-_14674_14742@39.58.20,trnalookalike8_GluTTC_5_-_93905172_93905240@39.58.20\|ATTGGTCGTGGTTGTAGTCC | >trnaMT_GluTTC_MT_-_14674_14742 | 39 | 58 | 20 | ATTGGTCGTGGTTGTAGTCC | 1607.0 | 8.6 | 1.1 | 8.1 | 4.5E-16 | 3.0E-15 | BT20/MB468 | MITO |
| >hg38_21_+_8433172_8446622_RNA45SN1_45S_50ntFlanks@10512.10529.18\|TCCGGAGTGGCGGAGATG | >hg38_21_+_8433172_8446622_RNA45SN1_45S_50ntFlanks | 10512 | 10529 | 18 | TCCGGAGTGGCGGAGATG | 253.3 | -11.8 | 1.5 | -8.1 | 4.1E-16 | 3.0E-15 | MB231/MB468 | MP |
| >hg38_21_+_8433172_8446622_RNA45SN1_45S_50ntFlanks@10411.10441.31\|CAGTCGGTCCTGAGAGATGGGCGAGCGCCGT | >hg38_21_+_8433172_8446622_RNA45SN1_45S_50ntFlanks | 10411 | 10441 | 31 | CAGTCGGTCCTGAGAGATGGGCGAGCGCCGT | 436.0 | -12.6 | 1.6 | -8.1 | 4.1E-16 | 3.0E-15 | MB231/MB468 | MP |
| >hg38_21_+_8433172_8446622_RNA45SN1_45S_50ntFlanks@10735.10752.18\|TGTAAATCTCGCGCCGGG | >hg38_21_+_8433172_8446622_RNA45SN1_45S_50ntFlanks | 10735 | 10752 | 18 | TGTAAATCTCGCGCCGGG | 293.0 | -12.0 | 1.5 | -8.1 | 6.8E-16 | 3.1E-15 | MB231/MB468 | MITO |
| >hg38_21_+_8433172_8446622_RNA45SN1_45S_50ntFlanks@9129.9149.21\|CGGGTCGGGGGGCGGGGCGGA | >hg38_21_+_8433172_8446622_RNA45SN1_45S_50ntFlanks | 9129 | 9149 | 21 | CGGGTCGGGGGGCGGGGCGGA | 307.4 | -12.0 | 1.5 | -8.1 | 6.8E-16 | 3.1E-15 | MB231/MB468 | MITO |
| >hg38_21_+_8433172_8446622_RNA45SN1_45S_50ntFlanks@5472.5501.30\|ACGGCCCTGGCGGAGCGCTGAGAAGACGGT | >hg38_21_+_8433172_8446622_RNA45SN1_45S_50ntFlanks | 5472 | 5501 | 30 | ACGGCCCTGGCGGAGCGCTGAGAAGACGGT | 7766.9 | 6.9 | 0.8 | 8.1 | 4.4E-16 | 3.1E-15 | MB231/MB468 | CYTO |
| >hg38_21_+_8433172_8446622_RNA45SN1_45S_50ntFlanks@11888.11905.18\|AGCTTGACTCTAGTCTGG | >hg38_21_+_8433172_8446622_RNA45SN1_45S_50ntFlanks | 11888 | 11905 | 18 | AGCTTGACTCTAGTCTGG | 223.7 | -11.7 | 1.4 | -8.1 | 4.1E-16 | 3.1E-15 | MB231/MB468 | MP |
| >hg38_21_+_8433172_8446622_RNA45SN1_45S_50ntFlanks@12712.12751.40\|CGGAGCCTCGGTTGGCCTCGGATAGCCGGTCCCCCGCCTG | >hg38_21_+_8433172_8446622_RNA45SN1_45S_50ntFlanks | 12712 | 12751 | 40 | CGGAGCCTCGGTTGGCCTCGGATAGCCGGTCCCCCGCCTG | 1136.1 | -14.0 | 1.7 | -8.1 | 4.1E-16 | 3.1E-15 | MB231/MB468 | MP |
| >MI0001445\|hsa-mir-423&WithFlank&17\|+\|30117073\|30117178@23.46.24&[MIMAT0004748&hsa-miR-423-5p&offsets\|0\|+1;m-141&17\|+\|30117095\|30117117&offsets\|0\|+1]\|TGAGGGGCAGAGAGCGAGACTTTT | >MI0001445\|hsa-mir-423&WithFlank&17\|+\|30117073\|30117178 | 23 | 46 | 24 | TGAGGGGCAGAGAGCGAGACTTTT | 695.2 | 5.7 | 0.7 | 8.1 | 5.1E-16 | 3.1E-15 | MB231/MB468 | NUC |
| >hg38_21_+_8433172_8446622_RNA45SN1_45S_50ntFlanks@5472.5511.40\|ACGGCCCTGGCGGAGCGCTGAGAAGACGGTCGAACTTGAC | >hg38_21_+_8433172_8446622_RNA45SN1_45S_50ntFlanks | 5472 | 5511 | 40 | ACGGCCCTGGCGGAGCGCTGAGAAGACGGTCGAACTTGAC | 507.0 | 7.5 | 0.9 | 8.1 | 5.1E-16 | 3.1E-15 | MB231/MB468 | NUC |
| >hg38_21_+_8433172_8446622_RNA45SN1_45S_50ntFlanks@4044.4063.20\|CCCATTCGAACGTCTGCCCT | >hg38_21_+_8433172_8446622_RNA45SN1_45S_50ntFlanks | 4044 | 4063 | 20 | CCCATTCGAACGTCTGCCCT | 306.6 | -11.2 | 1.4 | -8.1 | 5.1E-16 | 3.1E-15 | MB231/MB468 | NUC |
| >MI0000087\|hsa-mir-29a&WithFlank&7\|-\|130876741\|130876816@48.69.22&[MIMAT0000086&hsa-miR-29a-3p&offsets\|0\|0;m-26&7\|-\|130876749\|130876769&offsets\|0\|+1]\|TAGCACCATCTGAAATCGGTTA | >MI0000087\|hsa-mir-29a&WithFlank&7\|-\|130876741\|130876816 | 48 | 69 | 22 | TAGCACCATCTGAAATCGGTTA | 16590.1 | 9.0 | 1.1 | 8.1 | 6.8E-16 | 3.1E-15 | MB231/MB468 | MITO |
| >MI0000299\|hsa-mir-222&WithFlank&X\|-\|45747009\|45747130@75.98.24(+1G)&[MIMAT0000279&hsa-miR-222-3p&offsets\|0\|+3(+1G);m-140&X\|-\|45747033\|45747056&offsets\|0\|0(+1G)]\|AGCTACATCTGGCTACTGGGTCTCG | >MI0000299\|hsa-mir-222&WithFlank&X\|-\|45747009\|45747130 | 75 | 98 | 24(+1G) | AGCTACATCTGGCTACTGGGTCTCG | 1435.5 | 6.8 | 0.8 | 8.1 | 4.5E-16 | 3.1E-15 | MB231/MB468 | CYTO |
| >hg38_21_+_8433172_8446622_RNA45SN1_45S_50ntFlanks@4449.4470.22\|CCTCTCGGCGCCCCCTCGATGC | >hg38_21_+_8433172_8446622_RNA45SN1_45S_50ntFlanks | 4449 | 4470 | 22 | CCTCTCGGCGCCCCCTCGATGC | 326.4 | -12.1 | 1.5 | -8.1 | 6.8E-16 | 3.1E-15 | MB231/MB468 | MITO |
| >hg38_21_+_8433172_8446622_RNA45SN1_45S_50ntFlanks@5215.5233.19\|TCCGGGGCTGCACGCGCGC | >hg38_21_+_8433172_8446622_RNA45SN1_45S_50ntFlanks | 5215 | 5233 | 19 | TCCGGGGCTGCACGCGCGC | 756.9 | 11.2 | 1.4 | 8.1 | 4.5E-16 | 3.1E-15 | MB231/MB468 | CYTO |
| >MI0000542\|hsa-mir-320a&WithFlank&8\|-\|22244960\|22245043@48.69.22&[MIMAT0000510&hsa-miR-320a-3p&offsets\|0\|0;m-60&8\|-\|22244975\|22244996&offsets\|0\|0]\|AAAAGCTGGGTTGAGAGGGCGA | >MI0000542\|hsa-mir-320a&WithFlank&8\|-\|22244960\|22245043 | 48 | 69 | 22 | AAAAGCTGGGTTGAGAGGGCGA | 3357.1 | 7.0 | 0.9 | 8.1 | 4.5E-16 | 3.1E-15 | MB231/MB468 | CYTO |
| >hg38_21_+_8433172_8446622_RNA45SN1_45S_50ntFlanks@11567.11584.18\|AGCAGCCGACTTAGAACT | >hg38_21_+_8433172_8446622_RNA45SN1_45S_50ntFlanks | 11567 | 11584 | 18 | AGCAGCCGACTTAGAACT | 4045.1 | 5.8 | 0.7 | 8.1 | 4.5E-16 | 3.1E-15 | MB231/MB468 | CYTO |
| >hg38_21_+_8433172_8446622_RNA45SN1_45S_50ntFlanks@10350.10383.34\|ACGAGAACTTTGAAGGCCGAAGTGGAGAAGGGTT | >hg38_21_+_8433172_8446622_RNA45SN1_45S_50ntFlanks | 10350 | 10383 | 34 | ACGAGAACTTTGAAGGCCGAAGTGGAGAAGGGTT | 6088.5 | -4.0 | 0.5 | -8.1 | 6.4E-16 | 3.1E-15 | BT20/MB468 | CYTO |
| >hg38_21_+_8433172_8446622_RNA45SN1_45S_50ntFlanks@485.505.21\|GATGTGGTGACGTCGTGCTCT | >hg38_21_+_8433172_8446622_RNA45SN1_45S_50ntFlanks | 485 | 505 | 21 | GATGTGGTGACGTCGTGCTCT | 24572.7 | 15.9 | 2.0 | 8.1 | 5.1E-16 | 3.1E-15 | MB231/MB468 | NUC |
| >MI0000298\|hsa-mir-221&WithFlank&X\|-\|45746151\|45746272@71.93.23(+1G)&[MIMAT0000278&hsa-miR-221-3p&offsets\|0\|0(+1G);m-91&X\|-\|45746181\|45746202&offsets\|0\|+1(+1G)]\|AGCTACATTGTCTGCTGGGTTTCG | >MI0000298\|hsa-mir-221&WithFlank&X\|-\|45746151\|45746272 | 71 | 93 | 23(+1G) | AGCTACATTGTCTGCTGGGTTTCG | 543.3 | 5.1 | 0.6 | 8.1 | 6.4E-16 | 3.1E-15 | BT20/MB468 | CYTO |
| >hg38_21_+_8433172_8446622_RNA45SN1_45S_50ntFlanks@10349.10369.21\|AACGAGAACTTTGAAGGCCGA | >hg38_21_+_8433172_8446622_RNA45SN1_45S_50ntFlanks | 10349 | 10369 | 21 | AACGAGAACTTTGAAGGCCGA | 500.7 | -3.2 | 0.4 | -8.0 | 1.0E-15 | 3.1E-15 | BT20/MB468 | NUC |
| >hg38_21_+_8433172_8446622_RNA45SN1_45S_50ntFlanks@6746.6771.26\|CACTTCGAACGCACTTGCGGCCCCGG | >hg38_21_+_8433172_8446622_RNA45SN1_45S_50ntFlanks | 6746 | 6771 | 26 | CACTTCGAACGCACTTGCGGCCCCGG | 544.2 | -7.9 | 1.0 | -8.1 | 6.5E-16 | 3.1E-15 | BT20/MB468 | CYTO |
| >hg38_21_+_8433172_8446622_RNA45SN1_45S_50ntFlanks@11209.11238.30\|CGGCGACTCTGGACGCGAGCCGGGCCCTTC | >hg38_21_+_8433172_8446622_RNA45SN1_45S_50ntFlanks | 11209 | 11238 | 30 | CGGCGACTCTGGACGCGAGCCGGGCCCTTC | 284.7 | -12.0 | 1.5 | -8.1 | 4.2E-16 | 3.1E-15 | MB231/MB468 | MP |

<table>
<tr><td>>hg38_21_+_8433172_8446622_RNA45SN1_45S_50ntFlanks@12642.12659.18|CATCTGTGGGATTATGAC</td><td>>hg38_21_+_8433172_8446622_RNA45SN1_45S_50ntFlanks</td><td>12642</td><td>12659</td><td>18</td><td>CATCTGTGGGATTATGAC</td><td>422.7</td><td>-12.6</td><td>1.5</td><td>-8.1</td><td>4.2E-16</td><td>3.1E-15</td><td>MB231/MB468</td><td>MP</td></tr>
<tr><td>>trna116_GluCTC_1_-_145399233_145399304@3.35.33,trna59_GluCTC_1_+_249168447_249168518@3.35.33,trna71_GluCTC_1_-_161439189_161439260@3.35.33,trna74_GluCTC_1_-_161431809_161431880@3.35.33,trna77_GluCTC_1_-_161424398_161424469@3.35.33,trna77_GluCTC_6_+_28949976_28950047@3.35.33,trna80_GluCTC_1_-_161417018_161417089@3.35.33,trna87_GluCTC_6_-_126101393_126101464@3.35.33|CCTGGTGGTCTAGTGGTTAGGATTCGGCGCTCT</td><td>>trna116_GluCTC_1_-_145399233_145399304</td><td>3</td><td>35</td><td>33</td><td>CCTGGTGGTCTAGTGGTTAGGATTCGGCGCTCT</td><td>642.6</td><td>6.0</td><td>0.7</td><td>8.1</td><td>4.7E-16</td><td>3.1E-15</td><td>BT20/MB468</td><td>MITO</td></tr>
<tr><td>>hg38_21_+_8433172_8446622_RNA45SN1_45S_50ntFlanks@10351.10374.24|CGAGAACTTTGAAGGCCGAAGTGG</td><td>>hg38_21_+_8433172_8446622_RNA45SN1_45S_50ntFlanks</td><td>10351</td><td>10374</td><td>24</td><td>CGAGAACTTTGAAGGCCGAAGTGG</td><td>1989.5</td><td>-5.2</td><td>0.6</td><td>-8.1</td><td>6.5E-16</td><td>3.2E-15</td><td>BT20/MB468</td><td>CYTO</td></tr>
<tr><td>>hg38_21_+_8433172_8446622_RNA45SN1_45S_50ntFlanks@12488.12509.22|AACGTGAGCTGGGTTTAGACCG</td><td>>hg38_21_+_8433172_8446622_RNA45SN1_45S_50ntFlanks</td><td>12488</td><td>12509</td><td>22</td><td>AACGTGAGCTGGGTTTAGACCG</td><td>480.4</td><td>-4.0</td><td>0.5</td><td>-8.1</td><td>4.7E-16</td><td>3.2E-15</td><td>BT20/MB468</td><td>MITO</td></tr>
<tr><td>>MI0000650|hsa-mir-200c&WithFlank&12|+|6963693|6963772@49.72.24&[MIMAT0000617&hsa-miR-200c-3p&offsets|-1|0;m-54&12|+|6963742|6963764&offsets|-1|0]|CTAATACTGCCGGGTAATGATGGA</td><td>>MI0000650|hsa-mir-200c&WithFlank&12|+|6963693|6963772</td><td>49</td><td>72</td><td>24</td><td>CTAATACTGCCGGGTAATGATGGA</td><td>571.7</td><td>9.3</td><td>1.1</td><td>8.4</td><td>4.0E-17</td><td>3.2E-15</td><td>BT20/MB231</td><td>CYTO</td></tr>
<tr><td>>hg38_21_+_8433172_8446622_RNA45SN1_45S_50ntFlanks@12073.12110.38|CCGAGGGGCTCTCGCTTCTGGCGCCAAGCGCCCGGCCG</td><td>>hg38_21_+_8433172_8446622_RNA45SN1_45S_50ntFlanks</td><td>12073</td><td>12110</td><td>38</td><td>CCGAGGGGCTCTCGCTTCTGGCGCCAAGCGCCCGGCCG</td><td>425.6</td><td>-12.6</td><td>1.5</td><td>-8.1</td><td>4.3E-16</td><td>3.2E-15</td><td>MB231/MB468</td><td>MP</td></tr>
<tr><td>>MI0000746|hsa-mir-99b&WithFlank&19|+|51692606|51692687@51.72.22(+1C)&[MIMAT0004678&hsa-miR-99b-3p&offsets|0|0(+1C);m-151&19|+|51692656|51692678&offsets|0|-1(+1C)]|CAAGCTCGTGTCTGTGGGTCCGC</td><td>>MI0000746|hsa-mir-99b&WithFlank&19|+|51692606|51692687</td><td>51</td><td>72</td><td>22(+1C)</td><td>CAAGCTCGTGTCTGTGGGTCCGC</td><td>513.1</td><td>9.0</td><td>1.1</td><td>8.1</td><td>6.6E-16</td><td>3.2E-15</td><td>BT20/MB468</td><td>CYTO</td></tr>
<tr><td>>MI0003599|hsa-mir-589&WithFlank&7|-|5495813|5495923@30.52.23&[MIMAT0004799&hsa-miR-589-5p&offsets|0|+1;m-162&7|-|5495874|5495894&offsets|0|+2]|TGAGAACCACGTCTGCTCTGAGC</td><td>>MI0003599|hsa-mir-589&WithFlank&7|-|5495813|5495923</td><td>30</td><td>52</td><td>23</td><td>TGAGAACCACGTCTGCTCTGAGC</td><td>2327.7</td><td>10.5</td><td>1.3</td><td>8.1</td><td>6.6E-16</td><td>3.2E-15</td><td>BT20/MB468</td><td>CYTO</td></tr>
<tr><td>>hg38_21_+_8433172_8446622_RNA45SN1_45S_50ntFlanks@11667.11704.38|CTGCCCAGTGCTCTGAATGTCAAAGTGAAGAAATTCAA</td><td>>hg38_21_+_8433172_8446622_RNA45SN1_45S_50ntFlanks</td><td>11667</td><td>11704</td><td>38</td><td>CTGCCCAGTGCTCTGAATGTCAAAGTGAAGAAATTCAA</td><td>198.3</td><td>8.5</td><td>1.0</td><td>8.1</td><td>4.6E-16</td><td>3.2E-15</td><td>MB231/MB468</td><td>CYTO</td></tr>
<tr><td>>am_MI0016746|hsa-mir-548o-2&WithFlank&20|+|38516557|38516638@51.71.21(+1U);MI0006402|hsa-mir-548o&WithFlank&7|-|102405736|102405861@93.113.21(+1U)&[MIMAT0005919&hsa-miR-548o-3p&offsets|0|-1(+1U);m-431&20|+|38516607|38516628&offsets|0|-1(+1U)];[MIMAT0005919_1&hsa-miR-548o-3p&offsets|0|-1(+1U);m-430&7|-|102405748|102405769&offsets|0|-1(+1U)]|CCAAAACTGCAGTTACTTTTGT</td><td>>am_MI0016746|hsa-mir-548o-2&WithFlank&20|+|38516557|38516638</td><td>51</td><td>71</td><td>21(+1U)</td><td>CCAAAACTGCAGTTACTTTTGT</td><td>616.9</td><td>9.3</td><td>1.1</td><td>8.1</td><td>6.6E-16</td><td>3.2E-15</td><td>BT20/MB468</td><td>CYTO</td></tr>
<tr><td>>hg38_21_+_8433172_8446622_RNA45SN1_45S_50ntFlanks@11561.11580.20|GCGCCTAGCAGCCGACTTAG</td><td>>hg38_21_+_8433172_8446622_RNA45SN1_45S_50ntFlanks</td><td>11561</td><td>11580</td><td>20</td><td>GCGCCTAGCAGCCGACTTAG</td><td>340.7</td><td>-12.2</td><td>1.5</td><td>-8.1</td><td>7.1E-16</td><td>3.2E-15</td><td>MB231/MB468</td><td>MITO</td></tr>
<tr><td>>MI0003137|hsa-mir-193b&WithFlank&16|+|14303961|14304055@57.78.22(+1A)&[MIMAT0002819&hsa-miR-193b-3p&offsets|0|0(+1A);m-166&16|+|14304017|14304038&offsets|0|0(+1A)]|AACTGGCCCTCAAAGTCCCGCTA</td><td>>MI0003137|hsa-mir-193b&WithFlank&16|+|14303961|14304055</td><td>57</td><td>78</td><td>22(+1A)</td><td>AACTGGCCCTCAAAGTCCCGCTA</td><td>63.6</td><td>8.8</td><td>1.1</td><td>8.0</td><td>1.0E-15</td><td>3.2E-15</td><td>BT20/MB468</td><td>NUC</td></tr>
<tr><td>>MI0000263|hsa-mir-7-1&WithFlank&9|-|83969742|83969863@30.55.26&[MIMAT0000252_2&hsa-miR-7-5p&offsets|0|+2;m-401&9|-|83969811|83969834&offsets|0|+2]|TGGAAGACTAGTGATTTTGTTGTTTT</td><td>>MI0000263|hsa-mir-7-1&WithFlank&9|-|83969742|83969863</td><td>30</td><td>55</td><td>26</td><td>TGGAAGACTAGTGATTTTGTTGTTTT</td><td>214.2</td><td>5.8</td><td>0.7</td><td>8.1</td><td>5.3E-16</td><td>3.2E-15</td><td>MB231/MB468</td><td>NUC</td></tr>
<tr><td>>hg38_21_+_8433172_8446622_RNA45SN1_45S_50ntFlanks@10830.10858.29|CAAGCCGGATCCGTAACTTCGGGATAAGG</td><td>>hg38_21_+_8433172_8446622_RNA45SN1_45S_50ntFlanks</td><td>10830</td><td>10858</td><td>29</td><td>CAAGCCGGATCCGTAACTTCGGGATAAGG</td><td>928.6</td><td>-9.5</td><td>1.2</td><td>-8.1</td><td>7.1E-16</td><td>3.2E-15</td><td>MB231/MB468</td><td>MITO</td></tr>
<tr><td>>hg38_21_+_8433172_8446622_RNA45SN1_45S_50ntFlanks@4760.4779.20|TCAGATACCGTCGTAGTTCC</td><td>>hg38_21_+_8433172_8446622_RNA45SN1_45S_50ntFlanks</td><td>4760</td><td>4779</td><td>20</td><td>TCAGATACCGTCGTAGTTCC</td><td>386.7</td><td>-12.4</td><td>1.5</td><td>-8.1</td><td>7.1E-16</td><td>3.2E-15</td><td>MB231/MB468</td><td>MITO</td></tr>
<tr><td>>hg38_21_+_8433172_8446622_RNA45SN1_45S_50ntFlanks@11562.11591.30|CGCCTAGCAGCCGACTTAGAACTGGTGCGG</td><td>>hg38_21_+_8433172_8446622_RNA45SN1_45S_50ntFlanks</td><td>11562</td><td>11591</td><td>30</td><td>CGCCTAGCAGCCGACTTAGAACTGGTGCGG</td><td>574.1</td><td>-13.0</td><td>1.6</td><td>-8.1</td><td>4.3E-16</td><td>3.2E-15</td><td>MB231/MB468</td><td>MP</td></tr>
<tr><td>>hg38_21_+_8433172_8446622_RNA45SN1_45S_50ntFlanks@12615.12653.39|GCTGAGGAGCCAATGGGGCGAAGCTACCATCTGTGGGAT</td><td>>hg38_21_+_8433172_8446622_RNA45SN1_45S_50ntFlanks</td><td>12615</td><td>12653</td><td>39</td><td>GCTGAGGAGCCAATGGGGCGAAGCTACCATCTGTGGGAT</td><td>1935.6</td><td>-10.7</td><td>1.3</td><td>-8.1</td><td>5.3E-16</td><td>3.2E-15</td><td>MB231/MB468</td><td>NUC</td></tr>
<tr><td>>hg38_21_+_8433172_8446622_RNA45SN1_45S_50ntFlanks@11579.11603.25|AGAACTGGTGCGGACCAGGGGAATC</td><td>>hg38_21_+_8433172_8446622_RNA45SN1_45S_50ntFlanks</td><td>11579</td><td>11603</td><td>25</td><td>AGAACTGGTGCGGACCAGGGGAATC</td><td>14165.4</td><td>-10.6</td><td>1.3</td><td>-8.1</td><td>4.3E-16</td><td>3.2E-15</td><td>MB231/MB468</td><td>MP</td></tr>
<tr><td>>hg38_21_+_8433172_8446622_RNA45SN1_45S_50ntFlanks@10794.10820.27|CTGGCATGTTGGAACAATGTAGGTAAG</td><td>>hg38_21_+_8433172_8446622_RNA45SN1_45S_50ntFlanks</td><td>10794</td><td>10820</td><td>27</td><td>CTGGCATGTTGGAACAATGTAGGTAAG</td><td>230.4</td><td>-11.7</td><td>1.4</td><td>-8.1</td><td>4.3E-16</td><td>3.2E-15</td><td>MB231/MB468</td><td>MP</td></tr>
<tr><td>>hg38_21_+_8433172_8446622_RNA45SN1_45S_50ntFlanks@485.518.34|GATGTGGTGACGTCGTGCTCTCCCGGGCCGGGTC</td><td>>hg38_21_+_8433172_8446622_RNA45SN1_45S_50ntFlanks</td><td>485</td><td>518</td><td>34</td><td>GATGTGGTGACGTCGTGCTCTCCCGGGCCGGGTC</td><td>2739.1</td><td>29.4</td><td>3.6</td><td>8.1</td><td>5.4E-16</td><td>3.2E-15</td><td>MB231/MB468</td><td>NUC</td></tr>
<tr><td>>hg38_21_+_8433172_8446622_RNA45SN1_45S_50ntFlanks@485.527.43|GATGTGGTGACGTCGTGCTCTCCCGGGCCGGGTCCGAGCCGCG</td><td>>hg38_21_+_8433172_8446622_RNA45SN1_45S_50ntFlanks</td><td>485</td><td>527</td><td>43</td><td>GATGTGGTGACGTCGTGCTCTCCCGGGCCGGGTCCGAGCCGCG</td><td>3231.9</td><td>29.4</td><td>3.6</td><td>8.1</td><td>5.4E-16</td><td>3.2E-15</td><td>MB231/MB468</td><td>NUC</td></tr>
<tr><td>>hg38_21_+_8433172_8446622_RNA45SN1_45S_50ntFlanks@10845.10863.19|ACTTCGGGATAAGGATTGG</td><td>>hg38_21_+_8433172_8446622_RNA45SN1_45S_50ntFlanks</td><td>10845</td><td>10863</td><td>19</td><td>ACTTCGGGATAAGGATTGG</td><td>469.5</td><td>-11.8</td><td>1.5</td><td>-8.1</td><td>5.3E-16</td><td>3.2E-15</td><td>MB231/MB468</td><td>NUC</td></tr>
<tr><td>>hg38_21_+_8433172_8446622_RNA45SN1_45S_50ntFlanks@9737.9754.18|CGAAACGATCTCAACCTA</td><td>>hg38_21_+_8433172_8446622_RNA45SN1_45S_50ntFlanks</td><td>9737</td><td>9754</td><td>18</td><td>CGAAACGATCTCAACCTA</td><td>242.8</td><td>-11.8</td><td>1.5</td><td>-8.1</td><td>4.3E-16</td><td>3.2E-15</td><td>MB231/MB468</td><td>MP</td></tr>
<tr><td>>hg38_21_+_8433172_8446622_RNA45SN1_45S_50ntFlanks@10680.10712.33|GTTCCGGCGGCGTCCGGTGAGCTCTCGCTGGCC</td><td>>hg38_21_+_8433172_8446622_RNA45SN1_45S_50ntFlanks</td><td>10680</td><td>10712</td><td>33</td><td>GTTCCGGCGGCGTCCGGTGAGCTCTCGCTGGCC</td><td>5522.5</td><td>-5.6</td><td>0.7</td><td>-8.1</td><td>4.7E-16</td><td>3.2E-15</td><td>MB231/MB468</td><td>CYTO</td></tr>
<tr><td>>hg38_21_+_8433172_8446622_RNA45SN1_45S_50ntFlanks@10729.10755.27|AGAGGGTGTAAATCTCGCGCCGGGCCG</td><td>>hg38_21_+_8433172_8446622_RNA45SN1_45S_50ntFlanks</td><td>10729</td><td>10755</td><td>27</td><td>AGAGGGTGTAAATCTCGCGCCGGGCCG</td><td>861.9</td><td>-10.0</td><td>1.2</td><td>-8.1</td><td>5.4E-16</td><td>3.2E-15</td><td>MB231/MB468</td><td>NUC</td></tr>
<tr><td>>hg38_1_-_228634819_228635039_RNA5S12_5S_50ntFlanks@153.171.19|ACCGGGTGCTGTAGGCTTT</td><td>>hg38_1_-_228634819_228635039_RNA5S12_5S_50ntFlanks</td><td>153</td><td>171</td><td>19</td><td>ACCGGGTGCTGTAGGCTTT</td><td>19408.1</td><td>6.8</td><td>0.8</td><td>8.1</td><td>6.7E-16</td><td>3.2E-15</td><td>BT20/MB468</td><td>CYTO</td></tr>
<tr><td>>hg38_21_+_8433172_8446622_RNA45SN1_45S_50ntFlanks@5477.5516.40|CCTGGCGGAGCGCTGAGAAGACGGTCGAACTTGACTATCT</td><td>>hg38_21_+_8433172_8446622_RNA45SN1_45S_50ntFlanks</td><td>5477</td><td>5516</td><td>40</td><td>CCTGGCGGAGCGCTGAGAAGACGGTCGAACTTGACTATCT</td><td>2366.0</td><td>11.2</td><td>1.4</td><td>8.1</td><td>6.7E-16</td><td>3.2E-15</td><td>BT20/MB468</td><td>CYTO</td></tr>
<tr><td>>hg38_21_+_8433172_8446622_RNA45SN1_45S_50ntFlanks@8636.8658.23|CGGCGACCGGCCGCCGCCGGGCG</td><td>>hg38_21_+_8433172_8446622_RNA45SN1_45S_50ntFlanks</td><td>8636</td><td>8658</td><td>23</td><td>CGGCGACCGGCCGCCGCCGGGCG</td><td>1024.6</td><td>-13.8</td><td>1.7</td><td>-8.1</td><td>7.2E-16</td><td>3.2E-15</td><td>MB231/MB468</td><td>MITO</td></tr>
<tr><td>>hg38_21_+_8433172_8446622_RNA45SN1_45S_50ntFlanks@11618.11640.23|AAACAAAGCATCGCGAAGGCCCG</td><td>>hg38_21_+_8433172_8446622_RNA45SN1_45S_50ntFlanks</td><td>11618</td><td>11640</td><td>23</td><td>AAACAAAGCATCGCGAAGGCCCG</td><td>674.9</td><td>-11.5</td><td>1.4</td><td>-8.1</td><td>4.8E-16</td><td>3.2E-15</td><td>BT20/MB468</td><td>MITO</td></tr>
<tr><td>>hg38_21_+_8433172_8446622_RNA45SN1_45S_50ntFlanks@12991.13015.25|CTCCCTCGCTGCGATCTATTGAAAG</td><td>>hg38_21_+_8433172_8446622_RNA45SN1_45S_50ntFlanks</td><td>12991</td><td>13015</td><td>25</td><td>CTCCCTCGCTGCGATCTATTGAAAG</td><td>3789.5</td><td>-6.2</td><td>0.8</td><td>-8.1</td><td>4.8E-16</td><td>3.2E-15</td><td>BT20/MB468</td><td>MITO</td></tr>
<tr><td>>hg38_21_+_8433172_8446622_RNA45SN1_45S_50ntFlanks@11891.11910.20|TTGACTCTAGTCTGGCACGG</td><td>>hg38_21_+_8433172_8446622_RNA45SN1_45S_50ntFlanks</td><td>11891</td><td>11910</td><td>20</td><td>TTGACTCTAGTCTGGCACGG</td><td>187.9</td><td>-11.4</td><td>1.4</td><td>-8.1</td><td>4.4E-16</td><td>3.2E-15</td><td>MB231/MB468</td><td>MP</td></tr>
<tr><td>>hg38_21_+_8433172_8446622_RNA45SN1_45S_50ntFlanks@12934.12955.22|CTAAACCATTCGTAGACGACCT</td><td>>hg38_21_+_8433172_8446622_RNA45SN1_45S_50ntFlanks</td><td>12934</td><td>12955</td><td>22</td><td>CTAAACCATTCGTAGACGACCT</td><td>213.5</td><td>-11.6</td><td>1.4</td><td>-8.1</td><td>4.4E-16</td><td>3.2E-15</td><td>MB231/MB468</td><td>MP</td></tr>
<tr><td>>hg38_21_+_8433172_8446622_RNA45SN1_45S_50ntFlanks@8191.8215.25|CAGCCCGTGGACGGTGTGAGGCCGG</td><td>>hg38_21_+_8433172_8446622_RNA45SN1_45S_50ntFlanks</td><td>8191</td><td>8215</td><td>25</td><td>CAGCCCGTGGACGGTGTGAGGCCGG</td><td>1429.8</td><td>-10.0</td><td>1.2</td><td>-8.1</td><td>4.4E-16</td><td>3.2E-15</td><td>MB231/MB468</td><td>MP</td></tr>
<tr><td>>hg38_21_+_8433172_8446622_RNA45SN1_45S_50ntFlanks@11234.11254.21|CCTTCCCGTGGATCGCCCCAG</td><td>>hg38_21_+_8433172_8446622_RNA45SN1_45S_50ntFlanks</td><td>11234</td><td>11254</td><td>21</td><td>CCTTCCCGTGGATCGCCCCAG</td><td>405.4</td><td>-11.4</td><td>1.4</td><td>-8.1</td><td>4.8E-16</td><td>3.3E-15</td><td>MB231/MB468</td><td>CYTO</td></tr>
<tr><td>>MI0000783|hsa-mir-375&WithFlank&2|-|219001639|219001714@46.67.22&[MIMAT0000728&hsa-miR-375-3p&offsets|0|0;m-92&2|-|219001648|219001669&offsets|0|0]|TTTGTTCGTTCGGCTCGCGTGA</td><td>>MI0000783|hsa-mir-375&WithFlank&2|-|219001639|219001714</td><td>46</td><td>67</td><td>22</td><td>TTTGTTCGTTCGGCTCGCGTGA</td><td>271.1</td><td>-11.9</td><td>1.5</td><td>-8.1</td><td>4.4E-16</td><td>3.3E-15</td><td>MB231/MB468</td><td>MP</td></tr>
<tr><td>>hg38_21_+_8433172_8446622_RNA45SN1_45S_50ntFlanks@10463.10484.22|CTCCGTTGCCCTCGGCCGATCG</td><td>>hg38_21_+_8433172_8446622_RNA45SN1_45S_50ntFlanks</td><td>10463</td><td>10484</td><td>22</td><td>CTCCGTTGCCCTCGGCCGATCG</td><td>175.1</td><td>-5.9</td><td>0.7</td><td>-8.1</td><td>4.8E-16</td><td>3.3E-15</td><td>MB231/MB468</td><td>CYTO</td></tr>
<tr><td>>am_trna11_ProAGG_16_+_3241989_3242060@1.28.28,trna12_ProAGG_6_+_26555498_26555569@1.28.28,trna12_ProTGG_11_-_75946869_75946940@1.28.28,trna14_ProTGG_5_-_180615854_180615925@1.28.28,trna22_ProAGG_14_-_21081560_21081631@1.28.28,trna23_ProAGG_14_-_21077495_21077566@1.28.28,trna28_ProTGG_16_-_3234133_3234204@1.28.28,trna29_ProAGG_16_-_3232635_3232706@1.28.28,trna2_ProAGG_7_+_128423504_128423575@1.28.28,trna30_ProCGG_6_+_27059521_27059592@1.28.28,trna37_ProCGG_17_-_8126151_8126222@1.28.28,trna3_ProTGG_16_+_3208923_3208994@1.28.28,trna52_ProCGG_1_+_167683962_167684033@1.28.28,trna65_ProAGG_1_-_167684725_167684796@1.28.28,trna6_ProCGG_16_+_3222049_3222120@1.28.28,trna6_ProTGG_14_+_21152175_21152246@1.28.28,trna8_ProTGG_16_+_3238094_3238165@1.28.28,trna9_ProAGG_11_+_75946557_75946628@1.28.28,trna9_ProAGG_16_+_3239634_3239705@1.28.28|GGCTCGTTGGTCTAGGGGTATGATTCTC</td><td>>am_trna11_ProAGG_16_+_3241989_3242060</td><td>1</td><td>28</td><td>28</td><td>GGCTCGTTGGTCTAGGGGTATGATTCTC</td><td>5515.1</td><td>-12.6</td><td>1.6</td><td>-8.0</td><td>1.1E-15</td><td>3.3E-15</td><td>BT20/MB468</td><td>NUC</td></tr>
<tr><td>>hg38_21_+_8433172_8446622_RNA45SN1_45S_50ntFlanks@6730.6771.42|ACACATTGATCATCGACACTTCGAACGCACTTGCGGCCCCGG</td><td>>hg38_21_+_8433172_8446622_RNA45SN1_45S_50ntFlanks</td><td>6730</td><td>6771</td><td>42</td><td>ACACATTGATCATCGACACTTCGAACGCACTTGCGGCCCCGG</td><td>1602.7</td><td>-8.2</td><td>1.0</td><td>-8.1</td><td>7.4E-16</td><td>3.3E-15</td><td>MB231/MB468</td><td>MITO</td></tr>
<tr><td>>hg38_21_+_8433172_8446622_RNA45SN1_45S_50ntFlanks@8528.8551.24|CTTCCCCCGCCGCCCCTCCTCCTC</td><td>>hg38_21_+_8433172_8446622_RNA45SN1_45S_50ntFlanks</td><td>8528</td><td>8551</td><td>24</td><td>CTTCCCCCGCCGCCCCTCCTCCTC</td><td>407.3</td><td>-12.5</td><td>1.5</td><td>-8.1</td><td>7.4E-16</td><td>3.3E-15</td><td>MB231/MB468</td><td>MITO</td></tr>
<tr><td>>hg38_21_+_8433172_8446622_RNA45SN1_45S_50ntFlanks@12457.12478.22|CAAGCGTTGGATTGTTCACCCA</td><td>>hg38_21_+_8433172_8446622_RNA45SN1_45S_50ntFlanks</td><td>12457</td><td>12478</td><td>22</td><td>CAAGCGTTGGATTGTTCACCCA</td><td>1405.5</td><td>-4.8</td><td>0.6</td><td>-8.1</td><td>5.0E-16</td><td>3.3E-15</td><td>BT20/MB468</td><td>MITO</td></tr>
<tr><td>>MI0000266|hsa-mir-10a&WithFlank&17|-|48579832|48579953@29.48.20(+1C)&[MIMAT0000253&hsa-miR-10a-5p&offsets|+1|-2(+1C);m-18&17|-|48579905|48579926&offsets|+1|-1(+1C)]|ACCCTGTAGATCCGAATTTGC</td><td>>MI0000266|hsa-mir-10a&WithFlank&17|-|48579832|48579953</td><td>29</td><td>48</td><td>20(+1C)</td><td>ACCCTGTAGATCCGAATTTGC</td><td>57.9</td><td>8.7</td><td>1.1</td><td>8.0</td><td>1.1E-15</td><td>3.3E-15</td><td>BT20/MB468</td><td>NUC</td></tr>
</table>

| | | | | | | | | | | | | | |
|---|---|---|---|---|---|---|---|---|---|---|---|---|---|
| >hg38_21_+_8433172_8446622_RNA45SN1_45S_50ntFlanks@8107.8140.34\|CGGCGGGGCGCGGGACATGTGGCGTACGGAAGAC | >hg38_21_+_8433172_8446622_RNA45SN1_45S_50ntFlanks | 8107 | 8140 | 34 | CGGCGGGGCGCGGGACATGTGGCGTACGGAAGAC | 388.4 | -12.4 | 1.5 | -8.1 | 7.4E-16 | 3.3E-15 | MB231/MB468 | MITO |
| >hg38_21_+_8433172_8446622_RNA45SN1_45S_50ntFlanks@4444.4471.28\|CCTTGCCTCTCGGCGCCCCCTCGATGCT | >hg38_21_+_8433172_8446622_RNA45SN1_45S_50ntFlanks | 4444 | 4471 | 28 | CCTTGCCTCTCGGCGCCCCCTCGATGCT | 403.3 | -12.5 | 1.5 | -8.1 | 4.5E-16 | 3.3E-15 | MB231/MB468 | MP |
| >hg38_21_+_8433172_8446622_RNA45SN1_45S_50ntFlanks@8154.8178.25\|CCGCTCGTGGGGGGCCCAAGTCCTT | >hg38_21_+_8433172_8446622_RNA45SN1_45S_50ntFlanks | 8154 | 8178 | 25 | CCGCTCGTGGGGGGCCCAAGTCCTT | 420.2 | -12.6 | 1.5 | -8.1 | 4.5E-16 | 3.3E-15 | MB231/MB468 | MP |
| >hg38_21_+_8433172_8446622_RNA45SN1_45S_50ntFlanks@11660.11685.26\|GTGATTTCTGCCCAGTGCTCTGAATG | >hg38_21_+_8433172_8446622_RNA45SN1_45S_50ntFlanks | 11660 | 11685 | 26 | GTGATTTCTGCCCAGTGCTCTGAATG | 4912.5 | -9.2 | 1.1 | -8.1 | 7.4E-16 | 3.3E-15 | MB231/MB468 | MITO |
| >hg38_21_+_8433172_8446622_RNA45SN1_45S_50ntFlanks@9573.9595.23\|AAATCGGTCGTCCGACCTGGGTA | >hg38_21_+_8433172_8446622_RNA45SN1_45S_50ntFlanks | 9573 | 9595 | 23 | AAATCGGTCGTCCGACCTGGGTA | 5015.5 | -9.9 | 1.2 | -8.1 | 4.5E-16 | 3.3E-15 | MB231/MB468 | MP |
| >hg38_21_+_8433172_8446622_RNA45SN1_45S_50ntFlanks@5467.5491.25\|GGCCCACGGCCCTGGCGGAGCGCTG | >hg38_21_+_8433172_8446622_RNA45SN1_45S_50ntFlanks | 5467 | 5491 | 25 | GGCCCACGGCCCTGGCGGAGCGCTG | 260.2 | -11.0 | 1.4 | -8.1 | 5.6E-16 | 3.3E-15 | MB231/MB468 | NUC |
| >hg38_21_+_8433172_8446622_RNA45SN1_45S_50ntFlanks@6739.6771.33\|TCATCGACACTTCGAACGCACTTGCGGCCCCGG | >hg38_21_+_8433172_8446622_RNA45SN1_45S_50ntFlanks | 6739 | 6771 | 33 | TCATCGACACTTCGAACGCACTTGCGGCCCCGG | 496.2 | -12.8 | 1.6 | -8.1 | 4.5E-16 | 3.3E-15 | MB231/MB468 | MP |
| >hg38_21_+_8433172_8446622_RNA45SN1_45S_50ntFlanks@7978.8018.41\|GACCTCAGATCAGACGTGGCGACCCGCTGAATTTAAGCATA | >hg38_21_+_8433172_8446622_RNA45SN1_45S_50ntFlanks | 7978 | 8018 | 41 | GACCTCAGATCAGACGTGGCGACCCGCTGAATTTAAGCATA | 270.8 | 12.1 | 1.5 | 8.1 | 4.9E-16 | 3.3E-15 | MB231/MB468 | CYTO |
| >hg38_21_+_8433172_8446622_RNA45SN1_45S_50ntFlanks@8845.8867.23\|CGGCGGCGGCGGCGGCGGCGGGA | >hg38_21_+_8433172_8446622_RNA45SN1_45S_50ntFlanks | 8845 | 8867 | 23 | CGGCGGCGGCGGCGGCGGCGGGA | 998.7 | -12.1 | 1.5 | -8.1 | 5.0E-16 | 3.4E-15 | BT20/MB468 | MITO |
| >hg38_21_+_8433172_8446622_RNA45SN1_45S_50ntFlanks@4759.4786.28\|ATCAGATACCGTCGTAGTTCCGACCATA | >hg38_21_+_8433172_8446622_RNA45SN1_45S_50ntFlanks | 4759 | 4786 | 28 | ATCAGATACCGTCGTAGTTCCGACCATA | 871.1 | -12.8 | 1.5 | -8.5 | 1.4E-17 | 3.4E-15 | BT20/MB468 | MP |
| >hg38_21_+_8433172_8446622_RNA45SN1_45S_50ntFlanks@10186.10209.24\|TCCTCCCCTCCTCCCGCCCACGCC | >hg38_21_+_8433172_8446622_RNA45SN1_45S_50ntFlanks | 10186 | 10209 | 24 | TCCTCCCCTCCTCCCGCCCACGCC | 393.1 | -11.6 | 1.4 | -8.1 | 5.6E-16 | 3.4E-15 | MB231/MB468 | NUC |
| >am_trna106_HisGTG_1_-_149155828_149155899@7.28.22,trna111_HisGTG_1_-_147774845_147774916@7.28.22,trna118_HisGTG_1_-_145396881_145396952@7.28.22,trna16_HisGTG_1_+_146544773_146544844@7.28.22,trna1_HisGTG_15_+_45493349_45493420@7.28.22,trna21_HisGTG_1_+_147753471_147753542@7.28.22,trna33_HisGTG_6_+_27125906_27125977@7.28.22,trna7_HisGTG_9_-_14433938_14434009@7.28.22,trna8_HisGTG_15_-_45492611_45492682@7.28.22,trna9_HisGTG_15_-_45490804_45490875@7.28.22\|ATCGTATAGTGGTTAGTACTCT | >am_trna106_HisGTG_1_-_149155828_149155899 | 7 | 28 | 22 | ATCGTATAGTGGTTAGTACTCT | 908.8 | -12.5 | 1.5 | -8.1 | 4.9E-16 | 3.4E-15 | MB231/MB468 | CYTO |
| >trna14_LysTTT_11_-_59327808_59327880@1.33.33,trna2_LysTTT_17_+_8022473_8022545@1.33.33,trna54_LysTTT_1_+_204475655_204475727@1.33.33,trna5_LysTTT_11_+_59323902_59323974@1.33.33,trna62_LysTTT_1_-_204476158_204476230@1.33.33,trna76_LysTTT_6_+_28918806_28918878@1.33.33\|GCCCGGATAGCTCAGTCGGTAGAGCATCAGACT | >trna14_LysTTT_11_-_59327808_59327880 | 1 | 33 | 33 | GCCCGGATAGCTCAGTCGGTAGAGCATCAGACT | 4582.0 | 7.3 | 0.9 | 8.1 | 5.0E-16 | 3.4E-15 | MB231/MB468 | CYTO |
| >hg38_21_+_8433172_8446622_RNA45SN1_45S_50ntFlanks@11580.11605.26\|GAACTGGTGCGGACCAGGGGAATCCG | >hg38_21_+_8433172_8446622_RNA45SN1_45S_50ntFlanks | 11580 | 11605 | 26 | GAACTGGTGCGGACCAGGGGAATCCG | 2026.5 | -9.8 | 1.2 | -8.1 | 7.6E-16 | 3.4E-15 | MB231/MB468 | MITO |
| >hg38_21_+_8433172_8446622_RNA45SN1_45S_50ntFlanks@3858.3898.41\|TGGTAATTCTAGAGCTAATACATGCCGACGGGCGCTGACCC | >hg38_21_+_8433172_8446622_RNA45SN1_45S_50ntFlanks | 3858 | 3898 | 41 | TGGTAATTCTAGAGCTAATACATGCCGACGGGCGCTGACCC | 473.8 | -13.3 | 1.7 | -8.0 | 1.1E-15 | 3.4E-15 | BT20/MB468 | NUC |
| >trna10_MetCAT_8_-_124169470_124169542@31.51.21,trna22_MetCAT_16_-_87417628_87417700@31.51.21\|TCTCATAATCTGAAGGTCGTG | >trna10_MetCAT_8_-_124169470_124169542 | 31 | 51 | 21 | TCTCATAATCTGAAGGTCGTG | 1200.9 | -9.0 | 1.1 | -8.1 | 5.0E-16 | 3.4E-15 | MB231/MB468 | CYTO |
| >hg38_21_+_8433172_8446622_RNA45SN1_45S_50ntFlanks@12126.12150.25\|CGACCCGCTCCGGGGACAGTGCCAG | >hg38_21_+_8433172_8446622_RNA45SN1_45S_50ntFlanks | 12126 | 12150 | 25 | CGACCCGCTCCGGGGACAGTGCCAG | 650.4 | -8.6 | 1.1 | -8.1 | 5.0E-16 | 3.4E-15 | MB231/MB468 | CYTO |
| >am_trna128_GlyGCC_6_-_27870686_27870756@1.36.36,trna18_GlyGCC_16_+_70822597_70822667@1.36.36,trna19_GlyGCC_16_+_70823410_70823480@1.36.36,trna19_GlyGCC_2_-_157257659_157257729@1.36.36,trna24_GlyGCC_16_-_70812942_70813012@1.36.36,trna25_GlyGCC_16_-_70812114_70812184@1.36.36,trna5_GlyGCC_17_+_8029064_8029134@1.36.36,trna68_GlyGCC_1_-_161493637_161493707@1.36.36\|GCATTGGTGGTTCAGTGGTAGAATTCTCGCCTGCCA | >am_trna128_GlyGCC_6_-_27870686_27870756 | 1 | 36 | 36 | GCATTGGTGGTTCAGTGGTAGAATTCTCGCCTGCCA | 1315.5 | 5.2 | 0.6 | 8.1 | 5.0E-16 | 3.4E-15 | MB231/MB468 | CYTO |
| >MI0000477\|hsa-mir-146a&WithFlank&5\|+\|160485346\|160485456@27.49.23&[MIMAT0000449&hsa-miR-146a-5p&offsets\|0\|+1;m-129&5\|+\|160485372\|160485393&offsets\|0\|+1]\|TGAGAACTGAATTCCATGGGTTG | >MI0000477\|hsa-mir-146a&WithFlank&5\|+\|160485346\|160485456 | 27 | 49 | 23 | TGAGAACTGAATTCCATGGGTTG | 1529.4 | -13.7 | 1.6 | -8.3 | 7.7E-17 | 3.4E-15 | BT20/MB231 | NUC |
| >hg38_21_+_8433172_8446622_RNA45SN1_45S_50ntFlanks@10107.10125.19\|CGTGTGGTGTGCGTCGGAG | >hg38_21_+_8433172_8446622_RNA45SN1_45S_50ntFlanks | 10107 | 10125 | 19 | CGTGTGGTGTGCGTCGGAG | 458.5 | -11.8 | 1.5 | -8.1 | 5.7E-16 | 3.4E-15 | MB231/MB468 | NUC |
| >hg38_21_+_8433172_8446622_RNA45SN1_45S_50ntFlanks@474.505.32\|TCTTCGTGATCGATGTGGTGACGTCGTGCTCT | >hg38_21_+_8433172_8446622_RNA45SN1_45S_50ntFlanks | 474 | 505 | 32 | TCTTCGTGATCGATGTGGTGACGTCGTGCTCT | 361.6 | 9.1 | 1.1 | 8.1 | 7.1E-16 | 3.4E-15 | BT20/MB468 | CYTO |
| >hg38_21_+_8433172_8446622_RNA45SN1_45S_50ntFlanks@11579.11606.28\|AGAACTGGTGCGGACCAGGGGAATCCGA | >hg38_21_+_8433172_8446622_RNA45SN1_45S_50ntFlanks | 11579 | 11606 | 28 | AGAACTGGTGCGGACCAGGGGAATCCGA | 342.6 | -12.3 | 1.5 | -8.1 | 4.6E-16 | 3.4E-15 | MB231/MB468 | MP |
| >hg38_21_+_8433172_8446622_RNA45SN1_45S_50ntFlanks@9313.9334.22\|TCTAACACGTGCGCGAGTCGGG | >hg38_21_+_8433172_8446622_RNA45SN1_45S_50ntFlanks | 9313 | 9334 | 22 | TCTAACACGTGCGCGAGTCGGG | 492.2 | -12.8 | 1.6 | -8.1 | 4.6E-16 | 3.4E-15 | MB231/MB468 | MP |
| >hg38_21_+_8433172_8446622_RNA45SN1_45S_50ntFlanks@8106.8139.34\|GCGGCGGGGCGCGGGACATGTGGCGTACGGAAGA | >hg38_21_+_8433172_8446622_RNA45SN1_45S_50ntFlanks | 8106 | 8139 | 34 | GCGGCGGGGCGCGGGACATGTGGCGTACGGAAGA | 244.1 | -11.8 | 1.5 | -8.1 | 4.6E-16 | 3.4E-15 | MB231/MB468 | MP |
| >hg38_21_+_8433172_8446622_RNA45SN1_45S_50ntFlanks@4729.4752.24\|TCAAGAACGAAAGTCGGAGGTTCG | >hg38_21_+_8433172_8446622_RNA45SN1_45S_50ntFlanks | 4729 | 4752 | 24 | TCAAGAACGAAAGTCGGAGGTTCG | 263.0 | -11.8 | 1.5 | -8.1 | 7.7E-16 | 3.4E-15 | MB231/MB468 | MITO |
| >hg38_21_+_8433172_8446622_RNA45SN1_45S_50ntFlanks@7988.8012.25\|CAGACGTGGCGACCCGCTGAATTTA | >hg38_21_+_8433172_8446622_RNA45SN1_45S_50ntFlanks | 7988 | 8012 | 25 | CAGACGTGGCGACCCGCTGAATTTA | 1888.3 | -6.1 | 0.8 | -8.0 | 1.1E-15 | 3.4E-15 | BT20/MB468 | NUC |
| >MI0000440\|hsa-mir-27b&WithFlank&9\|+\|95085439\|95085547@67.87.21(+1G)&[MIMAT0000419&hsa-miR-27b-3p&offsets\|0\|0(+1G);m-39&9\|+\|95085505\|95085525&offsets\|0\|0(+1G)]\|TTCACAGTGGCTAAGTTCTGCG | >MI0000440\|hsa-mir-27b&WithFlank&9\|+\|95085439\|95085547 | 67 | 87 | 21(+1G) | TTCACAGTGGCTAAGTTCTGCG | 182.8 | 6.1 | 0.8 | 8.1 | 5.8E-16 | 3.4E-15 | MB231/MB468 | NUC |
| >hg38_21_+_8433172_8446622_RNA45SN1_45S_50ntFlanks@6719.6736.18\|TGAATTGCAGGACACATT | >hg38_21_+_8433172_8446622_RNA45SN1_45S_50ntFlanks | 6719 | 6736 | 18 | TGAATTGCAGGACACATT | 10900.7 | -10.6 | 1.3 | -8.0 | 1.1E-15 | 3.5E-15 | BT20/MB468 | NUC |
| >hg38_21_+_8433172_8446622_RNA45SN1_45S_50ntFlanks@12415.12445.31\|CTTCGATGTCGGCTCTTCCTATCATTGTGAA | >hg38_21_+_8433172_8446622_RNA45SN1_45S_50ntFlanks | 12415 | 12445 | 31 | CTTCGATGTCGGCTCTTCCTATCATTGTGAA | 864.1 | -4.8 | 0.6 | -8.0 | 1.1E-15 | 3.5E-15 | BT20/MB468 | NUC |
| >MI0000279\|hsa-mir-196a-2&WithFlank&12\|+\|53991732\|53991853@31.51.21;MI0000238\|hsa-mir-196a-1&WithFlank&17\|-\|48632484\|48632565@13.33.21&[MIMAT0000226&hsa-miR-196a-5p&offsets\|0\|-1;m-261&12\|+\|53991762\|53991783&offsets\|0\|-1];[MIMAT0000226_1&hsa-miR-196a-5p&offsets\|0\|-1;m-268&17\|-\|48632532\|48632553&offsets\|0\|-1]\|TAGGTAGTTTCATGTTGTTGG | >MI0000279\|hsa-mir-196a-2&WithFlank&12\|+\|53991732\|53991853 | 31 | 51 | 21 | TAGGTAGTTTCATGTTGTTGG | 272.7 | 11.7 | 1.4 | 8.4 | 6.2E-17 | 3.5E-15 | BT20/MB231 | MITO |
| >MI0000285\|hsa-mir-205&WithFlank&1\|+\|209432127\|209432248@40.60.21(+1U)&[MIMAT0000266&hsa-miR-205-5p&offsets\|0\|-1(+1U);m-351&1\|+\|209432166\|209432187&offsets\|0\|-1(+1U)]\|TCCTTCATTCCACCGGAGTCTT | >MI0000285\|hsa-mir-205&WithFlank&1\|+\|209432127\|209432248 | 40 | 60 | 21(+1U) | TCCTTCATTCCACCGGAGTCTT | 460.7 | 12.5 | 1.5 | 8.4 | 6.3E-17 | 3.5E-15 | BT20/MB231 | MITO |
| >hg38_21_+_8433172_8446622_RNA45SN1_45S_50ntFlanks@4370.4391.22\|TTAAAAAGCTCGTAGTTGGATC | >hg38_21_+_8433172_8446622_RNA45SN1_45S_50ntFlanks | 4370 | 4391 | 22 | TTAAAAAGCTCGTAGTTGGATC | 350.3 | -12.9 | 1.6 | -8.0 | 1.1E-15 | 3.5E-15 | BT20/MB468 | NUC |
| >hg38_21_+_8433172_8446622_RNA45SN1_45S_50ntFlanks@974.992.19\|CGCGTGGGTCCTGAGGGAG | >hg38_21_+_8433172_8446622_RNA45SN1_45S_50ntFlanks | 974 | 992 | 19 | CGCGTGGGTCCTGAGGGAG | 301.1 | -11.2 | 1.4 | -8.1 | 5.9E-16 | 3.5E-15 | MB231/MB468 | NUC |
| >hg38_21_+_8433172_8446622_RNA45SN1_45S_50ntFlanks@4003.4022.20\|ATAACCTCGGGCCGATCGCA | >hg38_21_+_8433172_8446622_RNA45SN1_45S_50ntFlanks | 4003 | 4022 | 20 | ATAACCTCGGGCCGATCGCA | 207.0 | -11.6 | 1.4 | -8.1 | 4.8E-16 | 3.5E-15 | MB231/MB468 | MP |
| >hg38_21_+_8433172_8446622_RNA45SN1_45S_50ntFlanks@10679.10706.28\|GGTTCCGGCGGCGTCCGGTGAGCTCTCG | >hg38_21_+_8433172_8446622_RNA45SN1_45S_50ntFlanks | 10679 | 10706 | 28 | GGTTCCGGCGGCGTCCGGTGAGCTCTCG | 4896.6 | -8.3 | 1.0 | -8.1 | 5.2E-16 | 3.5E-15 | MB231/MB468 | CYTO |
| >hg38_21_+_8433172_8446622_RNA45SN1_45S_50ntFlanks@4729.4758.30\|TCAAGAACGAAAGTCGGAGGTTCGAAGACG | >hg38_21_+_8433172_8446622_RNA45SN1_45S_50ntFlanks | 4729 | 4758 | 30 | TCAAGAACGAAAGTCGGAGGTTCGAAGACG | 4499.5 | -5.8 | 0.7 | -8.1 | 4.8E-16 | 3.5E-15 | MB231/MB468 | MP |
| >hg38_21_+_8433172_8446622_RNA45SN1_45S_50ntFlanks@10826.10858.33\|TCGGCAAGCCGGATCCGTAACTTCGGGATAAGG | >hg38_21_+_8433172_8446622_RNA45SN1_45S_50ntFlanks | 10826 | 10858 | 33 | TCGGCAAGCCGGATCCGTAACTTCGGGATAAGG | 305.8 | -12.0 | 1.5 | -8.1 | 7.9E-16 | 3.6E-15 | MB231/MB468 | MITO |
| >hg38_21_+_8433172_8446622_RNA45SN1_45S_50ntFlanks@4723.4755.33\|CATTAATCAAGAACGAAAGTCGGAGGTTCGAAG | >hg38_21_+_8433172_8446622_RNA45SN1_45S_50ntFlanks | 4723 | 4755 | 33 | CATTAATCAAGAACGAAAGTCGGAGGTTCGAAG | 284.4 | -11.9 | 1.5 | -8.1 | 8.0E-16 | 3.6E-15 | MB231/MB468 | MITO |
| >MI0000433\|hsa-let-7g&WithFlank&3\|-\|52268272\|52268367@11.31.21(+1A)&[MIMAT0000414&hsa-let-7g-5p&offsets\|0\|-1(+1A);m-58&3\|-\|52268336\|52268357&offsets\|0\|-1(+1A)]\|TGAGGTAGTAGTTTGTACAGTA | >MI0000433\|hsa-let-7g&WithFlank&3\|-\|52268272\|52268367 | 11 | 31 | 21(+1A) | TGAGGTAGTAGTTTGTACAGTA | 2219.2 | 5.2 | 0.6 | 8.1 | 6.0E-16 | 3.6E-15 | MB231/MB468 | NUC |
| >hg38_21_+_8433172_8446622_RNA45SN1_45S_50ntFlanks@12991.13008.18\|CTCCCTCGCTGCGATCTA | >hg38_21_+_8433172_8446622_RNA45SN1_45S_50ntFlanks | 12991 | 13008 | 18 | CTCCCTCGCTGCGATCTA | 661.0 | -12.3 | 1.5 | -8.1 | 6.0E-16 | 3.6E-15 | MB231/MB468 | NUC |
| >hg38_21_+_8433172_8446622_RNA45SN1_45S_50ntFlanks@12414.12442.29\|CCTTCGATGTCGGCTCTTCCTATCATTGT | >hg38_21_+_8433172_8446622_RNA45SN1_45S_50ntFlanks | 12414 | 12442 | 29 | CCTTCGATGTCGGCTCTTCCTATCATTGT | 396.4 | 7.0 | 0.9 | 8.1 | 5.3E-16 | 3.6E-15 | MB231/MB468 | CYTO |
| >hg38_21_+_8433172_8446622_RNA45SN1_45S_50ntFlanks@8356.8377.22\|GAAAAGAACTTTGAAGAGAGAG | >hg38_21_+_8433172_8446622_RNA45SN1_45S_50ntFlanks | 8356 | 8377 | 22 | GAAAAGAACTTTGAAGAGAGAG | 2447.4 | -9.1 | 1.1 | -8.1 | 4.9E-16 | 3.6E-15 | MB231/MB468 | MP |

<table>
<tr><td>>MI0000065|hsa-let-7d&WithFlank&9|+|94178828|94178926@14.34.21(+1A)&[MIMAT0000065&hsa-let-7d-5p&offsets|0|-1(+1A);m-90&9|+|94178841|94178862&offsets|0|-1(+1A)]|AGAGGTAGTAGGTTGCATAGTA</td><td>>MI0000065|hsa-let-7d&WithFlank&9|+|94178828|94178926</td><td>14</td><td>34</td><td>21(+1A)</td><td>AGAGGTAGTAGGTTGCATAGTA</td><td>963.8</td><td>9.0</td><td>1.1</td><td>8.1</td><td>7.5E-16</td><td>3.6E-15</td><td>BT20/MB468</td><td>CYTO</td></tr>
<tr><td>>hg38_21_+_8433172_8446622_RNA45SN1_45S_50ntFlanks@11999.12035.37|CCCTGCGGGCCGCCGGTGAAATACCACTACTCTGATC</td><td>>hg38_21_+_8433172_8446622_RNA45SN1_45S_50ntFlanks</td><td>11999</td><td>12035</td><td>37</td><td>CCCTGCGGGCCGCCGGTGAAATACCACTACTCTGATC</td><td>1310.7</td><td>-10.4</td><td>1.3</td><td>-8.1</td><td>8.0E-16</td><td>3.6E-15</td><td>MB231/MB468</td><td>MITO</td></tr>
<tr><td>>hg38_21_+_8433172_8446622_RNA45SN1_45S_50ntFlanks@11318.11336.19|CCCCACGTCTCGTCGCGCG</td><td>>hg38_21_+_8433172_8446622_RNA45SN1_45S_50ntFlanks</td><td>11318</td><td>11336</td><td>19</td><td>CCCCACGTCTCGTCGCGCG</td><td>796.7</td><td>-9.1</td><td>1.1</td><td>-8.1</td><td>5.3E-16</td><td>3.6E-15</td><td>MB231/MB468</td><td>CYTO</td></tr>
<tr><td>>hg38_21_+_8433172_8446622_RNA45SN1_45S_50ntFlanks@10346.10368.23|TCAAACGAGAACTTTGAAGGCCG</td><td>>hg38_21_+_8433172_8446622_RNA45SN1_45S_50ntFlanks</td><td>10346</td><td>10368</td><td>23</td><td>TCAAACGAGAACTTTGAAGGCCG</td><td>4415.5</td><td>-8.9</td><td>1.1</td><td>-8.1</td><td>4.9E-16</td><td>3.6E-15</td><td>MB231/MB468</td><td>MP</td></tr>
<tr><td>>hg38_21_+_8433172_8446622_RNA45SN1_45S_50ntFlanks@10408.10429.22|GGTCAGTCGGTCCTGAGAGATG</td><td>>hg38_21_+_8433172_8446622_RNA45SN1_45S_50ntFlanks</td><td>10408</td><td>10429</td><td>22</td><td>GGTCAGTCGGTCCTGAGAGATG</td><td>13806.0</td><td>-12.8</td><td>1.6</td><td>-8.0</td><td>1.2E-15</td><td>3.6E-15</td><td>BT20/MB468</td><td>NUC</td></tr>
<tr><td>>hg38_21_+_8433172_8446622_RNA45SN1_45S_50ntFlanks@12632.12654.23|GCGAAGCTACCATCTGTGGGATT</td><td>>hg38_21_+_8433172_8446622_RNA45SN1_45S_50ntFlanks</td><td>12632</td><td>12654</td><td>23</td><td>GCGAAGCTACCATCTGTGGGATT</td><td>2469.6</td><td>-9.1</td><td>1.1</td><td>-8.0</td><td>1.2E-15</td><td>3.6E-15</td><td>BT20/MB468</td><td>NUC</td></tr>
<tr><td>>hg38_21_+_8433172_8446622_RNA45SN1_45S_50ntFlanks@4724.4751.28|ATTAATCAAGAACGAAAGTCGGAGGTTC</td><td>>hg38_21_+_8433172_8446622_RNA45SN1_45S_50ntFlanks</td><td>4724</td><td>4751</td><td>28</td><td>ATTAATCAAGAACGAAAGTCGGAGGTTC</td><td>1422.3</td><td>-10.7</td><td>1.3</td><td>-8.0</td><td>1.2E-15</td><td>3.6E-15</td><td>BT20/MB468</td><td>NUC</td></tr>
<tr><td>>hg38_21_+_8433172_8446622_RNA45SN1_45S_50ntFlanks@7981.8008.28|CTCAGATCAGACGTGGCGACCCGCTGAA</td><td>>hg38_21_+_8433172_8446622_RNA45SN1_45S_50ntFlanks</td><td>7981</td><td>8008</td><td>28</td><td>CTCAGATCAGACGTGGCGACCCGCTGAA</td><td>1122.6</td><td>-5.8</td><td>0.7</td><td>-8.0</td><td>1.2E-15</td><td>3.6E-15</td><td>BT20/MB468</td><td>NUC</td></tr>
<tr><td>>hg38_21_+_8433172_8446622_RNA45SN1_45S_50ntFlanks@12526.12546.21|TTTTACCCTACTGATGATGTG</td><td>>hg38_21_+_8433172_8446622_RNA45SN1_45S_50ntFlanks</td><td>12526</td><td>12546</td><td>21</td><td>TTTTACCCTACTGATGATGTG</td><td>421.5</td><td>-12.6</td><td>1.6</td><td>-8.1</td><td>4.9E-16</td><td>3.6E-15</td><td>MB231/MB468</td><td>MP</td></tr>
<tr><td>>hg38_21_+_8433172_8446622_RNA45SN1_45S_50ntFlanks@11581.11601.21|AACTGGTGCGGACCAGGGGAA</td><td>>hg38_21_+_8433172_8446622_RNA45SN1_45S_50ntFlanks</td><td>11581</td><td>11601</td><td>21</td><td>AACTGGTGCGGACCAGGGGAA</td><td>2011.4</td><td>-6.5</td><td>0.8</td><td>-8.1</td><td>5.3E-16</td><td>3.6E-15</td><td>MB231/MB468</td><td>CYTO</td></tr>
<tr><td>>hg38_21_+_8433172_8446622_RNA45SN1_45S_50ntFlanks@10794.10822.29|CTGGCATGTTGGAACAATGTAGGTAAGGG</td><td>>hg38_21_+_8433172_8446622_RNA45SN1_45S_50ntFlanks</td><td>10794</td><td>10822</td><td>29</td><td>CTGGCATGTTGGAACAATGTAGGTAAGGG</td><td>668.5</td><td>-9.4</td><td>1.2</td><td>-8.1</td><td>8.2E-16</td><td>3.6E-15</td><td>MB231/MB468</td><td>MITO</td></tr>
<tr><td>>hg38_21_+_8433172_8446622_RNA45SN1_45S_50ntFlanks@4103.4121.19|CCACGGGTGACGGGGAATC</td><td>>hg38_21_+_8433172_8446622_RNA45SN1_45S_50ntFlanks</td><td>4103</td><td>4121</td><td>19</td><td>CCACGGGTGACGGGGAATC</td><td>821.4</td><td>-13.5</td><td>1.7</td><td>-8.1</td><td>8.2E-16</td><td>3.6E-15</td><td>MB231/MB468</td><td>MITO</td></tr>
<tr><td>>hg38_21_+_8433172_8446622_RNA45SN1_45S_50ntFlanks@10574.10596.23|AAGCCGGCGGGAGCCCCGGGGAG</td><td>>hg38_21_+_8433172_8446622_RNA45SN1_45S_50ntFlanks</td><td>10574</td><td>10596</td><td>23</td><td>AAGCCGGCGGGAGCCCCGGGGAG</td><td>274.2</td><td>-11.9</td><td>1.5</td><td>-8.1</td><td>8.2E-16</td><td>3.6E-15</td><td>MB231/MB468</td><td>MITO</td></tr>
<tr><td>>MI0000299|hsa-mir-222&WithFlank&X|-|45747009|45747130@75.99.25(+4U)&[MIMAT0000279&hsa-miR-222-3p&offsets|0|+4(+4U);m-140&X|-|45747033|45747056&offsets|0|+1(+4U)]|AGCTACATCTGGCTACTGGGTCTCTTTTT</td><td>>MI0000299|hsa-mir-222&WithFlank&X|-|45747009|45747130</td><td>75</td><td>99</td><td>25(+4U)</td><td>AGCTACATCTGGCTACTGGGTCTCTTTTT</td><td>751.5</td><td>10.7</td><td>1.3</td><td>8.1</td><td>5.4E-16</td><td>3.7E-15</td><td>MB231/MB468</td><td>CYTO</td></tr>
<tr><td>>hg38_21_+_8433172_8446622_RNA45SN1_45S_50ntFlanks@11594.11612.19|CAGGGGAATCCGACTGTTT</td><td>>hg38_21_+_8433172_8446622_RNA45SN1_45S_50ntFlanks</td><td>11594</td><td>11612</td><td>19</td><td>CAGGGGAATCCGACTGTTT</td><td>783.9</td><td>-4.3</td><td>0.5</td><td>-8.1</td><td>6.1E-16</td><td>3.7E-15</td><td>MB231/MB468</td><td>NUC</td></tr>
<tr><td>>hg38_21_+_8433172_8446622_RNA45SN1_45S_50ntFlanks@4259.4278.20|ATTGGAATGAGTCCACTTTA</td><td>>hg38_21_+_8433172_8446622_RNA45SN1_45S_50ntFlanks</td><td>4259</td><td>4278</td><td>20</td><td>ATTGGAATGAGTCCACTTTA</td><td>300.9</td><td>-12.1</td><td>1.5</td><td>-8.1</td><td>5.0E-16</td><td>3.7E-15</td><td>MB231/MB468</td><td>MP</td></tr>
<tr><td>>hg38_21_+_8433172_8446622_RNA45SN1_45S_50ntFlanks@11580.11611.32|GAACTGGTGCGGACCAGGGGAATCCGACTGTT</td><td>>hg38_21_+_8433172_8446622_RNA45SN1_45S_50ntFlanks</td><td>11580</td><td>11611</td><td>32</td><td>GAACTGGTGCGGACCAGGGGAATCCGACTGTT</td><td>3289.4</td><td>-6.1</td><td>0.7</td><td>-8.1</td><td>5.0E-16</td><td>3.7E-15</td><td>MB231/MB468</td><td>MP</td></tr>
<tr><td>>hg38_21_+_8433172_8446622_RNA45SN1_45S_50ntFlanks@10727.10750.24|GGAGAGGGTGTAAATCTCGCGCCG</td><td>>hg38_21_+_8433172_8446622_RNA45SN1_45S_50ntFlanks</td><td>10727</td><td>10750</td><td>24</td><td>GGAGAGGGTGTAAATCTCGCGCCG</td><td>442.2</td><td>-12.7</td><td>1.6</td><td>-8.1</td><td>5.0E-16</td><td>3.7E-15</td><td>MB231/MB468</td><td>MP</td></tr>
<tr><td>>am_trna119_LysCTT_1_-_145395522_145395594@7.35.29,trna11_LysCTT_5_-_180648979_180649051@7.35.29,trna128_LysCTT_1_-_55423542_55423614@7.35.29,trna13_LysCTT_6_+_26556774_26556846@7.35.29,trna6_LysCTT_19_-_52425396_52425468@7.35.29,trna7_LysCTT_16_+_3225692_3225764@7.35.29,trna9_LysCTT_5_+_180634755_180634827@7.35.29|CTAGCTCAGTCGGTAGAGCATGAGACTCT</td><td>>am_trna119_LysCTT_1_-_145395522_145395594</td><td>7</td><td>35</td><td>29</td><td>CTAGCTCAGTCGGTAGAGCATGAGACTCT</td><td>211.8</td><td>11.6</td><td>1.4</td><td>8.1</td><td>6.2E-16</td><td>3.7E-15</td><td>MB231/MB468</td><td>NUC</td></tr>
<tr><td>>hg38_21_+_8433172_8446622_RNA45SN1_45S_50ntFlanks@11200.11227.28|CGGCGGCGGCGGCGACTCTGGACGCGAG</td><td>>hg38_21_+_8433172_8446622_RNA45SN1_45S_50ntFlanks</td><td>11200</td><td>11227</td><td>28</td><td>CGGCGGCGGCGGCGACTCTGGACGCGAG</td><td>246.8</td><td>-11.8</td><td>1.5</td><td>-8.1</td><td>5.0E-16</td><td>3.7E-15</td><td>MB231/MB468</td><td>MP</td></tr>
<tr><td>>hg38_21_+_8433172_8446622_RNA45SN1_45S_50ntFlanks@12957.12980.24|CTTCTGGGTCGGGGTTTCGTACGT</td><td>>hg38_21_+_8433172_8446622_RNA45SN1_45S_50ntFlanks</td><td>12957</td><td>12980</td><td>24</td><td>CTTCTGGGTCGGGGTTTCGTACGT</td><td>358.9</td><td>-12.3</td><td>1.5</td><td>-8.0</td><td>8.3E-16</td><td>3.7E-15</td><td>MB231/MB468</td><td>MITO</td></tr>
<tr><td>>hg38_21_+_8433172_8446622_RNA45SN1_45S_50ntFlanks@5079.5112.34|GATAACGAACGAGACTCTGGCATGCTAACTAGTT</td><td>>hg38_21_+_8433172_8446622_RNA45SN1_45S_50ntFlanks</td><td>5079</td><td>5112</td><td>34</td><td>GATAACGAACGAGACTCTGGCATGCTAACTAGTT</td><td>436.6</td><td>7.1</td><td>0.9</td><td>8.1</td><td>5.5E-16</td><td>3.7E-15</td><td>MB231/MB468</td><td>CYTO</td></tr>
<tr><td>>hg38_21_+_8433172_8446622_RNA45SN1_45S_50ntFlanks@12686.12708.23|CCAGGCGGAACGATACGGCAGCG</td><td>>hg38_21_+_8433172_8446622_RNA45SN1_45S_50ntFlanks</td><td>12686</td><td>12708</td><td>23</td><td>CCAGGCGGAACGATACGGCAGCG</td><td>183.4</td><td>-11.4</td><td>1.4</td><td>-8.1</td><td>5.0E-16</td><td>3.7E-15</td><td>MB231/MB468</td><td>MP</td></tr>
<tr><td>>hg38_21_+_8433172_8446622_RNA45SN1_45S_50ntFlanks@12357.12376.20|CCACAGGGATAACTGGCTTG</td><td>>hg38_21_+_8433172_8446622_RNA45SN1_45S_50ntFlanks</td><td>12357</td><td>12376</td><td>20</td><td>CCACAGGGATAACTGGCTTG</td><td>909.9</td><td>-12.9</td><td>1.5</td><td>-8.5</td><td>1.5E-17</td><td>3.7E-15</td><td>BT20/MB468</td><td>MP</td></tr>
<tr><td>>hg38_21_+_8433172_8446622_RNA45SN1_45S_50ntFlanks@7984.8005.22|AGATCAGACGTGGCGACCCGCT</td><td>>hg38_21_+_8433172_8446622_RNA45SN1_45S_50ntFlanks</td><td>7984</td><td>8005</td><td>22</td><td>AGATCAGACGTGGCGACCCGCT</td><td>1151.1</td><td>-8.2</td><td>1.0</td><td>-8.0</td><td>8.3E-16</td><td>3.7E-15</td><td>MB231/MB468</td><td>MITO</td></tr>
<tr><td>>hg38_21_+_8433172_8446622_RNA45SN1_45S_50ntFlanks@5487.5518.32|CGCTGAGAAGACGGTCGAACTTGACTATCTAG</td><td>>hg38_21_+_8433172_8446622_RNA45SN1_45S_50ntFlanks</td><td>5487</td><td>5518</td><td>32</td><td>CGCTGAGAAGACGGTCGAACTTGACTATCTAG</td><td>264.6</td><td>-11.8</td><td>1.5</td><td>-8.0</td><td>8.4E-16</td><td>3.7E-15</td><td>MB231/MB468</td><td>MITO</td></tr>
<tr><td>>hg38_21_+_8433172_8446622_RNA45SN1_45S_50ntFlanks@10683.10705.23|CCGGCGGCGTCCGGTGAGCTCTC</td><td>>hg38_21_+_8433172_8446622_RNA45SN1_45S_50ntFlanks</td><td>10683</td><td>10705</td><td>23</td><td>CCGGCGGCGTCCGGTGAGCTCTC</td><td>1063.7</td><td>-4.7</td><td>0.6</td><td>-8.1</td><td>6.3E-16</td><td>3.7E-15</td><td>MB231/MB468</td><td>NUC</td></tr>
<tr><td>>hg38_21_+_8433172_8446622_RNA45SN1_45S_50ntFlanks@6772.6797.26|GTTCCTCCCGGGGCTACGCCTGTCTG</td><td>>hg38_21_+_8433172_8446622_RNA45SN1_45S_50ntFlanks</td><td>6772</td><td>6797</td><td>26</td><td>GTTCCTCCCGGGGCTACGCCTGTCTG</td><td>210.1</td><td>-10.4</td><td>1.3</td><td>-8.1</td><td>5.5E-16</td><td>3.7E-15</td><td>MB231/MB468</td><td>CYTO</td></tr>
<tr><td>>hg38_21_+_8433172_8446622_RNA45SN1_45S_50ntFlanks@9742.9765.24|CGATCTCAACCTATTCTCAAACTT</td><td>>hg38_21_+_8433172_8446622_RNA45SN1_45S_50ntFlanks</td><td>9742</td><td>9765</td><td>24</td><td>CGATCTCAACCTATTCTCAAACTT</td><td>876.1</td><td>-13.6</td><td>1.7</td><td>-8.0</td><td>8.4E-16</td><td>3.7E-15</td><td>MB231/MB468</td><td>MITO</td></tr>
<tr><td>>hg38_21_+_8433172_8446622_RNA45SN1_45S_50ntFlanks@3853.3881.29|AACTGTGGTAATTCTAGAGCTAATACATG</td><td>>hg38_21_+_8433172_8446622_RNA45SN1_45S_50ntFlanks</td><td>3853</td><td>3881</td><td>29</td><td>AACTGTGGTAATTCTAGAGCTAATACATG</td><td>1195.8</td><td>-8.2</td><td>1.0</td><td>-8.0</td><td>8.4E-16</td><td>3.7E-15</td><td>MB231/MB468</td><td>MITO</td></tr>
<tr><td>>hg38_21_+_8433172_8446622_RNA45SN1_45S_50ntFlanks@11795.11821.27|AACGAGATTCCCACTGTCCCTACCTAC</td><td>>hg38_21_+_8433172_8446622_RNA45SN1_45S_50ntFlanks</td><td>11795</td><td>11821</td><td>27</td><td>AACGAGATTCCCACTGTCCCTACCTAC</td><td>523.0</td><td>-12.8</td><td>1.6</td><td>-8.0</td><td>8.4E-16</td><td>3.7E-15</td><td>MB231/MB468</td><td>MITO</td></tr>
<tr><td>>hg38_21_+_8433172_8446622_RNA45SN1_45S_50ntFlanks@10921.10939.19|GCTGGACGAGGCGCCGCCG</td><td>>hg38_21_+_8433172_8446622_RNA45SN1_45S_50ntFlanks</td><td>10921</td><td>10939</td><td>19</td><td>GCTGGACGAGGCGCCGCCG</td><td>330.9</td><td>-12.2</td><td>1.5</td><td>-8.1</td><td>5.1E-16</td><td>3.7E-15</td><td>MB231/MB468</td><td>MP</td></tr>
<tr><td>>hg38_21_+_8433172_8446622_RNA45SN1_45S_50ntFlanks@4099.4117.19|GTGACCACGGGTGACGGGG</td><td>>hg38_21_+_8433172_8446622_RNA45SN1_45S_50ntFlanks</td><td>4099</td><td>4117</td><td>19</td><td>GTGACCACGGGTGACGGGG</td><td>209.4</td><td>-10.7</td><td>1.3</td><td>-8.1</td><td>6.3E-16</td><td>3.8E-15</td><td>MB231/MB468</td><td>NUC</td></tr>
<tr><td>>hg38_21_+_8433172_8446622_RNA45SN1_45S_50ntFlanks@11874.11910.37|AGAAGACCCTGTTGAGCTTGACTCTAGTCTGGCACGG</td><td>>hg38_21_+_8433172_8446622_RNA45SN1_45S_50ntFlanks</td><td>11874</td><td>11910</td><td>37</td><td>AGAAGACCCTGTTGAGCTTGACTCTAGTCTGGCACGG</td><td>287.4</td><td>-12.0</td><td>1.5</td><td>-8.1</td><td>5.1E-16</td><td>3.8E-15</td><td>MB231/MB468</td><td>MP</td></tr>
<tr><td>>MI0000115|hsa-mir-16-2&WithFlank&3|+|160404739|160404831@58.79.22&[MIMAT0004518&hsa-miR-16-2-3p&offsets|-1|-1;m-252&3|+|160404796|160404816&offsets|0|+1]|ACCAATATTACTGTGCTGCTTT</td><td>>MI0000115|hsa-mir-16-2&WithFlank&3|+|160404739|160404831</td><td>58</td><td>79</td><td>22</td><td>ACCAATATTACTGTGCTGCTTT</td><td>2030.3</td><td>6.6</td><td>0.8</td><td>8.1</td><td>7.9E-16</td><td>3.8E-15</td><td>BT20/MB468</td><td>CYTO</td></tr>
<tr><td>>hg38_21_+_8433172_8446622_RNA45SN1_45S_50ntFlanks@12433.12455.23|CTATCATTGTGAAGCAGAATTCA</td><td>>hg38_21_+_8433172_8446622_RNA45SN1_45S_50ntFlanks</td><td>12433</td><td>12455</td><td>23</td><td>CTATCATTGTGAAGCAGAATTCA</td><td>375.2</td><td>-12.3</td><td>1.5</td><td>-8.0</td><td>8.5E-16</td><td>3.8E-15</td><td>MB231/MB468</td><td>MITO</td></tr>
<tr><td>>hg38_21_+_8433172_8446622_RNA45SN1_45S_50ntFlanks@6667.6699.33|ATCACTCGGCTCGTGCGTCGATGAAGAACGCAG</td><td>>hg38_21_+_8433172_8446622_RNA45SN1_45S_50ntFlanks</td><td>6667</td><td>6699</td><td>33</td><td>ATCACTCGGCTCGTGCGTCGATGAAGAACGCAG</td><td>327.5</td><td>-12.2</td><td>1.5</td><td>-8.1</td><td>5.1E-16</td><td>3.8E-15</td><td>MB231/MB468</td><td>MP</td></tr>
<tr><td>>MI0000079|hsa-mir-23a&WithFlank&19|-|13836581|13836665@54.73.20(+1U)&[MIMAT0000078&hsa-miR-23a-3p&offsets|+3|+2(+1U);m-31&19|-|13836595|13836615&offsets|+3|+2(+1U)]|ACATTGCCAGGGATTTCCAAT</td><td>>MI0000079|hsa-mir-23a&WithFlank&19|-|13836581|13836665</td><td>54</td><td>73</td><td>20(+1U)</td><td>ACATTGCCAGGGATTTCCAAT</td><td>2599.3</td><td>29.3</td><td>3.6</td><td>8.1</td><td>6.4E-16</td><td>3.8E-15</td><td>MB231/MB468</td><td>NUC</td></tr>
<tr><td>>hg38_21_+_8433172_8446622_RNA45SN1_45S_50ntFlanks@9575.9596.22|ATCGGTCGTCCGACCTGGGTAT</td><td>>hg38_21_+_8433172_8446622_RNA45SN1_45S_50ntFlanks</td><td>9575</td><td>9596</td><td>22</td><td>ATCGGTCGTCCGACCTGGGTAT</td><td>257.9</td><td>-11.0</td><td>1.4</td><td>-8.1</td><td>6.4E-16</td><td>3.8E-15</td><td>MB231/MB468</td><td>NUC</td></tr>
<tr><td>>hg38_21_+_8433172_8446622_RNA45SN1_45S_50ntFlanks@13007.13039.33|TATTGAAAGTCAGCCCTCGACACAAGGGTTTGT</td><td>>hg38_21_+_8433172_8446622_RNA45SN1_45S_50ntFlanks</td><td>13007</td><td>13039</td><td>33</td><td>TATTGAAAGTCAGCCCTCGACACAAGGGTTTGT</td><td>6857.5</td><td>6.8</td><td>0.8</td><td>8.1</td><td>5.7E-16</td><td>3.8E-15</td><td>BT20/MB468</td><td>MITO</td></tr>
<tr><td>>hg38_21_+_8433172_8446622_RNA45SN1_45S_50ntFlanks@12539.12560.22|ATGATGTGTTGTTGCCATGGTA</td><td>>hg38_21_+_8433172_8446622_RNA45SN1_45S_50ntFlanks</td><td>12539</td><td>12560</td><td>22</td><td>ATGATGTGTTGTTGCCATGGTA</td><td>366.6</td><td>-12.4</td><td>1.5</td><td>-8.1</td><td>5.2E-16</td><td>3.8E-15</td><td>MB231/MB468</td><td>MP</td></tr>
<tr><td>>hg38_21_+_8433172_8446622_RNA45SN1_45S_50ntFlanks@4722.4758.37|TCATTAATCAAGAACGAAAGTCGGAGGTTCGAAGACG</td><td>>hg38_21_+_8433172_8446622_RNA45SN1_45S_50ntFlanks</td><td>4722</td><td>4758</td><td>37</td><td>TCATTAATCAAGAACGAAAGTCGGAGGTTCGAAGACG</td><td>8693.7</td><td>-10.4</td><td>1.3</td><td>-8.1</td><td>5.2E-16</td><td>3.8E-15</td><td>MB231/MB468</td><td>MP</td></tr>
<tr><td>>hg38_21_+_8433172_8446622_RNA45SN1_45S_50ntFlanks@10794.10814.21|CTGGCATGTTGGAACAATGTA</td><td>>hg38_21_+_8433172_8446622_RNA45SN1_45S_50ntFlanks</td><td>10794</td><td>10814</td><td>21</td><td>CTGGCATGTTGGAACAATGTA</td><td>178.5</td><td>-11.3</td><td>1.4</td><td>-8.1</td><td>5.2E-16</td><td>3.8E-15</td><td>MB231/MB468</td><td>MP</td></tr>
<tr><td>>hg38_21_+_8433172_8446622_RNA45SN1_45S_50ntFlanks@6651.6689.39|CGACTCTTAGCGGTGGATCACTCGGCTCGTGCGTCGATG</td><td>>hg38_21_+_8433172_8446622_RNA45SN1_45S_50ntFlanks</td><td>6651</td><td>6689</td><td>39</td><td>CGACTCTTAGCGGTGGATCACTCGGCTCGTGCGTCGATG</td><td>283798.9</td><td>-4.9</td><td>0.6</td><td>-8.0</td><td>8.5E-16</td><td>3.8E-15</td><td>MB231/MB468</td><td>MITO</td></tr>
<tr><td>>hg38_MT_+_1621_3279_RNR2_16S_50ntFlanks@622.652.31|TATAACTGAACTCCTCACACCCAATTGGACC</td><td>>hg38_MT_+_1621_3279_RNR2_16S_50ntFlanks</td><td>622</td><td>652</td><td>31</td><td>TATAACTGAACTCCTCACACCCAATTGGACC</td><td>771.0</td><td>9.0</td><td>1.1</td><td>8.1</td><td>5.7E-16</td><td>3.8E-15</td><td>BT20/MB468</td><td>MITO</td></tr>
<tr><td>>hg38_21_+_8433172_8446622_RNA45SN1_45S_50ntFlanks@10320.10341.22|CAGATCTTGGTGGTAGTAGCAA</td><td>>hg38_21_+_8433172_8446622_RNA45SN1_45S_50ntFlanks</td><td>10320</td><td>10341</td><td>22</td><td>CAGATCTTGGTGGTAGTAGCAA</td><td>298.7</td><td>-12.1</td><td>1.5</td><td>-8.1</td><td>5.2E-16</td><td>3.8E-15</td><td>MB231/MB468</td><td>MP</td></tr>
<tr><td>>hg38_21_+_8433172_8446622_RNA45SN1_45S_50ntFlanks@4729.4758.30|TCAAGAACGAAAGTCGGAGGTTCGAAGACG</td><td>>hg38_21_+_8433172_8446622_RNA45SN1_45S_50ntFlanks</td><td>4729</td><td>4758</td><td>30</td><td>TCAAGAACGAAAGTCGGAGGTTCGAAGACG</td><td>3308.5</td><td>-6.6</td><td>0.8</td><td>-8.0</td><td>8.6E-16</td><td>3.8E-15</td><td>MB231/MB468</td><td>MITO</td></tr>
<tr><td>>hg38_1_-_228634819_228635039_RNA5S12_5S_50ntFlanks@135.171.37|GGGAGACCGCCTGGGAATACCGGGTGCTGTAGGCTTT</td><td>>hg38_1_-_228634819_228635039_RNA5S12_5S_50ntFlanks</td><td>135</td><td>171</td><td>37</td><td>GGGAGACCGCCTGGGAATACCGGGTGCTGTAGGCTTT</td><td>10367.4</td><td>5.4</td><td>0.7</td><td>8.1</td><td>6.4E-16</td><td>3.8E-15</td><td>MB231/MB468</td><td>NUC</td></tr>
<tr><td>>hg38_21_+_8433172_8446622_RNA45SN1_45S_50ntFlanks@5420.5444.25|CCGATTGGATGGTTTAGTGAGGCCC</td><td>>hg38_21_+_8433172_8446622_RNA45SN1_45S_50ntFlanks</td><td>5420</td><td>5444</td><td>25</td><td>CCGATTGGATGGTTTAGTGAGGCCC</td><td>499.5</td><td>-12.8</td><td>1.6</td><td>-8.1</td><td>5.2E-16</td><td>3.8E-15</td><td>MB231/MB468</td><td>MP</td></tr>
</table>

<table>
<tr><td>>MI0000289|hsa-mir-181a-1&WithFlank&1|-|198859038|198859159@30.51.22;MI0000269|hsa-mir-181a-2&WithFlank&9|+|124692436|124692557@45.66.22&[MIMAT0000256&hsa-miR-181a-5p&offsets|0|-1;m-32&1|-|198859108|198859130&offsets|0|-1];[MIMAT0000256_1&hsa-miR-181a-5p&offsets|0|-1;m-32&9|+|124692480|124692502&offsets|0|-1]|AACATTCAACGCTGTCGGTGAG</td><td>>MI0000289|hsa-mir-181a-1&WithFlank&1|-|198859038|198859159</td><td>30</td><td>51</td><td>22</td><td>AACATTCAACGCTGTCGGTGAG</td><td>431.4</td><td>4.2</td><td>0.5</td><td>8.1</td><td>8.0E-16</td><td>3.8E-15</td><td>BT20/MB468</td><td>CYTO</td></tr>
<tr><td>>MI0000283|hsa-mir-203a&WithFlank&14|+|104117399|104117520@71.93.23&[MIMAT0000264&hsa-miR-203a-3p&offsets|0|+1;m-79&14|+|104117469|104117490&offsets|0|+1]|GTGAAATGTTTAGGACCACTAGA</td><td>>MI0000283|hsa-mir-203a&WithFlank&14|+|104117399|104117520</td><td>71</td><td>93</td><td>23</td><td>GTGAAATGTTTAGGACCACTAGA</td><td>274.1</td><td>11.9</td><td>1.4</td><td>8.3</td><td>8.8E-17</td><td>3.8E-15</td><td>BT20/MB231</td><td>NUC</td></tr>
<tr><td>>hg38_21_+_8433172_8446622_RNA45SN1_45S_50ntFlanks@10408.10437.30|GGTCAGTCGGTCCTGAGAGATGGGCGAGCG</td><td>>hg38_21_+_8433172_8446622_RNA45SN1_45S_50ntFlanks</td><td>10408</td><td>10437</td><td>30</td><td>GGTCAGTCGGTCCTGAGAGATGGGCGAGCG</td><td>256.4</td><td>-11.9</td><td>1.5</td><td>-8.1</td><td>5.3E-16</td><td>3.9E-15</td><td>MB231/MB468</td><td>MP</td></tr>
<tr><td>>hg38_21_+_8433172_8446622_RNA45SN1_45S_50ntFlanks@12161.12182.22|TGACTGGGGCGGTACACCTGTC</td><td>>hg38_21_+_8433172_8446622_RNA45SN1_45S_50ntFlanks</td><td>12161</td><td>12182</td><td>22</td><td>TGACTGGGGCGGTACACCTGTC</td><td>358.8</td><td>-12.4</td><td>1.5</td><td>-8.1</td><td>5.3E-16</td><td>3.9E-15</td><td>MB231/MB468</td><td>MP</td></tr>
<tr><td>>hg38_21_+_8433172_8446622_RNA45SN1_45S_50ntFlanks@5472.5501.30|ACGGCCCTGGCGGAGCGCTGAGAAGACGGT</td><td>>hg38_21_+_8433172_8446622_RNA45SN1_45S_50ntFlanks</td><td>5472</td><td>5501</td><td>30</td><td>ACGGCCCTGGCGGAGCGCTGAGAAGACGGT</td><td>2269.9</td><td>7.7</td><td>1.0</td><td>8.1</td><td>5.9E-16</td><td>3.9E-15</td><td>BT20/MB468</td><td>MITO</td></tr>
<tr><td>>MI0000234|hsa-mir-192&WithFlank&11|-|64891131|64891252@30.50.21&[MIMAT0000222&hsa-miR-192-5p&offsets|0|0;m-59&11|-|64891203|64891223&offsets|0|0]|CTGACCTATGAATTGACAGCC</td><td>>MI0000234|hsa-mir-192&WithFlank&11|-|64891131|64891252</td><td>30</td><td>50</td><td>21</td><td>CTGACCTATGAATTGACAGCC</td><td>1036.0</td><td>6.2</td><td>0.8</td><td>8.1</td><td>8.2E-16</td><td>3.9E-15</td><td>BT20/MB468</td><td>CYTO</td></tr>
<tr><td>>hg38_21_+_8433172_8446622_RNA45SN1_45S_50ntFlanks@9031.9076.46|TCCTCGTGGGGGGGCCGGGCCACCCCTCCCACGGCGCGACCGCTCT</td><td>>hg38_21_+_8433172_8446622_RNA45SN1_45S_50ntFlanks</td><td>9031</td><td>9076</td><td>46</td><td>TCCTCGTGGGGGGGCCGGGCCACCCCTCCCACGGCGCGACCGCTCT</td><td>562.0</td><td>-12.9</td><td>1.6</td><td>-8.0</td><td>8.8E-16</td><td>3.9E-15</td><td>MB231/MB468</td><td>MITO</td></tr>
<tr><td>>hg38_21_+_8433172_8446622_RNA45SN1_45S_50ntFlanks@12934.12969.36|CTAAACCATTCGTAGACGACCTGCTTCTGGGTCGGG</td><td>>hg38_21_+_8433172_8446622_RNA45SN1_45S_50ntFlanks</td><td>12934</td><td>12969</td><td>36</td><td>CTAAACCATTCGTAGACGACCTGCTTCTGGGTCGGG</td><td>678.8</td><td>-7.4</td><td>0.9</td><td>-8.1</td><td>5.8E-16</td><td>3.9E-15</td><td>MB231/MB468</td><td>CYTO</td></tr>
<tr><td>>hg38_21_+_8433172_8446622_RNA45SN1_45S_50ntFlanks@11641.11678.38|CGGCGGGTGTTGACGCGATGTGATTTCTGCCCAGTGCT</td><td>>hg38_21_+_8433172_8446622_RNA45SN1_45S_50ntFlanks</td><td>11641</td><td>11678</td><td>38</td><td>CGGCGGGTGTTGACGCGATGTGATTTCTGCCCAGTGCT</td><td>288.5</td><td>-12.0</td><td>1.5</td><td>-8.1</td><td>5.4E-16</td><td>3.9E-15</td><td>MB231/MB468</td><td>MP</td></tr>
<tr><td>>trnaMT_ValTAC_MT_+_1602_1670@43.72.30|GATTTCAACTTAACTTGACCGCTCTGACCA</td><td>>trnaMT_ValTAC_MT_+_1602_1670</td><td>43</td><td>72</td><td>30</td><td>GATTTCAACTTAACTTGACCGCTCTGACCA</td><td>533.9</td><td>5.3</td><td>0.7</td><td>8.1</td><td>6.6E-16</td><td>3.9E-15</td><td>MB231/MB468</td><td>NUC</td></tr>
<tr><td>>hg38_21_+_8433172_8446622_RNA45SN1_45S_50ntFlanks@12415.12436.22|CTTCGATGTCGGCTCTTCCTAT</td><td>>hg38_21_+_8433172_8446622_RNA45SN1_45S_50ntFlanks</td><td>12415</td><td>12436</td><td>22</td><td>CTTCGATGTCGGCTCTTCCTAT</td><td>11643.0</td><td>-5.9</td><td>0.7</td><td>-8.1</td><td>6.6E-16</td><td>3.9E-15</td><td>MB231/MB468</td><td>NUC</td></tr>
<tr><td>>hg38_21_+_8433172_8446622_RNA45SN1_45S_50ntFlanks@476.497.22|TTCGTGATCGATGTGGTGACGT</td><td>>hg38_21_+_8433172_8446622_RNA45SN1_45S_50ntFlanks</td><td>476</td><td>497</td><td>22</td><td>TTCGTGATCGATGTGGTGACGT</td><td>3691.4</td><td>6.8</td><td>0.8</td><td>8.1</td><td>6.7E-16</td><td>3.9E-15</td><td>MB231/MB468</td><td>NUC</td></tr>
<tr><td>>hg38_21_+_8433172_8446622_RNA45SN1_45S_50ntFlanks@4728.4752.25|ATCAAGAACGAAAGTCGGAGGTTCG</td><td>>hg38_21_+_8433172_8446622_RNA45SN1_45S_50ntFlanks</td><td>4728</td><td>4752</td><td>25</td><td>ATCAAGAACGAAAGTCGGAGGTTCG</td><td>403.4</td><td>-11.0</td><td>1.4</td><td>-8.1</td><td>8.2E-16</td><td>3.9E-15</td><td>BT20/MB468</td><td>CYTO</td></tr>
<tr><td>>hg38_21_+_8433172_8446622_RNA45SN1_45S_50ntFlanks@8179.8197.19|CTGATCGAGGCCCAGCCCG</td><td>>hg38_21_+_8433172_8446622_RNA45SN1_45S_50ntFlanks</td><td>8179</td><td>8197</td><td>19</td><td>CTGATCGAGGCCCAGCCCG</td><td>209.1</td><td>-11.6</td><td>1.4</td><td>-8.1</td><td>5.4E-16</td><td>3.9E-15</td><td>MB231/MB468</td><td>MP</td></tr>
<tr><td>>hg38_21_+_8433172_8446622_RNA45SN1_45S_50ntFlanks@10408.10425.18|GGTCAGTCGGTCCTGAGA</td><td>>hg38_21_+_8433172_8446622_RNA45SN1_45S_50ntFlanks</td><td>10408</td><td>10425</td><td>18</td><td>GGTCAGTCGGTCCTGAGA</td><td>355.5</td><td>-11.4</td><td>1.4</td><td>-8.1</td><td>6.7E-16</td><td>3.9E-15</td><td>MB231/MB468</td><td>NUC</td></tr>
<tr><td>>hg38_21_+_8433172_8446622_RNA45SN1_45S_50ntFlanks@3773.3792.20|CGGCCGGTACAGTGAAACTG</td><td>>hg38_21_+_8433172_8446622_RNA45SN1_45S_50ntFlanks</td><td>3773</td><td>3792</td><td>20</td><td>CGGCCGGTACAGTGAAACTG</td><td>420.4</td><td>-12.6</td><td>1.6</td><td>-8.1</td><td>5.4E-16</td><td>4.0E-15</td><td>MB231/MB468</td><td>MP</td></tr>
<tr><td>>hg38_21_+_8433172_8446622_RNA45SN1_45S_50ntFlanks@4657.4684.28|CTAGAGGTGAAATTCTTGGACCGGCGCA</td><td>>hg38_21_+_8433172_8446622_RNA45SN1_45S_50ntFlanks</td><td>4657</td><td>4684</td><td>28</td><td>CTAGAGGTGAAATTCTTGGACCGGCGCA</td><td>365.0</td><td>-12.4</td><td>1.5</td><td>-8.1</td><td>5.4E-16</td><td>4.0E-15</td><td>MB231/MB468</td><td>MP</td></tr>
<tr><td>>hg38_21_+_8433172_8446622_RNA45SN1_45S_50ntFlanks@10771.10797.27|CAGGTCTCCAAGGTGAACAGCCTCTGG</td><td>>hg38_21_+_8433172_8446622_RNA45SN1_45S_50ntFlanks</td><td>10771</td><td>10797</td><td>27</td><td>CAGGTCTCCAAGGTGAACAGCCTCTGG</td><td>280.5</td><td>-11.9</td><td>1.5</td><td>-8.0</td><td>8.9E-16</td><td>4.0E-15</td><td>MB231/MB468</td><td>MITO</td></tr>
<tr><td>>hg38_21_+_8433172_8446622_RNA45SN1_45S_50ntFlanks@8512.8531.20|CCTCCACCCGCCCTCCCTTC</td><td>>hg38_21_+_8433172_8446622_RNA45SN1_45S_50ntFlanks</td><td>8512</td><td>8531</td><td>20</td><td>CCTCCACCCGCCCTCCCTTC</td><td>349.2</td><td>-12.2</td><td>1.5</td><td>-8.0</td><td>9.0E-16</td><td>4.0E-15</td><td>MB231/MB468</td><td>MITO</td></tr>
<tr><td>>hg38_21_+_8433172_8446622_RNA45SN1_45S_50ntFlanks@12081.12109.29|CTCTCGCTTCTGGCGCCAAGCGCCCGGCC</td><td>>hg38_21_+_8433172_8446622_RNA45SN1_45S_50ntFlanks</td><td>12081</td><td>12109</td><td>29</td><td>CTCTCGCTTCTGGCGCCAAGCGCCCGGCC</td><td>296.2</td><td>-12.0</td><td>1.5</td><td>-8.0</td><td>9.0E-16</td><td>4.0E-15</td><td>MB231/MB468</td><td>MITO</td></tr>
<tr><td>>hg38_21_+_8433172_8446622_RNA45SN1_45S_50ntFlanks@6738.6771.34|ATCATCGACACTTCGAACGCACTTGCGGCCCCGG</td><td>>hg38_21_+_8433172_8446622_RNA45SN1_45S_50ntFlanks</td><td>6738</td><td>6771</td><td>34</td><td>ATCATCGACACTTCGAACGCACTTGCGGCCCCGG</td><td>2796.9</td><td>-14.5</td><td>1.7</td><td>-8.5</td><td>1.7E-17</td><td>4.0E-15</td><td>BT20/MB468</td><td>MP</td></tr>
<tr><td>>hg38_21_+_8433172_8446622_RNA45SN1_45S_50ntFlanks@485.505.21|GATGTGGTGACGTCGTGCTCT</td><td>>hg38_21_+_8433172_8446622_RNA45SN1_45S_50ntFlanks</td><td>485</td><td>505</td><td>21</td><td>GATGTGGTGACGTCGTGCTCT</td><td>3679.0</td><td>29.2</td><td>3.6</td><td>8.0</td><td>9.0E-16</td><td>4.0E-15</td><td>MB231/MB468</td><td>MITO</td></tr>
<tr><td>>hg38_21_+_8433172_8446622_RNA45SN1_45S_50ntFlanks@6651.6695.45|CGACTCTTAGCGGTGGATCACTCGGCTCGTGCGTCGATGAAGAAC</td><td>>hg38_21_+_8433172_8446622_RNA45SN1_45S_50ntFlanks</td><td>6651</td><td>6695</td><td>45</td><td>CGACTCTTAGCGGTGGATCACTCGGCTCGTGCGTCGATGAAGAAC</td><td>4116.4</td><td>7.3</td><td>0.9</td><td>8.1</td><td>5.9E-16</td><td>4.0E-15</td><td>MB231/MB468</td><td>CYTO</td></tr>
<tr><td>>hg38_21_+_8433172_8446622_RNA45SN1_45S_50ntFlanks@12676.12703.28|AGAATCCCGCCCAGGCGGAACGATACGG</td><td>>hg38_21_+_8433172_8446622_RNA45SN1_45S_50ntFlanks</td><td>12676</td><td>12703</td><td>28</td><td>AGAATCCCGCCCAGGCGGAACGATACGG</td><td>427.5</td><td>-11.5</td><td>1.4</td><td>-8.1</td><td>6.0E-16</td><td>4.0E-15</td><td>MB231/MB468</td><td>CYTO</td></tr>
<tr><td>>MI0000270|hsa-mir-181b-1&WithFlank&1|-|198858867|198858988@42.65.24;MI0000683|hsa-mir-181b-2&WithFlank&9|+|124693704|124693804@22.45.24&[MIMAT0000257&hsa-miR-181b-5p&offsets|0|+1;m-73&1|-|198858925|198858947&offsets|0|+1];[MIMAT0000257_1&hsa-miR-181b-5p&offsets|0|+1;m-72&9|+|124693725|124693747&offsets|0|+1]|AACATTCATTGCTGTCGGTGGGTT</td><td>>MI0000270|hsa-mir-181b-1&WithFlank&1|-|198858867|198858988</td><td>42</td><td>65</td><td>24</td><td>AACATTCATTGCTGTCGGTGGGTT</td><td>1023.7</td><td>6.0</td><td>0.7</td><td>8.1</td><td>6.0E-16</td><td>4.0E-15</td><td>BT20/MB468</td><td>MITO</td></tr>
<tr><td>>hg38_21_+_8433172_8446622_RNA45SN1_45S_50ntFlanks@11534.11551.18|GTTCCGCGCGGCGCCTCG</td><td>>hg38_21_+_8433172_8446622_RNA45SN1_45S_50ntFlanks</td><td>11534</td><td>11551</td><td>18</td><td>GTTCCGCGCGGCGCCTCG</td><td>9053.7</td><td>-12.7</td><td>1.6</td><td>-8.0</td><td>1.3E-15</td><td>4.0E-15</td><td>BT20/MB468</td><td>NUC</td></tr>
<tr><td>>MI0000811|hsa-mir-148b&WithFlank&12|+|54337210|54337320@69.90.22(+3U)&[MIMAT0000759&hsa-miR-148b-3p&offsets|0|0(+3U);m-77&12|+|54337278|54337299&offsets|0|0(+3U)]|TCAGTGCATCACAGAACTTTGTTTT</td><td>>MI0000811|hsa-mir-148b&WithFlank&12|+|54337210|54337320</td><td>69</td><td>90</td><td>22(+3U)</td><td>TCAGTGCATCACAGAACTTTGTTTT</td><td>206.4</td><td>5.2</td><td>0.6</td><td>8.1</td><td>6.9E-16</td><td>4.0E-15</td><td>MB231/MB468</td><td>NUC</td></tr>
<tr><td>>hg38_21_+_8433172_8446622_RNA45SN1_45S_50ntFlanks@5073.5113.41|AATTCCGATAACGAACGAGACTCTGGCATGCTAACTAGTTA</td><td>>hg38_21_+_8433172_8446622_RNA45SN1_45S_50ntFlanks</td><td>5073</td><td>5113</td><td>41</td><td>AATTCCGATAACGAACGAGACTCTGGCATGCTAACTAGTTA</td><td>87.5</td><td>-23.6</td><td>2.9</td><td>-8.1</td><td>5.6E-16</td><td>4.0E-15</td><td>MB231/MB468</td><td>MP</td></tr>
<tr><td>>hg38_21_+_8433172_8446622_RNA45SN1_45S_50ntFlanks@10819.10839.21|AGGGAAGTCGGCAAGCCGGAT</td><td>>hg38_21_+_8433172_8446622_RNA45SN1_45S_50ntFlanks</td><td>10819</td><td>10839</td><td>21</td><td>AGGGAAGTCGGCAAGCCGGAT</td><td>1149.7</td><td>-8.6</td><td>1.1</td><td>-8.1</td><td>5.6E-16</td><td>4.1E-15</td><td>MB231/MB468</td><td>MP</td></tr>
<tr><td>>hg38_21_+_8433172_8446622_RNA45SN1_45S_50ntFlanks@9240.9273.34|GCGGAGCGAGCGCACGGGGTCGGCGGCGACGTCG</td><td>>hg38_21_+_8433172_8446622_RNA45SN1_45S_50ntFlanks</td><td>9240</td><td>9273</td><td>34</td><td>GCGGAGCGAGCGCACGGGGTCGGCGGCGACGTCG</td><td>278.8</td><td>-12.0</td><td>1.5</td><td>-8.1</td><td>5.6E-16</td><td>4.1E-15</td><td>MB231/MB468</td><td>MP</td></tr>
<tr><td>>hg38_21_+_8433172_8446622_RNA45SN1_45S_50ntFlanks@4046.4064.19|CATTCGAACGTCTGCCCTA</td><td>>hg38_21_+_8433172_8446622_RNA45SN1_45S_50ntFlanks</td><td>4046</td><td>4064</td><td>19</td><td>CATTCGAACGTCTGCCCTA</td><td>190.7</td><td>-11.4</td><td>1.4</td><td>-8.1</td><td>5.6E-16</td><td>4.1E-15</td><td>MB231/MB468</td><td>MP</td></tr>
<tr><td>>hg38_21_+_8433172_8446622_RNA45SN1_45S_50ntFlanks@5035.5067.33|CATGGCCGTTCTTAGTTGGTGGAGCGATTTGTC</td><td>>hg38_21_+_8433172_8446622_RNA45SN1_45S_50ntFlanks</td><td>5035</td><td>5067</td><td>33</td><td>CATGGCCGTTCTTAGTTGGTGGAGCGATTTGTC</td><td>1168.4</td><td>-14.0</td><td>1.7</td><td>-8.0</td><td>9.2E-16</td><td>4.1E-15</td><td>MB231/MB468</td><td>MITO</td></tr>
<tr><td>>hg38_21_+_8433172_8446622_RNA45SN1_45S_50ntFlanks@4445.4470.26|CTTGCCTCTCGGCGCCCCCTCGATGC</td><td>>hg38_21_+_8433172_8446622_RNA45SN1_45S_50ntFlanks</td><td>4445</td><td>4470</td><td>26</td><td>CTTGCCTCTCGGCGCCCCCTCGATGC</td><td>311.7</td><td>-12.1</td><td>1.5</td><td>-8.0</td><td>9.2E-16</td><td>4.1E-15</td><td>MB231/MB468</td><td>MITO</td></tr>
<tr><td>>hg38_21_+_8433172_8446622_RNA45SN1_45S_50ntFlanks@4759.4784.26|ATCAGATACCGTCGTAGTTCCGACCA</td><td>>hg38_21_+_8433172_8446622_RNA45SN1_45S_50ntFlanks</td><td>4759</td><td>4784</td><td>26</td><td>ATCAGATACCGTCGTAGTTCCGACCA</td><td>270.1</td><td>-11.9</td><td>1.5</td><td>-8.1</td><td>5.6E-16</td><td>4.1E-15</td><td>MB231/MB468</td><td>MP</td></tr>
<tr><td>>hg38_21_+_8433172_8446622_RNA45SN1_45S_50ntFlanks@4538.4559.22|CCGAGCCGCCTGGATACCGCAG</td><td>>hg38_21_+_8433172_8446622_RNA45SN1_45S_50ntFlanks</td><td>4538</td><td>4559</td><td>22</td><td>CCGAGCCGCCTGGATACCGCAG</td><td>310.4</td><td>-12.1</td><td>1.5</td><td>-8.0</td><td>9.2E-16</td><td>4.1E-15</td><td>MB231/MB468</td><td>MITO</td></tr>
<tr><td>>hg38_21_+_8433172_8446622_RNA45SN1_45S_50ntFlanks@10878.10902.25|TCGGTCGGGCTGGGGCGCGAAGCGG</td><td>>hg38_21_+_8433172_8446622_RNA45SN1_45S_50ntFlanks</td><td>10878</td><td>10902</td><td>25</td><td>TCGGTCGGGCTGGGGCGCGAAGCGG</td><td>266.6</td><td>-11.8</td><td>1.5</td><td>-8.0</td><td>9.2E-16</td><td>4.1E-15</td><td>MB231/MB468</td><td>MITO</td></tr>
<tr><td>>hg38_21_+_8433172_8446622_RNA45SN1_45S_50ntFlanks@10805.10840.36|GAACAATGTAGGTAAGGGAAGTCGGCAAGCCGGATC</td><td>>hg38_21_+_8433172_8446622_RNA45SN1_45S_50ntFlanks</td><td>10805</td><td>10840</td><td>36</td><td>GAACAATGTAGGTAAGGGAAGTCGGCAAGCCGGATC</td><td>244.3</td><td>-11.8</td><td>1.5</td><td>-8.1</td><td>5.6E-16</td><td>4.1E-15</td><td>MB231/MB468</td><td>MP</td></tr>
<tr><td>>hg38_21_+_8433172_8446622_RNA45SN1_45S_50ntFlanks@12323.12346.24|TTTGGGTTTTAAGCAGGAGGTGTC</td><td>>hg38_21_+_8433172_8446622_RNA45SN1_45S_50ntFlanks</td><td>12323</td><td>12346</td><td>24</td><td>TTTGGGTTTTAAGCAGGAGGTGTC</td><td>190.0</td><td>-11.4</td><td>1.4</td><td>-8.1</td><td>5.6E-16</td><td>4.1E-15</td><td>MB231/MB468</td><td>MP</td></tr>
<tr><td>>hg38_21_+_8433172_8446622_RNA45SN1_45S_50ntFlanks@11452.11475.24|CTCCTCGGCGCGCGGCGGCGGCGG</td><td>>hg38_21_+_8433172_8446622_RNA45SN1_45S_50ntFlanks</td><td>11452</td><td>11475</td><td>24</td><td>CTCCTCGGCGCGCGGCGGCGGCGG</td><td>227.2</td><td>-10.5</td><td>1.3</td><td>-8.1</td><td>6.1E-16</td><td>4.1E-15</td><td>MB231/MB468</td><td>CYTO</td></tr>
<tr><td>>hg38_21_+_8433172_8446622_RNA45SN1_45S_50ntFlanks@9467.9503.37|GTGGAGCACGAGCGCACGTGTTAGGACCCGAAAGATG</td><td>>hg38_21_+_8433172_8446622_RNA45SN1_45S_50ntFlanks</td><td>9467</td><td>9503</td><td>37</td><td>GTGGAGCACGAGCGCACGTGTTAGGACCCGAAAGATG</td><td>598.2</td><td>-13.1</td><td>1.6</td><td>-8.1</td><td>5.6E-16</td><td>4.1E-15</td><td>MB231/MB468</td><td>MP</td></tr>
<tr><td>>hg38_21_+_8433172_8446622_RNA45SN1_45S_50ntFlanks@8170.8211.42|CAAGTCCTTCTGATCGAGGCCCAGCCCGTGGACGGTGTGAGG</td><td>>hg38_21_+_8433172_8446622_RNA45SN1_45S_50ntFlanks</td><td>8170</td><td>8211</td><td>42</td><td>CAAGTCCTTCTGATCGAGGCCCAGCCCGTGGACGGTGTGAGG</td><td>565.3</td><td>-12.9</td><td>1.6</td><td>-8.0</td><td>9.3E-16</td><td>4.1E-15</td><td>MB231/MB468</td><td>MITO</td></tr>
<tr><td>>hg38_21_+_8433172_8446622_RNA45SN1_45S_50ntFlanks@3835.3860.26|CTCCTCTCCTACTTGGATAACTGTGG</td><td>>hg38_21_+_8433172_8446622_RNA45SN1_45S_50ntFlanks</td><td>3835</td><td>3860</td><td>26</td><td>CTCCTCTCCTACTTGGATAACTGTGG</td><td>709.4</td><td>-13.3</td><td>1.6</td><td>-8.1</td><td>5.6E-16</td><td>4.1E-15</td><td>MB231/MB468</td><td>MP</td></tr>
<tr><td>>hg38_21_+_8433172_8446622_RNA45SN1_45S_50ntFlanks@12074.12095.22|CGAGGGGCTCTCGCTTCTGGCG</td><td>>hg38_21_+_8433172_8446622_RNA45SN1_45S_50ntFlanks</td><td>12074</td><td>12095</td><td>22</td><td>CGAGGGGCTCTCGCTTCTGGCG</td><td>1461.3</td><td>-7.8</td><td>1.0</td><td>-8.0</td><td>9.3E-16</td><td>4.1E-15</td><td>MB231/MB468</td><td>MITO</td></tr>
<tr><td>>hg38_1_-_228634819_228635039_RNA5S12_5S_50ntFlanks@149.170.22|GAATACCGGGTGCTGTAGGCTT</td><td>>hg38_1_-_228634819_228635039_RNA5S12_5S_50ntFlanks</td><td>149</td><td>170</td><td>22</td><td>GAATACCGGGTGCTGTAGGCTT</td><td>5093.1</td><td>5.9</td><td>0.7</td><td>8.1</td><td>6.1E-16</td><td>4.1E-15</td><td>MB231/MB468</td><td>CYTO</td></tr>
<tr><td>>MI0000081|hsa-mir-24-2&WithFlank&19|-|13836281|13836365@56.77.22(+1C);MI0000080|hsa-mir-24-1&WithFlank&9|+|95086015|95086094@50.71.22(+1C)&[MIMAT0000080&hsa-miR-24-3p&offsets|0|0(+1C);m-36&19|-|13836288|13836310&offsets|0|-1(+1C)];[MIMAT0000080_1&hsa-miR-24-3p&offsets|0|0(+1C);m-35&9|+|95086064|95086086&offsets|0|-1(+1C)]|TGGCTCAGTTCAGCAGGAACAGC</td><td>>MI0000081|hsa-mir-24-2&WithFlank&19|-|13836281|13836365</td><td>56</td><td>77</td><td>22(+1C)</td><td>TGGCTCAGTTCAGCAGGAACAGC</td><td>627.3</td><td>6.4</td><td>0.8</td><td>8.0</td><td>8.6E-16</td><td>4.1E-15</td><td>BT20/MB468</td><td>CYTO</td></tr>
<tr><td>>hg38_21_+_8433172_8446622_RNA45SN1_45S_50ntFlanks@10391.10409.19|AACAGCAGTTGAACATGGG</td><td>>hg38_21_+_8433172_8446622_RNA45SN1_45S_50ntFlanks</td><td>10391</td><td>10409</td><td>19</td><td>AACAGCAGTTGAACATGGG</td><td>344.2</td><td>-12.2</td><td>1.5</td><td>-8.0</td><td>9.4E-16</td><td>4.1E-15</td><td>MB231/MB468</td><td>MITO</td></tr>
<tr><td>>trnaMT_ValTAC_MT_+_1602_1670@53.71.19|TAACTTGACCGCTCTGACC</td><td>>trnaMT_ValTAC_MT_+_1602_1670</td><td>53</td><td>71</td><td>19</td><td>TAACTTGACCGCTCTGACC</td><td>1058.9</td><td>14.3</td><td>1.8</td><td>8.0</td><td>8.7E-16</td><td>4.2E-15</td><td>BT20/MB468</td><td>CYTO</td></tr>
<tr><td>>hg38_21_+_8433172_8446622_RNA45SN1_45S_50ntFlanks@9545.9570.26|GTGGAGGTCCGTAGCGGTCCTGACGT</td><td>>hg38_21_+_8433172_8446622_RNA45SN1_45S_50ntFlanks</td><td>9545</td><td>9570</td><td>26</td><td>GTGGAGGTCCGTAGCGGTCCTGACGT</td><td>863.2</td><td>-5.9</td><td>0.7</td><td>-8.1</td><td>5.7E-16</td><td>4.2E-15</td><td>MB231/MB468</td><td>MP</td></tr>
<tr><td>>hg38_21_+_8433172_8446622_RNA45SN1_45S_50ntFlanks@6731.6770.40|CACATTGATCATCGACACTTCGAACGCACTTGCGGCCCCG</td><td>>hg38_21_+_8433172_8446622_RNA45SN1_45S_50ntFlanks</td><td>6731</td><td>6770</td><td>40</td><td>CACATTGATCATCGACACTTCGAACGCACTTGCGGCCCCG</td><td>1989.0</td><td>-6.6</td><td>0.8</td><td>-8.1</td><td>7.1E-16</td><td>4.2E-15</td><td>MB231/MB468</td><td>NUC</td></tr>
<tr><td>>hg38_21_+_8433172_8446622_RNA45SN1_45S_50ntFlanks@4260.4278.19|TTGGAATGAGTCCACTTTA</td><td>>hg38_21_+_8433172_8446622_RNA45SN1_45S_50ntFlanks</td><td>4260</td><td>4278</td><td>19</td><td>TTGGAATGAGTCCACTTTA</td><td>211.1</td><td>-11.6</td><td>1.4</td><td>-8.1</td><td>5.8E-16</td><td>4.2E-15</td><td>MB231/MB468</td><td>MP</td></tr>
</table>

<table>
<tr><td>>MI0000272|hsa-mir-182&WithFlank&7|-|129770377|129770498@29.54.26(+1U)&[MIMAT0000259&hsa-miR-182-5p&offsets|0|+2(+1U);m-22&7|-|129770449|129770470&offsets|0|+4(+1U)]|TTTGGCAATGGTAGAACTCACACTGGT</td><td>>MI0000272|hsa-mir-182&WithFlank&7|-|129770377|129770498</td><td>29</td><td>54</td><td>26(+1U)</td><td>TTTGGCAATGGTAGAACTCACACTGGT</td><td>1347.6</td><td>8.3</td><td>1.0</td><td>8.0</td><td>8.8E-16</td><td>4.2E-15</td><td>BT20/MB468</td><td>CYTO</td></tr>
<tr><td>>hg38_21_+_8433172_8446622_RNA45SN1_45S_50ntFlanks@12004.12031.28|CGGGCCGCCGGTGAAATACCACTACTCT</td><td>>hg38_21_+_8433172_8446622_RNA45SN1_45S_50ntFlanks</td><td>12004</td><td>12031</td><td>28</td><td>CGGGCCGCCGGTGAAATACCACTACTCT</td><td>207.3</td><td>-11.6</td><td>1.4</td><td>-8.1</td><td>5.8E-16</td><td>4.2E-15</td><td>MB231/MB468</td><td>MP</td></tr>
<tr><td>>hg38_21_+_8433172_8446622_RNA45SN1_45S_50ntFlanks@4753.4782.30|AAGACGATCAGATACCGTCGTAGTTCCGAC</td><td>>hg38_21_+_8433172_8446622_RNA45SN1_45S_50ntFlanks</td><td>4753</td><td>4782</td><td>30</td><td>AAGACGATCAGATACCGTCGTAGTTCCGAC</td><td>552.4</td><td>-13.0</td><td>1.6</td><td>-8.1</td><td>5.8E-16</td><td>4.2E-15</td><td>MB231/MB468</td><td>MP</td></tr>
<tr><td>>hg38_21_+_8433172_8446622_RNA45SN1_45S_50ntFlanks@11214.11234.21|ACTCTGGACGCGAGCCGGGCC</td><td>>hg38_21_+_8433172_8446622_RNA45SN1_45S_50ntFlanks</td><td>11214</td><td>11234</td><td>21</td><td>ACTCTGGACGCGAGCCGGGCC</td><td>237.7</td><td>-10.6</td><td>1.3</td><td>-8.1</td><td>6.3E-16</td><td>4.2E-15</td><td>MB231/MB468</td><td>CYTO</td></tr>
<tr><td>>hg38_21_+_8433172_8446622_RNA45SN1_45S_50ntFlanks@4069.4095.27|CTTTCGATGGTAGTCGCCGTGCCTACC</td><td>>hg38_21_+_8433172_8446622_RNA45SN1_45S_50ntFlanks</td><td>4069</td><td>4095</td><td>27</td><td>CTTTCGATGGTAGTCGCCGTGCCTACC</td><td>949.6</td><td>-8.1</td><td>1.0</td><td>-8.1</td><td>5.8E-16</td><td>4.2E-15</td><td>MB231/MB468</td><td>MP</td></tr>
<tr><td>>hg38_21_+_8433172_8446622_RNA45SN1_45S_50ntFlanks@9391.9415.25|CCGAGGTGGGATCCCGAGGCCTCTC</td><td>>hg38_21_+_8433172_8446622_RNA45SN1_45S_50ntFlanks</td><td>9391</td><td>9415</td><td>25</td><td>CCGAGGTGGGATCCCGAGGCCTCTC</td><td>267.0</td><td>-11.9</td><td>1.5</td><td>-8.1</td><td>5.8E-16</td><td>4.2E-15</td><td>MB231/MB468</td><td>MP</td></tr>
<tr><td>>hg38_21_+_8433172_8446622_RNA45SN1_45S_50ntFlanks@11846.11872.27|GAACGGGCTTGGCGGAATCAGCGGGGA</td><td>>hg38_21_+_8433172_8446622_RNA45SN1_45S_50ntFlanks</td><td>11846</td><td>11872</td><td>27</td><td>GAACGGGCTTGGCGGAATCAGCGGGGA</td><td>280.3</td><td>-11.9</td><td>1.5</td><td>-8.0</td><td>9.6E-16</td><td>4.2E-15</td><td>MB231/MB468</td><td>MITO</td></tr>
<tr><td>>hg38_21_+_8433172_8446622_RNA45SN1_45S_50ntFlanks@3940.3959.20|ACCCGGTCAGCCCCTCTCCG</td><td>>hg38_21_+_8433172_8446622_RNA45SN1_45S_50ntFlanks</td><td>3940</td><td>3959</td><td>20</td><td>ACCCGGTCAGCCCCTCTCCG</td><td>338.5</td><td>-11.1</td><td>1.4</td><td>-8.1</td><td>6.3E-16</td><td>4.2E-15</td><td>MB231/MB468</td><td>CYTO</td></tr>
<tr><td>>hg38_21_+_8433172_8446622_RNA45SN1_45S_50ntFlanks@11210.11231.22|GGCGACTCTGGACGCGAGCCGG</td><td>>hg38_21_+_8433172_8446622_RNA45SN1_45S_50ntFlanks</td><td>11210</td><td>11231</td><td>22</td><td>GGCGACTCTGGACGCGAGCCGG</td><td>285.2</td><td>-12.0</td><td>1.5</td><td>-8.1</td><td>5.9E-16</td><td>4.2E-15</td><td>MB231/MB468</td><td>MP</td></tr>
<tr><td>>hg38_21_+_8433172_8446622_RNA45SN1_45S_50ntFlanks@9166.9187.22|CCGGGCGGGTCGCGCCGTCGGG</td><td>>hg38_21_+_8433172_8446622_RNA45SN1_45S_50ntFlanks</td><td>9166</td><td>9187</td><td>22</td><td>CCGGGCGGGTCGCGCCGTCGGG</td><td>771.5</td><td>-13.5</td><td>1.7</td><td>-8.1</td><td>5.9E-16</td><td>4.2E-15</td><td>MB231/MB468</td><td>MP</td></tr>
<tr><td>>MI0000457|hsa-mir-141&WithFlank&12|+|6964091|6964197@65.85.21&[MIMAT0000432&hsa-miR-141-3p&offsets|0|-1;m-134&12|+|6964155|6964175&offsets|0|0]|TAACACTGTCTGGTAAAGATG</td><td>>MI0000457|hsa-mir-141&WithFlank&12|+|6964091|6964197</td><td>65</td><td>85</td><td>21</td><td>TAACACTGTCTGGTAAAGATG</td><td>544.3</td><td>12.9</td><td>1.5</td><td>8.3</td><td>9.9E-17</td><td>4.2E-15</td><td>BT20/MB231</td><td>NUC</td></tr>
<tr><td>>MI0000079|hsa-mir-23a&WithFlank&19|-|13836581|13836665@54.73.20&[MIMAT0000078&hsa-miR-23a-3p&offsets|+3|+2;m-31&19|-|13836595|13836615&offsets|+3|+2]|ACATTGCCAGGGATTTCCAA</td><td>>MI0000079|hsa-mir-23a&WithFlank&19|-|13836581|13836665</td><td>54</td><td>73</td><td>20</td><td>ACATTGCCAGGGATTTCCAA</td><td>3196.8</td><td>29.3</td><td>3.6</td><td>8.1</td><td>7.2E-16</td><td>4.2E-15</td><td>MB231/MB468</td><td>NUC</td></tr>
<tr><td>>am_trnaMT_PheGAA_MT_+_577_647@41.62.22|GTTTAGACGGGCTCACATCACC</td><td>>am_trnaMT_PheGAA_MT_+_577_647</td><td>41</td><td>62</td><td>22</td><td>GTTTAGACGGGCTCACATCACC</td><td>2285.6</td><td>8.2</td><td>1.0</td><td>8.1</td><td>6.4E-16</td><td>4.3E-15</td><td>BT20/MB468</td><td>MITO</td></tr>
<tr><td>>hg38_21_+_8433172_8446622_RNA45SN1_45S_50ntFlanks@10793.10816.24|TCTGGCATGTTGGAACAATGTAGG</td><td>>hg38_21_+_8433172_8446622_RNA45SN1_45S_50ntFlanks</td><td>10793</td><td>10816</td><td>24</td><td>TCTGGCATGTTGGAACAATGTAGG</td><td>198.8</td><td>-11.5</td><td>1.4</td><td>-8.1</td><td>5.9E-16</td><td>4.3E-15</td><td>MB231/MB468</td><td>MP</td></tr>
<tr><td>>hg38_21_+_8433172_8446622_RNA45SN1_45S_50ntFlanks@9534.9561.28|AGGAAACTCTGGTGGAGGTCCGTAGCGG</td><td>>hg38_21_+_8433172_8446622_RNA45SN1_45S_50ntFlanks</td><td>9534</td><td>9561</td><td>28</td><td>AGGAAACTCTGGTGGAGGTCCGTAGCGG</td><td>3764.7</td><td>-10.8</td><td>1.3</td><td>-8.1</td><td>5.9E-16</td><td>4.3E-15</td><td>MB231/MB468</td><td>MP</td></tr>
<tr><td>>hg38_21_+_8433172_8446622_RNA45SN1_45S_50ntFlanks@12991.13027.37|CTCCCTCGCTGCGATCTATTGAAAGTCAGCCCTCGAC</td><td>>hg38_21_+_8433172_8446622_RNA45SN1_45S_50ntFlanks</td><td>12991</td><td>13027</td><td>37</td><td>CTCCCTCGCTGCGATCTATTGAAAGTCAGCCCTCGAC</td><td>669.4</td><td>-11.5</td><td>1.4</td><td>-8.1</td><td>6.4E-16</td><td>4.3E-15</td><td>BT20/MB468</td><td>MITO</td></tr>
<tr><td>>hg38_21_+_8433172_8446622_RNA45SN1_45S_50ntFlanks@11580.11600.21|GAACTGGTGCGGACCAGGGGA</td><td>>hg38_21_+_8433172_8446622_RNA45SN1_45S_50ntFlanks</td><td>11580</td><td>11600</td><td>21</td><td>GAACTGGTGCGGACCAGGGGA</td><td>659.1</td><td>-13.2</td><td>1.6</td><td>-8.1</td><td>5.9E-16</td><td>4.3E-15</td><td>MB231/MB468</td><td>MP</td></tr>
<tr><td>>hg38_21_+_8433172_8446622_RNA45SN1_45S_50ntFlanks@4752.4782.31|GAAGACGATCAGATACCGTCGTAGTTCCGAC</td><td>>hg38_21_+_8433172_8446622_RNA45SN1_45S_50ntFlanks</td><td>4752</td><td>4782</td><td>31</td><td>GAAGACGATCAGATACCGTCGTAGTTCCGAC</td><td>1140.5</td><td>-13.9</td><td>1.7</td><td>-8.0</td><td>9.7E-16</td><td>4.3E-15</td><td>MB231/MB468</td><td>MITO</td></tr>
<tr><td>>MI0000289|hsa-mir-181a-1&WithFlank&1|-|198859038|198859159@70.91.22&[MIMAT0000270&hsa-miR-181a-3p&offsets|0|0;m-125&1|-|198859069|198859090&offsets|0|0]|ACCATCGACCGTTGATTGTACC</td><td>>MI0000289|hsa-mir-181a-1&WithFlank&1|-|198859038|198859159</td><td>70</td><td>91</td><td>22</td><td>ACCATCGACCGTTGATTGTACC</td><td>2744.3</td><td>6.7</td><td>0.8</td><td>8.1</td><td>6.4E-16</td><td>4.3E-15</td><td>MB231/MB468</td><td>CYTO</td></tr>
<tr><td>>hg38_21_+_8433172_8446622_RNA45SN1_45S_50ntFlanks@12995.13035.41|CTCGCTGCGATCTATTGAAAGTCAGCCCTCGACACAAGGGT</td><td>>hg38_21_+_8433172_8446622_RNA45SN1_45S_50ntFlanks</td><td>12995</td><td>13035</td><td>41</td><td>CTCGCTGCGATCTATTGAAAGTCAGCCCTCGACACAAGGGT</td><td>837.3</td><td>10.4</td><td>1.3</td><td>8.1</td><td>6.4E-16</td><td>4.3E-15</td><td>MB231/MB468</td><td>CYTO</td></tr>
<tr><td>>hg38_21_+_8433172_8446622_RNA45SN1_45S_50ntFlanks@8357.8378.22|AAAAGAACTTTGAAGAGAGAGT</td><td>>hg38_21_+_8433172_8446622_RNA45SN1_45S_50ntFlanks</td><td>8357</td><td>8378</td><td>22</td><td>AAAAGAACTTTGAAGAGAGAGT</td><td>5252.4</td><td>-6.5</td><td>0.8</td><td>-8.0</td><td>9.8E-16</td><td>4.3E-15</td><td>MB231/MB468</td><td>MITO</td></tr>
<tr><td>>hg38_21_+_8433172_8446622_RNA45SN1_45S_50ntFlanks@12634.12657.24|GAAGCTACCATCTGTGGGATTATG</td><td>>hg38_21_+_8433172_8446622_RNA45SN1_45S_50ntFlanks</td><td>12634</td><td>12657</td><td>24</td><td>GAAGCTACCATCTGTGGGATTATG</td><td>224.4</td><td>-11.7</td><td>1.4</td><td>-8.1</td><td>5.9E-16</td><td>4.3E-15</td><td>MB231/MB468</td><td>MP</td></tr>
<tr><td>>hg38_21_+_8433172_8446622_RNA45SN1_45S_50ntFlanks@12033.12053.21|ATCGTTTTTTCACTGACCCGG</td><td>>hg38_21_+_8433172_8446622_RNA45SN1_45S_50ntFlanks</td><td>12033</td><td>12053</td><td>21</td><td>ATCGTTTTTTCACTGACCCGG</td><td>241.5</td><td>-10.9</td><td>1.3</td><td>-8.1</td><td>7.3E-16</td><td>4.3E-15</td><td>MB231/MB468</td><td>NUC</td></tr>
<tr><td>>hg38_21_+_8433172_8446622_RNA45SN1_45S_50ntFlanks@12154.12180.27|GGGAGTTTGACTGGGGCGGTACACCTG</td><td>>hg38_21_+_8433172_8446622_RNA45SN1_45S_50ntFlanks</td><td>12154</td><td>12180</td><td>27</td><td>GGGAGTTTGACTGGGGCGGTACACCTG</td><td>265.4</td><td>-11.8</td><td>1.5</td><td>-8.0</td><td>9.8E-16</td><td>4.3E-15</td><td>MB231/MB468</td><td>MITO</td></tr>
<tr><td>>hg38_21_+_8433172_8446622_RNA45SN1_45S_50ntFlanks@3875.3898.24|ATACATGCCGACGGGCGCTGACCC</td><td>>hg38_21_+_8433172_8446622_RNA45SN1_45S_50ntFlanks</td><td>3875</td><td>3898</td><td>24</td><td>ATACATGCCGACGGGCGCTGACCC</td><td>4596.6</td><td>-7.3</td><td>0.9</td><td>-8.1</td><td>7.3E-16</td><td>4.3E-15</td><td>MB231/MB468</td><td>NUC</td></tr>
<tr><td>>hg38_21_+_8433172_8446622_RNA45SN1_45S_50ntFlanks@7976.8010.35|GCGACCTCAGATCAGACGTGGCGACCCGCTGAATT</td><td>>hg38_21_+_8433172_8446622_RNA45SN1_45S_50ntFlanks</td><td>7976</td><td>8010</td><td>35</td><td>GCGACCTCAGATCAGACGTGGCGACCCGCTGAATT</td><td>1212.8</td><td>9.6</td><td>1.2</td><td>8.1</td><td>6.5E-16</td><td>4.3E-15</td><td>MB231/MB468</td><td>CYTO</td></tr>
<tr><td>>MI0000079|hsa-mir-23a&WithFlank&19|-|13836581|13836665@54.71.18&[MIMAT0000078&hsa-miR-23a-3p&offsets|+3|0;m-31&19|-|13836595|13836615&offsets|+3|0]|ACATTGCCAGGGATTTCC</td><td>>MI0000079|hsa-mir-23a&WithFlank&19|-|13836581|13836665</td><td>54</td><td>71</td><td>18</td><td>ACATTGCCAGGGATTTCC</td><td>3577.7</td><td>29.2</td><td>3.6</td><td>8.0</td><td>9.8E-16</td><td>4.3E-15</td><td>MB231/MB468</td><td>MITO</td></tr>
<tr><td>>hg38_21_+_8433172_8446622_RNA45SN1_45S_50ntFlanks@6721.6740.20|AATTGCAGGACACATTGATC</td><td>>hg38_21_+_8433172_8446622_RNA45SN1_45S_50ntFlanks</td><td>6721</td><td>6740</td><td>20</td><td>AATTGCAGGACACATTGATC</td><td>7020.0</td><td>-9.0</td><td>1.1</td><td>-8.1</td><td>6.0E-16</td><td>4.3E-15</td><td>MB231/MB468</td><td>MP</td></tr>
<tr><td>>MI0000079|hsa-mir-23a&WithFlank&19|-|13836581|13836665@54.73.20(+1U)&[MIMAT0000078&hsa-miR-23a-3p&offsets|+3|+2(+1U);m-31&19|-|13836595|13836615&offsets|+3|+2(+1U)]|ACATTGCCAGGGATTTCCAAT</td><td>>MI0000079|hsa-mir-23a&WithFlank&19|-|13836581|13836665</td><td>54</td><td>73</td><td>20(+1U)</td><td>ACATTGCCAGGGATTTCCAAT</td><td>3844.4</td><td>29.1</td><td>3.6</td><td>8.0</td><td>9.8E-16</td><td>4.3E-15</td><td>MB231/MB468</td><td>MITO</td></tr>
<tr><td>>hg38_21_+_8433172_8446622_RNA45SN1_45S_50ntFlanks@9029.9058.30|TCTCCTCGTGGGGGGGCCGGGCCACCCCTC</td><td>>hg38_21_+_8433172_8446622_RNA45SN1_45S_50ntFlanks</td><td>9029</td><td>9058</td><td>30</td><td>TCTCCTCGTGGGGGGGCCGGGCCACCCCTC</td><td>361.2</td><td>-12.3</td><td>1.5</td><td>-8.0</td><td>9.9E-16</td><td>4.3E-15</td><td>MB231/MB468</td><td>MITO</td></tr>
<tr><td>>hg38_21_+_8433172_8446622_RNA45SN1_45S_50ntFlanks@12635.12654.20|AAGCTACCATCTGTGGGATT</td><td>>hg38_21_+_8433172_8446622_RNA45SN1_45S_50ntFlanks</td><td>12635</td><td>12654</td><td>20</td><td>AAGCTACCATCTGTGGGATT</td><td>17290.3</td><td>-10.0</td><td>1.2</td><td>-8.5</td><td>1.8E-17</td><td>4.3E-15</td><td>BT20/MB468</td><td>MP</td></tr>
<tr><td>>hg38_21_+_8433172_8446622_RNA45SN1_45S_50ntFlanks@4003.4022.20|ATAACCTCGGGCCGATCGCA</td><td>>hg38_21_+_8433172_8446622_RNA45SN1_45S_50ntFlanks</td><td>4003</td><td>4022</td><td>20</td><td>ATAACCTCGGGCCGATCGCA</td><td>1353.3</td><td>-9.9</td><td>1.2</td><td>-8.0</td><td>9.9E-16</td><td>4.3E-15</td><td>MB231/MB468</td><td>MITO</td></tr>
<tr><td>>hg38_21_+_8433172_8446622_RNA45SN1_45S_50ntFlanks@11573.11605.33|CGACTTAGAACTGGTGCGGACCAGGGGAATCCG</td><td>>hg38_21_+_8433172_8446622_RNA45SN1_45S_50ntFlanks</td><td>11573</td><td>11605</td><td>33</td><td>CGACTTAGAACTGGTGCGGACCAGGGGAATCCG</td><td>689.9</td><td>-4.8</td><td>0.6</td><td>-8.1</td><td>7.4E-16</td><td>4.3E-15</td><td>MB231/MB468</td><td>NUC</td></tr>
<tr><td>>hg38_21_+_8433172_8446622_RNA45SN1_45S_50ntFlanks@11576.11605.30|CTTAGAACTGGTGCGGACCAGGGGAATCCG</td><td>>hg38_21_+_8433172_8446622_RNA45SN1_45S_50ntFlanks</td><td>11576</td><td>11605</td><td>30</td><td>CTTAGAACTGGTGCGGACCAGGGGAATCCG</td><td>1064.0</td><td>-9.6</td><td>1.2</td><td>-8.0</td><td>9.9E-16</td><td>4.3E-15</td><td>MB231/MB468</td><td>MITO</td></tr>
<tr><td>>hg38_21_+_8433172_8446622_RNA45SN1_45S_50ntFlanks@9567.9596.30|ACGTGCAAATCGGTCGTCCGACCTGGGTAT</td><td>>hg38_21_+_8433172_8446622_RNA45SN1_45S_50ntFlanks</td><td>9567</td><td>9596</td><td>30</td><td>ACGTGCAAATCGGTCGTCCGACCTGGGTAT</td><td>5829.3</td><td>-8.1</td><td>1.0</td><td>-8.1</td><td>6.5E-16</td><td>4.3E-15</td><td>MB231/MB468</td><td>CYTO</td></tr>
<tr><td>>hg38_21_+_8433172_8446622_RNA45SN1_45S_50ntFlanks@10572.10608.37|AGAAGCCGGCGGGAGCCCCGGGGAGAGTTCTCTTTTC</td><td>>hg38_21_+_8433172_8446622_RNA45SN1_45S_50ntFlanks</td><td>10572</td><td>10608</td><td>37</td><td>AGAAGCCGGCGGGAGCCCCGGGGAGAGTTCTCTTTTC</td><td>1338.6</td><td>-7.3</td><td>0.9</td><td>-8.1</td><td>6.5E-16</td><td>4.3E-15</td><td>MB231/MB468</td><td>CYTO</td></tr>
<tr><td>>hg38_21_+_8433172_8446622_RNA45SN1_45S_50ntFlanks@13006.13028.23|CTATTGAAAGTCAGCCCTCGACA</td><td>>hg38_21_+_8433172_8446622_RNA45SN1_45S_50ntFlanks</td><td>13006</td><td>13028</td><td>23</td><td>CTATTGAAAGTCAGCCCTCGACA</td><td>297.4</td><td>-12.0</td><td>1.5</td><td>-8.0</td><td>1.0E-15</td><td>4.3E-15</td><td>MB231/MB468</td><td>MITO</td></tr>
<tr><td>>hg38_21_+_8433172_8446622_RNA45SN1_45S_50ntFlanks@4005.4042.38|AACCTCGGGCCGATCGCACGCCCCCCGTGGCGGCGACG</td><td>>hg38_21_+_8433172_8446622_RNA45SN1_45S_50ntFlanks</td><td>4005</td><td>4042</td><td>38</td><td>AACCTCGGGCCGATCGCACGCCCCCCGTGGCGGCGACG</td><td>551.7</td><td>-5.8</td><td>0.7</td><td>-8.0</td><td>1.0E-15</td><td>4.3E-15</td><td>MB231/MB468</td><td>MITO</td></tr>
<tr><td>>hg38_21_+_8433172_8446622_RNA45SN1_45S_50ntFlanks@12405.12423.19|CTTTTTGATCCTTCGATGT</td><td>>hg38_21_+_8433172_8446622_RNA45SN1_45S_50ntFlanks</td><td>12405</td><td>12423</td><td>19</td><td>CTTTTTGATCCTTCGATGT</td><td>222.9</td><td>-11.7</td><td>1.4</td><td>-8.1</td><td>6.1E-16</td><td>4.4E-15</td><td>MB231/MB468</td><td>MP</td></tr>
<tr><td>>hg38_21_+_8433172_8446622_RNA45SN1_45S_50ntFlanks@10916.10936.21|CCGCGGCTGGACGAGGCGCCG</td><td>>hg38_21_+_8433172_8446622_RNA45SN1_45S_50ntFlanks</td><td>10916</td><td>10936</td><td>21</td><td>CCGCGGCTGGACGAGGCGCCG</td><td>435.9</td><td>-11.5</td><td>1.4</td><td>-8.1</td><td>6.6E-16</td><td>4.4E-15</td><td>MB231/MB468</td><td>CYTO</td></tr>
<tr><td>>hg38_21_+_8433172_8446622_RNA45SN1_45S_50ntFlanks@12437.12472.36|CATTGTGAAGCAGAATTCACCAAGCGTTGGATTGTT</td><td>>hg38_21_+_8433172_8446622_RNA45SN1_45S_50ntFlanks</td><td>12437</td><td>12472</td><td>36</td><td>CATTGTGAAGCAGAATTCACCAAGCGTTGGATTGTT</td><td>360.3</td><td>-5.6</td><td>0.7</td><td>-8.0</td><td>1.4E-15</td><td>4.4E-15</td><td>BT20/MB468</td><td>NUC</td></tr>
<tr><td>>hg38_21_+_8433172_8446622_RNA45SN1_45S_50ntFlanks@12288.12323.36|ACCGTGAAAGCGGGGCCTCACGATCCTTCTGACCTT</td><td>>hg38_21_+_8433172_8446622_RNA45SN1_45S_50ntFlanks</td><td>12288</td><td>12323</td><td>36</td><td>ACCGTGAAAGCGGGGCCTCACGATCCTTCTGACCTT</td><td>2394.8</td><td>-9.1</td><td>1.1</td><td>-8.0</td><td>1.0E-15</td><td>4.4E-15</td><td>MB231/MB468</td><td>MITO</td></tr>
<tr><td>>hg38_21_+_8433172_8446622_RNA45SN1_45S_50ntFlanks@6699.6717.19|GCTAGCTGCGAGAATTAAT</td><td>>hg38_21_+_8433172_8446622_RNA45SN1_45S_50ntFlanks</td><td>6699</td><td>6717</td><td>19</td><td>GCTAGCTGCGAGAATTAAT</td><td>852.3</td><td>-7.8</td><td>1.0</td><td>-8.0</td><td>1.0E-15</td><td>4.4E-15</td><td>MB231/MB468</td><td>MITO</td></tr>
<tr><td>>hg38_21_+_8433172_8446622_RNA45SN1_45S_50ntFlanks@8176.8197.22|CTTCTGATCGAGGCCCAGCCCG</td><td>>hg38_21_+_8433172_8446622_RNA45SN1_45S_50ntFlanks</td><td>8176</td><td>8197</td><td>22</td><td>CTTCTGATCGAGGCCCAGCCCG</td><td>810.5</td><td>-3.6</td><td>0.4</td><td>-8.1</td><td>7.5E-16</td><td>4.4E-15</td><td>MB231/MB468</td><td>NUC</td></tr>
<tr><td>>MI0000079|hsa-mir-23a&WithFlank&19|-|13836581|13836665@54.72.19&[MIMAT0000078&hsa-miR-23a-3p&offsets|+3|+1;m-31&19|-|13836595|13836615&offsets|+3|+1]|ACATTGCCAGGGATTTCCA</td><td>>MI0000079|hsa-mir-23a&WithFlank&19|-|13836581|13836665</td><td>54</td><td>72</td><td>19</td><td>ACATTGCCAGGGATTTCCA</td><td>4976.3</td><td>29.1</td><td>3.6</td><td>8.0</td><td>1.0E-15</td><td>4.4E-15</td><td>MB231/MB468</td><td>MITO</td></tr>
<tr><td>>trna10_LysCTT_16_+_3241501_3241573@1.35.35,trna13_LysCTT_14_-_58706613_58706685@1.35.35,trna2_LysCTT_15_+_79152904_79152976@1.35.35|GCCCGGCTAGCTCAGTCGGTAGAGCATGGGACTCT</td><td>>trna10_LysCTT_16_+_3241501_3241573</td><td>1</td><td>35</td><td>35</td><td>GCCCGGCTAGCTCAGTCGGTAGAGCATGGGACTCT</td><td>23144.8</td><td>8.6</td><td>1.1</td><td>8.1</td><td>6.6E-16</td><td>4.4E-15</td><td>MB231/MB468</td><td>CYTO</td></tr>
<tr><td>>hg38_21_+_8433172_8446622_RNA45SN1_45S_50ntFlanks@4472.4492.21|CTTAGCTGAGTGTCCCGCGGG</td><td>>hg38_21_+_8433172_8446622_RNA45SN1_45S_50ntFlanks</td><td>4472</td><td>4492</td><td>21</td><td>CTTAGCTGAGTGTCCCGCGGG</td><td>525.4</td><td>-12.8</td><td>1.6</td><td>-8.0</td><td>1.0E-15</td><td>4.4E-15</td><td>MB231/MB468</td><td>MITO</td></tr>
<tr><td>>hg38_21_+_8433172_8446622_RNA45SN1_45S_50ntFlanks@4472.4494.23|CTTAGCTGAGTGTCCCGCGGGGC</td><td>>hg38_21_+_8433172_8446622_RNA45SN1_45S_50ntFlanks</td><td>4472</td><td>4494</td><td>23</td><td>CTTAGCTGAGTGTCCCGCGGGGC</td><td>532.1</td><td>-11.1</td><td>1.4</td><td>-8.1</td><td>6.6E-16</td><td>4.4E-15</td><td>BT20/MB468</td><td>MITO</td></tr>
<tr><td>>hg38_21_+_8433172_8446622_RNA45SN1_45S_50ntFlanks@11568.11600.33|GCAGCCGACTTAGAACTGGTGCGGACCAGGGGA</td><td>>hg38_21_+_8433172_8446622_RNA45SN1_45S_50ntFlanks</td><td>11568</td><td>11600</td><td>33</td><td>GCAGCCGACTTAGAACTGGTGCGGACCAGGGGA</td><td>977.3</td><td>-4.0</td><td>0.5</td><td>-8.1</td><td>7.6E-16</td><td>4.4E-15</td><td>MB231/MB468</td><td>NUC</td></tr>
<tr><td>>hg38_21_+_8433172_8446622_RNA45SN1_45S_50ntFlanks@9572.9593.22|CAAATCGGTCGTCCGACCTGGG</td><td>>hg38_21_+_8433172_8446622_RNA45SN1_45S_50ntFlanks</td><td>9572</td><td>9593</td><td>22</td><td>CAAATCGGTCGTCCGACCTGGG</td><td>349.9</td><td>-12.3</td><td>1.5</td><td>-8.1</td><td>6.1E-16</td><td>4.4E-15</td><td>MB231/MB468</td><td>MP</td></tr>
<tr><td>>hg38_21_+_8433172_8446622_RNA45SN1_45S_50ntFlanks@10349.10390.42|AACGAGAACTTTGAAGGCCGAAGTGGAGAAGGGTTCCATGTG</td><td>>hg38_21_+_8433172_8446622_RNA45SN1_45S_50ntFlanks</td><td>10349</td><td>10390</td><td>42</td><td>AACGAGAACTTTGAAGGCCGAAGTGGAGAAGGGTTCCATGTG</td><td>3512.8</td><td>-7.2</td><td>0.9</td><td>-8.0</td><td>1.5E-15</td><td>4.4E-15</td><td>BT20/MB468</td><td>NUC</td></tr>
</table>

<table>
<tr><td></td><td></td><td></td><td></td><td></td><td></td><td></td><td></td><td></td><td></td><td></td><td></td><td></td><td></td></tr>
<tr><td>>hg38_21_+_8433172_8446622_RNA45SN1_45S_50ntFlanks@12627.12651.25|ATGGGGCGAAGCTACCATCTGTGGG</td><td>>hg38_21_+_8433172_8446622_RNA45SN1_45S_50ntFlanks</td><td>12627</td><td>12651</td><td>25</td><td>ATGGGGCGAAGCTACCATCTGTGGG</td><td>325.0</td><td>-12.1</td><td>1.5</td><td>-8.0</td><td>1.0E-15</td><td>4.5E-15</td><td>MB231/MB468</td><td>MITO</td></tr>
<tr><td>>hg38_21_+_8433172_8446622_RNA45SN1_45S_50ntFlanks@12616.12661.46|CTGAGGAGCCAATGGGGCGAAGCTACCATCTGTGGGATTATGACTG</td><td>>hg38_21_+_8433172_8446622_RNA45SN1_45S_50ntFlanks</td><td>12616</td><td>12661</td><td>46</td><td>CTGAGGAGCCAATGGGGCGAAGCTACCATCTGTGGGATTATGACTG</td><td>526.6</td><td>-13.5</td><td>1.7</td><td>-8.0</td><td>1.5E-15</td><td>4.5E-15</td><td>BT20/MB468</td><td>NUC</td></tr>
<tr><td>>hg38_21_+_8433172_8446622_RNA45SN1_45S_50ntFlanks@11768.11794.27|CTAATTAGTGACGCGCATGAATGGATG</td><td>>hg38_21_+_8433172_8446622_RNA45SN1_45S_50ntFlanks</td><td>11768</td><td>11794</td><td>27</td><td>CTAATTAGTGACGCGCATGAATGGATG</td><td>2981.7</td><td>-8.6</td><td>1.1</td><td>-8.1</td><td>6.2E-16</td><td>4.5E-15</td><td>MB231/MB468</td><td>MP</td></tr>
<tr><td>>hg38_21_+_8433172_8446622_RNA45SN1_45S_50ntFlanks@11769.11791.23|TAATTAGTGACGCGCATGAATGG</td><td>>hg38_21_+_8433172_8446622_RNA45SN1_45S_50ntFlanks</td><td>11769</td><td>11791</td><td>23</td><td>TAATTAGTGACGCGCATGAATGG</td><td>3989.6</td><td>-11.6</td><td>1.5</td><td>-8.0</td><td>1.5E-15</td><td>4.5E-15</td><td>BT20/MB468</td><td>NUC</td></tr>
<tr><td>>MI0000298|hsa-mir-221&WithFlank&X|-|45746151|45746272@71.93.23(+1C)&[MIMAT0000278&hsa-miR-221-3p&offsets|0|0(+1C);m-91&X|-|45746181|45746202&offsets|0|+1(+1C)]|AGCTACATTGTCTGCTGGGTTTCC</td><td>>MI0000298|hsa-mir-221&WithFlank&X|-|45746151|45746272</td><td>71</td><td>93</td><td>23(+1C)</td><td>AGCTACATTGTCTGCTGGGTTTCC</td><td>611.3</td><td>5.9</td><td>0.7</td><td>8.1</td><td>6.8E-16</td><td>4.5E-15</td><td>BT20/MB468</td><td>MITO</td></tr>
<tr><td>>hg38_21_+_8433172_8446622_RNA45SN1_45S_50ntFlanks@10873.10901.29|CTGGGTCGGTCGGGCTGGGGCGCGAAGCG</td><td>>hg38_21_+_8433172_8446622_RNA45SN1_45S_50ntFlanks</td><td>10873</td><td>10901</td><td>29</td><td>CTGGGTCGGTCGGGCTGGGGCGCGAAGCG</td><td>254.2</td><td>-11.8</td><td>1.5</td><td>-8.0</td><td>1.0E-15</td><td>4.5E-15</td><td>MB231/MB468</td><td>MITO</td></tr>
<tr><td>>MI0000465|hsa-mir-191&WithFlank&3|-|49020612|49020715@22.45.24&[MIMAT0000440&hsa-miR-191-5p&offsets|0|+1;m-48&3|-|49020672|49020694&offsets|0|+1]|CAACGGAATCCCAAAAGCAGCTGT</td><td>>MI0000465|hsa-mir-191&WithFlank&3|-|49020612|49020715</td><td>22</td><td>45</td><td>24</td><td>CAACGGAATCCCAAAAGCAGCTGT</td><td>4659.4</td><td>6.2</td><td>0.8</td><td>8.1</td><td>7.8E-16</td><td>4.5E-15</td><td>MB231/MB468</td><td>NUC</td></tr>
<tr><td>>hg38_21_+_8433172_8446622_RNA45SN1_45S_50ntFlanks@12418.12446.29|CGATGTCGGCTCTTCCTATCATTGTGAAG</td><td>>hg38_21_+_8433172_8446622_RNA45SN1_45S_50ntFlanks</td><td>12418</td><td>12446</td><td>29</td><td>CGATGTCGGCTCTTCCTATCATTGTGAAG</td><td>3926.0</td><td>-4.8</td><td>0.6</td><td>-8.1</td><td>7.8E-16</td><td>4.5E-15</td><td>MB231/MB468</td><td>NUC</td></tr>
<tr><td>>hg38_21_+_8433172_8446622_RNA45SN1_45S_50ntFlanks@6745.6772.28|ACACTTCGAACGCACTTGCGGCCCCGGG</td><td>>hg38_21_+_8433172_8446622_RNA45SN1_45S_50ntFlanks</td><td>6745</td><td>6772</td><td>28</td><td>ACACTTCGAACGCACTTGCGGCCCCGGG</td><td>249.3</td><td>-10.9</td><td>1.4</td><td>-8.1</td><td>7.8E-16</td><td>4.5E-15</td><td>MB231/MB468</td><td>NUC</td></tr>
<tr><td>>hg38_21_+_8433172_8446622_RNA45SN1_45S_50ntFlanks@10103.10125.23|CGCGCGTGTGGTGTGCGTCGGAG</td><td>>hg38_21_+_8433172_8446622_RNA45SN1_45S_50ntFlanks</td><td>10103</td><td>10125</td><td>23</td><td>CGCGCGTGTGGTGTGCGTCGGAG</td><td>239.4</td><td>-11.8</td><td>1.5</td><td>-8.1</td><td>6.3E-16</td><td>4.5E-15</td><td>MB231/MB468</td><td>MP</td></tr>
<tr><td>>hg38_21_+_8433172_8446622_RNA45SN1_45S_50ntFlanks@4007.4026.20|CCTCGGGCCGATCGCACGCC</td><td>>hg38_21_+_8433172_8446622_RNA45SN1_45S_50ntFlanks</td><td>4007</td><td>4026</td><td>20</td><td>CCTCGGGCCGATCGCACGCC</td><td>305.3</td><td>-12.1</td><td>1.5</td><td>-8.1</td><td>6.3E-16</td><td>4.5E-15</td><td>MB231/MB468</td><td>MP</td></tr>
<tr><td>>MI0000270|hsa-mir-181b-1&WithFlank&1|-|198858867|198858988@42.65.24;MI0000683|hsa-mir-181b-2&WithFlank&9|+|124693704|124693804@22.45.24&[MIMAT0000257&hsa-miR-181b-5p&offsets|0|+1;m-73&1|-|198858925|198858947&offsets|0|+1];[MIMAT0000257_1&hsa-miR-181b-5p&offsets|0|+1;m-72&9|+|124693725|124693747&offsets|0|+1]|AACATTCATTGCTGTCGGTGGGTT</td><td>>MI0000270|hsa-mir-181b-1&WithFlank&1|-|198858867|198858988</td><td>42</td><td>65</td><td>24</td><td>AACATTCATTGCTGTCGGTGGGTT</td><td>1910.8</td><td>6.2</td><td>0.8</td><td>8.1</td><td>7.8E-16</td><td>4.6E-15</td><td>MB231/MB468</td><td>NUC</td></tr>
<tr><td>>hg38_21_+_8433172_8446622_RNA45SN1_45S_50ntFlanks@6733.6771.39|CATTGATCATCGACACTTCGAACGCACTTGCGGCCCCGG</td><td>>hg38_21_+_8433172_8446622_RNA45SN1_45S_50ntFlanks</td><td>6733</td><td>6771</td><td>39</td><td>CATTGATCATCGACACTTCGAACGCACTTGCGGCCCCGG</td><td>1163.6</td><td>-14.0</td><td>1.7</td><td>-8.1</td><td>6.4E-16</td><td>4.6E-15</td><td>MB231/MB468</td><td>MP</td></tr>
<tr><td>>hg38_21_+_8433172_8446622_RNA45SN1_45S_50ntFlanks@7982.8004.23|TCAGATCAGACGTGGCGACCCGC</td><td>>hg38_21_+_8433172_8446622_RNA45SN1_45S_50ntFlanks</td><td>7982</td><td>8004</td><td>23</td><td>TCAGATCAGACGTGGCGACCCGC</td><td>302.8</td><td>-12.1</td><td>1.5</td><td>-8.1</td><td>6.4E-16</td><td>4.6E-15</td><td>MB231/MB468</td><td>MP</td></tr>
<tr><td>>hg38_21_+_8433172_8446622_RNA45SN1_45S_50ntFlanks@11570.11602.33|AGCCGACTTAGAACTGGTGCGGACCAGGGGAAT</td><td>>hg38_21_+_8433172_8446622_RNA45SN1_45S_50ntFlanks</td><td>11570</td><td>11602</td><td>33</td><td>AGCCGACTTAGAACTGGTGCGGACCAGGGGAAT</td><td>1048.2</td><td>-5.6</td><td>0.7</td><td>-8.1</td><td>6.4E-16</td><td>4.6E-15</td><td>MB231/MB468</td><td>MP</td></tr>
<tr><td>>hg38_21_+_8433172_8446622_RNA45SN1_45S_50ntFlanks@7983.8003.21|CAGATCAGACGTGGCGACCCG</td><td>>hg38_21_+_8433172_8446622_RNA45SN1_45S_50ntFlanks</td><td>7983</td><td>8003</td><td>21</td><td>CAGATCAGACGTGGCGACCCG</td><td>1221.9</td><td>-10.0</td><td>1.2</td><td>-8.1</td><td>6.4E-16</td><td>4.6E-15</td><td>MB231/MB468</td><td>MP</td></tr>
<tr><td>>trna12_TrpCCA_17_+_19411494_19411565@30.52.23,trna168_TrpCCA_6_-_26331672_26331743@30.52.23,trna170_TrpCCA_6_-_26319330_26319401@30.52.23,trna1_TrpCCA_7_+_99067307_99067378@30.52.23,trna39_TrpCCA_17_-_8124187_8124258@30.52.23,trna6_TrpCCA_17_+_8089676_8089747@30.52.23|ACTCCAGATCAGAAGGTTGCGTG</td><td>>trna12_TrpCCA_17_+_19411494_19411565</td><td>30</td><td>52</td><td>23</td><td>ACTCCAGATCAGAAGGTTGCGTG</td><td>518.3</td><td>-12.0</td><td>1.5</td><td>-8.1</td><td>7.9E-16</td><td>4.6E-15</td><td>MB231/MB468</td><td>NUC</td></tr>
<tr><td>>hg38_1_-_228634819_228635039_RNA5S12_5S_50ntFlanks@63.89.27|ACCACCCTGAACGCGCCCGATCTCGTC</td><td>>hg38_1_-_228634819_228635039_RNA5S12_5S_50ntFlanks</td><td>63</td><td>89</td><td>27</td><td>ACCACCCTGAACGCGCCCGATCTCGTC</td><td>508.3</td><td>9.7</td><td>1.2</td><td>8.1</td><td>6.9E-16</td><td>4.6E-15</td><td>MB231/MB468</td><td>CYTO</td></tr>
<tr><td>>hg38_21_+_8433172_8446622_RNA45SN1_45S_50ntFlanks@11319.11344.26|CCCACGTCTCGTCGCGCGCGCGTCCG</td><td>>hg38_21_+_8433172_8446622_RNA45SN1_45S_50ntFlanks</td><td>11319</td><td>11344</td><td>26</td><td>CCCACGTCTCGTCGCGCGCGCGTCCG</td><td>3003.4</td><td>-7.0</td><td>0.9</td><td>-8.1</td><td>6.9E-16</td><td>4.6E-15</td><td>MB231/MB468</td><td>CYTO</td></tr>
<tr><td>>hg38_21_+_8433172_8446622_RNA45SN1_45S_50ntFlanks@10587.10607.21|CCCCGGGGAGAGTTCTCTTTT</td><td>>hg38_21_+_8433172_8446622_RNA45SN1_45S_50ntFlanks</td><td>10587</td><td>10607</td><td>21</td><td>CCCCGGGGAGAGTTCTCTTTT</td><td>310.5</td><td>-12.1</td><td>1.5</td><td>-8.0</td><td>1.1E-15</td><td>4.6E-15</td><td>MB231/MB468</td><td>MITO</td></tr>
<tr><td>>hg38_21_+_8433172_8446622_RNA45SN1_45S_50ntFlanks@8380.8399.20|CAAGAGGGCGTGAAACCGTT</td><td>>hg38_21_+_8433172_8446622_RNA45SN1_45S_50ntFlanks</td><td>8380</td><td>8399</td><td>20</td><td>CAAGAGGGCGTGAAACCGTT</td><td>886.5</td><td>-9.7</td><td>1.2</td><td>-8.0</td><td>1.1E-15</td><td>4.6E-15</td><td>MB231/MB468</td><td>MITO</td></tr>
<tr><td>>trna2_GlyGCC_21_-_18827107_18827177@1.29.29,trna35_GlyGCC_1_+_161413094_161413164@1.29.29,trna37_GlyGCC_1_+_161420467_161420537@1.29.29,trna39_GlyGCC_1_+_161427898_161427968@1.29.29,trna41_GlyGCC_1_+_161435258_161435328@1.29.29|GCATGGGTGGTTCAGTGGTAGAATTCTCG</td><td>>trna2_GlyGCC_21_-_18827107_18827177</td><td>1</td><td>29</td><td>29</td><td>GCATGGGTGGTTCAGTGGTAGAATTCTCG</td><td>32987.7</td><td>-8.3</td><td>1.0</td><td>-8.0</td><td>1.5E-15</td><td>4.6E-15</td><td>BT20/MB468</td><td>NUC</td></tr>
<tr><td>>MI0000079|hsa-mir-23a&WithFlank&19|-|13836581|13836665@54.71.18&[MIMAT0000078&hsa-miR-23a-3p&offsets|+3|0;m-31&19|-|13836595|13836615&offsets|+3|0]|ACATTGCCAGGGATTTCC</td><td>>MI0000079|hsa-mir-23a&WithFlank&19|-|13836581|13836665</td><td>54</td><td>71</td><td>18</td><td>ACATTGCCAGGGATTTCC</td><td>4102.0</td><td>29.3</td><td>3.6</td><td>8.1</td><td>6.4E-16</td><td>4.6E-15</td><td>MB231/MB468</td><td>MP</td></tr>
<tr><td>>hg38_21_+_8433172_8446622_RNA45SN1_45S_50ntFlanks@11592.11613.22|ACCAGGGGAATCCGACTGTTTA</td><td>>hg38_21_+_8433172_8446622_RNA45SN1_45S_50ntFlanks</td><td>11592</td><td>11613</td><td>22</td><td>ACCAGGGGAATCCGACTGTTTA</td><td>263.6</td><td>-11.9</td><td>1.5</td><td>-8.1</td><td>6.4E-16</td><td>4.6E-15</td><td>MB231/MB468</td><td>MP</td></tr>
<tr><td>>hg38_21_+_8433172_8446622_RNA45SN1_45S_50ntFlanks@10074.10092.19|ACGGGAGCGGCGGGGGCGG</td><td>>hg38_21_+_8433172_8446622_RNA45SN1_45S_50ntFlanks</td><td>10074</td><td>10092</td><td>19</td><td>ACGGGAGCGGCGGGGGCGG</td><td>376.2</td><td>-12.4</td><td>1.5</td><td>-8.1</td><td>6.4E-16</td><td>4.6E-15</td><td>MB231/MB468</td><td>MP</td></tr>
<tr><td>>hg38_21_+_8433172_8446622_RNA45SN1_45S_50ntFlanks@6746.6772.27|CACTTCGAACGCACTTGCGGCCCCGGG</td><td>>hg38_21_+_8433172_8446622_RNA45SN1_45S_50ntFlanks</td><td>6746</td><td>6772</td><td>27</td><td>CACTTCGAACGCACTTGCGGCCCCGGG</td><td>353.7</td><td>-12.3</td><td>1.5</td><td>-8.1</td><td>6.4E-16</td><td>4.6E-15</td><td>MB231/MB468</td><td>MP</td></tr>
<tr><td>>hg38_21_+_8433172_8446622_RNA45SN1_45S_50ntFlanks@10391.10419.29|AACAGCAGTTGAACATGGGTCAGTCGGTC</td><td>>hg38_21_+_8433172_8446622_RNA45SN1_45S_50ntFlanks</td><td>10391</td><td>10419</td><td>29</td><td>AACAGCAGTTGAACATGGGTCAGTCGGTC</td><td>303.3</td><td>-12.0</td><td>1.5</td><td>-8.0</td><td>1.1E-15</td><td>4.6E-15</td><td>MB231/MB468</td><td>MITO</td></tr>
<tr><td>>hg38_21_+_8433172_8446622_RNA45SN1_45S_50ntFlanks@6734.6777.44|ATTGATCATCGACACTTCGAACGCACTTGCGGCCCCGGGTTCCT</td><td>>hg38_21_+_8433172_8446622_RNA45SN1_45S_50ntFlanks</td><td>6734</td><td>6777</td><td>44</td><td>ATTGATCATCGACACTTCGAACGCACTTGCGGCCCCGGGTTCCT</td><td>166.1</td><td>11.4</td><td>1.4</td><td>8.1</td><td>7.0E-16</td><td>4.6E-15</td><td>MB231/MB468</td><td>CYTO</td></tr>
<tr><td>>hg38_21_+_8433172_8446622_RNA45SN1_45S_50ntFlanks@4201.4219.19|CCGACCCGGGGAGGTAGTG</td><td>>hg38_21_+_8433172_8446622_RNA45SN1_45S_50ntFlanks</td><td>4201</td><td>4219</td><td>19</td><td>CCGACCCGGGGAGGTAGTG</td><td>377.3</td><td>-12.3</td><td>1.5</td><td>-8.0</td><td>1.1E-15</td><td>4.6E-15</td><td>MB231/MB468</td><td>MITO</td></tr>
<tr><td>>am_trna106_PheGAA_6_-_28775610_28775682@31.52.22,trna109_PheGAA_6_-_28758499_28758571@31.52.22,trna11_PheGAA_12_-_125412389_125412461@31.52.22,trna13_PheGAA_11_-_59333853_59333925@31.52.22,trna14_PheGAA_19_-_1383361_1383433@31.52.22,trna15_PheGAA_11_-_59324970_59325042@31.52.22,trna1_PheGAA_13_-_95201904_95201976@31.52.22,trna72_PheGAA_6_+_28732164_28732236@31.52.22,trna96_PheGAA_6_-_28949449_28949521@31.52.22|ACTGAAGATCTAAAGGTCCCTG</td><td>>am_trna106_PheGAA_6_-_28775610_28775682</td><td>31</td><td>52</td><td>22</td><td>ACTGAAGATCTAAAGGTCCCTG</td><td>174.6</td><td>-11.3</td><td>1.4</td><td>-8.1</td><td>6.5E-16</td><td>4.6E-15</td><td>MB231/MB468</td><td>MP</td></tr>
<tr><td>>hg38_21_+_8433172_8446622_RNA45SN1_45S_50ntFlanks@7979.8005.27|ACCTCAGATCAGACGTGGCGACCCGCT</td><td>>hg38_21_+_8433172_8446622_RNA45SN1_45S_50ntFlanks</td><td>7979</td><td>8005</td><td>27</td><td>ACCTCAGATCAGACGTGGCGACCCGCT</td><td>670.0</td><td>-4.3</td><td>0.5</td><td>-8.0</td><td>1.5E-15</td><td>4.6E-15</td><td>BT20/MB468</td><td>NUC</td></tr>
<tr><td>>hg38_21_+_8433172_8446622_RNA45SN1_45S_50ntFlanks@5420.5443.24|CCGATTGGATGGTTTAGTGAGGCC</td><td>>hg38_21_+_8433172_8446622_RNA45SN1_45S_50ntFlanks</td><td>5420</td><td>5443</td><td>24</td><td>CCGATTGGATGGTTTAGTGAGGCC</td><td>3383.6</td><td>-11.5</td><td>1.4</td><td>-8.0</td><td>1.5E-15</td><td>4.6E-15</td><td>BT20/MB468</td><td>NUC</td></tr>
<tr><td>>MI0000299|hsa-mir-222&WithFlank&X|-|45747009|45747130@75.96.22&[MIMAT0000279&hsa-miR-222-3p&offsets|0|+1;m-140&X|-|45747033|45747056&offsets|0|-2]|AGCTACATCTGGCTACTGGGTC</td><td>>MI0000299|hsa-mir-222&WithFlank&X|-|45747009|45747130</td><td>75</td><td>96</td><td>22</td><td>AGCTACATCTGGCTACTGGGTC</td><td>1929.7</td><td>5.5</td><td>0.7</td><td>8.1</td><td>8.0E-16</td><td>4.6E-15</td><td>MB231/MB468</td><td>NUC</td></tr>
<tr><td>>hg38_21_+_8433172_8446622_RNA45SN1_45S_50ntFlanks@5487.5510.24|CGCTGAGAAGACGGTCGAACTTGA</td><td>>hg38_21_+_8433172_8446622_RNA45SN1_45S_50ntFlanks</td><td>5487</td><td>5510</td><td>24</td><td>CGCTGAGAAGACGGTCGAACTTGA</td><td>284.9</td><td>-11.9</td><td>1.5</td><td>-8.0</td><td>1.1E-15</td><td>4.6E-15</td><td>MB231/MB468</td><td>MITO</td></tr>
<tr><td>>Antisense_hg38_21_-_8433172_8446622_RNA45SN1_45S_50ntFlanks@7674.7692.19|CTCTCTCTCTCTCTCCCCC</td><td>>Antisense_hg38_21_-_8433172_8446622_RNA45SN1_45S_50ntFlanks</td><td>7674</td><td>7692</td><td>19</td><td>CTCTCTCTCTCTCTCCCCC</td><td>306.7</td><td>-12.0</td><td>1.5</td><td>-8.0</td><td>1.1E-15</td><td>4.6E-15</td><td>MB231/MB468</td><td>MITO</td></tr>
<tr><td>>hg38_21_+_8433172_8446622_RNA45SN1_45S_50ntFlanks@11027.11045.19|CTCTCTCTCTCTCTCCCCC</td><td>>hg38_21_+_8433172_8446622_RNA45SN1_45S_50ntFlanks</td><td>11027</td><td>11045</td><td>19</td><td>CTCTCTCTCTCTCTCCCCC</td><td>306.7</td><td>-12.0</td><td>1.5</td><td>-8.0</td><td>1.1E-15</td><td>4.6E-15</td><td>MB231/MB468</td><td>MITO</td></tr>
<tr><td>>hg38_21_+_8433172_8446622_RNA45SN1_45S_50ntFlanks@9736.9759.24|CCGAAACGATCTCAACCTATTCTC</td><td>>hg38_21_+_8433172_8446622_RNA45SN1_45S_50ntFlanks</td><td>9736</td><td>9759</td><td>24</td><td>CCGAAACGATCTCAACCTATTCTC</td><td>1112.7</td><td>-13.9</td><td>1.7</td><td>-8.0</td><td>1.1E-15</td><td>4.7E-15</td><td>MB231/MB468</td><td>MITO</td></tr>
<tr><td>>MI0000737|hsa-mir-200a&WithFlank&1|+|1167857|1167958@23.43.21&[MIMAT0001620&hsa-miR-200a-5p&offsets|+1|0;m-154&1|+|1167878|1167899&offsets|+1|0]|ATCTTACCGGACAGTGCTGGA</td><td>>MI0000737|hsa-mir-200a&WithFlank&1|+|1167857|1167958</td><td>23</td><td>43</td><td>21</td><td>ATCTTACCGGACAGTGCTGGA</td><td>246.5</td><td>-11.8</td><td>1.5</td><td>-8.1</td><td>6.5E-16</td><td>4.7E-15</td><td>MB231/MB468</td><td>MP</td></tr>
<tr><td>>hg38_21_+_8433172_8446622_RNA45SN1_45S_50ntFlanks@9569.9592.24|GTGCAAATCGGTCGTCCGACCTGG</td><td>>hg38_21_+_8433172_8446622_RNA45SN1_45S_50ntFlanks</td><td>9569</td><td>9592</td><td>24</td><td>GTGCAAATCGGTCGTCCGACCTGG</td><td>211.7</td><td>-11.6</td><td>1.4</td><td>-8.1</td><td>6.5E-16</td><td>4.7E-15</td><td>MB231/MB468</td><td>MP</td></tr>
<tr><td>>MI0000283|hsa-mir-203a&WithFlank&14|+|104117399|104117520@72.91.20(+1A)&[MIMAT0000264&hsa-miR-203a-3p&offsets|+1|-1(+1A);m-79&14|+|104117469|104117490&offsets|+1|-1(+1A)]|TGAAATGTTTAGGACCACTAA</td><td>>MI0000283|hsa-mir-203a&WithFlank&14|+|104117399|104117520</td><td>72</td><td>91</td><td>20(+1A)</td><td>TGAAATGTTTAGGACCACTAA</td><td>392.4</td><td>12.0</td><td>1.4</td><td>8.4</td><td>6.0E-17</td><td>4.7E-15</td><td>BT20/MB231</td><td>CYTO</td></tr>
<tr><td>>hg38_21_+_8433172_8446622_RNA45SN1_45S_50ntFlanks@11874.11899.26|AGAAGACCCTGTTGAGCTTGACTCTA</td><td>>hg38_21_+_8433172_8446622_RNA45SN1_45S_50ntFlanks</td><td>11874</td><td>11899</td><td>26</td><td>AGAAGACCCTGTTGAGCTTGACTCTA</td><td>548.3</td><td>-4.6</td><td>0.6</td><td>-8.1</td><td>7.1E-16</td><td>4.7E-15</td><td>MB231/MB468</td><td>CYTO</td></tr>
<tr><td>>hg38_21_+_8433172_8446622_RNA45SN1_45S_50ntFlanks@12412.12443.32|ATCCTTCGATGTCGGCTCTTCCTATCATTGTG</td><td>>hg38_21_+_8433172_8446622_RNA45SN1_45S_50ntFlanks</td><td>12412</td><td>12443</td><td>32</td><td>ATCCTTCGATGTCGGCTCTTCCTATCATTGTG</td><td>479.6</td><td>-11.0</td><td>1.4</td><td>-8.1</td><td>7.1E-16</td><td>4.7E-15</td><td>BT20/MB468</td><td>MITO</td></tr>
<tr><td>>hg38_21_+_8433172_8446622_RNA45SN1_45S_50ntFlanks@12894.12933.40|CTCGCCCGTCACGCACCGCACGTTCGTGGGGAACCTGGCG</td><td>>hg38_21_+_8433172_8446622_RNA45SN1_45S_50ntFlanks</td><td>12894</td><td>12933</td><td>40</td><td>CTCGCCCGTCACGCACCGCACGTTCGTGGGGAACCTGGCG</td><td>325.7</td><td>-12.1</td><td>1.5</td><td>-8.0</td><td>1.1E-15</td><td>4.7E-15</td><td>MB231/MB468</td><td>MITO</td></tr>
<tr><td>>hg38_21_+_8433172_8446622_RNA45SN1_45S_50ntFlanks@4043.4063.21|ACCCATTCGAACGTCTGCCCT</td><td>>hg38_21_+_8433172_8446622_RNA45SN1_45S_50ntFlanks</td><td>4043</td><td>4063</td><td>21</td><td>ACCCATTCGAACGTCTGCCCT</td><td>591.2</td><td>-13.0</td><td>1.6</td><td>-8.0</td><td>1.1E-15</td><td>4.7E-15</td><td>MB231/MB468</td><td>MITO</td></tr>
<tr><td>>hg38_21_+_8433172_8446622_RNA45SN1_45S_50ntFlanks@9911.9928.18|CCAGAAAAGGTGTTGGTT</td><td>>hg38_21_+_8433172_8446622_RNA45SN1_45S_50ntFlanks</td><td>9911</td><td>9928</td><td>18</td><td>CCAGAAAAGGTGTTGGTT</td><td>400.7</td><td>-12.5</td><td>1.5</td><td>-8.1</td><td>6.6E-16</td><td>4.7E-15</td><td>MB231/MB468</td><td>MP</td></tr>
<tr><td>>hg38_21_+_8433172_8446622_RNA45SN1_45S_50ntFlanks@4045.4070.26|CCATTCGAACGTCTGCCCTATCAACT</td><td>>hg38_21_+_8433172_8446622_RNA45SN1_45S_50ntFlanks</td><td>4045</td><td>4070</td><td>26</td><td>CCATTCGAACGTCTGCCCTATCAACT</td><td>141.7</td><td>7.4</td><td>0.9</td><td>8.1</td><td>7.1E-16</td><td>4.7E-15</td><td>MB231/MB468</td><td>CYTO</td></tr>
</table>

| | | | | | | | | | | | | | |
|---|---|---|---|---|---|---|---|---|---|---|---|---|---|
| >MI0000434\|hsa-let-7i&WithFlank&12\|+\|62603680\|62603775@12.32.21(+1G)&[MIMAT0000415&hsa-let-7i-5p&offsets\|0\|-1(+1G);m-49&12\|+\|62603691\|62603712&offsets\|0\|-1(+1G)]\|TGAGGTAGTAGTTTGTGCTGTG | >MI0000434\|hsa-let-7i&WithFlank&12\|+\|62603680\|62603775 | 12 | 32 | 21(+1G) | TGAGGTAGTAGTTTGTGCTGTG | 651.4 | 6.3 | 0.8 | 8.1 | 7.1E-16 | 4.7E-15 | BT20/MB468 | MITO |
| >MI0000266\|hsa-mir-10a&WithFlank&17\|-\|48579832\|48579953@29.51.23&[MIMAT0000253&hsa-miR-10a-5p&offsets\|+1\|+1;m-18&17\|-\|48579905\|48579926&offsets\|+1\|+2]\|ACCCTGTAGATCCGAATTTGTGT | >MI0000266\|hsa-mir-10a&WithFlank&17\|-\|48579832\|48579953 | 29 | 51 | 23 | ACCCTGTAGATCCGAATTTGTGT | 278.9 | 11.3 | 1.4 | 8.1 | 6.7E-16 | 4.8E-15 | MB231/MB468 | MP |
| >hg38_21_+_8433172_8446622_RNA45SN1_45S_50ntFlanks@12488.12521.34\|AACGTGAGCTGGGTTTAGACCGTCGTGAGACAGG | >hg38_21_+_8433172_8446622_RNA45SN1_45S_50ntFlanks | 12488 | 12521 | 34 | AACGTGAGCTGGGTTTAGACCGTCGTGAGACAGG | 438.3 | -7.8 | 1.0 | -8.1 | 7.3E-16 | 4.8E-15 | MB231/MB468 | CYTO |
| >hg38_21_+_8433172_8446622_RNA45SN1_45S_50ntFlanks@4590.4613.24\|ATTTTGTTGGTTTTCGGAACTGAG | >hg38_21_+_8433172_8446622_RNA45SN1_45S_50ntFlanks | 4590 | 4613 | 24 | ATTTTGTTGGTTTTCGGAACTGAG | 238.6 | -11.8 | 1.5 | -8.1 | 6.8E-16 | 4.8E-15 | MB231/MB468 | MP |
| >hg38_21_+_8433172_8446622_RNA45SN1_45S_50ntFlanks@9624.9648.25\|AGTAGCTGGTTCCCTCCGAAGTTTC | >hg38_21_+_8433172_8446622_RNA45SN1_45S_50ntFlanks | 9624 | 9648 | 25 | AGTAGCTGGTTCCCTCCGAAGTTTC | 360.5 | -12.3 | 1.5 | -8.0 | 1.1E-15 | 4.8E-15 | MB231/MB468 | MITO |
| >hg38_21_+_8433172_8446622_RNA45SN1_45S_50ntFlanks@8836.8864.29\|CGGCGGTGGCGGCGGCGGCGGCGGCGGCG | >hg38_21_+_8433172_8446622_RNA45SN1_45S_50ntFlanks | 8836 | 8864 | 29 | CGGCGGTGGCGGCGGCGGCGGCGGCGGCG | 340.3 | -12.3 | 1.5 | -8.1 | 6.8E-16 | 4.8E-15 | MB231/MB468 | MP |
| >hg38_21_+_8433172_8446622_RNA45SN1_45S_50ntFlanks@11316.11335.20\|CACCCCACGTCTCGTCGCGC | >hg38_21_+_8433172_8446622_RNA45SN1_45S_50ntFlanks | 11316 | 11335 | 20 | CACCCCACGTCTCGTCGCGC | 333.0 | -12.2 | 1.5 | -8.0 | 1.1E-15 | 4.8E-15 | MB231/MB468 | MITO |
| >MI0000102\|hsa-mir-100&WithFlank&11\|-\|122152223\|122152314@19.39.21&[MIMAT0000098&hsa-miR-100-5p&offsets\|0\|-1;m-27&11\|-\|122152276\|122152296&offsets\|0\|0]\|AACCCGTAGATCCGAACTTGT | >MI0000102\|hsa-mir-100&WithFlank&11\|-\|122152223\|122152314 | 19 | 39 | 21 | AACCCGTAGATCCGAACTTGT | 159905.4 | 11.2 | 1.4 | 8.1 | 6.8E-16 | 4.8E-15 | MB231/MB468 | MP |
| >hg38_21_+_8433172_8446622_RNA45SN1_45S_50ntFlanks@12407.12435.29\|TTTTGATCCTTCGATGTCGGCTCTTCCTA | >hg38_21_+_8433172_8446622_RNA45SN1_45S_50ntFlanks | 12407 | 12435 | 29 | TTTTGATCCTTCGATGTCGGCTCTTCCTA | 203.4 | -11.5 | 1.4 | -8.1 | 6.8E-16 | 4.8E-15 | MB231/MB468 | MP |
| >hg38_21_+_8433172_8446622_RNA45SN1_45S_50ntFlanks@6721.6747.27\|AATTGCAGGACACATTGATCATCGACA | >hg38_21_+_8433172_8446622_RNA45SN1_45S_50ntFlanks | 6721 | 6747 | 27 | AATTGCAGGACACATTGATCATCGACA | 1741.1 | -8.3 | 1.0 | -8.1 | 6.8E-16 | 4.8E-15 | MB231/MB468 | MP |
| >hg38_21_+_8433172_8446622_RNA45SN1_45S_50ntFlanks@8182.8223.42\|ATCGAGGCCCAGCCCGTGGACGGTGTGAGGCCGGTAGCGGCC | >hg38_21_+_8433172_8446622_RNA45SN1_45S_50ntFlanks | 8182 | 8223 | 42 | ATCGAGGCCCAGCCCGTGGACGGTGTGAGGCCGGTAGCGGCC | 321.4 | -12.1 | 1.5 | -8.0 | 1.1E-15 | 4.8E-15 | MB231/MB468 | MITO |
| >hg38_21_+_8433172_8446622_RNA45SN1_45S_50ntFlanks@6747.6771.25\|ACTTCGAACGCACTTGCGGCCCCGG | >hg38_21_+_8433172_8446622_RNA45SN1_45S_50ntFlanks | 6747 | 6771 | 25 | ACTTCGAACGCACTTGCGGCCCCGG | 254.0 | -10.3 | 1.3 | -8.0 | 1.0E-15 | 4.8E-15 | BT20/MB468 | CYTO |
| >hg38_21_+_8433172_8446622_RNA45SN1_45S_50ntFlanks@11118.11145.28\|CAGGGGCCGGCGGCGGCCCGCCGCGGGG | >hg38_21_+_8433172_8446622_RNA45SN1_45S_50ntFlanks | 11118 | 11145 | 28 | CAGGGGCCGGCGGCGGCCCGCCGCGGGG | 331.8 | -12.2 | 1.5 | -8.0 | 1.1E-15 | 4.8E-15 | MB231/MB468 | MITO |
| >MI0000111\|hsa-mir-105-1&WithFlank&X\|-\|152392213\|152392305@19.40.22;MI0000112\|hsa-mir-105-2&WithFlank&X\|-\|152394406\|152394498@19.40.22&[MIMAT0000102&hsa-miR-105-5p&offsets\|0\|-1;m-735&X\|-\|152392266\|152392287&offsets\|0\|0];[MIMAT0000102_1&hsa-miR-105-5p&offsets\|0\|-1;m-735&X\|-\|152394459\|152394480&offsets\|0\|0]\|TCAAATGCTCAGACTCCTGTGG | >MI0000111\|hsa-mir-105-1&WithFlank&X\|-\|152392213\|152392305 | 19 | 40 | 22 | TCAAATGCTCAGACTCCTGTGG | 268.3 | 11.8 | 1.4 | 8.3 | 1.2E-16 | 4.9E-15 | BT20/MB231 | NUC |
| >MI0000650\|hsa-mir-200c&WithFlank&12\|+\|6963693\|6963772@49.70.22&[MIMAT0000617&hsa-miR-200c-3p&offsets\|-1\|-2;m-54&12\|+\|6963742\|6963764&offsets\|-1\|-2]\|CTAATACTGCCGGGTAATGATG | >MI0000650\|hsa-mir-200c&WithFlank&12\|+\|6963693\|6963772 | 49 | 70 | 22 | CTAATACTGCCGGGTAATGATG | 778.6 | 10.8 | 1.3 | 8.0 | 1.0E-15 | 4.9E-15 | BT20/MB468 | CYTO |
| >hg38_21_+_8433172_8446622_RNA45SN1_45S_50ntFlanks@12008.12041.34\|CCGCCGGTGAAATACCACTACTCTGATCGTTTTT | >hg38_21_+_8433172_8446622_RNA45SN1_45S_50ntFlanks | 12008 | 12041 | 34 | CCGCCGGTGAAATACCACTACTCTGATCGTTTTT | 626.9 | -12.3 | 1.5 | -8.0 | 8.4E-16 | 4.9E-15 | MB231/MB468 | NUC |
| >hg38_21_+_8433172_8446622_RNA45SN1_45S_50ntFlanks@4043.4063.21\|ACCCATTCGAACGTCTGCCCT | >hg38_21_+_8433172_8446622_RNA45SN1_45S_50ntFlanks | 4043 | 4063 | 21 | ACCCATTCGAACGTCTGCCCT | 571.2 | -12.1 | 1.5 | -8.0 | 8.4E-16 | 4.9E-15 | MB231/MB468 | NUC |
| >hg38_21_+_8433172_8446622_RNA45SN1_45S_50ntFlanks@4731.4767.37\|AAGAACGAAAGTCGGAGGTTCGAAGACGATCAGATAC | >hg38_21_+_8433172_8446622_RNA45SN1_45S_50ntFlanks | 4731 | 4767 | 37 | AAGAACGAAAGTCGGAGGTTCGAAGACGATCAGATAC | 4056.0 | 10.5 | 1.3 | 8.1 | 7.4E-16 | 4.9E-15 | MB231/MB468 | CYTO |
| >hg38_21_+_8433172_8446622_RNA45SN1_45S_50ntFlanks@8357.8378.22\|AAAAGAACTTTGAAGAGAGAGT | >hg38_21_+_8433172_8446622_RNA45SN1_45S_50ntFlanks | 8357 | 8378 | 22 | AAAAGAACTTTGAAGAGAGAGT | 5191.9 | -5.7 | 0.7 | -8.1 | 7.4E-16 | 4.9E-15 | MB231/MB468 | CYTO |
| >trna10_AspGTC_12_-_125424193_125424264@31.72.42,trna12_AspGTC_12_-_125411891_125411962@31.72.42,trna38_AspGTC_17_-_8125556_8125627@31.72.42,trna45_AspGTC_6_+_27447453_27447524@31.72.42,trna48_AspGTC_6_+_27471523_27471594@31.72.42,trna4_AspGTC_12_+_96429799_96429870@31.72.42,trna69_AspGTC_1_-_161440205_161440276@31.72.42,trna72_AspGTC_1_-_161432824_161432895@31.72.42,trna75_AspGTC_1_-_161425414_161425485@31.72.42,trna78_AspGTC_1_-_161418033_161418104@31.72.42,trna81_AspGTC_1_-_161410615_161410686@31.72.42\|CCTGTCACGCGGGAGACCGGGGTTCGATTCCCCGACGGGGAG | >trna10_AspGTC_12_-_125424193_125424264 | 31 | 72 | 42 | CCTGTCACGCGGGAGACCGGGGTTCGATTCCCCGACGGGGAG | 750.5 | -12.5 | 1.6 | -8.0 | 8.5E-16 | 4.9E-15 | MB231/MB468 | NUC |
| >MI0006444\|hsa-mir-1307&WithFlank&10\|-\|103394247\|103394407@86.107.22(+1U)&[MIMAT0005951&hsa-miR-1307-3p&offsets\|0\|0(+1U);m-66&10\|-\|103394300\|103394322&offsets\|0\|-1(+1U)]\|ACTCGGCGTGGCGTCGGTCGTGT | >MI0006444\|hsa-mir-1307&WithFlank&10\|-\|103394247\|103394407 | 86 | 107 | 22(+1U) | ACTCGGCGTGGCGTCGGTCGTGT | 991.4 | 6.4 | 0.8 | 8.1 | 7.5E-16 | 4.9E-15 | MB231/MB468 | CYTO |
| >MI0000542\|hsa-mir-320a&WithFlank&8\|-\|22244960\|22245043@48.71.24(+1U)&[MIMAT0000510&hsa-miR-320a-3p&offsets\|0\|+2(+1U);m-60&8\|-\|22244975\|22244996&offsets\|0\|+2(+1U)]\|AAAAGCTGGGTTGAGAGGGCGAAAT | >MI0000542\|hsa-mir-320a&WithFlank&8\|-\|22244960\|22245043 | 48 | 71 | 24(+1U) | AAAAGCTGGGTTGAGAGGGCGAAAT | 2745.7 | 12.0 | 1.5 | 8.0 | 1.0E-15 | 4.9E-15 | BT20/MB468 | CYTO |
| >hg38_21_+_8433172_8446622_RNA45SN1_45S_50ntFlanks@6773.6807.35\|TTCCTCCCGGGGCTACGCCTGTCTGAGCGTCGCTT | >hg38_21_+_8433172_8446622_RNA45SN1_45S_50ntFlanks | 6773 | 6807 | 35 | TTCCTCCCGGGGCTACGCCTGTCTGAGCGTCGCTT | 2985.8 | -4.5 | 0.6 | -8.0 | 1.1E-15 | 4.9E-15 | MB231/MB468 | MITO |
| >hg38_21_+_8433172_8446622_RNA45SN1_45S_50ntFlanks@10807.10826.20\|ACAATGTAGGTAAGGGAAGT | >hg38_21_+_8433172_8446622_RNA45SN1_45S_50ntFlanks | 10807 | 10826 | 20 | ACAATGTAGGTAAGGGAAGT | 290.8 | -6.4 | 0.8 | -8.1 | 7.5E-16 | 4.9E-15 | MB231/MB468 | CYTO |
| >hg38_21_+_8433172_8446622_RNA45SN1_45S_50ntFlanks@3771.3792.22\|CACGGCCGGTACAGTGAAACTG | >hg38_21_+_8433172_8446622_RNA45SN1_45S_50ntFlanks | 3771 | 3792 | 22 | CACGGCCGGTACAGTGAAACTG | 488.1 | -12.8 | 1.6 | -8.1 | 6.9E-16 | 4.9E-15 | MB231/MB468 | MP |
| >hg38_21_+_8433172_8446622_RNA45SN1_45S_50ntFlanks@4069.4098.30\|CTTTCGATGGTAGTCGCCGTGCCTACCATG | >hg38_21_+_8433172_8446622_RNA45SN1_45S_50ntFlanks | 4069 | 4098 | 30 | CTTTCGATGGTAGTCGCCGTGCCTACCATG | 180.9 | -11.4 | 1.4 | -8.1 | 6.9E-16 | 4.9E-15 | MB231/MB468 | MP |
| >hg38_21_+_8433172_8446622_RNA45SN1_45S_50ntFlanks@10404.10435.32\|CATGGGTCAGTCGGTCCTGAGAGATGGGCGAG | >hg38_21_+_8433172_8446622_RNA45SN1_45S_50ntFlanks | 10404 | 10435 | 32 | CATGGGTCAGTCGGTCCTGAGAGATGGGCGAG | 251.9 | -11.8 | 1.5 | -8.1 | 7.0E-16 | 4.9E-15 | MB231/MB468 | MP |
| >hg38_21_+_8433172_8446622_RNA45SN1_45S_50ntFlanks@10787.10822.36\|ACAGCCTCTGGCATGTTGGAACAATGTAGGTAAGGG | >hg38_21_+_8433172_8446622_RNA45SN1_45S_50ntFlanks | 10787 | 10822 | 36 | ACAGCCTCTGGCATGTTGGAACAATGTAGGTAAGGG | 258.7 | -11.9 | 1.5 | -8.1 | 7.0E-16 | 5.0E-15 | MB231/MB468 | MP |
| >MI0000542\|hsa-mir-320a&WithFlank&8\|-\|22244960\|22245043@48.68.21(+1G)&[MIMAT0000510&hsa-miR-320a-3p&offsets\|0\|-1(+1G);m-60&8\|-\|22244975\|22244996&offsets\|0\|-1(+1G)]\|AAAAGCTGGGTTGAGAGGGCGG | >MI0000542\|hsa-mir-320a&WithFlank&8\|-\|22244960\|22245043 | 48 | 68 | 21(+1G) | AAAAGCTGGGTTGAGAGGGCGG | 3374.6 | 10.9 | 1.4 | 8.0 | 1.0E-15 | 5.0E-15 | BT20/MB468 | CYTO |
| >hg38_21_+_8433172_8446622_RNA45SN1_45S_50ntFlanks@10678.10711.34\|CGGTTCCGGCGGCGTCCGGTGAGCTCTCGCTGGC | >hg38_21_+_8433172_8446622_RNA45SN1_45S_50ntFlanks | 10678 | 10711 | 34 | CGGTTCCGGCGGCGTCCGGTGAGCTCTCGCTGGC | 193903.7 | -10.3 | 1.3 | -8.0 | 1.6E-15 | 5.0E-15 | BT20/MB468 | NUC |
| >hg38_21_+_8433172_8446622_RNA45SN1_45S_50ntFlanks@8358.8379.22\|AAAGAACTTTGAAGAGAGAGTT | >hg38_21_+_8433172_8446622_RNA45SN1_45S_50ntFlanks | 8358 | 8379 | 22 | AAAGAACTTTGAAGAGAGAGTT | 1914.1 | -7.4 | 0.9 | -8.0 | 1.6E-15 | 5.0E-15 | BT20/MB468 | NUC |
| >hg38_21_+_8433172_8446622_RNA45SN1_45S_50ntFlanks@5446.5465.20\|CGGATCGGCCCCGCCGGGGT | >hg38_21_+_8433172_8446622_RNA45SN1_45S_50ntFlanks | 5446 | 5465 | 20 | CGGATCGGCCCCGCCGGGGT | 305.5 | -12.0 | 1.5 | -8.0 | 1.2E-15 | 5.0E-15 | MB231/MB468 | MITO |
| >hg38_21_+_8433172_8446622_RNA45SN1_45S_50ntFlanks@3899.3938.40\|CCTTCGCGGGGGGGATGCGTGCATTTATCAGATCAAAACC | >hg38_21_+_8433172_8446622_RNA45SN1_45S_50ntFlanks | 3899 | 3938 | 40 | CCTTCGCGGGGGGGATGCGTGCATTTATCAGATCAAAACC | 1345.0 | -14.2 | 1.8 | -8.0 | 1.2E-15 | 5.0E-15 | MB231/MB468 | MITO |
| >hg38_21_+_8433172_8446622_RNA45SN1_45S_50ntFlanks@12405.12430.26\|CTTTTTGATCCTTCGATGTCGGCTCT | >hg38_21_+_8433172_8446622_RNA45SN1_45S_50ntFlanks | 12405 | 12430 | 26 | CTTTTTGATCCTTCGATGTCGGCTCT | 1824.9 | -4.9 | 0.6 | -8.0 | 1.1E-15 | 5.0E-15 | BT20/MB468 | CYTO |
| >hg38_21_+_8433172_8446622_RNA45SN1_45S_50ntFlanks@10873.10892.20\|CTGGGTCGGTCGGGCTGGGG | >hg38_21_+_8433172_8446622_RNA45SN1_45S_50ntFlanks | 10873 | 10892 | 20 | CTGGGTCGGTCGGGCTGGGG | 935.3 | -7.1 | 0.9 | -8.1 | 7.6E-16 | 5.0E-15 | MB231/MB468 | CYTO |
| >hg38_21_+_8433172_8446622_RNA45SN1_45S_50ntFlanks@9544.9566.23\|GGTGGAGGTCCGTAGCGGTCCTG | >hg38_21_+_8433172_8446622_RNA45SN1_45S_50ntFlanks | 9544 | 9566 | 23 | GGTGGAGGTCCGTAGCGGTCCTG | 216.2 | -10.7 | 1.3 | -8.0 | 8.6E-16 | 5.0E-15 | MB231/MB468 | NUC |
| >hg38_21_+_8433172_8446622_RNA45SN1_45S_50ntFlanks@6663.6686.24\|GTGGATCACTCGGCTCGTGCGTCG | >hg38_21_+_8433172_8446622_RNA45SN1_45S_50ntFlanks | 6663 | 6686 | 24 | GTGGATCACTCGGCTCGTGCGTCG | 1942.9 | -7.7 | 1.0 | -8.0 | 1.2E-15 | 5.0E-15 | MB231/MB468 | MITO |
| >hg38_21_+_8433172_8446622_RNA45SN1_45S_50ntFlanks@8357.8387.31\|AAAAGAACTTTGAAGAGAGAGTTCAAGAGGG | >hg38_21_+_8433172_8446622_RNA45SN1_45S_50ntFlanks | 8357 | 8387 | 31 | AAAAGAACTTTGAAGAGAGAGTTCAAGAGGG | 336.0 | -11.4 | 1.4 | -8.0 | 8.7E-16 | 5.0E-15 | MB231/MB468 | NUC |
| >hg38_21_+_8433172_8446622_RNA45SN1_45S_50ntFlanks@10347.10383.37\|CAAACGAGAACTTTGAAGGCCGAAGTGGAGAAGGGTT | >hg38_21_+_8433172_8446622_RNA45SN1_45S_50ntFlanks | 10347 | 10383 | 37 | CAAACGAGAACTTTGAAGGCCGAAGTGGAGAAGGGTT | 555.4 | -5.5 | 0.7 | -8.0 | 1.7E-15 | 5.0E-15 | BT20/MB468 | NUC |
| >hg38_21_+_8433172_8446622_RNA45SN1_45S_50ntFlanks@4729.4767.39\|TCAAGAACGAAAGTCGGAGGTTCGAAGACGATCAGATAC | >hg38_21_+_8433172_8446622_RNA45SN1_45S_50ntFlanks | 4729 | 4767 | 39 | TCAAGAACGAAAGTCGGAGGTTCGAAGACGATCAGATAC | 326.1 | 10.3 | 1.3 | 8.0 | 1.1E-15 | 5.0E-15 | BT20/MB468 | CYTO |
| >hg38_21_+_8433172_8446622_RNA45SN1_45S_50ntFlanks@12414.12447.34\|CCTTCGATGTCGGCTCTTCCTATCATTGTGAAGC | >hg38_21_+_8433172_8446622_RNA45SN1_45S_50ntFlanks | 12414 | 12447 | 34 | CCTTCGATGTCGGCTCTTCCTATCATTGTGAAGC | 366.4 | 9.2 | 1.1 | 8.1 | 7.7E-16 | 5.0E-15 | MB231/MB468 | CYTO |
| >hg38_21_+_8433172_8446622_RNA45SN1_45S_50ntFlanks@6661.6685.25\|CGGTGGATCACTCGGCTCGTGCGTC | >hg38_21_+_8433172_8446622_RNA45SN1_45S_50ntFlanks | 6661 | 6685 | 25 | CGGTGGATCACTCGGCTCGTGCGTC | 746.1 | 5.9 | 0.7 | 8.0 | 1.1E-15 | 5.0E-15 | BT20/MB468 | CYTO |
| >MI0000088\|hsa-mir-30a&WithFlank&6\|-\|71403545\|71403627@53.70.18(+1U)&[MIMAT0000088&hsa-miR-30a-3p&offsets\|0\|-4(+1U);m-38&6\|-\|71403554\|71403575&offsets\|0\|-4(+1U)]\|CTTTCAGTCGGATGTTTGT | >MI0000088\|hsa-mir-30a&WithFlank&6\|-\|71403545\|71403627 | 53 | 70 | 18(+1U) | CTTTCAGTCGGATGTTTGT | 414.9 | 7.5 | 0.9 | 8.0 | 1.1E-15 | 5.0E-15 | BT20/MB468 | CYTO |
| >hg38_21_+_8433172_8446622_RNA45SN1_45S_50ntFlanks@9622.9648.27\|CTAGTAGCTGGTTCCCTCCGAAGTTTC | >hg38_21_+_8433172_8446622_RNA45SN1_45S_50ntFlanks | 9622 | 9648 | 27 | CTAGTAGCTGGTTCCCTCCGAAGTTTC | 557.2 | -13.0 | 1.6 | -8.1 | 7.1E-16 | 5.1E-15 | MB231/MB468 | MP |
| >hg38_21_+_8433172_8446622_RNA45SN1_45S_50ntFlanks@4160.4178.19\|CCACATCCAAGGAAGGCAG | >hg38_21_+_8433172_8446622_RNA45SN1_45S_50ntFlanks | 4160 | 4178 | 19 | CCACATCCAAGGAAGGCAG | 299.5 | -12.1 | 1.5 | -8.1 | 7.1E-16 | 5.1E-15 | MB231/MB468 | MP |

| | | | | | | | | | | | | | |
|---|---|---|---|---|---|---|---|---|---|---|---|---|---|
| >hg38_21_+_8433172_8446622_RNA45SN1_45S_50ntFlanks@5057.5098.42\|AGCGATTTGTCTGGTTAATTCCGATAACGAACGAGACTCTGG | >hg38_21_+_8433172_8446622_RNA45SN1_45S_50ntFlanks | 5057 | 5098 | 42 | AGCGATTTGTCTGGTTAATTCCGATAACGAACGAGACTCTGG | 85.4 | -23.6 | 2.9 | -8.1 | 7.2E-16 | 5.1E-15 | MB231/MB468 | MP |
| >hg38_21_+_8433172_8446622_RNA45SN1_45S_50ntFlanks@4472.4494.23\|CTTAGCTGAGTGTCCCGCGGGGC | >hg38_21_+_8433172_8446622_RNA45SN1_45S_50ntFlanks | 4472 | 4494 | 23 | CTTAGCTGAGTGTCCCGCGGGGC | 575.7 | -9.5 | 1.2 | -8.0 | 8.8E-16 | 5.1E-15 | MB231/MB468 | NUC |
| >hg38_21_+_8433172_8446622_RNA45SN1_45S_50ntFlanks@9713.9737.25\|GCGAATGATTAGAGGTCTTGGGGCC | >hg38_21_+_8433172_8446622_RNA45SN1_45S_50ntFlanks | 9713 | 9737 | 25 | GCGAATGATTAGAGGTCTTGGGGCC | 596.4 | -13.1 | 1.6 | -8.1 | 7.2E-16 | 5.1E-15 | MB231/MB468 | MP |
| >hg38_21_+_8433172_8446622_RNA45SN1_45S_50ntFlanks@12934.12970.37\|CTAAACCATTCGTAGACGACCTGCTTCTGGGTCGGGG | >hg38_21_+_8433172_8446622_RNA45SN1_45S_50ntFlanks | 12934 | 12970 | 37 | CTAAACCATTCGTAGACGACCTGCTTCTGGGTCGGGG | 264.2 | -11.8 | 1.5 | -8.0 | 1.2E-15 | 5.1E-15 | MB231/MB468 | MITO |
| >hg38_21_+_8433172_8446622_RNA45SN1_45S_50ntFlanks@10681.10711.31\|TTCCGGCGGCGTCCGGTGAGCTCTCGCTGGC | >hg38_21_+_8433172_8446622_RNA45SN1_45S_50ntFlanks | 10681 | 10711 | 31 | TTCCGGCGGCGTCCGGTGAGCTCTCGCTGGC | 701.2 | -3.9 | 0.5 | -8.1 | 7.8E-16 | 5.1E-15 | BT20/MB468 | MITO |
| >hg38_21_+_8433172_8446622_RNA45SN1_45S_50ntFlanks@7989.8006.18\|AGACGTGGCGACCCGCTG | >hg38_21_+_8433172_8446622_RNA45SN1_45S_50ntFlanks | 7989 | 8006 | 18 | AGACGTGGCGACCCGCTG | 4821.3 | -5.1 | 0.6 | -8.1 | 7.8E-16 | 5.1E-15 | BT20/MB468 | MITO |
| >hg38_21_+_8433172_8446622_RNA45SN1_45S_50ntFlanks@12141.12162.22\|ACAGTGCCAGGTGGGGAGTTTG | >hg38_21_+_8433172_8446622_RNA45SN1_45S_50ntFlanks | 12141 | 12162 | 22 | ACAGTGCCAGGTGGGGAGTTTG | 348.6 | -12.2 | 1.5 | -8.0 | 1.2E-15 | 5.1E-15 | MB231/MB468 | MITO |
| >hg38_21_+_8433172_8446622_RNA45SN1_45S_50ntFlanks@8079.8101.23\|AAGAGCCCAGCGCCGAATCCCCG | >hg38_21_+_8433172_8446622_RNA45SN1_45S_50ntFlanks | 8079 | 8101 | 23 | AAGAGCCCAGCGCCGAATCCCCG | 305.5 | -12.0 | 1.5 | -8.0 | 1.2E-15 | 5.1E-15 | MB231/MB468 | MITO |
| >hg38_21_+_8433172_8446622_RNA45SN1_45S_50ntFlanks@10402.10420.19\|AACATGGGTCAGTCGGTCC | >hg38_21_+_8433172_8446622_RNA45SN1_45S_50ntFlanks | 10402 | 10420 | 19 | AACATGGGTCAGTCGGTCC | 1062.7 | -4.5 | 0.6 | -8.1 | 7.8E-16 | 5.1E-15 | BT20/MB468 | MITO |
| >hg38_21_+_8433172_8446622_RNA45SN1_45S_50ntFlanks@3785.3812.28\|TGAAACTGCGAATGGCTCATTAAATCAG | >hg38_21_+_8433172_8446622_RNA45SN1_45S_50ntFlanks | 3785 | 3812 | 28 | TGAAACTGCGAATGGCTCATTAAATCAG | 831.8 | -12.7 | 1.6 | -8.0 | 8.9E-16 | 5.1E-15 | MB231/MB468 | NUC |
| >hg38_21_+_8433172_8446622_RNA45SN1_45S_50ntFlanks@8498.8526.29\|CGTTCCTCCCGACCCCTCCACCCGCCCTC | >hg38_21_+_8433172_8446622_RNA45SN1_45S_50ntFlanks | 8498 | 8526 | 29 | CGTTCCTCCCGACCCCTCCACCCGCCCTC | 254.6 | -11.8 | 1.5 | -8.0 | 1.2E-15 | 5.2E-15 | MB231/MB468 | MITO |
| >hg38_21_+_8433172_8446622_RNA45SN1_45S_50ntFlanks@11534.11555.22\|GTTCCGCGCGGCGCCTCGCCTC | >hg38_21_+_8433172_8446622_RNA45SN1_45S_50ntFlanks | 11534 | 11555 | 22 | GTTCCGCGCGGCGCCTCGCCTC | 446.7 | -8.7 | 1.1 | -8.1 | 7.9E-16 | 5.2E-15 | MB231/MB468 | CYTO |
| >hg38_21_+_8433172_8446622_RNA45SN1_45S_50ntFlanks@10411.10441.31\|CAGTCGGTCCTGAGAGATGGGCGAGCGCCGT | >hg38_21_+_8433172_8446622_RNA45SN1_45S_50ntFlanks | 10411 | 10441 | 31 | CAGTCGGTCCTGAGAGATGGGCGAGCGCCGT | 278.3 | -11.9 | 1.5 | -8.0 | 1.2E-15 | 5.2E-15 | MB231/MB468 | MITO |
| >hg38_1_-_228634819_228635039_RNA5S12_5S_50ntFlanks@153.170.18\|ACCGGGTGCTGTAGGCTT | >hg38_1_-_228634819_228635039_RNA5S12_5S_50ntFlanks | 153 | 170 | 18 | ACCGGGTGCTGTAGGCTT | 34541.8 | 7.1 | 0.9 | 8.1 | 7.9E-16 | 5.2E-15 | MB231/MB468 | CYTO |
| >hg38_21_+_8433172_8446622_RNA45SN1_45S_50ntFlanks@9122.9140.19\|CCGCGCGCGGGTCGGGGGG | >hg38_21_+_8433172_8446622_RNA45SN1_45S_50ntFlanks | 9122 | 9140 | 19 | CCGCGCGCGGGTCGGGGGG | 365.2 | -12.4 | 1.5 | -8.1 | 7.3E-16 | 5.2E-15 | MB231/MB468 | MP |
| >trnaMT_GluTTC_MT_-_14674_14742@40.58.19,trnalookalike8_GluTTC_5_-_93905172_93905240@40.58.19\|TTGGTCGTGGTTGTAGTCC | >trnaMT_GluTTC_MT_-_14674_14742 | 40 | 58 | 19 | TTGGTCGTGGTTGTAGTCC | 763.5 | 7.9 | 1.0 | 8.0 | 1.1E-15 | 5.2E-15 | BT20/MB468 | CYTO |
| >hg38_21_+_8433172_8446622_RNA45SN1_45S_50ntFlanks@12655.12673.19\|ATGACTGAACGCCTCTAAG | >hg38_21_+_8433172_8446622_RNA45SN1_45S_50ntFlanks | 12655 | 12673 | 19 | ATGACTGAACGCCTCTAAG | 271.0 | -5.9 | 0.7 | -8.1 | 8.0E-16 | 5.2E-15 | MB231/MB468 | CYTO |
| >hg38_21_+_8433172_8446622_RNA45SN1_45S_50ntFlanks@10392.10435.44\|ACAGCAGTTGAACATGGGTCAGTCGGTCCTGAGAGATGGGCGAG | >hg38_21_+_8433172_8446622_RNA45SN1_45S_50ntFlanks | 10392 | 10435 | 44 | ACAGCAGTTGAACATGGGTCAGTCGGTCCTGAGAGATGGGCGAG | 1444.3 | -13.5 | 1.7 | -8.0 | 9.1E-16 | 5.2E-15 | MB231/MB468 | NUC |
| >hg38_21_+_8433172_8446622_RNA45SN1_45S_50ntFlanks@6745.6762.18\|ACACTTCGAACGCACTTG | >hg38_21_+_8433172_8446622_RNA45SN1_45S_50ntFlanks | 6745 | 6762 | 18 | ACACTTCGAACGCACTTG | 204.3 | -11.5 | 1.4 | -8.1 | 7.4E-16 | 5.2E-15 | MB231/MB468 | MP |
| >MI0000440\|hsa-mir-27b&WithFlank&9\|+\|95085439\|95085547@67.88.22(+1U)&[MIMAT0000419&hsa-miR-27b-3p&offsets\|0\|+1(+1U);m-39&9\|+\|95085505\|95085525&offsets\|0\|+1(+1U)]\|TTCACAGTGGCTAAGTTCTGCAT | >MI0000440\|hsa-mir-27b&WithFlank&9\|+\|95085439\|95085547 | 67 | 88 | 22(+1U) | TTCACAGTGGCTAAGTTCTGCAT | 91.1 | 3.2 | 0.4 | 8.0 | 1.7E-15 | 5.3E-15 | BT20/MB468 | NUC |
| >hg38_21_+_8433172_8446622_RNA45SN1_45S_50ntFlanks@10411.10443.33\|CAGTCGGTCCTGAGAGATGGGCGAGCGCCGTTC | >hg38_21_+_8433172_8446622_RNA45SN1_45S_50ntFlanks | 10411 | 10443 | 33 | CAGTCGGTCCTGAGAGATGGGCGAGCGCCGTTC | 2664.5 | -10.5 | 1.3 | -8.0 | 1.2E-15 | 5.3E-15 | MB231/MB468 | MITO |
| >MI0000477\|hsa-mir-146a&WithFlank&5\|+\|160485346\|160485456@27.48.22(+1U)&[MIMAT0000449&hsa-miR-146a-5p&offsets\|0\|0(+1U);m-129&5\|+\|160485372\|160485393&offsets\|0\|0(+1U)]\|TGAGAACTGAATTCCATGGGTTT | >MI0000477\|hsa-mir-146a&WithFlank&5\|+\|160485346\|160485456 | 27 | 48 | 22(+1U) | TGAGAACTGAATTCCATGGGTTT | 1244.4 | 13.5 | 1.7 | 8.0 | 1.2E-15 | 5.3E-15 | MB231/MB468 | MITO |
| >hg38_21_+_8433172_8446622_RNA45SN1_45S_50ntFlanks@11315.11340.26\|CCACCCCACGTCTCGTCGCGCGCGCG | >hg38_21_+_8433172_8446622_RNA45SN1_45S_50ntFlanks | 11315 | 11340 | 26 | CCACCCCACGTCTCGTCGCGCGCGCG | 590.2 | -13.0 | 1.6 | -8.0 | 1.2E-15 | 5.3E-15 | MB231/MB468 | MITO |
| >hg38_21_+_8433172_8446622_RNA45SN1_45S_50ntFlanks@8194.8214.21\|CCCGTGGACGGTGTGAGGCCG | >hg38_21_+_8433172_8446622_RNA45SN1_45S_50ntFlanks | 8194 | 8214 | 21 | CCCGTGGACGGTGTGAGGCCG | 1574.3 | -8.9 | 1.1 | -8.1 | 7.5E-16 | 5.3E-15 | MB231/MB468 | MP |
| >hg38_21_+_8433172_8446622_RNA45SN1_45S_50ntFlanks@10408.10432.25\|GGTCAGTCGGTCCTGAGAGATGGGC | >hg38_21_+_8433172_8446622_RNA45SN1_45S_50ntFlanks | 10408 | 10432 | 25 | GGTCAGTCGGTCCTGAGAGATGGGC | 439.3 | -12.6 | 1.6 | -8.1 | 7.5E-16 | 5.3E-15 | MB231/MB468 | MP |
| >hg38_21_+_8433172_8446622_RNA45SN1_45S_50ntFlanks@4202.4219.18\|CGACCCGGGGAGGTAGTG | >hg38_21_+_8433172_8446622_RNA45SN1_45S_50ntFlanks | 4202 | 4219 | 18 | CGACCCGGGGAGGTAGTG | 396.3 | -12.4 | 1.6 | -8.0 | 1.2E-15 | 5.3E-15 | MB231/MB468 | MITO |
| >hg38_21_+_8433172_8446622_RNA45SN1_45S_50ntFlanks@9572.9596.25\|CAAATCGGTCGTCCGACCTGGGTAT | >hg38_21_+_8433172_8446622_RNA45SN1_45S_50ntFlanks | 9572 | 9596 | 25 | CAAATCGGTCGTCCGACCTGGGTAT | 17645.1 | -10.6 | 1.2 | -8.5 | 2.3E-17 | 5.3E-15 | BT20/MB468 | MP |
| >hg38_21_+_8433172_8446622_RNA45SN1_45S_50ntFlanks@11318.11340.23\|CCCCACGTCTCGTCGCGCGCGCG | >hg38_21_+_8433172_8446622_RNA45SN1_45S_50ntFlanks | 11318 | 11340 | 23 | CCCCACGTCTCGTCGCGCGCGCG | 462.3 | -12.6 | 1.6 | -8.0 | 1.2E-15 | 5.3E-15 | MB231/MB468 | MITO |
| >hg38_21_+_8433172_8446622_RNA45SN1_45S_50ntFlanks@12717.12751.35\|CCTCGGTTGGCCTCGGATAGCCGGTCCCCCGCCTG | >hg38_21_+_8433172_8446622_RNA45SN1_45S_50ntFlanks | 12717 | 12751 | 35 | CCTCGGTTGGCCTCGGATAGCCGGTCCCCCGCCTG | 281.8 | -11.9 | 1.5 | -8.0 | 1.2E-15 | 5.3E-15 | MB231/MB468 | MITO |
| >hg38_21_+_8433172_8446622_RNA45SN1_45S_50ntFlanks@12625.12654.30\|CAATGGGGCGAAGCTACCATCTGTGGGATT | >hg38_21_+_8433172_8446622_RNA45SN1_45S_50ntFlanks | 12625 | 12654 | 30 | CAATGGGGCGAAGCTACCATCTGTGGGATT | 3078.4 | -4.3 | 0.5 | -8.1 | 8.2E-16 | 5.3E-15 | MB231/MB468 | CYTO |
| >hg38_21_+_8433172_8446622_RNA45SN1_45S_50ntFlanks@4657.4682.26\|CTAGAGGTGAAATTCTTGGACCGGCG | >hg38_21_+_8433172_8446622_RNA45SN1_45S_50ntFlanks | 4657 | 4682 | 26 | CTAGAGGTGAAATTCTTGGACCGGCG | 297.1 | -12.1 | 1.5 | -8.1 | 7.5E-16 | 5.3E-15 | MB231/MB468 | MP |
| >hg38_21_+_8433172_8446622_RNA45SN1_45S_50ntFlanks@4449.4472.24\|CCTCTCGGCGCCCCCTCGATGCTC | >hg38_21_+_8433172_8446622_RNA45SN1_45S_50ntFlanks | 4449 | 4472 | 24 | CCTCTCGGCGCCCCCTCGATGCTC | 1174.2 | -12.3 | 1.5 | -8.1 | 8.1E-16 | 5.3E-15 | BT20/MB468 | MITO |
| >hg38_21_+_8433172_8446622_RNA45SN1_45S_50ntFlanks@12539.12565.27\|ATGATGTGTTGTTGCCATGGTAATCCT | >hg38_21_+_8433172_8446622_RNA45SN1_45S_50ntFlanks | 12539 | 12565 | 27 | ATGATGTGTTGTTGCCATGGTAATCCT | 216.1 | -10.5 | 1.3 | -8.1 | 8.2E-16 | 5.4E-15 | MB231/MB468 | CYTO |
| >hg38_21_+_8433172_8446622_RNA45SN1_45S_50ntFlanks@4005.4024.20\|AACCTCGGGCCGATCGCACG | >hg38_21_+_8433172_8446622_RNA45SN1_45S_50ntFlanks | 4005 | 4024 | 20 | AACCTCGGGCCGATCGCACG | 1131.8 | -5.9 | 0.7 | -8.0 | 1.3E-15 | 5.4E-15 | MB231/MB468 | MITO |
| >hg38_21_+_8433172_8446622_RNA45SN1_45S_50ntFlanks@9533.9555.23\|GAGGAAACTCTGGTGGAGGTCCG | >hg38_21_+_8433172_8446622_RNA45SN1_45S_50ntFlanks | 9533 | 9555 | 23 | GAGGAAACTCTGGTGGAGGTCCG | 85.4 | -23.6 | 2.9 | -8.1 | 7.6E-16 | 5.4E-15 | MB231/MB468 | MP |
| >MI0003513\|hsa-mir-455&WithFlank&9\|+\|114209428\|114209535@22.43.22&[MIMAT0003150&hsa-miR-455-5p&offsets\|0\|0;m-232&9\|+\|114209449\|114209470&offsets\|0\|0]\|TATGTGCCTTTGGACTACATCG | >MI0003513\|hsa-mir-455&WithFlank&9\|+\|114209428\|114209535 | 22 | 43 | 22 | TATGTGCCTTTGGACTACATCG | 303.8 | 11.5 | 1.4 | 8.1 | 7.6E-16 | 5.4E-15 | MB231/MB468 | MP |
| >MI0003560\|hsa-mir-92b&WithFlank&1\|+\|155195171\|155195278@67.87.21&[MIMAT0003218&hsa-miR-92b-3p&offsets\|0\|-1;m-115&1\|+\|155195237\|155195258&offsets\|0\|-1]\|TATTGCACTCGTCCCGGCCTC | >MI0003560\|hsa-mir-92b&WithFlank&1\|+\|155195171\|155195278 | 67 | 87 | 21 | TATTGCACTCGTCCCGGCCTC | 785.8 | 5.8 | 0.7 | 8.0 | 9.3E-16 | 5.4E-15 | MB231/MB468 | NUC |
| >hg38_21_+_8433172_8446622_RNA45SN1_45S_50ntFlanks@12589.12610.22\|GTTCAGACATTTGGTGTATGTG | >hg38_21_+_8433172_8446622_RNA45SN1_45S_50ntFlanks | 12589 | 12610 | 22 | GTTCAGACATTTGGTGTATGTG | 385.2 | -12.5 | 1.5 | -8.1 | 7.6E-16 | 5.4E-15 | MB231/MB468 | MP |
| >hg38_21_+_8433172_8446622_RNA45SN1_45S_50ntFlanks@11205.11254.50\|GCGGCGGCGACTCTGGACGCGAGCCGGGCCCTTCCCGTGGATCGCCCCAG | >hg38_21_+_8433172_8446622_RNA45SN1_45S_50ntFlanks | 11205 | 11254 | 50 | GCGGCGGCGACTCTGGACGCGAGCCGGGCCCTTCCCGTGGATCGCCCCAG | 650.4 | -13.8 | 1.7 | -8.0 | 1.8E-15 | 5.4E-15 | BT20/MB468 | NUC |
| >MI0000266\|hsa-mir-10a&WithFlank&17\|-\|48579832\|48579953@29.51.23&[MIMAT0000253&hsa-miR-10a-5p&offsets\|+1\|+1;m-18&17\|-\|48579905\|48579926&offsets\|+1\|+2]\|ACCCTGTAGATCCGAATTTGTGT | >MI0000266\|hsa-mir-10a&WithFlank&17\|-\|48579832\|48579953 | 29 | 51 | 23 | ACCCTGTAGATCCGAATTTGTGT | 692.6 | 12.7 | 1.6 | 8.0 | 1.3E-15 | 5.4E-15 | MB231/MB468 | MITO |
| >hg38_21_+_8433172_8446622_RNA45SN1_45S_50ntFlanks@8636.8658.23\|CGGCGACCGGCCGCCGCCGGGCG | >hg38_21_+_8433172_8446622_RNA45SN1_45S_50ntFlanks | 8636 | 8658 | 23 | CGGCGACCGGCCGCCGCCGGGCG | 370.6 | -12.4 | 1.5 | -8.1 | 7.6E-16 | 5.4E-15 | MB231/MB468 | MP |
| >hg38_21_+_8433172_8446622_RNA45SN1_45S_50ntFlanks@9304.9337.34\|ACCAAGGAGTCTAACACGTGCGCGAGTCGGGGGC | >hg38_21_+_8433172_8446622_RNA45SN1_45S_50ntFlanks | 9304 | 9337 | 34 | ACCAAGGAGTCTAACACGTGCGCGAGTCGGGGGC | 337.3 | -12.3 | 1.5 | -8.1 | 7.6E-16 | 5.4E-15 | MB231/MB468 | MP |
| >hg38_21_+_8433172_8446622_RNA45SN1_45S_50ntFlanks@10468.10486.19\|TTGCCCTCGGCCGATCGAA | >hg38_21_+_8433172_8446622_RNA45SN1_45S_50ntFlanks | 10468 | 10486 | 19 | TTGCCCTCGGCCGATCGAA | 597.3 | -5.8 | 0.7 | -8.0 | 1.8E-15 | 5.4E-15 | BT20/MB468 | NUC |
| >hg38_21_+_8433172_8446622_RNA45SN1_45S_50ntFlanks@9534.9561.28\|AGGAAACTCTGGTGGAGGTCCGTAGCGG | >hg38_21_+_8433172_8446622_RNA45SN1_45S_50ntFlanks | 9534 | 9561 | 28 | AGGAAACTCTGGTGGAGGTCCGTAGCGG | 6382.3 | -9.6 | 1.2 | -8.0 | 9.4E-16 | 5.4E-15 | MB231/MB468 | NUC |
| >hg38_21_+_8433172_8446622_RNA45SN1_45S_50ntFlanks@11641.11663.23\|CGGCGGGTGTTGACGCGATGTGA | >hg38_21_+_8433172_8446622_RNA45SN1_45S_50ntFlanks | 11641 | 11663 | 23 | CGGCGGGTGTTGACGCGATGTGA | 408.4 | -11.6 | 1.4 | -8.0 | 9.4E-16 | 5.4E-15 | MB231/MB468 | NUC |
| >hg38_21_+_8433172_8446622_RNA45SN1_45S_50ntFlanks@10806.10825.20\|AACAATGTAGGTAAGGGAAG | >hg38_21_+_8433172_8446622_RNA45SN1_45S_50ntFlanks | 10806 | 10825 | 20 | AACAATGTAGGTAAGGGAAG | 920.6 | -9.5 | 1.2 | -8.0 | 1.3E-15 | 5.4E-15 | MB231/MB468 | MITO |
| >hg38_21_+_8433172_8446622_RNA45SN1_45S_50ntFlanks@10805.10826.22\|GAACAATGTAGGTAAGGGAAGT | >hg38_21_+_8433172_8446622_RNA45SN1_45S_50ntFlanks | 10805 | 10826 | 22 | GAACAATGTAGGTAAGGGAAGT | 190.7 | -11.4 | 1.4 | -8.1 | 7.7E-16 | 5.4E-15 | MB231/MB468 | MP |
| >hg38_21_+_8433172_8446622_RNA45SN1_45S_50ntFlanks@12274.12293.20\|CAGTACGAATACAGACCGTG | >hg38_21_+_8433172_8446622_RNA45SN1_45S_50ntFlanks | 12274 | 12293 | 20 | CAGTACGAATACAGACCGTG | 426.8 | -12.5 | 1.6 | -8.0 | 1.3E-15 | 5.4E-15 | MB231/MB468 | MITO |
| >hg38_21_+_8433172_8446622_RNA45SN1_45S_50ntFlanks@9580.9597.18\|TCGTCCGACCTGGGTATA | >hg38_21_+_8433172_8446622_RNA45SN1_45S_50ntFlanks | 9580 | 9597 | 18 | TCGTCCGACCTGGGTATA | 297.8 | -12.0 | 1.5 | -8.0 | 1.3E-15 | 5.4E-15 | MB231/MB468 | MITO |
| >hg38_21_+_8433172_8446622_RNA45SN1_45S_50ntFlanks@11669.11693.25\|GCCCAGTGCTCTGAATGTCAAAGTG | >hg38_21_+_8433172_8446622_RNA45SN1_45S_50ntFlanks | 11669 | 11693 | 25 | GCCCAGTGCTCTGAATGTCAAAGTG | 644.5 | 6.0 | 0.7 | 8.0 | 8.3E-16 | 5.4E-15 | MB231/MB468 | CYTO |
| >hg38_21_+_8433172_8446622_RNA45SN1_45S_50ntFlanks@8170.8218.49\|CAAGTCCTTCTGATCGAGGCCCAGCCCGTGGACGGTGTGAGGCCGGTAG | >hg38_21_+_8433172_8446622_RNA45SN1_45S_50ntFlanks | 8170 | 8218 | 49 | CAAGTCCTTCTGATCGAGGCCCAGCCCGTGGACGGTGTGAGGCCGGTAG | 525.0 | -13.5 | 1.7 | -8.0 | 1.8E-15 | 5.4E-15 | BT20/MB468 | NUC |
| >MI0001445\|hsa-mir-423&WithFlank&17\|+\|30117073\|30117178@58.81.24&[MIMAT0001340&hsa-miR-423-3p&offsets\|-1\|0;m-100&17\|+\|30117131\|30117153&offsets\|-1\|0]\|AAGCTCGGTCTGAGGCCCCTCAGT | >MI0001445\|hsa-mir-423&WithFlank&17\|+\|30117073\|30117178 | 58 | 81 | 24 | AAGCTCGGTCTGAGGCCCCTCAGT | 2050.6 | 6.7 | 0.8 | 8.0 | 8.4E-16 | 5.4E-15 | MB231/MB468 | CYTO |
| >hg38_21_+_8433172_8446622_RNA45SN1_45S_50ntFlanks@10689.10708.20\|GCGTCCGGTGAGCTCTCGCT | >hg38_21_+_8433172_8446622_RNA45SN1_45S_50ntFlanks | 10689 | 10708 | 20 | GCGTCCGGTGAGCTCTCGCT | 1428.0 | -6.8 | 0.8 | -8.0 | 8.4E-16 | 5.4E-15 | MB231/MB468 | CYTO |

<table>
<tr><td>>hg38_21_+_8433172_8446622_RNA45SN1_45S_50ntFlanks@9625.9650.26|GTAGCTGGTTCCCTCCGAAGTTTCCC</td><td>>hg38_21_+_8433172_8446622_RNA45SN1_45S_50ntFlanks</td><td>9625</td><td>9650</td><td>26</td><td>GTAGCTGGTTCCCTCCGAAGTTTCCC</td><td>284.5</td><td>-11.9</td><td>1.5</td><td>-8.0</td><td>1.3E-15</td><td>5.5E-15</td><td>MB231/MB468</td><td>MITO</td></tr>
<tr><td>>hg38_1_-_228634819_228635039_RNA5S12_5S_50ntFlanks@134.171.38|TGGGAGACCGCCTGGGAATACCGGGTGCTGTAGGCTTT</td><td>>hg38_1_-_228634819_228635039_RNA5S12_5S_50ntFlanks</td><td>134</td><td>171</td><td>38</td><td>TGGGAGACCGCCTGGGAATACCGGGTGCTGTAGGCTTT</td><td>14289.6</td><td>6.6</td><td>0.8</td><td>8.0</td><td>1.2E-15</td><td>5.5E-15</td><td>BT20/MB468</td><td>CYTO</td></tr>
<tr><td>>hg38_21_+_8433172_8446622_RNA45SN1_45S_50ntFlanks@11930.11948.19|TAGAATAAGTGGGAGGCCC</td><td>>hg38_21_+_8433172_8446622_RNA45SN1_45S_50ntFlanks</td><td>11930</td><td>11948</td><td>19</td><td>TAGAATAAGTGGGAGGCCC</td><td>254.4</td><td>-11.8</td><td>1.5</td><td>-8.0</td><td>1.3E-15</td><td>5.5E-15</td><td>MB231/MB468</td><td>MITO</td></tr>
<tr><td>>hg38_21_+_8433172_8446622_RNA45SN1_45S_50ntFlanks@4027.4059.33|CCCCGTGGCGGCGACGACCCATTCGAACGTCTG</td><td>>hg38_21_+_8433172_8446622_RNA45SN1_45S_50ntFlanks</td><td>4027</td><td>4059</td><td>33</td><td>CCCCGTGGCGGCGACGACCCATTCGAACGTCTG</td><td>251.2</td><td>-11.8</td><td>1.5</td><td>-8.0</td><td>1.3E-15</td><td>5.5E-15</td><td>MB231/MB468</td><td>MITO</td></tr>
<tr><td>>hg38_21_+_8433172_8446622_RNA45SN1_45S_50ntFlanks@10101.10127.27|CGCGCGCGTGTGGTGTGCGTCGGAGGG</td><td>>hg38_21_+_8433172_8446622_RNA45SN1_45S_50ntFlanks</td><td>10101</td><td>10127</td><td>27</td><td>CGCGCGCGTGTGGTGTGCGTCGGAGGG</td><td>767.4</td><td>-12.5</td><td>1.6</td><td>-8.0</td><td>9.6E-16</td><td>5.5E-15</td><td>MB231/MB468</td><td>NUC</td></tr>
<tr><td>>hg38_21_+_8433172_8446622_RNA45SN1_45S_50ntFlanks@13004.13023.20|ATCTATTGAAAGTCAGCCCT</td><td>>hg38_21_+_8433172_8446622_RNA45SN1_45S_50ntFlanks</td><td>13004</td><td>13023</td><td>20</td><td>ATCTATTGAAAGTCAGCCCT</td><td>202.0</td><td>-10.6</td><td>1.3</td><td>-8.0</td><td>9.6E-16</td><td>5.5E-15</td><td>MB231/MB468</td><td>NUC</td></tr>
<tr><td>>hg38_21_+_8433172_8446622_RNA45SN1_45S_50ntFlanks@10391.10410.20|AACAGCAGTTGAACATGGGT</td><td>>hg38_21_+_8433172_8446622_RNA45SN1_45S_50ntFlanks</td><td>10391</td><td>10410</td><td>20</td><td>AACAGCAGTTGAACATGGGT</td><td>4548.5</td><td>-7.6</td><td>0.9</td><td>-8.0</td><td>8.5E-16</td><td>5.5E-15</td><td>MB231/MB468</td><td>CYTO</td></tr>
<tr><td>>hg38_21_+_8433172_8446622_RNA45SN1_45S_50ntFlanks@8357.8389.33|AAAAGAACTTTGAAGAGAGAGTTCAAGAGGGCG</td><td>>hg38_21_+_8433172_8446622_RNA45SN1_45S_50ntFlanks</td><td>8357</td><td>8389</td><td>33</td><td>AAAAGAACTTTGAAGAGAGAGTTCAAGAGGGCG</td><td>322.0</td><td>-12.2</td><td>1.5</td><td>-8.1</td><td>7.9E-16</td><td>5.5E-15</td><td>MB231/MB468</td><td>MP</td></tr>
<tr><td>>hg38_21_+_8433172_8446622_RNA45SN1_45S_50ntFlanks@4731.4771.41|AAGAACGAAAGTCGGAGGTTCGAAGACGATCAGATACCGTC</td><td>>hg38_21_+_8433172_8446622_RNA45SN1_45S_50ntFlanks</td><td>4731</td><td>4771</td><td>41</td><td>AAGAACGAAAGTCGGAGGTTCGAAGACGATCAGATACCGTC</td><td>406.8</td><td>7.0</td><td>0.9</td><td>8.0</td><td>8.6E-16</td><td>5.6E-15</td><td>MB231/MB468</td><td>CYTO</td></tr>
<tr><td>>hg38_1_-_228634819_228635039_RNA5S12_5S_50ntFlanks@151.171.21|ATACCGGGTGCTGTAGGCTTT</td><td>>hg38_1_-_228634819_228635039_RNA5S12_5S_50ntFlanks</td><td>151</td><td>171</td><td>21</td><td>ATACCGGGTGCTGTAGGCTTT</td><td>2914.1</td><td>5.2</td><td>0.6</td><td>8.0</td><td>1.2E-15</td><td>5.6E-15</td><td>BT20/MB468</td><td>CYTO</td></tr>
<tr><td>>hg38_21_+_8433172_8446622_RNA45SN1_45S_50ntFlanks@4724.4752.29|ATTAATCAAGAACGAAAGTCGGAGGTTCG</td><td>>hg38_21_+_8433172_8446622_RNA45SN1_45S_50ntFlanks</td><td>4724</td><td>4752</td><td>29</td><td>ATTAATCAAGAACGAAAGTCGGAGGTTCG</td><td>927.6</td><td>-11.9</td><td>1.5</td><td>-8.0</td><td>8.5E-16</td><td>5.6E-15</td><td>BT20/MB468</td><td>MITO</td></tr>
<tr><td>>MI0000734|hsa-mir-106b&WithFlank&7|-|100093987|100094080@18.39.22&[MIMAT0000680&hsa-miR-106b-5p&offsets|0|+1;m-103&7|-|100094043|100094063&offsets|0|+1]|TAAAGTGCTGACAGTGCAGATA</td><td>>MI0000734|hsa-mir-106b&WithFlank&7|-|100093987|100094080</td><td>18</td><td>39</td><td>22</td><td>TAAAGTGCTGACAGTGCAGATA</td><td>1250.0</td><td>5.7</td><td>0.7</td><td>8.0</td><td>9.7E-16</td><td>5.6E-15</td><td>MB231/MB468</td><td>NUC</td></tr>
<tr><td>>hg38_21_+_8433172_8446622_RNA45SN1_45S_50ntFlanks@5472.5496.25|ACGGCCCTGGCGGAGCGCTGAGAAG</td><td>>hg38_21_+_8433172_8446622_RNA45SN1_45S_50ntFlanks</td><td>5472</td><td>5496</td><td>25</td><td>ACGGCCCTGGCGGAGCGCTGAGAAG</td><td>475.1</td><td>6.9</td><td>0.9</td><td>8.0</td><td>8.6E-16</td><td>5.6E-15</td><td>MB231/MB468</td><td>CYTO</td></tr>
<tr><td>>hg38_21_+_8433172_8446622_RNA45SN1_45S_50ntFlanks@6719.6736.18|TGAATTGCAGGACACATT</td><td>>hg38_21_+_8433172_8446622_RNA45SN1_45S_50ntFlanks</td><td>6719</td><td>6736</td><td>18</td><td>TGAATTGCAGGACACATT</td><td>4996.3</td><td>-6.9</td><td>0.9</td><td>-8.0</td><td>9.8E-16</td><td>5.6E-15</td><td>MB231/MB468</td><td>NUC</td></tr>
<tr><td>>MI0000285|hsa-mir-205&WithFlank&1|+|209432127|209432248@40.61.22(+3A)&[MIMAT0000266&hsa-miR-205-5p&offsets|0|0(+3A);m-351&1|+|209432166|209432187&offsets|0|0(+3A)]|TCCTTCATTCCACCGGAGTCTGAAA</td><td>>MI0000285|hsa-mir-205&WithFlank&1|+|209432127|209432248</td><td>40</td><td>61</td><td>22(+3A)</td><td>TCCTTCATTCCACCGGAGTCTGAAA</td><td>494.4</td><td>12.3</td><td>1.5</td><td>8.3</td><td>7.4E-17</td><td>5.6E-15</td><td>BT20/MB231</td><td>CYTO</td></tr>
<tr><td>>hg38_21_+_8433172_8446622_RNA45SN1_45S_50ntFlanks@5473.5512.40|CGGCCCTGGCGGAGCGCTGAGAAGACGGTCGAACTTGACT</td><td>>hg38_21_+_8433172_8446622_RNA45SN1_45S_50ntFlanks</td><td>5473</td><td>5512</td><td>40</td><td>CGGCCCTGGCGGAGCGCTGAGAAGACGGTCGAACTTGACT</td><td>2074.7</td><td>7.2</td><td>0.9</td><td>8.0</td><td>1.2E-15</td><td>5.6E-15</td><td>BT20/MB468</td><td>CYTO</td></tr>
<tr><td>>hg38_21_+_8433172_8446622_RNA45SN1_45S_50ntFlanks@12418.12455.38|CGATGTCGGCTCTTCCTATCATTGTGAAGCAGAATTCA</td><td>>hg38_21_+_8433172_8446622_RNA45SN1_45S_50ntFlanks</td><td>12418</td><td>12455</td><td>38</td><td>CGATGTCGGCTCTTCCTATCATTGTGAAGCAGAATTCA</td><td>11081.0</td><td>-6.2</td><td>0.8</td><td>-8.0</td><td>1.3E-15</td><td>5.6E-15</td><td>MB231/MB468</td><td>MITO</td></tr>
<tr><td>>hg38_21_+_8433172_8446622_RNA45SN1_45S_50ntFlanks@4003.4026.24|ATAACCTCGGGCCGATCGCACGCC</td><td>>hg38_21_+_8433172_8446622_RNA45SN1_45S_50ntFlanks</td><td>4003</td><td>4026</td><td>24</td><td>ATAACCTCGGGCCGATCGCACGCC</td><td>858.9</td><td>-11.8</td><td>1.5</td><td>-8.0</td><td>8.6E-16</td><td>5.6E-15</td><td>BT20/MB468</td><td>MITO</td></tr>
<tr><td>>hg38_21_+_8433172_8446622_RNA45SN1_45S_50ntFlanks@5473.5501.29|CGGCCCTGGCGGAGCGCTGAGAAGACGGT</td><td>>hg38_21_+_8433172_8446622_RNA45SN1_45S_50ntFlanks</td><td>5473</td><td>5501</td><td>29</td><td>CGGCCCTGGCGGAGCGCTGAGAAGACGGT</td><td>7495.8</td><td>8.9</td><td>1.1</td><td>8.0</td><td>1.2E-15</td><td>5.6E-15</td><td>BT20/MB468</td><td>CYTO</td></tr>
<tr><td>>hg38_21_+_8433172_8446622_RNA45SN1_45S_50ntFlanks@12405.12434.30|CTTTTTGATCCTTCGATGTCGGCTCTTCCT</td><td>>hg38_21_+_8433172_8446622_RNA45SN1_45S_50ntFlanks</td><td>12405</td><td>12434</td><td>30</td><td>CTTTTTGATCCTTCGATGTCGGCTCTTCCT</td><td>12723.4</td><td>-6.4</td><td>0.8</td><td>-8.0</td><td>1.2E-15</td><td>5.6E-15</td><td>BT20/MB468</td><td>CYTO</td></tr>
<tr><td>>hg38_21_+_8433172_8446622_RNA45SN1_45S_50ntFlanks@12940.12965.26|CATTCGTAGACGACCTGCTTCTGGGT</td><td>>hg38_21_+_8433172_8446622_RNA45SN1_45S_50ntFlanks</td><td>12940</td><td>12965</td><td>26</td><td>CATTCGTAGACGACCTGCTTCTGGGT</td><td>191.6</td><td>-11.4</td><td>1.4</td><td>-8.1</td><td>8.0E-16</td><td>5.6E-15</td><td>MB231/MB468</td><td>MP</td></tr>
<tr><td>>hg38_21_+_8433172_8446622_RNA45SN1_45S_50ntFlanks@475.506.32|CTTCGTGATCGATGTGGTGACGTCGTGCTCTC</td><td>>hg38_21_+_8433172_8446622_RNA45SN1_45S_50ntFlanks</td><td>475</td><td>506</td><td>32</td><td>CTTCGTGATCGATGTGGTGACGTCGTGCTCTC</td><td>7108.5</td><td>9.2</td><td>1.1</td><td>8.0</td><td>1.2E-15</td><td>5.6E-15</td><td>BT20/MB468</td><td>CYTO</td></tr>
<tr><td>>hg38_21_+_8433172_8446622_RNA45SN1_45S_50ntFlanks@12128.12146.19|ACCCGCTCCGGGGACAGTG</td><td>>hg38_21_+_8433172_8446622_RNA45SN1_45S_50ntFlanks</td><td>12128</td><td>12146</td><td>19</td><td>ACCCGCTCCGGGGACAGTG</td><td>247.0</td><td>-8.1</td><td>1.0</td><td>-8.0</td><td>8.7E-16</td><td>5.6E-15</td><td>MB231/MB468</td><td>CYTO</td></tr>
<tr><td>>hg38_21_+_8433172_8446622_RNA45SN1_45S_50ntFlanks@10351.10375.25|CGAGAACTTTGAAGGCCGAAGTGGA</td><td>>hg38_21_+_8433172_8446622_RNA45SN1_45S_50ntFlanks</td><td>10351</td><td>10375</td><td>25</td><td>CGAGAACTTTGAAGGCCGAAGTGGA</td><td>948.1</td><td>-13.8</td><td>1.7</td><td>-8.1</td><td>8.0E-16</td><td>5.7E-15</td><td>MB231/MB468</td><td>MP</td></tr>
<tr><td>>MI0000066|hsa-let-7e&WithFlank&19|+|51692780|51692870@14.35.22(+1C)&[MIMAT0000066&hsa-let-7e-5p&offsets|0|0(+1C);m-50&19|+|51692793|51692814&offsets|0|0(+1C)]|TGAGGTAGGAGGTTGTATAGTTC</td><td>>MI0000066|hsa-let-7e&WithFlank&19|+|51692780|51692870</td><td>14</td><td>35</td><td>22(+1C)</td><td>TGAGGTAGGAGGTTGTATAGTTC</td><td>285.1</td><td>10.1</td><td>1.3</td><td>8.0</td><td>1.2E-15</td><td>5.7E-15</td><td>BT20/MB468</td><td>CYTO</td></tr>
<tr><td>>hg38_21_+_8433172_8446622_RNA45SN1_45S_50ntFlanks@11190.11208.19|CCCGGGGGGCCGGCGGCGG</td><td>>hg38_21_+_8433172_8446622_RNA45SN1_45S_50ntFlanks</td><td>11190</td><td>11208</td><td>19</td><td>CCCGGGGGGCCGGCGGCGG</td><td>482.7</td><td>-12.8</td><td>1.6</td><td>-8.1</td><td>8.1E-16</td><td>5.7E-15</td><td>MB231/MB468</td><td>MP</td></tr>
<tr><td>>hg38_21_+_8433172_8446622_RNA45SN1_45S_50ntFlanks@12605.12623.19|TATGTGCTTGGCTGAGGAG</td><td>>hg38_21_+_8433172_8446622_RNA45SN1_45S_50ntFlanks</td><td>12605</td><td>12623</td><td>19</td><td>TATGTGCTTGGCTGAGGAG</td><td>261.7</td><td>-11.8</td><td>1.5</td><td>-8.0</td><td>1.3E-15</td><td>5.7E-15</td><td>MB231/MB468</td><td>MITO</td></tr>
<tr><td>>hg38_21_+_8433172_8446622_RNA45SN1_45S_50ntFlanks@10362.10407.46|AAGGCCGAAGTGGAGAAGGGTTCCATGTGAACAGCAGTTGAACATG</td><td>>hg38_21_+_8433172_8446622_RNA45SN1_45S_50ntFlanks</td><td>10362</td><td>10407</td><td>46</td><td>AAGGCCGAAGTGGAGAAGGGTTCCATGTGAACAGCAGTTGAACATG</td><td>623.2</td><td>-13.7</td><td>1.7</td><td>-7.9</td><td>1.9E-15</td><td>5.7E-15</td><td>BT20/MB468</td><td>NUC</td></tr>
<tr><td>>hg38_21_+_8433172_8446622_RNA45SN1_45S_50ntFlanks@12451.12477.27|ATTCACCAAGCGTTGGATTGTTCACCC</td><td>>hg38_21_+_8433172_8446622_RNA45SN1_45S_50ntFlanks</td><td>12451</td><td>12477</td><td>27</td><td>ATTCACCAAGCGTTGGATTGTTCACCC</td><td>353.4</td><td>5.6</td><td>0.7</td><td>8.0</td><td>8.9E-16</td><td>5.7E-15</td><td>MB231/MB468</td><td>CYTO</td></tr>
<tr><td>>trna119_LysCTT_1_-_145395522_145395594@1.33.33,trna11_LysCTT_5_-_180648979_180649051@1.33.33,trna13_LysCTT_6_+_26556774_26556846@1.33.33,trna7_LysCTT_16_+_3225692_3225764@1.33.33,trna9_LysCTT_5_+_180634755_180634827@1.33.33|GCCCGGCTAGCTCAGTCGGTAGAGCATGAGACT</td><td>>trna119_LysCTT_1_-_145395522_145395594</td><td>1</td><td>33</td><td>33</td><td>GCCCGGCTAGCTCAGTCGGTAGAGCATGAGACT</td><td>57401.5</td><td>8.5</td><td>1.1</td><td>8.0</td><td>8.9E-16</td><td>5.7E-15</td><td>MB231/MB468</td><td>CYTO</td></tr>
<tr><td>>hg38_21_+_8433172_8446622_RNA45SN1_45S_50ntFlanks@5444.5465.22|CTCGGATCGGCCCCGCCGGGGT</td><td>>hg38_21_+_8433172_8446622_RNA45SN1_45S_50ntFlanks</td><td>5444</td><td>5465</td><td>22</td><td>CTCGGATCGGCCCCGCCGGGGT</td><td>206.0</td><td>-11.5</td><td>1.4</td><td>-8.1</td><td>8.2E-16</td><td>5.7E-15</td><td>MB231/MB468</td><td>MP</td></tr>
<tr><td>>hg38_21_+_8433172_8446622_RNA45SN1_45S_50ntFlanks@7991.8009.19|ACGTGGCGACCCGCTGAAT</td><td>>hg38_21_+_8433172_8446622_RNA45SN1_45S_50ntFlanks</td><td>7991</td><td>8009</td><td>19</td><td>ACGTGGCGACCCGCTGAAT</td><td>13040.6</td><td>-6.2</td><td>0.8</td><td>-8.1</td><td>8.2E-16</td><td>5.7E-15</td><td>MB231/MB468</td><td>MP</td></tr>
<tr><td>>hg38_21_+_8433172_8446622_RNA45SN1_45S_50ntFlanks@10466.10495.30|CGTTGCCCTCGGCCGATCGAAAGGGAGTCG</td><td>>hg38_21_+_8433172_8446622_RNA45SN1_45S_50ntFlanks</td><td>10466</td><td>10495</td><td>30</td><td>CGTTGCCCTCGGCCGATCGAAAGGGAGTCG</td><td>224.5</td><td>-11.7</td><td>1.4</td><td>-8.1</td><td>8.2E-16</td><td>5.7E-15</td><td>MB231/MB468</td><td>MP</td></tr>
<tr><td>>hg38_21_+_8433172_8446622_RNA45SN1_45S_50ntFlanks@9564.9592.29|CTGACGTGCAAATCGGTCGTCCGACCTGG</td><td>>hg38_21_+_8433172_8446622_RNA45SN1_45S_50ntFlanks</td><td>9564</td><td>9592</td><td>29</td><td>CTGACGTGCAAATCGGTCGTCCGACCTGG</td><td>2935.5</td><td>-5.1</td><td>0.6</td><td>-8.0</td><td>8.9E-16</td><td>5.8E-15</td><td>MB231/MB468</td><td>CYTO</td></tr>
<tr><td>>hg38_21_+_8433172_8446622_RNA45SN1_45S_50ntFlanks@12415.12437.23|CTTCGATGTCGGCTCTTCCTATC</td><td>>hg38_21_+_8433172_8446622_RNA45SN1_45S_50ntFlanks</td><td>12415</td><td>12437</td><td>23</td><td>CTTCGATGTCGGCTCTTCCTATC</td><td>6694.7</td><td>-5.4</td><td>0.7</td><td>-7.9</td><td>1.9E-15</td><td>5.8E-15</td><td>BT20/MB468</td><td>NUC</td></tr>
<tr><td>>MI0000477|hsa-mir-146a&WithFlank&5|+|160485346|160485456@27.50.24&[MIMAT0000449&hsa-miR-146a-5p&offsets|0|+2;m-129&5|+|160485372|160485393&offsets|0|+2]|TGAGAACTGAATTCCATGGGTTGT</td><td>>MI0000477|hsa-mir-146a&WithFlank&5|+|160485346|160485456</td><td>27</td><td>50</td><td>24</td><td>TGAGAACTGAATTCCATGGGTTGT</td><td>2092.1</td><td>-14.4</td><td>1.7</td><td>-8.5</td><td>1.3E-17</td><td>5.8E-15</td><td>BT20/MB231</td><td>MP</td></tr>
<tr><td>>hg38_21_+_8433172_8446622_RNA45SN1_45S_50ntFlanks@10731.10757.27|AGGGTGTAAATCTCGCGCCGGGCCGTA</td><td>>hg38_21_+_8433172_8446622_RNA45SN1_45S_50ntFlanks</td><td>10731</td><td>10757</td><td>27</td><td>AGGGTGTAAATCTCGCGCCGGGCCGTA</td><td>192.3</td><td>-11.5</td><td>1.4</td><td>-8.1</td><td>8.3E-16</td><td>5.8E-15</td><td>MB231/MB468</td><td>MP</td></tr>
<tr><td>>MI0003776|hsa-mir-320b-1&WithFlank&1|+|116671740|116671823@48.68.21;MI0003839|hsa-mir-320b-2&WithFlank&1|-|224257034|224257116@47.67.21&[MIMAT0005792&hsa-miR-320b&offsets|0|-1;m-183&1|+|116671787|116671806&offsets|0|+1];[MIMAT0005792_1&hsa-miR-320b&offsets|0|-1;m-183&1|-|224257051|224257070&offsets|0|+1]|AAAAGCTGGGTTGAGAGGGCA</td><td>>MI0003776|hsa-mir-320b-1&WithFlank&1|+|116671740|116671823</td><td>48</td><td>68</td><td>21</td><td>AAAAGCTGGGTTGAGAGGGCA</td><td>971.4</td><td>11.2</td><td>1.4</td><td>8.0</td><td>8.8E-16</td><td>5.8E-15</td><td>BT20/MB468</td><td>MITO</td></tr>
<tr><td>>MI0000067|hsa-let-7f-1&WithFlank&9|+|94176341|94176439@13.34.22(+1C);MI0000068|hsa-let-7f-2&WithFlank&X|-|53557186|53557280@14.35.22(+1C)&[MIMAT0000067&hsa-let-7f-5p&offsets|0|0(+1C);m-20&9|+|94176353|94176374&offsets|0|0(+1C)];[MIMAT0000067_1&hsa-let-7f-5p&offsets|0|0(+1C);m-19&X|-|53557246|53557267&offsets|0|0(+1C)]|TGAGGTAGTAGATTGTATAGTTC</td><td>>MI0000067|hsa-let-7f-1&WithFlank&9|+|94176341|94176439</td><td>13</td><td>34</td><td>22(+1C)</td><td>TGAGGTAGTAGATTGTATAGTTC</td><td>530.9</td><td>6.4</td><td>0.8</td><td>8.0</td><td>1.2E-15</td><td>5.8E-15</td><td>BT20/MB468</td><td>CYTO</td></tr>
<tr><td>>MI0000286|hsa-mir-210&WithFlank&11|-|568083|568204@72.93.22(+2A)&[MIMAT0000267&hsa-miR-210-3p&offsets|0|0(+2A);m-102&11|-|568112|568133&offsets|0|0(+2A)]|CTGTGCGTGTGACAGCGGCTGAAA</td><td>>MI0000286|hsa-mir-210&WithFlank&11|-|568083|568204</td><td>72</td><td>93</td><td>22(+2A)</td><td>CTGTGCGTGTGACAGCGGCTGAAA</td><td>478.7</td><td>6.4</td><td>0.8</td><td>8.0</td><td>1.0E-15</td><td>5.8E-15</td><td>MB231/MB468</td><td>NUC</td></tr>
<tr><td>>trna152_ValCAC_6_-_27248049_27248121@1.32.32|GCTTCTGTAGTGTAGTGGTTATCACGTTCGCC</td><td>>trna152_ValCAC_6_-_27248049_27248121</td><td>1</td><td>32</td><td>32</td><td>GCTTCTGTAGTGTAGTGGTTATCACGTTCGCC</td><td>1697.6</td><td>7.5</td><td>0.9</td><td>8.0</td><td>1.4E-15</td><td>5.8E-15</td><td>MB231/MB468</td><td>MITO</td></tr>
<tr><td>>hg38_21_+_8433172_8446622_RNA45SN1_45S_50ntFlanks@11322.11342.21|ACGTCTCGTCGCGCGCGCGTC</td><td>>hg38_21_+_8433172_8446622_RNA45SN1_45S_50ntFlanks</td><td>11322</td><td>11342</td><td>21</td><td>ACGTCTCGTCGCGCGCGCGTC</td><td>2425.1</td><td>-7.0</td><td>0.9</td><td>-8.0</td><td>1.4E-15</td><td>5.8E-15</td><td>MB231/MB468</td><td>MITO</td></tr>
<tr><td>>hg38_21_+_8433172_8446622_RNA45SN1_45S_50ntFlanks@6745.6766.22|ACACTTCGAACGCACTTGCGGC</td><td>>hg38_21_+_8433172_8446622_RNA45SN1_45S_50ntFlanks</td><td>6745</td><td>6766</td><td>22</td><td>ACACTTCGAACGCACTTGCGGC</td><td>317.3</td><td>-12.2</td><td>1.5</td><td>-8.0</td><td>8.3E-16</td><td>5.8E-15</td><td>MB231/MB468</td><td>MP</td></tr>
<tr><td>>MI0000081|hsa-mir-24-2&WithFlank&19|-|13836281|13836365@56.76.21;MI0000080|hsa-mir-24-1&WithFlank&9|+|95086015|95086094@50.70.21&[MIMAT0000080&hsa-miR-24-3p&offsets|0|-1;m-36&19|-|13836288|13836310&offsets|0|-2];[MIMAT0000080_1&hsa-miR-24-3p&offsets|0|-1;m-35&9|+|95086064|95086086&offsets|0|-2]|TGGCTCAGTTCAGCAGGAACA</td><td>>MI0000081|hsa-mir-24-2&WithFlank&19|-|13836281|13836365</td><td>56</td><td>76</td><td>21</td><td>TGGCTCAGTTCAGCAGGAACA</td><td>3035.7</td><td>5.3</td><td>0.7</td><td>8.0</td><td>1.2E-15</td><td>5.8E-15</td><td>BT20/MB468</td><td>CYTO</td></tr>
<tr><td>>trnaMT_PheGAA_MT_+_577_647@15.35.21|CCTCCTCAAAGCAATACACTG</td><td>>trnaMT_PheGAA_MT_+_577_647</td><td>15</td><td>35</td><td>21</td><td>CCTCCTCAAAGCAATACACTG</td><td>488.0</td><td>-13.4</td><td>1.7</td><td>-7.9</td><td>1.9E-15</td><td>5.8E-15</td><td>BT20/MB468</td><td>NUC</td></tr>
</table>

| | | | | | | | | | | | | | |
|---|---|---|---|---|---|---|---|---|---|---|---|---|---|
| >trna116_GluCTC_1_-_145399233_145399304@32.71.40,trna71_GluCTC_1_-_161439189_161439260@32.71.40,trna74_GluCTC_1_-_161431809_161431880@32.71.40,trna77_GluCTC_1_-_161424398_161424469@32.71.40,trna77_GluCTC_6_+_28949976_28950047@32.71.40,trna80_GluCTC_1_-_161417018_161417089@32.71.40,trna87_GluCTC_6_-_126101393_126101464@32.71.40\|CTCTCACCGCCGCGGCCCGGGTTCGATTCCCGGTCAGGGA | >trna116_GluCTC_1_-_145399233_145399304 | 32 | 71 | 40 | CTCTCACCGCCGCGGCCCGGGTTCGATTCCCGGTCAGGGA | 712.9 | -13.3 | 1.7 | -8.0 | 1.4E-15 | 5.8E-15 | MB231/MB468 | MITO |
| >MI0000061\|hsa-let-7a-2&WithFlank&11\|-\|122146516\|122146599@11.31.21(+1G);MI0000062\|hsa-let-7a-3&WithFlank&22\|+\|46112743\|46112828@10.30.21(+1G);MI0000060\|hsa-let-7a-1&WithFlank&9\|+\|94175951\|94176042@12.32.21(+1G)&[MIMAT0000062&hsa-let-7a-5p&offsets\|0\|-1(+1G);m-6&11\|-\|122146568\|122146589&offsets\|0\|-1(+1G)];[MIMAT0000062_1&hsa-let-7a-5p&offsets\|0\|-1(+1G);m-5&22\|+\|46112752\|46112773&offsets\|0\|-1(+1G)];[MIMAT0000062_2&hsa-let-7a-5p&offsets\|0\|-1(+1G);m-4&9\|+\|94175962\|94175983&offsets\|0\|-1(+1G)]\|TGAGGTAGTAGGTTGTATAGTG | >MI0000061\|hsa-let-7a-2&WithFlank&11\|-\|122146516\|122146599 | 11 | 31 | 21(+1G) | TGAGGTAGTAGGTTGTATAGTG | 319.1 | 5.9 | 0.7 | 8.0 | 8.9E-16 | 5.8E-15 | BT20/MB468 | MITO |
| >am_trna128_GlyGCC_6_-_27870686_27870756@1.32.32,trna133_GlyCCC_1_-_16872434_16872504@1.32.32,trna18_GlyGCC_16_+_70822597_70822667@1.32.32,trna19_GlyGCC_16_+_70823410_70823480@1.32.32,trna19_GlyGCC_2_-_157257659_157257729@1.32.32,trna24_GlyGCC_16_-_70812942_70813012@1.32.32,trna25_GlyGCC_16_-_70812114_70812184@1.32.32,trna4_GlyCCC_1_+_17188416_17188486@1.32.32,trna5_GlyGCC_17_+_8029064_8029134@1.32.32,trna68_GlyGCC_1_-_161493637_161493707@1.32.32\|GCATTGGTGGTTCAGTGGTAGAATTCTCGCCT | >am_trna128_GlyGCC_6_-_27870686_27870756 | 1 | 32 | 32 | GCATTGGTGGTTCAGTGGTAGAATTCTCGCCT | 20350.9 | 9.1 | 1.1 | 8.0 | 9.0E-16 | 5.9E-15 | BT20/MB468 | MITO |
| >hg38_21_+_8433172_8446622_RNA45SN1_45S_50ntFlanks@475.508.34\|CTTCGTGATCGATGTGGTGACGTCGTGCTCTCCC | >hg38_21_+_8433172_8446622_RNA45SN1_45S_50ntFlanks | 475 | 508 | 34 | CTTCGTGATCGATGTGGTGACGTCGTGCTCTCCC | 1284.7 | 7.4 | 0.9 | 8.0 | 9.0E-16 | 5.9E-15 | BT20/MB468 | MITO |
| >MI0000439\|hsa-mir-23b&WithFlank&9\|+\|95085202\|95085310@67.84.18&[MIMAT0000418&hsa-miR-23b-3p&offsets\|+3\|-2;m-42&9\|+\|95085265\|95085285&offsets\|+3\|0]\|ACATTGCCAGGGATTACC | >MI0000439\|hsa-mir-23b&WithFlank&9\|+\|95085202\|95085310 | 67 | 84 | 18 | ACATTGCCAGGGATTACC | 2118.3 | 29.2 | 3.6 | 8.0 | 9.2E-16 | 5.9E-15 | MB231/MB468 | CYTO |
| >MI0000750\|hsa-mir-26a-2&WithFlank&12\|-\|57824603\|57824698@20.39.20(+1U);MI0000083\|hsa-mir-26a-1&WithFlank&3\|+\|37969398\|37969486@16.35.20(+1U)&[MIMAT0000082&hsa-miR-26a-5p&offsets\|0\|-2(+1U);m-33&12\|-\|57824658\|57824679&offsets\|0\|-2(+1U)];[MIMAT0000082_1&hsa-miR-26a-5p&offsets\|0\|-2(+1U);m-34&3\|+\|37969413\|37969434&offsets\|0\|-2(+1U)]\|TTCAAGTAATCCAGGATAGGT | >MI0000750\|hsa-mir-26a-2&WithFlank&12\|-\|57824603\|57824698 | 20 | 39 | 20(+1U) | TTCAAGTAATCCAGGATAGGT | 291.9 | 5.6 | 0.7 | 8.0 | 1.0E-15 | 5.9E-15 | MB231/MB468 | NUC |
| >hg38_21_+_8433172_8446622_RNA45SN1_45S_50ntFlanks@11115.11132.18\|CGGCAGGGGCCGGCGGCG | >hg38_21_+_8433172_8446622_RNA45SN1_45S_50ntFlanks | 11115 | 11132 | 18 | CGGCAGGGGCCGGCGGCG | 450.9 | -11.8 | 1.5 | -8.0 | 1.0E-15 | 5.9E-15 | MB231/MB468 | NUC |
| >hg38_21_+_8433172_8446622_RNA45SN1_45S_50ntFlanks@10873.10894.22\|CTGGGTCGGTCGGGCTGGGGCG | >hg38_21_+_8433172_8446622_RNA45SN1_45S_50ntFlanks | 10873 | 10894 | 22 | CTGGGTCGGTCGGGCTGGGGCG | 412.8 | -6.1 | 0.8 | -8.0 | 9.2E-16 | 5.9E-15 | MB231/MB468 | CYTO |
| >hg38_21_+_8433172_8446622_RNA45SN1_45S_50ntFlanks@11452.11472.21\|CTCCTCGGCGCGCGGCGGCGG | >hg38_21_+_8433172_8446622_RNA45SN1_45S_50ntFlanks | 11452 | 11472 | 21 | CTCCTCGGCGCGCGGCGGCGG | 1232.6 | -13.2 | 1.6 | -8.0 | 1.0E-15 | 5.9E-15 | MB231/MB468 | NUC |
| >hg38_21_+_8433172_8446622_RNA45SN1_45S_50ntFlanks@3979.4001.23\|CGCCGGCGGCTTTGGTGACTCTA | >hg38_21_+_8433172_8446622_RNA45SN1_45S_50ntFlanks | 3979 | 4001 | 23 | CGCCGGCGGCTTTGGTGACTCTA | 728.9 | -13.3 | 1.7 | -8.0 | 1.4E-15 | 5.9E-15 | MB231/MB468 | MITO |
| >MI0003628\|hsa-mir-615&WithFlank&12\|+\|54033944\|54034051@67.87.21(+1C)&[MIMAT0003283&hsa-miR-615-3p&offsets\|0\|-1(+1C);m-495&12\|+\|54034010\|54034030&offsets\|0\|0(+1C)]\|TCCGAGCCTGGGTCTCCCTCTC | >MI0003628\|hsa-mir-615&WithFlank&12\|+\|54033944\|54034051 | 67 | 87 | 21(+1C) | TCCGAGCCTGGGTCTCCCTCTC | 246.3 | 11.6 | 1.4 | 8.3 | 1.1E-16 | 5.9E-15 | BT20/MB231 | MITO |
| >am_trna2_GlyGCC_21_-_18827107_18827177@1.26.26,trna35_GlyGCC_1_+_161413094_161413164@1.26.26,trna37_GlyGCC_1_+_161420467_161420537@1.26.26,trna39_GlyGCC_1_+_161427898_161427968@1.26.26,trna41_GlyGCC_1_+_161435258_161435328@1.26.26\|GCATGGGTGGTTCAGTGGTAGAATTC | >am_trna2_GlyGCC_21_-_18827107_18827177 | 1 | 26 | 26 | GCATGGGTGGTTCAGTGGTAGAATTC | 446.4 | 12.4 | 1.5 | 8.3 | 1.1E-16 | 5.9E-15 | BT20/MB231 | MITO |
| >hg38_21_+_8433172_8446622_RNA45SN1_45S_50ntFlanks@12956.12982.27\|GCTTCTGGGTCGGGGTTTCGTACGTAG | >hg38_21_+_8433172_8446622_RNA45SN1_45S_50ntFlanks | 12956 | 12982 | 27 | GCTTCTGGGTCGGGGTTTCGTACGTAG | 427.9 | -11.7 | 1.5 | -8.0 | 1.0E-15 | 5.9E-15 | MB231/MB468 | NUC |
| >hg38_21_+_8433172_8446622_RNA45SN1_45S_50ntFlanks@10883.10902.20\|CGGGCTGGGGCGCGAAGCGG | >hg38_21_+_8433172_8446622_RNA45SN1_45S_50ntFlanks | 10883 | 10902 | 20 | CGGGCTGGGGCGCGAAGCGG | 242.6 | -11.8 | 1.5 | -8.0 | 8.5E-16 | 5.9E-15 | MB231/MB468 | MP |
| >hg38_1_-_228634819_228635039_RNA5S12_5S_50ntFlanks@146.169.24\|TGGGAATACCGGGTGCTGTAGGCT | >hg38_1_-_228634819_228635039_RNA5S12_5S_50ntFlanks | 146 | 169 | 24 | TGGGAATACCGGGTGCTGTAGGCT | 515.3 | 5.3 | 0.7 | 8.0 | 9.1E-16 | 5.9E-15 | BT20/MB468 | MITO |
| >hg38_21_+_8433172_8446622_RNA45SN1_45S_50ntFlanks@4241.4258.18\|CTCTTTCGAGGCCCTGTA | >hg38_21_+_8433172_8446622_RNA45SN1_45S_50ntFlanks | 4241 | 4258 | 18 | CTCTTTCGAGGCCCTGTA | 1616.5 | -5.7 | 0.7 | -8.0 | 9.3E-16 | 6.0E-15 | MB231/MB468 | CYTO |
| >hg38_21_+_8433172_8446622_RNA45SN1_45S_50ntFlanks@8472.8497.26\|CGGCCCGGCGGATCTTTCCCGCCCCC | >hg38_21_+_8433172_8446622_RNA45SN1_45S_50ntFlanks | 8472 | 8497 | 26 | CGGCCCGGCGGATCTTTCCCGCCCCC | 396.6 | -12.5 | 1.6 | -8.0 | 8.5E-16 | 6.0E-15 | MB231/MB468 | MP |
| >trna111_HisGTG_1_-_147774845_147774916@1.31.31,trna118_HisGTG_1_-_145396881_145396952@1.31.31,trna16_HisGTG_1_+_146544773_146544844@1.31.31,trna1_HisGTG_15_+_45493349_45493420@1.31.31,trna21_HisGTG_1_+_147753471_147753542@1.31.31,trna33_HisGTG_6_+_27125906_27125977@1.31.31,trna7_HisGTG_9_-_14433938_14434009@1.31.31,trna8_HisGTG_15_-_45492611_45492682@1.31.31,trna9_HisGTG_15_-_45490804_45490875@1.31.31\|GCCGTGATCGTATAGTGGTTAGTACTCTGCG | >trna111_HisGTG_1_-_147774845_147774916 | 1 | 31 | 31 | GCCGTGATCGTATAGTGGTTAGTACTCTGCG | 28084.6 | -7.4 | 0.9 | -7.9 | 2.0E-15 | 6.0E-15 | BT20/MB468 | NUC |
| >hg38_21_+_8433172_8446622_RNA45SN1_45S_50ntFlanks@11847.11873.27\|AACGGGCTTGGCGGAATCAGCGGGGAA | >hg38_21_+_8433172_8446622_RNA45SN1_45S_50ntFlanks | 11847 | 11873 | 27 | AACGGGCTTGGCGGAATCAGCGGGGAA | 265.6 | -11.8 | 1.5 | -8.0 | 1.4E-15 | 6.0E-15 | MB231/MB468 | MITO |
| >MI0005756\|hsa-mir-934&WithFlank&X\|+\|136550872\|136550966@21.42.22&[MIMAT0004977&hsa-miR-934&offsets\|0\|0;m-943&X\|+\|136550892\|136550913&offsets\|0\|0]\|TGTCTACTACTGGAGACACTGG | >MI0005756\|hsa-mir-934&WithFlank&X\|+\|136550872\|136550966 | 21 | 42 | 22 | TGTCTACTACTGGAGACACTGG | 1406.4 | -14.3 | 1.8 | -8.0 | 8.6E-16 | 6.0E-15 | MB231/MB468 | MP |
| >hg38_21_+_8433172_8446622_RNA45SN1_45S_50ntFlanks@10351.10384.34\|CGAGAACTTTGAAGGCCGAAGTGGAGAAGGGTTC | >hg38_21_+_8433172_8446622_RNA45SN1_45S_50ntFlanks | 10351 | 10384 | 34 | CGAGAACTTTGAAGGCCGAAGTGGAGAAGGGTTC | 4388.2 | -8.8 | 1.1 | -8.0 | 1.4E-15 | 6.0E-15 | MB231/MB468 | MITO |
| >hg38_21_+_8433172_8446622_RNA45SN1_45S_50ntFlanks@9720.9748.29\|ATTAGAGGTCTTGGGGCCGAAACGATCTC | >hg38_21_+_8433172_8446622_RNA45SN1_45S_50ntFlanks | 9720 | 9748 | 29 | ATTAGAGGTCTTGGGGCCGAAACGATCTC | 303.8 | -12.0 | 1.5 | -8.0 | 1.4E-15 | 6.0E-15 | MB231/MB468 | MITO |
| >trna10_AlaCGC_6_+_26553731_26553802@54.75.22,trna119_AlaCGC_6_-_28641613_28641684@54.75.22,trna13_AlaTGC_12_-_125406301_125406372@54.75.22,trna66_AlaTGC_6_+_28611222_28611293@54.75.22,trna8_AlaTGC_5_+_180633868_180633939@54.75.22\|TCGATCCCCGGCATCTCCACCA | >trna10_AlaCGC_6_+_26553731_26553802 | 54 | 75 | 22 | TCGATCCCCGGCATCTCCACCA | 772.3 | 11.0 | 1.4 | 8.0 | 1.3E-15 | 6.0E-15 | BT20/MB468 | CYTO |
| >hg38_21_+_8433172_8446622_RNA45SN1_45S_50ntFlanks@485.510.26\|GATGTGGTGACGTCGTGCTCTCCCGG | >hg38_21_+_8433172_8446622_RNA45SN1_45S_50ntFlanks | 485 | 510 | 26 | GATGTGGTGACGTCGTGCTCTCCCGG | 3363.4 | 29.1 | 3.6 | 8.0 | 1.1E-15 | 6.0E-15 | MB231/MB468 | NUC |
| >hg38_21_+_8433172_8446622_RNA45SN1_45S_50ntFlanks@3899.3921.23\|CCTTCGCGGGGGGGATGCGTGCA | >hg38_21_+_8433172_8446622_RNA45SN1_45S_50ntFlanks | 3899 | 3921 | 23 | CCTTCGCGGGGGGGATGCGTGCA | 334.4 | -12.2 | 1.5 | -8.0 | 8.6E-16 | 6.0E-15 | MB231/MB468 | MP |
| >hg38_21_+_8433172_8446622_RNA45SN1_45S_50ntFlanks@11792.11820.29\|ATGAACGAGATTCCCACTGTCCCTACCTA | >hg38_21_+_8433172_8446622_RNA45SN1_45S_50ntFlanks | 11792 | 11820 | 29 | ATGAACGAGATTCCCACTGTCCCTACCTA | 306.5 | -12.1 | 1.5 | -8.0 | 8.7E-16 | 6.0E-15 | MB231/MB468 | MP |
| >hg38_21_+_8433172_8446622_RNA45SN1_45S_50ntFlanks@9566.9592.27\|GACGTGCAAATCGGTCGTCCGACCTGG | >hg38_21_+_8433172_8446622_RNA45SN1_45S_50ntFlanks | 9566 | 9592 | 27 | GACGTGCAAATCGGTCGTCCGACCTGG | 198.9 | -11.5 | 1.4 | -8.0 | 8.7E-16 | 6.1E-15 | MB231/MB468 | MP |
| >hg38_21_+_8433172_8446622_RNA45SN1_45S_50ntFlanks@9738.9755.18\|GAAACGATCTCAACCTAT | >hg38_21_+_8433172_8446622_RNA45SN1_45S_50ntFlanks | 9738 | 9755 | 18 | GAAACGATCTCAACCTAT | 866.2 | -8.1 | 1.0 | -8.0 | 9.4E-16 | 6.1E-15 | MB231/MB468 | CYTO |
| >hg38_21_+_8433172_8446622_RNA45SN1_45S_50ntFlanks@6732.6770.39\|ACATTGATCATCGACACTTCGAACGCACTTGCGGCCCCG | >hg38_21_+_8433172_8446622_RNA45SN1_45S_50ntFlanks | 6732 | 6770 | 39 | ACATTGATCATCGACACTTCGAACGCACTTGCGGCCCCG | 3507.5 | -5.9 | 0.7 | -8.0 | 1.4E-15 | 6.1E-15 | MB231/MB468 | MITO |
| >hg38_21_+_8433172_8446622_RNA45SN1_45S_50ntFlanks@11641.11671.31\|CGGCGGGTGTTGACGCGATGTGATTTCTGCC | >hg38_21_+_8433172_8446622_RNA45SN1_45S_50ntFlanks | 11641 | 11671 | 31 | CGGCGGGTGTTGACGCGATGTGATTTCTGCC | 394.8 | -12.5 | 1.6 | -8.0 | 8.7E-16 | 6.1E-15 | MB231/MB468 | MP |
| >MI0000266\|hsa-mir-10a&WithFlank&17\|-\|48579832\|48579953@28.51.24&[MIMAT0000253&hsa-miR-10a-5p&offsets\|0\|+1;m-18&17\|-\|48579905\|48579926&offsets\|0\|+2]\|TACCCTGTAGATCCGAATTTGTGT | >MI0000266\|hsa-mir-10a&WithFlank&17\|-\|48579832\|48579953 | 28 | 51 | 24 | TACCCTGTAGATCCGAATTTGTGT | 710.9 | 12.7 | 1.6 | 8.0 | 1.4E-15 | 6.1E-15 | MB231/MB468 | MITO |
| >hg38_21_+_8433172_8446622_RNA45SN1_45S_50ntFlanks@10768.10785.18\|CAGCAGGTCTCCAAGGTG | >hg38_21_+_8433172_8446622_RNA45SN1_45S_50ntFlanks | 10768 | 10785 | 18 | CAGCAGGTCTCCAAGGTG | 456.5 | -10.9 | 1.4 | -8.0 | 9.4E-16 | 6.1E-15 | BT20/MB468 | MITO |
| >hg38_21_+_8433172_8446622_RNA45SN1_45S_50ntFlanks@9240.9272.33\|GCGGAGCGAGCGCACGGGGTCGGCGGCGACGTC | >hg38_21_+_8433172_8446622_RNA45SN1_45S_50ntFlanks | 9240 | 9272 | 33 | GCGGAGCGAGCGCACGGGGTCGGCGGCGACGTC | 208.0 | -11.6 | 1.4 | -8.0 | 8.8E-16 | 6.2E-15 | MB231/MB468 | MP |
| >hg38_21_+_8433172_8446622_RNA45SN1_45S_50ntFlanks@11781.11803.23\|CGCATGAATGGATGAACGAGATT | >hg38_21_+_8433172_8446622_RNA45SN1_45S_50ntFlanks | 11781 | 11803 | 23 | CGCATGAATGGATGAACGAGATT | 6371.1 | -12.0 | 1.5 | -7.9 | 2.0E-15 | 6.2E-15 | BT20/MB468 | NUC |
| >hg38_21_+_8433172_8446622_RNA45SN1_45S_50ntFlanks@12931.12956.26\|GCGCTAAACCATTCGTAGACGACCTG | >hg38_21_+_8433172_8446622_RNA45SN1_45S_50ntFlanks | 12931 | 12956 | 26 | GCGCTAAACCATTCGTAGACGACCTG | 2333.9 | -8.9 | 1.1 | -7.9 | 2.0E-15 | 6.2E-15 | BT20/MB468 | NUC |
| >trna111_HisGTG_1_-_147774845_147774916@-1G.33.34,trna118_HisGTG_1_-_145396881_145396952@-1G.33.34,trna16_HisGTG_1_+_146544773_146544844@-1G.33.34,trna1_HisGTG_15_+_45493349_45493420@-1G.33.34,trna21_HisGTG_1_+_147753471_147753542@-1G.33.34,trna33_HisGTG_6_+_27125906_27125977@-1G.33.34,trna7_HisGTG_9_-_14433938_14434009@-1G.33.34,trna8_HisGTG_15_-_45492611_45492682@-1G.33.34,trna9_HisGTG_15_-_45490804_45490875@-1G.33.34\|GGCCGTGATCGTATAGTGGTTAGTACTCTGCGTT | >trna111_HisGTG_1_-_147774845_147774916 | -1G | 33 | 34 | GGCCGTGATCGTATAGTGGTTAGTACTCTGCGTT | 17095.1 | 8.3 | 1.0 | 8.0 | 1.5E-15 | 6.2E-15 | MB231/MB468 | MITO |

<table>
<tr><td>>hg38_21_+_8433172_8446622_RNA45SN1_45S_50ntFlanks@9736.9769.34|CCGAAACGATCTCAACCTATTCTCAAACTTTAAA</td><td>>hg38_21_+_8433172_8446622_RNA45SN1_45S_50ntFlanks</td><td>9736</td><td>9769</td><td>34</td><td>CCGAAACGATCTCAACCTATTCTCAAACTTTAAA</td><td>256.6</td><td>-11.8</td><td>1.5</td><td>-8.0</td><td>1.5E-15</td><td>6.2E-15</td><td>MB231/MB468</td><td>MITO</td></tr>
<tr><td>>trna10_LysCTT_16_+_3241501_3241573@3.35.33,trna13_LysCTT_14_-_58706613_58706685@3.35.33,trna2_LysCTT_15_+_79152904_79152976@3.35.33|CCGGCTAGCTCAGTCGGTAGAGCATGGGACTCT</td><td>>trna10_LysCTT_16_+_3241501_3241573</td><td>3</td><td>35</td><td>33</td><td>CCGGCTAGCTCAGTCGGTAGAGCATGGGACTCT</td><td>654.0</td><td>11.0</td><td>1.4</td><td>8.0</td><td>9.6E-16</td><td>6.2E-15</td><td>MB231/MB468</td><td>CYTO</td></tr>
<tr><td>>hg38_21_+_8433172_8446622_RNA45SN1_45S_50ntFlanks@9339.9371.33|CGCACGAAAGCCGCCGTGGCGCAATGAAGGTGA</td><td>>hg38_21_+_8433172_8446622_RNA45SN1_45S_50ntFlanks</td><td>9339</td><td>9371</td><td>33</td><td>CGCACGAAAGCCGCCGTGGCGCAATGAAGGTGA</td><td>494.5</td><td>-12.8</td><td>1.6</td><td>-8.0</td><td>8.8E-16</td><td>6.2E-15</td><td>MB231/MB468</td><td>MP</td></tr>
<tr><td>>hg38_21_+_8433172_8446622_RNA45SN1_45S_50ntFlanks@10369.10407.39|AAGTGGAGAAGGGTTCCATGTGAACAGCAGTTGAACATG</td><td>>hg38_21_+_8433172_8446622_RNA45SN1_45S_50ntFlanks</td><td>10369</td><td>10407</td><td>39</td><td>AAGTGGAGAAGGGTTCCATGTGAACAGCAGTTGAACATG</td><td>305.3</td><td>-12.0</td><td>1.5</td><td>-8.0</td><td>1.5E-15</td><td>6.2E-15</td><td>MB231/MB468</td><td>MITO</td></tr>
<tr><td>>hg38_21_+_8433172_8446622_RNA45SN1_45S_50ntFlanks@11104.11123.20|AAGGGTCGGGGCGGCAGGGG</td><td>>hg38_21_+_8433172_8446622_RNA45SN1_45S_50ntFlanks</td><td>11104</td><td>11123</td><td>20</td><td>AAGGGTCGGGGCGGCAGGGG</td><td>540.8</td><td>-8.8</td><td>1.1</td><td>-8.0</td><td>9.7E-16</td><td>6.2E-15</td><td>MB231/MB468</td><td>CYTO</td></tr>
<tr><td>>hg38_21_+_8433172_8446622_RNA45SN1_45S_50ntFlanks@4289.4317.29|CGAGGATCCATTGGAGGGCAAGTCTGGTG</td><td>>hg38_21_+_8433172_8446622_RNA45SN1_45S_50ntFlanks</td><td>4289</td><td>4317</td><td>29</td><td>CGAGGATCCATTGGAGGGCAAGTCTGGTG</td><td>381.3</td><td>-12.4</td><td>1.5</td><td>-8.0</td><td>8.9E-16</td><td>6.2E-15</td><td>MB231/MB468</td><td>MP</td></tr>
<tr><td>>hg38_21_+_8433172_8446622_RNA45SN1_45S_50ntFlanks@11586.11605.20|GTGCGGACCAGGGGAATCCG</td><td>>hg38_21_+_8433172_8446622_RNA45SN1_45S_50ntFlanks</td><td>11586</td><td>11605</td><td>20</td><td>GTGCGGACCAGGGGAATCCG</td><td>275.2</td><td>-11.1</td><td>1.4</td><td>-8.0</td><td>1.1E-15</td><td>6.2E-15</td><td>MB231/MB468</td><td>NUC</td></tr>
<tr><td>>hg38_21_+_8433172_8446622_RNA45SN1_45S_50ntFlanks@12456.12482.27|CCAAGCGTTGGATTGTTCACCCACTAA</td><td>>hg38_21_+_8433172_8446622_RNA45SN1_45S_50ntFlanks</td><td>12456</td><td>12482</td><td>27</td><td>CCAAGCGTTGGATTGTTCACCCACTAA</td><td>257.8</td><td>-11.8</td><td>1.5</td><td>-8.0</td><td>1.5E-15</td><td>6.2E-15</td><td>MB231/MB468</td><td>MITO</td></tr>
<tr><td>>hg38_21_+_8433172_8446622_RNA45SN1_45S_50ntFlanks@10992.11010.19|CGCGCGCCGCTCGCTCCCT</td><td>>hg38_21_+_8433172_8446622_RNA45SN1_45S_50ntFlanks</td><td>10992</td><td>11010</td><td>19</td><td>CGCGCGCCGCTCGCTCCCT</td><td>227.2</td><td>-10.5</td><td>1.3</td><td>-8.0</td><td>9.7E-16</td><td>6.2E-15</td><td>MB231/MB468</td><td>CYTO</td></tr>
<tr><td>>MI0000815|hsa-mir-339&WithFlank&7|-|1022927|1023032@21.43.23(+1A)&[MIMAT0000764&hsa-miR-339-5p&offsets|0|0(+1A);m-163&7|-|1022990|1023012&offsets|0|0(+1A)]|TCCCTGTCCTCCAGGAGCTCACGA</td><td>>MI0000815|hsa-mir-339&WithFlank&7|-|1022927|1023032</td><td>21</td><td>43</td><td>23(+1A)</td><td>TCCCTGTCCTCCAGGAGCTCACGA</td><td>395.4</td><td>8.8</td><td>1.1</td><td>8.0</td><td>1.3E-15</td><td>6.2E-15</td><td>BT20/MB468</td><td>CYTO</td></tr>
<tr><td>>hg38_21_+_8433172_8446622_RNA45SN1_45S_50ntFlanks@11561.11601.41|GCGCCTAGCAGCCGACTTAGAACTGGTGCGGACCAGGGGAA</td><td>>hg38_21_+_8433172_8446622_RNA45SN1_45S_50ntFlanks</td><td>11561</td><td>11601</td><td>41</td><td>GCGCCTAGCAGCCGACTTAGAACTGGTGCGGACCAGGGGAA</td><td>1848.2</td><td>-14.7</td><td>1.8</td><td>-8.0</td><td>9.0E-16</td><td>6.2E-15</td><td>MB231/MB468</td><td>MP</td></tr>
<tr><td>>hg38_21_+_8433172_8446622_RNA45SN1_45S_50ntFlanks@12156.12179.24|GAGTTTGACTGGGGCGGTACACCT</td><td>>hg38_21_+_8433172_8446622_RNA45SN1_45S_50ntFlanks</td><td>12156</td><td>12179</td><td>24</td><td>GAGTTTGACTGGGGCGGTACACCT</td><td>314.5</td><td>-12.2</td><td>1.5</td><td>-8.0</td><td>9.0E-16</td><td>6.3E-15</td><td>MB231/MB468</td><td>MP</td></tr>
<tr><td>>hg38_21_+_8433172_8446622_RNA45SN1_45S_50ntFlanks@8381.8401.21|AAGAGGGCGTGAAACCGTTAA</td><td>>hg38_21_+_8433172_8446622_RNA45SN1_45S_50ntFlanks</td><td>8381</td><td>8401</td><td>21</td><td>AAGAGGGCGTGAAACCGTTAA</td><td>370.1</td><td>-12.4</td><td>1.5</td><td>-8.0</td><td>9.0E-16</td><td>6.3E-15</td><td>MB231/MB468</td><td>MP</td></tr>
<tr><td>>hg38_21_+_8433172_8446622_RNA45SN1_45S_50ntFlanks@4000.4024.25|TAGATAACCTCGGGCCGATCGCACG</td><td>>hg38_21_+_8433172_8446622_RNA45SN1_45S_50ntFlanks</td><td>4000</td><td>4024</td><td>25</td><td>TAGATAACCTCGGGCCGATCGCACG</td><td>319.0</td><td>-12.2</td><td>1.5</td><td>-8.0</td><td>9.0E-16</td><td>6.3E-15</td><td>MB231/MB468</td><td>MP</td></tr>
<tr><td>>MI0000736|hsa-mir-30c-1&WithFlank&1|+|40757278|40757378@23.45.23(+1C);MI0000254|hsa-mir-30c-2&WithFlank&6|-|71376954|71377037@13.35.23(+1C)&[MIMAT0000244&hsa-miR-30c-5p&offsets|0|0(+1C);m-57&1|+|40757300|40757323&offsets|0|-1(+1C)];[MIMAT0000244_1&hsa-miR-30c-5p&offsets|0|0(+1C);m-56&6|-|71377002|71377025&offsets|0|-1(+1C)]|TGTAAACATCCTACACTCTCAGCC</td><td>>MI0000736|hsa-mir-30c-1&WithFlank&1|+|40757278|40757378</td><td>23</td><td>45</td><td>23(+1C)</td><td>TGTAAACATCCTACACTCTCAGCC</td><td>2304.9</td><td>5.6</td><td>0.7</td><td>8.0</td><td>1.3E-15</td><td>6.3E-15</td><td>BT20/MB468</td><td>CYTO</td></tr>
<tr><td>>hg38_21_+_8433172_8446622_RNA45SN1_45S_50ntFlanks@10347.10368.22|CAAACGAGAACTTTGAAGGCCG</td><td>>hg38_21_+_8433172_8446622_RNA45SN1_45S_50ntFlanks</td><td>10347</td><td>10368</td><td>22</td><td>CAAACGAGAACTTTGAAGGCCG</td><td>515.8</td><td>-3.2</td><td>0.4</td><td>-8.0</td><td>1.3E-15</td><td>6.3E-15</td><td>BT20/MB468</td><td>CYTO</td></tr>
<tr><td>>hg38_21_+_8433172_8446622_RNA45SN1_45S_50ntFlanks@10845.10868.24|ACTTCGGGATAAGGATTGGCTCTA</td><td>>hg38_21_+_8433172_8446622_RNA45SN1_45S_50ntFlanks</td><td>10845</td><td>10868</td><td>24</td><td>ACTTCGGGATAAGGATTGGCTCTA</td><td>3526.5</td><td>-11.6</td><td>1.5</td><td>-7.9</td><td>2.1E-15</td><td>6.3E-15</td><td>BT20/MB468</td><td>NUC</td></tr>
<tr><td>>hg38_21_+_8433172_8446622_RNA45SN1_45S_50ntFlanks@11217.11246.30|CTGGACGCGAGCCGGGCCCTTCCCGTGGAT</td><td>>hg38_21_+_8433172_8446622_RNA45SN1_45S_50ntFlanks</td><td>11217</td><td>11246</td><td>30</td><td>CTGGACGCGAGCCGGGCCCTTCCCGTGGAT</td><td>659.8</td><td>7.2</td><td>0.9</td><td>8.0</td><td>9.7E-16</td><td>6.3E-15</td><td>BT20/MB468</td><td>MITO</td></tr>
<tr><td>>hg38_21_+_8433172_8446622_RNA45SN1_45S_50ntFlanks@11211.11232.22|GCGACTCTGGACGCGAGCCGGG</td><td>>hg38_21_+_8433172_8446622_RNA45SN1_45S_50ntFlanks</td><td>11211</td><td>11232</td><td>22</td><td>GCGACTCTGGACGCGAGCCGGG</td><td>1314.5</td><td>-7.5</td><td>0.9</td><td>-8.0</td><td>9.9E-16</td><td>6.3E-15</td><td>MB231/MB468</td><td>CYTO</td></tr>
<tr><td>>hg38_21_+_8433172_8446622_RNA45SN1_45S_50ntFlanks@10361.10388.28|GAAGGCCGAAGTGGAGAAGGGTTCCATG</td><td>>hg38_21_+_8433172_8446622_RNA45SN1_45S_50ntFlanks</td><td>10361</td><td>10388</td><td>28</td><td>GAAGGCCGAAGTGGAGAAGGGTTCCATG</td><td>391.2</td><td>-12.4</td><td>1.6</td><td>-8.0</td><td>1.5E-15</td><td>6.3E-15</td><td>MB231/MB468</td><td>MITO</td></tr>
<tr><td>>hg38_21_+_8433172_8446622_RNA45SN1_45S_50ntFlanks@10794.10828.35|CTGGCATGTTGGAACAATGTAGGTAAGGGAAGTCG</td><td>>hg38_21_+_8433172_8446622_RNA45SN1_45S_50ntFlanks</td><td>10794</td><td>10828</td><td>35</td><td>CTGGCATGTTGGAACAATGTAGGTAAGGGAAGTCG</td><td>231.2</td><td>-11.7</td><td>1.5</td><td>-8.0</td><td>9.1E-16</td><td>6.3E-15</td><td>MB231/MB468</td><td>MP</td></tr>
<tr><td>>hg38_21_+_8433172_8446622_RNA45SN1_45S_50ntFlanks@4240.4258.19|ACTCTTTCGAGGCCCTGTA</td><td>>hg38_21_+_8433172_8446622_RNA45SN1_45S_50ntFlanks</td><td>4240</td><td>4258</td><td>19</td><td>ACTCTTTCGAGGCCCTGTA</td><td>255.1</td><td>-11.9</td><td>1.5</td><td>-8.0</td><td>9.1E-16</td><td>6.3E-15</td><td>MB231/MB468</td><td>MP</td></tr>
<tr><td>>hg38_21_+_8433172_8446622_RNA45SN1_45S_50ntFlanks@3975.4005.31|CGGGCGCCGGCGGCTTTGGTGACTCTAGATA</td><td>>hg38_21_+_8433172_8446622_RNA45SN1_45S_50ntFlanks</td><td>3975</td><td>4005</td><td>31</td><td>CGGGCGCCGGCGGCTTTGGTGACTCTAGATA</td><td>513.8</td><td>-12.8</td><td>1.6</td><td>-8.0</td><td>1.5E-15</td><td>6.3E-15</td><td>MB231/MB468</td><td>MITO</td></tr>
<tr><td>>hg38_21_+_8433172_8446622_RNA45SN1_45S_50ntFlanks@11661.11681.21|TGATTTCTGCCCAGTGCTCTG</td><td>>hg38_21_+_8433172_8446622_RNA45SN1_45S_50ntFlanks</td><td>11661</td><td>11681</td><td>21</td><td>TGATTTCTGCCCAGTGCTCTG</td><td>3759.0</td><td>-8.6</td><td>1.1</td><td>-8.0</td><td>1.5E-15</td><td>6.4E-15</td><td>MB231/MB468</td><td>MITO</td></tr>
<tr><td>>hg38_21_+_8433172_8446622_RNA45SN1_45S_50ntFlanks@12499.12518.20|GGTTTAGACCGTCGTGAGAC</td><td>>hg38_21_+_8433172_8446622_RNA45SN1_45S_50ntFlanks</td><td>12499</td><td>12518</td><td>20</td><td>GGTTTAGACCGTCGTGAGAC</td><td>287.4</td><td>-12.0</td><td>1.5</td><td>-8.0</td><td>9.2E-16</td><td>6.4E-15</td><td>MB231/MB468</td><td>MP</td></tr>
<tr><td>>hg38_21_+_8433172_8446622_RNA45SN1_45S_50ntFlanks@10806.10829.24|AACAATGTAGGTAAGGGAAGTCGG</td><td>>hg38_21_+_8433172_8446622_RNA45SN1_45S_50ntFlanks</td><td>10806</td><td>10829</td><td>24</td><td>AACAATGTAGGTAAGGGAAGTCGG</td><td>245.7</td><td>-11.8</td><td>1.5</td><td>-8.0</td><td>9.2E-16</td><td>6.4E-15</td><td>MB231/MB468</td><td>MP</td></tr>
<tr><td>>hg38_21_+_8433172_8446622_RNA45SN1_45S_50ntFlanks@4723.4759.37|CATTAATCAAGAACGAAAGTCGGAGGTTCGAAGACGA</td><td>>hg38_21_+_8433172_8446622_RNA45SN1_45S_50ntFlanks</td><td>4723</td><td>4759</td><td>37</td><td>CATTAATCAAGAACGAAAGTCGGAGGTTCGAAGACGA</td><td>1824.5</td><td>-9.0</td><td>1.1</td><td>-8.0</td><td>9.2E-16</td><td>6.4E-15</td><td>MB231/MB468</td><td>MP</td></tr>
<tr><td>>hg38_21_+_8433172_8446622_RNA45SN1_45S_50ntFlanks@10914.10931.18|CGCCGCGGCTGGACGAGG</td><td>>hg38_21_+_8433172_8446622_RNA45SN1_45S_50ntFlanks</td><td>10914</td><td>10931</td><td>18</td><td>CGCCGCGGCTGGACGAGG</td><td>957.0</td><td>-9.6</td><td>1.2</td><td>-8.0</td><td>1.5E-15</td><td>6.4E-15</td><td>MB231/MB468</td><td>MITO</td></tr>
<tr><td>>hg38_21_+_8433172_8446622_RNA45SN1_45S_50ntFlanks@11320.11338.19|CCACGTCTCGTCGCGCGCG</td><td>>hg38_21_+_8433172_8446622_RNA45SN1_45S_50ntFlanks</td><td>11320</td><td>11338</td><td>19</td><td>CCACGTCTCGTCGCGCGCG</td><td>323.4</td><td>-12.1</td><td>1.5</td><td>-8.0</td><td>1.5E-15</td><td>6.4E-15</td><td>MB231/MB468</td><td>MITO</td></tr>
<tr><td>>hg38_21_+_8433172_8446622_RNA45SN1_45S_50ntFlanks@4733.4766.34|GAACGAAAGTCGGAGGTTCGAAGACGATCAGATA</td><td>>hg38_21_+_8433172_8446622_RNA45SN1_45S_50ntFlanks</td><td>4733</td><td>4766</td><td>34</td><td>GAACGAAAGTCGGAGGTTCGAAGACGATCAGATA</td><td>1165.9</td><td>8.5</td><td>1.1</td><td>8.0</td><td>1.4E-15</td><td>6.4E-15</td><td>BT20/MB468</td><td>CYTO</td></tr>
<tr><td>>MI0000102|hsa-mir-100&WithFlank&11|-|122152223|122152314@19.38.20(+2G)&[MIMAT0000098&hsa-miR-100-5p&offsets|0|-2(+2G);m-27&11|-|122152276|122152296&offsets|0|-1(+2G)]|AACCCGTAGATCCGAACTTGGG</td><td>>MI0000102|hsa-mir-100&WithFlank&11|-|122152223|122152314</td><td>19</td><td>38</td><td>20(+2G)</td><td>AACCCGTAGATCCGAACTTGGG</td><td>1041.0</td><td>13.2</td><td>1.7</td><td>8.0</td><td>1.5E-15</td><td>6.4E-15</td><td>MB231/MB468</td><td>MITO</td></tr>
<tr><td>>am_trna117_GlyTCC_1_-_145397864_145397935@1.30.30,trna45_GlyTCC_1_+_161500903_161500974@1.30.30,trna70_GlyTCC_1_-_161439547_161439618@1.30.30,trna73_GlyTCC_1_-_161432166_161432237@1.30.30,trna76_GlyTCC_1_-_161424756_161424827@1.30.30,trna79_GlyTCC_1_-_161417375_161417446@1.30.30|GCGTTGGTGGTATAGTGGTGAGCATAGCTG</td><td>>am_trna117_GlyTCC_1_-_145397864_145397935</td><td>1</td><td>30</td><td>30</td><td>GCGTTGGTGGTATAGTGGTGAGCATAGCTG</td><td>6167.6</td><td>-7.1</td><td>0.9</td><td>-8.0</td><td>1.0E-15</td><td>6.4E-15</td><td>MB231/MB468</td><td>CYTO</td></tr>
<tr><td>>MI0000079|hsa-mir-23a&WithFlank&19|-|13836581|13836665@54.72.19(+1U)&[MIMAT0000078&hsa-miR-23a-3p&offsets|+3|+1(+1U);m-31&19|-|13836595|13836615&offsets|+3|+1(+1U)]|ACATTGCCAGGGATTTCCAT</td><td>>MI0000079|hsa-mir-23a&WithFlank&19|-|13836581|13836665</td><td>54</td><td>72</td><td>19(+1U)</td><td>ACATTGCCAGGGATTTCCAT</td><td>2050.6</td><td>29.1</td><td>3.6</td><td>8.0</td><td>1.0E-15</td><td>6.5E-15</td><td>MB231/MB468</td><td>CYTO</td></tr>
<tr><td>>hg38_21_+_8433172_8446622_RNA45SN1_45S_50ntFlanks@4919.4937.19|CCACCAGGAGTGGAGCCTG</td><td>>hg38_21_+_8433172_8446622_RNA45SN1_45S_50ntFlanks</td><td>4919</td><td>4937</td><td>19</td><td>CCACCAGGAGTGGAGCCTG</td><td>1223.5</td><td>-14.0</td><td>1.8</td><td>-8.0</td><td>1.5E-15</td><td>6.5E-15</td><td>MB231/MB468</td><td>MITO</td></tr>
<tr><td>>trnaMT_SerGCT_MT_+_12207_12265@8.38.31|CTCACAAGAACTGCTAACTCATGCCCCCATG</td><td>>trnaMT_SerGCT_MT_+_12207_12265</td><td>8</td><td>38</td><td>31</td><td>CTCACAAGAACTGCTAACTCATGCCCCCATG</td><td>1386.4</td><td>-8.6</td><td>1.1</td><td>-8.0</td><td>1.5E-15</td><td>6.5E-15</td><td>MB231/MB468</td><td>MITO</td></tr>
<tr><td>>hg38_21_+_8433172_8446622_RNA45SN1_45S_50ntFlanks@6737.6756.20|GATCATCGACACTTCGAACG</td><td>>hg38_21_+_8433172_8446622_RNA45SN1_45S_50ntFlanks</td><td>6737</td><td>6756</td><td>20</td><td>GATCATCGACACTTCGAACG</td><td>207.7</td><td>-11.6</td><td>1.4</td><td>-8.0</td><td>9.4E-16</td><td>6.5E-15</td><td>MB231/MB468</td><td>MP</td></tr>
<tr><td>>MI0000434|hsa-let-7i&WithFlank&12|+|62603680|62603775@12.33.22&[MIMAT0000415&hsa-let-7i-5p&offsets|0|0;m-49&12|+|62603691|62603712&offsets|0|0]|TGAGGTAGTAGTTTGTGCTGTT</td><td>>MI0000434|hsa-let-7i&WithFlank&12|+|62603680|62603775</td><td>12</td><td>33</td><td>22</td><td>TGAGGTAGTAGTTTGTGCTGTT</td><td>214804.7</td><td>5.9</td><td>0.7</td><td>8.0</td><td>1.1E-15</td><td>6.5E-15</td><td>MB231/MB468</td><td>NUC</td></tr>
<tr><td>>hg38_21_+_8433172_8446622_RNA45SN1_45S_50ntFlanks@8106.8138.33|GCGGCGGGGCGCGGGACATGTGGCGTACGGAAG</td><td>>hg38_21_+_8433172_8446622_RNA45SN1_45S_50ntFlanks</td><td>8106</td><td>8138</td><td>33</td><td>GCGGCGGGGCGCGGGACATGTGGCGTACGGAAG</td><td>366.8</td><td>-12.3</td><td>1.5</td><td>-8.0</td><td>1.6E-15</td><td>6.5E-15</td><td>MB231/MB468</td><td>MITO</td></tr>
<tr><td>>hg38_21_+_8433172_8446622_RNA45SN1_45S_50ntFlanks@9858.9899.42|CGGGATGAACCGAACGCCGGGTTAAGGCGCCCGATGCCGACG</td><td>>hg38_21_+_8433172_8446622_RNA45SN1_45S_50ntFlanks</td><td>9858</td><td>9899</td><td>42</td><td>CGGGATGAACCGAACGCCGGGTTAAGGCGCCCGATGCCGACG</td><td>342.6</td><td>-12.3</td><td>1.5</td><td>-8.0</td><td>9.4E-16</td><td>6.5E-15</td><td>MB231/MB468</td><td>MP</td></tr>
<tr><td>>hg38_21_+_8433172_8446622_RNA45SN1_45S_50ntFlanks@5467.5497.31|GGCCCACGGCCCTGGCGGAGCGCTGAGAAGA</td><td>>hg38_21_+_8433172_8446622_RNA45SN1_45S_50ntFlanks</td><td>5467</td><td>5497</td><td>31</td><td>GGCCCACGGCCCTGGCGGAGCGCTGAGAAGA</td><td>201.6</td><td>-10.6</td><td>1.3</td><td>-8.0</td><td>1.2E-15</td><td>6.5E-15</td><td>MB231/MB468</td><td>NUC</td></tr>
<tr><td>>hg38_21_+_8433172_8446622_RNA45SN1_45S_50ntFlanks@10351.10376.26|CGAGAACTTTGAAGGCCGAAGTGGAG</td><td>>hg38_21_+_8433172_8446622_RNA45SN1_45S_50ntFlanks</td><td>10351</td><td>10376</td><td>26</td><td>CGAGAACTTTGAAGGCCGAAGTGGAG</td><td>1859.5</td><td>-10.2</td><td>1.3</td><td>-8.0</td><td>1.6E-15</td><td>6.6E-15</td><td>MB231/MB468</td><td>MITO</td></tr>
<tr><td>>hg38_21_+_8433172_8446622_RNA45SN1_45S_50ntFlanks@6680.6718.39|TGCGTCGATGAAGAACGCAGCTAGCTGCGAGAATTAATG</td><td>>hg38_21_+_8433172_8446622_RNA45SN1_45S_50ntFlanks</td><td>6680</td><td>6718</td><td>39</td><td>TGCGTCGATGAAGAACGCAGCTAGCTGCGAGAATTAATG</td><td>451.5</td><td>-12.6</td><td>1.6</td><td>-8.0</td><td>1.6E-15</td><td>6.6E-15</td><td>MB231/MB468</td><td>MITO</td></tr>
<tr><td>>hg38_21_+_8433172_8446622_RNA45SN1_45S_50ntFlanks@5491.5516.26|GAGAAGACGGTCGAACTTGACTATCT</td><td>>hg38_21_+_8433172_8446622_RNA45SN1_45S_50ntFlanks</td><td>5491</td><td>5516</td><td>26</td><td>GAGAAGACGGTCGAACTTGACTATCT</td><td>26661.0</td><td>9.7</td><td>1.2</td><td>8.0</td><td>1.0E-15</td><td>6.6E-15</td><td>BT20/MB468</td><td>MITO</td></tr>
<tr><td>>hg38_21_+_8433172_8446622_RNA45SN1_45S_50ntFlanks@11200.11225.26|CGGCGGCGGCGGCGACTCTGGACGCG</td><td>>hg38_21_+_8433172_8446622_RNA45SN1_45S_50ntFlanks</td><td>11200</td><td>11225</td><td>26</td><td>CGGCGGCGGCGGCGACTCTGGACGCG</td><td>1099.7</td><td>-9.5</td><td>1.2</td><td>-8.0</td><td>1.6E-15</td><td>6.6E-15</td><td>MB231/MB468</td><td>MITO</td></tr>
<tr><td>>am_trnaMT_PheGAA_MT_+_577_647@41.58.18|GTTTAGACGGGCTCACAT</td><td>>am_trnaMT_PheGAA_MT_+_577_647</td><td>41</td><td>58</td><td>18</td><td>GTTTAGACGGGCTCACAT</td><td>546.7</td><td>6.0</td><td>0.7</td><td>8.0</td><td>1.2E-15</td><td>6.6E-15</td><td>MB231/MB468</td><td>NUC</td></tr>
<tr><td>>hg38_21_+_8433172_8446622_RNA45SN1_45S_50ntFlanks@10128.10148.21|CGGCGGCGGCGGCGGCGGCGG</td><td>>hg38_21_+_8433172_8446622_RNA45SN1_45S_50ntFlanks</td><td>10128</td><td>10148</td><td>21</td><td>CGGCGGCGGCGGCGGCGGCGG</td><td>325.7</td><td>-12.2</td><td>1.5</td><td>-8.0</td><td>9.6E-16</td><td>6.6E-15</td><td>MB231/MB468</td><td>MP</td></tr>
<tr><td>>hg38_21_+_8433172_8446622_RNA45SN1_45S_50ntFlanks@8845.8865.21|CGGCGGCGGCGGCGGCGGCGG</td><td>>hg38_21_+_8433172_8446622_RNA45SN1_45S_50ntFlanks</td><td>8845</td><td>8865</td><td>21</td><td>CGGCGGCGGCGGCGGCGGCGG</td><td>325.7</td><td>-12.2</td><td>1.5</td><td>-8.0</td><td>9.6E-16</td><td>6.6E-15</td><td>MB231/MB468</td><td>MP</td></tr>
<tr><td>>hg38_21_+_8433172_8446622_RNA45SN1_45S_50ntFlanks@6751.6770.20|CGAACGCACTTGCGGCCCCG</td><td>>hg38_21_+_8433172_8446622_RNA45SN1_45S_50ntFlanks</td><td>6751</td><td>6770</td><td>20</td><td>CGAACGCACTTGCGGCCCCG</td><td>264.2</td><td>-11.8</td><td>1.5</td><td>-8.0</td><td>1.6E-15</td><td>6.7E-15</td><td>MB231/MB468</td><td>MITO</td></tr>
<tr><td>>hg38_21_+_8433172_8446622_RNA45SN1_45S_50ntFlanks@12570.12587.18|AGTACGAGAGGAACCGCA</td><td>>hg38_21_+_8433172_8446622_RNA45SN1_45S_50ntFlanks</td><td>12570</td><td>12587</td><td>18</td><td>AGTACGAGAGGAACCGCA</td><td>2062.0</td><td>-10.4</td><td>1.3</td><td>-8.0</td><td>9.7E-16</td><td>6.7E-15</td><td>MB231/MB468</td><td>MP</td></tr>
<tr><td>>hg38_21_+_8433172_8446622_RNA45SN1_45S_50ntFlanks@11661.11679.19|TGATTTCTGCCCAGTGCTC</td><td>>hg38_21_+_8433172_8446622_RNA45SN1_45S_50ntFlanks</td><td>11661</td><td>11679</td><td>19</td><td>TGATTTCTGCCCAGTGCTC</td><td>284.9</td><td>-12.0</td><td>1.5</td><td>-8.0</td><td>9.7E-16</td><td>6.7E-15</td><td>MB231/MB468</td><td>MP</td></tr>
</table>

| | | | | | | | | | | | | | |
|---|---|---|---|---|---|---|---|---|---|---|---|---|---|
| >MI0001445\|hsa-mir-423&WithFlank&17\|+\|30117073\|30117178@59.81.23&[MIMAT0001340&hsa-miR-423-3p&offsets\|0\|0;m-100&17\|+\|30117131\|30117153&offsets\|0\|0]\|AGCTCGGTCTGAGGCCCCTCAGT | >MI0001445\|hsa-mir-423&WithFlank&17\|+\|30117073\|30117178 | 59 | 81 | 23 | AGCTCGGTCTGAGGCCCCTCAGT | 15424.4 | 7.0 | 0.9 | 8.0 | 1.1E-15 | 6.7E-15 | MB231/MB468 | CYTO |
| >MI0000440\|hsa-mir-27b&WithFlank&9\|+\|95085439\|95085547@67.88.22(+1A)&[MIMAT0000419&hsa-miR-27b-3p&offsets\|0\|+1(+1A);m-39&9\|+\|95085505\|95085525&offsets\|0\|+1(+1A)]\|TTCACAGTGGCTAAGTTCTGCAA | >MI0000440\|hsa-mir-27b&WithFlank&9\|+\|95085439\|95085547 | 67 | 88 | 22(+1A) | TTCACAGTGGCTAAGTTCTGCAA | 149.5 | 2.1 | 0.3 | 7.9 | 2.2E-15 | 6.8E-15 | BT20/MB468 | NUC |
| >hg38_21_+_8433172_8446622_RNA45SN1_45S_50ntFlanks@12435.12455.21\|ATCATTGTGAAGCAGAATTCA | >hg38_21_+_8433172_8446622_RNA45SN1_45S_50ntFlanks | 12435 | 12455 | 21 | ATCATTGTGAAGCAGAATTCA | 499.5 | -11.1 | 1.4 | -8.0 | 1.0E-15 | 6.8E-15 | BT20/MB468 | MITO |
| >MI0000084\|hsa-mir-26b&WithFlank&2\|+\|218402640\|218402728@18.38.21(+1C)&[MIMAT0000083&hsa-miR-26b-5p&offsets\|0\|0(+1C);m-53&2\|+\|218402657\|218402678&offsets\|0\|-1(+1C)]\|TTCAAGTAATTCAGGATAGGTC | >MI0000084\|hsa-mir-26b&WithFlank&2\|+\|218402640\|218402728 | 18 | 38 | 21(+1C) | TTCAAGTAATTCAGGATAGGTC | 348.9 | 3.1 | 0.4 | 7.9 | 2.2E-15 | 6.8E-15 | BT20/MB468 | NUC |
| >hg38_21_+_8433172_8446622_RNA45SN1_45S_50ntFlanks@11235.11253.19\|CTTCCCGTGGATCGCCCCA | >hg38_21_+_8433172_8446622_RNA45SN1_45S_50ntFlanks | 11235 | 11253 | 19 | CTTCCCGTGGATCGCCCCA | 1990.9 | -9.0 | 1.1 | -7.9 | 2.2E-15 | 6.8E-15 | BT20/MB468 | NUC |
| >MI0000070\|hsa-mir-16-1&WithFlank&13\|-\|50048967\|50049067@20.41.22(+1A);MI0000115\|hsa-mir-16-2&WithFlank&3\|+\|160404739\|160404831@16.37.22(+1A)&[MIMAT0000069&hsa-miR-16-5p&offsets\|0\|0(+1A);m-62&13\|-\|50049027\|50049048&offsets\|0\|0(+1A)];[MIMAT0000069_1&hsa-miR-16-5p&offsets\|0\|0(+1A);m-61&3\|+\|160404754\|160404775&offsets\|0\|0(+1A)]\|TAGCAGCACGTAAATATTGGCGA | >MI0000070\|hsa-mir-16-1&WithFlank&13\|-\|50048967\|50049067 | 20 | 41 | 22(+1A) | TAGCAGCACGTAAATATTGGCGA | 318.8 | 12.6 | 1.6 | 8.0 | 1.5E-15 | 6.8E-15 | BT20/MB468 | CYTO |
| >hg38_21_+_8433172_8446622_RNA45SN1_45S_50ntFlanks@11670.11702.33\|CCCAGTGCTCTGAATGTCAAAGTGAAGAAATTC | >hg38_21_+_8433172_8446622_RNA45SN1_45S_50ntFlanks | 11670 | 11702 | 33 | CCCAGTGCTCTGAATGTCAAAGTGAAGAAATTC | 15280.7 | -10.8 | 1.4 | -7.9 | 2.3E-15 | 6.8E-15 | BT20/MB468 | NUC |
| >hg38_21_+_8433172_8446622_RNA45SN1_45S_50ntFlanks@12635.12653.19\|AAGCTACCATCTGTGGGAT | >hg38_21_+_8433172_8446622_RNA45SN1_45S_50ntFlanks | 12635 | 12653 | 19 | AAGCTACCATCTGTGGGAT | 215.2 | -10.7 | 1.3 | -8.0 | 1.2E-15 | 6.8E-15 | MB231/MB468 | NUC |
| >hg38_21_+_8433172_8446622_RNA45SN1_45S_50ntFlanks@12459.12476.18\|AGCGTTGGATTGTTCACC | >hg38_21_+_8433172_8446622_RNA45SN1_45S_50ntFlanks | 12459 | 12476 | 18 | AGCGTTGGATTGTTCACC | 619.7 | -4.0 | 0.5 | -8.0 | 1.1E-15 | 6.8E-15 | BT20/MB468 | MITO |
| >hg38_21_+_8433172_8446622_RNA45SN1_45S_50ntFlanks@12397.12417.21\|CGACGTCGCTTTTTGATCCTT | >hg38_21_+_8433172_8446622_RNA45SN1_45S_50ntFlanks | 12397 | 12417 | 21 | CGACGTCGCTTTTTGATCCTT | 342.5 | -12.3 | 1.5 | -8.0 | 9.9E-16 | 6.8E-15 | MB231/MB468 | MP |
| >hg38_21_+_8433172_8446622_RNA45SN1_45S_50ntFlanks@9562.9597.36\|TCCTGACGTGCAAATCGGTCGTCCGACCTGGGTATA | >hg38_21_+_8433172_8446622_RNA45SN1_45S_50ntFlanks | 9562 | 9597 | 36 | TCCTGACGTGCAAATCGGTCGTCCGACCTGGGTATA | 659.7 | -7.4 | 0.9 | -8.0 | 1.1E-15 | 6.9E-15 | MB231/MB468 | CYTO |
| >hg38_21_+_8433172_8446622_RNA45SN1_45S_50ntFlanks@10794.10813.20\|CTGGCATGTTGGAACAATGT | >hg38_21_+_8433172_8446622_RNA45SN1_45S_50ntFlanks | 10794 | 10813 | 20 | CTGGCATGTTGGAACAATGT | 185.3 | -11.4 | 1.4 | -8.0 | 1.0E-15 | 6.9E-15 | MB231/MB468 | MP |
| >hg38_21_+_8433172_8446622_RNA45SN1_45S_50ntFlanks@485.529.45\|GATGTGGTGACGTCGTGCTCTCCCGGGCCGGGTCCGAGCCGCGAC | >hg38_21_+_8433172_8446622_RNA45SN1_45S_50ntFlanks | 485 | 529 | 45 | GATGTGGTGACGTCGTGCTCTCCCGGGCCGGGTCCGAGCCGCGAC | 2100.8 | 29.1 | 3.6 | 8.0 | 1.2E-15 | 6.9E-15 | MB231/MB468 | NUC |
| >hg38_21_+_8433172_8446622_RNA45SN1_45S_50ntFlanks@12954.12973.20\|CTGCTTCTGGGTCGGGGTTT | >hg38_21_+_8433172_8446622_RNA45SN1_45S_50ntFlanks | 12954 | 12973 | 20 | CTGCTTCTGGGTCGGGGTTT | 226.8 | -11.7 | 1.5 | -8.0 | 1.0E-15 | 6.9E-15 | MB231/MB468 | MP |
| >hg38_21_+_8433172_8446622_RNA45SN1_45S_50ntFlanks@9202.9226.25\|TCTCGGGGCCACGCGCGCGTCCCCC | >hg38_21_+_8433172_8446622_RNA45SN1_45S_50ntFlanks | 9202 | 9226 | 25 | TCTCGGGGCCACGCGCGCGTCCCCC | 332.4 | -12.2 | 1.5 | -8.0 | 1.0E-15 | 6.9E-15 | MB231/MB468 | MP |
| >hg38_21_+_8433172_8446622_RNA45SN1_45S_50ntFlanks@12249.12269.21\|CAAAAGCTCGCTTGATCTTGA | >hg38_21_+_8433172_8446622_RNA45SN1_45S_50ntFlanks | 12249 | 12269 | 21 | CAAAAGCTCGCTTGATCTTGA | 956.9 | -13.7 | 1.7 | -8.0 | 1.7E-15 | 6.9E-15 | MB231/MB468 | MITO |
| >hg38_21_+_8433172_8446622_RNA45SN1_45S_50ntFlanks@10808.10839.32\|CAATGTAGGTAAGGGAAGTCGGCAAGCCGGAT | >hg38_21_+_8433172_8446622_RNA45SN1_45S_50ntFlanks | 10808 | 10839 | 32 | CAATGTAGGTAAGGGAAGTCGGCAAGCCGGAT | 269.2 | -11.9 | 1.5 | -8.0 | 1.7E-15 | 6.9E-15 | MB231/MB468 | MITO |
| >hg38_21_+_8433172_8446622_RNA45SN1_45S_50ntFlanks@6652.6696.45\|GACTCTTAGCGGTGGATCACTCGGCTCGTGCGTCGATGAAGAACG | >hg38_21_+_8433172_8446622_RNA45SN1_45S_50ntFlanks | 6652 | 6696 | 45 | GACTCTTAGCGGTGGATCACTCGGCTCGTGCGTCGATGAAGAACG | 1492.1 | 5.6 | 0.7 | 8.0 | 1.1E-15 | 7.0E-15 | MB231/MB468 | CYTO |
| >MI0000482\|hsa-mir-185&WithFlank&22\|+\|20033133\|20033226@21.43.23&[MIMAT0000455&hsa-miR-185-5p&offsets\|0\|+1;m-145&22\|+\|20033153\|20033174&offsets\|0\|+1]\|TGGAGAGAAAGGCAGTTCCTGAT | >MI0000482\|hsa-mir-185&WithFlank&22\|+\|20033133\|20033226 | 21 | 43 | 23 | TGGAGAGAAAGGCAGTTCCTGAT | 507.0 | 7.5 | 0.9 | 8.0 | 1.1E-15 | 7.0E-15 | BT20/MB468 | MITO |
| >hg38_21_+_8433172_8446622_RNA45SN1_45S_50ntFlanks@11569.11605.37\|CAGCCGACTTAGAACTGGTGCGGACCAGGGGAATCCG | >hg38_21_+_8433172_8446622_RNA45SN1_45S_50ntFlanks | 11569 | 11605 | 37 | CAGCCGACTTAGAACTGGTGCGGACCAGGGGAATCCG | 3914.1 | -10.8 | 1.4 | -8.0 | 1.7E-15 | 7.0E-15 | MB231/MB468 | MITO |
| >hg38_21_+_8433172_8446622_RNA45SN1_45S_50ntFlanks@4003.4020.18\|ATAACCTCGGGCCGATCG | >hg38_21_+_8433172_8446622_RNA45SN1_45S_50ntFlanks | 4003 | 4020 | 18 | ATAACCTCGGGCCGATCG | 1764.0 | -10.0 | 1.3 | -8.0 | 1.7E-15 | 7.0E-15 | MB231/MB468 | MITO |
| >hg38_21_+_8433172_8446622_RNA45SN1_45S_50ntFlanks@8179.8204.26\|CTGATCGAGGCCCAGCCCGTGGACGG | >hg38_21_+_8433172_8446622_RNA45SN1_45S_50ntFlanks | 8179 | 8204 | 26 | CTGATCGAGGCCCAGCCCGTGGACGG | 260.9 | -11.9 | 1.5 | -8.0 | 1.0E-15 | 7.0E-15 | MB231/MB468 | MP |
| >MI0000433\|hsa-let-7g&WithFlank&3\|-\|52268272\|52268367@11.33.23&[MIMAT0000414&hsa-let-7g-5p&offsets\|0\|+1;m-58&3\|-\|52268336\|52268357&offsets\|0\|+1]\|TGAGGTAGTAGTTTGTACAGTTT | >MI0000433\|hsa-let-7g&WithFlank&3\|-\|52268272\|52268367 | 11 | 33 | 23 | TGAGGTAGTAGTTTGTACAGTTT | 1796.6 | 6.9 | 0.9 | 8.0 | 1.5E-15 | 7.0E-15 | BT20/MB468 | CYTO |
| >hg38_21_+_8433172_8446622_RNA45SN1_45S_50ntFlanks@10786.10821.36\|AACAGCCTCTGGCATGTTGGAACAATGTAGGTAAGG | >hg38_21_+_8433172_8446622_RNA45SN1_45S_50ntFlanks | 10786 | 10821 | 36 | AACAGCCTCTGGCATGTTGGAACAATGTAGGTAAGG | 540.8 | -5.2 | 0.6 | -8.0 | 1.0E-15 | 7.0E-15 | MB231/MB468 | MP |
| >hg38_21_+_8433172_8446622_RNA45SN1_45S_50ntFlanks@9167.9187.21\|CGGGCGGGTCGCGCCGTCGGG | >hg38_21_+_8433172_8446622_RNA45SN1_45S_50ntFlanks | 9167 | 9187 | 21 | CGGGCGGGTCGCGCCGTCGGG | 459.6 | -12.6 | 1.6 | -8.0 | 1.7E-15 | 7.0E-15 | MB231/MB468 | MITO |
| >hg38_21_+_8433172_8446622_RNA45SN1_45S_50ntFlanks@11573.11601.29\|CGACTTAGAACTGGTGCGGACCAGGGGAA | >hg38_21_+_8433172_8446622_RNA45SN1_45S_50ntFlanks | 11573 | 11601 | 29 | CGACTTAGAACTGGTGCGGACCAGGGGAA | 4438.4 | -9.5 | 1.2 | -7.9 | 2.3E-15 | 7.0E-15 | BT20/MB468 | NUC |
| >hg38_21_+_8433172_8446622_RNA45SN1_45S_50ntFlanks@6645.6669.25\|CTCGTACGACTCTTAGCGGTGGATC | >hg38_21_+_8433172_8446622_RNA45SN1_45S_50ntFlanks | 6645 | 6669 | 25 | CTCGTACGACTCTTAGCGGTGGATC | 35383.1 | 4.4 | 0.5 | 8.0 | 1.1E-15 | 7.0E-15 | BT20/MB468 | MITO |
| >hg38_21_+_8433172_8446622_RNA45SN1_45S_50ntFlanks@474.497.24\|TCTTCGTGATCGATGTGGTGACGT | >hg38_21_+_8433172_8446622_RNA45SN1_45S_50ntFlanks | 474 | 497 | 24 | TCTTCGTGATCGATGTGGTGACGT | 3205.5 | 6.9 | 0.9 | 8.0 | 1.2E-15 | 7.0E-15 | MB231/MB468 | NUC |
| >MI0000483\|hsa-mir-186&WithFlank&1\|-\|71067625\|71067722@21.40.20&[MIMAT0000456&hsa-miR-186-5p&offsets\|0\|-2;m-84&1\|-\|71067681\|71067702&offsets\|0\|-2]\|CAAAGAATTCTCCTTTTGGG | >MI0000483\|hsa-mir-186&WithFlank&1\|-\|71067625\|71067722 | 21 | 40 | 20 | CAAAGAATTCTCCTTTTGGG | 323.2 | 4.7 | 0.6 | 8.0 | 1.3E-15 | 7.1E-15 | MB231/MB468 | NUC |
| >hg38_21_+_8433172_8446622_RNA45SN1_45S_50ntFlanks@9031.9059.29\|TCCTCGTGGGGGGGCCGGGCCACCCCTCC | >hg38_21_+_8433172_8446622_RNA45SN1_45S_50ntFlanks | 9031 | 9059 | 29 | TCCTCGTGGGGGGGCCGGGCCACCCCTCC | 234.3 | -11.7 | 1.5 | -8.0 | 1.0E-15 | 7.1E-15 | MB231/MB468 | MP |
| >hg38_21_+_8433172_8446622_RNA45SN1_45S_50ntFlanks@8191.8212.22\|CAGCCCGTGGACGGTGTGAGGC | >hg38_21_+_8433172_8446622_RNA45SN1_45S_50ntFlanks | 8191 | 8212 | 22 | CAGCCCGTGGACGGTGTGAGGC | 1364.3 | -8.6 | 1.1 | -7.9 | 2.4E-15 | 7.1E-15 | BT20/MB468 | NUC |
| >hg38_21_+_8433172_8446622_RNA45SN1_45S_50ntFlanks@9457.9488.32\|CGCCGGGGAGGTGGAGCACGAGCGCACGTGTT | >hg38_21_+_8433172_8446622_RNA45SN1_45S_50ntFlanks | 9457 | 9488 | 32 | CGCCGGGGAGGTGGAGCACGAGCGCACGTGTT | 281.8 | -12.0 | 1.5 | -8.0 | 1.0E-15 | 7.1E-15 | MB231/MB468 | MP |
| >MI0000650\|hsa-mir-200c&WithFlank&12\|+\|6963693\|6963772@50.71.22&[MIMAT0000617&hsa-miR-200c-3p&offsets\|0\|-1;m-54&12\|+\|6963742\|6963764&offsets\|0\|-1]\|TAATACTGCCGGGTAATGATGG | >MI0000650\|hsa-mir-200c&WithFlank&12\|+\|6963693\|6963772 | 50 | 71 | 22 | TAATACTGCCGGGTAATGATGG | 6044.3 | 3.6 | 0.4 | 7.9 | 2.4E-15 | 7.1E-15 | BT20/MB468 | NUC |
| >hg38_21_+_8433172_8446622_RNA45SN1_45S_50ntFlanks@12288.12318.31\|ACCGTGAAAGCGGGGCCTCACGATCCTTCTG | >hg38_21_+_8433172_8446622_RNA45SN1_45S_50ntFlanks | 12288 | 12318 | 31 | ACCGTGAAAGCGGGGCCTCACGATCCTTCTG | 2667.1 | -10.9 | 1.4 | -8.0 | 1.7E-15 | 7.2E-15 | MB231/MB468 | MITO |
| >MI0000542\|hsa-mir-320a&WithFlank&8\|-\|22244960\|22245043@48.70.23(+1C)&[MIMAT0000510&hsa-miR-320a-3p&offsets\|0\|+1(+1C);m-60&8\|-\|22244975\|22244996&offsets\|0\|+1(+1C)]\|AAAAGCTGGGTTGAGAGGGCGAAC | >MI0000542\|hsa-mir-320a&WithFlank&8\|-\|22244960\|22245043 | 48 | 70 | 23(+1C) | AAAAGCTGGGTTGAGAGGGCGAAC | 176.5 | 3.9 | 0.5 | 7.9 | 2.4E-15 | 7.2E-15 | BT20/MB468 | NUC |
| >hg38_21_+_8433172_8446622_RNA45SN1_45S_50ntFlanks@6671.6698.28\|CTCGGCTCGTGCGTCGATGAAGAACGCA | >hg38_21_+_8433172_8446622_RNA45SN1_45S_50ntFlanks | 6671 | 6698 | 28 | CTCGGCTCGTGCGTCGATGAAGAACGCA | 20753.7 | -10.8 | 1.4 | -8.0 | 1.7E-15 | 7.2E-15 | MB231/MB468 | MITO |
| >hg38_21_+_8433172_8446622_RNA45SN1_45S_50ntFlanks@6650.6668.19\|ACGACTCTTAGCGGTGGAT | >hg38_21_+_8433172_8446622_RNA45SN1_45S_50ntFlanks | 6650 | 6668 | 19 | ACGACTCTTAGCGGTGGAT | 10058.9 | 6.7 | 0.8 | 8.0 | 1.1E-15 | 7.2E-15 | BT20/MB468 | MITO |
| >trna139_ValAAC_6_-_27618707_27618779@36.56.21\|CACGCGAAAGGTCCCTGGATC | >trna139_ValAAC_6_-_27618707_27618779 | 36 | 56 | 21 | CACGCGAAAGGTCCCTGGATC | 389.8 | 12.0 | 1.4 | 8.3 | 9.7E-17 | 7.2E-15 | BT20/MB231 | CYTO |
| >hg38_21_+_8433172_8446622_RNA45SN1_45S_50ntFlanks@10905.10941.37\|CTGGGCGCGCGCCGCGGCTGGACGAGGCGCCGCCGCC | >hg38_21_+_8433172_8446622_RNA45SN1_45S_50ntFlanks | 10905 | 10941 | 37 | CTGGGCGCGCGCCGCGGCTGGACGAGGCGCCGCCGCC | 311.7 | -12.1 | 1.5 | -8.0 | 1.7E-15 | 7.2E-15 | MB231/MB468 | MITO |
| >hg38_21_+_8433172_8446622_RNA45SN1_45S_50ntFlanks@7986.8016.31\|ATCAGACGTGGCGACCCGCTGAATTTAAGCA | >hg38_21_+_8433172_8446622_RNA45SN1_45S_50ntFlanks | 7986 | 8016 | 31 | ATCAGACGTGGCGACCCGCTGAATTTAAGCA | 696.8 | -11.5 | 1.4 | -8.0 | 1.1E-15 | 7.2E-15 | BT20/MB468 | MITO |
| >MI0000736\|hsa-mir-30c-1&WithFlank&1\|+\|40757278\|40757378@23.46.24;MI0000254\|hsa-mir-30c-2&WithFlank&6\|-\|71376954\|71377037@13.36.24&[MIMAT0000244&hsa-miR-30c-5p&offsets\|0\|+1;m-57&1\|+\|40757300\|40757323&offsets\|0\|0];[MIMAT0000244_1&hsa-miR-30c-5p&offsets\|0\|+1;m-56&6\|-\|71377002\|71377025&offsets\|0\|0]\|TGTAAACATCCTACACTCTCAGCT | >MI0000736\|hsa-mir-30c-1&WithFlank&1\|+\|40757278\|40757378 | 23 | 46 | 24 | TGTAAACATCCTACACTCTCAGCT | 16133.3 | 5.6 | 0.7 | 8.0 | 1.5E-15 | 7.2E-15 | BT20/MB468 | CYTO |
| >hg38_21_+_8433172_8446622_RNA45SN1_45S_50ntFlanks@11921.11949.29\|TGAGAGGTGTAGAATAAGTGGGAGGCCCC | >hg38_21_+_8433172_8446622_RNA45SN1_45S_50ntFlanks | 11921 | 11949 | 29 | TGAGAGGTGTAGAATAAGTGGGAGGCCCC | 378.1 | -12.4 | 1.5 | -8.0 | 1.1E-15 | 7.2E-15 | MB231/MB468 | MP |
| >MI0003581\|hsa-mir-574&WithFlank&4\|+\|38868026\|38868133@67.88.22&[MIMAT0003239&hsa-miR-574-3p&offsets\|0\|0;m-149&4\|+\|38868092\|38868113&offsets\|0\|0]\|CACGCTCATGCACACACCCACA | >MI0003581\|hsa-mir-574&WithFlank&4\|+\|38868026\|38868133 | 67 | 88 | 22 | CACGCTCATGCACACACCCACA | 386.5 | 4.9 | 0.6 | 8.0 | 1.1E-15 | 7.3E-15 | BT20/MB468 | MITO |
| >hg38_21_+_8433172_8446622_RNA45SN1_45S_50ntFlanks@9570.9592.23\|TGCAAATCGGTCGTCCGACCTGG | >hg38_21_+_8433172_8446622_RNA45SN1_45S_50ntFlanks | 9570 | 9592 | 23 | TGCAAATCGGTCGTCCGACCTGG | 453.3 | -12.7 | 1.6 | -8.0 | 1.1E-15 | 7.3E-15 | MB231/MB468 | MP |
| >hg38_21_+_8433172_8446622_RNA45SN1_45S_50ntFlanks@11197.11225.29\|GGCCGGCGGCGGCGGCGACTCTGGACGCG | >hg38_21_+_8433172_8446622_RNA45SN1_45S_50ntFlanks | 11197 | 11225 | 29 | GGCCGGCGGCGGCGGCGACTCTGGACGCG | 404.0 | -12.5 | 1.6 | -8.0 | 1.1E-15 | 7.3E-15 | MB231/MB468 | MP |
| >hg38_21_+_8433172_8446622_RNA45SN1_45S_50ntFlanks@11203.11225.23\|CGGCGGCGGCGACTCTGGACGCG | >hg38_21_+_8433172_8446622_RNA45SN1_45S_50ntFlanks | 11203 | 11225 | 23 | CGGCGGCGGCGACTCTGGACGCG | 311.6 | -6.1 | 0.8 | -8.0 | 1.1E-15 | 7.3E-15 | MB231/MB468 | CYTO |
| >hg38_21_+_8433172_8446622_RNA45SN1_45S_50ntFlanks@11199.11220.22\|CCGGCGGCGGCGGCGACTCTGG | >hg38_21_+_8433172_8446622_RNA45SN1_45S_50ntFlanks | 11199 | 11220 | 22 | CCGGCGGCGGCGGCGACTCTGG | 297.9 | -12.1 | 1.5 | -8.0 | 1.1E-15 | 7.3E-15 | MB231/MB468 | MP |
| >hg38_21_+_8433172_8446622_RNA45SN1_45S_50ntFlanks@12420.12455.36\|ATGTCGGCTCTTCCTATCATTGTGAAGCAGAATTCA | >hg38_21_+_8433172_8446622_RNA45SN1_45S_50ntFlanks | 12420 | 12455 | 36 | ATGTCGGCTCTTCCTATCATTGTGAAGCAGAATTCA | 1112.5 | -6.0 | 0.8 | -8.0 | 1.1E-15 | 7.3E-15 | MB231/MB468 | MP |

<table>
<tr><th></th><th></th><th></th><th></th><th></th><th></th><th></th><th></th><th></th><th></th><th></th><th></th><th></th><th></th></tr>
<tr><td>>MI0000434|hsa-let-7i&WithFlank&12|+|62603680|62603775@12.33.22&[MIMAT0000415&hsa-let-7i-5p&offsets|0|0;m-49&12|+|62603691|62603712&offsets|0|0]|TGAGGTAGTAGTTTGTGCTGTT</td><td>>MI0000434|hsa-let-7i&WithFlank&12|+|62603680|62603775</td><td>12</td><td>33</td><td>22</td><td>TGAGGTAGTAGTTTGTGCTGTT</td><td>75129.5</td><td>5.5</td><td>0.7</td><td>8.0</td><td>1.2E-15</td><td>7.3E-15</td><td>MB231/MB468</td><td>CYTO</td></tr>
<tr><td>>hg38_21_+_8433172_8446622_RNA45SN1_45S_50ntFlanks@9574.9592.19|AATCGGTCGTCCGACCTGG</td><td>>hg38_21_+_8433172_8446622_RNA45SN1_45S_50ntFlanks</td><td>9574</td><td>9592</td><td>19</td><td>AATCGGTCGTCCGACCTGG</td><td>190.8</td><td>-6.1</td><td>0.8</td><td>-8.0</td><td>1.2E-15</td><td>7.3E-15</td><td>MB231/MB468</td><td>CYTO</td></tr>
<tr><td>>hg38_21_+_8433172_8446622_RNA45SN1_45S_50ntFlanks@10922.10947.26|CTGGACGAGGCGCCGCCGCCCCCCCC</td><td>>hg38_21_+_8433172_8446622_RNA45SN1_45S_50ntFlanks</td><td>10922</td><td>10947</td><td>26</td><td>CTGGACGAGGCGCCGCCGCCCCCCCC</td><td>307.2</td><td>-11.0</td><td>1.4</td><td>-8.0</td><td>1.2E-15</td><td>7.3E-15</td><td>MB231/MB468</td><td>CYTO</td></tr>
<tr><td>>MI0001445|hsa-mir-423&WithFlank&17|+|30117073|30117178@59.81.23(+1U)&[MIMAT0001340&hsa-miR-423-3p&offsets|0|0(+1U);m-100&17|+|30117131|30117153&offsets|0|0(+1U)]|AGCTCGGTCTGAGGCCCCTCAGTT</td><td>>MI0001445|hsa-mir-423&WithFlank&17|+|30117073|30117178</td><td>59</td><td>81</td><td>23(+1U)</td><td>AGCTCGGTCTGAGGCCCCTCAGTT</td><td>1162.1</td><td>6.7</td><td>0.8</td><td>8.0</td><td>1.2E-15</td><td>7.3E-15</td><td>MB231/MB468</td><td>CYTO</td></tr>
<tr><td>>MI0000650|hsa-mir-200c&WithFlank&12|+|6963693|6963772@50.70.21(+1U)&[MIMAT0000617&hsa-miR-200c-3p&offsets|0|-2(+1U);m-54&12|+|6963742|6963764&offsets|0|-2(+1U)]|TAATACTGCCGGGTAATGATGT</td><td>>MI0000650|hsa-mir-200c&WithFlank&12|+|6963693|6963772</td><td>50</td><td>70</td><td>21(+1U)</td><td>TAATACTGCCGGGTAATGATGT</td><td>239.1</td><td>11.5</td><td>1.4</td><td>8.3</td><td>1.4E-16</td><td>7.3E-15</td><td>BT20/MB231</td><td>MITO</td></tr>
<tr><td>>MI0005757|hsa-mir-935&WithFlank&19|+|53982301|53982403@63.84.22&[MIMAT0004978&hsa-miR-935&offsets|+1|0;m-549&19|+|53982363|53982384&offsets|0|0]|CAGTTACCGCTTCCGCTACCGC</td><td>>MI0005757|hsa-mir-935&WithFlank&19|+|53982301|53982403</td><td>63</td><td>84</td><td>22</td><td>CAGTTACCGCTTCCGCTACCGC</td><td>232.6</td><td>11.5</td><td>1.4</td><td>8.3</td><td>1.4E-16</td><td>7.3E-15</td><td>BT20/MB231</td><td>MITO</td></tr>
<tr><td>>hg38_21_+_8433172_8446622_RNA45SN1_45S_50ntFlanks@11729.11747.19|TAACTATGACTCTCTTAAG</td><td>>hg38_21_+_8433172_8446622_RNA45SN1_45S_50ntFlanks</td><td>11729</td><td>11747</td><td>19</td><td>TAACTATGACTCTCTTAAG</td><td>16432.9</td><td>-10.8</td><td>1.4</td><td>-7.9</td><td>2.4E-15</td><td>7.3E-15</td><td>BT20/MB468</td><td>NUC</td></tr>
<tr><td>>hg38_21_+_8433172_8446622_RNA45SN1_45S_50ntFlanks@4724.4755.32|ATTAATCAAGAACGAAAGTCGGAGGTTCGAAG</td><td>>hg38_21_+_8433172_8446622_RNA45SN1_45S_50ntFlanks</td><td>4724</td><td>4755</td><td>32</td><td>ATTAATCAAGAACGAAAGTCGGAGGTTCGAAG</td><td>232.7</td><td>-11.7</td><td>1.5</td><td>-8.0</td><td>1.1E-15</td><td>7.3E-15</td><td>MB231/MB468</td><td>MP</td></tr>
<tr><td>>hg38_21_+_8433172_8446622_RNA45SN1_45S_50ntFlanks@11999.12038.40|CCCTGCGGGCCGCCGGTGAAATACCACTACTCTGATCGTT</td><td>>hg38_21_+_8433172_8446622_RNA45SN1_45S_50ntFlanks</td><td>11999</td><td>12038</td><td>40</td><td>CCCTGCGGGCCGCCGGTGAAATACCACTACTCTGATCGTT</td><td>449.6</td><td>-12.6</td><td>1.6</td><td>-8.0</td><td>1.8E-15</td><td>7.3E-15</td><td>MB231/MB468</td><td>MITO</td></tr>
<tr><td>>hg38_21_+_8433172_8446622_RNA45SN1_45S_50ntFlanks@10866.10904.39|CTAAGGGCTGGGTCGGTCGGGCTGGGGCGCGAAGCGGGG</td><td>>hg38_21_+_8433172_8446622_RNA45SN1_45S_50ntFlanks</td><td>10866</td><td>10904</td><td>39</td><td>CTAAGGGCTGGGTCGGTCGGGCTGGGGCGCGAAGCGGGG</td><td>349.1</td><td>-11.2</td><td>1.4</td><td>-8.0</td><td>1.2E-15</td><td>7.4E-15</td><td>MB231/MB468</td><td>CYTO</td></tr>
<tr><td>>hg38_21_+_8433172_8446622_RNA45SN1_45S_50ntFlanks@10866.10890.25|CTAAGGGCTGGGTCGGTCGGGCTGG</td><td>>hg38_21_+_8433172_8446622_RNA45SN1_45S_50ntFlanks</td><td>10866</td><td>10890</td><td>25</td><td>CTAAGGGCTGGGTCGGTCGGGCTGG</td><td>883.3</td><td>-9.6</td><td>1.2</td><td>-8.0</td><td>1.8E-15</td><td>7.4E-15</td><td>MB231/MB468</td><td>MITO</td></tr>
<tr><td>>hg38_21_+_8433172_8446622_RNA45SN1_45S_50ntFlanks@10099.10117.19|CGCGCGCGCGTGTGGTGTG</td><td>>hg38_21_+_8433172_8446622_RNA45SN1_45S_50ntFlanks</td><td>10099</td><td>10117</td><td>19</td><td>CGCGCGCGCGTGTGGTGTG</td><td>280.4</td><td>-11.1</td><td>1.4</td><td>-8.0</td><td>1.3E-15</td><td>7.4E-15</td><td>MB231/MB468</td><td>NUC</td></tr>
<tr><td>>hg38_21_+_8433172_8446622_RNA45SN1_45S_50ntFlanks@11641.11670.30|CGGCGGGTGTTGACGCGATGTGATTTCTGC</td><td>>hg38_21_+_8433172_8446622_RNA45SN1_45S_50ntFlanks</td><td>11641</td><td>11670</td><td>30</td><td>CGGCGGGTGTTGACGCGATGTGATTTCTGC</td><td>7577.3</td><td>-8.0</td><td>1.0</td><td>-8.0</td><td>1.1E-15</td><td>7.4E-15</td><td>BT20/MB468</td><td>MITO</td></tr>
<tr><td>>trna10_AspGTC_12_-_125424193_125424264@32.73.42,trna12_AspGTC_12_-_125411891_125411962@32.73.42,trna144_AspGTC_6_-_27551236_27551307@32.73.42,trna38_AspGTC_17_-_8125556_8125627@32.73.42,trna45_AspGTC_6_+_27447453_27447524@32.73.42,trna48_AspGTC_6_+_27471523_27471594@32.73.42,trna4_AspGTC_12_+_96429799_96429870@32.73.42,trna69_AspGTC_1_-_161440205_161440276@32.73.42,trna72_AspGTC_1_-_161432824_161432895@32.73.42,trna75_AspGTC_1_-_161425414_161425485@32.73.42,trna78_AspGTC_1_-_161418033_161418104@32.73.42,trna81_AspGTC_1_-_161410615_161410686@32.73.42|CTGTCACGCGGGAGACCGGGGTTCGATTCCCCGACGGGGAGC</td><td>>trna10_AspGTC_12_-_125424193_125424264</td><td>32</td><td>73</td><td>42</td><td>CTGTCACGCGGGAGACCGGGGTTCGATTCCCCGACGGGGAGC</td><td>336.8</td><td>-11.4</td><td>1.4</td><td>-8.0</td><td>1.3E-15</td><td>7.4E-15</td><td>MB231/MB468</td><td>NUC</td></tr>
<tr><td>>am_trnaMT_PheGAA_MT_+_577_647@41.60.20|GTTTAGACGGGCTCACATCA</td><td>>am_trnaMT_PheGAA_MT_+_577_647</td><td>41</td><td>60</td><td>20</td><td>GTTTAGACGGGCTCACATCA</td><td>243.1</td><td>12.2</td><td>1.5</td><td>8.0</td><td>1.6E-15</td><td>7.4E-15</td><td>BT20/MB468</td><td>CYTO</td></tr>
<tr><td>>hg38_21_+_8433172_8446622_RNA45SN1_45S_50ntFlanks@12156.12180.25|GAGTTTGACTGGGGCGGTACACCTG</td><td>>hg38_21_+_8433172_8446622_RNA45SN1_45S_50ntFlanks</td><td>12156</td><td>12180</td><td>25</td><td>GAGTTTGACTGGGGCGGTACACCTG</td><td>253.9</td><td>-11.8</td><td>1.5</td><td>-8.0</td><td>1.8E-15</td><td>7.4E-15</td><td>MB231/MB468</td><td>MITO</td></tr>
<tr><td>>hg38_21_+_8433172_8446622_RNA45SN1_45S_50ntFlanks@6652.6689.38|GACTCTTAGCGGTGGATCACTCGGCTCGTGCGTCGATG</td><td>>hg38_21_+_8433172_8446622_RNA45SN1_45S_50ntFlanks</td><td>6652</td><td>6689</td><td>38</td><td>GACTCTTAGCGGTGGATCACTCGGCTCGTGCGTCGATG</td><td>20191.4</td><td>-3.7</td><td>0.5</td><td>-7.9</td><td>2.5E-15</td><td>7.4E-15</td><td>BT20/MB468</td><td>NUC</td></tr>
<tr><td>>hg38_21_+_8433172_8446622_RNA45SN1_45S_50ntFlanks@6651.6668.18|CGACTCTTAGCGGTGGAT</td><td>>hg38_21_+_8433172_8446622_RNA45SN1_45S_50ntFlanks</td><td>6651</td><td>6668</td><td>18</td><td>CGACTCTTAGCGGTGGAT</td><td>94528.9</td><td>6.4</td><td>0.8</td><td>8.0</td><td>1.2E-15</td><td>7.5E-15</td><td>BT20/MB468</td><td>MITO</td></tr>
<tr><td>>hg38_21_+_8433172_8446622_RNA45SN1_45S_50ntFlanks@11035.11060.26|CTCTCTCCCCCGCTCCCCGTCCTCCC</td><td>>hg38_21_+_8433172_8446622_RNA45SN1_45S_50ntFlanks</td><td>11035</td><td>11060</td><td>26</td><td>CTCTCTCCCCCGCTCCCCGTCCTCCC</td><td>279.1</td><td>-11.9</td><td>1.5</td><td>-8.0</td><td>1.8E-15</td><td>7.5E-15</td><td>MB231/MB468</td><td>MITO</td></tr>
<tr><td>>MI0000085|hsa-mir-27a&WithFlank&19|-|13836434|13836523@57.74.18;MI0000440|hsa-mir-27b&WithFlank&9|+|95085439|95085547@67.84.18&[MIMAT0000084&hsa-miR-27a-3p&offsets|0|-3;m-43&19|-|13836447|13836467&offsets|0|-3];[MIMAT0000419&hsa-miR-27b-3p&offsets|0|-3;m-39&9|+|95085505|95085525&offsets|0|-3]|TTCACAGTGGCTAAGTTC</td><td>>MI0000085|hsa-mir-27a&WithFlank&19|-|13836434|13836523</td><td>57</td><td>74</td><td>18</td><td>TTCACAGTGGCTAAGTTC</td><td>593.6</td><td>5.9</td><td>0.7</td><td>8.0</td><td>1.3E-15</td><td>7.5E-15</td><td>MB231/MB468</td><td>NUC</td></tr>
<tr><td>>hg38_21_+_8433172_8446622_RNA45SN1_45S_50ntFlanks@12459.12476.18|AGCGTTGGATTGTTCACC</td><td>>hg38_21_+_8433172_8446622_RNA45SN1_45S_50ntFlanks</td><td>12459</td><td>12476</td><td>18</td><td>AGCGTTGGATTGTTCACC</td><td>1447.8</td><td>-6.0</td><td>0.8</td><td>-8.0</td><td>1.8E-15</td><td>7.5E-15</td><td>MB231/MB468</td><td>MITO</td></tr>
<tr><td>>hg38_21_+_8433172_8446622_RNA45SN1_45S_50ntFlanks@12456.12485.30|CCAAGCGTTGGATTGTTCACCCACTAATAG</td><td>>hg38_21_+_8433172_8446622_RNA45SN1_45S_50ntFlanks</td><td>12456</td><td>12485</td><td>30</td><td>CCAAGCGTTGGATTGTTCACCCACTAATAG</td><td>2415.6</td><td>-7.1</td><td>0.9</td><td>-8.0</td><td>1.2E-15</td><td>7.6E-15</td><td>MB231/MB468</td><td>CYTO</td></tr>
<tr><td>>hg38_21_+_8433172_8446622_RNA45SN1_45S_50ntFlanks@10410.10433.24|TCAGTCGGTCCTGAGAGATGGGCG</td><td>>hg38_21_+_8433172_8446622_RNA45SN1_45S_50ntFlanks</td><td>10410</td><td>10433</td><td>24</td><td>TCAGTCGGTCCTGAGAGATGGGCG</td><td>1186.4</td><td>-9.8</td><td>1.2</td><td>-8.0</td><td>1.8E-15</td><td>7.6E-15</td><td>MB231/MB468</td><td>MITO</td></tr>
<tr><td>>hg38_21_+_8433172_8446622_RNA45SN1_45S_50ntFlanks@11320.11338.19|CCACGTCTCGTCGCGCGCG</td><td>>hg38_21_+_8433172_8446622_RNA45SN1_45S_50ntFlanks</td><td>11320</td><td>11338</td><td>19</td><td>CCACGTCTCGTCGCGCGCG</td><td>276.2</td><td>-10.8</td><td>1.4</td><td>-8.0</td><td>1.2E-15</td><td>7.6E-15</td><td>MB231/MB468</td><td>CYTO</td></tr>
<tr><td>>hg38_21_+_8433172_8446622_RNA45SN1_45S_50ntFlanks@11873.11898.26|AAGAAGACCCTGTTGAGCTTGACTCT</td><td>>hg38_21_+_8433172_8446622_RNA45SN1_45S_50ntFlanks</td><td>11873</td><td>11898</td><td>26</td><td>AAGAAGACCCTGTTGAGCTTGACTCT</td><td>1077.6</td><td>-3.7</td><td>0.5</td><td>-8.0</td><td>1.6E-15</td><td>7.6E-15</td><td>BT20/MB468</td><td>CYTO</td></tr>
<tr><td>>hg38_21_+_8433172_8446622_RNA45SN1_45S_50ntFlanks@10797.10817.21|GCATGTTGGAACAATGTAGGT</td><td>>hg38_21_+_8433172_8446622_RNA45SN1_45S_50ntFlanks</td><td>10797</td><td>10817</td><td>21</td><td>GCATGTTGGAACAATGTAGGT</td><td>948.5</td><td>-7.7</td><td>1.0</td><td>-8.0</td><td>1.1E-15</td><td>7.6E-15</td><td>MB231/MB468</td><td>MP</td></tr>
<tr><td>>hg38_21_+_8433172_8446622_RNA45SN1_45S_50ntFlanks@11578.11605.28|TAGAACTGGTGCGGACCAGGGGAATCCG</td><td>>hg38_21_+_8433172_8446622_RNA45SN1_45S_50ntFlanks</td><td>11578</td><td>11605</td><td>28</td><td>TAGAACTGGTGCGGACCAGGGGAATCCG</td><td>556.0</td><td>-8.9</td><td>1.1</td><td>-8.0</td><td>1.4E-15</td><td>7.6E-15</td><td>MB231/MB468</td><td>NUC</td></tr>
<tr><td>>hg38_21_+_8433172_8446622_RNA45SN1_45S_50ntFlanks@12407.12433.27|TTTTGATCCTTCGATGTCGGCTCTTCC</td><td>>hg38_21_+_8433172_8446622_RNA45SN1_45S_50ntFlanks</td><td>12407</td><td>12433</td><td>27</td><td>TTTTGATCCTTCGATGTCGGCTCTTCC</td><td>2728.0</td><td>-5.3</td><td>0.7</td><td>-7.9</td><td>2.5E-15</td><td>7.6E-15</td><td>BT20/MB468</td><td>NUC</td></tr>
<tr><td>>hg38_21_+_8433172_8446622_RNA45SN1_45S_50ntFlanks@9454.9486.33|CCGCGCCGGGGAGGTGGAGCACGAGCGCACGTG</td><td>>hg38_21_+_8433172_8446622_RNA45SN1_45S_50ntFlanks</td><td>9454</td><td>9486</td><td>33</td><td>CCGCGCCGGGGAGGTGGAGCACGAGCGCACGTG</td><td>286.4</td><td>-11.9</td><td>1.5</td><td>-8.0</td><td>1.9E-15</td><td>7.7E-15</td><td>MB231/MB468</td><td>MITO</td></tr>
<tr><td>>MI0000087|hsa-mir-29a&WithFlank&7|-|130876741|130876816@48.69.22&[MIMAT0000086&hsa-miR-29a-3p&offsets|0|0;m-26&7|-|130876749|130876769&offsets|0|+1]|TAGCACCATCTGAAATCGGTTA</td><td>>MI0000087|hsa-mir-29a&WithFlank&7|-|130876741|130876816</td><td>48</td><td>69</td><td>22</td><td>TAGCACCATCTGAAATCGGTTA</td><td>2001.9</td><td>7.7</td><td>1.0</td><td>8.0</td><td>1.2E-15</td><td>7.7E-15</td><td>MB231/MB468</td><td>CYTO</td></tr>
<tr><td>>MI0000434|hsa-let-7i&WithFlank&12|+|62603680|62603775@12.33.22(+1U)&[MIMAT0000415&hsa-let-7i-5p&offsets|0|0(+1U);m-49&12|+|62603691|62603712&offsets|0|0(+1U)]|TGAGGTAGTAGTTTGTGCTGTTT</td><td>>MI0000434|hsa-let-7i&WithFlank&12|+|62603680|62603775</td><td>12</td><td>33</td><td>22(+1U)</td><td>TGAGGTAGTAGTTTGTGCTGTTT</td><td>3242.7</td><td>6.0</td><td>0.7</td><td>8.0</td><td>1.2E-15</td><td>7.7E-15</td><td>MB231/MB468</td><td>CYTO</td></tr>
<tr><td>>hg38_21_+_8433172_8446622_RNA45SN1_45S_50ntFlanks@11572.11590.19|CCGACTTAGAACTGGTGCG</td><td>>hg38_21_+_8433172_8446622_RNA45SN1_45S_50ntFlanks</td><td>11572</td><td>11590</td><td>19</td><td>CCGACTTAGAACTGGTGCG</td><td>1086.0</td><td>-5.0</td><td>0.6</td><td>-7.9</td><td>1.9E-15</td><td>7.7E-15</td><td>MB231/MB468</td><td>MITO</td></tr>
<tr><td>>hg38_21_+_8433172_8446622_RNA45SN1_45S_50ntFlanks@9940.9968.29|CAGGACGGTGGCCATGGAAGTCGGAATCC</td><td>>hg38_21_+_8433172_8446622_RNA45SN1_45S_50ntFlanks</td><td>9940</td><td>9968</td><td>29</td><td>CAGGACGGTGGCCATGGAAGTCGGAATCC</td><td>663.0</td><td>5.3</td><td>0.7</td><td>8.0</td><td>1.2E-15</td><td>7.7E-15</td><td>BT20/MB468</td><td>MITO</td></tr>
<tr><td>>hg38_21_+_8433172_8446622_RNA45SN1_45S_50ntFlanks@11641.11681.41|CGGCGGGTGTTGACGCGATGTGATTTCTGCCCAGTGCTCTG</td><td>>hg38_21_+_8433172_8446622_RNA45SN1_45S_50ntFlanks</td><td>11641</td><td>11681</td><td>41</td><td>CGGCGGGTGTTGACGCGATGTGATTTCTGCCCAGTGCTCTG</td><td>1894.9</td><td>-10.4</td><td>1.3</td><td>-7.9</td><td>1.9E-15</td><td>7.7E-15</td><td>MB231/MB468</td><td>MITO</td></tr>
<tr><td>>hg38_21_+_8433172_8446622_RNA45SN1_45S_50ntFlanks@11705.11728.24|TGAAGCGCGGGTAAACGGCGGGAG</td><td>>hg38_21_+_8433172_8446622_RNA45SN1_45S_50ntFlanks</td><td>11705</td><td>11728</td><td>24</td><td>TGAAGCGCGGGTAAACGGCGGGAG</td><td>452.3</td><td>-10.9</td><td>1.4</td><td>-8.0</td><td>1.2E-15</td><td>7.8E-15</td><td>BT20/MB468</td><td>MITO</td></tr>
<tr><td>>hg38_21_+_8433172_8446622_RNA45SN1_45S_50ntFlanks@10389.10408.20|TGAACAGCAGTTGAACATGG</td><td>>hg38_21_+_8433172_8446622_RNA45SN1_45S_50ntFlanks</td><td>10389</td><td>10408</td><td>20</td><td>TGAACAGCAGTTGAACATGG</td><td>286.9</td><td>-12.0</td><td>1.5</td><td>-8.0</td><td>1.1E-15</td><td>7.8E-15</td><td>MB231/MB468</td><td>MP</td></tr>
<tr><td>>hg38_21_+_8433172_8446622_RNA45SN1_45S_50ntFlanks@11768.11786.19|CTAATTAGTGACGCGCATG</td><td>>hg38_21_+_8433172_8446622_RNA45SN1_45S_50ntFlanks</td><td>11768</td><td>11786</td><td>19</td><td>CTAATTAGTGACGCGCATG</td><td>1300.8</td><td>-7.4</td><td>0.9</td><td>-8.0</td><td>1.2E-15</td><td>7.8E-15</td><td>MB231/MB468</td><td>CYTO</td></tr>
<tr><td>>hg38_21_+_8433172_8446622_RNA45SN1_45S_50ntFlanks@3875.3898.24|ATACATGCCGACGGGCGCTGACCC</td><td>>hg38_21_+_8433172_8446622_RNA45SN1_45S_50ntFlanks</td><td>3875</td><td>3898</td><td>24</td><td>ATACATGCCGACGGGCGCTGACCC</td><td>8954.2</td><td>-10.6</td><td>1.3</td><td>-7.9</td><td>2.6E-15</td><td>7.8E-15</td><td>BT20/MB468</td><td>NUC</td></tr>
<tr><td>>hg38_21_+_8433172_8446622_RNA45SN1_45S_50ntFlanks@10914.10932.19|CGCCGCGGCTGGACGAGGC</td><td>>hg38_21_+_8433172_8446622_RNA45SN1_45S_50ntFlanks</td><td>10914</td><td>10932</td><td>19</td><td>CGCCGCGGCTGGACGAGGC</td><td>885.6</td><td>-12.9</td><td>1.5</td><td>-8.4</td><td>3.5E-17</td><td>7.8E-15</td><td>BT20/MB468</td><td>MP</td></tr>
<tr><td>>hg38_21_+_8433172_8446622_RNA45SN1_45S_50ntFlanks@12712.12745.34|CGGAGCCTCGGTTGGCCTCGGATAGCCGGTCCCC</td><td>>hg38_21_+_8433172_8446622_RNA45SN1_45S_50ntFlanks</td><td>12712</td><td>12745</td><td>34</td><td>CGGAGCCTCGGTTGGCCTCGGATAGCCGGTCCCC</td><td>1679.4</td><td>-13.8</td><td>1.6</td><td>-8.4</td><td>3.5E-17</td><td>7.8E-15</td><td>BT20/MB468</td><td>MP</td></tr>
<tr><td>>hg38_21_+_8433172_8446622_RNA45SN1_45S_50ntFlanks@10128.10149.22|CGGCGGCGGCGGCGGCGGCGGG</td><td>>hg38_21_+_8433172_8446622_RNA45SN1_45S_50ntFlanks</td><td>10128</td><td>10149</td><td>22</td><td>CGGCGGCGGCGGCGGCGGCGGG</td><td>1165.2</td><td>-12.3</td><td>1.5</td><td>-8.0</td><td>1.2E-15</td><td>7.8E-15</td><td>BT20/MB468</td><td>MITO</td></tr>
<tr><td>>hg38_21_+_8433172_8446622_RNA45SN1_45S_50ntFlanks@8845.8866.22|CGGCGGCGGCGGCGGCGGCGGG</td><td>>hg38_21_+_8433172_8446622_RNA45SN1_45S_50ntFlanks</td><td>8845</td><td>8866</td><td>22</td><td>CGGCGGCGGCGGCGGCGGCGGG</td><td>1165.2</td><td>-12.3</td><td>1.5</td><td>-8.0</td><td>1.2E-15</td><td>7.8E-15</td><td>BT20/MB468</td><td>MITO</td></tr>
<tr><td>>hg38_21_+_8433172_8446622_RNA45SN1_45S_50ntFlanks@5500.5526.27|GTCGAACTTGACTATCTAGAGGAAGTA</td><td>>hg38_21_+_8433172_8446622_RNA45SN1_45S_50ntFlanks</td><td>5500</td><td>5526</td><td>27</td><td>GTCGAACTTGACTATCTAGAGGAAGTA</td><td>172.5</td><td>-11.3</td><td>1.4</td><td>-8.0</td><td>1.1E-15</td><td>7.8E-15</td><td>MB231/MB468</td><td>MP</td></tr>
<tr><td>>hg38_21_+_8433172_8446622_RNA45SN1_45S_50ntFlanks@4496.4524.29|CGAAGCGTTTACTTTGAAAAAATTAGAGT</td><td>>hg38_21_+_8433172_8446622_RNA45SN1_45S_50ntFlanks</td><td>4496</td><td>4524</td><td>29</td><td>CGAAGCGTTTACTTTGAAAAAATTAGAGT</td><td>345.6</td><td>5.8</td><td>0.7</td><td>8.0</td><td>1.7E-15</td><td>7.8E-15</td><td>BT20/MB468</td><td>CYTO</td></tr>
<tr><td>>hg38_21_+_8433172_8446622_RNA45SN1_45S_50ntFlanks@11235.11268.34|CTTCCCGTGGATCGCCCCAGCTGCGGCGGGCGTC</td><td>>hg38_21_+_8433172_8446622_RNA45SN1_45S_50ntFlanks</td><td>11235</td><td>11268</td><td>34</td><td>CTTCCCGTGGATCGCCCCAGCTGCGGCGGGCGTC</td><td>178.3</td><td>-7.2</td><td>0.9</td><td>-8.0</td><td>1.2E-15</td><td>7.8E-15</td><td>MB231/MB468</td><td>CYTO</td></tr>
<tr><td>>hg38_21_+_8433172_8446622_RNA45SN1_45S_50ntFlanks@9572.9596.25|CAAATCGGTCGTCCGACCTGGGTAT</td><td>>hg38_21_+_8433172_8446622_RNA45SN1_45S_50ntFlanks</td><td>9572</td><td>9596</td><td>25</td><td>CAAATCGGTCGTCCGACCTGGGTAT</td><td>2669.8</td><td>-11.3</td><td>1.4</td><td>-7.9</td><td>2.6E-15</td><td>7.8E-15</td><td>BT20/MB468</td><td>NUC</td></tr>
<tr><td>>hg38_21_+_8433172_8446622_RNA45SN1_45S_50ntFlanks@10396.10424.29|CAGTTGAACATGGGTCAGTCGGTCCTGAG</td><td>>hg38_21_+_8433172_8446622_RNA45SN1_45S_50ntFlanks</td><td>10396</td><td>10424</td><td>29</td><td>CAGTTGAACATGGGTCAGTCGGTCCTGAG</td><td>3855.3</td><td>4.8</td><td>0.6</td><td>8.0</td><td>1.7E-15</td><td>7.9E-15</td><td>BT20/MB468</td><td>CYTO</td></tr>
</table>

<table>
<tr><th></th><th></th><th></th><th></th><th></th><th></th><th></th><th></th><th></th><th></th><th></th><th></th><th></th><th></th></tr>
<tr><td>>MI0000736|hsa-mir-30c-1&WithFlank&1|+|40757278|40757378@23.44.22(+1A);MI0000254|hsa-mir-30c-2&WithFlank&6|-|71376954|71377037@13.34.22(+1A)&[MIMAT0000244&hsa-miR-30c-5p&offsets|0|-1(+1A);m-57&1|+|40757300|40757323&offsets|0|-2(+1A)];[MIMAT0000244_1&hsa-miR-30c-5p&offsets|0|-1(+1A);m-56&6|-|71377002|71377025&offsets|0|-2(+1A)]|TGTAAACATCCTACACTCTCAGA</td><td>>MI0000736|hsa-mir-30c-1&WithFlank&1|+|40757278|40757378</td><td>23</td><td>44</td><td>22(+1A)</td><td>TGTAAACATCCTACACTCTCAGA</td><td>1364.1</td><td>5.3</td><td>0.7</td><td>8.0</td><td>1.4E-15</td><td>7.9E-15</td><td>MB231/MB468</td><td>NUC</td></tr>
<tr><td>>MI0000743|hsa-mir-34c&WithFlank&11|+|111513433|111513521@19.41.23&[MIMAT0000686&hsa-miR-34c-5p&offsets|0|0;m-336&11|+|111513451|111513473&offsets|0|0]|AGGCAGTGTAGTTAGCTGATTGC</td><td>>MI0000743|hsa-mir-34c&WithFlank&11|+|111513433|111513521</td><td>19</td><td>41</td><td>23</td><td>AGGCAGTGTAGTTAGCTGATTGC</td><td>1835.6</td><td>-14.2</td><td>1.7</td><td>-8.5</td><td>1.8E-17</td><td>7.9E-15</td><td>BT20/MB231</td><td>MP</td></tr>
<tr><td>>MI0006442|hsa-mir-664a&WithFlank&1|-|220200532|220200625@55.77.23&[MIMAT0005949&hsa-miR-664a-3p&offsets|0|0;m-210&1|-|220200549|220200571&offsets|0|0]|TATTCATTTATCCCCAGCCTACA</td><td>>MI0006442|hsa-mir-664a&WithFlank&1|-|220200532|220200625</td><td>55</td><td>77</td><td>23</td><td>TATTCATTTATCCCCAGCCTACA</td><td>306.0</td><td>9.9</td><td>1.2</td><td>8.0</td><td>1.7E-15</td><td>7.9E-15</td><td>BT20/MB468</td><td>CYTO</td></tr>
<tr><td>>hg38_21_+_8433172_8446622_RNA45SN1_45S_50ntFlanks@3870.3898.29|AGCTAATACATGCCGACGGGCGCTGACCC</td><td>>hg38_21_+_8433172_8446622_RNA45SN1_45S_50ntFlanks</td><td>3870</td><td>3898</td><td>29</td><td>AGCTAATACATGCCGACGGGCGCTGACCC</td><td>522.0</td><td>-11.1</td><td>1.4</td><td>-8.0</td><td>1.2E-15</td><td>7.9E-15</td><td>BT20/MB468</td><td>MITO</td></tr>
<tr><td>>MI0000108|hsa-mir-103a-2&WithFlank&20|+|3917488|3917577@54.75.22(+1G);MI0000109|hsa-mir-103a-1&WithFlank&5|-|168560890|168560979@54.75.22(+1G)&[MIMAT0000101&hsa-miR-103a-3p&offsets|0|-1(+1G);m-12&20|+|3917541|3917563&offsets|0|-1(+1G)];[MIMAT0000101_1&hsa-miR-103a-3p&offsets|0|-1(+1G);m-13&5|-|168560904|168560926&offsets|0|-1(+1G);m-44220&5|-|168560908|168560926&offsets|0|+3(+1G)]|AGCAGCATTGTACAGGGCTATGG</td><td>>MI0000108|hsa-mir-103a-2&WithFlank&20|+|3917488|3917577</td><td>54</td><td>75</td><td>22(+1G)</td><td>AGCAGCATTGTACAGGGCTATGG</td><td>897.7</td><td>6.2</td><td>0.8</td><td>8.0</td><td>1.7E-15</td><td>7.9E-15</td><td>BT20/MB468</td><td>CYTO</td></tr>
<tr><td>>hg38_21_+_8433172_8446622_RNA45SN1_45S_50ntFlanks@12418.12443.26|CGATGTCGGCTCTTCCTATCATTGTG</td><td>>hg38_21_+_8433172_8446622_RNA45SN1_45S_50ntFlanks</td><td>12418</td><td>12443</td><td>26</td><td>CGATGTCGGCTCTTCCTATCATTGTG</td><td>8877.8</td><td>-4.7</td><td>0.6</td><td>-8.0</td><td>1.7E-15</td><td>7.9E-15</td><td>BT20/MB468</td><td>CYTO</td></tr>
<tr><td>>MI0000085|hsa-mir-27a&WithFlank&19|-|13836434|13836523@57.76.20(+2U)&[MIMAT0000084&hsa-miR-27a-3p&offsets|0|-1(+2U);m-43&19|-|13836447|13836467&offsets|0|-1(+2U)]|TTCACAGTGGCTAAGTTCCGTT</td><td>>MI0000085|hsa-mir-27a&WithFlank&19|-|13836434|13836523</td><td>57</td><td>76</td><td>20(+2U)</td><td>TTCACAGTGGCTAAGTTCCGTT</td><td>1874.3</td><td>5.4</td><td>0.7</td><td>8.0</td><td>1.4E-15</td><td>7.9E-15</td><td>MB231/MB468</td><td>NUC</td></tr>
<tr><td>>hg38_21_+_8433172_8446622_RNA45SN1_45S_50ntFlanks@12044.12068.25|ACTGACCCGGTGAGGCGGGGGGGCG</td><td>>hg38_21_+_8433172_8446622_RNA45SN1_45S_50ntFlanks</td><td>12044</td><td>12068</td><td>25</td><td>ACTGACCCGGTGAGGCGGGGGGGCG</td><td>258.2</td><td>-10.7</td><td>1.3</td><td>-8.0</td><td>1.3E-15</td><td>8.0E-15</td><td>MB231/MB468</td><td>CYTO</td></tr>
<tr><td>>hg38_21_+_8433172_8446622_RNA45SN1_45S_50ntFlanks@8498.8515.18|CGTTCCTCCCGACCCCTC</td><td>>hg38_21_+_8433172_8446622_RNA45SN1_45S_50ntFlanks</td><td>8498</td><td>8515</td><td>18</td><td>CGTTCCTCCCGACCCCTC</td><td>316.6</td><td>-12.1</td><td>1.5</td><td>-7.9</td><td>1.9E-15</td><td>8.0E-15</td><td>MB231/MB468</td><td>MITO</td></tr>
<tr><td>>MI0000746|hsa-mir-99b&WithFlank&19|+|51692606|51692687@13.32.20(+1U)&[MIMAT0000689&hsa-miR-99b-5p&offsets|0|-2(+1U);m-10&19|+|51692618|51692639&offsets|0|-2(+1U)]|CACCCGTAGAACCGACCTTGT</td><td>>MI0000746|hsa-mir-99b&WithFlank&19|+|51692606|51692687</td><td>13</td><td>32</td><td>20(+1U)</td><td>CACCCGTAGAACCGACCTTGT</td><td>1874.4</td><td>6.6</td><td>0.8</td><td>8.0</td><td>1.3E-15</td><td>8.0E-15</td><td>MB231/MB468</td><td>CYTO</td></tr>
<tr><td>>hg38_21_+_8433172_8446622_RNA45SN1_45S_50ntFlanks@10794.10817.24|CTGGCATGTTGGAACAATGTAGGT</td><td>>hg38_21_+_8433172_8446622_RNA45SN1_45S_50ntFlanks</td><td>10794</td><td>10817</td><td>24</td><td>CTGGCATGTTGGAACAATGTAGGT</td><td>776.9</td><td>-7.7</td><td>1.0</td><td>-7.9</td><td>1.9E-15</td><td>8.0E-15</td><td>MB231/MB468</td><td>MITO</td></tr>
<tr><td>>am_MI0005763|hsa-mir-941-1&WithFlank&20|+|63919443|63919526@53.74.22(+1A);MI0005764|hsa-mir-941-2&WithFlank&20|+|63919499|63919582@53.74.22(+1A);MI0005765|hsa-mir-941-3&WithFlank&20|+|63919555|63919638@53.74.22(+1A);MI0005766|hsa-mir-941-4&WithFlank&20|+|63919750|63919833@53.74.22(+1A);MI0031520|hsa-mir-941-5&WithFlank&20|+|63919862|63919945@53.74.22(+1A)&[MIMAT0004984&hsa-miR-941&offsets|0|-1(+1A);m-263&20|+|63919495|63919517&offsets|0|-1(+1A)];[MIMAT0004984_1&hsa-miR-941&offsets|0|-1(+1A);m-263&20|+|63919551|63919573&offsets|0|-1(+1A)];[MIMAT0004984_2&hsa-miR-941&offsets|0|-1(+1A);m-263&20|+|63919607|63919629&offsets|0|-1(+1A)];[MIMAT0004984_3&hsa-miR-941&offsets|0|-1(+1A);m-263&20|+|63919802|63919824&offsets|0|-1(+1A)];[MIMAT0004984_4&hsa-miR-941&offsets|0|-1(+1A);m-263&20|+|63919914|63919936&offsets|0|-1(+1A)]|CACCCGGCTGTGTGCACATGTGA</td><td>>am_MI0005763|hsa-mir-941-1&WithFlank&20|+|63919443|63919526</td><td>53</td><td>74</td><td>22(+1A)</td><td>CACCCGGCTGTGTGCACATGTGA</td><td>547.4</td><td>5.5</td><td>0.7</td><td>8.0</td><td>1.4E-15</td><td>8.0E-15</td><td>MB231/MB468</td><td>NUC</td></tr>
<tr><td>>hg38_21_+_8433172_8446622_RNA45SN1_45S_50ntFlanks@9341.9370.30|CACGAAAGCCGCCGTGGCGCAATGAAGGTG</td><td>>hg38_21_+_8433172_8446622_RNA45SN1_45S_50ntFlanks</td><td>9341</td><td>9370</td><td>30</td><td>CACGAAAGCCGCCGTGGCGCAATGAAGGTG</td><td>691.0</td><td>-13.3</td><td>1.7</td><td>-8.0</td><td>1.2E-15</td><td>8.0E-15</td><td>MB231/MB468</td><td>MP</td></tr>
<tr><td>>hg38_21_+_8433172_8446622_RNA45SN1_45S_50ntFlanks@4099.4127.29|GTGACCACGGGTGACGGGGAATCAGGGTT</td><td>>hg38_21_+_8433172_8446622_RNA45SN1_45S_50ntFlanks</td><td>4099</td><td>4127</td><td>29</td><td>GTGACCACGGGTGACGGGGAATCAGGGTT</td><td>231.2</td><td>-10.8</td><td>1.4</td><td>-8.0</td><td>1.4E-15</td><td>8.1E-15</td><td>MB231/MB468</td><td>NUC</td></tr>
<tr><td>>MI0003628|hsa-mir-615&WithFlank&12|+|54033944|54034051@67.86.20(+1A)&[MIMAT0003283&hsa-miR-615-3p&offsets|0|-2(+1A);m-495&12|+|54034010|54034030&offsets|0|-1(+1A)]|TCCGAGCCTGGGTCTCCCTCA</td><td>>MI0003628|hsa-mir-615&WithFlank&12|+|54033944|54034051</td><td>67</td><td>86</td><td>20(+1A)</td><td>TCCGAGCCTGGGTCTCCCTCA</td><td>640.8</td><td>8.0</td><td>1.0</td><td>8.0</td><td>1.3E-15</td><td>8.1E-15</td><td>BT20/MB468</td><td>MITO</td></tr>
<tr><td>>hg38_21_+_8433172_8446622_RNA45SN1_45S_50ntFlanks@12712.12757.46|CGGAGCCTCGGTTGGCCTCGGATAGCCGGTCCCCCGCCTGTCCCCG</td><td>>hg38_21_+_8433172_8446622_RNA45SN1_45S_50ntFlanks</td><td>12712</td><td>12757</td><td>46</td><td>CGGAGCCTCGGTTGGCCTCGGATAGCCGGTCCCCCGCCTGTCCCCG</td><td>430.3</td><td>-12.5</td><td>1.6</td><td>-7.9</td><td>2.0E-15</td><td>8.1E-15</td><td>MB231/MB468</td><td>MITO</td></tr>
<tr><td>>hg38_21_+_8433172_8446622_RNA45SN1_45S_50ntFlanks@11906.11929.24|CACGGTGAAGAGACATGAGAGGTG</td><td>>hg38_21_+_8433172_8446622_RNA45SN1_45S_50ntFlanks</td><td>11906</td><td>11929</td><td>24</td><td>CACGGTGAAGAGACATGAGAGGTG</td><td>346.2</td><td>-12.3</td><td>1.5</td><td>-8.0</td><td>1.2E-15</td><td>8.1E-15</td><td>MB231/MB468</td><td>MP</td></tr>
<tr><td>>trna119_LysCTT_1_-_145395522_145395594@1.34.34,trna11_LysCTT_5_-_180648979_180649051@1.34.34,trna13_LysCTT_6_+_26556774_26556846@1.34.34,trna7_LysCTT_16_+_3225692_3225764@1.34.34,trna9_LysCTT_5_+_180634755_180634827@1.34.34|GCCCGGCTAGCTCAGTCGGTAGAGCATGAGACTC</td><td>>trna119_LysCTT_1_-_145395522_145395594</td><td>1</td><td>34</td><td>34</td><td>GCCCGGCTAGCTCAGTCGGTAGAGCATGAGACTC</td><td>21638.9</td><td>9.4</td><td>1.2</td><td>8.0</td><td>1.3E-15</td><td>8.1E-15</td><td>MB231/MB468</td><td>CYTO</td></tr>
<tr><td>>hg38_1_-_228634819_228635039_RNA5S12_5S_50ntFlanks@150.169.20|AATACCGGGTGCTGTAGGCT</td><td>>hg38_1_-_228634819_228635039_RNA5S12_5S_50ntFlanks</td><td>150</td><td>169</td><td>20</td><td>AATACCGGGTGCTGTAGGCT</td><td>2106.5</td><td>5.2</td><td>0.7</td><td>8.0</td><td>1.7E-15</td><td>8.1E-15</td><td>BT20/MB468</td><td>CYTO</td></tr>
<tr><td>>hg38_21_+_8433172_8446622_RNA45SN1_45S_50ntFlanks@3854.3873.20|ACTGTGGTAATTCTAGAGCT</td><td>>hg38_21_+_8433172_8446622_RNA45SN1_45S_50ntFlanks</td><td>3854</td><td>3873</td><td>20</td><td>ACTGTGGTAATTCTAGAGCT</td><td>1164.9</td><td>-13.3</td><td>1.6</td><td>-8.4</td><td>3.7E-17</td><td>8.1E-15</td><td>BT20/MB468</td><td>MP</td></tr>
<tr><td>>MI0000088|hsa-mir-30a&WithFlank&6|-|71403545|71403627@12.34.23(+1G)&[MIMAT0000087&hsa-miR-30a-5p&offsets|0|+1(+1G);m-15&6|-|71403595|71403616&offsets|0|+1(+1G)]|TGTAAACATCCTCGACTGGAAGCG</td><td>>MI0000088|hsa-mir-30a&WithFlank&6|-|71403545|71403627</td><td>12</td><td>34</td><td>23(+1G)</td><td>TGTAAACATCCTCGACTGGAAGCG</td><td>2681.9</td><td>7.4</td><td>0.9</td><td>8.0</td><td>1.4E-15</td><td>8.1E-15</td><td>MB231/MB468</td><td>NUC</td></tr>
<tr><td>>hg38_21_+_8433172_8446622_RNA45SN1_45S_50ntFlanks@8182.8203.22|ATCGAGGCCCAGCCCGTGGACG</td><td>>hg38_21_+_8433172_8446622_RNA45SN1_45S_50ntFlanks</td><td>8182</td><td>8203</td><td>22</td><td>ATCGAGGCCCAGCCCGTGGACG</td><td>332.5</td><td>-10.7</td><td>1.3</td><td>-8.0</td><td>1.8E-15</td><td>8.1E-15</td><td>BT20/MB468</td><td>CYTO</td></tr>
<tr><td>>hg38_21_+_8433172_8446622_RNA45SN1_45S_50ntFlanks@12365.12383.19|ATAACTGGCTTGTGGCGGC</td><td>>hg38_21_+_8433172_8446622_RNA45SN1_45S_50ntFlanks</td><td>12365</td><td>12383</td><td>19</td><td>ATAACTGGCTTGTGGCGGC</td><td>278.2</td><td>-11.9</td><td>1.5</td><td>-7.9</td><td>2.0E-15</td><td>8.1E-15</td><td>MB231/MB468</td><td>MITO</td></tr>
<tr><td>>hg38_21_+_8433172_8446622_RNA45SN1_45S_50ntFlanks@12077.12095.19|GGGGCTCTCGCTTCTGGCG</td><td>>hg38_21_+_8433172_8446622_RNA45SN1_45S_50ntFlanks</td><td>12077</td><td>12095</td><td>19</td><td>GGGGCTCTCGCTTCTGGCG</td><td>725.7</td><td>-5.2</td><td>0.6</td><td>-8.0</td><td>1.5E-15</td><td>8.1E-15</td><td>MB231/MB468</td><td>NUC</td></tr>
<tr><td>>hg38_21_+_8433172_8446622_RNA45SN1_45S_50ntFlanks@12081.12106.26|CTCTCGCTTCTGGCGCCAAGCGCCCG</td><td>>hg38_21_+_8433172_8446622_RNA45SN1_45S_50ntFlanks</td><td>12081</td><td>12106</td><td>26</td><td>CTCTCGCTTCTGGCGCCAAGCGCCCG</td><td>1230.9</td><td>-7.8</td><td>1.0</td><td>-7.9</td><td>2.0E-15</td><td>8.1E-15</td><td>MB231/MB468</td><td>MITO</td></tr>
<tr><td>>am_MI0000809|hsa-mir-151a&WithFlank&8|-|140732558|140732659@53.70.18&[MIMAT0000757&hsa-miR-151a-3p&offsets|0|-3;m-30&8|-|140732587|140732607&offsets|0|-3]|CTAGACTGAAGCTCCTTG</td><td>>am_MI0000809|hsa-mir-151a&WithFlank&8|-|140732558|140732659</td><td>53</td><td>70</td><td>18</td><td>CTAGACTGAAGCTCCTTG</td><td>753.2</td><td>5.9</td><td>0.7</td><td>8.0</td><td>1.5E-15</td><td>8.2E-15</td><td>MB231/MB468</td><td>NUC</td></tr>
<tr><td>>hg38_21_+_8433172_8446622_RNA45SN1_45S_50ntFlanks@485.511.27|GATGTGGTGACGTCGTGCTCTCCCGGG</td><td>>hg38_21_+_8433172_8446622_RNA45SN1_45S_50ntFlanks</td><td>485</td><td>511</td><td>27</td><td>GATGTGGTGACGTCGTGCTCTCCCGGG</td><td>1982.9</td><td>29.0</td><td>3.6</td><td>8.0</td><td>1.5E-15</td><td>8.2E-15</td><td>MB231/MB468</td><td>NUC</td></tr>
<tr><td>>trna111_HisGTG_1_-_147774845_147774916@3.30.28,trna118_HisGTG_1_-_145396881_145396952@3.30.28,trna16_HisGTG_1_+_146544773_146544844@3.30.28,trna1_HisGTG_15_+_45493349_45493420@3.30.28,trna21_HisGTG_1_+_147753471_147753542@3.30.28,trna33_HisGTG_6_+_27125906_27125977@3.30.28,trna7_HisGTG_9_-_14433938_14434009@3.30.28,trna8_HisGTG_15_-_45492611_45492682@3.30.28,trna9_HisGTG_15_-_45490804_45490875@3.30.28|CGTGATCGTATAGTGGTTAGTACTCTGC</td><td>>trna111_HisGTG_1_-_147774845_147774916</td><td>3</td><td>30</td><td>28</td><td>CGTGATCGTATAGTGGTTAGTACTCTGC</td><td>927.5</td><td>5.0</td><td>0.6</td><td>8.0</td><td>1.8E-15</td><td>8.2E-15</td><td>BT20/MB468</td><td>CYTO</td></tr>
<tr><td>>hg38_21_+_8433172_8446622_RNA45SN1_45S_50ntFlanks@12626.12657.32|AATGGGGCGAAGCTACCATCTGTGGGATTATG</td><td>>hg38_21_+_8433172_8446622_RNA45SN1_45S_50ntFlanks</td><td>12626</td><td>12657</td><td>32</td><td>AATGGGGCGAAGCTACCATCTGTGGGATTATG</td><td>2735.6</td><td>-8.7</td><td>1.1</td><td>-7.9</td><td>2.0E-15</td><td>8.2E-15</td><td>MB231/MB468</td><td>MITO</td></tr>
<tr><td>>MI0000746|hsa-mir-99b&WithFlank&19|+|51692606|51692687@13.32.20(+1U)&[MIMAT0000689&hsa-miR-99b-5p&offsets|0|-2(+1U);m-10&19|+|51692618|51692639&offsets|0|-2(+1U)]|CACCCGTAGAACCGACCTTGT</td><td>>MI0000746|hsa-mir-99b&WithFlank&19|+|51692606|51692687</td><td>13</td><td>32</td><td>20(+1U)</td><td>CACCCGTAGAACCGACCTTGT</td><td>7618.4</td><td>7.1</td><td>0.8</td><td>8.4</td><td>3.8E-17</td><td>8.2E-15</td><td>BT20/MB468</td><td>MP</td></tr>
<tr><td>>MI0003628|hsa-mir-615&WithFlank&12|+|54033944|54034051@67.86.20(+1C)&[MIMAT0003283&hsa-miR-615-3p&offsets|0|-2(+1C);m-495&12|+|54034010|54034030&offsets|0|-1(+1C)]|TCCGAGCCTGGGTCTCCCTCC</td><td>>MI0003628|hsa-mir-615&WithFlank&12|+|54033944|54034051</td><td>67</td><td>86</td><td>20(+1C)</td><td>TCCGAGCCTGGGTCTCCCTCC</td><td>512.2</td><td>5.8</td><td>0.7</td><td>8.2</td><td>1.6E-16</td><td>8.3E-15</td><td>BT20/MB231</td><td>MITO</td></tr>
<tr><td>>hg38_21_+_8433172_8446622_RNA45SN1_45S_50ntFlanks@4599.4616.18|GTTTTCGGAACTGAGGCC</td><td>>hg38_21_+_8433172_8446622_RNA45SN1_45S_50ntFlanks</td><td>4599</td><td>4616</td><td>18</td><td>GTTTTCGGAACTGAGGCC</td><td>295.9</td><td>-12.1</td><td>1.5</td><td>-8.0</td><td>1.2E-15</td><td>8.3E-15</td><td>MB231/MB468</td><td>MP</td></tr>
<tr><td>>MI0000299|hsa-mir-222&WithFlank&X|-|45747009|45747130@75.99.25(+1U)&[MIMAT0000279&hsa-miR-222-3p&offsets|0|+4(+1U);m-140&X|-|45747033|45747056&offsets|0|+1(+1U)]|AGCTACATCTGGCTACTGGGTCTCTT</td><td>>MI0000299|hsa-mir-222&WithFlank&X|-|45747009|45747130</td><td>75</td><td>99</td><td>25(+1U)</td><td>AGCTACATCTGGCTACTGGGTCTCTT</td><td>3636.7</td><td>6.3</td><td>0.8</td><td>8.0</td><td>1.5E-15</td><td>8.3E-15</td><td>MB231/MB468</td><td>NUC</td></tr>
<tr><td>>hg38_21_+_8433172_8446622_RNA45SN1_45S_50ntFlanks@6719.6739.21|TGAATTGCAGGACACATTGAT</td><td>>hg38_21_+_8433172_8446622_RNA45SN1_45S_50ntFlanks</td><td>6719</td><td>6739</td><td>21</td><td>TGAATTGCAGGACACATTGAT</td><td>1365.6</td><td>-5.0</td><td>0.6</td><td>-8.0</td><td>1.5E-15</td><td>8.3E-15</td><td>MB231/MB468</td><td>NUC</td></tr>
<tr><td>>hg38_21_+_8433172_8446622_RNA45SN1_45S_50ntFlanks@12290.12318.29|CGTGAAAGCGGGGCCTCACGATCCTTCTG</td><td>>hg38_21_+_8433172_8446622_RNA45SN1_45S_50ntFlanks</td><td>12290</td><td>12318</td><td>29</td><td>CGTGAAAGCGGGGCCTCACGATCCTTCTG</td><td>799.9</td><td>-13.4</td><td>1.7</td><td>-7.9</td><td>2.0E-15</td><td>8.3E-15</td><td>MB231/MB468</td><td>MITO</td></tr>
<tr><td>>hg38_21_+_8433172_8446622_RNA45SN1_45S_50ntFlanks@9562.9596.35|TCCTGACGTGCAAATCGGTCGTCCGACCTGGGTAT</td><td>>hg38_21_+_8433172_8446622_RNA45SN1_45S_50ntFlanks</td><td>9562</td><td>9596</td><td>35</td><td>TCCTGACGTGCAAATCGGTCGTCCGACCTGGGTAT</td><td>439.2</td><td>-12.6</td><td>1.6</td><td>-8.0</td><td>1.2E-15</td><td>8.3E-15</td><td>MB231/MB468</td><td>MP</td></tr>
<tr><td>>hg38_21_+_8433172_8446622_RNA45SN1_45S_50ntFlanks@6646.6663.18|TCGTACGACTCTTAGCGG</td><td>>hg38_21_+_8433172_8446622_RNA45SN1_45S_50ntFlanks</td><td>6646</td><td>6663</td><td>18</td><td>TCGTACGACTCTTAGCGG</td><td>74301.1</td><td>4.8</td><td>0.6</td><td>8.0</td><td>1.3E-15</td><td>8.3E-15</td><td>BT20/MB468</td><td>MITO</td></tr>
<tr><td>>MI0001445|hsa-mir-423&WithFlank&17|+|30117073|30117178@58.81.24&[MIMAT0001340&hsa-miR-423-3p&offsets|-1|0;m-100&17|+|30117131|30117153&offsets|-1|0]|AAGCTCGGTCTGAGGCCCCTCAGT</td><td>>MI0001445|hsa-mir-423&WithFlank&17|+|30117073|30117178</td><td>58</td><td>81</td><td>24</td><td>AAGCTCGGTCTGAGGCCCCTCAGT</td><td>1816.6</td><td>5.7</td><td>0.7</td><td>8.0</td><td>1.5E-15</td><td>8.3E-15</td><td>MB231/MB468</td><td>NUC</td></tr>
</table>

<table>
<tr><td>>hg38_21_+_8433172_8446622_RNA45SN1_45S_50ntFlanks@7986.8017.32|ATCAGACGTGGCGACCCGCTGAATTTAAGCAT</td><td>>hg38_21_+_8433172_8446622_RNA45SN1_45S_50ntFlanks</td><td>7986</td><td>8017</td><td>32</td><td>ATCAGACGTGGCGACCCGCTGAATTTAAGCAT</td><td>873.8</td><td>-11.9</td><td>1.5</td><td>-8.0</td><td>1.3E-15</td><td>8.3E-15</td><td>BT20/MB468</td><td>MITO</td></tr>
<tr><td>>hg38_21_+_8433172_8446622_RNA45SN1_45S_50ntFlanks@12570.12587.18|AGTACGAGAGGAACCGCA</td><td>>hg38_21_+_8433172_8446622_RNA45SN1_45S_50ntFlanks</td><td>12570</td><td>12587</td><td>18</td><td>AGTACGAGAGGAACCGCA</td><td>626.2</td><td>-6.5</td><td>0.8</td><td>-8.0</td><td>1.3E-15</td><td>8.4E-15</td><td>MB231/MB468</td><td>CYTO</td></tr>
<tr><td>>hg38_21_+_8433172_8446622_RNA45SN1_45S_50ntFlanks@5484.5509.26|GAGCGCTGAGAAGACGGTCGAACTTG</td><td>>hg38_21_+_8433172_8446622_RNA45SN1_45S_50ntFlanks</td><td>5484</td><td>5509</td><td>26</td><td>GAGCGCTGAGAAGACGGTCGAACTTG</td><td>1418.2</td><td>-9.9</td><td>1.2</td><td>-8.0</td><td>1.2E-15</td><td>8.4E-15</td><td>MB231/MB468</td><td>MP</td></tr>
<tr><td>>trna17_ValTAC_11_-_59318102_59318174@1.30.30,trna4_ValTAC_X_-_18693029_18693101@1.30.30|GGTTCCATAGTGTAGTGGTTATCACGTCTG</td><td>>trna17_ValTAC_11_-_59318102_59318174</td><td>1</td><td>30</td><td>30</td><td>GGTTCCATAGTGTAGTGGTTATCACGTCTG</td><td>843.5</td><td>-8.7</td><td>1.1</td><td>-8.0</td><td>1.3E-15</td><td>8.4E-15</td><td>MB231/MB468</td><td>CYTO</td></tr>
<tr><td>>MI0000440|hsa-mir-27b&WithFlank&9|+|95085439|95085547@67.87.21&[MIMAT0000419&hsa-miR-27b-3p&offsets|0|0;m-39&9|+|95085505|95085525&offsets|0|0]|TTCACAGTGGCTAAGTTCTGC</td><td>>MI0000440|hsa-mir-27b&WithFlank&9|+|95085439|95085547</td><td>67</td><td>87</td><td>21</td><td>TTCACAGTGGCTAAGTTCTGC</td><td>12651.7</td><td>3.6</td><td>0.5</td><td>7.9</td><td>2.8E-15</td><td>8.4E-15</td><td>BT20/MB468</td><td>NUC</td></tr>
<tr><td>>hg38_21_+_8433172_8446622_RNA45SN1_45S_50ntFlanks@11614.11640.27|ATTAAAACAAAGCATCGCGAAGGCCCG</td><td>>hg38_21_+_8433172_8446622_RNA45SN1_45S_50ntFlanks</td><td>11614</td><td>11640</td><td>27</td><td>ATTAAAACAAAGCATCGCGAAGGCCCG</td><td>591.0</td><td>-11.3</td><td>1.4</td><td>-8.0</td><td>1.3E-15</td><td>8.4E-15</td><td>BT20/MB468</td><td>MITO</td></tr>
<tr><td>>hg38_21_+_8433172_8446622_RNA45SN1_45S_50ntFlanks@9201.9225.25|CTCTCGGGGCCACGCGCGCGTCCCC</td><td>>hg38_21_+_8433172_8446622_RNA45SN1_45S_50ntFlanks</td><td>9201</td><td>9225</td><td>25</td><td>CTCTCGGGGCCACGCGCGCGTCCCC</td><td>442.6</td><td>-12.6</td><td>1.6</td><td>-7.9</td><td>2.0E-15</td><td>8.4E-15</td><td>MB231/MB468</td><td>MITO</td></tr>
<tr><td>>hg38_21_+_8433172_8446622_RNA45SN1_45S_50ntFlanks@12033.12052.20|ATCGTTTTTTCACTGACCCG</td><td>>hg38_21_+_8433172_8446622_RNA45SN1_45S_50ntFlanks</td><td>12033</td><td>12052</td><td>20</td><td>ATCGTTTTTTCACTGACCCG</td><td>308.9</td><td>-12.1</td><td>1.5</td><td>-7.9</td><td>2.0E-15</td><td>8.4E-15</td><td>MB231/MB468</td><td>MITO</td></tr>
<tr><td>>hg38_21_+_8433172_8446622_RNA45SN1_45S_50ntFlanks@11199.11227.29|CCGGCGGCGGCGGCGACTCTGGACGCGAG</td><td>>hg38_21_+_8433172_8446622_RNA45SN1_45S_50ntFlanks</td><td>11199</td><td>11227</td><td>29</td><td>CCGGCGGCGGCGGCGACTCTGGACGCGAG</td><td>333.2</td><td>-12.2</td><td>1.5</td><td>-8.0</td><td>1.2E-15</td><td>8.4E-15</td><td>MB231/MB468</td><td>MP</td></tr>
<tr><td>>hg38_21_+_8433172_8446622_RNA45SN1_45S_50ntFlanks@12457.12484.28|CAAGCGTTGGATTGTTCACCCACTAATA</td><td>>hg38_21_+_8433172_8446622_RNA45SN1_45S_50ntFlanks</td><td>12457</td><td>12484</td><td>28</td><td>CAAGCGTTGGATTGTTCACCCACTAATA</td><td>272.0</td><td>-12.0</td><td>1.5</td><td>-8.0</td><td>1.2E-15</td><td>8.4E-15</td><td>MB231/MB468</td><td>MP</td></tr>
<tr><td>>hg38_21_+_8433172_8446622_RNA45SN1_45S_50ntFlanks@10905.10933.29|CTGGGCGCGCGCCGCGGCTGGACGAGGCG</td><td>>hg38_21_+_8433172_8446622_RNA45SN1_45S_50ntFlanks</td><td>10905</td><td>10933</td><td>29</td><td>CTGGGCGCGCGCCGCGGCTGGACGAGGCG</td><td>481.0</td><td>-12.7</td><td>1.6</td><td>-7.9</td><td>2.1E-15</td><td>8.4E-15</td><td>MB231/MB468</td><td>MITO</td></tr>
<tr><td>>hg38_21_+_8433172_8446622_RNA45SN1_45S_50ntFlanks@6698.6720.23|AGCTAGCTGCGAGAATTAATGTG</td><td>>hg38_21_+_8433172_8446622_RNA45SN1_45S_50ntFlanks</td><td>6698</td><td>6720</td><td>23</td><td>AGCTAGCTGCGAGAATTAATGTG</td><td>15589.4</td><td>-7.0</td><td>0.9</td><td>-7.9</td><td>2.1E-15</td><td>8.5E-15</td><td>MB231/MB468</td><td>MITO</td></tr>
<tr><td>>hg38_21_+_8433172_8446622_RNA45SN1_45S_50ntFlanks@476.507.32|TTCGTGATCGATGTGGTGACGTCGTGCTCTCC</td><td>>hg38_21_+_8433172_8446622_RNA45SN1_45S_50ntFlanks</td><td>476</td><td>507</td><td>32</td><td>TTCGTGATCGATGTGGTGACGTCGTGCTCTCC</td><td>894.3</td><td>7.1</td><td>0.9</td><td>8.0</td><td>1.3E-15</td><td>8.5E-15</td><td>BT20/MB468</td><td>MITO</td></tr>
<tr><td>>hg38_21_+_8433172_8446622_RNA45SN1_45S_50ntFlanks@9564.9597.34|CTGACGTGCAAATCGGTCGTCCGACCTGGGTATA</td><td>>hg38_21_+_8433172_8446622_RNA45SN1_45S_50ntFlanks</td><td>9564</td><td>9597</td><td>34</td><td>CTGACGTGCAAATCGGTCGTCCGACCTGGGTATA</td><td>7367.7</td><td>-5.3</td><td>0.7</td><td>-8.0</td><td>1.4E-15</td><td>8.5E-15</td><td>MB231/MB468</td><td>CYTO</td></tr>
<tr><td>>hg38_21_+_8433172_8446622_RNA45SN1_45S_50ntFlanks@4445.4471.27|CTTGCCTCTCGGCGCCCCCTCGATGCT</td><td>>hg38_21_+_8433172_8446622_RNA45SN1_45S_50ntFlanks</td><td>4445</td><td>4471</td><td>27</td><td>CTTGCCTCTCGGCGCCCCCTCGATGCT</td><td>504.5</td><td>-11.1</td><td>1.4</td><td>-8.0</td><td>1.3E-15</td><td>8.5E-15</td><td>BT20/MB468</td><td>MITO</td></tr>
<tr><td>>hg38_21_+_8433172_8446622_RNA45SN1_45S_50ntFlanks@11886.11905.20|TGAGCTTGACTCTAGTCTGG</td><td>>hg38_21_+_8433172_8446622_RNA45SN1_45S_50ntFlanks</td><td>11886</td><td>11905</td><td>20</td><td>TGAGCTTGACTCTAGTCTGG</td><td>179.4</td><td>-10.5</td><td>1.3</td><td>-8.0</td><td>1.5E-15</td><td>8.6E-15</td><td>MB231/MB468</td><td>NUC</td></tr>
<tr><td>>hg38_21_+_8433172_8446622_RNA45SN1_45S_50ntFlanks@11209.11227.19|CGGCGACTCTGGACGCGAG</td><td>>hg38_21_+_8433172_8446622_RNA45SN1_45S_50ntFlanks</td><td>11209</td><td>11227</td><td>19</td><td>CGGCGACTCTGGACGCGAG</td><td>2117.8</td><td>-8.1</td><td>1.0</td><td>-8.0</td><td>1.4E-15</td><td>8.6E-15</td><td>MB231/MB468</td><td>CYTO</td></tr>
<tr><td>>MI0000114|hsa-mir-107&WithFlank&10|-|89592741|89592833@56.77.22&[MIMAT0000104&hsa-miR-107&offsets|0|-1;m-78&10|-|89592758|89592778&offsets|0|+1]|AGCAGCATTGTACAGGGCTATC</td><td>>MI0000114|hsa-mir-107&WithFlank&10|-|89592741|89592833</td><td>56</td><td>77</td><td>22</td><td>AGCAGCATTGTACAGGGCTATC</td><td>694.9</td><td>5.7</td><td>0.7</td><td>8.0</td><td>1.9E-15</td><td>8.6E-15</td><td>BT20/MB468</td><td>CYTO</td></tr>
<tr><td>>hg38_21_+_8433172_8446622_RNA45SN1_45S_50ntFlanks@7986.8005.20|ATCAGACGTGGCGACCCGCT</td><td>>hg38_21_+_8433172_8446622_RNA45SN1_45S_50ntFlanks</td><td>7986</td><td>8005</td><td>20</td><td>ATCAGACGTGGCGACCCGCT</td><td>325.8</td><td>-10.7</td><td>1.3</td><td>-8.0</td><td>1.9E-15</td><td>8.6E-15</td><td>BT20/MB468</td><td>CYTO</td></tr>
<tr><td>>hg38_21_+_8433172_8446622_RNA45SN1_45S_50ntFlanks@11452.11471.20|CTCCTCGGCGCGCGGCGGCG</td><td>>hg38_21_+_8433172_8446622_RNA45SN1_45S_50ntFlanks</td><td>11452</td><td>11471</td><td>20</td><td>CTCCTCGGCGCGCGGCGGCG</td><td>321.5</td><td>-12.1</td><td>1.5</td><td>-7.9</td><td>2.1E-15</td><td>8.6E-15</td><td>MB231/MB468</td><td>MITO</td></tr>
<tr><td>>hg38_21_+_8433172_8446622_RNA45SN1_45S_50ntFlanks@9027.9060.34|CGTCTCCTCGTGGGGGGGCCGGGCCACCCCTCCC</td><td>>hg38_21_+_8433172_8446622_RNA45SN1_45S_50ntFlanks</td><td>9027</td><td>9060</td><td>34</td><td>CGTCTCCTCGTGGGGGGGCCGGGCCACCCCTCCC</td><td>306.9</td><td>-12.0</td><td>1.5</td><td>-7.9</td><td>2.1E-15</td><td>8.6E-15</td><td>MB231/MB468</td><td>MITO</td></tr>
<tr><td>>hg38_21_+_8433172_8446622_RNA45SN1_45S_50ntFlanks@6661.6709.49|CGGTGGATCACTCGGCTCGTGCGTCGATGAAGAACGCAGCTAGCTGCGA</td><td>>hg38_21_+_8433172_8446622_RNA45SN1_45S_50ntFlanks</td><td>6661</td><td>6709</td><td>49</td><td>CGGTGGATCACTCGGCTCGTGCGTCGATGAAGAACGCAGCTAGCTGCGA</td><td>2840.1</td><td>9.7</td><td>1.2</td><td>8.0</td><td>1.4E-15</td><td>8.6E-15</td><td>BT20/MB468</td><td>MITO</td></tr>
<tr><td>>hg38_21_+_8433172_8446622_RNA45SN1_45S_50ntFlanks@3899.3921.23|CCTTCGCGGGGGGGATGCGTGCA</td><td>>hg38_21_+_8433172_8446622_RNA45SN1_45S_50ntFlanks</td><td>3899</td><td>3921</td><td>23</td><td>CCTTCGCGGGGGGGATGCGTGCA</td><td>361.2</td><td>-12.3</td><td>1.5</td><td>-7.9</td><td>2.1E-15</td><td>8.6E-15</td><td>MB231/MB468</td><td>MITO</td></tr>
<tr><td>>hg38_21_+_8433172_8446622_RNA45SN1_45S_50ntFlanks@4047.4068.22|ATTCGAACGTCTGCCCTATCAA</td><td>>hg38_21_+_8433172_8446622_RNA45SN1_45S_50ntFlanks</td><td>4047</td><td>4068</td><td>22</td><td>ATTCGAACGTCTGCCCTATCAA</td><td>581.0</td><td>-13.0</td><td>1.6</td><td>-7.9</td><td>2.1E-15</td><td>8.6E-15</td><td>MB231/MB468</td><td>MITO</td></tr>
<tr><td>>hg38_21_+_8433172_8446622_RNA45SN1_45S_50ntFlanks@475.509.35|CTTCGTGATCGATGTGGTGACGTCGTGCTCTCCCG</td><td>>hg38_21_+_8433172_8446622_RNA45SN1_45S_50ntFlanks</td><td>475</td><td>509</td><td>35</td><td>CTTCGTGATCGATGTGGTGACGTCGTGCTCTCCCG</td><td>1274.2</td><td>8.1</td><td>1.0</td><td>8.0</td><td>1.4E-15</td><td>8.6E-15</td><td>MB231/MB468</td><td>CYTO</td></tr>
<tr><td>>hg38_21_+_8433172_8446622_RNA45SN1_45S_50ntFlanks@9504.9524.21|GTGAACTATGCCTGGGCAGGG</td><td>>hg38_21_+_8433172_8446622_RNA45SN1_45S_50ntFlanks</td><td>9504</td><td>9524</td><td>21</td><td>GTGAACTATGCCTGGGCAGGG</td><td>673.7</td><td>-8.7</td><td>1.1</td><td>-8.0</td><td>1.4E-15</td><td>8.6E-15</td><td>MB231/MB468</td><td>CYTO</td></tr>
<tr><td>>hg38_21_+_8433172_8446622_RNA45SN1_45S_50ntFlanks@8570.8593.24|CTCCGGCGGGTGCGGGGGTGGGCG</td><td>>hg38_21_+_8433172_8446622_RNA45SN1_45S_50ntFlanks</td><td>8570</td><td>8593</td><td>24</td><td>CTCCGGCGGGTGCGGGGGTGGGCG</td><td>181.4</td><td>-10.2</td><td>1.3</td><td>-8.0</td><td>1.4E-15</td><td>8.6E-15</td><td>MB231/MB468</td><td>CYTO</td></tr>
<tr><td>>hg38_21_+_8433172_8446622_RNA45SN1_45S_50ntFlanks@5423.5443.21|ATTGGATGGTTTAGTGAGGCC</td><td>>hg38_21_+_8433172_8446622_RNA45SN1_45S_50ntFlanks</td><td>5423</td><td>5443</td><td>21</td><td>ATTGGATGGTTTAGTGAGGCC</td><td>572.9</td><td>-13.0</td><td>1.6</td><td>-8.0</td><td>1.3E-15</td><td>8.6E-15</td><td>MB231/MB468</td><td>MP</td></tr>
<tr><td>>hg38_21_+_8433172_8446622_RNA45SN1_45S_50ntFlanks@5077.5098.22|CCGATAACGAACGAGACTCTGG</td><td>>hg38_21_+_8433172_8446622_RNA45SN1_45S_50ntFlanks</td><td>5077</td><td>5098</td><td>22</td><td>CCGATAACGAACGAGACTCTGG</td><td>838.5</td><td>-13.6</td><td>1.7</td><td>-8.0</td><td>1.3E-15</td><td>8.6E-15</td><td>MB231/MB468</td><td>MP</td></tr>
<tr><td>>hg38_21_+_8433172_8446622_RNA45SN1_45S_50ntFlanks@11212.11233.22|CGACTCTGGACGCGAGCCGGGC</td><td>>hg38_21_+_8433172_8446622_RNA45SN1_45S_50ntFlanks</td><td>11212</td><td>11233</td><td>22</td><td>CGACTCTGGACGCGAGCCGGGC</td><td>252.8</td><td>-10.7</td><td>1.3</td><td>-8.0</td><td>1.4E-15</td><td>8.6E-15</td><td>MB231/MB468</td><td>CYTO</td></tr>
<tr><td>>hg38_21_+_8433172_8446622_RNA45SN1_45S_50ntFlanks@11845.11872.28|GGAACGGGCTTGGCGGAATCAGCGGGGA</td><td>>hg38_21_+_8433172_8446622_RNA45SN1_45S_50ntFlanks</td><td>11845</td><td>11872</td><td>28</td><td>GGAACGGGCTTGGCGGAATCAGCGGGGA</td><td>232.1</td><td>-5.6</td><td>0.7</td><td>-8.0</td><td>1.4E-15</td><td>8.6E-15</td><td>MB231/MB468</td><td>CYTO</td></tr>
<tr><td>>trnaMT_PheGAA_MT_+_577_647@36.56.21|AAAATGTTTAGACGGGCTCAC</td><td>>trnaMT_PheGAA_MT_+_577_647</td><td>36</td><td>56</td><td>21</td><td>AAAATGTTTAGACGGGCTCAC</td><td>277.5</td><td>10.1</td><td>1.3</td><td>7.9</td><td>1.9E-15</td><td>8.6E-15</td><td>BT20/MB468</td><td>CYTO</td></tr>
<tr><td>>hg38_21_+_8433172_8446622_RNA45SN1_45S_50ntFlanks@9035.9058.24|CGTGGGGGGGCCGGGCCACCCCTC</td><td>>hg38_21_+_8433172_8446622_RNA45SN1_45S_50ntFlanks</td><td>9035</td><td>9058</td><td>24</td><td>CGTGGGGGGGCCGGGCCACCCCTC</td><td>464.8</td><td>-12.7</td><td>1.6</td><td>-8.0</td><td>1.3E-15</td><td>8.7E-15</td><td>MB231/MB468</td><td>MP</td></tr>
<tr><td>>hg38_21_+_8433172_8446622_RNA45SN1_45S_50ntFlanks@4759.4783.25|ATCAGATACCGTCGTAGTTCCGACC</td><td>>hg38_21_+_8433172_8446622_RNA45SN1_45S_50ntFlanks</td><td>4759</td><td>4783</td><td>25</td><td>ATCAGATACCGTCGTAGTTCCGACC</td><td>1525.8</td><td>-13.7</td><td>1.6</td><td>-8.4</td><td>4.1E-17</td><td>8.7E-15</td><td>BT20/MB468</td><td>MP</td></tr>
<tr><td>>hg38_21_+_8433172_8446622_RNA45SN1_45S_50ntFlanks@12586.12604.19|CAGGTTCAGACATTTGGTG</td><td>>hg38_21_+_8433172_8446622_RNA45SN1_45S_50ntFlanks</td><td>12586</td><td>12604</td><td>19</td><td>CAGGTTCAGACATTTGGTG</td><td>171.8</td><td>-11.3</td><td>1.4</td><td>-8.0</td><td>1.3E-15</td><td>8.7E-15</td><td>MB231/MB468</td><td>MP</td></tr>
<tr><td>>hg38_21_+_8433172_8446622_RNA45SN1_45S_50ntFlanks@9032.9056.25|CCTCGTGGGGGGGCCGGGCCACCCC</td><td>>hg38_21_+_8433172_8446622_RNA45SN1_45S_50ntFlanks</td><td>9032</td><td>9056</td><td>25</td><td>CCTCGTGGGGGGGCCGGGCCACCCC</td><td>222.1</td><td>-11.7</td><td>1.5</td><td>-8.0</td><td>1.3E-15</td><td>8.7E-15</td><td>MB231/MB468</td><td>MP</td></tr>
<tr><td>>hg38_21_+_8433172_8446622_RNA45SN1_45S_50ntFlanks@9572.9592.21|CAAATCGGTCGTCCGACCTGG</td><td>>hg38_21_+_8433172_8446622_RNA45SN1_45S_50ntFlanks</td><td>9572</td><td>9592</td><td>21</td><td>CAAATCGGTCGTCCGACCTGG</td><td>328.6</td><td>-11.1</td><td>1.4</td><td>-8.0</td><td>1.4E-15</td><td>8.7E-15</td><td>MB231/MB468</td><td>CYTO</td></tr>
<tr><td>>hg38_21_+_8433172_8446622_RNA45SN1_45S_50ntFlanks@12822.12856.35|CCGGGACCGGGGTCCGGTGCGGAGTGCCCTTCGTC</td><td>>hg38_21_+_8433172_8446622_RNA45SN1_45S_50ntFlanks</td><td>12822</td><td>12856</td><td>35</td><td>CCGGGACCGGGGTCCGGTGCGGAGTGCCCTTCGTC</td><td>215.7</td><td>-11.6</td><td>1.5</td><td>-8.0</td><td>1.3E-15</td><td>8.7E-15</td><td>MB231/MB468</td><td>MP</td></tr>
<tr><td>>hg38_21_+_8433172_8446622_RNA45SN1_45S_50ntFlanks@11209.11254.46|CGGCGACTCTGGACGCGAGCCGGGCCCTTCCCGTGGATCGCCCCAG</td><td>>hg38_21_+_8433172_8446622_RNA45SN1_45S_50ntFlanks</td><td>11209</td><td>11254</td><td>46</td><td>CGGCGACTCTGGACGCGAGCCGGGCCCTTCCCGTGGATCGCCCCAG</td><td>83879.0</td><td>-12.1</td><td>1.5</td><td>-7.9</td><td>2.9E-15</td><td>8.7E-15</td><td>BT20/MB468</td><td>NUC</td></tr>
<tr><td>>MI0000283|hsa-mir-203a&WithFlank&14|+|104117399|104117520@71.91.21(+1A)&[MIMAT0000264&hsa-miR-203a-3p&offsets|0|-1(+1A);m-79&14|+|104117469|104117490&offsets|0|-1(+1A)]|GTGAAATGTTTAGGACCACTAA</td><td>>MI0000283|hsa-mir-203a&WithFlank&14|+|104117399|104117520</td><td>71</td><td>91</td><td>21(+1A)</td><td>GTGAAATGTTTAGGACCACTAA</td><td>327.9</td><td>11.7</td><td>1.4</td><td>8.3</td><td>1.2E-16</td><td>8.8E-15</td><td>BT20/MB231</td><td>CYTO</td></tr>
<tr><td>>hg38_21_+_8433172_8446622_RNA45SN1_45S_50ntFlanks@10772.10805.34|AGGTCTCCAAGGTGAACAGCCTCTGGCATGTTGG</td><td>>hg38_21_+_8433172_8446622_RNA45SN1_45S_50ntFlanks</td><td>10772</td><td>10805</td><td>34</td><td>AGGTCTCCAAGGTGAACAGCCTCTGGCATGTTGG</td><td>273.1</td><td>-11.9</td><td>1.5</td><td>-7.9</td><td>2.2E-15</td><td>8.8E-15</td><td>MB231/MB468</td><td>MITO</td></tr>
<tr><td>>hg38_21_+_8433172_8446622_RNA45SN1_45S_50ntFlanks@8079.8101.23|AAGAGCCCAGCGCCGAATCCCCG</td><td>>hg38_21_+_8433172_8446622_RNA45SN1_45S_50ntFlanks</td><td>8079</td><td>8101</td><td>23</td><td>AAGAGCCCAGCGCCGAATCCCCG</td><td>751.9</td><td>-13.4</td><td>1.7</td><td>-8.0</td><td>1.3E-15</td><td>8.8E-15</td><td>MB231/MB468</td><td>MP</td></tr>
<tr><td>>hg38_21_+_8433172_8446622_RNA45SN1_45S_50ntFlanks@4734.4761.28|AACGAAAGTCGGAGGTTCGAAGACGATC</td><td>>hg38_21_+_8433172_8446622_RNA45SN1_45S_50ntFlanks</td><td>4734</td><td>4761</td><td>28</td><td>AACGAAAGTCGGAGGTTCGAAGACGATC</td><td>888.1</td><td>5.2</td><td>0.7</td><td>7.9</td><td>1.9E-15</td><td>8.9E-15</td><td>BT20/MB468</td><td>CYTO</td></tr>
<tr><td>>hg38_21_+_8433172_8446622_RNA45SN1_45S_50ntFlanks@7986.8003.18|ATCAGACGTGGCGACCCG</td><td>>hg38_21_+_8433172_8446622_RNA45SN1_45S_50ntFlanks</td><td>7986</td><td>8003</td><td>18</td><td>ATCAGACGTGGCGACCCG</td><td>3190.0</td><td>-10.2</td><td>1.3</td><td>-7.9</td><td>2.2E-15</td><td>8.9E-15</td><td>MB231/MB468</td><td>MITO</td></tr>
<tr><td>>hg38_21_+_8433172_8446622_RNA45SN1_45S_50ntFlanks@6671.6699.29|CTCGGCTCGTGCGTCGATGAAGAACGCAG</td><td>>hg38_21_+_8433172_8446622_RNA45SN1_45S_50ntFlanks</td><td>6671</td><td>6699</td><td>29</td><td>CTCGGCTCGTGCGTCGATGAAGAACGCAG</td><td>1643.9</td><td>-4.9</td><td>0.6</td><td>-8.0</td><td>1.6E-15</td><td>8.9E-15</td><td>MB231/MB468</td><td>NUC</td></tr>
<tr><td>>hg38_21_+_8433172_8446622_RNA45SN1_45S_50ntFlanks@13006.13023.18|CTATTGAAAGTCAGCCCT</td><td>>hg38_21_+_8433172_8446622_RNA45SN1_45S_50ntFlanks</td><td>13006</td><td>13023</td><td>18</td><td>CTATTGAAAGTCAGCCCT</td><td>215.0</td><td>-10.5</td><td>1.3</td><td>-8.0</td><td>1.4E-15</td><td>8.9E-15</td><td>MB231/MB468</td><td>CYTO</td></tr>
<tr><td>>hg38_21_+_8433172_8446622_RNA45SN1_45S_50ntFlanks@8153.8178.26|GCCGCTCGTGGGGGGCCCAAGTCCTT</td><td>>hg38_21_+_8433172_8446622_RNA45SN1_45S_50ntFlanks</td><td>8153</td><td>8178</td><td>26</td><td>GCCGCTCGTGGGGGGCCCAAGTCCTT</td><td>589.9</td><td>-13.0</td><td>1.6</td><td>-7.9</td><td>2.2E-15</td><td>8.9E-15</td><td>MB231/MB468</td><td>MITO</td></tr>
<tr><td>>hg38_21_+_8433172_8446622_RNA45SN1_45S_50ntFlanks@3843.3860.18|CTACTTGGATAACTGTGG</td><td>>hg38_21_+_8433172_8446622_RNA45SN1_45S_50ntFlanks</td><td>3843</td><td>3860</td><td>18</td><td>CTACTTGGATAACTGTGG</td><td>516.6</td><td>-12.9</td><td>1.6</td><td>-8.0</td><td>1.3E-15</td><td>8.9E-15</td><td>MB231/MB468</td><td>MP</td></tr>
<tr><td>>hg38_21_+_8433172_8446622_RNA45SN1_45S_50ntFlanks@11575.11599.25|ACTTAGAACTGGTGCGGACCAGGGG</td><td>>hg38_21_+_8433172_8446622_RNA45SN1_45S_50ntFlanks</td><td>11575</td><td>11599</td><td>25</td><td>ACTTAGAACTGGTGCGGACCAGGGG</td><td>350.8</td><td>-11.2</td><td>1.4</td><td>-8.0</td><td>1.4E-15</td><td>8.9E-15</td><td>MB231/MB468</td><td>CYTO</td></tr>
<tr><td>>MI0000299|hsa-mir-222&WithFlank&X|-|45747009|45747130@75.99.25(+1C)&[MIMAT0000279&hsa-miR-222-3p&offsets|0|+4(+1C);m-140&X|-|45747033|45747056&offsets|0|+1(+1C)]|AGCTACATCTGGCTACTGGGTCTCTC</td><td>>MI0000299|hsa-mir-222&WithFlank&X|-|45747009|45747130</td><td>75</td><td>99</td><td>25(+1C)</td><td>AGCTACATCTGGCTACTGGGTCTCTC</td><td>302.5</td><td>8.5</td><td>1.1</td><td>7.9</td><td>1.9E-15</td><td>9.0E-15</td><td>BT20/MB468</td><td>CYTO</td></tr>
<tr><td>>MI0003560|hsa-mir-92b&WithFlank&1|+|155195171|155195278@67.86.20&[MIMAT0003218&hsa-miR-92b-3p&offsets|0|-2;m-115&1|+|155195237|155195258&offsets|0|-2]|TATTGCACTCGTCCCGGCCT</td><td>>MI0003560|hsa-mir-92b&WithFlank&1|+|155195171|155195278</td><td>67</td><td>86</td><td>20</td><td>TATTGCACTCGTCCCGGCCT</td><td>1043.4</td><td>11.0</td><td>1.4</td><td>7.9</td><td>1.9E-15</td><td>9.0E-15</td><td>BT20/MB468</td><td>CYTO</td></tr>
<tr><td>>hg38_21_+_8433172_8446622_RNA45SN1_45S_50ntFlanks@4322.4339.18|CAGCCGCGGTAATTCCAG</td><td>>hg38_21_+_8433172_8446622_RNA45SN1_45S_50ntFlanks</td><td>4322</td><td>4339</td><td>18</td><td>CAGCCGCGGTAATTCCAG</td><td>292.5</td><td>-12.0</td><td>1.5</td><td>-7.9</td><td>2.2E-15</td><td>9.0E-15</td><td>MB231/MB468</td><td>MITO</td></tr>
<tr><td>>MI0000285|hsa-mir-205&WithFlank&1|+|209432127|209432248@40.59.20&[MIMAT0000266&hsa-miR-205-5p&offsets|0|-2;m-351&1|+|209432166|209432187&offsets|0|-2]|TCCTTCATTCCACCGGAGTC</td><td>>MI0000285|hsa-mir-205&WithFlank&1|+|209432127|209432248</td><td>40</td><td>59</td><td>20</td><td>TCCTTCATTCCACCGGAGTC</td><td>532.1</td><td>12.4</td><td>1.5</td><td>8.3</td><td>1.3E-16</td><td>9.0E-15</td><td>BT20/MB231</td><td>CYTO</td></tr>
</table>

| | | | | | | | | | | | | | |
|---|---|---|---|---|---|---|---|---|---|---|---|---|---|
| >MI0003628\|hsa-mir-615&WithFlank&12\|+\|54033944\|54034051@67.87.21&[MIMAT0003283&hsa-miR-615-3p&offsets\|0\|-1;m-495&12\|+\|54034010\|54034030&offsets\|0\|0]\|TCCGAGCCTGGGTCTCCCTCT | >MI0003628\|hsa-mir-615&WithFlank&12\|+\|54033944\|54034051 | 67 | 87 | 21 | TCCGAGCCTGGGTCTCCCTCT | 1902.7 | 5.9 | 0.7 | 8.3 | 1.3E-16 | 9.0E-15 | BT20/MB231 | CYTO |
| >hg38_21_+_8433172_8446622_RNA45SN1_45S_50ntFlanks@11671.11692.22\|CCAGTGCTCTGAATGTCAAAGT | >hg38_21_+_8433172_8446622_RNA45SN1_45S_50ntFlanks | 11671 | 11692 | 22 | CCAGTGCTCTGAATGTCAAAGT | 584.4 | 4.8 | 0.6 | 8.0 | 1.4E-15 | 9.0E-15 | BT20/MB468 | MITO |
| >Antisense_hg38_21_-_8433172_8446622_RNA45SN1_45S_50ntFlanks@7674.7693.20\|CTCTCTCTCTCTCTCCCCCG | >Antisense_hg38_21_-_8433172_8446622_RNA45SN1_45S_50ntFlanks | 7674 | 7693 | 20 | CTCTCTCTCTCTCTCCCCCG | 1770.7 | -13.8 | 1.7 | -8.0 | 1.6E-15 | 9.0E-15 | MB231/MB468 | NUC |
| >hg38_21_+_8433172_8446622_RNA45SN1_45S_50ntFlanks@11027.11046.20\|CTCTCTCTCTCTCTCCCCCG | >hg38_21_+_8433172_8446622_RNA45SN1_45S_50ntFlanks | 11027 | 11046 | 20 | CTCTCTCTCTCTCTCCCCCG | 1770.7 | -13.8 | 1.7 | -8.0 | 1.6E-15 | 9.0E-15 | MB231/MB468 | NUC |
| >MI0000477\|hsa-mir-146a&WithFlank&5\|+\|160485346\|160485456@27.46.20&[MIMAT0000449&hsa-miR-146a-5p&offsets\|0\|-2;m-129&5\|+\|160485372\|160485393&offsets\|0\|-2]\|TGAGAACTGAATTCCATGGG | >MI0000477\|hsa-mir-146a&WithFlank&5\|+\|160485346\|160485456 | 27 | 46 | 20 | TGAGAACTGAATTCCATGGG | 504.9 | 12.2 | 1.5 | 8.0 | 1.3E-15 | 9.0E-15 | MB231/MB468 | MP |
| >hg38_21_+_8433172_8446622_RNA45SN1_45S_50ntFlanks@11576.11605.30\|CTTAGAACTGGTGCGGACCAGGGGAATCCG | >hg38_21_+_8433172_8446622_RNA45SN1_45S_50ntFlanks | 11576 | 11605 | 30 | CTTAGAACTGGTGCGGACCAGGGGAATCCG | 1558.8 | -9.7 | 1.2 | -8.0 | 1.6E-15 | 9.0E-15 | MB231/MB468 | NUC |
| >hg38_21_+_8433172_8446622_RNA45SN1_45S_50ntFlanks@4657.4686.30\|CTAGAGGTGAAATTCTTGGACCGGCGCAAG | >hg38_21_+_8433172_8446622_RNA45SN1_45S_50ntFlanks | 4657 | 4686 | 30 | CTAGAGGTGAAATTCTTGGACCGGCGCAAG | 237.7 | -10.9 | 1.4 | -8.0 | 1.6E-15 | 9.0E-15 | MB231/MB468 | NUC |
| >hg38_21_+_8433172_8446622_RNA45SN1_45S_50ntFlanks@9339.9377.39\|CGCACGAAAGCCGCCGTGGCGCAATGAAGGTGAAGGCCG | >hg38_21_+_8433172_8446622_RNA45SN1_45S_50ntFlanks | 9339 | 9377 | 39 | CGCACGAAAGCCGCCGTGGCGCAATGAAGGTGAAGGCCG | 405.9 | -12.5 | 1.6 | -8.0 | 1.3E-15 | 9.1E-15 | MB231/MB468 | MP |
| >MI0003643\|hsa-mir-629&WithFlank&15\|-\|70079366\|70079474@28.48.21(+1C)&[MIMAT0004810&hsa-miR-629-5p&offsets\|0\|0(+1C);m-150&15\|-\|70079427\|70079447&offsets\|0\|0(+1C)]\|TGGGTTTACGTTGGGAGAACTC | >MI0003643\|hsa-mir-629&WithFlank&15\|-\|70079366\|70079474 | 28 | 48 | 21(+1C) | TGGGTTTACGTTGGGAGAACTC | 62.0 | 8.8 | 1.1 | 7.9 | 3.0E-15 | 9.1E-15 | BT20/MB468 | NUC |
| >hg38_1_-_228634819_228635039_RNA5S12_5S_50ntFlanks@134.169.36\|TGGGAGACCGCCTGGGAATACCGGGTGCTGTAGGCT | >hg38_1_-_228634819_228635039_RNA5S12_5S_50ntFlanks | 134 | 169 | 36 | TGGGAGACCGCCTGGGAATACCGGGTGCTGTAGGCT | 18698.1 | 7.3 | 0.9 | 8.0 | 1.5E-15 | 9.1E-15 | MB231/MB468 | CYTO |
| >hg38_21_+_8433172_8446622_RNA45SN1_45S_50ntFlanks@12419.12442.24\|GATGTCGGCTCTTCCTATCATTGT | >hg38_21_+_8433172_8446622_RNA45SN1_45S_50ntFlanks | 12419 | 12442 | 24 | GATGTCGGCTCTTCCTATCATTGT | 1167.0 | 5.0 | 0.6 | 7.9 | 2.0E-15 | 9.1E-15 | BT20/MB468 | CYTO |
| >hg38_21_+_8433172_8446622_RNA45SN1_45S_50ntFlanks@10794.10812.19\|CTGGCATGTTGGAACAATG | >hg38_21_+_8433172_8446622_RNA45SN1_45S_50ntFlanks | 10794 | 10812 | 19 | CTGGCATGTTGGAACAATG | 184.4 | -11.4 | 1.4 | -8.0 | 1.3E-15 | 9.1E-15 | MB231/MB468 | MP |
| >hg38_21_+_8433172_8446622_RNA45SN1_45S_50ntFlanks@6698.6720.23\|AGCTAGCTGCGAGAATTAATGTG | >hg38_21_+_8433172_8446622_RNA45SN1_45S_50ntFlanks | 6698 | 6720 | 23 | AGCTAGCTGCGAGAATTAATGTG | 14831.1 | -4.0 | 0.5 | -7.9 | 2.0E-15 | 9.1E-15 | BT20/MB468 | CYTO |
| >hg38_21_+_8433172_8446622_RNA45SN1_45S_50ntFlanks@10412.10431.20\|AGTCGGTCCTGAGAGATGGG | >hg38_21_+_8433172_8446622_RNA45SN1_45S_50ntFlanks | 10412 | 10431 | 20 | AGTCGGTCCTGAGAGATGGG | 180.5 | -7.7 | 1.0 | -8.0 | 1.5E-15 | 9.1E-15 | MB231/MB468 | CYTO |
| >hg38_21_+_8433172_8446622_RNA45SN1_45S_50ntFlanks@11343.11362.20\|CGCTGGGGGCGGGGAGCGGT | >hg38_21_+_8433172_8446622_RNA45SN1_45S_50ntFlanks | 11343 | 11362 | 20 | CGCTGGGGGCGGGGAGCGGT | 493.2 | -12.8 | 1.6 | -8.0 | 1.3E-15 | 9.2E-15 | MB231/MB468 | MP |
| >hg38_21_+_8433172_8446622_RNA45SN1_45S_50ntFlanks@6745.6770.26\|ACACTTCGAACGCACTTGCGGCCCCG | >hg38_21_+_8433172_8446622_RNA45SN1_45S_50ntFlanks | 6745 | 6770 | 26 | ACACTTCGAACGCACTTGCGGCCCCG | 5021.1 | -11.8 | 1.5 | -7.9 | 3.1E-15 | 9.2E-15 | BT20/MB468 | NUC |
| >hg38_21_+_8433172_8446622_RNA45SN1_45S_50ntFlanks@10819.10865.47\|AGGGAAGTCGGCAAGCCGGATCCGTAACTTCGGGATAAGGATTGGCT | >hg38_21_+_8433172_8446622_RNA45SN1_45S_50ntFlanks | 10819 | 10865 | 47 | AGGGAAGTCGGCAAGCCGGATCCGTAACTTCGGGATAAGGATTGGCT | 1116.5 | -14.6 | 1.8 | -7.9 | 3.1E-15 | 9.2E-15 | BT20/MB468 | NUC |
| >hg38_21_+_8433172_8446622_RNA45SN1_45S_50ntFlanks@12578.12604.27\|AGGAACCGCAGGTTCAGACATTTGGTG | >hg38_21_+_8433172_8446622_RNA45SN1_45S_50ntFlanks | 12578 | 12604 | 27 | AGGAACCGCAGGTTCAGACATTTGGTG | 707.5 | -12.5 | 1.5 | -8.4 | 4.4E-17 | 9.2E-15 | BT20/MB468 | MP |
| >hg38_21_+_8433172_8446622_RNA45SN1_45S_50ntFlanks@12628.12657.30\|TGGGGCGAAGCTACCATCTGTGGGATTATG | >hg38_21_+_8433172_8446622_RNA45SN1_45S_50ntFlanks | 12628 | 12657 | 30 | TGGGGCGAAGCTACCATCTGTGGGATTATG | 383.2 | -12.4 | 1.6 | -8.0 | 1.4E-15 | 9.2E-15 | MB231/MB468 | MP |
| >hg38_21_+_8433172_8446622_RNA45SN1_45S_50ntFlanks@9545.9564.20\|GTGGAGGTCCGTAGCGGTCC | >hg38_21_+_8433172_8446622_RNA45SN1_45S_50ntFlanks | 9545 | 9564 | 20 | GTGGAGGTCCGTAGCGGTCC | 266.0 | -11.8 | 1.5 | -7.9 | 2.3E-15 | 9.2E-15 | MB231/MB468 | MITO |
| >hg38_21_+_8433172_8446622_RNA45SN1_45S_50ntFlanks@12003.12031.29\|GCGGGCCGCCGGTGAAATACCACTACTCT | >hg38_21_+_8433172_8446622_RNA45SN1_45S_50ntFlanks | 12003 | 12031 | 29 | GCGGGCCGCCGGTGAAATACCACTACTCT | 1359.5 | -5.2 | 0.7 | -7.9 | 2.3E-15 | 9.3E-15 | MB231/MB468 | MITO |
| >hg38_21_+_8433172_8446622_RNA45SN1_45S_50ntFlanks@10350.10376.27\|ACGAGAACTTTGAAGGCCGAAGTGGAG | >hg38_21_+_8433172_8446622_RNA45SN1_45S_50ntFlanks | 10350 | 10376 | 27 | ACGAGAACTTTGAAGGCCGAAGTGGAG | 4646.8 | -10.1 | 1.3 | -8.0 | 1.4E-15 | 9.3E-15 | MB231/MB468 | MP |
| >hg38_21_+_8433172_8446622_RNA45SN1_45S_50ntFlanks@11663.11692.30\|ATTTCTGCCCAGTGCTCTGAATGTCAAAGT | >hg38_21_+_8433172_8446622_RNA45SN1_45S_50ntFlanks | 11663 | 11692 | 30 | ATTTCTGCCCAGTGCTCTGAATGTCAAAGT | 625.0 | 8.0 | 1.0 | 8.0 | 1.5E-15 | 9.3E-15 | BT20/MB468 | MITO |
| >hg38_21_+_8433172_8446622_RNA45SN1_45S_50ntFlanks@3844.3866.23\|TACTTGGATAACTGTGGTAATTC | >hg38_21_+_8433172_8446622_RNA45SN1_45S_50ntFlanks | 3844 | 3866 | 23 | TACTTGGATAACTGTGGTAATTC | 283.9 | 4.8 | 0.6 | 8.0 | 1.5E-15 | 9.3E-15 | MB231/MB468 | CYTO |
| >trna11_GluTTC_15_-_26327381_26327452@1.32.32,trna3_GluTTC_13_-_45492062_45492133@1.32.32\|TCCCACATGGTCTAGCGGTTAGGATTCCTGGT | >trna11_GluTTC_15_-_26327381_26327452 | 1 | 32 | 32 | TCCCACATGGTCTAGCGGTTAGGATTCCTGGT | 1207.1 | 9.1 | 1.1 | 8.0 | 1.5E-15 | 9.3E-15 | MB231/MB468 | CYTO |
| >hg38_21_+_8433172_8446622_RNA45SN1_45S_50ntFlanks@11874.11897.24\|AGAAGACCCTGTTGAGCTTGACTC | >hg38_21_+_8433172_8446622_RNA45SN1_45S_50ntFlanks | 11874 | 11897 | 24 | AGAAGACCCTGTTGAGCTTGACTC | 784.5 | -4.0 | 0.5 | -7.9 | 2.0E-15 | 9.3E-15 | BT20/MB468 | CYTO |
| >MI0000079\|hsa-mir-23a&WithFlank&19\|-\|13836581\|13836665@54.72.19(+2U)&[MIMAT0000078&hsa-miR-23a-3p&offsets\|+3\|+1(+2U);m-31&19\|-\|13836595\|13836615&offsets\|+3\|+1(+2U)]\|ACATTGCCAGGGATTTCCATT | >MI0000079\|hsa-mir-23a&WithFlank&19\|-\|13836581\|13836665 | 54 | 72 | 19(+2U) | ACATTGCCAGGGATTTCCATT | 1798.2 | 29.0 | 3.6 | 8.0 | 1.5E-15 | 9.3E-15 | MB231/MB468 | CYTO |
| >hg38_21_+_8433172_8446622_RNA45SN1_45S_50ntFlanks@7986.8004.19\|ATCAGACGTGGCGACCCGC | >hg38_21_+_8433172_8446622_RNA45SN1_45S_50ntFlanks | 7986 | 8004 | 19 | ATCAGACGTGGCGACCCGC | 617.5 | -11.4 | 1.4 | -8.0 | 1.5E-15 | 9.4E-15 | BT20/MB468 | MITO |
| >hg38_21_+_8433172_8446622_RNA45SN1_45S_50ntFlanks@10737.10755.19\|TAAATCTCGCGCCGGGCCG | >hg38_21_+_8433172_8446622_RNA45SN1_45S_50ntFlanks | 10737 | 10755 | 19 | TAAATCTCGCGCCGGGCCG | 194.6 | -11.5 | 1.4 | -8.0 | 1.4E-15 | 9.4E-15 | MB231/MB468 | MP |
| >hg38_21_+_8433172_8446622_RNA45SN1_45S_50ntFlanks@10571.10596.26\|GAGAAGCCGGCGGGAGCCCCGGGGAG | >hg38_21_+_8433172_8446622_RNA45SN1_45S_50ntFlanks | 10571 | 10596 | 26 | GAGAAGCCGGCGGGAGCCCCGGGGAG | 243.7 | -11.7 | 1.5 | -7.9 | 2.3E-15 | 9.4E-15 | MB231/MB468 | MITO |
| >hg38_21_+_8433172_8446622_RNA45SN1_45S_50ntFlanks@10878.10902.25\|TCGGTCGGGCTGGGGCGCGAAGCGG | >hg38_21_+_8433172_8446622_RNA45SN1_45S_50ntFlanks | 10878 | 10902 | 25 | TCGGTCGGGCTGGGGCGCGAAGCGG | 458.2 | -6.7 | 0.8 | -8.0 | 1.5E-15 | 9.4E-15 | MB231/MB468 | CYTO |
| >hg38_21_+_8433172_8446622_RNA45SN1_45S_50ntFlanks@12841.12862.22\|CGGAGTGCCCTTCGTCCTGGGA | >hg38_21_+_8433172_8446622_RNA45SN1_45S_50ntFlanks | 12841 | 12862 | 22 | CGGAGTGCCCTTCGTCCTGGGA | 1131.9 | -14.0 | 1.8 | -8.0 | 1.4E-15 | 9.4E-15 | MB231/MB468 | MP |
| >hg38_21_+_8433172_8446622_RNA45SN1_45S_50ntFlanks@9034.9059.26\|TCGTGGGGGGGCCGGGCCACCCCTCC | >hg38_21_+_8433172_8446622_RNA45SN1_45S_50ntFlanks | 9034 | 9059 | 26 | TCGTGGGGGGGCCGGGCCACCCCTCC | 338.4 | -12.3 | 1.5 | -8.0 | 1.4E-15 | 9.4E-15 | MB231/MB468 | MP |
| >hg38_21_+_8433172_8446622_RNA45SN1_45S_50ntFlanks@11882.11905.24\|CTGTTGAGCTTGACTCTAGTCTGG | >hg38_21_+_8433172_8446622_RNA45SN1_45S_50ntFlanks | 11882 | 11905 | 24 | CTGTTGAGCTTGACTCTAGTCTGG | 187.2 | -11.4 | 1.4 | -8.0 | 1.4E-15 | 9.5E-15 | MB231/MB468 | MP |
| >hg38_21_+_8433172_8446622_RNA45SN1_45S_50ntFlanks@12991.13025.35\|CTCCCTCGCTGCGATCTATTGAAAGTCAGCCCTCG | >hg38_21_+_8433172_8446622_RNA45SN1_45S_50ntFlanks | 12991 | 13025 | 35 | CTCCCTCGCTGCGATCTATTGAAAGTCAGCCCTCG | 276.5 | -11.9 | 1.5 | -7.9 | 2.3E-15 | 9.5E-15 | MB231/MB468 | MITO |
| >hg38_1_-_228634819_228635039_RNA5S12_5S_50ntFlanks@142.170.29\|CGCCTGGGAATACCGGGTGCTGTAGGCTT | >hg38_1_-_228634819_228635039_RNA5S12_5S_50ntFlanks | 142 | 170 | 29 | CGCCTGGGAATACCGGGTGCTGTAGGCTT | 1760.6 | 5.5 | 0.7 | 7.9 | 2.1E-15 | 9.5E-15 | BT20/MB468 | CYTO |
| >hg38_21_+_8433172_8446622_RNA45SN1_45S_50ntFlanks@10781.10802.22\|AGGTGAACAGCCTCTGGCATGT | >hg38_21_+_8433172_8446622_RNA45SN1_45S_50ntFlanks | 10781 | 10802 | 22 | AGGTGAACAGCCTCTGGCATGT | 235.2 | -11.7 | 1.5 | -8.0 | 1.4E-15 | 9.5E-15 | MB231/MB468 | MP |
| >hg38_21_+_8433172_8446622_RNA45SN1_45S_50ntFlanks@11233.11250.18\|CCCTTCCCGTGGATCGCC | >hg38_21_+_8433172_8446622_RNA45SN1_45S_50ntFlanks | 11233 | 11250 | 18 | CCCTTCCCGTGGATCGCC | 245.2 | -11.8 | 1.5 | -8.0 | 1.4E-15 | 9.5E-15 | MB231/MB468 | MP |
| >hg38_21_+_8433172_8446622_RNA45SN1_45S_50ntFlanks@10735.10757.23\|TGTAAATCTCGCGCCGGGCCGTA | >hg38_21_+_8433172_8446622_RNA45SN1_45S_50ntFlanks | 10735 | 10757 | 23 | TGTAAATCTCGCGCCGGGCCGTA | 299.5 | -12.0 | 1.5 | -7.9 | 2.3E-15 | 9.5E-15 | MB231/MB468 | MITO |
| >hg38_21_+_8433172_8446622_RNA45SN1_45S_50ntFlanks@12632.12653.22\|GCGAAGCTACCATCTGTGGGAT | >hg38_21_+_8433172_8446622_RNA45SN1_45S_50ntFlanks | 12632 | 12653 | 22 | GCGAAGCTACCATCTGTGGGAT | 477.8 | -6.7 | 0.8 | -8.0 | 1.5E-15 | 9.5E-15 | MB231/MB468 | CYTO |
| >hg38_21_+_8433172_8446622_RNA45SN1_45S_50ntFlanks@9004.9025.22\|TCTCCCCCGCGGGGGCGCGCCG | >hg38_21_+_8433172_8446622_RNA45SN1_45S_50ntFlanks | 9004 | 9025 | 22 | TCTCCCCCGCGGGGGCGCGCCG | 280.6 | -11.9 | 1.5 | -7.9 | 2.3E-15 | 9.5E-15 | MB231/MB468 | MITO |
| >MI0000093\|hsa-mir-92a-1&WithFlank&13\|+\|91351308\|91351397@54.75.22;MI0000094\|hsa-mir-92a-2&WithFlank&X\|-\|134169532\|134169618@54.75.22&[MIMAT0000092&hsa-miR-92a-3p&offsets\|0\|0;m-9&13\|+\|91351361\|91351382&offsets\|0\|0];[MIMAT0000092_1&hsa-miR-92a-3p&offsets\|0\|0;m-11&X\|-\|134169544\|134169565&offsets\|0\|0]\|TATTGCACTTGTCCCGGCCTGT | >MI0000093\|hsa-mir-92a-1&WithFlank&13\|+\|91351308\|91351397 | 54 | 75 | 22 | TATTGCACTTGTCCCGGCCTGT | 80833.5 | 6.7 | 0.8 | 8.0 | 1.6E-15 | 9.5E-15 | MB231/MB468 | CYTO |
| >hg38_21_+_8433172_8446622_RNA45SN1_45S_50ntFlanks@475.518.44\|CTTCGTGATCGATGTGGTGACGTCGTGCTCTCCCGGGCCGGGTC | >hg38_21_+_8433172_8446622_RNA45SN1_45S_50ntFlanks | 475 | 518 | 44 | CTTCGTGATCGATGTGGTGACGTCGTGCTCTCCCGGGCCGGGTC | 698.5 | 10.6 | 1.3 | 8.0 | 1.6E-15 | 9.5E-15 | MB231/MB468 | CYTO |
| >MI0000441\|hsa-mir-30b&WithFlank&8\|-\|134800514\|134800613@23.43.21(+1C)&[MIMAT0000420&hsa-miR-30b-5p&offsets\|0\|-1(+1C);m-76&8\|-\|134800570\|134800591&offsets\|0\|-1(+1C)]\|TGTAAACATCCTACACTCAGCC | >MI0000441\|hsa-mir-30b&WithFlank&8\|-\|134800514\|134800613 | 23 | 43 | 21(+1C) | TGTAAACATCCTACACTCAGCC | 650.8 | 8.2 | 1.0 | 7.9 | 2.1E-15 | 9.5E-15 | BT20/MB468 | CYTO |
| >hg38_21_+_8433172_8446622_RNA45SN1_45S_50ntFlanks@11612.11640.29\|TAATTAAAACAAAGCATCGCGAAGGCCCG | >hg38_21_+_8433172_8446622_RNA45SN1_45S_50ntFlanks | 11612 | 11640 | 29 | TAATTAAAACAAAGCATCGCGAAGGCCCG | 906.5 | -12.9 | 1.5 | -8.4 | 4.6E-17 | 9.6E-15 | BT20/MB468 | MP |
| >hg38_21_+_8433172_8446622_RNA45SN1_45S_50ntFlanks@11535.11552.18\|TTCCGCGCGGCGCCTCGC | >hg38_21_+_8433172_8446622_RNA45SN1_45S_50ntFlanks | 11535 | 11552 | 18 | TTCCGCGCGGCGCCTCGC | 583.0 | -12.2 | 1.5 | -8.0 | 1.7E-15 | 9.6E-15 | MB231/MB468 | NUC |
| >hg38_21_+_8433172_8446622_RNA45SN1_45S_50ntFlanks@12437.12473.37\|CATTGTGAAGCAGAATTCACCAAGCGTTGGATTGTTC | >hg38_21_+_8433172_8446622_RNA45SN1_45S_50ntFlanks | 12437 | 12473 | 37 | CATTGTGAAGCAGAATTCACCAAGCGTTGGATTGTTC | 5249.8 | -5.6 | 0.7 | -7.9 | 2.1E-15 | 9.6E-15 | BT20/MB468 | CYTO |
| >hg38_21_+_8433172_8446622_RNA45SN1_45S_50ntFlanks@11890.11910.21\|CTTGACTCTAGTCTGGCACGG | >hg38_21_+_8433172_8446622_RNA45SN1_45S_50ntFlanks | 11890 | 11910 | 21 | CTTGACTCTAGTCTGGCACGG | 189.3 | -11.4 | 1.4 | -8.0 | 1.4E-15 | 9.6E-15 | MB231/MB468 | MP |
| >MI0000446\|hsa-mir-125b-1&WithFlank&11\|-\|122099751\|122099850@21.40.20;MI0000470\|hsa-mir-125b-2&WithFlank&21\|+\|16590231\|16590331@23.42.20&[MIMAT0000423&hsa-miR-125b-5p&offsets\|0\|-2;m-67&11\|-\|122099809\|122099830&offsets\|0\|-2];[MIMAT0000423_1&hsa-miR-125b-5p&offsets\|0\|-2;m-68&21\|+\|16590253\|16590274&offsets\|0\|-2]\|TCCCTGAGACCCTAACTTGT | >MI0000446\|hsa-mir-125b-1&WithFlank&11\|-\|122099751\|122099850 | 21 | 40 | 20 | TCCCTGAGACCCTAACTTGT | 2071.5 | 7.6 | 1.0 | 7.9 | 2.4E-15 | 9.6E-15 | MB231/MB468 | MITO |

<table>
<tr><td>>hg38_21_+_8433172_8446622_RNA45SN1_45S_50ntFlanks@9623.9648.26|TAGTAGCTGGTTCCCTCCGAAGTTTC</td><td>>hg38_21_+_8433172_8446622_RNA45SN1_45S_50ntFlanks</td><td>9623</td><td>9648</td><td>26</td><td>TAGTAGCTGGTTCCCTCCGAAGTTTC</td><td>437.7</td><td>-12.6</td><td>1.6</td><td>-8.0</td><td>1.4E-15</td><td>9.6E-15</td><td>MB231/MB468</td><td>MP</td></tr>
<tr><td>>hg38_21_+_8433172_8446622_RNA45SN1_45S_50ntFlanks@4588.4616.29|CTATTTTGTTGGTTTTCGGAACTGAGGCC</td><td>>hg38_21_+_8433172_8446622_RNA45SN1_45S_50ntFlanks</td><td>4588</td><td>4616</td><td>29</td><td>CTATTTTGTTGGTTTTCGGAACTGAGGCC</td><td>4216.9</td><td>-11.2</td><td>1.4</td><td>-7.9</td><td>3.2E-15</td><td>9.6E-15</td><td>BT20/MB468</td><td>NUC</td></tr>
<tr><td>>hg38_21_+_8433172_8446622_RNA45SN1_45S_50ntFlanks@4472.4492.21|CTTAGCTGAGTGTCCCGCGGG</td><td>>hg38_21_+_8433172_8446622_RNA45SN1_45S_50ntFlanks</td><td>4472</td><td>4492</td><td>21</td><td>CTTAGCTGAGTGTCCCGCGGG</td><td>233.4</td><td>-10.8</td><td>1.4</td><td>-8.0</td><td>1.7E-15</td><td>9.6E-15</td><td>MB231/MB468</td><td>NUC</td></tr>
<tr><td>>hg38_21_+_8433172_8446622_RNA45SN1_45S_50ntFlanks@12141.12158.18|ACAGTGCCAGGTGGGGAG</td><td>>hg38_21_+_8433172_8446622_RNA45SN1_45S_50ntFlanks</td><td>12141</td><td>12158</td><td>18</td><td>ACAGTGCCAGGTGGGGAG</td><td>220.3</td><td>-10.7</td><td>1.4</td><td>-8.0</td><td>1.7E-15</td><td>9.6E-15</td><td>MB231/MB468</td><td>NUC</td></tr>
<tr><td>>hg38_21_+_8433172_8446622_RNA45SN1_45S_50ntFlanks@11781.11803.23|CGCATGAATGGATGAACGAGATT</td><td>>hg38_21_+_8433172_8446622_RNA45SN1_45S_50ntFlanks</td><td>11781</td><td>11803</td><td>23</td><td>CGCATGAATGGATGAACGAGATT</td><td>704.9</td><td>-7.4</td><td>0.9</td><td>-7.9</td><td>2.4E-15</td><td>9.6E-15</td><td>MB231/MB468</td><td>MITO</td></tr>
<tr><td>>hg38_21_+_8433172_8446622_RNA45SN1_45S_50ntFlanks@7982.8005.24|TCAGATCAGACGTGGCGACCCGCT</td><td>>hg38_21_+_8433172_8446622_RNA45SN1_45S_50ntFlanks</td><td>7982</td><td>8005</td><td>24</td><td>TCAGATCAGACGTGGCGACCCGCT</td><td>696.5</td><td>-5.5</td><td>0.7</td><td>-7.9</td><td>2.4E-15</td><td>9.6E-15</td><td>MB231/MB468</td><td>MITO</td></tr>
<tr><td>>hg38_21_+_8433172_8446622_RNA45SN1_45S_50ntFlanks@12161.12178.18|TGACTGGGGCGGTACACC</td><td>>hg38_21_+_8433172_8446622_RNA45SN1_45S_50ntFlanks</td><td>12161</td><td>12178</td><td>18</td><td>TGACTGGGGCGGTACACC</td><td>242.4</td><td>-11.8</td><td>1.5</td><td>-8.0</td><td>1.4E-15</td><td>9.7E-15</td><td>MB231/MB468</td><td>MP</td></tr>
<tr><td>>hg38_21_+_8433172_8446622_RNA45SN1_45S_50ntFlanks@12605.12623.19|TATGTGCTTGGCTGAGGAG</td><td>>hg38_21_+_8433172_8446622_RNA45SN1_45S_50ntFlanks</td><td>12605</td><td>12623</td><td>19</td><td>TATGTGCTTGGCTGAGGAG</td><td>1071.5</td><td>-7.6</td><td>1.0</td><td>-8.0</td><td>1.8E-15</td><td>9.7E-15</td><td>MB231/MB468</td><td>NUC</td></tr>
<tr><td>>hg38_21_+_8433172_8446622_RNA45SN1_45S_50ntFlanks@11576.11600.25|CTTAGAACTGGTGCGGACCAGGGGA</td><td>>hg38_21_+_8433172_8446622_RNA45SN1_45S_50ntFlanks</td><td>11576</td><td>11600</td><td>25</td><td>CTTAGAACTGGTGCGGACCAGGGGA</td><td>241.3</td><td>-11.7</td><td>1.5</td><td>-7.9</td><td>2.4E-15</td><td>9.7E-15</td><td>MB231/MB468</td><td>MITO</td></tr>
<tr><td>>hg38_21_+_8433172_8446622_RNA45SN1_45S_50ntFlanks@12631.12654.24|GGCGAAGCTACCATCTGTGGGATT</td><td>>hg38_21_+_8433172_8446622_RNA45SN1_45S_50ntFlanks</td><td>12631</td><td>12654</td><td>24</td><td>GGCGAAGCTACCATCTGTGGGATT</td><td>236.6</td><td>-5.3</td><td>0.7</td><td>-8.0</td><td>1.6E-15</td><td>9.8E-15</td><td>MB231/MB468</td><td>CYTO</td></tr>
<tr><td>>hg38_21_+_8433172_8446622_RNA45SN1_45S_50ntFlanks@12151.12182.32|GTGGGGAGTTTGACTGGGGCGGTACACCTGTC</td><td>>hg38_21_+_8433172_8446622_RNA45SN1_45S_50ntFlanks</td><td>12151</td><td>12182</td><td>32</td><td>GTGGGGAGTTTGACTGGGGCGGTACACCTGTC</td><td>389.3</td><td>-11.6</td><td>1.5</td><td>-8.0</td><td>1.8E-15</td><td>9.8E-15</td><td>MB231/MB468</td><td>NUC</td></tr>
<tr><td>>hg38_21_+_8433172_8446622_RNA45SN1_45S_50ntFlanks@10866.10886.21|CTAAGGGCTGGGTCGGTCGGG</td><td>>hg38_21_+_8433172_8446622_RNA45SN1_45S_50ntFlanks</td><td>10866</td><td>10886</td><td>21</td><td>CTAAGGGCTGGGTCGGTCGGG</td><td>331.5</td><td>-12.2</td><td>1.5</td><td>-7.9</td><td>2.4E-15</td><td>9.8E-15</td><td>MB231/MB468</td><td>MITO</td></tr>
<tr><td>>hg38_21_+_8433172_8446622_RNA45SN1_45S_50ntFlanks@12045.12071.27|CTGACCCGGTGAGGCGGGGGGGCGAGC</td><td>>hg38_21_+_8433172_8446622_RNA45SN1_45S_50ntFlanks</td><td>12045</td><td>12071</td><td>27</td><td>CTGACCCGGTGAGGCGGGGGGGCGAGC</td><td>275.9</td><td>-12.0</td><td>1.5</td><td>-8.0</td><td>1.5E-15</td><td>9.8E-15</td><td>MB231/MB468</td><td>MP</td></tr>
<tr><td>>MI0003763|hsa-mir-767&WithFlank&X|-|152393415|152393535@33.55.23&[MIMAT0003882&hsa-miR-767-5p&offsets|0|0;m-788&X|-|152393481|152393503&offsets|0|0]|TGCACCATGGTTGTCTGAGCATG</td><td>>MI0003763|hsa-mir-767&WithFlank&X|-|152393415|152393535</td><td>33</td><td>55</td><td>23</td><td>TGCACCATGGTTGTCTGAGCATG</td><td>332.5</td><td>11.8</td><td>1.4</td><td>8.3</td><td>1.4E-16</td><td>9.8E-15</td><td>BT20/MB231</td><td>CYTO</td></tr>
<tr><td>>MI0003591|hsa-mir-584&WithFlank&5|-|149062307|149062415@22.41.20(+1U)&[MIMAT0003249&hsa-miR-584-5p&offsets|0|-2(+1U);m-167&5|-|149062374|149062394&offsets|0|-1(+1U)]|TTATGGTTTGCCTGGGACTGT</td><td>>MI0003591|hsa-mir-584&WithFlank&5|-|149062307|149062415</td><td>22</td><td>41</td><td>20(+1U)</td><td>TTATGGTTTGCCTGGGACTGT</td><td>1147.6</td><td>11.2</td><td>1.4</td><td>8.0</td><td>1.6E-15</td><td>9.8E-15</td><td>BT20/MB468</td><td>MITO</td></tr>
<tr><td>>hg38_21_+_8433172_8446622_RNA45SN1_45S_50ntFlanks@12988.13014.27|CAGCTCCCTCGCTGCGATCTATTGAAA</td><td>>hg38_21_+_8433172_8446622_RNA45SN1_45S_50ntFlanks</td><td>12988</td><td>13014</td><td>27</td><td>CAGCTCCCTCGCTGCGATCTATTGAAA</td><td>246.1</td><td>4.0</td><td>0.5</td><td>8.0</td><td>1.6E-15</td><td>9.9E-15</td><td>MB231/MB468</td><td>CYTO</td></tr>
<tr><td>>hg38_21_+_8433172_8446622_RNA45SN1_45S_50ntFlanks@8168.8215.48|CCCAAGTCCTTCTGATCGAGGCCCAGCCCGTGGACGGTGTGAGGCCGG</td><td>>hg38_21_+_8433172_8446622_RNA45SN1_45S_50ntFlanks</td><td>8168</td><td>8215</td><td>48</td><td>CCCAAGTCCTTCTGATCGAGGCCCAGCCCGTGGACGGTGTGAGGCCGG</td><td>456.8</td><td>-12.6</td><td>1.6</td><td>-7.9</td><td>2.4E-15</td><td>9.9E-15</td><td>MB231/MB468</td><td>MITO</td></tr>
<tr><td>>MI0000736|hsa-mir-30c-1&WithFlank&1|+|40757278|40757378@23.44.22(+1U);MI0000254|hsa-mir-30c-2&WithFlank&6|-|71376954|71377037@13.34.22(+1U)&[MIMAT0000244&hsa-miR-30c-5p&offsets|0|-1(+1U);m-57&1|+|40757300|40757323&offsets|0|-2(+1U)];[MIMAT0000244_1&hsa-miR-30c-5p&offsets|0|-1(+1U);m-56&6|-|71377002|71377025&offsets|0|-2(+1U)]|TGTAAACATCCTACACTCTCAGT</td><td>>MI0000736|hsa-mir-30c-1&WithFlank&1|+|40757278|40757378</td><td>23</td><td>44</td><td>22(+1U)</td><td>TGTAAACATCCTACACTCTCAGT</td><td>6417.7</td><td>5.0</td><td>0.6</td><td>8.0</td><td>1.8E-15</td><td>9.9E-15</td><td>MB231/MB468</td><td>NUC</td></tr>
<tr><td>>hg38_21_+_8433172_8446622_RNA45SN1_45S_50ntFlanks@11576.11602.27|CTTAGAACTGGTGCGGACCAGGGGAAT</td><td>>hg38_21_+_8433172_8446622_RNA45SN1_45S_50ntFlanks</td><td>11576</td><td>11602</td><td>27</td><td>CTTAGAACTGGTGCGGACCAGGGGAAT</td><td>914.7</td><td>-7.4</td><td>0.9</td><td>-8.0</td><td>1.8E-15</td><td>9.9E-15</td><td>MB231/MB468</td><td>NUC</td></tr>
<tr><td>>hg38_21_+_8433172_8446622_RNA45SN1_45S_50ntFlanks@11783.11806.24|CATGAATGGATGAACGAGATTCCC</td><td>>hg38_21_+_8433172_8446622_RNA45SN1_45S_50ntFlanks</td><td>11783</td><td>11806</td><td>24</td><td>CATGAATGGATGAACGAGATTCCC</td><td>4203.7</td><td>-8.6</td><td>1.1</td><td>-8.0</td><td>1.5E-15</td><td>9.9E-15</td><td>MB231/MB468</td><td>MP</td></tr>
<tr><td>>MI0000078|hsa-mir-22&WithFlank&17|-|1713897|1713993@59.79.21(+1G)&[MIMAT0000077&hsa-miR-22-3p&offsets|0|-1(+1G);m-2&17|-|1713914|1713935&offsets|0|-1(+1G)]|AAGCTGCCAGTTGAAGAACTGG</td><td>>MI0000078|hsa-mir-22&WithFlank&17|-|1713897|1713993</td><td>59</td><td>79</td><td>21(+1G)</td><td>AAGCTGCCAGTTGAAGAACTGG</td><td>71.8</td><td>9.0</td><td>1.1</td><td>7.9</td><td>3.3E-15</td><td>9.9E-15</td><td>BT20/MB468</td><td>NUC</td></tr>
<tr><td>>hg38_21_+_8433172_8446622_RNA45SN1_45S_50ntFlanks@11255.11281.27|CTGCGGCGGGCGTCGCGGCCGCCCCCG</td><td>>hg38_21_+_8433172_8446622_RNA45SN1_45S_50ntFlanks</td><td>11255</td><td>11281</td><td>27</td><td>CTGCGGCGGGCGTCGCGGCCGCCCCCG</td><td>270.5</td><td>-11.9</td><td>1.5</td><td>-7.9</td><td>2.5E-15</td><td>9.9E-15</td><td>MB231/MB468</td><td>MITO</td></tr>
<tr><td>>hg38_21_+_8433172_8446622_RNA45SN1_45S_50ntFlanks@10771.10805.35|CAGGTCTCCAAGGTGAACAGCCTCTGGCATGTTGG</td><td>>hg38_21_+_8433172_8446622_RNA45SN1_45S_50ntFlanks</td><td>10771</td><td>10805</td><td>35</td><td>CAGGTCTCCAAGGTGAACAGCCTCTGGCATGTTGG</td><td>705.2</td><td>-4.2</td><td>0.5</td><td>-8.0</td><td>1.6E-15</td><td>1.0E-14</td><td>BT20/MB468</td><td>MITO</td></tr>
<tr><td>>hg38_21_+_8433172_8446622_RNA45SN1_45S_50ntFlanks@11317.11334.18|ACCCCACGTCTCGTCGCG</td><td>>hg38_21_+_8433172_8446622_RNA45SN1_45S_50ntFlanks</td><td>11317</td><td>11334</td><td>18</td><td>ACCCCACGTCTCGTCGCG</td><td>689.7</td><td>-12.1</td><td>1.5</td><td>-8.0</td><td>1.6E-15</td><td>1.0E-14</td><td>MB231/MB468</td><td>CYTO</td></tr>
<tr><td>>hg38_21_+_8433172_8446622_RNA45SN1_45S_50ntFlanks@11853.11872.20|CTTGGCGGAATCAGCGGGGA</td><td>>hg38_21_+_8433172_8446622_RNA45SN1_45S_50ntFlanks</td><td>11853</td><td>11872</td><td>20</td><td>CTTGGCGGAATCAGCGGGGA</td><td>333.6</td><td>-12.2</td><td>1.5</td><td>-7.9</td><td>2.5E-15</td><td>1.0E-14</td><td>MB231/MB468</td><td>MITO</td></tr>
<tr><td>>hg38_21_+_8433172_8446622_RNA45SN1_45S_50ntFlanks@12418.12440.23|CGATGTCGGCTCTTCCTATCATT</td><td>>hg38_21_+_8433172_8446622_RNA45SN1_45S_50ntFlanks</td><td>12418</td><td>12440</td><td>23</td><td>CGATGTCGGCTCTTCCTATCATT</td><td>1155.9</td><td>-5.1</td><td>0.7</td><td>-7.9</td><td>3.3E-15</td><td>1.0E-14</td><td>BT20/MB468</td><td>NUC</td></tr>
<tr><td>>hg38_21_+_8433172_8446622_RNA45SN1_45S_50ntFlanks@10355.10388.34|AACTTTGAAGGCCGAAGTGGAGAAGGGTTCCATG</td><td>>hg38_21_+_8433172_8446622_RNA45SN1_45S_50ntFlanks</td><td>10355</td><td>10388</td><td>34</td><td>AACTTTGAAGGCCGAAGTGGAGAAGGGTTCCATG</td><td>496.9</td><td>-12.8</td><td>1.6</td><td>-8.0</td><td>1.5E-15</td><td>1.0E-14</td><td>MB231/MB468</td><td>MP</td></tr>
<tr><td>>hg38_21_+_8433172_8446622_RNA45SN1_45S_50ntFlanks@6656.6680.25|CTTAGCGGTGGATCACTCGGCTCGT</td><td>>hg38_21_+_8433172_8446622_RNA45SN1_45S_50ntFlanks</td><td>6656</td><td>6680</td><td>25</td><td>CTTAGCGGTGGATCACTCGGCTCGT</td><td>279.7</td><td>-12.0</td><td>1.5</td><td>-8.0</td><td>1.5E-15</td><td>1.0E-14</td><td>MB231/MB468</td><td>MP</td></tr>
<tr><td>>hg38_21_+_8433172_8446622_RNA45SN1_45S_50ntFlanks@12420.12445.26|ATGTCGGCTCTTCCTATCATTGTGAA</td><td>>hg38_21_+_8433172_8446622_RNA45SN1_45S_50ntFlanks</td><td>12420</td><td>12445</td><td>26</td><td>ATGTCGGCTCTTCCTATCATTGTGAA</td><td>3064.7</td><td>-4.1</td><td>0.5</td><td>-8.0</td><td>1.8E-15</td><td>1.0E-14</td><td>MB231/MB468</td><td>NUC</td></tr>
<tr><td>>MI0000456|hsa-mir-140&WithFlank&16|+|69933075|69933186@69.90.22&[MIMAT0004597&hsa-miR-140-3p&offsets|+1|+2;m-45&16|+|69933143|69933164&offsets|0|0]|ACCACAGGGTAGAACCACGGAC</td><td>>MI0000456|hsa-mir-140&WithFlank&16|+|69933075|69933186</td><td>69</td><td>90</td><td>22</td><td>ACCACAGGGTAGAACCACGGAC</td><td>976.7</td><td>13.2</td><td>1.7</td><td>7.9</td><td>2.5E-15</td><td>1.0E-14</td><td>MB231/MB468</td><td>MITO</td></tr>
<tr><td>>hg38_21_+_8433172_8446622_RNA45SN1_45S_50ntFlanks@11104.11131.28|AAGGGTCGGGGCGGCAGGGGCCGGCGGC</td><td>>hg38_21_+_8433172_8446622_RNA45SN1_45S_50ntFlanks</td><td>11104</td><td>11131</td><td>28</td><td>AAGGGTCGGGGCGGCAGGGGCCGGCGGC</td><td>241.8</td><td>-10.9</td><td>1.4</td><td>-8.0</td><td>1.8E-15</td><td>1.0E-14</td><td>MB231/MB468</td><td>NUC</td></tr>
<tr><td>>hg38_21_+_8433172_8446622_RNA45SN1_45S_50ntFlanks@10823.10842.20|AAGTCGGCAAGCCGGATCCG</td><td>>hg38_21_+_8433172_8446622_RNA45SN1_45S_50ntFlanks</td><td>10823</td><td>10842</td><td>20</td><td>AAGTCGGCAAGCCGGATCCG</td><td>200.0</td><td>-11.5</td><td>1.4</td><td>-8.0</td><td>1.5E-15</td><td>1.0E-14</td><td>MB231/MB468</td><td>MP</td></tr>
<tr><td>>hg38_21_+_8433172_8446622_RNA45SN1_45S_50ntFlanks@4761.4780.20|CAGATACCGTCGTAGTTCCG</td><td>>hg38_21_+_8433172_8446622_RNA45SN1_45S_50ntFlanks</td><td>4761</td><td>4780</td><td>20</td><td>CAGATACCGTCGTAGTTCCG</td><td>273.8</td><td>-12.0</td><td>1.5</td><td>-8.0</td><td>1.5E-15</td><td>1.0E-14</td><td>MB231/MB468</td><td>MP</td></tr>
<tr><td>>hg38_21_+_8433172_8446622_RNA45SN1_45S_50ntFlanks@9534.9570.37|AGGAAACTCTGGTGGAGGTCCGTAGCGGTCCTGACGT</td><td>>hg38_21_+_8433172_8446622_RNA45SN1_45S_50ntFlanks</td><td>9534</td><td>9570</td><td>37</td><td>AGGAAACTCTGGTGGAGGTCCGTAGCGGTCCTGACGT</td><td>1198.7</td><td>-6.4</td><td>0.8</td><td>-8.0</td><td>1.5E-15</td><td>1.0E-14</td><td>MB231/MB468</td><td>MP</td></tr>
<tr><td>>trnaMT_PheGAA_MT_+_577_647@37.56.20|AAATGTTTAGACGGGCTCAC</td><td>>trnaMT_PheGAA_MT_+_577_647</td><td>37</td><td>56</td><td>20</td><td>AAATGTTTAGACGGGCTCAC</td><td>1026.1</td><td>10.1</td><td>1.3</td><td>8.0</td><td>1.6E-15</td><td>1.0E-14</td><td>BT20/MB468</td><td>MITO</td></tr>
<tr><td>>hg38_21_+_8433172_8446622_RNA45SN1_45S_50ntFlanks@12947.12990.44|AGACGACCTGCTTCTGGGTCGGGGTTTCGTACGTAGCAGAGCAG</td><td>>hg38_21_+_8433172_8446622_RNA45SN1_45S_50ntFlanks</td><td>12947</td><td>12990</td><td>44</td><td>AGACGACCTGCTTCTGGGTCGGGGTTTCGTACGTAGCAGAGCAG</td><td>515.2</td><td>-13.4</td><td>1.7</td><td>-7.9</td><td>3.4E-15</td><td>1.0E-14</td><td>BT20/MB468</td><td>NUC</td></tr>
<tr><td>>hg38_21_+_8433172_8446622_RNA45SN1_45S_50ntFlanks@3975.4002.28|CGGGCGCCGGCGGCTTTGGTGACTCTAG</td><td>>hg38_21_+_8433172_8446622_RNA45SN1_45S_50ntFlanks</td><td>3975</td><td>4002</td><td>28</td><td>CGGGCGCCGGCGGCTTTGGTGACTCTAG</td><td>408.8</td><td>-12.5</td><td>1.6</td><td>-7.9</td><td>2.6E-15</td><td>1.0E-14</td><td>MB231/MB468</td><td>MITO</td></tr>
<tr><td>>hg38_21_+_8433172_8446622_RNA45SN1_45S_50ntFlanks@12960.12990.31|CTGGGTCGGGGTTTCGTACGTAGCAGAGCAG</td><td>>hg38_21_+_8433172_8446622_RNA45SN1_45S_50ntFlanks</td><td>12960</td><td>12990</td><td>31</td><td>CTGGGTCGGGGTTTCGTACGTAGCAGAGCAG</td><td>525.2</td><td>-7.2</td><td>0.9</td><td>-7.9</td><td>1.9E-15</td><td>1.0E-14</td><td>MB231/MB468</td><td>NUC</td></tr>
<tr><td>>hg38_21_+_8433172_8446622_RNA45SN1_45S_50ntFlanks@4160.4182.23|CCACATCCAAGGAAGGCAGCAGG</td><td>>hg38_21_+_8433172_8446622_RNA45SN1_45S_50ntFlanks</td><td>4160</td><td>4182</td><td>23</td><td>CCACATCCAAGGAAGGCAGCAGG</td><td>154.3</td><td>-5.4</td><td>0.7</td><td>-8.0</td><td>1.7E-15</td><td>1.0E-14</td><td>MB231/MB468</td><td>CYTO</td></tr>
<tr><td>>hg38_21_+_8433172_8446622_RNA45SN1_45S_50ntFlanks@11188.11207.20|CACCCGGGGGGCCGGCGGCG</td><td>>hg38_21_+_8433172_8446622_RNA45SN1_45S_50ntFlanks</td><td>11188</td><td>11207</td><td>20</td><td>CACCCGGGGGGCCGGCGGCG</td><td>309.2</td><td>-12.1</td><td>1.5</td><td>-7.9</td><td>2.6E-15</td><td>1.0E-14</td><td>MB231/MB468</td><td>MITO</td></tr>
<tr><td>>hg38_MT_+_598_1651_RNR1_12S_50ntFlanks@369.404.36|AATAAAGCTAAAACTCACCTGAGTTGTAAAAAACTC</td><td>>hg38_MT_+_598_1651_RNR1_12S_50ntFlanks</td><td>369</td><td>404</td><td>36</td><td>AATAAAGCTAAAACTCACCTGAGTTGTAAAAAACTC</td><td>2101.0</td><td>5.7</td><td>0.7</td><td>8.0</td><td>1.6E-15</td><td>1.0E-14</td><td>BT20/MB468</td><td>MITO</td></tr>
<tr><td>>hg38_21_+_8433172_8446622_RNA45SN1_45S_50ntFlanks@10678.10710.33|CGGTTCCGGCGGCGTCCGGTGAGCTCTCGCTGG</td><td>>hg38_21_+_8433172_8446622_RNA45SN1_45S_50ntFlanks</td><td>10678</td><td>10710</td><td>33</td><td>CGGTTCCGGCGGCGTCCGGTGAGCTCTCGCTGG</td><td>774595.6</td><td>-10.4</td><td>1.3</td><td>-7.9</td><td>3.5E-15</td><td>1.0E-14</td><td>BT20/MB468</td><td>NUC</td></tr>
<tr><td>>hg38_21_+_8433172_8446622_RNA45SN1_45S_50ntFlanks@11919.11950.32|CATGAGAGGTGTAGAATAAGTGGGAGGCCCCC</td><td>>hg38_21_+_8433172_8446622_RNA45SN1_45S_50ntFlanks</td><td>11919</td><td>11950</td><td>32</td><td>CATGAGAGGTGTAGAATAAGTGGGAGGCCCCC</td><td>482.2</td><td>-12.8</td><td>1.6</td><td>-8.0</td><td>1.5E-15</td><td>1.0E-14</td><td>MB231/MB468</td><td>MP</td></tr>
<tr><td>>hg38_21_+_8433172_8446622_RNA45SN1_45S_50ntFlanks@10425.10456.32|AGATGGGCGAGCGCCGTTCCGAAGGGACGGGC</td><td>>hg38_21_+_8433172_8446622_RNA45SN1_45S_50ntFlanks</td><td>10425</td><td>10456</td><td>32</td><td>AGATGGGCGAGCGCCGTTCCGAAGGGACGGGC</td><td>627.4</td><td>-5.2</td><td>0.7</td><td>-7.9</td><td>1.9E-15</td><td>1.0E-14</td><td>MB231/MB468</td><td>NUC</td></tr>
<tr><td>>hg38_21_+_8433172_8446622_RNA45SN1_45S_50ntFlanks@6647.6669.23|CGTACGACTCTTAGCGGTGGATC</td><td>>hg38_21_+_8433172_8446622_RNA45SN1_45S_50ntFlanks</td><td>6647</td><td>6669</td><td>23</td><td>CGTACGACTCTTAGCGGTGGATC</td><td>5098.0</td><td>4.4</td><td>0.6</td><td>7.9</td><td>2.3E-15</td><td>1.0E-14</td><td>BT20/MB468</td><td>CYTO</td></tr>
<tr><td>>hg38_21_+_8433172_8446622_RNA45SN1_45S_50ntFlanks@10351.10376.26|CGAGAACTTTGAAGGCCGAAGTGGAG</td><td>>hg38_21_+_8433172_8446622_RNA45SN1_45S_50ntFlanks</td><td>10351</td><td>10376</td><td>26</td><td>CGAGAACTTTGAAGGCCGAAGTGGAG</td><td>1982.0</td><td>-8.1</td><td>1.0</td><td>-7.9</td><td>1.9E-15</td><td>1.0E-14</td><td>MB231/MB468</td><td>NUC</td></tr>
<tr><td>>hg38_21_+_8433172_8446622_RNA45SN1_45S_50ntFlanks@8176.8203.28|CTTCTGATCGAGGCCCAGCCCGTGGACG</td><td>>hg38_21_+_8433172_8446622_RNA45SN1_45S_50ntFlanks</td><td>8176</td><td>8203</td><td>28</td><td>CTTCTGATCGAGGCCCAGCCCGTGGACG</td><td>11261.5</td><td>-7.4</td><td>0.9</td><td>-8.0</td><td>1.6E-15</td><td>1.0E-14</td><td>MB231/MB468</td><td>MP</td></tr>
<tr><td>>hg38_21_+_8433172_8446622_RNA45SN1_45S_50ntFlanks@11320.11344.25|CCACGTCTCGTCGCGCGCGCGTCCG</td><td>>hg38_21_+_8433172_8446622_RNA45SN1_45S_50ntFlanks</td><td>11320</td><td>11344</td><td>25</td><td>CCACGTCTCGTCGCGCGCGCGTCCG</td><td>4793.6</td><td>-8.3</td><td>1.0</td><td>-8.0</td><td>1.7E-15</td><td>1.0E-14</td><td>MB231/MB468</td><td>CYTO</td></tr>
<tr><td>>hg38_21_+_8433172_8446622_RNA45SN1_45S_50ntFlanks@4727.4761.35|AATCAAGAACGAAAGTCGGAGGTTCGAAGACGATC</td><td>>hg38_21_+_8433172_8446622_RNA45SN1_45S_50ntFlanks</td><td>4727</td><td>4761</td><td>35</td><td>AATCAAGAACGAAAGTCGGAGGTTCGAAGACGATC</td><td>1255.8</td><td>5.3</td><td>0.7</td><td>8.0</td><td>1.7E-15</td><td>1.0E-14</td><td>MB231/MB468</td><td>CYTO</td></tr>
<tr><td>>hg38_21_+_8433172_8446622_RNA45SN1_45S_50ntFlanks@11027.11061.35|CTCTCTCTCTCTCTCCCCCGCTCCCCGTCCTCCCC</td><td>>hg38_21_+_8433172_8446622_RNA45SN1_45S_50ntFlanks</td><td>11027</td><td>11061</td><td>35</td><td>CTCTCTCTCTCTCTCCCCCGCTCCCCGTCCTCCCC</td><td>816.8</td><td>-12.6</td><td>1.6</td><td>-7.9</td><td>1.9E-15</td><td>1.0E-14</td><td>MB231/MB468</td><td>NUC</td></tr>
</table>

<table>
<tr><td>>am_trna162_MetCAT_6_-_26758550_26758622@31.53.23,trna164_MetCAT_6_-_26735574_26735646@31.53.23,trna20_MetCAT_16_+_71460396_71460468@31.53.23,trna21_MetCAT_6_+_26701712_26701784@31.53.23,trna27_MetCAT_6_+_26766444_26766516@31.53.23,trna75_MetCAT_6_+_28912352_28912424@31.53.23,trna97_MetCAT_6_-_28921042_28921114@31.53.23|TCTCATAATCTGAAGGTCCTGAG</td><td>>am_trna162_MetCAT_6_-_26758550_26758622</td><td>31</td><td>53</td><td>23</td><td>TCTCATAATCTGAAGGTCCTGAG</td><td>248.3</td><td>-11.8</td><td>1.5</td><td>-8.0</td><td>1.6E-15</td><td>1.0E-14</td><td>MB231/MB468</td><td>MP</td></tr>
<tr><td>>MI0000286|hsa-mir-210&WithFlank&11|-|568083|568204@72.93.22(+1A)&[MIMAT0000267&hsa-miR-210-3p&offsets|0|0(+1A);m-102&11|-|568112|568133&offsets|0|0(+1A)]|CTGTGCGTGTGACAGCGGCTGAA</td><td>>MI0000286|hsa-mir-210&WithFlank&11|-|568083|568204</td><td>72</td><td>93</td><td>22(+1A)</td><td>CTGTGCGTGTGACAGCGGCTGAA</td><td>1058.4</td><td>11.2</td><td>1.4</td><td>8.0</td><td>1.7E-15</td><td>1.1E-14</td><td>BT20/MB468</td><td>MITO</td></tr>
<tr><td>>hg38_21_+_8433172_8446622_RNA45SN1_45S_50ntFlanks@11190.11207.18|CCCGGGGGGCCGGCGGCG</td><td>>hg38_21_+_8433172_8446622_RNA45SN1_45S_50ntFlanks</td><td>11190</td><td>11207</td><td>18</td><td>CCCGGGGGGCCGGCGGCG</td><td>1052.8</td><td>-10.3</td><td>1.3</td><td>-7.9</td><td>1.9E-15</td><td>1.1E-14</td><td>MB231/MB468</td><td>NUC</td></tr>
<tr><td>>hg38_21_+_8433172_8446622_RNA45SN1_45S_50ntFlanks@11570.11603.34|AGCCGACTTAGAACTGGTGCGGACCAGGGGAATC</td><td>>hg38_21_+_8433172_8446622_RNA45SN1_45S_50ntFlanks</td><td>11570</td><td>11603</td><td>34</td><td>AGCCGACTTAGAACTGGTGCGGACCAGGGGAATC</td><td>4208.6</td><td>-8.4</td><td>1.1</td><td>-7.9</td><td>2.6E-15</td><td>1.1E-14</td><td>MB231/MB468</td><td>MITO</td></tr>
<tr><td>>hg38_1_-_228634819_228635039_RNA5S12_5S_50ntFlanks@135.171.37|GGGAGACCGCCTGGGAATACCGGGTGCTGTAGGCTTT</td><td>>hg38_1_-_228634819_228635039_RNA5S12_5S_50ntFlanks</td><td>135</td><td>171</td><td>37</td><td>GGGAGACCGCCTGGGAATACCGGGTGCTGTAGGCTTT</td><td>15088.8</td><td>6.7</td><td>0.8</td><td>7.9</td><td>2.3E-15</td><td>1.1E-14</td><td>BT20/MB468</td><td>CYTO</td></tr>
<tr><td>>MI0003643|hsa-mir-629&WithFlank&15|-|70079366|70079474@28.48.21(+1C)&[MIMAT0004810&hsa-miR-629-5p&offsets|0|0(+1C);m-150&15|-|70079427|70079447&offsets|0|0(+1C)]|TGGGTTTACGTTGGGAGAACTC</td><td>>MI0003643|hsa-mir-629&WithFlank&15|-|70079366|70079474</td><td>28</td><td>48</td><td>21(+1C)</td><td>TGGGTTTACGTTGGGAGAACTC</td><td>396.0</td><td>11.9</td><td>1.5</td><td>8.0</td><td>1.6E-15</td><td>1.1E-14</td><td>MB231/MB468</td><td>MP</td></tr>
<tr><td>>hg38_21_+_8433172_8446622_RNA45SN1_45S_50ntFlanks@4045.4062.18|CCATTCGAACGTCTGCCC</td><td>>hg38_21_+_8433172_8446622_RNA45SN1_45S_50ntFlanks</td><td>4045</td><td>4062</td><td>18</td><td>CCATTCGAACGTCTGCCC</td><td>636.4</td><td>-5.4</td><td>0.7</td><td>-7.9</td><td>1.9E-15</td><td>1.1E-14</td><td>MB231/MB468</td><td>NUC</td></tr>
<tr><td>>hg38_21_+_8433172_8446622_RNA45SN1_45S_50ntFlanks@10402.10420.19|AACATGGGTCAGTCGGTCC</td><td>>hg38_21_+_8433172_8446622_RNA45SN1_45S_50ntFlanks</td><td>10402</td><td>10420</td><td>19</td><td>AACATGGGTCAGTCGGTCC</td><td>1758.9</td><td>-10.8</td><td>1.4</td><td>-7.9</td><td>3.5E-15</td><td>1.1E-14</td><td>BT20/MB468</td><td>NUC</td></tr>
<tr><td>>hg38_21_+_8433172_8446622_RNA45SN1_45S_50ntFlanks@12655.12675.21|ATGACTGAACGCCTCTAAGTC</td><td>>hg38_21_+_8433172_8446622_RNA45SN1_45S_50ntFlanks</td><td>12655</td><td>12675</td><td>21</td><td>ATGACTGAACGCCTCTAAGTC</td><td>17212.4</td><td>-7.5</td><td>0.9</td><td>-8.0</td><td>1.7E-15</td><td>1.1E-14</td><td>MB231/MB468</td><td>CYTO</td></tr>
<tr><td>>hg38_21_+_8433172_8446622_RNA45SN1_45S_50ntFlanks@9034.9059.26|TCGTGGGGGGGCCGGGCCACCCCTCC</td><td>>hg38_21_+_8433172_8446622_RNA45SN1_45S_50ntFlanks</td><td>9034</td><td>9059</td><td>26</td><td>TCGTGGGGGGGCCGGGCCACCCCTCC</td><td>446.8</td><td>-12.6</td><td>1.6</td><td>-7.9</td><td>2.6E-15</td><td>1.1E-14</td><td>MB231/MB468</td><td>MITO</td></tr>
<tr><td>>hg38_21_+_8433172_8446622_RNA45SN1_45S_50ntFlanks@6750.6770.21|TCGAACGCACTTGCGGCCCCG</td><td>>hg38_21_+_8433172_8446622_RNA45SN1_45S_50ntFlanks</td><td>6750</td><td>6770</td><td>21</td><td>TCGAACGCACTTGCGGCCCCG</td><td>263.4</td><td>-11.9</td><td>1.5</td><td>-8.0</td><td>1.6E-15</td><td>1.1E-14</td><td>MB231/MB468</td><td>MP</td></tr>
<tr><td>>trna116_GluCTC_1_-_145399233_145399304@33.70.38,trna71_GluCTC_1_-_161439189_161439260@33.70.38,trna74_GluCTC_1_-_161431809_161431880@33.70.38,trna77_GluCTC_1_-_161424398_161424469@33.70.38,trna77_GluCTC_6_+_28949976_28950047@33.70.38,trna80_GluCTC_1_-_161417018_161417089@33.70.38,trna87_GluCTC_6_-_126101393_126101464@33.70.38|TCTCACCGCCGCGGCCCGGGTTCGATTCCCGGTCAGGG</td><td>>trna116_GluCTC_1_-_145399233_145399304</td><td>33</td><td>70</td><td>38</td><td>TCTCACCGCCGCGGCCCGGGTTCGATTCCCGGTCAGGG</td><td>212.3</td><td>-10.4</td><td>1.3</td><td>-8.0</td><td>1.7E-15</td><td>1.1E-14</td><td>MB231/MB468</td><td>CYTO</td></tr>
<tr><td>>hg38_21_+_8433172_8446622_RNA45SN1_45S_50ntFlanks@9202.9224.23|TCTCGGGGCCACGCGCGCGTCCC</td><td>>hg38_21_+_8433172_8446622_RNA45SN1_45S_50ntFlanks</td><td>9202</td><td>9224</td><td>23</td><td>TCTCGGGGCCACGCGCGCGTCCC</td><td>341.1</td><td>-12.3</td><td>1.5</td><td>-8.0</td><td>1.6E-15</td><td>1.1E-14</td><td>MB231/MB468</td><td>MP</td></tr>
<tr><td>>hg38_21_+_8433172_8446622_RNA45SN1_45S_50ntFlanks@6788.6806.19|CGCCTGTCTGAGCGTCGCT</td><td>>hg38_21_+_8433172_8446622_RNA45SN1_45S_50ntFlanks</td><td>6788</td><td>6806</td><td>19</td><td>CGCCTGTCTGAGCGTCGCT</td><td>4751.6</td><td>4.8</td><td>0.6</td><td>7.9</td><td>1.9E-15</td><td>1.1E-14</td><td>MB231/MB468</td><td>NUC</td></tr>
<tr><td>>hg38_21_+_8433172_8446622_RNA45SN1_45S_50ntFlanks@9141.9165.25|CGGGGCGGACTGTCCCCAGTGCGCC</td><td>>hg38_21_+_8433172_8446622_RNA45SN1_45S_50ntFlanks</td><td>9141</td><td>9165</td><td>25</td><td>CGGGGCGGACTGTCCCCAGTGCGCC</td><td>510.3</td><td>-12.9</td><td>1.6</td><td>-8.0</td><td>1.6E-15</td><td>1.1E-14</td><td>MB231/MB468</td><td>MP</td></tr>
<tr><td>>hg38_21_+_8433172_8446622_RNA45SN1_45S_50ntFlanks@6721.6738.18|AATTGCAGGACACATTGA</td><td>>hg38_21_+_8433172_8446622_RNA45SN1_45S_50ntFlanks</td><td>6721</td><td>6738</td><td>18</td><td>AATTGCAGGACACATTGA</td><td>284.2</td><td>-11.1</td><td>1.4</td><td>-7.9</td><td>2.0E-15</td><td>1.1E-14</td><td>MB231/MB468</td><td>NUC</td></tr>
<tr><td>>hg38_21_+_8433172_8446622_RNA45SN1_45S_50ntFlanks@11564.11591.28|CCTAGCAGCCGACTTAGAACTGGTGCGG</td><td>>hg38_21_+_8433172_8446622_RNA45SN1_45S_50ntFlanks</td><td>11564</td><td>11591</td><td>28</td><td>CCTAGCAGCCGACTTAGAACTGGTGCGG</td><td>2290.8</td><td>-7.1</td><td>0.9</td><td>-8.4</td><td>5.2E-17</td><td>1.1E-14</td><td>BT20/MB468</td><td>MP</td></tr>
<tr><td>>hg38_21_+_8433172_8446622_RNA45SN1_45S_50ntFlanks@4259.4277.19|ATTGGAATGAGTCCACTTT</td><td>>hg38_21_+_8433172_8446622_RNA45SN1_45S_50ntFlanks</td><td>4259</td><td>4277</td><td>19</td><td>ATTGGAATGAGTCCACTTT</td><td>341.9</td><td>-12.3</td><td>1.5</td><td>-8.0</td><td>1.6E-15</td><td>1.1E-14</td><td>MB231/MB468</td><td>MP</td></tr>
<tr><td>>hg38_21_+_8433172_8446622_RNA45SN1_45S_50ntFlanks@12365.12383.19|ATAACTGGCTTGTGGCGGC</td><td>>hg38_21_+_8433172_8446622_RNA45SN1_45S_50ntFlanks</td><td>12365</td><td>12383</td><td>19</td><td>ATAACTGGCTTGTGGCGGC</td><td>200.5</td><td>-11.5</td><td>1.4</td><td>-8.0</td><td>1.6E-15</td><td>1.1E-14</td><td>MB231/MB468</td><td>MP</td></tr>
<tr><td>>hg38_21_+_8433172_8446622_RNA45SN1_45S_50ntFlanks@6645.6678.34|CTCGTACGACTCTTAGCGGTGGATCACTCGGCTC</td><td>>hg38_21_+_8433172_8446622_RNA45SN1_45S_50ntFlanks</td><td>6645</td><td>6678</td><td>34</td><td>CTCGTACGACTCTTAGCGGTGGATCACTCGGCTC</td><td>437.8</td><td>5.1</td><td>0.6</td><td>7.9</td><td>2.3E-15</td><td>1.1E-14</td><td>BT20/MB468</td><td>CYTO</td></tr>
<tr><td>>hg38_21_+_8433172_8446622_RNA45SN1_45S_50ntFlanks@4473.4494.22|TTAGCTGAGTGTCCCGCGGGGC</td><td>>hg38_21_+_8433172_8446622_RNA45SN1_45S_50ntFlanks</td><td>4473</td><td>4494</td><td>22</td><td>TTAGCTGAGTGTCCCGCGGGGC</td><td>258.1</td><td>-10.7</td><td>1.3</td><td>-8.0</td><td>1.8E-15</td><td>1.1E-14</td><td>MB231/MB468</td><td>CYTO</td></tr>
<tr><td>>hg38_21_+_8433172_8446622_RNA45SN1_45S_50ntFlanks@8323.8354.32|ACCGATAGTCAACAAGTACCGTAAGGGAAAGT</td><td>>hg38_21_+_8433172_8446622_RNA45SN1_45S_50ntFlanks</td><td>8323</td><td>8354</td><td>32</td><td>ACCGATAGTCAACAAGTACCGTAAGGGAAAGT</td><td>172.3</td><td>-7.1</td><td>0.9</td><td>-8.0</td><td>1.8E-15</td><td>1.1E-14</td><td>MB231/MB468</td><td>CYTO</td></tr>
<tr><td>>MI0000089|hsa-mir-31&WithFlank&9|-|21512109|21512191@14.35.22(+1A)&[MIMAT0000089&hsa-miR-31-5p&offsets|0|+1(+1A);m-360&9|-|21512157|21512178&offsets|0|0(+1A)]|AGGCAAGATGCTGGCATAGCTGA</td><td>>MI0000089|hsa-mir-31&WithFlank&9|-|21512109|21512191</td><td>14</td><td>35</td><td>22(+1A)</td><td>AGGCAAGATGCTGGCATAGCTGA</td><td>568.0</td><td>9.5</td><td>1.2</td><td>7.9</td><td>2.3E-15</td><td>1.1E-14</td><td>BT20/MB468</td><td>CYTO</td></tr>
<tr><td>>MI0000471|hsa-mir-126&WithFlank&9|+|136670596|136670692@21.41.21&[MIMAT0000444&hsa-miR-126-5p&offsets|0|0;m-114&9|+|136670616|136670636&offsets|0|0]|CATTATTACTTTTGGTACGCG</td><td>>MI0000471|hsa-mir-126&WithFlank&9|+|136670596|136670692</td><td>21</td><td>41</td><td>21</td><td>CATTATTACTTTTGGTACGCG</td><td>176.8</td><td>11.4</td><td>1.4</td><td>7.9</td><td>2.0E-15</td><td>1.1E-14</td><td>MB231/MB468</td><td>NUC</td></tr>
<tr><td>>hg38_21_+_8433172_8446622_RNA45SN1_45S_50ntFlanks@9534.9563.30|AGGAAACTCTGGTGGAGGTCCGTAGCGGTC</td><td>>hg38_21_+_8433172_8446622_RNA45SN1_45S_50ntFlanks</td><td>9534</td><td>9563</td><td>30</td><td>AGGAAACTCTGGTGGAGGTCCGTAGCGGTC</td><td>20427.3</td><td>-6.1</td><td>0.8</td><td>-8.0</td><td>1.8E-15</td><td>1.1E-14</td><td>MB231/MB468</td><td>CYTO</td></tr>
<tr><td>>MI0005756|hsa-mir-934&WithFlank&X|+|136550872|136550966@21.42.22&[MIMAT0004977&hsa-miR-934&offsets|0|0;m-943&X|+|136550892|136550913&offsets|0|0]|TGTCTACTACTGGAGACACTGG</td><td>>MI0005756|hsa-mir-934&WithFlank&X|+|136550872|136550966</td><td>21</td><td>42</td><td>22</td><td>TGTCTACTACTGGAGACACTGG</td><td>2992.5</td><td>7.5</td><td>0.9</td><td>8.0</td><td>1.7E-15</td><td>1.1E-14</td><td>BT20/MB468</td><td>MITO</td></tr>
<tr><td>>am_trna2_GlyGCC_21_-_18827107_18827177@1.26.26,trna35_GlyGCC_1_+_161413094_161413164@1.26.26,trna37_GlyGCC_1_+_161420467_161420537@1.26.26,trna39_GlyGCC_1_+_161427898_161427968@1.26.26,trna41_GlyGCC_1_+_161435258_161435328@1.26.26|GCATGGGTGGTTCAGTGGTAGAATTC</td><td>>am_trna2_GlyGCC_21_-_18827107_18827177</td><td>1</td><td>26</td><td>26</td><td>GCATGGGTGGTTCAGTGGTAGAATTC</td><td>482.1</td><td>7.5</td><td>0.9</td><td>8.2</td><td>1.6E-16</td><td>1.1E-14</td><td>BT20/MB231</td><td>CYTO</td></tr>
<tr><td>>hg38_21_+_8433172_8446622_RNA45SN1_45S_50ntFlanks@4723.4751.29|CATTAATCAAGAACGAAAGTCGGAGGTTC</td><td>>hg38_21_+_8433172_8446622_RNA45SN1_45S_50ntFlanks</td><td>4723</td><td>4751</td><td>29</td><td>CATTAATCAAGAACGAAAGTCGGAGGTTC</td><td>1348.1</td><td>-9.8</td><td>1.2</td><td>-8.0</td><td>1.6E-15</td><td>1.1E-14</td><td>MB231/MB468</td><td>MP</td></tr>
<tr><td>>hg38_21_+_8433172_8446622_RNA45SN1_45S_50ntFlanks@3782.3814.33|CAGTGAAACTGCGAATGGCTCATTAAATCAGTT</td><td>>hg38_21_+_8433172_8446622_RNA45SN1_45S_50ntFlanks</td><td>3782</td><td>3814</td><td>33</td><td>CAGTGAAACTGCGAATGGCTCATTAAATCAGTT</td><td>394.2</td><td>-12.4</td><td>1.6</td><td>-7.9</td><td>2.7E-15</td><td>1.1E-14</td><td>MB231/MB468</td><td>MITO</td></tr>
<tr><td>>hg38_21_+_8433172_8446622_RNA45SN1_45S_50ntFlanks@12540.12569.30|TGATGTGTTGTTGCCATGGTAATCCTGCTC</td><td>>hg38_21_+_8433172_8446622_RNA45SN1_45S_50ntFlanks</td><td>12540</td><td>12569</td><td>30</td><td>TGATGTGTTGTTGCCATGGTAATCCTGCTC</td><td>326.8</td><td>-12.1</td><td>1.5</td><td>-7.9</td><td>2.7E-15</td><td>1.1E-14</td><td>MB231/MB468</td><td>MITO</td></tr>
<tr><td>>MI0000081|hsa-mir-24-2&WithFlank&19|-|13836281|13836365@56.77.22(+1A);MI0000080|hsa-mir-24-1&WithFlank&9|+|95086015|95086094@50.71.22(+1A)&[MIMAT0000080&hsa-miR-24-3p&offsets|0|0(+1A);m-36&19|-|13836288|13836310&offsets|0|-1(+1A)];[MIMAT0000080_1&hsa-miR-24-3p&offsets|0|0(+1A);m-35&9|+|95086064|95086086&offsets|0|-1(+1A)]|TGGCTCAGTTCAGCAGGAACAGA</td><td>>MI0000081|hsa-mir-24-2&WithFlank&19|-|13836281|13836365</td><td>56</td><td>77</td><td>22(+1A)</td><td>TGGCTCAGTTCAGCAGGAACAGA</td><td>679.8</td><td>6.4</td><td>0.8</td><td>7.9</td><td>2.4E-15</td><td>1.1E-14</td><td>BT20/MB468</td><td>CYTO</td></tr>
<tr><td>>hg38_21_+_8433172_8446622_RNA45SN1_45S_50ntFlanks@10128.10153.26|CGGCGGCGGCGGCGGCGGCGGGGGTG</td><td>>hg38_21_+_8433172_8446622_RNA45SN1_45S_50ntFlanks</td><td>10128</td><td>10153</td><td>26</td><td>CGGCGGCGGCGGCGGCGGCGGGGGTG</td><td>320.1</td><td>-12.2</td><td>1.5</td><td>-8.0</td><td>1.6E-15</td><td>1.1E-14</td><td>MB231/MB468</td><td>MP</td></tr>
<tr><td>>hg38_21_+_8433172_8446622_RNA45SN1_45S_50ntFlanks@4733.4765.33|GAACGAAAGTCGGAGGTTCGAAGACGATCAGAT</td><td>>hg38_21_+_8433172_8446622_RNA45SN1_45S_50ntFlanks</td><td>4733</td><td>4765</td><td>33</td><td>GAACGAAAGTCGGAGGTTCGAAGACGATCAGAT</td><td>816.6</td><td>7.1</td><td>0.9</td><td>8.0</td><td>1.8E-15</td><td>1.1E-14</td><td>MB231/MB468</td><td>CYTO</td></tr>
<tr><td>>hg38_21_+_8433172_8446622_RNA45SN1_45S_50ntFlanks@10411.10456.46|CAGTCGGTCCTGAGAGATGGGCGAGCGCCGTTCCGAAGGGACGGGC</td><td>>hg38_21_+_8433172_8446622_RNA45SN1_45S_50ntFlanks</td><td>10411</td><td>10456</td><td>46</td><td>CAGTCGGTCCTGAGAGATGGGCGAGCGCCGTTCCGAAGGGACGGGC</td><td>610.0</td><td>-13.7</td><td>1.7</td><td>-7.9</td><td>3.7E-15</td><td>1.1E-14</td><td>BT20/MB468</td><td>NUC</td></tr>
<tr><td>>hg38_21_+_8433172_8446622_RNA45SN1_45S_50ntFlanks@12937.12959.23|AACCATTCGTAGACGACCTGCTT</td><td>>hg38_21_+_8433172_8446622_RNA45SN1_45S_50ntFlanks</td><td>12937</td><td>12959</td><td>23</td><td>AACCATTCGTAGACGACCTGCTT</td><td>848.2</td><td>-4.3</td><td>0.5</td><td>-7.9</td><td>2.0E-15</td><td>1.1E-14</td><td>MB231/MB468</td><td>NUC</td></tr>
<tr><td>>hg38_21_+_8433172_8446622_RNA45SN1_45S_50ntFlanks@10410.10433.24|TCAGTCGGTCCTGAGAGATGGGCG</td><td>>hg38_21_+_8433172_8446622_RNA45SN1_45S_50ntFlanks</td><td>10410</td><td>10433</td><td>24</td><td>TCAGTCGGTCCTGAGAGATGGGCG</td><td>1234.2</td><td>-7.1</td><td>0.9</td><td>-8.0</td><td>1.8E-15</td><td>1.1E-14</td><td>MB231/MB468</td><td>CYTO</td></tr>
<tr><td>>hg38_21_+_8433172_8446622_RNA45SN1_45S_50ntFlanks@10457.10497.41|GATGGCCTCCGTTGCCCTCGGCCGATCGAAAGGGAGTCGGG</td><td>>hg38_21_+_8433172_8446622_RNA45SN1_45S_50ntFlanks</td><td>10457</td><td>10497</td><td>41</td><td>GATGGCCTCCGTTGCCCTCGGCCGATCGAAAGGGAGTCGGG</td><td>324.1</td><td>-12.1</td><td>1.5</td><td>-7.9</td><td>2.7E-15</td><td>1.1E-14</td><td>MB231/MB468</td><td>MITO</td></tr>
<tr><td>>hg38_21_+_8433172_8446622_RNA45SN1_45S_50ntFlanks@12938.12956.19|ACCATTCGTAGACGACCTG</td><td>>hg38_21_+_8433172_8446622_RNA45SN1_45S_50ntFlanks</td><td>12938</td><td>12956</td><td>19</td><td>ACCATTCGTAGACGACCTG</td><td>1372.6</td><td>-7.6</td><td>1.0</td><td>-7.9</td><td>2.7E-15</td><td>1.1E-14</td><td>MB231/MB468</td><td>MITO</td></tr>
<tr><td>>hg38_21_+_8433172_8446622_RNA45SN1_45S_50ntFlanks@9930.9957.28|ATATAGACAGCAGGACGGTGGCCATGGA</td><td>>hg38_21_+_8433172_8446622_RNA45SN1_45S_50ntFlanks</td><td>9930</td><td>9957</td><td>28</td><td>ATATAGACAGCAGGACGGTGGCCATGGA</td><td>772.8</td><td>-13.4</td><td>1.7</td><td>-7.9</td><td>2.7E-15</td><td>1.1E-14</td><td>MB231/MB468</td><td>MITO</td></tr>
<tr><td>>hg38_21_+_8433172_8446622_RNA45SN1_45S_50ntFlanks@10807.10825.19|ACAATGTAGGTAAGGGAAG</td><td>>hg38_21_+_8433172_8446622_RNA45SN1_45S_50ntFlanks</td><td>10807</td><td>10825</td><td>19</td><td>ACAATGTAGGTAAGGGAAG</td><td>208.0</td><td>-11.6</td><td>1.5</td><td>-8.0</td><td>1.6E-15</td><td>1.1E-14</td><td>MB231/MB468</td><td>MP</td></tr>
<tr><td>>hg38_21_+_8433172_8446622_RNA45SN1_45S_50ntFlanks@9534.9569.36|AGGAAACTCTGGTGGAGGTCCGTAGCGGTCCTGACG</td><td>>hg38_21_+_8433172_8446622_RNA45SN1_45S_50ntFlanks</td><td>9534</td><td>9569</td><td>36</td><td>AGGAAACTCTGGTGGAGGTCCGTAGCGGTCCTGACG</td><td>18094.4</td><td>-11.0</td><td>1.4</td><td>-7.9</td><td>3.7E-15</td><td>1.1E-14</td><td>BT20/MB468</td><td>NUC</td></tr>
<tr><td>>hg38_21_+_8433172_8446622_RNA45SN1_45S_50ntFlanks@12934.12960.27|CTAAACCATTCGTAGACGACCTGCTTC</td><td>>hg38_21_+_8433172_8446622_RNA45SN1_45S_50ntFlanks</td><td>12934</td><td>12960</td><td>27</td><td>CTAAACCATTCGTAGACGACCTGCTTC</td><td>566.9</td><td>-5.2</td><td>0.7</td><td>-7.9</td><td>2.7E-15</td><td>1.1E-14</td><td>MB231/MB468</td><td>MITO</td></tr>
<tr><td>>hg38_21_+_8433172_8446622_RNA45SN1_45S_50ntFlanks@11233.11252.20|CCCTTCCCGTGGATCGCCCC</td><td>>hg38_21_+_8433172_8446622_RNA45SN1_45S_50ntFlanks</td><td>11233</td><td>11252</td><td>20</td><td>CCCTTCCCGTGGATCGCCCC</td><td>305.5</td><td>-12.1</td><td>1.5</td><td>-8.0</td><td>1.6E-15</td><td>1.1E-14</td><td>MB231/MB468</td><td>MP</td></tr>
<tr><td>>MI0000081|hsa-mir-24-2&WithFlank&19|-|13836281|13836365@56.74.19(+1U);MI0000080|hsa-mir-24-1&WithFlank&9|+|95086015|95086094@50.68.19(+1U)&[MIMAT0000080&hsa-miR-24-3p&offsets|0|-3(+1U);m-36&19|-|13836288|13836310&offsets|0|-4(+1U)];[MIMAT0000080_1&hsa-miR-24-3p&offsets|0|-3(+1U);m-35&9|+|95086064|95086086&offsets|0|-4(+1U)]|TGGCTCAGTTCAGCAGGAAT</td><td>>MI0000081|hsa-mir-24-2&WithFlank&19|-|13836281|13836365</td><td>56</td><td>74</td><td>19(+1U)</td><td>TGGCTCAGTTCAGCAGGAAT</td><td>1739.1</td><td>7.2</td><td>0.9</td><td>7.9</td><td>2.4E-15</td><td>1.1E-14</td><td>BT20/MB468</td><td>CYTO</td></tr>
<tr><td>>hg38_21_+_8433172_8446622_RNA45SN1_45S_50ntFlanks@11316.11334.19|CACCCCACGTCTCGTCGCG</td><td>>hg38_21_+_8433172_8446622_RNA45SN1_45S_50ntFlanks</td><td>11316</td><td>11334</td><td>19</td><td>CACCCCACGTCTCGTCGCG</td><td>1760.8</td><td>-7.6</td><td>1.0</td><td>-8.0</td><td>1.8E-15</td><td>1.1E-14</td><td>MB231/MB468</td><td>CYTO</td></tr>
</table>

<table>
<tr><td>>hg38_21_+_8433172_8446622_RNA45SN1_45S_50ntFlanks@4399.4423.25|CGGGCGGGCGGTCCGCCGCGAGGCG</td><td>>hg38_21_+_8433172_8446622_RNA45SN1_45S_50ntFlanks</td><td>4399</td><td>4423</td><td>25</td><td>CGGGCGGGCGGTCCGCCGCGAGGCG</td><td>211.3</td><td>-11.6</td><td>1.5</td><td>-8.0</td><td>1.7E-15</td><td>1.1E-14</td><td>MB231/MB468</td><td>MP</td></tr>
<tr><td>>hg38_21_+_8433172_8446622_RNA45SN1_45S_50ntFlanks@11076.11103.28|CGCCGCGTGGGGGCGGCGGCGGGGGGAG</td><td>>hg38_21_+_8433172_8446622_RNA45SN1_45S_50ntFlanks</td><td>11076</td><td>11103</td><td>28</td><td>CGCCGCGTGGGGGCGGCGGCGGGGGGAG</td><td>576.0</td><td>-13.0</td><td>1.6</td><td>-7.9</td><td>2.8E-15</td><td>1.1E-14</td><td>MB231/MB468</td><td>MITO</td></tr>
<tr><td>>hg38_21_+_8433172_8446622_RNA45SN1_45S_50ntFlanks@10498.10516.19|TTCAGATCCCCGAATCCGG</td><td>>hg38_21_+_8433172_8446622_RNA45SN1_45S_50ntFlanks</td><td>10498</td><td>10516</td><td>19</td><td>TTCAGATCCCCGAATCCGG</td><td>269.0</td><td>-11.9</td><td>1.5</td><td>-8.0</td><td>1.7E-15</td><td>1.1E-14</td><td>MB231/MB468</td><td>MP</td></tr>
<tr><td>>hg38_21_+_8433172_8446622_RNA45SN1_45S_50ntFlanks@12157.12179.23|AGTTTGACTGGGGCGGTACACCT</td><td>>hg38_21_+_8433172_8446622_RNA45SN1_45S_50ntFlanks</td><td>12157</td><td>12179</td><td>23</td><td>AGTTTGACTGGGGCGGTACACCT</td><td>266.7</td><td>-11.0</td><td>1.4</td><td>-7.9</td><td>2.0E-15</td><td>1.1E-14</td><td>MB231/MB468</td><td>NUC</td></tr>
<tr><td>>hg38_21_+_8433172_8446622_RNA45SN1_45S_50ntFlanks@9554.9572.19|CGTAGCGGTCCTGACGTGC</td><td>>hg38_21_+_8433172_8446622_RNA45SN1_45S_50ntFlanks</td><td>9554</td><td>9572</td><td>19</td><td>CGTAGCGGTCCTGACGTGC</td><td>14079.2</td><td>-4.7</td><td>0.6</td><td>-7.9</td><td>2.0E-15</td><td>1.1E-14</td><td>MB231/MB468</td><td>NUC</td></tr>
<tr><td>>hg38_21_+_8433172_8446622_RNA45SN1_45S_50ntFlanks@7974.8004.31|ACGCGACCTCAGATCAGACGTGGCGACCCGC</td><td>>hg38_21_+_8433172_8446622_RNA45SN1_45S_50ntFlanks</td><td>7974</td><td>8004</td><td>31</td><td>ACGCGACCTCAGATCAGACGTGGCGACCCGC</td><td>9623.8</td><td>6.0</td><td>0.8</td><td>8.0</td><td>1.8E-15</td><td>1.1E-14</td><td>BT20/MB468</td><td>MITO</td></tr>
<tr><td>>hg38_21_+_8433172_8446622_RNA45SN1_45S_50ntFlanks@9504.9532.29|GTGAACTATGCCTGGGCAGGGCGAAGCCA</td><td>>hg38_21_+_8433172_8446622_RNA45SN1_45S_50ntFlanks</td><td>9504</td><td>9532</td><td>29</td><td>GTGAACTATGCCTGGGCAGGGCGAAGCCA</td><td>522.7</td><td>-12.9</td><td>1.6</td><td>-8.0</td><td>1.7E-15</td><td>1.1E-14</td><td>MB231/MB468</td><td>MP</td></tr>
<tr><td>>hg38_21_+_8433172_8446622_RNA45SN1_45S_50ntFlanks@4064.4098.35|ATCAACTTTCGATGGTAGTCGCCGTGCCTACCATG</td><td>>hg38_21_+_8433172_8446622_RNA45SN1_45S_50ntFlanks</td><td>4064</td><td>4098</td><td>35</td><td>ATCAACTTTCGATGGTAGTCGCCGTGCCTACCATG</td><td>287.5</td><td>-10.9</td><td>1.4</td><td>-8.0</td><td>1.9E-15</td><td>1.1E-14</td><td>MB231/MB468</td><td>CYTO</td></tr>
<tr><td>>MI0000746|hsa-mir-99b&WithFlank&19|+|51692606|51692687@51.71.21&[MIMAT0004678&hsa-miR-99b-3p&offsets|0|-1;m-151&19|+|51692656|51692678&offsets|0|-2]|CAAGCTCGTGTCTGTGGGTCC</td><td>>MI0000746|hsa-mir-99b&WithFlank&19|+|51692606|51692687</td><td>51</td><td>71</td><td>21</td><td>CAAGCTCGTGTCTGTGGGTCC</td><td>351.7</td><td>7.5</td><td>0.9</td><td>8.0</td><td>1.8E-15</td><td>1.1E-14</td><td>BT20/MB468</td><td>MITO</td></tr>
<tr><td>>hg38_21_+_8433172_8446622_RNA45SN1_45S_50ntFlanks@485.506.22|GATGTGGTGACGTCGTGCTCTC</td><td>>hg38_21_+_8433172_8446622_RNA45SN1_45S_50ntFlanks</td><td>485</td><td>506</td><td>22</td><td>GATGTGGTGACGTCGTGCTCTC</td><td>16471.3</td><td>15.7</td><td>2.0</td><td>7.9</td><td>2.1E-15</td><td>1.1E-14</td><td>MB231/MB468</td><td>NUC</td></tr>
<tr><td>>hg38_21_+_8433172_8446622_RNA45SN1_45S_50ntFlanks@10794.10818.25|CTGGCATGTTGGAACAATGTAGGTA</td><td>>hg38_21_+_8433172_8446622_RNA45SN1_45S_50ntFlanks</td><td>10794</td><td>10818</td><td>25</td><td>CTGGCATGTTGGAACAATGTAGGTA</td><td>1888.3</td><td>-11.1</td><td>1.4</td><td>-7.9</td><td>3.8E-15</td><td>1.1E-14</td><td>BT20/MB468</td><td>NUC</td></tr>
<tr><td>>hg38_21_+_8433172_8446622_RNA45SN1_45S_50ntFlanks@3988.4005.18|CTTTGGTGACTCTAGATA</td><td>>hg38_21_+_8433172_8446622_RNA45SN1_45S_50ntFlanks</td><td>3988</td><td>4005</td><td>18</td><td>CTTTGGTGACTCTAGATA</td><td>439.0</td><td>-4.9</td><td>0.6</td><td>-8.0</td><td>1.7E-15</td><td>1.1E-14</td><td>MB231/MB468</td><td>MP</td></tr>
<tr><td>>hg38_21_+_8433172_8446622_RNA45SN1_45S_50ntFlanks@12712.12747.36|CGGAGCCTCGGTTGGCCTCGGATAGCCGGTCCCCCG</td><td>>hg38_21_+_8433172_8446622_RNA45SN1_45S_50ntFlanks</td><td>12712</td><td>12747</td><td>36</td><td>CGGAGCCTCGGTTGGCCTCGGATAGCCGGTCCCCCG</td><td>1377.0</td><td>-13.5</td><td>1.6</td><td>-8.4</td><td>5.6E-17</td><td>1.1E-14</td><td>BT20/MB468</td><td>MP</td></tr>
<tr><td>>hg38_21_+_8433172_8446622_RNA45SN1_45S_50ntFlanks@4732.4762.31|AGAACGAAAGTCGGAGGTTCGAAGACGATCA</td><td>>hg38_21_+_8433172_8446622_RNA45SN1_45S_50ntFlanks</td><td>4732</td><td>4762</td><td>31</td><td>AGAACGAAAGTCGGAGGTTCGAAGACGATCA</td><td>645.0</td><td>8.2</td><td>1.0</td><td>7.9</td><td>2.5E-15</td><td>1.1E-14</td><td>BT20/MB468</td><td>CYTO</td></tr>
<tr><td>>hg38_21_+_8433172_8446622_RNA45SN1_45S_50ntFlanks@11884.11904.21|GTTGAGCTTGACTCTAGTCTG</td><td>>hg38_21_+_8433172_8446622_RNA45SN1_45S_50ntFlanks</td><td>11884</td><td>11904</td><td>21</td><td>GTTGAGCTTGACTCTAGTCTG</td><td>278.5</td><td>-12.0</td><td>1.5</td><td>-8.0</td><td>1.7E-15</td><td>1.1E-14</td><td>MB231/MB468</td><td>MP</td></tr>
<tr><td>>hg38_21_+_8433172_8446622_RNA45SN1_45S_50ntFlanks@4722.4753.32|TCATTAATCAAGAACGAAAGTCGGAGGTTCGA</td><td>>hg38_21_+_8433172_8446622_RNA45SN1_45S_50ntFlanks</td><td>4722</td><td>4753</td><td>32</td><td>TCATTAATCAAGAACGAAAGTCGGAGGTTCGA</td><td>307.5</td><td>-12.0</td><td>1.5</td><td>-7.9</td><td>2.8E-15</td><td>1.1E-14</td><td>MB231/MB468</td><td>MITO</td></tr>
<tr><td>>hg38_21_+_8433172_8446622_RNA45SN1_45S_50ntFlanks@12625.12654.30|CAATGGGGCGAAGCTACCATCTGTGGGATT</td><td>>hg38_21_+_8433172_8446622_RNA45SN1_45S_50ntFlanks</td><td>12625</td><td>12654</td><td>30</td><td>CAATGGGGCGAAGCTACCATCTGTGGGATT</td><td>6057.8</td><td>-4.9</td><td>0.6</td><td>-8.0</td><td>1.8E-15</td><td>1.1E-14</td><td>BT20/MB468</td><td>MITO</td></tr>
<tr><td>>hg38_21_+_8433172_8446622_RNA45SN1_45S_50ntFlanks@6738.6772.35|ATCATCGACACTTCGAACGCACTTGCGGCCCCGGG</td><td>>hg38_21_+_8433172_8446622_RNA45SN1_45S_50ntFlanks</td><td>6738</td><td>6772</td><td>35</td><td>ATCATCGACACTTCGAACGCACTTGCGGCCCCGGG</td><td>997.1</td><td>-13.8</td><td>1.7</td><td>-8.0</td><td>1.7E-15</td><td>1.1E-14</td><td>MB231/MB468</td><td>MP</td></tr>
<tr><td>>hg38_21_+_8433172_8446622_RNA45SN1_45S_50ntFlanks@10402.10430.29|AACATGGGTCAGTCGGTCCTGAGAGATGG</td><td>>hg38_21_+_8433172_8446622_RNA45SN1_45S_50ntFlanks</td><td>10402</td><td>10430</td><td>29</td><td>AACATGGGTCAGTCGGTCCTGAGAGATGG</td><td>2322.4</td><td>-8.1</td><td>1.0</td><td>-7.9</td><td>2.8E-15</td><td>1.1E-14</td><td>MB231/MB468</td><td>MITO</td></tr>
<tr><td>>hg38_21_+_8433172_8446622_RNA45SN1_45S_50ntFlanks@10873.10904.32|CTGGGTCGGTCGGGCTGGGGCGCGAAGCGGGG</td><td>>hg38_21_+_8433172_8446622_RNA45SN1_45S_50ntFlanks</td><td>10873</td><td>10904</td><td>32</td><td>CTGGGTCGGTCGGGCTGGGGCGCGAAGCGGGG</td><td>243.2</td><td>-11.8</td><td>1.5</td><td>-8.0</td><td>1.7E-15</td><td>1.1E-14</td><td>MB231/MB468</td><td>MP</td></tr>
<tr><td>>MI0000812|hsa-mir-331&WithFlank&12|+|95308414|95308519@67.86.20&[MIMAT0000760&hsa-miR-331-3p&offsets|0|-1;m-192&12|+|95308480|95308501&offsets|0|-2]|GCCCCTGGGCCTATCCTAGA</td><td>>MI0000812|hsa-mir-331&WithFlank&12|+|95308414|95308519</td><td>67</td><td>86</td><td>20</td><td>GCCCCTGGGCCTATCCTAGA</td><td>72.4</td><td>9.0</td><td>1.1</td><td>7.9</td><td>3.8E-15</td><td>1.1E-14</td><td>BT20/MB468</td><td>NUC</td></tr>
<tr><td>>MI0000764|hsa-mir-363&WithFlank&X|-|134169372|134169458@56.77.22&[MIMAT0000707&hsa-miR-363-3p&offsets|0|0;m-178&X|-|134169381|134169403&offsets|0|-1]|AATTGCACGGTATCCATCTGTA</td><td>>MI0000764|hsa-mir-363&WithFlank&X|-|134169372|134169458</td><td>56</td><td>77</td><td>22</td><td>AATTGCACGGTATCCATCTGTA</td><td>1518.0</td><td>5.5</td><td>0.7</td><td>7.9</td><td>2.5E-15</td><td>1.1E-14</td><td>BT20/MB468</td><td>CYTO</td></tr>
<tr><td>>hg38_21_+_8433172_8446622_RNA45SN1_45S_50ntFlanks@9736.9753.18|CCGAAACGATCTCAACCT</td><td>>hg38_21_+_8433172_8446622_RNA45SN1_45S_50ntFlanks</td><td>9736</td><td>9753</td><td>18</td><td>CCGAAACGATCTCAACCT</td><td>185.7</td><td>-11.4</td><td>1.4</td><td>-8.0</td><td>1.7E-15</td><td>1.1E-14</td><td>MB231/MB468</td><td>MP</td></tr>
<tr><td>>hg38_21_+_8433172_8446622_RNA45SN1_45S_50ntFlanks@4128.4146.19|CGATTCCGGAGAGGGAGCC</td><td>>hg38_21_+_8433172_8446622_RNA45SN1_45S_50ntFlanks</td><td>4128</td><td>4146</td><td>19</td><td>CGATTCCGGAGAGGGAGCC</td><td>186.3</td><td>-11.4</td><td>1.4</td><td>-8.0</td><td>1.7E-15</td><td>1.2E-14</td><td>MB231/MB468</td><td>MP</td></tr>
<tr><td>>hg38_21_+_8433172_8446622_RNA45SN1_45S_50ntFlanks@12345.12365.21|TCAGAAAAGTTACCACAGGGA</td><td>>hg38_21_+_8433172_8446622_RNA45SN1_45S_50ntFlanks</td><td>12345</td><td>12365</td><td>21</td><td>TCAGAAAAGTTACCACAGGGA</td><td>206.7</td><td>-10.4</td><td>1.3</td><td>-7.9</td><td>1.9E-15</td><td>1.2E-14</td><td>MB231/MB468</td><td>CYTO</td></tr>
<tr><td>>hg38_21_+_8433172_8446622_RNA45SN1_45S_50ntFlanks@13004.13025.22|ATCTATTGAAAGTCAGCCCTCG</td><td>>hg38_21_+_8433172_8446622_RNA45SN1_45S_50ntFlanks</td><td>13004</td><td>13025</td><td>22</td><td>ATCTATTGAAAGTCAGCCCTCG</td><td>1438.8</td><td>-8.9</td><td>1.1</td><td>-7.9</td><td>1.9E-15</td><td>1.2E-14</td><td>MB231/MB468</td><td>CYTO</td></tr>
<tr><td>>hg38_21_+_8433172_8446622_RNA45SN1_45S_50ntFlanks@4453.4474.22|TCGGCGCCCCCTCGATGCTCTT</td><td>>hg38_21_+_8433172_8446622_RNA45SN1_45S_50ntFlanks</td><td>4453</td><td>4474</td><td>22</td><td>TCGGCGCCCCCTCGATGCTCTT</td><td>269.7</td><td>12.1</td><td>1.5</td><td>7.9</td><td>1.9E-15</td><td>1.2E-14</td><td>MB231/MB468</td><td>CYTO</td></tr>
<tr><td>>hg38_21_+_8433172_8446622_RNA45SN1_45S_50ntFlanks@9166.9195.30|CCGGGCGGGTCGCGCCGTCGGGCCCGGGGG</td><td>>hg38_21_+_8433172_8446622_RNA45SN1_45S_50ntFlanks</td><td>9166</td><td>9195</td><td>30</td><td>CCGGGCGGGTCGCGCCGTCGGGCCCGGGGG</td><td>600.5</td><td>-13.0</td><td>1.6</td><td>-7.9</td><td>2.9E-15</td><td>1.2E-14</td><td>MB231/MB468</td><td>MITO</td></tr>
<tr><td>>hg38_21_+_8433172_8446622_RNA45SN1_45S_50ntFlanks@11452.11477.26|CTCCTCGGCGCGCGGCGGCGGCGGCG</td><td>>hg38_21_+_8433172_8446622_RNA45SN1_45S_50ntFlanks</td><td>11452</td><td>11477</td><td>26</td><td>CTCCTCGGCGCGCGGCGGCGGCGGCG</td><td>209.4</td><td>-11.6</td><td>1.5</td><td>-8.0</td><td>1.7E-15</td><td>1.2E-14</td><td>MB231/MB468</td><td>MP</td></tr>
<tr><td>>hg38_21_+_8433172_8446622_RNA45SN1_45S_50ntFlanks@11208.11232.25|GCGGCGACTCTGGACGCGAGCCGGG</td><td>>hg38_21_+_8433172_8446622_RNA45SN1_45S_50ntFlanks</td><td>11208</td><td>11232</td><td>25</td><td>GCGGCGACTCTGGACGCGAGCCGGG</td><td>18924.9</td><td>-10.0</td><td>1.3</td><td>-7.9</td><td>2.1E-15</td><td>1.2E-14</td><td>MB231/MB468</td><td>NUC</td></tr>
<tr><td>>MI0000477|hsa-mir-146a&WithFlank&5|+|160485346|160485456@27.48.22(+1A)&[MIMAT0000449&hsa-miR-146a-5p&offsets|0|0(+1A);m-129&5|+|160485372|160485393&offsets|0|0(+1A)]|TGAGAACTGAATTCCATGGGTTA</td><td>>MI0000477|hsa-mir-146a&WithFlank&5|+|160485346|160485456</td><td>27</td><td>48</td><td>22(+1A)</td><td>TGAGAACTGAATTCCATGGGTTA</td><td>561.2</td><td>-12.4</td><td>1.5</td><td>-8.2</td><td>2.3E-16</td><td>1.2E-14</td><td>BT20/MB231</td><td>MITO</td></tr>
<tr><td>>MI0000477|hsa-mir-146a&WithFlank&5|+|160485346|160485456@27.50.24&[MIMAT0000449&hsa-miR-146a-5p&offsets|0|+2;m-129&5|+|160485372|160485393&offsets|0|+2]|TGAGAACTGAATTCCATGGGTTGT</td><td>>MI0000477|hsa-mir-146a&WithFlank&5|+|160485346|160485456</td><td>27</td><td>50</td><td>24</td><td>TGAGAACTGAATTCCATGGGTTGT</td><td>524.3</td><td>-12.3</td><td>1.5</td><td>-8.2</td><td>2.4E-16</td><td>1.2E-14</td><td>BT20/MB231</td><td>MITO</td></tr>
<tr><td>>hg38_21_+_8433172_8446622_RNA45SN1_45S_50ntFlanks@10687.10705.19|CGGCGTCCGGTGAGCTCTC</td><td>>hg38_21_+_8433172_8446622_RNA45SN1_45S_50ntFlanks</td><td>10687</td><td>10705</td><td>19</td><td>CGGCGTCCGGTGAGCTCTC</td><td>243.3</td><td>-11.8</td><td>1.5</td><td>-8.0</td><td>1.8E-15</td><td>1.2E-14</td><td>MB231/MB468</td><td>MP</td></tr>
<tr><td>>hg38_21_+_8433172_8446622_RNA45SN1_45S_50ntFlanks@6671.6697.27|CTCGGCTCGTGCGTCGATGAAGAACGC</td><td>>hg38_21_+_8433172_8446622_RNA45SN1_45S_50ntFlanks</td><td>6671</td><td>6697</td><td>27</td><td>CTCGGCTCGTGCGTCGATGAAGAACGC</td><td>7602.1</td><td>-8.1</td><td>1.0</td><td>-7.9</td><td>2.9E-15</td><td>1.2E-14</td><td>MB231/MB468</td><td>MITO</td></tr>
<tr><td>>MI0003628|hsa-mir-615&WithFlank&12|+|54033944|54034051@67.88.22(+1U)&[MIMAT0003283&hsa-miR-615-3p&offsets|0|0(+1U);m-495&12|+|54034010|54034030&offsets|0|+1(+1U)]|TCCGAGCCTGGGTCTCCCTCTTT</td><td>>MI0003628|hsa-mir-615&WithFlank&12|+|54033944|54034051</td><td>67</td><td>88</td><td>22(+1U)</td><td>TCCGAGCCTGGGTCTCCCTCTTT</td><td>256.5</td><td>9.6</td><td>1.2</td><td>7.9</td><td>2.6E-15</td><td>1.2E-14</td><td>BT20/MB468</td><td>CYTO</td></tr>
<tr><td>>hg38_21_+_8433172_8446622_RNA45SN1_45S_50ntFlanks@5489.5514.26|CTGAGAAGACGGTCGAACTTGACTAT</td><td>>hg38_21_+_8433172_8446622_RNA45SN1_45S_50ntFlanks</td><td>5489</td><td>5514</td><td>26</td><td>CTGAGAAGACGGTCGAACTTGACTAT</td><td>1674.8</td><td>-9.6</td><td>1.2</td><td>-7.9</td><td>2.9E-15</td><td>1.2E-14</td><td>MB231/MB468</td><td>MITO</td></tr>
<tr><td>>MI0000462|hsa-mir-152&WithFlank&17|-|48037155|48037253@60.81.22&[MIMAT0000438&hsa-miR-152-3p&offsets|0|+1;m-97&17|-|48037174|48037194&offsets|0|+1]|TCAGTGCATGACAGAACTTGGG</td><td>>MI0000462|hsa-mir-152&WithFlank&17|-|48037155|48037253</td><td>60</td><td>81</td><td>22</td><td>TCAGTGCATGACAGAACTTGGG</td><td>461.8</td><td>9.2</td><td>1.2</td><td>7.9</td><td>2.2E-15</td><td>1.2E-14</td><td>MB231/MB468</td><td>NUC</td></tr>
<tr><td>>MI0000734|hsa-mir-106b&WithFlank&7|-|100093987|100094080@58.78.21(+1U)&[MIMAT0004672&hsa-miR-106b-3p&offsets|0|-1(+1U);m-87&7|-|100094003|100094023&offsets|0|0(+1U)]|CCGCACTGTGGGTACTTGCTGT</td><td>>MI0000734|hsa-mir-106b&WithFlank&7|-|100093987|100094080</td><td>58</td><td>78</td><td>21(+1U)</td><td>CCGCACTGTGGGTACTTGCTGT</td><td>1539.4</td><td>4.8</td><td>0.6</td><td>7.9</td><td>2.2E-15</td><td>1.2E-14</td><td>MB231/MB468</td><td>NUC</td></tr>
<tr><td>>MI0000062|hsa-let-7a-3&WithFlank&22|+|46112743|46112828@58.79.22;MI0000060|hsa-let-7a-1&WithFlank&9|+|94175951|94176042@63.84.22&[MIMAT0004481&hsa-let-7a-3p&offsets|0|+1;m-180&22|+|46112800|46112821&offsets|0|0];[MIMAT0004481_1&hsa-let-7a-3p&offsets|0|+1;m-181&9|+|94176013|94176034&offsets|0|0]|CTATACAATCTACTGTCTTTCC</td><td>>MI0000062|hsa-let-7a-3&WithFlank&22|+|46112743|46112828</td><td>58</td><td>79</td><td>22</td><td>CTATACAATCTACTGTCTTTCC</td><td>269.8</td><td>5.1</td><td>0.6</td><td>7.9</td><td>2.2E-15</td><td>1.2E-14</td><td>MB231/MB468</td><td>NUC</td></tr>
<tr><td>>hg38_21_+_8433172_8446622_RNA45SN1_45S_50ntFlanks@11581.11612.32|AACTGGTGCGGACCAGGGGAATCCGACTGTTT</td><td>>hg38_21_+_8433172_8446622_RNA45SN1_45S_50ntFlanks</td><td>11581</td><td>11612</td><td>32</td><td>AACTGGTGCGGACCAGGGGAATCCGACTGTTT</td><td>5129.1</td><td>-5.4</td><td>0.7</td><td>-7.9</td><td>2.9E-15</td><td>1.2E-14</td><td>MB231/MB468</td><td>MITO</td></tr>
<tr><td>>hg38_21_+_8433172_8446622_RNA45SN1_45S_50ntFlanks@12685.12703.19|CCCAGGCGGAACGATACGG</td><td>>hg38_21_+_8433172_8446622_RNA45SN1_45S_50ntFlanks</td><td>12685</td><td>12703</td><td>19</td><td>CCCAGGCGGAACGATACGG</td><td>358.7</td><td>-12.3</td><td>1.6</td><td>-7.9</td><td>2.9E-15</td><td>1.2E-14</td><td>MB231/MB468</td><td>MITO</td></tr>
<tr><td>>MI0005756|hsa-mir-934&WithFlank&X|+|136550872|136550966@21.43.23&[MIMAT0004977&hsa-miR-934&offsets|0|+1;m-943&X|+|136550892|136550913&offsets|0|+1]|TGTCTACTACTGGAGACACTGGT</td><td>>MI0005756|hsa-mir-934&WithFlank&X|+|136550872|136550966</td><td>21</td><td>43</td><td>23</td><td>TGTCTACTACTGGAGACACTGGT</td><td>3592.6</td><td>7.3</td><td>0.9</td><td>7.9</td><td>1.9E-15</td><td>1.2E-14</td><td>BT20/MB468</td><td>MITO</td></tr>
<tr><td>>hg38_21_+_8433172_8446622_RNA45SN1_45S_50ntFlanks@3854.3873.20|ACTGTGGTAATTCTAGAGCT</td><td>>hg38_21_+_8433172_8446622_RNA45SN1_45S_50ntFlanks</td><td>3854</td><td>3873</td><td>20</td><td>ACTGTGGTAATTCTAGAGCT</td><td>1683.6</td><td>-8.2</td><td>1.0</td><td>-8.0</td><td>1.8E-15</td><td>1.2E-14</td><td>MB231/MB468</td><td>MP</td></tr>
<tr><td>>hg38_21_+_8433172_8446622_RNA45SN1_45S_50ntFlanks@10686.10706.21|GCGGCGTCCGGTGAGCTCTCG</td><td>>hg38_21_+_8433172_8446622_RNA45SN1_45S_50ntFlanks</td><td>10686</td><td>10706</td><td>21</td><td>GCGGCGTCCGGTGAGCTCTCG</td><td>849.6</td><td>-6.7</td><td>0.8</td><td>-7.9</td><td>2.0E-15</td><td>1.2E-14</td><td>MB231/MB468</td><td>CYTO</td></tr>
<tr><td>>hg38_21_+_8433172_8446622_RNA45SN1_45S_50ntFlanks@8342.8378.37|CGTAAGGGAAAGTTGAAAAGAACTTTGAAGAGAGAGT</td><td>>hg38_21_+_8433172_8446622_RNA45SN1_45S_50ntFlanks</td><td>8342</td><td>8378</td><td>37</td><td>CGTAAGGGAAAGTTGAAAAGAACTTTGAAGAGAGAGT</td><td>2394.7</td><td>8.7</td><td>1.1</td><td>7.9</td><td>2.0E-15</td><td>1.2E-14</td><td>MB231/MB468</td><td>CYTO</td></tr>
<tr><td>>MI0000542|hsa-mir-320a&WithFlank&8|-|22244960|22245043@48.70.23(+1G)&[MIMAT0000510&hsa-miR-320a-3p&offsets|0|+1(+1G);m-60&8|-|22244975|22244996&offsets|0|+1(+1G)]|AAAAGCTGGGTTGAGAGGGCGAAG</td><td>>MI0000542|hsa-mir-320a&WithFlank&8|-|22244960|22245043</td><td>48</td><td>70</td><td>23(+1G)</td><td>AAAAGCTGGGTTGAGAGGGCGAAG</td><td>3060.6</td><td>12.0</td><td>1.5</td><td>7.9</td><td>2.6E-15</td><td>1.2E-14</td><td>BT20/MB468</td><td>CYTO</td></tr>
<tr><td>>MI0000086|hsa-mir-28&WithFlank&3|+|188688775|188688872@60.82.23&[MIMAT0004502&hsa-miR-28-3p&offsets|0|+1;m-28&3|+|188688834|188688855&offsets|0|+1]|CACTAGATTGTGAGCTCCTGGAG</td><td>>MI0000086|hsa-mir-28&WithFlank&3|+|188688775|188688872</td><td>60</td><td>82</td><td>23</td><td>CACTAGATTGTGAGCTCCTGGAG</td><td>132.0</td><td>4.3</td><td>0.6</td><td>7.9</td><td>4.0E-15</td><td>1.2E-14</td><td>BT20/MB468</td><td>NUC</td></tr>
<tr><td>>hg38_21_+_8433172_8446622_RNA45SN1_45S_50ntFlanks@10350.10376.27|ACGAGAACTTTGAAGGCCGAAGTGGAG</td><td>>hg38_21_+_8433172_8446622_RNA45SN1_45S_50ntFlanks</td><td>10350</td><td>10376</td><td>27</td><td>ACGAGAACTTTGAAGGCCGAAGTGGAG</td><td>4913.0</td><td>-11.7</td><td>1.5</td><td>-7.9</td><td>4.0E-15</td><td>1.2E-14</td><td>BT20/MB468</td><td>NUC</td></tr>
<tr><td>>hg38_1_-_228634819_228635039_RNA5S12_5S_50ntFlanks@145.169.25|CTGGGAATACCGGGTGCTGTAGGCT</td><td>>hg38_1_-_228634819_228635039_RNA5S12_5S_50ntFlanks</td><td>145</td><td>169</td><td>25</td><td>CTGGGAATACCGGGTGCTGTAGGCT</td><td>1016.3</td><td>6.3</td><td>0.8</td><td>7.9</td><td>2.0E-15</td><td>1.2E-14</td><td>MB231/MB468</td><td>CYTO</td></tr>
<tr><td>>hg38_21_+_8433172_8446622_RNA45SN1_45S_50ntFlanks@11881.11900.20|CCTGTTGAGCTTGACTCTAG</td><td>>hg38_21_+_8433172_8446622_RNA45SN1_45S_50ntFlanks</td><td>11881</td><td>11900</td><td>20</td><td>CCTGTTGAGCTTGACTCTAG</td><td>351.4</td><td>-12.3</td><td>1.5</td><td>-8.0</td><td>1.8E-15</td><td>1.2E-14</td><td>MB231/MB468</td><td>MP</td></tr>
</table>

<table>
<tr><th></th><th></th><th></th><th></th><th></th><th></th><th></th><th></th><th></th><th></th><th></th><th></th><th></th><th></th></tr>
<tr><td>>hg38_21_+_8433172_8446622_RNA45SN1_45S_50ntFlanks@3841.3862.22|TCCTACTTGGATAACTGTGGTA</td><td>>hg38_21_+_8433172_8446622_RNA45SN1_45S_50ntFlanks</td><td>3841</td><td>3862</td><td>22</td><td>TCCTACTTGGATAACTGTGGTA</td><td>570.3</td><td>5.7</td><td>0.7</td><td>7.9</td><td>2.6E-15</td><td>1.2E-14</td><td>BT20/MB468</td><td>CYTO</td></tr>
<tr><td>>hg38_21_+_8433172_8446622_RNA45SN1_45S_50ntFlanks@7990.8009.20|GACGTGGCGACCCGCTGAAT</td><td>>hg38_21_+_8433172_8446622_RNA45SN1_45S_50ntFlanks</td><td>7990</td><td>8009</td><td>20</td><td>GACGTGGCGACCCGCTGAAT</td><td>367.2</td><td>5.0</td><td>0.6</td><td>7.9</td><td>2.6E-15</td><td>1.2E-14</td><td>BT20/MB468</td><td>CYTO</td></tr>
<tr><td>>MI0000273|hsa-mir-183&WithFlank&7|-|129774899|129775020@33.53.21(+1A)&[MIMAT0000261&hsa-miR-183-5p&offsets|0|-1(+1A);m-41&7|-|129774966|129774987&offsets|-1|-2(+1A)]|TATGGCACTGGTAGAATTCACA</td><td>>MI0000273|hsa-mir-183&WithFlank&7|-|129774899|129775020</td><td>33</td><td>53</td><td>21(+1A)</td><td>TATGGCACTGGTAGAATTCACA</td><td>2002.6</td><td>8.6</td><td>1.1</td><td>7.9</td><td>2.6E-15</td><td>1.2E-14</td><td>BT20/MB468</td><td>CYTO</td></tr>
<tr><td>>MI0000809|hsa-mir-151a&WithFlank&8|-|140732558|140732659@53.74.22(+2A)&[MIMAT0000757&hsa-miR-151a-3p&offsets|0|+1(+2A);m-30&8|-|140732587|140732607&offsets|0|+1(+2A)]|CTAGACTGAAGCTCCTTGAGGAAA</td><td>>MI0000809|hsa-mir-151a&WithFlank&8|-|140732558|140732659</td><td>53</td><td>74</td><td>22(+2A)</td><td>CTAGACTGAAGCTCCTTGAGGAAA</td><td>369.4</td><td>4.3</td><td>0.5</td><td>7.9</td><td>2.2E-15</td><td>1.2E-14</td><td>MB231/MB468</td><td>NUC</td></tr>
<tr><td>>hg38_21_+_8433172_8446622_RNA45SN1_45S_50ntFlanks@10903.10939.37|GGCTGGGCGCGCGCCGCGGCTGGACGAGGCGCCGCCG</td><td>>hg38_21_+_8433172_8446622_RNA45SN1_45S_50ntFlanks</td><td>10903</td><td>10939</td><td>37</td><td>GGCTGGGCGCGCGCCGCGGCTGGACGAGGCGCCGCCG</td><td>278.5</td><td>-11.9</td><td>1.5</td><td>-7.9</td><td>3.0E-15</td><td>1.2E-14</td><td>MB231/MB468</td><td>MITO</td></tr>
<tr><td>>MI0001445|hsa-mir-423&WithFlank&17|+|30117073|30117178@59.80.22(+1C)&[MIMAT0001340&hsa-miR-423-3p&offsets|0|-1(+1C);m-100&17|+|30117131|30117153&offsets|0|-1(+1C)]|AGCTCGGTCTGAGGCCCCTCAGC</td><td>>MI0001445|hsa-mir-423&WithFlank&17|+|30117073|30117178</td><td>59</td><td>80</td><td>22(+1C)</td><td>AGCTCGGTCTGAGGCCCCTCAGC</td><td>1648.5</td><td>7.0</td><td>0.9</td><td>7.9</td><td>2.0E-15</td><td>1.2E-14</td><td>MB231/MB468</td><td>CYTO</td></tr>
<tr><td>>hg38_21_+_8433172_8446622_RNA45SN1_45S_50ntFlanks@8229.8252.24|CGCGCCGGGCCCGGGTCTTCCCGG</td><td>>hg38_21_+_8433172_8446622_RNA45SN1_45S_50ntFlanks</td><td>8229</td><td>8252</td><td>24</td><td>CGCGCCGGGCCCGGGTCTTCCCGG</td><td>398.1</td><td>12.6</td><td>1.6</td><td>7.9</td><td>2.0E-15</td><td>1.2E-14</td><td>MB231/MB468</td><td>CYTO</td></tr>
<tr><td>>hg38_21_+_8433172_8446622_RNA45SN1_45S_50ntFlanks@10349.10379.31|AACGAGAACTTTGAAGGCCGAAGTGGAGAAG</td><td>>hg38_21_+_8433172_8446622_RNA45SN1_45S_50ntFlanks</td><td>10349</td><td>10379</td><td>31</td><td>AACGAGAACTTTGAAGGCCGAAGTGGAGAAG</td><td>633.6</td><td>-4.6</td><td>0.6</td><td>-7.9</td><td>4.0E-15</td><td>1.2E-14</td><td>BT20/MB468</td><td>NUC</td></tr>
<tr><td>>MI0000270|hsa-mir-181b-1&WithFlank&1|-|198858867|198858988@42.64.23;MI0000683|hsa-mir-181b-2&WithFlank&9|+|124693704|124693804@22.44.23&[MIMAT0000257&hsa-miR-181b-5p&offsets|0|0;m-73&1|-|198858925|198858947&offsets|0|0];[MIMAT0000257_1&hsa-miR-181b-5p&offsets|0|0;m-72&9|+|124693725|124693747&offsets|0|0]|AACATTCATTGCTGTCGGTGGGT</td><td>>MI0000270|hsa-mir-181b-1&WithFlank&1|-|198858867|198858988</td><td>42</td><td>64</td><td>23</td><td>AACATTCATTGCTGTCGGTGGGT</td><td>2394.5</td><td>5.4</td><td>0.7</td><td>7.9</td><td>2.7E-15</td><td>1.2E-14</td><td>BT20/MB468</td><td>CYTO</td></tr>
<tr><td>>hg38_21_+_8433172_8446622_RNA45SN1_45S_50ntFlanks@9033.9058.26|CTCGTGGGGGGGCCGGGCCACCCCTC</td><td>>hg38_21_+_8433172_8446622_RNA45SN1_45S_50ntFlanks</td><td>9033</td><td>9058</td><td>26</td><td>CTCGTGGGGGGGCCGGGCCACCCCTC</td><td>691.5</td><td>-11.5</td><td>1.5</td><td>-7.9</td><td>1.9E-15</td><td>1.2E-14</td><td>BT20/MB468</td><td>MITO</td></tr>
<tr><td>>hg38_21_+_8433172_8446622_RNA45SN1_45S_50ntFlanks@8182.8203.22|ATCGAGGCCCAGCCCGTGGACG</td><td>>hg38_21_+_8433172_8446622_RNA45SN1_45S_50ntFlanks</td><td>8182</td><td>8203</td><td>22</td><td>ATCGAGGCCCAGCCCGTGGACG</td><td>359.9</td><td>-10.6</td><td>1.3</td><td>-7.9</td><td>1.9E-15</td><td>1.2E-14</td><td>BT20/MB468</td><td>MITO</td></tr>
<tr><td>>am_MI0003776|hsa-mir-320b-1&WithFlank&1|+|116671740|116671823@48.65.18;MI0003839|hsa-mir-320b-2&WithFlank&1|-|224257034|224257116@47.64.18;MI0008190|hsa-mir-320d-1&WithFlank&13|-|40727810|40727893@48.65.18;MI0003778|hsa-mir-320c-1&WithFlank&18|+|21683512|21683595@48.65.18;MI0008191|hsa-mir-320c-2&WithFlank&18|+|24321669|24321752@48.65.18;MI0000542|hsa-mir-320a&WithFlank&8|-|22244960|22245043@48.65.18;MI0008192|hsa-mir-320d-2&WithFlank&X|-|140926154|140926237@48.65.18&[MIMAT0005792&hsa-miR-320b&offsets|0|-4;m-183&1|+|116671787|116671806&offsets|0|-2];[MIMAT0005792_1&hsa-miR-320b&offsets|0|-4;m-183&1|-|224257051|224257070&offsets|0|-2];[MIMAT0006764&hsa-miR-320d&offsets|0|-1;m-246&13|-|40727829|40727846&offsets|0|0];[MIMAT0005793&hsa-miR-320c&offsets|0|-2;m-222&18|+|21683559|21683578&offsets|0|-2];[MIMAT0005793_1&hsa-miR-320c&offsets|0|-2;m-219&18|+|24321716|24321735&offsets|0|-2];[MIMAT0000510&hsa-miR-320a-3p&offsets|0|-4;m-60&8|-|22244975|22244996&offsets|0|-4];[MIMAT0006764_1&hsa-miR-320d&offsets|0|-1;m-253&X|-|140926173|140926190&offsets|0|0]|AAAAGCTGGGTTGAGAGG</td><td>>am_MI0003776|hsa-mir-320b-1&WithFlank&1|+|116671740|116671823</td><td>48</td><td>65</td><td>18</td><td>AAAAGCTGGGTTGAGAGG</td><td>106.1</td><td>3.2</td><td>0.4</td><td>7.9</td><td>4.1E-15</td><td>1.2E-14</td><td>BT20/MB468</td><td>NUC</td></tr>
<tr><td>>hg38_21_+_8433172_8446622_RNA45SN1_45S_50ntFlanks@4047.4066.20|ATTCGAACGTCTGCCCTATC</td><td>>hg38_21_+_8433172_8446622_RNA45SN1_45S_50ntFlanks</td><td>4047</td><td>4066</td><td>20</td><td>ATTCGAACGTCTGCCCTATC</td><td>1545.5</td><td>-7.8</td><td>1.0</td><td>-8.0</td><td>1.8E-15</td><td>1.2E-14</td><td>MB231/MB468</td><td>MP</td></tr>
<tr><td>>hg38_21_+_8433172_8446622_RNA45SN1_45S_50ntFlanks@10351.10381.31|CGAGAACTTTGAAGGCCGAAGTGGAGAAGGG</td><td>>hg38_21_+_8433172_8446622_RNA45SN1_45S_50ntFlanks</td><td>10351</td><td>10381</td><td>31</td><td>CGAGAACTTTGAAGGCCGAAGTGGAGAAGGG</td><td>2218.2</td><td>-8.6</td><td>1.1</td><td>-8.0</td><td>1.8E-15</td><td>1.2E-14</td><td>MB231/MB468</td><td>MP</td></tr>
<tr><td>>hg38_21_+_8433172_8446622_RNA45SN1_45S_50ntFlanks@12081.12107.27|CTCTCGCTTCTGGCGCCAAGCGCCCGG</td><td>>hg38_21_+_8433172_8446622_RNA45SN1_45S_50ntFlanks</td><td>12081</td><td>12107</td><td>27</td><td>CTCTCGCTTCTGGCGCCAAGCGCCCGG</td><td>624.1</td><td>-6.9</td><td>0.9</td><td>-7.9</td><td>2.0E-15</td><td>1.2E-14</td><td>MB231/MB468</td><td>CYTO</td></tr>
<tr><td>>hg38_21_+_8433172_8446622_RNA45SN1_45S_50ntFlanks@11918.11948.31|ACATGAGAGGTGTAGAATAAGTGGGAGGCCC</td><td>>hg38_21_+_8433172_8446622_RNA45SN1_45S_50ntFlanks</td><td>11918</td><td>11948</td><td>31</td><td>ACATGAGAGGTGTAGAATAAGTGGGAGGCCC</td><td>296.3</td><td>-12.0</td><td>1.5</td><td>-7.9</td><td>3.0E-15</td><td>1.2E-14</td><td>MB231/MB468</td><td>MITO</td></tr>
<tr><td>>hg38_MT_+_1621_3279_RNR2_16S_50ntFlanks@493.517.25|GCTCTTTGGACACTAGGAAAAAACC</td><td>>hg38_MT_+_1621_3279_RNR2_16S_50ntFlanks</td><td>493</td><td>517</td><td>25</td><td>GCTCTTTGGACACTAGGAAAAAACC</td><td>367.0</td><td>7.3</td><td>0.9</td><td>7.9</td><td>2.3E-15</td><td>1.2E-14</td><td>MB231/MB468</td><td>NUC</td></tr>
<tr><td>>hg38_21_+_8433172_8446622_RNA45SN1_45S_50ntFlanks@12363.12382.20|GGATAACTGGCTTGTGGCGG</td><td>>hg38_21_+_8433172_8446622_RNA45SN1_45S_50ntFlanks</td><td>12363</td><td>12382</td><td>20</td><td>GGATAACTGGCTTGTGGCGG</td><td>464.0</td><td>-11.6</td><td>1.5</td><td>-7.9</td><td>2.0E-15</td><td>1.2E-14</td><td>MB231/MB468</td><td>CYTO</td></tr>
<tr><td>>MI0000285|hsa-mir-205&WithFlank&1|+|209432127|209432248@40.60.21&[MIMAT0000266&hsa-miR-205-5p&offsets|0|-1;m-351&1|+|209432166|209432187&offsets|0|-1]|TCCTTCATTCCACCGGAGTCT</td><td>>MI0000285|hsa-mir-205&WithFlank&1|+|209432127|209432248</td><td>40</td><td>60</td><td>21</td><td>TCCTTCATTCCACCGGAGTCT</td><td>3282.9</td><td>15.0</td><td>1.8</td><td>8.4</td><td>3.4E-17</td><td>1.2E-14</td><td>BT20/MB231</td><td>MP</td></tr>
<tr><td>>MI0000650|hsa-mir-200c&WithFlank&12|+|6963693|6963772@50.69.20&[MIMAT0000617&hsa-miR-200c-3p&offsets|0|-3;m-54&12|+|6963742|6963764&offsets|0|-3]|TAATACTGCCGGGTAATGAT</td><td>>MI0000650|hsa-mir-200c&WithFlank&12|+|6963693|6963772</td><td>50</td><td>69</td><td>20</td><td>TAATACTGCCGGGTAATGAT</td><td>1172.8</td><td>13.5</td><td>1.6</td><td>8.4</td><td>3.0E-17</td><td>1.2E-14</td><td>BT20/MB231</td><td>MP</td></tr>
<tr><td>>MI0000764|hsa-mir-363&WithFlank&X|-|134169372|134169458@56.77.22&[MIMAT0000707&hsa-miR-363-3p&offsets|0|0;m-178&X|-|134169381|134169403&offsets|0|-1]|AATTGCACGGTATCCATCTGTA</td><td>>MI0000764|hsa-mir-363&WithFlank&X|-|134169372|134169458</td><td>56</td><td>77</td><td>22</td><td>AATTGCACGGTATCCATCTGTA</td><td>1198.2</td><td>13.6</td><td>1.6</td><td>8.4</td><td>3.2E-17</td><td>1.2E-14</td><td>BT20/MB231</td><td>MP</td></tr>
<tr><td>>MI0005756|hsa-mir-934&WithFlank&X|+|136550872|136550966@21.41.21&[MIMAT0004977&hsa-miR-934&offsets|0|-1;m-943&X|+|136550892|136550913&offsets|0|-1]|TGTCTACTACTGGAGACACTG</td><td>>MI0005756|hsa-mir-934&WithFlank&X|+|136550872|136550966</td><td>21</td><td>41</td><td>21</td><td>TGTCTACTACTGGAGACACTG</td><td>4054.0</td><td>15.3</td><td>1.8</td><td>8.4</td><td>3.4E-17</td><td>1.2E-14</td><td>BT20/MB231</td><td>MP</td></tr>
<tr><td>>MI0000108|hsa-mir-103a-2&WithFlank&20|+|3917488|3917577@54.76.23(+1U);MI0000109|hsa-mir-103a-1&WithFlank&5|-|168560890|168560979@54.76.23(+1U)&[MIMAT0000101&hsa-miR-103a-3p&offsets|0|0(+1U);m-12&20|+|3917541|3917563&offsets|0|0(+1U)];[MIMAT0000101_1&hsa-miR-103a-3p&offsets|0|0(+1U);m-13&5|-|168560904|168560926&offsets|0|0(+1U);m-44220&5|-|168560908|168560926&offsets|0|+4(+1U)]|AGCAGCATTGTACAGGGCTATGAT</td><td>>MI0000108|hsa-mir-103a-2&WithFlank&20|+|3917488|3917577</td><td>54</td><td>76</td><td>23(+1U)</td><td>AGCAGCATTGTACAGGGCTATGAT</td><td>1792.9</td><td>6.4</td><td>0.8</td><td>7.9</td><td>2.0E-15</td><td>1.2E-14</td><td>BT20/MB468</td><td>MITO</td></tr>
<tr><td>>hg38_21_+_8433172_8446622_RNA45SN1_45S_50ntFlanks@5420.5444.25|CCGATTGGATGGTTTAGTGAGGCCC</td><td>>hg38_21_+_8433172_8446622_RNA45SN1_45S_50ntFlanks</td><td>5420</td><td>5444</td><td>25</td><td>CCGATTGGATGGTTTAGTGAGGCCC</td><td>388.4</td><td>-12.4</td><td>1.6</td><td>-7.9</td><td>3.1E-15</td><td>1.2E-14</td><td>MB231/MB468</td><td>MITO</td></tr>
<tr><td>>trna10_LysCTT_16_+_3241501_3241573@3.35.33,trna13_LysCTT_14_-_58706613_58706685@3.35.33,trna2_LysCTT_15_+_79152904_79152976@3.35.33|CCGGCTAGCTCAGTCGGTAGAGCATGGGACTCT</td><td>>trna10_LysCTT_16_+_3241501_3241573</td><td>3</td><td>35</td><td>33</td><td>CCGGCTAGCTCAGTCGGTAGAGCATGGGACTCT</td><td>328.4</td><td>8.0</td><td>1.0</td><td>7.9</td><td>2.3E-15</td><td>1.2E-14</td><td>MB231/MB468</td><td>NUC</td></tr>
<tr><td>>hg38_21_+_8433172_8446622_RNA45SN1_45S_50ntFlanks@10827.10865.39|CGGCAAGCCGGATCCGTAACTTCGGGATAAGGATTGGCT</td><td>>hg38_21_+_8433172_8446622_RNA45SN1_45S_50ntFlanks</td><td>10827</td><td>10865</td><td>39</td><td>CGGCAAGCCGGATCCGTAACTTCGGGATAAGGATTGGCT</td><td>488.7</td><td>-11.7</td><td>1.5</td><td>-7.9</td><td>2.0E-15</td><td>1.2E-14</td><td>MB231/MB468</td><td>CYTO</td></tr>
<tr><td>>hg38_21_+_8433172_8446622_RNA45SN1_45S_50ntFlanks@10095.10117.23|CGCGCGCGCGCGCGTGTGGTGTG</td><td>>hg38_21_+_8433172_8446622_RNA45SN1_45S_50ntFlanks</td><td>10095</td><td>10117</td><td>23</td><td>CGCGCGCGCGCGCGTGTGGTGTG</td><td>199.2</td><td>-11.5</td><td>1.4</td><td>-8.0</td><td>1.9E-15</td><td>1.2E-14</td><td>MB231/MB468</td><td>MP</td></tr>
<tr><td>>hg38_21_+_8433172_8446622_RNA45SN1_45S_50ntFlanks@9454.9495.42|CCGCGCCGGGGAGGTGGAGCACGAGCGCACGTGTTAGGACCC</td><td>>hg38_21_+_8433172_8446622_RNA45SN1_45S_50ntFlanks</td><td>9454</td><td>9495</td><td>42</td><td>CCGCGCCGGGGAGGTGGAGCACGAGCGCACGTGTTAGGACCC</td><td>332.4</td><td>-12.2</td><td>1.5</td><td>-7.9</td><td>3.1E-15</td><td>1.2E-14</td><td>MB231/MB468</td><td>MITO</td></tr>
<tr><td>>hg38_21_+_8433172_8446622_RNA45SN1_45S_50ntFlanks@12927.12953.27|CCTGGCGCTAAACCATTCGTAGACGAC</td><td>>hg38_21_+_8433172_8446622_RNA45SN1_45S_50ntFlanks</td><td>12927</td><td>12953</td><td>27</td><td>CCTGGCGCTAAACCATTCGTAGACGAC</td><td>267.5</td><td>-11.9</td><td>1.5</td><td>-8.0</td><td>1.9E-15</td><td>1.2E-14</td><td>MB231/MB468</td><td>MP</td></tr>
<tr><td>>MI0000102|hsa-mir-100&WithFlank&11|-|122152223|122152314@19.40.22&[MIMAT0000098&hsa-miR-100-5p&offsets|0|0;m-27&11|-|122152276|122152296&offsets|0|+1]|AACCCGTAGATCCGAACTTGTG</td><td>>MI0000102|hsa-mir-100&WithFlank&11|-|122152223|122152314</td><td>19</td><td>40</td><td>22</td><td>AACCCGTAGATCCGAACTTGTG</td><td>627702.2</td><td>12.9</td><td>1.6</td><td>7.9</td><td>3.1E-15</td><td>1.2E-14</td><td>MB231/MB468</td><td>MITO</td></tr>
<tr><td>>hg38_21_+_8433172_8446622_RNA45SN1_45S_50ntFlanks@10349.10367.19|AACGAGAACTTTGAAGGCC</td><td>>hg38_21_+_8433172_8446622_RNA45SN1_45S_50ntFlanks</td><td>10349</td><td>10367</td><td>19</td><td>AACGAGAACTTTGAAGGCC</td><td>950.3</td><td>-4.9</td><td>0.6</td><td>-7.8</td><td>4.2E-15</td><td>1.2E-14</td><td>BT20/MB468</td><td>NUC</td></tr>
<tr><td>>hg38_21_+_8433172_8446622_RNA45SN1_45S_50ntFlanks@10516.10533.18|GAGTGGCGGAGATGGGCG</td><td>>hg38_21_+_8433172_8446622_RNA45SN1_45S_50ntFlanks</td><td>10516</td><td>10533</td><td>18</td><td>GAGTGGCGGAGATGGGCG</td><td>515.7</td><td>-12.0</td><td>1.5</td><td>-7.9</td><td>2.3E-15</td><td>1.2E-14</td><td>MB231/MB468</td><td>NUC</td></tr>
<tr><td>>MI0000063|hsa-let-7b&WithFlank&22|+|46113680|46113774@12.34.23(+1U)&[MIMAT0000063&hsa-let-7b-5p&offsets|0|+1(+1U);m-14&22|+|46113691|46113712&offsets|0|+1(+1U)]|TGAGGTAGTAGGTTGTGTGGTTTT</td><td>>MI0000063|hsa-let-7b&WithFlank&22|+|46113680|46113774</td><td>12</td><td>34</td><td>23(+1U)</td><td>TGAGGTAGTAGGTTGTGTGGTTTT</td><td>361.3</td><td>6.0</td><td>0.8</td><td>7.9</td><td>2.1E-15</td><td>1.2E-14</td><td>MB231/MB468</td><td>CYTO</td></tr>
<tr><td>>hg38_21_+_8433172_8446622_RNA45SN1_45S_50ntFlanks@12415.12448.34|CTTCGATGTCGGCTCTTCCTATCATTGTGAAGCA</td><td>>hg38_21_+_8433172_8446622_RNA45SN1_45S_50ntFlanks</td><td>12415</td><td>12448</td><td>34</td><td>CTTCGATGTCGGCTCTTCCTATCATTGTGAAGCA</td><td>2443.2</td><td>5.4</td><td>0.7</td><td>7.9</td><td>2.1E-15</td><td>1.2E-14</td><td>MB231/MB468</td><td>CYTO</td></tr>
<tr><td>>hg38_21_+_8433172_8446622_RNA45SN1_45S_50ntFlanks@11666.11685.20|TCTGCCCAGTGCTCTGAATG</td><td>>hg38_21_+_8433172_8446622_RNA45SN1_45S_50ntFlanks</td><td>11666</td><td>11685</td><td>20</td><td>TCTGCCCAGTGCTCTGAATG</td><td>3787.6</td><td>-6.3</td><td>0.8</td><td>-7.9</td><td>1.9E-15</td><td>1.2E-14</td><td>MB231/MB468</td><td>MP</td></tr>
<tr><td>>hg38_21_+_8433172_8446622_RNA45SN1_45S_50ntFlanks@10099.10127.29|CGCGCGCGCGTGTGGTGTGCGTCGGAGGG</td><td>>hg38_21_+_8433172_8446622_RNA45SN1_45S_50ntFlanks</td><td>10099</td><td>10127</td><td>29</td><td>CGCGCGCGCGTGTGGTGTGCGTCGGAGGG</td><td>292.2</td><td>-12.1</td><td>1.5</td><td>-7.9</td><td>1.9E-15</td><td>1.2E-14</td><td>MB231/MB468</td><td>MP</td></tr>
<tr><td>>hg38_21_+_8433172_8446622_RNA45SN1_45S_50ntFlanks@10713.10736.24|CTTGAAAATCCGGGGGAGAGGGTG</td><td>>hg38_21_+_8433172_8446622_RNA45SN1_45S_50ntFlanks</td><td>10713</td><td>10736</td><td>24</td><td>CTTGAAAATCCGGGGGAGAGGGTG</td><td>869.1</td><td>-7.9</td><td>1.0</td><td>-7.9</td><td>1.9E-15</td><td>1.2E-14</td><td>MB231/MB468</td><td>MP</td></tr>
<tr><td>>hg38_21_+_8433172_8446622_RNA45SN1_45S_50ntFlanks@7992.8012.21|CGTGGCGACCCGCTGAATTTA</td><td>>hg38_21_+_8433172_8446622_RNA45SN1_45S_50ntFlanks</td><td>7992</td><td>8012</td><td>21</td><td>CGTGGCGACCCGCTGAATTTA</td><td>273.6</td><td>-4.1</td><td>0.5</td><td>-7.8</td><td>4.2E-15</td><td>1.2E-14</td><td>BT20/MB468</td><td>NUC</td></tr>
<tr><td>>MI0003776|hsa-mir-320b-1&WithFlank&1|+|116671740|116671823@48.67.20;MI0003839|hsa-mir-320b-2&WithFlank&1|-|224257034|224257116@47.66.20;MI0000542|hsa-mir-320a&WithFlank&8|-|22244960|22245043@48.67.20&[MIMAT0005792&hsa-miR-320b&offsets|0|-2;m-183&1|+|116671787|116671806&offsets|0|0];[MIMAT0005792_1&hsa-miR-320b&offsets|0|-2;m-183&1|-|224257051|224257070&offsets|0|0];[MIMAT0000510&hsa-miR-320a-3p&offsets|0|-2;m-60&8|-|22244975|22244996&offsets|0|-2]|AAAAGCTGGGTTGAGAGGGC</td><td>>MI0003776|hsa-mir-320b-1&WithFlank&1|+|116671740|116671823</td><td>48</td><td>67</td><td>20</td><td>AAAAGCTGGGTTGAGAGGGC</td><td>7256.0</td><td>11.4</td><td>1.4</td><td>7.9</td><td>2.8E-15</td><td>1.3E-14</td><td>BT20/MB468</td><td>CYTO</td></tr>
</table>

| | | | | | | | | | | | | | |
|---|---|---|---|---|---|---|---|---|---|---|---|---|---|
| >hg38_21_+_8433172_8446622_RNA45SN1_45S_50ntFlanks@8176.8214.39\|CTTCTGATCGAGGCCCAGCCCGTGGACGGTGTGAGGCCG | >hg38_21_+_8433172_8446622_RNA45SN1_45S_50ntFlanks | 8176 | 8214 | 39 | CTTCTGATCGAGGCCCAGCCCGTGGACGGTGTGAGGCCG | 777.1 | -5.3 | 0.7 | -7.9 | 3.1E-15 | 1.3E-14 | MB231/MB468 | MITO |
| >hg38_21_+_8433172_8446622_RNA45SN1_45S_50ntFlanks@11664.11685.22\|TTTCTGCCCAGTGCTCTGAATG | >hg38_21_+_8433172_8446622_RNA45SN1_45S_50ntFlanks | 11664 | 11685 | 22 | TTTCTGCCCAGTGCTCTGAATG | 410.0 | -12.5 | 1.6 | -7.9 | 1.9E-15 | 1.3E-14 | MB231/MB468 | MP |
| >hg38_21_+_8433172_8446622_RNA45SN1_45S_50ntFlanks@4554.4573.20\|CCGCAGCTAGGAATAATGGA | >hg38_21_+_8433172_8446622_RNA45SN1_45S_50ntFlanks | 4554 | 4573 | 20 | CCGCAGCTAGGAATAATGGA | 249.2 | -11.8 | 1.5 | -7.9 | 1.9E-15 | 1.3E-14 | MB231/MB468 | MP |
| >hg38_21_+_8433172_8446622_RNA45SN1_45S_50ntFlanks@10409.10430.22\|GTCAGTCGGTCCTGAGAGATGG | >hg38_21_+_8433172_8446622_RNA45SN1_45S_50ntFlanks | 10409 | 10430 | 22 | GTCAGTCGGTCCTGAGAGATGG | 319.3 | -12.2 | 1.5 | -7.9 | 1.9E-15 | 1.3E-14 | MB231/MB468 | MP |
| >hg38_21_+_8433172_8446622_RNA45SN1_45S_50ntFlanks@9545.9574.30\|GTGGAGGTCCGTAGCGGTCCTGACGTGCAA | >hg38_21_+_8433172_8446622_RNA45SN1_45S_50ntFlanks | 9545 | 9574 | 30 | GTGGAGGTCCGTAGCGGTCCTGACGTGCAA | 2341.4 | -5.4 | 0.7 | -7.9 | 2.4E-15 | 1.3E-14 | MB231/MB468 | NUC |
| >hg38_21_+_8433172_8446622_RNA45SN1_45S_50ntFlanks@5525.5550.26\|TAAAAGTCGTAACAAGGTTTCCGTAG | >hg38_21_+_8433172_8446622_RNA45SN1_45S_50ntFlanks | 5525 | 5550 | 26 | TAAAAGTCGTAACAAGGTTTCCGTAG | 281.2 | -12.0 | 1.5 | -7.9 | 1.9E-15 | 1.3E-14 | MB231/MB468 | MP |
| >hg38_21_+_8433172_8446622_RNA45SN1_45S_50ntFlanks@11729.11767.39\|TAACTATGACTCTCTTAAGGTAGCCAAATGCCTCGTCAT | >hg38_21_+_8433172_8446622_RNA45SN1_45S_50ntFlanks | 11729 | 11767 | 39 | TAACTATGACTCTCTTAAGGTAGCCAAATGCCTCGTCAT | 1888.6 | -8.7 | 1.1 | -7.9 | 2.4E-15 | 1.3E-14 | MB231/MB468 | NUC |
| >hg38_21_+_8433172_8446622_RNA45SN1_45S_50ntFlanks@11786.11804.19\|GAATGGATGAACGAGATTC | >hg38_21_+_8433172_8446622_RNA45SN1_45S_50ntFlanks | 11786 | 11804 | 19 | GAATGGATGAACGAGATTC | 1308.3 | -5.8 | 0.7 | -7.9 | 2.4E-15 | 1.3E-14 | MB231/MB468 | NUC |
| >hg38_21_+_8433172_8446622_RNA45SN1_45S_50ntFlanks@11924.11948.25\|GAGGTGTAGAATAAGTGGGAGGCCC | >hg38_21_+_8433172_8446622_RNA45SN1_45S_50ntFlanks | 11924 | 11948 | 25 | GAGGTGTAGAATAAGTGGGAGGCCC | 159.9 | -11.2 | 1.4 | -7.9 | 1.9E-15 | 1.3E-14 | MB231/MB468 | MP |
| >hg38_21_+_8433172_8446622_RNA45SN1_45S_50ntFlanks@4264.4282.19\|AATGAGTCCACTTTAAATC | >hg38_21_+_8433172_8446622_RNA45SN1_45S_50ntFlanks | 4264 | 4282 | 19 | AATGAGTCCACTTTAAATC | 172.2 | -11.3 | 1.4 | -7.9 | 1.9E-15 | 1.3E-14 | MB231/MB468 | MP |
| >MI0003686\|hsa-mir-542&WithFlank&X\|-\|134541335\|134541443@59.80.22&[MIMAT0003389&hsa-miR-542-3p&offsets\|0\|0;m-136&X\|-\|134541364\|134541385&offsets\|0\|0]\|TGTGACAGATTGATAACTGAAA | >MI0003686\|hsa-mir-542&WithFlank&X\|-\|134541335\|134541443 | 59 | 80 | 22 | TGTGACAGATTGATAACTGAAA | 62.0 | 8.8 | 1.1 | 7.8 | 4.3E-15 | 1.3E-14 | BT20/MB468 | NUC |
| >MI0000746\|hsa-mir-99b&WithFlank&19\|+\|51692606\|51692687@51.73.23(+1U)&[MIMAT0004678&hsa-miR-99b-3p&offsets\|0\|+1(+1U);m-151&19\|+\|51692656\|51692678&offsets\|0\|0(+1U)]\|CAAGCTCGTGTCTGTGGGTCCGTT | >MI0000746\|hsa-mir-99b&WithFlank&19\|+\|51692606\|51692687 | 51 | 73 | 23(+1U) | CAAGCTCGTGTCTGTGGGTCCGTT | 62.4 | 2.7 | 0.4 | 7.8 | 4.3E-15 | 1.3E-14 | BT20/MB468 | NUC |
| >hg38_21_+_8433172_8446622_RNA45SN1_45S_50ntFlanks@12458.12496.39\|AAGCGTTGGATTGTTCACCCACTAATAGGGAACGTGAGC | >hg38_21_+_8433172_8446622_RNA45SN1_45S_50ntFlanks | 12458 | 12496 | 39 | AAGCGTTGGATTGTTCACCCACTAATAGGGAACGTGAGC | 620.3 | 5.2 | 0.7 | 7.9 | 2.2E-15 | 1.3E-14 | MB231/MB468 | CYTO |
| >hg38_21_+_8433172_8446622_RNA45SN1_45S_50ntFlanks@10841.10858.18\|CGTAACTTCGGGATAAGG | >hg38_21_+_8433172_8446622_RNA45SN1_45S_50ntFlanks | 10841 | 10858 | 18 | CGTAACTTCGGGATAAGG | 313.5 | -12.1 | 1.5 | -7.9 | 3.2E-15 | 1.3E-14 | MB231/MB468 | MITO |
| >hg38_21_+_8433172_8446622_RNA45SN1_45S_50ntFlanks@4003.4039.37\|ATAACCTCGGGCCGATCGCACGCCCCCGTGGCGGCG | >hg38_21_+_8433172_8446622_RNA45SN1_45S_50ntFlanks | 4003 | 4039 | 37 | ATAACCTCGGGCCGATCGCACGCCCCCGTGGCGGCG | 242.6 | -11.7 | 1.5 | -7.9 | 3.2E-15 | 1.3E-14 | MB231/MB468 | MITO |
| >hg38_21_+_8433172_8446622_RNA45SN1_45S_50ntFlanks@10806.10829.24\|AACAATGTAGGTAAGGGAAGTCGG | >hg38_21_+_8433172_8446622_RNA45SN1_45S_50ntFlanks | 10806 | 10829 | 24 | AACAATGTAGGTAAGGGAAGTCGG | 322.3 | -11.3 | 1.4 | -7.9 | 2.4E-15 | 1.3E-14 | MB231/MB468 | NUC |
| >MI0000239\|hsa-mir-197&WithFlank&1\|+\|109598887\|109598973@54.74.21&[MIMAT0000227&hsa-miR-197-3p&offsets\|0\|-1;m-99&1\|+\|109598940\|109598961&offsets\|0\|-1]\|TTCACCACCTTCTCCACCCAG | >MI0000239\|hsa-mir-197&WithFlank&1\|+\|109598887\|109598973 | 54 | 74 | 21 | TTCACCACCTTCTCCACCCAG | 433.6 | 10.2 | 1.3 | 7.9 | 2.1E-15 | 1.3E-14 | BT20/MB468 | MITO |
| >hg38_21_+_8433172_8446622_RNA45SN1_45S_50ntFlanks@12954.12982.29\|CTGCTTCTGGGTCGGGGTTTCGTACGTAG | >hg38_21_+_8433172_8446622_RNA45SN1_45S_50ntFlanks | 12954 | 12982 | 29 | CTGCTTCTGGGTCGGGGTTTCGTACGTAG | 184.8 | -5.3 | 0.7 | -7.9 | 2.2E-15 | 1.3E-14 | MB231/MB468 | CYTO |
| >MI0000483\|hsa-mir-186&WithFlank&1\|-\|71067625\|71067722@21.41.21&[MIMAT0000456&hsa-miR-186-5p&offsets\|0\|-1;m-84&1\|-\|71067681\|71067702&offsets\|0\|-1]\|CAAAGAATTCTCCTTTTGGGC | >MI0000483\|hsa-mir-186&WithFlank&1\|-\|71067625\|71067722 | 21 | 41 | 21 | CAAAGAATTCTCCTTTTGGGC | 858.0 | 7.0 | 0.9 | 7.9 | 2.9E-15 | 1.3E-14 | BT20/MB468 | CYTO |
| >hg38_21_+_8433172_8446622_RNA45SN1_45S_50ntFlanks@3855.3881.27\|CTGTGGTAATTCTAGAGCTAATACATG | >hg38_21_+_8433172_8446622_RNA45SN1_45S_50ntFlanks | 3855 | 3881 | 27 | CTGTGGTAATTCTAGAGCTAATACATG | 231.8 | -11.7 | 1.5 | -7.9 | 2.0E-15 | 1.3E-14 | MB231/MB468 | MP |
| >hg38_21_+_8433172_8446622_RNA45SN1_45S_50ntFlanks@11314.11342.29\|CCCACCCCACGTCTCGTCGCGCGCGCGTC | >hg38_21_+_8433172_8446622_RNA45SN1_45S_50ntFlanks | 11314 | 11342 | 29 | CCCACCCCACGTCTCGTCGCGCGCGCGTC | 307.7 | -12.0 | 1.5 | -7.9 | 3.3E-15 | 1.3E-14 | MB231/MB468 | MITO |
| >MI0003591\|hsa-mir-584&WithFlank&5\|-\|149062307\|149062415@22.42.21(+1A)&[MIMAT0003249&hsa-miR-584-5p&offsets\|0\|-1(+1A);m-167&5\|-\|149062374\|149062394&offsets\|0\|0(+1A)]\|TTATGGTTTGCCTGGGACTGAA | >MI0003591\|hsa-mir-584&WithFlank&5\|-\|149062307\|149062415 | 22 | 42 | 21(+1A) | TTATGGTTTGCCTGGGACTGAA | 264.9 | 9.7 | 1.2 | 7.9 | 2.9E-15 | 1.3E-14 | BT20/MB468 | CYTO |
| >MI0003188\|hsa-mir-503&WithFlank&X\|-\|134546322\|134546404@12.31.20&[MIMAT0002874&hsa-miR-503-5p&offsets\|0\|-3;m-297&X\|-\|134546371\|134546393&offsets\|0\|-3]\|TAGCAGCGGGAACAGTTCTG | >MI0003188\|hsa-mir-503&WithFlank&X\|-\|134546322\|134546404 | 12 | 31 | 20 | TAGCAGCGGGAACAGTTCTG | 300.2 | 11.5 | 1.4 | 7.9 | 2.0E-15 | 1.3E-14 | MB231/MB468 | MP |
| >hg38_21_+_8433172_8446622_RNA45SN1_45S_50ntFlanks@4001.4023.23\|AGATAACCTCGGGCCGATCGCAC | >hg38_21_+_8433172_8446622_RNA45SN1_45S_50ntFlanks | 4001 | 4023 | 23 | AGATAACCTCGGGCCGATCGCAC | 508.6 | 5.4 | 0.7 | 7.9 | 2.2E-15 | 1.3E-14 | MB231/MB468 | CYTO |
| >hg38_21_+_8433172_8446622_RNA45SN1_45S_50ntFlanks@4454.4473.20\|CGGCGCCCCCTCGATGCTCT | >hg38_21_+_8433172_8446622_RNA45SN1_45S_50ntFlanks | 4454 | 4473 | 20 | CGGCGCCCCCTCGATGCTCT | 316.1 | 12.3 | 1.6 | 7.9 | 2.2E-15 | 1.3E-14 | MB231/MB468 | CYTO |
| >hg38_21_+_8433172_8446622_RNA45SN1_45S_50ntFlanks@9545.9563.19\|GTGGAGGTCCGTAGCGGTC | >hg38_21_+_8433172_8446622_RNA45SN1_45S_50ntFlanks | 9545 | 9563 | 19 | GTGGAGGTCCGTAGCGGTC | 3153.8 | -11.2 | 1.4 | -7.9 | 2.0E-15 | 1.3E-14 | MB231/MB468 | MP |
| >hg38_21_+_8433172_8446622_RNA45SN1_45S_50ntFlanks@4660.4689.30\|GAGGTGAAATTCTTGGACCGGCGCAAGACG | >hg38_21_+_8433172_8446622_RNA45SN1_45S_50ntFlanks | 4660 | 4689 | 30 | GAGGTGAAATTCTTGGACCGGCGCAAGACG | 546.7 | -13.0 | 1.6 | -7.9 | 2.0E-15 | 1.3E-14 | MB231/MB468 | MP |
| >hg38_21_+_8433172_8446622_RNA45SN1_45S_50ntFlanks@11199.11224.26\|CCGGCGGCGGCGGCGACTCTGGACGC | >hg38_21_+_8433172_8446622_RNA45SN1_45S_50ntFlanks | 11199 | 11224 | 26 | CCGGCGGCGGCGGCGACTCTGGACGC | 260.6 | -7.5 | 0.9 | -7.9 | 2.2E-15 | 1.3E-14 | MB231/MB468 | CYTO |
| >MI0000067\|hsa-let-7f-1&WithFlank&9\|+\|94176341\|94176439@13.34.22(+1A);MI0000068\|hsa-let-7f-2&WithFlank&X\|-\|53557186\|53557280@14.35.22(+1A)&[MIMAT0000067&hsa-let-7f-5p&offsets\|0\|0(+1A);m-20&9\|+\|94176353\|94176374&offsets\|0\|0(+1A)];[MIMAT0000067_1&hsa-let-7f-5p&offsets\|0\|0(+1A);m-19&X\|-\|53557246\|53557267&offsets\|0\|0(+1A)]\|TGAGGTAGTAGATTGTATAGTTA | >MI0000067\|hsa-let-7f-1&WithFlank&9\|+\|94176341\|94176439 | 13 | 34 | 22(+1A) | TGAGGTAGTAGATTGTATAGTTA | 1346.0 | 3.6 | 0.5 | 7.9 | 2.5E-15 | 1.3E-14 | MB231/MB468 | NUC |
| >hg38_21_+_8433172_8446622_RNA45SN1_45S_50ntFlanks@11326.11345.20\|CTCGTCGCGCGCGCGTCCGC | >hg38_21_+_8433172_8446622_RNA45SN1_45S_50ntFlanks | 11326 | 11345 | 20 | CTCGTCGCGCGCGCGTCCGC | 2033.0 | 10.5 | 1.3 | 7.9 | 2.5E-15 | 1.3E-14 | MB231/MB468 | NUC |
| >hg38_21_+_8433172_8446622_RNA45SN1_45S_50ntFlanks@4726.4752.27\|TAATCAAGAACGAAAGTCGGAGGTTCG | >hg38_21_+_8433172_8446622_RNA45SN1_45S_50ntFlanks | 4726 | 4752 | 27 | TAATCAAGAACGAAAGTCGGAGGTTCG | 304.4 | -8.6 | 1.1 | -7.9 | 2.5E-15 | 1.3E-14 | MB231/MB468 | NUC |
| >MI0000465\|hsa-mir-191&WithFlank&3\|-\|49020612\|49020715@22.44.23(+1A)&[MIMAT0000440&hsa-miR-191-5p&offsets\|0\|0(+1A);m-48&3\|-\|49020672\|49020694&offsets\|0\|0(+1A)]\|CAACGGAATCCCAAAAGCAGCTGA | >MI0000465\|hsa-mir-191&WithFlank&3\|-\|49020612\|49020715 | 22 | 44 | 23(+1A) | CAACGGAATCCCAAAAGCAGCTGA | 764.6 | 7.2 | 0.9 | 7.9 | 2.2E-15 | 1.3E-14 | BT20/MB468 | MITO |
| >hg38_21_+_8433172_8446622_RNA45SN1_45S_50ntFlanks@4680.4720.41\|GCGCAAGACGGACCAGAGCGAAAGCATTTGCCAAGAATGTT | >hg38_21_+_8433172_8446622_RNA45SN1_45S_50ntFlanks | 4680 | 4720 | 41 | GCGCAAGACGGACCAGAGCGAAAGCATTTGCCAAGAATGTT | 352.4 | 10.1 | 1.3 | 7.9 | 2.3E-15 | 1.3E-14 | MB231/MB468 | CYTO |
| >hg38_21_+_8433172_8446622_RNA45SN1_45S_50ntFlanks@11569.11605.37\|CAGCCGACTTAGAACTGGTGCGGACCAGGGGAATCCG | >hg38_21_+_8433172_8446622_RNA45SN1_45S_50ntFlanks | 11569 | 11605 | 37 | CAGCCGACTTAGAACTGGTGCGGACCAGGGGAATCCG | 2979.1 | -11.3 | 1.4 | -7.8 | 4.6E-15 | 1.4E-14 | BT20/MB468 | NUC |
| >hg38_21_+_8433172_8446622_RNA45SN1_45S_50ntFlanks@11210.11232.23\|GGCGACTCTGGACGCGAGCCGGG | >hg38_21_+_8433172_8446622_RNA45SN1_45S_50ntFlanks | 11210 | 11232 | 23 | GGCGACTCTGGACGCGAGCCGGG | 1232.6 | -10.0 | 1.3 | -7.9 | 3.4E-15 | 1.4E-14 | MB231/MB468 | MITO |
| >am_MI0005763\|hsa-mir-941-1&WithFlank&20\|+\|63919443\|63919526@53.74.22;MI0005764\|hsa-mir-941-2&WithFlank&20\|+\|63919499\|63919582@53.74.22;MI0005765\|hsa-mir-941-3&WithFlank&20\|+\|63919555\|63919638@53.74.22;MI0005766\|hsa-mir-941-4&WithFlank&20\|+\|63919750\|63919833@53.74.22;MI0031520\|hsa-mir-941-5&WithFlank&20\|+\|63919862\|63919945@53.74.22&[MIMAT0004984&hsa-miR-941&offsets\|0\|-1;m-263&20\|+\|63919495\|63919517&offsets\|0\|-1];[MIMAT0004984_1&hsa-miR-941&offsets\|0\|-1;m-263&20\|+\|63919551\|63919573&offsets\|0\|-1];[MIMAT0004984_2&hsa-miR-941&offsets\|0\|-1;m-263&20\|+\|63919607\|63919629&offsets\|0\|-1];[MIMAT0004984_3&hsa-miR-941&offsets\|0\|-1;m-263&20\|+\|63919802\|63919824&offsets\|0\|-1];[MIMAT0004984_4&hsa-miR-941&offsets\|0\|-1;m-263&20\|+\|63919914\|63919936&offsets\|0\|-1]\|CACCCGGCTGTGTGCACATGTG | >am_MI0005763\|hsa-mir-941-1&WithFlank&20\|+\|63919443\|63919526 | 53 | 74 | 22 | CACCCGGCTGTGTGCACATGTG | 138.2 | 2.8 | 0.4 | 7.8 | 4.6E-15 | 1.4E-14 | BT20/MB468 | NUC |
| >hg38_21_+_8433172_8446622_RNA45SN1_45S_50ntFlanks@12717.12745.29\|CCTCGGTTGGCCTCGGATAGCCGGTCCCC | >hg38_21_+_8433172_8446622_RNA45SN1_45S_50ntFlanks | 12717 | 12745 | 29 | CCTCGGTTGGCCTCGGATAGCCGGTCCCC | 288.5 | -12.0 | 1.5 | -7.9 | 3.4E-15 | 1.4E-14 | MB231/MB468 | MITO |
| >hg38_21_+_8433172_8446622_RNA45SN1_45S_50ntFlanks@8020.8042.23\|TAGTCAGCGGAGGAGAAGAAACT | >hg38_21_+_8433172_8446622_RNA45SN1_45S_50ntFlanks | 8020 | 8042 | 23 | TAGTCAGCGGAGGAGAAGAAACT | 391.1 | 10.3 | 1.3 | 7.9 | 2.3E-15 | 1.4E-14 | MB231/MB468 | CYTO |
| >hg38_21_+_8433172_8446622_RNA45SN1_45S_50ntFlanks@10116.10134.19\|TGCGTCGGAGGGCGGCGGC | >hg38_21_+_8433172_8446622_RNA45SN1_45S_50ntFlanks | 10116 | 10134 | 19 | TGCGTCGGAGGGCGGCGGC | 214.6 | -10.5 | 1.3 | -7.9 | 2.3E-15 | 1.4E-14 | MB231/MB468 | CYTO |
| >hg38_21_+_8433172_8446622_RNA45SN1_45S_50ntFlanks@12635.12654.20\|AAGCTACCATCTGTGGGATT | >hg38_21_+_8433172_8446622_RNA45SN1_45S_50ntFlanks | 12635 | 12654 | 20 | AAGCTACCATCTGTGGGATT | 8381.4 | -7.1 | 0.9 | -7.9 | 2.2E-15 | 1.4E-14 | BT20/MB468 | MITO |
| >hg38_21_+_8433172_8446622_RNA45SN1_45S_50ntFlanks@4047.4073.27\|ATTCGAACGTCTGCCCTATCAACTTTC | >hg38_21_+_8433172_8446622_RNA45SN1_45S_50ntFlanks | 4047 | 4073 | 27 | ATTCGAACGTCTGCCCTATCAACTTTC | 166.1 | -11.2 | 1.4 | -7.9 | 2.1E-15 | 1.4E-14 | MB231/MB468 | MP |
| >hg38_21_+_8433172_8446622_RNA45SN1_45S_50ntFlanks@4731.4767.37\|AAGAACGAAAGTCGGAGGTTCGAAGACGATCAGATAC | >hg38_21_+_8433172_8446622_RNA45SN1_45S_50ntFlanks | 4731 | 4767 | 37 | AAGAACGAAAGTCGGAGGTTCGAAGACGATCAGATAC | 5589.6 | 11.3 | 1.4 | 7.9 | 3.0E-15 | 1.4E-14 | BT20/MB468 | CYTO |
| >hg38_21_+_8433172_8446622_RNA45SN1_45S_50ntFlanks@11974.11998.25\|AGGGGCCCGGGGCGGGGTCCGCCGG | >hg38_21_+_8433172_8446622_RNA45SN1_45S_50ntFlanks | 11974 | 11998 | 25 | AGGGGCCCGGGGCGGGGTCCGCCGG | 348.6 | -12.3 | 1.6 | -7.9 | 2.1E-15 | 1.4E-14 | MB231/MB468 | MP |
| >hg38_21_+_8433172_8446622_RNA45SN1_45S_50ntFlanks@10843.10865.23\|TAACTTCGGGATAAGGATTGGCT | >hg38_21_+_8433172_8446622_RNA45SN1_45S_50ntFlanks | 10843 | 10865 | 23 | TAACTTCGGGATAAGGATTGGCT | 2129.2 | -11.0 | 1.4 | -7.8 | 4.6E-15 | 1.4E-14 | BT20/MB468 | NUC |
| >hg38_21_+_8433172_8446622_RNA45SN1_45S_50ntFlanks@9505.9526.22\|TGAACTATGCCTGGGCAGGGCG | >hg38_21_+_8433172_8446622_RNA45SN1_45S_50ntFlanks | 9505 | 9526 | 22 | TGAACTATGCCTGGGCAGGGCG | 235.7 | -10.2 | 1.3 | -7.9 | 3.0E-15 | 1.4E-14 | BT20/MB468 | CYTO |
| >hg38_21_+_8433172_8446622_RNA45SN1_45S_50ntFlanks@12420.12446.27\|ATGTCGGCTCTTCCTATCATTGTGAAG | >hg38_21_+_8433172_8446622_RNA45SN1_45S_50ntFlanks | 12420 | 12446 | 27 | ATGTCGGCTCTTCCTATCATTGTGAAG | 24619.0 | -10.9 | 1.4 | -7.8 | 4.6E-15 | 1.4E-14 | BT20/MB468 | NUC |

<table>
<tr><th></th><th></th><th></th><th></th><th></th><th></th><th></th><th></th><th></th><th></th><th></th><th></th><th></th><th></th></tr>
<tr><td>>hg38_21_+_8433172_8446622_RNA45SN1_45S_50ntFlanks@9739.9760.22|AAACGATCTCAACCTATTCTCA</td><td>>hg38_21_+_8433172_8446622_RNA45SN1_45S_50ntFlanks</td><td>9739</td><td>9760</td><td>22</td><td>AAACGATCTCAACCTATTCTCA</td><td>169.3</td><td>-11.3</td><td>1.4</td><td>-7.9</td><td>2.1E-15</td><td>1.4E-14</td><td>MB231/MB468</td><td>MP</td></tr>
<tr><td>>trna116_GluCTC_1_-_145399233_145399304@32.51.20,trna59_GluCTC_1_+_249168447_249168518@32.51.20,trna71_GluCTC_1_-_161439189_161439260@32.51.20,trna74_GluCTC_1_-_161431809_161431880@32.51.20,trna77_GluCTC_1_-_161424398_161424469@32.51.20,trna77_GluCTC_6_+_28949976_28950047@32.51.20,trna80_GluCTC_1_-_161417018_161417089@32.51.20,trna87_GluCTC_6_-_126101393_126101464@32.51.20|CTCTCACCGCCGCGGCCCGG</td><td>>trna116_GluCTC_1_-_145399233_145399304</td><td>32</td><td>51</td><td>20</td><td>CTCTCACCGCCGCGGCCCGG</td><td>872.6</td><td>-14.2</td><td>1.8</td><td>-7.8</td><td>4.7E-15</td><td>1.4E-14</td><td>BT20/MB468</td><td>NUC</td></tr>
<tr><td>>hg38_21_+_8433172_8446622_RNA45SN1_45S_50ntFlanks@3993.4013.21|GTGACTCTAGATAACCTCGGG</td><td>>hg38_21_+_8433172_8446622_RNA45SN1_45S_50ntFlanks</td><td>3993</td><td>4013</td><td>21</td><td>GTGACTCTAGATAACCTCGGG</td><td>1157.5</td><td>-4.5</td><td>0.6</td><td>-7.9</td><td>2.6E-15</td><td>1.4E-14</td><td>MB231/MB468</td><td>NUC</td></tr>
<tr><td>>hg38_21_+_8433172_8446622_RNA45SN1_45S_50ntFlanks@10392.10409.18|ACAGCAGTTGAACATGGG</td><td>>hg38_21_+_8433172_8446622_RNA45SN1_45S_50ntFlanks</td><td>10392</td><td>10409</td><td>18</td><td>ACAGCAGTTGAACATGGG</td><td>3242.5</td><td>-7.8</td><td>1.0</td><td>-7.9</td><td>2.3E-15</td><td>1.4E-14</td><td>MB231/MB468</td><td>CYTO</td></tr>
<tr><td>>hg38_21_+_8433172_8446622_RNA45SN1_45S_50ntFlanks@4040.4059.20|ACGACCCATTCGAACGTCTG</td><td>>hg38_21_+_8433172_8446622_RNA45SN1_45S_50ntFlanks</td><td>4040</td><td>4059</td><td>20</td><td>ACGACCCATTCGAACGTCTG</td><td>247.0</td><td>-11.8</td><td>1.5</td><td>-7.9</td><td>2.1E-15</td><td>1.4E-14</td><td>MB231/MB468</td><td>MP</td></tr>
<tr><td>>hg38_21_+_8433172_8446622_RNA45SN1_45S_50ntFlanks@10713.10736.24|CTTGAAAATCCGGGGGAGAGGGTG</td><td>>hg38_21_+_8433172_8446622_RNA45SN1_45S_50ntFlanks</td><td>10713</td><td>10736</td><td>24</td><td>CTTGAAAATCCGGGGGAGAGGGTG</td><td>269.8</td><td>-11.0</td><td>1.4</td><td>-7.9</td><td>2.6E-15</td><td>1.4E-14</td><td>MB231/MB468</td><td>NUC</td></tr>
<tr><td>>hg38_21_+_8433172_8446622_RNA45SN1_45S_50ntFlanks@12360.12382.23|CAGGGATAACTGGCTTGTGGCGG</td><td>>hg38_21_+_8433172_8446622_RNA45SN1_45S_50ntFlanks</td><td>12360</td><td>12382</td><td>23</td><td>CAGGGATAACTGGCTTGTGGCGG</td><td>218.9</td><td>-10.5</td><td>1.3</td><td>-7.9</td><td>2.3E-15</td><td>1.4E-14</td><td>MB231/MB468</td><td>CYTO</td></tr>
<tr><td>>hg38_21_+_8433172_8446622_RNA45SN1_45S_50ntFlanks@10375.10407.33|AGAAGGGTTCCATGTGAACAGCAGTTGAACATG</td><td>>hg38_21_+_8433172_8446622_RNA45SN1_45S_50ntFlanks</td><td>10375</td><td>10407</td><td>33</td><td>AGAAGGGTTCCATGTGAACAGCAGTTGAACATG</td><td>264.3</td><td>-11.9</td><td>1.5</td><td>-7.9</td><td>2.1E-15</td><td>1.4E-14</td><td>MB231/MB468</td><td>MP</td></tr>
<tr><td>>MI0000650|hsa-mir-200c&WithFlank&12|+|6963693|6963772@49.70.22&[MIMAT0000617&hsa-miR-200c-3p&offsets|-1|-2;m-54&12|+|6963742|6963764&offsets|-1|-2]|CTAATACTGCCGGGTAATGATG</td><td>>MI0000650|hsa-mir-200c&WithFlank&12|+|6963693|6963772</td><td>49</td><td>70</td><td>22</td><td>CTAATACTGCCGGGTAATGATG</td><td>237.0</td><td>11.5</td><td>1.4</td><td>8.2</td><td>2.9E-16</td><td>1.4E-14</td><td>BT20/MB231</td><td>MITO</td></tr>
<tr><td>>hg38_21_+_8433172_8446622_RNA45SN1_45S_50ntFlanks@11568.11600.33|GCAGCCGACTTAGAACTGGTGCGGACCAGGGGA</td><td>>hg38_21_+_8433172_8446622_RNA45SN1_45S_50ntFlanks</td><td>11568</td><td>11600</td><td>33</td><td>GCAGCCGACTTAGAACTGGTGCGGACCAGGGGA</td><td>706.8</td><td>-5.2</td><td>0.7</td><td>-7.9</td><td>2.1E-15</td><td>1.4E-14</td><td>MB231/MB468</td><td>MP</td></tr>
<tr><td>>hg38_21_+_8433172_8446622_RNA45SN1_45S_50ntFlanks@10781.10822.42|AGGTGAACAGCCTCTGGCATGTTGGAACAATGTAGGTAAGGG</td><td>>hg38_21_+_8433172_8446622_RNA45SN1_45S_50ntFlanks</td><td>10781</td><td>10822</td><td>42</td><td>AGGTGAACAGCCTCTGGCATGTTGGAACAATGTAGGTAAGGG</td><td>380.2</td><td>-12.4</td><td>1.6</td><td>-7.9</td><td>3.5E-15</td><td>1.4E-14</td><td>MB231/MB468</td><td>MITO</td></tr>
<tr><td>>hg38_21_+_8433172_8446622_RNA45SN1_45S_50ntFlanks@12202.12222.21|CTAAGGCGAGCTCAGGGAGGA</td><td>>hg38_21_+_8433172_8446622_RNA45SN1_45S_50ntFlanks</td><td>12202</td><td>12222</td><td>21</td><td>CTAAGGCGAGCTCAGGGAGGA</td><td>748.0</td><td>-13.3</td><td>1.7</td><td>-7.9</td><td>3.5E-15</td><td>1.4E-14</td><td>MB231/MB468</td><td>MITO</td></tr>
<tr><td>>hg38_21_+_8433172_8446622_RNA45SN1_45S_50ntFlanks@10351.10377.27|CGAGAACTTTGAAGGCCGAAGTGGAGA</td><td>>hg38_21_+_8433172_8446622_RNA45SN1_45S_50ntFlanks</td><td>10351</td><td>10377</td><td>27</td><td>CGAGAACTTTGAAGGCCGAAGTGGAGA</td><td>5552.4</td><td>-8.4</td><td>1.1</td><td>-7.9</td><td>3.5E-15</td><td>1.4E-14</td><td>MB231/MB468</td><td>MITO</td></tr>
<tr><td>>hg38_21_+_8433172_8446622_RNA45SN1_45S_50ntFlanks@4367.4394.28|CAGTTAAAAAGCTCGTAGTTGGATCTTG</td><td>>hg38_21_+_8433172_8446622_RNA45SN1_45S_50ntFlanks</td><td>4367</td><td>4394</td><td>28</td><td>CAGTTAAAAAGCTCGTAGTTGGATCTTG</td><td>237.2</td><td>-11.7</td><td>1.5</td><td>-7.9</td><td>3.5E-15</td><td>1.4E-14</td><td>MB231/MB468</td><td>MITO</td></tr>
<tr><td>>hg38_21_+_8433172_8446622_RNA45SN1_45S_50ntFlanks@6661.6686.26|CGGTGGATCACTCGGCTCGTGCGTCG</td><td>>hg38_21_+_8433172_8446622_RNA45SN1_45S_50ntFlanks</td><td>6661</td><td>6686</td><td>26</td><td>CGGTGGATCACTCGGCTCGTGCGTCG</td><td>913.9</td><td>-4.8</td><td>0.6</td><td>-7.9</td><td>2.1E-15</td><td>1.4E-14</td><td>MB231/MB468</td><td>MP</td></tr>
<tr><td>>hg38_21_+_8433172_8446622_RNA45SN1_45S_50ntFlanks@11561.11581.21|GCGCCTAGCAGCCGACTTAGA</td><td>>hg38_21_+_8433172_8446622_RNA45SN1_45S_50ntFlanks</td><td>11561</td><td>11581</td><td>21</td><td>GCGCCTAGCAGCCGACTTAGA</td><td>313.2</td><td>-12.1</td><td>1.5</td><td>-7.9</td><td>3.5E-15</td><td>1.4E-14</td><td>MB231/MB468</td><td>MITO</td></tr>
<tr><td>>hg38_21_+_8433172_8446622_RNA45SN1_45S_50ntFlanks@3776.3801.26|CCGGTACAGTGAAACTGCGAATGGCT</td><td>>hg38_21_+_8433172_8446622_RNA45SN1_45S_50ntFlanks</td><td>3776</td><td>3801</td><td>26</td><td>CCGGTACAGTGAAACTGCGAATGGCT</td><td>1151.7</td><td>-9.8</td><td>1.2</td><td>-7.9</td><td>2.1E-15</td><td>1.4E-14</td><td>MB231/MB468</td><td>MP</td></tr>
<tr><td>>MI0000298|hsa-mir-221&WithFlank&X|-|45746151|45746272@71.92.22&[MIMAT0000278&hsa-miR-221-3p&offsets|0|-1;m-91&X|-|45746181|45746202&offsets|0|0]|AGCTACATTGTCTGCTGGGTTT</td><td>>MI0000298|hsa-mir-221&WithFlank&X|-|45746151|45746272</td><td>71</td><td>92</td><td>22</td><td>AGCTACATTGTCTGCTGGGTTT</td><td>11243.5</td><td>6.8</td><td>0.9</td><td>7.9</td><td>2.3E-15</td><td>1.4E-14</td><td>BT20/MB468</td><td>MITO</td></tr>
<tr><td>>hg38_21_+_8433172_8446622_RNA45SN1_45S_50ntFlanks@11320.11340.21|CCACGTCTCGTCGCGCGCGCG</td><td>>hg38_21_+_8433172_8446622_RNA45SN1_45S_50ntFlanks</td><td>11320</td><td>11340</td><td>21</td><td>CCACGTCTCGTCGCGCGCGCG</td><td>382.9</td><td>-12.4</td><td>1.6</td><td>-7.9</td><td>3.6E-15</td><td>1.4E-14</td><td>MB231/MB468</td><td>MITO</td></tr>
<tr><td>>MI0000447|hsa-mir-128-1&WithFlank&2|+|135665391|135665484@56.75.20;MI0000727|hsa-mir-128-2&WithFlank&3|+|35744470|35744565@58.77.20&[MIMAT0000424&hsa-miR-128-3p&offsets|0|-1;m-108&2|+|135665446|135665466&offsets|0|-1];[MIMAT0000424_1&hsa-miR-128-3p&offsets|0|-1;m-117&3|+|35744527|35744547&offsets|0|-1]|TCACAGTGAACCGGTCTCTT</td><td>>MI0000447|hsa-mir-128-1&WithFlank&2|+|135665391|135665484</td><td>56</td><td>75</td><td>20</td><td>TCACAGTGAACCGGTCTCTT</td><td>1122.7</td><td>5.4</td><td>0.7</td><td>7.9</td><td>2.3E-15</td><td>1.4E-14</td><td>BT20/MB468</td><td>MITO</td></tr>
<tr><td>>hg38_21_+_8433172_8446622_RNA45SN1_45S_50ntFlanks@5099.5136.38|CATGCTAACTAGTTACGCGACCCCCGAGCGGTCGGCGT</td><td>>hg38_21_+_8433172_8446622_RNA45SN1_45S_50ntFlanks</td><td>5099</td><td>5136</td><td>38</td><td>CATGCTAACTAGTTACGCGACCCCCGAGCGGTCGGCGT</td><td>135.5</td><td>-22.7</td><td>2.9</td><td>-7.9</td><td>2.2E-15</td><td>1.4E-14</td><td>MB231/MB468</td><td>MP</td></tr>
<tr><td>>hg38_21_+_8433172_8446622_RNA45SN1_45S_50ntFlanks@12942.12972.31|TTCGTAGACGACCTGCTTCTGGGTCGGGGTT</td><td>>hg38_21_+_8433172_8446622_RNA45SN1_45S_50ntFlanks</td><td>12942</td><td>12972</td><td>31</td><td>TTCGTAGACGACCTGCTTCTGGGTCGGGGTT</td><td>1045.3</td><td>-13.1</td><td>1.6</td><td>-8.3</td><td>7.2E-17</td><td>1.4E-14</td><td>BT20/MB468</td><td>MP</td></tr>
<tr><td>>hg38_21_+_8433172_8446622_RNA45SN1_45S_50ntFlanks@5421.5443.23|CGATTGGATGGTTTAGTGAGGCC</td><td>>hg38_21_+_8433172_8446622_RNA45SN1_45S_50ntFlanks</td><td>5421</td><td>5443</td><td>23</td><td>CGATTGGATGGTTTAGTGAGGCC</td><td>852.6</td><td>-10.0</td><td>1.3</td><td>-7.9</td><td>2.7E-15</td><td>1.4E-14</td><td>MB231/MB468</td><td>NUC</td></tr>
<tr><td>>MI0000298|hsa-mir-221&WithFlank&X|-|45746151|45746272@71.92.22(+1U)&[MIMAT0000278&hsa-miR-221-3p&offsets|0|-1(+1U);m-91&X|-|45746181|45746202&offsets|0|0(+1U)]|AGCTACATTGTCTGCTGGGTTTT</td><td>>MI0000298|hsa-mir-221&WithFlank&X|-|45746151|45746272</td><td>71</td><td>92</td><td>22(+1U)</td><td>AGCTACATTGTCTGCTGGGTTTT</td><td>2999.3</td><td>6.4</td><td>0.8</td><td>7.9</td><td>3.1E-15</td><td>1.4E-14</td><td>BT20/MB468</td><td>CYTO</td></tr>
<tr><td>>hg38_21_+_8433172_8446622_RNA45SN1_45S_50ntFlanks@9313.9338.26|TCTAACACGTGCGCGAGTCGGGGGCT</td><td>>hg38_21_+_8433172_8446622_RNA45SN1_45S_50ntFlanks</td><td>9313</td><td>9338</td><td>26</td><td>TCTAACACGTGCGCGAGTCGGGGGCT</td><td>881.4</td><td>-9.5</td><td>1.2</td><td>-7.9</td><td>3.6E-15</td><td>1.4E-14</td><td>MB231/MB468</td><td>MITO</td></tr>
<tr><td>>hg38_21_+_8433172_8446622_RNA45SN1_45S_50ntFlanks@11104.11132.29|AAGGGTCGGGGCGGCAGGGGCCGGCGGCG</td><td>>hg38_21_+_8433172_8446622_RNA45SN1_45S_50ntFlanks</td><td>11104</td><td>11132</td><td>29</td><td>AAGGGTCGGGGCGGCAGGGGCCGGCGGCG</td><td>208.1</td><td>-10.7</td><td>1.3</td><td>-7.9</td><td>2.7E-15</td><td>1.4E-14</td><td>MB231/MB468</td><td>NUC</td></tr>
<tr><td>>MI0000061|hsa-let-7a-2&WithFlank&11|-|122146516|122146599@11.30.20(+1C);MI0000062|hsa-let-7a-3&WithFlank&22|+|46112743|46112828@10.29.20(+1C);MI0000060|hsa-let-7a-1&WithFlank&9|+|94175951|94176042@12.31.20(+1C)&[MIMAT0000062&hsa-let-7a-5p&offsets|0|-2(+1C);m-6&11|-|122146568|122146589&offsets|0|-2(+1C)];[MIMAT0000062_1&hsa-let-7a-5p&offsets|0|-2(+1C);m-5&22|+|46112752|46112773&offsets|0|-2(+1C)];[MIMAT0000062_2&hsa-let-7a-5p&offsets|0|-2(+1C);m-4&9|+|94175962|94175983&offsets|0|-2(+1C)]|TGAGGTAGTAGGTTGTATAGC</td><td>>MI0000061|hsa-let-7a-2&WithFlank&11|-|122146516|122146599</td><td>11</td><td>30</td><td>20(+1C)</td><td>TGAGGTAGTAGGTTGTATAGC</td><td>785.5</td><td>6.3</td><td>0.8</td><td>7.9</td><td>2.4E-15</td><td>1.4E-14</td><td>MB231/MB468</td><td>CYTO</td></tr>
<tr><td>>hg38_21_+_8433172_8446622_RNA45SN1_45S_50ntFlanks@5420.5438.19|CCGATTGGATGGTTTAGTG</td><td>>hg38_21_+_8433172_8446622_RNA45SN1_45S_50ntFlanks</td><td>5420</td><td>5438</td><td>19</td><td>CCGATTGGATGGTTTAGTG</td><td>267.9</td><td>-6.9</td><td>0.9</td><td>-7.9</td><td>2.4E-15</td><td>1.4E-14</td><td>MB231/MB468</td><td>CYTO</td></tr>
<tr><td>>hg38_21_+_8433172_8446622_RNA45SN1_45S_50ntFlanks@6671.6689.19|CTCGGCTCGTGCGTCGATG</td><td>>hg38_21_+_8433172_8446622_RNA45SN1_45S_50ntFlanks</td><td>6671</td><td>6689</td><td>19</td><td>CTCGGCTCGTGCGTCGATG</td><td>358.4</td><td>-12.3</td><td>1.6</td><td>-7.9</td><td>3.6E-15</td><td>1.4E-14</td><td>MB231/MB468</td><td>MITO</td></tr>
<tr><td>>MI0000299|hsa-mir-222&WithFlank&X|-|45747009|45747130@75.99.25(+3U)&[MIMAT0000279&hsa-miR-222-3p&offsets|0|+4(+3U);m-140&X|-|45747033|45747056&offsets|0|+1(+3U)]|AGCTACATCTGGCTACTGGGTCTCTTTT</td><td>>MI0000299|hsa-mir-222&WithFlank&X|-|45747009|45747130</td><td>75</td><td>99</td><td>25(+3U)</td><td>AGCTACATCTGGCTACTGGGTCTCTTTT</td><td>2503.8</td><td>7.7</td><td>1.0</td><td>7.9</td><td>3.6E-15</td><td>1.4E-14</td><td>MB231/MB468</td><td>MITO</td></tr>
<tr><td>>hg38_21_+_8433172_8446622_RNA45SN1_45S_50ntFlanks@9459.9503.45|CCGGGGAGGTGGAGCACGAGCGCACGTGTTAGGACCCGAAAGATG</td><td>>hg38_21_+_8433172_8446622_RNA45SN1_45S_50ntFlanks</td><td>9459</td><td>9503</td><td>45</td><td>CCGGGGAGGTGGAGCACGAGCGCACGTGTTAGGACCCGAAAGATG</td><td>1120.5</td><td>-14.0</td><td>1.8</td><td>-7.9</td><td>2.2E-15</td><td>1.4E-14</td><td>MB231/MB468</td><td>MP</td></tr>
<tr><td>>hg38_21_+_8433172_8446622_RNA45SN1_45S_50ntFlanks@10682.10710.29|TCCGGCGGCGTCCGGTGAGCTCTCGCTGG</td><td>>hg38_21_+_8433172_8446622_RNA45SN1_45S_50ntFlanks</td><td>10682</td><td>10710</td><td>29</td><td>TCCGGCGGCGTCCGGTGAGCTCTCGCTGG</td><td>184.7</td><td>-11.4</td><td>1.4</td><td>-7.9</td><td>2.2E-15</td><td>1.4E-14</td><td>MB231/MB468</td><td>MP</td></tr>
<tr><td>>hg38_21_+_8433172_8446622_RNA45SN1_45S_50ntFlanks@12934.12958.25|CTAAACCATTCGTAGACGACCTGCT</td><td>>hg38_21_+_8433172_8446622_RNA45SN1_45S_50ntFlanks</td><td>12934</td><td>12958</td><td>25</td><td>CTAAACCATTCGTAGACGACCTGCT</td><td>176.7</td><td>-11.3</td><td>1.4</td><td>-7.9</td><td>2.2E-15</td><td>1.4E-14</td><td>MB231/MB468</td><td>MP</td></tr>
<tr><td>>hg38_21_+_8433172_8446622_RNA45SN1_45S_50ntFlanks@11614.11640.27|ATTAAAACAAAGCATCGCGAAGGCCCG</td><td>>hg38_21_+_8433172_8446622_RNA45SN1_45S_50ntFlanks</td><td>11614</td><td>11640</td><td>27</td><td>ATTAAAACAAAGCATCGCGAAGGCCCG</td><td>774.8</td><td>-12.7</td><td>1.5</td><td>-8.3</td><td>7.4E-17</td><td>1.4E-14</td><td>BT20/MB468</td><td>MP</td></tr>
<tr><td>>trna10_AspGTC_12_-_125424193_125424264@32.71.40,trna12_AspGTC_12_-_125411891_125411962@32.71.40,trna144_AspGTC_6_-_27551236_27551307@32.71.40,trna38_AspGTC_17_-_8125556_8125627@32.71.40,trna45_AspGTC_6_+_27447453_27447524@32.71.40,trna48_AspGTC_6_+_27471523_27471594@32.71.40,trna4_AspGTC_12_+_96429799_96429870@32.71.40,trna69_AspGTC_1_-_161440205_161440276@32.71.40,trna72_AspGTC_1_-_161432824_161432895@32.71.40,trna75_AspGTC_1_-_161425414_161425485@32.71.40,trna78_AspGTC_1_-_161418033_161418104@32.71.40,trna81_AspGTC_1_-_161410615_161410686@32.71.40|CTGTCACGCGGGAGACCGGGGTTCGATTCCCCGACGGGGA</td><td>>trna10_AspGTC_12_-_125424193_125424264</td><td>32</td><td>71</td><td>40</td><td>CTGTCACGCGGGAGACCGGGGTTCGATTCCCCGACGGGGA</td><td>261.9</td><td>-10.8</td><td>1.4</td><td>-7.9</td><td>2.5E-15</td><td>1.4E-14</td><td>MB231/MB468</td><td>CYTO</td></tr>
<tr><td>>hg38_21_+_8433172_8446622_RNA45SN1_45S_50ntFlanks@5485.5513.29|AGCGCTGAGAAGACGGTCGAACTTGACTA</td><td>>hg38_21_+_8433172_8446622_RNA45SN1_45S_50ntFlanks</td><td>5485</td><td>5513</td><td>29</td><td>AGCGCTGAGAAGACGGTCGAACTTGACTA</td><td>727.9</td><td>-9.1</td><td>1.2</td><td>-7.9</td><td>3.7E-15</td><td>1.4E-14</td><td>MB231/MB468</td><td>MITO</td></tr>
<tr><td>>hg38_21_+_8433172_8446622_RNA45SN1_45S_50ntFlanks@8850.8867.18|GCGGCGGCGGCGGCGGGA</td><td>>hg38_21_+_8433172_8446622_RNA45SN1_45S_50ntFlanks</td><td>8850</td><td>8867</td><td>18</td><td>GCGGCGGCGGCGGCGGGA</td><td>496.1</td><td>-12.7</td><td>1.6</td><td>-7.9</td><td>3.7E-15</td><td>1.4E-14</td><td>MB231/MB468</td><td>MITO</td></tr>
<tr><td>>hg38_21_+_8433172_8446622_RNA45SN1_45S_50ntFlanks@11208.11226.19|GCGGCGACTCTGGACGCGA</td><td>>hg38_21_+_8433172_8446622_RNA45SN1_45S_50ntFlanks</td><td>11208</td><td>11226</td><td>19</td><td>GCGGCGACTCTGGACGCGA</td><td>212.0</td><td>-10.4</td><td>1.3</td><td>-7.9</td><td>2.5E-15</td><td>1.4E-14</td><td>MB231/MB468</td><td>CYTO</td></tr>
<tr><td>>hg38_21_+_8433172_8446622_RNA45SN1_45S_50ntFlanks@4751.4768.18|CGAAGACGATCAGATACC</td><td>>hg38_21_+_8433172_8446622_RNA45SN1_45S_50ntFlanks</td><td>4751</td><td>4768</td><td>18</td><td>CGAAGACGATCAGATACC</td><td>151.3</td><td>8.6</td><td>1.1</td><td>7.9</td><td>2.5E-15</td><td>1.4E-14</td><td>MB231/MB468</td><td>CYTO</td></tr>
<tr><td>>hg38_21_+_8433172_8446622_RNA45SN1_45S_50ntFlanks@8552.8579.28|CTCCCCGGAGGGGGCGGGCTCCGGCGGG</td><td>>hg38_21_+_8433172_8446622_RNA45SN1_45S_50ntFlanks</td><td>8552</td><td>8579</td><td>28</td><td>CTCCCCGGAGGGGGCGGGCTCCGGCGGG</td><td>172.7</td><td>-10.2</td><td>1.3</td><td>-7.9</td><td>2.5E-15</td><td>1.4E-14</td><td>MB231/MB468</td><td>CYTO</td></tr>
<tr><td>>MI0000266|hsa-mir-10a&WithFlank&17|-|48579832|48579953@28.50.23(+1A)&[MIMAT0000253&hsa-miR-10a-5p&offsets|0|0(+1A);m-18&17|-|48579905|48579926&offsets|0|+1(+1A)]|TACCCTGTAGATCCGAATTTGTGA</td><td>>MI0000266|hsa-mir-10a&WithFlank&17|-|48579832|48579953</td><td>28</td><td>50</td><td>23(+1A)</td><td>TACCCTGTAGATCCGAATTTGTGA</td><td>472.5</td><td>12.1</td><td>1.5</td><td>7.9</td><td>3.7E-15</td><td>1.5E-14</td><td>MB231/MB468</td><td>MITO</td></tr>
<tr><td>>hg38_21_+_8433172_8446622_RNA45SN1_45S_50ntFlanks@12164.12182.19|CTGGGGCGGTACACCTGTC</td><td>>hg38_21_+_8433172_8446622_RNA45SN1_45S_50ntFlanks</td><td>12164</td><td>12182</td><td>19</td><td>CTGGGGCGGTACACCTGTC</td><td>464.4</td><td>-12.6</td><td>1.6</td><td>-7.9</td><td>3.7E-15</td><td>1.5E-14</td><td>MB231/MB468</td><td>MITO</td></tr>
</table>

<table>
<tr><td>>MI0000285|hsa-mir-205&WithFlank&1|+|209432127|209432248@40.62.23&[MIMAT0000266&hsa-miR-205-5p&offsets|0|+1;m-351&1|+|209432166|209432187&offsets|0|+1]|TCCTTCATTCCACCGGAGTCTGT</td><td>>MI0000285|hsa-mir-205&WithFlank&1|+|209432127|209432248</td><td>40</td><td>62</td><td>23</td><td>TCCTTCATTCCACCGGAGTCTGT</td><td>3658.7</td><td>-11.1</td><td>1.4</td><td>-7.9</td><td>2.2E-15</td><td>1.5E-14</td><td>MB231/MB468</td><td>MP</td></tr>
<tr><td>>hg38_21_+_8433172_8446622_RNA45SN1_45S_50ntFlanks@4615.4633.19|CCATGATTAAGAGGGACGG</td><td>>hg38_21_+_8433172_8446622_RNA45SN1_45S_50ntFlanks</td><td>4615</td><td>4633</td><td>19</td><td>CCATGATTAAGAGGGACGG</td><td>228.8</td><td>-11.7</td><td>1.5</td><td>-7.9</td><td>2.2E-15</td><td>1.5E-14</td><td>MB231/MB468</td><td>MP</td></tr>
<tr><td>>hg38_21_+_8433172_8446622_RNA45SN1_45S_50ntFlanks@9381.9409.29|CGCTCGCCGGCCGAGGTGGGATCCCGAGG</td><td>>hg38_21_+_8433172_8446622_RNA45SN1_45S_50ntFlanks</td><td>9381</td><td>9409</td><td>29</td><td>CGCTCGCCGGCCGAGGTGGGATCCCGAGG</td><td>598.8</td><td>-13.1</td><td>1.7</td><td>-7.9</td><td>2.2E-15</td><td>1.5E-14</td><td>MB231/MB468</td><td>MP</td></tr>
<tr><td>>hg38_21_+_8433172_8446622_RNA45SN1_45S_50ntFlanks@4853.4871.19|AAGTCTTTGGGTTCCGGGG</td><td>>hg38_21_+_8433172_8446622_RNA45SN1_45S_50ntFlanks</td><td>4853</td><td>4871</td><td>19</td><td>AAGTCTTTGGGTTCCGGGG</td><td>223.1</td><td>-11.7</td><td>1.5</td><td>-7.9</td><td>2.2E-15</td><td>1.5E-14</td><td>MB231/MB468</td><td>MP</td></tr>
<tr><td>>hg38_21_+_8433172_8446622_RNA45SN1_45S_50ntFlanks@4759.4781.23|ATCAGATACCGTCGTAGTTCCGA</td><td>>hg38_21_+_8433172_8446622_RNA45SN1_45S_50ntFlanks</td><td>4759</td><td>4781</td><td>23</td><td>ATCAGATACCGTCGTAGTTCCGA</td><td>356.9</td><td>-12.3</td><td>1.6</td><td>-7.9</td><td>3.7E-15</td><td>1.5E-14</td><td>MB231/MB468</td><td>MITO</td></tr>
<tr><td>>hg38_21_+_8433172_8446622_RNA45SN1_45S_50ntFlanks@12991.13016.26|CTCCCTCGCTGCGATCTATTGAAAGT</td><td>>hg38_21_+_8433172_8446622_RNA45SN1_45S_50ntFlanks</td><td>12991</td><td>13016</td><td>26</td><td>CTCCCTCGCTGCGATCTATTGAAAGT</td><td>534.6</td><td>-5.1</td><td>0.7</td><td>-7.9</td><td>3.7E-15</td><td>1.5E-14</td><td>MB231/MB468</td><td>MITO</td></tr>
<tr><td>>hg38_21_+_8433172_8446622_RNA45SN1_45S_50ntFlanks@4683.4707.25|CAAGACGGACCAGAGCGAAAGCATT</td><td>>hg38_21_+_8433172_8446622_RNA45SN1_45S_50ntFlanks</td><td>4683</td><td>4707</td><td>25</td><td>CAAGACGGACCAGAGCGAAAGCATT</td><td>1262.0</td><td>-7.8</td><td>1.0</td><td>-7.9</td><td>3.7E-15</td><td>1.5E-14</td><td>MB231/MB468</td><td>MITO</td></tr>
<tr><td>>hg38_21_+_8433172_8446622_RNA45SN1_45S_50ntFlanks@9621.9655.35|TCTAGTAGCTGGTTCCCTCCGAAGTTTCCCTCAGG</td><td>>hg38_21_+_8433172_8446622_RNA45SN1_45S_50ntFlanks</td><td>9621</td><td>9655</td><td>35</td><td>TCTAGTAGCTGGTTCCCTCCGAAGTTTCCCTCAGG</td><td>265.2</td><td>-11.8</td><td>1.5</td><td>-7.9</td><td>3.7E-15</td><td>1.5E-14</td><td>MB231/MB468</td><td>MITO</td></tr>
<tr><td>>hg38_21_+_8433172_8446622_RNA45SN1_45S_50ntFlanks@11206.11225.20|CGGCGGCGACTCTGGACGCG</td><td>>hg38_21_+_8433172_8446622_RNA45SN1_45S_50ntFlanks</td><td>11206</td><td>11225</td><td>20</td><td>CGGCGGCGACTCTGGACGCG</td><td>286.3</td><td>-7.8</td><td>1.0</td><td>-7.9</td><td>2.5E-15</td><td>1.5E-14</td><td>MB231/MB468</td><td>CYTO</td></tr>
<tr><td>>hg38_21_+_8433172_8446622_RNA45SN1_45S_50ntFlanks@11593.11611.19|CCAGGGGAATCCGACTGTT</td><td>>hg38_21_+_8433172_8446622_RNA45SN1_45S_50ntFlanks</td><td>11593</td><td>11611</td><td>19</td><td>CCAGGGGAATCCGACTGTT</td><td>4260.6</td><td>-6.9</td><td>0.9</td><td>-7.9</td><td>3.7E-15</td><td>1.5E-14</td><td>MB231/MB468</td><td>MITO</td></tr>
<tr><td>>trna111_HisGTG_1_-_147774845_147774916@-1A.23.24,trna118_HisGTG_1_-_145396881_145396952@-1A.23.24,trna16_HisGTG_1_+_146544773_146544844@-1A.23.24,trna1_HisGTG_15_+_45493349_45493420@-1A.23.24,trna21_HisGTG_1_+_147753471_147753542@-1A.23.24,trna33_HisGTG_6_+_27125906_27125977@-1A.23.24,trna7_HisGTG_9_-_14433938_14434009@-1A.23.24,trna8_HisGTG_15_-_45492611_45492682@-1A.23.24,trna9_HisGTG_15_-_45490804_45490875@-1A.23.24|AGCCGTGATCGTATAGTGGTTAGT</td><td>>trna111_HisGTG_1_-_147774845_147774916</td><td>-1A</td><td>23</td><td>24</td><td>AGCCGTGATCGTATAGTGGTTAGT</td><td>600.3</td><td>12.5</td><td>1.6</td><td>7.9</td><td>3.7E-15</td><td>1.5E-14</td><td>MB231/MB468</td><td>MITO</td></tr>
<tr><td>>hg38_21_+_8433172_8446622_RNA45SN1_45S_50ntFlanks@10905.10925.21|CTGGGCGCGCGCCGCGGCTGG</td><td>>hg38_21_+_8433172_8446622_RNA45SN1_45S_50ntFlanks</td><td>10905</td><td>10925</td><td>21</td><td>CTGGGCGCGCGCCGCGGCTGG</td><td>444.3</td><td>-12.7</td><td>1.6</td><td>-7.9</td><td>2.3E-15</td><td>1.5E-14</td><td>MB231/MB468</td><td>MP</td></tr>
<tr><td>>hg38_21_+_8433172_8446622_RNA45SN1_45S_50ntFlanks@7979.7996.18|ACCTCAGATCAGACGTGG</td><td>>hg38_21_+_8433172_8446622_RNA45SN1_45S_50ntFlanks</td><td>7979</td><td>7996</td><td>18</td><td>ACCTCAGATCAGACGTGG</td><td>252.7</td><td>-11.8</td><td>1.5</td><td>-7.9</td><td>2.3E-15</td><td>1.5E-14</td><td>MB231/MB468</td><td>MP</td></tr>
<tr><td>>MI0000299|hsa-mir-222&WithFlank&X|-|45747009|45747130@75.98.24(+1A)&[MIMAT0000279&hsa-miR-222-3p&offsets|0|+3(+1A);m-140&X|-|45747033|45747056&offsets|0|0(+1A)]|AGCTACATCTGGCTACTGGGTCTCA</td><td>>MI0000299|hsa-mir-222&WithFlank&X|-|45747009|45747130</td><td>75</td><td>98</td><td>24(+1A)</td><td>AGCTACATCTGGCTACTGGGTCTCA</td><td>1806.8</td><td>6.4</td><td>0.8</td><td>7.9</td><td>2.4E-15</td><td>1.5E-14</td><td>BT20/MB468</td><td>MITO</td></tr>
<tr><td>>hg38_21_+_8433172_8446622_RNA45SN1_45S_50ntFlanks@11189.11225.37|ACCCGGGGGGCCGGCGGCGGCGGCGACTCTGGACGCG</td><td>>hg38_21_+_8433172_8446622_RNA45SN1_45S_50ntFlanks</td><td>11189</td><td>11225</td><td>37</td><td>ACCCGGGGGGCCGGCGGCGGCGGCGACTCTGGACGCG</td><td>625.3</td><td>-12.3</td><td>1.6</td><td>-7.9</td><td>2.8E-15</td><td>1.5E-14</td><td>MB231/MB468</td><td>NUC</td></tr>
<tr><td>>trna111_HisGTG_1_-_147774845_147774916@-1T.32.33,trna118_HisGTG_1_-_145396881_145396952@-1T.32.33,trna16_HisGTG_1_+_146544773_146544844@-1T.32.33,trna1_HisGTG_15_+_45493349_45493420@-1T.32.33,trna21_HisGTG_1_+_147753471_147753542@-1T.32.33,trna33_HisGTG_6_+_27125906_27125977@-1T.32.33,trna7_HisGTG_9_-_14433938_14434009@-1T.32.33,trna8_HisGTG_15_-_45492611_45492682@-1T.32.33,trna9_HisGTG_15_-_45490804_45490875@-1T.32.33|TGCCGTGATCGTATAGTGGTTAGTACTCTGCGT</td><td>>trna111_HisGTG_1_-_147774845_147774916</td><td>-1T</td><td>32</td><td>33</td><td>TGCCGTGATCGTATAGTGGTTAGTACTCTGCGT</td><td>923.0</td><td>9.8</td><td>1.2</td><td>7.9</td><td>2.5E-15</td><td>1.5E-14</td><td>MB231/MB468</td><td>CYTO</td></tr>
<tr><td>>hg38_21_+_8433172_8446622_RNA45SN1_45S_50ntFlanks@4570.4587.18|TGGAATAGGACCGCGGTT</td><td>>hg38_21_+_8433172_8446622_RNA45SN1_45S_50ntFlanks</td><td>4570</td><td>4587</td><td>18</td><td>TGGAATAGGACCGCGGTT</td><td>248.9</td><td>-11.8</td><td>1.5</td><td>-7.9</td><td>2.3E-15</td><td>1.5E-14</td><td>MB231/MB468</td><td>MP</td></tr>
<tr><td>>hg38_21_+_8433172_8446622_RNA45SN1_45S_50ntFlanks@12418.12456.39|CGATGTCGGCTCTTCCTATCATTGTGAAGCAGAATTCAC</td><td>>hg38_21_+_8433172_8446622_RNA45SN1_45S_50ntFlanks</td><td>12418</td><td>12456</td><td>39</td><td>CGATGTCGGCTCTTCCTATCATTGTGAAGCAGAATTCAC</td><td>4874.5</td><td>-10.3</td><td>1.3</td><td>-7.8</td><td>5.0E-15</td><td>1.5E-14</td><td>BT20/MB468</td><td>NUC</td></tr>
<tr><td>>hg38_21_+_8433172_8446622_RNA45SN1_45S_50ntFlanks@12526.12549.24|TTTTACCCTACTGATGATGTGTTG</td><td>>hg38_21_+_8433172_8446622_RNA45SN1_45S_50ntFlanks</td><td>12526</td><td>12549</td><td>24</td><td>TTTTACCCTACTGATGATGTGTTG</td><td>298.8</td><td>-12.1</td><td>1.5</td><td>-7.9</td><td>2.3E-15</td><td>1.5E-14</td><td>MB231/MB468</td><td>MP</td></tr>
<tr><td>>hg38_21_+_8433172_8446622_RNA45SN1_45S_50ntFlanks@4724.4753.30|ATTAATCAAGAACGAAAGTCGGAGGTTCGA</td><td>>hg38_21_+_8433172_8446622_RNA45SN1_45S_50ntFlanks</td><td>4724</td><td>4753</td><td>30</td><td>ATTAATCAAGAACGAAAGTCGGAGGTTCGA</td><td>199.1</td><td>-11.5</td><td>1.5</td><td>-7.9</td><td>2.3E-15</td><td>1.5E-14</td><td>MB231/MB468</td><td>MP</td></tr>
<tr><td>>MI0000446|hsa-mir-125b-1&WithFlank&11|-|122099751|122099850@61.82.22&[MIMAT0004592&hsa-miR-125b-1-3p&offsets|0|0;m-485&11|-|122099769|122099790&offsets|0|0]|ACGGGTTAGGCTCTTGGGAGCT</td><td>>MI0000446|hsa-mir-125b-1&WithFlank&11|-|122099751|122099850</td><td>61</td><td>82</td><td>22</td><td>ACGGGTTAGGCTCTTGGGAGCT</td><td>423.3</td><td>11.9</td><td>1.5</td><td>7.9</td><td>3.8E-15</td><td>1.5E-14</td><td>MB231/MB468</td><td>MITO</td></tr>
<tr><td>>hg38_21_+_8433172_8446622_RNA45SN1_45S_50ntFlanks@10795.10822.28|TGGCATGTTGGAACAATGTAGGTAAGGG</td><td>>hg38_21_+_8433172_8446622_RNA45SN1_45S_50ntFlanks</td><td>10795</td><td>10822</td><td>28</td><td>TGGCATGTTGGAACAATGTAGGTAAGGG</td><td>189.7</td><td>-11.4</td><td>1.4</td><td>-7.9</td><td>2.3E-15</td><td>1.5E-14</td><td>MB231/MB468</td><td>MP</td></tr>
<tr><td>>hg38_21_+_8433172_8446622_RNA45SN1_45S_50ntFlanks@12576.12604.29|AGAGGAACCGCAGGTTCAGACATTTGGTG</td><td>>hg38_21_+_8433172_8446622_RNA45SN1_45S_50ntFlanks</td><td>12576</td><td>12604</td><td>29</td><td>AGAGGAACCGCAGGTTCAGACATTTGGTG</td><td>195.1</td><td>-11.5</td><td>1.4</td><td>-7.9</td><td>2.3E-15</td><td>1.5E-14</td><td>MB231/MB468</td><td>MP</td></tr>
<tr><td>>hg38_21_+_8433172_8446622_RNA45SN1_45S_50ntFlanks@10727.10752.26|GGAGAGGGTGTAAATCTCGCGCCGGG</td><td>>hg38_21_+_8433172_8446622_RNA45SN1_45S_50ntFlanks</td><td>10727</td><td>10752</td><td>26</td><td>GGAGAGGGTGTAAATCTCGCGCCGGG</td><td>185.7</td><td>-10.3</td><td>1.3</td><td>-7.9</td><td>2.6E-15</td><td>1.5E-14</td><td>MB231/MB468</td><td>CYTO</td></tr>
<tr><td>>hg38_21_+_8433172_8446622_RNA45SN1_45S_50ntFlanks@12008.12034.27|CCGCCGGTGAAATACCACTACTCTGAT</td><td>>hg38_21_+_8433172_8446622_RNA45SN1_45S_50ntFlanks</td><td>12008</td><td>12034</td><td>27</td><td>CCGCCGGTGAAATACCACTACTCTGAT</td><td>603.2</td><td>-8.0</td><td>1.0</td><td>-7.9</td><td>2.6E-15</td><td>1.5E-14</td><td>MB231/MB468</td><td>CYTO</td></tr>
<tr><td>>hg38_21_+_8433172_8446622_RNA45SN1_45S_50ntFlanks@11207.11233.27|GGCGGCGACTCTGGACGCGAGCCGGGC</td><td>>hg38_21_+_8433172_8446622_RNA45SN1_45S_50ntFlanks</td><td>11207</td><td>11233</td><td>27</td><td>GGCGGCGACTCTGGACGCGAGCCGGGC</td><td>223.0</td><td>-7.7</td><td>1.0</td><td>-7.9</td><td>2.6E-15</td><td>1.5E-14</td><td>MB231/MB468</td><td>CYTO</td></tr>
<tr><td>>hg38_21_+_8433172_8446622_RNA45SN1_45S_50ntFlanks@5446.5467.22|CGGATCGGCCCCGCCGGGGTCG</td><td>>hg38_21_+_8433172_8446622_RNA45SN1_45S_50ntFlanks</td><td>5446</td><td>5467</td><td>22</td><td>CGGATCGGCCCCGCCGGGGTCG</td><td>207.7</td><td>-10.7</td><td>1.3</td><td>-7.9</td><td>2.8E-15</td><td>1.5E-14</td><td>MB231/MB468</td><td>NUC</td></tr>
<tr><td>>hg38_21_+_8433172_8446622_RNA45SN1_45S_50ntFlanks@10425.10457.33|AGATGGGCGAGCGCCGTTCCGAAGGGACGGGCG</td><td>>hg38_21_+_8433172_8446622_RNA45SN1_45S_50ntFlanks</td><td>10425</td><td>10457</td><td>33</td><td>AGATGGGCGAGCGCCGTTCCGAAGGGACGGGCG</td><td>482.2</td><td>-8.2</td><td>1.0</td><td>-7.9</td><td>3.3E-15</td><td>1.5E-14</td><td>BT20/MB468</td><td>CYTO</td></tr>
<tr><td>>hg38_21_+_8433172_8446622_RNA45SN1_45S_50ntFlanks@6748.6771.24|CTTCGAACGCACTTGCGGCCCCGG</td><td>>hg38_21_+_8433172_8446622_RNA45SN1_45S_50ntFlanks</td><td>6748</td><td>6771</td><td>24</td><td>CTTCGAACGCACTTGCGGCCCCGG</td><td>130.6</td><td>-6.6</td><td>0.8</td><td>-7.9</td><td>2.6E-15</td><td>1.5E-14</td><td>MB231/MB468</td><td>CYTO</td></tr>
<tr><td>>hg38_21_+_8433172_8446622_RNA45SN1_45S_50ntFlanks@4367.4384.18|CAGTTAAAAAGCTCGTAG</td><td>>hg38_21_+_8433172_8446622_RNA45SN1_45S_50ntFlanks</td><td>4367</td><td>4384</td><td>18</td><td>CAGTTAAAAAGCTCGTAG</td><td>191.9</td><td>-11.4</td><td>1.4</td><td>-7.9</td><td>2.3E-15</td><td>1.5E-14</td><td>MB231/MB468</td><td>MP</td></tr>
<tr><td>>hg38_21_+_8433172_8446622_RNA45SN1_45S_50ntFlanks@6671.6692.22|CTCGGCTCGTGCGTCGATGAAG</td><td>>hg38_21_+_8433172_8446622_RNA45SN1_45S_50ntFlanks</td><td>6671</td><td>6692</td><td>22</td><td>CTCGGCTCGTGCGTCGATGAAG</td><td>459.5</td><td>-12.6</td><td>1.6</td><td>-7.9</td><td>3.9E-15</td><td>1.5E-14</td><td>MB231/MB468</td><td>MITO</td></tr>
<tr><td>>hg38_21_+_8433172_8446622_RNA45SN1_45S_50ntFlanks@8176.8206.31|CTTCTGATCGAGGCCCAGCCCGTGGACGGTG</td><td>>hg38_21_+_8433172_8446622_RNA45SN1_45S_50ntFlanks</td><td>8176</td><td>8206</td><td>31</td><td>CTTCTGATCGAGGCCCAGCCCGTGGACGGTG</td><td>760.0</td><td>-5.3</td><td>0.7</td><td>-7.9</td><td>3.9E-15</td><td>1.5E-14</td><td>MB231/MB468</td><td>MITO</td></tr>
<tr><td>>hg38_21_+_8433172_8446622_RNA45SN1_45S_50ntFlanks@10458.10497.40|ATGGCCTCCGTTGCCCTCGGCCGATCGAAAGGGAGTCGGG</td><td>>hg38_21_+_8433172_8446622_RNA45SN1_45S_50ntFlanks</td><td>10458</td><td>10497</td><td>40</td><td>ATGGCCTCCGTTGCCCTCGGCCGATCGAAAGGGAGTCGGG</td><td>894.4</td><td>-11.9</td><td>1.5</td><td>-7.9</td><td>2.4E-15</td><td>1.5E-14</td><td>BT20/MB468</td><td>MITO</td></tr>
<tr><td>>hg38_21_+_8433172_8446622_RNA45SN1_45S_50ntFlanks@12418.12437.20|CGATGTCGGCTCTTCCTATC</td><td>>hg38_21_+_8433172_8446622_RNA45SN1_45S_50ntFlanks</td><td>12418</td><td>12437</td><td>20</td><td>CGATGTCGGCTCTTCCTATC</td><td>10041.6</td><td>-5.6</td><td>0.7</td><td>-7.9</td><td>3.9E-15</td><td>1.5E-14</td><td>MB231/MB468</td><td>MITO</td></tr>
<tr><td>>hg38_21_+_8433172_8446622_RNA45SN1_45S_50ntFlanks@4975.5000.26|CCCGGACACGGACAGGATTGACAGAT</td><td>>hg38_21_+_8433172_8446622_RNA45SN1_45S_50ntFlanks</td><td>4975</td><td>5000</td><td>26</td><td>CCCGGACACGGACAGGATTGACAGAT</td><td>200.5</td><td>-11.5</td><td>1.5</td><td>-7.9</td><td>2.4E-15</td><td>1.5E-14</td><td>MB231/MB468</td><td>MP</td></tr>
<tr><td>>hg38_21_+_8433172_8446622_RNA45SN1_45S_50ntFlanks@12416.12448.33|TTCGATGTCGGCTCTTCCTATCATTGTGAAGCA</td><td>>hg38_21_+_8433172_8446622_RNA45SN1_45S_50ntFlanks</td><td>12416</td><td>12448</td><td>33</td><td>TTCGATGTCGGCTCTTCCTATCATTGTGAAGCA</td><td>653.4</td><td>4.3</td><td>0.5</td><td>7.9</td><td>2.5E-15</td><td>1.5E-14</td><td>BT20/MB468</td><td>MITO</td></tr>
<tr><td>>hg38_21_+_8433172_8446622_RNA45SN1_45S_50ntFlanks@4759.4779.21|ATCAGATACCGTCGTAGTTCC</td><td>>hg38_21_+_8433172_8446622_RNA45SN1_45S_50ntFlanks</td><td>4759</td><td>4779</td><td>21</td><td>ATCAGATACCGTCGTAGTTCC</td><td>519.3</td><td>-12.9</td><td>1.6</td><td>-7.9</td><td>2.4E-15</td><td>1.5E-14</td><td>MB231/MB468</td><td>MP</td></tr>
<tr><td>>hg38_21_+_8433172_8446622_RNA45SN1_45S_50ntFlanks@11783.11821.39|CATGAATGGATGAACGAGATTCCCACTGTCCCTACCTAC</td><td>>hg38_21_+_8433172_8446622_RNA45SN1_45S_50ntFlanks</td><td>11783</td><td>11821</td><td>39</td><td>CATGAATGGATGAACGAGATTCCCACTGTCCCTACCTAC</td><td>317.3</td><td>-12.1</td><td>1.5</td><td>-7.9</td><td>3.9E-15</td><td>1.5E-14</td><td>MB231/MB468</td><td>MITO</td></tr>
<tr><td>>hg38_21_+_8433172_8446622_RNA45SN1_45S_50ntFlanks@10731.10752.22|AGGGTGTAAATCTCGCGCCGGG</td><td>>hg38_21_+_8433172_8446622_RNA45SN1_45S_50ntFlanks</td><td>10731</td><td>10752</td><td>22</td><td>AGGGTGTAAATCTCGCGCCGGG</td><td>259.9</td><td>-11.8</td><td>1.5</td><td>-7.9</td><td>3.9E-15</td><td>1.5E-14</td><td>MB231/MB468</td><td>MITO</td></tr>
<tr><td>>hg38_21_+_8433172_8446622_RNA45SN1_45S_50ntFlanks@11273.11293.21|CCGCCCCCGGGGAGCCCGGCG</td><td>>hg38_21_+_8433172_8446622_RNA45SN1_45S_50ntFlanks</td><td>11273</td><td>11293</td><td>21</td><td>CCGCCCCCGGGGAGCCCGGCG</td><td>282.7</td><td>-11.9</td><td>1.5</td><td>-7.9</td><td>3.9E-15</td><td>1.5E-14</td><td>MB231/MB468</td><td>MITO</td></tr>
<tr><td>>hg38_21_+_8433172_8446622_RNA45SN1_45S_50ntFlanks@12630.12654.25|GGGCGAAGCTACCATCTGTGGGATT</td><td>>hg38_21_+_8433172_8446622_RNA45SN1_45S_50ntFlanks</td><td>12630</td><td>12654</td><td>25</td><td>GGGCGAAGCTACCATCTGTGGGATT</td><td>156.3</td><td>-4.8</td><td>0.6</td><td>-7.9</td><td>2.6E-15</td><td>1.5E-14</td><td>MB231/MB468</td><td>CYTO</td></tr>
<tr><td>>MI0000650|hsa-mir-200c&WithFlank&12|+|6963693|6963772@50.69.20(+1A)&[MIMAT0000617&hsa-miR-200c-3p&offsets|0|-3(+1A);m-54&12|+|6963742|6963764&offsets|0|-3(+1A)]|TAATACTGCCGGGTAATGATA</td><td>>MI0000650|hsa-mir-200c&WithFlank&12|+|6963693|6963772</td><td>50</td><td>69</td><td>20(+1A)</td><td>TAATACTGCCGGGTAATGATA</td><td>116.7</td><td>4.3</td><td>0.6</td><td>7.8</td><td>5.2E-15</td><td>1.5E-14</td><td>BT20/MB468</td><td>NUC</td></tr>
<tr><td>>hg38_21_+_8433172_8446622_RNA45SN1_45S_50ntFlanks@9308.9335.28|AGGAGTCTAACACGTGCGCGAGTCGGGG</td><td>>hg38_21_+_8433172_8446622_RNA45SN1_45S_50ntFlanks</td><td>9308</td><td>9335</td><td>28</td><td>AGGAGTCTAACACGTGCGCGAGTCGGGG</td><td>268.6</td><td>-11.9</td><td>1.5</td><td>-7.9</td><td>4.0E-15</td><td>1.5E-14</td><td>MB231/MB468</td><td>MITO</td></tr>
<tr><td>>hg38_21_+_8433172_8446622_RNA45SN1_45S_50ntFlanks@12932.12953.22|CGCTAAACCATTCGTAGACGAC</td><td>>hg38_21_+_8433172_8446622_RNA45SN1_45S_50ntFlanks</td><td>12932</td><td>12953</td><td>22</td><td>CGCTAAACCATTCGTAGACGAC</td><td>213.7</td><td>-11.6</td><td>1.5</td><td>-7.9</td><td>2.4E-15</td><td>1.5E-14</td><td>MB231/MB468</td><td>MP</td></tr>
<tr><td>>hg38_21_+_8433172_8446622_RNA45SN1_45S_50ntFlanks@12158.12180.23|GTTTGACTGGGGCGGTACACCTG</td><td>>hg38_21_+_8433172_8446622_RNA45SN1_45S_50ntFlanks</td><td>12158</td><td>12180</td><td>23</td><td>GTTTGACTGGGGCGGTACACCTG</td><td>306.8</td><td>-12.0</td><td>1.5</td><td>-7.9</td><td>4.0E-15</td><td>1.5E-14</td><td>MB231/MB468</td><td>MITO</td></tr>
<tr><td>>hg38_21_+_8433172_8446622_RNA45SN1_45S_50ntFlanks@12420.12455.36|ATGTCGGCTCTTCCTATCATTGTGAAGCAGAATTCA</td><td>>hg38_21_+_8433172_8446622_RNA45SN1_45S_50ntFlanks</td><td>12420</td><td>12455</td><td>36</td><td>ATGTCGGCTCTTCCTATCATTGTGAAGCAGAATTCA</td><td>9916.6</td><td>-10.9</td><td>1.4</td><td>-7.8</td><td>5.2E-15</td><td>1.5E-14</td><td>BT20/MB468</td><td>NUC</td></tr>
<tr><td>>hg38_21_+_8433172_8446622_RNA45SN1_45S_50ntFlanks@12358.12376.19|CACAGGGATAACTGGCTTG</td><td>>hg38_21_+_8433172_8446622_RNA45SN1_45S_50ntFlanks</td><td>12358</td><td>12376</td><td>19</td><td>CACAGGGATAACTGGCTTG</td><td>238.4</td><td>-11.8</td><td>1.5</td><td>-7.9</td><td>2.4E-15</td><td>1.5E-14</td><td>MB231/MB468</td><td>MP</td></tr>
</table>

<table>
<tr><td>>hg38_21_+_8433172_8446622_RNA45SN1_45S_50ntFlanks@4003.4022.20|ATAACCTCGGGCCGATCGCA</td><td>>hg38_21_+_8433172_8446622_RNA45SN1_45S_50ntFlanks</td><td>4003</td><td>4022</td><td>20</td><td>ATAACCTCGGGCCGATCGCA</td><td>2044.2</td><td>-8.7</td><td>1.1</td><td>-7.9</td><td>2.9E-15</td><td>1.5E-14</td><td>MB231/MB468</td><td>NUC</td></tr>
<tr><td>>am_MI0000809|hsa-mir-151a&WithFlank&8|-|140732558|140732659@53.70.18(+3G)&[MIMAT0000757&hsa-miR-151a-3p&offsets|0|-3(+3G);m-30&8|-|140732587|140732607&offsets|0|-3(+3G)]|CTAGACTGAAGCTCCTTGGGG</td><td>>am_MI0000809|hsa-mir-151a&WithFlank&8|-|140732558|140732659</td><td>53</td><td>70</td><td>18(+3G)</td><td>CTAGACTGAAGCTCCTTGGGG</td><td>788.7</td><td>9.2</td><td>1.2</td><td>7.9</td><td>3.4E-15</td><td>1.5E-14</td><td>BT20/MB468</td><td>CYTO</td></tr>
<tr><td>>MI0000071|hsa-mir-17&WithFlank&13|+|91350599|91350694@20.40.21&[MIMAT0000070&hsa-miR-17-5p&offsets|0|-2;m-88&13|+|91350618|91350640&offsets|0|-2]|CAAAGTGCTTACAGTGCAGGT</td><td>>MI0000071|hsa-mir-17&WithFlank&13|+|91350599|91350694</td><td>20</td><td>40</td><td>21</td><td>CAAAGTGCTTACAGTGCAGGT</td><td>900.9</td><td>5.1</td><td>0.6</td><td>7.9</td><td>2.9E-15</td><td>1.6E-14</td><td>MB231/MB468</td><td>NUC</td></tr>
<tr><td>>hg38_21_+_8433172_8446622_RNA45SN1_45S_50ntFlanks@12159.12178.20|TTTGACTGGGGCGGTACACC</td><td>>hg38_21_+_8433172_8446622_RNA45SN1_45S_50ntFlanks</td><td>12159</td><td>12178</td><td>20</td><td>TTTGACTGGGGCGGTACACC</td><td>651.0</td><td>-9.1</td><td>1.2</td><td>-7.9</td><td>4.0E-15</td><td>1.6E-14</td><td>MB231/MB468</td><td>MITO</td></tr>
<tr><td>>hg38_21_+_8433172_8446622_RNA45SN1_45S_50ntFlanks@3776.3801.26|CCGGTACAGTGAAACTGCGAATGGCT</td><td>>hg38_21_+_8433172_8446622_RNA45SN1_45S_50ntFlanks</td><td>3776</td><td>3801</td><td>26</td><td>CCGGTACAGTGAAACTGCGAATGGCT</td><td>862.9</td><td>-9.6</td><td>1.2</td><td>-7.9</td><td>4.0E-15</td><td>1.6E-14</td><td>MB231/MB468</td><td>MITO</td></tr>
<tr><td>>hg38_21_+_8433172_8446622_RNA45SN1_45S_50ntFlanks@9821.9838.18|CCTAGTGGGCCACTTTTG</td><td>>hg38_21_+_8433172_8446622_RNA45SN1_45S_50ntFlanks</td><td>9821</td><td>9838</td><td>18</td><td>CCTAGTGGGCCACTTTTG</td><td>446.5</td><td>-12.7</td><td>1.6</td><td>-7.9</td><td>2.4E-15</td><td>1.6E-14</td><td>MB231/MB468</td><td>MP</td></tr>
<tr><td>>hg38_21_+_8433172_8446622_RNA45SN1_45S_50ntFlanks@12477.12498.22|CACTAATAGGGAACGTGAGCTG</td><td>>hg38_21_+_8433172_8446622_RNA45SN1_45S_50ntFlanks</td><td>12477</td><td>12498</td><td>22</td><td>CACTAATAGGGAACGTGAGCTG</td><td>1291.3</td><td>-5.2</td><td>0.7</td><td>-7.9</td><td>2.9E-15</td><td>1.6E-14</td><td>MB231/MB468</td><td>NUC</td></tr>
<tr><td>>hg38_21_+_8433172_8446622_RNA45SN1_45S_50ntFlanks@7984.8007.24|AGATCAGACGTGGCGACCCGCTGA</td><td>>hg38_21_+_8433172_8446622_RNA45SN1_45S_50ntFlanks</td><td>7984</td><td>8007</td><td>24</td><td>AGATCAGACGTGGCGACCCGCTGA</td><td>1083.4</td><td>-9.7</td><td>1.2</td><td>-7.9</td><td>4.0E-15</td><td>1.6E-14</td><td>MB231/MB468</td><td>MITO</td></tr>
<tr><td>>hg38_21_+_8433172_8446622_RNA45SN1_45S_50ntFlanks@11575.11611.37|ACTTAGAACTGGTGCGGACCAGGGGAATCCGACTGTT</td><td>>hg38_21_+_8433172_8446622_RNA45SN1_45S_50ntFlanks</td><td>11575</td><td>11611</td><td>37</td><td>ACTTAGAACTGGTGCGGACCAGGGGAATCCGACTGTT</td><td>2668.7</td><td>-6.2</td><td>0.8</td><td>-7.9</td><td>4.0E-15</td><td>1.6E-14</td><td>MB231/MB468</td><td>MITO</td></tr>
<tr><td>>hg38_21_+_8433172_8446622_RNA45SN1_45S_50ntFlanks@11564.11594.31|CCTAGCAGCCGACTTAGAACTGGTGCGGACC</td><td>>hg38_21_+_8433172_8446622_RNA45SN1_45S_50ntFlanks</td><td>11564</td><td>11594</td><td>31</td><td>CCTAGCAGCCGACTTAGAACTGGTGCGGACC</td><td>220.7</td><td>-11.6</td><td>1.5</td><td>-7.9</td><td>2.4E-15</td><td>1.6E-14</td><td>MB231/MB468</td><td>MP</td></tr>
<tr><td>>hg38_21_+_8433172_8446622_RNA45SN1_45S_50ntFlanks@10571.10608.38|GAGAAGCCGGCGGGAGCCCCGGGGAGAGTTCTCTTTTC</td><td>>hg38_21_+_8433172_8446622_RNA45SN1_45S_50ntFlanks</td><td>10571</td><td>10608</td><td>38</td><td>GAGAAGCCGGCGGGAGCCCCGGGGAGAGTTCTCTTTTC</td><td>374.8</td><td>-11.5</td><td>1.5</td><td>-7.9</td><td>2.9E-15</td><td>1.6E-14</td><td>MB231/MB468</td><td>NUC</td></tr>
<tr><td>>hg38_21_+_8433172_8446622_RNA45SN1_45S_50ntFlanks@11233.11253.21|CCCTTCCCGTGGATCGCCCCA</td><td>>hg38_21_+_8433172_8446622_RNA45SN1_45S_50ntFlanks</td><td>11233</td><td>11253</td><td>21</td><td>CCCTTCCCGTGGATCGCCCCA</td><td>405.4</td><td>-11.6</td><td>1.5</td><td>-7.9</td><td>3.0E-15</td><td>1.6E-14</td><td>MB231/MB468</td><td>NUC</td></tr>
<tr><td>>hg38_21_+_8433172_8446622_RNA45SN1_45S_50ntFlanks@11702.11728.27|CAATGAAGCGCGGGTAAACGGCGGGAG</td><td>>hg38_21_+_8433172_8446622_RNA45SN1_45S_50ntFlanks</td><td>11702</td><td>11728</td><td>27</td><td>CAATGAAGCGCGGGTAAACGGCGGGAG</td><td>246.5</td><td>-11.7</td><td>1.5</td><td>-7.9</td><td>4.0E-15</td><td>1.6E-14</td><td>MB231/MB468</td><td>MITO</td></tr>
<tr><td>>MI0000650|hsa-mir-200c&WithFlank&12|+|6963693|6963772@50.72.23(+2U)&[MIMAT0000617&hsa-miR-200c-3p&offsets|0|0(+2U);m-54&12|+|6963742|6963764&offsets|0|0(+2U)]|TAATACTGCCGGGTAATGATGGATT</td><td>>MI0000650|hsa-mir-200c&WithFlank&12|+|6963693|6963772</td><td>50</td><td>72</td><td>23(+2U)</td><td>TAATACTGCCGGGTAATGATGGATT</td><td>545.1</td><td>9.8</td><td>1.2</td><td>8.2</td><td>2.4E-16</td><td>1.6E-14</td><td>BT20/MB231</td><td>CYTO</td></tr>
<tr><td>>hg38_21_+_8433172_8446622_RNA45SN1_45S_50ntFlanks@4917.4940.24|CACCACCAGGAGTGGAGCCTGCGG</td><td>>hg38_21_+_8433172_8446622_RNA45SN1_45S_50ntFlanks</td><td>4917</td><td>4940</td><td>24</td><td>CACCACCAGGAGTGGAGCCTGCGG</td><td>540.0</td><td>-12.9</td><td>1.6</td><td>-7.9</td><td>4.0E-15</td><td>1.6E-14</td><td>MB231/MB468</td><td>MITO</td></tr>
<tr><td>>hg38_21_+_8433172_8446622_RNA45SN1_45S_50ntFlanks@10101.10126.26|CGCGCGCGTGTGGTGTGCGTCGGAGG</td><td>>hg38_21_+_8433172_8446622_RNA45SN1_45S_50ntFlanks</td><td>10101</td><td>10126</td><td>26</td><td>CGCGCGCGTGTGGTGTGCGTCGGAGG</td><td>352.3</td><td>-11.2</td><td>1.4</td><td>-7.9</td><td>2.7E-15</td><td>1.6E-14</td><td>MB231/MB468</td><td>CYTO</td></tr>
<tr><td>>hg38_21_+_8433172_8446622_RNA45SN1_45S_50ntFlanks@12686.12708.23|CCAGGCGGAACGATACGGCAGCG</td><td>>hg38_21_+_8433172_8446622_RNA45SN1_45S_50ntFlanks</td><td>12686</td><td>12708</td><td>23</td><td>CCAGGCGGAACGATACGGCAGCG</td><td>292.1</td><td>-12.0</td><td>1.5</td><td>-7.9</td><td>4.1E-15</td><td>1.6E-14</td><td>MB231/MB468</td><td>MITO</td></tr>
<tr><td>>hg38_21_+_8433172_8446622_RNA45SN1_45S_50ntFlanks@3787.3812.26|AAACTGCGAATGGCTCATTAAATCAG</td><td>>hg38_21_+_8433172_8446622_RNA45SN1_45S_50ntFlanks</td><td>3787</td><td>3812</td><td>26</td><td>AAACTGCGAATGGCTCATTAAATCAG</td><td>575.6</td><td>-12.1</td><td>1.5</td><td>-7.9</td><td>3.0E-15</td><td>1.6E-14</td><td>MB231/MB468</td><td>NUC</td></tr>
<tr><td>>hg38_21_+_8433172_8446622_RNA45SN1_45S_50ntFlanks@11710.11728.19|CGCGGGTAAACGGCGGGAG</td><td>>hg38_21_+_8433172_8446622_RNA45SN1_45S_50ntFlanks</td><td>11710</td><td>11728</td><td>19</td><td>CGCGGGTAAACGGCGGGAG</td><td>576.4</td><td>-6.2</td><td>0.8</td><td>-7.9</td><td>2.7E-15</td><td>1.6E-14</td><td>MB231/MB468</td><td>CYTO</td></tr>
<tr><td>>hg38_21_+_8433172_8446622_RNA45SN1_45S_50ntFlanks@13004.13028.25|ATCTATTGAAAGTCAGCCCTCGACA</td><td>>hg38_21_+_8433172_8446622_RNA45SN1_45S_50ntFlanks</td><td>13004</td><td>13028</td><td>25</td><td>ATCTATTGAAAGTCAGCCCTCGACA</td><td>186.9</td><td>-11.4</td><td>1.4</td><td>-7.9</td><td>2.5E-15</td><td>1.6E-14</td><td>MB231/MB468</td><td>MP</td></tr>
<tr><td>>hg38_21_+_8433172_8446622_RNA45SN1_45S_50ntFlanks@8472.8490.19|CGGCCCGGCGGATCTTTCC</td><td>>hg38_21_+_8433172_8446622_RNA45SN1_45S_50ntFlanks</td><td>8472</td><td>8490</td><td>19</td><td>CGGCCCGGCGGATCTTTCC</td><td>608.9</td><td>-12.2</td><td>1.5</td><td>-7.9</td><td>3.0E-15</td><td>1.6E-14</td><td>MB231/MB468</td><td>NUC</td></tr>
<tr><td>>hg38_21_+_8433172_8446622_RNA45SN1_45S_50ntFlanks@10849.10868.20|CGGGATAAGGATTGGCTCTA</td><td>>hg38_21_+_8433172_8446622_RNA45SN1_45S_50ntFlanks</td><td>10849</td><td>10868</td><td>20</td><td>CGGGATAAGGATTGGCTCTA</td><td>1488.7</td><td>-10.7</td><td>1.4</td><td>-7.8</td><td>5.4E-15</td><td>1.6E-14</td><td>BT20/MB468</td><td>NUC</td></tr>
<tr><td>>hg38_21_+_8433172_8446622_RNA45SN1_45S_50ntFlanks@12405.12424.20|CTTTTTGATCCTTCGATGTC</td><td>>hg38_21_+_8433172_8446622_RNA45SN1_45S_50ntFlanks</td><td>12405</td><td>12424</td><td>20</td><td>CTTTTTGATCCTTCGATGTC</td><td>2794.9</td><td>-8.0</td><td>1.0</td><td>-7.9</td><td>2.8E-15</td><td>1.6E-14</td><td>MB231/MB468</td><td>CYTO</td></tr>
<tr><td>>hg38_21_+_8433172_8446622_RNA45SN1_45S_50ntFlanks@4823.4843.21|ATGACCCGCCGGGCAGCTTCC</td><td>>hg38_21_+_8433172_8446622_RNA45SN1_45S_50ntFlanks</td><td>4823</td><td>4843</td><td>21</td><td>ATGACCCGCCGGGCAGCTTCC</td><td>395.5</td><td>8.5</td><td>1.1</td><td>7.9</td><td>2.8E-15</td><td>1.6E-14</td><td>MB231/MB468</td><td>CYTO</td></tr>
<tr><td>>hg38_21_+_8433172_8446622_RNA45SN1_45S_50ntFlanks@10846.10865.20|CTTCGGGATAAGGATTGGCT</td><td>>hg38_21_+_8433172_8446622_RNA45SN1_45S_50ntFlanks</td><td>10846</td><td>10865</td><td>20</td><td>CTTCGGGATAAGGATTGGCT</td><td>6267.8</td><td>-6.0</td><td>0.8</td><td>-7.9</td><td>2.8E-15</td><td>1.6E-14</td><td>MB231/MB468</td><td>CYTO</td></tr>
<tr><td>>hg38_21_+_8433172_8446622_RNA45SN1_45S_50ntFlanks@10500.10533.34|CAGATCCCCGAATCCGGAGTGGCGGAGATGGGCG</td><td>>hg38_21_+_8433172_8446622_RNA45SN1_45S_50ntFlanks</td><td>10500</td><td>10533</td><td>34</td><td>CAGATCCCCGAATCCGGAGTGGCGGAGATGGGCG</td><td>236.2</td><td>-11.7</td><td>1.5</td><td>-7.9</td><td>2.5E-15</td><td>1.6E-14</td><td>MB231/MB468</td><td>MP</td></tr>
<tr><td>>hg38_21_+_8433172_8446622_RNA45SN1_45S_50ntFlanks@11316.11337.22|CACCCCACGTCTCGTCGCGCGC</td><td>>hg38_21_+_8433172_8446622_RNA45SN1_45S_50ntFlanks</td><td>11316</td><td>11337</td><td>22</td><td>CACCCCACGTCTCGTCGCGCGC</td><td>265.6</td><td>-11.8</td><td>1.5</td><td>-7.9</td><td>4.1E-15</td><td>1.6E-14</td><td>MB231/MB468</td><td>MITO</td></tr>
<tr><td>>hg38_21_+_8433172_8446622_RNA45SN1_45S_50ntFlanks@10351.10373.23|CGAGAACTTTGAAGGCCGAAGTG</td><td>>hg38_21_+_8433172_8446622_RNA45SN1_45S_50ntFlanks</td><td>10351</td><td>10373</td><td>23</td><td>CGAGAACTTTGAAGGCCGAAGTG</td><td>765.8</td><td>-3.6</td><td>0.5</td><td>-7.9</td><td>3.6E-15</td><td>1.6E-14</td><td>BT20/MB468</td><td>CYTO</td></tr>
<tr><td>>hg38_21_+_8433172_8446622_RNA45SN1_45S_50ntFlanks@12717.12742.26|CCTCGGTTGGCCTCGGATAGCCGGTC</td><td>>hg38_21_+_8433172_8446622_RNA45SN1_45S_50ntFlanks</td><td>12717</td><td>12742</td><td>26</td><td>CCTCGGTTGGCCTCGGATAGCCGGTC</td><td>1133.8</td><td>-13.2</td><td>1.6</td><td>-8.3</td><td>8.4E-17</td><td>1.6E-14</td><td>BT20/MB468</td><td>MP</td></tr>
<tr><td>>MI0001448|hsa-mir-425&WithFlank&3|-|49020142|49020240@20.44.25&[MIMAT0003393&hsa-miR-425-5p&offsets|0|+2;m-95&3|-|49020200|49020221&offsets|0|+3]|AATGACACGATCACTCCCGTTGAGT</td><td>>MI0001448|hsa-mir-425&WithFlank&3|-|49020142|49020240</td><td>20</td><td>44</td><td>25</td><td>AATGACACGATCACTCCCGTTGAGT</td><td>1924.2</td><td>6.3</td><td>0.8</td><td>7.9</td><td>2.6E-15</td><td>1.6E-14</td><td>BT20/MB468</td><td>MITO</td></tr>
<tr><td>>hg38_21_+_8433172_8446622_RNA45SN1_45S_50ntFlanks@6645.6662.18|CTCGTACGACTCTTAGCG</td><td>>hg38_21_+_8433172_8446622_RNA45SN1_45S_50ntFlanks</td><td>6645</td><td>6662</td><td>18</td><td>CTCGTACGACTCTTAGCG</td><td>12816.7</td><td>5.1</td><td>0.6</td><td>7.9</td><td>2.6E-15</td><td>1.6E-14</td><td>BT20/MB468</td><td>MITO</td></tr>
<tr><td>>MI0000103|hsa-mir-101-1&WithFlank&1|-|65058428|65058514@53.72.20;MI0000739|hsa-mir-101-2&WithFlank&9|+|4850291|4850381@55.74.20&[MIMAT0000099&hsa-miR-101-3p&offsets|0|-1;m-17&1|-|65058443|65058462&offsets|0|0];[MIMAT0000099_1&hsa-miR-101-3p&offsets|0|-1;m-16&9|+|4850345|4850364&offsets|0|0]|TACAGTACTGTGATAACTGA</td><td>>MI0000103|hsa-mir-101-1&WithFlank&1|-|65058428|65058514</td><td>53</td><td>72</td><td>20</td><td>TACAGTACTGTGATAACTGA</td><td>893.1</td><td>6.8</td><td>0.9</td><td>7.9</td><td>3.6E-15</td><td>1.6E-14</td><td>BT20/MB468</td><td>CYTO</td></tr>
<tr><td>>MI0000254|hsa-mir-30c-2&WithFlank&6|-|71376954|71377037@53.74.22&[MIMAT0004550&hsa-miR-30c-2-3p&offsets|0|0;m-206&6|-|71376964|71376985&offsets|0|0]|CTGGGAGAAGGCTGTTTACTCT</td><td>>MI0000254|hsa-mir-30c-2&WithFlank&6|-|71376954|71377037</td><td>53</td><td>74</td><td>22</td><td>CTGGGAGAAGGCTGTTTACTCT</td><td>2811.6</td><td>6.7</td><td>0.9</td><td>7.9</td><td>2.8E-15</td><td>1.6E-14</td><td>MB231/MB468</td><td>CYTO</td></tr>
<tr><td>>hg38_21_+_8433172_8446622_RNA45SN1_45S_50ntFlanks@4367.4398.32|CAGTTAAAAAGCTCGTAGTTGGATCTTGGGAG</td><td>>hg38_21_+_8433172_8446622_RNA45SN1_45S_50ntFlanks</td><td>4367</td><td>4398</td><td>32</td><td>CAGTTAAAAAGCTCGTAGTTGGATCTTGGGAG</td><td>327.5</td><td>-10.4</td><td>1.3</td><td>-7.9</td><td>2.7E-15</td><td>1.6E-14</td><td>BT20/MB468</td><td>MITO</td></tr>
<tr><td>>MI0000086|hsa-mir-28&WithFlank&3|+|188688775|188688872@60.81.22(+1U)&[MIMAT0004502&hsa-miR-28-3p&offsets|0|0(+1U);m-28&3|+|188688834|188688855&offsets|0|0(+1U)]|CACTAGATTGTGAGCTCCTGGAT</td><td>>MI0000086|hsa-mir-28&WithFlank&3|+|188688775|188688872</td><td>60</td><td>81</td><td>22(+1U)</td><td>CACTAGATTGTGAGCTCCTGGAT</td><td>107.5</td><td>3.4</td><td>0.4</td><td>7.8</td><td>5.6E-15</td><td>1.6E-14</td><td>BT20/MB468</td><td>NUC</td></tr>
<tr><td>>hg38_21_+_8433172_8446622_RNA45SN1_45S_50ntFlanks@6663.6686.24|GTGGATCACTCGGCTCGTGCGTCG</td><td>>hg38_21_+_8433172_8446622_RNA45SN1_45S_50ntFlanks</td><td>6663</td><td>6686</td><td>24</td><td>GTGGATCACTCGGCTCGTGCGTCG</td><td>1349.9</td><td>-10.0</td><td>1.3</td><td>-7.9</td><td>2.5E-15</td><td>1.6E-14</td><td>MB231/MB468</td><td>MP</td></tr>
<tr><td>>trna152_ValCAC_6_-_27248049_27248121@1.36.36|GCTTCTGTAGTGTAGTGGTTATCACGTTCGCCTCAC</td><td>>trna152_ValCAC_6_-_27248049_27248121</td><td>1</td><td>36</td><td>36</td><td>GCTTCTGTAGTGTAGTGGTTATCACGTTCGCCTCAC</td><td>791.8</td><td>7.7</td><td>1.0</td><td>7.9</td><td>3.1E-15</td><td>1.6E-14</td><td>MB231/MB468</td><td>NUC</td></tr>
<tr><td>>hg38_21_+_8433172_8446622_RNA45SN1_45S_50ntFlanks@12574.12604.31|CGAGAGGAACCGCAGGTTCAGACATTTGGTG</td><td>>hg38_21_+_8433172_8446622_RNA45SN1_45S_50ntFlanks</td><td>12574</td><td>12604</td><td>31</td><td>CGAGAGGAACCGCAGGTTCAGACATTTGGTG</td><td>663.9</td><td>-12.5</td><td>1.5</td><td>-8.3</td><td>8.7E-17</td><td>1.7E-14</td><td>BT20/MB468</td><td>MP</td></tr>
<tr><td>>hg38_21_+_8433172_8446622_RNA45SN1_45S_50ntFlanks@10786.10825.40|AACAGCCTCTGGCATGTTGGAACAATGTAGGTAAGGGAAG</td><td>>hg38_21_+_8433172_8446622_RNA45SN1_45S_50ntFlanks</td><td>10786</td><td>10825</td><td>40</td><td>AACAGCCTCTGGCATGTTGGAACAATGTAGGTAAGGGAAG</td><td>553.0</td><td>-13.0</td><td>1.6</td><td>-7.9</td><td>2.6E-15</td><td>1.7E-14</td><td>MB231/MB468</td><td>MP</td></tr>
<tr><td>>am_trna162_MetCAT_6_-_26758550_26758622@31.51.21,trna164_MetCAT_6_-_26735574_26735646@31.51.21,trna20_MetCAT_16_+_71460396_71460468@31.51.21,trna21_MetCAT_6_+_26701712_26701784@31.51.21,trna27_MetCAT_6_+_26766444_26766516@31.51.21,trna75_MetCAT_6_+_28912352_28912424@31.51.21,trna97_MetCAT_6_-_28921042_28921114@31.51.21|TCTCATAATCTGAAGGTCCTG</td><td>>am_trna162_MetCAT_6_-_26758550_26758622</td><td>31</td><td>51</td><td>21</td><td>TCTCATAATCTGAAGGTCCTG</td><td>265.3</td><td>-11.8</td><td>1.5</td><td>-7.8</td><td>4.3E-15</td><td>1.7E-14</td><td>MB231/MB468</td><td>MITO</td></tr>
<tr><td>>hg38_21_+_8433172_8446622_RNA45SN1_45S_50ntFlanks@4161.4182.22|CACATCCAAGGAAGGCAGCAGG</td><td>>hg38_21_+_8433172_8446622_RNA45SN1_45S_50ntFlanks</td><td>4161</td><td>4182</td><td>22</td><td>CACATCCAAGGAAGGCAGCAGG</td><td>187.4</td><td>-10.5</td><td>1.3</td><td>-7.9</td><td>3.1E-15</td><td>1.7E-14</td><td>MB231/MB468</td><td>NUC</td></tr>
<tr><td>>hg38_21_+_8433172_8446622_RNA45SN1_45S_50ntFlanks@8799.8823.25|CCTCGGGAGGGCGCGCGGGTCGGGG</td><td>>hg38_21_+_8433172_8446622_RNA45SN1_45S_50ntFlanks</td><td>8799</td><td>8823</td><td>25</td><td>CCTCGGGAGGGCGCGCGGGTCGGGG</td><td>390.1</td><td>-11.6</td><td>1.5</td><td>-7.9</td><td>3.1E-15</td><td>1.7E-14</td><td>MB231/MB468</td><td>NUC</td></tr>
<tr><td>>hg38_21_+_8433172_8446622_RNA45SN1_45S_50ntFlanks@11316.11337.22|CACCCCACGTCTCGTCGCGCGC</td><td>>hg38_21_+_8433172_8446622_RNA45SN1_45S_50ntFlanks</td><td>11316</td><td>11337</td><td>22</td><td>CACCCCACGTCTCGTCGCGCGC</td><td>178.6</td><td>-10.2</td><td>1.3</td><td>-7.9</td><td>2.9E-15</td><td>1.7E-14</td><td>MB231/MB468</td><td>CYTO</td></tr>
<tr><td>>hg38_21_+_8433172_8446622_RNA45SN1_45S_50ntFlanks@12539.12573.35|ATGATGTGTTGTTGCCATGGTAATCCTGCTCAGTA</td><td>>hg38_21_+_8433172_8446622_RNA45SN1_45S_50ntFlanks</td><td>12539</td><td>12573</td><td>35</td><td>ATGATGTGTTGTTGCCATGGTAATCCTGCTCAGTA</td><td>210.8</td><td>-10.4</td><td>1.3</td><td>-7.9</td><td>2.9E-15</td><td>1.7E-14</td><td>MB231/MB468</td><td>CYTO</td></tr>
<tr><td>>hg38_21_+_8433172_8446622_RNA45SN1_45S_50ntFlanks@11319.11338.20|CCCACGTCTCGTCGCGCGCG</td><td>>hg38_21_+_8433172_8446622_RNA45SN1_45S_50ntFlanks</td><td>11319</td><td>11338</td><td>20</td><td>CCCACGTCTCGTCGCGCGCG</td><td>271.4</td><td>-10.8</td><td>1.4</td><td>-7.9</td><td>2.9E-15</td><td>1.7E-14</td><td>MB231/MB468</td><td>CYTO</td></tr>
<tr><td>>hg38_21_+_8433172_8446622_RNA45SN1_45S_50ntFlanks@5188.5226.39|ATAACAGGTCTGTGATGCCCTTAGATGTCCGGGGCTGCA</td><td>>hg38_21_+_8433172_8446622_RNA45SN1_45S_50ntFlanks</td><td>5188</td><td>5226</td><td>39</td><td>ATAACAGGTCTGTGATGCCCTTAGATGTCCGGGGCTGCA</td><td>805.2</td><td>-13.4</td><td>1.7</td><td>-7.8</td><td>4.3E-15</td><td>1.7E-14</td><td>MB231/MB468</td><td>MITO</td></tr>
<tr><td>>hg38_21_+_8433172_8446622_RNA45SN1_45S_50ntFlanks@5498.5526.29|CGGTCGAACTTGACTATCTAGAGGAAGTA</td><td>>hg38_21_+_8433172_8446622_RNA45SN1_45S_50ntFlanks</td><td>5498</td><td>5526</td><td>29</td><td>CGGTCGAACTTGACTATCTAGAGGAAGTA</td><td>762.1</td><td>-13.4</td><td>1.7</td><td>-7.8</td><td>4.3E-15</td><td>1.7E-14</td><td>MB231/MB468</td><td>MITO</td></tr>
<tr><td>>hg38_21_+_8433172_8446622_RNA45SN1_45S_50ntFlanks@6773.6797.25|TTCCTCCCGGGGCTACGCCTGTCTG</td><td>>hg38_21_+_8433172_8446622_RNA45SN1_45S_50ntFlanks</td><td>6773</td><td>6797</td><td>25</td><td>TTCCTCCCGGGGCTACGCCTGTCTG</td><td>250.3</td><td>-10.7</td><td>1.4</td><td>-7.9</td><td>2.9E-15</td><td>1.7E-14</td><td>MB231/MB468</td><td>CYTO</td></tr>
<tr><td>>hg38_21_+_8433172_8446622_RNA45SN1_45S_50ntFlanks@12405.12444.40|CTTTTTGATCCTTCGATGTCGGCTCTTCCTATCATTGTGA</td><td>>hg38_21_+_8433172_8446622_RNA45SN1_45S_50ntFlanks</td><td>12405</td><td>12444</td><td>40</td><td>CTTTTTGATCCTTCGATGTCGGCTCTTCCTATCATTGTGA</td><td>6961.4</td><td>-7.1</td><td>0.9</td><td>-7.9</td><td>3.2E-15</td><td>1.7E-14</td><td>MB231/MB468</td><td>NUC</td></tr>
<tr><td>>hg38_21_+_8433172_8446622_RNA45SN1_45S_50ntFlanks@11665.11702.38|TTCTGCCCAGTGCTCTGAATGTCAAAGTGAAGAAATTC</td><td>>hg38_21_+_8433172_8446622_RNA45SN1_45S_50ntFlanks</td><td>11665</td><td>11702</td><td>38</td><td>TTCTGCCCAGTGCTCTGAATGTCAAAGTGAAGAAATTC</td><td>388.6</td><td>4.4</td><td>0.6</td><td>7.9</td><td>3.7E-15</td><td>1.7E-14</td><td>BT20/MB468</td><td>CYTO</td></tr>
</table>

<table>
<tr><td>>MI0003780|hsa-mir-1296&WithFlank&10|-|63372951|63373054@22.43.22&[MIMAT0005794&hsa-miR-1296-5p&offsets|0|0;m-304&10|-|63373012|63373033&offsets|0|0]|TTAGGGCCCTGGCTCCATCTCC</td><td>>MI0003780|hsa-mir-1296&WithFlank&10|-|63372951|63373054</td><td>22</td><td>43</td><td>22</td><td>TTAGGGCCCTGGCTCCATCTCC</td><td>69.8</td><td>4.2</td><td>0.5</td><td>7.8</td><td>5.7E-15</td><td>1.7E-14</td><td>BT20/MB468</td><td>NUC</td></tr>
<tr><td>>hg38_21_+_8433172_8446622_RNA45SN1_45S_50ntFlanks@12044.12066.23|ACTGACCCGGTGAGGCGGGGGGG</td><td>>hg38_21_+_8433172_8446622_RNA45SN1_45S_50ntFlanks</td><td>12044</td><td>12066</td><td>23</td><td>ACTGACCCGGTGAGGCGGGGGGG</td><td>412.7</td><td>-12.5</td><td>1.6</td><td>-7.8</td><td>4.3E-15</td><td>1.7E-14</td><td>MB231/MB468</td><td>MITO</td></tr>
<tr><td>>hg38_21_+_8433172_8446622_RNA45SN1_45S_50ntFlanks@12366.12383.18|TAACTGGCTTGTGGCGGC</td><td>>hg38_21_+_8433172_8446622_RNA45SN1_45S_50ntFlanks</td><td>12366</td><td>12383</td><td>18</td><td>TAACTGGCTTGTGGCGGC</td><td>511.0</td><td>-11.1</td><td>1.4</td><td>-7.9</td><td>2.7E-15</td><td>1.7E-14</td><td>BT20/MB468</td><td>MITO</td></tr>
<tr><td>>hg38_21_+_8433172_8446622_RNA45SN1_45S_50ntFlanks@12476.12497.22|CCACTAATAGGGAACGTGAGCT</td><td>>hg38_21_+_8433172_8446622_RNA45SN1_45S_50ntFlanks</td><td>12476</td><td>12497</td><td>22</td><td>CCACTAATAGGGAACGTGAGCT</td><td>1259.0</td><td>-5.2</td><td>0.7</td><td>-7.8</td><td>4.4E-15</td><td>1.7E-14</td><td>MB231/MB468</td><td>MITO</td></tr>
<tr><td>>hg38_21_+_8433172_8446622_RNA45SN1_45S_50ntFlanks@9575.9595.21|ATCGGTCGTCCGACCTGGGTA</td><td>>hg38_21_+_8433172_8446622_RNA45SN1_45S_50ntFlanks</td><td>9575</td><td>9595</td><td>21</td><td>ATCGGTCGTCCGACCTGGGTA</td><td>3366.6</td><td>-14.8</td><td>1.8</td><td>-8.3</td><td>9.0E-17</td><td>1.7E-14</td><td>BT20/MB468</td><td>MP</td></tr>
<tr><td>>hg38_1_-_228634819_228635039_RNA5S12_5S_50ntFlanks@134.173.40|TGGGAGACCGCCTGGGAATACCGGGTGCTGTAGGCTTTTT</td><td>>hg38_1_-_228634819_228635039_RNA5S12_5S_50ntFlanks</td><td>134</td><td>173</td><td>40</td><td>TGGGAGACCGCCTGGGAATACCGGGTGCTGTAGGCTTTTT</td><td>628.3</td><td>13.3</td><td>1.7</td><td>7.9</td><td>2.9E-15</td><td>1.7E-14</td><td>MB231/MB468</td><td>CYTO</td></tr>
<tr><td>>hg38_21_+_8433172_8446622_RNA45SN1_45S_50ntFlanks@11322.11347.26|ACGTCTCGTCGCGCGCGCGTCCGCTG</td><td>>hg38_21_+_8433172_8446622_RNA45SN1_45S_50ntFlanks</td><td>11322</td><td>11347</td><td>26</td><td>ACGTCTCGTCGCGCGCGCGTCCGCTG</td><td>3376.9</td><td>9.0</td><td>1.1</td><td>7.9</td><td>2.7E-15</td><td>1.7E-14</td><td>BT20/MB468</td><td>MITO</td></tr>
<tr><td>>hg38_21_+_8433172_8446622_RNA45SN1_45S_50ntFlanks@12054.12071.18|TGAGGCGGGGGGGCGAGC</td><td>>hg38_21_+_8433172_8446622_RNA45SN1_45S_50ntFlanks</td><td>12054</td><td>12071</td><td>18</td><td>TGAGGCGGGGGGGCGAGC</td><td>295.5</td><td>-12.0</td><td>1.5</td><td>-7.8</td><td>4.4E-15</td><td>1.7E-14</td><td>MB231/MB468</td><td>MITO</td></tr>
<tr><td>>hg38_21_+_8433172_8446622_RNA45SN1_45S_50ntFlanks@9548.9571.24|GAGGTCCGTAGCGGTCCTGACGTG</td><td>>hg38_21_+_8433172_8446622_RNA45SN1_45S_50ntFlanks</td><td>9548</td><td>9571</td><td>24</td><td>GAGGTCCGTAGCGGTCCTGACGTG</td><td>782.3</td><td>-4.6</td><td>0.6</td><td>-7.8</td><td>4.4E-15</td><td>1.7E-14</td><td>MB231/MB468</td><td>MITO</td></tr>
<tr><td>>hg38_21_+_8433172_8446622_RNA45SN1_45S_50ntFlanks@12072.12089.18|CCCGAGGGGCTCTCGCTT</td><td>>hg38_21_+_8433172_8446622_RNA45SN1_45S_50ntFlanks</td><td>12072</td><td>12089</td><td>18</td><td>CCCGAGGGGCTCTCGCTT</td><td>265.7</td><td>-11.8</td><td>1.5</td><td>-7.8</td><td>4.4E-15</td><td>1.7E-14</td><td>MB231/MB468</td><td>MITO</td></tr>
<tr><td>>MI0003560|hsa-mir-92b&WithFlank&1|+|155195171|155195278@67.87.21(+1A)&[MIMAT0003218&hsa-miR-92b-3p&offsets|0|-1(+1A);m-115&1|+|155195237|155195258&offsets|0|-1(+1A)]|TATTGCACTCGTCCCGGCCTCA</td><td>>MI0003560|hsa-mir-92b&WithFlank&1|+|155195171|155195278</td><td>67</td><td>87</td><td>21(+1A)</td><td>TATTGCACTCGTCCCGGCCTCA</td><td>524.4</td><td>5.4</td><td>0.7</td><td>7.9</td><td>3.2E-15</td><td>1.7E-14</td><td>MB231/MB468</td><td>NUC</td></tr>
<tr><td>>hg38_21_+_8433172_8446622_RNA45SN1_45S_50ntFlanks@12676.12703.28|AGAATCCCGCCCAGGCGGAACGATACGG</td><td>>hg38_21_+_8433172_8446622_RNA45SN1_45S_50ntFlanks</td><td>12676</td><td>12703</td><td>28</td><td>AGAATCCCGCCCAGGCGGAACGATACGG</td><td>3597.8</td><td>-10.8</td><td>1.4</td><td>-7.9</td><td>2.6E-15</td><td>1.7E-14</td><td>MB231/MB468</td><td>MP</td></tr>
<tr><td>>hg38_21_+_8433172_8446622_RNA45SN1_45S_50ntFlanks@6741.6772.32|ATCGACACTTCGAACGCACTTGCGGCCCCGGG</td><td>>hg38_21_+_8433172_8446622_RNA45SN1_45S_50ntFlanks</td><td>6741</td><td>6772</td><td>32</td><td>ATCGACACTTCGAACGCACTTGCGGCCCCGGG</td><td>528.7</td><td>-12.9</td><td>1.6</td><td>-7.9</td><td>2.6E-15</td><td>1.7E-14</td><td>MB231/MB468</td><td>MP</td></tr>
<tr><td>>hg38_21_+_8433172_8446622_RNA45SN1_45S_50ntFlanks@6646.6671.26|TCGTACGACTCTTAGCGGTGGATCAC</td><td>>hg38_21_+_8433172_8446622_RNA45SN1_45S_50ntFlanks</td><td>6646</td><td>6671</td><td>26</td><td>TCGTACGACTCTTAGCGGTGGATCAC</td><td>950.2</td><td>5.7</td><td>0.7</td><td>7.9</td><td>3.8E-15</td><td>1.7E-14</td><td>BT20/MB468</td><td>CYTO</td></tr>
<tr><td>>hg38_21_+_8433172_8446622_RNA45SN1_45S_50ntFlanks@6719.6752.34|TGAATTGCAGGACACATTGATCATCGACACTTCG</td><td>>hg38_21_+_8433172_8446622_RNA45SN1_45S_50ntFlanks</td><td>6719</td><td>6752</td><td>34</td><td>TGAATTGCAGGACACATTGATCATCGACACTTCG</td><td>580.6</td><td>-9.1</td><td>1.2</td><td>-7.8</td><td>4.4E-15</td><td>1.7E-14</td><td>MB231/MB468</td><td>MITO</td></tr>
<tr><td>>hg38_21_+_8433172_8446622_RNA45SN1_45S_50ntFlanks@12324.12345.22|TTGGGTTTTAAGCAGGAGGTGT</td><td>>hg38_21_+_8433172_8446622_RNA45SN1_45S_50ntFlanks</td><td>12324</td><td>12345</td><td>22</td><td>TTGGGTTTTAAGCAGGAGGTGT</td><td>335.0</td><td>-11.4</td><td>1.4</td><td>-7.9</td><td>3.2E-15</td><td>1.7E-14</td><td>MB231/MB468</td><td>NUC</td></tr>
<tr><td>>MI0000088|hsa-mir-30a&WithFlank&6|-|71403545|71403627@54.73.20(+2U)&[MIMAT0000088&hsa-miR-30a-3p&offsets|+1|-1(+2U);m-38&6|-|71403554|71403575&offsets|+1|-1(+2U)]|TTTCAGTCGGATGTTTGCAGTT</td><td>>MI0000088|hsa-mir-30a&WithFlank&6|-|71403545|71403627</td><td>54</td><td>73</td><td>20(+2U)</td><td>TTTCAGTCGGATGTTTGCAGTT</td><td>499.4</td><td>6.4</td><td>0.8</td><td>7.9</td><td>3.0E-15</td><td>1.7E-14</td><td>MB231/MB468</td><td>CYTO</td></tr>
<tr><td>>hg38_21_+_8433172_8446622_RNA45SN1_45S_50ntFlanks@9308.9338.31|AGGAGTCTAACACGTGCGCGAGTCGGGGGCT</td><td>>hg38_21_+_8433172_8446622_RNA45SN1_45S_50ntFlanks</td><td>9308</td><td>9338</td><td>31</td><td>AGGAGTCTAACACGTGCGCGAGTCGGGGGCT</td><td>4880.1</td><td>-10.8</td><td>1.4</td><td>-7.9</td><td>2.6E-15</td><td>1.7E-14</td><td>MB231/MB468</td><td>MP</td></tr>
<tr><td>>hg38_21_+_8433172_8446622_RNA45SN1_45S_50ntFlanks@6651.6682.32|CGACTCTTAGCGGTGGATCACTCGGCTCGTGC</td><td>>hg38_21_+_8433172_8446622_RNA45SN1_45S_50ntFlanks</td><td>6651</td><td>6682</td><td>32</td><td>CGACTCTTAGCGGTGGATCACTCGGCTCGTGC</td><td>16109.9</td><td>5.6</td><td>0.7</td><td>7.9</td><td>2.8E-15</td><td>1.7E-14</td><td>BT20/MB468</td><td>MITO</td></tr>
<tr><td>>hg38_21_+_8433172_8446622_RNA45SN1_45S_50ntFlanks@11455.11477.23|CTCGGCGCGCGGCGGCGGCGGCG</td><td>>hg38_21_+_8433172_8446622_RNA45SN1_45S_50ntFlanks</td><td>11455</td><td>11477</td><td>23</td><td>CTCGGCGCGCGGCGGCGGCGGCG</td><td>367.1</td><td>-12.4</td><td>1.6</td><td>-7.9</td><td>2.6E-15</td><td>1.7E-14</td><td>MB231/MB468</td><td>MP</td></tr>
<tr><td>>hg38_21_+_8433172_8446622_RNA45SN1_45S_50ntFlanks@12395.12414.20|AGCGACGTCGCTTTTTGATC</td><td>>hg38_21_+_8433172_8446622_RNA45SN1_45S_50ntFlanks</td><td>12395</td><td>12414</td><td>20</td><td>AGCGACGTCGCTTTTTGATC</td><td>770.7</td><td>4.0</td><td>0.5</td><td>7.9</td><td>3.2E-15</td><td>1.7E-14</td><td>MB231/MB468</td><td>NUC</td></tr>
<tr><td>>hg38_21_+_8433172_8446622_RNA45SN1_45S_50ntFlanks@10510.10533.24|AATCCGGAGTGGCGGAGATGGGCG</td><td>>hg38_21_+_8433172_8446622_RNA45SN1_45S_50ntFlanks</td><td>10510</td><td>10533</td><td>24</td><td>AATCCGGAGTGGCGGAGATGGGCG</td><td>12738.1</td><td>-11.7</td><td>1.5</td><td>-7.9</td><td>2.6E-15</td><td>1.7E-14</td><td>MB231/MB468</td><td>MP</td></tr>
<tr><td>>hg38_21_+_8433172_8446622_RNA45SN1_45S_50ntFlanks@11874.11904.31|AGAAGACCCTGTTGAGCTTGACTCTAGTCTG</td><td>>hg38_21_+_8433172_8446622_RNA45SN1_45S_50ntFlanks</td><td>11874</td><td>11904</td><td>31</td><td>AGAAGACCCTGTTGAGCTTGACTCTAGTCTG</td><td>263.0</td><td>-10.8</td><td>1.4</td><td>-7.9</td><td>3.0E-15</td><td>1.7E-14</td><td>MB231/MB468</td><td>CYTO</td></tr>
<tr><td>>hg38_21_+_8433172_8446622_RNA45SN1_45S_50ntFlanks@6651.6669.19|CGACTCTTAGCGGTGGATC</td><td>>hg38_21_+_8433172_8446622_RNA45SN1_45S_50ntFlanks</td><td>6651</td><td>6669</td><td>19</td><td>CGACTCTTAGCGGTGGATC</td><td>13002.3</td><td>3.9</td><td>0.5</td><td>7.9</td><td>3.8E-15</td><td>1.7E-14</td><td>BT20/MB468</td><td>CYTO</td></tr>
<tr><td>>hg38_21_+_8433172_8446622_RNA45SN1_45S_50ntFlanks@7975.8015.41|CGCGACCTCAGATCAGACGTGGCGACCCGCTGAATTTAAGC</td><td>>hg38_21_+_8433172_8446622_RNA45SN1_45S_50ntFlanks</td><td>7975</td><td>8015</td><td>41</td><td>CGCGACCTCAGATCAGACGTGGCGACCCGCTGAATTTAAGC</td><td>6637.1</td><td>7.5</td><td>0.9</td><td>7.9</td><td>3.8E-15</td><td>1.7E-14</td><td>BT20/MB468</td><td>CYTO</td></tr>
<tr><td>>MI0000285|hsa-mir-205&WithFlank&1|+|209432127|209432248@40.61.22(+1C)&[MIMAT0000266&hsa-miR-205-5p&offsets|0|0(+1C);m-351&1|+|209432166|209432187&offsets|0|0(+1C)]|TCCTTCATTCCACCGGAGTCTGC</td><td>>MI0000285|hsa-mir-205&WithFlank&1|+|209432127|209432248</td><td>40</td><td>61</td><td>22(+1C)</td><td>TCCTTCATTCCACCGGAGTCTGC</td><td>488.2</td><td>-12.8</td><td>1.6</td><td>-7.9</td><td>2.7E-15</td><td>1.7E-14</td><td>MB231/MB468</td><td>MP</td></tr>
<tr><td>>hg38_21_+_8433172_8446622_RNA45SN1_45S_50ntFlanks@8381.8405.25|AAGAGGGCGTGAAACCGTTAAGAGG</td><td>>hg38_21_+_8433172_8446622_RNA45SN1_45S_50ntFlanks</td><td>8381</td><td>8405</td><td>25</td><td>AAGAGGGCGTGAAACCGTTAAGAGG</td><td>3319.9</td><td>-6.8</td><td>0.9</td><td>-7.9</td><td>2.7E-15</td><td>1.7E-14</td><td>MB231/MB468</td><td>MP</td></tr>
<tr><td>>hg38_21_+_8433172_8446622_RNA45SN1_45S_50ntFlanks@11568.11591.24|GCAGCCGACTTAGAACTGGTGCGG</td><td>>hg38_21_+_8433172_8446622_RNA45SN1_45S_50ntFlanks</td><td>11568</td><td>11591</td><td>24</td><td>GCAGCCGACTTAGAACTGGTGCGG</td><td>734.4</td><td>-13.4</td><td>1.7</td><td>-7.9</td><td>2.7E-15</td><td>1.7E-14</td><td>MB231/MB468</td><td>MP</td></tr>
<tr><td>>hg38_21_+_8433172_8446622_RNA45SN1_45S_50ntFlanks@9157.9187.31|CAGTGCGCCCCGGGCGGGTCGCGCCGTCGGG</td><td>>hg38_21_+_8433172_8446622_RNA45SN1_45S_50ntFlanks</td><td>9157</td><td>9187</td><td>31</td><td>CAGTGCGCCCCGGGCGGGTCGCGCCGTCGGG</td><td>222.9</td><td>-11.7</td><td>1.5</td><td>-7.9</td><td>2.7E-15</td><td>1.7E-14</td><td>MB231/MB468</td><td>MP</td></tr>
<tr><td>>hg38_21_+_8433172_8446622_RNA45SN1_45S_50ntFlanks@3846.3871.26|CTTGGATAACTGTGGTAATTCTAGAG</td><td>>hg38_21_+_8433172_8446622_RNA45SN1_45S_50ntFlanks</td><td>3846</td><td>3871</td><td>26</td><td>CTTGGATAACTGTGGTAATTCTAGAG</td><td>173.1</td><td>-11.3</td><td>1.4</td><td>-7.9</td><td>2.7E-15</td><td>1.7E-14</td><td>MB231/MB468</td><td>MP</td></tr>
<tr><td>>hg38_21_+_8433172_8446622_RNA45SN1_45S_50ntFlanks@12712.12739.28|CGGAGCCTCGGTTGGCCTCGGATAGCCG</td><td>>hg38_21_+_8433172_8446622_RNA45SN1_45S_50ntFlanks</td><td>12712</td><td>12739</td><td>28</td><td>CGGAGCCTCGGTTGGCCTCGGATAGCCG</td><td>221.1</td><td>-11.7</td><td>1.5</td><td>-7.9</td><td>2.7E-15</td><td>1.7E-14</td><td>MB231/MB468</td><td>MP</td></tr>
<tr><td>>hg38_21_+_8433172_8446622_RNA45SN1_45S_50ntFlanks@11622.11640.19|AAAGCATCGCGAAGGCCCG</td><td>>hg38_21_+_8433172_8446622_RNA45SN1_45S_50ntFlanks</td><td>11622</td><td>11640</td><td>19</td><td>AAAGCATCGCGAAGGCCCG</td><td>1027.0</td><td>-13.1</td><td>1.6</td><td>-8.3</td><td>9.3E-17</td><td>1.7E-14</td><td>BT20/MB468</td><td>MP</td></tr>
<tr><td>>hg38_21_+_8433172_8446622_RNA45SN1_45S_50ntFlanks@11572.11593.22|CCGACTTAGAACTGGTGCGGAC</td><td>>hg38_21_+_8433172_8446622_RNA45SN1_45S_50ntFlanks</td><td>11572</td><td>11593</td><td>22</td><td>CCGACTTAGAACTGGTGCGGAC</td><td>829.8</td><td>-6.4</td><td>0.8</td><td>-7.8</td><td>4.5E-15</td><td>1.7E-14</td><td>MB231/MB468</td><td>MITO</td></tr>
<tr><td>>hg38_1_-_228634819_228635039_RNA5S12_5S_50ntFlanks@139.170.32|GACCGCCTGGGAATACCGGGTGCTGTAGGCTT</td><td>>hg38_1_-_228634819_228635039_RNA5S12_5S_50ntFlanks</td><td>139</td><td>170</td><td>32</td><td>GACCGCCTGGGAATACCGGGTGCTGTAGGCTT</td><td>803.4</td><td>4.5</td><td>0.6</td><td>7.9</td><td>2.8E-15</td><td>1.7E-14</td><td>BT20/MB468</td><td>MITO</td></tr>
<tr><td>>hg38_21_+_8433172_8446622_RNA45SN1_45S_50ntFlanks@5527.5550.24|AAAGTCGTAACAAGGTTTCCGTAG</td><td>>hg38_21_+_8433172_8446622_RNA45SN1_45S_50ntFlanks</td><td>5527</td><td>5550</td><td>24</td><td>AAAGTCGTAACAAGGTTTCCGTAG</td><td>2010.8</td><td>-9.7</td><td>1.2</td><td>-7.9</td><td>3.3E-15</td><td>1.7E-14</td><td>MB231/MB468</td><td>NUC</td></tr>
<tr><td>>hg38_21_+_8433172_8446622_RNA45SN1_45S_50ntFlanks@12419.12440.22|GATGTCGGCTCTTCCTATCATT</td><td>>hg38_21_+_8433172_8446622_RNA45SN1_45S_50ntFlanks</td><td>12419</td><td>12440</td><td>22</td><td>GATGTCGGCTCTTCCTATCATT</td><td>1789.6</td><td>5.2</td><td>0.7</td><td>7.9</td><td>3.0E-15</td><td>1.7E-14</td><td>MB231/MB468</td><td>CYTO</td></tr>
<tr><td>>hg38_21_+_8433172_8446622_RNA45SN1_45S_50ntFlanks@6732.6771.40|ACATTGATCATCGACACTTCGAACGCACTTGCGGCCCCGG</td><td>>hg38_21_+_8433172_8446622_RNA45SN1_45S_50ntFlanks</td><td>6732</td><td>6771</td><td>40</td><td>ACATTGATCATCGACACTTCGAACGCACTTGCGGCCCCGG</td><td>4609.6</td><td>-5.9</td><td>0.8</td><td>-7.8</td><td>4.5E-15</td><td>1.7E-14</td><td>MB231/MB468</td><td>MITO</td></tr>
<tr><td>>MI0000301|hsa-mir-224&WithFlank&X|-|151958572|151958664@14.34.21(+1C)&[MIMAT0000281&hsa-miR-224-5p&offsets|+1|-3(+1C);m-238&X|-|151958630|151958651&offsets|0|-1(+1C)]|CAAGTCACTAGTGGTTCCGTTC</td><td>>MI0000301|hsa-mir-224&WithFlank&X|-|151958572|151958664</td><td>14</td><td>34</td><td>21(+1C)</td><td>CAAGTCACTAGTGGTTCCGTTC</td><td>277.7</td><td>8.6</td><td>1.1</td><td>7.8</td><td>5.9E-15</td><td>1.7E-14</td><td>BT20/MB468</td><td>NUC</td></tr>
<tr><td>>hg38_21_+_8433172_8446622_RNA45SN1_45S_50ntFlanks@9252.9274.23|CACGGGGTCGGCGGCGACGTCGG</td><td>>hg38_21_+_8433172_8446622_RNA45SN1_45S_50ntFlanks</td><td>9252</td><td>9274</td><td>23</td><td>CACGGGGTCGGCGGCGACGTCGG</td><td>379.3</td><td>-12.3</td><td>1.6</td><td>-7.8</td><td>4.5E-15</td><td>1.7E-14</td><td>MB231/MB468</td><td>MITO</td></tr>
<tr><td>>hg38_21_+_8433172_8446622_RNA45SN1_45S_50ntFlanks@11603.11623.21|CCGACTGTTTAATTAAAACAA</td><td>>hg38_21_+_8433172_8446622_RNA45SN1_45S_50ntFlanks</td><td>11603</td><td>11623</td><td>21</td><td>CCGACTGTTTAATTAAAACAA</td><td>3042.5</td><td>6.1</td><td>0.8</td><td>7.9</td><td>2.8E-15</td><td>1.7E-14</td><td>BT20/MB468</td><td>MITO</td></tr>
<tr><td>>hg38_21_+_8433172_8446622_RNA45SN1_45S_50ntFlanks@10914.10939.26|CGCCGCGGCTGGACGAGGCGCCGCCG</td><td>>hg38_21_+_8433172_8446622_RNA45SN1_45S_50ntFlanks</td><td>10914</td><td>10939</td><td>26</td><td>CGCCGCGGCTGGACGAGGCGCCGCCG</td><td>245.7</td><td>-11.7</td><td>1.5</td><td>-7.8</td><td>4.5E-15</td><td>1.7E-14</td><td>MB231/MB468</td><td>MITO</td></tr>
<tr><td>>hg38_21_+_8433172_8446622_RNA45SN1_45S_50ntFlanks@11630.11660.31|GCGAAGGCCCGCGGCGGGTGTTGACGCGATG</td><td>>hg38_21_+_8433172_8446622_RNA45SN1_45S_50ntFlanks</td><td>11630</td><td>11660</td><td>31</td><td>GCGAAGGCCCGCGGCGGGTGTTGACGCGATG</td><td>365.1</td><td>-12.4</td><td>1.6</td><td>-7.9</td><td>2.7E-15</td><td>1.7E-14</td><td>MB231/MB468</td><td>MP</td></tr>
<tr><td>>hg38_21_+_8433172_8446622_RNA45SN1_45S_50ntFlanks@10727.10759.33|GGAGAGGGTGTAAATCTCGCGCCGGGCCGTACC</td><td>>hg38_21_+_8433172_8446622_RNA45SN1_45S_50ntFlanks</td><td>10727</td><td>10759</td><td>33</td><td>GGAGAGGGTGTAAATCTCGCGCCGGGCCGTACC</td><td>245.8</td><td>-10.9</td><td>1.4</td><td>-7.9</td><td>3.3E-15</td><td>1.7E-14</td><td>MB231/MB468</td><td>NUC</td></tr>
<tr><td>>MI0000084|hsa-mir-26b&WithFlank&2|+|218402640|218402728@18.38.21(+1C)&[MIMAT0000083&hsa-miR-26b-5p&offsets|0|0(+1C);m-53&2|+|218402657|218402678&offsets|0|-1(+1C)]|TTCAAGTAATTCAGGATAGGTC</td><td>>MI0000084|hsa-mir-26b&WithFlank&2|+|218402640|218402728</td><td>18</td><td>38</td><td>21(+1C)</td><td>TTCAAGTAATTCAGGATAGGTC</td><td>1689.1</td><td>6.7</td><td>0.9</td><td>7.9</td><td>3.9E-15</td><td>1.7E-14</td><td>BT20/MB468</td><td>CYTO</td></tr>
<tr><td>>hg38_21_+_8433172_8446622_RNA45SN1_45S_50ntFlanks@11574.11603.30|GACTTAGAACTGGTGCGGACCAGGGGAATC</td><td>>hg38_21_+_8433172_8446622_RNA45SN1_45S_50ntFlanks</td><td>11574</td><td>11603</td><td>30</td><td>GACTTAGAACTGGTGCGGACCAGGGGAATC</td><td>180.3</td><td>-11.4</td><td>1.4</td><td>-7.9</td><td>2.7E-15</td><td>1.8E-14</td><td>MB231/MB468</td><td>MP</td></tr>
<tr><td>>MI0000234|hsa-mir-192&WithFlank&11|-|64891131|64891252@31.53.23&[MIMAT0000222&hsa-miR-192-5p&offsets|+1|+3;m-59&11|-|64891203|64891223&offsets|+1|+3]|TGACCTATGAATTGACAGCCAGT</td><td>>MI0000234|hsa-mir-192&WithFlank&11|-|64891131|64891252</td><td>31</td><td>53</td><td>23</td><td>TGACCTATGAATTGACAGCCAGT</td><td>436.4</td><td>4.9</td><td>0.6</td><td>7.9</td><td>3.4E-15</td><td>1.8E-14</td><td>MB231/MB468</td><td>NUC</td></tr>
<tr><td>>hg38_21_+_8433172_8446622_RNA45SN1_45S_50ntFlanks@10375.10407.33|AGAAGGGTTCCATGTGAACAGCAGTTGAACATG</td><td>>hg38_21_+_8433172_8446622_RNA45SN1_45S_50ntFlanks</td><td>10375</td><td>10407</td><td>33</td><td>AGAAGGGTTCCATGTGAACAGCAGTTGAACATG</td><td>439.0</td><td>-9.1</td><td>1.2</td><td>-7.9</td><td>3.4E-15</td><td>1.8E-14</td><td>MB231/MB468</td><td>NUC</td></tr>
</table>

| | | | | | | | | | | | | | |
|---|---|---|---|---|---|---|---|---|---|---|---|---|---|
| >hg38_21_+_8433172_8446622_RNA45SN1_45S_50ntFlanks@3815.3845.31\|ATGGTTCCTTTGGTCGCTCGCTCCTCTCCTA | >hg38_21_+_8433172_8446622_RNA45SN1_45S_50ntFlanks | 3815 | 3845 | 31 | ATGGTTCCTTTGGTCGCTCGCTCCTCTCCTA | 239.3 | -10.9 | 1.4 | -7.9 | 3.4E-15 | 1.8E-14 | MB231/MB468 | NUC |
| >hg38_21_+_8433172_8446622_RNA45SN1_45S_50ntFlanks@10820.10840.21\|GGGAAGTCGGCAAGCCGGATC | >hg38_21_+_8433172_8446622_RNA45SN1_45S_50ntFlanks | 10820 | 10840 | 21 | GGGAAGTCGGCAAGCCGGATC | 3433.6 | -9.3 | 1.2 | -7.8 | 6.0E-15 | 1.8E-14 | BT20/MB468 | NUC |
| >hg38_21_+_8433172_8446622_RNA45SN1_45S_50ntFlanks@9534.9562.29\|AGGAAACTCTGGTGGAGGTCCGTAGCGGT | >hg38_21_+_8433172_8446622_RNA45SN1_45S_50ntFlanks | 9534 | 9562 | 29 | AGGAAACTCTGGTGGAGGTCCGTAGCGGT | 2155.8 | -4.5 | 0.6 | -7.9 | 3.1E-15 | 1.8E-14 | MB231/MB468 | CYTO |
| >MI0000471\|hsa-mir-126&WithFlank&9\|+\|136670596\|136670692@58.78.21&[MIMAT0000445&hsa-miR-126-3p&offsets\|0\|-1;m-40&9\|+\|136670653\|136670674&offsets\|0\|-1]\|TCGTACCGTGAGTAATAATGC | >MI0000471\|hsa-mir-126&WithFlank&9\|+\|136670596\|136670692 | 58 | 78 | 21 | TCGTACCGTGAGTAATAATGC | 862.2 | 8.1 | 1.0 | 7.9 | 3.1E-15 | 1.8E-14 | MB231/MB468 | CYTO |
| >hg38_21_+_8433172_8446622_RNA45SN1_45S_50ntFlanks@12718.12742.25\|CTCGGTTGGCCTCGGATAGCCGGTC | >hg38_21_+_8433172_8446622_RNA45SN1_45S_50ntFlanks | 12718 | 12742 | 25 | CTCGGTTGGCCTCGGATAGCCGGTC | 307.1 | -12.1 | 1.5 | -7.9 | 2.7E-15 | 1.8E-14 | MB231/MB468 | MP |
| >hg38_21_+_8433172_8446622_RNA45SN1_45S_50ntFlanks@10678.10696.19\|CGGTTCCGGCGGCGTCCGG | >hg38_21_+_8433172_8446622_RNA45SN1_45S_50ntFlanks | 10678 | 10696 | 19 | CGGTTCCGGCGGCGTCCGG | 236.7 | -10.6 | 1.3 | -7.9 | 3.1E-15 | 1.8E-14 | MB231/MB468 | CYTO |
| >hg38_21_+_8433172_8446622_RNA45SN1_45S_50ntFlanks@12710.12747.38\|CGCGGAGCCTCGGTTGGCCTCGGATAGCCGGTCCCCCG | >hg38_21_+_8433172_8446622_RNA45SN1_45S_50ntFlanks | 12710 | 12747 | 38 | CGCGGAGCCTCGGTTGGCCTCGGATAGCCGGTCCCCCG | 252.4 | -11.8 | 1.5 | -7.9 | 2.8E-15 | 1.8E-14 | MB231/MB468 | MP |
| >hg38_21_+_8433172_8446622_RNA45SN1_45S_50ntFlanks@13006.13027.22\|CTATTGAAAGTCAGCCCTCGAC | >hg38_21_+_8433172_8446622_RNA45SN1_45S_50ntFlanks | 13006 | 13027 | 22 | CTATTGAAAGTCAGCCCTCGAC | 176.7 | -11.3 | 1.4 | -7.9 | 2.8E-15 | 1.8E-14 | MB231/MB468 | MP |
| >hg38_21_+_8433172_8446622_RNA45SN1_45S_50ntFlanks@12416.12448.33\|TTCGATGTCGGCTCTTCCTATCATTGTGAAGCA | >hg38_21_+_8433172_8446622_RNA45SN1_45S_50ntFlanks | 12416 | 12448 | 33 | TTCGATGTCGGCTCTTCCTATCATTGTGAAGCA | 5705.7 | 7.1 | 0.9 | 7.9 | 3.1E-15 | 1.8E-14 | MB231/MB468 | CYTO |
| >hg38_21_+_8433172_8446622_RNA45SN1_45S_50ntFlanks@4443.4471.29\|CCCTTGCCTCTCGGCGCCCCCTCGATGCT | >hg38_21_+_8433172_8446622_RNA45SN1_45S_50ntFlanks | 4443 | 4471 | 29 | CCCTTGCCTCTCGGCGCCCCCTCGATGCT | 1528.5 | -9.0 | 1.1 | -7.9 | 3.1E-15 | 1.8E-14 | MB231/MB468 | CYTO |
| >hg38_21_+_8433172_8446622_RNA45SN1_45S_50ntFlanks@12436.12455.20\|TCATTGTGAAGCAGAATTCA | >hg38_21_+_8433172_8446622_RNA45SN1_45S_50ntFlanks | 12436 | 12455 | 20 | TCATTGTGAAGCAGAATTCA | 3919.0 | -11.2 | 1.3 | -8.3 | 9.8E-17 | 1.8E-14 | BT20/MB468 | MP |
| >MI0000098\|hsa-mir-96&WithFlank&7\|-\|129774686\|129774775@15.36.22(+1A)&[MIMAT0000095&hsa-miR-96-5p&offsets\|0\|-1(+1A);m-248&7\|-\|129774739\|129774761&offsets\|0\|-1(+1A)]\|TTTGGCACTAGCACATTTTTGCA | >MI0000098\|hsa-mir-96&WithFlank&7\|-\|129774686\|129774775 | 15 | 36 | 22(+1A) | TTTGGCACTAGCACATTTTTGCA | 720.1 | 7.4 | 0.9 | 7.9 | 4.0E-15 | 1.8E-14 | BT20/MB468 | CYTO |
| >hg38_21_+_8433172_8446622_RNA45SN1_45S_50ntFlanks@9459.9494.36\|CCGGGGAGGTGGAGCACGAGCGCACGTGTTAGGACC | >hg38_21_+_8433172_8446622_RNA45SN1_45S_50ntFlanks | 9459 | 9494 | 36 | CCGGGGAGGTGGAGCACGAGCGCACGTGTTAGGACC | 374.2 | -12.4 | 1.6 | -7.9 | 2.8E-15 | 1.8E-14 | MB231/MB468 | MP |
| >hg38_21_+_8433172_8446622_RNA45SN1_45S_50ntFlanks@12642.12661.20\|CATCTGTGGGATTATGACTG | >hg38_21_+_8433172_8446622_RNA45SN1_45S_50ntFlanks | 12642 | 12661 | 20 | CATCTGTGGGATTATGACTG | 953.7 | -9.6 | 1.2 | -7.9 | 2.8E-15 | 1.8E-14 | MB231/MB468 | MP |
| >hg38_21_+_8433172_8446622_RNA45SN1_45S_50ntFlanks@11876.11904.29\|AAGACCCTGTTGAGCTTGACTCTAGTCTG | >hg38_21_+_8433172_8446622_RNA45SN1_45S_50ntFlanks | 11876 | 11904 | 29 | AAGACCCTGTTGAGCTTGACTCTAGTCTG | 440.4 | -12.6 | 1.6 | -7.8 | 4.7E-15 | 1.8E-14 | MB231/MB468 | MITO |
| >hg38_21_+_8433172_8446622_RNA45SN1_45S_50ntFlanks@10349.10383.35\|AACGAGAACTTTGAAGGCCGAAGTGGAGAAGGGTT | >hg38_21_+_8433172_8446622_RNA45SN1_45S_50ntFlanks | 10349 | 10383 | 35 | AACGAGAACTTTGAAGGCCGAAGTGGAGAAGGGTT | 10624.5 | -5.7 | 0.7 | -7.9 | 3.4E-15 | 1.8E-14 | MB231/MB468 | NUC |
| >MI0000093\|hsa-mir-92a-1&WithFlank&13\|+\|91351308\|91351397@54.75.22(+1C);MI0000094\|hsa-mir-92a-2&WithFlank&X\|-\|134169532\|134169618@54.75.22(+1C)&[MIMAT0000092&hsa-miR-92a-3p&offsets\|0\|0(+1C);m-9&13\|+\|91351361\|91351382&offsets\|0\|0(+1C)];[MIMAT0000092_1&hsa-miR-92a-3p&offsets\|0\|0(+1C);m-11&X\|-\|134169544\|134169565&offsets\|0\|0(+1C)]\|TATTGCACTTGTCCCGGCCTGTC | >MI0000093\|hsa-mir-92a-1&WithFlank&13\|+\|91351308\|91351397 | 54 | 75 | 22(+1C) | TATTGCACTTGTCCCGGCCTGTC | 876.1 | 6.6 | 0.8 | 7.9 | 3.1E-15 | 1.8E-14 | MB231/MB468 | CYTO |
| >hg38_21_+_8433172_8446622_RNA45SN1_45S_50ntFlanks@9554.9584.31\|CGTAGCGGTCCTGACGTGCAAATCGGTCGTC | >hg38_21_+_8433172_8446622_RNA45SN1_45S_50ntFlanks | 9554 | 9584 | 31 | CGTAGCGGTCCTGACGTGCAAATCGGTCGTC | 1953.8 | -5.9 | 0.8 | -7.8 | 4.7E-15 | 1.8E-14 | MB231/MB468 | MITO |
| >hg38_21_+_8433172_8446622_RNA45SN1_45S_50ntFlanks@12957.12975.19\|CTTCTGGGTCGGGGTTTCG | >hg38_21_+_8433172_8446622_RNA45SN1_45S_50ntFlanks | 12957 | 12975 | 19 | CTTCTGGGTCGGGGTTTCG | 308.8 | -12.1 | 1.5 | -7.8 | 4.7E-15 | 1.8E-14 | MB231/MB468 | MITO |
| >hg38_21_+_8433172_8446622_RNA45SN1_45S_50ntFlanks@9203.9223.21\|CTCGGGGCCACGCGCGCGTCC | >hg38_21_+_8433172_8446622_RNA45SN1_45S_50ntFlanks | 9203 | 9223 | 21 | CTCGGGGCCACGCGCGCGTCC | 308.0 | -12.0 | 1.5 | -7.8 | 4.7E-15 | 1.8E-14 | MB231/MB468 | MITO |
| >hg38_21_+_8433172_8446622_RNA45SN1_45S_50ntFlanks@10431.10456.26\|GCGAGCGCCGTTCCGAAGGGACGGGC | >hg38_21_+_8433172_8446622_RNA45SN1_45S_50ntFlanks | 10431 | 10456 | 26 | GCGAGCGCCGTTCCGAAGGGACGGGC | 292.4 | -12.1 | 1.5 | -7.9 | 2.8E-15 | 1.8E-14 | MB231/MB468 | MP |
| >hg38_21_+_8433172_8446622_RNA45SN1_45S_50ntFlanks@12709.12745.37\|CCGCGGAGCCTCGGTTGGCCTCGGATAGCCGGTCCCC | >hg38_21_+_8433172_8446622_RNA45SN1_45S_50ntFlanks | 12709 | 12745 | 37 | CCGCGGAGCCTCGGTTGGCCTCGGATAGCCGGTCCCC | 242.3 | -11.8 | 1.5 | -7.9 | 2.8E-15 | 1.8E-14 | MB231/MB468 | MP |
| >hg38_21_+_8433172_8446622_RNA45SN1_45S_50ntFlanks@12973.12991.19\|TCGTACGTAGCAGAGCAGC | >hg38_21_+_8433172_8446622_RNA45SN1_45S_50ntFlanks | 12973 | 12991 | 19 | TCGTACGTAGCAGAGCAGC | 6270.0 | -5.5 | 0.7 | -7.9 | 3.2E-15 | 1.8E-14 | MB231/MB468 | CYTO |
| >hg38_21_+_8433172_8446622_RNA45SN1_45S_50ntFlanks@10781.10807.27\|AGGTGAACAGCCTCTGGCATGTTGGAA | >hg38_21_+_8433172_8446622_RNA45SN1_45S_50ntFlanks | 10781 | 10807 | 27 | AGGTGAACAGCCTCTGGCATGTTGGAA | 293.8 | -12.1 | 1.5 | -7.9 | 2.8E-15 | 1.8E-14 | MB231/MB468 | MP |
| >hg38_21_+_8433172_8446622_RNA45SN1_45S_50ntFlanks@11576.11601.26\|CTTAGAACTGGTGCGGACCAGGGGAA | >hg38_21_+_8433172_8446622_RNA45SN1_45S_50ntFlanks | 11576 | 11601 | 26 | CTTAGAACTGGTGCGGACCAGGGGAA | 1795.6 | -9.7 | 1.2 | -7.9 | 3.5E-15 | 1.8E-14 | MB231/MB468 | NUC |
| >hg38_21_+_8433172_8446622_RNA45SN1_45S_50ntFlanks@475.498.24\|CTTCGTGATCGATGTGGTGACGTC | >hg38_21_+_8433172_8446622_RNA45SN1_45S_50ntFlanks | 475 | 498 | 24 | CTTCGTGATCGATGTGGTGACGTC | 2312.6 | 8.6 | 1.1 | 7.9 | 3.2E-15 | 1.8E-14 | MB231/MB468 | CYTO |
| >hg38_21_+_8433172_8446622_RNA45SN1_45S_50ntFlanks@10427.10453.27\|ATGGGCGAGCGCCGTTCCGAAGGGACG | >hg38_21_+_8433172_8446622_RNA45SN1_45S_50ntFlanks | 10427 | 10453 | 27 | ATGGGCGAGCGCCGTTCCGAAGGGACG | 930.3 | -3.9 | 0.5 | -7.9 | 4.1E-15 | 1.8E-14 | BT20/MB468 | CYTO |
| >hg38_21_+_8433172_8446622_RNA45SN1_45S_50ntFlanks@12405.12428.24\|CTTTTTGATCCTTCGATGTCGGCT | >hg38_21_+_8433172_8446622_RNA45SN1_45S_50ntFlanks | 12405 | 12428 | 24 | CTTTTTGATCCTTCGATGTCGGCT | 284.1 | -12.0 | 1.5 | -7.9 | 2.9E-15 | 1.8E-14 | MB231/MB468 | MP |
| >hg38_21_+_8433172_8446622_RNA45SN1_45S_50ntFlanks@7988.8012.25\|CAGACGTGGCGACCCGCTGAATTTA | >hg38_21_+_8433172_8446622_RNA45SN1_45S_50ntFlanks | 7988 | 8012 | 25 | CAGACGTGGCGACCCGCTGAATTTA | 5494.1 | -5.0 | 0.6 | -7.9 | 3.0E-15 | 1.8E-14 | BT20/MB468 | MITO |
| >hg38_21_+_8433172_8446622_RNA45SN1_45S_50ntFlanks@5494.5514.21\|AAGACGGTCGAACTTGACTAT | >hg38_21_+_8433172_8446622_RNA45SN1_45S_50ntFlanks | 5494 | 5514 | 21 | AAGACGGTCGAACTTGACTAT | 3093.3 | -10.6 | 1.3 | -7.9 | 2.9E-15 | 1.8E-14 | MB231/MB468 | MP |
| >hg38_21_+_8433172_8446622_RNA45SN1_45S_50ntFlanks@10320.10341.22\|CAGATCTTGGTGGTAGTAGCAA | >hg38_21_+_8433172_8446622_RNA45SN1_45S_50ntFlanks | 10320 | 10341 | 22 | CAGATCTTGGTGGTAGTAGCAA | 307.1 | -12.0 | 1.5 | -7.8 | 4.8E-15 | 1.8E-14 | MB231/MB468 | MITO |
| >MI0000095\|hsa-mir-93&WithFlank&7\|-\|100093762\|100093853@17.39.23&[MIMAT0000093&hsa-miR-93-5p&offsets\|0\|0;m-29&7\|-\|100093815\|100093837&offsets\|0\|0]\|CAAAGTGCTGTTCGTGCAGGTAG | >MI0000095\|hsa-mir-93&WithFlank&7\|-\|100093762\|100093853 | 17 | 39 | 23 | CAAAGTGCTGTTCGTGCAGGTAG | 7114.1 | 5.8 | 0.7 | 7.9 | 3.0E-15 | 1.8E-14 | BT20/MB468 | MITO |
| >hg38_21_+_8433172_8446622_RNA45SN1_45S_50ntFlanks@10844.10865.22\|AACTTCGGGATAAGGATTGGCT | >hg38_21_+_8433172_8446622_RNA45SN1_45S_50ntFlanks | 10844 | 10865 | 22 | AACTTCGGGATAAGGATTGGCT | 238.2 | -11.8 | 1.5 | -7.9 | 2.9E-15 | 1.8E-14 | MB231/MB468 | MP |
| >hg38_21_+_8433172_8446622_RNA45SN1_45S_50ntFlanks@9027.9064.38\|CGTCTCCTCGTGGGGGGGCCGGGCCACCCCTCCCACGG | >hg38_21_+_8433172_8446622_RNA45SN1_45S_50ntFlanks | 9027 | 9064 | 38 | CGTCTCCTCGTGGGGGGGCCGGGCCACCCCTCCCACGG | 229.7 | -10.8 | 1.4 | -7.9 | 3.5E-15 | 1.8E-14 | MB231/MB468 | NUC |
| >hg38_21_+_8433172_8446622_RNA45SN1_45S_50ntFlanks@12942.12973.32\|TTCGTAGACGACCTGCTTCTGGGTCGGGGTTT | >hg38_21_+_8433172_8446622_RNA45SN1_45S_50ntFlanks | 12942 | 12973 | 32 | TTCGTAGACGACCTGCTTCTGGGTCGGGGTTT | 235.3 | -11.7 | 1.5 | -7.8 | 4.8E-15 | 1.8E-14 | MB231/MB468 | MITO |
| >hg38_21_+_8433172_8446622_RNA45SN1_45S_50ntFlanks@11573.11601.29\|CGACTTAGAACTGGTGCGGACCAGGGGAA | >hg38_21_+_8433172_8446622_RNA45SN1_45S_50ntFlanks | 11573 | 11601 | 29 | CGACTTAGAACTGGTGCGGACCAGGGGAA | 2269.3 | -8.5 | 1.1 | -7.9 | 2.9E-15 | 1.9E-14 | MB231/MB468 | MP |
| >hg38_21_+_8433172_8446622_RNA45SN1_45S_50ntFlanks@10806.10830.25\|AACAATGTAGGTAAGGGAAGTCGGC | >hg38_21_+_8433172_8446622_RNA45SN1_45S_50ntFlanks | 10806 | 10830 | 25 | AACAATGTAGGTAAGGGAAGTCGGC | 211.6 | -11.6 | 1.5 | -7.9 | 2.9E-15 | 1.9E-14 | MB231/MB468 | MP |
| >hg38_21_+_8433172_8446622_RNA45SN1_45S_50ntFlanks@3987.4026.40\|GCTTTGGTGACTCTAGATAACCTCGGGCCGATCGCACGCC | >hg38_21_+_8433172_8446622_RNA45SN1_45S_50ntFlanks | 3987 | 4026 | 40 | GCTTTGGTGACTCTAGATAACCTCGGGCCGATCGCACGCC | 1438.6 | -14.4 | 1.8 | -7.9 | 2.9E-15 | 1.9E-14 | MB231/MB468 | MP |
| >hg38_21_+_8433172_8446622_RNA45SN1_45S_50ntFlanks@4726.4759.34\|TAATCAAGAACGAAAGTCGGAGGTTCGAAGACGA | >hg38_21_+_8433172_8446622_RNA45SN1_45S_50ntFlanks | 4726 | 4759 | 34 | TAATCAAGAACGAAAGTCGGAGGTTCGAAGACGA | 2158.9 | -10.1 | 1.3 | -7.8 | 4.8E-15 | 1.9E-14 | MB231/MB468 | MITO |
| >MI0006444\|hsa-mir-1307&WithFlank&10\|-\|103394247\|103394407@86.109.24&[MIMAT0005951&hsa-miR-1307-3p&offsets\|0\|+2;m-66&10\|-\|103394300\|103394322&offsets\|0\|+1]\|ACTCGGCGTGGCGTCGGTCGTGGT | >MI0006444\|hsa-mir-1307&WithFlank&10\|-\|103394247\|103394407 | 86 | 109 | 24 | ACTCGGCGTGGCGTCGGTCGTGGT | 953.9 | 8.3 | 1.1 | 7.9 | 4.2E-15 | 1.9E-14 | BT20/MB468 | CYTO |
| >hg38_21_+_8433172_8446622_RNA45SN1_45S_50ntFlanks@10781.10804.24\|AGGTGAACAGCCTCTGGCATGTTG | >hg38_21_+_8433172_8446622_RNA45SN1_45S_50ntFlanks | 10781 | 10804 | 24 | AGGTGAACAGCCTCTGGCATGTTG | 259.7 | -11.8 | 1.5 | -7.8 | 4.9E-15 | 1.9E-14 | MB231/MB468 | MITO |
| >hg38_21_+_8433172_8446622_RNA45SN1_45S_50ntFlanks@8796.8825.30\|CTCCCTCGGGAGGGCGCGCGGGTCGGGGCG | >hg38_21_+_8433172_8446622_RNA45SN1_45S_50ntFlanks | 8796 | 8825 | 30 | CTCCCTCGGGAGGGCGCGCGGGTCGGGGCG | 1768.6 | 15.3 | 1.9 | 7.9 | 3.0E-15 | 1.9E-14 | BT20/MB468 | MITO |
| >MI0000070\|hsa-mir-16-1&WithFlank&13\|-\|50048967\|50049067@20.42.23;MI0000115\|hsa-mir-16-2&WithFlank&3\|+\|160404739\|160404831@16.38.23&[MIMAT0000069&hsa-miR-16-5p&offsets\|0\|+1;m-62&13\|-\|50049027\|50049048&offsets\|0\|+1];[MIMAT0000069_1&hsa-miR-16-5p&offsets\|0\|+1;m-61&3\|+\|160404754\|160404775&offsets\|0\|+1]\|TAGCAGCACGTAAATATTGGCGT | >MI0000070\|hsa-mir-16-1&WithFlank&13\|-\|50048967\|50049067 | 20 | 42 | 23 | TAGCAGCACGTAAATATTGGCGT | 1491.9 | 6.2 | 0.8 | 7.9 | 3.6E-15 | 1.9E-14 | MB231/MB468 | NUC |
| >hg38_21_+_8433172_8446622_RNA45SN1_45S_50ntFlanks@10792.10815.24\|CTCTGGCATGTTGGAACAATGTAG | >hg38_21_+_8433172_8446622_RNA45SN1_45S_50ntFlanks | 10792 | 10815 | 24 | CTCTGGCATGTTGGAACAATGTAG | 203.2 | -11.5 | 1.5 | -7.9 | 2.9E-15 | 1.9E-14 | MB231/MB468 | MP |
| >hg38_21_+_8433172_8446622_RNA45SN1_45S_50ntFlanks@11326.11345.20\|CTCGTCGCGCGCGCGTCCGC | >hg38_21_+_8433172_8446622_RNA45SN1_45S_50ntFlanks | 11326 | 11345 | 20 | CTCGTCGCGCGCGCGTCCGC | 2424.2 | 12.9 | 1.6 | 7.9 | 3.3E-15 | 1.9E-14 | MB231/MB468 | CYTO |

<table>
<tr><td>>trna116_GluCTC_1_-_145399233_145399304@1.39.39,trna59_GluCTC_1_+_249168447_249168518@1.39.39,trna71_GluCTC_1_-_161439189_161439260@1.39.39,trna74_GluCTC_1_-_161431809_161431880@1.39.39,trna77_GluCTC_1_-_161424398_161424469@1.39.39,trna77_GluCTC_6_+_28949976_28950047@1.39.39,trna80_GluCTC_1_-_161417018_161417089@1.39.39,trna87_GluCTC_6_-_126101393_126101464@1.39.39|TCCCTGGTGGTCTAGTGGTTAGGATTCGGCGCTCTCACC</td><td>>trna116_GluCTC_1_-_145399233_145399304</td><td>1</td><td>39</td><td>39</td><td>TCCCTGGTGGTCTAGTGGTTAGGATTCGGCGCTCTCACC</td><td>32399.4</td><td>7.2</td><td>0.9</td><td>7.9</td><td>3.3E-15</td><td>1.9E-14</td><td>MB231/MB468</td><td>CYTO</td></tr>
<tr><td>>hg38_21_+_8433172_8446622_RNA45SN1_45S_50ntFlanks@5423.5466.44|ATTGGATGGTTTAGTGAGGCCCTCGGATCGGCCCCGCCGGGGTC</td><td>>hg38_21_+_8433172_8446622_RNA45SN1_45S_50ntFlanks</td><td>5423</td><td>5466</td><td>44</td><td>ATTGGATGGTTTAGTGAGGCCCTCGGATCGGCCCCGCCGGGGTC</td><td>498.8</td><td>-12.7</td><td>1.6</td><td>-7.8</td><td>4.9E-15</td><td>1.9E-14</td><td>MB231/MB468</td><td>MITO</td></tr>
<tr><td>>hg38_1_-_228634819_228635039_RNA5S12_5S_50ntFlanks@131.170.40|GGATGGGAGACCGCCTGGGAATACCGGGTGCTGTAGGCTT</td><td>>hg38_1_-_228634819_228635039_RNA5S12_5S_50ntFlanks</td><td>131</td><td>170</td><td>40</td><td>GGATGGGAGACCGCCTGGGAATACCGGGTGCTGTAGGCTT</td><td>2446.4</td><td>7.5</td><td>1.0</td><td>7.9</td><td>3.1E-15</td><td>1.9E-14</td><td>BT20/MB468</td><td>MITO</td></tr>
<tr><td>>hg38_21_+_8433172_8446622_RNA45SN1_45S_50ntFlanks@6645.6673.29|CTCGTACGACTCTTAGCGGTGGATCACTC</td><td>>hg38_21_+_8433172_8446622_RNA45SN1_45S_50ntFlanks</td><td>6645</td><td>6673</td><td>29</td><td>CTCGTACGACTCTTAGCGGTGGATCACTC</td><td>1361.8</td><td>5.5</td><td>0.7</td><td>7.9</td><td>3.1E-15</td><td>1.9E-14</td><td>BT20/MB468</td><td>MITO</td></tr>
<tr><td>>hg38_21_+_8433172_8446622_RNA45SN1_45S_50ntFlanks@9352.9370.19|CCGTGGCGCAATGAAGGTG</td><td>>hg38_21_+_8433172_8446622_RNA45SN1_45S_50ntFlanks</td><td>9352</td><td>9370</td><td>19</td><td>CCGTGGCGCAATGAAGGTG</td><td>595.5</td><td>-9.0</td><td>1.1</td><td>-7.9</td><td>3.6E-15</td><td>1.9E-14</td><td>MB231/MB468</td><td>NUC</td></tr>
<tr><td>>hg38_21_+_8433172_8446622_RNA45SN1_45S_50ntFlanks@6771.6807.37|GGTTCCTCCCGGGGCTACGCCTGTCTGAGCGTCGCTT</td><td>>hg38_21_+_8433172_8446622_RNA45SN1_45S_50ntFlanks</td><td>6771</td><td>6807</td><td>37</td><td>GGTTCCTCCCGGGGCTACGCCTGTCTGAGCGTCGCTT</td><td>1133.5</td><td>-5.1</td><td>0.7</td><td>-7.8</td><td>4.9E-15</td><td>1.9E-14</td><td>MB231/MB468</td><td>MITO</td></tr>
<tr><td>>hg38_21_+_8433172_8446622_RNA45SN1_45S_50ntFlanks@8338.8375.38|GTACCGTAAGGGAAAGTTGAAAAGAACTTTGAAGAGAG</td><td>>hg38_21_+_8433172_8446622_RNA45SN1_45S_50ntFlanks</td><td>8338</td><td>8375</td><td>38</td><td>GTACCGTAAGGGAAAGTTGAAAAGAACTTTGAAGAGAG</td><td>702.7</td><td>13.2</td><td>1.6</td><td>8.3</td><td>1.1E-16</td><td>1.9E-14</td><td>BT20/MB468</td><td>MP</td></tr>
<tr><td>>hg38_21_+_8433172_8446622_RNA45SN1_45S_50ntFlanks@8107.8136.30|CGGCGGGGCGCGGGACATGTGGCGTACGGA</td><td>>hg38_21_+_8433172_8446622_RNA45SN1_45S_50ntFlanks</td><td>8107</td><td>8136</td><td>30</td><td>CGGCGGGGCGCGGGACATGTGGCGTACGGA</td><td>311.1</td><td>-12.1</td><td>1.5</td><td>-7.9</td><td>3.0E-15</td><td>1.9E-14</td><td>MB231/MB468</td><td>MP</td></tr>
<tr><td>>hg38_21_+_8433172_8446622_RNA45SN1_45S_50ntFlanks@11968.11985.18|CCCGCGAGGGGCCCGGGG</td><td>>hg38_21_+_8433172_8446622_RNA45SN1_45S_50ntFlanks</td><td>11968</td><td>11985</td><td>18</td><td>CCCGCGAGGGGCCCGGGG</td><td>250.8</td><td>-11.8</td><td>1.5</td><td>-7.8</td><td>5.0E-15</td><td>1.9E-14</td><td>MB231/MB468</td><td>MITO</td></tr>
<tr><td>>hg38_21_+_8433172_8446622_RNA45SN1_45S_50ntFlanks@475.498.24|CTTCGTGATCGATGTGGTGACGTC</td><td>>hg38_21_+_8433172_8446622_RNA45SN1_45S_50ntFlanks</td><td>475</td><td>498</td><td>24</td><td>CTTCGTGATCGATGTGGTGACGTC</td><td>21133.2</td><td>8.5</td><td>1.1</td><td>7.9</td><td>3.1E-15</td><td>1.9E-14</td><td>BT20/MB468</td><td>MITO</td></tr>
<tr><td>>hg38_21_+_8433172_8446622_RNA45SN1_45S_50ntFlanks@12958.12982.25|TTCTGGGTCGGGGTTTCGTACGTAG</td><td>>hg38_21_+_8433172_8446622_RNA45SN1_45S_50ntFlanks</td><td>12958</td><td>12982</td><td>25</td><td>TTCTGGGTCGGGGTTTCGTACGTAG</td><td>258.7</td><td>-11.0</td><td>1.4</td><td>-7.9</td><td>3.6E-15</td><td>1.9E-14</td><td>MB231/MB468</td><td>NUC</td></tr>
<tr><td>>hg38_21_+_8433172_8446622_RNA45SN1_45S_50ntFlanks@12893.12927.35|CCTCGCCCGTCACGCACCGCACGTTCGTGGGGAAC</td><td>>hg38_21_+_8433172_8446622_RNA45SN1_45S_50ntFlanks</td><td>12893</td><td>12927</td><td>35</td><td>CCTCGCCCGTCACGCACCGCACGTTCGTGGGGAAC</td><td>285.8</td><td>-11.9</td><td>1.5</td><td>-7.8</td><td>5.0E-15</td><td>1.9E-14</td><td>MB231/MB468</td><td>MITO</td></tr>
<tr><td>>hg38_21_+_8433172_8446622_RNA45SN1_45S_50ntFlanks@12321.12346.26|CTTTTGGGTTTTAAGCAGGAGGTGTC</td><td>>hg38_21_+_8433172_8446622_RNA45SN1_45S_50ntFlanks</td><td>12321</td><td>12346</td><td>26</td><td>CTTTTGGGTTTTAAGCAGGAGGTGTC</td><td>171.4</td><td>-11.3</td><td>1.4</td><td>-7.9</td><td>3.0E-15</td><td>1.9E-14</td><td>MB231/MB468</td><td>MP</td></tr>
<tr><td>>hg38_21_+_8433172_8446622_RNA45SN1_45S_50ntFlanks@9454.9494.41|CCGCGCCGGGGAGGTGGAGCACGAGCGCACGTGTTAGGACC</td><td>>hg38_21_+_8433172_8446622_RNA45SN1_45S_50ntFlanks</td><td>9454</td><td>9494</td><td>41</td><td>CCGCGCCGGGGAGGTGGAGCACGAGCGCACGTGTTAGGACC</td><td>524.1</td><td>-12.9</td><td>1.6</td><td>-7.9</td><td>3.0E-15</td><td>1.9E-14</td><td>MB231/MB468</td><td>MP</td></tr>
<tr><td>>hg38_21_+_8433172_8446622_RNA45SN1_45S_50ntFlanks@7977.7994.18|CGACCTCAGATCAGACGT</td><td>>hg38_21_+_8433172_8446622_RNA45SN1_45S_50ntFlanks</td><td>7977</td><td>7994</td><td>18</td><td>CGACCTCAGATCAGACGT</td><td>542.1</td><td>-4.0</td><td>0.5</td><td>-7.8</td><td>6.5E-15</td><td>1.9E-14</td><td>BT20/MB468</td><td>NUC</td></tr>
<tr><td>>MI0000434|hsa-let-7i&WithFlank&12|+|62603680|62603775@12.33.22(+1A)&[MIMAT0000415&hsa-let-7i-5p&offsets|0|0(+1A);m-49&12|+|62603691|62603712&offsets|0|0(+1A)]|TGAGGTAGTAGTTTGTGCTGTTA</td><td>>MI0000434|hsa-let-7i&WithFlank&12|+|62603680|62603775</td><td>12</td><td>33</td><td>22(+1A)</td><td>TGAGGTAGTAGTTTGTGCTGTTA</td><td>5395.9</td><td>5.3</td><td>0.7</td><td>7.9</td><td>3.7E-15</td><td>1.9E-14</td><td>MB231/MB468</td><td>NUC</td></tr>
<tr><td>>MI0000266|hsa-mir-10a&WithFlank&17|-|48579832|48579953@29.49.21(+1A)&[MIMAT0000253&hsa-miR-10a-5p&offsets|+1|-1(+1A);m-18&17|-|48579905|48579926&offsets|+1|0(+1A)]|ACCCTGTAGATCCGAATTTGTA</td><td>>MI0000266|hsa-mir-10a&WithFlank&17|-|48579832|48579953</td><td>29</td><td>49</td><td>21(+1A)</td><td>ACCCTGTAGATCCGAATTTGTA</td><td>268.9</td><td>11.3</td><td>1.4</td><td>7.9</td><td>3.0E-15</td><td>1.9E-14</td><td>MB231/MB468</td><td>MP</td></tr>
<tr><td>>hg38_21_+_8433172_8446622_RNA45SN1_45S_50ntFlanks@11845.11868.24|GGAACGGGCTTGGCGGAATCAGCG</td><td>>hg38_21_+_8433172_8446622_RNA45SN1_45S_50ntFlanks</td><td>11845</td><td>11868</td><td>24</td><td>GGAACGGGCTTGGCGGAATCAGCG</td><td>218.0</td><td>-11.6</td><td>1.5</td><td>-7.9</td><td>3.0E-15</td><td>1.9E-14</td><td>MB231/MB468</td><td>MP</td></tr>
<tr><td>>hg38_21_+_8433172_8446622_RNA45SN1_45S_50ntFlanks@11792.11820.29|ATGAACGAGATTCCCACTGTCCCTACCTA</td><td>>hg38_21_+_8433172_8446622_RNA45SN1_45S_50ntFlanks</td><td>11792</td><td>11820</td><td>29</td><td>ATGAACGAGATTCCCACTGTCCCTACCTA</td><td>231.2</td><td>-10.8</td><td>1.4</td><td>-7.9</td><td>3.7E-15</td><td>1.9E-14</td><td>MB231/MB468</td><td>NUC</td></tr>
<tr><td>>hg38_21_+_8433172_8446622_RNA45SN1_45S_50ntFlanks@12960.12990.31|CTGGGTCGGGGTTTCGTACGTAGCAGAGCAG</td><td>>hg38_21_+_8433172_8446622_RNA45SN1_45S_50ntFlanks</td><td>12960</td><td>12990</td><td>31</td><td>CTGGGTCGGGGTTTCGTACGTAGCAGAGCAG</td><td>1079.9</td><td>-10.3</td><td>1.3</td><td>-7.8</td><td>6.5E-15</td><td>1.9E-14</td><td>BT20/MB468</td><td>NUC</td></tr>
<tr><td>>MI0000299|hsa-mir-222&WithFlank&X|-|45747009|45747130@75.99.25&[MIMAT0000279&hsa-miR-222-3p&offsets|0|+4;m-140&X|-|45747033|45747056&offsets|0|+1]|AGCTACATCTGGCTACTGGGTCTCT</td><td>>MI0000299|hsa-mir-222&WithFlank&X|-|45747009|45747130</td><td>75</td><td>99</td><td>25</td><td>AGCTACATCTGGCTACTGGGTCTCT</td><td>34475.5</td><td>6.8</td><td>0.9</td><td>7.8</td><td>5.0E-15</td><td>1.9E-14</td><td>MB231/MB468</td><td>MITO</td></tr>
<tr><td>>MI0006444|hsa-mir-1307&WithFlank&10|-|103394247|103394407@87.107.21(+1U)&[MIMAT0005951&hsa-miR-1307-3p&offsets|+1|0(+1U);m-66&10|-|103394300|103394322&offsets|+1|-1(+1U)]|CTCGGCGTGGCGTCGGTCGTGT</td><td>>MI0006444|hsa-mir-1307&WithFlank&10|-|103394247|103394407</td><td>87</td><td>107</td><td>21(+1U)</td><td>CTCGGCGTGGCGTCGGTCGTGT</td><td>351.6</td><td>5.4</td><td>0.7</td><td>7.9</td><td>3.7E-15</td><td>1.9E-14</td><td>MB231/MB468</td><td>NUC</td></tr>
<tr><td>>MI0000750|hsa-mir-26a-2&WithFlank&12|-|57824603|57824698@20.41.22(+1U);MI0000083|hsa-mir-26a-1&WithFlank&3|+|37969398|37969486@16.37.22(+1U)&[MIMAT0000082&hsa-miR-26a-5p&offsets|0|0(+1U);m-33&12|-|57824658|57824679&offsets|0|0(+1U)];[MIMAT0000082_1&hsa-miR-26a-5p&offsets|0|0(+1U);m-34&3|+|37969413|37969434&offsets|0|0(+1U)]|TTCAAGTAATCCAGGATAGGCTT</td><td>>MI0000750|hsa-mir-26a-2&WithFlank&12|-|57824603|57824698</td><td>20</td><td>41</td><td>22(+1U)</td><td>TTCAAGTAATCCAGGATAGGCTT</td><td>3750.4</td><td>6.1</td><td>0.8</td><td>7.9</td><td>3.7E-15</td><td>1.9E-14</td><td>MB231/MB468</td><td>NUC</td></tr>
<tr><td>>hg38_21_+_8433172_8446622_RNA45SN1_45S_50ntFlanks@10912.10932.21|CGCGCCGCGGCTGGACGAGGC</td><td>>hg38_21_+_8433172_8446622_RNA45SN1_45S_50ntFlanks</td><td>10912</td><td>10932</td><td>21</td><td>CGCGCCGCGGCTGGACGAGGC</td><td>240.4</td><td>-11.7</td><td>1.5</td><td>-7.8</td><td>5.1E-15</td><td>1.9E-14</td><td>MB231/MB468</td><td>MITO</td></tr>
<tr><td>>hg38_21_+_8433172_8446622_RNA45SN1_45S_50ntFlanks@8336.8375.40|AAGTACCGTAAGGGAAAGTTGAAAAGAACTTTGAAGAGAG</td><td>>hg38_21_+_8433172_8446622_RNA45SN1_45S_50ntFlanks</td><td>8336</td><td>8375</td><td>40</td><td>AAGTACCGTAAGGGAAAGTTGAAAAGAACTTTGAAGAGAG</td><td>660.1</td><td>13.1</td><td>1.6</td><td>8.3</td><td>1.1E-16</td><td>1.9E-14</td><td>BT20/MB468</td><td>MP</td></tr>
<tr><td>>hg38_21_+_8433172_8446622_RNA45SN1_45S_50ntFlanks@13008.13028.21|ATTGAAAGTCAGCCCTCGACA</td><td>>hg38_21_+_8433172_8446622_RNA45SN1_45S_50ntFlanks</td><td>13008</td><td>13028</td><td>21</td><td>ATTGAAAGTCAGCCCTCGACA</td><td>289.6</td><td>-12.0</td><td>1.5</td><td>-7.9</td><td>3.0E-15</td><td>1.9E-14</td><td>MB231/MB468</td><td>MP</td></tr>
<tr><td>>MI0000283|hsa-mir-203a&WithFlank&14|+|104117399|104117520@71.90.20&[MIMAT0000264&hsa-miR-203a-3p&offsets|0|-2;m-79&14|+|104117469|104117490&offsets|0|-2]|GTGAAATGTTTAGGACCACT</td><td>>MI0000283|hsa-mir-203a&WithFlank&14|+|104117399|104117520</td><td>71</td><td>90</td><td>20</td><td>GTGAAATGTTTAGGACCACT</td><td>210.0</td><td>11.3</td><td>1.4</td><td>8.1</td><td>4.2E-16</td><td>1.9E-14</td><td>BT20/MB231</td><td>MITO</td></tr>
<tr><td>>MI0000283|hsa-mir-203a&WithFlank&14|+|104117399|104117520@72.90.19&[MIMAT0000264&hsa-miR-203a-3p&offsets|+1|-2;m-79&14|+|104117469|104117490&offsets|+1|-2]|TGAAATGTTTAGGACCACT</td><td>>MI0000283|hsa-mir-203a&WithFlank&14|+|104117399|104117520</td><td>72</td><td>90</td><td>19</td><td>TGAAATGTTTAGGACCACT</td><td>211.3</td><td>11.3</td><td>1.4</td><td>8.1</td><td>4.1E-16</td><td>1.9E-14</td><td>BT20/MB231</td><td>MITO</td></tr>
<tr><td>>MI0000294|hsa-mir-218-1&WithFlank&4|+|20528269|20528390@31.51.21;MI0000295|hsa-mir-218-2&WithFlank&5|-|168768140|168768261@31.51.21&[MIMAT0000275&hsa-miR-218-5p&offsets|0|0;m-185&4|+|20528299|20528319&offsets|0|0];[MIMAT0000275_1&hsa-miR-218-5p&offsets|0|0;m-187&5|-|168768211|168768231&offsets|0|0]|TTGTGCTTGATCTAACCATGT</td><td>>MI0000294|hsa-mir-218-1&WithFlank&4|+|20528269|20528390</td><td>31</td><td>51</td><td>21</td><td>TTGTGCTTGATCTAACCATGT</td><td>415.2</td><td>-11.9</td><td>1.5</td><td>-8.1</td><td>4.2E-16</td><td>1.9E-14</td><td>BT20/MB231</td><td>MITO</td></tr>
<tr><td>>hg38_21_+_8433172_8446622_RNA45SN1_45S_50ntFlanks@10458.10498.41|ATGGCCTCCGTTGCCCTCGGCCGATCGAAAGGGAGTCGGGT</td><td>>hg38_21_+_8433172_8446622_RNA45SN1_45S_50ntFlanks</td><td>10458</td><td>10498</td><td>41</td><td>ATGGCCTCCGTTGCCCTCGGCCGATCGAAAGGGAGTCGGGT</td><td>318.1</td><td>-12.1</td><td>1.5</td><td>-7.8</td><td>5.1E-15</td><td>1.9E-14</td><td>MB231/MB468</td><td>MITO</td></tr>
<tr><td>>hg38_21_+_8433172_8446622_RNA45SN1_45S_50ntFlanks@10537.10564.28|CGAGGCGTCCAGTGCGGTAACGCGACCG</td><td>>hg38_21_+_8433172_8446622_RNA45SN1_45S_50ntFlanks</td><td>10537</td><td>10564</td><td>28</td><td>CGAGGCGTCCAGTGCGGTAACGCGACCG</td><td>234.8</td><td>-11.7</td><td>1.5</td><td>-7.9</td><td>3.1E-15</td><td>1.9E-14</td><td>MB231/MB468</td><td>MP</td></tr>
<tr><td>>hg38_21_+_8433172_8446622_RNA45SN1_45S_50ntFlanks@4047.4070.24|ATTCGAACGTCTGCCCTATCAACT</td><td>>hg38_21_+_8433172_8446622_RNA45SN1_45S_50ntFlanks</td><td>4047</td><td>4070</td><td>24</td><td>ATTCGAACGTCTGCCCTATCAACT</td><td>163.1</td><td>4.1</td><td>0.5</td><td>7.9</td><td>3.4E-15</td><td>1.9E-14</td><td>MB231/MB468</td><td>CYTO</td></tr>
<tr><td>>hg38_21_+_8433172_8446622_RNA45SN1_45S_50ntFlanks@476.509.34|TTCGTGATCGATGTGGTGACGTCGTGCTCTCCCG</td><td>>hg38_21_+_8433172_8446622_RNA45SN1_45S_50ntFlanks</td><td>476</td><td>509</td><td>34</td><td>TTCGTGATCGATGTGGTGACGTCGTGCTCTCCCG</td><td>546.9</td><td>7.7</td><td>1.0</td><td>7.9</td><td>3.4E-15</td><td>1.9E-14</td><td>MB231/MB468</td><td>CYTO</td></tr>
<tr><td>>hg38_21_+_8433172_8446622_RNA45SN1_45S_50ntFlanks@10781.10807.27|AGGTGAACAGCCTCTGGCATGTTGGAA</td><td>>hg38_21_+_8433172_8446622_RNA45SN1_45S_50ntFlanks</td><td>10781</td><td>10807</td><td>27</td><td>AGGTGAACAGCCTCTGGCATGTTGGAA</td><td>257.7</td><td>-11.8</td><td>1.5</td><td>-7.8</td><td>5.1E-15</td><td>2.0E-14</td><td>MB231/MB468</td><td>MITO</td></tr>
<tr><td>>hg38_21_+_8433172_8446622_RNA45SN1_45S_50ntFlanks@5471.5499.29|CACGGCCCTGGCGGAGCGCTGAGAAGACG</td><td>>hg38_21_+_8433172_8446622_RNA45SN1_45S_50ntFlanks</td><td>5471</td><td>5499</td><td>29</td><td>CACGGCCCTGGCGGAGCGCTGAGAAGACG</td><td>2824.2</td><td>-5.6</td><td>0.7</td><td>-7.9</td><td>3.8E-15</td><td>2.0E-14</td><td>MB231/MB468</td><td>NUC</td></tr>
<tr><td>>hg38_21_+_8433172_8446622_RNA45SN1_45S_50ntFlanks@10404.10435.32|CATGGGTCAGTCGGTCCTGAGAGATGGGCGAG</td><td>>hg38_21_+_8433172_8446622_RNA45SN1_45S_50ntFlanks</td><td>10404</td><td>10435</td><td>32</td><td>CATGGGTCAGTCGGTCCTGAGAGATGGGCGAG</td><td>205.9</td><td>-5.7</td><td>0.7</td><td>-7.9</td><td>3.4E-15</td><td>2.0E-14</td><td>MB231/MB468</td><td>CYTO</td></tr>
<tr><td>>trnaMT_GluTTC_MT_-_14674_14742@26.61.36,trnalookalike8_GluTTC_5_-_93905172_93905240@26.61.36|GGTTTTTCATATCATTGGTCGTGGTTGTAGTCCGTG</td><td>>trnaMT_GluTTC_MT_-_14674_14742</td><td>26</td><td>61</td><td>36</td><td>GGTTTTTCATATCATTGGTCGTGGTTGTAGTCCGTG</td><td>954.0</td><td>7.2</td><td>0.9</td><td>7.9</td><td>3.2E-15</td><td>2.0E-14</td><td>BT20/MB468</td><td>MITO</td></tr>
<tr><td>>MI0000471|hsa-mir-126&WithFlank&9|+|136670596|136670692@58.77.20(+1U)&[MIMAT0000445&hsa-miR-126-3p&offsets|0|-2(+1U);m-40&9|+|136670653|136670674&offsets|0|-2(+1U)]|TCGTACCGTGAGTAATAATGT</td><td>>MI0000471|hsa-mir-126&WithFlank&9|+|136670596|136670692</td><td>58</td><td>77</td><td>20(+1U)</td><td>TCGTACCGTGAGTAATAATGT</td><td>558.3</td><td>12.3</td><td>1.6</td><td>7.8</td><td>5.2E-15</td><td>2.0E-14</td><td>MB231/MB468</td><td>MITO</td></tr>
<tr><td>>hg38_21_+_8433172_8446622_RNA45SN1_45S_50ntFlanks@11581.11603.23|AACTGGTGCGGACCAGGGGAATC</td><td>>hg38_21_+_8433172_8446622_RNA45SN1_45S_50ntFlanks</td><td>11581</td><td>11603</td><td>23</td><td>AACTGGTGCGGACCAGGGGAATC</td><td>2005.5</td><td>-4.4</td><td>0.6</td><td>-7.9</td><td>3.2E-15</td><td>2.0E-14</td><td>BT20/MB468</td><td>MITO</td></tr>
<tr><td>>hg38_21_+_8433172_8446622_RNA45SN1_45S_50ntFlanks@10362.10391.30|AAGGCCGAAGTGGAGAAGGGTTCCATGTGA</td><td>>hg38_21_+_8433172_8446622_RNA45SN1_45S_50ntFlanks</td><td>10362</td><td>10391</td><td>30</td><td>AAGGCCGAAGTGGAGAAGGGTTCCATGTGA</td><td>220.4</td><td>-11.6</td><td>1.5</td><td>-7.9</td><td>3.1E-15</td><td>2.0E-14</td><td>MB231/MB468</td><td>MP</td></tr>
<tr><td>>MI0000734|hsa-mir-106b&WithFlank&7|-|100093987|100094080@18.39.22&[MIMAT0000680&hsa-miR-106b-5p&offsets|0|+1;m-103&7|-|100094043|100094063&offsets|0|+1]|TAAAGTGCTGACAGTGCAGATA</td><td>>MI0000734|hsa-mir-106b&WithFlank&7|-|100093987|100094080</td><td>18</td><td>39</td><td>22</td><td>TAAAGTGCTGACAGTGCAGATA</td><td>481.6</td><td>4.8</td><td>0.6</td><td>7.9</td><td>3.2E-15</td><td>2.0E-14</td><td>BT20/MB468</td><td>MITO</td></tr>
<tr><td>>hg38_21_+_8433172_8446622_RNA45SN1_45S_50ntFlanks@3593.3610.18|CCCGTCCGTCCGTCCGTC</td><td>>hg38_21_+_8433172_8446622_RNA45SN1_45S_50ntFlanks</td><td>3593</td><td>3610</td><td>18</td><td>CCCGTCCGTCCGTCCGTC</td><td>381.4</td><td>-12.4</td><td>1.6</td><td>-7.9</td><td>3.1E-15</td><td>2.0E-14</td><td>MB231/MB468</td><td>MP</td></tr>
<tr><td>>hg38_21_+_8433172_8446622_RNA45SN1_45S_50ntFlanks@8737.8754.18|CCCGTCCGTCCGTCCGTC</td><td>>hg38_21_+_8433172_8446622_RNA45SN1_45S_50ntFlanks</td><td>8737</td><td>8754</td><td>18</td><td>CCCGTCCGTCCGTCCGTC</td><td>381.4</td><td>-12.4</td><td>1.6</td><td>-7.9</td><td>3.1E-15</td><td>2.0E-14</td><td>MB231/MB468</td><td>MP</td></tr>
<tr><td>>hg38_21_+_8433172_8446622_RNA45SN1_45S_50ntFlanks@9031.9058.28|TCCTCGTGGGGGGGCCGGGCCACCCCTC</td><td>>hg38_21_+_8433172_8446622_RNA45SN1_45S_50ntFlanks</td><td>9031</td><td>9058</td><td>28</td><td>TCCTCGTGGGGGGGCCGGGCCACCCCTC</td><td>266.4</td><td>-11.8</td><td>1.5</td><td>-7.8</td><td>5.2E-15</td><td>2.0E-14</td><td>MB231/MB468</td><td>MITO</td></tr>
</table>

| | | | | | | | | | | | | | |
|---|---|---|---|---|---|---|---|---|---|---|---|---|---|
| >hg38_21_+_8433172_8446622_RNA45SN1_45S_50ntFlanks@9738.9760.23\|GAAACGATCTCAACCTATTCTCA | >hg38_21_+_8433172_8446622_RNA45SN1_45S_50ntFlanks | 9738 | 9760 | 23 | GAAACGATCTCAACCTATTCTCA | 159.8 | -11.2 | 1.4 | -7.9 | 3.1E-15 | 2.0E-14 | MB231/MB468 | MP |
| >MI0000093\|hsa-mir-92a-1&WithFlank&13\|+\|91351308\|91351397@54.74.21;MI0000094\|hsa-mir-92a-2&WithFlank&X\|-\|134169532\|134169618@54.74.21&[MIMAT0000092&hsa-miR-92a-3p&offsets\|0\|-1;m-9&13\|+\|91351361\|91351382&offsets\|0\|-1];[MIMAT0000092_1&hsa-miR-92a-3p&offsets\|0\|-1;m-11&X\|-\|134169544\|134169565&offsets\|0\|-1]\|TATTGCACTTGTCCCGGCCTG | >MI0000093\|hsa-mir-92a-1&WithFlank&13\|+\|91351308\|91351397 | 54 | 74 | 21 | TATTGCACTTGTCCCGGCCTG | 386.3 | 5.2 | 0.7 | 7.9 | 3.3E-15 | 2.0E-14 | BT20/MB468 | MITO |
| >hg38_21_+_8433172_8446622_RNA45SN1_45S_50ntFlanks@13002.13030.29\|CGATCTATTGAAAGTCAGCCCTCGACACA | >hg38_21_+_8433172_8446622_RNA45SN1_45S_50ntFlanks | 13002 | 13030 | 29 | CGATCTATTGAAAGTCAGCCCTCGACACA | 239.5 | -10.9 | 1.4 | -7.9 | 3.8E-15 | 2.0E-14 | MB231/MB468 | NUC |
| >MI0000087\|hsa-mir-29a&WithFlank&7\|-\|130876741\|130876816@47.68.22(+1C)&[MIMAT0000086&hsa-miR-29a-3p&offsets\|-1\|-1(+1C);m-26&7\|-\|130876749\|130876769&offsets\|-1\|0(+1C)]\|CTAGCACCATCTGAAATCGGTTC | >MI0000087\|hsa-mir-29a&WithFlank&7\|-\|130876741\|130876816 | 47 | 68 | 22(+1C) | CTAGCACCATCTGAAATCGGTTC | 508.7 | 12.2 | 1.5 | 7.9 | 3.1E-15 | 2.0E-14 | MB231/MB468 | MP |
| >MI0000077\|hsa-mir-21&WithFlank&17\|+\|59841260\|59841343@52.72.21&[MIMAT0004494&hsa-miR-21-3p&offsets\|0\|0;m-52&17\|+\|59841311\|59841332&offsets\|0\|-1]\|CAACACCAGTCGATGGGCTGT | >MI0000077\|hsa-mir-21&WithFlank&17\|+\|59841260\|59841343 | 52 | 72 | 21 | CAACACCAGTCGATGGGCTGT | 528.2 | 10.7 | 1.4 | 7.8 | 4.4E-15 | 2.0E-14 | BT20/MB468 | CYTO |
| >MI0000078\|hsa-mir-22&WithFlank&17\|-\|1713897\|1713993@59.80.22&[MIMAT0000077&hsa-miR-22-3p&offsets\|0\|0;m-2&17\|-\|1713914\|1713935&offsets\|0\|0]\|AAGCTGCCAGTTGAAGAACTGT | >MI0000078\|hsa-mir-22&WithFlank&17\|-\|1713897\|1713993 | 59 | 80 | 22 | AAGCTGCCAGTTGAAGAACTGT | 3169.6 | 5.0 | 0.6 | 7.8 | 6.8E-15 | 2.0E-14 | BT20/MB468 | NUC |
| >trna111_HisGTG_1_-_147774845_147774916@-1A.31.32,trna118_HisGTG_1_-_145396881_145396952@-1A.31.32,trna16_HisGTG_1_+_146544773_146544844@-1A.31.32,trna1_HisGTG_15_+_45493349_45493420@-1A.31.32,trna21_HisGTG_1_+_147753471_147753542@-1A.31.32,trna33_HisGTG_6_+_27125906_27125977@-1A.31.32,trna7_HisGTG_9_-_14433938_14434009@-1A.31.32,trna8_HisGTG_15_-_45492611_45492682@-1A.31.32,trna9_HisGTG_15_-_45490804_45490875@-1A.31.32\|AGCCGTGATCGTATAGTGGTTAGTACTCTGCG | >trna111_HisGTG_1_-_147774845_147774916 | -1A | 31 | 32 | AGCCGTGATCGTATAGTGGTTAGTACTCTGCG | 3717.6 | -9.3 | 1.2 | -7.8 | 6.8E-15 | 2.0E-14 | BT20/MB468 | NUC |
| >hg38_21_+_8433172_8446622_RNA45SN1_45S_50ntFlanks@6671.6699.29\|CTCGGCTCGTGCGTCGATGAAGAACGCAG | >hg38_21_+_8433172_8446622_RNA45SN1_45S_50ntFlanks | 6671 | 6699 | 29 | CTCGGCTCGTGCGTCGATGAAGAACGCAG | 3453.2 | -9.4 | 1.2 | -7.8 | 6.8E-15 | 2.0E-14 | BT20/MB468 | NUC |
| >MI0000650\|hsa-mir-200c&WithFlank&12\|+\|6963693\|6963772@50.71.22(+1C)&[MIMAT0000617&hsa-miR-200c-3p&offsets\|0\|-1(+1C);m-54&12\|+\|6963742\|6963764&offsets\|0\|-1(+1C)]\|TAATACTGCCGGGTAATGATGGC | >MI0000650\|hsa-mir-200c&WithFlank&12\|+\|6963693\|6963772 | 50 | 71 | 22(+1C) | TAATACTGCCGGGTAATGATGGC | 312.0 | 11.7 | 1.4 | 8.2 | 3.1E-16 | 2.0E-14 | BT20/MB231 | CYTO |
| >hg38_21_+_8433172_8446622_RNA45SN1_45S_50ntFlanks@4759.4786.28\|ATCAGATACCGTCGTAGTTCCGACCATA | >hg38_21_+_8433172_8446622_RNA45SN1_45S_50ntFlanks | 4759 | 4786 | 28 | ATCAGATACCGTCGTAGTTCCGACCATA | 515.6 | -11.3 | 1.4 | -7.8 | 4.5E-15 | 2.0E-14 | BT20/MB468 | CYTO |
| >MI0000061\|hsa-let-7a-2&WithFlank&11\|-\|122146516\|122146599@11.32.22(+1G);MI0000062\|hsa-let-7a-3&WithFlank&22\|+\|46112743\|46112828@10.31.22(+1G);MI0000060\|hsa-let-7a-1&WithFlank&9\|+\|94175951\|94176042@12.33.22(+1G)&[MIMAT0000062&hsa-let-7a-5p&offsets\|0\|0(+1G);m-6&11\|-\|122146568\|122146589&offsets\|0\|0(+1G)];[MIMAT0000062_1&hsa-let-7a-5p&offsets\|0\|0(+1G);m-5&22\|+\|46112752\|46112773&offsets\|0\|0(+1G)];[MIMAT0000062_2&hsa-let-7a-5p&offsets\|0\|0(+1G);m-4&9\|+\|94175962\|94175983&offsets\|0\|0(+1G)]\|TGAGGTAGTAGGTTGTATAGTTG | >MI0000061\|hsa-let-7a-2&WithFlank&11\|-\|122146516\|122146599 | 11 | 32 | 22(+1G) | TGAGGTAGTAGGTTGTATAGTTG | 396.4 | 7.6 | 1.0 | 7.8 | 4.5E-15 | 2.0E-14 | BT20/MB468 | CYTO |
| >hg38_21_+_8433172_8446622_RNA45SN1_45S_50ntFlanks@10797.10822.26\|GCATGTTGGAACAATGTAGGTAAGGG | >hg38_21_+_8433172_8446622_RNA45SN1_45S_50ntFlanks | 10797 | 10822 | 26 | GCATGTTGGAACAATGTAGGTAAGGG | 989.0 | -7.7 | 1.0 | -7.9 | 3.2E-15 | 2.0E-14 | MB231/MB468 | MP |
| >hg38_21_+_8433172_8446622_RNA45SN1_45S_50ntFlanks@8848.8867.20\|CGGCGGCGGCGGCGGCGGGA | >hg38_21_+_8433172_8446622_RNA45SN1_45S_50ntFlanks | 8848 | 8867 | 20 | CGGCGGCGGCGGCGGCGGGA | 341.3 | -12.3 | 1.6 | -7.9 | 3.2E-15 | 2.0E-14 | MB231/MB468 | MP |
| >hg38_21_+_8433172_8446622_RNA45SN1_45S_50ntFlanks@12456.12483.28\|CCAAGCGTTGGATTGTTCACCCACTAAT | >hg38_21_+_8433172_8446622_RNA45SN1_45S_50ntFlanks | 12456 | 12483 | 28 | CCAAGCGTTGGATTGTTCACCCACTAAT | 1324.6 | -9.7 | 1.2 | -7.9 | 3.2E-15 | 2.0E-14 | MB231/MB468 | MP |
| >hg38_21_+_8433172_8446622_RNA45SN1_45S_50ntFlanks@11593.11612.20\|CCAGGGGAATCCGACTGTTT | >hg38_21_+_8433172_8446622_RNA45SN1_45S_50ntFlanks | 11593 | 11612 | 20 | CCAGGGGAATCCGACTGTTT | 1595.1 | -8.3 | 1.1 | -7.9 | 3.2E-15 | 2.0E-14 | MB231/MB468 | MP |
| >hg38_21_+_8433172_8446622_RNA45SN1_45S_50ntFlanks@5494.5514.21\|AAGACGGTCGAACTTGACTAT | >hg38_21_+_8433172_8446622_RNA45SN1_45S_50ntFlanks | 5494 | 5514 | 21 | AAGACGGTCGAACTTGACTAT | 368.5 | -12.3 | 1.6 | -7.8 | 5.3E-15 | 2.0E-14 | MB231/MB468 | MITO |
| >MI0000650\|hsa-mir-200c&WithFlank&12\|+\|6963693\|6963772@50.72.23(+2U)&[MIMAT0000617&hsa-miR-200c-3p&offsets\|0\|0(+2U);m-54&12\|+\|6963742\|6963764&offsets\|0\|0(+2U)]\|TAATACTGCCGGGTAATGATGGATT | >MI0000650\|hsa-mir-200c&WithFlank&12\|+\|6963693\|6963772 | 50 | 72 | 23(+2U) | TAATACTGCCGGGTAATGATGGATT | 314.3 | 12.1 | 1.5 | 8.1 | 4.9E-16 | 2.0E-14 | BT20/MB231 | NUC |
| >hg38_21_+_8433172_8446622_RNA45SN1_45S_50ntFlanks@8552.8577.26\|CTCCCCGGAGGGGGCGGGCTCCGGCG | >hg38_21_+_8433172_8446622_RNA45SN1_45S_50ntFlanks | 8552 | 8577 | 26 | CTCCCCGGAGGGGGCGGGCTCCGGCG | 522.7 | -12.9 | 1.6 | -7.9 | 3.2E-15 | 2.0E-14 | MB231/MB468 | MP |
| >hg38_21_+_8433172_8446622_RNA45SN1_45S_50ntFlanks@9032.9057.26\|CCTCGTGGGGGGGCCGGGCCACCCCT | >hg38_21_+_8433172_8446622_RNA45SN1_45S_50ntFlanks | 9032 | 9057 | 26 | CCTCGTGGGGGGGCCGGGCCACCCCT | 233.4 | -11.7 | 1.5 | -7.9 | 3.2E-15 | 2.0E-14 | MB231/MB468 | MP |
| >hg38_21_+_8433172_8446622_RNA45SN1_45S_50ntFlanks@10781.10818.38\|AGGTGAACAGCCTCTGGCATGTTGGAACAATGTAGGTA | >hg38_21_+_8433172_8446622_RNA45SN1_45S_50ntFlanks | 10781 | 10818 | 38 | AGGTGAACAGCCTCTGGCATGTTGGAACAATGTAGGTA | 1283.3 | -4.7 | 0.6 | -7.9 | 3.4E-15 | 2.1E-14 | BT20/MB468 | MITO |
| >hg38_21_+_8433172_8446622_RNA45SN1_45S_50ntFlanks@3843.3868.26\|CTACTTGGATAACTGTGGTAATTCTA | >hg38_21_+_8433172_8446622_RNA45SN1_45S_50ntFlanks | 3843 | 3868 | 26 | CTACTTGGATAACTGTGGTAATTCTA | 604.5 | -4.8 | 0.6 | -7.8 | 7.0E-15 | 2.1E-14 | BT20/MB468 | NUC |
| >trna152_ValCAC_6_-_27248049_27248121@1.34.34\|GCTTCTGTAGTGTAGTGGTTATCACGTTCGCCTC | >trna152_ValCAC_6_-_27248049_27248121 | 1 | 34 | 34 | GCTTCTGTAGTGTAGTGGTTATCACGTTCGCCTC | 1200.4 | 7.5 | 1.0 | 7.8 | 5.4E-15 | 2.1E-14 | MB231/MB468 | MITO |
| >hg38_21_+_8433172_8446622_RNA45SN1_45S_50ntFlanks@12043.12070.28\|CACTGACCCGGTGAGGCGGGGGGGCGAG | >hg38_21_+_8433172_8446622_RNA45SN1_45S_50ntFlanks | 12043 | 12070 | 28 | CACTGACCCGGTGAGGCGGGGGGGCGAG | 303.2 | -12.0 | 1.5 | -7.8 | 5.4E-15 | 2.1E-14 | MB231/MB468 | MITO |
| >hg38_21_+_8433172_8446622_RNA45SN1_45S_50ntFlanks@10840.10868.29\|CCGTAACTTCGGGATAAGGATTGGCTCTA | >hg38_21_+_8433172_8446622_RNA45SN1_45S_50ntFlanks | 10840 | 10868 | 29 | CCGTAACTTCGGGATAAGGATTGGCTCTA | 558.8 | -7.2 | 0.9 | -7.8 | 5.4E-15 | 2.1E-14 | MB231/MB468 | MITO |
| >hg38_21_+_8433172_8446622_RNA45SN1_45S_50ntFlanks@10786.10813.28\|AACAGCCTCTGGCATGTTGGAACAATGT | >hg38_21_+_8433172_8446622_RNA45SN1_45S_50ntFlanks | 10786 | 10813 | 28 | AACAGCCTCTGGCATGTTGGAACAATGT | 285.5 | -12.0 | 1.5 | -7.9 | 3.3E-15 | 2.1E-14 | MB231/MB468 | MP |
| >hg38_21_+_8433172_8446622_RNA45SN1_45S_50ntFlanks@10537.10567.31\|CGAGGCGTCCAGTGCGGTAACGCGACCGATC | >hg38_21_+_8433172_8446622_RNA45SN1_45S_50ntFlanks | 10537 | 10567 | 31 | CGAGGCGTCCAGTGCGGTAACGCGACCGATC | 199.7 | -11.5 | 1.5 | -7.9 | 3.3E-15 | 2.1E-14 | MB231/MB468 | MP |
| >hg38_21_+_8433172_8446622_RNA45SN1_45S_50ntFlanks@11452.11478.27\|CTCCTCGGCGCGCGGCGGCGGCGGCGG | >hg38_21_+_8433172_8446622_RNA45SN1_45S_50ntFlanks | 11452 | 11478 | 27 | CTCCTCGGCGCGCGGCGGCGGCGGCGG | 785.6 | -12.7 | 1.5 | -8.3 | 1.2E-16 | 2.1E-14 | BT20/MB468 | MP |
| >hg38_21_+_8433172_8446622_RNA45SN1_45S_50ntFlanks@10458.10487.30\|ATGGCCTCCGTTGCCCTCGGCCGATCGAAA | >hg38_21_+_8433172_8446622_RNA45SN1_45S_50ntFlanks | 10458 | 10487 | 30 | ATGGCCTCCGTTGCCCTCGGCCGATCGAAA | 153.8 | -11.1 | 1.4 | -7.9 | 3.3E-15 | 2.1E-14 | MB231/MB468 | MP |
| >MI0000298\|hsa-mir-221&WithFlank&X\|-\|45746151\|45746272@71.93.23(+1U)&[MIMAT0000278&hsa-miR-221-3p&offsets\|0\|0(+1U);m-91&X\|-\|45746181\|45746202&offsets\|0\|+1(+1U)]\|AGCTACATTGTCTGCTGGGTTTCT | >MI0000298\|hsa-mir-221&WithFlank&X\|-\|45746151\|45746272 | 71 | 93 | 23(+1U) | AGCTACATTGTCTGCTGGGTTTCT | 15588.7 | 6.5 | 0.8 | 7.9 | 3.6E-15 | 2.1E-14 | MB231/MB468 | CYTO |
| >hg38_21_+_8433172_8446622_RNA45SN1_45S_50ntFlanks@9033.9057.25\|CTCGTGGGGGGGCCGGGCCACCCCT | >hg38_21_+_8433172_8446622_RNA45SN1_45S_50ntFlanks | 9033 | 9057 | 25 | CTCGTGGGGGGGCCGGGCCACCCCT | 387.1 | -12.4 | 1.6 | -7.8 | 5.5E-15 | 2.1E-14 | MB231/MB468 | MITO |
| >hg38_21_+_8433172_8446622_RNA45SN1_45S_50ntFlanks@11886.11905.20\|TGAGCTTGACTCTAGTCTGG | >hg38_21_+_8433172_8446622_RNA45SN1_45S_50ntFlanks | 11886 | 11905 | 20 | TGAGCTTGACTCTAGTCTGG | 394.7 | -12.4 | 1.6 | -7.8 | 5.5E-15 | 2.1E-14 | MB231/MB468 | MITO |
| >hg38_21_+_8433172_8446622_RNA45SN1_45S_50ntFlanks@10905.10926.22\|CTGGGCGCGCGCCGCGGCTGGA | >hg38_21_+_8433172_8446622_RNA45SN1_45S_50ntFlanks | 10905 | 10926 | 22 | CTGGGCGCGCGCCGCGGCTGGA | 314.6 | -11.0 | 1.4 | -7.9 | 3.7E-15 | 2.1E-14 | MB231/MB468 | CYTO |
| >MI0000079\|hsa-mir-23a&WithFlank&19\|-\|13836581\|13836665@54.72.19&[MIMAT0000078&hsa-miR-23a-3p&offsets\|+3\|+1;m-31&19\|-\|13836595\|13836615&offsets\|+3\|+1]\|ACATTGCCAGGGATTTCCA | >MI0000079\|hsa-mir-23a&WithFlank&19\|-\|13836581\|13836665 | 54 | 72 | 19 | ACATTGCCAGGGATTTCCA | 1466.5 | 14.3 | 1.7 | 8.3 | 1.2E-16 | 2.1E-14 | BT20/MB468 | MP |
| >am_MI0003639\|hsa-mir-625&WithFlank&14\|+\|65471096\|65471192@58.80.23&[MIMAT0004808&hsa-miR-625-3p&offsets\|0\|+1;m-137&14\|+\|65471153\|65471174&offsets\|0\|+1]\|GACTATAGAACTTTCCCCCTCAT | >am_MI0003639\|hsa-mir-625&WithFlank&14\|+\|65471096\|65471192 | 58 | 80 | 23 | GACTATAGAACTTTCCCCCTCAT | 811.1 | 10.2 | 1.3 | 7.9 | 3.7E-15 | 2.1E-14 | MB231/MB468 | CYTO |
| >trna10_AspGTC_12_-_125424193_125424264@36.73.38,trna12_AspGTC_12_-_125411891_125411962@36.73.38,trna144_AspGTC_6_-_27551236_27551307@36.73.38,trna38_AspGTC_17_-_8125556_8125627@36.73.38,trna45_AspGTC_6_+_27447453_27447524@36.73.38,trna48_AspGTC_6_+_27471523_27471594@36.73.38,trna4_AspGTC_12_+_96429799_96429870@36.73.38,trna69_AspGTC_1_-_161440205_161440276@36.73.38,trna72_AspGTC_1_-_161432824_161432895@36.73.38,trna75_AspGTC_1_-_161425414_161425485@36.73.38,trna78_AspGTC_1_-_161418033_161418104@36.73.38,trna81_AspGTC_1_-_161410615_161410686@36.73.38\|CACGCGGGAGACCGGGGTTCGATTCCCCGACGGGGAGC | >trna10_AspGTC_12_-_125424193_125424264 | 36 | 73 | 38 | CACGCGGGAGACCGGGGTTCGATTCCCCGACGGGGAGC | 8314.7 | 8.2 | 1.0 | 7.9 | 3.7E-15 | 2.1E-14 | MB231/MB468 | CYTO |
| >trna70_AlaCGC_6_+_28697092_28697163@58.75.18\|CCCCCGGCTCCTCCACCA | >trna70_AlaCGC_6_+_28697092_28697163 | 58 | 75 | 18 | CCCCCGGCTCCTCCACCA | 881.0 | 9.8 | 1.2 | 7.8 | 4.7E-15 | 2.1E-14 | BT20/MB468 | CYTO |
| >hg38_21_+_8433172_8446622_RNA45SN1_45S_50ntFlanks@11783.11816.34\|CATGAATGGATGAACGAGATTCCCACTGTCCCTA | >hg38_21_+_8433172_8446622_RNA45SN1_45S_50ntFlanks | 11783 | 11816 | 34 | CATGAATGGATGAACGAGATTCCCACTGTCCCTA | 352.8 | -12.3 | 1.6 | -7.9 | 3.3E-15 | 2.1E-14 | MB231/MB468 | MP |
| >hg38_21_+_8433172_8446622_RNA45SN1_45S_50ntFlanks@6660.6689.30\|GCGGTGGATCACTCGGCTCGTGCGTCGATG | >hg38_21_+_8433172_8446622_RNA45SN1_45S_50ntFlanks | 6660 | 6689 | 30 | GCGGTGGATCACTCGGCTCGTGCGTCGATG | 2223.0 | -5.8 | 0.7 | -7.8 | 5.5E-15 | 2.1E-14 | MB231/MB468 | MITO |

<table>
<tr><td></td><td></td><td></td><td></td><td></td><td></td><td></td><td></td><td></td><td></td><td></td><td></td><td></td><td></td></tr>
<tr><td>>MI0000446|hsa-mir-125b-1&WithFlank&11|-|122099751|122099850@21.41.21;MI0000470|hsa-mir-125b-2&WithFlank&21|+|16590231|16590331@23.43.21&[MIMAT0000423&hsa-miR-125b-5p&offsets|0|-1;m-67&11|-|122099809|122099830&offsets|0|-1];[MIMAT0000423_1&hsa-miR-125b-5p&offsets|0|-1;m-68&21|+|16590253|16590274&offsets|0|-1]|TCCCTGAGACCCTAACTTGTG</td><td>>MI0000446|hsa-mir-125b-1&WithFlank&11|-|122099751|122099850</td><td>21</td><td>41</td><td>21</td><td>TCCCTGAGACCCTAACTTGTG</td><td>9218.0</td><td>7.2</td><td>0.9</td><td>7.8</td><td>5.5E-15</td><td>2.1E-14</td><td>MB231/MB468</td><td>MITO</td></tr>
<tr><td>>hg38_21_+_8433172_8446622_RNA45SN1_45S_50ntFlanks@10905.10925.21|CTGGGCGCGCGCCGCGGCTGG</td><td>>hg38_21_+_8433172_8446622_RNA45SN1_45S_50ntFlanks</td><td>10905</td><td>10925</td><td>21</td><td>CTGGGCGCGCGCCGCGGCTGG</td><td>261.0</td><td>-11.8</td><td>1.5</td><td>-7.8</td><td>5.5E-15</td><td>2.1E-14</td><td>MB231/MB468</td><td>MITO</td></tr>
<tr><td>>hg38_21_+_8433172_8446622_RNA45SN1_45S_50ntFlanks@10783.10818.36|GTGAACAGCCTCTGGCATGTTGGAACAATGTAGGTA</td><td>>hg38_21_+_8433172_8446622_RNA45SN1_45S_50ntFlanks</td><td>10783</td><td>10818</td><td>36</td><td>GTGAACAGCCTCTGGCATGTTGGAACAATGTAGGTA</td><td>2079.3</td><td>-5.8</td><td>0.7</td><td>-7.9</td><td>3.7E-15</td><td>2.1E-14</td><td>MB231/MB468</td><td>CYTO</td></tr>
<tr><td>>hg38_21_+_8433172_8446622_RNA45SN1_45S_50ntFlanks@10905.10935.31|CTGGGCGCGCGCCGCGGCTGGACGAGGCGCC</td><td>>hg38_21_+_8433172_8446622_RNA45SN1_45S_50ntFlanks</td><td>10905</td><td>10935</td><td>31</td><td>CTGGGCGCGCGCCGCGGCTGGACGAGGCGCC</td><td>195.2</td><td>-11.5</td><td>1.5</td><td>-7.9</td><td>3.3E-15</td><td>2.1E-14</td><td>MB231/MB468</td><td>MP</td></tr>
<tr><td>>hg38_21_+_8433172_8446622_RNA45SN1_45S_50ntFlanks@12632.12652.21|GCGAAGCTACCATCTGTGGGA</td><td>>hg38_21_+_8433172_8446622_RNA45SN1_45S_50ntFlanks</td><td>12632</td><td>12652</td><td>21</td><td>GCGAAGCTACCATCTGTGGGA</td><td>256.8</td><td>-11.9</td><td>1.5</td><td>-7.9</td><td>3.3E-15</td><td>2.1E-14</td><td>MB231/MB468</td><td>MP</td></tr>
<tr><td>>hg38_21_+_8433172_8446622_RNA45SN1_45S_50ntFlanks@9804.9838.35|GGCGTGGAATGCGAGTGCCTAGTGGGCCACTTTTG</td><td>>hg38_21_+_8433172_8446622_RNA45SN1_45S_50ntFlanks</td><td>9804</td><td>9838</td><td>35</td><td>GGCGTGGAATGCGAGTGCCTAGTGGGCCACTTTTG</td><td>227.3</td><td>-11.7</td><td>1.5</td><td>-7.9</td><td>3.4E-15</td><td>2.1E-14</td><td>MB231/MB468</td><td>MP</td></tr>
<tr><td>>hg38_21_+_8433172_8446622_RNA45SN1_45S_50ntFlanks@12417.12434.18|TCGATGTCGGCTCTTCCT</td><td>>hg38_21_+_8433172_8446622_RNA45SN1_45S_50ntFlanks</td><td>12417</td><td>12434</td><td>18</td><td>TCGATGTCGGCTCTTCCT</td><td>2991.5</td><td>-4.6</td><td>0.6</td><td>-7.8</td><td>5.6E-15</td><td>2.1E-14</td><td>MB231/MB468</td><td>MITO</td></tr>
<tr><td>>hg38_21_+_8433172_8446622_RNA45SN1_45S_50ntFlanks@3874.3898.25|AATACATGCCGACGGGCGCTGACCC</td><td>>hg38_21_+_8433172_8446622_RNA45SN1_45S_50ntFlanks</td><td>3874</td><td>3898</td><td>25</td><td>AATACATGCCGACGGGCGCTGACCC</td><td>6297.3</td><td>-10.5</td><td>1.3</td><td>-7.8</td><td>5.6E-15</td><td>2.1E-14</td><td>MB231/MB468</td><td>MITO</td></tr>
<tr><td>>MI0000266|hsa-mir-10a&WithFlank&17|-|48579832|48579953@29.50.22(+1A)&[MIMAT0000253&hsa-miR-10a-5p&offsets|+1|0(+1A);m-18&17|-|48579905|48579926&offsets|+1|+1(+1A)]|ACCCTGTAGATCCGAATTTGTGA</td><td>>MI0000266|hsa-mir-10a&WithFlank&17|-|48579832|48579953</td><td>29</td><td>50</td><td>22(+1A)</td><td>ACCCTGTAGATCCGAATTTGTGA</td><td>420.3</td><td>11.9</td><td>1.5</td><td>7.8</td><td>5.6E-15</td><td>2.1E-14</td><td>MB231/MB468</td><td>MITO</td></tr>
<tr><td>>MI0003591|hsa-mir-584&WithFlank&5|-|149062307|149062415@22.40.19&[MIMAT0003249&hsa-miR-584-5p&offsets|0|-3;m-167&5|-|149062374|149062394&offsets|0|-2]|TTATGGTTTGCCTGGGACT</td><td>>MI0003591|hsa-mir-584&WithFlank&5|-|149062307|149062415</td><td>22</td><td>40</td><td>19</td><td>TTATGGTTTGCCTGGGACT</td><td>723.2</td><td>9.3</td><td>1.2</td><td>7.8</td><td>4.8E-15</td><td>2.1E-14</td><td>BT20/MB468</td><td>CYTO</td></tr>
<tr><td>>MI0000088|hsa-mir-30a&WithFlank&6|-|71403545|71403627@12.35.24(+3U)&[MIMAT0000087&hsa-miR-30a-5p&offsets|0|+2(+3U);m-15&6|-|71403595|71403616&offsets|0|+2(+3U)]|TGTAAACATCCTCGACTGGAAGCTTTT</td><td>>MI0000088|hsa-mir-30a&WithFlank&6|-|71403545|71403627</td><td>12</td><td>35</td><td>24(+3U)</td><td>TGTAAACATCCTCGACTGGAAGCTTTT</td><td>202.9</td><td>11.6</td><td>1.5</td><td>7.9</td><td>4.1E-15</td><td>2.1E-14</td><td>MB231/MB468</td><td>NUC</td></tr>
<tr><td>>hg38_21_+_8433172_8446622_RNA45SN1_45S_50ntFlanks@11918.11950.33|ACATGAGAGGTGTAGAATAAGTGGGAGGCCCCC</td><td>>hg38_21_+_8433172_8446622_RNA45SN1_45S_50ntFlanks</td><td>11918</td><td>11950</td><td>33</td><td>ACATGAGAGGTGTAGAATAAGTGGGAGGCCCCC</td><td>1640.8</td><td>-5.1</td><td>0.7</td><td>-7.8</td><td>4.8E-15</td><td>2.1E-14</td><td>BT20/MB468</td><td>CYTO</td></tr>
<tr><td>>hg38_21_+_8433172_8446622_RNA45SN1_45S_50ntFlanks@11829.11856.28|CGAAACCACAGCCAAGGGAACGGGCTTG</td><td>>hg38_21_+_8433172_8446622_RNA45SN1_45S_50ntFlanks</td><td>11829</td><td>11856</td><td>28</td><td>CGAAACCACAGCCAAGGGAACGGGCTTG</td><td>296.5</td><td>-12.1</td><td>1.5</td><td>-7.9</td><td>3.4E-15</td><td>2.2E-14</td><td>MB231/MB468</td><td>MP</td></tr>
<tr><td>>hg38_21_+_8433172_8446622_RNA45SN1_45S_50ntFlanks@11874.11920.47|AGAAGACCCTGTTGAGCTTGACTCTAGTCTGGCACGGTGAAGAGACA</td><td>>hg38_21_+_8433172_8446622_RNA45SN1_45S_50ntFlanks</td><td>11874</td><td>11920</td><td>47</td><td>AGAAGACCCTGTTGAGCTTGACTCTAGTCTGGCACGGTGAAGAGACA</td><td>504.2</td><td>-12.8</td><td>1.6</td><td>-7.8</td><td>5.7E-15</td><td>2.2E-14</td><td>MB231/MB468</td><td>MITO</td></tr>
<tr><td>>hg38_21_+_8433172_8446622_RNA45SN1_45S_50ntFlanks@10508.10533.26|CGAATCCGGAGTGGCGGAGATGGGCG</td><td>>hg38_21_+_8433172_8446622_RNA45SN1_45S_50ntFlanks</td><td>10508</td><td>10533</td><td>26</td><td>CGAATCCGGAGTGGCGGAGATGGGCG</td><td>1748.9</td><td>-10.2</td><td>1.3</td><td>-7.9</td><td>3.4E-15</td><td>2.2E-14</td><td>MB231/MB468</td><td>MP</td></tr>
<tr><td>>hg38_21_+_8433172_8446622_RNA45SN1_45S_50ntFlanks@11784.11803.20|ATGAATGGATGAACGAGATT</td><td>>hg38_21_+_8433172_8446622_RNA45SN1_45S_50ntFlanks</td><td>11784</td><td>11803</td><td>20</td><td>ATGAATGGATGAACGAGATT</td><td>2442.9</td><td>-8.2</td><td>1.0</td><td>-7.9</td><td>3.4E-15</td><td>2.2E-14</td><td>MB231/MB468</td><td>MP</td></tr>
<tr><td>>trna13_ValCAC_19_-_4724647_4724719@1.34.34|GTTTCCGTAGTGTAGCGGTTATCACATTCGCCTC</td><td>>trna13_ValCAC_19_-_4724647_4724719</td><td>1</td><td>34</td><td>34</td><td>GTTTCCGTAGTGTAGCGGTTATCACATTCGCCTC</td><td>1628.7</td><td>10.7</td><td>1.4</td><td>7.9</td><td>3.8E-15</td><td>2.2E-14</td><td>MB231/MB468</td><td>CYTO</td></tr>
<tr><td>>MI0003591|hsa-mir-584&WithFlank&5|-|149062307|149062415@22.40.19&[MIMAT0003249&hsa-miR-584-5p&offsets|0|-3;m-167&5|-|149062374|149062394&offsets|0|-2]|TTATGGTTTGCCTGGGACT</td><td>>MI0003591|hsa-mir-584&WithFlank&5|-|149062307|149062415</td><td>22</td><td>40</td><td>19</td><td>TTATGGTTTGCCTGGGACT</td><td>767.7</td><td>9.7</td><td>1.2</td><td>7.9</td><td>3.6E-15</td><td>2.2E-14</td><td>BT20/MB468</td><td>MITO</td></tr>
<tr><td>>MI0000650|hsa-mir-200c&WithFlank&12|+|6963693|6963772@50.71.22(+1G)&[MIMAT0000617&hsa-miR-200c-3p&offsets|0|-1(+1G);m-54&12|+|6963742|6963764&offsets|0|-1(+1G)]|TAATACTGCCGGGTAATGATGGG</td><td>>MI0000650|hsa-mir-200c&WithFlank&12|+|6963693|6963772</td><td>50</td><td>71</td><td>22(+1G)</td><td>TAATACTGCCGGGTAATGATGGG</td><td>1390.8</td><td>4.1</td><td>0.5</td><td>7.8</td><td>7.4E-15</td><td>2.2E-14</td><td>BT20/MB468</td><td>NUC</td></tr>
<tr><td>>hg38_21_+_8433172_8446622_RNA45SN1_45S_50ntFlanks@10727.10756.30|GGAGAGGGTGTAAATCTCGCGCCGGGCCGT</td><td>>hg38_21_+_8433172_8446622_RNA45SN1_45S_50ntFlanks</td><td>10727</td><td>10756</td><td>30</td><td>GGAGAGGGTGTAAATCTCGCGCCGGGCCGT</td><td>335.8</td><td>-11.4</td><td>1.4</td><td>-7.8</td><td>4.2E-15</td><td>2.2E-14</td><td>MB231/MB468</td><td>NUC</td></tr>
<tr><td>>hg38_21_+_8433172_8446622_RNA45SN1_45S_50ntFlanks@12539.12565.27|ATGATGTGTTGTTGCCATGGTAATCCT</td><td>>hg38_21_+_8433172_8446622_RNA45SN1_45S_50ntFlanks</td><td>12539</td><td>12565</td><td>27</td><td>ATGATGTGTTGTTGCCATGGTAATCCT</td><td>467.3</td><td>-11.8</td><td>1.5</td><td>-7.8</td><td>4.2E-15</td><td>2.2E-14</td><td>MB231/MB468</td><td>NUC</td></tr>
<tr><td>>hg38_21_+_8433172_8446622_RNA45SN1_45S_50ntFlanks@12008.12043.36|CCGCCGGTGAAATACCACTACTCTGATCGTTTTTTC</td><td>>hg38_21_+_8433172_8446622_RNA45SN1_45S_50ntFlanks</td><td>12008</td><td>12043</td><td>36</td><td>CCGCCGGTGAAATACCACTACTCTGATCGTTTTTTC</td><td>11483.3</td><td>-10.2</td><td>1.3</td><td>-7.8</td><td>4.2E-15</td><td>2.2E-14</td><td>MB231/MB468</td><td>NUC</td></tr>
<tr><td>>hg38_21_+_8433172_8446622_RNA45SN1_45S_50ntFlanks@12971.12991.21|TTTCGTACGTAGCAGAGCAGC</td><td>>hg38_21_+_8433172_8446622_RNA45SN1_45S_50ntFlanks</td><td>12971</td><td>12991</td><td>21</td><td>TTTCGTACGTAGCAGAGCAGC</td><td>2112.1</td><td>-10.8</td><td>1.4</td><td>-7.8</td><td>7.4E-15</td><td>2.2E-14</td><td>BT20/MB468</td><td>NUC</td></tr>
<tr><td>>am_MI0000809|hsa-mir-151a&WithFlank&8|-|140732558|140732659@53.73.21(+2U)&[MIMAT0000757&hsa-miR-151a-3p&offsets|0|0(+2U);m-30&8|-|140732587|140732607&offsets|0|0(+2U)]|CTAGACTGAAGCTCCTTGAGGTT</td><td>>am_MI0000809|hsa-mir-151a&WithFlank&8|-|140732558|140732659</td><td>53</td><td>73</td><td>21(+2U)</td><td>CTAGACTGAAGCTCCTTGAGGTT</td><td>882.4</td><td>7.6</td><td>1.0</td><td>7.8</td><td>5.8E-15</td><td>2.2E-14</td><td>MB231/MB468</td><td>MITO</td></tr>
<tr><td>>hg38_21_+_8433172_8446622_RNA45SN1_45S_50ntFlanks@9564.9592.29|CTGACGTGCAAATCGGTCGTCCGACCTGG</td><td>>hg38_21_+_8433172_8446622_RNA45SN1_45S_50ntFlanks</td><td>9564</td><td>9592</td><td>29</td><td>CTGACGTGCAAATCGGTCGTCCGACCTGG</td><td>2810.9</td><td>-9.0</td><td>1.2</td><td>-7.8</td><td>7.5E-15</td><td>2.2E-14</td><td>BT20/MB468</td><td>NUC</td></tr>
<tr><td>>hg38_21_+_8433172_8446622_RNA45SN1_45S_50ntFlanks@12996.13036.41|TCGCTGCGATCTATTGAAAGTCAGCCCTCGACACAAGGGTT</td><td>>hg38_21_+_8433172_8446622_RNA45SN1_45S_50ntFlanks</td><td>12996</td><td>13036</td><td>41</td><td>TCGCTGCGATCTATTGAAAGTCAGCCCTCGACACAAGGGTT</td><td>5840.0</td><td>8.5</td><td>1.1</td><td>7.8</td><td>4.9E-15</td><td>2.2E-14</td><td>BT20/MB468</td><td>CYTO</td></tr>
<tr><td>>hg38_21_+_8433172_8446622_RNA45SN1_45S_50ntFlanks@10690.10709.20|CGTCCGGTGAGCTCTCGCTG</td><td>>hg38_21_+_8433172_8446622_RNA45SN1_45S_50ntFlanks</td><td>10690</td><td>10709</td><td>20</td><td>CGTCCGGTGAGCTCTCGCTG</td><td>2092.1</td><td>-7.1</td><td>0.9</td><td>-7.8</td><td>4.9E-15</td><td>2.2E-14</td><td>BT20/MB468</td><td>CYTO</td></tr>
<tr><td>>hg38_21_+_8433172_8446622_RNA45SN1_45S_50ntFlanks@10679.10712.34|GGTTCCGGCGGCGTCCGGTGAGCTCTCGCTGGCC</td><td>>hg38_21_+_8433172_8446622_RNA45SN1_45S_50ntFlanks</td><td>10679</td><td>10712</td><td>34</td><td>GGTTCCGGCGGCGTCCGGTGAGCTCTCGCTGGCC</td><td>541.1</td><td>-12.0</td><td>1.5</td><td>-7.8</td><td>4.2E-15</td><td>2.2E-14</td><td>MB231/MB468</td><td>NUC</td></tr>
<tr><td>>hg38_21_+_8433172_8446622_RNA45SN1_45S_50ntFlanks@11872.11893.22|AAAGAAGACCCTGTTGAGCTTG</td><td>>hg38_21_+_8433172_8446622_RNA45SN1_45S_50ntFlanks</td><td>11872</td><td>11893</td><td>22</td><td>AAAGAAGACCCTGTTGAGCTTG</td><td>243.0</td><td>-11.7</td><td>1.5</td><td>-7.8</td><td>5.8E-15</td><td>2.2E-14</td><td>MB231/MB468</td><td>MITO</td></tr>
<tr><td>>hg38_21_+_8433172_8446622_RNA45SN1_45S_50ntFlanks@10805.10825.21|GAACAATGTAGGTAAGGGAAG</td><td>>hg38_21_+_8433172_8446622_RNA45SN1_45S_50ntFlanks</td><td>10805</td><td>10825</td><td>21</td><td>GAACAATGTAGGTAAGGGAAG</td><td>167.8</td><td>-11.3</td><td>1.4</td><td>-7.9</td><td>3.5E-15</td><td>2.2E-14</td><td>MB231/MB468</td><td>MP</td></tr>
<tr><td>>hg38_21_+_8433172_8446622_RNA45SN1_45S_50ntFlanks@8112.8136.25|GGGCGCGGGACATGTGGCGTACGGA</td><td>>hg38_21_+_8433172_8446622_RNA45SN1_45S_50ntFlanks</td><td>8112</td><td>8136</td><td>25</td><td>GGGCGCGGGACATGTGGCGTACGGA</td><td>565.3</td><td>-13.0</td><td>1.7</td><td>-7.9</td><td>3.5E-15</td><td>2.2E-14</td><td>MB231/MB468</td><td>MP</td></tr>
<tr><td>>hg38_21_+_8433172_8446622_RNA45SN1_45S_50ntFlanks@11535.11553.19|TTCCGCGCGGCGCCTCGCC</td><td>>hg38_21_+_8433172_8446622_RNA45SN1_45S_50ntFlanks</td><td>11535</td><td>11553</td><td>19</td><td>TTCCGCGCGGCGCCTCGCC</td><td>230.2</td><td>-8.0</td><td>1.0</td><td>-7.9</td><td>3.9E-15</td><td>2.2E-14</td><td>MB231/MB468</td><td>CYTO</td></tr>
<tr><td>>MI0000088|hsa-mir-30a&WithFlank&6|-|71403545|71403627@12.34.23(+1G)&[MIMAT0000087&hsa-miR-30a-5p&offsets|0|+1(+1G);m-15&6|-|71403595|71403616&offsets|0|+1(+1G)]|TGTAAACATCCTCGACTGGAAGCG</td><td>>MI0000088|hsa-mir-30a&WithFlank&6|-|71403545|71403627</td><td>12</td><td>34</td><td>23(+1G)</td><td>TGTAAACATCCTCGACTGGAAGCG</td><td>494.8</td><td>5.1</td><td>0.6</td><td>7.9</td><td>3.6E-15</td><td>2.2E-14</td><td>BT20/MB468</td><td>MITO</td></tr>
<tr><td>>hg38_21_+_8433172_8446622_RNA45SN1_45S_50ntFlanks@12836.12861.26|CGGTGCGGAGTGCCCTTCGTCCTGGG</td><td>>hg38_21_+_8433172_8446622_RNA45SN1_45S_50ntFlanks</td><td>12836</td><td>12861</td><td>26</td><td>CGGTGCGGAGTGCCCTTCGTCCTGGG</td><td>253.7</td><td>-11.9</td><td>1.5</td><td>-7.9</td><td>3.5E-15</td><td>2.2E-14</td><td>MB231/MB468</td><td>MP</td></tr>
<tr><td>>hg38_21_+_8433172_8446622_RNA45SN1_45S_50ntFlanks@485.505.21|GATGTGGTGACGTCGTGCTCT</td><td>>hg38_21_+_8433172_8446622_RNA45SN1_45S_50ntFlanks</td><td>485</td><td>505</td><td>21</td><td>GATGTGGTGACGTCGTGCTCT</td><td>2644.7</td><td>28.6</td><td>3.6</td><td>7.9</td><td>3.5E-15</td><td>2.2E-14</td><td>MB231/MB468</td><td>MP</td></tr>
<tr><td>>hg38_21_+_8433172_8446622_RNA45SN1_45S_50ntFlanks@11766.11791.26|ATCTAATTAGTGACGCGCATGAATGG</td><td>>hg38_21_+_8433172_8446622_RNA45SN1_45S_50ntFlanks</td><td>11766</td><td>11791</td><td>26</td><td>ATCTAATTAGTGACGCGCATGAATGG</td><td>178.6</td><td>-11.3</td><td>1.4</td><td>-7.9</td><td>3.5E-15</td><td>2.2E-14</td><td>MB231/MB468</td><td>MP</td></tr>
<tr><td>>trna2_LeuTAA_4_-_156384978_156385052@1.18.18|GTTAAGATGGCAGAGCCT</td><td>>trna2_LeuTAA_4_-_156384978_156385052</td><td>1</td><td>18</td><td>18</td><td>GTTAAGATGGCAGAGCCT</td><td>295.0</td><td>12.5</td><td>1.6</td><td>7.8</td><td>5.0E-15</td><td>2.2E-14</td><td>BT20/MB468</td><td>CYTO</td></tr>
<tr><td>>hg38_21_+_8433172_8446622_RNA45SN1_45S_50ntFlanks@12624.12657.34|CCAATGGGGCGAAGCTACCATCTGTGGGATTATG</td><td>>hg38_21_+_8433172_8446622_RNA45SN1_45S_50ntFlanks</td><td>12624</td><td>12657</td><td>34</td><td>CCAATGGGGCGAAGCTACCATCTGTGGGATTATG</td><td>1570.3</td><td>-9.9</td><td>1.3</td><td>-7.9</td><td>3.5E-15</td><td>2.2E-14</td><td>MB231/MB468</td><td>MP</td></tr>
<tr><td>>MI0000471|hsa-mir-126&WithFlank&9|+|136670596|136670692@59.79.21&[MIMAT0000445&hsa-miR-126-3p&offsets|+1|0;m-40&9|+|136670653|136670674&offsets|+1|0]|CGTACCGTGAGTAATAATGCG</td><td>>MI0000471|hsa-mir-126&WithFlank&9|+|136670596|136670692</td><td>59</td><td>79</td><td>21</td><td>CGTACCGTGAGTAATAATGCG</td><td>397.1</td><td>11.9</td><td>1.5</td><td>7.9</td><td>3.5E-15</td><td>2.2E-14</td><td>MB231/MB468</td><td>MP</td></tr>
<tr><td>>hg38_21_+_8433172_8446622_RNA45SN1_45S_50ntFlanks@7981.8008.28|CTCAGATCAGACGTGGCGACCCGCTGAA</td><td>>hg38_21_+_8433172_8446622_RNA45SN1_45S_50ntFlanks</td><td>7981</td><td>8008</td><td>28</td><td>CTCAGATCAGACGTGGCGACCCGCTGAA</td><td>1755.4</td><td>-9.9</td><td>1.3</td><td>-7.9</td><td>3.5E-15</td><td>2.2E-14</td><td>MB231/MB468</td><td>MP</td></tr>
<tr><td>>hg38_21_+_8433172_8446622_RNA45SN1_45S_50ntFlanks@12957.12987.31|CTTCTGGGTCGGGGTTTCGTACGTAGCAGAG</td><td>>hg38_21_+_8433172_8446622_RNA45SN1_45S_50ntFlanks</td><td>12957</td><td>12987</td><td>31</td><td>CTTCTGGGTCGGGGTTTCGTACGTAGCAGAG</td><td>294.3</td><td>-5.6</td><td>0.7</td><td>-7.9</td><td>3.9E-15</td><td>2.2E-14</td><td>MB231/MB468</td><td>CYTO</td></tr>
<tr><td>>hg38_21_+_8433172_8446622_RNA45SN1_45S_50ntFlanks@6780.6799.20|CGGGGCTACGCCTGTCTGAG</td><td>>hg38_21_+_8433172_8446622_RNA45SN1_45S_50ntFlanks</td><td>6780</td><td>6799</td><td>20</td><td>CGGGGCTACGCCTGTCTGAG</td><td>222.9</td><td>-11.7</td><td>1.5</td><td>-7.9</td><td>3.5E-15</td><td>2.2E-14</td><td>MB231/MB468</td><td>MP</td></tr>
<tr><td>>trna2_GlyGCC_21_-_18827107_18827177@1.32.32,trna35_GlyGCC_1_+_161413094_161413164@1.32.32,trna37_GlyGCC_1_+_161420467_161420537@1.32.32,trna39_GlyGCC_1_+_161427898_161427968@1.32.32,trna41_GlyGCC_1_+_161435258_161435328@1.32.32|GCATGGGTGGTTCAGTGGTAGAATTCTCGCCT</td><td>>trna2_GlyGCC_21_-_18827107_18827177</td><td>1</td><td>32</td><td>32</td><td>GCATGGGTGGTTCAGTGGTAGAATTCTCGCCT</td><td>29220.7</td><td>10.0</td><td>1.3</td><td>7.9</td><td>3.7E-15</td><td>2.2E-14</td><td>BT20/MB468</td><td>MITO</td></tr>
<tr><td>>hg38_21_+_8433172_8446622_RNA45SN1_45S_50ntFlanks@10846.10865.20|CTTCGGGATAAGGATTGGCT</td><td>>hg38_21_+_8433172_8446622_RNA45SN1_45S_50ntFlanks</td><td>10846</td><td>10865</td><td>20</td><td>CTTCGGGATAAGGATTGGCT</td><td>3256.0</td><td>-10.3</td><td>1.3</td><td>-7.8</td><td>4.3E-15</td><td>2.2E-14</td><td>MB231/MB468</td><td>NUC</td></tr>
<tr><td>>hg38_21_+_8433172_8446622_RNA45SN1_45S_50ntFlanks@8115.8143.29|CGCGGGACATGTGGCGTACGGAAGACCCG</td><td>>hg38_21_+_8433172_8446622_RNA45SN1_45S_50ntFlanks</td><td>8115</td><td>8143</td><td>29</td><td>CGCGGGACATGTGGCGTACGGAAGACCCG</td><td>6091.4</td><td>-5.8</td><td>0.7</td><td>-7.9</td><td>3.7E-15</td><td>2.2E-14</td><td>BT20/MB468</td><td>MITO</td></tr>
<tr><td>>hg38_21_+_8433172_8446622_RNA45SN1_45S_50ntFlanks@11118.11145.28|CAGGGGCCGGCGGCGGCCCGCCGCGGGG</td><td>>hg38_21_+_8433172_8446622_RNA45SN1_45S_50ntFlanks</td><td>11118</td><td>11145</td><td>28</td><td>CAGGGGCCGGCGGCGGCCCGCCGCGGGG</td><td>387.8</td><td>-11.6</td><td>1.5</td><td>-7.8</td><td>4.3E-15</td><td>2.2E-14</td><td>MB231/MB468</td><td>NUC</td></tr>
</table>

<table>
<tr><td>>am_trna11_ProAGG_16_+_3241989_3242060@1.28.28,trna12_ProAGG_6_+_26555498_26555569@1.28.28,trna12_ProTGG_11_-_75946869_75946940@1.28.28,trna14_ProTGG_5_-_180615854_180615925@1.28.28,trna22_ProAGG_14_-_21081560_21081631@1.28.28,trna23_ProAGG_14_-_21077495_21077566@1.28.28,trna28_ProTGG_16_-_3234133_3234204@1.28.28,trna29_ProAGG_16_-_3232635_3232706@1.28.28,trna2_ProAGG_7_+_128423504_128423575@1.28.28,trna30_ProCGG_6_+_27059521_27059592@1.28.28,trna37_ProCGG_17_-_8126151_8126222@1.28.28,trna3_ProTGG_16_+_3208923_3208994@1.28.28,trna52_ProCGG_1_+_167683962_167684033@1.28.28,trna65_ProAGG_1_-_167684725_167684796@1.28.28,trna6_ProCGG_16_+_3222049_3222120@1.28.28,trna6_ProTGG_14_+_21152175_21152246@1.28.28,trna8_ProTGG_16_+_3238094_3238165@1.28.28,trna9_ProAGG_11_+_75946557_75946628@1.28.28,trna9_ProAGG_16_+_3239634_3239705@1.28.28|GGCTCGTTGGTCTAGGGGTATGATTCTC</td><td>>am_trna11_ProAGG_16_+_3241989_3242060</td><td>1</td><td>28</td><td>28</td><td>GGCTCGTTGGTCTAGGGGTATGATTCTC</td><td>2857.7</td><td>-11.2</td><td>1.4</td><td>-7.8</td><td>4.3E-15</td><td>2.2E-14</td><td>MB231/MB468</td><td>NUC</td></tr>
<tr><td>>am_trna2_GlyGCC_21_-_18827107_18827177@1.18.18,trna35_GlyGCC_1_+_161413094_161413164@1.18.18,trna37_GlyGCC_1_+_161420467_161420537@1.18.18,trna39_GlyGCC_1_+_161427898_161427968@1.18.18,trna41_GlyGCC_1_+_161435258_161435328@1.18.18|GCATGGGTGGTTCAGTGG</td><td>>am_trna2_GlyGCC_21_-_18827107_18827177</td><td>1</td><td>18</td><td>18</td><td>GCATGGGTGGTTCAGTGG</td><td>307.5</td><td>11.6</td><td>1.4</td><td>8.2</td><td>3.5E-16</td><td>2.2E-14</td><td>BT20/MB231</td><td>CYTO</td></tr>
<tr><td>>hg38_21_+_8433172_8446622_RNA45SN1_45S_50ntFlanks@10410.10443.34|TCAGTCGGTCCTGAGAGATGGGCGAGCGCCGTTC</td><td>>hg38_21_+_8433172_8446622_RNA45SN1_45S_50ntFlanks</td><td>10410</td><td>10443</td><td>34</td><td>TCAGTCGGTCCTGAGAGATGGGCGAGCGCCGTTC</td><td>477.8</td><td>-5.9</td><td>0.8</td><td>-7.8</td><td>5.0E-15</td><td>2.2E-14</td><td>BT20/MB468</td><td>CYTO</td></tr>
<tr><td>>hg38_21_+_8433172_8446622_RNA45SN1_45S_50ntFlanks@6730.6770.41|ACACATTGATCATCGACACTTCGAACGCACTTGCGGCCCCG</td><td>>hg38_21_+_8433172_8446622_RNA45SN1_45S_50ntFlanks</td><td>6730</td><td>6770</td><td>41</td><td>ACACATTGATCATCGACACTTCGAACGCACTTGCGGCCCCG</td><td>1305.2</td><td>-7.7</td><td>1.0</td><td>-7.8</td><td>5.9E-15</td><td>2.2E-14</td><td>MB231/MB468</td><td>MITO</td></tr>
<tr><td>>MI0000471|hsa-mir-126&WithFlank&9|+|136670596|136670692@57.79.23&[MIMAT0000445&hsa-miR-126-3p&offsets|-1|0;m-40&9|+|136670653|136670674&offsets|-1|0]|CTCGTACCGTGAGTAATAATGCG</td><td>>MI0000471|hsa-mir-126&WithFlank&9|+|136670596|136670692</td><td>57</td><td>79</td><td>23</td><td>CTCGTACCGTGAGTAATAATGCG</td><td>400.5</td><td>11.9</td><td>1.5</td><td>7.8</td><td>6.0E-15</td><td>2.2E-14</td><td>MB231/MB468</td><td>MITO</td></tr>
<tr><td>>hg38_21_+_8433172_8446622_RNA45SN1_45S_50ntFlanks@8513.8531.19|CTCCACCCGCCCTCCCTTC</td><td>>hg38_21_+_8433172_8446622_RNA45SN1_45S_50ntFlanks</td><td>8513</td><td>8531</td><td>19</td><td>CTCCACCCGCCCTCCCTTC</td><td>253.4</td><td>-11.8</td><td>1.5</td><td>-7.8</td><td>6.0E-15</td><td>2.2E-14</td><td>MB231/MB468</td><td>MITO</td></tr>
<tr><td>>hg38_21_+_8433172_8446622_RNA45SN1_45S_50ntFlanks@12934.12961.28|CTAAACCATTCGTAGACGACCTGCTTCT</td><td>>hg38_21_+_8433172_8446622_RNA45SN1_45S_50ntFlanks</td><td>12934</td><td>12961</td><td>28</td><td>CTAAACCATTCGTAGACGACCTGCTTCT</td><td>174.7</td><td>-11.3</td><td>1.4</td><td>-7.9</td><td>3.6E-15</td><td>2.2E-14</td><td>MB231/MB468</td><td>MP</td></tr>
<tr><td>>trna6_TrpCCA_12_+_98898030_98898101@30.52.23|ACTCCAGATCAGAAGGCTGCGTG</td><td>>trna6_TrpCCA_12_+_98898030_98898101</td><td>30</td><td>52</td><td>23</td><td>ACTCCAGATCAGAAGGCTGCGTG</td><td>429.4</td><td>-11.7</td><td>1.5</td><td>-7.8</td><td>4.4E-15</td><td>2.2E-14</td><td>MB231/MB468</td><td>NUC</td></tr>
<tr><td>>hg38_21_+_8433172_8446622_RNA45SN1_45S_50ntFlanks@9352.9369.18|CCGTGGCGCAATGAAGGT</td><td>>hg38_21_+_8433172_8446622_RNA45SN1_45S_50ntFlanks</td><td>9352</td><td>9369</td><td>18</td><td>CCGTGGCGCAATGAAGGT</td><td>283.9</td><td>-12.0</td><td>1.5</td><td>-7.9</td><td>3.6E-15</td><td>2.3E-14</td><td>MB231/MB468</td><td>MP</td></tr>
<tr><td>>trna130_GlnTTG_6_-_27763640_27763711@1.29.29,trna173_GlnTTG_6_-_26311975_26312046@1.29.29,trna174_GlnTTG_6_-_26311424_26311495@1.29.29|GGCCCCATGGTGTAATGGTTAGCACTCTG</td><td>>trna130_GlnTTG_6_-_27763640_27763711</td><td>1</td><td>29</td><td>29</td><td>GGCCCCATGGTGTAATGGTTAGCACTCTG</td><td>300.7</td><td>-12.0</td><td>1.5</td><td>-7.8</td><td>6.0E-15</td><td>2.3E-14</td><td>MB231/MB468</td><td>MITO</td></tr>
<tr><td>>am_trnaMT_HisGTG_MT_+_12138_12206@18.51.34|AAAACATCAGATTGTGAATCTGACAACAGAGGCT</td><td>>am_trnaMT_HisGTG_MT_+_12138_12206</td><td>18</td><td>51</td><td>34</td><td>AAAACATCAGATTGTGAATCTGACAACAGAGGCT</td><td>1331.9</td><td>10.5</td><td>1.3</td><td>7.9</td><td>3.7E-15</td><td>2.3E-14</td><td>BT20/MB468</td><td>MITO</td></tr>
<tr><td>>hg38_21_+_8433172_8446622_RNA45SN1_45S_50ntFlanks@10101.10124.24|CGCGCGCGTGTGGTGTGCGTCGGA</td><td>>hg38_21_+_8433172_8446622_RNA45SN1_45S_50ntFlanks</td><td>10101</td><td>10124</td><td>24</td><td>CGCGCGCGTGTGGTGTGCGTCGGA</td><td>333.6</td><td>-12.2</td><td>1.6</td><td>-7.9</td><td>3.6E-15</td><td>2.3E-14</td><td>MB231/MB468</td><td>MP</td></tr>
<tr><td>>hg38_21_+_8433172_8446622_RNA45SN1_45S_50ntFlanks@10771.10806.36|CAGGTCTCCAAGGTGAACAGCCTCTGGCATGTTGGA</td><td>>hg38_21_+_8433172_8446622_RNA45SN1_45S_50ntFlanks</td><td>10771</td><td>10806</td><td>36</td><td>CAGGTCTCCAAGGTGAACAGCCTCTGGCATGTTGGA</td><td>681.6</td><td>-8.8</td><td>1.1</td><td>-7.9</td><td>4.0E-15</td><td>2.3E-14</td><td>MB231/MB468</td><td>CYTO</td></tr>
<tr><td>>MI0000272|hsa-mir-182&WithFlank&7|-|129770377|129770498@29.50.22(+1U)&[MIMAT0000259&hsa-miR-182-5p&offsets|0|-2(+1U);m-22&7|-|129770449|129770470&offsets|0|0(+1U)]|TTTGGCAATGGTAGAACTCACAT</td><td>>MI0000272|hsa-mir-182&WithFlank&7|-|129770377|129770498</td><td>29</td><td>50</td><td>22(+1U)</td><td>TTTGGCAATGGTAGAACTCACAT</td><td>1486.2</td><td>11.7</td><td>1.5</td><td>7.8</td><td>5.1E-15</td><td>2.3E-14</td><td>BT20/MB468</td><td>CYTO</td></tr>
<tr><td>>MI0000063|hsa-let-7b&WithFlank&22|+|46113680|46113774@12.32.21(+1C)&[MIMAT0000063&hsa-let-7b-5p&offsets|0|-1(+1C);m-14&22|+|46113691|46113712&offsets|0|-1(+1C)]|TGAGGTAGTAGGTTGTGTGGTC</td><td>>MI0000063|hsa-let-7b&WithFlank&22|+|46113680|46113774</td><td>12</td><td>32</td><td>21(+1C)</td><td>TGAGGTAGTAGGTTGTGTGGTC</td><td>2263.4</td><td>3.2</td><td>0.4</td><td>7.8</td><td>7.7E-15</td><td>2.3E-14</td><td>BT20/MB468</td><td>NUC</td></tr>
<tr><td>>hg38_21_+_8433172_8446622_RNA45SN1_45S_50ntFlanks@10127.10145.19|GCGGCGGCGGCGGCGGCGG</td><td>>hg38_21_+_8433172_8446622_RNA45SN1_45S_50ntFlanks</td><td>10127</td><td>10145</td><td>19</td><td>GCGGCGGCGGCGGCGGCGG</td><td>429.3</td><td>-12.5</td><td>1.6</td><td>-7.8</td><td>6.0E-15</td><td>2.3E-14</td><td>MB231/MB468</td><td>MITO</td></tr>
<tr><td>>hg38_21_+_8433172_8446622_RNA45SN1_45S_50ntFlanks@10130.10148.19|GCGGCGGCGGCGGCGGCGG</td><td>>hg38_21_+_8433172_8446622_RNA45SN1_45S_50ntFlanks</td><td>10130</td><td>10148</td><td>19</td><td>GCGGCGGCGGCGGCGGCGG</td><td>429.3</td><td>-12.5</td><td>1.6</td><td>-7.8</td><td>6.0E-15</td><td>2.3E-14</td><td>MB231/MB468</td><td>MITO</td></tr>
<tr><td>>hg38_21_+_8433172_8446622_RNA45SN1_45S_50ntFlanks@8823.8841.19|GCGGCGGCGGCGGCGGCGG</td><td>>hg38_21_+_8433172_8446622_RNA45SN1_45S_50ntFlanks</td><td>8823</td><td>8841</td><td>19</td><td>GCGGCGGCGGCGGCGGCGG</td><td>429.3</td><td>-12.5</td><td>1.6</td><td>-7.8</td><td>6.0E-15</td><td>2.3E-14</td><td>MB231/MB468</td><td>MITO</td></tr>
<tr><td>>hg38_21_+_8433172_8446622_RNA45SN1_45S_50ntFlanks@8844.8862.19|GCGGCGGCGGCGGCGGCGG</td><td>>hg38_21_+_8433172_8446622_RNA45SN1_45S_50ntFlanks</td><td>8844</td><td>8862</td><td>19</td><td>GCGGCGGCGGCGGCGGCGG</td><td>429.3</td><td>-12.5</td><td>1.6</td><td>-7.8</td><td>6.0E-15</td><td>2.3E-14</td><td>MB231/MB468</td><td>MITO</td></tr>
<tr><td>>hg38_21_+_8433172_8446622_RNA45SN1_45S_50ntFlanks@8847.8865.19|GCGGCGGCGGCGGCGGCGG</td><td>>hg38_21_+_8433172_8446622_RNA45SN1_45S_50ntFlanks</td><td>8847</td><td>8865</td><td>19</td><td>GCGGCGGCGGCGGCGGCGG</td><td>429.3</td><td>-12.5</td><td>1.6</td><td>-7.8</td><td>6.0E-15</td><td>2.3E-14</td><td>MB231/MB468</td><td>MITO</td></tr>
<tr><td>>hg38_21_+_8433172_8446622_RNA45SN1_45S_50ntFlanks@9739.9757.19|AAACGATCTCAACCTATTC</td><td>>hg38_21_+_8433172_8446622_RNA45SN1_45S_50ntFlanks</td><td>9739</td><td>9757</td><td>19</td><td>AAACGATCTCAACCTATTC</td><td>6386.4</td><td>-11.2</td><td>1.4</td><td>-7.8</td><td>6.0E-15</td><td>2.3E-14</td><td>MB231/MB468</td><td>MITO</td></tr>
<tr><td>>hg38_21_+_8433172_8446622_RNA45SN1_45S_50ntFlanks@10547.10570.24|AGTGCGGTAACGCGACCGATCCCG</td><td>>hg38_21_+_8433172_8446622_RNA45SN1_45S_50ntFlanks</td><td>10547</td><td>10570</td><td>24</td><td>AGTGCGGTAACGCGACCGATCCCG</td><td>223.6</td><td>-11.7</td><td>1.5</td><td>-7.9</td><td>3.6E-15</td><td>2.3E-14</td><td>MB231/MB468</td><td>MP</td></tr>
<tr><td>>hg38_21_+_8433172_8446622_RNA45SN1_45S_50ntFlanks@10572.10607.36|AGAAGCCGGCGGGAGCCCCGGGGAGAGTTCTCTTTT</td><td>>hg38_21_+_8433172_8446622_RNA45SN1_45S_50ntFlanks</td><td>10572</td><td>10607</td><td>36</td><td>AGAAGCCGGCGGGAGCCCCGGGGAGAGTTCTCTTTT</td><td>1578.0</td><td>-10.8</td><td>1.4</td><td>-7.8</td><td>7.8E-15</td><td>2.3E-14</td><td>BT20/MB468</td><td>NUC</td></tr>
<tr><td>>hg38_21_+_8433172_8446622_RNA45SN1_45S_50ntFlanks@4899.4916.18|AAGGAATTGACGGAAGGG</td><td>>hg38_21_+_8433172_8446622_RNA45SN1_45S_50ntFlanks</td><td>4899</td><td>4916</td><td>18</td><td>AAGGAATTGACGGAAGGG</td><td>277.3</td><td>-12.0</td><td>1.5</td><td>-7.9</td><td>3.6E-15</td><td>2.3E-14</td><td>MB231/MB468</td><td>MP</td></tr>
<tr><td>>MI0000289|hsa-mir-181a-1&WithFlank&1|-|198859038|198859159@70.90.21(+1U)&[MIMAT0000270&hsa-miR-181a-3p&offsets|0|-1(+1U);m-125&1|-|198859069|198859090&offsets|0|-1(+1U)]|ACCATCGACCGTTGATTGTACT</td><td>>MI0000289|hsa-mir-181a-1&WithFlank&1|-|198859038|198859159</td><td>70</td><td>90</td><td>21(+1U)</td><td>ACCATCGACCGTTGATTGTACT</td><td>470.9</td><td>-12.1</td><td>1.5</td><td>-8.1</td><td>5.0E-16</td><td>2.3E-14</td><td>BT20/MB231</td><td>MITO</td></tr>
<tr><td>>hg38_21_+_8433172_8446622_RNA45SN1_45S_50ntFlanks@4006.4026.21|ACCTCGGGCCGATCGCACGCC</td><td>>hg38_21_+_8433172_8446622_RNA45SN1_45S_50ntFlanks</td><td>4006</td><td>4026</td><td>21</td><td>ACCTCGGGCCGATCGCACGCC</td><td>229.8</td><td>-11.7</td><td>1.5</td><td>-7.9</td><td>3.6E-15</td><td>2.3E-14</td><td>MB231/MB468</td><td>MP</td></tr>
<tr><td>>hg38_21_+_8433172_8446622_RNA45SN1_45S_50ntFlanks@9477.9497.21|AGCGCACGTGTTAGGACCCGA</td><td>>hg38_21_+_8433172_8446622_RNA45SN1_45S_50ntFlanks</td><td>9477</td><td>9497</td><td>21</td><td>AGCGCACGTGTTAGGACCCGA</td><td>2018.2</td><td>10.9</td><td>1.4</td><td>7.9</td><td>4.1E-15</td><td>2.3E-14</td><td>MB231/MB468</td><td>CYTO</td></tr>
<tr><td>>hg38_21_+_8433172_8446622_RNA45SN1_45S_50ntFlanks@7975.8019.45|CGCGACCTCAGATCAGACGTGGCGACCCGCTGAATTTAAGCATAT</td><td>>hg38_21_+_8433172_8446622_RNA45SN1_45S_50ntFlanks</td><td>7975</td><td>8019</td><td>45</td><td>CGCGACCTCAGATCAGACGTGGCGACCCGCTGAATTTAAGCATAT</td><td>338.1</td><td>9.1</td><td>1.2</td><td>7.9</td><td>4.1E-15</td><td>2.3E-14</td><td>MB231/MB468</td><td>CYTO</td></tr>
<tr><td>>hg38_21_+_8433172_8446622_RNA45SN1_45S_50ntFlanks@11027.11050.24|CTCTCTCTCTCTCTCCCCCGCTCC</td><td>>hg38_21_+_8433172_8446622_RNA45SN1_45S_50ntFlanks</td><td>11027</td><td>11050</td><td>24</td><td>CTCTCTCTCTCTCTCCCCCGCTCC</td><td>1602.4</td><td>-8.9</td><td>1.1</td><td>-7.9</td><td>4.1E-15</td><td>2.3E-14</td><td>MB231/MB468</td><td>CYTO</td></tr>
<tr><td>>hg38_21_+_8433172_8446622_RNA45SN1_45S_50ntFlanks@12935.12961.27|TAAACCATTCGTAGACGACCTGCTTCT</td><td>>hg38_21_+_8433172_8446622_RNA45SN1_45S_50ntFlanks</td><td>12935</td><td>12961</td><td>27</td><td>TAAACCATTCGTAGACGACCTGCTTCT</td><td>434.4</td><td>-4.6</td><td>0.6</td><td>-7.8</td><td>4.5E-15</td><td>2.3E-14</td><td>MB231/MB468</td><td>NUC</td></tr>
<tr><td>>hg38_21_+_8433172_8446622_RNA45SN1_45S_50ntFlanks@11562.11586.25|CGCCTAGCAGCCGACTTAGAACTGG</td><td>>hg38_21_+_8433172_8446622_RNA45SN1_45S_50ntFlanks</td><td>11562</td><td>11586</td><td>25</td><td>CGCCTAGCAGCCGACTTAGAACTGG</td><td>189.8</td><td>-11.4</td><td>1.5</td><td>-7.9</td><td>3.7E-15</td><td>2.3E-14</td><td>MB231/MB468</td><td>MP</td></tr>
<tr><td>>hg38_21_+_8433172_8446622_RNA45SN1_45S_50ntFlanks@10412.10429.18|AGTCGGTCCTGAGAGATG</td><td>>hg38_21_+_8433172_8446622_RNA45SN1_45S_50ntFlanks</td><td>10412</td><td>10429</td><td>18</td><td>AGTCGGTCCTGAGAGATG</td><td>286.0</td><td>-12.0</td><td>1.5</td><td>-7.9</td><td>3.7E-15</td><td>2.3E-14</td><td>MB231/MB468</td><td>MP</td></tr>
<tr><td>>hg38_21_+_8433172_8446622_RNA45SN1_45S_50ntFlanks@4064.4101.38|ATCAACTTTCGATGGTAGTCGCCGTGCCTACCATGGTG</td><td>>hg38_21_+_8433172_8446622_RNA45SN1_45S_50ntFlanks</td><td>4064</td><td>4101</td><td>38</td><td>ATCAACTTTCGATGGTAGTCGCCGTGCCTACCATGGTG</td><td>253.3</td><td>-11.8</td><td>1.5</td><td>-7.8</td><td>6.1E-15</td><td>2.3E-14</td><td>MB231/MB468</td><td>MITO</td></tr>
<tr><td>>MI0000264|hsa-mir-7-2&WithFlank&15|+|88611819|88611940@38.59.22;MI0000265|hsa-mir-7-3&WithFlank&19|+|4770664|4770785@37.58.22;MI0000263|hsa-mir-7-1&WithFlank&9|-|83969742|83969863@30.51.22&[MIMAT0000252&hsa-miR-7-5p&offsets|0|-2;m-400&15|+|88611856|88611879&offsets|0|-2];[MIMAT0000252_1&hsa-miR-7-5p&offsets|0|-2;m-402&19|+|4770700|4770723&offsets|0|-2];[MIMAT0000252_2&hsa-miR-7-5p&offsets|0|-2;m-401&9|-|83969811|83969834&offsets|0|-2]|TGGAAGACTAGTGATTTTGTTG</td><td>>MI0000264|hsa-mir-7-2&WithFlank&15|+|88611819|88611940</td><td>38</td><td>59</td><td>22</td><td>TGGAAGACTAGTGATTTTGTTG</td><td>1537.8</td><td>7.6</td><td>1.0</td><td>7.8</td><td>5.2E-15</td><td>2.3E-14</td><td>BT20/MB468</td><td>CYTO</td></tr>
<tr><td>>hg38_21_+_8433172_8446622_RNA45SN1_45S_50ntFlanks@10807.10826.20|ACAATGTAGGTAAGGGAAGT</td><td>>hg38_21_+_8433172_8446622_RNA45SN1_45S_50ntFlanks</td><td>10807</td><td>10826</td><td>20</td><td>ACAATGTAGGTAAGGGAAGT</td><td>794.2</td><td>-9.2</td><td>1.2</td><td>-7.8</td><td>6.2E-15</td><td>2.3E-14</td><td>MB231/MB468</td><td>MITO</td></tr>
<tr><td>>hg38_21_+_8433172_8446622_RNA45SN1_45S_50ntFlanks@10351.10376.26|CGAGAACTTTGAAGGCCGAAGTGGAG</td><td>>hg38_21_+_8433172_8446622_RNA45SN1_45S_50ntFlanks</td><td>10351</td><td>10376</td><td>26</td><td>CGAGAACTTTGAAGGCCGAAGTGGAG</td><td>4269.5</td><td>-11.7</td><td>1.5</td><td>-7.8</td><td>7.9E-15</td><td>2.3E-14</td><td>BT20/MB468</td><td>NUC</td></tr>
<tr><td>>hg38_21_+_8433172_8446622_RNA45SN1_45S_50ntFlanks@9739.9758.20|AAACGATCTCAACCTATTCT</td><td>>hg38_21_+_8433172_8446622_RNA45SN1_45S_50ntFlanks</td><td>9739</td><td>9758</td><td>20</td><td>AAACGATCTCAACCTATTCT</td><td>2008.6</td><td>-10.0</td><td>1.3</td><td>-7.9</td><td>3.7E-15</td><td>2.3E-14</td><td>MB231/MB468</td><td>MP</td></tr>
<tr><td>>hg38_21_+_8433172_8446622_RNA45SN1_45S_50ntFlanks@12365.12382.18|ATAACTGGCTTGTGGCGG</td><td>>hg38_21_+_8433172_8446622_RNA45SN1_45S_50ntFlanks</td><td>12365</td><td>12382</td><td>18</td><td>ATAACTGGCTTGTGGCGG</td><td>453.8</td><td>-8.7</td><td>1.1</td><td>-7.8</td><td>4.5E-15</td><td>2.3E-14</td><td>MB231/MB468</td><td>NUC</td></tr>
<tr><td>>hg38_21_+_8433172_8446622_RNA45SN1_45S_50ntFlanks@6772.6799.28|GTTCCTCCCGGGGCTACGCCTGTCTGAG</td><td>>hg38_21_+_8433172_8446622_RNA45SN1_45S_50ntFlanks</td><td>6772</td><td>6799</td><td>28</td><td>GTTCCTCCCGGGGCTACGCCTGTCTGAG</td><td>777.6</td><td>-9.0</td><td>1.2</td><td>-7.8</td><td>4.5E-15</td><td>2.3E-14</td><td>MB231/MB468</td><td>NUC</td></tr>
<tr><td>>MI0001445|hsa-mir-423&WithFlank&17|+|30117073|30117178@58.80.23(+1C)&[MIMAT0001340&hsa-miR-423-3p&offsets|-1|-1(+1C);m-100&17|+|30117131|30117153&offsets|-1|-1(+1C)]|AAGCTCGGTCTGAGGCCCCTCAGC</td><td>>MI0001445|hsa-mir-423&WithFlank&17|+|30117073|30117178</td><td>58</td><td>80</td><td>23(+1C)</td><td>AAGCTCGGTCTGAGGCCCCTCAGC</td><td>98.3</td><td>4.1</td><td>0.5</td><td>7.8</td><td>8.0E-15</td><td>2.3E-14</td><td>BT20/MB468</td><td>NUC</td></tr>
<tr><td>>hg38_21_+_8433172_8446622_RNA45SN1_45S_50ntFlanks@11783.11804.22|CATGAATGGATGAACGAGATTC</td><td>>hg38_21_+_8433172_8446622_RNA45SN1_45S_50ntFlanks</td><td>11783</td><td>11804</td><td>22</td><td>CATGAATGGATGAACGAGATTC</td><td>20108.5</td><td>-6.1</td><td>0.8</td><td>-7.9</td><td>4.1E-15</td><td>2.3E-14</td><td>MB231/MB468</td><td>CYTO</td></tr>
<tr><td>>hg38_21_+_8433172_8446622_RNA45SN1_45S_50ntFlanks@12626.12649.24|AATGGGGCGAAGCTACCATCTGTG</td><td>>hg38_21_+_8433172_8446622_RNA45SN1_45S_50ntFlanks</td><td>12626</td><td>12649</td><td>24</td><td>AATGGGGCGAAGCTACCATCTGTG</td><td>329.0</td><td>-12.1</td><td>1.6</td><td>-7.8</td><td>6.2E-15</td><td>2.3E-14</td><td>MB231/MB468</td><td>MITO</td></tr>
<tr><td>>hg38_21_+_8433172_8446622_RNA45SN1_45S_50ntFlanks@6731.6773.43|CACATTGATCATCGACACTTCGAACGCACTTGCGGCCCCGGGT</td><td>>hg38_21_+_8433172_8446622_RNA45SN1_45S_50ntFlanks</td><td>6731</td><td>6773</td><td>43</td><td>CACATTGATCATCGACACTTCGAACGCACTTGCGGCCCCGGGT</td><td>352.8</td><td>5.4</td><td>0.7</td><td>7.9</td><td>3.9E-15</td><td>2.3E-14</td><td>BT20/MB468</td><td>MITO</td></tr>
<tr><td>>hg38_21_+_8433172_8446622_RNA45SN1_45S_50ntFlanks@10389.10410.22|TGAACAGCAGTTGAACATGGGT</td><td>>hg38_21_+_8433172_8446622_RNA45SN1_45S_50ntFlanks</td><td>10389</td><td>10410</td><td>22</td><td>TGAACAGCAGTTGAACATGGGT</td><td>3906.1</td><td>-8.6</td><td>1.1</td><td>-7.8</td><td>4.6E-15</td><td>2.3E-14</td><td>MB231/MB468</td><td>NUC</td></tr>
<tr><td>>MI0000481|hsa-mir-184&WithFlank&15|+|79209782|79209877@59.80.22&[MIMAT0000454&hsa-miR-184&offsets|0|0;m-460&15|+|79209840|79209861&offsets|0|0]|TGGACGGAGAACTGATAAGGGT</td><td>>MI0000481|hsa-mir-184&WithFlank&15|+|79209782|79209877</td><td>59</td><td>80</td><td>22</td><td>TGGACGGAGAACTGATAAGGGT</td><td>265.4</td><td>7.5</td><td>0.9</td><td>8.1</td><td>3.7E-16</td><td>2.3E-14</td><td>BT20/MB231</td><td>CYTO</td></tr>
<tr><td>>hg38_21_+_8433172_8446622_RNA45SN1_45S_50ntFlanks@11190.11225.36|CCCGGGGGGCCGGCGGCGGCGGCGACTCTGGACGCG</td><td>>hg38_21_+_8433172_8446622_RNA45SN1_45S_50ntFlanks</td><td>11190</td><td>11225</td><td>36</td><td>CCCGGGGGGCCGGCGGCGGCGGCGACTCTGGACGCG</td><td>924.5</td><td>-13.7</td><td>1.7</td><td>-7.9</td><td>3.8E-15</td><td>2.3E-14</td><td>MB231/MB468</td><td>MP</td></tr>
</table>

<table>
<tr><th></th><th></th><th></th><th></th><th></th><th></th><th></th><th></th><th></th><th></th><th></th><th></th><th></th><th></th></tr>
<tr><td>>hg38_21_+_8433172_8446622_RNA45SN1_45S_50ntFlanks@12475.12498.24|CCCACTAATAGGGAACGTGAGCTG</td><td>>hg38_21_+_8433172_8446622_RNA45SN1_45S_50ntFlanks</td><td>12475</td><td>12498</td><td>24</td><td>CCCACTAATAGGGAACGTGAGCTG</td><td>652.1</td><td>-7.4</td><td>1.0</td><td>-7.8</td><td>6.3E-15</td><td>2.3E-14</td><td>MB231/MB468</td><td>MITO</td></tr>
<tr><td>>hg38_21_+_8433172_8446622_RNA45SN1_45S_50ntFlanks@9352.9379.28|CCGTGGCGCAATGAAGGTGAAGGCCGGC</td><td>>hg38_21_+_8433172_8446622_RNA45SN1_45S_50ntFlanks</td><td>9352</td><td>9379</td><td>28</td><td>CCGTGGCGCAATGAAGGTGAAGGCCGGC</td><td>289.5</td><td>-12.0</td><td>1.5</td><td>-7.8</td><td>6.3E-15</td><td>2.3E-14</td><td>MB231/MB468</td><td>MITO</td></tr>
<tr><td>>hg38_21_+_8433172_8446622_RNA45SN1_45S_50ntFlanks@10914.10945.32|CGCCGCGGCTGGACGAGGCGCCGCCGCCCCCC</td><td>>hg38_21_+_8433172_8446622_RNA45SN1_45S_50ntFlanks</td><td>10914</td><td>10945</td><td>32</td><td>CGCCGCGGCTGGACGAGGCGCCGCCGCCCCCC</td><td>234.2</td><td>-11.7</td><td>1.5</td><td>-7.9</td><td>3.8E-15</td><td>2.3E-14</td><td>MB231/MB468</td><td>MP</td></tr>
<tr><td>>hg38_21_+_8433172_8446622_RNA45SN1_45S_50ntFlanks@11580.11607.28|GAACTGGTGCGGACCAGGGGAATCCGAC</td><td>>hg38_21_+_8433172_8446622_RNA45SN1_45S_50ntFlanks</td><td>11580</td><td>11607</td><td>28</td><td>GAACTGGTGCGGACCAGGGGAATCCGAC</td><td>450.1</td><td>-4.4</td><td>0.6</td><td>-7.8</td><td>8.1E-15</td><td>2.4E-14</td><td>BT20/MB468</td><td>NUC</td></tr>
<tr><td>>hg38_21_+_8433172_8446622_RNA45SN1_45S_50ntFlanks@5526.5550.25|AAAAGTCGTAACAAGGTTTCCGTAG</td><td>>hg38_21_+_8433172_8446622_RNA45SN1_45S_50ntFlanks</td><td>5526</td><td>5550</td><td>25</td><td>AAAAGTCGTAACAAGGTTTCCGTAG</td><td>237.5</td><td>-11.8</td><td>1.5</td><td>-7.9</td><td>3.8E-15</td><td>2.4E-14</td><td>MB231/MB468</td><td>MP</td></tr>
<tr><td>>hg38_21_+_8433172_8446622_RNA45SN1_45S_50ntFlanks@11881.11906.26|CCTGTTGAGCTTGACTCTAGTCTGGC</td><td>>hg38_21_+_8433172_8446622_RNA45SN1_45S_50ntFlanks</td><td>11881</td><td>11906</td><td>26</td><td>CCTGTTGAGCTTGACTCTAGTCTGGC</td><td>162.1</td><td>-11.2</td><td>1.4</td><td>-7.9</td><td>3.8E-15</td><td>2.4E-14</td><td>MB231/MB468</td><td>MP</td></tr>
<tr><td>>trnaMT_MetCAT_MT_+_4402_4469@38.71.34|CGAAAATGTTGGTTATACCCTTCCCGTACTACCA</td><td>>trnaMT_MetCAT_MT_+_4402_4469</td><td>38</td><td>71</td><td>34</td><td>CGAAAATGTTGGTTATACCCTTCCCGTACTACCA</td><td>766.8</td><td>6.4</td><td>0.8</td><td>7.9</td><td>3.9E-15</td><td>2.4E-14</td><td>BT20/MB468</td><td>MITO</td></tr>
<tr><td>>hg38_21_+_8433172_8446622_RNA45SN1_45S_50ntFlanks@5083.5100.18|ACGAACGAGACTCTGGCA</td><td>>hg38_21_+_8433172_8446622_RNA45SN1_45S_50ntFlanks</td><td>5083</td><td>5100</td><td>18</td><td>ACGAACGAGACTCTGGCA</td><td>581.5</td><td>7.3</td><td>0.9</td><td>7.8</td><td>5.3E-15</td><td>2.4E-14</td><td>BT20/MB468</td><td>CYTO</td></tr>
<tr><td>>hg38_21_+_8433172_8446622_RNA45SN1_45S_50ntFlanks@6773.6797.25|TTCCTCCCGGGGCTACGCCTGTCTG</td><td>>hg38_21_+_8433172_8446622_RNA45SN1_45S_50ntFlanks</td><td>6773</td><td>6797</td><td>25</td><td>TTCCTCCCGGGGCTACGCCTGTCTG</td><td>226.1</td><td>-10.8</td><td>1.4</td><td>-7.8</td><td>4.6E-15</td><td>2.4E-14</td><td>MB231/MB468</td><td>NUC</td></tr>
<tr><td>>hg38_21_+_8433172_8446622_RNA45SN1_45S_50ntFlanks@476.503.28|TTCGTGATCGATGTGGTGACGTCGTGCT</td><td>>hg38_21_+_8433172_8446622_RNA45SN1_45S_50ntFlanks</td><td>476</td><td>503</td><td>28</td><td>TTCGTGATCGATGTGGTGACGTCGTGCT</td><td>1060.8</td><td>7.2</td><td>0.9</td><td>7.9</td><td>3.9E-15</td><td>2.4E-14</td><td>BT20/MB468</td><td>MITO</td></tr>
<tr><td>>hg38_21_+_8433172_8446622_RNA45SN1_45S_50ntFlanks@4005.4024.20|AACCTCGGGCCGATCGCACG</td><td>>hg38_21_+_8433172_8446622_RNA45SN1_45S_50ntFlanks</td><td>4005</td><td>4024</td><td>20</td><td>AACCTCGGGCCGATCGCACG</td><td>610.6</td><td>-7.4</td><td>0.9</td><td>-7.9</td><td>3.8E-15</td><td>2.4E-14</td><td>MB231/MB468</td><td>MP</td></tr>
<tr><td>>hg38_21_+_8433172_8446622_RNA45SN1_45S_50ntFlanks@4918.4941.24|ACCACCAGGAGTGGAGCCTGCGGC</td><td>>hg38_21_+_8433172_8446622_RNA45SN1_45S_50ntFlanks</td><td>4918</td><td>4941</td><td>24</td><td>ACCACCAGGAGTGGAGCCTGCGGC</td><td>258.7</td><td>-11.9</td><td>1.5</td><td>-7.9</td><td>3.8E-15</td><td>2.4E-14</td><td>MB231/MB468</td><td>MP</td></tr>
<tr><td>>hg38_21_+_8433172_8446622_RNA45SN1_45S_50ntFlanks@5498.5526.29|CGGTCGAACTTGACTATCTAGAGGAAGTA</td><td>>hg38_21_+_8433172_8446622_RNA45SN1_45S_50ntFlanks</td><td>5498</td><td>5526</td><td>29</td><td>CGGTCGAACTTGACTATCTAGAGGAAGTA</td><td>265.1</td><td>-11.9</td><td>1.5</td><td>-7.9</td><td>3.8E-15</td><td>2.4E-14</td><td>MB231/MB468</td><td>MP</td></tr>
<tr><td>>MI0000266|hsa-mir-10a&WithFlank&17|-|48579832|48579953@29.50.22(+1A)&[MIMAT0000253&hsa-miR-10a-5p&offsets|+1|0(+1A);m-18&17|-|48579905|48579926&offsets|+1|+1(+1A)]|ACCCTGTAGATCCGAATTTGTGA</td><td>>MI0000266|hsa-mir-10a&WithFlank&17|-|48579832|48579953</td><td>29</td><td>50</td><td>22(+1A)</td><td>ACCCTGTAGATCCGAATTTGTGA</td><td>233.3</td><td>11.1</td><td>1.4</td><td>7.9</td><td>3.8E-15</td><td>2.4E-14</td><td>MB231/MB468</td><td>MP</td></tr>
<tr><td>>hg38_21_+_8433172_8446622_RNA45SN1_45S_50ntFlanks@12628.12652.25|TGGGGCGAAGCTACCATCTGTGGGA</td><td>>hg38_21_+_8433172_8446622_RNA45SN1_45S_50ntFlanks</td><td>12628</td><td>12652</td><td>25</td><td>TGGGGCGAAGCTACCATCTGTGGGA</td><td>307.7</td><td>-12.1</td><td>1.5</td><td>-7.9</td><td>3.8E-15</td><td>2.4E-14</td><td>MB231/MB468</td><td>MP</td></tr>
<tr><td>>hg38_21_+_8433172_8446622_RNA45SN1_45S_50ntFlanks@5444.5467.24|CTCGGATCGGCCCCGCCGGGGTCG</td><td>>hg38_21_+_8433172_8446622_RNA45SN1_45S_50ntFlanks</td><td>5444</td><td>5467</td><td>24</td><td>CTCGGATCGGCCCCGCCGGGGTCG</td><td>4450.6</td><td>-10.6</td><td>1.3</td><td>-7.9</td><td>3.8E-15</td><td>2.4E-14</td><td>MB231/MB468</td><td>MP</td></tr>
<tr><td>>hg38_21_+_8433172_8446622_RNA45SN1_45S_50ntFlanks@8117.8139.23|CGGGACATGTGGCGTACGGAAGA</td><td>>hg38_21_+_8433172_8446622_RNA45SN1_45S_50ntFlanks</td><td>8117</td><td>8139</td><td>23</td><td>CGGGACATGTGGCGTACGGAAGA</td><td>262.3</td><td>-11.9</td><td>1.5</td><td>-7.9</td><td>3.8E-15</td><td>2.4E-14</td><td>MB231/MB468</td><td>MP</td></tr>
<tr><td>>hg38_21_+_8433172_8446622_RNA45SN1_45S_50ntFlanks@10366.10388.23|CCGAAGTGGAGAAGGGTTCCATG</td><td>>hg38_21_+_8433172_8446622_RNA45SN1_45S_50ntFlanks</td><td>10366</td><td>10388</td><td>23</td><td>CCGAAGTGGAGAAGGGTTCCATG</td><td>169.7</td><td>-11.3</td><td>1.4</td><td>-7.9</td><td>3.8E-15</td><td>2.4E-14</td><td>MB231/MB468</td><td>MP</td></tr>
<tr><td>>MI0000093|hsa-mir-92a-1&WithFlank&13|+|91351308|91351397@55.75.21;MI0000094|hsa-mir-92a-2&WithFlank&X|-|134169532|134169618@55.75.21&[MIMAT0000092&hsa-miR-92a-3p&offsets|+1|0;m-9&13|+|91351361|91351382&offsets|+1|0];[MIMAT0000092_1&hsa-miR-92a-3p&offsets|+1|0;m-11&X|-|134169544|134169565&offsets|+1|0]|ATTGCACTTGTCCCGGCCTGT</td><td>>MI0000093|hsa-mir-92a-1&WithFlank&13|+|91351308|91351397</td><td>55</td><td>75</td><td>21</td><td>ATTGCACTTGTCCCGGCCTGT</td><td>982.1</td><td>5.4</td><td>0.7</td><td>7.9</td><td>3.9E-15</td><td>2.4E-14</td><td>BT20/MB468</td><td>MITO</td></tr>
<tr><td>>hg38_21_+_8433172_8446622_RNA45SN1_45S_50ntFlanks@9806.9830.25|CGTGGAATGCGAGTGCCTAGTGGGC</td><td>>hg38_21_+_8433172_8446622_RNA45SN1_45S_50ntFlanks</td><td>9806</td><td>9830</td><td>25</td><td>CGTGGAATGCGAGTGCCTAGTGGGC</td><td>1128.9</td><td>-8.3</td><td>1.1</td><td>-7.9</td><td>3.8E-15</td><td>2.4E-14</td><td>MB231/MB468</td><td>MP</td></tr>
<tr><td>>MI0000299|hsa-mir-222&WithFlank&X|-|45747009|45747130@75.99.25(+1A)&[MIMAT0000279&hsa-miR-222-3p&offsets|0|+4(+1A);m-140&X|-|45747033|45747056&offsets|0|+1(+1A)]|AGCTACATCTGGCTACTGGGTCTCTA</td><td>>MI0000299|hsa-mir-222&WithFlank&X|-|45747009|45747130</td><td>75</td><td>99</td><td>25(+1A)</td><td>AGCTACATCTGGCTACTGGGTCTCTA</td><td>239.8</td><td>6.2</td><td>0.8</td><td>7.8</td><td>4.7E-15</td><td>2.4E-14</td><td>MB231/MB468</td><td>NUC</td></tr>
<tr><td>>hg38_21_+_8433172_8446622_RNA45SN1_45S_50ntFlanks@10679.10706.28|GGTTCCGGCGGCGTCCGGTGAGCTCTCG</td><td>>hg38_21_+_8433172_8446622_RNA45SN1_45S_50ntFlanks</td><td>10679</td><td>10706</td><td>28</td><td>GGTTCCGGCGGCGTCCGGTGAGCTCTCG</td><td>3415.1</td><td>-10.2</td><td>1.3</td><td>-7.8</td><td>4.7E-15</td><td>2.4E-14</td><td>MB231/MB468</td><td>NUC</td></tr>
<tr><td>>hg38_21_+_8433172_8446622_RNA45SN1_45S_50ntFlanks@8212.8230.19|CCGGTAGCGGCCCCCGGCG</td><td>>hg38_21_+_8433172_8446622_RNA45SN1_45S_50ntFlanks</td><td>8212</td><td>8230</td><td>19</td><td>CCGGTAGCGGCCCCCGGCG</td><td>169.0</td><td>-11.3</td><td>1.4</td><td>-7.9</td><td>3.9E-15</td><td>2.4E-14</td><td>MB231/MB468</td><td>MP</td></tr>
<tr><td>>hg38_21_+_8433172_8446622_RNA45SN1_45S_50ntFlanks@12990.13017.28|GCTCCCTCGCTGCGATCTATTGAAAGTC</td><td>>hg38_21_+_8433172_8446622_RNA45SN1_45S_50ntFlanks</td><td>12990</td><td>13017</td><td>28</td><td>GCTCCCTCGCTGCGATCTATTGAAAGTC</td><td>532.7</td><td>5.5</td><td>0.7</td><td>7.8</td><td>4.3E-15</td><td>2.4E-14</td><td>MB231/MB468</td><td>CYTO</td></tr>
<tr><td>>hg38_21_+_8433172_8446622_RNA45SN1_45S_50ntFlanks@11027.11049.23|CTCTCTCTCTCTCTCCCCCGCTC</td><td>>hg38_21_+_8433172_8446622_RNA45SN1_45S_50ntFlanks</td><td>11027</td><td>11049</td><td>23</td><td>CTCTCTCTCTCTCTCCCCCGCTC</td><td>1287.3</td><td>-10.5</td><td>1.3</td><td>-7.8</td><td>4.7E-15</td><td>2.4E-14</td><td>MB231/MB468</td><td>NUC</td></tr>
<tr><td>>hg38_21_+_8433172_8446622_RNA45SN1_45S_50ntFlanks@10116.10136.21|TGCGTCGGAGGGCGGCGGCGG</td><td>>hg38_21_+_8433172_8446622_RNA45SN1_45S_50ntFlanks</td><td>10116</td><td>10136</td><td>21</td><td>TGCGTCGGAGGGCGGCGGCGG</td><td>372.4</td><td>-11.5</td><td>1.5</td><td>-7.8</td><td>4.7E-15</td><td>2.4E-14</td><td>MB231/MB468</td><td>NUC</td></tr>
<tr><td>>hg38_21_+_8433172_8446622_RNA45SN1_45S_50ntFlanks@9718.9737.20|TGATTAGAGGTCTTGGGGCC</td><td>>hg38_21_+_8433172_8446622_RNA45SN1_45S_50ntFlanks</td><td>9718</td><td>9737</td><td>20</td><td>TGATTAGAGGTCTTGGGGCC</td><td>183.9</td><td>-11.4</td><td>1.4</td><td>-7.9</td><td>3.9E-15</td><td>2.4E-14</td><td>MB231/MB468</td><td>MP</td></tr>
<tr><td>>hg38_21_+_8433172_8446622_RNA45SN1_45S_50ntFlanks@4497.4520.24|GAAGCGTTTACTTTGAAAAAATTA</td><td>>hg38_21_+_8433172_8446622_RNA45SN1_45S_50ntFlanks</td><td>4497</td><td>4520</td><td>24</td><td>GAAGCGTTTACTTTGAAAAAATTA</td><td>449.2</td><td>5.9</td><td>0.8</td><td>7.8</td><td>5.5E-15</td><td>2.4E-14</td><td>BT20/MB468</td><td>CYTO</td></tr>
<tr><td>>MI0000298|hsa-mir-221&WithFlank&X|-|45746151|45746272@71.93.23(+1C)&[MIMAT0000278&hsa-miR-221-3p&offsets|0|0(+1C);m-91&X|-|45746181|45746202&offsets|0|+1(+1C)]|AGCTACATTGTCTGCTGGGTTTCC</td><td>>MI0000298|hsa-mir-221&WithFlank&X|-|45746151|45746272</td><td>71</td><td>93</td><td>23(+1C)</td><td>AGCTACATTGTCTGCTGGGTTTCC</td><td>1996.0</td><td>6.4</td><td>0.8</td><td>7.8</td><td>4.3E-15</td><td>2.4E-14</td><td>MB231/MB468</td><td>CYTO</td></tr>
<tr><td>>hg38_21_+_8433172_8446622_RNA45SN1_45S_50ntFlanks@7989.8017.29|AGACGTGGCGACCCGCTGAATTTAAGCAT</td><td>>hg38_21_+_8433172_8446622_RNA45SN1_45S_50ntFlanks</td><td>7989</td><td>8017</td><td>29</td><td>AGACGTGGCGACCCGCTGAATTTAAGCAT</td><td>642.8</td><td>5.9</td><td>0.7</td><td>7.8</td><td>4.3E-15</td><td>2.4E-14</td><td>MB231/MB468</td><td>CYTO</td></tr>
<tr><td>>hg38_21_+_8433172_8446622_RNA45SN1_45S_50ntFlanks@12324.12342.19|TTGGGTTTTAAGCAGGAGG</td><td>>hg38_21_+_8433172_8446622_RNA45SN1_45S_50ntFlanks</td><td>12324</td><td>12342</td><td>19</td><td>TTGGGTTTTAAGCAGGAGG</td><td>471.6</td><td>-8.6</td><td>1.1</td><td>-7.8</td><td>4.3E-15</td><td>2.4E-14</td><td>MB231/MB468</td><td>CYTO</td></tr>
<tr><td>>hg38_1_-_228634819_228635039_RNA5S12_5S_50ntFlanks@51.71.21|GTCTACGGCCATACCACCCTG</td><td>>hg38_1_-_228634819_228635039_RNA5S12_5S_50ntFlanks</td><td>51</td><td>71</td><td>21</td><td>GTCTACGGCCATACCACCCTG</td><td>2225.9</td><td>-5.7</td><td>0.7</td><td>-7.8</td><td>8.3E-15</td><td>2.4E-14</td><td>BT20/MB468</td><td>NUC</td></tr>
<tr><td>>hg38_21_+_8433172_8446622_RNA45SN1_45S_50ntFlanks@9202.9225.24|TCTCGGGGCCACGCGCGCGTCCCC</td><td>>hg38_21_+_8433172_8446622_RNA45SN1_45S_50ntFlanks</td><td>9202</td><td>9225</td><td>24</td><td>TCTCGGGGCCACGCGCGCGTCCCC</td><td>278.8</td><td>-11.9</td><td>1.5</td><td>-7.8</td><td>6.5E-15</td><td>2.4E-14</td><td>MB231/MB468</td><td>MITO</td></tr>
<tr><td>>hg38_21_+_8433172_8446622_RNA45SN1_45S_50ntFlanks@3775.3794.20|GCCGGTACAGTGAAACTGCG</td><td>>hg38_21_+_8433172_8446622_RNA45SN1_45S_50ntFlanks</td><td>3775</td><td>3794</td><td>20</td><td>GCCGGTACAGTGAAACTGCG</td><td>410.3</td><td>-12.5</td><td>1.6</td><td>-7.9</td><td>3.9E-15</td><td>2.4E-14</td><td>MB231/MB468</td><td>MP</td></tr>
<tr><td>>hg38_21_+_8433172_8446622_RNA45SN1_45S_50ntFlanks@8198.8225.28|TGGACGGTGTGAGGCCGGTAGCGGCCCC</td><td>>hg38_21_+_8433172_8446622_RNA45SN1_45S_50ntFlanks</td><td>8198</td><td>8225</td><td>28</td><td>TGGACGGTGTGAGGCCGGTAGCGGCCCC</td><td>648.1</td><td>-6.1</td><td>0.8</td><td>-7.8</td><td>5.5E-15</td><td>2.4E-14</td><td>BT20/MB468</td><td>CYTO</td></tr>
<tr><td>>hg38_21_+_8433172_8446622_RNA45SN1_45S_50ntFlanks@11667.11702.36|CTGCCCAGTGCTCTGAATGTCAAAGTGAAGAAATTC</td><td>>hg38_21_+_8433172_8446622_RNA45SN1_45S_50ntFlanks</td><td>11667</td><td>11702</td><td>36</td><td>CTGCCCAGTGCTCTGAATGTCAAAGTGAAGAAATTC</td><td>489.2</td><td>4.7</td><td>0.6</td><td>7.9</td><td>4.1E-15</td><td>2.4E-14</td><td>BT20/MB468</td><td>MITO</td></tr>
<tr><td>>hg38_21_+_8433172_8446622_RNA45SN1_45S_50ntFlanks@4378.4398.21|CTCGTAGTTGGATCTTGGGAG</td><td>>hg38_21_+_8433172_8446622_RNA45SN1_45S_50ntFlanks</td><td>4378</td><td>4398</td><td>21</td><td>CTCGTAGTTGGATCTTGGGAG</td><td>232.6</td><td>-11.7</td><td>1.5</td><td>-7.9</td><td>4.0E-15</td><td>2.5E-14</td><td>MB231/MB468</td><td>MP</td></tr>
<tr><td>>hg38_21_+_8433172_8446622_RNA45SN1_45S_50ntFlanks@10926.10946.21|ACGAGGCGCCGCCGCCCCCCC</td><td>>hg38_21_+_8433172_8446622_RNA45SN1_45S_50ntFlanks</td><td>10926</td><td>10946</td><td>21</td><td>ACGAGGCGCCGCCGCCCCCCC</td><td>264.2</td><td>-10.8</td><td>1.4</td><td>-7.8</td><td>4.4E-15</td><td>2.5E-14</td><td>MB231/MB468</td><td>CYTO</td></tr>
<tr><td>>MI0000477|hsa-mir-146a&WithFlank&5|+|160485346|160485456@27.48.22(+2U)&[MIMAT0000449&hsa-miR-146a-5p&offsets|0|0(+2U);m-129&5|+|160485372|160485393&offsets|0|0(+2U)]|TGAGAACTGAATTCCATGGGTTTT</td><td>>MI0000477|hsa-mir-146a&WithFlank&5|+|160485346|160485456</td><td>27</td><td>48</td><td>22(+2U)</td><td>TGAGAACTGAATTCCATGGGTTTT</td><td>1762.4</td><td>-14.2</td><td>1.7</td><td>-8.3</td><td>7.2E-17</td><td>2.5E-14</td><td>BT20/MB231</td><td>MP</td></tr>
<tr><td>>hg38_21_+_8433172_8446622_RNA45SN1_45S_50ntFlanks@11452.11475.24|CTCCTCGGCGCGCGGCGGCGGCGG</td><td>>hg38_21_+_8433172_8446622_RNA45SN1_45S_50ntFlanks</td><td>11452</td><td>11475</td><td>24</td><td>CTCCTCGGCGCGCGGCGGCGGCGG</td><td>242.5</td><td>-11.7</td><td>1.5</td><td>-7.8</td><td>6.6E-15</td><td>2.5E-14</td><td>MB231/MB468</td><td>MITO</td></tr>
<tr><td>>hg38_21_+_8433172_8446622_RNA45SN1_45S_50ntFlanks@10681.10706.26|TTCCGGCGGCGTCCGGTGAGCTCTCG</td><td>>hg38_21_+_8433172_8446622_RNA45SN1_45S_50ntFlanks</td><td>10681</td><td>10706</td><td>26</td><td>TTCCGGCGGCGTCCGGTGAGCTCTCG</td><td>3141.1</td><td>-6.4</td><td>0.8</td><td>-7.8</td><td>4.4E-15</td><td>2.5E-14</td><td>MB231/MB468</td><td>CYTO</td></tr>
<tr><td>>hg38_21_+_8433172_8446622_RNA45SN1_45S_50ntFlanks@11027.11075.49|CTCTCTCTCTCTCTCCCCCGCTCCCCGTCCTCCCCCCTCCCCGGGGGAG</td><td>>hg38_21_+_8433172_8446622_RNA45SN1_45S_50ntFlanks</td><td>11027</td><td>11075</td><td>49</td><td>CTCTCTCTCTCTCTCCCCCGCTCCCCGTCCTCCCCCCTCCCCGGGGGAG</td><td>708.6</td><td>-13.3</td><td>1.7</td><td>-7.8</td><td>6.6E-15</td><td>2.5E-14</td><td>MB231/MB468</td><td>MITO</td></tr>
<tr><td>>hg38_21_+_8433172_8446622_RNA45SN1_45S_50ntFlanks@6721.6740.20|AATTGCAGGACACATTGATC</td><td>>hg38_21_+_8433172_8446622_RNA45SN1_45S_50ntFlanks</td><td>6721</td><td>6740</td><td>20</td><td>AATTGCAGGACACATTGATC</td><td>8247.0</td><td>-6.4</td><td>0.8</td><td>-7.8</td><td>4.5E-15</td><td>2.5E-14</td><td>MB231/MB468</td><td>CYTO</td></tr>
<tr><td>>hg38_21_+_8433172_8446622_RNA45SN1_45S_50ntFlanks@8388.8405.18|CGTGAAACCGTTAAGAGG</td><td>>hg38_21_+_8433172_8446622_RNA45SN1_45S_50ntFlanks</td><td>8388</td><td>8405</td><td>18</td><td>CGTGAAACCGTTAAGAGG</td><td>257.0</td><td>-11.8</td><td>1.5</td><td>-7.8</td><td>6.7E-15</td><td>2.5E-14</td><td>MB231/MB468</td><td>MITO</td></tr>
<tr><td>>hg38_21_+_8433172_8446622_RNA45SN1_45S_50ntFlanks@9308.9337.30|AGGAGTCTAACACGTGCGCGAGTCGGGGGC</td><td>>hg38_21_+_8433172_8446622_RNA45SN1_45S_50ntFlanks</td><td>9308</td><td>9337</td><td>30</td><td>AGGAGTCTAACACGTGCGCGAGTCGGGGGC</td><td>305.3</td><td>-12.0</td><td>1.5</td><td>-7.8</td><td>6.7E-15</td><td>2.5E-14</td><td>MB231/MB468</td><td>MITO</td></tr>
<tr><td>>hg38_21_+_8433172_8446622_RNA45SN1_45S_50ntFlanks@5446.5464.19|CGGATCGGCCCCGCCGGGG</td><td>>hg38_21_+_8433172_8446622_RNA45SN1_45S_50ntFlanks</td><td>5446</td><td>5464</td><td>19</td><td>CGGATCGGCCCCGCCGGGG</td><td>193.1</td><td>-11.5</td><td>1.5</td><td>-7.9</td><td>4.0E-15</td><td>2.5E-14</td><td>MB231/MB468</td><td>MP</td></tr>
<tr><td>>hg38_21_+_8433172_8446622_RNA45SN1_45S_50ntFlanks@3870.3898.29|AGCTAATACATGCCGACGGGCGCTGACCC</td><td>>hg38_21_+_8433172_8446622_RNA45SN1_45S_50ntFlanks</td><td>3870</td><td>3898</td><td>29</td><td>AGCTAATACATGCCGACGGGCGCTGACCC</td><td>402.6</td><td>-7.2</td><td>0.9</td><td>-7.8</td><td>4.5E-15</td><td>2.5E-14</td><td>MB231/MB468</td><td>CYTO</td></tr>
<tr><td>>hg38_21_+_8433172_8446622_RNA45SN1_45S_50ntFlanks@11208.11233.26|GCGGCGACTCTGGACGCGAGCCGGGC</td><td>>hg38_21_+_8433172_8446622_RNA45SN1_45S_50ntFlanks</td><td>11208</td><td>11233</td><td>26</td><td>GCGGCGACTCTGGACGCGAGCCGGGC</td><td>2693.5</td><td>-8.6</td><td>1.1</td><td>-7.8</td><td>6.7E-15</td><td>2.5E-14</td><td>MB231/MB468</td><td>MITO</td></tr>
<tr><td>>hg38_21_+_8433172_8446622_RNA45SN1_45S_50ntFlanks@7988.8005.18|CAGACGTGGCGACCCGCT</td><td>>hg38_21_+_8433172_8446622_RNA45SN1_45S_50ntFlanks</td><td>7988</td><td>8005</td><td>18</td><td>CAGACGTGGCGACCCGCT</td><td>313.3</td><td>-12.1</td><td>1.5</td><td>-7.8</td><td>6.7E-15</td><td>2.5E-14</td><td>MB231/MB468</td><td>MITO</td></tr>
<tr><td>>hg38_21_+_8433172_8446622_RNA45SN1_45S_50ntFlanks@12162.12179.18|GACTGGGGCGGTACACCT</td><td>>hg38_21_+_8433172_8446622_RNA45SN1_45S_50ntFlanks</td><td>12162</td><td>12179</td><td>18</td><td>GACTGGGGCGGTACACCT</td><td>658.2</td><td>-13.1</td><td>1.7</td><td>-7.8</td><td>6.7E-15</td><td>2.5E-14</td><td>MB231/MB468</td><td>MITO</td></tr>
<tr><td>>hg38_MT_+_598_1651_RNR1_12S_50ntFlanks@570.611.42|AAGGACCTGGCGGTGCTTCATATCCCTCTAGAGGAGCCTGTT</td><td>>hg38_MT_+_598_1651_RNR1_12S_50ntFlanks</td><td>570</td><td>611</td><td>42</td><td>AAGGACCTGGCGGTGCTTCATATCCCTCTAGAGGAGCCTGTT</td><td>268.8</td><td>-11.3</td><td>1.4</td><td>-8.1</td><td>5.6E-16</td><td>2.5E-14</td><td>BT20/MB231</td><td>MITO</td></tr>
<tr><td>>hg38_21_+_8433172_8446622_RNA45SN1_45S_50ntFlanks@5482.5516.35|CGGAGCGCTGAGAAGACGGTCGAACTTGACTATCT</td><td>>hg38_21_+_8433172_8446622_RNA45SN1_45S_50ntFlanks</td><td>5482</td><td>5516</td><td>35</td><td>CGGAGCGCTGAGAAGACGGTCGAACTTGACTATCT</td><td>292.8</td><td>5.5</td><td>0.7</td><td>7.8</td><td>5.7E-15</td><td>2.5E-14</td><td>BT20/MB468</td><td>CYTO</td></tr>
<tr><td>>hg38_21_+_8433172_8446622_RNA45SN1_45S_50ntFlanks@9352.9379.28|CCGTGGCGCAATGAAGGTGAAGGCCGGC</td><td>>hg38_21_+_8433172_8446622_RNA45SN1_45S_50ntFlanks</td><td>9352</td><td>9379</td><td>28</td><td>CCGTGGCGCAATGAAGGTGAAGGCCGGC</td><td>290.8</td><td>-12.0</td><td>1.5</td><td>-7.9</td><td>4.1E-15</td><td>2.5E-14</td><td>MB231/MB468</td><td>MP</td></tr>
</table>

| | | | | | | | | | | | | | |
|---|---|---|---|---|---|---|---|---|---|---|---|---|---|
| >hg38_21_+_8433172_8446622_RNA45SN1_45S_50ntFlanks@9565.9596.32\|TGACGTGCAAATCGGTCGTCCGACCTGGGTAT | >hg38_21_+_8433172_8446622_RNA45SN1_45S_50ntFlanks | 9565 | 9596 | 32 | TGACGTGCAAATCGGTCGTCCGACCTGGGTAT | 22433.2 | -10.7 | 1.4 | -7.8 | 8.6E-15 | 2.5E-14 | BT20/MB468 | NUC |
| >hg38_21_+_8433172_8446622_RNA45SN1_45S_50ntFlanks@10806.10827.22\|AACAATGTAGGTAAGGGAAGTC | >hg38_21_+_8433172_8446622_RNA45SN1_45S_50ntFlanks | 10806 | 10827 | 22 | AACAATGTAGGTAAGGGAAGTC | 222.0 | -11.7 | 1.5 | -7.9 | 4.1E-15 | 2.5E-14 | MB231/MB468 | MP |
| >hg38_21_+_8433172_8446622_RNA45SN1_45S_50ntFlanks@11322.11342.21\|ACGTCTCGTCGCGCGCGTC | >hg38_21_+_8433172_8446622_RNA45SN1_45S_50ntFlanks | 11322 | 11342 | 21 | ACGTCTCGTCGCGCGCGTC | 930.0 | -9.4 | 1.2 | -7.9 | 4.1E-15 | 2.5E-14 | MB231/MB468 | MP |
| >hg38_21_+_8433172_8446622_RNA45SN1_45S_50ntFlanks@10127.10147.21\|GCGGCGGCGGCGGCGGCGGCG | >hg38_21_+_8433172_8446622_RNA45SN1_45S_50ntFlanks | 10127 | 10147 | 21 | GCGGCGGCGGCGGCGGCGGCG | 288.6 | -12.0 | 1.5 | -7.9 | 4.1E-15 | 2.5E-14 | MB231/MB468 | MP |
| >hg38_21_+_8433172_8446622_RNA45SN1_45S_50ntFlanks@10127.10149.23\|GCGGCGGCGGCGGCGGCGGCGGG | >hg38_21_+_8433172_8446622_RNA45SN1_45S_50ntFlanks | 10127 | 10149 | 23 | GCGGCGGCGGCGGCGGCGGCGGG | 342.8 | -12.3 | 1.6 | -7.9 | 4.1E-15 | 2.5E-14 | MB231/MB468 | MP |
| >hg38_21_+_8433172_8446622_RNA45SN1_45S_50ntFlanks@8844.8864.21\|GCGGCGGCGGCGGCGGCGGCG | >hg38_21_+_8433172_8446622_RNA45SN1_45S_50ntFlanks | 8844 | 8864 | 21 | GCGGCGGCGGCGGCGGCGGCG | 288.6 | -12.0 | 1.5 | -7.9 | 4.1E-15 | 2.5E-14 | MB231/MB468 | MP |
| >hg38_21_+_8433172_8446622_RNA45SN1_45S_50ntFlanks@8844.8866.23\|GCGGCGGCGGCGGCGGCGGCGGG | >hg38_21_+_8433172_8446622_RNA45SN1_45S_50ntFlanks | 8844 | 8866 | 23 | GCGGCGGCGGCGGCGGCGGCGGG | 342.8 | -12.3 | 1.6 | -7.9 | 4.1E-15 | 2.5E-14 | MB231/MB468 | MP |
| >hg38_21_+_8433172_8446622_RNA45SN1_45S_50ntFlanks@10684.10711.28\|CGGCGGCGTCCGGTGAGCTCTCGCTGGC | >hg38_21_+_8433172_8446622_RNA45SN1_45S_50ntFlanks | 10684 | 10711 | 28 | CGGCGGCGTCCGGTGAGCTCTCGCTGGC | 3476.6 | -4.1 | 0.5 | -7.8 | 4.2E-15 | 2.5E-14 | BT20/MB468 | MITO |
| >trna111_HisGTG_1_-_147774845_147774916@1.33.33,trna118_HisGTG_1_-_145396881_145396952@1.33.33,trna16_HisGTG_1_+_146544773_146544844@1.33.33,trna1_HisGTG_15_+_45493349_45493420@1.33.33,trna21_HisGTG_1_+_147753471_147753542@1.33.33,trna33_HisGTG_6_+_27125906_27125977@1.33.33,trna7_HisGTG_9_-_14433938_14434009@1.33.33,trna8_HisGTG_15_-_45492611_45492682@1.33.33,trna9_HisGTG_15_-_45490804_45490875@1.33.33\|GCCGTGATCGTATAGTGGTTAGTACTCTGCGTT | >trna111_HisGTG_1_-_147774845_147774916 | 1 | 33 | 33 | GCCGTGATCGTATAGTGGTTAGTACTCTGCGTT | 8099.4 | 7.2 | 0.9 | 7.8 | 4.6E-15 | 2.5E-14 | MB231/MB468 | CYTO |
| >MI0000736\|hsa-mir-30c-1&WithFlank&1\|+\|40757278\|40757378@23.46.24(+1U);MI0000254\|hsa-mir-30c-2&WithFlank&6\|-\|71376954\|71377037@13.36.24(+1U)&[MIMAT0000244&hsa-miR-30c-5p&offsets\|0\|+1(+1U);m-57&1\|+\|40757300\|40757323&offsets\|0\|0(+1U)];[MIMAT0000244_1&hsa-miR-30c-5p&offsets\|0\|+1(+1U);m-56&6\|-\|71377002\|71377025&offsets\|0\|0(+1U)]\|TGTAAACATCCTACACTCTCAGCTT | >MI0000736\|hsa-mir-30c-1&WithFlank&1\|+\|40757278\|40757378 | 23 | 46 | 24(+1U) | TGTAAACATCCTACACTCTCAGCTT | 1284.8 | 4.8 | 0.6 | 7.8 | 5.0E-15 | 2.5E-14 | MB231/MB468 | NUC |
| >hg38_21_+_8433172_8446622_RNA45SN1_45S_50ntFlanks@6779.6806.28\|CCGGGGCTACGCCTGTCTGAGCGTCGCT | >hg38_21_+_8433172_8446622_RNA45SN1_45S_50ntFlanks | 6779 | 6806 | 28 | CCGGGGCTACGCCTGTCTGAGCGTCGCT | 708.2 | 4.9 | 0.6 | 7.8 | 5.8E-15 | 2.5E-14 | BT20/MB468 | CYTO |
| >MI0000066\|hsa-let-7e&WithFlank&19\|+\|51692780\|51692870@14.34.21(+1A)&[MIMAT0000066&hsa-let-7e-5p&offsets\|0\|-1(+1A);m-50&19\|+\|51692793\|51692814&offsets\|0\|-1(+1A)]\|TGAGGTAGGAGGTTGTATAGTA | >MI0000066\|hsa-let-7e&WithFlank&19\|+\|51692780\|51692870 | 14 | 34 | 21(+1A) | TGAGGTAGGAGGTTGTATAGTA | 2185.5 | 9.4 | 1.2 | 7.8 | 5.8E-15 | 2.5E-14 | BT20/MB468 | CYTO |
| >hg38_21_+_8433172_8446622_RNA45SN1_45S_50ntFlanks@11212.11230.19\|CGACTCTGGACGCGAGCCG | >hg38_21_+_8433172_8446622_RNA45SN1_45S_50ntFlanks | 11212 | 11230 | 19 | CGACTCTGGACGCGAGCCG | 283.5 | -10.9 | 1.4 | -7.8 | 4.6E-15 | 2.5E-14 | MB231/MB468 | CYTO |
| >MI0000809\|hsa-mir-151a&WithFlank&8\|-\|140732558\|140732659@51.72.22(+1A)&[MIMAT0000757&hsa-miR-151a-3p&offsets\|-2\|-1(+1A);m-30&8\|-\|140732587\|140732607&offsets\|-2\|-1(+1A)]\|TACTAGACTGAAGCTCCTTGAGA | >MI0000809\|hsa-mir-151a&WithFlank&8\|-\|140732558\|140732659 | 51 | 72 | 22(+1A) | TACTAGACTGAAGCTCCTTGAGA | 131.8 | 2.1 | 0.3 | 7.8 | 8.7E-15 | 2.5E-14 | BT20/MB468 | NUC |
| >MI0003137\|hsa-mir-193b&WithFlank&16\|+\|14303961\|14304055@57.77.21(+1C)&[MIMAT0002819&hsa-miR-193b-3p&offsets\|0\|-1(+1C);m-166&16\|+\|14304017\|14304038&offsets\|0\|-1(+1C)]\|AACTGGCCCTCAAAGTCCCGCC | >MI0003137\|hsa-mir-193b&WithFlank&16\|+\|14303961\|14304055 | 57 | 77 | 21(+1C) | AACTGGCCCTCAAAGTCCCGCC | 259.7 | 6.1 | 0.8 | 7.8 | 8.8E-15 | 2.6E-14 | BT20/MB468 | NUC |
| >hg38_21_+_8433172_8446622_RNA45SN1_45S_50ntFlanks@13004.13024.21\|ATCTATTGAAAGTCAGCCCTC | >hg38_21_+_8433172_8446622_RNA45SN1_45S_50ntFlanks | 13004 | 13024 | 21 | ATCTATTGAAAGTCAGCCCTC | 167.3 | -10.1 | 1.3 | -7.8 | 4.6E-15 | 2.6E-14 | MB231/MB468 | CYTO |
| >hg38_21_+_8433172_8446622_RNA45SN1_45S_50ntFlanks@11561.11593.33\|GCGCCTAGCAGCCGACTTAGAACTGGTGCGGAC | >hg38_21_+_8433172_8446622_RNA45SN1_45S_50ntFlanks | 11561 | 11593 | 33 | GCGCCTAGCAGCCGACTTAGAACTGGTGCGGAC | 496.1 | -6.5 | 0.8 | -7.8 | 4.6E-15 | 2.6E-14 | MB231/MB468 | CYTO |
| >hg38_21_+_8433172_8446622_RNA45SN1_45S_50ntFlanks@10128.10147.20\|CGGCGGCGGCGGCGGCGGCG | >hg38_21_+_8433172_8446622_RNA45SN1_45S_50ntFlanks | 10128 | 10147 | 20 | CGGCGGCGGCGGCGGCGGCG | 200.4 | -10.6 | 1.4 | -7.8 | 5.0E-15 | 2.6E-14 | MB231/MB468 | NUC |
| >hg38_21_+_8433172_8446622_RNA45SN1_45S_50ntFlanks@8845.8864.20\|CGGCGGCGGCGGCGGCGGCG | >hg38_21_+_8433172_8446622_RNA45SN1_45S_50ntFlanks | 8845 | 8864 | 20 | CGGCGGCGGCGGCGGCGGCG | 200.4 | -10.6 | 1.4 | -7.8 | 5.0E-15 | 2.6E-14 | MB231/MB468 | NUC |
| >hg38_21_+_8433172_8446622_RNA45SN1_45S_50ntFlanks@12125.12146.22\|GCGACCCGCTCCGGGGACAGTG | >hg38_21_+_8433172_8446622_RNA45SN1_45S_50ntFlanks | 12125 | 12146 | 22 | GCGACCCGCTCCGGGGACAGTG | 390.8 | -12.5 | 1.6 | -7.9 | 4.2E-15 | 2.6E-14 | MB231/MB468 | MP |
| >hg38_21_+_8433172_8446622_RNA45SN1_45S_50ntFlanks@8238.8262.25\|CCCGGGTCTTCCCGGAGTCGGGTTG | >hg38_21_+_8433172_8446622_RNA45SN1_45S_50ntFlanks | 8238 | 8262 | 25 | CCCGGGTCTTCCCGGAGTCGGGTTG | 298.9 | -12.1 | 1.5 | -7.9 | 4.2E-15 | 2.6E-14 | MB231/MB468 | MP |
| >hg38_21_+_8433172_8446622_RNA45SN1_45S_50ntFlanks@485.504.20\|GATGTGGTGACGTCGTGCTC | >hg38_21_+_8433172_8446622_RNA45SN1_45S_50ntFlanks | 485 | 504 | 20 | GATGTGGTGACGTCGTGCTC | 1993.5 | 14.9 | 1.9 | 7.8 | 5.0E-15 | 2.6E-14 | MB231/MB468 | NUC |
| >MI0000439\|hsa-mir-23b&WithFlank&9\|+\|95085202\|95085310@67.85.19(+1U)&[MIMAT0000418&hsa-miR-23b-3p&offsets\|+3\|-1(+1U);m-42&9\|+\|95085265\|95085285&offsets\|+3\|+1(+1U)]\|ACATTGCCAGGGATTACCAT | >MI0000439\|hsa-mir-23b&WithFlank&9\|+\|95085202\|95085310 | 67 | 85 | 19(+1U) | ACATTGCCAGGGATTACCAT | 1246.2 | 28.5 | 3.6 | 7.8 | 4.6E-15 | 2.6E-14 | MB231/MB468 | CYTO |
| >hg38_21_+_8433172_8446622_RNA45SN1_45S_50ntFlanks@11670.11698.29\|CCCAGTGCTCTGAATGTCAAAGTGAAGAA | >hg38_21_+_8433172_8446622_RNA45SN1_45S_50ntFlanks | 11670 | 11698 | 29 | CCCAGTGCTCTGAATGTCAAAGTGAAGAA | 178.4 | -11.3 | 1.4 | -7.8 | 4.2E-15 | 2.6E-14 | MB231/MB468 | MP |
| >hg38_21_+_8433172_8446622_RNA45SN1_45S_50ntFlanks@10574.10607.34\|AAGCCGGCGGGAGCCCCGGGGAGAGTTCTCTTTT | >hg38_21_+_8433172_8446622_RNA45SN1_45S_50ntFlanks | 10574 | 10607 | 34 | AAGCCGGCGGGAGCCCCGGGGAGAGTTCTCTTTT | 253.3 | -11.8 | 1.5 | -7.8 | 6.9E-15 | 2.6E-14 | MB231/MB468 | MITO |
| >hg38_21_+_8433172_8446622_RNA45SN1_45S_50ntFlanks@4027.4044.18\|CCCCGTGGCGGCGACGAC | >hg38_21_+_8433172_8446622_RNA45SN1_45S_50ntFlanks | 4027 | 4044 | 18 | CCCCGTGGCGGCGACGAC | 376.4 | -12.3 | 1.6 | -7.8 | 6.9E-15 | 2.6E-14 | MB231/MB468 | MITO |
| >hg38_21_+_8433172_8446622_RNA45SN1_45S_50ntFlanks@10914.10932.19\|CGCCGCGGCTGGACGAGGC | >hg38_21_+_8433172_8446622_RNA45SN1_45S_50ntFlanks | 10914 | 10932 | 19 | CGCCGCGGCTGGACGAGGC | 658.3 | -12.1 | 1.5 | -7.8 | 4.7E-15 | 2.6E-14 | MB231/MB468 | CYTO |
| >hg38_21_+_8433172_8446622_RNA45SN1_45S_50ntFlanks@12687.12708.22\|CAGGCGGAACGATACGGCAGCG | >hg38_21_+_8433172_8446622_RNA45SN1_45S_50ntFlanks | 12687 | 12708 | 22 | CAGGCGGAACGATACGGCAGCG | 223.0 | -11.7 | 1.5 | -7.8 | 4.2E-15 | 2.6E-14 | MB231/MB468 | MP |
| >hg38_21_+_8433172_8446622_RNA45SN1_45S_50ntFlanks@4729.4766.38\|TCAAGAACGAAAGTCGGAGGTTCGAAGACGATCAGATA | >hg38_21_+_8433172_8446622_RNA45SN1_45S_50ntFlanks | 4729 | 4766 | 38 | TCAAGAACGAAAGTCGGAGGTTCGAAGACGATCAGATA | 265.6 | 6.2 | 0.8 | 7.8 | 4.7E-15 | 2.6E-14 | MB231/MB468 | CYTO |
| >hg38_21_+_8433172_8446622_RNA45SN1_45S_50ntFlanks@12631.12654.24\|GGCGAAGCTACCATCTGTGGGATT | >hg38_21_+_8433172_8446622_RNA45SN1_45S_50ntFlanks | 12631 | 12654 | 24 | GGCGAAGCTACCATCTGTGGGATT | 1266.7 | -9.6 | 1.2 | -7.8 | 7.0E-15 | 2.6E-14 | MB231/MB468 | MITO |
| >hg38_21_+_8433172_8446622_RNA45SN1_45S_50ntFlanks@11675.11702.28\|TGCTCTGAATGTCAAAGTGAAGAAATTC | >hg38_21_+_8433172_8446622_RNA45SN1_45S_50ntFlanks | 11675 | 11702 | 28 | TGCTCTGAATGTCAAAGTGAAGAAATTC | 728.3 | 2.4 | 0.3 | 7.8 | 5.9E-15 | 2.6E-14 | BT20/MB468 | CYTO |
| >hg38_21_+_8433172_8446622_RNA45SN1_45S_50ntFlanks@9568.9596.29\|CGTGCAAATCGGTCGTCCGACCTGGGTAT | >hg38_21_+_8433172_8446622_RNA45SN1_45S_50ntFlanks | 9568 | 9596 | 29 | CGTGCAAATCGGTCGTCCGACCTGGGTAT | 2879.5 | -8.3 | 1.1 | -7.8 | 5.1E-15 | 2.6E-14 | MB231/MB468 | NUC |
| >MI0000446\|hsa-mir-125b-1&WithFlank&11\|-\|122099751\|122099850@21.43.23(+1U)&[MIMAT0000423&hsa-miR-125b-5p&offsets\|0\|+1(+1U);m-67&11\|-\|122099809\|122099830&offsets\|0\|+1(+1U)]\|TCCCTGAGACCCTAACTTGTGATT | >MI0000446\|hsa-mir-125b-1&WithFlank&11\|-\|122099751\|122099850 | 21 | 43 | 23(+1U) | TCCCTGAGACCCTAACTTGTGATT | 430.3 | -12.0 | 1.5 | -8.1 | 5.9E-16 | 2.6E-14 | BT20/MB231 | MITO |
| >am_trna128_GlyGCC_6_-_27870686_27870756@1.35.35,trna18_GlyGCC_16_+_70822597_70822667@1.35.35,trna19_GlyGCC_16_+_70823410_70823480@1.35.35,trna19_GlyGCC_2_-_157257659_157257729@1.35.35,trna24_GlyGCC_16_-_70812942_70813012@1.35.35,trna25_GlyGCC_16_-_70812114_70812184@1.35.35,trna5_GlyGCC_17_+_8029064_8029134@1.35.35,trna68_GlyGCC_1_-_161493637_161493707@1.35.35\|GCATTGGTGGTTCAGTGGTAGAATTCTCGCCTGCC | >am_trna128_GlyGCC_6_-_27870686_27870756 | 1 | 35 | 35 | GCATTGGTGGTTCAGTGGTAGAATTCTCGCCTGCC | 15923.1 | 7.9 | 1.0 | 7.8 | 4.7E-15 | 2.6E-14 | MB231/MB468 | CYTO |
| >hg38_21_+_8433172_8446622_RNA45SN1_45S_50ntFlanks@11208.11250.43\|GCGGCGACTCTGGACGCGAGCCGGGCCCTTCCCGTGGATCGCC | >hg38_21_+_8433172_8446622_RNA45SN1_45S_50ntFlanks | 11208 | 11250 | 43 | GCGGCGACTCTGGACGCGAGCCGGGCCCTTCCCGTGGATCGCC | 294.5 | -12.0 | 1.5 | -7.8 | 7.1E-15 | 2.6E-14 | MB231/MB468 | MITO |
| >hg38_21_+_8433172_8446622_RNA45SN1_45S_50ntFlanks@10391.10430.40\|AACAGCAGTTGAACATGGGTCAGTCGGTCCTGAGAGATGG | >hg38_21_+_8433172_8446622_RNA45SN1_45S_50ntFlanks | 10391 | 10430 | 40 | AACAGCAGTTGAACATGGGTCAGTCGGTCCTGAGAGATGG | 1135.1 | -10.4 | 1.3 | -7.8 | 5.2E-15 | 2.6E-14 | MB231/MB468 | NUC |
| >hg38_21_+_8433172_8446622_RNA45SN1_45S_50ntFlanks@6652.6681.30\|GACTCTTAGCGGTGGATCACTCGGCTCGTG | >hg38_21_+_8433172_8446622_RNA45SN1_45S_50ntFlanks | 6652 | 6681 | 30 | GACTCTTAGCGGTGGATCACTCGGCTCGTG | 11744.9 | -4.1 | 0.5 | -7.8 | 9.0E-15 | 2.6E-14 | BT20/MB468 | NUC |
| >hg38_21_+_8433172_8446622_RNA45SN1_45S_50ntFlanks@9122.9152.31\|CCGCGCGCGGGTCGGGGGGCGGGGCGGACTG | >hg38_21_+_8433172_8446622_RNA45SN1_45S_50ntFlanks | 9122 | 9152 | 31 | CCGCGCGCGGGTCGGGGGGCGGGGCGGACTG | 468.2 | -12.7 | 1.6 | -7.8 | 4.3E-15 | 2.6E-14 | MB231/MB468 | MP |
| >hg38_21_+_8433172_8446622_RNA45SN1_45S_50ntFlanks@6671.6699.29\|CTCGGCTCGTGCGTCGATGAAGAACGCAG | >hg38_21_+_8433172_8446622_RNA45SN1_45S_50ntFlanks | 6671 | 6699 | 29 | CTCGGCTCGTGCGTCGATGAAGAACGCAG | 15346.0 | -7.0 | 0.9 | -7.8 | 4.4E-15 | 2.6E-14 | BT20/MB468 | MITO |
| >hg38_21_+_8433172_8446622_RNA45SN1_45S_50ntFlanks@12991.13013.23\|CTCCCTCGCTGCGATCTATTGAA | >hg38_21_+_8433172_8446622_RNA45SN1_45S_50ntFlanks | 12991 | 13013 | 23 | CTCCCTCGCTGCGATCTATTGAA | 1731.4 | -9.9 | 1.3 | -7.8 | 4.3E-15 | 2.6E-14 | MB231/MB468 | MP |
| >MI0000067\|hsa-let-7f-1&WithFlank&9\|+\|94176341\|94176439@13.30.18;MI0000068\|hsa-let-7f-2&WithFlank&X\|-\|53557186\|53557280@14.31.18&[MIMAT0000067&hsa-let-7f-5p&offsets\|0\|-4;m-20&9\|+\|94176353\|94176374&offsets\|0\|-4];[MIMAT0000067_1&hsa-let-7f-5p&offsets\|0\|-4;m-19&X\|-\|53557246\|53557267&offsets\|0\|-4]\|TGAGGTAGTAGATTGTAT | >MI0000067\|hsa-let-7f-1&WithFlank&9\|+\|94176341\|94176439 | 13 | 30 | 18 | TGAGGTAGTAGATTGTAT | 4314.4 | 4.7 | 0.6 | 7.8 | 5.2E-15 | 2.6E-14 | MB231/MB468 | NUC |
| >hg38_21_+_8433172_8446622_RNA45SN1_45S_50ntFlanks@12633.12660.28\|CGAAGCTACCATCTGTGGGATTATGACT | >hg38_21_+_8433172_8446622_RNA45SN1_45S_50ntFlanks | 12633 | 12660 | 28 | CGAAGCTACCATCTGTGGGATTATGACT | 244.1 | -11.8 | 1.5 | -7.8 | 4.3E-15 | 2.6E-14 | MB231/MB468 | MP |

<table>
<tr><th></th><th></th><th></th><th></th><th></th><th></th><th></th><th></th><th></th><th></th><th></th><th></th><th></th><th></th></tr>
<tr><td>>hg38_21_+_8433172_8446622_RNA45SN1_45S_50ntFlanks@12412.12431.20|ATCCTTCGATGTCGGCTCTT</td><td>>hg38_21_+_8433172_8446622_RNA45SN1_45S_50ntFlanks</td><td>12412</td><td>12431</td><td>20</td><td>ATCCTTCGATGTCGGCTCTT</td><td>519.0</td><td>-11.3</td><td>1.5</td><td>-7.8</td><td>6.0E-15</td><td>2.6E-14</td><td>BT20/MB468</td><td>CYTO</td></tr>
<tr><td>>hg38_21_+_8433172_8446622_RNA45SN1_45S_50ntFlanks@9736.9755.20|CCGAAACGATCTCAACCTAT</td><td>>hg38_21_+_8433172_8446622_RNA45SN1_45S_50ntFlanks</td><td>9736</td><td>9755</td><td>20</td><td>CCGAAACGATCTCAACCTAT</td><td>160.9</td><td>-11.2</td><td>1.4</td><td>-7.8</td><td>4.3E-15</td><td>2.6E-14</td><td>MB231/MB468</td><td>MP</td></tr>
<tr><td>>MI0000734|hsa-mir-106b&WithFlank&7|-|100093987|100094080@58.78.21(+1A)&[MIMAT0004672&hsa-miR-106b-3p&offsets|0|-1(+1A);m-87&7|-|100094003|100094023&offsets|0|0(+1A)]|CCGCACTGTGGGTACTTGCTGA</td><td>>MI0000734|hsa-mir-106b&WithFlank&7|-|100093987|100094080</td><td>58</td><td>78</td><td>21(+1A)</td><td>CCGCACTGTGGGTACTTGCTGA</td><td>300.5</td><td>4.7</td><td>0.6</td><td>7.8</td><td>5.2E-15</td><td>2.6E-14</td><td>MB231/MB468</td><td>NUC</td></tr>
<tr><td>>hg38_21_+_8433172_8446622_RNA45SN1_45S_50ntFlanks@11821.11856.36|CTATCCAGCGAAACCACAGCCAAGGGAACGGGCTTG</td><td>>hg38_21_+_8433172_8446622_RNA45SN1_45S_50ntFlanks</td><td>11821</td><td>11856</td><td>36</td><td>CTATCCAGCGAAACCACAGCCAAGGGAACGGGCTTG</td><td>192.3</td><td>-10.5</td><td>1.3</td><td>-7.8</td><td>5.2E-15</td><td>2.6E-14</td><td>MB231/MB468</td><td>NUC</td></tr>
<tr><td>>MI0000086|hsa-mir-28&WithFlank&3|+|188688775|188688872@60.81.22(+1A)&[MIMAT0004502&hsa-miR-28-3p&offsets|0|0(+1A);m-28&3|+|188688834|188688855&offsets|0|0(+1A)]|CACTAGATTGTGAGCTCCTGGAA</td><td>>MI0000086|hsa-mir-28&WithFlank&3|+|188688775|188688872</td><td>60</td><td>81</td><td>22(+1A)</td><td>CACTAGATTGTGAGCTCCTGGAA</td><td>958.4</td><td>8.4</td><td>1.1</td><td>7.8</td><td>4.8E-15</td><td>2.7E-14</td><td>MB231/MB468</td><td>CYTO</td></tr>
<tr><td>>hg38_21_+_8433172_8446622_RNA45SN1_45S_50ntFlanks@3979.4002.24|CGCCGGCGGCTTTGGTGACTCTAG</td><td>>hg38_21_+_8433172_8446622_RNA45SN1_45S_50ntFlanks</td><td>3979</td><td>4002</td><td>24</td><td>CGCCGGCGGCTTTGGTGACTCTAG</td><td>524.3</td><td>-12.8</td><td>1.6</td><td>-7.8</td><td>7.1E-15</td><td>2.7E-14</td><td>MB231/MB468</td><td>MITO</td></tr>
<tr><td>>trnaMT_ValTAC_MT_+_1602_1670@48.71.24|CAACTTAACTTGACCGCTCTGACC</td><td>>trnaMT_ValTAC_MT_+_1602_1670</td><td>48</td><td>71</td><td>24</td><td>CAACTTAACTTGACCGCTCTGACC</td><td>854.6</td><td>8.9</td><td>1.1</td><td>7.8</td><td>4.8E-15</td><td>2.7E-14</td><td>MB231/MB468</td><td>CYTO</td></tr>
<tr><td>>hg38_21_+_8433172_8446622_RNA45SN1_45S_50ntFlanks@11452.11471.20|CTCCTCGGCGCGCGGCGGCG</td><td>>hg38_21_+_8433172_8446622_RNA45SN1_45S_50ntFlanks</td><td>11452</td><td>11471</td><td>20</td><td>CTCCTCGGCGCGCGGCGGCG</td><td>612.3</td><td>-12.2</td><td>1.6</td><td>-7.8</td><td>5.3E-15</td><td>2.7E-14</td><td>MB231/MB468</td><td>NUC</td></tr>
<tr><td>>hg38_21_+_8433172_8446622_RNA45SN1_45S_50ntFlanks@9567.9597.31|ACGTGCAAATCGGTCGTCCGACCTGGGTATA</td><td>>hg38_21_+_8433172_8446622_RNA45SN1_45S_50ntFlanks</td><td>9567</td><td>9597</td><td>31</td><td>ACGTGCAAATCGGTCGTCCGACCTGGGTATA</td><td>5793.6</td><td>-11.8</td><td>1.5</td><td>-7.7</td><td>9.2E-15</td><td>2.7E-14</td><td>BT20/MB468</td><td>NUC</td></tr>
<tr><td>>hg38_21_+_8433172_8446622_RNA45SN1_45S_50ntFlanks@6644.6664.21|CCTCGTACGACTCTTAGCGGT</td><td>>hg38_21_+_8433172_8446622_RNA45SN1_45S_50ntFlanks</td><td>6644</td><td>6664</td><td>21</td><td>CCTCGTACGACTCTTAGCGGT</td><td>191.7</td><td>4.0</td><td>0.5</td><td>7.8</td><td>5.3E-15</td><td>2.7E-14</td><td>MB231/MB468</td><td>NUC</td></tr>
<tr><td>>hg38_21_+_8433172_8446622_RNA45SN1_45S_50ntFlanks@6645.6679.35|CTCGTACGACTCTTAGCGGTGGATCACTCGGCTCG</td><td>>hg38_21_+_8433172_8446622_RNA45SN1_45S_50ntFlanks</td><td>6645</td><td>6679</td><td>35</td><td>CTCGTACGACTCTTAGCGGTGGATCACTCGGCTCG</td><td>2064.7</td><td>-3.6</td><td>0.5</td><td>-7.7</td><td>9.3E-15</td><td>2.7E-14</td><td>BT20/MB468</td><td>NUC</td></tr>
<tr><td>>hg38_21_+_8433172_8446622_RNA45SN1_45S_50ntFlanks@10420.10443.24|CTGAGAGATGGGCGAGCGCCGTTC</td><td>>hg38_21_+_8433172_8446622_RNA45SN1_45S_50ntFlanks</td><td>10420</td><td>10443</td><td>24</td><td>CTGAGAGATGGGCGAGCGCCGTTC</td><td>249.6</td><td>-11.7</td><td>1.5</td><td>-7.8</td><td>7.3E-15</td><td>2.7E-14</td><td>MB231/MB468</td><td>MITO</td></tr>
<tr><td>>hg38_21_+_8433172_8446622_RNA45SN1_45S_50ntFlanks@10350.10390.41|ACGAGAACTTTGAAGGCCGAAGTGGAGAAGGGTTCCATGTG</td><td>>hg38_21_+_8433172_8446622_RNA45SN1_45S_50ntFlanks</td><td>10350</td><td>10390</td><td>41</td><td>ACGAGAACTTTGAAGGCCGAAGTGGAGAAGGGTTCCATGTG</td><td>6168.0</td><td>-6.6</td><td>0.8</td><td>-7.8</td><td>7.3E-15</td><td>2.7E-14</td><td>MB231/MB468</td><td>MITO</td></tr>
<tr><td>>hg38_21_+_8433172_8446622_RNA45SN1_45S_50ntFlanks@5446.5467.22|CGGATCGGCCCCGCCGGGGTCG</td><td>>hg38_21_+_8433172_8446622_RNA45SN1_45S_50ntFlanks</td><td>5446</td><td>5467</td><td>22</td><td>CGGATCGGCCCCGCCGGGGTCG</td><td>1334.6</td><td>-13.5</td><td>1.6</td><td>-8.2</td><td>1.6E-16</td><td>2.7E-14</td><td>BT20/MB468</td><td>MP</td></tr>
<tr><td>>hg38_21_+_8433172_8446622_RNA45SN1_45S_50ntFlanks@10681.10708.28|TTCCGGCGGCGTCCGGTGAGCTCTCGCT</td><td>>hg38_21_+_8433172_8446622_RNA45SN1_45S_50ntFlanks</td><td>10681</td><td>10708</td><td>28</td><td>TTCCGGCGGCGTCCGGTGAGCTCTCGCT</td><td>176.8</td><td>-11.3</td><td>1.4</td><td>-7.8</td><td>4.4E-15</td><td>2.7E-14</td><td>MB231/MB468</td><td>MP</td></tr>
<tr><td>>hg38_21_+_8433172_8446622_RNA45SN1_45S_50ntFlanks@5483.5514.32|GGAGCGCTGAGAAGACGGTCGAACTTGACTAT</td><td>>hg38_21_+_8433172_8446622_RNA45SN1_45S_50ntFlanks</td><td>5483</td><td>5514</td><td>32</td><td>GGAGCGCTGAGAAGACGGTCGAACTTGACTAT</td><td>344.1</td><td>-12.3</td><td>1.6</td><td>-7.8</td><td>4.4E-15</td><td>2.7E-14</td><td>MB231/MB468</td><td>MP</td></tr>
<tr><td>>hg38_21_+_8433172_8446622_RNA45SN1_45S_50ntFlanks@11783.11803.21|CATGAATGGATGAACGAGATT</td><td>>hg38_21_+_8433172_8446622_RNA45SN1_45S_50ntFlanks</td><td>11783</td><td>11803</td><td>21</td><td>CATGAATGGATGAACGAGATT</td><td>5983.0</td><td>-6.9</td><td>0.8</td><td>-8.2</td><td>1.6E-16</td><td>2.7E-14</td><td>BT20/MB468</td><td>MP</td></tr>
<tr><td>>hg38_21_+_8433172_8446622_RNA45SN1_45S_50ntFlanks@11199.11227.29|CCGGCGGCGGCGGCGACTCTGGACGCGAG</td><td>>hg38_21_+_8433172_8446622_RNA45SN1_45S_50ntFlanks</td><td>11199</td><td>11227</td><td>29</td><td>CCGGCGGCGGCGGCGACTCTGGACGCGAG</td><td>1086.7</td><td>-7.1</td><td>0.9</td><td>-7.8</td><td>4.9E-15</td><td>2.7E-14</td><td>MB231/MB468</td><td>CYTO</td></tr>
<tr><td>>MI0003763|hsa-mir-767&WithFlank&X|-|152393415|152393535@33.56.24&[MIMAT0003882&hsa-miR-767-5p&offsets|0|+1;m-788&X|-|152393481|152393503&offsets|0|+1]|TGCACCATGGTTGTCTGAGCATGC</td><td>>MI0003763|hsa-mir-767&WithFlank&X|-|152393415|152393535</td><td>33</td><td>56</td><td>24</td><td>TGCACCATGGTTGTCTGAGCATGC</td><td>267.4</td><td>11.8</td><td>1.5</td><td>8.1</td><td>6.7E-16</td><td>2.7E-14</td><td>BT20/MB231</td><td>NUC</td></tr>
<tr><td>>hg38_21_+_8433172_8446622_RNA45SN1_45S_50ntFlanks@9033.9056.24|CTCGTGGGGGGGCCGGGCCACCCC</td><td>>hg38_21_+_8433172_8446622_RNA45SN1_45S_50ntFlanks</td><td>9033</td><td>9056</td><td>24</td><td>CTCGTGGGGGGGCCGGGCCACCCC</td><td>877.4</td><td>-12.9</td><td>1.6</td><td>-8.2</td><td>1.6E-16</td><td>2.7E-14</td><td>BT20/MB468</td><td>MP</td></tr>
<tr><td>>hg38_21_+_8433172_8446622_RNA45SN1_45S_50ntFlanks@6667.6696.30|ATCACTCGGCTCGTGCGTCGATGAAGAACG</td><td>>hg38_21_+_8433172_8446622_RNA45SN1_45S_50ntFlanks</td><td>6667</td><td>6696</td><td>30</td><td>ATCACTCGGCTCGTGCGTCGATGAAGAACG</td><td>163.2</td><td>-11.2</td><td>1.4</td><td>-7.8</td><td>4.5E-15</td><td>2.7E-14</td><td>MB231/MB468</td><td>MP</td></tr>
<tr><td>>hg38_21_+_8433172_8446622_RNA45SN1_45S_50ntFlanks@10823.10840.18|AAGTCGGCAAGCCGGATC</td><td>>hg38_21_+_8433172_8446622_RNA45SN1_45S_50ntFlanks</td><td>10823</td><td>10840</td><td>18</td><td>AAGTCGGCAAGCCGGATC</td><td>677.0</td><td>-13.3</td><td>1.7</td><td>-7.8</td><td>4.5E-15</td><td>2.7E-14</td><td>MB231/MB468</td><td>MP</td></tr>
<tr><td>>hg38_21_+_8433172_8446622_RNA45SN1_45S_50ntFlanks@3846.3869.24|CTTGGATAACTGTGGTAATTCTAG</td><td>>hg38_21_+_8433172_8446622_RNA45SN1_45S_50ntFlanks</td><td>3846</td><td>3869</td><td>24</td><td>CTTGGATAACTGTGGTAATTCTAG</td><td>701.6</td><td>-7.8</td><td>1.0</td><td>-7.7</td><td>9.4E-15</td><td>2.7E-14</td><td>BT20/MB468</td><td>NUC</td></tr>
<tr><td>>hg38_21_+_8433172_8446622_RNA45SN1_45S_50ntFlanks@7973.7997.25|GACGCGACCTCAGATCAGACGTGGC</td><td>>hg38_21_+_8433172_8446622_RNA45SN1_45S_50ntFlanks</td><td>7973</td><td>7997</td><td>25</td><td>GACGCGACCTCAGATCAGACGTGGC</td><td>264.7</td><td>4.1</td><td>0.5</td><td>7.8</td><td>5.4E-15</td><td>2.7E-14</td><td>MB231/MB468</td><td>NUC</td></tr>
<tr><td>>hg38_21_+_8433172_8446622_RNA45SN1_45S_50ntFlanks@10420.10443.24|CTGAGAGATGGGCGAGCGCCGTTC</td><td>>hg38_21_+_8433172_8446622_RNA45SN1_45S_50ntFlanks</td><td>10420</td><td>10443</td><td>24</td><td>CTGAGAGATGGGCGAGCGCCGTTC</td><td>1086.1</td><td>-13.9</td><td>1.8</td><td>-7.8</td><td>4.5E-15</td><td>2.8E-14</td><td>MB231/MB468</td><td>MP</td></tr>
<tr><td>>hg38_21_+_8433172_8446622_RNA45SN1_45S_50ntFlanks@5417.5445.29|CTACCGATTGGATGGTTTAGTGAGGCCCT</td><td>>hg38_21_+_8433172_8446622_RNA45SN1_45S_50ntFlanks</td><td>5417</td><td>5445</td><td>29</td><td>CTACCGATTGGATGGTTTAGTGAGGCCCT</td><td>332.9</td><td>-12.2</td><td>1.6</td><td>-7.8</td><td>4.5E-15</td><td>2.8E-14</td><td>MB231/MB468</td><td>MP</td></tr>
<tr><td>>hg38_21_+_8433172_8446622_RNA45SN1_45S_50ntFlanks@5485.5514.30|AGCGCTGAGAAGACGGTCGAACTTGACTAT</td><td>>hg38_21_+_8433172_8446622_RNA45SN1_45S_50ntFlanks</td><td>5485</td><td>5514</td><td>30</td><td>AGCGCTGAGAAGACGGTCGAACTTGACTAT</td><td>5466.4</td><td>-5.9</td><td>0.8</td><td>-7.8</td><td>5.0E-15</td><td>2.8E-14</td><td>MB231/MB468</td><td>CYTO</td></tr>
<tr><td>>trna6_ValTAC_10_-_5895674_5895746@1.30.30|GGTTCCATAGTGTAGTGGTTATCACATCTG</td><td>>trna6_ValTAC_10_-_5895674_5895746</td><td>1</td><td>30</td><td>30</td><td>GGTTCCATAGTGTAGTGGTTATCACATCTG</td><td>189.5</td><td>-10.3</td><td>1.3</td><td>-7.8</td><td>5.0E-15</td><td>2.8E-14</td><td>MB231/MB468</td><td>CYTO</td></tr>
<tr><td>>hg38_21_+_8433172_8446622_RNA45SN1_45S_50ntFlanks@11873.11904.32|AAGAAGACCCTGTTGAGCTTGACTCTAGTCTG</td><td>>hg38_21_+_8433172_8446622_RNA45SN1_45S_50ntFlanks</td><td>11873</td><td>11904</td><td>32</td><td>AAGAAGACCCTGTTGAGCTTGACTCTAGTCTG</td><td>354.6</td><td>-8.2</td><td>1.0</td><td>-7.8</td><td>5.0E-15</td><td>2.8E-14</td><td>MB231/MB468</td><td>CYTO</td></tr>
<tr><td>>MI0000102|hsa-mir-100&WithFlank&11|-|122152223|122152314@19.40.22(+3A)&[MIMAT0000098&hsa-miR-100-5p&offsets|0|0(+3A);m-27&11|-|122152276|122152296&offsets|0|+1(+3A)]|AACCCGTAGATCCGAACTTGTGAAA</td><td>>MI0000102|hsa-mir-100&WithFlank&11|-|122152223|122152314</td><td>19</td><td>40</td><td>22(+3A)</td><td>AACCCGTAGATCCGAACTTGTGAAA</td><td>275.9</td><td>-11.4</td><td>1.4</td><td>-8.1</td><td>6.3E-16</td><td>2.8E-14</td><td>BT20/MB231</td><td>MITO</td></tr>
<tr><td>>hg38_21_+_8433172_8446622_RNA45SN1_45S_50ntFlanks@11617.11640.24|AAAACAAAGCATCGCGAAGGCCCG</td><td>>hg38_21_+_8433172_8446622_RNA45SN1_45S_50ntFlanks</td><td>11617</td><td>11640</td><td>24</td><td>AAAACAAAGCATCGCGAAGGCCCG</td><td>844.1</td><td>-12.8</td><td>1.6</td><td>-8.2</td><td>1.7E-16</td><td>2.8E-14</td><td>BT20/MB468</td><td>MP</td></tr>
<tr><td>>hg38_21_+_8433172_8446622_RNA45SN1_45S_50ntFlanks@11783.11806.24|CATGAATGGATGAACGAGATTCCC</td><td>>hg38_21_+_8433172_8446622_RNA45SN1_45S_50ntFlanks</td><td>11783</td><td>11806</td><td>24</td><td>CATGAATGGATGAACGAGATTCCC</td><td>2932.5</td><td>-6.6</td><td>0.8</td><td>-8.2</td><td>1.7E-16</td><td>2.8E-14</td><td>BT20/MB468</td><td>MP</td></tr>
<tr><td>>MI0000299|hsa-mir-222&WithFlank&X|-|45747009|45747130@75.96.22(+1C)&[MIMAT0000279&hsa-miR-222-3p&offsets|0|+1(+1C);m-140&X|-|45747033|45747056&offsets|0|-2(+1C)]|AGCTACATCTGGCTACTGGGTCC</td><td>>MI0000299|hsa-mir-222&WithFlank&X|-|45747009|45747130</td><td>75</td><td>96</td><td>22(+1C)</td><td>AGCTACATCTGGCTACTGGGTCC</td><td>710.9</td><td>7.0</td><td>0.9</td><td>7.8</td><td>5.0E-15</td><td>2.8E-14</td><td>MB231/MB468</td><td>CYTO</td></tr>
<tr><td>>hg38_21_+_8433172_8446622_RNA45SN1_45S_50ntFlanks@7979.8008.30|ACCTCAGATCAGACGTGGCGACCCGCTGAA</td><td>>hg38_21_+_8433172_8446622_RNA45SN1_45S_50ntFlanks</td><td>7979</td><td>8008</td><td>30</td><td>ACCTCAGATCAGACGTGGCGACCCGCTGAA</td><td>4781.0</td><td>-4.4</td><td>0.6</td><td>-7.8</td><td>7.5E-15</td><td>2.8E-14</td><td>MB231/MB468</td><td>MITO</td></tr>
<tr><td>>hg38_21_+_8433172_8446622_RNA45SN1_45S_50ntFlanks@11455.11478.24|CTCGGCGCGCGGCGGCGGCGGCGG</td><td>>hg38_21_+_8433172_8446622_RNA45SN1_45S_50ntFlanks</td><td>11455</td><td>11478</td><td>24</td><td>CTCGGCGCGCGGCGGCGGCGGCGG</td><td>219.7</td><td>-10.5</td><td>1.3</td><td>-7.8</td><td>5.1E-15</td><td>2.8E-14</td><td>MB231/MB468</td><td>CYTO</td></tr>
<tr><td>>MI0003591|hsa-mir-584&WithFlank&5|-|149062307|149062415@22.42.21(+1A)&[MIMAT0003249&hsa-miR-584-5p&offsets|0|-1(+1A);m-167&5|-|149062374|149062394&offsets|0|0(+1A)]|TTATGGTTTGCCTGGGACTGAA</td><td>>MI0003591|hsa-mir-584&WithFlank&5|-|149062307|149062415</td><td>22</td><td>42</td><td>21(+1A)</td><td>TTATGGTTTGCCTGGGACTGAA</td><td>529.7</td><td>9.4</td><td>1.2</td><td>7.8</td><td>4.6E-15</td><td>2.8E-14</td><td>BT20/MB468</td><td>MITO</td></tr>
<tr><td>>hg38_21_+_8433172_8446622_RNA45SN1_45S_50ntFlanks@11768.11806.39|CTAATTAGTGACGCGCATGAATGGATGAACGAGATTCCC</td><td>>hg38_21_+_8433172_8446622_RNA45SN1_45S_50ntFlanks</td><td>11768</td><td>11806</td><td>39</td><td>CTAATTAGTGACGCGCATGAATGGATGAACGAGATTCCC</td><td>425.3</td><td>-11.7</td><td>1.5</td><td>-7.8</td><td>5.5E-15</td><td>2.8E-14</td><td>MB231/MB468</td><td>NUC</td></tr>
<tr><td>>hg38_21_+_8433172_8446622_RNA45SN1_45S_50ntFlanks@12285.12317.33|CAGACCGTGAAAGCGGGGCCTCACGATCCTTCT</td><td>>hg38_21_+_8433172_8446622_RNA45SN1_45S_50ntFlanks</td><td>12285</td><td>12317</td><td>33</td><td>CAGACCGTGAAAGCGGGGCCTCACGATCCTTCT</td><td>479.2</td><td>-12.7</td><td>1.6</td><td>-7.8</td><td>7.5E-15</td><td>2.8E-14</td><td>MB231/MB468</td><td>MITO</td></tr>
<tr><td>>hg38_21_+_8433172_8446622_RNA45SN1_45S_50ntFlanks@5379.5410.32|ATTAAGTCCCTGCCCTTTGTACACACCGCCCG</td><td>>hg38_21_+_8433172_8446622_RNA45SN1_45S_50ntFlanks</td><td>5379</td><td>5410</td><td>32</td><td>ATTAAGTCCCTGCCCTTTGTACACACCGCCCG</td><td>635.4</td><td>-13.1</td><td>1.7</td><td>-7.8</td><td>7.6E-15</td><td>2.8E-14</td><td>MB231/MB468</td><td>MITO</td></tr>
<tr><td>>hg38_21_+_8433172_8446622_RNA45SN1_45S_50ntFlanks@8472.8496.25|CGGCCCGGCGGATCTTTCCCGCCCC</td><td>>hg38_21_+_8433172_8446622_RNA45SN1_45S_50ntFlanks</td><td>8472</td><td>8496</td><td>25</td><td>CGGCCCGGCGGATCTTTCCCGCCCC</td><td>277.2</td><td>-11.9</td><td>1.5</td><td>-7.8</td><td>7.6E-15</td><td>2.8E-14</td><td>MB231/MB468</td><td>MITO</td></tr>
<tr><td>>hg38_21_+_8433172_8446622_RNA45SN1_45S_50ntFlanks@12624.12653.30|CCAATGGGGCGAAGCTACCATCTGTGGGAT</td><td>>hg38_21_+_8433172_8446622_RNA45SN1_45S_50ntFlanks</td><td>12624</td><td>12653</td><td>30</td><td>CCAATGGGGCGAAGCTACCATCTGTGGGAT</td><td>291.4</td><td>-10.9</td><td>1.4</td><td>-7.8</td><td>5.1E-15</td><td>2.8E-14</td><td>MB231/MB468</td><td>CYTO</td></tr>
<tr><td>>hg38_1_-_228634819_228635039_RNA5S12_5S_50ntFlanks@51.88.38|GTCTACGGCCATACCACCCTGAACGCGCCCGATCTCGT</td><td>>hg38_1_-_228634819_228635039_RNA5S12_5S_50ntFlanks</td><td>51</td><td>88</td><td>38</td><td>GTCTACGGCCATACCACCCTGAACGCGCCCGATCTCGT</td><td>7556.9</td><td>9.6</td><td>1.2</td><td>7.8</td><td>6.4E-15</td><td>2.8E-14</td><td>BT20/MB468</td><td>CYTO</td></tr>
<tr><td>>hg38_21_+_8433172_8446622_RNA45SN1_45S_50ntFlanks@9833.9850.18|CTTTTGGTAAGCAGAACT</td><td>>hg38_21_+_8433172_8446622_RNA45SN1_45S_50ntFlanks</td><td>9833</td><td>9850</td><td>18</td><td>CTTTTGGTAAGCAGAACT</td><td>215.6</td><td>-11.6</td><td>1.5</td><td>-7.8</td><td>4.6E-15</td><td>2.8E-14</td><td>MB231/MB468</td><td>MP</td></tr>
<tr><td>>hg38_21_+_8433172_8446622_RNA45SN1_45S_50ntFlanks@9201.9224.24|CTCTCGGGGCCACGCGCGCGTCCC</td><td>>hg38_21_+_8433172_8446622_RNA45SN1_45S_50ntFlanks</td><td>9201</td><td>9224</td><td>24</td><td>CTCTCGGGGCCACGCGCGCGTCCC</td><td>257.3</td><td>-11.9</td><td>1.5</td><td>-7.8</td><td>4.6E-15</td><td>2.8E-14</td><td>MB231/MB468</td><td>MP</td></tr>
<tr><td>>hg38_21_+_8433172_8446622_RNA45SN1_45S_50ntFlanks@12456.12477.22|CCAAGCGTTGGATTGTTCACCC</td><td>>hg38_21_+_8433172_8446622_RNA45SN1_45S_50ntFlanks</td><td>12456</td><td>12477</td><td>22</td><td>CCAAGCGTTGGATTGTTCACCC</td><td>8375.8</td><td>-5.0</td><td>0.6</td><td>-7.8</td><td>5.2E-15</td><td>2.8E-14</td><td>MB231/MB468</td><td>CYTO</td></tr>
<tr><td>>hg38_21_+_8433172_8446622_RNA45SN1_45S_50ntFlanks@11561.11591.31|GCGCCTAGCAGCCGACTTAGAACTGGTGCGG</td><td>>hg38_21_+_8433172_8446622_RNA45SN1_45S_50ntFlanks</td><td>11561</td><td>11591</td><td>31</td><td>GCGCCTAGCAGCCGACTTAGAACTGGTGCGG</td><td>945.5</td><td>-7.3</td><td>0.9</td><td>-7.8</td><td>5.2E-15</td><td>2.8E-14</td><td>MB231/MB468</td><td>CYTO</td></tr>
<tr><td>>MI0000809|hsa-mir-151a&WithFlank&8|-|140732558|140732659@51.73.23(+1U)&[MIMAT0000757&hsa-miR-151a-3p&offsets|-2|0(+1U);m-30&8|-|140732587|140732607&offsets|-2|0(+1U)]|TACTAGACTGAAGCTCCTTGAGGT</td><td>>MI0000809|hsa-mir-151a&WithFlank&8|-|140732558|140732659</td><td>51</td><td>73</td><td>23(+1U)</td><td>TACTAGACTGAAGCTCCTTGAGGT</td><td>639.6</td><td>7.3</td><td>0.9</td><td>7.8</td><td>6.4E-15</td><td>2.8E-14</td><td>BT20/MB468</td><td>CYTO</td></tr>
<tr><td>>hg38_21_+_8433172_8446622_RNA45SN1_45S_50ntFlanks@10841.10865.25|CGTAACTTCGGGATAAGGATTGGCT</td><td>>hg38_21_+_8433172_8446622_RNA45SN1_45S_50ntFlanks</td><td>10841</td><td>10865</td><td>25</td><td>CGTAACTTCGGGATAAGGATTGGCT</td><td>74408.5</td><td>-6.5</td><td>0.8</td><td>-7.8</td><td>6.4E-15</td><td>2.8E-14</td><td>BT20/MB468</td><td>CYTO</td></tr>
<tr><td>>hg38_21_+_8433172_8446622_RNA45SN1_45S_50ntFlanks@3939.3956.18|AACCCGGTCAGCCCCTCT</td><td>>hg38_21_+_8433172_8446622_RNA45SN1_45S_50ntFlanks</td><td>3939</td><td>3956</td><td>18</td><td>AACCCGGTCAGCCCCTCT</td><td>221.2</td><td>-11.6</td><td>1.5</td><td>-7.8</td><td>7.7E-15</td><td>2.8E-14</td><td>MB231/MB468</td><td>MITO</td></tr>
<tr><td>>hg38_21_+_8433172_8446622_RNA45SN1_45S_50ntFlanks@3899.3920.22|CCTTCGCGGGGGGGATGCGTGC</td><td>>hg38_21_+_8433172_8446622_RNA45SN1_45S_50ntFlanks</td><td>3899</td><td>3920</td><td>22</td><td>CCTTCGCGGGGGGGATGCGTGC</td><td>977.9</td><td>-12.9</td><td>1.7</td><td>-7.8</td><td>5.6E-15</td><td>2.8E-14</td><td>MB231/MB468</td><td>NUC</td></tr>
</table>

<table>
<tr><td>>hg38_21_+_8433172_8446622_RNA45SN1_45S_50ntFlanks@9623.9655.33|TAGTAGCTGGTTCCCTCCGAAGTTTCCCTCAGG</td><td>>hg38_21_+_8433172_8446622_RNA45SN1_45S_50ntFlanks</td><td>9623</td><td>9655</td><td>33</td><td>TAGTAGCTGGTTCCCTCCGAAGTTTCCCTCAGG</td><td>697.2</td><td>-9.1</td><td>1.2</td><td>-7.8</td><td>7.7E-15</td><td>2.8E-14</td><td>MB231/MB468</td><td>MITO</td></tr>
<tr><td>>hg38_21_+_8433172_8446622_RNA45SN1_45S_50ntFlanks@12654.12674.21|TATGACTGAACGCCTCTAAGT</td><td>>hg38_21_+_8433172_8446622_RNA45SN1_45S_50ntFlanks</td><td>12654</td><td>12674</td><td>21</td><td>TATGACTGAACGCCTCTAAGT</td><td>167.1</td><td>-11.2</td><td>1.4</td><td>-7.8</td><td>4.6E-15</td><td>2.8E-14</td><td>MB231/MB468</td><td>MP</td></tr>
<tr><td>>hg38_21_+_8433172_8446622_RNA45SN1_45S_50ntFlanks@482.529.48|ATCGATGTGGTGACGTCGTGCTCTCCCGGGCCGGGTCCGAGCCGCGAC</td><td>>hg38_21_+_8433172_8446622_RNA45SN1_45S_50ntFlanks</td><td>482</td><td>529</td><td>48</td><td>ATCGATGTGGTGACGTCGTGCTCTCCCGGGCCGGGTCCGAGCCGCGAC</td><td>1817.6</td><td>-15.3</td><td>2.0</td><td>-7.7</td><td>9.8E-15</td><td>2.8E-14</td><td>BT20/MB468</td><td>NUC</td></tr>
<tr><td>>hg38_21_+_8433172_8446622_RNA45SN1_45S_50ntFlanks@12632.12654.23|GCGAAGCTACCATCTGTGGGATT</td><td>>hg38_21_+_8433172_8446622_RNA45SN1_45S_50ntFlanks</td><td>12632</td><td>12654</td><td>23</td><td>GCGAAGCTACCATCTGTGGGATT</td><td>1194.5</td><td>-5.2</td><td>0.7</td><td>-7.8</td><td>5.7E-15</td><td>2.8E-14</td><td>MB231/MB468</td><td>NUC</td></tr>
<tr><td>>hg38_21_+_8433172_8446622_RNA45SN1_45S_50ntFlanks@9578.9605.28|GGTCGTCCGACCTGGGTATAGGGGCGAA</td><td>>hg38_21_+_8433172_8446622_RNA45SN1_45S_50ntFlanks</td><td>9578</td><td>9605</td><td>28</td><td>GGTCGTCCGACCTGGGTATAGGGGCGAA</td><td>481.8</td><td>12.7</td><td>1.5</td><td>8.2</td><td>1.8E-16</td><td>2.8E-14</td><td>BT20/MB468</td><td>MP</td></tr>
<tr><td>>hg38_21_+_8433172_8446622_RNA45SN1_45S_50ntFlanks@6734.6770.37|ATTGATCATCGACACTTCGAACGCACTTGCGGCCCCG</td><td>>hg38_21_+_8433172_8446622_RNA45SN1_45S_50ntFlanks</td><td>6734</td><td>6770</td><td>37</td><td>ATTGATCATCGACACTTCGAACGCACTTGCGGCCCCG</td><td>2596.2</td><td>-5.4</td><td>0.7</td><td>-7.8</td><td>7.7E-15</td><td>2.9E-14</td><td>MB231/MB468</td><td>MITO</td></tr>
<tr><td>>hg38_21_+_8433172_8446622_RNA45SN1_45S_50ntFlanks@3776.3801.26|CCGGTACAGTGAAACTGCGAATGGCT</td><td>>hg38_21_+_8433172_8446622_RNA45SN1_45S_50ntFlanks</td><td>3776</td><td>3801</td><td>26</td><td>CCGGTACAGTGAAACTGCGAATGGCT</td><td>402.8</td><td>-5.0</td><td>0.6</td><td>-7.8</td><td>5.7E-15</td><td>2.9E-14</td><td>MB231/MB468</td><td>NUC</td></tr>
<tr><td>>MI0003137|hsa-mir-193b&WithFlank&16|+|14303961|14304055@57.79.23&[MIMAT0002819&hsa-miR-193b-3p&offsets|0|+1;m-166&16|+|14304017|14304038&offsets|0|+1]|AACTGGCCCTCAAAGTCCCGCTT</td><td>>MI0003137|hsa-mir-193b&WithFlank&16|+|14303961|14304055</td><td>57</td><td>79</td><td>23</td><td>AACTGGCCCTCAAAGTCCCGCTT</td><td>64.8</td><td>8.9</td><td>1.1</td><td>7.7</td><td>9.8E-15</td><td>2.9E-14</td><td>BT20/MB468</td><td>NUC</td></tr>
<tr><td>>hg38_21_+_8433172_8446622_RNA45SN1_45S_50ntFlanks@11781.11800.20|CGCATGAATGGATGAACGAG</td><td>>hg38_21_+_8433172_8446622_RNA45SN1_45S_50ntFlanks</td><td>11781</td><td>11800</td><td>20</td><td>CGCATGAATGGATGAACGAG</td><td>170.6</td><td>-11.3</td><td>1.4</td><td>-7.8</td><td>4.7E-15</td><td>2.9E-14</td><td>MB231/MB468</td><td>MP</td></tr>
<tr><td>>hg38_21_+_8433172_8446622_RNA45SN1_45S_50ntFlanks@5074.5098.25|ATTCCGATAACGAACGAGACTCTGG</td><td>>hg38_21_+_8433172_8446622_RNA45SN1_45S_50ntFlanks</td><td>5074</td><td>5098</td><td>25</td><td>ATTCCGATAACGAACGAGACTCTGG</td><td>839.8</td><td>-13.5</td><td>1.7</td><td>-7.8</td><td>7.8E-15</td><td>2.9E-14</td><td>MB231/MB468</td><td>MITO</td></tr>
<tr><td>>hg38_21_+_8433172_8446622_RNA45SN1_45S_50ntFlanks@9568.9592.25|CGTGCAAATCGGTCGTCCGACCTGG</td><td>>hg38_21_+_8433172_8446622_RNA45SN1_45S_50ntFlanks</td><td>9568</td><td>9592</td><td>25</td><td>CGTGCAAATCGGTCGTCCGACCTGG</td><td>1129.5</td><td>-7.2</td><td>0.9</td><td>-7.8</td><td>5.3E-15</td><td>2.9E-14</td><td>MB231/MB468</td><td>CYTO</td></tr>
<tr><td>>hg38_21_+_8433172_8446622_RNA45SN1_45S_50ntFlanks@8552.8569.18|CTCCCCGGAGGGGGCGGG</td><td>>hg38_21_+_8433172_8446622_RNA45SN1_45S_50ntFlanks</td><td>8552</td><td>8569</td><td>18</td><td>CTCCCCGGAGGGGGCGGG</td><td>1930.7</td><td>-14.0</td><td>1.7</td><td>-8.2</td><td>1.8E-16</td><td>2.9E-14</td><td>BT20/MB468</td><td>MP</td></tr>
<tr><td>>MI0000439|hsa-mir-23b&WithFlank&9|+|95085202|95085310@67.86.20(+1C)&[MIMAT0000418&hsa-miR-23b-3p&offsets|+3|0(+1C);m-42&9|+|95085265|95085285&offsets|+3|+2(+1C)]|ACATTGCCAGGGATTACCACC</td><td>>MI0000439|hsa-mir-23b&WithFlank&9|+|95085202|95085310</td><td>67</td><td>86</td><td>20(+1C)</td><td>ACATTGCCAGGGATTACCACC</td><td>1193.4</td><td>28.4</td><td>3.6</td><td>7.8</td><td>5.3E-15</td><td>2.9E-14</td><td>MB231/MB468</td><td>CYTO</td></tr>
<tr><td>>MI0000477|hsa-mir-146a&WithFlank&5|+|160485346|160485456@27.48.22(+1U)&[MIMAT0000449&hsa-miR-146a-5p&offsets|0|0(+1U);m-129&5|+|160485372|160485393&offsets|0|0(+1U)]|TGAGAACTGAATTCCATGGGTTT</td><td>>MI0000477|hsa-mir-146a&WithFlank&5|+|160485346|160485456</td><td>27</td><td>48</td><td>22(+1U)</td><td>TGAGAACTGAATTCCATGGGTTT</td><td>273.4</td><td>12.1</td><td>1.5</td><td>7.8</td><td>5.3E-15</td><td>2.9E-14</td><td>MB231/MB468</td><td>CYTO</td></tr>
<tr><td>>hg38_21_+_8433172_8446622_RNA45SN1_45S_50ntFlanks@5472.5502.31|ACGGCCCTGGCGGAGCGCTGAGAAGACGGTC</td><td>>hg38_21_+_8433172_8446622_RNA45SN1_45S_50ntFlanks</td><td>5472</td><td>5502</td><td>31</td><td>ACGGCCCTGGCGGAGCGCTGAGAAGACGGTC</td><td>2056.7</td><td>6.0</td><td>0.8</td><td>7.8</td><td>5.3E-15</td><td>2.9E-14</td><td>MB231/MB468</td><td>CYTO</td></tr>
<tr><td>>hg38_21_+_8433172_8446622_RNA45SN1_45S_50ntFlanks@4069.4098.30|CTTTCGATGGTAGTCGCCGTGCCTACCATG</td><td>>hg38_21_+_8433172_8446622_RNA45SN1_45S_50ntFlanks</td><td>4069</td><td>4098</td><td>30</td><td>CTTTCGATGGTAGTCGCCGTGCCTACCATG</td><td>465.1</td><td>-8.6</td><td>1.1</td><td>-7.8</td><td>5.3E-15</td><td>2.9E-14</td><td>MB231/MB468</td><td>CYTO</td></tr>
<tr><td>>hg38_21_+_8433172_8446622_RNA45SN1_45S_50ntFlanks@11672.11707.36|CAGTGCTCTGAATGTCAAAGTGAAGAAATTCAATGA</td><td>>hg38_21_+_8433172_8446622_RNA45SN1_45S_50ntFlanks</td><td>11672</td><td>11707</td><td>36</td><td>CAGTGCTCTGAATGTCAAAGTGAAGAAATTCAATGA</td><td>544.1</td><td>-4.7</td><td>0.6</td><td>-7.7</td><td>9.9E-15</td><td>2.9E-14</td><td>BT20/MB468</td><td>NUC</td></tr>
<tr><td>>MI0000234|hsa-mir-192&WithFlank&11|-|64891131|64891252@30.50.21&[MIMAT0000222&hsa-miR-192-5p&offsets|0|0;m-59&11|-|64891203|64891223&offsets|0|0]|CTGACCTATGAATTGACAGCC</td><td>>MI0000234|hsa-mir-192&WithFlank&11|-|64891131|64891252</td><td>30</td><td>50</td><td>21</td><td>CTGACCTATGAATTGACAGCC</td><td>703.0</td><td>5.1</td><td>0.7</td><td>7.8</td><td>4.8E-15</td><td>2.9E-14</td><td>BT20/MB468</td><td>MITO</td></tr>
<tr><td>>hg38_21_+_8433172_8446622_RNA45SN1_45S_50ntFlanks@6667.6686.20|ATCACTCGGCTCGTGCGTCG</td><td>>hg38_21_+_8433172_8446622_RNA45SN1_45S_50ntFlanks</td><td>6667</td><td>6686</td><td>20</td><td>ATCACTCGGCTCGTGCGTCG</td><td>505.5</td><td>-11.1</td><td>1.4</td><td>-7.8</td><td>4.8E-15</td><td>2.9E-14</td><td>BT20/MB468</td><td>MITO</td></tr>
<tr><td>>hg38_21_+_8433172_8446622_RNA45SN1_45S_50ntFlanks@10101.10127.27|CGCGCGCGTGTGGTGTGCGTCGGAGGG</td><td>>hg38_21_+_8433172_8446622_RNA45SN1_45S_50ntFlanks</td><td>10101</td><td>10127</td><td>27</td><td>CGCGCGCGTGTGGTGTGCGTCGGAGGG</td><td>712.7</td><td>-8.6</td><td>1.1</td><td>-7.8</td><td>5.3E-15</td><td>2.9E-14</td><td>MB231/MB468</td><td>CYTO</td></tr>
<tr><td>>hg38_21_+_8433172_8446622_RNA45SN1_45S_50ntFlanks@11452.11469.18|CTCCTCGGCGCGCGGCGG</td><td>>hg38_21_+_8433172_8446622_RNA45SN1_45S_50ntFlanks</td><td>11452</td><td>11469</td><td>18</td><td>CTCCTCGGCGCGCGGCGG</td><td>156.7</td><td>-10.0</td><td>1.3</td><td>-7.8</td><td>5.3E-15</td><td>2.9E-14</td><td>MB231/MB468</td><td>CYTO</td></tr>
<tr><td>>hg38_21_+_8433172_8446622_RNA45SN1_45S_50ntFlanks@10411.10433.23|CAGTCGGTCCTGAGAGATGGGCG</td><td>>hg38_21_+_8433172_8446622_RNA45SN1_45S_50ntFlanks</td><td>10411</td><td>10433</td><td>23</td><td>CAGTCGGTCCTGAGAGATGGGCG</td><td>1346.4</td><td>-7.6</td><td>1.0</td><td>-7.8</td><td>5.3E-15</td><td>2.9E-14</td><td>MB231/MB468</td><td>CYTO</td></tr>
<tr><td>>hg38_21_+_8433172_8446622_RNA45SN1_45S_50ntFlanks@6780.6806.27|CGGGGCTACGCCTGTCTGAGCGTCGCT</td><td>>hg38_21_+_8433172_8446622_RNA45SN1_45S_50ntFlanks</td><td>6780</td><td>6806</td><td>27</td><td>CGGGGCTACGCCTGTCTGAGCGTCGCT</td><td>271.7</td><td>4.9</td><td>0.6</td><td>7.8</td><td>5.8E-15</td><td>2.9E-14</td><td>MB231/MB468</td><td>NUC</td></tr>
<tr><td>>hg38_21_+_8433172_8446622_RNA45SN1_45S_50ntFlanks@10840.10864.25|CCGTAACTTCGGGATAAGGATTGGC</td><td>>hg38_21_+_8433172_8446622_RNA45SN1_45S_50ntFlanks</td><td>10840</td><td>10864</td><td>25</td><td>CCGTAACTTCGGGATAAGGATTGGC</td><td>326.1</td><td>-12.2</td><td>1.6</td><td>-7.8</td><td>4.8E-15</td><td>2.9E-14</td><td>MB231/MB468</td><td>MP</td></tr>
<tr><td>>hg38_21_+_8433172_8446622_RNA45SN1_45S_50ntFlanks@8194.8211.18|CCCGTGGACGGTGTGAGG</td><td>>hg38_21_+_8433172_8446622_RNA45SN1_45S_50ntFlanks</td><td>8194</td><td>8211</td><td>18</td><td>CCCGTGGACGGTGTGAGG</td><td>973.7</td><td>-9.7</td><td>1.3</td><td>-7.8</td><td>7.9E-15</td><td>2.9E-14</td><td>MB231/MB468</td><td>MITO</td></tr>
<tr><td>>hg38_21_+_8433172_8446622_RNA45SN1_45S_50ntFlanks@8380.8415.36|CAAGAGGGCGTGAAACCGTTAAGAGGTAAACGGGTG</td><td>>hg38_21_+_8433172_8446622_RNA45SN1_45S_50ntFlanks</td><td>8380</td><td>8415</td><td>36</td><td>CAAGAGGGCGTGAAACCGTTAAGAGGTAAACGGGTG</td><td>322.4</td><td>-12.2</td><td>1.6</td><td>-7.8</td><td>4.8E-15</td><td>2.9E-14</td><td>MB231/MB468</td><td>MP</td></tr>
<tr><td>>hg38_21_+_8433172_8446622_RNA45SN1_45S_50ntFlanks@5485.5523.39|AGCGCTGAGAAGACGGTCGAACTTGACTATCTAGAGGAA</td><td>>hg38_21_+_8433172_8446622_RNA45SN1_45S_50ntFlanks</td><td>5485</td><td>5523</td><td>39</td><td>AGCGCTGAGAAGACGGTCGAACTTGACTATCTAGAGGAA</td><td>236.5</td><td>-11.7</td><td>1.5</td><td>-7.8</td><td>8.0E-15</td><td>2.9E-14</td><td>MB231/MB468</td><td>MITO</td></tr>
<tr><td>>hg38_21_+_8433172_8446622_RNA45SN1_45S_50ntFlanks@12367.12384.18|AACTGGCTTGTGGCGGCC</td><td>>hg38_21_+_8433172_8446622_RNA45SN1_45S_50ntFlanks</td><td>12367</td><td>12384</td><td>18</td><td>AACTGGCTTGTGGCGGCC</td><td>311.3</td><td>-12.1</td><td>1.6</td><td>-7.8</td><td>8.0E-15</td><td>2.9E-14</td><td>MB231/MB468</td><td>MITO</td></tr>
<tr><td>>hg38_21_+_8433172_8446622_RNA45SN1_45S_50ntFlanks@8174.8202.29|TCCTTCTGATCGAGGCCCAGCCCGTGGAC</td><td>>hg38_21_+_8433172_8446622_RNA45SN1_45S_50ntFlanks</td><td>8174</td><td>8202</td><td>29</td><td>TCCTTCTGATCGAGGCCCAGCCCGTGGAC</td><td>386.1</td><td>6.2</td><td>0.8</td><td>7.8</td><td>5.4E-15</td><td>2.9E-14</td><td>MB231/MB468</td><td>CYTO</td></tr>
<tr><td>>hg38_21_+_8433172_8446622_RNA45SN1_45S_50ntFlanks@11208.11231.24|GCGGCGACTCTGGACGCGAGCCGG</td><td>>hg38_21_+_8433172_8446622_RNA45SN1_45S_50ntFlanks</td><td>11208</td><td>11231</td><td>24</td><td>GCGGCGACTCTGGACGCGAGCCGG</td><td>1054.5</td><td>-7.5</td><td>1.0</td><td>-7.8</td><td>8.0E-15</td><td>3.0E-14</td><td>MB231/MB468</td><td>MITO</td></tr>
<tr><td>>am_trnaMT_GluTTC_MT_-_14674_14742@27.52.26,trnalookalike8_GluTTC_5_-_93905172_93905240@27.52.26|GTTTTTCATATCATTGGTCGTGGTTG</td><td>>am_trnaMT_GluTTC_MT_-_14674_14742</td><td>27</td><td>52</td><td>26</td><td>GTTTTTCATATCATTGGTCGTGGTTG</td><td>213.6</td><td>-10.7</td><td>1.4</td><td>-7.8</td><td>5.9E-15</td><td>3.0E-14</td><td>MB231/MB468</td><td>NUC</td></tr>
<tr><td>>hg38_21_+_8433172_8446622_RNA45SN1_45S_50ntFlanks@6700.6718.19|CTAGCTGCGAGAATTAATG</td><td>>hg38_21_+_8433172_8446622_RNA45SN1_45S_50ntFlanks</td><td>6700</td><td>6718</td><td>19</td><td>CTAGCTGCGAGAATTAATG</td><td>10737.8</td><td>-5.1</td><td>0.7</td><td>-7.8</td><td>5.5E-15</td><td>3.0E-14</td><td>MB231/MB468</td><td>CYTO</td></tr>
<tr><td>>hg38_21_+_8433172_8446622_RNA45SN1_45S_50ntFlanks@13002.13026.25|CGATCTATTGAAAGTCAGCCCTCGA</td><td>>hg38_21_+_8433172_8446622_RNA45SN1_45S_50ntFlanks</td><td>13002</td><td>13026</td><td>25</td><td>CGATCTATTGAAAGTCAGCCCTCGA</td><td>312.8</td><td>-10.6</td><td>1.4</td><td>-7.8</td><td>6.7E-15</td><td>3.0E-14</td><td>BT20/MB468</td><td>CYTO</td></tr>
<tr><td>>hg38_21_+_8433172_8446622_RNA45SN1_45S_50ntFlanks@11534.11552.19|GTTCCGCGCGGCGCCTCGC</td><td>>hg38_21_+_8433172_8446622_RNA45SN1_45S_50ntFlanks</td><td>11534</td><td>11552</td><td>19</td><td>GTTCCGCGCGGCGCCTCGC</td><td>2107.5</td><td>-11.2</td><td>1.4</td><td>-7.8</td><td>5.9E-15</td><td>3.0E-14</td><td>MB231/MB468</td><td>NUC</td></tr>
<tr><td>>MI0000088|hsa-mir-30a&WithFlank&6|-|71403545|71403627@12.35.24(+1U)&[MIMAT0000087&hsa-miR-30a-5p&offsets|0|+2(+1U);m-15&6|-|71403595|71403616&offsets|0|+2(+1U)]|TGTAAACATCCTCGACTGGAAGCTT</td><td>>MI0000088|hsa-mir-30a&WithFlank&6|-|71403545|71403627</td><td>12</td><td>35</td><td>24(+1U)</td><td>TGTAAACATCCTCGACTGGAAGCTT</td><td>8855.8</td><td>8.3</td><td>1.1</td><td>7.8</td><td>8.1E-15</td><td>3.0E-14</td><td>MB231/MB468</td><td>MITO</td></tr>
<tr><td>>hg38_21_+_8433172_8446622_RNA45SN1_45S_50ntFlanks@11323.11341.19|CGTCTCGTCGCGCGCGCGT</td><td>>hg38_21_+_8433172_8446622_RNA45SN1_45S_50ntFlanks</td><td>11323</td><td>11341</td><td>19</td><td>CGTCTCGTCGCGCGCGCGT</td><td>197.1</td><td>-10.3</td><td>1.3</td><td>-7.8</td><td>5.5E-15</td><td>3.0E-14</td><td>MB231/MB468</td><td>CYTO</td></tr>
<tr><td>>hg38_21_+_8433172_8446622_RNA45SN1_45S_50ntFlanks@11769.11792.24|TAATTAGTGACGCGCATGAATGGA</td><td>>hg38_21_+_8433172_8446622_RNA45SN1_45S_50ntFlanks</td><td>11769</td><td>11792</td><td>24</td><td>TAATTAGTGACGCGCATGAATGGA</td><td>254.0</td><td>-11.9</td><td>1.5</td><td>-7.8</td><td>4.9E-15</td><td>3.0E-14</td><td>MB231/MB468</td><td>MP</td></tr>
<tr><td>>hg38_21_+_8433172_8446622_RNA45SN1_45S_50ntFlanks@10346.10369.24|TCAAACGAGAACTTTGAAGGCCGA</td><td>>hg38_21_+_8433172_8446622_RNA45SN1_45S_50ntFlanks</td><td>10346</td><td>10369</td><td>24</td><td>TCAAACGAGAACTTTGAAGGCCGA</td><td>748.7</td><td>-5.4</td><td>0.7</td><td>-7.7</td><td>1.0E-14</td><td>3.0E-14</td><td>BT20/MB468</td><td>NUC</td></tr>
<tr><td>>MI0000301|hsa-mir-224&WithFlank&X|-|151958572|151958664@14.37.24&[MIMAT0000281&hsa-miR-224-5p&offsets|+1|0;m-238&X|-|151958630|151958651&offsets|0|+2]|CAAGTCACTAGTGGTTCCGTTTAG</td><td>>MI0000301|hsa-mir-224&WithFlank&X|-|151958572|151958664</td><td>14</td><td>37</td><td>24</td><td>CAAGTCACTAGTGGTTCCGTTTAG</td><td>1662.0</td><td>8.4</td><td>1.1</td><td>7.8</td><td>6.0E-15</td><td>3.0E-14</td><td>MB231/MB468</td><td>NUC</td></tr>
<tr><td>>hg38_21_+_8433172_8446622_RNA45SN1_45S_50ntFlanks@12646.12667.22|TGTGGGATTATGACTGAACGCC</td><td>>hg38_21_+_8433172_8446622_RNA45SN1_45S_50ntFlanks</td><td>12646</td><td>12667</td><td>22</td><td>TGTGGGATTATGACTGAACGCC</td><td>412.2</td><td>9.8</td><td>1.3</td><td>7.8</td><td>5.5E-15</td><td>3.0E-14</td><td>MB231/MB468</td><td>CYTO</td></tr>
<tr><td>>hg38_21_+_8433172_8446622_RNA45SN1_45S_50ntFlanks@4364.4391.28|CTGCAGTTAAAAAGCTCGTAGTTGGATC</td><td>>hg38_21_+_8433172_8446622_RNA45SN1_45S_50ntFlanks</td><td>4364</td><td>4391</td><td>28</td><td>CTGCAGTTAAAAAGCTCGTAGTTGGATC</td><td>183.6</td><td>-11.4</td><td>1.5</td><td>-7.8</td><td>4.9E-15</td><td>3.0E-14</td><td>MB231/MB468</td><td>MP</td></tr>
<tr><td>>hg38_21_+_8433172_8446622_RNA45SN1_45S_50ntFlanks@10678.10695.18|CGGTTCCGGCGGCGTCCG</td><td>>hg38_21_+_8433172_8446622_RNA45SN1_45S_50ntFlanks</td><td>10678</td><td>10695</td><td>18</td><td>CGGTTCCGGCGGCGTCCG</td><td>150.6</td><td>-10.0</td><td>1.3</td><td>-7.8</td><td>5.5E-15</td><td>3.0E-14</td><td>MB231/MB468</td><td>CYTO</td></tr>
<tr><td>>MI0000542|hsa-mir-320a&WithFlank&8|-|22244960|22245043@48.69.22&[MIMAT0000510&hsa-miR-320a-3p&offsets|0|0;m-60&8|-|22244975|22244996&offsets|0|0]|AAAAGCTGGGTTGAGAGGGCGA</td><td>>MI0000542|hsa-mir-320a&WithFlank&8|-|22244960|22245043</td><td>48</td><td>69</td><td>22</td><td>AAAAGCTGGGTTGAGAGGGCGA</td><td>88553.1</td><td>8.2</td><td>1.0</td><td>8.2</td><td>1.9E-16</td><td>3.0E-14</td><td>BT20/MB468</td><td>MP</td></tr>
<tr><td>>hg38_21_+_8433172_8446622_RNA45SN1_45S_50ntFlanks@5468.5514.47|GCCCACGGCCCTGGCGGAGCGCTGAGAAGACGGTCGAACTTGACTAT</td><td>>hg38_21_+_8433172_8446622_RNA45SN1_45S_50ntFlanks</td><td>5468</td><td>5514</td><td>47</td><td>GCCCACGGCCCTGGCGGAGCGCTGAGAAGACGGTCGAACTTGACTAT</td><td>451.8</td><td>-13.2</td><td>1.7</td><td>-7.7</td><td>1.0E-14</td><td>3.0E-14</td><td>BT20/MB468</td><td>NUC</td></tr>
<tr><td>>hg38_21_+_8433172_8446622_RNA45SN1_45S_50ntFlanks@3813.3842.30|TTATGGTTCCTTTGGTCGCTCGCTCCTCTC</td><td>>hg38_21_+_8433172_8446622_RNA45SN1_45S_50ntFlanks</td><td>3813</td><td>3842</td><td>30</td><td>TTATGGTTCCTTTGGTCGCTCGCTCCTCTC</td><td>361.6</td><td>-11.5</td><td>1.5</td><td>-7.8</td><td>6.0E-15</td><td>3.0E-14</td><td>MB231/MB468</td><td>NUC</td></tr>
<tr><td>>hg38_21_+_8433172_8446622_RNA45SN1_45S_50ntFlanks@10408.10433.26|GGTCAGTCGGTCCTGAGAGATGGGCG</td><td>>hg38_21_+_8433172_8446622_RNA45SN1_45S_50ntFlanks</td><td>10408</td><td>10433</td><td>26</td><td>GGTCAGTCGGTCCTGAGAGATGGGCG</td><td>4363.7</td><td>-11.9</td><td>1.5</td><td>-7.7</td><td>1.0E-14</td><td>3.0E-14</td><td>BT20/MB468</td><td>NUC</td></tr>
<tr><td>>hg38_21_+_8433172_8446622_RNA45SN1_45S_50ntFlanks@11843.11872.30|AGGGAACGGGCTTGGCGGAATCAGCGGGGA</td><td>>hg38_21_+_8433172_8446622_RNA45SN1_45S_50ntFlanks</td><td>11843</td><td>11872</td><td>30</td><td>AGGGAACGGGCTTGGCGGAATCAGCGGGGA</td><td>228.0</td><td>-8.0</td><td>1.0</td><td>-7.8</td><td>5.6E-15</td><td>3.0E-14</td><td>MB231/MB468</td><td>CYTO</td></tr>
<tr><td>>hg38_21_+_8433172_8446622_RNA45SN1_45S_50ntFlanks@12133.12150.18|CTCCGGGGACAGTGCCAG</td><td>>hg38_21_+_8433172_8446622_RNA45SN1_45S_50ntFlanks</td><td>12133</td><td>12150</td><td>18</td><td>CTCCGGGGACAGTGCCAG</td><td>2098.2</td><td>-9.4</td><td>1.2</td><td>-7.8</td><td>6.0E-15</td><td>3.0E-14</td><td>MB231/MB468</td><td>NUC</td></tr>
<tr><td>>hg38_21_+_8433172_8446622_RNA45SN1_45S_50ntFlanks@10514.10533.20|CGGAGTGGCGGAGATGGGCG</td><td>>hg38_21_+_8433172_8446622_RNA45SN1_45S_50ntFlanks</td><td>10514</td><td>10533</td><td>20</td><td>CGGAGTGGCGGAGATGGGCG</td><td>3422.8</td><td>-10.7</td><td>1.4</td><td>-7.8</td><td>4.9E-15</td><td>3.0E-14</td><td>MB231/MB468</td><td>MP</td></tr>
<tr><td>>hg38_21_+_8433172_8446622_RNA45SN1_45S_50ntFlanks@9568.9586.19|CGTGCAAATCGGTCGTCCG</td><td>>hg38_21_+_8433172_8446622_RNA45SN1_45S_50ntFlanks</td><td>9568</td><td>9586</td><td>19</td><td>CGTGCAAATCGGTCGTCCG</td><td>170.9</td><td>-11.3</td><td>1.4</td><td>-7.8</td><td>5.0E-15</td><td>3.0E-14</td><td>MB231/MB468</td><td>MP</td></tr>
<tr><td>>MI0000477|hsa-mir-146a&WithFlank&5|+|160485346|160485456@27.50.24&[MIMAT0000449&hsa-miR-146a-5p&offsets|0|+2;m-129&5|+|160485372|160485393&offsets|0|+2]|TGAGAACTGAATTCCATGGGTTGT</td><td>>MI0000477|hsa-mir-146a&WithFlank&5|+|160485346|160485456</td><td>27</td><td>50</td><td>24</td><td>TGAGAACTGAATTCCATGGGTTGT</td><td>247.6</td><td>12.0</td><td>1.5</td><td>7.8</td><td>5.6E-15</td><td>3.0E-14</td><td>MB231/MB468</td><td>CYTO</td></tr>
<tr><td>>MI0000077|hsa-mir-21&WithFlank&17|+|59841260|59841343@14.36.23(+1C)&[MIMAT0000076&hsa-miR-21-5p&offsets|0|+1(+1C);m-1&17|+|59841273|59841294&offsets|0|+1(+1C)]|TAGCTTATCAGACTGATGTTGACC</td><td>>MI0000077|hsa-mir-21&WithFlank&17|+|59841260|59841343</td><td>14</td><td>36</td><td>23(+1C)</td><td>TAGCTTATCAGACTGATGTTGACC</td><td>35871.9</td><td>7.0</td><td>0.9</td><td>7.8</td><td>8.3E-15</td><td>3.0E-14</td><td>MB231/MB468</td><td>MITO</td></tr>
</table>

<table>
<tr><td>>hg38_21_+_8433172_8446622_RNA45SN1_45S_50ntFlanks@11534.11560.27|GTTCCGCGCGGCGCCTCGCCTCGGCCG</td><td>>hg38_21_+_8433172_8446622_RNA45SN1_45S_50ntFlanks</td><td>11534</td><td>11560</td><td>27</td><td>GTTCCGCGCGGCGCCTCGCCTCGGCCG</td><td>265.2</td><td>-11.9</td><td>1.5</td><td>-7.8</td><td>5.0E-15</td><td>3.1E-14</td><td>MB231/MB468</td><td>MP</td></tr>
<tr><td>>hg38_21_+_8433172_8446622_RNA45SN1_45S_50ntFlanks@11781.11798.18|CGCATGAATGGATGAACG</td><td>>hg38_21_+_8433172_8446622_RNA45SN1_45S_50ntFlanks</td><td>11781</td><td>11798</td><td>18</td><td>CGCATGAATGGATGAACG</td><td>898.3</td><td>-7.4</td><td>0.9</td><td>-7.8</td><td>5.7E-15</td><td>3.1E-14</td><td>MB231/MB468</td><td>CYTO</td></tr>
<tr><td>>hg38_21_+_8433172_8446622_RNA45SN1_45S_50ntFlanks@9714.9738.25|CGAATGATTAGAGGTCTTGGGGCCG</td><td>>hg38_21_+_8433172_8446622_RNA45SN1_45S_50ntFlanks</td><td>9714</td><td>9738</td><td>25</td><td>CGAATGATTAGAGGTCTTGGGGCCG</td><td>190.4</td><td>-11.4</td><td>1.5</td><td>-7.8</td><td>5.0E-15</td><td>3.1E-14</td><td>MB231/MB468</td><td>MP</td></tr>
<tr><td>>am_trna10_LysCTT_16_+_3241501_3241573@31.51.21,trna119_LysCTT_1_-_145395522_145395594@31.51.21,trna11_LysCTT_5_-_180648979_180649051@31.51.21,trna13_LysCTT_16_+_3246153_3246225@31.51.21,trna13_LysCTT_6_+_26556774_26556846@31.51.21,trna30_LysCTT_16_-_3230555_3230627@31.51.21,trna5_LysCTT_16_+_3214939_3215011@31.51.21,trna7_LysCTT_16_+_3225692_3225764@31.51.21,trna8_LysCTT_1_+_39970195_39970267@31.51.21,trna9_LysCTT_5_+_180634755_180634827@31.51.21|ACTCTTAATCTCAGGGTCGTG</td><td>>am_trna10_LysCTT_16_+_3241501_3241573</td><td>31</td><td>51</td><td>21</td><td>ACTCTTAATCTCAGGGTCGTG</td><td>930.0</td><td>-8.6</td><td>1.1</td><td>-7.8</td><td>5.7E-15</td><td>3.1E-14</td><td>MB231/MB468</td><td>CYTO</td></tr>
<tr><td>>hg38_MT_+_598_1651_RNR1_12S_50ntFlanks@970.1004.35|ATGGTAAGTGTACTGGAAAGTGCACTTGGACGAAC</td><td>>hg38_MT_+_598_1651_RNR1_12S_50ntFlanks</td><td>970</td><td>1004</td><td>35</td><td>ATGGTAAGTGTACTGGAAAGTGCACTTGGACGAAC</td><td>175.8</td><td>11.4</td><td>1.5</td><td>7.8</td><td>6.1E-15</td><td>3.1E-14</td><td>MB231/MB468</td><td>NUC</td></tr>
<tr><td>>MI0001448|hsa-mir-425&WithFlank&3|-|49020142|49020240@20.43.24(+1A)&[MIMAT0003393&hsa-miR-425-5p&offsets|0|+1(+1A);m-95&3|-|49020200|49020221&offsets|0|+2(+1A)]|AATGACACGATCACTCCCGTTGAGA</td><td>>MI0001448|hsa-mir-425&WithFlank&3|-|49020142|49020240</td><td>20</td><td>43</td><td>24(+1A)</td><td>AATGACACGATCACTCCCGTTGAGA</td><td>614.3</td><td>9.9</td><td>1.3</td><td>7.8</td><td>7.0E-15</td><td>3.1E-14</td><td>BT20/MB468</td><td>CYTO</td></tr>
<tr><td>>hg38_21_+_8433172_8446622_RNA45SN1_45S_50ntFlanks@9339.9365.27|CGCACGAAAGCCGCCGTGGCGCAATGA</td><td>>hg38_21_+_8433172_8446622_RNA45SN1_45S_50ntFlanks</td><td>9339</td><td>9365</td><td>27</td><td>CGCACGAAAGCCGCCGTGGCGCAATGA</td><td>221.2</td><td>-11.7</td><td>1.5</td><td>-7.8</td><td>5.0E-15</td><td>3.1E-14</td><td>MB231/MB468</td><td>MP</td></tr>
<tr><td>>hg38_21_+_8433172_8446622_RNA45SN1_45S_50ntFlanks@11584.11623.40|TGGTGCGGACCAGGGGAATCCGACTGTTTAATTAAAACAA</td><td>>hg38_21_+_8433172_8446622_RNA45SN1_45S_50ntFlanks</td><td>11584</td><td>11623</td><td>40</td><td>TGGTGCGGACCAGGGGAATCCGACTGTTTAATTAAAACAA</td><td>397.1</td><td>12.9</td><td>1.7</td><td>7.8</td><td>7.0E-15</td><td>3.1E-14</td><td>BT20/MB468</td><td>CYTO</td></tr>
<tr><td>>hg38_21_+_8433172_8446622_RNA45SN1_45S_50ntFlanks@11923.11951.29|AGAGGTGTAGAATAAGTGGGAGGCCCCCG</td><td>>hg38_21_+_8433172_8446622_RNA45SN1_45S_50ntFlanks</td><td>11923</td><td>11951</td><td>29</td><td>AGAGGTGTAGAATAAGTGGGAGGCCCCCG</td><td>178.7</td><td>-11.3</td><td>1.4</td><td>-7.8</td><td>5.1E-15</td><td>3.1E-14</td><td>MB231/MB468</td><td>MP</td></tr>
<tr><td>>hg38_21_+_8433172_8446622_RNA45SN1_45S_50ntFlanks@6749.6770.22|TTCGAACGCACTTGCGGCCCCG</td><td>>hg38_21_+_8433172_8446622_RNA45SN1_45S_50ntFlanks</td><td>6749</td><td>6770</td><td>22</td><td>TTCGAACGCACTTGCGGCCCCG</td><td>210.0</td><td>-11.6</td><td>1.5</td><td>-7.8</td><td>5.1E-15</td><td>3.1E-14</td><td>MB231/MB468</td><td>MP</td></tr>
<tr><td>>MI0000439|hsa-mir-23b&WithFlank&9|+|95085202|95085310@67.85.19(+2U)&[MIMAT0000418&hsa-miR-23b-3p&offsets|+3|-1(+2U);m-42&9|+|95085265|95085285&offsets|+3|+1(+2U)]|ACATTGCCAGGGATTACCATT</td><td>>MI0000439|hsa-mir-23b&WithFlank&9|+|95085202|95085310</td><td>67</td><td>85</td><td>19(+2U)</td><td>ACATTGCCAGGGATTACCATT</td><td>1298.1</td><td>28.4</td><td>3.6</td><td>7.8</td><td>5.7E-15</td><td>3.1E-14</td><td>MB231/MB468</td><td>CYTO</td></tr>
<tr><td>>MI0000074|hsa-mir-19b-1&WithFlank&13|+|91351186|91351284@60.82.23;MI0000075|hsa-mir-19b-2&WithFlank&X|-|134169665|134169772@68.90.23&[MIMAT0000074&hsa-miR-19b-3p&offsets|0|0;m-122&13|+|91351245|91351267&offsets|0|0];[MIMAT0000074_1&hsa-miR-19b-3p&offsets|0|0;m-121&X|-|134169683|134169705&offsets|0|0]|TGTGCAAATCCATGCAAAACTGA</td><td>>MI0000074|hsa-mir-19b-1&WithFlank&13|+|91351186|91351284</td><td>60</td><td>82</td><td>23</td><td>TGTGCAAATCCATGCAAAACTGA</td><td>581.1</td><td>6.3</td><td>0.8</td><td>7.8</td><td>6.2E-15</td><td>3.1E-14</td><td>MB231/MB468</td><td>NUC</td></tr>
<tr><td>>trnaMT_LeuTAA_MT_+_3230_3304@11.44.34|CAGAGCCCGGTAATCGCATAAAACTTAAAACTTT</td><td>>trnaMT_LeuTAA_MT_+_3230_3304</td><td>11</td><td>44</td><td>34</td><td>CAGAGCCCGGTAATCGCATAAAACTTAAAACTTT</td><td>329.1</td><td>-12.8</td><td>1.7</td><td>-7.7</td><td>1.1E-14</td><td>3.1E-14</td><td>BT20/MB468</td><td>NUC</td></tr>
<tr><td>>hg38_21_+_8433172_8446622_RNA45SN1_45S_50ntFlanks@8158.8178.21|TCGTGGGGGGCCCAAGTCCTT</td><td>>hg38_21_+_8433172_8446622_RNA45SN1_45S_50ntFlanks</td><td>8158</td><td>8178</td><td>21</td><td>TCGTGGGGGGCCCAAGTCCTT</td><td>645.6</td><td>-13.1</td><td>1.7</td><td>-7.8</td><td>8.4E-15</td><td>3.1E-14</td><td>MB231/MB468</td><td>MITO</td></tr>
<tr><td>>hg38_21_+_8433172_8446622_RNA45SN1_45S_50ntFlanks@10432.10453.22|CGAGCGCCGTTCCGAAGGGACG</td><td>>hg38_21_+_8433172_8446622_RNA45SN1_45S_50ntFlanks</td><td>10432</td><td>10453</td><td>22</td><td>CGAGCGCCGTTCCGAAGGGACG</td><td>311.9</td><td>-12.1</td><td>1.6</td><td>-7.8</td><td>8.4E-15</td><td>3.1E-14</td><td>MB231/MB468</td><td>MITO</td></tr>
<tr><td>>hg38_21_+_8433172_8446622_RNA45SN1_45S_50ntFlanks@12902.12930.29|TCACGCACCGCACGTTCGTGGGGAACCTG</td><td>>hg38_21_+_8433172_8446622_RNA45SN1_45S_50ntFlanks</td><td>12902</td><td>12930</td><td>29</td><td>TCACGCACCGCACGTTCGTGGGGAACCTG</td><td>193.2</td><td>-11.5</td><td>1.5</td><td>-7.8</td><td>5.1E-15</td><td>3.1E-14</td><td>MB231/MB468</td><td>MP</td></tr>
<tr><td>>hg38_21_+_8433172_8446622_RNA45SN1_45S_50ntFlanks@11233.11253.21|CCCTTCCCGTGGATCGCCCCA</td><td>>hg38_21_+_8433172_8446622_RNA45SN1_45S_50ntFlanks</td><td>11233</td><td>11253</td><td>21</td><td>CCCTTCCCGTGGATCGCCCCA</td><td>415.7</td><td>-11.4</td><td>1.5</td><td>-7.8</td><td>5.8E-15</td><td>3.1E-14</td><td>MB231/MB468</td><td>CYTO</td></tr>
<tr><td>>hg38_21_+_8433172_8446622_RNA45SN1_45S_50ntFlanks@12415.12444.30|CTTCGATGTCGGCTCTTCCTATCATTGTGA</td><td>>hg38_21_+_8433172_8446622_RNA45SN1_45S_50ntFlanks</td><td>12415</td><td>12444</td><td>30</td><td>CTTCGATGTCGGCTCTTCCTATCATTGTGA</td><td>2273.0</td><td>-4.8</td><td>0.6</td><td>-7.7</td><td>1.1E-14</td><td>3.1E-14</td><td>BT20/MB468</td><td>NUC</td></tr>
<tr><td>>hg38_21_+_8433172_8446622_RNA45SN1_45S_50ntFlanks@12893.12925.33|CCTCGCCCGTCACGCACCGCACGTTCGTGGGGA</td><td>>hg38_21_+_8433172_8446622_RNA45SN1_45S_50ntFlanks</td><td>12893</td><td>12925</td><td>33</td><td>CCTCGCCCGTCACGCACCGCACGTTCGTGGGGA</td><td>282.5</td><td>-11.9</td><td>1.5</td><td>-7.8</td><td>8.4E-15</td><td>3.1E-14</td><td>MB231/MB468</td><td>MITO</td></tr>
<tr><td>>hg38_21_+_8433172_8446622_RNA45SN1_45S_50ntFlanks@10807.10825.19|ACAATGTAGGTAAGGGAAG</td><td>>hg38_21_+_8433172_8446622_RNA45SN1_45S_50ntFlanks</td><td>10807</td><td>10825</td><td>19</td><td>ACAATGTAGGTAAGGGAAG</td><td>363.7</td><td>-11.2</td><td>1.4</td><td>-7.8</td><td>5.8E-15</td><td>3.1E-14</td><td>MB231/MB468</td><td>CYTO</td></tr>
<tr><td>>MI0000542|hsa-mir-320a&WithFlank&8|-|22244960|22245043@49.70.22&[MIMAT0000510&hsa-miR-320a-3p&offsets|+1|+1;m-60&8|-|22244975|22244996&offsets|+1|+1]|AAAGCTGGGTTGAGAGGGCGAA</td><td>>MI0000542|hsa-mir-320a&WithFlank&8|-|22244960|22245043</td><td>49</td><td>70</td><td>22</td><td>AAAGCTGGGTTGAGAGGGCGAA</td><td>1552.5</td><td>11.3</td><td>1.5</td><td>7.8</td><td>7.1E-15</td><td>3.1E-14</td><td>BT20/MB468</td><td>CYTO</td></tr>
<tr><td>>MI0000088|hsa-mir-30a&WithFlank&6|-|71403545|71403627@12.35.24(+1C)&[MIMAT0000087&hsa-miR-30a-5p&offsets|0|+2(+1C);m-15&6|-|71403595|71403616&offsets|0|+2(+1C)]|TGTAAACATCCTCGACTGGAAGCTC</td><td>>MI0000088|hsa-mir-30a&WithFlank&6|-|71403545|71403627</td><td>12</td><td>35</td><td>24(+1C)</td><td>TGTAAACATCCTCGACTGGAAGCTC</td><td>1295.8</td><td>13.6</td><td>1.7</td><td>7.8</td><td>8.4E-15</td><td>3.1E-14</td><td>MB231/MB468</td><td>MITO</td></tr>
<tr><td>>trnaMT_ValTAC_MT_+_1602_1670@42.72.31|AGATTTCAACTTAACTTGACCGCTCTGACCA</td><td>>trnaMT_ValTAC_MT_+_1602_1670</td><td>42</td><td>72</td><td>31</td><td>AGATTTCAACTTAACTTGACCGCTCTGACCA</td><td>304.2</td><td>8.5</td><td>1.1</td><td>7.8</td><td>7.1E-15</td><td>3.1E-14</td><td>BT20/MB468</td><td>CYTO</td></tr>
<tr><td>>hg38_21_+_8433172_8446622_RNA45SN1_45S_50ntFlanks@10687.10708.22|CGGCGTCCGGTGAGCTCTCGCT</td><td>>hg38_21_+_8433172_8446622_RNA45SN1_45S_50ntFlanks</td><td>10687</td><td>10708</td><td>22</td><td>CGGCGTCCGGTGAGCTCTCGCT</td><td>2751.3</td><td>-5.4</td><td>0.7</td><td>-7.8</td><td>8.5E-15</td><td>3.1E-14</td><td>MB231/MB468</td><td>MITO</td></tr>
<tr><td>>hg38_21_+_8433172_8446622_RNA45SN1_45S_50ntFlanks@12636.12657.22|AGCTACCATCTGTGGGATTATG</td><td>>hg38_21_+_8433172_8446622_RNA45SN1_45S_50ntFlanks</td><td>12636</td><td>12657</td><td>22</td><td>AGCTACCATCTGTGGGATTATG</td><td>1209.8</td><td>-10.2</td><td>1.3</td><td>-7.8</td><td>8.5E-15</td><td>3.1E-14</td><td>MB231/MB468</td><td>MITO</td></tr>
<tr><td>>hg38_21_+_8433172_8446622_RNA45SN1_45S_50ntFlanks@11920.11968.49|ATGAGAGGTGTAGAATAAGTGGGAGGCCCCCGGCGCCCCCCCGGTGTCC</td><td>>hg38_21_+_8433172_8446622_RNA45SN1_45S_50ntFlanks</td><td>11920</td><td>11968</td><td>49</td><td>ATGAGAGGTGTAGAATAAGTGGGAGGCCCCCGGCGCCCCCCCGGTGTCC</td><td>533.2</td><td>-13.5</td><td>1.7</td><td>-7.7</td><td>1.1E-14</td><td>3.1E-14</td><td>BT20/MB468</td><td>NUC</td></tr>
<tr><td>>hg38_21_+_8433172_8446622_RNA45SN1_45S_50ntFlanks@12928.12951.24|CTGGCGCTAAACCATTCGTAGACG</td><td>>hg38_21_+_8433172_8446622_RNA45SN1_45S_50ntFlanks</td><td>12928</td><td>12951</td><td>24</td><td>CTGGCGCTAAACCATTCGTAGACG</td><td>205.9</td><td>-11.5</td><td>1.5</td><td>-7.8</td><td>5.1E-15</td><td>3.1E-14</td><td>MB231/MB468</td><td>MP</td></tr>
<tr><td>>hg38_21_+_8433172_8446622_RNA45SN1_45S_50ntFlanks@10572.10592.21|AGAAGCCGGCGGGAGCCCCGG</td><td>>hg38_21_+_8433172_8446622_RNA45SN1_45S_50ntFlanks</td><td>10572</td><td>10592</td><td>21</td><td>AGAAGCCGGCGGGAGCCCCGG</td><td>173.3</td><td>-11.3</td><td>1.4</td><td>-7.8</td><td>5.1E-15</td><td>3.1E-14</td><td>MB231/MB468</td><td>MP</td></tr>
<tr><td>>hg38_21_+_8433172_8446622_RNA45SN1_45S_50ntFlanks@11703.11740.38|AATGAAGCGCGGGTAAACGGCGGGAGTAACTATGACTC</td><td>>hg38_21_+_8433172_8446622_RNA45SN1_45S_50ntFlanks</td><td>11703</td><td>11740</td><td>38</td><td>AATGAAGCGCGGGTAAACGGCGGGAGTAACTATGACTC</td><td>972.9</td><td>-8.3</td><td>1.1</td><td>-7.7</td><td>1.1E-14</td><td>3.1E-14</td><td>BT20/MB468</td><td>NUC</td></tr>
<tr><td>>hg38_21_+_8433172_8446622_RNA45SN1_45S_50ntFlanks@5074.5102.29|ATTCCGATAACGAACGAGACTCTGGCATG</td><td>>hg38_21_+_8433172_8446622_RNA45SN1_45S_50ntFlanks</td><td>5074</td><td>5102</td><td>29</td><td>ATTCCGATAACGAACGAGACTCTGGCATG</td><td>772.4</td><td>-13.4</td><td>1.7</td><td>-7.8</td><td>8.6E-15</td><td>3.1E-14</td><td>MB231/MB468</td><td>MITO</td></tr>
<tr><td>>hg38_21_+_8433172_8446622_RNA45SN1_45S_50ntFlanks@8179.8211.33|CTGATCGAGGCCCAGCCCGTGGACGGTGTGAGG</td><td>>hg38_21_+_8433172_8446622_RNA45SN1_45S_50ntFlanks</td><td>8179</td><td>8211</td><td>33</td><td>CTGATCGAGGCCCAGCCCGTGGACGGTGTGAGG</td><td>272.0</td><td>-11.9</td><td>1.5</td><td>-7.8</td><td>8.6E-15</td><td>3.1E-14</td><td>MB231/MB468</td><td>MITO</td></tr>
<tr><td>>hg38_21_+_8433172_8446622_RNA45SN1_45S_50ntFlanks@9352.9377.26|CCGTGGCGCAATGAAGGTGAAGGCCG</td><td>>hg38_21_+_8433172_8446622_RNA45SN1_45S_50ntFlanks</td><td>9352</td><td>9377</td><td>26</td><td>CCGTGGCGCAATGAAGGTGAAGGCCG</td><td>416.7</td><td>-12.6</td><td>1.6</td><td>-7.8</td><td>5.2E-15</td><td>3.1E-14</td><td>MB231/MB468</td><td>MP</td></tr>
<tr><td>>hg38_21_+_8433172_8446622_RNA45SN1_45S_50ntFlanks@8104.8143.40|CCGCGGCGGGGCGCGGGACATGTGGCGTACGGAAGACCCG</td><td>>hg38_21_+_8433172_8446622_RNA45SN1_45S_50ntFlanks</td><td>8104</td><td>8143</td><td>40</td><td>CCGCGGCGGGGCGCGGGACATGTGGCGTACGGAAGACCCG</td><td>33327.5</td><td>-11.9</td><td>1.5</td><td>-7.8</td><td>5.2E-15</td><td>3.1E-14</td><td>MB231/MB468</td><td>MP</td></tr>
<tr><td>>hg38_21_+_8433172_8446622_RNA45SN1_45S_50ntFlanks@11564.11593.30|CCTAGCAGCCGACTTAGAACTGGTGCGGAC</td><td>>hg38_21_+_8433172_8446622_RNA45SN1_45S_50ntFlanks</td><td>11564</td><td>11593</td><td>30</td><td>CCTAGCAGCCGACTTAGAACTGGTGCGGAC</td><td>382.7</td><td>-12.4</td><td>1.6</td><td>-7.8</td><td>5.2E-15</td><td>3.2E-14</td><td>MB231/MB468</td><td>MP</td></tr>
<tr><td>>hg38_21_+_8433172_8446622_RNA45SN1_45S_50ntFlanks@9534.9555.22|AGGAAACTCTGGTGGAGGTCCG</td><td>>hg38_21_+_8433172_8446622_RNA45SN1_45S_50ntFlanks</td><td>9534</td><td>9555</td><td>22</td><td>AGGAAACTCTGGTGGAGGTCCG</td><td>874.0</td><td>-13.6</td><td>1.7</td><td>-7.8</td><td>5.2E-15</td><td>3.2E-14</td><td>MB231/MB468</td><td>MP</td></tr>
<tr><td>>hg38_21_+_8433172_8446622_RNA45SN1_45S_50ntFlanks@10713.10734.22|CTTGAAAATCCGGGGGAGAGGG</td><td>>hg38_21_+_8433172_8446622_RNA45SN1_45S_50ntFlanks</td><td>10713</td><td>10734</td><td>22</td><td>CTTGAAAATCCGGGGGAGAGGG</td><td>213.6</td><td>-11.6</td><td>1.5</td><td>-7.8</td><td>5.2E-15</td><td>3.2E-14</td><td>MB231/MB468</td><td>MP</td></tr>
<tr><td>>hg38_21_+_8433172_8446622_RNA45SN1_45S_50ntFlanks@10795.10826.32|TGGCATGTTGGAACAATGTAGGTAAGGGAAGT</td><td>>hg38_21_+_8433172_8446622_RNA45SN1_45S_50ntFlanks</td><td>10795</td><td>10826</td><td>32</td><td>TGGCATGTTGGAACAATGTAGGTAAGGGAAGT</td><td>177.2</td><td>-11.3</td><td>1.4</td><td>-7.8</td><td>5.3E-15</td><td>3.2E-14</td><td>MB231/MB468</td><td>MP</td></tr>
<tr><td>>hg38_21_+_8433172_8446622_RNA45SN1_45S_50ntFlanks@10927.10946.20|CGAGGCGCCGCCGCCCCCCC</td><td>>hg38_21_+_8433172_8446622_RNA45SN1_45S_50ntFlanks</td><td>10927</td><td>10946</td><td>20</td><td>CGAGGCGCCGCCGCCCCCCC</td><td>248.8</td><td>-10.7</td><td>1.4</td><td>-7.8</td><td>5.9E-15</td><td>3.2E-14</td><td>MB231/MB468</td><td>CYTO</td></tr>
<tr><td>>hg38_21_+_8433172_8446622_RNA45SN1_45S_50ntFlanks@11918.11950.33|ACATGAGAGGTGTAGAATAAGTGGGAGGCCCCC</td><td>>hg38_21_+_8433172_8446622_RNA45SN1_45S_50ntFlanks</td><td>11918</td><td>11950</td><td>33</td><td>ACATGAGAGGTGTAGAATAAGTGGGAGGCCCCC</td><td>371.9</td><td>-11.5</td><td>1.5</td><td>-7.8</td><td>6.4E-15</td><td>3.2E-14</td><td>MB231/MB468</td><td>NUC</td></tr>
<tr><td>>hg38_21_+_8433172_8446622_RNA45SN1_45S_50ntFlanks@9539.9571.33|ACTCTGGTGGAGGTCCGTAGCGGTCCTGACGTG</td><td>>hg38_21_+_8433172_8446622_RNA45SN1_45S_50ntFlanks</td><td>9539</td><td>9571</td><td>33</td><td>ACTCTGGTGGAGGTCCGTAGCGGTCCTGACGTG</td><td>644.7</td><td>-5.3</td><td>0.7</td><td>-7.8</td><td>8.8E-15</td><td>3.2E-14</td><td>MB231/MB468</td><td>MITO</td></tr>
<tr><td>>hg38_21_+_8433172_8446622_RNA45SN1_45S_50ntFlanks@4730.4759.30|CAAGAACGAAAGTCGGAGGTTCGAAGACGA</td><td>>hg38_21_+_8433172_8446622_RNA45SN1_45S_50ntFlanks</td><td>4730</td><td>4759</td><td>30</td><td>CAAGAACGAAAGTCGGAGGTTCGAAGACGA</td><td>1176.4</td><td>-6.4</td><td>0.8</td><td>-7.8</td><td>8.8E-15</td><td>3.2E-14</td><td>MB231/MB468</td><td>MITO</td></tr>
<tr><td>>hg38_21_+_8433172_8446622_RNA45SN1_45S_50ntFlanks@8170.8194.25|CAAGTCCTTCTGATCGAGGCCCAGC</td><td>>hg38_21_+_8433172_8446622_RNA45SN1_45S_50ntFlanks</td><td>8170</td><td>8194</td><td>25</td><td>CAAGTCCTTCTGATCGAGGCCCAGC</td><td>246.5</td><td>-11.8</td><td>1.5</td><td>-7.8</td><td>5.3E-15</td><td>3.2E-14</td><td>MB231/MB468</td><td>MP</td></tr>
<tr><td>>hg38_21_+_8433172_8446622_RNA45SN1_45S_50ntFlanks@11874.11917.44|AGAAGACCCTGTTGAGCTTGACTCTAGTCTGGCACGGTGAAGAG</td><td>>hg38_21_+_8433172_8446622_RNA45SN1_45S_50ntFlanks</td><td>11874</td><td>11917</td><td>44</td><td>AGAAGACCCTGTTGAGCTTGACTCTAGTCTGGCACGGTGAAGAG</td><td>475.4</td><td>-12.7</td><td>1.6</td><td>-7.8</td><td>8.8E-15</td><td>3.2E-14</td><td>MB231/MB468</td><td>MITO</td></tr>
<tr><td>>MI0000283|hsa-mir-203a&WithFlank&14|+|104117399|104117520@72.93.22&[MIMAT0000264&hsa-miR-203a-3p&offsets|+1|+1;m-79&14|+|104117469|104117490&offsets|+1|+1]|TGAAATGTTTAGGACCACTAGA</td><td>>MI0000283|hsa-mir-203a&WithFlank&14|+|104117399|104117520</td><td>72</td><td>93</td><td>22</td><td>TGAAATGTTTAGGACCACTAGA</td><td>211.6</td><td>11.3</td><td>1.4</td><td>8.1</td><td>7.5E-16</td><td>3.2E-14</td><td>BT20/MB231</td><td>MITO</td></tr>
<tr><td>>hg38_21_+_8433172_8446622_RNA45SN1_45S_50ntFlanks@11190.11208.19|CCCGGGGGGCCGGCGGCGG</td><td>>hg38_21_+_8433172_8446622_RNA45SN1_45S_50ntFlanks</td><td>11190</td><td>11208</td><td>19</td><td>CCCGGGGGGCCGGCGGCGG</td><td>623.5</td><td>-12.2</td><td>1.6</td><td>-7.8</td><td>6.5E-15</td><td>3.2E-14</td><td>MB231/MB468</td><td>NUC</td></tr>
<tr><td>>hg38_21_+_8433172_8446622_RNA45SN1_45S_50ntFlanks@10788.10815.28|CAGCCTCTGGCATGTTGGAACAATGTAG</td><td>>hg38_21_+_8433172_8446622_RNA45SN1_45S_50ntFlanks</td><td>10788</td><td>10815</td><td>28</td><td>CAGCCTCTGGCATGTTGGAACAATGTAG</td><td>1495.4</td><td>-8.7</td><td>1.1</td><td>-7.7</td><td>1.1E-14</td><td>3.2E-14</td><td>BT20/MB468</td><td>NUC</td></tr>
</table>

| | | | | | | | | | | | | | |
|---|---|---|---|---|---|---|---|---|---|---|---|---|---|
| >hg38_21_+_8433172_8446622_RNA45SN1_45S_50ntFlanks@12418.12443.26\|CGATGTCGGCTCTTCCTATCATTGTG | >hg38_21_+_8433172_8446622_RNA45SN1_45S_50ntFlanks | 12418 | 12443 | 26 | CGATGTCGGCTCTTCCTATCATTGTG | 15275.8 | -5.6 | 0.7 | -7.8 | 6.5E-15 | 3.2E-14 | MB231/MB468 | NUC |
| >hg38_21_+_8433172_8446622_RNA45SN1_45S_50ntFlanks@10905.10926.22\|CTGGGCGCGCGCCGCGGCTGGA | >hg38_21_+_8433172_8446622_RNA45SN1_45S_50ntFlanks | 10905 | 10926 | 22 | CTGGGCGCGCGCCGCGGCTGGA | 381.3 | -12.4 | 1.6 | -7.8 | 5.3E-15 | 3.2E-14 | MB231/MB468 | MP |
| >hg38_21_+_8433172_8446622_RNA45SN1_45S_50ntFlanks@8104.8142.39\|CCGCGGCGGGGCGCGGGACATGTGGCGTACGGAAGACCC | >hg38_21_+_8433172_8446622_RNA45SN1_45S_50ntFlanks | 8104 | 8142 | 39 | CCGCGGCGGGGCGCGGGACATGTGGCGTACGGAAGACCC | 969.4 | -13.8 | 1.8 | -7.8 | 5.3E-15 | 3.2E-14 | MB231/MB468 | MP |
| >hg38_21_+_8433172_8446622_RNA45SN1_45S_50ntFlanks@9143.9163.21\|GGGCGGACTGTCCCCAGTGCG | >hg38_21_+_8433172_8446622_RNA45SN1_45S_50ntFlanks | 9143 | 9163 | 21 | GGGCGGACTGTCCCCAGTGCG | 301.0 | -12.1 | 1.5 | -7.8 | 5.3E-15 | 3.2E-14 | MB231/MB468 | MP |
| >hg38_21_+_8433172_8446622_RNA45SN1_45S_50ntFlanks@485.528.44\|GATGTGGTGACGTCGTGCTCTCCCGGGCCGGGTCCGAGCCGCGA | >hg38_21_+_8433172_8446622_RNA45SN1_45S_50ntFlanks | 485 | 528 | 44 | GATGTGGTGACGTCGTGCTCTCCCGGGCCGGGTCCGAGCCGCGA | 1216.2 | 28.3 | 3.6 | 7.8 | 6.5E-15 | 3.2E-14 | MB231/MB468 | NUC |
| >hg38_21_+_8433172_8446622_RNA45SN1_45S_50ntFlanks@12957.12989.33\|CTTCTGGGTCGGGGTTTCGTACGTAGCAGAGCA | >hg38_21_+_8433172_8446622_RNA45SN1_45S_50ntFlanks | 12957 | 12989 | 33 | CTTCTGGGTCGGGGTTTCGTACGTAGCAGAGCA | 631.8 | -9.1 | 1.2 | -7.8 | 6.5E-15 | 3.2E-14 | MB231/MB468 | NUC |
| >trnaMT_GluTTC_MT_-_14674_14742@40.63.24,trnalookalike8_GluTTC_5_-_93905172_93905240@40.63.24\|TTGGTCGTGGTTGTAGTCCGTGCG | >trnaMT_GluTTC_MT_-_14674_14742 | 40 | 63 | 24 | TTGGTCGTGGTTGTAGTCCGTGCG | 1354.3 | 9.1 | 1.2 | 7.8 | 5.4E-15 | 3.2E-14 | BT20/MB468 | MITO |
| >hg38_21_+_8433172_8446622_RNA45SN1_45S_50ntFlanks@485.506.22\|GATGTGGTGACGTCGTGCTCTC | >hg38_21_+_8433172_8446622_RNA45SN1_45S_50ntFlanks | 485 | 506 | 22 | GATGTGGTGACGTCGTGCTCTC | 1728.1 | 28.2 | 3.6 | 7.8 | 8.8E-15 | 3.2E-14 | MB231/MB468 | MITO |
| >hg38_21_+_8433172_8446622_RNA45SN1_45S_50ntFlanks@10421.10456.36\|TGAGAGATGGGCGAGCGCCGTTCCGAAGGGACGGGC | >hg38_21_+_8433172_8446622_RNA45SN1_45S_50ntFlanks | 10421 | 10456 | 36 | TGAGAGATGGGCGAGCGCCGTTCCGAAGGGACGGGC | 3881.9 | -11.5 | 1.5 | -7.7 | 1.1E-14 | 3.2E-14 | BT20/MB468 | NUC |
| >hg38_21_+_8433172_8446622_RNA45SN1_45S_50ntFlanks@3848.3866.19\|TGGATAACTGTGGTAATTC | >hg38_21_+_8433172_8446622_RNA45SN1_45S_50ntFlanks | 3848 | 3866 | 19 | TGGATAACTGTGGTAATTC | 368.7 | 5.5 | 0.7 | 7.8 | 6.1E-15 | 3.3E-14 | MB231/MB468 | CYTO |
| >hg38_21_+_8433172_8446622_RNA45SN1_45S_50ntFlanks@9740.9757.18\|AACGATCTCAACCTATTC | >hg38_21_+_8433172_8446622_RNA45SN1_45S_50ntFlanks | 9740 | 9757 | 18 | AACGATCTCAACCTATTC | 187.3 | -5.8 | 0.7 | -7.8 | 6.1E-15 | 3.3E-14 | MB231/MB468 | CYTO |
| >hg38_21_+_8433172_8446622_RNA45SN1_45S_50ntFlanks@10404.10430.27\|CATGGGTCAGTCGGTCCTGAGAGATGG | >hg38_21_+_8433172_8446622_RNA45SN1_45S_50ntFlanks | 10404 | 10430 | 27 | CATGGGTCAGTCGGTCCTGAGAGATGG | 1050.9 | -7.6 | 1.0 | -7.8 | 6.6E-15 | 3.3E-14 | MB231/MB468 | NUC |
| >hg38_21_+_8433172_8446622_RNA45SN1_45S_50ntFlanks@12124.12146.23\|CGCGACCCGCTCCGGGGACAGTG | >hg38_21_+_8433172_8446622_RNA45SN1_45S_50ntFlanks | 12124 | 12146 | 23 | CGCGACCCGCTCCGGGGACAGTG | 339.8 | -12.3 | 1.6 | -7.8 | 5.4E-15 | 3.3E-14 | MB231/MB468 | MP |
| >hg38_21_+_8433172_8446622_RNA45SN1_45S_50ntFlanks@4759.4778.20\|ATCAGATACCGTCGTAGTTC | >hg38_21_+_8433172_8446622_RNA45SN1_45S_50ntFlanks | 4759 | 4778 | 20 | ATCAGATACCGTCGTAGTTC | 773.9 | -11.7 | 1.5 | -7.8 | 5.5E-15 | 3.3E-14 | BT20/MB468 | MITO |
| >hg38_21_+_8433172_8446622_RNA45SN1_45S_50ntFlanks@9567.9595.29\|ACGTGCAAATCGGTCGTCCGACCTGGGTA | >hg38_21_+_8433172_8446622_RNA45SN1_45S_50ntFlanks | 9567 | 9595 | 29 | ACGTGCAAATCGGTCGTCCGACCTGGGTA | 2214.2 | -10.2 | 1.3 | -7.8 | 5.4E-15 | 3.3E-14 | MB231/MB468 | MP |
| >hg38_21_+_8433172_8446622_RNA45SN1_45S_50ntFlanks@12141.12162.22\|ACAGTGCCAGGTGGGGAGTTTG | >hg38_21_+_8433172_8446622_RNA45SN1_45S_50ntFlanks | 12141 | 12162 | 22 | ACAGTGCCAGGTGGGGAGTTTG | 207.1 | -10.7 | 1.4 | -7.8 | 6.6E-15 | 3.3E-14 | MB231/MB468 | NUC |
| >hg38_21_+_8433172_8446622_RNA45SN1_45S_50ntFlanks@10351.10388.38\|CGAGAACTTTGAAGGCCGAAGTGGAGAAGGGTTCCATG | >hg38_21_+_8433172_8446622_RNA45SN1_45S_50ntFlanks | 10351 | 10388 | 38 | CGAGAACTTTGAAGGCCGAAGTGGAGAAGGGTTCCATG | 7837.5 | -6.7 | 0.9 | -7.8 | 6.1E-15 | 3.3E-14 | MB231/MB468 | CYTO |
| >hg38_21_+_8433172_8446622_RNA45SN1_45S_50ntFlanks@3782.3801.20\|CAGTGAAACTGCGAATGGCT | >hg38_21_+_8433172_8446622_RNA45SN1_45S_50ntFlanks | 3782 | 3801 | 20 | CAGTGAAACTGCGAATGGCT | 1183.8 | -4.4 | 0.6 | -7.8 | 6.1E-15 | 3.3E-14 | MB231/MB468 | CYTO |
| >hg38_21_+_8433172_8446622_RNA45SN1_45S_50ntFlanks@3927.3959.33\|CAGATCAAAACCAACCCGGTCAGCCCCTCTCCG | >hg38_21_+_8433172_8446622_RNA45SN1_45S_50ntFlanks | 3927 | 3959 | 33 | CAGATCAAAACCAACCCGGTCAGCCCCTCTCCG | 167.6 | -5.3 | 0.7 | -7.8 | 6.1E-15 | 3.3E-14 | MB231/MB468 | CYTO |
| >hg38_21_+_8433172_8446622_RNA45SN1_45S_50ntFlanks@9454.9494.41\|CCGCGCCGGGGAGGTGGAGCACGAGCGCACGTGTTAGGACC | >hg38_21_+_8433172_8446622_RNA45SN1_45S_50ntFlanks | 9454 | 9494 | 41 | CCGCGCCGGGGAGGTGGAGCACGAGCGCACGTGTTAGGACC | 159.3 | -10.3 | 1.3 | -7.8 | 6.6E-15 | 3.3E-14 | MB231/MB468 | NUC |
| >hg38_21_+_8433172_8446622_RNA45SN1_45S_50ntFlanks@12570.12587.18\|AGTACGAGAGGAACCGCA | >hg38_21_+_8433172_8446622_RNA45SN1_45S_50ntFlanks | 12570 | 12587 | 18 | AGTACGAGAGGAACCGCA | 580.9 | -4.8 | 0.6 | -7.8 | 5.5E-15 | 3.3E-14 | BT20/MB468 | MITO |
| >MI0000272\|hsa-mir-182&WithFlank&7\|-\|129770377\|129770498@29.53.25(+1C)&[MIMAT0000259&hsa-miR-182-5p&offsets\|0\|+1(+1C);m-22&7\|-\|129770449\|129770470&offsets\|0\|+3(+1C)]\|TTTGGCAATGGTAGAACTCACACTGC | >MI0000272\|hsa-mir-182&WithFlank&7\|-\|129770377\|129770498 | 29 | 53 | 25(+1C) | TTTGGCAATGGTAGAACTCACACTGC | 707.4 | 7.9 | 1.0 | 7.8 | 7.5E-15 | 3.3E-14 | BT20/MB468 | CYTO |
| >hg38_21_+_8433172_8446622_RNA45SN1_45S_50ntFlanks@9575.9596.22\|ATCGGTCGTCCGACCTGGGTAT | >hg38_21_+_8433172_8446622_RNA45SN1_45S_50ntFlanks | 9575 | 9596 | 22 | ATCGGTCGTCCGACCTGGGTAT | 5145.9 | -10.6 | 1.4 | -7.8 | 9.0E-15 | 3.3E-14 | MB231/MB468 | MITO |
| >hg38_21_+_8433172_8446622_RNA45SN1_45S_50ntFlanks@12233.12258.26\|CCGTGGAGCAGAAGGGCAAAAGCTCG | >hg38_21_+_8433172_8446622_RNA45SN1_45S_50ntFlanks | 12233 | 12258 | 26 | CCGTGGAGCAGAAGGGCAAAAGCTCG | 227.1 | -11.7 | 1.5 | -7.8 | 5.5E-15 | 3.3E-14 | MB231/MB468 | MP |
| >hg38_21_+_8433172_8446622_RNA45SN1_45S_50ntFlanks@7986.8018.33\|ATCAGACGTGGCGACCCGCTGAATTTAAGCATA | >hg38_21_+_8433172_8446622_RNA45SN1_45S_50ntFlanks | 7986 | 8018 | 33 | ATCAGACGTGGCGACCCGCTGAATTTAAGCATA | 643.6 | -11.4 | 1.5 | -7.8 | 5.6E-15 | 3.3E-14 | BT20/MB468 | MITO |
| >hg38_21_+_8433172_8446622_RNA45SN1_45S_50ntFlanks@10539.10571.33\|AGGCGTCCAGTGCGGTAACGCGACCGATCCCGG | >hg38_21_+_8433172_8446622_RNA45SN1_45S_50ntFlanks | 10539 | 10571 | 33 | AGGCGTCCAGTGCGGTAACGCGACCGATCCCGG | 241.0 | -11.8 | 1.5 | -7.8 | 5.5E-15 | 3.3E-14 | MB231/MB468 | MP |
| >hg38_21_+_8433172_8446622_RNA45SN1_45S_50ntFlanks@10786.10826.41\|AACAGCCTCTGGCATGTTGGAACAATGTAGGTAAGGGAAGT | >hg38_21_+_8433172_8446622_RNA45SN1_45S_50ntFlanks | 10786 | 10826 | 41 | AACAGCCTCTGGCATGTTGGAACAATGTAGGTAAGGGAAGT | 4310.9 | -8.8 | 1.1 | -7.8 | 9.1E-15 | 3.3E-14 | MB231/MB468 | MITO |
| >hg38_21_+_8433172_8446622_RNA45SN1_45S_50ntFlanks@3899.3938.40\|CCTTCGCGGGGGGGATGCGTGCATTTATCAGATCAAAACC | >hg38_21_+_8433172_8446622_RNA45SN1_45S_50ntFlanks | 3899 | 3938 | 40 | CCTTCGCGGGGGGGATGCGTGCATTTATCAGATCAAAACC | 553.4 | -13.0 | 1.7 | -7.8 | 5.5E-15 | 3.3E-14 | MB231/MB468 | MP |
| >MI0000079\|hsa-mir-23a&WithFlank&19\|-\|13836581\|13836665@54.72.19&[MIMAT0000078&hsa-miR-23a-3p&offsets\|+3\|+1;m-31&19\|-\|13836595\|13836615&offsets\|+3\|+1]\|ACATTGCCAGGGATTTCCA | >MI0000079\|hsa-mir-23a&WithFlank&19\|-\|13836581\|13836665 | 54 | 72 | 19 | ACATTGCCAGGGATTTCCA | 2285.9 | 28.4 | 3.6 | 7.8 | 5.5E-15 | 3.3E-14 | MB231/MB468 | MP |
| >MI0003591\|hsa-mir-584&WithFlank&5\|-\|149062307\|149062415@22.40.19&[MIMAT0003249&hsa-miR-584-5p&offsets\|0\|-3;m-167&5\|-\|149062374\|149062394&offsets\|0\|-2]\|TTATGGTTTGCCTGGGACT | >MI0003591\|hsa-mir-584&WithFlank&5\|-\|149062307\|149062415 | 22 | 40 | 19 | TTATGGTTTGCCTGGGACT | 119.4 | 3.0 | 0.4 | 7.7 | 1.2E-14 | 3.3E-14 | BT20/MB468 | NUC |
| >hg38_21_+_8433172_8446622_RNA45SN1_45S_50ntFlanks@7989.8012.24\|AGACGTGGCGACCCGCTGAATTTA | >hg38_21_+_8433172_8446622_RNA45SN1_45S_50ntFlanks | 7989 | 8012 | 24 | AGACGTGGCGACCCGCTGAATTTA | 8627.1 | -5.6 | 0.7 | -7.8 | 9.1E-15 | 3.3E-14 | MB231/MB468 | MITO |
| >hg38_21_+_8433172_8446622_RNA45SN1_45S_50ntFlanks@12324.12355.32\|TTGGGTTTTAAGCAGGAGGTGTCAGAAAAGTT | >hg38_21_+_8433172_8446622_RNA45SN1_45S_50ntFlanks | 12324 | 12355 | 32 | TTGGGTTTTAAGCAGGAGGTGTCAGAAAAGTT | 208.2 | -10.4 | 1.3 | -7.8 | 6.2E-15 | 3.3E-14 | MB231/MB468 | CYTO |
| >MI0000093\|hsa-mir-92a-1&WithFlank&13\|+\|91351308\|91351397@54.74.21(+1G);MI0000094\|hsa-mir-92a-2&WithFlank&X\|-\|134169532\|134169618@54.74.21(+1G)&[MIMAT0000092&hsa-miR-92a-3p&offsets\|0\|-1(+1G);m-9&13\|+\|91351361\|91351382&offsets\|0\|-1(+1G)];[MIMAT0000092_1&hsa-miR-92a-3p&offsets\|0\|-1(+1G);m-11&X\|-\|134169544\|134169565&offsets\|0\|-1(+1G)]\|TATTGCACTTGTCCCGGCCTGG | >MI0000093\|hsa-mir-92a-1&WithFlank&13\|+\|91351308\|91351397 | 54 | 74 | 21(+1G) | TATTGCACTTGTCCCGGCCTGG | 1893.0 | 5.5 | 0.7 | 7.8 | 7.6E-15 | 3.3E-14 | BT20/MB468 | CYTO |
| >hg38_21_+_8433172_8446622_RNA45SN1_45S_50ntFlanks@7986.8016.31\|ATCAGACGTGGCGACCCGCTGAATTTAAGCA | >hg38_21_+_8433172_8446622_RNA45SN1_45S_50ntFlanks | 7986 | 8016 | 31 | ATCAGACGTGGCGACCCGCTGAATTTAAGCA | 659.9 | -12.4 | 1.5 | -8.2 | 2.2E-16 | 3.3E-14 | BT20/MB468 | MP |
| >hg38_21_+_8433172_8446622_RNA45SN1_45S_50ntFlanks@12998.13017.20\|GCTGCGATCTATTGAAAGTC | >hg38_21_+_8433172_8446622_RNA45SN1_45S_50ntFlanks | 12998 | 13017 | 20 | GCTGCGATCTATTGAAAGTC | 240.7 | 6.9 | 0.9 | 7.8 | 6.8E-15 | 3.4E-14 | MB231/MB468 | NUC |
| >hg38_21_+_8433172_8446622_RNA45SN1_45S_50ntFlanks@12151.12179.29\|GTGGGGAGTTTGACTGGGGCGGTACACCT | >hg38_21_+_8433172_8446622_RNA45SN1_45S_50ntFlanks | 12151 | 12179 | 29 | GTGGGGAGTTTGACTGGGGCGGTACACCT | 263.8 | -11.0 | 1.4 | -7.8 | 6.8E-15 | 3.4E-14 | MB231/MB468 | NUC |
| >hg38_21_+_8433172_8446622_RNA45SN1_45S_50ntFlanks@8357.8375.19\|AAAAGAACTTTGAAGAGAG | >hg38_21_+_8433172_8446622_RNA45SN1_45S_50ntFlanks | 8357 | 8375 | 19 | AAAAGAACTTTGAAGAGAG | 2554.3 | -5.2 | 0.7 | -7.8 | 6.8E-15 | 3.4E-14 | MB231/MB468 | NUC |
| >hg38_21_+_8433172_8446622_RNA45SN1_45S_50ntFlanks@11890.11909.20\|CTTGACTCTAGTCTGGCACG | >hg38_21_+_8433172_8446622_RNA45SN1_45S_50ntFlanks | 11890 | 11909 | 20 | CTTGACTCTAGTCTGGCACG | 254.3 | -11.9 | 1.5 | -7.8 | 5.6E-15 | 3.4E-14 | MB231/MB468 | MP |
| >MI0003684\|hsa-mir-660&WithFlank&X\|+\|50013235\|50013343@22.44.23&[MIMAT0003338&hsa-miR-660-5p&offsets\|0\|+1;m-157&X\|+\|50013256\|50013278&offsets\|0\|0]\|TACCCATTGCATATCGGAGTTGT | >MI0003684\|hsa-mir-660&WithFlank&X\|+\|50013235\|50013343 | 22 | 44 | 23 | TACCCATTGCATATCGGAGTTGT | 176.8 | 4.6 | 0.6 | 7.8 | 6.8E-15 | 3.4E-14 | MB231/MB468 | NUC |
| >hg38_21_+_8433172_8446622_RNA45SN1_45S_50ntFlanks@10905.10925.21\|CTGGGCGCGCGCCGCGGCTGG | >hg38_21_+_8433172_8446622_RNA45SN1_45S_50ntFlanks | 10905 | 10925 | 21 | CTGGGCGCGCGCCGCGGCTGG | 348.5 | -11.2 | 1.4 | -7.8 | 6.3E-15 | 3.4E-14 | MB231/MB468 | CYTO |
| >hg38_21_+_8433172_8446622_RNA45SN1_45S_50ntFlanks@11188.11207.20\|CACCCGGGGGGCCGGCGGCG | >hg38_21_+_8433172_8446622_RNA45SN1_45S_50ntFlanks | 11188 | 11207 | 20 | CACCCGGGGGGCCGGCGGCG | 190.7 | -10.3 | 1.3 | -7.8 | 6.3E-15 | 3.4E-14 | MB231/MB468 | CYTO |
| >hg38_21_+_8433172_8446622_RNA45SN1_45S_50ntFlanks@12289.12318.30\|CCGTGAAAGCGGGGCCTCACGATCCTTCTG | >hg38_21_+_8433172_8446622_RNA45SN1_45S_50ntFlanks | 12289 | 12318 | 30 | CCGTGAAAGCGGGGCCTCACGATCCTTCTG | 2518.6 | -13.4 | 1.7 | -7.8 | 5.7E-15 | 3.4E-14 | BT20/MB468 | MITO |
| >hg38_21_+_8433172_8446622_RNA45SN1_45S_50ntFlanks@12437.12454.18\|CATTGTGAAGCAGAATTC | >hg38_21_+_8433172_8446622_RNA45SN1_45S_50ntFlanks | 12437 | 12454 | 18 | CATTGTGAAGCAGAATTC | 6467.2 | -9.2 | 1.2 | -7.8 | 5.7E-15 | 3.4E-14 | MB231/MB468 | MP |
| >MI0000301\|hsa-mir-224&WithFlank&X\|-\|151958572\|151958664@14.38.25&[MIMAT0000281&hsa-miR-224-5p&offsets\|+1\|+1;m-238&X\|-\|151958630\|151958651&offsets\|0\|+3]\|CAAGTCACTAGTGGTTCCGTTTAGT | >MI0000301\|hsa-mir-224&WithFlank&X\|-\|151958572\|151958664 | 14 | 38 | 25 | CAAGTCACTAGTGGTTCCGTTTAGT | 620.4 | 12.5 | 1.6 | 7.7 | 9.3E-15 | 3.4E-14 | MB231/MB468 | MITO |
| >hg38_21_+_8433172_8446622_RNA45SN1_45S_50ntFlanks@8738.8758.21\|CCGTCCGTCCGTCCGTCCGTC | >hg38_21_+_8433172_8446622_RNA45SN1_45S_50ntFlanks | 8738 | 8758 | 21 | CCGTCCGTCCGTCCGTCCGTC | 226.9 | -11.6 | 1.5 | -7.7 | 9.4E-15 | 3.4E-14 | MB231/MB468 | MITO |
| >hg38_21_+_8433172_8446622_RNA45SN1_45S_50ntFlanks@11784.11804.21\|ATGAATGGATGAACGAGATTC | >hg38_21_+_8433172_8446622_RNA45SN1_45S_50ntFlanks | 11784 | 11804 | 21 | ATGAATGGATGAACGAGATTC | 9287.5 | -9.0 | 1.2 | -7.8 | 5.7E-15 | 3.4E-14 | MB231/MB468 | MP |
| >hg38_21_+_8433172_8446622_RNA45SN1_45S_50ntFlanks@6731.6758.28\|CACATTGATCATCGACACTTCGAACGCA | >hg38_21_+_8433172_8446622_RNA45SN1_45S_50ntFlanks | 6731 | 6758 | 28 | CACATTGATCATCGACACTTCGAACGCA | 206.5 | -11.6 | 1.5 | -7.8 | 5.7E-15 | 3.4E-14 | MB231/MB468 | MP |
| >hg38_21_+_8433172_8446622_RNA45SN1_45S_50ntFlanks@8106.8143.38\|GCGGCGGGGCGCGGGACATGTGGCGTACGGAAGACCCG | >hg38_21_+_8433172_8446622_RNA45SN1_45S_50ntFlanks | 8106 | 8143 | 38 | GCGGCGGGGCGCGGGACATGTGGCGTACGGAAGACCCG | 185.6 | -10.5 | 1.3 | -7.8 | 6.9E-15 | 3.4E-14 | MB231/MB468 | NUC |
| >hg38_21_+_8433172_8446622_RNA45SN1_45S_50ntFlanks@4452.4471.20\|CTCGGCGCCCCCTCGATGCT | >hg38_21_+_8433172_8446622_RNA45SN1_45S_50ntFlanks | 4452 | 4471 | 20 | CTCGGCGCCCCCTCGATGCT | 245.8 | -11.8 | 1.5 | -7.8 | 5.7E-15 | 3.4E-14 | MB231/MB468 | MP |
| >MI0000255\|hsa-mir-30d&WithFlank&8\|-\|134804870\|134804951@12.31.20&[MIMAT0000245&hsa-miR-30d-5p&offsets\|0\|-2;m-21&8\|-\|134804919\|134804940&offsets\|0\|-2]\|TGTAAACATCCCCGACTGGA | >MI0000255\|hsa-mir-30d&WithFlank&8\|-\|134804870\|134804951 | 12 | 31 | 20 | TGTAAACATCCCCGACTGGA | 473.2 | 3.4 | 0.4 | 7.7 | 1.2E-14 | 3.4E-14 | BT20/MB468 | NUC |
| >MI0003137\|hsa-mir-193b&WithFlank&16\|+\|14303961\|14304055@57.77.21&[MIMAT0002819&hsa-miR-193b-3p&offsets\|0\|-1;m-166&16\|+\|14304017\|14304038&offsets\|0\|-1]\|AACTGGCCCTCAAAGTCCCGC | >MI0003137\|hsa-mir-193b&WithFlank&16\|+\|14303961\|14304055 | 57 | 77 | 21 | AACTGGCCCTCAAAGTCCCGC | 58.7 | 8.7 | 1.1 | 7.7 | 1.2E-14 | 3.4E-14 | BT20/MB468 | NUC |
| >hg38_21_+_8433172_8446622_RNA45SN1_45S_50ntFlanks@10727.10754.28\|GGAGAGGGTGTAAATCTCGCGCCGGGCC | >hg38_21_+_8433172_8446622_RNA45SN1_45S_50ntFlanks | 10727 | 10754 | 28 | GGAGAGGGTGTAAATCTCGCGCCGGGCC | 195.2 | -11.5 | 1.5 | -7.8 | 5.7E-15 | 3.4E-14 | MB231/MB468 | MP |
| >hg38_21_+_8433172_8446622_RNA45SN1_45S_50ntFlanks@11781.11798.18\|CGCATGAATGGATGAACG | >hg38_21_+_8433172_8446622_RNA45SN1_45S_50ntFlanks | 11781 | 11798 | 18 | CGCATGAATGGATGAACG | 854.6 | -4.6 | 0.6 | -7.8 | 7.0E-15 | 3.5E-14 | MB231/MB468 | NUC |

| | | | | | | | | | | | | | |
|---|---|---|---|---|---|---|---|---|---|---|---|---|---|
| >hg38_21_+_8433172_8446622_RNA45SN1_45S_50ntFlanks@10871.10894.24\|GGCTGGGTCGGTCGGGCTGGGGCG | >hg38_21_+_8433172_8446622_RNA45SN1_45S_50ntFlanks | 10871 | 10894 | 24 | GGCTGGGTCGGTCGGGCTGGGGCG | 232.8 | -11.7 | 1.5 | -7.8 | 5.8E-15 | 3.5E-14 | MB231/MB468 | MP |
| >hg38_21_+_8433172_8446622_RNA45SN1_45S_50ntFlanks@9712.9729.18\|AGCGAATGATTAGAGGTC | >hg38_21_+_8433172_8446622_RNA45SN1_45S_50ntFlanks | 9712 | 9729 | 18 | AGCGAATGATTAGAGGTC | 671.7 | -13.2 | 1.7 | -7.7 | 9.5E-15 | 3.5E-14 | MB231/MB468 | MITO |
| >hg38_21_+_8433172_8446622_RNA45SN1_45S_50ntFlanks@3775.3792.18\|GCCGGTACAGTGAAACTG | >hg38_21_+_8433172_8446622_RNA45SN1_45S_50ntFlanks | 3775 | 3792 | 18 | GCCGGTACAGTGAAACTG | 4576.2 | -10.8 | 1.4 | -7.7 | 9.5E-15 | 3.5E-14 | MB231/MB468 | MITO |
| >hg38_21_+_8433172_8446622_RNA45SN1_45S_50ntFlanks@8381.8404.24\|AAGAGGGCGTGAAACCGTTAAGAG | >hg38_21_+_8433172_8446622_RNA45SN1_45S_50ntFlanks | 8381 | 8404 | 24 | AAGAGGGCGTGAAACCGTTAAGAG | 447.3 | -12.6 | 1.6 | -7.7 | 9.5E-15 | 3.5E-14 | MB231/MB468 | MITO |
| >trna2_GlyGCC_21_-_18827107_18827177@1.29.29,trna35_GlyGCC_1_+_161413094_161413164@1.29.29,trna37_GlyGCC_1_+_161420467_161420537@1.29.29,trna39_GlyGCC_1_+_161427898_161427968@1.29.29,trna41_GlyGCC_1_+_161435258_161435328@1.29.29\|GCATGGGTGGTTCAGTGGTAGAATTCTCG | >trna2_GlyGCC_21_-_18827107_18827177 | 1 | 29 | 29 | GCATGGGTGGTTCAGTGGTAGAATTCTCG | 45296.2 | -7.4 | 0.9 | -7.8 | 6.5E-15 | 3.5E-14 | MB231/MB468 | CYTO |
| >hg38_21_+_8433172_8446622_RNA45SN1_45S_50ntFlanks@6745.6765.21\|ACACTTCGAACGCACTTGCGG | >hg38_21_+_8433172_8446622_RNA45SN1_45S_50ntFlanks | 6745 | 6765 | 21 | ACACTTCGAACGCACTTGCGG | 321.2 | -12.2 | 1.6 | -7.8 | 5.8E-15 | 3.5E-14 | MB231/MB468 | MP |
| >hg38_21_+_8433172_8446622_RNA45SN1_45S_50ntFlanks@4390.4408.19\|TCTTGGGAGCGGGCGGGCG | >hg38_21_+_8433172_8446622_RNA45SN1_45S_50ntFlanks | 4390 | 4408 | 19 | TCTTGGGAGCGGGCGGGCG | 286.0 | -12.0 | 1.5 | -7.8 | 5.8E-15 | 3.5E-14 | MB231/MB468 | MP |
| >hg38_21_+_8433172_8446622_RNA45SN1_45S_50ntFlanks@12490.12521.32\|CGTGAGCTGGGTTTAGACCGTCGTGAGACAGG | >hg38_21_+_8433172_8446622_RNA45SN1_45S_50ntFlanks | 12490 | 12521 | 32 | CGTGAGCTGGGTTTAGACCGTCGTGAGACAGG | 347.8 | -12.2 | 1.6 | -7.7 | 9.6E-15 | 3.5E-14 | MB231/MB468 | MITO |
| >hg38_21_+_8433172_8446622_RNA45SN1_45S_50ntFlanks@10410.10435.26\|TCAGTCGGTCCTGAGAGATGGGCGAG | >hg38_21_+_8433172_8446622_RNA45SN1_45S_50ntFlanks | 10410 | 10435 | 26 | TCAGTCGGTCCTGAGAGATGGGCGAG | 1345.1 | -13.5 | 1.6 | -8.2 | 2.3E-16 | 3.5E-14 | BT20/MB468 | MP |
| >MI0000061\|hsa-let-7a-2&WithFlank&11\|-\|122146516\|122146599@11.33.23;MI0000062\|hsa-let-7a-3&WithFlank&22\|+\|46112743\|46112828@10.32.23;MI0000060\|hsa-let-7a-1&WithFlank&9\|+\|94175951\|94176042@12.34.23&[MIMAT0000062&hsa-let-7a-5p&offsets\|0\|+1;m-6&11\|-\|122146568\|122146589&offsets\|0\|+1];[MIMAT0000062_1&hsa-let-7a-5p&offsets\|0\|+1;m-5&22\|+\|46112752\|46112773&offsets\|0\|+1];[MIMAT0000062_2&hsa-let-7a-5p&offsets\|0\|+1;m-4&9\|+\|94175962\|94175983&offsets\|0\|+1]\|TGAGGTAGTAGGTTGTATAGTTT | >MI0000061\|hsa-let-7a-2&WithFlank&11\|-\|122146516\|122146599 | 11 | 33 | 23 | TGAGGTAGTAGGTTGTATAGTTT | 2770.0 | 6.9 | 0.9 | 7.7 | 9.6E-15 | 3.5E-14 | MB231/MB468 | MITO |
| >hg38_21_+_8433172_8446622_RNA45SN1_45S_50ntFlanks@8380.8400.21\|CAAGAGGGCGTGAAACCGTTA | >hg38_21_+_8433172_8446622_RNA45SN1_45S_50ntFlanks | 8380 | 8400 | 21 | CAAGAGGGCGTGAAACCGTTA | 554.0 | -8.9 | 1.1 | -7.8 | 7.1E-15 | 3.5E-14 | MB231/MB468 | NUC |
| >hg38_21_+_8433172_8446622_RNA45SN1_45S_50ntFlanks@4725.4759.35\|TTAATCAAGAACGAAAGTCGGAGGTTCGAAGACGA | >hg38_21_+_8433172_8446622_RNA45SN1_45S_50ntFlanks | 4725 | 4759 | 35 | TTAATCAAGAACGAAAGTCGGAGGTTCGAAGACGA | 568.2 | -4.7 | 0.6 | -7.8 | 7.1E-15 | 3.5E-14 | MB231/MB468 | NUC |
| >hg38_21_+_8433172_8446622_RNA45SN1_45S_50ntFlanks@10391.10410.20\|AACAGCAGTTGAACATGGGT | >hg38_21_+_8433172_8446622_RNA45SN1_45S_50ntFlanks | 10391 | 10410 | 20 | AACAGCAGTTGAACATGGGT | 2632.6 | -10.3 | 1.3 | -7.8 | 5.9E-15 | 3.5E-14 | MB231/MB468 | MP |
| >hg38_21_+_8433172_8446622_RNA45SN1_45S_50ntFlanks@11670.11704.35\|CCCAGTGCTCTGAATGTCAAAGTGAAGAAATTCAA | >hg38_21_+_8433172_8446622_RNA45SN1_45S_50ntFlanks | 11670 | 11704 | 35 | CCCAGTGCTCTGAATGTCAAAGTGAAGAAATTCAA | 749.9 | -4.2 | 0.5 | -7.8 | 7.1E-15 | 3.5E-14 | MB231/MB468 | NUC |
| >MI0000299\|hsa-mir-222&WithFlank&X\|-\|45747009\|45747130@75.99.25(+1U)&[MIMAT0000279&hsa-miR-222-3p&offsets\|0\|+4(+1U);m-140&X\|-\|45747033\|45747056&offsets\|0\|+1(+1U)]\|AGCTACATCTGGCTACTGGGTCTCTT | >MI0000299\|hsa-mir-222&WithFlank&X\|-\|45747009\|45747130 | 75 | 99 | 25(+1U) | AGCTACATCTGGCTACTGGGTCTCTT | 4517.0 | 7.1 | 0.9 | 7.7 | 9.7E-15 | 3.5E-14 | MB231/MB468 | MITO |
| >hg38_21_+_8433172_8446622_RNA45SN1_45S_50ntFlanks@12420.12446.27\|ATGTCGGCTCTTCCTATCATTGTGAAG | >hg38_21_+_8433172_8446622_RNA45SN1_45S_50ntFlanks | 12420 | 12446 | 27 | ATGTCGGCTCTTCCTATCATTGTGAAG | 8317.4 | -4.9 | 0.6 | -7.8 | 8.1E-15 | 3.5E-14 | BT20/MB468 | CYTO |
| >MI0000077\|hsa-mir-21&WithFlank&17\|+\|59841260\|59841343@14.35.22&[MIMAT0000076&hsa-miR-21-5p&offsets\|0\|0;m-1&17\|+\|59841273\|59841294&offsets\|0\|0]\|TAGCTTATCAGACTGATGTTGA | >MI0000077\|hsa-mir-21&WithFlank&17\|+\|59841260\|59841343 | 14 | 35 | 22 | TAGCTTATCAGACTGATGTTGA | 36924.9 | 4.6 | 0.6 | 7.7 | 1.2E-14 | 3.6E-14 | BT20/MB468 | NUC |
| >hg38_21_+_8433172_8446622_RNA45SN1_45S_50ntFlanks@10826.10865.40\|TCGGCAAGCCGGATCCGTAACTTCGGGATAAGGATTGGCT | >hg38_21_+_8433172_8446622_RNA45SN1_45S_50ntFlanks | 10826 | 10865 | 40 | TCGGCAAGCCGGATCCGTAACTTCGGGATAAGGATTGGCT | 226.6 | -11.6 | 1.5 | -7.7 | 9.8E-15 | 3.6E-14 | MB231/MB468 | MITO |
| >hg38_21_+_8433172_8446622_RNA45SN1_45S_50ntFlanks@9621.9656.36\|TCTAGTAGCTGGTTCCCTCCGAAGTTTCCCTCAGGA | >hg38_21_+_8433172_8446622_RNA45SN1_45S_50ntFlanks | 9621 | 9656 | 36 | TCTAGTAGCTGGTTCCCTCCGAAGTTTCCCTCAGGA | 227.1 | -11.6 | 1.5 | -7.7 | 9.8E-15 | 3.6E-14 | MB231/MB468 | MITO |
| >hg38_21_+_8433172_8446622_RNA45SN1_45S_50ntFlanks@8423.8470.48\|CGCAGTCCGCCCGGAGGATTCAACCCGGCGGCGGGTCCGGCCGTGTCG | >hg38_21_+_8433172_8446622_RNA45SN1_45S_50ntFlanks | 8423 | 8470 | 48 | CGCAGTCCGCCCGGAGGATTCAACCCGGCGGCGGGTCCGGCCGTGTCG | 423.2 | -12.5 | 1.6 | -7.7 | 9.8E-15 | 3.6E-14 | MB231/MB468 | MITO |
| >hg38_21_+_8433172_8446622_RNA45SN1_45S_50ntFlanks@8191.8215.25\|CAGCCCGTGGACGGTGTGAGGCCGG | >hg38_21_+_8433172_8446622_RNA45SN1_45S_50ntFlanks | 8191 | 8215 | 25 | CAGCCCGTGGACGGTGTGAGGCCGG | 1637.9 | -6.3 | 0.8 | -7.7 | 9.8E-15 | 3.6E-14 | MB231/MB468 | MITO |
| >MI0000084\|hsa-mir-26b&WithFlank&2\|+\|218402640\|218402728@18.38.21(+1C)&[MIMAT0000083&hsa-miR-26b-5p&offsets\|0\|0(+1C);m-53&2\|+\|218402657\|218402678&offsets\|0\|-1(+1C)]\|TTCAAGTAATTCAGGATAGGTC | >MI0000084\|hsa-mir-26b&WithFlank&2\|+\|218402640\|218402728 | 18 | 38 | 21(+1C) | TTCAAGTAATTCAGGATAGGTC | 470.5 | 4.7 | 0.6 | 7.8 | 7.3E-15 | 3.6E-14 | MB231/MB468 | NUC |
| >hg38_21_+_8433172_8446622_RNA45SN1_45S_50ntFlanks@4002.4026.25\|GATAACCTCGGGCCGATCGCACGCC | >hg38_21_+_8433172_8446622_RNA45SN1_45S_50ntFlanks | 4002 | 4026 | 25 | GATAACCTCGGGCCGATCGCACGCC | 396.8 | -11.6 | 1.5 | -7.8 | 7.3E-15 | 3.6E-14 | MB231/MB468 | NUC |
| >hg38_21_+_8433172_8446622_RNA45SN1_45S_50ntFlanks@6644.6661.18\|CCTCGTACGACTCTTAGC | >hg38_21_+_8433172_8446622_RNA45SN1_45S_50ntFlanks | 6644 | 6661 | 18 | CCTCGTACGACTCTTAGC | 325.0 | 4.4 | 0.6 | 7.8 | 7.3E-15 | 3.6E-14 | MB231/MB468 | NUC |
| >hg38_21_+_8433172_8446622_RNA45SN1_45S_50ntFlanks@8381.8402.22\|AAGAGGGCGTGAAACCGTTAAG | >hg38_21_+_8433172_8446622_RNA45SN1_45S_50ntFlanks | 8381 | 8402 | 22 | AAGAGGGCGTGAAACCGTTAAG | 3299.2 | -10.3 | 1.3 | -7.7 | 9.9E-15 | 3.6E-14 | MB231/MB468 | MITO |
| >hg38_21_+_8433172_8446622_RNA45SN1_45S_50ntFlanks@11234.11252.19\|CCTTCCCGTGGATCGCCCC | >hg38_21_+_8433172_8446622_RNA45SN1_45S_50ntFlanks | 11234 | 11252 | 19 | CCTTCCCGTGGATCGCCCC | 275.8 | -11.9 | 1.5 | -7.7 | 9.9E-15 | 3.6E-14 | MB231/MB468 | MITO |
| >hg38_21_+_8433172_8446622_RNA45SN1_45S_50ntFlanks@12624.12661.38\|CCAATGGGGCGAAGCTACCATCTGTGGGATTATGACTG | >hg38_21_+_8433172_8446622_RNA45SN1_45S_50ntFlanks | 12624 | 12661 | 38 | CCAATGGGGCGAAGCTACCATCTGTGGGATTATGACTG | 13493.5 | -9.5 | 1.2 | -7.7 | 9.9E-15 | 3.6E-14 | MB231/MB468 | MITO |
| >hg38_21_+_8433172_8446622_RNA45SN1_45S_50ntFlanks@7986.8013.28\|ATCAGACGTGGCGACCCGCTGAATTTAA | >hg38_21_+_8433172_8446622_RNA45SN1_45S_50ntFlanks | 7986 | 8013 | 28 | ATCAGACGTGGCGACCCGCTGAATTTAA | 181.7 | -11.4 | 1.5 | -7.8 | 6.0E-15 | 3.6E-14 | MB231/MB468 | MP |
| >hg38_21_+_8433172_8446622_RNA45SN1_45S_50ntFlanks@12624.12653.30\|CCAATGGGGCGAAGCTACCATCTGTGGGAT | >hg38_21_+_8433172_8446622_RNA45SN1_45S_50ntFlanks | 12624 | 12653 | 30 | CCAATGGGGCGAAGCTACCATCTGTGGGAT | 1361.4 | -13.5 | 1.6 | -8.2 | 2.4E-16 | 3.6E-14 | BT20/MB468 | MP |
| >MI0000269\|hsa-mir-181a-2&WithFlank&9\|+\|124692436\|124692557@83.103.21(+1U)&[MIMAT0004558&hsa-miR-181a-2-3p&offsets\|0\|-1(+1U);m-71&9\|+\|124692518\|124692539&offsets\|0\|-1(+1U)]\|ACCACTGACCGTTGACTGTACT | >MI0000269\|hsa-mir-181a-2&WithFlank&9\|+\|124692436\|124692557 | 83 | 103 | 21(+1U) | ACCACTGACCGTTGACTGTACT | 894.7 | 9.4 | 1.2 | 7.8 | 6.7E-15 | 3.6E-14 | MB231/MB468 | CYTO |
| >hg38_21_+_8433172_8446622_RNA45SN1_45S_50ntFlanks@4554.4572.19\|CCGCAGCTAGGAATAATGG | >hg38_21_+_8433172_8446622_RNA45SN1_45S_50ntFlanks | 4554 | 4572 | 19 | CCGCAGCTAGGAATAATGG | 215.4 | -11.6 | 1.5 | -7.8 | 6.0E-15 | 3.6E-14 | MB231/MB468 | MP |
| >hg38_MT_+_598_1651_RNR1_12S_50ntFlanks@370.404.35\|ATAAAGCTAAAACTCACCTGAGTTGTAAAAAACTC | >hg38_MT_+_598_1651_RNR1_12S_50ntFlanks | 370 | 404 | 35 | ATAAAGCTAAAACTCACCTGAGTTGTAAAAAACTC | 1428.2 | 5.7 | 0.7 | 7.8 | 6.0E-15 | 3.6E-14 | BT20/MB468 | MITO |
| >hg38_21_+_8433172_8446622_RNA45SN1_45S_50ntFlanks@11188.11225.38\|CACCCGGGGGGCCGGCGGCGGCGGCGACTCTGGACGCG | >hg38_21_+_8433172_8446622_RNA45SN1_45S_50ntFlanks | 11188 | 11225 | 38 | CACCCGGGGGGCCGGCGGCGGCGGCGACTCTGGACGCG | 392.5 | -11.6 | 1.5 | -7.8 | 7.3E-15 | 3.6E-14 | MB231/MB468 | NUC |
| >hg38_21_+_8433172_8446622_RNA45SN1_45S_50ntFlanks@9738.9764.27\|GAAACGATCTCAACCTATTCTCAAACT | >hg38_21_+_8433172_8446622_RNA45SN1_45S_50ntFlanks | 9738 | 9764 | 27 | GAAACGATCTCAACCTATTCTCAAACT | 804.2 | -5.6 | 0.7 | -7.7 | 1.0E-14 | 3.6E-14 | MB231/MB468 | MITO |
| >hg38_21_+_8433172_8446622_RNA45SN1_45S_50ntFlanks@10727.10757.31\|GGAGAGGGTGTAAATCTCGCGCCGGGCCGTA | >hg38_21_+_8433172_8446622_RNA45SN1_45S_50ntFlanks | 10727 | 10757 | 31 | GGAGAGGGTGTAAATCTCGCGCCGGGCCGTA | 1832.4 | -10.8 | 1.4 | -7.7 | 1.2E-14 | 3.6E-14 | BT20/MB468 | NUC |
| >hg38_21_+_8433172_8446622_RNA45SN1_45S_50ntFlanks@5485.5526.42\|AGCGCTGAGAAGACGGTCGAACTTGACTATCTAGAGGAAGTA | >hg38_21_+_8433172_8446622_RNA45SN1_45S_50ntFlanks | 5485 | 5526 | 42 | AGCGCTGAGAAGACGGTCGAACTTGACTATCTAGAGGAAGTA | 687.8 | -12.4 | 1.6 | -7.8 | 7.4E-15 | 3.6E-14 | MB231/MB468 | NUC |
| >hg38_21_+_8433172_8446622_RNA45SN1_45S_50ntFlanks@12072.12097.26\|CCCGAGGGGCTCTCGCTTCTGGCGCC | >hg38_21_+_8433172_8446622_RNA45SN1_45S_50ntFlanks | 12072 | 12097 | 26 | CCCGAGGGGCTCTCGCTTCTGGCGCC | 635.9 | -4.9 | 0.6 | -7.8 | 7.4E-15 | 3.6E-14 | MB231/MB468 | NUC |
| >MI0000301\|hsa-mir-224&WithFlank&X\|-\|151958572\|151958664@14.38.25&[MIMAT0000281&hsa-miR-224-5p&offsets\|+1\|+1;m-238&X\|-\|151958630\|151958651&offsets\|0\|+3]\|CAAGTCACTAGTGGTTCCGTTTAGT | >MI0000301\|hsa-mir-224&WithFlank&X\|-\|151958572\|151958664 | 14 | 38 | 25 | CAAGTCACTAGTGGTTCCGTTTAGT | 569.0 | 10.8 | 1.4 | 7.8 | 6.8E-15 | 3.6E-14 | MB231/MB468 | CYTO |
| >hg38_21_+_8433172_8446622_RNA45SN1_45S_50ntFlanks@10410.10435.26\|TCAGTCGGTCCTGAGAGATGGGCGAG | >hg38_21_+_8433172_8446622_RNA45SN1_45S_50ntFlanks | 10410 | 10435 | 26 | TCAGTCGGTCCTGAGAGATGGGCGAG | 275.7 | -10.8 | 1.4 | -7.8 | 6.8E-15 | 3.6E-14 | MB231/MB468 | CYTO |
| >hg38_21_+_8433172_8446622_RNA45SN1_45S_50ntFlanks@8737.8756.20\|CCCGTCCGTCCGTCCGTCCG | >hg38_21_+_8433172_8446622_RNA45SN1_45S_50ntFlanks | 8737 | 8756 | 20 | CCCGTCCGTCCGTCCGTCCG | 237.8 | -10.6 | 1.4 | -7.8 | 6.8E-15 | 3.6E-14 | MB231/MB468 | CYTO |
| >trna116_GluCTC_1_-_145399233_145399304@32.72.41,trna71_GluCTC_1_-_161439189_161439260@32.72.41,trna74_GluCTC_1_-_161431809_161431880@32.72.41,trna77_GluCTC_1_-_161424398_161424469@32.72.41,trna77_GluCTC_6_+_28949976_28950047@32.72.41,trna80_GluCTC_1_-_161417018_161417089@32.72.41,trna87_GluCTC_6_-_126101393_126101464@32.72.41\|CTCTCACCGCCGCGGCCCGGGTTCGATTCCCGGTCAGGGAA | >trna116_GluCTC_1_-_145399233_145399304 | 32 | 72 | 41 | CTCTCACCGCCGCGGCCCGGGTTCGATTCCCGGTCAGGGAA | 308.9 | -11.2 | 1.4 | -7.8 | 7.4E-15 | 3.6E-14 | MB231/MB468 | NUC |
| >hg38_21_+_8433172_8446622_RNA45SN1_45S_50ntFlanks@11573.11605.33\|CGACTTAGAACTGGTGCGGACCAGGGGAATCCG | >hg38_21_+_8433172_8446622_RNA45SN1_45S_50ntFlanks | 11573 | 11605 | 33 | CGACTTAGAACTGGTGCGGACCAGGGGAATCCG | 405.9 | -3.4 | 0.4 | -7.8 | 8.3E-15 | 3.6E-14 | BT20/MB468 | CYTO |
| >MI0000266\|hsa-mir-10a&WithFlank&17\|-\|48579832\|48579953@28.48.21(+1G)&[MIMAT0000253&hsa-miR-10a-5p&offsets\|0\|-2(+1G);m-18&17\|-\|48579905\|48579926&offsets\|0\|-1(+1G)]\|TACCCTGTAGATCCGAATTTGG | >MI0000266\|hsa-mir-10a&WithFlank&17\|-\|48579832\|48579953 | 28 | 48 | 21(+1G) | TACCCTGTAGATCCGAATTTGG | 79.7 | 9.2 | 1.2 | 7.7 | 1.3E-14 | 3.6E-14 | BT20/MB468 | NUC |
| >hg38_21_+_8433172_8446622_RNA45SN1_45S_50ntFlanks@10664.10710.47\|CTTGGAAAGCGTCGCGGTTCCGGCGGCGTCCGGTGAGCTCTCGCTGG | >hg38_21_+_8433172_8446622_RNA45SN1_45S_50ntFlanks | 10664 | 10710 | 47 | CTTGGAAAGCGTCGCGGTTCCGGCGGCGTCCGGTGAGCTCTCGCTGG | 1234.7 | -14.7 | 1.9 | -7.7 | 1.3E-14 | 3.7E-14 | BT20/MB468 | NUC |
| >hg38_21_+_8433172_8446622_RNA45SN1_45S_50ntFlanks@8357.8380.24\|AAAAGAACTTTGAAGAGAGAGTTC | >hg38_21_+_8433172_8446622_RNA45SN1_45S_50ntFlanks | 8357 | 8380 | 24 | AAAAGAACTTTGAAGAGAGAGTTC | 226.8 | -11.7 | 1.5 | -7.8 | 6.1E-15 | 3.7E-14 | MB231/MB468 | MP |
| >hg38_21_+_8433172_8446622_RNA45SN1_45S_50ntFlanks@12045.12072.28\|CTGACCCGGTGAGGCGGGGGGGCGAGCC | >hg38_21_+_8433172_8446622_RNA45SN1_45S_50ntFlanks | 12045 | 12072 | 28 | CTGACCCGGTGAGGCGGGGGGGCGAGCC | 221.0 | -11.7 | 1.5 | -7.8 | 6.1E-15 | 3.7E-14 | MB231/MB468 | MP |

| | | | | | | | | | | | | | |
|---|---|---|---|---|---|---|---|---|---|---|---|---|---|
| >hg38_21_+_8433172_8446622_RNA45SN1_45S_50ntFlanks@10391.10426.36\|AACAGCAGTTGAACATGGGTCAGTCGGTCCTGAGAG | >hg38_21_+_8433172_8446622_RNA45SN1_45S_50ntFlanks | 10391 | 10426 | 36 | AACAGCAGTTGAACATGGGTCAGTCGGTCCTGAGAG | 314.2 | -12.2 | 1.6 | -7.8 | 6.1E-15 | 3.7E-14 | MB231/MB468 | MP |
| >hg38_21_+_8433172_8446622_RNA45SN1_45S_50ntFlanks@11321.11344.24\|CACGTCTCGTCGCGCGCGTCCG | >hg38_21_+_8433172_8446622_RNA45SN1_45S_50ntFlanks | 11321 | 11344 | 24 | CACGTCTCGTCGCGCGCGTCCG | 7574.2 | -7.4 | 0.9 | -7.8 | 7.0E-15 | 3.7E-14 | MB231/MB468 | CYTO |
| >hg38_21_+_8433172_8446622_RNA45SN1_45S_50ntFlanks@10782.10818.37\|GGTGAACAGCCTCTGGCATGTTGGAACAATGTAGGTA | >hg38_21_+_8433172_8446622_RNA45SN1_45S_50ntFlanks | 10782 | 10818 | 37 | GGTGAACAGCCTCTGGCATGTTGGAACAATGTAGGTA | 276.2 | -12.0 | 1.5 | -7.8 | 6.2E-15 | 3.7E-14 | MB231/MB468 | MP |
| >hg38_21_+_8433172_8446622_RNA45SN1_45S_50ntFlanks@12147.12179.33\|CCAGGTGGGGAGTTTGACTGGGGCGGTACACCT | >hg38_21_+_8433172_8446622_RNA45SN1_45S_50ntFlanks | 12147 | 12179 | 33 | CCAGGTGGGGAGTTTGACTGGGGCGGTACACCT | 148.6 | -11.1 | 1.4 | -7.8 | 6.2E-15 | 3.7E-14 | MB231/MB468 | MP |
| >hg38_21_+_8433172_8446622_RNA45SN1_45S_50ntFlanks@11535.11552.18\|TTCCGCGCGGCGCCTCGC | >hg38_21_+_8433172_8446622_RNA45SN1_45S_50ntFlanks | 11535 | 11552 | 18 | TTCCGCGCGGCGCCTCGC | 184.0 | -6.4 | 0.8 | -7.8 | 7.0E-15 | 3.7E-14 | MB231/MB468 | CYTO |
| >hg38_21_+_8433172_8446622_RNA45SN1_45S_50ntFlanks@5420.5443.24\|CCGATTGGATGGTTTAGTGAGGCC | >hg38_21_+_8433172_8446622_RNA45SN1_45S_50ntFlanks | 5420 | 5443 | 24 | CCGATTGGATGGTTTAGTGAGGCC | 3777.2 | -7.6 | 1.0 | -7.8 | 8.5E-15 | 3.7E-14 | BT20/MB468 | CYTO |
| >hg38_21_+_8433172_8446622_RNA45SN1_45S_50ntFlanks@12156.12182.27\|GAGTTTGACTGGGGCGGTACACCTGTC | >hg38_21_+_8433172_8446622_RNA45SN1_45S_50ntFlanks | 12156 | 12182 | 27 | GAGTTTGACTGGGGCGGTACACCTGTC | 373.5 | -12.4 | 1.6 | -7.8 | 6.2E-15 | 3.7E-14 | MB231/MB468 | MP |
| >hg38_21_+_8433172_8446622_RNA45SN1_45S_50ntFlanks@10574.10608.35\|AAGCCGGCGGGAGCCCCGGGGAGAGTTCTCTTTTC | >hg38_21_+_8433172_8446622_RNA45SN1_45S_50ntFlanks | 10574 | 10608 | 35 | AAGCCGGCGGGAGCCCCGGGGAGAGTTCTCTTTTC | 266.9 | -11.9 | 1.5 | -7.8 | 6.2E-15 | 3.7E-14 | MB231/MB468 | MP |
| >hg38_21_+_8433172_8446622_RNA45SN1_45S_50ntFlanks@6735.6770.36\|TTGATCATCGACACTTCGAACGCACTTGCGGCCCCG | >hg38_21_+_8433172_8446622_RNA45SN1_45S_50ntFlanks | 6735 | 6770 | 36 | TTGATCATCGACACTTCGAACGCACTTGCGGCCCCG | 1002.0 | -7.3 | 0.9 | -7.7 | 1.0E-14 | 3.7E-14 | MB231/MB468 | MITO |
| >hg38_21_+_8433172_8446622_RNA45SN1_45S_50ntFlanks@12289.12322.34\|CCGTGAAAGCGGGGCCTCACGATCCTTCTGACCT | >hg38_21_+_8433172_8446622_RNA45SN1_45S_50ntFlanks | 12289 | 12322 | 34 | CCGTGAAAGCGGGGCCTCACGATCCTTCTGACCT | 847.9 | -13.5 | 1.7 | -7.7 | 1.0E-14 | 3.7E-14 | MB231/MB468 | MITO |
| >hg38_21_+_8433172_8446622_RNA45SN1_45S_50ntFlanks@12685.12708.24\|CCCAGGCGGAACGATACGGCAGCG | >hg38_21_+_8433172_8446622_RNA45SN1_45S_50ntFlanks | 12685 | 12708 | 24 | CCCAGGCGGAACGATACGGCAGCG | 327.7 | -12.1 | 1.6 | -7.7 | 1.0E-14 | 3.7E-14 | MB231/MB468 | MITO |
| >MI0000439\|hsa-mir-23b&WithFlank&9\|+\|95085202\|95085310@65.86.22(+1U)&[MIMAT0000418&hsa-miR-23b-3p&offsets\|+1\|0(+1U);m-42&9\|+\|95085265\|95085285&offsets\|+1\|+2(+1U)]\|TCACATTGCCAGGGATTACCACT | >MI0000439\|hsa-mir-23b&WithFlank&9\|+\|95085202\|95085310 | 65 | 86 | 22(+1U) | TCACATTGCCAGGGATTACCACT | 270.8 | 5.9 | 0.8 | 7.8 | 7.6E-15 | 3.7E-14 | MB231/MB468 | NUC |
| >hg38_21_+_8433172_8446622_RNA45SN1_45S_50ntFlanks@9821.9850.30\|CCTAGTGGGCCACTTTTGGTAAGCAGAACT | >hg38_21_+_8433172_8446622_RNA45SN1_45S_50ntFlanks | 9821 | 9850 | 30 | CCTAGTGGGCCACTTTTGGTAAGCAGAACT | 3220.4 | -8.7 | 1.1 | -7.7 | 1.0E-14 | 3.7E-14 | MB231/MB468 | MITO |
| >MI0000471\|hsa-mir-126&WithFlank&9\|+\|136670596\|136670692@58.78.21&[MIMAT0000445&hsa-miR-126-3p&offsets\|0\|-1;m-40&9\|+\|136670653\|136670674&offsets\|0\|-1]\|TCGTACCGTGAGTAATAATGC | >MI0000471\|hsa-mir-126&WithFlank&9\|+\|136670596\|136670692 | 58 | 78 | 21 | TCGTACCGTGAGTAATAATGC | 3894.2 | 10.6 | 1.4 | 7.8 | 8.6E-15 | 3.7E-14 | BT20/MB468 | CYTO |
| >hg38_21_+_8433172_8446622_RNA45SN1_45S_50ntFlanks@12914.12935.22\|CGTTCGTGGGGAACCTGGCGCT | >hg38_21_+_8433172_8446622_RNA45SN1_45S_50ntFlanks | 12914 | 12935 | 22 | CGTTCGTGGGGAACCTGGCGCT | 240.8 | -11.8 | 1.5 | -7.8 | 6.3E-15 | 3.7E-14 | MB231/MB468 | MP |
| >hg38_21_+_8433172_8446622_RNA45SN1_45S_50ntFlanks@11704.11728.25\|ATGAAGCGCGGGTAAACGGCGGGAG | >hg38_21_+_8433172_8446622_RNA45SN1_45S_50ntFlanks | 11704 | 11728 | 25 | ATGAAGCGCGGGTAAACGGCGGGAG | 738.4 | -8.1 | 1.0 | -7.8 | 7.1E-15 | 3.8E-14 | MB231/MB468 | CYTO |
| >hg38_21_+_8433172_8446622_RNA45SN1_45S_50ntFlanks@11163.11189.27\|CGGTCCCCCGCGAGGGGGGCCCGGGCA | >hg38_21_+_8433172_8446622_RNA45SN1_45S_50ntFlanks | 11163 | 11189 | 27 | CGGTCCCCCGCGAGGGGGGCCCGGGCA | 327.8 | -12.1 | 1.6 | -7.7 | 1.0E-14 | 3.8E-14 | MB231/MB468 | MITO |
| >am_trnaMT_ValTAC_MT_+_1602_1670@29.52.24\|ACTTACACTTAGGAGATTTCAACT | >am_trnaMT_ValTAC_MT_+_1602_1670 | 29 | 52 | 24 | ACTTACACTTAGGAGATTTCAACT | 651.3 | 11.0 | 1.4 | 7.8 | 8.6E-15 | 3.8E-14 | BT20/MB468 | CYTO |
| >hg38_21_+_8433172_8446622_RNA45SN1_45S_50ntFlanks@11567.11584.18\|AGCAGCCGACTTAGAACT | >hg38_21_+_8433172_8446622_RNA45SN1_45S_50ntFlanks | 11567 | 11584 | 18 | AGCAGCCGACTTAGAACT | 2578.8 | 5.4 | 0.7 | 7.8 | 8.6E-15 | 3.8E-14 | BT20/MB468 | CYTO |
| >trna10_LysCTT_16_+_3241501_3241573@3.33.31,trna13_LysCTT_14_-_58706613_58706685@3.33.31,trna2_LysCTT_15_+_79152904_79152976@3.33.31\|CCGGCTAGCTCAGTCGGTAGAGCATGGGACT | >trna10_LysCTT_16_+_3241501_3241573 | 3 | 33 | 31 | CCGGCTAGCTCAGTCGGTAGAGCATGGGACT | 1627.9 | 6.7 | 0.9 | 7.7 | 1.0E-14 | 3.8E-14 | MB231/MB468 | MITO |
| >MI0000650\|hsa-mir-200c&WithFlank&12\|+\|6963693\|6963772@50.72.23(+1A)&[MIMAT0000617&hsa-miR-200c-3p&offsets\|0\|0(+1A);m-54&12\|+\|6963742\|6963764&offsets\|0\|0(+1A)]\|TAATACTGCCGGGTAATGATGGAA | >MI0000650\|hsa-mir-200c&WithFlank&12\|+\|6963693\|6963772 | 50 | 72 | 23(+1A) | TAATACTGCCGGGTAATGATGGAA | 210.7 | -11.6 | 1.5 | -7.8 | 6.3E-15 | 3.8E-14 | MB231/MB468 | MP |
| >MI0003815\|hsa-mir-1301&WithFlank&2\|-\|25328634\|25328727@54.77.24&[MIMAT0005797&hsa-miR-1301-3p&offsets\|0\|0;m-256&2\|-\|25328650\|25328674&offsets\|0\|-1]\|TTGCAGCTGCCTGGGAGTGACTTC | >MI0003815\|hsa-mir-1301&WithFlank&2\|-\|25328634\|25328727 | 54 | 77 | 24 | TTGCAGCTGCCTGGGAGTGACTTC | 617.8 | 10.5 | 1.3 | 7.8 | 6.3E-15 | 3.8E-14 | BT20/MB468 | MITO |
| >hg38_21_+_8433172_8446622_RNA45SN1_45S_50ntFlanks@11571.11603.33\|GCCGACTTAGAACTGGTGCGGACCAGGGGAATC | >hg38_21_+_8433172_8446622_RNA45SN1_45S_50ntFlanks | 11571 | 11603 | 33 | GCCGACTTAGAACTGGTGCGGACCAGGGGAATC | 689.4 | -5.5 | 0.7 | -7.8 | 6.3E-15 | 3.8E-14 | MB231/MB468 | MP |
| >hg38_21_+_8433172_8446622_RNA45SN1_45S_50ntFlanks@10786.10819.34\|AACAGCCTCTGGCATGTTGGAACAATGTAGGTAA | >hg38_21_+_8433172_8446622_RNA45SN1_45S_50ntFlanks | 10786 | 10819 | 34 | AACAGCCTCTGGCATGTTGGAACAATGTAGGTAA | 196.0 | -11.5 | 1.5 | -7.8 | 6.3E-15 | 3.8E-14 | MB231/MB468 | MP |
| >hg38_21_+_8433172_8446622_RNA45SN1_45S_50ntFlanks@9622.9658.37\|CTAGTAGCTGGTTCCCTCCGAAGTTTCCCTCAGGATA | >hg38_21_+_8433172_8446622_RNA45SN1_45S_50ntFlanks | 9622 | 9658 | 37 | CTAGTAGCTGGTTCCCTCCGAAGTTTCCCTCAGGATA | 278.7 | -12.0 | 1.5 | -7.8 | 6.3E-15 | 3.8E-14 | MB231/MB468 | MP |
| >hg38_21_+_8433172_8446622_RNA45SN1_45S_50ntFlanks@11027.11049.23\|CTCTCTCTCTCTCTCCCCCGCTC | >hg38_21_+_8433172_8446622_RNA45SN1_45S_50ntFlanks | 11027 | 11049 | 23 | CTCTCTCTCTCTCTCCCCCGCTC | 926.4 | -8.5 | 1.1 | -7.8 | 7.1E-15 | 3.8E-14 | MB231/MB468 | CYTO |
| >hg38_21_+_8433172_8446622_RNA45SN1_45S_50ntFlanks@10587.10607.21\|CCCCGGGGAGAGTTCTCTTTT | >hg38_21_+_8433172_8446622_RNA45SN1_45S_50ntFlanks | 10587 | 10607 | 21 | CCCCGGGGAGAGTTCTCTTTT | 284.3 | -8.5 | 1.1 | -7.8 | 7.8E-15 | 3.8E-14 | MB231/MB468 | NUC |
| >MI0000650\|hsa-mir-200c&WithFlank&12\|+\|6963693\|6963772@50.70.21(+1A)&[MIMAT0000617&hsa-miR-200c-3p&offsets\|0\|-2(+1A);m-54&12\|+\|6963742\|6963764&offsets\|0\|-2(+1A)]\|TAATACTGCCGGGTAATGATGA | >MI0000650\|hsa-mir-200c&WithFlank&12\|+\|6963693\|6963772 | 50 | 70 | 21(+1A) | TAATACTGCCGGGTAATGATGA | 941.8 | 4.3 | 0.6 | 7.7 | 1.3E-14 | 3.8E-14 | BT20/MB468 | NUC |
| >hg38_21_+_8433172_8446622_RNA45SN1_45S_50ntFlanks@10684.10711.28\|CGGCGGCGTCCGGTGAGCTCTCGCTGGC | >hg38_21_+_8433172_8446622_RNA45SN1_45S_50ntFlanks | 10684 | 10711 | 28 | CGGCGGCGTCCGGTGAGCTCTCGCTGGC | 5148.2 | -5.2 | 0.7 | -7.8 | 7.2E-15 | 3.8E-14 | MB231/MB468 | CYTO |
| >hg38_21_+_8433172_8446622_RNA45SN1_45S_50ntFlanks@11564.11600.37\|CCTAGCAGCCGACTTAGAACTGGTGCGGACCAGGGGA | >hg38_21_+_8433172_8446622_RNA45SN1_45S_50ntFlanks | 11564 | 11600 | 37 | CCTAGCAGCCGACTTAGAACTGGTGCGGACCAGGGGA | 1538.4 | -5.2 | 0.7 | -7.8 | 7.8E-15 | 3.8E-14 | MB231/MB468 | NUC |
| >hg38_21_+_8433172_8446622_RNA45SN1_45S_50ntFlanks@4588.4608.21\|CTATTTTGTTGGTTTTCGGAA | >hg38_21_+_8433172_8446622_RNA45SN1_45S_50ntFlanks | 4588 | 4608 | 21 | CTATTTTGTTGGTTTTCGGAA | 188.1 | -11.4 | 1.5 | -7.8 | 6.4E-15 | 3.8E-14 | MB231/MB468 | MP |
| >hg38_21_+_8433172_8446622_RNA45SN1_45S_50ntFlanks@13008.13027.20\|ATTGAAAGTCAGCCCTCGAC | >hg38_21_+_8433172_8446622_RNA45SN1_45S_50ntFlanks | 13008 | 13027 | 20 | ATTGAAAGTCAGCCCTCGAC | 219.7 | -11.6 | 1.5 | -7.8 | 6.4E-15 | 3.8E-14 | MB231/MB468 | MP |
| >hg38_21_+_8433172_8446622_RNA45SN1_45S_50ntFlanks@10351.10384.34\|CGAGAACTTTGAAGGCCGAAGTGGAGAAGGGTTC | >hg38_21_+_8433172_8446622_RNA45SN1_45S_50ntFlanks | 10351 | 10384 | 34 | CGAGAACTTTGAAGGCCGAAGTGGAGAAGGGTTC | 10698.0 | -10.5 | 1.3 | -7.8 | 6.5E-15 | 3.8E-14 | MB231/MB468 | MP |
| >hg38_21_+_8433172_8446622_RNA45SN1_45S_50ntFlanks@12002.12029.28\|TGCGGGCCGCCGGTGAAATACCACTACT | >hg38_21_+_8433172_8446622_RNA45SN1_45S_50ntFlanks | 12002 | 12029 | 28 | TGCGGGCCGCCGGTGAAATACCACTACT | 217.9 | -11.6 | 1.5 | -7.8 | 6.5E-15 | 3.8E-14 | MB231/MB468 | MP |
| >hg38_21_+_8433172_8446622_RNA45SN1_45S_50ntFlanks@9353.9370.18\|CGTGGCGCAATGAAGGTG | >hg38_21_+_8433172_8446622_RNA45SN1_45S_50ntFlanks | 9353 | 9370 | 18 | CGTGGCGCAATGAAGGTG | 1485.1 | -10.3 | 1.3 | -7.8 | 6.5E-15 | 3.8E-14 | MB231/MB468 | MP |
| >hg38_21_+_8433172_8446622_RNA45SN1_45S_50ntFlanks@6721.6753.33\|AATTGCAGGACACATTGATCATCGACACTTCGA | >hg38_21_+_8433172_8446622_RNA45SN1_45S_50ntFlanks | 6721 | 6753 | 33 | AATTGCAGGACACATTGATCATCGACACTTCGA | 356.3 | -12.3 | 1.6 | -7.8 | 6.5E-15 | 3.9E-14 | MB231/MB468 | MP |
| >hg38_21_+_8433172_8446622_RNA45SN1_45S_50ntFlanks@12417.12454.38\|TCGATGTCGGCTCTTCCTATCATTGTGAAGCAGAATTC | >hg38_21_+_8433172_8446622_RNA45SN1_45S_50ntFlanks | 12417 | 12454 | 38 | TCGATGTCGGCTCTTCCTATCATTGTGAAGCAGAATTC | 268.1 | -5.5 | 0.7 | -7.7 | 1.3E-14 | 3.9E-14 | BT20/MB468 | NUC |
| >hg38_21_+_8433172_8446622_RNA45SN1_45S_50ntFlanks@11575.11605.31\|ACTTAGAACTGGTGCGGACCAGGGGAATCCG | >hg38_21_+_8433172_8446622_RNA45SN1_45S_50ntFlanks | 11575 | 11605 | 31 | ACTTAGAACTGGTGCGGACCAGGGGAATCCG | 2001.1 | -9.8 | 1.3 | -7.8 | 7.9E-15 | 3.9E-14 | MB231/MB468 | NUC |
| >am_trna10_LysCTT_16_+_3241501_3241573@32.51.20,trna119_LysCTT_1_-_145395522_145395594@32.51.20,trna11_LysCTT_5_-_180648979_180649051@32.51.20,trna13_LysCTT_16_+_3246153_3246225@32.51.20,trna13_LysCTT_6_+_26556774_26556846@32.51.20,trna30_LysCTT_16_-_3230555_3230627@32.51.20,trna5_LysCTT_16_+_3214939_3215011@32.51.20,trna7_LysCTT_16_+_3225692_3225764@32.51.20,trna8_LysCTT_1_+_39970195_39970267@32.51.20,trna9_LysCTT_5_+_180634755_180634827@32.51.20\|CTCTTAATCTCAGGGTCGTG | >am_trna10_LysCTT_16_+_3241501_3241573 | 32 | 51 | 20 | CTCTTAATCTCAGGGTCGTG | 236.4 | -11.7 | 1.5 | -7.8 | 6.5E-15 | 3.9E-14 | MB231/MB468 | MP |
| >hg38_21_+_8433172_8446622_RNA45SN1_45S_50ntFlanks@12073.12093.21\|CCGAGGGGCTCTCGCTTCTGG | >hg38_21_+_8433172_8446622_RNA45SN1_45S_50ntFlanks | 12073 | 12093 | 21 | CCGAGGGGCTCTCGCTTCTGG | 328.9 | -12.2 | 1.6 | -7.8 | 6.5E-15 | 3.9E-14 | MB231/MB468 | MP |
| >trna10_AspGTC_12_-_125424193_125424264@32.72.41,trna12_AspGTC_12_-_125411891_125411962@32.72.41,trna144_AspGTC_6_-_27551236_27551307@32.72.41,trna38_AspGTC_17_-_8125556_8125627@32.72.41,trna45_AspGTC_6_+_27447453_27447524@32.72.41,trna48_AspGTC_6_+_27471523_27471594@32.72.41,trna4_AspGTC_12_+_96429799_96429870@32.72.41,trna69_AspGTC_1_-_161440205_161440276@32.72.41,trna72_AspGTC_1_-_161432824_161432895@32.72.41,trna75_AspGTC_1_-_161425414_161425485@32.72.41,trna78_AspGTC_1_-_161418033_161418104@32.72.41,trna81_AspGTC_1_-_161410615_161410686@32.72.41\|CTGTCACGCGGGAGACCGGGGTTCGATTCCCCGACGGGGAG | >trna10_AspGTC_12_-_125424193_125424264 | 32 | 72 | 41 | CTGTCACGCGGGAGACCGGGGTTCGATTCCCCGACGGGGAG | 364.9 | -12.4 | 1.6 | -7.8 | 6.5E-15 | 3.9E-14 | MB231/MB468 | MP |
| >MI0003778\|hsa-mir-320c-1&WithFlank&18\|+\|21683512\|21683595@48.67.20;MI0008191\|hsa-mir-320c-2&WithFlank&18\|+\|24321669\|24321752@48.67.20&[MIMAT0005793&hsa-miR-320c&offsets\|0\|0;m-222&18\|+\|21683559\|21683578&offsets\|0\|0];[MIMAT0005793_1&hsa-miR-320c&offsets\|0\|0;m-219&18\|+\|24321716\|24321735&offsets\|0\|0]\|AAAAGCTGGGTTGAGAGGGT | >MI0003778\|hsa-mir-320c-1&WithFlank&18\|+\|21683512\|21683595 | 48 | 67 | 20 | AAAAGCTGGGTTGAGAGGGT | 146.1 | 4.0 | 0.5 | 7.7 | 1.4E-14 | 3.9E-14 | BT20/MB468 | NUC |
| >hg38_21_+_8433172_8446622_RNA45SN1_45S_50ntFlanks@11881.11905.25\|CCTGTTGAGCTTGACTCTAGTCTGG | >hg38_21_+_8433172_8446622_RNA45SN1_45S_50ntFlanks | 11881 | 11905 | 25 | CCTGTTGAGCTTGACTCTAGTCTGG | 244.3 | -10.7 | 1.4 | -7.8 | 7.3E-15 | 3.9E-14 | MB231/MB468 | CYTO |
| >hg38_21_+_8433172_8446622_RNA45SN1_45S_50ntFlanks@6648.6673.26\|GTACGACTCTTAGCGGTGGATCACTC | >hg38_21_+_8433172_8446622_RNA45SN1_45S_50ntFlanks | 6648 | 6673 | 26 | GTACGACTCTTAGCGGTGGATCACTC | 1302.9 | 5.4 | 0.7 | 7.8 | 6.6E-15 | 3.9E-14 | BT20/MB468 | MITO |

| | | | | | | | | | | | | | |
|---|---|---|---|---|---|---|---|---|---|---|---|---|---|
| >hg38_21_+_8433172_8446622_RNA45SN1_45S_50ntFlanks@10806.10830.25\|AACAATGTAGGTAAGGGAAGTCGGC | >hg38_21_+_8433172_8446622_RNA45SN1_45S_50ntFlanks | 10806 | 10830 | 25 | AACAATGTAGGTAAGGGAAGTCGGC | 169.2 | -4.9 | 0.6 | -7.8 | 7.4E-15 | 3.9E-14 | MB231/MB468 | CYTO |
| >hg38_21_+_8433172_8446622_RNA45SN1_45S_50ntFlanks@3776.3800.25\|CCGGTACAGTGAAACTGCGAATGGC | >hg38_21_+_8433172_8446622_RNA45SN1_45S_50ntFlanks | 3776 | 3800 | 25 | CCGGTACAGTGAAACTGCGAATGGC | 268.9 | -11.9 | 1.5 | -7.8 | 6.6E-15 | 3.9E-14 | MB231/MB468 | MP |
| >hg38_21_+_8433172_8446622_RNA45SN1_45S_50ntFlanks@9504.9522.19\|GTGAACTATGCCTGGGCAG | >hg38_21_+_8433172_8446622_RNA45SN1_45S_50ntFlanks | 9504 | 9522 | 19 | GTGAACTATGCCTGGGCAG | 908.5 | -7.4 | 1.0 | -7.7 | 1.1E-14 | 3.9E-14 | MB231/MB468 | MITO |
| >MI0003599\|hsa-mir-589&WithFlank&7\|-\|5495813\|5495923@30.52.23&[MIMAT0004799&hsa-miR-589-5p&offsets\|0\|+1;m-162&7\|-\|5495874\|5495894&offsets\|0\|+2]\|TGAGAACCACGTCTGCTCTGAGC | >MI0003599\|hsa-mir-589&WithFlank&7\|-\|5495813\|5495923 | 30 | 52 | 23 | TGAGAACCACGTCTGCTCTGAGC | 829.3 | 6.4 | 0.8 | 7.8 | 8.1E-15 | 3.9E-14 | MB231/MB468 | NUC |
| >hg38_21_+_8433172_8446622_RNA45SN1_45S_50ntFlanks@6662.6688.27\|GGTGGATCACTCGGCTCGTGCGTCGAT | >hg38_21_+_8433172_8446622_RNA45SN1_45S_50ntFlanks | 6662 | 6688 | 27 | GGTGGATCACTCGGCTCGTGCGTCGAT | 388.6 | 4.5 | 0.6 | 7.8 | 6.6E-15 | 3.9E-14 | BT20/MB468 | MITO |
| >hg38_21_+_8433172_8446622_RNA45SN1_45S_50ntFlanks@10546.10567.22\|CAGTGCGGTAACGCGACCGATC | >hg38_21_+_8433172_8446622_RNA45SN1_45S_50ntFlanks | 10546 | 10567 | 22 | CAGTGCGGTAACGCGACCGATC | 987.6 | -6.2 | 0.8 | -7.8 | 6.6E-15 | 3.9E-14 | MB231/MB468 | MP |
| >hg38_21_+_8433172_8446622_RNA45SN1_45S_50ntFlanks@10727.10751.25\|GGAGAGGGTGTAAATCTCGCGCCGG | >hg38_21_+_8433172_8446622_RNA45SN1_45S_50ntFlanks | 10727 | 10751 | 25 | GGAGAGGGTGTAAATCTCGCGCCGG | 340.2 | -12.3 | 1.6 | -7.8 | 6.6E-15 | 3.9E-14 | MB231/MB468 | MP |
| >hg38_21_+_8433172_8446622_RNA45SN1_45S_50ntFlanks@4003.4025.23\|ATAACCTCGGGCCGATCGCACGC | >hg38_21_+_8433172_8446622_RNA45SN1_45S_50ntFlanks | 4003 | 4025 | 23 | ATAACCTCGGGCCGATCGCACGC | 184.8 | -11.4 | 1.5 | -7.8 | 6.6E-15 | 3.9E-14 | MB231/MB468 | MP |
| >trna111_HisGTG_1_-_147774845_147774916@-1T.26.27,trna118_HisGTG_1_-_145396881_145396952@-1T.26.27,trna16_HisGTG_1_+_146544773_146544844@-1T.26.27,trna1_HisGTG_15_+_45493349_45493420@-1T.26.27,trna21_HisGTG_1_+_147753471_147753542@-1T.26.27,trna33_HisGTG_6_+_27125906_27125977@-1T.26.27,trna7_HisGTG_9_-_14433938_14434009@-1T.26.27,trna8_HisGTG_15_-_45492611_45492682@-1T.26.27,trna9_HisGTG_15_-_45490804_45490875@-1T.26.27\|TGCCGTGATCGTATAGTGGTTAGTACT | >trna111_HisGTG_1_-_147774845_147774916 | -1T | 26 | 27 | TGCCGTGATCGTATAGTGGTTAGTACT | 3265.6 | 6.8 | 0.9 | 7.7 | 1.1E-14 | 3.9E-14 | MB231/MB468 | MITO |
| >MI0000542\|hsa-mir-320a&WithFlank&8\|-\|22244960\|22245043@49.69.21&[MIMAT0000510&hsa-miR-320a-3p&offsets\|+1\|0;m-60&8\|-\|22244975\|22244996&offsets\|+1\|0]\|AAAGCTGGGTTGAGAGGGCGA | >MI0000542\|hsa-mir-320a&WithFlank&8\|-\|22244960\|22245043 | 49 | 69 | 21 | AAAGCTGGGTTGAGAGGGCGA | 172.9 | 3.2 | 0.4 | 7.7 | 1.4E-14 | 3.9E-14 | BT20/MB468 | NUC |
| >hg38_21_+_8433172_8446622_RNA45SN1_45S_50ntFlanks@9127.9152.26\|CGCGGGTCGGGGGGCGGGGCGGACTG | >hg38_21_+_8433172_8446622_RNA45SN1_45S_50ntFlanks | 9127 | 9152 | 26 | CGCGGGTCGGGGGGCGGGGCGGACTG | 303.1 | -12.1 | 1.6 | -7.8 | 6.7E-15 | 4.0E-14 | MB231/MB468 | MP |
| >hg38_21_+_8433172_8446622_RNA45SN1_45S_50ntFlanks@12405.12422.18\|CTTTTTGATCCTTCGATG | >hg38_21_+_8433172_8446622_RNA45SN1_45S_50ntFlanks | 12405 | 12422 | 18 | CTTTTTGATCCTTCGATG | 197.1 | -10.6 | 1.4 | -7.8 | 8.1E-15 | 4.0E-14 | MB231/MB468 | NUC |
| >hg38_21_+_8433172_8446622_RNA45SN1_45S_50ntFlanks@11319.11342.24\|CCCACGTCTCGTCGCGCGCGCGTC | >hg38_21_+_8433172_8446622_RNA45SN1_45S_50ntFlanks | 11319 | 11342 | 24 | CCCACGTCTCGTCGCGCGCGCGTC | 913.9 | -7.1 | 0.9 | -7.8 | 7.5E-15 | 4.0E-14 | MB231/MB468 | CYTO |
| >MI0000065\|hsa-let-7d&WithFlank&9\|+\|94178828\|94178926@68.88.21(+1C)&[MIMAT0004484&hsa-let-7d-3p&offsets\|0\|-1(+1C);m-98&9\|+\|94178895\|94178916&offsets\|0\|-1(+1C)]\|CTATACGACCTGCTGCCTTTCC | >MI0000065\|hsa-let-7d&WithFlank&9\|+\|94178828\|94178926 | 68 | 88 | 21(+1C) | CTATACGACCTGCTGCCTTTCC | 866.8 | 8.0 | 1.0 | 7.8 | 9.1E-15 | 4.0E-14 | BT20/MB468 | CYTO |
| >MI0000456\|hsa-mir-140&WithFlank&16\|+\|69933075\|69933186@69.89.21&[MIMAT0004597&hsa-miR-140-3p&offsets\|+1\|+1;m-45&16\|+\|69933143\|69933164&offsets\|0\|-1]\|ACCACAGGGTAGAACCACGGA | >MI0000456\|hsa-mir-140&WithFlank&16\|+\|69933075\|69933186 | 69 | 89 | 21 | ACCACAGGGTAGAACCACGGA | 845.2 | 12.9 | 1.7 | 7.7 | 1.1E-14 | 4.0E-14 | MB231/MB468 | MITO |
| >MI0000077\|hsa-mir-21&WithFlank&17\|+\|59841260\|59841343@14.33.20(+1A)&[MIMAT0000076&hsa-miR-21-5p&offsets\|0\|-2(+1A);m-1&17\|+\|59841273\|59841294&offsets\|0\|-2(+1A)]\|TAGCTTATCAGACTGATGTTA | >MI0000077\|hsa-mir-21&WithFlank&17\|+\|59841260\|59841343 | 14 | 33 | 20(+1A) | TAGCTTATCAGACTGATGTTA | 493.2 | 5.0 | 0.7 | 7.7 | 1.4E-14 | 4.0E-14 | BT20/MB468 | NUC |
| >hg38_21_+_8433172_8446622_RNA45SN1_45S_50ntFlanks@485.505.21\|GATGTGGTGACGTCGTGCTCT | >hg38_21_+_8433172_8446622_RNA45SN1_45S_50ntFlanks | 485 | 505 | 21 | GATGTGGTGACGTCGTGCTCT | 607.9 | 13.2 | 1.7 | 7.8 | 7.5E-15 | 4.0E-14 | MB231/MB468 | CYTO |
| >hg38_21_+_8433172_8446622_RNA45SN1_45S_50ntFlanks@4443.4472.30\|CCCTTGCCTCTCGGCGCCCCCTCGATGCTC | >hg38_21_+_8433172_8446622_RNA45SN1_45S_50ntFlanks | 4443 | 4472 | 30 | CCCTTGCCTCTCGGCGCCCCCTCGATGCTC | 765.9 | -9.3 | 1.2 | -7.8 | 7.5E-15 | 4.0E-14 | MB231/MB468 | CYTO |
| >hg38_21_+_8433172_8446622_RNA45SN1_45S_50ntFlanks@6682.6718.37\|CGTCGATGAAGAACGCAGCTAGCTGCGAGAATTAATG | >hg38_21_+_8433172_8446622_RNA45SN1_45S_50ntFlanks | 6682 | 6718 | 37 | CGTCGATGAAGAACGCAGCTAGCTGCGAGAATTAATG | 153.7 | -10.0 | 1.3 | -7.8 | 7.5E-15 | 4.0E-14 | MB231/MB468 | CYTO |
| >hg38_21_+_8433172_8446622_RNA45SN1_45S_50ntFlanks@11208.11234.27\|GCGGCGACTCTGGACGCGAGCCGGGCC | >hg38_21_+_8433172_8446622_RNA45SN1_45S_50ntFlanks | 11208 | 11234 | 27 | GCGGCGACTCTGGACGCGAGCCGGGCC | 2849.0 | -8.5 | 1.1 | -7.8 | 8.2E-15 | 4.0E-14 | MB231/MB468 | NUC |
| >hg38_21_+_8433172_8446622_RNA45SN1_45S_50ntFlanks@11561.11600.40\|GCGCCTAGCAGCCGACTTAGAACTGGTGCGGACCAGGGGA | >hg38_21_+_8433172_8446622_RNA45SN1_45S_50ntFlanks | 11561 | 11600 | 40 | GCGCCTAGCAGCCGACTTAGAACTGGTGCGGACCAGGGGA | 759.6 | -13.4 | 1.7 | -7.8 | 6.7E-15 | 4.0E-14 | MB231/MB468 | MP |
| >hg38_21_+_8433172_8446622_RNA45SN1_45S_50ntFlanks@8107.8142.36\|CGGCGGGGCGCGGGACATGTGGCGTACGGAAGACCC | >hg38_21_+_8433172_8446622_RNA45SN1_45S_50ntFlanks | 8107 | 8142 | 36 | CGGCGGGGCGCGGGACATGTGGCGTACGGAAGACCC | 211.8 | -11.6 | 1.5 | -7.8 | 6.7E-15 | 4.0E-14 | MB231/MB468 | MP |
| >hg38_21_+_8433172_8446622_RNA45SN1_45S_50ntFlanks@4472.4496.25\|CTTAGCTGAGTGTCCCGCGGGGCCC | >hg38_21_+_8433172_8446622_RNA45SN1_45S_50ntFlanks | 4472 | 4496 | 25 | CTTAGCTGAGTGTCCCGCGGGGCCC | 249.3 | -11.8 | 1.5 | -7.8 | 6.7E-15 | 4.0E-14 | MB231/MB468 | MP |
| >hg38_21_+_8433172_8446622_RNA45SN1_45S_50ntFlanks@12640.12659.20\|ACCATCTGTGGGATTATGAC | >hg38_21_+_8433172_8446622_RNA45SN1_45S_50ntFlanks | 12640 | 12659 | 20 | ACCATCTGTGGGATTATGAC | 2401.2 | -6.7 | 0.9 | -7.7 | 1.1E-14 | 4.0E-14 | MB231/MB468 | MITO |
| >hg38_21_+_8433172_8446622_RNA45SN1_45S_50ntFlanks@11576.11602.27\|CTTAGAACTGGTGCGGACCAGGGGAAT | >hg38_21_+_8433172_8446622_RNA45SN1_45S_50ntFlanks | 11576 | 11602 | 27 | CTTAGAACTGGTGCGGACCAGGGGAAT | 506.0 | -9.0 | 1.2 | -7.7 | 1.1E-14 | 4.0E-14 | MB231/MB468 | MITO |
| >MI0000102\|hsa-mir-100&WithFlank&11\|-\|122152223\|122152314@19.38.20&[MIMAT0000098&hsa-miR-100-5p&offsets\|0\|-2;m-27&11\|-\|122152276\|122152296&offsets\|0\|-1]\|AACCCGTAGATCCGAACTTG | >MI0000102\|hsa-mir-100&WithFlank&11\|-\|122152223\|122152314 | 19 | 38 | 20 | AACCCGTAGATCCGAACTTG | 2800.0 | 15.7 | 2.0 | 7.8 | 9.2E-15 | 4.0E-14 | BT20/MB468 | CYTO |
| >hg38_21_+_8433172_8446622_RNA45SN1_45S_50ntFlanks@10343.10382.40\|TATTCAAACGAGAACTTTGAAGGCCGAAGTGGAGAAGGGT | >hg38_21_+_8433172_8446622_RNA45SN1_45S_50ntFlanks | 10343 | 10382 | 40 | TATTCAAACGAGAACTTTGAAGGCCGAAGTGGAGAAGGGT | 270.1 | 8.1 | 1.0 | 7.8 | 7.6E-15 | 4.0E-14 | MB231/MB468 | CYTO |
| >hg38_21_+_8433172_8446622_RNA45SN1_45S_50ntFlanks@10534.10568.35\|CCGCGAGGCGTCCAGTGCGGTAACGCGACCGATCC | >hg38_21_+_8433172_8446622_RNA45SN1_45S_50ntFlanks | 10534 | 10568 | 35 | CCGCGAGGCGTCCAGTGCGGTAACGCGACCGATCC | 187.7 | -11.4 | 1.5 | -7.8 | 6.8E-15 | 4.0E-14 | MB231/MB468 | MP |
| >hg38_21_+_8433172_8446622_RNA45SN1_45S_50ntFlanks@12994.13036.43\|CCTCGCTGCGATCTATTGAAAGTCAGCCCTCGACACAAGGGTT | >hg38_21_+_8433172_8446622_RNA45SN1_45S_50ntFlanks | 12994 | 13036 | 43 | CCTCGCTGCGATCTATTGAAAGTCAGCCCTCGACACAAGGGTT | 427.8 | 5.6 | 0.7 | 7.8 | 7.6E-15 | 4.0E-14 | MB231/MB468 | CYTO |
| >hg38_21_+_8433172_8446622_RNA45SN1_45S_50ntFlanks@10829.10853.25\|GCAAGCCGGATCCGTAACTTCGGGA | >hg38_21_+_8433172_8446622_RNA45SN1_45S_50ntFlanks | 10829 | 10853 | 25 | GCAAGCCGGATCCGTAACTTCGGGA | 195.4 | -11.5 | 1.5 | -7.8 | 6.8E-15 | 4.0E-14 | MB231/MB468 | MP |
| >hg38_21_+_8433172_8446622_RNA45SN1_45S_50ntFlanks@4103.4121.19\|CCACGGGTGACGGGGAATC | >hg38_21_+_8433172_8446622_RNA45SN1_45S_50ntFlanks | 4103 | 4121 | 19 | CCACGGGTGACGGGGAATC | 1602.7 | -10.3 | 1.3 | -7.8 | 6.8E-15 | 4.0E-14 | MB231/MB468 | MP |
| >hg38_21_+_8433172_8446622_RNA45SN1_45S_50ntFlanks@9565.9596.32\|TGACGTGCAAATCGGTCGTCCGACCTGGGTAT | >hg38_21_+_8433172_8446622_RNA45SN1_45S_50ntFlanks | 9565 | 9596 | 32 | TGACGTGCAAATCGGTCGTCCGACCTGGGTAT | 8469.7 | -5.1 | 0.7 | -7.8 | 6.8E-15 | 4.0E-14 | BT20/MB468 | MITO |
| >hg38_21_+_8433172_8446622_RNA45SN1_45S_50ntFlanks@11316.11339.24\|CACCCCACGTCTCGTCGCGCGCGC | >hg38_21_+_8433172_8446622_RNA45SN1_45S_50ntFlanks | 11316 | 11339 | 24 | CACCCCACGTCTCGTCGCGCGCGC | 229.4 | -11.6 | 1.5 | -7.7 | 1.1E-14 | 4.0E-14 | MB231/MB468 | MITO |
| >hg38_21_+_8433172_8446622_RNA45SN1_45S_50ntFlanks@4220.4243.24\|ACGAAAAATAACAATACAGGACTC | >hg38_21_+_8433172_8446622_RNA45SN1_45S_50ntFlanks | 4220 | 4243 | 24 | ACGAAAAATAACAATACAGGACTC | 266.0 | -11.8 | 1.5 | -7.7 | 1.1E-14 | 4.0E-14 | MB231/MB468 | MITO |
| >hg38_21_+_8433172_8446622_RNA45SN1_45S_50ntFlanks@12627.12652.26\|ATGGGGCGAAGCTACCATCTGTGGGA | >hg38_21_+_8433172_8446622_RNA45SN1_45S_50ntFlanks | 12627 | 12652 | 26 | ATGGGGCGAAGCTACCATCTGTGGGA | 200.3 | -10.6 | 1.4 | -7.8 | 8.3E-15 | 4.0E-14 | MB231/MB468 | NUC |
| >MI0000483\|hsa-mir-186&WithFlank&1\|-\|71067625\|71067722@21.42.22&[MIMAT0000456&hsa-miR-186-5p&offsets\|0\|0;m-84&1\|-\|71067681\|71067702&offsets\|0\|0]\|CAAAGAATTCTCCTTTTGGGCT | >MI0000483\|hsa-mir-186&WithFlank&1\|-\|71067625\|71067722 | 21 | 42 | 22 | CAAAGAATTCTCCTTTTGGGCT | 585.0 | 3.2 | 0.4 | 7.7 | 1.4E-14 | 4.0E-14 | BT20/MB468 | NUC |
| >hg38_21_+_8433172_8446622_RNA45SN1_45S_50ntFlanks@476.504.29\|TTCGTGATCGATGTGGTGACGTCGTGCTC | >hg38_21_+_8433172_8446622_RNA45SN1_45S_50ntFlanks | 476 | 504 | 29 | TTCGTGATCGATGTGGTGACGTCGTGCTC | 1941.3 | 7.9 | 1.0 | 7.8 | 6.8E-15 | 4.0E-14 | BT20/MB468 | MITO |
| >hg38_21_+_8433172_8446622_RNA45SN1_45S_50ntFlanks@10926.10946.21\|ACGAGGCGCCGCCGCCCCCCC | >hg38_21_+_8433172_8446622_RNA45SN1_45S_50ntFlanks | 10926 | 10946 | 21 | ACGAGGCGCCGCCGCCCCCCC | 2092.9 | -14.9 | 1.9 | -7.8 | 6.8E-15 | 4.0E-14 | MB231/MB468 | MP |
| >hg38_21_+_8433172_8446622_RNA45SN1_45S_50ntFlanks@6772.6797.26\|GTTCCTCCCGGGGCTACGCCTGTCTG | >hg38_21_+_8433172_8446622_RNA45SN1_45S_50ntFlanks | 6772 | 6797 | 26 | GTTCCTCCCGGGGCTACGCCTGTCTG | 257.9 | -11.9 | 1.5 | -7.8 | 6.9E-15 | 4.1E-14 | MB231/MB468 | MP |
| >hg38_21_+_8433172_8446622_RNA45SN1_45S_50ntFlanks@4099.4121.23\|GTGACCACGGGTGACGGGGAATC | >hg38_21_+_8433172_8446622_RNA45SN1_45S_50ntFlanks | 4099 | 4121 | 23 | GTGACCACGGGTGACGGGGAATC | 414.3 | -11.7 | 1.5 | -7.8 | 8.4E-15 | 4.1E-14 | MB231/MB468 | NUC |
| >hg38_21_+_8433172_8446622_RNA45SN1_45S_50ntFlanks@12405.12440.36\|CTTTTTGATCCTTCGATGTCGGCTCTTCCTATCATT | >hg38_21_+_8433172_8446622_RNA45SN1_45S_50ntFlanks | 12405 | 12440 | 36 | CTTTTTGATCCTTCGATGTCGGCTCTTCCTATCATT | 4493.1 | -9.6 | 1.2 | -7.7 | 1.4E-14 | 4.1E-14 | BT20/MB468 | NUC |
| >MI0000285\|hsa-mir-205&WithFlank&1\|+\|209432127\|209432248@40.60.21&[MIMAT0000266&hsa-miR-205-5p&offsets\|0\|-1;m-351&1\|+\|209432166\|209432187&offsets\|0\|-1]\|TCCTTCATTCCACCGGAGTCT | >MI0000285\|hsa-mir-205&WithFlank&1\|+\|209432127\|209432248 | 40 | 60 | 21 | TCCTTCATTCCACCGGAGTCT | 257.5 | -11.8 | 1.5 | -7.7 | 1.1E-14 | 4.1E-14 | MB231/MB468 | MITO |
| >hg38_21_+_8433172_8446622_RNA45SN1_45S_50ntFlanks@13002.13020.19\|CGATCTATTGAAAGTCAGC | >hg38_21_+_8433172_8446622_RNA45SN1_45S_50ntFlanks | 13002 | 13020 | 19 | CGATCTATTGAAAGTCAGC | 378.5 | -8.0 | 1.0 | -7.8 | 7.8E-15 | 4.1E-14 | MB231/MB468 | CYTO |
| >am_trna11_ProAGG_16_+_3241989_3242060@1.29.29,trna12_ProAGG_6_+_26555498_26555569@1.29.29,trna12_ProTGG_11_-_75946869_75946940@1.29.29,trna14_ProTGG_5_-_180615854_180615925@1.29.29,trna22_ProAGG_14_-_21081560_21081631@1.29.29,trna23_ProAGG_14_-_21077495_21077566@1.29.29,trna28_ProTGG_16_-_3234133_3234204@1.29.29,trna29_ProAGG_16_-_3232635_3232706@1.29.29,trna2_ProAGG_7_+_128423504_128423575@1.29.29,trna30_ProCGG_6_+_27059521_27059592@1.29.29,trna37_ProCGG_17_-_8126151_8126222@1.29.29,trna3_ProTGG_16_+_3208923_3208994@1.29.29,trna52_ProCGG_1_+_167683962_167684033@1.29.29,trna65_ProAGG_1_-_167684725_167684796@1.29.29,trna6_ProCGG_16_+_3222049_3222120@1.29.29,trna6_ProTGG_14_+_21152175_21152246@1.29.29,trna8_ProTGG_16_+_3238094_3238165@1.29.29,trna9_ProAGG_11_+_75946557_75946628@1.29.29,trna9_ProAGG_16_+_3239634_3239705@1.29.29\|GGCTCGTTGGTCTAGGGGTATGATTCTCG | >am_trna11_ProAGG_16_+_3241989_3242060 | 1 | 29 | 29 | GGCTCGTTGGTCTAGGGGTATGATTCTCG | 2863.4 | -10.1 | 1.3 | -7.8 | 8.5E-15 | 4.1E-14 | MB231/MB468 | NUC |

<table>
<tr><td>>MI0000273|hsa-mir-183&WithFlank&7|-|129774899|129775020@34.55.22(+1C)&[MIMAT0000261&hsa-miR-183-5p&offsets|+1|+1(+1C);m-41&7|-|129774966|129774987&offsets|0|0(+1C)]|ATGGCACTGGTAGAATTCACTGC</td><td>>MI0000273|hsa-mir-183&WithFlank&7|-|129774899|129775020</td><td>34</td><td>55</td><td>22(+1C)</td><td>ATGGCACTGGTAGAATTCACTGC</td><td>488.6</td><td>9.0</td><td>1.2</td><td>7.7</td><td>9.4E-15</td><td>4.1E-14</td><td>BT20/MB468</td><td>CYTO</td></tr>
<tr><td>>hg38_21_+_8433172_8446622_RNA45SN1_45S_50ntFlanks@12456.12482.27|CCAAGCGTTGGATTGTTCACCCACTAA</td><td>>hg38_21_+_8433172_8446622_RNA45SN1_45S_50ntFlanks</td><td>12456</td><td>12482</td><td>27</td><td>CCAAGCGTTGGATTGTTCACCCACTAA</td><td>297.4</td><td>-12.1</td><td>1.6</td><td>-7.8</td><td>7.0E-15</td><td>4.1E-14</td><td>MB231/MB468</td><td>MP</td></tr>
<tr><td>>hg38_21_+_8433172_8446622_RNA45SN1_45S_50ntFlanks@11190.11208.19|CCCGGGGGGCCGGCGGCGG</td><td>>hg38_21_+_8433172_8446622_RNA45SN1_45S_50ntFlanks</td><td>11190</td><td>11208</td><td>19</td><td>CCCGGGGGGCCGGCGGCGG</td><td>227.6</td><td>-10.5</td><td>1.4</td><td>-7.8</td><td>7.9E-15</td><td>4.1E-14</td><td>MB231/MB468</td><td>CYTO</td></tr>
<tr><td>>hg38_21_+_8433172_8446622_RNA45SN1_45S_50ntFlanks@4730.4761.32|CAAGAACGAAAGTCGGAGGTTCGAAGACGATC</td><td>>hg38_21_+_8433172_8446622_RNA45SN1_45S_50ntFlanks</td><td>4730</td><td>4761</td><td>32</td><td>CAAGAACGAAAGTCGGAGGTTCGAAGACGATC</td><td>855.7</td><td>5.6</td><td>0.7</td><td>7.7</td><td>9.5E-15</td><td>4.1E-14</td><td>BT20/MB468</td><td>CYTO</td></tr>
<tr><td>>hg38_21_+_8433172_8446622_RNA45SN1_45S_50ntFlanks@7986.8004.19|ATCAGACGTGGCGACCCGC</td><td>>hg38_21_+_8433172_8446622_RNA45SN1_45S_50ntFlanks</td><td>7986</td><td>8004</td><td>19</td><td>ATCAGACGTGGCGACCCGC</td><td>880.9</td><td>-12.9</td><td>1.6</td><td>-8.2</td><td>2.8E-16</td><td>4.1E-14</td><td>BT20/MB468</td><td>MP</td></tr>
<tr><td>>hg38_21_+_8433172_8446622_RNA45SN1_45S_50ntFlanks@11766.11789.24|ATCTAATTAGTGACGCGCATGAAT</td><td>>hg38_21_+_8433172_8446622_RNA45SN1_45S_50ntFlanks</td><td>11766</td><td>11789</td><td>24</td><td>ATCTAATTAGTGACGCGCATGAAT</td><td>293.6</td><td>-11.2</td><td>1.4</td><td>-7.8</td><td>8.5E-15</td><td>4.1E-14</td><td>MB231/MB468</td><td>NUC</td></tr>
<tr><td>>hg38_21_+_8433172_8446622_RNA45SN1_45S_50ntFlanks@4452.4471.20|CTCGGCGCCCCCTCGATGCT</td><td>>hg38_21_+_8433172_8446622_RNA45SN1_45S_50ntFlanks</td><td>4452</td><td>4471</td><td>20</td><td>CTCGGCGCCCCCTCGATGCT</td><td>252.1</td><td>-11.8</td><td>1.5</td><td>-7.7</td><td>1.2E-14</td><td>4.1E-14</td><td>MB231/MB468</td><td>MITO</td></tr>
<tr><td>>hg38_21_+_8433172_8446622_RNA45SN1_45S_50ntFlanks@12513.12538.26|TGAGACAGGTTAGTTTTACCCTACTG</td><td>>hg38_21_+_8433172_8446622_RNA45SN1_45S_50ntFlanks</td><td>12513</td><td>12538</td><td>26</td><td>TGAGACAGGTTAGTTTTACCCTACTG</td><td>295.3</td><td>-12.1</td><td>1.6</td><td>-7.8</td><td>7.0E-15</td><td>4.1E-14</td><td>MB231/MB468</td><td>MP</td></tr>
<tr><td>>hg38_21_+_8433172_8446622_RNA45SN1_45S_50ntFlanks@11783.11804.22|CATGAATGGATGAACGAGATTC</td><td>>hg38_21_+_8433172_8446622_RNA45SN1_45S_50ntFlanks</td><td>11783</td><td>11804</td><td>22</td><td>CATGAATGGATGAACGAGATTC</td><td>17081.6</td><td>-6.5</td><td>0.8</td><td>-7.7</td><td>9.5E-15</td><td>4.1E-14</td><td>BT20/MB468</td><td>CYTO</td></tr>
<tr><td>>hg38_21_+_8433172_8446622_RNA45SN1_45S_50ntFlanks@11027.11060.34|CTCTCTCTCTCTCTCCCCCGCTCCCCGTCCTCCC</td><td>>hg38_21_+_8433172_8446622_RNA45SN1_45S_50ntFlanks</td><td>11027</td><td>11060</td><td>34</td><td>CTCTCTCTCTCTCTCCCCCGCTCCCCGTCCTCCC</td><td>1211.2</td><td>-8.8</td><td>1.1</td><td>-7.8</td><td>7.9E-15</td><td>4.1E-14</td><td>MB231/MB468</td><td>CYTO</td></tr>
<tr><td>>hg38_21_+_8433172_8446622_RNA45SN1_45S_50ntFlanks@9565.9597.33|TGACGTGCAAATCGGTCGTCCGACCTGGGTATA</td><td>>hg38_21_+_8433172_8446622_RNA45SN1_45S_50ntFlanks</td><td>9565</td><td>9597</td><td>33</td><td>TGACGTGCAAATCGGTCGTCCGACCTGGGTATA</td><td>9485.4</td><td>-6.5</td><td>0.8</td><td>-7.7</td><td>1.2E-14</td><td>4.1E-14</td><td>MB231/MB468</td><td>MITO</td></tr>
<tr><td>>hg38_21_+_8433172_8446622_RNA45SN1_45S_50ntFlanks@11669.11703.35|GCCCAGTGCTCTGAATGTCAAAGTGAAGAAATTCA</td><td>>hg38_21_+_8433172_8446622_RNA45SN1_45S_50ntFlanks</td><td>11669</td><td>11703</td><td>35</td><td>GCCCAGTGCTCTGAATGTCAAAGTGAAGAAATTCA</td><td>466.4</td><td>4.9</td><td>0.6</td><td>7.7</td><td>9.6E-15</td><td>4.2E-14</td><td>BT20/MB468</td><td>CYTO</td></tr>
<tr><td>>hg38_21_+_8433172_8446622_RNA45SN1_45S_50ntFlanks@6655.6685.31|TCTTAGCGGTGGATCACTCGGCTCGTGCGTC</td><td>>hg38_21_+_8433172_8446622_RNA45SN1_45S_50ntFlanks</td><td>6655</td><td>6685</td><td>31</td><td>TCTTAGCGGTGGATCACTCGGCTCGTGCGTC</td><td>362.4</td><td>3.9</td><td>0.5</td><td>7.8</td><td>7.0E-15</td><td>4.2E-14</td><td>BT20/MB468</td><td>MITO</td></tr>
<tr><td>>MI0000089|hsa-mir-31&WithFlank&9|-|21512109|21512191@14.35.22(+1A)&[MIMAT0000089&hsa-miR-31-5p&offsets|0|+1(+1A);m-360&9|-|21512157|21512178&offsets|0|0(+1A)]|AGGCAAGATGCTGGCATAGCTGA</td><td>>MI0000089|hsa-mir-31&WithFlank&9|-|21512109|21512191</td><td>14</td><td>35</td><td>22(+1A)</td><td>AGGCAAGATGCTGGCATAGCTGA</td><td>299.6</td><td>12.0</td><td>1.5</td><td>8.0</td><td>1.0E-15</td><td>4.2E-14</td><td>BT20/MB231</td><td>NUC</td></tr>
<tr><td>>hg38_21_+_8433172_8446622_RNA45SN1_45S_50ntFlanks@11581.11613.33|AACTGGTGCGGACCAGGGGAATCCGACTGTTTA</td><td>>hg38_21_+_8433172_8446622_RNA45SN1_45S_50ntFlanks</td><td>11581</td><td>11613</td><td>33</td><td>AACTGGTGCGGACCAGGGGAATCCGACTGTTTA</td><td>506.0</td><td>-4.3</td><td>0.6</td><td>-7.7</td><td>1.5E-14</td><td>4.2E-14</td><td>BT20/MB468</td><td>NUC</td></tr>
<tr><td>>hg38_21_+_8433172_8446622_RNA45SN1_45S_50ntFlanks@6742.6770.29|TCGACACTTCGAACGCACTTGCGGCCCCG</td><td>>hg38_21_+_8433172_8446622_RNA45SN1_45S_50ntFlanks</td><td>6742</td><td>6770</td><td>29</td><td>TCGACACTTCGAACGCACTTGCGGCCCCG</td><td>677.3</td><td>-9.1</td><td>1.2</td><td>-7.7</td><td>1.2E-14</td><td>4.2E-14</td><td>MB231/MB468</td><td>MITO</td></tr>
<tr><td>>hg38_21_+_8433172_8446622_RNA45SN1_45S_50ntFlanks@12928.12972.45|CTGGCGCTAAACCATTCGTAGACGACCTGCTTCTGGGTCGGGGTT</td><td>>hg38_21_+_8433172_8446622_RNA45SN1_45S_50ntFlanks</td><td>12928</td><td>12972</td><td>45</td><td>CTGGCGCTAAACCATTCGTAGACGACCTGCTTCTGGGTCGGGGTT</td><td>456.1</td><td>-12.6</td><td>1.6</td><td>-7.7</td><td>1.2E-14</td><td>4.2E-14</td><td>MB231/MB468</td><td>MITO</td></tr>
<tr><td>>MI0000299|hsa-mir-222&WithFlank&X|-|45747009|45747130@75.95.21&[MIMAT0000279&hsa-miR-222-3p&offsets|0|0;m-140&X|-|45747033|45747056&offsets|0|-3]|AGCTACATCTGGCTACTGGGT</td><td>>MI0000299|hsa-mir-222&WithFlank&X|-|45747009|45747130</td><td>75</td><td>95</td><td>21</td><td>AGCTACATCTGGCTACTGGGT</td><td>1201.0</td><td>6.2</td><td>0.8</td><td>7.8</td><td>8.0E-15</td><td>4.2E-14</td><td>MB231/MB468</td><td>CYTO</td></tr>
<tr><td>>hg38_21_+_8433172_8446622_RNA45SN1_45S_50ntFlanks@7973.8013.41|GACGCGACCTCAGATCAGACGTGGCGACCCGCTGAATTTAA</td><td>>hg38_21_+_8433172_8446622_RNA45SN1_45S_50ntFlanks</td><td>7973</td><td>8013</td><td>41</td><td>GACGCGACCTCAGATCAGACGTGGCGACCCGCTGAATTTAA</td><td>291.2</td><td>9.3</td><td>1.2</td><td>7.8</td><td>8.0E-15</td><td>4.2E-14</td><td>MB231/MB468</td><td>CYTO</td></tr>
<tr><td>>hg38_21_+_8433172_8446622_RNA45SN1_45S_50ntFlanks@11576.11602.27|CTTAGAACTGGTGCGGACCAGGGGAAT</td><td>>hg38_21_+_8433172_8446622_RNA45SN1_45S_50ntFlanks</td><td>11576</td><td>11602</td><td>27</td><td>CTTAGAACTGGTGCGGACCAGGGGAAT</td><td>1669.9</td><td>-5.7</td><td>0.7</td><td>-7.8</td><td>8.0E-15</td><td>4.2E-14</td><td>MB231/MB468</td><td>CYTO</td></tr>
<tr><td>>hg38_1_-_228634819_228635039_RNA5S12_5S_50ntFlanks@84.112.29|CTCGTCTGATCTCGGAAGCTAAGCAGGGT</td><td>>hg38_1_-_228634819_228635039_RNA5S12_5S_50ntFlanks</td><td>84</td><td>112</td><td>29</td><td>CTCGTCTGATCTCGGAAGCTAAGCAGGGT</td><td>149.0</td><td>-11.1</td><td>1.4</td><td>-7.8</td><td>7.1E-15</td><td>4.2E-14</td><td>MB231/MB468</td><td>MP</td></tr>
<tr><td>>hg38_21_+_8433172_8446622_RNA45SN1_45S_50ntFlanks@11577.11603.27|TTAGAACTGGTGCGGACCAGGGGAATC</td><td>>hg38_21_+_8433172_8446622_RNA45SN1_45S_50ntFlanks</td><td>11577</td><td>11603</td><td>27</td><td>TTAGAACTGGTGCGGACCAGGGGAATC</td><td>904.0</td><td>-9.5</td><td>1.2</td><td>-7.7</td><td>1.2E-14</td><td>4.2E-14</td><td>MB231/MB468</td><td>MITO</td></tr>
<tr><td>>hg38_21_+_8433172_8446622_RNA45SN1_45S_50ntFlanks@11575.11600.26|ACTTAGAACTGGTGCGGACCAGGGGA</td><td>>hg38_21_+_8433172_8446622_RNA45SN1_45S_50ntFlanks</td><td>11575</td><td>11600</td><td>26</td><td>ACTTAGAACTGGTGCGGACCAGGGGA</td><td>948.5</td><td>-6.0</td><td>0.8</td><td>-7.8</td><td>8.0E-15</td><td>4.2E-14</td><td>MB231/MB468</td><td>CYTO</td></tr>
<tr><td>>hg38_21_+_8433172_8446622_RNA45SN1_45S_50ntFlanks@12053.12070.18|GTGAGGCGGGGGGGCGAG</td><td>>hg38_21_+_8433172_8446622_RNA45SN1_45S_50ntFlanks</td><td>12053</td><td>12070</td><td>18</td><td>GTGAGGCGGGGGGGCGAG</td><td>385.9</td><td>-11.6</td><td>1.5</td><td>-7.8</td><td>8.7E-15</td><td>4.2E-14</td><td>MB231/MB468</td><td>NUC</td></tr>
<tr><td>>hg38_21_+_8433172_8446622_RNA45SN1_45S_50ntFlanks@12054.12072.19|TGAGGCGGGGGGGCGAGCC</td><td>>hg38_21_+_8433172_8446622_RNA45SN1_45S_50ntFlanks</td><td>12054</td><td>12072</td><td>19</td><td>TGAGGCGGGGGGGCGAGCC</td><td>365.4</td><td>-11.5</td><td>1.5</td><td>-7.8</td><td>8.7E-15</td><td>4.2E-14</td><td>MB231/MB468</td><td>NUC</td></tr>
<tr><td>>hg38_21_+_8433172_8446622_RNA45SN1_45S_50ntFlanks@10351.10390.40|CGAGAACTTTGAAGGCCGAAGTGGAGAAGGGTTCCATGTG</td><td>>hg38_21_+_8433172_8446622_RNA45SN1_45S_50ntFlanks</td><td>10351</td><td>10390</td><td>40</td><td>CGAGAACTTTGAAGGCCGAAGTGGAGAAGGGTTCCATGTG</td><td>8292.0</td><td>-5.3</td><td>0.7</td><td>-7.7</td><td>9.7E-15</td><td>4.2E-14</td><td>BT20/MB468</td><td>CYTO</td></tr>
<tr><td>>hg38_21_+_8433172_8446622_RNA45SN1_45S_50ntFlanks@11787.11805.19|AATGGATGAACGAGATTCC</td><td>>hg38_21_+_8433172_8446622_RNA45SN1_45S_50ntFlanks</td><td>11787</td><td>11805</td><td>19</td><td>AATGGATGAACGAGATTCC</td><td>185.6</td><td>-11.4</td><td>1.5</td><td>-7.8</td><td>7.2E-15</td><td>4.2E-14</td><td>MB231/MB468</td><td>MP</td></tr>
<tr><td>>hg38_21_+_8433172_8446622_RNA45SN1_45S_50ntFlanks@12635.12667.33|AAGCTACCATCTGTGGGATTATGACTGAACGCC</td><td>>hg38_21_+_8433172_8446622_RNA45SN1_45S_50ntFlanks</td><td>12635</td><td>12667</td><td>33</td><td>AAGCTACCATCTGTGGGATTATGACTGAACGCC</td><td>416.2</td><td>4.9</td><td>0.6</td><td>7.8</td><td>7.2E-15</td><td>4.2E-14</td><td>BT20/MB468</td><td>MITO</td></tr>
<tr><td>>MI0000102|hsa-mir-100&WithFlank&11|-|122152223|122152314@19.40.22(+1A)&[MIMAT0000098&hsa-miR-100-5p&offsets|0|0(+1A);m-27&11|-|122152276|122152296&offsets|0|+1(+1A)]|AACCCGTAGATCCGAACTTGTGA</td><td>>MI0000102|hsa-mir-100&WithFlank&11|-|122152223|122152314</td><td>19</td><td>40</td><td>22(+1A)</td><td>AACCCGTAGATCCGAACTTGTGA</td><td>34232.8</td><td>13.1</td><td>1.7</td><td>7.7</td><td>1.2E-14</td><td>4.2E-14</td><td>MB231/MB468</td><td>MITO</td></tr>
<tr><td>>hg38_21_+_8433172_8446622_RNA45SN1_45S_50ntFlanks@10466.10484.19|CGTTGCCCTCGGCCGATCG</td><td>>hg38_21_+_8433172_8446622_RNA45SN1_45S_50ntFlanks</td><td>10466</td><td>10484</td><td>19</td><td>CGTTGCCCTCGGCCGATCG</td><td>137.7</td><td>-5.2</td><td>0.7</td><td>-7.8</td><td>8.2E-15</td><td>4.3E-14</td><td>MB231/MB468</td><td>CYTO</td></tr>
<tr><td>>hg38_21_+_8433172_8446622_RNA45SN1_45S_50ntFlanks@5445.5466.22|TCGGATCGGCCCCGCCGGGGTC</td><td>>hg38_21_+_8433172_8446622_RNA45SN1_45S_50ntFlanks</td><td>5445</td><td>5466</td><td>22</td><td>TCGGATCGGCCCCGCCGGGGTC</td><td>5808.3</td><td>-5.8</td><td>0.8</td><td>-7.8</td><td>8.2E-15</td><td>4.3E-14</td><td>MB231/MB468</td><td>CYTO</td></tr>
<tr><td>>hg38_21_+_8433172_8446622_RNA45SN1_45S_50ntFlanks@10760.10785.26|CATATCCGCAGCAGGTCTCCAAGGTG</td><td>>hg38_21_+_8433172_8446622_RNA45SN1_45S_50ntFlanks</td><td>10760</td><td>10785</td><td>26</td><td>CATATCCGCAGCAGGTCTCCAAGGTG</td><td>278.0</td><td>-12.0</td><td>1.5</td><td>-7.8</td><td>7.2E-15</td><td>4.3E-14</td><td>MB231/MB468</td><td>MP</td></tr>
<tr><td>>hg38_21_+_8433172_8446622_RNA45SN1_45S_50ntFlanks@12128.12146.19|ACCCGCTCCGGGGACAGTG</td><td>>hg38_21_+_8433172_8446622_RNA45SN1_45S_50ntFlanks</td><td>12128</td><td>12146</td><td>19</td><td>ACCCGCTCCGGGGACAGTG</td><td>331.8</td><td>-12.2</td><td>1.6</td><td>-7.7</td><td>1.2E-14</td><td>4.3E-14</td><td>MB231/MB468</td><td>MITO</td></tr>
<tr><td>>hg38_21_+_8433172_8446622_RNA45SN1_45S_50ntFlanks@12640.12661.22|ACCATCTGTGGGATTATGACTG</td><td>>hg38_21_+_8433172_8446622_RNA45SN1_45S_50ntFlanks</td><td>12640</td><td>12661</td><td>22</td><td>ACCATCTGTGGGATTATGACTG</td><td>3738.9</td><td>-11.0</td><td>1.4</td><td>-7.8</td><td>7.3E-15</td><td>4.3E-14</td><td>MB231/MB468</td><td>MP</td></tr>
<tr><td>>MI0000079|hsa-mir-23a&WithFlank&19|-|13836581|13836665@54.73.20(+2U)&[MIMAT0000078&hsa-miR-23a-3p&offsets|+3|+2(+2U);m-31&19|-|13836595|13836615&offsets|+3|+2(+2U)]|ACATTGCCAGGGATTTCCAATT</td><td>>MI0000079|hsa-mir-23a&WithFlank&19|-|13836581|13836665</td><td>54</td><td>73</td><td>20(+2U)</td><td>ACATTGCCAGGGATTTCCAATT</td><td>1029.1</td><td>28.2</td><td>3.6</td><td>7.8</td><td>8.2E-15</td><td>4.3E-14</td><td>MB231/MB468</td><td>CYTO</td></tr>
<tr><td>>hg38_21_+_8433172_8446622_RNA45SN1_45S_50ntFlanks@10686.10711.26|GCGGCGTCCGGTGAGCTCTCGCTGGC</td><td>>hg38_21_+_8433172_8446622_RNA45SN1_45S_50ntFlanks</td><td>10686</td><td>10711</td><td>26</td><td>GCGGCGTCCGGTGAGCTCTCGCTGGC</td><td>1080.3</td><td>-4.7</td><td>0.6</td><td>-7.7</td><td>1.2E-14</td><td>4.3E-14</td><td>MB231/MB468</td><td>MITO</td></tr>
<tr><td>>hg38_21_+_8433172_8446622_RNA45SN1_45S_50ntFlanks@12949.12990.42|ACGACCTGCTTCTGGGTCGGGGTTTCGTACGTAGCAGAGCAG</td><td>>hg38_21_+_8433172_8446622_RNA45SN1_45S_50ntFlanks</td><td>12949</td><td>12990</td><td>42</td><td>ACGACCTGCTTCTGGGTCGGGGTTTCGTACGTAGCAGAGCAG</td><td>297.1</td><td>-12.0</td><td>1.6</td><td>-7.7</td><td>1.2E-14</td><td>4.3E-14</td><td>MB231/MB468</td><td>MITO</td></tr>
<tr><td>>hg38_21_+_8433172_8446622_RNA45SN1_45S_50ntFlanks@8552.8569.18|CTCCCCGGAGGGGGCGGG</td><td>>hg38_21_+_8433172_8446622_RNA45SN1_45S_50ntFlanks</td><td>8552</td><td>8569</td><td>18</td><td>CTCCCCGGAGGGGGCGGG</td><td>262.5</td><td>-11.0</td><td>1.4</td><td>-7.8</td><td>8.9E-15</td><td>4.3E-14</td><td>MB231/MB468</td><td>NUC</td></tr>
<tr><td>>hg38_21_+_8433172_8446622_RNA45SN1_45S_50ntFlanks@4728.4747.20|ATCAAGAACGAAAGTCGGAG</td><td>>hg38_21_+_8433172_8446622_RNA45SN1_45S_50ntFlanks</td><td>4728</td><td>4747</td><td>20</td><td>ATCAAGAACGAAAGTCGGAG</td><td>260.7</td><td>-11.9</td><td>1.5</td><td>-7.8</td><td>7.3E-15</td><td>4.3E-14</td><td>MB231/MB468</td><td>MP</td></tr>
<tr><td>>hg38_21_+_8433172_8446622_RNA45SN1_45S_50ntFlanks@11315.11338.24|CCACCCCACGTCTCGTCGCGCGCG</td><td>>hg38_21_+_8433172_8446622_RNA45SN1_45S_50ntFlanks</td><td>11315</td><td>11338</td><td>24</td><td>CCACCCCACGTCTCGTCGCGCGCG</td><td>174.3</td><td>-11.3</td><td>1.5</td><td>-7.8</td><td>7.3E-15</td><td>4.3E-14</td><td>MB231/MB468</td><td>MP</td></tr>
<tr><td>>MI0000077|hsa-mir-21&WithFlank&17|+|59841260|59841343@14.35.22(+1A)&[MIMAT0000076&hsa-miR-21-5p&offsets|0|0(+1A);m-1&17|+|59841273|59841294&offsets|0|0(+1A)]|TAGCTTATCAGACTGATGTTGAA</td><td>>MI0000077|hsa-mir-21&WithFlank&17|+|59841260|59841343</td><td>14</td><td>35</td><td>22(+1A)</td><td>TAGCTTATCAGACTGATGTTGAA</td><td>4474.5</td><td>4.2</td><td>0.5</td><td>7.7</td><td>1.5E-14</td><td>4.3E-14</td><td>BT20/MB468</td><td>NUC</td></tr>
<tr><td>>hg38_21_+_8433172_8446622_RNA45SN1_45S_50ntFlanks@476.498.23|TTCGTGATCGATGTGGTGACGTC</td><td>>hg38_21_+_8433172_8446622_RNA45SN1_45S_50ntFlanks</td><td>476</td><td>498</td><td>23</td><td>TTCGTGATCGATGTGGTGACGTC</td><td>10491.0</td><td>6.7</td><td>0.9</td><td>7.8</td><td>8.9E-15</td><td>4.3E-14</td><td>MB231/MB468</td><td>NUC</td></tr>
<tr><td>>MI0015866|hsa-mir-4326&WithFlank&20|+|63286802|63286872@17.39.23&[MIMAT0016888&hsa-miR-4326&offsets|0|+3;m-339&20|+|63286818|63286841&offsets|0|-1]|TGTTCCTCTGTCTCCCAGACTCT</td><td>>MI0015866|hsa-mir-4326&WithFlank&20|+|63286802|63286872</td><td>17</td><td>39</td><td>23</td><td>TGTTCCTCTGTCTCCCAGACTCT</td><td>310.2</td><td>9.6</td><td>1.2</td><td>7.8</td><td>8.3E-15</td><td>4.3E-14</td><td>MB231/MB468</td><td>CYTO</td></tr>
<tr><td>>trnaMT_GluTTC_MT_-_14674_14742@40.58.19,trnalookalike8_GluTTC_5_-_93905172_93905240@40.58.19|TTGGTCGTGGTTGTAGTCC</td><td>>trnaMT_GluTTC_MT_-_14674_14742</td><td>40</td><td>58</td><td>19</td><td>TTGGTCGTGGTTGTAGTCC</td><td>793.3</td><td>7.7</td><td>1.0</td><td>7.8</td><td>8.3E-15</td><td>4.3E-14</td><td>MB231/MB468</td><td>CYTO</td></tr>
<tr><td>>trna16_ValTAC_11_-_59318460_59318532@1.30.30|GGTTCCATAGTGTAGCGGTTATCACGTCTG</td><td>>trna16_ValTAC_11_-_59318460_59318532</td><td>1</td><td>30</td><td>30</td><td>GGTTCCATAGTGTAGCGGTTATCACGTCTG</td><td>318.9</td><td>-7.6</td><td>1.0</td><td>-7.8</td><td>8.3E-15</td><td>4.3E-14</td><td>MB231/MB468</td><td>CYTO</td></tr>
<tr><td>>MI0000342|hsa-mir-200b&WithFlank&1|+|1167098|1167204@63.84.22(+1A)&[MIMAT0000318&hsa-miR-200b-3p&offsets|0|0(+1A);m-93&1|+|1167160|1167182&offsets|0|-1(+1A)]|TAATACTGCCTGGTAATGATGAA</td><td>>MI0000342|hsa-mir-200b&WithFlank&1|+|1167098|1167204</td><td>63</td><td>84</td><td>22(+1A)</td><td>TAATACTGCCTGGTAATGATGAA</td><td>238.2</td><td>-11.8</td><td>1.5</td><td>-7.8</td><td>7.4E-15</td><td>4.3E-14</td><td>MB231/MB468</td><td>MP</td></tr>
<tr><td>>hg38_21_+_8433172_8446622_RNA45SN1_45S_50ntFlanks@3853.3884.32|AACTGTGGTAATTCTAGAGCTAATACATGCCG</td><td>>hg38_21_+_8433172_8446622_RNA45SN1_45S_50ntFlanks</td><td>3853</td><td>3884</td><td>32</td><td>AACTGTGGTAATTCTAGAGCTAATACATGCCG</td><td>416.9</td><td>-4.2</td><td>0.5</td><td>-7.7</td><td>1.5E-14</td><td>4.3E-14</td><td>BT20/MB468</td><td>NUC</td></tr>
<tr><td>>MI0000239|hsa-mir-197&WithFlank&1|+|109598887|109598973@54.74.21(+1U)&[MIMAT0000227&hsa-miR-197-3p&offsets|0|-1(+1U);m-99&1|+|109598940|109598961&offsets|0|-1(+1U)]|TTCACCACCTTCTCCACCCAGT</td><td>>MI0000239|hsa-mir-197&WithFlank&1|+|109598887|109598973</td><td>54</td><td>74</td><td>21(+1U)</td><td>TTCACCACCTTCTCCACCCAGT</td><td>829.8</td><td>3.7</td><td>0.5</td><td>7.7</td><td>1.5E-14</td><td>4.4E-14</td><td>BT20/MB468</td><td>NUC</td></tr>
<tr><td>>hg38_21_+_8433172_8446622_RNA45SN1_45S_50ntFlanks@12678.12704.27|AATCCCGCCCAGGCGGAACGATACGGC</td><td>>hg38_21_+_8433172_8446622_RNA45SN1_45S_50ntFlanks</td><td>12678</td><td>12704</td><td>27</td><td>AATCCCGCCCAGGCGGAACGATACGGC</td><td>324.9</td><td>-12.2</td><td>1.6</td><td>-7.8</td><td>7.4E-15</td><td>4.4E-14</td><td>MB231/MB468</td><td>MP</td></tr>
<tr><td>>hg38_21_+_8433172_8446622_RNA45SN1_45S_50ntFlanks@3843.3869.27|CTACTTGGATAACTGTGGTAATTCTAG</td><td>>hg38_21_+_8433172_8446622_RNA45SN1_45S_50ntFlanks</td><td>3843</td><td>3869</td><td>27</td><td>CTACTTGGATAACTGTGGTAATTCTAG</td><td>286.0</td><td>-12.0</td><td>1.5</td><td>-7.8</td><td>7.4E-15</td><td>4.4E-14</td><td>MB231/MB468</td><td>MP</td></tr>
</table>

<table>
<tr><td>>hg38_21_+_8433172_8446622_RNA45SN1_45S_50ntFlanks@8246.8268.23|TTCCCGGAGTCGGGTTGCTTGGG</td><td>>hg38_21_+_8433172_8446622_RNA45SN1_45S_50ntFlanks</td><td>8246</td><td>8268</td><td>23</td><td>TTCCCGGAGTCGGGTTGCTTGGG</td><td>164.2</td><td>-11.2</td><td>1.4</td><td>-7.8</td><td>7.4E-15</td><td>4.4E-14</td><td>MB231/MB468</td><td>MP</td></tr>
<tr><td>>hg38_21_+_8433172_8446622_RNA45SN1_45S_50ntFlanks@9027.9065.39|CGTCTCCTCGTGGGGGGGCCGGGCCACCCCTCCCACGGC</td><td>>hg38_21_+_8433172_8446622_RNA45SN1_45S_50ntFlanks</td><td>9027</td><td>9065</td><td>39</td><td>CGTCTCCTCGTGGGGGGGCCGGGCCACCCCTCCCACGGC</td><td>176.3</td><td>-10.4</td><td>1.3</td><td>-7.8</td><td>9.0E-15</td><td>4.4E-14</td><td>MB231/MB468</td><td>NUC</td></tr>
<tr><td>>hg38_21_+_8433172_8446622_RNA45SN1_45S_50ntFlanks@12407.12436.30|TTTTGATCCTTCGATGTCGGCTCTTCCTAT</td><td>>hg38_21_+_8433172_8446622_RNA45SN1_45S_50ntFlanks</td><td>12407</td><td>12436</td><td>30</td><td>TTTTGATCCTTCGATGTCGGCTCTTCCTAT</td><td>3831.0</td><td>-8.2</td><td>1.1</td><td>-7.7</td><td>1.2E-14</td><td>4.4E-14</td><td>MB231/MB468</td><td>MITO</td></tr>
<tr><td>>hg38_21_+_8433172_8446622_RNA45SN1_45S_50ntFlanks@3738.3760.23|TCTCAAAGATTAAGCCATGCATG</td><td>>hg38_21_+_8433172_8446622_RNA45SN1_45S_50ntFlanks</td><td>3738</td><td>3760</td><td>23</td><td>TCTCAAAGATTAAGCCATGCATG</td><td>264.6</td><td>-11.9</td><td>1.5</td><td>-7.8</td><td>7.4E-15</td><td>4.4E-14</td><td>MB231/MB468</td><td>MP</td></tr>
<tr><td>>hg38_21_+_8433172_8446622_RNA45SN1_45S_50ntFlanks@7975.8015.41|CGCGACCTCAGATCAGACGTGGCGACCCGCTGAATTTAAGC</td><td>>hg38_21_+_8433172_8446622_RNA45SN1_45S_50ntFlanks</td><td>7975</td><td>8015</td><td>41</td><td>CGCGACCTCAGATCAGACGTGGCGACCCGCTGAATTTAAGC</td><td>78196.9</td><td>6.8</td><td>0.9</td><td>7.8</td><td>7.4E-15</td><td>4.4E-14</td><td>BT20/MB468</td><td>MITO</td></tr>
<tr><td>>MI0003628|hsa-mir-615&WithFlank&12|+|54033944|54034051@67.87.21&[MIMAT0003283&hsa-miR-615-3p&offsets|0|-1;m-495&12|+|54034010|54034030&offsets|0|0]|TCCGAGCCTGGGTCTCCCTCT</td><td>>MI0003628|hsa-mir-615&WithFlank&12|+|54033944|54034051</td><td>67</td><td>87</td><td>21</td><td>TCCGAGCCTGGGTCTCCCTCT</td><td>1721.8</td><td>4.9</td><td>0.6</td><td>8.0</td><td>1.1E-15</td><td>4.4E-14</td><td>BT20/MB231</td><td>NUC</td></tr>
<tr><td>>MI0000446|hsa-mir-125b-1&WithFlank&11|-|122099751|122099850@21.39.19;MI0000470|hsa-mir-125b-2&WithFlank&21|+|16590231|16590331@23.41.19&[MIMAT0000423&hsa-miR-125b-5p&offsets|0|-3;m-67&11|-|122099809|122099830&offsets|0|-3];[MIMAT0000423_1&hsa-miR-125b-5p&offsets|0|-3;m-68&21|+|16590253|16590274&offsets|0|-3]|TCCCTGAGACCCTAACTTG</td><td>>MI0000446|hsa-mir-125b-1&WithFlank&11|-|122099751|122099850</td><td>21</td><td>39</td><td>19</td><td>TCCCTGAGACCCTAACTTG</td><td>1600.9</td><td>9.2</td><td>1.2</td><td>7.8</td><td>9.2E-15</td><td>4.4E-14</td><td>MB231/MB468</td><td>NUC</td></tr>
<tr><td>>trnaMT_GluTTC_MT_-_14674_14742@27.65.39,trnalookalike8_GluTTC_5_-_93905172_93905240@27.65.39|GTTTTTCATATCATTGGTCGTGGTTGTAGTCCGTGCGAG</td><td>>trnaMT_GluTTC_MT_-_14674_14742</td><td>27</td><td>65</td><td>39</td><td>GTTTTTCATATCATTGGTCGTGGTTGTAGTCCGTGCGAG</td><td>484.9</td><td>-13.3</td><td>1.7</td><td>-7.7</td><td>1.5E-14</td><td>4.4E-14</td><td>BT20/MB468</td><td>NUC</td></tr>
<tr><td>>hg38_21_+_8433172_8446622_RNA45SN1_45S_50ntFlanks@4047.4081.35|ATTCGAACGTCTGCCCTATCAACTTTCGATGGTAG</td><td>>hg38_21_+_8433172_8446622_RNA45SN1_45S_50ntFlanks</td><td>4047</td><td>4081</td><td>35</td><td>ATTCGAACGTCTGCCCTATCAACTTTCGATGGTAG</td><td>539.8</td><td>-11.2</td><td>1.4</td><td>-7.8</td><td>7.5E-15</td><td>4.4E-14</td><td>BT20/MB468</td><td>MITO</td></tr>
<tr><td>>am_trnaMT_PheGAA_MT_+_577_647@41.62.22|GTTTAGACGGGCTCACATCACC</td><td>>am_trnaMT_PheGAA_MT_+_577_647</td><td>41</td><td>62</td><td>22</td><td>GTTTAGACGGGCTCACATCACC</td><td>673.2</td><td>5.2</td><td>0.7</td><td>7.7</td><td>1.2E-14</td><td>4.4E-14</td><td>MB231/MB468</td><td>MITO</td></tr>
<tr><td>>hg38_21_+_8433172_8446622_RNA45SN1_45S_50ntFlanks@475.514.40|CTTCGTGATCGATGTGGTGACGTCGTGCTCTCCCGGGCCG</td><td>>hg38_21_+_8433172_8446622_RNA45SN1_45S_50ntFlanks</td><td>475</td><td>514</td><td>40</td><td>CTTCGTGATCGATGTGGTGACGTCGTGCTCTCCCGGGCCG</td><td>527.8</td><td>7.9</td><td>1.0</td><td>7.8</td><td>8.5E-15</td><td>4.4E-14</td><td>MB231/MB468</td><td>CYTO</td></tr>
<tr><td>>hg38_21_+_8433172_8446622_RNA45SN1_45S_50ntFlanks@11323.11341.19|CGTCTCGTCGCGCGCGCGT</td><td>>hg38_21_+_8433172_8446622_RNA45SN1_45S_50ntFlanks</td><td>11323</td><td>11341</td><td>19</td><td>CGTCTCGTCGCGCGCGCGT</td><td>230.0</td><td>-11.6</td><td>1.5</td><td>-7.7</td><td>1.2E-14</td><td>4.4E-14</td><td>MB231/MB468</td><td>MITO</td></tr>
<tr><td>>hg38_21_+_8433172_8446622_RNA45SN1_45S_50ntFlanks@6646.6690.45|TCGTACGACTCTTAGCGGTGGATCACTCGGCTCGTGCGTCGATGA</td><td>>hg38_21_+_8433172_8446622_RNA45SN1_45S_50ntFlanks</td><td>6646</td><td>6690</td><td>45</td><td>TCGTACGACTCTTAGCGGTGGATCACTCGGCTCGTGCGTCGATGA</td><td>1727.1</td><td>4.0</td><td>0.5</td><td>7.8</td><td>7.5E-15</td><td>4.4E-14</td><td>BT20/MB468</td><td>MITO</td></tr>
<tr><td>>MI0000279|hsa-mir-196a-2&WithFlank&12|+|53991732|53991853@31.52.22;MI0000238|hsa-mir-196a-1&WithFlank&17|-|48632484|48632565@13.34.22&[MIMAT0000226&hsa-miR-196a-5p&offsets|0|0;m-261&12|+|53991762|53991783&offsets|0|0];[MIMAT0000226_1&hsa-miR-196a-5p&offsets|0|0;m-268&17|-|48632532|48632553&offsets|0|0]|TAGGTAGTTTCATGTTGTTGGG</td><td>>MI0000279|hsa-mir-196a-2&WithFlank&12|+|53991732|53991853</td><td>31</td><td>52</td><td>22</td><td>TAGGTAGTTTCATGTTGTTGGG</td><td>235.5</td><td>-11.7</td><td>1.5</td><td>-7.8</td><td>7.6E-15</td><td>4.4E-14</td><td>MB231/MB468</td><td>MP</td></tr>
<tr><td>>hg38_21_+_8433172_8446622_RNA45SN1_45S_50ntFlanks@12994.13015.22|CCTCGCTGCGATCTATTGAAAG</td><td>>hg38_21_+_8433172_8446622_RNA45SN1_45S_50ntFlanks</td><td>12994</td><td>13015</td><td>22</td><td>CCTCGCTGCGATCTATTGAAAG</td><td>2208.3</td><td>-8.7</td><td>1.1</td><td>-7.7</td><td>1.6E-14</td><td>4.4E-14</td><td>BT20/MB468</td><td>NUC</td></tr>
<tr><td>>hg38_21_+_8433172_8446622_RNA45SN1_45S_50ntFlanks@12488.12519.32|AACGTGAGCTGGGTTTAGACCGTCGTGAGACA</td><td>>hg38_21_+_8433172_8446622_RNA45SN1_45S_50ntFlanks</td><td>12488</td><td>12519</td><td>32</td><td>AACGTGAGCTGGGTTTAGACCGTCGTGAGACA</td><td>175.7</td><td>-11.3</td><td>1.5</td><td>-7.8</td><td>7.6E-15</td><td>4.5E-14</td><td>MB231/MB468</td><td>MP</td></tr>
<tr><td>>hg38_21_+_8433172_8446622_RNA45SN1_45S_50ntFlanks@7975.8009.35|CGCGACCTCAGATCAGACGTGGCGACCCGCTGAAT</td><td>>hg38_21_+_8433172_8446622_RNA45SN1_45S_50ntFlanks</td><td>7975</td><td>8009</td><td>35</td><td>CGCGACCTCAGATCAGACGTGGCGACCCGCTGAAT</td><td>858753.6</td><td>5.9</td><td>0.8</td><td>7.8</td><td>7.6E-15</td><td>4.5E-14</td><td>BT20/MB468</td><td>MITO</td></tr>
<tr><td>>hg38_21_+_8433172_8446622_RNA45SN1_45S_50ntFlanks@10362.10390.29|AAGGCCGAAGTGGAGAAGGGTTCCATGTG</td><td>>hg38_21_+_8433172_8446622_RNA45SN1_45S_50ntFlanks</td><td>10362</td><td>10390</td><td>29</td><td>AAGGCCGAAGTGGAGAAGGGTTCCATGTG</td><td>5570.3</td><td>-12.0</td><td>1.6</td><td>-7.7</td><td>1.6E-14</td><td>4.5E-14</td><td>BT20/MB468</td><td>NUC</td></tr>
<tr><td>>hg38_21_+_8433172_8446622_RNA45SN1_45S_50ntFlanks@5446.5467.22|CGGATCGGCCCCGCCGGGGTCG</td><td>>hg38_21_+_8433172_8446622_RNA45SN1_45S_50ntFlanks</td><td>5446</td><td>5467</td><td>22</td><td>CGGATCGGCCCCGCCGGGGTCG</td><td>369.2</td><td>-8.3</td><td>1.1</td><td>-7.8</td><td>8.6E-15</td><td>4.5E-14</td><td>MB231/MB468</td><td>CYTO</td></tr>
<tr><td>>MI0003137|hsa-mir-193b&WithFlank&16|+|14303961|14304055@57.78.22&[MIMAT0002819&hsa-miR-193b-3p&offsets|0|0;m-166&16|+|14304017|14304038&offsets|0|0]|AACTGGCCCTCAAAGTCCCGCT</td><td>>MI0003137|hsa-mir-193b&WithFlank&16|+|14303961|14304055</td><td>57</td><td>78</td><td>22</td><td>AACTGGCCCTCAAAGTCCCGCT</td><td>5191.5</td><td>5.3</td><td>0.7</td><td>8.0</td><td>1.1E-15</td><td>4.5E-14</td><td>BT20/MB231</td><td>NUC</td></tr>
<tr><td>>hg38_21_+_8433172_8446622_RNA45SN1_45S_50ntFlanks@11234.11254.21|CCTTCCCGTGGATCGCCCCAG</td><td>>hg38_21_+_8433172_8446622_RNA45SN1_45S_50ntFlanks</td><td>11234</td><td>11254</td><td>21</td><td>CCTTCCCGTGGATCGCCCCAG</td><td>277.8</td><td>-12.0</td><td>1.5</td><td>-7.8</td><td>7.6E-15</td><td>4.5E-14</td><td>MB231/MB468</td><td>MP</td></tr>
<tr><td>>MI0000285|hsa-mir-205&WithFlank&1|+|209432127|209432248@40.59.20(+1A)&[MIMAT0000266&hsa-miR-205-5p&offsets|0|-2(+1A);m-351&1|+|209432166|209432187&offsets|0|-2(+1A)]|TCCTTCATTCCACCGGAGTCA</td><td>>MI0000285|hsa-mir-205&WithFlank&1|+|209432127|209432248</td><td>40</td><td>59</td><td>20(+1A)</td><td>TCCTTCATTCCACCGGAGTCA</td><td>274.4</td><td>11.7</td><td>1.5</td><td>8.0</td><td>1.1E-15</td><td>4.5E-14</td><td>BT20/MB231</td><td>MITO</td></tr>
<tr><td>>hg38_21_+_8433172_8446622_RNA45SN1_45S_50ntFlanks@12640.12659.20|ACCATCTGTGGGATTATGAC</td><td>>hg38_21_+_8433172_8446622_RNA45SN1_45S_50ntFlanks</td><td>12640</td><td>12659</td><td>20</td><td>ACCATCTGTGGGATTATGAC</td><td>1827.3</td><td>-9.1</td><td>1.2</td><td>-7.8</td><td>7.7E-15</td><td>4.5E-14</td><td>MB231/MB468</td><td>MP</td></tr>
<tr><td>>hg38_21_+_8433172_8446622_RNA45SN1_45S_50ntFlanks@12448.12480.33|AGAATTCACCAAGCGTTGGATTGTTCACCCACT</td><td>>hg38_21_+_8433172_8446622_RNA45SN1_45S_50ntFlanks</td><td>12448</td><td>12480</td><td>33</td><td>AGAATTCACCAAGCGTTGGATTGTTCACCCACT</td><td>565.7</td><td>7.6</td><td>1.0</td><td>7.8</td><td>8.7E-15</td><td>4.5E-14</td><td>MB231/MB468</td><td>CYTO</td></tr>
<tr><td>>hg38_21_+_8433172_8446622_RNA45SN1_45S_50ntFlanks@11668.11707.40|TGCCCAGTGCTCTGAATGTCAAAGTGAAGAAATTCAATGA</td><td>>hg38_21_+_8433172_8446622_RNA45SN1_45S_50ntFlanks</td><td>11668</td><td>11707</td><td>40</td><td>TGCCCAGTGCTCTGAATGTCAAAGTGAAGAAATTCAATGA</td><td>447.8</td><td>7.7</td><td>1.0</td><td>7.7</td><td>1.0E-14</td><td>4.5E-14</td><td>BT20/MB468</td><td>CYTO</td></tr>
<tr><td>>MI0000471|hsa-mir-126&WithFlank&9|+|136670596|136670692@59.79.21&[MIMAT0000445&hsa-miR-126-3p&offsets|+1|0;m-40&9|+|136670653|136670674&offsets|+1|0]|CGTACCGTGAGTAATAATGCG</td><td>>MI0000471|hsa-mir-126&WithFlank&9|+|136670596|136670692</td><td>59</td><td>79</td><td>21</td><td>CGTACCGTGAGTAATAATGCG</td><td>178.5</td><td>9.4</td><td>1.2</td><td>7.7</td><td>9.4E-15</td><td>4.5E-14</td><td>MB231/MB468</td><td>NUC</td></tr>
<tr><td>>hg38_21_+_8433172_8446622_RNA45SN1_45S_50ntFlanks@11670.11703.34|CCCAGTGCTCTGAATGTCAAAGTGAAGAAATTCA</td><td>>hg38_21_+_8433172_8446622_RNA45SN1_45S_50ntFlanks</td><td>11670</td><td>11703</td><td>34</td><td>CCCAGTGCTCTGAATGTCAAAGTGAAGAAATTCA</td><td>1677.2</td><td>-4.7</td><td>0.6</td><td>-7.7</td><td>9.3E-15</td><td>4.5E-14</td><td>MB231/MB468</td><td>NUC</td></tr>
<tr><td>>hg38_21_+_8433172_8446622_RNA45SN1_45S_50ntFlanks@11883.11904.22|TGTTGAGCTTGACTCTAGTCTG</td><td>>hg38_21_+_8433172_8446622_RNA45SN1_45S_50ntFlanks</td><td>11883</td><td>11904</td><td>22</td><td>TGTTGAGCTTGACTCTAGTCTG</td><td>183.8</td><td>-11.4</td><td>1.5</td><td>-7.8</td><td>7.7E-15</td><td>4.5E-14</td><td>MB231/MB468</td><td>MP</td></tr>
<tr><td>>hg38_21_+_8433172_8446622_RNA45SN1_45S_50ntFlanks@8317.8356.40|CACGAGACCGATAGTCAACAAGTACCGTAAGGGAAAGTTG</td><td>>hg38_21_+_8433172_8446622_RNA45SN1_45S_50ntFlanks</td><td>8317</td><td>8356</td><td>40</td><td>CACGAGACCGATAGTCAACAAGTACCGTAAGGGAAAGTTG</td><td>1378.6</td><td>-10.6</td><td>1.4</td><td>-7.7</td><td>1.6E-14</td><td>4.5E-14</td><td>BT20/MB468</td><td>NUC</td></tr>
<tr><td>>hg38_21_+_8433172_8446622_RNA45SN1_45S_50ntFlanks@10843.10862.20|TAACTTCGGGATAAGGATTG</td><td>>hg38_21_+_8433172_8446622_RNA45SN1_45S_50ntFlanks</td><td>10843</td><td>10862</td><td>20</td><td>TAACTTCGGGATAAGGATTG</td><td>249.7</td><td>-11.8</td><td>1.5</td><td>-7.8</td><td>7.7E-15</td><td>4.5E-14</td><td>MB231/MB468</td><td>MP</td></tr>
<tr><td>>MI0000298|hsa-mir-221&WithFlank&X|-|45746151|45746272@71.92.22&[MIMAT0000278&hsa-miR-221-3p&offsets|0|-1;m-91&X|-|45746181|45746202&offsets|0|0]|AGCTACATTGTCTGCTGGGTTT</td><td>>MI0000298|hsa-mir-221&WithFlank&X|-|45746151|45746272</td><td>71</td><td>92</td><td>22</td><td>AGCTACATTGTCTGCTGGGTTT</td><td>32439.1</td><td>7.1</td><td>0.9</td><td>7.7</td><td>1.3E-14</td><td>4.5E-14</td><td>MB231/MB468</td><td>MITO</td></tr>
<tr><td>>trnaMT_ValTAC_MT_+_1602_1670@49.72.24|AACTTAACTTGACCGCTCTGACCA</td><td>>trnaMT_ValTAC_MT_+_1602_1670</td><td>49</td><td>72</td><td>24</td><td>AACTTAACTTGACCGCTCTGACCA</td><td>1668.7</td><td>8.7</td><td>1.1</td><td>7.7</td><td>1.0E-14</td><td>4.5E-14</td><td>BT20/MB468</td><td>CYTO</td></tr>
<tr><td>>hg38_21_+_8433172_8446622_RNA45SN1_45S_50ntFlanks@12942.12966.25|TTCGTAGACGACCTGCTTCTGGGTC</td><td>>hg38_21_+_8433172_8446622_RNA45SN1_45S_50ntFlanks</td><td>12942</td><td>12966</td><td>25</td><td>TTCGTAGACGACCTGCTTCTGGGTC</td><td>1177.6</td><td>-7.3</td><td>0.9</td><td>-7.8</td><td>8.8E-15</td><td>4.6E-14</td><td>MB231/MB468</td><td>CYTO</td></tr>
<tr><td>>hg38_21_+_8433172_8446622_RNA45SN1_45S_50ntFlanks@8170.8188.19|CAAGTCCTTCTGATCGAGG</td><td>>hg38_21_+_8433172_8446622_RNA45SN1_45S_50ntFlanks</td><td>8170</td><td>8188</td><td>19</td><td>CAAGTCCTTCTGATCGAGG</td><td>618.0</td><td>-12.4</td><td>1.5</td><td>-8.2</td><td>3.1E-16</td><td>4.6E-14</td><td>BT20/MB468</td><td>MP</td></tr>
<tr><td>>MI0000285|hsa-mir-205&WithFlank&1|+|209432127|209432248@40.59.20&[MIMAT0000266&hsa-miR-205-5p&offsets|0|-2;m-351&1|+|209432166|209432187&offsets|0|-2]|TCCTTCATTCCACCGGAGTC</td><td>>MI0000285|hsa-mir-205&WithFlank&1|+|209432127|209432248</td><td>40</td><td>59</td><td>20</td><td>TCCTTCATTCCACCGGAGTC</td><td>488.7</td><td>12.7</td><td>1.6</td><td>8.0</td><td>1.2E-15</td><td>4.6E-14</td><td>BT20/MB231</td><td>NUC</td></tr>
<tr><td>>hg38_21_+_8433172_8446622_RNA45SN1_45S_50ntFlanks@12435.12455.21|ATCATTGTGAAGCAGAATTCA</td><td>>hg38_21_+_8433172_8446622_RNA45SN1_45S_50ntFlanks</td><td>12435</td><td>12455</td><td>21</td><td>ATCATTGTGAAGCAGAATTCA</td><td>1653.8</td><td>-10.0</td><td>1.3</td><td>-7.8</td><td>8.8E-15</td><td>4.6E-14</td><td>MB231/MB468</td><td>CYTO</td></tr>
<tr><td>>hg38_21_+_8433172_8446622_RNA45SN1_45S_50ntFlanks@8115.8141.27|CGCGGGACATGTGGCGTACGGAAGACC</td><td>>hg38_21_+_8433172_8446622_RNA45SN1_45S_50ntFlanks</td><td>8115</td><td>8141</td><td>27</td><td>CGCGGGACATGTGGCGTACGGAAGACC</td><td>466.3</td><td>12.5</td><td>1.6</td><td>8.0</td><td>1.1E-15</td><td>4.6E-14</td><td>BT20/MB231</td><td>MITO</td></tr>
<tr><td>>hg38_21_+_8433172_8446622_RNA45SN1_45S_50ntFlanks@8107.8141.35|CGGCGGGGCGCGGGACATGTGGCGTACGGAAGACC</td><td>>hg38_21_+_8433172_8446622_RNA45SN1_45S_50ntFlanks</td><td>8107</td><td>8141</td><td>35</td><td>CGGCGGGGCGCGGGACATGTGGCGTACGGAAGACC</td><td>392.6</td><td>-12.4</td><td>1.6</td><td>-7.7</td><td>1.3E-14</td><td>4.6E-14</td><td>MB231/MB468</td><td>MITO</td></tr>
<tr><td>>hg38_21_+_8433172_8446622_RNA45SN1_45S_50ntFlanks@7986.8005.20|ATCAGACGTGGCGACCCGCT</td><td>>hg38_21_+_8433172_8446622_RNA45SN1_45S_50ntFlanks</td><td>7986</td><td>8005</td><td>20</td><td>ATCAGACGTGGCGACCCGCT</td><td>3095.0</td><td>-10.3</td><td>1.3</td><td>-7.8</td><td>7.9E-15</td><td>4.6E-14</td><td>MB231/MB468</td><td>MP</td></tr>
<tr><td>>hg38_21_+_8433172_8446622_RNA45SN1_45S_50ntFlanks@9572.9603.32|CAAATCGGTCGTCCGACCTGGGTATAGGGGCG</td><td>>hg38_21_+_8433172_8446622_RNA45SN1_45S_50ntFlanks</td><td>9572</td><td>9603</td><td>32</td><td>CAAATCGGTCGTCCGACCTGGGTATAGGGGCG</td><td>7697.8</td><td>-7.5</td><td>1.0</td><td>-7.8</td><td>7.9E-15</td><td>4.6E-14</td><td>MB231/MB468</td><td>MP</td></tr>
<tr><td>>MI0000462|hsa-mir-152&WithFlank&17|-|48037155|48037253@60.80.21(+1U)&[MIMAT0000438&hsa-miR-152-3p&offsets|0|0(+1U);m-97&17|-|48037174|48037194&offsets|0|0(+1U)]|TCAGTGCATGACAGAACTTGGT</td><td>>MI0000462|hsa-mir-152&WithFlank&17|-|48037155|48037253</td><td>60</td><td>80</td><td>21(+1U)</td><td>TCAGTGCATGACAGAACTTGGT</td><td>260.1</td><td>10.0</td><td>1.3</td><td>7.7</td><td>9.6E-15</td><td>4.6E-14</td><td>MB231/MB468</td><td>NUC</td></tr>
<tr><td>>MI0005756|hsa-mir-934&WithFlank&X|+|136550872|136550966@21.41.21(+1A)&[MIMAT0004977&hsa-miR-934&offsets|0|-1(+1A);m-943&X|+|136550892|136550913&offsets|0|-1(+1A)]|TGTCTACTACTGGAGACACTGA</td><td>>MI0005756|hsa-mir-934&WithFlank&X|+|136550872|136550966</td><td>21</td><td>41</td><td>21(+1A)</td><td>TGTCTACTACTGGAGACACTGA</td><td>412.8</td><td>12.5</td><td>1.6</td><td>8.0</td><td>1.2E-15</td><td>4.6E-14</td><td>BT20/MB231</td><td>NUC</td></tr>
<tr><td>>hg38_21_+_8433172_8446622_RNA45SN1_45S_50ntFlanks@6740.6772.33|CATCGACACTTCGAACGCACTTGCGGCCCCGGG</td><td>>hg38_21_+_8433172_8446622_RNA45SN1_45S_50ntFlanks</td><td>6740</td><td>6772</td><td>33</td><td>CATCGACACTTCGAACGCACTTGCGGCCCCGGG</td><td>487.4</td><td>-12.8</td><td>1.6</td><td>-7.8</td><td>7.9E-15</td><td>4.6E-14</td><td>MB231/MB468</td><td>MP</td></tr>
<tr><td>>hg38_21_+_8433172_8446622_RNA45SN1_45S_50ntFlanks@8472.8492.21|CGGCCCGGCGGATCTTTCCCG</td><td>>hg38_21_+_8433172_8446622_RNA45SN1_45S_50ntFlanks</td><td>8472</td><td>8492</td><td>21</td><td>CGGCCCGGCGGATCTTTCCCG</td><td>166.3</td><td>-11.2</td><td>1.4</td><td>-7.8</td><td>7.9E-15</td><td>4.6E-14</td><td>MB231/MB468</td><td>MP</td></tr>
<tr><td>>hg38_21_+_8433172_8446622_RNA45SN1_45S_50ntFlanks@11583.11601.19|CTGGTGCGGACCAGGGGAA</td><td>>hg38_21_+_8433172_8446622_RNA45SN1_45S_50ntFlanks</td><td>11583</td><td>11601</td><td>19</td><td>CTGGTGCGGACCAGGGGAA</td><td>739.9</td><td>-7.3</td><td>1.0</td><td>-7.7</td><td>1.3E-14</td><td>4.6E-14</td><td>MB231/MB468</td><td>MITO</td></tr>
<tr><td>>hg38_21_+_8433172_8446622_RNA45SN1_45S_50ntFlanks@11235.11253.19|CTTCCCGTGGATCGCCCCA</td><td>>hg38_21_+_8433172_8446622_RNA45SN1_45S_50ntFlanks</td><td>11235</td><td>11253</td><td>19</td><td>CTTCCCGTGGATCGCCCCA</td><td>1409.1</td><td>-5.7</td><td>0.7</td><td>-7.7</td><td>1.1E-14</td><td>4.6E-14</td><td>BT20/MB468</td><td>CYTO</td></tr>
<tr><td>>hg38_21_+_8433172_8446622_RNA45SN1_45S_50ntFlanks@12258.12284.27|GCTTGATCTTGATTTTCAGTACGAATA</td><td>>hg38_21_+_8433172_8446622_RNA45SN1_45S_50ntFlanks</td><td>12258</td><td>12284</td><td>27</td><td>GCTTGATCTTGATTTTCAGTACGAATA</td><td>722.4</td><td>-4.6</td><td>0.6</td><td>-7.7</td><td>9.7E-15</td><td>4.6E-14</td><td>MB231/MB468</td><td>NUC</td></tr>
<tr><td>>hg38_21_+_8433172_8446622_RNA45SN1_45S_50ntFlanks@5164.5185.22|CGTTCAGCCACCCGAGATTGAG</td><td>>hg38_21_+_8433172_8446622_RNA45SN1_45S_50ntFlanks</td><td>5164</td><td>5185</td><td>22</td><td>CGTTCAGCCACCCGAGATTGAG</td><td>331.1</td><td>-12.2</td><td>1.6</td><td>-7.8</td><td>8.0E-15</td><td>4.7E-14</td><td>MB231/MB468</td><td>MP</td></tr>
<tr><td>>hg38_21_+_8433172_8446622_RNA45SN1_45S_50ntFlanks@11641.11674.34|CGGCGGGTGTTGACGCGATGTGATTTCTGCCCAG</td><td>>hg38_21_+_8433172_8446622_RNA45SN1_45S_50ntFlanks</td><td>11641</td><td>11674</td><td>34</td><td>CGGCGGGTGTTGACGCGATGTGATTTCTGCCCAG</td><td>265.4</td><td>-11.8</td><td>1.5</td><td>-7.7</td><td>1.3E-14</td><td>4.7E-14</td><td>MB231/MB468</td><td>MITO</td></tr>
<tr><td>>hg38_21_+_8433172_8446622_RNA45SN1_45S_50ntFlanks@12685.12711.27|CCCAGGCGGAACGATACGGCAGCGCCG</td><td>>hg38_21_+_8433172_8446622_RNA45SN1_45S_50ntFlanks</td><td>12685</td><td>12711</td><td>27</td><td>CCCAGGCGGAACGATACGGCAGCGCCG</td><td>1074.1</td><td>-8.2</td><td>1.1</td><td>-7.7</td><td>1.3E-14</td><td>4.7E-14</td><td>MB231/MB468</td><td>MITO</td></tr>
</table>

<table>
<tr><td>>am_MI0031829|hsa-mir-9901&WithFlank&16|+|34163572|34163679@25.50.26&[MIMAT0039321&hsa-miR-9901&offsets|-5|0;TJU_CMC_MD2.ID01155.5p-miR&offsets|0|+3;m-2778&16|+|34163596|34163618&offsets|0|+3]|CGTCCCGGTCGCCGCGGTTCGCCGCC</td><td>>am_MI0031829|hsa-mir-9901&WithFlank&16|+|34163572|34163679</td><td>25</td><td>50</td><td>26</td><td>CGTCCCGGTCGCCGCGGTTCGCCGCC</td><td>242.2</td><td>-10.9</td><td>1.4</td><td>-7.7</td><td>9.7E-15</td><td>4.7E-14</td><td>MB231/MB468</td><td>NUC</td></tr>
<tr><td>>hg38_21_+_8433172_8446622_RNA45SN1_45S_50ntFlanks@7340.7365.26|CGTCCCGGTCGCCGCGGTTCGCCGCC</td><td>>hg38_21_+_8433172_8446622_RNA45SN1_45S_50ntFlanks</td><td>7340</td><td>7365</td><td>26</td><td>CGTCCCGGTCGCCGCGGTTCGCCGCC</td><td>242.2</td><td>-10.9</td><td>1.4</td><td>-7.7</td><td>9.7E-15</td><td>4.7E-14</td><td>MB231/MB468</td><td>NUC</td></tr>
<tr><td>>hg38_21_+_8433172_8446622_RNA45SN1_45S_50ntFlanks@9129.9151.23|CGGGTCGGGGGGCGGGGCGGACT</td><td>>hg38_21_+_8433172_8446622_RNA45SN1_45S_50ntFlanks</td><td>9129</td><td>9151</td><td>23</td><td>CGGGTCGGGGGGCGGGGCGGACT</td><td>309.1</td><td>-12.1</td><td>1.6</td><td>-7.8</td><td>8.0E-15</td><td>4.7E-14</td><td>MB231/MB468</td><td>MP</td></tr>
<tr><td>>hg38_21_+_8433172_8446622_RNA45SN1_45S_50ntFlanks@10846.10864.19|CTTCGGGATAAGGATTGGC</td><td>>hg38_21_+_8433172_8446622_RNA45SN1_45S_50ntFlanks</td><td>10846</td><td>10864</td><td>19</td><td>CTTCGGGATAAGGATTGGC</td><td>982.7</td><td>-10.1</td><td>1.3</td><td>-7.7</td><td>1.6E-14</td><td>4.7E-14</td><td>BT20/MB468</td><td>NUC</td></tr>
<tr><td>>MI0000234|hsa-mir-192&WithFlank&11|-|64891131|64891252@30.51.22(+1U)&[MIMAT0000222&hsa-miR-192-5p&offsets|0|+1(+1U);m-59&11|-|64891203|64891223&offsets|0|+1(+1U)]|CTGACCTATGAATTGACAGCCAT</td><td>>MI0000234|hsa-mir-192&WithFlank&11|-|64891131|64891252</td><td>30</td><td>51</td><td>22(+1U)</td><td>CTGACCTATGAATTGACAGCCAT</td><td>458.9</td><td>5.2</td><td>0.7</td><td>7.8</td><td>8.0E-15</td><td>4.7E-14</td><td>BT20/MB468</td><td>MITO</td></tr>
<tr><td>>hg38_21_+_8433172_8446622_RNA45SN1_45S_50ntFlanks@10423.10453.31|AGAGATGGGCGAGCGCCGTTCCGAAGGGACG</td><td>>hg38_21_+_8433172_8446622_RNA45SN1_45S_50ntFlanks</td><td>10423</td><td>10453</td><td>31</td><td>AGAGATGGGCGAGCGCCGTTCCGAAGGGACG</td><td>1422.6</td><td>-10.5</td><td>1.4</td><td>-7.7</td><td>1.6E-14</td><td>4.7E-14</td><td>BT20/MB468</td><td>NUC</td></tr>
<tr><td>>MI0000263|hsa-mir-7-1&WithFlank&9|-|83969742|83969863@30.54.25&[MIMAT0000252_2&hsa-miR-7-5p&offsets|0|+1;m-401&9|-|83969811|83969834&offsets|0|+1]|TGGAAGACTAGTGATTTTGTTGTTT</td><td>>MI0000263|hsa-mir-7-1&WithFlank&9|-|83969742|83969863</td><td>30</td><td>54</td><td>25</td><td>TGGAAGACTAGTGATTTTGTTGTTT</td><td>967.6</td><td>6.6</td><td>0.8</td><td>7.8</td><td>8.0E-15</td><td>4.7E-14</td><td>BT20/MB468</td><td>MITO</td></tr>
<tr><td>>trnaMT_PheGAA_MT_+_577_647@36.62.27|AAAATGTTTAGACGGGCTCACATCACC</td><td>>trnaMT_PheGAA_MT_+_577_647</td><td>36</td><td>62</td><td>27</td><td>AAAATGTTTAGACGGGCTCACATCACC</td><td>1477.7</td><td>9.4</td><td>1.2</td><td>7.8</td><td>8.0E-15</td><td>4.7E-14</td><td>BT20/MB468</td><td>MITO</td></tr>
<tr><td>>hg38_21_+_8433172_8446622_RNA45SN1_45S_50ntFlanks@6772.6799.28|GTTCCTCCCGGGGCTACGCCTGTCTGAG</td><td>>hg38_21_+_8433172_8446622_RNA45SN1_45S_50ntFlanks</td><td>6772</td><td>6799</td><td>28</td><td>GTTCCTCCCGGGGCTACGCCTGTCTGAG</td><td>694.5</td><td>-6.8</td><td>0.9</td><td>-7.8</td><td>9.1E-15</td><td>4.7E-14</td><td>MB231/MB468</td><td>CYTO</td></tr>
<tr><td>>MI0000272|hsa-mir-182&WithFlank&7|-|129770377|129770498@29.51.23(+1C)&[MIMAT0000259&hsa-miR-182-5p&offsets|0|-1(+1C);m-22&7|-|129770449|129770470&offsets|0|+1(+1C)]|TTTGGCAATGGTAGAACTCACACC</td><td>>MI0000272|hsa-mir-182&WithFlank&7|-|129770377|129770498</td><td>29</td><td>51</td><td>23(+1C)</td><td>TTTGGCAATGGTAGAACTCACACC</td><td>7443.8</td><td>3.9</td><td>0.5</td><td>8.0</td><td>1.3E-15</td><td>4.7E-14</td><td>BT20/MB231</td><td>NUC</td></tr>
<tr><td>>hg38_21_+_8433172_8446622_RNA45SN1_45S_50ntFlanks@10099.10117.19|CGCGCGCGCGTGTGGTGTG</td><td>>hg38_21_+_8433172_8446622_RNA45SN1_45S_50ntFlanks</td><td>10099</td><td>10117</td><td>19</td><td>CGCGCGCGCGTGTGGTGTG</td><td>402.7</td><td>-7.9</td><td>1.0</td><td>-7.8</td><td>9.1E-15</td><td>4.7E-14</td><td>MB231/MB468</td><td>CYTO</td></tr>
<tr><td>>hg38_21_+_8433172_8446622_RNA45SN1_45S_50ntFlanks@12385.12414.30|AAGCGTTCATAGCGACGTCGCTTTTTGATC</td><td>>hg38_21_+_8433172_8446622_RNA45SN1_45S_50ntFlanks</td><td>12385</td><td>12414</td><td>30</td><td>AAGCGTTCATAGCGACGTCGCTTTTTGATC</td><td>528.2</td><td>-12.8</td><td>1.7</td><td>-7.7</td><td>1.3E-14</td><td>4.7E-14</td><td>MB231/MB468</td><td>MITO</td></tr>
<tr><td>>hg38_21_+_8433172_8446622_RNA45SN1_45S_50ntFlanks@11799.11820.22|AGATTCCCACTGTCCCTACCTA</td><td>>hg38_21_+_8433172_8446622_RNA45SN1_45S_50ntFlanks</td><td>11799</td><td>11820</td><td>22</td><td>AGATTCCCACTGTCCCTACCTA</td><td>197.2</td><td>-10.6</td><td>1.4</td><td>-7.7</td><td>9.8E-15</td><td>4.7E-14</td><td>MB231/MB468</td><td>NUC</td></tr>
<tr><td>>hg38_21_+_8433172_8446622_RNA45SN1_45S_50ntFlanks@12221.12241.21|GACAGAAACCTCCCGTGGAGC</td><td>>hg38_21_+_8433172_8446622_RNA45SN1_45S_50ntFlanks</td><td>12221</td><td>12241</td><td>21</td><td>GACAGAAACCTCCCGTGGAGC</td><td>194.7</td><td>-11.5</td><td>1.5</td><td>-7.8</td><td>8.1E-15</td><td>4.7E-14</td><td>MB231/MB468</td><td>MP</td></tr>
<tr><td>>hg38_21_+_8433172_8446622_RNA45SN1_45S_50ntFlanks@6743.6770.28|CGACACTTCGAACGCACTTGCGGCCCCG</td><td>>hg38_21_+_8433172_8446622_RNA45SN1_45S_50ntFlanks</td><td>6743</td><td>6770</td><td>28</td><td>CGACACTTCGAACGCACTTGCGGCCCCG</td><td>3089.8</td><td>-9.9</td><td>1.3</td><td>-7.7</td><td>1.3E-14</td><td>4.7E-14</td><td>MB231/MB468</td><td>MITO</td></tr>
<tr><td>>hg38_21_+_8433172_8446622_RNA45SN1_45S_50ntFlanks@10412.10430.19|AGTCGGTCCTGAGAGATGG</td><td>>hg38_21_+_8433172_8446622_RNA45SN1_45S_50ntFlanks</td><td>10412</td><td>10430</td><td>19</td><td>AGTCGGTCCTGAGAGATGG</td><td>348.2</td><td>-7.8</td><td>1.0</td><td>-7.8</td><td>9.2E-15</td><td>4.7E-14</td><td>MB231/MB468</td><td>CYTO</td></tr>
<tr><td>>hg38_21_+_8433172_8446622_RNA45SN1_45S_50ntFlanks@6706.6732.27|GCGAGAATTAATGTGAATTGCAGGACA</td><td>>hg38_21_+_8433172_8446622_RNA45SN1_45S_50ntFlanks</td><td>6706</td><td>6732</td><td>27</td><td>GCGAGAATTAATGTGAATTGCAGGACA</td><td>203.8</td><td>8.4</td><td>1.1</td><td>7.8</td><td>9.2E-15</td><td>4.7E-14</td><td>MB231/MB468</td><td>CYTO</td></tr>
<tr><td>>hg38_21_+_8433172_8446622_RNA45SN1_45S_50ntFlanks@12071.12095.25|CCCCGAGGGGCTCTCGCTTCTGGCG</td><td>>hg38_21_+_8433172_8446622_RNA45SN1_45S_50ntFlanks</td><td>12071</td><td>12095</td><td>25</td><td>CCCCGAGGGGCTCTCGCTTCTGGCG</td><td>2873.8</td><td>-6.0</td><td>0.8</td><td>-7.7</td><td>1.1E-14</td><td>4.7E-14</td><td>BT20/MB468</td><td>CYTO</td></tr>
<tr><td>>MI0000093|hsa-mir-92a-1&WithFlank&13|+|91351308|91351397@54.74.21(+1C);MI0000094|hsa-mir-92a-2&WithFlank&X|-|134169532|134169618@54.74.21(+1C)&[MIMAT0000092&hsa-miR-92a-3p&offsets|0|-1(+1C);m-9&13|+|91351361|91351382&offsets|0|-1(+1C)];[MIMAT0000092_1&hsa-miR-92a-3p&offsets|0|-1(+1C);m-11&X|-|134169544|134169565&offsets|0|-1(+1C)]|TATTGCACTTGTCCCGGCCTGC</td><td>>MI0000093|hsa-mir-92a-1&WithFlank&13|+|91351308|91351397</td><td>54</td><td>74</td><td>21(+1C)</td><td>TATTGCACTTGTCCCGGCCTGC</td><td>9145.7</td><td>6.6</td><td>0.8</td><td>7.7</td><td>9.2E-15</td><td>4.8E-14</td><td>MB231/MB468</td><td>CYTO</td></tr>
<tr><td>>hg38_21_+_8433172_8446622_RNA45SN1_45S_50ntFlanks@12944.12972.29|CGTAGACGACCTGCTTCTGGGTCGGGGTT</td><td>>hg38_21_+_8433172_8446622_RNA45SN1_45S_50ntFlanks</td><td>12944</td><td>12972</td><td>29</td><td>CGTAGACGACCTGCTTCTGGGTCGGGGTT</td><td>257.7</td><td>-11.8</td><td>1.5</td><td>-7.7</td><td>1.3E-14</td><td>4.8E-14</td><td>MB231/MB468</td><td>MITO</td></tr>
<tr><td>>hg38_21_+_8433172_8446622_RNA45SN1_45S_50ntFlanks@12202.12240.39|CTAAGGCGAGCTCAGGGAGGACAGAAACCTCCCGTGGAG</td><td>>hg38_21_+_8433172_8446622_RNA45SN1_45S_50ntFlanks</td><td>12202</td><td>12240</td><td>39</td><td>CTAAGGCGAGCTCAGGGAGGACAGAAACCTCCCGTGGAG</td><td>512.3</td><td>-12.8</td><td>1.7</td><td>-7.7</td><td>1.3E-14</td><td>4.8E-14</td><td>MB231/MB468</td><td>MITO</td></tr>
<tr><td>>hg38_21_+_8433172_8446622_RNA45SN1_45S_50ntFlanks@12071.12110.40|CCCCGAGGGGCTCTCGCTTCTGGCGCCAAGCGCCCGGCCG</td><td>>hg38_21_+_8433172_8446622_RNA45SN1_45S_50ntFlanks</td><td>12071</td><td>12110</td><td>40</td><td>CCCCGAGGGGCTCTCGCTTCTGGCGCCAAGCGCCCGGCCG</td><td>663.5</td><td>-9.2</td><td>1.2</td><td>-7.7</td><td>1.3E-14</td><td>4.8E-14</td><td>MB231/MB468</td><td>MITO</td></tr>
<tr><td>>hg38_21_+_8433172_8446622_RNA45SN1_45S_50ntFlanks@10404.10429.26|CATGGGTCAGTCGGTCCTGAGAGATG</td><td>>hg38_21_+_8433172_8446622_RNA45SN1_45S_50ntFlanks</td><td>10404</td><td>10429</td><td>26</td><td>CATGGGTCAGTCGGTCCTGAGAGATG</td><td>1729.7</td><td>-5.4</td><td>0.7</td><td>-7.7</td><td>1.4E-14</td><td>4.8E-14</td><td>MB231/MB468</td><td>MITO</td></tr>
<tr><td>>hg38_21_+_8433172_8446622_RNA45SN1_45S_50ntFlanks@10689.10710.22|GCGTCCGGTGAGCTCTCGCTGG</td><td>>hg38_21_+_8433172_8446622_RNA45SN1_45S_50ntFlanks</td><td>10689</td><td>10710</td><td>22</td><td>GCGTCCGGTGAGCTCTCGCTGG</td><td>274.7</td><td>-12.0</td><td>1.5</td><td>-7.8</td><td>8.2E-15</td><td>4.8E-14</td><td>MB231/MB468</td><td>MP</td></tr>
<tr><td>>hg38_21_+_8433172_8446622_RNA45SN1_45S_50ntFlanks@10391.10411.21|AACAGCAGTTGAACATGGGTC</td><td>>hg38_21_+_8433172_8446622_RNA45SN1_45S_50ntFlanks</td><td>10391</td><td>10411</td><td>21</td><td>AACAGCAGTTGAACATGGGTC</td><td>261.8</td><td>-3.8</td><td>0.5</td><td>-7.7</td><td>1.0E-14</td><td>4.8E-14</td><td>MB231/MB468</td><td>NUC</td></tr>
<tr><td>>hg38_21_+_8433172_8446622_RNA45SN1_45S_50ntFlanks@4725.4759.35|TTAATCAAGAACGAAAGTCGGAGGTTCGAAGACGA</td><td>>hg38_21_+_8433172_8446622_RNA45SN1_45S_50ntFlanks</td><td>4725</td><td>4759</td><td>35</td><td>TTAATCAAGAACGAAAGTCGGAGGTTCGAAGACGA</td><td>672.1</td><td>-7.6</td><td>1.0</td><td>-7.7</td><td>1.4E-14</td><td>4.8E-14</td><td>MB231/MB468</td><td>MITO</td></tr>
<tr><td>>MI0000102|hsa-mir-100&WithFlank&11|-|122152223|122152314@19.38.20(+1C)&[MIMAT0000098&hsa-miR-100-5p&offsets|0|-2(+1C);m-27&11|-|122152276|122152296&offsets|0|-1(+1C)]|AACCCGTAGATCCGAACTTGC</td><td>>MI0000102|hsa-mir-100&WithFlank&11|-|122152223|122152314</td><td>19</td><td>38</td><td>20(+1C)</td><td>AACCCGTAGATCCGAACTTGC</td><td>2731.7</td><td>15.7</td><td>2.0</td><td>7.7</td><td>1.1E-14</td><td>4.8E-14</td><td>BT20/MB468</td><td>CYTO</td></tr>
<tr><td>>hg38_21_+_8433172_8446622_RNA45SN1_45S_50ntFlanks@11592.11609.18|ACCAGGGGAATCCGACTG</td><td>>hg38_21_+_8433172_8446622_RNA45SN1_45S_50ntFlanks</td><td>11592</td><td>11609</td><td>18</td><td>ACCAGGGGAATCCGACTG</td><td>256.3</td><td>-11.9</td><td>1.5</td><td>-7.8</td><td>8.2E-15</td><td>4.8E-14</td><td>MB231/MB468</td><td>MP</td></tr>
<tr><td>>hg38_21_+_8433172_8446622_RNA45SN1_45S_50ntFlanks@11718.11737.20|AACGGCGGGAGTAACTATGA</td><td>>hg38_21_+_8433172_8446622_RNA45SN1_45S_50ntFlanks</td><td>11718</td><td>11737</td><td>20</td><td>AACGGCGGGAGTAACTATGA</td><td>201.6</td><td>-11.5</td><td>1.5</td><td>-7.8</td><td>8.3E-15</td><td>4.8E-14</td><td>MB231/MB468</td><td>MP</td></tr>
<tr><td>>am_MI0000786|hsa-mir-378a&WithFlank&5|+|149732819|149732896@50.69.20(+1U)&[MIMAT0000732&hsa-miR-378a-3p&offsets|+1|-1(+1U);m-44&5|+|149732867|149732888&offsets|+1|-1(+1U)]|CTGGACTTGGAGTCAGAAGGT</td><td>>am_MI0000786|hsa-mir-378a&WithFlank&5|+|149732819|149732896</td><td>50</td><td>69</td><td>20(+1U)</td><td>CTGGACTTGGAGTCAGAAGGT</td><td>409.4</td><td>5.8</td><td>0.7</td><td>7.8</td><td>8.2E-15</td><td>4.8E-14</td><td>BT20/MB468</td><td>MITO</td></tr>
<tr><td>>hg38_21_+_8433172_8446622_RNA45SN1_45S_50ntFlanks@10802.10828.27|TTGGAACAATGTAGGTAAGGGAAGTCG</td><td>>hg38_21_+_8433172_8446622_RNA45SN1_45S_50ntFlanks</td><td>10802</td><td>10828</td><td>27</td><td>TTGGAACAATGTAGGTAAGGGAAGTCG</td><td>194.2</td><td>-11.5</td><td>1.5</td><td>-7.8</td><td>8.3E-15</td><td>4.8E-14</td><td>MB231/MB468</td><td>MP</td></tr>
<tr><td>>hg38_21_+_8433172_8446622_RNA45SN1_45S_50ntFlanks@12926.12956.31|ACCTGGCGCTAAACCATTCGTAGACGACCTG</td><td>>hg38_21_+_8433172_8446622_RNA45SN1_45S_50ntFlanks</td><td>12926</td><td>12956</td><td>31</td><td>ACCTGGCGCTAAACCATTCGTAGACGACCTG</td><td>980.6</td><td>-9.3</td><td>1.2</td><td>-7.7</td><td>1.4E-14</td><td>4.9E-14</td><td>MB231/MB468</td><td>MITO</td></tr>
<tr><td>>hg38_21_+_8433172_8446622_RNA45SN1_45S_50ntFlanks@8113.8143.31|GGCGCGGGACATGTGGCGTACGGAAGACCCG</td><td>>hg38_21_+_8433172_8446622_RNA45SN1_45S_50ntFlanks</td><td>8113</td><td>8143</td><td>31</td><td>GGCGCGGGACATGTGGCGTACGGAAGACCCG</td><td>459.0</td><td>-10.9</td><td>1.4</td><td>-7.8</td><td>8.3E-15</td><td>4.9E-14</td><td>BT20/MB468</td><td>MITO</td></tr>
<tr><td>>hg38_21_+_8433172_8446622_RNA45SN1_45S_50ntFlanks@6653.6681.29|ACTCTTAGCGGTGGATCACTCGGCTCGTG</td><td>>hg38_21_+_8433172_8446622_RNA45SN1_45S_50ntFlanks</td><td>6653</td><td>6681</td><td>29</td><td>ACTCTTAGCGGTGGATCACTCGGCTCGTG</td><td>4007.4</td><td>-3.7</td><td>0.5</td><td>-7.7</td><td>1.7E-14</td><td>4.9E-14</td><td>BT20/MB468</td><td>NUC</td></tr>
<tr><td>>trnaMT_ValTAC_MT_+_1602_1670@54.71.18|AACTTGACCGCTCTGACC</td><td>>trnaMT_ValTAC_MT_+_1602_1670</td><td>54</td><td>71</td><td>18</td><td>AACTTGACCGCTCTGACC</td><td>586.8</td><td>9.3</td><td>1.2</td><td>7.7</td><td>9.5E-15</td><td>4.9E-14</td><td>MB231/MB468</td><td>CYTO</td></tr>
<tr><td>>hg38_21_+_8433172_8446622_RNA45SN1_45S_50ntFlanks@4064.4081.18|ATCAACTTTCGATGGTAG</td><td>>hg38_21_+_8433172_8446622_RNA45SN1_45S_50ntFlanks</td><td>4064</td><td>4081</td><td>18</td><td>ATCAACTTTCGATGGTAG</td><td>660.0</td><td>-12.1</td><td>1.6</td><td>-7.7</td><td>9.5E-15</td><td>4.9E-14</td><td>MB231/MB468</td><td>CYTO</td></tr>
<tr><td>>MI0000743|hsa-mir-34c&WithFlank&11|+|111513433|111513521@19.41.23&[MIMAT0000686&hsa-miR-34c-5p&offsets|0|0;m-336&11|+|111513451|111513473&offsets|0|0]|AGGCAGTGTAGTTAGCTGATTGC</td><td>>MI0000743|hsa-mir-34c&WithFlank&11|+|111513433|111513521</td><td>19</td><td>41</td><td>23</td><td>AGGCAGTGTAGTTAGCTGATTGC</td><td>831.8</td><td>12.9</td><td>1.7</td><td>7.7</td><td>1.4E-14</td><td>4.9E-14</td><td>MB231/MB468</td><td>MITO</td></tr>
<tr><td>>hg38_21_+_8433172_8446622_RNA45SN1_45S_50ntFlanks@10463.10495.33|CTCCGTTGCCCTCGGCCGATCGAAAGGGAGTCG</td><td>>hg38_21_+_8433172_8446622_RNA45SN1_45S_50ntFlanks</td><td>10463</td><td>10495</td><td>33</td><td>CTCCGTTGCCCTCGGCCGATCGAAAGGGAGTCG</td><td>154.0</td><td>-11.1</td><td>1.4</td><td>-7.8</td><td>8.4E-15</td><td>4.9E-14</td><td>MB231/MB468</td><td>MP</td></tr>
<tr><td>>hg38_21_+_8433172_8446622_RNA45SN1_45S_50ntFlanks@3793.3810.18|CGAATGGCTCATTAAATC</td><td>>hg38_21_+_8433172_8446622_RNA45SN1_45S_50ntFlanks</td><td>3793</td><td>3810</td><td>18</td><td>CGAATGGCTCATTAAATC</td><td>233.0</td><td>-11.7</td><td>1.5</td><td>-7.8</td><td>8.4E-15</td><td>4.9E-14</td><td>MB231/MB468</td><td>MP</td></tr>
<tr><td>>MI0003834|hsa-mir-769&WithFlank&19|+|46018926|46019055@36.56.21&[MIMAT0003886&hsa-miR-769-5p&offsets|0|-1;m-216&19|+|46018961|46018982&offsets|0|-1]|TGAGACCTCTGGGTTCTGAGC</td><td>>MI0003834|hsa-mir-769&WithFlank&19|+|46018926|46019055</td><td>36</td><td>56</td><td>21</td><td>TGAGACCTCTGGGTTCTGAGC</td><td>78.8</td><td>2.8</td><td>0.4</td><td>7.7</td><td>1.7E-14</td><td>4.9E-14</td><td>BT20/MB468</td><td>NUC</td></tr>
<tr><td>>am_trna117_GlyTCC_1_-_145397864_145397935@1.29.29,trna45_GlyTCC_1_+_161500903_161500974@1.29.29,trna70_GlyTCC_1_-_161439547_161439618@1.29.29,trna73_GlyTCC_1_-_161432166_161432237@1.29.29,trna76_GlyTCC_1_-_161424756_161424827@1.29.29,trna79_GlyTCC_1_-_161417375_161417446@1.29.29|GCGTTGGTGGTATAGTGGTGAGCATAGCT</td><td>>am_trna117_GlyTCC_1_-_145397864_145397935</td><td>1</td><td>29</td><td>29</td><td>GCGTTGGTGGTATAGTGGTGAGCATAGCT</td><td>805.3</td><td>-6.3</td><td>0.8</td><td>-7.7</td><td>1.4E-14</td><td>4.9E-14</td><td>MB231/MB468</td><td>MITO</td></tr>
<tr><td>>hg38_21_+_8433172_8446622_RNA45SN1_45S_50ntFlanks@9471.9500.30|AGCACGAGCGCACGTGTTAGGACCCGAAAG</td><td>>hg38_21_+_8433172_8446622_RNA45SN1_45S_50ntFlanks</td><td>9471</td><td>9500</td><td>30</td><td>AGCACGAGCGCACGTGTTAGGACCCGAAAG</td><td>154.0</td><td>-11.1</td><td>1.4</td><td>-7.8</td><td>8.5E-15</td><td>4.9E-14</td><td>MB231/MB468</td><td>MP</td></tr>
<tr><td>>hg38_21_+_8433172_8446622_RNA45SN1_45S_50ntFlanks@9738.9767.30|GAAACGATCTCAACCTATTCTCAAACTTTA</td><td>>hg38_21_+_8433172_8446622_RNA45SN1_45S_50ntFlanks</td><td>9738</td><td>9767</td><td>30</td><td>GAAACGATCTCAACCTATTCTCAAACTTTA</td><td>1180.7</td><td>-4.2</td><td>0.5</td><td>-7.7</td><td>1.0E-14</td><td>4.9E-14</td><td>MB231/MB468</td><td>NUC</td></tr>
<tr><td>>hg38_21_+_8433172_8446622_RNA45SN1_45S_50ntFlanks@6740.6772.33|CATCGACACTTCGAACGCACTTGCGGCCCCGGG</td><td>>hg38_21_+_8433172_8446622_RNA45SN1_45S_50ntFlanks</td><td>6740</td><td>6772</td><td>33</td><td>CATCGACACTTCGAACGCACTTGCGGCCCCGGG</td><td>2836.5</td><td>-9.9</td><td>1.3</td><td>-7.7</td><td>1.4E-14</td><td>4.9E-14</td><td>MB231/MB468</td><td>MITO</td></tr>
<tr><td>>trna119_LysCTT_1_-_145395522_145395594@4.35.32,trna11_LysCTT_5_-_180648979_180649051@4.35.32,trna13_LysCTT_6_+_26556774_26556846@4.35.32,trna7_LysCTT_16_+_3225692_3225764@4.35.32,trna9_LysCTT_5_+_180634755_180634827@4.35.32|CGGCTAGCTCAGTCGGTAGAGCATGAGACTCT</td><td>>trna119_LysCTT_1_-_145395522_145395594</td><td>4</td><td>35</td><td>32</td><td>CGGCTAGCTCAGTCGGTAGAGCATGAGACTCT</td><td>4917.5</td><td>6.9</td><td>0.9</td><td>7.7</td><td>1.4E-14</td><td>5.0E-14</td><td>MB231/MB468</td><td>MITO</td></tr>
</table>

<table>
<tr><td></td><td></td><td></td><td></td><td></td><td></td><td></td><td></td><td></td><td></td><td></td><td></td><td></td><td></td></tr>
<tr><td>>hg38_21_+_8433172_8446622_RNA45SN1_45S_50ntFlanks@10458.10497.40|ATGGCCTCCGTTGCCCTCGGCCGATCGAAAGGGAGTCGGG</td><td>>hg38_21_+_8433172_8446622_RNA45SN1_45S_50ntFlanks</td><td>10458</td><td>10497</td><td>40</td><td>ATGGCCTCCGTTGCCCTCGGCCGATCGAAAGGGAGTCGGG</td><td>430.9</td><td>-11.7</td><td>1.5</td><td>-7.7</td><td>1.0E-14</td><td>5.0E-14</td><td>MB231/MB468</td><td>NUC</td></tr>
<tr><td>>MI0000298|hsa-mir-221&WithFlank&X|-|45746151|45746272@31.53.23&[MIMAT0004568&hsa-miR-221-5p&offsets|0|+1;m-405&X|-|45746220|45746242&offsets|0|0]|ACCTGGCATACAATGTAGATTTC</td><td>>MI0000298|hsa-mir-221&WithFlank&X|-|45746151|45746272</td><td>31</td><td>53</td><td>23</td><td>ACCTGGCATACAATGTAGATTTC</td><td>615.5</td><td>5.2</td><td>0.7</td><td>7.7</td><td>1.0E-14</td><td>5.0E-14</td><td>MB231/MB468</td><td>NUC</td></tr>
<tr><td>>hg38_21_+_8433172_8446622_RNA45SN1_45S_50ntFlanks@10366.10390.25|CCGAAGTGGAGAAGGGTTCCATGTG</td><td>>hg38_21_+_8433172_8446622_RNA45SN1_45S_50ntFlanks</td><td>10366</td><td>10390</td><td>25</td><td>CCGAAGTGGAGAAGGGTTCCATGTG</td><td>143.1</td><td>-11.0</td><td>1.4</td><td>-7.8</td><td>8.6E-15</td><td>5.0E-14</td><td>MB231/MB468</td><td>MP</td></tr>
<tr><td>>hg38_21_+_8433172_8446622_RNA45SN1_45S_50ntFlanks@8170.8188.19|CAAGTCCTTCTGATCGAGG</td><td>>hg38_21_+_8433172_8446622_RNA45SN1_45S_50ntFlanks</td><td>8170</td><td>8188</td><td>19</td><td>CAAGTCCTTCTGATCGAGG</td><td>635.5</td><td>-12.0</td><td>1.6</td><td>-7.7</td><td>9.8E-15</td><td>5.0E-14</td><td>MB231/MB468</td><td>CYTO</td></tr>
<tr><td>>hg38_21_+_8433172_8446622_RNA45SN1_45S_50ntFlanks@12160.12182.23|TTGACTGGGGCGGTACACCTGTC</td><td>>hg38_21_+_8433172_8446622_RNA45SN1_45S_50ntFlanks</td><td>12160</td><td>12182</td><td>23</td><td>TTGACTGGGGCGGTACACCTGTC</td><td>274.2</td><td>-12.0</td><td>1.5</td><td>-7.8</td><td>8.7E-15</td><td>5.0E-14</td><td>MB231/MB468</td><td>MP</td></tr>
<tr><td>>MI0000299|hsa-mir-222&WithFlank&X|-|45747009|45747130@75.99.25(+2U)&[MIMAT0000279&hsa-miR-222-3p&offsets|0|+4(+2U);m-140&X|-|45747033|45747056&offsets|0|+1(+2U)]|AGCTACATCTGGCTACTGGGTCTCTTT</td><td>>MI0000299|hsa-mir-222&WithFlank&X|-|45747009|45747130</td><td>75</td><td>99</td><td>25(+2U)</td><td>AGCTACATCTGGCTACTGGGTCTCTTT</td><td>2332.7</td><td>6.5</td><td>0.8</td><td>7.7</td><td>1.1E-14</td><td>5.0E-14</td><td>MB231/MB468</td><td>NUC</td></tr>
<tr><td>>hg38_21_+_8433172_8446622_RNA45SN1_45S_50ntFlanks@12419.12449.31|GATGTCGGCTCTTCCTATCATTGTGAAGCAG</td><td>>hg38_21_+_8433172_8446622_RNA45SN1_45S_50ntFlanks</td><td>12419</td><td>12449</td><td>31</td><td>GATGTCGGCTCTTCCTATCATTGTGAAGCAG</td><td>2787.7</td><td>4.9</td><td>0.6</td><td>7.7</td><td>9.9E-15</td><td>5.1E-14</td><td>MB231/MB468</td><td>CYTO</td></tr>
<tr><td>>hg38_21_+_8433172_8446622_RNA45SN1_45S_50ntFlanks@3785.3812.28|TGAAACTGCGAATGGCTCATTAAATCAG</td><td>>hg38_21_+_8433172_8446622_RNA45SN1_45S_50ntFlanks</td><td>3785</td><td>3812</td><td>28</td><td>TGAAACTGCGAATGGCTCATTAAATCAG</td><td>189.0</td><td>-5.5</td><td>0.7</td><td>-7.7</td><td>9.9E-15</td><td>5.1E-14</td><td>MB231/MB468</td><td>CYTO</td></tr>
<tr><td>>hg38_21_+_8433172_8446622_RNA45SN1_45S_50ntFlanks@11766.11794.29|ATCTAATTAGTGACGCGCATGAATGGATG</td><td>>hg38_21_+_8433172_8446622_RNA45SN1_45S_50ntFlanks</td><td>11766</td><td>11794</td><td>29</td><td>ATCTAATTAGTGACGCGCATGAATGGATG</td><td>245.9</td><td>-10.7</td><td>1.4</td><td>-7.7</td><td>9.9E-15</td><td>5.1E-14</td><td>MB231/MB468</td><td>CYTO</td></tr>
<tr><td>>hg38_21_+_8433172_8446622_RNA45SN1_45S_50ntFlanks@10807.10840.34|ACAATGTAGGTAAGGGAAGTCGGCAAGCCGGATC</td><td>>hg38_21_+_8433172_8446622_RNA45SN1_45S_50ntFlanks</td><td>10807</td><td>10840</td><td>34</td><td>ACAATGTAGGTAAGGGAAGTCGGCAAGCCGGATC</td><td>1160.2</td><td>-7.6</td><td>1.0</td><td>-7.7</td><td>1.4E-14</td><td>5.1E-14</td><td>MB231/MB468</td><td>MITO</td></tr>
<tr><td>>hg38_21_+_8433172_8446622_RNA45SN1_45S_50ntFlanks@5080.5098.19|ATAACGAACGAGACTCTGG</td><td>>hg38_21_+_8433172_8446622_RNA45SN1_45S_50ntFlanks</td><td>5080</td><td>5098</td><td>19</td><td>ATAACGAACGAGACTCTGG</td><td>568.0</td><td>-13.0</td><td>1.7</td><td>-7.8</td><td>8.8E-15</td><td>5.1E-14</td><td>MB231/MB468</td><td>MP</td></tr>
<tr><td>>hg38_21_+_8433172_8446622_RNA45SN1_45S_50ntFlanks@5186.5206.21|CAATAACAGGTCTGTGATGCC</td><td>>hg38_21_+_8433172_8446622_RNA45SN1_45S_50ntFlanks</td><td>5186</td><td>5206</td><td>21</td><td>CAATAACAGGTCTGTGATGCC</td><td>1012.1</td><td>-13.8</td><td>1.8</td><td>-7.7</td><td>1.4E-14</td><td>5.1E-14</td><td>MB231/MB468</td><td>MITO</td></tr>
<tr><td>>MI0000084|hsa-mir-26b&WithFlank&2|+|218402640|218402728@18.38.21&[MIMAT0000083&hsa-miR-26b-5p&offsets|0|0;m-53&2|+|218402657|218402678&offsets|0|-1]|TTCAAGTAATTCAGGATAGGT</td><td>>MI0000084|hsa-mir-26b&WithFlank&2|+|218402640|218402728</td><td>18</td><td>38</td><td>21</td><td>TTCAAGTAATTCAGGATAGGT</td><td>1971.5</td><td>6.8</td><td>0.9</td><td>7.7</td><td>1.2E-14</td><td>5.1E-14</td><td>BT20/MB468</td><td>CYTO</td></tr>
<tr><td>>hg38_21_+_8433172_8446622_RNA45SN1_45S_50ntFlanks@10843.10867.25|TAACTTCGGGATAAGGATTGGCTCT</td><td>>hg38_21_+_8433172_8446622_RNA45SN1_45S_50ntFlanks</td><td>10843</td><td>10867</td><td>25</td><td>TAACTTCGGGATAAGGATTGGCTCT</td><td>359.1</td><td>-4.3</td><td>0.6</td><td>-7.7</td><td>1.1E-14</td><td>5.1E-14</td><td>MB231/MB468</td><td>NUC</td></tr>
<tr><td>>hg38_21_+_8433172_8446622_RNA45SN1_45S_50ntFlanks@12324.12344.21|TTGGGTTTTAAGCAGGAGGTG</td><td>>hg38_21_+_8433172_8446622_RNA45SN1_45S_50ntFlanks</td><td>12324</td><td>12344</td><td>21</td><td>TTGGGTTTTAAGCAGGAGGTG</td><td>513.4</td><td>-11.1</td><td>1.4</td><td>-7.8</td><td>8.8E-15</td><td>5.1E-14</td><td>BT20/MB468</td><td>MITO</td></tr>
<tr><td>>hg38_21_+_8433172_8446622_RNA45SN1_45S_50ntFlanks@9554.9571.18|CGTAGCGGTCCTGACGTG</td><td>>hg38_21_+_8433172_8446622_RNA45SN1_45S_50ntFlanks</td><td>9554</td><td>9571</td><td>18</td><td>CGTAGCGGTCCTGACGTG</td><td>36928.4</td><td>-4.8</td><td>0.6</td><td>-7.8</td><td>8.8E-15</td><td>5.2E-14</td><td>BT20/MB468</td><td>MITO</td></tr>
<tr><td>>hg38_21_+_8433172_8446622_RNA45SN1_45S_50ntFlanks@12958.12993.36|TTCTGGGTCGGGGTTTCGTACGTAGCAGAGCAGCTC</td><td>>hg38_21_+_8433172_8446622_RNA45SN1_45S_50ntFlanks</td><td>12958</td><td>12993</td><td>36</td><td>TTCTGGGTCGGGGTTTCGTACGTAGCAGAGCAGCTC</td><td>1474.5</td><td>8.6</td><td>1.1</td><td>7.7</td><td>1.0E-14</td><td>5.2E-14</td><td>MB231/MB468</td><td>CYTO</td></tr>
<tr><td>>hg38_1_-_228634819_228635039_RNA5S12_5S_50ntFlanks@142.169.28|CGCCTGGGAATACCGGGTGCTGTAGGCT</td><td>>hg38_1_-_228634819_228635039_RNA5S12_5S_50ntFlanks</td><td>142</td><td>169</td><td>28</td><td>CGCCTGGGAATACCGGGTGCTGTAGGCT</td><td>923.2</td><td>5.9</td><td>0.8</td><td>7.7</td><td>1.2E-14</td><td>5.2E-14</td><td>BT20/MB468</td><td>CYTO</td></tr>
<tr><td>>hg38_21_+_8433172_8446622_RNA45SN1_45S_50ntFlanks@4103.4127.25|CCACGGGTGACGGGGAATCAGGGTT</td><td>>hg38_21_+_8433172_8446622_RNA45SN1_45S_50ntFlanks</td><td>4103</td><td>4127</td><td>25</td><td>CCACGGGTGACGGGGAATCAGGGTT</td><td>290.2</td><td>-11.1</td><td>1.4</td><td>-7.7</td><td>1.1E-14</td><td>5.2E-14</td><td>MB231/MB468</td><td>NUC</td></tr>
<tr><td>>MI0000086|hsa-mir-28&WithFlank&3|+|188688775|188688872@60.81.22&[MIMAT0004502&hsa-miR-28-3p&offsets|0|0;m-28&3|+|188688834|188688855&offsets|0|0]|CACTAGATTGTGAGCTCCTGGA</td><td>>MI0000086|hsa-mir-28&WithFlank&3|+|188688775|188688872</td><td>60</td><td>81</td><td>22</td><td>CACTAGATTGTGAGCTCCTGGA</td><td>2659.6</td><td>7.5</td><td>1.0</td><td>7.7</td><td>1.5E-14</td><td>5.2E-14</td><td>MB231/MB468</td><td>MITO</td></tr>
<tr><td>>hg38_21_+_8433172_8446622_RNA45SN1_45S_50ntFlanks@11210.11234.25|GGCGACTCTGGACGCGAGCCGGGCC</td><td>>hg38_21_+_8433172_8446622_RNA45SN1_45S_50ntFlanks</td><td>11210</td><td>11234</td><td>25</td><td>GGCGACTCTGGACGCGAGCCGGGCC</td><td>204.2</td><td>-11.5</td><td>1.5</td><td>-7.8</td><td>8.9E-15</td><td>5.2E-14</td><td>MB231/MB468</td><td>MP</td></tr>
<tr><td>>hg38_21_+_8433172_8446622_RNA45SN1_45S_50ntFlanks@4390.4408.19|TCTTGGGAGCGGGCGGGCG</td><td>>hg38_21_+_8433172_8446622_RNA45SN1_45S_50ntFlanks</td><td>4390</td><td>4408</td><td>19</td><td>TCTTGGGAGCGGGCGGGCG</td><td>192.5</td><td>-10.6</td><td>1.4</td><td>-7.7</td><td>1.1E-14</td><td>5.2E-14</td><td>MB231/MB468</td><td>NUC</td></tr>
<tr><td>>MI0000061|hsa-let-7a-2&WithFlank&11|-|122146516|122146599@11.31.21;MI0000062|hsa-let-7a-3&WithFlank&22|+|46112743|46112828@10.30.21;MI0000060|hsa-let-7a-1&WithFlank&9|+|94175951|94176042@12.32.21&[MIMAT0000062&hsa-let-7a-5p&offsets|0|-1;m-6&11|-|122146568|122146589&offsets|0|-1];[MIMAT0000062_1&hsa-let-7a-5p&offsets|0|-1;m-5&22|+|46112752|46112773&offsets|0|-1];[MIMAT0000062_2&hsa-let-7a-5p&offsets|0|-1;m-4&9|+|94175962|94175983&offsets|0|-1]|TGAGGTAGTAGGTTGTATAGT</td><td>>MI0000061|hsa-let-7a-2&WithFlank&11|-|122146516|122146599</td><td>11</td><td>31</td><td>21</td><td>TGAGGTAGTAGGTTGTATAGT</td><td>8131.8</td><td>6.6</td><td>0.9</td><td>7.7</td><td>1.5E-14</td><td>5.2E-14</td><td>MB231/MB468</td><td>MITO</td></tr>
<tr><td>>hg38_21_+_8433172_8446622_RNA45SN1_45S_50ntFlanks@4547.4565.19|CTGGATACCGCAGCTAGGA</td><td>>hg38_21_+_8433172_8446622_RNA45SN1_45S_50ntFlanks</td><td>4547</td><td>4565</td><td>19</td><td>CTGGATACCGCAGCTAGGA</td><td>272.5</td><td>-11.9</td><td>1.5</td><td>-7.7</td><td>1.5E-14</td><td>5.2E-14</td><td>MB231/MB468</td><td>MITO</td></tr>
<tr><td>>hg38_21_+_8433172_8446622_RNA45SN1_45S_50ntFlanks@11197.11225.29|GGCCGGCGGCGGCGGCGACTCTGGACGCG</td><td>>hg38_21_+_8433172_8446622_RNA45SN1_45S_50ntFlanks</td><td>11197</td><td>11225</td><td>29</td><td>GGCCGGCGGCGGCGGCGACTCTGGACGCG</td><td>181.4</td><td>-10.2</td><td>1.3</td><td>-7.7</td><td>1.0E-14</td><td>5.2E-14</td><td>MB231/MB468</td><td>CYTO</td></tr>
<tr><td>>hg38_21_+_8433172_8446622_RNA45SN1_45S_50ntFlanks@4064.4081.18|ATCAACTTTCGATGGTAG</td><td>>hg38_21_+_8433172_8446622_RNA45SN1_45S_50ntFlanks</td><td>4064</td><td>4081</td><td>18</td><td>ATCAACTTTCGATGGTAG</td><td>818.1</td><td>-13.5</td><td>1.8</td><td>-7.7</td><td>1.5E-14</td><td>5.2E-14</td><td>MB231/MB468</td><td>MITO</td></tr>
<tr><td>>MI0000082|hsa-mir-25&WithFlank&7|-|100093554|100093649@58.79.22(+1U)&[MIMAT0000081&hsa-miR-25-3p&offsets|0|0(+1U);m-23&7|-|100093571|100093592&offsets|0|0(+1U)]|CATTGCACTTGTCTCGGTCTGAT</td><td>>MI0000082|hsa-mir-25&WithFlank&7|-|100093554|100093649</td><td>58</td><td>79</td><td>22(+1U)</td><td>CATTGCACTTGTCTCGGTCTGAT</td><td>1416.1</td><td>6.2</td><td>0.8</td><td>7.7</td><td>1.0E-14</td><td>5.2E-14</td><td>MB231/MB468</td><td>CYTO</td></tr>
<tr><td>>am_trna162_MetCAT_6_-_26758550_26758622@31.51.21,trna164_MetCAT_6_-_26735574_26735646@31.51.21,trna20_MetCAT_16_+_71460396_71460468@31.51.21,trna21_MetCAT_6_+_26701712_26701784@31.51.21,trna27_MetCAT_6_+_26766444_26766516@31.51.21,trna75_MetCAT_6_+_28912352_28912424@31.51.21,trna97_MetCAT_6_-_28921042_28921114@31.51.21|TCTCATAATCTGAAGGTCCTG</td><td>>am_trna162_MetCAT_6_-_26758550_26758622</td><td>31</td><td>51</td><td>21</td><td>TCTCATAATCTGAAGGTCCTG</td><td>227.2</td><td>-10.8</td><td>1.4</td><td>-7.7</td><td>1.1E-14</td><td>5.2E-14</td><td>MB231/MB468</td><td>NUC</td></tr>
<tr><td>>am_trna2_GlyGCC_21_-_18827107_18827177@1.25.25,trna35_GlyGCC_1_+_161413094_161413164@1.25.25,trna37_GlyGCC_1_+_161420467_161420537@1.25.25,trna39_GlyGCC_1_+_161427898_161427968@1.25.25,trna41_GlyGCC_1_+_161435258_161435328@1.25.25|GCATGGGTGGTTCAGTGGTAGAATT</td><td>>am_trna2_GlyGCC_21_-_18827107_18827177</td><td>1</td><td>25</td><td>25</td><td>GCATGGGTGGTTCAGTGGTAGAATT</td><td>259.5</td><td>11.6</td><td>1.5</td><td>8.0</td><td>1.3E-15</td><td>5.2E-14</td><td>BT20/MB231</td><td>MITO</td></tr>
<tr><td>>hg38_21_+_8433172_8446622_RNA45SN1_45S_50ntFlanks@10402.10424.23|AACATGGGTCAGTCGGTCCTGAG</td><td>>hg38_21_+_8433172_8446622_RNA45SN1_45S_50ntFlanks</td><td>10402</td><td>10424</td><td>23</td><td>AACATGGGTCAGTCGGTCCTGAG</td><td>20520.6</td><td>4.0</td><td>0.5</td><td>7.8</td><td>9.0E-15</td><td>5.2E-14</td><td>BT20/MB468</td><td>MITO</td></tr>
<tr><td>>MI0000298|hsa-mir-221&WithFlank&X|-|45746151|45746272@71.91.21(+1C)&[MIMAT0000278&hsa-miR-221-3p&offsets|0|-2(+1C);m-91&X|-|45746181|45746202&offsets|0|-1(+1C)]|AGCTACATTGTCTGCTGGGTTC</td><td>>MI0000298|hsa-mir-221&WithFlank&X|-|45746151|45746272</td><td>71</td><td>91</td><td>21(+1C)</td><td>AGCTACATTGTCTGCTGGGTTC</td><td>1408.3</td><td>6.8</td><td>0.9</td><td>7.8</td><td>9.0E-15</td><td>5.2E-14</td><td>BT20/MB468</td><td>MITO</td></tr>
<tr><td>>hg38_21_+_8433172_8446622_RNA45SN1_45S_50ntFlanks@9201.9218.18|CTCTCGGGGCCACGCGCG</td><td>>hg38_21_+_8433172_8446622_RNA45SN1_45S_50ntFlanks</td><td>9201</td><td>9218</td><td>18</td><td>CTCTCGGGGCCACGCGCG</td><td>733.3</td><td>-12.6</td><td>1.5</td><td>-8.2</td><td>3.6E-16</td><td>5.2E-14</td><td>BT20/MB468</td><td>MP</td></tr>
<tr><td>>MI0000269|hsa-mir-181a-2&WithFlank&9|+|124692436|124692557@83.103.21(+1U)&[MIMAT0004558&hsa-miR-181a-2-3p&offsets|0|-1(+1U);m-71&9|+|124692518|124692539&offsets|0|-1(+1U)]|ACCACTGACCGTTGACTGTACT</td><td>>MI0000269|hsa-mir-181a-2&WithFlank&9|+|124692436|124692557</td><td>83</td><td>103</td><td>21(+1U)</td><td>ACCACTGACCGTTGACTGTACT</td><td>400.8</td><td>5.4</td><td>0.7</td><td>7.7</td><td>1.1E-14</td><td>5.2E-14</td><td>MB231/MB468</td><td>NUC</td></tr>
<tr><td>>hg38_21_+_8433172_8446622_RNA45SN1_45S_50ntFlanks@10369.10407.39|AAGTGGAGAAGGGTTCCATGTGAACAGCAGTTGAACATG</td><td>>hg38_21_+_8433172_8446622_RNA45SN1_45S_50ntFlanks</td><td>10369</td><td>10407</td><td>39</td><td>AAGTGGAGAAGGGTTCCATGTGAACAGCAGTTGAACATG</td><td>1148.3</td><td>-10.4</td><td>1.3</td><td>-7.7</td><td>1.1E-14</td><td>5.2E-14</td><td>MB231/MB468</td><td>NUC</td></tr>
<tr><td>>hg38_21_+_8433172_8446622_RNA45SN1_45S_50ntFlanks@10404.10433.30|CATGGGTCAGTCGGTCCTGAGAGATGGGCG</td><td>>hg38_21_+_8433172_8446622_RNA45SN1_45S_50ntFlanks</td><td>10404</td><td>10433</td><td>30</td><td>CATGGGTCAGTCGGTCCTGAGAGATGGGCG</td><td>207.2</td><td>-11.6</td><td>1.5</td><td>-7.8</td><td>9.0E-15</td><td>5.3E-14</td><td>MB231/MB468</td><td>MP</td></tr>
<tr><td>>hg38_21_+_8433172_8446622_RNA45SN1_45S_50ntFlanks@476.499.24|TTCGTGATCGATGTGGTGACGTCG</td><td>>hg38_21_+_8433172_8446622_RNA45SN1_45S_50ntFlanks</td><td>476</td><td>499</td><td>24</td><td>TTCGTGATCGATGTGGTGACGTCG</td><td>848.4</td><td>8.0</td><td>1.0</td><td>7.7</td><td>1.2E-14</td><td>5.3E-14</td><td>BT20/MB468</td><td>CYTO</td></tr>
<tr><td>>hg38_21_+_8433172_8446622_RNA45SN1_45S_50ntFlanks@11455.11478.24|CTCGGCGCGCGGCGGCGGCGGCGG</td><td>>hg38_21_+_8433172_8446622_RNA45SN1_45S_50ntFlanks</td><td>11455</td><td>11478</td><td>24</td><td>CTCGGCGCGCGGCGGCGGCGGCGG</td><td>286.2</td><td>-12.0</td><td>1.6</td><td>-7.8</td><td>9.1E-15</td><td>5.3E-14</td><td>MB231/MB468</td><td>MP</td></tr>
<tr><td>>trna3_AlaAGC_2_+_27274082_27274154@59.76.18,trna6_AlaAGC_8_+_67026424_67026496@59.76.18|TGCCCGCATCCTCCACCA</td><td>>trna3_AlaAGC_2_+_27274082_27274154</td><td>59</td><td>76</td><td>18</td><td>TGCCCGCATCCTCCACCA</td><td>368.6</td><td>9.2</td><td>1.2</td><td>7.7</td><td>1.2E-14</td><td>5.3E-14</td><td>BT20/MB468</td><td>CYTO</td></tr>
<tr><td>>hg38_21_+_8433172_8446622_RNA45SN1_45S_50ntFlanks@10481.10498.18|ATCGAAAGGGAGTCGGGT</td><td>>hg38_21_+_8433172_8446622_RNA45SN1_45S_50ntFlanks</td><td>10481</td><td>10498</td><td>18</td><td>ATCGAAAGGGAGTCGGGT</td><td>280.0</td><td>-10.4</td><td>1.4</td><td>-7.7</td><td>1.2E-14</td><td>5.3E-14</td><td>BT20/MB468</td><td>CYTO</td></tr>
<tr><td>>hg38_21_+_8433172_8446622_RNA45SN1_45S_50ntFlanks@13002.13020.19|CGATCTATTGAAAGTCAGC</td><td>>hg38_21_+_8433172_8446622_RNA45SN1_45S_50ntFlanks</td><td>13002</td><td>13020</td><td>19</td><td>CGATCTATTGAAAGTCAGC</td><td>318.3</td><td>-10.6</td><td>1.4</td><td>-7.7</td><td>1.2E-14</td><td>5.3E-14</td><td>BT20/MB468</td><td>CYTO</td></tr>
<tr><td>>MI0000439|hsa-mir-23b&WithFlank&9|+|95085202|95085310@67.85.19&[MIMAT0000418&hsa-miR-23b-3p&offsets|+3|-1;m-42&9|+|95085265|95085285&offsets|+3|+1]|ACATTGCCAGGGATTACCA</td><td>>MI0000439|hsa-mir-23b&WithFlank&9|+|95085202|95085310</td><td>67</td><td>85</td><td>19</td><td>ACATTGCCAGGGATTACCA</td><td>949.5</td><td>28.1</td><td>3.6</td><td>7.7</td><td>1.0E-14</td><td>5.3E-14</td><td>MB231/MB468</td><td>CYTO</td></tr>
<tr><td>>hg38_21_+_8433172_8446622_RNA45SN1_45S_50ntFlanks@12678.12703.26|AATCCCGCCCAGGCGGAACGATACGG</td><td>>hg38_21_+_8433172_8446622_RNA45SN1_45S_50ntFlanks</td><td>12678</td><td>12703</td><td>26</td><td>AATCCCGCCCAGGCGGAACGATACGG</td><td>338.3</td><td>-11.1</td><td>1.4</td><td>-7.7</td><td>1.0E-14</td><td>5.3E-14</td><td>MB231/MB468</td><td>CYTO</td></tr>
<tr><td>>hg38_21_+_8433172_8446622_RNA45SN1_45S_50ntFlanks@8193.8211.19|GCCCGTGGACGGTGTGAGG</td><td>>hg38_21_+_8433172_8446622_RNA45SN1_45S_50ntFlanks</td><td>8193</td><td>8211</td><td>19</td><td>GCCCGTGGACGGTGTGAGG</td><td>334.2</td><td>-12.2</td><td>1.6</td><td>-7.8</td><td>9.1E-15</td><td>5.3E-14</td><td>MB231/MB468</td><td>MP</td></tr>
<tr><td>>hg38_21_+_8433172_8446622_RNA45SN1_45S_50ntFlanks@10761.10785.25|ATATCCGCAGCAGGTCTCCAAGGTG</td><td>>hg38_21_+_8433172_8446622_RNA45SN1_45S_50ntFlanks</td><td>10761</td><td>10785</td><td>25</td><td>ATATCCGCAGCAGGTCTCCAAGGTG</td><td>192.7</td><td>-11.5</td><td>1.5</td><td>-7.8</td><td>9.1E-15</td><td>5.3E-14</td><td>MB231/MB468</td><td>MP</td></tr>
<tr><td>>MI0000279|hsa-mir-196a-2&WithFlank&12|+|53991732|53991853@31.53.23&[MIMAT0000226&hsa-miR-196a-5p&offsets|0|+1;m-261&12|+|53991762|53991783&offsets|0|+1]|TAGGTAGTTTCATGTTGTTGGGA</td><td>>MI0000279|hsa-mir-196a-2&WithFlank&12|+|53991732|53991853</td><td>31</td><td>53</td><td>23</td><td>TAGGTAGTTTCATGTTGTTGGGA</td><td>636.5</td><td>8.0</td><td>1.0</td><td>7.8</td><td>9.1E-15</td><td>5.3E-14</td><td>BT20/MB468</td><td>MITO</td></tr>
<tr><td>>hg38_21_+_8433172_8446622_RNA45SN1_45S_50ntFlanks@4068.4089.22|ACTTTCGATGGTAGTCGCCGTG</td><td>>hg38_21_+_8433172_8446622_RNA45SN1_45S_50ntFlanks</td><td>4068</td><td>4089</td><td>22</td><td>ACTTTCGATGGTAGTCGCCGTG</td><td>1093.3</td><td>-13.9</td><td>1.8</td><td>-7.7</td><td>1.5E-14</td><td>5.3E-14</td><td>MB231/MB468</td><td>MITO</td></tr>
<tr><td>>hg38_21_+_8433172_8446622_RNA45SN1_45S_50ntFlanks@8194.8212.19|CCCGTGGACGGTGTGAGGC</td><td>>hg38_21_+_8433172_8446622_RNA45SN1_45S_50ntFlanks</td><td>8194</td><td>8212</td><td>19</td><td>CCCGTGGACGGTGTGAGGC</td><td>1839.1</td><td>-6.2</td><td>0.8</td><td>-7.7</td><td>1.1E-14</td><td>5.3E-14</td><td>MB231/MB468</td><td>NUC</td></tr>
<tr><td>>MI0003560|hsa-mir-92b&WithFlank&1|+|155195171|155195278@67.86.20&[MIMAT0003218&hsa-miR-92b-3p&offsets|0|-2;m-115&1|+|155195237|155195258&offsets|0|-2]|TATTGCACTCGTCCCGGCCT</td><td>>MI0003560|hsa-mir-92b&WithFlank&1|+|155195171|155195278</td><td>67</td><td>86</td><td>20</td><td>TATTGCACTCGTCCCGGCCT</td><td>838.9</td><td>10.4</td><td>1.3</td><td>7.7</td><td>1.0E-14</td><td>5.3E-14</td><td>MB231/MB468</td><td>CYTO</td></tr>
<tr><td>>MI0000077|hsa-mir-21&WithFlank&17|+|59841260|59841343@14.36.23&[MIMAT0000076&hsa-miR-21-5p&offsets|0|+1;m-1&17|+|59841273|59841294&offsets|0|+1]|TAGCTTATCAGACTGATGTTGAC</td><td>>MI0000077|hsa-mir-21&WithFlank&17|+|59841260|59841343</td><td>14</td><td>36</td><td>23</td><td>TAGCTTATCAGACTGATGTTGAC</td><td>176065.9</td><td>3.3</td><td>0.4</td><td>7.7</td><td>1.9E-14</td><td>5.3E-14</td><td>BT20/MB468</td><td>NUC</td></tr>
</table>

<table>
<tr><th></th><th></th><th></th><th></th><th></th><th></th><th></th><th></th><th></th><th></th><th></th><th></th><th></th><th></th></tr>
<tr><td>>hg38_21_+_8433172_8446622_RNA45SN1_45S_50ntFlanks@10408.10435.28|GGTCAGTCGGTCCTGAGAGATGGGCGAG</td><td>>hg38_21_+_8433172_8446622_RNA45SN1_45S_50ntFlanks</td><td>10408</td><td>10435</td><td>28</td><td>GGTCAGTCGGTCCTGAGAGATGGGCGAG</td><td>2208.0</td><td>-7.8</td><td>1.0</td><td>-7.7</td><td>1.0E-14</td><td>5.3E-14</td><td>MB231/MB468</td><td>CYTO</td></tr>
<tr><td>>hg38_21_+_8433172_8446622_RNA45SN1_45S_50ntFlanks@11317.11342.26|ACCCCACGTCTCGTCGCGCGCGCGTC</td><td>>hg38_21_+_8433172_8446622_RNA45SN1_45S_50ntFlanks</td><td>11317</td><td>11342</td><td>26</td><td>ACCCCACGTCTCGTCGCGCGCGCGTC</td><td>254.2</td><td>-11.9</td><td>1.5</td><td>-7.8</td><td>9.2E-15</td><td>5.3E-14</td><td>MB231/MB468</td><td>MP</td></tr>
<tr><td>>hg38_21_+_8433172_8446622_RNA45SN1_45S_50ntFlanks@11163.11188.26|CGGTCCCCCGCGAGGGGGGCCCGGGC</td><td>>hg38_21_+_8433172_8446622_RNA45SN1_45S_50ntFlanks</td><td>11163</td><td>11188</td><td>26</td><td>CGGTCCCCCGCGAGGGGGGCCCGGGC</td><td>296.1</td><td>-12.0</td><td>1.6</td><td>-7.7</td><td>1.5E-14</td><td>5.3E-14</td><td>MB231/MB468</td><td>MITO</td></tr>
<tr><td>>hg38_21_+_8433172_8446622_RNA45SN1_45S_50ntFlanks@11694.11728.35|AAGAAATTCAATGAAGCGCGGGTAAACGGCGGGAG</td><td>>hg38_21_+_8433172_8446622_RNA45SN1_45S_50ntFlanks</td><td>11694</td><td>11728</td><td>35</td><td>AAGAAATTCAATGAAGCGCGGGTAAACGGCGGGAG</td><td>409.8</td><td>-11.6</td><td>1.5</td><td>-7.7</td><td>1.1E-14</td><td>5.3E-14</td><td>MB231/MB468</td><td>NUC</td></tr>
<tr><td>>hg38_21_+_8433172_8446622_RNA45SN1_45S_50ntFlanks@9033.9060.28|CTCGTGGGGGGGCCGGGCCACCCCTCCC</td><td>>hg38_21_+_8433172_8446622_RNA45SN1_45S_50ntFlanks</td><td>9033</td><td>9060</td><td>28</td><td>CTCGTGGGGGGGCCGGGCCACCCCTCCC</td><td>232.3</td><td>-11.7</td><td>1.5</td><td>-7.7</td><td>9.2E-15</td><td>5.3E-14</td><td>MB231/MB468</td><td>MP</td></tr>
<tr><td>>MI0000061|hsa-let-7a-2&WithFlank&11|-|122146516|122146599@11.31.21(+1G);MI0000062|hsa-let-7a-3&WithFlank&22|+|46112743|46112828@10.30.21(+1G);MI0000060|hsa-let-7a-1&WithFlank&9|+|94175951|94176042@12.32.21(+1G)&[MIMAT0000062&hsa-let-7a-5p&offsets|0|-1(+1G);m-6&11|-|122146568|122146589&offsets|0|-1(+1G)];[MIMAT0000062_1&hsa-let-7a-5p&offsets|0|-1(+1G);m-5&22|+|46112752|46112773&offsets|0|-1(+1G)];[MIMAT0000062_2&hsa-let-7a-5p&offsets|0|-1(+1G);m-4&9|+|94175962|94175983&offsets|0|-1(+1G)]|TGAGGTAGTAGGTTGTATAGTG</td><td>>MI0000061|hsa-let-7a-2&WithFlank&11|-|122146516|122146599</td><td>11</td><td>31</td><td>21(+1G)</td><td>TGAGGTAGTAGGTTGTATAGTG</td><td>2429.1</td><td>6.4</td><td>0.8</td><td>7.7</td><td>1.2E-14</td><td>5.3E-14</td><td>BT20/MB468</td><td>CYTO</td></tr>
<tr><td>>hg38_21_+_8433172_8446622_RNA45SN1_45S_50ntFlanks@4045.4062.18|CCATTCGAACGTCTGCCC</td><td>>hg38_21_+_8433172_8446622_RNA45SN1_45S_50ntFlanks</td><td>4045</td><td>4062</td><td>18</td><td>CCATTCGAACGTCTGCCC</td><td>654.4</td><td>-3.4</td><td>0.4</td><td>-7.7</td><td>1.2E-14</td><td>5.3E-14</td><td>BT20/MB468</td><td>CYTO</td></tr>
<tr><td>>hg38_21_+_8433172_8446622_RNA45SN1_45S_50ntFlanks@7977.8018.42|CGACCTCAGATCAGACGTGGCGACCCGCTGAATTTAAGCATA</td><td>>hg38_21_+_8433172_8446622_RNA45SN1_45S_50ntFlanks</td><td>7977</td><td>8018</td><td>42</td><td>CGACCTCAGATCAGACGTGGCGACCCGCTGAATTTAAGCATA</td><td>507.7</td><td>10.7</td><td>1.4</td><td>7.7</td><td>1.0E-14</td><td>5.3E-14</td><td>MB231/MB468</td><td>CYTO</td></tr>
<tr><td>>hg38_21_+_8433172_8446622_RNA45SN1_45S_50ntFlanks@11565.11588.24|CTAGCAGCCGACTTAGAACTGGTG</td><td>>hg38_21_+_8433172_8446622_RNA45SN1_45S_50ntFlanks</td><td>11565</td><td>11588</td><td>24</td><td>CTAGCAGCCGACTTAGAACTGGTG</td><td>174.7</td><td>-11.3</td><td>1.5</td><td>-7.7</td><td>9.2E-15</td><td>5.3E-14</td><td>MB231/MB468</td><td>MP</td></tr>
<tr><td>>hg38_21_+_8433172_8446622_RNA45SN1_45S_50ntFlanks@10805.10826.22|GAACAATGTAGGTAAGGGAAGT</td><td>>hg38_21_+_8433172_8446622_RNA45SN1_45S_50ntFlanks</td><td>10805</td><td>10826</td><td>22</td><td>GAACAATGTAGGTAAGGGAAGT</td><td>262.3</td><td>-11.0</td><td>1.4</td><td>-7.7</td><td>1.1E-14</td><td>5.3E-14</td><td>MB231/MB468</td><td>NUC</td></tr>
<tr><td>>hg38_21_+_8433172_8446622_RNA45SN1_45S_50ntFlanks@6722.6740.19|ATTGCAGGACACATTGATC</td><td>>hg38_21_+_8433172_8446622_RNA45SN1_45S_50ntFlanks</td><td>6722</td><td>6740</td><td>19</td><td>ATTGCAGGACACATTGATC</td><td>1152.6</td><td>-5.1</td><td>0.7</td><td>-7.7</td><td>1.1E-14</td><td>5.3E-14</td><td>MB231/MB468</td><td>NUC</td></tr>
<tr><td>>hg38_21_+_8433172_8446622_RNA45SN1_45S_50ntFlanks@3985.4003.19|CGGCTTTGGTGACTCTAGA</td><td>>hg38_21_+_8433172_8446622_RNA45SN1_45S_50ntFlanks</td><td>3985</td><td>4003</td><td>19</td><td>CGGCTTTGGTGACTCTAGA</td><td>433.9</td><td>-12.5</td><td>1.6</td><td>-7.7</td><td>1.5E-14</td><td>5.4E-14</td><td>MB231/MB468</td><td>MITO</td></tr>
<tr><td>>hg38_21_+_8433172_8446622_RNA45SN1_45S_50ntFlanks@7989.8013.25|AGACGTGGCGACCCGCTGAATTTAA</td><td>>hg38_21_+_8433172_8446622_RNA45SN1_45S_50ntFlanks</td><td>7989</td><td>8013</td><td>25</td><td>AGACGTGGCGACCCGCTGAATTTAA</td><td>445.7</td><td>-5.0</td><td>0.6</td><td>-7.7</td><td>9.3E-15</td><td>5.4E-14</td><td>MB231/MB468</td><td>MP</td></tr>
<tr><td>>MI0000478|hsa-mir-149&WithFlank&2|+|240455995|240456095@21.42.22(+1A)&[MIMAT0000450&hsa-miR-149-5p&offsets|0|-1(+1A);m-195&2|+|240456015|240456037&offsets|0|-1(+1A)]|TCTGGCTCCGTGTCTTCACTCCA</td><td>>MI0000478|hsa-mir-149&WithFlank&2|+|240455995|240456095</td><td>21</td><td>42</td><td>22(+1A)</td><td>TCTGGCTCCGTGTCTTCACTCCA</td><td>113.7</td><td>9.7</td><td>1.3</td><td>7.7</td><td>1.9E-14</td><td>5.4E-14</td><td>BT20/MB468</td><td>NUC</td></tr>
<tr><td>>hg38_21_+_8433172_8446622_RNA45SN1_45S_50ntFlanks@12182.12211.30|CAAACGGTAACGCAGGTGTCCTAAGGCGAG</td><td>>hg38_21_+_8433172_8446622_RNA45SN1_45S_50ntFlanks</td><td>12182</td><td>12211</td><td>30</td><td>CAAACGGTAACGCAGGTGTCCTAAGGCGAG</td><td>212.6</td><td>-11.5</td><td>1.5</td><td>-7.7</td><td>1.5E-14</td><td>5.4E-14</td><td>MB231/MB468</td><td>MITO</td></tr>
<tr><td>>hg38_21_+_8433172_8446622_RNA45SN1_45S_50ntFlanks@11569.11603.35|CAGCCGACTTAGAACTGGTGCGGACCAGGGGAATC</td><td>>hg38_21_+_8433172_8446622_RNA45SN1_45S_50ntFlanks</td><td>11569</td><td>11603</td><td>35</td><td>CAGCCGACTTAGAACTGGTGCGGACCAGGGGAATC</td><td>6302.0</td><td>-11.1</td><td>1.4</td><td>-7.7</td><td>9.3E-15</td><td>5.4E-14</td><td>MB231/MB468</td><td>MP</td></tr>
<tr><td>>hg38_21_+_8433172_8446622_RNA45SN1_45S_50ntFlanks@13006.13028.23|CTATTGAAAGTCAGCCCTCGACA</td><td>>hg38_21_+_8433172_8446622_RNA45SN1_45S_50ntFlanks</td><td>13006</td><td>13028</td><td>23</td><td>CTATTGAAAGTCAGCCCTCGACA</td><td>172.3</td><td>-11.3</td><td>1.5</td><td>-7.7</td><td>9.3E-15</td><td>5.4E-14</td><td>MB231/MB468</td><td>MP</td></tr>
<tr><td>>MI0000298|hsa-mir-221&WithFlank&X|-|45746151|45746272@31.54.24&[MIMAT0004568&hsa-miR-221-5p&offsets|0|+2;m-405&X|-|45746220|45746242&offsets|0|+1]|ACCTGGCATACAATGTAGATTTCT</td><td>>MI0000298|hsa-mir-221&WithFlank&X|-|45746151|45746272</td><td>31</td><td>54</td><td>24</td><td>ACCTGGCATACAATGTAGATTTCT</td><td>2234.4</td><td>6.8</td><td>0.9</td><td>7.7</td><td>1.1E-14</td><td>5.4E-14</td><td>MB231/MB468</td><td>CYTO</td></tr>
<tr><td>>hg38_21_+_8433172_8446622_RNA45SN1_45S_50ntFlanks@12007.12043.37|GCCGCCGGTGAAATACCACTACTCTGATCGTTTTTTC</td><td>>hg38_21_+_8433172_8446622_RNA45SN1_45S_50ntFlanks</td><td>12007</td><td>12043</td><td>37</td><td>GCCGCCGGTGAAATACCACTACTCTGATCGTTTTTTC</td><td>1456.2</td><td>-14.4</td><td>1.9</td><td>-7.7</td><td>9.4E-15</td><td>5.4E-14</td><td>MB231/MB468</td><td>MP</td></tr>
<tr><td>>hg38_21_+_8433172_8446622_RNA45SN1_45S_50ntFlanks@8177.8203.27|TTCTGATCGAGGCCCAGCCCGTGGACG</td><td>>hg38_21_+_8433172_8446622_RNA45SN1_45S_50ntFlanks</td><td>8177</td><td>8203</td><td>27</td><td>TTCTGATCGAGGCCCAGCCCGTGGACG</td><td>1264.0</td><td>-4.8</td><td>0.6</td><td>-7.7</td><td>1.9E-14</td><td>5.4E-14</td><td>BT20/MB468</td><td>NUC</td></tr>
<tr><td>>hg38_21_+_8433172_8446622_RNA45SN1_45S_50ntFlanks@7975.8000.26|CGCGACCTCAGATCAGACGTGGCGAC</td><td>>hg38_21_+_8433172_8446622_RNA45SN1_45S_50ntFlanks</td><td>7975</td><td>8000</td><td>26</td><td>CGCGACCTCAGATCAGACGTGGCGAC</td><td>520.2</td><td>4.5</td><td>0.6</td><td>7.7</td><td>1.9E-14</td><td>5.4E-14</td><td>BT20/MB468</td><td>NUC</td></tr>
<tr><td>>hg38_21_+_8433172_8446622_RNA45SN1_45S_50ntFlanks@11882.11900.19|CTGTTGAGCTTGACTCTAG</td><td>>hg38_21_+_8433172_8446622_RNA45SN1_45S_50ntFlanks</td><td>11882</td><td>11900</td><td>19</td><td>CTGTTGAGCTTGACTCTAG</td><td>252.0</td><td>-11.8</td><td>1.5</td><td>-7.7</td><td>9.5E-15</td><td>5.5E-14</td><td>MB231/MB468</td><td>MP</td></tr>
<tr><td>>hg38_21_+_8433172_8446622_RNA45SN1_45S_50ntFlanks@3711.3737.27|GTTGATCCTGCCAGTAGCATATGCTTG</td><td>>hg38_21_+_8433172_8446622_RNA45SN1_45S_50ntFlanks</td><td>3711</td><td>3737</td><td>27</td><td>GTTGATCCTGCCAGTAGCATATGCTTG</td><td>447.2</td><td>-12.7</td><td>1.6</td><td>-7.7</td><td>9.5E-15</td><td>5.5E-14</td><td>MB231/MB468</td><td>MP</td></tr>
<tr><td>>hg38_21_+_8433172_8446622_RNA45SN1_45S_50ntFlanks@9471.9494.24|AGCACGAGCGCACGTGTTAGGACC</td><td>>hg38_21_+_8433172_8446622_RNA45SN1_45S_50ntFlanks</td><td>9471</td><td>9494</td><td>24</td><td>AGCACGAGCGCACGTGTTAGGACC</td><td>197.1</td><td>-11.5</td><td>1.5</td><td>-7.7</td><td>9.4E-15</td><td>5.5E-14</td><td>MB231/MB468</td><td>MP</td></tr>
<tr><td>>hg38_21_+_8433172_8446622_RNA45SN1_45S_50ntFlanks@6735.6756.22|TTGATCATCGACACTTCGAACG</td><td>>hg38_21_+_8433172_8446622_RNA45SN1_45S_50ntFlanks</td><td>6735</td><td>6756</td><td>22</td><td>TTGATCATCGACACTTCGAACG</td><td>170.2</td><td>-11.3</td><td>1.5</td><td>-7.7</td><td>9.5E-15</td><td>5.5E-14</td><td>MB231/MB468</td><td>MP</td></tr>
<tr><td>>hg38_21_+_8433172_8446622_RNA45SN1_45S_50ntFlanks@11208.11225.18|GCGGCGACTCTGGACGCG</td><td>>hg38_21_+_8433172_8446622_RNA45SN1_45S_50ntFlanks</td><td>11208</td><td>11225</td><td>18</td><td>GCGGCGACTCTGGACGCG</td><td>813.0</td><td>-9.1</td><td>1.2</td><td>-7.7</td><td>1.2E-14</td><td>5.5E-14</td><td>MB231/MB468</td><td>NUC</td></tr>
<tr><td>>hg38_21_+_8433172_8446622_RNA45SN1_45S_50ntFlanks@4000.4024.25|TAGATAACCTCGGGCCGATCGCACG</td><td>>hg38_21_+_8433172_8446622_RNA45SN1_45S_50ntFlanks</td><td>4000</td><td>4024</td><td>25</td><td>TAGATAACCTCGGGCCGATCGCACG</td><td>1443.1</td><td>-10.5</td><td>1.4</td><td>-7.7</td><td>1.9E-14</td><td>5.5E-14</td><td>BT20/MB468</td><td>NUC</td></tr>
<tr><td>>hg38_21_+_8433172_8446622_RNA45SN1_45S_50ntFlanks@9176.9197.22|CGCGCCGTCGGGCCCGGGGGAG</td><td>>hg38_21_+_8433172_8446622_RNA45SN1_45S_50ntFlanks</td><td>9176</td><td>9197</td><td>22</td><td>CGCGCCGTCGGGCCCGGGGGAG</td><td>468.4</td><td>-12.7</td><td>1.6</td><td>-7.7</td><td>1.6E-14</td><td>5.5E-14</td><td>MB231/MB468</td><td>MITO</td></tr>
<tr><td>>hg38_21_+_8433172_8446622_RNA45SN1_45S_50ntFlanks@8109.8140.32|GCGGGGCGCGGGACATGTGGCGTACGGAAGAC</td><td>>hg38_21_+_8433172_8446622_RNA45SN1_45S_50ntFlanks</td><td>8109</td><td>8140</td><td>32</td><td>GCGGGGCGCGGGACATGTGGCGTACGGAAGAC</td><td>425.9</td><td>-12.6</td><td>1.6</td><td>-7.7</td><td>9.5E-15</td><td>5.5E-14</td><td>MB231/MB468</td><td>MP</td></tr>
<tr><td>>MI0000734|hsa-mir-106b&WithFlank&7|-|100093987|100094080@58.77.20&[MIMAT0004672&hsa-miR-106b-3p&offsets|0|-2;m-87&7|-|100094003|100094023&offsets|0|-1]|CCGCACTGTGGGTACTTGCT</td><td>>MI0000734|hsa-mir-106b&WithFlank&7|-|100093987|100094080</td><td>58</td><td>77</td><td>20</td><td>CCGCACTGTGGGTACTTGCT</td><td>1035.8</td><td>5.2</td><td>0.7</td><td>7.7</td><td>1.2E-14</td><td>5.5E-14</td><td>MB231/MB468</td><td>NUC</td></tr>
<tr><td>>hg38_21_+_8433172_8446622_RNA45SN1_45S_50ntFlanks@11849.11871.23|CGGGCTTGGCGGAATCAGCGGGG</td><td>>hg38_21_+_8433172_8446622_RNA45SN1_45S_50ntFlanks</td><td>11849</td><td>11871</td><td>23</td><td>CGGGCTTGGCGGAATCAGCGGGG</td><td>225.1</td><td>-11.6</td><td>1.5</td><td>-7.7</td><td>1.6E-14</td><td>5.5E-14</td><td>MB231/MB468</td><td>MITO</td></tr>
<tr><td>>hg38_21_+_8433172_8446622_RNA45SN1_45S_50ntFlanks@10764.10785.22|TCCGCAGCAGGTCTCCAAGGTG</td><td>>hg38_21_+_8433172_8446622_RNA45SN1_45S_50ntFlanks</td><td>10764</td><td>10785</td><td>22</td><td>TCCGCAGCAGGTCTCCAAGGTG</td><td>177.8</td><td>-10.4</td><td>1.4</td><td>-7.7</td><td>1.2E-14</td><td>5.5E-14</td><td>MB231/MB468</td><td>NUC</td></tr>
<tr><td>>hg38_21_+_8433172_8446622_RNA45SN1_45S_50ntFlanks@12991.13009.19|CTCCCTCGCTGCGATCTAT</td><td>>hg38_21_+_8433172_8446622_RNA45SN1_45S_50ntFlanks</td><td>12991</td><td>13009</td><td>19</td><td>CTCCCTCGCTGCGATCTAT</td><td>741.5</td><td>-12.5</td><td>1.6</td><td>-7.7</td><td>1.2E-14</td><td>5.5E-14</td><td>MB231/MB468</td><td>NUC</td></tr>
<tr><td>>hg38_21_+_8433172_8446622_RNA45SN1_45S_50ntFlanks@11535.11560.26|TTCCGCGCGGCGCCTCGCCTCGGCCG</td><td>>hg38_21_+_8433172_8446622_RNA45SN1_45S_50ntFlanks</td><td>11535</td><td>11560</td><td>26</td><td>TTCCGCGCGGCGCCTCGCCTCGGCCG</td><td>228.9</td><td>-11.6</td><td>1.5</td><td>-7.7</td><td>1.6E-14</td><td>5.5E-14</td><td>MB231/MB468</td><td>MITO</td></tr>
<tr><td>>MI0000077|hsa-mir-21&WithFlank&17|+|59841260|59841343@14.34.21(+1U)&[MIMAT0000076&hsa-miR-21-5p&offsets|0|-1(+1U);m-1&17|+|59841273|59841294&offsets|0|-1(+1U)]|TAGCTTATCAGACTGATGTTGT</td><td>>MI0000077|hsa-mir-21&WithFlank&17|+|59841260|59841343</td><td>14</td><td>34</td><td>21(+1U)</td><td>TAGCTTATCAGACTGATGTTGT</td><td>692.0</td><td>5.6</td><td>0.7</td><td>7.7</td><td>1.9E-14</td><td>5.5E-14</td><td>BT20/MB468</td><td>NUC</td></tr>
<tr><td>>hg38_21_+_8433172_8446622_RNA45SN1_45S_50ntFlanks@11321.11340.20|CACGTCTCGTCGCGCGCGCG</td><td>>hg38_21_+_8433172_8446622_RNA45SN1_45S_50ntFlanks</td><td>11321</td><td>11340</td><td>20</td><td>CACGTCTCGTCGCGCGCGCG</td><td>147.8</td><td>-11.1</td><td>1.4</td><td>-7.7</td><td>9.6E-15</td><td>5.5E-14</td><td>MB231/MB468</td><td>MP</td></tr>
<tr><td>>hg38_21_+_8433172_8446622_RNA45SN1_45S_50ntFlanks@11641.11679.39|CGGCGGGTGTTGACGCGATGTGATTTCTGCCCAGTGCTC</td><td>>hg38_21_+_8433172_8446622_RNA45SN1_45S_50ntFlanks</td><td>11641</td><td>11679</td><td>39</td><td>CGGCGGGTGTTGACGCGATGTGATTTCTGCCCAGTGCTC</td><td>493.5</td><td>-12.8</td><td>1.7</td><td>-7.7</td><td>9.6E-15</td><td>5.5E-14</td><td>MB231/MB468</td><td>MP</td></tr>
<tr><td>>hg38_21_+_8433172_8446622_RNA45SN1_45S_50ntFlanks@10922.10947.26|CTGGACGAGGCGCCGCCGCCCCCCCC</td><td>>hg38_21_+_8433172_8446622_RNA45SN1_45S_50ntFlanks</td><td>10922</td><td>10947</td><td>26</td><td>CTGGACGAGGCGCCGCCGCCCCCCCC</td><td>276.1</td><td>-11.9</td><td>1.5</td><td>-7.7</td><td>1.6E-14</td><td>5.5E-14</td><td>MB231/MB468</td><td>MITO</td></tr>
<tr><td>>hg38_21_+_8433172_8446622_RNA45SN1_45S_50ntFlanks@5467.5499.33|GGCCCACGGCCCTGGCGGAGCGCTGAGAAGACG</td><td>>hg38_21_+_8433172_8446622_RNA45SN1_45S_50ntFlanks</td><td>5467</td><td>5499</td><td>33</td><td>GGCCCACGGCCCTGGCGGAGCGCTGAGAAGACG</td><td>7633.0</td><td>-9.5</td><td>1.2</td><td>-7.7</td><td>1.2E-14</td><td>5.5E-14</td><td>MB231/MB468</td><td>NUC</td></tr>
<tr><td>>hg38_21_+_8433172_8446622_RNA45SN1_45S_50ntFlanks@12201.12220.20|CCTAAGGCGAGCTCAGGGAG</td><td>>hg38_21_+_8433172_8446622_RNA45SN1_45S_50ntFlanks</td><td>12201</td><td>12220</td><td>20</td><td>CCTAAGGCGAGCTCAGGGAG</td><td>162.1</td><td>-11.2</td><td>1.4</td><td>-7.7</td><td>9.6E-15</td><td>5.6E-14</td><td>MB231/MB468</td><td>MP</td></tr>
<tr><td>>am_trna134_GluTTC_1_-_16861774_16861845@1.30.30,trna5_GluTTC_1_+_17199078_17199149@1.30.30,trna84_GluTTC_1_-_161391883_161391954@1.30.30,trna94_GluTTC_1_-_149664355_149664427@1.30.30|TCCCTGGTGGTCTAGTGGCTAGGATTCGGC</td><td>>am_trna134_GluTTC_1_-_16861774_16861845</td><td>1</td><td>30</td><td>30</td><td>TCCCTGGTGGTCTAGTGGCTAGGATTCGGC</td><td>291.0</td><td>-12.0</td><td>1.6</td><td>-7.7</td><td>9.7E-15</td><td>5.6E-14</td><td>MB231/MB468</td><td>MP</td></tr>
<tr><td>>hg38_21_+_8433172_8446622_RNA45SN1_45S_50ntFlanks@11577.11601.25|TTAGAACTGGTGCGGACCAGGGGAA</td><td>>hg38_21_+_8433172_8446622_RNA45SN1_45S_50ntFlanks</td><td>11577</td><td>11601</td><td>25</td><td>TTAGAACTGGTGCGGACCAGGGGAA</td><td>414.5</td><td>-8.6</td><td>1.1</td><td>-7.7</td><td>1.1E-14</td><td>5.6E-14</td><td>MB231/MB468</td><td>CYTO</td></tr>
<tr><td>>trna12_TrpCCA_17_+_19411494_19411565@30.52.23,trna168_TrpCCA_6_-_26331672_26331743@30.52.23,trna170_TrpCCA_6_-_26319330_26319401@30.52.23,trna1_TrpCCA_7_+_99067307_99067378@30.52.23,trna39_TrpCCA_17_-_8124187_8124258@30.52.23,trna6_TrpCCA_17_+_8089676_8089747@30.52.23|ACTCCAGATCAGAAGGTTGCGTG</td><td>>trna12_TrpCCA_17_+_19411494_19411565</td><td>30</td><td>52</td><td>23</td><td>ACTCCAGATCAGAAGGTTGCGTG</td><td>143.0</td><td>-9.9</td><td>1.3</td><td>-7.7</td><td>1.1E-14</td><td>5.6E-14</td><td>MB231/MB468</td><td>CYTO</td></tr>
<tr><td>>MI0015866|hsa-mir-4326&WithFlank&20|+|63286802|63286872@17.40.24(+1U)&[MIMAT0016888&hsa-miR-4326&offsets|0|+4(+1U);m-339&20|+|63286818|63286841&offsets|0|0(+1U)]|TGTTCCTCTGTCTCCCAGACTCTGT</td><td>>MI0015866|hsa-mir-4326&WithFlank&20|+|63286802|63286872</td><td>17</td><td>40</td><td>24(+1U)</td><td>TGTTCCTCTGTCTCCCAGACTCTGT</td><td>279.1</td><td>9.8</td><td>1.3</td><td>7.7</td><td>1.3E-14</td><td>5.6E-14</td><td>BT20/MB468</td><td>CYTO</td></tr>
<tr><td>>hg38_21_+_8433172_8446622_RNA45SN1_45S_50ntFlanks@4759.4782.24|ATCAGATACCGTCGTAGTTCCGAC</td><td>>hg38_21_+_8433172_8446622_RNA45SN1_45S_50ntFlanks</td><td>4759</td><td>4782</td><td>24</td><td>ATCAGATACCGTCGTAGTTCCGAC</td><td>2769.6</td><td>-13.5</td><td>1.7</td><td>-7.7</td><td>9.6E-15</td><td>5.6E-14</td><td>BT20/MB468</td><td>MITO</td></tr>
<tr><td>>MI0000809|hsa-mir-151a&WithFlank&8|-|140732558|140732659@53.74.22(+2U)&[MIMAT0000757&hsa-miR-151a-3p&offsets|0|+1(+2U);m-30&8|-|140732587|140732607&offsets|0|+1(+2U)]|CTAGACTGAAGCTCCTTGAGGATT</td><td>>MI0000809|hsa-mir-151a&WithFlank&8|-|140732558|140732659</td><td>53</td><td>74</td><td>22(+2U)</td><td>CTAGACTGAAGCTCCTTGAGGATT</td><td>534.3</td><td>8.2</td><td>1.1</td><td>7.7</td><td>9.6E-15</td><td>5.6E-14</td><td>BT20/MB468</td><td>MITO</td></tr>
</table>

<table>
<tr><td></td><td></td><td></td><td></td><td></td><td></td><td></td><td></td><td></td><td></td><td></td><td></td><td></td><td></td></tr>
<tr><td>>am_trna11_ProAGG_16_+_3241989_3242060@1.29.29,trna12_ProAGG_6_+_26555498_26555569@1.29.29,trna12_ProTGG_11_-_75946869_75946940@1.29.29,trna14_ProTGG_5_-_180615854_180615925@1.29.29,trna22_ProAGG_14_-_21081560_21081631@1.29.29,trna23_ProAGG_14_-_21077495_21077566@1.29.29,trna28_ProTGG_16_-_3234133_3234204@1.29.29,trna29_ProAGG_16_-_3232635_3232706@1.29.29,trna2_ProAGG_7_+_128423504_128423575@1.29.29,trna30_ProCGG_6_+_27059521_27059592@1.29.29,trna37_ProCGG_17_-_8126151_8126222@1.29.29,trna3_ProTGG_16_+_3208923_3208994@1.29.29,trna52_ProCGG_1_+_167683962_167684033@1.29.29,trna65_ProAGG_1_-_167684725_167684796@1.29.29,trna6_ProCGG_16_+_3222049_3222120@1.29.29,trna6_ProTGG_14_+_21152175_21152246@1.29.29,trna8_ProTGG_16_+_3238094_3238165@1.29.29,trna9_ProAGG_11_+_75946557_75946628@1.29.29,trna9_ProAGG_16_+_3239634_3239705@1.29.29|GGCTCGTTGGTCTAGGGGTATGATTCTCG</td><td>>am_trna11_ProAGG_16_+_3241989_3242060</td><td>1</td><td>29</td><td>29</td><td>GGCTCGTTGGTCTAGGGGTATGATTCTCG</td><td>7404.0</td><td>-6.6</td><td>0.9</td><td>-7.7</td><td>1.1E-14</td><td>5.6E-14</td><td>MB231/MB468</td><td>CYTO</td></tr>
<tr><td>>hg38_21_+_8433172_8446622_RNA45SN1_45S_50ntFlanks@10412.10431.20|AGTCGGTCCTGAGAGATGGG</td><td>>hg38_21_+_8433172_8446622_RNA45SN1_45S_50ntFlanks</td><td>10412</td><td>10431</td><td>20</td><td>AGTCGGTCCTGAGAGATGGG</td><td>185.4</td><td>-11.4</td><td>1.5</td><td>-7.7</td><td>9.8E-15</td><td>5.6E-14</td><td>MB231/MB468</td><td>MP</td></tr>
<tr><td>>hg38_21_+_8433172_8446622_RNA45SN1_45S_50ntFlanks@9027.9044.18|CGTCTCCTCGTGGGGGGG</td><td>>hg38_21_+_8433172_8446622_RNA45SN1_45S_50ntFlanks</td><td>9027</td><td>9044</td><td>18</td><td>CGTCTCCTCGTGGGGGGG</td><td>262.2</td><td>-11.8</td><td>1.5</td><td>-7.7</td><td>1.6E-14</td><td>5.6E-14</td><td>MB231/MB468</td><td>MITO</td></tr>
<tr><td>>hg38_21_+_8433172_8446622_RNA45SN1_45S_50ntFlanks@9554.9596.43|CGTAGCGGTCCTGACGTGCAAATCGGTCGTCCGACCTGGGTAT</td><td>>hg38_21_+_8433172_8446622_RNA45SN1_45S_50ntFlanks</td><td>9554</td><td>9596</td><td>43</td><td>CGTAGCGGTCCTGACGTGCAAATCGGTCGTCCGACCTGGGTAT</td><td>1469.5</td><td>-14.9</td><td>2.0</td><td>-7.7</td><td>2.0E-14</td><td>5.7E-14</td><td>BT20/MB468</td><td>NUC</td></tr>
<tr><td>>hg38_21_+_8433172_8446622_RNA45SN1_45S_50ntFlanks@10344.10382.39|ATTCAAACGAGAACTTTGAAGGCCGAAGTGGAGAAGGGT</td><td>>hg38_21_+_8433172_8446622_RNA45SN1_45S_50ntFlanks</td><td>10344</td><td>10382</td><td>39</td><td>ATTCAAACGAGAACTTTGAAGGCCGAAGTGGAGAAGGGT</td><td>314.2</td><td>9.2</td><td>1.2</td><td>7.7</td><td>1.1E-14</td><td>5.7E-14</td><td>MB231/MB468</td><td>CYTO</td></tr>
<tr><td>>hg38_21_+_8433172_8446622_RNA45SN1_45S_50ntFlanks@4557.4587.31|CAGCTAGGAATAATGGAATAGGACCGCGGTT</td><td>>hg38_21_+_8433172_8446622_RNA45SN1_45S_50ntFlanks</td><td>4557</td><td>4587</td><td>31</td><td>CAGCTAGGAATAATGGAATAGGACCGCGGTT</td><td>239.5</td><td>-11.8</td><td>1.5</td><td>-7.7</td><td>9.9E-15</td><td>5.7E-14</td><td>MB231/MB468</td><td>MP</td></tr>
<tr><td>>hg38_21_+_8433172_8446622_RNA45SN1_45S_50ntFlanks@10829.10852.24|GCAAGCCGGATCCGTAACTTCGGG</td><td>>hg38_21_+_8433172_8446622_RNA45SN1_45S_50ntFlanks</td><td>10829</td><td>10852</td><td>24</td><td>GCAAGCCGGATCCGTAACTTCGGG</td><td>246.7</td><td>-11.8</td><td>1.5</td><td>-7.7</td><td>9.9E-15</td><td>5.7E-14</td><td>MB231/MB468</td><td>MP</td></tr>
<tr><td>>hg38_21_+_8433172_8446622_RNA45SN1_45S_50ntFlanks@11873.11898.26|AAGAAGACCCTGTTGAGCTTGACTCT</td><td>>hg38_21_+_8433172_8446622_RNA45SN1_45S_50ntFlanks</td><td>11873</td><td>11898</td><td>26</td><td>AAGAAGACCCTGTTGAGCTTGACTCT</td><td>1260.0</td><td>-4.2</td><td>0.5</td><td>-7.7</td><td>1.1E-14</td><td>5.7E-14</td><td>MB231/MB468</td><td>CYTO</td></tr>
<tr><td>>hg38_21_+_8433172_8446622_RNA45SN1_45S_50ntFlanks@9930.9960.31|ATATAGACAGCAGGACGGTGGCCATGGAAGT</td><td>>hg38_21_+_8433172_8446622_RNA45SN1_45S_50ntFlanks</td><td>9930</td><td>9960</td><td>31</td><td>ATATAGACAGCAGGACGGTGGCCATGGAAGT</td><td>142.0</td><td>-11.0</td><td>1.4</td><td>-7.7</td><td>9.9E-15</td><td>5.7E-14</td><td>MB231/MB468</td><td>MP</td></tr>
<tr><td>>hg38_21_+_8433172_8446622_RNA45SN1_45S_50ntFlanks@11561.11583.23|GCGCCTAGCAGCCGACTTAGAAC</td><td>>hg38_21_+_8433172_8446622_RNA45SN1_45S_50ntFlanks</td><td>11561</td><td>11583</td><td>23</td><td>GCGCCTAGCAGCCGACTTAGAAC</td><td>223.4</td><td>-11.7</td><td>1.5</td><td>-7.7</td><td>9.9E-15</td><td>5.7E-14</td><td>MB231/MB468</td><td>MP</td></tr>
<tr><td>>hg38_21_+_8433172_8446622_RNA45SN1_45S_50ntFlanks@4298.4317.20|ATTGGAGGGCAAGTCTGGTG</td><td>>hg38_21_+_8433172_8446622_RNA45SN1_45S_50ntFlanks</td><td>4298</td><td>4317</td><td>20</td><td>ATTGGAGGGCAAGTCTGGTG</td><td>240.9</td><td>-11.8</td><td>1.5</td><td>-7.7</td><td>9.9E-15</td><td>5.7E-14</td><td>MB231/MB468</td><td>MP</td></tr>
<tr><td>>hg38_21_+_8433172_8446622_RNA45SN1_45S_50ntFlanks@4976.4993.18|CCGGACACGGACAGGATT</td><td>>hg38_21_+_8433172_8446622_RNA45SN1_45S_50ntFlanks</td><td>4976</td><td>4993</td><td>18</td><td>CCGGACACGGACAGGATT</td><td>648.2</td><td>9.1</td><td>1.2</td><td>7.7</td><td>1.3E-14</td><td>5.7E-14</td><td>BT20/MB468</td><td>CYTO</td></tr>
<tr><td>>hg38_21_+_8433172_8446622_RNA45SN1_45S_50ntFlanks@12151.12181.31|GTGGGGAGTTTGACTGGGGCGGTACACCTGT</td><td>>hg38_21_+_8433172_8446622_RNA45SN1_45S_50ntFlanks</td><td>12151</td><td>12181</td><td>31</td><td>GTGGGGAGTTTGACTGGGGCGGTACACCTGT</td><td>236.9</td><td>-10.9</td><td>1.4</td><td>-7.7</td><td>1.2E-14</td><td>5.7E-14</td><td>MB231/MB468</td><td>NUC</td></tr>
<tr><td>>MI0000088|hsa-mir-30a&WithFlank&6|-|71403545|71403627@54.74.21&[MIMAT0000088&hsa-miR-30a-3p&offsets|+1|0;m-38&6|-|71403554|71403575&offsets|+1|0]|TTTCAGTCGGATGTTTGCAGC</td><td>>MI0000088|hsa-mir-30a&WithFlank&6|-|71403545|71403627</td><td>54</td><td>74</td><td>21</td><td>TTTCAGTCGGATGTTTGCAGC</td><td>2927.2</td><td>7.0</td><td>0.9</td><td>7.7</td><td>1.6E-14</td><td>5.7E-14</td><td>MB231/MB468</td><td>MITO</td></tr>
<tr><td>>MI0000272|hsa-mir-182&WithFlank&7|-|129770377|129770498@29.52.24(+1A)&[MIMAT0000259&hsa-miR-182-5p&offsets|0|0(+1A);m-22&7|-|129770449|129770470&offsets|0|+2(+1A)]|TTTGGCAATGGTAGAACTCACACTA</td><td>>MI0000272|hsa-mir-182&WithFlank&7|-|129770377|129770498</td><td>29</td><td>52</td><td>24(+1A)</td><td>TTTGGCAATGGTAGAACTCACACTA</td><td>396.1</td><td>3.4</td><td>0.4</td><td>7.7</td><td>2.0E-14</td><td>5.7E-14</td><td>BT20/MB468</td><td>NUC</td></tr>
<tr><td>>trna11_GluTTC_15_-_26327381_26327452@1.32.32,trna3_GluTTC_13_-_45492062_45492133@1.32.32|TCCCACATGGTCTAGCGGTTAGGATTCCTGGT</td><td>>trna11_GluTTC_15_-_26327381_26327452</td><td>1</td><td>32</td><td>32</td><td>TCCCACATGGTCTAGCGGTTAGGATTCCTGGT</td><td>617.9</td><td>12.5</td><td>1.6</td><td>7.7</td><td>1.6E-14</td><td>5.7E-14</td><td>MB231/MB468</td><td>MITO</td></tr>
<tr><td>>hg38_21_+_8433172_8446622_RNA45SN1_45S_50ntFlanks@9711.9737.27|AAGCGAATGATTAGAGGTCTTGGGGCC</td><td>>hg38_21_+_8433172_8446622_RNA45SN1_45S_50ntFlanks</td><td>9711</td><td>9737</td><td>27</td><td>AAGCGAATGATTAGAGGTCTTGGGGCC</td><td>437.8</td><td>-12.6</td><td>1.6</td><td>-7.7</td><td>1.6E-14</td><td>5.7E-14</td><td>MB231/MB468</td><td>MITO</td></tr>
<tr><td>>hg38_21_+_8433172_8446622_RNA45SN1_45S_50ntFlanks@12894.12932.39|CTCGCCCGTCACGCACCGCACGTTCGTGGGGAACCTGGC</td><td>>hg38_21_+_8433172_8446622_RNA45SN1_45S_50ntFlanks</td><td>12894</td><td>12932</td><td>39</td><td>CTCGCCCGTCACGCACCGCACGTTCGTGGGGAACCTGGC</td><td>691.5</td><td>-5.4</td><td>0.7</td><td>-7.7</td><td>1.6E-14</td><td>5.7E-14</td><td>MB231/MB468</td><td>MITO</td></tr>
<tr><td>>hg38_21_+_8433172_8446622_RNA45SN1_45S_50ntFlanks@9454.9503.50|CCGCGCCGGGGAGGTGGAGCACGAGCGCACGTGTTAGGACCCGAAAGATG</td><td>>hg38_21_+_8433172_8446622_RNA45SN1_45S_50ntFlanks</td><td>9454</td><td>9503</td><td>50</td><td>CCGCGCCGGGGAGGTGGAGCACGAGCGCACGTGTTAGGACCCGAAAGATG</td><td>2575.2</td><td>-15.2</td><td>2.0</td><td>-7.7</td><td>1.0E-14</td><td>5.7E-14</td><td>MB231/MB468</td><td>MP</td></tr>
<tr><td>>hg38_21_+_8433172_8446622_RNA45SN1_45S_50ntFlanks@12081.12110.30|CTCTCGCTTCTGGCGCCAAGCGCCCGGCCG</td><td>>hg38_21_+_8433172_8446622_RNA45SN1_45S_50ntFlanks</td><td>12081</td><td>12110</td><td>30</td><td>CTCTCGCTTCTGGCGCCAAGCGCCCGGCCG</td><td>2334.5</td><td>-9.7</td><td>1.3</td><td>-7.7</td><td>1.6E-14</td><td>5.7E-14</td><td>MB231/MB468</td><td>MITO</td></tr>
<tr><td>>trna116_GluCTC_1_-_145399233_145399304@14.31.18,trna25_GluCTC_2_-_71273488_71273560@14.31.18,trna59_GluCTC_1_+_249168447_249168518@14.31.18,trna71_GluCTC_1_-_161439189_161439260@14.31.18,trna74_GluCTC_1_-_161431809_161431880@14.31.18,trna77_GluCTC_1_-_161424398_161424469@14.31.18,trna77_GluCTC_6_+_28949976_28950047@14.31.18,trna80_GluCTC_1_-_161417018_161417089@14.31.18,trna87_GluCTC_6_-_126101393_126101464@14.31.18|AGTGGTTAGGATTCGGCG</td><td>>trna116_GluCTC_1_-_145399233_145399304</td><td>14</td><td>31</td><td>18</td><td>AGTGGTTAGGATTCGGCG</td><td>900.2</td><td>5.0</td><td>0.6</td><td>7.7</td><td>1.3E-14</td><td>5.7E-14</td><td>BT20/MB468</td><td>CYTO</td></tr>
<tr><td>>hg38_21_+_8433172_8446622_RNA45SN1_45S_50ntFlanks@11314.11338.25|CCCACCCCACGTCTCGTCGCGCGCG</td><td>>hg38_21_+_8433172_8446622_RNA45SN1_45S_50ntFlanks</td><td>11314</td><td>11338</td><td>25</td><td>CCCACCCCACGTCTCGTCGCGCGCG</td><td>294.4</td><td>-11.2</td><td>1.4</td><td>-7.7</td><td>1.2E-14</td><td>5.7E-14</td><td>MB231/MB468</td><td>NUC</td></tr>
<tr><td>>hg38_21_+_8433172_8446622_RNA45SN1_45S_50ntFlanks@5421.5445.25|CGATTGGATGGTTTAGTGAGGCCCT</td><td>>hg38_21_+_8433172_8446622_RNA45SN1_45S_50ntFlanks</td><td>5421</td><td>5445</td><td>25</td><td>CGATTGGATGGTTTAGTGAGGCCCT</td><td>654.0</td><td>-12.3</td><td>1.6</td><td>-7.7</td><td>1.2E-14</td><td>5.7E-14</td><td>MB231/MB468</td><td>NUC</td></tr>
<tr><td>>hg38_21_+_8433172_8446622_RNA45SN1_45S_50ntFlanks@11561.11603.43|GCGCCTAGCAGCCGACTTAGAACTGGTGCGGACCAGGGGAATC</td><td>>hg38_21_+_8433172_8446622_RNA45SN1_45S_50ntFlanks</td><td>11561</td><td>11603</td><td>43</td><td>GCGCCTAGCAGCCGACTTAGAACTGGTGCGGACCAGGGGAATC</td><td>1161.4</td><td>-14.0</td><td>1.8</td><td>-7.7</td><td>1.0E-14</td><td>5.7E-14</td><td>MB231/MB468</td><td>MP</td></tr>
<tr><td>>hg38_21_+_8433172_8446622_RNA45SN1_45S_50ntFlanks@10873.10904.32|CTGGGTCGGTCGGGCTGGGGCGCGAAGCGGGG</td><td>>hg38_21_+_8433172_8446622_RNA45SN1_45S_50ntFlanks</td><td>10873</td><td>10904</td><td>32</td><td>CTGGGTCGGTCGGGCTGGGGCGCGAAGCGGGG</td><td>338.7</td><td>-7.7</td><td>1.0</td><td>-7.7</td><td>1.1E-14</td><td>5.8E-14</td><td>MB231/MB468</td><td>CYTO</td></tr>
<tr><td>>MI0000439|hsa-mir-23b&WithFlank&9|+|95085202|95085310@67.86.20(+1A)&[MIMAT0000418&hsa-miR-23b-3p&offsets|+3|0(+1A);m-42&9|+|95085265|95085285&offsets|+3|+2(+1A)]|ACATTGCCAGGGATTACCACA</td><td>>MI0000439|hsa-mir-23b&WithFlank&9|+|95085202|95085310</td><td>67</td><td>86</td><td>20(+1A)</td><td>ACATTGCCAGGGATTACCACA</td><td>921.5</td><td>28.0</td><td>3.6</td><td>7.7</td><td>1.1E-14</td><td>5.8E-14</td><td>MB231/MB468</td><td>CYTO</td></tr>
<tr><td>>hg38_21_+_8433172_8446622_RNA45SN1_45S_50ntFlanks@6684.6706.23|TCGATGAAGAACGCAGCTAGCTG</td><td>>hg38_21_+_8433172_8446622_RNA45SN1_45S_50ntFlanks</td><td>6684</td><td>6706</td><td>23</td><td>TCGATGAAGAACGCAGCTAGCTG</td><td>189.3</td><td>-11.4</td><td>1.5</td><td>-7.7</td><td>1.0E-14</td><td>5.8E-14</td><td>MB231/MB468</td><td>MP</td></tr>
<tr><td>>hg38_21_+_8433172_8446622_RNA45SN1_45S_50ntFlanks@8324.8356.33|CCGATAGTCAACAAGTACCGTAAGGGAAAGTTG</td><td>>hg38_21_+_8433172_8446622_RNA45SN1_45S_50ntFlanks</td><td>8324</td><td>8356</td><td>33</td><td>CCGATAGTCAACAAGTACCGTAAGGGAAAGTTG</td><td>809.7</td><td>-8.3</td><td>1.1</td><td>-7.7</td><td>1.1E-14</td><td>5.8E-14</td><td>MB231/MB468</td><td>CYTO</td></tr>
<tr><td>>hg38_21_+_8433172_8446622_RNA45SN1_45S_50ntFlanks@4128.4162.35|CGATTCCGGAGAGGGAGCCTGAGAAACGGCTACCA</td><td>>hg38_21_+_8433172_8446622_RNA45SN1_45S_50ntFlanks</td><td>4128</td><td>4162</td><td>35</td><td>CGATTCCGGAGAGGGAGCCTGAGAAACGGCTACCA</td><td>537.8</td><td>-6.2</td><td>0.8</td><td>-7.7</td><td>1.1E-14</td><td>5.8E-14</td><td>MB231/MB468</td><td>CYTO</td></tr>
<tr><td>>hg38_21_+_8433172_8446622_RNA45SN1_45S_50ntFlanks@10808.10828.21|CAATGTAGGTAAGGGAAGTCG</td><td>>hg38_21_+_8433172_8446622_RNA45SN1_45S_50ntFlanks</td><td>10808</td><td>10828</td><td>21</td><td>CAATGTAGGTAAGGGAAGTCG</td><td>2024.1</td><td>-11.2</td><td>1.5</td><td>-7.6</td><td>2.0E-14</td><td>5.8E-14</td><td>BT20/MB468</td><td>NUC</td></tr>
<tr><td>>hg38_21_+_8433172_8446622_RNA45SN1_45S_50ntFlanks@7985.8008.24|GATCAGACGTGGCGACCCGCTGAA</td><td>>hg38_21_+_8433172_8446622_RNA45SN1_45S_50ntFlanks</td><td>7985</td><td>8008</td><td>24</td><td>GATCAGACGTGGCGACCCGCTGAA</td><td>2418.6</td><td>-10.2</td><td>1.3</td><td>-7.7</td><td>1.0E-14</td><td>5.8E-14</td><td>MB231/MB468</td><td>MP</td></tr>
<tr><td>>hg38_21_+_8433172_8446622_RNA45SN1_45S_50ntFlanks@12081.12106.26|CTCTCGCTTCTGGCGCCAAGCGCCCG</td><td>>hg38_21_+_8433172_8446622_RNA45SN1_45S_50ntFlanks</td><td>12081</td><td>12106</td><td>26</td><td>CTCTCGCTTCTGGCGCCAAGCGCCCG</td><td>655.7</td><td>-7.1</td><td>0.9</td><td>-7.7</td><td>1.2E-14</td><td>5.8E-14</td><td>MB231/MB468</td><td>CYTO</td></tr>
<tr><td>>MI0000746|hsa-mir-99b&WithFlank&19|+|51692606|51692687@13.32.20(+1G)&[MIMAT0000689&hsa-miR-99b-5p&offsets|0|-2(+1G);m-10&19|+|51692618|51692639&offsets|0|-2(+1G)]|CACCCGTAGAACCGACCTTGG</td><td>>MI0000746|hsa-mir-99b&WithFlank&19|+|51692606|51692687</td><td>13</td><td>32</td><td>20(+1G)</td><td>CACCCGTAGAACCGACCTTGG</td><td>63.1</td><td>6.0</td><td>0.8</td><td>7.6</td><td>2.1E-14</td><td>5.8E-14</td><td>BT20/MB468</td><td>NUC</td></tr>
<tr><td>>hg38_21_+_8433172_8446622_RNA45SN1_45S_50ntFlanks@12160.12179.20|TTGACTGGGGCGGTACACCT</td><td>>hg38_21_+_8433172_8446622_RNA45SN1_45S_50ntFlanks</td><td>12160</td><td>12179</td><td>20</td><td>TTGACTGGGGCGGTACACCT</td><td>358.5</td><td>-12.3</td><td>1.6</td><td>-7.7</td><td>1.7E-14</td><td>5.8E-14</td><td>MB231/MB468</td><td>MITO</td></tr>
<tr><td>>hg38_21_+_8433172_8446622_RNA45SN1_45S_50ntFlanks@10808.10839.32|CAATGTAGGTAAGGGAAGTCGGCAAGCCGGAT</td><td>>hg38_21_+_8433172_8446622_RNA45SN1_45S_50ntFlanks</td><td>10808</td><td>10839</td><td>32</td><td>CAATGTAGGTAAGGGAAGTCGGCAAGCCGGAT</td><td>219.2</td><td>-11.6</td><td>1.5</td><td>-7.7</td><td>1.0E-14</td><td>5.8E-14</td><td>MB231/MB468</td><td>MP</td></tr>
<tr><td>>hg38_21_+_8433172_8446622_RNA45SN1_45S_50ntFlanks@10869.10891.23|AGGGCTGGGTCGGTCGGGCTGGG</td><td>>hg38_21_+_8433172_8446622_RNA45SN1_45S_50ntFlanks</td><td>10869</td><td>10891</td><td>23</td><td>AGGGCTGGGTCGGTCGGGCTGGG</td><td>1578.2</td><td>-8.3</td><td>1.1</td><td>-7.7</td><td>1.0E-14</td><td>5.8E-14</td><td>MB231/MB468</td><td>MP</td></tr>
<tr><td>>hg38_21_+_8433172_8446622_RNA45SN1_45S_50ntFlanks@11569.11603.35|CAGCCGACTTAGAACTGGTGCGGACCAGGGGAATC</td><td>>hg38_21_+_8433172_8446622_RNA45SN1_45S_50ntFlanks</td><td>11569</td><td>11603</td><td>35</td><td>CAGCCGACTTAGAACTGGTGCGGACCAGGGGAATC</td><td>7667.7</td><td>-4.7</td><td>0.6</td><td>-7.7</td><td>1.0E-14</td><td>5.8E-14</td><td>BT20/MB468</td><td>MITO</td></tr>
<tr><td>>hg38_21_+_8433172_8446622_RNA45SN1_45S_50ntFlanks@6689.6720.32|GAAGAACGCAGCTAGCTGCGAGAATTAATGTG</td><td>>hg38_21_+_8433172_8446622_RNA45SN1_45S_50ntFlanks</td><td>6689</td><td>6720</td><td>32</td><td>GAAGAACGCAGCTAGCTGCGAGAATTAATGTG</td><td>226.1</td><td>-11.6</td><td>1.5</td><td>-7.7</td><td>1.7E-14</td><td>5.9E-14</td><td>MB231/MB468</td><td>MITO</td></tr>
<tr><td>>hg38_21_+_8433172_8446622_RNA45SN1_45S_50ntFlanks@11875.11906.32|GAAGACCCTGTTGAGCTTGACTCTAGTCTGGC</td><td>>hg38_21_+_8433172_8446622_RNA45SN1_45S_50ntFlanks</td><td>11875</td><td>11906</td><td>32</td><td>GAAGACCCTGTTGAGCTTGACTCTAGTCTGGC</td><td>143.9</td><td>-11.0</td><td>1.4</td><td>-7.7</td><td>1.0E-14</td><td>5.9E-14</td><td>MB231/MB468</td><td>MP</td></tr>
<tr><td>>MI0000298|hsa-mir-221&WithFlank&X|-|45746151|45746272@31.54.24&[MIMAT0004568&hsa-miR-221-5p&offsets|0|+2;m-405&X|-|45746220|45746242&offsets|0|+1]|ACCTGGCATACAATGTAGATTTCT</td><td>>MI0000298|hsa-mir-221&WithFlank&X|-|45746151|45746272</td><td>31</td><td>54</td><td>24</td><td>ACCTGGCATACAATGTAGATTTCT</td><td>745.9</td><td>5.5</td><td>0.7</td><td>7.7</td><td>1.4E-14</td><td>5.9E-14</td><td>BT20/MB468</td><td>CYTO</td></tr>
<tr><td>>hg38_21_+_8433172_8446622_RNA45SN1_45S_50ntFlanks@8686.8706.21|CCGGCTCCGGGACGGCTGGGA</td><td>>hg38_21_+_8433172_8446622_RNA45SN1_45S_50ntFlanks</td><td>8686</td><td>8706</td><td>21</td><td>CCGGCTCCGGGACGGCTGGGA</td><td>1575.7</td><td>10.8</td><td>1.4</td><td>7.7</td><td>1.2E-14</td><td>5.9E-14</td><td>MB231/MB468</td><td>CYTO</td></tr>
<tr><td>>hg38_21_+_8433172_8446622_RNA45SN1_45S_50ntFlanks@9504.9522.19|GTGAACTATGCCTGGGCAG</td><td>>hg38_21_+_8433172_8446622_RNA45SN1_45S_50ntFlanks</td><td>9504</td><td>9522</td><td>19</td><td>GTGAACTATGCCTGGGCAG</td><td>1307.6</td><td>-6.9</td><td>0.9</td><td>-7.7</td><td>1.2E-14</td><td>5.9E-14</td><td>MB231/MB468</td><td>CYTO</td></tr>
</table>

<table>
<tr><td>>hg38_21_+_8433172_8446622_RNA45SN1_45S_50ntFlanks@12141.12180.40|ACAGTGCCAGGTGGGGAGTTTGACTGGGGCGGTACACCTG</td><td>>hg38_21_+_8433172_8446622_RNA45SN1_45S_50ntFlanks</td><td>12141</td><td>12180</td><td>40</td><td>ACAGTGCCAGGTGGGGAGTTTGACTGGGGCGGTACACCTG</td><td>259.9</td><td>-11.8</td><td>1.5</td><td>-7.7</td><td>1.7E-14</td><td>5.9E-14</td><td>MB231/MB468</td><td>MITO</td></tr>
<tr><td>>hg38_21_+_8433172_8446622_RNA45SN1_45S_50ntFlanks@10393.10410.18|CAGCAGTTGAACATGGGT</td><td>>hg38_21_+_8433172_8446622_RNA45SN1_45S_50ntFlanks</td><td>10393</td><td>10410</td><td>18</td><td>CAGCAGTTGAACATGGGT</td><td>3222.0</td><td>-8.3</td><td>1.1</td><td>-7.7</td><td>1.3E-14</td><td>5.9E-14</td><td>MB231/MB468</td><td>NUC</td></tr>
<tr><td>>hg38_21_+_8433172_8446622_RNA45SN1_45S_50ntFlanks@12159.12182.24|TTTGACTGGGGCGGTACACCTGTC</td><td>>hg38_21_+_8433172_8446622_RNA45SN1_45S_50ntFlanks</td><td>12159</td><td>12182</td><td>24</td><td>TTTGACTGGGGCGGTACACCTGTC</td><td>144.7</td><td>-10.1</td><td>1.3</td><td>-7.7</td><td>1.3E-14</td><td>5.9E-14</td><td>MB231/MB468</td><td>NUC</td></tr>
<tr><td>>hg38_21_+_8433172_8446622_RNA45SN1_45S_50ntFlanks@11875.11893.19|GAAGACCCTGTTGAGCTTG</td><td>>hg38_21_+_8433172_8446622_RNA45SN1_45S_50ntFlanks</td><td>11875</td><td>11893</td><td>19</td><td>GAAGACCCTGTTGAGCTTG</td><td>1251.5</td><td>-9.8</td><td>1.3</td><td>-7.7</td><td>1.0E-14</td><td>5.9E-14</td><td>MB231/MB468</td><td>MP</td></tr>
<tr><td>>hg38_21_+_8433172_8446622_RNA45SN1_45S_50ntFlanks@9736.9766.31|CCGAAACGATCTCAACCTATTCTCAAACTTT</td><td>>hg38_21_+_8433172_8446622_RNA45SN1_45S_50ntFlanks</td><td>9736</td><td>9766</td><td>31</td><td>CCGAAACGATCTCAACCTATTCTCAAACTTT</td><td>365.7</td><td>-12.3</td><td>1.6</td><td>-7.7</td><td>1.7E-14</td><td>5.9E-14</td><td>MB231/MB468</td><td>MITO</td></tr>
<tr><td>>hg38_21_+_8433172_8446622_RNA45SN1_45S_50ntFlanks@6671.6689.19|CTCGGCTCGTGCGTCGATG</td><td>>hg38_21_+_8433172_8446622_RNA45SN1_45S_50ntFlanks</td><td>6671</td><td>6689</td><td>19</td><td>CTCGGCTCGTGCGTCGATG</td><td>991.2</td><td>-13.8</td><td>1.8</td><td>-7.7</td><td>1.0E-14</td><td>5.9E-14</td><td>MB231/MB468</td><td>MP</td></tr>
<tr><td>>hg38_21_+_8433172_8446622_RNA45SN1_45S_50ntFlanks@4588.4622.35|CTATTTTGTTGGTTTTCGGAACTGAGGCCATGATT</td><td>>hg38_21_+_8433172_8446622_RNA45SN1_45S_50ntFlanks</td><td>4588</td><td>4622</td><td>35</td><td>CTATTTTGTTGGTTTTCGGAACTGAGGCCATGATT</td><td>448.4</td><td>-7.2</td><td>0.9</td><td>-7.7</td><td>1.7E-14</td><td>5.9E-14</td><td>MB231/MB468</td><td>MITO</td></tr>
<tr><td>>hg38_21_+_8433172_8446622_RNA45SN1_45S_50ntFlanks@6786.6807.22|TACGCCTGTCTGAGCGTCGCTT</td><td>>hg38_21_+_8433172_8446622_RNA45SN1_45S_50ntFlanks</td><td>6786</td><td>6807</td><td>22</td><td>TACGCCTGTCTGAGCGTCGCTT</td><td>230.5</td><td>4.4</td><td>0.6</td><td>7.7</td><td>1.3E-14</td><td>6.0E-14</td><td>MB231/MB468</td><td>NUC</td></tr>
<tr><td>>hg38_21_+_8433172_8446622_RNA45SN1_45S_50ntFlanks@6649.6695.47|TACGACTCTTAGCGGTGGATCACTCGGCTCGTGCGTCGATGAAGAAC</td><td>>hg38_21_+_8433172_8446622_RNA45SN1_45S_50ntFlanks</td><td>6649</td><td>6695</td><td>47</td><td>TACGACTCTTAGCGGTGGATCACTCGGCTCGTGCGTCGATGAAGAAC</td><td>198.5</td><td>8.7</td><td>1.1</td><td>7.7</td><td>1.2E-14</td><td>6.0E-14</td><td>MB231/MB468</td><td>CYTO</td></tr>
<tr><td>>hg38_21_+_8433172_8446622_RNA45SN1_45S_50ntFlanks@11452.11474.23|CTCCTCGGCGCGCGGCGGCGGCG</td><td>>hg38_21_+_8433172_8446622_RNA45SN1_45S_50ntFlanks</td><td>11452</td><td>11474</td><td>23</td><td>CTCCTCGGCGCGCGGCGGCGGCG</td><td>136.7</td><td>-9.8</td><td>1.3</td><td>-7.7</td><td>1.2E-14</td><td>6.0E-14</td><td>MB231/MB468</td><td>CYTO</td></tr>
<tr><td>>hg38_21_+_8433172_8446622_RNA45SN1_45S_50ntFlanks@10408.10429.22|GGTCAGTCGGTCCTGAGAGATG</td><td>>hg38_21_+_8433172_8446622_RNA45SN1_45S_50ntFlanks</td><td>10408</td><td>10429</td><td>22</td><td>GGTCAGTCGGTCCTGAGAGATG</td><td>2433.3</td><td>-11.0</td><td>1.4</td><td>-7.7</td><td>1.0E-14</td><td>6.0E-14</td><td>MB231/MB468</td><td>MP</td></tr>
<tr><td>>hg38_21_+_8433172_8446622_RNA45SN1_45S_50ntFlanks@12605.12623.19|TATGTGCTTGGCTGAGGAG</td><td>>hg38_21_+_8433172_8446622_RNA45SN1_45S_50ntFlanks</td><td>12605</td><td>12623</td><td>19</td><td>TATGTGCTTGGCTGAGGAG</td><td>171.3</td><td>-11.3</td><td>1.5</td><td>-7.7</td><td>1.0E-14</td><td>6.0E-14</td><td>MB231/MB468</td><td>MP</td></tr>
<tr><td>>hg38_21_+_8433172_8446622_RNA45SN1_45S_50ntFlanks@6663.6698.36|GTGGATCACTCGGCTCGTGCGTCGATGAAGAACGCA</td><td>>hg38_21_+_8433172_8446622_RNA45SN1_45S_50ntFlanks</td><td>6663</td><td>6698</td><td>36</td><td>GTGGATCACTCGGCTCGTGCGTCGATGAAGAACGCA</td><td>223.5</td><td>-11.7</td><td>1.5</td><td>-7.7</td><td>1.0E-14</td><td>6.0E-14</td><td>MB231/MB468</td><td>MP</td></tr>
<tr><td>>hg38_21_+_8433172_8446622_RNA45SN1_45S_50ntFlanks@8437.8470.34|AGGATTCAACCCGGCGGCGGGTCCGGCCGTGTCG</td><td>>hg38_21_+_8433172_8446622_RNA45SN1_45S_50ntFlanks</td><td>8437</td><td>8470</td><td>34</td><td>AGGATTCAACCCGGCGGCGGGTCCGGCCGTGTCG</td><td>157.2</td><td>-11.2</td><td>1.4</td><td>-7.7</td><td>1.0E-14</td><td>6.0E-14</td><td>MB231/MB468</td><td>MP</td></tr>
<tr><td>>hg38_21_+_8433172_8446622_RNA45SN1_45S_50ntFlanks@4722.4751.30|TCATTAATCAAGAACGAAAGTCGGAGGTTC</td><td>>hg38_21_+_8433172_8446622_RNA45SN1_45S_50ntFlanks</td><td>4722</td><td>4751</td><td>30</td><td>TCATTAATCAAGAACGAAAGTCGGAGGTTC</td><td>380.5</td><td>-10.7</td><td>1.4</td><td>-7.7</td><td>1.0E-14</td><td>6.0E-14</td><td>BT20/MB468</td><td>MITO</td></tr>
<tr><td>>hg38_21_+_8433172_8446622_RNA45SN1_45S_50ntFlanks@12510.12527.18|TCGTGAGACAGGTTAGTT</td><td>>hg38_21_+_8433172_8446622_RNA45SN1_45S_50ntFlanks</td><td>12510</td><td>12527</td><td>18</td><td>TCGTGAGACAGGTTAGTT</td><td>1371.3</td><td>-7.9</td><td>1.0</td><td>-7.7</td><td>1.2E-14</td><td>6.0E-14</td><td>MB231/MB468</td><td>CYTO</td></tr>
<tr><td>>hg38_21_+_8433172_8446622_RNA45SN1_45S_50ntFlanks@3870.3898.29|AGCTAATACATGCCGACGGGCGCTGACCC</td><td>>hg38_21_+_8433172_8446622_RNA45SN1_45S_50ntFlanks</td><td>3870</td><td>3898</td><td>29</td><td>AGCTAATACATGCCGACGGGCGCTGACCC</td><td>348.0</td><td>-12.3</td><td>1.6</td><td>-7.7</td><td>1.1E-14</td><td>6.0E-14</td><td>MB231/MB468</td><td>MP</td></tr>
<tr><td>>hg38_21_+_8433172_8446622_RNA45SN1_45S_50ntFlanks@12568.12587.20|TCAGTACGAGAGGAACCGCA</td><td>>hg38_21_+_8433172_8446622_RNA45SN1_45S_50ntFlanks</td><td>12568</td><td>12587</td><td>20</td><td>TCAGTACGAGAGGAACCGCA</td><td>206.1</td><td>-11.6</td><td>1.5</td><td>-7.7</td><td>1.1E-14</td><td>6.0E-14</td><td>MB231/MB468</td><td>MP</td></tr>
<tr><td>>hg38_21_+_8433172_8446622_RNA45SN1_45S_50ntFlanks@4884.4918.35|CAAAGCTGAAACTTAAAGGAATTGACGGAAGGGCA</td><td>>hg38_21_+_8433172_8446622_RNA45SN1_45S_50ntFlanks</td><td>4884</td><td>4918</td><td>35</td><td>CAAAGCTGAAACTTAAAGGAATTGACGGAAGGGCA</td><td>261.2</td><td>-11.9</td><td>1.5</td><td>-7.7</td><td>1.1E-14</td><td>6.0E-14</td><td>MB231/MB468</td><td>MP</td></tr>
<tr><td>>MI0000283|hsa-mir-203a&WithFlank&14|+|104117399|104117520@72.92.21&[MIMAT0000264&hsa-miR-203a-3p&offsets|+1|0;m-79&14|+|104117469|104117490&offsets|+1|0]|TGAAATGTTTAGGACCACTAG</td><td>>MI0000283|hsa-mir-203a&WithFlank&14|+|104117399|104117520</td><td>72</td><td>92</td><td>21</td><td>TGAAATGTTTAGGACCACTAG</td><td>770.6</td><td>9.3</td><td>1.2</td><td>7.7</td><td>1.0E-14</td><td>6.0E-14</td><td>BT20/MB468</td><td>MITO</td></tr>
<tr><td>>hg38_21_+_8433172_8446622_RNA45SN1_45S_50ntFlanks@12624.12661.38|CCAATGGGGCGAAGCTACCATCTGTGGGATTATGACTG</td><td>>hg38_21_+_8433172_8446622_RNA45SN1_45S_50ntFlanks</td><td>12624</td><td>12661</td><td>38</td><td>CCAATGGGGCGAAGCTACCATCTGTGGGATTATGACTG</td><td>2521.1</td><td>-11.0</td><td>1.4</td><td>-7.7</td><td>1.1E-14</td><td>6.0E-14</td><td>MB231/MB468</td><td>MP</td></tr>
<tr><td>>hg38_21_+_8433172_8446622_RNA45SN1_45S_50ntFlanks@3813.3837.25|TTATGGTTCCTTTGGTCGCTCGCTC</td><td>>hg38_21_+_8433172_8446622_RNA45SN1_45S_50ntFlanks</td><td>3813</td><td>3837</td><td>25</td><td>TTATGGTTCCTTTGGTCGCTCGCTC</td><td>182.2</td><td>-11.4</td><td>1.5</td><td>-7.7</td><td>1.1E-14</td><td>6.0E-14</td><td>MB231/MB468</td><td>MP</td></tr>
<tr><td>>MI0003137|hsa-mir-193b&WithFlank&16|+|14303961|14304055@57.77.21(+1G)&[MIMAT0002819&hsa-miR-193b-3p&offsets|0|-1(+1G);m-166&16|+|14304017|14304038&offsets|0|-1(+1G)]|AACTGGCCCTCAAAGTCCCGCG</td><td>>MI0003137|hsa-mir-193b&WithFlank&16|+|14303961|14304055</td><td>57</td><td>77</td><td>21(+1G)</td><td>AACTGGCCCTCAAAGTCCCGCG</td><td>51.2</td><td>8.5</td><td>1.1</td><td>7.6</td><td>2.1E-14</td><td>6.0E-14</td><td>BT20/MB468</td><td>NUC</td></tr>
<tr><td>>hg38_21_+_8433172_8446622_RNA45SN1_45S_50ntFlanks@12685.12708.24|CCCAGGCGGAACGATACGGCAGCG</td><td>>hg38_21_+_8433172_8446622_RNA45SN1_45S_50ntFlanks</td><td>12685</td><td>12708</td><td>24</td><td>CCCAGGCGGAACGATACGGCAGCG</td><td>203.3</td><td>-10.6</td><td>1.4</td><td>-7.7</td><td>1.3E-14</td><td>6.0E-14</td><td>MB231/MB468</td><td>NUC</td></tr>
<tr><td>>hg38_21_+_8433172_8446622_RNA45SN1_45S_50ntFlanks@4044.4063.20|CCCATTCGAACGTCTGCCCT</td><td>>hg38_21_+_8433172_8446622_RNA45SN1_45S_50ntFlanks</td><td>4044</td><td>4063</td><td>20</td><td>CCCATTCGAACGTCTGCCCT</td><td>339.4</td><td>-12.2</td><td>1.6</td><td>-7.7</td><td>1.7E-14</td><td>6.0E-14</td><td>MB231/MB468</td><td>MITO</td></tr>
<tr><td>>hg38_21_+_8433172_8446622_RNA45SN1_45S_50ntFlanks@11873.11897.25|AAGAAGACCCTGTTGAGCTTGACTC</td><td>>hg38_21_+_8433172_8446622_RNA45SN1_45S_50ntFlanks</td><td>11873</td><td>11897</td><td>25</td><td>AAGAAGACCCTGTTGAGCTTGACTC</td><td>1498.4</td><td>-8.4</td><td>1.1</td><td>-7.6</td><td>2.1E-14</td><td>6.1E-14</td><td>BT20/MB468</td><td>NUC</td></tr>
<tr><td>>hg38_21_+_8433172_8446622_RNA45SN1_45S_50ntFlanks@12822.12871.50|CCGGGACCGGGGTCCGGTGCGGAGTGCCCTTCGTCCTGGGAAACGGGGCG</td><td>>hg38_21_+_8433172_8446622_RNA45SN1_45S_50ntFlanks</td><td>12822</td><td>12871</td><td>50</td><td>CCGGGACCGGGGTCCGGTGCGGAGTGCCCTTCGTCCTGGGAAACGGGGCG</td><td>426.3</td><td>-12.5</td><td>1.6</td><td>-7.7</td><td>1.7E-14</td><td>6.1E-14</td><td>MB231/MB468</td><td>MITO</td></tr>
<tr><td>>hg38_21_+_8433172_8446622_RNA45SN1_45S_50ntFlanks@10351.10368.18|CGAGAACTTTGAAGGCCG</td><td>>hg38_21_+_8433172_8446622_RNA45SN1_45S_50ntFlanks</td><td>10351</td><td>10368</td><td>18</td><td>CGAGAACTTTGAAGGCCG</td><td>13319.8</td><td>-6.8</td><td>0.9</td><td>-7.7</td><td>1.4E-14</td><td>6.1E-14</td><td>BT20/MB468</td><td>CYTO</td></tr>
<tr><td>>hg38_21_+_8433172_8446622_RNA45SN1_45S_50ntFlanks@8737.8758.22|CCCGTCCGTCCGTCCGTCCGTC</td><td>>hg38_21_+_8433172_8446622_RNA45SN1_45S_50ntFlanks</td><td>8737</td><td>8758</td><td>22</td><td>CCCGTCCGTCCGTCCGTCCGTC</td><td>385.8</td><td>-12.5</td><td>1.6</td><td>-7.7</td><td>1.1E-14</td><td>6.1E-14</td><td>MB231/MB468</td><td>MP</td></tr>
<tr><td>>hg38_21_+_8433172_8446622_RNA45SN1_45S_50ntFlanks@9156.9187.32|CCAGTGCGCCCCGGGCGGGTCGCGCCGTCGGG</td><td>>hg38_21_+_8433172_8446622_RNA45SN1_45S_50ntFlanks</td><td>9156</td><td>9187</td><td>32</td><td>CCAGTGCGCCCCGGGCGGGTCGCGCCGTCGGG</td><td>154.0</td><td>-11.1</td><td>1.4</td><td>-7.7</td><td>1.1E-14</td><td>6.1E-14</td><td>MB231/MB468</td><td>MP</td></tr>
<tr><td>>hg38_21_+_8433172_8446622_RNA45SN1_45S_50ntFlanks@13005.13025.21|TCTATTGAAAGTCAGCCCTCG</td><td>>hg38_21_+_8433172_8446622_RNA45SN1_45S_50ntFlanks</td><td>13005</td><td>13025</td><td>21</td><td>TCTATTGAAAGTCAGCCCTCG</td><td>360.7</td><td>-6.9</td><td>0.9</td><td>-7.7</td><td>1.2E-14</td><td>6.1E-14</td><td>MB231/MB468</td><td>CYTO</td></tr>
<tr><td>>hg38_21_+_8433172_8446622_RNA45SN1_45S_50ntFlanks@10798.10828.31|CATGTTGGAACAATGTAGGTAAGGGAAGTCG</td><td>>hg38_21_+_8433172_8446622_RNA45SN1_45S_50ntFlanks</td><td>10798</td><td>10828</td><td>31</td><td>CATGTTGGAACAATGTAGGTAAGGGAAGTCG</td><td>161.6</td><td>-11.2</td><td>1.4</td><td>-7.7</td><td>1.1E-14</td><td>6.1E-14</td><td>MB231/MB468</td><td>MP</td></tr>
<tr><td>>hg38_21_+_8433172_8446622_RNA45SN1_45S_50ntFlanks@12324.12345.22|TTGGGTTTTAAGCAGGAGGTGT</td><td>>hg38_21_+_8433172_8446622_RNA45SN1_45S_50ntFlanks</td><td>12324</td><td>12345</td><td>22</td><td>TTGGGTTTTAAGCAGGAGGTGT</td><td>688.1</td><td>-9.5</td><td>1.2</td><td>-7.7</td><td>1.4E-14</td><td>6.1E-14</td><td>BT20/MB468</td><td>CYTO</td></tr>
<tr><td>>hg38_21_+_8433172_8446622_RNA45SN1_45S_50ntFlanks@9786.9820.35|CTCGCTGGCGTGGAGCCGGGCGTGGAATGCGAGTG</td><td>>hg38_21_+_8433172_8446622_RNA45SN1_45S_50ntFlanks</td><td>9786</td><td>9820</td><td>35</td><td>CTCGCTGGCGTGGAGCCGGGCGTGGAATGCGAGTG</td><td>170.5</td><td>-11.3</td><td>1.5</td><td>-7.7</td><td>1.1E-14</td><td>6.1E-14</td><td>MB231/MB468</td><td>MP</td></tr>
<tr><td>>MI0003643|hsa-mir-629&WithFlank&15|-|70079366|70079474@28.49.22&[MIMAT0004810&hsa-miR-629-5p&offsets|0|+1;m-150&15|-|70079427|70079447&offsets|0|+1]|TGGGTTTACGTTGGGAGAACTT</td><td>>MI0003643|hsa-mir-629&WithFlank&15|-|70079366|70079474</td><td>28</td><td>49</td><td>22</td><td>TGGGTTTACGTTGGGAGAACTT</td><td>450.5</td><td>12.0</td><td>1.6</td><td>7.7</td><td>1.8E-14</td><td>6.1E-14</td><td>MB231/MB468</td><td>MITO</td></tr>
<tr><td>>hg38_21_+_8433172_8446622_RNA45SN1_45S_50ntFlanks@11208.11230.23|GCGGCGACTCTGGACGCGAGCCG</td><td>>hg38_21_+_8433172_8446622_RNA45SN1_45S_50ntFlanks</td><td>11208</td><td>11230</td><td>23</td><td>GCGGCGACTCTGGACGCGAGCCG</td><td>965.4</td><td>-7.3</td><td>1.0</td><td>-7.7</td><td>1.8E-14</td><td>6.1E-14</td><td>MB231/MB468</td><td>MITO</td></tr>
<tr><td>>am_trna2_GlyGCC_21_-_18827107_18827177@1.26.26,trna35_GlyGCC_1_+_161413094_161413164@1.26.26,trna37_GlyGCC_1_+_161420467_161420537@1.26.26,trna39_GlyGCC_1_+_161427898_161427968@1.26.26,trna41_GlyGCC_1_+_161435258_161435328@1.26.26|GCATGGGTGGTTCAGTGGTAGAATTC</td><td>>am_trna2_GlyGCC_21_-_18827107_18827177</td><td>1</td><td>26</td><td>26</td><td>GCATGGGTGGTTCAGTGGTAGAATTC</td><td>785.9</td><td>7.2</td><td>0.9</td><td>7.7</td><td>1.1E-14</td><td>6.1E-14</td><td>BT20/MB468</td><td>MITO</td></tr>
<tr><td>>hg38_21_+_8433172_8446622_RNA45SN1_45S_50ntFlanks@11821.11844.24|CTATCCAGCGAAACCACAGCCAAG</td><td>>hg38_21_+_8433172_8446622_RNA45SN1_45S_50ntFlanks</td><td>11821</td><td>11844</td><td>24</td><td>CTATCCAGCGAAACCACAGCCAAG</td><td>182.5</td><td>-7.0</td><td>0.9</td><td>-7.7</td><td>1.2E-14</td><td>6.1E-14</td><td>MB231/MB468</td><td>CYTO</td></tr>
<tr><td>>hg38_21_+_8433172_8446622_RNA45SN1_45S_50ntFlanks@3979.4001.23|CGCCGGCGGCTTTGGTGACTCTA</td><td>>hg38_21_+_8433172_8446622_RNA45SN1_45S_50ntFlanks</td><td>3979</td><td>4001</td><td>23</td><td>CGCCGGCGGCTTTGGTGACTCTA</td><td>386.3</td><td>-11.6</td><td>1.5</td><td>-7.7</td><td>1.3E-14</td><td>6.1E-14</td><td>MB231/MB468</td><td>NUC</td></tr>
<tr><td>>hg38_21_+_8433172_8446622_RNA45SN1_45S_50ntFlanks@7969.8009.41|CCGAGACGCGACCTCAGATCAGACGTGGCGACCCGCTGAAT</td><td>>hg38_21_+_8433172_8446622_RNA45SN1_45S_50ntFlanks</td><td>7969</td><td>8009</td><td>41</td><td>CCGAGACGCGACCTCAGATCAGACGTGGCGACCCGCTGAAT</td><td>187.0</td><td>5.3</td><td>0.7</td><td>7.7</td><td>1.3E-14</td><td>6.1E-14</td><td>MB231/MB468</td><td>NUC</td></tr>
<tr><td>>hg38_21_+_8433172_8446622_RNA45SN1_45S_50ntFlanks@6734.6774.41|ATTGATCATCGACACTTCGAACGCACTTGCGGCCCCGGGTT</td><td>>hg38_21_+_8433172_8446622_RNA45SN1_45S_50ntFlanks</td><td>6734</td><td>6774</td><td>41</td><td>ATTGATCATCGACACTTCGAACGCACTTGCGGCCCCGGGTT</td><td>454.6</td><td>8.5</td><td>1.1</td><td>7.7</td><td>1.2E-14</td><td>6.1E-14</td><td>MB231/MB468</td><td>CYTO</td></tr>
<tr><td>>hg38_21_+_8433172_8446622_RNA45SN1_45S_50ntFlanks@4731.4761.31|AAGAACGAAAGTCGGAGGTTCGAAGACGATC</td><td>>hg38_21_+_8433172_8446622_RNA45SN1_45S_50ntFlanks</td><td>4731</td><td>4761</td><td>31</td><td>AAGAACGAAAGTCGGAGGTTCGAAGACGATC</td><td>3662.4</td><td>5.2</td><td>0.7</td><td>7.7</td><td>1.4E-14</td><td>6.1E-14</td><td>BT20/MB468</td><td>CYTO</td></tr>
<tr><td>>hg38_21_+_8433172_8446622_RNA45SN1_45S_50ntFlanks@8497.8514.18|CCGTTCCTCCCGACCCCT</td><td>>hg38_21_+_8433172_8446622_RNA45SN1_45S_50ntFlanks</td><td>8497</td><td>8514</td><td>18</td><td>CCGTTCCTCCCGACCCCT</td><td>259.1</td><td>-11.8</td><td>1.5</td><td>-7.7</td><td>1.8E-14</td><td>6.1E-14</td><td>MB231/MB468</td><td>MITO</td></tr>
<tr><td>>hg38_21_+_8433172_8446622_RNA45SN1_45S_50ntFlanks@12147.12166.20|CCAGGTGGGGAGTTTGACTG</td><td>>hg38_21_+_8433172_8446622_RNA45SN1_45S_50ntFlanks</td><td>12147</td><td>12166</td><td>20</td><td>CCAGGTGGGGAGTTTGACTG</td><td>231.0</td><td>-11.7</td><td>1.5</td><td>-7.7</td><td>1.1E-14</td><td>6.1E-14</td><td>MB231/MB468</td><td>MP</td></tr>
<tr><td>>MI0000077|hsa-mir-21&WithFlank&17|+|59841260|59841343@16.36.21&[MIMAT0000076&hsa-miR-21-5p&offsets|+2|+1;m-1&17|+|59841273|59841294&offsets|+2|+1]|GCTTATCAGACTGATGTTGAC</td><td>>MI0000077|hsa-mir-21&WithFlank&17|+|59841260|59841343</td><td>16</td><td>36</td><td>21</td><td>GCTTATCAGACTGATGTTGAC</td><td>257.8</td><td>10.0</td><td>1.3</td><td>7.7</td><td>1.3E-14</td><td>6.2E-14</td><td>MB231/MB468</td><td>NUC</td></tr>
<tr><td>>hg38_21_+_8433172_8446622_RNA45SN1_45S_50ntFlanks@10663.10710.48|CCTTGGAAAGCGTCGCGGTTCCGGCGGCGTCCGGTGAGCTCTCGCTGG</td><td>>hg38_21_+_8433172_8446622_RNA45SN1_45S_50ntFlanks</td><td>10663</td><td>10710</td><td>48</td><td>CCTTGGAAAGCGTCGCGGTTCCGGCGGCGTCCGGTGAGCTCTCGCTGG</td><td>1515.1</td><td>-15.0</td><td>2.0</td><td>-7.6</td><td>2.2E-14</td><td>6.2E-14</td><td>BT20/MB468</td><td>NUC</td></tr>
<tr><td>>hg38_21_+_8433172_8446622_RNA45SN1_45S_50ntFlanks@3900.3932.33|CTTCGCGGGGGGGATGCGTGCATTTATCAGATC</td><td>>hg38_21_+_8433172_8446622_RNA45SN1_45S_50ntFlanks</td><td>3900</td><td>3932</td><td>33</td><td>CTTCGCGGGGGGGATGCGTGCATTTATCAGATC</td><td>283.3</td><td>-12.0</td><td>1.6</td><td>-7.7</td><td>1.1E-14</td><td>6.2E-14</td><td>MB231/MB468</td><td>MP</td></tr>
<tr><td>>hg38_21_+_8433172_8446622_RNA45SN1_45S_50ntFlanks@9623.9658.36|TAGTAGCTGGTTCCCTCCGAAGTTTCCCTCAGGATA</td><td>>hg38_21_+_8433172_8446622_RNA45SN1_45S_50ntFlanks</td><td>9623</td><td>9658</td><td>36</td><td>TAGTAGCTGGTTCCCTCCGAAGTTTCCCTCAGGATA</td><td>343.9</td><td>-12.3</td><td>1.6</td><td>-7.7</td><td>1.1E-14</td><td>6.2E-14</td><td>MB231/MB468</td><td>MP</td></tr>
<tr><td>>hg38_21_+_8433172_8446622_RNA45SN1_45S_50ntFlanks@6733.6754.22|CATTGATCATCGACACTTCGAA</td><td>>hg38_21_+_8433172_8446622_RNA45SN1_45S_50ntFlanks</td><td>6733</td><td>6754</td><td>22</td><td>CATTGATCATCGACACTTCGAA</td><td>238.0</td><td>-11.8</td><td>1.5</td><td>-7.7</td><td>1.1E-14</td><td>6.2E-14</td><td>MB231/MB468</td><td>MP</td></tr>
<tr><td>>hg38_21_+_8433172_8446622_RNA45SN1_45S_50ntFlanks@4922.4941.20|CCAGGAGTGGAGCCTGCGGC</td><td>>hg38_21_+_8433172_8446622_RNA45SN1_45S_50ntFlanks</td><td>4922</td><td>4941</td><td>20</td><td>CCAGGAGTGGAGCCTGCGGC</td><td>515.2</td><td>-5.3</td><td>0.7</td><td>-7.7</td><td>1.1E-14</td><td>6.2E-14</td><td>MB231/MB468</td><td>MP</td></tr>
<tr><td>>MI0000102|hsa-mir-100&WithFlank&11|-|122152223|122152314@19.38.20(+1C)&[MIMAT0000098&hsa-miR-100-5p&offsets|0|-2(+1C);m-27&11|-|122152276|122152296&offsets|0|-1(+1C)]|AACCCGTAGATCCGAACTTGC</td><td>>MI0000102|hsa-mir-100&WithFlank&11|-|122152223|122152314</td><td>19</td><td>38</td><td>20(+1C)</td><td>AACCCGTAGATCCGAACTTGC</td><td>26597.1</td><td>11.9</td><td>1.5</td><td>7.7</td><td>1.1E-14</td><td>6.2E-14</td><td>MB231/MB468</td><td>MP</td></tr>
</table>

<table>
<tr><td>>hg38_21_+_8433172_8446622_RNA45SN1_45S_50ntFlanks@10869.10892.24|AGGGCTGGGTCGGTCGGGCTGGGG</td><td>>hg38_21_+_8433172_8446622_RNA45SN1_45S_50ntFlanks</td><td>10869</td><td>10892</td><td>24</td><td>AGGGCTGGGTCGGTCGGGCTGGGG</td><td>697.1</td><td>-12.2</td><td>1.6</td><td>-7.7</td><td>1.2E-14</td><td>6.2E-14</td><td>MB231/MB468</td><td>CYTO</td></tr>
<tr><td>>MI0000767|hsa-mir-365a&WithFlank&16|+|14309279|14309377@62.81.20(+2U);MI0000769|hsa-mir-365b&WithFlank&17|+|31575405|31575527@74.93.20(+2U)&[MIMAT0000710&hsa-miR-365a-3p&offsets|0|-2(+2U);m-176&16|+|14309340|14309361&offsets|0|-2(+2U)];[MIMAT0022834&hsa-miR-365b-3p&offsets|0|-2(+2U);m-175&17|+|31575478|31575499&offsets|0|-2(+2U)]|TAATGCCCCTAAAAATCCTTTT</td><td>>MI0000767|hsa-mir-365a&WithFlank&16|+|14309279|14309377</td><td>62</td><td>81</td><td>20(+2U)</td><td>TAATGCCCCTAAAAATCCTTTT</td><td>865.5</td><td>11.0</td><td>1.4</td><td>7.7</td><td>1.5E-14</td><td>6.2E-14</td><td>BT20/MB468</td><td>CYTO</td></tr>
<tr><td>>MI0016005|hsa-mir-3615&WithFlank&17|+|74748607|74748705@57.79.23&[MIMAT0017994&hsa-miR-3615&offsets|0|+2;m-271&17|+|74748663|74748684&offsets|0|+1]|TCTCTCGGCTCCTCGCGGCTCGC</td><td>>MI0016005|hsa-mir-3615&WithFlank&17|+|74748607|74748705</td><td>57</td><td>79</td><td>23</td><td>TCTCTCGGCTCCTCGCGGCTCGC</td><td>477.3</td><td>5.9</td><td>0.8</td><td>7.7</td><td>1.2E-14</td><td>6.2E-14</td><td>MB231/MB468</td><td>CYTO</td></tr>
<tr><td>>MI0000750|hsa-mir-26a-2&WithFlank&12|-|57824603|57824698@20.41.22(+1C);MI0000083|hsa-mir-26a-1&WithFlank&3|+|37969398|37969486@16.37.22(+1C)&[MIMAT0000082&hsa-miR-26a-5p&offsets|0|0(+1C);m-33&12|-|57824658|57824679&offsets|0|0(+1C)];[MIMAT0000082_1&hsa-miR-26a-5p&offsets|0|0(+1C);m-34&3|+|37969413|37969434&offsets|0|0(+1C)]|TTCAAGTAATCCAGGATAGGCTC</td><td>>MI0000750|hsa-mir-26a-2&WithFlank&12|-|57824603|57824698</td><td>20</td><td>41</td><td>22(+1C)</td><td>TTCAAGTAATCCAGGATAGGCTC</td><td>429.7</td><td>6.2</td><td>0.8</td><td>7.7</td><td>1.3E-14</td><td>6.2E-14</td><td>MB231/MB468</td><td>NUC</td></tr>
<tr><td>>hg38_21_+_8433172_8446622_RNA45SN1_45S_50ntFlanks@11661.11679.19|TGATTTCTGCCCAGTGCTC</td><td>>hg38_21_+_8433172_8446622_RNA45SN1_45S_50ntFlanks</td><td>11661</td><td>11679</td><td>19</td><td>TGATTTCTGCCCAGTGCTC</td><td>3586.0</td><td>-8.6</td><td>1.1</td><td>-7.7</td><td>1.2E-14</td><td>6.2E-14</td><td>MB231/MB468</td><td>CYTO</td></tr>
<tr><td>>hg38_21_+_8433172_8446622_RNA45SN1_45S_50ntFlanks@12412.12432.21|ATCCTTCGATGTCGGCTCTTC</td><td>>hg38_21_+_8433172_8446622_RNA45SN1_45S_50ntFlanks</td><td>12412</td><td>12432</td><td>21</td><td>ATCCTTCGATGTCGGCTCTTC</td><td>353.4</td><td>-10.6</td><td>1.4</td><td>-7.7</td><td>1.1E-14</td><td>6.2E-14</td><td>BT20/MB468</td><td>MITO</td></tr>
<tr><td>>hg38_21_+_8433172_8446622_RNA45SN1_45S_50ntFlanks@10128.10149.22|CGGCGGCGGCGGCGGCGGCGGG</td><td>>hg38_21_+_8433172_8446622_RNA45SN1_45S_50ntFlanks</td><td>10128</td><td>10149</td><td>22</td><td>CGGCGGCGGCGGCGGCGGCGGG</td><td>150.2</td><td>-10.2</td><td>1.3</td><td>-7.7</td><td>1.3E-14</td><td>6.2E-14</td><td>MB231/MB468</td><td>NUC</td></tr>
<tr><td>>hg38_21_+_8433172_8446622_RNA45SN1_45S_50ntFlanks@8845.8866.22|CGGCGGCGGCGGCGGCGGCGGG</td><td>>hg38_21_+_8433172_8446622_RNA45SN1_45S_50ntFlanks</td><td>8845</td><td>8866</td><td>22</td><td>CGGCGGCGGCGGCGGCGGCGGG</td><td>150.2</td><td>-10.2</td><td>1.3</td><td>-7.7</td><td>1.3E-14</td><td>6.2E-14</td><td>MB231/MB468</td><td>NUC</td></tr>
<tr><td>>MI0000087|hsa-mir-29a&WithFlank&7|-|130876741|130876816@47.68.22&[MIMAT0000086&hsa-miR-29a-3p&offsets|-1|-1;m-26&7|-|130876749|130876769&offsets|-1|0]|CTAGCACCATCTGAAATCGGTT</td><td>>MI0000087|hsa-mir-29a&WithFlank&7|-|130876741|130876816</td><td>47</td><td>68</td><td>22</td><td>CTAGCACCATCTGAAATCGGTT</td><td>21554.8</td><td>5.8</td><td>0.8</td><td>7.7</td><td>1.1E-14</td><td>6.2E-14</td><td>MB231/MB468</td><td>MP</td></tr>
<tr><td>>hg38_21_+_8433172_8446622_RNA45SN1_45S_50ntFlanks@11759.11777.19|CCTCGTCATCTAATTAGTG</td><td>>hg38_21_+_8433172_8446622_RNA45SN1_45S_50ntFlanks</td><td>11759</td><td>11777</td><td>19</td><td>CCTCGTCATCTAATTAGTG</td><td>150.9</td><td>-11.1</td><td>1.4</td><td>-7.7</td><td>1.1E-14</td><td>6.2E-14</td><td>MB231/MB468</td><td>MP</td></tr>
<tr><td>>hg38_21_+_8433172_8446622_RNA45SN1_45S_50ntFlanks@11534.11555.22|GTTCCGCGCGGCGCCTCGCCTC</td><td>>hg38_21_+_8433172_8446622_RNA45SN1_45S_50ntFlanks</td><td>11534</td><td>11555</td><td>22</td><td>GTTCCGCGCGGCGCCTCGCCTC</td><td>364.9</td><td>-11.5</td><td>1.5</td><td>-7.7</td><td>1.3E-14</td><td>6.2E-14</td><td>MB231/MB468</td><td>NUC</td></tr>
<tr><td>>hg38_21_+_8433172_8446622_RNA45SN1_45S_50ntFlanks@8179.8211.33|CTGATCGAGGCCCAGCCCGTGGACGGTGTGAGG</td><td>>hg38_21_+_8433172_8446622_RNA45SN1_45S_50ntFlanks</td><td>8179</td><td>8211</td><td>33</td><td>CTGATCGAGGCCCAGCCCGTGGACGGTGTGAGG</td><td>244.1</td><td>-10.9</td><td>1.4</td><td>-7.7</td><td>1.3E-14</td><td>6.2E-14</td><td>MB231/MB468</td><td>NUC</td></tr>
<tr><td>>MI0000298|hsa-mir-221&WithFlank&X|-|45746151|45746272@71.90.20&[MIMAT0000278&hsa-miR-221-3p&offsets|0|-3;m-91&X|-|45746181|45746202&offsets|0|-2]|AGCTACATTGTCTGCTGGGT</td><td>>MI0000298|hsa-mir-221&WithFlank&X|-|45746151|45746272</td><td>71</td><td>90</td><td>20</td><td>AGCTACATTGTCTGCTGGGT</td><td>608.9</td><td>7.5</td><td>1.0</td><td>7.7</td><td>1.5E-14</td><td>6.3E-14</td><td>BT20/MB468</td><td>CYTO</td></tr>
<tr><td>>hg38_21_+_8433172_8446622_RNA45SN1_45S_50ntFlanks@5003.5034.32|ATAGCTCTTTCTCGATTCCGTGGGTGGTGGTG</td><td>>hg38_21_+_8433172_8446622_RNA45SN1_45S_50ntFlanks</td><td>5003</td><td>5034</td><td>32</td><td>ATAGCTCTTTCTCGATTCCGTGGGTGGTGGTG</td><td>854.2</td><td>-13.5</td><td>1.8</td><td>-7.7</td><td>1.8E-14</td><td>6.3E-14</td><td>MB231/MB468</td><td>MITO</td></tr>
<tr><td>>MI0000074|hsa-mir-19b-1&WithFlank&13|+|91351186|91351284@60.81.22;MI0000075|hsa-mir-19b-2&WithFlank&X|-|134169665|134169772@68.89.22&[MIMAT0000074&hsa-miR-19b-3p&offsets|0|-1;m-122&13|+|91351245|91351267&offsets|0|-1];[MIMAT0000074_1&hsa-miR-19b-3p&offsets|0|-1;m-121&X|-|134169683|134169705&offsets|0|-1]|TGTGCAAATCCATGCAAAACTG</td><td>>MI0000074|hsa-mir-19b-1&WithFlank&13|+|91351186|91351284</td><td>60</td><td>81</td><td>22</td><td>TGTGCAAATCCATGCAAAACTG</td><td>276.7</td><td>5.9</td><td>0.8</td><td>7.7</td><td>1.3E-14</td><td>6.3E-14</td><td>MB231/MB468</td><td>NUC</td></tr>
<tr><td>>hg38_21_+_8433172_8446622_RNA45SN1_45S_50ntFlanks@11934.11967.34|ATAAGTGGGAGGCCCCCGGCGCCCCCCCGGTGTC</td><td>>hg38_21_+_8433172_8446622_RNA45SN1_45S_50ntFlanks</td><td>11934</td><td>11967</td><td>34</td><td>ATAAGTGGGAGGCCCCCGGCGCCCCCCCGGTGTC</td><td>182.6</td><td>-11.4</td><td>1.5</td><td>-7.7</td><td>1.1E-14</td><td>6.3E-14</td><td>MB231/MB468</td><td>MP</td></tr>
<tr><td>>hg38_21_+_8433172_8446622_RNA45SN1_45S_50ntFlanks@12670.12703.34|TAAGTCAGAATCCCGCCCAGGCGGAACGATACGG</td><td>>hg38_21_+_8433172_8446622_RNA45SN1_45S_50ntFlanks</td><td>12670</td><td>12703</td><td>34</td><td>TAAGTCAGAATCCCGCCCAGGCGGAACGATACGG</td><td>220.7</td><td>-11.7</td><td>1.5</td><td>-7.7</td><td>1.1E-14</td><td>6.3E-14</td><td>MB231/MB468</td><td>MP</td></tr>
<tr><td>>hg38_21_+_8433172_8446622_RNA45SN1_45S_50ntFlanks@4069.4093.25|CTTTCGATGGTAGTCGCCGTGCCTA</td><td>>hg38_21_+_8433172_8446622_RNA45SN1_45S_50ntFlanks</td><td>4069</td><td>4093</td><td>25</td><td>CTTTCGATGGTAGTCGCCGTGCCTA</td><td>220.8</td><td>-11.6</td><td>1.5</td><td>-7.7</td><td>1.1E-14</td><td>6.3E-14</td><td>MB231/MB468</td><td>MP</td></tr>
<tr><td>>MI0000457|hsa-mir-141&WithFlank&12|+|6964091|6964197@65.85.21&[MIMAT0000432&hsa-miR-141-3p&offsets|0|-1;m-134&12|+|6964155|6964175&offsets|0|0]|TAACACTGTCTGGTAAAGATG</td><td>>MI0000457|hsa-mir-141&WithFlank&12|+|6964091|6964197</td><td>65</td><td>85</td><td>21</td><td>TAACACTGTCTGGTAAAGATG</td><td>396.0</td><td>12.0</td><td>1.5</td><td>8.0</td><td>1.0E-15</td><td>6.3E-14</td><td>BT20/MB231</td><td>CYTO</td></tr>
<tr><td>>MI0003628|hsa-mir-615&WithFlank&12|+|54033944|54034051@67.86.20(+1C)&[MIMAT0003283&hsa-miR-615-3p&offsets|0|-2(+1C);m-495&12|+|54034010|54034030&offsets|0|-1(+1C)]|TCCGAGCCTGGGTCTCCCTCC</td><td>>MI0003628|hsa-mir-615&WithFlank&12|+|54033944|54034051</td><td>67</td><td>86</td><td>20(+1C)</td><td>TCCGAGCCTGGGTCTCCCTCC</td><td>264.4</td><td>7.0</td><td>0.9</td><td>8.0</td><td>1.1E-15</td><td>6.3E-14</td><td>BT20/MB231</td><td>CYTO</td></tr>
<tr><td>>am_trna2_GlyGCC_21_-_18827107_18827177@1.25.25,trna35_GlyGCC_1_+_161413094_161413164@1.25.25,trna37_GlyGCC_1_+_161420467_161420537@1.25.25,trna39_GlyGCC_1_+_161427898_161427968@1.25.25,trna41_GlyGCC_1_+_161435258_161435328@1.25.25|GCATGGGTGGTTCAGTGGTAGAATT</td><td>>am_trna2_GlyGCC_21_-_18827107_18827177</td><td>1</td><td>25</td><td>25</td><td>GCATGGGTGGTTCAGTGGTAGAATT</td><td>1378.8</td><td>6.2</td><td>0.8</td><td>8.0</td><td>1.0E-15</td><td>6.3E-14</td><td>BT20/MB231</td><td>CYTO</td></tr>
<tr><td>>hg38_21_+_8433172_8446622_RNA45SN1_45S_50ntFlanks@12405.12426.22|CTTTTTGATCCTTCGATGTCGG</td><td>>hg38_21_+_8433172_8446622_RNA45SN1_45S_50ntFlanks</td><td>12405</td><td>12426</td><td>22</td><td>CTTTTTGATCCTTCGATGTCGG</td><td>6629.8</td><td>-8.5</td><td>1.1</td><td>-7.7</td><td>1.3E-14</td><td>6.4E-14</td><td>MB231/MB468</td><td>CYTO</td></tr>
<tr><td>>hg38_21_+_8433172_8446622_RNA45SN1_45S_50ntFlanks@4732.4766.35|AGAACGAAAGTCGGAGGTTCGAAGACGATCAGATA</td><td>>hg38_21_+_8433172_8446622_RNA45SN1_45S_50ntFlanks</td><td>4732</td><td>4766</td><td>35</td><td>AGAACGAAAGTCGGAGGTTCGAAGACGATCAGATA</td><td>2263.3</td><td>8.8</td><td>1.2</td><td>7.7</td><td>1.5E-14</td><td>6.4E-14</td><td>BT20/MB468</td><td>CYTO</td></tr>
<tr><td>>MI0000482|hsa-mir-185&WithFlank&22|+|20033133|20033226@21.42.22(+1A)&[MIMAT0000455&hsa-miR-185-5p&offsets|0|0(+1A);m-145&22|+|20033153|20033174&offsets|0|0(+1A)]|TGGAGAGAAAGGCAGTTCCTGAA</td><td>>MI0000482|hsa-mir-185&WithFlank&22|+|20033133|20033226</td><td>21</td><td>42</td><td>22(+1A)</td><td>TGGAGAGAAAGGCAGTTCCTGAA</td><td>693.0</td><td>8.0</td><td>1.0</td><td>7.7</td><td>1.5E-14</td><td>6.4E-14</td><td>BT20/MB468</td><td>CYTO</td></tr>
<tr><td>>MI0000095|hsa-mir-93&WithFlank&7|-|100093762|100093853@17.38.22(+1A)&[MIMAT0000093&hsa-miR-93-5p&offsets|0|-1(+1A);m-29&7|-|100093815|100093837&offsets|0|-1(+1A)]|CAAAGTGCTGTTCGTGCAGGTAA</td><td>>MI0000095|hsa-mir-93&WithFlank&7|-|100093762|100093853</td><td>17</td><td>38</td><td>22(+1A)</td><td>CAAAGTGCTGTTCGTGCAGGTAA</td><td>621.5</td><td>5.1</td><td>0.7</td><td>7.7</td><td>1.4E-14</td><td>6.4E-14</td><td>MB231/MB468</td><td>NUC</td></tr>
<tr><td>>hg38_21_+_8433172_8446622_RNA45SN1_45S_50ntFlanks@6646.6678.33|TCGTACGACTCTTAGCGGTGGATCACTCGGCTC</td><td>>hg38_21_+_8433172_8446622_RNA45SN1_45S_50ntFlanks</td><td>6646</td><td>6678</td><td>33</td><td>TCGTACGACTCTTAGCGGTGGATCACTCGGCTC</td><td>2341.3</td><td>5.1</td><td>0.7</td><td>7.7</td><td>1.5E-14</td><td>6.4E-14</td><td>BT20/MB468</td><td>CYTO</td></tr>
<tr><td>>hg38_21_+_8433172_8446622_RNA45SN1_45S_50ntFlanks@11661.11682.22|TGATTTCTGCCCAGTGCTCTGA</td><td>>hg38_21_+_8433172_8446622_RNA45SN1_45S_50ntFlanks</td><td>11661</td><td>11682</td><td>22</td><td>TGATTTCTGCCCAGTGCTCTGA</td><td>657.2</td><td>-4.8</td><td>0.6</td><td>-7.7</td><td>1.5E-14</td><td>6.4E-14</td><td>BT20/MB468</td><td>CYTO</td></tr>
<tr><td>>hg38_21_+_8433172_8446622_RNA45SN1_45S_50ntFlanks@4728.4759.32|ATCAAGAACGAAAGTCGGAGGTTCGAAGACGA</td><td>>hg38_21_+_8433172_8446622_RNA45SN1_45S_50ntFlanks</td><td>4728</td><td>4759</td><td>32</td><td>ATCAAGAACGAAAGTCGGAGGTTCGAAGACGA</td><td>310.7</td><td>-10.6</td><td>1.4</td><td>-7.7</td><td>1.5E-14</td><td>6.4E-14</td><td>BT20/MB468</td><td>CYTO</td></tr>
<tr><td>>hg38_21_+_8433172_8446622_RNA45SN1_45S_50ntFlanks@10689.10711.23|GCGTCCGGTGAGCTCTCGCTGGC</td><td>>hg38_21_+_8433172_8446622_RNA45SN1_45S_50ntFlanks</td><td>10689</td><td>10711</td><td>23</td><td>GCGTCCGGTGAGCTCTCGCTGGC</td><td>417.6</td><td>-12.6</td><td>1.6</td><td>-7.7</td><td>1.1E-14</td><td>6.4E-14</td><td>MB231/MB468</td><td>MP</td></tr>
<tr><td>>am_MI0000809|hsa-mir-151a&WithFlank&8|-|140732558|140732659@53.72.20(+2U)&[MIMAT0000757&hsa-miR-151a-3p&offsets|0|-1(+2U);m-30&8|-|140732587|140732607&offsets|0|-1(+2U)]|CTAGACTGAAGCTCCTTGAGTT</td><td>>am_MI0000809|hsa-mir-151a&WithFlank&8|-|140732558|140732659</td><td>53</td><td>72</td><td>20(+2U)</td><td>CTAGACTGAAGCTCCTTGAGTT</td><td>198.7</td><td>7.2</td><td>0.9</td><td>7.7</td><td>1.4E-14</td><td>6.4E-14</td><td>MB231/MB468</td><td>NUC</td></tr>
<tr><td>>hg38_21_+_8433172_8446622_RNA45SN1_45S_50ntFlanks@9574.9596.23|AATCGGTCGTCCGACCTGGGTAT</td><td>>hg38_21_+_8433172_8446622_RNA45SN1_45S_50ntFlanks</td><td>9574</td><td>9596</td><td>23</td><td>AATCGGTCGTCCGACCTGGGTAT</td><td>888.8</td><td>-4.3</td><td>0.6</td><td>-7.7</td><td>1.4E-14</td><td>6.4E-14</td><td>MB231/MB468</td><td>NUC</td></tr>
<tr><td>>hg38_21_+_8433172_8446622_RNA45SN1_45S_50ntFlanks@12008.12039.32|CCGCCGGTGAAATACCACTACTCTGATCGTTT</td><td>>hg38_21_+_8433172_8446622_RNA45SN1_45S_50ntFlanks</td><td>12008</td><td>12039</td><td>32</td><td>CCGCCGGTGAAATACCACTACTCTGATCGTTT</td><td>459.8</td><td>-12.7</td><td>1.6</td><td>-7.7</td><td>1.1E-14</td><td>6.4E-14</td><td>MB231/MB468</td><td>MP</td></tr>
<tr><td>>hg38_21_+_8433172_8446622_RNA45SN1_45S_50ntFlanks@12954.12982.29|CTGCTTCTGGGTCGGGGTTTCGTACGTAG</td><td>>hg38_21_+_8433172_8446622_RNA45SN1_45S_50ntFlanks</td><td>12954</td><td>12982</td><td>29</td><td>CTGCTTCTGGGTCGGGGTTTCGTACGTAG</td><td>158.7</td><td>-11.2</td><td>1.4</td><td>-7.7</td><td>1.1E-14</td><td>6.4E-14</td><td>MB231/MB468</td><td>MP</td></tr>
<tr><td>>MI0000760|hsa-mir-361&WithFlank&X|-|85903630|85903713@51.74.24&[MIMAT0004682&hsa-miR-361-3p&offsets|0|+1;m-124&X|-|85903642|85903663&offsets|0|+2]|TCCCCCAGGTGTGATTCTGATTTG</td><td>>MI0000760|hsa-mir-361&WithFlank&X|-|85903630|85903713</td><td>51</td><td>74</td><td>24</td><td>TCCCCCAGGTGTGATTCTGATTTG</td><td>320.5</td><td>5.1</td><td>0.7</td><td>7.7</td><td>1.4E-14</td><td>6.4E-14</td><td>MB231/MB468</td><td>NUC</td></tr>
<tr><td>>hg38_21_+_8433172_8446622_RNA45SN1_45S_50ntFlanks@6662.6681.20|GGTGGATCACTCGGCTCGTG</td><td>>hg38_21_+_8433172_8446622_RNA45SN1_45S_50ntFlanks</td><td>6662</td><td>6681</td><td>20</td><td>GGTGGATCACTCGGCTCGTG</td><td>227.2</td><td>-11.7</td><td>1.5</td><td>-7.7</td><td>1.1E-14</td><td>6.4E-14</td><td>MB231/MB468</td><td>MP</td></tr>
<tr><td>>hg38_21_+_8433172_8446622_RNA45SN1_45S_50ntFlanks@12957.12981.25|CTTCTGGGTCGGGGTTTCGTACGTA</td><td>>hg38_21_+_8433172_8446622_RNA45SN1_45S_50ntFlanks</td><td>12957</td><td>12981</td><td>25</td><td>CTTCTGGGTCGGGGTTTCGTACGTA</td><td>195.3</td><td>-11.5</td><td>1.5</td><td>-7.7</td><td>1.1E-14</td><td>6.4E-14</td><td>MB231/MB468</td><td>MP</td></tr>
<tr><td>>hg38_21_+_8433172_8446622_RNA45SN1_45S_50ntFlanks@10681.10709.29|TTCCGGCGGCGTCCGGTGAGCTCTCGCTG</td><td>>hg38_21_+_8433172_8446622_RNA45SN1_45S_50ntFlanks</td><td>10681</td><td>10709</td><td>29</td><td>TTCCGGCGGCGTCCGGTGAGCTCTCGCTG</td><td>237.6</td><td>-10.6</td><td>1.4</td><td>-7.7</td><td>1.3E-14</td><td>6.5E-14</td><td>MB231/MB468</td><td>CYTO</td></tr>
<tr><td>>hg38_21_+_8433172_8446622_RNA45SN1_45S_50ntFlanks@5527.5549.23|AAAGTCGTAACAAGGTTTCCGTA</td><td>>hg38_21_+_8433172_8446622_RNA45SN1_45S_50ntFlanks</td><td>5527</td><td>5549</td><td>23</td><td>AAAGTCGTAACAAGGTTTCCGTA</td><td>257.3</td><td>-11.8</td><td>1.5</td><td>-7.7</td><td>1.9E-14</td><td>6.5E-14</td><td>MB231/MB468</td><td>MITO</td></tr>
<tr><td>>hg38_21_+_8433172_8446622_RNA45SN1_45S_50ntFlanks@11076.11103.28|CGCCGCGTGGGGGCGGCGGCGGGGGGAG</td><td>>hg38_21_+_8433172_8446622_RNA45SN1_45S_50ntFlanks</td><td>11076</td><td>11103</td><td>28</td><td>CGCCGCGTGGGGGCGGCGGCGGGGGGAG</td><td>202.9</td><td>-10.4</td><td>1.3</td><td>-7.7</td><td>1.3E-14</td><td>6.5E-14</td><td>MB231/MB468</td><td>CYTO</td></tr>
<tr><td>>am_MI0000809|hsa-mir-151a&WithFlank&8|-|140732558|140732659@52.73.22&[MIMAT0000757&hsa-miR-151a-3p&offsets|-1|0;m-30&8|-|140732587|140732607&offsets|-1|0]|ACTAGACTGAAGCTCCTTGAGG</td><td>>am_MI0000809|hsa-mir-151a&WithFlank&8|-|140732558|140732659</td><td>52</td><td>73</td><td>22</td><td>ACTAGACTGAAGCTCCTTGAGG</td><td>877.2</td><td>5.9</td><td>0.8</td><td>7.7</td><td>1.3E-14</td><td>6.5E-14</td><td>MB231/MB468</td><td>CYTO</td></tr>
<tr><td>>hg38_21_+_8433172_8446622_RNA45SN1_45S_50ntFlanks@10922.10947.26|CTGGACGAGGCGCCGCCGCCCCCCCC</td><td>>hg38_21_+_8433172_8446622_RNA45SN1_45S_50ntFlanks</td><td>10922</td><td>10947</td><td>26</td><td>CTGGACGAGGCGCCGCCGCCCCCCCC</td><td>343.1</td><td>-12.3</td><td>1.6</td><td>-7.7</td><td>1.1E-14</td><td>6.5E-14</td><td>MB231/MB468</td><td>MP</td></tr>
<tr><td>>MI0000299|hsa-mir-222&WithFlank&X|-|45747009|45747130@75.95.21(+1U)&[MIMAT0000279&hsa-miR-222-3p&offsets|0|0(+1U);m-140&X|-|45747033|45747056&offsets|0|-3(+1U)]|AGCTACATCTGGCTACTGGGTT</td><td>>MI0000299|hsa-mir-222&WithFlank&X|-|45747009|45747130</td><td>75</td><td>95</td><td>21(+1U)</td><td>AGCTACATCTGGCTACTGGGTT</td><td>249.6</td><td>9.6</td><td>1.2</td><td>7.7</td><td>1.3E-14</td><td>6.5E-14</td><td>MB231/MB468</td><td>CYTO</td></tr>
<tr><td>>hg38_21_+_8433172_8446622_RNA45SN1_45S_50ntFlanks@6652.6701.50|GACTCTTAGCGGTGGATCACTCGGCTCGTGCGTCGATGAAGAACGCAGCT</td><td>>hg38_21_+_8433172_8446622_RNA45SN1_45S_50ntFlanks</td><td>6652</td><td>6701</td><td>50</td><td>GACTCTTAGCGGTGGATCACTCGGCTCGTGCGTCGATGAAGAACGCAGCT</td><td>6023.7</td><td>8.3</td><td>1.1</td><td>7.7</td><td>1.1E-14</td><td>6.5E-14</td><td>BT20/MB468</td><td>MITO</td></tr>
</table>

<table>
<tr><td></td><td></td><td></td><td></td><td></td><td></td><td></td><td></td><td></td><td></td><td></td><td></td><td></td><td></td></tr>
<tr><td>>MI0000067|hsa-let-7f-1&WithFlank&9|+|94176341|94176439@13.32.20(+1A);MI0000068|hsa-let-7f-2&WithFlank&X|-|53557186|53557280@14.33.20(+1A)&[MIMAT0000067&hsa-let-7f-5p&offsets|0|-2(+1A);m-20&9|+|94176353|94176374&offsets|0|-2(+1A)];[MIMAT0000067_1&hsa-let-7f-5p&offsets|0|-2(+1A);m-19&X|-|53557246|53557267&offsets|0|-2(+1A)]|TGAGGTAGTAGATTGTATAGA</td><td>>MI0000067|hsa-let-7f-1&WithFlank&9|+|94176341|94176439</td><td>13</td><td>32</td><td>20(+1A)</td><td>TGAGGTAGTAGATTGTATAGA</td><td>479.5</td><td>9.5</td><td>1.2</td><td>7.7</td><td>1.5E-14</td><td>6.5E-14</td><td>BT20/MB468</td><td>CYTO</td></tr>
<tr><td>>hg38_21_+_8433172_8446622_RNA45SN1_45S_50ntFlanks@3866.3898.33|CTAGAGCTAATACATGCCGACGGGCGCTGACCC</td><td>>hg38_21_+_8433172_8446622_RNA45SN1_45S_50ntFlanks</td><td>3866</td><td>3898</td><td>33</td><td>CTAGAGCTAATACATGCCGACGGGCGCTGACCC</td><td>449.5</td><td>-12.7</td><td>1.6</td><td>-7.7</td><td>1.2E-14</td><td>6.5E-14</td><td>MB231/MB468</td><td>MP</td></tr>
<tr><td>>trnaMT_PheGAA_MT_+_577_647@39.57.19|ATGTTTAGACGGGCTCACA</td><td>>trnaMT_PheGAA_MT_+_577_647</td><td>39</td><td>57</td><td>19</td><td>ATGTTTAGACGGGCTCACA</td><td>705.0</td><td>10.9</td><td>1.4</td><td>7.7</td><td>1.5E-14</td><td>6.5E-14</td><td>BT20/MB468</td><td>CYTO</td></tr>
<tr><td>>MI0000809|hsa-mir-151a&WithFlank&8|-|140732558|140732659@53.74.22(+1A)&[MIMAT0000757&hsa-miR-151a-3p&offsets|0|+1(+1A);m-30&8|-|140732587|140732607&offsets|0|+1(+1A)]|CTAGACTGAAGCTCCTTGAGGAA</td><td>>MI0000809|hsa-mir-151a&WithFlank&8|-|140732558|140732659</td><td>53</td><td>74</td><td>22(+1A)</td><td>CTAGACTGAAGCTCCTTGAGGAA</td><td>1601.9</td><td>5.5</td><td>0.7</td><td>7.7</td><td>1.3E-14</td><td>6.5E-14</td><td>MB231/MB468</td><td>CYTO</td></tr>
<tr><td>>hg38_21_+_8433172_8446622_RNA45SN1_45S_50ntFlanks@12163.12182.20|ACTGGGGCGGTACACCTGTC</td><td>>hg38_21_+_8433172_8446622_RNA45SN1_45S_50ntFlanks</td><td>12163</td><td>12182</td><td>20</td><td>ACTGGGGCGGTACACCTGTC</td><td>8587.6</td><td>-6.4</td><td>0.8</td><td>-8.1</td><td>4.5E-16</td><td>6.5E-14</td><td>BT20/MB468</td><td>MP</td></tr>
<tr><td>>trnaMT_GluTTC_MT_-_14674_14742@35.66.32,trnalookalike8_GluTTC_5_-_93905172_93905240@35.66.32|TATCATTGGTCGTGGTTGTAGTCCGTGCGAGA</td><td>>trnaMT_GluTTC_MT_-_14674_14742</td><td>35</td><td>66</td><td>32</td><td>TATCATTGGTCGTGGTTGTAGTCCGTGCGAGA</td><td>691.0</td><td>8.4</td><td>1.1</td><td>7.7</td><td>1.1E-14</td><td>6.5E-14</td><td>BT20/MB468</td><td>MITO</td></tr>
<tr><td>>hg38_21_+_8433172_8446622_RNA45SN1_45S_50ntFlanks@12631.12659.29|GGCGAAGCTACCATCTGTGGGATTATGAC</td><td>>hg38_21_+_8433172_8446622_RNA45SN1_45S_50ntFlanks</td><td>12631</td><td>12659</td><td>29</td><td>GGCGAAGCTACCATCTGTGGGATTATGAC</td><td>539.0</td><td>-12.9</td><td>1.7</td><td>-7.7</td><td>1.2E-14</td><td>6.5E-14</td><td>MB231/MB468</td><td>MP</td></tr>
<tr><td>>hg38_21_+_8433172_8446622_RNA45SN1_45S_50ntFlanks@11836.11872.37|ACAGCCAAGGGAACGGGCTTGGCGGAATCAGCGGGGA</td><td>>hg38_21_+_8433172_8446622_RNA45SN1_45S_50ntFlanks</td><td>11836</td><td>11872</td><td>37</td><td>ACAGCCAAGGGAACGGGCTTGGCGGAATCAGCGGGGA</td><td>216.4</td><td>-11.6</td><td>1.5</td><td>-7.7</td><td>1.2E-14</td><td>6.5E-14</td><td>MB231/MB468</td><td>MP</td></tr>
<tr><td>>hg38_21_+_8433172_8446622_RNA45SN1_45S_50ntFlanks@9029.9064.36|TCTCCTCGTGGGGGGGCCGGGCCACCCCTCCCACGG</td><td>>hg38_21_+_8433172_8446622_RNA45SN1_45S_50ntFlanks</td><td>9029</td><td>9064</td><td>36</td><td>TCTCCTCGTGGGGGGGCCGGGCCACCCCTCCCACGG</td><td>370.7</td><td>-12.3</td><td>1.6</td><td>-7.7</td><td>1.9E-14</td><td>6.6E-14</td><td>MB231/MB468</td><td>MITO</td></tr>
<tr><td>>MI0000298|hsa-mir-221&WithFlank&X|-|45746151|45746272@71.92.22(+1U)&[MIMAT0000278&hsa-miR-221-3p&offsets|0|-1(+1U);m-91&X|-|45746181|45746202&offsets|0|0(+1U)]|AGCTACATTGTCTGCTGGGTTTT</td><td>>MI0000298|hsa-mir-221&WithFlank&X|-|45746151|45746272</td><td>71</td><td>92</td><td>22(+1U)</td><td>AGCTACATTGTCTGCTGGGTTTT</td><td>1760.8</td><td>6.2</td><td>0.8</td><td>7.7</td><td>1.1E-14</td><td>6.6E-14</td><td>BT20/MB468</td><td>MITO</td></tr>
<tr><td>>hg38_21_+_8433172_8446622_RNA45SN1_45S_50ntFlanks@12894.12930.37|CTCGCCCGTCACGCACCGCACGTTCGTGGGGAACCTG</td><td>>hg38_21_+_8433172_8446622_RNA45SN1_45S_50ntFlanks</td><td>12894</td><td>12930</td><td>37</td><td>CTCGCCCGTCACGCACCGCACGTTCGTGGGGAACCTG</td><td>313.4</td><td>-12.2</td><td>1.6</td><td>-7.7</td><td>1.2E-14</td><td>6.6E-14</td><td>MB231/MB468</td><td>MP</td></tr>
<tr><td>>hg38_21_+_8433172_8446622_RNA45SN1_45S_50ntFlanks@12407.12435.29|TTTTGATCCTTCGATGTCGGCTCTTCCTA</td><td>>hg38_21_+_8433172_8446622_RNA45SN1_45S_50ntFlanks</td><td>12407</td><td>12435</td><td>29</td><td>TTTTGATCCTTCGATGTCGGCTCTTCCTA</td><td>1063.4</td><td>-7.5</td><td>1.0</td><td>-7.7</td><td>1.9E-14</td><td>6.6E-14</td><td>MB231/MB468</td><td>MITO</td></tr>
<tr><td>>hg38_21_+_8433172_8446622_RNA45SN1_45S_50ntFlanks@11161.11188.28|CACGGTCCCCCGCGAGGGGGGCCCGGGC</td><td>>hg38_21_+_8433172_8446622_RNA45SN1_45S_50ntFlanks</td><td>11161</td><td>11188</td><td>28</td><td>CACGGTCCCCCGCGAGGGGGGCCCGGGC</td><td>625.6</td><td>-11.4</td><td>1.5</td><td>-7.7</td><td>1.2E-14</td><td>6.6E-14</td><td>BT20/MB468</td><td>MITO</td></tr>
<tr><td>>hg38_21_+_8433172_8446622_RNA45SN1_45S_50ntFlanks@4728.4759.32|ATCAAGAACGAAAGTCGGAGGTTCGAAGACGA</td><td>>hg38_21_+_8433172_8446622_RNA45SN1_45S_50ntFlanks</td><td>4728</td><td>4759</td><td>32</td><td>ATCAAGAACGAAAGTCGGAGGTTCGAAGACGA</td><td>816.3</td><td>-12.8</td><td>1.6</td><td>-8.1</td><td>4.6E-16</td><td>6.7E-14</td><td>BT20/MB468</td><td>MP</td></tr>
<tr><td>>hg38_21_+_8433172_8446622_RNA45SN1_45S_50ntFlanks@7974.7992.19|ACGCGACCTCAGATCAGAC</td><td>>hg38_21_+_8433172_8446622_RNA45SN1_45S_50ntFlanks</td><td>7974</td><td>7992</td><td>19</td><td>ACGCGACCTCAGATCAGAC</td><td>23586.7</td><td>6.0</td><td>0.8</td><td>7.7</td><td>1.2E-14</td><td>6.7E-14</td><td>BT20/MB468</td><td>MITO</td></tr>
<tr><td>>hg38_21_+_8433172_8446622_RNA45SN1_45S_50ntFlanks@475.502.28|CTTCGTGATCGATGTGGTGACGTCGTGC</td><td>>hg38_21_+_8433172_8446622_RNA45SN1_45S_50ntFlanks</td><td>475</td><td>502</td><td>28</td><td>CTTCGTGATCGATGTGGTGACGTCGTGC</td><td>1202.2</td><td>7.8</td><td>1.0</td><td>7.7</td><td>1.2E-14</td><td>6.7E-14</td><td>BT20/MB468</td><td>MITO</td></tr>
<tr><td>>hg38_21_+_8433172_8446622_RNA45SN1_45S_50ntFlanks@476.502.27|TTCGTGATCGATGTGGTGACGTCGTGC</td><td>>hg38_21_+_8433172_8446622_RNA45SN1_45S_50ntFlanks</td><td>476</td><td>502</td><td>27</td><td>TTCGTGATCGATGTGGTGACGTCGTGC</td><td>2708.5</td><td>4.8</td><td>0.6</td><td>7.7</td><td>1.4E-14</td><td>6.7E-14</td><td>MB231/MB468</td><td>NUC</td></tr>
<tr><td>>trna5_AspGTC_12_+_98897281_98897352@36.73.38|CACGCGGGAGACCGGGGTTCAATTCCCCGACGGGGAGC</td><td>>trna5_AspGTC_12_+_98897281_98897352</td><td>36</td><td>73</td><td>38</td><td>CACGCGGGAGACCGGGGTTCAATTCCCCGACGGGGAGC</td><td>1097.1</td><td>10.1</td><td>1.3</td><td>7.7</td><td>1.4E-14</td><td>6.7E-14</td><td>MB231/MB468</td><td>CYTO</td></tr>
<tr><td>>hg38_21_+_8433172_8446622_RNA45SN1_45S_50ntFlanks@12366.12384.19|TAACTGGCTTGTGGCGGCC</td><td>>hg38_21_+_8433172_8446622_RNA45SN1_45S_50ntFlanks</td><td>12366</td><td>12384</td><td>19</td><td>TAACTGGCTTGTGGCGGCC</td><td>772.6</td><td>-9.2</td><td>1.2</td><td>-7.7</td><td>1.6E-14</td><td>6.7E-14</td><td>BT20/MB468</td><td>CYTO</td></tr>
<tr><td>>hg38_21_+_8433172_8446622_RNA45SN1_45S_50ntFlanks@8570.8593.24|CTCCGGCGGGTGCGGGGGTGGGCG</td><td>>hg38_21_+_8433172_8446622_RNA45SN1_45S_50ntFlanks</td><td>8570</td><td>8593</td><td>24</td><td>CTCCGGCGGGTGCGGGGGTGGGCG</td><td>408.7</td><td>-11.6</td><td>1.5</td><td>-7.7</td><td>1.5E-14</td><td>6.7E-14</td><td>MB231/MB468</td><td>NUC</td></tr>
<tr><td>>hg38_21_+_8433172_8446622_RNA45SN1_45S_50ntFlanks@9568.9594.27|CGTGCAAATCGGTCGTCCGACCTGGGT</td><td>>hg38_21_+_8433172_8446622_RNA45SN1_45S_50ntFlanks</td><td>9568</td><td>9594</td><td>27</td><td>CGTGCAAATCGGTCGTCCGACCTGGGT</td><td>1182.2</td><td>-9.3</td><td>1.2</td><td>-7.7</td><td>2.0E-14</td><td>6.8E-14</td><td>MB231/MB468</td><td>MITO</td></tr>
<tr><td>>hg38_21_+_8433172_8446622_RNA45SN1_45S_50ntFlanks@11669.11702.34|GCCCAGTGCTCTGAATGTCAAAGTGAAGAAATTC</td><td>>hg38_21_+_8433172_8446622_RNA45SN1_45S_50ntFlanks</td><td>11669</td><td>11702</td><td>34</td><td>GCCCAGTGCTCTGAATGTCAAAGTGAAGAAATTC</td><td>5324.8</td><td>6.4</td><td>0.8</td><td>7.7</td><td>1.4E-14</td><td>6.8E-14</td><td>MB231/MB468</td><td>CYTO</td></tr>
<tr><td>>hg38_21_+_8433172_8446622_RNA45SN1_45S_50ntFlanks@6668.6686.19|TCACTCGGCTCGTGCGTCG</td><td>>hg38_21_+_8433172_8446622_RNA45SN1_45S_50ntFlanks</td><td>6668</td><td>6686</td><td>19</td><td>TCACTCGGCTCGTGCGTCG</td><td>157.5</td><td>-11.2</td><td>1.4</td><td>-7.7</td><td>1.2E-14</td><td>6.8E-14</td><td>MB231/MB468</td><td>MP</td></tr>
<tr><td>>hg38_21_+_8433172_8446622_RNA45SN1_45S_50ntFlanks@12126.12149.24|CGACCCGCTCCGGGGACAGTGCCA</td><td>>hg38_21_+_8433172_8446622_RNA45SN1_45S_50ntFlanks</td><td>12126</td><td>12149</td><td>24</td><td>CGACCCGCTCCGGGGACAGTGCCA</td><td>256.4</td><td>-11.8</td><td>1.5</td><td>-7.7</td><td>2.0E-14</td><td>6.8E-14</td><td>MB231/MB468</td><td>MITO</td></tr>
<tr><td>>hg38_21_+_8433172_8446622_RNA45SN1_45S_50ntFlanks@11235.11255.21|CTTCCCGTGGATCGCCCCAGC</td><td>>hg38_21_+_8433172_8446622_RNA45SN1_45S_50ntFlanks</td><td>11235</td><td>11255</td><td>21</td><td>CTTCCCGTGGATCGCCCCAGC</td><td>228.6</td><td>-11.7</td><td>1.5</td><td>-7.7</td><td>1.2E-14</td><td>6.8E-14</td><td>MB231/MB468</td><td>MP</td></tr>
<tr><td>>MI0000750|hsa-mir-26a-2&WithFlank&12|-|57824603|57824698@20.40.21(+1G);MI0000083|hsa-mir-26a-1&WithFlank&3|+|37969398|37969486@16.36.21(+1G)&[MIMAT0000082&hsa-miR-26a-5p&offsets|0|-1(+1G);m-33&12|-|57824658|57824679&offsets|0|-1(+1G)];[MIMAT0000082_1&hsa-miR-26a-5p&offsets|0|-1(+1G);m-34&3|+|37969413|37969434&offsets|0|-1(+1G)]|TTCAAGTAATCCAGGATAGGCG</td><td>>MI0000750|hsa-mir-26a-2&WithFlank&12|-|57824603|57824698</td><td>20</td><td>40</td><td>21(+1G)</td><td>TTCAAGTAATCCAGGATAGGCG</td><td>3378.8</td><td>5.0</td><td>0.7</td><td>7.7</td><td>1.6E-14</td><td>6.8E-14</td><td>BT20/MB468</td><td>CYTO</td></tr>
<tr><td>>hg38_21_+_8433172_8446622_RNA45SN1_45S_50ntFlanks@12141.12158.18|ACAGTGCCAGGTGGGGAG</td><td>>hg38_21_+_8433172_8446622_RNA45SN1_45S_50ntFlanks</td><td>12141</td><td>12158</td><td>18</td><td>ACAGTGCCAGGTGGGGAG</td><td>174.1</td><td>-11.3</td><td>1.5</td><td>-7.7</td><td>1.2E-14</td><td>6.8E-14</td><td>MB231/MB468</td><td>MP</td></tr>
<tr><td>>Antisense_hg38_21_-_8433172_8446622_RNA45SN1_45S_50ntFlanks@7123.7140.18|GCGGCGGCGGCGGCGGCG</td><td>>Antisense_hg38_21_-_8433172_8446622_RNA45SN1_45S_50ntFlanks</td><td>7123</td><td>7140</td><td>18</td><td>GCGGCGGCGGCGGCGGCG</td><td>216.9</td><td>-11.6</td><td>1.5</td><td>-7.7</td><td>1.2E-14</td><td>6.8E-14</td><td>MB231/MB468</td><td>MP</td></tr>
<tr><td>>hg38_21_+_8433172_8446622_RNA45SN1_45S_50ntFlanks@10127.10144.18|GCGGCGGCGGCGGCGGCG</td><td>>hg38_21_+_8433172_8446622_RNA45SN1_45S_50ntFlanks</td><td>10127</td><td>10144</td><td>18</td><td>GCGGCGGCGGCGGCGGCG</td><td>216.9</td><td>-11.6</td><td>1.5</td><td>-7.7</td><td>1.2E-14</td><td>6.8E-14</td><td>MB231/MB468</td><td>MP</td></tr>
<tr><td>>hg38_21_+_8433172_8446622_RNA45SN1_45S_50ntFlanks@10130.10147.18|GCGGCGGCGGCGGCGGCG</td><td>>hg38_21_+_8433172_8446622_RNA45SN1_45S_50ntFlanks</td><td>10130</td><td>10147</td><td>18</td><td>GCGGCGGCGGCGGCGGCG</td><td>216.9</td><td>-11.6</td><td>1.5</td><td>-7.7</td><td>1.2E-14</td><td>6.8E-14</td><td>MB231/MB468</td><td>MP</td></tr>
<tr><td>>hg38_21_+_8433172_8446622_RNA45SN1_45S_50ntFlanks@8823.8840.18|GCGGCGGCGGCGGCGGCG</td><td>>hg38_21_+_8433172_8446622_RNA45SN1_45S_50ntFlanks</td><td>8823</td><td>8840</td><td>18</td><td>GCGGCGGCGGCGGCGGCG</td><td>216.9</td><td>-11.6</td><td>1.5</td><td>-7.7</td><td>1.2E-14</td><td>6.8E-14</td><td>MB231/MB468</td><td>MP</td></tr>
<tr><td>>hg38_21_+_8433172_8446622_RNA45SN1_45S_50ntFlanks@8844.8861.18|GCGGCGGCGGCGGCGGCG</td><td>>hg38_21_+_8433172_8446622_RNA45SN1_45S_50ntFlanks</td><td>8844</td><td>8861</td><td>18</td><td>GCGGCGGCGGCGGCGGCG</td><td>216.9</td><td>-11.6</td><td>1.5</td><td>-7.7</td><td>1.2E-14</td><td>6.8E-14</td><td>MB231/MB468</td><td>MP</td></tr>
<tr><td>>hg38_21_+_8433172_8446622_RNA45SN1_45S_50ntFlanks@8847.8864.18|GCGGCGGCGGCGGCGGCG</td><td>>hg38_21_+_8433172_8446622_RNA45SN1_45S_50ntFlanks</td><td>8847</td><td>8864</td><td>18</td><td>GCGGCGGCGGCGGCGGCG</td><td>216.9</td><td>-11.6</td><td>1.5</td><td>-7.7</td><td>1.2E-14</td><td>6.8E-14</td><td>MB231/MB468</td><td>MP</td></tr>
<tr><td>>hg38_21_+_8433172_8446622_RNA45SN1_45S_50ntFlanks@4731.4750.20|AAGAACGAAAGTCGGAGGTT</td><td>>hg38_21_+_8433172_8446622_RNA45SN1_45S_50ntFlanks</td><td>4731</td><td>4750</td><td>20</td><td>AAGAACGAAAGTCGGAGGTT</td><td>401.1</td><td>-4.4</td><td>0.6</td><td>-7.6</td><td>2.4E-14</td><td>6.9E-14</td><td>BT20/MB468</td><td>NUC</td></tr>
<tr><td>>MI0000477|hsa-mir-146a&WithFlank&5|+|160485346|160485456@27.46.20(+1C)&[MIMAT0000449&hsa-miR-146a-5p&offsets|0|-2(+1C);m-129&5|+|160485372|160485393&offsets|0|-2(+1C)]|TGAGAACTGAATTCCATGGGC</td><td>>MI0000477|hsa-mir-146a&WithFlank&5|+|160485346|160485456</td><td>27</td><td>46</td><td>20(+1C)</td><td>TGAGAACTGAATTCCATGGGC</td><td>295.3</td><td>12.1</td><td>1.6</td><td>7.7</td><td>1.5E-14</td><td>6.9E-14</td><td>MB231/MB468</td><td>NUC</td></tr>
<tr><td>>trna119_LysCTT_1_-_145395522_145395594@2.33.32,trna11_LysCTT_5_-_180648979_180649051@2.33.32,trna13_LysCTT_6_+_26556774_26556846@2.33.32,trna7_LysCTT_16_+_3225692_3225764@2.33.32,trna9_LysCTT_5_+_180634755_180634827@2.33.32|CCCGGCTAGCTCAGTCGGTAGAGCATGAGACT</td><td>>trna119_LysCTT_1_-_145395522_145395594</td><td>2</td><td>33</td><td>32</td><td>CCCGGCTAGCTCAGTCGGTAGAGCATGAGACT</td><td>552.9</td><td>13.1</td><td>1.7</td><td>7.7</td><td>1.4E-14</td><td>6.9E-14</td><td>MB231/MB468</td><td>CYTO</td></tr>
<tr><td>>hg38_21_+_8433172_8446622_RNA45SN1_45S_50ntFlanks@12347.12364.18|AGAAAAGTTACCACAGGG</td><td>>hg38_21_+_8433172_8446622_RNA45SN1_45S_50ntFlanks</td><td>12347</td><td>12364</td><td>18</td><td>AGAAAAGTTACCACAGGG</td><td>182.7</td><td>-10.2</td><td>1.3</td><td>-7.7</td><td>1.4E-14</td><td>6.9E-14</td><td>MB231/MB468</td><td>CYTO</td></tr>
<tr><td>>trnaMT_GluTTC_MT_-_14674_14742@39.63.25,trnalookalike8_GluTTC_5_-_93905172_93905240@39.63.25|ATTGGTCGTGGTTGTAGTCCGTGCG</td><td>>trnaMT_GluTTC_MT_-_14674_14742</td><td>39</td><td>63</td><td>25</td><td>ATTGGTCGTGGTTGTAGTCCGTGCG</td><td>1778.3</td><td>9.1</td><td>1.2</td><td>7.7</td><td>1.2E-14</td><td>6.9E-14</td><td>BT20/MB468</td><td>MITO</td></tr>
<tr><td>>hg38_21_+_8433172_8446622_RNA45SN1_45S_50ntFlanks@12936.12955.20|AAACCATTCGTAGACGACCT</td><td>>hg38_21_+_8433172_8446622_RNA45SN1_45S_50ntFlanks</td><td>12936</td><td>12955</td><td>20</td><td>AAACCATTCGTAGACGACCT</td><td>161.0</td><td>-11.2</td><td>1.5</td><td>-7.7</td><td>1.2E-14</td><td>6.9E-14</td><td>MB231/MB468</td><td>MP</td></tr>
<tr><td>>hg38_21_+_8433172_8446622_RNA45SN1_45S_50ntFlanks@11052.11075.24|CGTCCTCCCCCCTCCCCGGGGGAG</td><td>>hg38_21_+_8433172_8446622_RNA45SN1_45S_50ntFlanks</td><td>11052</td><td>11075</td><td>24</td><td>CGTCCTCCCCCCTCCCCGGGGGAG</td><td>213.4</td><td>-10.0</td><td>1.3</td><td>-7.7</td><td>1.6E-14</td><td>6.9E-14</td><td>BT20/MB468</td><td>CYTO</td></tr>
<tr><td>>hg38_21_+_8433172_8446622_RNA45SN1_45S_50ntFlanks@11190.11225.36|CCCGGGGGGGCCGGCGGCGGCGGCGACTCTGGACGCG</td><td>>hg38_21_+_8433172_8446622_RNA45SN1_45S_50ntFlanks</td><td>11190</td><td>11225</td><td>36</td><td>CCCGGGGGGGCCGGCGGCGGCGGCGACTCTGGACGCG</td><td>1092.5</td><td>-13.9</td><td>1.8</td><td>-7.7</td><td>2.0E-14</td><td>6.9E-14</td><td>MB231/MB468</td><td>MITO</td></tr>
<tr><td>>trna111_HisGTG_1_-_147774845_147774916@-1T.29.30,trna118_HisGTG_1_-_145396881_145396952@-1T.29.30,trna16_HisGTG_1_+_146544773_146544844@-1T.29.30,trna1_HisGTG_15_+_45493349_45493420@-1T.29.30,trna21_HisGTG_1_+_147753471_147753542@-1T.29.30,trna33_HisGTG_6_+_27125906_27125977@-1T.29.30,trna7_HisGTG_9_-_14433938_14434009@-1T.29.30,trna8_HisGTG_15_-_45492611_45492682@-1T.29.30,trna9_HisGTG_15_-_45490804_45490875@-1T.29.30|TGCCGTGATCGTATAGTGGTTAGTACTCTG</td><td>>trna111_HisGTG_1_-_147774845_147774916</td><td>-1T</td><td>29</td><td>30</td><td>TGCCGTGATCGTATAGTGGTTAGTACTCTG</td><td>11252.6</td><td>-5.9</td><td>0.8</td><td>-7.7</td><td>1.4E-14</td><td>6.9E-14</td><td>MB231/MB468</td><td>CYTO</td></tr>
<tr><td>>hg38_21_+_8433172_8446622_RNA45SN1_45S_50ntFlanks@12927.12951.25|CCTGGCGCTAAACCATTCGTAGACG</td><td>>hg38_21_+_8433172_8446622_RNA45SN1_45S_50ntFlanks</td><td>12927</td><td>12951</td><td>25</td><td>CCTGGCGCTAAACCATTCGTAGACG</td><td>260.6</td><td>-11.9</td><td>1.5</td><td>-7.7</td><td>1.2E-14</td><td>6.9E-14</td><td>MB231/MB468</td><td>MP</td></tr>
<tr><td>>hg38_21_+_8433172_8446622_RNA45SN1_45S_50ntFlanks@4759.4786.28|ATCAGATACCGTCGTAGTTCCGACCATA</td><td>>hg38_21_+_8433172_8446622_RNA45SN1_45S_50ntFlanks</td><td>4759</td><td>4786</td><td>28</td><td>ATCAGATACCGTCGTAGTTCCGACCATA</td><td>871.8</td><td>-11.9</td><td>1.5</td><td>-7.7</td><td>1.2E-14</td><td>6.9E-14</td><td>BT20/MB468</td><td>MITO</td></tr>
<tr><td>>hg38_21_+_8433172_8446622_RNA45SN1_45S_50ntFlanks@12944.12964.21|CGTAGACGACCTGCTTCTGGG</td><td>>hg38_21_+_8433172_8446622_RNA45SN1_45S_50ntFlanks</td><td>12944</td><td>12964</td><td>21</td><td>CGTAGACGACCTGCTTCTGGG</td><td>315.1</td><td>-12.2</td><td>1.6</td><td>-7.7</td><td>1.2E-14</td><td>6.9E-14</td><td>MB231/MB468</td><td>MP</td></tr>
<tr><td>>hg38_21_+_8433172_8446622_RNA45SN1_45S_50ntFlanks@11668.11688.21|TGCCCAGTGCTCTGAATGTCA</td><td>>hg38_21_+_8433172_8446622_RNA45SN1_45S_50ntFlanks</td><td>11668</td><td>11688</td><td>21</td><td>TGCCCAGTGCTCTGAATGTCA</td><td>260.8</td><td>4.9</td><td>0.6</td><td>7.7</td><td>1.6E-14</td><td>6.9E-14</td><td>BT20/MB468</td><td>CYTO</td></tr>
</table>

<table>
<tr><td>>MI0000082|hsa-mir-25&WithFlank&7|-|100093554|100093649@58.79.22&[MIMAT0000081&hsa-miR-25-3p&offsets|0|0;m-23&7|-|100093571|100093592&offsets|0|0]|CATTGCACTTGTCTCGGTCTGA</td><td>>MI0000082|hsa-mir-25&WithFlank&7|-|100093554|100093649</td><td>58</td><td>79</td><td>22</td><td>CATTGCACTTGTCTCGGTCTGA</td><td>29801.9</td><td>6.0</td><td>0.8</td><td>7.7</td><td>1.4E-14</td><td>6.9E-14</td><td>MB231/MB468</td><td>CYTO</td></tr>
<tr><td>>hg38_21_+_8433172_8446622_RNA45SN1_45S_50ntFlanks@8113.8137.25|GGCGCGGGACATGTGGCGTACGGAA</td><td>>hg38_21_+_8433172_8446622_RNA45SN1_45S_50ntFlanks</td><td>8113</td><td>8137</td><td>25</td><td>GGCGCGGGACATGTGGCGTACGGAA</td><td>282.2</td><td>-12.0</td><td>1.6</td><td>-7.7</td><td>1.2E-14</td><td>7.0E-14</td><td>MB231/MB468</td><td>MP</td></tr>
<tr><td>>hg38_21_+_8433172_8446622_RNA45SN1_45S_50ntFlanks@12407.12434.28|TTTTGATCCTTCGATGTCGGCTCTTCCT</td><td>>hg38_21_+_8433172_8446622_RNA45SN1_45S_50ntFlanks</td><td>12407</td><td>12434</td><td>28</td><td>TTTTGATCCTTCGATGTCGGCTCTTCCT</td><td>8759.1</td><td>-6.0</td><td>0.8</td><td>-7.6</td><td>2.5E-14</td><td>7.0E-14</td><td>BT20/MB468</td><td>NUC</td></tr>
<tr><td>>hg38_21_+_8433172_8446622_RNA45SN1_45S_50ntFlanks@9534.9559.26|AGGAAACTCTGGTGGAGGTCCGTAGC</td><td>>hg38_21_+_8433172_8446622_RNA45SN1_45S_50ntFlanks</td><td>9534</td><td>9559</td><td>26</td><td>AGGAAACTCTGGTGGAGGTCCGTAGC</td><td>1513.0</td><td>-8.1</td><td>1.1</td><td>-7.7</td><td>1.2E-14</td><td>7.0E-14</td><td>MB231/MB468</td><td>MP</td></tr>
<tr><td>>MI0001448|hsa-mir-425&WithFlank&3|-|49020142|49020240@20.44.25&[MIMAT0003393&hsa-miR-425-5p&offsets|0|+2;m-95&3|-|49020200|49020221&offsets|0|+3]|AATGACACGATCACTCCCGTTGAGT</td><td>>MI0001448|hsa-mir-425&WithFlank&3|-|49020142|49020240</td><td>20</td><td>44</td><td>25</td><td>AATGACACGATCACTCCCGTTGAGT</td><td>3697.5</td><td>6.3</td><td>0.8</td><td>7.7</td><td>1.4E-14</td><td>7.0E-14</td><td>MB231/MB468</td><td>CYTO</td></tr>
<tr><td>>hg38_21_+_8433172_8446622_RNA45SN1_45S_50ntFlanks@10404.10431.28|CATGGGTCAGTCGGTCCTGAGAGATGGG</td><td>>hg38_21_+_8433172_8446622_RNA45SN1_45S_50ntFlanks</td><td>10404</td><td>10431</td><td>28</td><td>CATGGGTCAGTCGGTCCTGAGAGATGGG</td><td>206.3</td><td>-11.6</td><td>1.5</td><td>-7.7</td><td>1.3E-14</td><td>7.0E-14</td><td>MB231/MB468</td><td>MP</td></tr>
<tr><td>>hg38_21_+_8433172_8446622_RNA45SN1_45S_50ntFlanks@8169.8187.19|CCAAGTCCTTCTGATCGAG</td><td>>hg38_21_+_8433172_8446622_RNA45SN1_45S_50ntFlanks</td><td>8169</td><td>8187</td><td>19</td><td>CCAAGTCCTTCTGATCGAG</td><td>170.0</td><td>-11.3</td><td>1.5</td><td>-7.7</td><td>1.3E-14</td><td>7.0E-14</td><td>MB231/MB468</td><td>MP</td></tr>
<tr><td>>hg38_21_+_8433172_8446622_RNA45SN1_45S_50ntFlanks@11207.11234.28|GGCGGCGACTCTGGACGCGAGCCGGGCC</td><td>>hg38_21_+_8433172_8446622_RNA45SN1_45S_50ntFlanks</td><td>11207</td><td>11234</td><td>28</td><td>GGCGGCGACTCTGGACGCGAGCCGGGCC</td><td>244.6</td><td>-11.8</td><td>1.5</td><td>-7.7</td><td>1.3E-14</td><td>7.0E-14</td><td>MB231/MB468</td><td>MP</td></tr>
<tr><td>>hg38_21_+_8433172_8446622_RNA45SN1_45S_50ntFlanks@6737.6770.34|GATCATCGACACTTCGAACGCACTTGCGGCCCCG</td><td>>hg38_21_+_8433172_8446622_RNA45SN1_45S_50ntFlanks</td><td>6737</td><td>6770</td><td>34</td><td>GATCATCGACACTTCGAACGCACTTGCGGCCCCG</td><td>273.1</td><td>-10.4</td><td>1.4</td><td>-7.7</td><td>1.7E-14</td><td>7.0E-14</td><td>BT20/MB468</td><td>CYTO</td></tr>
<tr><td>>hg38_21_+_8433172_8446622_RNA45SN1_45S_50ntFlanks@10922.10944.23|CTGGACGAGGCGCCGCCGCCCCC</td><td>>hg38_21_+_8433172_8446622_RNA45SN1_45S_50ntFlanks</td><td>10922</td><td>10944</td><td>23</td><td>CTGGACGAGGCGCCGCCGCCCCC</td><td>283.7</td><td>-12.0</td><td>1.6</td><td>-7.7</td><td>1.3E-14</td><td>7.0E-14</td><td>MB231/MB468</td><td>MP</td></tr>
<tr><td>>hg38_21_+_8433172_8446622_RNA45SN1_45S_50ntFlanks@12526.12557.32|TTTTACCCTACTGATGATGTGTTGTTGCCATG</td><td>>hg38_21_+_8433172_8446622_RNA45SN1_45S_50ntFlanks</td><td>12526</td><td>12557</td><td>32</td><td>TTTTACCCTACTGATGATGTGTTGTTGCCATG</td><td>242.6</td><td>-11.8</td><td>1.5</td><td>-7.7</td><td>1.3E-14</td><td>7.1E-14</td><td>MB231/MB468</td><td>MP</td></tr>
<tr><td>>hg38_21_+_8433172_8446622_RNA45SN1_45S_50ntFlanks@12202.12221.20|CTAAGGCGAGCTCAGGGAGG</td><td>>hg38_21_+_8433172_8446622_RNA45SN1_45S_50ntFlanks</td><td>12202</td><td>12221</td><td>20</td><td>CTAAGGCGAGCTCAGGGAGG</td><td>1129.5</td><td>-10.2</td><td>1.3</td><td>-7.6</td><td>2.0E-14</td><td>7.1E-14</td><td>MB231/MB468</td><td>MITO</td></tr>
<tr><td>>hg38_21_+_8433172_8446622_RNA45SN1_45S_50ntFlanks@10872.10894.23|GCTGGGTCGGTCGGGCTGGGGCG</td><td>>hg38_21_+_8433172_8446622_RNA45SN1_45S_50ntFlanks</td><td>10872</td><td>10894</td><td>23</td><td>GCTGGGTCGGTCGGGCTGGGGCG</td><td>265.2</td><td>-11.9</td><td>1.5</td><td>-7.7</td><td>1.3E-14</td><td>7.1E-14</td><td>MB231/MB468</td><td>MP</td></tr>
<tr><td>>hg38_21_+_8433172_8446622_RNA45SN1_45S_50ntFlanks@12163.12182.20|ACTGGGGCGGTACACCTGTC</td><td>>hg38_21_+_8433172_8446622_RNA45SN1_45S_50ntFlanks</td><td>12163</td><td>12182</td><td>20</td><td>ACTGGGGCGGTACACCTGTC</td><td>693.9</td><td>-5.2</td><td>0.7</td><td>-7.7</td><td>1.5E-14</td><td>7.1E-14</td><td>MB231/MB468</td><td>NUC</td></tr>
<tr><td>>hg38_21_+_8433172_8446622_RNA45SN1_45S_50ntFlanks@10513.10534.22|CCGGAGTGGCGGAGATGGGCGC</td><td>>hg38_21_+_8433172_8446622_RNA45SN1_45S_50ntFlanks</td><td>10513</td><td>10534</td><td>22</td><td>CCGGAGTGGCGGAGATGGGCGC</td><td>370.2</td><td>-11.5</td><td>1.5</td><td>-7.7</td><td>1.5E-14</td><td>7.1E-14</td><td>MB231/MB468</td><td>NUC</td></tr>
<tr><td>>hg38_21_+_8433172_8446622_RNA45SN1_45S_50ntFlanks@8189.8215.27|CCCAGCCCGTGGACGGTGTGAGGCCGG</td><td>>hg38_21_+_8433172_8446622_RNA45SN1_45S_50ntFlanks</td><td>8189</td><td>8215</td><td>27</td><td>CCCAGCCCGTGGACGGTGTGAGGCCGG</td><td>203.1</td><td>-11.5</td><td>1.5</td><td>-7.7</td><td>1.3E-14</td><td>7.1E-14</td><td>MB231/MB468</td><td>MP</td></tr>
<tr><td>>hg38_21_+_8433172_8446622_RNA45SN1_45S_50ntFlanks@11452.11478.27|CTCCTCGGCGCGCGGCGGCGGCGGCGG</td><td>>hg38_21_+_8433172_8446622_RNA45SN1_45S_50ntFlanks</td><td>11452</td><td>11478</td><td>27</td><td>CTCCTCGGCGCGCGGCGGCGGCGGCGG</td><td>1018.0</td><td>-8.4</td><td>1.1</td><td>-7.7</td><td>1.4E-14</td><td>7.1E-14</td><td>MB231/MB468</td><td>CYTO</td></tr>
<tr><td>>hg38_21_+_8433172_8446622_RNA45SN1_45S_50ntFlanks@10787.10818.32|ACAGCCTCTGGCATGTTGGAACAATGTAGGTA</td><td>>hg38_21_+_8433172_8446622_RNA45SN1_45S_50ntFlanks</td><td>10787</td><td>10818</td><td>32</td><td>ACAGCCTCTGGCATGTTGGAACAATGTAGGTA</td><td>2638.4</td><td>-8.1</td><td>1.1</td><td>-7.6</td><td>2.1E-14</td><td>7.1E-14</td><td>MB231/MB468</td><td>MITO</td></tr>
<tr><td>>hg38_21_+_8433172_8446622_RNA45SN1_45S_50ntFlanks@9563.9596.34|CCTGACGTGCAAATCGGTCGTCCGACCTGGGTAT</td><td>>hg38_21_+_8433172_8446622_RNA45SN1_45S_50ntFlanks</td><td>9563</td><td>9596</td><td>34</td><td>CCTGACGTGCAAATCGGTCGTCCGACCTGGGTAT</td><td>7940.4</td><td>-8.5</td><td>1.1</td><td>-7.6</td><td>2.1E-14</td><td>7.1E-14</td><td>MB231/MB468</td><td>MITO</td></tr>
<tr><td>>hg38_21_+_8433172_8446622_RNA45SN1_45S_50ntFlanks@4728.4752.25|ATCAAGAACGAAAGTCGGAGGTTCG</td><td>>hg38_21_+_8433172_8446622_RNA45SN1_45S_50ntFlanks</td><td>4728</td><td>4752</td><td>25</td><td>ATCAAGAACGAAAGTCGGAGGTTCG</td><td>617.4</td><td>-12.4</td><td>1.5</td><td>-8.1</td><td>5.0E-16</td><td>7.1E-14</td><td>BT20/MB468</td><td>MP</td></tr>
<tr><td>>am_MI0000809|hsa-mir-151a&WithFlank&8|-|140732558|140732659@53.73.21&[MIMAT0000757&hsa-miR-151a-3p&offsets|0|0;m-30&8|-|140732587|140732607&offsets|0|0]|CTAGACTGAAGCTCCTTGAGG</td><td>>am_MI0000809|hsa-mir-151a&WithFlank&8|-|140732558|140732659</td><td>53</td><td>73</td><td>21</td><td>CTAGACTGAAGCTCCTTGAGG</td><td>43894.6</td><td>6.8</td><td>0.9</td><td>7.7</td><td>1.4E-14</td><td>7.1E-14</td><td>MB231/MB468</td><td>CYTO</td></tr>
<tr><td>>hg38_21_+_8433172_8446622_RNA45SN1_45S_50ntFlanks@3705.3735.31|TACCTGGTTGATCCTGCCAGTAGCATATGCT</td><td>>hg38_21_+_8433172_8446622_RNA45SN1_45S_50ntFlanks</td><td>3705</td><td>3735</td><td>31</td><td>TACCTGGTTGATCCTGCCAGTAGCATATGCT</td><td>111.6</td><td>3.7</td><td>0.5</td><td>7.6</td><td>2.5E-14</td><td>7.1E-14</td><td>BT20/MB468</td><td>NUC</td></tr>
<tr><td>>MI0000750|hsa-mir-26a-2&WithFlank&12|-|57824603|57824698@23.41.19;MI0000083|hsa-mir-26a-1&WithFlank&3|+|37969398|37969486@19.37.19&[MIMAT0000082&hsa-miR-26a-5p&offsets|+3|0;m-33&12|-|57824658|57824679&offsets|+3|0];[MIMAT0000082_1&hsa-miR-26a-5p&offsets|+3|0;m-34&3|+|37969413|37969434&offsets|+3|0]|AAGTAATCCAGGATAGGCT</td><td>>MI0000750|hsa-mir-26a-2&WithFlank&12|-|57824603|57824698</td><td>23</td><td>41</td><td>19</td><td>AAGTAATCCAGGATAGGCT</td><td>1371.7</td><td>27.8</td><td>3.6</td><td>7.6</td><td>2.1E-14</td><td>7.2E-14</td><td>MB231/MB468</td><td>MITO</td></tr>
<tr><td>>hg38_21_+_8433172_8446622_RNA45SN1_45S_50ntFlanks@11124.11146.23|CCGGCGGCGGCCCGCCGCGGGGC</td><td>>hg38_21_+_8433172_8446622_RNA45SN1_45S_50ntFlanks</td><td>11124</td><td>11146</td><td>23</td><td>CCGGCGGCGGCCCGCCGCGGGGC</td><td>182.2</td><td>-10.2</td><td>1.3</td><td>-7.7</td><td>1.5E-14</td><td>7.2E-14</td><td>MB231/MB468</td><td>CYTO</td></tr>
<tr><td>>hg38_1_-_228634819_228635039_RNA5S12_5S_50ntFlanks@141.171.31|CCGCCTGGGAATACCGGGTGCTGTAGGCTTT</td><td>>hg38_1_-_228634819_228635039_RNA5S12_5S_50ntFlanks</td><td>141</td><td>171</td><td>31</td><td>CCGCCTGGGAATACCGGGTGCTGTAGGCTTT</td><td>4115.3</td><td>5.7</td><td>0.7</td><td>7.7</td><td>1.7E-14</td><td>7.2E-14</td><td>BT20/MB468</td><td>CYTO</td></tr>
<tr><td>>hg38_21_+_8433172_8446622_RNA45SN1_45S_50ntFlanks@485.531.47|GATGTGGTGACGTCGTGCTCTCCCGGGCCGGGTCCGAGCCGCGACGG</td><td>>hg38_21_+_8433172_8446622_RNA45SN1_45S_50ntFlanks</td><td>485</td><td>531</td><td>47</td><td>GATGTGGTGACGTCGTGCTCTCCCGGGCCGGGTCCGAGCCGCGACGG</td><td>902.5</td><td>27.9</td><td>3.6</td><td>7.7</td><td>1.6E-14</td><td>7.2E-14</td><td>MB231/MB468</td><td>NUC</td></tr>
<tr><td>>hg38_21_+_8433172_8446622_RNA45SN1_45S_50ntFlanks@11115.11133.19|CGGCAGGGGCCGGCGGCGG</td><td>>hg38_21_+_8433172_8446622_RNA45SN1_45S_50ntFlanks</td><td>11115</td><td>11133</td><td>19</td><td>CGGCAGGGGCCGGCGGCGG</td><td>302.1</td><td>-12.1</td><td>1.6</td><td>-7.7</td><td>1.3E-14</td><td>7.2E-14</td><td>MB231/MB468</td><td>MP</td></tr>
<tr><td>>hg38_21_+_8433172_8446622_RNA45SN1_45S_50ntFlanks@11207.11227.21|GGCGGCGACTCTGGACGCGAG</td><td>>hg38_21_+_8433172_8446622_RNA45SN1_45S_50ntFlanks</td><td>11207</td><td>11227</td><td>21</td><td>GGCGGCGACTCTGGACGCGAG</td><td>193.3</td><td>-10.3</td><td>1.3</td><td>-7.7</td><td>1.5E-14</td><td>7.2E-14</td><td>MB231/MB468</td><td>CYTO</td></tr>
<tr><td>>MI0000086|hsa-mir-28&WithFlank&3|+|188688775|188688872@20.41.22(+1A)&[MIMAT0000085&hsa-miR-28-5p&offsets|0|0(+1A);m-133&3|+|188688794|188688815&offsets|0|0(+1A)]|AAGGAGCTCACAGTCTATTGAGA</td><td>>MI0000086|hsa-mir-28&WithFlank&3|+|188688775|188688872</td><td>20</td><td>41</td><td>22(+1A)</td><td>AAGGAGCTCACAGTCTATTGAGA</td><td>92.7</td><td>6.7</td><td>0.9</td><td>7.6</td><td>2.5E-14</td><td>7.2E-14</td><td>BT20/MB468</td><td>NUC</td></tr>
<tr><td>>MI0000650|hsa-mir-200c&WithFlank&12|+|6963693|6963772@50.72.23(+1C)&[MIMAT0000617&hsa-miR-200c-3p&offsets|0|0(+1C);m-54&12|+|6963742|6963764&offsets|0|0(+1C)]|TAATACTGCCGGGTAATGATGGAC</td><td>>MI0000650|hsa-mir-200c&WithFlank&12|+|6963693|6963772</td><td>50</td><td>72</td><td>23(+1C)</td><td>TAATACTGCCGGGTAATGATGGAC</td><td>396.7</td><td>3.2</td><td>0.4</td><td>7.6</td><td>2.5E-14</td><td>7.2E-14</td><td>BT20/MB468</td><td>NUC</td></tr>
<tr><td>>hg38_21_+_8433172_8446622_RNA45SN1_45S_50ntFlanks@6663.6681.19|GTGGATCACTCGGCTCGTG</td><td>>hg38_21_+_8433172_8446622_RNA45SN1_45S_50ntFlanks</td><td>6663</td><td>6681</td><td>19</td><td>GTGGATCACTCGGCTCGTG</td><td>3439.3</td><td>-9.9</td><td>1.3</td><td>-7.6</td><td>2.1E-14</td><td>7.2E-14</td><td>MB231/MB468</td><td>MITO</td></tr>
<tr><td>>hg38_21_+_8433172_8446622_RNA45SN1_45S_50ntFlanks@10481.10498.18|ATCGAAAGGGAGTCGGGT</td><td>>hg38_21_+_8433172_8446622_RNA45SN1_45S_50ntFlanks</td><td>10481</td><td>10498</td><td>18</td><td>ATCGAAAGGGAGTCGGGT</td><td>257.1</td><td>-11.9</td><td>1.5</td><td>-7.7</td><td>1.3E-14</td><td>7.2E-14</td><td>MB231/MB468</td><td>MP</td></tr>
<tr><td>>hg38_21_+_8433172_8446622_RNA45SN1_45S_50ntFlanks@6666.6689.24|GATCACTCGGCTCGTGCGTCGATG</td><td>>hg38_21_+_8433172_8446622_RNA45SN1_45S_50ntFlanks</td><td>6666</td><td>6689</td><td>24</td><td>GATCACTCGGCTCGTGCGTCGATG</td><td>833.7</td><td>-12.8</td><td>1.6</td><td>-8.1</td><td>5.1E-16</td><td>7.2E-14</td><td>BT20/MB468</td><td>MP</td></tr>
<tr><td>>hg38_21_+_8433172_8446622_RNA45SN1_45S_50ntFlanks@7977.8006.30|CGACCTCAGATCAGACGTGGCGACCCGCTG</td><td>>hg38_21_+_8433172_8446622_RNA45SN1_45S_50ntFlanks</td><td>7977</td><td>8006</td><td>30</td><td>CGACCTCAGATCAGACGTGGCGACCCGCTG</td><td>4738.7</td><td>-6.2</td><td>0.8</td><td>-7.6</td><td>2.1E-14</td><td>7.2E-14</td><td>MB231/MB468</td><td>MITO</td></tr>
<tr><td>>hg38_21_+_8433172_8446622_RNA45SN1_45S_50ntFlanks@12582.12604.23|ACCGCAGGTTCAGACATTTGGTG</td><td>>hg38_21_+_8433172_8446622_RNA45SN1_45S_50ntFlanks</td><td>12582</td><td>12604</td><td>23</td><td>ACCGCAGGTTCAGACATTTGGTG</td><td>146.4</td><td>-11.1</td><td>1.4</td><td>-7.7</td><td>1.3E-14</td><td>7.2E-14</td><td>MB231/MB468</td><td>MP</td></tr>
<tr><td>>MI0000077|hsa-mir-21&WithFlank&17|+|59841260|59841343@14.37.24&[MIMAT0000076&hsa-miR-21-5p&offsets|0|+2;m-1&17|+|59841273|59841294&offsets|0|+2]|TAGCTTATCAGACTGATGTTGACT</td><td>>MI0000077|hsa-mir-21&WithFlank&17|+|59841260|59841343</td><td>14</td><td>37</td><td>24</td><td>TAGCTTATCAGACTGATGTTGACT</td><td>176384.4</td><td>6.8</td><td>0.9</td><td>7.6</td><td>2.1E-14</td><td>7.2E-14</td><td>MB231/MB468</td><td>MITO</td></tr>
<tr><td>>hg38_21_+_8433172_8446622_RNA45SN1_45S_50ntFlanks@10116.10139.24|TGCGTCGGAGGGCGGCGGCGGCGG</td><td>>hg38_21_+_8433172_8446622_RNA45SN1_45S_50ntFlanks</td><td>10116</td><td>10139</td><td>24</td><td>TGCGTCGGAGGGCGGCGGCGGCGG</td><td>298.0</td><td>-10.9</td><td>1.4</td><td>-7.7</td><td>1.5E-14</td><td>7.2E-14</td><td>MB231/MB468</td><td>CYTO</td></tr>
<tr><td>>hg38_21_+_8433172_8446622_RNA45SN1_45S_50ntFlanks@5071.5098.28|TTAATTCCGATAACGAACGAGACTCTGG</td><td>>hg38_21_+_8433172_8446622_RNA45SN1_45S_50ntFlanks</td><td>5071</td><td>5098</td><td>28</td><td>TTAATTCCGATAACGAACGAGACTCTGG</td><td>2444.4</td><td>-15.1</td><td>2.0</td><td>-7.7</td><td>1.3E-14</td><td>7.2E-14</td><td>MB231/MB468</td><td>MP</td></tr>
<tr><td>>hg38_21_+_8433172_8446622_RNA45SN1_45S_50ntFlanks@11320.11342.23|CCACGTCTCGTCGCGCGCGCGTC</td><td>>hg38_21_+_8433172_8446622_RNA45SN1_45S_50ntFlanks</td><td>11320</td><td>11342</td><td>23</td><td>CCACGTCTCGTCGCGCGCGCGTC</td><td>1792.5</td><td>-7.6</td><td>1.0</td><td>-7.7</td><td>1.5E-14</td><td>7.3E-14</td><td>MB231/MB468</td><td>CYTO</td></tr>
<tr><td>>am_MI0000809|hsa-mir-151a&WithFlank&8|-|140732558|140732659@53.71.19(+1A)&[MIMAT0000757&hsa-miR-151a-3p&offsets|0|-2(+1A);m-30&8|-|140732587|140732607&offsets|0|-2(+1A)]|CTAGACTGAAGCTCCTTGAA</td><td>>am_MI0000809|hsa-mir-151a&WithFlank&8|-|140732558|140732659</td><td>53</td><td>71</td><td>19(+1A)</td><td>CTAGACTGAAGCTCCTTGAA</td><td>389.3</td><td>10.1</td><td>1.3</td><td>7.7</td><td>1.3E-14</td><td>7.3E-14</td><td>BT20/MB468</td><td>MITO</td></tr>
<tr><td>>hg38_21_+_8433172_8446622_RNA45SN1_45S_50ntFlanks@4722.4751.30|TCATTAATCAAGAACGAAAGTCGGAGGTTC</td><td>>hg38_21_+_8433172_8446622_RNA45SN1_45S_50ntFlanks</td><td>4722</td><td>4751</td><td>30</td><td>TCATTAATCAAGAACGAAAGTCGGAGGTTC</td><td>615.6</td><td>-7.0</td><td>0.9</td><td>-7.7</td><td>1.6E-14</td><td>7.3E-14</td><td>MB231/MB468</td><td>NUC</td></tr>
<tr><td>>hg38_21_+_8433172_8446622_RNA45SN1_45S_50ntFlanks@8170.8188.19|CAAGTCCTTCTGATCGAGG</td><td>>hg38_21_+_8433172_8446622_RNA45SN1_45S_50ntFlanks</td><td>8170</td><td>8188</td><td>19</td><td>CAAGTCCTTCTGATCGAGG</td><td>406.9</td><td>-12.5</td><td>1.6</td><td>-7.6</td><td>2.1E-14</td><td>7.3E-14</td><td>MB231/MB468</td><td>MITO</td></tr>
<tr><td>>hg38_21_+_8433172_8446622_RNA45SN1_45S_50ntFlanks@4727.4768.42|AATCAAGAACGAAAGTCGGAGGTTCGAAGACGATCAGATACC</td><td>>hg38_21_+_8433172_8446622_RNA45SN1_45S_50ntFlanks</td><td>4727</td><td>4768</td><td>42</td><td>AATCAAGAACGAAAGTCGGAGGTTCGAAGACGATCAGATACC</td><td>2833.5</td><td>11.1</td><td>1.4</td><td>7.7</td><td>1.5E-14</td><td>7.3E-14</td><td>MB231/MB468</td><td>CYTO</td></tr>
<tr><td>>trnaMT_LysTTT_MT_+_8295_8364@1.38.38,trnalookalike7_LysTTT_1_+_568844_568913@1.38.38|CACTGTAAAGCTAACTTAGCATTAACCTTTTAAGTTAA</td><td>>trnaMT_LysTTT_MT_+_8295_8364</td><td>1</td><td>38</td><td>38</td><td>CACTGTAAAGCTAACTTAGCATTAACCTTTTAAGTTAA</td><td>1210.9</td><td>7.0</td><td>0.9</td><td>7.7</td><td>1.3E-14</td><td>7.3E-14</td><td>BT20/MB468</td><td>MITO</td></tr>
<tr><td>>hg38_21_+_8433172_8446622_RNA45SN1_45S_50ntFlanks@11918.11954.37|ACATGAGAGGTGTAGAATAAGTGGGAGGCCCCCGGCG</td><td>>hg38_21_+_8433172_8446622_RNA45SN1_45S_50ntFlanks</td><td>11918</td><td>11954</td><td>37</td><td>ACATGAGAGGTGTAGAATAAGTGGGAGGCCCCCGGCG</td><td>224.0</td><td>-10.8</td><td>1.4</td><td>-7.7</td><td>1.6E-14</td><td>7.3E-14</td><td>MB231/MB468</td><td>NUC</td></tr>
<tr><td>>hg38_21_+_8433172_8446622_RNA45SN1_45S_50ntFlanks@12259.12288.30|CTTGATCTTGATTTTCAGTACGAATACAGA</td><td>>hg38_21_+_8433172_8446622_RNA45SN1_45S_50ntFlanks</td><td>12259</td><td>12288</td><td>30</td><td>CTTGATCTTGATTTTCAGTACGAATACAGA</td><td>313.3</td><td>-12.2</td><td>1.6</td><td>-7.7</td><td>1.3E-14</td><td>7.3E-14</td><td>MB231/MB468</td><td>MP</td></tr>
<tr><td>>trna5_AspGTC_12_+_98897281_98897352@32.72.41|CTGTCACGCGGGAGACCGGGGTTCAATTCCCCGACGGGGAG</td><td>>trna5_AspGTC_12_+_98897281_98897352</td><td>32</td><td>72</td><td>41</td><td>CTGTCACGCGGGAGACCGGGGTTCAATTCCCCGACGGGGAG</td><td>169.0</td><td>-10.1</td><td>1.3</td><td>-7.7</td><td>1.5E-14</td><td>7.3E-14</td><td>MB231/MB468</td><td>CYTO</td></tr>
<tr><td>>hg38_21_+_8433172_8446622_RNA45SN1_45S_50ntFlanks@12499.12516.18|GGTTTAGACCGTCGTGAG</td><td>>hg38_21_+_8433172_8446622_RNA45SN1_45S_50ntFlanks</td><td>12499</td><td>12516</td><td>18</td><td>GGTTTAGACCGTCGTGAG</td><td>170.9</td><td>-6.9</td><td>0.9</td><td>-7.7</td><td>1.5E-14</td><td>7.3E-14</td><td>MB231/MB468</td><td>CYTO</td></tr>
<tr><td>>hg38_21_+_8433172_8446622_RNA45SN1_45S_50ntFlanks@9575.9592.18|ATCGGTCGTCCGACCTGG</td><td>>hg38_21_+_8433172_8446622_RNA45SN1_45S_50ntFlanks</td><td>9575</td><td>9592</td><td>18</td><td>ATCGGTCGTCCGACCTGG</td><td>491.2</td><td>-12.8</td><td>1.7</td><td>-7.7</td><td>1.3E-14</td><td>7.3E-14</td><td>MB231/MB468</td><td>MP</td></tr>
<tr><td>>hg38_1_-_228634819_228635039_RNA5S12_5S_50ntFlanks@51.91.41|GTCTACGGCCATACCACCCTGAACGCGCCCGATCTCGTCTG</td><td>>hg38_1_-_228634819_228635039_RNA5S12_5S_50ntFlanks</td><td>51</td><td>91</td><td>41</td><td>GTCTACGGCCATACCACCCTGAACGCGCCCGATCTCGTCTG</td><td>1298.0</td><td>6.1</td><td>0.8</td><td>7.7</td><td>1.5E-14</td><td>7.3E-14</td><td>MB231/MB468</td><td>CYTO</td></tr>
</table>

<table>
<tr><td>>MI0000434|hsa-let-7i&WithFlank&12|+|62603680|62603775@12.33.22(+1C)&[MIMAT0000415&hsa-let-7i-5p&offsets|0|0(+1C);m-49&12|+|62603691|62603712&offsets|0|0(+1C)]|TGAGGTAGTAGTTTGTGCTGTTC</td><td>>MI0000434|hsa-let-7i&WithFlank&12|+|62603680|62603775</td><td>12</td><td>33</td><td>22(+1C)</td><td>TGAGGTAGTAGTTTGTGCTGTTC</td><td>118.1</td><td>2.5</td><td>0.3</td><td>7.6</td><td>2.6E-14</td><td>7.3E-14</td><td>BT20/MB468</td><td>NUC</td></tr>
<tr><td>>hg38_21_+_8433172_8446622_RNA45SN1_45S_50ntFlanks@12405.12441.37|CTTTTTGATCCTTCGATGTCGGCTCTTCCTATCATTG</td><td>>hg38_21_+_8433172_8446622_RNA45SN1_45S_50ntFlanks</td><td>12405</td><td>12441</td><td>37</td><td>CTTTTTGATCCTTCGATGTCGGCTCTTCCTATCATTG</td><td>40403.6</td><td>-13.5</td><td>1.8</td><td>-7.6</td><td>2.6E-14</td><td>7.3E-14</td><td>BT20/MB468</td><td>NUC</td></tr>
<tr><td>>hg38_21_+_8433172_8446622_RNA45SN1_45S_50ntFlanks@8116.8138.23|GCGGGACATGTGGCGTACGGAAG</td><td>>hg38_21_+_8433172_8446622_RNA45SN1_45S_50ntFlanks</td><td>8116</td><td>8138</td><td>23</td><td>GCGGGACATGTGGCGTACGGAAG</td><td>289.7</td><td>-12.0</td><td>1.6</td><td>-7.7</td><td>1.3E-14</td><td>7.3E-14</td><td>MB231/MB468</td><td>MP</td></tr>
<tr><td>>hg38_21_+_8433172_8446622_RNA45SN1_45S_50ntFlanks@9462.9503.42|GGGAGGTGGAGCACGAGCGCACGTGTTAGGACCCGAAAGATG</td><td>>hg38_21_+_8433172_8446622_RNA45SN1_45S_50ntFlanks</td><td>9462</td><td>9503</td><td>42</td><td>GGGAGGTGGAGCACGAGCGCACGTGTTAGGACCCGAAAGATG</td><td>354.1</td><td>-12.3</td><td>1.6</td><td>-7.6</td><td>2.1E-14</td><td>7.4E-14</td><td>MB231/MB468</td><td>MITO</td></tr>
<tr><td>>hg38_21_+_8433172_8446622_RNA45SN1_45S_50ntFlanks@12539.12572.34|ATGATGTGTTGTTGCCATGGTAATCCTGCTCAGT</td><td>>hg38_21_+_8433172_8446622_RNA45SN1_45S_50ntFlanks</td><td>12539</td><td>12572</td><td>34</td><td>ATGATGTGTTGTTGCCATGGTAATCCTGCTCAGT</td><td>169.9</td><td>-11.3</td><td>1.5</td><td>-7.7</td><td>1.3E-14</td><td>7.4E-14</td><td>MB231/MB468</td><td>MP</td></tr>
<tr><td>>hg38_21_+_8433172_8446622_RNA45SN1_45S_50ntFlanks@13004.13023.20|ATCTATTGAAAGTCAGCCCT</td><td>>hg38_21_+_8433172_8446622_RNA45SN1_45S_50ntFlanks</td><td>13004</td><td>13023</td><td>20</td><td>ATCTATTGAAAGTCAGCCCT</td><td>253.3</td><td>-10.3</td><td>1.3</td><td>-7.7</td><td>1.7E-14</td><td>7.4E-14</td><td>BT20/MB468</td><td>CYTO</td></tr>
<tr><td>>hg38_21_+_8433172_8446622_RNA45SN1_45S_50ntFlanks@8844.8867.24|GCGGCGGCGGCGGCGGCGGCGGGA</td><td>>hg38_21_+_8433172_8446622_RNA45SN1_45S_50ntFlanks</td><td>8844</td><td>8867</td><td>24</td><td>GCGGCGGCGGCGGCGGCGGCGGGA</td><td>270.2</td><td>-11.9</td><td>1.6</td><td>-7.7</td><td>1.3E-14</td><td>7.4E-14</td><td>MB231/MB468</td><td>MP</td></tr>
<tr><td>>hg38_21_+_8433172_8446622_RNA45SN1_45S_50ntFlanks@4005.4042.38|AACCTCGGGCCGATCGCACGCCCCCCGTGGCGGCGACG</td><td>>hg38_21_+_8433172_8446622_RNA45SN1_45S_50ntFlanks</td><td>4005</td><td>4042</td><td>38</td><td>AACCTCGGGCCGATCGCACGCCCCCCGTGGCGGCGACG</td><td>562.7</td><td>-5.0</td><td>0.7</td><td>-7.7</td><td>1.8E-14</td><td>7.4E-14</td><td>BT20/MB468</td><td>CYTO</td></tr>
<tr><td>>hg38_21_+_8433172_8446622_RNA45SN1_45S_50ntFlanks@10879.10901.23|CGGTCGGGCTGGGGCGCGAAGCG</td><td>>hg38_21_+_8433172_8446622_RNA45SN1_45S_50ntFlanks</td><td>10879</td><td>10901</td><td>23</td><td>CGGTCGGGCTGGGGCGCGAAGCG</td><td>213.5</td><td>-11.6</td><td>1.5</td><td>-7.7</td><td>1.3E-14</td><td>7.4E-14</td><td>MB231/MB468</td><td>MP</td></tr>
<tr><td>>hg38_21_+_8433172_8446622_RNA45SN1_45S_50ntFlanks@4733.4765.33|GAACGAAAGTCGGAGGTTCGAAGACGATCAGAT</td><td>>hg38_21_+_8433172_8446622_RNA45SN1_45S_50ntFlanks</td><td>4733</td><td>4765</td><td>33</td><td>GAACGAAAGTCGGAGGTTCGAAGACGATCAGAT</td><td>1129.5</td><td>7.8</td><td>1.0</td><td>7.7</td><td>1.8E-14</td><td>7.4E-14</td><td>BT20/MB468</td><td>CYTO</td></tr>
<tr><td>>hg38_21_+_8433172_8446622_RNA45SN1_45S_50ntFlanks@12073.12096.24|CCGAGGGGCTCTCGCTTCTGGCGC</td><td>>hg38_21_+_8433172_8446622_RNA45SN1_45S_50ntFlanks</td><td>12073</td><td>12096</td><td>24</td><td>CCGAGGGGCTCTCGCTTCTGGCGC</td><td>1232.8</td><td>-9.4</td><td>1.2</td><td>-7.6</td><td>2.2E-14</td><td>7.4E-14</td><td>MB231/MB468</td><td>MITO</td></tr>
<tr><td>>MI0000299|hsa-mir-222&WithFlank&X|-|45747009|45747130@75.95.21(+1U)&[MIMAT0000279&hsa-miR-222-3p&offsets|0|0(+1U);m-140&X|-|45747033|45747056&offsets|0|-3(+1U)]|AGCTACATCTGGCTACTGGGTT</td><td>>MI0000299|hsa-mir-222&WithFlank&X|-|45747009|45747130</td><td>75</td><td>95</td><td>21(+1U)</td><td>AGCTACATCTGGCTACTGGGTT</td><td>261.6</td><td>5.6</td><td>0.7</td><td>7.7</td><td>1.6E-14</td><td>7.5E-14</td><td>MB231/MB468</td><td>NUC</td></tr>
<tr><td>>hg38_21_+_8433172_8446622_RNA45SN1_45S_50ntFlanks@10574.10607.34|AAGCCGGCGGGAGCCCCGGGGAGAGTTCTCTTTT</td><td>>hg38_21_+_8433172_8446622_RNA45SN1_45S_50ntFlanks</td><td>10574</td><td>10607</td><td>34</td><td>AAGCCGGCGGGAGCCCCGGGGAGAGTTCTCTTTT</td><td>450.7</td><td>-7.0</td><td>0.9</td><td>-7.7</td><td>1.5E-14</td><td>7.5E-14</td><td>MB231/MB468</td><td>CYTO</td></tr>
<tr><td>>hg38_21_+_8433172_8446622_RNA45SN1_45S_50ntFlanks@12054.12071.18|TGAGGCGGGGGGGCGAGC</td><td>>hg38_21_+_8433172_8446622_RNA45SN1_45S_50ntFlanks</td><td>12054</td><td>12071</td><td>18</td><td>TGAGGCGGGGGGGCGAGC</td><td>257.9</td><td>-11.0</td><td>1.4</td><td>-7.7</td><td>1.6E-14</td><td>7.5E-14</td><td>MB231/MB468</td><td>NUC</td></tr>
<tr><td>>hg38_21_+_8433172_8446622_RNA45SN1_45S_50ntFlanks@8116.8140.25|GCGGGACATGTGGCGTACGGAAGAC</td><td>>hg38_21_+_8433172_8446622_RNA45SN1_45S_50ntFlanks</td><td>8116</td><td>8140</td><td>25</td><td>GCGGGACATGTGGCGTACGGAAGAC</td><td>212.2</td><td>-11.6</td><td>1.5</td><td>-7.7</td><td>1.3E-14</td><td>7.5E-14</td><td>MB231/MB468</td><td>MP</td></tr>
<tr><td>>hg38_21_+_8433172_8446622_RNA45SN1_45S_50ntFlanks@10410.10440.31|TCAGTCGGTCCTGAGAGATGGGCGAGCGCCG</td><td>>hg38_21_+_8433172_8446622_RNA45SN1_45S_50ntFlanks</td><td>10410</td><td>10440</td><td>31</td><td>TCAGTCGGTCCTGAGAGATGGGCGAGCGCCG</td><td>156.1</td><td>-11.2</td><td>1.4</td><td>-7.7</td><td>1.4E-14</td><td>7.5E-14</td><td>MB231/MB468</td><td>MP</td></tr>
<tr><td>>MI0000095|hsa-mir-93&WithFlank&7|-|100093762|100093853@17.39.23(+1A)&[MIMAT0000093&hsa-miR-93-5p&offsets|0|0(+1A);m-29&7|-|100093815|100093837&offsets|0|0(+1A)]|CAAAGTGCTGTTCGTGCAGGTAGA</td><td>>MI0000095|hsa-mir-93&WithFlank&7|-|100093762|100093853</td><td>17</td><td>39</td><td>23(+1A)</td><td>CAAAGTGCTGTTCGTGCAGGTAGA</td><td>1725.3</td><td>6.0</td><td>0.8</td><td>7.7</td><td>1.3E-14</td><td>7.5E-14</td><td>BT20/MB468</td><td>MITO</td></tr>
<tr><td>>hg38_21_+_8433172_8446622_RNA45SN1_45S_50ntFlanks@5494.5516.23|AAGACGGTCGAACTTGACTATCT</td><td>>hg38_21_+_8433172_8446622_RNA45SN1_45S_50ntFlanks</td><td>5494</td><td>5516</td><td>23</td><td>AAGACGGTCGAACTTGACTATCT</td><td>2954.7</td><td>6.0</td><td>0.8</td><td>7.7</td><td>1.3E-14</td><td>7.5E-14</td><td>BT20/MB468</td><td>MITO</td></tr>
<tr><td>>hg38_21_+_8433172_8446622_RNA45SN1_45S_50ntFlanks@12141.12158.18|ACAGTGCCAGGTGGGGAG</td><td>>hg38_21_+_8433172_8446622_RNA45SN1_45S_50ntFlanks</td><td>12141</td><td>12158</td><td>18</td><td>ACAGTGCCAGGTGGGGAG</td><td>155.1</td><td>-10.0</td><td>1.3</td><td>-7.7</td><td>1.5E-14</td><td>7.5E-14</td><td>MB231/MB468</td><td>CYTO</td></tr>
<tr><td>>hg38_21_+_8433172_8446622_RNA45SN1_45S_50ntFlanks@11602.11619.18|TCCGACTGTTTAATTAAA</td><td>>hg38_21_+_8433172_8446622_RNA45SN1_45S_50ntFlanks</td><td>11602</td><td>11619</td><td>18</td><td>TCCGACTGTTTAATTAAA</td><td>918.7</td><td>-5.2</td><td>0.7</td><td>-7.6</td><td>2.2E-14</td><td>7.5E-14</td><td>MB231/MB468</td><td>MITO</td></tr>
<tr><td>>hg38_21_+_8433172_8446622_RNA45SN1_45S_50ntFlanks@6719.6740.22|TGAATTGCAGGACACATTGATC</td><td>>hg38_21_+_8433172_8446622_RNA45SN1_45S_50ntFlanks</td><td>6719</td><td>6740</td><td>22</td><td>TGAATTGCAGGACACATTGATC</td><td>14192.5</td><td>-12.5</td><td>1.6</td><td>-7.6</td><td>2.7E-14</td><td>7.5E-14</td><td>BT20/MB468</td><td>NUC</td></tr>
<tr><td>>hg38_21_+_8433172_8446622_RNA45SN1_45S_50ntFlanks@3813.3834.22|TTATGGTTCCTTTGGTCGCTCG</td><td>>hg38_21_+_8433172_8446622_RNA45SN1_45S_50ntFlanks</td><td>3813</td><td>3834</td><td>22</td><td>TTATGGTTCCTTTGGTCGCTCG</td><td>156.9</td><td>-11.2</td><td>1.4</td><td>-7.7</td><td>1.4E-14</td><td>7.5E-14</td><td>MB231/MB468</td><td>MP</td></tr>
<tr><td>>hg38_1_-_228634819_228635039_RNA5S12_5S_50ntFlanks@147.169.23|GGGAATACCGGGTGCTGTAGGCT</td><td>>hg38_1_-_228634819_228635039_RNA5S12_5S_50ntFlanks</td><td>147</td><td>169</td><td>23</td><td>GGGAATACCGGGTGCTGTAGGCT</td><td>1478.5</td><td>5.0</td><td>0.7</td><td>7.7</td><td>1.5E-14</td><td>7.6E-14</td><td>MB231/MB468</td><td>CYTO</td></tr>
<tr><td>>hg38_21_+_8433172_8446622_RNA45SN1_45S_50ntFlanks@9564.9595.32|CTGACGTGCAAATCGGTCGTCCGACCTGGGTA</td><td>>hg38_21_+_8433172_8446622_RNA45SN1_45S_50ntFlanks</td><td>9564</td><td>9595</td><td>32</td><td>CTGACGTGCAAATCGGTCGTCCGACCTGGGTA</td><td>1038.9</td><td>-9.4</td><td>1.2</td><td>-7.7</td><td>1.4E-14</td><td>7.6E-14</td><td>MB231/MB468</td><td>MP</td></tr>
<tr><td>>MI0000061|hsa-let-7a-2&WithFlank&11|-|122146516|122146599@11.31.21(+1G);MI0000062|hsa-let-7a-3&WithFlank&22|+|46112743|46112828@10.30.21(+1G);MI0000060|hsa-let-7a-1&WithFlank&9|+|94175951|94176042@12.32.21(+1G)&[MIMAT0000062&hsa-let-7a-5p&offsets|0|-1(+1G);m-6&11|-|122146568|122146589&offsets|0|-1(+1G)];[MIMAT0000062_1&hsa-let-7a-5p&offsets|0|-1(+1G);m-5&22|+|46112752|46112773&offsets|0|-1(+1G)];[MIMAT0000062_2&hsa-let-7a-5p&offsets|0|-1(+1G);m-4&9|+|94175962|94175983&offsets|0|-1(+1G)]|TGAGGTAGTAGGTTGTATAGTG</td><td>>MI0000061|hsa-let-7a-2&WithFlank&11|-|122146516|122146599</td><td>11</td><td>31</td><td>21(+1G)</td><td>TGAGGTAGTAGGTTGTATAGTG</td><td>1002.2</td><td>5.1</td><td>0.7</td><td>7.7</td><td>1.6E-14</td><td>7.6E-14</td><td>MB231/MB468</td><td>NUC</td></tr>
<tr><td>>am_trna2_GlyGCC_21_-_18827107_18827177@57.74.18,trna35_GlyGCC_1_+_161413094_161413164@57.74.18,trna37_GlyGCC_1_+_161420467_161420537@57.74.18,trna39_GlyGCC_1_+_161427898_161427968@57.74.18,trna41_GlyGCC_1_+_161435258_161435328@57.74.18|TTCCCGGCCCATGCACCA</td><td>>am_trna2_GlyGCC_21_-_18827107_18827177</td><td>57</td><td>74</td><td>18</td><td>TTCCCGGCCCATGCACCA</td><td>205.1</td><td>11.4</td><td>1.4</td><td>7.9</td><td>2.1E-15</td><td>7.6E-14</td><td>BT20/MB231</td><td>NUC</td></tr>
<tr><td>>hg38_21_+_8433172_8446622_RNA45SN1_45S_50ntFlanks@10728.10755.28|GAGAGGGTGTAAATCTCGCGCCGGGCCG</td><td>>hg38_21_+_8433172_8446622_RNA45SN1_45S_50ntFlanks</td><td>10728</td><td>10755</td><td>28</td><td>GAGAGGGTGTAAATCTCGCGCCGGGCCG</td><td>180.6</td><td>-10.2</td><td>1.3</td><td>-7.7</td><td>1.5E-14</td><td>7.6E-14</td><td>MB231/MB468</td><td>CYTO</td></tr>
<tr><td>>hg38_21_+_8433172_8446622_RNA45SN1_45S_50ntFlanks@12202.12232.31|CTAAGGCGAGCTCAGGGAGGACAGAAACCTC</td><td>>hg38_21_+_8433172_8446622_RNA45SN1_45S_50ntFlanks</td><td>12202</td><td>12232</td><td>31</td><td>CTAAGGCGAGCTCAGGGAGGACAGAAACCTC</td><td>754.9</td><td>-13.3</td><td>1.7</td><td>-7.6</td><td>2.2E-14</td><td>7.6E-14</td><td>MB231/MB468</td><td>MITO</td></tr>
<tr><td>>MI0000084|hsa-mir-26b&WithFlank&2|+|218402640|218402728@18.39.22&[MIMAT0000083&hsa-miR-26b-5p&offsets|0|+1;m-53&2|+|218402657|218402678&offsets|0|0]|TTCAAGTAATTCAGGATAGGTT</td><td>>MI0000084|hsa-mir-26b&WithFlank&2|+|218402640|218402728</td><td>18</td><td>39</td><td>22</td><td>TTCAAGTAATTCAGGATAGGTT</td><td>4747.7</td><td>4.7</td><td>0.6</td><td>7.7</td><td>1.6E-14</td><td>7.6E-14</td><td>MB231/MB468</td><td>NUC</td></tr>
<tr><td>>hg38_21_+_8433172_8446622_RNA45SN1_45S_50ntFlanks@3854.3872.19|ACTGTGGTAATTCTAGAGC</td><td>>hg38_21_+_8433172_8446622_RNA45SN1_45S_50ntFlanks</td><td>3854</td><td>3872</td><td>19</td><td>ACTGTGGTAATTCTAGAGC</td><td>1270.7</td><td>-8.2</td><td>1.1</td><td>-7.7</td><td>1.4E-14</td><td>7.6E-14</td><td>MB231/MB468</td><td>MP</td></tr>
<tr><td>>hg38_21_+_8433172_8446622_RNA45SN1_45S_50ntFlanks@8194.8223.30|CCCGTGGACGGTGTGAGGCCGGTAGCGGCC</td><td>>hg38_21_+_8433172_8446622_RNA45SN1_45S_50ntFlanks</td><td>8194</td><td>8223</td><td>30</td><td>CCCGTGGACGGTGTGAGGCCGGTAGCGGCC</td><td>149.9</td><td>-11.1</td><td>1.4</td><td>-7.7</td><td>1.4E-14</td><td>7.6E-14</td><td>MB231/MB468</td><td>MP</td></tr>
<tr><td>>hg38_21_+_8433172_8446622_RNA45SN1_45S_50ntFlanks@12155.12181.27|GGAGTTTGACTGGGGCGGTACACCTGT</td><td>>hg38_21_+_8433172_8446622_RNA45SN1_45S_50ntFlanks</td><td>12155</td><td>12181</td><td>27</td><td>GGAGTTTGACTGGGGCGGTACACCTGT</td><td>167.6</td><td>-11.3</td><td>1.5</td><td>-7.7</td><td>1.4E-14</td><td>7.6E-14</td><td>MB231/MB468</td><td>MP</td></tr>
<tr><td>>hg38_21_+_8433172_8446622_RNA45SN1_45S_50ntFlanks@9454.9493.40|CCGCGCCGGGGAGGTGGAGCACGAGCGCACGTGTTAGGAC</td><td>>hg38_21_+_8433172_8446622_RNA45SN1_45S_50ntFlanks</td><td>9454</td><td>9493</td><td>40</td><td>CCGCGCCGGGGAGGTGGAGCACGAGCGCACGTGTTAGGAC</td><td>147.6</td><td>-10.2</td><td>1.3</td><td>-7.7</td><td>1.7E-14</td><td>7.6E-14</td><td>MB231/MB468</td><td>NUC</td></tr>
<tr><td>>hg38_21_+_8433172_8446622_RNA45SN1_45S_50ntFlanks@5489.5512.24|CTGAGAAGACGGTCGAACTTGACT</td><td>>hg38_21_+_8433172_8446622_RNA45SN1_45S_50ntFlanks</td><td>5489</td><td>5512</td><td>24</td><td>CTGAGAAGACGGTCGAACTTGACT</td><td>1271.3</td><td>5.8</td><td>0.8</td><td>7.7</td><td>1.8E-14</td><td>7.6E-14</td><td>BT20/MB468</td><td>CYTO</td></tr>
<tr><td>>MI0000102|hsa-mir-100&WithFlank&11|-|122152223|122152314@22.40.19&[MIMAT0000098&hsa-miR-100-5p&offsets|+3|0;m-27&11|-|122152276|122152296&offsets|+3|+1]|CCGTAGATCCGAACTTGTG</td><td>>MI0000102|hsa-mir-100&WithFlank&11|-|122152223|122152314</td><td>22</td><td>40</td><td>19</td><td>CCGTAGATCCGAACTTGTG</td><td>808.6</td><td>13.6</td><td>1.8</td><td>7.7</td><td>1.7E-14</td><td>7.6E-14</td><td>MB231/MB468</td><td>NUC</td></tr>
<tr><td>>am_trnaMT_MetCAT_MT_+_4402_4469@19.36.18|AGCTATCGGGCCCATACC</td><td>>am_trnaMT_MetCAT_MT_+_4402_4469</td><td>19</td><td>36</td><td>18</td><td>AGCTATCGGGCCCATACC</td><td>328.0</td><td>9.8</td><td>1.3</td><td>7.7</td><td>1.8E-14</td><td>7.6E-14</td><td>BT20/MB468</td><td>CYTO</td></tr>
<tr><td>>hg38_21_+_8433172_8446622_RNA45SN1_45S_50ntFlanks@8357.8379.23|AAAAGAACTTTGAAGAGAGAGTT</td><td>>hg38_21_+_8433172_8446622_RNA45SN1_45S_50ntFlanks</td><td>8357</td><td>8379</td><td>23</td><td>AAAAGAACTTTGAAGAGAGAGTT</td><td>24712.0</td><td>-7.6</td><td>1.0</td><td>-7.7</td><td>1.8E-14</td><td>7.6E-14</td><td>BT20/MB468</td><td>CYTO</td></tr>
<tr><td>>hg38_21_+_8433172_8446622_RNA45SN1_45S_50ntFlanks@4762.4783.22|AGATACCGTCGTAGTTCCGACC</td><td>>hg38_21_+_8433172_8446622_RNA45SN1_45S_50ntFlanks</td><td>4762</td><td>4783</td><td>22</td><td>AGATACCGTCGTAGTTCCGACC</td><td>242.2</td><td>5.3</td><td>0.7</td><td>7.7</td><td>1.8E-14</td><td>7.7E-14</td><td>BT20/MB468</td><td>CYTO</td></tr>
<tr><td>>hg38_21_+_8433172_8446622_RNA45SN1_45S_50ntFlanks@9566.9597.32|GACGTGCAAATCGGTCGTCCGACCTGGGTATA</td><td>>hg38_21_+_8433172_8446622_RNA45SN1_45S_50ntFlanks</td><td>9566</td><td>9597</td><td>32</td><td>GACGTGCAAATCGGTCGTCCGACCTGGGTATA</td><td>1032.4</td><td>-4.6</td><td>0.6</td><td>-7.7</td><td>1.8E-14</td><td>7.7E-14</td><td>BT20/MB468</td><td>CYTO</td></tr>
<tr><td>>hg38_21_+_8433172_8446622_RNA45SN1_45S_50ntFlanks@12936.12960.25|AAACCATTCGTAGACGACCTGCTTC</td><td>>hg38_21_+_8433172_8446622_RNA45SN1_45S_50ntFlanks</td><td>12936</td><td>12960</td><td>25</td><td>AAACCATTCGTAGACGACCTGCTTC</td><td>166.7</td><td>-11.2</td><td>1.5</td><td>-7.7</td><td>1.4E-14</td><td>7.7E-14</td><td>MB231/MB468</td><td>MP</td></tr>
<tr><td>>hg38_21_+_8433172_8446622_RNA45SN1_45S_50ntFlanks@12639.12661.23|TACCATCTGTGGGATTATGACTG</td><td>>hg38_21_+_8433172_8446622_RNA45SN1_45S_50ntFlanks</td><td>12639</td><td>12661</td><td>23</td><td>TACCATCTGTGGGATTATGACTG</td><td>1110.1</td><td>-6.0</td><td>0.8</td><td>-7.6</td><td>2.2E-14</td><td>7.7E-14</td><td>MB231/MB468</td><td>MITO</td></tr>
<tr><td>>hg38_21_+_8433172_8446622_RNA45SN1_45S_50ntFlanks@9241.9274.34|CGGAGCGAGCGCACGGGGTCGGCGGCGACGTCGG</td><td>>hg38_21_+_8433172_8446622_RNA45SN1_45S_50ntFlanks</td><td>9241</td><td>9274</td><td>34</td><td>CGGAGCGAGCGCACGGGGTCGGCGGCGACGTCGG</td><td>206.3</td><td>-11.6</td><td>1.5</td><td>-7.7</td><td>1.4E-14</td><td>7.7E-14</td><td>MB231/MB468</td><td>MP</td></tr>
<tr><td>>hg38_21_+_8433172_8446622_RNA45SN1_45S_50ntFlanks@9303.9336.34|GACCAAGGAGTCTAACACGTGCGCGAGTCGGGGG</td><td>>hg38_21_+_8433172_8446622_RNA45SN1_45S_50ntFlanks</td><td>9303</td><td>9336</td><td>34</td><td>GACCAAGGAGTCTAACACGTGCGCGAGTCGGGGG</td><td>135.5</td><td>-10.9</td><td>1.4</td><td>-7.7</td><td>1.4E-14</td><td>7.7E-14</td><td>MB231/MB468</td><td>MP</td></tr>
<tr><td>>hg38_21_+_8433172_8446622_RNA45SN1_45S_50ntFlanks@10729.10750.22|AGAGGGTGTAAATCTCGCGCCG</td><td>>hg38_21_+_8433172_8446622_RNA45SN1_45S_50ntFlanks</td><td>10729</td><td>10750</td><td>22</td><td>AGAGGGTGTAAATCTCGCGCCG</td><td>239.4</td><td>-11.8</td><td>1.5</td><td>-7.7</td><td>1.4E-14</td><td>7.7E-14</td><td>MB231/MB468</td><td>MP</td></tr>
<tr><td>>hg38_21_+_8433172_8446622_RNA45SN1_45S_50ntFlanks@12907.12935.29|CACCGCACGTTCGTGGGGAACCTGGCGCT</td><td>>hg38_21_+_8433172_8446622_RNA45SN1_45S_50ntFlanks</td><td>12907</td><td>12935</td><td>29</td><td>CACCGCACGTTCGTGGGGAACCTGGCGCT</td><td>238.6</td><td>-11.8</td><td>1.5</td><td>-7.7</td><td>1.4E-14</td><td>7.8E-14</td><td>MB231/MB468</td><td>MP</td></tr>
<tr><td>>hg38_21_+_8433172_8446622_RNA45SN1_45S_50ntFlanks@6645.6664.20|CTCGTACGACTCTTAGCGGT</td><td>>hg38_21_+_8433172_8446622_RNA45SN1_45S_50ntFlanks</td><td>6645</td><td>6664</td><td>20</td><td>CTCGTACGACTCTTAGCGGT</td><td>4492.2</td><td>5.7</td><td>0.7</td><td>7.7</td><td>1.9E-14</td><td>7.8E-14</td><td>BT20/MB468</td><td>CYTO</td></tr>
<tr><td>>MI0000255|hsa-mir-30d&WithFlank&8|-|134804870|134804951@12.34.23(+1G)&[MIMAT0000245&hsa-miR-30d-5p&offsets|0|+1(+1G);m-21&8|-|134804919|134804940&offsets|0|+1(+1G)]|TGTAAACATCCCCGACTGGAAGCG</td><td>>MI0000255|hsa-mir-30d&WithFlank&8|-|134804870|134804951</td><td>12</td><td>34</td><td>23(+1G)</td><td>TGTAAACATCCCCGACTGGAAGCG</td><td>1328.3</td><td>6.1</td><td>0.8</td><td>7.7</td><td>1.7E-14</td><td>7.8E-14</td><td>MB231/MB468</td><td>NUC</td></tr>
</table>

<table>
<tr><td></td><td></td><td></td><td></td><td></td><td></td><td></td><td></td><td></td><td></td><td></td><td></td><td></td><td></td></tr>
<tr><td>>hg38_21_+_8433172_8446622_RNA45SN1_45S_50ntFlanks@12636.12654.19|AGCTACCATCTGTGGGATT</td><td>>hg38_21_+_8433172_8446622_RNA45SN1_45S_50ntFlanks</td><td>12636</td><td>12654</td><td>19</td><td>AGCTACCATCTGTGGGATT</td><td>262.8</td><td>-4.0</td><td>0.5</td><td>-7.7</td><td>1.7E-14</td><td>7.8E-14</td><td>MB231/MB468</td><td>NUC</td></tr>
<tr><td>>hg38_21_+_8433172_8446622_RNA45SN1_45S_50ntFlanks@6787.6806.20|ACGCCTGTCTGAGCGTCGCT</td><td>>hg38_21_+_8433172_8446622_RNA45SN1_45S_50ntFlanks</td><td>6787</td><td>6806</td><td>20</td><td>ACGCCTGTCTGAGCGTCGCT</td><td>2280.9</td><td>5.3</td><td>0.7</td><td>7.7</td><td>1.6E-14</td><td>7.8E-14</td><td>MB231/MB468</td><td>CYTO</td></tr>
<tr><td>>hg38_21_+_8433172_8446622_RNA45SN1_45S_50ntFlanks@7987.8005.19|TCAGACGTGGCGACCCGCT</td><td>>hg38_21_+_8433172_8446622_RNA45SN1_45S_50ntFlanks</td><td>7987</td><td>8005</td><td>19</td><td>TCAGACGTGGCGACCCGCT</td><td>174.8</td><td>-11.3</td><td>1.5</td><td>-7.7</td><td>1.4E-14</td><td>7.8E-14</td><td>MB231/MB468</td><td>MP</td></tr>
<tr><td>>hg38_21_+_8433172_8446622_RNA45SN1_45S_50ntFlanks@11792.11819.28|ATGAACGAGATTCCCACTGTCCCTACCT</td><td>>hg38_21_+_8433172_8446622_RNA45SN1_45S_50ntFlanks</td><td>11792</td><td>11819</td><td>28</td><td>ATGAACGAGATTCCCACTGTCCCTACCT</td><td>221.5</td><td>-11.7</td><td>1.5</td><td>-7.7</td><td>1.4E-14</td><td>7.8E-14</td><td>MB231/MB468</td><td>MP</td></tr>
<tr><td>>hg38_21_+_8433172_8446622_RNA45SN1_45S_50ntFlanks@5486.5514.29|GCGCTGAGAAGACGGTCGAACTTGACTAT</td><td>>hg38_21_+_8433172_8446622_RNA45SN1_45S_50ntFlanks</td><td>5486</td><td>5514</td><td>29</td><td>GCGCTGAGAAGACGGTCGAACTTGACTAT</td><td>2003.1</td><td>-9.6</td><td>1.3</td><td>-7.6</td><td>2.3E-14</td><td>7.8E-14</td><td>MB231/MB468</td><td>MITO</td></tr>
<tr><td>>hg38_21_+_8433172_8446622_RNA45SN1_45S_50ntFlanks@12836.12870.35|CGGTGCGGAGTGCCCTTCGTCCTGGGAAACGGGGC</td><td>>hg38_21_+_8433172_8446622_RNA45SN1_45S_50ntFlanks</td><td>12836</td><td>12870</td><td>35</td><td>CGGTGCGGAGTGCCCTTCGTCCTGGGAAACGGGGC</td><td>168.5</td><td>-11.3</td><td>1.5</td><td>-7.7</td><td>1.4E-14</td><td>7.9E-14</td><td>MB231/MB468</td><td>MP</td></tr>
<tr><td>>hg38_21_+_8433172_8446622_RNA45SN1_45S_50ntFlanks@9545.9571.27|GTGGAGGTCCGTAGCGGTCCTGACGTG</td><td>>hg38_21_+_8433172_8446622_RNA45SN1_45S_50ntFlanks</td><td>9545</td><td>9571</td><td>27</td><td>GTGGAGGTCCGTAGCGGTCCTGACGTG</td><td>3817.7</td><td>-3.3</td><td>0.4</td><td>-7.7</td><td>1.4E-14</td><td>7.9E-14</td><td>BT20/MB468</td><td>MITO</td></tr>
<tr><td>>hg38_21_+_8433172_8446622_RNA45SN1_45S_50ntFlanks@4160.4182.23|CCACATCCAAGGAAGGCAGCAGG</td><td>>hg38_21_+_8433172_8446622_RNA45SN1_45S_50ntFlanks</td><td>4160</td><td>4182</td><td>23</td><td>CCACATCCAAGGAAGGCAGCAGG</td><td>503.9</td><td>-12.8</td><td>1.7</td><td>-7.7</td><td>1.4E-14</td><td>7.9E-14</td><td>MB231/MB468</td><td>MP</td></tr>
<tr><td>>hg38_21_+_8433172_8446622_RNA45SN1_45S_50ntFlanks@6699.6719.21|GCTAGCTGCGAGAATTAATGT</td><td>>hg38_21_+_8433172_8446622_RNA45SN1_45S_50ntFlanks</td><td>6699</td><td>6719</td><td>21</td><td>GCTAGCTGCGAGAATTAATGT</td><td>6302.5</td><td>-11.7</td><td>1.5</td><td>-7.6</td><td>2.8E-14</td><td>7.9E-14</td><td>BT20/MB468</td><td>NUC</td></tr>
<tr><td>>hg38_21_+_8433172_8446622_RNA45SN1_45S_50ntFlanks@4884.4916.33|CAAAGCTGAAACTTAAAGGAATTGACGGAAGGG</td><td>>hg38_21_+_8433172_8446622_RNA45SN1_45S_50ntFlanks</td><td>4884</td><td>4916</td><td>33</td><td>CAAAGCTGAAACTTAAAGGAATTGACGGAAGGG</td><td>187.2</td><td>-11.4</td><td>1.5</td><td>-7.7</td><td>1.4E-14</td><td>7.9E-14</td><td>MB231/MB468</td><td>MP</td></tr>
<tr><td>>hg38_21_+_8433172_8446622_RNA45SN1_45S_50ntFlanks@11564.11602.39|CCTAGCAGCCGACTTAGAACTGGTGCGGACCAGGGGAAT</td><td>>hg38_21_+_8433172_8446622_RNA45SN1_45S_50ntFlanks</td><td>11564</td><td>11602</td><td>39</td><td>CCTAGCAGCCGACTTAGAACTGGTGCGGACCAGGGGAAT</td><td>2583.0</td><td>-5.3</td><td>0.7</td><td>-7.7</td><td>1.7E-14</td><td>8.0E-14</td><td>MB231/MB468</td><td>NUC</td></tr>
<tr><td>>MI0000089|hsa-mir-31&WithFlank&9|-|21512109|21512191@14.35.22(+1A)&[MIMAT0000089&hsa-miR-31-5p&offsets|0|+1(+1A);m-360&9|-|21512157|21512178&offsets|0|0(+1A)]|AGGCAAGATGCTGGCATAGCTGA</td><td>>MI0000089|hsa-mir-31&WithFlank&9|-|21512109|21512191</td><td>14</td><td>35</td><td>22(+1A)</td><td>AGGCAAGATGCTGGCATAGCTGA</td><td>325.9</td><td>11.7</td><td>1.5</td><td>8.0</td><td>1.4E-15</td><td>8.0E-14</td><td>BT20/MB231</td><td>CYTO</td></tr>
<tr><td>>hg38_21_+_8433172_8446622_RNA45SN1_45S_50ntFlanks@5467.5484.18|GGCCCACGGCCCTGGCGG</td><td>>hg38_21_+_8433172_8446622_RNA45SN1_45S_50ntFlanks</td><td>5467</td><td>5484</td><td>18</td><td>GGCCCACGGCCCTGGCGG</td><td>314.5</td><td>-11.3</td><td>1.5</td><td>-7.7</td><td>1.7E-14</td><td>8.0E-14</td><td>MB231/MB468</td><td>NUC</td></tr>
<tr><td>>MI0000108|hsa-mir-103a-2&WithFlank&20|+|3917488|3917577@54.75.22(+1G);MI0000109|hsa-mir-103a-1&WithFlank&5|-|168560890|168560979@54.75.22(+1G)&[MIMAT0000101&hsa-miR-103a-3p&offsets|0|-1(+1G);m-12&20|+|3917541|3917563&offsets|0|-1(+1G)];[MIMAT0000101_1&hsa-miR-103a-3p&offsets|0|-1(+1G);m-13&5|-|168560904|168560926&offsets|0|-1(+1G);m-44220&5|-|168560908|168560926&offsets|0|+3(+1G)]|AGCAGCATTGTACAGGGCTATGG</td><td>>MI0000108|hsa-mir-103a-2&WithFlank&20|+|3917488|3917577</td><td>54</td><td>75</td><td>22(+1G)</td><td>AGCAGCATTGTACAGGGCTATGG</td><td>404.9</td><td>5.2</td><td>0.7</td><td>7.7</td><td>1.7E-14</td><td>8.0E-14</td><td>MB231/MB468</td><td>NUC</td></tr>
<tr><td>>hg38_21_+_8433172_8446622_RNA45SN1_45S_50ntFlanks@11315.11338.24|CCACCCCACGTCTCGTCGCGCGCG</td><td>>hg38_21_+_8433172_8446622_RNA45SN1_45S_50ntFlanks</td><td>11315</td><td>11338</td><td>24</td><td>CCACCCCACGTCTCGTCGCGCGCG</td><td>1394.5</td><td>-10.2</td><td>1.3</td><td>-7.7</td><td>1.7E-14</td><td>8.0E-14</td><td>MB231/MB468</td><td>NUC</td></tr>
<tr><td>>hg38_21_+_8433172_8446622_RNA45SN1_45S_50ntFlanks@8109.8138.30|GCGGGGCGCGGGACATGTGGCGTACGGAAG</td><td>>hg38_21_+_8433172_8446622_RNA45SN1_45S_50ntFlanks</td><td>8109</td><td>8138</td><td>30</td><td>GCGGGGCGCGGGACATGTGGCGTACGGAAG</td><td>463.5</td><td>-12.7</td><td>1.7</td><td>-7.7</td><td>1.5E-14</td><td>8.0E-14</td><td>MB231/MB468</td><td>MP</td></tr>
<tr><td>>hg38_21_+_8433172_8446622_RNA45SN1_45S_50ntFlanks@11669.11702.34|GCCCAGTGCTCTGAATGTCAAAGTGAAGAAATTC</td><td>>hg38_21_+_8433172_8446622_RNA45SN1_45S_50ntFlanks</td><td>11669</td><td>11702</td><td>34</td><td>GCCCAGTGCTCTGAATGTCAAAGTGAAGAAATTC</td><td>517.4</td><td>4.2</td><td>0.5</td><td>7.7</td><td>1.4E-14</td><td>8.0E-14</td><td>BT20/MB468</td><td>MITO</td></tr>
<tr><td>>trnaMT_PheGAA_MT_+_577_647@39.62.24|ATGTTTAGACGGGCTCACATCACC</td><td>>trnaMT_PheGAA_MT_+_577_647</td><td>39</td><td>62</td><td>24</td><td>ATGTTTAGACGGGCTCACATCACC</td><td>1574.8</td><td>7.3</td><td>0.9</td><td>7.7</td><td>1.4E-14</td><td>8.0E-14</td><td>BT20/MB468</td><td>MITO</td></tr>
<tr><td>>hg38_21_+_8433172_8446622_RNA45SN1_45S_50ntFlanks@4759.4783.25|ATCAGATACCGTCGTAGTTCCGACC</td><td>>hg38_21_+_8433172_8446622_RNA45SN1_45S_50ntFlanks</td><td>4759</td><td>4783</td><td>25</td><td>ATCAGATACCGTCGTAGTTCCGACC</td><td>1718.3</td><td>-12.8</td><td>1.7</td><td>-7.7</td><td>1.4E-14</td><td>8.0E-14</td><td>BT20/MB468</td><td>MITO</td></tr>
<tr><td>>hg38_21_+_8433172_8446622_RNA45SN1_45S_50ntFlanks@11729.11754.26|TAACTATGACTCTCTTAAGGTAGCCA</td><td>>hg38_21_+_8433172_8446622_RNA45SN1_45S_50ntFlanks</td><td>11729</td><td>11754</td><td>26</td><td>TAACTATGACTCTCTTAAGGTAGCCA</td><td>401.2</td><td>-12.5</td><td>1.6</td><td>-7.7</td><td>1.5E-14</td><td>8.1E-14</td><td>MB231/MB468</td><td>MP</td></tr>
<tr><td>>am_MI0003583|hsa-mir-576&WithFlank&4|+|109488692|109488801@61.83.23&[MIMAT0004796&hsa-miR-576-3p&offsets|0|+1;m-524&4|+|109488752|109488772&offsets|0|+2]|AAGATGTGGAAAAATTGGAATCC</td><td>>am_MI0003583|hsa-mir-576&WithFlank&4|+|109488692|109488801</td><td>61</td><td>83</td><td>23</td><td>AAGATGTGGAAAAATTGGAATCC</td><td>465.7</td><td>9.3</td><td>1.2</td><td>7.7</td><td>1.9E-14</td><td>8.1E-14</td><td>BT20/MB468</td><td>CYTO</td></tr>
<tr><td>>hg38_21_+_8433172_8446622_RNA45SN1_45S_50ntFlanks@8169.8215.47|CCAAGTCCTTCTGATCGAGGCCCAGCCCGTGGACGGTGTGAGGCCGG</td><td>>hg38_21_+_8433172_8446622_RNA45SN1_45S_50ntFlanks</td><td>8169</td><td>8215</td><td>47</td><td>CCAAGTCCTTCTGATCGAGGCCCAGCCCGTGGACGGTGTGAGGCCGG</td><td>407.5</td><td>-12.5</td><td>1.6</td><td>-7.6</td><td>2.4E-14</td><td>8.1E-14</td><td>MB231/MB468</td><td>MITO</td></tr>
<tr><td>>hg38_21_+_8433172_8446622_RNA45SN1_45S_50ntFlanks@11701.11728.28|TCAATGAAGCGCGGGTAAACGGCGGGAG</td><td>>hg38_21_+_8433172_8446622_RNA45SN1_45S_50ntFlanks</td><td>11701</td><td>11728</td><td>28</td><td>TCAATGAAGCGCGGGTAAACGGCGGGAG</td><td>222.5</td><td>-11.6</td><td>1.5</td><td>-7.6</td><td>2.4E-14</td><td>8.1E-14</td><td>MB231/MB468</td><td>MITO</td></tr>
<tr><td>>MI0000746|hsa-mir-99b&WithFlank&19|+|51692606|51692687@13.34.22(+1U)&[MIMAT0000689&hsa-miR-99b-5p&offsets|0|0(+1U);m-10&19|+|51692618|51692639&offsets|0|0(+1U)]|CACCCGTAGAACCGACCTTGCGT</td><td>>MI0000746|hsa-mir-99b&WithFlank&19|+|51692606|51692687</td><td>13</td><td>34</td><td>22(+1U)</td><td>CACCCGTAGAACCGACCTTGCGT</td><td>3544.7</td><td>6.6</td><td>0.9</td><td>7.7</td><td>1.7E-14</td><td>8.1E-14</td><td>MB231/MB468</td><td>CYTO</td></tr>
<tr><td>>hg38_21_+_8433172_8446622_RNA45SN1_45S_50ntFlanks@8198.8224.27|TGGACGGTGTGAGGCCGGTAGCGGCCC</td><td>>hg38_21_+_8433172_8446622_RNA45SN1_45S_50ntFlanks</td><td>8198</td><td>8224</td><td>27</td><td>TGGACGGTGTGAGGCCGGTAGCGGCCC</td><td>825.6</td><td>-9.9</td><td>1.3</td><td>-7.6</td><td>2.9E-14</td><td>8.1E-14</td><td>BT20/MB468</td><td>NUC</td></tr>
<tr><td>>hg38_21_+_8433172_8446622_RNA45SN1_45S_50ntFlanks@11913.11948.36|AAGAGACATGAGAGGTGTAGAATAAGTGGGAGGCCC</td><td>>hg38_21_+_8433172_8446622_RNA45SN1_45S_50ntFlanks</td><td>11913</td><td>11948</td><td>36</td><td>AAGAGACATGAGAGGTGTAGAATAAGTGGGAGGCCC</td><td>409.3</td><td>-12.5</td><td>1.6</td><td>-7.7</td><td>1.5E-14</td><td>8.1E-14</td><td>MB231/MB468</td><td>MP</td></tr>
<tr><td>>hg38_21_+_8433172_8446622_RNA45SN1_45S_50ntFlanks@9308.9337.30|AGGAGTCTAACACGTGCGCGAGTCGGGGGC</td><td>>hg38_21_+_8433172_8446622_RNA45SN1_45S_50ntFlanks</td><td>9308</td><td>9337</td><td>30</td><td>AGGAGTCTAACACGTGCGCGAGTCGGGGGC</td><td>1278.1</td><td>-10.0</td><td>1.3</td><td>-7.7</td><td>1.5E-14</td><td>8.1E-14</td><td>MB231/MB468</td><td>MP</td></tr>
<tr><td>>hg38_21_+_8433172_8446622_RNA45SN1_45S_50ntFlanks@12934.12957.24|CTAAACCATTCGTAGACGACCTGC</td><td>>hg38_21_+_8433172_8446622_RNA45SN1_45S_50ntFlanks</td><td>12934</td><td>12957</td><td>24</td><td>CTAAACCATTCGTAGACGACCTGC</td><td>1231.5</td><td>-10.2</td><td>1.3</td><td>-7.6</td><td>2.9E-14</td><td>8.2E-14</td><td>BT20/MB468</td><td>NUC</td></tr>
<tr><td>>hg38_21_+_8433172_8446622_RNA45SN1_45S_50ntFlanks@13002.13031.30|CGATCTATTGAAAGTCAGCCCTCGACACAA</td><td>>hg38_21_+_8433172_8446622_RNA45SN1_45S_50ntFlanks</td><td>13002</td><td>13031</td><td>30</td><td>CGATCTATTGAAAGTCAGCCCTCGACACAA</td><td>185.5</td><td>-10.5</td><td>1.4</td><td>-7.7</td><td>1.8E-14</td><td>8.2E-14</td><td>MB231/MB468</td><td>NUC</td></tr>
<tr><td>>trnaMT_GluTTC_MT_-_14674_14742@40.72.33,trnalookalike8_GluTTC_5_-_93905172_93905240@40.72.33|TTGGTCGTGGTTGTAGTCCGTGCGAGAATACCA</td><td>>trnaMT_GluTTC_MT_-_14674_14742</td><td>40</td><td>72</td><td>33</td><td>TTGGTCGTGGTTGTAGTCCGTGCGAGAATACCA</td><td>17896.3</td><td>7.7</td><td>1.0</td><td>7.7</td><td>1.7E-14</td><td>8.2E-14</td><td>MB231/MB468</td><td>CYTO</td></tr>
<tr><td>>hg38_21_+_8433172_8446622_RNA45SN1_45S_50ntFlanks@9554.9573.20|CGTAGCGGTCCTGACGTGCA</td><td>>hg38_21_+_8433172_8446622_RNA45SN1_45S_50ntFlanks</td><td>9554</td><td>9573</td><td>20</td><td>CGTAGCGGTCCTGACGTGCA</td><td>8595.8</td><td>-5.0</td><td>0.7</td><td>-7.6</td><td>2.4E-14</td><td>8.2E-14</td><td>MB231/MB468</td><td>MITO</td></tr>
<tr><td>>hg38_21_+_8433172_8446622_RNA45SN1_45S_50ntFlanks@11925.11951.27|AGGTGTAGAATAAGTGGGAGGCCCCCG</td><td>>hg38_21_+_8433172_8446622_RNA45SN1_45S_50ntFlanks</td><td>11925</td><td>11951</td><td>27</td><td>AGGTGTAGAATAAGTGGGAGGCCCCCG</td><td>213.0</td><td>-11.6</td><td>1.5</td><td>-7.7</td><td>1.5E-14</td><td>8.2E-14</td><td>MB231/MB468</td><td>MP</td></tr>
<tr><td>>hg38_21_+_8433172_8446622_RNA45SN1_45S_50ntFlanks@9535.9553.19|GGAAACTCTGGTGGAGGTC</td><td>>hg38_21_+_8433172_8446622_RNA45SN1_45S_50ntFlanks</td><td>9535</td><td>9553</td><td>19</td><td>GGAAACTCTGGTGGAGGTC</td><td>146.3</td><td>-10.2</td><td>1.3</td><td>-7.7</td><td>1.8E-14</td><td>8.2E-14</td><td>MB231/MB468</td><td>NUC</td></tr>
<tr><td>>hg38_21_+_8433172_8446622_RNA45SN1_45S_50ntFlanks@8346.8373.28|AGGGAAAGTTGAAAAGAACTTTGAAGAG</td><td>>hg38_21_+_8433172_8446622_RNA45SN1_45S_50ntFlanks</td><td>8346</td><td>8373</td><td>28</td><td>AGGGAAAGTTGAAAAGAACTTTGAAGAG</td><td>658.7</td><td>13.1</td><td>1.6</td><td>8.1</td><td>6.0E-16</td><td>8.2E-14</td><td>BT20/MB468</td><td>MP</td></tr>
<tr><td>>hg38_21_+_8433172_8446622_RNA45SN1_45S_50ntFlanks@9575.9594.20|ATCGGTCGTCCGACCTGGGT</td><td>>hg38_21_+_8433172_8446622_RNA45SN1_45S_50ntFlanks</td><td>9575</td><td>9594</td><td>20</td><td>ATCGGTCGTCCGACCTGGGT</td><td>1582.3</td><td>-13.7</td><td>1.7</td><td>-8.1</td><td>5.9E-16</td><td>8.2E-14</td><td>BT20/MB468</td><td>MP</td></tr>
<tr><td>>hg38_21_+_8433172_8446622_RNA45SN1_45S_50ntFlanks@12415.12448.34|CTTCGATGTCGGCTCTTCCTATCATTGTGAAGCA</td><td>>hg38_21_+_8433172_8446622_RNA45SN1_45S_50ntFlanks</td><td>12415</td><td>12448</td><td>34</td><td>CTTCGATGTCGGCTCTTCCTATCATTGTGAAGCA</td><td>1113.3</td><td>4.5</td><td>0.6</td><td>7.7</td><td>2.0E-14</td><td>8.3E-14</td><td>BT20/MB468</td><td>CYTO</td></tr>
<tr><td>>am_MI0000809|hsa-mir-151a&WithFlank&8|-|140732558|140732659@53.73.21(+1U)&[MIMAT0000757&hsa-miR-151a-3p&offsets|0|0(+1U);m-30&8|-|140732587|140732607&offsets|0|0(+1U)]|CTAGACTGAAGCTCCTTGAGGT</td><td>>am_MI0000809|hsa-mir-151a&WithFlank&8|-|140732558|140732659</td><td>53</td><td>73</td><td>21(+1U)</td><td>CTAGACTGAAGCTCCTTGAGGT</td><td>6126.6</td><td>6.9</td><td>0.9</td><td>7.6</td><td>2.4E-14</td><td>8.3E-14</td><td>MB231/MB468</td><td>MITO</td></tr>
<tr><td>>hg38_21_+_8433172_8446622_RNA45SN1_45S_50ntFlanks@11781.11798.18|CGCATGAATGGATGAACG</td><td>>hg38_21_+_8433172_8446622_RNA45SN1_45S_50ntFlanks</td><td>11781</td><td>11798</td><td>18</td><td>CGCATGAATGGATGAACG</td><td>1698.1</td><td>-10.7</td><td>1.4</td><td>-7.6</td><td>2.9E-14</td><td>8.3E-14</td><td>BT20/MB468</td><td>NUC</td></tr>
<tr><td>>hg38_21_+_8433172_8446622_RNA45SN1_45S_50ntFlanks@12632.12651.20|GCGAAGCTACCATCTGTGGG</td><td>>hg38_21_+_8433172_8446622_RNA45SN1_45S_50ntFlanks</td><td>12632</td><td>12651</td><td>20</td><td>GCGAAGCTACCATCTGTGGG</td><td>194.1</td><td>-11.5</td><td>1.5</td><td>-7.7</td><td>1.5E-14</td><td>8.3E-14</td><td>MB231/MB468</td><td>MP</td></tr>
<tr><td>>hg38_1_-_228634819_228635039_RNA5S12_5S_50ntFlanks@151.171.21|ATACCGGGTGCTGTAGGCTTT</td><td>>hg38_1_-_228634819_228635039_RNA5S12_5S_50ntFlanks</td><td>151</td><td>171</td><td>21</td><td>ATACCGGGTGCTGTAGGCTTT</td><td>5395.5</td><td>5.8</td><td>0.8</td><td>7.7</td><td>1.7E-14</td><td>8.3E-14</td><td>MB231/MB468</td><td>CYTO</td></tr>
<tr><td>>hg38_21_+_8433172_8446622_RNA45SN1_45S_50ntFlanks@4733.4771.39|GAACGAAAGTCGGAGGTTCGAAGACGATCAGATACCGTC</td><td>>hg38_21_+_8433172_8446622_RNA45SN1_45S_50ntFlanks</td><td>4733</td><td>4771</td><td>39</td><td>GAACGAAAGTCGGAGGTTCGAAGACGATCAGATACCGTC</td><td>356.2</td><td>8.1</td><td>1.1</td><td>7.7</td><td>1.7E-14</td><td>8.3E-14</td><td>MB231/MB468</td><td>CYTO</td></tr>
<tr><td>>hg38_21_+_8433172_8446622_RNA45SN1_45S_50ntFlanks@476.506.31|TTCGTGATCGATGTGGTGACGTCGTGCTCTC</td><td>>hg38_21_+_8433172_8446622_RNA45SN1_45S_50ntFlanks</td><td>476</td><td>506</td><td>31</td><td>TTCGTGATCGATGTGGTGACGTCGTGCTCTC</td><td>955.2</td><td>7.7</td><td>1.0</td><td>7.7</td><td>1.7E-14</td><td>8.3E-14</td><td>MB231/MB468</td><td>CYTO</td></tr>
<tr><td>>MI0003628|hsa-mir-615&WithFlank&12|+|54033944|54034051@67.86.20(+1C)&[MIMAT0003283&hsa-miR-615-3p&offsets|0|-2(+1C);m-495&12|+|54034010|54034030&offsets|0|-1(+1C)]|TCCGAGCCTGGGTCTCCCTCC</td><td>>MI0003628|hsa-mir-615&WithFlank&12|+|54033944|54034051</td><td>67</td><td>86</td><td>20(+1C)</td><td>TCCGAGCCTGGGTCTCCCTCC</td><td>856.2</td><td>8.0</td><td>1.0</td><td>7.7</td><td>1.5E-14</td><td>8.3E-14</td><td>BT20/MB468</td><td>MITO</td></tr>
<tr><td>>hg38_21_+_8433172_8446622_RNA45SN1_45S_50ntFlanks@4657.4688.32|CTAGAGGTGAAATTCTTGGACCGGCGCAAGAC</td><td>>hg38_21_+_8433172_8446622_RNA45SN1_45S_50ntFlanks</td><td>4657</td><td>4688</td><td>32</td><td>CTAGAGGTGAAATTCTTGGACCGGCGCAAGAC</td><td>1055.3</td><td>-12.1</td><td>1.6</td><td>-7.7</td><td>1.5E-14</td><td>8.3E-14</td><td>BT20/MB468</td><td>MITO</td></tr>
<tr><td>>hg38_21_+_8433172_8446622_RNA45SN1_45S_50ntFlanks@6651.6672.22|CGACTCTTAGCGGTGGATCACT</td><td>>hg38_21_+_8433172_8446622_RNA45SN1_45S_50ntFlanks</td><td>6651</td><td>6672</td><td>22</td><td>CGACTCTTAGCGGTGGATCACT</td><td>39901.7</td><td>6.4</td><td>0.8</td><td>7.7</td><td>1.5E-14</td><td>8.3E-14</td><td>BT20/MB468</td><td>MITO</td></tr>
<tr><td>>hg38_21_+_8433172_8446622_RNA45SN1_45S_50ntFlanks@11641.11660.20|CGGCGGGTGTTGACGCGATG</td><td>>hg38_21_+_8433172_8446622_RNA45SN1_45S_50ntFlanks</td><td>11641</td><td>11660</td><td>20</td><td>CGGCGGGTGTTGACGCGATG</td><td>164682.1</td><td>-11.5</td><td>1.4</td><td>-8.1</td><td>6.1E-16</td><td>8.3E-14</td><td>BT20/MB468</td><td>MP</td></tr>
<tr><td>>hg38_21_+_8433172_8446622_RNA45SN1_45S_50ntFlanks@7974.8012.39|ACGCGACCTCAGATCAGACGTGGCGACCCGCTGAATTTA</td><td>>hg38_21_+_8433172_8446622_RNA45SN1_45S_50ntFlanks</td><td>7974</td><td>8012</td><td>39</td><td>ACGCGACCTCAGATCAGACGTGGCGACCCGCTGAATTTA</td><td>2599.1</td><td>4.5</td><td>0.6</td><td>7.7</td><td>1.7E-14</td><td>8.3E-14</td><td>MB231/MB468</td><td>CYTO</td></tr>
<tr><td>>trna2_GlyGCC_21_-_18827107_18827177@1.28.28,trna35_GlyGCC_1_+_161413094_161413164@1.28.28,trna37_GlyGCC_1_+_161420467_161420537@1.28.28,trna39_GlyGCC_1_+_161427898_161427968@1.28.28,trna41_GlyGCC_1_+_161435258_161435328@1.28.28|GCATGGGTGGTTCAGTGGTAGAATTCTC</td><td>>trna2_GlyGCC_21_-_18827107_18827177</td><td>1</td><td>28</td><td>28</td><td>GCATGGGTGGTTCAGTGGTAGAATTCTC</td><td>47039.6</td><td>-6.9</td><td>0.9</td><td>-7.7</td><td>1.7E-14</td><td>8.3E-14</td><td>MB231/MB468</td><td>CYTO</td></tr>
<tr><td>>MI0000088|hsa-mir-30a&WithFlank&6|-|71403545|71403627@12.34.23(+1C)&[MIMAT0000087&hsa-miR-30a-5p&offsets|0|+1(+1C);m-15&6|-|71403595|71403616&offsets|0|+1(+1C)]|TGTAAACATCCTCGACTGGAAGCC</td><td>>MI0000088|hsa-mir-30a&WithFlank&6|-|71403545|71403627</td><td>12</td><td>34</td><td>23(+1C)</td><td>TGTAAACATCCTCGACTGGAAGCC</td><td>4281.7</td><td>6.6</td><td>0.9</td><td>7.7</td><td>2.0E-14</td><td>8.3E-14</td><td>BT20/MB468</td><td>CYTO</td></tr>
<tr><td>>hg38_21_+_8433172_8446622_RNA45SN1_45S_50ntFlanks@12940.12972.33|CATTCGTAGACGACCTGCTTCTGGGTCGGGGTT</td><td>>hg38_21_+_8433172_8446622_RNA45SN1_45S_50ntFlanks</td><td>12940</td><td>12972</td><td>33</td><td>CATTCGTAGACGACCTGCTTCTGGGTCGGGGTT</td><td>211.5</td><td>-11.5</td><td>1.5</td><td>-7.6</td><td>2.4E-14</td><td>8.4E-14</td><td>MB231/MB468</td><td>MITO</td></tr>
</table>

<table>
<tr><td>>hg38_21_+_8433172_8446622_RNA45SN1_45S_50ntFlanks@9739.9769.31|AAACGATCTCAACCTATTCTCAAACTTTAAA</td><td>>hg38_21_+_8433172_8446622_RNA45SN1_45S_50ntFlanks</td><td>9739</td><td>9769</td><td>31</td><td>AAACGATCTCAACCTATTCTCAAACTTTAAA</td><td>649.4</td><td>-13.2</td><td>1.7</td><td>-7.7</td><td>1.5E-14</td><td>8.4E-14</td><td>MB231/MB468</td><td>MP</td></tr>
<tr><td>>hg38_21_+_8433172_8446622_RNA45SN1_45S_50ntFlanks@12627.12649.23|ATGGGGCGAAGCTACCATCTGTG</td><td>>hg38_21_+_8433172_8446622_RNA45SN1_45S_50ntFlanks</td><td>12627</td><td>12649</td><td>23</td><td>ATGGGGCGAAGCTACCATCTGTG</td><td>152.5</td><td>-11.1</td><td>1.4</td><td>-7.7</td><td>1.5E-14</td><td>8.4E-14</td><td>MB231/MB468</td><td>MP</td></tr>
<tr><td>>hg38_21_+_8433172_8446622_RNA45SN1_45S_50ntFlanks@4100.4121.22|TGACCACGGGTGACGGGGAATC</td><td>>hg38_21_+_8433172_8446622_RNA45SN1_45S_50ntFlanks</td><td>4100</td><td>4121</td><td>22</td><td>TGACCACGGGTGACGGGGAATC</td><td>187.8</td><td>-11.4</td><td>1.5</td><td>-7.7</td><td>1.5E-14</td><td>8.4E-14</td><td>MB231/MB468</td><td>MP</td></tr>
<tr><td>>am_MI0000809|hsa-mir-151a&WithFlank&8|-|140732558|140732659@53.72.20(+1U)&[MIMAT0000757&hsa-miR-151a-3p&offsets|0|-1(+1U);m-30&8|-|140732587|140732607&offsets|0|-1(+1U)]|CTAGACTGAAGCTCCTTGAGT</td><td>>am_MI0000809|hsa-mir-151a&WithFlank&8|-|140732558|140732659</td><td>53</td><td>72</td><td>20(+1U)</td><td>CTAGACTGAAGCTCCTTGAGT</td><td>2767.8</td><td>7.1</td><td>0.9</td><td>7.6</td><td>2.4E-14</td><td>8.4E-14</td><td>MB231/MB468</td><td>MITO</td></tr>
<tr><td>>hg38_21_+_8433172_8446622_RNA45SN1_45S_50ntFlanks@10350.10368.19|ACGAGAACTTTGAAGGCCG</td><td>>hg38_21_+_8433172_8446622_RNA45SN1_45S_50ntFlanks</td><td>10350</td><td>10368</td><td>19</td><td>ACGAGAACTTTGAAGGCCG</td><td>4524.9</td><td>-5.8</td><td>0.8</td><td>-7.7</td><td>2.0E-14</td><td>8.4E-14</td><td>BT20/MB468</td><td>CYTO</td></tr>
<tr><td>>hg38_21_+_8433172_8446622_RNA45SN1_45S_50ntFlanks@12677.12708.32|GAATCCCGCCCAGGCGGAACGATACGGCAGCG</td><td>>hg38_21_+_8433172_8446622_RNA45SN1_45S_50ntFlanks</td><td>12677</td><td>12708</td><td>32</td><td>GAATCCCGCCCAGGCGGAACGATACGGCAGCG</td><td>143.8</td><td>-11.0</td><td>1.4</td><td>-7.7</td><td>1.5E-14</td><td>8.4E-14</td><td>MB231/MB468</td><td>MP</td></tr>
<tr><td>>hg38_21_+_8433172_8446622_RNA45SN1_45S_50ntFlanks@8017.8046.30|TATTAGTCAGCGGAGGAGAAGAAACTAACC</td><td>>hg38_21_+_8433172_8446622_RNA45SN1_45S_50ntFlanks</td><td>8017</td><td>8046</td><td>30</td><td>TATTAGTCAGCGGAGGAGAAGAAACTAACC</td><td>443.6</td><td>9.9</td><td>1.3</td><td>7.7</td><td>1.7E-14</td><td>8.4E-14</td><td>MB231/MB468</td><td>CYTO</td></tr>
<tr><td>>hg38_21_+_8433172_8446622_RNA45SN1_45S_50ntFlanks@11534.11552.19|GTTCCGCGCGGCGCCTCGC</td><td>>hg38_21_+_8433172_8446622_RNA45SN1_45S_50ntFlanks</td><td>11534</td><td>11552</td><td>19</td><td>GTTCCGCGCGGCGCCTCGC</td><td>741.9</td><td>-6.8</td><td>0.9</td><td>-7.7</td><td>1.7E-14</td><td>8.4E-14</td><td>MB231/MB468</td><td>CYTO</td></tr>
<tr><td>>hg38_21_+_8433172_8446622_RNA45SN1_45S_50ntFlanks@4550.4573.24|GATACCGCAGCTAGGAATAATGGA</td><td>>hg38_21_+_8433172_8446622_RNA45SN1_45S_50ntFlanks</td><td>4550</td><td>4573</td><td>24</td><td>GATACCGCAGCTAGGAATAATGGA</td><td>210.5</td><td>-11.6</td><td>1.5</td><td>-7.7</td><td>1.5E-14</td><td>8.4E-14</td><td>MB231/MB468</td><td>MP</td></tr>
<tr><td>>hg38_21_+_8433172_8446622_RNA45SN1_45S_50ntFlanks@11882.11903.22|CTGTTGAGCTTGACTCTAGTCT</td><td>>hg38_21_+_8433172_8446622_RNA45SN1_45S_50ntFlanks</td><td>11882</td><td>11903</td><td>22</td><td>CTGTTGAGCTTGACTCTAGTCT</td><td>172.0</td><td>-11.3</td><td>1.5</td><td>-7.7</td><td>1.5E-14</td><td>8.4E-14</td><td>MB231/MB468</td><td>MP</td></tr>
<tr><td>>hg38_21_+_8433172_8446622_RNA45SN1_45S_50ntFlanks@12691.12708.18|CGGAACGATACGGCAGCG</td><td>>hg38_21_+_8433172_8446622_RNA45SN1_45S_50ntFlanks</td><td>12691</td><td>12708</td><td>18</td><td>CGGAACGATACGGCAGCG</td><td>343.1</td><td>-12.3</td><td>1.6</td><td>-7.7</td><td>1.5E-14</td><td>8.5E-14</td><td>MB231/MB468</td><td>MP</td></tr>
<tr><td>>hg38_21_+_8433172_8446622_RNA45SN1_45S_50ntFlanks@10534.10569.36|CCGCGAGGCGTCCAGTGCGGTAACGCGACCGATCCC</td><td>>hg38_21_+_8433172_8446622_RNA45SN1_45S_50ntFlanks</td><td>10534</td><td>10569</td><td>36</td><td>CCGCGAGGCGTCCAGTGCGGTAACGCGACCGATCCC</td><td>225.4</td><td>-11.7</td><td>1.5</td><td>-7.7</td><td>1.5E-14</td><td>8.5E-14</td><td>MB231/MB468</td><td>MP</td></tr>
<tr><td>>am_MI0000809|hsa-mir-151a&WithFlank&8|-|140732558|140732659@53.73.21&[MIMAT0000757&hsa-miR-151a-3p&offsets|0|0;m-30&8|-|140732587|140732607&offsets|0|0]|CTAGACTGAAGCTCCTTGAGG</td><td>>am_MI0000809|hsa-mir-151a&WithFlank&8|-|140732558|140732659</td><td>53</td><td>73</td><td>21</td><td>CTAGACTGAAGCTCCTTGAGG</td><td>25872.1</td><td>5.8</td><td>0.8</td><td>7.6</td><td>2.5E-14</td><td>8.5E-14</td><td>MB231/MB468</td><td>MITO</td></tr>
<tr><td>>trnaMT_MetCAT_MT_+_4402_4469@35.71.37|CCCCGAAAATGTTGGTTATACCCTTCCCGTACTACCA</td><td>>trnaMT_MetCAT_MT_+_4402_4469</td><td>35</td><td>71</td><td>37</td><td>CCCCGAAAATGTTGGTTATACCCTTCCCGTACTACCA</td><td>548.2</td><td>8.1</td><td>1.1</td><td>7.7</td><td>1.9E-14</td><td>8.5E-14</td><td>MB231/MB468</td><td>NUC</td></tr>
<tr><td>>hg38_21_+_8433172_8446622_RNA45SN1_45S_50ntFlanks@10475.10497.23|CGGCCGATCGAAAGGGAGTCGGG</td><td>>hg38_21_+_8433172_8446622_RNA45SN1_45S_50ntFlanks</td><td>10475</td><td>10497</td><td>23</td><td>CGGCCGATCGAAAGGGAGTCGGG</td><td>344.4</td><td>-12.3</td><td>1.6</td><td>-7.7</td><td>1.6E-14</td><td>8.5E-14</td><td>MB231/MB468</td><td>MP</td></tr>
<tr><td>>hg38_21_+_8433172_8446622_RNA45SN1_45S_50ntFlanks@9134.9152.19|CGGGGGGCGGGGCGGACTG</td><td>>hg38_21_+_8433172_8446622_RNA45SN1_45S_50ntFlanks</td><td>9134</td><td>9152</td><td>19</td><td>CGGGGGGCGGGGCGGACTG</td><td>324.6</td><td>-12.2</td><td>1.6</td><td>-7.7</td><td>1.6E-14</td><td>8.5E-14</td><td>MB231/MB468</td><td>MP</td></tr>
<tr><td>>hg38_21_+_8433172_8446622_RNA45SN1_45S_50ntFlanks@12676.12706.31|AGAATCCCGCCCAGGCGGAACGATACGGCAG</td><td>>hg38_21_+_8433172_8446622_RNA45SN1_45S_50ntFlanks</td><td>12676</td><td>12706</td><td>31</td><td>AGAATCCCGCCCAGGCGGAACGATACGGCAG</td><td>145.4</td><td>-11.0</td><td>1.4</td><td>-7.7</td><td>1.6E-14</td><td>8.5E-14</td><td>MB231/MB468</td><td>MP</td></tr>
<tr><td>>trna152_ValCAC_6_-_27248049_27248121@1.29.29|GCTTCTGTAGTGTAGTGGTTATCACGTTC</td><td>>trna152_ValCAC_6_-_27248049_27248121</td><td>1</td><td>29</td><td>29</td><td>GCTTCTGTAGTGTAGTGGTTATCACGTTC</td><td>13207.3</td><td>-10.1</td><td>1.3</td><td>-7.6</td><td>3.0E-14</td><td>8.5E-14</td><td>BT20/MB468</td><td>NUC</td></tr>
<tr><td>>hg38_21_+_8433172_8446622_RNA45SN1_45S_50ntFlanks@10903.10939.37|GGCTGGGCGCGCGCCGCGGCTGGACGAGGCGCCGCCG</td><td>>hg38_21_+_8433172_8446622_RNA45SN1_45S_50ntFlanks</td><td>10903</td><td>10939</td><td>37</td><td>GGCTGGGCGCGCGCCGCGGCTGGACGAGGCGCCGCCG</td><td>157.7</td><td>-11.2</td><td>1.5</td><td>-7.7</td><td>1.6E-14</td><td>8.5E-14</td><td>MB231/MB468</td><td>MP</td></tr>
<tr><td>>hg38_21_+_8433172_8446622_RNA45SN1_45S_50ntFlanks@11569.11592.24|CAGCCGACTTAGAACTGGTGCGGA</td><td>>hg38_21_+_8433172_8446622_RNA45SN1_45S_50ntFlanks</td><td>11569</td><td>11592</td><td>24</td><td>CAGCCGACTTAGAACTGGTGCGGA</td><td>415.5</td><td>-7.0</td><td>0.9</td><td>-7.7</td><td>1.8E-14</td><td>8.5E-14</td><td>MB231/MB468</td><td>CYTO</td></tr>
<tr><td>>hg38_21_+_8433172_8446622_RNA45SN1_45S_50ntFlanks@12635.12652.18|AAGCTACCATCTGTGGGA</td><td>>hg38_21_+_8433172_8446622_RNA45SN1_45S_50ntFlanks</td><td>12635</td><td>12652</td><td>18</td><td>AAGCTACCATCTGTGGGA</td><td>146.3</td><td>-7.4</td><td>1.0</td><td>-7.7</td><td>1.8E-14</td><td>8.5E-14</td><td>MB231/MB468</td><td>CYTO</td></tr>
<tr><td>>hg38_21_+_8433172_8446622_RNA45SN1_45S_50ntFlanks@10384.10429.46|CCATGTGAACAGCAGTTGAACATGGGTCAGTCGGTCCTGAGAGATG</td><td>>hg38_21_+_8433172_8446622_RNA45SN1_45S_50ntFlanks</td><td>10384</td><td>10429</td><td>46</td><td>CCATGTGAACAGCAGTTGAACATGGGTCAGTCGGTCCTGAGAGATG</td><td>1092.1</td><td>-14.5</td><td>1.9</td><td>-7.6</td><td>3.0E-14</td><td>8.5E-14</td><td>BT20/MB468</td><td>NUC</td></tr>
<tr><td>>hg38_21_+_8433172_8446622_RNA45SN1_45S_50ntFlanks@12456.12486.31|CCAAGCGTTGGATTGTTCACCCACTAATAGG</td><td>>hg38_21_+_8433172_8446622_RNA45SN1_45S_50ntFlanks</td><td>12456</td><td>12486</td><td>31</td><td>CCAAGCGTTGGATTGTTCACCCACTAATAGG</td><td>5284.2</td><td>-6.3</td><td>0.8</td><td>-7.6</td><td>2.0E-14</td><td>8.6E-14</td><td>BT20/MB468</td><td>CYTO</td></tr>
<tr><td>>hg38_21_+_8433172_8446622_RNA45SN1_45S_50ntFlanks@6663.6688.26|GTGGATCACTCGGCTCGTGCGTCGAT</td><td>>hg38_21_+_8433172_8446622_RNA45SN1_45S_50ntFlanks</td><td>6663</td><td>6688</td><td>26</td><td>GTGGATCACTCGGCTCGTGCGTCGAT</td><td>994.6</td><td>-7.2</td><td>0.9</td><td>-7.6</td><td>2.5E-14</td><td>8.6E-14</td><td>MB231/MB468</td><td>MITO</td></tr>
<tr><td>>hg38_21_+_8433172_8446622_RNA45SN1_45S_50ntFlanks@11660.11685.26|GTGATTTCTGCCCAGTGCTCTGAATG</td><td>>hg38_21_+_8433172_8446622_RNA45SN1_45S_50ntFlanks</td><td>11660</td><td>11685</td><td>26</td><td>GTGATTTCTGCCCAGTGCTCTGAATG</td><td>873.2</td><td>-9.4</td><td>1.2</td><td>-7.7</td><td>1.6E-14</td><td>8.6E-14</td><td>MB231/MB468</td><td>MP</td></tr>
<tr><td>>hg38_21_+_8433172_8446622_RNA45SN1_45S_50ntFlanks@6687.6706.20|ATGAAGAACGCAGCTAGCTG</td><td>>hg38_21_+_8433172_8446622_RNA45SN1_45S_50ntFlanks</td><td>6687</td><td>6706</td><td>20</td><td>ATGAAGAACGCAGCTAGCTG</td><td>186.2</td><td>-11.4</td><td>1.5</td><td>-7.7</td><td>1.6E-14</td><td>8.6E-14</td><td>MB231/MB468</td><td>MP</td></tr>
<tr><td>>hg38_21_+_8433172_8446622_RNA45SN1_45S_50ntFlanks@12939.12966.28|CCATTCGTAGACGACCTGCTTCTGGGTC</td><td>>hg38_21_+_8433172_8446622_RNA45SN1_45S_50ntFlanks</td><td>12939</td><td>12966</td><td>28</td><td>CCATTCGTAGACGACCTGCTTCTGGGTC</td><td>306.1</td><td>-6.4</td><td>0.8</td><td>-7.7</td><td>1.8E-14</td><td>8.6E-14</td><td>MB231/MB468</td><td>CYTO</td></tr>
<tr><td>>hg38_21_+_8433172_8446622_RNA45SN1_45S_50ntFlanks@11783.11820.38|CATGAATGGATGAACGAGATTCCCACTGTCCCTACCTA</td><td>>hg38_21_+_8433172_8446622_RNA45SN1_45S_50ntFlanks</td><td>11783</td><td>11820</td><td>38</td><td>CATGAATGGATGAACGAGATTCCCACTGTCCCTACCTA</td><td>323.2</td><td>-12.2</td><td>1.6</td><td>-7.7</td><td>1.6E-14</td><td>8.6E-14</td><td>MB231/MB468</td><td>MP</td></tr>
<tr><td>>hg38_21_+_8433172_8446622_RNA45SN1_45S_50ntFlanks@6661.6683.23|CGGTGGATCACTCGGCTCGTGCG</td><td>>hg38_21_+_8433172_8446622_RNA45SN1_45S_50ntFlanks</td><td>6661</td><td>6683</td><td>23</td><td>CGGTGGATCACTCGGCTCGTGCG</td><td>8145.9</td><td>5.4</td><td>0.7</td><td>7.7</td><td>1.9E-14</td><td>8.6E-14</td><td>MB231/MB468</td><td>NUC</td></tr>
<tr><td>>hg38_21_+_8433172_8446622_RNA45SN1_45S_50ntFlanks@6661.6681.21|CGGTGGATCACTCGGCTCGTG</td><td>>hg38_21_+_8433172_8446622_RNA45SN1_45S_50ntFlanks</td><td>6661</td><td>6681</td><td>21</td><td>CGGTGGATCACTCGGCTCGTG</td><td>4085.6</td><td>-6.2</td><td>0.8</td><td>-7.7</td><td>1.6E-14</td><td>8.6E-14</td><td>MB231/MB468</td><td>MP</td></tr>
<tr><td>>hg38_21_+_8433172_8446622_RNA45SN1_45S_50ntFlanks@9343.9370.28|CGAAAGCCGCCGTGGCGCAATGAAGGTG</td><td>>hg38_21_+_8433172_8446622_RNA45SN1_45S_50ntFlanks</td><td>9343</td><td>9370</td><td>28</td><td>CGAAAGCCGCCGTGGCGCAATGAAGGTG</td><td>620.1</td><td>-5.0</td><td>0.7</td><td>-7.7</td><td>1.5E-14</td><td>8.6E-14</td><td>BT20/MB468</td><td>MITO</td></tr>
<tr><td>>hg38_21_+_8433172_8446622_RNA45SN1_45S_50ntFlanks@12717.12757.41|CCTCGGTTGGCCTCGGATAGCCGGTCCCCCGCCTGTCCCCG</td><td>>hg38_21_+_8433172_8446622_RNA45SN1_45S_50ntFlanks</td><td>12717</td><td>12757</td><td>41</td><td>CCTCGGTTGGCCTCGGATAGCCGGTCCCCCGCCTGTCCCCG</td><td>328.3</td><td>-11.3</td><td>1.5</td><td>-7.7</td><td>1.9E-14</td><td>8.6E-14</td><td>MB231/MB468</td><td>NUC</td></tr>
<tr><td>>hg38_21_+_8433172_8446622_RNA45SN1_45S_50ntFlanks@7973.8011.39|GACGCGACCTCAGATCAGACGTGGCGACCCGCTGAATTT</td><td>>hg38_21_+_8433172_8446622_RNA45SN1_45S_50ntFlanks</td><td>7973</td><td>8011</td><td>39</td><td>GACGCGACCTCAGATCAGACGTGGCGACCCGCTGAATTT</td><td>720.9</td><td>6.4</td><td>0.8</td><td>7.6</td><td>2.1E-14</td><td>8.6E-14</td><td>BT20/MB468</td><td>CYTO</td></tr>
<tr><td>>hg38_21_+_8433172_8446622_RNA45SN1_45S_50ntFlanks@11662.11685.24|GATTTCTGCCCAGTGCTCTGAATG</td><td>>hg38_21_+_8433172_8446622_RNA45SN1_45S_50ntFlanks</td><td>11662</td><td>11685</td><td>24</td><td>GATTTCTGCCCAGTGCTCTGAATG</td><td>334.4</td><td>-4.5</td><td>0.6</td><td>-7.6</td><td>3.1E-14</td><td>8.6E-14</td><td>BT20/MB468</td><td>NUC</td></tr>
<tr><td>>trna111_HisGTG_1_-_147774845_147774916@-1T.20.21,trna118_HisGTG_1_-_145396881_145396952@-1T.20.21,trna16_HisGTG_1_+_146544773_146544844@-1T.20.21,trna1_HisGTG_15_+_45493349_45493420@-1T.20.21,trna21_HisGTG_1_+_147753471_147753542@-1T.20.21,trna33_HisGTG_6_+_27125906_27125977@-1T.20.21,trna7_HisGTG_9_-_14433938_14434009@-1T.20.21,trna8_HisGTG_15_-_45492611_45492682@-1T.20.21,trna9_HisGTG_15_-_45490804_45490875@-1T.20.21|TGCCGTGATCGTATAGTGGTT</td><td>>trna111_HisGTG_1_-_147774845_147774916</td><td>-1T</td><td>20</td><td>21</td><td>TGCCGTGATCGTATAGTGGTT</td><td>219.1</td><td>11.7</td><td>1.5</td><td>7.7</td><td>1.9E-14</td><td>8.7E-14</td><td>MB231/MB468</td><td>NUC</td></tr>
<tr><td>>hg38_21_+_8433172_8446622_RNA45SN1_45S_50ntFlanks@7977.8017.41|CGACCTCAGATCAGACGTGGCGACCCGCTGAATTTAAGCAT</td><td>>hg38_21_+_8433172_8446622_RNA45SN1_45S_50ntFlanks</td><td>7977</td><td>8017</td><td>41</td><td>CGACCTCAGATCAGACGTGGCGACCCGCTGAATTTAAGCAT</td><td>769.0</td><td>8.1</td><td>1.1</td><td>7.7</td><td>1.8E-14</td><td>8.7E-14</td><td>MB231/MB468</td><td>CYTO</td></tr>
<tr><td>>MI0000456|hsa-mir-140&WithFlank&16|+|69933075|69933186@69.90.22(+1U)&[MIMAT0004597&hsa-miR-140-3p&offsets|+1|+2(+1U);m-45&16|+|69933143|69933164&offsets|0|0(+1U)]|ACCACAGGGTAGAACCACGGACT</td><td>>MI0000456|hsa-mir-140&WithFlank&16|+|69933075|69933186</td><td>69</td><td>90</td><td>22(+1U)</td><td>ACCACAGGGTAGAACCACGGACT</td><td>717.6</td><td>12.7</td><td>1.7</td><td>7.6</td><td>2.5E-14</td><td>8.7E-14</td><td>MB231/MB468</td><td>MITO</td></tr>
<tr><td>>hg38_21_+_8433172_8446622_RNA45SN1_45S_50ntFlanks@4008.4025.18|CTCGGGCCGATCGCACGC</td><td>>hg38_21_+_8433172_8446622_RNA45SN1_45S_50ntFlanks</td><td>4008</td><td>4025</td><td>18</td><td>CTCGGGCCGATCGCACGC</td><td>210.9</td><td>-11.6</td><td>1.5</td><td>-7.7</td><td>1.6E-14</td><td>8.7E-14</td><td>MB231/MB468</td><td>MP</td></tr>
<tr><td>>am_trna128_GlyGCC_6_-_27870686_27870756@1.28.28,trna133_GlyCCC_1_-_16872434_16872504@1.28.28,trna18_GlyGCC_16_+_70822597_70822667@1.28.28,trna19_GlyGCC_16_+_70823410_70823480@1.28.28,trna19_GlyGCC_2_-_157257659_157257729@1.28.28,trna24_GlyGCC_16_-_70812942_70813012@1.28.28,trna25_GlyGCC_16_-_70812114_70812184@1.28.28,trna4_GlyCCC_1_+_17188416_17188486@1.28.28,trna5_GlyGCC_17_+_8029064_8029134@1.28.28,trna68_GlyGCC_1_-_161493637_161493707@1.28.28|GCATTGGTGGTTCAGTGGTAGAATTCTC</td><td>>am_trna128_GlyGCC_6_-_27870686_27870756</td><td>1</td><td>28</td><td>28</td><td>GCATTGGTGGTTCAGTGGTAGAATTCTC</td><td>52842.2</td><td>-6.6</td><td>0.9</td><td>-7.7</td><td>1.9E-14</td><td>8.7E-14</td><td>MB231/MB468</td><td>NUC</td></tr>
<tr><td>>hg38_21_+_8433172_8446622_RNA45SN1_45S_50ntFlanks@11319.11338.20|CCCACGTCTCGTCGCGCGCG</td><td>>hg38_21_+_8433172_8446622_RNA45SN1_45S_50ntFlanks</td><td>11319</td><td>11338</td><td>20</td><td>CCCACGTCTCGTCGCGCGCG</td><td>264.9</td><td>-11.8</td><td>1.6</td><td>-7.6</td><td>2.6E-14</td><td>8.8E-14</td><td>MB231/MB468</td><td>MITO</td></tr>
<tr><td>>am_trna2_GlyGCC_21_-_18827107_18827177@1.20.20,trna35_GlyGCC_1_+_161413094_161413164@1.20.20,trna37_GlyGCC_1_+_161420467_161420537@1.20.20,trna39_GlyGCC_1_+_161427898_161427968@1.20.20,trna41_GlyGCC_1_+_161435258_161435328@1.20.20|GCATGGGTGGTTCAGTGGTA</td><td>>am_trna2_GlyGCC_21_-_18827107_18827177</td><td>1</td><td>20</td><td>20</td><td>GCATGGGTGGTTCAGTGGTA</td><td>747.0</td><td>9.7</td><td>1.2</td><td>8.0</td><td>1.5E-15</td><td>8.8E-14</td><td>BT20/MB231</td><td>CYTO</td></tr>
<tr><td>>MI0000061|hsa-let-7a-2&WithFlank&11|-|122146516|122146599@11.31.21(+1C);MI0000062|hsa-let-7a-3&WithFlank&22|+|46112743|46112828@10.30.21(+1C);MI0000060|hsa-let-7a-1&WithFlank&9|+|94175951|94176042@12.32.21(+1C)&[MIMAT0000062&hsa-let-7a-5p&offsets|0|-1(+1C);m-6&11|-|122146568|122146589&offsets|0|-1(+1C)];[MIMAT0000062_1&hsa-let-7a-5p&offsets|0|-1(+1C);m-5&22|+|46112752|46112773&offsets|0|-1(+1C)];[MIMAT0000062_2&hsa-let-7a-5p&offsets|0|-1(+1C);m-4&9|+|94175962|94175983&offsets|0|-1(+1C)]|TGAGGTAGTAGGTTGTATAGTC</td><td>>MI0000061|hsa-let-7a-2&WithFlank&11|-|122146516|122146599</td><td>11</td><td>31</td><td>21(+1C)</td><td>TGAGGTAGTAGGTTGTATAGTC</td><td>8634.6</td><td>6.1</td><td>0.8</td><td>7.7</td><td>1.8E-14</td><td>8.8E-14</td><td>MB231/MB468</td><td>CYTO</td></tr>
<tr><td>>trnaMT_GluTTC_MT_-_14674_14742@38.58.21,trnalookalike8_GluTTC_5_-_93905172_93905240@38.58.21|CATTGGTCGTGGTTGTAGTCC</td><td>>trnaMT_GluTTC_MT_-_14674_14742</td><td>38</td><td>58</td><td>21</td><td>CATTGGTCGTGGTTGTAGTCC</td><td>425.0</td><td>7.7</td><td>1.0</td><td>7.7</td><td>1.6E-14</td><td>8.9E-14</td><td>BT20/MB468</td><td>MITO</td></tr>
</table>

| | | | | | | | | | | | | | |
|---|---|---|---|---|---|---|---|---|---|---|---|---|---|
| >trna112_GlnCTG_1_-_147737382_147737453@32.50.19,trna131_GlnCTG_6_-_27759135_27759206@32.50.19,trna146_GlnCTG_6_-_27515531_27515602@32.50.19,trna15_GlnCTG_1_+_145963304_145963375@32.50.19,trna19_GlnCTG_1_+_147505038_147505109@32.50.19,trna1_GlnCTG_6_+_18836402_18836473@32.50.19,trna28_GlnCTG_1_+_149186125_149186196@32.50.19,trna3_GlnCTG_17_+_8023070_8023141@32.50.19,trna49_GlnCTG_6_+_27487308_27487379@32.50.19,trna7_GlnCTG_15_-_66161400_66161471@32.50.19,trna99_GlnCTG_6_-_28909378_28909449@32.50.19\|CTCTGAATCCAGCGATCCG | >trna112_GlnCTG_1_-_147737382_147737453 | 32 | 50 | 19 | CTCTGAATCCAGCGATCCG | 158.2 | -11.2 | 1.5 | -7.7 | 1.6E-14 | 8.9E-14 | MB231/MB468 | MP |
| >MI0000477\|hsa-mir-146a&WithFlank&5\|+\|160485346\|160485456@27.46.20(+1C)&[MIMAT0000449&hsa-miR-146a-5p&offsets\|0\|-2(+1C);m-129&5\|+\|160485372\|160485393&offsets\|0\|-2(+1C)]\|TGAGAACTGAATTCCATGGGC | >MI0000477\|hsa-mir-146a&WithFlank&5\|+\|160485346\|160485456 | 27 | 46 | 20(+1C) | TGAGAACTGAATTCCATGGGC | 2475.1 | -14.7 | 1.8 | -8.2 | 2.7E-16 | 8.9E-14 | BT20/MB231 | MP |
| >hg38_21_+_8433172_8446622_RNA45SN1_45S_50ntFlanks@11933.11968.36\|AATAAGTGGGAGGCCCCCGGCGCCCCCCCGGTGTCC | >hg38_21_+_8433172_8446622_RNA45SN1_45S_50ntFlanks | 11933 | 11968 | 36 | AATAAGTGGGAGGCCCCCGGCGCCCCCCCGGTGTCC | 209.9 | -11.6 | 1.5 | -7.7 | 1.6E-14 | 8.9E-14 | MB231/MB468 | MP |
| >hg38_21_+_8433172_8446622_RNA45SN1_45S_50ntFlanks@3899.3932.34\|CCTTCGCGGGGGGGATGCGTGCATTTATCAGATC | >hg38_21_+_8433172_8446622_RNA45SN1_45S_50ntFlanks | 3899 | 3932 | 34 | CCTTCGCGGGGGGGATGCGTGCATTTATCAGATC | 279.8 | -12.0 | 1.6 | -7.7 | 1.6E-14 | 8.9E-14 | MB231/MB468 | MP |
| >hg38_21_+_8433172_8446622_RNA45SN1_45S_50ntFlanks@7986.8011.26\|ATCAGACGTGGCGACCCGCTGAATTT | >hg38_21_+_8433172_8446622_RNA45SN1_45S_50ntFlanks | 7986 | 8011 | 26 | ATCAGACGTGGCGACCCGCTGAATTT | 483.1 | -11.0 | 1.4 | -7.7 | 1.6E-14 | 8.9E-14 | BT20/MB468 | MITO |
| >hg38_21_+_8433172_8446622_RNA45SN1_45S_50ntFlanks@12655.12675.21\|ATGACTGAACGCCTCTAAGTC | >hg38_21_+_8433172_8446622_RNA45SN1_45S_50ntFlanks | 12655 | 12675 | 21 | ATGACTGAACGCCTCTAAGTC | 3850.4 | -5.2 | 0.7 | -7.7 | 1.6E-14 | 8.9E-14 | BT20/MB468 | MITO |
| >hg38_21_+_8433172_8446622_RNA45SN1_45S_50ntFlanks@5446.5466.21\|CGGATCGGCCCCGCCGGGGTC | >hg38_21_+_8433172_8446622_RNA45SN1_45S_50ntFlanks | 5446 | 5466 | 21 | CGGATCGGCCCCGCCGGGGTC | 7718.6 | -5.3 | 0.7 | -7.6 | 2.1E-14 | 9.0E-14 | BT20/MB468 | CYTO |
| >MI0000279\|hsa-mir-196a-2&WithFlank&12\|+\|53991732\|53991853@31.51.21(+1A);MI0000238\|hsa-mir-196a-1&WithFlank&17\|-\|48632484\|48632565@13.33.21(+1A)&[MIMAT0000226&hsa-miR-196a-5p&offsets\|0\|-1(+1A);m-261&12\|+\|53991762\|53991783&offsets\|0\|-1(+1A)];[MIMAT0000226_1&hsa-miR-196a-5p&offsets\|0\|-1(+1A);m-268&17\|-\|48632532\|48632553&offsets\|0\|-1(+1A)]\|TAGGTAGTTTCATGTTGTTGGA | >MI0000279\|hsa-mir-196a-2&WithFlank&12\|+\|53991732\|53991853 | 31 | 51 | 21(+1A) | TAGGTAGTTTCATGTTGTTGGA | 209.6 | 3.3 | 0.4 | 7.6 | 3.2E-14 | 9.0E-14 | BT20/MB468 | NUC |
| >hg38_21_+_8433172_8446622_RNA45SN1_45S_50ntFlanks@12619.12661.43\|AGGAGCCAATGGGGCGAAGCTACCATCTGTGGGATTATGACTG | >hg38_21_+_8433172_8446622_RNA45SN1_45S_50ntFlanks | 12619 | 12661 | 43 | AGGAGCCAATGGGGCGAAGCTACCATCTGTGGGATTATGACTG | 449.6 | -12.6 | 1.7 | -7.6 | 2.6E-14 | 9.0E-14 | MB231/MB468 | MITO |
| >hg38_21_+_8433172_8446622_RNA45SN1_45S_50ntFlanks@9122.9152.31\|CCGCGCGCGGGTCGGGGGGCGGGGCGGACTG | >hg38_21_+_8433172_8446622_RNA45SN1_45S_50ntFlanks | 9122 | 9152 | 31 | CCGCGCGCGGGTCGGGGGGCGGGGCGGACTG | 429.2 | -12.5 | 1.6 | -7.6 | 2.6E-14 | 9.0E-14 | MB231/MB468 | MITO |
| >hg38_21_+_8433172_8446622_RNA45SN1_45S_50ntFlanks@8021.8042.22\|AGTCAGCGGAGGAGAAGAAACT | >hg38_21_+_8433172_8446622_RNA45SN1_45S_50ntFlanks | 8021 | 8042 | 22 | AGTCAGCGGAGGAGAAGAAACT | 212.1 | 8.4 | 1.1 | 7.7 | 1.9E-14 | 9.0E-14 | MB231/MB468 | CYTO |
| >trnaMT_LeuTAA_MT_+_3230_3304@16.45.30\|CCCGGTAATCGCATAAAACTTAAAACTTTA | >trnaMT_LeuTAA_MT_+_3230_3304 | 16 | 45 | 30 | CCCGGTAATCGCATAAAACTTAAAACTTTA | 277.0 | -11.1 | 1.4 | -7.7 | 2.0E-14 | 9.0E-14 | MB231/MB468 | NUC |
| >hg38_21_+_8433172_8446622_RNA45SN1_45S_50ntFlanks@8169.8186.18\|CCAAGTCCTTCTGATCGA | >hg38_21_+_8433172_8446622_RNA45SN1_45S_50ntFlanks | 8169 | 8186 | 18 | CCAAGTCCTTCTGATCGA | 623.0 | -13.1 | 1.7 | -7.7 | 1.7E-14 | 9.0E-14 | MB231/MB468 | MP |
| >hg38_21_+_8433172_8446622_RNA45SN1_45S_50ntFlanks@12414.12433.20\|CCTTCGATGTCGGCTCTTCC | >hg38_21_+_8433172_8446622_RNA45SN1_45S_50ntFlanks | 12414 | 12433 | 20 | CCTTCGATGTCGGCTCTTCC | 661.1 | 5.4 | 0.7 | 7.7 | 1.9E-14 | 9.0E-14 | MB231/MB468 | CYTO |
| >hg38_21_+_8433172_8446622_RNA45SN1_45S_50ntFlanks@476.505.30\|TTCGTGATCGATGTGGTGACGTCGTGCTCT | >hg38_21_+_8433172_8446622_RNA45SN1_45S_50ntFlanks | 476 | 505 | 30 | TTCGTGATCGATGTGGTGACGTCGTGCTCT | 989.2 | 7.8 | 1.0 | 7.7 | 1.9E-14 | 9.0E-14 | MB231/MB468 | CYTO |
| >hg38_21_+_8433172_8446622_RNA45SN1_45S_50ntFlanks@6742.6770.29\|TCGACACTTCGAACGCACTTGCGGCCCCG | >hg38_21_+_8433172_8446622_RNA45SN1_45S_50ntFlanks | 6742 | 6770 | 29 | TCGACACTTCGAACGCACTTGCGGCCCCG | 567.8 | -6.9 | 0.9 | -7.7 | 1.9E-14 | 9.0E-14 | MB231/MB468 | CYTO |
| >hg38_21_+_8433172_8446622_RNA45SN1_45S_50ntFlanks@9507.9526.20\|AACTATGCCTGGGCAGGGCG | >hg38_21_+_8433172_8446622_RNA45SN1_45S_50ntFlanks | 9507 | 9526 | 20 | AACTATGCCTGGGCAGGGCG | 177.9 | -10.2 | 1.3 | -7.7 | 1.9E-14 | 9.0E-14 | MB231/MB468 | CYTO |
| >MI0003583\|hsa-mir-576&WithFlank&4\|+\|109488692\|109488801@22.43.22&[MIMAT0003241&hsa-miR-576-5p&offsets\|0\|0;m-264&4\|+\|109488713\|109488735&offsets\|0\|-1]\|ATTCTAATTTCTCCACGTCTTT | >MI0003583\|hsa-mir-576&WithFlank&4\|+\|109488692\|109488801 | 22 | 43 | 22 | ATTCTAATTTCTCCACGTCTTT | 426.1 | 9.4 | 1.2 | 7.6 | 2.2E-14 | 9.1E-14 | BT20/MB468 | CYTO |
| >hg38_21_+_8433172_8446622_RNA45SN1_45S_50ntFlanks@4003.4025.23\|ATAACCTCGGGCCGATCGCACGC | >hg38_21_+_8433172_8446622_RNA45SN1_45S_50ntFlanks | 4003 | 4025 | 23 | ATAACCTCGGGCCGATCGCACGC | 1667.2 | -8.2 | 1.1 | -7.7 | 2.0E-14 | 9.1E-14 | MB231/MB468 | NUC |
| >hg38_21_+_8433172_8446622_RNA45SN1_45S_50ntFlanks@10905.10936.32\|CTGGGCGCGCGCCGCGGCTGGACGAGGCGCCG | >hg38_21_+_8433172_8446622_RNA45SN1_45S_50ntFlanks | 10905 | 10936 | 32 | CTGGGCGCGCGCCGCGGCTGGACGAGGCGCCG | 441.0 | -11.5 | 1.5 | -7.7 | 1.9E-14 | 9.1E-14 | MB231/MB468 | CYTO |
| >hg38_21_+_8433172_8446622_RNA45SN1_45S_50ntFlanks@10829.10851.23\|GCAAGCCGGATCCGTAACTTCGG | >hg38_21_+_8433172_8446622_RNA45SN1_45S_50ntFlanks | 10829 | 10851 | 23 | GCAAGCCGGATCCGTAACTTCGG | 287.3 | -4.8 | 0.6 | -7.6 | 3.2E-14 | 9.1E-14 | BT20/MB468 | NUC |
| >hg38_21_+_8433172_8446622_RNA45SN1_45S_50ntFlanks@12936.12966.31\|AAACCATTCGTAGACGACCTGCTTCTGGGTC | >hg38_21_+_8433172_8446622_RNA45SN1_45S_50ntFlanks | 12936 | 12966 | 31 | AAACCATTCGTAGACGACCTGCTTCTGGGTC | 627.4 | -6.8 | 0.9 | -7.7 | 1.9E-14 | 9.1E-14 | MB231/MB468 | CYTO |
| >hg38_21_+_8433172_8446622_RNA45SN1_45S_50ntFlanks@12588.12614.27\|GGTTCAGACATTTGGTGTATGTGCTTG | >hg38_21_+_8433172_8446622_RNA45SN1_45S_50ntFlanks | 12588 | 12614 | 27 | GGTTCAGACATTTGGTGTATGTGCTTG | 228.3 | -11.6 | 1.5 | -7.6 | 2.7E-14 | 9.1E-14 | MB231/MB468 | MITO |
| >hg38_21_+_8433172_8446622_RNA45SN1_45S_50ntFlanks@4725.4758.34\|TTAATCAAGAACGAAAGTCGGAGGTTCGAAGACG | >hg38_21_+_8433172_8446622_RNA45SN1_45S_50ntFlanks | 4725 | 4758 | 34 | TTAATCAAGAACGAAAGTCGGAGGTTCGAAGACG | 3800.0 | -9.2 | 1.2 | -7.6 | 3.2E-14 | 9.1E-14 | BT20/MB468 | NUC |
| >hg38_21_+_8433172_8446622_RNA45SN1_45S_50ntFlanks@4590.4616.27\|ATTTTGTTGGTTTTCGGAACTGAGGCC | >hg38_21_+_8433172_8446622_RNA45SN1_45S_50ntFlanks | 4590 | 4616 | 27 | ATTTTGTTGGTTTTCGGAACTGAGGCC | 868.7 | -7.7 | 1.0 | -7.7 | 1.7E-14 | 9.1E-14 | MB231/MB468 | MP |
| >hg38_21_+_8433172_8446622_RNA45SN1_45S_50ntFlanks@3930.3959.30\|ATCAAAACCAACCCGGTCAGCCCCTCTCCG | >hg38_21_+_8433172_8446622_RNA45SN1_45S_50ntFlanks | 3930 | 3959 | 30 | ATCAAAACCAACCCGGTCAGCCCCTCTCCG | 138.4 | -11.0 | 1.4 | -7.7 | 1.7E-14 | 9.1E-14 | MB231/MB468 | MP |
| >hg38_21_+_8433172_8446622_RNA45SN1_45S_50ntFlanks@7987.8008.22\|TCAGACGTGGCGACCCGCTGAA | >hg38_21_+_8433172_8446622_RNA45SN1_45S_50ntFlanks | 7987 | 8008 | 22 | TCAGACGTGGCGACCCGCTGAA | 620.5 | -6.3 | 0.8 | -7.7 | 1.7E-14 | 9.1E-14 | MB231/MB468 | MP |
| >MI0000095\|hsa-mir-93&WithFlank&7\|-\|100093762\|100093853@17.39.23&[MIMAT0000093&hsa-miR-93-5p&offsets\|0\|0;m-29&7\|-\|100093815\|100093837&offsets\|0\|0]\|CAAAGTGCTGTTCGTGCAGGTAG | >MI0000095\|hsa-mir-93&WithFlank&7\|-\|100093762\|100093853 | 17 | 39 | 23 | CAAAGTGCTGTTCGTGCAGGTAG | 9356.0 | 4.8 | 0.6 | 7.6 | 2.0E-14 | 9.1E-14 | MB231/MB468 | NUC |
| >MI0000734\|hsa-mir-106b&WithFlank&7\|-\|100093987\|100094080@58.78.21&[MIMAT0004672&hsa-miR-106b-3p&offsets\|0\|-1;m-87&7\|-\|100094003\|100094023&offsets\|0\|0]\|CCGCACTGTGGGTACTTGCTG | >MI0000734\|hsa-mir-106b&WithFlank&7\|-\|100093987\|100094080 | 58 | 78 | 21 | CCGCACTGTGGGTACTTGCTG | 1742.3 | 5.0 | 0.7 | 7.6 | 2.0E-14 | 9.1E-14 | MB231/MB468 | NUC |
| >hg38_21_+_8433172_8446622_RNA45SN1_45S_50ntFlanks@12928.12953.26\|CTGGCGCTAAACCATTCGTAGACGAC | >hg38_21_+_8433172_8446622_RNA45SN1_45S_50ntFlanks | 12928 | 12953 | 26 | CTGGCGCTAAACCATTCGTAGACGAC | 150.7 | -11.1 | 1.4 | -7.7 | 1.7E-14 | 9.2E-14 | MB231/MB468 | MP |
| >hg38_21_+_8433172_8446622_RNA45SN1_45S_50ntFlanks@3822.3852.31\|CTTTGGTCGCTCGCTCCTCTCCTACTTGGAT | >hg38_21_+_8433172_8446622_RNA45SN1_45S_50ntFlanks | 3822 | 3852 | 31 | CTTTGGTCGCTCGCTCCTCTCCTACTTGGAT | 342.9 | -12.2 | 1.6 | -7.6 | 2.7E-14 | 9.2E-14 | MB231/MB468 | MITO |
| >trna2_GlyGCC_21_-_18827107_18827177@1.29.29,trna35_GlyGCC_1_+_161413094_161413164@1.29.29,trna37_GlyGCC_1_+_161420467_161420537@1.29.29,trna39_GlyGCC_1_+_161427898_161427968@1.29.29,trna41_GlyGCC_1_+_161435258_161435328@1.29.29\|GCATGGGTGGTTCAGTGGTAGAATTCTCG | >trna2_GlyGCC_21_-_18827107_18827177 | 1 | 29 | 29 | GCATGGGTGGTTCAGTGGTAGAATTCTCG | 17043.9 | -10.0 | 1.3 | -7.7 | 1.7E-14 | 9.2E-14 | MB231/MB468 | MP |
| >hg38_21_+_8433172_8446622_RNA45SN1_45S_50ntFlanks@11576.11603.28\|CTTAGAACTGGTGCGGACCAGGGGAATC | >hg38_21_+_8433172_8446622_RNA45SN1_45S_50ntFlanks | 11576 | 11603 | 28 | CTTAGAACTGGTGCGGACCAGGGGAATC | 1027.6 | -4.0 | 0.5 | -7.7 | 1.6E-14 | 9.2E-14 | BT20/MB468 | MITO |
| >MI0000650\|hsa-mir-200c&WithFlank&12\|+\|6963693\|6963772@50.71.22(+1G)&[MIMAT0000617&hsa-miR-200c-3p&offsets\|0\|-1(+1G);m-54&12\|+\|6963742\|6963764&offsets\|0\|-1(+1G)]\|TAATACTGCCGGGTAATGATGGG | >MI0000650\|hsa-mir-200c&WithFlank&12\|+\|6963693\|6963772 | 50 | 71 | 22(+1G) | TAATACTGCCGGGTAATGATGGG | 1290.4 | 13.7 | 1.7 | 8.2 | 2.9E-16 | 9.2E-14 | BT20/MB231 | MP |
| >trnaMT_PheGAA_MT_+_577_647@35.53.19\|GAAAATGTTTAGACGGGCT | >trnaMT_PheGAA_MT_+_577_647 | 35 | 53 | 19 | GAAAATGTTTAGACGGGCT | 824.8 | 6.2 | 0.8 | 7.7 | 1.6E-14 | 9.2E-14 | BT20/MB468 | MITO |
| >MI0000285\|hsa-mir-205&WithFlank&1\|+\|209432127\|209432248@40.63.24&[MIMAT0000266&hsa-miR-205-5p&offsets\|0\|+2;m-351&1\|+\|209432166\|209432187&offsets\|0\|+2]\|TCCTTCATTCCACCGGAGTCTGTC | >MI0000285\|hsa-mir-205&WithFlank&1\|+\|209432127\|209432248 | 40 | 63 | 24 | TCCTTCATTCCACCGGAGTCTGTC | 349.3 | 11.8 | 1.5 | 8.0 | 1.6E-15 | 9.2E-14 | BT20/MB231 | CYTO |
| >hg38_21_+_8433172_8446622_RNA45SN1_45S_50ntFlanks@12912.12930.19\|CACGTTCGTGGGGAACCTG | >hg38_21_+_8433172_8446622_RNA45SN1_45S_50ntFlanks | 12912 | 12930 | 19 | CACGTTCGTGGGGAACCTG | 312.4 | -12.2 | 1.6 | -7.7 | 1.7E-14 | 9.2E-14 | MB231/MB468 | MP |
| >hg38_21_+_8433172_8446622_RNA45SN1_45S_50ntFlanks@11104.11121.18\|AAGGGTCGGGGCGGCAGG | >hg38_21_+_8433172_8446622_RNA45SN1_45S_50ntFlanks | 11104 | 11121 | 18 | AAGGGTCGGGGCGGCAGG | 226.7 | -11.7 | 1.5 | -7.7 | 1.7E-14 | 9.2E-14 | MB231/MB468 | MP |
| >hg38_21_+_8433172_8446622_RNA45SN1_45S_50ntFlanks@12589.12614.26\|GTTCAGACATTTGGTGTATGTGCTTG | >hg38_21_+_8433172_8446622_RNA45SN1_45S_50ntFlanks | 12589 | 12614 | 26 | GTTCAGACATTTGGTGTATGTGCTTG | 642.1 | -12.4 | 1.5 | -8.1 | 6.8E-16 | 9.2E-14 | BT20/MB468 | MP |
| >hg38_21_+_8433172_8446622_RNA45SN1_45S_50ntFlanks@4003.4026.24\|ATAACCTCGGGCCGATCGCACGCC | >hg38_21_+_8433172_8446622_RNA45SN1_45S_50ntFlanks | 4003 | 4026 | 24 | ATAACCTCGGGCCGATCGCACGCC | 841.1 | -6.6 | 0.9 | -7.7 | 1.9E-14 | 9.2E-14 | MB231/MB468 | CYTO |
| >MI0000087\|hsa-mir-29a&WithFlank&7\|-\|130876741\|130876816@48.68.21&[MIMAT0000086&hsa-miR-29a-3p&offsets\|0\|-1;m-26&7\|-\|130876749\|130876769&offsets\|0\|0]\|TAGCACCATCTGAAATCGGTT | >MI0000087\|hsa-mir-29a&WithFlank&7\|-\|130876741\|130876816 | 48 | 68 | 21 | TAGCACCATCTGAAATCGGTT | 4298.0 | 7.8 | 1.0 | 7.7 | 1.9E-14 | 9.2E-14 | MB231/MB468 | CYTO |
| >hg38_21_+_8433172_8446622_RNA45SN1_45S_50ntFlanks@11834.11852.19\|CCACAGCCAAGGGAACGGG | >hg38_21_+_8433172_8446622_RNA45SN1_45S_50ntFlanks | 11834 | 11852 | 19 | CCACAGCCAAGGGAACGGG | 147.6 | -11.1 | 1.4 | -7.7 | 1.7E-14 | 9.2E-14 | MB231/MB468 | MP |
| >hg38_21_+_8433172_8446622_RNA45SN1_45S_50ntFlanks@6656.6686.31\|CTTAGCGGTGGATCACTCGGCTCGTGCGTCG | >hg38_21_+_8433172_8446622_RNA45SN1_45S_50ntFlanks | 6656 | 6686 | 31 | CTTAGCGGTGGATCACTCGGCTCGTGCGTCG | 240.2 | -11.8 | 1.5 | -7.7 | 1.7E-14 | 9.3E-14 | MB231/MB468 | MP |
| >hg38_21_+_8433172_8446622_RNA45SN1_45S_50ntFlanks@11727.11744.18\|AGTAACTATGACTCTCTT | >hg38_21_+_8433172_8446622_RNA45SN1_45S_50ntFlanks | 11727 | 11744 | 18 | AGTAACTATGACTCTCTT | 164.3 | -11.2 | 1.5 | -7.7 | 1.7E-14 | 9.3E-14 | MB231/MB468 | MP |
| >hg38_21_+_8433172_8446622_RNA45SN1_45S_50ntFlanks@12249.12268.20\|CAAAAGCTCGCTTGATCTTG | >hg38_21_+_8433172_8446622_RNA45SN1_45S_50ntFlanks | 12249 | 12268 | 20 | CAAAAGCTCGCTTGATCTTG | 200.7 | -11.5 | 1.5 | -7.7 | 1.7E-14 | 9.3E-14 | MB231/MB468 | MP |

| | | | | | | | | | | | | | |
|---|---|---|---|---|---|---|---|---|---|---|---|---|---|
| >MI0000483\|hsa-mir-186&WithFlank&1\|-\|71067625\|71067722@21.41.21(+1C)&[MIMAT0000456&hsa-miR-186-5p&offsets\|0\|-1(+1C);m-84&1\|-\|71067681\|71067702&offsets\|0\|-1(+1C)]\|CAAAGAATTCTCCTTTTGGGCC | >MI0000483\|hsa-mir-186&WithFlank&1\|-\|71067625\|71067722 | 21 | 41 | 21(+1C) | CAAAGAATTCTCCTTTTGGGCC | 85.7 | 3.7 | 0.5 | 7.6 | 3.3E-14 | 9.3E-14 | BT20/MB468 | NUC |
| >MI0000061\|hsa-let-7a-2&WithFlank&11\|-\|122146516\|122146599@11.30.20;MI0000062\|hsa-let-7a-3&WithFlank&22\|+\|46112743\|46112828@10.29.20;MI0000060\|hsa-let-7a-1&WithFlank&9\|+\|94175951\|94176042@12.31.20&[MIMAT0000062&hsa-let-7a-5p&offsets\|0\|-2;m-6&11\|-\|122146568\|122146589&offsets\|0\|-2];[MIMAT0000062_1&hsa-let-7a-5p&offsets\|0\|-2;m-5&22\|+\|46112752\|46112773&offsets\|0\|-2];[MIMAT0000062_2&hsa-let-7a-5p&offsets\|0\|-2;m-4&9\|+\|94175962\|94175983&offsets\|0\|-2]\|TGAGGTAGTAGGTTGTATAG | >MI0000061\|hsa-let-7a-2&WithFlank&11\|-\|122146516\|122146599 | 11 | 30 | 20 | TGAGGTAGTAGGTTGTATAG | 280.9 | 5.8 | 0.8 | 7.6 | 2.1E-14 | 9.3E-14 | MB231/MB468 | NUC |
| >hg38_21_+_8433172_8446622_RNA45SN1_45S_50ntFlanks@4727.4761.35\|AATCAAGAACGAAAGTCGGAGGTTCGAAGACGATC | >hg38_21_+_8433172_8446622_RNA45SN1_45S_50ntFlanks | 4727 | 4761 | 35 | AATCAAGAACGAAAGTCGGAGGTTCGAAGACGATC | 1286.3 | 5.6 | 0.7 | 7.6 | 2.2E-14 | 9.3E-14 | BT20/MB468 | CYTO |
| >hg38_21_+_8433172_8446622_RNA45SN1_45S_50ntFlanks@11564.11592.29\|CCTAGCAGCCGACTTAGAACTGGTGCGGA | >hg38_21_+_8433172_8446622_RNA45SN1_45S_50ntFlanks | 11564 | 11592 | 29 | CCTAGCAGCCGACTTAGAACTGGTGCGGA | 445.3 | -6.6 | 0.9 | -7.7 | 1.9E-14 | 9.3E-14 | MB231/MB468 | CYTO |
| >hg38_21_+_8433172_8446622_RNA45SN1_45S_50ntFlanks@4759.4793.35\|ATCAGATACCGTCGTAGTTCCGACCATAAACGATG | >hg38_21_+_8433172_8446622_RNA45SN1_45S_50ntFlanks | 4759 | 4793 | 35 | ATCAGATACCGTCGTAGTTCCGACCATAAACGATG | 377.8 | -10.7 | 1.4 | -7.7 | 1.7E-14 | 9.4E-14 | BT20/MB468 | MITO |
| >hg38_21_+_8433172_8446622_RNA45SN1_45S_50ntFlanks@11200.11223.24\|CGGCGGCGGCGGCGACTCTGGACG | >hg38_21_+_8433172_8446622_RNA45SN1_45S_50ntFlanks | 11200 | 11223 | 24 | CGGCGGCGGCGGCGACTCTGGACG | 352.0 | -7.8 | 1.0 | -7.7 | 2.0E-14 | 9.4E-14 | MB231/MB468 | CYTO |
| >hg38_21_+_8433172_8446622_RNA45SN1_45S_50ntFlanks@12347.12366.20\|AGAAAAGTTACCACAGGGAT | >hg38_21_+_8433172_8446622_RNA45SN1_45S_50ntFlanks | 12347 | 12366 | 20 | AGAAAAGTTACCACAGGGAT | 178.9 | -10.2 | 1.3 | -7.7 | 2.0E-14 | 9.4E-14 | MB231/MB468 | CYTO |
| >hg38_21_+_8433172_8446622_RNA45SN1_45S_50ntFlanks@11320.11338.19\|CCACGTCTCGTCGCGCGCG | >hg38_21_+_8433172_8446622_RNA45SN1_45S_50ntFlanks | 11320 | 11338 | 19 | CCACGTCTCGTCGCGCGCG | 190.1 | -11.4 | 1.5 | -7.7 | 1.7E-14 | 9.4E-14 | MB231/MB468 | MP |
| >hg38_21_+_8433172_8446622_RNA45SN1_45S_50ntFlanks@11562.11588.27\|CGCCTAGCAGCCGACTTAGAACTGGTG | >hg38_21_+_8433172_8446622_RNA45SN1_45S_50ntFlanks | 11562 | 11588 | 27 | CGCCTAGCAGCCGACTTAGAACTGGTG | 845.1 | -9.4 | 1.2 | -7.7 | 1.7E-14 | 9.4E-14 | MB231/MB468 | MP |
| >hg38_21_+_8433172_8446622_RNA45SN1_45S_50ntFlanks@11933.11952.20\|AATAAGTGGGAGGCCCCCGG | >hg38_21_+_8433172_8446622_RNA45SN1_45S_50ntFlanks | 11933 | 11952 | 20 | AATAAGTGGGAGGCCCCCGG | 247.1 | -11.8 | 1.5 | -7.7 | 1.7E-14 | 9.4E-14 | MB231/MB468 | MP |
| >hg38_21_+_8433172_8446622_RNA45SN1_45S_50ntFlanks@10408.10434.27\|GGTCAGTCGGTCCTGAGAGATGGGCGA | >hg38_21_+_8433172_8446622_RNA45SN1_45S_50ntFlanks | 10408 | 10434 | 27 | GGTCAGTCGGTCCTGAGAGATGGGCGA | 254.2 | -7.2 | 0.9 | -7.7 | 2.0E-14 | 9.4E-14 | MB231/MB468 | CYTO |
| >hg38_21_+_8433172_8446622_RNA45SN1_45S_50ntFlanks@10423.10453.31\|AGAGATGGGCGAGCGCCGTTCCGAAGGGACG | >hg38_21_+_8433172_8446622_RNA45SN1_45S_50ntFlanks | 10423 | 10453 | 31 | AGAGATGGGCGAGCGCCGTTCCGAAGGGACG | 714.1 | -4.9 | 0.6 | -7.6 | 2.1E-14 | 9.4E-14 | MB231/MB468 | NUC |
| >hg38_21_+_8433172_8446622_RNA45SN1_45S_50ntFlanks@10389.10433.45\|TGAACAGCAGTTGAACATGGGTCAGTCGGTCCTGAGAGATGGGCG | >hg38_21_+_8433172_8446622_RNA45SN1_45S_50ntFlanks | 10389 | 10433 | 45 | TGAACAGCAGTTGAACATGGGTCAGTCGGTCCTGAGAGATGGGCG | 560.7 | -13.6 | 1.8 | -7.6 | 3.4E-14 | 9.4E-14 | BT20/MB468 | NUC |
| >MI0000767\|hsa-mir-365a&WithFlank&16\|+\|14309279\|14309377@62.83.22;MI0000769\|hsa-mir-365b&WithFlank&17\|+\|31575405\|31575527@74.95.22&[MIMAT0000710&hsa-miR-365a-3p&offsets\|0\|0;m-176&16\|+\|14309340\|14309361&offsets\|0\|0];[MIMAT0022834&hsa-miR-365b-3p&offsets\|0\|0;m-175&17\|+\|31575478\|31575499&offsets\|0\|0]\|TAATGCCCCTAAAAATCCTTAT | >MI0000767\|hsa-mir-365a&WithFlank&16\|+\|14309279\|14309377 | 62 | 83 | 22 | TAATGCCCCTAAAAATCCTTAT | 1837.9 | 4.3 | 0.5 | 7.9 | 2.6E-15 | 9.4E-14 | BT20/MB231 | NUC |
| >hg38_21_+_8433172_8446622_RNA45SN1_45S_50ntFlanks@5485.5514.30\|AGCGCTGAGAAGACGGTCGAACTTGACTAT | >hg38_21_+_8433172_8446622_RNA45SN1_45S_50ntFlanks | 5485 | 5514 | 30 | AGCGCTGAGAAGACGGTCGAACTTGACTAT | 4165.0 | -5.3 | 0.7 | -7.7 | 1.7E-14 | 9.5E-14 | BT20/MB468 | MITO |
| >hg38_21_+_8433172_8446622_RNA45SN1_45S_50ntFlanks@12386.12414.29\|AGCGTTCATAGCGACGTCGCTTTTTGATC | >hg38_21_+_8433172_8446622_RNA45SN1_45S_50ntFlanks | 12386 | 12414 | 29 | AGCGTTCATAGCGACGTCGCTTTTTGATC | 443.0 | -6.7 | 0.9 | -7.7 | 2.0E-14 | 9.5E-14 | MB231/MB468 | CYTO |
| >hg38_21_+_8433172_8446622_RNA45SN1_45S_50ntFlanks@10479.10497.19\|CGATCGAAAGGGAGTCGGG | >hg38_21_+_8433172_8446622_RNA45SN1_45S_50ntFlanks | 10479 | 10497 | 19 | CGATCGAAAGGGAGTCGGG | 220.0 | -10.7 | 1.4 | -7.6 | 2.1E-14 | 9.5E-14 | MB231/MB468 | NUC |
| >hg38_21_+_8433172_8446622_RNA45SN1_45S_50ntFlanks@11593.11611.19\|CCAGGGGAATCCGACTGTT | >hg38_21_+_8433172_8446622_RNA45SN1_45S_50ntFlanks | 11593 | 11611 | 19 | CCAGGGGAATCCGACTGTT | 2408.7 | -5.7 | 0.7 | -7.6 | 2.1E-14 | 9.5E-14 | MB231/MB468 | NUC |
| >hg38_21_+_8433172_8446622_RNA45SN1_45S_50ntFlanks@11315.11343.29\|CCACCCCACGTCTCGTCGCGCGCGCGTCC | >hg38_21_+_8433172_8446622_RNA45SN1_45S_50ntFlanks | 11315 | 11343 | 29 | CCACCCCACGTCTCGTCGCGCGCGCGTCC | 188.0 | -10.3 | 1.3 | -7.7 | 2.0E-14 | 9.5E-14 | MB231/MB468 | CYTO |
| >hg38_21_+_8433172_8446622_RNA45SN1_45S_50ntFlanks@9125.9152.28\|CGCGCGGGTCGGGGGGCGGGGCGGACTG | >hg38_21_+_8433172_8446622_RNA45SN1_45S_50ntFlanks | 9125 | 9152 | 28 | CGCGCGGGTCGGGGGGCGGGGCGGACTG | 321.0 | -12.1 | 1.6 | -7.6 | 2.8E-14 | 9.5E-14 | MB231/MB468 | MITO |
| >hg38_21_+_8433172_8446622_RNA45SN1_45S_50ntFlanks@10689.10710.22\|GCGTCCGGTGAGCTCTCGCTGG | >hg38_21_+_8433172_8446622_RNA45SN1_45S_50ntFlanks | 10689 | 10710 | 22 | GCGTCCGGTGAGCTCTCGCTGG | 243.4 | -10.6 | 1.4 | -7.7 | 2.0E-14 | 9.5E-14 | MB231/MB468 | CYTO |
| >hg38_21_+_8433172_8446622_RNA45SN1_45S_50ntFlanks@5472.5501.30\|ACGGCCCTGGCGGAGCGCTGAGAAGACGGT | >hg38_21_+_8433172_8446622_RNA45SN1_45S_50ntFlanks | 5472 | 5501 | 30 | ACGGCCCTGGCGGAGCGCTGAGAAGACGGT | 23240.9 | 8.7 | 1.1 | 7.6 | 2.3E-14 | 9.5E-14 | BT20/MB468 | CYTO |
| >am_trna106_HisGTG_1_-_149155828_149155899@9.29.21,trna111_HisGTG_1_-_147774845_147774916@9.29.21,trna118_HisGTG_1_-_145396881_145396952@9.29.21,trna16_HisGTG_1_+_146544773_146544844@9.29.21,trna1_HisGTG_15_+_45493349_45493420@9.29.21,trna21_HisGTG_1_+_147753471_147753542@9.29.21,trna33_HisGTG_6_+_27125906_27125977@9.29.21,trna7_HisGTG_9_-_14433938_14434009@9.29.21,trna8_HisGTG_15_-_45492611_45492682@9.29.21,trna9_HisGTG_15_-_45490804_45490875@9.29.21\|CGTATAGTGGTTAGTACTCTG | >am_trna106_HisGTG_1_-_149155828_149155899 | 9 | 29 | 21 | CGTATAGTGGTTAGTACTCTG | 155.4 | -11.1 | 1.5 | -7.7 | 1.8E-14 | 9.5E-14 | MB231/MB468 | MP |
| >hg38_21_+_8433172_8446622_RNA45SN1_45S_50ntFlanks@4390.4423.34\|TCTTGGGAGCGGGCGGGCGGTCCGCCGCGAGGCG | >hg38_21_+_8433172_8446622_RNA45SN1_45S_50ntFlanks | 4390 | 4423 | 34 | TCTTGGGAGCGGGCGGGCGGTCCGCCGCGAGGCG | 244.1 | -8.1 | 1.1 | -7.7 | 2.0E-14 | 9.5E-14 | MB231/MB468 | CYTO |
| >hg38_21_+_8433172_8446622_RNA45SN1_45S_50ntFlanks@9574.9597.24\|AATCGGTCGTCCGACCTGGGTATA | >hg38_21_+_8433172_8446622_RNA45SN1_45S_50ntFlanks | 9574 | 9597 | 24 | AATCGGTCGTCCGACCTGGGTATA | 3401.9 | -5.8 | 0.8 | -7.6 | 2.3E-14 | 9.5E-14 | BT20/MB468 | CYTO |
| >hg38_21_+_8433172_8446622_RNA45SN1_45S_50ntFlanks@11453.11478.26\|TCCTCGGCGCGCGGCGGCGGCGGCGG | >hg38_21_+_8433172_8446622_RNA45SN1_45S_50ntFlanks | 11453 | 11478 | 26 | TCCTCGGCGCGCGGCGGCGGCGGCGG | 170.0 | -10.1 | 1.3 | -7.6 | 2.0E-14 | 9.6E-14 | MB231/MB468 | CYTO |
| >hg38_21_+_8433172_8446622_RNA45SN1_45S_50ntFlanks@9736.9758.23\|CCGAAACGATCTCAACCTATTCT | >hg38_21_+_8433172_8446622_RNA45SN1_45S_50ntFlanks | 9736 | 9758 | 23 | CCGAAACGATCTCAACCTATTCT | 211.0 | -10.4 | 1.4 | -7.6 | 2.0E-14 | 9.6E-14 | MB231/MB468 | CYTO |
| >hg38_21_+_8433172_8446622_RNA45SN1_45S_50ntFlanks@11876.11893.18\|AAGACCCTGTTGAGCTTG | >hg38_21_+_8433172_8446622_RNA45SN1_45S_50ntFlanks | 11876 | 11893 | 18 | AAGACCCTGTTGAGCTTG | 3549.6 | -4.8 | 0.6 | -7.6 | 2.3E-14 | 9.6E-14 | BT20/MB468 | CYTO |
| >hg38_21_+_8433172_8446622_RNA45SN1_45S_50ntFlanks@11667.11706.40\|CTGCCCAGTGCTCTGAATGTCAAAGTGAAGAAATTCAATG | >hg38_21_+_8433172_8446622_RNA45SN1_45S_50ntFlanks | 11667 | 11706 | 40 | CTGCCCAGTGCTCTGAATGTCAAAGTGAAGAAATTCAATG | 1079.7 | 9.4 | 1.2 | 7.6 | 2.0E-14 | 9.6E-14 | MB231/MB468 | CYTO |
| >hg38_21_+_8433172_8446622_RNA45SN1_45S_50ntFlanks@10350.10376.27\|ACGAGAACTTTGAAGGCCGAAGTGGAG | >hg38_21_+_8433172_8446622_RNA45SN1_45S_50ntFlanks | 10350 | 10376 | 27 | ACGAGAACTTTGAAGGCCGAAGTGGAG | 1223.6 | -4.2 | 0.5 | -7.6 | 2.0E-14 | 9.6E-14 | MB231/MB468 | CYTO |
| >hg38_21_+_8433172_8446622_RNA45SN1_45S_50ntFlanks@10392.10420.29\|ACAGCAGTTGAACATGGGTCAGTCGGTCC | >hg38_21_+_8433172_8446622_RNA45SN1_45S_50ntFlanks | 10392 | 10420 | 29 | ACAGCAGTTGAACATGGGTCAGTCGGTCC | 430.4 | -6.0 | 0.8 | -7.6 | 2.0E-14 | 9.6E-14 | MB231/MB468 | CYTO |
| >hg38_21_+_8433172_8446622_RNA45SN1_45S_50ntFlanks@10767.10785.19\|GCAGCAGGTCTCCAAGGTG | >hg38_21_+_8433172_8446622_RNA45SN1_45S_50ntFlanks | 10767 | 10785 | 19 | GCAGCAGGTCTCCAAGGTG | 298.3 | -12.1 | 1.6 | -7.7 | 1.8E-14 | 9.7E-14 | MB231/MB468 | MP |
| >hg38_21_+_8433172_8446622_RNA45SN1_45S_50ntFlanks@4732.4768.37\|AGAACGAAAGTCGGAGGTTCGAAGACGATCAGATACC | >hg38_21_+_8433172_8446622_RNA45SN1_45S_50ntFlanks | 4732 | 4768 | 37 | AGAACGAAAGTCGGAGGTTCGAAGACGATCAGATACC | 1567.5 | 10.1 | 1.3 | 7.6 | 2.3E-14 | 9.7E-14 | BT20/MB468 | CYTO |
| >hg38_21_+_8433172_8446622_RNA45SN1_45S_50ntFlanks@12540.12569.30\|TGATGTGTTGTTGCCATGGTAATCCTGCTC | >hg38_21_+_8433172_8446622_RNA45SN1_45S_50ntFlanks | 12540 | 12569 | 30 | TGATGTGTTGTTGCCATGGTAATCCTGCTC | 152.1 | -11.1 | 1.4 | -7.7 | 1.8E-14 | 9.7E-14 | MB231/MB468 | MP |
| >hg38_21_+_8433172_8446622_RNA45SN1_45S_50ntFlanks@11115.11133.19\|CGGCAGGGGCCGGCGGCGG | >hg38_21_+_8433172_8446622_RNA45SN1_45S_50ntFlanks | 11115 | 11133 | 19 | CGGCAGGGGCCGGCGGCGG | 275.0 | -11.9 | 1.6 | -7.6 | 2.9E-14 | 9.7E-14 | MB231/MB468 | MITO |
| >hg38_21_+_8433172_8446622_RNA45SN1_45S_50ntFlanks@12300.12317.18\|GGGCCTCACGATCCTTCT | >hg38_21_+_8433172_8446622_RNA45SN1_45S_50ntFlanks | 12300 | 12317 | 18 | GGGCCTCACGATCCTTCT | 536.2 | -12.8 | 1.7 | -7.6 | 2.9E-14 | 9.7E-14 | MB231/MB468 | MITO |
| >hg38_21_+_8433172_8446622_RNA45SN1_45S_50ntFlanks@4730.4766.37\|CAAGAACGAAAGTCGGAGGTTCGAAGACGATCAGATA | >hg38_21_+_8433172_8446622_RNA45SN1_45S_50ntFlanks | 4730 | 4766 | 37 | CAAGAACGAAAGTCGGAGGTTCGAAGACGATCAGATA | 432.6 | 6.8 | 0.9 | 7.6 | 2.1E-14 | 9.7E-14 | MB231/MB468 | CYTO |
| >MI0000542\|hsa-mir-320a&WithFlank&8\|-\|22244960\|22245043@49.69.21&[MIMAT0000510&hsa-miR-320a-3p&offsets\|+1\|0;m-60&8\|-\|22244975\|22244996&offsets\|+1\|0]\|AAAGCTGGGTTGAGAGGGCGA | >MI0000542\|hsa-mir-320a&WithFlank&8\|-\|22244960\|22245043 | 49 | 69 | 21 | AAAGCTGGGTTGAGAGGGCGA | 579.9 | 10.4 | 1.4 | 7.7 | 1.7E-14 | 9.8E-14 | BT20/MB468 | MITO |
| >hg38_21_+_8433172_8446622_RNA45SN1_45S_50ntFlanks@9539.9566.28\|ACTCTGGTGGAGGTCCGTAGCGGTCCTG | >hg38_21_+_8433172_8446622_RNA45SN1_45S_50ntFlanks | 9539 | 9566 | 28 | ACTCTGGTGGAGGTCCGTAGCGGTCCTG | 190.0 | -10.5 | 1.4 | -7.6 | 2.2E-14 | 9.8E-14 | MB231/MB468 | NUC |
| >hg38_21_+_8433172_8446622_RNA45SN1_45S_50ntFlanks@12407.12444.38\|TTTTGATCCTTCGATGTCGGCTCTTCCTATCATTGTGA | >hg38_21_+_8433172_8446622_RNA45SN1_45S_50ntFlanks | 12407 | 12444 | 38 | TTTTGATCCTTCGATGTCGGCTCTTCCTATCATTGTGA | 1797.4 | -6.2 | 0.8 | -7.6 | 3.5E-14 | 9.8E-14 | BT20/MB468 | NUC |
| >MI0002470\|hsa-mir-486-1&WithFlank&8\|-\|41660435\|41660514@10.31.22(+1U);MI0023622\|hsa-mir-486-2&WithFlank&8\|+\|41660438\|41660513@7.28.22(+1U)&[MIMAT0002177&hsa-miR-486-5p&offsets\|0\|0(+1U);m-107&8\|-\|41660484\|41660505&offsets\|0\|0(+1U)];[MIMAT0002177_1&hsa-miR-486-5p&offsets\|0\|0(+1U);TJU_CMC_MD2.ID03138.5p-miR&offsets\|0\|-2(+1U);m-106&8\|+\|41660444\|41660465&offsets\|0\|0(+1U)]\|TCCTGTACTGAGCTGCCCCGAGT | >MI0002470\|hsa-mir-486-1&WithFlank&8\|-\|41660435\|41660514 | 10 | 31 | 22(+1U) | TCCTGTACTGAGCTGCCCCGAGT | 267.2 | -11.3 | 1.4 | -7.9 | 2.4E-15 | 9.8E-14 | BT20/MB231 | MITO |
| >hg38_21_+_8433172_8446622_RNA45SN1_45S_50ntFlanks@11188.11207.20\|CACCCGGGGGGCCGGCGGCG | >hg38_21_+_8433172_8446622_RNA45SN1_45S_50ntFlanks | 11188 | 11207 | 20 | CACCCGGGGGGCCGGCGGCG | 330.1 | -12.2 | 1.6 | -7.7 | 1.8E-14 | 9.8E-14 | MB231/MB468 | MP |
| >hg38_21_+_8433172_8446622_RNA45SN1_45S_50ntFlanks@12395.12414.20\|AGCGACGTCGCTTTTTGATC | >hg38_21_+_8433172_8446622_RNA45SN1_45S_50ntFlanks | 12395 | 12414 | 20 | AGCGACGTCGCTTTTTGATC | 337.3 | 3.6 | 0.5 | 7.6 | 2.4E-14 | 9.8E-14 | BT20/MB468 | CYTO |
| >hg38_21_+_8433172_8446622_RNA45SN1_45S_50ntFlanks@10403.10430.28\|ACATGGGTCAGTCGGTCCTGAGAGATGG | >hg38_21_+_8433172_8446622_RNA45SN1_45S_50ntFlanks | 10403 | 10430 | 28 | ACATGGGTCAGTCGGTCCTGAGAGATGG | 203.2 | -11.5 | 1.5 | -7.7 | 1.8E-14 | 9.8E-14 | MB231/MB468 | MP |

<table>
<tr><td>>hg38_21_+_8433172_8446622_RNA45SN1_45S_50ntFlanks@12934.12962.29|CTAAACCATTCGTAGACGACCTGCTTCTG</td><td>>hg38_21_+_8433172_8446622_RNA45SN1_45S_50ntFlanks</td><td>12934</td><td>12962</td><td>29</td><td>CTAAACCATTCGTAGACGACCTGCTTCTG</td><td>1833.6</td><td>-10.6</td><td>1.4</td><td>-7.6</td><td>3.5E-14</td><td>9.9E-14</td><td>BT20/MB468</td><td>NUC</td></tr>
<tr><td>>hg38_21_+_8433172_8446622_RNA45SN1_45S_50ntFlanks@9739.9769.31|AAACGATCTCAACCTATTCTCAAACTTTAAA</td><td>>hg38_21_+_8433172_8446622_RNA45SN1_45S_50ntFlanks</td><td>9739</td><td>9769</td><td>31</td><td>AAACGATCTCAACCTATTCTCAAACTTTAAA</td><td>2729.0</td><td>-8.0</td><td>1.0</td><td>-7.6</td><td>2.9E-14</td><td>9.9E-14</td><td>MB231/MB468</td><td>MITO</td></tr>
<tr><td>>hg38_21_+_8433172_8446622_RNA45SN1_45S_50ntFlanks@485.507.23|GATGTGGTGACGTCGTGCTCTCC</td><td>>hg38_21_+_8433172_8446622_RNA45SN1_45S_50ntFlanks</td><td>485</td><td>507</td><td>23</td><td>GATGTGGTGACGTCGTGCTCTCC</td><td>747.5</td><td>27.8</td><td>3.6</td><td>7.6</td><td>2.1E-14</td><td>9.9E-14</td><td>MB231/MB468</td><td>CYTO</td></tr>
<tr><td>>hg38_21_+_8433172_8446622_RNA45SN1_45S_50ntFlanks@11564.11602.39|CCTAGCAGCCGACTTAGAACTGGTGCGGACCAGGGGAAT</td><td>>hg38_21_+_8433172_8446622_RNA45SN1_45S_50ntFlanks</td><td>11564</td><td>11602</td><td>39</td><td>CCTAGCAGCCGACTTAGAACTGGTGCGGACCAGGGGAAT</td><td>5210.7</td><td>-9.4</td><td>1.2</td><td>-7.6</td><td>3.5E-14</td><td>9.9E-14</td><td>BT20/MB468</td><td>NUC</td></tr>
<tr><td>>hg38_21_+_8433172_8446622_RNA45SN1_45S_50ntFlanks@11205.11227.23|GCGGCGGCGACTCTGGACGCGAG</td><td>>hg38_21_+_8433172_8446622_RNA45SN1_45S_50ntFlanks</td><td>11205</td><td>11227</td><td>23</td><td>GCGGCGGCGACTCTGGACGCGAG</td><td>167.1</td><td>-11.2</td><td>1.5</td><td>-7.7</td><td>1.8E-14</td><td>9.9E-14</td><td>MB231/MB468</td><td>MP</td></tr>
<tr><td>>hg38_21_+_8433172_8446622_RNA45SN1_45S_50ntFlanks@485.530.46|GATGTGGTGACGTCGTGCTCTCCCGGGCCGGGTCCGAGCCGCGACG</td><td>>hg38_21_+_8433172_8446622_RNA45SN1_45S_50ntFlanks</td><td>485</td><td>530</td><td>46</td><td>GATGTGGTGACGTCGTGCTCTCCCGGGCCGGGTCCGAGCCGCGACG</td><td>933.8</td><td>27.7</td><td>3.6</td><td>7.6</td><td>2.2E-14</td><td>9.9E-14</td><td>MB231/MB468</td><td>NUC</td></tr>
<tr><td>>hg38_21_+_8433172_8446622_RNA45SN1_45S_50ntFlanks@12008.12032.25|CCGCCGGTGAAATACCACTACTCTG</td><td>>hg38_21_+_8433172_8446622_RNA45SN1_45S_50ntFlanks</td><td>12008</td><td>12032</td><td>25</td><td>CCGCCGGTGAAATACCACTACTCTG</td><td>47350.2</td><td>-12.7</td><td>1.6</td><td>-8.1</td><td>7.4E-16</td><td>9.9E-14</td><td>BT20/MB468</td><td>MP</td></tr>
<tr><td>>hg38_21_+_8433172_8446622_RNA45SN1_45S_50ntFlanks@3984.4004.21|GCGGCTTTGGTGACTCTAGAT</td><td>>hg38_21_+_8433172_8446622_RNA45SN1_45S_50ntFlanks</td><td>3984</td><td>4004</td><td>21</td><td>GCGGCTTTGGTGACTCTAGAT</td><td>435.9</td><td>-12.6</td><td>1.7</td><td>-7.6</td><td>2.9E-14</td><td>9.9E-14</td><td>MB231/MB468</td><td>MITO</td></tr>
<tr><td>>hg38_21_+_8433172_8446622_RNA45SN1_45S_50ntFlanks@10905.10926.22|CTGGGCGCGCGCCGCGGCTGGA</td><td>>hg38_21_+_8433172_8446622_RNA45SN1_45S_50ntFlanks</td><td>10905</td><td>10926</td><td>22</td><td>CTGGGCGCGCGCCGCGGCTGGA</td><td>232.8</td><td>-11.6</td><td>1.5</td><td>-7.6</td><td>2.9E-14</td><td>9.9E-14</td><td>MB231/MB468</td><td>MITO</td></tr>
<tr><td>>MI0000456|hsa-mir-140&WithFlank&16|+|69933075|69933186@69.90.22(+1U)&[MIMAT0004597&hsa-miR-140-3p&offsets|+1|+2(+1U);m-45&16|+|69933143|69933164&offsets|0|0(+1U)]|ACCACAGGGTAGAACCACGGACT</td><td>>MI0000456|hsa-mir-140&WithFlank&16|+|69933075|69933186</td><td>69</td><td>90</td><td>22(+1U)</td><td>ACCACAGGGTAGAACCACGGACT</td><td>1308.9</td><td>11.1</td><td>1.4</td><td>7.6</td><td>2.1E-14</td><td>9.9E-14</td><td>MB231/MB468</td><td>CYTO</td></tr>
<tr><td>>MI0000077|hsa-mir-21&WithFlank&17|+|59841260|59841343@15.37.23&[MIMAT0000076&hsa-miR-21-5p&offsets|+1|+2;m-1&17|+|59841273|59841294&offsets|+1|+2]|AGCTTATCAGACTGATGTTGACT</td><td>>MI0000077|hsa-mir-21&WithFlank&17|+|59841260|59841343</td><td>15</td><td>37</td><td>23</td><td>AGCTTATCAGACTGATGTTGACT</td><td>210.4</td><td>6.9</td><td>0.9</td><td>7.6</td><td>2.2E-14</td><td>1.0E-13</td><td>MB231/MB468</td><td>NUC</td></tr>
<tr><td>>hg38_21_+_8433172_8446622_RNA45SN1_45S_50ntFlanks@10387.10424.38|TGTGAACAGCAGTTGAACATGGGTCAGTCGGTCCTGAG</td><td>>hg38_21_+_8433172_8446622_RNA45SN1_45S_50ntFlanks</td><td>10387</td><td>10424</td><td>38</td><td>TGTGAACAGCAGTTGAACATGGGTCAGTCGGTCCTGAG</td><td>1804.4</td><td>7.6</td><td>1.0</td><td>7.6</td><td>2.1E-14</td><td>1.0E-13</td><td>MB231/MB468</td><td>CYTO</td></tr>
<tr><td>>hg38_21_+_8433172_8446622_RNA45SN1_45S_50ntFlanks@10798.10822.25|CATGTTGGAACAATGTAGGTAAGGG</td><td>>hg38_21_+_8433172_8446622_RNA45SN1_45S_50ntFlanks</td><td>10798</td><td>10822</td><td>25</td><td>CATGTTGGAACAATGTAGGTAAGGG</td><td>474.2</td><td>-7.2</td><td>1.0</td><td>-7.6</td><td>3.0E-14</td><td>1.0E-13</td><td>MB231/MB468</td><td>MITO</td></tr>
<tr><td>>hg38_21_+_8433172_8446622_RNA45SN1_45S_50ntFlanks@12942.12964.23|TTCGTAGACGACCTGCTTCTGGG</td><td>>hg38_21_+_8433172_8446622_RNA45SN1_45S_50ntFlanks</td><td>12942</td><td>12964</td><td>23</td><td>TTCGTAGACGACCTGCTTCTGGG</td><td>208.6</td><td>-11.6</td><td>1.5</td><td>-7.7</td><td>1.9E-14</td><td>1.0E-13</td><td>MB231/MB468</td><td>MP</td></tr>
<tr><td>>hg38_21_+_8433172_8446622_RNA45SN1_45S_50ntFlanks@11315.11342.28|CCACCCCACGTCTCGTCGCGCGCGCGTC</td><td>>hg38_21_+_8433172_8446622_RNA45SN1_45S_50ntFlanks</td><td>11315</td><td>11342</td><td>28</td><td>CCACCCCACGTCTCGTCGCGCGCGCGTC</td><td>302.0</td><td>-12.1</td><td>1.6</td><td>-7.7</td><td>1.9E-14</td><td>1.0E-13</td><td>MB231/MB468</td><td>MP</td></tr>
<tr><td>>hg38_21_+_8433172_8446622_RNA45SN1_45S_50ntFlanks@10912.10930.19|CGCGCCGCGGCTGGACGAG</td><td>>hg38_21_+_8433172_8446622_RNA45SN1_45S_50ntFlanks</td><td>10912</td><td>10930</td><td>19</td><td>CGCGCCGCGGCTGGACGAG</td><td>270.2</td><td>-11.9</td><td>1.6</td><td>-7.7</td><td>1.9E-14</td><td>1.0E-13</td><td>MB231/MB468</td><td>MP</td></tr>
<tr><td>>hg38_21_+_8433172_8446622_RNA45SN1_45S_50ntFlanks@9310.9335.26|GAGTCTAACACGTGCGCGAGTCGGGG</td><td>>hg38_21_+_8433172_8446622_RNA45SN1_45S_50ntFlanks</td><td>9310</td><td>9335</td><td>26</td><td>GAGTCTAACACGTGCGCGAGTCGGGG</td><td>376.4</td><td>-12.4</td><td>1.6</td><td>-7.7</td><td>1.9E-14</td><td>1.0E-13</td><td>MB231/MB468</td><td>MP</td></tr>
<tr><td>>hg38_21_+_8433172_8446622_RNA45SN1_45S_50ntFlanks@11853.11872.20|CTTGGCGGAATCAGCGGGGA</td><td>>hg38_21_+_8433172_8446622_RNA45SN1_45S_50ntFlanks</td><td>11853</td><td>11872</td><td>20</td><td>CTTGGCGGAATCAGCGGGGA</td><td>213.8</td><td>-10.7</td><td>1.4</td><td>-7.6</td><td>2.2E-14</td><td>1.0E-13</td><td>MB231/MB468</td><td>NUC</td></tr>
<tr><td>>hg38_21_+_8433172_8446622_RNA45SN1_45S_50ntFlanks@11894.11912.19|ACTCTAGTCTGGCACGGTG</td><td>>hg38_21_+_8433172_8446622_RNA45SN1_45S_50ntFlanks</td><td>11894</td><td>11912</td><td>19</td><td>ACTCTAGTCTGGCACGGTG</td><td>1295.4</td><td>-9.4</td><td>1.2</td><td>-7.6</td><td>2.2E-14</td><td>1.0E-13</td><td>MB231/MB468</td><td>NUC</td></tr>
<tr><td>>MI0000082|hsa-mir-25&WithFlank&7|-|100093554|100093649@58.79.22&[MIMAT0000081&hsa-miR-25-3p&offsets|0|0;m-23&7|-|100093571|100093592&offsets|0|0]|CATTGCACTTGTCTCGGTCTGA</td><td>>MI0000082|hsa-mir-25&WithFlank&7|-|100093554|100093649</td><td>58</td><td>79</td><td>22</td><td>CATTGCACTTGTCTCGGTCTGA</td><td>53383.5</td><td>4.6</td><td>0.6</td><td>7.6</td><td>2.2E-14</td><td>1.0E-13</td><td>MB231/MB468</td><td>NUC</td></tr>
<tr><td>>hg38_21_+_8433172_8446622_RNA45SN1_45S_50ntFlanks@10116.10133.18|TGCGTCGGAGGGCGGCGG</td><td>>hg38_21_+_8433172_8446622_RNA45SN1_45S_50ntFlanks</td><td>10116</td><td>10133</td><td>18</td><td>TGCGTCGGAGGGCGGCGG</td><td>181.5</td><td>-11.4</td><td>1.5</td><td>-7.7</td><td>1.9E-14</td><td>1.0E-13</td><td>MB231/MB468</td><td>MP</td></tr>
<tr><td>>hg38_21_+_8433172_8446622_RNA45SN1_45S_50ntFlanks@10383.10429.47|TCCATGTGAACAGCAGTTGAACATGGGTCAGTCGGTCCTGAGAGATG</td><td>>hg38_21_+_8433172_8446622_RNA45SN1_45S_50ntFlanks</td><td>10383</td><td>10429</td><td>47</td><td>TCCATGTGAACAGCAGTTGAACATGGGTCAGTCGGTCCTGAGAGATG</td><td>358.3</td><td>9.2</td><td>1.2</td><td>7.6</td><td>2.1E-14</td><td>1.0E-13</td><td>MB231/MB468</td><td>CYTO</td></tr>
<tr><td>>hg38_21_+_8433172_8446622_RNA45SN1_45S_50ntFlanks@11873.11899.27|AAGAAGACCCTGTTGAGCTTGACTCTA</td><td>>hg38_21_+_8433172_8446622_RNA45SN1_45S_50ntFlanks</td><td>11873</td><td>11899</td><td>27</td><td>AAGAAGACCCTGTTGAGCTTGACTCTA</td><td>320.3</td><td>-3.3</td><td>0.4</td><td>-7.6</td><td>2.4E-14</td><td>1.0E-13</td><td>BT20/MB468</td><td>CYTO</td></tr>
<tr><td>>hg38_21_+_8433172_8446622_RNA45SN1_45S_50ntFlanks@10350.10368.19|ACGAGAACTTTGAAGGCCG</td><td>>hg38_21_+_8433172_8446622_RNA45SN1_45S_50ntFlanks</td><td>10350</td><td>10368</td><td>19</td><td>ACGAGAACTTTGAAGGCCG</td><td>11717.3</td><td>-9.2</td><td>1.2</td><td>-7.7</td><td>1.9E-14</td><td>1.0E-13</td><td>MB231/MB468</td><td>MP</td></tr>
<tr><td>>hg38_21_+_8433172_8446622_RNA45SN1_45S_50ntFlanks@11575.11601.27|ACTTAGAACTGGTGCGGACCAGGGGAA</td><td>>hg38_21_+_8433172_8446622_RNA45SN1_45S_50ntFlanks</td><td>11575</td><td>11601</td><td>27</td><td>ACTTAGAACTGGTGCGGACCAGGGGAA</td><td>2145.3</td><td>-9.7</td><td>1.3</td><td>-7.6</td><td>2.2E-14</td><td>1.0E-13</td><td>MB231/MB468</td><td>NUC</td></tr>
<tr><td>>MI0000079|hsa-mir-23a&WithFlank&19|-|13836581|13836665@54.73.20&[MIMAT0000078&hsa-miR-23a-3p&offsets|+3|+2;m-31&19|-|13836595|13836615&offsets|+3|+2]|ACATTGCCAGGGATTTCCAA</td><td>>MI0000079|hsa-mir-23a&WithFlank&19|-|13836581|13836665</td><td>54</td><td>73</td><td>20</td><td>ACATTGCCAGGGATTTCCAA</td><td>1476.9</td><td>27.6</td><td>3.6</td><td>7.6</td><td>3.0E-14</td><td>1.0E-13</td><td>MB231/MB468</td><td>MITO</td></tr>
<tr><td>>hg38_21_+_8433172_8446622_RNA45SN1_45S_50ntFlanks@8104.8136.33|CCGCGGCGGGGCGCGGGACATGTGGCGTACGGA</td><td>>hg38_21_+_8433172_8446622_RNA45SN1_45S_50ntFlanks</td><td>8104</td><td>8136</td><td>33</td><td>CCGCGGCGGGGCGCGGGACATGTGGCGTACGGA</td><td>483.9</td><td>-12.7</td><td>1.7</td><td>-7.6</td><td>3.0E-14</td><td>1.0E-13</td><td>MB231/MB468</td><td>MITO</td></tr>
<tr><td>>hg38_21_+_8433172_8446622_RNA45SN1_45S_50ntFlanks@3705.3726.22|TACCTGGTTGATCCTGCCAGTA</td><td>>hg38_21_+_8433172_8446622_RNA45SN1_45S_50ntFlanks</td><td>3705</td><td>3726</td><td>22</td><td>TACCTGGTTGATCCTGCCAGTA</td><td>1405.5</td><td>10.1</td><td>1.3</td><td>7.6</td><td>2.5E-14</td><td>1.0E-13</td><td>BT20/MB468</td><td>CYTO</td></tr>
<tr><td>>hg38_21_+_8433172_8446622_RNA45SN1_45S_50ntFlanks@8356.8380.25|GAAAAGAACTTTGAAGAGAGAGTTC</td><td>>hg38_21_+_8433172_8446622_RNA45SN1_45S_50ntFlanks</td><td>8356</td><td>8380</td><td>25</td><td>GAAAAGAACTTTGAAGAGAGAGTTC</td><td>254.1</td><td>-11.9</td><td>1.5</td><td>-7.7</td><td>1.9E-14</td><td>1.0E-13</td><td>MB231/MB468</td><td>MP</td></tr>
<tr><td>>hg38_1_-_228634819_228635039_RNA5S12_5S_50ntFlanks@124.147.24|AGTACTTGGATGGGAGACCGCCTG</td><td>>hg38_1_-_228634819_228635039_RNA5S12_5S_50ntFlanks</td><td>124</td><td>147</td><td>24</td><td>AGTACTTGGATGGGAGACCGCCTG</td><td>368.5</td><td>-12.4</td><td>1.6</td><td>-7.7</td><td>1.9E-14</td><td>1.0E-13</td><td>MB231/MB468</td><td>MP</td></tr>
<tr><td>>hg38_21_+_8433172_8446622_RNA45SN1_45S_50ntFlanks@5489.5510.22|CTGAGAAGACGGTCGAACTTGA</td><td>>hg38_21_+_8433172_8446622_RNA45SN1_45S_50ntFlanks</td><td>5489</td><td>5510</td><td>22</td><td>CTGAGAAGACGGTCGAACTTGA</td><td>1145.3</td><td>-9.5</td><td>1.2</td><td>-7.7</td><td>1.9E-14</td><td>1.0E-13</td><td>MB231/MB468</td><td>MP</td></tr>
<tr><td>>hg38_21_+_8433172_8446622_RNA45SN1_45S_50ntFlanks@10474.10497.24|TCGGCCGATCGAAAGGGAGTCGGG</td><td>>hg38_21_+_8433172_8446622_RNA45SN1_45S_50ntFlanks</td><td>10474</td><td>10497</td><td>24</td><td>TCGGCCGATCGAAAGGGAGTCGGG</td><td>238.9</td><td>-11.8</td><td>1.5</td><td>-7.7</td><td>1.9E-14</td><td>1.0E-13</td><td>MB231/MB468</td><td>MP</td></tr>
<tr><td>>hg38_21_+_8433172_8446622_RNA45SN1_45S_50ntFlanks@10351.10368.18|CGAGAACTTTGAAGGCCG</td><td>>hg38_21_+_8433172_8446622_RNA45SN1_45S_50ntFlanks</td><td>10351</td><td>10368</td><td>18</td><td>CGAGAACTTTGAAGGCCG</td><td>7495.3</td><td>-5.6</td><td>0.7</td><td>-7.7</td><td>1.8E-14</td><td>1.0E-13</td><td>BT20/MB468</td><td>MITO</td></tr>
<tr><td>>hg38_21_+_8433172_8446622_RNA45SN1_45S_50ntFlanks@5485.5508.24|AGCGCTGAGAAGACGGTCGAACTT</td><td>>hg38_21_+_8433172_8446622_RNA45SN1_45S_50ntFlanks</td><td>5485</td><td>5508</td><td>24</td><td>AGCGCTGAGAAGACGGTCGAACTT</td><td>136.7</td><td>-11.0</td><td>1.4</td><td>-7.7</td><td>1.9E-14</td><td>1.0E-13</td><td>MB231/MB468</td><td>MP</td></tr>
<tr><td>>MI0001448|hsa-mir-425&WithFlank&3|-|49020142|49020240@20.43.24&[MIMAT0003393&hsa-miR-425-5p&offsets|0|+1;m-95&3|-|49020200|49020221&offsets|0|+2]|AATGACACGATCACTCCCGTTGAG</td><td>>MI0001448|hsa-mir-425&WithFlank&3|-|49020142|49020240</td><td>20</td><td>43</td><td>24</td><td>AATGACACGATCACTCCCGTTGAG</td><td>909.1</td><td>6.6</td><td>0.9</td><td>7.7</td><td>1.8E-14</td><td>1.0E-13</td><td>BT20/MB468</td><td>MITO</td></tr>
<tr><td>>am_trna106_HisGTG_1_-_149155828_149155899@11.29.19,trna111_HisGTG_1_-_147774845_147774916@11.29.19,trna118_HisGTG_1_-_145396881_145396952@11.29.19,trna16_HisGTG_1_+_146544773_146544844@11.29.19,trna1_HisGTG_15_+_45493349_45493420@11.29.19,trna21_HisGTG_1_+_147753471_147753542@11.29.19,trna33_HisGTG_6_+_27125906_27125977@11.29.19,trna7_HisGTG_9_-_14433938_14434009@11.29.19,trna8_HisGTG_15_-_45492611_45492682@11.29.19,trna9_HisGTG_15_-_45490804_45490875@11.29.19|TATAGTGGTTAGTACTCTG</td><td>>am_trna106_HisGTG_1_-_149155828_149155899</td><td>11</td><td>29</td><td>19</td><td>TATAGTGGTTAGTACTCTG</td><td>145.9</td><td>-11.1</td><td>1.4</td><td>-7.7</td><td>1.9E-14</td><td>1.0E-13</td><td>MB231/MB468</td><td>MP</td></tr>
<tr><td>>hg38_21_+_8433172_8446622_RNA45SN1_45S_50ntFlanks@11452.11472.21|CTCCTCGGCGCGCGGCGGCGG</td><td>>hg38_21_+_8433172_8446622_RNA45SN1_45S_50ntFlanks</td><td>11452</td><td>11472</td><td>21</td><td>CTCCTCGGCGCGCGGCGGCGG</td><td>219.1</td><td>-11.6</td><td>1.5</td><td>-7.7</td><td>1.9E-14</td><td>1.0E-13</td><td>MB231/MB468</td><td>MP</td></tr>
<tr><td>>hg38_21_+_8433172_8446622_RNA45SN1_45S_50ntFlanks@9572.9594.23|CAAATCGGTCGTCCGACCTGGGT</td><td>>hg38_21_+_8433172_8446622_RNA45SN1_45S_50ntFlanks</td><td>9572</td><td>9594</td><td>23</td><td>CAAATCGGTCGTCCGACCTGGGT</td><td>2025.6</td><td>-7.5</td><td>1.0</td><td>-7.6</td><td>3.0E-14</td><td>1.0E-13</td><td>MB231/MB468</td><td>MITO</td></tr>
<tr><td>>hg38_21_+_8433172_8446622_RNA45SN1_45S_50ntFlanks@11573.11600.28|CGACTTAGAACTGGTGCGGACCAGGGGA</td><td>>hg38_21_+_8433172_8446622_RNA45SN1_45S_50ntFlanks</td><td>11573</td><td>11600</td><td>28</td><td>CGACTTAGAACTGGTGCGGACCAGGGGA</td><td>767.2</td><td>-4.2</td><td>0.5</td><td>-7.6</td><td>2.3E-14</td><td>1.0E-13</td><td>MB231/MB468</td><td>NUC</td></tr>
<tr><td>>hg38_21_+_8433172_8446622_RNA45SN1_45S_50ntFlanks@11321.11340.20|CACGTCTCGTCGCGCGCGCG</td><td>>hg38_21_+_8433172_8446622_RNA45SN1_45S_50ntFlanks</td><td>11321</td><td>11340</td><td>20</td><td>CACGTCTCGTCGCGCGCGCG</td><td>365.8</td><td>-6.6</td><td>0.9</td><td>-7.6</td><td>2.2E-14</td><td>1.0E-13</td><td>MB231/MB468</td><td>CYTO</td></tr>
<tr><td>>MI0000542|hsa-mir-320a&WithFlank&8|-|22244960|22245043@49.68.20&[MIMAT0000510&hsa-miR-320a-3p&offsets|+1|-1;m-60&8|-|22244975|22244996&offsets|+1|-1]|AAAGCTGGGTTGAGAGGGCG</td><td>>MI0000542|hsa-mir-320a&WithFlank&8|-|22244960|22245043</td><td>49</td><td>68</td><td>20</td><td>AAAGCTGGGTTGAGAGGGCG</td><td>71.3</td><td>2.1</td><td>0.3</td><td>7.6</td><td>3.7E-14</td><td>1.0E-13</td><td>BT20/MB468</td><td>NUC</td></tr>
<tr><td>>hg38_21_+_8433172_8446622_RNA45SN1_45S_50ntFlanks@8168.8203.36|CCCAAGTCCTTCTGATCGAGGCCCAGCCCGTGGACG</td><td>>hg38_21_+_8433172_8446622_RNA45SN1_45S_50ntFlanks</td><td>8168</td><td>8203</td><td>36</td><td>CCCAAGTCCTTCTGATCGAGGCCCAGCCCGTGGACG</td><td>374.0</td><td>-12.3</td><td>1.6</td><td>-7.6</td><td>3.1E-14</td><td>1.0E-13</td><td>MB231/MB468</td><td>MITO</td></tr>
<tr><td>>MI0000456|hsa-mir-140&WithFlank&16|+|69933075|69933186@69.90.22(+1U)&[MIMAT0004597&hsa-miR-140-3p&offsets|+1|+2(+1U);m-45&16|+|69933143|69933164&offsets|0|0(+1U)]|ACCACAGGGTAGAACCACGGACT</td><td>>MI0000456|hsa-mir-140&WithFlank&16|+|69933075|69933186</td><td>69</td><td>90</td><td>22(+1U)</td><td>ACCACAGGGTAGAACCACGGACT</td><td>645.2</td><td>7.9</td><td>1.0</td><td>7.6</td><td>2.3E-14</td><td>1.0E-13</td><td>MB231/MB468</td><td>NUC</td></tr>
<tr><td>>hg38_21_+_8433172_8446622_RNA45SN1_45S_50ntFlanks@12183.12211.29|AAACGGTAACGCAGGTGTCCTAAGGCGAG</td><td>>hg38_21_+_8433172_8446622_RNA45SN1_45S_50ntFlanks</td><td>12183</td><td>12211</td><td>29</td><td>AAACGGTAACGCAGGTGTCCTAAGGCGAG</td><td>221.2</td><td>-11.6</td><td>1.5</td><td>-7.6</td><td>3.1E-14</td><td>1.0E-13</td><td>MB231/MB468</td><td>MITO</td></tr>
<tr><td>>hg38_21_+_8433172_8446622_RNA45SN1_45S_50ntFlanks@7989.8020.32|AGACGTGGCGACCCGCTGAATTTAAGCATATT</td><td>>hg38_21_+_8433172_8446622_RNA45SN1_45S_50ntFlanks</td><td>7989</td><td>8020</td><td>32</td><td>AGACGTGGCGACCCGCTGAATTTAAGCATATT</td><td>173.9</td><td>7.8</td><td>1.0</td><td>7.6</td><td>2.2E-14</td><td>1.0E-13</td><td>MB231/MB468</td><td>CYTO</td></tr>
<tr><td>>hg38_21_+_8433172_8446622_RNA45SN1_45S_50ntFlanks@5472.5489.18|ACGGCCCTGGCGGAGCGC</td><td>>hg38_21_+_8433172_8446622_RNA45SN1_45S_50ntFlanks</td><td>5472</td><td>5489</td><td>18</td><td>ACGGCCCTGGCGGAGCGC</td><td>3217.4</td><td>10.9</td><td>1.4</td><td>7.6</td><td>2.5E-14</td><td>1.0E-13</td><td>BT20/MB468</td><td>CYTO</td></tr>
<tr><td>>hg38_21_+_8433172_8446622_RNA45SN1_45S_50ntFlanks@12286.12318.33|AGACCGTGAAAGCGGGGCCTCACGATCCTTCTG</td><td>>hg38_21_+_8433172_8446622_RNA45SN1_45S_50ntFlanks</td><td>12286</td><td>12318</td><td>33</td><td>AGACCGTGAAAGCGGGGCCTCACGATCCTTCTG</td><td>470.5</td><td>-12.7</td><td>1.7</td><td>-7.6</td><td>3.1E-14</td><td>1.0E-13</td><td>MB231/MB468</td><td>MITO</td></tr>
<tr><td>>hg38_21_+_8433172_8446622_RNA45SN1_45S_50ntFlanks@10427.10457.31|ATGGGCGAGCGCCGTTCCGAAGGGACGGGCG</td><td>>hg38_21_+_8433172_8446622_RNA45SN1_45S_50ntFlanks</td><td>10427</td><td>10457</td><td>31</td><td>ATGGGCGAGCGCCGTTCCGAAGGGACGGGCG</td><td>174.2</td><td>-11.3</td><td>1.5</td><td>-7.7</td><td>1.9E-14</td><td>1.0E-13</td><td>MB231/MB468</td><td>MP</td></tr>
</table>

| | | | | | | | | | | | | | |
|---|---|---|---|---|---|---|---|---|---|---|---|---|---|
| >hg38_21_+_8433172_8446622_RNA45SN1_45S_50ntFlanks@8117.8142.26\|CGGGACATGTGGCGTACGGAAGACCC | >hg38_21_+_8433172_8446622_RNA45SN1_45S_50ntFlanks | 8117 | 8142 | 26 | CGGGACATGTGGCGTACGGAAGACCC | 181.8 | -11.4 | 1.5 | -7.7 | 1.9E-14 | 1.0E-13 | MB231/MB468 | MP |
| >hg38_21_+_8433172_8446622_RNA45SN1_45S_50ntFlanks@4560.4579.20\|CTAGGAATAATGGAATAGGA | >hg38_21_+_8433172_8446622_RNA45SN1_45S_50ntFlanks | 4560 | 4579 | 20 | CTAGGAATAATGGAATAGGA | 377.4 | -4.0 | 0.5 | -7.6 | 2.5E-14 | 1.0E-13 | BT20/MB468 | CYTO |
| >hg38_1_-_228634819_228635039_RNA5S12_5S_50ntFlanks@135.171.37\|GGGAGACCGCCTGGGAATACCGGGTGCTGTAGGCTTT | >hg38_1_-_228634819_228635039_RNA5S12_5S_50ntFlanks | 135 | 171 | 37 | GGGAGACCGCCTGGGAATACCGGGTGCTGTAGGCTTT | 16311.8 | 6.6 | 0.9 | 7.6 | 2.2E-14 | 1.0E-13 | MB231/MB468 | CYTO |
| >hg38_21_+_8433172_8446622_RNA45SN1_45S_50ntFlanks@10571.10596.26\|GAGAAGCCGGCGGGAGCCCCGGGGAG | >hg38_21_+_8433172_8446622_RNA45SN1_45S_50ntFlanks | 10571 | 10596 | 26 | GAGAAGCCGGCGGGAGCCCCGGGGAG | 191.4 | -10.3 | 1.3 | -7.6 | 2.2E-14 | 1.1E-13 | MB231/MB468 | CYTO |
| >hg38_1_-_228634819_228635039_RNA5S12_5S_50ntFlanks@141.170.30\|CCGCCTGGGAATACCGGGTGCTGTAGGCTT | >hg38_1_-_228634819_228635039_RNA5S12_5S_50ntFlanks | 141 | 170 | 30 | CCGCCTGGGAATACCGGGTGCTGTAGGCTT | 4290.3 | 5.7 | 0.7 | 7.6 | 2.5E-14 | 1.1E-13 | BT20/MB468 | CYTO |
| >hg38_21_+_8433172_8446622_RNA45SN1_45S_50ntFlanks@12625.12653.29\|CAATGGGGCGAAGCTACCATCTGTGGGAT | >hg38_21_+_8433172_8446622_RNA45SN1_45S_50ntFlanks | 12625 | 12653 | 29 | CAATGGGGCGAAGCTACCATCTGTGGGAT | 1136.4 | -6.2 | 0.8 | -7.6 | 2.2E-14 | 1.1E-13 | MB231/MB468 | CYTO |
| >MI0000087\|hsa-mir-29a&WithFlank&7\|-\|130876741\|130876816@47.67.21(+1A)&[MIMAT0000086&hsa-miR-29a-3p&offsets\|-1\|-2(+1A);m-26&7\|-\|130876749\|130876769&offsets\|-1\|-1(+1A)]\|CTAGCACCATCTGAAATCGGTA | >MI0000087\|hsa-mir-29a&WithFlank&7\|-\|130876741\|130876816 | 47 | 67 | 21(+1A) | CTAGCACCATCTGAAATCGGTA | 283.1 | 8.4 | 1.1 | 7.6 | 2.3E-14 | 1.1E-13 | MB231/MB468 | NUC |
| >am_MI0000809\|hsa-mir-151a&WithFlank&8\|-\|140732558\|140732659@53.73.21(+1G)&[MIMAT0000757&hsa-miR-151a-3p&offsets\|0\|0(+1G);m-30&8\|-\|140732587\|140732607&offsets\|0\|0(+1G)]\|CTAGACTGAAGCTCCTTGAGGG | >am_MI0000809\|hsa-mir-151a&WithFlank&8\|-\|140732558\|140732659 | 53 | 73 | 21(+1G) | CTAGACTGAAGCTCCTTGAGGG | 293.2 | 5.4 | 0.7 | 7.6 | 2.4E-14 | 1.1E-13 | MB231/MB468 | NUC |
| >hg38_21_+_8433172_8446622_RNA45SN1_45S_50ntFlanks@3899.3918.20\|CCTTCGCGGGGGGGATGCGT | >hg38_21_+_8433172_8446622_RNA45SN1_45S_50ntFlanks | 3899 | 3918 | 20 | CCTTCGCGGGGGGGATGCGT | 276.5 | -12.0 | 1.6 | -7.7 | 2.0E-14 | 1.1E-13 | MB231/MB468 | MP |
| >hg38_21_+_8433172_8446622_RNA45SN1_45S_50ntFlanks@4443.4460.18\|CCCTTGCCTCTCGGCGCC | >hg38_21_+_8433172_8446622_RNA45SN1_45S_50ntFlanks | 4443 | 4460 | 18 | CCCTTGCCTCTCGGCGCC | 257.5 | -11.9 | 1.6 | -7.7 | 2.0E-14 | 1.1E-13 | MB231/MB468 | MP |
| >am_trna10_LysCTT_16_+_3241501_3241573@31.51.21,trna119_LysCTT_1_-_145395522_145395594@31.51.21,trna11_LysCTT_5_-_180648979_180649051@31.51.21,trna13_LysCTT_16_+_3246153_3246225@31.51.21,trna13_LysCTT_6_+_26556774_26556846@31.51.21,trna30_LysCTT_16_-_3230555_3230627@31.51.21,trna5_LysCTT_16_+_3214939_3215011@31.51.21,trna7_LysCTT_16_+_3225692_3225764@31.51.21,trna8_LysCTT_1_+_39970195_39970267@31.51.21,trna9_LysCTT_5_+_180634755_180634827@31.51.21\|ACTCTTAATCTCAGGGTCGTG | >am_trna10_LysCTT_16_+_3241501_3241573 | 31 | 51 | 21 | ACTCTTAATCTCAGGGTCGTG | 207.7 | -11.5 | 1.5 | -7.6 | 3.1E-14 | 1.1E-13 | MB231/MB468 | MITO |
| >hg38_21_+_8433172_8446622_RNA45SN1_45S_50ntFlanks@12488.12521.34\|AACGTGAGCTGGGTTTAGACCGTCGTGAGACAGG | >hg38_21_+_8433172_8446622_RNA45SN1_45S_50ntFlanks | 12488 | 12521 | 34 | AACGTGAGCTGGGTTTAGACCGTCGTGAGACAGG | 1017.0 | -9.5 | 1.3 | -7.6 | 3.1E-14 | 1.1E-13 | MB231/MB468 | MITO |
| >trnaMT_GluTTC_MT_-_14674_14742@32.72.41,trnalookalike8_GluTTC_5_-_93905172_93905240@32.72.41\|TCATATCATTGGTCGTGGTTGTAGTCCGTGCGAGAATACCA | >trnaMT_GluTTC_MT_-_14674_14742 | 32 | 72 | 41 | TCATATCATTGGTCGTGGTTGTAGTCCGTGCGAGAATACCA | 1414.2 | 5.8 | 0.8 | 7.7 | 1.9E-14 | 1.1E-13 | BT20/MB468 | MITO |
| >hg38_21_+_8433172_8446622_RNA45SN1_45S_50ntFlanks@4560.4587.28\|CTAGGAATAATGGAATAGGACCGCGGTT | >hg38_21_+_8433172_8446622_RNA45SN1_45S_50ntFlanks | 4560 | 4587 | 28 | CTAGGAATAATGGAATAGGACCGCGGTT | 723.0 | -8.9 | 1.2 | -7.6 | 2.4E-14 | 1.1E-13 | MB231/MB468 | NUC |
| >hg38_21_+_8433172_8446622_RNA45SN1_45S_50ntFlanks@10544.10564.21\|TCCAGTGCGGTAACGCGACCG | >hg38_21_+_8433172_8446622_RNA45SN1_45S_50ntFlanks | 10544 | 10564 | 21 | TCCAGTGCGGTAACGCGACCG | 167.1 | -11.2 | 1.5 | -7.7 | 2.0E-14 | 1.1E-13 | MB231/MB468 | MP |
| >hg38_21_+_8433172_8446622_RNA45SN1_45S_50ntFlanks@10905.10941.37\|CTGGGCGCGCGCCGCGGCTGGACGAGGCGCCGCCGCC | >hg38_21_+_8433172_8446622_RNA45SN1_45S_50ntFlanks | 10905 | 10941 | 37 | CTGGGCGCGCGCCGCGGCTGGACGAGGCGCCGCCGCC | 390.2 | -11.3 | 1.5 | -7.6 | 2.3E-14 | 1.1E-13 | MB231/MB468 | CYTO |
| >hg38_21_+_8433172_8446622_RNA45SN1_45S_50ntFlanks@12126.12146.21\|CGACCCGCTCCGGGGACAGTG | >hg38_21_+_8433172_8446622_RNA45SN1_45S_50ntFlanks | 12126 | 12146 | 21 | CGACCCGCTCCGGGGACAGTG | 147.6 | -10.2 | 1.3 | -7.6 | 2.4E-14 | 1.1E-13 | MB231/MB468 | NUC |
| >hg38_21_+_8433172_8446622_RNA45SN1_45S_50ntFlanks@12073.12091.19\|CCGAGGGGCTCTCGCTTCT | >hg38_21_+_8433172_8446622_RNA45SN1_45S_50ntFlanks | 12073 | 12091 | 19 | CCGAGGGGCTCTCGCTTCT | 144.7 | -11.0 | 1.4 | -7.7 | 2.0E-14 | 1.1E-13 | MB231/MB468 | MP |
| >hg38_21_+_8433172_8446622_RNA45SN1_45S_50ntFlanks@9567.9597.31\|ACGTGCAAATCGGTCGTCCGACCTGGGTATA | >hg38_21_+_8433172_8446622_RNA45SN1_45S_50ntFlanks | 9567 | 9597 | 31 | ACGTGCAAATCGGTCGTCCGACCTGGGTATA | 4218.7 | -10.5 | 1.4 | -7.7 | 2.0E-14 | 1.1E-13 | MB231/MB468 | MP |
| >hg38_21_+_8433172_8446622_RNA45SN1_45S_50ntFlanks@11453.11471.19\|TCCTCGGCGCGCGGCGGCG | >hg38_21_+_8433172_8446622_RNA45SN1_45S_50ntFlanks | 11453 | 11471 | 19 | TCCTCGGCGCGCGGCGGCG | 198.8 | -7.1 | 0.9 | -7.6 | 2.3E-14 | 1.1E-13 | MB231/MB468 | CYTO |
| >MI0003581\|hsa-mir-574&WithFlank&4\|+\|38868026\|38868133@67.88.22&[MIMAT0003239&hsa-miR-574-3p&offsets\|0\|0;m-149&4\|+\|38868092\|38868113&offsets\|0\|0]\|CACGCTCATGCACACACCCACA | >MI0003581\|hsa-mir-574&WithFlank&4\|+\|38868026\|38868133 | 67 | 88 | 22 | CACGCTCATGCACACACCCACA | 6323.5 | 8.0 | 1.0 | 7.6 | 2.3E-14 | 1.1E-13 | MB231/MB468 | CYTO |
| >trna119_LysCTT_1_-_145395522_145395594@1.30.30,trna11_LysCTT_5_-_180648979_180649051@1.30.30,trna13_LysCTT_6_+_26556774_26556846@1.30.30,trna32_LysCTT_16_-_3207406_3207478@1.30.30,trna7_LysCTT_16_+_3225692_3225764@1.30.30,trna9_LysCTT_5_+_180634755_180634827@1.30.30\|GCCCGGCTAGCTCAGTCGGTAGAGCATGAG | >trna119_LysCTT_1_-_145395522_145395594 | 1 | 30 | 30 | GCCCGGCTAGCTCAGTCGGTAGAGCATGAG | 11236.9 | -5.8 | 0.8 | -7.6 | 3.2E-14 | 1.1E-13 | MB231/MB468 | MITO |
| >hg38_21_+_8433172_8446622_RNA45SN1_45S_50ntFlanks@9572.9603.32\|CAAATCGGTCGTCCGACCTGGGTATAGGGGCG | >hg38_21_+_8433172_8446622_RNA45SN1_45S_50ntFlanks | 9572 | 9603 | 32 | CAAATCGGTCGTCCGACCTGGGTATAGGGGCG | 10216.5 | -6.4 | 0.8 | -7.6 | 3.2E-14 | 1.1E-13 | MB231/MB468 | MITO |
| >am_trnaMT_ValTAC_MT_+_1602_1670@50.68.19\|ACTTAACTTGACCGCTCTG | >am_trnaMT_ValTAC_MT_+_1602_1670 | 50 | 68 | 19 | ACTTAACTTGACCGCTCTG | 3874.7 | -8.3 | 1.1 | -7.7 | 2.0E-14 | 1.1E-13 | MB231/MB468 | MP |
| >hg38_21_+_8433172_8446622_RNA45SN1_45S_50ntFlanks@12001.12030.30\|CTGCGGGCCGCCGGTGAAATACCACTACTC | >hg38_21_+_8433172_8446622_RNA45SN1_45S_50ntFlanks | 12001 | 12030 | 30 | CTGCGGGCCGCCGGTGAAATACCACTACTC | 155.7 | -11.1 | 1.5 | -7.7 | 2.0E-14 | 1.1E-13 | MB231/MB468 | MP |
| >hg38_21_+_8433172_8446622_RNA45SN1_45S_50ntFlanks@11319.11344.26\|CCCACGTCTCGTCGCGCGCGCGTCCG | >hg38_21_+_8433172_8446622_RNA45SN1_45S_50ntFlanks | 11319 | 11344 | 26 | CCCACGTCTCGTCGCGCGCGCGTCCG | 182.9 | -11.4 | 1.5 | -7.6 | 2.0E-14 | 1.1E-13 | MB231/MB468 | MP |
| >hg38_21_+_8433172_8446622_RNA45SN1_45S_50ntFlanks@10808.10827.20\|CAATGTAGGTAAGGGAAGTC | >hg38_21_+_8433172_8446622_RNA45SN1_45S_50ntFlanks | 10808 | 10827 | 20 | CAATGTAGGTAAGGGAAGTC | 489.8 | -11.0 | 1.4 | -7.7 | 1.9E-14 | 1.1E-13 | BT20/MB468 | MITO |
| >hg38_21_+_8433172_8446622_RNA45SN1_45S_50ntFlanks@10571.10596.26\|GAGAAGCCGGCGGGAGCCCCGGGGAG | >hg38_21_+_8433172_8446622_RNA45SN1_45S_50ntFlanks | 10571 | 10596 | 26 | GAGAAGCCGGCGGGAGCCCCGGGGAG | 241.0 | -11.8 | 1.5 | -7.6 | 2.0E-14 | 1.1E-13 | MB231/MB468 | MP |
| >MI0000477\|hsa-mir-146a&WithFlank&5\|+\|160485346\|160485456@27.47.21(+1A)&[MIMAT0000449&hsa-miR-146a-5p&offsets\|0\|-1(+1A);m-129&5\|+\|160485372\|160485393&offsets\|0\|-1(+1A)]\|TGAGAACTGAATTCCATGGGTA | >MI0000477\|hsa-mir-146a&WithFlank&5\|+\|160485346\|160485456 | 27 | 47 | 21(+1A) | TGAGAACTGAATTCCATGGGTA | 582.7 | 13.2 | 1.7 | 7.6 | 2.3E-14 | 1.1E-13 | MB231/MB468 | CYTO |
| >hg38_21_+_8433172_8446622_RNA45SN1_45S_50ntFlanks@4735.4766.32\|ACGAAAGTCGGAGGTTCGAAGACGATCAGATA | >hg38_21_+_8433172_8446622_RNA45SN1_45S_50ntFlanks | 4735 | 4766 | 32 | ACGAAAGTCGGAGGTTCGAAGACGATCAGATA | 388.3 | 6.8 | 0.9 | 7.6 | 2.3E-14 | 1.1E-13 | MB231/MB468 | CYTO |
| >hg38_21_+_8433172_8446622_RNA45SN1_45S_50ntFlanks@8213.8230.18\|CGGTAGCGGCCCCCGGCG | >hg38_21_+_8433172_8446622_RNA45SN1_45S_50ntFlanks | 8213 | 8230 | 18 | CGGTAGCGGCCCCCGGCG | 160.7 | -11.2 | 1.5 | -7.6 | 2.0E-14 | 1.1E-13 | MB231/MB468 | MP |
| >hg38_21_+_8433172_8446622_RNA45SN1_45S_50ntFlanks@6773.6793.21\|TTCCTCCCGGGGCTACGCCTG | >hg38_21_+_8433172_8446622_RNA45SN1_45S_50ntFlanks | 6773 | 6793 | 21 | TTCCTCCCGGGGCTACGCCTG | 420.9 | -10.8 | 1.4 | -7.7 | 1.9E-14 | 1.1E-13 | BT20/MB468 | MITO |
| >hg38_21_+_8433172_8446622_RNA45SN1_45S_50ntFlanks@3789.3812.24\|ACTGCGAATGGCTCATTAAATCAG | >hg38_21_+_8433172_8446622_RNA45SN1_45S_50ntFlanks | 3789 | 3812 | 24 | ACTGCGAATGGCTCATTAAATCAG | 785.9 | -13.4 | 1.8 | -7.6 | 3.2E-14 | 1.1E-13 | MB231/MB468 | MITO |
| >hg38_21_+_8433172_8446622_RNA45SN1_45S_50ntFlanks@12640.12667.28\|ACCATCTGTGGGATTATGACTGAACGCC | >hg38_21_+_8433172_8446622_RNA45SN1_45S_50ntFlanks | 12640 | 12667 | 28 | ACCATCTGTGGGATTATGACTGAACGCC | 536.8 | 7.7 | 1.0 | 7.6 | 2.3E-14 | 1.1E-13 | MB231/MB468 | CYTO |
| >MI0003820\|hsa-mir-454&WithFlank&17\|-\|59137752\|59137878@70.92.23&[MIMAT0003885&hsa-miR-454-3p&offsets\|0\|0;m-272&17\|-\|59137787\|59137809&offsets\|0\|0]\|TAGTGCAATATTGCTTATAGGGT | >MI0003820\|hsa-mir-454&WithFlank&17\|-\|59137752\|59137878 | 70 | 92 | 23 | TAGTGCAATATTGCTTATAGGGT | 546.3 | 8.1 | 1.1 | 7.6 | 2.6E-14 | 1.1E-13 | BT20/MB468 | CYTO |
| >hg38_21_+_8433172_8446622_RNA45SN1_45S_50ntFlanks@10866.10904.39\|CTAAGGGCTGGGTCGGTCGGGCTGGGGCGCGAAGCGGGG | >hg38_21_+_8433172_8446622_RNA45SN1_45S_50ntFlanks | 10866 | 10904 | 39 | CTAAGGGCTGGGTCGGTCGGGCTGGGGCGCGAAGCGGGG | 215.8 | -11.6 | 1.5 | -7.6 | 2.0E-14 | 1.1E-13 | MB231/MB468 | MP |
| >hg38_21_+_8433172_8446622_RNA45SN1_45S_50ntFlanks@12630.12654.25\|GGGCGAAGCTACCATCTGTGGGATT | >hg38_21_+_8433172_8446622_RNA45SN1_45S_50ntFlanks | 12630 | 12654 | 25 | GGGCGAAGCTACCATCTGTGGGATT | 208.1 | -10.7 | 1.4 | -7.6 | 2.4E-14 | 1.1E-13 | MB231/MB468 | NUC |
| >hg38_21_+_8433172_8446622_RNA45SN1_45S_50ntFlanks@12674.12708.35\|TCAGAATCCCGCCCAGGCGGAACGATACGGCAGCG | >hg38_21_+_8433172_8446622_RNA45SN1_45S_50ntFlanks | 12674 | 12708 | 35 | TCAGAATCCCGCCCAGGCGGAACGATACGGCAGCG | 153.6 | -11.1 | 1.5 | -7.6 | 2.0E-14 | 1.1E-13 | MB231/MB468 | MP |
| >hg38_21_+_8433172_8446622_RNA45SN1_45S_50ntFlanks@12409.12443.35\|TTGATCCTTCGATGTCGGCTCTTCCTATCATTGTG | >hg38_21_+_8433172_8446622_RNA45SN1_45S_50ntFlanks | 12409 | 12443 | 35 | TTGATCCTTCGATGTCGGCTCTTCCTATCATTGTG | 939.7 | -5.4 | 0.7 | -7.6 | 3.9E-14 | 1.1E-13 | BT20/MB468 | NUC |
| >hg38_21_+_8433172_8446622_RNA45SN1_45S_50ntFlanks@12435.12455.21\|ATCATTGTGAAGCAGAATTCA | >hg38_21_+_8433172_8446622_RNA45SN1_45S_50ntFlanks | 12435 | 12455 | 21 | ATCATTGTGAAGCAGAATTCA | 1411.5 | -12.8 | 1.7 | -7.6 | 2.6E-14 | 1.1E-13 | BT20/MB468 | CYTO |
| >hg38_21_+_8433172_8446622_RNA45SN1_45S_50ntFlanks@4724.4747.24\|ATTAATCAAGAACGAAAGTCGGAG | >hg38_21_+_8433172_8446622_RNA45SN1_45S_50ntFlanks | 4724 | 4747 | 24 | ATTAATCAAGAACGAAAGTCGGAG | 187.5 | -11.4 | 1.5 | -7.6 | 2.0E-14 | 1.1E-13 | MB231/MB468 | MP |
| >hg38_21_+_8433172_8446622_RNA45SN1_45S_50ntFlanks@8378.8400.23\|TTCAAGAGGGCGTGAAACCGTTA | >hg38_21_+_8433172_8446622_RNA45SN1_45S_50ntFlanks | 8378 | 8400 | 23 | TTCAAGAGGGCGTGAAACCGTTA | 232.5 | -11.7 | 1.5 | -7.6 | 2.0E-14 | 1.1E-13 | MB231/MB468 | MP |
| >hg38_21_+_8433172_8446622_RNA45SN1_45S_50ntFlanks@12499.12518.20\|GGTTTAGACCGTCGTGAGAC | >hg38_21_+_8433172_8446622_RNA45SN1_45S_50ntFlanks | 12499 | 12518 | 20 | GGTTTAGACCGTCGTGAGAC | 310.5 | -12.1 | 1.6 | -7.6 | 3.2E-14 | 1.1E-13 | MB231/MB468 | MITO |
| >trna10_GlyTCC_17_+_8124866_8124937@1.30.30\|GCGTTGGTGGTATAGTGGTAAGCATAGCTG | >trna10_GlyTCC_17_+_8124866_8124937 | 1 | 30 | 30 | GCGTTGGTGGTATAGTGGTAAGCATAGCTG | 305.5 | -12.0 | 1.6 | -7.6 | 3.3E-14 | 1.1E-13 | MB231/MB468 | MITO |
| >hg38_21_+_8433172_8446622_RNA45SN1_45S_50ntFlanks@12289.12316.28\|CCGTGAAAGCGGGGCCTCACGATCCTTC | >hg38_21_+_8433172_8446622_RNA45SN1_45S_50ntFlanks | 12289 | 12316 | 28 | CCGTGAAAGCGGGGCCTCACGATCCTTC | 641.2 | -13.1 | 1.7 | -7.6 | 3.3E-14 | 1.1E-13 | MB231/MB468 | MITO |
| >hg38_21_+_8433172_8446622_RNA45SN1_45S_50ntFlanks@10727.10752.26\|GGAGAGGGTGTAAATCTCGCGCCGGG | >hg38_21_+_8433172_8446622_RNA45SN1_45S_50ntFlanks | 10727 | 10752 | 26 | GGAGAGGGTGTAAATCTCGCGCCGGG | 695.0 | -12.5 | 1.6 | -8.0 | 8.3E-16 | 1.1E-13 | BT20/MB468 | MP |
| >hg38_21_+_8433172_8446622_RNA45SN1_45S_50ntFlanks@9736.9754.19\|CCGAAACGATCTCAACCTA | >hg38_21_+_8433172_8446622_RNA45SN1_45S_50ntFlanks | 9736 | 9754 | 19 | CCGAAACGATCTCAACCTA | 1128.9 | -9.6 | 1.3 | -7.6 | 3.3E-14 | 1.1E-13 | MB231/MB468 | MITO |
| >hg38_21_+_8433172_8446622_RNA45SN1_45S_50ntFlanks@8319.8356.38\|CGAGACCGATAGTCAACAAGTACCGTAAGGGAAAGTTG | >hg38_21_+_8433172_8446622_RNA45SN1_45S_50ntFlanks | 8319 | 8356 | 38 | CGAGACCGATAGTCAACAAGTACCGTAAGGGAAAGTTG | 345.3 | -12.3 | 1.6 | -7.6 | 2.1E-14 | 1.1E-13 | MB231/MB468 | MP |
| >hg38_21_+_8433172_8446622_RNA45SN1_45S_50ntFlanks@4550.4573.24\|GATACCGCAGCTAGGAATAATGGA | >hg38_21_+_8433172_8446622_RNA45SN1_45S_50ntFlanks | 4550 | 4573 | 24 | GATACCGCAGCTAGGAATAATGGA | 157.5 | -10.3 | 1.3 | -7.6 | 2.5E-14 | 1.1E-13 | MB231/MB468 | NUC |

<table>
<tr><td></td><td></td><td></td><td></td><td></td><td></td><td></td><td></td><td></td><td></td><td></td><td></td><td></td><td></td><td></td></tr>
<tr><td>>hg38_21_+_8433172_8446622_RNA45SN1_45S_50ntFlanks@10843.10865.23|TAACTTCGGGATAAGGATTGGCT</td><td>>hg38_21_+_8433172_8446622_RNA45SN1_45S_50ntFlanks</td><td>10843</td><td>10865</td><td>23</td><td>TAACTTCGGGATAAGGATTGGCT</td><td>1060.3</td><td>-9.4</td><td>1.2</td><td>-7.6</td><td>2.5E-14</td><td>1.1E-13</td><td>MB231/MB468</td><td>NUC</td></tr>
<tr><td>>hg38_21_+_8433172_8446622_RNA45SN1_45S_50ntFlanks@10808.10829.22|CAATGTAGGTAAGGGAAGTCGG</td><td>>hg38_21_+_8433172_8446622_RNA45SN1_45S_50ntFlanks</td><td>10808</td><td>10829</td><td>22</td><td>CAATGTAGGTAAGGGAAGTCGG</td><td>465.5</td><td>-11.0</td><td>1.4</td><td>-7.7</td><td>2.0E-14</td><td>1.1E-13</td><td>BT20/MB468</td><td>MITO</td></tr>
<tr><td>>hg38_21_+_8433172_8446622_RNA45SN1_45S_50ntFlanks@4759.4778.20|ATCAGATACCGTCGTAGTTC</td><td>>hg38_21_+_8433172_8446622_RNA45SN1_45S_50ntFlanks</td><td>4759</td><td>4778</td><td>20</td><td>ATCAGATACCGTCGTAGTTC</td><td>1871.5</td><td>-10.1</td><td>1.3</td><td>-7.6</td><td>2.1E-14</td><td>1.1E-13</td><td>MB231/MB468</td><td>MP</td></tr>
<tr><td>>hg38_21_+_8433172_8446622_RNA45SN1_45S_50ntFlanks@7984.8014.31|AGATCAGACGTGGCGACCCGCTGAATTTAAG</td><td>>hg38_21_+_8433172_8446622_RNA45SN1_45S_50ntFlanks</td><td>7984</td><td>8014</td><td>31</td><td>AGATCAGACGTGGCGACCCGCTGAATTTAAG</td><td>132.9</td><td>-10.9</td><td>1.4</td><td>-7.6</td><td>2.1E-14</td><td>1.1E-13</td><td>MB231/MB468</td><td>MP</td></tr>
<tr><td>>hg38_21_+_8433172_8446622_RNA45SN1_45S_50ntFlanks@12717.12747.31|CCTCGGTTGGCCTCGGATAGCCGGTCCCCCG</td><td>>hg38_21_+_8433172_8446622_RNA45SN1_45S_50ntFlanks</td><td>12717</td><td>12747</td><td>31</td><td>CCTCGGTTGGCCTCGGATAGCCGGTCCCCCG</td><td>413.7</td><td>-13.1</td><td>1.7</td><td>-7.6</td><td>3.9E-14</td><td>1.1E-13</td><td>BT20/MB468</td><td>NUC</td></tr>
<tr><td>>hg38_21_+_8433172_8446622_RNA45SN1_45S_50ntFlanks@6690.6719.30|AAGAACGCAGCTAGCTGCGAGAATTAATGT</td><td>>hg38_21_+_8433172_8446622_RNA45SN1_45S_50ntFlanks</td><td>6690</td><td>6719</td><td>30</td><td>AAGAACGCAGCTAGCTGCGAGAATTAATGT</td><td>244.8</td><td>-11.7</td><td>1.5</td><td>-7.6</td><td>3.3E-14</td><td>1.1E-13</td><td>MB231/MB468</td><td>MITO</td></tr>
<tr><td>>hg38_21_+_8433172_8446622_RNA45SN1_45S_50ntFlanks@3705.3730.26|TACCTGGTTGATCCTGCCAGTAGCAT</td><td>>hg38_21_+_8433172_8446622_RNA45SN1_45S_50ntFlanks</td><td>3705</td><td>3730</td><td>26</td><td>TACCTGGTTGATCCTGCCAGTAGCAT</td><td>376.6</td><td>10.4</td><td>1.4</td><td>7.6</td><td>2.7E-14</td><td>1.1E-13</td><td>BT20/MB468</td><td>CYTO</td></tr>
<tr><td>>hg38_21_+_8433172_8446622_RNA45SN1_45S_50ntFlanks@4588.4619.32|CTATTTTGTTGGTTTTCGGAACTGAGGCCATG</td><td>>hg38_21_+_8433172_8446622_RNA45SN1_45S_50ntFlanks</td><td>4588</td><td>4619</td><td>32</td><td>CTATTTTGTTGGTTTTCGGAACTGAGGCCATG</td><td>517.0</td><td>-7.1</td><td>0.9</td><td>-7.6</td><td>3.3E-14</td><td>1.1E-13</td><td>MB231/MB468</td><td>MITO</td></tr>
<tr><td>>MI0000088|hsa-mir-30a&WithFlank&6|-|71403545|71403627@12.32.21&[MIMAT0000087&hsa-miR-30a-5p&offsets|0|-1;m-15&6|-|71403595|71403616&offsets|0|-1]|TGTAAACATCCTCGACTGGAA</td><td>>MI0000088|hsa-mir-30a&WithFlank&6|-|71403545|71403627</td><td>12</td><td>32</td><td>21</td><td>TGTAAACATCCTCGACTGGAA</td><td>859.4</td><td>6.8</td><td>0.9</td><td>7.6</td><td>2.5E-14</td><td>1.1E-13</td><td>MB231/MB468</td><td>NUC</td></tr>
<tr><td>>hg38_21_+_8433172_8446622_RNA45SN1_45S_50ntFlanks@8327.8356.30|ATAGTCAACAAGTACCGTAAGGGAAAGTTG</td><td>>hg38_21_+_8433172_8446622_RNA45SN1_45S_50ntFlanks</td><td>8327</td><td>8356</td><td>30</td><td>ATAGTCAACAAGTACCGTAAGGGAAAGTTG</td><td>130.8</td><td>-10.9</td><td>1.4</td><td>-7.6</td><td>2.1E-14</td><td>1.1E-13</td><td>MB231/MB468</td><td>MP</td></tr>
<tr><td>>MI0000299|hsa-mir-222&WithFlank&X|-|45747009|45747130@75.97.23(+1A)&[MIMAT0000279&hsa-miR-222-3p&offsets|0|+2(+1A);m-140&X|-|45747033|45747056&offsets|0|-1(+1A)]|AGCTACATCTGGCTACTGGGTCTA</td><td>>MI0000299|hsa-mir-222&WithFlank&X|-|45747009|45747130</td><td>75</td><td>97</td><td>23(+1A)</td><td>AGCTACATCTGGCTACTGGGTCTA</td><td>1951.5</td><td>8.4</td><td>1.1</td><td>7.6</td><td>2.4E-14</td><td>1.1E-13</td><td>MB231/MB468</td><td>CYTO</td></tr>
<tr><td>>hg38_21_+_8433172_8446622_RNA45SN1_45S_50ntFlanks@12004.12029.26|CGGGCCGCCGGTGAAATACCACTACT</td><td>>hg38_21_+_8433172_8446622_RNA45SN1_45S_50ntFlanks</td><td>12004</td><td>12029</td><td>26</td><td>CGGGCCGCCGGTGAAATACCACTACT</td><td>529.4</td><td>4.4</td><td>0.6</td><td>7.7</td><td>2.0E-14</td><td>1.1E-13</td><td>BT20/MB468</td><td>MITO</td></tr>
<tr><td>>hg38_21_+_8433172_8446622_RNA45SN1_45S_50ntFlanks@5472.5517.46|ACGGCCCTGGCGGAGCGCTGAGAAGACGGTCGAACTTGACTATCTA</td><td>>hg38_21_+_8433172_8446622_RNA45SN1_45S_50ntFlanks</td><td>5472</td><td>5517</td><td>46</td><td>ACGGCCCTGGCGGAGCGCTGAGAAGACGGTCGAACTTGACTATCTA</td><td>274.1</td><td>9.2</td><td>1.2</td><td>7.6</td><td>2.4E-14</td><td>1.1E-13</td><td>MB231/MB468</td><td>CYTO</td></tr>
<tr><td>>hg38_21_+_8433172_8446622_RNA45SN1_45S_50ntFlanks@11209.11253.45|CGGCGACTCTGGACGCGAGCCGGGCCCTTCCCGTGGATCGCCCCA</td><td>>hg38_21_+_8433172_8446622_RNA45SN1_45S_50ntFlanks</td><td>11209</td><td>11253</td><td>45</td><td>CGGCGACTCTGGACGCGAGCCGGGCCCTTCCCGTGGATCGCCCCA</td><td>320.5</td><td>-12.1</td><td>1.6</td><td>-7.6</td><td>3.3E-14</td><td>1.1E-13</td><td>MB231/MB468</td><td>MITO</td></tr>
<tr><td>>hg38_21_+_8433172_8446622_RNA45SN1_45S_50ntFlanks@11633.11660.28|AAGGCCCGCGGCGGGTGTTGACGCGATG</td><td>>hg38_21_+_8433172_8446622_RNA45SN1_45S_50ntFlanks</td><td>11633</td><td>11660</td><td>28</td><td>AAGGCCCGCGGCGGGTGTTGACGCGATG</td><td>601.7</td><td>-13.0</td><td>1.7</td><td>-7.6</td><td>3.3E-14</td><td>1.1E-13</td><td>MB231/MB468</td><td>MITO</td></tr>
<tr><td>>hg38_21_+_8433172_8446622_RNA45SN1_45S_50ntFlanks@11483.11502.20|CGGAGGGGCCGCGGGCCGGT</td><td>>hg38_21_+_8433172_8446622_RNA45SN1_45S_50ntFlanks</td><td>11483</td><td>11502</td><td>20</td><td>CGGAGGGGCCGCGGGCCGGT</td><td>129.4</td><td>-7.2</td><td>0.9</td><td>-7.6</td><td>2.4E-14</td><td>1.1E-13</td><td>MB231/MB468</td><td>CYTO</td></tr>
<tr><td>>hg38_21_+_8433172_8446622_RNA45SN1_45S_50ntFlanks@12643.12661.19|ATCTGTGGGATTATGACTG</td><td>>hg38_21_+_8433172_8446622_RNA45SN1_45S_50ntFlanks</td><td>12643</td><td>12661</td><td>19</td><td>ATCTGTGGGATTATGACTG</td><td>662.5</td><td>-12.5</td><td>1.5</td><td>-8.0</td><td>8.6E-16</td><td>1.1E-13</td><td>BT20/MB468</td><td>MP</td></tr>
<tr><td>>hg38_21_+_8433172_8446622_RNA45SN1_45S_50ntFlanks@11569.11597.29|CAGCCGACTTAGAACTGGTGCGGACCAGG</td><td>>hg38_21_+_8433172_8446622_RNA45SN1_45S_50ntFlanks</td><td>11569</td><td>11597</td><td>29</td><td>CAGCCGACTTAGAACTGGTGCGGACCAGG</td><td>571.8</td><td>-5.5</td><td>0.7</td><td>-7.6</td><td>4.0E-14</td><td>1.1E-13</td><td>BT20/MB468</td><td>NUC</td></tr>
<tr><td>>hg38_21_+_8433172_8446622_RNA45SN1_45S_50ntFlanks@4760.4793.34|TCAGATACCGTCGTAGTTCCGACCATAAACGATG</td><td>>hg38_21_+_8433172_8446622_RNA45SN1_45S_50ntFlanks</td><td>4760</td><td>4793</td><td>34</td><td>TCAGATACCGTCGTAGTTCCGACCATAAACGATG</td><td>233.2</td><td>-11.7</td><td>1.5</td><td>-7.6</td><td>2.1E-14</td><td>1.1E-13</td><td>MB231/MB468</td><td>MP</td></tr>
<tr><td>>MI0000088|hsa-mir-30a&WithFlank&6|-|71403545|71403627@12.35.24(+1U)&[MIMAT0000087&hsa-miR-30a-5p&offsets|0|+2(+1U);m-15&6|-|71403595|71403616&offsets|0|+2(+1U)]|TGTAAACATCCTCGACTGGAAGCTT</td><td>>MI0000088|hsa-mir-30a&WithFlank&6|-|71403545|71403627</td><td>12</td><td>35</td><td>24(+1U)</td><td>TGTAAACATCCTCGACTGGAAGCTT</td><td>623.2</td><td>5.7</td><td>0.7</td><td>7.6</td><td>2.0E-14</td><td>1.1E-13</td><td>BT20/MB468</td><td>MITO</td></tr>
<tr><td>>hg38_21_+_8433172_8446622_RNA45SN1_45S_50ntFlanks@11200.11227.28|CGGCGGCGGCGGCGACTCTGGACGCGAG</td><td>>hg38_21_+_8433172_8446622_RNA45SN1_45S_50ntFlanks</td><td>11200</td><td>11227</td><td>28</td><td>CGGCGGCGGCGGCGACTCTGGACGCGAG</td><td>224.5</td><td>-11.6</td><td>1.5</td><td>-7.6</td><td>3.4E-14</td><td>1.1E-13</td><td>MB231/MB468</td><td>MITO</td></tr>
<tr><td>>hg38_21_+_8433172_8446622_RNA45SN1_45S_50ntFlanks@9720.9737.18|ATTAGAGGTCTTGGGGCC</td><td>>hg38_21_+_8433172_8446622_RNA45SN1_45S_50ntFlanks</td><td>9720</td><td>9737</td><td>18</td><td>ATTAGAGGTCTTGGGGCC</td><td>922.5</td><td>-12.9</td><td>1.6</td><td>-8.0</td><td>8.7E-16</td><td>1.1E-13</td><td>BT20/MB468</td><td>MP</td></tr>
<tr><td>>hg38_21_+_8433172_8446622_RNA45SN1_45S_50ntFlanks@4102.4129.28|ACCACGGGTGACGGGGAATCAGGGTTCG</td><td>>hg38_21_+_8433172_8446622_RNA45SN1_45S_50ntFlanks</td><td>4102</td><td>4129</td><td>28</td><td>ACCACGGGTGACGGGGAATCAGGGTTCG</td><td>236.4</td><td>-10.8</td><td>1.4</td><td>-7.6</td><td>2.5E-14</td><td>1.1E-13</td><td>MB231/MB468</td><td>NUC</td></tr>
<tr><td>>hg38_21_+_8433172_8446622_RNA45SN1_45S_50ntFlanks@11874.11909.36|AGAAGACCCTGTTGAGCTTGACTCTAGTCTGGCACG</td><td>>hg38_21_+_8433172_8446622_RNA45SN1_45S_50ntFlanks</td><td>11874</td><td>11909</td><td>36</td><td>AGAAGACCCTGTTGAGCTTGACTCTAGTCTGGCACG</td><td>214.2</td><td>-11.6</td><td>1.5</td><td>-7.6</td><td>2.1E-14</td><td>1.1E-13</td><td>MB231/MB468</td><td>MP</td></tr>
<tr><td>>hg38_21_+_8433172_8446622_RNA45SN1_45S_50ntFlanks@5515.5535.21|CTAGAGGAAGTAAAAGTCGTA</td><td>>hg38_21_+_8433172_8446622_RNA45SN1_45S_50ntFlanks</td><td>5515</td><td>5535</td><td>21</td><td>CTAGAGGAAGTAAAAGTCGTA</td><td>932.1</td><td>-9.3</td><td>1.2</td><td>-7.6</td><td>2.1E-14</td><td>1.1E-13</td><td>MB231/MB468</td><td>MP</td></tr>
<tr><td>>hg38_21_+_8433172_8446622_RNA45SN1_45S_50ntFlanks@12444.12480.37|AAGCAGAATTCACCAAGCGTTGGATTGTTCACCCACT</td><td>>hg38_21_+_8433172_8446622_RNA45SN1_45S_50ntFlanks</td><td>12444</td><td>12480</td><td>37</td><td>AAGCAGAATTCACCAAGCGTTGGATTGTTCACCCACT</td><td>750.6</td><td>-3.6</td><td>0.5</td><td>-7.6</td><td>4.1E-14</td><td>1.1E-13</td><td>BT20/MB468</td><td>NUC</td></tr>
<tr><td>>hg38_21_+_8433172_8446622_RNA45SN1_45S_50ntFlanks@10410.10430.21|TCAGTCGGTCCTGAGAGATGG</td><td>>hg38_21_+_8433172_8446622_RNA45SN1_45S_50ntFlanks</td><td>10410</td><td>10430</td><td>21</td><td>TCAGTCGGTCCTGAGAGATGG</td><td>1214.9</td><td>-4.4</td><td>0.6</td><td>-7.6</td><td>2.0E-14</td><td>1.1E-13</td><td>BT20/MB468</td><td>MITO</td></tr>
<tr><td>>hg38_21_+_8433172_8446622_RNA45SN1_45S_50ntFlanks@9569.9597.29|GTGCAAATCGGTCGTCCGACCTGGGTATA</td><td>>hg38_21_+_8433172_8446622_RNA45SN1_45S_50ntFlanks</td><td>9569</td><td>9597</td><td>29</td><td>GTGCAAATCGGTCGTCCGACCTGGGTATA</td><td>2495.8</td><td>-5.3</td><td>0.7</td><td>-7.6</td><td>3.4E-14</td><td>1.1E-13</td><td>MB231/MB468</td><td>MITO</td></tr>
<tr><td>>hg38_21_+_8433172_8446622_RNA45SN1_45S_50ntFlanks@10690.10707.18|CGTCCGGTGAGCTCTCGC</td><td>>hg38_21_+_8433172_8446622_RNA45SN1_45S_50ntFlanks</td><td>10690</td><td>10707</td><td>18</td><td>CGTCCGGTGAGCTCTCGC</td><td>154.8</td><td>-6.9</td><td>0.9</td><td>-7.6</td><td>2.4E-14</td><td>1.1E-13</td><td>MB231/MB468</td><td>CYTO</td></tr>
<tr><td>>MI0000462|hsa-mir-152&WithFlank&17|-|48037155|48037253@60.80.21&[MIMAT0000438&hsa-miR-152-3p&offsets|0|0;m-97&17|-|48037174|48037194&offsets|0|0]|TCAGTGCATGACAGAACTTGG</td><td>>MI0000462|hsa-mir-152&WithFlank&17|-|48037155|48037253</td><td>60</td><td>80</td><td>21</td><td>TCAGTGCATGACAGAACTTGG</td><td>364.3</td><td>12.5</td><td>1.6</td><td>7.6</td><td>2.4E-14</td><td>1.1E-13</td><td>MB231/MB468</td><td>CYTO</td></tr>
<tr><td>>hg38_21_+_8433172_8446622_RNA45SN1_45S_50ntFlanks@7980.8004.25|CCTCAGATCAGACGTGGCGACCCGC</td><td>>hg38_21_+_8433172_8446622_RNA45SN1_45S_50ntFlanks</td><td>7980</td><td>8004</td><td>25</td><td>CCTCAGATCAGACGTGGCGACCCGC</td><td>178.9</td><td>-11.3</td><td>1.5</td><td>-7.6</td><td>2.1E-14</td><td>1.1E-13</td><td>MB231/MB468</td><td>MP</td></tr>
<tr><td>>MI0000068|hsa-let-7f-2&WithFlank&X|-|53557186|53557280@14.36.23&[MIMAT0000067_1&hsa-let-7f-5p&offsets|0|+1;m-19&X|-|53557246|53557267&offsets|0|+1]|TGAGGTAGTAGATTGTATAGTTT</td><td>>MI0000068|hsa-let-7f-2&WithFlank&X|-|53557186|53557280</td><td>14</td><td>36</td><td>23</td><td>TGAGGTAGTAGATTGTATAGTTT</td><td>4307.0</td><td>6.0</td><td>0.8</td><td>7.6</td><td>2.8E-14</td><td>1.1E-13</td><td>BT20/MB468</td><td>CYTO</td></tr>
<tr><td>>hg38_21_+_8433172_8446622_RNA45SN1_45S_50ntFlanks@11583.11600.18|CTGGTGCGGACCAGGGGA</td><td>>hg38_21_+_8433172_8446622_RNA45SN1_45S_50ntFlanks</td><td>11583</td><td>11600</td><td>18</td><td>CTGGTGCGGACCAGGGGA</td><td>331.8</td><td>-3.6</td><td>0.5</td><td>-7.6</td><td>2.6E-14</td><td>1.1E-13</td><td>MB231/MB468</td><td>NUC</td></tr>
<tr><td>>MI0000103|hsa-mir-101-1&WithFlank&1|-|65058428|65058514@52.73.22;MI0000739|hsa-mir-101-2&WithFlank&9|+|4850291|4850381@54.75.22&[MIMAT0000099&hsa-miR-101-3p&offsets|-1|0;m-17&1|-|65058443|65058462&offsets|-1|+1];[MIMAT0000099_1&hsa-miR-101-3p&offsets|-1|0;m-16&9|+|4850345|4850364&offsets|-1|+1]|GTACAGTACTGTGATAACTGAA</td><td>>MI0000103|hsa-mir-101-1&WithFlank&1|-|65058428|65058514</td><td>52</td><td>73</td><td>22</td><td>GTACAGTACTGTGATAACTGAA</td><td>1217.7</td><td>5.6</td><td>0.7</td><td>7.6</td><td>2.6E-14</td><td>1.1E-13</td><td>MB231/MB468</td><td>NUC</td></tr>
<tr><td>>hg38_21_+_8433172_8446622_RNA45SN1_45S_50ntFlanks@9243.9274.32|GAGCGAGCGCACGGGGTCGGCGGCGACGTCGG</td><td>>hg38_21_+_8433172_8446622_RNA45SN1_45S_50ntFlanks</td><td>9243</td><td>9274</td><td>32</td><td>GAGCGAGCGCACGGGGTCGGCGGCGACGTCGG</td><td>248.3</td><td>-11.8</td><td>1.5</td><td>-7.6</td><td>2.2E-14</td><td>1.2E-13</td><td>MB231/MB468</td><td>MP</td></tr>
<tr><td>>hg38_21_+_8433172_8446622_RNA45SN1_45S_50ntFlanks@12008.12044.37|CCGCCGGTGAAATACCACTACTCTGATCGTTTTTTCA</td><td>>hg38_21_+_8433172_8446622_RNA45SN1_45S_50ntFlanks</td><td>12008</td><td>12044</td><td>37</td><td>CCGCCGGTGAAATACCACTACTCTGATCGTTTTTTCA</td><td>351.7</td><td>-12.2</td><td>1.6</td><td>-7.6</td><td>3.4E-14</td><td>1.2E-13</td><td>MB231/MB468</td><td>MITO</td></tr>
<tr><td>>hg38_21_+_8433172_8446622_RNA45SN1_45S_50ntFlanks@13002.13023.22|CGATCTATTGAAAGTCAGCCCT</td><td>>hg38_21_+_8433172_8446622_RNA45SN1_45S_50ntFlanks</td><td>13002</td><td>13023</td><td>22</td><td>CGATCTATTGAAAGTCAGCCCT</td><td>731.9</td><td>-12.6</td><td>1.6</td><td>-8.0</td><td>9.0E-16</td><td>1.2E-13</td><td>BT20/MB468</td><td>MP</td></tr>
<tr><td>>MI0000734|hsa-mir-106b&WithFlank&7|-|100093987|100094080@58.76.19&[MIMAT0004672&hsa-miR-106b-3p&offsets|0|-3;m-87&7|-|100094003|100094023&offsets|0|-2]|CCGCACTGTGGGTACTTGC</td><td>>MI0000734|hsa-mir-106b&WithFlank&7|-|100093987|100094080</td><td>58</td><td>76</td><td>19</td><td>CCGCACTGTGGGTACTTGC</td><td>570.0</td><td>9.0</td><td>1.2</td><td>7.6</td><td>2.8E-14</td><td>1.2E-13</td><td>BT20/MB468</td><td>CYTO</td></tr>
<tr><td>>hg38_21_+_8433172_8446622_RNA45SN1_45S_50ntFlanks@6663.6689.27|GTGGATCACTCGGCTCGTGCGTCGATG</td><td>>hg38_21_+_8433172_8446622_RNA45SN1_45S_50ntFlanks</td><td>6663</td><td>6689</td><td>27</td><td>GTGGATCACTCGGCTCGTGCGTCGATG</td><td>1051.3</td><td>-4.5</td><td>0.6</td><td>-7.6</td><td>2.6E-14</td><td>1.2E-13</td><td>MB231/MB468</td><td>NUC</td></tr>
<tr><td>>hg38_21_+_8433172_8446622_RNA45SN1_45S_50ntFlanks@4735.4766.32|ACGAAAGTCGGAGGTTCGAAGACGATCAGATA</td><td>>hg38_21_+_8433172_8446622_RNA45SN1_45S_50ntFlanks</td><td>4735</td><td>4766</td><td>32</td><td>ACGAAAGTCGGAGGTTCGAAGACGATCAGATA</td><td>725.0</td><td>8.0</td><td>1.0</td><td>7.6</td><td>2.8E-14</td><td>1.2E-13</td><td>BT20/MB468</td><td>CYTO</td></tr>
<tr><td>>hg38_21_+_8433172_8446622_RNA45SN1_45S_50ntFlanks@10458.10497.40|ATGGCCTCCGTTGCCCTCGGCCGATCGAAAGGGAGTCGGG</td><td>>hg38_21_+_8433172_8446622_RNA45SN1_45S_50ntFlanks</td><td>10458</td><td>10497</td><td>40</td><td>ATGGCCTCCGTTGCCCTCGGCCGATCGAAAGGGAGTCGGG</td><td>524.4</td><td>-8.5</td><td>1.1</td><td>-7.6</td><td>2.5E-14</td><td>1.2E-13</td><td>MB231/MB468</td><td>CYTO</td></tr>
<tr><td>>hg38_21_+_8433172_8446622_RNA45SN1_45S_50ntFlanks@5400.5419.20|CACACCGCCCGTCGCTACTA</td><td>>hg38_21_+_8433172_8446622_RNA45SN1_45S_50ntFlanks</td><td>5400</td><td>5419</td><td>20</td><td>CACACCGCCCGTCGCTACTA</td><td>347.1</td><td>-12.2</td><td>1.6</td><td>-7.6</td><td>3.5E-14</td><td>1.2E-13</td><td>MB231/MB468</td><td>MITO</td></tr>
<tr><td>>MI0000081|hsa-mir-24-2&WithFlank&19|-|13836281|13836365@18.41.24&[MIMAT0004497&hsa-miR-24-2-5p&offsets|-1|+1;m-226&19|-|13836326|13836348&offsets|0|+1]|GTGCCTACTGAGCTGAAACACAGT</td><td>>MI0000081|hsa-mir-24-2&WithFlank&19|-|13836281|13836365</td><td>18</td><td>41</td><td>24</td><td>GTGCCTACTGAGCTGAAACACAGT</td><td>344.8</td><td>5.2</td><td>0.7</td><td>7.6</td><td>2.6E-14</td><td>1.2E-13</td><td>MB231/MB468</td><td>NUC</td></tr>
<tr><td>>hg38_21_+_8433172_8446622_RNA45SN1_45S_50ntFlanks@10408.10433.26|GGTCAGTCGGTCCTGAGAGATGGGCG</td><td>>hg38_21_+_8433172_8446622_RNA45SN1_45S_50ntFlanks</td><td>10408</td><td>10433</td><td>26</td><td>GGTCAGTCGGTCCTGAGAGATGGGCG</td><td>2151.7</td><td>-6.8</td><td>0.9</td><td>-7.6</td><td>2.5E-14</td><td>1.2E-13</td><td>MB231/MB468</td><td>CYTO</td></tr>
<tr><td>>hg38_21_+_8433172_8446622_RNA45SN1_45S_50ntFlanks@4443.4462.20|CCCTTGCCTCTCGGCGCCCC</td><td>>hg38_21_+_8433172_8446622_RNA45SN1_45S_50ntFlanks</td><td>4443</td><td>4462</td><td>20</td><td>CCCTTGCCTCTCGGCGCCCC</td><td>209.7</td><td>-11.6</td><td>1.5</td><td>-7.6</td><td>2.2E-14</td><td>1.2E-13</td><td>MB231/MB468</td><td>MP</td></tr>
<tr><td>>hg38_21_+_8433172_8446622_RNA45SN1_45S_50ntFlanks@12456.12480.25|CCAAGCGTTGGATTGTTCACCCACT</td><td>>hg38_21_+_8433172_8446622_RNA45SN1_45S_50ntFlanks</td><td>12456</td><td>12480</td><td>25</td><td>CCAAGCGTTGGATTGTTCACCCACT</td><td>2780.8</td><td>-5.3</td><td>0.7</td><td>-7.6</td><td>2.5E-14</td><td>1.2E-13</td><td>MB231/MB468</td><td>CYTO</td></tr>
<tr><td>>hg38_21_+_8433172_8446622_RNA45SN1_45S_50ntFlanks@6745.6772.28|ACACTTCGAACGCACTTGCGGCCCCGGG</td><td>>hg38_21_+_8433172_8446622_RNA45SN1_45S_50ntFlanks</td><td>6745</td><td>6772</td><td>28</td><td>ACACTTCGAACGCACTTGCGGCCCCGGG</td><td>310.3</td><td>-7.3</td><td>1.0</td><td>-7.6</td><td>2.5E-14</td><td>1.2E-13</td><td>MB231/MB468</td><td>CYTO</td></tr>
<tr><td>>trna12_TrpCCA_17_+_19411494_19411565@30.52.23,trna168_TrpCCA_6_-_26331672_26331743@30.52.23,trna170_TrpCCA_6_-_26319330_26319401@30.52.23,trna1_TrpCCA_7_+_99067307_99067378@30.52.23,trna39_TrpCCA_17_-_8124187_8124258@30.52.23,trna6_TrpCCA_17_+_8089676_8089747@30.52.23|ACTCCAGATCAGAAGGTTGCGTG</td><td>>trna12_TrpCCA_17_+_19411494_19411565</td><td>30</td><td>52</td><td>23</td><td>ACTCCAGATCAGAAGGTTGCGTG</td><td>160.2</td><td>-11.2</td><td>1.5</td><td>-7.6</td><td>2.2E-14</td><td>1.2E-13</td><td>MB231/MB468</td><td>MP</td></tr>
<tr><td>>hg38_21_+_8433172_8446622_RNA45SN1_45S_50ntFlanks@4657.4682.26|CTAGAGGTGAAATTCTTGGACCGGCG</td><td>>hg38_21_+_8433172_8446622_RNA45SN1_45S_50ntFlanks</td><td>4657</td><td>4682</td><td>26</td><td>CTAGAGGTGAAATTCTTGGACCGGCG</td><td>574.0</td><td>-11.3</td><td>1.5</td><td>-7.6</td><td>2.1E-14</td><td>1.2E-13</td><td>BT20/MB468</td><td>MITO</td></tr>
</table>

<table>
<tr><td>>am_trna11_ProAGG_16_+_3241989_3242060@1.29.29,trna12_ProAGG_6_+_26555498_26555569@1.29.29,trna12_ProTGG_11_-_75946869_75946940@1.29.29,trna14_ProTGG_5_-_180615854_180615925@1.29.29,trna22_ProAGG_14_-_21081560_21081631@1.29.29,trna23_ProAGG_14_-_21077495_21077566@1.29.29,trna28_ProTGG_16_-_3234133_3234204@1.29.29,trna29_ProAGG_16_-_3232635_3232706@1.29.29,trna2_ProAGG_7_+_128423504_128423575@1.29.29,trna30_ProCGG_6_+_27059521_27059592@1.29.29,trna37_ProCGG_17_-_8126151_8126222@1.29.29,trna3_ProTGG_16_+_3208923_3208994@1.29.29,trna52_ProCGG_1_+_167683962_167684033@1.29.29,trna65_ProAGG_1_-_167684725_167684796@1.29.29,trna6_ProCGG_16_+_3222049_3222120@1.29.29,trna6_ProTGG_14_+_21152175_21152246@1.29.29,trna8_ProTGG_16_+_3238094_3238165@1.29.29,trna9_ProAGG_11_+_75946557_75946628@1.29.29,trna9_ProAGG_16_+_3239634_3239705@1.29.29|GGCTCGTTGGTCTAGGGGTATGATTCTCG</td><td>>am_trna11_ProAGG_16_+_3241989_3242060</td><td>1</td><td>29</td><td>29</td><td>GGCTCGTTGGTCTAGGGGTATGATTCTCG</td><td>348.0</td><td>-12.3</td><td>1.6</td><td>-7.6</td><td>2.2E-14</td><td>1.2E-13</td><td>MB231/MB468</td><td>MP</td></tr>
<tr><td>>MI0003560|hsa-mir-92b&WithFlank&1|+|155195171|155195278@67.88.22(+1U)&[MIMAT0003218&hsa-miR-92b-3p&offsets|0|0(+1U);m-115&1|+|155195237|155195258&offsets|0|0(+1U)]|TATTGCACTCGTCCCGGCCTCCT</td><td>>MI0003560|hsa-mir-92b&WithFlank&1|+|155195171|155195278</td><td>67</td><td>88</td><td>22(+1U)</td><td>TATTGCACTCGTCCCGGCCTCCT</td><td>677.7</td><td>9.0</td><td>1.2</td><td>7.6</td><td>2.5E-14</td><td>1.2E-13</td><td>MB231/MB468</td><td>CYTO</td></tr>
<tr><td>>hg38_21_+_8433172_8446622_RNA45SN1_45S_50ntFlanks@6743.6772.30|CGACACTTCGAACGCACTTGCGGCCCCGGG</td><td>>hg38_21_+_8433172_8446622_RNA45SN1_45S_50ntFlanks</td><td>6743</td><td>6772</td><td>30</td><td>CGACACTTCGAACGCACTTGCGGCCCCGGG</td><td>547.8</td><td>-13.0</td><td>1.7</td><td>-7.6</td><td>2.2E-14</td><td>1.2E-13</td><td>MB231/MB468</td><td>MP</td></tr>
<tr><td>>hg38_21_+_8433172_8446622_RNA45SN1_45S_50ntFlanks@12957.12987.31|CTTCTGGGTCGGGGTTTCGTACGTAGCAGAG</td><td>>hg38_21_+_8433172_8446622_RNA45SN1_45S_50ntFlanks</td><td>12957</td><td>12987</td><td>31</td><td>CTTCTGGGTCGGGGTTTCGTACGTAGCAGAG</td><td>1013.9</td><td>-7.7</td><td>1.0</td><td>-7.6</td><td>2.7E-14</td><td>1.2E-13</td><td>MB231/MB468</td><td>NUC</td></tr>
<tr><td>>hg38_21_+_8433172_8446622_RNA45SN1_45S_50ntFlanks@9571.9596.26|GCAAATCGGTCGTCCGACCTGGGTAT</td><td>>hg38_21_+_8433172_8446622_RNA45SN1_45S_50ntFlanks</td><td>9571</td><td>9596</td><td>26</td><td>GCAAATCGGTCGTCCGACCTGGGTAT</td><td>756.6</td><td>-9.2</td><td>1.2</td><td>-7.6</td><td>3.5E-14</td><td>1.2E-13</td><td>MB231/MB468</td><td>MITO</td></tr>
<tr><td>>hg38_21_+_8433172_8446622_RNA45SN1_45S_50ntFlanks@4130.4162.33|ATTCCGGAGAGGGAGCCTGAGAAACGGCTACCA</td><td>>hg38_21_+_8433172_8446622_RNA45SN1_45S_50ntFlanks</td><td>4130</td><td>4162</td><td>33</td><td>ATTCCGGAGAGGGAGCCTGAGAAACGGCTACCA</td><td>663.8</td><td>-4.8</td><td>0.6</td><td>-7.6</td><td>2.9E-14</td><td>1.2E-13</td><td>BT20/MB468</td><td>CYTO</td></tr>
<tr><td>>hg38_21_+_8433172_8446622_RNA45SN1_45S_50ntFlanks@10798.10815.18|CATGTTGGAACAATGTAG</td><td>>hg38_21_+_8433172_8446622_RNA45SN1_45S_50ntFlanks</td><td>10798</td><td>10815</td><td>18</td><td>CATGTTGGAACAATGTAG</td><td>139.6</td><td>-11.0</td><td>1.4</td><td>-7.6</td><td>2.2E-14</td><td>1.2E-13</td><td>MB231/MB468</td><td>MP</td></tr>
<tr><td>>hg38_21_+_8433172_8446622_RNA45SN1_45S_50ntFlanks@8845.8867.23|CGGCGGCGGCGGCGGCGGCGGGA</td><td>>hg38_21_+_8433172_8446622_RNA45SN1_45S_50ntFlanks</td><td>8845</td><td>8867</td><td>23</td><td>CGGCGGCGGCGGCGGCGGCGGGA</td><td>243.2</td><td>-10.9</td><td>1.4</td><td>-7.6</td><td>2.7E-14</td><td>1.2E-13</td><td>MB231/MB468</td><td>NUC</td></tr>
<tr><td>>hg38_21_+_8433172_8446622_RNA45SN1_45S_50ntFlanks@12894.12924.31|CTCGCCCGTCACGCACCGCACGTTCGTGGGG</td><td>>hg38_21_+_8433172_8446622_RNA45SN1_45S_50ntFlanks</td><td>12894</td><td>12924</td><td>31</td><td>CTCGCCCGTCACGCACCGCACGTTCGTGGGG</td><td>140.9</td><td>-11.0</td><td>1.4</td><td>-7.6</td><td>2.2E-14</td><td>1.2E-13</td><td>MB231/MB468</td><td>MP</td></tr>
<tr><td>>hg38_21_+_8433172_8446622_RNA45SN1_45S_50ntFlanks@7985.8014.30|GATCAGACGTGGCGACCCGCTGAATTTAAG</td><td>>hg38_21_+_8433172_8446622_RNA45SN1_45S_50ntFlanks</td><td>7985</td><td>8014</td><td>30</td><td>GATCAGACGTGGCGACCCGCTGAATTTAAG</td><td>132.5</td><td>-10.9</td><td>1.4</td><td>-7.6</td><td>2.2E-14</td><td>1.2E-13</td><td>MB231/MB468</td><td>MP</td></tr>
<tr><td>>hg38_21_+_8433172_8446622_RNA45SN1_45S_50ntFlanks@9143.9161.19|GGGCGGACTGTCCCCAGTG</td><td>>hg38_21_+_8433172_8446622_RNA45SN1_45S_50ntFlanks</td><td>9143</td><td>9161</td><td>19</td><td>GGGCGGACTGTCCCCAGTG</td><td>168.6</td><td>-11.3</td><td>1.5</td><td>-7.6</td><td>2.2E-14</td><td>1.2E-13</td><td>MB231/MB468</td><td>MP</td></tr>
<tr><td>>MI0000093|hsa-mir-92a-1&WithFlank&13|+|91351308|91351397@54.75.22(+2A);MI0000094|hsa-mir-92a-2&WithFlank&X|-|134169532|134169618@54.75.22(+2A)&[MIMAT0000092&hsa-miR-92a-3p&offsets|0|0(+2A);m-9&13|+|91351361|91351382&offsets|0|0(+2A)];[MIMAT0000092_1&hsa-miR-92a-3p&offsets|0|0(+2A);m-11&X|-|134169544|134169565&offsets|0|0(+2A)]|TATTGCACTTGTCCCGGCCTGTAA</td><td>>MI0000093|hsa-mir-92a-1&WithFlank&13|+|91351308|91351397</td><td>54</td><td>75</td><td>22(+2A)</td><td>TATTGCACTTGTCCCGGCCTGTAA</td><td>655.2</td><td>5.3</td><td>0.7</td><td>7.6</td><td>2.1E-14</td><td>1.2E-13</td><td>BT20/MB468</td><td>MITO</td></tr>
<tr><td>>hg38_21_+_8433172_8446622_RNA45SN1_45S_50ntFlanks@5472.5505.34|ACGGCCCTGGCGGAGCGCTGAGAAGACGGTCGAA</td><td>>hg38_21_+_8433172_8446622_RNA45SN1_45S_50ntFlanks</td><td>5472</td><td>5505</td><td>34</td><td>ACGGCCCTGGCGGAGCGCTGAGAAGACGGTCGAA</td><td>506.7</td><td>7.2</td><td>0.9</td><td>7.6</td><td>2.6E-14</td><td>1.2E-13</td><td>MB231/MB468</td><td>CYTO</td></tr>
<tr><td>>hg38_21_+_8433172_8446622_RNA45SN1_45S_50ntFlanks@5421.5438.18|CGATTGGATGGTTTAGTG</td><td>>hg38_21_+_8433172_8446622_RNA45SN1_45S_50ntFlanks</td><td>5421</td><td>5438</td><td>18</td><td>CGATTGGATGGTTTAGTG</td><td>258.0</td><td>-11.9</td><td>1.6</td><td>-7.6</td><td>2.3E-14</td><td>1.2E-13</td><td>MB231/MB468</td><td>MP</td></tr>
<tr><td>>hg38_21_+_8433172_8446622_RNA45SN1_45S_50ntFlanks@3927.3959.33|CAGATCAAAACCAACCCGGTCAGCCCCTCTCCG</td><td>>hg38_21_+_8433172_8446622_RNA45SN1_45S_50ntFlanks</td><td>3927</td><td>3959</td><td>33</td><td>CAGATCAAAACCAACCCGGTCAGCCCCTCTCCG</td><td>219.2</td><td>-10.7</td><td>1.4</td><td>-7.6</td><td>2.7E-14</td><td>1.2E-13</td><td>MB231/MB468</td><td>NUC</td></tr>
<tr><td>>hg38_21_+_8433172_8446622_RNA45SN1_45S_50ntFlanks@5472.5495.24|ACGGCCCTGGCGGAGCGCTGAGAA</td><td>>hg38_21_+_8433172_8446622_RNA45SN1_45S_50ntFlanks</td><td>5472</td><td>5495</td><td>24</td><td>ACGGCCCTGGCGGAGCGCTGAGAA</td><td>253.3</td><td>6.1</td><td>0.8</td><td>7.6</td><td>2.6E-14</td><td>1.2E-13</td><td>MB231/MB468</td><td>CYTO</td></tr>
<tr><td>>hg38_21_+_8433172_8446622_RNA45SN1_45S_50ntFlanks@3937.3959.23|CCAACCCGGTCAGCCCCTCTCCG</td><td>>hg38_21_+_8433172_8446622_RNA45SN1_45S_50ntFlanks</td><td>3937</td><td>3959</td><td>23</td><td>CCAACCCGGTCAGCCCCTCTCCG</td><td>430.0</td><td>-12.5</td><td>1.7</td><td>-7.6</td><td>3.6E-14</td><td>1.2E-13</td><td>MB231/MB468</td><td>MITO</td></tr>
<tr><td>>hg38_21_+_8433172_8446622_RNA45SN1_45S_50ntFlanks@12627.12654.28|ATGGGGCGAAGCTACCATCTGTGGGATT</td><td>>hg38_21_+_8433172_8446622_RNA45SN1_45S_50ntFlanks</td><td>12627</td><td>12654</td><td>28</td><td>ATGGGGCGAAGCTACCATCTGTGGGATT</td><td>15497.4</td><td>-12.4</td><td>1.6</td><td>-7.6</td><td>4.3E-14</td><td>1.2E-13</td><td>BT20/MB468</td><td>NUC</td></tr>
<tr><td>>hg38_21_+_8433172_8446622_RNA45SN1_45S_50ntFlanks@10664.10706.43|CTTGGAAAGCGTCGCGGTTCCGGCGGCGTCCGGTGAGCTCTCG</td><td>>hg38_21_+_8433172_8446622_RNA45SN1_45S_50ntFlanks</td><td>10664</td><td>10706</td><td>43</td><td>CTTGGAAAGCGTCGCGGTTCCGGCGGCGTCCGGTGAGCTCTCG</td><td>1537.9</td><td>-9.8</td><td>1.3</td><td>-7.6</td><td>3.6E-14</td><td>1.2E-13</td><td>MB231/MB468</td><td>MITO</td></tr>
<tr><td>>trnaMT_MetCAT_MT_+_4402_4469@37.71.35|CCGAAAATGTTGGTTATACCCTTCCCGTACTACCA</td><td>>trnaMT_MetCAT_MT_+_4402_4469</td><td>37</td><td>71</td><td>35</td><td>CCGAAAATGTTGGTTATACCCTTCCCGTACTACCA</td><td>1283.1</td><td>6.8</td><td>0.9</td><td>7.6</td><td>2.2E-14</td><td>1.2E-13</td><td>BT20/MB468</td><td>MITO</td></tr>
<tr><td>>hg38_21_+_8433172_8446622_RNA45SN1_45S_50ntFlanks@485.507.23|GATGTGGTGACGTCGTGCTCTCC</td><td>>hg38_21_+_8433172_8446622_RNA45SN1_45S_50ntFlanks</td><td>485</td><td>507</td><td>23</td><td>GATGTGGTGACGTCGTGCTCTCC</td><td>1076.5</td><td>27.5</td><td>3.6</td><td>7.6</td><td>3.6E-14</td><td>1.2E-13</td><td>MB231/MB468</td><td>MITO</td></tr>
<tr><td>>hg38_21_+_8433172_8446622_RNA45SN1_45S_50ntFlanks@4657.4688.32|CTAGAGGTGAAATTCTTGGACCGGCGCAAGAC</td><td>>hg38_21_+_8433172_8446622_RNA45SN1_45S_50ntFlanks</td><td>4657</td><td>4688</td><td>32</td><td>CTAGAGGTGAAATTCTTGGACCGGCGCAAGAC</td><td>267.8</td><td>-11.9</td><td>1.6</td><td>-7.6</td><td>2.3E-14</td><td>1.2E-13</td><td>MB231/MB468</td><td>MP</td></tr>
<tr><td>>MI0003667|hsa-mir-652&WithFlank&X|+|110055323|110055432@67.87.21&[MIMAT0003322&hsa-miR-652-3p&offsets|0|0;m-148&X|+|110055389|110055410&offsets|0|-1]|AATGGCGCCACTAGGGTTGTG</td><td>>MI0003667|hsa-mir-652&WithFlank&X|+|110055323|110055432</td><td>67</td><td>87</td><td>21</td><td>AATGGCGCCACTAGGGTTGTG</td><td>69.5</td><td>3.0</td><td>0.4</td><td>7.6</td><td>4.3E-14</td><td>1.2E-13</td><td>BT20/MB468</td><td>NUC</td></tr>
<tr><td>>am_MI0000809|hsa-mir-151a&WithFlank&8|-|140732558|140732659@53.73.21(+1C)&[MIMAT0000757&hsa-miR-151a-3p&offsets|0|0(+1C);m-30&8|-|140732587|140732607&offsets|0|0(+1C)]|CTAGACTGAAGCTCCTTGAGGC</td><td>>am_MI0000809|hsa-mir-151a&WithFlank&8|-|140732558|140732659</td><td>53</td><td>73</td><td>21(+1C)</td><td>CTAGACTGAAGCTCCTTGAGGC</td><td>744.2</td><td>7.5</td><td>1.0</td><td>7.6</td><td>2.6E-14</td><td>1.2E-13</td><td>MB231/MB468</td><td>CYTO</td></tr>
<tr><td>>hg38_21_+_8433172_8446622_RNA45SN1_45S_50ntFlanks@12623.12654.32|GCCAATGGGGCGAAGCTACCATCTGTGGGATT</td><td>>hg38_21_+_8433172_8446622_RNA45SN1_45S_50ntFlanks</td><td>12623</td><td>12654</td><td>32</td><td>GCCAATGGGGCGAAGCTACCATCTGTGGGATT</td><td>147.7</td><td>-5.4</td><td>0.7</td><td>-7.6</td><td>2.6E-14</td><td>1.2E-13</td><td>MB231/MB468</td><td>CYTO</td></tr>
<tr><td>>MI0000071|hsa-mir-17&WithFlank&13|+|91350599|91350694@20.42.23(+1A)&[MIMAT0000070&hsa-miR-17-5p&offsets|0|0(+1A);m-88&13|+|91350618|91350640&offsets|0|0(+1A)]|CAAAGTGCTTACAGTGCAGGTAGA</td><td>>MI0000071|hsa-mir-17&WithFlank&13|+|91350599|91350694</td><td>20</td><td>42</td><td>23(+1A)</td><td>CAAAGTGCTTACAGTGCAGGTAGA</td><td>3137.8</td><td>4.7</td><td>0.6</td><td>7.6</td><td>2.7E-14</td><td>1.2E-13</td><td>MB231/MB468</td><td>NUC</td></tr>
<tr><td>>MI0000254|hsa-mir-30c-2&WithFlank&6|-|71376954|71377037@53.73.21(+1C)&[MIMAT0004550&hsa-miR-30c-2-3p&offsets|0|-1(+1C);m-206&6|-|71376964|71376985&offsets|0|-1(+1C)]|CTGGGAGAAGGCTGTTTACTCC</td><td>>MI0000254|hsa-mir-30c-2&WithFlank&6|-|71376954|71377037</td><td>53</td><td>73</td><td>21(+1C)</td><td>CTGGGAGAAGGCTGTTTACTCC</td><td>463.2</td><td>7.4</td><td>1.0</td><td>7.6</td><td>2.9E-14</td><td>1.2E-13</td><td>BT20/MB468</td><td>CYTO</td></tr>
<tr><td>>MI0000456|hsa-mir-140&WithFlank&16|+|69933075|69933186@69.88.20&[MIMAT0004597&hsa-miR-140-3p&offsets|+1|0;m-45&16|+|69933143|69933164&offsets|0|-2]|ACCACAGGGTAGAACCACGG</td><td>>MI0000456|hsa-mir-140&WithFlank&16|+|69933075|69933186</td><td>69</td><td>88</td><td>20</td><td>ACCACAGGGTAGAACCACGG</td><td>225.3</td><td>9.8</td><td>1.3</td><td>7.6</td><td>2.8E-14</td><td>1.2E-13</td><td>MB231/MB468</td><td>NUC</td></tr>
<tr><td>>hg38_21_+_8433172_8446622_RNA45SN1_45S_50ntFlanks@10574.10596.23|AAGCCGGCGGGAGCCCCGGGGAG</td><td>>hg38_21_+_8433172_8446622_RNA45SN1_45S_50ntFlanks</td><td>10574</td><td>10596</td><td>23</td><td>AAGCCGGCGGGAGCCCCGGGGAG</td><td>291.9</td><td>-7.9</td><td>1.0</td><td>-7.6</td><td>2.6E-14</td><td>1.2E-13</td><td>MB231/MB468</td><td>CYTO</td></tr>
<tr><td>>hg38_21_+_8433172_8446622_RNA45SN1_45S_50ntFlanks@13002.13025.24|CGATCTATTGAAAGTCAGCCCTCG</td><td>>hg38_21_+_8433172_8446622_RNA45SN1_45S_50ntFlanks</td><td>13002</td><td>13025</td><td>24</td><td>CGATCTATTGAAAGTCAGCCCTCG</td><td>657.2</td><td>-8.1</td><td>1.1</td><td>-7.6</td><td>2.6E-14</td><td>1.2E-13</td><td>MB231/MB468</td><td>CYTO</td></tr>
<tr><td>>hg38_21_+_8433172_8446622_RNA45SN1_45S_50ntFlanks@5003.5022.20|ATAGCTCTTTCTCGATTCCG</td><td>>hg38_21_+_8433172_8446622_RNA45SN1_45S_50ntFlanks</td><td>5003</td><td>5022</td><td>20</td><td>ATAGCTCTTTCTCGATTCCG</td><td>498.6</td><td>-12.7</td><td>1.7</td><td>-7.6</td><td>3.7E-14</td><td>1.2E-13</td><td>MB231/MB468</td><td>MITO</td></tr>
<tr><td>>hg38_21_+_8433172_8446622_RNA45SN1_45S_50ntFlanks@10136.10158.23|GCGGCGGCGGCGGGGGTGTGGGG</td><td>>hg38_21_+_8433172_8446622_RNA45SN1_45S_50ntFlanks</td><td>10136</td><td>10158</td><td>23</td><td>GCGGCGGCGGCGGGGGTGTGGGG</td><td>277.3</td><td>-12.0</td><td>1.6</td><td>-7.6</td><td>2.3E-14</td><td>1.2E-13</td><td>MB231/MB468</td><td>MP</td></tr>
<tr><td>>hg38_21_+_8433172_8446622_RNA45SN1_45S_50ntFlanks@10409.10426.18|GTCAGTCGGTCCTGAGAG</td><td>>hg38_21_+_8433172_8446622_RNA45SN1_45S_50ntFlanks</td><td>10409</td><td>10426</td><td>18</td><td>GTCAGTCGGTCCTGAGAG</td><td>162.6</td><td>-11.2</td><td>1.5</td><td>-7.6</td><td>2.3E-14</td><td>1.2E-13</td><td>MB231/MB468</td><td>MP</td></tr>
<tr><td>>hg38_21_+_8433172_8446622_RNA45SN1_45S_50ntFlanks@12437.12473.37|CATTGTGAAGCAGAATTCACCAAGCGTTGGATTGTTC</td><td>>hg38_21_+_8433172_8446622_RNA45SN1_45S_50ntFlanks</td><td>12437</td><td>12473</td><td>37</td><td>CATTGTGAAGCAGAATTCACCAAGCGTTGGATTGTTC</td><td>2069.6</td><td>-5.5</td><td>0.7</td><td>-7.6</td><td>2.8E-14</td><td>1.2E-13</td><td>MB231/MB468</td><td>NUC</td></tr>
<tr><td>>hg38_21_+_8433172_8446622_RNA45SN1_45S_50ntFlanks@12499.12517.19|GGTTTAGACCGTCGTGAGA</td><td>>hg38_21_+_8433172_8446622_RNA45SN1_45S_50ntFlanks</td><td>12499</td><td>12517</td><td>19</td><td>GGTTTAGACCGTCGTGAGA</td><td>196.6</td><td>-10.3</td><td>1.4</td><td>-7.6</td><td>2.6E-14</td><td>1.2E-13</td><td>MB231/MB468</td><td>CYTO</td></tr>
<tr><td>>hg38_21_+_8433172_8446622_RNA45SN1_45S_50ntFlanks@3835.3859.25|CTCCTCTCCTACTTGGATAACTGTG</td><td>>hg38_21_+_8433172_8446622_RNA45SN1_45S_50ntFlanks</td><td>3835</td><td>3859</td><td>25</td><td>CTCCTCTCCTACTTGGATAACTGTG</td><td>568.4</td><td>-12.9</td><td>1.7</td><td>-7.6</td><td>3.7E-14</td><td>1.2E-13</td><td>MB231/MB468</td><td>MITO</td></tr>
<tr><td>>trnaMT_GluTTC_MT_-_14674_14742@35.72.38,trnalookalike8_GluTTC_5_-_93905172_93905240@35.72.38|TATCATTGGTCGTGGTTGTAGTCCGTGCGAGAATACCA</td><td>>trnaMT_GluTTC_MT_-_14674_14742</td><td>35</td><td>72</td><td>38</td><td>TATCATTGGTCGTGGTTGTAGTCCGTGCGAGAATACCA</td><td>6580.4</td><td>6.5</td><td>0.8</td><td>7.6</td><td>2.2E-14</td><td>1.2E-13</td><td>BT20/MB468</td><td>MITO</td></tr>
<tr><td>>hg38_21_+_8433172_8446622_RNA45SN1_45S_50ntFlanks@6733.6771.39|CATTGATCATCGACACTTCGAACGCACTTGCGGCCCCGG</td><td>>hg38_21_+_8433172_8446622_RNA45SN1_45S_50ntFlanks</td><td>6733</td><td>6771</td><td>39</td><td>CATTGATCATCGACACTTCGAACGCACTTGCGGCCCCGG</td><td>3610.6</td><td>-4.4</td><td>0.6</td><td>-7.6</td><td>2.2E-14</td><td>1.2E-13</td><td>BT20/MB468</td><td>MITO</td></tr>
<tr><td>>MI0000299|hsa-mir-222&WithFlank&X|-|45747009|45747130@75.99.25&[MIMAT0000279&hsa-miR-222-3p&offsets|0|+4;m-140&X|-|45747033|45747056&offsets|0|+1]|AGCTACATCTGGCTACTGGGTCTCT</td><td>>MI0000299|hsa-mir-222&WithFlank&X|-|45747009|45747130</td><td>75</td><td>99</td><td>25</td><td>AGCTACATCTGGCTACTGGGTCTCT</td><td>31679.3</td><td>5.9</td><td>0.8</td><td>7.6</td><td>2.8E-14</td><td>1.2E-13</td><td>MB231/MB468</td><td>NUC</td></tr>
<tr><td>>hg38_21_+_8433172_8446622_RNA45SN1_45S_50ntFlanks@9029.9065.37|TCTCCTCGTGGGGGGGCCGGGCCACCCCTCCCACGGC</td><td>>hg38_21_+_8433172_8446622_RNA45SN1_45S_50ntFlanks</td><td>9029</td><td>9065</td><td>37</td><td>TCTCCTCGTGGGGGGGCCGGGCCACCCCTCCCACGGC</td><td>316.4</td><td>-12.1</td><td>1.6</td><td>-7.6</td><td>3.7E-14</td><td>1.2E-13</td><td>MB231/MB468</td><td>MITO</td></tr>
<tr><td>>hg38_21_+_8433172_8446622_RNA45SN1_45S_50ntFlanks@10914.10946.33|CGCCGCGGCTGGACGAGGCGCCGCCGCCCCCCC</td><td>>hg38_21_+_8433172_8446622_RNA45SN1_45S_50ntFlanks</td><td>10914</td><td>10946</td><td>33</td><td>CGCCGCGGCTGGACGAGGCGCCGCCGCCCCCCC</td><td>388.6</td><td>-11.6</td><td>1.5</td><td>-7.6</td><td>2.8E-14</td><td>1.2E-13</td><td>MB231/MB468</td><td>NUC</td></tr>
<tr><td>>hg38_21_+_8433172_8446622_RNA45SN1_45S_50ntFlanks@9033.9055.23|CTCGTGGGGGGGCCGGGCCACCC</td><td>>hg38_21_+_8433172_8446622_RNA45SN1_45S_50ntFlanks</td><td>9033</td><td>9055</td><td>23</td><td>CTCGTGGGGGGGCCGGGCCACCC</td><td>343.4</td><td>-12.2</td><td>1.6</td><td>-7.6</td><td>3.7E-14</td><td>1.2E-13</td><td>MB231/MB468</td><td>MITO</td></tr>
<tr><td>>hg38_21_+_8433172_8446622_RNA45SN1_45S_50ntFlanks@11315.11332.18|CCACCCCACGTCTCGTCG</td><td>>hg38_21_+_8433172_8446622_RNA45SN1_45S_50ntFlanks</td><td>11315</td><td>11332</td><td>18</td><td>CCACCCCACGTCTCGTCG</td><td>1290.9</td><td>-13.3</td><td>1.7</td><td>-7.6</td><td>2.8E-14</td><td>1.2E-13</td><td>MB231/MB468</td><td>NUC</td></tr>
</table>

| | | | | | | | | | | | | | |
|---|---|---|---|---|---|---|---|---|---|---|---|---|---|
| >MI0000466\|hsa-mir-9-1&WithFlank&1\|-\|156420335\|156420435@22.44.23;MI0000468\|hsa-mir-9-3&WithFlank&15\|+\|89368011\|89368112@22.44.23;MI0000467\|hsa-mir-9-2&WithFlank&5\|-\|88666847\|88666945@22.44.23&[MIMAT0000441&hsa-miR-9-5p&offsets\|0\|0;m-104&1\|-\|156420392\|156420414&offsets\|0\|0];[MIMAT0000441_1&hsa-miR-9-5p&offsets\|0\|0;m-105&15\|+\|89368032\|89368054&offsets\|0\|0];[MIMAT0000441_2&hsa-miR-9-5p&offsets\|0\|0;m-104&5\|-\|88666902\|88666924&offsets\|0\|0]\|TCTTTGGTTATCTAGCTGTATGA | >MI0000466\|hsa-mir-9-1&WithFlank&1\|-\|156420335\|156420435 | 22 | 44 | 23 | TCTTTGGTTATCTAGCTGTATGA | 347.4 | 6.3 | 0.8 | 7.6 | 2.8E-14 | 1.2E-13 | MB231/MB468 | NUC |
| >hg38_21_+_8433172_8446622_RNA45SN1_45S_50ntFlanks@10375.10406.32\|AGAAGGGTTCCATGTGAACAGCAGTTGAACAT | >hg38_21_+_8433172_8446622_RNA45SN1_45S_50ntFlanks | 10375 | 10406 | 32 | AGAAGGGTTCCATGTGAACAGCAGTTGAACAT | 214.0 | -11.6 | 1.5 | -7.6 | 2.3E-14 | 1.2E-13 | MB231/MB468 | MP |
| >hg38_21_+_8433172_8446622_RNA45SN1_45S_50ntFlanks@12073.12097.25\|CCGAGGGGCTCTCGCTTCTGGCGCC | >hg38_21_+_8433172_8446622_RNA45SN1_45S_50ntFlanks | 12073 | 12097 | 25 | CCGAGGGGCTCTCGCTTCTGGCGCC | 3696.9 | -8.0 | 1.1 | -7.6 | 3.7E-14 | 1.2E-13 | MB231/MB468 | MITO |
| >hg38_21_+_8433172_8446622_RNA45SN1_45S_50ntFlanks@10128.10147.20\|CGGCGGCGGCGGCGGCGGCG | >hg38_21_+_8433172_8446622_RNA45SN1_45S_50ntFlanks | 10128 | 10147 | 20 | CGGCGGCGGCGGCGGCGGCG | 143.9 | -9.9 | 1.3 | -7.6 | 2.7E-14 | 1.2E-13 | MB231/MB468 | CYTO |
| >hg38_21_+_8433172_8446622_RNA45SN1_45S_50ntFlanks@8845.8864.20\|CGGCGGCGGCGGCGGCGGCG | >hg38_21_+_8433172_8446622_RNA45SN1_45S_50ntFlanks | 8845 | 8864 | 20 | CGGCGGCGGCGGCGGCGGCG | 143.9 | -9.9 | 1.3 | -7.6 | 2.7E-14 | 1.2E-13 | MB231/MB468 | CYTO |
| >hg38_21_+_8433172_8446622_RNA45SN1_45S_50ntFlanks@10534.10566.33\|CCGCGAGGCGTCCAGTGCGGTAACGCGACCGAT | >hg38_21_+_8433172_8446622_RNA45SN1_45S_50ntFlanks | 10534 | 10566 | 33 | CCGCGAGGCGTCCAGTGCGGTAACGCGACCGAT | 151.9 | -11.1 | 1.5 | -7.6 | 2.4E-14 | 1.2E-13 | MB231/MB468 | MP |
| >hg38_21_+_8433172_8446622_RNA45SN1_45S_50ntFlanks@12404.12431.28\|GCTTTTTGATCCTTCGATGTCGGCTCTT | >hg38_21_+_8433172_8446622_RNA45SN1_45S_50ntFlanks | 12404 | 12431 | 28 | GCTTTTTGATCCTTCGATGTCGGCTCTT | 416.0 | -3.8 | 0.5 | -7.6 | 3.0E-14 | 1.3E-13 | BT20/MB468 | CYTO |
| >hg38_21_+_8433172_8446622_RNA45SN1_45S_50ntFlanks@12588.12616.29\|GGTTCAGACATTTGGTGTATGTGCTTGGC | >hg38_21_+_8433172_8446622_RNA45SN1_45S_50ntFlanks | 12588 | 12616 | 29 | GGTTCAGACATTTGGTGTATGTGCTTGGC | 187.4 | -11.4 | 1.5 | -7.6 | 2.4E-14 | 1.3E-13 | MB231/MB468 | MP |
| >hg38_21_+_8433172_8446622_RNA45SN1_45S_50ntFlanks@3985.4025.41\|CGGCTTTGGTGACTCTAGATAACCTCGGGCCGATCGCACGC | >hg38_21_+_8433172_8446622_RNA45SN1_45S_50ntFlanks | 3985 | 4025 | 41 | CGGCTTTGGTGACTCTAGATAACCTCGGGCCGATCGCACGC | 357.4 | -12.3 | 1.6 | -7.6 | 3.8E-14 | 1.3E-13 | MB231/MB468 | MITO |
| >hg38_21_+_8433172_8446622_RNA45SN1_45S_50ntFlanks@11190.11208.19\|CCCGGGGGGCCGGCGGCGG | >hg38_21_+_8433172_8446622_RNA45SN1_45S_50ntFlanks | 11190 | 11208 | 19 | CCCGGGGGGCCGGCGGCGG | 374.3 | -12.3 | 1.6 | -7.6 | 3.8E-14 | 1.3E-13 | MB231/MB468 | MITO |
| >hg38_21_+_8433172_8446622_RNA45SN1_45S_50ntFlanks@10805.10825.21\|GAACAATGTAGGTAAGGGAAG | >hg38_21_+_8433172_8446622_RNA45SN1_45S_50ntFlanks | 10805 | 10825 | 21 | GAACAATGTAGGTAAGGGAAG | 212.1 | -7.9 | 1.0 | -7.6 | 2.7E-14 | 1.3E-13 | MB231/MB468 | CYTO |
| >hg38_21_+_8433172_8446622_RNA45SN1_45S_50ntFlanks@12184.12211.28\|AACGGTAACGCAGGTGTCCTAAGGCGAG | >hg38_21_+_8433172_8446622_RNA45SN1_45S_50ntFlanks | 12184 | 12211 | 28 | AACGGTAACGCAGGTGTCCTAAGGCGAG | 176.1 | -11.3 | 1.5 | -7.6 | 2.4E-14 | 1.3E-13 | MB231/MB468 | MP |
| >hg38_21_+_8433172_8446622_RNA45SN1_45S_50ntFlanks@12615.12657.43\|GCTGAGGAGCCAATGGGGCGAAGCTACCATCTGTGGGATTATG | >hg38_21_+_8433172_8446622_RNA45SN1_45S_50ntFlanks | 12615 | 12657 | 43 | GCTGAGGAGCCAATGGGGCGAAGCTACCATCTGTGGGATTATG | 1250.2 | -8.3 | 1.1 | -7.6 | 3.8E-14 | 1.3E-13 | MB231/MB468 | MITO |
| >hg38_21_+_8433172_8446622_RNA45SN1_45S_50ntFlanks@12934.12953.20\|CTAAACCATTCGTAGACGAC | >hg38_21_+_8433172_8446622_RNA45SN1_45S_50ntFlanks | 12934 | 12953 | 20 | CTAAACCATTCGTAGACGAC | 673.9 | -8.9 | 1.2 | -7.6 | 3.8E-14 | 1.3E-13 | MB231/MB468 | MITO |
| >MI0000108\|hsa-mir-103a-2&WithFlank&20\|+\|3917488\|3917577@54.75.22;MI0000109\|hsa-mir-103a-1&WithFlank&5\|-\|168560890\|168560979@54.75.22&[MIMAT0000101&hsa-miR-103a-3p&offsets\|0\|-1;m-12&20\|+\|3917541\|3917563&offsets\|0\|-1];[MIMAT0000101_1&hsa-miR-103a-3p&offsets\|0\|-1;m-13&5\|-\|168560904\|168560926&offsets\|0\|-1;m-44220&5\|-\|168560908\|168560926&offsets\|0\|+3]\|AGCAGCATTGTACAGGGCTATG | >MI0000108\|hsa-mir-103a-2&WithFlank&20\|+\|3917488\|3917577 | 54 | 75 | 22 | AGCAGCATTGTACAGGGCTATG | 8451.9 | 7.0 | 0.9 | 7.6 | 2.3E-14 | 1.3E-13 | BT20/MB468 | MITO |
| >hg38_21_+_8433172_8446622_RNA45SN1_45S_50ntFlanks@11323.11340.18\|CGTCTCGTCGCGCGCGCG | >hg38_21_+_8433172_8446622_RNA45SN1_45S_50ntFlanks | 11323 | 11340 | 18 | CGTCTCGTCGCGCGCGCG | 1013.7 | -7.1 | 0.9 | -7.6 | 2.7E-14 | 1.3E-13 | MB231/MB468 | CYTO |
| >hg38_21_+_8433172_8446622_RNA45SN1_45S_50ntFlanks@10771.10797.27\|CAGGTCTCCAAGGTGAACAGCCTCTGG | >hg38_21_+_8433172_8446622_RNA45SN1_45S_50ntFlanks | 10771 | 10797 | 27 | CAGGTCTCCAAGGTGAACAGCCTCTGG | 647.3 | -12.4 | 1.5 | -8.0 | 9.9E-16 | 1.3E-13 | BT20/MB468 | MP |
| >hg38_21_+_8433172_8446622_RNA45SN1_45S_50ntFlanks@11661.11679.19\|TGATTTCTGCCCAGTGCTC | >hg38_21_+_8433172_8446622_RNA45SN1_45S_50ntFlanks | 11661 | 11679 | 19 | TGATTTCTGCCCAGTGCTC | 2307.6 | -8.5 | 1.1 | -7.6 | 2.9E-14 | 1.3E-13 | MB231/MB468 | NUC |
| >hg38_21_+_8433172_8446622_RNA45SN1_45S_50ntFlanks@8115.8140.26\|CGCGGGACATGTGGCGTACGGAAGAC | >hg38_21_+_8433172_8446622_RNA45SN1_45S_50ntFlanks | 8115 | 8140 | 26 | CGCGGGACATGTGGCGTACGGAAGAC | 221.6 | -10.8 | 1.4 | -7.6 | 2.9E-14 | 1.3E-13 | MB231/MB468 | NUC |
| >hg38_21_+_8433172_8446622_RNA45SN1_45S_50ntFlanks@12406.12443.38\|TTTTTGATCCTTCGATGTCGGCTCTTCCTATCATTGTG | >hg38_21_+_8433172_8446622_RNA45SN1_45S_50ntFlanks | 12406 | 12443 | 38 | TTTTTGATCCTTCGATGTCGGCTCTTCCTATCATTGTG | 3863.6 | -9.6 | 1.3 | -7.5 | 4.5E-14 | 1.3E-13 | BT20/MB468 | NUC |
| >hg38_21_+_8433172_8446622_RNA45SN1_45S_50ntFlanks@11617.11640.24\|AAAACAAAGCATCGCGAAGGCCCG | >hg38_21_+_8433172_8446622_RNA45SN1_45S_50ntFlanks | 11617 | 11640 | 24 | AAAACAAAGCATCGCGAAGGCCCG | 386.1 | -10.7 | 1.4 | -7.6 | 2.3E-14 | 1.3E-13 | BT20/MB468 | MITO |
| >hg38_21_+_8433172_8446622_RNA45SN1_45S_50ntFlanks@11729.11766.38\|TAACTATGACTCTCTTAAGGTAGCCAAATGCCTCGTCA | >hg38_21_+_8433172_8446622_RNA45SN1_45S_50ntFlanks | 11729 | 11766 | 38 | TAACTATGACTCTCTTAAGGTAGCCAAATGCCTCGTCA | 323.0 | -12.1 | 1.6 | -7.6 | 3.8E-14 | 1.3E-13 | MB231/MB468 | MITO |
| >hg38_21_+_8433172_8446622_RNA45SN1_45S_50ntFlanks@9598.9616.19\|GGGGCGAAAGACTAATCGA | >hg38_21_+_8433172_8446622_RNA45SN1_45S_50ntFlanks | 9598 | 9616 | 19 | GGGGCGAAAGACTAATCGA | 177.4 | -11.3 | 1.5 | -7.6 | 2.4E-14 | 1.3E-13 | MB231/MB468 | MP |
| >MI0000289\|hsa-mir-181a-1&WithFlank&1\|-\|198859038\|198859159@30.52.23;MI0000269\|hsa-mir-181a-2&WithFlank&9\|+\|124692436\|124692557@45.67.23&[MIMAT0000256&hsa-miR-181a-5p&offsets\|0\|0;m-32&1\|-\|198859108\|198859130&offsets\|0\|0];[MIMAT0000256_1&hsa-miR-181a-5p&offsets\|0\|0;m-32&9\|+\|124692480\|124692502&offsets\|0\|0]\|AACATTCAACGCTGTCGGTGAGT | >MI0000289\|hsa-mir-181a-1&WithFlank&1\|-\|198859038\|198859159 | 30 | 52 | 23 | AACATTCAACGCTGTCGGTGAGT | 1592.6 | 4.8 | 0.6 | 7.6 | 2.3E-14 | 1.3E-13 | BT20/MB468 | MITO |
| >MI0006444\|hsa-mir-1307&WithFlank&10\|-\|103394247\|103394407@86.107.22&[MIMAT0005951&hsa-miR-1307-3p&offsets\|0\|0;m-66&10\|-\|103394300\|103394322&offsets\|0\|-1]\|ACTCGGCGTGGCGTCGGTCGTG | >MI0006444\|hsa-mir-1307&WithFlank&10\|-\|103394247\|103394407 | 86 | 107 | 22 | ACTCGGCGTGGCGTCGGTCGTG | 766.1 | 5.7 | 0.8 | 7.6 | 2.7E-14 | 1.3E-13 | MB231/MB468 | CYTO |
| >hg38_21_+_8433172_8446622_RNA45SN1_45S_50ntFlanks@11561.11592.32\|GCGCCTAGCAGCCGACTTAGAACTGGTGCGGA | >hg38_21_+_8433172_8446622_RNA45SN1_45S_50ntFlanks | 11561 | 11592 | 32 | GCGCCTAGCAGCCGACTTAGAACTGGTGCGGA | 1319.2 | -8.5 | 1.1 | -7.6 | 2.7E-14 | 1.3E-13 | MB231/MB468 | CYTO |
| >Antisense_hg38_21_-_8433172_8446622_RNA45SN1_45S_50ntFlanks@7674.7693.20\|CTCTCTCTCTCTCTCCCCCG | >Antisense_hg38_21_-_8433172_8446622_RNA45SN1_45S_50ntFlanks | 7674 | 7693 | 20 | CTCTCTCTCTCTCTCCCCCG | 1020.5 | -8.9 | 1.2 | -7.6 | 2.8E-14 | 1.3E-13 | MB231/MB468 | CYTO |
| >hg38_21_+_8433172_8446622_RNA45SN1_45S_50ntFlanks@11027.11046.20\|CTCTCTCTCTCTCTCCCCCG | >hg38_21_+_8433172_8446622_RNA45SN1_45S_50ntFlanks | 11027 | 11046 | 20 | CTCTCTCTCTCTCTCCCCCG | 1020.5 | -8.9 | 1.2 | -7.6 | 2.8E-14 | 1.3E-13 | MB231/MB468 | CYTO |
| >MI0000816\|hsa-mir-335&WithFlank&7\|+\|130496105\|130496210@58.78.21(+1A)&[MIMAT0004703&hsa-miR-335-3p&offsets\|0\|-1(+1A);m-159&7\|+\|130496162\|130496183&offsets\|0\|-1(+1A)]\|TTTTTCATTATTGCTCCTGACA | >MI0000816\|hsa-mir-335&WithFlank&7\|+\|130496105\|130496210 | 58 | 78 | 21(+1A) | TTTTTCATTATTGCTCCTGACA | 741.6 | 7.9 | 1.0 | 7.6 | 3.1E-14 | 1.3E-13 | BT20/MB468 | CYTO |
| >hg38_21_+_8433172_8446622_RNA45SN1_45S_50ntFlanks@12676.12707.32\|AGAATCCCGCCCAGGCGGAACGATACGGCAGC | >hg38_21_+_8433172_8446622_RNA45SN1_45S_50ntFlanks | 12676 | 12707 | 32 | AGAATCCCGCCCAGGCGGAACGATACGGCAGC | 132.4 | -10.9 | 1.4 | -7.6 | 2.4E-14 | 1.3E-13 | MB231/MB468 | MP |
| >hg38_21_+_8433172_8446622_RNA45SN1_45S_50ntFlanks@10458.10484.27\|ATGGCCTCCGTTGCCCTCGGCCGATCG | >hg38_21_+_8433172_8446622_RNA45SN1_45S_50ntFlanks | 10458 | 10484 | 27 | ATGGCCTCCGTTGCCCTCGGCCGATCG | 286.9 | -12.0 | 1.6 | -7.6 | 2.4E-14 | 1.3E-13 | MB231/MB468 | MP |
| >hg38_21_+_8433172_8446622_RNA45SN1_45S_50ntFlanks@10408.10425.18\|GGTCAGTCGGTCCTGAGA | >hg38_21_+_8433172_8446622_RNA45SN1_45S_50ntFlanks | 10408 | 10425 | 18 | GGTCAGTCGGTCCTGAGA | 211.6 | -11.6 | 1.5 | -7.6 | 2.4E-14 | 1.3E-13 | MB231/MB468 | MP |
| >MI0000477\|hsa-mir-146a&WithFlank&5\|+\|160485346\|160485456@27.47.21(+1G)&[MIMAT0000449&hsa-miR-146a-5p&offsets\|0\|-1(+1G);m-129&5\|+\|160485372\|160485393&offsets\|0\|-1(+1G)]\|TGAGAACTGAATTCCATGGGTG | >MI0000477\|hsa-mir-146a&WithFlank&5\|+\|160485346\|160485456 | 27 | 47 | 21(+1G) | TGAGAACTGAATTCCATGGGTG | 367.4 | 12.4 | 1.6 | 7.6 | 2.9E-14 | 1.3E-13 | MB231/MB468 | NUC |
| >hg38_21_+_8433172_8446622_RNA45SN1_45S_50ntFlanks@4732.4759.28\|AGAACGAAAGTCGGAGGTTCGAAGACGA | >hg38_21_+_8433172_8446622_RNA45SN1_45S_50ntFlanks | 4732 | 4759 | 28 | AGAACGAAAGTCGGAGGTTCGAAGACGA | 1674.1 | -5.2 | 0.7 | -7.6 | 2.4E-14 | 1.3E-13 | MB231/MB468 | MP |
| >hg38_21_+_8433172_8446622_RNA45SN1_45S_50ntFlanks@12258.12284.27\|GCTTGATCTTGATTTTCAGTACGAATA | >hg38_21_+_8433172_8446622_RNA45SN1_45S_50ntFlanks | 12258 | 12284 | 27 | GCTTGATCTTGATTTTCAGTACGAATA | 635.3 | -13.2 | 1.7 | -7.6 | 2.4E-14 | 1.3E-13 | MB231/MB468 | MP |
| >hg38_21_+_8433172_8446622_RNA45SN1_45S_50ntFlanks@11874.11895.22\|AGAAGACCCTGTTGAGCTTGAC | >hg38_21_+_8433172_8446622_RNA45SN1_45S_50ntFlanks | 11874 | 11895 | 22 | AGAAGACCCTGTTGAGCTTGAC | 162.2 | -11.2 | 1.5 | -7.6 | 2.5E-14 | 1.3E-13 | MB231/MB468 | MP |
| >MI0000077\|hsa-mir-21&WithFlank&17\|+\|59841260\|59841343@13.34.22&[MIMAT0000076&hsa-miR-21-5p&offsets\|-1\|-1;m-1&17\|+\|59841273\|59841294&offsets\|-1\|-1]\|GTAGCTTATCAGACTGATGTTG | >MI0000077\|hsa-mir-21&WithFlank&17\|+\|59841260\|59841343 | 13 | 34 | 22 | GTAGCTTATCAGACTGATGTTG | 405.2 | 7.0 | 0.9 | 7.6 | 3.2E-14 | 1.3E-13 | BT20/MB468 | CYTO |
| >hg38_21_+_8433172_8446622_RNA45SN1_45S_50ntFlanks@9554.9596.43\|CGTAGCGGTCCTGACGTGCAAATCGGTCGTCCGACCTGGGTAT | >hg38_21_+_8433172_8446622_RNA45SN1_45S_50ntFlanks | 9554 | 9596 | 43 | CGTAGCGGTCCTGACGTGCAAATCGGTCGTCCGACCTGGGTAT | 2888.4 | -9.4 | 1.2 | -7.6 | 3.9E-14 | 1.3E-13 | MB231/MB468 | MITO |
| >hg38_21_+_8433172_8446622_RNA45SN1_45S_50ntFlanks@6699.6718.20\|GCTAGCTGCGAGAATTAATG | >hg38_21_+_8433172_8446622_RNA45SN1_45S_50ntFlanks | 6699 | 6718 | 20 | GCTAGCTGCGAGAATTAATG | 42869.7 | -10.2 | 1.3 | -7.6 | 2.5E-14 | 1.3E-13 | MB231/MB468 | MP |
| >hg38_21_+_8433172_8446622_RNA45SN1_45S_50ntFlanks@11491.11516.26\|CCGCGGGCCGGTCCCCCCCGCCGGGT | >hg38_21_+_8433172_8446622_RNA45SN1_45S_50ntFlanks | 11491 | 11516 | 26 | CCGCGGGCCGGTCCCCCCCGCCGGGT | 140.4 | -9.9 | 1.3 | -7.6 | 2.8E-14 | 1.3E-13 | MB231/MB468 | CYTO |
| >hg38_21_+_8433172_8446622_RNA45SN1_45S_50ntFlanks@12526.12552.27\|TTTTACCCTACTGATGATGTGTTGTTG | >hg38_21_+_8433172_8446622_RNA45SN1_45S_50ntFlanks | 12526 | 12552 | 27 | TTTTACCCTACTGATGATGTGTTGTTG | 182.8 | -11.4 | 1.5 | -7.6 | 2.5E-14 | 1.3E-13 | MB231/MB468 | MP |
| >hg38_21_+_8433172_8446622_RNA45SN1_45S_50ntFlanks@4725.4752.28\|TTAATCAAGAACGAAAGTCGGAGGTTCG | >hg38_21_+_8433172_8446622_RNA45SN1_45S_50ntFlanks | 4725 | 4752 | 28 | TTAATCAAGAACGAAAGTCGGAGGTTCG | 152.8 | -11.1 | 1.5 | -7.6 | 2.5E-14 | 1.3E-13 | MB231/MB468 | MP |
| >hg38_21_+_8433172_8446622_RNA45SN1_45S_50ntFlanks@4370.4389.20\|TTAAAAAGCTCGTAGTTGGA | >hg38_21_+_8433172_8446622_RNA45SN1_45S_50ntFlanks | 4370 | 4389 | 20 | TTAAAAAGCTCGTAGTTGGA | 294.1 | -8.1 | 1.1 | -7.6 | 2.8E-14 | 1.3E-13 | MB231/MB468 | CYTO |
| >hg38_21_+_8433172_8446622_RNA45SN1_45S_50ntFlanks@12626.12661.36\|AATGGGGCGAAGCTACCATCTGTGGGATTATGACTG | >hg38_21_+_8433172_8446622_RNA45SN1_45S_50ntFlanks | 12626 | 12661 | 36 | AATGGGGCGAAGCTACCATCTGTGGGATTATGACTG | 4015.4 | -8.9 | 1.2 | -7.6 | 2.5E-14 | 1.3E-13 | MB231/MB468 | MP |
| >hg38_21_+_8433172_8446622_RNA45SN1_45S_50ntFlanks@13002.13032.31\|CGATCTATTGAAAGTCAGCCCTCGACACAAG | >hg38_21_+_8433172_8446622_RNA45SN1_45S_50ntFlanks | 13002 | 13032 | 31 | CGATCTATTGAAAGTCAGCCCTCGACACAAG | 200.6 | -10.6 | 1.4 | -7.6 | 3.0E-14 | 1.3E-13 | MB231/MB468 | NUC |
| >hg38_21_+_8433172_8446622_RNA45SN1_45S_50ntFlanks@10905.10939.35\|CTGGGCGCGCGCCGCGGCTGGACGAGGCGCCGCCG | >hg38_21_+_8433172_8446622_RNA45SN1_45S_50ntFlanks | 10905 | 10939 | 35 | CTGGGCGCGCGCCGCGGCTGGACGAGGCGCCGCCG | 228.4 | -11.6 | 1.5 | -7.6 | 3.9E-14 | 1.3E-13 | MB231/MB468 | MITO |
| >hg38_21_+_8433172_8446622_RNA45SN1_45S_50ntFlanks@10873.10891.19\|CTGGGTCGGTCGGGCTGGG | >hg38_21_+_8433172_8446622_RNA45SN1_45S_50ntFlanks | 10873 | 10891 | 19 | CTGGGTCGGTCGGGCTGGG | 858.6 | -7.1 | 0.9 | -7.6 | 2.9E-14 | 1.3E-13 | MB231/MB468 | CYTO |

<table>
<tr><th></th><th></th><th></th><th></th><th></th><th></th><th></th><th></th><th></th><th></th><th></th><th></th><th></th><th></th></tr>
<tr><td>>hg38_21_+_8433172_8446622_RNA45SN1_45S_50ntFlanks@12626.12657.32|AATGGGGCGAAGCTACCATCTGTGGGATTATG</td><td>>hg38_21_+_8433172_8446622_RNA45SN1_45S_50ntFlanks</td><td>12626</td><td>12657</td><td>32</td><td>AATGGGGCGAAGCTACCATCTGTGGGATTATG</td><td>585.8</td><td>-4.9</td><td>0.6</td><td>-7.6</td><td>3.0E-14</td><td>1.3E-13</td><td>MB231/MB468</td><td>NUC</td></tr>
<tr><td>>hg38_21_+_8433172_8446622_RNA45SN1_45S_50ntFlanks@12357.12375.19|CCACAGGGATAACTGGCTT</td><td>>hg38_21_+_8433172_8446622_RNA45SN1_45S_50ntFlanks</td><td>12357</td><td>12375</td><td>19</td><td>CCACAGGGATAACTGGCTT</td><td>255.7</td><td>-8.3</td><td>1.1</td><td>-7.6</td><td>3.0E-14</td><td>1.3E-13</td><td>MB231/MB468</td><td>NUC</td></tr>
<tr><td>>hg38_21_+_8433172_8446622_RNA45SN1_45S_50ntFlanks@9125.9152.28|CGCGCGGGTCGGGGGGCGGGGCGGACTG</td><td>>hg38_21_+_8433172_8446622_RNA45SN1_45S_50ntFlanks</td><td>9125</td><td>9152</td><td>28</td><td>CGCGCGGGTCGGGGGGCGGGGCGGACTG</td><td>324.2</td><td>-12.2</td><td>1.6</td><td>-7.6</td><td>2.5E-14</td><td>1.3E-13</td><td>MB231/MB468</td><td>MP</td></tr>
<tr><td>>hg38_21_+_8433172_8446622_RNA45SN1_45S_50ntFlanks@9739.9765.27|AAACGATCTCAACCTATTCTCAAACTT</td><td>>hg38_21_+_8433172_8446622_RNA45SN1_45S_50ntFlanks</td><td>9739</td><td>9765</td><td>27</td><td>AAACGATCTCAACCTATTCTCAAACTT</td><td>5112.4</td><td>-9.4</td><td>1.2</td><td>-7.5</td><td>4.7E-14</td><td>1.3E-13</td><td>BT20/MB468</td><td>NUC</td></tr>
<tr><td>>hg38_21_+_8433172_8446622_RNA45SN1_45S_50ntFlanks@11658.11685.28|ATGTGATTTCTGCCCAGTGCTCTGAATG</td><td>>hg38_21_+_8433172_8446622_RNA45SN1_45S_50ntFlanks</td><td>11658</td><td>11685</td><td>28</td><td>ATGTGATTTCTGCCCAGTGCTCTGAATG</td><td>195.8</td><td>-11.5</td><td>1.5</td><td>-7.6</td><td>2.5E-14</td><td>1.3E-13</td><td>MB231/MB468</td><td>MP</td></tr>
<tr><td>>hg38_21_+_8433172_8446622_RNA45SN1_45S_50ntFlanks@8155.8178.24|CGCTCGTGGGGGGCCCAAGTCCTT</td><td>>hg38_21_+_8433172_8446622_RNA45SN1_45S_50ntFlanks</td><td>8155</td><td>8178</td><td>24</td><td>CGCTCGTGGGGGGCCCAAGTCCTT</td><td>164.3</td><td>-11.2</td><td>1.5</td><td>-7.6</td><td>2.5E-14</td><td>1.3E-13</td><td>MB231/MB468</td><td>MP</td></tr>
<tr><td>>hg38_21_+_8433172_8446622_RNA45SN1_45S_50ntFlanks@11843.11861.19|AGGGAACGGGCTTGGCGGA</td><td>>hg38_21_+_8433172_8446622_RNA45SN1_45S_50ntFlanks</td><td>11843</td><td>11861</td><td>19</td><td>AGGGAACGGGCTTGGCGGA</td><td>273.5</td><td>-12.0</td><td>1.6</td><td>-7.6</td><td>2.5E-14</td><td>1.3E-13</td><td>MB231/MB468</td><td>MP</td></tr>
<tr><td>>hg38_21_+_8433172_8446622_RNA45SN1_45S_50ntFlanks@12632.12654.23|GCGAAGCTACCATCTGTGGGATT</td><td>>hg38_21_+_8433172_8446622_RNA45SN1_45S_50ntFlanks</td><td>12632</td><td>12654</td><td>23</td><td>GCGAAGCTACCATCTGTGGGATT</td><td>2732.1</td><td>-4.2</td><td>0.6</td><td>-7.6</td><td>2.4E-14</td><td>1.3E-13</td><td>BT20/MB468</td><td>MITO</td></tr>
<tr><td>>hg38_21_+_8433172_8446622_RNA45SN1_45S_50ntFlanks@11534.11551.18|GTTCCGCGCGGCGCCTCG</td><td>>hg38_21_+_8433172_8446622_RNA45SN1_45S_50ntFlanks</td><td>11534</td><td>11551</td><td>18</td><td>GTTCCGCGCGGCGCCTCG</td><td>2499.7</td><td>-8.1</td><td>1.1</td><td>-7.6</td><td>4.0E-14</td><td>1.3E-13</td><td>MB231/MB468</td><td>MITO</td></tr>
<tr><td>>hg38_21_+_8433172_8446622_RNA45SN1_45S_50ntFlanks@7984.8012.29|AGATCAGACGTGGCGACCCGCTGAATTTA</td><td>>hg38_21_+_8433172_8446622_RNA45SN1_45S_50ntFlanks</td><td>7984</td><td>8012</td><td>29</td><td>AGATCAGACGTGGCGACCCGCTGAATTTA</td><td>944.3</td><td>-9.4</td><td>1.2</td><td>-7.6</td><td>4.0E-14</td><td>1.3E-13</td><td>MB231/MB468</td><td>MITO</td></tr>
<tr><td>>hg38_21_+_8433172_8446622_RNA45SN1_45S_50ntFlanks@4724.4753.30|ATTAATCAAGAACGAAAGTCGGAGGTTCGA</td><td>>hg38_21_+_8433172_8446622_RNA45SN1_45S_50ntFlanks</td><td>4724</td><td>4753</td><td>30</td><td>ATTAATCAAGAACGAAAGTCGGAGGTTCGA</td><td>369.0</td><td>-12.3</td><td>1.6</td><td>-7.6</td><td>4.0E-14</td><td>1.3E-13</td><td>MB231/MB468</td><td>MITO</td></tr>
<tr><td>>am_trna143_LysTTT_6_-_27559593_27559665@7.24.18,trna14_LysTTT_11_-_59327808_59327880@7.24.18,trna2_LysTTT_17_+_8022473_8022545@7.24.18,trna54_LysTTT_1_+_204475655_204475727@7.24.18,trna5_LysTTT_11_+_59323902_59323974@7.24.18,trna62_LysTTT_1_-_204476158_204476230@7.24.18,trna76_LysTTT_6_+_28918806_28918878@7.24.18|ATAGCTCAGTCGGTAGAG</td><td>>am_trna143_LysTTT_6_-_27559593_27559665</td><td>7</td><td>24</td><td>18</td><td>ATAGCTCAGTCGGTAGAG</td><td>180.7</td><td>-11.4</td><td>1.5</td><td>-7.6</td><td>2.5E-14</td><td>1.3E-13</td><td>MB231/MB468</td><td>MP</td></tr>
<tr><td>>hg38_21_+_8433172_8446622_RNA45SN1_45S_50ntFlanks@11729.11750.22|TAACTATGACTCTCTTAAGGTA</td><td>>hg38_21_+_8433172_8446622_RNA45SN1_45S_50ntFlanks</td><td>11729</td><td>11750</td><td>22</td><td>TAACTATGACTCTCTTAAGGTA</td><td>1567.5</td><td>-4.8</td><td>0.6</td><td>-7.6</td><td>3.0E-14</td><td>1.3E-13</td><td>MB231/MB468</td><td>NUC</td></tr>
<tr><td>>hg38_21_+_8433172_8446622_RNA45SN1_45S_50ntFlanks@485.506.22|GATGTGGTGACGTCGTGCTCTC</td><td>>hg38_21_+_8433172_8446622_RNA45SN1_45S_50ntFlanks</td><td>485</td><td>506</td><td>22</td><td>GATGTGGTGACGTCGTGCTCTC</td><td>793.0</td><td>13.4</td><td>1.7</td><td>8.0</td><td>1.1E-15</td><td>1.3E-13</td><td>BT20/MB468</td><td>MP</td></tr>
<tr><td>>hg38_21_+_8433172_8446622_RNA45SN1_45S_50ntFlanks@13006.13027.22|CTATTGAAAGTCAGCCCTCGAC</td><td>>hg38_21_+_8433172_8446622_RNA45SN1_45S_50ntFlanks</td><td>13006</td><td>13027</td><td>22</td><td>CTATTGAAAGTCAGCCCTCGAC</td><td>1592.5</td><td>-7.8</td><td>1.0</td><td>-7.6</td><td>2.9E-14</td><td>1.3E-13</td><td>MB231/MB468</td><td>CYTO</td></tr>
<tr><td>>hg38_21_+_8433172_8446622_RNA45SN1_45S_50ntFlanks@10781.10823.43|AGGTGAACAGCCTCTGGCATGTTGGAACAATGTAGGTAAGGGA</td><td>>hg38_21_+_8433172_8446622_RNA45SN1_45S_50ntFlanks</td><td>10781</td><td>10823</td><td>43</td><td>AGGTGAACAGCCTCTGGCATGTTGGAACAATGTAGGTAAGGGA</td><td>374.5</td><td>-12.3</td><td>1.6</td><td>-7.6</td><td>4.0E-14</td><td>1.3E-13</td><td>MB231/MB468</td><td>MITO</td></tr>
<tr><td>>MI0003591|hsa-mir-584&WithFlank&5|-|149062307|149062415@22.41.20&[MIMAT0003249&hsa-miR-584-5p&offsets|0|-2;m-167&5|-|149062374|149062394&offsets|0|-1]|TTATGGTTTGCCTGGGACTG</td><td>>MI0003591|hsa-mir-584&WithFlank&5|-|149062307|149062415</td><td>22</td><td>41</td><td>20</td><td>TTATGGTTTGCCTGGGACTG</td><td>88.0</td><td>2.8</td><td>0.4</td><td>7.5</td><td>4.8E-14</td><td>1.3E-13</td><td>BT20/MB468</td><td>NUC</td></tr>
<tr><td>>hg38_21_+_8433172_8446622_RNA45SN1_45S_50ntFlanks@11210.11233.24|GGCGACTCTGGACGCGAGCCGGGC</td><td>>hg38_21_+_8433172_8446622_RNA45SN1_45S_50ntFlanks</td><td>11210</td><td>11233</td><td>24</td><td>GGCGACTCTGGACGCGAGCCGGGC</td><td>507.0</td><td>-8.5</td><td>1.1</td><td>-7.6</td><td>2.9E-14</td><td>1.3E-13</td><td>MB231/MB468</td><td>CYTO</td></tr>
<tr><td>>hg38_21_+_8433172_8446622_RNA45SN1_45S_50ntFlanks@12044.12070.27|ACTGACCCGGTGAGGCGGGGGGGCGAG</td><td>>hg38_21_+_8433172_8446622_RNA45SN1_45S_50ntFlanks</td><td>12044</td><td>12070</td><td>27</td><td>ACTGACCCGGTGAGGCGGGGGGGCGAG</td><td>2956.2</td><td>-11.1</td><td>1.5</td><td>-7.5</td><td>4.8E-14</td><td>1.3E-13</td><td>BT20/MB468</td><td>NUC</td></tr>
<tr><td>>hg38_21_+_8433172_8446622_RNA45SN1_45S_50ntFlanks@12008.12035.28|CCGCCGGTGAAATACCACTACTCTGATC</td><td>>hg38_21_+_8433172_8446622_RNA45SN1_45S_50ntFlanks</td><td>12008</td><td>12035</td><td>28</td><td>CCGCCGGTGAAATACCACTACTCTGATC</td><td>248.7</td><td>-5.5</td><td>0.7</td><td>-7.6</td><td>2.9E-14</td><td>1.3E-13</td><td>MB231/MB468</td><td>CYTO</td></tr>
<tr><td>>hg38_21_+_8433172_8446622_RNA45SN1_45S_50ntFlanks@12457.12487.31|CAAGCGTTGGATTGTTCACCCACTAATAGGG</td><td>>hg38_21_+_8433172_8446622_RNA45SN1_45S_50ntFlanks</td><td>12457</td><td>12487</td><td>31</td><td>CAAGCGTTGGATTGTTCACCCACTAATAGGG</td><td>1872.2</td><td>-7.0</td><td>0.9</td><td>-7.6</td><td>2.9E-14</td><td>1.3E-13</td><td>MB231/MB468</td><td>CYTO</td></tr>
<tr><td>>MI0000285|hsa-mir-205&WithFlank&1|+|209432127|209432248@40.59.20(+1G)&[MIMAT0000266&hsa-miR-205-5p&offsets|0|-2(+1G);m-351&1|+|209432166|209432187&offsets|0|-2(+1G)]|TCCTTCATTCCACCGGAGTCG</td><td>>MI0000285|hsa-mir-205&WithFlank&1|+|209432127|209432248</td><td>40</td><td>59</td><td>20(+1G)</td><td>TCCTTCATTCCACCGGAGTCG</td><td>231.7</td><td>11.2</td><td>1.4</td><td>7.9</td><td>2.4E-15</td><td>1.3E-13</td><td>BT20/MB231</td><td>CYTO</td></tr>
<tr><td>>hg38_21_+_8433172_8446622_RNA45SN1_45S_50ntFlanks@11670.11687.18|CCCAGTGCTCTGAATGTC</td><td>>hg38_21_+_8433172_8446622_RNA45SN1_45S_50ntFlanks</td><td>11670</td><td>11687</td><td>18</td><td>CCCAGTGCTCTGAATGTC</td><td>801.4</td><td>-5.1</td><td>0.7</td><td>-7.6</td><td>2.6E-14</td><td>1.3E-13</td><td>MB231/MB468</td><td>MP</td></tr>
<tr><td>>hg38_21_+_8433172_8446622_RNA45SN1_45S_50ntFlanks@6719.6751.33|TGAATTGCAGGACACATTGATCATCGACACTTC</td><td>>hg38_21_+_8433172_8446622_RNA45SN1_45S_50ntFlanks</td><td>6719</td><td>6751</td><td>33</td><td>TGAATTGCAGGACACATTGATCATCGACACTTC</td><td>526.3</td><td>-4.6</td><td>0.6</td><td>-7.6</td><td>3.1E-14</td><td>1.3E-13</td><td>MB231/MB468</td><td>NUC</td></tr>
<tr><td>>hg38_21_+_8433172_8446622_RNA45SN1_45S_50ntFlanks@11768.11804.37|CTAATTAGTGACGCGCATGAATGGATGAACGAGATTC</td><td>>hg38_21_+_8433172_8446622_RNA45SN1_45S_50ntFlanks</td><td>11768</td><td>11804</td><td>37</td><td>CTAATTAGTGACGCGCATGAATGGATGAACGAGATTC</td><td>895.2</td><td>-7.6</td><td>1.0</td><td>-7.6</td><td>3.1E-14</td><td>1.3E-13</td><td>MB231/MB468</td><td>NUC</td></tr>
<tr><td>>hg38_21_+_8433172_8446622_RNA45SN1_45S_50ntFlanks@10407.10429.23|GGGTCAGTCGGTCCTGAGAGATG</td><td>>hg38_21_+_8433172_8446622_RNA45SN1_45S_50ntFlanks</td><td>10407</td><td>10429</td><td>23</td><td>GGGTCAGTCGGTCCTGAGAGATG</td><td>387.4</td><td>-12.5</td><td>1.6</td><td>-7.6</td><td>2.6E-14</td><td>1.3E-13</td><td>MB231/MB468</td><td>MP</td></tr>
<tr><td>>hg38_21_+_8433172_8446622_RNA45SN1_45S_50ntFlanks@11211.11231.21|GCGACTCTGGACGCGAGCCGG</td><td>>hg38_21_+_8433172_8446622_RNA45SN1_45S_50ntFlanks</td><td>11211</td><td>11231</td><td>21</td><td>GCGACTCTGGACGCGAGCCGG</td><td>151.2</td><td>-11.1</td><td>1.5</td><td>-7.6</td><td>2.6E-14</td><td>1.3E-13</td><td>MB231/MB468</td><td>MP</td></tr>
<tr><td>>MI0000279|hsa-mir-196a-2&WithFlank&12|+|53991732|53991853@31.51.21(+1U);MI0000238|hsa-mir-196a-1&WithFlank&17|-|48632484|48632565@13.33.21(+1U)&[MIMAT0000226&hsa-miR-196a-5p&offsets|0|-1(+1U);m-261&12|+|53991762|53991783&offsets|0|-1(+1U)];[MIMAT0000226_1&hsa-miR-196a-5p&offsets|0|-1(+1U);m-268&17|-|48632532|48632553&offsets|0|-1(+1U)]|TAGGTAGTTTCATGTTGTTGGT</td><td>>MI0000279|hsa-mir-196a-2&WithFlank&12|+|53991732|53991853</td><td>31</td><td>51</td><td>21(+1U)</td><td>TAGGTAGTTTCATGTTGTTGGT</td><td>399.7</td><td>10.6</td><td>1.4</td><td>7.6</td><td>3.3E-14</td><td>1.3E-13</td><td>BT20/MB468</td><td>CYTO</td></tr>
<tr><td>>hg38_21_+_8433172_8446622_RNA45SN1_45S_50ntFlanks@12636.12653.18|AGCTACCATCTGTGGGAT</td><td>>hg38_21_+_8433172_8446622_RNA45SN1_45S_50ntFlanks</td><td>12636</td><td>12653</td><td>18</td><td>AGCTACCATCTGTGGGAT</td><td>204.2</td><td>-10.0</td><td>1.3</td><td>-7.6</td><td>3.3E-14</td><td>1.3E-13</td><td>BT20/MB468</td><td>CYTO</td></tr>
<tr><td>>hg38_21_+_8433172_8446622_RNA45SN1_45S_50ntFlanks@10786.10816.31|AACAGCCTCTGGCATGTTGGAACAATGTAGG</td><td>>hg38_21_+_8433172_8446622_RNA45SN1_45S_50ntFlanks</td><td>10786</td><td>10816</td><td>31</td><td>AACAGCCTCTGGCATGTTGGAACAATGTAGG</td><td>2595.7</td><td>-5.3</td><td>0.7</td><td>-7.6</td><td>3.0E-14</td><td>1.4E-13</td><td>MB231/MB468</td><td>CYTO</td></tr>
<tr><td>>hg38_21_+_8433172_8446622_RNA45SN1_45S_50ntFlanks@10794.10818.25|CTGGCATGTTGGAACAATGTAGGTA</td><td>>hg38_21_+_8433172_8446622_RNA45SN1_45S_50ntFlanks</td><td>10794</td><td>10818</td><td>25</td><td>CTGGCATGTTGGAACAATGTAGGTA</td><td>1888.7</td><td>-4.8</td><td>0.6</td><td>-7.6</td><td>3.3E-14</td><td>1.4E-13</td><td>BT20/MB468</td><td>CYTO</td></tr>
<tr><td>>hg38_21_+_8433172_8446622_RNA45SN1_45S_50ntFlanks@10107.10124.18|CGTGTGGTGTGCGTCGGA</td><td>>hg38_21_+_8433172_8446622_RNA45SN1_45S_50ntFlanks</td><td>10107</td><td>10124</td><td>18</td><td>CGTGTGGTGTGCGTCGGA</td><td>299.6</td><td>-12.1</td><td>1.6</td><td>-7.6</td><td>2.6E-14</td><td>1.4E-13</td><td>MB231/MB468</td><td>MP</td></tr>
<tr><td>>hg38_21_+_8433172_8446622_RNA45SN1_45S_50ntFlanks@10805.10823.19|GAACAATGTAGGTAAGGGA</td><td>>hg38_21_+_8433172_8446622_RNA45SN1_45S_50ntFlanks</td><td>10805</td><td>10823</td><td>19</td><td>GAACAATGTAGGTAAGGGA</td><td>236.8</td><td>-10.8</td><td>1.4</td><td>-7.6</td><td>3.1E-14</td><td>1.4E-13</td><td>MB231/MB468</td><td>NUC</td></tr>
<tr><td>>hg38_21_+_8433172_8446622_RNA45SN1_45S_50ntFlanks@6661.6682.22|CGGTGGATCACTCGGCTCGTGC</td><td>>hg38_21_+_8433172_8446622_RNA45SN1_45S_50ntFlanks</td><td>6661</td><td>6682</td><td>22</td><td>CGGTGGATCACTCGGCTCGTGC</td><td>723.0</td><td>6.6</td><td>0.9</td><td>7.6</td><td>2.5E-14</td><td>1.4E-13</td><td>BT20/MB468</td><td>MITO</td></tr>
<tr><td>>hg38_21_+_8433172_8446622_RNA45SN1_45S_50ntFlanks@10402.10419.18|AACATGGGTCAGTCGGTC</td><td>>hg38_21_+_8433172_8446622_RNA45SN1_45S_50ntFlanks</td><td>10402</td><td>10419</td><td>18</td><td>AACATGGGTCAGTCGGTC</td><td>2989.8</td><td>-11.0</td><td>1.5</td><td>-7.5</td><td>4.9E-14</td><td>1.4E-13</td><td>BT20/MB468</td><td>NUC</td></tr>
<tr><td>>hg38_21_+_8433172_8446622_RNA45SN1_45S_50ntFlanks@11315.11338.24|CCACCCCACGTCTCGTCGCGCGCG</td><td>>hg38_21_+_8433172_8446622_RNA45SN1_45S_50ntFlanks</td><td>11315</td><td>11338</td><td>24</td><td>CCACCCCACGTCTCGTCGCGCGCG</td><td>637.6</td><td>-8.4</td><td>1.1</td><td>-7.6</td><td>3.0E-14</td><td>1.4E-13</td><td>MB231/MB468</td><td>CYTO</td></tr>
<tr><td>>hg38_21_+_8433172_8446622_RNA45SN1_45S_50ntFlanks@12934.12968.35|CTAAACCATTCGTAGACGACCTGCTTCTGGGTCGG</td><td>>hg38_21_+_8433172_8446622_RNA45SN1_45S_50ntFlanks</td><td>12934</td><td>12968</td><td>35</td><td>CTAAACCATTCGTAGACGACCTGCTTCTGGGTCGG</td><td>164.5</td><td>-10.1</td><td>1.3</td><td>-7.6</td><td>3.0E-14</td><td>1.4E-13</td><td>MB231/MB468</td><td>CYTO</td></tr>
<tr><td>>hg38_21_+_8433172_8446622_RNA45SN1_45S_50ntFlanks@10771.10807.37|CAGGTCTCCAAGGTGAACAGCCTCTGGCATGTTGGAA</td><td>>hg38_21_+_8433172_8446622_RNA45SN1_45S_50ntFlanks</td><td>10771</td><td>10807</td><td>37</td><td>CAGGTCTCCAAGGTGAACAGCCTCTGGCATGTTGGAA</td><td>226.9</td><td>-11.6</td><td>1.5</td><td>-7.6</td><td>4.1E-14</td><td>1.4E-13</td><td>MB231/MB468</td><td>MITO</td></tr>
<tr><td>>hg38_21_+_8433172_8446622_RNA45SN1_45S_50ntFlanks@10421.10457.37|TGAGAGATGGGCGAGCGCCGTTCCGAAGGGACGGGCG</td><td>>hg38_21_+_8433172_8446622_RNA45SN1_45S_50ntFlanks</td><td>10421</td><td>10457</td><td>37</td><td>TGAGAGATGGGCGAGCGCCGTTCCGAAGGGACGGGCG</td><td>805.8</td><td>-6.3</td><td>0.8</td><td>-7.6</td><td>3.0E-14</td><td>1.4E-13</td><td>MB231/MB468</td><td>CYTO</td></tr>
<tr><td>>hg38_21_+_8433172_8446622_RNA45SN1_45S_50ntFlanks@7976.8009.34|GCGACCTCAGATCAGACGTGGCGACCCGCTGAAT</td><td>>hg38_21_+_8433172_8446622_RNA45SN1_45S_50ntFlanks</td><td>7976</td><td>8009</td><td>34</td><td>GCGACCTCAGATCAGACGTGGCGACCCGCTGAAT</td><td>1855.8</td><td>9.8</td><td>1.3</td><td>7.6</td><td>3.0E-14</td><td>1.4E-13</td><td>MB231/MB468</td><td>CYTO</td></tr>
<tr><td>>hg38_21_+_8433172_8446622_RNA45SN1_45S_50ntFlanks@12655.12674.20|ATGACTGAACGCCTCTAAGT</td><td>>hg38_21_+_8433172_8446622_RNA45SN1_45S_50ntFlanks</td><td>12655</td><td>12674</td><td>20</td><td>ATGACTGAACGCCTCTAAGT</td><td>3155.8</td><td>-4.9</td><td>0.6</td><td>-7.6</td><td>3.0E-14</td><td>1.4E-13</td><td>MB231/MB468</td><td>CYTO</td></tr>
<tr><td>>trna119_LysCTT_1_-_145395522_145395594@4.34.31,trna11_LysCTT_5_-_180648979_180649051@4.34.31,trna13_LysCTT_6_+_26556774_26556846@4.34.31,trna7_LysCTT_16_+_3225692_3225764@4.34.31,trna9_LysCTT_5_+_180634755_180634827@4.34.31|CGGCTAGCTCAGTCGGTAGAGCATGAGACTC</td><td>>trna119_LysCTT_1_-_145395522_145395594</td><td>4</td><td>34</td><td>31</td><td>CGGCTAGCTCAGTCGGTAGAGCATGAGACTC</td><td>441.6</td><td>8.2</td><td>1.1</td><td>7.6</td><td>3.1E-14</td><td>1.4E-13</td><td>MB231/MB468</td><td>NUC</td></tr>
<tr><td>>hg38_21_+_8433172_8446622_RNA45SN1_45S_50ntFlanks@11663.11691.29|ATTTCTGCCCAGTGCTCTGAATGTCAAAG</td><td>>hg38_21_+_8433172_8446622_RNA45SN1_45S_50ntFlanks</td><td>11663</td><td>11691</td><td>29</td><td>ATTTCTGCCCAGTGCTCTGAATGTCAAAG</td><td>963.4</td><td>5.3</td><td>0.7</td><td>7.6</td><td>2.5E-14</td><td>1.4E-13</td><td>BT20/MB468</td><td>MITO</td></tr>
<tr><td>>hg38_21_+_8433172_8446622_RNA45SN1_45S_50ntFlanks@12540.12566.27|TGATGTGTTGTTGCCATGGTAATCCTG</td><td>>hg38_21_+_8433172_8446622_RNA45SN1_45S_50ntFlanks</td><td>12540</td><td>12566</td><td>27</td><td>TGATGTGTTGTTGCCATGGTAATCCTG</td><td>156.4</td><td>-11.2</td><td>1.5</td><td>-7.6</td><td>2.6E-14</td><td>1.4E-13</td><td>MB231/MB468</td><td>MP</td></tr>
<tr><td>>hg38_21_+_8433172_8446622_RNA45SN1_45S_50ntFlanks@11211.11230.20|GCGACTCTGGACGCGAGCCG</td><td>>hg38_21_+_8433172_8446622_RNA45SN1_45S_50ntFlanks</td><td>11211</td><td>11230</td><td>20</td><td>GCGACTCTGGACGCGAGCCG</td><td>147.7</td><td>-11.1</td><td>1.5</td><td>-7.6</td><td>2.6E-14</td><td>1.4E-13</td><td>MB231/MB468</td><td>MP</td></tr>
<tr><td>>am_trna134_GluTTC_1_-_16861774_16861845@1.27.27,trna5_GluTTC_1_+_17199078_17199149@1.27.27,trna84_GluTTC_1_-_161391883_161391954@1.27.27,trna94_GluTTC_1_-_149664355_149664427@1.27.27|TCCCTGGTGGTCTAGTGGCTAGGATTC</td><td>>am_trna134_GluTTC_1_-_16861774_16861845</td><td>1</td><td>27</td><td>27</td><td>TCCCTGGTGGTCTAGTGGCTAGGATTC</td><td>214.9</td><td>-6.0</td><td>0.8</td><td>-7.6</td><td>3.0E-14</td><td>1.4E-13</td><td>MB231/MB468</td><td>CYTO</td></tr>
<tr><td>>hg38_21_+_8433172_8446622_RNA45SN1_45S_50ntFlanks@10571.10608.38|GAGAAGCCGGCGGGAGCCCCGGGGAGAGTTCTCTTTTC</td><td>>hg38_21_+_8433172_8446622_RNA45SN1_45S_50ntFlanks</td><td>10571</td><td>10608</td><td>38</td><td>GAGAAGCCGGCGGGAGCCCCGGGGAGAGTTCTCTTTTC</td><td>329.3</td><td>-12.1</td><td>1.6</td><td>-7.6</td><td>4.2E-14</td><td>1.4E-13</td><td>MB231/MB468</td><td>MITO</td></tr>
<tr><td>>hg38_21_+_8433172_8446622_RNA45SN1_45S_50ntFlanks@13004.13028.25|ATCTATTGAAAGTCAGCCCTCGACA</td><td>>hg38_21_+_8433172_8446622_RNA45SN1_45S_50ntFlanks</td><td>13004</td><td>13028</td><td>25</td><td>ATCTATTGAAAGTCAGCCCTCGACA</td><td>1603.3</td><td>-9.0</td><td>1.2</td><td>-7.6</td><td>3.0E-14</td><td>1.4E-13</td><td>MB231/MB468</td><td>CYTO</td></tr>
</table>

<table>
<tr><td>>hg38_21_+_8433172_8446622_RNA45SN1_45S_50ntFlanks@4130.4162.33|ATTCCGGAGAGGGAGCCTGAGAAACGGCTACCA</td><td>>hg38_21_+_8433172_8446622_RNA45SN1_45S_50ntFlanks</td><td>4130</td><td>4162</td><td>33</td><td>ATTCCGGAGAGGGAGCCTGAGAAACGGCTACCA</td><td>1014.0</td><td>-9.4</td><td>1.2</td><td>-7.6</td><td>4.2E-14</td><td>1.4E-13</td><td>MB231/MB468</td><td>MITO</td></tr>
<tr><td>>hg38_21_+_8433172_8446622_RNA45SN1_45S_50ntFlanks@8115.8136.22|CGCGGGACATGTGGCGTACGGA</td><td>>hg38_21_+_8433172_8446622_RNA45SN1_45S_50ntFlanks</td><td>8115</td><td>8136</td><td>22</td><td>CGCGGGACATGTGGCGTACGGA</td><td>314.6</td><td>-12.1</td><td>1.6</td><td>-7.6</td><td>4.2E-14</td><td>1.4E-13</td><td>MB231/MB468</td><td>MITO</td></tr>
<tr><td>>MI0000088|hsa-mir-30a&WithFlank&6|-|71403545|71403627@54.73.20(+1A)&[MIMAT0000088&hsa-miR-30a-3p&offsets|+1|-1(+1A);m-38&6|-|71403554|71403575&offsets|+1|-1(+1A)]|TTTCAGTCGGATGTTTGCAGA</td><td>>MI0000088|hsa-mir-30a&WithFlank&6|-|71403545|71403627</td><td>54</td><td>73</td><td>20(+1A)</td><td>TTTCAGTCGGATGTTTGCAGA</td><td>398.3</td><td>7.9</td><td>1.0</td><td>7.6</td><td>3.4E-14</td><td>1.4E-13</td><td>BT20/MB468</td><td>CYTO</td></tr>
<tr><td>>hg38_21_+_8433172_8446622_RNA45SN1_45S_50ntFlanks@10866.10901.36|CTAAGGGCTGGGTCGGTCGGGCTGGGGCGCGAAGCG</td><td>>hg38_21_+_8433172_8446622_RNA45SN1_45S_50ntFlanks</td><td>10866</td><td>10901</td><td>36</td><td>CTAAGGGCTGGGTCGGTCGGGCTGGGGCGCGAAGCG</td><td>654.4</td><td>-6.9</td><td>0.9</td><td>-7.6</td><td>3.0E-14</td><td>1.4E-13</td><td>MB231/MB468</td><td>CYTO</td></tr>
<tr><td>>MI0003591|hsa-mir-584&WithFlank&5|-|149062307|149062415@22.42.21&[MIMAT0003249&hsa-miR-584-5p&offsets|0|-1;m-167&5|-|149062374|149062394&offsets|0|0]|TTATGGTTTGCCTGGGACTGA</td><td>>MI0003591|hsa-mir-584&WithFlank&5|-|149062307|149062415</td><td>22</td><td>42</td><td>21</td><td>TTATGGTTTGCCTGGGACTGA</td><td>1607.4</td><td>6.6</td><td>0.9</td><td>7.6</td><td>4.2E-14</td><td>1.4E-13</td><td>MB231/MB468</td><td>MITO</td></tr>
<tr><td>>hg38_21_+_8433172_8446622_RNA45SN1_45S_50ntFlanks@10731.10751.21|AGGGTGTAAATCTCGCGCCGG</td><td>>hg38_21_+_8433172_8446622_RNA45SN1_45S_50ntFlanks</td><td>10731</td><td>10751</td><td>21</td><td>AGGGTGTAAATCTCGCGCCGG</td><td>203.7</td><td>-11.5</td><td>1.5</td><td>-7.6</td><td>2.6E-14</td><td>1.4E-13</td><td>MB231/MB468</td><td>MP</td></tr>
<tr><td>>hg38_21_+_8433172_8446622_RNA45SN1_45S_50ntFlanks@7973.7991.19|GACGCGACCTCAGATCAGA</td><td>>hg38_21_+_8433172_8446622_RNA45SN1_45S_50ntFlanks</td><td>7973</td><td>7991</td><td>19</td><td>GACGCGACCTCAGATCAGA</td><td>3744.9</td><td>6.1</td><td>0.8</td><td>7.6</td><td>2.5E-14</td><td>1.4E-13</td><td>BT20/MB468</td><td>MITO</td></tr>
<tr><td>>hg38_21_+_8433172_8446622_RNA45SN1_45S_50ntFlanks@11540.11560.21|CGCGGCGCCTCGCCTCGGCCG</td><td>>hg38_21_+_8433172_8446622_RNA45SN1_45S_50ntFlanks</td><td>11540</td><td>11560</td><td>21</td><td>CGCGGCGCCTCGCCTCGGCCG</td><td>302.9</td><td>-11.0</td><td>1.4</td><td>-7.6</td><td>3.1E-14</td><td>1.4E-13</td><td>MB231/MB468</td><td>CYTO</td></tr>
<tr><td>>MI0003776|hsa-mir-320b-1&WithFlank&1|+|116671740|116671823@48.67.20;MI0003839|hsa-mir-320b-2&WithFlank&1|-|224257034|224257116@47.66.20;MI0000542|hsa-mir-320a&WithFlank&8|-|22244960|22245043@48.67.20&[MIMAT0005792&hsa-miR-320b&offsets|0|-2;m-183&1|+|116671787|116671806&offsets|0|0];[MIMAT0005792_1&hsa-miR-320b&offsets|0|-2;m-183&1|-|224257051|224257070&offsets|0|0];[MIMAT0000510&hsa-miR-320a-3p&offsets|0|-2;m-60&8|-|22244975|22244996&offsets|0|-2]|AAAAGCTGGGTTGAGAGGGC</td><td>>MI0003776|hsa-mir-320b-1&WithFlank&1|+|116671740|116671823</td><td>48</td><td>67</td><td>20</td><td>AAAAGCTGGGTTGAGAGGGC</td><td>12305.7</td><td>8.1</td><td>1.0</td><td>8.0</td><td>1.1E-15</td><td>1.4E-13</td><td>BT20/MB468</td><td>MP</td></tr>
<tr><td>>hg38_21_+_8433172_8446622_RNA45SN1_45S_50ntFlanks@10869.10903.35|AGGGCTGGGTCGGTCGGGCTGGGGCGCGAAGCGGG</td><td>>hg38_21_+_8433172_8446622_RNA45SN1_45S_50ntFlanks</td><td>10869</td><td>10903</td><td>35</td><td>AGGGCTGGGTCGGTCGGGCTGGGGCGCGAAGCGGG</td><td>240.2</td><td>-11.8</td><td>1.5</td><td>-7.6</td><td>2.7E-14</td><td>1.4E-13</td><td>MB231/MB468</td><td>MP</td></tr>
<tr><td>>am_trna2_GlyTCC_19_+_4724082_4724153@1.30.30|GCGTTGGTGGTATAGTGGTTAGCATAGCTG</td><td>>am_trna2_GlyTCC_19_+_4724082_4724153</td><td>1</td><td>30</td><td>30</td><td>GCGTTGGTGGTATAGTGGTTAGCATAGCTG</td><td>264.9</td><td>-11.8</td><td>1.6</td><td>-7.6</td><td>4.2E-14</td><td>1.4E-13</td><td>MB231/MB468</td><td>MITO</td></tr>
<tr><td>>MI0000439|hsa-mir-23b&WithFlank&9|+|95085202|95085310@67.86.20(+1U)&[MIMAT0000418&hsa-miR-23b-3p&offsets|+3|0(+1U);m-42&9|+|95085265|95085285&offsets|+3|+2(+1U)]|ACATTGCCAGGGATTACCACT</td><td>>MI0000439|hsa-mir-23b&WithFlank&9|+|95085202|95085310</td><td>67</td><td>86</td><td>20(+1U)</td><td>ACATTGCCAGGGATTACCACT</td><td>1255.7</td><td>27.6</td><td>3.6</td><td>7.6</td><td>2.7E-14</td><td>1.4E-13</td><td>MB231/MB468</td><td>MP</td></tr>
<tr><td>>hg38_21_+_8433172_8446622_RNA45SN1_45S_50ntFlanks@13009.13027.19|TTGAAAGTCAGCCCTCGAC</td><td>>hg38_21_+_8433172_8446622_RNA45SN1_45S_50ntFlanks</td><td>13009</td><td>13027</td><td>19</td><td>TTGAAAGTCAGCCCTCGAC</td><td>169.6</td><td>-10.4</td><td>1.4</td><td>-7.6</td><td>3.2E-14</td><td>1.4E-13</td><td>MB231/MB468</td><td>NUC</td></tr>
<tr><td>>hg38_21_+_8433172_8446622_RNA45SN1_45S_50ntFlanks@4760.4793.34|TCAGATACCGTCGTAGTTCCGACCATAAACGATG</td><td>>hg38_21_+_8433172_8446622_RNA45SN1_45S_50ntFlanks</td><td>4760</td><td>4793</td><td>34</td><td>TCAGATACCGTCGTAGTTCCGACCATAAACGATG</td><td>442.7</td><td>-4.9</td><td>0.6</td><td>-7.6</td><td>3.2E-14</td><td>1.4E-13</td><td>MB231/MB468</td><td>NUC</td></tr>
<tr><td>>hg38_21_+_8433172_8446622_RNA45SN1_45S_50ntFlanks@8170.8212.43|CAAGTCCTTCTGATCGAGGCCCAGCCCGTGGACGGTGTGAGGC</td><td>>hg38_21_+_8433172_8446622_RNA45SN1_45S_50ntFlanks</td><td>8170</td><td>8212</td><td>43</td><td>CAAGTCCTTCTGATCGAGGCCCAGCCCGTGGACGGTGTGAGGC</td><td>4508.1</td><td>-5.3</td><td>0.7</td><td>-7.6</td><td>2.5E-14</td><td>1.4E-13</td><td>BT20/MB468</td><td>MITO</td></tr>
<tr><td>>hg38_21_+_8433172_8446622_RNA45SN1_45S_50ntFlanks@11873.11908.36|AAGAAGACCCTGTTGAGCTTGACTCTAGTCTGGCAC</td><td>>hg38_21_+_8433172_8446622_RNA45SN1_45S_50ntFlanks</td><td>11873</td><td>11908</td><td>36</td><td>AAGAAGACCCTGTTGAGCTTGACTCTAGTCTGGCAC</td><td>319.3</td><td>-7.5</td><td>1.0</td><td>-7.6</td><td>3.1E-14</td><td>1.4E-13</td><td>MB231/MB468</td><td>CYTO</td></tr>
<tr><td>>hg38_21_+_8433172_8446622_RNA45SN1_45S_50ntFlanks@10574.10591.18|AAGCCGGCGGGAGCCCCG</td><td>>hg38_21_+_8433172_8446622_RNA45SN1_45S_50ntFlanks</td><td>10574</td><td>10591</td><td>18</td><td>AAGCCGGCGGGAGCCCCG</td><td>202.8</td><td>-10.6</td><td>1.4</td><td>-7.6</td><td>3.2E-14</td><td>1.4E-13</td><td>MB231/MB468</td><td>NUC</td></tr>
<tr><td>>hg38_21_+_8433172_8446622_RNA45SN1_45S_50ntFlanks@10127.10149.23|GCGGCGGCGGCGGCGGCGGCGGG</td><td>>hg38_21_+_8433172_8446622_RNA45SN1_45S_50ntFlanks</td><td>10127</td><td>10149</td><td>23</td><td>GCGGCGGCGGCGGCGGCGGCGGG</td><td>365.7</td><td>-12.3</td><td>1.6</td><td>-7.6</td><td>4.3E-14</td><td>1.4E-13</td><td>MB231/MB468</td><td>MITO</td></tr>
<tr><td>>hg38_21_+_8433172_8446622_RNA45SN1_45S_50ntFlanks@8844.8866.23|GCGGCGGCGGCGGCGGCGGCGGG</td><td>>hg38_21_+_8433172_8446622_RNA45SN1_45S_50ntFlanks</td><td>8844</td><td>8866</td><td>23</td><td>GCGGCGGCGGCGGCGGCGGCGGG</td><td>365.7</td><td>-12.3</td><td>1.6</td><td>-7.6</td><td>4.3E-14</td><td>1.4E-13</td><td>MB231/MB468</td><td>MITO</td></tr>
<tr><td>>hg38_21_+_8433172_8446622_RNA45SN1_45S_50ntFlanks@6650.6678.29|ACGACTCTTAGCGGTGGATCACTCGGCTC</td><td>>hg38_21_+_8433172_8446622_RNA45SN1_45S_50ntFlanks</td><td>6650</td><td>6678</td><td>29</td><td>ACGACTCTTAGCGGTGGATCACTCGGCTC</td><td>6708.1</td><td>4.8</td><td>0.6</td><td>7.6</td><td>2.6E-14</td><td>1.4E-13</td><td>BT20/MB468</td><td>MITO</td></tr>
<tr><td>>hg38_21_+_8433172_8446622_RNA45SN1_45S_50ntFlanks@10808.10840.33|CAATGTAGGTAAGGGAAGTCGGCAAGCCGGATC</td><td>>hg38_21_+_8433172_8446622_RNA45SN1_45S_50ntFlanks</td><td>10808</td><td>10840</td><td>33</td><td>CAATGTAGGTAAGGGAAGTCGGCAAGCCGGATC</td><td>1556.3</td><td>-9.7</td><td>1.3</td><td>-7.6</td><td>3.2E-14</td><td>1.4E-13</td><td>MB231/MB468</td><td>NUC</td></tr>
<tr><td>>hg38_21_+_8433172_8446622_RNA45SN1_45S_50ntFlanks@9568.9593.26|CGTGCAAATCGGTCGTCCGACCTGGG</td><td>>hg38_21_+_8433172_8446622_RNA45SN1_45S_50ntFlanks</td><td>9568</td><td>9593</td><td>26</td><td>CGTGCAAATCGGTCGTCCGACCTGGG</td><td>135.1</td><td>-10.9</td><td>1.4</td><td>-7.6</td><td>2.7E-14</td><td>1.4E-13</td><td>MB231/MB468</td><td>MP</td></tr>
<tr><td>>hg38_21_+_8433172_8446622_RNA45SN1_45S_50ntFlanks@10786.10824.39|AACAGCCTCTGGCATGTTGGAACAATGTAGGTAAGGGAA</td><td>>hg38_21_+_8433172_8446622_RNA45SN1_45S_50ntFlanks</td><td>10786</td><td>10824</td><td>39</td><td>AACAGCCTCTGGCATGTTGGAACAATGTAGGTAAGGGAA</td><td>260.6</td><td>-11.8</td><td>1.6</td><td>-7.6</td><td>4.3E-14</td><td>1.4E-13</td><td>MB231/MB468</td><td>MITO</td></tr>
<tr><td>>trnaMT_SerGCT_MT_+_12207_12265@1.22.22|GAGAAAGCTCACAAGAACTGCT</td><td>>trnaMT_SerGCT_MT_+_12207_12265</td><td>1</td><td>22</td><td>22</td><td>GAGAAAGCTCACAAGAACTGCT</td><td>1086.2</td><td>6.4</td><td>0.8</td><td>7.6</td><td>3.5E-14</td><td>1.4E-13</td><td>BT20/MB468</td><td>CYTO</td></tr>
<tr><td>>hg38_21_+_8433172_8446622_RNA45SN1_45S_50ntFlanks@9390.9415.26|GCCGAGGTGGGATCCCGAGGCCTCTC</td><td>>hg38_21_+_8433172_8446622_RNA45SN1_45S_50ntFlanks</td><td>9390</td><td>9415</td><td>26</td><td>GCCGAGGTGGGATCCCGAGGCCTCTC</td><td>196.4</td><td>-11.5</td><td>1.5</td><td>-7.6</td><td>2.7E-14</td><td>1.4E-13</td><td>MB231/MB468</td><td>MP</td></tr>
<tr><td>>MI0003643|hsa-mir-629&WithFlank&15|-|70079366|70079474@28.48.21(+1C)&[MIMAT0004810&hsa-miR-629-5p&offsets|0|0(+1C);m-150&15|-|70079427|70079447&offsets|0|0(+1C)]|TGGGTTTACGTTGGGAGAACTC</td><td>>MI0003643|hsa-mir-629&WithFlank&15|-|70079366|70079474</td><td>28</td><td>48</td><td>21(+1C)</td><td>TGGGTTTACGTTGGGAGAACTC</td><td>1772.1</td><td>14.5</td><td>1.8</td><td>8.0</td><td>1.2E-15</td><td>1.4E-13</td><td>BT20/MB468</td><td>MP</td></tr>
<tr><td>>hg38_21_+_8433172_8446622_RNA45SN1_45S_50ntFlanks@6776.6799.24|CTCCCGGGGCTACGCCTGTCTGAG</td><td>>hg38_21_+_8433172_8446622_RNA45SN1_45S_50ntFlanks</td><td>6776</td><td>6799</td><td>24</td><td>CTCCCGGGGCTACGCCTGTCTGAG</td><td>164.1</td><td>-11.2</td><td>1.5</td><td>-7.6</td><td>2.7E-14</td><td>1.4E-13</td><td>MB231/MB468</td><td>MP</td></tr>
<tr><td>>hg38_21_+_8433172_8446622_RNA45SN1_45S_50ntFlanks@10348.10374.27|AAACGAGAACTTTGAAGGCCGAAGTGG</td><td>>hg38_21_+_8433172_8446622_RNA45SN1_45S_50ntFlanks</td><td>10348</td><td>10374</td><td>27</td><td>AAACGAGAACTTTGAAGGCCGAAGTGG</td><td>1078.3</td><td>-4.5</td><td>0.6</td><td>-7.6</td><td>3.3E-14</td><td>1.4E-13</td><td>MB231/MB468</td><td>NUC</td></tr>
<tr><td>>hg38_21_+_8433172_8446622_RNA45SN1_45S_50ntFlanks@11894.11912.19|ACTCTAGTCTGGCACGGTG</td><td>>hg38_21_+_8433172_8446622_RNA45SN1_45S_50ntFlanks</td><td>11894</td><td>11912</td><td>19</td><td>ACTCTAGTCTGGCACGGTG</td><td>1755.9</td><td>-7.3</td><td>1.0</td><td>-7.6</td><td>3.1E-14</td><td>1.4E-13</td><td>MB231/MB468</td><td>CYTO</td></tr>
<tr><td>>hg38_21_+_8433172_8446622_RNA45SN1_45S_50ntFlanks@11934.11951.18|ATAAGTGGGAGGCCCCCG</td><td>>hg38_21_+_8433172_8446622_RNA45SN1_45S_50ntFlanks</td><td>11934</td><td>11951</td><td>18</td><td>ATAAGTGGGAGGCCCCCG</td><td>248.9</td><td>-11.8</td><td>1.6</td><td>-7.6</td><td>2.7E-14</td><td>1.4E-13</td><td>MB231/MB468</td><td>MP</td></tr>
<tr><td>>hg38_21_+_8433172_8446622_RNA45SN1_45S_50ntFlanks@10069.10088.20|AGTGGACGGGAGCGGCGGGG</td><td>>hg38_21_+_8433172_8446622_RNA45SN1_45S_50ntFlanks</td><td>10069</td><td>10088</td><td>20</td><td>AGTGGACGGGAGCGGCGGGG</td><td>243.1</td><td>-11.8</td><td>1.5</td><td>-7.6</td><td>2.7E-14</td><td>1.4E-13</td><td>MB231/MB468</td><td>MP</td></tr>
<tr><td>>hg38_21_+_8433172_8446622_RNA45SN1_45S_50ntFlanks@10534.10571.38|CCGCGAGGCGTCCAGTGCGGTAACGCGACCGATCCCGG</td><td>>hg38_21_+_8433172_8446622_RNA45SN1_45S_50ntFlanks</td><td>10534</td><td>10571</td><td>38</td><td>CCGCGAGGCGTCCAGTGCGGTAACGCGACCGATCCCGG</td><td>159.4</td><td>-11.2</td><td>1.5</td><td>-7.6</td><td>2.7E-14</td><td>1.4E-13</td><td>MB231/MB468</td><td>MP</td></tr>
<tr><td>>hg38_21_+_8433172_8446622_RNA45SN1_45S_50ntFlanks@3991.4026.36|TGGTGACTCTAGATAACCTCGGGCCGATCGCACGCC</td><td>>hg38_21_+_8433172_8446622_RNA45SN1_45S_50ntFlanks</td><td>3991</td><td>4026</td><td>36</td><td>TGGTGACTCTAGATAACCTCGGGCCGATCGCACGCC</td><td>305.9</td><td>-12.1</td><td>1.6</td><td>-7.6</td><td>2.7E-14</td><td>1.4E-13</td><td>MB231/MB468</td><td>MP</td></tr>
<tr><td>>hg38_21_+_8433172_8446622_RNA45SN1_45S_50ntFlanks@9738.9769.32|GAAACGATCTCAACCTATTCTCAAACTTTAAA</td><td>>hg38_21_+_8433172_8446622_RNA45SN1_45S_50ntFlanks</td><td>9738</td><td>9769</td><td>32</td><td>GAAACGATCTCAACCTATTCTCAAACTTTAAA</td><td>2166.3</td><td>-7.8</td><td>1.0</td><td>-7.5</td><td>4.4E-14</td><td>1.4E-13</td><td>MB231/MB468</td><td>MITO</td></tr>
<tr><td>>MI0000108|hsa-mir-103a-2&WithFlank&20|+|3917488|3917577@54.77.24(+1U);MI0000109|hsa-mir-103a-1&WithFlank&5|-|168560890|168560979@54.77.24(+1U)&[MIMAT0000101&hsa-miR-103a-3p&offsets|0|+1(+1U);m-12&20|+|3917541|3917563&offsets|0|+1(+1U)];[MIMAT0000101_1&hsa-miR-103a-3p&offsets|0|+1(+1U);m-13&5|-|168560904|168560926&offsets|0|+1(+1U);m-44220&5|-|168560908|168560926&offsets|0|+5(+1U)]|AGCAGCATTGTACAGGGCTATGAAT</td><td>>MI0000108|hsa-mir-103a-2&WithFlank&20|+|3917488|3917577</td><td>54</td><td>77</td><td>24(+1U)</td><td>AGCAGCATTGTACAGGGCTATGAAT</td><td>433.3</td><td>5.9</td><td>0.8</td><td>7.6</td><td>3.3E-14</td><td>1.4E-13</td><td>MB231/MB468</td><td>NUC</td></tr>
<tr><td>>trna30_ProCGG_6_+_27059521_27059592@1.33.33,trna37_ProCGG_17_-_8126151_8126222@1.33.33,trna52_ProCGG_1_+_167683962_167684033@1.33.33,trna6_ProCGG_16_+_3222049_3222120@1.33.33|GGCTCGTTGGTCTAGGGGTATGATTCTCGCTTC</td><td>>trna30_ProCGG_6_+_27059521_27059592</td><td>1</td><td>33</td><td>33</td><td>GGCTCGTTGGTCTAGGGGTATGATTCTCGCTTC</td><td>413.1</td><td>12.7</td><td>1.7</td><td>7.6</td><td>3.2E-14</td><td>1.4E-13</td><td>MB231/MB468</td><td>CYTO</td></tr>
<tr><td>>MI0001445|hsa-mir-423&WithFlank&17|+|30117073|30117178@59.80.22&[MIMAT0001340&hsa-miR-423-3p&offsets|0|-1;m-100&17|+|30117131|30117153&offsets|0|-1]|AGCTCGGTCTGAGGCCCCTCAG</td><td>>MI0001445|hsa-mir-423&WithFlank&17|+|30117073|30117178</td><td>59</td><td>80</td><td>22</td><td>AGCTCGGTCTGAGGCCCCTCAG</td><td>1259.2</td><td>6.5</td><td>0.9</td><td>7.6</td><td>3.2E-14</td><td>1.4E-13</td><td>MB231/MB468</td><td>CYTO</td></tr>
<tr><td>>hg38_21_+_8433172_8446622_RNA45SN1_45S_50ntFlanks@8839.8864.26|CGGTGGCGGCGGCGGCGGCGGCGGCG</td><td>>hg38_21_+_8433172_8446622_RNA45SN1_45S_50ntFlanks</td><td>8839</td><td>8864</td><td>26</td><td>CGGTGGCGGCGGCGGCGGCGGCGGCG</td><td>265.0</td><td>-11.9</td><td>1.6</td><td>-7.6</td><td>2.8E-14</td><td>1.4E-13</td><td>MB231/MB468</td><td>MP</td></tr>
<tr><td>>hg38_21_+_8433172_8446622_RNA45SN1_45S_50ntFlanks@12150.12180.31|GGTGGGGAGTTTGACTGGGGCGGTACACCTG</td><td>>hg38_21_+_8433172_8446622_RNA45SN1_45S_50ntFlanks</td><td>12150</td><td>12180</td><td>31</td><td>GGTGGGGAGTTTGACTGGGGCGGTACACCTG</td><td>170.7</td><td>-11.3</td><td>1.5</td><td>-7.6</td><td>2.8E-14</td><td>1.4E-13</td><td>MB231/MB468</td><td>MP</td></tr>
<tr><td>>MI0000478|hsa-mir-149&WithFlank&2|+|240455995|240456095@21.42.22(+1U)&[MIMAT0000450&hsa-miR-149-5p&offsets|0|-1(+1U);m-195&2|+|240456015|240456037&offsets|0|-1(+1U)]|TCTGGCTCCGTGTCTTCACTCCT</td><td>>MI0000478|hsa-mir-149&WithFlank&2|+|240455995|240456095</td><td>21</td><td>42</td><td>22(+1U)</td><td>TCTGGCTCCGTGTCTTCACTCCT</td><td>322.0</td><td>7.8</td><td>1.0</td><td>7.5</td><td>5.2E-14</td><td>1.5E-13</td><td>BT20/MB468</td><td>NUC</td></tr>
<tr><td>>MI0000095|hsa-mir-93&WithFlank&7|-|100093762|100093853@17.38.22&[MIMAT0000093&hsa-miR-93-5p&offsets|0|-1;m-29&7|-|100093815|100093837&offsets|0|-1]|CAAAGTGCTGTTCGTGCAGGTA</td><td>>MI0000095|hsa-mir-93&WithFlank&7|-|100093762|100093853</td><td>17</td><td>38</td><td>22</td><td>CAAAGTGCTGTTCGTGCAGGTA</td><td>457.0</td><td>5.3</td><td>0.7</td><td>7.6</td><td>3.3E-14</td><td>1.5E-13</td><td>MB231/MB468</td><td>NUC</td></tr>
<tr><td>>hg38_21_+_8433172_8446622_RNA45SN1_45S_50ntFlanks@4661.4682.22|AGGTGAAATTCTTGGACCGGCG</td><td>>hg38_21_+_8433172_8446622_RNA45SN1_45S_50ntFlanks</td><td>4661</td><td>4682</td><td>22</td><td>AGGTGAAATTCTTGGACCGGCG</td><td>134.5</td><td>-10.0</td><td>1.3</td><td>-7.6</td><td>3.3E-14</td><td>1.5E-13</td><td>MB231/MB468</td><td>NUC</td></tr>
<tr><td>>hg38_21_+_8433172_8446622_RNA45SN1_45S_50ntFlanks@4264.4282.19|AATGAGTCCACTTTAAATC</td><td>>hg38_21_+_8433172_8446622_RNA45SN1_45S_50ntFlanks</td><td>4264</td><td>4282</td><td>19</td><td>AATGAGTCCACTTTAAATC</td><td>364.9</td><td>-12.3</td><td>1.6</td><td>-7.5</td><td>4.4E-14</td><td>1.5E-13</td><td>MB231/MB468</td><td>MITO</td></tr>
<tr><td>>hg38_21_+_8433172_8446622_RNA45SN1_45S_50ntFlanks@9787.9821.35|TCGCTGGCGTGGAGCCGGGCGTGGAATGCGAGTGC</td><td>>hg38_21_+_8433172_8446622_RNA45SN1_45S_50ntFlanks</td><td>9787</td><td>9821</td><td>35</td><td>TCGCTGGCGTGGAGCCGGGCGTGGAATGCGAGTGC</td><td>142.1</td><td>-11.0</td><td>1.4</td><td>-7.6</td><td>2.8E-14</td><td>1.5E-13</td><td>MB231/MB468</td><td>MP</td></tr>
<tr><td>>hg38_21_+_8433172_8446622_RNA45SN1_45S_50ntFlanks@11668.11709.42|TGCCCAGTGCTCTGAATGTCAAAGTGAAGAAATTCAATGAAG</td><td>>hg38_21_+_8433172_8446622_RNA45SN1_45S_50ntFlanks</td><td>11668</td><td>11709</td><td>42</td><td>TGCCCAGTGCTCTGAATGTCAAAGTGAAGAAATTCAATGAAG</td><td>968.6</td><td>10.2</td><td>1.3</td><td>7.6</td><td>3.2E-14</td><td>1.5E-13</td><td>MB231/MB468</td><td>CYTO</td></tr>
<tr><td>>hg38_21_+_8433172_8446622_RNA45SN1_45S_50ntFlanks@10794.10818.25|CTGGCATGTTGGAACAATGTAGGTA</td><td>>hg38_21_+_8433172_8446622_RNA45SN1_45S_50ntFlanks</td><td>10794</td><td>10818</td><td>25</td><td>CTGGCATGTTGGAACAATGTAGGTA</td><td>884.4</td><td>-7.5</td><td>1.0</td><td>-7.6</td><td>3.4E-14</td><td>1.5E-13</td><td>MB231/MB468</td><td>NUC</td></tr>
<tr><td>>hg38_21_+_8433172_8446622_RNA45SN1_45S_50ntFlanks@10101.10125.25|CGCGCGCGTGTGGTGTGCGTCGGAG</td><td>>hg38_21_+_8433172_8446622_RNA45SN1_45S_50ntFlanks</td><td>10101</td><td>10125</td><td>25</td><td>CGCGCGCGTGTGGTGTGCGTCGGAG</td><td>603.8</td><td>-11.3</td><td>1.5</td><td>-7.6</td><td>2.7E-14</td><td>1.5E-13</td><td>BT20/MB468</td><td>MITO</td></tr>
<tr><td>>hg38_21_+_8433172_8446622_RNA45SN1_45S_50ntFlanks@12490.12521.32|CGTGAGCTGGGTTTAGACCGTCGTGAGACAGG</td><td>>hg38_21_+_8433172_8446622_RNA45SN1_45S_50ntFlanks</td><td>12490</td><td>12521</td><td>32</td><td>CGTGAGCTGGGTTTAGACCGTCGTGAGACAGG</td><td>119.9</td><td>-7.1</td><td>0.9</td><td>-7.6</td><td>3.2E-14</td><td>1.5E-13</td><td>MB231/MB468</td><td>CYTO</td></tr>
</table>

<table>
<tr><td></td><td></td><td></td><td></td><td></td><td></td><td></td><td></td><td></td><td></td><td></td><td></td><td></td><td></td></tr>
<tr><td>>hg38_21_+_8433172_8446622_RNA45SN1_45S_50ntFlanks@12655.12674.20|ATGACTGAACGCCTCTAAGT</td><td>>hg38_21_+_8433172_8446622_RNA45SN1_45S_50ntFlanks</td><td>12655</td><td>12674</td><td>20</td><td>ATGACTGAACGCCTCTAAGT</td><td>9597.6</td><td>-6.1</td><td>0.8</td><td>-7.6</td><td>3.4E-14</td><td>1.5E-13</td><td>MB231/MB468</td><td>NUC</td></tr>
<tr><td>>hg38_21_+_8433172_8446622_RNA45SN1_45S_50ntFlanks@12108.12125.18|CCGCGCGCCGGCCGGGCG</td><td>>hg38_21_+_8433172_8446622_RNA45SN1_45S_50ntFlanks</td><td>12108</td><td>12125</td><td>18</td><td>CCGCGCGCCGGCCGGGCG</td><td>277.8</td><td>-11.9</td><td>1.6</td><td>-7.5</td><td>4.5E-14</td><td>1.5E-13</td><td>MB231/MB468</td><td>MITO</td></tr>
<tr><td>>MI0000446|hsa-mir-125b-1&WithFlank&11|-|122099751|122099850@22.42.21;MI0000470|hsa-mir-125b-2&WithFlank&21|+|16590231|16590331@24.44.21&[MIMAT0000423&hsa-miR-125b-5p&offsets|+1|0;m-67&11|-|122099809|122099830&offsets|+1|0];[MIMAT0000423_1&hsa-miR-125b-5p&offsets|+1|0;m-68&21|+|16590253|16590274&offsets|+1|0]|CCCTGAGACCCTAACTTGTGA</td><td>>MI0000446|hsa-mir-125b-1&WithFlank&11|-|122099751|122099850</td><td>22</td><td>42</td><td>21</td><td>CCCTGAGACCCTAACTTGTGA</td><td>298.5</td><td>-11.5</td><td>1.5</td><td>-7.9</td><td>3.7E-15</td><td>1.5E-13</td><td>BT20/MB231</td><td>MITO</td></tr>
<tr><td>>hg38_21_+_8433172_8446622_RNA45SN1_45S_50ntFlanks@6772.6793.22|GTTCCTCCCGGGGCTACGCCTG</td><td>>hg38_21_+_8433172_8446622_RNA45SN1_45S_50ntFlanks</td><td>6772</td><td>6793</td><td>22</td><td>GTTCCTCCCGGGGCTACGCCTG</td><td>293.3</td><td>-11.2</td><td>1.5</td><td>-7.6</td><td>3.4E-14</td><td>1.5E-13</td><td>MB231/MB468</td><td>NUC</td></tr>
<tr><td>>hg38_21_+_8433172_8446622_RNA45SN1_45S_50ntFlanks@3835.3859.25|CTCCTCTCCTACTTGGATAACTGTG</td><td>>hg38_21_+_8433172_8446622_RNA45SN1_45S_50ntFlanks</td><td>3835</td><td>3859</td><td>25</td><td>CTCCTCTCCTACTTGGATAACTGTG</td><td>175.6</td><td>-10.4</td><td>1.4</td><td>-7.6</td><td>3.4E-14</td><td>1.5E-13</td><td>MB231/MB468</td><td>NUC</td></tr>
<tr><td>>hg38_21_+_8433172_8446622_RNA45SN1_45S_50ntFlanks@12417.12436.20|TCGATGTCGGCTCTTCCTAT</td><td>>hg38_21_+_8433172_8446622_RNA45SN1_45S_50ntFlanks</td><td>12417</td><td>12436</td><td>20</td><td>TCGATGTCGGCTCTTCCTAT</td><td>9175.9</td><td>-4.4</td><td>0.6</td><td>-7.6</td><td>3.3E-14</td><td>1.5E-13</td><td>MB231/MB468</td><td>CYTO</td></tr>
<tr><td>>hg38_21_+_8433172_8446622_RNA45SN1_45S_50ntFlanks@9539.9571.33|ACTCTGGTGGAGGTCCGTAGCGGTCCTGACGTG</td><td>>hg38_21_+_8433172_8446622_RNA45SN1_45S_50ntFlanks</td><td>9539</td><td>9571</td><td>33</td><td>ACTCTGGTGGAGGTCCGTAGCGGTCCTGACGTG</td><td>345.9</td><td>-12.3</td><td>1.6</td><td>-7.6</td><td>2.8E-14</td><td>1.5E-13</td><td>MB231/MB468</td><td>MP</td></tr>
<tr><td>>hg38_21_+_8433172_8446622_RNA45SN1_45S_50ntFlanks@9565.9596.32|TGACGTGCAAATCGGTCGTCCGACCTGGGTAT</td><td>>hg38_21_+_8433172_8446622_RNA45SN1_45S_50ntFlanks</td><td>9565</td><td>9596</td><td>32</td><td>TGACGTGCAAATCGGTCGTCCGACCTGGGTAT</td><td>3190.8</td><td>-8.4</td><td>1.1</td><td>-7.6</td><td>2.8E-14</td><td>1.5E-13</td><td>MB231/MB468</td><td>MP</td></tr>
<tr><td>>hg38_21_+_8433172_8446622_RNA45SN1_45S_50ntFlanks@13004.13025.22|ATCTATTGAAAGTCAGCCCTCG</td><td>>hg38_21_+_8433172_8446622_RNA45SN1_45S_50ntFlanks</td><td>13004</td><td>13025</td><td>22</td><td>ATCTATTGAAAGTCAGCCCTCG</td><td>155.1</td><td>-11.1</td><td>1.5</td><td>-7.6</td><td>2.9E-14</td><td>1.5E-13</td><td>MB231/MB468</td><td>MP</td></tr>
<tr><td>>hg38_21_+_8433172_8446622_RNA45SN1_45S_50ntFlanks@9623.9659.37|TAGTAGCTGGTTCCCTCCGAAGTTTCCCTCAGGATAG</td><td>>hg38_21_+_8433172_8446622_RNA45SN1_45S_50ntFlanks</td><td>9623</td><td>9659</td><td>37</td><td>TAGTAGCTGGTTCCCTCCGAAGTTTCCCTCAGGATAG</td><td>219.4</td><td>-11.6</td><td>1.5</td><td>-7.5</td><td>4.5E-14</td><td>1.5E-13</td><td>MB231/MB468</td><td>MITO</td></tr>
<tr><td>>hg38_21_+_8433172_8446622_RNA45SN1_45S_50ntFlanks@9573.9596.24|AAATCGGTCGTCCGACCTGGGTAT</td><td>>hg38_21_+_8433172_8446622_RNA45SN1_45S_50ntFlanks</td><td>9573</td><td>9596</td><td>24</td><td>AAATCGGTCGTCCGACCTGGGTAT</td><td>7521.9</td><td>-7.6</td><td>0.9</td><td>-8.0</td><td>1.2E-15</td><td>1.5E-13</td><td>BT20/MB468</td><td>MP</td></tr>
<tr><td>>MI0000077|hsa-mir-21&WithFlank&17|+|59841260|59841343@14.34.21&[MIMAT0000076&hsa-miR-21-5p&offsets|0|-1;m-1&17|+|59841273|59841294&offsets|0|-1]|TAGCTTATCAGACTGATGTTG</td><td>>MI0000077|hsa-mir-21&WithFlank&17|+|59841260|59841343</td><td>14</td><td>34</td><td>21</td><td>TAGCTTATCAGACTGATGTTG</td><td>3935.7</td><td>4.3</td><td>0.6</td><td>7.5</td><td>5.4E-14</td><td>1.5E-13</td><td>BT20/MB468</td><td>NUC</td></tr>
<tr><td>>hg38_21_+_8433172_8446622_RNA45SN1_45S_50ntFlanks@10130.10149.20|GCGGCGGCGGCGGCGGCGGG</td><td>>hg38_21_+_8433172_8446622_RNA45SN1_45S_50ntFlanks</td><td>10130</td><td>10149</td><td>20</td><td>GCGGCGGCGGCGGCGGCGGG</td><td>250.6</td><td>-11.8</td><td>1.6</td><td>-7.6</td><td>2.9E-14</td><td>1.5E-13</td><td>MB231/MB468</td><td>MP</td></tr>
<tr><td>>hg38_21_+_8433172_8446622_RNA45SN1_45S_50ntFlanks@8847.8866.20|GCGGCGGCGGCGGCGGCGGG</td><td>>hg38_21_+_8433172_8446622_RNA45SN1_45S_50ntFlanks</td><td>8847</td><td>8866</td><td>20</td><td>GCGGCGGCGGCGGCGGCGGG</td><td>250.6</td><td>-11.8</td><td>1.6</td><td>-7.6</td><td>2.9E-14</td><td>1.5E-13</td><td>MB231/MB468</td><td>MP</td></tr>
<tr><td>>hg38_21_+_8433172_8446622_RNA45SN1_45S_50ntFlanks@12133.12150.18|CTCCGGGGACAGTGCCAG</td><td>>hg38_21_+_8433172_8446622_RNA45SN1_45S_50ntFlanks</td><td>12133</td><td>12150</td><td>18</td><td>CTCCGGGGACAGTGCCAG</td><td>1773.2</td><td>-7.2</td><td>1.0</td><td>-7.6</td><td>3.3E-14</td><td>1.5E-13</td><td>MB231/MB468</td><td>CYTO</td></tr>
<tr><td>>hg38_21_+_8433172_8446622_RNA45SN1_45S_50ntFlanks@5078.5105.28|CGATAACGAACGAGACTCTGGCATGCTA</td><td>>hg38_21_+_8433172_8446622_RNA45SN1_45S_50ntFlanks</td><td>5078</td><td>5105</td><td>28</td><td>CGATAACGAACGAGACTCTGGCATGCTA</td><td>553.0</td><td>-12.9</td><td>1.7</td><td>-7.5</td><td>4.6E-14</td><td>1.5E-13</td><td>MB231/MB468</td><td>MITO</td></tr>
<tr><td>>hg38_21_+_8433172_8446622_RNA45SN1_45S_50ntFlanks@5472.5498.27|ACGGCCCTGGCGGAGCGCTGAGAAGAC</td><td>>hg38_21_+_8433172_8446622_RNA45SN1_45S_50ntFlanks</td><td>5472</td><td>5498</td><td>27</td><td>ACGGCCCTGGCGGAGCGCTGAGAAGAC</td><td>1230.1</td><td>4.7</td><td>0.6</td><td>7.6</td><td>3.3E-14</td><td>1.5E-13</td><td>MB231/MB468</td><td>CYTO</td></tr>
<tr><td>>hg38_21_+_8433172_8446622_RNA45SN1_45S_50ntFlanks@11454.11477.24|CCTCGGCGCGCGGCGGCGGCGG</td><td>>hg38_21_+_8433172_8446622_RNA45SN1_45S_50ntFlanks</td><td>11454</td><td>11477</td><td>24</td><td>CCTCGGCGCGCGGCGGCGGCGGCG</td><td>181.7</td><td>-10.2</td><td>1.3</td><td>-7.6</td><td>3.3E-14</td><td>1.5E-13</td><td>MB231/MB468</td><td>CYTO</td></tr>
<tr><td>>hg38_21_+_8433172_8446622_RNA45SN1_45S_50ntFlanks@4102.4121.20|ACCACGGGTGACGGGGAATC</td><td>>hg38_21_+_8433172_8446622_RNA45SN1_45S_50ntFlanks</td><td>4102</td><td>4121</td><td>20</td><td>ACCACGGGTGACGGGGAATC</td><td>1557.8</td><td>-4.7</td><td>0.6</td><td>-7.6</td><td>3.4E-14</td><td>1.5E-13</td><td>MB231/MB468</td><td>CYTO</td></tr>
<tr><td>>hg38_21_+_8433172_8446622_RNA45SN1_45S_50ntFlanks@10597.10617.21|AGTTCTCTTTTCTTTGTGAAG</td><td>>hg38_21_+_8433172_8446622_RNA45SN1_45S_50ntFlanks</td><td>10597</td><td>10617</td><td>21</td><td>AGTTCTCTTTTCTTTGTGAAG</td><td>154.1</td><td>-11.1</td><td>1.5</td><td>-7.6</td><td>2.9E-14</td><td>1.5E-13</td><td>MB231/MB468</td><td>MP</td></tr>
<tr><td>>hg38_21_+_8433172_8446622_RNA45SN1_45S_50ntFlanks@5073.5097.25|AATTCCGATAACGAACGAGACTCTG</td><td>>hg38_21_+_8433172_8446622_RNA45SN1_45S_50ntFlanks</td><td>5073</td><td>5097</td><td>25</td><td>AATTCCGATAACGAACGAGACTCTG</td><td>1733.5</td><td>-14.6</td><td>1.9</td><td>-7.6</td><td>2.9E-14</td><td>1.5E-13</td><td>MB231/MB468</td><td>MP</td></tr>
<tr><td>>hg38_21_+_8433172_8446622_RNA45SN1_45S_50ntFlanks@5494.5526.33|AAGACGGTCGAACTTGACTATCTAGAGGAAGTA</td><td>>hg38_21_+_8433172_8446622_RNA45SN1_45S_50ntFlanks</td><td>5494</td><td>5526</td><td>33</td><td>AAGACGGTCGAACTTGACTATCTAGAGGAAGTA</td><td>1004.3</td><td>-7.8</td><td>1.0</td><td>-7.6</td><td>2.9E-14</td><td>1.5E-13</td><td>MB231/MB468</td><td>MP</td></tr>
<tr><td>>hg38_21_+_8433172_8446622_RNA45SN1_45S_50ntFlanks@12160.12180.21|TTGACTGGGGCGGTACACCTG</td><td>>hg38_21_+_8433172_8446622_RNA45SN1_45S_50ntFlanks</td><td>12160</td><td>12180</td><td>21</td><td>TTGACTGGGGCGGTACACCTG</td><td>290.7</td><td>-12.0</td><td>1.6</td><td>-7.5</td><td>4.6E-14</td><td>1.5E-13</td><td>MB231/MB468</td><td>MITO</td></tr>
<tr><td>>hg38_21_+_8433172_8446622_RNA45SN1_45S_50ntFlanks@4477.4494.18|CTGAGTGTCCCGCGGGGC</td><td>>hg38_21_+_8433172_8446622_RNA45SN1_45S_50ntFlanks</td><td>4477</td><td>4494</td><td>18</td><td>CTGAGTGTCCCGCGGGGC</td><td>249.1</td><td>-11.8</td><td>1.6</td><td>-7.6</td><td>2.9E-14</td><td>1.5E-13</td><td>MB231/MB468</td><td>MP</td></tr>
<tr><td>>hg38_21_+_8433172_8446622_RNA45SN1_45S_50ntFlanks@7983.8008.26|CAGATCAGACGTGGCGACCCGCTGAA</td><td>>hg38_21_+_8433172_8446622_RNA45SN1_45S_50ntFlanks</td><td>7983</td><td>8008</td><td>26</td><td>CAGATCAGACGTGGCGACCCGCTGAA</td><td>5606.3</td><td>-8.8</td><td>1.2</td><td>-7.6</td><td>2.9E-14</td><td>1.5E-13</td><td>MB231/MB468</td><td>MP</td></tr>
<tr><td>>MI0000449|hsa-mir-132&WithFlank&17|-|2049902|2050014@65.86.22(+1U)&[MIMAT0000426&hsa-miR-132-3p&offsets|0|0(+1U);m-138&17|-|2049929|2049950&offsets|0|0(+1U)]|TAACAGTCTACAGCCATGGTCGT</td><td>>MI0000449|hsa-mir-132&WithFlank&17|-|2049902|2050014</td><td>65</td><td>86</td><td>22(+1U)</td><td>TAACAGTCTACAGCCATGGTCGT</td><td>442.9</td><td>10.5</td><td>1.4</td><td>7.6</td><td>3.4E-14</td><td>1.5E-13</td><td>MB231/MB468</td><td>CYTO</td></tr>
<tr><td>>hg38_21_+_8433172_8446622_RNA45SN1_45S_50ntFlanks@11197.11225.29|GGCCGGCGGCGGCGGCGACTCTGGACGCG</td><td>>hg38_21_+_8433172_8446622_RNA45SN1_45S_50ntFlanks</td><td>11197</td><td>11225</td><td>29</td><td>GGCCGGCGGCGGCGGCGACTCTGGACGCG</td><td>163.7</td><td>-10.3</td><td>1.4</td><td>-7.6</td><td>3.5E-14</td><td>1.5E-13</td><td>MB231/MB468</td><td>NUC</td></tr>
<tr><td>>hg38_21_+_8433172_8446622_RNA45SN1_45S_50ntFlanks@11768.11791.24|CTAATTAGTGACGCGCATGAATGG</td><td>>hg38_21_+_8433172_8446622_RNA45SN1_45S_50ntFlanks</td><td>11768</td><td>11791</td><td>24</td><td>CTAATTAGTGACGCGCATGAATGG</td><td>1288.0</td><td>-9.7</td><td>1.3</td><td>-7.6</td><td>3.5E-14</td><td>1.5E-13</td><td>MB231/MB468</td><td>NUC</td></tr>
<tr><td>>MI0000299|hsa-mir-222&WithFlank&X|-|45747009|45747130@75.96.22(+1A)&[MIMAT0000279&hsa-miR-222-3p&offsets|0|+1(+1A);m-140&X|-|45747033|45747056&offsets|0|-2(+1A)]|AGCTACATCTGGCTACTGGGTCA</td><td>>MI0000299|hsa-mir-222&WithFlank&X|-|45747009|45747130</td><td>75</td><td>96</td><td>22(+1A)</td><td>AGCTACATCTGGCTACTGGGTCA</td><td>594.7</td><td>7.3</td><td>1.0</td><td>7.6</td><td>3.4E-14</td><td>1.5E-13</td><td>MB231/MB468</td><td>CYTO</td></tr>
<tr><td>>hg38_21_+_8433172_8446622_RNA45SN1_45S_50ntFlanks@6721.6756.36|AATTGCAGGACACATTGATCATCGACACTTCGAACG</td><td>>hg38_21_+_8433172_8446622_RNA45SN1_45S_50ntFlanks</td><td>6721</td><td>6756</td><td>36</td><td>AATTGCAGGACACATTGATCATCGACACTTCGAACG</td><td>768.6</td><td>-4.7</td><td>0.6</td><td>-7.6</td><td>3.5E-14</td><td>1.5E-13</td><td>MB231/MB468</td><td>NUC</td></tr>
<tr><td>>hg38_21_+_8433172_8446622_RNA45SN1_45S_50ntFlanks@6659.6680.22|AGCGGTGGATCACTCGGCTCGT</td><td>>hg38_21_+_8433172_8446622_RNA45SN1_45S_50ntFlanks</td><td>6659</td><td>6680</td><td>22</td><td>AGCGGTGGATCACTCGGCTCGT</td><td>228.0</td><td>-11.7</td><td>1.5</td><td>-7.6</td><td>2.9E-14</td><td>1.5E-13</td><td>MB231/MB468</td><td>MP</td></tr>
<tr><td>>hg38_21_+_8433172_8446622_RNA45SN1_45S_50ntFlanks@11671.11702.32|CCAGTGCTCTGAATGTCAAAGTGAAGAAATTC</td><td>>hg38_21_+_8433172_8446622_RNA45SN1_45S_50ntFlanks</td><td>11671</td><td>11702</td><td>32</td><td>CCAGTGCTCTGAATGTCAAAGTGAAGAAATTC</td><td>1086.3</td><td>-5.2</td><td>0.7</td><td>-7.5</td><td>4.6E-14</td><td>1.5E-13</td><td>MB231/MB468</td><td>MITO</td></tr>
<tr><td>>hg38_21_+_8433172_8446622_RNA45SN1_45S_50ntFlanks@4160.4186.27|CCACATCCAAGGAAGGCAGCAGGCGCG</td><td>>hg38_21_+_8433172_8446622_RNA45SN1_45S_50ntFlanks</td><td>4160</td><td>4186</td><td>27</td><td>CCACATCCAAGGAAGGCAGCAGGCGCG</td><td>173.9</td><td>-10.4</td><td>1.4</td><td>-7.6</td><td>3.5E-14</td><td>1.5E-13</td><td>MB231/MB468</td><td>NUC</td></tr>
<tr><td>>hg38_21_+_8433172_8446622_RNA45SN1_45S_50ntFlanks@7987.8012.26|TCAGACGTGGCGACCCGCTGAATTTA</td><td>>hg38_21_+_8433172_8446622_RNA45SN1_45S_50ntFlanks</td><td>7987</td><td>8012</td><td>26</td><td>TCAGACGTGGCGACCCGCTGAATTTA</td><td>170.3</td><td>-11.3</td><td>1.5</td><td>-7.6</td><td>3.0E-14</td><td>1.5E-13</td><td>MB231/MB468</td><td>MP</td></tr>
<tr><td>>hg38_21_+_8433172_8446622_RNA45SN1_45S_50ntFlanks@9623.9652.30|TAGTAGCTGGTTCCCTCCGAAGTTTCCCTC</td><td>>hg38_21_+_8433172_8446622_RNA45SN1_45S_50ntFlanks</td><td>9623</td><td>9652</td><td>30</td><td>TAGTAGCTGGTTCCCTCCGAAGTTTCCCTC</td><td>1361.6</td><td>-9.3</td><td>1.2</td><td>-7.5</td><td>4.7E-14</td><td>1.5E-13</td><td>MB231/MB468</td><td>MITO</td></tr>
<tr><td>>hg38_21_+_8433172_8446622_RNA45SN1_45S_50ntFlanks@4003.4023.21|ATAACCTCGGGCCGATCGCAC</td><td>>hg38_21_+_8433172_8446622_RNA45SN1_45S_50ntFlanks</td><td>4003</td><td>4023</td><td>21</td><td>ATAACCTCGGGCCGATCGCAC</td><td>141.7</td><td>-11.0</td><td>1.4</td><td>-7.6</td><td>3.0E-14</td><td>1.5E-13</td><td>MB231/MB468</td><td>MP</td></tr>
<tr><td>>hg38_21_+_8433172_8446622_RNA45SN1_45S_50ntFlanks@9862.9899.38|ATGAACCGAACGCCGGGTTAAGGCGCCCGATGCCGACG</td><td>>hg38_21_+_8433172_8446622_RNA45SN1_45S_50ntFlanks</td><td>9862</td><td>9899</td><td>38</td><td>ATGAACCGAACGCCGGGTTAAGGCGCCCGATGCCGACG</td><td>490.4</td><td>-12.8</td><td>1.7</td><td>-7.6</td><td>3.0E-14</td><td>1.5E-13</td><td>MB231/MB468</td><td>MP</td></tr>
<tr><td>>hg38_21_+_8433172_8446622_RNA45SN1_45S_50ntFlanks@11641.11662.22|CGGCGGGTGTTGACGCGATGTG</td><td>>hg38_21_+_8433172_8446622_RNA45SN1_45S_50ntFlanks</td><td>11641</td><td>11662</td><td>22</td><td>CGGCGGGTGTTGACGCGATGTG</td><td>514.2</td><td>-6.2</td><td>0.8</td><td>-7.6</td><td>3.4E-14</td><td>1.5E-13</td><td>MB231/MB468</td><td>CYTO</td></tr>
<tr><td>>hg38_21_+_8433172_8446622_RNA45SN1_45S_50ntFlanks@10727.10755.29|GGAGAGGGTGTAAATCTCGCGCCGGGCCG</td><td>>hg38_21_+_8433172_8446622_RNA45SN1_45S_50ntFlanks</td><td>10727</td><td>10755</td><td>29</td><td>GGAGAGGGTGTAAATCTCGCGCCGGGCCG</td><td>359.5</td><td>-10.6</td><td>1.4</td><td>-7.6</td><td>2.8E-14</td><td>1.5E-13</td><td>BT20/MB468</td><td>MITO</td></tr>
<tr><td>>hg38_21_+_8433172_8446622_RNA45SN1_45S_50ntFlanks@10916.10935.20|CCGCGGCTGGACGAGGCGCC</td><td>>hg38_21_+_8433172_8446622_RNA45SN1_45S_50ntFlanks</td><td>10916</td><td>10935</td><td>20</td><td>CCGCGGCTGGACGAGGCGCC</td><td>195.2</td><td>-11.5</td><td>1.5</td><td>-7.6</td><td>3.0E-14</td><td>1.6E-13</td><td>MB231/MB468</td><td>MP</td></tr>
<tr><td>>hg38_21_+_8433172_8446622_RNA45SN1_45S_50ntFlanks@10539.10570.32|AGGCGTCCAGTGCGGTAACGCGACCGATCCCG</td><td>>hg38_21_+_8433172_8446622_RNA45SN1_45S_50ntFlanks</td><td>10539</td><td>10570</td><td>32</td><td>AGGCGTCCAGTGCGGTAACGCGACCGATCCCG</td><td>136.9</td><td>-11.0</td><td>1.4</td><td>-7.6</td><td>3.0E-14</td><td>1.6E-13</td><td>MB231/MB468</td><td>MP</td></tr>
<tr><td>>hg38_21_+_8433172_8446622_RNA45SN1_45S_50ntFlanks@7973.8010.38|GACGCGACCTCAGATCAGACGTGGCGACCCGCTGAATT</td><td>>hg38_21_+_8433172_8446622_RNA45SN1_45S_50ntFlanks</td><td>7973</td><td>8010</td><td>38</td><td>GACGCGACCTCAGATCAGACGTGGCGACCCGCTGAATT</td><td>2125.6</td><td>4.7</td><td>0.6</td><td>7.6</td><td>3.6E-14</td><td>1.6E-13</td><td>MB231/MB468</td><td>NUC</td></tr>
<tr><td>>hg38_21_+_8433172_8446622_RNA45SN1_45S_50ntFlanks@12423.12458.36|TCGGCTCTTCCTATCATTGTGAAGCAGAATTCACCA</td><td>>hg38_21_+_8433172_8446622_RNA45SN1_45S_50ntFlanks</td><td>12423</td><td>12458</td><td>36</td><td>TCGGCTCTTCCTATCATTGTGAAGCAGAATTCACCA</td><td>229.9</td><td>4.2</td><td>0.5</td><td>7.6</td><td>3.5E-14</td><td>1.6E-13</td><td>MB231/MB468</td><td>CYTO</td></tr>
<tr><td>>hg38_21_+_8433172_8446622_RNA45SN1_45S_50ntFlanks@4489.4511.23|CGGGGCCCGAAGCGTTTACTTTG</td><td>>hg38_21_+_8433172_8446622_RNA45SN1_45S_50ntFlanks</td><td>4489</td><td>4511</td><td>23</td><td>CGGGGCCCGAAGCGTTTACTTTG</td><td>478.7</td><td>-12.7</td><td>1.7</td><td>-7.5</td><td>4.7E-14</td><td>1.6E-13</td><td>MB231/MB468</td><td>MITO</td></tr>
<tr><td>>hg38_21_+_8433172_8446622_RNA45SN1_45S_50ntFlanks@4723.4751.29|CATTAATCAAGAACGAAAGTCGGAGGTTC</td><td>>hg38_21_+_8433172_8446622_RNA45SN1_45S_50ntFlanks</td><td>4723</td><td>4751</td><td>29</td><td>CATTAATCAAGAACGAAAGTCGGAGGTTC</td><td>899.1</td><td>-9.3</td><td>1.2</td><td>-7.5</td><td>4.7E-14</td><td>1.6E-13</td><td>MB231/MB468</td><td>MITO</td></tr>
<tr><td>>hg38_21_+_8433172_8446622_RNA45SN1_45S_50ntFlanks@3900.3927.28|CTTCGCGGGGGGGATGCGTGCATTTATC</td><td>>hg38_21_+_8433172_8446622_RNA45SN1_45S_50ntFlanks</td><td>3900</td><td>3927</td><td>28</td><td>CTTCGCGGGGGGGATGCGTGCATTTATC</td><td>263.2</td><td>-11.9</td><td>1.6</td><td>-7.6</td><td>3.0E-14</td><td>1.6E-13</td><td>MB231/MB468</td><td>MP</td></tr>
<tr><td>>hg38_21_+_8433172_8446622_RNA45SN1_45S_50ntFlanks@8155.8182.28|CGCTCGTGGGGGGCCCAAGTCCTTCTGA</td><td>>hg38_21_+_8433172_8446622_RNA45SN1_45S_50ntFlanks</td><td>8155</td><td>8182</td><td>28</td><td>CGCTCGTGGGGGGCCCAAGTCCTTCTGA</td><td>292.6</td><td>-12.0</td><td>1.6</td><td>-7.5</td><td>4.7E-14</td><td>1.6E-13</td><td>MB231/MB468</td><td>MITO</td></tr>
<tr><td>>MI0000446|hsa-mir-125b-1&WithFlank&11|-|122099751|122099850@61.81.21&[MIMAT0004592&hsa-miR-125b-1-3p&offsets|0|-1;m-485&11|-|122099769|122099790&offsets|0|-1]|ACGGGTTAGGCTCTTGGGAGC</td><td>>MI0000446|hsa-mir-125b-1&WithFlank&11|-|122099751|122099850</td><td>61</td><td>81</td><td>21</td><td>ACGGGTTAGGCTCTTGGGAGC</td><td>304.1</td><td>11.5</td><td>1.5</td><td>7.6</td><td>3.0E-14</td><td>1.6E-13</td><td>MB231/MB468</td><td>MP</td></tr>
<tr><td>>MI0000734|hsa-mir-106b&WithFlank&7|-|100093987|100094080@58.79.22&[MIMAT0004672&hsa-miR-106b-3p&offsets|0|0;m-87&7|-|100094003|100094023&offsets|0|+1]|CCGCACTGTGGGTACTTGCTGC</td><td>>MI0000734|hsa-mir-106b&WithFlank&7|-|100093987|100094080</td><td>58</td><td>79</td><td>22</td><td>CCGCACTGTGGGTACTTGCTGC</td><td>7084.3</td><td>5.8</td><td>0.8</td><td>7.6</td><td>3.5E-14</td><td>1.6E-13</td><td>MB231/MB468</td><td>CYTO</td></tr>
<tr><td>>hg38_21_+_8433172_8446622_RNA45SN1_45S_50ntFlanks@12691.12711.21|CGGAACGATACGGCAGCGCCG</td><td>>hg38_21_+_8433172_8446622_RNA45SN1_45S_50ntFlanks</td><td>12691</td><td>12711</td><td>21</td><td>CGGAACGATACGGCAGCGCCG</td><td>186.9</td><td>-11.4</td><td>1.5</td><td>-7.6</td><td>3.0E-14</td><td>1.6E-13</td><td>MB231/MB468</td><td>MP</td></tr>
</table>

| | | | | | | | | | | | | | |
|---|---|---|---|---|---|---|---|---|---|---|---|---|---|
| >hg38_21_+_8433172_8446622_RNA45SN1_45S_50ntFlanks@8356.8379.24\|GAAAAGAACTTTGAAGAGAGAGTT | >hg38_21_+_8433172_8446622_RNA45SN1_45S_50ntFlanks | 8356 | 8379 | 24 | GAAAAGAACTTTGAAGAGAGAGTT | 6159.4 | -8.0 | 1.1 | -7.6 | 3.5E-14 | 1.6E-13 | MB231/MB468 | CYTO |
| >trna10_AspGTC_12_-_125424193_125424264@31.73.43,trna12_AspGTC_12_-_125411891_125411962@31.73.43,trna38_AspGTC_17_-_8125556_8125627@31.73.43,trna45_AspGTC_6_+_27447453_27447524@31.73.43,trna48_AspGTC_6_+_27471523_27471594@31.73.43,trna4_AspGTC_12_+_96429799_96429870@31.73.43,trna69_AspGTC_1_-_161440205_161440276@31.73.43,trna72_AspGTC_1_-_161432824_161432895@31.73.43,trna75_AspGTC_1_-_161425414_161425485@31.73.43,trna78_AspGTC_1_-_161418033_161418104@31.73.43,trna81_AspGTC_1_-_161410615_161410686@31.73.43\|CCTGTCACGCGGGAGACCGGGGTTCGATTCCCCGACGGGGAGC | >trna10_AspGTC_12_-_125424193_125424264 | 31 | 73 | 43 | CCTGTCACGCGGGAGACCGGGGTTCGATTCCCCGACGGGGAGC | 230.1 | -10.6 | 1.4 | -7.6 | 3.5E-14 | 1.6E-13 | MB231/MB468 | CYTO |
| >hg38_21_+_8433172_8446622_RNA45SN1_45S_50ntFlanks@12635.12662.28\|AAGCTACCATCTGTGGGATTATGACTGA | >hg38_21_+_8433172_8446622_RNA45SN1_45S_50ntFlanks | 12635 | 12662 | 28 | AAGCTACCATCTGTGGGATTATGACTGA | 195.7 | -11.5 | 1.5 | -7.6 | 3.0E-14 | 1.6E-13 | MB231/MB468 | MP |
| >hg38_21_+_8433172_8446622_RNA45SN1_45S_50ntFlanks@11561.11585.25\|GCGCCTAGCAGCCGACTTAGAACTG | >hg38_21_+_8433172_8446622_RNA45SN1_45S_50ntFlanks | 11561 | 11585 | 25 | GCGCCTAGCAGCCGACTTAGAACTG | 1294.1 | -6.5 | 0.9 | -7.6 | 3.5E-14 | 1.6E-13 | MB231/MB468 | CYTO |
| >hg38_21_+_8433172_8446622_RNA45SN1_45S_50ntFlanks@11628.11660.33\|TCGCGAAGGCCCGCGGCGGGTGTTGACGCGATG | >hg38_21_+_8433172_8446622_RNA45SN1_45S_50ntFlanks | 11628 | 11660 | 33 | TCGCGAAGGCCCGCGGCGGGTGTTGACGCGATG | 1128.5 | -6.7 | 0.9 | -7.6 | 3.0E-14 | 1.6E-13 | MB231/MB468 | MP |
| >hg38_21_+_8433172_8446622_RNA45SN1_45S_50ntFlanks@9545.9564.20\|GTGGAGGTCCGTAGCGGTCC | >hg38_21_+_8433172_8446622_RNA45SN1_45S_50ntFlanks | 9545 | 9564 | 20 | GTGGAGGTCCGTAGCGGTCC | 364.9 | -12.4 | 1.6 | -7.6 | 3.0E-14 | 1.6E-13 | MB231/MB468 | MP |
| >hg38_21_+_8433172_8446622_RNA45SN1_45S_50ntFlanks@11564.11586.23\|CCTAGCAGCCGACTTAGAACTGG | >hg38_21_+_8433172_8446622_RNA45SN1_45S_50ntFlanks | 11564 | 11586 | 23 | CCTAGCAGCCGACTTAGAACTGG | 671.3 | -3.7 | 0.5 | -7.6 | 3.6E-14 | 1.6E-13 | MB231/MB468 | NUC |
| >hg38_21_+_8433172_8446622_RNA45SN1_45S_50ntFlanks@10781.10825.45\|AGGTGAACAGCCTCTGGCATGTTGGAACAATGTAGGTAAGGGAAG | >hg38_21_+_8433172_8446622_RNA45SN1_45S_50ntFlanks | 10781 | 10825 | 45 | AGGTGAACAGCCTCTGGCATGTTGGAACAATGTAGGTAAGGGAAG | 350.9 | -12.2 | 1.6 | -7.5 | 4.8E-14 | 1.6E-13 | MB231/MB468 | MITO |
| >hg38_21_+_8433172_8446622_RNA45SN1_45S_50ntFlanks@4047.4071.25\|ATTCGAACGTCTGCCCTATCAACTT | >hg38_21_+_8433172_8446622_RNA45SN1_45S_50ntFlanks | 4047 | 4071 | 25 | ATTCGAACGTCTGCCCTATCAACTT | 507.3 | -12.8 | 1.7 | -7.5 | 4.8E-14 | 1.6E-13 | MB231/MB468 | MITO |
| >hg38_21_+_8433172_8446622_RNA45SN1_45S_50ntFlanks@12476.12500.25\|CCACTAATAGGGAACGTGAGCTGGG | >hg38_21_+_8433172_8446622_RNA45SN1_45S_50ntFlanks | 12476 | 12500 | 25 | CCACTAATAGGGAACGTGAGCTGGG | 1012.0 | -9.1 | 1.2 | -7.6 | 3.6E-14 | 1.6E-13 | MB231/MB468 | NUC |
| >hg38_21_+_8433172_8446622_RNA45SN1_45S_50ntFlanks@10516.10533.18\|GAGTGGCGGAGATGGGCG | >hg38_21_+_8433172_8446622_RNA45SN1_45S_50ntFlanks | 10516 | 10533 | 18 | GAGTGGCGGAGATGGGCG | 296.4 | -6.6 | 0.9 | -7.6 | 3.5E-14 | 1.6E-13 | MB231/MB468 | CYTO |
| >hg38_21_+_8433172_8446622_RNA45SN1_45S_50ntFlanks@7991.8018.28\|ACGTGGCGACCCGCTGAATTTAAGCATA | >hg38_21_+_8433172_8446622_RNA45SN1_45S_50ntFlanks | 7991 | 8018 | 28 | ACGTGGCGACCCGCTGAATTTAAGCATA | 649.3 | -3.7 | 0.5 | -7.6 | 2.9E-14 | 1.6E-13 | BT20/MB468 | MITO |
| >hg38_21_+_8433172_8446622_RNA45SN1_45S_50ntFlanks@7976.8016.41\|GCGACCTCAGATCAGACGTGGCGACCCGCTGAATTTAAGCA | >hg38_21_+_8433172_8446622_RNA45SN1_45S_50ntFlanks | 7976 | 8016 | 41 | GCGACCTCAGATCAGACGTGGCGACCCGCTGAATTTAAGCA | 568.8 | 5.7 | 0.7 | 7.6 | 2.9E-14 | 1.6E-13 | BT20/MB468 | MITO |
| >hg38_21_+_8433172_8446622_RNA45SN1_45S_50ntFlanks@4288.4317.30\|ACGAGGATCCATTGGAGGGCAAGTCTGGTG | >hg38_21_+_8433172_8446622_RNA45SN1_45S_50ntFlanks | 4288 | 4317 | 30 | ACGAGGATCCATTGGAGGGCAAGTCTGGTG | 353.2 | -12.3 | 1.6 | -7.6 | 3.1E-14 | 1.6E-13 | MB231/MB468 | MP |
| >hg38_21_+_8433172_8446622_RNA45SN1_45S_50ntFlanks@9566.9597.32\|GACGTGCAAATCGGTCGTCCGACCTGGGTATA | >hg38_21_+_8433172_8446622_RNA45SN1_45S_50ntFlanks | 9566 | 9597 | 32 | GACGTGCAAATCGGTCGTCCGACCTGGGTATA | 4133.8 | -5.7 | 0.8 | -7.5 | 4.8E-14 | 1.6E-13 | MB231/MB468 | MITO |
| >hg38_21_+_8433172_8446622_RNA45SN1_45S_50ntFlanks@8170.8201.32\|CAAGTCCTTCTGATCGAGGCCCAGCCCGTGGA | >hg38_21_+_8433172_8446622_RNA45SN1_45S_50ntFlanks | 8170 | 8201 | 32 | CAAGTCCTTCTGATCGAGGCCCAGCCCGTGGA | 218.1 | -11.6 | 1.5 | -7.6 | 3.1E-14 | 1.6E-13 | MB231/MB468 | MP |
| >hg38_21_+_8433172_8446622_RNA45SN1_45S_50ntFlanks@6722.6770.49\|ATTGCAGGACACATTGATCATCGACACTTCGAACGCACTTGCGGCCCCG | >hg38_21_+_8433172_8446622_RNA45SN1_45S_50ntFlanks | 6722 | 6770 | 49 | ATTGCAGGACACATTGATCATCGACACTTCGAACGCACTTGCGGCCCCG | 87.9 | -22.1 | 2.9 | -7.6 | 3.1E-14 | 1.6E-13 | MB231/MB468 | MP |
| >hg38_21_+_8433172_8446622_RNA45SN1_45S_50ntFlanks@12787.12805.19\|CCGCGCGCGCGGGAGGGCG | >hg38_21_+_8433172_8446622_RNA45SN1_45S_50ntFlanks | 12787 | 12805 | 19 | CCGCGCGCGCGGGAGGGCG | 222.0 | -11.7 | 1.5 | -7.6 | 3.1E-14 | 1.6E-13 | MB231/MB468 | MP |
| >trna3_ArgTCT_11_+_59318767_59318852@1.29.29,trna4_ArgTCT_17_+_8024243_8024330@1.29.29,trna4_ArgTCT_9_-_131102355_131102445@1.29.29,trna52_ArgTCT_6_+_27529963_27530049@1.29.29\|GGCTCTGTGGCGCAATGGATAGCGCATTG | >trna3_ArgTCT_11_+_59318767_59318852 | 1 | 29 | 29 | GGCTCTGTGGCGCAATGGATAGCGCATTG | 167.0 | -10.1 | 1.3 | -7.6 | 3.6E-14 | 1.6E-13 | MB231/MB468 | CYTO |
| >hg38_21_+_8433172_8446622_RNA45SN1_45S_50ntFlanks@6738.6758.21\|ATCATCGACACTTCGAACGCA | >hg38_21_+_8433172_8446622_RNA45SN1_45S_50ntFlanks | 6738 | 6758 | 21 | ATCATCGACACTTCGAACGCA | 174.1 | -11.3 | 1.5 | -7.6 | 3.1E-14 | 1.6E-13 | MB231/MB468 | MP |
| >MI0000440\|hsa-mir-27b&WithFlank&9\|+\|95085439\|95085547@67.85.19&[MIMAT0000419&hsa-miR-27b-3p&offsets\|0\|-2;m-39&9\|+\|95085505\|95085525&offsets\|0\|-2]\|TTCACAGTGGCTAAGTTCT | >MI0000440\|hsa-mir-27b&WithFlank&9\|+\|95085439\|95085547 | 67 | 85 | 19 | TTCACAGTGGCTAAGTTCT | 889.5 | 5.6 | 0.7 | 7.6 | 3.6E-14 | 1.6E-13 | MB231/MB468 | CYTO |
| >hg38_21_+_8433172_8446622_RNA45SN1_45S_50ntFlanks@6781.6799.19\|GGGGCTACGCCTGTCTGAG | >hg38_21_+_8433172_8446622_RNA45SN1_45S_50ntFlanks | 6781 | 6799 | 19 | GGGGCTACGCCTGTCTGAG | 136.3 | -11.0 | 1.4 | -7.6 | 3.1E-14 | 1.6E-13 | MB231/MB468 | MP |
| >hg38_1_-_228634819_228635039_RNA5S12_5S_50ntFlanks@51.89.39\|GTCTACGGCCATACCACCCTGAACGCGCCCGATCTCGTC | >hg38_1_-_228634819_228635039_RNA5S12_5S_50ntFlanks | 51 | 89 | 39 | GTCTACGGCCATACCACCCTGAACGCGCCCGATCTCGTC | 25484.9 | 6.2 | 0.8 | 7.6 | 2.9E-14 | 1.6E-13 | BT20/MB468 | MITO |
| >hg38_21_+_8433172_8446622_RNA45SN1_45S_50ntFlanks@11316.11340.25\|CACCCCACGTCTCGTCGCGCGCGCG | >hg38_21_+_8433172_8446622_RNA45SN1_45S_50ntFlanks | 11316 | 11340 | 25 | CACCCCACGTCTCGTCGCGCGCGCG | 202.6 | -11.5 | 1.5 | -7.6 | 3.1E-14 | 1.6E-13 | MB231/MB468 | MP |
| >hg38_21_+_8433172_8446622_RNA45SN1_45S_50ntFlanks@11452.11477.26\|CTCCTCGGCGCGCGGCGGCGGCGGCG | >hg38_21_+_8433172_8446622_RNA45SN1_45S_50ntFlanks | 11452 | 11477 | 26 | CTCCTCGGCGCGCGGCGGCGGCGGCG | 218.7 | -10.7 | 1.4 | -7.6 | 3.7E-14 | 1.6E-13 | MB231/MB468 | NUC |
| >hg38_21_+_8433172_8446622_RNA45SN1_45S_50ntFlanks@5445.5467.23\|TCGGATCGGCCCCGCCGGGGTCG | >hg38_21_+_8433172_8446622_RNA45SN1_45S_50ntFlanks | 5445 | 5467 | 23 | TCGGATCGGCCCCGCCGGGGTCG | 736.5 | -12.6 | 1.6 | -8.0 | 1.3E-15 | 1.6E-13 | BT20/MB468 | MP |
| >hg38_21_+_8433172_8446622_RNA45SN1_45S_50ntFlanks@4103.4120.18\|CCACGGGTGACGGGGAAT | >hg38_21_+_8433172_8446622_RNA45SN1_45S_50ntFlanks | 4103 | 4120 | 18 | CCACGGGTGACGGGGAAT | 209.8 | -11.6 | 1.5 | -7.6 | 3.1E-14 | 1.6E-13 | MB231/MB468 | MP |
| >hg38_21_+_8433172_8446622_RNA45SN1_45S_50ntFlanks@12413.12432.20\|TCCTTCGATGTCGGCTCTTC | >hg38_21_+_8433172_8446622_RNA45SN1_45S_50ntFlanks | 12413 | 12432 | 20 | TCCTTCGATGTCGGCTCTTC | 192.3 | -11.5 | 1.5 | -7.6 | 3.1E-14 | 1.6E-13 | MB231/MB468 | MP |
| >hg38_21_+_8433172_8446622_RNA45SN1_45S_50ntFlanks@12633.12651.19\|CGAAGCTACCATCTGTGGG | >hg38_21_+_8433172_8446622_RNA45SN1_45S_50ntFlanks | 12633 | 12651 | 19 | CGAAGCTACCATCTGTGGG | 1069.5 | -7.4 | 1.0 | -7.5 | 4.9E-14 | 1.6E-13 | MB231/MB468 | MITO |
| >hg38_21_+_8433172_8446622_RNA45SN1_45S_50ntFlanks@6661.6689.29\|CGGTGGATCACTCGGCTCGTGCGTCGATG | >hg38_21_+_8433172_8446622_RNA45SN1_45S_50ntFlanks | 6661 | 6689 | 29 | CGGTGGATCACTCGGCTCGTGCGTCGATG | 23541.7 | -5.5 | 0.7 | -7.5 | 4.9E-14 | 1.6E-13 | MB231/MB468 | MITO |
| >hg38_21_+_8433172_8446622_RNA45SN1_45S_50ntFlanks@11785.11803.19\|TGAATGGATGAACGAGATT | >hg38_21_+_8433172_8446622_RNA45SN1_45S_50ntFlanks | 11785 | 11803 | 19 | TGAATGGATGAACGAGATT | 595.0 | -5.5 | 0.7 | -7.5 | 4.9E-14 | 1.6E-13 | MB231/MB468 | MITO |
| >hg38_21_+_8433172_8446622_RNA45SN1_45S_50ntFlanks@11580.11605.26\|GAACTGGTGCGGACCAGGGGAATCCG | >hg38_21_+_8433172_8446622_RNA45SN1_45S_50ntFlanks | 11580 | 11605 | 26 | GAACTGGTGCGGACCAGGGGAATCCG | 900.8 | -4.3 | 0.6 | -7.6 | 2.9E-14 | 1.6E-13 | BT20/MB468 | MITO |
| >MI0000108\|hsa-mir-103a-2&WithFlank&20\|+\|3917488\|3917577@54.75.22(+1U);MI0000109\|hsa-mir-103a-1&WithFlank&5\|-\|168560890\|168560979@54.75.22(+1U)&[MIMAT0000101&hsa-miR-103a-3p&offsets\|0\|-1(+1U);m-12&20\|+\|3917541\|3917563&offsets\|0\|-1(+1U)];[MIMAT0000101_1&hsa-miR-103a-3p&offsets\|0\|-1(+1U);m-13&5\|-\|168560904\|168560926&offsets\|0\|-1(+1U);m-44220&5\|-\|168560908\|168560926&offsets\|0\|+3(+1U)]\|AGCAGCATTGTACAGGGCTATGT | >MI0000108\|hsa-mir-103a-2&WithFlank&20\|+\|3917488\|3917577 | 54 | 75 | 22(+1U) | AGCAGCATTGTACAGGGCTATGT | 872.1 | 6.5 | 0.9 | 7.6 | 2.9E-14 | 1.6E-13 | BT20/MB468 | MITO |
| >hg38_21_+_8433172_8446622_RNA45SN1_45S_50ntFlanks@9804.9821.18\|GGCGTGGAATGCGAGTGC | >hg38_21_+_8433172_8446622_RNA45SN1_45S_50ntFlanks | 9804 | 9821 | 18 | GGCGTGGAATGCGAGTGC | 213.0 | -11.6 | 1.5 | -7.6 | 3.1E-14 | 1.6E-13 | MB231/MB468 | MP |
| >hg38_21_+_8433172_8446622_RNA45SN1_45S_50ntFlanks@9505.9523.19\|TGAACTATGCCTGGGCAGG | >hg38_21_+_8433172_8446622_RNA45SN1_45S_50ntFlanks | 9505 | 9523 | 19 | TGAACTATGCCTGGGCAGG | 220.5 | -7.5 | 1.0 | -7.6 | 3.6E-14 | 1.6E-13 | MB231/MB468 | CYTO |
| >hg38_21_+_8433172_8446622_RNA45SN1_45S_50ntFlanks@11821.11842.22\|CTATCCAGCGAAACCACAGCCA | >hg38_21_+_8433172_8446622_RNA45SN1_45S_50ntFlanks | 11821 | 11842 | 22 | CTATCCAGCGAAACCACAGCCA | 174.4 | -10.2 | 1.3 | -7.6 | 3.6E-14 | 1.6E-13 | MB231/MB468 | CYTO |
| >hg38_21_+_8433172_8446622_RNA45SN1_45S_50ntFlanks@8323.8356.34\|ACCGATAGTCAACAAGTACCGTAAGGGAAAGTTG | >hg38_21_+_8433172_8446622_RNA45SN1_45S_50ntFlanks | 8323 | 8356 | 34 | ACCGATAGTCAACAAGTACCGTAAGGGAAAGTTG | 166.3 | -11.2 | 1.5 | -7.6 | 3.2E-14 | 1.6E-13 | MB231/MB468 | MP |
| >hg38_21_+_8433172_8446622_RNA45SN1_45S_50ntFlanks@12957.12991.35\|CTTCTGGGTCGGGGTTTCGTACGTAGCAGAGCAGC | >hg38_21_+_8433172_8446622_RNA45SN1_45S_50ntFlanks | 12957 | 12991 | 35 | CTTCTGGGTCGGGGTTTCGTACGTAGCAGAGCAGC | 389.5 | -8.9 | 1.2 | -7.6 | 3.8E-14 | 1.6E-13 | MB231/MB468 | NUC |
| >hg38_21_+_8433172_8446622_RNA45SN1_45S_50ntFlanks@12405.12446.42\|CTTTTTGATCCTTCGATGTCGGCTCTTCCTATCATTGTGAAG | >hg38_21_+_8433172_8446622_RNA45SN1_45S_50ntFlanks | 12405 | 12446 | 42 | CTTTTTGATCCTTCGATGTCGGCTCTTCCTATCATTGTGAAG | 2812.2 | -6.1 | 0.8 | -7.5 | 5.0E-14 | 1.6E-13 | MB231/MB468 | MITO |
| >hg38_21_+_8433172_8446622_RNA45SN1_45S_50ntFlanks@8176.8208.33\|CTTCTGATCGAGGCCCAGCCCGTGGACGGTGTG | >hg38_21_+_8433172_8446622_RNA45SN1_45S_50ntFlanks | 8176 | 8208 | 33 | CTTCTGATCGAGGCCCAGCCCGTGGACGGTGTG | 1125.4 | -4.0 | 0.5 | -7.6 | 3.8E-14 | 1.6E-13 | MB231/MB468 | NUC |
| >hg38_21_+_8433172_8446622_RNA45SN1_45S_50ntFlanks@3899.3917.19\|CCTTCGCGGGGGGGATGCG | >hg38_21_+_8433172_8446622_RNA45SN1_45S_50ntFlanks | 3899 | 3917 | 19 | CCTTCGCGGGGGGGATGCG | 234.9 | -11.7 | 1.5 | -7.6 | 3.2E-14 | 1.6E-13 | MB231/MB468 | MP |
| >hg38_21_+_8433172_8446622_RNA45SN1_45S_50ntFlanks@11316.11345.30\|CACCCCACGTCTCGTCGCGCGCGCGTCCGC | >hg38_21_+_8433172_8446622_RNA45SN1_45S_50ntFlanks | 11316 | 11345 | 30 | CACCCCACGTCTCGTCGCGCGCGCGTCCGC | 265.1 | -11.8 | 1.6 | -7.5 | 5.0E-14 | 1.6E-13 | MB231/MB468 | MITO |
| >hg38_21_+_8433172_8446622_RNA45SN1_45S_50ntFlanks@11092.11114.23\|CGGCGGGGGGAGAAGGGTCGGGG | >hg38_21_+_8433172_8446622_RNA45SN1_45S_50ntFlanks | 11092 | 11114 | 23 | CGGCGGGGGGAGAAGGGTCGGGG | 172.8 | -10.4 | 1.4 | -7.6 | 3.8E-14 | 1.6E-13 | MB231/MB468 | NUC |
| >hg38_21_+_8433172_8446622_RNA45SN1_45S_50ntFlanks@11873.11897.25\|AAGAAGACCCTGTTGAGCTTGACTC | >hg38_21_+_8433172_8446622_RNA45SN1_45S_50ntFlanks | 11873 | 11897 | 25 | AAGAAGACCCTGTTGAGCTTGACTC | 683.9 | -4.8 | 0.6 | -7.6 | 3.7E-14 | 1.6E-13 | MB231/MB468 | CYTO |
| >hg38_21_+_8433172_8446622_RNA45SN1_45S_50ntFlanks@11564.11582.19\|CCTAGCAGCCGACTTAGAA | >hg38_21_+_8433172_8446622_RNA45SN1_45S_50ntFlanks | 11564 | 11582 | 19 | CCTAGCAGCCGACTTAGAA | 1664.4 | -4.6 | 0.6 | -7.6 | 3.0E-14 | 1.6E-13 | BT20/MB468 | MITO |
| >hg38_21_+_8433172_8446622_RNA45SN1_45S_50ntFlanks@993.1027.35\|CTCGTCGGTGTGGGGTTCGAGGCGGTTTGAGTGAG | >hg38_21_+_8433172_8446622_RNA45SN1_45S_50ntFlanks | 993 | 1027 | 35 | CTCGTCGGTGTGGGGTTCGAGGCGGTTTGAGTGAG | 146.2 | -10.2 | 1.3 | -7.6 | 3.8E-14 | 1.6E-13 | MB231/MB468 | NUC |
| >hg38_21_+_8433172_8446622_RNA45SN1_45S_50ntFlanks@4444.4471.28\|CCTTGCCTCTCGGCGCCCCCTCGATGCT | >hg38_21_+_8433172_8446622_RNA45SN1_45S_50ntFlanks | 4444 | 4471 | 28 | CCTTGCCTCTCGGCGCCCCCTCGATGCT | 417.5 | -11.7 | 1.5 | -7.6 | 3.8E-14 | 1.6E-13 | MB231/MB468 | NUC |
| >hg38_21_+_8433172_8446622_RNA45SN1_45S_50ntFlanks@11702.11728.27\|CAATGAAGCGCGGGTAAACGGCGGGAG | >hg38_21_+_8433172_8446622_RNA45SN1_45S_50ntFlanks | 11702 | 11728 | 27 | CAATGAAGCGCGGGTAAACGGCGGGAG | 238.7 | -11.8 | 1.5 | -7.6 | 3.2E-14 | 1.6E-13 | MB231/MB468 | MP |

<table>
<tr><th></th><th></th><th></th><th></th><th></th><th></th><th></th><th></th><th></th><th></th><th></th><th></th><th></th><th></th></tr>
<tr><td>>hg38_21_+_8433172_8446622_RNA45SN1_45S_50ntFlanks@485.527.43|GATGTGGTGACGTCGTGCTCTCCCGGGCCGGGTCCGAGCCGCG</td><td>>hg38_21_+_8433172_8446622_RNA45SN1_45S_50ntFlanks</td><td>485</td><td>527</td><td>43</td><td>GATGTGGTGACGTCGTGCTCTCCCGGGCCGGGTCCGAGCCGCG</td><td>591.5</td><td>12.0</td><td>1.6</td><td>7.5</td><td>5.9E-14</td><td>1.6E-13</td><td>BT20/MB468</td><td>NUC</td></tr>
<tr><td>>hg38_21_+_8433172_8446622_RNA45SN1_45S_50ntFlanks@4047.4068.22|ATTCGAACGTCTGCCCTATCAA</td><td>>hg38_21_+_8433172_8446622_RNA45SN1_45S_50ntFlanks</td><td>4047</td><td>4068</td><td>22</td><td>ATTCGAACGTCTGCCCTATCAA</td><td>254.7</td><td>-11.0</td><td>1.4</td><td>-7.6</td><td>3.8E-14</td><td>1.6E-13</td><td>MB231/MB468</td><td>NUC</td></tr>
<tr><td>>hg38_21_+_8433172_8446622_RNA45SN1_45S_50ntFlanks@9104.9121.18|CCGGCGACGGGGGGGGTG</td><td>>hg38_21_+_8433172_8446622_RNA45SN1_45S_50ntFlanks</td><td>9104</td><td>9121</td><td>18</td><td>CCGGCGACGGGGGGGGTG</td><td>212.8</td><td>-11.6</td><td>1.5</td><td>-7.6</td><td>3.2E-14</td><td>1.6E-13</td><td>MB231/MB468</td><td>MP</td></tr>
<tr><td>>hg38_21_+_8433172_8446622_RNA45SN1_45S_50ntFlanks@12329.12346.18|TTTTAAGCAGGAGGTGTC</td><td>>hg38_21_+_8433172_8446622_RNA45SN1_45S_50ntFlanks</td><td>12329</td><td>12346</td><td>18</td><td>TTTTAAGCAGGAGGTGTC</td><td>193.2</td><td>-11.5</td><td>1.5</td><td>-7.6</td><td>3.2E-14</td><td>1.6E-13</td><td>MB231/MB468</td><td>MP</td></tr>
<tr><td>>hg38_21_+_8433172_8446622_RNA45SN1_45S_50ntFlanks@11575.11605.31|ACTTAGAACTGGTGCGGACCAGGGGAATCCG</td><td>>hg38_21_+_8433172_8446622_RNA45SN1_45S_50ntFlanks</td><td>11575</td><td>11605</td><td>31</td><td>ACTTAGAACTGGTGCGGACCAGGGGAATCCG</td><td>2606.2</td><td>-6.9</td><td>0.9</td><td>-7.6</td><td>3.7E-14</td><td>1.6E-13</td><td>MB231/MB468</td><td>CYTO</td></tr>
<tr><td>>hg38_21_+_8433172_8446622_RNA45SN1_45S_50ntFlanks@10379.10424.46|GGGTTCCATGTGAACAGCAGTTGAACATGGGTCAGTCGGTCCTGAG</td><td>>hg38_21_+_8433172_8446622_RNA45SN1_45S_50ntFlanks</td><td>10379</td><td>10424</td><td>46</td><td>GGGTTCCATGTGAACAGCAGTTGAACATGGGTCAGTCGGTCCTGAG</td><td>2198.0</td><td>7.4</td><td>1.0</td><td>7.6</td><td>3.0E-14</td><td>1.6E-13</td><td>BT20/MB468</td><td>MITO</td></tr>
<tr><td>>hg38_21_+_8433172_8446622_RNA45SN1_45S_50ntFlanks@11572.11603.32|CCGACTTAGAACTGGTGCGGACCAGGGGAATC</td><td>>hg38_21_+_8433172_8446622_RNA45SN1_45S_50ntFlanks</td><td>11572</td><td>11603</td><td>32</td><td>CCGACTTAGAACTGGTGCGGACCAGGGGAATC</td><td>8596.9</td><td>-4.7</td><td>0.6</td><td>-7.6</td><td>3.0E-14</td><td>1.6E-13</td><td>BT20/MB468</td><td>MITO</td></tr>
<tr><td>>hg38_21_+_8433172_8446622_RNA45SN1_45S_50ntFlanks@3916.3938.23|CGTGCATTTATCAGATCAAAACC</td><td>>hg38_21_+_8433172_8446622_RNA45SN1_45S_50ntFlanks</td><td>3916</td><td>3938</td><td>23</td><td>CGTGCATTTATCAGATCAAAACC</td><td>200.9</td><td>-11.5</td><td>1.5</td><td>-7.6</td><td>3.2E-14</td><td>1.6E-13</td><td>MB231/MB468</td><td>MP</td></tr>
<tr><td>>hg38_21_+_8433172_8446622_RNA45SN1_45S_50ntFlanks@12405.12441.37|CTTTTTGATCCTTCGATGTCGGCTCTTCCTATCATTG</td><td>>hg38_21_+_8433172_8446622_RNA45SN1_45S_50ntFlanks</td><td>12405</td><td>12441</td><td>37</td><td>CTTTTTGATCCTTCGATGTCGGCTCTTCCTATCATTG</td><td>474.0</td><td>-12.8</td><td>1.7</td><td>-7.6</td><td>3.2E-14</td><td>1.6E-13</td><td>MB231/MB468</td><td>MP</td></tr>
<tr><td>>MI0000061|hsa-let-7a-2&WithFlank&11|-|122146516|122146599@11.30.20(+1C);MI0000062|hsa-let-7a-3&WithFlank&22|+|46112743|46112828@10.29.20(+1C);MI0000060|hsa-let-7a-1&WithFlank&9|+|94175951|94176042@12.31.20(+1C)&[MIMAT0000062&hsa-let-7a-5p&offsets|0|-2(+1C);m-6&11|-|122146568|122146589&offsets|0|-2(+1C)];[MIMAT0000062_1&hsa-let-7a-5p&offsets|0|-2(+1C);m-5&22|+|46112752|46112773&offsets|0|-2(+1C)];[MIMAT0000062_2&hsa-let-7a-5p&offsets|0|-2(+1C);m-4&9|+|94175962|94175983&offsets|0|-2(+1C)]|TGAGGTAGTAGGTTGTATAGC</td><td>>MI0000061|hsa-let-7a-2&WithFlank&11|-|122146516|122146599</td><td>11</td><td>30</td><td>20(+1C)</td><td>TGAGGTAGTAGGTTGTATAGC</td><td>1073.7</td><td>7.2</td><td>1.0</td><td>7.5</td><td>5.0E-14</td><td>1.7E-13</td><td>MB231/MB468</td><td>MITO</td></tr>
<tr><td>>trna111_HisGTG_1_-_147774845_147774916@-1G.36.37,trna118_HisGTG_1_-_145396881_145396952@-1G.36.37,trna16_HisGTG_1_+_146544773_146544844@-1G.36.37,trna1_HisGTG_15_+_45493349_45493420@-1G.36.37,trna21_HisGTG_1_+_147753471_147753542@-1G.36.37,trna33_HisGTG_6_+_27125906_27125977@-1G.36.37,trna7_HisGTG_9_-_14433938_14434009@-1G.36.37,trna8_HisGTG_15_-_45492611_45492682@-1G.36.37,trna9_HisGTG_15_-_45490804_45490875@-1G.36.37|GGCCGTGATCGTATAGTGGTTAGTACTCTGCGTTGTG</td><td>>trna111_HisGTG_1_-_147774845_147774916</td><td>-1G</td><td>36</td><td>37</td><td>GGCCGTGATCGTATAGTGGTTAGTACTCTGCGTTGTG</td><td>222.0</td><td>8.2</td><td>1.1</td><td>7.6</td><td>3.8E-14</td><td>1.7E-13</td><td>MB231/MB468</td><td>NUC</td></tr>
<tr><td>>MI0000446|hsa-mir-125b-1&WithFlank&11|-|122099751|122099850@21.42.22(+2A);MI0000470|hsa-mir-125b-2&WithFlank&21|+|16590231|16590331@23.44.22(+2A)&[MIMAT0000423&hsa-miR-125b-5p&offsets|0|0(+2A);m-67&11|-|122099809|122099830&offsets|0|0(+2A)];[MIMAT0000423_1&hsa-miR-125b-5p&offsets|0|0(+2A);m-68&21|+|16590253|16590274&offsets|0|0(+2A)]|TCCCTGAGACCCTAACTTGTGAAA</td><td>>MI0000446|hsa-mir-125b-1&WithFlank&11|-|122099751|122099850</td><td>21</td><td>42</td><td>22(+2A)</td><td>TCCCTGAGACCCTAACTTGTGAAA</td><td>1122.6</td><td>10.0</td><td>1.3</td><td>7.6</td><td>3.7E-14</td><td>1.7E-13</td><td>MB231/MB468</td><td>CYTO</td></tr>
<tr><td>>hg38_21_+_8433172_8446622_RNA45SN1_45S_50ntFlanks@10689.10711.23|GCGTCCGGTGAGCTCTCGCTGGC</td><td>>hg38_21_+_8433172_8446622_RNA45SN1_45S_50ntFlanks</td><td>10689</td><td>10711</td><td>23</td><td>GCGTCCGGTGAGCTCTCGCTGGC</td><td>266.7</td><td>-10.8</td><td>1.4</td><td>-7.6</td><td>3.7E-14</td><td>1.7E-13</td><td>MB231/MB468</td><td>CYTO</td></tr>
<tr><td>>hg38_21_+_8433172_8446622_RNA45SN1_45S_50ntFlanks@6719.6742.24|TGAATTGCAGGACACATTGATCAT</td><td>>hg38_21_+_8433172_8446622_RNA45SN1_45S_50ntFlanks</td><td>6719</td><td>6742</td><td>24</td><td>TGAATTGCAGGACACATTGATCAT</td><td>205.2</td><td>-11.5</td><td>1.5</td><td>-7.6</td><td>3.2E-14</td><td>1.7E-13</td><td>MB231/MB468</td><td>MP</td></tr>
<tr><td>>hg38_21_+_8433172_8446622_RNA45SN1_45S_50ntFlanks@4006.4026.21|ACCTCGGGCCGATCGCACGCC</td><td>>hg38_21_+_8433172_8446622_RNA45SN1_45S_50ntFlanks</td><td>4006</td><td>4026</td><td>21</td><td>ACCTCGGGCCGATCGCACGCC</td><td>349.4</td><td>-11.4</td><td>1.5</td><td>-7.6</td><td>3.9E-14</td><td>1.7E-13</td><td>MB231/MB468</td><td>NUC</td></tr>
<tr><td>>hg38_21_+_8433172_8446622_RNA45SN1_45S_50ntFlanks@11668.11687.20|TGCCCAGTGCTCTGAATGTC</td><td>>hg38_21_+_8433172_8446622_RNA45SN1_45S_50ntFlanks</td><td>11668</td><td>11687</td><td>20</td><td>TGCCCAGTGCTCTGAATGTC</td><td>927.0</td><td>5.2</td><td>0.7</td><td>7.6</td><td>4.0E-14</td><td>1.7E-13</td><td>BT20/MB468</td><td>CYTO</td></tr>
<tr><td>>hg38_21_+_8433172_8446622_RNA45SN1_45S_50ntFlanks@11572.11593.22|CCGACTTAGAACTGGTGCGGAC</td><td>>hg38_21_+_8433172_8446622_RNA45SN1_45S_50ntFlanks</td><td>11572</td><td>11593</td><td>22</td><td>CCGACTTAGAACTGGTGCGGAC</td><td>1507.0</td><td>-7.9</td><td>1.0</td><td>-7.6</td><td>3.2E-14</td><td>1.7E-13</td><td>MB231/MB468</td><td>MP</td></tr>
<tr><td>>hg38_21_+_8433172_8446622_RNA45SN1_45S_50ntFlanks@11663.11687.25|ATTTCTGCCCAGTGCTCTGAATGTC</td><td>>hg38_21_+_8433172_8446622_RNA45SN1_45S_50ntFlanks</td><td>11663</td><td>11687</td><td>25</td><td>ATTTCTGCCCAGTGCTCTGAATGTC</td><td>314.6</td><td>5.7</td><td>0.8</td><td>7.6</td><td>3.1E-14</td><td>1.7E-13</td><td>BT20/MB468</td><td>MITO</td></tr>
<tr><td>>hg38_21_+_8433172_8446622_RNA45SN1_45S_50ntFlanks@485.512.28|GATGTGGTGACGTCGTGCTCTCCCGGGC</td><td>>hg38_21_+_8433172_8446622_RNA45SN1_45S_50ntFlanks</td><td>485</td><td>512</td><td>28</td><td>GATGTGGTGACGTCGTGCTCTCCCGGGC</td><td>61.9</td><td>8.8</td><td>1.2</td><td>7.5</td><td>6.0E-14</td><td>1.7E-13</td><td>BT20/MB468</td><td>NUC</td></tr>
<tr><td>>hg38_21_+_8433172_8446622_RNA45SN1_45S_50ntFlanks@12003.12030.28|GCGGGCCGCCGGTGAAATACCACTACTC</td><td>>hg38_21_+_8433172_8446622_RNA45SN1_45S_50ntFlanks</td><td>12003</td><td>12030</td><td>28</td><td>GCGGGCCGCCGGTGAAATACCACTACTC</td><td>712.5</td><td>-5.1</td><td>0.7</td><td>-7.5</td><td>5.1E-14</td><td>1.7E-13</td><td>MB231/MB468</td><td>MITO</td></tr>
<tr><td>>MI0000266|hsa-mir-10a&WithFlank&17|-|48579832|48579953@28.49.22(+1A)&[MIMAT0000253&hsa-miR-10a-5p&offsets|0|-1(+1A);m-18&17|-|48579905|48579926&offsets|0|0(+1A)]|TACCCTGTAGATCCGAATTTGTA</td><td>>MI0000266|hsa-mir-10a&WithFlank&17|-|48579832|48579953</td><td>28</td><td>49</td><td>22(+1A)</td><td>TACCCTGTAGATCCGAATTTGTA</td><td>507.8</td><td>12.2</td><td>1.6</td><td>7.5</td><td>5.1E-14</td><td>1.7E-13</td><td>MB231/MB468</td><td>MITO</td></tr>
<tr><td>>hg38_21_+_8433172_8446622_RNA45SN1_45S_50ntFlanks@4147.4175.29|TGAGAAACGGCTACCACATCCAAGGAAGG</td><td>>hg38_21_+_8433172_8446622_RNA45SN1_45S_50ntFlanks</td><td>4147</td><td>4175</td><td>29</td><td>TGAGAAACGGCTACCACATCCAAGGAAGG</td><td>235.4</td><td>-10.8</td><td>1.4</td><td>-7.6</td><td>3.9E-14</td><td>1.7E-13</td><td>MB231/MB468</td><td>NUC</td></tr>
<tr><td>>am_trna162_MetCAT_6_-_26758550_26758622@31.53.23,trna164_MetCAT_6_-_26735574_26735646@31.53.23,trna20_MetCAT_16_+_71460396_71460468@31.53.23,trna21_MetCAT_6_+_26701712_26701784@31.53.23,trna27_MetCAT_6_+_26766444_26766516@31.53.23,trna75_MetCAT_6_+_28912352_28912424@31.53.23,trna97_MetCAT_6_-_28921042_28921114@31.53.23|TCTCATAATCTGAAGGTCCTGAG</td><td>>am_trna162_MetCAT_6_-_26758550_26758622</td><td>31</td><td>53</td><td>23</td><td>TCTCATAATCTGAAGGTCCTGAG</td><td>215.8</td><td>-11.5</td><td>1.5</td><td>-7.5</td><td>5.1E-14</td><td>1.7E-13</td><td>MB231/MB468</td><td>MITO</td></tr>
<tr><td>>MI0000286|hsa-mir-210&WithFlank&11|-|568083|568204@72.94.23&[MIMAT0000267&hsa-miR-210-3p&offsets|0|+1;m-102&11|-|568112|568133&offsets|0|+1]|CTGTGCGTGTGACAGCGGCTGAT</td><td>>MI0000286|hsa-mir-210&WithFlank&11|-|568083|568204</td><td>72</td><td>94</td><td>23</td><td>CTGTGCGTGTGACAGCGGCTGAT</td><td>99.2</td><td>9.5</td><td>1.3</td><td>7.5</td><td>6.0E-14</td><td>1.7E-13</td><td>BT20/MB468</td><td>NUC</td></tr>
<tr><td>>MI0000477|hsa-mir-146a&WithFlank&5|+|160485346|160485456@27.48.22(+1C)&[MIMAT0000449&hsa-miR-146a-5p&offsets|0|0(+1C);m-129&5|+|160485372|160485393&offsets|0|0(+1C)]|TGAGAACTGAATTCCATGGGTTC</td><td>>MI0000477|hsa-mir-146a&WithFlank&5|+|160485346|160485456</td><td>27</td><td>48</td><td>22(+1C)</td><td>TGAGAACTGAATTCCATGGGTTC</td><td>2152.9</td><td>-14.5</td><td>1.8</td><td>-8.1</td><td>5.5E-16</td><td>1.7E-13</td><td>BT20/MB231</td><td>MP</td></tr>
<tr><td>>hg38_21_+_8433172_8446622_RNA45SN1_45S_50ntFlanks@11572.11599.28|CCGACTTAGAACTGGTGCGGACCAGGGG</td><td>>hg38_21_+_8433172_8446622_RNA45SN1_45S_50ntFlanks</td><td>11572</td><td>11599</td><td>28</td><td>CCGACTTAGAACTGGTGCGGACCAGGGG</td><td>674.1</td><td>-3.4</td><td>0.4</td><td>-7.6</td><td>4.1E-14</td><td>1.7E-13</td><td>BT20/MB468</td><td>CYTO</td></tr>
<tr><td>>hg38_21_+_8433172_8446622_RNA45SN1_45S_50ntFlanks@5420.5444.25|CCGATTGGATGGTTTAGTGAGGCCC</td><td>>hg38_21_+_8433172_8446622_RNA45SN1_45S_50ntFlanks</td><td>5420</td><td>5444</td><td>25</td><td>CCGATTGGATGGTTTAGTGAGGCCC</td><td>711.3</td><td>-6.0</td><td>0.8</td><td>-7.6</td><td>4.1E-14</td><td>1.7E-13</td><td>BT20/MB468</td><td>CYTO</td></tr>
<tr><td>>MI0000264|hsa-mir-7-2&WithFlank&15|+|88611819|88611940@38.59.22(+1G);MI0000265|hsa-mir-7-3&WithFlank&19|+|4770664|4770785@37.58.22(+1G);MI0000263|hsa-mir-7-1&WithFlank&9|-|83969742|83969863@30.51.22(+1G)&[MIMAT0000252&hsa-miR-7-5p&offsets|0|-2(+1G);m-400&15|+|88611856|88611879&offsets|0|-2(+1G)];[MIMAT0000252_1&hsa-miR-7-5p&offsets|0|-2(+1G);m-402&19|+|4770700|4770723&offsets|0|-2(+1G)];[MIMAT0000252_2&hsa-miR-7-5p&offsets|0|-2(+1G);m-401&9|-|83969811|83969834&offsets|0|-2(+1G)]|TGGAAGACTAGTGATTTTGTTGG</td><td>>MI0000264|hsa-mir-7-2&WithFlank&15|+|88611819|88611940</td><td>38</td><td>59</td><td>22(+1G)</td><td>TGGAAGACTAGTGATTTTGTTGG</td><td>61.0</td><td>3.8</td><td>0.5</td><td>7.5</td><td>6.0E-14</td><td>1.7E-13</td><td>BT20/MB468</td><td>NUC</td></tr>
<tr><td>>hg38_21_+_8433172_8446622_RNA45SN1_45S_50ntFlanks@7985.8012.28|GATCAGACGTGGCGACCCGCTGAATTTA</td><td>>hg38_21_+_8433172_8446622_RNA45SN1_45S_50ntFlanks</td><td>7985</td><td>8012</td><td>28</td><td>GATCAGACGTGGCGACCCGCTGAATTTA</td><td>508.2</td><td>-11.1</td><td>1.5</td><td>-7.6</td><td>3.1E-14</td><td>1.7E-13</td><td>BT20/MB468</td><td>MITO</td></tr>
<tr><td>>hg38_21_+_8433172_8446622_RNA45SN1_45S_50ntFlanks@11877.11904.28|AGACCCTGTTGAGCTTGACTCTAGTCTG</td><td>>hg38_21_+_8433172_8446622_RNA45SN1_45S_50ntFlanks</td><td>11877</td><td>11904</td><td>28</td><td>AGACCCTGTTGAGCTTGACTCTAGTCTG</td><td>216.4</td><td>-11.6</td><td>1.5</td><td>-7.6</td><td>3.3E-14</td><td>1.7E-13</td><td>MB231/MB468</td><td>MP</td></tr>
<tr><td>>hg38_21_+_8433172_8446622_RNA45SN1_45S_50ntFlanks@3830.3860.31|GCTCGCTCCTCTCCTACTTGGATAACTGTGG</td><td>>hg38_21_+_8433172_8446622_RNA45SN1_45S_50ntFlanks</td><td>3830</td><td>3860</td><td>31</td><td>GCTCGCTCCTCTCCTACTTGGATAACTGTGG</td><td>775.8</td><td>-5.2</td><td>0.7</td><td>-7.6</td><td>3.9E-14</td><td>1.7E-13</td><td>MB231/MB468</td><td>NUC</td></tr>
<tr><td>>hg38_21_+_8433172_8446622_RNA45SN1_45S_50ntFlanks@7985.8005.21|GATCAGACGTGGCGACCCGCT</td><td>>hg38_21_+_8433172_8446622_RNA45SN1_45S_50ntFlanks</td><td>7985</td><td>8005</td><td>21</td><td>GATCAGACGTGGCGACCCGCT</td><td>544.2</td><td>-12.2</td><td>1.5</td><td>-8.0</td><td>1.4E-15</td><td>1.7E-13</td><td>BT20/MB468</td><td>MP</td></tr>
<tr><td>>MI0000299|hsa-mir-222&WithFlank&X|-|45747009|45747130@75.98.24&[MIMAT0000279&hsa-miR-222-3p&offsets|0|+3;m-140&X|-|45747033|45747056&offsets|0|0]|AGCTACATCTGGCTACTGGGTCTC</td><td>>MI0000299|hsa-mir-222&WithFlank&X|-|45747009|45747130</td><td>75</td><td>98</td><td>24</td><td>AGCTACATCTGGCTACTGGGTCTC</td><td>16722.1</td><td>6.1</td><td>0.8</td><td>7.5</td><td>5.2E-14</td><td>1.7E-13</td><td>MB231/MB468</td><td>MITO</td></tr>
<tr><td>>hg38_21_+_8433172_8446622_RNA45SN1_45S_50ntFlanks@10791.10822.32|CCTCTGGCATGTTGGAACAATGTAGGTAAGGG</td><td>>hg38_21_+_8433172_8446622_RNA45SN1_45S_50ntFlanks</td><td>10791</td><td>10822</td><td>32</td><td>CCTCTGGCATGTTGGAACAATGTAGGTAAGGG</td><td>134.7</td><td>-10.9</td><td>1.4</td><td>-7.6</td><td>3.3E-14</td><td>1.7E-13</td><td>MB231/MB468</td><td>MP</td></tr>
<tr><td>>hg38_21_+_8433172_8446622_RNA45SN1_45S_50ntFlanks@10877.10902.26|GTCGGTCGGGCTGGGGCGCGAAGCGG</td><td>>hg38_21_+_8433172_8446622_RNA45SN1_45S_50ntFlanks</td><td>10877</td><td>10902</td><td>26</td><td>GTCGGTCGGGCTGGGGCGCGAAGCGG</td><td>194.6</td><td>-11.5</td><td>1.5</td><td>-7.6</td><td>3.3E-14</td><td>1.7E-13</td><td>MB231/MB468</td><td>MP</td></tr>
<tr><td>>hg38_21_+_8433172_8446622_RNA45SN1_45S_50ntFlanks@11270.11295.26|CGGCCGCCCCCGGGGAGCCCGGCGGG</td><td>>hg38_21_+_8433172_8446622_RNA45SN1_45S_50ntFlanks</td><td>11270</td><td>11295</td><td>26</td><td>CGGCCGCCCCCGGGGAGCCCGGCGGG</td><td>987.8</td><td>-13.0</td><td>1.6</td><td>-8.0</td><td>1.4E-15</td><td>1.7E-13</td><td>BT20/MB468</td><td>MP</td></tr>
<tr><td>>hg38_21_+_8433172_8446622_RNA45SN1_45S_50ntFlanks@11314.11342.29|CCCACCCCACGTCTCGTCGCGCGCGCGTC</td><td>>hg38_21_+_8433172_8446622_RNA45SN1_45S_50ntFlanks</td><td>11314</td><td>11342</td><td>29</td><td>CCCACCCCACGTCTCGTCGCGCGCGCGTC</td><td>444.8</td><td>-9.1</td><td>1.2</td><td>-7.6</td><td>4.0E-14</td><td>1.7E-13</td><td>MB231/MB468</td><td>NUC</td></tr>
<tr><td>>hg38_21_+_8433172_8446622_RNA45SN1_45S_50ntFlanks@10348.10384.37|AAACGAGAACTTTGAAGGCCGAAGTGGAGAAGGGTTC</td><td>>hg38_21_+_8433172_8446622_RNA45SN1_45S_50ntFlanks</td><td>10348</td><td>10384</td><td>37</td><td>AAACGAGAACTTTGAAGGCCGAAGTGGAGAAGGGTTC</td><td>775.3</td><td>-6.0</td><td>0.8</td><td>-7.5</td><td>6.1E-14</td><td>1.7E-13</td><td>BT20/MB468</td><td>NUC</td></tr>
<tr><td>>MI0000299|hsa-mir-222&WithFlank&X|-|45747009|45747130@75.99.25(+2U)&[MIMAT0000279&hsa-miR-222-3p&offsets|0|+4(+2U);m-140&X|-|45747033|45747056&offsets|0|+1(+2U)]|AGCTACATCTGGCTACTGGGTCTCTTT</td><td>>MI0000299|hsa-mir-222&WithFlank&X|-|45747009|45747130</td><td>75</td><td>99</td><td>25(+2U)</td><td>AGCTACATCTGGCTACTGGGTCTCTTT</td><td>2563.5</td><td>7.3</td><td>1.0</td><td>7.5</td><td>5.2E-14</td><td>1.7E-13</td><td>MB231/MB468</td><td>MITO</td></tr>
</table>

| | | | | | | | | | | | | | |
|---|---|---|---|---|---|---|---|---|---|---|---|---|---|
| >hg38_21_+_8433172_8446622_RNA45SN1_45S_50ntFlanks@9310.9334.25\|GAGTCTAACACGTGCGCGAGTCGGG | >hg38_21_+_8433172_8446622_RNA45SN1_45S_50ntFlanks | 9310 | 9334 | 25 | GAGTCTAACACGTGCGCGAGTCGGG | 320.4 | -12.2 | 1.6 | -7.6 | 3.3E-14 | 1.7E-13 | MB231/MB468 | MP |
| >hg38_21_+_8433172_8446622_RNA45SN1_45S_50ntFlanks@10841.10863.23\|CGTAACTTCGGGATAAGGATTGG | >hg38_21_+_8433172_8446622_RNA45SN1_45S_50ntFlanks | 10841 | 10863 | 23 | CGTAACTTCGGGATAAGGATTGG | 2489.8 | -8.2 | 1.1 | -7.6 | 4.0E-14 | 1.7E-13 | MB231/MB468 | NUC |
| >hg38_21_+_8433172_8446622_RNA45SN1_45S_50ntFlanks@10712.10736.25\|CCTTGAAAATCCGGGGGAGAGGGTG | >hg38_21_+_8433172_8446622_RNA45SN1_45S_50ntFlanks | 10712 | 10736 | 25 | CCTTGAAAATCCGGGGGAGAGGGTG | 173.2 | -11.3 | 1.5 | -7.6 | 3.3E-14 | 1.7E-13 | MB231/MB468 | MP |
| >hg38_21_+_8433172_8446622_RNA45SN1_45S_50ntFlanks@7986.8013.28\|ATCAGACGTGGCGACCCGCTGAATTTAA | >hg38_21_+_8433172_8446622_RNA45SN1_45S_50ntFlanks | 7986 | 8013 | 28 | ATCAGACGTGGCGACCCGCTGAATTTAA | 241.3 | -11.7 | 1.6 | -7.5 | 5.2E-14 | 1.7E-13 | MB231/MB468 | MITO |
| >hg38_21_+_8433172_8446622_RNA45SN1_45S_50ntFlanks@10826.10844.19\|TCGGCAAGCCGGATCCGTA | >hg38_21_+_8433172_8446622_RNA45SN1_45S_50ntFlanks | 10826 | 10844 | 19 | TCGGCAAGCCGGATCCGTA | 158.6 | -11.2 | 1.5 | -7.6 | 3.3E-14 | 1.7E-13 | MB231/MB468 | MP |
| >hg38_21_+_8433172_8446622_RNA45SN1_45S_50ntFlanks@11792.11810.19\|ATGAACGAGATTCCCACTG | >hg38_21_+_8433172_8446622_RNA45SN1_45S_50ntFlanks | 11792 | 11810 | 19 | ATGAACGAGATTCCCACTG | 157.6 | -10.3 | 1.4 | -7.6 | 4.0E-14 | 1.7E-13 | MB231/MB468 | NUC |
| >hg38_21_+_8433172_8446622_RNA45SN1_45S_50ntFlanks@12241.12272.32\|CAGAAGGGCAAAAGCTCGCTTGATCTTGATTT | >hg38_21_+_8433172_8446622_RNA45SN1_45S_50ntFlanks | 12241 | 12272 | 32 | CAGAAGGGCAAAAGCTCGCTTGATCTTGATTT | 2415.6 | -6.1 | 0.8 | -7.5 | 5.2E-14 | 1.7E-13 | MB231/MB468 | MITO |
| >MI0000650\|hsa-mir-200c&WithFlank&12\|+\|6963693\|6963772@50.72.23&[MIMAT0000617&hsa-miR-200c-3p&offsets\|0\|0;m-54&12\|+\|6963742\|6963764&offsets\|0\|0]\|TAATACTGCCGGGTAATGATGGA | >MI0000650\|hsa-mir-200c&WithFlank&12\|+\|6963693\|6963772 | 50 | 72 | 23 | TAATACTGCCGGGTAATGATGGA | 47547.3 | 3.5 | 0.5 | 7.5 | 6.2E-14 | 1.7E-13 | BT20/MB468 | NUC |
| >am_trna116_GluCTC_1_-_145399233_145399304@1.25.25,trna59_GluCTC_1_+_249168447_249168518@1.25.25,trna71_GluCTC_1_-_161439189_161439260@1.25.25,trna74_GluCTC_1_-_161431809_161431880@1.25.25,trna77_GluCTC_1_-_161424398_161424469@1.25.25,trna77_GluCTC_6_+_28949976_28950047@1.25.25,trna80_GluCTC_1_-_161417018_161417089@1.25.25,trna87_GluCTC_6_-_126101393_126101464@1.25.25\|TCCCTGGTGGTCTAGTGGTTAGGAT | >am_trna116_GluCTC_1_-_145399233_145399304 | 1 | 25 | 25 | TCCCTGGTGGTCTAGTGGTTAGGAT | 619.8 | 4.1 | 0.5 | 7.6 | 4.2E-14 | 1.7E-13 | BT20/MB468 | CYTO |
| >hg38_21_+_8433172_8446622_RNA45SN1_45S_50ntFlanks@11873.11906.34\|AAGAAGACCCTGTTGAGCTTGACTCTAGTCTGGC | >hg38_21_+_8433172_8446622_RNA45SN1_45S_50ntFlanks | 11873 | 11906 | 34 | AAGAAGACCCTGTTGAGCTTGACTCTAGTCTGGC | 1404.9 | -4.0 | 0.5 | -7.6 | 3.1E-14 | 1.7E-13 | BT20/MB468 | MITO |
| >hg38_21_+_8433172_8446622_RNA45SN1_45S_50ntFlanks@3863.3898.36\|ATTCTAGAGCTAATACATGCCGACGGGCGCTGACCC | >hg38_21_+_8433172_8446622_RNA45SN1_45S_50ntFlanks | 3863 | 3898 | 36 | ATTCTAGAGCTAATACATGCCGACGGGCGCTGACCC | 596.7 | -8.8 | 1.2 | -7.5 | 5.3E-14 | 1.7E-13 | MB231/MB468 | MITO |
| >hg38_21_+_8433172_8446622_RNA45SN1_45S_50ntFlanks@10458.10489.32\|ATGGCCTCCGTTGCCCTCGGCCGATCGAAAGG | >hg38_21_+_8433172_8446622_RNA45SN1_45S_50ntFlanks | 10458 | 10489 | 32 | ATGGCCTCCGTTGCCCTCGGCCGATCGAAAGG | 135.4 | -10.9 | 1.4 | -7.6 | 3.4E-14 | 1.7E-13 | MB231/MB468 | MP |
| >hg38_21_+_8433172_8446622_RNA45SN1_45S_50ntFlanks@10690.10711.22\|CGTCCGGTGAGCTCTCGCTGGC | >hg38_21_+_8433172_8446622_RNA45SN1_45S_50ntFlanks | 10690 | 10711 | 22 | CGTCCGGTGAGCTCTCGCTGGC | 490.9 | -12.8 | 1.7 | -7.6 | 3.4E-14 | 1.7E-13 | MB231/MB468 | MP |
| >hg38_21_+_8433172_8446622_RNA45SN1_45S_50ntFlanks@6644.6664.21\|CCTCGTACGACTCTTAGCGGT | >hg38_21_+_8433172_8446622_RNA45SN1_45S_50ntFlanks | 6644 | 6664 | 21 | CCTCGTACGACTCTTAGCGGT | 494.9 | 5.5 | 0.7 | 7.6 | 3.2E-14 | 1.7E-13 | BT20/MB468 | MITO |
| >hg38_21_+_8433172_8446622_RNA45SN1_45S_50ntFlanks@12942.12990.49\|TTCGTAGACGACCTGCTTCTGGGTCGGGGTTTCGTACGTAGCAGAGCAG | >hg38_21_+_8433172_8446622_RNA45SN1_45S_50ntFlanks | 12942 | 12990 | 49 | TTCGTAGACGACCTGCTTCTGGGTCGGGGTTTCGTACGTAGCAGAGCAG | 546.9 | -13.5 | 1.8 | -7.5 | 6.2E-14 | 1.7E-13 | BT20/MB468 | NUC |
| >hg38_21_+_8433172_8446622_RNA45SN1_45S_50ntFlanks@11564.11585.22\|CCTAGCAGCCGACTTAGAACTG | >hg38_21_+_8433172_8446622_RNA45SN1_45S_50ntFlanks | 11564 | 11585 | 22 | CCTAGCAGCCGACTTAGAACTG | 5506.0 | -11.5 | 1.5 | -7.5 | 6.2E-14 | 1.7E-13 | BT20/MB468 | NUC |
| >hg38_21_+_8433172_8446622_RNA45SN1_45S_50ntFlanks@10819.10838.20\|AGGGAAGTCGGCAAGCCGGA | >hg38_21_+_8433172_8446622_RNA45SN1_45S_50ntFlanks | 10819 | 10838 | 20 | AGGGAAGTCGGCAAGCCGGA | 915.2 | -7.8 | 1.0 | -7.6 | 3.4E-14 | 1.7E-13 | MB231/MB468 | MP |
| >hg38_21_+_8433172_8446622_RNA45SN1_45S_50ntFlanks@12637.12661.25\|GCTACCATCTGTGGGATTATGACTG | >hg38_21_+_8433172_8446622_RNA45SN1_45S_50ntFlanks | 12637 | 12661 | 25 | GCTACCATCTGTGGGATTATGACTG | 3451.0 | -10.9 | 1.4 | -7.6 | 3.4E-14 | 1.7E-13 | MB231/MB468 | MP |
| >hg38_21_+_8433172_8446622_RNA45SN1_45S_50ntFlanks@11930.11952.23\|TAGAATAAGTGGGAGGCCCCCGG | >hg38_21_+_8433172_8446622_RNA45SN1_45S_50ntFlanks | 11930 | 11952 | 23 | TAGAATAAGTGGGAGGCCCCCGG | 239.6 | -11.8 | 1.6 | -7.6 | 3.4E-14 | 1.7E-13 | MB231/MB468 | MP |
| >hg38_21_+_8433172_8446622_RNA45SN1_45S_50ntFlanks@3853.3881.29\|AACTGTGGTAATTCTAGAGCTAATACATG | >hg38_21_+_8433172_8446622_RNA45SN1_45S_50ntFlanks | 3853 | 3881 | 29 | AACTGTGGTAATTCTAGAGCTAATACATG | 497.6 | -11.0 | 1.5 | -7.6 | 3.2E-14 | 1.7E-13 | BT20/MB468 | MITO |
| >am_trna143_LysTTT_6_-_27559593_27559665@6.24.19,trna14_LysTTT_11_-_59327808_59327880@6.24.19,trna2_LysTTT_17_+_8022473_8022545@6.24.19,trna54_LysTTT_1_+_204475655_204475727@6.24.19,trna5_LysTTT_11_+_59323902_59323974@6.24.19,trna62_LysTTT_1_-_204476158_204476230@6.24.19,trna76_LysTTT_6_+_28918806_28918878@6.24.19\|GATAGCTCAGTCGGTAGAG | >am_trna143_LysTTT_6_-_27559593_27559665 | 6 | 24 | 19 | GATAGCTCAGTCGGTAGAG | 211.3 | -11.6 | 1.5 | -7.6 | 3.4E-14 | 1.7E-13 | MB231/MB468 | MP |
| >hg38_21_+_8433172_8446622_RNA45SN1_45S_50ntFlanks@8238.8261.24\|CCCGGGTCTTCCCGGAGTCGGGTT | >hg38_21_+_8433172_8446622_RNA45SN1_45S_50ntFlanks | 8238 | 8261 | 24 | CCCGGGTCTTCCCGGAGTCGGGTT | 180.5 | -11.4 | 1.5 | -7.6 | 3.4E-14 | 1.7E-13 | MB231/MB468 | MP |
| >MI0000086\|hsa-mir-28&WithFlank&3\|+\|188688775\|188688872@60.81.22(+1U)&[MIMAT0004502&hsa-miR-28-3p&offsets\|0\|0(+1U);m-28&3\|+\|188688834\|188688855&offsets\|0\|0(+1U)]\|CACTAGATTGTGAGCTCCTGGAT | >MI0000086\|hsa-mir-28&WithFlank&3\|+\|188688775\|188688872 | 60 | 81 | 22(+1U) | CACTAGATTGTGAGCTCCTGGAT | 528.1 | 12.3 | 1.6 | 7.5 | 5.4E-14 | 1.7E-13 | MB231/MB468 | MITO |
| >MI0000079\|hsa-mir-23a&WithFlank&19\|-\|13836581\|13836665@54.74.21&[MIMAT0000078&hsa-miR-23a-3p&offsets\|+3\|+3;m-31&19\|-\|13836595\|13836615&offsets\|+3\|+3]\|ACATTGCCAGGGATTTCCAAC | >MI0000079\|hsa-mir-23a&WithFlank&19\|-\|13836581\|13836665 | 54 | 74 | 21 | ACATTGCCAGGGATTTCCAAC | 1037.7 | 27.5 | 3.6 | 7.6 | 3.9E-14 | 1.8E-13 | MB231/MB468 | CYTO |
| >trna6_TrpCCA_12_+_98898030_98898101@30.52.23\|ACTCCAGATCAGAAGGCTGCGTG | >trna6_TrpCCA_12_+_98898030_98898101 | 30 | 52 | 23 | ACTCCAGATCAGAAGGCTGCGTG | 189.9 | -11.4 | 1.5 | -7.6 | 3.4E-14 | 1.8E-13 | MB231/MB468 | MP |
| >hg38_21_+_8433172_8446622_RNA45SN1_45S_50ntFlanks@11886.11904.19\|TGAGCTTGACTCTAGTCTG | >hg38_21_+_8433172_8446622_RNA45SN1_45S_50ntFlanks | 11886 | 11904 | 19 | TGAGCTTGACTCTAGTCTG | 1289.4 | -9.7 | 1.3 | -7.6 | 4.3E-14 | 1.8E-13 | BT20/MB468 | CYTO |
| >hg38_21_+_8433172_8446622_RNA45SN1_45S_50ntFlanks@12397.12414.18\|CGACGTCGCTTTTTGATC | >hg38_21_+_8433172_8446622_RNA45SN1_45S_50ntFlanks | 12397 | 12414 | 18 | CGACGTCGCTTTTTGATC | 1718.1 | -6.4 | 0.8 | -7.6 | 3.4E-14 | 1.8E-13 | MB231/MB468 | MP |
| >hg38_21_+_8433172_8446622_RNA45SN1_45S_50ntFlanks@12715.12745.31\|AGCCTCGGTTGGCCTCGGATAGCCGGTCCCC | >hg38_21_+_8433172_8446622_RNA45SN1_45S_50ntFlanks | 12715 | 12745 | 31 | AGCCTCGGTTGGCCTCGGATAGCCGGTCCCC | 220.0 | -11.6 | 1.5 | -7.6 | 3.4E-14 | 1.8E-13 | MB231/MB468 | MP |
| >hg38_21_+_8433172_8446622_RNA45SN1_45S_50ntFlanks@12223.12241.19\|CAGAAACCTCCCGTGGAGC | >hg38_21_+_8433172_8446622_RNA45SN1_45S_50ntFlanks | 12223 | 12241 | 19 | CAGAAACCTCCCGTGGAGC | 190.9 | -11.4 | 1.5 | -7.6 | 3.4E-14 | 1.8E-13 | MB231/MB468 | MP |
| >hg38_21_+_8433172_8446622_RNA45SN1_45S_50ntFlanks@5423.5445.23\|ATTGGATGGTTTAGTGAGGCCCT | >hg38_21_+_8433172_8446622_RNA45SN1_45S_50ntFlanks | 5423 | 5445 | 23 | ATTGGATGGTTTAGTGAGGCCCT | 351.5 | -12.3 | 1.6 | -7.6 | 3.4E-14 | 1.8E-13 | MB231/MB468 | MP |
| >MI0000088\|hsa-mir-30a&WithFlank&6\|-\|71403545\|71403627@53.70.18(+1U)&[MIMAT0000088&hsa-miR-30a-3p&offsets\|0\|-4(+1U);m-38&6\|-\|71403554\|71403575&offsets\|0\|-4(+1U)]\|CTTTCAGTCGGATGTTTGT | >MI0000088\|hsa-mir-30a&WithFlank&6\|-\|71403545\|71403627 | 53 | 70 | 18(+1U) | CTTTCAGTCGGATGTTTGT | 556.2 | 7.6 | 1.0 | 7.6 | 4.0E-14 | 1.8E-13 | MB231/MB468 | CYTO |
| >hg38_21_+_8433172_8446622_RNA45SN1_45S_50ntFlanks@11027.11052.26\|CTCTCTCTCTCTCTCCCCCGCTCCCC | >hg38_21_+_8433172_8446622_RNA45SN1_45S_50ntFlanks | 11027 | 11052 | 26 | CTCTCTCTCTCTCTCCCCCGCTCCCC | 158.3 | -10.0 | 1.3 | -7.6 | 4.0E-14 | 1.8E-13 | MB231/MB468 | CYTO |
| >hg38_MT_+_1621_3279_RNR2_16S_50ntFlanks@713.740.28\|GCATAAGCCTGCGTCAGATTAAAACACT | >hg38_MT_+_1621_3279_RNR2_16S_50ntFlanks | 713 | 740 | 28 | GCATAAGCCTGCGTCAGATTAAAACACT | 239.3 | 11.1 | 1.5 | 7.5 | 5.4E-14 | 1.8E-13 | MB231/MB468 | MITO |
| >hg38_21_+_8433172_8446622_RNA45SN1_45S_50ntFlanks@11886.11903.18\|TGAGCTTGACTCTAGTCT | >hg38_21_+_8433172_8446622_RNA45SN1_45S_50ntFlanks | 11886 | 11903 | 18 | TGAGCTTGACTCTAGTCT | 349.0 | -11.2 | 1.5 | -7.6 | 4.0E-14 | 1.8E-13 | MB231/MB468 | CYTO |
| >hg38_21_+_8433172_8446622_RNA45SN1_45S_50ntFlanks@4760.4782.23\|TCAGATACCGTCGTAGTTCCGAC | >hg38_21_+_8433172_8446622_RNA45SN1_45S_50ntFlanks | 4760 | 4782 | 23 | TCAGATACCGTCGTAGTTCCGAC | 463.0 | -7.2 | 1.0 | -7.6 | 4.2E-14 | 1.8E-13 | MB231/MB468 | NUC |
| >hg38_21_+_8433172_8446622_RNA45SN1_45S_50ntFlanks@12126.12148.23\|CGACCCGCTCCGGGGACAGTGCC | >hg38_21_+_8433172_8446622_RNA45SN1_45S_50ntFlanks | 12126 | 12148 | 23 | CGACCCGCTCCGGGGACAGTGCC | 192.6 | -10.3 | 1.4 | -7.6 | 4.0E-14 | 1.8E-13 | MB231/MB468 | CYTO |
| >hg38_21_+_8433172_8446622_RNA45SN1_45S_50ntFlanks@9304.9323.20\|ACCAAGGAGTCTAACACGTG | >hg38_21_+_8433172_8446622_RNA45SN1_45S_50ntFlanks | 9304 | 9323 | 20 | ACCAAGGAGTCTAACACGTG | 219.2 | -11.6 | 1.5 | -7.5 | 5.5E-14 | 1.8E-13 | MB231/MB468 | MITO |
| >hg38_21_+_8433172_8446622_RNA45SN1_45S_50ntFlanks@12289.12319.31\|CCGTGAAAGCGGGGCCTCACGATCCTTCTGA | >hg38_21_+_8433172_8446622_RNA45SN1_45S_50ntFlanks | 12289 | 12319 | 31 | CCGTGAAAGCGGGGCCTCACGATCCTTCTGA | 1973.9 | -10.8 | 1.4 | -7.5 | 5.5E-14 | 1.8E-13 | MB231/MB468 | MITO |
| >hg38_21_+_8433172_8446622_RNA45SN1_45S_50ntFlanks@11202.11232.31\|GCGGCGGCGGCGACTCTGGACGCGAGCCGGG | >hg38_21_+_8433172_8446622_RNA45SN1_45S_50ntFlanks | 11202 | 11232 | 31 | GCGGCGGCGGCGACTCTGGACGCGAGCCGGG | 154.4 | -11.1 | 1.5 | -7.6 | 3.5E-14 | 1.8E-13 | MB231/MB468 | MP |
| >hg38_21_+_8433172_8446622_RNA45SN1_45S_50ntFlanks@9551.9568.18\|GTCCGTAGCGGTCCTGAC | >hg38_21_+_8433172_8446622_RNA45SN1_45S_50ntFlanks | 9551 | 9568 | 18 | GTCCGTAGCGGTCCTGAC | 351.1 | 5.1 | 0.7 | 7.6 | 4.0E-14 | 1.8E-13 | MB231/MB468 | CYTO |
| >hg38_21_+_8433172_8446622_RNA45SN1_45S_50ntFlanks@12412.12434.23\|ATCCTTCGATGTCGGCTCTTCCT | >hg38_21_+_8433172_8446622_RNA45SN1_45S_50ntFlanks | 12412 | 12434 | 23 | ATCCTTCGATGTCGGCTCTTCCT | 563.5 | -11.4 | 1.5 | -7.5 | 4.4E-14 | 1.8E-13 | BT20/MB468 | CYTO |
| >hg38_21_+_8433172_8446622_RNA45SN1_45S_50ntFlanks@11314.11338.25\|CCCACCCCACGTCTCGTCGCGCGCG | >hg38_21_+_8433172_8446622_RNA45SN1_45S_50ntFlanks | 11314 | 11338 | 25 | CCCACCCCACGTCTCGTCGCGCGCG | 271.1 | -11.9 | 1.6 | -7.5 | 5.5E-14 | 1.8E-13 | MB231/MB468 | MITO |
| >hg38_21_+_8433172_8446622_RNA45SN1_45S_50ntFlanks@6773.6802.30\|TTCCTCCCGGGGCTACGCCTGTCTGAGCGT | >hg38_21_+_8433172_8446622_RNA45SN1_45S_50ntFlanks | 6773 | 6802 | 30 | TTCCTCCCGGGGCTACGCCTGTCTGAGCGT | 496.0 | -4.1 | 0.5 | -7.5 | 6.5E-14 | 1.8E-13 | BT20/MB468 | NUC |
| >MI0000088\|hsa-mir-30a&WithFlank&6\|-\|71403545\|71403627@54.71.18&[MIMAT0000088&hsa-miR-30a-3p&offsets\|+1\|-3;m-38&6\|-\|71403554\|71403575&offsets\|+1\|-3]\|TTTCAGTCGGATGTTTGC | >MI0000088\|hsa-mir-30a&WithFlank&6\|-\|71403545\|71403627 | 54 | 71 | 18 | TTTCAGTCGGATGTTTGC | 905.6 | 7.6 | 1.0 | 7.6 | 4.1E-14 | 1.8E-13 | MB231/MB468 | CYTO |
| >hg38_21_+_8433172_8446622_RNA45SN1_45S_50ntFlanks@10766.10797.32\|CGCAGCAGGTCTCCAAGGTGAACAGCCTCTGG | >hg38_21_+_8433172_8446622_RNA45SN1_45S_50ntFlanks | 10766 | 10797 | 32 | CGCAGCAGGTCTCCAAGGTGAACAGCCTCTGG | 147.4 | -5.3 | 0.7 | -7.6 | 4.1E-14 | 1.8E-13 | MB231/MB468 | CYTO |
| >hg38_21_+_8433172_8446622_RNA45SN1_45S_50ntFlanks@10866.10890.25\|CTAAGGGCTGGGTCGGTCGGGCTGG | >hg38_21_+_8433172_8446622_RNA45SN1_45S_50ntFlanks | 10866 | 10890 | 25 | CTAAGGGCTGGGTCGGTCGGGCTGG | 872.4 | -7.4 | 1.0 | -7.6 | 4.2E-14 | 1.8E-13 | MB231/MB468 | NUC |
| >hg38_21_+_8433172_8446622_RNA45SN1_45S_50ntFlanks@10534.10563.30\|CCGCGAGGCGTCCAGTGCGGTAACGCGACC | >hg38_21_+_8433172_8446622_RNA45SN1_45S_50ntFlanks | 10534 | 10563 | 30 | CCGCGAGGCGTCCAGTGCGGTAACGCGACC | 161.5 | -11.2 | 1.5 | -7.6 | 3.6E-14 | 1.8E-13 | MB231/MB468 | MP |
| >hg38_21_+_8433172_8446622_RNA45SN1_45S_50ntFlanks@4736.4758.23\|CGAAAGTCGGAGGTTCGAAGACG | >hg38_21_+_8433172_8446622_RNA45SN1_45S_50ntFlanks | 4736 | 4758 | 23 | CGAAAGTCGGAGGTTCGAAGACG | 1414.6 | -8.4 | 1.1 | -7.5 | 6.6E-14 | 1.8E-13 | BT20/MB468 | NUC |
| >hg38_21_+_8433172_8446622_RNA45SN1_45S_50ntFlanks@11315.11340.26\|CCACCCCACGTCTCGTCGCGCGCGCG | >hg38_21_+_8433172_8446622_RNA45SN1_45S_50ntFlanks | 11315 | 11340 | 26 | CCACCCCACGTCTCGTCGCGCGCGCG | 262.4 | -11.0 | 1.5 | -7.6 | 4.3E-14 | 1.8E-13 | MB231/MB468 | NUC |
| >hg38_21_+_8433172_8446622_RNA45SN1_45S_50ntFlanks@11849.11868.20\|CGGGCTTGGCGGAATCAGCG | >hg38_21_+_8433172_8446622_RNA45SN1_45S_50ntFlanks | 11849 | 11868 | 20 | CGGGCTTGGCGGAATCAGCG | 209.8 | -11.6 | 1.5 | -7.6 | 3.6E-14 | 1.8E-13 | MB231/MB468 | MP |

| | | | | | | | | | | | | | |
|---|---|---|---|---|---|---|---|---|---|---|---|---|---|
| >MI0000077\|hsa-mir-21&WithFlank&17\|+\|59841260\|59841343@13.36.24&[MIMAT0000076&hsa-miR-21-5p&offsets\|-1\|+1;m-1&17\|+\|59841273\|59841294&offsets\|-1\|+1]\|GTAGCTTATCAGACTGATGTTGAC | >MI0000077\|hsa-mir-21&WithFlank&17\|+\|59841260\|59841343 | 13 | 36 | 24 | GTAGCTTATCAGACTGATGTTGAC | 468.9 | 6.8 | 0.9 | 7.5 | 4.5E-14 | 1.8E-13 | BT20/MB468 | CYTO |
| >am_trnaMT_SerGCT_MT_+_12207_12265@14.35.22\|AGAACTGCTAACTCATGCCCCC | >am_trnaMT_SerGCT_MT_+_12207_12265 | 14 | 35 | 22 | AGAACTGCTAACTCATGCCCCC | 369.7 | 9.9 | 1.3 | 7.5 | 4.5E-14 | 1.8E-13 | BT20/MB468 | CYTO |
| >hg38_21_+_8433172_8446622_RNA45SN1_45S_50ntFlanks@11319.11336.18\|CCCACGTCTCGTCGCGCG | >hg38_21_+_8433172_8446622_RNA45SN1_45S_50ntFlanks | 11319 | 11336 | 18 | CCCACGTCTCGTCGCGCG | 274.7 | -10.8 | 1.4 | -7.6 | 4.1E-14 | 1.8E-13 | MB231/MB468 | CYTO |
| >hg38_21_+_8433172_8446622_RNA45SN1_45S_50ntFlanks@4616.4633.18\|CATGATTAAGAGGGACGG | >hg38_21_+_8433172_8446622_RNA45SN1_45S_50ntFlanks | 4616 | 4633 | 18 | CATGATTAAGAGGGACGG | 188.5 | -11.4 | 1.5 | -7.6 | 3.6E-14 | 1.8E-13 | MB231/MB468 | MP |
| >hg38_21_+_8433172_8446622_RNA45SN1_45S_50ntFlanks@11882.11904.23\|CTGTTGAGCTTGACTCTAGTCTG | >hg38_21_+_8433172_8446622_RNA45SN1_45S_50ntFlanks | 11882 | 11904 | 23 | CTGTTGAGCTTGACTCTAGTCTG | 1077.4 | -7.0 | 0.9 | -7.6 | 4.1E-14 | 1.8E-13 | MB231/MB468 | CYTO |
| >hg38_21_+_8433172_8446622_RNA45SN1_45S_50ntFlanks@12676.12708.33\|AGAATCCCGCCCAGGCGGAACGATACGGCAGCG | >hg38_21_+_8433172_8446622_RNA45SN1_45S_50ntFlanks | 12676 | 12708 | 33 | AGAATCCCGCCCAGGCGGAACGATACGGCAGCG | 231.0 | -10.2 | 1.3 | -7.5 | 4.5E-14 | 1.8E-13 | BT20/MB468 | CYTO |
| >hg38_21_+_8433172_8446622_RNA45SN1_45S_50ntFlanks@12233.12262.30\|CCGTGGAGCAGAAGGGCAAAAGCTCGCTTG | >hg38_21_+_8433172_8446622_RNA45SN1_45S_50ntFlanks | 12233 | 12262 | 30 | CCGTGGAGCAGAAGGGCAAAAGCTCGCTTG | 156.6 | -11.2 | 1.5 | -7.6 | 3.6E-14 | 1.8E-13 | MB231/MB468 | MP |
| >hg38_21_+_8433172_8446622_RNA45SN1_45S_50ntFlanks@12657.12674.18\|GACTGAACGCCTCTAAGT | >hg38_21_+_8433172_8446622_RNA45SN1_45S_50ntFlanks | 12657 | 12674 | 18 | GACTGAACGCCTCTAAGT | 405.3 | -3.9 | 0.5 | -7.6 | 4.1E-14 | 1.8E-13 | MB231/MB468 | CYTO |
| >hg38_21_+_8433172_8446622_RNA45SN1_45S_50ntFlanks@6652.6694.43\|GACTCTTAGCGGTGGATCACTCGGCTCGTGCGTCGATGAAGAA | >hg38_21_+_8433172_8446622_RNA45SN1_45S_50ntFlanks | 6652 | 6694 | 43 | GACTCTTAGCGGTGGATCACTCGGCTCGTGCGTCGATGAAGAA | 1935.9 | 4.7 | 0.6 | 7.6 | 3.4E-14 | 1.8E-13 | BT20/MB468 | MITO |
| >hg38_21_+_8433172_8446622_RNA45SN1_45S_50ntFlanks@11318.11342.25\|CCCCACGTCTCGTCGCGCGCGCGTC | >hg38_21_+_8433172_8446622_RNA45SN1_45S_50ntFlanks | 11318 | 11342 | 25 | CCCCACGTCTCGTCGCGCGCGCGTC | 179.0 | -11.3 | 1.5 | -7.6 | 3.6E-14 | 1.8E-13 | MB231/MB468 | MP |
| >hg38_21_+_8433172_8446622_RNA45SN1_45S_50ntFlanks@3705.3736.32\|TACCTGGTTGATCCTGCCAGTAGCATATGCTT | >hg38_21_+_8433172_8446622_RNA45SN1_45S_50ntFlanks | 3705 | 3736 | 32 | TACCTGGTTGATCCTGCCAGTAGCATATGCTT | 802.2 | 5.1 | 0.7 | 7.6 | 4.2E-14 | 1.8E-13 | MB231/MB468 | CYTO |
| >hg38_21_+_8433172_8446622_RNA45SN1_45S_50ntFlanks@5491.5525.35\|GAGAAGACGGTCGAACTTGACTATCTAGAGGAAGT | >hg38_21_+_8433172_8446622_RNA45SN1_45S_50ntFlanks | 5491 | 5525 | 35 | GAGAAGACGGTCGAACTTGACTATCTAGAGGAAGT | 206.8 | 7.2 | 1.0 | 7.6 | 4.2E-14 | 1.8E-13 | MB231/MB468 | CYTO |
| >hg38_21_+_8433172_8446622_RNA45SN1_45S_50ntFlanks@11999.12029.31\|CCCTGCGGGCCGCCGGTGAAATACCACTACT | >hg38_21_+_8433172_8446622_RNA45SN1_45S_50ntFlanks | 11999 | 12029 | 31 | CCCTGCGGGCCGCCGGTGAAATACCACTACT | 1350.0 | -10.4 | 1.4 | -7.5 | 6.7E-14 | 1.8E-13 | BT20/MB468 | NUC |
| >hg38_21_+_8433172_8446622_RNA45SN1_45S_50ntFlanks@10234.10261.28\|CGGACGCTACGCCGCGACGAGTAGGAGG | >hg38_21_+_8433172_8446622_RNA45SN1_45S_50ntFlanks | 10234 | 10261 | 28 | CGGACGCTACGCCGCGACGAGTAGGAGG | 144.9 | -11.0 | 1.5 | -7.6 | 3.6E-14 | 1.8E-13 | MB231/MB468 | MP |
| >hg38_21_+_8433172_8446622_RNA45SN1_45S_50ntFlanks@12405.12429.25\|CTTTTTGATCCTTCGATGTCGGCTC | >hg38_21_+_8433172_8446622_RNA45SN1_45S_50ntFlanks | 12405 | 12429 | 25 | CTTTTTGATCCTTCGATGTCGGCTC | 14885.4 | -12.4 | 1.6 | -7.5 | 6.7E-14 | 1.8E-13 | BT20/MB468 | NUC |
| >hg38_21_+_8433172_8446622_RNA45SN1_45S_50ntFlanks@11922.11950.29\|GAGAGGTGTAGAATAAGTGGGAGGCCCCC | >hg38_21_+_8433172_8446622_RNA45SN1_45S_50ntFlanks | 11922 | 11950 | 29 | GAGAGGTGTAGAATAAGTGGGAGGCCCCC | 3281.5 | 7.1 | 0.9 | 7.6 | 4.2E-14 | 1.8E-13 | MB231/MB468 | CYTO |
| >hg38_21_+_8433172_8446622_RNA45SN1_45S_50ntFlanks@12418.12441.24\|CGATGTCGGCTCTTCCTATCATTG | >hg38_21_+_8433172_8446622_RNA45SN1_45S_50ntFlanks | 12418 | 12441 | 24 | CGATGTCGGCTCTTCCTATCATTG | 2958.3 | -6.5 | 0.9 | -7.5 | 5.7E-14 | 1.8E-13 | MB231/MB468 | MITO |
| >hg38_21_+_8433172_8446622_RNA45SN1_45S_50ntFlanks@6646.6684.39\|TCGTACGACTCTTAGCGGTGGATCACTCGGCTCGTGCGT | >hg38_21_+_8433172_8446622_RNA45SN1_45S_50ntFlanks | 6646 | 6684 | 39 | TCGTACGACTCTTAGCGGTGGATCACTCGGCTCGTGCGT | 58.3 | 5.3 | 0.7 | 7.5 | 6.7E-14 | 1.9E-13 | BT20/MB468 | NUC |
| >hg38_21_+_8433172_8446622_RNA45SN1_45S_50ntFlanks@11207.11234.28\|GGCGGCGACTCTGGACGCGAGCCGGGCC | >hg38_21_+_8433172_8446622_RNA45SN1_45S_50ntFlanks | 11207 | 11234 | 28 | GGCGGCGACTCTGGACGCGAGCCGGGCC | 308.1 | -11.0 | 1.5 | -7.6 | 4.2E-14 | 1.9E-13 | MB231/MB468 | CYTO |
| >hg38_21_+_8433172_8446622_RNA45SN1_45S_50ntFlanks@11208.11225.18\|GCGGCGACTCTGGACGCG | >hg38_21_+_8433172_8446622_RNA45SN1_45S_50ntFlanks | 11208 | 11225 | 18 | GCGGCGACTCTGGACGCG | 7331.3 | -9.8 | 1.3 | -7.6 | 4.2E-14 | 1.9E-13 | MB231/MB468 | CYTO |
| >MI0000542\|hsa-mir-320a&WithFlank&8\|-\|22244960\|22245043@48.71.24&[MIMAT0000510&hsa-miR-320a-3p&offsets\|0\|+2;m-60&8\|-\|22244975\|22244996&offsets\|0\|+2]\|AAAAGCTGGGTTGAGAGGGCGAAA | >MI0000542\|hsa-mir-320a&WithFlank&8\|-\|22244960\|22245043 | 48 | 71 | 24 | AAAAGCTGGGTTGAGAGGGCGAAA | 582.4 | 3.5 | 0.5 | 7.5 | 6.7E-14 | 1.9E-13 | BT20/MB468 | NUC |
| >hg38_21_+_8433172_8446622_RNA45SN1_45S_50ntFlanks@11197.11232.36\|GGCCGGCGGCGGCGGCGACTCTGGACGCGAGCCGGG | >hg38_21_+_8433172_8446622_RNA45SN1_45S_50ntFlanks | 11197 | 11232 | 36 | GGCCGGCGGCGGCGGCGACTCTGGACGCGAGCCGGG | 205.7 | -10.4 | 1.4 | -7.6 | 4.2E-14 | 1.9E-13 | MB231/MB468 | CYTO |
| >am_trna128_GlyGCC_6_-_27870686_27870756@57.74.18,trna133_GlyCCC_1_-_16872434_16872504@57.74.18,trna19_GlyGCC_16_+_70823410_70823480@57.74.18,trna19_GlyGCC_2_-_157257659_157257729@57.74.18,trna24_GlyGCC_16_-_70812942_70813012@57.74.18,trna2_GlyCCC_1_+_17053780_17053850@57.74.18,trna4_GlyCCC_1_+_17188416_17188486@57.74.18,trna5_GlyGCC_17_+_8029064_8029134@57.74.18,trna68_GlyGCC_1_-_161493637_161493707@57.74.18\|TTCCCGGCCAATGCACCA | >am_trna128_GlyGCC_6_-_27870686_27870756 | 57 | 74 | 18 | TTCCCGGCCAATGCACCA | 356.8 | 8.6 | 1.1 | 7.5 | 4.6E-14 | 1.9E-13 | BT20/MB468 | CYTO |
| >hg38_21_+_8433172_8446622_RNA45SN1_45S_50ntFlanks@9391.9411.21\|CCGAGGTGGGATCCCGAGGCC | >hg38_21_+_8433172_8446622_RNA45SN1_45S_50ntFlanks | 9391 | 9411 | 21 | CCGAGGTGGGATCCCGAGGCC | 171.9 | -11.3 | 1.5 | -7.6 | 3.7E-14 | 1.9E-13 | MB231/MB468 | MP |
| >hg38_21_+_8433172_8446622_RNA45SN1_45S_50ntFlanks@9788.9818.31\|CGCTGGCGTGGAGCCGGGCGTGGAATGCGAG | >hg38_21_+_8433172_8446622_RNA45SN1_45S_50ntFlanks | 9788 | 9818 | 31 | CGCTGGCGTGGAGCCGGGCGTGGAATGCGAG | 186.1 | -11.4 | 1.5 | -7.6 | 3.7E-14 | 1.9E-13 | MB231/MB468 | MP |
| >hg38_21_+_8433172_8446622_RNA45SN1_45S_50ntFlanks@11876.11912.37\|AAGACCCTGTTGAGCTTGACTCTAGTCTGGCACGGTG | >hg38_21_+_8433172_8446622_RNA45SN1_45S_50ntFlanks | 11876 | 11912 | 37 | AAGACCCTGTTGAGCTTGACTCTAGTCTGGCACGGTG | 231.9 | -11.7 | 1.5 | -7.6 | 3.7E-14 | 1.9E-13 | MB231/MB468 | MP |
| >hg38_21_+_8433172_8446622_RNA45SN1_45S_50ntFlanks@11572.11590.19\|CCGACTTAGAACTGGTGCG | >hg38_21_+_8433172_8446622_RNA45SN1_45S_50ntFlanks | 11572 | 11590 | 19 | CCGACTTAGAACTGGTGCG | 2890.4 | -4.2 | 0.6 | -7.5 | 4.4E-14 | 1.9E-13 | MB231/MB468 | NUC |
| >hg38_21_+_8433172_8446622_RNA45SN1_45S_50ntFlanks@12676.12711.36\|AGAATCCCGCCCAGGCGGAACGATACGGCAGCGCCG | >hg38_21_+_8433172_8446622_RNA45SN1_45S_50ntFlanks | 12676 | 12711 | 36 | AGAATCCCGCCCAGGCGGAACGATACGGCAGCGCCG | 161.4 | -11.2 | 1.5 | -7.6 | 3.7E-14 | 1.9E-13 | MB231/MB468 | MP |
| >hg38_21_+_8433172_8446622_RNA45SN1_45S_50ntFlanks@11569.11600.32\|CAGCCGACTTAGAACTGGTGCGGACCAGGGGA | >hg38_21_+_8433172_8446622_RNA45SN1_45S_50ntFlanks | 11569 | 11600 | 32 | CAGCCGACTTAGAACTGGTGCGGACCAGGGGA | 4042.4 | -8.5 | 1.1 | -7.6 | 3.7E-14 | 1.9E-13 | MB231/MB468 | MP |
| >hg38_21_+_8433172_8446622_RNA45SN1_45S_50ntFlanks@7975.8016.42\|CGCGACCTCAGATCAGACGTGGCGACCCGCTGAATTTAAGCA | >hg38_21_+_8433172_8446622_RNA45SN1_45S_50ntFlanks | 7975 | 8016 | 42 | CGCGACCTCAGATCAGACGTGGCGACCCGCTGAATTTAAGCA | 101794.9 | 5.3 | 0.7 | 7.6 | 3.4E-14 | 1.9E-13 | BT20/MB468 | MITO |
| >hg38_21_+_8433172_8446622_RNA45SN1_45S_50ntFlanks@5496.5514.19\|GACGGTCGAACTTGACTAT | >hg38_21_+_8433172_8446622_RNA45SN1_45S_50ntFlanks | 5496 | 5514 | 19 | GACGGTCGAACTTGACTAT | 330.8 | -12.2 | 1.6 | -7.6 | 3.7E-14 | 1.9E-13 | MB231/MB468 | MP |
| >MI0003599\|hsa-mir-589&WithFlank&7\|-\|5495813\|5495923@30.50.21&[MIMAT0004799&hsa-miR-589-5p&offsets\|0\|-1;m-162&7\|-\|5495874\|5495894&offsets\|0\|0]\|TGAGAACCACGTCTGCTCTGA | >MI0003599\|hsa-mir-589&WithFlank&7\|-\|5495813\|5495923 | 30 | 50 | 21 | TGAGAACCACGTCTGCTCTGA | 413.2 | 11.9 | 1.6 | 7.5 | 5.8E-14 | 1.9E-13 | MB231/MB468 | MITO |
| >hg38_21_+_8433172_8446622_RNA45SN1_45S_50ntFlanks@4588.4608.21\|CTATTTTGTTGGTTTTCGGAA | >hg38_21_+_8433172_8446622_RNA45SN1_45S_50ntFlanks | 4588 | 4608 | 21 | CTATTTTGTTGGTTTTCGGAA | 198.3 | -11.4 | 1.5 | -7.5 | 5.8E-14 | 1.9E-13 | MB231/MB468 | MITO |
| >hg38_21_+_8433172_8446622_RNA45SN1_45S_50ntFlanks@11918.11951.34\|ACATGAGAGGTGTAGAATAAGTGGGAGGCCCCCG | >hg38_21_+_8433172_8446622_RNA45SN1_45S_50ntFlanks | 11918 | 11951 | 34 | ACATGAGAGGTGTAGAATAAGTGGGAGGCCCCCG | 306.6 | -8.6 | 1.1 | -7.5 | 4.4E-14 | 1.9E-13 | MB231/MB468 | NUC |
| >hg38_21_+_8433172_8446622_RNA45SN1_45S_50ntFlanks@11846.11871.26\|GAACGGGCTTGGCGGAATCAGCGGGG | >hg38_21_+_8433172_8446622_RNA45SN1_45S_50ntFlanks | 11846 | 11871 | 26 | GAACGGGCTTGGCGGAATCAGCGGGG | 148.8 | -11.1 | 1.5 | -7.6 | 3.7E-14 | 1.9E-13 | MB231/MB468 | MP |
| >hg38_21_+_8433172_8446622_RNA45SN1_45S_50ntFlanks@12894.12933.40\|CTCGCCCGTCACGCACCGCACGTTCGTGGGGAACCTGGCG | >hg38_21_+_8433172_8446622_RNA45SN1_45S_50ntFlanks | 12894 | 12933 | 40 | CTCGCCCGTCACGCACCGCACGTTCGTGGGGAACCTGGCG | 213.8 | -11.6 | 1.5 | -7.6 | 3.7E-14 | 1.9E-13 | MB231/MB468 | MP |
| >hg38_21_+_8433172_8446622_RNA45SN1_45S_50ntFlanks@12937.12955.19\|AACCATTCGTAGACGACCT | >hg38_21_+_8433172_8446622_RNA45SN1_45S_50ntFlanks | 12937 | 12955 | 19 | AACCATTCGTAGACGACCT | 144.4 | -11.0 | 1.5 | -7.6 | 3.7E-14 | 1.9E-13 | MB231/MB468 | MP |
| >hg38_21_+_8433172_8446622_RNA45SN1_45S_50ntFlanks@8157.8175.19\|CTCGTGGGGGGCCCAAGTC | >hg38_21_+_8433172_8446622_RNA45SN1_45S_50ntFlanks | 8157 | 8175 | 19 | CTCGTGGGGGGCCCAAGTC | 210.3 | -11.6 | 1.5 | -7.6 | 3.7E-14 | 1.9E-13 | MB231/MB468 | MP |
| >hg38_21_+_8433172_8446622_RNA45SN1_45S_50ntFlanks@10840.10863.24\|CCGTAACTTCGGGATAAGGATTGG | >hg38_21_+_8433172_8446622_RNA45SN1_45S_50ntFlanks | 10840 | 10863 | 24 | CCGTAACTTCGGGATAAGGATTGG | 237.0 | -11.8 | 1.6 | -7.6 | 3.7E-14 | 1.9E-13 | MB231/MB468 | MP |
| >hg38_21_+_8433172_8446622_RNA45SN1_45S_50ntFlanks@12634.12654.21\|GAAGCTACCATCTGTGGGATT | >hg38_21_+_8433172_8446622_RNA45SN1_45S_50ntFlanks | 12634 | 12654 | 21 | GAAGCTACCATCTGTGGGATT | 887.4 | -9.8 | 1.3 | -7.5 | 5.8E-14 | 1.9E-13 | MB231/MB468 | MITO |
| >MI0003834\|hsa-mir-769&WithFlank&19\|+\|46018926\|46019055@36.56.21&[MIMAT0003886&hsa-miR-769-5p&offsets\|0\|-1;m-216&19\|+\|46018961\|46018982&offsets\|0\|-1]\|TGAGACCTCTGGGTTCTGAGC | >MI0003834\|hsa-mir-769&WithFlank&19\|+\|46018926\|46019055 | 36 | 56 | 21 | TGAGACCTCTGGGTTCTGAGC | 252.0 | 5.8 | 0.8 | 7.5 | 4.4E-14 | 1.9E-13 | MB231/MB468 | NUC |
| >hg38_21_+_8433172_8446622_RNA45SN1_45S_50ntFlanks@12628.12661.34\|TGGGGCGAAGCTACCATCTGTGGGATTATGACTG | >hg38_21_+_8433172_8446622_RNA45SN1_45S_50ntFlanks | 12628 | 12661 | 34 | TGGGGCGAAGCTACCATCTGTGGGATTATGACTG | 7996.4 | -11.4 | 1.5 | -7.5 | 5.8E-14 | 1.9E-13 | MB231/MB468 | MITO |
| >hg38_21_+_8433172_8446622_RNA45SN1_45S_50ntFlanks@4726.4759.34\|TAATCAAGAACGAAAGTCGGAGGTTCGAAGACGA | >hg38_21_+_8433172_8446622_RNA45SN1_45S_50ntFlanks | 4726 | 4759 | 34 | TAATCAAGAACGAAAGTCGGAGGTTCGAAGACGA | 3980.3 | -11.3 | 1.5 | -7.5 | 6.9E-14 | 1.9E-13 | BT20/MB468 | NUC |
| >hg38_21_+_8433172_8446622_RNA45SN1_45S_50ntFlanks@12423.12441.19\|TCGGCTCTTCCTATCATTG | >hg38_21_+_8433172_8446622_RNA45SN1_45S_50ntFlanks | 12423 | 12441 | 19 | TCGGCTCTTCCTATCATTG | 774.1 | -9.8 | 1.3 | -7.5 | 6.9E-14 | 1.9E-13 | BT20/MB468 | NUC |
| >hg38_21_+_8433172_8446622_RNA45SN1_45S_50ntFlanks@3775.3799.25\|GCCGGTACAGTGAAACTGCGAATGG | >hg38_21_+_8433172_8446622_RNA45SN1_45S_50ntFlanks | 3775 | 3799 | 25 | GCCGGTACAGTGAAACTGCGAATGG | 338.9 | -12.2 | 1.6 | -7.5 | 5.9E-14 | 1.9E-13 | MB231/MB468 | MITO |
| >MI0000062\|hsa-let-7a-3&WithFlank&22\|+\|46112743\|46112828@9.31.23&[MIMAT0000062_1&hsa-let-7a-5p&offsets\|-1\|0;m-5&22\|+\|46112752\|46112773&offsets\|-1\|0]\|GTGAGGTAGTAGGTTGTATAGTT | >MI0000062\|hsa-let-7a-3&WithFlank&22\|+\|46112743\|46112828 | 9 | 31 | 23 | GTGAGGTAGTAGGTTGTATAGTT | 442.2 | 7.1 | 0.9 | 7.5 | 4.7E-14 | 1.9E-13 | BT20/MB468 | CYTO |
| >hg38_21_+_8433172_8446622_RNA45SN1_45S_50ntFlanks@11580.11609.30\|GAACTGGTGCGGACCAGGGGAATCCGACTG | >hg38_21_+_8433172_8446622_RNA45SN1_45S_50ntFlanks | 11580 | 11609 | 30 | GAACTGGTGCGGACCAGGGGAATCCGACTG | 1056.0 | -4.8 | 0.6 | -7.5 | 6.9E-14 | 1.9E-13 | BT20/MB468 | NUC |
| >hg38_21_+_8433172_8446622_RNA45SN1_45S_50ntFlanks@9339.9366.28\|CGCACGAAAGCCGCCGTGGCGCAATGAA | >hg38_21_+_8433172_8446622_RNA45SN1_45S_50ntFlanks | 9339 | 9366 | 28 | CGCACGAAAGCCGCCGTGGCGCAATGAA | 368.2 | -12.3 | 1.6 | -7.5 | 5.9E-14 | 1.9E-13 | MB231/MB468 | MITO |
| >hg38_21_+_8433172_8446622_RNA45SN1_45S_50ntFlanks@8194.8225.32\|CCCGTGGACGGTGTGAGGCCGGTAGCGGCCCC | >hg38_21_+_8433172_8446622_RNA45SN1_45S_50ntFlanks | 8194 | 8225 | 32 | CCCGTGGACGGTGTGAGGCCGGTAGCGGCCCC | 401.7 | -8.3 | 1.1 | -7.5 | 4.7E-14 | 1.9E-13 | BT20/MB468 | CYTO |
| >hg38_21_+_8433172_8446622_RNA45SN1_45S_50ntFlanks@8196.8214.19\|CGTGGACGGTGTGAGGCCG | >hg38_21_+_8433172_8446622_RNA45SN1_45S_50ntFlanks | 8196 | 8214 | 19 | CGTGGACGGTGTGAGGCCG | 1013.6 | -4.5 | 0.6 | -7.5 | 4.5E-14 | 1.9E-13 | MB231/MB468 | NUC |
| >hg38_21_+_8433172_8446622_RNA45SN1_45S_50ntFlanks@10408.10431.24\|GGTCAGTCGGTCCTGAGAGATGGG | >hg38_21_+_8433172_8446622_RNA45SN1_45S_50ntFlanks | 10408 | 10431 | 24 | GGTCAGTCGGTCCTGAGAGATGGG | 960.5 | -6.5 | 0.9 | -7.5 | 4.4E-14 | 1.9E-13 | MB231/MB468 | CYTO |
| >hg38_21_+_8433172_8446622_RNA45SN1_45S_50ntFlanks@12634.12669.36\|GAAGCTACCATCTGTGGGATTATGACTGAACGCCTC | >hg38_21_+_8433172_8446622_RNA45SN1_45S_50ntFlanks | 12634 | 12669 | 36 | GAAGCTACCATCTGTGGGATTATGACTGAACGCCTC | 570.9 | 4.5 | 0.6 | 7.6 | 3.5E-14 | 1.9E-13 | BT20/MB468 | MITO |
| >hg38_21_+_8433172_8446622_RNA45SN1_45S_50ntFlanks@11575.11601.27\|ACTTAGAACTGGTGCGGACCAGGGGAA | >hg38_21_+_8433172_8446622_RNA45SN1_45S_50ntFlanks | 11575 | 11601 | 27 | ACTTAGAACTGGTGCGGACCAGGGGAA | 4425.3 | -11.3 | 1.5 | -7.5 | 7.0E-14 | 1.9E-13 | BT20/MB468 | NUC |

| | | | | | | | | | | | | | |
|---|---|---|---|---|---|---|---|---|---|---|---|---|---|
| >hg38_21_+_8433172_8446622_RNA45SN1_45S_50ntFlanks@6656.6686.31\|CTTAGCGGTGGATCACTCGGCTCGTGCGTCG | >hg38_21_+_8433172_8446622_RNA45SN1_45S_50ntFlanks | 6656 | 6686 | 31 | CTTAGCGGTGGATCACTCGGCTCGTGCGTCG | 335.1 | -4.4 | 0.6 | -7.5 | 7.0E-14 | 1.9E-13 | BT20/MB468 | NUC |
| >hg38_21_+_8433172_8446622_RNA45SN1_45S_50ntFlanks@4569.4587.19\|ATGGAATAGGACCGCGGTT | >hg38_21_+_8433172_8446622_RNA45SN1_45S_50ntFlanks | 4569 | 4587 | 19 | ATGGAATAGGACCGCGGTT | 1150.5 | -10.4 | 1.4 | -7.5 | 7.0E-14 | 1.9E-13 | BT20/MB468 | NUC |
| >hg38_21_+_8433172_8446622_RNA45SN1_45S_50ntFlanks@10400.10424.25\|TGAACATGGGTCAGTCGGTCCTGAG | >hg38_21_+_8433172_8446622_RNA45SN1_45S_50ntFlanks | 10400 | 10424 | 25 | TGAACATGGGTCAGTCGGTCCTGAG | 4845.0 | 7.6 | 0.9 | 8.0 | 1.6E-15 | 1.9E-13 | BT20/MB468 | MP |
| >hg38_21_+_8433172_8446622_RNA45SN1_45S_50ntFlanks@8110.8139.30\|CGGGGCGCGGGACATGTGGCGTACGGAAGA | >hg38_21_+_8433172_8446622_RNA45SN1_45S_50ntFlanks | 8110 | 8139 | 30 | CGGGGCGCGGGACATGTGGCGTACGGAAGA | 147.8 | -11.1 | 1.5 | -7.6 | 3.8E-14 | 1.9E-13 | MB231/MB468 | MP |
| >MI0000071\|hsa-mir-17&WithFlank&13\|+\|91350599\|91350694@20.42.23(+1C)&[MIMAT0000070&hsa-miR-17-5p&offsets\|0\|0(+1C);m-88&13\|+\|91350618\|91350640&offsets\|0\|0(+1C)]\|CAAAGTGCTTACAGTGCAGGTAGC | >MI0000071\|hsa-mir-17&WithFlank&13\|+\|91350599\|91350694 | 20 | 42 | 23(+1C) | CAAAGTGCTTACAGTGCAGGTAGC | 253.4 | 5.4 | 0.7 | 7.5 | 4.5E-14 | 1.9E-13 | MB231/MB468 | NUC |
| >hg38_21_+_8433172_8446622_RNA45SN1_45S_50ntFlanks@10795.10823.29\|TGGCATGTTGGAACAATGTAGGTAAGGGA | >hg38_21_+_8433172_8446622_RNA45SN1_45S_50ntFlanks | 10795 | 10823 | 29 | TGGCATGTTGGAACAATGTAGGTAAGGGA | 140.1 | -11.0 | 1.5 | -7.6 | 3.8E-14 | 1.9E-13 | MB231/MB468 | MP |
| >hg38_21_+_8433172_8446622_RNA45SN1_45S_50ntFlanks@11845.11864.20\|GGAACGGGCTTGGCGGAATC | >hg38_21_+_8433172_8446622_RNA45SN1_45S_50ntFlanks | 11845 | 11864 | 20 | GGAACGGGCTTGGCGGAATC | 221.3 | -11.7 | 1.5 | -7.6 | 3.8E-14 | 1.9E-13 | MB231/MB468 | MP |
| >hg38_21_+_8433172_8446622_RNA45SN1_45S_50ntFlanks@10105.10125.21\|CGCGTGTGGTGTGCGTCGGAG | >hg38_21_+_8433172_8446622_RNA45SN1_45S_50ntFlanks | 10105 | 10125 | 21 | CGCGTGTGGTGTGCGTCGGAG | 179.2 | -11.3 | 1.5 | -7.6 | 3.8E-14 | 1.9E-13 | MB231/MB468 | MP |
| >hg38_21_+_8433172_8446622_RNA45SN1_45S_50ntFlanks@10369.10406.38\|AAGTGGAGAAGGGTTCCATGTGAACAGCAGTTGAACAT | >hg38_21_+_8433172_8446622_RNA45SN1_45S_50ntFlanks | 10369 | 10406 | 38 | AAGTGGAGAAGGGTTCCATGTGAACAGCAGTTGAACAT | 335.3 | -11.4 | 1.5 | -7.5 | 4.6E-14 | 1.9E-13 | MB231/MB468 | NUC |
| >hg38_21_+_8433172_8446622_RNA45SN1_45S_50ntFlanks@9203.9226.24\|CTCGGGGCCACGCGCGCGTCCCCC | >hg38_21_+_8433172_8446622_RNA45SN1_45S_50ntFlanks | 9203 | 9226 | 24 | CTCGGGGCCACGCGCGCGTCCCCC | 226.8 | -11.7 | 1.5 | -7.6 | 3.9E-14 | 2.0E-13 | MB231/MB468 | MP |
| >hg38_21_+_8433172_8446622_RNA45SN1_45S_50ntFlanks@10069.10098.30\|AGTGGACGGGAGCGGCGGGGGCGGCGCGCG | >hg38_21_+_8433172_8446622_RNA45SN1_45S_50ntFlanks | 10069 | 10098 | 30 | AGTGGACGGGAGCGGCGGGGGCGGCGCGCG | 244.3 | -11.8 | 1.6 | -7.6 | 3.9E-14 | 2.0E-13 | MB231/MB468 | MP |
| >hg38_21_+_8433172_8446622_RNA45SN1_45S_50ntFlanks@4128.4162.35\|CGATTCCGGAGAGGGAGCCTGAGAAACGGCTACCA | >hg38_21_+_8433172_8446622_RNA45SN1_45S_50ntFlanks | 4128 | 4162 | 35 | CGATTCCGGAGAGGGAGCCTGAGAAACGGCTACCA | 1331.7 | -10.6 | 1.4 | -7.5 | 7.1E-14 | 2.0E-13 | BT20/MB468 | NUC |
| >hg38_21_+_8433172_8446622_RNA45SN1_45S_50ntFlanks@4723.4751.29\|CATTAATCAAGAACGAAAGTCGGAGGTTC | >hg38_21_+_8433172_8446622_RNA45SN1_45S_50ntFlanks | 4723 | 4751 | 29 | CATTAATCAAGAACGAAAGTCGGAGGTTC | 377.9 | -10.7 | 1.4 | -7.6 | 3.6E-14 | 2.0E-13 | BT20/MB468 | MITO |
| >hg38_21_+_8433172_8446622_RNA45SN1_45S_50ntFlanks@11848.11872.25\|ACGGGCTTGGCGGAATCAGCGGGGA | >hg38_21_+_8433172_8446622_RNA45SN1_45S_50ntFlanks | 11848 | 11872 | 25 | ACGGGCTTGGCGGAATCAGCGGGGA | 138.0 | -5.2 | 0.7 | -7.5 | 4.5E-14 | 2.0E-13 | MB231/MB468 | CYTO |
| >hg38_21_+_8433172_8446622_RNA45SN1_45S_50ntFlanks@11452.11477.26\|CTCCTCGGCGCGCGGCGGCGGCGGCG | >hg38_21_+_8433172_8446622_RNA45SN1_45S_50ntFlanks | 11452 | 11477 | 26 | CTCCTCGGCGCGCGGCGGCGGCGGCG | 247.3 | -7.7 | 1.0 | -7.5 | 4.5E-14 | 2.0E-13 | MB231/MB468 | CYTO |
| >hg38_21_+_8433172_8446622_RNA45SN1_45S_50ntFlanks@10788.10823.36\|CAGCCTCTGGCATGTTGGAACAATGTAGGTAAGGGA | >hg38_21_+_8433172_8446622_RNA45SN1_45S_50ntFlanks | 10788 | 10823 | 36 | CAGCCTCTGGCATGTTGGAACAATGTAGGTAAGGGA | 192.1 | -11.5 | 1.5 | -7.6 | 3.9E-14 | 2.0E-13 | MB231/MB468 | MP |
| >MI0000102\|hsa-mir-100&WithFlank&11\|-\|122152223\|122152314@19.38.20(+1C)&[MIMAT0000098&hsa-miR-100-5p&offsets\|0\|-2(+1C);m-27&11\|-\|122152276\|122152296&offsets\|0\|-1(+1C)]\|AACCCGTAGATCCGAACTTGC | >MI0000102\|hsa-mir-100&WithFlank&11\|-\|122152223\|122152314 | 19 | 38 | 20(+1C) | AACCCGTAGATCCGAACTTGC | 20878.6 | 12.7 | 1.7 | 7.5 | 6.1E-14 | 2.0E-13 | MB231/MB468 | MITO |
| >hg38_21_+_8433172_8446622_RNA45SN1_45S_50ntFlanks@6690.6719.30\|AAGAACGCAGCTAGCTGCGAGAATTAATGT | >hg38_21_+_8433172_8446622_RNA45SN1_45S_50ntFlanks | 6690 | 6719 | 30 | AAGAACGCAGCTAGCTGCGAGAATTAATGT | 168.1 | -7.0 | 0.9 | -7.5 | 4.5E-14 | 2.0E-13 | MB231/MB468 | CYTO |
| >hg38_21_+_8433172_8446622_RNA45SN1_45S_50ntFlanks@12414.12448.35\|CCTTCGATGTCGGCTCTTCCTATCATTGTGAAGCA | >hg38_21_+_8433172_8446622_RNA45SN1_45S_50ntFlanks | 12414 | 12448 | 35 | CCTTCGATGTCGGCTCTTCCTATCATTGTGAAGCA | 866.1 | 7.7 | 1.0 | 7.5 | 4.5E-14 | 2.0E-13 | MB231/MB468 | CYTO |
| >hg38_21_+_8433172_8446622_RNA45SN1_45S_50ntFlanks@12488.12518.31\|AACGTGAGCTGGGTTTAGACCGTCGTGAGAC | >hg38_21_+_8433172_8446622_RNA45SN1_45S_50ntFlanks | 12488 | 12518 | 31 | AACGTGAGCTGGGTTTAGACCGTCGTGAGAC | 248.2 | -5.5 | 0.7 | -7.5 | 4.5E-14 | 2.0E-13 | MB231/MB468 | CYTO |
| >hg38_21_+_8433172_8446622_RNA45SN1_45S_50ntFlanks@10482.10511.30\|TCGAAAGGGAGTCGGGTTCAGATCCCCGAA | >hg38_21_+_8433172_8446622_RNA45SN1_45S_50ntFlanks | 10482 | 10511 | 30 | TCGAAAGGGAGTCGGGTTCAGATCCCCGAA | 486.4 | -8.9 | 1.2 | -7.5 | 4.5E-14 | 2.0E-13 | MB231/MB468 | CYTO |
| >hg38_21_+_8433172_8446622_RNA45SN1_45S_50ntFlanks@10921.10946.26\|GCTGGACGAGGCGCCGCCGCCCCCC | >hg38_21_+_8433172_8446622_RNA45SN1_45S_50ntFlanks | 10921 | 10946 | 26 | GCTGGACGAGGCGCCGCCGCCCCCC | 374.1 | -7.9 | 1.0 | -7.5 | 4.5E-14 | 2.0E-13 | MB231/MB468 | CYTO |
| >hg38_21_+_8433172_8446622_RNA45SN1_45S_50ntFlanks@9572.9605.34\|CAAATCGGTCGTCCGACCTGGGTATAGGGGCGAA | >hg38_21_+_8433172_8446622_RNA45SN1_45S_50ntFlanks | 9572 | 9605 | 34 | CAAATCGGTCGTCCGACCTGGGTATAGGGGCGAA | 1886.2 | -5.9 | 0.8 | -7.5 | 6.1E-14 | 2.0E-13 | MB231/MB468 | MITO |
| >MI0000077\|hsa-mir-21&WithFlank&17\|+\|59841260\|59841343@14.33.20(+1A)&[MIMAT0000076&hsa-miR-21-5p&offsets\|0\|-2(+1A);m-1&17\|+\|59841273\|59841294&offsets\|0\|-2(+1A)]\|TAGCTTATCAGACTGATGTTA | >MI0000077\|hsa-mir-21&WithFlank&17\|+\|59841260\|59841343 | 14 | 33 | 20(+1A) | TAGCTTATCAGACTGATGTTA | 5244.0 | 8.7 | 1.2 | 7.5 | 4.9E-14 | 2.0E-13 | BT20/MB468 | CYTO |
| >MI0000746\|hsa-mir-99b&WithFlank&19\|+\|51692606\|51692687@51.70.20(+1A)&[MIMAT0004678&hsa-miR-99b-3p&offsets\|0\|-2(+1A);m-151&19\|+\|51692656\|51692678&offsets\|0\|-3(+1A)]\|CAAGCTCGTGTCTGTGGGTCA | >MI0000746\|hsa-mir-99b&WithFlank&19\|+\|51692606\|51692687 | 51 | 70 | 20(+1A) | CAAGCTCGTGTCTGTGGGTCA | 429.1 | 10.4 | 1.4 | 7.5 | 4.9E-14 | 2.0E-13 | BT20/MB468 | CYTO |
| >MI0000471\|hsa-mir-126&WithFlank&9\|+\|136670596\|136670692@58.79.22&[MIMAT0000445&hsa-miR-126-3p&offsets\|0\|0;m-40&9\|+\|136670653\|136670674&offsets\|0\|0]\|TCGTACCGTGAGTAATAATGCG | >MI0000471\|hsa-mir-126&WithFlank&9\|+\|136670596\|136670692 | 58 | 79 | 22 | TCGTACCGTGAGTAATAATGCG | 2539.8 | 8.7 | 1.2 | 7.5 | 6.1E-14 | 2.0E-13 | MB231/MB468 | MITO |
| >hg38_21_+_8433172_8446622_RNA45SN1_45S_50ntFlanks@8104.8143.40\|CCGCGGCGGGGCGCGGGACATGTGGCGTACGGAAGACCCG | >hg38_21_+_8433172_8446622_RNA45SN1_45S_50ntFlanks | 8104 | 8143 | 40 | CCGCGGCGGGGCGCGGGACATGTGGCGTACGGAAGACCCG | 6591.8 | -9.1 | 1.2 | -7.5 | 4.7E-14 | 2.0E-13 | MB231/MB468 | NUC |
| >hg38_21_+_8433172_8446622_RNA45SN1_45S_50ntFlanks@11670.11699.30\|CCCAGTGCTCTGAATGTCAAAGTGAAGAAA | >hg38_21_+_8433172_8446622_RNA45SN1_45S_50ntFlanks | 11670 | 11699 | 30 | CCCAGTGCTCTGAATGTCAAAGTGAAGAAA | 166.0 | -11.2 | 1.5 | -7.6 | 3.9E-14 | 2.0E-13 | MB231/MB468 | MP |
| >hg38_21_+_8433172_8446622_RNA45SN1_45S_50ntFlanks@11661.11679.19\|TGATTTCTGCCCAGTGCTC | >hg38_21_+_8433172_8446622_RNA45SN1_45S_50ntFlanks | 11661 | 11679 | 19 | TGATTTCTGCCCAGTGCTC | 436.1 | -3.8 | 0.5 | -7.6 | 3.7E-14 | 2.0E-13 | BT20/MB468 | MITO |
| >MI0000239\|hsa-mir-197&WithFlank&1\|+\|109598887\|109598973@54.74.21(+1A)&[MIMAT0000227&hsa-miR-197-3p&offsets\|0\|-1(+1A);m-99&1\|+\|109598940\|109598961&offsets\|0\|-1(+1A)]\|TTCACCACCTTCTCCACCCAGA | >MI0000239\|hsa-mir-197&WithFlank&1\|+\|109598887\|109598973 | 54 | 74 | 21(+1A) | TTCACCACCTTCTCCACCCAGA | 2936.2 | 12.1 | 1.6 | 7.5 | 4.9E-14 | 2.0E-13 | BT20/MB468 | CYTO |
| >hg38_21_+_8433172_8446622_RNA45SN1_45S_50ntFlanks@3899.3918.20\|CCTTCGCGGGGGGGATGCGT | >hg38_21_+_8433172_8446622_RNA45SN1_45S_50ntFlanks | 3899 | 3918 | 20 | CCTTCGCGGGGGGGATGCGT | 271.9 | -11.9 | 1.6 | -7.5 | 6.2E-14 | 2.0E-13 | MB231/MB468 | MITO |
| >hg38_21_+_8433172_8446622_RNA45SN1_45S_50ntFlanks@12643.12661.19\|ATCTGTGGGATTATGACTG | >hg38_21_+_8433172_8446622_RNA45SN1_45S_50ntFlanks | 12643 | 12661 | 19 | ATCTGTGGGATTATGACTG | 669.6 | -11.5 | 1.5 | -7.6 | 3.7E-14 | 2.0E-13 | BT20/MB468 | MITO |
| >trnaMT_PheGAA_MT_+_577_647@35.54.20\|GAAAATGTTTAGACGGGCTC | >trnaMT_PheGAA_MT_+_577_647 | 35 | 54 | 20 | GAAAATGTTTAGACGGGCTC | 361.1 | 7.7 | 1.0 | 7.5 | 4.7E-14 | 2.0E-13 | MB231/MB468 | NUC |
| >MI0000477\|hsa-mir-146a&WithFlank&5\|+\|160485346\|160485456@27.47.21(+1A)&[MIMAT0000449&hsa-miR-146a-5p&offsets\|0\|-1(+1A);m-129&5\|+\|160485372\|160485393&offsets\|0\|-1(+1A)]\|TGAGAACTGAATTCCATGGGTA | >MI0000477\|hsa-mir-146a&WithFlank&5\|+\|160485346\|160485456 | 27 | 47 | 21(+1A) | TGAGAACTGAATTCCATGGGTA | 791.1 | 12.9 | 1.7 | 7.5 | 6.2E-14 | 2.0E-13 | MB231/MB468 | MITO |
| >hg38_21_+_8433172_8446622_RNA45SN1_45S_50ntFlanks@12480.12497.18\|TAATAGGGAACGTGAGCT | >hg38_21_+_8433172_8446622_RNA45SN1_45S_50ntFlanks | 12480 | 12497 | 18 | TAATAGGGAACGTGAGCT | 1915.4 | 4.2 | 0.6 | 7.5 | 4.6E-14 | 2.0E-13 | MB231/MB468 | CYTO |
| >MI0000477\|hsa-mir-146a&WithFlank&5\|+\|160485346\|160485456@27.48.22(+1U)&[MIMAT0000449&hsa-miR-146a-5p&offsets\|0\|0(+1U);m-129&5\|+\|160485372\|160485393&offsets\|0\|0(+1U)]\|TGAGAACTGAATTCCATGGGTTT | >MI0000477\|hsa-mir-146a&WithFlank&5\|+\|160485346\|160485456 | 27 | 48 | 22(+1U) | TGAGAACTGAATTCCATGGGTTT | 961.3 | -13.0 | 1.7 | -7.8 | 5.6E-15 | 2.0E-13 | BT20/MB231 | NUC |
| >hg38_21_+_8433172_8446622_RNA45SN1_45S_50ntFlanks@9505.9523.19\|TGAACTATGCCTGGGCAGG | >hg38_21_+_8433172_8446622_RNA45SN1_45S_50ntFlanks | 9505 | 9523 | 19 | TGAACTATGCCTGGGCAGG | 285.4 | -12.0 | 1.6 | -7.6 | 4.0E-14 | 2.0E-13 | MB231/MB468 | MP |
| >hg38_21_+_8433172_8446622_RNA45SN1_45S_50ntFlanks@7984.8008.25\|AGATCAGACGTGGCGACCCGCTGAA | >hg38_21_+_8433172_8446622_RNA45SN1_45S_50ntFlanks | 7984 | 8008 | 25 | AGATCAGACGTGGCGACCCGCTGAA | 6167.3 | -6.8 | 0.9 | -7.5 | 6.3E-14 | 2.0E-13 | MB231/MB468 | MITO |
| >MI0000439\|hsa-mir-23b&WithFlank&9\|+\|95085202\|95085310@67.86.20(+1C)&[MIMAT0000418&hsa-miR-23b-3p&offsets\|+3\|0(+1C);m-42&9\|+\|95085265\|95085285&offsets\|+3\|+2(+1C)]\|ACATTGCCAGGGATTACCACC | >MI0000439\|hsa-mir-23b&WithFlank&9\|+\|95085202\|95085310 | 67 | 86 | 20(+1C) | ACATTGCCAGGGATTACCACC | 888.6 | 27.2 | 3.6 | 7.5 | 6.3E-14 | 2.0E-13 | MB231/MB468 | MITO |
| >hg38_21_+_8433172_8446622_RNA45SN1_45S_50ntFlanks@8170.8214.45\|CAAGTCCTTCTGATCGAGGCCCAGCCCGTGGACGGTGTGAGGCCG | >hg38_21_+_8433172_8446622_RNA45SN1_45S_50ntFlanks | 8170 | 8214 | 45 | CAAGTCCTTCTGATCGAGGCCCAGCCCGTGGACGGTGTGAGGCCG | 6506.1 | -12.0 | 1.6 | -7.5 | 7.4E-14 | 2.0E-13 | BT20/MB468 | NUC |
| >hg38_21_+_8433172_8446622_RNA45SN1_45S_50ntFlanks@11873.11890.18\|AAGAAGACCCTGTTGAGC | >hg38_21_+_8433172_8446622_RNA45SN1_45S_50ntFlanks | 11873 | 11890 | 18 | AAGAAGACCCTGTTGAGC | 1534.3 | -7.7 | 1.0 | -7.5 | 4.8E-14 | 2.0E-13 | MB231/MB468 | NUC |
| >hg38_21_+_8433172_8446622_RNA45SN1_45S_50ntFlanks@9740.9765.26\|AACGATCTCAACCTATTCTCAAACTT | >hg38_21_+_8433172_8446622_RNA45SN1_45S_50ntFlanks | 9740 | 9765 | 26 | AACGATCTCAACCTATTCTCAAACTT | 3353.8 | -6.4 | 0.9 | -7.5 | 6.3E-14 | 2.0E-13 | MB231/MB468 | MITO |
| >hg38_21_+_8433172_8446622_RNA45SN1_45S_50ntFlanks@12008.12038.31\|CCGCCGGTGAAATACCACTACTCTGATCGTT | >hg38_21_+_8433172_8446622_RNA45SN1_45S_50ntFlanks | 12008 | 12038 | 31 | CCGCCGGTGAAATACCACTACTCTGATCGTT | 256.5 | -11.0 | 1.5 | -7.5 | 4.8E-14 | 2.0E-13 | MB231/MB468 | NUC |
| >hg38_21_+_8433172_8446622_RNA45SN1_45S_50ntFlanks@3991.4026.36\|TGGTGACTCTAGATAACCTCGGGCCGATCGCACGCC | >hg38_21_+_8433172_8446622_RNA45SN1_45S_50ntFlanks | 3991 | 4026 | 36 | TGGTGACTCTAGATAACCTCGGGCCGATCGCACGCC | 312.7 | -3.6 | 0.5 | -7.6 | 3.8E-14 | 2.0E-13 | BT20/MB468 | MITO |
| >MI0000085\|hsa-mir-27a&WithFlank&19\|-\|13836434\|13836523@57.74.18;MI0000440\|hsa-mir-27b&WithFlank&9\|+\|95085439\|95085547@67.84.18&[MIMAT0000084&hsa-miR-27a-3p&offsets\|0\|-3;m-43&19\|-\|13836447\|13836467&offsets\|0\|-3];[MIMAT0000419&hsa-miR-27b-3p&offsets\|0\|-3;m-39&9\|+\|95085505\|95085525&offsets\|0\|-3]\|TTCACAGTGGCTAAGTTC | >MI0000085\|hsa-mir-27a&WithFlank&19\|-\|13836434\|13836523 | 57 | 74 | 18 | TTCACAGTGGCTAAGTTC | 669.5 | 6.7 | 0.9 | 7.5 | 5.0E-14 | 2.0E-13 | BT20/MB468 | CYTO |
| >MI0001446\|hsa-mir-424&WithFlank&X\|-\|134546608\|134546717@54.74.21&[MIMAT0004749&hsa-miR-424-3p&offsets\|0\|0;m-305&X\|-\|134546644\|134546664&offsets\|0\|0]\|CAAAACGTGAGGCGCTGCTAT | >MI0001446\|hsa-mir-424&WithFlank&X\|-\|134546608\|134546717 | 54 | 74 | 21 | CAAAACGTGAGGCGCTGCTAT | 432.8 | 10.4 | 1.4 | 7.5 | 4.7E-14 | 2.0E-13 | MB231/MB468 | CYTO |
| >hg38_21_+_8433172_8446622_RNA45SN1_45S_50ntFlanks@10574.10596.23\|AAGCCGGCGGGAGCCCCGGGGAG | >hg38_21_+_8433172_8446622_RNA45SN1_45S_50ntFlanks | 10574 | 10596 | 23 | AAGCCGGCGGGAGCCCCGGGGAG | 170.8 | -11.3 | 1.5 | -7.6 | 4.0E-14 | 2.0E-13 | MB231/MB468 | MP |
| >hg38_21_+_8433172_8446622_RNA45SN1_45S_50ntFlanks@4390.4408.19\|TCTTGGGAGCGGGCGGGCG | >hg38_21_+_8433172_8446622_RNA45SN1_45S_50ntFlanks | 4390 | 4408 | 19 | TCTTGGGAGCGGGCGGGCG | 229.6 | -10.6 | 1.4 | -7.5 | 4.7E-14 | 2.0E-13 | MB231/MB468 | CYTO |

<table>
<tr><td>>hg38_21_+_8433172_8446622_RNA45SN1_45S_50ntFlanks@4732.4768.37|AGAACGAAAGTCGGAGGTTCGAAGACGATCAGATACC</td><td>>hg38_21_+_8433172_8446622_RNA45SN1_45S_50ntFlanks</td><td>4732</td><td>4768</td><td>37</td><td>AGAACGAAAGTCGGAGGTTCGAAGACGATCAGATACC</td><td>1234.1</td><td>9.5</td><td>1.3</td><td>7.5</td><td>4.7E-14</td><td>2.0E-13</td><td>MB231/MB468</td><td>CYTO</td></tr>
<tr><td>>MI0000471|hsa-mir-126&WithFlank&9|+|136670596|136670692@58.77.20(+1U)&[MIMAT0000445&hsa-miR-126-3p&offsets|0|-2(+1U);m-40&9|+|136670653|136670674&offsets|0|-2(+1U)]|TCGTACCGTGAGTAATAATGT</td><td>>MI0000471|hsa-mir-126&WithFlank&9|+|136670596|136670692</td><td>58</td><td>77</td><td>20(+1U)</td><td>TCGTACCGTGAGTAATAATGT</td><td>159.5</td><td>10.2</td><td>1.4</td><td>7.5</td><td>7.5E-14</td><td>2.1E-13</td><td>BT20/MB468</td><td>NUC</td></tr>
<tr><td>>hg38_21_+_8433172_8446622_RNA45SN1_45S_50ntFlanks@4006.4025.20|ACCTCGGGCCGATCGCACGC</td><td>>hg38_21_+_8433172_8446622_RNA45SN1_45S_50ntFlanks</td><td>4006</td><td>4025</td><td>20</td><td>ACCTCGGGCCGATCGCACGC</td><td>461.5</td><td>-4.0</td><td>0.5</td><td>-7.5</td><td>4.7E-14</td><td>2.1E-13</td><td>MB231/MB468</td><td>CYTO</td></tr>
<tr><td>>MI0000066|hsa-let-7e&WithFlank&19|+|51692780|51692870@14.34.21&[MIMAT0000066&hsa-let-7e-5p&offsets|0|-1;m-50&19|+|51692793|51692814&offsets|0|-1]|TGAGGTAGGAGGTTGTATAGT</td><td>>MI0000066|hsa-let-7e&WithFlank&19|+|51692780|51692870</td><td>14</td><td>34</td><td>21</td><td>TGAGGTAGGAGGTTGTATAGT</td><td>2614.9</td><td>6.7</td><td>0.9</td><td>7.5</td><td>4.7E-14</td><td>2.1E-13</td><td>MB231/MB468</td><td>CYTO</td></tr>
<tr><td>>hg38_21_+_8433172_8446622_RNA45SN1_45S_50ntFlanks@7977.8016.40|CGACCTCAGATCAGACGTGGCGACCCGCTGAATTTAAGCA</td><td>>hg38_21_+_8433172_8446622_RNA45SN1_45S_50ntFlanks</td><td>7977</td><td>8016</td><td>40</td><td>CGACCTCAGATCAGACGTGGCGACCCGCTGAATTTAAGCA</td><td>2460.1</td><td>9.0</td><td>1.2</td><td>7.5</td><td>4.7E-14</td><td>2.1E-13</td><td>MB231/MB468</td><td>CYTO</td></tr>
<tr><td>>hg38_21_+_8433172_8446622_RNA45SN1_45S_50ntFlanks@10509.10533.25|GAATCCGGAGTGGCGGAGATGGGCG</td><td>>hg38_21_+_8433172_8446622_RNA45SN1_45S_50ntFlanks</td><td>10509</td><td>10533</td><td>25</td><td>GAATCCGGAGTGGCGGAGATGGGCG</td><td>1719.6</td><td>-10.4</td><td>1.4</td><td>-7.6</td><td>4.1E-14</td><td>2.1E-13</td><td>MB231/MB468</td><td>MP</td></tr>
<tr><td>>hg38_21_+_8433172_8446622_RNA45SN1_45S_50ntFlanks@9576.9595.20|TCGGTCGTCCGACCTGGGTA</td><td>>hg38_21_+_8433172_8446622_RNA45SN1_45S_50ntFlanks</td><td>9576</td><td>9595</td><td>20</td><td>TCGGTCGTCCGACCTGGGTA</td><td>328.1</td><td>-12.2</td><td>1.6</td><td>-7.6</td><td>4.1E-14</td><td>2.1E-13</td><td>MB231/MB468</td><td>MP</td></tr>
<tr><td>>hg38_21_+_8433172_8446622_RNA45SN1_45S_50ntFlanks@6644.6681.38|CCTCGTACGACTCTTAGCGGTGGATCACTCGGCTCGTG</td><td>>hg38_21_+_8433172_8446622_RNA45SN1_45S_50ntFlanks</td><td>6644</td><td>6681</td><td>38</td><td>CCTCGTACGACTCTTAGCGGTGGATCACTCGGCTCGTG</td><td>365.4</td><td>-3.5</td><td>0.5</td><td>-7.5</td><td>7.5E-14</td><td>2.1E-13</td><td>BT20/MB468</td><td>NUC</td></tr>
<tr><td>>hg38_21_+_8433172_8446622_RNA45SN1_45S_50ntFlanks@11620.11660.41|ACAAAGCATCGCGAAGGCCCGCGGCGGGTGTTGACGCGATG</td><td>>hg38_21_+_8433172_8446622_RNA45SN1_45S_50ntFlanks</td><td>11620</td><td>11660</td><td>41</td><td>ACAAAGCATCGCGAAGGCCCGCGGCGGGTGTTGACGCGATG</td><td>410.2</td><td>-12.5</td><td>1.7</td><td>-7.5</td><td>6.4E-14</td><td>2.1E-13</td><td>MB231/MB468</td><td>MITO</td></tr>
<tr><td>>hg38_21_+_8433172_8446622_RNA45SN1_45S_50ntFlanks@11769.11792.24|TAATTAGTGACGCGCATGAATGGA</td><td>>hg38_21_+_8433172_8446622_RNA45SN1_45S_50ntFlanks</td><td>11769</td><td>11792</td><td>24</td><td>TAATTAGTGACGCGCATGAATGGA</td><td>1485.8</td><td>-7.5</td><td>1.0</td><td>-7.5</td><td>4.9E-14</td><td>2.1E-13</td><td>MB231/MB468</td><td>NUC</td></tr>
<tr><td>>hg38_21_+_8433172_8446622_RNA45SN1_45S_50ntFlanks@12474.12498.25|ACCCACTAATAGGGAACGTGAGCTG</td><td>>hg38_21_+_8433172_8446622_RNA45SN1_45S_50ntFlanks</td><td>12474</td><td>12498</td><td>25</td><td>ACCCACTAATAGGGAACGTGAGCTG</td><td>313.6</td><td>-3.4</td><td>0.5</td><td>-7.5</td><td>4.9E-14</td><td>2.1E-13</td><td>MB231/MB468</td><td>NUC</td></tr>
<tr><td>>hg38_21_+_8433172_8446622_RNA45SN1_45S_50ntFlanks@6785.6806.22|CTACGCCTGTCTGAGCGTCGCT</td><td>>hg38_21_+_8433172_8446622_RNA45SN1_45S_50ntFlanks</td><td>6785</td><td>6806</td><td>22</td><td>CTACGCCTGTCTGAGCGTCGCT</td><td>875.0</td><td>4.4</td><td>0.6</td><td>7.5</td><td>4.9E-14</td><td>2.1E-13</td><td>MB231/MB468</td><td>NUC</td></tr>
<tr><td>>hg38_21_+_8433172_8446622_RNA45SN1_45S_50ntFlanks@10457.10497.41|GATGGCCTCCGTTGCCCTCGGCCGATCGAAAGGGAGTCGGG</td><td>>hg38_21_+_8433172_8446622_RNA45SN1_45S_50ntFlanks</td><td>10457</td><td>10497</td><td>41</td><td>GATGGCCTCCGTTGCCCTCGGCCGATCGAAAGGGAGTCGGG</td><td>173.2</td><td>-11.3</td><td>1.5</td><td>-7.6</td><td>4.1E-14</td><td>2.1E-13</td><td>MB231/MB468</td><td>MP</td></tr>
<tr><td>>hg38_21_+_8433172_8446622_RNA45SN1_45S_50ntFlanks@12432.12455.24|CCTATCATTGTGAAGCAGAATTCA</td><td>>hg38_21_+_8433172_8446622_RNA45SN1_45S_50ntFlanks</td><td>12432</td><td>12455</td><td>24</td><td>CCTATCATTGTGAAGCAGAATTCA</td><td>557.9</td><td>-9.0</td><td>1.2</td><td>-7.5</td><td>6.4E-14</td><td>2.1E-13</td><td>MB231/MB468</td><td>MITO</td></tr>
<tr><td>>hg38_21_+_8433172_8446622_RNA45SN1_45S_50ntFlanks@11270.11293.24|CGGCCGCCCCCGGGGAGCCCGGCG</td><td>>hg38_21_+_8433172_8446622_RNA45SN1_45S_50ntFlanks</td><td>11270</td><td>11293</td><td>24</td><td>CGGCCGCCCCCGGGGAGCCCGGCG</td><td>402.7</td><td>-10.7</td><td>1.4</td><td>-7.6</td><td>3.9E-14</td><td>2.1E-13</td><td>BT20/MB468</td><td>MITO</td></tr>
<tr><td>>hg38_21_+_8433172_8446622_RNA45SN1_45S_50ntFlanks@8190.8212.23|CCAGCCCGTGGACGGTGTGAGGC</td><td>>hg38_21_+_8433172_8446622_RNA45SN1_45S_50ntFlanks</td><td>8190</td><td>8212</td><td>23</td><td>CCAGCCCGTGGACGGTGTGAGGC</td><td>1132.7</td><td>-10.3</td><td>1.4</td><td>-7.5</td><td>7.6E-14</td><td>2.1E-13</td><td>BT20/MB468</td><td>NUC</td></tr>
<tr><td>>hg38_21_+_8433172_8446622_RNA45SN1_45S_50ntFlanks@12490.12520.31|CGTGAGCTGGGTTTAGACCGTCGTGAGACAG</td><td>>hg38_21_+_8433172_8446622_RNA45SN1_45S_50ntFlanks</td><td>12490</td><td>12520</td><td>31</td><td>CGTGAGCTGGGTTTAGACCGTCGTGAGACAG</td><td>165.2</td><td>-11.2</td><td>1.5</td><td>-7.6</td><td>4.2E-14</td><td>2.1E-13</td><td>MB231/MB468</td><td>MP</td></tr>
<tr><td>>hg38_21_+_8433172_8446622_RNA45SN1_45S_50ntFlanks@11118.11145.28|CAGGGGCCGGCGGCGGCCCGCCGCGGGG</td><td>>hg38_21_+_8433172_8446622_RNA45SN1_45S_50ntFlanks</td><td>11118</td><td>11145</td><td>28</td><td>CAGGGGCCGGCGGCGGCCCGCCGCGGGG</td><td>310.6</td><td>-11.0</td><td>1.5</td><td>-7.5</td><td>4.8E-14</td><td>2.1E-13</td><td>MB231/MB468</td><td>CYTO</td></tr>
<tr><td>>MI0000261|hsa-mir-139&WithFlank&11|-|72615057|72615136@13.35.23&[MIMAT0000250&hsa-miR-139-5p&offsets|0|0;m-186&11|-|72615102|72615124&offsets|0|0]|TCTACAGTGCACGTGTCTCCAGT</td><td>>MI0000261|hsa-mir-139&WithFlank&11|-|72615057|72615136</td><td>13</td><td>35</td><td>23</td><td>TCTACAGTGCACGTGTCTCCAGT</td><td>103.2</td><td>9.5</td><td>1.3</td><td>7.5</td><td>7.6E-14</td><td>2.1E-13</td><td>BT20/MB468</td><td>NUC</td></tr>
<tr><td>>hg38_21_+_8433172_8446622_RNA45SN1_45S_50ntFlanks@10768.10785.18|CAGCAGGTCTCCAAGGTG</td><td>>hg38_21_+_8433172_8446622_RNA45SN1_45S_50ntFlanks</td><td>10768</td><td>10785</td><td>18</td><td>CAGCAGGTCTCCAAGGTG</td><td>1072.7</td><td>-9.2</td><td>1.2</td><td>-7.5</td><td>6.5E-14</td><td>2.1E-13</td><td>MB231/MB468</td><td>MITO</td></tr>
<tr><td>>MI0000102|hsa-mir-100&WithFlank&11|-|122152223|122152314@19.39.21(+1A)&[MIMAT0000098&hsa-miR-100-5p&offsets|0|-1(+1A);m-27&11|-|122152276|122152296&offsets|0|0(+1A)]|AACCCGTAGATCCGAACTTGTA</td><td>>MI0000102|hsa-mir-100&WithFlank&11|-|122152223|122152314</td><td>19</td><td>39</td><td>21(+1A)</td><td>AACCCGTAGATCCGAACTTGTA</td><td>69047.3</td><td>11.0</td><td>1.5</td><td>7.6</td><td>4.2E-14</td><td>2.1E-13</td><td>MB231/MB468</td><td>MP</td></tr>
<tr><td>>hg38_21_+_8433172_8446622_RNA45SN1_45S_50ntFlanks@11759.11780.22|CCTCGTCATCTAATTAGTGACG</td><td>>hg38_21_+_8433172_8446622_RNA45SN1_45S_50ntFlanks</td><td>11759</td><td>11780</td><td>22</td><td>CCTCGTCATCTAATTAGTGACG</td><td>275.9</td><td>-10.8</td><td>1.4</td><td>-7.5</td><td>4.9E-14</td><td>2.1E-13</td><td>MB231/MB468</td><td>CYTO</td></tr>
<tr><td>>MI0000342|hsa-mir-200b&WithFlank&1|+|1167098|1167204@63.85.23&[MIMAT0000318&hsa-miR-200b-3p&offsets|0|+1;m-93&1|+|1167160|1167182&offsets|0|0]|TAATACTGCCTGGTAATGATGAC</td><td>>MI0000342|hsa-mir-200b&WithFlank&1|+|1167098|1167204</td><td>63</td><td>85</td><td>23</td><td>TAATACTGCCTGGTAATGATGAC</td><td>4481.7</td><td>-4.5</td><td>0.6</td><td>-7.5</td><td>5.0E-14</td><td>2.1E-13</td><td>MB231/MB468</td><td>NUC</td></tr>
<tr><td>>hg38_21_+_8433172_8446622_RNA45SN1_45S_50ntFlanks@12711.12745.35|GCGGAGCCTCGGTTGGCCTCGGATAGCCGGTCCCC</td><td>>hg38_21_+_8433172_8446622_RNA45SN1_45S_50ntFlanks</td><td>12711</td><td>12745</td><td>35</td><td>GCGGAGCCTCGGTTGGCCTCGGATAGCCGGTCCCC</td><td>184.4</td><td>-11.4</td><td>1.5</td><td>-7.6</td><td>4.2E-14</td><td>2.1E-13</td><td>MB231/MB468</td><td>MP</td></tr>
<tr><td>>MI0000299|hsa-mir-222&WithFlank&X|-|45747009|45747130@75.99.25(+3U)&[MIMAT0000279&hsa-miR-222-3p&offsets|0|+4(+3U);m-140&X|-|45747033|45747056&offsets|0|+1(+3U)]|AGCTACATCTGGCTACTGGGTCTCTTTT</td><td>>MI0000299|hsa-mir-222&WithFlank&X|-|45747009|45747130</td><td>75</td><td>99</td><td>25(+3U)</td><td>AGCTACATCTGGCTACTGGGTCTCTTTT</td><td>1677.4</td><td>6.6</td><td>0.9</td><td>7.5</td><td>5.0E-14</td><td>2.1E-13</td><td>MB231/MB468</td><td>NUC</td></tr>
<tr><td>>hg38_21_+_8433172_8446622_RNA45SN1_45S_50ntFlanks@10351.10390.40|CGAGAACTTTGAAGGCCGAAGTGGAGAAGGGTTCCATGTG</td><td>>hg38_21_+_8433172_8446622_RNA45SN1_45S_50ntFlanks</td><td>10351</td><td>10390</td><td>40</td><td>CGAGAACTTTGAAGGCCGAAGTGGAGAAGGGTTCCATGTG</td><td>33282.5</td><td>-10.8</td><td>1.4</td><td>-7.5</td><td>7.7E-14</td><td>2.1E-13</td><td>BT20/MB468</td><td>NUC</td></tr>
<tr><td>>hg38_21_+_8433172_8446622_RNA45SN1_45S_50ntFlanks@10786.10817.32|AACAGCCTCTGGCATGTTGGAACAATGTAGGT</td><td>>hg38_21_+_8433172_8446622_RNA45SN1_45S_50ntFlanks</td><td>10786</td><td>10817</td><td>32</td><td>AACAGCCTCTGGCATGTTGGAACAATGTAGGT</td><td>479.7</td><td>-3.9</td><td>0.5</td><td>-7.5</td><td>5.0E-14</td><td>2.1E-13</td><td>MB231/MB468</td><td>NUC</td></tr>
<tr><td>>hg38_21_+_8433172_8446622_RNA45SN1_45S_50ntFlanks@4683.4722.40|CAAGACGGACCAGAGCGAAAGCATTTGCCAAGAATGTTTT</td><td>>hg38_21_+_8433172_8446622_RNA45SN1_45S_50ntFlanks</td><td>4683</td><td>4722</td><td>40</td><td>CAAGACGGACCAGAGCGAAAGCATTTGCCAAGAATGTTTT</td><td>991.5</td><td>-8.0</td><td>1.1</td><td>-7.5</td><td>6.6E-14</td><td>2.1E-13</td><td>MB231/MB468</td><td>MITO</td></tr>
<tr><td>>hg38_21_+_8433172_8446622_RNA45SN1_45S_50ntFlanks@12657.12674.18|GACTGAACGCCTCTAAGT</td><td>>hg38_21_+_8433172_8446622_RNA45SN1_45S_50ntFlanks</td><td>12657</td><td>12674</td><td>18</td><td>GACTGAACGCCTCTAAGT</td><td>242.7</td><td>-11.8</td><td>1.6</td><td>-7.6</td><td>4.2E-14</td><td>2.1E-13</td><td>MB231/MB468</td><td>MP</td></tr>
<tr><td>>MI0003815|hsa-mir-1301&WithFlank&2|-|25328634|25328727@54.77.24&[MIMAT0005797&hsa-miR-1301-3p&offsets|0|0;m-256&2|-|25328650|25328674&offsets|0|-1]|TTGCAGCTGCCTGGGAGTGACTTC</td><td>>MI0003815|hsa-mir-1301&WithFlank&2|-|25328634|25328727</td><td>54</td><td>77</td><td>24</td><td>TTGCAGCTGCCTGGGAGTGACTTC</td><td>268.7</td><td>8.2</td><td>1.1</td><td>7.5</td><td>5.3E-14</td><td>2.1E-13</td><td>BT20/MB468</td><td>CYTO</td></tr>
<tr><td>>hg38_21_+_8433172_8446622_RNA45SN1_45S_50ntFlanks@12405.12424.20|CTTTTTGATCCTTCGATGTC</td><td>>hg38_21_+_8433172_8446622_RNA45SN1_45S_50ntFlanks</td><td>12405</td><td>12424</td><td>20</td><td>CTTTTTGATCCTTCGATGTC</td><td>1026.3</td><td>-9.7</td><td>1.3</td><td>-7.5</td><td>6.6E-14</td><td>2.1E-13</td><td>MB231/MB468</td><td>MITO</td></tr>
<tr><td>>hg38_21_+_8433172_8446622_RNA45SN1_45S_50ntFlanks@12894.12927.34|CTCGCCCGTCACGCACCGCACGTTCGTGGGGAAC</td><td>>hg38_21_+_8433172_8446622_RNA45SN1_45S_50ntFlanks</td><td>12894</td><td>12927</td><td>34</td><td>CTCGCCCGTCACGCACCGCACGTTCGTGGGGAAC</td><td>148.5</td><td>-11.1</td><td>1.5</td><td>-7.6</td><td>4.3E-14</td><td>2.1E-13</td><td>MB231/MB468</td><td>MP</td></tr>
<tr><td>>hg38_21_+_8433172_8446622_RNA45SN1_45S_50ntFlanks@3900.3917.18|CTTCGCGGGGGGGATGCG</td><td>>hg38_21_+_8433172_8446622_RNA45SN1_45S_50ntFlanks</td><td>3900</td><td>3917</td><td>18</td><td>CTTCGCGGGGGGGATGCG</td><td>218.3</td><td>-11.6</td><td>1.5</td><td>-7.6</td><td>4.3E-14</td><td>2.1E-13</td><td>MB231/MB468</td><td>MP</td></tr>
<tr><td>>hg38_21_+_8433172_8446622_RNA45SN1_45S_50ntFlanks@12949.12982.34|ACGACCTGCTTCTGGGTCGGGGTTTCGTACGTAG</td><td>>hg38_21_+_8433172_8446622_RNA45SN1_45S_50ntFlanks</td><td>12949</td><td>12982</td><td>34</td><td>ACGACCTGCTTCTGGGTCGGGGTTTCGTACGTAG</td><td>206.9</td><td>-10.7</td><td>1.4</td><td>-7.5</td><td>5.1E-14</td><td>2.1E-13</td><td>MB231/MB468</td><td>NUC</td></tr>
<tr><td>>hg38_21_+_8433172_8446622_RNA45SN1_45S_50ntFlanks@13004.13029.26|ATCTATTGAAAGTCAGCCCTCGACAC</td><td>>hg38_21_+_8433172_8446622_RNA45SN1_45S_50ntFlanks</td><td>13004</td><td>13029</td><td>26</td><td>ATCTATTGAAAGTCAGCCCTCGACAC</td><td>171.1</td><td>-10.4</td><td>1.4</td><td>-7.5</td><td>5.1E-14</td><td>2.1E-13</td><td>MB231/MB468</td><td>NUC</td></tr>
<tr><td>>hg38_21_+_8433172_8446622_RNA45SN1_45S_50ntFlanks@11925.11950.26|AGGTGTAGAATAAGTGGGAGGCCCCC</td><td>>hg38_21_+_8433172_8446622_RNA45SN1_45S_50ntFlanks</td><td>11925</td><td>11950</td><td>26</td><td>AGGTGTAGAATAAGTGGGAGGCCCCC</td><td>176.3</td><td>-11.3</td><td>1.5</td><td>-7.6</td><td>4.3E-14</td><td>2.2E-13</td><td>MB231/MB468</td><td>MP</td></tr>
<tr><td>>MI0000298|hsa-mir-221&WithFlank&X|-|45746151|45746272@71.92.22(+1U)&[MIMAT0000278&hsa-miR-221-3p&offsets|0|-1(+1U);m-91&X|-|45746181|45746202&offsets|0|0(+1U)]|AGCTACATTGTCTGCTGGGTTTT</td><td>>MI0000298|hsa-mir-221&WithFlank&X|-|45746151|45746272</td><td>71</td><td>92</td><td>22(+1U)</td><td>AGCTACATTGTCTGCTGGGTTTT</td><td>5912.2</td><td>6.7</td><td>0.9</td><td>7.5</td><td>6.7E-14</td><td>2.2E-13</td><td>MB231/MB468</td><td>MITO</td></tr>
<tr><td>>hg38_21_+_8433172_8446622_RNA45SN1_45S_50ntFlanks@5059.5082.24|CGATTTGTCTGGTTAATTCCGATA</td><td>>hg38_21_+_8433172_8446622_RNA45SN1_45S_50ntFlanks</td><td>5059</td><td>5082</td><td>24</td><td>CGATTTGTCTGGTTAATTCCGATA</td><td>502.9</td><td>-12.8</td><td>1.7</td><td>-7.6</td><td>4.3E-14</td><td>2.2E-13</td><td>MB231/MB468</td><td>MP</td></tr>
<tr><td>>hg38_21_+_8433172_8446622_RNA45SN1_45S_50ntFlanks@11846.11873.28|GAACGGGCTTGGCGGAATCAGCGGGGAA</td><td>>hg38_21_+_8433172_8446622_RNA45SN1_45S_50ntFlanks</td><td>11846</td><td>11873</td><td>28</td><td>GAACGGGCTTGGCGGAATCAGCGGGGAA</td><td>153.6</td><td>-11.1</td><td>1.5</td><td>-7.6</td><td>4.3E-14</td><td>2.2E-13</td><td>MB231/MB468</td><td>MP</td></tr>
<tr><td>>MI0000255|hsa-mir-30d&WithFlank&8|-|134804870|134804951@12.35.24(+1C)&[MIMAT0000245&hsa-miR-30d-5p&offsets|0|+2(+1C);m-21&8|-|134804919|134804940&offsets|0|+2(+1C)]|TGTAAACATCCCCGACTGGAAGCTC</td><td>>MI0000255|hsa-mir-30d&WithFlank&8|-|134804870|134804951</td><td>12</td><td>35</td><td>24(+1C)</td><td>TGTAAACATCCCCGACTGGAAGCTC</td><td>303.6</td><td>7.1</td><td>0.9</td><td>7.5</td><td>5.2E-14</td><td>2.2E-13</td><td>MB231/MB468</td><td>NUC</td></tr>
<tr><td>>hg38_21_+_8433172_8446622_RNA45SN1_45S_50ntFlanks@3990.4024.35|TTGGTGACTCTAGATAACCTCGGGCCGATCGCACG</td><td>>hg38_21_+_8433172_8446622_RNA45SN1_45S_50ntFlanks</td><td>3990</td><td>4024</td><td>35</td><td>TTGGTGACTCTAGATAACCTCGGGCCGATCGCACG</td><td>1170.0</td><td>-5.9</td><td>0.8</td><td>-7.5</td><td>8.0E-14</td><td>2.2E-13</td><td>BT20/MB468</td><td>NUC</td></tr>
<tr><td>>hg38_21_+_8433172_8446622_RNA45SN1_45S_50ntFlanks@12627.12657.31|ATGGGGCGAAGCTACCATCTGTGGGATTATG</td><td>>hg38_21_+_8433172_8446622_RNA45SN1_45S_50ntFlanks</td><td>12627</td><td>12657</td><td>31</td><td>ATGGGGCGAAGCTACCATCTGTGGGATTATG</td><td>792.2</td><td>-7.4</td><td>1.0</td><td>-7.5</td><td>5.2E-14</td><td>2.2E-13</td><td>MB231/MB468</td><td>NUC</td></tr>
<tr><td>>am_trna11_ProAGG_16_+_3241989_3242060@1.29.29,trna12_ProAGG_6_+_26555498_26555569@1.29.29,trna12_ProTGG_11_-_75946869_75946940@1.29.29,trna14_ProTGG_5_-_180615854_180615925@1.29.29,trna22_ProAGG_14_-_21081560_21081631@1.29.29,trna23_ProAGG_14_-_21077495_21077566@1.29.29,trna28_ProTGG_16_-_3234133_3234204@1.29.29,trna29_ProAGG_16_-_3232635_3232706@1.29.29,trna2_ProAGG_7_+_128423504_128423575@1.29.29,trna30_ProCGG_6_+_27059521_27059592@1.29.29,trna37_ProCGG_17_-_8126151_8126222@1.29.29,trna3_ProTGG_16_+_3208923_3208994@1.29.29,trna52_ProCGG_1_+_167683962_167684033@1.29.29,trna65_ProAGG_1_-_167684725_167684796@1.29.29,trna6_ProCGG_16_+_3222049_3222120@1.29.29,trna6_ProTGG_14_+_21152175_21152246@1.29.29,trna8_ProTGG_16_+_3238094_3238165@1.29.29,trna9_ProAGG_11_+_75946557_75946628@1.29.29,trna9_ProAGG_16_+_3239634_3239705@1.29.29|GGCTCGTTGGTCTAGGGGTATGATTCTCG</td><td>>am_trna11_ProAGG_16_+_3241989_3242060</td><td>1</td><td>29</td><td>29</td><td>GGCTCGTTGGTCTAGGGGTATGATTCTCG</td><td>817.4</td><td>-9.3</td><td>1.2</td><td>-7.5</td><td>6.8E-14</td><td>2.2E-13</td><td>MB231/MB468</td><td>MITO</td></tr>
<tr><td>>hg38_21_+_8433172_8446622_RNA45SN1_45S_50ntFlanks@7981.8007.27|CTCAGATCAGACGTGGCGACCCGCTGA</td><td>>hg38_21_+_8433172_8446622_RNA45SN1_45S_50ntFlanks</td><td>7981</td><td>8007</td><td>27</td><td>CTCAGATCAGACGTGGCGACCCGCTGA</td><td>642.3</td><td>-7.2</td><td>1.0</td><td>-7.5</td><td>6.8E-14</td><td>2.2E-13</td><td>MB231/MB468</td><td>MITO</td></tr>
<tr><td>>hg38_21_+_8433172_8446622_RNA45SN1_45S_50ntFlanks@10408.10429.22|GGTCAGTCGGTCCTGAGAGATG</td><td>>hg38_21_+_8433172_8446622_RNA45SN1_45S_50ntFlanks</td><td>10408</td><td>10429</td><td>22</td><td>GGTCAGTCGGTCCTGAGAGATG</td><td>1511.9</td><td>-10.0</td><td>1.3</td><td>-7.5</td><td>6.8E-14</td><td>2.2E-13</td><td>MB231/MB468</td><td>MITO</td></tr>
</table>

<table>
<tr><th></th><th></th><th></th><th></th><th></th><th></th><th></th><th></th><th></th><th></th><th></th><th></th><th></th><th></th></tr>
<tr><td>>hg38_21_+_8433172_8446622_RNA45SN1_45S_50ntFlanks@4759.4781.23|ATCAGATACCGTCGTAGTTCCGA</td><td>>hg38_21_+_8433172_8446622_RNA45SN1_45S_50ntFlanks</td><td>4759</td><td>4781</td><td>23</td><td>ATCAGATACCGTCGTAGTTCCGA</td><td>432.7</td><td>-11.1</td><td>1.5</td><td>-7.5</td><td>5.4E-14</td><td>2.2E-13</td><td>BT20/MB468</td><td>CYTO</td></tr>
<tr><td>>MI0000542|hsa-mir-320a&WithFlank&8|-|22244960|22245043@48.70.23&[MIMAT0000510&hsa-miR-320a-3p&offsets|0|+1;m-60&8|-|22244975|22244996&offsets|0|+1]|AAAAGCTGGGTTGAGAGGGCGAA</td><td>>MI0000542|hsa-mir-320a&WithFlank&8|-|22244960|22245043</td><td>48</td><td>70</td><td>23</td><td>AAAAGCTGGGTTGAGAGGGCGAA</td><td>11546.6</td><td>7.1</td><td>0.9</td><td>7.9</td><td>1.9E-15</td><td>2.2E-13</td><td>BT20/MB468</td><td>MP</td></tr>
<tr><td>>hg38_21_+_8433172_8446622_RNA45SN1_45S_50ntFlanks@4046.4068.23|CATTCGAACGTCTGCCCTATCAA</td><td>>hg38_21_+_8433172_8446622_RNA45SN1_45S_50ntFlanks</td><td>4046</td><td>4068</td><td>23</td><td>CATTCGAACGTCTGCCCTATCAA</td><td>172.3</td><td>-10.4</td><td>1.4</td><td>-7.5</td><td>5.3E-14</td><td>2.2E-13</td><td>MB231/MB468</td><td>NUC</td></tr>
<tr><td>>hg38_21_+_8433172_8446622_RNA45SN1_45S_50ntFlanks@12637.12659.23|GCTACCATCTGTGGGATTATGAC</td><td>>hg38_21_+_8433172_8446622_RNA45SN1_45S_50ntFlanks</td><td>12637</td><td>12659</td><td>23</td><td>GCTACCATCTGTGGGATTATGAC</td><td>2401.5</td><td>-10.5</td><td>1.4</td><td>-7.5</td><td>4.4E-14</td><td>2.2E-13</td><td>MB231/MB468</td><td>MP</td></tr>
<tr><td>>trna116_GluCTC_1_-_145399233_145399304@3.33.31,trna59_GluCTC_1_+_249168447_249168518@3.33.31,trna71_GluCTC_1_-_161439189_161439260@3.33.31,trna74_GluCTC_1_-_161431809_161431880@3.33.31,trna77_GluCTC_1_-_161424398_161424469@3.33.31,trna77_GluCTC_6_+_28949976_28950047@3.33.31,trna80_GluCTC_1_-_161417018_161417089@3.33.31,trna87_GluCTC_6_-_126101393_126101464@3.33.31|CCTGGTGGTCTAGTGGTTAGGATTCGGCGCT</td><td>>trna116_GluCTC_1_-_145399233_145399304</td><td>3</td><td>33</td><td>31</td><td>CCTGGTGGTCTAGTGGTTAGGATTCGGCGCT</td><td>817.8</td><td>6.3</td><td>0.8</td><td>7.6</td><td>4.1E-14</td><td>2.2E-13</td><td>BT20/MB468</td><td>MITO</td></tr>
<tr><td>>hg38_21_+_8433172_8446622_RNA45SN1_45S_50ntFlanks@11579.11603.25|AGAACTGGTGCGGACCAGGGGAATC</td><td>>hg38_21_+_8433172_8446622_RNA45SN1_45S_50ntFlanks</td><td>11579</td><td>11603</td><td>25</td><td>AGAACTGGTGCGGACCAGGGGAATC</td><td>4891.2</td><td>-4.6</td><td>0.6</td><td>-7.5</td><td>5.5E-14</td><td>2.2E-13</td><td>BT20/MB468</td><td>CYTO</td></tr>
<tr><td>>MI0000086|hsa-mir-28&WithFlank&3|+|188688775|188688872@60.80.21&[MIMAT0004502&hsa-miR-28-3p&offsets|0|-1;m-28&3|+|188688834|188688855&offsets|0|-1]|CACTAGATTGTGAGCTCCTGG</td><td>>MI0000086|hsa-mir-28&WithFlank&3|+|188688775|188688872</td><td>60</td><td>80</td><td>21</td><td>CACTAGATTGTGAGCTCCTGG</td><td>1453.0</td><td>8.5</td><td>1.1</td><td>7.5</td><td>5.1E-14</td><td>2.2E-13</td><td>MB231/MB468</td><td>CYTO</td></tr>
<tr><td>>hg38_21_+_8433172_8446622_RNA45SN1_45S_50ntFlanks@9035.9055.21|CGTGGGGGGGCCGGGCCACCC</td><td>>hg38_21_+_8433172_8446622_RNA45SN1_45S_50ntFlanks</td><td>9035</td><td>9055</td><td>21</td><td>CGTGGGGGGGCCGGGCCACCC</td><td>294.4</td><td>-12.1</td><td>1.6</td><td>-7.5</td><td>4.4E-14</td><td>2.2E-13</td><td>MB231/MB468</td><td>MP</td></tr>
<tr><td>>hg38_21_+_8433172_8446622_RNA45SN1_45S_50ntFlanks@10587.10608.22|CCCCGGGGAGAGTTCTCTTTTC</td><td>>hg38_21_+_8433172_8446622_RNA45SN1_45S_50ntFlanks</td><td>10587</td><td>10608</td><td>22</td><td>CCCCGGGGAGAGTTCTCTTTTC</td><td>187.7</td><td>-10.5</td><td>1.4</td><td>-7.5</td><td>5.3E-14</td><td>2.2E-13</td><td>MB231/MB468</td><td>NUC</td></tr>
<tr><td>>hg38_21_+_8433172_8446622_RNA45SN1_45S_50ntFlanks@11577.11611.35|TTAGAACTGGTGCGGACCAGGGGAATCCGACTGTT</td><td>>hg38_21_+_8433172_8446622_RNA45SN1_45S_50ntFlanks</td><td>11577</td><td>11611</td><td>35</td><td>TTAGAACTGGTGCGGACCAGGGGAATCCGACTGTT</td><td>728.7</td><td>-5.2</td><td>0.7</td><td>-7.5</td><td>6.9E-14</td><td>2.2E-13</td><td>MB231/MB468</td><td>MITO</td></tr>
<tr><td>>hg38_1_-_228634819_228635039_RNA5S12_5S_50ntFlanks@134.170.37|TGGGAGACCGCCTGGGAATACCGGGTGCTGTAGGCTT</td><td>>hg38_1_-_228634819_228635039_RNA5S12_5S_50ntFlanks</td><td>134</td><td>170</td><td>37</td><td>TGGGAGACCGCCTGGGAATACCGGGTGCTGTAGGCTT</td><td>88934.1</td><td>7.6</td><td>1.0</td><td>7.6</td><td>4.1E-14</td><td>2.2E-13</td><td>BT20/MB468</td><td>MITO</td></tr>
<tr><td>>hg38_21_+_8433172_8446622_RNA45SN1_45S_50ntFlanks@5472.5502.31|ACGGCCCTGGCGGAGCGCTGAGAAGACGGTC</td><td>>hg38_21_+_8433172_8446622_RNA45SN1_45S_50ntFlanks</td><td>5472</td><td>5502</td><td>31</td><td>ACGGCCCTGGCGGAGCGCTGAGAAGACGGTC</td><td>777.3</td><td>6.7</td><td>0.9</td><td>7.6</td><td>4.1E-14</td><td>2.2E-13</td><td>BT20/MB468</td><td>MITO</td></tr>
<tr><td>>hg38_MT_+_598_1651_RNR1_12S_50ntFlanks@915.946.32|AAAGGACATTTAACTAAAACCCCTACGCATTT</td><td>>hg38_MT_+_598_1651_RNR1_12S_50ntFlanks</td><td>915</td><td>946</td><td>32</td><td>AAAGGACATTTAACTAAAACCCCTACGCATTT</td><td>671.0</td><td>6.9</td><td>0.9</td><td>7.6</td><td>4.1E-14</td><td>2.2E-13</td><td>BT20/MB468</td><td>MITO</td></tr>
<tr><td>>hg38_21_+_8433172_8446622_RNA45SN1_45S_50ntFlanks@12416.12458.43|TTCGATGTCGGCTCTTCCTATCATTGTGAAGCAGAATTCACCA</td><td>>hg38_21_+_8433172_8446622_RNA45SN1_45S_50ntFlanks</td><td>12416</td><td>12458</td><td>43</td><td>TTCGATGTCGGCTCTTCCTATCATTGTGAAGCAGAATTCACCA</td><td>776.9</td><td>8.1</td><td>1.1</td><td>7.5</td><td>5.1E-14</td><td>2.2E-13</td><td>MB231/MB468</td><td>CYTO</td></tr>
<tr><td>>trna119_LysCTT_1_-_145395522_145395594@1.35.35,trna11_LysCTT_5_-_180648979_180649051@1.35.35,trna13_LysCTT_6_+_26556774_26556846@1.35.35,trna7_LysCTT_16_+_3225692_3225764@1.35.35,trna9_LysCTT_5_+_180634755_180634827@1.35.35|GCCCGGCTAGCTCAGTCGGTAGAGCATGAGACTCT</td><td>>trna119_LysCTT_1_-_145395522_145395594</td><td>1</td><td>35</td><td>35</td><td>GCCCGGCTAGCTCAGTCGGTAGAGCATGAGACTCT</td><td>1287.6</td><td>6.4</td><td>0.9</td><td>7.5</td><td>5.3E-14</td><td>2.2E-13</td><td>MB231/MB468</td><td>NUC</td></tr>
<tr><td>>trna6_TrpCCA_12_+_98898030_98898101@30.52.23|ACTCCAGATCAGAAGGCTGCGTG</td><td>>trna6_TrpCCA_12_+_98898030_98898101</td><td>30</td><td>52</td><td>23</td><td>ACTCCAGATCAGAAGGCTGCGTG</td><td>128.5</td><td>-9.7</td><td>1.3</td><td>-7.5</td><td>5.1E-14</td><td>2.2E-13</td><td>MB231/MB468</td><td>CYTO</td></tr>
<tr><td>>hg38_21_+_8433172_8446622_RNA45SN1_45S_50ntFlanks@5475.5516.42|GCCCTGGCGGAGCGCTGAGAAGACGGTCGAACTTGACTATCT</td><td>>hg38_21_+_8433172_8446622_RNA45SN1_45S_50ntFlanks</td><td>5475</td><td>5516</td><td>42</td><td>GCCCTGGCGGAGCGCTGAGAAGACGGTCGAACTTGACTATCT</td><td>5460.3</td><td>13.6</td><td>1.8</td><td>7.6</td><td>4.1E-14</td><td>2.2E-13</td><td>BT20/MB468</td><td>MITO</td></tr>
<tr><td>>hg38_21_+_8433172_8446622_RNA45SN1_45S_50ntFlanks@3994.4013.20|TGACTCTAGATAACCTCGGG</td><td>>hg38_21_+_8433172_8446622_RNA45SN1_45S_50ntFlanks</td><td>3994</td><td>4013</td><td>20</td><td>TGACTCTAGATAACCTCGGG</td><td>145.1</td><td>-11.0</td><td>1.5</td><td>-7.5</td><td>4.5E-14</td><td>2.2E-13</td><td>MB231/MB468</td><td>MP</td></tr>
<tr><td>>hg38_21_+_8433172_8446622_RNA45SN1_45S_50ntFlanks@6738.6769.32|ATCATCGACACTTCGAACGCACTTGCGGCCCC</td><td>>hg38_21_+_8433172_8446622_RNA45SN1_45S_50ntFlanks</td><td>6738</td><td>6769</td><td>32</td><td>ATCATCGACACTTCGAACGCACTTGCGGCCCC</td><td>221.1</td><td>-11.7</td><td>1.5</td><td>-7.5</td><td>4.5E-14</td><td>2.2E-13</td><td>MB231/MB468</td><td>MP</td></tr>
<tr><td>>hg38_21_+_8433172_8446622_RNA45SN1_45S_50ntFlanks@12944.12965.22|CGTAGACGACCTGCTTCTGGGT</td><td>>hg38_21_+_8433172_8446622_RNA45SN1_45S_50ntFlanks</td><td>12944</td><td>12965</td><td>22</td><td>CGTAGACGACCTGCTTCTGGGT</td><td>182.4</td><td>-11.4</td><td>1.5</td><td>-7.5</td><td>4.5E-14</td><td>2.2E-13</td><td>MB231/MB468</td><td>MP</td></tr>
<tr><td>>hg38_21_+_8433172_8446622_RNA45SN1_45S_50ntFlanks@11568.11605.38|GCAGCCGACTTAGAACTGGTGCGGACCAGGGGAATCCG</td><td>>hg38_21_+_8433172_8446622_RNA45SN1_45S_50ntFlanks</td><td>11568</td><td>11605</td><td>38</td><td>GCAGCCGACTTAGAACTGGTGCGGACCAGGGGAATCCG</td><td>307.7</td><td>-12.1</td><td>1.6</td><td>-7.5</td><td>4.5E-14</td><td>2.2E-13</td><td>MB231/MB468</td><td>MP</td></tr>
<tr><td>>hg38_21_+_8433172_8446622_RNA45SN1_45S_50ntFlanks@4047.4074.28|ATTCGAACGTCTGCCCTATCAACTTTCG</td><td>>hg38_21_+_8433172_8446622_RNA45SN1_45S_50ntFlanks</td><td>4047</td><td>4074</td><td>28</td><td>ATTCGAACGTCTGCCCTATCAACTTTCG</td><td>127.2</td><td>-10.9</td><td>1.4</td><td>-7.5</td><td>4.5E-14</td><td>2.2E-13</td><td>MB231/MB468</td><td>MP</td></tr>
<tr><td>>hg38_21_+_8433172_8446622_RNA45SN1_45S_50ntFlanks@9739.9757.19|AAACGATCTCAACCTATTC</td><td>>hg38_21_+_8433172_8446622_RNA45SN1_45S_50ntFlanks</td><td>9739</td><td>9757</td><td>19</td><td>AAACGATCTCAACCTATTC</td><td>600.4</td><td>-4.6</td><td>0.6</td><td>-7.5</td><td>5.4E-14</td><td>2.2E-13</td><td>MB231/MB468</td><td>NUC</td></tr>
<tr><td>>hg38_21_+_8433172_8446622_RNA45SN1_45S_50ntFlanks@4025.4045.21|CCCCCCGTGGCGGCGACGACC</td><td>>hg38_21_+_8433172_8446622_RNA45SN1_45S_50ntFlanks</td><td>4025</td><td>4045</td><td>21</td><td>CCCCCCGTGGCGGCGACGACC</td><td>329.0</td><td>-12.1</td><td>1.6</td><td>-7.5</td><td>7.0E-14</td><td>2.2E-13</td><td>MB231/MB468</td><td>MITO</td></tr>
<tr><td>>hg38_21_+_8433172_8446622_RNA45SN1_45S_50ntFlanks@4367.4406.40|CAGTTAAAAAGCTCGTAGTTGGATCTTGGGAGCGGGCGGG</td><td>>hg38_21_+_8433172_8446622_RNA45SN1_45S_50ntFlanks</td><td>4367</td><td>4406</td><td>40</td><td>CAGTTAAAAAGCTCGTAGTTGGATCTTGGGAGCGGGCGGG</td><td>255.6</td><td>-11.8</td><td>1.6</td><td>-7.5</td><td>7.0E-14</td><td>2.2E-13</td><td>MB231/MB468</td><td>MITO</td></tr>
<tr><td>>trna111_HisGTG_1_-_147774845_147774916@-1T.23.24,trna118_HisGTG_1_-_145396881_145396952@-1T.23.24,trna16_HisGTG_1_+_146544773_146544844@-1T.23.24,trna1_HisGTG_15_+_45493349_45493420@-1T.23.24,trna21_HisGTG_1_+_147753471_147753542@-1T.23.24,trna33_HisGTG_6_+_27125906_27125977@-1T.23.24,trna7_HisGTG_9_-_14433938_14434009@-1T.23.24,trna8_HisGTG_15_-_45492611_45492682@-1T.23.24,trna9_HisGTG_15_-_45490804_45490875@-1T.23.24|TGCCGTGATCGTATAGTGGTTAGT</td><td>>trna111_HisGTG_1_-_147774845_147774916</td><td>-1T</td><td>23</td><td>24</td><td>TGCCGTGATCGTATAGTGGTTAGT</td><td>4510.7</td><td>9.1</td><td>1.2</td><td>7.5</td><td>7.0E-14</td><td>2.3E-13</td><td>MB231/MB468</td><td>MITO</td></tr>
<tr><td>>MI0000088|hsa-mir-30a&WithFlank&6|-|71403545|71403627@54.73.20(+1A)&[MIMAT0000088&hsa-miR-30a-3p&offsets|+1|-1(+1A);m-38&6|-|71403554|71403575&offsets|+1|-1(+1A)]|TTTCAGTCGGATGTTTGCAGA</td><td>>MI0000088|hsa-mir-30a&WithFlank&6|-|71403545|71403627</td><td>54</td><td>73</td><td>20(+1A)</td><td>TTTCAGTCGGATGTTTGCAGA</td><td>205.0</td><td>7.9</td><td>1.1</td><td>7.5</td><td>5.4E-14</td><td>2.3E-13</td><td>MB231/MB468</td><td>NUC</td></tr>
<tr><td>>hg38_1_-_228634819_228635039_RNA5S12_5S_50ntFlanks@126.171.46|TACTTGGATGGGAGACCGCCTGGGAATACCGGGTGCTGTAGGCTTT</td><td>>hg38_1_-_228634819_228635039_RNA5S12_5S_50ntFlanks</td><td>126</td><td>171</td><td>46</td><td>TACTTGGATGGGAGACCGCCTGGGAATACCGGGTGCTGTAGGCTTT</td><td>3215.5</td><td>7.4</td><td>1.0</td><td>7.5</td><td>5.2E-14</td><td>2.3E-13</td><td>MB231/MB468</td><td>CYTO</td></tr>
<tr><td>>hg38_21_+_8433172_8446622_RNA45SN1_45S_50ntFlanks@10548.10574.27|GTGCGGTAACGCGACCGATCCCGGAGA</td><td>>hg38_21_+_8433172_8446622_RNA45SN1_45S_50ntFlanks</td><td>10548</td><td>10574</td><td>27</td><td>GTGCGGTAACGCGACCGATCCCGGAGA</td><td>408.2</td><td>6.9</td><td>0.9</td><td>7.6</td><td>4.2E-14</td><td>2.3E-13</td><td>BT20/MB468</td><td>MITO</td></tr>
<tr><td>>hg38_21_+_8433172_8446622_RNA45SN1_45S_50ntFlanks@12625.12659.35|CAATGGGGCGAAGCTACCATCTGTGGGATTATGAC</td><td>>hg38_21_+_8433172_8446622_RNA45SN1_45S_50ntFlanks</td><td>12625</td><td>12659</td><td>35</td><td>CAATGGGGCGAAGCTACCATCTGTGGGATTATGAC</td><td>388.8</td><td>-12.5</td><td>1.7</td><td>-7.5</td><td>4.5E-14</td><td>2.3E-13</td><td>MB231/MB468</td><td>MP</td></tr>
<tr><td>>hg38_21_+_8433172_8446622_RNA45SN1_45S_50ntFlanks@3984.4004.21|GCGGCTTTGGTGACTCTAGAT</td><td>>hg38_21_+_8433172_8446622_RNA45SN1_45S_50ntFlanks</td><td>3984</td><td>4004</td><td>21</td><td>GCGGCTTTGGTGACTCTAGAT</td><td>194.4</td><td>-11.5</td><td>1.5</td><td>-7.5</td><td>4.5E-14</td><td>2.3E-13</td><td>MB231/MB468</td><td>MP</td></tr>
<tr><td>>MI0000102|hsa-mir-100&WithFlank&11|-|122152223|122152314@20.39.20&[MIMAT0000098&hsa-miR-100-5p&offsets|+1|-1;m-27&11|-|122152276|122152296&offsets|+1|0]|ACCCGTAGATCCGAACTTGT</td><td>>MI0000102|hsa-mir-100&WithFlank&11|-|122152223|122152314</td><td>20</td><td>39</td><td>20</td><td>ACCCGTAGATCCGAACTTGT</td><td>252.5</td><td>-11.2</td><td>1.4</td><td>-7.8</td><td>5.7E-15</td><td>2.3E-13</td><td>BT20/MB231</td><td>MITO</td></tr>
<tr><td>>MI0000076|hsa-mir-20a&WithFlank&13|+|91351059|91351141@14.34.21&[MIMAT0000075&hsa-miR-20a-5p&offsets|0|-2;m-96&13|+|91351072|91351094&offsets|0|-2]|TAAAGTGCTTATAGTGCAGGT</td><td>>MI0000076|hsa-mir-20a&WithFlank&13|+|91351059|91351141</td><td>14</td><td>34</td><td>21</td><td>TAAAGTGCTTATAGTGCAGGT</td><td>1013.4</td><td>6.0</td><td>0.8</td><td>7.5</td><td>5.4E-14</td><td>2.3E-13</td><td>MB231/MB468</td><td>NUC</td></tr>
<tr><td>>MI0000071|hsa-mir-17&WithFlank&13|+|91350599|91350694@20.41.22&[MIMAT0000070&hsa-miR-17-5p&offsets|0|-1;m-88&13|+|91350618|91350640&offsets|0|-1]|CAAAGTGCTTACAGTGCAGGTA</td><td>>MI0000071|hsa-mir-17&WithFlank&13|+|91350599|91350694</td><td>20</td><td>41</td><td>22</td><td>CAAAGTGCTTACAGTGCAGGTA</td><td>410.4</td><td>4.8</td><td>0.6</td><td>7.5</td><td>5.6E-14</td><td>2.3E-13</td><td>BT20/MB468</td><td>CYTO</td></tr>
<tr><td>>MI0016887|hsa-mir-4521&WithFlank&17|+|8186939|8187010@14.35.22&[MIMAT0019058&hsa-miR-4521&offsets|+4|+4;m-765&17|+|8186948|8186969&offsets|+4|+4]|AGGAAGTCCTGTGCTCAGTTTT</td><td>>MI0016887|hsa-mir-4521&WithFlank&17|+|8186939|8187010</td><td>14</td><td>35</td><td>22</td><td>AGGAAGTCCTGTGCTCAGTTTT</td><td>570.2</td><td>9.6</td><td>1.3</td><td>7.5</td><td>5.6E-14</td><td>2.3E-13</td><td>BT20/MB468</td><td>CYTO</td></tr>
<tr><td>>hg38_21_+_8433172_8446622_RNA45SN1_45S_50ntFlanks@11919.11967.49|CATGAGAGGTGTAGAATAAGTGGGAGGCCCCCGGCGCCCCCCCGGTGTC</td><td>>hg38_21_+_8433172_8446622_RNA45SN1_45S_50ntFlanks</td><td>11919</td><td>11967</td><td>49</td><td>CATGAGAGGTGTAGAATAAGTGGGAGGCCCCCGGCGCCCCCCCGGTGTC</td><td>841.1</td><td>-14.1</td><td>1.9</td><td>-7.5</td><td>8.3E-14</td><td>2.3E-13</td><td>BT20/MB468</td><td>NUC</td></tr>
<tr><td>>am_trna11_ProAGG_16_+_3241989_3242060@1.32.32,trna12_ProAGG_6_+_26555498_26555569@1.32.32,trna14_ProTGG_5_-_180615854_180615925@1.32.32,trna22_ProAGG_14_-_21081560_21081631@1.32.32,trna23_ProAGG_14_-_21077495_21077566@1.32.32,trna28_ProTGG_16_-_3234133_3234204@1.32.32,trna29_ProAGG_16_-_3232635_3232706@1.32.32,trna2_ProAGG_7_+_128423504_128423575@1.32.32,trna30_ProCGG_6_+_27059521_27059592@1.32.32,trna37_ProCGG_17_-_8126151_8126222@1.32.32,trna3_ProTGG_16_+_3208923_3208994@1.32.32,trna52_ProCGG_1_+_167683962_167684033@1.32.32,trna65_ProAGG_1_-_167684725_167684796@1.32.32,trna6_ProCGG_16_+_3222049_3222120@1.32.32,trna6_ProTGG_14_+_21152175_21152246@1.32.32,trna8_ProTGG_16_+_3238094_3238165@1.32.32,trna9_ProAGG_11_+_75946557_75946628@1.32.32,trna9_ProAGG_16_+_3239634_3239705@1.32.32|GGCTCGTTGGTCTAGGGGTATGATTCTCGCTT</td><td>>am_trna11_ProAGG_16_+_3241989_3242060</td><td>1</td><td>32</td><td>32</td><td>GGCTCGTTGGTCTAGGGGTATGATTCTCGCTT</td><td>278.5</td><td>10.1</td><td>1.3</td><td>7.5</td><td>5.6E-14</td><td>2.3E-13</td><td>BT20/MB468</td><td>CYTO</td></tr>
<tr><td>>hg38_21_+_8433172_8446622_RNA45SN1_45S_50ntFlanks@6648.6672.25|GTACGACTCTTAGCGGTGGATCACT</td><td>>hg38_21_+_8433172_8446622_RNA45SN1_45S_50ntFlanks</td><td>6648</td><td>6672</td><td>25</td><td>GTACGACTCTTAGCGGTGGATCACT</td><td>3152.2</td><td>6.4</td><td>0.8</td><td>7.6</td><td>4.3E-14</td><td>2.3E-13</td><td>BT20/MB468</td><td>MITO</td></tr>
<tr><td>>hg38_21_+_8433172_8446622_RNA45SN1_45S_50ntFlanks@7991.8018.28|ACGTGGCGACCCGCTGAATTTAAGCATA</td><td>>hg38_21_+_8433172_8446622_RNA45SN1_45S_50ntFlanks</td><td>7991</td><td>8018</td><td>28</td><td>ACGTGGCGACCCGCTGAATTTAAGCATA</td><td>1486.6</td><td>-5.1</td><td>0.7</td><td>-7.5</td><td>7.1E-14</td><td>2.3E-13</td><td>MB231/MB468</td><td>MITO</td></tr>
<tr><td>>hg38_21_+_8433172_8446622_RNA45SN1_45S_50ntFlanks@11665.11687.23|TTCTGCCCAGTGCTCTGAATGTC</td><td>>hg38_21_+_8433172_8446622_RNA45SN1_45S_50ntFlanks</td><td>11665</td><td>11687</td><td>23</td><td>TTCTGCCCAGTGCTCTGAATGTC</td><td>353.5</td><td>4.6</td><td>0.6</td><td>7.5</td><td>5.7E-14</td><td>2.3E-13</td><td>BT20/MB468</td><td>CYTO</td></tr>
</table>

| | | | | | | | | | | | | | |
|---|---|---|---|---|---|---|---|---|---|---|---|---|---|
| >hg38_21_+_8433172_8446622_RNA45SN1_45S_50ntFlanks@12991.13010.20\|CTCCCTCGCTGCGATCTATT | >hg38_21_+_8433172_8446622_RNA45SN1_45S_50ntFlanks | 12991 | 13010 | 20 | CTCCCTCGCTGCGATCTATT | 922.9 | -9.6 | 1.3 | -7.5 | 5.5E-14 | 2.3E-13 | MB231/MB468 | NUC |
| >hg38_21_+_8433172_8446622_RNA45SN1_45S_50ntFlanks@11161.11189.29\|CACGGTCCCCCGCGAGGGGGGCCCGGGCA | >hg38_21_+_8433172_8446622_RNA45SN1_45S_50ntFlanks | 11161 | 11189 | 29 | CACGGTCCCCCGCGAGGGGGGCCCGGGCA | 294.1 | -10.9 | 1.5 | -7.5 | 5.3E-14 | 2.3E-13 | MB231/MB468 | CYTO |
| >hg38_21_+_8433172_8446622_RNA45SN1_45S_50ntFlanks@9736.9765.30\|CCGAAACGATCTCAACCTATTCTCAAACTT | >hg38_21_+_8433172_8446622_RNA45SN1_45S_50ntFlanks | 9736 | 9765 | 30 | CCGAAACGATCTCAACCTATTCTCAAACTT | 146.8 | -11.1 | 1.5 | -7.5 | 4.6E-14 | 2.3E-13 | MB231/MB468 | MP |
| >MI0000088\|hsa-mir-30a&WithFlank&6\|-\|71403545\|71403627@53.73.21(+1A)&[MIMAT0000088&hsa-miR-30a-3p&offsets\|0\|-1(+1A);m-38&6\|-\|71403554\|71403575&offsets\|0\|-1(+1A)]\|CTTTCAGTCGGATGTTTGCAGA | >MI0000088\|hsa-mir-30a&WithFlank&6\|-\|71403545\|71403627 | 53 | 73 | 21(+1A) | CTTTCAGTCGGATGTTTGCAGA | 1159.6 | 8.1 | 1.1 | 7.5 | 5.7E-14 | 2.3E-13 | BT20/MB468 | CYTO |
| >hg38_1_-_228634819_228635039_RNA5S12_5S_50ntFlanks@51.75.25\|GTCTACGGCCATACCACCCTGAACG | >hg38_1_-_228634819_228635039_RNA5S12_5S_50ntFlanks | 51 | 75 | 25 | GTCTACGGCCATACCACCCTGAACG | 1605.1 | 5.1 | 0.7 | 7.5 | 5.7E-14 | 2.3E-13 | BT20/MB468 | CYTO |
| >hg38_21_+_8433172_8446622_RNA45SN1_45S_50ntFlanks@10678.10695.18\|CGGTTCCGGCGGCGTCCG | >hg38_21_+_8433172_8446622_RNA45SN1_45S_50ntFlanks | 10678 | 10695 | 18 | CGGTTCCGGCGGCGTCCG | 194.8 | -11.4 | 1.5 | -7.5 | 7.2E-14 | 2.3E-13 | MB231/MB468 | MITO |
| >MI0000088\|hsa-mir-30a&WithFlank&6\|-\|71403545\|71403627@54.73.20(+2U)&[MIMAT0000088&hsa-miR-30a-3p&offsets\|+1\|-1(+2U);m-38&6\|-\|71403554\|71403575&offsets\|+1\|-1(+2U)]\|TTTCAGTCGGATGTTTGCAGTT | >MI0000088\|hsa-mir-30a&WithFlank&6\|-\|71403545\|71403627 | 54 | 73 | 20(+2U) | TTTCAGTCGGATGTTTGCAGTT | 597.9 | 6.0 | 0.8 | 7.5 | 5.5E-14 | 2.3E-13 | MB231/MB468 | NUC |
| >hg38_21_+_8433172_8446622_RNA45SN1_45S_50ntFlanks@10410.10432.23\|TCAGTCGGTCCTGAGAGATGGGC | >hg38_21_+_8433172_8446622_RNA45SN1_45S_50ntFlanks | 10410 | 10432 | 23 | TCAGTCGGTCCTGAGAGATGGGC | 720.0 | -7.1 | 0.9 | -7.5 | 7.2E-14 | 2.3E-13 | MB231/MB468 | MITO |
| >MI0000087\|hsa-mir-29a&WithFlank&7\|-\|130876741\|130876816@47.69.23&[MIMAT0000086&hsa-miR-29a-3p&offsets\|-1\|0;m-26&7\|-\|130876749\|130876769&offsets\|-1\|+1]\|CTAGCACCATCTGAAATCGGTTA | >MI0000087\|hsa-mir-29a&WithFlank&7\|-\|130876741\|130876816 | 47 | 69 | 23 | CTAGCACCATCTGAAATCGGTTA | 937.8 | 8.8 | 1.2 | 7.6 | 4.3E-14 | 2.3E-13 | BT20/MB468 | MITO |
| >hg38_21_+_8433172_8446622_RNA45SN1_45S_50ntFlanks@11646.11670.25\|GGTGTTGACGCGATGTGATTTCTGC | >hg38_21_+_8433172_8446622_RNA45SN1_45S_50ntFlanks | 11646 | 11670 | 25 | GGTGTTGACGCGATGTGATTTCTGC | 402.2 | -12.5 | 1.7 | -7.5 | 4.6E-14 | 2.3E-13 | MB231/MB468 | MP |
| >hg38_21_+_8433172_8446622_RNA45SN1_45S_50ntFlanks@12625.12651.27\|CAATGGGGCGAAGCTACCATCTGTGGG | >hg38_21_+_8433172_8446622_RNA45SN1_45S_50ntFlanks | 12625 | 12651 | 27 | CAATGGGGCGAAGCTACCATCTGTGGG | 150.5 | -11.1 | 1.5 | -7.5 | 4.6E-14 | 2.3E-13 | MB231/MB468 | MP |
| >hg38_21_+_8433172_8446622_RNA45SN1_45S_50ntFlanks@11783.11807.25\|CATGAATGGATGAACGAGATTCCCA | >hg38_21_+_8433172_8446622_RNA45SN1_45S_50ntFlanks | 11783 | 11807 | 25 | CATGAATGGATGAACGAGATTCCCA | 249.8 | -11.7 | 1.6 | -7.5 | 7.2E-14 | 2.3E-13 | MB231/MB468 | MITO |
| >hg38_21_+_8433172_8446622_RNA45SN1_45S_50ntFlanks@9858.9899.42\|CGGGATGAACCGAACGCCGGGTTAAGGCGCCCGATGCCGACG | >hg38_21_+_8433172_8446622_RNA45SN1_45S_50ntFlanks | 9858 | 9899 | 42 | CGGGATGAACCGAACGCCGGGTTAAGGCGCCCGATGCCGACG | 3119.5 | -11.7 | 1.6 | -7.5 | 7.2E-14 | 2.3E-13 | MB231/MB468 | MITO |
| >MI0000279\|hsa-mir-196a-2&WithFlank&12\|+\|53991732\|53991853@31.50.20(+1A);MI0000238\|hsa-mir-196a-1&WithFlank&17\|-\|48632484\|48632565@13.32.20(+1A)&[MIMAT0000226&hsa-miR-196a-5p&offsets\|0\|-2(+1A);m-261&12\|+\|53991762\|53991783&offsets\|0\|-2(+1A)];[MIMAT0000226_1&hsa-miR-196a-5p&offsets\|0\|-2(+1A);m-268&17\|-\|48632532\|48632553&offsets\|0\|-2(+1A)]\|TAGGTAGTTTCATGTTGTTGA | >MI0000279\|hsa-mir-196a-2&WithFlank&12\|+\|53991732\|53991853 | 31 | 50 | 20(+1A) | TAGGTAGTTTCATGTTGTTGA | 313.4 | 11.7 | 1.5 | 7.8 | 4.2E-15 | 2.3E-13 | BT20/MB231 | CYTO |
| >hg38_21_+_8433172_8446622_RNA45SN1_45S_50ntFlanks@11572.11601.30\|CCGACTTAGAACTGGTGCGGACCAGGGGAA | >hg38_21_+_8433172_8446622_RNA45SN1_45S_50ntFlanks | 11572 | 11601 | 30 | CCGACTTAGAACTGGTGCGGACCAGGGGAA | 11176.4 | -5.6 | 0.7 | -7.5 | 5.8E-14 | 2.3E-13 | BT20/MB468 | CYTO |
| >hg38_21_+_8433172_8446622_RNA45SN1_45S_50ntFlanks@10684.10702.19\|CGGCGGCGTCCGGTGAGCT | >hg38_21_+_8433172_8446622_RNA45SN1_45S_50ntFlanks | 10684 | 10702 | 19 | CGGCGGCGTCCGGTGAGCT | 219.5 | -11.6 | 1.5 | -7.5 | 4.7E-14 | 2.3E-13 | MB231/MB468 | MP |
| >hg38_21_+_8433172_8446622_RNA45SN1_45S_50ntFlanks@4731.4751.21\|AAGAACGAAAGTCGGAGGTTC | >hg38_21_+_8433172_8446622_RNA45SN1_45S_50ntFlanks | 4731 | 4751 | 21 | AAGAACGAAAGTCGGAGGTTC | 3309.5 | -4.9 | 0.7 | -7.5 | 4.7E-14 | 2.3E-13 | MB231/MB468 | MP |
| >hg38_21_+_8433172_8446622_RNA45SN1_45S_50ntFlanks@10574.10612.39\|AAGCCGGCGGGAGCCCCGGGGAGAGTTCTCTTTTCTTTG | >hg38_21_+_8433172_8446622_RNA45SN1_45S_50ntFlanks | 10574 | 10612 | 39 | AAGCCGGCGGGAGCCCCGGGGAGAGTTCTCTTTTCTTTG | 274.5 | -11.1 | 1.5 | -7.5 | 5.6E-14 | 2.3E-13 | MB231/MB468 | NUC |
| >hg38_21_+_8433172_8446622_RNA45SN1_45S_50ntFlanks@3854.3871.18\|ACTGTGGTAATTCTAGAG | >hg38_21_+_8433172_8446622_RNA45SN1_45S_50ntFlanks | 3854 | 3871 | 18 | ACTGTGGTAATTCTAGAG | 501.7 | -12.1 | 1.5 | -7.9 | 2.0E-15 | 2.3E-13 | BT20/MB468 | MP |
| >hg38_21_+_8433172_8446622_RNA45SN1_45S_50ntFlanks@8153.8182.30\|GCCGCTCGTGGGGGGCCCAAGTCCTTCTGA | >hg38_21_+_8433172_8446622_RNA45SN1_45S_50ntFlanks | 8153 | 8182 | 30 | GCCGCTCGTGGGGGGCCCAAGTCCTTCTGA | 1432.6 | -10.3 | 1.4 | -7.5 | 7.3E-14 | 2.3E-13 | MB231/MB468 | MITO |
| >hg38_21_+_8433172_8446622_RNA45SN1_45S_50ntFlanks@3992.4023.32\|GGTGACTCTAGATAACCTCGGGCCGATCGCAC | >hg38_21_+_8433172_8446622_RNA45SN1_45S_50ntFlanks | 3992 | 4023 | 32 | GGTGACTCTAGATAACCTCGGGCCGATCGCAC | 583.8 | 5.4 | 0.7 | 7.5 | 5.4E-14 | 2.4E-13 | MB231/MB468 | CYTO |
| >trna119_LysCTT_1_-_145395522_145395594@1.30.30,trna11_LysCTT_5_-_180648979_180649051@1.30.30,trna13_LysCTT_6_+_26556774_26556846@1.30.30,trna32_LysCTT_16_-_3207406_3207478@1.30.30,trna7_LysCTT_16_+_3225692_3225764@1.30.30,trna9_LysCTT_5_+_180634755_180634827@1.30.30\|GCCCGGCTAGCTCAGTCGGTAGAGCATGAG | >trna119_LysCTT_1_-_145395522_145395594 | 1 | 30 | 30 | GCCCGGCTAGCTCAGTCGGTAGAGCATGAG | 37064.1 | -6.3 | 0.8 | -7.5 | 5.5E-14 | 2.4E-13 | MB231/MB468 | CYTO |
| >hg38_21_+_8433172_8446622_RNA45SN1_45S_50ntFlanks@4130.4162.33\|ATTCCGGAGAGGGAGCCTGAGAAACGGCTACCA | >hg38_21_+_8433172_8446622_RNA45SN1_45S_50ntFlanks | 4130 | 4162 | 33 | ATTCCGGAGAGGGAGCCTGAGAAACGGCTACCA | 421.5 | -10.8 | 1.4 | -7.5 | 4.4E-14 | 2.4E-13 | BT20/MB468 | MITO |
| >MI0003628\|hsa-mir-615&WithFlank&12\|+\|54033944\|54034051@67.86.20(+1A)&[MIMAT0003283&hsa-miR-615-3p&offsets\|0\|-2(+1A);m-495&12\|+\|54034010\|54034030&offsets\|0\|-1(+1A)]\|TCCGAGCCTGGGTCTCCCTCA | >MI0003628\|hsa-mir-615&WithFlank&12\|+\|54033944\|54034051 | 67 | 86 | 20(+1A) | TCCGAGCCTGGGTCTCCCTCA | 299.7 | 8.2 | 1.0 | 7.8 | 4.3E-15 | 2.4E-13 | BT20/MB231 | CYTO |
| >hg38_21_+_8433172_8446622_RNA45SN1_45S_50ntFlanks@12709.12742.34\|CCGCGGAGCCTCGGTTGGCCTCGGATAGCCGGTC | >hg38_21_+_8433172_8446622_RNA45SN1_45S_50ntFlanks | 12709 | 12742 | 34 | CCGCGGAGCCTCGGTTGGCCTCGGATAGCCGGTC | 591.3 | -12.3 | 1.5 | -7.9 | 2.1E-15 | 2.4E-13 | BT20/MB468 | MP |
| >hg38_21_+_8433172_8446622_RNA45SN1_45S_50ntFlanks@5524.5550.27\|GTAAAAGTCGTAACAAGGTTTCCGTAG | >hg38_21_+_8433172_8446622_RNA45SN1_45S_50ntFlanks | 5524 | 5550 | 27 | GTAAAAGTCGTAACAAGGTTTCCGTAG | 192.7 | -11.5 | 1.5 | -7.5 | 4.8E-14 | 2.4E-13 | MB231/MB468 | MP |
| >hg38_21_+_8433172_8446622_RNA45SN1_45S_50ntFlanks@12605.12624.20\|TATGTGCTTGGCTGAGGAGC | >hg38_21_+_8433172_8446622_RNA45SN1_45S_50ntFlanks | 12605 | 12624 | 20 | TATGTGCTTGGCTGAGGAGC | 668.2 | -5.1 | 0.7 | -7.5 | 4.8E-14 | 2.4E-13 | MB231/MB468 | MP |
| >hg38_21_+_8433172_8446622_RNA45SN1_45S_50ntFlanks@6718.6739.22\|GTGAATTGCAGGACACATTGAT | >hg38_21_+_8433172_8446622_RNA45SN1_45S_50ntFlanks | 6718 | 6739 | 22 | GTGAATTGCAGGACACATTGAT | 300.8 | 5.0 | 0.7 | 7.5 | 5.7E-14 | 2.4E-13 | MB231/MB468 | NUC |
| >hg38_21_+_8433172_8446622_RNA45SN1_45S_50ntFlanks@3854.3874.21\|ACTGTGGTAATTCTAGAGCTA | >hg38_21_+_8433172_8446622_RNA45SN1_45S_50ntFlanks | 3854 | 3874 | 21 | ACTGTGGTAATTCTAGAGCTA | 485.2 | -12.0 | 1.5 | -7.9 | 2.1E-15 | 2.4E-13 | BT20/MB468 | MP |
| >hg38_21_+_8433172_8446622_RNA45SN1_45S_50ntFlanks@11564.11586.23\|CCTAGCAGCCGACTTAGAACTGG | >hg38_21_+_8433172_8446622_RNA45SN1_45S_50ntFlanks | 11564 | 11586 | 23 | CCTAGCAGCCGACTTAGAACTGG | 1326.5 | -8.2 | 1.1 | -7.5 | 8.7E-14 | 2.4E-13 | BT20/MB468 | NUC |
| >hg38_21_+_8433172_8446622_RNA45SN1_45S_50ntFlanks@11215.11233.19\|CTCTGGACGCGAGCCGGGC | >hg38_21_+_8433172_8446622_RNA45SN1_45S_50ntFlanks | 11215 | 11233 | 19 | CTCTGGACGCGAGCCGGGC | 142.2 | -9.9 | 1.3 | -7.5 | 5.5E-14 | 2.4E-13 | MB231/MB468 | CYTO |
| >hg38_21_+_8433172_8446622_RNA45SN1_45S_50ntFlanks@12418.12460.43\|CGATGTCGGCTCTTCCTATCATTGTGAAGCAGAATTCACCAAG | >hg38_21_+_8433172_8446622_RNA45SN1_45S_50ntFlanks | 12418 | 12460 | 43 | CGATGTCGGCTCTTCCTATCATTGTGAAGCAGAATTCACCAAG | 1144.7 | -14.6 | 2.0 | -7.5 | 8.7E-14 | 2.4E-13 | BT20/MB468 | NUC |
| >MI0000434\|hsa-let-7i&WithFlank&12\|+\|62603680\|62603775@12.31.20(+1C)&[MIMAT0000415&hsa-let-7i-5p&offsets\|0\|-2(+1C);m-49&12\|+\|62603691\|62603712&offsets\|0\|-2(+1C)]\|TGAGGTAGTAGTTTGTGCTGC | >MI0000434\|hsa-let-7i&WithFlank&12\|+\|62603680\|62603775 | 12 | 31 | 20(+1C) | TGAGGTAGTAGTTTGTGCTGC | 454.1 | 6.3 | 0.8 | 7.5 | 5.5E-14 | 2.4E-13 | MB231/MB468 | CYTO |
| >hg38_21_+_8433172_8446622_RNA45SN1_45S_50ntFlanks@9470.9500.31\|GAGCACGAGCGCACGTGTTAGGACCCGAAAG | >hg38_21_+_8433172_8446622_RNA45SN1_45S_50ntFlanks | 9470 | 9500 | 31 | GAGCACGAGCGCACGTGTTAGGACCCGAAAG | 182.8 | -11.4 | 1.5 | -7.5 | 4.8E-14 | 2.4E-13 | MB231/MB468 | MP |
| >am_MI0000809\|hsa-mir-151a&WithFlank&8\|-\|140732558\|140732659@53.72.20(+1A)&[MIMAT0000757&hsa-miR-151a-3p&offsets\|0\|-1(+1A);m-30&8\|-\|140732587\|140732607&offsets\|0\|-1(+1A)]\|CTAGACTGAAGCTCCTTGAGA | >am_MI0000809\|hsa-mir-151a&WithFlank&8\|-\|140732558\|140732659 | 53 | 72 | 20(+1A) | CTAGACTGAAGCTCCTTGAGA | 3840.3 | 6.2 | 0.8 | 7.5 | 7.5E-14 | 2.4E-13 | MB231/MB468 | MITO |
| >MI0000063\|hsa-let-7b&WithFlank&22\|+\|46113680\|46113774@12.33.22(+1C)&[MIMAT0000063&hsa-let-7b-5p&offsets\|0\|0(+1C);m-14&22\|+\|46113691\|46113712&offsets\|0\|0(+1C)]\|TGAGGTAGTAGGTTGTGTGGTTC | >MI0000063\|hsa-let-7b&WithFlank&22\|+\|46113680\|46113774 | 12 | 33 | 22(+1C) | TGAGGTAGTAGGTTGTGTGGTTC | 700.1 | 9.4 | 1.2 | 7.5 | 4.5E-14 | 2.4E-13 | BT20/MB468 | MITO |
| >hg38_21_+_8433172_8446622_RNA45SN1_45S_50ntFlanks@10905.10928.24\|CTGGGCGCGCGCCGCGGCTGGACG | >hg38_21_+_8433172_8446622_RNA45SN1_45S_50ntFlanks | 10905 | 10928 | 24 | CTGGGCGCGCGCCGCGGCTGGACG | 172.5 | -11.3 | 1.5 | -7.5 | 4.8E-14 | 2.4E-13 | MB231/MB468 | MP |
| >hg38_21_+_8433172_8446622_RNA45SN1_45S_50ntFlanks@7990.8012.23\|GACGTGGCGACCCGCTGAATTTA | >hg38_21_+_8433172_8446622_RNA45SN1_45S_50ntFlanks | 7990 | 8012 | 23 | GACGTGGCGACCCGCTGAATTTA | 1126.9 | -5.4 | 0.7 | -7.5 | 4.8E-14 | 2.4E-13 | MB231/MB468 | MP |
| >hg38_21_+_8433172_8446622_RNA45SN1_45S_50ntFlanks@11886.11912.27\|TGAGCTTGACTCTAGTCTGGCACGGTG | >hg38_21_+_8433172_8446622_RNA45SN1_45S_50ntFlanks | 11886 | 11912 | 27 | TGAGCTTGACTCTAGTCTGGCACGGTG | 283.2 | -12.0 | 1.6 | -7.5 | 4.8E-14 | 2.4E-13 | MB231/MB468 | MP |
| >hg38_1_-_228634819_228635039_RNA5S12_5S_50ntFlanks@84.112.29\|CTCGTCTGATCTCGGAAGCTAAGCAGGGT | >hg38_1_-_228634819_228635039_RNA5S12_5S_50ntFlanks | 84 | 112 | 29 | CTCGTCTGATCTCGGAAGCTAAGCAGGGT | 197.0 | -11.4 | 1.5 | -7.5 | 7.6E-14 | 2.4E-13 | MB231/MB468 | MITO |
| >hg38_21_+_8433172_8446622_RNA45SN1_45S_50ntFlanks@11209.11227.19\|CGGCGACTCTGGACGCGAG | >hg38_21_+_8433172_8446622_RNA45SN1_45S_50ntFlanks | 11209 | 11227 | 19 | CGGCGACTCTGGACGCGAG | 864.9 | -9.0 | 1.2 | -7.5 | 7.6E-14 | 2.4E-13 | MB231/MB468 | MITO |
| >hg38_21_+_8433172_8446622_RNA45SN1_45S_50ntFlanks@5379.5413.35\|ATTAAGTCCCTGCCCTTTGTACACACCGCCCGTCG | >hg38_21_+_8433172_8446622_RNA45SN1_45S_50ntFlanks | 5379 | 5413 | 35 | ATTAAGTCCCTGCCCTTTGTACACACCGCCCGTCG | 458.9 | -12.6 | 1.7 | -7.5 | 7.6E-14 | 2.4E-13 | MB231/MB468 | MITO |
| >MI0003205\|hsa-mir-532&WithFlank&X\|+\|50003142\|50003244@26.46.21(+1A)&[MIMAT0002888&hsa-miR-532-5p&offsets\|0\|-1(+1A);m-65&X\|+\|50003167\|50003188&offsets\|0\|-1(+1A)]\|CATGCCTTGAGTGTAGGACCGA | >MI0003205\|hsa-mir-532&WithFlank&X\|+\|50003142\|50003244 | 26 | 46 | 21(+1A) | CATGCCTTGAGTGTAGGACCGA | 350.1 | 7.0 | 0.9 | 7.5 | 6.0E-14 | 2.4E-13 | BT20/MB468 | CYTO |
| >hg38_21_+_8433172_8446622_RNA45SN1_45S_50ntFlanks@10409.10435.27\|GTCAGTCGGTCCTGAGAGATGGGCGAG | >hg38_21_+_8433172_8446622_RNA45SN1_45S_50ntFlanks | 10409 | 10435 | 27 | GTCAGTCGGTCCTGAGAGATGGGCGAG | 218.9 | -7.2 | 1.0 | -7.5 | 5.6E-14 | 2.4E-13 | MB231/MB468 | CYTO |
| >hg38_21_+_8433172_8446622_RNA45SN1_45S_50ntFlanks@10130.10149.20\|GCGGCGGCGGCGGCGGCGGG | >hg38_21_+_8433172_8446622_RNA45SN1_45S_50ntFlanks | 10130 | 10149 | 20 | GCGGCGGCGGCGGCGGCGGG | 484.3 | -12.7 | 1.7 | -7.5 | 7.6E-14 | 2.4E-13 | MB231/MB468 | MITO |
| >hg38_21_+_8433172_8446622_RNA45SN1_45S_50ntFlanks@8847.8866.20\|GCGGCGGCGGCGGCGGCGGG | >hg38_21_+_8433172_8446622_RNA45SN1_45S_50ntFlanks | 8847 | 8866 | 20 | GCGGCGGCGGCGGCGGCGGG | 484.3 | -12.7 | 1.7 | -7.5 | 7.6E-14 | 2.4E-13 | MB231/MB468 | MITO |

| | | | | | | | | | | | | | |
|---|---|---|---|---|---|---|---|---|---|---|---|---|---|
| >MI0000298\|hsa-mir-221&WithFlank&X\|-\|45746151\|45746272@71.91.21(+1C)&[MIMAT0000278&hsa-miR-221-3p&offsets\|0\|-2(+1C);m-91&X\|-\|45746181\|45746202&offsets\|0\|-1(+1C)]\|AGCTACATTGTCTGCTGGGTTC | >MI0000298\|hsa-mir-221&WithFlank&X\|-\|45746151\|45746272 | 71 | 91 | 21(+1C) | AGCTACATTGTCTGCTGGGTTC | 4856.5 | 7.3 | 1.0 | 7.5 | 7.6E-14 | 2.4E-13 | MB231/MB468 | MITO |
| >hg38_21_+_8433172_8446622_RNA45SN1_45S_50ntFlanks@10475.10492.18\|CGGCCGATCGAAAGGGAG | >hg38_21_+_8433172_8446622_RNA45SN1_45S_50ntFlanks | 10475 | 10492 | 18 | CGGCCGATCGAAAGGGAG | 170.7 | -11.3 | 1.5 | -7.5 | 4.9E-14 | 2.4E-13 | MB231/MB468 | MP |
| >hg38_21_+_8433172_8446622_RNA45SN1_45S_50ntFlanks@12405.12446.42\|CTTTTTGATCCTTCGATGTCGGCTCTTCCTATCATTGTGAAG | >hg38_21_+_8433172_8446622_RNA45SN1_45S_50ntFlanks | 12405 | 12446 | 42 | CTTTTTGATCCTTCGATGTCGGCTCTTCCTATCATTGTGAAG | 7874.9 | -7.4 | 1.0 | -7.5 | 5.8E-14 | 2.4E-13 | MB231/MB468 | NUC |
| >hg38_21_+_8433172_8446622_RNA45SN1_45S_50ntFlanks@10663.10712.50\|CCTTGGAAAGCGTCGCGGTTCCGGCGGCGTCCGGTGAGCTCTCGCTGGCC | >hg38_21_+_8433172_8446622_RNA45SN1_45S_50ntFlanks | 10663 | 10712 | 50 | CCTTGGAAAGCGTCGCGGTTCCGGCGGCGTCCGGTGAGCTCTCGCTGGCC | 290.2 | -12.0 | 1.6 | -7.5 | 7.7E-14 | 2.4E-13 | MB231/MB468 | MITO |
| >hg38_21_+_8433172_8446622_RNA45SN1_45S_50ntFlanks@9862.9899.38\|ATGAACCGAACGCCGGGTTAAGGCGCCCGATGCCGACG | >hg38_21_+_8433172_8446622_RNA45SN1_45S_50ntFlanks | 9862 | 9899 | 38 | ATGAACCGAACGCCGGGTTAAGGCGCCCGATGCCGACG | 765.4 | -13.4 | 1.8 | -7.5 | 7.7E-14 | 2.4E-13 | MB231/MB468 | MITO |
| >hg38_21_+_8433172_8446622_RNA45SN1_45S_50ntFlanks@10802.10823.22\|TTGGAACAATGTAGGTAAGGGA | >hg38_21_+_8433172_8446622_RNA45SN1_45S_50ntFlanks | 10802 | 10823 | 22 | TTGGAACAATGTAGGTAAGGGA | 142.0 | -9.9 | 1.3 | -7.5 | 5.7E-14 | 2.4E-13 | MB231/MB468 | CYTO |
| >hg38_21_+_8433172_8446622_RNA45SN1_45S_50ntFlanks@3705.3748.44\|TACCTGGTTGATCCTGCCAGTAGCATATGCTTGTCTCAAAGATT | >hg38_21_+_8433172_8446622_RNA45SN1_45S_50ntFlanks | 3705 | 3748 | 44 | TACCTGGTTGATCCTGCCAGTAGCATATGCTTGTCTCAAAGATT | 2190.6 | 5.8 | 0.8 | 7.5 | 5.7E-14 | 2.5E-13 | MB231/MB468 | CYTO |
| >hg38_21_+_8433172_8446622_RNA45SN1_45S_50ntFlanks@3985.4004.20\|CGGCTTTGGTGACTCTAGAT | >hg38_21_+_8433172_8446622_RNA45SN1_45S_50ntFlanks | 3985 | 4004 | 20 | CGGCTTTGGTGACTCTAGAT | 420.7 | -9.0 | 1.2 | -7.5 | 5.9E-14 | 2.5E-13 | MB231/MB468 | NUC |
| >hg38_21_+_8433172_8446622_RNA45SN1_45S_50ntFlanks@9459.9494.36\|CCGGGGAGGTGGAGCACGAGCGCACGTGTTAGGACC | >hg38_21_+_8433172_8446622_RNA45SN1_45S_50ntFlanks | 9459 | 9494 | 36 | CCGGGGAGGTGGAGCACGAGCGCACGTGTTAGGACC | 242.9 | -11.7 | 1.6 | -7.5 | 7.7E-14 | 2.5E-13 | MB231/MB468 | MITO |
| >am_MI0005763\|hsa-mir-941-1&WithFlank&20\|+\|63919443\|63919526@53.75.23(+1U);MI0005764\|hsa-mir-941-2&WithFlank&20\|+\|63919499\|63919582@53.75.23(+1U);MI0005765\|hsa-mir-941-3&WithFlank&20\|+\|63919555\|63919638@53.75.23(+1U);MI0005766\|hsa-mir-941-4&WithFlank&20\|+\|63919750\|63919833@53.75.23(+1U);MI0031520\|hsa-mir-941-5&WithFlank&20\|+\|63919862\|63919945@53.75.23(+1U)&[MIMAT0004984&hsa-miR-941&offsets\|0\|0(+1U);m-263&20\|+\|63919495\|63919517&offsets\|0\|0(+1U)];[MIMAT0004984_1&hsa-miR-941&offsets\|0\|0(+1U);m-263&20\|+\|63919551\|63919573&offsets\|0\|0(+1U)];[MIMAT0004984_2&hsa-miR-941&offsets\|0\|0(+1U);m-263&20\|+\|63919607\|63919629&offsets\|0\|0(+1U)];[MIMAT0004984_3&hsa-miR-941&offsets\|0\|0(+1U);m-263&20\|+\|63919802\|63919824&offsets\|0\|0(+1U)];[MIMAT0004984_4&hsa-miR-941&offsets\|0\|0(+1U);m-263&20\|+\|63919914\|63919936&offsets\|0\|0(+1U)]\|CACCCGGCTGTGTGCACATGTGCT | >am_MI0005763\|hsa-mir-941-1&WithFlank&20\|+\|63919443\|63919526 | 53 | 75 | 23(+1U) | CACCCGGCTGTGTGCACATGTGCT | 397.3 | 5.7 | 0.8 | 7.5 | 5.9E-14 | 2.5E-13 | MB231/MB468 | NUC |
| >hg38_21_+_8433172_8446622_RNA45SN1_45S_50ntFlanks@5225.5244.20\|CACGCGCGCTACACTGACTG | >hg38_21_+_8433172_8446622_RNA45SN1_45S_50ntFlanks | 5225 | 5244 | 20 | CACGCGCGCTACACTGACTG | 1316.2 | -14.1 | 1.9 | -7.5 | 7.8E-14 | 2.5E-13 | MB231/MB468 | MITO |
| >am_MI0003639\|hsa-mir-625&WithFlank&14\|+\|65471096\|65471192@58.80.23&[MIMAT0004808&hsa-miR-625-3p&offsets\|0\|+1;m-137&14\|+\|65471153\|65471174&offsets\|0\|+1]\|GACTATAGAACTTTCCCCCTCAT | >am_MI0003639\|hsa-mir-625&WithFlank&14\|+\|65471096\|65471192 | 58 | 80 | 23 | GACTATAGAACTTTCCCCCTCAT | 322.5 | 11.6 | 1.5 | 7.5 | 7.8E-14 | 2.5E-13 | MB231/MB468 | MITO |
| >hg38_21_+_8433172_8446622_RNA45SN1_45S_50ntFlanks@10408.10457.50\|GGTCAGTCGGTCCTGAGAGATGGGCGAGCGCCGTTCCGAAGGGACGGGCG | >hg38_21_+_8433172_8446622_RNA45SN1_45S_50ntFlanks | 10408 | 10457 | 50 | GGTCAGTCGGTCCTGAGAGATGGGCGAGCGCCGTTCCGAAGGGACGGGCG | 429.0 | -13.2 | 1.8 | -7.5 | 9.0E-14 | 2.5E-13 | BT20/MB468 | NUC |
| >MI0000299\|hsa-mir-222&WithFlank&X\|-\|45747009\|45747130@75.99.25(+1C)&[MIMAT0000279&hsa-miR-222-3p&offsets\|0\|+4(+1C);m-140&X\|-\|45747033\|45747056&offsets\|0\|+1(+1C)]\|AGCTACATCTGGCTACTGGGTCTCTC | >MI0000299\|hsa-mir-222&WithFlank&X\|-\|45747009\|45747130 | 75 | 99 | 25(+1C) | AGCTACATCTGGCTACTGGGTCTCTC | 359.6 | 6.1 | 0.8 | 7.5 | 5.9E-14 | 2.5E-13 | MB231/MB468 | NUC |
| >trna10_AspGTC_12_-_125424193_125424264@31.71.41,trna12_AspGTC_12_-_125411891_125411962@31.71.41,trna38_AspGTC_17_-_8125556_8125627@31.71.41,trna45_AspGTC_6_+_27447453_27447524@31.71.41,trna48_AspGTC_6_+_27471523_27471594@31.71.41,trna4_AspGTC_12_+_96429799_96429870@31.71.41,trna69_AspGTC_1_-_161440205_161440276@31.71.41,trna72_AspGTC_1_-_161432824_161432895@31.71.41,trna75_AspGTC_1_-_161425414_161425485@31.71.41,trna78_AspGTC_1_-_161418033_161418104@31.71.41,trna81_AspGTC_1_-_161410615_161410686@31.71.41\|CCTGTCACGCGGGAGACCGGGGTTCGATTCCCCGACGGGGA | >trna10_AspGTC_12_-_125424193_125424264 | 31 | 71 | 41 | CCTGTCACGCGGGAGACCGGGGTTCGATTCCCCGACGGGGA | 237.8 | -10.6 | 1.4 | -7.5 | 5.8E-14 | 2.5E-13 | MB231/MB468 | CYTO |
| >hg38_21_+_8433172_8446622_RNA45SN1_45S_50ntFlanks@11641.11681.41\|CGGCGGGTGTTGACGCGATGTGATTTCTGCCCAGTGCTCTG | >hg38_21_+_8433172_8446622_RNA45SN1_45S_50ntFlanks | 11641 | 11681 | 41 | CGGCGGGTGTTGACGCGATGTGATTTCTGCCCAGTGCTCTG | 411.9 | -12.6 | 1.7 | -7.5 | 5.0E-14 | 2.5E-13 | MB231/MB468 | MP |
| >hg38_21_+_8433172_8446622_RNA45SN1_45S_50ntFlanks@9545.9562.18\|GTGGAGGTCCGTAGCGGT | >hg38_21_+_8433172_8446622_RNA45SN1_45S_50ntFlanks | 9545 | 9562 | 18 | GTGGAGGTCCGTAGCGGT | 529.7 | -7.1 | 0.9 | -7.5 | 6.0E-14 | 2.5E-13 | MB231/MB468 | NUC |
| >hg38_21_+_8433172_8446622_RNA45SN1_45S_50ntFlanks@10410.10440.31\|TCAGTCGGTCCTGAGAGATGGGCGAGCGCCG | >hg38_21_+_8433172_8446622_RNA45SN1_45S_50ntFlanks | 10410 | 10440 | 31 | TCAGTCGGTCCTGAGAGATGGGCGAGCGCCG | 227.0 | -5.1 | 0.7 | -7.5 | 5.8E-14 | 2.5E-13 | MB231/MB468 | CYTO |
| >hg38_21_+_8433172_8446622_RNA45SN1_45S_50ntFlanks@8839.8859.21\|CGGTGGCGGCGGCGGCGGCGG | >hg38_21_+_8433172_8446622_RNA45SN1_45S_50ntFlanks | 8839 | 8859 | 21 | CGGTGGCGGCGGCGGCGGCGG | 242.9 | -11.8 | 1.6 | -7.5 | 5.0E-14 | 2.5E-13 | MB231/MB468 | MP |
| >hg38_21_+_8433172_8446622_RNA45SN1_45S_50ntFlanks@7981.7998.18\|CTCAGATCAGACGTGGCG | >hg38_21_+_8433172_8446622_RNA45SN1_45S_50ntFlanks | 7981 | 7998 | 18 | CTCAGATCAGACGTGGCG | 128.0 | -10.9 | 1.4 | -7.5 | 5.0E-14 | 2.5E-13 | MB231/MB468 | MP |
| >hg38_21_+_8433172_8446622_RNA45SN1_45S_50ntFlanks@11104.11131.28\|AAGGGTCGGGGCGGCAGGGGCCGGCGGC | >hg38_21_+_8433172_8446622_RNA45SN1_45S_50ntFlanks | 11104 | 11131 | 28 | AAGGGTCGGGGCGGCAGGGGCCGGCGGC | 237.2 | -11.8 | 1.6 | -7.5 | 5.0E-14 | 2.5E-13 | MB231/MB468 | MP |
| >hg38_21_+_8433172_8446622_RNA45SN1_45S_50ntFlanks@9546.9573.28\|TGGAGGTCCGTAGCGGTCCTGACGTGCA | >hg38_21_+_8433172_8446622_RNA45SN1_45S_50ntFlanks | 9546 | 9573 | 28 | TGGAGGTCCGTAGCGGTCCTGACGTGCA | 385.3 | -4.2 | 0.6 | -7.5 | 9.1E-14 | 2.5E-13 | BT20/MB468 | NUC |
| >hg38_21_+_8433172_8446622_RNA45SN1_45S_50ntFlanks@12938.12956.19\|ACCATTCGTAGACGACCTG | >hg38_21_+_8433172_8446622_RNA45SN1_45S_50ntFlanks | 12938 | 12956 | 19 | ACCATTCGTAGACGACCTG | 1317.2 | -9.5 | 1.3 | -7.5 | 5.0E-14 | 2.5E-13 | MB231/MB468 | MP |
| >hg38_21_+_8433172_8446622_RNA45SN1_45S_50ntFlanks@10663.10711.49\|CCTTGGAAAGCGTCGCGGTTCCGGCGGCGTCCGGTGAGCTCTCGCTGGC | >hg38_21_+_8433172_8446622_RNA45SN1_45S_50ntFlanks | 10663 | 10711 | 49 | CCTTGGAAAGCGTCGCGGTTCCGGCGGCGTCCGGTGAGCTCTCGCTGGC | 1672.4 | -15.1 | 2.0 | -7.5 | 9.2E-14 | 2.5E-13 | BT20/MB468 | NUC |
| >hg38_21_+_8433172_8446622_RNA45SN1_45S_50ntFlanks@10805.10822.18\|GAACAATGTAGGTAAGGG | >hg38_21_+_8433172_8446622_RNA45SN1_45S_50ntFlanks | 10805 | 10822 | 18 | GAACAATGTAGGTAAGGG | 225.4 | -10.8 | 1.4 | -7.5 | 6.0E-14 | 2.5E-13 | MB231/MB468 | NUC |
| >MI0000269\|hsa-mir-181a-2&WithFlank&9\|+\|124692436\|124692557@83.104.22&[MIMAT0004558&hsa-miR-181a-2-3p&offsets\|0\|0;m-71&9\|+\|124692518\|124692539&offsets\|0\|0]\|ACCACTGACCGTTGACTGTACC | >MI0000269\|hsa-mir-181a-2&WithFlank&9\|+\|124692436\|124692557 | 83 | 104 | 22 | ACCACTGACCGTTGACTGTACC | 1015.8 | 6.5 | 0.9 | 7.5 | 7.9E-14 | 2.5E-13 | MB231/MB468 | MITO |
| >hg38_21_+_8433172_8446622_RNA45SN1_45S_50ntFlanks@10871.10892.22\|GGCTGGGTCGGTCGGGCTGGGG | >hg38_21_+_8433172_8446622_RNA45SN1_45S_50ntFlanks | 10871 | 10892 | 22 | GGCTGGGTCGGTCGGGCTGGGG | 170.0 | -10.1 | 1.3 | -7.5 | 5.8E-14 | 2.5E-13 | MB231/MB468 | CYTO |
| >hg38_21_+_8433172_8446622_RNA45SN1_45S_50ntFlanks@12456.12475.20\|CCAAGCGTTGGATTGTTCAC | >hg38_21_+_8433172_8446622_RNA45SN1_45S_50ntFlanks | 12456 | 12475 | 20 | CCAAGCGTTGGATTGTTCAC | 13775.7 | -11.4 | 1.5 | -7.5 | 5.0E-14 | 2.5E-13 | MB231/MB468 | MP |
| >hg38_21_+_8433172_8446622_RNA45SN1_45S_50ntFlanks@4241.4258.18\|CTCTTTCGAGGCCCTGTA | >hg38_21_+_8433172_8446622_RNA45SN1_45S_50ntFlanks | 4241 | 4258 | 18 | CTCTTTCGAGGCCCTGTA | 755.3 | -9.8 | 1.3 | -7.5 | 9.2E-14 | 2.5E-13 | BT20/MB468 | NUC |
| >hg38_21_+_8433172_8446622_RNA45SN1_45S_50ntFlanks@8104.8139.36\|CCGCGGCGGGGCGCGGGACATGTGGCGTACGGAAGA | >hg38_21_+_8433172_8446622_RNA45SN1_45S_50ntFlanks | 8104 | 8139 | 36 | CCGCGGCGGGGCGCGGGACATGTGGCGTACGGAAGA | 537.8 | -9.1 | 1.2 | -7.5 | 7.9E-14 | 2.5E-13 | MB231/MB468 | MITO |
| >hg38_21_+_8433172_8446622_RNA45SN1_45S_50ntFlanks@7974.8010.37\|ACGCGACCTCAGATCAGACGTGGCGACCCGCTGAATT | >hg38_21_+_8433172_8446622_RNA45SN1_45S_50ntFlanks | 7974 | 8010 | 37 | ACGCGACCTCAGATCAGACGTGGCGACCCGCTGAATT | 9966.8 | 4.4 | 0.6 | 7.5 | 6.1E-14 | 2.5E-13 | MB231/MB468 | NUC |
| >hg38_21_+_8433172_8446622_RNA45SN1_45S_50ntFlanks@10349.10377.29\|AACGAGAACTTTGAAGGCCGAAGTGGAGA | >hg38_21_+_8433172_8446622_RNA45SN1_45S_50ntFlanks | 10349 | 10377 | 29 | AACGAGAACTTTGAAGGCCGAAGTGGAGA | 1910.1 | -4.2 | 0.6 | -7.5 | 6.1E-14 | 2.5E-13 | MB231/MB468 | NUC |
| >hg38_21_+_8433172_8446622_RNA45SN1_45S_50ntFlanks@3992.4026.35\|GGTGACTCTAGATAACCTCGGGCCGATCGCACGCC | >hg38_21_+_8433172_8446622_RNA45SN1_45S_50ntFlanks | 3992 | 4026 | 35 | GGTGACTCTAGATAACCTCGGGCCGATCGCACGCC | 246.2 | -11.8 | 1.6 | -7.5 | 5.1E-14 | 2.5E-13 | MB231/MB468 | MP |
| >hg38_21_+_8433172_8446622_RNA45SN1_45S_50ntFlanks@4772.4793.22\|GTAGTTCCGACCATAAACGATG | >hg38_21_+_8433172_8446622_RNA45SN1_45S_50ntFlanks | 4772 | 4793 | 22 | GTAGTTCCGACCATAAACGATG | 235.7 | -11.7 | 1.6 | -7.5 | 5.1E-14 | 2.5E-13 | MB231/MB468 | MP |
| >MI0000294\|hsa-mir-218-1&WithFlank&4\|+\|20528269\|20528390@31.51.21;MI0000295\|hsa-mir-218-2&WithFlank&5\|-\|168768140\|168768261@31.51.21&[MIMAT0000275&hsa-miR-218-5p&offsets\|0\|0;m-185&4\|+\|20528299\|20528319&offsets\|0\|0];[MIMAT0000275_1&hsa-miR-218-5p&offsets\|0\|0;m-187&5\|-\|168768211\|168768231&offsets\|0\|0]\|TTGTGCTTGATCTAACCATGT | >MI0000294\|hsa-mir-218-1&WithFlank&4\|+\|20528269\|20528390 | 31 | 51 | 21 | TTGTGCTTGATCTAACCATGT | 312.1 | -11.4 | 1.5 | -7.8 | 7.2E-15 | 2.5E-13 | BT20/MB231 | NUC |
| >MI0000477\|hsa-mir-146a&WithFlank&5\|+\|160485346\|160485456@27.48.22&[MIMAT0000449&hsa-miR-146a-5p&offsets\|0\|0;m-129&5\|+\|160485372\|160485393&offsets\|0\|0]\|TGAGAACTGAATTCCATGGGTT | >MI0000477\|hsa-mir-146a&WithFlank&5\|+\|160485346\|160485456 | 27 | 48 | 22 | TGAGAACTGAATTCCATGGGTT | 19380.2 | -10.4 | 1.3 | -7.8 | 7.2E-15 | 2.5E-13 | BT20/MB231 | NUC |
| >hg38_21_+_8433172_8446622_RNA45SN1_45S_50ntFlanks@3920.3952.33\|CATTTATCAGATCAAAACCAACCCGGTCAGCCC | >hg38_21_+_8433172_8446622_RNA45SN1_45S_50ntFlanks | 3920 | 3952 | 33 | CATTTATCAGATCAAAACCAACCCGGTCAGCCC | 247.4 | -11.7 | 1.6 | -7.5 | 8.0E-14 | 2.5E-13 | MB231/MB468 | MITO |
| >hg38_21_+_8433172_8446622_RNA45SN1_45S_50ntFlanks@13002.13025.24\|CGATCTATTGAAAGTCAGCCCTCG | >hg38_21_+_8433172_8446622_RNA45SN1_45S_50ntFlanks | 13002 | 13025 | 24 | CGATCTATTGAAAGTCAGCCCTCG | 198.7 | -11.4 | 1.5 | -7.5 | 8.0E-14 | 2.5E-13 | MB231/MB468 | MITO |
| >hg38_21_+_8433172_8446622_RNA45SN1_45S_50ntFlanks@12456.12491.36\|CCAAGCGTTGGATTGTTCACCCACTAATAGGGAACG | >hg38_21_+_8433172_8446622_RNA45SN1_45S_50ntFlanks | 12456 | 12491 | 36 | CCAAGCGTTGGATTGTTCACCCACTAATAGGGAACG | 333.2 | -12.2 | 1.6 | -7.5 | 5.1E-14 | 2.5E-13 | MB231/MB468 | MP |
| >MI0000078\|hsa-mir-22&WithFlank&17\|-\|1713897\|1713993@59.80.22&[MIMAT0000077&hsa-miR-22-3p&offsets\|0\|0;m-2&17\|-\|1713914\|1713935&offsets\|0\|0]\|AAGCTGCCAGTTGAAGAACTGT | >MI0000078\|hsa-mir-22&WithFlank&17\|-\|1713897\|1713993 | 59 | 80 | 22 | AAGCTGCCAGTTGAAGAACTGT | 12734.5 | 6.9 | 0.9 | 7.5 | 8.0E-14 | 2.5E-13 | MB231/MB468 | MITO |
| >hg38_21_+_8433172_8446622_RNA45SN1_45S_50ntFlanks@12274.12305.32\|CAGTACGAATACAGACCGTGAAAGCGGGGCCT | >hg38_21_+_8433172_8446622_RNA45SN1_45S_50ntFlanks | 12274 | 12305 | 32 | CAGTACGAATACAGACCGTGAAAGCGGGGCCT | 459.2 | -12.6 | 1.7 | -7.5 | 8.0E-14 | 2.5E-13 | MB231/MB468 | MITO |
| >hg38_21_+_8433172_8446622_RNA45SN1_45S_50ntFlanks@11575.11602.28\|ACTTAGAACTGGTGCGGACCAGGGGAAT | >hg38_21_+_8433172_8446622_RNA45SN1_45S_50ntFlanks | 11575 | 11602 | 28 | ACTTAGAACTGGTGCGGACCAGGGGAAT | 2527.8 | -8.8 | 1.2 | -7.5 | 9.3E-14 | 2.5E-13 | BT20/MB468 | NUC |

<table>
<tr><th></th><th></th><th></th><th></th><th></th><th></th><th></th><th></th><th></th><th></th><th></th><th></th><th></th><th></th></tr>
<tr><td>>hg38_21_+_8433172_8446622_RNA45SN1_45S_50ntFlanks@10866.10890.25|CTAAGGGCTGGGTCGGTCGGGCTGG</td><td>>hg38_21_+_8433172_8446622_RNA45SN1_45S_50ntFlanks</td><td>10866</td><td>10890</td><td>25</td><td>CTAAGGGCTGGGTCGGTCGGGCTGG</td><td>800.7</td><td>-9.4</td><td>1.2</td><td>-7.5</td><td>5.1E-14</td><td>2.5E-13</td><td>MB231/MB468</td><td>MP</td></tr>
<tr><td>>hg38_21_+_8433172_8446622_RNA45SN1_45S_50ntFlanks@12841.12861.21|CGGAGTGCCCTTCGTCCTGGG</td><td>>hg38_21_+_8433172_8446622_RNA45SN1_45S_50ntFlanks</td><td>12841</td><td>12861</td><td>21</td><td>CGGAGTGCCCTTCGTCCTGGG</td><td>580.8</td><td>-13.0</td><td>1.7</td><td>-7.5</td><td>5.1E-14</td><td>2.5E-13</td><td>MB231/MB468</td><td>MP</td></tr>
<tr><td>>hg38_21_+_8433172_8446622_RNA45SN1_45S_50ntFlanks@5420.5445.26|CCGATTGGATGGTTTAGTGAGGCCCT</td><td>>hg38_21_+_8433172_8446622_RNA45SN1_45S_50ntFlanks</td><td>5420</td><td>5445</td><td>26</td><td>CCGATTGGATGGTTTAGTGAGGCCCT</td><td>622.1</td><td>-11.4</td><td>1.5</td><td>-7.5</td><td>4.8E-14</td><td>2.5E-13</td><td>BT20/MB468</td><td>MITO</td></tr>
<tr><td>>MI0000086|hsa-mir-28&WithFlank&3|+|188688775|188688872@20.41.22(+1A)&[MIMAT0000085&hsa-miR-28-5p&offsets|0|0(+1A);m-133&3|+|188688794|188688815&offsets|0|0(+1A)]|AAGGAGCTCACAGTCTATTGAGA</td><td>>MI0000086|hsa-mir-28&WithFlank&3|+|188688775|188688872</td><td>20</td><td>41</td><td>22(+1A)</td><td>AAGGAGCTCACAGTCTATTGAGA</td><td>419.5</td><td>10.4</td><td>1.4</td><td>7.5</td><td>6.3E-14</td><td>2.6E-13</td><td>BT20/MB468</td><td>CYTO</td></tr>
<tr><td>>hg38_21_+_8433172_8446622_RNA45SN1_45S_50ntFlanks@12306.12323.18|CACGATCCTTCTGACCTT</td><td>>hg38_21_+_8433172_8446622_RNA45SN1_45S_50ntFlanks</td><td>12306</td><td>12323</td><td>18</td><td>CACGATCCTTCTGACCTT</td><td>241.0</td><td>-11.8</td><td>1.6</td><td>-7.5</td><td>5.2E-14</td><td>2.6E-13</td><td>MB231/MB468</td><td>MP</td></tr>
<tr><td>>hg38_21_+_8433172_8446622_RNA45SN1_45S_50ntFlanks@12624.12656.33|CCAATGGGGCGAAGCTACCATCTGTGGGATTAT</td><td>>hg38_21_+_8433172_8446622_RNA45SN1_45S_50ntFlanks</td><td>12624</td><td>12656</td><td>33</td><td>CCAATGGGGCGAAGCTACCATCTGTGGGATTAT</td><td>574.8</td><td>-9.0</td><td>1.2</td><td>-7.5</td><td>8.1E-14</td><td>2.6E-13</td><td>MB231/MB468</td><td>MITO</td></tr>
<tr><td>>MI0005756|hsa-mir-934&WithFlank&X|+|136550872|136550966@21.43.23&[MIMAT0004977&hsa-miR-934&offsets|0|+1;m-943&X|+|136550892|136550913&offsets|0|+1]|TGTCTACTACTGGAGACACTGGT</td><td>>MI0005756|hsa-mir-934&WithFlank&X|+|136550872|136550966</td><td>21</td><td>43</td><td>23</td><td>TGTCTACTACTGGAGACACTGGT</td><td>486.8</td><td>-12.8</td><td>1.7</td><td>-7.5</td><td>5.2E-14</td><td>2.6E-13</td><td>MB231/MB468</td><td>MP</td></tr>
<tr><td>>hg38_21_+_8433172_8446622_RNA45SN1_45S_50ntFlanks@12633.12661.29|CGAAGCTACCATCTGTGGGATTATGACTG</td><td>>hg38_21_+_8433172_8446622_RNA45SN1_45S_50ntFlanks</td><td>12633</td><td>12661</td><td>29</td><td>CGAAGCTACCATCTGTGGGATTATGACTG</td><td>2954.8</td><td>-8.9</td><td>1.2</td><td>-7.4</td><td>9.4E-14</td><td>2.6E-13</td><td>BT20/MB468</td><td>NUC</td></tr>
<tr><td>>hg38_21_+_8433172_8446622_RNA45SN1_45S_50ntFlanks@5035.5052.18|CATGGCCGTTCTTAGTTG</td><td>>hg38_21_+_8433172_8446622_RNA45SN1_45S_50ntFlanks</td><td>5035</td><td>5052</td><td>18</td><td>CATGGCCGTTCTTAGTTG</td><td>1592.8</td><td>-14.4</td><td>1.9</td><td>-7.5</td><td>8.1E-14</td><td>2.6E-13</td><td>MB231/MB468</td><td>MITO</td></tr>
<tr><td>>MI0000102|hsa-mir-100&WithFlank&11|-|122152223|122152314@19.40.22(+1A)&[MIMAT0000098&hsa-miR-100-5p&offsets|0|0(+1A);m-27&11|-|122152276|122152296&offsets|0|+1(+1A)]|AACCCGTAGATCCGAACTTGTGA</td><td>>MI0000102|hsa-mir-100&WithFlank&11|-|122152223|122152314</td><td>19</td><td>40</td><td>22(+1A)</td><td>AACCCGTAGATCCGAACTTGTGA</td><td>39196.1</td><td>-6.8</td><td>0.9</td><td>-7.8</td><td>6.6E-15</td><td>2.6E-13</td><td>BT20/MB231</td><td>MITO</td></tr>
<tr><td>>hg38_21_+_8433172_8446622_RNA45SN1_45S_50ntFlanks@8198.8215.18|TGGACGGTGTGAGGCCGG</td><td>>hg38_21_+_8433172_8446622_RNA45SN1_45S_50ntFlanks</td><td>8198</td><td>8215</td><td>18</td><td>TGGACGGTGTGAGGCCGG</td><td>750.8</td><td>-9.5</td><td>1.3</td><td>-7.5</td><td>8.1E-14</td><td>2.6E-13</td><td>MB231/MB468</td><td>MITO</td></tr>
<tr><td>>hg38_21_+_8433172_8446622_RNA45SN1_45S_50ntFlanks@9162.9187.26|CGCCCCGGGCGGGTCGCGCCGTCGGG</td><td>>hg38_21_+_8433172_8446622_RNA45SN1_45S_50ntFlanks</td><td>9162</td><td>9187</td><td>26</td><td>CGCCCCGGGCGGGTCGCGCCGTCGGG</td><td>179.2</td><td>-11.3</td><td>1.5</td><td>-7.5</td><td>5.2E-14</td><td>2.6E-13</td><td>MB231/MB468</td><td>MP</td></tr>
<tr><td>>hg38_21_+_8433172_8446622_RNA45SN1_45S_50ntFlanks@10731.10750.20|AGGGTGTAAATCTCGCGCCG</td><td>>hg38_21_+_8433172_8446622_RNA45SN1_45S_50ntFlanks</td><td>10731</td><td>10750</td><td>20</td><td>AGGGTGTAAATCTCGCGCCG</td><td>226.3</td><td>-11.7</td><td>1.6</td><td>-7.5</td><td>5.2E-14</td><td>2.6E-13</td><td>MB231/MB468</td><td>MP</td></tr>
<tr><td>>hg38_21_+_8433172_8446622_RNA45SN1_45S_50ntFlanks@8388.8405.18|CGTGAAACCGTTAAGAGG</td><td>>hg38_21_+_8433172_8446622_RNA45SN1_45S_50ntFlanks</td><td>8388</td><td>8405</td><td>18</td><td>CGTGAAACCGTTAAGAGG</td><td>461.8</td><td>-8.4</td><td>1.1</td><td>-7.5</td><td>6.0E-14</td><td>2.6E-13</td><td>MB231/MB468</td><td>CYTO</td></tr>
<tr><td>>trna20_GluTTC_2_-_131094701_131094772@1.33.33,trna5_GluTTC_13_-_41634874_41634945@1.33.33|TCCCATATGGTCTAGCGGTTAGGATTCCTGGTT</td><td>>trna20_GluTTC_2_-_131094701_131094772</td><td>1</td><td>33</td><td>33</td><td>TCCCATATGGTCTAGCGGTTAGGATTCCTGGTT</td><td>825.0</td><td>10.0</td><td>1.3</td><td>7.5</td><td>6.2E-14</td><td>2.6E-13</td><td>MB231/MB468</td><td>NUC</td></tr>
<tr><td>>hg38_21_+_8433172_8446622_RNA45SN1_45S_50ntFlanks@11207.11225.19|GGCGGCGACTCTGGACGCG</td><td>>hg38_21_+_8433172_8446622_RNA45SN1_45S_50ntFlanks</td><td>11207</td><td>11225</td><td>19</td><td>GGCGGCGACTCTGGACGCG</td><td>1289.9</td><td>-9.5</td><td>1.3</td><td>-7.5</td><td>5.2E-14</td><td>2.6E-13</td><td>MB231/MB468</td><td>MP</td></tr>
<tr><td>>hg38_21_+_8433172_8446622_RNA45SN1_45S_50ntFlanks@10411.10445.35|CAGTCGGTCCTGAGAGATGGGCGAGCGCCGTTCCG</td><td>>hg38_21_+_8433172_8446622_RNA45SN1_45S_50ntFlanks</td><td>10411</td><td>10445</td><td>35</td><td>CAGTCGGTCCTGAGAGATGGGCGAGCGCCGTTCCG</td><td>173.0</td><td>-11.3</td><td>1.5</td><td>-7.5</td><td>5.2E-14</td><td>2.6E-13</td><td>MB231/MB468</td><td>MP</td></tr>
<tr><td>>hg38_21_+_8433172_8446622_RNA45SN1_45S_50ntFlanks@10872.10902.31|GCTGGGTCGGTCGGGCTGGGGCGCGAAGCGG</td><td>>hg38_21_+_8433172_8446622_RNA45SN1_45S_50ntFlanks</td><td>10872</td><td>10902</td><td>31</td><td>GCTGGGTCGGTCGGGCTGGGGCGCGAAGCGG</td><td>183.9</td><td>-11.4</td><td>1.5</td><td>-7.5</td><td>5.3E-14</td><td>2.6E-13</td><td>MB231/MB468</td><td>MP</td></tr>
<tr><td>>hg38_21_+_8433172_8446622_RNA45SN1_45S_50ntFlanks@6700.6717.18|CTAGCTGCGAGAATTAAT</td><td>>hg38_21_+_8433172_8446622_RNA45SN1_45S_50ntFlanks</td><td>6700</td><td>6717</td><td>18</td><td>CTAGCTGCGAGAATTAAT</td><td>327.7</td><td>-12.2</td><td>1.6</td><td>-7.5</td><td>5.3E-14</td><td>2.6E-13</td><td>MB231/MB468</td><td>MP</td></tr>
<tr><td>>hg38_21_+_8433172_8446622_RNA45SN1_45S_50ntFlanks@11273.11293.21|CCGCCCCCGGGGAGCCCGGCG</td><td>>hg38_21_+_8433172_8446622_RNA45SN1_45S_50ntFlanks</td><td>11273</td><td>11293</td><td>21</td><td>CCGCCCCCGGGGAGCCCGGCG</td><td>317.7</td><td>-7.6</td><td>1.0</td><td>-7.5</td><td>6.1E-14</td><td>2.6E-13</td><td>MB231/MB468</td><td>CYTO</td></tr>
<tr><td>>hg38_21_+_8433172_8446622_RNA45SN1_45S_50ntFlanks@9573.9598.26|AAATCGGTCGTCCGACCTGGGTATAG</td><td>>hg38_21_+_8433172_8446622_RNA45SN1_45S_50ntFlanks</td><td>9573</td><td>9598</td><td>26</td><td>AAATCGGTCGTCCGACCTGGGTATAG</td><td>139.2</td><td>-11.0</td><td>1.5</td><td>-7.5</td><td>5.3E-14</td><td>2.6E-13</td><td>MB231/MB468</td><td>MP</td></tr>
<tr><td>>hg38_21_+_8433172_8446622_RNA45SN1_45S_50ntFlanks@5084.5113.30|CGAACGAGACTCTGGCATGCTAACTAGTTA</td><td>>hg38_21_+_8433172_8446622_RNA45SN1_45S_50ntFlanks</td><td>5084</td><td>5113</td><td>30</td><td>CGAACGAGACTCTGGCATGCTAACTAGTTA</td><td>286.1</td><td>-12.0</td><td>1.6</td><td>-7.5</td><td>5.3E-14</td><td>2.6E-13</td><td>MB231/MB468</td><td>MP</td></tr>
<tr><td>>hg38_21_+_8433172_8446622_RNA45SN1_45S_50ntFlanks@10927.10946.20|CGAGGCGCCGCCGCCCCCCC</td><td>>hg38_21_+_8433172_8446622_RNA45SN1_45S_50ntFlanks</td><td>10927</td><td>10946</td><td>20</td><td>CGAGGCGCCGCCGCCCCCCC</td><td>1260.2</td><td>-14.2</td><td>1.9</td><td>-7.5</td><td>5.3E-14</td><td>2.6E-13</td><td>MB231/MB468</td><td>MP</td></tr>
<tr><td>>hg38_21_+_8433172_8446622_RNA45SN1_45S_50ntFlanks@3705.3735.31|TACCTGGTTGATCCTGCCAGTAGCATATGCT</td><td>>hg38_21_+_8433172_8446622_RNA45SN1_45S_50ntFlanks</td><td>3705</td><td>3735</td><td>31</td><td>TACCTGGTTGATCCTGCCAGTAGCATATGCT</td><td>473.6</td><td>5.5</td><td>0.7</td><td>7.5</td><td>6.1E-14</td><td>2.6E-13</td><td>MB231/MB468</td><td>CYTO</td></tr>
<tr><td>>hg38_21_+_8433172_8446622_RNA45SN1_45S_50ntFlanks@12418.12445.28|CGATGTCGGCTCTTCCTATCATTGTGAA</td><td>>hg38_21_+_8433172_8446622_RNA45SN1_45S_50ntFlanks</td><td>12418</td><td>12445</td><td>28</td><td>CGATGTCGGCTCTTCCTATCATTGTGAA</td><td>1778.4</td><td>-3.7</td><td>0.5</td><td>-7.5</td><td>6.3E-14</td><td>2.6E-13</td><td>MB231/MB468</td><td>NUC</td></tr>
<tr><td>>hg38_21_+_8433172_8446622_RNA45SN1_45S_50ntFlanks@11569.11592.24|CAGCCGACTTAGAACTGGTGCGGA</td><td>>hg38_21_+_8433172_8446622_RNA45SN1_45S_50ntFlanks</td><td>11569</td><td>11592</td><td>24</td><td>CAGCCGACTTAGAACTGGTGCGGA</td><td>5285.8</td><td>-11.9</td><td>1.6</td><td>-7.5</td><td>5.3E-14</td><td>2.6E-13</td><td>MB231/MB468</td><td>MP</td></tr>
<tr><td>>hg38_21_+_8433172_8446622_RNA45SN1_45S_50ntFlanks@12827.12847.21|ACCGGGGTCCGGTGCGGAGTG</td><td>>hg38_21_+_8433172_8446622_RNA45SN1_45S_50ntFlanks</td><td>12827</td><td>12847</td><td>21</td><td>ACCGGGGTCCGGTGCGGAGTG</td><td>257.0</td><td>-11.9</td><td>1.6</td><td>-7.5</td><td>5.3E-14</td><td>2.6E-13</td><td>MB231/MB468</td><td>MP</td></tr>
<tr><td>>trnaMT_GluTTC_MT_-_14674_14742@39.71.33,trnalookalike8_GluTTC_5_-_93905172_93905240@39.71.33|ATTGGTCGTGGTTGTAGTCCGTGCGAGAATACC</td><td>>trnaMT_GluTTC_MT_-_14674_14742</td><td>39</td><td>71</td><td>33</td><td>ATTGGTCGTGGTTGTAGTCCGTGCGAGAATACC</td><td>3522.6</td><td>10.6</td><td>1.4</td><td>7.5</td><td>6.1E-14</td><td>2.6E-13</td><td>MB231/MB468</td><td>CYTO</td></tr>
<tr><td>>hg38_21_+_8433172_8446622_RNA45SN1_45S_50ntFlanks@10871.10891.21|GGCTGGGTCGGTCGGGCTGGG</td><td>>hg38_21_+_8433172_8446622_RNA45SN1_45S_50ntFlanks</td><td>10871</td><td>10891</td><td>21</td><td>GGCTGGGTCGGTCGGGCTGGG</td><td>214.2</td><td>-11.6</td><td>1.5</td><td>-7.5</td><td>5.3E-14</td><td>2.6E-13</td><td>MB231/MB468</td><td>MP</td></tr>
<tr><td>>MI0000438|hsa-mir-15b&WithFlank&3|+|160404582|160404691@26.47.22&[MIMAT0000417&hsa-miR-15b-5p&offsets|0|0;m-110&3|+|160404607|160404628&offsets|0|0]|TAGCAGCACATCATGGTTTACA</td><td>>MI0000438|hsa-mir-15b&WithFlank&3|+|160404582|160404691</td><td>26</td><td>47</td><td>22</td><td>TAGCAGCACATCATGGTTTACA</td><td>1000.6</td><td>5.8</td><td>0.8</td><td>7.5</td><td>4.9E-14</td><td>2.6E-13</td><td>BT20/MB468</td><td>MITO</td></tr>
<tr><td>>MI0000061|hsa-let-7a-2&WithFlank&11|-|122146516|122146599@11.32.22(+1C);MI0000062|hsa-let-7a-3&WithFlank&22|+|46112743|46112828@10.31.22(+1C);MI0000060|hsa-let-7a-1&WithFlank&9|+|94175951|94176042@12.33.22(+1C)&[MIMAT0000062&hsa-let-7a-5p&offsets|0|0(+1C);m-6&11|-|122146568|122146589&offsets|0|0(+1C)];[MIMAT0000062_1&hsa-let-7a-5p&offsets|0|0(+1C);m-5&22|+|46112752|46112773&offsets|0|0(+1C)];[MIMAT0000062_2&hsa-let-7a-5p&offsets|0|0(+1C);m-4&9|+|94175962|94175983&offsets|0|0(+1C)]|TGAGGTAGTAGGTTGTATAGTTC</td><td>>MI0000061|hsa-let-7a-2&WithFlank&11|-|122146516|122146599</td><td>11</td><td>32</td><td>22(+1C)</td><td>TGAGGTAGTAGGTTGTATAGTTC</td><td>488.6</td><td>7.4</td><td>1.0</td><td>7.5</td><td>6.6E-14</td><td>2.6E-13</td><td>BT20/MB468</td><td>CYTO</td></tr>
<tr><td>>hg38_21_+_8433172_8446622_RNA45SN1_45S_50ntFlanks@10798.10829.32|CATGTTGGAACAATGTAGGTAAGGGAAGTCGG</td><td>>hg38_21_+_8433172_8446622_RNA45SN1_45S_50ntFlanks</td><td>10798</td><td>10829</td><td>32</td><td>CATGTTGGAACAATGTAGGTAAGGGAAGTCGG</td><td>129.2</td><td>-10.9</td><td>1.4</td><td>-7.5</td><td>5.4E-14</td><td>2.6E-13</td><td>MB231/MB468</td><td>MP</td></tr>
<tr><td>>hg38_21_+_8433172_8446622_RNA45SN1_45S_50ntFlanks@9313.9335.23|TCTAACACGTGCGCGAGTCGGGG</td><td>>hg38_21_+_8433172_8446622_RNA45SN1_45S_50ntFlanks</td><td>9313</td><td>9335</td><td>23</td><td>TCTAACACGTGCGCGAGTCGGGG</td><td>227.7</td><td>-11.6</td><td>1.6</td><td>-7.5</td><td>8.4E-14</td><td>2.6E-13</td><td>MB231/MB468</td><td>MITO</td></tr>
<tr><td>>hg38_21_+_8433172_8446622_RNA45SN1_45S_50ntFlanks@5074.5098.25|ATTCCGATAACGAACGAGACTCTGG</td><td>>hg38_21_+_8433172_8446622_RNA45SN1_45S_50ntFlanks</td><td>5074</td><td>5098</td><td>25</td><td>ATTCCGATAACGAACGAGACTCTGG</td><td>1282.6</td><td>-14.2</td><td>1.9</td><td>-7.5</td><td>5.4E-14</td><td>2.7E-13</td><td>MB231/MB468</td><td>MP</td></tr>
<tr><td>>hg38_21_+_8433172_8446622_RNA45SN1_45S_50ntFlanks@8380.8397.18|CAAGAGGGCGTGAAACCG</td><td>>hg38_21_+_8433172_8446622_RNA45SN1_45S_50ntFlanks</td><td>8380</td><td>8397</td><td>18</td><td>CAAGAGGGCGTGAAACCG</td><td>966.2</td><td>-6.5</td><td>0.9</td><td>-7.5</td><td>6.2E-14</td><td>2.7E-13</td><td>MB231/MB468</td><td>CYTO</td></tr>
<tr><td>>hg38_21_+_8433172_8446622_RNA45SN1_45S_50ntFlanks@6722.6740.19|ATTGCAGGACACATTGATC</td><td>>hg38_21_+_8433172_8446622_RNA45SN1_45S_50ntFlanks</td><td>6722</td><td>6740</td><td>19</td><td>ATTGCAGGACACATTGATC</td><td>2451.7</td><td>-9.1</td><td>1.2</td><td>-7.4</td><td>9.8E-14</td><td>2.7E-13</td><td>BT20/MB468</td><td>NUC</td></tr>
<tr><td>>hg38_21_+_8433172_8446622_RNA45SN1_45S_50ntFlanks@5473.5502.30|CGGCCCTGGCGGAGCGCTGAGAAGACGGTC</td><td>>hg38_21_+_8433172_8446622_RNA45SN1_45S_50ntFlanks</td><td>5473</td><td>5502</td><td>30</td><td>CGGCCCTGGCGGAGCGCTGAGAAGACGGTC</td><td>2694.4</td><td>7.8</td><td>1.0</td><td>7.5</td><td>6.6E-14</td><td>2.7E-13</td><td>BT20/MB468</td><td>CYTO</td></tr>
<tr><td>>hg38_21_+_8433172_8446622_RNA45SN1_45S_50ntFlanks@12347.12376.30|AGAAAAGTTACCACAGGGATAACTGGCTTG</td><td>>hg38_21_+_8433172_8446622_RNA45SN1_45S_50ntFlanks</td><td>12347</td><td>12376</td><td>30</td><td>AGAAAAGTTACCACAGGGATAACTGGCTTG</td><td>338.0</td><td>-12.3</td><td>1.6</td><td>-7.5</td><td>5.4E-14</td><td>2.7E-13</td><td>MB231/MB468</td><td>MP</td></tr>
<tr><td>>trnaMT_ValTAC_MT_+_1602_1670@51.71.21|CTTAACTTGACCGCTCTGACC</td><td>>trnaMT_ValTAC_MT_+_1602_1670</td><td>51</td><td>71</td><td>21</td><td>CTTAACTTGACCGCTCTGACC</td><td>679.1</td><td>-3.9</td><td>0.5</td><td>-7.5</td><td>5.4E-14</td><td>2.7E-13</td><td>MB231/MB468</td><td>MP</td></tr>
<tr><td>>hg38_21_+_8433172_8446622_RNA45SN1_45S_50ntFlanks@4047.4065.19|ATTCGAACGTCTGCCCTAT</td><td>>hg38_21_+_8433172_8446622_RNA45SN1_45S_50ntFlanks</td><td>4047</td><td>4065</td><td>19</td><td>ATTCGAACGTCTGCCCTAT</td><td>137.4</td><td>-11.0</td><td>1.5</td><td>-7.5</td><td>5.4E-14</td><td>2.7E-13</td><td>MB231/MB468</td><td>MP</td></tr>
<tr><td>>MI0000483|hsa-mir-186&WithFlank&1|-|71067625|71067722@21.43.23&[MIMAT0000456&hsa-miR-186-5p&offsets|0|+1;m-84&1|-|71067681|71067702&offsets|0|+1]|CAAAGAATTCTCCTTTTGGGCTT</td><td>>MI0000483|hsa-mir-186&WithFlank&1|-|71067625|71067722</td><td>21</td><td>43</td><td>23</td><td>CAAAGAATTCTCCTTTTGGGCTT</td><td>417.7</td><td>5.4</td><td>0.7</td><td>7.5</td><td>6.5E-14</td><td>2.7E-13</td><td>MB231/MB468</td><td>NUC</td></tr>
<tr><td>>hg38_21_+_8433172_8446622_RNA45SN1_45S_50ntFlanks@9201.9226.26|CTCTCGGGGCCACGCGCGCGTCCCCC</td><td>>hg38_21_+_8433172_8446622_RNA45SN1_45S_50ntFlanks</td><td>9201</td><td>9226</td><td>26</td><td>CTCTCGGGGCCACGCGCGCGTCCCCC</td><td>170.2</td><td>-11.3</td><td>1.5</td><td>-7.5</td><td>5.4E-14</td><td>2.7E-13</td><td>MB231/MB468</td><td>MP</td></tr>
<tr><td>>hg38_21_+_8433172_8446622_RNA45SN1_45S_50ntFlanks@12249.12273.25|CAAAAGCTCGCTTGATCTTGATTTT</td><td>>hg38_21_+_8433172_8446622_RNA45SN1_45S_50ntFlanks</td><td>12249</td><td>12273</td><td>25</td><td>CAAAAGCTCGCTTGATCTTGATTTT</td><td>153.7</td><td>-11.1</td><td>1.5</td><td>-7.5</td><td>5.4E-14</td><td>2.7E-13</td><td>MB231/MB468</td><td>MP</td></tr>
<tr><td>>hg38_21_+_8433172_8446622_RNA45SN1_45S_50ntFlanks@4099.4121.23|GTGACCACGGGTGACGGGGAATC</td><td>>hg38_21_+_8433172_8446622_RNA45SN1_45S_50ntFlanks</td><td>4099</td><td>4121</td><td>23</td><td>GTGACCACGGGTGACGGGGAATC</td><td>203.1</td><td>-5.5</td><td>0.7</td><td>-7.5</td><td>6.3E-14</td><td>2.7E-13</td><td>MB231/MB468</td><td>CYTO</td></tr>
<tr><td>>hg38_21_+_8433172_8446622_RNA45SN1_45S_50ntFlanks@11208.11254.47|GCGGCGACTCTGGACGCGAGCCGGGCCCTTCCCGTGGATCGCCCCAG</td><td>>hg38_21_+_8433172_8446622_RNA45SN1_45S_50ntFlanks</td><td>11208</td><td>11254</td><td>47</td><td>GCGGCGACTCTGGACGCGAGCCGGGCCCTTCCCGTGGATCGCCCCAG</td><td>837.3</td><td>-13.6</td><td>1.8</td><td>-7.5</td><td>5.5E-14</td><td>2.7E-13</td><td>MB231/MB468</td><td>MP</td></tr>
<tr><td>>hg38_21_+_8433172_8446622_RNA45SN1_45S_50ntFlanks@11233.11255.23|CCCTTCCCGTGGATCGCCCCAGC</td><td>>hg38_21_+_8433172_8446622_RNA45SN1_45S_50ntFlanks</td><td>11233</td><td>11255</td><td>23</td><td>CCCTTCCCGTGGATCGCCCCAGC</td><td>217.3</td><td>-11.5</td><td>1.5</td><td>-7.5</td><td>8.5E-14</td><td>2.7E-13</td><td>MB231/MB468</td><td>MITO</td></tr>
<tr><td>>hg38_21_+_8433172_8446622_RNA45SN1_45S_50ntFlanks@3831.3852.22|CTCGCTCCTCTCCTACTTGGAT</td><td>>hg38_21_+_8433172_8446622_RNA45SN1_45S_50ntFlanks</td><td>3831</td><td>3852</td><td>22</td><td>CTCGCTCCTCTCCTACTTGGAT</td><td>285.0</td><td>-11.1</td><td>1.5</td><td>-7.5</td><td>6.5E-14</td><td>2.7E-13</td><td>MB231/MB468</td><td>NUC</td></tr>
<tr><td>>hg38_21_+_8433172_8446622_RNA45SN1_45S_50ntFlanks@10686.10709.24|GCGGCGTCCGGTGAGCTCTCGCTG</td><td>>hg38_21_+_8433172_8446622_RNA45SN1_45S_50ntFlanks</td><td>10686</td><td>10709</td><td>24</td><td>GCGGCGTCCGGTGAGCTCTCGCTG</td><td>834.0</td><td>-4.2</td><td>0.6</td><td>-7.5</td><td>6.3E-14</td><td>2.7E-13</td><td>MB231/MB468</td><td>CYTO</td></tr>
<tr><td>>hg38_21_+_8433172_8446622_RNA45SN1_45S_50ntFlanks@11876.11906.31|AAGACCCTGTTGAGCTTGACTCTAGTCTGGC</td><td>>hg38_21_+_8433172_8446622_RNA45SN1_45S_50ntFlanks</td><td>11876</td><td>11906</td><td>31</td><td>AAGACCCTGTTGAGCTTGACTCTAGTCTGGC</td><td>389.3</td><td>-7.0</td><td>0.9</td><td>-7.5</td><td>6.3E-14</td><td>2.7E-13</td><td>MB231/MB468</td><td>CYTO</td></tr>
<tr><td>>hg38_21_+_8433172_8446622_RNA45SN1_45S_50ntFlanks@12459.12476.18|AGCGTTGGATTGTTCACC</td><td>>hg38_21_+_8433172_8446622_RNA45SN1_45S_50ntFlanks</td><td>12459</td><td>12476</td><td>18</td><td>AGCGTTGGATTGTTCACC</td><td>3000.7</td><td>-4.7</td><td>0.6</td><td>-7.5</td><td>6.8E-14</td><td>2.7E-13</td><td>BT20/MB468</td><td>CYTO</td></tr>
<tr><td>>hg38_21_+_8433172_8446622_RNA45SN1_45S_50ntFlanks@12488.12505.18|AACGTGAGCTGGGTTTAG</td><td>>hg38_21_+_8433172_8446622_RNA45SN1_45S_50ntFlanks</td><td>12488</td><td>12505</td><td>18</td><td>AACGTGAGCTGGGTTTAG</td><td>404.1</td><td>-11.6</td><td>1.6</td><td>-7.5</td><td>6.6E-14</td><td>2.7E-13</td><td>MB231/MB468</td><td>NUC</td></tr>
<tr><td>>hg38_21_+_8433172_8446622_RNA45SN1_45S_50ntFlanks@11641.11670.30|CGGCGGGTGTTGACGCGATGTGATTTCTGC</td><td>>hg38_21_+_8433172_8446622_RNA45SN1_45S_50ntFlanks</td><td>11641</td><td>11670</td><td>30</td><td>CGGCGGGTGTTGACGCGATGTGATTTCTGC</td><td>4391.9</td><td>-10.7</td><td>1.4</td><td>-7.5</td><td>6.6E-14</td><td>2.7E-13</td><td>MB231/MB468</td><td>NUC</td></tr>
<tr><td>>MI0000285|hsa-mir-205&WithFlank&1|+|209432127|209432248@40.61.22&[MIMAT0000266&hsa-miR-205-5p&offsets|0|0;m-351&1|+|209432166|209432187&offsets|0|0]|TCCTTCATTCCACCGGAGTCTG</td><td>>MI0000285|hsa-mir-205&WithFlank&1|+|209432127|209432248</td><td>40</td><td>61</td><td>22</td><td>TCCTTCATTCCACCGGAGTCTG</td><td>22616.3</td><td>13.0</td><td>1.6</td><td>8.0</td><td>9.3E-16</td><td>2.7E-13</td><td>BT20/MB231</td><td>MP</td></tr>
</table>

<table>
<tr><th></th><th></th><th></th><th></th><th></th><th></th><th></th><th></th><th></th><th></th><th></th><th></th><th></th><th></th><th></th></tr>
<tr><td>>hg38_21_+_8433172_8446622_RNA45SN1_45S_50ntFlanks@4472.4492.21|CTTAGCTGAGTGTCCCGCGGG</td><td>>hg38_21_+_8433172_8446622_RNA45SN1_45S_50ntFlanks</td><td>4472</td><td>4492</td><td>21</td><td>CTTAGCTGAGTGTCCCGCGGG</td><td>306.4</td><td>-5.6</td><td>0.8</td><td>-7.5</td><td>6.4E-14</td><td>2.7E-13</td><td>MB231/MB468</td><td>CYTO</td><td></td></tr>
<tr><td>>hg38_21_+_8433172_8446622_RNA45SN1_45S_50ntFlanks@9570.9592.23|TGCAAATCGGTCGTCCGACCTGG</td><td>>hg38_21_+_8433172_8446622_RNA45SN1_45S_50ntFlanks</td><td>9570</td><td>9592</td><td>23</td><td>TGCAAATCGGTCGTCCGACCTGG</td><td>211.6</td><td>-7.4</td><td>1.0</td><td>-7.5</td><td>6.4E-14</td><td>2.7E-13</td><td>MB231/MB468</td><td>CYTO</td><td></td></tr>
<tr><td>>hg38_21_+_8433172_8446622_RNA45SN1_45S_50ntFlanks@12191.12211.21|ACGCAGGTGTCCTAAGGCGAG</td><td>>hg38_21_+_8433172_8446622_RNA45SN1_45S_50ntFlanks</td><td>12191</td><td>12211</td><td>21</td><td>ACGCAGGTGTCCTAAGGCGAG</td><td>193.5</td><td>-11.5</td><td>1.5</td><td>-7.5</td><td>5.6E-14</td><td>2.7E-13</td><td>MB231/MB468</td><td>MP</td><td></td></tr>
<tr><td>>hg38_21_+_8433172_8446622_RNA45SN1_45S_50ntFlanks@12357.12376.20|CCACAGGGATAACTGGCTTG</td><td>>hg38_21_+_8433172_8446622_RNA45SN1_45S_50ntFlanks</td><td>12357</td><td>12376</td><td>20</td><td>CCACAGGGATAACTGGCTTG</td><td>339.8</td><td>-8.2</td><td>1.1</td><td>-7.5</td><td>6.6E-14</td><td>2.7E-13</td><td>MB231/MB468</td><td>NUC</td><td></td></tr>
<tr><td>>hg38_21_+_8433172_8446622_RNA45SN1_45S_50ntFlanks@12412.12441.30|ATCCTTCGATGTCGGCTCTTCCTATCATTG</td><td>>hg38_21_+_8433172_8446622_RNA45SN1_45S_50ntFlanks</td><td>12412</td><td>12441</td><td>30</td><td>ATCCTTCGATGTCGGCTCTTCCTATCATTG</td><td>222.7</td><td>-10.1</td><td>1.3</td><td>-7.5</td><td>6.8E-14</td><td>2.7E-13</td><td>BT20/MB468</td><td>CYTO</td><td></td></tr>
<tr><td>>hg38_21_+_8433172_8446622_RNA45SN1_45S_50ntFlanks@9035.9056.22|CGTGGGGGGGCCGGGCCACCCC</td><td>>hg38_21_+_8433172_8446622_RNA45SN1_45S_50ntFlanks</td><td>9035</td><td>9056</td><td>22</td><td>CGTGGGGGGGCCGGGCCACCCC</td><td>357.9</td><td>-12.3</td><td>1.6</td><td>-7.5</td><td>8.7E-14</td><td>2.8E-13</td><td>MB231/MB468</td><td>MITO</td><td></td></tr>
<tr><td>>hg38_21_+_8433172_8446622_RNA45SN1_45S_50ntFlanks@10921.10945.25|GCTGGACGAGGCGCCGCCGCCCCC</td><td>>hg38_21_+_8433172_8446622_RNA45SN1_45S_50ntFlanks</td><td>10921</td><td>10945</td><td>25</td><td>GCTGGACGAGGCGCCGCCGCCCCC</td><td>206.9</td><td>-6.1</td><td>0.8</td><td>-7.5</td><td>6.5E-14</td><td>2.8E-13</td><td>MB231/MB468</td><td>CYTO</td><td></td></tr>
<tr><td>>hg38_21_+_8433172_8446622_RNA45SN1_45S_50ntFlanks@12159.12182.24|TTTGACTGGGGCGGTACACCTGTC</td><td>>hg38_21_+_8433172_8446622_RNA45SN1_45S_50ntFlanks</td><td>12159</td><td>12182</td><td>24</td><td>TTTGACTGGGGCGGTACACCTGTC</td><td>261.8</td><td>-10.8</td><td>1.4</td><td>-7.5</td><td>6.5E-14</td><td>2.8E-13</td><td>MB231/MB468</td><td>CYTO</td><td></td></tr>
<tr><td>>hg38_21_+_8433172_8446622_RNA45SN1_45S_50ntFlanks@10678.10707.30|CGGTTCCGGCGGCGTCCGGTGAGCTCTCGC</td><td>>hg38_21_+_8433172_8446622_RNA45SN1_45S_50ntFlanks</td><td>10678</td><td>10707</td><td>30</td><td>CGGTTCCGGCGGCGTCCGGTGAGCTCTCGC</td><td>1733.3</td><td>-6.4</td><td>0.9</td><td>-7.5</td><td>8.7E-14</td><td>2.8E-13</td><td>MB231/MB468</td><td>MITO</td><td></td></tr>
<tr><td>>hg38_21_+_8433172_8446622_RNA45SN1_45S_50ntFlanks@11629.11665.37|CGCGAAGGCCCGCGGCGGGTGTTGACGCGATGTGATT</td><td>>hg38_21_+_8433172_8446622_RNA45SN1_45S_50ntFlanks</td><td>11629</td><td>11665</td><td>37</td><td>CGCGAAGGCCCGCGGCGGGTGTTGACGCGATGTGATT</td><td>193.7</td><td>-11.5</td><td>1.5</td><td>-7.5</td><td>5.6E-14</td><td>2.8E-13</td><td>MB231/MB468</td><td>MP</td><td></td></tr>
<tr><td>>hg38_21_+_8433172_8446622_RNA45SN1_45S_50ntFlanks@6788.6807.20|CGCCTGTCTGAGCGTCGCTT</td><td>>hg38_21_+_8433172_8446622_RNA45SN1_45S_50ntFlanks</td><td>6788</td><td>6807</td><td>20</td><td>CGCCTGTCTGAGCGTCGCTT</td><td>1138.3</td><td>5.0</td><td>0.7</td><td>7.5</td><td>6.9E-14</td><td>2.8E-13</td><td>BT20/MB468</td><td>CYTO</td><td></td></tr>
<tr><td>>hg38_21_+_8433172_8446622_RNA45SN1_45S_50ntFlanks@10539.10567.29|AGGCGTCCAGTGCGGTAACGCGACCGATC</td><td>>hg38_21_+_8433172_8446622_RNA45SN1_45S_50ntFlanks</td><td>10539</td><td>10567</td><td>29</td><td>AGGCGTCCAGTGCGGTAACGCGACCGATC</td><td>192.7</td><td>-11.5</td><td>1.5</td><td>-7.5</td><td>5.7E-14</td><td>2.8E-13</td><td>MB231/MB468</td><td>MP</td><td></td></tr>
<tr><td>>hg38_21_+_8433172_8446622_RNA45SN1_45S_50ntFlanks@8194.8225.32|CCCGTGGACGGTGTGAGGCCGGTAGCGGCCCC</td><td>>hg38_21_+_8433172_8446622_RNA45SN1_45S_50ntFlanks</td><td>8194</td><td>8225</td><td>32</td><td>CCCGTGGACGGTGTGAGGCCGGTAGCGGCCCC</td><td>223.6</td><td>-11.7</td><td>1.6</td><td>-7.5</td><td>5.7E-14</td><td>2.8E-13</td><td>MB231/MB468</td><td>MP</td><td></td></tr>
<tr><td>>MI0000448|hsa-mir-130a&WithFlank&11|+|57641192|57641292@61.83.23&[MIMAT0000425&hsa-miR-130a-3p&offsets|0|+1;m-139&11|+|57641252|57641273&offsets|0|+1]|CAGTGCAATGTTAAAAGGGCATT</td><td>>MI0000448|hsa-mir-130a&WithFlank&11|+|57641192|57641292</td><td>61</td><td>83</td><td>23</td><td>CAGTGCAATGTTAAAAGGGCATT</td><td>325.3</td><td>7.6</td><td>1.0</td><td>7.5</td><td>6.8E-14</td><td>2.8E-13</td><td>MB231/MB468</td><td>NUC</td><td></td></tr>
<tr><td>>MI0000087|hsa-mir-29a&WithFlank&7|-|130876741|130876816@48.67.20&[MIMAT0000086&hsa-miR-29a-3p&offsets|0|-2;m-26&7|-|130876749|130876769&offsets|0|-1]|TAGCACCATCTGAAATCGGT</td><td>>MI0000087|hsa-mir-29a&WithFlank&7|-|130876741|130876816</td><td>48</td><td>67</td><td>20</td><td>TAGCACCATCTGAAATCGGT</td><td>275.3</td><td>9.7</td><td>1.3</td><td>7.5</td><td>6.8E-14</td><td>2.8E-13</td><td>MB231/MB468</td><td>NUC</td><td></td></tr>
<tr><td>>hg38_21_+_8433172_8446622_RNA45SN1_45S_50ntFlanks@12957.12986.30|CTTCTGGGTCGGGGTTTCGTACGTAGCAGA</td><td>>hg38_21_+_8433172_8446622_RNA45SN1_45S_50ntFlanks</td><td>12957</td><td>12986</td><td>30</td><td>CTTCTGGGTCGGGGTTTCGTACGTAGCAGA</td><td>758.2</td><td>-4.9</td><td>0.7</td><td>-7.5</td><td>6.8E-14</td><td>2.8E-13</td><td>MB231/MB468</td><td>NUC</td><td></td></tr>
<tr><td>>hg38_21_+_8433172_8446622_RNA45SN1_45S_50ntFlanks@11199.11233.35|CCGGCGGCGGCGGCGACTCTGGACGCGAGCCGGGC</td><td>>hg38_21_+_8433172_8446622_RNA45SN1_45S_50ntFlanks</td><td>11199</td><td>11233</td><td>35</td><td>CCGGCGGCGGCGGCGACTCTGGACGCGAGCCGGGC</td><td>277.3</td><td>-3.7</td><td>0.5</td><td>-7.5</td><td>7.0E-14</td><td>2.8E-13</td><td>BT20/MB468</td><td>CYTO</td><td></td></tr>
<tr><td>>hg38_21_+_8433172_8446622_RNA45SN1_45S_50ntFlanks@11882.11905.24|CTGTTGAGCTTGACTCTAGTCTGG</td><td>>hg38_21_+_8433172_8446622_RNA45SN1_45S_50ntFlanks</td><td>11882</td><td>11905</td><td>24</td><td>CTGTTGAGCTTGACTCTAGTCTGG</td><td>134.9</td><td>-9.8</td><td>1.3</td><td>-7.5</td><td>6.6E-14</td><td>2.8E-13</td><td>MB231/MB468</td><td>CYTO</td><td></td></tr>
<tr><td>>hg38_1_-_228634819_228635039_RNA5S12_5S_50ntFlanks@127.170.44|ACTTGGATGGGAGACCGCCTGGGAATACCGGGTGCTGTAGGCTT</td><td>>hg38_1_-_228634819_228635039_RNA5S12_5S_50ntFlanks</td><td>127</td><td>170</td><td>44</td><td>ACTTGGATGGGAGACCGCCTGGGAATACCGGGTGCTGTAGGCTT</td><td>4094.2</td><td>10.4</td><td>1.4</td><td>7.5</td><td>6.6E-14</td><td>2.8E-13</td><td>MB231/MB468</td><td>CYTO</td><td></td></tr>
<tr><td>>hg38_21_+_8433172_8446622_RNA45SN1_45S_50ntFlanks@10391.10408.18|AACAGCAGTTGAACATGG</td><td>>hg38_21_+_8433172_8446622_RNA45SN1_45S_50ntFlanks</td><td>10391</td><td>10408</td><td>18</td><td>AACAGCAGTTGAACATGG</td><td>558.3</td><td>-6.8</td><td>0.9</td><td>-7.5</td><td>6.6E-14</td><td>2.8E-13</td><td>MB231/MB468</td><td>CYTO</td><td></td></tr>
<tr><td>>hg38_21_+_8433172_8446622_RNA45SN1_45S_50ntFlanks@12223.12248.26|CAGAAACCTCCCGTGGAGCAGAAGGG</td><td>>hg38_21_+_8433172_8446622_RNA45SN1_45S_50ntFlanks</td><td>12223</td><td>12248</td><td>26</td><td>CAGAAACCTCCCGTGGAGCAGAAGGG</td><td>163.0</td><td>-11.2</td><td>1.5</td><td>-7.5</td><td>5.7E-14</td><td>2.8E-13</td><td>MB231/MB468</td><td>MP</td><td></td></tr>
<tr><td>>MI0000298|hsa-mir-221&WithFlank&X|-|45746151|45746272@71.91.21&[MIMAT0000278&hsa-miR-221-3p&offsets|0|-2;m-91&X|-|45746181|45746202&offsets|0|-1]|AGCTACATTGTCTGCTGGGTT</td><td>>MI0000298|hsa-mir-221&WithFlank&X|-|45746151|45746272</td><td>71</td><td>91</td><td>21</td><td>AGCTACATTGTCTGCTGGGTT</td><td>1194.6</td><td>6.8</td><td>0.9</td><td>7.5</td><td>6.6E-14</td><td>2.8E-13</td><td>MB231/MB468</td><td>CYTO</td><td></td></tr>
<tr><td>>hg38_21_+_8433172_8446622_RNA45SN1_45S_50ntFlanks@9553.9576.24|CCGTAGCGGTCCTGACGTGCAAAT</td><td>>hg38_21_+_8433172_8446622_RNA45SN1_45S_50ntFlanks</td><td>9553</td><td>9576</td><td>24</td><td>CCGTAGCGGTCCTGACGTGCAAAT</td><td>251.4</td><td>6.2</td><td>0.8</td><td>7.5</td><td>6.6E-14</td><td>2.8E-13</td><td>MB231/MB468</td><td>CYTO</td><td></td></tr>
<tr><td>>hg38_21_+_8433172_8446622_RNA45SN1_45S_50ntFlanks@12993.13015.23|CCCTCGCTGCGATCTATTGAAAG</td><td>>hg38_21_+_8433172_8446622_RNA45SN1_45S_50ntFlanks</td><td>12993</td><td>13015</td><td>23</td><td>CCCTCGCTGCGATCTATTGAAAG</td><td>692.2</td><td>-4.1</td><td>0.5</td><td>-7.5</td><td>6.8E-14</td><td>2.8E-13</td><td>MB231/MB468</td><td>NUC</td><td></td></tr>
<tr><td>>trnaMT_GluTTC_MT_-_14674_14742@34.72.39,trnalookalike8_GluTTC_5_-_93905172_93905240@34.72.39|ATATCATTGGTCGTGGTTGTAGTCCGTGCGAGAATACCA</td><td>>trnaMT_GluTTC_MT_-_14674_14742</td><td>34</td><td>72</td><td>39</td><td>ATATCATTGGTCGTGGTTGTAGTCCGTGCGAGAATACCA</td><td>4191.7</td><td>9.6</td><td>1.3</td><td>7.5</td><td>6.7E-14</td><td>2.8E-13</td><td>MB231/MB468</td><td>CYTO</td><td></td></tr>
<tr><td>>hg38_21_+_8433172_8446622_RNA45SN1_45S_50ntFlanks@10500.10533.34|CAGATCCCCGAATCCGGAGTGGCGGAGATGGGCG</td><td>>hg38_21_+_8433172_8446622_RNA45SN1_45S_50ntFlanks</td><td>10500</td><td>10533</td><td>34</td><td>CAGATCCCCGAATCCGGAGTGGCGGAGATGGGCG</td><td>193.8</td><td>-11.4</td><td>1.5</td><td>-7.5</td><td>8.9E-14</td><td>2.8E-13</td><td>MB231/MB468</td><td>MITO</td><td></td></tr>
<tr><td>>MI0000279|hsa-mir-196a-2&WithFlank&12|+|53991732|53991853@31.53.23&[MIMAT0000226&hsa-miR-196a-5p&offsets|0|+1;m-261&12|+|53991762|53991783&offsets|0|+1]|TAGGTAGTTTCATGTTGTTGGGA</td><td>>MI0000279|hsa-mir-196a-2&WithFlank&12|+|53991732|53991853</td><td>31</td><td>53</td><td>23</td><td>TAGGTAGTTTCATGTTGTTGGGA</td><td>491.0</td><td>3.3</td><td>0.4</td><td>7.4</td><td>1.0E-13</td><td>2.8E-13</td><td>BT20/MB468</td><td>NUC</td><td></td></tr>
<tr><td>>hg38_21_+_8433172_8446622_RNA45SN1_45S_50ntFlanks@9862.9890.29|ATGAACCGAACGCCGGGTTAAGGCGCCCG</td><td>>hg38_21_+_8433172_8446622_RNA45SN1_45S_50ntFlanks</td><td>9862</td><td>9890</td><td>29</td><td>ATGAACCGAACGCCGGGTTAAGGCGCCCG</td><td>285.8</td><td>-12.0</td><td>1.6</td><td>-7.5</td><td>5.8E-14</td><td>2.8E-13</td><td>MB231/MB468</td><td>MP</td><td></td></tr>
<tr><td>>hg38_21_+_8433172_8446622_RNA45SN1_45S_50ntFlanks@12960.12982.23|CTGGGTCGGGGTTTCGTACGTAG</td><td>>hg38_21_+_8433172_8446622_RNA45SN1_45S_50ntFlanks</td><td>12960</td><td>12982</td><td>23</td><td>CTGGGTCGGGGTTTCGTACGTAG</td><td>218.2</td><td>-11.6</td><td>1.5</td><td>-7.5</td><td>5.8E-14</td><td>2.8E-13</td><td>MB231/MB468</td><td>MP</td><td></td></tr>
<tr><td>>hg38_21_+_8433172_8446622_RNA45SN1_45S_50ntFlanks@11209.11230.22|CGGCGACTCTGGACGCGAGCCG</td><td>>hg38_21_+_8433172_8446622_RNA45SN1_45S_50ntFlanks</td><td>11209</td><td>11230</td><td>22</td><td>CGGCGACTCTGGACGCGAGCCG</td><td>1317.0</td><td>-7.4</td><td>1.0</td><td>-7.5</td><td>6.7E-14</td><td>2.9E-13</td><td>MB231/MB468</td><td>CYTO</td><td></td></tr>
<tr><td>>hg38_21_+_8433172_8446622_RNA45SN1_45S_50ntFlanks@10407.10430.24|GGGTCAGTCGGTCCTGAGAGATGG</td><td>>hg38_21_+_8433172_8446622_RNA45SN1_45S_50ntFlanks</td><td>10407</td><td>10430</td><td>24</td><td>GGGTCAGTCGGTCCTGAGAGATGG</td><td>184.0</td><td>-10.5</td><td>1.4</td><td>-7.5</td><td>7.0E-14</td><td>2.9E-13</td><td>MB231/MB468</td><td>NUC</td><td></td></tr>
<tr><td>>hg38_21_+_8433172_8446622_RNA45SN1_45S_50ntFlanks@8170.8201.32|CAAGTCCTTCTGATCGAGGCCCAGCCCGTGGA</td><td>>hg38_21_+_8433172_8446622_RNA45SN1_45S_50ntFlanks</td><td>8170</td><td>8201</td><td>32</td><td>CAAGTCCTTCTGATCGAGGCCCAGCCCGTGGA</td><td>502.3</td><td>-11.9</td><td>1.6</td><td>-7.5</td><td>7.0E-14</td><td>2.9E-13</td><td>MB231/MB468</td><td>NUC</td><td></td></tr>
<tr><td>>hg38_21_+_8433172_8446622_RNA45SN1_45S_50ntFlanks@8105.8141.37|CGCGGCGGGGCGCGGGACATGTGGCGTACGGAAGACC</td><td>>hg38_21_+_8433172_8446622_RNA45SN1_45S_50ntFlanks</td><td>8105</td><td>8141</td><td>37</td><td>CGCGGCGGGGCGCGGGACATGTGGCGTACGGAAGACC</td><td>5396.2</td><td>-7.7</td><td>1.0</td><td>-7.5</td><td>5.8E-14</td><td>2.9E-13</td><td>MB231/MB468</td><td>MP</td><td></td></tr>
<tr><td>>hg38_21_+_8433172_8446622_RNA45SN1_45S_50ntFlanks@6643.6681.39|ACCTCGTACGACTCTTAGCGGTGGATCACTCGGCTCGTG</td><td>>hg38_21_+_8433172_8446622_RNA45SN1_45S_50ntFlanks</td><td>6643</td><td>6681</td><td>39</td><td>ACCTCGTACGACTCTTAGCGGTGGATCACTCGGCTCGTG</td><td>550.5</td><td>5.6</td><td>0.7</td><td>7.5</td><td>7.0E-14</td><td>2.9E-13</td><td>MB231/MB468</td><td>NUC</td><td></td></tr>
<tr><td>>hg38_21_+_8433172_8446622_RNA45SN1_45S_50ntFlanks@7986.8017.32|ATCAGACGTGGCGACCCGCTGAATTTAAGCAT</td><td>>hg38_21_+_8433172_8446622_RNA45SN1_45S_50ntFlanks</td><td>7986</td><td>8017</td><td>32</td><td>ATCAGACGTGGCGACCCGCTGAATTTAAGCAT</td><td>701.1</td><td>-12.5</td><td>1.6</td><td>-7.9</td><td>2.5E-15</td><td>2.9E-13</td><td>BT20/MB468</td><td>MP</td><td></td></tr>
<tr><td>>hg38_21_+_8433172_8446622_RNA45SN1_45S_50ntFlanks@10797.10818.22|GCATGTTGGAACAATGTAGGTA</td><td>>hg38_21_+_8433172_8446622_RNA45SN1_45S_50ntFlanks</td><td>10797</td><td>10818</td><td>22</td><td>GCATGTTGGAACAATGTAGGTA</td><td>3439.3</td><td>-5.2</td><td>0.7</td><td>-7.5</td><td>6.8E-14</td><td>2.9E-13</td><td>MB231/MB468</td><td>CYTO</td><td></td></tr>
<tr><td>>hg38_21_+_8433172_8446622_RNA45SN1_45S_50ntFlanks@4982.5016.35|ACGGACAGGATTGACAGATTGATAGCTCTTTCTCG</td><td>>hg38_21_+_8433172_8446622_RNA45SN1_45S_50ntFlanks</td><td>4982</td><td>5016</td><td>35</td><td>ACGGACAGGATTGACAGATTGATAGCTCTTTCTCG</td><td>670.0</td><td>-13.2</td><td>1.8</td><td>-7.5</td><td>9.1E-14</td><td>2.9E-13</td><td>MB231/MB468</td><td>MITO</td><td></td></tr>
<tr><td>>hg38_21_+_8433172_8446622_RNA45SN1_45S_50ntFlanks@10351.10378.28|CGAGAACTTTGAAGGCCGAAGTGGAGAA</td><td>>hg38_21_+_8433172_8446622_RNA45SN1_45S_50ntFlanks</td><td>10351</td><td>10378</td><td>28</td><td>CGAGAACTTTGAAGGCCGAAGTGGAGAA</td><td>929.1</td><td>-7.3</td><td>1.0</td><td>-7.5</td><td>9.1E-14</td><td>2.9E-13</td><td>MB231/MB468</td><td>MITO</td><td></td></tr>
<tr><td>>hg38_21_+_8433172_8446622_RNA45SN1_45S_50ntFlanks@12033.12053.21|ATCGTTTTTTCACTGACCCGG</td><td>>hg38_21_+_8433172_8446622_RNA45SN1_45S_50ntFlanks</td><td>12033</td><td>12053</td><td>21</td><td>ATCGTTTTTTCACTGACCCGG</td><td>372.6</td><td>-10.9</td><td>1.4</td><td>-7.5</td><td>7.2E-14</td><td>2.9E-13</td><td>BT20/MB468</td><td>CYTO</td><td></td></tr>
<tr><td>>hg38_21_+_8433172_8446622_RNA45SN1_45S_50ntFlanks@12073.12109.37|CCGAGGGGCTCTCGCTTCTGGCGCCAAGCGCCCGGCC</td><td>>hg38_21_+_8433172_8446622_RNA45SN1_45S_50ntFlanks</td><td>12073</td><td>12109</td><td>37</td><td>CCGAGGGGCTCTCGCTTCTGGCGCCAAGCGCCCGGCC</td><td>158.5</td><td>-11.2</td><td>1.5</td><td>-7.5</td><td>5.9E-14</td><td>2.9E-13</td><td>MB231/MB468</td><td>MP</td><td></td></tr>
<tr><td>>hg38_21_+_8433172_8446622_RNA45SN1_45S_50ntFlanks@6725.6747.23|GCAGGACACATTGATCATCGACA</td><td>>hg38_21_+_8433172_8446622_RNA45SN1_45S_50ntFlanks</td><td>6725</td><td>6747</td><td>23</td><td>GCAGGACACATTGATCATCGACA</td><td>659.2</td><td>5.7</td><td>0.8</td><td>7.5</td><td>7.2E-14</td><td>2.9E-13</td><td>BT20/MB468</td><td>CYTO</td><td></td></tr>
<tr><td>>hg38_21_+_8433172_8446622_RNA45SN1_45S_50ntFlanks@4759.4780.22|ATCAGATACCGTCGTAGTTCCG</td><td>>hg38_21_+_8433172_8446622_RNA45SN1_45S_50ntFlanks</td><td>4759</td><td>4780</td><td>22</td><td>ATCAGATACCGTCGTAGTTCCG</td><td>12640.8</td><td>-11.1</td><td>1.5</td><td>-7.5</td><td>9.2E-14</td><td>2.9E-13</td><td>MB231/MB468</td><td>MITO</td><td></td></tr>
<tr><td>>hg38_1_-_228634819_228635039_RNA5S12_5S_50ntFlanks@130.171.42|TGGATGGGAGACCGCCTGGGAATACCGGGTGCTGTAGGCTTT</td><td>>hg38_1_-_228634819_228635039_RNA5S12_5S_50ntFlanks</td><td>130</td><td>171</td><td>42</td><td>TGGATGGGAGACCGCCTGGGAATACCGGGTGCTGTAGGCTTT</td><td>4189.6</td><td>9.4</td><td>1.3</td><td>7.5</td><td>7.3E-14</td><td>2.9E-13</td><td>BT20/MB468</td><td>CYTO</td><td></td></tr>
<tr><td>>MI0000750|hsa-mir-26a-2&WithFlank&12|-|57824603|57824698@20.40.21;MI0000083|hsa-mir-26a-1&WithFlank&3|+|37969398|37969486@16.36.21&[MIMAT0000082&hsa-miR-26a-5p&offsets|0|-1;m-33&12|-|57824658|57824679&offsets|0|-1];[MIMAT0000082_1&hsa-miR-26a-5p&offsets|0|-1;m-34&3|+|37969413|37969434&offsets|0|-1]|TTCAAGTAATCCAGGATAGGC</td><td>>MI0000750|hsa-mir-26a-2&WithFlank&12|-|57824603|57824698</td><td>20</td><td>40</td><td>21</td><td>TTCAAGTAATCCAGGATAGGC</td><td>517.7</td><td>5.6</td><td>0.7</td><td>7.5</td><td>5.4E-14</td><td>2.9E-13</td><td>BT20/MB468</td><td>MITO</td><td></td></tr>
<tr><td>>hg38_21_+_8433172_8446622_RNA45SN1_45S_50ntFlanks@4472.4493.22|CTTAGCTGAGTGTCCCGCGGGG</td><td>>hg38_21_+_8433172_8446622_RNA45SN1_45S_50ntFlanks</td><td>4472</td><td>4493</td><td>22</td><td>CTTAGCTGAGTGTCCCGCGGGG</td><td>258.2</td><td>-8.3</td><td>1.1</td><td>-7.5</td><td>7.1E-14</td><td>2.9E-13</td><td>MB231/MB468</td><td>NUC</td><td></td></tr>
<tr><td>>hg38_21_+_8433172_8446622_RNA45SN1_45S_50ntFlanks@4764.4781.18|ATACCGTCGTAGTTCCGA</td><td>>hg38_21_+_8433172_8446622_RNA45SN1_45S_50ntFlanks</td><td>4764</td><td>4781</td><td>18</td><td>ATACCGTCGTAGTTCCGA</td><td>222.4</td><td>-5.8</td><td>0.8</td><td>-7.5</td><td>6.9E-14</td><td>2.9E-13</td><td>MB231/MB468</td><td>CYTO</td><td></td></tr>
<tr><td>>trna11_GluTTC_15_-_26327381_26327452@1.32.32,trna3_GluTTC_13_-_45492062_45492133@1.32.32|TCCCACATGGTCTAGCGGTTAGGATTCCTGGT</td><td>>trna11_GluTTC_15_-_26327381_26327452</td><td>1</td><td>32</td><td>32</td><td>TCCCACATGGTCTAGCGGTTAGGATTCCTGGT</td><td>225.0</td><td>11.0</td><td>1.5</td><td>7.5</td><td>5.9E-14</td><td>2.9E-13</td><td>MB231/MB468</td><td>MP</td><td></td></tr>
<tr><td>>hg38_21_+_8433172_8446622_RNA45SN1_45S_50ntFlanks@8170.8204.35|CAAGTCCTTCTGATCGAGGCCCAGCCCGTGGACGG</td><td>>hg38_21_+_8433172_8446622_RNA45SN1_45S_50ntFlanks</td><td>8170</td><td>8204</td><td>35</td><td>CAAGTCCTTCTGATCGAGGCCCAGCCCGTGGACGG</td><td>362.0</td><td>-12.4</td><td>1.6</td><td>-7.5</td><td>6.0E-14</td><td>2.9E-13</td><td>MB231/MB468</td><td>MP</td><td></td></tr>
</table>

<table>
<tr><td>>hg38_1_-_228634819_228635039_RNA5S12_5S_50ntFlanks@51.91.41|GTCTACGGCCATACCACCCTGAACGCGCCCGATCTCGTCTG</td><td>>hg38_1_-_228634819_228635039_RNA5S12_5S_50ntFlanks</td><td>51</td><td>91</td><td>41</td><td>GTCTACGGCCATACCACCCTGAACGCGCCCGATCTCGTCTG</td><td>6949.7</td><td>8.8</td><td>1.2</td><td>7.5</td><td>7.3E-14</td><td>2.9E-13</td><td>BT20/MB468</td><td>CYTO</td></tr>
<tr><td>>MI0000103|hsa-mir-101-1&WithFlank&1|-|65058428|65058514@53.72.20;MI0000739|hsa-mir-101-2&WithFlank&9|+|4850291|4850381@55.74.20&[MIMAT0000099&hsa-miR-101-3p&offsets|0|-1;m-17&1|-|65058443|65058462&offsets|0|0];[MIMAT0000099_1&hsa-miR-101-3p&offsets|0|-1;m-16&9|+|4850345|4850364&offsets|0|0]|TACAGTACTGTGATAACTGA</td><td>>MI0000103|hsa-mir-101-1&WithFlank&1|-|65058428|65058514</td><td>53</td><td>72</td><td>20</td><td>TACAGTACTGTGATAACTGA</td><td>1090.4</td><td>5.8</td><td>0.8</td><td>7.5</td><td>7.1E-14</td><td>2.9E-13</td><td>MB231/MB468</td><td>NUC</td></tr>
<tr><td>>MI0000737|hsa-mir-200a&WithFlank&1|+|1167857|1167958@22.43.22(+1A)&[MIMAT0001620&hsa-miR-200a-5p&offsets|0|0(+1A);m-154&1|+|1167878|1167899&offsets|0|0(+1A)]|CATCTTACCGGACAGTGCTGGAA</td><td>>MI0000737|hsa-mir-200a&WithFlank&1|+|1167857|1167958</td><td>22</td><td>43</td><td>22(+1A)</td><td>CATCTTACCGGACAGTGCTGGAA</td><td>145.7</td><td>-11.1</td><td>1.5</td><td>-7.5</td><td>6.0E-14</td><td>2.9E-13</td><td>MB231/MB468</td><td>MP</td></tr>
<tr><td>>hg38_21_+_8433172_8446622_RNA45SN1_45S_50ntFlanks@10099.10116.18|CGCGCGCGCGTGTGGTGT</td><td>>hg38_21_+_8433172_8446622_RNA45SN1_45S_50ntFlanks</td><td>10099</td><td>10116</td><td>18</td><td>CGCGCGCGCGTGTGGTGT</td><td>145.2</td><td>-11.0</td><td>1.5</td><td>-7.5</td><td>6.0E-14</td><td>2.9E-13</td><td>MB231/MB468</td><td>MP</td></tr>
<tr><td>>hg38_21_+_8433172_8446622_RNA45SN1_45S_50ntFlanks@482.531.50|ATCGATGTGGTGACGTCGTGCTCTCCCGGGCCGGGTCCGAGCCGCGACGG</td><td>>hg38_21_+_8433172_8446622_RNA45SN1_45S_50ntFlanks</td><td>482</td><td>531</td><td>50</td><td>ATCGATGTGGTGACGTCGTGCTCTCCCGGGCCGGGTCCGAGCCGCGACGG</td><td>2025.4</td><td>-15.4</td><td>2.1</td><td>-7.4</td><td>1.1E-13</td><td>2.9E-13</td><td>BT20/MB468</td><td>NUC</td></tr>
<tr><td>>hg38_21_+_8433172_8446622_RNA45SN1_45S_50ntFlanks@12994.13014.21|CCTCGCTGCGATCTATTGAAA</td><td>>hg38_21_+_8433172_8446622_RNA45SN1_45S_50ntFlanks</td><td>12994</td><td>13014</td><td>21</td><td>CCTCGCTGCGATCTATTGAAA</td><td>987.3</td><td>-3.8</td><td>0.5</td><td>-7.5</td><td>7.2E-14</td><td>2.9E-13</td><td>MB231/MB468</td><td>NUC</td></tr>
<tr><td>>hg38_21_+_8433172_8446622_RNA45SN1_45S_50ntFlanks@10808.10828.21|CAATGTAGGTAAGGGAAGTCG</td><td>>hg38_21_+_8433172_8446622_RNA45SN1_45S_50ntFlanks</td><td>10808</td><td>10828</td><td>21</td><td>CAATGTAGGTAAGGGAAGTCG</td><td>1144.3</td><td>-6.4</td><td>0.9</td><td>-7.5</td><td>6.9E-14</td><td>2.9E-13</td><td>MB231/MB468</td><td>CYTO</td></tr>
<tr><td>>hg38_21_+_8433172_8446622_RNA45SN1_45S_50ntFlanks@4064.4081.18|ATCAACTTTCGATGGTAG</td><td>>hg38_21_+_8433172_8446622_RNA45SN1_45S_50ntFlanks</td><td>4064</td><td>4081</td><td>18</td><td>ATCAACTTTCGATGGTAG</td><td>472.7</td><td>-12.0</td><td>1.5</td><td>-7.9</td><td>2.6E-15</td><td>2.9E-13</td><td>BT20/MB468</td><td>MP</td></tr>
<tr><td>>hg38_21_+_8433172_8446622_RNA45SN1_45S_50ntFlanks@10351.10371.21|CGAGAACTTTGAAGGCCGAAG</td><td>>hg38_21_+_8433172_8446622_RNA45SN1_45S_50ntFlanks</td><td>10351</td><td>10371</td><td>21</td><td>CGAGAACTTTGAAGGCCGAAG</td><td>1842.0</td><td>-5.6</td><td>0.7</td><td>-7.4</td><td>9.3E-14</td><td>2.9E-13</td><td>MB231/MB468</td><td>MITO</td></tr>
<tr><td>>hg38_21_+_8433172_8446622_RNA45SN1_45S_50ntFlanks@4380.4410.31|CGTAGTTGGATCTTGGGAGCGGGCGGGCGGT</td><td>>hg38_21_+_8433172_8446622_RNA45SN1_45S_50ntFlanks</td><td>4380</td><td>4410</td><td>31</td><td>CGTAGTTGGATCTTGGGAGCGGGCGGGCGGT</td><td>196.1</td><td>-11.5</td><td>1.5</td><td>-7.5</td><td>6.0E-14</td><td>3.0E-13</td><td>MB231/MB468</td><td>MP</td></tr>
<tr><td>>MI0000266|hsa-mir-10a&WithFlank&17|-|48579832|48579953@28.49.22&[MIMAT0000253&hsa-miR-10a-5p&offsets|0|-1;m-18&17|-|48579905|48579926&offsets|0|0]|TACCCTGTAGATCCGAATTTGT</td><td>>MI0000266|hsa-mir-10a&WithFlank&17|-|48579832|48579953</td><td>28</td><td>49</td><td>22</td><td>TACCCTGTAGATCCGAATTTGT</td><td>10180.0</td><td>12.9</td><td>1.7</td><td>7.5</td><td>5.6E-14</td><td>3.0E-13</td><td>BT20/MB468</td><td>MITO</td></tr>
<tr><td>>hg38_21_+_8433172_8446622_RNA45SN1_45S_50ntFlanks@10853.10875.23|ATAAGGATTGGCTCTAAGGGCTG</td><td>>hg38_21_+_8433172_8446622_RNA45SN1_45S_50ntFlanks</td><td>10853</td><td>10875</td><td>23</td><td>ATAAGGATTGGCTCTAAGGGCTG</td><td>160.8</td><td>-11.2</td><td>1.5</td><td>-7.5</td><td>6.1E-14</td><td>3.0E-13</td><td>MB231/MB468</td><td>MP</td></tr>
<tr><td>>hg38_21_+_8433172_8446622_RNA45SN1_45S_50ntFlanks@6664.6699.36|TGGATCACTCGGCTCGTGCGTCGATGAAGAACGCAG</td><td>>hg38_21_+_8433172_8446622_RNA45SN1_45S_50ntFlanks</td><td>6664</td><td>6699</td><td>36</td><td>TGGATCACTCGGCTCGTGCGTCGATGAAGAACGCAG</td><td>219.3</td><td>-11.6</td><td>1.6</td><td>-7.5</td><td>6.1E-14</td><td>3.0E-13</td><td>MB231/MB468</td><td>MP</td></tr>
<tr><td>>hg38_21_+_8433172_8446622_RNA45SN1_45S_50ntFlanks@10786.10823.38|AACAGCCTCTGGCATGTTGGAACAATGTAGGTAAGGGA</td><td>>hg38_21_+_8433172_8446622_RNA45SN1_45S_50ntFlanks</td><td>10786</td><td>10823</td><td>38</td><td>AACAGCCTCTGGCATGTTGGAACAATGTAGGTAAGGGA</td><td>2606.1</td><td>-9.8</td><td>1.3</td><td>-7.4</td><td>9.5E-14</td><td>3.0E-13</td><td>MB231/MB468</td><td>MITO</td></tr>
<tr><td>>hg38_21_+_8433172_8446622_RNA45SN1_45S_50ntFlanks@10388.10409.22|GTGAACAGCAGTTGAACATGGG</td><td>>hg38_21_+_8433172_8446622_RNA45SN1_45S_50ntFlanks</td><td>10388</td><td>10409</td><td>22</td><td>GTGAACAGCAGTTGAACATGGG</td><td>183.3</td><td>-5.6</td><td>0.7</td><td>-7.5</td><td>7.1E-14</td><td>3.0E-13</td><td>MB231/MB468</td><td>CYTO</td></tr>
<tr><td>>MI0000103|hsa-mir-101-1&WithFlank&1|-|65058428|65058514@53.73.21;MI0000739|hsa-mir-101-2&WithFlank&9|+|4850291|4850381@55.75.21&[MIMAT0000099&hsa-miR-101-3p&offsets|0|0;m-17&1|-|65058443|65058462&offsets|0|+1];[MIMAT0000099_1&hsa-miR-101-3p&offsets|0|0;m-16&9|+|4850345|4850364&offsets|0|+1]|TACAGTACTGTGATAACTGAA</td><td>>MI0000103|hsa-mir-101-1&WithFlank&1|-|65058428|65058514</td><td>53</td><td>73</td><td>21</td><td>TACAGTACTGTGATAACTGAA</td><td>393.9</td><td>5.9</td><td>0.8</td><td>7.5</td><td>7.5E-14</td><td>3.0E-13</td><td>BT20/MB468</td><td>CYTO</td></tr>
<tr><td>>hg38_21_+_8433172_8446622_RNA45SN1_45S_50ntFlanks@11233.11250.18|CCCTTCCCGTGGATCGCC</td><td>>hg38_21_+_8433172_8446622_RNA45SN1_45S_50ntFlanks</td><td>11233</td><td>11250</td><td>18</td><td>CCCTTCCCGTGGATCGCC</td><td>346.5</td><td>-10.5</td><td>1.4</td><td>-7.5</td><td>5.6E-14</td><td>3.0E-13</td><td>BT20/MB468</td><td>MITO</td></tr>
<tr><td>>MI0000102|hsa-mir-100&WithFlank&11|-|122152223|122152314@20.39.20&[MIMAT0000098&hsa-miR-100-5p&offsets|+1|-1;m-27&11|-|122152276|122152296&offsets|+1|0]|ACCCGTAGATCCGAACTTGT</td><td>>MI0000102|hsa-mir-100&WithFlank&11|-|122152223|122152314</td><td>20</td><td>39</td><td>20</td><td>ACCCGTAGATCCGAACTTGT</td><td>263.9</td><td>11.3</td><td>1.5</td><td>7.5</td><td>6.1E-14</td><td>3.0E-13</td><td>MB231/MB468</td><td>MP</td></tr>
<tr><td>>MI0000471|hsa-mir-126&WithFlank&9|+|136670596|136670692@59.78.20&[MIMAT0000445&hsa-miR-126-3p&offsets|+1|-1;m-40&9|+|136670653|136670674&offsets|+1|-1]|CGTACCGTGAGTAATAATGC</td><td>>MI0000471|hsa-mir-126&WithFlank&9|+|136670596|136670692</td><td>59</td><td>78</td><td>20</td><td>CGTACCGTGAGTAATAATGC</td><td>1441.4</td><td>12.4</td><td>1.7</td><td>7.5</td><td>7.5E-14</td><td>3.0E-13</td><td>BT20/MB468</td><td>CYTO</td></tr>
<tr><td>>hg38_21_+_8433172_8446622_RNA45SN1_45S_50ntFlanks@12126.12146.21|CGACCCGCTCCGGGGACAGTG</td><td>>hg38_21_+_8433172_8446622_RNA45SN1_45S_50ntFlanks</td><td>12126</td><td>12146</td><td>21</td><td>CGACCCGCTCCGGGGACAGTG</td><td>474.7</td><td>-8.5</td><td>1.1</td><td>-7.5</td><td>7.5E-14</td><td>3.0E-13</td><td>BT20/MB468</td><td>CYTO</td></tr>
<tr><td>>hg38_21_+_8433172_8446622_RNA45SN1_45S_50ntFlanks@10291.10310.20|CGCGGGCCCGGGTGGAGCCG</td><td>>hg38_21_+_8433172_8446622_RNA45SN1_45S_50ntFlanks</td><td>10291</td><td>10310</td><td>20</td><td>CGCGGGCCCGGGTGGAGCCG</td><td>194.0</td><td>-11.5</td><td>1.5</td><td>-7.5</td><td>6.1E-14</td><td>3.0E-13</td><td>MB231/MB468</td><td>MP</td></tr>
<tr><td>>hg38_21_+_8433172_8446622_RNA45SN1_45S_50ntFlanks@12676.12709.34|AGAATCCCGCCCAGGCGGAACGATACGGCAGCGC</td><td>>hg38_21_+_8433172_8446622_RNA45SN1_45S_50ntFlanks</td><td>12676</td><td>12709</td><td>34</td><td>AGAATCCCGCCCAGGCGGAACGATACGGCAGCGC</td><td>124.2</td><td>-10.8</td><td>1.4</td><td>-7.5</td><td>6.1E-14</td><td>3.0E-13</td><td>MB231/MB468</td><td>MP</td></tr>
<tr><td>>hg38_21_+_8433172_8446622_RNA45SN1_45S_50ntFlanks@12630.12653.24|GGGCGAAGCTACCATCTGTGGGAT</td><td>>hg38_21_+_8433172_8446622_RNA45SN1_45S_50ntFlanks</td><td>12630</td><td>12653</td><td>24</td><td>GGGCGAAGCTACCATCTGTGGGAT</td><td>181.3</td><td>-11.4</td><td>1.5</td><td>-7.5</td><td>6.2E-14</td><td>3.0E-13</td><td>MB231/MB468</td><td>MP</td></tr>
<tr><td>>MI0000065|hsa-let-7d&WithFlank&9|+|94178828|94178926@14.35.22&[MIMAT0000065&hsa-let-7d-5p&offsets|0|0;m-90&9|+|94178841|94178862&offsets|0|0]|AGAGGTAGTAGGTTGCATAGTT</td><td>>MI0000065|hsa-let-7d&WithFlank&9|+|94178828|94178926</td><td>14</td><td>35</td><td>22</td><td>AGAGGTAGTAGGTTGCATAGTT</td><td>5102.8</td><td>5.7</td><td>0.8</td><td>7.5</td><td>7.1E-14</td><td>3.0E-13</td><td>MB231/MB468</td><td>CYTO</td></tr>
<tr><td>>hg38_21_+_8433172_8446622_RNA45SN1_45S_50ntFlanks@4367.4388.22|CAGTTAAAAAGCTCGTAGTTGG</td><td>>hg38_21_+_8433172_8446622_RNA45SN1_45S_50ntFlanks</td><td>4367</td><td>4388</td><td>22</td><td>CAGTTAAAAAGCTCGTAGTTGG</td><td>170.6</td><td>-11.3</td><td>1.5</td><td>-7.5</td><td>6.2E-14</td><td>3.0E-13</td><td>MB231/MB468</td><td>MP</td></tr>
<tr><td>>hg38_21_+_8433172_8446622_RNA45SN1_45S_50ntFlanks@6771.6799.29|GGTTCCTCCCGGGGCTACGCCTGTCTGAG</td><td>>hg38_21_+_8433172_8446622_RNA45SN1_45S_50ntFlanks</td><td>6771</td><td>6799</td><td>29</td><td>GGTTCCTCCCGGGGCTACGCCTGTCTGAG</td><td>148.4</td><td>-9.9</td><td>1.3</td><td>-7.5</td><td>7.1E-14</td><td>3.0E-13</td><td>MB231/MB468</td><td>CYTO</td></tr>
<tr><td>>hg38_21_+_8433172_8446622_RNA45SN1_45S_50ntFlanks@10798.10818.21|CATGTTGGAACAATGTAGGTA</td><td>>hg38_21_+_8433172_8446622_RNA45SN1_45S_50ntFlanks</td><td>10798</td><td>10818</td><td>21</td><td>CATGTTGGAACAATGTAGGTA</td><td>4116.9</td><td>-5.8</td><td>0.8</td><td>-7.5</td><td>7.1E-14</td><td>3.0E-13</td><td>MB231/MB468</td><td>CYTO</td></tr>
<tr><td>>hg38_21_+_8433172_8446622_RNA45SN1_45S_50ntFlanks@10782.10818.37|GGTGAACAGCCTCTGGCATGTTGGAACAATGTAGGTA</td><td>>hg38_21_+_8433172_8446622_RNA45SN1_45S_50ntFlanks</td><td>10782</td><td>10818</td><td>37</td><td>GGTGAACAGCCTCTGGCATGTTGGAACAATGTAGGTA</td><td>420.8</td><td>-6.2</td><td>0.8</td><td>-7.5</td><td>7.2E-14</td><td>3.0E-13</td><td>MB231/MB468</td><td>CYTO</td></tr>
<tr><td>>hg38_21_+_8433172_8446622_RNA45SN1_45S_50ntFlanks@10362.10390.29|AAGGCCGAAGTGGAGAAGGGTTCCATGTG</td><td>>hg38_21_+_8433172_8446622_RNA45SN1_45S_50ntFlanks</td><td>10362</td><td>10390</td><td>29</td><td>AAGGCCGAAGTGGAGAAGGGTTCCATGTG</td><td>2852.0</td><td>-8.7</td><td>1.2</td><td>-7.5</td><td>7.4E-14</td><td>3.0E-13</td><td>MB231/MB468</td><td>NUC</td></tr>
<tr><td>>hg38_21_+_8433172_8446622_RNA45SN1_45S_50ntFlanks@11200.11232.33|CGGCGGCGGCGGCGACTCTGGACGCGAGCCGGG</td><td>>hg38_21_+_8433172_8446622_RNA45SN1_45S_50ntFlanks</td><td>11200</td><td>11232</td><td>33</td><td>CGGCGGCGGCGGCGACTCTGGACGCGAGCCGGG</td><td>2210.4</td><td>-8.6</td><td>1.1</td><td>-7.5</td><td>6.2E-14</td><td>3.0E-13</td><td>MB231/MB468</td><td>MP</td></tr>
<tr><td>>am_trna2_GlyGCC_21_-_18827107_18827177@1.24.24,trna35_GlyGCC_1_+_161413094_161413164@1.24.24,trna37_GlyGCC_1_+_161420467_161420537@1.24.24,trna39_GlyGCC_1_+_161427898_161427968@1.24.24,trna41_GlyGCC_1_+_161435258_161435328@1.24.24|GCATGGGTGGTTCAGTGGTAGAAT</td><td>>am_trna2_GlyGCC_21_-_18827107_18827177</td><td>1</td><td>24</td><td>24</td><td>GCATGGGTGGTTCAGTGGTAGAAT</td><td>231.4</td><td>11.2</td><td>1.4</td><td>7.8</td><td>5.6E-15</td><td>3.0E-13</td><td>BT20/MB231</td><td>CYTO</td></tr>
<tr><td>>hg38_21_+_8433172_8446622_RNA45SN1_45S_50ntFlanks@6644.6667.24|CCTCGTACGACTCTTAGCGGTGGA</td><td>>hg38_21_+_8433172_8446622_RNA45SN1_45S_50ntFlanks</td><td>6644</td><td>6667</td><td>24</td><td>CCTCGTACGACTCTTAGCGGTGGA</td><td>576.0</td><td>5.9</td><td>0.8</td><td>7.5</td><td>5.7E-14</td><td>3.0E-13</td><td>BT20/MB468</td><td>MITO</td></tr>
<tr><td>>am_MI0003639|hsa-mir-625&WithFlank&14|+|65471096|65471192@58.78.21(+1U)&[MIMAT0004808&hsa-miR-625-3p&offsets|0|-1(+1U);m-137&14|+|65471153|65471174&offsets|0|-1(+1U)]|GACTATAGAACTTTCCCCCTCT</td><td>>am_MI0003639|hsa-mir-625&WithFlank&14|+|65471096|65471192</td><td>58</td><td>78</td><td>21(+1U)</td><td>GACTATAGAACTTTCCCCCTCT</td><td>386.3</td><td>9.5</td><td>1.3</td><td>7.5</td><td>7.2E-14</td><td>3.0E-13</td><td>MB231/MB468</td><td>CYTO</td></tr>
<tr><td>>trnaMT_ValTAC_MT_+_1602_1670@50.72.23|ACTTAACTTGACCGCTCTGACCA</td><td>>trnaMT_ValTAC_MT_+_1602_1670</td><td>50</td><td>72</td><td>23</td><td>ACTTAACTTGACCGCTCTGACCA</td><td>3364.1</td><td>7.0</td><td>0.9</td><td>7.5</td><td>7.6E-14</td><td>3.0E-13</td><td>BT20/MB468</td><td>CYTO</td></tr>
<tr><td>>hg38_21_+_8433172_8446622_RNA45SN1_45S_50ntFlanks@10405.10424.20|ATGGGTCAGTCGGTCCTGAG</td><td>>hg38_21_+_8433172_8446622_RNA45SN1_45S_50ntFlanks</td><td>10405</td><td>10424</td><td>20</td><td>ATGGGTCAGTCGGTCCTGAG</td><td>1266.3</td><td>5.1</td><td>0.7</td><td>7.5</td><td>7.2E-14</td><td>3.0E-13</td><td>MB231/MB468</td><td>CYTO</td></tr>
<tr><td>>am_trna162_MetCAT_6_-_26758550_26758622@31.51.21,trna164_MetCAT_6_-_26735574_26735646@31.51.21,trna20_MetCAT_16_+_71460396_71460468@31.51.21,trna21_MetCAT_6_+_26701712_26701784@31.51.21,trna27_MetCAT_6_+_26766444_26766516@31.51.21,trna75_MetCAT_6_+_28912352_28912424@31.51.21,trna97_MetCAT_6_-_28921042_28921114@31.51.21|TCTCATAATCTGAAGGTCCTG</td><td>>am_trna162_MetCAT_6_-_26758550_26758622</td><td>31</td><td>51</td><td>21</td><td>TCTCATAATCTGAAGGTCCTG</td><td>349.6</td><td>-11.2</td><td>1.5</td><td>-7.5</td><td>7.2E-14</td><td>3.0E-13</td><td>MB231/MB468</td><td>CYTO</td></tr>
<tr><td>>hg38_21_+_8433172_8446622_RNA45SN1_45S_50ntFlanks@11886.11903.18|TGAGCTTGACTCTAGTCT</td><td>>hg38_21_+_8433172_8446622_RNA45SN1_45S_50ntFlanks</td><td>11886</td><td>11903</td><td>18</td><td>TGAGCTTGACTCTAGTCT</td><td>260.1</td><td>-11.8</td><td>1.6</td><td>-7.4</td><td>9.7E-14</td><td>3.0E-13</td><td>MB231/MB468</td><td>MITO</td></tr>
<tr><td>>hg38_21_+_8433172_8446622_RNA45SN1_45S_50ntFlanks@11786.11803.18|GAATGGATGAACGAGATT</td><td>>hg38_21_+_8433172_8446622_RNA45SN1_45S_50ntFlanks</td><td>11786</td><td>11803</td><td>18</td><td>GAATGGATGAACGAGATT</td><td>2013.6</td><td>-5.2</td><td>0.7</td><td>-7.5</td><td>7.4E-14</td><td>3.1E-13</td><td>MB231/MB468</td><td>NUC</td></tr>
<tr><td>>hg38_21_+_8433172_8446622_RNA45SN1_45S_50ntFlanks@4047.4064.18|ATTCGAACGTCTGCCCTA</td><td>>hg38_21_+_8433172_8446622_RNA45SN1_45S_50ntFlanks</td><td>4047</td><td>4064</td><td>18</td><td>ATTCGAACGTCTGCCCTA</td><td>343.8</td><td>-4.0</td><td>0.5</td><td>-7.5</td><td>7.7E-14</td><td>3.1E-13</td><td>BT20/MB468</td><td>CYTO</td></tr>
<tr><td>>MI0000079|hsa-mir-23a&WithFlank&19|-|13836581|13836665@54.72.19(+1U)&[MIMAT0000078&hsa-miR-23a-3p&offsets|+3|+1(+1U);m-31&19|-|13836595|13836615&offsets|+3|+1(+1U)]|ACATTGCCAGGGATTTCCAT</td><td>>MI0000079|hsa-mir-23a&WithFlank&19|-|13836581|13836665</td><td>54</td><td>72</td><td>19(+1U)</td><td>ACATTGCCAGGGATTTCCAT</td><td>898.9</td><td>27.0</td><td>3.6</td><td>7.4</td><td>9.7E-14</td><td>3.1E-13</td><td>MB231/MB468</td><td>MITO</td></tr>
<tr><td>>hg38_21_+_8433172_8446622_RNA45SN1_45S_50ntFlanks@9337.9370.34|CTCGCACGAAAGCCGCCGTGGCGCAATGAAGGTG</td><td>>hg38_21_+_8433172_8446622_RNA45SN1_45S_50ntFlanks</td><td>9337</td><td>9370</td><td>34</td><td>CTCGCACGAAAGCCGCCGTGGCGCAATGAAGGTG</td><td>287.4</td><td>-12.0</td><td>1.6</td><td>-7.4</td><td>9.7E-14</td><td>3.1E-13</td><td>MB231/MB468</td><td>MITO</td></tr>
<tr><td>>hg38_21_+_8433172_8446622_RNA45SN1_45S_50ntFlanks@12957.12980.24|CTTCTGGGTCGGGGTTTCGTACGT</td><td>>hg38_21_+_8433172_8446622_RNA45SN1_45S_50ntFlanks</td><td>12957</td><td>12980</td><td>24</td><td>CTTCTGGGTCGGGGTTTCGTACGT</td><td>172.4</td><td>-4.9</td><td>0.7</td><td>-7.5</td><td>7.3E-14</td><td>3.1E-13</td><td>MB231/MB468</td><td>CYTO</td></tr>
<tr><td>>hg38_21_+_8433172_8446622_RNA45SN1_45S_50ntFlanks@3987.4010.24|GCTTTGGTGACTCTAGATAACCTC</td><td>>hg38_21_+_8433172_8446622_RNA45SN1_45S_50ntFlanks</td><td>3987</td><td>4010</td><td>24</td><td>GCTTTGGTGACTCTAGATAACCTC</td><td>310.0</td><td>4.1</td><td>0.6</td><td>7.5</td><td>7.7E-14</td><td>3.1E-13</td><td>BT20/MB468</td><td>CYTO</td></tr>
<tr><td>>hg38_21_+_8433172_8446622_RNA45SN1_45S_50ntFlanks@10692.10709.18|TCCGGTGAGCTCTCGCTG</td><td>>hg38_21_+_8433172_8446622_RNA45SN1_45S_50ntFlanks</td><td>10692</td><td>10709</td><td>18</td><td>TCCGGTGAGCTCTCGCTG</td><td>167.8</td><td>-10.1</td><td>1.4</td><td>-7.5</td><td>7.3E-14</td><td>3.1E-13</td><td>MB231/MB468</td><td>CYTO</td></tr>
<tr><td>>trnaMT_PheGAA_MT_+_577_647@38.55.18|AATGTTTAGACGGGCTCA</td><td>>trnaMT_PheGAA_MT_+_577_647</td><td>38</td><td>55</td><td>18</td><td>AATGTTTAGACGGGCTCA</td><td>393.0</td><td>9.0</td><td>1.2</td><td>7.5</td><td>5.8E-14</td><td>3.1E-13</td><td>BT20/MB468</td><td>MITO</td></tr>
<tr><td>>hg38_21_+_8433172_8446622_RNA45SN1_45S_50ntFlanks@8380.8405.26|CAAGAGGGCGTGAAACCGTTAAGAGG</td><td>>hg38_21_+_8433172_8446622_RNA45SN1_45S_50ntFlanks</td><td>8380</td><td>8405</td><td>26</td><td>CAAGAGGGCGTGAAACCGTTAAGAGG</td><td>2147.1</td><td>-5.9</td><td>0.8</td><td>-7.5</td><td>5.8E-14</td><td>3.1E-13</td><td>BT20/MB468</td><td>MITO</td></tr>
<tr><td>>hg38_21_+_8433172_8446622_RNA45SN1_45S_50ntFlanks@10140.10157.18|CGGCGGCGGGGGTGTGGG</td><td>>hg38_21_+_8433172_8446622_RNA45SN1_45S_50ntFlanks</td><td>10140</td><td>10157</td><td>18</td><td>CGGCGGCGGGGGTGTGGG</td><td>210.0</td><td>-11.6</td><td>1.5</td><td>-7.5</td><td>6.3E-14</td><td>3.1E-13</td><td>MB231/MB468</td><td>MP</td></tr>
<tr><td>>hg38_21_+_8433172_8446622_RNA45SN1_45S_50ntFlanks@4163.4181.19|CATCCAAGGAAGGCAGCAG</td><td>>hg38_21_+_8433172_8446622_RNA45SN1_45S_50ntFlanks</td><td>4163</td><td>4181</td><td>19</td><td>CATCCAAGGAAGGCAGCAG</td><td>148.3</td><td>-10.2</td><td>1.4</td><td>-7.5</td><td>7.5E-14</td><td>3.1E-13</td><td>MB231/MB468</td><td>NUC</td></tr>
</table>

<table>
<tr><th></th><th></th><th></th><th></th><th></th><th></th><th></th><th></th><th></th><th></th><th></th><th></th><th></th><th></th></tr>
<tr><td>>MI0000067|hsa-let-7f-1&WithFlank&9|+|94176341|94176439@13.33.21;MI0000068|hsa-let-7f-2&WithFlank&X|-|53557186|53557280@14.34.21&[MIMAT0000067&hsa-let-7f-5p&offsets|0|-1;m-20&9|+|94176353|94176374&offsets|0|-1];[MIMAT0000067_1&hsa-let-7f-5p&offsets|0|-1;m-19&X|-|53557246|53557267&offsets|0|-1]|TGAGGTAGTAGATTGTATAGT</td><td>>MI0000067|hsa-let-7f-1&WithFlank&9|+|94176341|94176439</td><td>13</td><td>33</td><td>21</td><td>TGAGGTAGTAGATTGTATAGT</td><td>4686.2</td><td>4.8</td><td>0.6</td><td>7.5</td><td>7.4E-14</td><td>3.1E-13</td><td>MB231/MB468</td><td>CYTO</td></tr>
<tr><td>>hg38_21_+_8433172_8446622_RNA45SN1_45S_50ntFlanks@10797.10826.30|GCATGTTGGAACAATGTAGGTAAGGGAAGT</td><td>>hg38_21_+_8433172_8446622_RNA45SN1_45S_50ntFlanks</td><td>10797</td><td>10826</td><td>30</td><td>GCATGTTGGAACAATGTAGGTAAGGGAAGT</td><td>975.1</td><td>-7.4</td><td>1.0</td><td>-7.5</td><td>6.4E-14</td><td>3.1E-13</td><td>MB231/MB468</td><td>MP</td></tr>
<tr><td>>hg38_21_+_8433172_8446622_RNA45SN1_45S_50ntFlanks@11876.11908.33|AAGACCCTGTTGAGCTTGACTCTAGTCTGGCAC</td><td>>hg38_21_+_8433172_8446622_RNA45SN1_45S_50ntFlanks</td><td>11876</td><td>11908</td><td>33</td><td>AAGACCCTGTTGAGCTTGACTCTAGTCTGGCAC</td><td>773.9</td><td>-8.9</td><td>1.2</td><td>-7.4</td><td>9.8E-14</td><td>3.1E-13</td><td>MB231/MB468</td><td>MITO</td></tr>
<tr><td>>hg38_21_+_8433172_8446622_RNA45SN1_45S_50ntFlanks@4660.4682.23|GAGGTGAAATTCTTGGACCGGCG</td><td>>hg38_21_+_8433172_8446622_RNA45SN1_45S_50ntFlanks</td><td>4660</td><td>4682</td><td>23</td><td>GAGGTGAAATTCTTGGACCGGCG</td><td>202.5</td><td>-11.5</td><td>1.5</td><td>-7.5</td><td>6.4E-14</td><td>3.1E-13</td><td>MB231/MB468</td><td>MP</td></tr>
<tr><td>>hg38_21_+_8433172_8446622_RNA45SN1_45S_50ntFlanks@11783.11814.32|CATGAATGGATGAACGAGATTCCCACTGTCCC</td><td>>hg38_21_+_8433172_8446622_RNA45SN1_45S_50ntFlanks</td><td>11783</td><td>11814</td><td>32</td><td>CATGAATGGATGAACGAGATTCCCACTGTCCC</td><td>192.5</td><td>-11.5</td><td>1.5</td><td>-7.5</td><td>6.4E-14</td><td>3.1E-13</td><td>MB231/MB468</td><td>MP</td></tr>
<tr><td>>hg38_21_+_8433172_8446622_RNA45SN1_45S_50ntFlanks@12073.12108.36|CCGAGGGGCTCTCGCTTCTGGCGCCAAGCGCCCGGC</td><td>>hg38_21_+_8433172_8446622_RNA45SN1_45S_50ntFlanks</td><td>12073</td><td>12108</td><td>36</td><td>CCGAGGGGCTCTCGCTTCTGGCGCCAAGCGCCCGGC</td><td>617.5</td><td>-8.8</td><td>1.2</td><td>-7.4</td><td>9.9E-14</td><td>3.1E-13</td><td>MB231/MB468</td><td>MITO</td></tr>
<tr><td>>hg38_21_+_8433172_8446622_RNA45SN1_45S_50ntFlanks@11572.11591.20|CCGACTTAGAACTGGTGCGG</td><td>>hg38_21_+_8433172_8446622_RNA45SN1_45S_50ntFlanks</td><td>11572</td><td>11591</td><td>20</td><td>CCGACTTAGAACTGGTGCGG</td><td>6063.4</td><td>-6.9</td><td>0.9</td><td>-7.5</td><td>7.4E-14</td><td>3.1E-13</td><td>MB231/MB468</td><td>CYTO</td></tr>
<tr><td>>MI0000542|hsa-mir-320a&WithFlank&8|-|22244960|22245043@48.68.21(+2U)&[MIMAT0000510&hsa-miR-320a-3p&offsets|0|-1(+2U);m-60&8|-|22244975|22244996&offsets|0|-1(+2U)]|AAAAGCTGGGTTGAGAGGGCGTT</td><td>>MI0000542|hsa-mir-320a&WithFlank&8|-|22244960|22245043</td><td>48</td><td>68</td><td>21(+2U)</td><td>AAAAGCTGGGTTGAGAGGGCGTT</td><td>2233.4</td><td>11.5</td><td>1.5</td><td>7.5</td><td>5.9E-14</td><td>3.1E-13</td><td>BT20/MB468</td><td>MITO</td></tr>
<tr><td>>hg38_21_+_8433172_8446622_RNA45SN1_45S_50ntFlanks@6762.6806.45|GCGGCCCCGGGTTCCTCCCGGGGCTACGCCTGTCTGAGCGTCGCT</td><td>>hg38_21_+_8433172_8446622_RNA45SN1_45S_50ntFlanks</td><td>6762</td><td>6806</td><td>45</td><td>GCGGCCCCGGGTTCCTCCCGGGGCTACGCCTGTCTGAGCGTCGCT</td><td>450.3</td><td>10.5</td><td>1.4</td><td>7.5</td><td>7.5E-14</td><td>3.1E-13</td><td>MB231/MB468</td><td>CYTO</td></tr>
<tr><td>>hg38_21_+_8433172_8446622_RNA45SN1_45S_50ntFlanks@3873.3899.27|TAATACATGCCGACGGGCGCTGACCCC</td><td>>hg38_21_+_8433172_8446622_RNA45SN1_45S_50ntFlanks</td><td>3873</td><td>3899</td><td>27</td><td>TAATACATGCCGACGGGCGCTGACCCC</td><td>262.5</td><td>-10.1</td><td>1.3</td><td>-7.5</td><td>5.9E-14</td><td>3.1E-13</td><td>BT20/MB468</td><td>MITO</td></tr>
<tr><td>>hg38_21_+_8433172_8446622_RNA45SN1_45S_50ntFlanks@10571.10592.22|GAGAAGCCGGCGGGAGCCCCGG</td><td>>hg38_21_+_8433172_8446622_RNA45SN1_45S_50ntFlanks</td><td>10571</td><td>10592</td><td>22</td><td>GAGAAGCCGGCGGGAGCCCCGG</td><td>180.2</td><td>-11.4</td><td>1.5</td><td>-7.5</td><td>6.4E-14</td><td>3.1E-13</td><td>MB231/MB468</td><td>MP</td></tr>
<tr><td>>hg38_21_+_8433172_8446622_RNA45SN1_45S_50ntFlanks@11845.11869.25|GGAACGGGCTTGGCGGAATCAGCGG</td><td>>hg38_21_+_8433172_8446622_RNA45SN1_45S_50ntFlanks</td><td>11845</td><td>11869</td><td>25</td><td>GGAACGGGCTTGGCGGAATCAGCGG</td><td>166.0</td><td>-11.2</td><td>1.5</td><td>-7.5</td><td>6.4E-14</td><td>3.1E-13</td><td>MB231/MB468</td><td>MP</td></tr>
<tr><td>>hg38_21_+_8433172_8446622_RNA45SN1_45S_50ntFlanks@5379.5398.20|ATTAAGTCCCTGCCCTTTGT</td><td>>hg38_21_+_8433172_8446622_RNA45SN1_45S_50ntFlanks</td><td>5379</td><td>5398</td><td>20</td><td>ATTAAGTCCCTGCCCTTTGT</td><td>865.1</td><td>-13.5</td><td>1.8</td><td>-7.4</td><td>1.0E-13</td><td>3.1E-13</td><td>MB231/MB468</td><td>MITO</td></tr>
<tr><td>>MI0000285|hsa-mir-205&WithFlank&1|+|209432127|209432248@40.60.21(+1C)&[MIMAT0000266&hsa-miR-205-5p&offsets|0|-1(+1C);m-351&1|+|209432166|209432187&offsets|0|-1(+1C)]|TCCTTCATTCCACCGGAGTCTC</td><td>>MI0000285|hsa-mir-205&WithFlank&1|+|209432127|209432248</td><td>40</td><td>60</td><td>21(+1C)</td><td>TCCTTCATTCCACCGGAGTCTC</td><td>252.3</td><td>11.4</td><td>1.5</td><td>7.8</td><td>5.9E-15</td><td>3.1E-13</td><td>BT20/MB231</td><td>CYTO</td></tr>
<tr><td>>am_trna2_GlyGCC_21_-_18827107_18827177@1.27.27,trna35_GlyGCC_1_+_161413094_161413164@1.27.27,trna37_GlyGCC_1_+_161420467_161420537@1.27.27,trna39_GlyGCC_1_+_161427898_161427968@1.27.27,trna41_GlyGCC_1_+_161435258_161435328@1.27.27|GCATGGGTGGTTCAGTGGTAGAATTCT</td><td>>am_trna2_GlyGCC_21_-_18827107_18827177</td><td>1</td><td>27</td><td>27</td><td>GCATGGGTGGTTCAGTGGTAGAATTCT</td><td>1908.9</td><td>8.6</td><td>1.1</td><td>7.8</td><td>8.2E-15</td><td>3.2E-13</td><td>BT20/MB231</td><td>MITO</td></tr>
<tr><td>>hg38_21_+_8433172_8446622_RNA45SN1_45S_50ntFlanks@10508.10534.27|CGAATCCGGAGTGGCGGAGATGGGCGC</td><td>>hg38_21_+_8433172_8446622_RNA45SN1_45S_50ntFlanks</td><td>10508</td><td>10534</td><td>27</td><td>CGAATCCGGAGTGGCGGAGATGGGCGC</td><td>161.4</td><td>-11.2</td><td>1.5</td><td>-7.5</td><td>6.5E-14</td><td>3.2E-13</td><td>MB231/MB468</td><td>MP</td></tr>
<tr><td>>hg38_21_+_8433172_8446622_RNA45SN1_45S_50ntFlanks@10394.10429.36|AGCAGTTGAACATGGGTCAGTCGGTCCTGAGAGATG</td><td>>hg38_21_+_8433172_8446622_RNA45SN1_45S_50ntFlanks</td><td>10394</td><td>10429</td><td>36</td><td>AGCAGTTGAACATGGGTCAGTCGGTCCTGAGAGATG</td><td>730.8</td><td>-5.4</td><td>0.7</td><td>-7.4</td><td>1.2E-13</td><td>3.2E-13</td><td>BT20/MB468</td><td>NUC</td></tr>
<tr><td>>MI0001448|hsa-mir-425&WithFlank&3|-|49020142|49020240@21.44.24&[MIMAT0003393&hsa-miR-425-5p&offsets|+1|+2;m-95&3|-|49020200|49020221&offsets|+1|+3]|ATGACACGATCACTCCCGTTGAGT</td><td>>MI0001448|hsa-mir-425&WithFlank&3|-|49020142|49020240</td><td>21</td><td>44</td><td>24</td><td>ATGACACGATCACTCCCGTTGAGT</td><td>274.9</td><td>5.4</td><td>0.7</td><td>7.5</td><td>7.7E-14</td><td>3.2E-13</td><td>MB231/MB468</td><td>NUC</td></tr>
<tr><td>>MI0000061|hsa-let-7a-2&WithFlank&11|-|122146516|122146599@11.32.22(+1A);MI0000062|hsa-let-7a-3&WithFlank&22|+|46112743|46112828@10.31.22(+1A);MI0000060|hsa-let-7a-1&WithFlank&9|+|94175951|94176042@12.33.22(+1A)&[MIMAT0000062&hsa-let-7a-5p&offsets|0|0(+1A);m-6&11|-|122146568|122146589&offsets|0|0(+1A)];[MIMAT0000062_1&hsa-let-7a-5p&offsets|0|0(+1A);m-5&22|+|46112752|46112773&offsets|0|0(+1A)];[MIMAT0000062_2&hsa-let-7a-5p&offsets|0|0(+1A);m-4&9|+|94175962|94175983&offsets|0|0(+1A)]|TGAGGTAGTAGGTTGTATAGTTA</td><td>>MI0000061|hsa-let-7a-2&WithFlank&11|-|122146516|122146599</td><td>11</td><td>32</td><td>22(+1A)</td><td>TGAGGTAGTAGGTTGTATAGTTA</td><td>1269.6</td><td>4.4</td><td>0.6</td><td>7.5</td><td>7.7E-14</td><td>3.2E-13</td><td>MB231/MB468</td><td>NUC</td></tr>
<tr><td>>am_trna162_MetCAT_6_-_26758550_26758622@31.53.23,trna164_MetCAT_6_-_26735574_26735646@31.53.23,trna20_MetCAT_16_+_71460396_71460468@31.53.23,trna21_MetCAT_6_+_26701712_26701784@31.53.23,trna27_MetCAT_6_+_26766444_26766516@31.53.23,trna75_MetCAT_6_+_28912352_28912424@31.53.23,trna97_MetCAT_6_-_28921042_28921114@31.53.23|TCTCATAATCTGAAGGTCCTGAG</td><td>>am_trna162_MetCAT_6_-_26758550_26758622</td><td>31</td><td>53</td><td>23</td><td>TCTCATAATCTGAAGGTCCTGAG</td><td>269.8</td><td>-10.8</td><td>1.4</td><td>-7.5</td><td>7.6E-14</td><td>3.2E-13</td><td>MB231/MB468</td><td>CYTO</td></tr>
<tr><td>>hg38_21_+_8433172_8446622_RNA45SN1_45S_50ntFlanks@4367.4389.23|CAGTTAAAAAGCTCGTAGTTGGA</td><td>>hg38_21_+_8433172_8446622_RNA45SN1_45S_50ntFlanks</td><td>4367</td><td>4389</td><td>23</td><td>CAGTTAAAAAGCTCGTAGTTGGA</td><td>1655.7</td><td>-6.3</td><td>0.8</td><td>-7.5</td><td>7.6E-14</td><td>3.2E-13</td><td>MB231/MB468</td><td>CYTO</td></tr>
<tr><td>>hg38_21_+_8433172_8446622_RNA45SN1_45S_50ntFlanks@9736.9767.32|CCGAAACGATCTCAACCTATTCTCAAACTTTA</td><td>>hg38_21_+_8433172_8446622_RNA45SN1_45S_50ntFlanks</td><td>9736</td><td>9767</td><td>32</td><td>CCGAAACGATCTCAACCTATTCTCAAACTTTA</td><td>1208.3</td><td>-9.5</td><td>1.3</td><td>-7.4</td><td>1.0E-13</td><td>3.2E-13</td><td>MB231/MB468</td><td>MITO</td></tr>
<tr><td>>hg38_21_+_8433172_8446622_RNA45SN1_45S_50ntFlanks@11641.11669.29|CGGCGGGTGTTGACGCGATGTGATTTCTG</td><td>>hg38_21_+_8433172_8446622_RNA45SN1_45S_50ntFlanks</td><td>11641</td><td>11669</td><td>29</td><td>CGGCGGGTGTTGACGCGATGTGATTTCTG</td><td>2934.0</td><td>-7.7</td><td>1.0</td><td>-7.5</td><td>6.0E-14</td><td>3.2E-13</td><td>BT20/MB468</td><td>MITO</td></tr>
<tr><td>>hg38_1_-_228634819_228635039_RNA5S12_5S_50ntFlanks@84.102.19|CTCGTCTGATCTCGGAAGC</td><td>>hg38_1_-_228634819_228635039_RNA5S12_5S_50ntFlanks</td><td>84</td><td>102</td><td>19</td><td>CTCGTCTGATCTCGGAAGC</td><td>153.7</td><td>-11.1</td><td>1.5</td><td>-7.5</td><td>6.5E-14</td><td>3.2E-13</td><td>MB231/MB468</td><td>MP</td></tr>
<tr><td>>hg38_21_+_8433172_8446622_RNA45SN1_45S_50ntFlanks@12324.12355.32|TTGGGTTTTAAGCAGGAGGTGTCAGAAAAGTT</td><td>>hg38_21_+_8433172_8446622_RNA45SN1_45S_50ntFlanks</td><td>12324</td><td>12355</td><td>32</td><td>TTGGGTTTTAAGCAGGAGGTGTCAGAAAAGTT</td><td>227.9</td><td>-11.6</td><td>1.6</td><td>-7.4</td><td>1.0E-13</td><td>3.2E-13</td><td>MB231/MB468</td><td>MITO</td></tr>
<tr><td>>hg38_21_+_8433172_8446622_RNA45SN1_45S_50ntFlanks@11876.11903.28|AAGACCCTGTTGAGCTTGACTCTAGTCT</td><td>>hg38_21_+_8433172_8446622_RNA45SN1_45S_50ntFlanks</td><td>11876</td><td>11903</td><td>28</td><td>AAGACCCTGTTGAGCTTGACTCTAGTCT</td><td>165.4</td><td>-11.2</td><td>1.5</td><td>-7.5</td><td>6.6E-14</td><td>3.2E-13</td><td>MB231/MB468</td><td>MP</td></tr>
<tr><td>>hg38_21_+_8433172_8446622_RNA45SN1_45S_50ntFlanks@5077.5102.26|CCGATAACGAACGAGACTCTGGCATG</td><td>>hg38_21_+_8433172_8446622_RNA45SN1_45S_50ntFlanks</td><td>5077</td><td>5102</td><td>26</td><td>CCGATAACGAACGAGACTCTGGCATG</td><td>310.0</td><td>-12.1</td><td>1.6</td><td>-7.5</td><td>6.6E-14</td><td>3.2E-13</td><td>MB231/MB468</td><td>MP</td></tr>
<tr><td>>hg38_21_+_8433172_8446622_RNA45SN1_45S_50ntFlanks@10514.10533.20|CGGAGTGGCGGAGATGGGCG</td><td>>hg38_21_+_8433172_8446622_RNA45SN1_45S_50ntFlanks</td><td>10514</td><td>10533</td><td>20</td><td>CGGAGTGGCGGAGATGGGCG</td><td>1191.9</td><td>-9.8</td><td>1.3</td><td>-7.4</td><td>1.0E-13</td><td>3.2E-13</td><td>MB231/MB468</td><td>MITO</td></tr>
<tr><td>>hg38_21_+_8433172_8446622_RNA45SN1_45S_50ntFlanks@11209.11231.23|CGGCGACTCTGGACGCGAGCCGG</td><td>>hg38_21_+_8433172_8446622_RNA45SN1_45S_50ntFlanks</td><td>11209</td><td>11231</td><td>23</td><td>CGGCGACTCTGGACGCGAGCCGG</td><td>1105.6</td><td>-7.2</td><td>1.0</td><td>-7.4</td><td>1.0E-13</td><td>3.2E-13</td><td>MB231/MB468</td><td>MITO</td></tr>
<tr><td>>hg38_21_+_8433172_8446622_RNA45SN1_45S_50ntFlanks@10480.10497.18|GATCGAAAGGGAGTCGGG</td><td>>hg38_21_+_8433172_8446622_RNA45SN1_45S_50ntFlanks</td><td>10480</td><td>10497</td><td>18</td><td>GATCGAAAGGGAGTCGGG</td><td>274.7</td><td>-12.0</td><td>1.6</td><td>-7.5</td><td>6.6E-14</td><td>3.2E-13</td><td>MB231/MB468</td><td>MP</td></tr>
<tr><td>>hg38_21_+_8433172_8446622_RNA45SN1_45S_50ntFlanks@10807.10830.24|ACAATGTAGGTAAGGGAAGTCGGC</td><td>>hg38_21_+_8433172_8446622_RNA45SN1_45S_50ntFlanks</td><td>10807</td><td>10830</td><td>24</td><td>ACAATGTAGGTAAGGGAAGTCGGC</td><td>160.1</td><td>-11.2</td><td>1.5</td><td>-7.5</td><td>6.6E-14</td><td>3.2E-13</td><td>MB231/MB468</td><td>MP</td></tr>
<tr><td>>hg38_21_+_8433172_8446622_RNA45SN1_45S_50ntFlanks@10766.10797.32|CGCAGCAGGTCTCCAAGGTGAACAGCCTCTGG</td><td>>hg38_21_+_8433172_8446622_RNA45SN1_45S_50ntFlanks</td><td>10766</td><td>10797</td><td>32</td><td>CGCAGCAGGTCTCCAAGGTGAACAGCCTCTGG</td><td>223.4</td><td>-11.7</td><td>1.6</td><td>-7.5</td><td>6.6E-14</td><td>3.2E-13</td><td>MB231/MB468</td><td>MP</td></tr>
<tr><td>>hg38_21_+_8433172_8446622_RNA45SN1_45S_50ntFlanks@475.507.33|CTTCGTGATCGATGTGGTGACGTCGTGCTCTCC</td><td>>hg38_21_+_8433172_8446622_RNA45SN1_45S_50ntFlanks</td><td>475</td><td>507</td><td>33</td><td>CTTCGTGATCGATGTGGTGACGTCGTGCTCTCC</td><td>2855.9</td><td>7.4</td><td>1.0</td><td>7.5</td><td>6.1E-14</td><td>3.2E-13</td><td>BT20/MB468</td><td>MITO</td></tr>
<tr><td>>hg38_21_+_8433172_8446622_RNA45SN1_45S_50ntFlanks@7982.8006.25|TCAGATCAGACGTGGCGACCCGCTG</td><td>>hg38_21_+_8433172_8446622_RNA45SN1_45S_50ntFlanks</td><td>7982</td><td>8006</td><td>25</td><td>TCAGATCAGACGTGGCGACCCGCTG</td><td>4806.5</td><td>-4.8</td><td>0.6</td><td>-7.5</td><td>6.1E-14</td><td>3.2E-13</td><td>BT20/MB468</td><td>MITO</td></tr>
<tr><td>>hg38_21_+_8433172_8446622_RNA45SN1_45S_50ntFlanks@11199.11223.25|CCGGCGGCGGCGGCGACTCTGGACG</td><td>>hg38_21_+_8433172_8446622_RNA45SN1_45S_50ntFlanks</td><td>11199</td><td>11223</td><td>25</td><td>CCGGCGGCGGCGGCGACTCTGGACG</td><td>620.9</td><td>-7.9</td><td>1.1</td><td>-7.5</td><td>7.7E-14</td><td>3.2E-13</td><td>MB231/MB468</td><td>CYTO</td></tr>
<tr><td>>hg38_21_+_8433172_8446622_RNA45SN1_45S_50ntFlanks@10389.10419.31|TGAACAGCAGTTGAACATGGGTCAGTCGGTC</td><td>>hg38_21_+_8433172_8446622_RNA45SN1_45S_50ntFlanks</td><td>10389</td><td>10419</td><td>31</td><td>TGAACAGCAGTTGAACATGGGTCAGTCGGTC</td><td>124.6</td><td>-6.2</td><td>0.8</td><td>-7.5</td><td>7.7E-14</td><td>3.2E-13</td><td>MB231/MB468</td><td>CYTO</td></tr>
<tr><td>>hg38_21_+_8433172_8446622_RNA45SN1_45S_50ntFlanks@10391.10430.40|AACAGCAGTTGAACATGGGTCAGTCGGTCCTGAGAGATGG</td><td>>hg38_21_+_8433172_8446622_RNA45SN1_45S_50ntFlanks</td><td>10391</td><td>10430</td><td>40</td><td>AACAGCAGTTGAACATGGGTCAGTCGGTCCTGAGAGATGG</td><td>357.9</td><td>-12.3</td><td>1.6</td><td>-7.5</td><td>6.7E-14</td><td>3.2E-13</td><td>MB231/MB468</td><td>MP</td></tr>
<tr><td>>hg38_21_+_8433172_8446622_RNA45SN1_45S_50ntFlanks@13005.13027.23|TCTATTGAAAGTCAGCCCTCGAC</td><td>>hg38_21_+_8433172_8446622_RNA45SN1_45S_50ntFlanks</td><td>13005</td><td>13027</td><td>23</td><td>TCTATTGAAAGTCAGCCCTCGAC</td><td>193.2</td><td>-10.6</td><td>1.4</td><td>-7.5</td><td>7.9E-14</td><td>3.2E-13</td><td>MB231/MB468</td><td>NUC</td></tr>
<tr><td>>hg38_21_+_8433172_8446622_RNA45SN1_45S_50ntFlanks@8194.8224.31|CCCGTGGACGGTGTGAGGCCGGTAGCGGCCC</td><td>>hg38_21_+_8433172_8446622_RNA45SN1_45S_50ntFlanks</td><td>8194</td><td>8224</td><td>31</td><td>CCCGTGGACGGTGTGAGGCCGGTAGCGGCCC</td><td>161.0</td><td>-11.2</td><td>1.5</td><td>-7.5</td><td>6.7E-14</td><td>3.2E-13</td><td>MB231/MB468</td><td>MP</td></tr>
<tr><td>>hg38_21_+_8433172_8446622_RNA45SN1_45S_50ntFlanks@6666.6686.21|GATCACTCGGCTCGTGCGTCG</td><td>>hg38_21_+_8433172_8446622_RNA45SN1_45S_50ntFlanks</td><td>6666</td><td>6686</td><td>21</td><td>GATCACTCGGCTCGTGCGTCG</td><td>304.1</td><td>-10.3</td><td>1.4</td><td>-7.5</td><td>6.1E-14</td><td>3.2E-13</td><td>BT20/MB468</td><td>MITO</td></tr>
<tr><td>>hg38_21_+_8433172_8446622_RNA45SN1_45S_50ntFlanks@11781.11820.40|CGCATGAATGGATGAACGAGATTCCCACTGTCCCTACCTA</td><td>>hg38_21_+_8433172_8446622_RNA45SN1_45S_50ntFlanks</td><td>11781</td><td>11820</td><td>40</td><td>CGCATGAATGGATGAACGAGATTCCCACTGTCCCTACCTA</td><td>296.8</td><td>-12.0</td><td>1.6</td><td>-7.4</td><td>1.0E-13</td><td>3.3E-13</td><td>MB231/MB468</td><td>MITO</td></tr>
<tr><td>>hg38_21_+_8433172_8446622_RNA45SN1_45S_50ntFlanks@12934.12954.21|CTAAACCATTCGTAGACGACC</td><td>>hg38_21_+_8433172_8446622_RNA45SN1_45S_50ntFlanks</td><td>12934</td><td>12954</td><td>21</td><td>CTAAACCATTCGTAGACGACC</td><td>360.6</td><td>-3.9</td><td>0.5</td><td>-7.5</td><td>8.0E-14</td><td>3.3E-13</td><td>MB231/MB468</td><td>NUC</td></tr>
<tr><td>>hg38_21_+_8433172_8446622_RNA45SN1_45S_50ntFlanks@9534.9566.33|AGGAAACTCTGGTGGAGGTCCGTAGCGGTCCTG</td><td>>hg38_21_+_8433172_8446622_RNA45SN1_45S_50ntFlanks</td><td>9534</td><td>9566</td><td>33</td><td>AGGAAACTCTGGTGGAGGTCCGTAGCGGTCCTG</td><td>4521.5</td><td>-10.3</td><td>1.4</td><td>-7.5</td><td>8.0E-14</td><td>3.3E-13</td><td>MB231/MB468</td><td>NUC</td></tr>
<tr><td>>hg38_21_+_8433172_8446622_RNA45SN1_45S_50ntFlanks@12939.12972.34|CCATTCGTAGACGACCTGCTTCTGGGTCGGGGTT</td><td>>hg38_21_+_8433172_8446622_RNA45SN1_45S_50ntFlanks</td><td>12939</td><td>12972</td><td>34</td><td>CCATTCGTAGACGACCTGCTTCTGGGTCGGGGTT</td><td>187.3</td><td>-11.4</td><td>1.5</td><td>-7.5</td><td>6.7E-14</td><td>3.3E-13</td><td>MB231/MB468</td><td>MP</td></tr>
</table>

<table>
<tr><td>>MI0001445|hsa-mir-423&WithFlank&17|+|30117073|30117178@59.80.22(+1C)&[MIMAT0001340&hsa-miR-423-3p&offsets|0|-1(+1C);m-100&17|+|30117131|30117153&offsets|0|-1(+1C)]|AGCTCGGTCTGAGGCCCCTCAGC</td><td>>MI0001445|hsa-mir-423&WithFlank&17|+|30117073|30117178</td><td>59</td><td>80</td><td>22(+1C)</td><td>AGCTCGGTCTGAGGCCCCTCAGC</td><td>1448.8</td><td>5.7</td><td>0.8</td><td>7.5</td><td>8.0E-14</td><td>3.3E-13</td><td>MB231/MB468</td><td>NUC</td></tr>
<tr><td>>hg38_21_+_8433172_8446622_RNA45SN1_45S_50ntFlanks@11564.11607.44|CCTAGCAGCCGACTTAGAACTGGTGCGGACCAGGGGAATCCGAC</td><td>>hg38_21_+_8433172_8446622_RNA45SN1_45S_50ntFlanks</td><td>11564</td><td>11607</td><td>44</td><td>CCTAGCAGCCGACTTAGAACTGGTGCGGACCAGGGGAATCCGAC</td><td>313.2</td><td>-12.1</td><td>1.6</td><td>-7.4</td><td>1.0E-13</td><td>3.3E-13</td><td>MB231/MB468</td><td>MITO</td></tr>
<tr><td>>hg38_21_+_8433172_8446622_RNA45SN1_45S_50ntFlanks@9570.9595.26|TGCAAATCGGTCGTCCGACCTGGGTA</td><td>>hg38_21_+_8433172_8446622_RNA45SN1_45S_50ntFlanks</td><td>9570</td><td>9595</td><td>26</td><td>TGCAAATCGGTCGTCCGACCTGGGTA</td><td>4203.8</td><td>-8.4</td><td>1.1</td><td>-7.4</td><td>1.0E-13</td><td>3.3E-13</td><td>MB231/MB468</td><td>MITO</td></tr>
<tr><td>>hg38_21_+_8433172_8446622_RNA45SN1_45S_50ntFlanks@11534.11557.24|GTTCCGCGCGGCGCCTCGCCTCGG</td><td>>hg38_21_+_8433172_8446622_RNA45SN1_45S_50ntFlanks</td><td>11534</td><td>11557</td><td>24</td><td>GTTCCGCGCGGCGCCTCGCCTCGG</td><td>365.9</td><td>-11.2</td><td>1.5</td><td>-7.5</td><td>7.8E-14</td><td>3.3E-13</td><td>MB231/MB468</td><td>CYTO</td></tr>
<tr><td>>hg38_21_+_8433172_8446622_RNA45SN1_45S_50ntFlanks@4722.4748.27|TCATTAATCAAGAACGAAAGTCGGAGG</td><td>>hg38_21_+_8433172_8446622_RNA45SN1_45S_50ntFlanks</td><td>4722</td><td>4748</td><td>27</td><td>TCATTAATCAAGAACGAAAGTCGGAGG</td><td>815.6</td><td>-9.9</td><td>1.3</td><td>-7.4</td><td>1.2E-13</td><td>3.3E-13</td><td>BT20/MB468</td><td>NUC</td></tr>
<tr><td>>hg38_21_+_8433172_8446622_RNA45SN1_45S_50ntFlanks@9459.9493.35|CCGGGGAGGTGGAGCACGAGCGCACGTGTTAGGAC</td><td>>hg38_21_+_8433172_8446622_RNA45SN1_45S_50ntFlanks</td><td>9459</td><td>9493</td><td>35</td><td>CCGGGGAGGTGGAGCACGAGCGCACGTGTTAGGAC</td><td>145.7</td><td>-11.1</td><td>1.5</td><td>-7.5</td><td>6.8E-14</td><td>3.3E-13</td><td>MB231/MB468</td><td>MP</td></tr>
<tr><td>>hg38_21_+_8433172_8446622_RNA45SN1_45S_50ntFlanks@11767.11794.28|TCTAATTAGTGACGCGCATGAATGGATG</td><td>>hg38_21_+_8433172_8446622_RNA45SN1_45S_50ntFlanks</td><td>11767</td><td>11794</td><td>28</td><td>TCTAATTAGTGACGCGCATGAATGGATG</td><td>578.6</td><td>-5.1</td><td>0.7</td><td>-7.4</td><td>1.1E-13</td><td>3.3E-13</td><td>MB231/MB468</td><td>MITO</td></tr>
<tr><td>>MI0000815|hsa-mir-339&WithFlank&7|-|1022927|1023032@56.76.21&[MIMAT0004702&hsa-miR-339-3p&offsets|0|-2;m-235&7|-|1022957|1022977&offsets|0|0]|TGAGCGCCTCGACGACAGAGC</td><td>>MI0000815|hsa-mir-339&WithFlank&7|-|1022927|1023032</td><td>56</td><td>76</td><td>21</td><td>TGAGCGCCTCGACGACAGAGC</td><td>570.5</td><td>9.1</td><td>1.2</td><td>7.5</td><td>8.3E-14</td><td>3.3E-13</td><td>BT20/MB468</td><td>CYTO</td></tr>
<tr><td>>hg38_21_+_8433172_8446622_RNA45SN1_45S_50ntFlanks@6779.6806.28|CCGGGGCTACGCCTGTCTGAGCGTCGCT</td><td>>hg38_21_+_8433172_8446622_RNA45SN1_45S_50ntFlanks</td><td>6779</td><td>6806</td><td>28</td><td>CCGGGGCTACGCCTGTCTGAGCGTCGCT</td><td>597.3</td><td>4.4</td><td>0.6</td><td>7.5</td><td>7.9E-14</td><td>3.3E-13</td><td>MB231/MB468</td><td>CYTO</td></tr>
<tr><td>>hg38_21_+_8433172_8446622_RNA45SN1_45S_50ntFlanks@11318.11340.23|CCCCACGTCTCGTCGCGCGCGCG</td><td>>hg38_21_+_8433172_8446622_RNA45SN1_45S_50ntFlanks</td><td>11318</td><td>11340</td><td>23</td><td>CCCCACGTCTCGTCGCGCGCGCG</td><td>174.3</td><td>-10.4</td><td>1.4</td><td>-7.5</td><td>8.1E-14</td><td>3.3E-13</td><td>MB231/MB468</td><td>NUC</td></tr>
<tr><td>>hg38_21_+_8433172_8446622_RNA45SN1_45S_50ntFlanks@3900.3937.38|CTTCGCGGGGGGGATGCGTGCATTTATCAGATCAAAAC</td><td>>hg38_21_+_8433172_8446622_RNA45SN1_45S_50ntFlanks</td><td>3900</td><td>3937</td><td>38</td><td>CTTCGCGGGGGGGATGCGTGCATTTATCAGATCAAAAC</td><td>272.4</td><td>-11.9</td><td>1.6</td><td>-7.4</td><td>1.1E-13</td><td>3.3E-13</td><td>MB231/MB468</td><td>MITO</td></tr>
<tr><td>>hg38_21_+_8433172_8446622_RNA45SN1_45S_50ntFlanks@10571.10608.38|GAGAAGCCGGCGGGAGCCCCGGGGAGAGTTCTCTTTTC</td><td>>hg38_21_+_8433172_8446622_RNA45SN1_45S_50ntFlanks</td><td>10571</td><td>10608</td><td>38</td><td>GAGAAGCCGGCGGGAGCCCCGGGGAGAGTTCTCTTTTC</td><td>138.9</td><td>-11.0</td><td>1.5</td><td>-7.5</td><td>6.8E-14</td><td>3.3E-13</td><td>MB231/MB468</td><td>MP</td></tr>
<tr><td>>hg38_21_+_8433172_8446622_RNA45SN1_45S_50ntFlanks@11671.11709.39|CCAGTGCTCTGAATGTCAAAGTGAAGAAATTCAATGAAG</td><td>>hg38_21_+_8433172_8446622_RNA45SN1_45S_50ntFlanks</td><td>11671</td><td>11709</td><td>39</td><td>CCAGTGCTCTGAATGTCAAAGTGAAGAAATTCAATGAAG</td><td>1120.0</td><td>-8.2</td><td>1.1</td><td>-7.4</td><td>1.2E-13</td><td>3.3E-13</td><td>BT20/MB468</td><td>NUC</td></tr>
<tr><td>>hg38_21_+_8433172_8446622_RNA45SN1_45S_50ntFlanks@9623.9650.28|TAGTAGCTGGTTCCCTCCGAAGTTTCCC</td><td>>hg38_21_+_8433172_8446622_RNA45SN1_45S_50ntFlanks</td><td>9623</td><td>9650</td><td>28</td><td>TAGTAGCTGGTTCCCTCCGAAGTTTCCC</td><td>854.0</td><td>-4.5</td><td>0.6</td><td>-7.5</td><td>6.3E-14</td><td>3.3E-13</td><td>BT20/MB468</td><td>MITO</td></tr>
<tr><td>>hg38_21_+_8433172_8446622_RNA45SN1_45S_50ntFlanks@9454.9490.37|CCGCGCCGGGGAGGTGGAGCACGAGCGCACGTGTTAG</td><td>>hg38_21_+_8433172_8446622_RNA45SN1_45S_50ntFlanks</td><td>9454</td><td>9490</td><td>37</td><td>CCGCGCCGGGGAGGTGGAGCACGAGCGCACGTGTTAG</td><td>162.2</td><td>-11.2</td><td>1.5</td><td>-7.5</td><td>6.8E-14</td><td>3.3E-13</td><td>MB231/MB468</td><td>MP</td></tr>
<tr><td>>MI0000115|hsa-mir-16-2&WithFlank&3|+|160404739|160404831@58.79.22&[MIMAT0004518&hsa-miR-16-2-3p&offsets|-1|-1;m-252&3|+|160404796|160404816&offsets|0|+1]|ACCAATATTACTGTGCTGCTTT</td><td>>MI0000115|hsa-mir-16-2&WithFlank&3|+|160404739|160404831</td><td>58</td><td>79</td><td>22</td><td>ACCAATATTACTGTGCTGCTTT</td><td>1040.2</td><td>6.2</td><td>0.8</td><td>7.5</td><td>6.3E-14</td><td>3.3E-13</td><td>BT20/MB468</td><td>MITO</td></tr>
<tr><td>>hg38_21_+_8433172_8446622_RNA45SN1_45S_50ntFlanks@6744.6770.27|GACACTTCGAACGCACTTGCGGCCCCG</td><td>>hg38_21_+_8433172_8446622_RNA45SN1_45S_50ntFlanks</td><td>6744</td><td>6770</td><td>27</td><td>GACACTTCGAACGCACTTGCGGCCCCG</td><td>1232.4</td><td>-5.8</td><td>0.8</td><td>-7.5</td><td>8.2E-14</td><td>3.3E-13</td><td>MB231/MB468</td><td>NUC</td></tr>
<tr><td>>MI0000440|hsa-mir-27b&WithFlank&9|+|95085439|95085547@66.87.22&[MIMAT0000419&hsa-miR-27b-3p&offsets|-1|0;m-39&9|+|95085505|95085525&offsets|-1|0]|GTTCACAGTGGCTAAGTTCTGC</td><td>>MI0000440|hsa-mir-27b&WithFlank&9|+|95085439|95085547</td><td>66</td><td>87</td><td>22</td><td>GTTCACAGTGGCTAAGTTCTGC</td><td>306.2</td><td>5.6</td><td>0.8</td><td>7.5</td><td>8.2E-14</td><td>3.3E-13</td><td>MB231/MB468</td><td>NUC</td></tr>
<tr><td>>hg38_21_+_8433172_8446622_RNA45SN1_45S_50ntFlanks@4588.4617.30|CTATTTTGTTGGTTTTCGGAACTGAGGCCA</td><td>>hg38_21_+_8433172_8446622_RNA45SN1_45S_50ntFlanks</td><td>4588</td><td>4617</td><td>30</td><td>CTATTTTGTTGGTTTTCGGAACTGAGGCCA</td><td>222.6</td><td>-11.7</td><td>1.6</td><td>-7.5</td><td>6.9E-14</td><td>3.3E-13</td><td>MB231/MB468</td><td>MP</td></tr>
<tr><td>>hg38_21_+_8433172_8446622_RNA45SN1_45S_50ntFlanks@3996.4024.29|ACTCTAGATAACCTCGGGCCGATCGCACG</td><td>>hg38_21_+_8433172_8446622_RNA45SN1_45S_50ntFlanks</td><td>3996</td><td>4024</td><td>29</td><td>ACTCTAGATAACCTCGGGCCGATCGCACG</td><td>208.0</td><td>-11.6</td><td>1.5</td><td>-7.5</td><td>6.9E-14</td><td>3.3E-13</td><td>MB231/MB468</td><td>MP</td></tr>
<tr><td>>hg38_21_+_8433172_8446622_RNA45SN1_45S_50ntFlanks@12479.12500.22|CTAATAGGGAACGTGAGCTGGG</td><td>>hg38_21_+_8433172_8446622_RNA45SN1_45S_50ntFlanks</td><td>12479</td><td>12500</td><td>22</td><td>CTAATAGGGAACGTGAGCTGGG</td><td>563.6</td><td>-5.3</td><td>0.7</td><td>-7.5</td><td>8.2E-14</td><td>3.3E-13</td><td>MB231/MB468</td><td>NUC</td></tr>
<tr><td>>hg38_21_+_8433172_8446622_RNA45SN1_45S_50ntFlanks@6648.6668.21|GTACGACTCTTAGCGGTGGAT</td><td>>hg38_21_+_8433172_8446622_RNA45SN1_45S_50ntFlanks</td><td>6648</td><td>6668</td><td>21</td><td>GTACGACTCTTAGCGGTGGAT</td><td>8560.4</td><td>6.2</td><td>0.8</td><td>7.5</td><td>6.3E-14</td><td>3.3E-13</td><td>BT20/MB468</td><td>MITO</td></tr>
<tr><td>>hg38_21_+_8433172_8446622_RNA45SN1_45S_50ntFlanks@10436.10453.18|CGCCGTTCCGAAGGGACG</td><td>>hg38_21_+_8433172_8446622_RNA45SN1_45S_50ntFlanks</td><td>10436</td><td>10453</td><td>18</td><td>CGCCGTTCCGAAGGGACG</td><td>543.0</td><td>-11.4</td><td>1.5</td><td>-7.5</td><td>8.4E-14</td><td>3.3E-13</td><td>BT20/MB468</td><td>CYTO</td></tr>
<tr><td>>hg38_21_+_8433172_8446622_RNA45SN1_45S_50ntFlanks@9175.9197.23|TCGCGCCGTCGGGCCCGGGGGAG</td><td>>hg38_21_+_8433172_8446622_RNA45SN1_45S_50ntFlanks</td><td>9175</td><td>9197</td><td>23</td><td>TCGCGCCGTCGGGCCCGGGGGAG</td><td>250.1</td><td>-11.8</td><td>1.6</td><td>-7.5</td><td>6.9E-14</td><td>3.4E-13</td><td>MB231/MB468</td><td>MP</td></tr>
<tr><td>>MI0000087|hsa-mir-29a&WithFlank&7|-|130876741|130876816@47.67.21(+1C)&[MIMAT0000086&hsa-miR-29a-3p&offsets|-1|-2(+1C);m-26&7|-|130876749|130876769&offsets|-1|-1(+1C)]|CTAGCACCATCTGAAATCGGTC</td><td>>MI0000087|hsa-mir-29a&WithFlank&7|-|130876741|130876816</td><td>47</td><td>67</td><td>21(+1C)</td><td>CTAGCACCATCTGAAATCGGTC</td><td>421.8</td><td>5.5</td><td>0.7</td><td>7.4</td><td>1.2E-13</td><td>3.4E-13</td><td>BT20/MB468</td><td>NUC</td></tr>
<tr><td>>trna116_GluCTC_1_-_145399233_145399304@32.74.43,trna71_GluCTC_1_-_161439189_161439260@32.74.43,trna74_GluCTC_1_-_161431809_161431880@32.74.43,trna77_GluCTC_1_-_161424398_161424469@32.74.43,trna77_GluCTC_6_+_28949976_28950047@32.74.43,trna80_GluCTC_1_-_161417018_161417089@32.74.43,trna87_GluCTC_6_-_126101393_126101464@32.74.43|CTCTCACCGCCGCGGCCCGGGTTCGATTCCCGGTCAGGGAACC</td><td>>trna116_GluCTC_1_-_145399233_145399304</td><td>32</td><td>74</td><td>43</td><td>CTCTCACCGCCGCGGCCCGGGTTCGATTCCCGGTCAGGGAACC</td><td>250.6</td><td>-10.9</td><td>1.5</td><td>-7.5</td><td>8.3E-14</td><td>3.4E-13</td><td>MB231/MB468</td><td>NUC</td></tr>
<tr><td>>hg38_21_+_8433172_8446622_RNA45SN1_45S_50ntFlanks@11729.11750.22|TAACTATGACTCTCTTAAGGTA</td><td>>hg38_21_+_8433172_8446622_RNA45SN1_45S_50ntFlanks</td><td>11729</td><td>11750</td><td>22</td><td>TAACTATGACTCTCTTAAGGTA</td><td>3252.6</td><td>-11.1</td><td>1.5</td><td>-7.4</td><td>1.2E-13</td><td>3.4E-13</td><td>BT20/MB468</td><td>NUC</td></tr>
<tr><td>>hg38_21_+_8433172_8446622_RNA45SN1_45S_50ntFlanks@10683.10711.29|CCGGCGGCGTCCGGTGAGCTCTCGCTGGC</td><td>>hg38_21_+_8433172_8446622_RNA45SN1_45S_50ntFlanks</td><td>10683</td><td>10711</td><td>29</td><td>CCGGCGGCGTCCGGTGAGCTCTCGCTGGC</td><td>1833.0</td><td>-6.8</td><td>0.9</td><td>-7.5</td><td>8.1E-14</td><td>3.4E-13</td><td>MB231/MB468</td><td>CYTO</td></tr>
<tr><td>>hg38_21_+_8433172_8446622_RNA45SN1_45S_50ntFlanks@10914.10936.23|CGCCGCGGCTGGACGAGGCGCCG</td><td>>hg38_21_+_8433172_8446622_RNA45SN1_45S_50ntFlanks</td><td>10914</td><td>10936</td><td>23</td><td>CGCCGCGGCTGGACGAGGCGCCG</td><td>215.6</td><td>-10.7</td><td>1.4</td><td>-7.5</td><td>8.3E-14</td><td>3.4E-13</td><td>MB231/MB468</td><td>NUC</td></tr>
<tr><td>>MI0003137|hsa-mir-193b&WithFlank&16|+|14303961|14304055@57.77.21(+1C)&[MIMAT0002819&hsa-miR-193b-3p&offsets|0|-1(+1C);m-166&16|+|14304017|14304038&offsets|0|-1(+1C)]|AACTGGCCCTCAAAGTCCCGCC</td><td>>MI0003137|hsa-mir-193b&WithFlank&16|+|14303961|14304055</td><td>57</td><td>77</td><td>21(+1C)</td><td>AACTGGCCCTCAAAGTCCCGCC</td><td>902.1</td><td>5.5</td><td>0.7</td><td>7.7</td><td>9.8E-15</td><td>3.4E-13</td><td>BT20/MB231</td><td>NUC</td></tr>
<tr><td>>hg38_21_+_8433172_8446622_RNA45SN1_45S_50ntFlanks@12907.12933.27|CACCGCACGTTCGTGGGGAACCTGGCG</td><td>>hg38_21_+_8433172_8446622_RNA45SN1_45S_50ntFlanks</td><td>12907</td><td>12933</td><td>27</td><td>CACCGCACGTTCGTGGGGAACCTGGCG</td><td>2745.3</td><td>-8.0</td><td>1.1</td><td>-7.4</td><td>1.1E-13</td><td>3.4E-13</td><td>MB231/MB468</td><td>MITO</td></tr>
<tr><td>>hg38_21_+_8433172_8446622_RNA45SN1_45S_50ntFlanks@12126.12145.20|CGACCCGCTCCGGGGACAGT</td><td>>hg38_21_+_8433172_8446622_RNA45SN1_45S_50ntFlanks</td><td>12126</td><td>12145</td><td>20</td><td>CGACCCGCTCCGGGGACAGT</td><td>212.5</td><td>-11.6</td><td>1.5</td><td>-7.5</td><td>7.0E-14</td><td>3.4E-13</td><td>MB231/MB468</td><td>MP</td></tr>
<tr><td>>hg38_21_+_8433172_8446622_RNA45SN1_45S_50ntFlanks@10798.10818.21|CATGTTGGAACAATGTAGGTA</td><td>>hg38_21_+_8433172_8446622_RNA45SN1_45S_50ntFlanks</td><td>10798</td><td>10818</td><td>21</td><td>CATGTTGGAACAATGTAGGTA</td><td>549.6</td><td>-5.0</td><td>0.7</td><td>-7.5</td><td>8.3E-14</td><td>3.4E-13</td><td>MB231/MB468</td><td>NUC</td></tr>
<tr><td>>hg38_21_+_8433172_8446622_RNA45SN1_45S_50ntFlanks@6721.6739.19|AATTGCAGGACACATTGAT</td><td>>hg38_21_+_8433172_8446622_RNA45SN1_45S_50ntFlanks</td><td>6721</td><td>6739</td><td>19</td><td>AATTGCAGGACACATTGAT</td><td>712.9</td><td>-12.6</td><td>1.6</td><td>-7.9</td><td>3.0E-15</td><td>3.4E-13</td><td>BT20/MB468</td><td>MP</td></tr>
<tr><td>>MI0000299|hsa-mir-222&WithFlank&X|-|45747009|45747130@75.99.25(+1A)&[MIMAT0000279&hsa-miR-222-3p&offsets|0|+4(+1A);m-140&X|-|45747033|45747056&offsets|0|+1(+1A)]|AGCTACATCTGGCTACTGGGTCTCTA</td><td>>MI0000299|hsa-mir-222&WithFlank&X|-|45747009|45747130</td><td>75</td><td>99</td><td>25(+1A)</td><td>AGCTACATCTGGCTACTGGGTCTCTA</td><td>293.9</td><td>8.2</td><td>1.1</td><td>7.5</td><td>8.5E-14</td><td>3.4E-13</td><td>BT20/MB468</td><td>CYTO</td></tr>
<tr><td>>hg38_21_+_8433172_8446622_RNA45SN1_45S_50ntFlanks@8170.8211.42|CAAGTCCTTCTGATCGAGGCCCAGCCCGTGGACGGTGTGAGG</td><td>>hg38_21_+_8433172_8446622_RNA45SN1_45S_50ntFlanks</td><td>8170</td><td>8211</td><td>42</td><td>CAAGTCCTTCTGATCGAGGCCCAGCCCGTGGACGGTGTGAGG</td><td>302.9</td><td>-12.1</td><td>1.6</td><td>-7.5</td><td>7.0E-14</td><td>3.4E-13</td><td>MB231/MB468</td><td>MP</td></tr>
<tr><td>>hg38_21_+_8433172_8446622_RNA45SN1_45S_50ntFlanks@6737.6771.35|GATCATCGACACTTCGAACGCACTTGCGGCCCCGG</td><td>>hg38_21_+_8433172_8446622_RNA45SN1_45S_50ntFlanks</td><td>6737</td><td>6771</td><td>35</td><td>GATCATCGACACTTCGAACGCACTTGCGGCCCCGG</td><td>344.9</td><td>-12.3</td><td>1.6</td><td>-7.5</td><td>7.0E-14</td><td>3.4E-13</td><td>MB231/MB468</td><td>MP</td></tr>
<tr><td>>hg38_21_+_8433172_8446622_RNA45SN1_45S_50ntFlanks@4385.4408.24|TTGGATCTTGGGAGCGGGCGGGCG</td><td>>hg38_21_+_8433172_8446622_RNA45SN1_45S_50ntFlanks</td><td>4385</td><td>4408</td><td>24</td><td>TTGGATCTTGGGAGCGGGCGGGCG</td><td>219.2</td><td>-11.6</td><td>1.6</td><td>-7.5</td><td>7.0E-14</td><td>3.4E-13</td><td>MB231/MB468</td><td>MP</td></tr>
<tr><td>>hg38_21_+_8433172_8446622_RNA45SN1_45S_50ntFlanks@12306.12325.20|CACGATCCTTCTGACCTTTT</td><td>>hg38_21_+_8433172_8446622_RNA45SN1_45S_50ntFlanks</td><td>12306</td><td>12325</td><td>20</td><td>CACGATCCTTCTGACCTTTT</td><td>2270.3</td><td>-9.0</td><td>1.2</td><td>-7.4</td><td>1.1E-13</td><td>3.4E-13</td><td>MB231/MB468</td><td>MITO</td></tr>
<tr><td>>hg38_21_+_8433172_8446622_RNA45SN1_45S_50ntFlanks@12004.12029.26|CGGGCCGCCGGTGAAATACCACTACT</td><td>>hg38_21_+_8433172_8446622_RNA45SN1_45S_50ntFlanks</td><td>12004</td><td>12029</td><td>26</td><td>CGGGCCGCCGGTGAAATACCACTACT</td><td>1024.7</td><td>6.5</td><td>0.9</td><td>7.5</td><td>8.2E-14</td><td>3.4E-13</td><td>MB231/MB468</td><td>CYTO</td></tr>
<tr><td>>hg38_21_+_8433172_8446622_RNA45SN1_45S_50ntFlanks@5490.5525.36|TGAGAAGACGGTCGAACTTGACTATCTAGAGGAAGT</td><td>>hg38_21_+_8433172_8446622_RNA45SN1_45S_50ntFlanks</td><td>5490</td><td>5525</td><td>36</td><td>TGAGAAGACGGTCGAACTTGACTATCTAGAGGAAGT</td><td>555.8</td><td>10.7</td><td>1.4</td><td>7.5</td><td>8.6E-14</td><td>3.4E-13</td><td>BT20/MB468</td><td>CYTO</td></tr>
<tr><td>>hg38_21_+_8433172_8446622_RNA45SN1_45S_50ntFlanks@6649.6673.25|TACGACTCTTAGCGGTGGATCACTC</td><td>>hg38_21_+_8433172_8446622_RNA45SN1_45S_50ntFlanks</td><td>6649</td><td>6673</td><td>25</td><td>TACGACTCTTAGCGGTGGATCACTC</td><td>640.0</td><td>5.4</td><td>0.7</td><td>7.5</td><td>6.5E-14</td><td>3.4E-13</td><td>BT20/MB468</td><td>MITO</td></tr>
<tr><td>>hg38_21_+_8433172_8446622_RNA45SN1_45S_50ntFlanks@12626.12653.28|AATGGGGCGAAGCTACCATCTGTGGGAT</td><td>>hg38_21_+_8433172_8446622_RNA45SN1_45S_50ntFlanks</td><td>12626</td><td>12653</td><td>28</td><td>AATGGGGCGAAGCTACCATCTGTGGGAT</td><td>4496.2</td><td>-7.8</td><td>1.0</td><td>-7.5</td><td>6.5E-14</td><td>3.4E-13</td><td>BT20/MB468</td><td>MITO</td></tr>
<tr><td>>MI0000433|hsa-let-7g&WithFlank&3|-|52268272|52268367@11.30.20(+1C)&[MIMAT0000414&hsa-let-7g-5p&offsets|0|-2(+1C);m-58&3|-|52268336|52268357&offsets|0|-2(+1C)]|TGAGGTAGTAGTTTGTACAGC</td><td>>MI0000433|hsa-let-7g&WithFlank&3|-|52268272|52268367</td><td>11</td><td>30</td><td>20(+1C)</td><td>TGAGGTAGTAGTTTGTACAGC</td><td>445.3</td><td>7.9</td><td>1.1</td><td>7.5</td><td>6.5E-14</td><td>3.4E-13</td><td>BT20/MB468</td><td>MITO</td></tr>
<tr><td>>hg38_21_+_8433172_8446622_RNA45SN1_45S_50ntFlanks@4443.4472.30|CCCTTGCCTCTCGGCGCCCCCTCGATGCTC</td><td>>hg38_21_+_8433172_8446622_RNA45SN1_45S_50ntFlanks</td><td>4443</td><td>4472</td><td>30</td><td>CCCTTGCCTCTCGGCGCCCCCTCGATGCTC</td><td>902.2</td><td>-12.8</td><td>1.7</td><td>-7.5</td><td>8.5E-14</td><td>3.4E-13</td><td>MB231/MB468</td><td>NUC</td></tr>
<tr><td>>MI0015866|hsa-mir-4326&WithFlank&20|+|63286802|63286872@17.40.24&[MIMAT0016888&hsa-miR-4326&offsets|0|+4;m-339&20|+|63286818|63286841&offsets|0|0]|TGTTCCTCTGTCTCCCAGACTCTG</td><td>>MI0015866|hsa-mir-4326&WithFlank&20|+|63286802|63286872</td><td>17</td><td>40</td><td>24</td><td>TGTTCCTCTGTCTCCCAGACTCTG</td><td>385.9</td><td>8.6</td><td>1.2</td><td>7.5</td><td>8.2E-14</td><td>3.4E-13</td><td>MB231/MB468</td><td>CYTO</td></tr>
<tr><td>>hg38_21_+_8433172_8446622_RNA45SN1_45S_50ntFlanks@12996.13037.42|TCGCTGCGATCTATTGAAAGTCAGCCCTCGACACAAGGGTTT</td><td>>hg38_21_+_8433172_8446622_RNA45SN1_45S_50ntFlanks</td><td>12996</td><td>13037</td><td>42</td><td>TCGCTGCGATCTATTGAAAGTCAGCCCTCGACACAAGGGTTT</td><td>1333.8</td><td>10.4</td><td>1.4</td><td>7.5</td><td>8.2E-14</td><td>3.4E-13</td><td>MB231/MB468</td><td>CYTO</td></tr>
</table>

| | | | | | | | | | | | | | |
|---|---|---|---|---|---|---|---|---|---|---|---|---|---|
| >hg38_21_+_8433172_8446622_RNA45SN1_45S_50ntFlanks@12991.13029.39\|CTCCCTCGCTGCGATCTATTGAAAGTCAGCCCTCGACAC | >hg38_21_+_8433172_8446622_RNA45SN1_45S_50ntFlanks | 12991 | 13029 | 39 | CTCCCTCGCTGCGATCTATTGAAAGTCAGCCCTCGACAC | 241.1 | -11.7 | 1.6 | -7.4 | 1.1E-13 | 3.4E-13 | MB231/MB468 | MITO |
| >MI0003599\|hsa-mir-589&WithFlank&7\|-\|5495813\|5495923@30.52.23&[MIMAT0004799&hsa-miR-589-5p&offsets\|0\|+1;m-162&7\|-\|5495874\|5495894&offsets\|0\|+2]\|TGAGAACCACGTCTGCTCTGAGC | >MI0003599\|hsa-mir-589&WithFlank&7\|-\|5495813\|5495923 | 30 | 52 | 23 | TGAGAACCACGTCTGCTCTGAGC | 229.6 | 3.3 | 0.4 | 7.4 | 1.3E-13 | 3.5E-13 | BT20/MB468 | NUC |
| >MI0000812\|hsa-mir-331&WithFlank&12\|+\|95308414\|95308519@67.87.21(+1U)&[MIMAT0000760&hsa-miR-331-3p&offsets\|0\|0(+1U);m-192&12\|+\|95308480\|95308501&offsets\|0\|-1(+1U)]\|GCCCCTGGGCCTATCCTAGAAT | >MI0000812\|hsa-mir-331&WithFlank&12\|+\|95308414\|95308519 | 67 | 87 | 21(+1U) | GCCCCTGGGCCTATCCTAGAAT | 730.2 | 6.4 | 0.9 | 7.5 | 8.5E-14 | 3.5E-13 | MB231/MB468 | NUC |
| >hg38_21_+_8433172_8446622_RNA45SN1_45S_50ntFlanks@10678.10712.35\|CGGTTCCGGCGGCGTCCGGTGAGCTCTCGCTGGCC | >hg38_21_+_8433172_8446622_RNA45SN1_45S_50ntFlanks | 10678 | 10712 | 35 | CGGTTCCGGCGGCGTCCGGTGAGCTCTCGCTGGCC | 20494.8 | -6.2 | 0.8 | -7.5 | 6.6E-14 | 3.5E-13 | BT20/MB468 | MITO |
| >hg38_21_+_8433172_8446622_RNA45SN1_45S_50ntFlanks@6655.6695.41\|TCTTAGCGGTGGATCACTCGGCTCGTGCGTCGATGAAGAAC | >hg38_21_+_8433172_8446622_RNA45SN1_45S_50ntFlanks | 6655 | 6695 | 41 | TCTTAGCGGTGGATCACTCGGCTCGTGCGTCGATGAAGAAC | 467.1 | 7.1 | 0.9 | 7.5 | 6.6E-14 | 3.5E-13 | BT20/MB468 | MITO |
| >hg38_21_+_8433172_8446622_RNA45SN1_45S_50ntFlanks@4732.4767.36\|AGAACGAAAGTCGGAGGTTCGAAGACGATCAGATAC | >hg38_21_+_8433172_8446622_RNA45SN1_45S_50ntFlanks | 4732 | 4767 | 36 | AGAACGAAAGTCGGAGGTTCGAAGACGATCAGATAC | 2995.0 | 10.4 | 1.4 | 7.5 | 8.7E-14 | 3.5E-13 | BT20/MB468 | CYTO |
| >hg38_21_+_8433172_8446622_RNA45SN1_45S_50ntFlanks@9034.9058.25\|TCGTGGGGGGGCCGGGCCACCCCTC | >hg38_21_+_8433172_8446622_RNA45SN1_45S_50ntFlanks | 9034 | 9058 | 25 | TCGTGGGGGGGCCGGGCCACCCCTC | 283.2 | -12.0 | 1.6 | -7.5 | 7.2E-14 | 3.5E-13 | MB231/MB468 | MP |
| >MI0000298\|hsa-mir-221&WithFlank&X\|-\|45746151\|45746272@71.92.22(+1A)&[MIMAT0000278&hsa-miR-221-3p&offsets\|0\|-1(+1A);m-91&X\|-\|45746181\|45746202&offsets\|0\|0(+1A)]\|AGCTACATTGTCTGCTGGGTTTA | >MI0000298\|hsa-mir-221&WithFlank&X\|-\|45746151\|45746272 | 71 | 92 | 22(+1A) | AGCTACATTGTCTGCTGGGTTTA | 1300.1 | 6.4 | 0.9 | 7.5 | 8.8E-14 | 3.5E-13 | BT20/MB468 | CYTO |
| >MI0000299\|hsa-mir-222&WithFlank&X\|-\|45747009\|45747130@75.96.22(+1A)&[MIMAT0000279&hsa-miR-222-3p&offsets\|0\|+1(+1A);m-140&X\|-\|45747033\|45747056&offsets\|0\|-2(+1A)]\|AGCTACATCTGGCTACTGGGTCA | >MI0000299\|hsa-mir-222&WithFlank&X\|-\|45747009\|45747130 | 75 | 96 | 22(+1A) | AGCTACATCTGGCTACTGGGTCA | 548.0 | 5.8 | 0.8 | 7.5 | 8.6E-14 | 3.5E-13 | MB231/MB468 | NUC |
| >MI0000063\|hsa-let-7b&WithFlank&22\|+\|46113680\|46113774@12.34.23&[MIMAT0000063&hsa-let-7b-5p&offsets\|0\|+1;m-14&22\|+\|46113691\|46113712&offsets\|0\|+1]\|TGAGGTAGTAGGTTGTGTGGTTT | >MI0000063\|hsa-let-7b&WithFlank&22\|+\|46113680\|46113774 | 12 | 34 | 23 | TGAGGTAGTAGGTTGTGTGGTTT | 2446.0 | 5.8 | 0.8 | 7.5 | 8.4E-14 | 3.5E-13 | MB231/MB468 | CYTO |
| >hg38_21_+_8433172_8446622_RNA45SN1_45S_50ntFlanks@9504.9529.26\|GTGAACTATGCCTGGGCAGGGCGAAG | >hg38_21_+_8433172_8446622_RNA45SN1_45S_50ntFlanks | 9504 | 9529 | 26 | GTGAACTATGCCTGGGCAGGGCGAAG | 432.5 | -11.5 | 1.5 | -7.5 | 8.4E-14 | 3.5E-13 | MB231/MB468 | CYTO |
| >hg38_21_+_8433172_8446622_RNA45SN1_45S_50ntFlanks@12510.12538.29\|TCGTGAGACAGGTTAGTTTTACCCTACTG | >hg38_21_+_8433172_8446622_RNA45SN1_45S_50ntFlanks | 12510 | 12538 | 29 | TCGTGAGACAGGTTAGTTTTACCCTACTG | 235.8 | -11.7 | 1.6 | -7.5 | 7.2E-14 | 3.5E-13 | MB231/MB468 | MP |
| >hg38_21_+_8433172_8446622_RNA45SN1_45S_50ntFlanks@6773.6793.21\|TTCCTCCCGGGGCTACGCCTG | >hg38_21_+_8433172_8446622_RNA45SN1_45S_50ntFlanks | 6773 | 6793 | 21 | TTCCTCCCGGGGCTACGCCTG | 651.0 | -9.6 | 1.3 | -7.5 | 8.6E-14 | 3.5E-13 | MB231/MB468 | NUC |
| >MI0000542\|hsa-mir-320a&WithFlank&8\|-\|22244960\|22245043@48.68.21(+1U)&[MIMAT0000510&hsa-miR-320a-3p&offsets\|0\|-1(+1U);m-60&8\|-\|22244975\|22244996&offsets\|0\|-1(+1U)]\|AAAAGCTGGGTTGAGAGGGCGT | >MI0000542\|hsa-mir-320a&WithFlank&8\|-\|22244960\|22245043 | 48 | 68 | 21(+1U) | AAAAGCTGGGTTGAGAGGGCGT | 2906.6 | 10.7 | 1.4 | 7.5 | 6.7E-14 | 3.5E-13 | BT20/MB468 | MITO |
| >hg38_21_+_8433172_8446622_RNA45SN1_45S_50ntFlanks@12676.12703.28\|AGAATCCCGCCCAGGCGGAACGATACGG | >hg38_21_+_8433172_8446622_RNA45SN1_45S_50ntFlanks | 12676 | 12703 | 28 | AGAATCCCGCCCAGGCGGAACGATACGG | 353.2 | -10.6 | 1.4 | -7.5 | 6.7E-14 | 3.5E-13 | BT20/MB468 | MITO |
| >hg38_21_+_8433172_8446622_RNA45SN1_45S_50ntFlanks@11199.11233.35\|CCGGCGGCGGCGGCGACTCTGGACGCGAGCCGGGC | >hg38_21_+_8433172_8446622_RNA45SN1_45S_50ntFlanks | 11199 | 11233 | 35 | CCGGCGGCGGCGGCGACTCTGGACGCGAGCCGGGC | 226.5 | -11.7 | 1.6 | -7.5 | 7.3E-14 | 3.5E-13 | MB231/MB468 | MP |
| >hg38_21_+_8433172_8446622_RNA45SN1_45S_50ntFlanks@8153.8181.29\|GCCGCTCGTGGGGGGCCCAAGTCCTTCTG | >hg38_21_+_8433172_8446622_RNA45SN1_45S_50ntFlanks | 8153 | 8181 | 29 | GCCGCTCGTGGGGGGCCCAAGTCCTTCTG | 2453.8 | -8.3 | 1.1 | -7.4 | 1.1E-13 | 3.5E-13 | MB231/MB468 | MITO |
| >hg38_21_+_8433172_8446622_RNA45SN1_45S_50ntFlanks@12939.12956.18\|CCATTCGTAGACGACCTG | >hg38_21_+_8433172_8446622_RNA45SN1_45S_50ntFlanks | 12939 | 12956 | 18 | CCATTCGTAGACGACCTG | 2101.0 | -10.7 | 1.4 | -7.4 | 1.3E-13 | 3.5E-13 | BT20/MB468 | NUC |
| >MI0000285\|hsa-mir-205&WithFlank&1\|+\|209432127\|209432248@40.59.20(+1C)&[MIMAT0000266&hsa-miR-205-5p&offsets\|0\|-2(+1C);m-351&1\|+\|209432166\|209432187&offsets\|0\|-2(+1C)]\|TCCTTCATTCCACCGGAGTCC | >MI0000285\|hsa-mir-205&WithFlank&1\|+\|209432127\|209432248 | 40 | 59 | 20(+1C) | TCCTTCATTCCACCGGAGTCC | 213.6 | -11.6 | 1.6 | -7.5 | 7.3E-14 | 3.5E-13 | MB231/MB468 | MP |
| >hg38_21_+_8433172_8446622_RNA45SN1_45S_50ntFlanks@10508.10533.26\|CGAATCCGGAGTGGCGGAGATGGGCG | >hg38_21_+_8433172_8446622_RNA45SN1_45S_50ntFlanks | 10508 | 10533 | 26 | CGAATCCGGAGTGGCGGAGATGGGCG | 319.3 | -11.3 | 1.5 | -7.5 | 8.7E-14 | 3.5E-13 | MB231/MB468 | NUC |
| >trnaMT_GluTTC_MT_-_14674_14742@39.71.33,trnalookalike8_GluTTC_5_-_93905172_93905240@39.71.33\|ATTGGTCGTGGTTGTAGTCCGTGCGAGAATACC | >trnaMT_GluTTC_MT_-_14674_14742 | 39 | 71 | 33 | ATTGGTCGTGGTTGTAGTCCGTGCGAGAATACC | 2883.8 | 7.0 | 0.9 | 7.5 | 8.7E-14 | 3.5E-13 | MB231/MB468 | NUC |
| >hg38_21_+_8433172_8446622_RNA45SN1_45S_50ntFlanks@10402.10435.34\|AACATGGGTCAGTCGGTCCTGAGAGATGGGCGAG | >hg38_21_+_8433172_8446622_RNA45SN1_45S_50ntFlanks | 10402 | 10435 | 34 | AACATGGGTCAGTCGGTCCTGAGAGATGGGCGAG | 1836.2 | -9.7 | 1.3 | -7.5 | 8.7E-14 | 3.5E-13 | MB231/MB468 | NUC |
| >hg38_21_+_8433172_8446622_RNA45SN1_45S_50ntFlanks@10463.10490.28\|CTCCGTTGCCCTCGGCCGATCGAAAGGG | >hg38_21_+_8433172_8446622_RNA45SN1_45S_50ntFlanks | 10463 | 10490 | 28 | CTCCGTTGCCCTCGGCCGATCGAAAGGG | 129.1 | -10.9 | 1.5 | -7.5 | 7.3E-14 | 3.5E-13 | MB231/MB468 | MP |
| >hg38_21_+_8433172_8446622_RNA45SN1_45S_50ntFlanks@6753.6771.19\|AACGCACTTGCGGCCCCGG | >hg38_21_+_8433172_8446622_RNA45SN1_45S_50ntFlanks | 6753 | 6771 | 19 | AACGCACTTGCGGCCCCGG | 178.3 | -11.3 | 1.5 | -7.5 | 7.3E-14 | 3.5E-13 | MB231/MB468 | MP |
| >MI0000111\|hsa-mir-105-1&WithFlank&X\|-\|152392213\|152392305@19.40.22;MI0000112\|hsa-mir-105-2&WithFlank&X\|-\|152394406\|152394498@19.40.22&[MIMAT0000102&hsa-miR-105-5p&offsets\|0\|-1;m-735&X\|-\|152392266\|152392287&offsets\|0\|0];[MIMAT0000102_1&hsa-miR-105-5p&offsets\|0\|-1;m-735&X\|-\|152394459\|152394480&offsets\|0\|0]\|TCAAATGCTCAGACTCCTGTGG | >MI0000111\|hsa-mir-105-1&WithFlank&X\|-\|152392213\|152392305 | 19 | 40 | 22 | TCAAATGCTCAGACTCCTGTGG | 830.6 | 4.6 | 0.6 | 7.5 | 8.9E-14 | 3.5E-13 | BT20/MB468 | CYTO |
| >MI0000471\|hsa-mir-126&WithFlank&9\|+\|136670596\|136670692@59.78.20&[MIMAT0000445&hsa-miR-126-3p&offsets\|+1\|-1;m-40&9\|+\|136670653\|136670674&offsets\|+1\|-1]\|CGTACCGTGAGTAATAATGC | >MI0000471\|hsa-mir-126&WithFlank&9\|+\|136670596\|136670692 | 59 | 78 | 20 | CGTACCGTGAGTAATAATGC | 315.0 | 11.5 | 1.6 | 7.4 | 1.1E-13 | 3.5E-13 | MB231/MB468 | MITO |
| >trna129_MetCAT_6_-_27870271_27870342@31.51.21,trna142_MetCAT_6_-_27560600_27560671@31.51.21,trna150_MetCAT_6_-_27300764_27300835@31.51.21,trna169_MetCAT_6_-_26330529_26330600@31.51.21,trna171_MetCAT_6_-_26313352_26313423@31.51.21,trna20_MetCAT_17_-_80452597_80452668@31.51.21,trna2_MetCAT_6_+_26286754_26286825@31.51.21,trna32_MetCAT_1_+_153643726_153643797@31.51.21,trna61_MetCAT_6_+_27745664_27745735@31.51.21\|CCCATAACCCAGAGGTCGATG | >trna129_MetCAT_6_-_27870271_27870342 | 31 | 51 | 21 | CCCATAACCCAGAGGTCGATG | 629.0 | -9.3 | 1.3 | -7.5 | 8.9E-14 | 3.5E-13 | BT20/MB468 | CYTO |
| >hg38_21_+_8433172_8446622_RNA45SN1_45S_50ntFlanks@8104.8143.40\|CCGCGGCGGGGCGCGGGACATGTGGCGTACGGAAGACCCG | >hg38_21_+_8433172_8446622_RNA45SN1_45S_50ntFlanks | 8104 | 8143 | 40 | CCGCGGCGGGGCGCGGGACATGTGGCGTACGGAAGACCCG | 1163.4 | -7.0 | 0.9 | -7.5 | 8.5E-14 | 3.5E-13 | MB231/MB468 | CYTO |
| >hg38_21_+_8433172_8446622_RNA45SN1_45S_50ntFlanks@11577.11601.25\|TTAGAACTGGTGCGGACCAGGGGAA | >hg38_21_+_8433172_8446622_RNA45SN1_45S_50ntFlanks | 11577 | 11601 | 25 | TTAGAACTGGTGCGGACCAGGGGAA | 206.3 | -11.6 | 1.5 | -7.5 | 7.3E-14 | 3.5E-13 | MB231/MB468 | MP |
| >hg38_21_+_8433172_8446622_RNA45SN1_45S_50ntFlanks@7981.8004.24\|CTCAGATCAGACGTGGCGACCCGC | >hg38_21_+_8433172_8446622_RNA45SN1_45S_50ntFlanks | 7981 | 8004 | 24 | CTCAGATCAGACGTGGCGACCCGC | 206.0 | -11.6 | 1.5 | -7.5 | 7.3E-14 | 3.5E-13 | MB231/MB468 | MP |
| >MI0000079\|hsa-mir-23a&WithFlank&19\|-\|13836581\|13836665@54.72.19(+2U)&[MIMAT0000078&hsa-miR-23a-3p&offsets\|+3\|+1(+2U);m-31&19\|-\|13836595\|13836615&offsets\|+3\|+1(+2U)]\|ACATTGCCAGGGATTTCCATT | >MI0000079\|hsa-mir-23a&WithFlank&19\|-\|13836581\|13836665 | 54 | 72 | 19(+2U) | ACATTGCCAGGGATTTCCATT | 719.0 | 27.0 | 3.6 | 7.4 | 1.1E-13 | 3.6E-13 | MB231/MB468 | MITO |
| >trna10_AlaCGC_6_+_26553731_26553802@58.75.18,trna119_AlaCGC_6_-_28641613_28641684@58.75.18,trna13_AlaCGC_2_+_157257281_157257352@58.75.18,trna13_AlaTGC_12_-_125406301_125406372@58.75.18,trna66_AlaTGC_6_+_28611222_28611293@58.75.18,trna8_AlaTGC_12_+_125424512_125424583@58.75.18,trna8_AlaTGC_5_+_180633868_180633939@58.75.18\|TCCCCGGCATCTCCACCA | >trna10_AlaCGC_6_+_26553731_26553802 | 58 | 75 | 18 | TCCCCGGCATCTCCACCA | 9733.8 | 6.7 | 0.9 | 7.5 | 8.6E-14 | 3.6E-13 | MB231/MB468 | CYTO |
| >trnaMT_ValTAC_MT_+_1602_1670@51.72.22\|CTTAACTTGACCGCTCTGACCA | >trnaMT_ValTAC_MT_+_1602_1670 | 51 | 72 | 22 | CTTAACTTGACCGCTCTGACCA | 11917.3 | -7.6 | 1.0 | -7.4 | 1.3E-13 | 3.6E-13 | BT20/MB468 | NUC |
| >hg38_21_+_8433172_8446622_RNA45SN1_45S_50ntFlanks@9250.9274.25\|CGCACGGGGTCGGCGGCGACGTCGG | >hg38_21_+_8433172_8446622_RNA45SN1_45S_50ntFlanks | 9250 | 9274 | 25 | CGCACGGGGTCGGCGGCGACGTCGG | 192.1 | -11.4 | 1.5 | -7.5 | 7.4E-14 | 3.6E-13 | MB231/MB468 | MP |
| >hg38_21_+_8433172_8446622_RNA45SN1_45S_50ntFlanks@12397.12417.21\|CGACGTCGCTTTTTGATCCTT | >hg38_21_+_8433172_8446622_RNA45SN1_45S_50ntFlanks | 12397 | 12417 | 21 | CGACGTCGCTTTTTGATCCTT | 2861.0 | -6.6 | 0.9 | -7.4 | 1.1E-13 | 3.6E-13 | MB231/MB468 | MITO |
| >hg38_21_+_8433172_8446622_RNA45SN1_45S_50ntFlanks@12510.12530.21\|TCGTGAGACAGGTTAGTTTTA | >hg38_21_+_8433172_8446622_RNA45SN1_45S_50ntFlanks | 12510 | 12530 | 21 | TCGTGAGACAGGTTAGTTTTA | 137.0 | -5.0 | 0.7 | -7.5 | 8.6E-14 | 3.6E-13 | MB231/MB468 | CYTO |
| >hg38_21_+_8433172_8446622_RNA45SN1_45S_50ntFlanks@5051.5081.31\|TGGTGGAGCGATTTGTCTGGTTAATTCCGAT | >hg38_21_+_8433172_8446622_RNA45SN1_45S_50ntFlanks | 5051 | 5081 | 31 | TGGTGGAGCGATTTGTCTGGTTAATTCCGAT | 363.2 | 12.3 | 1.6 | 7.9 | 3.2E-15 | 3.6E-13 | BT20/MB468 | MP |
| >MI0000342\|hsa-mir-200b&WithFlank&1\|+\|1167098\|1167204@63.85.23(+1A)&[MIMAT0000318&hsa-miR-200b-3p&offsets\|0\|+1(+1A);m-93&1\|+\|1167160\|1167182&offsets\|0\|0(+1A)]\|TAATACTGCCTGGTAATGATGACA | >MI0000342\|hsa-mir-200b&WithFlank&1\|+\|1167098\|1167204 | 63 | 85 | 23(+1A) | TAATACTGCCTGGTAATGATGACA | 506.9 | -12.1 | 1.5 | -7.9 | 3.3E-15 | 3.6E-13 | BT20/MB468 | MP |
| >hg38_21_+_8433172_8446622_RNA45SN1_45S_50ntFlanks@10807.10826.20\|ACAATGTAGGTAAGGGAAGT | >hg38_21_+_8433172_8446622_RNA45SN1_45S_50ntFlanks | 10807 | 10826 | 20 | ACAATGTAGGTAAGGGAAGT | 278.8 | -11.1 | 1.5 | -7.5 | 8.9E-14 | 3.6E-13 | MB231/MB468 | NUC |
| >hg38_21_+_8433172_8446622_RNA45SN1_45S_50ntFlanks@475.498.24\|CTTCGTGATCGATGTGGTGACGTC | >hg38_21_+_8433172_8446622_RNA45SN1_45S_50ntFlanks | 475 | 498 | 24 | CTTCGTGATCGATGTGGTGACGTC | 2053.5 | 8.7 | 1.2 | 7.5 | 9.1E-14 | 3.6E-13 | BT20/MB468 | CYTO |
| >am_MI0000809\|hsa-mir-151a&WithFlank&8\|-\|140732558\|140732659@17.38.22&[MIMAT0004697&hsa-miR-151a-5p&offsets\|0\|+1;m-75&8\|-\|140732623\|140732643&offsets\|0\|+1]\|TCGAGGAGCTCACAGTCTAGTA | >am_MI0000809\|hsa-mir-151a&WithFlank&8\|-\|140732558\|140732659 | 17 | 38 | 22 | TCGAGGAGCTCACAGTCTAGTA | 1565.9 | 9.8 | 1.3 | 7.5 | 9.1E-14 | 3.6E-13 | BT20/MB468 | CYTO |
| >hg38_21_+_8433172_8446622_RNA45SN1_45S_50ntFlanks@12358.12375.18\|CACAGGGATAACTGGCTT | >hg38_21_+_8433172_8446622_RNA45SN1_45S_50ntFlanks | 12358 | 12375 | 18 | CACAGGGATAACTGGCTT | 123.7 | -10.8 | 1.4 | -7.5 | 7.5E-14 | 3.6E-13 | MB231/MB468 | MP |
| >hg38_21_+_8433172_8446622_RNA45SN1_45S_50ntFlanks@7977.8003.27\|CGACCTCAGATCAGACGTGGCGACCCG | >hg38_21_+_8433172_8446622_RNA45SN1_45S_50ntFlanks | 7977 | 8003 | 27 | CGACCTCAGATCAGACGTGGCGACCCG | 138.8 | -11.0 | 1.5 | -7.5 | 7.5E-14 | 3.6E-13 | MB231/MB468 | MP |

<table>
<tr><th></th><th></th><th></th><th></th><th></th><th></th><th></th><th></th><th></th><th></th><th></th><th></th><th></th><th></th></tr>
<tr><td>>hg38_21_+_8433172_8446622_RNA45SN1_45S_50ntFlanks@4854.4883.30|AGTCTTTGGGTTCCGGGGGGAGTATGGTTG</td><td>>hg38_21_+_8433172_8446622_RNA45SN1_45S_50ntFlanks</td><td>4854</td><td>4883</td><td>30</td><td>AGTCTTTGGGTTCCGGGGGGAGTATGGTTG</td><td>135.5</td><td>-10.9</td><td>1.5</td><td>-7.5</td><td>7.6E-14</td><td>3.6E-13</td><td>MB231/MB468</td><td>MP</td></tr>
<tr><td>>hg38_21_+_8433172_8446622_RNA45SN1_45S_50ntFlanks@4661.4682.22|AGGTGAAATTCTTGGACCGGCG</td><td>>hg38_21_+_8433172_8446622_RNA45SN1_45S_50ntFlanks</td><td>4661</td><td>4682</td><td>22</td><td>AGGTGAAATTCTTGGACCGGCG</td><td>267.2</td><td>-11.9</td><td>1.6</td><td>-7.5</td><td>7.6E-14</td><td>3.6E-13</td><td>MB231/MB468</td><td>MP</td></tr>
<tr><td>>hg38_21_+_8433172_8446622_RNA45SN1_45S_50ntFlanks@8552.8569.18|CTCCCCGGAGGGGGCGGG</td><td>>hg38_21_+_8433172_8446622_RNA45SN1_45S_50ntFlanks</td><td>8552</td><td>8569</td><td>18</td><td>CTCCCCGGAGGGGGCGGG</td><td>623.4</td><td>-11.4</td><td>1.5</td><td>-7.5</td><td>6.9E-14</td><td>3.6E-13</td><td>BT20/MB468</td><td>MITO</td></tr>
<tr><td>>hg38_21_+_8433172_8446622_RNA45SN1_45S_50ntFlanks@10425.10456.32|AGATGGGCGAGCGCCGTTCCGAAGGGACGGGC</td><td>>hg38_21_+_8433172_8446622_RNA45SN1_45S_50ntFlanks</td><td>10425</td><td>10456</td><td>32</td><td>AGATGGGCGAGCGCCGTTCCGAAGGGACGGGC</td><td>1241.8</td><td>-10.3</td><td>1.4</td><td>-7.4</td><td>1.3E-13</td><td>3.6E-13</td><td>BT20/MB468</td><td>NUC</td></tr>
<tr><td>>MI0006444|hsa-mir-1307&WithFlank&10|-|103394247|103394407@86.108.23(+1A)&[MIMAT0005951&hsa-miR-1307-3p&offsets|0|+1(+1A);m-66&10|-|103394300|103394322&offsets|0|0(+1A)]|ACTCGGCGTGGCGTCGGTCGTGGA</td><td>>MI0006444|hsa-mir-1307&WithFlank&10|-|103394247|103394407</td><td>86</td><td>108</td><td>23(+1A)</td><td>ACTCGGCGTGGCGTCGGTCGTGGA</td><td>519.1</td><td>7.9</td><td>1.1</td><td>7.5</td><td>7.0E-14</td><td>3.6E-13</td><td>BT20/MB468</td><td>MITO</td></tr>
<tr><td>>hg38_21_+_8433172_8446622_RNA45SN1_45S_50ntFlanks@11876.11899.24|AAGACCCTGTTGAGCTTGACTCTA</td><td>>hg38_21_+_8433172_8446622_RNA45SN1_45S_50ntFlanks</td><td>11876</td><td>11899</td><td>24</td><td>AAGACCCTGTTGAGCTTGACTCTA</td><td>164.8</td><td>-11.2</td><td>1.5</td><td>-7.5</td><td>7.6E-14</td><td>3.6E-13</td><td>MB231/MB468</td><td>MP</td></tr>
<tr><td>>hg38_21_+_8433172_8446622_RNA45SN1_45S_50ntFlanks@8196.8215.20|CGTGGACGGTGTGAGGCCGG</td><td>>hg38_21_+_8433172_8446622_RNA45SN1_45S_50ntFlanks</td><td>8196</td><td>8215</td><td>20</td><td>CGTGGACGGTGTGAGGCCGG</td><td>3596.8</td><td>-5.0</td><td>0.7</td><td>-7.5</td><td>9.2E-14</td><td>3.7E-13</td><td>BT20/MB468</td><td>CYTO</td></tr>
<tr><td>>hg38_21_+_8433172_8446622_RNA45SN1_45S_50ntFlanks@10826.10864.39|TCGGCAAGCCGGATCCGTAACTTCGGGATAAGGATTGGC</td><td>>hg38_21_+_8433172_8446622_RNA45SN1_45S_50ntFlanks</td><td>10826</td><td>10864</td><td>39</td><td>TCGGCAAGCCGGATCCGTAACTTCGGGATAAGGATTGGC</td><td>266.8</td><td>-10.8</td><td>1.4</td><td>-7.5</td><td>8.8E-14</td><td>3.7E-13</td><td>MB231/MB468</td><td>CYTO</td></tr>
<tr><td>>hg38_21_+_8433172_8446622_RNA45SN1_45S_50ntFlanks@11686.11717.32|TCAAAGTGAAGAAATTCAATGAAGCGCGGGTA</td><td>>hg38_21_+_8433172_8446622_RNA45SN1_45S_50ntFlanks</td><td>11686</td><td>11717</td><td>32</td><td>TCAAAGTGAAGAAATTCAATGAAGCGCGGGTA</td><td>199.4</td><td>-11.4</td><td>1.5</td><td>-7.4</td><td>1.2E-13</td><td>3.7E-13</td><td>MB231/MB468</td><td>MITO</td></tr>
<tr><td>>hg38_21_+_8433172_8446622_RNA45SN1_45S_50ntFlanks@6648.6670.23|GTACGACTCTTAGCGGTGGATCA</td><td>>hg38_21_+_8433172_8446622_RNA45SN1_45S_50ntFlanks</td><td>6648</td><td>6670</td><td>23</td><td>GTACGACTCTTAGCGGTGGATCA</td><td>203.6</td><td>-11.5</td><td>1.5</td><td>-7.5</td><td>7.6E-14</td><td>3.7E-13</td><td>MB231/MB468</td><td>MP</td></tr>
<tr><td>>hg38_21_+_8433172_8446622_RNA45SN1_45S_50ntFlanks@8158.8178.21|TCGTGGGGGGCCCAAGTCCTT</td><td>>hg38_21_+_8433172_8446622_RNA45SN1_45S_50ntFlanks</td><td>8158</td><td>8178</td><td>21</td><td>TCGTGGGGGGCCCAAGTCCTT</td><td>152.9</td><td>-11.1</td><td>1.5</td><td>-7.5</td><td>7.6E-14</td><td>3.7E-13</td><td>MB231/MB468</td><td>MP</td></tr>
<tr><td>>hg38_21_+_8433172_8446622_RNA45SN1_45S_50ntFlanks@11552.11574.23|CCTCGGCCGGCGCCTAGCAGCCG</td><td>>hg38_21_+_8433172_8446622_RNA45SN1_45S_50ntFlanks</td><td>11552</td><td>11574</td><td>23</td><td>CCTCGGCCGGCGCCTAGCAGCCG</td><td>155.8</td><td>-11.1</td><td>1.5</td><td>-7.5</td><td>7.6E-14</td><td>3.7E-13</td><td>MB231/MB468</td><td>MP</td></tr>
<tr><td>>hg38_21_+_8433172_8446622_RNA45SN1_45S_50ntFlanks@3940.3959.20|ACCCGGTCAGCCCCTCTCCG</td><td>>hg38_21_+_8433172_8446622_RNA45SN1_45S_50ntFlanks</td><td>3940</td><td>3959</td><td>20</td><td>ACCCGGTCAGCCCCTCTCCG</td><td>234.2</td><td>-10.8</td><td>1.5</td><td>-7.5</td><td>9.1E-14</td><td>3.7E-13</td><td>MB231/MB468</td><td>NUC</td></tr>
<tr><td>>MI0000750|hsa-mir-26a-2&WithFlank&12|-|57824603|57824698@20.40.21(+1A);MI0000083|hsa-mir-26a-1&WithFlank&3|+|37969398|37969486@16.36.21(+1A)&[MIMAT0000082&hsa-miR-26a-5p&offsets|0|-1(+1A);m-33&12|-|57824658|57824679&offsets|0|-1(+1A)];[MIMAT0000082_1&hsa-miR-26a-5p&offsets|0|-1(+1A);m-34&3|+|37969413|37969434&offsets|0|-1(+1A)]|TTCAAGTAATCCAGGATAGGCA</td><td>>MI0000750|hsa-mir-26a-2&WithFlank&12|-|57824603|57824698</td><td>20</td><td>40</td><td>21(+1A)</td><td>TTCAAGTAATCCAGGATAGGCA</td><td>5224.6</td><td>5.0</td><td>0.7</td><td>7.5</td><td>9.1E-14</td><td>3.7E-13</td><td>MB231/MB468</td><td>NUC</td></tr>
<tr><td>>hg38_21_+_8433172_8446622_RNA45SN1_45S_50ntFlanks@6744.6770.27|GACACTTCGAACGCACTTGCGGCCCCG</td><td>>hg38_21_+_8433172_8446622_RNA45SN1_45S_50ntFlanks</td><td>6744</td><td>6770</td><td>27</td><td>GACACTTCGAACGCACTTGCGGCCCCG</td><td>2471.1</td><td>-10.1</td><td>1.3</td><td>-7.5</td><td>7.7E-14</td><td>3.7E-13</td><td>MB231/MB468</td><td>MP</td></tr>
<tr><td>>hg38_21_+_8433172_8446622_RNA45SN1_45S_50ntFlanks@12405.12444.40|CTTTTTGATCCTTCGATGTCGGCTCTTCCTATCATTGTGA</td><td>>hg38_21_+_8433172_8446622_RNA45SN1_45S_50ntFlanks</td><td>12405</td><td>12444</td><td>40</td><td>CTTTTTGATCCTTCGATGTCGGCTCTTCCTATCATTGTGA</td><td>1516.3</td><td>-5.5</td><td>0.7</td><td>-7.4</td><td>1.2E-13</td><td>3.7E-13</td><td>MB231/MB468</td><td>MITO</td></tr>
<tr><td>>hg38_21_+_8433172_8446622_RNA45SN1_45S_50ntFlanks@6664.6689.26|TGGATCACTCGGCTCGTGCGTCGATG</td><td>>hg38_21_+_8433172_8446622_RNA45SN1_45S_50ntFlanks</td><td>6664</td><td>6689</td><td>26</td><td>TGGATCACTCGGCTCGTGCGTCGATG</td><td>1068.7</td><td>-7.3</td><td>1.0</td><td>-7.4</td><td>1.2E-13</td><td>3.7E-13</td><td>MB231/MB468</td><td>MITO</td></tr>
<tr><td>>MI0000446|hsa-mir-125b-1&WithFlank&11|-|122099751|122099850@61.82.22(+1U)&[MIMAT0004592&hsa-miR-125b-1-3p&offsets|0|0(+1U);m-485&11|-|122099769|122099790&offsets|0|0(+1U)]|ACGGGTTAGGCTCTTGGGAGCTT</td><td>>MI0000446|hsa-mir-125b-1&WithFlank&11|-|122099751|122099850</td><td>61</td><td>82</td><td>22(+1U)</td><td>ACGGGTTAGGCTCTTGGGAGCTT</td><td>231.4</td><td>11.1</td><td>1.5</td><td>7.5</td><td>7.7E-14</td><td>3.7E-13</td><td>MB231/MB468</td><td>MP</td></tr>
<tr><td>>hg38_21_+_8433172_8446622_RNA45SN1_45S_50ntFlanks@12457.12478.22|CAAGCGTTGGATTGTTCACCCA</td><td>>hg38_21_+_8433172_8446622_RNA45SN1_45S_50ntFlanks</td><td>12457</td><td>12478</td><td>22</td><td>CAAGCGTTGGATTGTTCACCCA</td><td>1422.5</td><td>-9.7</td><td>1.3</td><td>-7.5</td><td>7.7E-14</td><td>3.7E-13</td><td>MB231/MB468</td><td>MP</td></tr>
<tr><td>>hg38_21_+_8433172_8446622_RNA45SN1_45S_50ntFlanks@10664.10711.48|CTTGGAAAGCGTCGCGGTTCCGGCGGCGTCCGGTGAGCTCTCGCTGGC</td><td>>hg38_21_+_8433172_8446622_RNA45SN1_45S_50ntFlanks</td><td>10664</td><td>10711</td><td>48</td><td>CTTGGAAAGCGTCGCGGTTCCGGCGGCGTCCGGTGAGCTCTCGCTGGC</td><td>1644.3</td><td>-15.1</td><td>2.0</td><td>-7.4</td><td>1.4E-13</td><td>3.7E-13</td><td>BT20/MB468</td><td>NUC</td></tr>
<tr><td>>hg38_21_+_8433172_8446622_RNA45SN1_45S_50ntFlanks@6666.6699.34|GATCACTCGGCTCGTGCGTCGATGAAGAACGCAG</td><td>>hg38_21_+_8433172_8446622_RNA45SN1_45S_50ntFlanks</td><td>6666</td><td>6699</td><td>34</td><td>GATCACTCGGCTCGTGCGTCGATGAAGAACGCAG</td><td>243.0</td><td>-10.0</td><td>1.3</td><td>-7.5</td><td>7.1E-14</td><td>3.7E-13</td><td>BT20/MB468</td><td>MITO</td></tr>
<tr><td>>hg38_21_+_8433172_8446622_RNA45SN1_45S_50ntFlanks@10684.10706.23|CGGCGGCGTCCGGTGAGCTCTCG</td><td>>hg38_21_+_8433172_8446622_RNA45SN1_45S_50ntFlanks</td><td>10684</td><td>10706</td><td>23</td><td>CGGCGGCGTCCGGTGAGCTCTCG</td><td>11318.2</td><td>-9.1</td><td>1.2</td><td>-7.5</td><td>9.3E-14</td><td>3.7E-13</td><td>MB231/MB468</td><td>NUC</td></tr>
<tr><td>>hg38_21_+_8433172_8446622_RNA45SN1_45S_50ntFlanks@10342.10361.20|ATATTCAAACGAGAACTTTG</td><td>>hg38_21_+_8433172_8446622_RNA45SN1_45S_50ntFlanks</td><td>10342</td><td>10361</td><td>20</td><td>ATATTCAAACGAGAACTTTG</td><td>953.3</td><td>-5.7</td><td>0.8</td><td>-7.4</td><td>1.2E-13</td><td>3.7E-13</td><td>MB231/MB468</td><td>MITO</td></tr>
<tr><td>>hg38_21_+_8433172_8446622_RNA45SN1_45S_50ntFlanks@8552.8581.30|CTCCCCGGAGGGGGCGGGCTCCGGCGGGTG</td><td>>hg38_21_+_8433172_8446622_RNA45SN1_45S_50ntFlanks</td><td>8552</td><td>8581</td><td>30</td><td>CTCCCCGGAGGGGGCGGGCTCCGGCGGGTG</td><td>270.1</td><td>-11.0</td><td>1.5</td><td>-7.4</td><td>9.4E-14</td><td>3.8E-13</td><td>MB231/MB468</td><td>NUC</td></tr>
<tr><td>>hg38_21_+_8433172_8446622_RNA45SN1_45S_50ntFlanks@10465.10497.33|CCGTTGCCCTCGGCCGATCGAAAGGGAGTCGGG</td><td>>hg38_21_+_8433172_8446622_RNA45SN1_45S_50ntFlanks</td><td>10465</td><td>10497</td><td>33</td><td>CCGTTGCCCTCGGCCGATCGAAAGGGAGTCGGG</td><td>147.3</td><td>-11.1</td><td>1.5</td><td>-7.5</td><td>7.9E-14</td><td>3.8E-13</td><td>MB231/MB468</td><td>MP</td></tr>
<tr><td>>MI0000077|hsa-mir-21&WithFlank&17|+|59841260|59841343@14.34.21(+1U)&[MIMAT0000076&hsa-miR-21-5p&offsets|0|-1(+1U);m-1&17|+|59841273|59841294&offsets|0|-1(+1U)]|TAGCTTATCAGACTGATGTTGT</td><td>>MI0000077|hsa-mir-21&WithFlank&17|+|59841260|59841343</td><td>14</td><td>34</td><td>21(+1U)</td><td>TAGCTTATCAGACTGATGTTGT</td><td>5098.1</td><td>8.5</td><td>1.1</td><td>7.4</td><td>9.6E-14</td><td>3.8E-13</td><td>BT20/MB468</td><td>CYTO</td></tr>
<tr><td>>hg38_21_+_8433172_8446622_RNA45SN1_45S_50ntFlanks@11799.11820.22|AGATTCCCACTGTCCCTACCTA</td><td>>hg38_21_+_8433172_8446622_RNA45SN1_45S_50ntFlanks</td><td>11799</td><td>11820</td><td>22</td><td>AGATTCCCACTGTCCCTACCTA</td><td>225.7</td><td>-11.6</td><td>1.6</td><td>-7.4</td><td>1.2E-13</td><td>3.8E-13</td><td>MB231/MB468</td><td>MITO</td></tr>
<tr><td>>am_trna2_GlyGCC_21_-_18827107_18827177@1.19.19,trna35_GlyGCC_1_+_161413094_161413164@1.19.19,trna37_GlyGCC_1_+_161420467_161420537@1.19.19,trna39_GlyGCC_1_+_161427898_161427968@1.19.19,trna41_GlyGCC_1_+_161435258_161435328@1.19.19|GCATGGGTGGTTCAGTGGT</td><td>>am_trna2_GlyGCC_21_-_18827107_18827177</td><td>1</td><td>19</td><td>19</td><td>GCATGGGTGGTTCAGTGGT</td><td>282.4</td><td>11.5</td><td>1.5</td><td>7.8</td><td>7.3E-15</td><td>3.8E-13</td><td>BT20/MB231</td><td>CYTO</td></tr>
<tr><td>>hg38_21_+_8433172_8446622_RNA45SN1_45S_50ntFlanks@10683.10710.28|CCGGCGGCGTCCGGTGAGCTCTCGCTGG</td><td>>hg38_21_+_8433172_8446622_RNA45SN1_45S_50ntFlanks</td><td>10683</td><td>10710</td><td>28</td><td>CCGGCGGCGTCCGGTGAGCTCTCGCTGG</td><td>719.7</td><td>-9.0</td><td>1.2</td><td>-7.4</td><td>1.2E-13</td><td>3.8E-13</td><td>MB231/MB468</td><td>MITO</td></tr>
<tr><td>>hg38_21_+_8433172_8446622_RNA45SN1_45S_50ntFlanks@11873.11895.23|AAGAAGACCCTGTTGAGCTTGAC</td><td>>hg38_21_+_8433172_8446622_RNA45SN1_45S_50ntFlanks</td><td>11873</td><td>11895</td><td>23</td><td>AAGAAGACCCTGTTGAGCTTGAC</td><td>145.7</td><td>-11.1</td><td>1.5</td><td>-7.5</td><td>7.9E-14</td><td>3.8E-13</td><td>MB231/MB468</td><td>MP</td></tr>
<tr><td>>MI0000542|hsa-mir-320a&WithFlank&8|-|22244960|22245043@48.68.21(+1U)&[MIMAT0000510&hsa-miR-320a-3p&offsets|0|-1(+1U);m-60&8|-|22244975|22244996&offsets|0|-1(+1U)]|AAAAGCTGGGTTGAGAGGGCGT</td><td>>MI0000542|hsa-mir-320a&WithFlank&8|-|22244960|22245043</td><td>48</td><td>68</td><td>21(+1U)</td><td>AAAAGCTGGGTTGAGAGGGCGT</td><td>42143.5</td><td>8.5</td><td>1.1</td><td>7.9</td><td>3.5E-15</td><td>3.8E-13</td><td>BT20/MB468</td><td>MP</td></tr>
<tr><td>>MI0000737|hsa-mir-200a&WithFlank&1|+|1167857|1167958@60.81.22(+1C)&[MIMAT0000682&hsa-miR-200a-3p&offsets|0|0(+1C);m-142&1|+|1167916|1167938&offsets|0|-1(+1C)]|TAACACTGTCTGGTAACGATGTC</td><td>>MI0000737|hsa-mir-200a&WithFlank&1|+|1167857|1167958</td><td>60</td><td>81</td><td>22(+1C)</td><td>TAACACTGTCTGGTAACGATGTC</td><td>773.8</td><td>-12.7</td><td>1.6</td><td>-7.9</td><td>3.6E-15</td><td>3.8E-13</td><td>BT20/MB468</td><td>MP</td></tr>
<tr><td>>hg38_21_+_8433172_8446622_RNA45SN1_45S_50ntFlanks@12366.12384.19|TAACTGGCTTGTGGCGGCC</td><td>>hg38_21_+_8433172_8446622_RNA45SN1_45S_50ntFlanks</td><td>12366</td><td>12384</td><td>19</td><td>TAACTGGCTTGTGGCGGCC</td><td>625.8</td><td>-12.4</td><td>1.6</td><td>-7.9</td><td>3.6E-15</td><td>3.8E-13</td><td>BT20/MB468</td><td>MP</td></tr>
<tr><td>>hg38_21_+_8433172_8446622_RNA45SN1_45S_50ntFlanks@10509.10534.26|GAATCCGGAGTGGCGGAGATGGGCGC</td><td>>hg38_21_+_8433172_8446622_RNA45SN1_45S_50ntFlanks</td><td>10509</td><td>10534</td><td>26</td><td>GAATCCGGAGTGGCGGAGATGGGCGC</td><td>164.6</td><td>-11.2</td><td>1.5</td><td>-7.5</td><td>8.0E-14</td><td>3.8E-13</td><td>MB231/MB468</td><td>MP</td></tr>
<tr><td>>hg38_21_+_8433172_8446622_RNA45SN1_45S_50ntFlanks@11876.11894.19|AAGACCCTGTTGAGCTTGA</td><td>>hg38_21_+_8433172_8446622_RNA45SN1_45S_50ntFlanks</td><td>11876</td><td>11894</td><td>19</td><td>AAGACCCTGTTGAGCTTGA</td><td>118.5</td><td>-10.8</td><td>1.4</td><td>-7.5</td><td>8.0E-14</td><td>3.8E-13</td><td>MB231/MB468</td><td>MP</td></tr>
<tr><td>>hg38_21_+_8433172_8446622_RNA45SN1_45S_50ntFlanks@12157.12181.25|AGTTTGACTGGGGCGGTACACCTGT</td><td>>hg38_21_+_8433172_8446622_RNA45SN1_45S_50ntFlanks</td><td>12157</td><td>12181</td><td>25</td><td>AGTTTGACTGGGGCGGTACACCTGT</td><td>253.5</td><td>-7.1</td><td>1.0</td><td>-7.5</td><td>9.3E-14</td><td>3.8E-13</td><td>MB231/MB468</td><td>CYTO</td></tr>
<tr><td>>hg38_21_+_8433172_8446622_RNA45SN1_45S_50ntFlanks@10910.10931.22|CGCGCGCCGCGGCTGGACGAGG</td><td>>hg38_21_+_8433172_8446622_RNA45SN1_45S_50ntFlanks</td><td>10910</td><td>10931</td><td>22</td><td>CGCGCGCCGCGGCTGGACGAGG</td><td>171.4</td><td>-11.3</td><td>1.5</td><td>-7.5</td><td>8.0E-14</td><td>3.9E-13</td><td>MB231/MB468</td><td>MP</td></tr>
<tr><td>>hg38_21_+_8433172_8446622_RNA45SN1_45S_50ntFlanks@6721.6758.38|AATTGCAGGACACATTGATCATCGACACTTCGAACGCA</td><td>>hg38_21_+_8433172_8446622_RNA45SN1_45S_50ntFlanks</td><td>6721</td><td>6758</td><td>38</td><td>AATTGCAGGACACATTGATCATCGACACTTCGAACGCA</td><td>1315.7</td><td>-8.6</td><td>1.1</td><td>-7.5</td><td>8.0E-14</td><td>3.9E-13</td><td>MB231/MB468</td><td>MP</td></tr>
<tr><td>>hg38_21_+_8433172_8446622_RNA45SN1_45S_50ntFlanks@10116.10139.24|TGCGTCGGAGGGCGGCGGCGGCGG</td><td>>hg38_21_+_8433172_8446622_RNA45SN1_45S_50ntFlanks</td><td>10116</td><td>10139</td><td>24</td><td>TGCGTCGGAGGGCGGCGGCGGCGG</td><td>294.7</td><td>-11.2</td><td>1.5</td><td>-7.4</td><td>9.6E-14</td><td>3.9E-13</td><td>MB231/MB468</td><td>NUC</td></tr>
<tr><td>>hg38_21_+_8433172_8446622_RNA45SN1_45S_50ntFlanks@4005.4024.20|AACCTCGGGCCGATCGCACG</td><td>>hg38_21_+_8433172_8446622_RNA45SN1_45S_50ntFlanks</td><td>4005</td><td>4024</td><td>20</td><td>AACCTCGGGCCGATCGCACG</td><td>1956.0</td><td>-8.6</td><td>1.2</td><td>-7.4</td><td>1.4E-13</td><td>3.9E-13</td><td>BT20/MB468</td><td>NUC</td></tr>
<tr><td>>hg38_21_+_8433172_8446622_RNA45SN1_45S_50ntFlanks@12934.12969.36|CTAAACCATTCGTAGACGACCTGCTTCTGGGTCGGG</td><td>>hg38_21_+_8433172_8446622_RNA45SN1_45S_50ntFlanks</td><td>12934</td><td>12969</td><td>36</td><td>CTAAACCATTCGTAGACGACCTGCTTCTGGGTCGGG</td><td>297.9</td><td>-12.1</td><td>1.6</td><td>-7.5</td><td>8.1E-14</td><td>3.9E-13</td><td>MB231/MB468</td><td>MP</td></tr>
<tr><td>>hg38_21_+_8433172_8446622_RNA45SN1_45S_50ntFlanks@10689.10709.21|GCGTCCGGTGAGCTCTCGCTG</td><td>>hg38_21_+_8433172_8446622_RNA45SN1_45S_50ntFlanks</td><td>10689</td><td>10709</td><td>21</td><td>GCGTCCGGTGAGCTCTCGCTG</td><td>1799.5</td><td>-10.6</td><td>1.4</td><td>-7.4</td><td>1.4E-13</td><td>3.9E-13</td><td>BT20/MB468</td><td>NUC</td></tr>
<tr><td>>hg38_21_+_8433172_8446622_RNA45SN1_45S_50ntFlanks@3989.4023.35|TTTGGTGACTCTAGATAACCTCGGGCCGATCGCAC</td><td>>hg38_21_+_8433172_8446622_RNA45SN1_45S_50ntFlanks</td><td>3989</td><td>4023</td><td>35</td><td>TTTGGTGACTCTAGATAACCTCGGGCCGATCGCAC</td><td>335.4</td><td>4.4</td><td>0.6</td><td>7.4</td><td>9.4E-14</td><td>3.9E-13</td><td>MB231/MB468</td><td>CYTO</td></tr>
<tr><td>>hg38_21_+_8433172_8446622_RNA45SN1_45S_50ntFlanks@12934.12965.32|CTAAACCATTCGTAGACGACCTGCTTCTGGGT</td><td>>hg38_21_+_8433172_8446622_RNA45SN1_45S_50ntFlanks</td><td>12934</td><td>12965</td><td>32</td><td>CTAAACCATTCGTAGACGACCTGCTTCTGGGT</td><td>583.0</td><td>-6.9</td><td>0.9</td><td>-7.4</td><td>1.3E-13</td><td>3.9E-13</td><td>MB231/MB468</td><td>MITO</td></tr>
<tr><td>>hg38_1_-_228634819_228635039_RNA5S12_5S_50ntFlanks@128.170.43|CTTGGATGGGAGACCGCCTGGGAATACCGGGTGCTGTAGGCTT</td><td>>hg38_1_-_228634819_228635039_RNA5S12_5S_50ntFlanks</td><td>128</td><td>170</td><td>43</td><td>CTTGGATGGGAGACCGCCTGGGAATACCGGGTGCTGTAGGCTT</td><td>3749.8</td><td>8.9</td><td>1.2</td><td>7.4</td><td>9.9E-14</td><td>3.9E-13</td><td>BT20/MB468</td><td>CYTO</td></tr>
<tr><td>>hg38_21_+_8433172_8446622_RNA45SN1_45S_50ntFlanks@10350.10388.39|ACGAGAACTTTGAAGGCCGAAGTGGAGAAGGGTTCCATG</td><td>>hg38_21_+_8433172_8446622_RNA45SN1_45S_50ntFlanks</td><td>10350</td><td>10388</td><td>39</td><td>ACGAGAACTTTGAAGGCCGAAGTGGAGAAGGGTTCCATG</td><td>3411.3</td><td>-6.2</td><td>0.8</td><td>-7.4</td><td>9.8E-14</td><td>3.9E-13</td><td>MB231/MB468</td><td>NUC</td></tr>
<tr><td>>hg38_21_+_8433172_8446622_RNA45SN1_45S_50ntFlanks@12046.12070.25|TGACCCGGTGAGGCGGGGGGGCGAG</td><td>>hg38_21_+_8433172_8446622_RNA45SN1_45S_50ntFlanks</td><td>12046</td><td>12070</td><td>25</td><td>TGACCCGGTGAGGCGGGGGGGCGAG</td><td>409.2</td><td>-11.6</td><td>1.6</td><td>-7.4</td><td>9.8E-14</td><td>3.9E-13</td><td>MB231/MB468</td><td>NUC</td></tr>
<tr><td>>hg38_21_+_8433172_8446622_RNA45SN1_45S_50ntFlanks@4003.4020.18|ATAACCTCGGGCCGATCG</td><td>>hg38_21_+_8433172_8446622_RNA45SN1_45S_50ntFlanks</td><td>4003</td><td>4020</td><td>18</td><td>ATAACCTCGGGCCGATCG</td><td>3074.6</td><td>-8.4</td><td>1.1</td><td>-7.4</td><td>9.8E-14</td><td>3.9E-13</td><td>MB231/MB468</td><td>NUC</td></tr>
</table>

<table>
<tr><th></th><th></th><th></th><th></th><th></th><th></th><th></th><th></th><th></th><th></th><th></th><th></th><th></th><th></th></tr>
<tr><td>>hg38_21_+_8433172_8446622_RNA45SN1_45S_50ntFlanks@10832.10862.31|AGCCGGATCCGTAACTTCGGGATAAGGATTG</td><td>>hg38_21_+_8433172_8446622_RNA45SN1_45S_50ntFlanks</td><td>10832</td><td>10862</td><td>31</td><td>AGCCGGATCCGTAACTTCGGGATAAGGATTG</td><td>222.1</td><td>-11.7</td><td>1.6</td><td>-7.5</td><td>8.2E-14</td><td>3.9E-13</td><td>MB231/MB468</td><td>MP</td></tr>
<tr><td>>MI0000299|hsa-mir-222&WithFlank&X|-|45747009|45747130@75.97.23(+1A)&[MIMAT0000279&hsa-miR-222-3p&offsets|0|+2(+1A);m-140&X|-|45747033|45747056&offsets|0|-1(+1A)]|AGCTACATCTGGCTACTGGGTCTA</td><td>>MI0000299|hsa-mir-222&WithFlank&X|-|45747009|45747130</td><td>75</td><td>97</td><td>23(+1A)</td><td>AGCTACATCTGGCTACTGGGTCTA</td><td>1001.9</td><td>7.7</td><td>1.0</td><td>7.4</td><td>1.0E-13</td><td>3.9E-13</td><td>BT20/MB468</td><td>CYTO</td></tr>
<tr><td>>hg38_21_+_8433172_8446622_RNA45SN1_45S_50ntFlanks@12991.13027.37|CTCCCTCGCTGCGATCTATTGAAAGTCAGCCCTCGAC</td><td>>hg38_21_+_8433172_8446622_RNA45SN1_45S_50ntFlanks</td><td>12991</td><td>13027</td><td>37</td><td>CTCCCTCGCTGCGATCTATTGAAAGTCAGCCCTCGAC</td><td>413.3</td><td>-6.6</td><td>0.9</td><td>-7.4</td><td>9.5E-14</td><td>4.0E-13</td><td>MB231/MB468</td><td>CYTO</td></tr>
<tr><td>>hg38_21_+_8433172_8446622_RNA45SN1_45S_50ntFlanks@11768.11786.19|CTAATTAGTGACGCGCATG</td><td>>hg38_21_+_8433172_8446622_RNA45SN1_45S_50ntFlanks</td><td>11768</td><td>11786</td><td>19</td><td>CTAATTAGTGACGCGCATG</td><td>291.3</td><td>-10.3</td><td>1.4</td><td>-7.5</td><td>7.6E-14</td><td>4.0E-13</td><td>BT20/MB468</td><td>MITO</td></tr>
<tr><td>>hg38_21_+_8433172_8446622_RNA45SN1_45S_50ntFlanks@10024.10072.49|GCGCTGGAGCGTCGGGCCCATACCCGGCCGTCGCCGGCAGTCGAGAGTG</td><td>>hg38_21_+_8433172_8446622_RNA45SN1_45S_50ntFlanks</td><td>10024</td><td>10072</td><td>49</td><td>GCGCTGGAGCGTCGGGCCCATACCCGGCCGTCGCCGGCAGTCGAGAGTG</td><td>656.5</td><td>-13.1</td><td>1.8</td><td>-7.4</td><td>1.3E-13</td><td>4.0E-13</td><td>MB231/MB468</td><td>MITO</td></tr>
<tr><td>>hg38_21_+_8433172_8446622_RNA45SN1_45S_50ntFlanks@4099.4121.23|GTGACCACGGGTGACGGGGAATC</td><td>>hg38_21_+_8433172_8446622_RNA45SN1_45S_50ntFlanks</td><td>4099</td><td>4121</td><td>23</td><td>GTGACCACGGGTGACGGGGAATC</td><td>302.4</td><td>-12.0</td><td>1.6</td><td>-7.4</td><td>1.3E-13</td><td>4.0E-13</td><td>MB231/MB468</td><td>MITO</td></tr>
<tr><td>>hg38_21_+_8433172_8446622_RNA45SN1_45S_50ntFlanks@10186.10208.23|TCCTCCCCTCCTCCCGCCCACGC</td><td>>hg38_21_+_8433172_8446622_RNA45SN1_45S_50ntFlanks</td><td>10186</td><td>10208</td><td>23</td><td>TCCTCCCCTCCTCCCGCCCACGC</td><td>209.8</td><td>-11.5</td><td>1.6</td><td>-7.4</td><td>1.3E-13</td><td>4.0E-13</td><td>MB231/MB468</td><td>MITO</td></tr>
<tr><td>>hg38_21_+_8433172_8446622_RNA45SN1_45S_50ntFlanks@9345.9370.26|AAAGCCGCCGTGGCGCAATGAAGGTG</td><td>>hg38_21_+_8433172_8446622_RNA45SN1_45S_50ntFlanks</td><td>9345</td><td>9370</td><td>26</td><td>AAAGCCGCCGTGGCGCAATGAAGGTG</td><td>1069.4</td><td>-9.6</td><td>1.3</td><td>-7.5</td><td>8.3E-14</td><td>4.0E-13</td><td>MB231/MB468</td><td>MP</td></tr>
<tr><td>>hg38_21_+_8433172_8446622_RNA45SN1_45S_50ntFlanks@6647.6677.31|CGTACGACTCTTAGCGGTGGATCACTCGGCT</td><td>>hg38_21_+_8433172_8446622_RNA45SN1_45S_50ntFlanks</td><td>6647</td><td>6677</td><td>31</td><td>CGTACGACTCTTAGCGGTGGATCACTCGGCT</td><td>460.7</td><td>4.7</td><td>0.6</td><td>7.4</td><td>1.0E-13</td><td>4.0E-13</td><td>BT20/MB468</td><td>CYTO</td></tr>
<tr><td>>hg38_21_+_8433172_8446622_RNA45SN1_45S_50ntFlanks@12622.12653.32|AGCCAATGGGGCGAAGCTACCATCTGTGGGAT</td><td>>hg38_21_+_8433172_8446622_RNA45SN1_45S_50ntFlanks</td><td>12622</td><td>12653</td><td>32</td><td>AGCCAATGGGGCGAAGCTACCATCTGTGGGAT</td><td>134.1</td><td>-10.9</td><td>1.5</td><td>-7.5</td><td>8.3E-14</td><td>4.0E-13</td><td>MB231/MB468</td><td>MP</td></tr>
<tr><td>>hg38_21_+_8433172_8446622_RNA45SN1_45S_50ntFlanks@12412.12434.23|ATCCTTCGATGTCGGCTCTTCCT</td><td>>hg38_21_+_8433172_8446622_RNA45SN1_45S_50ntFlanks</td><td>12412</td><td>12434</td><td>23</td><td>ATCCTTCGATGTCGGCTCTTCCT</td><td>276.7</td><td>-10.2</td><td>1.4</td><td>-7.5</td><td>7.6E-14</td><td>4.0E-13</td><td>BT20/MB468</td><td>MITO</td></tr>
<tr><td>>am_trna10_ValCAC_5_-_180649395_180649467@1.33.33,trna12_ValAAC_5_-_180645270_180645342@1.33.33,trna132_ValAAC_6_-_27721179_27721251@1.33.33,trna136_ValAAC_6_-_27648885_27648957@1.33.33,trna139_ValAAC_6_-_27618707_27618779@1.33.33,trna18_ValCAC_5_-_180529253_180529325@1.33.33,trna2_ValAAC_3_+_169490018_169490090@1.33.33,trna2_ValCAC_5_+_180524070_180524142@1.33.33,trna4_ValAAC_5_+_180591154_180591226@1.33.33,trna5_ValAAC_5_+_180596610_180596682@1.33.33,trna6_ValCAC_5_+_180600650_180600722@1.33.33,trna85_ValCAC_1_-_161369490_161369562@1.33.33,trna90_ValCAC_1_-_149684088_149684161@1.33.33,trna98_ValCAC_1_-_149298555_149298627@1.33.33,trna9_ValCAC_6_+_26538282_26538354@1.33.33|GTTTCCGTAGTGTAGTGGTTATCACGTTCGCCT</td><td>>am_trna10_ValCAC_5_-_180649395_180649467</td><td>1</td><td>33</td><td>33</td><td>GTTTCCGTAGTGTAGTGGTTATCACGTTCGCCT</td><td>5835.4</td><td>9.7</td><td>1.3</td><td>7.5</td><td>7.7E-14</td><td>4.0E-13</td><td>BT20/MB468</td><td>MITO</td></tr>
<tr><td>>hg38_21_+_8433172_8446622_RNA45SN1_45S_50ntFlanks@3787.3810.24|AAACTGCGAATGGCTCATTAAATC</td><td>>hg38_21_+_8433172_8446622_RNA45SN1_45S_50ntFlanks</td><td>3787</td><td>3810</td><td>24</td><td>AAACTGCGAATGGCTCATTAAATC</td><td>2381.6</td><td>-8.6</td><td>1.2</td><td>-7.4</td><td>1.5E-13</td><td>4.0E-13</td><td>BT20/MB468</td><td>NUC</td></tr>
<tr><td>>MI0001445|hsa-mir-423&WithFlank&17|+|30117073|30117178@23.46.24(+1U)&[MIMAT0004748&hsa-miR-423-5p&offsets|0|+1(+1U);m-141&17|+|30117095|30117117&offsets|0|+1(+1U)]|TGAGGGGCAGAGAGCGAGACTTTTT</td><td>>MI0001445|hsa-mir-423&WithFlank&17|+|30117073|30117178</td><td>23</td><td>46</td><td>24(+1U)</td><td>TGAGGGGCAGAGAGCGAGACTTTTT</td><td>263.0</td><td>9.4</td><td>1.3</td><td>7.4</td><td>1.0E-13</td><td>4.0E-13</td><td>BT20/MB468</td><td>CYTO</td></tr>
<tr><td>>hg38_21_+_8433172_8446622_RNA45SN1_45S_50ntFlanks@5210.5240.31|AGATGTCCGGGGCTGCACGCGCGCTACACTG</td><td>>hg38_21_+_8433172_8446622_RNA45SN1_45S_50ntFlanks</td><td>5210</td><td>5240</td><td>31</td><td>AGATGTCCGGGGCTGCACGCGCGCTACACTG</td><td>997.2</td><td>-13.7</td><td>1.9</td><td>-7.4</td><td>1.3E-13</td><td>4.0E-13</td><td>MB231/MB468</td><td>MITO</td></tr>
<tr><td>>hg38_21_+_8433172_8446622_RNA45SN1_45S_50ntFlanks@11887.11904.18|GAGCTTGACTCTAGTCTG</td><td>>hg38_21_+_8433172_8446622_RNA45SN1_45S_50ntFlanks</td><td>11887</td><td>11904</td><td>18</td><td>GAGCTTGACTCTAGTCTG</td><td>131.1</td><td>-10.9</td><td>1.5</td><td>-7.5</td><td>8.4E-14</td><td>4.0E-13</td><td>MB231/MB468</td><td>MP</td></tr>
<tr><td>>hg38_21_+_8433172_8446622_RNA45SN1_45S_50ntFlanks@4002.4020.19|GATAACCTCGGGCCGATCG</td><td>>hg38_21_+_8433172_8446622_RNA45SN1_45S_50ntFlanks</td><td>4002</td><td>4020</td><td>19</td><td>GATAACCTCGGGCCGATCG</td><td>174.5</td><td>-11.3</td><td>1.5</td><td>-7.5</td><td>8.4E-14</td><td>4.0E-13</td><td>MB231/MB468</td><td>MP</td></tr>
<tr><td>>hg38_21_+_8433172_8446622_RNA45SN1_45S_50ntFlanks@12214.12232.19|CAGGGAGGACAGAAACCTC</td><td>>hg38_21_+_8433172_8446622_RNA45SN1_45S_50ntFlanks</td><td>12214</td><td>12232</td><td>19</td><td>CAGGGAGGACAGAAACCTC</td><td>120.0</td><td>-10.8</td><td>1.4</td><td>-7.5</td><td>8.4E-14</td><td>4.0E-13</td><td>MB231/MB468</td><td>MP</td></tr>
<tr><td>>hg38_21_+_8433172_8446622_RNA45SN1_45S_50ntFlanks@7991.8014.24|ACGTGGCGACCCGCTGAATTTAAG</td><td>>hg38_21_+_8433172_8446622_RNA45SN1_45S_50ntFlanks</td><td>7991</td><td>8014</td><td>24</td><td>ACGTGGCGACCCGCTGAATTTAAG</td><td>3896.7</td><td>-5.7</td><td>0.8</td><td>-7.4</td><td>1.3E-13</td><td>4.0E-13</td><td>MB231/MB468</td><td>MITO</td></tr>
<tr><td>>trna20_GluTTC_2_-_131094701_131094772@1.32.32,trna5_GluTTC_13_-_41634874_41634945@1.32.32|TCCCATATGGTCTAGCGGTTAGGATTCCTGGT</td><td>>trna20_GluTTC_2_-_131094701_131094772</td><td>1</td><td>32</td><td>32</td><td>TCCCATATGGTCTAGCGGTTAGGATTCCTGGT</td><td>245.6</td><td>11.2</td><td>1.5</td><td>7.5</td><td>8.4E-14</td><td>4.0E-13</td><td>MB231/MB468</td><td>MP</td></tr>
<tr><td>>hg38_21_+_8433172_8446622_RNA45SN1_45S_50ntFlanks@10826.10865.40|TCGGCAAGCCGGATCCGTAACTTCGGGATAAGGATTGGCT</td><td>>hg38_21_+_8433172_8446622_RNA45SN1_45S_50ntFlanks</td><td>10826</td><td>10865</td><td>40</td><td>TCGGCAAGCCGGATCCGTAACTTCGGGATAAGGATTGGCT</td><td>1039.1</td><td>-9.5</td><td>1.3</td><td>-7.4</td><td>9.8E-14</td><td>4.0E-13</td><td>MB231/MB468</td><td>CYTO</td></tr>
<tr><td>>MI0000471|hsa-mir-126&WithFlank&9|+|136670596|136670692@58.77.20(+1U)&[MIMAT0000445&hsa-miR-126-3p&offsets|0|-2(+1U);m-40&9|+|136670653|136670674&offsets|0|-2(+1U)]|TCGTACCGTGAGTAATAATGT</td><td>>MI0000471|hsa-mir-126&WithFlank&9|+|136670596|136670692</td><td>58</td><td>77</td><td>20(+1U)</td><td>TCGTACCGTGAGTAATAATGT</td><td>1120.1</td><td>11.8</td><td>1.6</td><td>7.4</td><td>1.0E-13</td><td>4.1E-13</td><td>BT20/MB468</td><td>CYTO</td></tr>
<tr><td>>hg38_21_+_8433172_8446622_RNA45SN1_45S_50ntFlanks@8238.8262.25|CCCGGGTCTTCCCGGAGTCGGGTTG</td><td>>hg38_21_+_8433172_8446622_RNA45SN1_45S_50ntFlanks</td><td>8238</td><td>8262</td><td>25</td><td>CCCGGGTCTTCCCGGAGTCGGGTTG</td><td>978.4</td><td>-9.3</td><td>1.3</td><td>-7.4</td><td>1.3E-13</td><td>4.1E-13</td><td>MB231/MB468</td><td>MITO</td></tr>
<tr><td>>hg38_21_+_8433172_8446622_RNA45SN1_45S_50ntFlanks@6745.6770.26|ACACTTCGAACGCACTTGCGGCCCCG</td><td>>hg38_21_+_8433172_8446622_RNA45SN1_45S_50ntFlanks</td><td>6745</td><td>6770</td><td>26</td><td>ACACTTCGAACGCACTTGCGGCCCCG</td><td>5449.9</td><td>-8.0</td><td>1.1</td><td>-7.5</td><td>7.8E-14</td><td>4.1E-13</td><td>BT20/MB468</td><td>MITO</td></tr>
<tr><td>>hg38_21_+_8433172_8446622_RNA45SN1_45S_50ntFlanks@6747.6772.26|ACTTCGAACGCACTTGCGGCCCCGGG</td><td>>hg38_21_+_8433172_8446622_RNA45SN1_45S_50ntFlanks</td><td>6747</td><td>6772</td><td>26</td><td>ACTTCGAACGCACTTGCGGCCCCGGG</td><td>214.9</td><td>-11.6</td><td>1.6</td><td>-7.5</td><td>8.5E-14</td><td>4.1E-13</td><td>MB231/MB468</td><td>MP</td></tr>
<tr><td>>hg38_21_+_8433172_8446622_RNA45SN1_45S_50ntFlanks@10105.10127.23|CGCGTGTGGTGTGCGTCGGAGGG</td><td>>hg38_21_+_8433172_8446622_RNA45SN1_45S_50ntFlanks</td><td>10105</td><td>10127</td><td>23</td><td>CGCGTGTGGTGTGCGTCGGAGGG</td><td>142.3</td><td>-11.0</td><td>1.5</td><td>-7.5</td><td>8.5E-14</td><td>4.1E-13</td><td>MB231/MB468</td><td>MP</td></tr>
<tr><td>>hg38_21_+_8433172_8446622_RNA45SN1_45S_50ntFlanks@5557.5580.24|CTGCGGAAGGATCATTAACGGAGC</td><td>>hg38_21_+_8433172_8446622_RNA45SN1_45S_50ntFlanks</td><td>5557</td><td>5580</td><td>24</td><td>CTGCGGAAGGATCATTAACGGAGC</td><td>223.9</td><td>7.3</td><td>1.0</td><td>7.4</td><td>9.9E-14</td><td>4.1E-13</td><td>MB231/MB468</td><td>CYTO</td></tr>
<tr><td>>hg38_21_+_8433172_8446622_RNA45SN1_45S_50ntFlanks@12076.12096.21|AGGGGCTCTCGCTTCTGGCGC</td><td>>hg38_21_+_8433172_8446622_RNA45SN1_45S_50ntFlanks</td><td>12076</td><td>12096</td><td>21</td><td>AGGGGCTCTCGCTTCTGGCGC</td><td>409.1</td><td>-4.7</td><td>0.6</td><td>-7.4</td><td>1.0E-13</td><td>4.1E-13</td><td>MB231/MB468</td><td>NUC</td></tr>
<tr><td>>hg38_21_+_8433172_8446622_RNA45SN1_45S_50ntFlanks@10771.10805.35|CAGGTCTCCAAGGTGAACAGCCTCTGGCATGTTGG</td><td>>hg38_21_+_8433172_8446622_RNA45SN1_45S_50ntFlanks</td><td>10771</td><td>10805</td><td>35</td><td>CAGGTCTCCAAGGTGAACAGCCTCTGGCATGTTGG</td><td>1255.1</td><td>-9.1</td><td>1.2</td><td>-7.4</td><td>1.0E-13</td><td>4.1E-13</td><td>MB231/MB468</td><td>NUC</td></tr>
<tr><td>>hg38_21_+_8433172_8446622_RNA45SN1_45S_50ntFlanks@8182.8206.25|ATCGAGGCCCAGCCCGTGGACGGTG</td><td>>hg38_21_+_8433172_8446622_RNA45SN1_45S_50ntFlanks</td><td>8182</td><td>8206</td><td>25</td><td>ATCGAGGCCCAGCCCGTGGACGGTG</td><td>241.3</td><td>-10.0</td><td>1.3</td><td>-7.5</td><td>7.9E-14</td><td>4.1E-13</td><td>BT20/MB468</td><td>MITO</td></tr>
<tr><td>>hg38_21_+_8433172_8446622_RNA45SN1_45S_50ntFlanks@4367.4389.23|CAGTTAAAAAGCTCGTAGTTGGA</td><td>>hg38_21_+_8433172_8446622_RNA45SN1_45S_50ntFlanks</td><td>4367</td><td>4389</td><td>23</td><td>CAGTTAAAAAGCTCGTAGTTGGA</td><td>341.1</td><td>-10.5</td><td>1.4</td><td>-7.5</td><td>7.9E-14</td><td>4.1E-13</td><td>BT20/MB468</td><td>MITO</td></tr>
<tr><td>>hg38_21_+_8433172_8446622_RNA45SN1_45S_50ntFlanks@6753.6770.18|AACGCACTTGCGGCCCCG</td><td>>hg38_21_+_8433172_8446622_RNA45SN1_45S_50ntFlanks</td><td>6753</td><td>6770</td><td>18</td><td>AACGCACTTGCGGCCCCG</td><td>202.6</td><td>-11.4</td><td>1.5</td><td>-7.4</td><td>1.3E-13</td><td>4.1E-13</td><td>MB231/MB468</td><td>MITO</td></tr>
<tr><td>>hg38_21_+_8433172_8446622_RNA45SN1_45S_50ntFlanks@9937.9955.19|CAGCAGGACGGTGGCCATG</td><td>>hg38_21_+_8433172_8446622_RNA45SN1_45S_50ntFlanks</td><td>9937</td><td>9955</td><td>19</td><td>CAGCAGGACGGTGGCCATG</td><td>760.1</td><td>13.8</td><td>1.9</td><td>7.4</td><td>1.0E-13</td><td>4.1E-13</td><td>BT20/MB468</td><td>CYTO</td></tr>
<tr><td>>hg38_1_-_228634819_228635039_RNA5S12_5S_50ntFlanks@51.90.40|GTCTACGGCCATACCACCCTGAACGCGCCCGATCTCGTCT</td><td>>hg38_1_-_228634819_228635039_RNA5S12_5S_50ntFlanks</td><td>51</td><td>90</td><td>40</td><td>GTCTACGGCCATACCACCCTGAACGCGCCCGATCTCGTCT</td><td>443.4</td><td>5.9</td><td>0.8</td><td>7.5</td><td>7.9E-14</td><td>4.1E-13</td><td>BT20/MB468</td><td>MITO</td></tr>
<tr><td>>hg38_21_+_8433172_8446622_RNA45SN1_45S_50ntFlanks@8321.8356.36|AGACCGATAGTCAACAAGTACCGTAAGGGAAAGTTG</td><td>>hg38_21_+_8433172_8446622_RNA45SN1_45S_50ntFlanks</td><td>8321</td><td>8356</td><td>36</td><td>AGACCGATAGTCAACAAGTACCGTAAGGGAAAGTTG</td><td>155.3</td><td>-11.1</td><td>1.5</td><td>-7.5</td><td>8.6E-14</td><td>4.1E-13</td><td>MB231/MB468</td><td>MP</td></tr>
<tr><td>>MI0000299|hsa-mir-222&WithFlank&X|-|45747009|45747130@75.99.25(+1A)&[MIMAT0000279&hsa-miR-222-3p&offsets|0|+4(+1A);m-140&X|-|45747033|45747056&offsets|0|+1(+1A)]|AGCTACATCTGGCTACTGGGTCTCTA</td><td>>MI0000299|hsa-mir-222&WithFlank&X|-|45747009|45747130</td><td>75</td><td>99</td><td>25(+1A)</td><td>AGCTACATCTGGCTACTGGGTCTCTA</td><td>607.7</td><td>9.0</td><td>1.2</td><td>7.4</td><td>1.0E-13</td><td>4.1E-13</td><td>MB231/MB468</td><td>CYTO</td></tr>
<tr><td>>hg38_21_+_8433172_8446622_RNA45SN1_45S_50ntFlanks@9575.9598.24|ATCGGTCGTCCGACCTGGGTATAG</td><td>>hg38_21_+_8433172_8446622_RNA45SN1_45S_50ntFlanks</td><td>9575</td><td>9598</td><td>24</td><td>ATCGGTCGTCCGACCTGGGTATAG</td><td>143.8</td><td>-11.0</td><td>1.5</td><td>-7.5</td><td>8.6E-14</td><td>4.1E-13</td><td>MB231/MB468</td><td>MP</td></tr>
<tr><td>>trna10_GlyTCC_17_+_8124866_8124937@1.29.29|GCGTTGGTGGTATAGTGGTAAGCATAGCT</td><td>>trna10_GlyTCC_17_+_8124866_8124937</td><td>1</td><td>29</td><td>29</td><td>GCGTTGGTGGTATAGTGGTAAGCATAGCT</td><td>138.5</td><td>-11.0</td><td>1.5</td><td>-7.5</td><td>8.7E-14</td><td>4.1E-13</td><td>MB231/MB468</td><td>MP</td></tr>
<tr><td>>MI0000255|hsa-mir-30d&WithFlank&8|-|134804870|134804951@12.34.23(+1C)&[MIMAT0000245&hsa-miR-30d-5p&offsets|0|+1(+1C);m-21&8|-|134804919|134804940&offsets|0|+1(+1C)]|TGTAAACATCCCCGACTGGAAGCC</td><td>>MI0000255|hsa-mir-30d&WithFlank&8|-|134804870|134804951</td><td>12</td><td>34</td><td>23(+1C)</td><td>TGTAAACATCCCCGACTGGAAGCC</td><td>3485.3</td><td>6.7</td><td>0.9</td><td>7.4</td><td>1.0E-13</td><td>4.1E-13</td><td>MB231/MB468</td><td>CYTO</td></tr>
<tr><td>>hg38_21_+_8433172_8446622_RNA45SN1_45S_50ntFlanks@9716.9737.22|AATGATTAGAGGTCTTGGGGCC</td><td>>hg38_21_+_8433172_8446622_RNA45SN1_45S_50ntFlanks</td><td>9716</td><td>9737</td><td>22</td><td>AATGATTAGAGGTCTTGGGGCC</td><td>331.9</td><td>-4.3</td><td>0.6</td><td>-7.4</td><td>1.0E-13</td><td>4.1E-13</td><td>MB231/MB468</td><td>NUC</td></tr>
<tr><td>>hg38_21_+_8433172_8446622_RNA45SN1_45S_50ntFlanks@12615.12659.45|GCTGAGGAGCCAATGGGGCGAAGCTACCATCTGTGGGATTATGAC</td><td>>hg38_21_+_8433172_8446622_RNA45SN1_45S_50ntFlanks</td><td>12615</td><td>12659</td><td>45</td><td>GCTGAGGAGCCAATGGGGCGAAGCTACCATCTGTGGGATTATGAC</td><td>1229.5</td><td>-10.1</td><td>1.4</td><td>-7.4</td><td>1.3E-13</td><td>4.2E-13</td><td>MB231/MB468</td><td>MITO</td></tr>
<tr><td>>hg38_21_+_8433172_8446622_RNA45SN1_45S_50ntFlanks@4982.5015.34|ACGGACAGGATTGACAGATTGATAGCTCTTTCTC</td><td>>hg38_21_+_8433172_8446622_RNA45SN1_45S_50ntFlanks</td><td>4982</td><td>5015</td><td>34</td><td>ACGGACAGGATTGACAGATTGATAGCTCTTTCTC</td><td>686.9</td><td>-13.2</td><td>1.8</td><td>-7.4</td><td>1.3E-13</td><td>4.2E-13</td><td>MB231/MB468</td><td>MITO</td></tr>
<tr><td>>hg38_21_+_8433172_8446622_RNA45SN1_45S_50ntFlanks@4555.4587.33|CGCAGCTAGGAATAATGGAATAGGACCGCGGTT</td><td>>hg38_21_+_8433172_8446622_RNA45SN1_45S_50ntFlanks</td><td>4555</td><td>4587</td><td>33</td><td>CGCAGCTAGGAATAATGGAATAGGACCGCGGTT</td><td>335.2</td><td>-8.2</td><td>1.1</td><td>-7.4</td><td>1.0E-13</td><td>4.2E-13</td><td>MB231/MB468</td><td>NUC</td></tr>
<tr><td>>MI0000070|hsa-mir-16-1&WithFlank&13|-|50048967|50049067@20.39.20;MI0000115|hsa-mir-16-2&WithFlank&3|+|160404739|160404831@16.35.20&[MIMAT0000069&hsa-miR-16-5p&offsets|0|-2;m-62&13|-|50049027|50049048&offsets|0|-2];[MIMAT0000069_1&hsa-miR-16-5p&offsets|0|-2;m-61&3|+|160404754|160404775&offsets|0|-2]|TAGCAGCACGTAAATATTGG</td><td>>MI0000070|hsa-mir-16-1&WithFlank&13|-|50048967|50049067</td><td>20</td><td>39</td><td>20</td><td>TAGCAGCACGTAAATATTGG</td><td>257.5</td><td>5.5</td><td>0.7</td><td>7.4</td><td>1.0E-13</td><td>4.2E-13</td><td>MB231/MB468</td><td>NUC</td></tr>
<tr><td>>hg38_21_+_8433172_8446622_RNA45SN1_45S_50ntFlanks@7979.7998.20|ACCTCAGATCAGACGTGGCG</td><td>>hg38_21_+_8433172_8446622_RNA45SN1_45S_50ntFlanks</td><td>7979</td><td>7998</td><td>20</td><td>ACCTCAGATCAGACGTGGCG</td><td>133.7</td><td>-10.9</td><td>1.5</td><td>-7.5</td><td>8.8E-14</td><td>4.2E-13</td><td>MB231/MB468</td><td>MP</td></tr>
<tr><td>>hg38_21_+_8433172_8446622_RNA45SN1_45S_50ntFlanks@10689.10706.18|GCGTCCGGTGAGCTCTCG</td><td>>hg38_21_+_8433172_8446622_RNA45SN1_45S_50ntFlanks</td><td>10689</td><td>10706</td><td>18</td><td>GCGTCCGGTGAGCTCTCG</td><td>1942.6</td><td>-8.0</td><td>1.1</td><td>-7.4</td><td>1.0E-13</td><td>4.2E-13</td><td>MB231/MB468</td><td>CYTO</td></tr>
</table>

<table>
<tr><th></th><th></th><th></th><th></th><th></th><th></th><th></th><th></th><th></th><th></th><th></th><th></th><th></th><th></th></tr>
<tr><td>>hg38_21_+_8433172_8446622_RNA45SN1_45S_50ntFlanks@4725.4758.34|TTAATCAAGAACGAAAGTCGGAGGTTCGAAGACG</td><td>>hg38_21_+_8433172_8446622_RNA45SN1_45S_50ntFlanks</td><td>4725</td><td>4758</td><td>34</td><td>TTAATCAAGAACGAAAGTCGGAGGTTCGAAGACG</td><td>2186.9</td><td>-7.9</td><td>1.1</td><td>-7.4</td><td>1.4E-13</td><td>4.2E-13</td><td>MB231/MB468</td><td>MITO</td></tr>
<tr><td>>hg38_21_+_8433172_8446622_RNA45SN1_45S_50ntFlanks@10103.10127.25|CGCGCGTGTGGTGTGCGTCGGAGGG</td><td>>hg38_21_+_8433172_8446622_RNA45SN1_45S_50ntFlanks</td><td>10103</td><td>10127</td><td>25</td><td>CGCGCGTGTGGTGTGCGTCGGAGGG</td><td>174.1</td><td>-11.3</td><td>1.5</td><td>-7.5</td><td>8.8E-14</td><td>4.2E-13</td><td>MB231/MB468</td><td>MP</td></tr>
<tr><td>>hg38_21_+_8433172_8446622_RNA45SN1_45S_50ntFlanks@6647.6679.33|CGTACGACTCTTAGCGGTGGATCACTCGGCTCG</td><td>>hg38_21_+_8433172_8446622_RNA45SN1_45S_50ntFlanks</td><td>6647</td><td>6679</td><td>33</td><td>CGTACGACTCTTAGCGGTGGATCACTCGGCTCG</td><td>903.8</td><td>-3.2</td><td>0.4</td><td>-7.4</td><td>1.5E-13</td><td>4.2E-13</td><td>BT20/MB468</td><td>NUC</td></tr>
<tr><td>>hg38_21_+_8433172_8446622_RNA45SN1_45S_50ntFlanks@10408.10433.26|GGTCAGTCGGTCCTGAGAGATGGGCG</td><td>>hg38_21_+_8433172_8446622_RNA45SN1_45S_50ntFlanks</td><td>10408</td><td>10433</td><td>26</td><td>GGTCAGTCGGTCCTGAGAGATGGGCG</td><td>2215.6</td><td>-10.2</td><td>1.4</td><td>-7.4</td><td>1.0E-13</td><td>4.2E-13</td><td>MB231/MB468</td><td>NUC</td></tr>
<tr><td>>hg38_1_-_228634819_228635039_RNA5S12_5S_50ntFlanks@137.170.34|GAGACCGCCTGGGAATACCGGGTGCTGTAGGCTT</td><td>>hg38_1_-_228634819_228635039_RNA5S12_5S_50ntFlanks</td><td>137</td><td>170</td><td>34</td><td>GAGACCGCCTGGGAATACCGGGTGCTGTAGGCTT</td><td>846.6</td><td>7.7</td><td>1.0</td><td>7.4</td><td>1.0E-13</td><td>4.2E-13</td><td>MB231/MB468</td><td>CYTO</td></tr>
<tr><td>>hg38_21_+_8433172_8446622_RNA45SN1_45S_50ntFlanks@12539.12573.35|ATGATGTGTTGTTGCCATGGTAATCCTGCTCAGTA</td><td>>hg38_21_+_8433172_8446622_RNA45SN1_45S_50ntFlanks</td><td>12539</td><td>12573</td><td>35</td><td>ATGATGTGTTGTTGCCATGGTAATCCTGCTCAGTA</td><td>609.1</td><td>-12.3</td><td>1.6</td><td>-7.9</td><td>3.9E-15</td><td>4.2E-13</td><td>BT20/MB468</td><td>MP</td></tr>
<tr><td>>hg38_21_+_8433172_8446622_RNA45SN1_45S_50ntFlanks@3899.3937.39|CCTTCGCGGGGGGGATGCGTGCATTTATCAGATCAAAAC</td><td>>hg38_21_+_8433172_8446622_RNA45SN1_45S_50ntFlanks</td><td>3899</td><td>3937</td><td>39</td><td>CCTTCGCGGGGGGGATGCGTGCATTTATCAGATCAAAAC</td><td>1014.4</td><td>-13.9</td><td>1.9</td><td>-7.5</td><td>8.8E-14</td><td>4.2E-13</td><td>MB231/MB468</td><td>MP</td></tr>
<tr><td>>Antisense_hg38_21_-_8433172_8446622_RNA45SN1_45S_50ntFlanks@7674.7691.18|CTCTCTCTCTCTCTCCCC</td><td>>Antisense_hg38_21_-_8433172_8446622_RNA45SN1_45S_50ntFlanks</td><td>7674</td><td>7691</td><td>18</td><td>CTCTCTCTCTCTCTCCCC</td><td>206.6</td><td>-10.4</td><td>1.4</td><td>-7.4</td><td>1.0E-13</td><td>4.2E-13</td><td>MB231/MB468</td><td>CYTO</td></tr>
<tr><td>>hg38_21_+_8433172_8446622_RNA45SN1_45S_50ntFlanks@11027.11044.18|CTCTCTCTCTCTCTCCCC</td><td>>hg38_21_+_8433172_8446622_RNA45SN1_45S_50ntFlanks</td><td>11027</td><td>11044</td><td>18</td><td>CTCTCTCTCTCTCTCCCC</td><td>206.6</td><td>-10.4</td><td>1.4</td><td>-7.4</td><td>1.0E-13</td><td>4.2E-13</td><td>MB231/MB468</td><td>CYTO</td></tr>
<tr><td>>hg38_21_+_8433172_8446622_RNA45SN1_45S_50ntFlanks@12416.12450.35|TTCGATGTCGGCTCTTCCTATCATTGTGAAGCAGA</td><td>>hg38_21_+_8433172_8446622_RNA45SN1_45S_50ntFlanks</td><td>12416</td><td>12450</td><td>35</td><td>TTCGATGTCGGCTCTTCCTATCATTGTGAAGCAGA</td><td>375.0</td><td>4.9</td><td>0.7</td><td>7.4</td><td>1.0E-13</td><td>4.2E-13</td><td>MB231/MB468</td><td>CYTO</td></tr>
<tr><td>>hg38_21_+_8433172_8446622_RNA45SN1_45S_50ntFlanks@3705.3746.42|TACCTGGTTGATCCTGCCAGTAGCATATGCTTGTCTCAAAGA</td><td>>hg38_21_+_8433172_8446622_RNA45SN1_45S_50ntFlanks</td><td>3705</td><td>3746</td><td>42</td><td>TACCTGGTTGATCCTGCCAGTAGCATATGCTTGTCTCAAAGA</td><td>1917.7</td><td>7.9</td><td>1.1</td><td>7.4</td><td>1.0E-13</td><td>4.2E-13</td><td>MB231/MB468</td><td>CYTO</td></tr>
<tr><td>>hg38_MT_+_1621_3279_RNR2_16S_50ntFlanks@493.518.26|GCTCTTTGGACACTAGGAAAAAACCT</td><td>>hg38_MT_+_1621_3279_RNR2_16S_50ntFlanks</td><td>493</td><td>518</td><td>26</td><td>GCTCTTTGGACACTAGGAAAAAACCT</td><td>382.1</td><td>7.5</td><td>1.0</td><td>7.4</td><td>1.1E-13</td><td>4.2E-13</td><td>MB231/MB468</td><td>NUC</td></tr>
<tr><td>>MI0000456|hsa-mir-140&WithFlank&16|+|69933075|69933186@69.89.21&[MIMAT0004597&hsa-miR-140-3p&offsets|+1|+1;m-45&16|+|69933143|69933164&offsets|0|-1]|ACCACAGGGTAGAACCACGGA</td><td>>MI0000456|hsa-mir-140&WithFlank&16|+|69933075|69933186</td><td>69</td><td>89</td><td>21</td><td>ACCACAGGGTAGAACCACGGA</td><td>189.9</td><td>6.7</td><td>0.9</td><td>7.4</td><td>1.6E-13</td><td>4.2E-13</td><td>BT20/MB468</td><td>NUC</td></tr>
<tr><td>>hg38_1_-_228634819_228635039_RNA5S12_5S_50ntFlanks@148.171.24|GGAATACCGGGTGCTGTAGGCTTT</td><td>>hg38_1_-_228634819_228635039_RNA5S12_5S_50ntFlanks</td><td>148</td><td>171</td><td>24</td><td>GGAATACCGGGTGCTGTAGGCTTT</td><td>20595.3</td><td>-6.0</td><td>0.8</td><td>-7.5</td><td>8.9E-14</td><td>4.2E-13</td><td>MB231/MB468</td><td>MP</td></tr>
<tr><td>>MI0000102|hsa-mir-100&WithFlank&11|-|122152223|122152314@19.40.22(+3U)&[MIMAT0000098&hsa-miR-100-5p&offsets|0|0(+3U);m-27&11|-|122152276|122152296&offsets|0|+1(+3U)]|AACCCGTAGATCCGAACTTGTGTTT</td><td>>MI0000102|hsa-mir-100&WithFlank&11|-|122152223|122152314</td><td>19</td><td>40</td><td>22(+3U)</td><td>AACCCGTAGATCCGAACTTGTGTTT</td><td>215.8</td><td>-11.0</td><td>1.4</td><td>-7.7</td><td>1.1E-14</td><td>4.2E-13</td><td>BT20/MB231</td><td>MITO</td></tr>
<tr><td>>hg38_21_+_8433172_8446622_RNA45SN1_45S_50ntFlanks@5186.5206.21|CAATAACAGGTCTGTGATGCC</td><td>>hg38_21_+_8433172_8446622_RNA45SN1_45S_50ntFlanks</td><td>5186</td><td>5206</td><td>21</td><td>CAATAACAGGTCTGTGATGCC</td><td>654.9</td><td>-13.2</td><td>1.8</td><td>-7.5</td><td>8.9E-14</td><td>4.2E-13</td><td>MB231/MB468</td><td>MP</td></tr>
<tr><td>>hg38_21_+_8433172_8446622_RNA45SN1_45S_50ntFlanks@11667.11704.38|CTGCCCAGTGCTCTGAATGTCAAAGTGAAGAAATTCAA</td><td>>hg38_21_+_8433172_8446622_RNA45SN1_45S_50ntFlanks</td><td>11667</td><td>11704</td><td>38</td><td>CTGCCCAGTGCTCTGAATGTCAAAGTGAAGAAATTCAA</td><td>359.0</td><td>7.6</td><td>1.0</td><td>7.5</td><td>8.2E-14</td><td>4.2E-13</td><td>BT20/MB468</td><td>MITO</td></tr>
<tr><td>>hg38_21_+_8433172_8446622_RNA45SN1_45S_50ntFlanks@4442.4471.30|CCCCTTGCCTCTCGGCGCCCCCTCGATGCT</td><td>>hg38_21_+_8433172_8446622_RNA45SN1_45S_50ntFlanks</td><td>4442</td><td>4471</td><td>30</td><td>CCCCTTGCCTCTCGGCGCCCCCTCGATGCT</td><td>166.0</td><td>-7.1</td><td>1.0</td><td>-7.4</td><td>1.0E-13</td><td>4.2E-13</td><td>MB231/MB468</td><td>CYTO</td></tr>
<tr><td>>MI0003185|hsa-mir-501&WithFlank&X|+|50009716|50009811@57.78.22&[MIMAT0004774&hsa-miR-501-3p&offsets|0|0;m-168&X|+|50009772|50009793&offsets|0|0]|AATGCACCCGGGCAAGGATTCT</td><td>>MI0003185|hsa-mir-501&WithFlank&X|+|50009716|50009811</td><td>57</td><td>78</td><td>22</td><td>AATGCACCCGGGCAAGGATTCT</td><td>424.2</td><td>6.7</td><td>0.9</td><td>7.4</td><td>1.1E-13</td><td>4.3E-13</td><td>BT20/MB468</td><td>CYTO</td></tr>
<tr><td>>hg38_21_+_8433172_8446622_RNA45SN1_45S_50ntFlanks@6721.6757.37|AATTGCAGGACACATTGATCATCGACACTTCGAACGC</td><td>>hg38_21_+_8433172_8446622_RNA45SN1_45S_50ntFlanks</td><td>6721</td><td>6757</td><td>37</td><td>AATTGCAGGACACATTGATCATCGACACTTCGAACGC</td><td>272.0</td><td>-12.0</td><td>1.6</td><td>-7.5</td><td>9.0E-14</td><td>4.3E-13</td><td>MB231/MB468</td><td>MP</td></tr>
<tr><td>>am_trna106_HisGTG_1_-_149155828_149155899@7.29.23,trna111_HisGTG_1_-_147774845_147774916@7.29.23,trna118_HisGTG_1_-_145396881_145396952@7.29.23,trna16_HisGTG_1_+_146544773_146544844@7.29.23,trna1_HisGTG_15_+_45493349_45493420@7.29.23,trna21_HisGTG_1_+_147753471_147753542@7.29.23,trna33_HisGTG_6_+_27125906_27125977@7.29.23,trna7_HisGTG_9_-_14433938_14434009@7.29.23,trna8_HisGTG_15_-_45492611_45492682@7.29.23,trna9_HisGTG_15_-_45490804_45490875@7.29.23|ATCGTATAGTGGTTAGTACTCTG</td><td>>am_trna106_HisGTG_1_-_149155828_149155899</td><td>7</td><td>29</td><td>23</td><td>ATCGTATAGTGGTTAGTACTCTG</td><td>505.4</td><td>-12.1</td><td>1.5</td><td>-7.9</td><td>4.1E-15</td><td>4.3E-13</td><td>BT20/MB468</td><td>MP</td></tr>
<tr><td>>hg38_21_+_8433172_8446622_RNA45SN1_45S_50ntFlanks@12435.12454.20|ATCATTGTGAAGCAGAATTC</td><td>>hg38_21_+_8433172_8446622_RNA45SN1_45S_50ntFlanks</td><td>12435</td><td>12454</td><td>20</td><td>ATCATTGTGAAGCAGAATTC</td><td>717.2</td><td>-12.6</td><td>1.6</td><td>-7.9</td><td>4.1E-15</td><td>4.3E-13</td><td>BT20/MB468</td><td>MP</td></tr>
<tr><td>>hg38_21_+_8433172_8446622_RNA45SN1_45S_50ntFlanks@12046.12070.25|TGACCCGGTGAGGCGGGGGGGCGAG</td><td>>hg38_21_+_8433172_8446622_RNA45SN1_45S_50ntFlanks</td><td>12046</td><td>12070</td><td>25</td><td>TGACCCGGTGAGGCGGGGGGGCGAG</td><td>561.0</td><td>-12.9</td><td>1.7</td><td>-7.4</td><td>1.4E-13</td><td>4.3E-13</td><td>MB231/MB468</td><td>MITO</td></tr>
<tr><td>>hg38_21_+_8433172_8446622_RNA45SN1_45S_50ntFlanks@5472.5502.31|ACGGCCCTGGCGGAGCGCTGAGAAGACGGTC</td><td>>hg38_21_+_8433172_8446622_RNA45SN1_45S_50ntFlanks</td><td>5472</td><td>5502</td><td>31</td><td>ACGGCCCTGGCGGAGCGCTGAGAAGACGGTC</td><td>6531.9</td><td>8.0</td><td>1.1</td><td>7.4</td><td>1.1E-13</td><td>4.3E-13</td><td>BT20/MB468</td><td>CYTO</td></tr>
<tr><td>>hg38_21_+_8433172_8446622_RNA45SN1_45S_50ntFlanks@10867.10902.36|TAAGGGCTGGGTCGGTCGGGCTGGGGCGCGAAGCGG</td><td>>hg38_21_+_8433172_8446622_RNA45SN1_45S_50ntFlanks</td><td>10867</td><td>10902</td><td>36</td><td>TAAGGGCTGGGTCGGTCGGGCTGGGGCGCGAAGCGG</td><td>189.9</td><td>-10.5</td><td>1.4</td><td>-7.4</td><td>1.1E-13</td><td>4.3E-13</td><td>MB231/MB468</td><td>NUC</td></tr>
<tr><td>>MI0000440|hsa-mir-27b&WithFlank&9|+|95085439|95085547@67.88.22(+1A)&[MIMAT0000419&hsa-miR-27b-3p&offsets|0|+1(+1A);m-39&9|+|95085505|95085525&offsets|0|+1(+1A)]|TTCACAGTGGCTAAGTTCTGCAA</td><td>>MI0000440|hsa-mir-27b&WithFlank&9|+|95085439|95085547</td><td>67</td><td>88</td><td>22(+1A)</td><td>TTCACAGTGGCTAAGTTCTGCAA</td><td>350.1</td><td>4.6</td><td>0.6</td><td>7.4</td><td>1.1E-13</td><td>4.3E-13</td><td>MB231/MB468</td><td>NUC</td></tr>
<tr><td>>hg38_21_+_8433172_8446622_RNA45SN1_45S_50ntFlanks@11205.11227.23|GCGGCGGCGACTCTGGACGCGAG</td><td>>hg38_21_+_8433172_8446622_RNA45SN1_45S_50ntFlanks</td><td>11205</td><td>11227</td><td>23</td><td>GCGGCGGCGACTCTGGACGCGAG</td><td>160.9</td><td>-6.3</td><td>0.8</td><td>-7.4</td><td>1.1E-13</td><td>4.3E-13</td><td>MB231/MB468</td><td>CYTO</td></tr>
<tr><td>>hg38_21_+_8433172_8446622_RNA45SN1_45S_50ntFlanks@10690.10711.22|CGTCCGGTGAGCTCTCGCTGGC</td><td>>hg38_21_+_8433172_8446622_RNA45SN1_45S_50ntFlanks</td><td>10690</td><td>10711</td><td>22</td><td>CGTCCGGTGAGCTCTCGCTGGC</td><td>265.4</td><td>-10.8</td><td>1.4</td><td>-7.4</td><td>1.1E-13</td><td>4.3E-13</td><td>MB231/MB468</td><td>CYTO</td></tr>
<tr><td>>MI0000086|hsa-mir-28&WithFlank&3|+|188688775|188688872@61.82.22&[MIMAT0004502&hsa-miR-28-3p&offsets|+1|+1;m-28&3|+|188688834|188688855&offsets|+1|+1]|ACTAGATTGTGAGCTCCTGGAG</td><td>>MI0000086|hsa-mir-28&WithFlank&3|+|188688775|188688872</td><td>61</td><td>82</td><td>22</td><td>ACTAGATTGTGAGCTCCTGGAG</td><td>108.3</td><td>3.7</td><td>0.5</td><td>7.4</td><td>1.6E-13</td><td>4.3E-13</td><td>BT20/MB468</td><td>NUC</td></tr>
<tr><td>>hg38_21_+_8433172_8446622_RNA45SN1_45S_50ntFlanks@5079.5108.30|GATAACGAACGAGACTCTGGCATGCTAACT</td><td>>hg38_21_+_8433172_8446622_RNA45SN1_45S_50ntFlanks</td><td>5079</td><td>5108</td><td>30</td><td>GATAACGAACGAGACTCTGGCATGCTAACT</td><td>246.6</td><td>6.5</td><td>0.9</td><td>7.4</td><td>1.1E-13</td><td>4.3E-13</td><td>MB231/MB468</td><td>CYTO</td></tr>
<tr><td>>trnaMT_ValTAC_MT_+_1602_1670@44.72.29|ATTTCAACTTAACTTGACCGCTCTGACCA</td><td>>trnaMT_ValTAC_MT_+_1602_1670</td><td>44</td><td>72</td><td>29</td><td>ATTTCAACTTAACTTGACCGCTCTGACCA</td><td>824.1</td><td>4.6</td><td>0.6</td><td>7.4</td><td>1.1E-13</td><td>4.4E-13</td><td>MB231/MB468</td><td>NUC</td></tr>
<tr><td>>hg38_21_+_8433172_8446622_RNA45SN1_45S_50ntFlanks@8381.8402.22|AAGAGGGCGTGAAACCGTTAAG</td><td>>hg38_21_+_8433172_8446622_RNA45SN1_45S_50ntFlanks</td><td>8381</td><td>8402</td><td>22</td><td>AAGAGGGCGTGAAACCGTTAAG</td><td>1389.7</td><td>-5.5</td><td>0.7</td><td>-7.5</td><td>8.4E-14</td><td>4.4E-13</td><td>BT20/MB468</td><td>MITO</td></tr>
<tr><td>>hg38_21_+_8433172_8446622_RNA45SN1_45S_50ntFlanks@12234.12257.24|CGTGGAGCAGAAGGGCAAAAGCTC</td><td>>hg38_21_+_8433172_8446622_RNA45SN1_45S_50ntFlanks</td><td>12234</td><td>12257</td><td>24</td><td>CGTGGAGCAGAAGGGCAAAAGCTC</td><td>245.2</td><td>-11.8</td><td>1.6</td><td>-7.5</td><td>9.2E-14</td><td>4.4E-13</td><td>MB231/MB468</td><td>MP</td></tr>
<tr><td>>hg38_21_+_8433172_8446622_RNA45SN1_45S_50ntFlanks@10136.10153.18|GCGGCGGCGGCGGGGGTG</td><td>>hg38_21_+_8433172_8446622_RNA45SN1_45S_50ntFlanks</td><td>10136</td><td>10153</td><td>18</td><td>GCGGCGGCGGCGGGGGTG</td><td>221.9</td><td>-11.7</td><td>1.6</td><td>-7.5</td><td>9.2E-14</td><td>4.4E-13</td><td>MB231/MB468</td><td>MP</td></tr>
<tr><td>>MI0000299|hsa-mir-222&WithFlank&X|-|45747009|45747130@75.98.24(+1C)&[MIMAT0000279&hsa-miR-222-3p&offsets|0|+3(+1C);m-140&X|-|45747033|45747056&offsets|0|0(+1C)]|AGCTACATCTGGCTACTGGGTCTCC</td><td>>MI0000299|hsa-mir-222&WithFlank&X|-|45747009|45747130</td><td>75</td><td>98</td><td>24(+1C)</td><td>AGCTACATCTGGCTACTGGGTCTCC</td><td>8410.2</td><td>6.5</td><td>0.9</td><td>7.4</td><td>1.4E-13</td><td>4.4E-13</td><td>MB231/MB468</td><td>MITO</td></tr>
<tr><td>>hg38_21_+_8433172_8446622_RNA45SN1_45S_50ntFlanks@11570.11605.36|AGCCGACTTAGAACTGGTGCGGACCAGGGGAATCCG</td><td>>hg38_21_+_8433172_8446622_RNA45SN1_45S_50ntFlanks</td><td>11570</td><td>11605</td><td>36</td><td>AGCCGACTTAGAACTGGTGCGGACCAGGGGAATCCG</td><td>273.4</td><td>-12.0</td><td>1.6</td><td>-7.5</td><td>9.2E-14</td><td>4.4E-13</td><td>MB231/MB468</td><td>MP</td></tr>
<tr><td>>hg38_21_+_8433172_8446622_RNA45SN1_45S_50ntFlanks@4729.4759.31|TCAAGAACGAAAGTCGGAGGTTCGAAGACGA</td><td>>hg38_21_+_8433172_8446622_RNA45SN1_45S_50ntFlanks</td><td>4729</td><td>4759</td><td>31</td><td>TCAAGAACGAAAGTCGGAGGTTCGAAGACGA</td><td>746.6</td><td>-5.7</td><td>0.8</td><td>-7.4</td><td>1.4E-13</td><td>4.4E-13</td><td>MB231/MB468</td><td>MITO</td></tr>
<tr><td>>hg38_21_+_8433172_8446622_RNA45SN1_45S_50ntFlanks@11314.11342.29|CCCACCCCACGTCTCGTCGCGCGCGCGTC</td><td>>hg38_21_+_8433172_8446622_RNA45SN1_45S_50ntFlanks</td><td>11314</td><td>11342</td><td>29</td><td>CCCACCCCACGTCTCGTCGCGCGCGCGTC</td><td>201.8</td><td>-10.4</td><td>1.4</td><td>-7.4</td><td>1.1E-13</td><td>4.4E-13</td><td>MB231/MB468</td><td>CYTO</td></tr>
<tr><td>>hg38_21_+_8433172_8446622_RNA45SN1_45S_50ntFlanks@11198.11225.28|GCCGGCGGCGGCGGCGACTCTGGACGCG</td><td>>hg38_21_+_8433172_8446622_RNA45SN1_45S_50ntFlanks</td><td>11198</td><td>11225</td><td>28</td><td>GCCGGCGGCGGCGGCGACTCTGGACGCG</td><td>132.3</td><td>-9.8</td><td>1.3</td><td>-7.4</td><td>1.1E-13</td><td>4.4E-13</td><td>MB231/MB468</td><td>CYTO</td></tr>
<tr><td>>hg38_21_+_8433172_8446622_RNA45SN1_45S_50ntFlanks@11564.11584.21|CCTAGCAGCCGACTTAGAACT</td><td>>hg38_21_+_8433172_8446622_RNA45SN1_45S_50ntFlanks</td><td>11564</td><td>11584</td><td>21</td><td>CCTAGCAGCCGACTTAGAACT</td><td>563.5</td><td>-4.2</td><td>0.6</td><td>-7.4</td><td>1.6E-13</td><td>4.4E-13</td><td>BT20/MB468</td><td>NUC</td></tr>
<tr><td>>am_trna134_GluTTC_1_-_16861774_16861845@1.28.28,trna5_GluTTC_1_+_17199078_17199149@1.28.28,trna84_GluTTC_1_-_161391883_161391954@1.28.28,trna94_GluTTC_1_-_149664355_149664427@1.28.28|TCCCTGGTGGTCTAGTGGCTAGGATTCG</td><td>>am_trna134_GluTTC_1_-_16861774_16861845</td><td>1</td><td>28</td><td>28</td><td>TCCCTGGTGGTCTAGTGGCTAGGATTCG</td><td>198.0</td><td>-11.5</td><td>1.5</td><td>-7.5</td><td>9.3E-14</td><td>4.4E-13</td><td>MB231/MB468</td><td>MP</td></tr>
<tr><td>>hg38_21_+_8433172_8446622_RNA45SN1_45S_50ntFlanks@10781.10808.28|AGGTGAACAGCCTCTGGCATGTTGGAAC</td><td>>hg38_21_+_8433172_8446622_RNA45SN1_45S_50ntFlanks</td><td>10781</td><td>10808</td><td>28</td><td>AGGTGAACAGCCTCTGGCATGTTGGAAC</td><td>160.5</td><td>-10.3</td><td>1.4</td><td>-7.4</td><td>1.1E-13</td><td>4.4E-13</td><td>MB231/MB468</td><td>NUC</td></tr>
<tr><td>>hg38_21_+_8433172_8446622_RNA45SN1_45S_50ntFlanks@12992.13014.23|TCCCTCGCTGCGATCTATTGAAA</td><td>>hg38_21_+_8433172_8446622_RNA45SN1_45S_50ntFlanks</td><td>12992</td><td>13014</td><td>23</td><td>TCCCTCGCTGCGATCTATTGAAA</td><td>1536.9</td><td>-4.9</td><td>0.7</td><td>-7.4</td><td>1.1E-13</td><td>4.4E-13</td><td>MB231/MB468</td><td>NUC</td></tr>
<tr><td>>MI0000089|hsa-mir-31&WithFlank&9|-|21512109|21512191@14.34.21&[MIMAT0000089&hsa-miR-31-5p&offsets|0|0;m-360&9|-|21512157|21512178&offsets|0|-1]|AGGCAAGATGCTGGCATAGCT</td><td>>MI0000089|hsa-mir-31&WithFlank&9|-|21512109|21512191</td><td>14</td><td>34</td><td>21</td><td>AGGCAAGATGCTGGCATAGCT</td><td>217.4</td><td>11.1</td><td>1.4</td><td>7.8</td><td>8.6E-15</td><td>4.4E-13</td><td>BT20/MB231</td><td>CYTO</td></tr>
<tr><td>>hg38_21_+_8433172_8446622_RNA45SN1_45S_50ntFlanks@12081.12102.22|CTCTCGCTTCTGGCGCCAAGCG</td><td>>hg38_21_+_8433172_8446622_RNA45SN1_45S_50ntFlanks</td><td>12081</td><td>12102</td><td>22</td><td>CTCTCGCTTCTGGCGCCAAGCG</td><td>233.6</td><td>-6.6</td><td>0.9</td><td>-7.4</td><td>1.1E-13</td><td>4.4E-13</td><td>MB231/MB468</td><td>CYTO</td></tr>
<tr><td>>hg38_21_+_8433172_8446622_RNA45SN1_45S_50ntFlanks@12488.12509.22|AACGTGAGCTGGGTTTAGACCG</td><td>>hg38_21_+_8433172_8446622_RNA45SN1_45S_50ntFlanks</td><td>12488</td><td>12509</td><td>22</td><td>AACGTGAGCTGGGTTTAGACCG</td><td>3928.4</td><td>-7.3</td><td>1.0</td><td>-7.4</td><td>1.1E-13</td><td>4.4E-13</td><td>MB231/MB468</td><td>CYTO</td></tr>
</table>

| | | | | | | | | | | | | | |
|---|---|---|---|---|---|---|---|---|---|---|---|---|---|
| >hg38_21_+_8433172_8446622_RNA45SN1_45S_50ntFlanks@10808.10840.33\|CAATGTAGGTAAGGGAAGTCGGCAAGCCGGATC | >hg38_21_+_8433172_8446622_RNA45SN1_45S_50ntFlanks | 10808 | 10840 | 33 | CAATGTAGGTAAGGGAAGTCGGCAAGCCGGATC | 1108.1 | -4.8 | 0.6 | -7.4 | 1.1E-13 | 4.4E-13 | BT20/MB468 | CYTO |
| >hg38_21_+_8433172_8446622_RNA45SN1_45S_50ntFlanks@9454.9487.34\|CCGCGCCGGGGAGGTGGAGCACGAGCGCACGTGT | >hg38_21_+_8433172_8446622_RNA45SN1_45S_50ntFlanks | 9454 | 9487 | 34 | CCGCGCCGGGGAGGTGGAGCACGAGCGCACGTGT | 137.7 | -11.0 | 1.5 | -7.4 | 9.4E-14 | 4.4E-13 | MB231/MB468 | MP |
| >hg38_21_+_8433172_8446622_RNA45SN1_45S_50ntFlanks@10512.10534.23\|TCCGGAGTGGCGGAGATGGGCGC | >hg38_21_+_8433172_8446622_RNA45SN1_45S_50ntFlanks | 10512 | 10534 | 23 | TCCGGAGTGGCGGAGATGGGCGC | 310.5 | -11.2 | 1.5 | -7.4 | 1.1E-13 | 4.4E-13 | MB231/MB468 | NUC |
| >hg38_21_+_8433172_8446622_RNA45SN1_45S_50ntFlanks@11189.11207.19\|ACCCGGGGGGCCGGCGGCG | >hg38_21_+_8433172_8446622_RNA45SN1_45S_50ntFlanks | 11189 | 11207 | 19 | ACCCGGGGGGCCGGCGGCG | 301.9 | -11.2 | 1.5 | -7.4 | 1.1E-13 | 4.5E-13 | MB231/MB468 | NUC |
| >hg38_21_+_8433172_8446622_RNA45SN1_45S_50ntFlanks@13008.13039.32\|ATTGAAAGTCAGCCCTCGACACAAGGGTTTGT | >hg38_21_+_8433172_8446622_RNA45SN1_45S_50ntFlanks | 13008 | 13039 | 32 | ATTGAAAGTCAGCCCTCGACACAAGGGTTTGT | 4835.1 | 6.0 | 0.8 | 7.4 | 1.1E-13 | 4.5E-13 | BT20/MB468 | CYTO |
| >hg38_21_+_8433172_8446622_RNA45SN1_45S_50ntFlanks@11455.11475.21\|CTCGGCGCGCGGCGGCGGCGG | >hg38_21_+_8433172_8446622_RNA45SN1_45S_50ntFlanks | 11455 | 11475 | 21 | CTCGGCGCGCGGCGGCGGCGG | 205.8 | -11.5 | 1.6 | -7.4 | 9.4E-14 | 4.5E-13 | MB231/MB468 | MP |
| >hg38_21_+_8433172_8446622_RNA45SN1_45S_50ntFlanks@8182.8206.25\|ATCGAGGCCCAGCCCGTGGACGGTG | >hg38_21_+_8433172_8446622_RNA45SN1_45S_50ntFlanks | 8182 | 8206 | 25 | ATCGAGGCCCAGCCCGTGGACGGTG | 190.0 | -9.9 | 1.3 | -7.4 | 1.1E-13 | 4.5E-13 | BT20/MB468 | CYTO |
| >hg38_21_+_8433172_8446622_RNA45SN1_45S_50ntFlanks@5017.5034.18\|ATTCCGTGGGTGGTGGTG | >hg38_21_+_8433172_8446622_RNA45SN1_45S_50ntFlanks | 5017 | 5034 | 18 | ATTCCGTGGGTGGTGGTG | 1759.1 | -14.6 | 2.0 | -7.4 | 1.4E-13 | 4.5E-13 | MB231/MB468 | MITO |
| >hg38_21_+_8433172_8446622_RNA45SN1_45S_50ntFlanks@10408.10434.27\|GGTCAGTCGGTCCTGAGAGATGGGCGA | >hg38_21_+_8433172_8446622_RNA45SN1_45S_50ntFlanks | 10408 | 10434 | 27 | GGTCAGTCGGTCCTGAGAGATGGGCGA | 213.2 | -7.3 | 1.0 | -7.4 | 1.1E-13 | 4.5E-13 | BT20/MB468 | CYTO |
| >MI0000446\|hsa-mir-125b-1&WithFlank&11\|-\|122099751\|122099850@21.41.21(+1G);MI0000470\|hsa-mir-125b-2&WithFlank&21\|+\|16590231\|16590331@23.43.21(+1G)&[MIMAT0000423&hsa-miR-125b-5p&offsets\|0\|-1(+1G);m-67&11\|-\|122099809\|122099830&offsets\|0\|-1(+1G)];[MIMAT0000423_1&hsa-miR-125b-5p&offsets\|0\|-1(+1G);m-68&21\|+\|16590253\|16590274&offsets\|0\|-1(+1G)]\|TCCCTGAGACCCTAACTTGTGG | >MI0000446\|hsa-mir-125b-1&WithFlank&11\|-\|122099751\|122099850 | 21 | 41 | 21(+1G) | TCCCTGAGACCCTAACTTGTGG | 346.3 | -11.7 | 1.5 | -7.7 | 1.2E-14 | 4.5E-13 | BT20/MB231 | MITO |
| >hg38_21_+_8433172_8446622_RNA45SN1_45S_50ntFlanks@11579.11601.23\|AGAACTGGTGCGGACCAGGGGAA | >hg38_21_+_8433172_8446622_RNA45SN1_45S_50ntFlanks | 11579 | 11601 | 23 | AGAACTGGTGCGGACCAGGGGAA | 526.6 | -8.8 | 1.2 | -7.4 | 1.5E-13 | 4.5E-13 | MB231/MB468 | MITO |
| >hg38_21_+_8433172_8446622_RNA45SN1_45S_50ntFlanks@9534.9561.28\|AGGAAACTCTGGTGGAGGTCCGTAGCGG | >hg38_21_+_8433172_8446622_RNA45SN1_45S_50ntFlanks | 9534 | 9561 | 28 | AGGAAACTCTGGTGGAGGTCCGTAGCGG | 2181.0 | -9.9 | 1.3 | -7.4 | 1.5E-13 | 4.5E-13 | MB231/MB468 | MITO |
| >hg38_21_+_8433172_8446622_RNA45SN1_45S_50ntFlanks@12785.12805.21\|CCCCGCGCGCGCGGGAGGGCG | >hg38_21_+_8433172_8446622_RNA45SN1_45S_50ntFlanks | 12785 | 12805 | 21 | CCCCGCGCGCGCGGGAGGGCG | 1962.0 | -9.8 | 1.3 | -7.4 | 1.1E-13 | 4.5E-13 | MB231/MB468 | NUC |
| >hg38_21_+_8433172_8446622_RNA45SN1_45S_50ntFlanks@4163.4184.22\|CATCCAAGGAAGGCAGCAGGCG | >hg38_21_+_8433172_8446622_RNA45SN1_45S_50ntFlanks | 4163 | 4184 | 22 | CATCCAAGGAAGGCAGCAGGCG | 622.1 | -4.3 | 0.6 | -7.4 | 1.1E-13 | 4.5E-13 | MB231/MB468 | CYTO |
| >hg38_21_+_8433172_8446622_RNA45SN1_45S_50ntFlanks@11821.11845.25\|CTATCCAGCGAAACCACAGCCAAGG | >hg38_21_+_8433172_8446622_RNA45SN1_45S_50ntFlanks | 11821 | 11845 | 25 | CTATCCAGCGAAACCACAGCCAAGG | 147.5 | -11.1 | 1.5 | -7.4 | 9.5E-14 | 4.5E-13 | MB231/MB468 | MP |
| >hg38_21_+_8433172_8446622_RNA45SN1_45S_50ntFlanks@4473.4494.22\|TTAGCTGAGTGTCCCGCGGGGC | >hg38_21_+_8433172_8446622_RNA45SN1_45S_50ntFlanks | 4473 | 4494 | 22 | TTAGCTGAGTGTCCCGCGGGGC | 146.3 | -11.1 | 1.5 | -7.4 | 9.5E-14 | 4.5E-13 | MB231/MB468 | MP |
| >hg38_21_+_8433172_8446622_RNA45SN1_45S_50ntFlanks@9535.9563.29\|GGAAACTCTGGTGGAGGTCCGTAGCGGTC | >hg38_21_+_8433172_8446622_RNA45SN1_45S_50ntFlanks | 9535 | 9563 | 29 | GGAAACTCTGGTGGAGGTCCGTAGCGGTC | 209.8 | -10.7 | 1.4 | -7.4 | 1.1E-13 | 4.5E-13 | MB231/MB468 | NUC |
| >hg38_21_+_8433172_8446622_RNA45SN1_45S_50ntFlanks@12035.12053.19\|CGTTTTTTCACTGACCCGG | >hg38_21_+_8433172_8446622_RNA45SN1_45S_50ntFlanks | 12035 | 12053 | 19 | CGTTTTTTCACTGACCCGG | 151.4 | -10.0 | 1.3 | -7.4 | 1.1E-13 | 4.5E-13 | MB231/MB468 | CYTO |
| >hg38_21_+_8433172_8446622_RNA45SN1_45S_50ntFlanks@12539.12560.22\|ATGATGTGTTGTTGCCATGGTA | >hg38_21_+_8433172_8446622_RNA45SN1_45S_50ntFlanks | 12539 | 12560 | 22 | ATGATGTGTTGTTGCCATGGTA | 139.6 | -6.6 | 0.9 | -7.4 | 1.1E-13 | 4.5E-13 | MB231/MB468 | CYTO |
| >hg38_21_+_8433172_8446622_RNA45SN1_45S_50ntFlanks@485.526.42\|GATGTGGTGACGTCGTGCTCTCCCGGGCCGGGTCCGAGCCGC | >hg38_21_+_8433172_8446622_RNA45SN1_45S_50ntFlanks | 485 | 526 | 42 | GATGTGGTGACGTCGTGCTCTCCCGGGCCGGGTCCGAGCCGC | 340.5 | 11.2 | 1.5 | 7.4 | 1.7E-13 | 4.5E-13 | BT20/MB468 | NUC |
| >hg38_21_+_8433172_8446622_RNA45SN1_45S_50ntFlanks@4733.4764.32\|GAACGAAAGTCGGAGGTTCGAAGACGATCAGA | >hg38_21_+_8433172_8446622_RNA45SN1_45S_50ntFlanks | 4733 | 4764 | 32 | GAACGAAAGTCGGAGGTTCGAAGACGATCAGA | 281.5 | 8.3 | 1.1 | 7.4 | 1.1E-13 | 4.5E-13 | MB231/MB468 | CYTO |
| >hg38_21_+_8433172_8446622_RNA45SN1_45S_50ntFlanks@11561.11582.22\|GCGCCTAGCAGCCGACTTAGAA | >hg38_21_+_8433172_8446622_RNA45SN1_45S_50ntFlanks | 11561 | 11582 | 22 | GCGCCTAGCAGCCGACTTAGAA | 151.0 | -11.1 | 1.5 | -7.4 | 9.6E-14 | 4.5E-13 | MB231/MB468 | MP |
| >hg38_21_+_8433172_8446622_RNA45SN1_45S_50ntFlanks@12419.12446.28\|GATGTCGGCTCTTCCTATCATTGTGAAG | >hg38_21_+_8433172_8446622_RNA45SN1_45S_50ntFlanks | 12419 | 12446 | 28 | GATGTCGGCTCTTCCTATCATTGTGAAG | 1620.3 | -6.5 | 0.9 | -7.4 | 1.7E-13 | 4.5E-13 | BT20/MB468 | NUC |
| >MI0000477\|hsa-mir-146a&WithFlank&5\|+\|160485346\|160485456@26.48.23&[MIMAT0000449&hsa-miR-146a-5p&offsets\|-1\|0;m-129&5\|+\|160485372\|160485393&offsets\|-1\|0]\|TTGAGAACTGAATTCCATGGGTT | >MI0000477\|hsa-mir-146a&WithFlank&5\|+\|160485346\|160485456 | 26 | 48 | 23 | TTGAGAACTGAATTCCATGGGTT | 322.8 | 11.6 | 1.6 | 7.4 | 9.6E-14 | 4.5E-13 | MB231/MB468 | MP |
| >hg38_21_+_8433172_8446622_RNA45SN1_45S_50ntFlanks@5446.5463.18\|CGGATCGGCCCCGCCGGG | >hg38_21_+_8433172_8446622_RNA45SN1_45S_50ntFlanks | 5446 | 5463 | 18 | CGGATCGGCCCCGCCGGG | 252.6 | -11.8 | 1.6 | -7.4 | 9.6E-14 | 4.5E-13 | MB231/MB468 | MP |
| >hg38_21_+_8433172_8446622_RNA45SN1_45S_50ntFlanks@9033.9054.22\|CTCGTGGGGGGGCCGGGCCACC | >hg38_21_+_8433172_8446622_RNA45SN1_45S_50ntFlanks | 9033 | 9054 | 22 | CTCGTGGGGGGGCCGGGCCACC | 147.6 | -11.1 | 1.5 | -7.4 | 9.6E-14 | 4.5E-13 | MB231/MB468 | MP |
| >hg38_21_+_8433172_8446622_RNA45SN1_45S_50ntFlanks@12691.12711.21\|CGGAACGATACGGCAGCGCCG | >hg38_21_+_8433172_8446622_RNA45SN1_45S_50ntFlanks | 12691 | 12711 | 21 | CGGAACGATACGGCAGCGCCG | 352.4 | -12.2 | 1.7 | -7.4 | 1.5E-13 | 4.6E-13 | MB231/MB468 | MITO |
| >hg38_21_+_8433172_8446622_RNA45SN1_45S_50ntFlanks@10402.10430.29\|AACATGGGTCAGTCGGTCCTGAGAGATGG | >hg38_21_+_8433172_8446622_RNA45SN1_45S_50ntFlanks | 10402 | 10430 | 29 | AACATGGGTCAGTCGGTCCTGAGAGATGG | 1035.3 | -4.3 | 0.6 | -7.5 | 8.8E-14 | 4.6E-13 | BT20/MB468 | MITO |
| >hg38_21_+_8433172_8446622_RNA45SN1_45S_50ntFlanks@10352.10372.21\|GAGAACTTTGAAGGCCGAAGT | >hg38_21_+_8433172_8446622_RNA45SN1_45S_50ntFlanks | 10352 | 10372 | 21 | GAGAACTTTGAAGGCCGAAGT | 361.8 | 5.6 | 0.8 | 7.4 | 1.2E-13 | 4.6E-13 | MB231/MB468 | NUC |
| >hg38_21_+_8433172_8446622_RNA45SN1_45S_50ntFlanks@11552.11574.23\|CCTCGGCCGGCGCCTAGCAGCCG | >hg38_21_+_8433172_8446622_RNA45SN1_45S_50ntFlanks | 11552 | 11574 | 23 | CCTCGGCCGGCGCCTAGCAGCCG | 242.2 | -5.5 | 0.7 | -7.4 | 1.1E-13 | 4.6E-13 | MB231/MB468 | CYTO |
| >am_trna128_GlyGCC_6_-_27870686_27870756@1.34.34,trna18_GlyGCC_16_+_70822597_70822667@1.34.34,trna19_GlyGCC_16_+_70823410_70823480@1.34.34,trna19_GlyGCC_2_-_157257659_157257729@1.34.34,trna24_GlyGCC_16_-_70812942_70813012@1.34.34,trna25_GlyGCC_16_-_70812114_70812184@1.34.34,trna5_GlyGCC_17_+_8029064_8029134@1.34.34,trna68_GlyGCC_1_-_161493637_161493707@1.34.34\|GCATTGGTGGTTCAGTGGTAGAATTCTCGCCTGC | >am_trna128_GlyGCC_6_-_27870686_27870756 | 1 | 34 | 34 | GCATTGGTGGTTCAGTGGTAGAATTCTCGCCTGC | 5805.3 | 7.8 | 1.1 | 7.4 | 1.1E-13 | 4.6E-13 | MB231/MB468 | CYTO |
| >hg38_21_+_8433172_8446622_RNA45SN1_45S_50ntFlanks@11999.12034.36\|CCCTGCGGGCCGCCGGTGAAATACCACTACTCTGAT | >hg38_21_+_8433172_8446622_RNA45SN1_45S_50ntFlanks | 11999 | 12034 | 36 | CCCTGCGGGCCGCCGGTGAAATACCACTACTCTGAT | 227.1 | -10.5 | 1.4 | -7.4 | 1.1E-13 | 4.6E-13 | MB231/MB468 | CYTO |
| >hg38_21_+_8433172_8446622_RNA45SN1_45S_50ntFlanks@8170.8187.18\|CAAGTCCTTCTGATCGAG | >hg38_21_+_8433172_8446622_RNA45SN1_45S_50ntFlanks | 8170 | 8187 | 18 | CAAGTCCTTCTGATCGAG | 659.5 | -4.8 | 0.6 | -7.4 | 1.2E-13 | 4.6E-13 | MB231/MB468 | NUC |
| >MI0000283\|hsa-mir-203a&WithFlank&14\|+\|104117399\|104117520@71.90.20&[MIMAT0000264&hsa-miR-203a-3p&offsets\|0\|-2;m-79&14\|+\|104117469\|104117490&offsets\|0\|-2]\|GTGAAATGTTTAGGACCACT | >MI0000283\|hsa-mir-203a&WithFlank&14\|+\|104117399\|104117520 | 71 | 90 | 20 | GTGAAATGTTTAGGACCACT | 1095.9 | 9.6 | 1.3 | 7.4 | 1.2E-13 | 4.6E-13 | BT20/MB468 | CYTO |
| >hg38_21_+_8433172_8446622_RNA45SN1_45S_50ntFlanks@13005.13028.24\|TCTATTGAAAGTCAGCCCTCGACA | >hg38_21_+_8433172_8446622_RNA45SN1_45S_50ntFlanks | 13005 | 13028 | 24 | TCTATTGAAAGTCAGCCCTCGACA | 386.4 | -8.6 | 1.2 | -7.4 | 1.1E-13 | 4.6E-13 | MB231/MB468 | CYTO |
| >hg38_21_+_8433172_8446622_RNA45SN1_45S_50ntFlanks@7986.8014.29\|ATCAGACGTGGCGACCCGCTGAATTTAAG | >hg38_21_+_8433172_8446622_RNA45SN1_45S_50ntFlanks | 7986 | 8014 | 29 | ATCAGACGTGGCGACCCGCTGAATTTAAG | 480.4 | -12.0 | 1.5 | -7.8 | 4.4E-15 | 4.6E-13 | BT20/MB468 | MP |
| >hg38_21_+_8433172_8446622_RNA45SN1_45S_50ntFlanks@12141.12162.22\|ACAGTGCCAGGTGGGGAGTTTG | >hg38_21_+_8433172_8446622_RNA45SN1_45S_50ntFlanks | 12141 | 12162 | 22 | ACAGTGCCAGGTGGGGAGTTTG | 123.2 | -9.7 | 1.3 | -7.4 | 1.1E-13 | 4.6E-13 | MB231/MB468 | CYTO |
| >hg38_21_+_8433172_8446622_RNA45SN1_45S_50ntFlanks@11919.11949.31\|CATGAGAGGTGTAGAATAAGTGGGAGGCCCC | >hg38_21_+_8433172_8446622_RNA45SN1_45S_50ntFlanks | 11919 | 11949 | 31 | CATGAGAGGTGTAGAATAAGTGGGAGGCCCC | 249.2 | -11.8 | 1.6 | -7.4 | 9.8E-14 | 4.6E-13 | MB231/MB468 | MP |
| >hg38_21_+_8433172_8446622_RNA45SN1_45S_50ntFlanks@10808.10838.31\|CAATGTAGGTAAGGGAAGTCGGCAAGCCGGA | >hg38_21_+_8433172_8446622_RNA45SN1_45S_50ntFlanks | 10808 | 10838 | 31 | CAATGTAGGTAAGGGAAGTCGGCAAGCCGGA | 180.8 | -11.4 | 1.5 | -7.4 | 9.8E-14 | 4.6E-13 | MB231/MB468 | MP |
| >hg38_21_+_8433172_8446622_RNA45SN1_45S_50ntFlanks@9569.9596.28\|GTGCAAATCGGTCGTCCGACCTGGGTAT | >hg38_21_+_8433172_8446622_RNA45SN1_45S_50ntFlanks | 9569 | 9596 | 28 | GTGCAAATCGGTCGTCCGACCTGGGTAT | 760.6 | -7.5 | 1.0 | -7.4 | 9.8E-14 | 4.6E-13 | MB231/MB468 | MP |
| >hg38_21_+_8433172_8446622_RNA45SN1_45S_50ntFlanks@10109.10127.19\|TGTGGTGTGCGTCGGAGGG | >hg38_21_+_8433172_8446622_RNA45SN1_45S_50ntFlanks | 10109 | 10127 | 19 | TGTGGTGTGCGTCGGAGGG | 176.3 | -10.2 | 1.4 | -7.4 | 1.1E-13 | 4.6E-13 | MB231/MB468 | CYTO |
| >hg38_21_+_8433172_8446622_RNA45SN1_45S_50ntFlanks@10916.10934.19\|CCGCGGCTGGACGAGGCGC | >hg38_21_+_8433172_8446622_RNA45SN1_45S_50ntFlanks | 10916 | 10934 | 19 | CCGCGGCTGGACGAGGCGC | 132.1 | -10.9 | 1.5 | -7.4 | 9.8E-14 | 4.6E-13 | MB231/MB468 | MP |
| >hg38_21_+_8433172_8446622_RNA45SN1_45S_50ntFlanks@11999.12035.37\|CCCTGCGGGCCGCCGGTGAAATACCACTACTCTGATC | >hg38_21_+_8433172_8446622_RNA45SN1_45S_50ntFlanks | 11999 | 12035 | 37 | CCCTGCGGGCCGCCGGTGAAATACCACTACTCTGATC | 174.9 | -11.3 | 1.5 | -7.4 | 9.8E-14 | 4.6E-13 | MB231/MB468 | MP |
| >am_trnaMT_GlyTCC_MT_+_9991_10058@28.48.21\|ACTTCCAATTAACTAGTTTTG | >am_trnaMT_GlyTCC_MT_+_9991_10058 | 28 | 48 | 21 | ACTTCCAATTAACTAGTTTTG | 164.4 | -11.2 | 1.5 | -7.4 | 9.8E-14 | 4.6E-13 | MB231/MB468 | MP |
| >hg38_21_+_8433172_8446622_RNA45SN1_45S_50ntFlanks@11873.11893.21\|AAGAAGACCCTGTTGAGCTTG | >hg38_21_+_8433172_8446622_RNA45SN1_45S_50ntFlanks | 11873 | 11893 | 21 | AAGAAGACCCTGTTGAGCTTG | 8231.5 | -9.7 | 1.3 | -7.4 | 1.7E-13 | 4.7E-13 | BT20/MB468 | NUC |
| >hg38_21_+_8433172_8446622_RNA45SN1_45S_50ntFlanks@12406.12435.30\|TTTTTGATCCTTCGATGTCGGCTCTTCCTA | >hg38_21_+_8433172_8446622_RNA45SN1_45S_50ntFlanks | 12406 | 12435 | 30 | TTTTTGATCCTTCGATGTCGGCTCTTCCTA | 1630.1 | -10.8 | 1.5 | -7.4 | 1.7E-13 | 4.7E-13 | BT20/MB468 | NUC |
| >MI0000299\|hsa-mir-222&WithFlank&X\|-\|45747009\|45747130@75.99.25(+1U)&[MIMAT0000279&hsa-miR-222-3p&offsets\|0\|+4(+1U);m-140&X\|-\|45747033\|45747056&offsets\|0\|+1(+1U)]\|AGCTACATCTGGCTACTGGGTCTCTT | >MI0000299\|hsa-mir-222&WithFlank&X\|-\|45747009\|45747130 | 75 | 99 | 25(+1U) | AGCTACATCTGGCTACTGGGTCTCTT | 1574.1 | 6.8 | 0.9 | 7.5 | 9.0E-14 | 4.7E-13 | BT20/MB468 | MITO |

<table>
<tr><td>>hg38_21_+_8433172_8446622_RNA45SN1_45S_50ntFlanks@11581.11599.19|AACTGGTGCGGACCAGGGG</td><td>>hg38_21_+_8433172_8446622_RNA45SN1_45S_50ntFlanks</td><td>11581</td><td>11599</td><td>19</td><td>AACTGGTGCGGACCAGGGG</td><td>225.0</td><td>-8.0</td><td>1.1</td><td>-7.4</td><td>1.1E-13</td><td>4.7E-13</td><td>MB231/MB468</td><td>CYTO</td></tr>
<tr><td>>MI0000086|hsa-mir-28&WithFlank&3|+|188688775|188688872@60.80.21(+1U)&[MIMAT0004502&hsa-miR-28-3p&offsets|0|-1(+1U);m-28&3|+|188688834|188688855&offsets|0|-1(+1U)]|CACTAGATTGTGAGCTCCTGGT</td><td>>MI0000086|hsa-mir-28&WithFlank&3|+|188688775|188688872</td><td>60</td><td>80</td><td>21(+1U)</td><td>CACTAGATTGTGAGCTCCTGGT</td><td>224.5</td><td>8.1</td><td>1.1</td><td>7.4</td><td>1.2E-13</td><td>4.7E-13</td><td>MB231/MB468</td><td>NUC</td></tr>
<tr><td>>trna111_HisGTG_1_-_147774845_147774916@-1T.29.30,trna118_HisGTG_1_-_145396881_145396952@-1T.29.30,trna16_HisGTG_1_+_146544773_146544844@-1T.29.30,trna1_HisGTG_15_+_45493349_45493420@-1T.29.30,trna21_HisGTG_1_+_147753471_147753542@-1T.29.30,trna33_HisGTG_6_+_27125906_27125977@-1T.29.30,trna7_HisGTG_9_-_14433938_14434009@-1T.29.30,trna8_HisGTG_15_-_45492611_45492682@-1T.29.30,trna9_HisGTG_15_-_45490804_45490875@-1T.29.30|TGCCGTGATCGTATAGTGGTTAGTACTCTG</td><td>>trna111_HisGTG_1_-_147774845_147774916</td><td>-1T</td><td>29</td><td>30</td><td>TGCCGTGATCGTATAGTGGTTAGTACTCTG</td><td>6439.5</td><td>-5.8</td><td>0.8</td><td>-7.4</td><td>1.2E-13</td><td>4.7E-13</td><td>MB231/MB468</td><td>NUC</td></tr>
<tr><td>>hg38_21_+_8433172_8446622_RNA45SN1_45S_50ntFlanks@10388.10409.22|GTGAACAGCAGTTGAACATGGG</td><td>>hg38_21_+_8433172_8446622_RNA45SN1_45S_50ntFlanks</td><td>10388</td><td>10409</td><td>22</td><td>GTGAACAGCAGTTGAACATGGG</td><td>184.5</td><td>-10.5</td><td>1.4</td><td>-7.4</td><td>1.2E-13</td><td>4.7E-13</td><td>MB231/MB468</td><td>NUC</td></tr>
<tr><td>>hg38_21_+_8433172_8446622_RNA45SN1_45S_50ntFlanks@10116.10138.23|TGCGTCGGAGGGCGGCGGCGGCG</td><td>>hg38_21_+_8433172_8446622_RNA45SN1_45S_50ntFlanks</td><td>10116</td><td>10138</td><td>23</td><td>TGCGTCGGAGGGCGGCGGCGGCG</td><td>198.8</td><td>-10.4</td><td>1.4</td><td>-7.4</td><td>1.2E-13</td><td>4.7E-13</td><td>MB231/MB468</td><td>CYTO</td></tr>
<tr><td>>MI0003666|hsa-mir-651&WithFlank&X|+|8126959|8127067@22.43.22&[MIMAT0003321&hsa-miR-651-5p&offsets|0|0;m-399&X|+|8126980|8127001&offsets|0|0]|TTTAGGATAAGCTTGACTTTTG</td><td>>MI0003666|hsa-mir-651&WithFlank&X|+|8126959|8127067</td><td>22</td><td>43</td><td>22</td><td>TTTAGGATAAGCTTGACTTTTG</td><td>373.7</td><td>10.4</td><td>1.4</td><td>7.4</td><td>1.2E-13</td><td>4.7E-13</td><td>BT20/MB468</td><td>CYTO</td></tr>
<tr><td>>hg38_21_+_8433172_8446622_RNA45SN1_45S_50ntFlanks@12157.12176.20|AGTTTGACTGGGGCGGTACA</td><td>>hg38_21_+_8433172_8446622_RNA45SN1_45S_50ntFlanks</td><td>12157</td><td>12176</td><td>20</td><td>AGTTTGACTGGGGCGGTACA</td><td>143.9</td><td>-11.0</td><td>1.5</td><td>-7.4</td><td>9.9E-14</td><td>4.7E-13</td><td>MB231/MB468</td><td>MP</td></tr>
<tr><td>>hg38_21_+_8433172_8446622_RNA45SN1_45S_50ntFlanks@9623.9655.33|TAGTAGCTGGTTCCCTCCGAAGTTTCCCTCAGG</td><td>>hg38_21_+_8433172_8446622_RNA45SN1_45S_50ntFlanks</td><td>9623</td><td>9655</td><td>33</td><td>TAGTAGCTGGTTCCCTCCGAAGTTTCCCTCAGG</td><td>179.7</td><td>-11.4</td><td>1.5</td><td>-7.4</td><td>9.9E-14</td><td>4.7E-13</td><td>MB231/MB468</td><td>MP</td></tr>
<tr><td>>hg38_21_+_8433172_8446622_RNA45SN1_45S_50ntFlanks@11200.11225.26|CGGCGGCGGCGGCGACTCTGGACGCG</td><td>>hg38_21_+_8433172_8446622_RNA45SN1_45S_50ntFlanks</td><td>11200</td><td>11225</td><td>26</td><td>CGGCGGCGGCGGCGACTCTGGACGCG</td><td>1576.6</td><td>-5.7</td><td>0.8</td><td>-7.4</td><td>1.2E-13</td><td>4.7E-13</td><td>MB231/MB468</td><td>CYTO</td></tr>
<tr><td>>MI0000063|hsa-let-7b&WithFlank&22|+|46113680|46113774@12.31.20(+1C)&[MIMAT0000063&hsa-let-7b-5p&offsets|0|-2(+1C);m-14&22|+|46113691|46113712&offsets|0|-2(+1C)]|TGAGGTAGTAGGTTGTGTGGC</td><td>>MI0000063|hsa-let-7b&WithFlank&22|+|46113680|46113774</td><td>12</td><td>31</td><td>20(+1C)</td><td>TGAGGTAGTAGGTTGTGTGGC</td><td>556.8</td><td>8.9</td><td>1.2</td><td>7.5</td><td>9.1E-14</td><td>4.7E-13</td><td>BT20/MB468</td><td>MITO</td></tr>
<tr><td>>hg38_21_+_8433172_8446622_RNA45SN1_45S_50ntFlanks@6656.6689.34|CTTAGCGGTGGATCACTCGGCTCGTGCGTCGATG</td><td>>hg38_21_+_8433172_8446622_RNA45SN1_45S_50ntFlanks</td><td>6656</td><td>6689</td><td>34</td><td>CTTAGCGGTGGATCACTCGGCTCGTGCGTCGATG</td><td>929.6</td><td>-8.0</td><td>1.1</td><td>-7.4</td><td>1.0E-13</td><td>4.7E-13</td><td>MB231/MB468</td><td>MP</td></tr>
<tr><td>>hg38_21_+_8433172_8446622_RNA45SN1_45S_50ntFlanks@6671.6690.20|CTCGGCTCGTGCGTCGATGA</td><td>>hg38_21_+_8433172_8446622_RNA45SN1_45S_50ntFlanks</td><td>6671</td><td>6690</td><td>20</td><td>CTCGGCTCGTGCGTCGATGA</td><td>152.0</td><td>-11.1</td><td>1.5</td><td>-7.4</td><td>1.0E-13</td><td>4.7E-13</td><td>MB231/MB468</td><td>MP</td></tr>
<tr><td>>hg38_21_+_8433172_8446622_RNA45SN1_45S_50ntFlanks@8323.8355.33|ACCGATAGTCAACAAGTACCGTAAGGGAAAGTT</td><td>>hg38_21_+_8433172_8446622_RNA45SN1_45S_50ntFlanks</td><td>8323</td><td>8355</td><td>33</td><td>ACCGATAGTCAACAAGTACCGTAAGGGAAAGTT</td><td>144.7</td><td>-11.0</td><td>1.5</td><td>-7.4</td><td>1.0E-13</td><td>4.7E-13</td><td>MB231/MB468</td><td>MP</td></tr>
<tr><td>>MI0000102|hsa-mir-100&WithFlank&11|-|122152223|122152314@19.40.22(+2U)&[MIMAT0000098&hsa-miR-100-5p&offsets|0|0(+2U);m-27&11|-|122152276|122152296&offsets|0|+1(+2U)]|AACCCGTAGATCCGAACTTGTGTT</td><td>>MI0000102|hsa-mir-100&WithFlank&11|-|122152223|122152314</td><td>19</td><td>40</td><td>22(+2U)</td><td>AACCCGTAGATCCGAACTTGTGTT</td><td>222.9</td><td>11.0</td><td>1.5</td><td>7.4</td><td>1.0E-13</td><td>4.7E-13</td><td>MB231/MB468</td><td>MP</td></tr>
<tr><td>>am_trnaMT_MetCAT_MT_+_4402_4469@1.21.21|AGTAAGGTCAGCTAAATAAGC</td><td>>am_trnaMT_MetCAT_MT_+_4402_4469</td><td>1</td><td>21</td><td>21</td><td>AGTAAGGTCAGCTAAATAAGC</td><td>639.0</td><td>4.7</td><td>0.6</td><td>7.4</td><td>1.2E-13</td><td>4.7E-13</td><td>MB231/MB468</td><td>NUC</td></tr>
<tr><td>>MI0000102|hsa-mir-100&WithFlank&11|-|122152223|122152314@19.36.18&[MIMAT0000098&hsa-miR-100-5p&offsets|0|-4;m-27&11|-|122152276|122152296&offsets|0|-3]|AACCCGTAGATCCGAACT</td><td>>MI0000102|hsa-mir-100&WithFlank&11|-|122152223|122152314</td><td>19</td><td>36</td><td>18</td><td>AACCCGTAGATCCGAACT</td><td>369.3</td><td>11.8</td><td>1.6</td><td>7.4</td><td>1.5E-13</td><td>4.7E-13</td><td>MB231/MB468</td><td>MITO</td></tr>
<tr><td>>hg38_21_+_8433172_8446622_RNA45SN1_45S_50ntFlanks@10116.10139.24|TGCGTCGGAGGGCGGCGGCGGCGG</td><td>>hg38_21_+_8433172_8446622_RNA45SN1_45S_50ntFlanks</td><td>10116</td><td>10139</td><td>24</td><td>TGCGTCGGAGGGCGGCGGCGGCGG</td><td>276.3</td><td>-12.0</td><td>1.6</td><td>-7.4</td><td>1.0E-13</td><td>4.7E-13</td><td>MB231/MB468</td><td>MP</td></tr>
<tr><td>>hg38_21_+_8433172_8446622_RNA45SN1_45S_50ntFlanks@3981.4001.21|CCGGCGGCTTTGGTGACTCTA</td><td>>hg38_21_+_8433172_8446622_RNA45SN1_45S_50ntFlanks</td><td>3981</td><td>4001</td><td>21</td><td>CCGGCGGCTTTGGTGACTCTA</td><td>331.5</td><td>-12.2</td><td>1.6</td><td>-7.4</td><td>1.5E-13</td><td>4.8E-13</td><td>MB231/MB468</td><td>MITO</td></tr>
<tr><td>>am_MI0000815|hsa-mir-339&WithFlank&7|-|1022927|1023032@21.42.22(+1A)&[MIMAT0000764&hsa-miR-339-5p&offsets|0|-1(+1A);m-163&7|-|1022990|1023012&offsets|0|-1(+1A)]|TCCCTGTCCTCCAGGAGCTCACA</td><td>>am_MI0000815|hsa-mir-339&WithFlank&7|-|1022927|1023032</td><td>21</td><td>42</td><td>22(+1A)</td><td>TCCCTGTCCTCCAGGAGCTCACA</td><td>723.9</td><td>8.1</td><td>1.1</td><td>7.4</td><td>1.2E-13</td><td>4.8E-13</td><td>BT20/MB468</td><td>CYTO</td></tr>
<tr><td>>trna11_GluTTC_15_-_26327381_26327452@1.34.34,trna3_GluTTC_13_-_45492062_45492133@1.34.34|TCCCACATGGTCTAGCGGTTAGGATTCCTGGTTT</td><td>>trna11_GluTTC_15_-_26327381_26327452</td><td>1</td><td>34</td><td>34</td><td>TCCCACATGGTCTAGCGGTTAGGATTCCTGGTTT</td><td>788.9</td><td>10.9</td><td>1.5</td><td>7.4</td><td>1.2E-13</td><td>4.8E-13</td><td>BT20/MB468</td><td>CYTO</td></tr>
<tr><td>>hg38_21_+_8433172_8446622_RNA45SN1_45S_50ntFlanks@7993.8012.20|GTGGCGACCCGCTGAATTTA</td><td>>hg38_21_+_8433172_8446622_RNA45SN1_45S_50ntFlanks</td><td>7993</td><td>8012</td><td>20</td><td>GTGGCGACCCGCTGAATTTA</td><td>168.2</td><td>-11.3</td><td>1.5</td><td>-7.4</td><td>1.0E-13</td><td>4.8E-13</td><td>MB231/MB468</td><td>MP</td></tr>
<tr><td>>hg38_21_+_8433172_8446622_RNA45SN1_45S_50ntFlanks@9132.9152.21|GTCGGGGGGCGGGGCGGACTG</td><td>>hg38_21_+_8433172_8446622_RNA45SN1_45S_50ntFlanks</td><td>9132</td><td>9152</td><td>21</td><td>GTCGGGGGGCGGGGCGGACTG</td><td>327.0</td><td>-12.2</td><td>1.6</td><td>-7.4</td><td>1.0E-13</td><td>4.8E-13</td><td>MB231/MB468</td><td>MP</td></tr>
<tr><td>>hg38_21_+_8433172_8446622_RNA45SN1_45S_50ntFlanks@4734.4761.28|AACGAAAGTCGGAGGTTCGAAGACGATC</td><td>>hg38_21_+_8433172_8446622_RNA45SN1_45S_50ntFlanks</td><td>4734</td><td>4761</td><td>28</td><td>AACGAAAGTCGGAGGTTCGAAGACGATC</td><td>636.0</td><td>4.4</td><td>0.6</td><td>7.4</td><td>1.2E-13</td><td>4.8E-13</td><td>MB231/MB468</td><td>CYTO</td></tr>
<tr><td>>MI0000102|hsa-mir-100&WithFlank&11|-|122152223|122152314@20.39.20&[MIMAT0000098&hsa-miR-100-5p&offsets|+1|-1;m-27&11|-|122152276|122152296&offsets|+1|0]|ACCCGTAGATCCGAACTTGT</td><td>>MI0000102|hsa-mir-100&WithFlank&11|-|122152223|122152314</td><td>20</td><td>39</td><td>20</td><td>ACCCGTAGATCCGAACTTGT</td><td>289.5</td><td>-11.3</td><td>1.5</td><td>-7.7</td><td>1.4E-14</td><td>4.8E-13</td><td>BT20/MB231</td><td>NUC</td></tr>
<tr><td>>hg38_21_+_8433172_8446622_RNA45SN1_45S_50ntFlanks@11999.12029.31|CCCTGCGGGCCGCCGGTGAAATACCACTACT</td><td>>hg38_21_+_8433172_8446622_RNA45SN1_45S_50ntFlanks</td><td>11999</td><td>12029</td><td>31</td><td>CCCTGCGGGCCGCCGGTGAAATACCACTACT</td><td>215.0</td><td>-11.5</td><td>1.6</td><td>-7.4</td><td>1.6E-13</td><td>4.8E-13</td><td>MB231/MB468</td><td>MITO</td></tr>
<tr><td>>hg38_21_+_8433172_8446622_RNA45SN1_45S_50ntFlanks@5073.5105.33|AATTCCGATAACGAACGAGACTCTGGCATGCTA</td><td>>hg38_21_+_8433172_8446622_RNA45SN1_45S_50ntFlanks</td><td>5073</td><td>5105</td><td>33</td><td>AATTCCGATAACGAACGAGACTCTGGCATGCTA</td><td>477.3</td><td>-12.7</td><td>1.7</td><td>-7.4</td><td>1.6E-13</td><td>4.8E-13</td><td>MB231/MB468</td><td>MITO</td></tr>
<tr><td>>MI0000102|hsa-mir-100&WithFlank&11|-|122152223|122152314@19.40.22(+1A)&[MIMAT0000098&hsa-miR-100-5p&offsets|0|0(+1A);m-27&11|-|122152276|122152296&offsets|0|+1(+1A)]|AACCCGTAGATCCGAACTTGTGA</td><td>>MI0000102|hsa-mir-100&WithFlank&11|-|122152223|122152314</td><td>19</td><td>40</td><td>22(+1A)</td><td>AACCCGTAGATCCGAACTTGTGA</td><td>3173.0</td><td>13.3</td><td>1.8</td><td>7.4</td><td>1.2E-13</td><td>4.8E-13</td><td>BT20/MB468</td><td>CYTO</td></tr>
<tr><td>>hg38_MT_+_1621_3279_RNR2_16S_50ntFlanks@1184.1215.32|AATTTCGGTTGGGGCGACCTCGGAGCAGAACC</td><td>>hg38_MT_+_1621_3279_RNR2_16S_50ntFlanks</td><td>1184</td><td>1215</td><td>32</td><td>AATTTCGGTTGGGGCGACCTCGGAGCAGAACC</td><td>1015.8</td><td>8.8</td><td>1.2</td><td>7.5</td><td>9.3E-14</td><td>4.8E-13</td><td>BT20/MB468</td><td>MITO</td></tr>
<tr><td>>hg38_21_+_8433172_8446622_RNA45SN1_45S_50ntFlanks@9473.9503.31|CACGAGCGCACGTGTTAGGACCCGAAAGATG</td><td>>hg38_21_+_8433172_8446622_RNA45SN1_45S_50ntFlanks</td><td>9473</td><td>9503</td><td>31</td><td>CACGAGCGCACGTGTTAGGACCCGAAAGATG</td><td>262.3</td><td>-11.9</td><td>1.6</td><td>-7.4</td><td>1.0E-13</td><td>4.8E-13</td><td>MB231/MB468</td><td>MP</td></tr>
<tr><td>>hg38_21_+_8433172_8446622_RNA45SN1_45S_50ntFlanks@11786.11804.19|GAATGGATGAACGAGATTC</td><td>>hg38_21_+_8433172_8446622_RNA45SN1_45S_50ntFlanks</td><td>11786</td><td>11804</td><td>19</td><td>GAATGGATGAACGAGATTC</td><td>2273.5</td><td>-8.1</td><td>1.1</td><td>-7.4</td><td>1.0E-13</td><td>4.8E-13</td><td>MB231/MB468</td><td>MP</td></tr>
<tr><td>>hg38_21_+_8433172_8446622_RNA45SN1_45S_50ntFlanks@10237.10260.24|ACGCTACGCCGCGACGAGTAGGAG</td><td>>hg38_21_+_8433172_8446622_RNA45SN1_45S_50ntFlanks</td><td>10237</td><td>10260</td><td>24</td><td>ACGCTACGCCGCGACGAGTAGGAG</td><td>142.7</td><td>-11.0</td><td>1.5</td><td>-7.4</td><td>1.0E-13</td><td>4.8E-13</td><td>MB231/MB468</td><td>MP</td></tr>
<tr><td>>hg38_21_+_8433172_8446622_RNA45SN1_45S_50ntFlanks@4588.4612.25|CTATTTTGTTGGTTTTCGGAACTGA</td><td>>hg38_21_+_8433172_8446622_RNA45SN1_45S_50ntFlanks</td><td>4588</td><td>4612</td><td>25</td><td>CTATTTTGTTGGTTTTCGGAACTGA</td><td>163.7</td><td>-11.2</td><td>1.5</td><td>-7.4</td><td>1.0E-13</td><td>4.8E-13</td><td>MB231/MB468</td><td>MP</td></tr>
<tr><td>>hg38_21_+_8433172_8446622_RNA45SN1_45S_50ntFlanks@9166.9195.30|CCGGGCGGGTCGCGCCGTCGGGCCCGGGGG</td><td>>hg38_21_+_8433172_8446622_RNA45SN1_45S_50ntFlanks</td><td>9166</td><td>9195</td><td>30</td><td>CCGGGCGGGTCGCGCCGTCGGGCCCGGGGG</td><td>296.1</td><td>-12.1</td><td>1.6</td><td>-7.4</td><td>1.0E-13</td><td>4.8E-13</td><td>MB231/MB468</td><td>MP</td></tr>
<tr><td>>hg38_21_+_8433172_8446622_RNA45SN1_45S_50ntFlanks@12365.12384.20|ATAACTGGCTTGTGGCGGCC</td><td>>hg38_21_+_8433172_8446622_RNA45SN1_45S_50ntFlanks</td><td>12365</td><td>12384</td><td>20</td><td>ATAACTGGCTTGTGGCGGCC</td><td>141.7</td><td>-9.9</td><td>1.3</td><td>-7.4</td><td>1.2E-13</td><td>4.8E-13</td><td>MB231/MB468</td><td>CYTO</td></tr>
<tr><td>>hg38_21_+_8433172_8446622_RNA45SN1_45S_50ntFlanks@3705.3722.18|TACCTGGTTGATCCTGCC</td><td>>hg38_21_+_8433172_8446622_RNA45SN1_45S_50ntFlanks</td><td>3705</td><td>3722</td><td>18</td><td>TACCTGGTTGATCCTGCC</td><td>22589.8</td><td>4.7</td><td>0.6</td><td>7.4</td><td>1.2E-13</td><td>4.8E-13</td><td>MB231/MB468</td><td>CYTO</td></tr>
<tr><td>>hg38_21_+_8433172_8446622_RNA45SN1_45S_50ntFlanks@4732.4767.36|AGAACGAAAGTCGGAGGTTCGAAGACGATCAGATAC</td><td>>hg38_21_+_8433172_8446622_RNA45SN1_45S_50ntFlanks</td><td>4732</td><td>4767</td><td>36</td><td>AGAACGAAAGTCGGAGGTTCGAAGACGATCAGATAC</td><td>2008.1</td><td>9.5</td><td>1.3</td><td>7.4</td><td>1.2E-13</td><td>4.8E-13</td><td>MB231/MB468</td><td>CYTO</td></tr>
<tr><td>>hg38_21_+_8433172_8446622_RNA45SN1_45S_50ntFlanks@6738.6756.19|ATCATCGACACTTCGAACG</td><td>>hg38_21_+_8433172_8446622_RNA45SN1_45S_50ntFlanks</td><td>6738</td><td>6756</td><td>19</td><td>ATCATCGACACTTCGAACG</td><td>624.9</td><td>-12.4</td><td>1.6</td><td>-7.8</td><td>4.7E-15</td><td>4.9E-13</td><td>BT20/MB468</td><td>MP</td></tr>
<tr><td>>hg38_21_+_8433172_8446622_RNA45SN1_45S_50ntFlanks@11211.11232.22|GCGACTCTGGACGCGAGCCGGG</td><td>>hg38_21_+_8433172_8446622_RNA45SN1_45S_50ntFlanks</td><td>11211</td><td>11232</td><td>22</td><td>GCGACTCTGGACGCGAGCCGGG</td><td>697.0</td><td>-9.1</td><td>1.2</td><td>-7.4</td><td>1.6E-13</td><td>4.9E-13</td><td>MB231/MB468</td><td>MITO</td></tr>
<tr><td>>hg38_21_+_8433172_8446622_RNA45SN1_45S_50ntFlanks@11575.11593.19|ACTTAGAACTGGTGCGGAC</td><td>>hg38_21_+_8433172_8446622_RNA45SN1_45S_50ntFlanks</td><td>11575</td><td>11593</td><td>19</td><td>ACTTAGAACTGGTGCGGAC</td><td>3397.0</td><td>-11.1</td><td>1.5</td><td>-7.4</td><td>1.8E-13</td><td>4.9E-13</td><td>BT20/MB468</td><td>NUC</td></tr>
<tr><td>>hg38_21_+_8433172_8446622_RNA45SN1_45S_50ntFlanks@8971.8991.21|CCCACCCCCGCGGGGAATCCC</td><td>>hg38_21_+_8433172_8446622_RNA45SN1_45S_50ntFlanks</td><td>8971</td><td>8991</td><td>21</td><td>CCCACCCCCGCGGGGAATCCC</td><td>258.3</td><td>-11.8</td><td>1.6</td><td>-7.4</td><td>1.6E-13</td><td>4.9E-13</td><td>MB231/MB468</td><td>MITO</td></tr>
<tr><td>>hg38_21_+_8433172_8446622_RNA45SN1_45S_50ntFlanks@12405.12427.23|CTTTTTGATCCTTCGATGTCGGC</td><td>>hg38_21_+_8433172_8446622_RNA45SN1_45S_50ntFlanks</td><td>12405</td><td>12427</td><td>23</td><td>CTTTTTGATCCTTCGATGTCGGC</td><td>158.0</td><td>-11.2</td><td>1.5</td><td>-7.4</td><td>1.0E-13</td><td>4.9E-13</td><td>MB231/MB468</td><td>MP</td></tr>
<tr><td>>hg38_21_+_8433172_8446622_RNA45SN1_45S_50ntFlanks@8114.8135.22|GCGCGGGACATGTGGCGTACGG</td><td>>hg38_21_+_8433172_8446622_RNA45SN1_45S_50ntFlanks</td><td>8114</td><td>8135</td><td>22</td><td>GCGCGGGACATGTGGCGTACGG</td><td>225.0</td><td>-11.7</td><td>1.6</td><td>-7.4</td><td>1.0E-13</td><td>4.9E-13</td><td>MB231/MB468</td><td>MP</td></tr>
<tr><td>>MI0000465|hsa-mir-191&WithFlank&3|-|49020612|49020715@22.45.24&[MIMAT0000440&hsa-miR-191-5p&offsets|0|+1;m-48&3|-|49020672|49020694&offsets|0|+1]|CAACGGAATCCCAAAAGCAGCTGT</td><td>>MI0000465|hsa-mir-191&WithFlank&3|-|49020612|49020715</td><td>22</td><td>45</td><td>24</td><td>CAACGGAATCCCAAAAGCAGCTGT</td><td>2849.1</td><td>7.4</td><td>1.0</td><td>7.4</td><td>9.4E-14</td><td>4.9E-13</td><td>BT20/MB468</td><td>MITO</td></tr>
<tr><td>>hg38_21_+_8433172_8446622_RNA45SN1_45S_50ntFlanks@4735.4764.30|ACGAAAGTCGGAGGTTCGAAGACGATCAGA</td><td>>hg38_21_+_8433172_8446622_RNA45SN1_45S_50ntFlanks</td><td>4735</td><td>4764</td><td>30</td><td>ACGAAAGTCGGAGGTTCGAAGACGATCAGA</td><td>427.9</td><td>10.3</td><td>1.4</td><td>7.4</td><td>1.2E-13</td><td>4.9E-13</td><td>BT20/MB468</td><td>CYTO</td></tr>
<tr><td>>hg38_21_+_8433172_8446622_RNA45SN1_45S_50ntFlanks@6721.6745.25|AATTGCAGGACACATTGATCATCGA</td><td>>hg38_21_+_8433172_8446622_RNA45SN1_45S_50ntFlanks</td><td>6721</td><td>6745</td><td>25</td><td>AATTGCAGGACACATTGATCATCGA</td><td>601.6</td><td>-8.9</td><td>1.2</td><td>-7.4</td><td>1.6E-13</td><td>4.9E-13</td><td>MB231/MB468</td><td>MITO</td></tr>
<tr><td>>hg38_21_+_8433172_8446622_RNA45SN1_45S_50ntFlanks@12937.12956.20|AACCATTCGTAGACGACCTG</td><td>>hg38_21_+_8433172_8446622_RNA45SN1_45S_50ntFlanks</td><td>12937</td><td>12956</td><td>20</td><td>AACCATTCGTAGACGACCTG</td><td>4400.1</td><td>-4.9</td><td>0.7</td><td>-7.4</td><td>1.3E-13</td><td>4.9E-13</td><td>BT20/MB468</td><td>CYTO</td></tr>
<tr><td>>hg38_21_+_8433172_8446622_RNA45SN1_45S_50ntFlanks@12044.12063.20|ACTGACCCGGTGAGGCGGGG</td><td>>hg38_21_+_8433172_8446622_RNA45SN1_45S_50ntFlanks</td><td>12044</td><td>12063</td><td>20</td><td>ACTGACCCGGTGAGGCGGGG</td><td>357.2</td><td>-12.3</td><td>1.7</td><td>-7.4</td><td>1.0E-13</td><td>4.9E-13</td><td>MB231/MB468</td><td>MP</td></tr>
<tr><td>>hg38_21_+_8433172_8446622_RNA45SN1_45S_50ntFlanks@12539.12566.28|ATGATGTGTTGTTGCCATGGTAATCCTG</td><td>>hg38_21_+_8433172_8446622_RNA45SN1_45S_50ntFlanks</td><td>12539</td><td>12566</td><td>28</td><td>ATGATGTGTTGTTGCCATGGTAATCCTG</td><td>1500.0</td><td>-9.4</td><td>1.3</td><td>-7.4</td><td>1.2E-13</td><td>4.9E-13</td><td>MB231/MB468</td><td>NUC</td></tr>
</table>

| | | | | | | | | | | | | |
|---|---|---|---|---|---|---|---|---|---|---|---|---|
| >hg38_21_+_8433172_8446622_RNA45SN1_45S_50ntFlanks@5035.5070.36\|CATGGCCGTTCTTAGTTGGTGGAGCGATTTGTCTGG | >hg38_21_+_8433172_8446622_RNA45SN1_45S_50ntFlanks | 5035 | 5070 | 36 | CATGGCCGTTCTTAGTTGGTGGAGCGATTTGTCTGG | 566.0 | -12.9 | 1.8 | -7.4 | 1.6E-13 | 4.9E-13 | MB231/MB468 | MITO |
| >hg38_21_+_8433172_8446622_RNA45SN1_45S_50ntFlanks@8176.8197.22\|CTTCTGATCGAGGCCCAGCCCG | >hg38_21_+_8433172_8446622_RNA45SN1_45S_50ntFlanks | 8176 | 8197 | 22 | CTTCTGATCGAGGCCCAGCCCG | 794.5 | -3.3 | 0.4 | -7.4 | 1.3E-13 | 4.9E-13 | BT20/MB468 | CYTO |
| >hg38_21_+_8433172_8446622_RNA45SN1_45S_50ntFlanks@4724.4748.25\|ATTAATCAAGAACGAAAGTCGGAGG | >hg38_21_+_8433172_8446622_RNA45SN1_45S_50ntFlanks | 4724 | 4748 | 25 | ATTAATCAAGAACGAAAGTCGGAGG | 799.7 | -9.8 | 1.3 | -7.4 | 1.8E-13 | 4.9E-13 | BT20/MB468 | NUC |
| >hg38_21_+_8433172_8446622_RNA45SN1_45S_50ntFlanks@12412.12436.25\|ATCCTTCGATGTCGGCTCTTCCTAT | >hg38_21_+_8433172_8446622_RNA45SN1_45S_50ntFlanks | 12412 | 12436 | 25 | ATCCTTCGATGTCGGCTCTTCCTAT | 568.2 | -11.2 | 1.5 | -7.4 | 9.6E-14 | 4.9E-13 | BT20/MB468 | MITO |
| >hg38_21_+_8433172_8446622_RNA45SN1_45S_50ntFlanks@9310.9336.27\|GAGTCTAACACGTGCGCGAGTCGGGGG | >hg38_21_+_8433172_8446622_RNA45SN1_45S_50ntFlanks | 9310 | 9336 | 27 | GAGTCTAACACGTGCGCGAGTCGGGGG | 222.7 | -11.7 | 1.6 | -7.4 | 1.1E-13 | 5.0E-13 | MB231/MB468 | MP |
| >hg38_21_+_8433172_8446622_RNA45SN1_45S_50ntFlanks@6721.6739.19\|AATTGCAGGACACATTGAT | >hg38_21_+_8433172_8446622_RNA45SN1_45S_50ntFlanks | 6721 | 6739 | 19 | AATTGCAGGACACATTGAT | 900.6 | -7.4 | 1.0 | -7.4 | 1.6E-13 | 5.0E-13 | MB231/MB468 | MITO |
| >hg38_21_+_8433172_8446622_RNA45SN1_45S_50ntFlanks@10853.10874.22\|ATAAGGATTGGCTCTAAGGGCT | >hg38_21_+_8433172_8446622_RNA45SN1_45S_50ntFlanks | 10853 | 10874 | 22 | ATAAGGATTGGCTCTAAGGGCT | 141.3 | -11.0 | 1.5 | -7.4 | 1.1E-13 | 5.0E-13 | MB231/MB468 | MP |
| >hg38_21_+_8433172_8446622_RNA45SN1_45S_50ntFlanks@9621.9649.29\|TCTAGTAGCTGGTTCCCTCCGAAGTTTCC | >hg38_21_+_8433172_8446622_RNA45SN1_45S_50ntFlanks | 9621 | 9649 | 29 | TCTAGTAGCTGGTTCCCTCCGAAGTTTCC | 544.0 | -8.7 | 1.2 | -7.4 | 1.6E-13 | 5.0E-13 | MB231/MB468 | MITO |
| >MI0001445\|hsa-mir-423&WithFlank&17\|+\|30117073\|30117178@23.44.22&[MIMAT0004748&hsa-miR-423-5p&offsets\|0\|-1;m-141&17\|+\|30117095\|30117117&offsets\|0\|-1]\|TGAGGGGCAGAGAGCGAGACTT | >MI0001445\|hsa-mir-423&WithFlank&17\|+\|30117073\|30117178 | 23 | 44 | 22 | TGAGGGGCAGAGAGCGAGACTT | 451.5 | 5.7 | 0.8 | 7.4 | 1.3E-13 | 5.0E-13 | MB231/MB468 | NUC |
| >hg38_21_+_8433172_8446622_RNA45SN1_45S_50ntFlanks@8463.8497.35\|CCGTGTCGGCGGCCCGGCGGATCTTTCCCGCCCCC | >hg38_21_+_8433172_8446622_RNA45SN1_45S_50ntFlanks | 8463 | 8497 | 35 | CCGTGTCGGCGGCCCGGCGGATCTTTCCCGCCCCC | 1068.9 | -10.1 | 1.4 | -7.4 | 1.6E-13 | 5.0E-13 | MB231/MB468 | MITO |
| >am_trna162_MetCAT_6_-_26758550_26758622@31.53.23,trna164_MetCAT_6_-_26735574_26735646@31.53.23,trna20_MetCAT_16_+_71460396_71460468@31.53.23,trna21_MetCAT_6_+_26701712_26701784@31.53.23,trna27_MetCAT_6_+_26766444_26766516@31.53.23,trna75_MetCAT_6_+_28912352_28912424@31.53.23,trna97_MetCAT_6_-_28921042_28921114@31.53.23\|TCTCATAATCTGAAGGTCCTGAG | >am_trna162_MetCAT_6_-_26758550_26758622 | 31 | 53 | 23 | TCTCATAATCTGAAGGTCCTGAG | 174.7 | -10.4 | 1.4 | -7.4 | 1.3E-13 | 5.0E-13 | MB231/MB468 | NUC |
| >MI0000093\|hsa-mir-92a-1&WithFlank&13\|+\|91351308\|91351397@54.74.21(+1A);MI0000094\|hsa-mir-92a-2&WithFlank&X\|-\|134169532\|134169618@54.74.21(+1A)&[MIMAT0000092&hsa-miR-92a-3p&offsets\|0\|-1(+1A);m-9&13\|+\|91351361\|91351382&offsets\|0\|-1(+1A)];[MIMAT0000092_1&hsa-miR-92a-3p&offsets\|0\|-1(+1A);m-11&X\|-\|134169544\|134169565&offsets\|0\|-1(+1A)]\|TATTGCACTTGTCCCGGCCTGA | >MI0000093\|hsa-mir-92a-1&WithFlank&13\|+\|91351308\|91351397 | 54 | 74 | 21(+1A) | TATTGCACTTGTCCCGGCCTGA | 2827.1 | 6.7 | 0.9 | 7.4 | 1.2E-13 | 5.0E-13 | MB231/MB468 | CYTO |
| >trnaMT_ProTGG_MT_-_15956_16023@40.58.19\|AATGGTGGAGTTAAAGACT | >trnaMT_ProTGG_MT_-_15956_16023 | 40 | 58 | 19 | AATGGTGGAGTTAAAGACT | 1430.7 | 6.0 | 0.8 | 7.4 | 9.7E-14 | 5.0E-13 | BT20/MB468 | MITO |
| >hg38_21_+_8433172_8446622_RNA45SN1_45S_50ntFlanks@11561.11602.42\|GCGCCTAGCAGCCGACTTAGAACTGGTGCGGACCAGGGGAAT | >hg38_21_+_8433172_8446622_RNA45SN1_45S_50ntFlanks | 11561 | 11602 | 42 | GCGCCTAGCAGCCGACTTAGAACTGGTGCGGACCAGGGGAAT | 954.5 | -13.8 | 1.9 | -7.4 | 1.1E-13 | 5.0E-13 | MB231/MB468 | MP |
| >hg38_21_+_8433172_8446622_RNA45SN1_45S_50ntFlanks@5492.5516.25\|AGAAGACGGTCGAACTTGACTATCT | >hg38_21_+_8433172_8446622_RNA45SN1_45S_50ntFlanks | 5492 | 5516 | 25 | AGAAGACGGTCGAACTTGACTATCT | 21817.0 | 7.4 | 1.0 | 7.4 | 9.7E-14 | 5.0E-13 | BT20/MB468 | MITO |
| >hg38_21_+_8433172_8446622_RNA45SN1_45S_50ntFlanks@12416.12433.18\|TTCGATGTCGGCTCTTCC | >hg38_21_+_8433172_8446622_RNA45SN1_45S_50ntFlanks | 12416 | 12433 | 18 | TTCGATGTCGGCTCTTCC | 9943.0 | 5.6 | 0.8 | 7.4 | 1.2E-13 | 5.0E-13 | MB231/MB468 | CYTO |
| >hg38_21_+_8433172_8446622_RNA45SN1_45S_50ntFlanks@10678.10706.29\|CGGTTCCGGCGGCGTCCGGTGAGCTCTCG | >hg38_21_+_8433172_8446622_RNA45SN1_45S_50ntFlanks | 10678 | 10706 | 29 | CGGTTCCGGCGGCGTCCGGTGAGCTCTCG | 26829.7 | -7.0 | 0.9 | -7.8 | 4.9E-15 | 5.0E-13 | BT20/MB468 | MP |
| >hg38_21_+_8433172_8446622_RNA45SN1_45S_50ntFlanks@7986.8009.24\|ATCAGACGTGGCGACCCGCTGAAT | >hg38_21_+_8433172_8446622_RNA45SN1_45S_50ntFlanks | 7986 | 8009 | 24 | ATCAGACGTGGCGACCCGCTGAAT | 914.8 | -12.9 | 1.7 | -7.8 | 4.9E-15 | 5.0E-13 | BT20/MB468 | MP |
| >MI0000067\|hsa-let-7f-1&WithFlank&9\|+\|94176341\|94176439@13.33.21(+1A);MI0000068\|hsa-let-7f-2&WithFlank&X\|-\|53557186\|53557280@14.34.21(+1A)&[MIMAT0000067&hsa-let-7f-5p&offsets\|0\|-1(+1A);m-20&9\|+\|94176353\|94176374&offsets\|0\|-1(+1A)];[MIMAT0000067_1&hsa-let-7f-5p&offsets\|0\|-1(+1A);m-19&X\|-\|53557246\|53557267&offsets\|0\|-1(+1A)]\|TGAGGTAGTAGATTGTATAGTA | >MI0000067\|hsa-let-7f-1&WithFlank&9\|+\|94176341\|94176439 | 13 | 33 | 21(+1A) | TGAGGTAGTAGATTGTATAGTA | 1994.6 | 4.6 | 0.6 | 7.4 | 1.3E-13 | 5.0E-13 | MB231/MB468 | NUC |
| >MI0000076\|hsa-mir-20a&WithFlank&13\|+\|91351059\|91351141@14.36.23(+1A)&[MIMAT0000075&hsa-miR-20a-5p&offsets\|0\|0(+1A);m-96&13\|+\|91351072\|91351094&offsets\|0\|0(+1A)]\|TAAAGTGCTTATAGTGCAGGTAGA | >MI0000076\|hsa-mir-20a&WithFlank&13\|+\|91351059\|91351141 | 14 | 36 | 23(+1A) | TAAAGTGCTTATAGTGCAGGTAGA | 2559.2 | 5.4 | 0.7 | 7.4 | 1.3E-13 | 5.0E-13 | MB231/MB468 | NUC |
| >hg38_21_+_8433172_8446622_RNA45SN1_45S_50ntFlanks@12447.12475.29\|CAGAATTCACCAAGCGTTGGATTGTTCAC | >hg38_21_+_8433172_8446622_RNA45SN1_45S_50ntFlanks | 12447 | 12475 | 29 | CAGAATTCACCAAGCGTTGGATTGTTCAC | 272.2 | -12.0 | 1.6 | -7.4 | 1.1E-13 | 5.0E-13 | MB231/MB468 | MP |
| >hg38_21_+_8433172_8446622_RNA45SN1_45S_50ntFlanks@9533.9553.21\|GAGGAAACTCTGGTGGAGGTC | >hg38_21_+_8433172_8446622_RNA45SN1_45S_50ntFlanks | 9533 | 9553 | 21 | GAGGAAACTCTGGTGGAGGTC | 14085.3 | -6.2 | 0.8 | -7.4 | 1.3E-13 | 5.0E-13 | MB231/MB468 | NUC |
| >hg38_21_+_8433172_8446622_RNA45SN1_45S_50ntFlanks@10819.10840.22\|AGGGAAGTCGGCAAGCCGGATC | >hg38_21_+_8433172_8446622_RNA45SN1_45S_50ntFlanks | 10819 | 10840 | 22 | AGGGAAGTCGGCAAGCCGGATC | 4348.5 | -4.6 | 0.6 | -7.4 | 1.3E-13 | 5.0E-13 | BT20/MB468 | CYTO |
| >am_trna10_LysCTT_16_+_3241501_3241573@31.51.21,trna119_LysCTT_1_-_145395522_145395594@31.51.21,trna11_LysCTT_5_-_180648979_180649051@31.51.21,trna13_LysCTT_16_+_3246153_3246225@31.51.21,trna13_LysCTT_6_+_26556774_26556846@31.51.21,trna30_LysCTT_16_-_3230555_3230627@31.51.21,trna5_LysCTT_16_+_3214939_3215011@31.51.21,trna7_LysCTT_16_+_3225692_3225764@31.51.21,trna8_LysCTT_1_+_39970195_39970267@31.51.21,trna9_LysCTT_5_+_180634755_180634827@31.51.21\|ACTCTTAATCTCAGGGTCGTG | >am_trna10_LysCTT_16_+_3241501_3241573 | 31 | 51 | 21 | ACTCTTAATCTCAGGGTCGTG | 288.2 | -11.1 | 1.5 | -7.4 | 1.3E-13 | 5.1E-13 | MB231/MB468 | NUC |
| >hg38_21_+_8433172_8446622_RNA45SN1_45S_50ntFlanks@8355.8375.21\|TGAAAAGAACTTTGAAGAGAG | >hg38_21_+_8433172_8446622_RNA45SN1_45S_50ntFlanks | 8355 | 8375 | 21 | TGAAAAGAACTTTGAAGAGAG | 5298.6 | 9.6 | 1.2 | 7.8 | 5.0E-15 | 5.1E-13 | BT20/MB468 | MP |
| >hg38_21_+_8433172_8446622_RNA45SN1_45S_50ntFlanks@7974.8014.41\|ACGCGACCTCAGATCAGACGTGGCGACCCGCTGAATTTAAG | >hg38_21_+_8433172_8446622_RNA45SN1_45S_50ntFlanks | 7974 | 8014 | 41 | ACGCGACCTCAGATCAGACGTGGCGACCCGCTGAATTTAAG | 3154.9 | 4.2 | 0.6 | 7.4 | 9.9E-14 | 5.1E-13 | BT20/MB468 | MITO |
| >hg38_21_+_8433172_8446622_RNA45SN1_45S_50ntFlanks@9534.9579.46\|AGGAAACTCTGGTGGAGGTCCGTAGCGGTCCTGACGTGCAAATCGG | >hg38_21_+_8433172_8446622_RNA45SN1_45S_50ntFlanks | 9534 | 9579 | 46 | AGGAAACTCTGGTGGAGGTCCGTAGCGGTCCTGACGTGCAAATCGG | 417.5 | -13.1 | 1.8 | -7.4 | 1.9E-13 | 5.1E-13 | BT20/MB468 | NUC |
| >hg38_21_+_8433172_8446622_RNA45SN1_45S_50ntFlanks@11873.11910.38\|AAGAAGACCCTGTTGAGCTTGACTCTAGTCTGGCACGG | >hg38_21_+_8433172_8446622_RNA45SN1_45S_50ntFlanks | 11873 | 11910 | 38 | AAGAAGACCCTGTTGAGCTTGACTCTAGTCTGGCACGG | 209.2 | -11.6 | 1.6 | -7.4 | 1.1E-13 | 5.1E-13 | MB231/MB468 | MP |
| >hg38_21_+_8433172_8446622_RNA45SN1_45S_50ntFlanks@12632.12659.28\|GCGAAGCTACCATCTGTGGGATTATGAC | >hg38_21_+_8433172_8446622_RNA45SN1_45S_50ntFlanks | 12632 | 12659 | 28 | GCGAAGCTACCATCTGTGGGATTATGAC | 1915.9 | -8.9 | 1.2 | -7.4 | 1.1E-13 | 5.1E-13 | MB231/MB468 | MP |
| >MI0000077\|hsa-mir-21&WithFlank&17\|+\|59841260\|59841343@14.36.23&[MIMAT0000076&hsa-miR-21-5p&offsets\|0\|+1;m-1&17\|+\|59841273\|59841294&offsets\|0\|+1]\|TAGCTTATCAGACTGATGTTGAC | >MI0000077\|hsa-mir-21&WithFlank&17\|+\|59841260\|59841343 | 14 | 36 | 23 | TAGCTTATCAGACTGATGTTGAC | 868152.1 | 6.6 | 0.9 | 7.4 | 1.7E-13 | 5.1E-13 | MB231/MB468 | MITO |
| >hg38_21_+_8433172_8446622_RNA45SN1_45S_50ntFlanks@12605.12654.50\|TATGTGCTTGGCTGAGGAGCCAATGGGGCGAAGCTACCATCTGTGGGATT | >hg38_21_+_8433172_8446622_RNA45SN1_45S_50ntFlanks | 12605 | 12654 | 50 | TATGTGCTTGGCTGAGGAGCCAATGGGGCGAAGCTACCATCTGTGGGATT | 307.2 | -12.0 | 1.6 | -7.4 | 1.7E-13 | 5.1E-13 | MB231/MB468 | MITO |
| >hg38_21_+_8433172_8446622_RNA45SN1_45S_50ntFlanks@9533.9572.40\|GAGGAAACTCTGGTGGAGGTCCGTAGCGGTCCTGACGTGC | >hg38_21_+_8433172_8446622_RNA45SN1_45S_50ntFlanks | 9533 | 9572 | 40 | GAGGAAACTCTGGTGGAGGTCCGTAGCGGTCCTGACGTGC | 2983.2 | -7.3 | 1.0 | -7.4 | 1.9E-13 | 5.1E-13 | BT20/MB468 | NUC |
| >hg38_21_+_8433172_8446622_RNA45SN1_45S_50ntFlanks@10798.10823.26\|CATGTTGGAACAATGTAGGTAAGGGA | >hg38_21_+_8433172_8446622_RNA45SN1_45S_50ntFlanks | 10798 | 10823 | 26 | CATGTTGGAACAATGTAGGTAAGGGA | 396.5 | -8.6 | 1.2 | -7.4 | 1.7E-13 | 5.1E-13 | MB231/MB468 | MITO |
| >hg38_21_+_8433172_8446622_RNA45SN1_45S_50ntFlanks@10866.10903.38\|CTAAGGGCTGGGTCGGTCGGGCTGGGGCGCGAAGCGGG | >hg38_21_+_8433172_8446622_RNA45SN1_45S_50ntFlanks | 10866 | 10903 | 38 | CTAAGGGCTGGGTCGGTCGGGCTGGGGCGCGAAGCGGG | 225.1 | -10.8 | 1.5 | -7.4 | 1.3E-13 | 5.1E-13 | MB231/MB468 | NUC |
| >hg38_21_+_8433172_8446622_RNA45SN1_45S_50ntFlanks@11888.11912.25\|AGCTTGACTCTAGTCTGGCACGGTG | >hg38_21_+_8433172_8446622_RNA45SN1_45S_50ntFlanks | 11888 | 11912 | 25 | AGCTTGACTCTAGTCTGGCACGGTG | 165.8 | -11.2 | 1.5 | -7.4 | 1.1E-13 | 5.1E-13 | MB231/MB468 | MP |
| >hg38_21_+_8433172_8446622_RNA45SN1_45S_50ntFlanks@8194.8215.22\|CCCGTGGACGGTGTGAGGCCGG | >hg38_21_+_8433172_8446622_RNA45SN1_45S_50ntFlanks | 8194 | 8215 | 22 | CCCGTGGACGGTGTGAGGCCGG | 2781.7 | -10.5 | 1.4 | -7.4 | 1.1E-13 | 5.1E-13 | MB231/MB468 | MP |
| >hg38_21_+_8433172_8446622_RNA45SN1_45S_50ntFlanks@12054.12071.18\|TGAGGCGGGGGGGCGAGC | >hg38_21_+_8433172_8446622_RNA45SN1_45S_50ntFlanks | 12054 | 12071 | 18 | TGAGGCGGGGGGGCGAGC | 159.5 | -11.2 | 1.5 | -7.4 | 1.1E-13 | 5.2E-13 | MB231/MB468 | MP |
| >hg38_21_+_8433172_8446622_RNA45SN1_45S_50ntFlanks@7966.7983.18\|CCTCCGAGACGCGACCTC | >hg38_21_+_8433172_8446622_RNA45SN1_45S_50ntFlanks | 7966 | 7983 | 18 | CCTCCGAGACGCGACCTC | 185.9 | 4.8 | 0.6 | 7.4 | 1.3E-13 | 5.2E-13 | MB231/MB468 | NUC |
| >hg38_21_+_8433172_8446622_RNA45SN1_45S_50ntFlanks@12628.12661.34\|TGGGGCGAAGCTACCATCTGTGGGATTATGACTG | >hg38_21_+_8433172_8446622_RNA45SN1_45S_50ntFlanks | 12628 | 12661 | 34 | TGGGGCGAAGCTACCATCTGTGGGATTATGACTG | 1140.2 | -9.5 | 1.3 | -7.4 | 1.3E-13 | 5.2E-13 | MB231/MB468 | NUC |
| >hg38_21_+_8433172_8446622_RNA45SN1_45S_50ntFlanks@11874.11912.39\|AGAAGACCCTGTTGAGCTTGACTCTAGTCTGGCACGGTG | >hg38_21_+_8433172_8446622_RNA45SN1_45S_50ntFlanks | 11874 | 11912 | 39 | AGAAGACCCTGTTGAGCTTGACTCTAGTCTGGCACGGTG | 216.1 | -10.5 | 1.4 | -7.4 | 1.3E-13 | 5.2E-13 | MB231/MB468 | CYTO |
| >hg38_21_+_8433172_8446622_RNA45SN1_45S_50ntFlanks@10392.10426.35\|ACAGCAGTTGAACATGGGTCAGTCGGTCCTGAGAG | >hg38_21_+_8433172_8446622_RNA45SN1_45S_50ntFlanks | 10392 | 10426 | 35 | ACAGCAGTTGAACATGGGTCAGTCGGTCCTGAGAG | 2131.6 | -11.0 | 1.5 | -7.4 | 1.9E-13 | 5.2E-13 | BT20/MB468 | NUC |
| >hg38_21_+_8433172_8446622_RNA45SN1_45S_50ntFlanks@9718.9737.20\|TGATTAGAGGTCTTGGGGCC | >hg38_21_+_8433172_8446622_RNA45SN1_45S_50ntFlanks | 9718 | 9737 | 20 | TGATTAGAGGTCTTGGGGCC | 365.1 | -12.3 | 1.7 | -7.4 | 1.7E-13 | 5.2E-13 | MB231/MB468 | MITO |
| >hg38_21_+_8433172_8446622_RNA45SN1_45S_50ntFlanks@10808.10840.33\|CAATGTAGGTAAGGGAAGTCGGCAAGCCGGATC | >hg38_21_+_8433172_8446622_RNA45SN1_45S_50ntFlanks | 10808 | 10840 | 33 | CAATGTAGGTAAGGGAAGTCGGCAAGCCGGATC | 3126.5 | -11.1 | 1.5 | -7.4 | 1.9E-13 | 5.2E-13 | BT20/MB468 | NUC |
| >MI0000082\|hsa-mir-25&WithFlank&7\|-\|100093554\|100093649@58.79.22(+1A)&[MIMAT0000081&hsa-miR-25-3p&offsets\|0\|0(+1A);m-23&7\|-\|100093571\|100093592&offsets\|0\|0(+1A)]\|CATTGCACTTGTCTCGGTCTGAA | >MI0000082\|hsa-mir-25&WithFlank&7\|-\|100093554\|100093649 | 58 | 79 | 22(+1A) | CATTGCACTTGTCTCGGTCTGAA | 721.6 | 5.7 | 0.8 | 7.4 | 1.3E-13 | 5.2E-13 | MB231/MB468 | CYTO |
| >hg38_21_+_8433172_8446622_RNA45SN1_45S_50ntFlanks@4296.4317.22\|CCATTGGAGGGCAAGTCTGGTG | >hg38_21_+_8433172_8446622_RNA45SN1_45S_50ntFlanks | 4296 | 4317 | 22 | CCATTGGAGGGCAAGTCTGGTG | 153.8 | -11.1 | 1.5 | -7.4 | 1.1E-13 | 5.2E-13 | MB231/MB468 | MP |

<table>
<tr><td></td><td></td><td></td><td></td><td></td><td></td><td></td><td></td><td></td><td></td><td></td><td></td><td></td><td></td></tr>
<tr><td>>hg38_21_+_8433172_8446622_RNA45SN1_45S_50ntFlanks@7985.8004.20|GATCAGACGTGGCGACCCGC</td><td>>hg38_21_+_8433172_8446622_RNA45SN1_45S_50ntFlanks</td><td>7985</td><td>8004</td><td>20</td><td>GATCAGACGTGGCGACCCGC</td><td>444.5</td><td>-10.9</td><td>1.5</td><td>-7.4</td><td>1.0E-13</td><td>5.2E-13</td><td>BT20/MB468</td><td>MITO</td></tr>
<tr><td>>MI0003560|hsa-mir-92b&WithFlank&1|+|155195171|155195278@67.87.21&[MIMAT0003218&hsa-miR-92b-3p&offsets|0|-1;m-115&1|+|155195237|155195258&offsets|0|-1]|TATTGCACTCGTCCCGGCCTC</td><td>>MI0003560|hsa-mir-92b&WithFlank&1|+|155195171|155195278</td><td>67</td><td>87</td><td>21</td><td>TATTGCACTCGTCCCGGCCTC</td><td>2155.1</td><td>9.9</td><td>1.3</td><td>7.4</td><td>1.3E-13</td><td>5.2E-13</td><td>BT20/MB468</td><td>CYTO</td></tr>
<tr><td>>hg38_21_+_8433172_8446622_RNA45SN1_45S_50ntFlanks@11920.11954.35|ATGAGAGGTGTAGAATAAGTGGGAGGCCCCCGGCG</td><td>>hg38_21_+_8433172_8446622_RNA45SN1_45S_50ntFlanks</td><td>11920</td><td>11954</td><td>35</td><td>ATGAGAGGTGTAGAATAAGTGGGAGGCCCCCGGCG</td><td>212.0</td><td>-11.6</td><td>1.6</td><td>-7.4</td><td>1.1E-13</td><td>5.2E-13</td><td>MB231/MB468</td><td>MP</td></tr>
<tr><td>>hg38_21_+_8433172_8446622_RNA45SN1_45S_50ntFlanks@11766.11794.29|ATCTAATTAGTGACGCGCATGAATGGATG</td><td>>hg38_21_+_8433172_8446622_RNA45SN1_45S_50ntFlanks</td><td>11766</td><td>11794</td><td>29</td><td>ATCTAATTAGTGACGCGCATGAATGGATG</td><td>138.0</td><td>-11.0</td><td>1.5</td><td>-7.4</td><td>1.1E-13</td><td>5.2E-13</td><td>MB231/MB468</td><td>MP</td></tr>
<tr><td>>hg38_21_+_8433172_8446622_RNA45SN1_45S_50ntFlanks@10351.10379.29|CGAGAACTTTGAAGGCCGAAGTGGAGAAG</td><td>>hg38_21_+_8433172_8446622_RNA45SN1_45S_50ntFlanks</td><td>10351</td><td>10379</td><td>29</td><td>CGAGAACTTTGAAGGCCGAAGTGGAGAAG</td><td>3021.7</td><td>-8.8</td><td>1.2</td><td>-7.4</td><td>1.1E-13</td><td>5.2E-13</td><td>MB231/MB468</td><td>MP</td></tr>
<tr><td>>hg38_21_+_8433172_8446622_RNA45SN1_45S_50ntFlanks@11729.11752.24|TAACTATGACTCTCTTAAGGTAGC</td><td>>hg38_21_+_8433172_8446622_RNA45SN1_45S_50ntFlanks</td><td>11729</td><td>11752</td><td>24</td><td>TAACTATGACTCTCTTAAGGTAGC</td><td>570.8</td><td>-3.3</td><td>0.4</td><td>-7.4</td><td>1.3E-13</td><td>5.2E-13</td><td>MB231/MB468</td><td>NUC</td></tr>
<tr><td>>hg38_21_+_8433172_8446622_RNA45SN1_45S_50ntFlanks@12151.12180.30|GTGGGGAGTTTGACTGGGGCGGTACACCTG</td><td>>hg38_21_+_8433172_8446622_RNA45SN1_45S_50ntFlanks</td><td>12151</td><td>12180</td><td>30</td><td>GTGGGGAGTTTGACTGGGGCGGTACACCTG</td><td>266.2</td><td>-10.1</td><td>1.4</td><td>-7.4</td><td>1.0E-13</td><td>5.3E-13</td><td>BT20/MB468</td><td>MITO</td></tr>
<tr><td>>hg38_21_+_8433172_8446622_RNA45SN1_45S_50ntFlanks@12892.12933.42|CCCTCGCCCGTCACGCACCGCACGTTCGTGGGGAACCTGGCG</td><td>>hg38_21_+_8433172_8446622_RNA45SN1_45S_50ntFlanks</td><td>12892</td><td>12933</td><td>42</td><td>CCCTCGCCCGTCACGCACCGCACGTTCGTGGGGAACCTGGCG</td><td>512.3</td><td>-12.9</td><td>1.7</td><td>-7.4</td><td>1.1E-13</td><td>5.3E-13</td><td>MB231/MB468</td><td>MP</td></tr>
<tr><td>>hg38_21_+_8433172_8446622_RNA45SN1_45S_50ntFlanks@7984.8016.33|AGATCAGACGTGGCGACCCGCTGAATTTAAGCA</td><td>>hg38_21_+_8433172_8446622_RNA45SN1_45S_50ntFlanks</td><td>7984</td><td>8016</td><td>33</td><td>AGATCAGACGTGGCGACCCGCTGAATTTAAGCA</td><td>141.2</td><td>-11.0</td><td>1.5</td><td>-7.4</td><td>1.1E-13</td><td>5.3E-13</td><td>MB231/MB468</td><td>MP</td></tr>
<tr><td>>hg38_21_+_8433172_8446622_RNA45SN1_45S_50ntFlanks@8182.8204.23|ATCGAGGCCCAGCCCGTGGACGG</td><td>>hg38_21_+_8433172_8446622_RNA45SN1_45S_50ntFlanks</td><td>8182</td><td>8204</td><td>23</td><td>ATCGAGGCCCAGCCCGTGGACGG</td><td>190.4</td><td>-9.9</td><td>1.3</td><td>-7.4</td><td>1.3E-13</td><td>5.3E-13</td><td>BT20/MB468</td><td>CYTO</td></tr>
<tr><td>>hg38_21_+_8433172_8446622_RNA45SN1_45S_50ntFlanks@8106.8136.31|GCGGCGGGGCGCGGGACATGTGGCGTACGGA</td><td>>hg38_21_+_8433172_8446622_RNA45SN1_45S_50ntFlanks</td><td>8106</td><td>8136</td><td>31</td><td>GCGGCGGGGCGCGGGACATGTGGCGTACGGA</td><td>253.9</td><td>-11.9</td><td>1.6</td><td>-7.4</td><td>1.1E-13</td><td>5.3E-13</td><td>MB231/MB468</td><td>MP</td></tr>
<tr><td>>hg38_21_+_8433172_8446622_RNA45SN1_45S_50ntFlanks@3838.3860.23|CTCTCCTACTTGGATAACTGTGG</td><td>>hg38_21_+_8433172_8446622_RNA45SN1_45S_50ntFlanks</td><td>3838</td><td>3860</td><td>23</td><td>CTCTCCTACTTGGATAACTGTGG</td><td>888.9</td><td>-13.7</td><td>1.8</td><td>-7.4</td><td>1.1E-13</td><td>5.3E-13</td><td>MB231/MB468</td><td>MP</td></tr>
<tr><td>>hg38_21_+_8433172_8446622_RNA45SN1_45S_50ntFlanks@11573.11603.31|CGACTTAGAACTGGTGCGGACCAGGGGAATC</td><td>>hg38_21_+_8433172_8446622_RNA45SN1_45S_50ntFlanks</td><td>11573</td><td>11603</td><td>31</td><td>CGACTTAGAACTGGTGCGGACCAGGGGAATC</td><td>1214.4</td><td>-7.9</td><td>1.1</td><td>-7.4</td><td>1.1E-13</td><td>5.3E-13</td><td>MB231/MB468</td><td>MP</td></tr>
<tr><td>>hg38_21_+_8433172_8446622_RNA45SN1_45S_50ntFlanks@7988.8017.30|CAGACGTGGCGACCCGCTGAATTTAAGCAT</td><td>>hg38_21_+_8433172_8446622_RNA45SN1_45S_50ntFlanks</td><td>7988</td><td>8017</td><td>30</td><td>CAGACGTGGCGACCCGCTGAATTTAAGCAT</td><td>989.1</td><td>-7.7</td><td>1.0</td><td>-7.4</td><td>1.1E-13</td><td>5.3E-13</td><td>MB231/MB468</td><td>MP</td></tr>
<tr><td>>hg38_21_+_8433172_8446622_RNA45SN1_45S_50ntFlanks@6719.6745.27|TGAATTGCAGGACACATTGATCATCGA</td><td>>hg38_21_+_8433172_8446622_RNA45SN1_45S_50ntFlanks</td><td>6719</td><td>6745</td><td>27</td><td>TGAATTGCAGGACACATTGATCATCGA</td><td>578.2</td><td>-8.6</td><td>1.2</td><td>-7.4</td><td>1.3E-13</td><td>5.3E-13</td><td>MB231/MB468</td><td>NUC</td></tr>
<tr><td>>hg38_21_+_8433172_8446622_RNA45SN1_45S_50ntFlanks@5073.5102.30|AATTCCGATAACGAACGAGACTCTGGCATG</td><td>>hg38_21_+_8433172_8446622_RNA45SN1_45S_50ntFlanks</td><td>5073</td><td>5102</td><td>30</td><td>AATTCCGATAACGAACGAGACTCTGGCATG</td><td>795.7</td><td>-13.5</td><td>1.8</td><td>-7.4</td><td>1.1E-13</td><td>5.3E-13</td><td>MB231/MB468</td><td>MP</td></tr>
<tr><td>>MI0000255|hsa-mir-30d&WithFlank&8|-|134804870|134804951@12.33.22(+2U)&[MIMAT0000245&hsa-miR-30d-5p&offsets|0|0(+2U);m-21&8|-|134804919|134804940&offsets|0|0(+2U)]|TGTAAACATCCCCGACTGGAAGTT</td><td>>MI0000255|hsa-mir-30d&WithFlank&8|-|134804870|134804951</td><td>12</td><td>33</td><td>22(+2U)</td><td>TGTAAACATCCCCGACTGGAAGTT</td><td>451.6</td><td>10.2</td><td>1.4</td><td>7.4</td><td>1.4E-13</td><td>5.3E-13</td><td>BT20/MB468</td><td>CYTO</td></tr>
<tr><td>>trnaMT_ValTAC_MT_+_1602_1670@52.72.21|TTAACTTGACCGCTCTGACCA</td><td>>trnaMT_ValTAC_MT_+_1602_1670</td><td>52</td><td>72</td><td>21</td><td>TTAACTTGACCGCTCTGACCA</td><td>692.9</td><td>10.6</td><td>1.4</td><td>7.4</td><td>1.4E-13</td><td>5.3E-13</td><td>BT20/MB468</td><td>CYTO</td></tr>
<tr><td>>hg38_21_+_8433172_8446622_RNA45SN1_45S_50ntFlanks@11783.11804.22|CATGAATGGATGAACGAGATTC</td><td>>hg38_21_+_8433172_8446622_RNA45SN1_45S_50ntFlanks</td><td>11783</td><td>11804</td><td>22</td><td>CATGAATGGATGAACGAGATTC</td><td>21079.1</td><td>-7.0</td><td>0.9</td><td>-7.8</td><td>5.3E-15</td><td>5.3E-13</td><td>BT20/MB468</td><td>MP</td></tr>
<tr><td>>hg38_21_+_8433172_8446622_RNA45SN1_45S_50ntFlanks@11050.11075.26|CCCGTCCTCCCCCCTCCCCGGGGGAG</td><td>>hg38_21_+_8433172_8446622_RNA45SN1_45S_50ntFlanks</td><td>11050</td><td>11075</td><td>26</td><td>CCCGTCCTCCCCCCTCCCCGGGGGAG</td><td>129.6</td><td>-9.7</td><td>1.3</td><td>-7.4</td><td>1.3E-13</td><td>5.3E-13</td><td>MB231/MB468</td><td>CYTO</td></tr>
<tr><td>>hg38_21_+_8433172_8446622_RNA45SN1_45S_50ntFlanks@12624.12654.31|CCAATGGGGCGAAGCTACCATCTGTGGGATT</td><td>>hg38_21_+_8433172_8446622_RNA45SN1_45S_50ntFlanks</td><td>12624</td><td>12654</td><td>31</td><td>CCAATGGGGCGAAGCTACCATCTGTGGGATT</td><td>3292.8</td><td>-11.2</td><td>1.5</td><td>-7.4</td><td>2.0E-13</td><td>5.3E-13</td><td>BT20/MB468</td><td>NUC</td></tr>
<tr><td>>hg38_21_+_8433172_8446622_RNA45SN1_45S_50ntFlanks@11029.11050.22|CTCTCTCTCTCTCCCCCGCTCC</td><td>>hg38_21_+_8433172_8446622_RNA45SN1_45S_50ntFlanks</td><td>11029</td><td>11050</td><td>22</td><td>CTCTCTCTCTCTCCCCCGCTCC</td><td>188.6</td><td>-10.3</td><td>1.4</td><td>-7.4</td><td>1.3E-13</td><td>5.3E-13</td><td>MB231/MB468</td><td>CYTO</td></tr>
<tr><td>>hg38_21_+_8433172_8446622_RNA45SN1_45S_50ntFlanks@12490.12521.32|CGTGAGCTGGGTTTAGACCGTCGTGAGACAGG</td><td>>hg38_21_+_8433172_8446622_RNA45SN1_45S_50ntFlanks</td><td>12490</td><td>12521</td><td>32</td><td>CGTGAGCTGGGTTTAGACCGTCGTGAGACAGG</td><td>173.4</td><td>-11.3</td><td>1.5</td><td>-7.4</td><td>1.1E-13</td><td>5.3E-13</td><td>MB231/MB468</td><td>MP</td></tr>
<tr><td>>MI0000239|hsa-mir-197&WithFlank&1|+|109598887|109598973@54.74.21(+1A)&[MIMAT0000227&hsa-miR-197-3p&offsets|0|-1(+1A);m-99&1|+|109598940|109598961&offsets|0|-1(+1A)]|TTCACCACCTTCTCCACCCAGA</td><td>>MI0000239|hsa-mir-197&WithFlank&1|+|109598887|109598973</td><td>54</td><td>74</td><td>21(+1A)</td><td>TTCACCACCTTCTCCACCCAGA</td><td>658.5</td><td>9.0</td><td>1.2</td><td>7.4</td><td>1.0E-13</td><td>5.3E-13</td><td>BT20/MB468</td><td>MITO</td></tr>
<tr><td>>MI0000086|hsa-mir-28&WithFlank&3|+|188688775|188688872@20.41.22&[MIMAT0000085&hsa-miR-28-5p&offsets|0|0;m-133&3|+|188688794|188688815&offsets|0|0]|AAGGAGCTCACAGTCTATTGAG</td><td>>MI0000086|hsa-mir-28&WithFlank&3|+|188688775|188688872</td><td>20</td><td>41</td><td>22</td><td>AAGGAGCTCACAGTCTATTGAG</td><td>543.1</td><td>9.0</td><td>1.2</td><td>7.4</td><td>1.0E-13</td><td>5.3E-13</td><td>BT20/MB468</td><td>MITO</td></tr>
<tr><td>>hg38_21_+_8433172_8446622_RNA45SN1_45S_50ntFlanks@4560.4578.19|CTAGGAATAATGGAATAGG</td><td>>hg38_21_+_8433172_8446622_RNA45SN1_45S_50ntFlanks</td><td>4560</td><td>4578</td><td>19</td><td>CTAGGAATAATGGAATAGG</td><td>157.4</td><td>-11.2</td><td>1.5</td><td>-7.4</td><td>1.1E-13</td><td>5.3E-13</td><td>MB231/MB468</td><td>MP</td></tr>
<tr><td>>MI0005566|hsa-mir-374b&WithFlank&X|-|74218541|74218624@17.38.22&[MIMAT0004955&hsa-miR-374b-5p&offsets|0|0;m-153&X|-|74218587|74218608&offsets|0|0]|ATATAATACAACCTGCTAAGTG</td><td>>MI0005566|hsa-mir-374b&WithFlank&X|-|74218541|74218624</td><td>17</td><td>38</td><td>22</td><td>ATATAATACAACCTGCTAAGTG</td><td>992.9</td><td>5.5</td><td>0.7</td><td>7.4</td><td>1.4E-13</td><td>5.3E-13</td><td>MB231/MB468</td><td>NUC</td></tr>
<tr><td>>hg38_21_+_8433172_8446622_RNA45SN1_45S_50ntFlanks@12457.12488.32|CAAGCGTTGGATTGTTCACCCACTAATAGGGA</td><td>>hg38_21_+_8433172_8446622_RNA45SN1_45S_50ntFlanks</td><td>12457</td><td>12488</td><td>32</td><td>CAAGCGTTGGATTGTTCACCCACTAATAGGGA</td><td>1017.3</td><td>-7.3</td><td>1.0</td><td>-7.4</td><td>1.4E-13</td><td>5.4E-13</td><td>MB231/MB468</td><td>NUC</td></tr>
<tr><td>>hg38_21_+_8433172_8446622_RNA45SN1_45S_50ntFlanks@12635.12659.25|AAGCTACCATCTGTGGGATTATGAC</td><td>>hg38_21_+_8433172_8446622_RNA45SN1_45S_50ntFlanks</td><td>12635</td><td>12659</td><td>25</td><td>AAGCTACCATCTGTGGGATTATGAC</td><td>1085.4</td><td>-5.1</td><td>0.7</td><td>-7.4</td><td>1.4E-13</td><td>5.4E-13</td><td>MB231/MB468</td><td>NUC</td></tr>
<tr><td>>hg38_21_+_8433172_8446622_RNA45SN1_45S_50ntFlanks@6719.6747.29|TGAATTGCAGGACACATTGATCATCGACA</td><td>>hg38_21_+_8433172_8446622_RNA45SN1_45S_50ntFlanks</td><td>6719</td><td>6747</td><td>29</td><td>TGAATTGCAGGACACATTGATCATCGACA</td><td>867.3</td><td>-5.2</td><td>0.7</td><td>-7.4</td><td>1.4E-13</td><td>5.4E-13</td><td>MB231/MB468</td><td>NUC</td></tr>
<tr><td>>hg38_21_+_8433172_8446622_RNA45SN1_45S_50ntFlanks@3881.3898.18|GCCGACGGGCGCTGACCC</td><td>>hg38_21_+_8433172_8446622_RNA45SN1_45S_50ntFlanks</td><td>3881</td><td>3898</td><td>18</td><td>GCCGACGGGCGCTGACCC</td><td>670.8</td><td>-9.4</td><td>1.3</td><td>-7.4</td><td>1.8E-13</td><td>5.4E-13</td><td>MB231/MB468</td><td>MITO</td></tr>
<tr><td>>hg38_21_+_8433172_8446622_RNA45SN1_45S_50ntFlanks@10914.10931.18|CGCCGCGGCTGGACGAGG</td><td>>hg38_21_+_8433172_8446622_RNA45SN1_45S_50ntFlanks</td><td>10914</td><td>10931</td><td>18</td><td>CGCCGCGGCTGGACGAGG</td><td>754.7</td><td>-7.6</td><td>1.0</td><td>-7.4</td><td>1.3E-13</td><td>5.4E-13</td><td>MB231/MB468</td><td>CYTO</td></tr>
<tr><td>>hg38_21_+_8433172_8446622_RNA45SN1_45S_50ntFlanks@9738.9755.18|GAAACGATCTCAACCTAT</td><td>>hg38_21_+_8433172_8446622_RNA45SN1_45S_50ntFlanks</td><td>9738</td><td>9755</td><td>18</td><td>GAAACGATCTCAACCTAT</td><td>618.0</td><td>-9.0</td><td>1.2</td><td>-7.4</td><td>1.4E-13</td><td>5.4E-13</td><td>MB231/MB468</td><td>NUC</td></tr>
<tr><td>>trnaMT_ProTGG_MT_-_15956_16023@42.67.26|TGGTGGAGTTAAAGACTTTTTCTCTG</td><td>>trnaMT_ProTGG_MT_-_15956_16023</td><td>42</td><td>67</td><td>26</td><td>TGGTGGAGTTAAAGACTTTTTCTCTG</td><td>121.7</td><td>-10.8</td><td>1.5</td><td>-7.4</td><td>1.2E-13</td><td>5.4E-13</td><td>MB231/MB468</td><td>MP</td></tr>
<tr><td>>hg38_21_+_8433172_8446622_RNA45SN1_45S_50ntFlanks@9027.9057.31|CGTCTCCTCGTGGGGGGGCCGGGCCACCCCT</td><td>>hg38_21_+_8433172_8446622_RNA45SN1_45S_50ntFlanks</td><td>9027</td><td>9057</td><td>31</td><td>CGTCTCCTCGTGGGGGGGCCGGGCCACCCCT</td><td>251.9</td><td>-10.1</td><td>1.4</td><td>-7.4</td><td>1.1E-13</td><td>5.4E-13</td><td>BT20/MB468</td><td>MITO</td></tr>
<tr><td>>hg38_21_+_8433172_8446622_RNA45SN1_45S_50ntFlanks@4047.4067.21|ATTCGAACGTCTGCCCTATCA</td><td>>hg38_21_+_8433172_8446622_RNA45SN1_45S_50ntFlanks</td><td>4047</td><td>4067</td><td>21</td><td>ATTCGAACGTCTGCCCTATCA</td><td>1154.9</td><td>-8.3</td><td>1.1</td><td>-7.4</td><td>1.8E-13</td><td>5.4E-13</td><td>MB231/MB468</td><td>MITO</td></tr>
<tr><td>>hg38_21_+_8433172_8446622_RNA45SN1_45S_50ntFlanks@6646.6683.38|TCGTACGACTCTTAGCGGTGGATCACTCGGCTCGTGCG</td><td>>hg38_21_+_8433172_8446622_RNA45SN1_45S_50ntFlanks</td><td>6646</td><td>6683</td><td>38</td><td>TCGTACGACTCTTAGCGGTGGATCACTCGGCTCGTGCG</td><td>1426.3</td><td>4.2</td><td>0.6</td><td>7.4</td><td>1.1E-13</td><td>5.4E-13</td><td>BT20/MB468</td><td>MITO</td></tr>
<tr><td>>MI0000253|hsa-mir-148a&WithFlank&7|-|25949913|25949992@50.71.22(+1A)&[MIMAT0000243&hsa-miR-148a-3p&offsets|0|0(+1A);m-7&7|-|25949922|25949943&offsets|0|0(+1A)]|TCAGTGCACTACAGAACTTTGTA</td><td>>MI0000253|hsa-mir-148a&WithFlank&7|-|25949913|25949992</td><td>50</td><td>71</td><td>22(+1A)</td><td>TCAGTGCACTACAGAACTTTGTA</td><td>257.2</td><td>8.2</td><td>1.1</td><td>7.7</td><td>1.1E-14</td><td>5.4E-13</td><td>BT20/MB231</td><td>CYTO</td></tr>
<tr><td>>hg38_21_+_8433172_8446622_RNA45SN1_45S_50ntFlanks@11641.11682.42|CGGCGGGTGTTGACGCGATGTGATTTCTGCCCAGTGCTCTGA</td><td>>hg38_21_+_8433172_8446622_RNA45SN1_45S_50ntFlanks</td><td>11641</td><td>11682</td><td>42</td><td>CGGCGGGTGTTGACGCGATGTGATTTCTGCCCAGTGCTCTGA</td><td>819.9</td><td>-9.7</td><td>1.3</td><td>-7.4</td><td>1.8E-13</td><td>5.4E-13</td><td>MB231/MB468</td><td>MITO</td></tr>
<tr><td>>hg38_21_+_8433172_8446622_RNA45SN1_45S_50ntFlanks@10572.10608.37|AGAAGCCGGCGGGAGCCCCGGGGAGAGTTCTCTTTTC</td><td>>hg38_21_+_8433172_8446622_RNA45SN1_45S_50ntFlanks</td><td>10572</td><td>10608</td><td>37</td><td>AGAAGCCGGCGGGAGCCCCGGGGAGAGTTCTCTTTTC</td><td>1330.1</td><td>-9.5</td><td>1.3</td><td>-7.4</td><td>1.4E-13</td><td>5.4E-13</td><td>MB231/MB468</td><td>NUC</td></tr>
<tr><td>>hg38_21_+_8433172_8446622_RNA45SN1_45S_50ntFlanks@11848.11872.25|ACGGGCTTGGCGGAATCAGCGGGGA</td><td>>hg38_21_+_8433172_8446622_RNA45SN1_45S_50ntFlanks</td><td>11848</td><td>11872</td><td>25</td><td>ACGGGCTTGGCGGAATCAGCGGGGA</td><td>624.6</td><td>-9.5</td><td>1.3</td><td>-7.3</td><td>2.0E-13</td><td>5.4E-13</td><td>BT20/MB468</td><td>NUC</td></tr>
<tr><td>>hg38_21_+_8433172_8446622_RNA45SN1_45S_50ntFlanks@3875.3898.24|ATACATGCCGACGGGCGCTGACCC</td><td>>hg38_21_+_8433172_8446622_RNA45SN1_45S_50ntFlanks</td><td>3875</td><td>3898</td><td>24</td><td>ATACATGCCGACGGGCGCTGACCC</td><td>4231.2</td><td>-4.9</td><td>0.7</td><td>-7.4</td><td>1.4E-13</td><td>5.4E-13</td><td>MB231/MB468</td><td>CYTO</td></tr>
<tr><td>>trna111_HisGTG_1_-_147774845_147774916@-1T.29.30,trna118_HisGTG_1_-_145396881_145396952@-1T.29.30,trna16_HisGTG_1_+_146544773_146544844@-1T.29.30,trna1_HisGTG_15_+_45493349_45493420@-1T.29.30,trna21_HisGTG_1_+_147753471_147753542@-1T.29.30,trna33_HisGTG_6_+_27125906_27125977@-1T.29.30,trna7_HisGTG_9_-_14433938_14434009@-1T.29.30,trna8_HisGTG_15_-_45492611_45492682@-1T.29.30,trna9_HisGTG_15_-_45490804_45490875@-1T.29.30|TGCCGTGATCGTATAGTGGTTAGTACTCTG</td><td>>trna111_HisGTG_1_-_147774845_147774916</td><td>-1T</td><td>29</td><td>30</td><td>TGCCGTGATCGTATAGTGGTTAGTACTCTG</td><td>2551.1</td><td>-5.4</td><td>0.7</td><td>-7.4</td><td>1.8E-13</td><td>5.5E-13</td><td>MB231/MB468</td><td>MITO</td></tr>
<tr><td>>hg38_21_+_8433172_8446622_RNA45SN1_45S_50ntFlanks@10830.10855.26|CAAGCCGGATCCGTAACTTCGGGATA</td><td>>hg38_21_+_8433172_8446622_RNA45SN1_45S_50ntFlanks</td><td>10830</td><td>10855</td><td>26</td><td>CAAGCCGGATCCGTAACTTCGGGATA</td><td>265.3</td><td>-11.9</td><td>1.6</td><td>-7.4</td><td>1.2E-13</td><td>5.5E-13</td><td>MB231/MB468</td><td>MP</td></tr>
<tr><td>>hg38_21_+_8433172_8446622_RNA45SN1_45S_50ntFlanks@4560.4589.30|CTAGGAATAATGGAATAGGACCGCGGTTCT</td><td>>hg38_21_+_8433172_8446622_RNA45SN1_45S_50ntFlanks</td><td>4560</td><td>4589</td><td>30</td><td>CTAGGAATAATGGAATAGGACCGCGGTTCT</td><td>259.6</td><td>-11.9</td><td>1.6</td><td>-7.4</td><td>1.2E-13</td><td>5.5E-13</td><td>MB231/MB468</td><td>MP</td></tr>
<tr><td>>hg38_21_+_8433172_8446622_RNA45SN1_45S_50ntFlanks@5472.5505.34|ACGGCCCTGGCGGAGCGCTGAGAAGACGGTCGAA</td><td>>hg38_21_+_8433172_8446622_RNA45SN1_45S_50ntFlanks</td><td>5472</td><td>5505</td><td>34</td><td>ACGGCCCTGGCGGAGCGCTGAGAAGACGGTCGAA</td><td>1426.6</td><td>9.0</td><td>1.2</td><td>7.4</td><td>1.4E-13</td><td>5.5E-13</td><td>BT20/MB468</td><td>CYTO</td></tr>
<tr><td>>hg38_21_+_8433172_8446622_RNA45SN1_45S_50ntFlanks@11921.11967.47|TGAGAGGTGTAGAATAAGTGGGAGGCCCCCGGCGCCCCCCCGGTGTC</td><td>>hg38_21_+_8433172_8446622_RNA45SN1_45S_50ntFlanks</td><td>11921</td><td>11967</td><td>47</td><td>TGAGAGGTGTAGAATAAGTGGGAGGCCCCCGGCGCCCCCCCGGTGTC</td><td>676.2</td><td>-12.4</td><td>1.7</td><td>-7.4</td><td>1.4E-13</td><td>5.5E-13</td><td>MB231/MB468</td><td>NUC</td></tr>
<tr><td>>MI0000272|hsa-mir-182&WithFlank&7|-|129770377|129770498@29.51.23&[MIMAT0000259&hsa-miR-182-5p&offsets|0|-1;m-22&7|-|129770449|129770470&offsets|0|+1]|TTTGGCAATGGTAGAACTCACAC</td><td>>MI0000272|hsa-mir-182&WithFlank&7|-|129770377|129770498</td><td>29</td><td>51</td><td>23</td><td>TTTGGCAATGGTAGAACTCACAC</td><td>3095.6</td><td>3.8</td><td>0.5</td><td>7.7</td><td>1.6E-14</td><td>5.5E-13</td><td>BT20/MB231</td><td>NUC</td></tr>
<tr><td>>hg38_21_+_8433172_8446622_RNA45SN1_45S_50ntFlanks@11888.11905.18|AGCTTGACTCTAGTCTGG</td><td>>hg38_21_+_8433172_8446622_RNA45SN1_45S_50ntFlanks</td><td>11888</td><td>11905</td><td>18</td><td>AGCTTGACTCTAGTCTGG</td><td>221.4</td><td>-10.5</td><td>1.4</td><td>-7.4</td><td>1.4E-13</td><td>5.5E-13</td><td>MB231/MB468</td><td>CYTO</td></tr>
<tr><td>>hg38_21_+_8433172_8446622_RNA45SN1_45S_50ntFlanks@485.514.30|GATGTGGTGACGTCGTGCTCTCCCGGGCCG</td><td>>hg38_21_+_8433172_8446622_RNA45SN1_45S_50ntFlanks</td><td>485</td><td>514</td><td>30</td><td>GATGTGGTGACGTCGTGCTCTCCCGGGCCG</td><td>3628.2</td><td>12.1</td><td>1.6</td><td>7.4</td><td>1.4E-13</td><td>5.5E-13</td><td>MB231/MB468</td><td>NUC</td></tr>
</table>

| | | | | | | | | | | | | | |
|---|---|---|---|---|---|---|---|---|---|---|---|---|---|
| >hg38_21_+_8433172_8446622_RNA45SN1_45S_50ntFlanks@10411.10443.33\|CAGTCGGTCCTGAGAGATGGGCGAGCGCCGTTC | >hg38_21_+_8433172_8446622_RNA45SN1_45S_50ntFlanks | 10411 | 10443 | 33 | CAGTCGGTCCTGAGAGATGGGCGAGCGCCGTTC | 879.1 | -7.4 | 1.0 | -7.4 | 1.4E-13 | 5.5E-13 | MB231/MB468 | NUC |
| >hg38_21_+_8433172_8446622_RNA45SN1_45S_50ntFlanks@8381.8398.18\|AAGAGGGCGTGAAACCGT | >hg38_21_+_8433172_8446622_RNA45SN1_45S_50ntFlanks | 8381 | 8398 | 18 | AAGAGGGCGTGAAACCGT | 170.4 | -11.3 | 1.5 | -7.4 | 1.2E-13 | 5.5E-13 | MB231/MB468 | MP |
| >hg38_21_+_8433172_8446622_RNA45SN1_45S_50ntFlanks@9252.9271.20\|CACGGGGTCGGCGGCGACGT | >hg38_21_+_8433172_8446622_RNA45SN1_45S_50ntFlanks | 9252 | 9271 | 20 | CACGGGGTCGGCGGCGACGT | 132.2 | -10.9 | 1.5 | -7.4 | 1.2E-13 | 5.5E-13 | MB231/MB468 | MP |
| >hg38_21_+_8433172_8446622_RNA45SN1_45S_50ntFlanks@10839.10868.30\|TCCGTAACTTCGGGATAAGGATTGGCTCTA | >hg38_21_+_8433172_8446622_RNA45SN1_45S_50ntFlanks | 10839 | 10868 | 30 | TCCGTAACTTCGGGATAAGGATTGGCTCTA | 759.1 | -9.8 | 1.3 | -7.3 | 2.1E-13 | 5.5E-13 | BT20/MB468 | NUC |
| >hg38_21_+_8433172_8446622_RNA45SN1_45S_50ntFlanks@11482.11502.21\|GCGGAGGGGCCGCGGGCCGGT | >hg38_21_+_8433172_8446622_RNA45SN1_45S_50ntFlanks | 11482 | 11502 | 21 | GCGGAGGGGCCGCGGGCCGGT | 145.2 | -5.1 | 0.7 | -7.4 | 1.4E-13 | 5.5E-13 | MB231/MB468 | CYTO |
| >MI0003591\|hsa-mir-584&WithFlank&5\|-\|149062307\|149062415@22.42.21(+1U)&[MIMAT0003249&hsa-miR-584-5p&offsets\|0\|-1(+1U);m-167&5\|-\|149062374\|149062394&offsets\|0\|0(+1U)]\|TTATGGTTTGCCTGGGACTGAT | >MI0003591\|hsa-mir-584&WithFlank&5\|-\|149062307\|149062415 | 22 | 42 | 21(+1U) | TTATGGTTTGCCTGGGACTGAT | 397.8 | 10.0 | 1.3 | 7.4 | 1.4E-13 | 5.5E-13 | BT20/MB468 | CYTO |
| >hg38_21_+_8433172_8446622_RNA45SN1_45S_50ntFlanks@6650.6681.32\|ACGACTCTTAGCGGTGGATCACTCGGCTCGTG | >hg38_21_+_8433172_8446622_RNA45SN1_45S_50ntFlanks | 6650 | 6681 | 32 | ACGACTCTTAGCGGTGGATCACTCGGCTCGTG | 6390.6 | -6.2 | 0.8 | -7.4 | 1.2E-13 | 5.5E-13 | MB231/MB468 | MP |
| >MI0000264\|hsa-mir-7-2&WithFlank&15\|+\|88611819\|88611940@38.59.22(+1A);MI0000265\|hsa-mir-7-3&WithFlank&19\|+\|4770664\|4770785@37.58.22(+1A);MI0000263\|hsa-mir-7-1&WithFlank&9\|-\|83969742\|83969863@30.51.22(+1A)&[MIMAT0000252&hsa-miR-7-5p&offsets\|0\|-2(+1A);m-400&15\|+\|88611856\|88611879&offsets\|0\|-2(+1A)];[MIMAT0000252_1&hsa-miR-7-5p&offsets\|0\|-2(+1A);m-402&19\|+\|4770700\|4770723&offsets\|0\|-2(+1A)];[MIMAT0000252_2&hsa-miR-7-5p&offsets\|0\|-2(+1A);m-401&9\|-\|83969811\|83969834&offsets\|0\|-2(+1A)]\|TGGAAGACTAGTGATTTTGTTGA | >MI0000264\|hsa-mir-7-2&WithFlank&15\|+\|88611819\|88611940 | 38 | 59 | 22(+1A) | TGGAAGACTAGTGATTTTGTTGA | 1290.0 | 9.8 | 1.3 | 7.4 | 1.4E-13 | 5.5E-13 | BT20/MB468 | CYTO |
| >hg38_21_+_8433172_8446622_RNA45SN1_45S_50ntFlanks@11584.11607.24\|TGGTGCGGACCAGGGGAATCCGAC | >hg38_21_+_8433172_8446622_RNA45SN1_45S_50ntFlanks | 11584 | 11607 | 24 | TGGTGCGGACCAGGGGAATCCGAC | 357.7 | 7.8 | 1.1 | 7.4 | 1.1E-13 | 5.5E-13 | BT20/MB468 | MITO |
| >hg38_21_+_8433172_8446622_RNA45SN1_45S_50ntFlanks@11564.11591.28\|CCTAGCAGCCGACTTAGAACTGGTGCGG | >hg38_21_+_8433172_8446622_RNA45SN1_45S_50ntFlanks | 11564 | 11591 | 28 | CCTAGCAGCCGACTTAGAACTGGTGCGG | 1164.9 | -7.2 | 1.0 | -7.4 | 1.4E-13 | 5.5E-13 | MB231/MB468 | CYTO |
| >hg38_21_+_8433172_8446622_RNA45SN1_45S_50ntFlanks@9575.9596.22\|ATCGGTCGTCCGACCTGGGTAT | >hg38_21_+_8433172_8446622_RNA45SN1_45S_50ntFlanks | 9575 | 9596 | 22 | ATCGGTCGTCCGACCTGGGTAT | 1491.0 | -8.6 | 1.2 | -7.4 | 1.4E-13 | 5.5E-13 | MB231/MB468 | CYTO |
| >hg38_21_+_8433172_8446622_RNA45SN1_45S_50ntFlanks@11567.11586.20\|AGCAGCCGACTTAGAACTGG | >hg38_21_+_8433172_8446622_RNA45SN1_45S_50ntFlanks | 11567 | 11586 | 20 | AGCAGCCGACTTAGAACTGG | 611.2 | -3.5 | 0.5 | -7.3 | 2.1E-13 | 5.5E-13 | BT20/MB468 | NUC |
| >hg38_21_+_8433172_8446622_RNA45SN1_45S_50ntFlanks@12581.12604.24\|AACCGCAGGTTCAGACATTTGGTG | >hg38_21_+_8433172_8446622_RNA45SN1_45S_50ntFlanks | 12581 | 12604 | 24 | AACCGCAGGTTCAGACATTTGGTG | 202.2 | -11.5 | 1.6 | -7.4 | 1.2E-13 | 5.5E-13 | MB231/MB468 | MP |
| >hg38_21_+_8433172_8446622_RNA45SN1_45S_50ntFlanks@3802.3834.33\|CATTAAATCAGTTATGGTTCCTTTGGTCGCTCG | >hg38_21_+_8433172_8446622_RNA45SN1_45S_50ntFlanks | 3802 | 3834 | 33 | CATTAAATCAGTTATGGTTCCTTTGGTCGCTCG | 547.3 | -4.0 | 0.5 | -7.4 | 1.4E-13 | 5.6E-13 | BT20/MB468 | CYTO |
| >hg38_21_+_8433172_8446622_RNA45SN1_45S_50ntFlanks@12126.12151.26\|CGACCCGCTCCGGGGACAGTGCCAGG | >hg38_21_+_8433172_8446622_RNA45SN1_45S_50ntFlanks | 12126 | 12151 | 26 | CGACCCGCTCCGGGGACAGTGCCAGG | 259.6 | -11.0 | 1.5 | -7.4 | 1.4E-13 | 5.6E-13 | MB231/MB468 | NUC |
| >hg38_21_+_8433172_8446622_RNA45SN1_45S_50ntFlanks@12202.12221.20\|CTAAGGCGAGCTCAGGGAGG | >hg38_21_+_8433172_8446622_RNA45SN1_45S_50ntFlanks | 12202 | 12221 | 20 | CTAAGGCGAGCTCAGGGAGG | 810.3 | -9.9 | 1.3 | -7.3 | 2.1E-13 | 5.6E-13 | BT20/MB468 | NUC |
| >hg38_21_+_8433172_8446622_RNA45SN1_45S_50ntFlanks@11577.11602.26\|TTAGAACTGGTGCGGACCAGGGGAAT | >hg38_21_+_8433172_8446622_RNA45SN1_45S_50ntFlanks | 11577 | 11602 | 26 | TTAGAACTGGTGCGGACCAGGGGAAT | 321.8 | -6.0 | 0.8 | -7.4 | 1.4E-13 | 5.6E-13 | MB231/MB468 | CYTO |
| >hg38_21_+_8433172_8446622_RNA45SN1_45S_50ntFlanks@13007.13025.19\|TATTGAAAGTCAGCCCTCG | >hg38_21_+_8433172_8446622_RNA45SN1_45S_50ntFlanks | 13007 | 13025 | 19 | TATTGAAAGTCAGCCCTCG | 148.6 | -11.1 | 1.5 | -7.4 | 1.2E-13 | 5.6E-13 | MB231/MB468 | MP |
| >hg38_21_+_8433172_8446622_RNA45SN1_45S_50ntFlanks@6723.6746.24\|TTGCAGGACACATTGATCATCGAC | >hg38_21_+_8433172_8446622_RNA45SN1_45S_50ntFlanks | 6723 | 6746 | 24 | TTGCAGGACACATTGATCATCGAC | 148.7 | -11.1 | 1.5 | -7.4 | 1.2E-13 | 5.6E-13 | MB231/MB468 | MP |
| >hg38_21_+_8433172_8446622_RNA45SN1_45S_50ntFlanks@9929.9959.31\|GATATAGACAGCAGGACGGTGGCCATGGAAG | >hg38_21_+_8433172_8446622_RNA45SN1_45S_50ntFlanks | 9929 | 9959 | 31 | GATATAGACAGCAGGACGGTGGCCATGGAAG | 368.1 | -12.4 | 1.7 | -7.4 | 1.2E-13 | 5.6E-13 | MB231/MB468 | MP |
| >trna119_LysCTT_1_-_145395522_145395594@1.30.30,trna11_LysCTT_5_-_180648979_180649051@1.30.30,trna13_LysCTT_6_+_26556774_26556846@1.30.30,trna32_LysCTT_16_-_3207406_3207478@1.30.30,trna7_LysCTT_16_+_3225692_3225764@1.30.30,trna9_LysCTT_5_+_180634755_180634827@1.30.30\|GCCCGGCTAGCTCAGTCGGTAGAGCATGAG | >trna119_LysCTT_1_-_145395522_145395594 | 1 | 30 | 30 | GCCCGGCTAGCTCAGTCGGTAGAGCATGAG | 30607.4 | -6.6 | 0.9 | -7.4 | 1.4E-13 | 5.6E-13 | BT20/MB468 | CYTO |
| >hg38_21_+_8433172_8446622_RNA45SN1_45S_50ntFlanks@12459.12487.29\|AGCGTTGGATTGTTCACCCACTAATAGGG | >hg38_21_+_8433172_8446622_RNA45SN1_45S_50ntFlanks | 12459 | 12487 | 29 | AGCGTTGGATTGTTCACCCACTAATAGGG | 439.2 | -7.8 | 1.1 | -7.4 | 1.4E-13 | 5.6E-13 | BT20/MB468 | CYTO |
| >hg38_21_+_8433172_8446622_RNA45SN1_45S_50ntFlanks@11199.11226.28\|CCGGCGGCGGCGGCGACTCTGGACGCGA | >hg38_21_+_8433172_8446622_RNA45SN1_45S_50ntFlanks | 11199 | 11226 | 28 | CCGGCGGCGGCGGCGACTCTGGACGCGA | 110.9 | -9.5 | 1.3 | -7.4 | 1.4E-13 | 5.6E-13 | MB231/MB468 | CYTO |
| >MI0000299\|hsa-mir-222&WithFlank&X\|-\|45747009\|45747130@75.97.23(+2U)&[MIMAT0000279&hsa-miR-222-3p&offsets\|0\|+2(+2U);m-140&X\|-\|45747033\|45747056&offsets\|0\|-1(+2U)]\|AGCTACATCTGGCTACTGGGTCTTT | >MI0000299\|hsa-mir-222&WithFlank&X\|-\|45747009\|45747130 | 75 | 97 | 23(+2U) | AGCTACATCTGGCTACTGGGTCTTT | 1216.0 | 6.8 | 0.9 | 7.4 | 1.8E-13 | 5.6E-13 | MB231/MB468 | MITO |
| >hg38_21_+_8433172_8446622_RNA45SN1_45S_50ntFlanks@12424.12457.34\|CGGCTCTTCCTATCATTGTGAAGCAGAATTCACC | >hg38_21_+_8433172_8446622_RNA45SN1_45S_50ntFlanks | 12424 | 12457 | 34 | CGGCTCTTCCTATCATTGTGAAGCAGAATTCACC | 271.6 | 7.5 | 1.0 | 7.4 | 1.4E-13 | 5.6E-13 | MB231/MB468 | CYTO |
| >hg38_21_+_8433172_8446622_RNA45SN1_45S_50ntFlanks@10690.10710.21\|CGTCCGGTGAGCTCTCGCTGG | >hg38_21_+_8433172_8446622_RNA45SN1_45S_50ntFlanks | 10690 | 10710 | 21 | CGTCCGGTGAGCTCTCGCTGG | 359.6 | -10.6 | 1.4 | -7.4 | 1.1E-13 | 5.6E-13 | BT20/MB468 | MITO |
| >hg38_1_-_228634819_228635039_RNA5S12_5S_50ntFlanks@139.171.33\|GACCGCCTGGGAATACCGGGTGCTGTAGGCTTT | >hg38_1_-_228634819_228635039_RNA5S12_5S_50ntFlanks | 139 | 171 | 33 | GACCGCCTGGGAATACCGGGTGCTGTAGGCTTT | 917.8 | 5.5 | 0.7 | 7.4 | 1.4E-13 | 5.6E-13 | MB231/MB468 | NUC |
| >hg38_21_+_8433172_8446622_RNA45SN1_45S_50ntFlanks@5525.5549.25\|TAAAAGTCGTAACAAGGTTTCCGTA | >hg38_21_+_8433172_8446622_RNA45SN1_45S_50ntFlanks | 5525 | 5549 | 25 | TAAAAGTCGTAACAAGGTTTCCGTA | 165.4 | -11.2 | 1.5 | -7.4 | 1.2E-13 | 5.6E-13 | MB231/MB468 | MP |
| >hg38_1_-_228634819_228635039_RNA5S12_5S_50ntFlanks@129.169.41\|TTGGATGGGAGACCGCCTGGGAATACCGGGTGCTGTAGGCT | >hg38_1_-_228634819_228635039_RNA5S12_5S_50ntFlanks | 129 | 169 | 41 | TTGGATGGGAGACCGCCTGGGAATACCGGGTGCTGTAGGCT | 1075.2 | 9.0 | 1.2 | 7.4 | 1.4E-13 | 5.6E-13 | MB231/MB468 | CYTO |
| >MI0000088\|hsa-mir-30a&WithFlank&6\|-\|71403545\|71403627@53.73.21(+1U)&[MIMAT0000088&hsa-miR-30a-3p&offsets\|0\|-1(+1U);m-38&6\|-\|71403554\|71403575&offsets\|0\|-1(+1U)]\|CTTTCAGTCGGATGTTTGCAGT | >MI0000088\|hsa-mir-30a&WithFlank&6\|-\|71403545\|71403627 | 53 | 73 | 21(+1U) | CTTTCAGTCGGATGTTTGCAGT | 6512.4 | 6.5 | 0.9 | 7.4 | 1.8E-13 | 5.6E-13 | MB231/MB468 | MITO |
| >hg38_21_+_8433172_8446622_RNA45SN1_45S_50ntFlanks@11760.11777.18\|CTCGTCATCTAATTAGTG | >hg38_21_+_8433172_8446622_RNA45SN1_45S_50ntFlanks | 11760 | 11777 | 18 | CTCGTCATCTAATTAGTG | 129.3 | -10.9 | 1.5 | -7.4 | 1.2E-13 | 5.6E-13 | MB231/MB468 | MP |
| >hg38_21_+_8433172_8446622_RNA45SN1_45S_50ntFlanks@12685.12703.19\|CCCAGGCGGAACGATACGG | >hg38_21_+_8433172_8446622_RNA45SN1_45S_50ntFlanks | 12685 | 12703 | 19 | CCCAGGCGGAACGATACGG | 221.2 | -11.7 | 1.6 | -7.4 | 1.2E-13 | 5.7E-13 | MB231/MB468 | MP |
| >hg38_21_+_8433172_8446622_RNA45SN1_45S_50ntFlanks@9571.9596.26\|GCAAATCGGTCGTCCGACCTGGGTAT | >hg38_21_+_8433172_8446622_RNA45SN1_45S_50ntFlanks | 9571 | 9596 | 26 | GCAAATCGGTCGTCCGACCTGGGTAT | 210.0 | -7.2 | 1.0 | -7.4 | 1.4E-13 | 5.7E-13 | MB231/MB468 | CYTO |
| >hg38_21_+_8433172_8446622_RNA45SN1_45S_50ntFlanks@11785.11803.19\|TGAATGGATGAACGAGATT | >hg38_21_+_8433172_8446622_RNA45SN1_45S_50ntFlanks | 11785 | 11803 | 19 | TGAATGGATGAACGAGATT | 2540.7 | -8.1 | 1.1 | -7.4 | 1.2E-13 | 5.7E-13 | MB231/MB468 | MP |
| >hg38_21_+_8433172_8446622_RNA45SN1_45S_50ntFlanks@11210.11230.21\|GGCGACTCTGGACGCGAGCCG | >hg38_21_+_8433172_8446622_RNA45SN1_45S_50ntFlanks | 11210 | 11230 | 21 | GGCGACTCTGGACGCGAGCCG | 194.9 | -11.5 | 1.5 | -7.4 | 1.2E-13 | 5.7E-13 | MB231/MB468 | MP |
| >hg38_21_+_8433172_8446622_RNA45SN1_45S_50ntFlanks@12973.13001.29\|TCGTACGTAGCAGAGCAGCTCCCTCGCTG | >hg38_21_+_8433172_8446622_RNA45SN1_45S_50ntFlanks | 12973 | 13001 | 29 | TCGTACGTAGCAGAGCAGCTCCCTCGCTG | 420.7 | -6.0 | 0.8 | -7.4 | 1.4E-13 | 5.7E-13 | MB231/MB468 | CYTO |
| >hg38_21_+_8433172_8446622_RNA45SN1_45S_50ntFlanks@12444.12467.24\|AAGCAGAATTCACCAAGCGTTGGA | >hg38_21_+_8433172_8446622_RNA45SN1_45S_50ntFlanks | 12444 | 12467 | 24 | AAGCAGAATTCACCAAGCGTTGGA | 169.6 | -11.3 | 1.5 | -7.4 | 1.2E-13 | 5.7E-13 | MB231/MB468 | MP |
| >hg38_21_+_8433172_8446622_RNA45SN1_45S_50ntFlanks@9141.9163.23\|CGGGGCGGACTGTCCCCAGTGCG | >hg38_21_+_8433172_8446622_RNA45SN1_45S_50ntFlanks | 9141 | 9163 | 23 | CGGGGCGGACTGTCCCCAGTGCG | 137.1 | -11.0 | 1.5 | -7.4 | 1.2E-13 | 5.7E-13 | MB231/MB468 | MP |
| >MI0000082\|hsa-mir-25&WithFlank&7\|-\|100093554\|100093649@58.79.22(+1U)&[MIMAT0000081&hsa-miR-25-3p&offsets\|0\|0(+1U);m-23&7\|-\|100093571\|100093592&offsets\|0\|0(+1U)]\|CATTGCACTTGTCTCGGTCTGAT | >MI0000082\|hsa-mir-25&WithFlank&7\|-\|100093554\|100093649 | 58 | 79 | 22(+1U) | CATTGCACTTGTCTCGGTCTGAT | 1037.1 | 5.1 | 0.7 | 7.4 | 1.5E-13 | 5.7E-13 | MB231/MB468 | NUC |
| >hg38_21_+_8433172_8446622_RNA45SN1_45S_50ntFlanks@5087.5104.18\|ACGAGACTCTGGCATGCT | >hg38_21_+_8433172_8446622_RNA45SN1_45S_50ntFlanks | 5087 | 5104 | 18 | ACGAGACTCTGGCATGCT | 258.3 | 8.3 | 1.1 | 7.4 | 1.5E-13 | 5.7E-13 | BT20/MB468 | CYTO |
| >hg38_1_-_228634819_228635039_RNA5S12_5S_50ntFlanks@144.169.26\|CCTGGGAATACCGGGTGCTGTAGGCT | >hg38_1_-_228634819_228635039_RNA5S12_5S_50ntFlanks | 144 | 169 | 26 | CCTGGGAATACCGGGTGCTGTAGGCT | 868.8 | 7.8 | 1.1 | 7.4 | 1.5E-13 | 5.7E-13 | BT20/MB468 | CYTO |
| >hg38_21_+_8433172_8446622_RNA45SN1_45S_50ntFlanks@9564.9594.31\|CTGACGTGCAAATCGGTCGTCCGACCTGGGT | >hg38_21_+_8433172_8446622_RNA45SN1_45S_50ntFlanks | 9564 | 9594 | 31 | CTGACGTGCAAATCGGTCGTCCGACCTGGGT | 1031.9 | -4.1 | 0.6 | -7.4 | 1.5E-13 | 5.7E-13 | MB231/MB468 | NUC |
| >hg38_21_+_8433172_8446622_RNA45SN1_45S_50ntFlanks@12002.12031.30\|TGCGGGCCGCCGGTGAAATACCACTACTCT | >hg38_21_+_8433172_8446622_RNA45SN1_45S_50ntFlanks | 12002 | 12031 | 30 | TGCGGGCCGCCGGTGAAATACCACTACTCT | 449.9 | -11.9 | 1.5 | -7.8 | 5.8E-15 | 5.7E-13 | BT20/MB468 | MP |
| >hg38_21_+_8433172_8446622_RNA45SN1_45S_50ntFlanks@11579.11607.29\|AGAACTGGTGCGGACCAGGGGAATCCGAC | >hg38_21_+_8433172_8446622_RNA45SN1_45S_50ntFlanks | 11579 | 11607 | 29 | AGAACTGGTGCGGACCAGGGGAATCCGAC | 730.7 | -5.9 | 0.8 | -7.4 | 1.2E-13 | 5.8E-13 | MB231/MB468 | MP |
| >hg38_21_+_8433172_8446622_RNA45SN1_45S_50ntFlanks@12456.12477.22\|CCAAGCGTTGGATTGTTCACCC | >hg38_21_+_8433172_8446622_RNA45SN1_45S_50ntFlanks | 12456 | 12477 | 22 | CCAAGCGTTGGATTGTTCACCC | 9576.0 | -9.3 | 1.3 | -7.4 | 1.2E-13 | 5.8E-13 | MB231/MB468 | MP |
| >hg38_21_+_8433172_8446622_RNA45SN1_45S_50ntFlanks@12626.12659.34\|AATGGGGCGAAGCTACCATCTGTGGGATTATGAC | >hg38_21_+_8433172_8446622_RNA45SN1_45S_50ntFlanks | 12626 | 12659 | 34 | AATGGGGCGAAGCTACCATCTGTGGGATTATGAC | 6632.1 | -9.5 | 1.3 | -7.4 | 1.9E-13 | 5.8E-13 | MB231/MB468 | MITO |
| >hg38_21_+_8433172_8446622_RNA45SN1_45S_50ntFlanks@12994.13012.19\|CCTCGCTGCGATCTATTGA | >hg38_21_+_8433172_8446622_RNA45SN1_45S_50ntFlanks | 12994 | 13012 | 19 | CCTCGCTGCGATCTATTGA | 675.6 | -3.3 | 0.5 | -7.4 | 1.5E-13 | 5.8E-13 | MB231/MB468 | NUC |

| | | | | | | | | | | | | | |
|---|---|---|---|---|---|---|---|---|---|---|---|---|---|
| >hg38_21_+_8433172_8446622_RNA45SN1_45S_50ntFlanks@4442.4471.30\|CCCCTTGCCTCTCGGCGCCCCCTCGATGCT | >hg38_21_+_8433172_8446622_RNA45SN1_45S_50ntFlanks | 4442 | 4471 | 30 | CCCCTTGCCTCTCGGCGCCCCCTCGATGCT | 241.0 | -10.0 | 1.3 | -7.4 | 1.1E-13 | 5.8E-13 | BT20/MB468 | MITO |
| >hg38_21_+_8433172_8446622_RNA45SN1_45S_50ntFlanks@11570.11603.34\|AGCCGACTTAGAACTGGTGCGGACCAGGGGAATC | >hg38_21_+_8433172_8446622_RNA45SN1_45S_50ntFlanks | 11570 | 11603 | 34 | AGCCGACTTAGAACTGGTGCGGACCAGGGGAATC | 1599.1 | -8.0 | 1.1 | -7.4 | 1.3E-13 | 5.8E-13 | MB231/MB468 | MP |
| >MI0000462\|hsa-mir-152&WithFlank&17\|-\|48037155\|48037253@60.79.20&[MIMAT0000438&hsa-miR-152-3p&offsets\|0\|-1;m-97&17\|-\|48037174\|48037194&offsets\|0\|-1]\|TCAGTGCATGACAGAACTTG | >MI0000462\|hsa-mir-152&WithFlank&17\|-\|48037155\|48037253 | 60 | 79 | 20 | TCAGTGCATGACAGAACTTG | 208.7 | 10.9 | 1.5 | 7.4 | 1.3E-13 | 5.8E-13 | MB231/MB468 | MP |
| >hg38_21_+_8433172_8446622_RNA45SN1_45S_50ntFlanks@7977.7994.18\|CGACCTCAGATCAGACGT | >hg38_21_+_8433172_8446622_RNA45SN1_45S_50ntFlanks | 7977 | 7994 | 18 | CGACCTCAGATCAGACGT | 471.8 | -4.9 | 0.7 | -7.4 | 1.3E-13 | 5.8E-13 | MB231/MB468 | MP |
| >hg38_21_+_8433172_8446622_RNA45SN1_45S_50ntFlanks@9720.9737.18\|ATTAGAGGTCTTGGGGCC | >hg38_21_+_8433172_8446622_RNA45SN1_45S_50ntFlanks | 9720 | 9737 | 18 | ATTAGAGGTCTTGGGGCC | 166.3 | -7.6 | 1.0 | -7.4 | 1.5E-13 | 5.8E-13 | MB231/MB468 | CYTO |
| >trnaMT_ValTAC_MT_+_1602_1670@42.72.31\|AGATTTCAACTTAACTTGACCGCTCTGACCA | >trnaMT_ValTAC_MT_+_1602_1670 | 42 | 72 | 31 | AGATTTCAACTTAACTTGACCGCTCTGACCA | 365.3 | 8.5 | 1.2 | 7.4 | 1.5E-13 | 5.8E-13 | MB231/MB468 | CYTO |
| >hg38_21_+_8433172_8446622_RNA45SN1_45S_50ntFlanks@8154.8175.22\|CCGCTCGTGGGGGGCCCAAGTC | >hg38_21_+_8433172_8446622_RNA45SN1_45S_50ntFlanks | 8154 | 8175 | 22 | CCGCTCGTGGGGGGCCCAAGTC | 260.0 | -11.9 | 1.6 | -7.4 | 1.3E-13 | 5.8E-13 | MB231/MB468 | MP |
| >hg38_21_+_8433172_8446622_RNA45SN1_45S_50ntFlanks@11027.11075.49\|CTCTCTCTCTCTCTCCCCCGCTCCCCGTCCTCCCCCCTCCCCGGGGGAG | >hg38_21_+_8433172_8446622_RNA45SN1_45S_50ntFlanks | 11027 | 11075 | 49 | CTCTCTCTCTCTCTCCCCCGCTCCCCGTCCTCCCCCCTCCCCGGGGGAG | 1227.4 | -13.2 | 1.8 | -7.4 | 1.5E-13 | 5.9E-13 | MB231/MB468 | NUC |
| >hg38_21_+_8433172_8446622_RNA45SN1_45S_50ntFlanks@7979.8010.32\|ACCTCAGATCAGACGTGGCGACCCGCTGAATT | >hg38_21_+_8433172_8446622_RNA45SN1_45S_50ntFlanks | 7979 | 8010 | 32 | ACCTCAGATCAGACGTGGCGACCCGCTGAATT | 219.5 | 3.9 | 0.5 | 7.4 | 1.5E-13 | 5.9E-13 | MB231/MB468 | CYTO |
| >hg38_21_+_8433172_8446622_RNA45SN1_45S_50ntFlanks@5189.5226.38\|TAACAGGTCTGTGATGCCCTTAGATGTCCGGGGCTGCA | >hg38_21_+_8433172_8446622_RNA45SN1_45S_50ntFlanks | 5189 | 5226 | 38 | TAACAGGTCTGTGATGCCCTTAGATGTCCGGGGCTGCA | 565.4 | -12.9 | 1.8 | -7.4 | 1.9E-13 | 5.9E-13 | MB231/MB468 | MITO |
| >hg38_21_+_8433172_8446622_RNA45SN1_45S_50ntFlanks@10116.10135.20\|TGCGTCGGAGGGCGGCGGCG | >hg38_21_+_8433172_8446622_RNA45SN1_45S_50ntFlanks | 10116 | 10135 | 20 | TGCGTCGGAGGGCGGCGGCG | 210.9 | -10.7 | 1.4 | -7.4 | 1.5E-13 | 5.9E-13 | MB231/MB468 | NUC |
| >hg38_21_+_8433172_8446622_RNA45SN1_45S_50ntFlanks@11834.11872.39\|CCACAGCCAAGGGAACGGGCTTGGCGGAATCAGCGGGGA | >hg38_21_+_8433172_8446622_RNA45SN1_45S_50ntFlanks | 11834 | 11872 | 39 | CCACAGCCAAGGGAACGGGCTTGGCGGAATCAGCGGGGA | 216.0 | -11.6 | 1.6 | -7.4 | 1.3E-13 | 5.9E-13 | MB231/MB468 | MP |
| >hg38_21_+_8433172_8446622_RNA45SN1_45S_50ntFlanks@4923.4940.18\|CAGGAGTGGAGCCTGCGG | >hg38_21_+_8433172_8446622_RNA45SN1_45S_50ntFlanks | 4923 | 4940 | 18 | CAGGAGTGGAGCCTGCGG | 231.7 | -11.7 | 1.6 | -7.4 | 1.3E-13 | 5.9E-13 | MB231/MB468 | MP |
| >trna70_AlaCGC_6_+_28697092_28697163@58.75.18\|CCCCCGGCTCCTCCACCA | >trna70_AlaCGC_6_+_28697092_28697163 | 58 | 75 | 18 | CCCCCGGCTCCTCCACCA | 346.2 | 8.1 | 1.1 | 7.4 | 1.5E-13 | 5.9E-13 | MB231/MB468 | CYTO |
| >hg38_21_+_8433172_8446622_RNA45SN1_45S_50ntFlanks@4562.4587.26\|AGGAATAATGGAATAGGACCGCGGTT | >hg38_21_+_8433172_8446622_RNA45SN1_45S_50ntFlanks | 4562 | 4587 | 26 | AGGAATAATGGAATAGGACCGCGGTT | 183.0 | -11.4 | 1.5 | -7.4 | 1.3E-13 | 5.9E-13 | MB231/MB468 | MP |
| >MI0005756\|hsa-mir-934&WithFlank&X\|+\|136550872\|136550966@21.41.21(+1A)&[MIMAT0004977&hsa-miR-934&offsets\|0\|-1(+1A);m-943&X\|+\|136550892\|136550913&offsets\|0\|-1(+1A)]\|TGTCTACTACTGGAGACACTGA | >MI0005756\|hsa-mir-934&WithFlank&X\|+\|136550872\|136550966 | 21 | 41 | 21(+1A) | TGTCTACTACTGGAGACACTGA | 206.9 | 11.1 | 1.4 | 7.7 | 1.2E-14 | 5.9E-13 | BT20/MB231 | CYTO |
| >hg38_21_+_8433172_8446622_RNA45SN1_45S_50ntFlanks@12679.12703.25\|ATCCCGCCCAGGCGGAACGATACGG | >hg38_21_+_8433172_8446622_RNA45SN1_45S_50ntFlanks | 12679 | 12703 | 25 | ATCCCGCCCAGGCGGAACGATACGG | 197.6 | -11.5 | 1.6 | -7.4 | 1.3E-13 | 5.9E-13 | MB231/MB468 | MP |
| >hg38_21_+_8433172_8446622_RNA45SN1_45S_50ntFlanks@11783.11803.21\|CATGAATGGATGAACGAGATT | >hg38_21_+_8433172_8446622_RNA45SN1_45S_50ntFlanks | 11783 | 11803 | 21 | CATGAATGGATGAACGAGATT | 35332.5 | -12.9 | 1.8 | -7.3 | 2.2E-13 | 5.9E-13 | BT20/MB468 | NUC |
| >hg38_21_+_8433172_8446622_RNA45SN1_45S_50ntFlanks@4730.4766.37\|CAAGAACGAAAGTCGGAGGTTCGAAGACGATCAGATA | >hg38_21_+_8433172_8446622_RNA45SN1_45S_50ntFlanks | 4730 | 4766 | 37 | CAAGAACGAAAGTCGGAGGTTCGAAGACGATCAGATA | 1288.9 | 8.6 | 1.2 | 7.4 | 1.5E-13 | 5.9E-13 | BT20/MB468 | CYTO |
| >hg38_21_+_8433172_8446622_RNA45SN1_45S_50ntFlanks@10423.10456.34\|AGAGATGGGCGAGCGCCGTTCCGAAGGGACGGGC | >hg38_21_+_8433172_8446622_RNA45SN1_45S_50ntFlanks | 10423 | 10456 | 34 | AGAGATGGGCGAGCGCCGTTCCGAAGGGACGGGC | 1064.6 | -4.2 | 0.6 | -7.4 | 1.5E-13 | 5.9E-13 | MB231/MB468 | NUC |
| >trna116_GluCTC_1_-_145399233_145399304@3.34.32,trna59_GluCTC_1_+_249168447_249168518@3.34.32,trna71_GluCTC_1_-_161439189_161439260@3.34.32,trna74_GluCTC_1_-_161431809_161431880@3.34.32,trna77_GluCTC_1_-_161424398_161424469@3.34.32,trna77_GluCTC_6_+_28949976_28950047@3.34.32,trna80_GluCTC_1_-_161417018_161417089@3.34.32,trna87_GluCTC_6_-_126101393_126101464@3.34.32\|CCTGGTGGTCTAGTGGTTAGGATTCGGCGCTC | >trna116_GluCTC_1_-_145399233_145399304 | 3 | 34 | 32 | CCTGGTGGTCTAGTGGTTAGGATTCGGCGCTC | 461.2 | 6.5 | 0.9 | 7.4 | 1.2E-13 | 5.9E-13 | BT20/MB468 | MITO |
| >hg38_21_+_8433172_8446622_RNA45SN1_45S_50ntFlanks@10683.10710.28\|CCGGCGGCGTCCGGTGAGCTCTCGCTGG | >hg38_21_+_8433172_8446622_RNA45SN1_45S_50ntFlanks | 10683 | 10710 | 28 | CCGGCGGCGTCCGGTGAGCTCTCGCTGG | 303.6 | -10.3 | 1.4 | -7.4 | 1.2E-13 | 5.9E-13 | BT20/MB468 | MITO |
| >hg38_21_+_8433172_8446622_RNA45SN1_45S_50ntFlanks@10392.10422.31\|ACAGCAGTTGAACATGGGTCAGTCGGTCCTG | >hg38_21_+_8433172_8446622_RNA45SN1_45S_50ntFlanks | 10392 | 10422 | 31 | ACAGCAGTTGAACATGGGTCAGTCGGTCCTG | 2398.9 | -10.8 | 1.5 | -7.3 | 2.2E-13 | 5.9E-13 | BT20/MB468 | NUC |
| >hg38_21_+_8433172_8446622_RNA45SN1_45S_50ntFlanks@12157.12179.23\|AGTTTGACTGGGGCGGTACACCT | >hg38_21_+_8433172_8446622_RNA45SN1_45S_50ntFlanks | 12157 | 12179 | 23 | AGTTTGACTGGGGCGGTACACCT | 307.1 | -7.0 | 1.0 | -7.4 | 1.5E-13 | 5.9E-13 | MB231/MB468 | CYTO |
| >hg38_21_+_8433172_8446622_RNA45SN1_45S_50ntFlanks@12437.12455.19\|CATTGTGAAGCAGAATTCA | >hg38_21_+_8433172_8446622_RNA45SN1_45S_50ntFlanks | 12437 | 12455 | 19 | CATTGTGAAGCAGAATTCA | 19217.3 | -9.5 | 1.3 | -7.4 | 1.5E-13 | 5.9E-13 | MB231/MB468 | CYTO |
| >am_trna2_ArgCCT_16_+_3202901_3202973@14.31.18,trna3_ArgCCT_7_+_139025446_139025518@14.31.18\|AATGGATAAGGCATTGGC | >am_trna2_ArgCCT_16_+_3202901_3202973 | 14 | 31 | 18 | AATGGATAAGGCATTGGC | 136.2 | -11.0 | 1.5 | -7.4 | 1.3E-13 | 6.0E-13 | MB231/MB468 | MP |
| >hg38_21_+_8433172_8446622_RNA45SN1_45S_50ntFlanks@6651.6684.34\|CGACTCTTAGCGGTGGATCACTCGGCTCGTGCGT | >hg38_21_+_8433172_8446622_RNA45SN1_45S_50ntFlanks | 6651 | 6684 | 34 | CGACTCTTAGCGGTGGATCACTCGGCTCGTGCGT | 10584.4 | 5.2 | 0.7 | 7.4 | 1.5E-13 | 6.0E-13 | MB231/MB468 | NUC |
| >hg38_21_+_8433172_8446622_RNA45SN1_45S_50ntFlanks@11104.11121.18\|AAGGGTCGGGGCGGCAGG | >hg38_21_+_8433172_8446622_RNA45SN1_45S_50ntFlanks | 11104 | 11121 | 18 | AAGGGTCGGGGCGGCAGG | 201.9 | -7.4 | 1.0 | -7.4 | 1.5E-13 | 6.0E-13 | MB231/MB468 | CYTO |
| >trna152_ValCAC_6_-_27248049_27248121@1.24.24\|GCTTCTGTAGTGTAGTGGTTATCA | >trna152_ValCAC_6_-_27248049_27248121 | 1 | 24 | 24 | GCTTCTGTAGTGTAGTGGTTATCA | 507.1 | -13.4 | 1.8 | -7.3 | 2.2E-13 | 6.0E-13 | BT20/MB468 | NUC |
| >hg38_21_+_8433172_8446622_RNA45SN1_45S_50ntFlanks@9738.9758.21\|GAAACGATCTCAACCTATTCT | >hg38_21_+_8433172_8446622_RNA45SN1_45S_50ntFlanks | 9738 | 9758 | 21 | GAAACGATCTCAACCTATTCT | 4338.9 | -6.0 | 0.8 | -7.4 | 1.2E-13 | 6.0E-13 | BT20/MB468 | MITO |
| >hg38_21_+_8433172_8446622_RNA45SN1_45S_50ntFlanks@9504.9529.26\|GTGAACTATGCCTGGGCAGGGCGAAG | >hg38_21_+_8433172_8446622_RNA45SN1_45S_50ntFlanks | 9504 | 9529 | 26 | GTGAACTATGCCTGGGCAGGGCGAAG | 786.6 | -12.7 | 1.6 | -7.8 | 6.0E-15 | 6.0E-13 | BT20/MB468 | MP |
| >hg38_21_+_8433172_8446622_RNA45SN1_45S_50ntFlanks@3988.4013.26\|CTTTGGTGACTCTAGATAACCTCGGG | >hg38_21_+_8433172_8446622_RNA45SN1_45S_50ntFlanks | 3988 | 4013 | 26 | CTTTGGTGACTCTAGATAACCTCGGG | 156.4 | -11.2 | 1.5 | -7.4 | 1.3E-13 | 6.0E-13 | MB231/MB468 | MP |
| >hg38_21_+_8433172_8446622_RNA45SN1_45S_50ntFlanks@10547.10567.21\|AGTGCGGTAACGCGACCGATC | >hg38_21_+_8433172_8446622_RNA45SN1_45S_50ntFlanks | 10547 | 10567 | 21 | AGTGCGGTAACGCGACCGATC | 257.3 | -11.9 | 1.6 | -7.4 | 1.3E-13 | 6.0E-13 | MB231/MB468 | MP |
| >hg38_21_+_8433172_8446622_RNA45SN1_45S_50ntFlanks@11092.11116.25\|CGGCGGGGGGAGAAGGGTCGGGGCG | >hg38_21_+_8433172_8446622_RNA45SN1_45S_50ntFlanks | 11092 | 11116 | 25 | CGGCGGGGGGAGAAGGGTCGGGGCG | 209.6 | -10.7 | 1.4 | -7.4 | 1.5E-13 | 6.0E-13 | MB231/MB468 | NUC |
| >hg38_21_+_8433172_8446622_RNA45SN1_45S_50ntFlanks@10845.10864.20\|ACTTCGGGATAAGGATTGGC | >hg38_21_+_8433172_8446622_RNA45SN1_45S_50ntFlanks | 10845 | 10864 | 20 | ACTTCGGGATAAGGATTGGC | 1460.1 | -10.3 | 1.4 | -7.3 | 2.2E-13 | 6.0E-13 | BT20/MB468 | NUC |
| >MI0001445\|hsa-mir-423&WithFlank&17\|+\|30117073\|30117178@23.43.21&[MIMAT0004748&hsa-miR-423-5p&offsets\|0\|-2;m-141&17\|+\|30117095\|30117117&offsets\|0\|-2]\|TGAGGGGCAGAGAGCGAGACT | >MI0001445\|hsa-mir-423&WithFlank&17\|+\|30117073\|30117178 | 23 | 43 | 21 | TGAGGGGCAGAGAGCGAGACT | 766.0 | 5.2 | 0.7 | 7.4 | 1.5E-13 | 6.0E-13 | MB231/MB468 | NUC |
| >hg38_21_+_8433172_8446622_RNA45SN1_45S_50ntFlanks@10771.10790.20\|CAGGTCTCCAAGGTGAACAG | >hg38_21_+_8433172_8446622_RNA45SN1_45S_50ntFlanks | 10771 | 10790 | 20 | CAGGTCTCCAAGGTGAACAG | 132.2 | -10.9 | 1.5 | -7.4 | 1.3E-13 | 6.0E-13 | MB231/MB468 | MP |
| >trna10_AspGTC_12_-_125424193_125424264@32.72.41,trna12_AspGTC_12_-_125411891_125411962@32.72.41,trna144_AspGTC_6_-_27551236_27551307@32.72.41,trna38_AspGTC_17_-_8125556_8125627@32.72.41,trna45_AspGTC_6_+_27447453_27447524@32.72.41,trna48_AspGTC_6_+_27471523_27471594@32.72.41,trna4_AspGTC_12_+_96429799_96429870@32.72.41,trna69_AspGTC_1_-_161440205_161440276@32.72.41,trna72_AspGTC_1_-_161432824_161432895@32.72.41,trna75_AspGTC_1_-_161425414_161425485@32.72.41,trna78_AspGTC_1_-_161418033_161418104@32.72.41,trna81_AspGTC_1_-_161410615_161410686@32.72.41\|CTGTCACGCGGGAGACCGGGGTTCGATTCCCCGACGGGGAG | >trna10_AspGTC_12_-_125424193_125424264 | 32 | 72 | 41 | CTGTCACGCGGGAGACCGGGGTTCGATTCCCCGACGGGGAG | 533.8 | -12.8 | 1.7 | -7.4 | 2.0E-13 | 6.0E-13 | MB231/MB468 | MITO |
| >hg38_21_+_8433172_8446622_RNA45SN1_45S_50ntFlanks@12616.12659.44\|CTGAGGAGCCAATGGGGCGAAGCTACCATCTGTGGGATTATGAC | >hg38_21_+_8433172_8446622_RNA45SN1_45S_50ntFlanks | 12616 | 12659 | 44 | CTGAGGAGCCAATGGGGCGAAGCTACCATCTGTGGGATTATGAC | 321.8 | -12.1 | 1.6 | -7.4 | 2.0E-13 | 6.0E-13 | MB231/MB468 | MITO |
| >MI0000255\|hsa-mir-30d&WithFlank&8\|-\|134804870\|134804951@12.35.24&[MIMAT0000245&hsa-miR-30d-5p&offsets\|0\|+2;m-21&8\|-\|134804919\|134804940&offsets\|0\|+2]\|TGTAAACATCCCCGACTGGAAGCT | >MI0000255\|hsa-mir-30d&WithFlank&8\|-\|134804870\|134804951 | 12 | 35 | 24 | TGTAAACATCCCCGACTGGAAGCT | 26092.9 | 6.7 | 0.9 | 7.4 | 1.5E-13 | 6.0E-13 | MB231/MB468 | CYTO |
| >hg38_21_+_8433172_8446622_RNA45SN1_45S_50ntFlanks@12419.12450.32\|GATGTCGGCTCTTCCTATCATTGTGAAGCAGA | >hg38_21_+_8433172_8446622_RNA45SN1_45S_50ntFlanks | 12419 | 12450 | 32 | GATGTCGGCTCTTCCTATCATTGTGAAGCAGA | 724.0 | 4.6 | 0.6 | 7.4 | 1.5E-13 | 6.0E-13 | MB231/MB468 | CYTO |
| >hg38_21_+_8433172_8446622_RNA45SN1_45S_50ntFlanks@11589.11607.19\|CGGACCAGGGGAATCCGAC | >hg38_21_+_8433172_8446622_RNA45SN1_45S_50ntFlanks | 11589 | 11607 | 19 | CGGACCAGGGGAATCCGAC | 681.4 | 4.2 | 0.6 | 7.4 | 1.2E-13 | 6.0E-13 | BT20/MB468 | MITO |
| >hg38_21_+_8433172_8446622_RNA45SN1_45S_50ntFlanks@12181.12220.40\|TCAAACGGTAACGCAGGTGTCCTAAGGCGAGCTCAGGGAG | >hg38_21_+_8433172_8446622_RNA45SN1_45S_50ntFlanks | 12181 | 12220 | 40 | TCAAACGGTAACGCAGGTGTCCTAAGGCGAGCTCAGGGAG | 198.9 | -11.4 | 1.6 | -7.4 | 2.0E-13 | 6.0E-13 | MB231/MB468 | MITO |
| >hg38_21_+_8433172_8446622_RNA45SN1_45S_50ntFlanks@10136.10157.22\|GCGGCGGCGGCGGGGGTGTGGG | >hg38_21_+_8433172_8446622_RNA45SN1_45S_50ntFlanks | 10136 | 10157 | 22 | GCGGCGGCGGCGGGGGTGTGGG | 236.9 | -11.8 | 1.6 | -7.4 | 1.3E-13 | 6.0E-13 | MB231/MB468 | MP |
| >hg38_21_+_8433172_8446622_RNA45SN1_45S_50ntFlanks@11317.11344.28\|ACCCCACGTCTCGTCGCGCGCGCGTCCG | >hg38_21_+_8433172_8446622_RNA45SN1_45S_50ntFlanks | 11317 | 11344 | 28 | ACCCCACGTCTCGTCGCGCGCGCGTCCG | 24153.4 | -9.5 | 1.3 | -7.4 | 1.6E-13 | 6.0E-13 | MB231/MB468 | NUC |
| >hg38_21_+_8433172_8446622_RNA45SN1_45S_50ntFlanks@11317.11336.20\|ACCCCACGTCTCGTCGCGCG | >hg38_21_+_8433172_8446622_RNA45SN1_45S_50ntFlanks | 11317 | 11336 | 20 | ACCCCACGTCTCGTCGCGCG | 1412.3 | -9.8 | 1.3 | -7.4 | 1.6E-13 | 6.0E-13 | MB231/MB468 | NUC |
| >hg38_21_+_8433172_8446622_RNA45SN1_45S_50ntFlanks@7989.8010.22\|AGACGTGGCGACCCGCTGAATT | >hg38_21_+_8433172_8446622_RNA45SN1_45S_50ntFlanks | 7989 | 8010 | 22 | AGACGTGGCGACCCGCTGAATT | 3024.1 | 7.2 | 1.0 | 7.4 | 1.5E-13 | 6.0E-13 | MB231/MB468 | CYTO |

<table>
<tr><td></td><td></td><td></td><td></td><td></td><td></td><td></td><td></td><td></td><td></td><td></td><td></td><td></td><td></td></tr>
<tr><td>>hg38_21_+_8433172_8446622_RNA45SN1_45S_50ntFlanks@7985.8005.21|GATCAGACGTGGCGACCCGCT</td><td>>hg38_21_+_8433172_8446622_RNA45SN1_45S_50ntFlanks</td><td>7985</td><td>8005</td><td>21</td><td>GATCAGACGTGGCGACCCGCT</td><td>341.7</td><td>-10.5</td><td>1.4</td><td>-7.4</td><td>1.2E-13</td><td>6.1E-13</td><td>BT20/MB468</td><td>MITO</td></tr>
<tr><td>>hg38_21_+_8433172_8446622_RNA45SN1_45S_50ntFlanks@9572.9593.22|CAAATCGGTCGTCCGACCTGGG</td><td>>hg38_21_+_8433172_8446622_RNA45SN1_45S_50ntFlanks</td><td>9572</td><td>9593</td><td>22</td><td>CAAATCGGTCGTCCGACCTGGG</td><td>144.3</td><td>-9.9</td><td>1.3</td><td>-7.4</td><td>1.5E-13</td><td>6.1E-13</td><td>MB231/MB468</td><td>CYTO</td></tr>
<tr><td>>hg38_21_+_8433172_8446622_RNA45SN1_45S_50ntFlanks@10458.10495.38|ATGGCCTCCGTTGCCCTCGGCCGATCGAAAGGGAGTCG</td><td>>hg38_21_+_8433172_8446622_RNA45SN1_45S_50ntFlanks</td><td>10458</td><td>10495</td><td>38</td><td>ATGGCCTCCGTTGCCCTCGGCCGATCGAAAGGGAGTCG</td><td>231.7</td><td>-9.9</td><td>1.3</td><td>-7.4</td><td>1.2E-13</td><td>6.1E-13</td><td>BT20/MB468</td><td>MITO</td></tr>
<tr><td>>hg38_21_+_8433172_8446622_RNA45SN1_45S_50ntFlanks@4163.4184.22|CATCCAAGGAAGGCAGCAGGCG</td><td>>hg38_21_+_8433172_8446622_RNA45SN1_45S_50ntFlanks</td><td>4163</td><td>4184</td><td>22</td><td>CATCCAAGGAAGGCAGCAGGCG</td><td>275.1</td><td>-10.2</td><td>1.4</td><td>-7.4</td><td>1.2E-13</td><td>6.1E-13</td><td>BT20/MB468</td><td>MITO</td></tr>
<tr><td>>MI0000456|hsa-mir-140&WithFlank&16|+|69933075|69933186@68.90.23(+1U)&[MIMAT0004597&hsa-miR-140-3p&offsets|0|+2(+1U);m-45&16|+|69933143|69933164&offsets|-1|0(+1U)]|TACCACAGGGTAGAACCACGGACT</td><td>>MI0000456|hsa-mir-140&WithFlank&16|+|69933075|69933186</td><td>68</td><td>90</td><td>23(+1U)</td><td>TACCACAGGGTAGAACCACGGACT</td><td>513.1</td><td>10.7</td><td>1.4</td><td>7.4</td><td>1.5E-13</td><td>6.1E-13</td><td>MB231/MB468</td><td>CYTO</td></tr>
<tr><td>>hg38_21_+_8433172_8446622_RNA45SN1_45S_50ntFlanks@3984.4026.43|GCGGCTTTGGTGACTCTAGATAACCTCGGGCCGATCGCACGCC</td><td>>hg38_21_+_8433172_8446622_RNA45SN1_45S_50ntFlanks</td><td>3984</td><td>4026</td><td>43</td><td>GCGGCTTTGGTGACTCTAGATAACCTCGGGCCGATCGCACGCC</td><td>198.0</td><td>-10.6</td><td>1.4</td><td>-7.4</td><td>1.6E-13</td><td>6.1E-13</td><td>MB231/MB468</td><td>NUC</td></tr>
<tr><td>>hg38_21_+_8433172_8446622_RNA45SN1_45S_50ntFlanks@8154.8175.22|CCGCTCGTGGGGGGCCCAAGTC</td><td>>hg38_21_+_8433172_8446622_RNA45SN1_45S_50ntFlanks</td><td>8154</td><td>8175</td><td>22</td><td>CCGCTCGTGGGGGGCCCAAGTC</td><td>365.3</td><td>-10.6</td><td>1.4</td><td>-7.4</td><td>1.2E-13</td><td>6.1E-13</td><td>BT20/MB468</td><td>MITO</td></tr>
<tr><td>>hg38_21_+_8433172_8446622_RNA45SN1_45S_50ntFlanks@4976.5001.26|CCGGACACGGACAGGATTGACAGATT</td><td>>hg38_21_+_8433172_8446622_RNA45SN1_45S_50ntFlanks</td><td>4976</td><td>5001</td><td>26</td><td>CCGGACACGGACAGGATTGACAGATT</td><td>672.1</td><td>9.1</td><td>1.2</td><td>7.4</td><td>1.6E-13</td><td>6.1E-13</td><td>BT20/MB468</td><td>CYTO</td></tr>
<tr><td>>MI0000095|hsa-mir-93&WithFlank&7|-|100093762|100093853@17.37.21&[MIMAT0000093&hsa-miR-93-5p&offsets|0|-2;m-29&7|-|100093815|100093837&offsets|0|-2]|CAAAGTGCTGTTCGTGCAGGT</td><td>>MI0000095|hsa-mir-93&WithFlank&7|-|100093762|100093853</td><td>17</td><td>37</td><td>21</td><td>CAAAGTGCTGTTCGTGCAGGT</td><td>620.4</td><td>5.5</td><td>0.7</td><td>7.4</td><td>1.2E-13</td><td>6.1E-13</td><td>BT20/MB468</td><td>MITO</td></tr>
<tr><td>>MI0000087|hsa-mir-29a&WithFlank&7|-|130876741|130876816@47.67.21(+1C)&[MIMAT0000086&hsa-miR-29a-3p&offsets|-1|-2(+1C);m-26&7|-|130876749|130876769&offsets|-1|-1(+1C)]|CTAGCACCATCTGAAATCGGTC</td><td>>MI0000087|hsa-mir-29a&WithFlank&7|-|130876741|130876816</td><td>47</td><td>67</td><td>21(+1C)</td><td>CTAGCACCATCTGAAATCGGTC</td><td>3490.8</td><td>6.3</td><td>0.8</td><td>7.4</td><td>1.3E-13</td><td>6.1E-13</td><td>MB231/MB468</td><td>MP</td></tr>
<tr><td>>MI0033423|hsa-mir-10399&WithFlank&7|-|139044094|139044163@43.63.21(+1U)&[MIMAT0041630&hsa-miR-10399-3p&offsets|-1|-1(+1U);TJU_CMC_MD2.ID03061.3p-miR&offsets|+3|+2(+1U);m-2657&7|-|139044103|139044124&offsets|+3|+2(+1U)]|TCTCTCGGACAAGCTGTAGGTT</td><td>>MI0033423|hsa-mir-10399&WithFlank&7|-|139044094|139044163</td><td>43</td><td>63</td><td>21(+1U)</td><td>TCTCTCGGACAAGCTGTAGGTT</td><td>382.6</td><td>9.7</td><td>1.3</td><td>7.4</td><td>1.5E-13</td><td>6.1E-13</td><td>MB231/MB468</td><td>CYTO</td></tr>
<tr><td>>hg38_21_+_8433172_8446622_RNA45SN1_45S_50ntFlanks@10402.10435.34|AACATGGGTCAGTCGGTCCTGAGAGATGGGCGAG</td><td>>hg38_21_+_8433172_8446622_RNA45SN1_45S_50ntFlanks</td><td>10402</td><td>10435</td><td>34</td><td>AACATGGGTCAGTCGGTCCTGAGAGATGGGCGAG</td><td>3712.2</td><td>-11.2</td><td>1.5</td><td>-7.3</td><td>2.3E-13</td><td>6.1E-13</td><td>BT20/MB468</td><td>NUC</td></tr>
<tr><td>>MI0000065|hsa-let-7d&WithFlank&9|+|94178828|94178926@14.34.21&[MIMAT0000065&hsa-let-7d-5p&offsets|0|-1;m-90&9|+|94178841|94178862&offsets|0|-1]|AGAGGTAGTAGGTTGCATAGT</td><td>>MI0000065|hsa-let-7d&WithFlank&9|+|94178828|94178926</td><td>14</td><td>34</td><td>21</td><td>AGAGGTAGTAGGTTGCATAGT</td><td>1247.8</td><td>5.0</td><td>0.7</td><td>7.4</td><td>1.6E-13</td><td>6.1E-13</td><td>MB231/MB468</td><td>NUC</td></tr>
<tr><td>>hg38_21_+_8433172_8446622_RNA45SN1_45S_50ntFlanks@11766.11790.25|ATCTAATTAGTGACGCGCATGAATG</td><td>>hg38_21_+_8433172_8446622_RNA45SN1_45S_50ntFlanks</td><td>11766</td><td>11790</td><td>25</td><td>ATCTAATTAGTGACGCGCATGAATG</td><td>300.6</td><td>-12.1</td><td>1.6</td><td>-7.4</td><td>1.3E-13</td><td>6.2E-13</td><td>MB231/MB468</td><td>MP</td></tr>
<tr><td>>hg38_21_+_8433172_8446622_RNA45SN1_45S_50ntFlanks@12001.12043.43|CTGCGGGCCGCCGGTGAAATACCACTACTCTGATCGTTTTTTC</td><td>>hg38_21_+_8433172_8446622_RNA45SN1_45S_50ntFlanks</td><td>12001</td><td>12043</td><td>43</td><td>CTGCGGGCCGCCGGTGAAATACCACTACTCTGATCGTTTTTTC</td><td>567.8</td><td>-12.1</td><td>1.6</td><td>-7.4</td><td>1.6E-13</td><td>6.2E-13</td><td>MB231/MB468</td><td>NUC</td></tr>
<tr><td>>hg38_21_+_8433172_8446622_RNA45SN1_45S_50ntFlanks@12927.12953.27|CCTGGCGCTAAACCATTCGTAGACGAC</td><td>>hg38_21_+_8433172_8446622_RNA45SN1_45S_50ntFlanks</td><td>12927</td><td>12953</td><td>27</td><td>CCTGGCGCTAAACCATTCGTAGACGAC</td><td>835.7</td><td>-9.2</td><td>1.3</td><td>-7.3</td><td>2.0E-13</td><td>6.2E-13</td><td>MB231/MB468</td><td>MITO</td></tr>
<tr><td>>hg38_21_+_8433172_8446622_RNA45SN1_45S_50ntFlanks@5471.5516.46|CACGGCCCTGGCGGAGCGCTGAGAAGACGGTCGAACTTGACTATCT</td><td>>hg38_21_+_8433172_8446622_RNA45SN1_45S_50ntFlanks</td><td>5471</td><td>5516</td><td>46</td><td>CACGGCCCTGGCGGAGCGCTGAGAAGACGGTCGAACTTGACTATCT</td><td>12575.8</td><td>7.5</td><td>1.0</td><td>7.4</td><td>1.6E-13</td><td>6.2E-13</td><td>MB231/MB468</td><td>CYTO</td></tr>
<tr><td>>hg38_21_+_8433172_8446622_RNA45SN1_45S_50ntFlanks@6721.6750.30|AATTGCAGGACACATTGATCATCGACACTT</td><td>>hg38_21_+_8433172_8446622_RNA45SN1_45S_50ntFlanks</td><td>6721</td><td>6750</td><td>30</td><td>AATTGCAGGACACATTGATCATCGACACTT</td><td>1039.6</td><td>-8.1</td><td>1.1</td><td>-7.3</td><td>2.3E-13</td><td>6.2E-13</td><td>BT20/MB468</td><td>NUC</td></tr>
<tr><td>>hg38_21_+_8433172_8446622_RNA45SN1_45S_50ntFlanks@8194.8213.20|CCCGTGGACGGTGTGAGGCC</td><td>>hg38_21_+_8433172_8446622_RNA45SN1_45S_50ntFlanks</td><td>8194</td><td>8213</td><td>20</td><td>CCCGTGGACGGTGTGAGGCC</td><td>4857.2</td><td>-9.1</td><td>1.2</td><td>-7.3</td><td>2.3E-13</td><td>6.2E-13</td><td>BT20/MB468</td><td>NUC</td></tr>
<tr><td>>hg38_21_+_8433172_8446622_RNA45SN1_45S_50ntFlanks@8012.8046.35|AAGCATATTAGTCAGCGGAGGAGAAGAAACTAACC</td><td>>hg38_21_+_8433172_8446622_RNA45SN1_45S_50ntFlanks</td><td>8012</td><td>8046</td><td>35</td><td>AAGCATATTAGTCAGCGGAGGAGAAGAAACTAACC</td><td>225.4</td><td>8.9</td><td>1.2</td><td>7.4</td><td>1.6E-13</td><td>6.2E-13</td><td>MB231/MB468</td><td>CYTO</td></tr>
<tr><td>>hg38_21_+_8433172_8446622_RNA45SN1_45S_50ntFlanks@4367.4389.23|CAGTTAAAAAGCTCGTAGTTGGA</td><td>>hg38_21_+_8433172_8446622_RNA45SN1_45S_50ntFlanks</td><td>4367</td><td>4389</td><td>23</td><td>CAGTTAAAAAGCTCGTAGTTGGA</td><td>841.3</td><td>-9.9</td><td>1.3</td><td>-7.4</td><td>1.6E-13</td><td>6.2E-13</td><td>MB231/MB468</td><td>NUC</td></tr>
<tr><td>>hg38_21_+_8433172_8446622_RNA45SN1_45S_50ntFlanks@12456.12485.30|CCAAGCGTTGGATTGTTCACCCACTAATAG</td><td>>hg38_21_+_8433172_8446622_RNA45SN1_45S_50ntFlanks</td><td>12456</td><td>12485</td><td>30</td><td>CCAAGCGTTGGATTGTTCACCCACTAATAG</td><td>5428.4</td><td>-8.9</td><td>1.2</td><td>-7.4</td><td>1.6E-13</td><td>6.2E-13</td><td>MB231/MB468</td><td>NUC</td></tr>
<tr><td>>hg38_21_+_8433172_8446622_RNA45SN1_45S_50ntFlanks@8113.8134.22|GGCGCGGGACATGTGGCGTACG</td><td>>hg38_21_+_8433172_8446622_RNA45SN1_45S_50ntFlanks</td><td>8113</td><td>8134</td><td>22</td><td>GGCGCGGGACATGTGGCGTACG</td><td>134.4</td><td>-10.9</td><td>1.5</td><td>-7.4</td><td>1.3E-13</td><td>6.2E-13</td><td>MB231/MB468</td><td>MP</td></tr>
<tr><td>>hg38_21_+_8433172_8446622_RNA45SN1_45S_50ntFlanks@4443.4470.28|CCCTTGCCTCTCGGCGCCCCCTCGATGC</td><td>>hg38_21_+_8433172_8446622_RNA45SN1_45S_50ntFlanks</td><td>4443</td><td>4470</td><td>28</td><td>CCCTTGCCTCTCGGCGCCCCCTCGATGC</td><td>271.6</td><td>-10.2</td><td>1.4</td><td>-7.4</td><td>1.2E-13</td><td>6.2E-13</td><td>BT20/MB468</td><td>MITO</td></tr>
<tr><td>>MI0000089|hsa-mir-31&WithFlank&9|-|21512109|21512191@14.34.21&[MIMAT0000089&hsa-miR-31-5p&offsets|0|0;m-360&9|-|21512157|21512178&offsets|0|-1]|AGGCAAGATGCTGGCATAGCT</td><td>>MI0000089|hsa-mir-31&WithFlank&9|-|21512109|21512191</td><td>14</td><td>34</td><td>21</td><td>AGGCAAGATGCTGGCATAGCT</td><td>382.3</td><td>10.0</td><td>1.4</td><td>7.4</td><td>1.6E-13</td><td>6.2E-13</td><td>BT20/MB468</td><td>CYTO</td></tr>
<tr><td>>hg38_21_+_8433172_8446622_RNA45SN1_45S_50ntFlanks@5472.5516.45|ACGGCCCTGGCGGAGCGCTGAGAAGACGGTCGAACTTGACTATCT</td><td>>hg38_21_+_8433172_8446622_RNA45SN1_45S_50ntFlanks</td><td>5472</td><td>5516</td><td>45</td><td>ACGGCCCTGGCGGAGCGCTGAGAAGACGGTCGAACTTGACTATCT</td><td>20720.7</td><td>10.9</td><td>1.5</td><td>7.4</td><td>1.2E-13</td><td>6.2E-13</td><td>BT20/MB468</td><td>MITO</td></tr>
<tr><td>>hg38_21_+_8433172_8446622_RNA45SN1_45S_50ntFlanks@10411.10442.32|CAGTCGGTCCTGAGAGATGGGCGAGCGCCGTT</td><td>>hg38_21_+_8433172_8446622_RNA45SN1_45S_50ntFlanks</td><td>10411</td><td>10442</td><td>32</td><td>CAGTCGGTCCTGAGAGATGGGCGAGCGCCGTT</td><td>1242.0</td><td>-6.5</td><td>0.9</td><td>-7.3</td><td>2.0E-13</td><td>6.2E-13</td><td>MB231/MB468</td><td>MITO</td></tr>
<tr><td>>hg38_21_+_8433172_8446622_RNA45SN1_45S_50ntFlanks@9566.9597.32|GACGTGCAAATCGGTCGTCCGACCTGGGTATA</td><td>>hg38_21_+_8433172_8446622_RNA45SN1_45S_50ntFlanks</td><td>9566</td><td>9597</td><td>32</td><td>GACGTGCAAATCGGTCGTCCGACCTGGGTATA</td><td>1834.4</td><td>-3.5</td><td>0.5</td><td>-7.4</td><td>1.2E-13</td><td>6.2E-13</td><td>BT20/MB468</td><td>MITO</td></tr>
<tr><td>>hg38_21_+_8433172_8446622_RNA45SN1_45S_50ntFlanks@6778.6806.29|CCCGGGGCTACGCCTGTCTGAGCGTCGCT</td><td>>hg38_21_+_8433172_8446622_RNA45SN1_45S_50ntFlanks</td><td>6778</td><td>6806</td><td>29</td><td>CCCGGGGCTACGCCTGTCTGAGCGTCGCT</td><td>7386.5</td><td>5.6</td><td>0.8</td><td>7.4</td><td>1.2E-13</td><td>6.2E-13</td><td>BT20/MB468</td><td>MITO</td></tr>
<tr><td>>hg38_21_+_8433172_8446622_RNA45SN1_45S_50ntFlanks@10390.10424.35|GAACAGCAGTTGAACATGGGTCAGTCGGTCCTGAG</td><td>>hg38_21_+_8433172_8446622_RNA45SN1_45S_50ntFlanks</td><td>10390</td><td>10424</td><td>35</td><td>GAACAGCAGTTGAACATGGGTCAGTCGGTCCTGAG</td><td>966.3</td><td>7.5</td><td>1.0</td><td>7.4</td><td>1.6E-13</td><td>6.2E-13</td><td>MB231/MB468</td><td>CYTO</td></tr>
<tr><td>>hg38_21_+_8433172_8446622_RNA45SN1_45S_50ntFlanks@10411.10435.25|CAGTCGGTCCTGAGAGATGGGCGAG</td><td>>hg38_21_+_8433172_8446622_RNA45SN1_45S_50ntFlanks</td><td>10411</td><td>10435</td><td>25</td><td>CAGTCGGTCCTGAGAGATGGGCGAG</td><td>703.4</td><td>-11.5</td><td>1.6</td><td>-7.4</td><td>1.2E-13</td><td>6.2E-13</td><td>BT20/MB468</td><td>MITO</td></tr>
<tr><td>>hg38_21_+_8433172_8446622_RNA45SN1_45S_50ntFlanks@8201.8223.23|ACGGTGTGAGGCCGGTAGCGGCC</td><td>>hg38_21_+_8433172_8446622_RNA45SN1_45S_50ntFlanks</td><td>8201</td><td>8223</td><td>23</td><td>ACGGTGTGAGGCCGGTAGCGGCC</td><td>132.3</td><td>-10.9</td><td>1.5</td><td>-7.4</td><td>1.4E-13</td><td>6.2E-13</td><td>MB231/MB468</td><td>MP</td></tr>
<tr><td>>hg38_21_+_8433172_8446622_RNA45SN1_45S_50ntFlanks@10116.10135.20|TGCGTCGGAGGGCGGCGGCG</td><td>>hg38_21_+_8433172_8446622_RNA45SN1_45S_50ntFlanks</td><td>10116</td><td>10135</td><td>20</td><td>TGCGTCGGAGGGCGGCGGCG</td><td>366.9</td><td>-12.4</td><td>1.7</td><td>-7.4</td><td>1.4E-13</td><td>6.3E-13</td><td>MB231/MB468</td><td>MP</td></tr>
<tr><td>>hg38_21_+_8433172_8446622_RNA45SN1_45S_50ntFlanks@8157.8181.25|CTCGTGGGGGGCCCAAGTCCTTCTG</td><td>>hg38_21_+_8433172_8446622_RNA45SN1_45S_50ntFlanks</td><td>8157</td><td>8181</td><td>25</td><td>CTCGTGGGGGGCCCAAGTCCTTCTG</td><td>1757.0</td><td>-9.6</td><td>1.3</td><td>-7.4</td><td>1.4E-13</td><td>6.3E-13</td><td>MB231/MB468</td><td>MP</td></tr>
<tr><td>>hg38_21_+_8433172_8446622_RNA45SN1_45S_50ntFlanks@6732.6774.43|ACATTGATCATCGACACTTCGAACGCACTTGCGGCCCCGGGTT</td><td>>hg38_21_+_8433172_8446622_RNA45SN1_45S_50ntFlanks</td><td>6732</td><td>6774</td><td>43</td><td>ACATTGATCATCGACACTTCGAACGCACTTGCGGCCCCGGGTT</td><td>472.6</td><td>9.4</td><td>1.3</td><td>7.4</td><td>1.6E-13</td><td>6.3E-13</td><td>MB231/MB468</td><td>CYTO</td></tr>
<tr><td>>hg38_21_+_8433172_8446622_RNA45SN1_45S_50ntFlanks@3830.3860.31|GCTCGCTCCTCTCCTACTTGGATAACTGTGG</td><td>>hg38_21_+_8433172_8446622_RNA45SN1_45S_50ntFlanks</td><td>3830</td><td>3860</td><td>31</td><td>GCTCGCTCCTCTCCTACTTGGATAACTGTGG</td><td>1537.9</td><td>-10.4</td><td>1.4</td><td>-7.3</td><td>2.3E-13</td><td>6.3E-13</td><td>BT20/MB468</td><td>NUC</td></tr>
<tr><td>>MI0000088|hsa-mir-30a&WithFlank&6|-|71403545|71403627@12.33.22(+1U)&[MIMAT0000087&hsa-miR-30a-5p&offsets|0|0(+1U);m-15&6|-|71403595|71403616&offsets|0|0(+1U)]|TGTAAACATCCTCGACTGGAAGT</td><td>>MI0000088|hsa-mir-30a&WithFlank&6|-|71403545|71403627</td><td>12</td><td>33</td><td>22(+1U)</td><td>TGTAAACATCCTCGACTGGAAGT</td><td>4560.3</td><td>7.6</td><td>1.0</td><td>7.4</td><td>1.6E-13</td><td>6.3E-13</td><td>BT20/MB468</td><td>CYTO</td></tr>
<tr><td>>hg38_21_+_8433172_8446622_RNA45SN1_45S_50ntFlanks@12365.12384.20|ATAACTGGCTTGTGGCGGCC</td><td>>hg38_21_+_8433172_8446622_RNA45SN1_45S_50ntFlanks</td><td>12365</td><td>12384</td><td>20</td><td>ATAACTGGCTTGTGGCGGCC</td><td>231.0</td><td>-11.7</td><td>1.6</td><td>-7.4</td><td>1.4E-13</td><td>6.3E-13</td><td>MB231/MB468</td><td>MP</td></tr>
<tr><td>>hg38_21_+_8433172_8446622_RNA45SN1_45S_50ntFlanks@11027.11062.36|CTCTCTCTCTCTCTCCCCCGCTCCCCGTCCTCCCCC</td><td>>hg38_21_+_8433172_8446622_RNA45SN1_45S_50ntFlanks</td><td>11027</td><td>11062</td><td>36</td><td>CTCTCTCTCTCTCTCCCCCGCTCCCCGTCCTCCCCC</td><td>353.6</td><td>-12.2</td><td>1.7</td><td>-7.3</td><td>2.1E-13</td><td>6.3E-13</td><td>MB231/MB468</td><td>MITO</td></tr>
<tr><td>>MI0000103|hsa-mir-101-1&WithFlank&1|-|65058428|65058514@52.72.21;MI0000739|hsa-mir-101-2&WithFlank&9|+|4850291|4850381@54.74.21&[MIMAT0000099&hsa-miR-101-3p&offsets|-1|-1;m-17&1|-|65058443|65058462&offsets|-1|0];[MIMAT0000099_1&hsa-miR-101-3p&offsets|-1|-1;m-16&9|+|4850345|4850364&offsets|-1|0]|GTACAGTACTGTGATAACTGA</td><td>>MI0000103|hsa-mir-101-1&WithFlank&1|-|65058428|65058514</td><td>52</td><td>72</td><td>21</td><td>GTACAGTACTGTGATAACTGA</td><td>375.2</td><td>5.7</td><td>0.8</td><td>7.4</td><td>1.6E-13</td><td>6.3E-13</td><td>BT20/MB468</td><td>CYTO</td></tr>
<tr><td>>hg38_21_+_8433172_8446622_RNA45SN1_45S_50ntFlanks@12676.12708.33|AGAATCCCGCCCAGGCGGAACGATACGGCAGCG</td><td>>hg38_21_+_8433172_8446622_RNA45SN1_45S_50ntFlanks</td><td>12676</td><td>12708</td><td>33</td><td>AGAATCCCGCCCAGGCGGAACGATACGGCAGCG</td><td>251.9</td><td>-10.1</td><td>1.4</td><td>-7.4</td><td>1.3E-13</td><td>6.3E-13</td><td>BT20/MB468</td><td>MITO</td></tr>
<tr><td>>hg38_21_+_8433172_8446622_RNA45SN1_45S_50ntFlanks@9027.9044.18|CGTCTCCTCGTGGGGGGG</td><td>>hg38_21_+_8433172_8446622_RNA45SN1_45S_50ntFlanks</td><td>9027</td><td>9044</td><td>18</td><td>CGTCTCCTCGTGGGGGGG</td><td>307.5</td><td>-12.1</td><td>1.6</td><td>-7.4</td><td>1.4E-13</td><td>6.3E-13</td><td>MB231/MB468</td><td>MP</td></tr>
<tr><td>>hg38_21_+_8433172_8446622_RNA45SN1_45S_50ntFlanks@10788.10807.20|CAGCCTCTGGCATGTTGGAA</td><td>>hg38_21_+_8433172_8446622_RNA45SN1_45S_50ntFlanks</td><td>10788</td><td>10807</td><td>20</td><td>CAGCCTCTGGCATGTTGGAA</td><td>127.2</td><td>-10.9</td><td>1.5</td><td>-7.4</td><td>1.4E-13</td><td>6.3E-13</td><td>MB231/MB468</td><td>MP</td></tr>
<tr><td>>hg38_21_+_8433172_8446622_RNA45SN1_45S_50ntFlanks@4297.4320.24|CATTGGAGGGCAAGTCTGGTGCCA</td><td>>hg38_21_+_8433172_8446622_RNA45SN1_45S_50ntFlanks</td><td>4297</td><td>4320</td><td>24</td><td>CATTGGAGGGCAAGTCTGGTGCCA</td><td>165.8</td><td>-11.2</td><td>1.5</td><td>-7.4</td><td>1.4E-13</td><td>6.3E-13</td><td>MB231/MB468</td><td>MP</td></tr>
<tr><td>>trna111_HisGTG_1_-_147774845_147774916@-1A.33.34,trna118_HisGTG_1_-_145396881_145396952@-1A.33.34,trna16_HisGTG_1_+_146544773_146544844@-1A.33.34,trna1_HisGTG_15_+_45493349_45493420@-1A.33.34,trna21_HisGTG_1_+_147753471_147753542@-1A.33.34,trna33_HisGTG_6_+_27125906_27125977@-1A.33.34,trna7_HisGTG_9_-_14433938_14434009@-1A.33.34,trna8_HisGTG_15_-_45492611_45492682@-1A.33.34,trna9_HisGTG_15_-_45490804_45490875@-1A.33.34|AGCCGTGATCGTATAGTGGTTAGTACTCTGCGTT</td><td>>trna111_HisGTG_1_-_147774845_147774916</td><td>-1A</td><td>33</td><td>34</td><td>AGCCGTGATCGTATAGTGGTTAGTACTCTGCGTT</td><td>1082.5</td><td>9.6</td><td>1.3</td><td>7.4</td><td>1.6E-13</td><td>6.3E-13</td><td>MB231/MB468</td><td>CYTO</td></tr>
</table>

<table>
<tr><td>>hg38_21_+_8433172_8446622_RNA45SN1_45S_50ntFlanks@12259.12287.29|CTTGATCTTGATTTTCAGTACGAATACAG</td><td>>hg38_21_+_8433172_8446622_RNA45SN1_45S_50ntFlanks</td><td>12259</td><td>12287</td><td>29</td><td>CTTGATCTTGATTTTCAGTACGAATACAG</td><td>157.3</td><td>-11.2</td><td>1.5</td><td>-7.4</td><td>1.4E-13</td><td>6.3E-13</td><td>MB231/MB468</td><td>MP</td></tr>
<tr><td>>hg38_21_+_8433172_8446622_RNA45SN1_45S_50ntFlanks@5087.5124.38|ACGAGACTCTGGCATGCTAACTAGTTACGCGACCCCCG</td><td>>hg38_21_+_8433172_8446622_RNA45SN1_45S_50ntFlanks</td><td>5087</td><td>5124</td><td>38</td><td>ACGAGACTCTGGCATGCTAACTAGTTACGCGACCCCCG</td><td>238.7</td><td>12.2</td><td>1.6</td><td>7.4</td><td>1.6E-13</td><td>6.4E-13</td><td>BT20/MB468</td><td>CYTO</td></tr>
<tr><td>>hg38_21_+_8433172_8446622_RNA45SN1_45S_50ntFlanks@6721.6739.19|AATTGCAGGACACATTGAT</td><td>>hg38_21_+_8433172_8446622_RNA45SN1_45S_50ntFlanks</td><td>6721</td><td>6739</td><td>19</td><td>AATTGCAGGACACATTGAT</td><td>1673.9</td><td>-7.9</td><td>1.1</td><td>-7.4</td><td>1.7E-13</td><td>6.4E-13</td><td>MB231/MB468</td><td>NUC</td></tr>
<tr><td>>hg38_21_+_8433172_8446622_RNA45SN1_45S_50ntFlanks@10348.10369.22|AAACGAGAACTTTGAAGGCCGA</td><td>>hg38_21_+_8433172_8446622_RNA45SN1_45S_50ntFlanks</td><td>10348</td><td>10369</td><td>22</td><td>AAACGAGAACTTTGAAGGCCGA</td><td>736.6</td><td>4.1</td><td>0.6</td><td>7.4</td><td>1.3E-13</td><td>6.4E-13</td><td>BT20/MB468</td><td>MITO</td></tr>
<tr><td>>hg38_21_+_8433172_8446622_RNA45SN1_45S_50ntFlanks@12367.12384.18|AACTGGCTTGTGGCGGCC</td><td>>hg38_21_+_8433172_8446622_RNA45SN1_45S_50ntFlanks</td><td>12367</td><td>12384</td><td>18</td><td>AACTGGCTTGTGGCGGCC</td><td>264.7</td><td>-5.8</td><td>0.8</td><td>-7.4</td><td>1.6E-13</td><td>6.4E-13</td><td>MB231/MB468</td><td>CYTO</td></tr>
<tr><td>>hg38_21_+_8433172_8446622_RNA45SN1_45S_50ntFlanks@12937.12966.30|AACCATTCGTAGACGACCTGCTTCTGGGTC</td><td>>hg38_21_+_8433172_8446622_RNA45SN1_45S_50ntFlanks</td><td>12937</td><td>12966</td><td>30</td><td>AACCATTCGTAGACGACCTGCTTCTGGGTC</td><td>198.6</td><td>-11.5</td><td>1.6</td><td>-7.4</td><td>1.4E-13</td><td>6.4E-13</td><td>MB231/MB468</td><td>MP</td></tr>
<tr><td>>hg38_21_+_8433172_8446622_RNA45SN1_45S_50ntFlanks@11212.11254.43|CGACTCTGGACGCGAGCCGGGCCCTTCCCGTGGATCGCCCCAG</td><td>>hg38_21_+_8433172_8446622_RNA45SN1_45S_50ntFlanks</td><td>11212</td><td>11254</td><td>43</td><td>CGACTCTGGACGCGAGCCGGGCCCTTCCCGTGGATCGCCCCAG</td><td>721.0</td><td>-12.5</td><td>1.7</td><td>-7.4</td><td>1.7E-13</td><td>6.4E-13</td><td>MB231/MB468</td><td>NUC</td></tr>
<tr><td>>hg38_21_+_8433172_8446622_RNA45SN1_45S_50ntFlanks@5469.5499.31|CCCACGGCCCTGGCGGAGCGCTGAGAAGACG</td><td>>hg38_21_+_8433172_8446622_RNA45SN1_45S_50ntFlanks</td><td>5469</td><td>5499</td><td>31</td><td>CCCACGGCCCTGGCGGAGCGCTGAGAAGACG</td><td>693.7</td><td>-8.8</td><td>1.2</td><td>-7.4</td><td>1.7E-13</td><td>6.4E-13</td><td>MB231/MB468</td><td>NUC</td></tr>
<tr><td>>hg38_21_+_8433172_8446622_RNA45SN1_45S_50ntFlanks@12084.12107.24|TCGCTTCTGGCGCCAAGCGCCCGG</td><td>>hg38_21_+_8433172_8446622_RNA45SN1_45S_50ntFlanks</td><td>12084</td><td>12107</td><td>24</td><td>TCGCTTCTGGCGCCAAGCGCCCGG</td><td>314.5</td><td>-12.2</td><td>1.6</td><td>-7.4</td><td>1.4E-13</td><td>6.4E-13</td><td>MB231/MB468</td><td>MP</td></tr>
<tr><td>>MI0000065|hsa-let-7d&WithFlank&9|+|94178828|94178926@14.34.21&[MIMAT0000065&hsa-let-7d-5p&offsets|0|-1;m-90&9|+|94178841|94178862&offsets|0|-1]|AGAGGTAGTAGGTTGCATAGT</td><td>>MI0000065|hsa-let-7d&WithFlank&9|+|94178828|94178926</td><td>14</td><td>34</td><td>21</td><td>AGAGGTAGTAGGTTGCATAGT</td><td>932.8</td><td>5.7</td><td>0.8</td><td>7.4</td><td>1.6E-13</td><td>6.4E-13</td><td>MB231/MB468</td><td>CYTO</td></tr>
<tr><td>>hg38_21_+_8433172_8446622_RNA45SN1_45S_50ntFlanks@5425.5450.26|TGGATGGTTTAGTGAGGCCCTCGGAT</td><td>>hg38_21_+_8433172_8446622_RNA45SN1_45S_50ntFlanks</td><td>5425</td><td>5450</td><td>26</td><td>TGGATGGTTTAGTGAGGCCCTCGGAT</td><td>333.9</td><td>9.3</td><td>1.3</td><td>7.4</td><td>1.6E-13</td><td>6.4E-13</td><td>MB231/MB468</td><td>CYTO</td></tr>
<tr><td>>hg38_21_+_8433172_8446622_RNA45SN1_45S_50ntFlanks@10758.10785.28|CCCATATCCGCAGCAGGTCTCCAAGGTG</td><td>>hg38_21_+_8433172_8446622_RNA45SN1_45S_50ntFlanks</td><td>10758</td><td>10785</td><td>28</td><td>CCCATATCCGCAGCAGGTCTCCAAGGTG</td><td>220.4</td><td>-11.6</td><td>1.6</td><td>-7.3</td><td>2.1E-13</td><td>6.4E-13</td><td>MB231/MB468</td><td>MITO</td></tr>
<tr><td>>hg38_21_+_8433172_8446622_RNA45SN1_45S_50ntFlanks@11694.11728.35|AAGAAATTCAATGAAGCGCGGGTAAACGGCGGGAG</td><td>>hg38_21_+_8433172_8446622_RNA45SN1_45S_50ntFlanks</td><td>11694</td><td>11728</td><td>35</td><td>AAGAAATTCAATGAAGCGCGGGTAAACGGCGGGAG</td><td>175.0</td><td>-11.3</td><td>1.5</td><td>-7.4</td><td>1.4E-13</td><td>6.5E-13</td><td>MB231/MB468</td><td>MP</td></tr>
<tr><td>>hg38_21_+_8433172_8446622_RNA45SN1_45S_50ntFlanks@5485.5517.33|AGCGCTGAGAAGACGGTCGAACTTGACTATCTA</td><td>>hg38_21_+_8433172_8446622_RNA45SN1_45S_50ntFlanks</td><td>5485</td><td>5517</td><td>33</td><td>AGCGCTGAGAAGACGGTCGAACTTGACTATCTA</td><td>138.4</td><td>-11.0</td><td>1.5</td><td>-7.4</td><td>1.4E-13</td><td>6.5E-13</td><td>MB231/MB468</td><td>MP</td></tr>
<tr><td>>hg38_21_+_8433172_8446622_RNA45SN1_45S_50ntFlanks@10807.10827.21|ACAATGTAGGTAAGGGAAGTC</td><td>>hg38_21_+_8433172_8446622_RNA45SN1_45S_50ntFlanks</td><td>10807</td><td>10827</td><td>21</td><td>ACAATGTAGGTAAGGGAAGTC</td><td>146.9</td><td>-11.1</td><td>1.5</td><td>-7.4</td><td>1.4E-13</td><td>6.5E-13</td><td>MB231/MB468</td><td>MP</td></tr>
<tr><td>>hg38_21_+_8433172_8446622_RNA45SN1_45S_50ntFlanks@9597.9616.20|AGGGGCGAAAGACTAATCGA</td><td>>hg38_21_+_8433172_8446622_RNA45SN1_45S_50ntFlanks</td><td>9597</td><td>9616</td><td>20</td><td>AGGGGCGAAAGACTAATCGA</td><td>166.8</td><td>-11.2</td><td>1.5</td><td>-7.4</td><td>1.4E-13</td><td>6.5E-13</td><td>MB231/MB468</td><td>MP</td></tr>
<tr><td>>hg38_21_+_8433172_8446622_RNA45SN1_45S_50ntFlanks@12078.12095.18|GGGCTCTCGCTTCTGGCG</td><td>>hg38_21_+_8433172_8446622_RNA45SN1_45S_50ntFlanks</td><td>12078</td><td>12095</td><td>18</td><td>GGGCTCTCGCTTCTGGCG</td><td>844.7</td><td>-9.1</td><td>1.2</td><td>-7.4</td><td>1.7E-13</td><td>6.5E-13</td><td>MB231/MB468</td><td>NUC</td></tr>
<tr><td>>hg38_21_+_8433172_8446622_RNA45SN1_45S_50ntFlanks@5446.5466.21|CGGATCGGCCCCGCCGGGGTC</td><td>>hg38_21_+_8433172_8446622_RNA45SN1_45S_50ntFlanks</td><td>5446</td><td>5466</td><td>21</td><td>CGGATCGGCCCCGCCGGGGTC</td><td>37175.3</td><td>-8.1</td><td>1.0</td><td>-7.8</td><td>6.6E-15</td><td>6.5E-13</td><td>BT20/MB468</td><td>MP</td></tr>
<tr><td>>hg38_21_+_8433172_8446622_RNA45SN1_45S_50ntFlanks@4390.4410.21|TCTTGGGAGCGGGCGGGCGGT</td><td>>hg38_21_+_8433172_8446622_RNA45SN1_45S_50ntFlanks</td><td>4390</td><td>4410</td><td>21</td><td>TCTTGGGAGCGGGCGGGCGGT</td><td>262.7</td><td>-11.9</td><td>1.6</td><td>-7.4</td><td>1.4E-13</td><td>6.5E-13</td><td>MB231/MB468</td><td>MP</td></tr>
<tr><td>>hg38_21_+_8433172_8446622_RNA45SN1_45S_50ntFlanks@11752.11780.29|CCAAATGCCTCGTCATCTAATTAGTGACG</td><td>>hg38_21_+_8433172_8446622_RNA45SN1_45S_50ntFlanks</td><td>11752</td><td>11780</td><td>29</td><td>CCAAATGCCTCGTCATCTAATTAGTGACG</td><td>393.1</td><td>-8.2</td><td>1.1</td><td>-7.4</td><td>1.7E-13</td><td>6.5E-13</td><td>BT20/MB468</td><td>CYTO</td></tr>
<tr><td>>hg38_21_+_8433172_8446622_RNA45SN1_45S_50ntFlanks@12437.12473.37|CATTGTGAAGCAGAATTCACCAAGCGTTGGATTGTTC</td><td>>hg38_21_+_8433172_8446622_RNA45SN1_45S_50ntFlanks</td><td>12437</td><td>12473</td><td>37</td><td>CATTGTGAAGCAGAATTCACCAAGCGTTGGATTGTTC</td><td>1234.3</td><td>-6.2</td><td>0.8</td><td>-7.3</td><td>2.2E-13</td><td>6.5E-13</td><td>MB231/MB468</td><td>MITO</td></tr>
<tr><td>>am_trnaMT_ValTAC_MT_+_1602_1670@49.68.20|AACTTAACTTGACCGCTCTG</td><td>>am_trnaMT_ValTAC_MT_+_1602_1670</td><td>49</td><td>68</td><td>20</td><td>AACTTAACTTGACCGCTCTG</td><td>7456.1</td><td>-8.0</td><td>1.1</td><td>-7.3</td><td>2.5E-13</td><td>6.5E-13</td><td>BT20/MB468</td><td>NUC</td></tr>
<tr><td>>MI0000272|hsa-mir-182&WithFlank&7|-|129770377|129770498@29.50.22(+1U)&[MIMAT0000259&hsa-miR-182-5p&offsets|0|-2(+1U);m-22&7|-|129770449|129770470&offsets|0|0(+1U)]|TTTGGCAATGGTAGAACTCACAT</td><td>>MI0000272|hsa-mir-182&WithFlank&7|-|129770377|129770498</td><td>29</td><td>50</td><td>22(+1U)</td><td>TTTGGCAATGGTAGAACTCACAT</td><td>587.8</td><td>4.6</td><td>0.6</td><td>7.7</td><td>2.0E-14</td><td>6.5E-13</td><td>BT20/MB231</td><td>NUC</td></tr>
<tr><td>>hg38_21_+_8433172_8446622_RNA45SN1_45S_50ntFlanks@3866.3898.33|CTAGAGCTAATACATGCCGACGGGCGCTGACCC</td><td>>hg38_21_+_8433172_8446622_RNA45SN1_45S_50ntFlanks</td><td>3866</td><td>3898</td><td>33</td><td>CTAGAGCTAATACATGCCGACGGGCGCTGACCC</td><td>380.2</td><td>-8.1</td><td>1.1</td><td>-7.4</td><td>1.7E-13</td><td>6.6E-13</td><td>MB231/MB468</td><td>CYTO</td></tr>
<tr><td>>MI0000466|hsa-mir-9-1&WithFlank&1|-|156420335|156420435@22.44.23;MI0000468|hsa-mir-9-3&WithFlank&15|+|89368011|89368112@22.44.23;MI0000467|hsa-mir-9-2&WithFlank&5|-|88666847|88666945@22.44.23&[MIMAT0000441&hsa-miR-9-5p&offsets|0|0;m-104&1|-|156420392|156420414&offsets|0|0];[MIMAT0000441_1&hsa-miR-9-5p&offsets|0|0;m-105&15|+|89368032|89368054&offsets|0|0];[MIMAT0000441_2&hsa-miR-9-5p&offsets|0|0;m-104&5|-|88666902|88666924&offsets|0|0]|TCTTTGGTTATCTAGCTGTATGA</td><td>>MI0000466|hsa-mir-9-1&WithFlank&1|-|156420335|156420435</td><td>22</td><td>44</td><td>23</td><td>TCTTTGGTTATCTAGCTGTATGA</td><td>485.2</td><td>5.7</td><td>0.8</td><td>7.3</td><td>2.5E-13</td><td>6.6E-13</td><td>BT20/MB468</td><td>NUC</td></tr>
<tr><td>>hg38_21_+_8433172_8446622_RNA45SN1_45S_50ntFlanks@12349.12376.28|AAAAGTTACCACAGGGATAACTGGCTTG</td><td>>hg38_21_+_8433172_8446622_RNA45SN1_45S_50ntFlanks</td><td>12349</td><td>12376</td><td>28</td><td>AAAAGTTACCACAGGGATAACTGGCTTG</td><td>195.7</td><td>-11.5</td><td>1.6</td><td>-7.4</td><td>1.4E-13</td><td>6.6E-13</td><td>MB231/MB468</td><td>MP</td></tr>
<tr><td>>hg38_21_+_8433172_8446622_RNA45SN1_45S_50ntFlanks@10683.10710.28|CCGGCGGCGTCCGGTGAGCTCTCGCTGG</td><td>>hg38_21_+_8433172_8446622_RNA45SN1_45S_50ntFlanks</td><td>10683</td><td>10710</td><td>28</td><td>CCGGCGGCGTCCGGTGAGCTCTCGCTGG</td><td>353.5</td><td>-11.2</td><td>1.5</td><td>-7.4</td><td>1.7E-13</td><td>6.6E-13</td><td>MB231/MB468</td><td>CYTO</td></tr>
<tr><td>>hg38_21_+_8433172_8446622_RNA45SN1_45S_50ntFlanks@8179.8211.33|CTGATCGAGGCCCAGCCCGTGGACGGTGTGAGG</td><td>>hg38_21_+_8433172_8446622_RNA45SN1_45S_50ntFlanks</td><td>8179</td><td>8211</td><td>33</td><td>CTGATCGAGGCCCAGCCCGTGGACGGTGTGAGG</td><td>290.2</td><td>-10.9</td><td>1.5</td><td>-7.4</td><td>1.7E-13</td><td>6.6E-13</td><td>MB231/MB468</td><td>CYTO</td></tr>
<tr><td>>hg38_21_+_8433172_8446622_RNA45SN1_45S_50ntFlanks@11874.11897.24|AGAAGACCCTGTTGAGCTTGACTC</td><td>>hg38_21_+_8433172_8446622_RNA45SN1_45S_50ntFlanks</td><td>11874</td><td>11897</td><td>24</td><td>AGAAGACCCTGTTGAGCTTGACTC</td><td>1901.6</td><td>-8.4</td><td>1.1</td><td>-7.3</td><td>2.5E-13</td><td>6.6E-13</td><td>BT20/MB468</td><td>NUC</td></tr>
<tr><td>>hg38_21_+_8433172_8446622_RNA45SN1_45S_50ntFlanks@9622.9655.34|CTAGTAGCTGGTTCCCTCCGAAGTTTCCCTCAGG</td><td>>hg38_21_+_8433172_8446622_RNA45SN1_45S_50ntFlanks</td><td>9622</td><td>9655</td><td>34</td><td>CTAGTAGCTGGTTCCCTCCGAAGTTTCCCTCAGG</td><td>146.0</td><td>-11.1</td><td>1.5</td><td>-7.4</td><td>1.4E-13</td><td>6.6E-13</td><td>MB231/MB468</td><td>MP</td></tr>
<tr><td>>hg38_21_+_8433172_8446622_RNA45SN1_45S_50ntFlanks@3991.4013.23|TGGTGACTCTAGATAACCTCGGG</td><td>>hg38_21_+_8433172_8446622_RNA45SN1_45S_50ntFlanks</td><td>3991</td><td>4013</td><td>23</td><td>TGGTGACTCTAGATAACCTCGGG</td><td>646.3</td><td>-9.5</td><td>1.3</td><td>-7.3</td><td>2.5E-13</td><td>6.6E-13</td><td>BT20/MB468</td><td>NUC</td></tr>
<tr><td>>hg38_21_+_8433172_8446622_RNA45SN1_45S_50ntFlanks@4760.4793.34|TCAGATACCGTCGTAGTTCCGACCATAAACGATG</td><td>>hg38_21_+_8433172_8446622_RNA45SN1_45S_50ntFlanks</td><td>4760</td><td>4793</td><td>34</td><td>TCAGATACCGTCGTAGTTCCGACCATAAACGATG</td><td>401.0</td><td>-3.5</td><td>0.5</td><td>-7.4</td><td>1.7E-13</td><td>6.6E-13</td><td>BT20/MB468</td><td>CYTO</td></tr>
<tr><td>>hg38_21_+_8433172_8446622_RNA45SN1_45S_50ntFlanks@6671.6690.20|CTCGGCTCGTGCGTCGATGA</td><td>>hg38_21_+_8433172_8446622_RNA45SN1_45S_50ntFlanks</td><td>6671</td><td>6690</td><td>20</td><td>CTCGGCTCGTGCGTCGATGA</td><td>903.5</td><td>-13.6</td><td>1.9</td><td>-7.3</td><td>2.2E-13</td><td>6.6E-13</td><td>MB231/MB468</td><td>MITO</td></tr>
<tr><td>>hg38_21_+_8433172_8446622_RNA45SN1_45S_50ntFlanks@10347.10374.28|CAAACGAGAACTTTGAAGGCCGAAGTGG</td><td>>hg38_21_+_8433172_8446622_RNA45SN1_45S_50ntFlanks</td><td>10347</td><td>10374</td><td>28</td><td>CAAACGAGAACTTTGAAGGCCGAAGTGG</td><td>362.1</td><td>-5.1</td><td>0.7</td><td>-7.3</td><td>2.2E-13</td><td>6.6E-13</td><td>MB231/MB468</td><td>MITO</td></tr>
<tr><td>>hg38_21_+_8433172_8446622_RNA45SN1_45S_50ntFlanks@11641.11660.20|CGGCGGGTGTTGACGCGATG</td><td>>hg38_21_+_8433172_8446622_RNA45SN1_45S_50ntFlanks</td><td>11641</td><td>11660</td><td>20</td><td>CGGCGGGTGTTGACGCGATG</td><td>5839.5</td><td>-11.7</td><td>1.6</td><td>-7.3</td><td>2.5E-13</td><td>6.6E-13</td><td>BT20/MB468</td><td>NUC</td></tr>
<tr><td>>hg38_21_+_8433172_8446622_RNA45SN1_45S_50ntFlanks@12621.12654.34|GAGCCAATGGGGCGAAGCTACCATCTGTGGGATT</td><td>>hg38_21_+_8433172_8446622_RNA45SN1_45S_50ntFlanks</td><td>12621</td><td>12654</td><td>34</td><td>GAGCCAATGGGGCGAAGCTACCATCTGTGGGATT</td><td>122.0</td><td>-5.0</td><td>0.7</td><td>-7.4</td><td>1.7E-13</td><td>6.6E-13</td><td>MB231/MB468</td><td>CYTO</td></tr>
<tr><td>>hg38_21_+_8433172_8446622_RNA45SN1_45S_50ntFlanks@12597.12614.18|ATTTGGTGTATGTGCTTG</td><td>>hg38_21_+_8433172_8446622_RNA45SN1_45S_50ntFlanks</td><td>12597</td><td>12614</td><td>18</td><td>ATTTGGTGTATGTGCTTG</td><td>306.2</td><td>-12.1</td><td>1.6</td><td>-7.4</td><td>1.5E-13</td><td>6.6E-13</td><td>MB231/MB468</td><td>MP</td></tr>
<tr><td>>hg38_21_+_8433172_8446622_RNA45SN1_45S_50ntFlanks@11661.11685.25|TGATTTCTGCCCAGTGCTCTGAATG</td><td>>hg38_21_+_8433172_8446622_RNA45SN1_45S_50ntFlanks</td><td>11661</td><td>11685</td><td>25</td><td>TGATTTCTGCCCAGTGCTCTGAATG</td><td>3741.2</td><td>-4.5</td><td>0.6</td><td>-7.4</td><td>1.3E-13</td><td>6.7E-13</td><td>BT20/MB468</td><td>MITO</td></tr>
<tr><td>>hg38_21_+_8433172_8446622_RNA45SN1_45S_50ntFlanks@11766.11791.26|ATCTAATTAGTGACGCGCATGAATGG</td><td>>hg38_21_+_8433172_8446622_RNA45SN1_45S_50ntFlanks</td><td>11766</td><td>11791</td><td>26</td><td>ATCTAATTAGTGACGCGCATGAATGG</td><td>249.6</td><td>-10.3</td><td>1.4</td><td>-7.4</td><td>1.7E-13</td><td>6.7E-13</td><td>BT20/MB468</td><td>CYTO</td></tr>
<tr><td>>hg38_21_+_8433172_8446622_RNA45SN1_45S_50ntFlanks@6663.6683.21|GTGGATCACTCGGCTCGTGCG</td><td>>hg38_21_+_8433172_8446622_RNA45SN1_45S_50ntFlanks</td><td>6663</td><td>6683</td><td>21</td><td>GTGGATCACTCGGCTCGTGCG</td><td>153.4</td><td>-11.1</td><td>1.5</td><td>-7.4</td><td>1.5E-13</td><td>6.7E-13</td><td>MB231/MB468</td><td>MP</td></tr>
<tr><td>>hg38_21_+_8433172_8446622_RNA45SN1_45S_50ntFlanks@12045.12070.26|CTGACCCGGTGAGGCGGGGGGGCGAG</td><td>>hg38_21_+_8433172_8446622_RNA45SN1_45S_50ntFlanks</td><td>12045</td><td>12070</td><td>26</td><td>CTGACCCGGTGAGGCGGGGGGGCGAG</td><td>512.4</td><td>-9.3</td><td>1.3</td><td>-7.4</td><td>1.7E-13</td><td>6.7E-13</td><td>MB231/MB468</td><td>NUC</td></tr>
<tr><td>>hg38_21_+_8433172_8446622_RNA45SN1_45S_50ntFlanks@6660.6707.48|GCGGTGGATCACTCGGCTCGTGCGTCGATGAAGAACGCAGCTAGCTGC</td><td>>hg38_21_+_8433172_8446622_RNA45SN1_45S_50ntFlanks</td><td>6660</td><td>6707</td><td>48</td><td>GCGGTGGATCACTCGGCTCGTGCGTCGATGAAGAACGCAGCTAGCTGC</td><td>1353.6</td><td>8.3</td><td>1.1</td><td>7.4</td><td>1.3E-13</td><td>6.7E-13</td><td>BT20/MB468</td><td>MITO</td></tr>
<tr><td>>hg38_21_+_8433172_8446622_RNA45SN1_45S_50ntFlanks@11669.11701.33|GCCCAGTGCTCTGAATGTCAAAGTGAAGAAATT</td><td>>hg38_21_+_8433172_8446622_RNA45SN1_45S_50ntFlanks</td><td>11669</td><td>11701</td><td>33</td><td>GCCCAGTGCTCTGAATGTCAAAGTGAAGAAATT</td><td>354.1</td><td>5.5</td><td>0.7</td><td>7.4</td><td>1.7E-13</td><td>6.7E-13</td><td>MB231/MB468</td><td>CYTO</td></tr>
<tr><td>>hg38_21_+_8433172_8446622_RNA45SN1_45S_50ntFlanks@5084.5117.34|CGAACGAGACTCTGGCATGCTAACTAGTTACGCG</td><td>>hg38_21_+_8433172_8446622_RNA45SN1_45S_50ntFlanks</td><td>5084</td><td>5117</td><td>34</td><td>CGAACGAGACTCTGGCATGCTAACTAGTTACGCG</td><td>565.1</td><td>-13.0</td><td>1.8</td><td>-7.4</td><td>1.5E-13</td><td>6.7E-13</td><td>MB231/MB468</td><td>MP</td></tr>
<tr><td>>hg38_21_+_8433172_8446622_RNA45SN1_45S_50ntFlanks@6732.6773.42|ACATTGATCATCGACACTTCGAACGCACTTGCGGCCCCGGGT</td><td>>hg38_21_+_8433172_8446622_RNA45SN1_45S_50ntFlanks</td><td>6732</td><td>6773</td><td>42</td><td>ACATTGATCATCGACACTTCGAACGCACTTGCGGCCCCGGGT</td><td>668.1</td><td>7.4</td><td>1.0</td><td>7.4</td><td>1.7E-13</td><td>6.7E-13</td><td>MB231/MB468</td><td>CYTO</td></tr>
<tr><td>>MI0000264|hsa-mir-7-2&WithFlank&15|+|88611819|88611940@38.60.23(+1A);MI0000265|hsa-mir-7-3&WithFlank&19|+|4770664|4770785@37.59.23(+1A);MI0000263|hsa-mir-7-1&WithFlank&9|-|83969742|83969863@30.52.23(+1A)&[MIMAT0000252&hsa-miR-7-5p&offsets|0|-1(+1A);m-400&15|+|88611856|88611879&offsets|0|-1(+1A)];[MIMAT0000252_1&hsa-miR-7-5p&offsets|0|-1(+1A);m-402&19|+|4770700|4770723&offsets|0|-1(+1A)];[MIMAT0000252_2&hsa-miR-7-5p&offsets|0|-1(+1A);m-401&9|-|83969811|83969834&offsets|0|-1(+1A)]|TGGAAGACTAGTGATTTTGTTGTA</td><td>>MI0000264|hsa-mir-7-2&WithFlank&15|+|88611819|88611940</td><td>38</td><td>60</td><td>23(+1A)</td><td>TGGAAGACTAGTGATTTTGTTGTA</td><td>642.5</td><td>4.1</td><td>0.6</td><td>7.3</td><td>2.5E-13</td><td>6.7E-13</td><td>BT20/MB468</td><td>NUC</td></tr>
<tr><td>>MI0000088|hsa-mir-30a&WithFlank&6|-|71403545|71403627@53.73.21(+2U)&[MIMAT0000088&hsa-miR-30a-3p&offsets|0|-1(+2U);m-38&6|-|71403554|71403575&offsets|0|-1(+2U)]|CTTTCAGTCGGATGTTTGCAGTT</td><td>>MI0000088|hsa-mir-30a&WithFlank&6|-|71403545|71403627</td><td>53</td><td>73</td><td>21(+2U)</td><td>CTTTCAGTCGGATGTTTGCAGTT</td><td>830.5</td><td>8.9</td><td>1.2</td><td>7.4</td><td>1.7E-13</td><td>6.7E-13</td><td>MB231/MB468</td><td>CYTO</td></tr>
<tr><td>>hg38_21_+_8433172_8446622_RNA45SN1_45S_50ntFlanks@5073.5109.37|AATTCCGATAACGAACGAGACTCTGGCATGCTAACTA</td><td>>hg38_21_+_8433172_8446622_RNA45SN1_45S_50ntFlanks</td><td>5073</td><td>5109</td><td>37</td><td>AATTCCGATAACGAACGAGACTCTGGCATGCTAACTA</td><td>359.1</td><td>-12.3</td><td>1.7</td><td>-7.3</td><td>2.2E-13</td><td>6.7E-13</td><td>MB231/MB468</td><td>MITO</td></tr>
</table>

<table>
<tr><td></td><td></td><td></td><td></td><td></td><td></td><td></td><td></td><td></td><td></td><td></td><td></td><td></td><td></td></tr>
<tr><td>>hg38_21_+_8433172_8446622_RNA45SN1_45S_50ntFlanks@4918.4941.24|ACCACCAGGAGTGGAGCCTGCGGC</td><td>>hg38_21_+_8433172_8446622_RNA45SN1_45S_50ntFlanks</td><td>4918</td><td>4941</td><td>24</td><td>ACCACCAGGAGTGGAGCCTGCGGC</td><td>552.5</td><td>-12.9</td><td>1.8</td><td>-7.3</td><td>2.2E-13</td><td>6.7E-13</td><td>MB231/MB468</td><td>MITO</td></tr>
<tr><td>>hg38_21_+_8433172_8446622_RNA45SN1_45S_50ntFlanks@11785.11802.18|TGAATGGATGAACGAGAT</td><td>>hg38_21_+_8433172_8446622_RNA45SN1_45S_50ntFlanks</td><td>11785</td><td>11802</td><td>18</td><td>TGAATGGATGAACGAGAT</td><td>539.0</td><td>-12.9</td><td>1.8</td><td>-7.4</td><td>1.5E-13</td><td>6.7E-13</td><td>MB231/MB468</td><td>MP</td></tr>
<tr><td>>hg38_21_+_8433172_8446622_RNA45SN1_45S_50ntFlanks@10389.10430.42|TGAACAGCAGTTGAACATGGGTCAGTCGGTCCTGAGAGATGG</td><td>>hg38_21_+_8433172_8446622_RNA45SN1_45S_50ntFlanks</td><td>10389</td><td>10430</td><td>42</td><td>TGAACAGCAGTTGAACATGGGTCAGTCGGTCCTGAGAGATGG</td><td>323.9</td><td>-12.1</td><td>1.7</td><td>-7.3</td><td>2.2E-13</td><td>6.7E-13</td><td>MB231/MB468</td><td>MITO</td></tr>
<tr><td>>hg38_1_-_228634819_228635039_RNA5S12_5S_50ntFlanks@152.169.18|TACCGGGTGCTGTAGGCT</td><td>>hg38_1_-_228634819_228635039_RNA5S12_5S_50ntFlanks</td><td>152</td><td>169</td><td>18</td><td>TACCGGGTGCTGTAGGCT</td><td>1226.3</td><td>5.5</td><td>0.7</td><td>7.4</td><td>1.7E-13</td><td>6.7E-13</td><td>MB231/MB468</td><td>CYTO</td></tr>
<tr><td>>hg38_21_+_8433172_8446622_RNA45SN1_45S_50ntFlanks@5420.5443.24|CCGATTGGATGGTTTAGTGAGGCC</td><td>>hg38_21_+_8433172_8446622_RNA45SN1_45S_50ntFlanks</td><td>5420</td><td>5443</td><td>24</td><td>CCGATTGGATGGTTTAGTGAGGCC</td><td>4564.1</td><td>-6.5</td><td>0.9</td><td>-7.4</td><td>1.7E-13</td><td>6.8E-13</td><td>MB231/MB468</td><td>CYTO</td></tr>
<tr><td>>hg38_21_+_8433172_8446622_RNA45SN1_45S_50ntFlanks@6773.6797.25|TTCCTCCCGGGGCTACGCCTGTCTG</td><td>>hg38_21_+_8433172_8446622_RNA45SN1_45S_50ntFlanks</td><td>6773</td><td>6797</td><td>25</td><td>TTCCTCCCGGGGCTACGCCTGTCTG</td><td>157.7</td><td>-11.2</td><td>1.5</td><td>-7.4</td><td>1.5E-13</td><td>6.8E-13</td><td>MB231/MB468</td><td>MP</td></tr>
<tr><td>>hg38_21_+_8433172_8446622_RNA45SN1_45S_50ntFlanks@9563.9597.35|CCTGACGTGCAAATCGGTCGTCCGACCTGGGTATA</td><td>>hg38_21_+_8433172_8446622_RNA45SN1_45S_50ntFlanks</td><td>9563</td><td>9597</td><td>35</td><td>CCTGACGTGCAAATCGGTCGTCCGACCTGGGTATA</td><td>263.3</td><td>-11.9</td><td>1.6</td><td>-7.4</td><td>1.5E-13</td><td>6.8E-13</td><td>MB231/MB468</td><td>MP</td></tr>
<tr><td>>hg38_21_+_8433172_8446622_RNA45SN1_45S_50ntFlanks@6719.6740.22|TGAATTGCAGGACACATTGATC</td><td>>hg38_21_+_8433172_8446622_RNA45SN1_45S_50ntFlanks</td><td>6719</td><td>6740</td><td>22</td><td>TGAATTGCAGGACACATTGATC</td><td>3443.5</td><td>-5.5</td><td>0.8</td><td>-7.4</td><td>1.7E-13</td><td>6.8E-13</td><td>MB231/MB468</td><td>CYTO</td></tr>
<tr><td>>hg38_21_+_8433172_8446622_RNA45SN1_45S_50ntFlanks@11210.11227.18|GGCGACTCTGGACGCGAG</td><td>>hg38_21_+_8433172_8446622_RNA45SN1_45S_50ntFlanks</td><td>11210</td><td>11227</td><td>18</td><td>GGCGACTCTGGACGCGAG</td><td>367.0</td><td>-11.2</td><td>1.5</td><td>-7.4</td><td>1.7E-13</td><td>6.8E-13</td><td>MB231/MB468</td><td>CYTO</td></tr>
<tr><td>>trnaMT_HisGTG_MT_+_12138_12206@28.66.39|ATTGTGAATCTGACAACAGAGGCTTACGACCCCTTATTT</td><td>>trnaMT_HisGTG_MT_+_12138_12206</td><td>28</td><td>66</td><td>39</td><td>ATTGTGAATCTGACAACAGAGGCTTACGACCCCTTATTT</td><td>330.1</td><td>-12.8</td><td>1.7</td><td>-7.3</td><td>2.6E-13</td><td>6.8E-13</td><td>BT20/MB468</td><td>NUC</td></tr>
<tr><td>>am_trna116_GluCTC_1_-_145399233_145399304@13.30.18,trna25_GluCTC_2_-_71273488_71273560@13.30.18,trna59_GluCTC_1_+_249168447_249168518@13.30.18,trna71_GluCTC_1_-_161439189_161439260@13.30.18,trna74_GluCTC_1_-_161431809_161431880@13.30.18,trna77_GluCTC_1_-_161424398_161424469@13.30.18,trna77_GluCTC_6_+_28949976_28950047@13.30.18,trna80_GluCTC_1_-_161417018_161417089@13.30.18,trna87_GluCTC_6_-_126101393_126101464@13.30.18|TAGTGGTTAGGATTCGGC</td><td>>am_trna116_GluCTC_1_-_145399233_145399304</td><td>13</td><td>30</td><td>18</td><td>TAGTGGTTAGGATTCGGC</td><td>2554.3</td><td>5.6</td><td>0.8</td><td>7.4</td><td>1.8E-13</td><td>6.8E-13</td><td>BT20/MB468</td><td>CYTO</td></tr>
<tr><td>>hg38_21_+_8433172_8446622_RNA45SN1_45S_50ntFlanks@10916.10933.18|CCGCGGCTGGACGAGGCG</td><td>>hg38_21_+_8433172_8446622_RNA45SN1_45S_50ntFlanks</td><td>10916</td><td>10933</td><td>18</td><td>CCGCGGCTGGACGAGGCG</td><td>468.4</td><td>-8.9</td><td>1.2</td><td>-7.3</td><td>2.3E-13</td><td>6.8E-13</td><td>MB231/MB468</td><td>MITO</td></tr>
<tr><td>>hg38_21_+_8433172_8446622_RNA45SN1_45S_50ntFlanks@3986.4024.39|GGCTTTGGTGACTCTAGATAACCTCGGGCCGATCGCACG</td><td>>hg38_21_+_8433172_8446622_RNA45SN1_45S_50ntFlanks</td><td>3986</td><td>4024</td><td>39</td><td>GGCTTTGGTGACTCTAGATAACCTCGGGCCGATCGCACG</td><td>758.4</td><td>-13.4</td><td>1.8</td><td>-7.4</td><td>1.5E-13</td><td>6.8E-13</td><td>MB231/MB468</td><td>MP</td></tr>
<tr><td>>hg38_21_+_8433172_8446622_RNA45SN1_45S_50ntFlanks@10590.10608.19|CGGGGAGAGTTCTCTTTTC</td><td>>hg38_21_+_8433172_8446622_RNA45SN1_45S_50ntFlanks</td><td>10590</td><td>10608</td><td>19</td><td>CGGGGAGAGTTCTCTTTTC</td><td>140.8</td><td>-11.0</td><td>1.5</td><td>-7.4</td><td>1.5E-13</td><td>6.8E-13</td><td>MB231/MB468</td><td>MP</td></tr>
<tr><td>>hg38_21_+_8433172_8446622_RNA45SN1_45S_50ntFlanks@4161.4184.24|CACATCCAAGGAAGGCAGCAGGCG</td><td>>hg38_21_+_8433172_8446622_RNA45SN1_45S_50ntFlanks</td><td>4161</td><td>4184</td><td>24</td><td>CACATCCAAGGAAGGCAGCAGGCG</td><td>201.7</td><td>-11.5</td><td>1.6</td><td>-7.4</td><td>1.5E-13</td><td>6.8E-13</td><td>MB231/MB468</td><td>MP</td></tr>
<tr><td>>hg38_21_+_8433172_8446622_RNA45SN1_45S_50ntFlanks@12893.12932.40|CCTCGCCCGTCACGCACCGCACGTTCGTGGGGAACCTGGC</td><td>>hg38_21_+_8433172_8446622_RNA45SN1_45S_50ntFlanks</td><td>12893</td><td>12932</td><td>40</td><td>CCTCGCCCGTCACGCACCGCACGTTCGTGGGGAACCTGGC</td><td>477.0</td><td>-4.8</td><td>0.7</td><td>-7.4</td><td>1.8E-13</td><td>6.9E-13</td><td>MB231/MB468</td><td>NUC</td></tr>
<tr><td>>hg38_21_+_8433172_8446622_RNA45SN1_45S_50ntFlanks@7994.8011.18|TGGCGACCCGCTGAATTT</td><td>>hg38_21_+_8433172_8446622_RNA45SN1_45S_50ntFlanks</td><td>7994</td><td>8011</td><td>18</td><td>TGGCGACCCGCTGAATTT</td><td>486.9</td><td>5.2</td><td>0.7</td><td>7.4</td><td>1.4E-13</td><td>6.9E-13</td><td>BT20/MB468</td><td>MITO</td></tr>
<tr><td>>hg38_21_+_8433172_8446622_RNA45SN1_45S_50ntFlanks@12418.12443.26|CGATGTCGGCTCTTCCTATCATTGTG</td><td>>hg38_21_+_8433172_8446622_RNA45SN1_45S_50ntFlanks</td><td>12418</td><td>12443</td><td>26</td><td>CGATGTCGGCTCTTCCTATCATTGTG</td><td>14640.6</td><td>-6.7</td><td>0.9</td><td>-7.3</td><td>2.3E-13</td><td>6.9E-13</td><td>MB231/MB468</td><td>MITO</td></tr>
<tr><td>>hg38_21_+_8433172_8446622_RNA45SN1_45S_50ntFlanks@10864.10892.29|CTCTAAGGGCTGGGTCGGTCGGGCTGGGG</td><td>>hg38_21_+_8433172_8446622_RNA45SN1_45S_50ntFlanks</td><td>10864</td><td>10892</td><td>29</td><td>CTCTAAGGGCTGGGTCGGTCGGGCTGGGG</td><td>209.7</td><td>-11.6</td><td>1.6</td><td>-7.4</td><td>1.5E-13</td><td>6.9E-13</td><td>MB231/MB468</td><td>MP</td></tr>
<tr><td>>hg38_21_+_8433172_8446622_RNA45SN1_45S_50ntFlanks@10821.10838.18|GGAAGTCGGCAAGCCGGA</td><td>>hg38_21_+_8433172_8446622_RNA45SN1_45S_50ntFlanks</td><td>10821</td><td>10838</td><td>18</td><td>GGAAGTCGGCAAGCCGGA</td><td>1372.8</td><td>-8.0</td><td>1.1</td><td>-7.4</td><td>1.5E-13</td><td>6.9E-13</td><td>MB231/MB468</td><td>MP</td></tr>
<tr><td>>MI0000114|hsa-mir-107&WithFlank&10|-|89592741|89592833@56.78.23&[MIMAT0000104&hsa-miR-107&offsets|0|0;m-78&10|-|89592758|89592778&offsets|0|+2]|AGCAGCATTGTACAGGGCTATCA</td><td>>MI0000114|hsa-mir-107&WithFlank&10|-|89592741|89592833</td><td>56</td><td>78</td><td>23</td><td>AGCAGCATTGTACAGGGCTATCA</td><td>747.6</td><td>5.0</td><td>0.7</td><td>7.4</td><td>1.8E-13</td><td>6.9E-13</td><td>BT20/MB468</td><td>CYTO</td></tr>
<tr><td>>hg38_21_+_8433172_8446622_RNA45SN1_45S_50ntFlanks@10410.10443.34|TCAGTCGGTCCTGAGAGATGGGCGAGCGCCGTTC</td><td>>hg38_21_+_8433172_8446622_RNA45SN1_45S_50ntFlanks</td><td>10410</td><td>10443</td><td>34</td><td>TCAGTCGGTCCTGAGAGATGGGCGAGCGCCGTTC</td><td>825.7</td><td>-9.9</td><td>1.4</td><td>-7.3</td><td>2.6E-13</td><td>6.9E-13</td><td>BT20/MB468</td><td>NUC</td></tr>
<tr><td>>hg38_21_+_8433172_8446622_RNA45SN1_45S_50ntFlanks@11876.11907.32|AAGACCCTGTTGAGCTTGACTCTAGTCTGGCA</td><td>>hg38_21_+_8433172_8446622_RNA45SN1_45S_50ntFlanks</td><td>11876</td><td>11907</td><td>32</td><td>AAGACCCTGTTGAGCTTGACTCTAGTCTGGCA</td><td>122.3</td><td>-10.8</td><td>1.5</td><td>-7.4</td><td>1.5E-13</td><td>6.9E-13</td><td>MB231/MB468</td><td>MP</td></tr>
<tr><td>>hg38_21_+_8433172_8446622_RNA45SN1_45S_50ntFlanks@10369.10407.39|AAGTGGAGAAGGGTTCCATGTGAACAGCAGTTGAACATG</td><td>>hg38_21_+_8433172_8446622_RNA45SN1_45S_50ntFlanks</td><td>10369</td><td>10407</td><td>39</td><td>AAGTGGAGAAGGGTTCCATGTGAACAGCAGTTGAACATG</td><td>193.3</td><td>-11.5</td><td>1.6</td><td>-7.4</td><td>1.5E-13</td><td>6.9E-13</td><td>MB231/MB468</td><td>MP</td></tr>
<tr><td>>hg38_21_+_8433172_8446622_RNA45SN1_45S_50ntFlanks@4995.5022.28|ACAGATTGATAGCTCTTTCTCGATTCCG</td><td>>hg38_21_+_8433172_8446622_RNA45SN1_45S_50ntFlanks</td><td>4995</td><td>5022</td><td>28</td><td>ACAGATTGATAGCTCTTTCTCGATTCCG</td><td>737.2</td><td>-13.4</td><td>1.8</td><td>-7.4</td><td>1.5E-13</td><td>6.9E-13</td><td>MB231/MB468</td><td>MP</td></tr>
<tr><td>>hg38_21_+_8433172_8446622_RNA45SN1_45S_50ntFlanks@11561.11590.30|GCGCCTAGCAGCCGACTTAGAACTGGTGCG</td><td>>hg38_21_+_8433172_8446622_RNA45SN1_45S_50ntFlanks</td><td>11561</td><td>11590</td><td>30</td><td>GCGCCTAGCAGCCGACTTAGAACTGGTGCG</td><td>510.3</td><td>-6.8</td><td>0.9</td><td>-7.3</td><td>2.3E-13</td><td>6.9E-13</td><td>MB231/MB468</td><td>MITO</td></tr>
<tr><td>>hg38_21_+_8433172_8446622_RNA45SN1_45S_50ntFlanks@9806.9850.45|CGTGGAATGCGAGTGCCTAGTGGGCCACTTTTGGTAAGCAGAACT</td><td>>hg38_21_+_8433172_8446622_RNA45SN1_45S_50ntFlanks</td><td>9806</td><td>9850</td><td>45</td><td>CGTGGAATGCGAGTGCCTAGTGGGCCACTTTTGGTAAGCAGAACT</td><td>325.7</td><td>-12.1</td><td>1.7</td><td>-7.3</td><td>2.3E-13</td><td>6.9E-13</td><td>MB231/MB468</td><td>MITO</td></tr>
<tr><td>>hg38_21_+_8433172_8446622_RNA45SN1_45S_50ntFlanks@11781.11806.26|CGCATGAATGGATGAACGAGATTCCC</td><td>>hg38_21_+_8433172_8446622_RNA45SN1_45S_50ntFlanks</td><td>11781</td><td>11806</td><td>26</td><td>CGCATGAATGGATGAACGAGATTCCC</td><td>609.4</td><td>-6.6</td><td>0.9</td><td>-7.4</td><td>1.8E-13</td><td>6.9E-13</td><td>MB231/MB468</td><td>CYTO</td></tr>
<tr><td>>MI0000301|hsa-mir-224&WithFlank&X|-|151958572|151958664@14.38.25&[MIMAT0000281&hsa-miR-224-5p&offsets|+1|+1;m-238&X|-|151958630|151958651&offsets|0|+3]|CAAGTCACTAGTGGTTCCGTTTAGT</td><td>>MI0000301|hsa-mir-224&WithFlank&X|-|151958572|151958664</td><td>14</td><td>38</td><td>25</td><td>CAAGTCACTAGTGGTTCCGTTTAGT</td><td>618.5</td><td>9.1</td><td>1.2</td><td>7.4</td><td>1.8E-13</td><td>6.9E-13</td><td>MB231/MB468</td><td>NUC</td></tr>
<tr><td>>MI0000087|hsa-mir-29a&WithFlank&7|-|130876741|130876816@47.68.22(+1U)&[MIMAT0000086&hsa-miR-29a-3p&offsets|-1|-1(+1U);m-26&7|-|130876749|130876769&offsets|-1|0(+1U)]|CTAGCACCATCTGAAATCGGTTT</td><td>>MI0000087|hsa-mir-29a&WithFlank&7|-|130876741|130876816</td><td>47</td><td>68</td><td>22(+1U)</td><td>CTAGCACCATCTGAAATCGGTTT</td><td>848.5</td><td>11.4</td><td>1.5</td><td>7.4</td><td>1.8E-13</td><td>7.0E-13</td><td>BT20/MB468</td><td>CYTO</td></tr>
<tr><td>>hg38_21_+_8433172_8446622_RNA45SN1_45S_50ntFlanks@13006.13025.20|CTATTGAAAGTCAGCCCTCG</td><td>>hg38_21_+_8433172_8446622_RNA45SN1_45S_50ntFlanks</td><td>13006</td><td>13025</td><td>20</td><td>CTATTGAAAGTCAGCCCTCG</td><td>1191.0</td><td>-7.0</td><td>1.0</td><td>-7.4</td><td>1.8E-13</td><td>7.0E-13</td><td>MB231/MB468</td><td>CYTO</td></tr>
<tr><td>>hg38_21_+_8433172_8446622_RNA45SN1_45S_50ntFlanks@10926.10945.20|ACGAGGCGCCGCCGCCCCCC</td><td>>hg38_21_+_8433172_8446622_RNA45SN1_45S_50ntFlanks</td><td>10926</td><td>10945</td><td>20</td><td>ACGAGGCGCCGCCGCCCCCC</td><td>951.3</td><td>-13.8</td><td>1.9</td><td>-7.4</td><td>1.5E-13</td><td>7.0E-13</td><td>MB231/MB468</td><td>MP</td></tr>
<tr><td>>hg38_21_+_8433172_8446622_RNA45SN1_45S_50ntFlanks@12542.12569.28|ATGTGTTGTTGCCATGGTAATCCTGCTC</td><td>>hg38_21_+_8433172_8446622_RNA45SN1_45S_50ntFlanks</td><td>12542</td><td>12569</td><td>28</td><td>ATGTGTTGTTGCCATGGTAATCCTGCTC</td><td>159.7</td><td>-11.2</td><td>1.5</td><td>-7.4</td><td>1.5E-13</td><td>7.0E-13</td><td>MB231/MB468</td><td>MP</td></tr>
<tr><td>>hg38_21_+_8433172_8446622_RNA45SN1_45S_50ntFlanks@5016.5034.19|GATTCCGTGGGTGGTGGTG</td><td>>hg38_21_+_8433172_8446622_RNA45SN1_45S_50ntFlanks</td><td>5016</td><td>5034</td><td>19</td><td>GATTCCGTGGGTGGTGGTG</td><td>1387.9</td><td>-14.2</td><td>1.9</td><td>-7.3</td><td>2.3E-13</td><td>7.0E-13</td><td>MB231/MB468</td><td>MITO</td></tr>
<tr><td>>MI0000299|hsa-mir-222&WithFlank&X|-|45747009|45747130@75.97.23(+2U)&[MIMAT0000279&hsa-miR-222-3p&offsets|0|+2(+2U);m-140&X|-|45747033|45747056&offsets|0|-1(+2U)]|AGCTACATCTGGCTACTGGGTCTTT</td><td>>MI0000299|hsa-mir-222&WithFlank&X|-|45747009|45747130</td><td>75</td><td>97</td><td>23(+2U)</td><td>AGCTACATCTGGCTACTGGGTCTTT</td><td>988.0</td><td>9.4</td><td>1.3</td><td>7.4</td><td>1.8E-13</td><td>7.0E-13</td><td>MB231/MB468</td><td>CYTO</td></tr>
<tr><td>>hg38_21_+_8433172_8446622_RNA45SN1_45S_50ntFlanks@12128.12146.19|ACCCGCTCCGGGGACAGTG</td><td>>hg38_21_+_8433172_8446622_RNA45SN1_45S_50ntFlanks</td><td>12128</td><td>12146</td><td>19</td><td>ACCCGCTCCGGGGACAGTG</td><td>137.5</td><td>-11.0</td><td>1.5</td><td>-7.4</td><td>1.5E-13</td><td>7.0E-13</td><td>MB231/MB468</td><td>MP</td></tr>
<tr><td>>hg38_21_+_8433172_8446622_RNA45SN1_45S_50ntFlanks@11874.11904.31|AGAAGACCCTGTTGAGCTTGACTCTAGTCTG</td><td>>hg38_21_+_8433172_8446622_RNA45SN1_45S_50ntFlanks</td><td>11874</td><td>11904</td><td>31</td><td>AGAAGACCCTGTTGAGCTTGACTCTAGTCTG</td><td>2229.5</td><td>-6.7</td><td>0.9</td><td>-7.8</td><td>7.2E-15</td><td>7.0E-13</td><td>BT20/MB468</td><td>MP</td></tr>
<tr><td>>hg38_21_+_8433172_8446622_RNA45SN1_45S_50ntFlanks@12539.12559.21|ATGATGTGTTGTTGCCATGGT</td><td>>hg38_21_+_8433172_8446622_RNA45SN1_45S_50ntFlanks</td><td>12539</td><td>12559</td><td>21</td><td>ATGATGTGTTGTTGCCATGGT</td><td>179.7</td><td>-11.4</td><td>1.5</td><td>-7.4</td><td>1.5E-13</td><td>7.0E-13</td><td>MB231/MB468</td><td>MP</td></tr>
<tr><td>>hg38_21_+_8433172_8446622_RNA45SN1_45S_50ntFlanks@9430.9453.24|CGCACCACCGGCCCGTCTCGCCCG</td><td>>hg38_21_+_8433172_8446622_RNA45SN1_45S_50ntFlanks</td><td>9430</td><td>9453</td><td>24</td><td>CGCACCACCGGCCCGTCTCGCCCG</td><td>426.3</td><td>-12.5</td><td>1.7</td><td>-7.3</td><td>2.3E-13</td><td>7.0E-13</td><td>MB231/MB468</td><td>MITO</td></tr>
<tr><td>>hg38_21_+_8433172_8446622_RNA45SN1_45S_50ntFlanks@9504.9521.18|GTGAACTATGCCTGGGCA</td><td>>hg38_21_+_8433172_8446622_RNA45SN1_45S_50ntFlanks</td><td>9504</td><td>9521</td><td>18</td><td>GTGAACTATGCCTGGGCA</td><td>1221.9</td><td>-7.3</td><td>1.0</td><td>-7.3</td><td>2.3E-13</td><td>7.0E-13</td><td>MB231/MB468</td><td>MITO</td></tr>
<tr><td>>hg38_21_+_8433172_8446622_RNA45SN1_45S_50ntFlanks@12432.12455.24|CCTATCATTGTGAAGCAGAATTCA</td><td>>hg38_21_+_8433172_8446622_RNA45SN1_45S_50ntFlanks</td><td>12432</td><td>12455</td><td>24</td><td>CCTATCATTGTGAAGCAGAATTCA</td><td>252.5</td><td>-6.3</td><td>0.9</td><td>-7.4</td><td>1.8E-13</td><td>7.0E-13</td><td>MB231/MB468</td><td>CYTO</td></tr>
<tr><td>>hg38_21_+_8433172_8446622_RNA45SN1_45S_50ntFlanks@4919.4941.23|CCACCAGGAGTGGAGCCTGCGGC</td><td>>hg38_21_+_8433172_8446622_RNA45SN1_45S_50ntFlanks</td><td>4919</td><td>4941</td><td>23</td><td>CCACCAGGAGTGGAGCCTGCGGC</td><td>2396.1</td><td>-11.3</td><td>1.5</td><td>-7.3</td><td>2.3E-13</td><td>7.0E-13</td><td>MB231/MB468</td><td>MITO</td></tr>
<tr><td>>hg38_21_+_8433172_8446622_RNA45SN1_45S_50ntFlanks@12076.12097.22|AGGGGCTCTCGCTTCTGGCGCC</td><td>>hg38_21_+_8433172_8446622_RNA45SN1_45S_50ntFlanks</td><td>12076</td><td>12097</td><td>22</td><td>AGGGGCTCTCGCTTCTGGCGCC</td><td>294.3</td><td>-4.1</td><td>0.6</td><td>-7.4</td><td>1.8E-13</td><td>7.0E-13</td><td>MB231/MB468</td><td>NUC</td></tr>
<tr><td>>MI0000447|hsa-mir-128-1&WithFlank&2|+|135665391|135665484@56.76.21(+2A);MI0000727|hsa-mir-128-2&WithFlank&3|+|35744470|35744565@58.78.21(+2A)&[MIMAT0000424&hsa-miR-128-3p&offsets|0|0(+2A);m-108&2|+|135665446|135665466&offsets|0|0(+2A)];[MIMAT0000424_1&hsa-miR-128-3p&offsets|0|0(+2A);m-117&3|+|35744527|35744547&offsets|0|0(+2A)]|TCACAGTGAACCGGTCTCTTTAA</td><td>>MI0000447|hsa-mir-128-1&WithFlank&2|+|135665391|135665484</td><td>56</td><td>76</td><td>21(+2A)</td><td>TCACAGTGAACCGGTCTCTTTAA</td><td>846.3</td><td>4.8</td><td>0.6</td><td>7.4</td><td>1.4E-13</td><td>7.0E-13</td><td>BT20/MB468</td><td>MITO</td></tr>
<tr><td>>hg38_21_+_8433172_8446622_RNA45SN1_45S_50ntFlanks@11575.11606.32|ACTTAGAACTGGTGCGGACCAGGGGAATCCGA</td><td>>hg38_21_+_8433172_8446622_RNA45SN1_45S_50ntFlanks</td><td>11575</td><td>11606</td><td>32</td><td>ACTTAGAACTGGTGCGGACCAGGGGAATCCGA</td><td>212.0</td><td>-11.6</td><td>1.6</td><td>-7.4</td><td>1.6E-13</td><td>7.1E-13</td><td>MB231/MB468</td><td>MP</td></tr>
<tr><td>>hg38_21_+_8433172_8446622_RNA45SN1_45S_50ntFlanks@6748.6772.25|CTTCGAACGCACTTGCGGCCCCGGG</td><td>>hg38_21_+_8433172_8446622_RNA45SN1_45S_50ntFlanks</td><td>6748</td><td>6772</td><td>25</td><td>CTTCGAACGCACTTGCGGCCCCGGG</td><td>176.4</td><td>-11.3</td><td>1.5</td><td>-7.4</td><td>1.6E-13</td><td>7.1E-13</td><td>MB231/MB468</td><td>MP</td></tr>
<tr><td>>hg38_21_+_8433172_8446622_RNA45SN1_45S_50ntFlanks@9738.9766.29|GAAACGATCTCAACCTATTCTCAAACTTT</td><td>>hg38_21_+_8433172_8446622_RNA45SN1_45S_50ntFlanks</td><td>9738</td><td>9766</td><td>29</td><td>GAAACGATCTCAACCTATTCTCAAACTTT</td><td>140.4</td><td>-11.0</td><td>1.5</td><td>-7.4</td><td>1.6E-13</td><td>7.1E-13</td><td>MB231/MB468</td><td>MP</td></tr>
<tr><td>>hg38_21_+_8433172_8446622_RNA45SN1_45S_50ntFlanks@11918.11954.37|ACATGAGAGGTGTAGAATAAGTGGGAGGCCCCCGGCG</td><td>>hg38_21_+_8433172_8446622_RNA45SN1_45S_50ntFlanks</td><td>11918</td><td>11954</td><td>37</td><td>ACATGAGAGGTGTAGAATAAGTGGGAGGCCCCCGGCG</td><td>237.1</td><td>-11.8</td><td>1.6</td><td>-7.4</td><td>1.6E-13</td><td>7.1E-13</td><td>MB231/MB468</td><td>MP</td></tr>
<tr><td>>hg38_21_+_8433172_8446622_RNA45SN1_45S_50ntFlanks@10101.10124.24|CGCGCGCGTGTGGTGTGCGTCGGA</td><td>>hg38_21_+_8433172_8446622_RNA45SN1_45S_50ntFlanks</td><td>10101</td><td>10124</td><td>24</td><td>CGCGCGCGTGTGGTGTGCGTCGGA</td><td>361.1</td><td>-12.3</td><td>1.7</td><td>-7.3</td><td>2.4E-13</td><td>7.1E-13</td><td>MB231/MB468</td><td>MITO</td></tr>
<tr><td>>hg38_21_+_8433172_8446622_RNA45SN1_45S_50ntFlanks@11233.11253.21|CCCTTCCCGTGGATCGCCCCA</td><td>>hg38_21_+_8433172_8446622_RNA45SN1_45S_50ntFlanks</td><td>11233</td><td>11253</td><td>21</td><td>CCCTTCCCGTGGATCGCCCCA</td><td>182.9</td><td>-11.4</td><td>1.5</td><td>-7.4</td><td>1.6E-13</td><td>7.1E-13</td><td>MB231/MB468</td><td>MP</td></tr>
</table>

| | | | | | | | | | | | | | |
|---|---|---|---|---|---|---|---|---|---|---|---|---|---|
| >hg38_21_+_8433172_8446622_RNA45SN1_45S_50ntFlanks@10728.10755.28\|GAGAGGGTGTAAATCTCGCGCCGGGCCG | >hg38_21_+_8433172_8446622_RNA45SN1_45S_50ntFlanks | 10728 | 10755 | 28 | GAGAGGGTGTAAATCTCGCGCCGGGCCG | 131.9 | -10.9 | 1.5 | -7.4 | 1.6E-13 | 7.1E-13 | MB231/MB468 | MP |
| >hg38_21_+_8433172_8446622_RNA45SN1_45S_50ntFlanks@4380.4398.19\|CGTAGTTGGATCTTGGGAG | >hg38_21_+_8433172_8446622_RNA45SN1_45S_50ntFlanks | 4380 | 4398 | 19 | CGTAGTTGGATCTTGGGAG | 181.8 | -11.4 | 1.5 | -7.4 | 1.6E-13 | 7.1E-13 | MB231/MB468 | MP |
| >hg38_21_+_8433172_8446622_RNA45SN1_45S_50ntFlanks@11646.11674.29\|GGTGTTGACGCGATGTGATTTCTGCCCAG | >hg38_21_+_8433172_8446622_RNA45SN1_45S_50ntFlanks | 11646 | 11674 | 29 | GGTGTTGACGCGATGTGATTTCTGCCCAG | 229.9 | -11.7 | 1.6 | -7.4 | 1.6E-13 | 7.1E-13 | MB231/MB468 | MP |
| >hg38_21_+_8433172_8446622_RNA45SN1_45S_50ntFlanks@10792.10817.26\|CTCTGGCATGTTGGAACAATGTAGGT | >hg38_21_+_8433172_8446622_RNA45SN1_45S_50ntFlanks | 10792 | 10817 | 26 | CTCTGGCATGTTGGAACAATGTAGGT | 154.3 | -11.1 | 1.5 | -7.4 | 1.6E-13 | 7.1E-13 | MB231/MB468 | MP |
| >hg38_21_+_8433172_8446622_RNA45SN1_45S_50ntFlanks@3813.3842.30\|TTATGGTTCCTTTGGTCGCTCGCTCCTCTC | >hg38_21_+_8433172_8446622_RNA45SN1_45S_50ntFlanks | 3813 | 3842 | 30 | TTATGGTTCCTTTGGTCGCTCGCTCCTCTC | 170.7 | -11.3 | 1.5 | -7.4 | 1.6E-13 | 7.1E-13 | MB231/MB468 | MP |
| >hg38_21_+_8433172_8446622_RNA45SN1_45S_50ntFlanks@13002.13026.25\|CGATCTATTGAAAGTCAGCCCTCGA | >hg38_21_+_8433172_8446622_RNA45SN1_45S_50ntFlanks | 13002 | 13026 | 25 | CGATCTATTGAAAGTCAGCCCTCGA | 157.8 | -10.3 | 1.4 | -7.4 | 1.9E-13 | 7.1E-13 | MB231/MB468 | NUC |
| >hg38_21_+_8433172_8446622_RNA45SN1_45S_50ntFlanks@6656.6681.26\|CTTAGCGGTGGATCACTCGGCTCGTG | >hg38_21_+_8433172_8446622_RNA45SN1_45S_50ntFlanks | 6656 | 6681 | 26 | CTTAGCGGTGGATCACTCGGCTCGTG | 2757.2 | -2.7 | 0.4 | -7.4 | 1.4E-13 | 7.1E-13 | BT20/MB468 | MITO |
| >hg38_21_+_8433172_8446622_RNA45SN1_45S_50ntFlanks@10826.10865.40\|TCGGCAAGCCGGATCCGTAACTTCGGGATAAGGATTGGCT | >hg38_21_+_8433172_8446622_RNA45SN1_45S_50ntFlanks | 10826 | 10865 | 40 | TCGGCAAGCCGGATCCGTAACTTCGGGATAAGGATTGGCT | 889.7 | -12.1 | 1.6 | -7.4 | 1.9E-13 | 7.2E-13 | BT20/MB468 | CYTO |
| >hg38_21_+_8433172_8446622_RNA45SN1_45S_50ntFlanks@4472.4496.25\|CTTAGCTGAGTGTCCCGCGGGGCCC | >hg38_21_+_8433172_8446622_RNA45SN1_45S_50ntFlanks | 4472 | 4496 | 25 | CTTAGCTGAGTGTCCCGCGGGGCCC | 232.8 | -10.0 | 1.3 | -7.4 | 1.4E-13 | 7.2E-13 | BT20/MB468 | MITO |
| >hg38_21_+_8433172_8446622_RNA45SN1_45S_50ntFlanks@12043.12060.18\|CACTGACCCGGTGAGGCG | >hg38_21_+_8433172_8446622_RNA45SN1_45S_50ntFlanks | 12043 | 12060 | 18 | CACTGACCCGGTGAGGCG | 156.0 | -11.1 | 1.5 | -7.4 | 1.6E-13 | 7.2E-13 | MB231/MB468 | MP |
| >trna32_LysCTT_16_-_3207406_3207478@1.33.33\|GCCCGGCTAGCTCAGTCGGTAGAGCATGAGACC | >trna32_LysCTT_16_-_3207406_3207478 | 1 | 33 | 33 | GCCCGGCTAGCTCAGTCGGTAGAGCATGAGACC | 1788.6 | 8.4 | 1.1 | 7.4 | 1.9E-13 | 7.2E-13 | MB231/MB468 | NUC |
| >hg38_21_+_8433172_8446622_RNA45SN1_45S_50ntFlanks@11027.11062.36\|CTCTCTCTCTCTCTCCCCCGCTCCCCGTCCTCCCCC | >hg38_21_+_8433172_8446622_RNA45SN1_45S_50ntFlanks | 11027 | 11062 | 36 | CTCTCTCTCTCTCTCCCCCGCTCCCCGTCCTCCCCC | 206.9 | -10.7 | 1.4 | -7.4 | 1.9E-13 | 7.2E-13 | MB231/MB468 | NUC |
| >hg38_21_+_8433172_8446622_RNA45SN1_45S_50ntFlanks@10014.10040.27\|AAAATGGATGGCGCTGGAGCGTCGGGC | >hg38_21_+_8433172_8446622_RNA45SN1_45S_50ntFlanks | 10014 | 10040 | 27 | AAAATGGATGGCGCTGGAGCGTCGGGC | 281.1 | -12.0 | 1.6 | -7.4 | 1.6E-13 | 7.2E-13 | MB231/MB468 | MP |
| >hg38_21_+_8433172_8446622_RNA45SN1_45S_50ntFlanks@9738.9769.32\|GAAACGATCTCAACCTATTCTCAAACTTTAAA | >hg38_21_+_8433172_8446622_RNA45SN1_45S_50ntFlanks | 9738 | 9769 | 32 | GAAACGATCTCAACCTATTCTCAAACTTTAAA | 371.2 | -12.4 | 1.7 | -7.4 | 1.6E-13 | 7.2E-13 | MB231/MB468 | MP |
| >hg38_21_+_8433172_8446622_RNA45SN1_45S_50ntFlanks@12412.12430.19\|ATCCTTCGATGTCGGCTCT | >hg38_21_+_8433172_8446622_RNA45SN1_45S_50ntFlanks | 12412 | 12430 | 19 | ATCCTTCGATGTCGGCTCT | 266.4 | -10.4 | 1.4 | -7.4 | 1.9E-13 | 7.2E-13 | BT20/MB468 | CYTO |
| >hg38_21_+_8433172_8446622_RNA45SN1_45S_50ntFlanks@11821.11844.24\|CTATCCAGCGAAACCACAGCCAAG | >hg38_21_+_8433172_8446622_RNA45SN1_45S_50ntFlanks | 11821 | 11844 | 24 | CTATCCAGCGAAACCACAGCCAAG | 273.8 | -10.2 | 1.4 | -7.4 | 1.4E-13 | 7.2E-13 | BT20/MB468 | MITO |
| >hg38_21_+_8433172_8446622_RNA45SN1_45S_50ntFlanks@10805.10826.22\|GAACAATGTAGGTAAGGGAAGT | >hg38_21_+_8433172_8446622_RNA45SN1_45S_50ntFlanks | 10805 | 10826 | 22 | GAACAATGTAGGTAAGGGAAGT | 224.8 | -9.9 | 1.3 | -7.4 | 1.4E-13 | 7.2E-13 | BT20/MB468 | MITO |
| >hg38_21_+_8433172_8446622_RNA45SN1_45S_50ntFlanks@12983.13001.19\|CAGAGCAGCTCCCTCGCTG | >hg38_21_+_8433172_8446622_RNA45SN1_45S_50ntFlanks | 12983 | 13001 | 19 | CAGAGCAGCTCCCTCGCTG | 171.6 | -10.1 | 1.4 | -7.4 | 1.8E-13 | 7.2E-13 | MB231/MB468 | CYTO |
| >hg38_21_+_8433172_8446622_RNA45SN1_45S_50ntFlanks@11783.11805.23\|CATGAATGGATGAACGAGATTCC | >hg38_21_+_8433172_8446622_RNA45SN1_45S_50ntFlanks | 11783 | 11805 | 23 | CATGAATGGATGAACGAGATTCC | 2080.7 | -4.5 | 0.6 | -7.4 | 1.9E-13 | 7.2E-13 | MB231/MB468 | NUC |
| >hg38_21_+_8433172_8446622_RNA45SN1_45S_50ntFlanks@7991.8013.23\|ACGTGGCGACCCGCTGAATTTAA | >hg38_21_+_8433172_8446622_RNA45SN1_45S_50ntFlanks | 7991 | 8013 | 23 | ACGTGGCGACCCGCTGAATTTAA | 1440.1 | -4.6 | 0.6 | -7.3 | 2.4E-13 | 7.2E-13 | MB231/MB468 | MITO |
| >hg38_21_+_8433172_8446622_RNA45SN1_45S_50ntFlanks@10681.10708.28\|TTCCGGCGGCGTCCGGTGAGCTCTCGCT | >hg38_21_+_8433172_8446622_RNA45SN1_45S_50ntFlanks | 10681 | 10708 | 28 | TTCCGGCGGCGTCCGGTGAGCTCTCGCT | 666.1 | -7.0 | 1.0 | -7.3 | 2.4E-13 | 7.2E-13 | MB231/MB468 | MITO |
| >hg38_21_+_8433172_8446622_RNA45SN1_45S_50ntFlanks@12934.12951.18\|CTAAACCATTCGTAGACG | >hg38_21_+_8433172_8446622_RNA45SN1_45S_50ntFlanks | 12934 | 12951 | 18 | CTAAACCATTCGTAGACG | 928.3 | -7.1 | 1.0 | -7.3 | 2.4E-13 | 7.2E-13 | MB231/MB468 | MITO |
| >hg38_21_+_8433172_8446622_RNA45SN1_45S_50ntFlanks@12804.12821.18\|CGCGTGCCCCGCCGCGCG | >hg38_21_+_8433172_8446622_RNA45SN1_45S_50ntFlanks | 12804 | 12821 | 18 | CGCGTGCCCCGCCGCGCG | 315.1 | -12.2 | 1.6 | -7.4 | 1.6E-13 | 7.2E-13 | MB231/MB468 | MP |
| >hg38_21_+_8433172_8446622_RNA45SN1_45S_50ntFlanks@4103.4121.19\|CCACGGGTGACGGGGAATC | >hg38_21_+_8433172_8446622_RNA45SN1_45S_50ntFlanks | 4103 | 4121 | 19 | CCACGGGTGACGGGGAATC | 1001.0 | -10.2 | 1.4 | -7.4 | 1.9E-13 | 7.3E-13 | MB231/MB468 | NUC |
| >hg38_21_+_8433172_8446622_RNA45SN1_45S_50ntFlanks@9567.9593.27\|ACGTGCAAATCGGTCGTCCGACCTGGG | >hg38_21_+_8433172_8446622_RNA45SN1_45S_50ntFlanks | 9567 | 9593 | 27 | ACGTGCAAATCGGTCGTCCGACCTGGG | 127.2 | -10.9 | 1.5 | -7.4 | 1.6E-13 | 7.3E-13 | MB231/MB468 | MP |
| >hg38_21_+_8433172_8446622_RNA45SN1_45S_50ntFlanks@7978.8006.29\|GACCTCAGATCAGACGTGGCGACCCGCTG | >hg38_21_+_8433172_8446622_RNA45SN1_45S_50ntFlanks | 7978 | 8006 | 29 | GACCTCAGATCAGACGTGGCGACCCGCTG | 430.6 | -3.9 | 0.5 | -7.4 | 1.9E-13 | 7.3E-13 | MB231/MB468 | NUC |
| >hg38_21_+_8433172_8446622_RNA45SN1_45S_50ntFlanks@12241.12273.33\|CAGAAGGGCAAAAGCTCGCTTGATCTTGATTTT | >hg38_21_+_8433172_8446622_RNA45SN1_45S_50ntFlanks | 12241 | 12273 | 33 | CAGAAGGGCAAAAGCTCGCTTGATCTTGATTTT | 827.4 | -4.3 | 0.6 | -7.4 | 1.5E-13 | 7.3E-13 | BT20/MB468 | MITO |
| >hg38_21_+_8433172_8446622_RNA45SN1_45S_50ntFlanks@11882.11904.23\|CTGTTGAGCTTGACTCTAGTCTG | >hg38_21_+_8433172_8446622_RNA45SN1_45S_50ntFlanks | 11882 | 11904 | 23 | CTGTTGAGCTTGACTCTAGTCTG | 1368.4 | -9.5 | 1.3 | -7.3 | 2.4E-13 | 7.3E-13 | MB231/MB468 | MITO |
| >hg38_21_+_8433172_8446622_RNA45SN1_45S_50ntFlanks@12386.12414.29\|AGCGTTCATAGCGACGTCGCTTTTTGATC | >hg38_21_+_8433172_8446622_RNA45SN1_45S_50ntFlanks | 12386 | 12414 | 29 | AGCGTTCATAGCGACGTCGCTTTTTGATC | 428.3 | -11.8 | 1.5 | -7.8 | 7.5E-15 | 7.3E-13 | BT20/MB468 | MP |
| >hg38_21_+_8433172_8446622_RNA45SN1_45S_50ntFlanks@10827.10858.32\|CGGCAAGCCGGATCCGTAACTTCGGGATAAGG | >hg38_21_+_8433172_8446622_RNA45SN1_45S_50ntFlanks | 10827 | 10858 | 32 | CGGCAAGCCGGATCCGTAACTTCGGGATAAGG | 341.9 | -12.3 | 1.7 | -7.4 | 1.6E-13 | 7.3E-13 | MB231/MB468 | MP |
| >hg38_21_+_8433172_8446622_RNA45SN1_45S_50ntFlanks@8176.8223.48\|CTTCTGATCGAGGCCCAGCCCGTGGACGGTGTGAGGCCGGTAGCGGCC | >hg38_21_+_8433172_8446622_RNA45SN1_45S_50ntFlanks | 8176 | 8223 | 48 | CTTCTGATCGAGGCCCAGCCCGTGGACGGTGTGAGGCCGGTAGCGGCC | 365.2 | -12.3 | 1.7 | -7.3 | 2.4E-13 | 7.3E-13 | MB231/MB468 | MITO |
| >hg38_21_+_8433172_8446622_RNA45SN1_45S_50ntFlanks@11104.11132.29\|AAGGGTCGGGGCGGCAGGGGCCGGCGGCG | >hg38_21_+_8433172_8446622_RNA45SN1_45S_50ntFlanks | 11104 | 11132 | 29 | AAGGGTCGGGGCGGCAGGGGCCGGCGGCG | 241.2 | -11.8 | 1.6 | -7.4 | 1.6E-13 | 7.3E-13 | MB231/MB468 | MP |
| >MI0000483\|hsa-mir-186&WithFlank&1\|-\|71067625\|71067722@21.40.20&[MIMAT0000456&hsa-miR-186-5p&offsets\|0\|-2;m-84&1\|-\|71067681\|71067702&offsets\|0\|-2]\|CAAAGAATTCTCCTTTTGGG | >MI0000483\|hsa-mir-186&WithFlank&1\|-\|71067625\|71067722 | 21 | 40 | 20 | CAAAGAATTCTCCTTTTGGG | 433.8 | 6.1 | 0.8 | 7.4 | 1.9E-13 | 7.3E-13 | BT20/MB468 | CYTO |
| >MI0003628\|hsa-mir-615&WithFlank&12\|+\|54033944\|54034051@67.86.20(+1A)&[MIMAT0003283&hsa-miR-615-3p&offsets\|0\|-2(+1A);m-495&12\|+\|54034010\|54034030&offsets\|0\|-1(+1A)]\|TCCGAGCCTGGGTCTCCCTCA | >MI0003628\|hsa-mir-615&WithFlank&12\|+\|54033944\|54034051 | 67 | 86 | 20(+1A) | TCCGAGCCTGGGTCTCCCTCA | 366.7 | 7.8 | 1.0 | 7.7 | 2.0E-14 | 7.3E-13 | BT20/MB231 | MITO |
| >hg38_21_+_8433172_8446622_RNA45SN1_45S_50ntFlanks@10806.10840.35\|AACAATGTAGGTAAGGGAAGTCGGCAAGCCGGATC | >hg38_21_+_8433172_8446622_RNA45SN1_45S_50ntFlanks | 10806 | 10840 | 35 | AACAATGTAGGTAAGGGAAGTCGGCAAGCCGGATC | 1670.6 | -7.4 | 1.0 | -7.3 | 2.4E-13 | 7.3E-13 | MB231/MB468 | MITO |
| >hg38_1_-_228634819_228635039_RNA5S12_5S_50ntFlanks@144.170.27\|CCTGGGAATACCGGGTGCTGTAGGCTT | >hg38_1_-_228634819_228635039_RNA5S12_5S_50ntFlanks | 144 | 170 | 27 | CCTGGGAATACCGGGTGCTGTAGGCTT | 4633.6 | 6.2 | 0.8 | 7.4 | 1.9E-13 | 7.4E-13 | MB231/MB468 | CYTO |
| >hg38_21_+_8433172_8446622_RNA45SN1_45S_50ntFlanks@483.520.38\|TCGATGTGGTGACGTCGTGCTCTCCCGGGCCGGGTCCG | >hg38_21_+_8433172_8446622_RNA45SN1_45S_50ntFlanks | 483 | 520 | 38 | TCGATGTGGTGACGTCGTGCTCTCCCGGGCCGGGTCCG | 537.7 | 5.3 | 0.7 | 7.4 | 1.9E-13 | 7.4E-13 | MB231/MB468 | NUC |
| >hg38_21_+_8433172_8446622_RNA45SN1_45S_50ntFlanks@11718.11738.21\|AACGGCGGGAGTAACTATGAC | >hg38_21_+_8433172_8446622_RNA45SN1_45S_50ntFlanks | 11718 | 11738 | 21 | AACGGCGGGAGTAACTATGAC | 1468.0 | -8.0 | 1.1 | -7.4 | 1.9E-13 | 7.4E-13 | MB231/MB468 | NUC |
| >hg38_21_+_8433172_8446622_RNA45SN1_45S_50ntFlanks@3835.3868.34\|CTCCTCTCCTACTTGGATAACTGTGGTAATTCTA | >hg38_21_+_8433172_8446622_RNA45SN1_45S_50ntFlanks | 3835 | 3868 | 34 | CTCCTCTCCTACTTGGATAACTGTGGTAATTCTA | 1137.4 | -10.1 | 1.4 | -7.3 | 2.8E-13 | 7.4E-13 | BT20/MB468 | NUC |
| >MI0000087\|hsa-mir-29a&WithFlank&7\|-\|130876741\|130876816@48.68.21&[MIMAT0000086&hsa-miR-29a-3p&offsets\|0\|-1;m-26&7\|-\|130876749\|130876769&offsets\|0\|0]\|TAGCACCATCTGAAATCGGTT | >MI0000087\|hsa-mir-29a&WithFlank&7\|-\|130876741\|130876816 | 48 | 68 | 21 | TAGCACCATCTGAAATCGGTT | 2514.5 | 6.7 | 0.9 | 7.3 | 2.8E-13 | 7.4E-13 | BT20/MB468 | NUC |
| >hg38_21_+_8433172_8446622_RNA45SN1_45S_50ntFlanks@9621.9658.38\|TCTAGTAGCTGGTTCCCTCCGAAGTTTCCCTCAGGATA | >hg38_21_+_8433172_8446622_RNA45SN1_45S_50ntFlanks | 9621 | 9658 | 38 | TCTAGTAGCTGGTTCCCTCCGAAGTTTCCCTCAGGATA | 1042.3 | -7.4 | 1.0 | -7.3 | 2.5E-13 | 7.4E-13 | MB231/MB468 | MITO |
| >hg38_21_+_8433172_8446622_RNA45SN1_45S_50ntFlanks@12418.12455.38\|CGATGTCGGCTCTTCCTATCATTGTGAAGCAGAATTCA | >hg38_21_+_8433172_8446622_RNA45SN1_45S_50ntFlanks | 12418 | 12455 | 38 | CGATGTCGGCTCTTCCTATCATTGTGAAGCAGAATTCA | 4946.5 | -5.4 | 0.7 | -7.4 | 1.5E-13 | 7.4E-13 | BT20/MB468 | MITO |
| >hg38_21_+_8433172_8446622_RNA45SN1_45S_50ntFlanks@9651.9682.32\|TCAGGATAGCTGGCGCTCTCGCAGACCCGACG | >hg38_21_+_8433172_8446622_RNA45SN1_45S_50ntFlanks | 9651 | 9682 | 32 | TCAGGATAGCTGGCGCTCTCGCAGACCCGACG | 108.7 | -10.6 | 1.4 | -7.4 | 1.6E-13 | 7.4E-13 | MB231/MB468 | MP |
| >hg38_21_+_8433172_8446622_RNA45SN1_45S_50ntFlanks@4128.4158.31\|CGATTCCGGAGAGGGAGCCTGAGAAACGGCT | >hg38_21_+_8433172_8446622_RNA45SN1_45S_50ntFlanks | 4128 | 4158 | 31 | CGATTCCGGAGAGGGAGCCTGAGAAACGGCT | 195.2 | -11.5 | 1.6 | -7.4 | 1.6E-13 | 7.4E-13 | MB231/MB468 | MP |
| >hg38_21_+_8433172_8446622_RNA45SN1_45S_50ntFlanks@8212.8230.19\|CCGGTAGCGGCCCCCGGCG | >hg38_21_+_8433172_8446622_RNA45SN1_45S_50ntFlanks | 8212 | 8230 | 19 | CCGGTAGCGGCCCCCGGCG | 155.6 | -5.1 | 0.7 | -7.4 | 1.9E-13 | 7.4E-13 | MB231/MB468 | CYTO |
| >hg38_21_+_8433172_8446622_RNA45SN1_45S_50ntFlanks@6661.6702.42\|CGGTGGATCACTCGGCTCGTGCGTCGATGAAGAACGCAGCTA | >hg38_21_+_8433172_8446622_RNA45SN1_45S_50ntFlanks | 6661 | 6702 | 42 | CGGTGGATCACTCGGCTCGTGCGTCGATGAAGAACGCAGCTA | 748.6 | 5.3 | 0.7 | 7.4 | 1.5E-13 | 7.4E-13 | BT20/MB468 | MITO |
| >hg38_21_+_8433172_8446622_RNA45SN1_45S_50ntFlanks@9240.9274.35\|GCGGAGCGAGCGCACGGGGTCGGCGGCGACGTCGG | >hg38_21_+_8433172_8446622_RNA45SN1_45S_50ntFlanks | 9240 | 9274 | 35 | GCGGAGCGAGCGCACGGGGTCGGCGGCGACGTCGG | 314.9 | -12.1 | 1.7 | -7.3 | 2.5E-13 | 7.4E-13 | MB231/MB468 | MITO |
| >hg38_21_+_8433172_8446622_RNA45SN1_45S_50ntFlanks@9129.9152.24\|CGGGTCGGGGGGCGGGGCGGACTG | >hg38_21_+_8433172_8446622_RNA45SN1_45S_50ntFlanks | 9129 | 9152 | 24 | CGGGTCGGGGGGCGGGGCGGACTG | 210.8 | -10.7 | 1.5 | -7.4 | 1.9E-13 | 7.4E-13 | MB231/MB468 | NUC |
| >MI0000477\|hsa-mir-146a&WithFlank&5\|+\|160485346\|160485456@27.47.21(+1G)&[MIMAT0000449&hsa-miR-146a-5p&offsets\|0\|-1(+1G);m-129&5\|+\|160485372\|160485393&offsets\|0\|-1(+1G)]\|TGAGAACTGAATTCCATGGGTG | >MI0000477\|hsa-mir-146a&WithFlank&5\|+\|160485346\|160485456 | 27 | 47 | 21(+1G) | TGAGAACTGAATTCCATGGGTG | 2252.5 | -14.5 | 1.8 | -7.9 | 2.6E-15 | 7.4E-13 | BT20/MB231 | MP |
| >am_trna2_GlyGCC_21_-_18827107_18827177@1.22.22,trna35_GlyGCC_1_+_161413094_161413164@1.22.22,trna37_GlyGCC_1_+_161420467_161420537@1.22.22,trna39_GlyGCC_1_+_161427898_161427968@1.22.22,trna41_GlyGCC_1_+_161435258_161435328@1.22.22\|GCATGGGTGGTTCAGTGGTAGA | >am_trna2_GlyGCC_21_-_18827107_18827177 | 1 | 22 | 22 | GCATGGGTGGTTCAGTGGTAGA | 366.3 | 8.5 | 1.1 | 7.7 | 1.5E-14 | 7.4E-13 | BT20/MB231 | CYTO |
| >hg38_21_+_8433172_8446622_RNA45SN1_45S_50ntFlanks@11161.11189.29\|CACGGTCCCCCGCGAGGGGGGCCCGGGCA | >hg38_21_+_8433172_8446622_RNA45SN1_45S_50ntFlanks | 11161 | 11189 | 29 | CACGGTCCCCCGCGAGGGGGGCCCGGGCA | 423.9 | -12.6 | 1.7 | -7.4 | 1.6E-13 | 7.4E-13 | MB231/MB468 | MP |
| >hg38_21_+_8433172_8446622_RNA45SN1_45S_50ntFlanks@7973.8010.38\|GACGCGACCTCAGATCAGACGTGGCGACCCGCTGAATT | >hg38_21_+_8433172_8446622_RNA45SN1_45S_50ntFlanks | 7973 | 8010 | 38 | GACGCGACCTCAGATCAGACGTGGCGACCCGCTGAATT | 1500.0 | 8.1 | 1.1 | 7.4 | 1.9E-13 | 7.4E-13 | BT20/MB468 | CYTO |
| >hg38_21_+_8433172_8446622_RNA45SN1_45S_50ntFlanks@12850.12867.18\|CTTCGTCCTGGGAAACGG | >hg38_21_+_8433172_8446622_RNA45SN1_45S_50ntFlanks | 12850 | 12867 | 18 | CTTCGTCCTGGGAAACGG | 183.1 | -11.4 | 1.5 | -7.4 | 1.6E-13 | 7.4E-13 | MB231/MB468 | MP |

<table>
<tr><td>>hg38_21_+_8433172_8446622_RNA45SN1_45S_50ntFlanks@11209.11232.24|CGGCGACTCTGGACGCGAGCCGGG</td><td>>hg38_21_+_8433172_8446622_RNA45SN1_45S_50ntFlanks</td><td>11209</td><td>11232</td><td>24</td><td>CGGCGACTCTGGACGCGAGCCGGG</td><td>24516.4</td><td>-7.6</td><td>1.0</td><td>-7.4</td><td>1.9E-13</td><td>7.4E-13</td><td>MB231/MB468</td><td>CYTO</td></tr>
<tr><td>>hg38_21_+_8433172_8446622_RNA45SN1_45S_50ntFlanks@3850.3881.32|GATAACTGTGGTAATTCTAGAGCTAATACATG</td><td>>hg38_21_+_8433172_8446622_RNA45SN1_45S_50ntFlanks</td><td>3850</td><td>3881</td><td>32</td><td>GATAACTGTGGTAATTCTAGAGCTAATACATG</td><td>235.4</td><td>-11.7</td><td>1.6</td><td>-7.4</td><td>1.7E-13</td><td>7.5E-13</td><td>MB231/MB468</td><td>MP</td></tr>
<tr><td>>hg38_21_+_8433172_8446622_RNA45SN1_45S_50ntFlanks@4726.4759.34|TAATCAAGAACGAAAGTCGGAGGTTCGAAGACGA</td><td>>hg38_21_+_8433172_8446622_RNA45SN1_45S_50ntFlanks</td><td>4726</td><td>4759</td><td>34</td><td>TAATCAAGAACGAAAGTCGGAGGTTCGAAGACGA</td><td>917.8</td><td>-4.5</td><td>0.6</td><td>-7.4</td><td>1.5E-13</td><td>7.5E-13</td><td>BT20/MB468</td><td>MITO</td></tr>
<tr><td>>hg38_21_+_8433172_8446622_RNA45SN1_45S_50ntFlanks@12399.12436.38|ACGTCGCTTTTTGATCCTTCGATGTCGGCTCTTCCTAT</td><td>>hg38_21_+_8433172_8446622_RNA45SN1_45S_50ntFlanks</td><td>12399</td><td>12436</td><td>38</td><td>ACGTCGCTTTTTGATCCTTCGATGTCGGCTCTTCCTAT</td><td>369.1</td><td>-10.6</td><td>1.4</td><td>-7.4</td><td>1.5E-13</td><td>7.5E-13</td><td>BT20/MB468</td><td>MITO</td></tr>
<tr><td>>hg38_21_+_8433172_8446622_RNA45SN1_45S_50ntFlanks@7981.7999.19|CTCAGATCAGACGTGGCGA</td><td>>hg38_21_+_8433172_8446622_RNA45SN1_45S_50ntFlanks</td><td>7981</td><td>7999</td><td>19</td><td>CTCAGATCAGACGTGGCGA</td><td>139.2</td><td>-11.0</td><td>1.5</td><td>-7.4</td><td>1.7E-13</td><td>7.5E-13</td><td>MB231/MB468</td><td>MP</td></tr>
<tr><td>>hg38_21_+_8433172_8446622_RNA45SN1_45S_50ntFlanks@11729.11749.21|TAACTATGACTCTCTTAAGGT</td><td>>hg38_21_+_8433172_8446622_RNA45SN1_45S_50ntFlanks</td><td>11729</td><td>11749</td><td>21</td><td>TAACTATGACTCTCTTAAGGT</td><td>2482.3</td><td>-6.1</td><td>0.8</td><td>-7.3</td><td>2.5E-13</td><td>7.5E-13</td><td>MB231/MB468</td><td>MITO</td></tr>
<tr><td>>am_trna10_ValCAC_5_-_180649395_180649467@1.32.32,trna12_ValAAC_5_-_180645270_180645342@1.32.32,trna132_ValAAC_6_-_27721179_27721251@1.32.32,trna136_ValAAC_6_-_27648885_27648957@1.32.32,trna139_ValAAC_6_-_27618707_27618779@1.32.32,trna18_ValCAC_5_-_180529253_180529325@1.32.32,trna2_ValAAC_3_+_169490018_169490090@1.32.32,trna2_ValCAC_5_+_180524070_180524142@1.32.32,trna4_ValAAC_5_+_180591154_180591226@1.32.32,trna5_ValAAC_5_+_180596610_180596682@1.32.32,trna6_ValCAC_5_+_180600650_180600722@1.32.32,trna85_ValCAC_1_-_161369490_161369562@1.32.32,trna90_ValCAC_1_-_149684088_149684161@1.32.32,trna98_ValCAC_1_-_149298555_149298627@1.32.32,trna9_ValCAC_6_+_26538282_26538354@1.32.32|GTTTCCGTAGTGTAGTGGTTATCACGTTCGCC</td><td>>am_trna10_ValCAC_5_-_180649395_180649467</td><td>1</td><td>32</td><td>32</td><td>GTTTCCGTAGTGTAGTGGTTATCACGTTCGCC</td><td>4690.3</td><td>6.5</td><td>0.9</td><td>7.3</td><td>2.5E-13</td><td>7.5E-13</td><td>MB231/MB468</td><td>MITO</td></tr>
<tr><td>>hg38_21_+_8433172_8446622_RNA45SN1_45S_50ntFlanks@4640.4662.23|GCATTCGTATTGCGCCGCTAGAG</td><td>>hg38_21_+_8433172_8446622_RNA45SN1_45S_50ntFlanks</td><td>4640</td><td>4662</td><td>23</td><td>GCATTCGTATTGCGCCGCTAGAG</td><td>114.1</td><td>-10.7</td><td>1.5</td><td>-7.4</td><td>1.7E-13</td><td>7.5E-13</td><td>MB231/MB468</td><td>MP</td></tr>
<tr><td>>am_trnaMT_ArgTCG_MT_+_10405_10469@28.48.21|ATTTCGACTCATTAAATTATG</td><td>>am_trnaMT_ArgTCG_MT_+_10405_10469</td><td>28</td><td>48</td><td>21</td><td>ATTTCGACTCATTAAATTATG</td><td>610.3</td><td>-7.2</td><td>1.0</td><td>-7.4</td><td>1.7E-13</td><td>7.5E-13</td><td>MB231/MB468</td><td>MP</td></tr>
<tr><td>>hg38_21_+_8433172_8446622_RNA45SN1_45S_50ntFlanks@10684.10706.23|CGGCGGCGTCCGGTGAGCTCTCG</td><td>>hg38_21_+_8433172_8446622_RNA45SN1_45S_50ntFlanks</td><td>10684</td><td>10706</td><td>23</td><td>CGGCGGCGTCCGGTGAGCTCTCG</td><td>11931.1</td><td>-10.6</td><td>1.4</td><td>-7.3</td><td>2.5E-13</td><td>7.5E-13</td><td>MB231/MB468</td><td>MITO</td></tr>
<tr><td>>hg38_21_+_8433172_8446622_RNA45SN1_45S_50ntFlanks@11209.11239.31|CGGCGACTCTGGACGCGAGCCGGGCCCTTCC</td><td>>hg38_21_+_8433172_8446622_RNA45SN1_45S_50ntFlanks</td><td>11209</td><td>11239</td><td>31</td><td>CGGCGACTCTGGACGCGAGCCGGGCCCTTCC</td><td>208.6</td><td>-7.4</td><td>1.0</td><td>-7.4</td><td>1.9E-13</td><td>7.5E-13</td><td>MB231/MB468</td><td>CYTO</td></tr>
<tr><td>>hg38_21_+_8433172_8446622_RNA45SN1_45S_50ntFlanks@12944.12972.29|CGTAGACGACCTGCTTCTGGGTCGGGGTT</td><td>>hg38_21_+_8433172_8446622_RNA45SN1_45S_50ntFlanks</td><td>12944</td><td>12972</td><td>29</td><td>CGTAGACGACCTGCTTCTGGGTCGGGGTT</td><td>545.5</td><td>-12.2</td><td>1.6</td><td>-7.8</td><td>7.9E-15</td><td>7.5E-13</td><td>BT20/MB468</td><td>MP</td></tr>
<tr><td>>hg38_21_+_8433172_8446622_RNA45SN1_45S_50ntFlanks@9166.9183.18|CCGGGCGGGTCGCGCCGT</td><td>>hg38_21_+_8433172_8446622_RNA45SN1_45S_50ntFlanks</td><td>9166</td><td>9183</td><td>18</td><td>CCGGGCGGGTCGCGCCGT</td><td>579.4</td><td>-12.3</td><td>1.6</td><td>-7.8</td><td>7.9E-15</td><td>7.5E-13</td><td>BT20/MB468</td><td>MP</td></tr>
<tr><td>>hg38_21_+_8433172_8446622_RNA45SN1_45S_50ntFlanks@12907.12933.27|CACCGCACGTTCGTGGGGAACCTGGCG</td><td>>hg38_21_+_8433172_8446622_RNA45SN1_45S_50ntFlanks</td><td>12907</td><td>12933</td><td>27</td><td>CACCGCACGTTCGTGGGGAACCTGGCG</td><td>1254.3</td><td>-3.6</td><td>0.5</td><td>-7.4</td><td>1.5E-13</td><td>7.5E-13</td><td>BT20/MB468</td><td>MITO</td></tr>
<tr><td>>hg38_21_+_8433172_8446622_RNA45SN1_45S_50ntFlanks@10905.10935.31|CTGGGCGCGCGCCGCGGCTGGACGAGGCGCC</td><td>>hg38_21_+_8433172_8446622_RNA45SN1_45S_50ntFlanks</td><td>10905</td><td>10935</td><td>31</td><td>CTGGGCGCGCGCCGCGGCTGGACGAGGCGCC</td><td>230.1</td><td>-10.6</td><td>1.4</td><td>-7.4</td><td>1.9E-13</td><td>7.6E-13</td><td>MB231/MB468</td><td>CYTO</td></tr>
<tr><td>>hg38_21_+_8433172_8446622_RNA45SN1_45S_50ntFlanks@12324.12356.33|TTGGGTTTTAAGCAGGAGGTGTCAGAAAAGTTA</td><td>>hg38_21_+_8433172_8446622_RNA45SN1_45S_50ntFlanks</td><td>12324</td><td>12356</td><td>33</td><td>TTGGGTTTTAAGCAGGAGGTGTCAGAAAAGTTA</td><td>238.5</td><td>-11.7</td><td>1.6</td><td>-7.3</td><td>2.5E-13</td><td>7.6E-13</td><td>MB231/MB468</td><td>MITO</td></tr>
<tr><td>>MI0000111|hsa-mir-105-1&WithFlank&X|-|152392213|152392305@19.40.22;MI0000112|hsa-mir-105-2&WithFlank&X|-|152394406|152394498@19.40.22&[MIMAT0000102&hsa-miR-105-5p&offsets|0|-1;m-735&X|-|152392266|152392287&offsets|0|0];[MIMAT0000102_1&hsa-miR-105-5p&offsets|0|-1;m-735&X|-|152394459|152394480&offsets|0|0]|TCAAATGCTCAGACTCCTGTGG</td><td>>MI0000111|hsa-mir-105-1&WithFlank&X|-|152392213|152392305</td><td>19</td><td>40</td><td>22</td><td>TCAAATGCTCAGACTCCTGTGG</td><td>463.0</td><td>8.4</td><td>1.1</td><td>7.7</td><td>1.6E-14</td><td>7.6E-13</td><td>BT20/MB231</td><td>CYTO</td></tr>
<tr><td>>trna111_HisGTG_1_-_147774845_147774916@-1G.35.36,trna118_HisGTG_1_-_145396881_145396952@-1G.35.36,trna16_HisGTG_1_+_146544773_146544844@-1G.35.36,trna1_HisGTG_15_+_45493349_45493420@-1G.35.36,trna21_HisGTG_1_+_147753471_147753542@-1G.35.36,trna33_HisGTG_6_+_27125906_27125977@-1G.35.36,trna7_HisGTG_9_-_14433938_14434009@-1G.35.36,trna8_HisGTG_15_-_45492611_45492682@-1G.35.36,trna9_HisGTG_15_-_45490804_45490875@-1G.35.36|GGCCGTGATCGTATAGTGGTTAGTACTCTGCGTTGT</td><td>>trna111_HisGTG_1_-_147774845_147774916</td><td>-1G</td><td>35</td><td>36</td><td>GGCCGTGATCGTATAGTGGTTAGTACTCTGCGTTGT</td><td>534.5</td><td>12.3</td><td>1.7</td><td>7.4</td><td>1.7E-13</td><td>7.6E-13</td><td>MB231/MB468</td><td>MP</td></tr>
<tr><td>>hg38_21_+_8433172_8446622_RNA45SN1_45S_50ntFlanks@9534.9564.31|AGGAAACTCTGGTGGAGGTCCGTAGCGGTCC</td><td>>hg38_21_+_8433172_8446622_RNA45SN1_45S_50ntFlanks</td><td>9534</td><td>9564</td><td>31</td><td>AGGAAACTCTGGTGGAGGTCCGTAGCGGTCC</td><td>11179.3</td><td>-12.1</td><td>1.7</td><td>-7.3</td><td>2.9E-13</td><td>7.6E-13</td><td>BT20/MB468</td><td>NUC</td></tr>
<tr><td>>hg38_MT_+_598_1651_RNR1_12S_50ntFlanks@561.605.45|TAAAACTCAAAGGACCTGGCGGTGCTTCATATCCCTCTAGAGGAG</td><td>>hg38_MT_+_598_1651_RNR1_12S_50ntFlanks</td><td>561</td><td>605</td><td>45</td><td>TAAAACTCAAAGGACCTGGCGGTGCTTCATATCCCTCTAGAGGAG</td><td>177.3</td><td>-10.7</td><td>1.4</td><td>-7.6</td><td>2.1E-14</td><td>7.6E-13</td><td>BT20/MB231</td><td>MITO</td></tr>
<tr><td>>hg38_21_+_8433172_8446622_RNA45SN1_45S_50ntFlanks@12126.12148.23|CGACCCGCTCCGGGGACAGTGCC</td><td>>hg38_21_+_8433172_8446622_RNA45SN1_45S_50ntFlanks</td><td>12126</td><td>12148</td><td>23</td><td>CGACCCGCTCCGGGGACAGTGCC</td><td>442.0</td><td>-10.9</td><td>1.5</td><td>-7.4</td><td>1.5E-13</td><td>7.6E-13</td><td>BT20/MB468</td><td>MITO</td></tr>
<tr><td>>trna111_HisGTG_1_-_147774845_147774916@1.35.35,trna118_HisGTG_1_-_145396881_145396952@1.35.35,trna16_HisGTG_1_+_146544773_146544844@1.35.35,trna1_HisGTG_15_+_45493349_45493420@1.35.35,trna21_HisGTG_1_+_147753471_147753542@1.35.35,trna33_HisGTG_6_+_27125906_27125977@1.35.35,trna7_HisGTG_9_-_14433938_14434009@1.35.35,trna8_HisGTG_15_-_45492611_45492682@1.35.35,trna9_HisGTG_15_-_45490804_45490875@1.35.35|GCCGTGATCGTATAGTGGTTAGTACTCTGCGTTGT</td><td>>trna111_HisGTG_1_-_147774845_147774916</td><td>1</td><td>35</td><td>35</td><td>GCCGTGATCGTATAGTGGTTAGTACTCTGCGTTGT</td><td>4256.8</td><td>12.0</td><td>1.6</td><td>7.3</td><td>2.0E-13</td><td>7.6E-13</td><td>BT20/MB468</td><td>CYTO</td></tr>
<tr><td>>hg38_21_+_8433172_8446622_RNA45SN1_45S_50ntFlanks@9574.9596.23|AATCGGTCGTCCGACCTGGGTAT</td><td>>hg38_21_+_8433172_8446622_RNA45SN1_45S_50ntFlanks</td><td>9574</td><td>9596</td><td>23</td><td>AATCGGTCGTCCGACCTGGGTAT</td><td>3907.6</td><td>-7.3</td><td>1.0</td><td>-7.4</td><td>2.0E-13</td><td>7.6E-13</td><td>MB231/MB468</td><td>CYTO</td></tr>
<tr><td>>hg38_21_+_8433172_8446622_RNA45SN1_45S_50ntFlanks@3990.4023.34|TTGGTGACTCTAGATAACCTCGGGCCGATCGCAC</td><td>>hg38_21_+_8433172_8446622_RNA45SN1_45S_50ntFlanks</td><td>3990</td><td>4023</td><td>34</td><td>TTGGTGACTCTAGATAACCTCGGGCCGATCGCAC</td><td>817.4</td><td>4.2</td><td>0.6</td><td>7.4</td><td>2.0E-13</td><td>7.6E-13</td><td>MB231/MB468</td><td>CYTO</td></tr>
<tr><td>>hg38_21_+_8433172_8446622_RNA45SN1_45S_50ntFlanks@4027.4045.19|CCCCGTGGCGGCGACGACC</td><td>>hg38_21_+_8433172_8446622_RNA45SN1_45S_50ntFlanks</td><td>4027</td><td>4045</td><td>19</td><td>CCCCGTGGCGGCGACGACC</td><td>168.4</td><td>-11.3</td><td>1.5</td><td>-7.4</td><td>1.7E-13</td><td>7.6E-13</td><td>MB231/MB468</td><td>MP</td></tr>
<tr><td>>hg38_21_+_8433172_8446622_RNA45SN1_45S_50ntFlanks@11594.11613.20|CAGGGGAATCCGACTGTTTA</td><td>>hg38_21_+_8433172_8446622_RNA45SN1_45S_50ntFlanks</td><td>11594</td><td>11613</td><td>20</td><td>CAGGGGAATCCGACTGTTTA</td><td>1019.1</td><td>-6.0</td><td>0.8</td><td>-7.3</td><td>2.5E-13</td><td>7.7E-13</td><td>MB231/MB468</td><td>MITO</td></tr>
<tr><td>>hg38_21_+_8433172_8446622_RNA45SN1_45S_50ntFlanks@9534.9555.22|AGGAAACTCTGGTGGAGGTCCG</td><td>>hg38_21_+_8433172_8446622_RNA45SN1_45S_50ntFlanks</td><td>9534</td><td>9555</td><td>22</td><td>AGGAAACTCTGGTGGAGGTCCG</td><td>263.5</td><td>-10.8</td><td>1.5</td><td>-7.4</td><td>2.0E-13</td><td>7.7E-13</td><td>MB231/MB468</td><td>CYTO</td></tr>
<tr><td>>hg38_21_+_8433172_8446622_RNA45SN1_45S_50ntFlanks@9475.9503.29|CGAGCGCACGTGTTAGGACCCGAAAGATG</td><td>>hg38_21_+_8433172_8446622_RNA45SN1_45S_50ntFlanks</td><td>9475</td><td>9503</td><td>29</td><td>CGAGCGCACGTGTTAGGACCCGAAAGATG</td><td>1227.2</td><td>-10.1</td><td>1.4</td><td>-7.3</td><td>2.6E-13</td><td>7.7E-13</td><td>MB231/MB468</td><td>MITO</td></tr>
<tr><td>>MI0000079|hsa-mir-23a&WithFlank&19|-|13836581|13836665@54.74.21&[MIMAT0000078&hsa-miR-23a-3p&offsets|+3|+3;m-31&19|-|13836595|13836615&offsets|+3|+3]|ACATTGCCAGGGATTTCCAAC</td><td>>MI0000079|hsa-mir-23a&WithFlank&19|-|13836581|13836665</td><td>54</td><td>74</td><td>21</td><td>ACATTGCCAGGGATTTCCAAC</td><td>541.4</td><td>26.6</td><td>3.6</td><td>7.3</td><td>2.6E-13</td><td>7.7E-13</td><td>MB231/MB468</td><td>MITO</td></tr>
<tr><td>>hg38_21_+_8433172_8446622_RNA45SN1_45S_50ntFlanks@9534.9571.38|AGGAAACTCTGGTGGAGGTCCGTAGCGGTCCTGACGTG</td><td>>hg38_21_+_8433172_8446622_RNA45SN1_45S_50ntFlanks</td><td>9534</td><td>9571</td><td>38</td><td>AGGAAACTCTGGTGGAGGTCCGTAGCGGTCCTGACGTG</td><td>16303.4</td><td>-9.5</td><td>1.3</td><td>-7.4</td><td>1.7E-13</td><td>7.7E-13</td><td>MB231/MB468</td><td>MP</td></tr>
<tr><td>>hg38_21_+_8433172_8446622_RNA45SN1_45S_50ntFlanks@8194.8217.24|CCCGTGGACGGTGTGAGGCCGGTA</td><td>>hg38_21_+_8433172_8446622_RNA45SN1_45S_50ntFlanks</td><td>8194</td><td>8217</td><td>24</td><td>CCCGTGGACGGTGTGAGGCCGGTA</td><td>143.9</td><td>-11.0</td><td>1.5</td><td>-7.4</td><td>1.7E-13</td><td>7.7E-13</td><td>MB231/MB468</td><td>MP</td></tr>
<tr><td>>hg38_21_+_8433172_8446622_RNA45SN1_45S_50ntFlanks@12933.12956.24|GCTAAACCATTCGTAGACGACCTG</td><td>>hg38_21_+_8433172_8446622_RNA45SN1_45S_50ntFlanks</td><td>12933</td><td>12956</td><td>24</td><td>GCTAAACCATTCGTAGACGACCTG</td><td>6480.3</td><td>-11.7</td><td>1.6</td><td>-7.3</td><td>2.9E-13</td><td>7.7E-13</td><td>BT20/MB468</td><td>NUC</td></tr>
<tr><td>>hg38_21_+_8433172_8446622_RNA45SN1_45S_50ntFlanks@11586.11609.24|GTGCGGACCAGGGGAATCCGACTG</td><td>>hg38_21_+_8433172_8446622_RNA45SN1_45S_50ntFlanks</td><td>11586</td><td>11609</td><td>24</td><td>GTGCGGACCAGGGGAATCCGACTG</td><td>682.0</td><td>-5.2</td><td>0.7</td><td>-7.3</td><td>2.9E-13</td><td>7.7E-13</td><td>BT20/MB468</td><td>NUC</td></tr>
<tr><td>>hg38_21_+_8433172_8446622_RNA45SN1_45S_50ntFlanks@10350.10374.25|ACGAGAACTTTGAAGGCCGAAGTGG</td><td>>hg38_21_+_8433172_8446622_RNA45SN1_45S_50ntFlanks</td><td>10350</td><td>10374</td><td>25</td><td>ACGAGAACTTTGAAGGCCGAAGTGG</td><td>7123.7</td><td>-9.2</td><td>1.3</td><td>-7.4</td><td>1.7E-13</td><td>7.7E-13</td><td>MB231/MB468</td><td>MP</td></tr>
<tr><td>>hg38_21_+_8433172_8446622_RNA45SN1_45S_50ntFlanks@6777.6806.30|TCCCGGGGCTACGCCTGTCTGAGCGTCGCT</td><td>>hg38_21_+_8433172_8446622_RNA45SN1_45S_50ntFlanks</td><td>6777</td><td>6806</td><td>30</td><td>TCCCGGGGCTACGCCTGTCTGAGCGTCGCT</td><td>3005.0</td><td>5.6</td><td>0.8</td><td>7.4</td><td>1.6E-13</td><td>7.8E-13</td><td>BT20/MB468</td><td>MITO</td></tr>
<tr><td>>hg38_21_+_8433172_8446622_RNA45SN1_45S_50ntFlanks@6645.6670.26|CTCGTACGACTCTTAGCGGTGGATCA</td><td>>hg38_21_+_8433172_8446622_RNA45SN1_45S_50ntFlanks</td><td>6645</td><td>6670</td><td>26</td><td>CTCGTACGACTCTTAGCGGTGGATCA</td><td>743.4</td><td>-7.2</td><td>1.0</td><td>-7.4</td><td>1.7E-13</td><td>7.8E-13</td><td>MB231/MB468</td><td>MP</td></tr>
<tr><td>>hg38_21_+_8433172_8446622_RNA45SN1_45S_50ntFlanks@4003.4021.19|ATAACCTCGGGCCGATCGC</td><td>>hg38_21_+_8433172_8446622_RNA45SN1_45S_50ntFlanks</td><td>4003</td><td>4021</td><td>19</td><td>ATAACCTCGGGCCGATCGC</td><td>224.8</td><td>-10.8</td><td>1.5</td><td>-7.3</td><td>2.0E-13</td><td>7.8E-13</td><td>MB231/MB468</td><td>NUC</td></tr>
<tr><td>>hg38_21_+_8433172_8446622_RNA45SN1_45S_50ntFlanks@12625.12654.30|CAATGGGGCGAAGCTACCATCTGTGGGATT</td><td>>hg38_21_+_8433172_8446622_RNA45SN1_45S_50ntFlanks</td><td>12625</td><td>12654</td><td>30</td><td>CAATGGGGCGAAGCTACCATCTGTGGGATT</td><td>10131.2</td><td>-9.6</td><td>1.3</td><td>-7.3</td><td>2.9E-13</td><td>7.8E-13</td><td>BT20/MB468</td><td>NUC</td></tr>
<tr><td>>MI0000077|hsa-mir-21&WithFlank&17|+|59841260|59841343@14.35.22(+2U)&[MIMAT0000076&hsa-miR-21-5p&offsets|0|0(+2U);m-1&17|+|59841273|59841294&offsets|0|0(+2U)]|TAGCTTATCAGACTGATGTTGATT</td><td>>MI0000077|hsa-mir-21&WithFlank&17|+|59841260|59841343</td><td>14</td><td>35</td><td>22(+2U)</td><td>TAGCTTATCAGACTGATGTTGATT</td><td>655.6</td><td>7.6</td><td>1.0</td><td>7.3</td><td>2.0E-13</td><td>7.8E-13</td><td>MB231/MB468</td><td>NUC</td></tr>
<tr><td>>hg38_21_+_8433172_8446622_RNA45SN1_45S_50ntFlanks@12202.12232.31|CTAAGGCGAGCTCAGGGAGGACAGAAACCTC</td><td>>hg38_21_+_8433172_8446622_RNA45SN1_45S_50ntFlanks</td><td>12202</td><td>12232</td><td>31</td><td>CTAAGGCGAGCTCAGGGAGGACAGAAACCTC</td><td>341.6</td><td>-12.3</td><td>1.7</td><td>-7.4</td><td>1.7E-13</td><td>7.8E-13</td><td>MB231/MB468</td><td>MP</td></tr>
<tr><td>>hg38_21_+_8433172_8446622_RNA45SN1_45S_50ntFlanks@4722.4767.46|TCATTAATCAAGAACGAAAGTCGGAGGTTCGAAGACGATCAGATAC</td><td>>hg38_21_+_8433172_8446622_RNA45SN1_45S_50ntFlanks</td><td>4722</td><td>4767</td><td>46</td><td>TCATTAATCAAGAACGAAAGTCGGAGGTTCGAAGACGATCAGATAC</td><td>280.3</td><td>9.2</td><td>1.3</td><td>7.3</td><td>2.0E-13</td><td>7.8E-13</td><td>MB231/MB468</td><td>CYTO</td></tr>
<tr><td>>hg38_21_+_8433172_8446622_RNA45SN1_45S_50ntFlanks@5487.5519.33|CGCTGAGAAGACGGTCGAACTTGACTATCTAGA</td><td>>hg38_21_+_8433172_8446622_RNA45SN1_45S_50ntFlanks</td><td>5487</td><td>5519</td><td>33</td><td>CGCTGAGAAGACGGTCGAACTTGACTATCTAGA</td><td>268.9</td><td>-11.9</td><td>1.6</td><td>-7.4</td><td>1.7E-13</td><td>7.8E-13</td><td>MB231/MB468</td><td>MP</td></tr>
</table>

<table>
<tr><td>>hg38_21_+_8433172_8446622_RNA45SN1_45S_50ntFlanks@4723.4755.33|CATTAATCAAGAACGAAAGTCGGAGGTTCGAAG</td><td>>hg38_21_+_8433172_8446622_RNA45SN1_45S_50ntFlanks</td><td>4723</td><td>4755</td><td>33</td><td>CATTAATCAAGAACGAAAGTCGGAGGTTCGAAG</td><td>185.4</td><td>-11.4</td><td>1.5</td><td>-7.4</td><td>1.7E-13</td><td>7.8E-13</td><td>MB231/MB468</td><td>MP</td></tr>
<tr><td>>hg38_21_+_8433172_8446622_RNA45SN1_45S_50ntFlanks@10407.10429.23|GGGTCAGTCGGTCCTGAGAGATG</td><td>>hg38_21_+_8433172_8446622_RNA45SN1_45S_50ntFlanks</td><td>10407</td><td>10429</td><td>23</td><td>GGGTCAGTCGGTCCTGAGAGATG</td><td>484.8</td><td>-9.2</td><td>1.2</td><td>-7.3</td><td>2.1E-13</td><td>7.8E-13</td><td>MB231/MB468</td><td>NUC</td></tr>
<tr><td>>hg38_21_+_8433172_8446622_RNA45SN1_45S_50ntFlanks@7981.8005.25|CTCAGATCAGACGTGGCGACCCGCT</td><td>>hg38_21_+_8433172_8446622_RNA45SN1_45S_50ntFlanks</td><td>7981</td><td>8005</td><td>25</td><td>CTCAGATCAGACGTGGCGACCCGCT</td><td>334.0</td><td>-4.0</td><td>0.5</td><td>-7.3</td><td>2.1E-13</td><td>7.8E-13</td><td>MB231/MB468</td><td>NUC</td></tr>
<tr><td>>hg38_21_+_8433172_8446622_RNA45SN1_45S_50ntFlanks@10808.10825.18|CAATGTAGGTAAGGGAAG</td><td>>hg38_21_+_8433172_8446622_RNA45SN1_45S_50ntFlanks</td><td>10808</td><td>10825</td><td>18</td><td>CAATGTAGGTAAGGGAAG</td><td>505.5</td><td>-12.1</td><td>1.6</td><td>-7.8</td><td>8.3E-15</td><td>7.9E-13</td><td>BT20/MB468</td><td>MP</td></tr>
<tr><td>>MI0000301|hsa-mir-224&WithFlank&X|-|151958572|151958664@14.34.21&[MIMAT0000281&hsa-miR-224-5p&offsets|+1|-3;m-238&X|-|151958630|151958651&offsets|0|-1]|CAAGTCACTAGTGGTTCCGTT</td><td>>MI0000301|hsa-mir-224&WithFlank&X|-|151958572|151958664</td><td>14</td><td>34</td><td>21</td><td>CAAGTCACTAGTGGTTCCGTT</td><td>672.7</td><td>13.1</td><td>1.7</td><td>7.8</td><td>8.4E-15</td><td>7.9E-13</td><td>BT20/MB468</td><td>MP</td></tr>
<tr><td>>hg38_21_+_8433172_8446622_RNA45SN1_45S_50ntFlanks@12407.12424.18|TTTTGATCCTTCGATGTC</td><td>>hg38_21_+_8433172_8446622_RNA45SN1_45S_50ntFlanks</td><td>12407</td><td>12424</td><td>18</td><td>TTTTGATCCTTCGATGTC</td><td>142.8</td><td>-11.0</td><td>1.5</td><td>-7.4</td><td>1.8E-13</td><td>7.9E-13</td><td>MB231/MB468</td><td>MP</td></tr>
<tr><td>>hg38_21_+_8433172_8446622_RNA45SN1_45S_50ntFlanks@9545.9574.30|GTGGAGGTCCGTAGCGGTCCTGACGTGCAA</td><td>>hg38_21_+_8433172_8446622_RNA45SN1_45S_50ntFlanks</td><td>9545</td><td>9574</td><td>30</td><td>GTGGAGGTCCGTAGCGGTCCTGACGTGCAA</td><td>4724.5</td><td>-11.3</td><td>1.5</td><td>-7.3</td><td>3.0E-13</td><td>7.9E-13</td><td>BT20/MB468</td><td>NUC</td></tr>
<tr><td>>am_MI0005763|hsa-mir-941-1&WithFlank&20|+|63919443|63919526@53.74.22(+1U);MI0005764|hsa-mir-941-2&WithFlank&20|+|63919499|63919582@53.74.22(+1U);MI0005765|hsa-mir-941-3&WithFlank&20|+|63919555|63919638@53.74.22(+1U);MI0005766|hsa-mir-941-4&WithFlank&20|+|63919750|63919833@53.74.22(+1U);MI0031520|hsa-mir-941-5&WithFlank&20|+|63919862|63919945@53.74.22(+1U)&[MIMAT0004984&hsa-miR-941&offsets|0|-1(+1U);m-263&20|+|63919495|63919517&offsets|0|-1(+1U)];[MIMAT0004984_1&hsa-miR-941&offsets|0|-1(+1U);m-263&20|+|63919551|63919573&offsets|0|-1(+1U)];[MIMAT0004984_2&hsa-miR-941&offsets|0|-1(+1U);m-263&20|+|63919607|63919629&offsets|0|-1(+1U)];[MIMAT0004984_3&hsa-miR-941&offsets|0|-1(+1U);m-263&20|+|63919802|63919824&offsets|0|-1(+1U)];[MIMAT0004984_4&hsa-miR-941&offsets|0|-1(+1U);m-263&20|+|63919914|63919936&offsets|0|-1(+1U)]|CACCCGGCTGTGTGCACATGTGT</td><td>>am_MI0005763|hsa-mir-941-1&WithFlank&20|+|63919443|63919526</td><td>53</td><td>74</td><td>22(+1U)</td><td>CACCCGGCTGTGTGCACATGTGT</td><td>679.6</td><td>2.2</td><td>0.3</td><td>7.3</td><td>3.0E-13</td><td>7.9E-13</td><td>BT20/MB468</td><td>NUC</td></tr>
<tr><td>>hg38_21_+_8433172_8446622_RNA45SN1_45S_50ntFlanks@9032.9056.25|CCTCGTGGGGGGGCCGGGCCACCCC</td><td>>hg38_21_+_8433172_8446622_RNA45SN1_45S_50ntFlanks</td><td>9032</td><td>9056</td><td>25</td><td>CCTCGTGGGGGGGCCGGGCCACCCC</td><td>321.8</td><td>-12.1</td><td>1.7</td><td>-7.3</td><td>2.6E-13</td><td>7.9E-13</td><td>MB231/MB468</td><td>MITO</td></tr>
<tr><td>>hg38_21_+_8433172_8446622_RNA45SN1_45S_50ntFlanks@12998.13036.39|GCTGCGATCTATTGAAAGTCAGCCCTCGACACAAGGGTT</td><td>>hg38_21_+_8433172_8446622_RNA45SN1_45S_50ntFlanks</td><td>12998</td><td>13036</td><td>39</td><td>GCTGCGATCTATTGAAAGTCAGCCCTCGACACAAGGGTT</td><td>305.6</td><td>12.5</td><td>1.7</td><td>7.3</td><td>2.1E-13</td><td>7.9E-13</td><td>BT20/MB468</td><td>CYTO</td></tr>
<tr><td>>hg38_21_+_8433172_8446622_RNA45SN1_45S_50ntFlanks@12488.12525.38|AACGTGAGCTGGGTTTAGACCGTCGTGAGACAGGTTAG</td><td>>hg38_21_+_8433172_8446622_RNA45SN1_45S_50ntFlanks</td><td>12488</td><td>12525</td><td>38</td><td>AACGTGAGCTGGGTTTAGACCGTCGTGAGACAGGTTAG</td><td>193.5</td><td>-7.8</td><td>1.1</td><td>-7.3</td><td>2.0E-13</td><td>7.9E-13</td><td>MB231/MB468</td><td>CYTO</td></tr>
<tr><td>>MI0000078|hsa-mir-22&WithFlank&17|-|1713897|1713993@21.41.21&[MIMAT0004495&hsa-miR-22-5p&offsets|0|-1;m-217&17|-|1713953|1713973&offsets|0|0]|AGTTCTTCAGTGGCAAGCTTT</td><td>>MI0000078|hsa-mir-22&WithFlank&17|-|1713897|1713993</td><td>21</td><td>41</td><td>21</td><td>AGTTCTTCAGTGGCAAGCTTT</td><td>656.2</td><td>8.1</td><td>1.1</td><td>7.4</td><td>1.6E-13</td><td>7.9E-13</td><td>BT20/MB468</td><td>MITO</td></tr>
<tr><td>>trna10_LysCTT_16_+_3241501_3241573@1.35.35,trna13_LysCTT_14_-_58706613_58706685@1.35.35,trna2_LysCTT_15_+_79152904_79152976@1.35.35|GCCCGGCTAGCTCAGTCGGTAGAGCATGGGACTCT</td><td>>trna10_LysCTT_16_+_3241501_3241573</td><td>1</td><td>35</td><td>35</td><td>GCCCGGCTAGCTCAGTCGGTAGAGCATGGGACTCT</td><td>1071.7</td><td>5.5</td><td>0.8</td><td>7.3</td><td>3.0E-13</td><td>7.9E-13</td><td>BT20/MB468</td><td>NUC</td></tr>
<tr><td>>hg38_21_+_8433172_8446622_RNA45SN1_45S_50ntFlanks@11701.11728.28|TCAATGAAGCGCGGGTAAACGGCGGGAG</td><td>>hg38_21_+_8433172_8446622_RNA45SN1_45S_50ntFlanks</td><td>11701</td><td>11728</td><td>28</td><td>TCAATGAAGCGCGGGTAAACGGCGGGAG</td><td>179.7</td><td>-11.4</td><td>1.5</td><td>-7.4</td><td>1.8E-13</td><td>7.9E-13</td><td>MB231/MB468</td><td>MP</td></tr>
<tr><td>>hg38_21_+_8433172_8446622_RNA45SN1_45S_50ntFlanks@10713.10733.21|CTTGAAAATCCGGGGGAGAGG</td><td>>hg38_21_+_8433172_8446622_RNA45SN1_45S_50ntFlanks</td><td>10713</td><td>10733</td><td>21</td><td>CTTGAAAATCCGGGGGAGAGG</td><td>208.8</td><td>-11.6</td><td>1.6</td><td>-7.4</td><td>1.8E-13</td><td>7.9E-13</td><td>MB231/MB468</td><td>MP</td></tr>
<tr><td>>hg38_21_+_8433172_8446622_RNA45SN1_45S_50ntFlanks@6742.6771.30|TCGACACTTCGAACGCACTTGCGGCCCCGG</td><td>>hg38_21_+_8433172_8446622_RNA45SN1_45S_50ntFlanks</td><td>6742</td><td>6771</td><td>30</td><td>TCGACACTTCGAACGCACTTGCGGCCCCGG</td><td>837.5</td><td>-8.8</td><td>1.2</td><td>-7.3</td><td>2.7E-13</td><td>7.9E-13</td><td>MB231/MB468</td><td>MITO</td></tr>
<tr><td>>hg38_21_+_8433172_8446622_RNA45SN1_45S_50ntFlanks@10866.10904.39|CTAAGGGCTGGGTCGGTCGGGCTGGGGCGCGAAGCGGGG</td><td>>hg38_21_+_8433172_8446622_RNA45SN1_45S_50ntFlanks</td><td>10866</td><td>10904</td><td>39</td><td>CTAAGGGCTGGGTCGGTCGGGCTGGGGCGCGAAGCGGGG</td><td>1789.8</td><td>-11.0</td><td>1.5</td><td>-7.3</td><td>3.0E-13</td><td>7.9E-13</td><td>BT20/MB468</td><td>NUC</td></tr>
<tr><td>>hg38_21_+_8433172_8446622_RNA45SN1_45S_50ntFlanks@12128.12146.19|ACCCGCTCCGGGGACAGTG</td><td>>hg38_21_+_8433172_8446622_RNA45SN1_45S_50ntFlanks</td><td>12128</td><td>12146</td><td>19</td><td>ACCCGCTCCGGGGACAGTG</td><td>208.9</td><td>-10.0</td><td>1.4</td><td>-7.3</td><td>2.1E-13</td><td>8.0E-13</td><td>BT20/MB468</td><td>CYTO</td></tr>
<tr><td>>hg38_21_+_8433172_8446622_RNA45SN1_45S_50ntFlanks@10410.10432.23|TCAGTCGGTCCTGAGAGATGGGC</td><td>>hg38_21_+_8433172_8446622_RNA45SN1_45S_50ntFlanks</td><td>10410</td><td>10432</td><td>23</td><td>TCAGTCGGTCCTGAGAGATGGGC</td><td>1027.1</td><td>-5.0</td><td>0.7</td><td>-7.3</td><td>2.1E-13</td><td>8.0E-13</td><td>MB231/MB468</td><td>CYTO</td></tr>
<tr><td>>hg38_21_+_8433172_8446622_RNA45SN1_45S_50ntFlanks@11453.11471.19|TCCTCGGCGCGCGGCGGCG</td><td>>hg38_21_+_8433172_8446622_RNA45SN1_45S_50ntFlanks</td><td>11453</td><td>11471</td><td>19</td><td>TCCTCGGCGCGCGGCGGCG</td><td>165.4</td><td>-9.7</td><td>1.3</td><td>-7.3</td><td>2.1E-13</td><td>8.0E-13</td><td>BT20/MB468</td><td>CYTO</td></tr>
<tr><td>>MI0006444|hsa-mir-1307&WithFlank&10|-|103394247|103394407@86.107.22(+1A)&[MIMAT0005951&hsa-miR-1307-3p&offsets|0|0(+1A);m-66&10|-|103394300|103394322&offsets|0|-1(+1A)]|ACTCGGCGTGGCGTCGGTCGTGA</td><td>>MI0006444|hsa-mir-1307&WithFlank&10|-|103394247|103394407</td><td>86</td><td>107</td><td>22(+1A)</td><td>ACTCGGCGTGGCGTCGGTCGTGA</td><td>449.2</td><td>5.1</td><td>0.7</td><td>7.3</td><td>2.1E-13</td><td>8.0E-13</td><td>MB231/MB468</td><td>NUC</td></tr>
<tr><td>>MI0016887|hsa-mir-4521&WithFlank&17|+|8186939|8187010@10.34.25&[MIMAT0019058&hsa-miR-4521&offsets|0|+3;m-765&17|+|8186948|8186969&offsets|0|+3]|GCTAAGGAAGTCCTGTGCTCAGTTT</td><td>>MI0016887|hsa-mir-4521&WithFlank&17|+|8186939|8187010</td><td>10</td><td>34</td><td>25</td><td>GCTAAGGAAGTCCTGTGCTCAGTTT</td><td>221.9</td><td>6.6</td><td>0.9</td><td>7.3</td><td>2.1E-13</td><td>8.0E-13</td><td>MB231/MB468</td><td>NUC</td></tr>
<tr><td>>MI0005561|hsa-mir-877&WithFlank&6|+|30584326|30584423@7.28.22&[MIMAT0004949&hsa-miR-877-5p&offsets|0|+2;m-382&6|+|30584332|30584354&offsets|0|-1]|GTAGAGGAGATGGCGCAGGGGA</td><td>>MI0005561|hsa-mir-877&WithFlank&6|+|30584326|30584423</td><td>7</td><td>28</td><td>22</td><td>GTAGAGGAGATGGCGCAGGGGA</td><td>557.3</td><td>9.1</td><td>1.2</td><td>7.3</td><td>2.1E-13</td><td>8.0E-13</td><td>BT20/MB468</td><td>CYTO</td></tr>
<tr><td>>am_trna2_GlyGCC_21_-_18827107_18827177@1.27.27,trna35_GlyGCC_1_+_161413094_161413164@1.27.27,trna37_GlyGCC_1_+_161420467_161420537@1.27.27,trna39_GlyGCC_1_+_161427898_161427968@1.27.27,trna41_GlyGCC_1_+_161435258_161435328@1.27.27|GCATGGGTGGTTCAGTGGTAGAATTCT</td><td>>am_trna2_GlyGCC_21_-_18827107_18827177</td><td>1</td><td>27</td><td>27</td><td>GCATGGGTGGTTCAGTGGTAGAATTCT</td><td>2322.0</td><td>6.6</td><td>0.9</td><td>7.7</td><td>1.7E-14</td><td>8.0E-13</td><td>BT20/MB231</td><td>CYTO</td></tr>
<tr><td>>MI0001448|hsa-mir-425&WithFlank&3|-|49020142|49020240@60.81.22&[MIMAT0001343&hsa-miR-425-3p&offsets|-1|-1;m-258&3|-|49020160|49020181&offsets|0|0]|CATCGGGAATGTCGTGTCCGCC</td><td>>MI0001448|hsa-mir-425&WithFlank&3|-|49020142|49020240</td><td>60</td><td>81</td><td>22</td><td>CATCGGGAATGTCGTGTCCGCC</td><td>405.4</td><td>8.7</td><td>1.2</td><td>7.3</td><td>2.1E-13</td><td>8.0E-13</td><td>BT20/MB468</td><td>CYTO</td></tr>
<tr><td>>hg38_21_+_8433172_8446622_RNA45SN1_45S_50ntFlanks@11315.11336.22|CCACCCCACGTCTCGTCGCGCG</td><td>>hg38_21_+_8433172_8446622_RNA45SN1_45S_50ntFlanks</td><td>11315</td><td>11336</td><td>22</td><td>CCACCCCACGTCTCGTCGCGCG</td><td>155.1</td><td>-11.1</td><td>1.5</td><td>-7.4</td><td>1.8E-13</td><td>8.0E-13</td><td>MB231/MB468</td><td>MP</td></tr>
<tr><td>>hg38_21_+_8433172_8446622_RNA45SN1_45S_50ntFlanks@6649.6674.26|TACGACTCTTAGCGGTGGATCACTCG</td><td>>hg38_21_+_8433172_8446622_RNA45SN1_45S_50ntFlanks</td><td>6649</td><td>6674</td><td>26</td><td>TACGACTCTTAGCGGTGGATCACTCG</td><td>1337.8</td><td>5.2</td><td>0.7</td><td>7.4</td><td>1.6E-13</td><td>8.0E-13</td><td>BT20/MB468</td><td>MITO</td></tr>
<tr><td>>MI0000061|hsa-let-7a-2&WithFlank&11|-|122146516|122146599@11.32.22;MI0000062|hsa-let-7a-3&WithFlank&22|+|46112743|46112828@10.31.22;MI0000060|hsa-let-7a-1&WithFlank&9|+|94175951|94176042@12.33.22&[MIMAT0000062&hsa-let-7a-5p&offsets|0|0;m-6&11|-|122146568|122146589&offsets|0|0];[MIMAT0000062_1&hsa-let-7a-5p&offsets|0|0;m-5&22|+|46112752|46112773&offsets|0|0];[MIMAT0000062_2&hsa-let-7a-5p&offsets|0|0;m-4&9|+|94175962|94175983&offsets|0|0]|TGAGGTAGTAGGTTGTATAGTT</td><td>>MI0000061|hsa-let-7a-2&WithFlank&11|-|122146516|122146599</td><td>11</td><td>32</td><td>22</td><td>TGAGGTAGTAGGTTGTATAGTT</td><td>85035.8</td><td>6.3</td><td>0.9</td><td>7.3</td><td>2.7E-13</td><td>8.0E-13</td><td>MB231/MB468</td><td>MITO</td></tr>
<tr><td>>hg38_21_+_8433172_8446622_RNA45SN1_45S_50ntFlanks@11320.11343.24|CCACGTCTCGTCGCGCGCGTCC</td><td>>hg38_21_+_8433172_8446622_RNA45SN1_45S_50ntFlanks</td><td>11320</td><td>11343</td><td>24</td><td>CCACGTCTCGTCGCGCGCGCGTCC</td><td>554.3</td><td>-8.7</td><td>1.2</td><td>-7.3</td><td>2.7E-13</td><td>8.0E-13</td><td>MB231/MB468</td><td>MITO</td></tr>
<tr><td>>hg38_21_+_8433172_8446622_RNA45SN1_45S_50ntFlanks@11876.11900.25|AAGACCCTGTTGAGCTTGACTCTAG</td><td>>hg38_21_+_8433172_8446622_RNA45SN1_45S_50ntFlanks</td><td>11876</td><td>11900</td><td>25</td><td>AAGACCCTGTTGAGCTTGACTCTAG</td><td>530.6</td><td>-6.9</td><td>0.9</td><td>-7.3</td><td>2.7E-13</td><td>8.0E-13</td><td>MB231/MB468</td><td>MITO</td></tr>
<tr><td>>MI0000093|hsa-mir-92a-1&WithFlank&13|+|91351308|91351397@54.76.23&[MIMAT0000092&hsa-miR-92a-3p&offsets|0|+1;m-9&13|+|91351361|91351382&offsets|0|+1]|TATTGCACTTGTCCCGGCCTGTT</td><td>>MI0000093|hsa-mir-92a-1&WithFlank&13|+|91351308|91351397</td><td>54</td><td>76</td><td>23</td><td>TATTGCACTTGTCCCGGCCTGTT</td><td>2524.7</td><td>5.3</td><td>0.7</td><td>7.4</td><td>1.6E-13</td><td>8.0E-13</td><td>BT20/MB468</td><td>MITO</td></tr>
<tr><td>>hg38_21_+_8433172_8446622_RNA45SN1_45S_50ntFlanks@9339.9370.32|CGCACGAAAGCCGCCGTGGCGCAATGAAGGTG</td><td>>hg38_21_+_8433172_8446622_RNA45SN1_45S_50ntFlanks</td><td>9339</td><td>9370</td><td>32</td><td>CGCACGAAAGCCGCCGTGGCGCAATGAAGGTG</td><td>488.5</td><td>-11.0</td><td>1.5</td><td>-7.4</td><td>1.6E-13</td><td>8.0E-13</td><td>BT20/MB468</td><td>MITO</td></tr>
<tr><td>>hg38_21_+_8433172_8446622_RNA45SN1_45S_50ntFlanks@4853.4873.21|AAGTCTTTGGGTTCCGGGGGG</td><td>>hg38_21_+_8433172_8446622_RNA45SN1_45S_50ntFlanks</td><td>4853</td><td>4873</td><td>21</td><td>AAGTCTTTGGGTTCCGGGGGG</td><td>137.6</td><td>-11.0</td><td>1.5</td><td>-7.4</td><td>1.8E-13</td><td>8.0E-13</td><td>MB231/MB468</td><td>MP</td></tr>
<tr><td>>hg38_21_+_8433172_8446622_RNA45SN1_45S_50ntFlanks@12626.12654.29|AATGGGGCGAAGCTACCATCTGTGGGATT</td><td>>hg38_21_+_8433172_8446622_RNA45SN1_45S_50ntFlanks</td><td>12626</td><td>12654</td><td>29</td><td>AATGGGGCGAAGCTACCATCTGTGGGATT</td><td>19229.2</td><td>-10.1</td><td>1.4</td><td>-7.3</td><td>3.0E-13</td><td>8.0E-13</td><td>BT20/MB468</td><td>NUC</td></tr>
<tr><td>>MI0006273|hsa-mir-1180&WithFlank&17|-|19344500|19344580@47.68.22&[MIMAT0005825&hsa-miR-1180-3p&offsets|0|0;m-214&17|-|19344513|19344534&offsets|0|0]|TTTCCGGCTCGCGTGGGTGTGT</td><td>>MI0006273|hsa-mir-1180&WithFlank&17|-|19344500|19344580</td><td>47</td><td>68</td><td>22</td><td>TTTCCGGCTCGCGTGGGTGTGT</td><td>628.7</td><td>5.9</td><td>0.8</td><td>7.3</td><td>2.1E-13</td><td>8.1E-13</td><td>MB231/MB468</td><td>CYTO</td></tr>
<tr><td>>hg38_21_+_8433172_8446622_RNA45SN1_45S_50ntFlanks@9304.9334.31|ACCAAGGAGTCTAACACGTGCGCGAGTCGGG</td><td>>hg38_21_+_8433172_8446622_RNA45SN1_45S_50ntFlanks</td><td>9304</td><td>9334</td><td>31</td><td>ACCAAGGAGTCTAACACGTGCGCGAGTCGGG</td><td>183.8</td><td>-11.4</td><td>1.5</td><td>-7.4</td><td>1.8E-13</td><td>8.1E-13</td><td>MB231/MB468</td><td>MP</td></tr>
<tr><td>>hg38_21_+_8433172_8446622_RNA45SN1_45S_50ntFlanks@12170.12188.19|CGGTACACCTGTCAAACGG</td><td>>hg38_21_+_8433172_8446622_RNA45SN1_45S_50ntFlanks</td><td>12170</td><td>12188</td><td>19</td><td>CGGTACACCTGTCAAACGG</td><td>930.6</td><td>-13.6</td><td>1.9</td><td>-7.3</td><td>2.7E-13</td><td>8.1E-13</td><td>MB231/MB468</td><td>MITO</td></tr>
<tr><td>>hg38_21_+_8433172_8446622_RNA45SN1_45S_50ntFlanks@5420.5444.25|CCGATTGGATGGTTTAGTGAGGCCC</td><td>>hg38_21_+_8433172_8446622_RNA45SN1_45S_50ntFlanks</td><td>5420</td><td>5444</td><td>25</td><td>CCGATTGGATGGTTTAGTGAGGCCC</td><td>579.0</td><td>-12.1</td><td>1.7</td><td>-7.3</td><td>2.1E-13</td><td>8.1E-13</td><td>MB231/MB468</td><td>NUC</td></tr>
<tr><td>>hg38_21_+_8433172_8446622_RNA45SN1_45S_50ntFlanks@12418.12435.18|CGATGTCGGCTCTTCCTA</td><td>>hg38_21_+_8433172_8446622_RNA45SN1_45S_50ntFlanks</td><td>12418</td><td>12435</td><td>18</td><td>CGATGTCGGCTCTTCCTA</td><td>14060.0</td><td>-5.2</td><td>0.7</td><td>-7.3</td><td>2.1E-13</td><td>8.1E-13</td><td>MB231/MB468</td><td>CYTO</td></tr>
<tr><td>>hg38_21_+_8433172_8446622_RNA45SN1_45S_50ntFlanks@3802.3830.29|CATTAAATCAGTTATGGTTCCTTTGGTCG</td><td>>hg38_21_+_8433172_8446622_RNA45SN1_45S_50ntFlanks</td><td>3802</td><td>3830</td><td>29</td><td>CATTAAATCAGTTATGGTTCCTTTGGTCG</td><td>124.0</td><td>-9.9</td><td>1.4</td><td>-7.3</td><td>2.1E-13</td><td>8.1E-13</td><td>MB231/MB468</td><td>NUC</td></tr>
<tr><td>>hg38_21_+_8433172_8446622_RNA45SN1_45S_50ntFlanks@11891.11912.22|TTGACTCTAGTCTGGCACGGTG</td><td>>hg38_21_+_8433172_8446622_RNA45SN1_45S_50ntFlanks</td><td>11891</td><td>11912</td><td>22</td><td>TTGACTCTAGTCTGGCACGGTG</td><td>1115.6</td><td>-9.1</td><td>1.2</td><td>-7.3</td><td>2.7E-13</td><td>8.1E-13</td><td>MB231/MB468</td><td>MITO</td></tr>
<tr><td>>hg38_21_+_8433172_8446622_RNA45SN1_45S_50ntFlanks@8154.8173.20|CCGCTCGTGGGGGGCCCAAG</td><td>>hg38_21_+_8433172_8446622_RNA45SN1_45S_50ntFlanks</td><td>8154</td><td>8173</td><td>20</td><td>CCGCTCGTGGGGGGCCCAAG</td><td>356.6</td><td>-12.3</td><td>1.7</td><td>-7.3</td><td>2.7E-13</td><td>8.2E-13</td><td>MB231/MB468</td><td>MITO</td></tr>
<tr><td>>hg38_21_+_8433172_8446622_RNA45SN1_45S_50ntFlanks@12002.12032.31|TGCGGGCCGCCGGTGAAATACCACTACTCTG</td><td>>hg38_21_+_8433172_8446622_RNA45SN1_45S_50ntFlanks</td><td>12002</td><td>12032</td><td>31</td><td>TGCGGGCCGCCGGTGAAATACCACTACTCTG</td><td>11784.1</td><td>-12.1</td><td>1.7</td><td>-7.3</td><td>3.1E-13</td><td>8.2E-13</td><td>BT20/MB468</td><td>NUC</td></tr>
<tr><td>>MI0000342|hsa-mir-200b&WithFlank&1|+|1167098|1167204@63.83.21&[MIMAT0000318&hsa-miR-200b-3p&offsets|0|-1;m-93&1|+|1167160|1167182&offsets|0|-2]|TAATACTGCCTGGTAATGATG</td><td>>MI0000342|hsa-mir-200b&WithFlank&1|+|1167098|1167204</td><td>63</td><td>83</td><td>21</td><td>TAATACTGCCTGGTAATGATG</td><td>1572.8</td><td>-5.7</td><td>0.8</td><td>-7.4</td><td>1.8E-13</td><td>8.2E-13</td><td>MB231/MB468</td><td>MP</td></tr>
</table>

| | | | | | | | | | | | | | |
|---|---|---|---|---|---|---|---|---|---|---|---|---|---|
| >hg38_21_+_8433172_8446622_RNA45SN1_45S_50ntFlanks@4002.4022.21\|GATAACCTCGGGCCGATCGCA | >hg38_21_+_8433172_8446622_RNA45SN1_45S_50ntFlanks | 4002 | 4022 | 21 | GATAACCTCGGGCCGATCGCA | 133.6 | -10.9 | 1.5 | -7.4 | 1.8E-13 | 8.2E-13 | MB231/MB468 | MP |
| >MI0000683\|hsa-mir-181b-2&WithFlank&9\|+\|124693704\|124693804@22.46.25(+1U)&[MIMAT0000257_1&hsa-miR-181b-5p&offsets\|0\|+2(+1U);m-72&9\|+\|124693725\|124693747&offsets\|0\|+2(+1U)]\|AACATTCATTGCTGTCGGTGGGTTTT | >MI0000683\|hsa-mir-181b-2&WithFlank&9\|+\|124693704\|124693804 | 22 | 46 | 25(+1U) | AACATTCATTGCTGTCGGTGGGTTTT | 339.2 | 8.9 | 1.2 | 7.3 | 2.2E-13 | 8.2E-13 | MB231/MB468 | NUC |
| >hg38_1_-_228634819_228635039_RNA5S12_5S_50ntFlanks@134.171.38\|TGGGAGACCGCCTGGGAATACCGGGTGCTGTAGGCTTT | >hg38_1_-_228634819_228635039_RNA5S12_5S_50ntFlanks | 134 | 171 | 38 | TGGGAGACCGCCTGGGAATACCGGGTGCTGTAGGCTTT | 37807.3 | 6.6 | 0.9 | 7.3 | 2.2E-13 | 8.2E-13 | MB231/MB468 | NUC |
| >hg38_21_+_8433172_8446622_RNA45SN1_45S_50ntFlanks@4731.4765.35\|AAGAACGAAAGTCGGAGGTTCGAAGACGATCAGAT | >hg38_21_+_8433172_8446622_RNA45SN1_45S_50ntFlanks | 4731 | 4765 | 35 | AAGAACGAAAGTCGGAGGTTCGAAGACGATCAGAT | 2550.4 | 6.1 | 0.8 | 7.3 | 2.1E-13 | 8.2E-13 | MB231/MB468 | CYTO |
| >hg38_21_+_8433172_8446622_RNA45SN1_45S_50ntFlanks@12937.12956.20\|AACCATTCGTAGACGACCTG | >hg38_21_+_8433172_8446622_RNA45SN1_45S_50ntFlanks | 12937 | 12956 | 20 | AACCATTCGTAGACGACCTG | 5300.5 | -4.4 | 0.6 | -7.3 | 2.1E-13 | 8.2E-13 | MB231/MB468 | CYTO |
| >hg38_21_+_8433172_8446622_RNA45SN1_45S_50ntFlanks@13007.13025.19\|TATTGAAAGTCAGCCCTCG | >hg38_21_+_8433172_8446622_RNA45SN1_45S_50ntFlanks | 13007 | 13025 | 19 | TATTGAAAGTCAGCCCTCG | 1629.3 | -10.5 | 1.4 | -7.3 | 3.1E-13 | 8.2E-13 | BT20/MB468 | NUC |
| >hg38_21_+_8433172_8446622_RNA45SN1_45S_50ntFlanks@11795.11820.26\|AACGAGATTCCCACTGTCCCTACCTA | >hg38_21_+_8433172_8446622_RNA45SN1_45S_50ntFlanks | 11795 | 11820 | 26 | AACGAGATTCCCACTGTCCCTACCTA | 315.6 | -8.6 | 1.2 | -7.3 | 2.2E-13 | 8.2E-13 | MB231/MB468 | NUC |
| >hg38_21_+_8433172_8446622_RNA45SN1_45S_50ntFlanks@10683.10703.21\|CCGGCGGCGTCCGGTGAGCTC | >hg38_21_+_8433172_8446622_RNA45SN1_45S_50ntFlanks | 10683 | 10703 | 21 | CCGGCGGCGTCCGGTGAGCTC | 138.2 | -11.0 | 1.5 | -7.4 | 1.8E-13 | 8.2E-13 | MB231/MB468 | MP |
| >MI0000087\|hsa-mir-29a&WithFlank&7\|-\|130876741\|130876816@48.68.21(+1U)&[MIMAT0000086&hsa-miR-29a-3p&offsets\|0\|-1(+1U);m-26&7\|-\|130876749\|130876769&offsets\|0\|0(+1U)]\|TAGCACCATCTGAAATCGGTTT | >MI0000087\|hsa-mir-29a&WithFlank&7\|-\|130876741\|130876816 | 48 | 68 | 21(+1U) | TAGCACCATCTGAAATCGGTTT | 718.9 | 12.7 | 1.7 | 7.3 | 2.8E-13 | 8.2E-13 | MB231/MB468 | MITO |
| >hg38_21_+_8433172_8446622_RNA45SN1_45S_50ntFlanks@3922.3959.38\|TTTATCAGATCAAAACCAACCCGGTCAGCCCCTCTCCG | >hg38_21_+_8433172_8446622_RNA45SN1_45S_50ntFlanks | 3922 | 3959 | 38 | TTTATCAGATCAAAACCAACCCGGTCAGCCCCTCTCCG | 352.3 | -11.4 | 1.6 | -7.3 | 2.2E-13 | 8.2E-13 | MB231/MB468 | NUC |
| >hg38_21_+_8433172_8446622_RNA45SN1_45S_50ntFlanks@12954.12972.19\|CTGCTTCTGGGTCGGGGTT | >hg38_21_+_8433172_8446622_RNA45SN1_45S_50ntFlanks | 12954 | 12972 | 19 | CTGCTTCTGGGTCGGGGTT | 533.6 | -5.9 | 0.8 | -7.3 | 2.1E-13 | 8.2E-13 | MB231/MB468 | CYTO |
| >hg38_21_+_8433172_8446622_RNA45SN1_45S_50ntFlanks@6646.6677.32\|TCGTACGACTCTTAGCGGTGGATCACTCGGCT | >hg38_21_+_8433172_8446622_RNA45SN1_45S_50ntFlanks | 6646 | 6677 | 32 | TCGTACGACTCTTAGCGGTGGATCACTCGGCT | 22895.1 | 5.1 | 0.7 | 7.4 | 1.7E-13 | 8.2E-13 | BT20/MB468 | MITO |
| >hg38_21_+_8433172_8446622_RNA45SN1_45S_50ntFlanks@10362.10407.46\|AAGGCCGAAGTGGAGAAGGGTTCCATGTGAACAGCAGTTGAACATG | >hg38_21_+_8433172_8446622_RNA45SN1_45S_50ntFlanks | 10362 | 10407 | 46 | AAGGCCGAAGTGGAGAAGGGTTCCATGTGAACAGCAGTTGAACATG | 272.8 | -11.9 | 1.6 | -7.3 | 2.8E-13 | 8.3E-13 | MB231/MB468 | MITO |
| >hg38_21_+_8433172_8446622_RNA45SN1_45S_50ntFlanks@12163.12188.26\|ACTGGGGCGGTACACCTGTCAAACGG | >hg38_21_+_8433172_8446622_RNA45SN1_45S_50ntFlanks | 12163 | 12188 | 26 | ACTGGGGCGGTACACCTGTCAAACGG | 523.8 | -12.8 | 1.8 | -7.3 | 2.8E-13 | 8.3E-13 | MB231/MB468 | MITO |
| >hg38_21_+_8433172_8446622_RNA45SN1_45S_50ntFlanks@12405.12423.19\|CTTTTTGATCCTTCGATGT | >hg38_21_+_8433172_8446622_RNA45SN1_45S_50ntFlanks | 12405 | 12423 | 19 | CTTTTTGATCCTTCGATGT | 466.4 | -8.5 | 1.2 | -7.3 | 2.2E-13 | 8.3E-13 | BT20/MB468 | CYTO |
| >hg38_21_+_8433172_8446622_RNA45SN1_45S_50ntFlanks@11576.11593.18\|CTTAGAACTGGTGCGGAC | >hg38_21_+_8433172_8446622_RNA45SN1_45S_50ntFlanks | 11576 | 11593 | 18 | CTTAGAACTGGTGCGGAC | 2551.5 | -10.9 | 1.5 | -7.3 | 3.1E-13 | 8.3E-13 | BT20/MB468 | NUC |
| >hg38_21_+_8433172_8446622_RNA45SN1_45S_50ntFlanks@9736.9759.24\|CCGAAACGATCTCAACCTATTCTC | >hg38_21_+_8433172_8446622_RNA45SN1_45S_50ntFlanks | 9736 | 9759 | 24 | CCGAAACGATCTCAACCTATTCTC | 125.0 | -10.8 | 1.5 | -7.4 | 1.9E-13 | 8.3E-13 | MB231/MB468 | MP |
| >hg38_21_+_8433172_8446622_RNA45SN1_45S_50ntFlanks@11564.11591.28\|CCTAGCAGCCGACTTAGAACTGGTGCGG | >hg38_21_+_8433172_8446622_RNA45SN1_45S_50ntFlanks | 11564 | 11591 | 28 | CCTAGCAGCCGACTTAGAACTGGTGCGG | 1475.2 | -4.8 | 0.7 | -7.4 | 1.7E-13 | 8.3E-13 | BT20/MB468 | MITO |
| >hg38_21_+_8433172_8446622_RNA45SN1_45S_50ntFlanks@4884.4904.21\|CAAAGCTGAAACTTAAAGGAA | >hg38_21_+_8433172_8446622_RNA45SN1_45S_50ntFlanks | 4884 | 4904 | 21 | CAAAGCTGAAACTTAAAGGAA | 144.4 | -11.0 | 1.5 | -7.4 | 1.9E-13 | 8.3E-13 | MB231/MB468 | MP |
| >MI0000289\|hsa-mir-181a-1&WithFlank&1\|-\|198859038\|198859159@70.90.21(+1U)&[MIMAT0000270&hsa-miR-181a-3p&offsets\|0\|-1(+1U);m-125&1\|-\|198859069\|198859090&offsets\|0\|-1(+1U)]\|ACCATCGACCGTTGATTGTACT | >MI0000289\|hsa-mir-181a-1&WithFlank&1\|-\|198859038\|198859159 | 70 | 90 | 21(+1U) | ACCATCGACCGTTGATTGTACT | 850.8 | 7.8 | 1.1 | 7.3 | 2.1E-13 | 8.3E-13 | MB231/MB468 | CYTO |
| >hg38_21_+_8433172_8446622_RNA45SN1_45S_50ntFlanks@11873.11900.28\|AAGAAGACCCTGTTGAGCTTGACTCTAG | >hg38_21_+_8433172_8446622_RNA45SN1_45S_50ntFlanks | 11873 | 11900 | 28 | AAGAAGACCCTGTTGAGCTTGACTCTAG | 1153.8 | -7.3 | 1.0 | -7.3 | 2.2E-13 | 8.3E-13 | MB231/MB468 | NUC |
| >am_trnaMT_ProTGG_MT_-_15956_16023@52.70.19\|AAAGACTTTTTCTCTGACC | >am_trnaMT_ProTGG_MT_-_15956_16023 | 52 | 70 | 19 | AAAGACTTTTTCTCTGACC | 401.9 | 10.0 | 1.4 | 7.3 | 2.2E-13 | 8.3E-13 | MB231/MB468 | CYTO |
| >hg38_21_+_8433172_8446622_RNA45SN1_45S_50ntFlanks@11729.11753.25\|TAACTATGACTCTCTTAAGGTAGCC | >hg38_21_+_8433172_8446622_RNA45SN1_45S_50ntFlanks | 11729 | 11753 | 25 | TAACTATGACTCTCTTAAGGTAGCC | 1945.5 | -9.3 | 1.3 | -7.3 | 2.2E-13 | 8.3E-13 | BT20/MB468 | CYTO |
| >hg38_21_+_8433172_8446622_RNA45SN1_45S_50ntFlanks@9534.9562.29\|AGGAAACTCTGGTGGAGGTCCGTAGCGGT | >hg38_21_+_8433172_8446622_RNA45SN1_45S_50ntFlanks | 9534 | 9562 | 29 | AGGAAACTCTGGTGGAGGTCCGTAGCGGT | 1850.6 | -4.0 | 0.6 | -7.3 | 2.2E-13 | 8.3E-13 | BT20/MB468 | CYTO |
| >hg38_21_+_8433172_8446622_RNA45SN1_45S_50ntFlanks@7975.7997.23\|CGCGACCTCAGATCAGACGTGGC | >hg38_21_+_8433172_8446622_RNA45SN1_45S_50ntFlanks | 7975 | 7997 | 23 | CGCGACCTCAGATCAGACGTGGC | 14652.0 | 5.6 | 0.8 | 7.3 | 2.2E-13 | 8.3E-13 | BT20/MB468 | CYTO |
| >hg38_21_+_8433172_8446622_RNA45SN1_45S_50ntFlanks@3773.3792.20\|CGGCCGGTACAGTGAAACTG | >hg38_21_+_8433172_8446622_RNA45SN1_45S_50ntFlanks | 3773 | 3792 | 20 | CGGCCGGTACAGTGAAACTG | 1988.1 | -10.8 | 1.5 | -7.3 | 2.8E-13 | 8.3E-13 | MB231/MB468 | MITO |
| >hg38_21_+_8433172_8446622_RNA45SN1_45S_50ntFlanks@10357.10374.18\|CTTTGAAGGCCGAAGTGG | >hg38_21_+_8433172_8446622_RNA45SN1_45S_50ntFlanks | 10357 | 10374 | 18 | CTTTGAAGGCCGAAGTGG | 186.2 | -11.4 | 1.6 | -7.4 | 1.9E-13 | 8.4E-13 | MB231/MB468 | MP |
| >hg38_21_+_8433172_8446622_RNA45SN1_45S_50ntFlanks@12044.12066.23\|ACTGACCCGGTGAGGCGGGGGGG | >hg38_21_+_8433172_8446622_RNA45SN1_45S_50ntFlanks | 12044 | 12066 | 23 | ACTGACCCGGTGAGGCGGGGGGG | 306.6 | -12.1 | 1.6 | -7.4 | 1.9E-13 | 8.4E-13 | MB231/MB468 | MP |
| >hg38_21_+_8433172_8446622_RNA45SN1_45S_50ntFlanks@9552.9569.18\|TCCGTAGCGGTCCTGACG | >hg38_21_+_8433172_8446622_RNA45SN1_45S_50ntFlanks | 9552 | 9569 | 18 | TCCGTAGCGGTCCTGACG | 891.5 | -5.5 | 0.8 | -7.3 | 3.2E-13 | 8.4E-13 | BT20/MB468 | NUC |
| >hg38_21_+_8433172_8446622_RNA45SN1_45S_50ntFlanks@4512.4537.26\|AAAAAATTAGAGTGTTCAAAGCAGGC | >hg38_21_+_8433172_8446622_RNA45SN1_45S_50ntFlanks | 4512 | 4537 | 26 | AAAAAATTAGAGTGTTCAAAGCAGGC | 165.4 | -11.2 | 1.5 | -7.4 | 1.9E-13 | 8.4E-13 | MB231/MB468 | MP |
| >hg38_21_+_8433172_8446622_RNA45SN1_45S_50ntFlanks@10678.10695.18\|CGGTTCCGGCGGCGTCCG | >hg38_21_+_8433172_8446622_RNA45SN1_45S_50ntFlanks | 10678 | 10695 | 18 | CGGTTCCGGCGGCGTCCG | 137.3 | -10.1 | 1.4 | -7.3 | 2.2E-13 | 8.4E-13 | MB231/MB468 | NUC |
| >hg38_21_+_8433172_8446622_RNA45SN1_45S_50ntFlanks@8388.8406.19\|CGTGAAACCGTTAAGAGGT | >hg38_21_+_8433172_8446622_RNA45SN1_45S_50ntFlanks | 8388 | 8406 | 19 | CGTGAAACCGTTAAGAGGT | 141.3 | -11.0 | 1.5 | -7.4 | 1.9E-13 | 8.4E-13 | MB231/MB468 | MP |
| >hg38_21_+_8433172_8446622_RNA45SN1_45S_50ntFlanks@11564.11585.22\|CCTAGCAGCCGACTTAGAACTG | >hg38_21_+_8433172_8446622_RNA45SN1_45S_50ntFlanks | 11564 | 11585 | 22 | CCTAGCAGCCGACTTAGAACTG | 3727.7 | -8.0 | 1.1 | -7.3 | 2.8E-13 | 8.4E-13 | MB231/MB468 | MITO |
| >hg38_21_+_8433172_8446622_RNA45SN1_45S_50ntFlanks@4823.4851.29\|ATGACCCGCCGGGCAGCTTCCGGGAAACC | >hg38_21_+_8433172_8446622_RNA45SN1_45S_50ntFlanks | 4823 | 4851 | 29 | ATGACCCGCCGGGCAGCTTCCGGGAAACC | 439.4 | 8.6 | 1.2 | 7.3 | 2.2E-13 | 8.4E-13 | BT20/MB468 | CYTO |
| >MI0000255\|hsa-mir-30d&WithFlank&8\|-\|134804870\|134804951@12.31.20&[MIMAT0000245&hsa-miR-30d-5p&offsets\|0\|-2;m-21&8\|-\|134804919\|134804940&offsets\|0\|-2]\|TGTAAACATCCCCGACTGGA | >MI0000255\|hsa-mir-30d&WithFlank&8\|-\|134804870\|134804951 | 12 | 31 | 20 | TGTAAACATCCCCGACTGGA | 1496.1 | 6.3 | 0.9 | 7.3 | 2.2E-13 | 8.4E-13 | MB231/MB468 | NUC |
| >hg38_21_+_8433172_8446622_RNA45SN1_45S_50ntFlanks@6653.6685.33\|ACTCTTAGCGGTGGATCACTCGGCTCGTGCGTC | >hg38_21_+_8433172_8446622_RNA45SN1_45S_50ntFlanks | 6653 | 6685 | 33 | ACTCTTAGCGGTGGATCACTCGGCTCGTGCGTC | 609.5 | 4.3 | 0.6 | 7.4 | 1.7E-13 | 8.5E-13 | BT20/MB468 | MITO |
| >hg38_21_+_8433172_8446622_RNA45SN1_45S_50ntFlanks@7986.8011.26\|ATCAGACGTGGCGACCCGCTGAATTT | >hg38_21_+_8433172_8446622_RNA45SN1_45S_50ntFlanks | 7986 | 8011 | 26 | ATCAGACGTGGCGACCCGCTGAATTT | 684.5 | -12.5 | 1.6 | -7.8 | 9.1E-15 | 8.5E-13 | BT20/MB468 | MP |
| >MI0000746\|hsa-mir-99b&WithFlank&19\|+\|51692606\|51692687@13.33.21&[MIMAT0000689&hsa-miR-99b-5p&offsets\|0\|-1;m-10&19\|+\|51692618\|51692639&offsets\|0\|-1]\|CACCCGTAGAACCGACCTTGC | >MI0000746\|hsa-mir-99b&WithFlank&19\|+\|51692606\|51692687 | 13 | 33 | 21 | CACCCGTAGAACCGACCTTGC | 10306.5 | 6.1 | 0.8 | 7.3 | 2.2E-13 | 8.5E-13 | MB231/MB468 | CYTO |
| >MI0000737\|hsa-mir-200a&WithFlank&1\|+\|1167857\|1167958@60.82.23&[MIMAT0000682&hsa-miR-200a-3p&offsets\|0\|+1;m-142&1\|+\|1167916\|1167938&offsets\|0\|0]\|TAACACTGTCTGGTAACGATGTT | >MI0000737\|hsa-mir-200a&WithFlank&1\|+\|1167857\|1167958 | 60 | 82 | 23 | TAACACTGTCTGGTAACGATGTT | 8390.2 | -6.2 | 0.8 | -7.4 | 1.9E-13 | 8.5E-13 | MB231/MB468 | MP |
| >hg38_21_+_8433172_8446622_RNA45SN1_45S_50ntFlanks@9573.9597.25\|AAATCGGTCGTCCGACCTGGGTATA | >hg38_21_+_8433172_8446622_RNA45SN1_45S_50ntFlanks | 9573 | 9597 | 25 | AAATCGGTCGTCCGACCTGGGTATA | 1706.7 | -5.7 | 0.8 | -7.3 | 2.2E-13 | 8.5E-13 | BT20/MB468 | CYTO |
| >hg38_21_+_8433172_8446622_RNA45SN1_45S_50ntFlanks@13005.13027.23\|TCTATTGAAAGTCAGCCCTCGAC | >hg38_21_+_8433172_8446622_RNA45SN1_45S_50ntFlanks | 13005 | 13027 | 23 | TCTATTGAAAGTCAGCCCTCGAC | 510.8 | -7.0 | 1.0 | -7.3 | 2.2E-13 | 8.5E-13 | MB231/MB468 | CYTO |
| >MI0000736\|hsa-mir-30c-1&WithFlank&1\|+\|40757278\|40757378@23.45.23(+1A);MI0000254\|hsa-mir-30c-2&WithFlank&6\|-\|71376954\|71377037@13.35.23(+1A)&[MIMAT0000244&hsa-miR-30c-5p&offsets\|0\|0(+1A);m-57&1\|+\|40757300\|40757323&offsets\|0\|-1(+1A)];[MIMAT0000244_1&hsa-miR-30c-5p&offsets\|0\|0(+1A);m-56&6\|-\|71377002\|71377025&offsets\|0\|-1(+1A)]\|TGTAAACATCCTACACTCTCAGCA | >MI0000736\|hsa-mir-30c-1&WithFlank&1\|+\|40757278\|40757378 | 23 | 45 | 23(+1A) | TGTAAACATCCTACACTCTCAGCA | 2263.2 | 4.3 | 0.6 | 7.3 | 2.3E-13 | 8.5E-13 | MB231/MB468 | NUC |
| >hg38_21_+_8433172_8446622_RNA45SN1_45S_50ntFlanks@5374.5395.22\|CGTTGATTAAGTCCCTGCCCTT | >hg38_21_+_8433172_8446622_RNA45SN1_45S_50ntFlanks | 5374 | 5395 | 22 | CGTTGATTAAGTCCCTGCCCTT | 605.0 | -13.0 | 1.8 | -7.3 | 2.9E-13 | 8.6E-13 | MB231/MB468 | MITO |
| >hg38_21_+_8433172_8446622_RNA45SN1_45S_50ntFlanks@10797.10818.22\|GCATGTTGGAACAATGTAGGTA | >hg38_21_+_8433172_8446622_RNA45SN1_45S_50ntFlanks | 10797 | 10818 | 22 | GCATGTTGGAACAATGTAGGTA | 2600.2 | -6.7 | 0.9 | -7.7 | 9.2E-15 | 8.6E-13 | BT20/MB468 | MP |
| >hg38_21_+_8433172_8446622_RNA45SN1_45S_50ntFlanks@6747.6771.25\|ACTTCGAACGCACTTGCGGCCCCGG | >hg38_21_+_8433172_8446622_RNA45SN1_45S_50ntFlanks | 6747 | 6771 | 25 | ACTTCGAACGCACTTGCGGCCCCGG | 304.0 | -7.1 | 1.0 | -7.3 | 2.2E-13 | 8.6E-13 | MB231/MB468 | CYTO |
| >trnaMT_ValTAC_MT_+_1602_1670@54.71.18\|AACTTGACCGCTCTGACC | >trnaMT_ValTAC_MT_+_1602_1670 | 54 | 71 | 18 | AACTTGACCGCTCTGACC | 1657.1 | 11.1 | 1.5 | 7.3 | 2.3E-13 | 8.6E-13 | BT20/MB468 | CYTO |
| >hg38_21_+_8433172_8446622_RNA45SN1_45S_50ntFlanks@7975.8006.32\|CGCGACCTCAGATCAGACGTGGCGACCCGCTG | >hg38_21_+_8433172_8446622_RNA45SN1_45S_50ntFlanks | 7975 | 8006 | 32 | CGCGACCTCAGATCAGACGTGGCGACCCGCTG | 24737.4 | -5.3 | 0.7 | -7.4 | 1.9E-13 | 8.6E-13 | MB231/MB468 | MP |
| >MI0000078\|hsa-mir-22&WithFlank&17\|-\|1713897\|1713993@59.81.23&[MIMAT0000077&hsa-miR-22-3p&offsets\|0\|+1;m-2&17\|-\|1713914\|1713935&offsets\|0\|+1]\|AAGCTGCCAGTTGAAGAACTGTT | >MI0000078\|hsa-mir-22&WithFlank&17\|-\|1713897\|1713993 | 59 | 81 | 23 | AAGCTGCCAGTTGAAGAACTGTT | 320.2 | 9.9 | 1.4 | 7.3 | 2.3E-13 | 8.6E-13 | MB231/MB468 | NUC |
| >trnaMT_GluTTC_MT_-_14674_14742@37.72.36,trnalookalike8_GluTTC_5_-_93905172_93905240@37.72.36\|TCATTGGTCGTGGTTGTAGTCCGTGCGAGAATACCA | >trnaMT_GluTTC_MT_-_14674_14742 | 37 | 72 | 36 | TCATTGGTCGTGGTTGTAGTCCGTGCGAGAATACCA | 8887.3 | 9.4 | 1.3 | 7.3 | 2.2E-13 | 8.6E-13 | MB231/MB468 | CYTO |
| >hg38_21_+_8433172_8446622_RNA45SN1_45S_50ntFlanks@12438.12455.18\|ATTGTGAAGCAGAATTCA | >hg38_21_+_8433172_8446622_RNA45SN1_45S_50ntFlanks | 12438 | 12455 | 18 | ATTGTGAAGCAGAATTCA | 3266.9 | -8.1 | 1.1 | -7.3 | 2.2E-13 | 8.6E-13 | MB231/MB468 | CYTO |
| >MI0000066\|hsa-let-7e&WithFlank&19\|+\|51692780\|51692870@14.33.20(+1C)&[MIMAT0000066&hsa-let-7e-5p&offsets\|0\|-2(+1C);m-50&19\|+\|51692793\|51692814&offsets\|0\|-2(+1C)]\|TGAGGTAGGAGGTTGTATAGC | >MI0000066\|hsa-let-7e&WithFlank&19\|+\|51692780\|51692870 | 14 | 33 | 20(+1C) | TGAGGTAGGAGGTTGTATAGC | 906.3 | 9.5 | 1.3 | 7.3 | 2.3E-13 | 8.7E-13 | BT20/MB468 | CYTO |

| | | | | | | | | | | | | | |
|---|---|---|---|---|---|---|---|---|---|---|---|---|---|
| >MI0000102\|hsa-mir-100&WithFlank&11\|-\|122152223\|122152314@19.40.22(+3A)&[MIMAT0000098&hsa-miR-100-5p&offsets\|0\|0(+3A);m-27&11\|-\|122152276\|122152296&offsets\|0\|+1(+3A)]\|AACCCGTAGATCCGAACTTGTGAAA | >MI0000102\|hsa-mir-100&WithFlank&11\|-\|122152223\|122152314 | 19 | 40 | 22(+3A) | AACCCGTAGATCCGAACTTGTGAAA | 256.9 | -11.1 | 1.5 | -7.6 | 2.7E-14 | 8.7E-13 | BT20/MB231 | NUC |
| >hg38_21_+_8433172_8446622_RNA45SN1_45S_50ntFlanks@6719.6738.20\|TGAATTGCAGGACACATTGA | >hg38_21_+_8433172_8446622_RNA45SN1_45S_50ntFlanks | 6719 | 6738 | 20 | TGAATTGCAGGACACATTGA | 153.0 | -11.1 | 1.5 | -7.4 | 1.9E-13 | 8.7E-13 | MB231/MB468 | MP |
| >hg38_21_+_8433172_8446622_RNA45SN1_45S_50ntFlanks@9470.9503.34\|GAGCACGAGCGCACGTGTTAGGACCCGAAAGATG | >hg38_21_+_8433172_8446622_RNA45SN1_45S_50ntFlanks | 9470 | 9503 | 34 | GAGCACGAGCGCACGTGTTAGGACCCGAAAGATG | 1448.7 | -9.3 | 1.3 | -7.3 | 2.9E-13 | 8.7E-13 | MB231/MB468 | MITO |
| >hg38_21_+_8433172_8446622_RNA45SN1_45S_50ntFlanks@4729.4758.30\|TCAAGAACGAAAGTCGGAGGTTCGAAGACG | >hg38_21_+_8433172_8446622_RNA45SN1_45S_50ntFlanks | 4729 | 4758 | 30 | TCAAGAACGAAAGTCGGAGGTTCGAAGACG | 4583.9 | -9.1 | 1.3 | -7.3 | 3.3E-13 | 8.7E-13 | BT20/MB468 | NUC |
| >hg38_21_+_8433172_8446622_RNA45SN1_45S_50ntFlanks@9598.9617.20\|GGGGCGAAAGACTAATCGAA | >hg38_21_+_8433172_8446622_RNA45SN1_45S_50ntFlanks | 9598 | 9617 | 20 | GGGGCGAAAGACTAATCGAA | 153.4 | -11.1 | 1.5 | -7.4 | 2.0E-13 | 8.7E-13 | MB231/MB468 | MP |
| >hg38_21_+_8433172_8446622_RNA45SN1_45S_50ntFlanks@12633.12650.18\|CGAAGCTACCATCTGTGG | >hg38_21_+_8433172_8446622_RNA45SN1_45S_50ntFlanks | 12633 | 12650 | 18 | CGAAGCTACCATCTGTGG | 160.2 | -11.2 | 1.5 | -7.4 | 2.0E-13 | 8.7E-13 | MB231/MB468 | MP |
| >hg38_21_+_8433172_8446622_RNA45SN1_45S_50ntFlanks@12635.12667.33\|AAGCTACCATCTGTGGGATTATGACTGAACGCC | >hg38_21_+_8433172_8446622_RNA45SN1_45S_50ntFlanks | 12635 | 12667 | 33 | AAGCTACCATCTGTGGGATTATGACTGAACGCC | 497.4 | 7.1 | 1.0 | 7.3 | 2.3E-13 | 8.7E-13 | MB231/MB468 | CYTO |
| >hg38_21_+_8433172_8446622_RNA45SN1_45S_50ntFlanks@12412.12444.33\|ATCCTTCGATGTCGGCTCTTCCTATCATTGTGA | >hg38_21_+_8433172_8446622_RNA45SN1_45S_50ntFlanks | 12412 | 12444 | 33 | ATCCTTCGATGTCGGCTCTTCCTATCATTGTGA | 170.6 | -9.7 | 1.3 | -7.3 | 2.3E-13 | 8.7E-13 | BT20/MB468 | CYTO |
| >hg38_21_+_8433172_8446622_RNA45SN1_45S_50ntFlanks@12004.12036.33\|CGGGCCGCCGGTGAAATACCACTACTCTGATCG | >hg38_21_+_8433172_8446622_RNA45SN1_45S_50ntFlanks | 12004 | 12036 | 33 | CGGGCCGCCGGTGAAATACCACTACTCTGATCG | 840.2 | 5.8 | 0.8 | 7.4 | 1.8E-13 | 8.7E-13 | BT20/MB468 | MITO |
| >hg38_21_+_8433172_8446622_RNA45SN1_45S_50ntFlanks@12033.12052.20\|ATCGTTTTTTCACTGACCCG | >hg38_21_+_8433172_8446622_RNA45SN1_45S_50ntFlanks | 12033 | 12052 | 20 | ATCGTTTTTTCACTGACCCG | 142.2 | -11.0 | 1.5 | -7.4 | 2.0E-13 | 8.8E-13 | MB231/MB468 | MP |
| >hg38_21_+_8433172_8446622_RNA45SN1_45S_50ntFlanks@9563.9592.30\|CCTGACGTGCAAATCGGTCGTCCGACCTGG | >hg38_21_+_8433172_8446622_RNA45SN1_45S_50ntFlanks | 9563 | 9592 | 30 | CCTGACGTGCAAATCGGTCGTCCGACCTGG | 120.0 | -10.8 | 1.5 | -7.4 | 2.0E-13 | 8.8E-13 | MB231/MB468 | MP |
| >hg38_21_+_8433172_8446622_RNA45SN1_45S_50ntFlanks@11562.11601.40\|CGCCTAGCAGCCGACTTAGAACTGGTGCGGACCAGGGGAA | >hg38_21_+_8433172_8446622_RNA45SN1_45S_50ntFlanks | 11562 | 11601 | 40 | CGCCTAGCAGCCGACTTAGAACTGGTGCGGACCAGGGGAA | 753.4 | -7.5 | 1.0 | -7.3 | 3.0E-13 | 8.8E-13 | MB231/MB468 | MITO |
| >hg38_21_+_8433172_8446622_RNA45SN1_45S_50ntFlanks@9572.9592.21\|CAAATCGGTCGTCCGACCTGG | >hg38_21_+_8433172_8446622_RNA45SN1_45S_50ntFlanks | 9572 | 9592 | 21 | CAAATCGGTCGTCCGACCTGG | 277.8 | -10.4 | 1.4 | -7.3 | 2.3E-13 | 8.8E-13 | BT20/MB468 | CYTO |
| >hg38_21_+_8433172_8446622_RNA45SN1_45S_50ntFlanks@12937.12960.24\|AACCATTCGTAGACGACCTGCTTC | >hg38_21_+_8433172_8446622_RNA45SN1_45S_50ntFlanks | 12937 | 12960 | 24 | AACCATTCGTAGACGACCTGCTTC | 126.3 | -10.8 | 1.5 | -7.4 | 2.0E-13 | 8.8E-13 | MB231/MB468 | MP |
| >hg38_21_+_8433172_8446622_RNA45SN1_45S_50ntFlanks@5473.5505.33\|CGGCCCTGGCGGAGCGCTGAGAAGACGGTCGAA | >hg38_21_+_8433172_8446622_RNA45SN1_45S_50ntFlanks | 5473 | 5505 | 33 | CGGCCCTGGCGGAGCGCTGAGAAGACGGTCGAA | 2356.5 | 9.1 | 1.2 | 7.3 | 2.3E-13 | 8.8E-13 | BT20/MB468 | CYTO |
| >hg38_21_+_8433172_8446622_RNA45SN1_45S_50ntFlanks@12935.12953.19\|TAAACCATTCGTAGACGAC | >hg38_21_+_8433172_8446622_RNA45SN1_45S_50ntFlanks | 12935 | 12953 | 19 | TAAACCATTCGTAGACGAC | 160.4 | -11.2 | 1.5 | -7.4 | 2.0E-13 | 8.8E-13 | MB231/MB468 | MP |
| >hg38_21_+_8433172_8446622_RNA45SN1_45S_50ntFlanks@5379.5397.19\|ATTAAGTCCCTGCCCTTTG | >hg38_21_+_8433172_8446622_RNA45SN1_45S_50ntFlanks | 5379 | 5397 | 19 | ATTAAGTCCCTGCCCTTTG | 2830.5 | -11.0 | 1.5 | -7.3 | 3.0E-13 | 8.9E-13 | MB231/MB468 | MITO |
| >hg38_21_+_8433172_8446622_RNA45SN1_45S_50ntFlanks@10461.10497.37\|GCCTCCGTTGCCCTCGGCCGATCGAAAGGGAGTCGGG | >hg38_21_+_8433172_8446622_RNA45SN1_45S_50ntFlanks | 10461 | 10497 | 37 | GCCTCCGTTGCCCTCGGCCGATCGAAAGGGAGTCGGG | 179.4 | -7.4 | 1.0 | -7.3 | 2.3E-13 | 8.9E-13 | MB231/MB468 | CYTO |
| >hg38_21_+_8433172_8446622_RNA45SN1_45S_50ntFlanks@11580.11601.22\|GAACTGGTGCGGACCAGGGGAA | >hg38_21_+_8433172_8446622_RNA45SN1_45S_50ntFlanks | 11580 | 11601 | 22 | GAACTGGTGCGGACCAGGGGAA | 1559.5 | -5.6 | 0.8 | -7.3 | 2.3E-13 | 8.9E-13 | MB231/MB468 | CYTO |
| >hg38_21_+_8433172_8446622_RNA45SN1_45S_50ntFlanks@12874.12892.19\|GCTGGAAAGGCGGCCGCCC | >hg38_21_+_8433172_8446622_RNA45SN1_45S_50ntFlanks | 12874 | 12892 | 19 | GCTGGAAAGGCGGCCGCCC | 176.7 | -7.2 | 1.0 | -7.3 | 2.3E-13 | 8.9E-13 | MB231/MB468 | CYTO |
| >hg38_21_+_8433172_8446622_RNA45SN1_45S_50ntFlanks@10787.10823.37\|ACAGCCTCTGGCATGTTGGAACAATGTAGGTAAGGGA | >hg38_21_+_8433172_8446622_RNA45SN1_45S_50ntFlanks | 10787 | 10823 | 37 | ACAGCCTCTGGCATGTTGGAACAATGTAGGTAAGGGA | 202.4 | -11.5 | 1.6 | -7.3 | 2.0E-13 | 8.9E-13 | MB231/MB468 | MP |
| >trna10_GlyTCC_17_+_8124866_8124937@1.30.30\|GCGTTGGTGGTATAGTGGTAAGCATAGCTG | >trna10_GlyTCC_17_+_8124866_8124937 | 1 | 30 | 30 | GCGTTGGTGGTATAGTGGTAAGCATAGCTG | 907.0 | -4.3 | 0.6 | -7.3 | 2.4E-13 | 8.9E-13 | BT20/MB468 | CYTO |
| >hg38_21_+_8433172_8446622_RNA45SN1_45S_50ntFlanks@10677.10706.30\|GCGGTTCCGGCGGCGTCCGGTGAGCTCTCG | >hg38_21_+_8433172_8446622_RNA45SN1_45S_50ntFlanks | 10677 | 10706 | 30 | GCGGTTCCGGCGGCGTCCGGTGAGCTCTCG | 232.3 | -11.7 | 1.6 | -7.3 | 2.0E-13 | 8.9E-13 | MB231/MB468 | MP |
| >hg38_21_+_8433172_8446622_RNA45SN1_45S_50ntFlanks@11576.11609.34\|CTTAGAACTGGTGCGGACCAGGGGAATCCGACTG | >hg38_21_+_8433172_8446622_RNA45SN1_45S_50ntFlanks | 11576 | 11609 | 34 | CTTAGAACTGGTGCGGACCAGGGGAATCCGACTG | 1083.3 | -8.3 | 1.1 | -7.3 | 2.0E-13 | 8.9E-13 | MB231/MB468 | MP |
| >hg38_21_+_8433172_8446622_RNA45SN1_45S_50ntFlanks@10681.10711.31\|TTCCGGCGGCGTCCGGTGAGCTCTCGCTGGC | >hg38_21_+_8433172_8446622_RNA45SN1_45S_50ntFlanks | 10681 | 10711 | 31 | TTCCGGCGGCGTCCGGTGAGCTCTCGCTGGC | 1560.1 | -7.5 | 1.0 | -7.3 | 3.0E-13 | 8.9E-13 | MB231/MB468 | MITO |
| >trna10_GlyTCC_17_+_8124866_8124937@1.29.29\|GCGTTGGTGGTATAGTGGTAAGCATAGCT | >trna10_GlyTCC_17_+_8124866_8124937 | 1 | 29 | 29 | GCGTTGGTGGTATAGTGGTAAGCATAGCT | 391.7 | -7.8 | 1.1 | -7.3 | 2.3E-13 | 9.0E-13 | MB231/MB468 | CYTO |
| >hg38_21_+_8433172_8446622_RNA45SN1_45S_50ntFlanks@3985.4014.30\|CGGCTTTGGTGACTCTAGATAACCTCGGGC | >hg38_21_+_8433172_8446622_RNA45SN1_45S_50ntFlanks | 3985 | 4014 | 30 | CGGCTTTGGTGACTCTAGATAACCTCGGGC | 160.7 | -11.2 | 1.5 | -7.3 | 2.0E-13 | 9.0E-13 | MB231/MB468 | MP |
| >MI0000815\|hsa-mir-339&WithFlank&7\|-\|1022927\|1023032@21.44.24&[MIMAT0000764&hsa-miR-339-5p&offsets\|0\|+1;m-163&7\|-\|1022990\|1023012&offsets\|0\|+1]\|TCCCTGTCCTCCAGGAGCTCACGT | >MI0000815\|hsa-mir-339&WithFlank&7\|-\|1022927\|1023032 | 21 | 44 | 24 | TCCCTGTCCTCCAGGAGCTCACGT | 271.2 | 7.1 | 1.0 | 7.3 | 2.4E-13 | 9.0E-13 | BT20/MB468 | CYTO |
| >hg38_21_+_8433172_8446622_RNA45SN1_45S_50ntFlanks@11641.11666.26\|CGGCGGGTGTTGACGCGATGTGATTT | >hg38_21_+_8433172_8446622_RNA45SN1_45S_50ntFlanks | 11641 | 11666 | 26 | CGGCGGGTGTTGACGCGATGTGATTT | 706.2 | -9.3 | 1.3 | -7.3 | 3.0E-13 | 9.0E-13 | MB231/MB468 | MITO |
| >MI0003628\|hsa-mir-615&WithFlank&12\|+\|54033944\|54034051@67.87.21(+1C)&[MIMAT0003283&hsa-miR-615-3p&offsets\|0\|-1(+1C);m-495&12\|+\|54034010\|54034030&offsets\|0\|0(+1C)]\|TCCGAGCCTGGGTCTCCCTCTC | >MI0003628\|hsa-mir-615&WithFlank&12\|+\|54033944\|54034051 | 67 | 87 | 21(+1C) | TCCGAGCCTGGGTCTCCCTCTC | 185.0 | 10.9 | 1.4 | 7.7 | 1.9E-14 | 9.0E-13 | BT20/MB231 | CYTO |
| >MI0000434\|hsa-let-7i&WithFlank&12\|+\|62603680\|62603775@12.34.23&[MIMAT0000415&hsa-let-7i-5p&offsets\|0\|+1;m-49&12\|+\|62603691\|62603712&offsets\|0\|+1]\|TGAGGTAGTAGTTTGTGCTGTTG | >MI0000434\|hsa-let-7i&WithFlank&12\|+\|62603680\|62603775 | 12 | 34 | 23 | TGAGGTAGTAGTTTGTGCTGTTG | 270.6 | 5.2 | 0.7 | 7.3 | 2.4E-13 | 9.0E-13 | MB231/MB468 | NUC |
| >hg38_21_+_8433172_8446622_RNA45SN1_45S_50ntFlanks@5527.5548.22\|AAAGTCGTAACAAGGTTTCCGT | >hg38_21_+_8433172_8446622_RNA45SN1_45S_50ntFlanks | 5527 | 5548 | 22 | AAAGTCGTAACAAGGTTTCCGT | 184.4 | -10.5 | 1.4 | -7.3 | 2.4E-13 | 9.0E-13 | MB231/MB468 | NUC |
| >trnaMT_SerGCT_MT_+_12207_12265@7.38.32\|GCTCACAAGAACTGCTAACTCATGCCCCCATG | >trnaMT_SerGCT_MT_+_12207_12265 | 7 | 38 | 32 | GCTCACAAGAACTGCTAACTCATGCCCCCATG | 460.8 | -9.1 | 1.2 | -7.3 | 2.4E-13 | 9.0E-13 | MB231/MB468 | NUC |
| >MI0000085\|hsa-mir-27a&WithFlank&19\|-\|13836434\|13836523@57.76.20(+1U)&[MIMAT0000084&hsa-miR-27a-3p&offsets\|0\|-1(+1U);m-43&19\|-\|13836447\|13836467&offsets\|0\|-1(+1U)]\|TTCACAGTGGCTAAGTTCCGT | >MI0000085\|hsa-mir-27a&WithFlank&19\|-\|13836434\|13836523 | 57 | 76 | 20(+1U) | TTCACAGTGGCTAAGTTCCGT | 6038.5 | 5.7 | 0.8 | 7.3 | 2.4E-13 | 9.0E-13 | MB231/MB468 | NUC |
| >hg38_21_+_8433172_8446622_RNA45SN1_45S_50ntFlanks@10402.10430.29\|AACATGGGTCAGTCGGTCCTGAGAGATGG | >hg38_21_+_8433172_8446622_RNA45SN1_45S_50ntFlanks | 10402 | 10430 | 29 | AACATGGGTCAGTCGGTCCTGAGAGATGG | 7312.1 | -11.6 | 1.6 | -7.3 | 3.4E-13 | 9.0E-13 | BT20/MB468 | NUC |
| >hg38_21_+_8433172_8446622_RNA45SN1_45S_50ntFlanks@10233.10260.28\|GCGGACGCTACGCCGCGACGAGTAGGAG | >hg38_21_+_8433172_8446622_RNA45SN1_45S_50ntFlanks | 10233 | 10260 | 28 | GCGGACGCTACGCCGCGACGAGTAGGAG | 141.2 | -11.0 | 1.5 | -7.3 | 2.0E-13 | 9.0E-13 | MB231/MB468 | MP |
| >hg38_21_+_8433172_8446622_RNA45SN1_45S_50ntFlanks@11707.11728.22\|AAGCGCGGGTAAACGGCGGGAG | >hg38_21_+_8433172_8446622_RNA45SN1_45S_50ntFlanks | 11707 | 11728 | 22 | AAGCGCGGGTAAACGGCGGGAG | 304.5 | -5.8 | 0.8 | -7.3 | 2.4E-13 | 9.0E-13 | MB231/MB468 | CYTO |
| >hg38_21_+_8433172_8446622_RNA45SN1_45S_50ntFlanks@10458.10478.21\|ATGGCCTCCGTTGCCCTCGGC | >hg38_21_+_8433172_8446622_RNA45SN1_45S_50ntFlanks | 10458 | 10478 | 21 | ATGGCCTCCGTTGCCCTCGGC | 4606.9 | -8.5 | 1.2 | -7.3 | 2.4E-13 | 9.0E-13 | MB231/MB468 | NUC |
| >hg38_21_+_8433172_8446622_RNA45SN1_45S_50ntFlanks@7979.8003.25\|ACCTCAGATCAGACGTGGCGACCCG | >hg38_21_+_8433172_8446622_RNA45SN1_45S_50ntFlanks | 7979 | 8003 | 25 | ACCTCAGATCAGACGTGGCGACCCG | 501.8 | -8.7 | 1.2 | -7.3 | 2.0E-13 | 9.0E-13 | MB231/MB468 | MP |
| >hg38_21_+_8433172_8446622_RNA45SN1_45S_50ntFlanks@10786.10828.43\|AACAGCCTCTGGCATGTTGGAACAATGTAGGTAAGGGAAGTCG | >hg38_21_+_8433172_8446622_RNA45SN1_45S_50ntFlanks | 10786 | 10828 | 43 | AACAGCCTCTGGCATGTTGGAACAATGTAGGTAAGGGAAGTCG | 1845.0 | -8.3 | 1.1 | -7.3 | 3.0E-13 | 9.0E-13 | MB231/MB468 | MITO |
| >am_trna106_PheGAA_6_-_28775610_28775682@31.52.22,trna109_PheGAA_6_-_28758499_28758571@31.52.22,trna11_PheGAA_12_-_125412389_125412461@31.52.22,trna13_PheGAA_11_-_59333853_59333925@31.52.22,trna14_PheGAA_19_-_1383361_1383433@31.52.22,trna15_PheGAA_11_-_59324970_59325042@31.52.22,trna1_PheGAA_13_-_95201904_95201976@31.52.22,trna72_PheGAA_6_+_28732164_28732236@31.52.22,trna96_PheGAA_6_-_28949449_28949521@31.52.22\|ACTGAAGATCTAAAGGTCCCTG | >am_trna106_PheGAA_6_-_28775610_28775682 | 31 | 52 | 22 | ACTGAAGATCTAAAGGTCCCTG | 227.0 | -10.8 | 1.5 | -7.3 | 2.4E-13 | 9.0E-13 | MB231/MB468 | NUC |
| >hg38_21_+_8433172_8446622_RNA45SN1_45S_50ntFlanks@3831.3853.23\|CTCGCTCCTCTCCTACTTGGATA | >hg38_21_+_8433172_8446622_RNA45SN1_45S_50ntFlanks | 3831 | 3853 | 23 | CTCGCTCCTCTCCTACTTGGATA | 168.5 | -10.4 | 1.4 | -7.3 | 2.4E-13 | 9.0E-13 | MB231/MB468 | NUC |
| >hg38_21_+_8433172_8446622_RNA45SN1_45S_50ntFlanks@11718.11751.34\|AACGGCGGGAGTAACTATGACTCTCTTAAGGTAG | >hg38_21_+_8433172_8446622_RNA45SN1_45S_50ntFlanks | 11718 | 11751 | 34 | AACGGCGGGAGTAACTATGACTCTCTTAAGGTAG | 529.5 | -7.0 | 1.0 | -7.3 | 2.4E-13 | 9.0E-13 | MB231/MB468 | NUC |
| >hg38_21_+_8433172_8446622_RNA45SN1_45S_50ntFlanks@12635.12661.27\|AAGCTACCATCTGTGGGATTATGACTG | >hg38_21_+_8433172_8446622_RNA45SN1_45S_50ntFlanks | 12635 | 12661 | 27 | AAGCTACCATCTGTGGGATTATGACTG | 2169.5 | -4.2 | 0.6 | -7.3 | 2.4E-13 | 9.0E-13 | MB231/MB468 | NUC |
| >hg38_21_+_8433172_8446622_RNA45SN1_45S_50ntFlanks@12478.12496.19\|ACTAATAGGGAACGTGAGC | >hg38_21_+_8433172_8446622_RNA45SN1_45S_50ntFlanks | 12478 | 12496 | 19 | ACTAATAGGGAACGTGAGC | 930.9 | -3.9 | 0.5 | -7.3 | 3.4E-13 | 9.1E-13 | BT20/MB468 | NUC |
| >hg38_21_+_8433172_8446622_RNA45SN1_45S_50ntFlanks@12489.12521.33\|ACGTGAGCTGGGTTTAGACCGTCGTGAGACAGG | >hg38_21_+_8433172_8446622_RNA45SN1_45S_50ntFlanks | 12489 | 12521 | 33 | ACGTGAGCTGGGTTTAGACCGTCGTGAGACAGG | 131.1 | -10.9 | 1.5 | -7.3 | 2.0E-13 | 9.1E-13 | MB231/MB468 | MP |
| >hg38_21_+_8433172_8446622_RNA45SN1_45S_50ntFlanks@11587.11605.19\|TGCGGACCAGGGGAATCCG | >hg38_21_+_8433172_8446622_RNA45SN1_45S_50ntFlanks | 11587 | 11605 | 19 | TGCGGACCAGGGGAATCCG | 342.9 | -12.3 | 1.7 | -7.3 | 2.0E-13 | 9.1E-13 | MB231/MB468 | MP |
| >hg38_21_+_8433172_8446622_RNA45SN1_45S_50ntFlanks@12405.12425.21\|CTTTTTGATCCTTCGATGTCG | >hg38_21_+_8433172_8446622_RNA45SN1_45S_50ntFlanks | 12405 | 12425 | 21 | CTTTTTGATCCTTCGATGTCG | 482.9 | -6.8 | 0.9 | -7.3 | 2.4E-13 | 9.1E-13 | MB231/MB468 | NUC |
| >hg38_21_+_8433172_8446622_RNA45SN1_45S_50ntFlanks@10805.10828.24\|GAACAATGTAGGTAAGGGAAGTCG | >hg38_21_+_8433172_8446622_RNA45SN1_45S_50ntFlanks | 10805 | 10828 | 24 | GAACAATGTAGGTAAGGGAAGTCG | 135.6 | -10.9 | 1.5 | -7.3 | 2.1E-13 | 9.1E-13 | MB231/MB468 | MP |
| >hg38_21_+_8433172_8446622_RNA45SN1_45S_50ntFlanks@4555.4575.21\|CGCAGCTAGGAATAATGGAAT | >hg38_21_+_8433172_8446622_RNA45SN1_45S_50ntFlanks | 4555 | 4575 | 21 | CGCAGCTAGGAATAATGGAAT | 134.8 | -10.9 | 1.5 | -7.3 | 2.1E-13 | 9.1E-13 | MB231/MB468 | MP |
| >hg38_21_+_8433172_8446622_RNA45SN1_45S_50ntFlanks@4773.4793.21\|TAGTTCCGACCATAAACGATG | >hg38_21_+_8433172_8446622_RNA45SN1_45S_50ntFlanks | 4773 | 4793 | 21 | TAGTTCCGACCATAAACGATG | 2278.0 | -6.0 | 0.8 | -7.3 | 2.5E-13 | 9.2E-13 | MB231/MB468 | NUC |

| | | | | | | | | | | | | | |
|---|---|---|---|---|---|---|---|---|---|---|---|---|---|
| >hg38_21_+_8433172_8446622_RNA45SN1_45S_50ntFlanks@11455.11474.20\|CTCGGCGCGCGGCGGCGGCG | >hg38_21_+_8433172_8446622_RNA45SN1_45S_50ntFlanks | 11455 | 11474 | 20 | CTCGGCGCGCGGCGGCGGCG | 199.2 | -11.5 | 1.6 | -7.3 | 2.1E-13 | 9.2E-13 | MB231/MB468 | MP |
| >hg38_21_+_8433172_8446622_RNA45SN1_45S_50ntFlanks@9196.9222.27\|AGGTTCTCTCGGGGCCACGCGCGCGTC | >hg38_21_+_8433172_8446622_RNA45SN1_45S_50ntFlanks | 9196 | 9222 | 27 | AGGTTCTCTCGGGGCCACGCGCGCGTC | 257.6 | -11.9 | 1.6 | -7.3 | 2.1E-13 | 9.2E-13 | MB231/MB468 | MP |
| >MI0000087\|hsa-mir-29a&WithFlank&7\|-\|130876741\|130876816@48.67.20(+1C)&[MIMAT0000086&hsa-miR-29a-3p&offsets\|0\|-2(+1C);m-26&7\|-\|130876749\|130876769&offsets\|0\|-1(+1C)]\|TAGCACCATCTGAAATCGGTC | >MI0000087\|hsa-mir-29a&WithFlank&7\|-\|130876741\|130876816 | 48 | 67 | 20(+1C) | TAGCACCATCTGAAATCGGTC | 4727.8 | 6.7 | 0.9 | 7.3 | 2.1E-13 | 9.3E-13 | MB231/MB468 | MP |
| >hg38_21_+_8433172_8446622_RNA45SN1_45S_50ntFlanks@11027.11059.33\|CTCTCTCTCTCTCTCCCCCGCTCCCCGTCCTCC | >hg38_21_+_8433172_8446622_RNA45SN1_45S_50ntFlanks | 11027 | 11059 | 33 | CTCTCTCTCTCTCTCCCCCGCTCCCCGTCCTCC | 521.8 | -12.0 | 1.6 | -7.3 | 2.5E-13 | 9.3E-13 | MB231/MB468 | NUC |
| >hg38_21_+_8433172_8446622_RNA45SN1_45S_50ntFlanks@12589.12614.26\|GTTCAGACATTTGGTGTATGTGCTTG | >hg38_21_+_8433172_8446622_RNA45SN1_45S_50ntFlanks | 12589 | 12614 | 26 | GTTCAGACATTTGGTGTATGTGCTTG | 326.9 | -11.1 | 1.5 | -7.3 | 2.4E-13 | 9.3E-13 | MB231/MB468 | CYTO |
| >trna116_GluCTC_1_-_145399233_145399304@37.74.38,trna134_GluTTC_1_-_16861774_16861845@37.74.38,trna71_GluCTC_1_-_161439189_161439260@37.74.38,trna74_GluCTC_1_-_161431809_161431880@37.74.38,trna77_GluCTC_1_-_161424398_161424469@37.74.38,trna77_GluCTC_6_+_28949976_28950047@37.74.38,trna80_GluCTC_1_-_161417018_161417089@37.74.38,trna84_GluTTC_1_-_161391883_161391954@37.74.38,trna87_GluCTC_6_-_126101393_126101464@37.74.38\|ACCGCCGCGGCCCGGGTTCGATTCCCGGTCAGGGAACC | >trna116_GluCTC_1_-_145399233_145399304 | 37 | 74 | 38 | ACCGCCGCGGCCCGGGTTCGATTCCCGGTCAGGGAACC | 5796.7 | 9.5 | 1.3 | 7.3 | 2.5E-13 | 9.3E-13 | MB231/MB468 | NUC |
| >hg38_21_+_8433172_8446622_RNA45SN1_45S_50ntFlanks@12457.12478.22\|CAAGCGTTGGATTGTTCACCCA | >hg38_21_+_8433172_8446622_RNA45SN1_45S_50ntFlanks | 12457 | 12478 | 22 | CAAGCGTTGGATTGTTCACCCA | 3374.8 | -8.1 | 1.1 | -7.3 | 3.1E-13 | 9.3E-13 | MB231/MB468 | MITO |
| >hg38_21_+_8433172_8446622_RNA45SN1_45S_50ntFlanks@9564.9597.34\|CTGACGTGCAAATCGGTCGTCCGACCTGGGTATA | >hg38_21_+_8433172_8446622_RNA45SN1_45S_50ntFlanks | 9564 | 9597 | 34 | CTGACGTGCAAATCGGTCGTCCGACCTGGGTATA | 26748.5 | -12.7 | 1.7 | -7.3 | 3.5E-13 | 9.3E-13 | BT20/MB468 | NUC |
| >MI0003628\|hsa-mir-615&WithFlank&12\|+\|54033944\|54034051@67.86.20&[MIMAT0003283&hsa-miR-615-3p&offsets\|0\|-2;m-495&12\|+\|54034010\|54034030&offsets\|0\|-1]\|TCCGAGCCTGGGTCTCCCTC | >MI0003628\|hsa-mir-615&WithFlank&12\|+\|54033944\|54034051 | 67 | 86 | 20 | TCCGAGCCTGGGTCTCCCTC | 857.1 | 5.3 | 0.7 | 7.6 | 2.6E-14 | 9.3E-13 | BT20/MB231 | MITO |
| >hg38_21_+_8433172_8446622_RNA45SN1_45S_50ntFlanks@10427.10453.27\|ATGGGCGAGCGCCGTTCCGAAGGGACG | >hg38_21_+_8433172_8446622_RNA45SN1_45S_50ntFlanks | 10427 | 10453 | 27 | ATGGGCGAGCGCCGTTCCGAAGGGACG | 1946.1 | -10.7 | 1.5 | -7.3 | 3.5E-13 | 9.3E-13 | BT20/MB468 | NUC |
| >hg38_21_+_8433172_8446622_RNA45SN1_45S_50ntFlanks@4283.4320.38\|CTTTAACGAGGATCCATTGGAGGGCAAGTCTGGTGCCA | >hg38_21_+_8433172_8446622_RNA45SN1_45S_50ntFlanks | 4283 | 4320 | 38 | CTTTAACGAGGATCCATTGGAGGGCAAGTCTGGTGCCA | 195.7 | -10.6 | 1.4 | -7.3 | 2.5E-13 | 9.3E-13 | MB231/MB468 | NUC |
| >hg38_21_+_8433172_8446622_RNA45SN1_45S_50ntFlanks@11873.11909.37\|AAGAAGACCCTGTTGAGCTTGACTCTAGTCTGGCACG | >hg38_21_+_8433172_8446622_RNA45SN1_45S_50ntFlanks | 11873 | 11909 | 37 | AAGAAGACCCTGTTGAGCTTGACTCTAGTCTGGCACG | 173.8 | -11.3 | 1.5 | -7.3 | 2.1E-13 | 9.3E-13 | MB231/MB468 | MP |
| >hg38_21_+_8433172_8446622_RNA45SN1_45S_50ntFlanks@10947.10967.21\|CACGCCCGGGGCACCCCCCTC | >hg38_21_+_8433172_8446622_RNA45SN1_45S_50ntFlanks | 10947 | 10967 | 21 | CACGCCCGGGGCACCCCCCTC | 148.4 | -9.9 | 1.4 | -7.3 | 2.4E-13 | 9.3E-13 | MB231/MB468 | CYTO |
| >hg38_21_+_8433172_8446622_RNA45SN1_45S_50ntFlanks@12627.12657.31\|ATGGGGCGAAGCTACCATCTGTGGGATTATG | >hg38_21_+_8433172_8446622_RNA45SN1_45S_50ntFlanks | 12627 | 12657 | 31 | ATGGGGCGAAGCTACCATCTGTGGGATTATG | 1631.3 | -10.5 | 1.4 | -7.3 | 3.6E-13 | 9.4E-13 | BT20/MB468 | NUC |
| >hg38_21_+_8433172_8446622_RNA45SN1_45S_50ntFlanks@10385.10409.25\|CATGTGAACAGCAGTTGAACATGGG | >hg38_21_+_8433172_8446622_RNA45SN1_45S_50ntFlanks | 10385 | 10409 | 25 | CATGTGAACAGCAGTTGAACATGGG | 261.8 | -11.9 | 1.6 | -7.3 | 2.1E-13 | 9.4E-13 | MB231/MB468 | MP |
| >MI0000736\|hsa-mir-30c-1&WithFlank&1\|+\|40757278\|40757378@23.46.24(+1U);MI0000254\|hsa-mir-30c-2&WithFlank&6\|-\|71376954\|71377037@13.36.24(+1U)&[MIMAT0000244&hsa-miR-30c-5p&offsets\|0\|+1(+1U);m-57&1\|+\|40757300\|40757323&offsets\|0\|0(+1U)];[MIMAT0000244_1&hsa-miR-30c-5p&offsets\|0\|+1(+1U);m-56&6\|-\|71377002\|71377025&offsets\|0\|0(+1U)]\|TGTAAACATCCTACACTCTCAGCTT | >MI0000736\|hsa-mir-30c-1&WithFlank&1\|+\|40757278\|40757378 | 23 | 46 | 24(+1U) | TGTAAACATCCTACACTCTCAGCTT | 457.2 | 5.5 | 0.8 | 7.3 | 2.5E-13 | 9.4E-13 | MB231/MB468 | CYTO |
| >trna20_GluTTC_2_-_131094701_131094772@1.33.33,trna5_GluTTC_13_-_41634874_41634945@1.33.33\|TCCCATATGGTCTAGCGGTTAGGATTCCTGGTT | >trna20_GluTTC_2_-_131094701_131094772 | 1 | 33 | 33 | TCCCATATGGTCTAGCGGTTAGGATTCCTGGTT | 1667.9 | 10.6 | 1.4 | 7.3 | 3.2E-13 | 9.4E-13 | MB231/MB468 | MITO |
| >hg38_21_+_8433172_8446622_RNA45SN1_45S_50ntFlanks@12991.13028.38\|CTCCCTCGCTGCGATCTATTGAAAGTCAGCCCTCGACA | >hg38_21_+_8433172_8446622_RNA45SN1_45S_50ntFlanks | 12991 | 13028 | 38 | CTCCCTCGCTGCGATCTATTGAAAGTCAGCCCTCGACA | 316.5 | -6.4 | 0.9 | -7.3 | 2.5E-13 | 9.5E-13 | MB231/MB468 | CYTO |
| >am_trna128_GlyGCC_6_-_27870686_27870756@3.35.33,trna18_GlyGCC_16_+_70822597_70822667@3.35.33,trna19_GlyGCC_16_+_70823410_70823480@3.35.33,trna19_GlyGCC_2_-_157257659_157257729@3.35.33,trna24_GlyGCC_16_-_70812942_70813012@3.35.33,trna25_GlyGCC_16_-_70812114_70812184@3.35.33,trna5_GlyGCC_17_+_8029064_8029134@3.35.33,trna68_GlyGCC_1_-_161493637_161493707@3.35.33\|ATTGGTGGTTCAGTGGTAGAATTCTCGCCTGCC | >am_trna128_GlyGCC_6_-_27870686_27870756 | 3 | 35 | 33 | ATTGGTGGTTCAGTGGTAGAATTCTCGCCTGCC | 552.1 | 10.3 | 1.4 | 7.3 | 2.5E-13 | 9.5E-13 | MB231/MB468 | NUC |
| >hg38_21_+_8433172_8446622_RNA45SN1_45S_50ntFlanks@7976.8008.33\|GCGACCTCAGATCAGACGTGGCGACCCGCTGAA | >hg38_21_+_8433172_8446622_RNA45SN1_45S_50ntFlanks | 7976 | 8008 | 33 | GCGACCTCAGATCAGACGTGGCGACCCGCTGAA | 1047.9 | 6.7 | 0.9 | 7.3 | 2.5E-13 | 9.5E-13 | MB231/MB468 | CYTO |
| >hg38_21_+_8433172_8446622_RNA45SN1_45S_50ntFlanks@10947.10981.35\|CACGCCCGGGGCACCCCCCTCGCGGCCCTCCCCCG | >hg38_21_+_8433172_8446622_RNA45SN1_45S_50ntFlanks | 10947 | 10981 | 35 | CACGCCCGGGGCACCCCCCTCGCGGCCCTCCCCCG | 124.9 | -6.7 | 0.9 | -7.3 | 2.5E-13 | 9.5E-13 | MB231/MB468 | CYTO |
| >hg38_21_+_8433172_8446622_RNA45SN1_45S_50ntFlanks@8168.8188.21\|CCCAAGTCCTTCTGATCGAGG | >hg38_21_+_8433172_8446622_RNA45SN1_45S_50ntFlanks | 8168 | 8188 | 21 | CCCAAGTCCTTCTGATCGAGG | 397.5 | -11.7 | 1.5 | -7.7 | 1.0E-14 | 9.5E-13 | BT20/MB468 | MP |
| >hg38_21_+_8433172_8446622_RNA45SN1_45S_50ntFlanks@3985.4026.42\|CGGCTTTGGTGACTCTAGATAACCTCGGGCCGATCGCACGCC | >hg38_21_+_8433172_8446622_RNA45SN1_45S_50ntFlanks | 3985 | 4026 | 42 | CGGCTTTGGTGACTCTAGATAACCTCGGGCCGATCGCACGCC | 918.7 | -9.2 | 1.3 | -7.3 | 2.5E-13 | 9.5E-13 | MB231/MB468 | NUC |
| >hg38_21_+_8433172_8446622_RNA45SN1_45S_50ntFlanks@12420.12455.36\|ATGTCGGCTCTTCCTATCATTGTGAAGCAGAATTCA | >hg38_21_+_8433172_8446622_RNA45SN1_45S_50ntFlanks | 12420 | 12455 | 36 | ATGTCGGCTCTTCCTATCATTGTGAAGCAGAATTCA | 2831.7 | -5.6 | 0.8 | -7.3 | 2.5E-13 | 9.5E-13 | BT20/MB468 | CYTO |
| >hg38_21_+_8433172_8446622_RNA45SN1_45S_50ntFlanks@11718.11740.23\|AACGGCGGGAGTAACTATGACTC | >hg38_21_+_8433172_8446622_RNA45SN1_45S_50ntFlanks | 11718 | 11740 | 23 | AACGGCGGGAGTAACTATGACTC | 1247.4 | -7.8 | 1.1 | -7.3 | 3.2E-13 | 9.5E-13 | MB231/MB468 | MITO |
| >hg38_21_+_8433172_8446622_RNA45SN1_45S_50ntFlanks@485.532.48\|GATGTGGTGACGTCGTGCTCTCCCGGGCCGGGTCCGAGCCGCGACGGG | >hg38_21_+_8433172_8446622_RNA45SN1_45S_50ntFlanks | 485 | 532 | 48 | GATGTGGTGACGTCGTGCTCTCCCGGGCCGGGTCCGAGCCGCGACGGG | 705.7 | 26.5 | 3.6 | 7.3 | 3.2E-13 | 9.5E-13 | MB231/MB468 | MITO |
| >hg38_21_+_8433172_8446622_RNA45SN1_45S_50ntFlanks@4494.4511.18\|CCCGAAGCGTTTACTTTG | >hg38_21_+_8433172_8446622_RNA45SN1_45S_50ntFlanks | 4494 | 4511 | 18 | CCCGAAGCGTTTACTTTG | 382.1 | -12.4 | 1.7 | -7.3 | 3.2E-13 | 9.5E-13 | MB231/MB468 | MITO |
| >hg38_21_+_8433172_8446622_RNA45SN1_45S_50ntFlanks@10805.10828.24\|GAACAATGTAGGTAAGGGAAGTCG | >hg38_21_+_8433172_8446622_RNA45SN1_45S_50ntFlanks | 10805 | 10828 | 24 | GAACAATGTAGGTAAGGGAAGTCG | 185.6 | -5.2 | 0.7 | -7.3 | 2.5E-13 | 9.6E-13 | MB231/MB468 | CYTO |
| >MI0000266\|hsa-mir-10a&WithFlank&17\|-\|48579832\|48579953@28.48.21(+1A)&[MIMAT0000253&hsa-miR-10a-5p&offsets\|0\|-2(+1A);m-18&17\|-\|48579905\|48579926&offsets\|0\|-1(+1A)]\|TACCCTGTAGATCCGAATTTGA | >MI0000266\|hsa-mir-10a&WithFlank&17\|-\|48579832\|48579953 | 28 | 48 | 21(+1A) | TACCCTGTAGATCCGAATTTGA | 205.3 | 10.9 | 1.5 | 7.3 | 2.2E-13 | 9.6E-13 | MB231/MB468 | MP |
| >hg38_21_+_8433172_8446622_RNA45SN1_45S_50ntFlanks@11668.11703.36\|TGCCCAGTGCTCTGAATGTCAAAGTGAAGAAATTCA | >hg38_21_+_8433172_8446622_RNA45SN1_45S_50ntFlanks | 11668 | 11703 | 36 | TGCCCAGTGCTCTGAATGTCAAAGTGAAGAAATTCA | 2144.9 | 5.1 | 0.7 | 7.3 | 2.5E-13 | 9.6E-13 | MB231/MB468 | CYTO |
| >hg38_21_+_8433172_8446622_RNA45SN1_45S_50ntFlanks@12151.12182.32\|GTGGGGAGTTTGACTGGGGCGGTACACCTGTC | >hg38_21_+_8433172_8446622_RNA45SN1_45S_50ntFlanks | 12151 | 12182 | 32 | GTGGGGAGTTTGACTGGGGCGGTACACCTGTC | 290.7 | -10.3 | 1.4 | -7.4 | 1.9E-13 | 9.6E-13 | BT20/MB468 | MITO |
| >hg38_21_+_8433172_8446622_RNA45SN1_45S_50ntFlanks@10682.10706.25\|TCCGGCGGCGTCCGGTGAGCTCTCG | >hg38_21_+_8433172_8446622_RNA45SN1_45S_50ntFlanks | 10682 | 10706 | 25 | TCCGGCGGCGTCCGGTGAGCTCTCG | 2185.5 | -5.7 | 0.8 | -7.3 | 2.5E-13 | 9.6E-13 | MB231/MB468 | CYTO |
| >hg38_21_+_8433172_8446622_RNA45SN1_45S_50ntFlanks@11876.11910.35\|AAGACCCTGTTGAGCTTGACTCTAGTCTGGCACGG | >hg38_21_+_8433172_8446622_RNA45SN1_45S_50ntFlanks | 11876 | 11910 | 35 | AAGACCCTGTTGAGCTTGACTCTAGTCTGGCACGG | 127.8 | -10.9 | 1.5 | -7.3 | 2.2E-13 | 9.6E-13 | MB231/MB468 | MP |
| >hg38_21_+_8433172_8446622_RNA45SN1_45S_50ntFlanks@10351.10391.41\|CGAGAACTTTGAAGGCCGAAGTGGAGAAGGGTTCCATGTGA | >hg38_21_+_8433172_8446622_RNA45SN1_45S_50ntFlanks | 10351 | 10391 | 41 | CGAGAACTTTGAAGGCCGAAGTGGAGAAGGGTTCCATGTGA | 2683.4 | -9.2 | 1.3 | -7.3 | 3.6E-13 | 9.6E-13 | BT20/MB468 | NUC |
| >hg38_21_+_8433172_8446622_RNA45SN1_45S_50ntFlanks@11905.11922.18\|GCACGGTGAAGAGACATG | >hg38_21_+_8433172_8446622_RNA45SN1_45S_50ntFlanks | 11905 | 11922 | 18 | GCACGGTGAAGAGACATG | 183.6 | -11.4 | 1.6 | -7.3 | 2.2E-13 | 9.6E-13 | MB231/MB468 | MP |
| >trna116_GluCTC_1_-_145399233_145399304@33.70.38,trna71_GluCTC_1_-_161439189_161439260@33.70.38,trna74_GluCTC_1_-_161431809_161431880@33.70.38,trna77_GluCTC_1_-_161424398_161424469@33.70.38,trna77_GluCTC_6_+_28949976_28950047@33.70.38,trna80_GluCTC_1_-_161417018_161417089@33.70.38,trna87_GluCTC_6_-_126101393_126101464@33.70.38\|TCTCACCGCCGCGGCCCGGGTTCGATTCCCGGTCAGGG | >trna116_GluCTC_1_-_145399233_145399304 | 33 | 70 | 38 | TCTCACCGCCGCGGCCCGGGTTCGATTCCCGGTCAGGG | 179.1 | -9.8 | 1.3 | -7.3 | 2.5E-13 | 9.6E-13 | BT20/MB468 | CYTO |
| >hg38_21_+_8433172_8446622_RNA45SN1_45S_50ntFlanks@10069.10100.32\|AGTGGACGGGAGCGGCGGGGGGCGGCGCGCGCG | >hg38_21_+_8433172_8446622_RNA45SN1_45S_50ntFlanks | 10069 | 10100 | 32 | AGTGGACGGGAGCGGCGGGGGGCGGCGCGCGCG | 204.9 | -11.5 | 1.6 | -7.3 | 2.2E-13 | 9.6E-13 | MB231/MB468 | MP |
| >trna11_GluTTC_15_-_26327381_26327452@1.36.36,trna3_GluTTC_13_-_45492062_45492133@1.36.36\|TCCCACATGGTCTAGCGGTTAGGATTCCTGGTTTTC | >trna11_GluTTC_15_-_26327381_26327452 | 1 | 36 | 36 | TCCCACATGGTCTAGCGGTTAGGATTCCTGGTTTTC | 2567.1 | 10.6 | 1.4 | 7.3 | 2.5E-13 | 9.6E-13 | MB231/MB468 | CYTO |
| >hg38_21_+_8433172_8446622_RNA45SN1_45S_50ntFlanks@11573.11602.30\|CGACTTAGAACTGGTGCGGACCAGGGGAAT | >hg38_21_+_8433172_8446622_RNA45SN1_45S_50ntFlanks | 11573 | 11602 | 30 | CGACTTAGAACTGGTGCGGACCAGGGGAAT | 1197.2 | -4.5 | 0.6 | -7.3 | 2.6E-13 | 9.6E-13 | MB231/MB468 | NUC |
| >hg38_21_+_8433172_8446622_RNA45SN1_45S_50ntFlanks@10726.10755.30\|GGGAGAGGGTGTAAATCTCGCGCCGGGCCG | >hg38_21_+_8433172_8446622_RNA45SN1_45S_50ntFlanks | 10726 | 10755 | 30 | GGGAGAGGGTGTAAATCTCGCGCCGGGCCG | 383.1 | -11.5 | 1.6 | -7.3 | 2.6E-13 | 9.7E-13 | MB231/MB468 | NUC |
| >hg38_21_+_8433172_8446622_RNA45SN1_45S_50ntFlanks@4283.4319.37\|CTTTAACGAGGATCCATTGGAGGGCAAGTCTGGTGCC | >hg38_21_+_8433172_8446622_RNA45SN1_45S_50ntFlanks | 4283 | 4319 | 37 | CTTTAACGAGGATCCATTGGAGGGCAAGTCTGGTGCC | 188.4 | -10.5 | 1.4 | -7.3 | 2.6E-13 | 9.7E-13 | MB231/MB468 | NUC |
| >hg38_21_+_8433172_8446622_RNA45SN1_45S_50ntFlanks@11568.11585.18\|GCAGCCGACTTAGAACTG | >hg38_21_+_8433172_8446622_RNA45SN1_45S_50ntFlanks | 11568 | 11585 | 18 | GCAGCCGACTTAGAACTG | 3086.6 | -5.7 | 0.8 | -7.3 | 3.7E-13 | 9.7E-13 | BT20/MB468 | NUC |

| | | | | | | | | | | | | | |
|---|---|---|---|---|---|---|---|---|---|---|---|---|---|
| >MI0000743\|hsa-mir-34c&WithFlank&11\|+\|111513433\|111513521@19.41.23&[MIMAT0000686&hsa-miR-34c-5p&offsets\|0\|0;m-336&11\|+\|111513451\|111513473&offsets\|0\|0]\|AGGCAGTGTAGTTAGCTGATTGC | >MI0000743\|hsa-mir-34c&WithFlank&11\|+\|111513433\|111513521 | 19 | 41 | 23 | AGGCAGTGTAGTTAGCTGATTGC | 227.5 | -10.9 | 1.4 | -7.6 | 3.0E-14 | 9.7E-13 | BT20/MB231 | NUC |
| >hg38_21_+_8433172_8446622_RNA45SN1_45S_50ntFlanks@3785.3810.26\|TGAAACTGCGAATGGCTCATTAAATC | >hg38_21_+_8433172_8446622_RNA45SN1_45S_50ntFlanks | 3785 | 3810 | 26 | TGAAACTGCGAATGGCTCATTAAATC | 568.8 | -4.9 | 0.7 | -7.3 | 2.6E-13 | 9.7E-13 | MB231/MB468 | NUC |
| >hg38_21_+_8433172_8446622_RNA45SN1_45S_50ntFlanks@8192.8211.20\|AGCCCGTGGACGGTGTGAGG | >hg38_21_+_8433172_8446622_RNA45SN1_45S_50ntFlanks | 8192 | 8211 | 20 | AGCCCGTGGACGGTGTGAGG | 708.2 | -5.7 | 0.8 | -7.3 | 2.2E-13 | 9.7E-13 | MB231/MB468 | MP |
| >MI0000077\|hsa-mir-21&WithFlank&17\|+\|59841260\|59841343@13.35.23&[MIMAT0000076&hsa-miR-21-5p&offsets\|-1\|0;m-1&17\|+\|59841273\|59841294&offsets\|-1\|0]\|GTAGCTTATCAGACTGATGTTGA | >MI0000077\|hsa-mir-21&WithFlank&17\|+\|59841260\|59841343 | 13 | 35 | 23 | GTAGCTTATCAGACTGATGTTGA | 180.8 | 3.0 | 0.4 | 7.3 | 3.7E-13 | 9.7E-13 | BT20/MB468 | NUC |
| >MI0005756\|hsa-mir-934&WithFlank&X\|+\|136550872\|136550966@21.42.22&[MIMAT0004977&hsa-miR-934&offsets\|0\|0;m-943&X\|+\|136550892\|136550913&offsets\|0\|0]\|TGTCTACTACTGGAGACACTGG | >MI0005756\|hsa-mir-934&WithFlank&X\|+\|136550872\|136550966 | 21 | 42 | 22 | TGTCTACTACTGGAGACACTGG | 720.5 | 4.8 | 0.7 | 7.3 | 3.7E-13 | 9.7E-13 | BT20/MB468 | NUC |
| >MI0003776\|hsa-mir-320b-1&WithFlank&1\|+\|116671740\|116671823@48.67.20(+1U);MI0003839\|hsa-mir-320b-2&WithFlank&1\|-\|224257034\|224257116@47.66.20(+1U);MI0000542\|hsa-mir-320a&WithFlank&8\|-\|22244960\|22245043@48.67.20(+1U)&[MIMAT0005792&hsa-miR-320b&offsets\|0\|-2(+1U);m-183&1\|+\|116671787\|116671806&offsets\|0\|0(+1U)];[MIMAT0005792_1&hsa-miR-320b&offsets\|0\|-2(+1U);m-183&1\|-\|224257051\|224257070&offsets\|0\|0(+1U)];[MIMAT0000510&hsa-miR-320a-3p&offsets\|0\|-2(+1U);m-60&8\|-\|22244975\|22244996&offsets\|0\|-2(+1U)]\|AAAAGCTGGGTTGAGAGGGCT | >MI0003776\|hsa-mir-320b-1&WithFlank&1\|+\|116671740\|116671823 | 48 | 67 | 20(+1U) | AAAAGCTGGGTTGAGAGGGCT | 2048.0 | 11.5 | 1.6 | 7.3 | 2.6E-13 | 9.7E-13 | BT20/MB468 | CYTO |
| >hg38_21_+_8433172_8446622_RNA45SN1_45S_50ntFlanks@9738.9763.26\|GAAACGATCTCAACCTATTCTCAAAC | >hg38_21_+_8433172_8446622_RNA45SN1_45S_50ntFlanks | 9738 | 9763 | 26 | GAAACGATCTCAACCTATTCTCAAAC | 167.3 | -11.3 | 1.5 | -7.3 | 2.2E-13 | 9.7E-13 | MB231/MB468 | MP |
| >hg38_21_+_8433172_8446622_RNA45SN1_45S_50ntFlanks@6731.6756.26\|CACATTGATCATCGACACTTCGAACG | >hg38_21_+_8433172_8446622_RNA45SN1_45S_50ntFlanks | 6731 | 6756 | 26 | CACATTGATCATCGACACTTCGAACG | 798.3 | -7.2 | 1.0 | -7.3 | 2.2E-13 | 9.7E-13 | MB231/MB468 | MP |
| >hg38_1_-_228634819_228635039_RNA5S12_5S_50ntFlanks@151.170.20\|ATACCGGGTGCTGTAGGCTT | >hg38_1_-_228634819_228635039_RNA5S12_5S_50ntFlanks | 151 | 170 | 20 | ATACCGGGTGCTGTAGGCTT | 4269.7 | 5.5 | 0.7 | 7.3 | 2.6E-13 | 9.7E-13 | MB231/MB468 | CYTO |
| >hg38_21_+_8433172_8446622_RNA45SN1_45S_50ntFlanks@8357.8400.44\|AAAAGAACTTTGAAGAGAGAGTTCAAGAGGGCGTGAAACCGTTA | >hg38_21_+_8433172_8446622_RNA45SN1_45S_50ntFlanks | 8357 | 8400 | 44 | AAAAGAACTTTGAAGAGAGAGTTCAAGAGGGCGTGAAACCGTTA | 327.5 | -12.1 | 1.7 | -7.3 | 3.3E-13 | 9.7E-13 | MB231/MB468 | MITO |
| >hg38_21_+_8433172_8446622_RNA45SN1_45S_50ntFlanks@11564.11588.25\|CCTAGCAGCCGACTTAGAACTGGTG | >hg38_21_+_8433172_8446622_RNA45SN1_45S_50ntFlanks | 11564 | 11588 | 25 | CCTAGCAGCCGACTTAGAACTGGTG | 1097.5 | -5.1 | 0.7 | -7.3 | 2.6E-13 | 9.8E-13 | MB231/MB468 | CYTO |
| >hg38_21_+_8433172_8446622_RNA45SN1_45S_50ntFlanks@11703.11728.26\|AATGAAGCGCGGGTAAACGGCGGGAG | >hg38_21_+_8433172_8446622_RNA45SN1_45S_50ntFlanks | 11703 | 11728 | 26 | AATGAAGCGCGGGTAAACGGCGGGAG | 1829.8 | -6.4 | 0.9 | -7.3 | 2.6E-13 | 9.8E-13 | MB231/MB468 | CYTO |
| >hg38_21_+_8433172_8446622_RNA45SN1_45S_50ntFlanks@9571.9597.27\|GCAAATCGGTCGTCCGACCTGGGTATA | >hg38_21_+_8433172_8446622_RNA45SN1_45S_50ntFlanks | 9571 | 9597 | 27 | GCAAATCGGTCGTCCGACCTGGGTATA | 196.2 | -5.9 | 0.8 | -7.3 | 2.6E-13 | 9.8E-13 | MB231/MB468 | CYTO |
| >trnaMT_ValTAC_MT_+_1602_1670@48.72.25\|CAACTTAACTTGACCGCTCTGACCA | >trnaMT_ValTAC_MT_+_1602_1670 | 48 | 72 | 25 | CAACTTAACTTGACCGCTCTGACCA | 776.8 | 7.9 | 1.1 | 7.3 | 2.6E-13 | 9.8E-13 | MB231/MB468 | CYTO |
| >am_trna106_HisGTG_1_-_149155828_149155899@7.29.23,trna111_HisGTG_1_-_147774845_147774916@7.29.23,trna118_HisGTG_1_-_145396881_145396952@7.29.23,trna16_HisGTG_1_+_146544773_146544844@7.29.23,trna1_HisGTG_15_+_45493349_45493420@7.29.23,trna21_HisGTG_1_+_147753471_147753542@7.29.23,trna33_HisGTG_6_+_27125906_27125977@7.29.23,trna7_HisGTG_9_-_14433938_14434009@7.29.23,trna8_HisGTG_15_-_45492611_45492682@7.29.23,trna9_HisGTG_15_-_45490804_45490875@7.29.23\|ATCGTATAGTGGTTAGTACTCTG | >am_trna106_HisGTG_1_-_149155828_149155899 | 7 | 29 | 23 | ATCGTATAGTGGTTAGTACTCTG | 246.0 | -10.0 | 1.4 | -7.4 | 2.0E-13 | 9.8E-13 | BT20/MB468 | MITO |
| >hg38_21_+_8433172_8446622_RNA45SN1_45S_50ntFlanks@11768.11806.39\|CTAATTAGTGACGCGCATGAATGGATGAACGAGATTCCC | >hg38_21_+_8433172_8446622_RNA45SN1_45S_50ntFlanks | 11768 | 11806 | 39 | CTAATTAGTGACGCGCATGAATGGATGAACGAGATTCCC | 225.3 | -5.3 | 0.7 | -7.3 | 2.6E-13 | 9.8E-13 | MB231/MB468 | CYTO |
| >hg38_21_+_8433172_8446622_RNA45SN1_45S_50ntFlanks@10801.10818.18\|GTTGGAACAATGTAGGTA | >hg38_21_+_8433172_8446622_RNA45SN1_45S_50ntFlanks | 10801 | 10818 | 18 | GTTGGAACAATGTAGGTA | 185.0 | -11.4 | 1.6 | -7.3 | 2.2E-13 | 9.8E-13 | MB231/MB468 | MP |
| >hg38_21_+_8433172_8446622_RNA45SN1_45S_50ntFlanks@6652.6700.49\|GACTCTTAGCGGTGGATCACTCGGCTCGTGCGTCGATGAAGAACGCAGC | >hg38_21_+_8433172_8446622_RNA45SN1_45S_50ntFlanks | 6652 | 6700 | 49 | GACTCTTAGCGGTGGATCACTCGGCTCGTGCGTCGATGAAGAACGCAGC | 7216.3 | 7.3 | 1.0 | 7.3 | 2.0E-13 | 9.8E-13 | BT20/MB468 | MITO |
| >hg38_21_+_8433172_8446622_RNA45SN1_45S_50ntFlanks@9352.9376.25\|CCGTGGCGCAATGAAGGTGAAGGCC | >hg38_21_+_8433172_8446622_RNA45SN1_45S_50ntFlanks | 9352 | 9376 | 25 | CCGTGGCGCAATGAAGGTGAAGGCC | 182.0 | -11.4 | 1.6 | -7.3 | 2.2E-13 | 9.8E-13 | MB231/MB468 | MP |
| >hg38_21_+_8433172_8446622_RNA45SN1_45S_50ntFlanks@9545.9572.28\|GTGGAGGTCCGTAGCGGTCCTGACGTGC | >hg38_21_+_8433172_8446622_RNA45SN1_45S_50ntFlanks | 9545 | 9572 | 28 | GTGGAGGTCCGTAGCGGTCCTGACGTGC | 2305.9 | -4.4 | 0.6 | -7.3 | 2.6E-13 | 9.8E-13 | MB231/MB468 | NUC |
| >hg38_21_+_8433172_8446622_RNA45SN1_45S_50ntFlanks@9806.9832.27\|CGTGGAATGCGAGTGCCTAGTGGGCCA | >hg38_21_+_8433172_8446622_RNA45SN1_45S_50ntFlanks | 9806 | 9832 | 27 | CGTGGAATGCGAGTGCCTAGTGGGCCA | 314.9 | -12.2 | 1.7 | -7.3 | 2.2E-13 | 9.8E-13 | MB231/MB468 | MP |
| >hg38_21_+_8433172_8446622_RNA45SN1_45S_50ntFlanks@10806.10825.20\|AACAATGTAGGTAAGGGAAG | >hg38_21_+_8433172_8446622_RNA45SN1_45S_50ntFlanks | 10806 | 10825 | 20 | AACAATGTAGGTAAGGGAAG | 427.9 | -6.7 | 0.9 | -7.3 | 2.6E-13 | 9.8E-13 | MB231/MB468 | CYTO |
| >hg38_21_+_8433172_8446622_RNA45SN1_45S_50ntFlanks@475.506.32\|CTTCGTGATCGATGTGGTGACGTCGTGCTCTC | >hg38_21_+_8433172_8446622_RNA45SN1_45S_50ntFlanks | 475 | 506 | 32 | CTTCGTGATCGATGTGGTGACGTCGTGCTCTC | 3868.7 | 8.0 | 1.1 | 7.3 | 2.6E-13 | 9.9E-13 | MB231/MB468 | CYTO |
| >hg38_21_+_8433172_8446622_RNA45SN1_45S_50ntFlanks@6721.6740.20\|AATTGCAGGACACATTGATC | >hg38_21_+_8433172_8446622_RNA45SN1_45S_50ntFlanks | 6721 | 6740 | 20 | AATTGCAGGACACATTGATC | 3108.7 | -7.8 | 1.1 | -7.3 | 3.3E-13 | 9.9E-13 | MB231/MB468 | MITO |
| >hg38_21_+_8433172_8446622_RNA45SN1_45S_50ntFlanks@11078.11103.26\|CCGCGTGGGGGCGGCGGCGGGGGGAG | >hg38_21_+_8433172_8446622_RNA45SN1_45S_50ntFlanks | 11078 | 11103 | 26 | CCGCGTGGGGGCGGCGGCGGGGGGAG | 201.1 | -10.6 | 1.5 | -7.3 | 2.7E-13 | 9.9E-13 | MB231/MB468 | NUC |
| >hg38_21_+_8433172_8446622_RNA45SN1_45S_50ntFlanks@11211.11234.24\|GCGACTCTGGACGCGAGCCGGGCC | >hg38_21_+_8433172_8446622_RNA45SN1_45S_50ntFlanks | 11211 | 11234 | 24 | GCGACTCTGGACGCGAGCCGGGCC | 138.5 | -11.0 | 1.5 | -7.3 | 2.2E-13 | 9.9E-13 | MB231/MB468 | MP |
| >hg38_21_+_8433172_8446622_RNA45SN1_45S_50ntFlanks@10687.10708.22\|CGGCGTCCGGTGAGCTCTCGCT | >hg38_21_+_8433172_8446622_RNA45SN1_45S_50ntFlanks | 10687 | 10708 | 22 | CGGCGTCCGGTGAGCTCTCGCT | 2374.3 | -4.4 | 0.6 | -7.3 | 2.6E-13 | 9.9E-13 | MB231/MB468 | CYTO |
| >hg38_21_+_8433172_8446622_RNA45SN1_45S_50ntFlanks@7978.8012.35\|GACCTCAGATCAGACGTGGCGACCCGCTGAATTTA | >hg38_21_+_8433172_8446622_RNA45SN1_45S_50ntFlanks | 7978 | 8012 | 35 | GACCTCAGATCAGACGTGGCGACCCGCTGAATTTA | 459.1 | 6.7 | 0.9 | 7.3 | 2.6E-13 | 9.9E-13 | MB231/MB468 | CYTO |
| >MI0000477\|hsa-mir-146a&WithFlank&5\|+\|160485346\|160485456@27.48.22(+1A)&[MIMAT0000449&hsa-miR-146a-5p&offsets\|0\|0(+1A);m-129&5\|+\|160485372\|160485393&offsets\|0\|0(+1A)]\|TGAGAACTGAATTCCATGGGTTA | >MI0000477\|hsa-mir-146a&WithFlank&5\|+\|160485346\|160485456 | 27 | 48 | 22(+1A) | TGAGAACTGAATTCCATGGGTTA | 466.8 | 12.1 | 1.7 | 7.3 | 3.4E-13 | 9.9E-13 | MB231/MB468 | MITO |
| >hg38_21_+_8433172_8446622_RNA45SN1_45S_50ntFlanks@10544.10570.27\|TCCAGTGCGGTAACGCGACCGATCCCG | >hg38_21_+_8433172_8446622_RNA45SN1_45S_50ntFlanks | 10544 | 10570 | 27 | TCCAGTGCGGTAACGCGACCGATCCCG | 114.5 | -10.7 | 1.5 | -7.3 | 2.3E-13 | 9.9E-13 | MB231/MB468 | MP |
| >MI0001448\|hsa-mir-425&WithFlank&3\|-\|49020142\|49020240@20.43.24&[MIMAT0003393&hsa-miR-425-5p&offsets\|0\|+1;m-95&3\|-\|49020200\|49020221&offsets\|0\|+2]\|AATGACACGATCACTCCCGTTGAG | >MI0001448\|hsa-mir-425&WithFlank&3\|-\|49020142\|49020240 | 20 | 43 | 24 | AATGACACGATCACTCCCGTTGAG | 1253.1 | 5.9 | 0.8 | 7.3 | 2.6E-13 | 9.9E-13 | MB231/MB468 | CYTO |
| >hg38_21_+_8433172_8446622_RNA45SN1_45S_50ntFlanks@10392.10420.29\|ACAGCAGTTGAACATGGGTCAGTCGGTCC | >hg38_21_+_8433172_8446622_RNA45SN1_45S_50ntFlanks | 10392 | 10420 | 29 | ACAGCAGTTGAACATGGGTCAGTCGGTCC | 583.4 | -8.9 | 1.2 | -7.3 | 3.4E-13 | 1.0E-12 | MB231/MB468 | MITO |
| >hg38_21_+_8433172_8446622_RNA45SN1_45S_50ntFlanks@12936.12956.21\|AAACCATTCGTAGACGACCTG | >hg38_21_+_8433172_8446622_RNA45SN1_45S_50ntFlanks | 12936 | 12956 | 21 | AAACCATTCGTAGACGACCTG | 2380.0 | -4.4 | 0.6 | -7.3 | 2.0E-13 | 1.0E-12 | BT20/MB468 | MITO |
| >hg38_21_+_8433172_8446622_RNA45SN1_45S_50ntFlanks@4983.5013.31\|CGGACAGGATTGACAGATTGATAGCTCTTTC | >hg38_21_+_8433172_8446622_RNA45SN1_45S_50ntFlanks | 4983 | 5013 | 31 | CGGACAGGATTGACAGATTGATAGCTCTTTC | 466.8 | -12.6 | 1.7 | -7.3 | 3.4E-13 | 1.0E-12 | MB231/MB468 | MITO |
| >hg38_21_+_8433172_8446622_RNA45SN1_45S_50ntFlanks@10781.10814.34\|AGGTGAACAGCCTCTGGCATGTTGGAACAATGTA | >hg38_21_+_8433172_8446622_RNA45SN1_45S_50ntFlanks | 10781 | 10814 | 34 | AGGTGAACAGCCTCTGGCATGTTGGAACAATGTA | 141.3 | -11.0 | 1.5 | -7.3 | 2.3E-13 | 1.0E-12 | MB231/MB468 | MP |
| >hg38_21_+_8433172_8446622_RNA45SN1_45S_50ntFlanks@8182.8211.30\|ATCGAGGCCCAGCCCGTGGACGGTGTGAGG | >hg38_21_+_8433172_8446622_RNA45SN1_45S_50ntFlanks | 8182 | 8211 | 30 | ATCGAGGCCCAGCCCGTGGACGGTGTGAGG | 179.4 | -9.8 | 1.3 | -7.3 | 2.6E-13 | 1.0E-12 | BT20/MB468 | CYTO |
| >hg38_21_+_8433172_8446622_RNA45SN1_45S_50ntFlanks@10819.10838.20\|AGGGAAGTCGGCAAGCCGGA | >hg38_21_+_8433172_8446622_RNA45SN1_45S_50ntFlanks | 10819 | 10838 | 20 | AGGGAAGTCGGCAAGCCGGA | 227.1 | -10.8 | 1.5 | -7.3 | 2.7E-13 | 1.0E-12 | MB231/MB468 | NUC |
| >MI0005756\|hsa-mir-934&WithFlank&X\|+\|136550872\|136550966@21.40.20(+1A)&[MIMAT0004977&hsa-miR-934&offsets\|0\|-2(+1A);m-943&X\|+\|136550892\|136550913&offsets\|0\|-2(+1A)]\|TGTCTACTACTGGAGACACTA | >MI0005756\|hsa-mir-934&WithFlank&X\|+\|136550872\|136550966 | 21 | 40 | 20(+1A) | TGTCTACTACTGGAGACACTA | 223.9 | 11.6 | 1.5 | 7.6 | 3.1E-14 | 1.0E-12 | BT20/MB231 | NUC |
| >am_trna106_HisGTG_1_-_149155828_149155899@7.28.22,trna111_HisGTG_1_-_147774845_147774916@7.28.22,trna118_HisGTG_1_-_145396881_145396952@7.28.22,trna16_HisGTG_1_+_146544773_146544844@7.28.22,trna1_HisGTG_15_+_45493349_45493420@7.28.22,trna21_HisGTG_1_+_147753471_147753542@7.28.22,trna33_HisGTG_6_+_27125906_27125977@7.28.22,trna7_HisGTG_9_-_14433938_14434009@7.28.22,trna8_HisGTG_15_-_45492611_45492682@7.28.22,trna9_HisGTG_15_-_45490804_45490875@7.28.22\|ATCGTATAGTGGTTAGTACTCT | >am_trna106_HisGTG_1_-_149155828_149155899 | 7 | 28 | 22 | ATCGTATAGTGGTTAGTACTCT | 772.3 | -11.9 | 1.6 | -7.3 | 2.7E-13 | 1.0E-12 | BT20/MB468 | CYTO |
| >hg38_21_+_8433172_8446622_RNA45SN1_45S_50ntFlanks@12957.12987.31\|CTTCTGGGTCGGGGTTTCGTACGTAGCAGAG | >hg38_21_+_8433172_8446622_RNA45SN1_45S_50ntFlanks | 12957 | 12987 | 31 | CTTCTGGGTCGGGGTTTCGTACGTAGCAGAG | 149.4 | -11.1 | 1.5 | -7.3 | 2.3E-13 | 1.0E-12 | MB231/MB468 | MP |
| >hg38_21_+_8433172_8446622_RNA45SN1_45S_50ntFlanks@12405.12427.23\|CTTTTTGATCCTTCGATGTCGGC | >hg38_21_+_8433172_8446622_RNA45SN1_45S_50ntFlanks | 12405 | 12427 | 23 | CTTTTTGATCCTTCGATGTCGGC | 226.3 | -7.8 | 1.1 | -7.3 | 2.7E-13 | 1.0E-12 | BT20/MB468 | CYTO |
| >hg38_21_+_8433172_8446622_RNA45SN1_45S_50ntFlanks@11641.11664.24\|CGGCGGGTGTTGACGCGATGTGAT | >hg38_21_+_8433172_8446622_RNA45SN1_45S_50ntFlanks | 11641 | 11664 | 24 | CGGCGGGTGTTGACGCGATGTGAT | 3498.9 | -10.2 | 1.4 | -7.3 | 3.4E-13 | 1.0E-12 | MB231/MB468 | MITO |
| >hg38_21_+_8433172_8446622_RNA45SN1_45S_50ntFlanks@4449.4471.23\|CCTCTCGGCGCCCCCTCGATGCT | >hg38_21_+_8433172_8446622_RNA45SN1_45S_50ntFlanks | 4449 | 4471 | 23 | CCTCTCGGCGCCCCCTCGATGCT | 1444.0 | -9.9 | 1.3 | -7.3 | 2.7E-13 | 1.0E-12 | MB231/MB468 | NUC |

<table>
<tr><td>>hg38_21_+_8433172_8446622_RNA45SN1_45S_50ntFlanks@12934.12966.33|CTAAACCATTCGTAGACGACCTGCTTCTGGGTC</td><td>>hg38_21_+_8433172_8446622_RNA45SN1_45S_50ntFlanks</td><td>12934</td><td>12966</td><td>33</td><td>CTAAACCATTCGTAGACGACCTGCTTCTGGGTC</td><td>1112.9</td><td>-8.7</td><td>1.2</td><td>-7.3</td><td>2.7E-13</td><td>1.0E-12</td><td>BT20/MB468</td><td>CYTO</td></tr>
<tr><td>>MI0006273|hsa-mir-1180&WithFlank&17|-|19344500|19344580@47.67.21(+1A)&[MIMAT0005825&hsa-miR-1180-3p&offsets|0|-1(+1A);m-214&17|-|19344513|19344534&offsets|0|-1(+1A)]|TTTCCGGCTCGCGTGGGTGTGA</td><td>>MI0006273|hsa-mir-1180&WithFlank&17|-|19344500|19344580</td><td>47</td><td>67</td><td>21(+1A)</td><td>TTTCCGGCTCGCGTGGGTGTGA</td><td>442.5</td><td>10.2</td><td>1.4</td><td>7.3</td><td>2.7E-13</td><td>1.0E-12</td><td>BT20/MB468</td><td>CYTO</td></tr>
<tr><td>>hg38_21_+_8433172_8446622_RNA45SN1_45S_50ntFlanks@4760.4780.21|TCAGATACCGTCGTAGTTCCG</td><td>>hg38_21_+_8433172_8446622_RNA45SN1_45S_50ntFlanks</td><td>4760</td><td>4780</td><td>21</td><td>TCAGATACCGTCGTAGTTCCG</td><td>4255.0</td><td>-7.7</td><td>1.1</td><td>-7.3</td><td>2.7E-13</td><td>1.0E-12</td><td>BT20/MB468</td><td>CYTO</td></tr>
<tr><td>>MI0003188|hsa-mir-503&WithFlank&X|-|134546322|134546404@12.31.20&[MIMAT0002874&hsa-miR-503-5p&offsets|0|-3;m-297&X|-|134546371|134546393&offsets|0|-3]|TAGCAGCGGGAACAGTTCTG</td><td>>MI0003188|hsa-mir-503&WithFlank&X|-|134546322|134546404</td><td>12</td><td>31</td><td>20</td><td>TAGCAGCGGGAACAGTTCTG</td><td>211.3</td><td>9.3</td><td>1.3</td><td>7.3</td><td>2.7E-13</td><td>1.0E-12</td><td>MB231/MB468</td><td>NUC</td></tr>
<tr><td>>hg38_21_+_8433172_8446622_RNA45SN1_45S_50ntFlanks@10678.10703.26|CGGTTCCGGCGGCGTCCGGTGAGCTC</td><td>>hg38_21_+_8433172_8446622_RNA45SN1_45S_50ntFlanks</td><td>10678</td><td>10703</td><td>26</td><td>CGGTTCCGGCGGCGTCCGGTGAGCTC</td><td>137.7</td><td>-11.0</td><td>1.5</td><td>-7.3</td><td>2.3E-13</td><td>1.0E-12</td><td>MB231/MB468</td><td>MP</td></tr>
<tr><td>>trna116_GluCTC_1_-_145399233_145399304@32.75.44,trna71_GluCTC_1_-_161439189_161439260@32.75.44,trna74_GluCTC_1_-_161431809_161431880@32.75.44,trna77_GluCTC_1_-_161424398_161424469@32.75.44,trna77_GluCTC_6_+_28949976_28950047@32.75.44,trna80_GluCTC_1_-_161417018_161417089@32.75.44,trna87_GluCTC_6_-_126101393_126101464@32.75.44|CTCTCACCGCCGCGGCCCGGGTTCGATTCCCGGTCAGGGAACCA</td><td>>trna116_GluCTC_1_-_145399233_145399304</td><td>32</td><td>75</td><td>44</td><td>CTCTCACCGCCGCGGCCCGGGTTCGATTCCCGGTCAGGGAACCA</td><td>318.7</td><td>-12.1</td><td>1.7</td><td>-7.3</td><td>3.4E-13</td><td>1.0E-12</td><td>MB231/MB468</td><td>MITO</td></tr>
<tr><td>>hg38_21_+_8433172_8446622_RNA45SN1_45S_50ntFlanks@10871.10902.32|GGCTGGGTCGGTCGGGCTGGGGCGCGAAGCGG</td><td>>hg38_21_+_8433172_8446622_RNA45SN1_45S_50ntFlanks</td><td>10871</td><td>10902</td><td>32</td><td>GGCTGGGTCGGTCGGGCTGGGGCGCGAAGCGG</td><td>176.3</td><td>-11.3</td><td>1.5</td><td>-7.3</td><td>2.3E-13</td><td>1.0E-12</td><td>MB231/MB468</td><td>MP</td></tr>
<tr><td>>hg38_21_+_8433172_8446622_RNA45SN1_45S_50ntFlanks@12992.13012.21|TCCCTCGCTGCGATCTATTGA</td><td>>hg38_21_+_8433172_8446622_RNA45SN1_45S_50ntFlanks</td><td>12992</td><td>13012</td><td>21</td><td>TCCCTCGCTGCGATCTATTGA</td><td>1302.7</td><td>-5.0</td><td>0.7</td><td>-7.3</td><td>2.7E-13</td><td>1.0E-12</td><td>MB231/MB468</td><td>NUC</td></tr>
<tr><td>>hg38_21_+_8433172_8446622_RNA45SN1_45S_50ntFlanks@10841.10862.22|CGTAACTTCGGGATAAGGATTG</td><td>>hg38_21_+_8433172_8446622_RNA45SN1_45S_50ntFlanks</td><td>10841</td><td>10862</td><td>22</td><td>CGTAACTTCGGGATAAGGATTG</td><td>2318.2</td><td>-7.8</td><td>1.1</td><td>-7.3</td><td>2.7E-13</td><td>1.0E-12</td><td>MB231/MB468</td><td>NUC</td></tr>
<tr><td>>hg38_21_+_8433172_8446622_RNA45SN1_45S_50ntFlanks@9459.9486.28|CCGGGGAGGTGGAGCACGAGCGCACGTG</td><td>>hg38_21_+_8433172_8446622_RNA45SN1_45S_50ntFlanks</td><td>9459</td><td>9486</td><td>28</td><td>CCGGGGAGGTGGAGCACGAGCGCACGTG</td><td>177.2</td><td>-11.3</td><td>1.5</td><td>-7.3</td><td>2.3E-13</td><td>1.0E-12</td><td>MB231/MB468</td><td>MP</td></tr>
<tr><td>>hg38_1_-_228634819_228635039_RNA5S12_5S_50ntFlanks@51.88.38|GTCTACGGCCATACCACCCTGAACGCGCCCGATCTCGT</td><td>>hg38_1_-_228634819_228635039_RNA5S12_5S_50ntFlanks</td><td>51</td><td>88</td><td>38</td><td>GTCTACGGCCATACCACCCTGAACGCGCCCGATCTCGT</td><td>8906.0</td><td>9.5</td><td>1.3</td><td>7.3</td><td>2.7E-13</td><td>1.0E-12</td><td>MB231/MB468</td><td>CYTO</td></tr>
<tr><td>>hg38_21_+_8433172_8446622_RNA45SN1_45S_50ntFlanks@12420.12463.44|ATGTCGGCTCTTCCTATCATTGTGAAGCAGAATTCACCAAGCGT</td><td>>hg38_21_+_8433172_8446622_RNA45SN1_45S_50ntFlanks</td><td>12420</td><td>12463</td><td>44</td><td>ATGTCGGCTCTTCCTATCATTGTGAAGCAGAATTCACCAAGCGT</td><td>348.1</td><td>7.0</td><td>1.0</td><td>7.3</td><td>2.7E-13</td><td>1.0E-12</td><td>MB231/MB468</td><td>CYTO</td></tr>
<tr><td>>hg38_21_+_8433172_8446622_RNA45SN1_45S_50ntFlanks@475.504.30|CTTCGTGATCGATGTGGTGACGTCGTGCTC</td><td>>hg38_21_+_8433172_8446622_RNA45SN1_45S_50ntFlanks</td><td>475</td><td>504</td><td>30</td><td>CTTCGTGATCGATGTGGTGACGTCGTGCTC</td><td>1824.3</td><td>7.7</td><td>1.0</td><td>7.3</td><td>2.7E-13</td><td>1.0E-12</td><td>MB231/MB468</td><td>CYTO</td></tr>
<tr><td>>hg38_21_+_8433172_8446622_RNA45SN1_45S_50ntFlanks@3875.3899.25|ATACATGCCGACGGGCGCTGACCCC</td><td>>hg38_21_+_8433172_8446622_RNA45SN1_45S_50ntFlanks</td><td>3875</td><td>3899</td><td>25</td><td>ATACATGCCGACGGGCGCTGACCCC</td><td>238.9</td><td>-11.8</td><td>1.6</td><td>-7.3</td><td>2.3E-13</td><td>1.0E-12</td><td>MB231/MB468</td><td>MP</td></tr>
<tr><td>>hg38_21_+_8433172_8446622_RNA45SN1_45S_50ntFlanks@10392.10429.38|ACAGCAGTTGAACATGGGTCAGTCGGTCCTGAGAGATG</td><td>>hg38_21_+_8433172_8446622_RNA45SN1_45S_50ntFlanks</td><td>10392</td><td>10429</td><td>38</td><td>ACAGCAGTTGAACATGGGTCAGTCGGTCCTGAGAGATG</td><td>8042.3</td><td>-9.7</td><td>1.3</td><td>-7.3</td><td>3.9E-13</td><td>1.0E-12</td><td>BT20/MB468</td><td>NUC</td></tr>
<tr><td>>hg38_21_+_8433172_8446622_RNA45SN1_45S_50ntFlanks@10402.10429.28|AACATGGGTCAGTCGGTCCTGAGAGATG</td><td>>hg38_21_+_8433172_8446622_RNA45SN1_45S_50ntFlanks</td><td>10402</td><td>10429</td><td>28</td><td>AACATGGGTCAGTCGGTCCTGAGAGATG</td><td>5797.2</td><td>-8.6</td><td>1.2</td><td>-7.3</td><td>2.3E-13</td><td>1.0E-12</td><td>MB231/MB468</td><td>MP</td></tr>
<tr><td>>hg38_21_+_8433172_8446622_RNA45SN1_45S_50ntFlanks@11320.11338.19|CCACGTCTCGTCGCGCGCG</td><td>>hg38_21_+_8433172_8446622_RNA45SN1_45S_50ntFlanks</td><td>11320</td><td>11338</td><td>19</td><td>CCACGTCTCGTCGCGCGCG</td><td>153.3</td><td>-10.2</td><td>1.4</td><td>-7.3</td><td>2.8E-13</td><td>1.0E-12</td><td>MB231/MB468</td><td>NUC</td></tr>
<tr><td>>trnaMT_GluTTC_MT_-_14674_14742@41.72.32,trnalookalike8_GluTTC_5_-_93905172_93905240@41.72.32|TGGTCGTGGTTGTAGTCCGTGCGAGAATACCA</td><td>>trnaMT_GluTTC_MT_-_14674_14742</td><td>41</td><td>72</td><td>32</td><td>TGGTCGTGGTTGTAGTCCGTGCGAGAATACCA</td><td>10284.4</td><td>7.7</td><td>1.0</td><td>7.3</td><td>2.7E-13</td><td>1.0E-12</td><td>MB231/MB468</td><td>CYTO</td></tr>
<tr><td>>hg38_21_+_8433172_8446622_RNA45SN1_45S_50ntFlanks@4325.4342.18|CCGCGGTAATTCCAGCTC</td><td>>hg38_21_+_8433172_8446622_RNA45SN1_45S_50ntFlanks</td><td>4325</td><td>4342</td><td>18</td><td>CCGCGGTAATTCCAGCTC</td><td>136.1</td><td>-11.0</td><td>1.5</td><td>-7.3</td><td>2.3E-13</td><td>1.0E-12</td><td>MB231/MB468</td><td>MP</td></tr>
<tr><td>>hg38_21_+_8433172_8446622_RNA45SN1_45S_50ntFlanks@10905.10935.31|CTGGGCGCGCGCCGCGGCTGGACGAGGCGCC</td><td>>hg38_21_+_8433172_8446622_RNA45SN1_45S_50ntFlanks</td><td>10905</td><td>10935</td><td>31</td><td>CTGGGCGCGCGCCGCGGCTGGACGAGGCGCC</td><td>183.4</td><td>-10.5</td><td>1.4</td><td>-7.3</td><td>2.8E-13</td><td>1.0E-12</td><td>MB231/MB468</td><td>NUC</td></tr>
<tr><td>>hg38_21_+_8433172_8446622_RNA45SN1_45S_50ntFlanks@6648.6679.32|GTACGACTCTTAGCGGTGGATCACTCGGCTCG</td><td>>hg38_21_+_8433172_8446622_RNA45SN1_45S_50ntFlanks</td><td>6648</td><td>6679</td><td>32</td><td>GTACGACTCTTAGCGGTGGATCACTCGGCTCG</td><td>2043.3</td><td>-3.6</td><td>0.5</td><td>-7.3</td><td>3.9E-13</td><td>1.0E-12</td><td>BT20/MB468</td><td>NUC</td></tr>
<tr><td>>hg38_21_+_8433172_8446622_RNA45SN1_45S_50ntFlanks@12711.12747.37|GCGGAGCCTCGGTTGGCCTCGGATAGCCGGTCCCCCG</td><td>>hg38_21_+_8433172_8446622_RNA45SN1_45S_50ntFlanks</td><td>12711</td><td>12747</td><td>37</td><td>GCGGAGCCTCGGTTGGCCTCGGATAGCCGGTCCCCCG</td><td>215.3</td><td>-11.6</td><td>1.6</td><td>-7.3</td><td>2.4E-13</td><td>1.0E-12</td><td>MB231/MB468</td><td>MP</td></tr>
<tr><td>>hg38_21_+_8433172_8446622_RNA45SN1_45S_50ntFlanks@11580.11605.26|GAACTGGTGCGGACCAGGGGAATCCG</td><td>>hg38_21_+_8433172_8446622_RNA45SN1_45S_50ntFlanks</td><td>11580</td><td>11605</td><td>26</td><td>GAACTGGTGCGGACCAGGGGAATCCG</td><td>1373.4</td><td>-7.4</td><td>1.0</td><td>-7.3</td><td>2.8E-13</td><td>1.0E-12</td><td>MB231/MB468</td><td>NUC</td></tr>
<tr><td>>MI0000085|hsa-mir-27a&WithFlank&19|-|13836434|13836523@57.76.20(+1A)&[MIMAT0000084&hsa-miR-27a-3p&offsets|0|-1(+1A);m-43&19|-|13836447|13836467&offsets|0|-1(+1A)]|TTCACAGTGGCTAAGTTCCGA</td><td>>MI0000085|hsa-mir-27a&WithFlank&19|-|13836434|13836523</td><td>57</td><td>76</td><td>20(+1A)</td><td>TTCACAGTGGCTAAGTTCCGA</td><td>2678.2</td><td>5.3</td><td>0.7</td><td>7.3</td><td>2.8E-13</td><td>1.0E-12</td><td>MB231/MB468</td><td>NUC</td></tr>
<tr><td>>hg38_21_+_8433172_8446622_RNA45SN1_45S_50ntFlanks@8115.8137.23|CGCGGGACATGTGGCGTACGGAA</td><td>>hg38_21_+_8433172_8446622_RNA45SN1_45S_50ntFlanks</td><td>8115</td><td>8137</td><td>23</td><td>CGCGGGACATGTGGCGTACGGAA</td><td>210.1</td><td>-11.6</td><td>1.6</td><td>-7.3</td><td>2.4E-13</td><td>1.0E-12</td><td>MB231/MB468</td><td>MP</td></tr>
<tr><td>>hg38_21_+_8433172_8446622_RNA45SN1_45S_50ntFlanks@6650.6670.21|ACGACTCTTAGCGGTGGATCA</td><td>>hg38_21_+_8433172_8446622_RNA45SN1_45S_50ntFlanks</td><td>6650</td><td>6670</td><td>21</td><td>ACGACTCTTAGCGGTGGATCA</td><td>154.5</td><td>-11.1</td><td>1.5</td><td>-7.3</td><td>2.4E-13</td><td>1.0E-12</td><td>MB231/MB468</td><td>MP</td></tr>
<tr><td>>hg38_21_+_8433172_8446622_RNA45SN1_45S_50ntFlanks@4128.4145.18|CGATTCCGGAGAGGGAGC</td><td>>hg38_21_+_8433172_8446622_RNA45SN1_45S_50ntFlanks</td><td>4128</td><td>4145</td><td>18</td><td>CGATTCCGGAGAGGGAGC</td><td>164.5</td><td>-11.2</td><td>1.5</td><td>-7.3</td><td>2.4E-13</td><td>1.0E-12</td><td>MB231/MB468</td><td>MP</td></tr>
<tr><td>>hg38_21_+_8433172_8446622_RNA45SN1_45S_50ntFlanks@10542.10567.26|CGTCCAGTGCGGTAACGCGACCGATC</td><td>>hg38_21_+_8433172_8446622_RNA45SN1_45S_50ntFlanks</td><td>10542</td><td>10567</td><td>26</td><td>CGTCCAGTGCGGTAACGCGACCGATC</td><td>163.7</td><td>-11.2</td><td>1.5</td><td>-7.3</td><td>2.4E-13</td><td>1.0E-12</td><td>MB231/MB468</td><td>MP</td></tr>
<tr><td>>hg38_21_+_8433172_8446622_RNA45SN1_45S_50ntFlanks@12053.12070.18|GTGAGGCGGGGGGGCGAG</td><td>>hg38_21_+_8433172_8446622_RNA45SN1_45S_50ntFlanks</td><td>12053</td><td>12070</td><td>18</td><td>GTGAGGCGGGGGGGCGAG</td><td>214.8</td><td>-11.6</td><td>1.6</td><td>-7.3</td><td>2.4E-13</td><td>1.0E-12</td><td>MB231/MB468</td><td>MP</td></tr>
<tr><td>>hg38_21_+_8433172_8446622_RNA45SN1_45S_50ntFlanks@12934.12964.31|CTAAACCATTCGTAGACGACCTGCTTCTGGG</td><td>>hg38_21_+_8433172_8446622_RNA45SN1_45S_50ntFlanks</td><td>12934</td><td>12964</td><td>31</td><td>CTAAACCATTCGTAGACGACCTGCTTCTGGG</td><td>123.2</td><td>-10.8</td><td>1.5</td><td>-7.3</td><td>2.4E-13</td><td>1.0E-12</td><td>MB231/MB468</td><td>MP</td></tr>
<tr><td>>hg38_21_+_8433172_8446622_RNA45SN1_45S_50ntFlanks@6734.6771.38|ATTGATCATCGACACTTCGAACGCACTTGCGGCCCCGG</td><td>>hg38_21_+_8433172_8446622_RNA45SN1_45S_50ntFlanks</td><td>6734</td><td>6771</td><td>38</td><td>ATTGATCATCGACACTTCGAACGCACTTGCGGCCCCGG</td><td>2725.7</td><td>-5.2</td><td>0.7</td><td>-7.3</td><td>3.5E-13</td><td>1.0E-12</td><td>MB231/MB468</td><td>MITO</td></tr>
<tr><td>>MI0000102|hsa-mir-100&WithFlank&11|-|122152223|122152314@19.38.20(+2G)&[MIMAT0000098&hsa-miR-100-5p&offsets|0|-2(+2G);m-27&11|-|122152276|122152296&offsets|0|-1(+2G)]|AACCCGTAGATCCGAACTTGGG</td><td>>MI0000102|hsa-mir-100&WithFlank&11|-|122152223|122152314</td><td>19</td><td>38</td><td>20(+2G)</td><td>AACCCGTAGATCCGAACTTGGG</td><td>832.7</td><td>-13.1</td><td>1.7</td><td>-7.9</td><td>3.8E-15</td><td>1.0E-12</td><td>BT20/MB231</td><td>MP</td></tr>
<tr><td>>MI0000093|hsa-mir-92a-1&WithFlank&13|+|91351308|91351397@54.73.20;MI0000094|hsa-mir-92a-2&WithFlank&X|-|134169532|134169618@54.73.20&[MIMAT0000092&hsa-miR-92a-3p&offsets|0|-2;m-9&13|+|91351361|91351382&offsets|0|-2];[MIMAT0000092_1&hsa-miR-92a-3p&offsets|0|-2;m-11&X|-|134169544|134169565&offsets|0|-2]|TATTGCACTTGTCCCGGCCT</td><td>>MI0000093|hsa-mir-92a-1&WithFlank&13|+|91351308|91351397</td><td>54</td><td>73</td><td>20</td><td>TATTGCACTTGTCCCGGCCT</td><td>248.8</td><td>5.0</td><td>0.7</td><td>7.3</td><td>2.8E-13</td><td>1.1E-12</td><td>MB231/MB468</td><td>NUC</td></tr>
<tr><td>>hg38_21_+_8433172_8446622_RNA45SN1_45S_50ntFlanks@13004.13029.26|ATCTATTGAAAGTCAGCCCTCGACAC</td><td>>hg38_21_+_8433172_8446622_RNA45SN1_45S_50ntFlanks</td><td>13004</td><td>13029</td><td>26</td><td>ATCTATTGAAAGTCAGCCCTCGACAC</td><td>247.3</td><td>-10.3</td><td>1.4</td><td>-7.3</td><td>2.8E-13</td><td>1.1E-12</td><td>BT20/MB468</td><td>CYTO</td></tr>
<tr><td>>hg38_21_+_8433172_8446622_RNA45SN1_45S_50ntFlanks@4827.4847.21|CCCGCCGGGCAGCTTCCGGGA</td><td>>hg38_21_+_8433172_8446622_RNA45SN1_45S_50ntFlanks</td><td>4827</td><td>4847</td><td>21</td><td>CCCGCCGGGCAGCTTCCGGGA</td><td>382.3</td><td>7.6</td><td>1.0</td><td>7.3</td><td>2.8E-13</td><td>1.1E-12</td><td>MB231/MB468</td><td>CYTO</td></tr>
<tr><td>>hg38_21_+_8433172_8446622_RNA45SN1_45S_50ntFlanks@11920.11967.48|ATGAGAGGTGTAGAATAAGTGGGAGGCCCCCGGCGCCCCCCCGGTGTC</td><td>>hg38_21_+_8433172_8446622_RNA45SN1_45S_50ntFlanks</td><td>11920</td><td>11967</td><td>48</td><td>ATGAGAGGTGTAGAATAAGTGGGAGGCCCCCGGCGCCCCCCCGGTGTC</td><td>739.1</td><td>-12.5</td><td>1.7</td><td>-7.3</td><td>2.9E-13</td><td>1.1E-12</td><td>MB231/MB468</td><td>NUC</td></tr>
<tr><td>>hg38_21_+_8433172_8446622_RNA45SN1_45S_50ntFlanks@476.510.35|TTCGTGATCGATGTGGTGACGTCGTGCTCTCCCGG</td><td>>hg38_21_+_8433172_8446622_RNA45SN1_45S_50ntFlanks</td><td>476</td><td>510</td><td>35</td><td>TTCGTGATCGATGTGGTGACGTCGTGCTCTCCCGG</td><td>655.6</td><td>9.0</td><td>1.2</td><td>7.3</td><td>2.8E-13</td><td>1.1E-12</td><td>BT20/MB468</td><td>CYTO</td></tr>
<tr><td>>trnaMT_MetCAT_MT_+_4402_4469@37.71.35|CCGAAAATGTTGGTTATACCCTTCCCGTACTACCA</td><td>>trnaMT_MetCAT_MT_+_4402_4469</td><td>37</td><td>71</td><td>35</td><td>CCGAAAATGTTGGTTATACCCTTCCCGTACTACCA</td><td>660.5</td><td>7.9</td><td>1.1</td><td>7.3</td><td>2.9E-13</td><td>1.1E-12</td><td>MB231/MB468</td><td>NUC</td></tr>
<tr><td>>hg38_21_+_8433172_8446622_RNA45SN1_45S_50ntFlanks@12944.12973.30|CGTAGACGACCTGCTTCTGGGTCGGGGTTT</td><td>>hg38_21_+_8433172_8446622_RNA45SN1_45S_50ntFlanks</td><td>12944</td><td>12973</td><td>30</td><td>CGTAGACGACCTGCTTCTGGGTCGGGGTTT</td><td>151.1</td><td>-11.1</td><td>1.5</td><td>-7.3</td><td>2.4E-13</td><td>1.1E-12</td><td>MB231/MB468</td><td>MP</td></tr>
<tr><td>>hg38_21_+_8433172_8446622_RNA45SN1_45S_50ntFlanks@12633.12662.30|CGAAGCTACCATCTGTGGGATTATGACTGA</td><td>>hg38_21_+_8433172_8446622_RNA45SN1_45S_50ntFlanks</td><td>12633</td><td>12662</td><td>30</td><td>CGAAGCTACCATCTGTGGGATTATGACTGA</td><td>125.1</td><td>-10.8</td><td>1.5</td><td>-7.3</td><td>2.4E-13</td><td>1.1E-12</td><td>MB231/MB468</td><td>MP</td></tr>
<tr><td>>hg38_21_+_8433172_8446622_RNA45SN1_45S_50ntFlanks@11905.11924.20|GCACGGTGAAGAGACATGAG</td><td>>hg38_21_+_8433172_8446622_RNA45SN1_45S_50ntFlanks</td><td>11905</td><td>11924</td><td>20</td><td>GCACGGTGAAGAGACATGAG</td><td>133.2</td><td>-10.9</td><td>1.5</td><td>-7.3</td><td>2.4E-13</td><td>1.1E-12</td><td>MB231/MB468</td><td>MP</td></tr>
<tr><td>>hg38_21_+_8433172_8446622_RNA45SN1_45S_50ntFlanks@7986.8012.27|ATCAGACGTGGCGACCCGCTGAATTTA</td><td>>hg38_21_+_8433172_8446622_RNA45SN1_45S_50ntFlanks</td><td>7986</td><td>8012</td><td>27</td><td>ATCAGACGTGGCGACCCGCTGAATTTA</td><td>133.6</td><td>-10.0</td><td>1.4</td><td>-7.3</td><td>2.9E-13</td><td>1.1E-12</td><td>MB231/MB468</td><td>NUC</td></tr>
<tr><td>>MI0000477|hsa-mir-146a&WithFlank&5|+|160485346|160485456@27.50.24&[MIMAT0000449&hsa-miR-146a-5p&offsets|0|+2;m-129&5|+|160485372|160485393&offsets|0|+2]|TGAGAACTGAATTCCATGGGTTGT</td><td>>MI0000477|hsa-mir-146a&WithFlank&5|+|160485346|160485456</td><td>27</td><td>50</td><td>24</td><td>TGAGAACTGAATTCCATGGGTTGT</td><td>436.4</td><td>12.0</td><td>1.6</td><td>7.3</td><td>3.6E-13</td><td>1.1E-12</td><td>MB231/MB468</td><td>MITO</td></tr>
<tr><td>>hg38_21_+_8433172_8446622_RNA45SN1_45S_50ntFlanks@11778.11795.18|ACGCGCATGAATGGATGA</td><td>>hg38_21_+_8433172_8446622_RNA45SN1_45S_50ntFlanks</td><td>11778</td><td>11795</td><td>18</td><td>ACGCGCATGAATGGATGA</td><td>144.1</td><td>-11.0</td><td>1.5</td><td>-7.3</td><td>2.4E-13</td><td>1.1E-12</td><td>MB231/MB468</td><td>MP</td></tr>
<tr><td>>MI0000471|hsa-mir-126&WithFlank&9|+|136670596|136670692@58.77.20(+1A)&[MIMAT0000445&hsa-miR-126-3p&offsets|0|-2(+1A);m-40&9|+|136670653|136670674&offsets|0|-2(+1A)]|TCGTACCGTGAGTAATAATGA</td><td>>MI0000471|hsa-mir-126&WithFlank&9|+|136670596|136670692</td><td>58</td><td>77</td><td>20(+1A)</td><td>TCGTACCGTGAGTAATAATGA</td><td>303.8</td><td>11.5</td><td>1.6</td><td>7.3</td><td>2.4E-13</td><td>1.1E-12</td><td>MB231/MB468</td><td>MP</td></tr>
<tr><td>>hg38_21_+_8433172_8446622_RNA45SN1_45S_50ntFlanks@4724.4757.34|ATTAATCAAGAACGAAAGTCGGAGGTTCGAAGAC</td><td>>hg38_21_+_8433172_8446622_RNA45SN1_45S_50ntFlanks</td><td>4724</td><td>4757</td><td>34</td><td>ATTAATCAAGAACGAAAGTCGGAGGTTCGAAGAC</td><td>281.1</td><td>-12.0</td><td>1.6</td><td>-7.3</td><td>2.4E-13</td><td>1.1E-12</td><td>MB231/MB468</td><td>MP</td></tr>
<tr><td>>hg38_21_+_8433172_8446622_RNA45SN1_45S_50ntFlanks@11739.11768.30|TCTCTTAAGGTAGCCAAATGCCTCGTCATC</td><td>>hg38_21_+_8433172_8446622_RNA45SN1_45S_50ntFlanks</td><td>11739</td><td>11768</td><td>30</td><td>TCTCTTAAGGTAGCCAAATGCCTCGTCATC</td><td>147.4</td><td>-11.1</td><td>1.5</td><td>-7.3</td><td>2.4E-13</td><td>1.1E-12</td><td>MB231/MB468</td><td>MP</td></tr>
</table>

<table>
<tr><td>>MI0000063|hsa-let-7b&WithFlank&22|+|46113680|46113774@12.34.23(+1U)&[MIMAT0000063&hsa-let-7b-5p&offsets|0|+1(+1U);m-14&22|+|46113691|46113712&offsets|0|+1(+1U)]|TGAGGTAGTAGGTTGTGTGGTTTT</td><td>>MI0000063|hsa-let-7b&WithFlank&22|+|46113680|46113774</td><td>12</td><td>34</td><td>23(+1U)</td><td>TGAGGTAGTAGGTTGTGTGGTTTT</td><td>490.0</td><td>7.8</td><td>1.1</td><td>7.3</td><td>2.2E-13</td><td>1.1E-12</td><td>BT20/MB468</td><td>MITO</td></tr>
<tr><td>>hg38_21_+_8433172_8446622_RNA45SN1_45S_50ntFlanks@12073.12090.18|CCGAGGGGCTCTCGCTTC</td><td>>hg38_21_+_8433172_8446622_RNA45SN1_45S_50ntFlanks</td><td>12073</td><td>12090</td><td>18</td><td>CCGAGGGGCTCTCGCTTC</td><td>468.7</td><td>-12.0</td><td>1.5</td><td>-7.7</td><td>1.2E-14</td><td>1.1E-12</td><td>BT20/MB468</td><td>MP</td></tr>
<tr><td>>hg38_21_+_8433172_8446622_RNA45SN1_45S_50ntFlanks@11847.11872.26|AACGGGCTTGGCGGAATCAGCGGGGA</td><td>>hg38_21_+_8433172_8446622_RNA45SN1_45S_50ntFlanks</td><td>11847</td><td>11872</td><td>26</td><td>AACGGGCTTGGCGGAATCAGCGGGGA</td><td>497.6</td><td>-8.4</td><td>1.1</td><td>-7.3</td><td>2.9E-13</td><td>1.1E-12</td><td>MB231/MB468</td><td>NUC</td></tr>
<tr><td>>hg38_21_+_8433172_8446622_RNA45SN1_45S_50ntFlanks@9839.9864.26|GTAAGCAGAACTGGCGCTGCGGGATG</td><td>>hg38_21_+_8433172_8446622_RNA45SN1_45S_50ntFlanks</td><td>9839</td><td>9864</td><td>26</td><td>GTAAGCAGAACTGGCGCTGCGGGATG</td><td>322.8</td><td>-12.2</td><td>1.7</td><td>-7.3</td><td>2.4E-13</td><td>1.1E-12</td><td>MB231/MB468</td><td>MP</td></tr>
<tr><td>>hg38_21_+_8433172_8446622_RNA45SN1_45S_50ntFlanks@8170.8206.37|CAAGTCCTTCTGATCGAGGCCCAGCCCGTGGACGGTG</td><td>>hg38_21_+_8433172_8446622_RNA45SN1_45S_50ntFlanks</td><td>8170</td><td>8206</td><td>37</td><td>CAAGTCCTTCTGATCGAGGCCCAGCCCGTGGACGGTG</td><td>337.3</td><td>-12.3</td><td>1.7</td><td>-7.3</td><td>2.4E-13</td><td>1.1E-12</td><td>MB231/MB468</td><td>MP</td></tr>
<tr><td>>hg38_21_+_8433172_8446622_RNA45SN1_45S_50ntFlanks@9605.9622.18|AAGACTAATCGAACCATC</td><td>>hg38_21_+_8433172_8446622_RNA45SN1_45S_50ntFlanks</td><td>9605</td><td>9622</td><td>18</td><td>AAGACTAATCGAACCATC</td><td>694.9</td><td>-4.1</td><td>0.6</td><td>-7.3</td><td>2.8E-13</td><td>1.1E-12</td><td>MB231/MB468</td><td>CYTO</td></tr>
<tr><td>>hg38_21_+_8433172_8446622_RNA45SN1_45S_50ntFlanks@12622.12654.33|AGCCAATGGGGCGAAGCTACCATCTGTGGGATT</td><td>>hg38_21_+_8433172_8446622_RNA45SN1_45S_50ntFlanks</td><td>12622</td><td>12654</td><td>33</td><td>AGCCAATGGGGCGAAGCTACCATCTGTGGGATT</td><td>214.4</td><td>-5.5</td><td>0.7</td><td>-7.3</td><td>2.8E-13</td><td>1.1E-12</td><td>MB231/MB468</td><td>CYTO</td></tr>
<tr><td>>hg38_21_+_8433172_8446622_RNA45SN1_45S_50ntFlanks@3984.4026.43|GCGGCTTTGGTGACTCTAGATAACCTCGGGCCGATCGCACGCC</td><td>>hg38_21_+_8433172_8446622_RNA45SN1_45S_50ntFlanks</td><td>3984</td><td>4026</td><td>43</td><td>GCGGCTTTGGTGACTCTAGATAACCTCGGGCCGATCGCACGCC</td><td>1403.6</td><td>-12.5</td><td>1.7</td><td>-7.3</td><td>2.2E-13</td><td>1.1E-12</td><td>BT20/MB468</td><td>MITO</td></tr>
<tr><td>>MI0000299|hsa-mir-222&WithFlank&X|-|45747009|45747130@75.98.24(+1A)&[MIMAT0000279&hsa-miR-222-3p&offsets|0|+3(+1A);m-140&X|-|45747033|45747056&offsets|0|0(+1A)]|AGCTACATCTGGCTACTGGGTCTCA</td><td>>MI0000299|hsa-mir-222&WithFlank&X|-|45747009|45747130</td><td>75</td><td>98</td><td>24(+1A)</td><td>AGCTACATCTGGCTACTGGGTCTCA</td><td>4432.0</td><td>6.5</td><td>0.9</td><td>7.3</td><td>3.6E-13</td><td>1.1E-12</td><td>MB231/MB468</td><td>MITO</td></tr>
<tr><td>>MI0000093|hsa-mir-92a-1&WithFlank&13|+|91351308|91351397@54.74.21;MI0000094|hsa-mir-92a-2&WithFlank&X|-|134169532|134169618@54.74.21&[MIMAT0000092&hsa-miR-92a-3p&offsets|0|-1;m-9&13|+|91351361|91351382&offsets|0|-1];[MIMAT0000092_1&hsa-miR-92a-3p&offsets|0|-1;m-11&X|-|134169544|134169565&offsets|0|-1]|TATTGCACTTGTCCCGGCCTG</td><td>>MI0000093|hsa-mir-92a-1&WithFlank&13|+|91351308|91351397</td><td>54</td><td>74</td><td>21</td><td>TATTGCACTTGTCCCGGCCTG</td><td>1190.8</td><td>6.1</td><td>0.8</td><td>7.3</td><td>2.9E-13</td><td>1.1E-12</td><td>MB231/MB468</td><td>CYTO</td></tr>
<tr><td>>hg38_21_+_8433172_8446622_RNA45SN1_45S_50ntFlanks@8357.8405.49|AAAAGAACTTTGAAGAGAGAGTTCAAGAGGGCGTGAAACCGTTAAGAGG</td><td>>hg38_21_+_8433172_8446622_RNA45SN1_45S_50ntFlanks</td><td>8357</td><td>8405</td><td>49</td><td>AAAAGAACTTTGAAGAGAGAGTTCAAGAGGGCGTGAAACCGTTAAGAGG</td><td>270.2</td><td>-11.9</td><td>1.6</td><td>-7.3</td><td>3.7E-13</td><td>1.1E-12</td><td>MB231/MB468</td><td>MITO</td></tr>
<tr><td>>am_trna106_HisGTG_1_-_149155828_149155899@7.29.23,trna111_HisGTG_1_-_147774845_147774916@7.29.23,trna118_HisGTG_1_-_145396881_145396952@7.29.23,trna16_HisGTG_1_+_146544773_146544844@7.29.23,trna1_HisGTG_15_+_45493349_45493420@7.29.23,trna21_HisGTG_1_+_147753471_147753542@7.29.23,trna33_HisGTG_6_+_27125906_27125977@7.29.23,trna7_HisGTG_9_-_14433938_14434009@7.29.23,trna8_HisGTG_15_-_45492611_45492682@7.29.23,trna9_HisGTG_15_-_45490804_45490875@7.29.23|ATCGTATAGTGGTTAGTACTCTG</td><td>>am_trna106_HisGTG_1_-_149155828_149155899</td><td>7</td><td>29</td><td>23</td><td>ATCGTATAGTGGTTAGTACTCTG</td><td>522.4</td><td>-11.3</td><td>1.6</td><td>-7.3</td><td>2.9E-13</td><td>1.1E-12</td><td>BT20/MB468</td><td>CYTO</td></tr>
<tr><td>>hg38_21_+_8433172_8446622_RNA45SN1_45S_50ntFlanks@8238.8266.29|CCCGGGTCTTCCCGGAGTCGGGTTGCTTG</td><td>>hg38_21_+_8433172_8446622_RNA45SN1_45S_50ntFlanks</td><td>8238</td><td>8266</td><td>29</td><td>CCCGGGTCTTCCCGGAGTCGGGTTGCTTG</td><td>154.5</td><td>-11.1</td><td>1.5</td><td>-7.3</td><td>2.5E-13</td><td>1.1E-12</td><td>MB231/MB468</td><td>MP</td></tr>
<tr><td>>hg38_21_+_8433172_8446622_RNA45SN1_45S_50ntFlanks@8317.8356.40|CACGAGACCGATAGTCAACAAGTACCGTAAGGGAAAGTTG</td><td>>hg38_21_+_8433172_8446622_RNA45SN1_45S_50ntFlanks</td><td>8317</td><td>8356</td><td>40</td><td>CACGAGACCGATAGTCAACAAGTACCGTAAGGGAAAGTTG</td><td>518.3</td><td>-12.9</td><td>1.8</td><td>-7.3</td><td>2.5E-13</td><td>1.1E-12</td><td>MB231/MB468</td><td>MP</td></tr>
<tr><td>>hg38_21_+_8433172_8446622_RNA45SN1_45S_50ntFlanks@6731.6771.41|CACATTGATCATCGACACTTCGAACGCACTTGCGGCCCCGG</td><td>>hg38_21_+_8433172_8446622_RNA45SN1_45S_50ntFlanks</td><td>6731</td><td>6771</td><td>41</td><td>CACATTGATCATCGACACTTCGAACGCACTTGCGGCCCCGG</td><td>1586.0</td><td>-6.9</td><td>0.9</td><td>-7.3</td><td>2.9E-13</td><td>1.1E-12</td><td>MB231/MB468</td><td>NUC</td></tr>
<tr><td>>MI0000086|hsa-mir-28&WithFlank&3|+|188688775|188688872@20.40.21&[MIMAT0000085&hsa-miR-28-5p&offsets|0|-1;m-133&3|+|188688794|188688815&offsets|0|-1]|AAGGAGCTCACAGTCTATTGA</td><td>>MI0000086|hsa-mir-28&WithFlank&3|+|188688775|188688872</td><td>20</td><td>40</td><td>21</td><td>AAGGAGCTCACAGTCTATTGA</td><td>201.7</td><td>5.9</td><td>0.8</td><td>7.3</td><td>2.9E-13</td><td>1.1E-12</td><td>MB231/MB468</td><td>NUC</td></tr>
<tr><td>>hg38_21_+_8433172_8446622_RNA45SN1_45S_50ntFlanks@8738.8758.21|CCGTCCGTCCGTCCGTCCGTC</td><td>>hg38_21_+_8433172_8446622_RNA45SN1_45S_50ntFlanks</td><td>8738</td><td>8758</td><td>21</td><td>CCGTCCGTCCGTCCGTCCGTC</td><td>143.2</td><td>-9.9</td><td>1.4</td><td>-7.3</td><td>2.9E-13</td><td>1.1E-12</td><td>MB231/MB468</td><td>CYTO</td></tr>
<tr><td>>trna134_GluTTC_1_-_16861774_16861845@1.32.32,trna5_GluTTC_1_+_17199078_17199149@1.32.32,trna84_GluTTC_1_-_161391883_161391954@1.32.32,trna94_GluTTC_1_-_149664355_149664427@1.32.32|TCCCTGGTGGTCTAGTGGCTAGGATTCGGCGC</td><td>>trna134_GluTTC_1_-_16861774_16861845</td><td>1</td><td>32</td><td>32</td><td>TCCCTGGTGGTCTAGTGGCTAGGATTCGGCGC</td><td>356.7</td><td>6.6</td><td>0.9</td><td>7.3</td><td>2.9E-13</td><td>1.1E-12</td><td>BT20/MB468</td><td>CYTO</td></tr>
<tr><td>>MI0000736|hsa-mir-30c-1&WithFlank&1|+|40757278|40757378@23.42.20;MI0000254|hsa-mir-30c-2&WithFlank&6|-|71376954|71377037@13.32.20&[MIMAT0000244&hsa-miR-30c-5p&offsets|0|-3;m-57&1|+|40757300|40757323&offsets|0|-4];[MIMAT0000244_1&hsa-miR-30c-5p&offsets|0|-3;m-56&6|-|71377002|71377025&offsets|0|-4]|TGTAAACATCCTACACTCTC</td><td>>MI0000736|hsa-mir-30c-1&WithFlank&1|+|40757278|40757378</td><td>23</td><td>42</td><td>20</td><td>TGTAAACATCCTACACTCTC</td><td>1126.2</td><td>6.7</td><td>0.9</td><td>7.3</td><td>2.9E-13</td><td>1.1E-12</td><td>BT20/MB468</td><td>CYTO</td></tr>
<tr><td>>hg38_21_+_8433172_8446622_RNA45SN1_45S_50ntFlanks@10919.10946.28|CGGCTGGACGAGGCGCCGCCGCCCCCCC</td><td>>hg38_21_+_8433172_8446622_RNA45SN1_45S_50ntFlanks</td><td>10919</td><td>10946</td><td>28</td><td>CGGCTGGACGAGGCGCCGCCGCCCCCCC</td><td>158.1</td><td>-10.3</td><td>1.4</td><td>-7.3</td><td>3.0E-13</td><td>1.1E-12</td><td>MB231/MB468</td><td>NUC</td></tr>
<tr><td>>hg38_21_+_8433172_8446622_RNA45SN1_45S_50ntFlanks@12126.12151.26|CGACCCGCTCCGGGGACAGTGCCAGG</td><td>>hg38_21_+_8433172_8446622_RNA45SN1_45S_50ntFlanks</td><td>12126</td><td>12151</td><td>26</td><td>CGACCCGCTCCGGGGACAGTGCCAGG</td><td>520.5</td><td>-11.1</td><td>1.5</td><td>-7.3</td><td>2.2E-13</td><td>1.1E-12</td><td>BT20/MB468</td><td>MITO</td></tr>
<tr><td>>hg38_21_+_8433172_8446622_RNA45SN1_45S_50ntFlanks@10392.10410.19|ACAGCAGTTGAACATGGGT</td><td>>hg38_21_+_8433172_8446622_RNA45SN1_45S_50ntFlanks</td><td>10392</td><td>10410</td><td>19</td><td>ACAGCAGTTGAACATGGGT</td><td>1726.0</td><td>-9.4</td><td>1.3</td><td>-7.3</td><td>3.7E-13</td><td>1.1E-12</td><td>MB231/MB468</td><td>MITO</td></tr>
<tr><td>>hg38_21_+_8433172_8446622_RNA45SN1_45S_50ntFlanks@11452.11475.24|CTCCTCGGCGCGCGGCGGCGGCGG</td><td>>hg38_21_+_8433172_8446622_RNA45SN1_45S_50ntFlanks</td><td>11452</td><td>11475</td><td>24</td><td>CTCCTCGGCGCGCGGCGGCGGCGG</td><td>235.4</td><td>-10.8</td><td>1.5</td><td>-7.3</td><td>3.0E-13</td><td>1.1E-12</td><td>MB231/MB468</td><td>NUC</td></tr>
<tr><td>>hg38_21_+_8433172_8446622_RNA45SN1_45S_50ntFlanks@12324.12355.32|TTGGGTTTTAAGCAGGAGGTGTCAGAAAAGTT</td><td>>hg38_21_+_8433172_8446622_RNA45SN1_45S_50ntFlanks</td><td>12324</td><td>12355</td><td>32</td><td>TTGGGTTTTAAGCAGGAGGTGTCAGAAAAGTT</td><td>223.9</td><td>-10.8</td><td>1.5</td><td>-7.3</td><td>3.0E-13</td><td>1.1E-12</td><td>MB231/MB468</td><td>NUC</td></tr>
<tr><td>>trnaMT_ValTAC_MT_+_1602_1670@45.72.28|TTTCAACTTAACTTGACCGCTCTGACCA</td><td>>trnaMT_ValTAC_MT_+_1602_1670</td><td>45</td><td>72</td><td>28</td><td>TTTCAACTTAACTTGACCGCTCTGACCA</td><td>868.1</td><td>6.1</td><td>0.8</td><td>7.3</td><td>3.0E-13</td><td>1.1E-12</td><td>MB231/MB468</td><td>NUC</td></tr>
<tr><td>>hg38_21_+_8433172_8446622_RNA45SN1_45S_50ntFlanks@4724.4757.34|ATTAATCAAGAACGAAAGTCGGAGGTTCGAAGAC</td><td>>hg38_21_+_8433172_8446622_RNA45SN1_45S_50ntFlanks</td><td>4724</td><td>4757</td><td>34</td><td>ATTAATCAAGAACGAAAGTCGGAGGTTCGAAGAC</td><td>630.7</td><td>-9.3</td><td>1.3</td><td>-7.3</td><td>3.7E-13</td><td>1.1E-12</td><td>MB231/MB468</td><td>MITO</td></tr>
<tr><td>>hg38_21_+_8433172_8446622_RNA45SN1_45S_50ntFlanks@4163.4186.24|CATCCAAGGAAGGCAGCAGGCGCG</td><td>>hg38_21_+_8433172_8446622_RNA45SN1_45S_50ntFlanks</td><td>4163</td><td>4186</td><td>24</td><td>CATCCAAGGAAGGCAGCAGGCGCG</td><td>608.3</td><td>-9.0</td><td>1.2</td><td>-7.3</td><td>3.7E-13</td><td>1.1E-12</td><td>MB231/MB468</td><td>MITO</td></tr>
<tr><td>>hg38_21_+_8433172_8446622_RNA45SN1_45S_50ntFlanks@11999.12039.41|CCCTGCGGGCCGCCGGTGAAATACCACTACTCTGATCGTTT</td><td>>hg38_21_+_8433172_8446622_RNA45SN1_45S_50ntFlanks</td><td>11999</td><td>12039</td><td>41</td><td>CCCTGCGGGCCGCCGGTGAAATACCACTACTCTGATCGTTT</td><td>319.9</td><td>-12.1</td><td>1.7</td><td>-7.3</td><td>3.7E-13</td><td>1.1E-12</td><td>MB231/MB468</td><td>MITO</td></tr>
<tr><td>>trna10_LysCTT_16_+_3241501_3241573@3.33.31,trna13_LysCTT_14_-_58706613_58706685@3.33.31,trna2_LysCTT_15_+_79152904_79152976@3.33.31|CCGGCTAGCTCAGTCGGTAGAGCATGGGACT</td><td>>trna10_LysCTT_16_+_3241501_3241573</td><td>3</td><td>33</td><td>31</td><td>CCGGCTAGCTCAGTCGGTAGAGCATGGGACT</td><td>705.8</td><td>6.8</td><td>0.9</td><td>7.3</td><td>2.2E-13</td><td>1.1E-12</td><td>BT20/MB468</td><td>MITO</td></tr>
<tr><td>>hg38_21_+_8433172_8446622_RNA45SN1_45S_50ntFlanks@10805.10825.21|GAACAATGTAGGTAAGGGAAG</td><td>>hg38_21_+_8433172_8446622_RNA45SN1_45S_50ntFlanks</td><td>10805</td><td>10825</td><td>21</td><td>GAACAATGTAGGTAAGGGAAG</td><td>265.3</td><td>-11.0</td><td>1.5</td><td>-7.3</td><td>3.0E-13</td><td>1.1E-12</td><td>MB231/MB468</td><td>NUC</td></tr>
<tr><td>>hg38_21_+_8433172_8446622_RNA45SN1_45S_50ntFlanks@10116.10138.23|TGCGTCGGAGGGCGGCGGCGGCG</td><td>>hg38_21_+_8433172_8446622_RNA45SN1_45S_50ntFlanks</td><td>10116</td><td>10138</td><td>23</td><td>TGCGTCGGAGGGCGGCGGCGGCG</td><td>305.3</td><td>-12.1</td><td>1.7</td><td>-7.3</td><td>2.5E-13</td><td>1.1E-12</td><td>MB231/MB468</td><td>MP</td></tr>
<tr><td>>hg38_21_+_8433172_8446622_RNA45SN1_45S_50ntFlanks@12995.13040.46|CTCGCTGCGATCTATTGAAAGTCAGCCCTCGACACAAGGGTTTGTC</td><td>>hg38_21_+_8433172_8446622_RNA45SN1_45S_50ntFlanks</td><td>12995</td><td>13040</td><td>46</td><td>CTCGCTGCGATCTATTGAAAGTCAGCCCTCGACACAAGGGTTTGTC</td><td>530.0</td><td>9.6</td><td>1.3</td><td>7.3</td><td>2.9E-13</td><td>1.1E-12</td><td>MB231/MB468</td><td>CYTO</td></tr>
<tr><td>>hg38_21_+_8433172_8446622_RNA45SN1_45S_50ntFlanks@9930.9959.30|ATATAGACAGCAGGACGGTGGCCATGGAAG</td><td>>hg38_21_+_8433172_8446622_RNA45SN1_45S_50ntFlanks</td><td>9930</td><td>9959</td><td>30</td><td>ATATAGACAGCAGGACGGTGGCCATGGAAG</td><td>371.0</td><td>-12.4</td><td>1.7</td><td>-7.3</td><td>2.5E-13</td><td>1.1E-12</td><td>MB231/MB468</td><td>MP</td></tr>
<tr><td>>hg38_21_+_8433172_8446622_RNA45SN1_45S_50ntFlanks@11561.11593.33|GCGCCTAGCAGCCGACTTAGAACTGGTGCGGAC</td><td>>hg38_21_+_8433172_8446622_RNA45SN1_45S_50ntFlanks</td><td>11561</td><td>11593</td><td>33</td><td>GCGCCTAGCAGCCGACTTAGAACTGGTGCGGAC</td><td>872.0</td><td>-8.9</td><td>1.2</td><td>-7.3</td><td>3.8E-13</td><td>1.1E-12</td><td>MB231/MB468</td><td>MITO</td></tr>
<tr><td>>hg38_21_+_8433172_8446622_RNA45SN1_45S_50ntFlanks@9906.9928.23|AGACCCCAGAAAAGGTGTTGGTT</td><td>>hg38_21_+_8433172_8446622_RNA45SN1_45S_50ntFlanks</td><td>9906</td><td>9928</td><td>23</td><td>AGACCCCAGAAAAGGTGTTGGTT</td><td>111.5</td><td>9.6</td><td>1.3</td><td>7.2</td><td>4.2E-13</td><td>1.1E-12</td><td>BT20/MB468</td><td>NUC</td></tr>
<tr><td>>hg38_21_+_8433172_8446622_RNA45SN1_45S_50ntFlanks@6650.6695.46|ACGACTCTTAGCGGTGGATCACTCGGCTCGTGCGTCGATGAAGAAC</td><td>>hg38_21_+_8433172_8446622_RNA45SN1_45S_50ntFlanks</td><td>6650</td><td>6695</td><td>46</td><td>ACGACTCTTAGCGGTGGATCACTCGGCTCGTGCGTCGATGAAGAAC</td><td>3392.0</td><td>6.8</td><td>0.9</td><td>7.3</td><td>2.3E-13</td><td>1.1E-12</td><td>BT20/MB468</td><td>MITO</td></tr>
<tr><td>>hg38_21_+_8433172_8446622_RNA45SN1_45S_50ntFlanks@10841.10868.28|CGTAACTTCGGGATAAGGATTGGCTCTA</td><td>>hg38_21_+_8433172_8446622_RNA45SN1_45S_50ntFlanks</td><td>10841</td><td>10868</td><td>28</td><td>CGTAACTTCGGGATAAGGATTGGCTCTA</td><td>13281.4</td><td>-6.5</td><td>0.9</td><td>-7.3</td><td>3.0E-13</td><td>1.1E-12</td><td>MB231/MB468</td><td>NUC</td></tr>
<tr><td>>hg38_21_+_8433172_8446622_RNA45SN1_45S_50ntFlanks@11999.12033.35|CCCTGCGGGCCGCCGGTGAAATACCACTACTCTGA</td><td>>hg38_21_+_8433172_8446622_RNA45SN1_45S_50ntFlanks</td><td>11999</td><td>12033</td><td>35</td><td>CCCTGCGGGCCGCCGGTGAAATACCACTACTCTGA</td><td>152.4</td><td>-11.1</td><td>1.5</td><td>-7.3</td><td>2.5E-13</td><td>1.1E-12</td><td>MB231/MB468</td><td>MP</td></tr>
<tr><td>>MI0000737|hsa-mir-200a&WithFlank&1|+|1167857|1167958@23.44.22&[MIMAT0001620&hsa-miR-200a-5p&offsets|+1|+1;m-154&1|+|1167878|1167899&offsets|+1|+1]|ATCTTACCGGACAGTGCTGGAT</td><td>>MI0000737|hsa-mir-200a&WithFlank&1|+|1167857|1167958</td><td>23</td><td>44</td><td>22</td><td>ATCTTACCGGACAGTGCTGGAT</td><td>134.4</td><td>-10.9</td><td>1.5</td><td>-7.3</td><td>2.6E-13</td><td>1.1E-12</td><td>MB231/MB468</td><td>MP</td></tr>
<tr><td>>MI0000299|hsa-mir-222&WithFlank&X|-|45747009|45747130@75.98.24&[MIMAT0000279&hsa-miR-222-3p&offsets|0|+3;m-140&X|-|45747033|45747056&offsets|0|0]|AGCTACATCTGGCTACTGGGTCTC</td><td>>MI0000299|hsa-mir-222&WithFlank&X|-|45747009|45747130</td><td>75</td><td>98</td><td>24</td><td>AGCTACATCTGGCTACTGGGTCTC</td><td>18641.4</td><td>5.6</td><td>0.8</td><td>7.3</td><td>3.0E-13</td><td>1.1E-12</td><td>MB231/MB468</td><td>NUC</td></tr>
<tr><td>>MI0000476|hsa-mir-138-1&WithFlank&3|+|44114206|44114316@29.52.24&[MIMAT0000430_1&hsa-miR-138-5p&offsets|0|+1;m-497&3|+|44114234|44114256&offsets|0|+1]|AGCTGGTGTTGTGAATCAGGCCGT</td><td>>MI0000476|hsa-mir-138-1&WithFlank&3|+|44114206|44114316</td><td>29</td><td>52</td><td>24</td><td>AGCTGGTGTTGTGAATCAGGCCGT</td><td>256.9</td><td>-11.3</td><td>1.5</td><td>-7.6</td><td>3.1E-14</td><td>1.1E-12</td><td>BT20/MB231</td><td>MITO</td></tr>
<tr><td>>MI0000111|hsa-mir-105-1&WithFlank&X|-|152392213|152392305@19.41.23;MI0000112|hsa-mir-105-2&WithFlank&X|-|152394406|152394498@19.41.23&[MIMAT0000102&hsa-miR-105-5p&offsets|0|0;m-735&X|-|152392266|152392287&offsets|0|+1];[MIMAT0000102_1&hsa-miR-105-5p&offsets|0|0;m-735&X|-|152394459|152394480&offsets|0|+1]|TCAAATGCTCAGACTCCTGTGGT</td><td>>MI0000111|hsa-mir-105-1&WithFlank&X|-|152392213|152392305</td><td>19</td><td>41</td><td>23</td><td>TCAAATGCTCAGACTCCTGTGGT</td><td>194.0</td><td>11.0</td><td>1.4</td><td>7.6</td><td>2.4E-14</td><td>1.1E-12</td><td>BT20/MB231</td><td>CYTO</td></tr>
<tr><td>>hg38_21_+_8433172_8446622_RNA45SN1_45S_50ntFlanks@10401.10424.24|GAACATGGGTCAGTCGGTCCTGAG</td><td>>hg38_21_+_8433172_8446622_RNA45SN1_45S_50ntFlanks</td><td>10401</td><td>10424</td><td>24</td><td>GAACATGGGTCAGTCGGTCCTGAG</td><td>2508.6</td><td>4.7</td><td>0.6</td><td>7.3</td><td>3.0E-13</td><td>1.1E-12</td><td>MB231/MB468</td><td>CYTO</td></tr>
<tr><td>>hg38_21_+_8433172_8446622_RNA45SN1_45S_50ntFlanks@9789.9820.32|GCTGGCGTGGAGCCGGGCGTGGAATGCGAGTG</td><td>>hg38_21_+_8433172_8446622_RNA45SN1_45S_50ntFlanks</td><td>9789</td><td>9820</td><td>32</td><td>GCTGGCGTGGAGCCGGGCGTGGAATGCGAGTG</td><td>161.6</td><td>-11.2</td><td>1.5</td><td>-7.3</td><td>2.6E-13</td><td>1.1E-12</td><td>MB231/MB468</td><td>MP</td></tr>
</table>

<table>
<tr><td>>MI0000299|hsa-mir-222&WithFlank&X|-|45747009|45747130@75.97.23(+1U)&[MIMAT0000279&hsa-miR-222-3p&offsets|0|+2(+1U);m-140&X|-|45747033|45747056&offsets|0|-1(+1U)]|AGCTACATCTGGCTACTGGGTCTT</td><td>>MI0000299|hsa-mir-222&WithFlank&X|-|45747009|45747130</td><td>75</td><td>97</td><td>23(+1U)</td><td>AGCTACATCTGGCTACTGGGTCTT</td><td>4114.8</td><td>6.3</td><td>0.9</td><td>7.3</td><td>3.8E-13</td><td>1.1E-12</td><td>MB231/MB468</td><td>MITO</td></tr>
<tr><td>>hg38_21_+_8433172_8446622_RNA45SN1_45S_50ntFlanks@10350.10388.39|ACGAGAACTTTGAAGGCCGAAGTGGAGAAGGGTTCCATG</td><td>>hg38_21_+_8433172_8446622_RNA45SN1_45S_50ntFlanks</td><td>10350</td><td>10388</td><td>39</td><td>ACGAGAACTTTGAAGGCCGAAGTGGAGAAGGGTTCCATG</td><td>6691.5</td><td>-9.8</td><td>1.3</td><td>-7.2</td><td>4.3E-13</td><td>1.1E-12</td><td>BT20/MB468</td><td>NUC</td></tr>
<tr><td>>hg38_21_+_8433172_8446622_RNA45SN1_45S_50ntFlanks@4099.4118.20|GTGACCACGGGTGACGGGGA</td><td>>hg38_21_+_8433172_8446622_RNA45SN1_45S_50ntFlanks</td><td>4099</td><td>4118</td><td>20</td><td>GTGACCACGGGTGACGGGGA</td><td>159.4</td><td>-11.2</td><td>1.5</td><td>-7.3</td><td>2.6E-13</td><td>1.1E-12</td><td>MB231/MB468</td><td>MP</td></tr>
<tr><td>>hg38_21_+_8433172_8446622_RNA45SN1_45S_50ntFlanks@12405.12440.36|CTTTTTGATCCTTCGATGTCGGCTCTTCCTATCATT</td><td>>hg38_21_+_8433172_8446622_RNA45SN1_45S_50ntFlanks</td><td>12405</td><td>12440</td><td>36</td><td>CTTTTTGATCCTTCGATGTCGGCTCTTCCTATCATT</td><td>2175.2</td><td>-5.0</td><td>0.7</td><td>-7.3</td><td>3.1E-13</td><td>1.1E-12</td><td>MB231/MB468</td><td>NUC</td></tr>
<tr><td>>hg38_21_+_8433172_8446622_RNA45SN1_45S_50ntFlanks@9759.9787.29|CAAACTTTAAATGGGTAAGAAGCCCGGCT</td><td>>hg38_21_+_8433172_8446622_RNA45SN1_45S_50ntFlanks</td><td>9759</td><td>9787</td><td>29</td><td>CAAACTTTAAATGGGTAAGAAGCCCGGCT</td><td>411.2</td><td>-12.5</td><td>1.7</td><td>-7.3</td><td>2.6E-13</td><td>1.1E-12</td><td>MB231/MB468</td><td>MP</td></tr>
<tr><td>>hg38_21_+_8433172_8446622_RNA45SN1_45S_50ntFlanks@10788.10815.28|CAGCCTCTGGCATGTTGGAACAATGTAG</td><td>>hg38_21_+_8433172_8446622_RNA45SN1_45S_50ntFlanks</td><td>10788</td><td>10815</td><td>28</td><td>CAGCCTCTGGCATGTTGGAACAATGTAG</td><td>194.9</td><td>-11.5</td><td>1.6</td><td>-7.3</td><td>2.6E-13</td><td>1.1E-12</td><td>MB231/MB468</td><td>MP</td></tr>
<tr><td>>hg38_21_+_8433172_8446622_RNA45SN1_45S_50ntFlanks@6671.6699.29|CTCGGCTCGTGCGTCGATGAAGAACGCAG</td><td>>hg38_21_+_8433172_8446622_RNA45SN1_45S_50ntFlanks</td><td>6671</td><td>6699</td><td>29</td><td>CTCGGCTCGTGCGTCGATGAAGAACGCAG</td><td>4925.0</td><td>-6.2</td><td>0.8</td><td>-7.7</td><td>1.3E-14</td><td>1.1E-12</td><td>BT20/MB468</td><td>MP</td></tr>
<tr><td>>hg38_21_+_8433172_8446622_RNA45SN1_45S_50ntFlanks@8182.8206.25|ATCGAGGCCCAGCCCGTGGACGGTG</td><td>>hg38_21_+_8433172_8446622_RNA45SN1_45S_50ntFlanks</td><td>8182</td><td>8206</td><td>25</td><td>ATCGAGGCCCAGCCCGTGGACGGTG</td><td>540.8</td><td>-12.2</td><td>1.6</td><td>-7.7</td><td>1.3E-14</td><td>1.1E-12</td><td>BT20/MB468</td><td>MP</td></tr>
<tr><td>>hg38_21_+_8433172_8446622_RNA45SN1_45S_50ntFlanks@7991.8017.27|ACGTGGCGACCCGCTGAATTTAAGCAT</td><td>>hg38_21_+_8433172_8446622_RNA45SN1_45S_50ntFlanks</td><td>7991</td><td>8017</td><td>27</td><td>ACGTGGCGACCCGCTGAATTTAAGCAT</td><td>1726.8</td><td>-4.2</td><td>0.6</td><td>-7.3</td><td>3.9E-13</td><td>1.1E-12</td><td>MB231/MB468</td><td>MITO</td></tr>
<tr><td>>hg38_21_+_8433172_8446622_RNA45SN1_45S_50ntFlanks@476.507.32|TTCGTGATCGATGTGGTGACGTCGTGCTCTCC</td><td>>hg38_21_+_8433172_8446622_RNA45SN1_45S_50ntFlanks</td><td>476</td><td>507</td><td>32</td><td>TTCGTGATCGATGTGGTGACGTCGTGCTCTCC</td><td>946.9</td><td>9.0</td><td>1.2</td><td>7.3</td><td>3.0E-13</td><td>1.1E-12</td><td>MB231/MB468</td><td>CYTO</td></tr>
<tr><td>>hg38_21_+_8433172_8446622_RNA45SN1_45S_50ntFlanks@10786.10816.31|AACAGCCTCTGGCATGTTGGAACAATGTAGG</td><td>>hg38_21_+_8433172_8446622_RNA45SN1_45S_50ntFlanks</td><td>10786</td><td>10816</td><td>31</td><td>AACAGCCTCTGGCATGTTGGAACAATGTAGG</td><td>959.3</td><td>-7.3</td><td>1.0</td><td>-7.3</td><td>3.9E-13</td><td>1.1E-12</td><td>MB231/MB468</td><td>MITO</td></tr>
<tr><td>>hg38_21_+_8433172_8446622_RNA45SN1_45S_50ntFlanks@3985.4003.19|CGGCTTTGGTGACTCTAGA</td><td>>hg38_21_+_8433172_8446622_RNA45SN1_45S_50ntFlanks</td><td>3985</td><td>4003</td><td>19</td><td>CGGCTTTGGTGACTCTAGA</td><td>256.9</td><td>-11.9</td><td>1.6</td><td>-7.3</td><td>2.6E-13</td><td>1.1E-12</td><td>MB231/MB468</td><td>MP</td></tr>
<tr><td>>hg38_21_+_8433172_8446622_RNA45SN1_45S_50ntFlanks@12526.12551.26|TTTTACCCTACTGATGATGTGTTGTT</td><td>>hg38_21_+_8433172_8446622_RNA45SN1_45S_50ntFlanks</td><td>12526</td><td>12551</td><td>26</td><td>TTTTACCCTACTGATGATGTGTTGTT</td><td>123.9</td><td>-10.8</td><td>1.5</td><td>-7.3</td><td>2.6E-13</td><td>1.1E-12</td><td>MB231/MB468</td><td>MP</td></tr>
<tr><td>>MI0000266|hsa-mir-10a&WithFlank&17|-|48579832|48579953@29.49.21(+1A)&[MIMAT0000253&hsa-miR-10a-5p&offsets|+1|-1(+1A);m-18&17|-|48579905|48579926&offsets|+1|0(+1A)]|ACCCTGTAGATCCGAATTTGTA</td><td>>MI0000266|hsa-mir-10a&WithFlank&17|-|48579832|48579953</td><td>29</td><td>49</td><td>21(+1A)</td><td>ACCCTGTAGATCCGAATTTGTA</td><td>378.8</td><td>11.8</td><td>1.6</td><td>7.3</td><td>3.9E-13</td><td>1.1E-12</td><td>MB231/MB468</td><td>MITO</td></tr>
<tr><td>>hg38_21_+_8433172_8446622_RNA45SN1_45S_50ntFlanks@9021.9039.19|CGCCGGCGTCTCCTCGTGG</td><td>>hg38_21_+_8433172_8446622_RNA45SN1_45S_50ntFlanks</td><td>9021</td><td>9039</td><td>19</td><td>CGCCGGCGTCTCCTCGTGG</td><td>184.2</td><td>-10.5</td><td>1.4</td><td>-7.3</td><td>3.1E-13</td><td>1.1E-12</td><td>MB231/MB468</td><td>NUC</td></tr>
<tr><td>>hg38_21_+_8433172_8446622_RNA45SN1_45S_50ntFlanks@10391.10435.45|AACAGCAGTTGAACATGGGTCAGTCGGTCCTGAGAGATGGGCGAG</td><td>>hg38_21_+_8433172_8446622_RNA45SN1_45S_50ntFlanks</td><td>10391</td><td>10435</td><td>45</td><td>AACAGCAGTTGAACATGGGTCAGTCGGTCCTGAGAGATGGGCGAG</td><td>851.6</td><td>-12.7</td><td>1.7</td><td>-7.3</td><td>3.1E-13</td><td>1.1E-12</td><td>MB231/MB468</td><td>NUC</td></tr>
<tr><td>>hg38_21_+_8433172_8446622_RNA45SN1_45S_50ntFlanks@12324.12365.42|TTGGGTTTTAAGCAGGAGGTGTCAGAAAAGTTACCACAGGGA</td><td>>hg38_21_+_8433172_8446622_RNA45SN1_45S_50ntFlanks</td><td>12324</td><td>12365</td><td>42</td><td>TTGGGTTTTAAGCAGGAGGTGTCAGAAAAGTTACCACAGGGA</td><td>309.0</td><td>-12.1</td><td>1.7</td><td>-7.3</td><td>3.9E-13</td><td>1.1E-12</td><td>MB231/MB468</td><td>MITO</td></tr>
<tr><td>>hg38_21_+_8433172_8446622_RNA45SN1_45S_50ntFlanks@11581.11605.25|AACTGGTGCGGACCAGGGGAATCCG</td><td>>hg38_21_+_8433172_8446622_RNA45SN1_45S_50ntFlanks</td><td>11581</td><td>11605</td><td>25</td><td>AACTGGTGCGGACCAGGGGAATCCG</td><td>3042.4</td><td>-8.3</td><td>1.1</td><td>-7.3</td><td>3.9E-13</td><td>1.2E-12</td><td>MB231/MB468</td><td>MITO</td></tr>
<tr><td>>hg38_21_+_8433172_8446622_RNA45SN1_45S_50ntFlanks@8738.8758.21|CCGTCCGTCCGTCCGTCCGTC</td><td>>hg38_21_+_8433172_8446622_RNA45SN1_45S_50ntFlanks</td><td>8738</td><td>8758</td><td>21</td><td>CCGTCCGTCCGTCCGTCCGTC</td><td>427.5</td><td>-12.6</td><td>1.7</td><td>-7.3</td><td>2.6E-13</td><td>1.2E-12</td><td>MB231/MB468</td><td>MP</td></tr>
<tr><td>>hg38_21_+_8433172_8446622_RNA45SN1_45S_50ntFlanks@12351.12376.26|AAGTTACCACAGGGATAACTGGCTTG</td><td>>hg38_21_+_8433172_8446622_RNA45SN1_45S_50ntFlanks</td><td>12351</td><td>12376</td><td>26</td><td>AAGTTACCACAGGGATAACTGGCTTG</td><td>143.5</td><td>-11.0</td><td>1.5</td><td>-7.3</td><td>2.6E-13</td><td>1.2E-12</td><td>MB231/MB468</td><td>MP</td></tr>
<tr><td>>hg38_21_+_8433172_8446622_RNA45SN1_45S_50ntFlanks@12539.12563.25|ATGATGTGTTGTTGCCATGGTAATC</td><td>>hg38_21_+_8433172_8446622_RNA45SN1_45S_50ntFlanks</td><td>12539</td><td>12563</td><td>25</td><td>ATGATGTGTTGTTGCCATGGTAATC</td><td>1552.4</td><td>-10.4</td><td>1.4</td><td>-7.2</td><td>4.4E-13</td><td>1.2E-12</td><td>BT20/MB468</td><td>NUC</td></tr>
<tr><td>>hg38_21_+_8433172_8446622_RNA45SN1_45S_50ntFlanks@11873.11912.40|AAGAAGACCCTGTTGAGCTTGACTCTAGTCTGGCACGGTG</td><td>>hg38_21_+_8433172_8446622_RNA45SN1_45S_50ntFlanks</td><td>11873</td><td>11912</td><td>40</td><td>AAGAAGACCCTGTTGAGCTTGACTCTAGTCTGGCACGGTG</td><td>194.3</td><td>-10.3</td><td>1.4</td><td>-7.3</td><td>3.1E-13</td><td>1.2E-12</td><td>MB231/MB468</td><td>CYTO</td></tr>
<tr><td>>MI0003591|hsa-mir-584&WithFlank&5|-|149062307|149062415@22.42.21&[MIMAT0003249&hsa-miR-584-5p&offsets|0|-1;m-167&5|-|149062374|149062394&offsets|0|0]|TTATGGTTTGCCTGGGACTGA</td><td>>MI0003591|hsa-mir-584&WithFlank&5|-|149062307|149062415</td><td>22</td><td>42</td><td>21</td><td>TTATGGTTTGCCTGGGACTGA</td><td>617.8</td><td>7.3</td><td>1.0</td><td>7.3</td><td>3.1E-13</td><td>1.2E-12</td><td>MB231/MB468</td><td>CYTO</td></tr>
<tr><td>>hg38_21_+_8433172_8446622_RNA45SN1_45S_50ntFlanks@6773.6803.31|TTCCTCCCGGGGCTACGCCTGTCTGAGCGTC</td><td>>hg38_21_+_8433172_8446622_RNA45SN1_45S_50ntFlanks</td><td>6773</td><td>6803</td><td>31</td><td>TTCCTCCCGGGGCTACGCCTGTCTGAGCGTC</td><td>1759.9</td><td>-8.4</td><td>1.2</td><td>-7.2</td><td>4.4E-13</td><td>1.2E-12</td><td>BT20/MB468</td><td>NUC</td></tr>
<tr><td>>hg38_21_+_8433172_8446622_RNA45SN1_45S_50ntFlanks@12285.12318.34|CAGACCGTGAAAGCGGGGCCTCACGATCCTTCTG</td><td>>hg38_21_+_8433172_8446622_RNA45SN1_45S_50ntFlanks</td><td>12285</td><td>12318</td><td>34</td><td>CAGACCGTGAAAGCGGGGCCTCACGATCCTTCTG</td><td>129.6</td><td>-10.9</td><td>1.5</td><td>-7.3</td><td>2.7E-13</td><td>1.2E-12</td><td>MB231/MB468</td><td>MP</td></tr>
<tr><td>>MI0000760|hsa-mir-361&WithFlank&X|-|85903630|85903713@51.74.24&[MIMAT0004682&hsa-miR-361-3p&offsets|0|+1;m-124&X|-|85903642|85903663&offsets|0|+2]|TCCCCCAGGTGTGATTCTGATTTG</td><td>>MI0000760|hsa-mir-361&WithFlank&X|-|85903630|85903713</td><td>51</td><td>74</td><td>24</td><td>TCCCCCAGGTGTGATTCTGATTTG</td><td>856.5</td><td>7.5</td><td>1.0</td><td>7.3</td><td>3.1E-13</td><td>1.2E-12</td><td>BT20/MB468</td><td>CYTO</td></tr>
<tr><td>>hg38_21_+_8433172_8446622_RNA45SN1_45S_50ntFlanks@4783.4803.21|CATAAACGATGCCGACCGGCG</td><td>>hg38_21_+_8433172_8446622_RNA45SN1_45S_50ntFlanks</td><td>4783</td><td>4803</td><td>21</td><td>CATAAACGATGCCGACCGGCG</td><td>148.6</td><td>-11.1</td><td>1.5</td><td>-7.3</td><td>2.7E-13</td><td>1.2E-12</td><td>MB231/MB468</td><td>MP</td></tr>
<tr><td>>hg38_21_+_8433172_8446622_RNA45SN1_45S_50ntFlanks@10731.10753.23|AGGGTGTAAATCTCGCGCCGGGC</td><td>>hg38_21_+_8433172_8446622_RNA45SN1_45S_50ntFlanks</td><td>10731</td><td>10753</td><td>23</td><td>AGGGTGTAAATCTCGCGCCGGGC</td><td>118.4</td><td>-10.8</td><td>1.5</td><td>-7.3</td><td>2.7E-13</td><td>1.2E-12</td><td>MB231/MB468</td><td>MP</td></tr>
<tr><td>>hg38_21_+_8433172_8446622_RNA45SN1_45S_50ntFlanks@12459.12476.18|AGCGTTGGATTGTTCACC</td><td>>hg38_21_+_8433172_8446622_RNA45SN1_45S_50ntFlanks</td><td>12459</td><td>12476</td><td>18</td><td>AGCGTTGGATTGTTCACC</td><td>2445.5</td><td>-8.2</td><td>1.1</td><td>-7.3</td><td>2.7E-13</td><td>1.2E-12</td><td>MB231/MB468</td><td>MP</td></tr>
<tr><td>>hg38_21_+_8433172_8446622_RNA45SN1_45S_50ntFlanks@12477.12497.21|CACTAATAGGGAACGTGAGCT</td><td>>hg38_21_+_8433172_8446622_RNA45SN1_45S_50ntFlanks</td><td>12477</td><td>12497</td><td>21</td><td>CACTAATAGGGAACGTGAGCT</td><td>639.7</td><td>-3.4</td><td>0.5</td><td>-7.2</td><td>4.4E-13</td><td>1.2E-12</td><td>BT20/MB468</td><td>NUC</td></tr>
<tr><td>>trna119_LysCTT_1_-_145395522_145395594@2.33.32,trna11_LysCTT_5_-_180648979_180649051@2.33.32,trna13_LysCTT_6_+_26556774_26556846@2.33.32,trna7_LysCTT_16_+_3225692_3225764@2.33.32,trna9_LysCTT_5_+_180634755_180634827@2.33.32|CCCGGCTAGCTCAGTCGGTAGAGCATGAGACT</td><td>>trna119_LysCTT_1_-_145395522_145395594</td><td>2</td><td>33</td><td>32</td><td>CCCGGCTAGCTCAGTCGGTAGAGCATGAGACT</td><td>328.2</td><td>11.6</td><td>1.6</td><td>7.3</td><td>4.0E-13</td><td>1.2E-12</td><td>MB231/MB468</td><td>MITO</td></tr>
<tr><td>>hg38_21_+_8433172_8446622_RNA45SN1_45S_50ntFlanks@5078.5112.35|CGATAACGAACGAGACTCTGGCATGCTAACTAGTT</td><td>>hg38_21_+_8433172_8446622_RNA45SN1_45S_50ntFlanks</td><td>5078</td><td>5112</td><td>35</td><td>CGATAACGAACGAGACTCTGGCATGCTAACTAGTT</td><td>206.0</td><td>3.2</td><td>0.4</td><td>7.3</td><td>3.1E-13</td><td>1.2E-12</td><td>MB231/MB468</td><td>CYTO</td></tr>
<tr><td>>MI0000809|hsa-mir-151a&WithFlank&8|-|140732558|140732659@51.73.23&[MIMAT0000757&hsa-miR-151a-3p&offsets|-2|0;m-30&8|-|140732587|140732607&offsets|-2|0]|TACTAGACTGAAGCTCCTTGAGG</td><td>>MI0000809|hsa-mir-151a&WithFlank&8|-|140732558|140732659</td><td>51</td><td>73</td><td>23</td><td>TACTAGACTGAAGCTCCTTGAGG</td><td>2722.7</td><td>6.1</td><td>0.8</td><td>7.3</td><td>4.0E-13</td><td>1.2E-12</td><td>MB231/MB468</td><td>MITO</td></tr>
<tr><td>>hg38_21_+_8433172_8446622_RNA45SN1_45S_50ntFlanks@10351.10379.29|CGAGAACTTTGAAGGCCGAAGTGGAGAAG</td><td>>hg38_21_+_8433172_8446622_RNA45SN1_45S_50ntFlanks</td><td>10351</td><td>10379</td><td>29</td><td>CGAGAACTTTGAAGGCCGAAGTGGAGAAG</td><td>2481.3</td><td>-8.5</td><td>1.2</td><td>-7.2</td><td>4.5E-13</td><td>1.2E-12</td><td>BT20/MB468</td><td>NUC</td></tr>
<tr><td>>MI0000285|hsa-mir-205&WithFlank&1|+|209432127|209432248@40.61.22(+1A)&[MIMAT0000266&hsa-miR-205-5p&offsets|0|0(+1A);m-351&1|+|209432166|209432187&offsets|0|0(+1A)]|TCCTTCATTCCACCGGAGTCTGA</td><td>>MI0000285|hsa-mir-205&WithFlank&1|+|209432127|209432248</td><td>40</td><td>61</td><td>22(+1A)</td><td>TCCTTCATTCCACCGGAGTCTGA</td><td>79039.8</td><td>6.4</td><td>0.9</td><td>7.3</td><td>2.4E-13</td><td>1.2E-12</td><td>BT20/MB468</td><td>MITO</td></tr>
<tr><td>>hg38_21_+_8433172_8446622_RNA45SN1_45S_50ntFlanks@11583.11601.19|CTGGTGCGGACCAGGGGAA</td><td>>hg38_21_+_8433172_8446622_RNA45SN1_45S_50ntFlanks</td><td>11583</td><td>11601</td><td>19</td><td>CTGGTGCGGACCAGGGGAA</td><td>1152.3</td><td>-9.5</td><td>1.3</td><td>-7.3</td><td>2.7E-13</td><td>1.2E-12</td><td>MB231/MB468</td><td>MP</td></tr>
<tr><td>>hg38_21_+_8433172_8446622_RNA45SN1_45S_50ntFlanks@8401.8418.18|AGAGGTAAACGGGTGGGG</td><td>>hg38_21_+_8433172_8446622_RNA45SN1_45S_50ntFlanks</td><td>8401</td><td>8418</td><td>18</td><td>AGAGGTAAACGGGTGGGG</td><td>167.1</td><td>-11.2</td><td>1.5</td><td>-7.3</td><td>2.7E-13</td><td>1.2E-12</td><td>MB231/MB468</td><td>MP</td></tr>
<tr><td>>hg38_21_+_8433172_8446622_RNA45SN1_45S_50ntFlanks@10910.10932.23|CGCGCGCCGCGGCTGGACGAGGC</td><td>>hg38_21_+_8433172_8446622_RNA45SN1_45S_50ntFlanks</td><td>10910</td><td>10932</td><td>23</td><td>CGCGCGCCGCGGCTGGACGAGGC</td><td>294.4</td><td>-10.9</td><td>1.5</td><td>-7.3</td><td>3.1E-13</td><td>1.2E-12</td><td>MB231/MB468</td><td>CYTO</td></tr>
<tr><td>>hg38_21_+_8433172_8446622_RNA45SN1_45S_50ntFlanks@5515.5545.31|CTAGAGGAAGTAAAAGTCGTAACAAGGTTTC</td><td>>hg38_21_+_8433172_8446622_RNA45SN1_45S_50ntFlanks</td><td>5515</td><td>5545</td><td>31</td><td>CTAGAGGAAGTAAAAGTCGTAACAAGGTTTC</td><td>166.8</td><td>-11.2</td><td>1.5</td><td>-7.3</td><td>2.7E-13</td><td>1.2E-12</td><td>MB231/MB468</td><td>MP</td></tr>
<tr><td>>hg38_21_+_8433172_8446622_RNA45SN1_45S_50ntFlanks@9309.9338.30|GGAGTCTAACACGTGCGCGAGTCGGGGGCT</td><td>>hg38_21_+_8433172_8446622_RNA45SN1_45S_50ntFlanks</td><td>9309</td><td>9338</td><td>30</td><td>GGAGTCTAACACGTGCGCGAGTCGGGGGCT</td><td>182.1</td><td>-11.4</td><td>1.6</td><td>-7.3</td><td>2.7E-13</td><td>1.2E-12</td><td>MB231/MB468</td><td>MP</td></tr>
<tr><td>>MI0000477|hsa-mir-146a&WithFlank&5|+|160485346|160485456@27.46.20(+1A)&[MIMAT0000449&hsa-miR-146a-5p&offsets|0|-2(+1A);m-129&5|+|160485372|160485393&offsets|0|-2(+1A)]|TGAGAACTGAATTCCATGGGA</td><td>>MI0000477|hsa-mir-146a&WithFlank&5|+|160485346|160485456</td><td>27</td><td>46</td><td>20(+1A)</td><td>TGAGAACTGAATTCCATGGGA</td><td>426.3</td><td>12.0</td><td>1.6</td><td>7.3</td><td>2.7E-13</td><td>1.2E-12</td><td>MB231/MB468</td><td>MP</td></tr>
<tr><td>>hg38_21_+_8433172_8446622_RNA45SN1_45S_50ntFlanks@11661.11678.18|TGATTTCTGCCCAGTGCT</td><td>>hg38_21_+_8433172_8446622_RNA45SN1_45S_50ntFlanks</td><td>11661</td><td>11678</td><td>18</td><td>TGATTTCTGCCCAGTGCT</td><td>427.7</td><td>-8.7</td><td>1.2</td><td>-7.3</td><td>3.2E-13</td><td>1.2E-12</td><td>MB231/MB468</td><td>CYTO</td></tr>
<tr><td>>hg38_21_+_8433172_8446622_RNA45SN1_45S_50ntFlanks@12461.12487.27|CGTTGGATTGTTCACCCACTAATAGGG</td><td>>hg38_21_+_8433172_8446622_RNA45SN1_45S_50ntFlanks</td><td>12461</td><td>12487</td><td>27</td><td>CGTTGGATTGTTCACCCACTAATAGGG</td><td>157.9</td><td>-10.3</td><td>1.4</td><td>-7.3</td><td>3.2E-13</td><td>1.2E-12</td><td>MB231/MB468</td><td>NUC</td></tr>
<tr><td>>hg38_21_+_8433172_8446622_RNA45SN1_45S_50ntFlanks@10733.10755.23|GGTGTAAATCTCGCGCCGGGCCG</td><td>>hg38_21_+_8433172_8446622_RNA45SN1_45S_50ntFlanks</td><td>10733</td><td>10755</td><td>23</td><td>GGTGTAAATCTCGCGCCGGGCCG</td><td>132.1</td><td>-10.9</td><td>1.5</td><td>-7.3</td><td>2.7E-13</td><td>1.2E-12</td><td>MB231/MB468</td><td>MP</td></tr>
<tr><td>>hg38_21_+_8433172_8446622_RNA45SN1_45S_50ntFlanks@6773.6799.27|TTCCTCCCGGGGCTACGCCTGTCTGAG</td><td>>hg38_21_+_8433172_8446622_RNA45SN1_45S_50ntFlanks</td><td>6773</td><td>6799</td><td>27</td><td>TTCCTCCCGGGGCTACGCCTGTCTGAG</td><td>361.5</td><td>-7.6</td><td>1.0</td><td>-7.3</td><td>3.2E-13</td><td>1.2E-12</td><td>MB231/MB468</td><td>CYTO</td></tr>
<tr><td>>hg38_21_+_8433172_8446622_RNA45SN1_45S_50ntFlanks@11785.11806.22|TGAATGGATGAACGAGATTCCC</td><td>>hg38_21_+_8433172_8446622_RNA45SN1_45S_50ntFlanks</td><td>11785</td><td>11806</td><td>22</td><td>TGAATGGATGAACGAGATTCCC</td><td>883.2</td><td>-5.9</td><td>0.8</td><td>-7.7</td><td>1.3E-14</td><td>1.2E-12</td><td>BT20/MB468</td><td>MP</td></tr>
<tr><td>>hg38_21_+_8433172_8446622_RNA45SN1_45S_50ntFlanks@10351.10387.37|CGAGAACTTTGAAGGCCGAAGTGGAGAAGGGTTCCAT</td><td>>hg38_21_+_8433172_8446622_RNA45SN1_45S_50ntFlanks</td><td>10351</td><td>10387</td><td>37</td><td>CGAGAACTTTGAAGGCCGAAGTGGAGAAGGGTTCCAT</td><td>199.2</td><td>-11.5</td><td>1.6</td><td>-7.3</td><td>2.7E-13</td><td>1.2E-12</td><td>MB231/MB468</td><td>MP</td></tr>
<tr><td>>MI0000286|hsa-mir-210&WithFlank&11|-|568083|568204@72.93.22(+2A)&[MIMAT0000267&hsa-miR-210-3p&offsets|0|0(+2A);m-102&11|-|568112|568133&offsets|0|0(+2A)]|CTGTGCGTGTGACAGCGGCTGAAA</td><td>>MI0000286|hsa-mir-210&WithFlank&11|-|568083|568204</td><td>72</td><td>93</td><td>22(+2A)</td><td>CTGTGCGTGTGACAGCGGCTGAAA</td><td>871.3</td><td>6.3</td><td>0.9</td><td>7.2</td><td>4.6E-13</td><td>1.2E-12</td><td>BT20/MB468</td><td>NUC</td></tr>
<tr><td>>hg38_21_+_8433172_8446622_RNA45SN1_45S_50ntFlanks@12354.12376.23|TTACCACAGGGATAACTGGCTTG</td><td>>hg38_21_+_8433172_8446622_RNA45SN1_45S_50ntFlanks</td><td>12354</td><td>12376</td><td>23</td><td>TTACCACAGGGATAACTGGCTTG</td><td>107.6</td><td>-10.6</td><td>1.5</td><td>-7.3</td><td>2.8E-13</td><td>1.2E-12</td><td>MB231/MB468</td><td>MP</td></tr>
</table>

| | | | | | | | | | | | | | |
|---|---|---|---|---|---|---|---|---|---|---|---|---|---|
| >hg38_21_+_8433172_8446622_RNA45SN1_45S_50ntFlanks@3900.3938.39\|CTTCGCGGGGGGGATGCGTGCATTTATCAGATCAAAACC | >hg38_21_+_8433172_8446622_RNA45SN1_45S_50ntFlanks | 3900 | 3938 | 39 | CTTCGCGGGGGGGATGCGTGCATTTATCAGATCAAAACC | 533.5 | -12.9 | 1.8 | -7.3 | 2.8E-13 | 1.2E-12 | MB231/MB468 | MP |
| >hg38_21_+_8433172_8446622_RNA45SN1_45S_50ntFlanks@11873.11892.20\|AAGAAGACCCTGTTGAGCTT | >hg38_21_+_8433172_8446622_RNA45SN1_45S_50ntFlanks | 11873 | 11892 | 20 | AAGAAGACCCTGTTGAGCTT | 554.4 | -4.4 | 0.6 | -7.3 | 3.2E-13 | 1.2E-12 | MB231/MB468 | CYTO |
| >MI0000746\|hsa-mir-99b&WithFlank&19\|+\|51692606\|51692687@51.70.20&[MIMAT0004678&hsa-miR-99b-3p&offsets\|0\|-2;m-151&19\|+\|51692656\|51692678&offsets\|0\|-3]\|CAAGCTCGTGTCTGTGGGTC | >MI0000746\|hsa-mir-99b&WithFlank&19\|+\|51692606\|51692687 | 51 | 70 | 20 | CAAGCTCGTGTCTGTGGGTC | 1131.4 | 7.8 | 1.1 | 7.3 | 3.2E-13 | 1.2E-12 | BT20/MB468 | CYTO |
| >hg38_21_+_8433172_8446622_RNA45SN1_45S_50ntFlanks@12617.12657.41\|TGAGGAGCCAATGGGGCGAAGCTACCATCTGTGGGATTATG | >hg38_21_+_8433172_8446622_RNA45SN1_45S_50ntFlanks | 12617 | 12657 | 41 | TGAGGAGCCAATGGGGCGAAGCTACCATCTGTGGGATTATG | 273.2 | -11.9 | 1.6 | -7.3 | 4.1E-13 | 1.2E-12 | MB231/MB468 | MITO |
| >hg38_21_+_8433172_8446622_RNA45SN1_45S_50ntFlanks@10392.10430.39\|ACAGCAGTTGAACATGGGTCAGTCGGTCCTGAGAGATGG | >hg38_21_+_8433172_8446622_RNA45SN1_45S_50ntFlanks | 10392 | 10430 | 39 | ACAGCAGTTGAACATGGGTCAGTCGGTCCTGAGAGATGG | 3380.0 | -11.2 | 1.5 | -7.2 | 4.6E-13 | 1.2E-12 | BT20/MB468 | NUC |
| >MI0001445\|hsa-mir-423&WithFlank&17\|+\|30117073\|30117178@23.44.22&[MIMAT0004748&hsa-miR-423-5p&offsets\|0\|-1;m-141&17\|+\|30117095\|30117117&offsets\|0\|-1]\|TGAGGGGCAGAGAGCGAGACTT | >MI0001445\|hsa-mir-423&WithFlank&17\|+\|30117073\|30117178 | 23 | 44 | 22 | TGAGGGGCAGAGAGCGAGACTT | 2528.2 | 10.1 | 1.4 | 7.3 | 3.2E-13 | 1.2E-12 | BT20/MB468 | CYTO |
| >hg38_21_+_8433172_8446622_RNA45SN1_45S_50ntFlanks@4064.4098.35\|ATCAACTTTCGATGGTAGTCGCCGTGCCTACCATG | >hg38_21_+_8433172_8446622_RNA45SN1_45S_50ntFlanks | 4064 | 4098 | 35 | ATCAACTTTCGATGGTAGTCGCCGTGCCTACCATG | 242.9 | -10.2 | 1.4 | -7.3 | 3.2E-13 | 1.2E-12 | BT20/MB468 | CYTO |
| >am_MI0005763\|hsa-mir-941-1&WithFlank&20\|+\|63919443\|63919526@53.74.22(+1U);MI0005764\|hsa-mir-941-2&WithFlank&20\|+\|63919499\|63919582@53.74.22(+1U);MI0005765\|hsa-mir-941-3&WithFlank&20\|+\|63919555\|63919638@53.74.22(+1U);MI0005766\|hsa-mir-941-4&WithFlank&20\|+\|63919750\|63919833@53.74.22(+1U);MI0031520\|hsa-mir-941-5&WithFlank&20\|+\|63919862\|63919945@53.74.22(+1U)&[MIMAT0004984&hsa-miR-941&offsets\|0\|-1(+1U);m-263&20\|+\|63919495\|63919517&offsets\|0\|-1(+1U)];[MIMAT0004984_1&hsa-miR-941&offsets\|0\|-1(+1U);m-263&20\|+\|63919551\|63919573&offsets\|0\|-1(+1U)];[MIMAT0004984_2&hsa-miR-941&offsets\|0\|-1(+1U);m-263&20\|+\|63919607\|63919629&offsets\|0\|-1(+1U)];[MIMAT0004984_3&hsa-miR-941&offsets\|0\|-1(+1U);m-263&20\|+\|63919802\|63919824&offsets\|0\|-1(+1U)];[MIMAT0004984_4&hsa-miR-941&offsets\|0\|-1(+1U);m-263&20\|+\|63919914\|63919936&offsets\|0\|-1(+1U)]\|CACCCGGCTGTGTGCACATGTGT | >am_MI0005763\|hsa-mir-941-1&WithFlank&20\|+\|63919443\|63919526 | 53 | 74 | 22(+1U) | CACCCGGCTGTGTGCACATGTGT | 2010.5 | 6.4 | 0.9 | 7.3 | 4.1E-13 | 1.2E-12 | MB231/MB468 | MITO |
| >hg38_21_+_8433172_8446622_RNA45SN1_45S_50ntFlanks@8182.8203.22\|ATCGAGGCCCAGCCCGTGGACG | >hg38_21_+_8433172_8446622_RNA45SN1_45S_50ntFlanks | 8182 | 8203 | 22 | ATCGAGGCCCAGCCCGTGGACG | 503.5 | -12.1 | 1.6 | -7.7 | 1.4E-14 | 1.2E-12 | BT20/MB468 | MP |
| >hg38_21_+_8433172_8446622_RNA45SN1_45S_50ntFlanks@11569.11603.35\|CAGCCGACTTAGAACTGGTGCGGACCAGGGGAATC | >hg38_21_+_8433172_8446622_RNA45SN1_45S_50ntFlanks | 11569 | 11603 | 35 | CAGCCGACTTAGAACTGGTGCGGACCAGGGGAATC | 1885.2 | -4.3 | 0.6 | -7.3 | 3.2E-13 | 1.2E-12 | BT20/MB468 | CYTO |
| >MI0000079\|hsa-mir-23a&WithFlank&19\|-\|13836581\|13836665@51.72.22&[MIMAT0000078&hsa-miR-23a-3p&offsets\|0\|+1;m-31&19\|-\|13836595\|13836615&offsets\|0\|+1]\|ATCACATTGCCAGGGATTTCCA | >MI0000079\|hsa-mir-23a&WithFlank&19\|-\|13836581\|13836665 | 51 | 72 | 22 | ATCACATTGCCAGGGATTTCCA | 178.7 | -9.8 | 1.3 | -7.3 | 3.2E-13 | 1.2E-12 | BT20/MB468 | CYTO |
| >trna10_LysCTT_16_+_3241501_3241573@1.35.35,trna13_LysCTT_14_-_58706613_58706685@1.35.35,trna2_LysCTT_15_+_79152904_79152976@1.35.35\|GCCCGGCTAGCTCAGTCGGTAGAGCATGGGACTCT | >trna10_LysCTT_16_+_3241501_3241573 | 1 | 35 | 35 | GCCCGGCTAGCTCAGTCGGTAGAGCATGGGACTCT | 1287.9 | 6.8 | 0.9 | 7.3 | 3.3E-13 | 1.2E-12 | MB231/MB468 | NUC |
| >trna20_GluTTC_2_-_131094701_131094772@1.36.36,trna5_GluTTC_13_-_41634874_41634945@1.36.36\|TCCCATATGGTCTAGCGGTTAGGATTCCTGGTTTTC | >trna20_GluTTC_2_-_131094701_131094772 | 1 | 36 | 36 | TCCCATATGGTCTAGCGGTTAGGATTCCTGGTTTTC | 396.0 | 11.9 | 1.6 | 7.3 | 4.1E-13 | 1.2E-12 | MB231/MB468 | MITO |
| >hg38_21_+_8433172_8446622_RNA45SN1_45S_50ntFlanks@12615.12634.20\|GCTGAGGAGCCAATGGGGCG | >hg38_21_+_8433172_8446622_RNA45SN1_45S_50ntFlanks | 12615 | 12634 | 20 | GCTGAGGAGCCAATGGGGCG | 136.7 | -11.0 | 1.5 | -7.3 | 2.8E-13 | 1.2E-12 | MB231/MB468 | MP |
| >hg38_21_+_8433172_8446622_RNA45SN1_45S_50ntFlanks@8155.8181.27\|CGCTCGTGGGGGGCCCAAGTCCTTCTG | >hg38_21_+_8433172_8446622_RNA45SN1_45S_50ntFlanks | 8155 | 8181 | 27 | CGCTCGTGGGGGGCCCAAGTCCTTCTG | 526.1 | -4.4 | 0.6 | -7.3 | 3.3E-13 | 1.2E-12 | MB231/MB468 | NUC |
| >trna111_HisGTG_1_-_147774845_147774916@-1G.35.36,trna118_HisGTG_1_-_145396881_145396952@-1G.35.36,trna16_HisGTG_1_+_146544773_146544844@-1G.35.36,trna1_HisGTG_15_+_45493349_45493420@-1G.35.36,trna21_HisGTG_1_+_147753471_147753542@-1G.35.36,trna33_HisGTG_6_+_27125906_27125977@-1G.35.36,trna7_HisGTG_9_-_14433938_14434009@-1G.35.36,trna8_HisGTG_15_-_45492611_45492682@-1G.35.36,trna9_HisGTG_15_-_45490804_45490875@-1G.35.36\|GGCCGTGATCGTATAGTGGTTAGTACTCTGCGTTGT | >trna111_HisGTG_1_-_147774845_147774916 | -1G | 35 | 36 | GGCCGTGATCGTATAGTGGTTAGTACTCTGCGTTGT | 690.2 | 11.0 | 1.5 | 7.3 | 3.3E-13 | 1.2E-12 | BT20/MB468 | CYTO |
| >hg38_21_+_8433172_8446622_RNA45SN1_45S_50ntFlanks@11571.11601.31\|GCCGACTTAGAACTGGTGCGGACCAGGGGAA | >hg38_21_+_8433172_8446622_RNA45SN1_45S_50ntFlanks | 11571 | 11601 | 31 | GCCGACTTAGAACTGGTGCGGACCAGGGGAA | 972.6 | -5.2 | 0.7 | -7.3 | 2.8E-13 | 1.2E-12 | MB231/MB468 | MP |
| >hg38_21_+_8433172_8446622_RNA45SN1_45S_50ntFlanks@4283.4302.20\|CTTTAACGAGGATCCATTGG | >hg38_21_+_8433172_8446622_RNA45SN1_45S_50ntFlanks | 4283 | 4302 | 20 | CTTTAACGAGGATCCATTGG | 135.6 | -10.9 | 1.5 | -7.3 | 2.8E-13 | 1.2E-12 | MB231/MB468 | MP |
| >hg38_21_+_8433172_8446622_RNA45SN1_45S_50ntFlanks@10572.10596.25\|AGAAGCCGGCGGGAGCCCCGGGGAG | >hg38_21_+_8433172_8446622_RNA45SN1_45S_50ntFlanks | 10572 | 10596 | 25 | AGAAGCCGGCGGGAGCCCCGGGGAG | 147.1 | -11.1 | 1.5 | -7.3 | 2.8E-13 | 1.2E-12 | MB231/MB468 | MP |
| >hg38_21_+_8433172_8446622_RNA45SN1_45S_50ntFlanks@10387.10424.38\|TGTGAACAGCAGTTGAACATGGGTCAGTCGGTCCTGAG | >hg38_21_+_8433172_8446622_RNA45SN1_45S_50ntFlanks | 10387 | 10424 | 38 | TGTGAACAGCAGTTGAACATGGGTCAGTCGGTCCTGAG | 2048.8 | 8.1 | 1.1 | 7.3 | 3.3E-13 | 1.2E-12 | BT20/MB468 | CYTO |
| >hg38_21_+_8433172_8446622_RNA45SN1_45S_50ntFlanks@11666.11685.20\|TCTGCCCAGTGCTCTGAATG | >hg38_21_+_8433172_8446622_RNA45SN1_45S_50ntFlanks | 11666 | 11685 | 20 | TCTGCCCAGTGCTCTGAATG | 4260.9 | -6.1 | 0.8 | -7.2 | 4.2E-13 | 1.2E-12 | MB231/MB468 | MITO |
| >am_trna18_ArgCCT_17_+_73030001_73030073@14.31.18,trna21_ArgCCT_17_-_73030526_73030598@14.31.18\|AATGGATAAGGCACTGGC | >am_trna18_ArgCCT_17_+_73030001_73030073 | 14 | 31 | 18 | AATGGATAAGGCACTGGC | 169.3 | -10.4 | 1.4 | -7.3 | 3.4E-13 | 1.2E-12 | MB231/MB468 | NUC |
| >hg38_21_+_8433172_8446622_RNA45SN1_45S_50ntFlanks@6788.6806.19\|CGCCTGTCTGAGCGTCGCT | >hg38_21_+_8433172_8446622_RNA45SN1_45S_50ntFlanks | 6788 | 6806 | 19 | CGCCTGTCTGAGCGTCGCT | 9337.5 | 5.5 | 0.8 | 7.3 | 3.3E-13 | 1.2E-12 | MB231/MB468 | CYTO |
| >hg38_21_+_8433172_8446622_RNA45SN1_45S_50ntFlanks@10473.10490.18\|CTCGGCCGATCGAAAGGG | >hg38_21_+_8433172_8446622_RNA45SN1_45S_50ntFlanks | 10473 | 10490 | 18 | CTCGGCCGATCGAAAGGG | 122.9 | -10.8 | 1.5 | -7.3 | 2.8E-13 | 1.2E-12 | MB231/MB468 | MP |
| >hg38_21_+_8433172_8446622_RNA45SN1_45S_50ntFlanks@10843.10863.21\|TAACTTCGGGATAAGGATTGG | >hg38_21_+_8433172_8446622_RNA45SN1_45S_50ntFlanks | 10843 | 10863 | 21 | TAACTTCGGGATAAGGATTGG | 186.5 | -11.4 | 1.6 | -7.3 | 2.8E-13 | 1.2E-12 | MB231/MB468 | MP |
| >hg38_21_+_8433172_8446622_RNA45SN1_45S_50ntFlanks@12605.12622.18\|TATGTGCTTGGCTGAGGA | >hg38_21_+_8433172_8446622_RNA45SN1_45S_50ntFlanks | 12605 | 12622 | 18 | TATGTGCTTGGCTGAGGA | 491.3 | -6.8 | 0.9 | -7.3 | 3.4E-13 | 1.2E-12 | MB231/MB468 | NUC |
| >hg38_21_+_8433172_8446622_RNA45SN1_45S_50ntFlanks@10808.10844.37\|CAATGTAGGTAAGGGAAGTCGGCAAGCCGGATCCGTA | >hg38_21_+_8433172_8446622_RNA45SN1_45S_50ntFlanks | 10808 | 10844 | 37 | CAATGTAGGTAAGGGAAGTCGGCAAGCCGGATCCGTA | 789.3 | -9.0 | 1.2 | -7.2 | 4.2E-13 | 1.2E-12 | MB231/MB468 | MITO |
| >hg38_21_+_8433172_8446622_RNA45SN1_45S_50ntFlanks@10116.10133.18\|TGCGTCGGAGGGCGGCGG | >hg38_21_+_8433172_8446622_RNA45SN1_45S_50ntFlanks | 10116 | 10133 | 18 | TGCGTCGGAGGGCGGCGG | 126.7 | -9.7 | 1.3 | -7.3 | 3.3E-13 | 1.2E-12 | MB231/MB468 | CYTO |
| >hg38_21_+_8433172_8446622_RNA45SN1_45S_50ntFlanks@11667.11687.21\|CTGCCCAGTGCTCTGAATGTC | >hg38_21_+_8433172_8446622_RNA45SN1_45S_50ntFlanks | 11667 | 11687 | 21 | CTGCCCAGTGCTCTGAATGTC | 1156.9 | 5.6 | 0.8 | 7.3 | 3.3E-13 | 1.2E-12 | MB231/MB468 | CYTO |
| >hg38_21_+_8433172_8446622_RNA45SN1_45S_50ntFlanks@11452.11471.20\|CTCCTCGGCGCGCGGCGGCG | >hg38_21_+_8433172_8446622_RNA45SN1_45S_50ntFlanks | 11452 | 11471 | 20 | CTCCTCGGCGCGCGGCGGCG | 177.0 | -11.3 | 1.6 | -7.3 | 2.8E-13 | 1.2E-12 | MB231/MB468 | MP |
| >hg38_21_+_8433172_8446622_RNA45SN1_45S_50ntFlanks@3852.3881.30\|TAACTGTGGTAATTCTAGAGCTAATACATG | >hg38_21_+_8433172_8446622_RNA45SN1_45S_50ntFlanks | 3852 | 3881 | 30 | TAACTGTGGTAATTCTAGAGCTAATACATG | 171.0 | -11.3 | 1.5 | -7.3 | 2.8E-13 | 1.2E-12 | MB231/MB468 | MP |
| >hg38_21_+_8433172_8446622_RNA45SN1_45S_50ntFlanks@10389.10430.42\|TGAACAGCAGTTGAACATGGGTCAGTCGGTCCTGAGAGATGG | >hg38_21_+_8433172_8446622_RNA45SN1_45S_50ntFlanks | 10389 | 10430 | 42 | TGAACAGCAGTTGAACATGGGTCAGTCGGTCCTGAGAGATGG | 525.9 | -12.0 | 1.6 | -7.3 | 3.4E-13 | 1.2E-12 | MB231/MB468 | NUC |
| >hg38_21_+_8433172_8446622_RNA45SN1_45S_50ntFlanks@11766.11794.29\|ATCTAATTAGTGACGCGCATGAATGGATG | >hg38_21_+_8433172_8446622_RNA45SN1_45S_50ntFlanks | 11766 | 11794 | 29 | ATCTAATTAGTGACGCGCATGAATGGATG | 151.1 | -10.2 | 1.4 | -7.3 | 3.4E-13 | 1.2E-12 | MB231/MB468 | NUC |
| >hg38_21_+_8433172_8446622_RNA45SN1_45S_50ntFlanks@4005.4026.22\|AACCTCGGGCCGATCGCACGCC | >hg38_21_+_8433172_8446622_RNA45SN1_45S_50ntFlanks | 4005 | 4026 | 22 | AACCTCGGGCCGATCGCACGCC | 161.9 | -11.2 | 1.5 | -7.3 | 2.8E-13 | 1.2E-12 | MB231/MB468 | MP |
| >hg38_21_+_8433172_8446622_RNA45SN1_45S_50ntFlanks@4297.4324.28\|CATTGGAGGGCAAGTCTGGTGCCAGCAG | >hg38_21_+_8433172_8446622_RNA45SN1_45S_50ntFlanks | 4297 | 4324 | 28 | CATTGGAGGGCAAGTCTGGTGCCAGCAG | 115.4 | -10.7 | 1.5 | -7.3 | 2.8E-13 | 1.2E-12 | MB231/MB468 | MP |
| >hg38_21_+_8433172_8446622_RNA45SN1_45S_50ntFlanks@10572.10608.37\|AGAAGCCGGCGGGAGCCCCGGGGAGAGTTCTCTTTTC | >hg38_21_+_8433172_8446622_RNA45SN1_45S_50ntFlanks | 10572 | 10608 | 37 | AGAAGCCGGCGGGAGCCCCGGGGAGAGTTCTCTTTTC | 217.9 | -11.6 | 1.6 | -7.3 | 2.8E-13 | 1.2E-12 | MB231/MB468 | MP |
| >hg38_21_+_8433172_8446622_RNA45SN1_45S_50ntFlanks@10391.10420.30\|AACAGCAGTTGAACATGGGTCAGTCGGTCC | >hg38_21_+_8433172_8446622_RNA45SN1_45S_50ntFlanks | 10391 | 10420 | 30 | AACAGCAGTTGAACATGGGTCAGTCGGTCC | 813.0 | -8.9 | 1.2 | -7.2 | 4.2E-13 | 1.2E-12 | MB231/MB468 | MITO |
| >hg38_21_+_8433172_8446622_RNA45SN1_45S_50ntFlanks@10408.10428.21\|GGTCAGTCGGTCCTGAGAGAT | >hg38_21_+_8433172_8446622_RNA45SN1_45S_50ntFlanks | 10408 | 10428 | 21 | GGTCAGTCGGTCCTGAGAGAT | 275.0 | -12.0 | 1.6 | -7.3 | 2.9E-13 | 1.2E-12 | MB231/MB468 | MP |
| >hg38_21_+_8433172_8446622_RNA45SN1_45S_50ntFlanks@12500.12518.19\|GTTTAGACCGTCGTGAGAC | >hg38_21_+_8433172_8446622_RNA45SN1_45S_50ntFlanks | 12500 | 12518 | 19 | GTTTAGACCGTCGTGAGAC | 145.3 | -11.0 | 1.5 | -7.3 | 2.9E-13 | 1.2E-12 | MB231/MB468 | MP |
| >hg38_21_+_8433172_8446622_RNA45SN1_45S_50ntFlanks@11718.11742.25\|AACGGCGGGAGTAACTATGACTCTC | >hg38_21_+_8433172_8446622_RNA45SN1_45S_50ntFlanks | 11718 | 11742 | 25 | AACGGCGGGAGTAACTATGACTCTC | 709.9 | -6.9 | 1.0 | -7.3 | 3.4E-13 | 1.2E-12 | MB231/MB468 | NUC |
| >hg38_21_+_8433172_8446622_RNA45SN1_45S_50ntFlanks@4099.4127.29\|GTGACCACGGGTGACGGGGAATCAGGGTT | >hg38_21_+_8433172_8446622_RNA45SN1_45S_50ntFlanks | 4099 | 4127 | 29 | GTGACCACGGGTGACGGGGAATCAGGGTT | 132.6 | -10.9 | 1.5 | -7.3 | 2.9E-13 | 1.2E-12 | MB231/MB468 | MP |
| >hg38_21_+_8433172_8446622_RNA45SN1_45S_50ntFlanks@6738.6772.35\|ATCATCGACACTTCGAACGCACTTGCGGCCCCGGG | >hg38_21_+_8433172_8446622_RNA45SN1_45S_50ntFlanks | 6738 | 6772 | 35 | ATCATCGACACTTCGAACGCACTTGCGGCCCCGGG | 205.6 | -10.0 | 1.4 | -7.3 | 3.3E-13 | 1.2E-12 | BT20/MB468 | CYTO |
| >hg38_21_+_8433172_8446622_RNA45SN1_45S_50ntFlanks@10362.10388.27\|AAGGCCGAAGTGGAGAAGGGTTCCATG | >hg38_21_+_8433172_8446622_RNA45SN1_45S_50ntFlanks | 10362 | 10388 | 27 | AAGGCCGAAGTGGAGAAGGGTTCCATG | 4720.1 | -8.7 | 1.2 | -7.3 | 3.3E-13 | 1.2E-12 | MB231/MB468 | CYTO |
| >hg38_21_+_8433172_8446622_RNA45SN1_45S_50ntFlanks@12996.13013.18\|TCGCTGCGATCTATTGAA | >hg38_21_+_8433172_8446622_RNA45SN1_45S_50ntFlanks | 12996 | 13013 | 18 | TCGCTGCGATCTATTGAA | 1566.3 | -3.6 | 0.5 | -7.2 | 4.7E-13 | 1.2E-12 | BT20/MB468 | NUC |

<table>
<tr><td></td><td></td><td></td><td></td><td></td><td></td><td></td><td></td><td></td><td></td><td></td><td></td><td></td><td></td></tr>
<tr><td>>MI0000077|hsa-mir-21&WithFlank&17|+|59841260|59841343@14.34.21(+2C)&[MIMAT0000076&hsa-miR-21-5p&offsets|0|-1(+2C);m-1&17|+|59841273|59841294&offsets|0|-1(+2C)]|TAGCTTATCAGACTGATGTTGCC</td><td>>MI0000077|hsa-mir-21&WithFlank&17|+|59841260|59841343</td><td>14</td><td>34</td><td>21(+2C)</td><td>TAGCTTATCAGACTGATGTTGCC</td><td>2487.4</td><td>7.8</td><td>1.1</td><td>7.2</td><td>4.2E-13</td><td>1.2E-12</td><td>MB231/MB468</td><td>MITO</td></tr>
<tr><td>>hg38_21_+_8433172_8446622_RNA45SN1_45S_50ntFlanks@8116.8141.26|GCGGGACATGTGGCGTACGGAAGACC</td><td>>hg38_21_+_8433172_8446622_RNA45SN1_45S_50ntFlanks</td><td>8116</td><td>8141</td><td>26</td><td>GCGGGACATGTGGCGTACGGAAGACC</td><td>161.1</td><td>-11.2</td><td>1.5</td><td>-7.3</td><td>2.9E-13</td><td>1.2E-12</td><td>MB231/MB468</td><td>MP</td></tr>
<tr><td>>hg38_21_+_8433172_8446622_RNA45SN1_45S_50ntFlanks@10534.10570.37|CCGCGAGGCGTCCAGTGCGGTAACGCGACCGATCCCG</td><td>>hg38_21_+_8433172_8446622_RNA45SN1_45S_50ntFlanks</td><td>10534</td><td>10570</td><td>37</td><td>CCGCGAGGCGTCCAGTGCGGTAACGCGACCGATCCCG</td><td>163.5</td><td>-10.3</td><td>1.4</td><td>-7.3</td><td>3.4E-13</td><td>1.2E-12</td><td>MB231/MB468</td><td>NUC</td></tr>
<tr><td>>hg38_21_+_8433172_8446622_RNA45SN1_45S_50ntFlanks@12836.12856.21|CGGTGCGGAGTGCCCTTCGTC</td><td>>hg38_21_+_8433172_8446622_RNA45SN1_45S_50ntFlanks</td><td>12836</td><td>12856</td><td>21</td><td>CGGTGCGGAGTGCCCTTCGTC</td><td>543.7</td><td>-8.8</td><td>1.2</td><td>-7.3</td><td>2.9E-13</td><td>1.2E-12</td><td>MB231/MB468</td><td>MP</td></tr>
<tr><td>>trna116_GluCTC_1_-_145399233_145399304@3.36.34,trna59_GluCTC_1_+_249168447_249168518@3.36.34,trna71_GluCTC_1_-_161439189_161439260@3.36.34,trna74_GluCTC_1_-_161431809_161431880@3.36.34,trna77_GluCTC_1_-_161424398_161424469@3.36.34,trna77_GluCTC_6_+_28949976_28950047@3.36.34,trna80_GluCTC_1_-_161417018_161417089@3.36.34,trna87_GluCTC_6_-_126101393_126101464@3.36.34|CCTGGTGGTCTAGTGGTTAGGATTCGGCGCTCTC</td><td>>trna116_GluCTC_1_-_145399233_145399304</td><td>3</td><td>36</td><td>34</td><td>CCTGGTGGTCTAGTGGTTAGGATTCGGCGCTCTC</td><td>508.9</td><td>5.6</td><td>0.8</td><td>7.3</td><td>2.5E-13</td><td>1.2E-12</td><td>BT20/MB468</td><td>MITO</td></tr>
<tr><td>>hg38_21_+_8433172_8446622_RNA45SN1_45S_50ntFlanks@9623.9652.30|TAGTAGCTGGTTCCCTCCGAAGTTTCCCTC</td><td>>hg38_21_+_8433172_8446622_RNA45SN1_45S_50ntFlanks</td><td>9623</td><td>9652</td><td>30</td><td>TAGTAGCTGGTTCCCTCCGAAGTTTCCCTC</td><td>620.7</td><td>-3.9</td><td>0.5</td><td>-7.3</td><td>2.5E-13</td><td>1.2E-12</td><td>BT20/MB468</td><td>MITO</td></tr>
<tr><td>>hg38_21_+_8433172_8446622_RNA45SN1_45S_50ntFlanks@11235.11252.18|CTTCCCGTGGATCGCCCC</td><td>>hg38_21_+_8433172_8446622_RNA45SN1_45S_50ntFlanks</td><td>11235</td><td>11252</td><td>18</td><td>CTTCCCGTGGATCGCCCC</td><td>2173.9</td><td>-8.1</td><td>1.1</td><td>-7.3</td><td>3.4E-13</td><td>1.2E-12</td><td>MB231/MB468</td><td>NUC</td></tr>
<tr><td>>hg38_21_+_8433172_8446622_RNA45SN1_45S_50ntFlanks@9545.9571.27|GTGGAGGTCCGTAGCGGTCCTGACGTG</td><td>>hg38_21_+_8433172_8446622_RNA45SN1_45S_50ntFlanks</td><td>9545</td><td>9571</td><td>27</td><td>GTGGAGGTCCGTAGCGGTCCTGACGTG</td><td>32403.0</td><td>-10.7</td><td>1.5</td><td>-7.2</td><td>4.8E-13</td><td>1.2E-12</td><td>BT20/MB468</td><td>NUC</td></tr>
<tr><td>>MI0000285|hsa-mir-205&WithFlank&1|+|209432127|209432248@40.60.21&[MIMAT0000266&hsa-miR-205-5p&offsets|0|-1;m-351&1|+|209432166|209432187&offsets|0|-1]|TCCTTCATTCCACCGGAGTCT</td><td>>MI0000285|hsa-mir-205&WithFlank&1|+|209432127|209432248</td><td>40</td><td>60</td><td>21</td><td>TCCTTCATTCCACCGGAGTCT</td><td>134.8</td><td>-10.0</td><td>1.4</td><td>-7.3</td><td>3.4E-13</td><td>1.2E-12</td><td>MB231/MB468</td><td>NUC</td></tr>
<tr><td>>hg38_21_+_8433172_8446622_RNA45SN1_45S_50ntFlanks@6661.6684.24|CGGTGGATCACTCGGCTCGTGCGT</td><td>>hg38_21_+_8433172_8446622_RNA45SN1_45S_50ntFlanks</td><td>6661</td><td>6684</td><td>24</td><td>CGGTGGATCACTCGGCTCGTGCGT</td><td>1058.5</td><td>5.3</td><td>0.7</td><td>7.3</td><td>2.6E-13</td><td>1.2E-12</td><td>BT20/MB468</td><td>MITO</td></tr>
<tr><td>>hg38_21_+_8433172_8446622_RNA45SN1_45S_50ntFlanks@8437.8459.23|AGGATTCAACCCGGCGGCGGGTC</td><td>>hg38_21_+_8433172_8446622_RNA45SN1_45S_50ntFlanks</td><td>8437</td><td>8459</td><td>23</td><td>AGGATTCAACCCGGCGGCGGGTC</td><td>278.2</td><td>-12.0</td><td>1.6</td><td>-7.3</td><td>2.9E-13</td><td>1.2E-12</td><td>MB231/MB468</td><td>MP</td></tr>
<tr><td>>hg38_21_+_8433172_8446622_RNA45SN1_45S_50ntFlanks@12476.12509.34|CCACTAATAGGGAACGTGAGCTGGGTTTAGACCG</td><td>>hg38_21_+_8433172_8446622_RNA45SN1_45S_50ntFlanks</td><td>12476</td><td>12509</td><td>34</td><td>CCACTAATAGGGAACGTGAGCTGGGTTTAGACCG</td><td>224.9</td><td>-10.8</td><td>1.5</td><td>-7.3</td><td>3.4E-13</td><td>1.2E-12</td><td>MB231/MB468</td><td>NUC</td></tr>
<tr><td>>hg38_21_+_8433172_8446622_RNA45SN1_45S_50ntFlanks@8176.8197.22|CTTCTGATCGAGGCCCAGCCCG</td><td>>hg38_21_+_8433172_8446622_RNA45SN1_45S_50ntFlanks</td><td>8176</td><td>8197</td><td>22</td><td>CTTCTGATCGAGGCCCAGCCCG</td><td>1169.7</td><td>-7.2</td><td>1.0</td><td>-7.2</td><td>4.3E-13</td><td>1.2E-12</td><td>MB231/MB468</td><td>MITO</td></tr>
<tr><td>>hg38_21_+_8433172_8446622_RNA45SN1_45S_50ntFlanks@10713.10737.25|CTTGAAAATCCGGGGGAGAGGGTGT</td><td>>hg38_21_+_8433172_8446622_RNA45SN1_45S_50ntFlanks</td><td>10713</td><td>10737</td><td>25</td><td>CTTGAAAATCCGGGGGAGAGGGTGT</td><td>125.1</td><td>-10.8</td><td>1.5</td><td>-7.3</td><td>2.9E-13</td><td>1.2E-12</td><td>MB231/MB468</td><td>MP</td></tr>
<tr><td>>hg38_21_+_8433172_8446622_RNA45SN1_45S_50ntFlanks@8405.8422.18|GTAAACGGGTGGGGTCCG</td><td>>hg38_21_+_8433172_8446622_RNA45SN1_45S_50ntFlanks</td><td>8405</td><td>8422</td><td>18</td><td>GTAAACGGGTGGGGTCCG</td><td>136.8</td><td>-11.0</td><td>1.5</td><td>-7.3</td><td>2.9E-13</td><td>1.3E-12</td><td>MB231/MB468</td><td>MP</td></tr>
<tr><td>>MI0000088|hsa-mir-30a&WithFlank&6|-|71403545|71403627@54.71.18&[MIMAT0000088&hsa-miR-30a-3p&offsets|+1|-3;m-38&6|-|71403554|71403575&offsets|+1|-3]|TTTCAGTCGGATGTTTGC</td><td>>MI0000088|hsa-mir-30a&WithFlank&6|-|71403545|71403627</td><td>54</td><td>71</td><td>18</td><td>TTTCAGTCGGATGTTTGC</td><td>696.8</td><td>7.5</td><td>1.0</td><td>7.3</td><td>3.4E-13</td><td>1.3E-12</td><td>BT20/MB468</td><td>CYTO</td></tr>
<tr><td>>hg38_21_+_8433172_8446622_RNA45SN1_45S_50ntFlanks@485.523.39|GATGTGGTGACGTCGTGCTCTCCCGGGCCGGGTCCGAGC</td><td>>hg38_21_+_8433172_8446622_RNA45SN1_45S_50ntFlanks</td><td>485</td><td>523</td><td>39</td><td>GATGTGGTGACGTCGTGCTCTCCCGGGCCGGGTCCGAGC</td><td>89.2</td><td>9.3</td><td>1.3</td><td>7.2</td><td>4.8E-13</td><td>1.3E-12</td><td>BT20/MB468</td><td>NUC</td></tr>
<tr><td>>hg38_21_+_8433172_8446622_RNA45SN1_45S_50ntFlanks@12996.13035.40|TCGCTGCGATCTATTGAAAGTCAGCCCTCGACACAAGGGT</td><td>>hg38_21_+_8433172_8446622_RNA45SN1_45S_50ntFlanks</td><td>12996</td><td>13035</td><td>40</td><td>TCGCTGCGATCTATTGAAAGTCAGCCCTCGACACAAGGGT</td><td>508.0</td><td>9.7</td><td>1.3</td><td>7.3</td><td>3.4E-13</td><td>1.3E-12</td><td>MB231/MB468</td><td>CYTO</td></tr>
<tr><td>>hg38_21_+_8433172_8446622_RNA45SN1_45S_50ntFlanks@10788.10823.36|CAGCCTCTGGCATGTTGGAACAATGTAGGTAAGGGA</td><td>>hg38_21_+_8433172_8446622_RNA45SN1_45S_50ntFlanks</td><td>10788</td><td>10823</td><td>36</td><td>CAGCCTCTGGCATGTTGGAACAATGTAGGTAAGGGA</td><td>390.7</td><td>-8.6</td><td>1.2</td><td>-7.2</td><td>4.3E-13</td><td>1.3E-12</td><td>MB231/MB468</td><td>MITO</td></tr>
<tr><td>>hg38_1_-_228634819_228635039_RNA5S12_5S_50ntFlanks@148.170.23|GGAATACCGGGTGCTGTAGGCTT</td><td>>hg38_1_-_228634819_228635039_RNA5S12_5S_50ntFlanks</td><td>148</td><td>170</td><td>23</td><td>GGAATACCGGGTGCTGTAGGCTT</td><td>17830.1</td><td>-5.5</td><td>0.8</td><td>-7.3</td><td>2.9E-13</td><td>1.3E-12</td><td>MB231/MB468</td><td>MP</td></tr>
<tr><td>>hg38_21_+_8433172_8446622_RNA45SN1_45S_50ntFlanks@11641.11677.37|CGGCGGGTGTTGACGCGATGTGATTTCTGCCCAGTGC</td><td>>hg38_21_+_8433172_8446622_RNA45SN1_45S_50ntFlanks</td><td>11641</td><td>11677</td><td>37</td><td>CGGCGGGTGTTGACGCGATGTGATTTCTGCCCAGTGC</td><td>366.4</td><td>-10.6</td><td>1.5</td><td>-7.3</td><td>2.6E-13</td><td>1.3E-12</td><td>BT20/MB468</td><td>MITO</td></tr>
<tr><td>>hg38_21_+_8433172_8446622_RNA45SN1_45S_50ntFlanks@11885.11912.28|TTGAGCTTGACTCTAGTCTGGCACGGTG</td><td>>hg38_21_+_8433172_8446622_RNA45SN1_45S_50ntFlanks</td><td>11885</td><td>11912</td><td>28</td><td>TTGAGCTTGACTCTAGTCTGGCACGGTG</td><td>147.0</td><td>-9.9</td><td>1.4</td><td>-7.3</td><td>3.4E-13</td><td>1.3E-12</td><td>MB231/MB468</td><td>CYTO</td></tr>
<tr><td>>MI0003188|hsa-mir-503&WithFlank&X|-|134546322|134546404@12.31.20&[MIMAT0002874&hsa-miR-503-5p&offsets|0|-3;m-297&X|-|134546371|134546393&offsets|0|-3]|TAGCAGCGGGAACAGTTCTG</td><td>>MI0003188|hsa-mir-503&WithFlank&X|-|134546322|134546404</td><td>12</td><td>31</td><td>20</td><td>TAGCAGCGGGAACAGTTCTG</td><td>907.3</td><td>13.6</td><td>1.8</td><td>7.7</td><td>1.4E-14</td><td>1.3E-12</td><td>BT20/MB468</td><td>MP</td></tr>
<tr><td>>hg38_21_+_8433172_8446622_RNA45SN1_45S_50ntFlanks@4046.4066.21|CATTCGAACGTCTGCCCTATC</td><td>>hg38_21_+_8433172_8446622_RNA45SN1_45S_50ntFlanks</td><td>4046</td><td>4066</td><td>21</td><td>CATTCGAACGTCTGCCCTATC</td><td>4220.6</td><td>-6.8</td><td>0.9</td><td>-7.2</td><td>4.3E-13</td><td>1.3E-12</td><td>MB231/MB468</td><td>MITO</td></tr>
<tr><td>>hg38_21_+_8433172_8446622_RNA45SN1_45S_50ntFlanks@9308.9335.28|AGGAGTCTAACACGTGCGCGAGTCGGGG</td><td>>hg38_21_+_8433172_8446622_RNA45SN1_45S_50ntFlanks</td><td>9308</td><td>9335</td><td>28</td><td>AGGAGTCTAACACGTGCGCGAGTCGGGG</td><td>1178.5</td><td>-9.7</td><td>1.3</td><td>-7.3</td><td>2.9E-13</td><td>1.3E-12</td><td>MB231/MB468</td><td>MP</td></tr>
<tr><td>>hg38_21_+_8433172_8446622_RNA45SN1_45S_50ntFlanks@12627.12653.27|ATGGGGCGAAGCTACCATCTGTGGGAT</td><td>>hg38_21_+_8433172_8446622_RNA45SN1_45S_50ntFlanks</td><td>12627</td><td>12653</td><td>27</td><td>ATGGGGCGAAGCTACCATCTGTGGGAT</td><td>4776.1</td><td>-9.4</td><td>1.3</td><td>-7.3</td><td>2.6E-13</td><td>1.3E-12</td><td>BT20/MB468</td><td>MITO</td></tr>
<tr><td>>hg38_21_+_8433172_8446622_RNA45SN1_45S_50ntFlanks@3875.3899.25|ATACATGCCGACGGGCGCTGACCCC</td><td>>hg38_21_+_8433172_8446622_RNA45SN1_45S_50ntFlanks</td><td>3875</td><td>3899</td><td>25</td><td>ATACATGCCGACGGGCGCTGACCCC</td><td>1612.3</td><td>-9.3</td><td>1.3</td><td>-7.2</td><td>4.3E-13</td><td>1.3E-12</td><td>MB231/MB468</td><td>MITO</td></tr>
<tr><td>>hg38_21_+_8433172_8446622_RNA45SN1_45S_50ntFlanks@12385.12414.30|AAGCGTTCATAGCGACGTCGCTTTTTGATC</td><td>>hg38_21_+_8433172_8446622_RNA45SN1_45S_50ntFlanks</td><td>12385</td><td>12414</td><td>30</td><td>AAGCGTTCATAGCGACGTCGCTTTTTGATC</td><td>130.5</td><td>-10.9</td><td>1.5</td><td>-7.3</td><td>2.9E-13</td><td>1.3E-12</td><td>MB231/MB468</td><td>MP</td></tr>
<tr><td>>hg38_21_+_8433172_8446622_RNA45SN1_45S_50ntFlanks@12539.12558.20|ATGATGTGTTGTTGCCATGG</td><td>>hg38_21_+_8433172_8446622_RNA45SN1_45S_50ntFlanks</td><td>12539</td><td>12558</td><td>20</td><td>ATGATGTGTTGTTGCCATGG</td><td>166.0</td><td>-11.2</td><td>1.5</td><td>-7.3</td><td>3.0E-13</td><td>1.3E-12</td><td>MB231/MB468</td><td>MP</td></tr>
<tr><td>>hg38_21_+_8433172_8446622_RNA45SN1_45S_50ntFlanks@8110.8136.27|CGGGGCGCGGGACATGTGGCGTACGGA</td><td>>hg38_21_+_8433172_8446622_RNA45SN1_45S_50ntFlanks</td><td>8110</td><td>8136</td><td>27</td><td>CGGGGCGCGGGACATGTGGCGTACGGA</td><td>157.7</td><td>-11.2</td><td>1.5</td><td>-7.3</td><td>3.0E-13</td><td>1.3E-12</td><td>MB231/MB468</td><td>MP</td></tr>
<tr><td>>hg38_21_+_8433172_8446622_RNA45SN1_45S_50ntFlanks@9597.9617.21|AGGGGCGAAAGACTAATCGAA</td><td>>hg38_21_+_8433172_8446622_RNA45SN1_45S_50ntFlanks</td><td>9597</td><td>9617</td><td>21</td><td>AGGGGCGAAAGACTAATCGAA</td><td>196.5</td><td>-11.5</td><td>1.6</td><td>-7.3</td><td>3.0E-13</td><td>1.3E-12</td><td>MB231/MB468</td><td>MP</td></tr>
<tr><td>>hg38_21_+_8433172_8446622_RNA45SN1_45S_50ntFlanks@8154.8178.25|CCGCTCGTGGGGGGCCCAAGTCCTT</td><td>>hg38_21_+_8433172_8446622_RNA45SN1_45S_50ntFlanks</td><td>8154</td><td>8178</td><td>25</td><td>CCGCTCGTGGGGGGCCCAAGTCCTT</td><td>350.7</td><td>-10.5</td><td>1.4</td><td>-7.3</td><td>2.6E-13</td><td>1.3E-12</td><td>BT20/MB468</td><td>MITO</td></tr>
<tr><td>>MI0000746|hsa-mir-99b&WithFlank&19|+|51692606|51692687@13.32.20(+2U)&[MIMAT0000689&hsa-miR-99b-5p&offsets|0|-2(+2U);m-10&19|+|51692618|51692639&offsets|0|-2(+2U)]|CACCCGTAGAACCGACCTTGTT</td><td>>MI0000746|hsa-mir-99b&WithFlank&19|+|51692606|51692687</td><td>13</td><td>32</td><td>20(+2U)</td><td>CACCCGTAGAACCGACCTTGTT</td><td>358.5</td><td>12.2</td><td>1.6</td><td>7.7</td><td>1.4E-14</td><td>1.3E-12</td><td>BT20/MB468</td><td>MP</td></tr>
<tr><td>>hg38_21_+_8433172_8446622_RNA45SN1_45S_50ntFlanks@12233.12257.25|CCGTGGAGCAGAAGGGCAAAAGCTC</td><td>>hg38_21_+_8433172_8446622_RNA45SN1_45S_50ntFlanks</td><td>12233</td><td>12257</td><td>25</td><td>CCGTGGAGCAGAAGGGCAAAAGCTC</td><td>154.4</td><td>-11.1</td><td>1.5</td><td>-7.3</td><td>3.0E-13</td><td>1.3E-12</td><td>MB231/MB468</td><td>MP</td></tr>
<tr><td>>hg38_21_+_8433172_8446622_RNA45SN1_45S_50ntFlanks@12997.13016.20|CGCTGCGATCTATTGAAAGT</td><td>>hg38_21_+_8433172_8446622_RNA45SN1_45S_50ntFlanks</td><td>12997</td><td>13016</td><td>20</td><td>CGCTGCGATCTATTGAAAGT</td><td>503.5</td><td>5.8</td><td>0.8</td><td>7.3</td><td>3.5E-13</td><td>1.3E-12</td><td>MB231/MB468</td><td>NUC</td></tr>
<tr><td>>hg38_21_+_8433172_8446622_RNA45SN1_45S_50ntFlanks@11668.11692.25|TGCCCAGTGCTCTGAATGTCAAAGT</td><td>>hg38_21_+_8433172_8446622_RNA45SN1_45S_50ntFlanks</td><td>11668</td><td>11692</td><td>25</td><td>TGCCCAGTGCTCTGAATGTCAAAGT</td><td>426.3</td><td>6.3</td><td>0.9</td><td>7.3</td><td>3.4E-13</td><td>1.3E-12</td><td>MB231/MB468</td><td>CYTO</td></tr>
<tr><td>>hg38_21_+_8433172_8446622_RNA45SN1_45S_50ntFlanks@11847.11868.22|AACGGGCTTGGCGGAATCAGCG</td><td>>hg38_21_+_8433172_8446622_RNA45SN1_45S_50ntFlanks</td><td>11847</td><td>11868</td><td>22</td><td>AACGGGCTTGGCGGAATCAGCG</td><td>187.2</td><td>-11.4</td><td>1.6</td><td>-7.3</td><td>3.0E-13</td><td>1.3E-12</td><td>MB231/MB468</td><td>MP</td></tr>
<tr><td>>hg38_21_+_8433172_8446622_RNA45SN1_45S_50ntFlanks@12628.12657.30|TGGGGCGAAGCTACCATCTGTGGGATTATG</td><td>>hg38_21_+_8433172_8446622_RNA45SN1_45S_50ntFlanks</td><td>12628</td><td>12657</td><td>30</td><td>TGGGGCGAAGCTACCATCTGTGGGATTATG</td><td>538.1</td><td>-11.2</td><td>1.5</td><td>-7.3</td><td>2.6E-13</td><td>1.3E-12</td><td>BT20/MB468</td><td>MITO</td></tr>
<tr><td>>MI0000456|hsa-mir-140&WithFlank&16|+|69933075|69933186@29.50.22&[MIMAT0000431&hsa-miR-140-5p&offsets|0|0;m-173&16|+|69933103|69933124&offsets|0|0]|CAGTGGTTTTACCCTATGGTAG</td><td>>MI0000456|hsa-mir-140&WithFlank&16|+|69933075|69933186</td><td>29</td><td>50</td><td>22</td><td>CAGTGGTTTTACCCTATGGTAG</td><td>301.3</td><td>9.5</td><td>1.3</td><td>7.3</td><td>3.5E-13</td><td>1.3E-12</td><td>MB231/MB468</td><td>NUC</td></tr>
<tr><td>>hg38_21_+_8433172_8446622_RNA45SN1_45S_50ntFlanks@12274.12293.20|CAGTACGAATACAGACCGTG</td><td>>hg38_21_+_8433172_8446622_RNA45SN1_45S_50ntFlanks</td><td>12274</td><td>12293</td><td>20</td><td>CAGTACGAATACAGACCGTG</td><td>785.5</td><td>-12.7</td><td>1.7</td><td>-7.7</td><td>1.5E-14</td><td>1.3E-12</td><td>BT20/MB468</td><td>MP</td></tr>
<tr><td>>MI0000087|hsa-mir-29a&WithFlank&7|-|130876741|130876816@48.68.21(+1C)&[MIMAT0000086&hsa-miR-29a-3p&offsets|0|-1(+1C);m-26&7|-|130876749|130876769&offsets|0|0(+1C)]|TAGCACCATCTGAAATCGGTTC</td><td>>MI0000087|hsa-mir-29a&WithFlank&7|-|130876741|130876816</td><td>48</td><td>68</td><td>21(+1C)</td><td>TAGCACCATCTGAAATCGGTTC</td><td>277.7</td><td>11.3</td><td>1.6</td><td>7.3</td><td>3.0E-13</td><td>1.3E-12</td><td>MB231/MB468</td><td>MP</td></tr>
<tr><td>>hg38_21_+_8433172_8446622_RNA45SN1_45S_50ntFlanks@6690.6718.29|AAGAACGCAGCTAGCTGCGAGAATTAATG</td><td>>hg38_21_+_8433172_8446622_RNA45SN1_45S_50ntFlanks</td><td>6690</td><td>6718</td><td>29</td><td>AAGAACGCAGCTAGCTGCGAGAATTAATG</td><td>175.7</td><td>-10.2</td><td>1.4</td><td>-7.3</td><td>3.5E-13</td><td>1.3E-12</td><td>MB231/MB468</td><td>CYTO</td></tr>
<tr><td>>am_trna128_GlyGCC_6_-_27870686_27870756@1.31.31,trna133_GlyCCC_1_-_16872434_16872504@1.31.31,trna18_GlyGCC_16_+_70822597_70822667@1.31.31,trna19_GlyGCC_16_+_70823410_70823480@1.31.31,trna19_GlyGCC_2_-_157257659_157257729@1.31.31,trna24_GlyGCC_16_-_70812942_70813012@1.31.31,trna25_GlyGCC_16_-_70812114_70812184@1.31.31,trna4_GlyCCC_1_+_17188416_17188486@1.31.31,trna5_GlyGCC_17_+_8029064_8029134@1.31.31,trna68_GlyGCC_1_-_161493637_161493707@1.31.31|GCATTGGTGGTTCAGTGGTAGAATTCTCGCC</td><td>>am_trna128_GlyGCC_6_-_27870686_27870756</td><td>1</td><td>31</td><td>31</td><td>GCATTGGTGGTTCAGTGGTAGAATTCTCGCC</td><td>27215.9</td><td>7.4</td><td>1.0</td><td>7.3</td><td>2.6E-13</td><td>1.3E-12</td><td>BT20/MB468</td><td>MITO</td></tr>
<tr><td>>hg38_21_+_8433172_8446622_RNA45SN1_45S_50ntFlanks@10461.10480.20|GCCTCCGTTGCCCTCGGCCG</td><td>>hg38_21_+_8433172_8446622_RNA45SN1_45S_50ntFlanks</td><td>10461</td><td>10480</td><td>20</td><td>GCCTCCGTTGCCCTCGGCCG</td><td>177.5</td><td>-10.4</td><td>1.4</td><td>-7.3</td><td>3.6E-13</td><td>1.3E-12</td><td>MB231/MB468</td><td>NUC</td></tr>
<tr><td>>hg38_21_+_8433172_8446622_RNA45SN1_45S_50ntFlanks@9120.9152.33|TGCCGCGCGCGGGTCGGGGGGCGGGGCGGACTG</td><td>>hg38_21_+_8433172_8446622_RNA45SN1_45S_50ntFlanks</td><td>9120</td><td>9152</td><td>33</td><td>TGCCGCGCGCGGGTCGGGGGGCGGGGCGGACTG</td><td>325.2</td><td>-12.2</td><td>1.7</td><td>-7.3</td><td>3.0E-13</td><td>1.3E-12</td><td>MB231/MB468</td><td>MP</td></tr>
</table>

| | | | | | | | | | | | | | |
|---|---|---|---|---|---|---|---|---|---|---|---|---|---|
| >hg38_21_+_8433172_8446622_RNA45SN1_45S_50ntFlanks@11211.11232.22\|GCGACTCTGGACGCGAGCCGGG | >hg38_21_+_8433172_8446622_RNA45SN1_45S_50ntFlanks | 11211 | 11232 | 22 | GCGACTCTGGACGCGAGCCGGG | 1142.4 | -4.7 | 0.6 | -7.3 | 3.5E-13 | 1.3E-12 | BT20/MB468 | CYTO |
| >hg38_21_+_8433172_8446622_RNA45SN1_45S_50ntFlanks@10392.10419.28\|ACAGCAGTTGAACATGGGTCAGTCGGTC | >hg38_21_+_8433172_8446622_RNA45SN1_45S_50ntFlanks | 10392 | 10419 | 28 | ACAGCAGTTGAACATGGGTCAGTCGGTC | 2701.6 | -10.9 | 1.5 | -7.2 | 4.9E-13 | 1.3E-12 | BT20/MB468 | NUC |
| >hg38_21_+_8433172_8446622_RNA45SN1_45S_50ntFlanks@10101.10126.26\|CGCGCGCGTGTGGTGTGCGTCGGAGG | >hg38_21_+_8433172_8446622_RNA45SN1_45S_50ntFlanks | 10101 | 10126 | 26 | CGCGCGCGTGTGGTGTGCGTCGGAGG | 276.4 | -11.1 | 1.5 | -7.3 | 3.6E-13 | 1.3E-12 | MB231/MB468 | NUC |
| >hg38_21_+_8433172_8446622_RNA45SN1_45S_50ntFlanks@9622.9649.28\|CTAGTAGCTGGTTCCCTCCGAAGTTTCC | >hg38_21_+_8433172_8446622_RNA45SN1_45S_50ntFlanks | 9622 | 9649 | 28 | CTAGTAGCTGGTTCCCTCCGAAGTTTCC | 545.0 | -8.6 | 1.2 | -7.2 | 4.4E-13 | 1.3E-12 | MB231/MB468 | MITO |
| >hg38_21_+_8433172_8446622_RNA45SN1_45S_50ntFlanks@6719.6739.21\|TGAATTGCAGGACACATTGAT | >hg38_21_+_8433172_8446622_RNA45SN1_45S_50ntFlanks | 6719 | 6739 | 21 | TGAATTGCAGGACACATTGAT | 2917.6 | -10.8 | 1.5 | -7.2 | 5.0E-13 | 1.3E-12 | BT20/MB468 | NUC |
| >hg38_21_+_8433172_8446622_RNA45SN1_45S_50ntFlanks@12008.12039.32\|CCGCCGGTGAAATACCACTACTCTGATCGTTT | >hg38_21_+_8433172_8446622_RNA45SN1_45S_50ntFlanks | 12008 | 12039 | 32 | CCGCCGGTGAAATACCACTACTCTGATCGTTT | 433.3 | -6.4 | 0.9 | -7.3 | 3.5E-13 | 1.3E-12 | MB231/MB468 | CYTO |
| >hg38_21_+_8433172_8446622_RNA45SN1_45S_50ntFlanks@8358.8375.18\|AAAGAACTTTGAAGAGAG | >hg38_21_+_8433172_8446622_RNA45SN1_45S_50ntFlanks | 8358 | 8375 | 18 | AAAGAACTTTGAAGAGAG | 1364.1 | 6.1 | 0.8 | 7.3 | 2.7E-13 | 1.3E-12 | BT20/MB468 | MITO |
| >hg38_21_+_8433172_8446622_RNA45SN1_45S_50ntFlanks@4657.4688.32\|CTAGAGGTGAAATTCTTGGACCGGCGCAAGAC | >hg38_21_+_8433172_8446622_RNA45SN1_45S_50ntFlanks | 4657 | 4688 | 32 | CTAGAGGTGAAATTCTTGGACCGGCGCAAGAC | 507.7 | -9.2 | 1.3 | -7.3 | 3.6E-13 | 1.3E-12 | MB231/MB468 | NUC |
| >hg38_21_+_8433172_8446622_RNA45SN1_45S_50ntFlanks@12838.12869.32\|GTGCGGAGTGCCCTTCGTCCTGGGAAACGGGG | >hg38_21_+_8433172_8446622_RNA45SN1_45S_50ntFlanks | 12838 | 12869 | 32 | GTGCGGAGTGCCCTTCGTCCTGGGAAACGGGG | 132.7 | -10.9 | 1.5 | -7.3 | 3.0E-13 | 1.3E-12 | MB231/MB468 | MP |
| >MI0003581\|hsa-mir-574&WithFlank&4\|+\|38868026\|38868133@67.87.21(+1U)&[MIMAT0003239&hsa-miR-574-3p&offsets\|0\|-1(+1U);m-149&4\|+\|38868092\|38868113&offsets\|0\|-1(+1U)]\|CACGCTCATGCACACACCCACT | >MI0003581\|hsa-mir-574&WithFlank&4\|+\|38868026\|38868133 | 67 | 87 | 21(+1U) | CACGCTCATGCACACACCCACT | 419.4 | 9.1 | 1.3 | 7.3 | 3.5E-13 | 1.3E-12 | MB231/MB468 | CYTO |
| >MI0001448\|hsa-mir-425&WithFlank&3\|-\|49020142\|49020240@20.41.22&[MIMAT0003393&hsa-miR-425-5p&offsets\|0\|-1;m-95&3\|-\|49020200\|49020221&offsets\|0\|0]\|AATGACACGATCACTCCCGTTG | >MI0001448\|hsa-mir-425&WithFlank&3\|-\|49020142\|49020240 | 20 | 41 | 22 | AATGACACGATCACTCCCGTTG | 804.3 | 8.3 | 1.1 | 7.3 | 3.5E-13 | 1.3E-12 | BT20/MB468 | CYTO |
| >hg38_1_-_228634819_228635039_RNA5S12_5S_50ntFlanks@129.170.42\|TTGGATGGGAGACCGCCTGGGAATACCGGGTGCTGTAGGCTT | >hg38_1_-_228634819_228635039_RNA5S12_5S_50ntFlanks | 129 | 170 | 42 | TTGGATGGGAGACCGCCTGGGAATACCGGGTGCTGTAGGCTT | 4816.9 | 10.1 | 1.4 | 7.3 | 3.5E-13 | 1.3E-12 | BT20/MB468 | CYTO |
| >MI0000093\|hsa-mir-92a-1&WithFlank&13\|+\|91351308\|91351397@55.75.21;MI0000094\|hsa-mir-92a-2&WithFlank&X\|-\|134169532\|134169618@55.75.21&[MIMAT0000092&hsa-miR-92a-3p&offsets\|+1\|0;m-9&13\|+\|91351361\|91351382&offsets\|+1\|0];[MIMAT0000092_1&hsa-miR-92a-3p&offsets\|+1\|0;m-11&X\|-\|134169544\|134169565&offsets\|+1\|0]\|ATTGCACTTGTCCCGGCCTGT | >MI0000093\|hsa-mir-92a-1&WithFlank&13\|+\|91351308\|91351397 | 55 | 75 | 21 | ATTGCACTTGTCCCGGCCTGT | 1232.1 | 6.2 | 0.9 | 7.3 | 3.5E-13 | 1.3E-12 | MB231/MB468 | CYTO |
| >MI0000750\|hsa-mir-26a-2&WithFlank&12\|-\|57824603\|57824698@20.39.20(+1U);MI0000083\|hsa-mir-26a-1&WithFlank&3\|+\|37969398\|37969486@16.35.20(+1U)&[MIMAT0000082&hsa-miR-26a-5p&offsets\|0\|-2(+1U);m-33&12\|-\|57824658\|57824679&offsets\|0\|-2(+1U)];[MIMAT0000082_1&hsa-miR-26a-5p&offsets\|0\|-2(+1U);m-34&3\|+\|37969413\|37969434&offsets\|0\|-2(+1U)]\|TTCAAGTAATCCAGGATAGGT | >MI0000750\|hsa-mir-26a-2&WithFlank&12\|-\|57824603\|57824698 | 20 | 39 | 20(+1U) | TTCAAGTAATCCAGGATAGGT | 656.0 | 7.9 | 1.1 | 7.3 | 3.5E-13 | 1.3E-12 | BT20/MB468 | CYTO |
| >MI0000746\|hsa-mir-99b&WithFlank&19\|+\|51692606\|51692687@13.34.22(+1U)&[MIMAT0000689&hsa-miR-99b-5p&offsets\|0\|0(+1U);m-10&19\|+\|51692618\|51692639&offsets\|0\|0(+1U)]\|CACCCGTAGAACCGACCTTGCGT | >MI0000746\|hsa-mir-99b&WithFlank&19\|+\|51692606\|51692687 | 13 | 34 | 22(+1U) | CACCCGTAGAACCGACCTTGCGT | 2205.2 | 5.1 | 0.7 | 7.3 | 3.6E-13 | 1.3E-12 | MB231/MB468 | NUC |
| >hg38_21_+_8433172_8446622_RNA45SN1_45S_50ntFlanks@11849.11869.21\|CGGGCTTGGCGGAATCAGCGG | >hg38_21_+_8433172_8446622_RNA45SN1_45S_50ntFlanks | 11849 | 11869 | 21 | CGGGCTTGGCGGAATCAGCGG | 130.3 | -10.9 | 1.5 | -7.3 | 3.0E-13 | 1.3E-12 | MB231/MB468 | MP |
| >hg38_21_+_8433172_8446622_RNA45SN1_45S_50ntFlanks@10390.10424.35\|GAACAGCAGTTGAACATGGGTCAGTCGGTCCTGAG | >hg38_21_+_8433172_8446622_RNA45SN1_45S_50ntFlanks | 10390 | 10424 | 35 | GAACAGCAGTTGAACATGGGTCAGTCGGTCCTGAG | 799.4 | 7.5 | 1.0 | 7.3 | 3.5E-13 | 1.3E-12 | BT20/MB468 | CYTO |
| >hg38_21_+_8433172_8446622_RNA45SN1_45S_50ntFlanks@9804.9830.27\|GGCGTGGAATGCGAGTGCCTAGTGGGC | >hg38_21_+_8433172_8446622_RNA45SN1_45S_50ntFlanks | 9804 | 9830 | 27 | GGCGTGGAATGCGAGTGCCTAGTGGGC | 199.9 | -11.5 | 1.6 | -7.3 | 3.1E-13 | 1.3E-12 | MB231/MB468 | MP |
| >hg38_21_+_8433172_8446622_RNA45SN1_45S_50ntFlanks@7985.8012.28\|GATCAGACGTGGCGACCCGCTGAATTTA | >hg38_21_+_8433172_8446622_RNA45SN1_45S_50ntFlanks | 7985 | 8012 | 28 | GATCAGACGTGGCGACCCGCTGAATTTA | 451.5 | -11.9 | 1.5 | -7.7 | 1.5E-14 | 1.3E-12 | BT20/MB468 | MP |
| >MI0000736\|hsa-mir-30c-1&WithFlank&1\|+\|40757278\|40757378@23.46.24(+3U);MI0000254\|hsa-mir-30c-2&WithFlank&6\|-\|71376954\|71377037@13.36.24(+3U)&[MIMAT0000244&hsa-miR-30c-5p&offsets\|0\|+1(+3U);m-57&1\|+\|40757300\|40757323&offsets\|0\|0(+3U)];[MIMAT0000244_1&hsa-miR-30c-5p&offsets\|0\|+1(+3U);m-56&6\|-\|71377002\|71377025&offsets\|0\|0(+3U)]\|TGTAAACATCCTACACTCTCAGCTTTT | >MI0000736\|hsa-mir-30c-1&WithFlank&1\|+\|40757278\|40757378 | 23 | 46 | 24(+3U) | TGTAAACATCCTACACTCTCAGCTTTT | 308.0 | -11.5 | 1.5 | -7.6 | 3.7E-14 | 1.3E-12 | BT20/MB231 | MITO |
| >hg38_21_+_8433172_8446622_RNA45SN1_45S_50ntFlanks@12203.12221.19\|TAAGGCGAGCTCAGGGAGG | >hg38_21_+_8433172_8446622_RNA45SN1_45S_50ntFlanks | 12203 | 12221 | 19 | TAAGGCGAGCTCAGGGAGG | 381.6 | -12.4 | 1.7 | -7.2 | 4.5E-13 | 1.3E-12 | MB231/MB468 | MITO |
| >MI0000078\|hsa-mir-22&WithFlank&17\|-\|1713897\|1713993@59.81.23&[MIMAT0000077&hsa-miR-22-3p&offsets\|0\|+1;m-2&17\|-\|1713914\|1713935&offsets\|0\|+1]\|AAGCTGCCAGTTGAAGAACTGTT | >MI0000078\|hsa-mir-22&WithFlank&17\|-\|1713897\|1713993 | 59 | 81 | 23 | AAGCTGCCAGTTGAAGAACTGTT | 359.2 | 10.4 | 1.4 | 7.3 | 3.6E-13 | 1.3E-12 | BT20/MB468 | CYTO |
| >hg38_21_+_8433172_8446622_RNA45SN1_45S_50ntFlanks@9033.9068.36\|CTCGTGGGGGGGCCGGGCCACCCCTCCCACGGCGCG | >hg38_21_+_8433172_8446622_RNA45SN1_45S_50ntFlanks | 9033 | 9068 | 36 | CTCGTGGGGGGGCCGGGCCACCCCTCCCACGGCGCG | 109.5 | -10.6 | 1.5 | -7.3 | 3.1E-13 | 1.3E-12 | MB231/MB468 | MP |
| >hg38_21_+_8433172_8446622_RNA45SN1_45S_50ntFlanks@9622.9649.28\|CTAGTAGCTGGTTCCCTCCGAAGTTTCC | >hg38_21_+_8433172_8446622_RNA45SN1_45S_50ntFlanks | 9622 | 9649 | 28 | CTAGTAGCTGGTTCCCTCCGAAGTTTCC | 115.3 | -10.7 | 1.5 | -7.3 | 3.1E-13 | 1.3E-12 | MB231/MB468 | MP |
| >hg38_21_+_8433172_8446622_RNA45SN1_45S_50ntFlanks@10411.10456.46\|CAGTCGGTCCTGAGAGATGGGCGAGCGCCGTTCCGAAGGGACGGGC | >hg38_21_+_8433172_8446622_RNA45SN1_45S_50ntFlanks | 10411 | 10456 | 46 | CAGTCGGTCCTGAGAGATGGGCGAGCGCCGTTCCGAAGGGACGGGC | 266.7 | -11.9 | 1.6 | -7.3 | 3.1E-13 | 1.3E-12 | MB231/MB468 | MP |
| >hg38_21_+_8433172_8446622_RNA45SN1_45S_50ntFlanks@4220.4243.24\|ACGAAAAATAACAATACAGGACTC | >hg38_21_+_8433172_8446622_RNA45SN1_45S_50ntFlanks | 4220 | 4243 | 24 | ACGAAAAATAACAATACAGGACTC | 504.6 | -7.1 | 1.0 | -7.3 | 3.1E-13 | 1.3E-12 | MB231/MB468 | MP |
| >hg38_21_+_8433172_8446622_RNA45SN1_45S_50ntFlanks@12992.13013.22\|TCCCTCGCTGCGATCTATTGAA | >hg38_21_+_8433172_8446622_RNA45SN1_45S_50ntFlanks | 12992 | 13013 | 22 | TCCCTCGCTGCGATCTATTGAA | 1205.1 | -7.2 | 1.0 | -7.2 | 4.5E-13 | 1.3E-12 | MB231/MB468 | MITO |
| >hg38_21_+_8433172_8446622_RNA45SN1_45S_50ntFlanks@10823.10844.22\|AAGTCGGCAAGCCGGATCCGTA | >hg38_21_+_8433172_8446622_RNA45SN1_45S_50ntFlanks | 10823 | 10844 | 22 | AAGTCGGCAAGCCGGATCCGTA | 164.5 | -11.2 | 1.5 | -7.3 | 3.1E-13 | 1.3E-12 | MB231/MB468 | MP |
| >MI0000063\|hsa-let-7b&WithFlank&22\|+\|46113680\|46113774@12.31.20(+1G)&[MIMAT0000063&hsa-let-7b-5p&offsets\|0\|-2(+1G);m-14&22\|+\|46113691\|46113712&offsets\|0\|-2(+1G)]\|TGAGGTAGTAGGTTGTGTGGG | >MI0000063\|hsa-let-7b&WithFlank&22\|+\|46113680\|46113774 | 12 | 31 | 20(+1G) | TGAGGTAGTAGGTTGTGTGGG | 395.9 | 7.7 | 1.1 | 7.3 | 3.6E-13 | 1.3E-12 | BT20/MB468 | CYTO |
| >hg38_21_+_8433172_8446622_RNA45SN1_45S_50ntFlanks@11604.11640.37\|CGACTGTTTAATTAAAACAAAGCATCGCGAAGGCCCG | >hg38_21_+_8433172_8446622_RNA45SN1_45S_50ntFlanks | 11604 | 11640 | 37 | CGACTGTTTAATTAAAACAAAGCATCGCGAAGGCCCG | 386.3 | -12.4 | 1.7 | -7.2 | 4.6E-13 | 1.3E-12 | MB231/MB468 | MITO |
| >hg38_21_+_8433172_8446622_RNA45SN1_45S_50ntFlanks@11570.11602.33\|AGCCGACTTAGAACTGGTGCGGACCAGGGGAAT | >hg38_21_+_8433172_8446622_RNA45SN1_45S_50ntFlanks | 11570 | 11602 | 33 | AGCCGACTTAGAACTGGTGCGGACCAGGGGAAT | 1387.9 | -5.7 | 0.8 | -7.2 | 4.6E-13 | 1.3E-12 | MB231/MB468 | MITO |
| >hg38_21_+_8433172_8446622_RNA45SN1_45S_50ntFlanks@10403.10429.27\|ACATGGGTCAGTCGGTCCTGAGAGATG | >hg38_21_+_8433172_8446622_RNA45SN1_45S_50ntFlanks | 10403 | 10429 | 27 | ACATGGGTCAGTCGGTCCTGAGAGATG | 949.5 | -3.8 | 0.5 | -7.3 | 3.7E-13 | 1.3E-12 | MB231/MB468 | NUC |
| >hg38_21_+_8433172_8446622_RNA45SN1_45S_50ntFlanks@10101.10125.25\|CGCGCGCGTGTGGTGTGCGTCGGAG | >hg38_21_+_8433172_8446622_RNA45SN1_45S_50ntFlanks | 10101 | 10125 | 25 | CGCGCGCGTGTGGTGTGCGTCGGAG | 285.1 | -10.9 | 1.5 | -7.3 | 3.6E-13 | 1.3E-12 | MB231/MB468 | CYTO |
| >hg38_21_+_8433172_8446622_RNA45SN1_45S_50ntFlanks@480.497.18\|TGATCGATGTGGTGACGT | >hg38_21_+_8433172_8446622_RNA45SN1_45S_50ntFlanks | 480 | 497 | 18 | TGATCGATGTGGTGACGT | 534.9 | 5.5 | 0.8 | 7.3 | 3.7E-13 | 1.3E-12 | MB231/MB468 | NUC |
| >hg38_21_+_8433172_8446622_RNA45SN1_45S_50ntFlanks@12996.13014.19\|TCGCTGCGATCTATTGAAA | >hg38_21_+_8433172_8446622_RNA45SN1_45S_50ntFlanks | 12996 | 13014 | 19 | TCGCTGCGATCTATTGAAA | 2686.1 | 4.6 | 0.6 | 7.3 | 3.6E-13 | 1.3E-12 | MB231/MB468 | CYTO |
| >hg38_21_+_8433172_8446622_RNA45SN1_45S_50ntFlanks@10846.10868.23\|CTTCGGGATAAGGATTGGCTCTA | >hg38_21_+_8433172_8446622_RNA45SN1_45S_50ntFlanks | 10846 | 10868 | 23 | CTTCGGGATAAGGATTGGCTCTA | 378.5 | -8.6 | 1.2 | -7.2 | 4.6E-13 | 1.3E-12 | MB231/MB468 | MITO |
| >hg38_21_+_8433172_8446622_RNA45SN1_45S_50ntFlanks@12244.12269.26\|AAGGGCAAAAGCTCGCTTGATCTTGA | >hg38_21_+_8433172_8446622_RNA45SN1_45S_50ntFlanks | 12244 | 12269 | 26 | AAGGGCAAAAGCTCGCTTGATCTTGA | 141.3 | -11.0 | 1.5 | -7.3 | 3.1E-13 | 1.3E-12 | MB231/MB468 | MP |
| >hg38_21_+_8433172_8446622_RNA45SN1_45S_50ntFlanks@7982.8003.22\|TCAGATCAGACGTGGCGACCCG | >hg38_21_+_8433172_8446622_RNA45SN1_45S_50ntFlanks | 7982 | 8003 | 22 | TCAGATCAGACGTGGCGACCCG | 162.4 | -11.2 | 1.5 | -7.3 | 3.1E-13 | 1.3E-12 | MB231/MB468 | MP |
| >hg38_21_+_8433172_8446622_RNA45SN1_45S_50ntFlanks@10756.10785.30\|TACCCATATCCGCAGCAGGTCTCCAAGGTG | >hg38_21_+_8433172_8446622_RNA45SN1_45S_50ntFlanks | 10756 | 10785 | 30 | TACCCATATCCGCAGCAGGTCTCCAAGGTG | 221.4 | -11.6 | 1.6 | -7.2 | 4.6E-13 | 1.3E-12 | MB231/MB468 | MITO |
| >hg38_21_+_8433172_8446622_RNA45SN1_45S_50ntFlanks@9504.9528.25\|GTGAACTATGCCTGGGCAGGGCGAA | >hg38_21_+_8433172_8446622_RNA45SN1_45S_50ntFlanks | 9504 | 9528 | 25 | GTGAACTATGCCTGGGCAGGGCGAA | 188.6 | -11.4 | 1.6 | -7.3 | 3.1E-13 | 1.3E-12 | MB231/MB468 | MP |
| >hg38_21_+_8433172_8446622_RNA45SN1_45S_50ntFlanks@10392.10429.38\|ACAGCAGTTGAACATGGGTCAGTCGGTCCTGAGAGATG | >hg38_21_+_8433172_8446622_RNA45SN1_45S_50ntFlanks | 10392 | 10429 | 38 | ACAGCAGTTGAACATGGGTCAGTCGGTCCTGAGAGATG | 2246.8 | -7.2 | 1.0 | -7.3 | 3.2E-13 | 1.3E-12 | MB231/MB468 | MP |
| >hg38_21_+_8433172_8446622_RNA45SN1_45S_50ntFlanks@11752.11777.26\|CCAAATGCCTCGTCATCTAATTAGTG | >hg38_21_+_8433172_8446622_RNA45SN1_45S_50ntFlanks | 11752 | 11777 | 26 | CCAAATGCCTCGTCATCTAATTAGTG | 147.0 | -10.2 | 1.4 | -7.3 | 3.7E-13 | 1.4E-12 | MB231/MB468 | NUC |
| >hg38_21_+_8433172_8446622_RNA45SN1_45S_50ntFlanks@3776.3799.24\|CCGGTACAGTGAAACTGCGAATGG | >hg38_21_+_8433172_8446622_RNA45SN1_45S_50ntFlanks | 3776 | 3799 | 24 | CCGGTACAGTGAAACTGCGAATGG | 136.4 | -11.0 | 1.5 | -7.3 | 3.2E-13 | 1.4E-12 | MB231/MB468 | MP |
| >hg38_21_+_8433172_8446622_RNA45SN1_45S_50ntFlanks@10391.10431.41\|AACAGCAGTTGAACATGGGTCAGTCGGTCCTGAGAGATGGG | >hg38_21_+_8433172_8446622_RNA45SN1_45S_50ntFlanks | 10391 | 10431 | 41 | AACAGCAGTTGAACATGGGTCAGTCGGTCCTGAGAGATGGG | 314.6 | -12.1 | 1.7 | -7.2 | 4.7E-13 | 1.4E-12 | MB231/MB468 | MITO |
| >hg38_21_+_8433172_8446622_RNA45SN1_45S_50ntFlanks@10539.10564.26\|AGGCGTCCAGTGCGGTAACGCGACCG | >hg38_21_+_8433172_8446622_RNA45SN1_45S_50ntFlanks | 10539 | 10564 | 26 | AGGCGTCCAGTGCGGTAACGCGACCG | 139.8 | -11.0 | 1.5 | -7.3 | 3.2E-13 | 1.4E-12 | MB231/MB468 | MP |
| >hg38_21_+_8433172_8446622_RNA45SN1_45S_50ntFlanks@3988.4026.39\|CTTTGGTGACTCTAGATAACCTCGGGCCGATCGCACGCC | >hg38_21_+_8433172_8446622_RNA45SN1_45S_50ntFlanks | 3988 | 4026 | 39 | CTTTGGTGACTCTAGATAACCTCGGGCCGATCGCACGCC | 199.8 | -11.4 | 1.6 | -7.2 | 4.7E-13 | 1.4E-12 | MB231/MB468 | MITO |

| | | | | | | | | | | | | | |
|---|---|---|---|---|---|---|---|---|---|---|---|---|---|
| >hg38_21_+_8433172_8446622_RNA45SN1_45S_50ntFlanks@10843.10865.23\|TAACTTCGGGATAAGGATTGGCT | >hg38_21_+_8433172_8446622_RNA45SN1_45S_50ntFlanks | 10843 | 10865 | 23 | TAACTTCGGGATAAGGATTGGCT | 2390.3 | -4.9 | 0.7 | -7.3 | 3.7E-13 | 1.4E-12 | MB231/MB468 | CYTO |
| >hg38_21_+_8433172_8446622_RNA45SN1_45S_50ntFlanks@11572.11594.23\|CCGACTTAGAACTGGTGCGGACC | >hg38_21_+_8433172_8446622_RNA45SN1_45S_50ntFlanks | 11572 | 11594 | 23 | CCGACTTAGAACTGGTGCGGACC | 617.5 | -4.8 | 0.7 | -7.3 | 3.2E-13 | 1.4E-12 | MB231/MB468 | MP |
| >hg38_21_+_8433172_8446622_RNA45SN1_45S_50ntFlanks@11703.11725.23\|AATGAAGCGCGGGTAAACGGCGG | >hg38_21_+_8433172_8446622_RNA45SN1_45S_50ntFlanks | 11703 | 11725 | 23 | AATGAAGCGCGGGTAAACGGCGG | 232.9 | -11.7 | 1.6 | -7.3 | 3.2E-13 | 1.4E-12 | MB231/MB468 | MP |
| >hg38_21_+_8433172_8446622_RNA45SN1_45S_50ntFlanks@12938.12966.29\|ACCATTCGTAGACGACCTGCTTCTGGGTC | >hg38_21_+_8433172_8446622_RNA45SN1_45S_50ntFlanks | 12938 | 12966 | 29 | ACCATTCGTAGACGACCTGCTTCTGGGTC | 283.9 | -6.1 | 0.8 | -7.3 | 3.7E-13 | 1.4E-12 | MB231/MB468 | CYTO |
| >hg38_21_+_8433172_8446622_RNA45SN1_45S_50ntFlanks@11766.11791.26\|ATCTAATTAGTGACGCGCATGAATGG | >hg38_21_+_8433172_8446622_RNA45SN1_45S_50ntFlanks | 11766 | 11791 | 26 | ATCTAATTAGTGACGCGCATGAATGG | 295.6 | -7.5 | 1.0 | -7.3 | 3.7E-13 | 1.4E-12 | MB231/MB468 | CYTO |
| >hg38_MT_+_598_1651_RNR1_12S_50ntFlanks@568.611.44\|CAAAGGACCTGGCGGTGCTTCATATCCCTCTAGAGGAGCCTGTT | >hg38_MT_+_598_1651_RNR1_12S_50ntFlanks | 568 | 611 | 44 | CAAAGGACCTGGCGGTGCTTCATATCCCTCTAGAGGAGCCTGTT | 188.0 | -10.8 | 1.4 | -7.6 | 3.9E-14 | 1.4E-12 | BT20/MB231 | MITO |
| >hg38_21_+_8433172_8446622_RNA45SN1_45S_50ntFlanks@5083.5105.23\|ACGAACGAGACTCTGGCATGCTA | >hg38_21_+_8433172_8446622_RNA45SN1_45S_50ntFlanks | 5083 | 5105 | 23 | ACGAACGAGACTCTGGCATGCTA | 471.5 | -12.7 | 1.8 | -7.2 | 4.7E-13 | 1.4E-12 | MB231/MB468 | MITO |
| >hg38_21_+_8433172_8446622_RNA45SN1_45S_50ntFlanks@12456.12490.35\|CCAAGCGTTGGATTGTTCACCCACTAATAGGGAAC | >hg38_21_+_8433172_8446622_RNA45SN1_45S_50ntFlanks | 12456 | 12490 | 35 | CCAAGCGTTGGATTGTTCACCCACTAATAGGGAAC | 4847.2 | -9.9 | 1.4 | -7.2 | 5.2E-13 | 1.4E-12 | BT20/MB468 | NUC |
| >hg38_21_+_8433172_8446622_RNA45SN1_45S_50ntFlanks@10458.10492.35\|ATGGCCTCCGTTGCCCTCGGCCGATCGAAAGGGAG | >hg38_21_+_8433172_8446622_RNA45SN1_45S_50ntFlanks | 10458 | 10492 | 35 | ATGGCCTCCGTTGCCCTCGGCCGATCGAAAGGGAG | 179.0 | -11.3 | 1.6 | -7.3 | 3.2E-13 | 1.4E-12 | MB231/MB468 | MP |
| >MI0000085\|hsa-mir-27a&WithFlank&19\|-\|13836434\|13836523@16.36.21&[MIMAT0004501&hsa-miR-27a-5p&offsets\|0\|-1;m-270&19\|-\|13836487\|13836508&offsets\|0\|-1]\|AGGGCTTAGCTGCTTGTGAGC | >MI0000085\|hsa-mir-27a&WithFlank&19\|-\|13836434\|13836523 | 16 | 36 | 21 | AGGGCTTAGCTGCTTGTGAGC | 1286.3 | 6.5 | 0.9 | 7.3 | 3.7E-13 | 1.4E-12 | MB231/MB468 | CYTO |
| >hg38_21_+_8433172_8446622_RNA45SN1_45S_50ntFlanks@11729.11748.20\|TAACTATGACTCTCTTAAGG | >hg38_21_+_8433172_8446622_RNA45SN1_45S_50ntFlanks | 11729 | 11748 | 20 | TAACTATGACTCTCTTAAGG | 7273.1 | -4.6 | 0.6 | -7.3 | 3.8E-13 | 1.4E-12 | MB231/MB468 | NUC |
| >hg38_21_+_8433172_8446622_RNA45SN1_45S_50ntFlanks@8443.8464.22\|CAACCCGGCGGCGGGTCCGGCC | >hg38_21_+_8433172_8446622_RNA45SN1_45S_50ntFlanks | 8443 | 8464 | 22 | CAACCCGGCGGCGGGTCCGGCC | 195.8 | -11.5 | 1.6 | -7.3 | 3.2E-13 | 1.4E-12 | MB231/MB468 | MP |
| >hg38_21_+_8433172_8446622_RNA45SN1_45S_50ntFlanks@10807.10825.19\|ACAATGTAGGTAAGGGAAG | >hg38_21_+_8433172_8446622_RNA45SN1_45S_50ntFlanks | 10807 | 10825 | 19 | ACAATGTAGGTAAGGGAAG | 347.1 | -11.4 | 1.6 | -7.3 | 3.8E-13 | 1.4E-12 | MB231/MB468 | NUC |
| >hg38_21_+_8433172_8446622_RNA45SN1_45S_50ntFlanks@11874.11893.20\|AGAAGACCCTGTTGAGCTTG | >hg38_21_+_8433172_8446622_RNA45SN1_45S_50ntFlanks | 11874 | 11893 | 20 | AGAAGACCCTGTTGAGCTTG | 4136.2 | -8.5 | 1.2 | -7.3 | 3.8E-13 | 1.4E-12 | MB231/MB468 | NUC |
| >hg38_21_+_8433172_8446622_RNA45SN1_45S_50ntFlanks@9572.9594.23\|CAAATCGGTCGTCCGACCTGGGT | >hg38_21_+_8433172_8446622_RNA45SN1_45S_50ntFlanks | 9572 | 9594 | 23 | CAAATCGGTCGTCCGACCTGGGT | 797.7 | -5.4 | 0.7 | -7.3 | 3.7E-13 | 1.4E-12 | MB231/MB468 | CYTO |
| >MI0000816\|hsa-mir-335&WithFlank&7\|+\|130496105\|130496210@58.78.21&[MIMAT0004703&hsa-miR-335-3p&offsets\|0\|-1;m-159&7\|+\|130496162\|130496183&offsets\|0\|-1]\|TTTTTCATTATTGCTCCTGAC | >MI0000816\|hsa-mir-335&WithFlank&7\|+\|130496105\|130496210 | 58 | 78 | 21 | TTTTTCATTATTGCTCCTGAC | 608.1 | 6.8 | 0.9 | 7.3 | 3.7E-13 | 1.4E-12 | MB231/MB468 | CYTO |
| >hg38_21_+_8433172_8446622_RNA45SN1_45S_50ntFlanks@11999.12042.44\|CCCTGCGGGCCGCCGGTGAAATACCACTACTCTGATCGTTTTTT | >hg38_21_+_8433172_8446622_RNA45SN1_45S_50ntFlanks | 11999 | 12042 | 44 | CCCTGCGGGCCGCCGGTGAAATACCACTACTCTGATCGTTTTTT | 695.3 | -13.9 | 1.9 | -7.2 | 5.3E-13 | 1.4E-12 | BT20/MB468 | NUC |
| >hg38_21_+_8433172_8446622_RNA45SN1_45S_50ntFlanks@10511.10533.23\|ATCCGGAGTGGCGGAGATGGGCG | >hg38_21_+_8433172_8446622_RNA45SN1_45S_50ntFlanks | 10511 | 10533 | 23 | ATCCGGAGTGGCGGAGATGGGCG | 420.8 | -11.7 | 1.6 | -7.3 | 3.8E-13 | 1.4E-12 | MB231/MB468 | NUC |
| >hg38_21_+_8433172_8446622_RNA45SN1_45S_50ntFlanks@12412.12429.18\|ATCCTTCGATGTCGGCTC | >hg38_21_+_8433172_8446622_RNA45SN1_45S_50ntFlanks | 12412 | 12429 | 18 | ATCCTTCGATGTCGGCTC | 260.1 | -10.3 | 1.4 | -7.3 | 3.7E-13 | 1.4E-12 | BT20/MB468 | CYTO |
| >MI0001445\|hsa-mir-423&WithFlank&17\|+\|30117073\|30117178@58.79.22&[MIMAT0001340&hsa-miR-423-3p&offsets\|-1\|-2;m-100&17\|+\|30117131\|30117153&offsets\|-1\|-2]\|AAGCTCGGTCTGAGGCCCCTCA | >MI0001445\|hsa-mir-423&WithFlank&17\|+\|30117073\|30117178 | 58 | 79 | 22 | AAGCTCGGTCTGAGGCCCCTCA | 800.1 | 8.9 | 1.2 | 7.3 | 3.7E-13 | 1.4E-12 | BT20/MB468 | CYTO |
| >trna10_AspGTC_12_-_125424193_125424264@31.53.23,trna12_AspGTC_12_-_125411891_125411962@31.53.23,trna38_AspGTC_17_-_8125556_8125627@31.53.23,trna45_AspGTC_6_+_27447453_27447524@31.53.23,trna48_AspGTC_6_+_27471523_27471594@31.53.23,trna4_AspGTC_12_+_96429799_96429870@31.53.23,trna5_AspGTC_12_+_98897281_98897352@31.53.23,trna69_AspGTC_1_-_161440205_161440276@31.53.23,trna6_AspGTC_9_-_77517990_77518061@31.53.23,trna72_AspGTC_1_-_161432824_161432895@31.53.23,trna75_AspGTC_1_-_161425414_161425485@31.53.23,trna78_AspGTC_1_-_161418033_161418104@31.53.23,trna81_AspGTC_1_-_161410615_161410686@31.53.23\|CCTGTCACGCGGGAGACCGGGGT | >trna10_AspGTC_12_-_125424193_125424264 | 31 | 53 | 23 | CCTGTCACGCGGGAGACCGGGGT | 1601.2 | 5.1 | 0.7 | 7.3 | 3.7E-13 | 1.4E-12 | BT20/MB468 | CYTO |
| >trna59_GluCTC_1_+_249168447_249168518@32.70.39\|CTCTCACCGCCGCGGCCCGGGTTCGATTCCCGGTCAGGA | >trna59_GluCTC_1_+_249168447_249168518 | 32 | 70 | 39 | CTCTCACCGCCGCGGCCCGGGTTCGATTCCCGGTCAGGA | 487.1 | -7.8 | 1.1 | -7.3 | 3.7E-13 | 1.4E-12 | BT20/MB468 | CYTO |
| >hg38_21_+_8433172_8446622_RNA45SN1_45S_50ntFlanks@4439.4471.33\|CCGCCCCTTGCCTCTCGGCGCCCCCTCGATGCT | >hg38_21_+_8433172_8446622_RNA45SN1_45S_50ntFlanks | 4439 | 4471 | 33 | CCGCCCCTTGCCTCTCGGCGCCCCCTCGATGCT | 257.6 | -11.0 | 1.5 | -7.3 | 3.9E-13 | 1.4E-12 | MB231/MB468 | NUC |
| >hg38_21_+_8433172_8446622_RNA45SN1_45S_50ntFlanks@12420.12447.28\|ATGTCGGCTCTTCCTATCATTGTGAAGC | >hg38_21_+_8433172_8446622_RNA45SN1_45S_50ntFlanks | 12420 | 12447 | 28 | ATGTCGGCTCTTCCTATCATTGTGAAGC | 733.4 | -5.5 | 0.8 | -7.2 | 5.3E-13 | 1.4E-12 | BT20/MB468 | NUC |
| >hg38_21_+_8433172_8446622_RNA45SN1_45S_50ntFlanks@12991.13025.35\|CTCCCTCGCTGCGATCTATTGAAAGTCAGCCCTCG | >hg38_21_+_8433172_8446622_RNA45SN1_45S_50ntFlanks | 12991 | 13025 | 35 | CTCCCTCGCTGCGATCTATTGAAAGTCAGCCCTCG | 187.3 | -10.5 | 1.4 | -7.3 | 3.9E-13 | 1.4E-12 | MB231/MB468 | NUC |
| >hg38_21_+_8433172_8446622_RNA45SN1_45S_50ntFlanks@10408.10432.25\|GGTCAGTCGGTCCTGAGAGATGGGC | >hg38_21_+_8433172_8446622_RNA45SN1_45S_50ntFlanks | 10408 | 10432 | 25 | GGTCAGTCGGTCCTGAGAGATGGGC | 2757.4 | -11.0 | 1.5 | -7.2 | 5.3E-13 | 1.4E-12 | BT20/MB468 | NUC |
| >hg38_21_+_8433172_8446622_RNA45SN1_45S_50ntFlanks@11874.11902.29\|AGAAGACCCTGTTGAGCTTGACTCTAGTC | >hg38_21_+_8433172_8446622_RNA45SN1_45S_50ntFlanks | 11874 | 11902 | 29 | AGAAGACCCTGTTGAGCTTGACTCTAGTC | 775.6 | -9.8 | 1.4 | -7.2 | 5.4E-13 | 1.4E-12 | BT20/MB468 | NUC |
| >MI0000746\|hsa-mir-99b&WithFlank&19\|+\|51692606\|51692687@51.71.21(+1A)&[MIMAT0004678&hsa-miR-99b-3p&offsets\|0\|-1(+1A);m-151&19\|+\|51692656\|51692678&offsets\|0\|-2(+1A)]\|CAAGCTCGTGTCTGTGGGTCCA | >MI0000746\|hsa-mir-99b&WithFlank&19\|+\|51692606\|51692687 | 51 | 71 | 21(+1A) | CAAGCTCGTGTCTGTGGGTCCA | 470.0 | 7.4 | 1.0 | 7.3 | 2.9E-13 | 1.4E-12 | BT20/MB468 | MITO |
| >hg38_21_+_8433172_8446622_RNA45SN1_45S_50ntFlanks@3931.3959.29\|TCAAAACCAACCCGGTCAGCCCCTCTCCG | >hg38_21_+_8433172_8446622_RNA45SN1_45S_50ntFlanks | 3931 | 3959 | 29 | TCAAAACCAACCCGGTCAGCCCCTCTCCG | 500.9 | -6.0 | 0.8 | -7.3 | 3.8E-13 | 1.4E-12 | MB231/MB468 | CYTO |
| >MI0000070\|hsa-mir-16-1&WithFlank&13\|-\|50048967\|50049067@20.41.22;MI0000115\|hsa-mir-16-2&WithFlank&3\|+\|160404739\|160404831@16.37.22&[MIMAT0000069&hsa-miR-16-5p&offsets\|0\|0;m-62&13\|-\|50049027\|50049048&offsets\|0\|0];[MIMAT0000069_1&hsa-miR-16-5p&offsets\|0\|0;m-61&3\|+\|160404754\|160404775&offsets\|0\|0]\|TAGCAGCACGTAAATATTGGCG | >MI0000070\|hsa-mir-16-1&WithFlank&13\|-\|50048967\|50049067 | 20 | 41 | 22 | TAGCAGCACGTAAATATTGGCG | 2448.4 | 5.4 | 0.7 | 7.3 | 3.9E-13 | 1.4E-12 | MB231/MB468 | NUC |
| >hg38_21_+_8433172_8446622_RNA45SN1_45S_50ntFlanks@9554.9581.28\|CGTAGCGGTCCTGACGTGCAAATCGGTC | >hg38_21_+_8433172_8446622_RNA45SN1_45S_50ntFlanks | 9554 | 9581 | 28 | CGTAGCGGTCCTGACGTGCAAATCGGTC | 1450.9 | -5.7 | 0.8 | -7.2 | 5.4E-13 | 1.4E-12 | BT20/MB468 | NUC |
| >hg38_21_+_8433172_8446622_RNA45SN1_45S_50ntFlanks@5527.5550.24\|AAAGTCGTAACAAGGTTTCCGTAG | >hg38_21_+_8433172_8446622_RNA45SN1_45S_50ntFlanks | 5527 | 5550 | 24 | AAAGTCGTAACAAGGTTTCCGTAG | 304.4 | -10.3 | 1.4 | -7.3 | 2.9E-13 | 1.4E-12 | BT20/MB468 | MITO |
| >hg38_21_+_8433172_8446622_RNA45SN1_45S_50ntFlanks@12072.12090.19\|CCCGAGGGGCTCTCGCTTC | >hg38_21_+_8433172_8446622_RNA45SN1_45S_50ntFlanks | 12072 | 12090 | 19 | CCCGAGGGGCTCTCGCTTC | 128.7 | -10.9 | 1.5 | -7.3 | 3.3E-13 | 1.4E-12 | MB231/MB468 | MP |
| >hg38_21_+_8433172_8446622_RNA45SN1_45S_50ntFlanks@4590.4611.22\|ATTTTGTTGGTTTTCGGAACTG | >hg38_21_+_8433172_8446622_RNA45SN1_45S_50ntFlanks | 4590 | 4611 | 22 | ATTTTGTTGGTTTTCGGAACTG | 169.5 | -11.3 | 1.5 | -7.3 | 3.3E-13 | 1.4E-12 | MB231/MB468 | MP |
| >hg38_21_+_8433172_8446622_RNA45SN1_45S_50ntFlanks@10404.10431.28\|CATGGGTCAGTCGGTCCTGAGAGATGGG | >hg38_21_+_8433172_8446622_RNA45SN1_45S_50ntFlanks | 10404 | 10431 | 28 | CATGGGTCAGTCGGTCCTGAGAGATGGG | 412.4 | -9.0 | 1.2 | -7.3 | 3.9E-13 | 1.4E-12 | MB231/MB468 | NUC |
| >hg38_21_+_8433172_8446622_RNA45SN1_45S_50ntFlanks@12456.12493.38\|CCAAGCGTTGGATTGTTCACCCACTAATAGGGAACGTG | >hg38_21_+_8433172_8446622_RNA45SN1_45S_50ntFlanks | 12456 | 12493 | 38 | CCAAGCGTTGGATTGTTCACCCACTAATAGGGAACGTG | 893.9 | -5.8 | 0.8 | -7.2 | 4.9E-13 | 1.4E-12 | MB231/MB468 | MITO |
| >hg38_21_+_8433172_8446622_RNA45SN1_45S_50ntFlanks@12149.12179.31\|AGGTGGGGAGTTTGACTGGGGCGGTACACCT | >hg38_21_+_8433172_8446622_RNA45SN1_45S_50ntFlanks | 12149 | 12179 | 31 | AGGTGGGGAGTTTGACTGGGGCGGTACACCT | 105.8 | -10.6 | 1.5 | -7.3 | 3.3E-13 | 1.4E-12 | MB231/MB468 | MP |
| >hg38_21_+_8433172_8446622_RNA45SN1_45S_50ntFlanks@12716.12742.27\|GCCTCGGTTGGCCTCGGATAGCCGGTC | >hg38_21_+_8433172_8446622_RNA45SN1_45S_50ntFlanks | 12716 | 12742 | 27 | GCCTCGGTTGGCCTCGGATAGCCGGTC | 131.0 | -10.9 | 1.5 | -7.3 | 3.3E-13 | 1.4E-12 | MB231/MB468 | MP |
| >hg38_21_+_8433172_8446622_RNA45SN1_45S_50ntFlanks@12071.12096.26\|CCCCGAGGGGCTCTCGCTTCTGGCGC | >hg38_21_+_8433172_8446622_RNA45SN1_45S_50ntFlanks | 12071 | 12096 | 26 | CCCCGAGGGGCTCTCGCTTCTGGCGC | 598.1 | -8.8 | 1.2 | -7.2 | 4.9E-13 | 1.4E-12 | MB231/MB468 | MITO |
| >hg38_21_+_8433172_8446622_RNA45SN1_45S_50ntFlanks@10841.10875.35\|CGTAACTTCGGGATAAGGATTGGCTCTAAGGGCTG | >hg38_21_+_8433172_8446622_RNA45SN1_45S_50ntFlanks | 10841 | 10875 | 35 | CGTAACTTCGGGATAAGGATTGGCTCTAAGGGCTG | 264.1 | -11.9 | 1.6 | -7.3 | 3.3E-13 | 1.4E-12 | MB231/MB468 | MP |
| >hg38_21_+_8433172_8446622_RNA45SN1_45S_50ntFlanks@12574.12604.31\|CGAGAGGAACCGCAGGTTCAGACATTTGGTG | >hg38_21_+_8433172_8446622_RNA45SN1_45S_50ntFlanks | 12574 | 12604 | 31 | CGAGAGGAACCGCAGGTTCAGACATTTGGTG | 193.0 | -9.7 | 1.3 | -7.3 | 2.9E-13 | 1.4E-12 | BT20/MB468 | MITO |
| >hg38_21_+_8433172_8446622_RNA45SN1_45S_50ntFlanks@11537.11560.24\|CCGCGCGGCGCCTCGCCTCGGCCG | >hg38_21_+_8433172_8446622_RNA45SN1_45S_50ntFlanks | 11537 | 11560 | 24 | CCGCGCGGCGCCTCGCCTCGGCCG | 157.7 | -11.2 | 1.5 | -7.3 | 3.3E-13 | 1.4E-12 | MB231/MB468 | MP |
| >hg38_21_+_8433172_8446622_RNA45SN1_45S_50ntFlanks@11874.11892.19\|AGAAGACCCTGTTGAGCTT | >hg38_21_+_8433172_8446622_RNA45SN1_45S_50ntFlanks | 11874 | 11892 | 19 | AGAAGACCCTGTTGAGCTT | 429.0 | -3.8 | 0.5 | -7.3 | 3.8E-13 | 1.4E-12 | BT20/MB468 | CYTO |
| >hg38_21_+_8433172_8446622_RNA45SN1_45S_50ntFlanks@11641.11668.28\|CGGCGGGTGTTGACGCGATGTGATTTCT | >hg38_21_+_8433172_8446622_RNA45SN1_45S_50ntFlanks | 11641 | 11668 | 28 | CGGCGGGTGTTGACGCGATGTGATTTCT | 1820.8 | -9.7 | 1.3 | -7.2 | 4.9E-13 | 1.4E-12 | MB231/MB468 | MITO |
| >hg38_21_+_8433172_8446622_RNA45SN1_45S_50ntFlanks@9204.9224.21\|TCGGGGCCACGCGCGCGTCCC | >hg38_21_+_8433172_8446622_RNA45SN1_45S_50ntFlanks | 9204 | 9224 | 21 | TCGGGGCCACGCGCGCGTCCC | 243.8 | -11.8 | 1.6 | -7.3 | 3.4E-13 | 1.4E-12 | MB231/MB468 | MP |
| >MI0000478\|hsa-mir-149&WithFlank&2\|+\|240455995\|240456095@21.41.21&[MIMAT0000450&hsa-miR-149-5p&offsets\|0\|-2;m-195&2\|+\|240456015\|240456037&offsets\|0\|-2]\|TCTGGCTCCGTGTCTTCACTC | >MI0000478\|hsa-mir-149&WithFlank&2\|+\|240455995\|240456095 | 21 | 41 | 21 | TCTGGCTCCGTGTCTTCACTC | 545.6 | 10.1 | 1.4 | 7.3 | 3.0E-13 | 1.4E-12 | BT20/MB468 | MITO |

<table>
<tr><td>>MI0000085|hsa-mir-27a&WithFlank&19|-|13836434|13836523@57.75.19(+1A)&[MIMAT0000084&hsa-miR-27a-3p&offsets|0|-2(+1A);m-43&19|-|13836447|13836467&offsets|0|-2(+1A)]|TTCACAGTGGCTAAGTTCCA</td><td>>MI0000085|hsa-mir-27a&WithFlank&19|-|13836434|13836523</td><td>57</td><td>75</td><td>19(+1A)</td><td>TTCACAGTGGCTAAGTTCCA</td><td>1519.3</td><td>5.9</td><td>0.8</td><td>7.3</td><td>3.9E-13</td><td>1.4E-12</td><td>BT20/MB468</td><td>CYTO</td></tr>
<tr><td>>hg38_21_+_8433172_8446622_RNA45SN1_45S_50ntFlanks@8171.8214.44|AAGTCCTTCTGATCGAGGCCCAGCCCGTGGACGGTGTGAGGCCG</td><td>>hg38_21_+_8433172_8446622_RNA45SN1_45S_50ntFlanks</td><td>8171</td><td>8214</td><td>44</td><td>AAGTCCTTCTGATCGAGGCCCAGCCCGTGGACGGTGTGAGGCCG</td><td>401.1</td><td>-12.5</td><td>1.7</td><td>-7.3</td><td>3.4E-13</td><td>1.4E-12</td><td>MB231/MB468</td><td>MP</td></tr>
<tr><td>>hg38_21_+_8433172_8446622_RNA45SN1_45S_50ntFlanks@4996.5013.18|CAGATTGATAGCTCTTTC</td><td>>hg38_21_+_8433172_8446622_RNA45SN1_45S_50ntFlanks</td><td>4996</td><td>5013</td><td>18</td><td>CAGATTGATAGCTCTTTC</td><td>279.8</td><td>-12.0</td><td>1.6</td><td>-7.3</td><td>3.4E-13</td><td>1.4E-12</td><td>MB231/MB468</td><td>MP</td></tr>
<tr><td>>hg38_21_+_8433172_8446622_RNA45SN1_45S_50ntFlanks@12616.12654.39|CTGAGGAGCCAATGGGGCGAAGCTACCATCTGTGGGATT</td><td>>hg38_21_+_8433172_8446622_RNA45SN1_45S_50ntFlanks</td><td>12616</td><td>12654</td><td>39</td><td>CTGAGGAGCCAATGGGGCGAAGCTACCATCTGTGGGATT</td><td>895.7</td><td>-9.5</td><td>1.3</td><td>-7.3</td><td>3.4E-13</td><td>1.4E-12</td><td>MB231/MB468</td><td>MP</td></tr>
<tr><td>>hg38_21_+_8433172_8446622_RNA45SN1_45S_50ntFlanks@12488.12522.35|AACGTGAGCTGGGTTTAGACCGTCGTGAGACAGGT</td><td>>hg38_21_+_8433172_8446622_RNA45SN1_45S_50ntFlanks</td><td>12488</td><td>12522</td><td>35</td><td>AACGTGAGCTGGGTTTAGACCGTCGTGAGACAGGT</td><td>150.2</td><td>-11.1</td><td>1.5</td><td>-7.3</td><td>3.4E-13</td><td>1.4E-12</td><td>MB231/MB468</td><td>MP</td></tr>
<tr><td>>hg38_21_+_8433172_8446622_RNA45SN1_45S_50ntFlanks@10402.10429.28|AACATGGGTCAGTCGGTCCTGAGAGATG</td><td>>hg38_21_+_8433172_8446622_RNA45SN1_45S_50ntFlanks</td><td>10402</td><td>10429</td><td>28</td><td>AACATGGGTCAGTCGGTCCTGAGAGATG</td><td>4069.0</td><td>-4.4</td><td>0.6</td><td>-7.3</td><td>3.9E-13</td><td>1.4E-12</td><td>MB231/MB468</td><td>CYTO</td></tr>
<tr><td>>hg38_21_+_8433172_8446622_RNA45SN1_45S_50ntFlanks@12983.13001.19|CAGAGCAGCTCCCTCGCTG</td><td>>hg38_21_+_8433172_8446622_RNA45SN1_45S_50ntFlanks</td><td>12983</td><td>13001</td><td>19</td><td>CAGAGCAGCTCCCTCGCTG</td><td>147.0</td><td>-11.1</td><td>1.5</td><td>-7.3</td><td>3.4E-13</td><td>1.4E-12</td><td>MB231/MB468</td><td>MP</td></tr>
<tr><td>>hg38_21_+_8433172_8446622_RNA45SN1_45S_50ntFlanks@4760.4784.25|TCAGATACCGTCGTAGTTCCGACCA</td><td>>hg38_21_+_8433172_8446622_RNA45SN1_45S_50ntFlanks</td><td>4760</td><td>4784</td><td>25</td><td>TCAGATACCGTCGTAGTTCCGACCA</td><td>134.2</td><td>-10.9</td><td>1.5</td><td>-7.3</td><td>3.4E-13</td><td>1.4E-12</td><td>MB231/MB468</td><td>MP</td></tr>
<tr><td>>hg38_21_+_8433172_8446622_RNA45SN1_45S_50ntFlanks@11661.11682.22|TGATTTCTGCCCAGTGCTCTGA</td><td>>hg38_21_+_8433172_8446622_RNA45SN1_45S_50ntFlanks</td><td>11661</td><td>11682</td><td>22</td><td>TGATTTCTGCCCAGTGCTCTGA</td><td>858.9</td><td>-8.9</td><td>1.2</td><td>-7.3</td><td>4.0E-13</td><td>1.4E-12</td><td>MB231/MB468</td><td>NUC</td></tr>
<tr><td>>hg38_21_+_8433172_8446622_RNA45SN1_45S_50ntFlanks@11452.11475.24|CTCCTCGGCGCGCGGCGGCGGCGG</td><td>>hg38_21_+_8433172_8446622_RNA45SN1_45S_50ntFlanks</td><td>11452</td><td>11475</td><td>24</td><td>CTCCTCGGCGCGCGGCGGCGGCGG</td><td>191.2</td><td>-7.6</td><td>1.0</td><td>-7.3</td><td>3.9E-13</td><td>1.4E-12</td><td>BT20/MB468</td><td>CYTO</td></tr>
<tr><td>>hg38_21_+_8433172_8446622_RNA45SN1_45S_50ntFlanks@6648.6668.21|GTACGACTCTTAGCGGTGGAT</td><td>>hg38_21_+_8433172_8446622_RNA45SN1_45S_50ntFlanks</td><td>6648</td><td>6668</td><td>21</td><td>GTACGACTCTTAGCGGTGGAT</td><td>1117.2</td><td>4.9</td><td>0.7</td><td>7.3</td><td>3.9E-13</td><td>1.4E-12</td><td>BT20/MB468</td><td>CYTO</td></tr>
<tr><td>>hg38_21_+_8433172_8446622_RNA45SN1_45S_50ntFlanks@10534.10570.37|CCGCGAGGCGTCCAGTGCGGTAACGCGACCGATCCCG</td><td>>hg38_21_+_8433172_8446622_RNA45SN1_45S_50ntFlanks</td><td>10534</td><td>10570</td><td>37</td><td>CCGCGAGGCGTCCAGTGCGGTAACGCGACCGATCCCG</td><td>241.6</td><td>-10.6</td><td>1.5</td><td>-7.3</td><td>3.9E-13</td><td>1.4E-12</td><td>MB231/MB468</td><td>CYTO</td></tr>
<tr><td>>hg38_21_+_8433172_8446622_RNA45SN1_45S_50ntFlanks@7972.8010.39|AGACGCGACCTCAGATCAGACGTGGCGACCCGCTGAATT</td><td>>hg38_21_+_8433172_8446622_RNA45SN1_45S_50ntFlanks</td><td>7972</td><td>8010</td><td>39</td><td>AGACGCGACCTCAGATCAGACGTGGCGACCCGCTGAATT</td><td>249.8</td><td>4.8</td><td>0.7</td><td>7.3</td><td>4.0E-13</td><td>1.4E-12</td><td>MB231/MB468</td><td>NUC</td></tr>
<tr><td>>hg38_21_+_8433172_8446622_RNA45SN1_45S_50ntFlanks@10827.10864.38|CGGCAAGCCGGATCCGTAACTTCGGGATAAGGATTGGC</td><td>>hg38_21_+_8433172_8446622_RNA45SN1_45S_50ntFlanks</td><td>10827</td><td>10864</td><td>38</td><td>CGGCAAGCCGGATCCGTAACTTCGGGATAAGGATTGGC</td><td>144.9</td><td>-9.9</td><td>1.4</td><td>-7.3</td><td>3.9E-13</td><td>1.5E-12</td><td>MB231/MB468</td><td>CYTO</td></tr>
<tr><td>>hg38_21_+_8433172_8446622_RNA45SN1_45S_50ntFlanks@10766.10785.20|CGCAGCAGGTCTCCAAGGTG</td><td>>hg38_21_+_8433172_8446622_RNA45SN1_45S_50ntFlanks</td><td>10766</td><td>10785</td><td>20</td><td>CGCAGCAGGTCTCCAAGGTG</td><td>279.2</td><td>-4.1</td><td>0.6</td><td>-7.3</td><td>4.1E-13</td><td>1.5E-12</td><td>MB231/MB468</td><td>NUC</td></tr>
<tr><td>>hg38_21_+_8433172_8446622_RNA45SN1_45S_50ntFlanks@9534.9573.40|AGGAAACTCTGGTGGAGGTCCGTAGCGGTCCTGACGTGCA</td><td>>hg38_21_+_8433172_8446622_RNA45SN1_45S_50ntFlanks</td><td>9534</td><td>9573</td><td>40</td><td>AGGAAACTCTGGTGGAGGTCCGTAGCGGTCCTGACGTGCA</td><td>3614.2</td><td>-5.5</td><td>0.8</td><td>-7.3</td><td>4.1E-13</td><td>1.5E-12</td><td>MB231/MB468</td><td>NUC</td></tr>
<tr><td>>hg38_21_+_8433172_8446622_RNA45SN1_45S_50ntFlanks@10391.10419.29|AACAGCAGTTGAACATGGGTCAGTCGGTC</td><td>>hg38_21_+_8433172_8446622_RNA45SN1_45S_50ntFlanks</td><td>10391</td><td>10419</td><td>29</td><td>AACAGCAGTTGAACATGGGTCAGTCGGTC</td><td>524.7</td><td>-6.0</td><td>0.8</td><td>-7.3</td><td>4.0E-13</td><td>1.5E-12</td><td>MB231/MB468</td><td>CYTO</td></tr>
<tr><td>>hg38_21_+_8433172_8446622_RNA45SN1_45S_50ntFlanks@7983.8000.18|CAGATCAGACGTGGCGAC</td><td>>hg38_21_+_8433172_8446622_RNA45SN1_45S_50ntFlanks</td><td>7983</td><td>8000</td><td>18</td><td>CAGATCAGACGTGGCGAC</td><td>138.7</td><td>-11.0</td><td>1.5</td><td>-7.3</td><td>3.4E-13</td><td>1.5E-12</td><td>MB231/MB468</td><td>MP</td></tr>
<tr><td>>hg38_21_+_8433172_8446622_RNA45SN1_45S_50ntFlanks@8423.8471.49|CGCAGTCCGCCCGGAGGATTCAACCCGGCGGCGGGTCCGGCCGTGTCGG</td><td>>hg38_21_+_8433172_8446622_RNA45SN1_45S_50ntFlanks</td><td>8423</td><td>8471</td><td>49</td><td>CGCAGTCCGCCCGGAGGATTCAACCCGGCGGCGGGTCCGGCCGTGTCGG</td><td>657.4</td><td>-11.5</td><td>1.6</td><td>-7.3</td><td>3.0E-13</td><td>1.5E-12</td><td>BT20/MB468</td><td>MITO</td></tr>
<tr><td>>hg38_21_+_8433172_8446622_RNA45SN1_45S_50ntFlanks@4296.4314.19|CCATTGGAGGGCAAGTCTG</td><td>>hg38_21_+_8433172_8446622_RNA45SN1_45S_50ntFlanks</td><td>4296</td><td>4314</td><td>19</td><td>CCATTGGAGGGCAAGTCTG</td><td>128.3</td><td>-10.9</td><td>1.5</td><td>-7.3</td><td>3.5E-13</td><td>1.5E-12</td><td>MB231/MB468</td><td>MP</td></tr>
<tr><td>>MI0003820|hsa-mir-454&WithFlank&17|-|59137752|59137878@70.92.23&[MIMAT0003885&hsa-miR-454-3p&offsets|0|0;m-272&17|-|59137787|59137809&offsets|0|0]|TAGTGCAATATTGCTTATAGGGT</td><td>>MI0003820|hsa-mir-454&WithFlank&17|-|59137752|59137878</td><td>70</td><td>92</td><td>23</td><td>TAGTGCAATATTGCTTATAGGGT</td><td>210.5</td><td>4.9</td><td>0.7</td><td>7.3</td><td>4.1E-13</td><td>1.5E-12</td><td>MB231/MB468</td><td>NUC</td></tr>
<tr><td>>am_MI0005763|hsa-mir-941-1&WithFlank&20|+|63919443|63919526@53.75.23;MI0005764|hsa-mir-941-2&WithFlank&20|+|63919499|63919582@53.75.23;MI0005765|hsa-mir-941-3&WithFlank&20|+|63919555|63919638@53.75.23;MI0005766|hsa-mir-941-4&WithFlank&20|+|63919750|63919833@53.75.23;MI0031520|hsa-mir-941-5&WithFlank&20|+|63919862|63919945@53.75.23&[MIMAT0004984&hsa-miR-941&offsets|0|0;m-263&20|+|63919495|63919517&offsets|0|0];[MIMAT0004984_1&hsa-miR-941&offsets|0|0;m-263&20|+|63919551|63919573&offsets|0|0];[MIMAT0004984_2&hsa-miR-941&offsets|0|0;m-263&20|+|63919607|63919629&offsets|0|0];[MIMAT0004984_3&hsa-miR-941&offsets|0|0;m-263&20|+|63919802|63919824&offsets|0|0];[MIMAT0004984_4&hsa-miR-941&offsets|0|0;m-263&20|+|63919914|63919936&offsets|0|0]|CACCCGGCTGTGTGCACATGTGC</td><td>>am_MI0005763|hsa-mir-941-1&WithFlank&20|+|63919443|63919526</td><td>53</td><td>75</td><td>23</td><td>CACCCGGCTGTGTGCACATGTGC</td><td>5750.1</td><td>5.4</td><td>0.8</td><td>7.2</td><td>5.1E-13</td><td>1.5E-12</td><td>MB231/MB468</td><td>MITO</td></tr>
<tr><td>>MI0000767|hsa-mir-365a&WithFlank&16|+|14309279|14309377@62.81.20(+2U);MI0000769|hsa-mir-365b&WithFlank&17|+|31575405|31575527@74.93.20(+2U)&[MIMAT0000710&hsa-miR-365a-3p&offsets|0|-2(+2U);m-176&16|+|14309340|14309361&offsets|0|-2(+2U)];[MIMAT0022834&hsa-miR-365b-3p&offsets|0|-2(+2U);m-175&17|+|31575478|31575499&offsets|0|-2(+2U)]|TAATGCCCCTAAAAATCCTTTT</td><td>>MI0000767|hsa-mir-365a&WithFlank&16|+|14309279|14309377</td><td>62</td><td>81</td><td>20(+2U)</td><td>TAATGCCCCTAAAAATCCTTTT</td><td>90.4</td><td>3.0</td><td>0.4</td><td>7.2</td><td>5.6E-13</td><td>1.5E-12</td><td>BT20/MB468</td><td>NUC</td></tr>
<tr><td>>hg38_21_+_8433172_8446622_RNA45SN1_45S_50ntFlanks@11255.11281.27|CTGCGGCGGGCGTCGCGGCCGCCCCCG</td><td>>hg38_21_+_8433172_8446622_RNA45SN1_45S_50ntFlanks</td><td>11255</td><td>11281</td><td>27</td><td>CTGCGGCGGGCGTCGCGGCCGCCCCCG</td><td>188.5</td><td>-10.3</td><td>1.4</td><td>-7.3</td><td>4.0E-13</td><td>1.5E-12</td><td>MB231/MB468</td><td>CYTO</td></tr>
<tr><td>>hg38_21_+_8433172_8446622_RNA45SN1_45S_50ntFlanks@12499.12518.20|GGTTTAGACCGTCGTGAGAC</td><td>>hg38_21_+_8433172_8446622_RNA45SN1_45S_50ntFlanks</td><td>12499</td><td>12518</td><td>20</td><td>GGTTTAGACCGTCGTGAGAC</td><td>217.2</td><td>-10.5</td><td>1.4</td><td>-7.3</td><td>4.0E-13</td><td>1.5E-12</td><td>MB231/MB468</td><td>CYTO</td></tr>
<tr><td>>hg38_21_+_8433172_8446622_RNA45SN1_45S_50ntFlanks@12163.12180.18|ACTGGGGCGGTACACCTG</td><td>>hg38_21_+_8433172_8446622_RNA45SN1_45S_50ntFlanks</td><td>12163</td><td>12180</td><td>18</td><td>ACTGGGGCGGTACACCTG</td><td>1023.3</td><td>-10.2</td><td>1.4</td><td>-7.2</td><td>5.7E-13</td><td>1.5E-12</td><td>BT20/MB468</td><td>NUC</td></tr>
<tr><td>>hg38_21_+_8433172_8446622_RNA45SN1_45S_50ntFlanks@4380.4402.23|CGTAGTTGGATCTTGGGAGCGGG</td><td>>hg38_21_+_8433172_8446622_RNA45SN1_45S_50ntFlanks</td><td>4380</td><td>4402</td><td>23</td><td>CGTAGTTGGATCTTGGGAGCGGG</td><td>146.7</td><td>-11.1</td><td>1.5</td><td>-7.3</td><td>3.5E-13</td><td>1.5E-12</td><td>MB231/MB468</td><td>MP</td></tr>
<tr><td>>MI0000456|hsa-mir-140&WithFlank&16|+|69933075|69933186@68.89.22&[MIMAT0004597&hsa-miR-140-3p&offsets|0|+1;m-45&16|+|69933143|69933164&offsets|-1|-1]|TACCACAGGGTAGAACCACGGA</td><td>>MI0000456|hsa-mir-140&WithFlank&16|+|69933075|69933186</td><td>68</td><td>89</td><td>22</td><td>TACCACAGGGTAGAACCACGGA</td><td>486.7</td><td>12.1</td><td>1.7</td><td>7.2</td><td>5.1E-13</td><td>1.5E-12</td><td>MB231/MB468</td><td>MITO</td></tr>
<tr><td>>hg38_21_+_8433172_8446622_RNA45SN1_45S_50ntFlanks@11729.11766.38|TAACTATGACTCTCTTAAGGTAGCCAAATGCCTCGTCA</td><td>>hg38_21_+_8433172_8446622_RNA45SN1_45S_50ntFlanks</td><td>11729</td><td>11766</td><td>38</td><td>TAACTATGACTCTCTTAAGGTAGCCAAATGCCTCGTCA</td><td>407.9</td><td>-11.6</td><td>1.6</td><td>-7.3</td><td>4.1E-13</td><td>1.5E-12</td><td>MB231/MB468</td><td>NUC</td></tr>
<tr><td>>hg38_21_+_8433172_8446622_RNA45SN1_45S_50ntFlanks@4367.4408.42|CAGTTAAAAAGCTCGTAGTTGGATCTTGGGAGCGGGCGGGCG</td><td>>hg38_21_+_8433172_8446622_RNA45SN1_45S_50ntFlanks</td><td>4367</td><td>4408</td><td>42</td><td>CAGTTAAAAAGCTCGTAGTTGGATCTTGGGAGCGGGCGGGCG</td><td>237.5</td><td>-11.7</td><td>1.6</td><td>-7.2</td><td>5.1E-13</td><td>1.5E-12</td><td>MB231/MB468</td><td>MITO</td></tr>
<tr><td>>hg38_21_+_8433172_8446622_RNA45SN1_45S_50ntFlanks@12415.12447.33|CTTCGATGTCGGCTCTTCCTATCATTGTGAAGC</td><td>>hg38_21_+_8433172_8446622_RNA45SN1_45S_50ntFlanks</td><td>12415</td><td>12447</td><td>33</td><td>CTTCGATGTCGGCTCTTCCTATCATTGTGAAGC</td><td>584.1</td><td>3.3</td><td>0.5</td><td>7.3</td><td>3.1E-13</td><td>1.5E-12</td><td>BT20/MB468</td><td>MITO</td></tr>
<tr><td>>hg38_21_+_8433172_8446622_RNA45SN1_45S_50ntFlanks@5487.5514.28|CGCTGAGAAGACGGTCGAACTTGACTAT</td><td>>hg38_21_+_8433172_8446622_RNA45SN1_45S_50ntFlanks</td><td>5487</td><td>5514</td><td>28</td><td>CGCTGAGAAGACGGTCGAACTTGACTAT</td><td>4176.6</td><td>-5.6</td><td>0.8</td><td>-7.3</td><td>4.0E-13</td><td>1.5E-12</td><td>MB231/MB468</td><td>CYTO</td></tr>
<tr><td>>hg38_21_+_8433172_8446622_RNA45SN1_45S_50ntFlanks@11570.11592.23|AGCCGACTTAGAACTGGTGCGGA</td><td>>hg38_21_+_8433172_8446622_RNA45SN1_45S_50ntFlanks</td><td>11570</td><td>11592</td><td>23</td><td>AGCCGACTTAGAACTGGTGCGGA</td><td>1417.2</td><td>-8.1</td><td>1.1</td><td>-7.3</td><td>3.5E-13</td><td>1.5E-12</td><td>MB231/MB468</td><td>MP</td></tr>
<tr><td>>hg38_21_+_8433172_8446622_RNA45SN1_45S_50ntFlanks@9739.9758.20|AAACGATCTCAACCTATTCT</td><td>>hg38_21_+_8433172_8446622_RNA45SN1_45S_50ntFlanks</td><td>9739</td><td>9758</td><td>20</td><td>AAACGATCTCAACCTATTCT</td><td>2082.1</td><td>-4.7</td><td>0.7</td><td>-7.3</td><td>4.0E-13</td><td>1.5E-12</td><td>BT20/MB468</td><td>CYTO</td></tr>
<tr><td>>hg38_21_+_8433172_8446622_RNA45SN1_45S_50ntFlanks@10808.10827.20|CAATGTAGGTAAGGGAAGTC</td><td>>hg38_21_+_8433172_8446622_RNA45SN1_45S_50ntFlanks</td><td>10808</td><td>10827</td><td>20</td><td>CAATGTAGGTAAGGGAAGTC</td><td>1214.2</td><td>-9.4</td><td>1.3</td><td>-7.2</td><td>5.1E-13</td><td>1.5E-12</td><td>MB231/MB468</td><td>MITO</td></tr>
<tr><td>>trna111_HisGTG_1_-_147774845_147774916@1.34.34,trna118_HisGTG_1_-_145396881_145396952@1.34.34,trna16_HisGTG_1_+_146544773_146544844@1.34.34,trna1_HisGTG_15_+_45493349_45493420@1.34.34,trna21_HisGTG_1_+_147753471_147753542@1.34.34,trna33_HisGTG_6_+_27125906_27125977@1.34.34,trna7_HisGTG_9_-_14433938_14434009@1.34.34,trna8_HisGTG_15_-_45492611_45492682@1.34.34,trna9_HisGTG_15_-_45490804_45490875@1.34.34|GCCGTGATCGTATAGTGGTTAGTACTCTGCGTTG</td><td>>trna111_HisGTG_1_-_147774845_147774916</td><td>1</td><td>34</td><td>34</td><td>GCCGTGATCGTATAGTGGTTAGTACTCTGCGTTG</td><td>3646.3</td><td>7.6</td><td>1.0</td><td>7.3</td><td>3.1E-13</td><td>1.5E-12</td><td>BT20/MB468</td><td>MITO</td></tr>
<tr><td>>hg38_1_-_228634819_228635039_RNA5S12_5S_50ntFlanks@125.170.46|GTACTTGGATGGGAGACCGCCTGGGAATACCGGGTGCTGTAGGCTT</td><td>>hg38_1_-_228634819_228635039_RNA5S12_5S_50ntFlanks</td><td>125</td><td>170</td><td>46</td><td>GTACTTGGATGGGAGACCGCCTGGGAATACCGGGTGCTGTAGGCTT</td><td>1237.7</td><td>9.7</td><td>1.3</td><td>7.3</td><td>4.0E-13</td><td>1.5E-12</td><td>MB231/MB468</td><td>CYTO</td></tr>
<tr><td>>hg38_21_+_8433172_8446622_RNA45SN1_45S_50ntFlanks@12157.12179.23|AGTTTGACTGGGGCGGTACACCT</td><td>>hg38_21_+_8433172_8446622_RNA45SN1_45S_50ntFlanks</td><td>12157</td><td>12179</td><td>23</td><td>AGTTTGACTGGGGCGGTACACCT</td><td>581.0</td><td>-4.7</td><td>0.6</td><td>-7.3</td><td>3.1E-13</td><td>1.5E-12</td><td>BT20/MB468</td><td>MITO</td></tr>
<tr><td>>MI0000071|hsa-mir-17&WithFlank&13|+|91350599|91350694@20.42.23&[MIMAT0000070&hsa-miR-17-5p&offsets|0|0;m-88&13|+|91350618|91350640&offsets|0|0]|CAAAGTGCTTACAGTGCAGGTAG</td><td>>MI0000071|hsa-mir-17&WithFlank&13|+|91350599|91350694</td><td>20</td><td>42</td><td>23</td><td>CAAAGTGCTTACAGTGCAGGTAG</td><td>5292.5</td><td>4.8</td><td>0.7</td><td>7.3</td><td>4.0E-13</td><td>1.5E-12</td><td>BT20/MB468</td><td>CYTO</td></tr>
<tr><td>>hg38_21_+_8433172_8446622_RNA45SN1_45S_50ntFlanks@10461.10497.37|GCCTCCGTTGCCCTCGGCCGATCGAAAGGGAGTCGGG</td><td>>hg38_21_+_8433172_8446622_RNA45SN1_45S_50ntFlanks</td><td>10461</td><td>10497</td><td>37</td><td>GCCTCCGTTGCCCTCGGCCGATCGAAAGGGAGTCGGG</td><td>230.5</td><td>-9.9</td><td>1.4</td><td>-7.3</td><td>3.1E-13</td><td>1.5E-12</td><td>BT20/MB468</td><td>MITO</td></tr>
<tr><td>>hg38_21_+_8433172_8446622_RNA45SN1_45S_50ntFlanks@12456.12487.32|CCAAGCGTTGGATTGTTCACCCACTAATAGGG</td><td>>hg38_21_+_8433172_8446622_RNA45SN1_45S_50ntFlanks</td><td>12456</td><td>12487</td><td>32</td><td>CCAAGCGTTGGATTGTTCACCCACTAATAGGG</td><td>9838.0</td><td>-8.1</td><td>1.1</td><td>-7.3</td><td>3.1E-13</td><td>1.5E-12</td><td>BT20/MB468</td><td>MITO</td></tr>
<tr><td>>hg38_21_+_8433172_8446622_RNA45SN1_45S_50ntFlanks@11668.11702.35|TGCCCAGTGCTCTGAATGTCAAAGTGAAGAAATTC</td><td>>hg38_21_+_8433172_8446622_RNA45SN1_45S_50ntFlanks</td><td>11668</td><td>11702</td><td>35</td><td>TGCCCAGTGCTCTGAATGTCAAAGTGAAGAAATTC</td><td>6475.4</td><td>-6.2</td><td>0.9</td><td>-7.2</td><td>5.8E-13</td><td>1.5E-12</td><td>BT20/MB468</td><td>NUC</td></tr>
<tr><td>>am_trna11_ArgACG_3_-_45730491_45730563@39.56.18,trna138_ArgACG_6_-_27638344_27638416@39.56.18,trna156_ArgACG_6_-_27181623_27181695@39.56.18,trna36_ArgACG_6_+_27182952_27183024@39.56.18|TCAGAAGATTCTAGGTTC</td><td>>am_trna11_ArgACG_3_-_45730491_45730563</td><td>39</td><td>56</td><td>18</td><td>TCAGAAGATTCTAGGTTC</td><td>147.5</td><td>-6.4</td><td>0.9</td><td>-7.3</td><td>4.1E-13</td><td>1.5E-12</td><td>MB231/MB468</td><td>CYTO</td></tr>
<tr><td>>hg38_21_+_8433172_8446622_RNA45SN1_45S_50ntFlanks@11452.11477.26|CTCCTCGGCGCGCGGCGGCGGCGGCG</td><td>>hg38_21_+_8433172_8446622_RNA45SN1_45S_50ntFlanks</td><td>11452</td><td>11477</td><td>26</td><td>CTCCTCGGCGCGCGGCGGCGGCGGCG</td><td>204.8</td><td>-10.0</td><td>1.4</td><td>-7.3</td><td>4.1E-13</td><td>1.5E-12</td><td>BT20/MB468</td><td>CYTO</td></tr>
</table>

<table>
<tr><td>>hg38_21_+_8433172_8446622_RNA45SN1_45S_50ntFlanks@9470.9494.25|GAGCACGAGCGCACGTGTTAGGACC</td><td>>hg38_21_+_8433172_8446622_RNA45SN1_45S_50ntFlanks</td><td>9470</td><td>9494</td><td>25</td><td>GAGCACGAGCGCACGTGTTAGGACC</td><td>156.0</td><td>-11.1</td><td>1.5</td><td>-7.3</td><td>3.5E-13</td><td>1.5E-12</td><td>MB231/MB468</td><td>MP</td></tr>
<tr><td>>MI0000061|hsa-let-7a-2&WithFlank&11|-|122146516|122146599@11.31.21(+1C);MI0000062|hsa-let-7a-3&WithFlank&22|+|46112743|46112828@10.30.21(+1C);MI0000060|hsa-let-7a-1&WithFlank&9|+|94175951|94176042@12.32.21(+1C)&[MIMAT0000062&hsa-let-7a-5p&offsets|0|-1(+1C);m-6&11|-|122146568|122146589&offsets|0|-1(+1C)];[MIMAT0000062_1&hsa-let-7a-5p&offsets|0|-1(+1C);m-5&22|+|46112752|46112773&offsets|0|-1(+1C)];[MIMAT0000062_2&hsa-let-7a-5p&offsets|0|-1(+1C);m-4&9|+|94175962|94175983&offsets|0|-1(+1C)]|TGAGGTAGTAGGTTGTATAGTC</td><td>>MI0000061|hsa-let-7a-2&WithFlank&11|-|122146516|122146599</td><td>11</td><td>31</td><td>21(+1C)</td><td>TGAGGTAGTAGGTTGTATAGTC</td><td>12087.5</td><td>6.6</td><td>0.9</td><td>7.2</td><td>5.2E-13</td><td>1.5E-12</td><td>MB231/MB468</td><td>MITO</td></tr>
<tr><td>>hg38_21_+_8433172_8446622_RNA45SN1_45S_50ntFlanks@11315.11332.18|CCACCCCACGTCTCGTCG</td><td>>hg38_21_+_8433172_8446622_RNA45SN1_45S_50ntFlanks</td><td>11315</td><td>11332</td><td>18</td><td>CCACCCCACGTCTCGTCG</td><td>122.1</td><td>-10.8</td><td>1.5</td><td>-7.3</td><td>3.5E-13</td><td>1.5E-12</td><td>MB231/MB468</td><td>MP</td></tr>
<tr><td>>hg38_21_+_8433172_8446622_RNA45SN1_45S_50ntFlanks@12718.12747.30|CTCGGTTGGCCTCGGATAGCCGGTCCCCCG</td><td>>hg38_21_+_8433172_8446622_RNA45SN1_45S_50ntFlanks</td><td>12718</td><td>12747</td><td>30</td><td>CTCGGTTGGCCTCGGATAGCCGGTCCCCCG</td><td>142.1</td><td>-11.0</td><td>1.5</td><td>-7.3</td><td>3.5E-13</td><td>1.5E-12</td><td>MB231/MB468</td><td>MP</td></tr>
<tr><td>>hg38_21_+_8433172_8446622_RNA45SN1_45S_50ntFlanks@11666.11702.37|TCTGCCCAGTGCTCTGAATGTCAAAGTGAAGAAATTC</td><td>>hg38_21_+_8433172_8446622_RNA45SN1_45S_50ntFlanks</td><td>11666</td><td>11702</td><td>37</td><td>TCTGCCCAGTGCTCTGAATGTCAAAGTGAAGAAATTC</td><td>1000.2</td><td>-5.6</td><td>0.8</td><td>-7.2</td><td>5.8E-13</td><td>1.5E-12</td><td>BT20/MB468</td><td>NUC</td></tr>
<tr><td>>hg38_21_+_8433172_8446622_RNA45SN1_45S_50ntFlanks@11620.11660.41|ACAAAGCATCGCGAAGGCCCGCGGCGGGTGTTGACGCGATG</td><td>>hg38_21_+_8433172_8446622_RNA45SN1_45S_50ntFlanks</td><td>11620</td><td>11660</td><td>41</td><td>ACAAAGCATCGCGAAGGCCCGCGGCGGGTGTTGACGCGATG</td><td>134.5</td><td>-10.9</td><td>1.5</td><td>-7.3</td><td>3.5E-13</td><td>1.5E-12</td><td>MB231/MB468</td><td>MP</td></tr>
<tr><td>>hg38_21_+_8433172_8446622_RNA45SN1_45S_50ntFlanks@11578.11601.24|TAGAACTGGTGCGGACCAGGGGAA</td><td>>hg38_21_+_8433172_8446622_RNA45SN1_45S_50ntFlanks</td><td>11578</td><td>11601</td><td>24</td><td>TAGAACTGGTGCGGACCAGGGGAA</td><td>288.1</td><td>-6.2</td><td>0.9</td><td>-7.3</td><td>4.1E-13</td><td>1.5E-12</td><td>MB231/MB468</td><td>CYTO</td></tr>
<tr><td>>hg38_21_+_8433172_8446622_RNA45SN1_45S_50ntFlanks@10400.10424.25|TGAACATGGGTCAGTCGGTCCTGAG</td><td>>hg38_21_+_8433172_8446622_RNA45SN1_45S_50ntFlanks</td><td>10400</td><td>10424</td><td>25</td><td>TGAACATGGGTCAGTCGGTCCTGAG</td><td>1697.4</td><td>5.3</td><td>0.7</td><td>7.2</td><td>4.2E-13</td><td>1.5E-12</td><td>MB231/MB468</td><td>NUC</td></tr>
<tr><td>>MI0003591|hsa-mir-584&WithFlank&5|-|149062307|149062415@22.42.21(+1U)&[MIMAT0003249&hsa-miR-584-5p&offsets|0|-1(+1U);m-167&5|-|149062374|149062394&offsets|0|0(+1U)]|TTATGGTTTGCCTGGGACTGAT</td><td>>MI0003591|hsa-mir-584&WithFlank&5|-|149062307|149062415</td><td>22</td><td>42</td><td>21(+1U)</td><td>TTATGGTTTGCCTGGGACTGAT</td><td>404.5</td><td>11.9</td><td>1.6</td><td>7.2</td><td>5.2E-13</td><td>1.5E-12</td><td>MB231/MB468</td><td>MITO</td></tr>
<tr><td>>hg38_21_+_8433172_8446622_RNA45SN1_45S_50ntFlanks@10393.10429.37|CAGCAGTTGAACATGGGTCAGTCGGTCCTGAGAGATG</td><td>>hg38_21_+_8433172_8446622_RNA45SN1_45S_50ntFlanks</td><td>10393</td><td>10429</td><td>37</td><td>CAGCAGTTGAACATGGGTCAGTCGGTCCTGAGAGATG</td><td>2453.2</td><td>-4.9</td><td>0.7</td><td>-7.2</td><td>4.2E-13</td><td>1.5E-12</td><td>MB231/MB468</td><td>NUC</td></tr>
<tr><td>>hg38_21_+_8433172_8446622_RNA45SN1_45S_50ntFlanks@9564.9595.32|CTGACGTGCAAATCGGTCGTCCGACCTGGGTA</td><td>>hg38_21_+_8433172_8446622_RNA45SN1_45S_50ntFlanks</td><td>9564</td><td>9595</td><td>32</td><td>CTGACGTGCAAATCGGTCGTCCGACCTGGGTA</td><td>2888.8</td><td>-4.6</td><td>0.6</td><td>-7.3</td><td>3.1E-13</td><td>1.5E-12</td><td>BT20/MB468</td><td>MITO</td></tr>
<tr><td>>hg38_21_+_8433172_8446622_RNA45SN1_45S_50ntFlanks@12084.12110.27|TCGCTTCTGGCGCCAAGCGCCCGGCCG</td><td>>hg38_21_+_8433172_8446622_RNA45SN1_45S_50ntFlanks</td><td>12084</td><td>12110</td><td>27</td><td>TCGCTTCTGGCGCCAAGCGCCCGGCCG</td><td>257.7</td><td>-11.9</td><td>1.6</td><td>-7.3</td><td>3.6E-13</td><td>1.5E-12</td><td>MB231/MB468</td><td>MP</td></tr>
<tr><td>>hg38_21_+_8433172_8446622_RNA45SN1_45S_50ntFlanks@10291.10319.29|CGCGGGCCCGGGTGGAGCCGCCGCAGGTG</td><td>>hg38_21_+_8433172_8446622_RNA45SN1_45S_50ntFlanks</td><td>10291</td><td>10319</td><td>29</td><td>CGCGGGCCCGGGTGGAGCCGCCGCAGGTG</td><td>1488.6</td><td>-10.0</td><td>1.4</td><td>-7.3</td><td>3.6E-13</td><td>1.5E-12</td><td>MB231/MB468</td><td>MP</td></tr>
<tr><td>>hg38_21_+_8433172_8446622_RNA45SN1_45S_50ntFlanks@6664.6683.20|TGGATCACTCGGCTCGTGCG</td><td>>hg38_21_+_8433172_8446622_RNA45SN1_45S_50ntFlanks</td><td>6664</td><td>6683</td><td>20</td><td>TGGATCACTCGGCTCGTGCG</td><td>132.0</td><td>-10.9</td><td>1.5</td><td>-7.3</td><td>3.6E-13</td><td>1.5E-12</td><td>MB231/MB468</td><td>MP</td></tr>
<tr><td>>hg38_21_+_8433172_8446622_RNA45SN1_45S_50ntFlanks@11881.11904.24|CCTGTTGAGCTTGACTCTAGTCTG</td><td>>hg38_21_+_8433172_8446622_RNA45SN1_45S_50ntFlanks</td><td>11881</td><td>11904</td><td>24</td><td>CCTGTTGAGCTTGACTCTAGTCTG</td><td>1860.6</td><td>-5.7</td><td>0.7</td><td>-7.7</td><td>1.8E-14</td><td>1.5E-12</td><td>BT20/MB468</td><td>MP</td></tr>
<tr><td>>hg38_21_+_8433172_8446622_RNA45SN1_45S_50ntFlanks@4259.4282.24|ATTGGAATGAGTCCACTTTAAATC</td><td>>hg38_21_+_8433172_8446622_RNA45SN1_45S_50ntFlanks</td><td>4259</td><td>4282</td><td>24</td><td>ATTGGAATGAGTCCACTTTAAATC</td><td>655.0</td><td>-12.4</td><td>1.6</td><td>-7.7</td><td>1.8E-14</td><td>1.5E-12</td><td>BT20/MB468</td><td>MP</td></tr>
<tr><td>>am_trna143_LysTTT_6_-_27559593_27559665@6.24.19,trna14_LysTTT_11_-_59327808_59327880@6.24.19,trna2_LysTTT_17_+_8022473_8022545@6.24.19,trna54_LysTTT_1_+_204475655_204475727@6.24.19,trna5_LysTTT_11_+_59323902_59323974@6.24.19,trna62_LysTTT_1_-_204476158_204476230@6.24.19,trna76_LysTTT_6_+_28918806_28918878@6.24.19|GATAGCTCAGTCGGTAGAG</td><td>>am_trna143_LysTTT_6_-_27559593_27559665</td><td>6</td><td>24</td><td>19</td><td>GATAGCTCAGTCGGTAGAG</td><td>513.5</td><td>-11.7</td><td>1.6</td><td>-7.3</td><td>4.1E-13</td><td>1.5E-12</td><td>MB231/MB468</td><td>CYTO</td></tr>
<tr><td>>hg38_21_+_8433172_8446622_RNA45SN1_45S_50ntFlanks@3991.4013.23|TGGTGACTCTAGATAACCTCGGG</td><td>>hg38_21_+_8433172_8446622_RNA45SN1_45S_50ntFlanks</td><td>3991</td><td>4013</td><td>23</td><td>TGGTGACTCTAGATAACCTCGGG</td><td>154.8</td><td>-11.1</td><td>1.5</td><td>-7.3</td><td>3.6E-13</td><td>1.5E-12</td><td>MB231/MB468</td><td>MP</td></tr>
<tr><td>>hg38_21_+_8433172_8446622_RNA45SN1_45S_50ntFlanks@4641.4662.22|CATTCGTATTGCGCCGCTAGAG</td><td>>hg38_21_+_8433172_8446622_RNA45SN1_45S_50ntFlanks</td><td>4641</td><td>4662</td><td>22</td><td>CATTCGTATTGCGCCGCTAGAG</td><td>218.4</td><td>-9.9</td><td>1.4</td><td>-7.3</td><td>3.2E-13</td><td>1.5E-12</td><td>BT20/MB468</td><td>MITO</td></tr>
<tr><td>>MI0000063|hsa-let-7b&WithFlank&22|+|46113680|46113774@12.33.22(+1C)&[MIMAT0000063&hsa-let-7b-5p&offsets|0|0(+1C);m-14&22|+|46113691|46113712&offsets|0|0(+1C)]|TGAGGTAGTAGGTTGTGTGGTTC</td><td>>MI0000063|hsa-let-7b&WithFlank&22|+|46113680|46113774</td><td>12</td><td>33</td><td>22(+1C)</td><td>TGAGGTAGTAGGTTGTGTGGTTC</td><td>246.8</td><td>3.3</td><td>0.5</td><td>7.2</td><td>5.9E-13</td><td>1.5E-12</td><td>BT20/MB468</td><td>NUC</td></tr>
<tr><td>>hg38_21_+_8433172_8446622_RNA45SN1_45S_50ntFlanks@10782.10801.20|GGTGAACAGCCTCTGGCATG</td><td>>hg38_21_+_8433172_8446622_RNA45SN1_45S_50ntFlanks</td><td>10782</td><td>10801</td><td>20</td><td>GGTGAACAGCCTCTGGCATG</td><td>603.0</td><td>-8.7</td><td>1.2</td><td>-7.2</td><td>5.3E-13</td><td>1.5E-12</td><td>MB231/MB468</td><td>MITO</td></tr>
<tr><td>>trna134_GluTTC_1_-_16861774_16861845@1.33.33,trna5_GluTTC_1_+_17199078_17199149@1.33.33,trna84_GluTTC_1_-_161391883_161391954@1.33.33,trna94_GluTTC_1_-_149664355_149664427@1.33.33|TCCCTGGTGGTCTAGTGGCTAGGATTCGGCGCT</td><td>>trna134_GluTTC_1_-_16861774_16861845</td><td>1</td><td>33</td><td>33</td><td>TCCCTGGTGGTCTAGTGGCTAGGATTCGGCGCT</td><td>2374.7</td><td>9.5</td><td>1.3</td><td>7.3</td><td>3.2E-13</td><td>1.5E-12</td><td>BT20/MB468</td><td>MITO</td></tr>
<tr><td>>hg38_21_+_8433172_8446622_RNA45SN1_45S_50ntFlanks@5084.5105.22|CGAACGAGACTCTGGCATGCTA</td><td>>hg38_21_+_8433172_8446622_RNA45SN1_45S_50ntFlanks</td><td>5084</td><td>5105</td><td>22</td><td>CGAACGAGACTCTGGCATGCTA</td><td>680.3</td><td>-12.4</td><td>1.7</td><td>-7.2</td><td>4.3E-13</td><td>1.5E-12</td><td>MB231/MB468</td><td>NUC</td></tr>
<tr><td>>MI0000070|hsa-mir-16-1&WithFlank&13|-|50048967|50049067@20.41.22(+1C);MI0000115|hsa-mir-16-2&WithFlank&3|+|160404739|160404831@16.37.22(+1C)&[MIMAT0000069&hsa-miR-16-5p&offsets|0|0(+1C);m-62&13|-|50049027|50049048&offsets|0|0(+1C)];[MIMAT0000069_1&hsa-miR-16-5p&offsets|0|0(+1C);m-61&3|+|160404754|160404775&offsets|0|0(+1C)]|TAGCAGCACGTAAATATTGGCGC</td><td>>MI0000070|hsa-mir-16-1&WithFlank&13|-|50048967|50049067</td><td>20</td><td>41</td><td>22(+1C)</td><td>TAGCAGCACGTAAATATTGGCGC</td><td>492.3</td><td>8.1</td><td>1.1</td><td>7.3</td><td>4.1E-13</td><td>1.5E-12</td><td>BT20/MB468</td><td>CYTO</td></tr>
<tr><td>>hg38_21_+_8433172_8446622_RNA45SN1_45S_50ntFlanks@10853.10873.21|ATAAGGATTGGCTCTAAGGGC</td><td>>hg38_21_+_8433172_8446622_RNA45SN1_45S_50ntFlanks</td><td>10853</td><td>10873</td><td>21</td><td>ATAAGGATTGGCTCTAAGGGC</td><td>235.0</td><td>-11.7</td><td>1.6</td><td>-7.3</td><td>3.6E-13</td><td>1.5E-12</td><td>MB231/MB468</td><td>MP</td></tr>
<tr><td>>hg38_21_+_8433172_8446622_RNA45SN1_45S_50ntFlanks@12710.12739.30|CGCGGAGCCTCGGTTGGCCTCGGATAGCCG</td><td>>hg38_21_+_8433172_8446622_RNA45SN1_45S_50ntFlanks</td><td>12710</td><td>12739</td><td>30</td><td>CGCGGAGCCTCGGTTGGCCTCGGATAGCCG</td><td>113.3</td><td>-10.7</td><td>1.5</td><td>-7.3</td><td>3.6E-13</td><td>1.5E-12</td><td>MB231/MB468</td><td>MP</td></tr>
<tr><td>>MI0000298|hsa-mir-221&WithFlank&X|-|45746151|45746272@71.91.21&[MIMAT0000278&hsa-miR-221-3p&offsets|0|-2;m-91&X|-|45746181|45746202&offsets|0|-1]|AGCTACATTGTCTGCTGGGTT</td><td>>MI0000298|hsa-mir-221&WithFlank&X|-|45746151|45746272</td><td>71</td><td>91</td><td>21</td><td>AGCTACATTGTCTGCTGGGTT</td><td>2281.8</td><td>6.4</td><td>0.9</td><td>7.2</td><td>5.3E-13</td><td>1.5E-12</td><td>MB231/MB468</td><td>MITO</td></tr>
<tr><td>>hg38_21_+_8433172_8446622_RNA45SN1_45S_50ntFlanks@11452.11471.20|CTCCTCGGCGCGCGGCGGCG</td><td>>hg38_21_+_8433172_8446622_RNA45SN1_45S_50ntFlanks</td><td>11452</td><td>11471</td><td>20</td><td>CTCCTCGGCGCGCGGCGGCG</td><td>319.9</td><td>-7.2</td><td>1.0</td><td>-7.3</td><td>4.2E-13</td><td>1.5E-12</td><td>MB231/MB468</td><td>CYTO</td></tr>
<tr><td>>hg38_21_+_8433172_8446622_RNA45SN1_45S_50ntFlanks@11027.11062.36|CTCTCTCTCTCTCTCCCCCGCTCCCCGTCCTCCCCC</td><td>>hg38_21_+_8433172_8446622_RNA45SN1_45S_50ntFlanks</td><td>11027</td><td>11062</td><td>36</td><td>CTCTCTCTCTCTCTCCCCCGCTCCCCGTCCTCCCCC</td><td>214.9</td><td>-10.5</td><td>1.4</td><td>-7.3</td><td>4.2E-13</td><td>1.5E-12</td><td>MB231/MB468</td><td>CYTO</td></tr>
<tr><td>>hg38_21_+_8433172_8446622_RNA45SN1_45S_50ntFlanks@4550.4572.23|GATACCGCAGCTAGGAATAATGG</td><td>>hg38_21_+_8433172_8446622_RNA45SN1_45S_50ntFlanks</td><td>4550</td><td>4572</td><td>23</td><td>GATACCGCAGCTAGGAATAATGG</td><td>150.2</td><td>-11.1</td><td>1.5</td><td>-7.3</td><td>3.6E-13</td><td>1.5E-12</td><td>MB231/MB468</td><td>MP</td></tr>
<tr><td>>hg38_21_+_8433172_8446622_RNA45SN1_45S_50ntFlanks@12234.12258.25|CGTGGAGCAGAAGGGCAAAAGCTCG</td><td>>hg38_21_+_8433172_8446622_RNA45SN1_45S_50ntFlanks</td><td>12234</td><td>12258</td><td>25</td><td>CGTGGAGCAGAAGGGCAAAAGCTCG</td><td>173.1</td><td>-11.3</td><td>1.6</td><td>-7.3</td><td>3.6E-13</td><td>1.5E-12</td><td>MB231/MB468</td><td>MP</td></tr>
<tr><td>>hg38_21_+_8433172_8446622_RNA45SN1_45S_50ntFlanks@10872.10892.21|GCTGGGTCGGTCGGGCTGGGG</td><td>>hg38_21_+_8433172_8446622_RNA45SN1_45S_50ntFlanks</td><td>10872</td><td>10892</td><td>21</td><td>GCTGGGTCGGTCGGGCTGGGG</td><td>127.7</td><td>-6.7</td><td>0.9</td><td>-7.2</td><td>4.2E-13</td><td>1.5E-12</td><td>MB231/MB468</td><td>CYTO</td></tr>
<tr><td>>hg38_21_+_8433172_8446622_RNA45SN1_45S_50ntFlanks@10684.10711.28|CGGCGGCGTCCGGTGAGCTCTCGCTGGC</td><td>>hg38_21_+_8433172_8446622_RNA45SN1_45S_50ntFlanks</td><td>10684</td><td>10711</td><td>28</td><td>CGGCGGCGTCCGGTGAGCTCTCGCTGGC</td><td>1756.4</td><td>-9.5</td><td>1.3</td><td>-7.3</td><td>3.6E-13</td><td>1.5E-12</td><td>MB231/MB468</td><td>MP</td></tr>
<tr><td>>hg38_21_+_8433172_8446622_RNA45SN1_45S_50ntFlanks@9738.9767.30|GAAACGATCTCAACCTATTCTCAAACTTTA</td><td>>hg38_21_+_8433172_8446622_RNA45SN1_45S_50ntFlanks</td><td>9738</td><td>9767</td><td>30</td><td>GAAACGATCTCAACCTATTCTCAAACTTTA</td><td>4702.2</td><td>-4.5</td><td>0.6</td><td>-7.3</td><td>3.2E-13</td><td>1.5E-12</td><td>BT20/MB468</td><td>MITO</td></tr>
<tr><td>>hg38_21_+_8433172_8446622_RNA45SN1_45S_50ntFlanks@3876.3898.23|TACATGCCGACGGGCGCTGACCC</td><td>>hg38_21_+_8433172_8446622_RNA45SN1_45S_50ntFlanks</td><td>3876</td><td>3898</td><td>23</td><td>TACATGCCGACGGGCGCTGACCC</td><td>902.4</td><td>-4.2</td><td>0.6</td><td>-7.3</td><td>3.2E-13</td><td>1.5E-12</td><td>BT20/MB468</td><td>MITO</td></tr>
<tr><td>>hg38_MT_+_1621_3279_RNR2_16S_50ntFlanks@51.79.29|GCTAAACCTAGCCCCAAACCCACTCCACC</td><td>>hg38_MT_+_1621_3279_RNR2_16S_50ntFlanks</td><td>51</td><td>79</td><td>29</td><td>GCTAAACCTAGCCCCAAACCCACTCCACC</td><td>326.5</td><td>9.5</td><td>1.3</td><td>7.2</td><td>4.2E-13</td><td>1.5E-12</td><td>MB231/MB468</td><td>CYTO</td></tr>
<tr><td>>hg38_21_+_8433172_8446622_RNA45SN1_45S_50ntFlanks@10838.10865.28|ATCCGTAACTTCGGGATAAGGATTGGCT</td><td>>hg38_21_+_8433172_8446622_RNA45SN1_45S_50ntFlanks</td><td>10838</td><td>10865</td><td>28</td><td>ATCCGTAACTTCGGGATAAGGATTGGCT</td><td>562.0</td><td>-11.4</td><td>1.6</td><td>-7.3</td><td>4.2E-13</td><td>1.5E-12</td><td>BT20/MB468</td><td>CYTO</td></tr>
<tr><td>>hg38_21_+_8433172_8446622_RNA45SN1_45S_50ntFlanks@6744.6772.29|GACACTTCGAACGCACTTGCGGCCCCGGG</td><td>>hg38_21_+_8433172_8446622_RNA45SN1_45S_50ntFlanks</td><td>6744</td><td>6772</td><td>29</td><td>GACACTTCGAACGCACTTGCGGCCCCGGG</td><td>406.2</td><td>-12.5</td><td>1.7</td><td>-7.3</td><td>3.6E-13</td><td>1.5E-12</td><td>MB231/MB468</td><td>MP</td></tr>
<tr><td>>trna10_GlyTCC_17_+_8124866_8124937@32.53.22,trna117_GlyTCC_1_-_145397864_145397935@32.53.22,trna2_GlyTCC_19_+_4724082_4724153@32.53.22,trna45_GlyTCC_1_+_161500903_161500974@32.53.22,trna70_GlyTCC_1_-_161439547_161439618@32.53.22,trna73_GlyTCC_1_-_161432166_161432237@32.53.22,trna76_GlyTCC_1_-_161424756_161424827@32.53.22,trna79_GlyTCC_1_-_161417375_161417446@32.53.22|CTTCCAAGCAGTTGACCCGGGT</td><td>>trna10_GlyTCC_17_+_8124866_8124937</td><td>32</td><td>53</td><td>22</td><td>CTTCCAAGCAGTTGACCCGGGT</td><td>264.2</td><td>5.2</td><td>0.7</td><td>7.3</td><td>4.2E-13</td><td>1.5E-12</td><td>BT20/MB468</td><td>CYTO</td></tr>
<tr><td>>MI0001445|hsa-mir-423&WithFlank&17|+|30117073|30117178@59.77.19(+1U)&[MIMAT0001340&hsa-miR-423-3p&offsets|0|-4(+1U);m-100&17|+|30117131|30117153&offsets|0|-4(+1U)]|AGCTCGGTCTGAGGCCCCTT</td><td>>MI0001445|hsa-mir-423&WithFlank&17|+|30117073|30117178</td><td>59</td><td>77</td><td>19(+1U)</td><td>AGCTCGGTCTGAGGCCCCTT</td><td>240.3</td><td>7.6</td><td>1.0</td><td>7.2</td><td>4.2E-13</td><td>1.5E-12</td><td>MB231/MB468</td><td>CYTO</td></tr>
<tr><td>>hg38_21_+_8433172_8446622_RNA45SN1_45S_50ntFlanks@11561.11586.26|GCGCCTAGCAGCCGACTTAGAACTGG</td><td>>hg38_21_+_8433172_8446622_RNA45SN1_45S_50ntFlanks</td><td>11561</td><td>11586</td><td>26</td><td>GCGCCTAGCAGCCGACTTAGAACTGG</td><td>769.1</td><td>-4.5</td><td>0.6</td><td>-7.2</td><td>4.3E-13</td><td>1.5E-12</td><td>MB231/MB468</td><td>NUC</td></tr>
<tr><td>>hg38_21_+_8433172_8446622_RNA45SN1_45S_50ntFlanks@9545.9574.30|GTGGAGGTCCGTAGCGGTCCTGACGTGCAA</td><td>>hg38_21_+_8433172_8446622_RNA45SN1_45S_50ntFlanks</td><td>9545</td><td>9574</td><td>30</td><td>GTGGAGGTCCGTAGCGGTCCTGACGTGCAA</td><td>921.1</td><td>-7.0</td><td>1.0</td><td>-7.2</td><td>5.3E-13</td><td>1.5E-12</td><td>MB231/MB468</td><td>MITO</td></tr>
</table>

<table>
<tr><td>>hg38_21_+_8433172_8446622_RNA45SN1_45S_50ntFlanks@12836.12862.27|CGGTGCGGAGTGCCCTTCGTCCTGGGA</td><td>>hg38_21_+_8433172_8446622_RNA45SN1_45S_50ntFlanks</td><td>12836</td><td>12862</td><td>27</td><td>CGGTGCGGAGTGCCCTTCGTCCTGGGA</td><td>261.8</td><td>-11.9</td><td>1.6</td><td>-7.3</td><td>3.6E-13</td><td>1.5E-12</td><td>MB231/MB468</td><td>MP</td></tr>
<tr><td>>hg38_21_+_8433172_8446622_RNA45SN1_45S_50ntFlanks@10384.10404.21|CCATGTGAACAGCAGTTGAAC</td><td>>hg38_21_+_8433172_8446622_RNA45SN1_45S_50ntFlanks</td><td>10384</td><td>10404</td><td>21</td><td>CCATGTGAACAGCAGTTGAAC</td><td>206.4</td><td>-11.6</td><td>1.6</td><td>-7.3</td><td>3.6E-13</td><td>1.5E-12</td><td>MB231/MB468</td><td>MP</td></tr>
<tr><td>>hg38_21_+_8433172_8446622_RNA45SN1_45S_50ntFlanks@12073.12096.24|CCGAGGGGCTCTCGCTTCTGGCGC</td><td>>hg38_21_+_8433172_8446622_RNA45SN1_45S_50ntFlanks</td><td>12073</td><td>12096</td><td>24</td><td>CCGAGGGGCTCTCGCTTCTGGCGC</td><td>2132.4</td><td>-5.1</td><td>0.7</td><td>-7.2</td><td>4.2E-13</td><td>1.5E-12</td><td>MB231/MB468</td><td>CYTO</td></tr>
<tr><td>>hg38_21_+_8433172_8446622_RNA45SN1_45S_50ntFlanks@9575.9595.21|ATCGGTCGTCCGACCTGGGTA</td><td>>hg38_21_+_8433172_8446622_RNA45SN1_45S_50ntFlanks</td><td>9575</td><td>9595</td><td>21</td><td>ATCGGTCGTCCGACCTGGGTA</td><td>603.9</td><td>-11.3</td><td>1.6</td><td>-7.3</td><td>3.2E-13</td><td>1.5E-12</td><td>BT20/MB468</td><td>MITO</td></tr>
<tr><td>>hg38_21_+_8433172_8446622_RNA45SN1_45S_50ntFlanks@11873.11899.27|AAGAAGACCCTGTTGAGCTTGACTCTA</td><td>>hg38_21_+_8433172_8446622_RNA45SN1_45S_50ntFlanks</td><td>11873</td><td>11899</td><td>27</td><td>AAGAAGACCCTGTTGAGCTTGACTCTA</td><td>538.3</td><td>-8.6</td><td>1.2</td><td>-7.2</td><td>5.4E-13</td><td>1.5E-12</td><td>MB231/MB468</td><td>MITO</td></tr>
<tr><td>>hg38_21_+_8433172_8446622_RNA45SN1_45S_50ntFlanks@9313.9337.25|TCTAACACGTGCGCGAGTCGGGGGC</td><td>>hg38_21_+_8433172_8446622_RNA45SN1_45S_50ntFlanks</td><td>9313</td><td>9337</td><td>25</td><td>TCTAACACGTGCGCGAGTCGGGGGC</td><td>229.2</td><td>-11.7</td><td>1.6</td><td>-7.3</td><td>3.7E-13</td><td>1.5E-12</td><td>MB231/MB468</td><td>MP</td></tr>
<tr><td>>hg38_21_+_8433172_8446622_RNA45SN1_45S_50ntFlanks@5186.5205.20|CAATAACAGGTCTGTGATGC</td><td>>hg38_21_+_8433172_8446622_RNA45SN1_45S_50ntFlanks</td><td>5186</td><td>5205</td><td>20</td><td>CAATAACAGGTCTGTGATGC</td><td>832.2</td><td>-13.6</td><td>1.9</td><td>-7.3</td><td>3.7E-13</td><td>1.5E-12</td><td>MB231/MB468</td><td>MP</td></tr>
<tr><td>>hg38_21_+_8433172_8446622_RNA45SN1_45S_50ntFlanks@4640.4663.24|GCATTCGTATTGCGCCGCTAGAGG</td><td>>hg38_21_+_8433172_8446622_RNA45SN1_45S_50ntFlanks</td><td>4640</td><td>4663</td><td>24</td><td>GCATTCGTATTGCGCCGCTAGAGG</td><td>110.5</td><td>-10.7</td><td>1.5</td><td>-7.3</td><td>3.7E-13</td><td>1.6E-12</td><td>MB231/MB468</td><td>MP</td></tr>
<tr><td>>MI0000279|hsa-mir-196a-2&WithFlank&12|+|53991732|53991853@31.51.21(+1U);MI0000238|hsa-mir-196a-1&WithFlank&17|-|48632484|48632565@13.33.21(+1U)&[MIMAT0000226&hsa-miR-196a-5p&offsets|0|-1(+1U);m-261&12|+|53991762|53991783&offsets|0|-1(+1U)];[MIMAT0000226_1&hsa-miR-196a-5p&offsets|0|-1(+1U);m-268&17|-|48632532|48632553&offsets|0|-1(+1U)]|TAGGTAGTTTCATGTTGTTGGT</td><td>>MI0000279|hsa-mir-196a-2&WithFlank&12|+|53991732|53991853</td><td>31</td><td>51</td><td>21(+1U)</td><td>TAGGTAGTTTCATGTTGTTGGT</td><td>215.0</td><td>11.1</td><td>1.5</td><td>7.6</td><td>3.4E-14</td><td>1.6E-12</td><td>BT20/MB231</td><td>CYTO</td></tr>
<tr><td>>trnaMT_GluTTC_MT_-_14674_14742@31.72.42,trnalookalike8_GluTTC_5_-_93905172_93905240@31.72.42|TTCATATCATTGGTCGTGGTTGTAGTCCGTGCGAGAATACCA</td><td>>trnaMT_GluTTC_MT_-_14674_14742</td><td>31</td><td>72</td><td>42</td><td>TTCATATCATTGGTCGTGGTTGTAGTCCGTGCGAGAATACCA</td><td>2958.6</td><td>9.0</td><td>1.2</td><td>7.2</td><td>4.3E-13</td><td>1.6E-12</td><td>MB231/MB468</td><td>CYTO</td></tr>
<tr><td>>hg38_21_+_8433172_8446622_RNA45SN1_45S_50ntFlanks@10369.10390.22|AAGTGGAGAAGGGTTCCATGTG</td><td>>hg38_21_+_8433172_8446622_RNA45SN1_45S_50ntFlanks</td><td>10369</td><td>10390</td><td>22</td><td>AAGTGGAGAAGGGTTCCATGTG</td><td>118.9</td><td>-10.8</td><td>1.5</td><td>-7.3</td><td>3.7E-13</td><td>1.6E-12</td><td>MB231/MB468</td><td>MP</td></tr>
<tr><td>>hg38_21_+_8433172_8446622_RNA45SN1_45S_50ntFlanks@10482.10509.28|TCGAAAGGGAGTCGGGTTCAGATCCCCG</td><td>>hg38_21_+_8433172_8446622_RNA45SN1_45S_50ntFlanks</td><td>10482</td><td>10509</td><td>28</td><td>TCGAAAGGGAGTCGGGTTCAGATCCCCG</td><td>204.3</td><td>-10.4</td><td>1.4</td><td>-7.2</td><td>4.3E-13</td><td>1.6E-12</td><td>MB231/MB468</td><td>CYTO</td></tr>
<tr><td>>hg38_21_+_8433172_8446622_RNA45SN1_45S_50ntFlanks@12033.12053.21|ATCGTTTTTTCACTGACCCGG</td><td>>hg38_21_+_8433172_8446622_RNA45SN1_45S_50ntFlanks</td><td>12033</td><td>12053</td><td>21</td><td>ATCGTTTTTTCACTGACCCGG</td><td>501.7</td><td>-12.1</td><td>1.6</td><td>-7.7</td><td>1.8E-14</td><td>1.6E-12</td><td>BT20/MB468</td><td>MP</td></tr>
<tr><td>>hg38_21_+_8433172_8446622_RNA45SN1_45S_50ntFlanks@6652.6695.44|GACTCTTAGCGGTGGATCACTCGGCTCGTGCGTCGATGAAGAAC</td><td>>hg38_21_+_8433172_8446622_RNA45SN1_45S_50ntFlanks</td><td>6652</td><td>6695</td><td>44</td><td>GACTCTTAGCGGTGGATCACTCGGCTCGTGCGTCGATGAAGAAC</td><td>1340.1</td><td>7.7</td><td>1.1</td><td>7.2</td><td>4.3E-13</td><td>1.6E-12</td><td>MB231/MB468</td><td>CYTO</td></tr>
<tr><td>>hg38_21_+_8433172_8446622_RNA45SN1_45S_50ntFlanks@11316.11338.23|CACCCCACGTCTCGTCGCGCGCG</td><td>>hg38_21_+_8433172_8446622_RNA45SN1_45S_50ntFlanks</td><td>11316</td><td>11338</td><td>23</td><td>CACCCCACGTCTCGTCGCGCGCG</td><td>2937.7</td><td>-8.7</td><td>1.2</td><td>-7.2</td><td>4.3E-13</td><td>1.6E-12</td><td>MB231/MB468</td><td>CYTO</td></tr>
<tr><td>>hg38_21_+_8433172_8446622_RNA45SN1_45S_50ntFlanks@8175.8216.42|CCTTCTGATCGAGGCCCAGCCCGTGGACGGTGTGAGGCCGGT</td><td>>hg38_21_+_8433172_8446622_RNA45SN1_45S_50ntFlanks</td><td>8175</td><td>8216</td><td>42</td><td>CCTTCTGATCGAGGCCCAGCCCGTGGACGGTGTGAGGCCGGT</td><td>275.7</td><td>5.2</td><td>0.7</td><td>7.2</td><td>4.3E-13</td><td>1.6E-12</td><td>MB231/MB468</td><td>CYTO</td></tr>
<tr><td>>hg38_21_+_8433172_8446622_RNA45SN1_45S_50ntFlanks@11666.11704.39|TCTGCCCAGTGCTCTGAATGTCAAAGTGAAGAAATTCAA</td><td>>hg38_21_+_8433172_8446622_RNA45SN1_45S_50ntFlanks</td><td>11666</td><td>11704</td><td>39</td><td>TCTGCCCAGTGCTCTGAATGTCAAAGTGAAGAAATTCAA</td><td>207.5</td><td>4.2</td><td>0.6</td><td>7.2</td><td>4.3E-13</td><td>1.6E-12</td><td>MB231/MB468</td><td>CYTO</td></tr>
<tr><td>>hg38_21_+_8433172_8446622_RNA45SN1_45S_50ntFlanks@12971.12991.21|TTTCGTACGTAGCAGAGCAGC</td><td>>hg38_21_+_8433172_8446622_RNA45SN1_45S_50ntFlanks</td><td>12971</td><td>12991</td><td>21</td><td>TTTCGTACGTAGCAGAGCAGC</td><td>596.2</td><td>-4.2</td><td>0.6</td><td>-7.2</td><td>4.3E-13</td><td>1.6E-12</td><td>MB231/MB468</td><td>CYTO</td></tr>
<tr><td>>hg38_21_+_8433172_8446622_RNA45SN1_45S_50ntFlanks@12712.12744.33|CGGAGCCTCGGTTGGCCTCGGATAGCCGGTCCC</td><td>>hg38_21_+_8433172_8446622_RNA45SN1_45S_50ntFlanks</td><td>12712</td><td>12744</td><td>33</td><td>CGGAGCCTCGGTTGGCCTCGGATAGCCGGTCCC</td><td>170.3</td><td>-11.3</td><td>1.6</td><td>-7.3</td><td>3.7E-13</td><td>1.6E-12</td><td>MB231/MB468</td><td>MP</td></tr>
<tr><td>>hg38_21_+_8433172_8446622_RNA45SN1_45S_50ntFlanks@9226.9249.24|CGAAGAGGGGGACGGCGGAGCGAG</td><td>>hg38_21_+_8433172_8446622_RNA45SN1_45S_50ntFlanks</td><td>9226</td><td>9249</td><td>24</td><td>CGAAGAGGGGGACGGCGGAGCGAG</td><td>122.6</td><td>-10.8</td><td>1.5</td><td>-7.3</td><td>3.7E-13</td><td>1.6E-12</td><td>MB231/MB468</td><td>MP</td></tr>
<tr><td>>hg38_21_+_8433172_8446622_RNA45SN1_45S_50ntFlanks@11702.11728.27|CAATGAAGCGCGGGTAAACGGCGGGAG</td><td>>hg38_21_+_8433172_8446622_RNA45SN1_45S_50ntFlanks</td><td>11702</td><td>11728</td><td>27</td><td>CAATGAAGCGCGGGTAAACGGCGGGAG</td><td>131.9</td><td>-6.3</td><td>0.9</td><td>-7.2</td><td>4.3E-13</td><td>1.6E-12</td><td>MB231/MB468</td><td>CYTO</td></tr>
<tr><td>>hg38_21_+_8433172_8446622_RNA45SN1_45S_50ntFlanks@12418.12437.20|CGATGTCGGCTCTTCCTATC</td><td>>hg38_21_+_8433172_8446622_RNA45SN1_45S_50ntFlanks</td><td>12418</td><td>12437</td><td>20</td><td>CGATGTCGGCTCTTCCTATC</td><td>9253.8</td><td>-3.8</td><td>0.5</td><td>-7.2</td><td>4.4E-13</td><td>1.6E-12</td><td>MB231/MB468</td><td>NUC</td></tr>
<tr><td>>hg38_21_+_8433172_8446622_RNA45SN1_45S_50ntFlanks@12990.13019.30|GCTCCCTCGCTGCGATCTATTGAAAGTCAG</td><td>>hg38_21_+_8433172_8446622_RNA45SN1_45S_50ntFlanks</td><td>12990</td><td>13019</td><td>30</td><td>GCTCCCTCGCTGCGATCTATTGAAAGTCAG</td><td>157.7</td><td>-10.3</td><td>1.4</td><td>-7.2</td><td>4.4E-13</td><td>1.6E-12</td><td>MB231/MB468</td><td>NUC</td></tr>
<tr><td>>hg38_21_+_8433172_8446622_RNA45SN1_45S_50ntFlanks@4128.4162.35|CGATTCCGGAGAGGGAGCCTGAGAAACGGCTACCA</td><td>>hg38_21_+_8433172_8446622_RNA45SN1_45S_50ntFlanks</td><td>4128</td><td>4162</td><td>35</td><td>CGATTCCGGAGAGGGAGCCTGAGAAACGGCTACCA</td><td>660.9</td><td>-9.1</td><td>1.3</td><td>-7.2</td><td>4.4E-13</td><td>1.6E-12</td><td>MB231/MB468</td><td>NUC</td></tr>
<tr><td>>hg38_21_+_8433172_8446622_RNA45SN1_45S_50ntFlanks@6650.6682.33|ACGACTCTTAGCGGTGGATCACTCGGCTCGTGC</td><td>>hg38_21_+_8433172_8446622_RNA45SN1_45S_50ntFlanks</td><td>6650</td><td>6682</td><td>33</td><td>ACGACTCTTAGCGGTGGATCACTCGGCTCGTGC</td><td>1062.5</td><td>5.6</td><td>0.8</td><td>7.3</td><td>3.3E-13</td><td>1.6E-12</td><td>BT20/MB468</td><td>MITO</td></tr>
<tr><td>>hg38_21_+_8433172_8446622_RNA45SN1_45S_50ntFlanks@10830.10848.19|CAAGCCGGATCCGTAACTT</td><td>>hg38_21_+_8433172_8446622_RNA45SN1_45S_50ntFlanks</td><td>10830</td><td>10848</td><td>19</td><td>CAAGCCGGATCCGTAACTT</td><td>131.1</td><td>-10.9</td><td>1.5</td><td>-7.3</td><td>3.8E-13</td><td>1.6E-12</td><td>MB231/MB468</td><td>MP</td></tr>
<tr><td>>hg38_21_+_8433172_8446622_RNA45SN1_45S_50ntFlanks@4899.4917.19|AAGGAATTGACGGAAGGGC</td><td>>hg38_21_+_8433172_8446622_RNA45SN1_45S_50ntFlanks</td><td>4899</td><td>4917</td><td>19</td><td>AAGGAATTGACGGAAGGGC</td><td>266.9</td><td>-11.9</td><td>1.6</td><td>-7.3</td><td>3.8E-13</td><td>1.6E-12</td><td>MB231/MB468</td><td>MP</td></tr>
<tr><td>>hg38_21_+_8433172_8446622_RNA45SN1_45S_50ntFlanks@9381.9411.31|CGCTCGCCGGCCGAGGTGGGATCCCGAGGCC</td><td>>hg38_21_+_8433172_8446622_RNA45SN1_45S_50ntFlanks</td><td>9381</td><td>9411</td><td>31</td><td>CGCTCGCCGGCCGAGGTGGGATCCCGAGGCC</td><td>386.1</td><td>-12.4</td><td>1.7</td><td>-7.2</td><td>5.5E-13</td><td>1.6E-12</td><td>MB231/MB468</td><td>MITO</td></tr>
<tr><td>>hg38_21_+_8433172_8446622_RNA45SN1_45S_50ntFlanks@11569.11609.41|CAGCCGACTTAGAACTGGTGCGGACCAGGGGAATCCGACTG</td><td>>hg38_21_+_8433172_8446622_RNA45SN1_45S_50ntFlanks</td><td>11569</td><td>11609</td><td>41</td><td>CAGCCGACTTAGAACTGGTGCGGACCAGGGGAATCCGACTG</td><td>1869.5</td><td>-5.9</td><td>0.8</td><td>-7.2</td><td>5.5E-13</td><td>1.6E-12</td><td>MB231/MB468</td><td>MITO</td></tr>
<tr><td>>MI0000107|hsa-mir-29b-2&WithFlank&1|-|207802437|207802529@58.80.23;MI0000105|hsa-mir-29b-1&WithFlank&7|-|130877453|130877545@57.79.23&[MIMAT0000100&hsa-miR-29b-3p&offsets|0|0;m-81&1|-|207802450|207802472&offsets|0|0];[MIMAT0000100_1&hsa-miR-29b-3p&offsets|0|0;m-82&7|-|130877467|130877489&offsets|0|0]|TAGCACCATTTGAAATCAGTGTT</td><td>>MI0000107|hsa-mir-29b-2&WithFlank&1|-|207802437|207802529</td><td>58</td><td>80</td><td>23</td><td>TAGCACCATTTGAAATCAGTGTT</td><td>663.9</td><td>12.6</td><td>1.7</td><td>7.2</td><td>5.5E-13</td><td>1.6E-12</td><td>MB231/MB468</td><td>MITO</td></tr>
<tr><td>>MI0000108|hsa-mir-103a-2&WithFlank&20|+|3917488|3917577@54.76.23(+1C);MI0000109|hsa-mir-103a-1&WithFlank&5|-|168560890|168560979@54.76.23(+1C)&[MIMAT0000101&hsa-miR-103a-3p&offsets|0|0(+1C);m-12&20|+|3917541|3917563&offsets|0|0(+1C)];[MIMAT0000101_1&hsa-miR-103a-3p&offsets|0|0(+1C);m-13&5|-|168560904|168560926&offsets|0|0(+1C);m-44220&5|-|168560908|168560926&offsets|0|+4(+1C)]|AGCAGCATTGTACAGGGCTATGAC</td><td>>MI0000108|hsa-mir-103a-2&WithFlank&20|+|3917488|3917577</td><td>54</td><td>76</td><td>23(+1C)</td><td>AGCAGCATTGTACAGGGCTATGAC</td><td>421.3</td><td>7.1</td><td>1.0</td><td>7.2</td><td>4.3E-13</td><td>1.6E-12</td><td>BT20/MB468</td><td>CYTO</td></tr>
<tr><td>>hg38_21_+_8433172_8446622_RNA45SN1_45S_50ntFlanks@10366.10383.18|CCGAAGTGGAGAAGGGTT</td><td>>hg38_21_+_8433172_8446622_RNA45SN1_45S_50ntFlanks</td><td>10366</td><td>10383</td><td>18</td><td>CCGAAGTGGAGAAGGGTT</td><td>112.4</td><td>-10.7</td><td>1.5</td><td>-7.3</td><td>3.8E-13</td><td>1.6E-12</td><td>MB231/MB468</td><td>MP</td></tr>
<tr><td>>hg38_21_+_8433172_8446622_RNA45SN1_45S_50ntFlanks@11324.11344.21|GTCTCGTCGCGCGCGCGTCCG</td><td>>hg38_21_+_8433172_8446622_RNA45SN1_45S_50ntFlanks</td><td>11324</td><td>11344</td><td>21</td><td>GTCTCGTCGCGCGCGCGTCCG</td><td>160.8</td><td>-11.2</td><td>1.5</td><td>-7.3</td><td>3.8E-13</td><td>1.6E-12</td><td>MB231/MB468</td><td>MP</td></tr>
<tr><td>>hg38_21_+_8433172_8446622_RNA45SN1_45S_50ntFlanks@12406.12431.26|TTTTTGATCCTTCGATGTCGGCTCTT</td><td>>hg38_21_+_8433172_8446622_RNA45SN1_45S_50ntFlanks</td><td>12406</td><td>12431</td><td>26</td><td>TTTTTGATCCTTCGATGTCGGCTCTT</td><td>681.6</td><td>-7.7</td><td>1.1</td><td>-7.2</td><td>6.2E-13</td><td>1.6E-12</td><td>BT20/MB468</td><td>NUC</td></tr>
<tr><td>>hg38_21_+_8433172_8446622_RNA45SN1_45S_50ntFlanks@11576.11603.28|CTTAGAACTGGTGCGGACCAGGGGAATC</td><td>>hg38_21_+_8433172_8446622_RNA45SN1_45S_50ntFlanks</td><td>11576</td><td>11603</td><td>28</td><td>CTTAGAACTGGTGCGGACCAGGGGAATC</td><td>2624.7</td><td>-6.5</td><td>0.9</td><td>-7.7</td><td>1.9E-14</td><td>1.6E-12</td><td>BT20/MB468</td><td>MP</td></tr>
<tr><td>>hg38_21_+_8433172_8446622_RNA45SN1_45S_50ntFlanks@12006.12033.28|GGCCGCCGGTGAAATACCACTACTCTGA</td><td>>hg38_21_+_8433172_8446622_RNA45SN1_45S_50ntFlanks</td><td>12006</td><td>12033</td><td>28</td><td>GGCCGCCGGTGAAATACCACTACTCTGA</td><td>472.3</td><td>5.9</td><td>0.8</td><td>7.2</td><td>4.4E-13</td><td>1.6E-12</td><td>BT20/MB468</td><td>CYTO</td></tr>
<tr><td>>hg38_21_+_8433172_8446622_RNA45SN1_45S_50ntFlanks@11873.11910.38|AAGAAGACCCTGTTGAGCTTGACTCTAGTCTGGCACGG</td><td>>hg38_21_+_8433172_8446622_RNA45SN1_45S_50ntFlanks</td><td>11873</td><td>11910</td><td>38</td><td>AAGAAGACCCTGTTGAGCTTGACTCTAGTCTGGCACGG</td><td>149.9</td><td>-6.9</td><td>1.0</td><td>-7.2</td><td>4.4E-13</td><td>1.6E-12</td><td>MB231/MB468</td><td>CYTO</td></tr>
<tr><td>>hg38_21_+_8433172_8446622_RNA45SN1_45S_50ntFlanks@12181.12211.31|TCAAACGGTAACGCAGGTGTCCTAAGGCGAG</td><td>>hg38_21_+_8433172_8446622_RNA45SN1_45S_50ntFlanks</td><td>12181</td><td>12211</td><td>31</td><td>TCAAACGGTAACGCAGGTGTCCTAAGGCGAG</td><td>119.4</td><td>-10.8</td><td>1.5</td><td>-7.3</td><td>3.8E-13</td><td>1.6E-12</td><td>MB231/MB468</td><td>MP</td></tr>
<tr><td>>hg38_21_+_8433172_8446622_RNA45SN1_45S_50ntFlanks@4546.4573.28|CCTGGATACCGCAGCTAGGAATAATGGA</td><td>>hg38_21_+_8433172_8446622_RNA45SN1_45S_50ntFlanks</td><td>4546</td><td>4573</td><td>28</td><td>CCTGGATACCGCAGCTAGGAATAATGGA</td><td>483.0</td><td>-12.8</td><td>1.8</td><td>-7.3</td><td>3.8E-13</td><td>1.6E-12</td><td>MB231/MB468</td><td>MP</td></tr>
<tr><td>>hg38_21_+_8433172_8446622_RNA45SN1_45S_50ntFlanks@11534.11551.18|GTTCCGCGCGGCGCCTCG</td><td>>hg38_21_+_8433172_8446622_RNA45SN1_45S_50ntFlanks</td><td>11534</td><td>11551</td><td>18</td><td>GTTCCGCGCGGCGCCTCG</td><td>145.8</td><td>-11.1</td><td>1.5</td><td>-7.3</td><td>3.8E-13</td><td>1.6E-12</td><td>MB231/MB468</td><td>MP</td></tr>
<tr><td>>hg38_21_+_8433172_8446622_RNA45SN1_45S_50ntFlanks@3838.3861.24|CTCTCCTACTTGGATAACTGTGGT</td><td>>hg38_21_+_8433172_8446622_RNA45SN1_45S_50ntFlanks</td><td>3838</td><td>3861</td><td>24</td><td>CTCTCCTACTTGGATAACTGTGGT</td><td>687.9</td><td>-5.1</td><td>0.7</td><td>-7.2</td><td>6.2E-13</td><td>1.6E-12</td><td>BT20/MB468</td><td>NUC</td></tr>
<tr><td>>hg38_21_+_8433172_8446622_RNA45SN1_45S_50ntFlanks@5487.5514.28|CGCTGAGAAGACGGTCGAACTTGACTAT</td><td>>hg38_21_+_8433172_8446622_RNA45SN1_45S_50ntFlanks</td><td>5487</td><td>5514</td><td>28</td><td>CGCTGAGAAGACGGTCGAACTTGACTAT</td><td>1104.6</td><td>-7.4</td><td>1.0</td><td>-7.2</td><td>4.5E-13</td><td>1.6E-12</td><td>MB231/MB468</td><td>NUC</td></tr>
<tr><td>>hg38_21_+_8433172_8446622_RNA45SN1_45S_50ntFlanks@9564.9592.29|CTGACGTGCAAATCGGTCGTCCGACCTGG</td><td>>hg38_21_+_8433172_8446622_RNA45SN1_45S_50ntFlanks</td><td>9564</td><td>9592</td><td>29</td><td>CTGACGTGCAAATCGGTCGTCCGACCTGG</td><td>1440.2</td><td>-7.4</td><td>1.0</td><td>-7.2</td><td>5.6E-13</td><td>1.6E-12</td><td>MB231/MB468</td><td>MITO</td></tr>
<tr><td>>hg38_21_+_8433172_8446622_RNA45SN1_45S_50ntFlanks@10411.10431.21|CAGTCGGTCCTGAGAGATGGG</td><td>>hg38_21_+_8433172_8446622_RNA45SN1_45S_50ntFlanks</td><td>10411</td><td>10431</td><td>21</td><td>CAGTCGGTCCTGAGAGATGGG</td><td>4892.3</td><td>-7.7</td><td>1.1</td><td>-7.2</td><td>4.4E-13</td><td>1.6E-12</td><td>BT20/MB468</td><td>CYTO</td></tr>
<tr><td>>hg38_21_+_8433172_8446622_RNA45SN1_45S_50ntFlanks@6788.6806.19|CGCCTGTCTGAGCGTCGCT</td><td>>hg38_21_+_8433172_8446622_RNA45SN1_45S_50ntFlanks</td><td>6788</td><td>6806</td><td>19</td><td>CGCCTGTCTGAGCGTCGCT</td><td>5052.7</td><td>4.9</td><td>0.7</td><td>7.2</td><td>4.4E-13</td><td>1.6E-12</td><td>BT20/MB468</td><td>CYTO</td></tr>
<tr><td>>hg38_21_+_8433172_8446622_RNA45SN1_45S_50ntFlanks@6738.6756.19|ATCATCGACACTTCGAACG</td><td>>hg38_21_+_8433172_8446622_RNA45SN1_45S_50ntFlanks</td><td>6738</td><td>6756</td><td>19</td><td>ATCATCGACACTTCGAACG</td><td>627.5</td><td>-11.6</td><td>1.6</td><td>-7.2</td><td>4.4E-13</td><td>1.6E-12</td><td>BT20/MB468</td><td>CYTO</td></tr>
<tr><td>>MI0000650|hsa-mir-200c&WithFlank&12|+|6963693|6963772@49.70.22&[MIMAT0000617&hsa-miR-200c-3p&offsets|-1|-2;m-54&12|+|6963742|6963764&offsets|-1|-2]|CTAATACTGCCGGGTAATGATG</td><td>>MI0000650|hsa-mir-200c&WithFlank&12|+|6963693|6963772</td><td>49</td><td>70</td><td>22</td><td>CTAATACTGCCGGGTAATGATG</td><td>379.1</td><td>9.8</td><td>1.3</td><td>7.3</td><td>3.4E-13</td><td>1.6E-12</td><td>BT20/MB468</td><td>MITO</td></tr>
<tr><td>>hg38_21_+_8433172_8446622_RNA45SN1_45S_50ntFlanks@10468.10487.20|TTGCCCTCGGCCGATCGAAA</td><td>>hg38_21_+_8433172_8446622_RNA45SN1_45S_50ntFlanks</td><td>10468</td><td>10487</td><td>20</td><td>TTGCCCTCGGCCGATCGAAA</td><td>952.3</td><td>-5.2</td><td>0.7</td><td>-7.2</td><td>6.3E-13</td><td>1.6E-12</td><td>BT20/MB468</td><td>NUC</td></tr>
<tr><td>>hg38_21_+_8433172_8446622_RNA45SN1_45S_50ntFlanks@12405.12443.39|CTTTTTGATCCTTCGATGTCGGCTCTTCCTATCATTGTG</td><td>>hg38_21_+_8433172_8446622_RNA45SN1_45S_50ntFlanks</td><td>12405</td><td>12443</td><td>39</td><td>CTTTTTGATCCTTCGATGTCGGCTCTTCCTATCATTGTG</td><td>4551.2</td><td>-6.2</td><td>0.9</td><td>-7.2</td><td>4.4E-13</td><td>1.6E-12</td><td>BT20/MB468</td><td>CYTO</td></tr>
</table>

<table>
<tr><td>>trna10_AspGTC_12_-_125424193_125424264@32.53.22,trna12_AspGTC_12_-_125411891_125411962@32.53.22,trna144_AspGTC_6_-_27551236_27551307@32.53.22,trna38_AspGTC_17_-_8125556_8125627@32.53.22,trna45_AspGTC_6_+_27447453_27447524@32.53.22,trna48_AspGTC_6_+_27471523_27471594@32.53.22,trna4_AspGTC_12_+_96429799_96429870@32.53.22,trna5_AspGTC_12_+_98897281_98897352@32.53.22,trna69_AspGTC_1_-_161440205_161440276@32.53.22,trna6_AspGTC_9_-_77517990_77518061@32.53.22,trna72_AspGTC_1_-_161432824_161432895@32.53.22,trna75_AspGTC_1_-_161425414_161425485@32.53.22,trna78_AspGTC_1_-_161418033_161418104@32.53.22,trna81_AspGTC_1_-_161410615_161410686@32.53.22|CTGTCACGCGGGAGACCGGGGT</td><td>>trna10_AspGTC_12_-_125424193_125424264</td><td>32</td><td>53</td><td>22</td><td>CTGTCACGCGGGAGACCGGGGT</td><td>740.6</td><td>4.9</td><td>0.7</td><td>7.2</td><td>4.4E-13</td><td>1.6E-12</td><td>BT20/MB468</td><td>CYTO</td></tr>
<tr><td>>hg38_21_+_8433172_8446622_RNA45SN1_45S_50ntFlanks@10386.10424.39|ATGTGAACAGCAGTTGAACATGGGTCAGTCGGTCCTGAG</td><td>>hg38_21_+_8433172_8446622_RNA45SN1_45S_50ntFlanks</td><td>10386</td><td>10424</td><td>39</td><td>ATGTGAACAGCAGTTGAACATGGGTCAGTCGGTCCTGAG</td><td>2087.0</td><td>7.3</td><td>1.0</td><td>7.2</td><td>4.5E-13</td><td>1.6E-12</td><td>MB231/MB468</td><td>CYTO</td></tr>
<tr><td>>hg38_21_+_8433172_8446622_RNA45SN1_45S_50ntFlanks@12459.12487.29|AGCGTTGGATTGTTCACCCACTAATAGGG</td><td>>hg38_21_+_8433172_8446622_RNA45SN1_45S_50ntFlanks</td><td>12459</td><td>12487</td><td>29</td><td>AGCGTTGGATTGTTCACCCACTAATAGGG</td><td>268.4</td><td>-11.9</td><td>1.6</td><td>-7.3</td><td>3.9E-13</td><td>1.6E-12</td><td>MB231/MB468</td><td>MP</td></tr>
<tr><td>>hg38_21_+_8433172_8446622_RNA45SN1_45S_50ntFlanks@9339.9366.28|CGCACGAAAGCCGCCGTGGCGCAATGAA</td><td>>hg38_21_+_8433172_8446622_RNA45SN1_45S_50ntFlanks</td><td>9339</td><td>9366</td><td>28</td><td>CGCACGAAAGCCGCCGTGGCGCAATGAA</td><td>214.7</td><td>-11.6</td><td>1.6</td><td>-7.3</td><td>3.9E-13</td><td>1.6E-12</td><td>MB231/MB468</td><td>MP</td></tr>
<tr><td>>hg38_21_+_8433172_8446622_RNA45SN1_45S_50ntFlanks@12233.12250.18|CCGTGGAGCAGAAGGGCA</td><td>>hg38_21_+_8433172_8446622_RNA45SN1_45S_50ntFlanks</td><td>12233</td><td>12250</td><td>18</td><td>CCGTGGAGCAGAAGGGCA</td><td>194.2</td><td>-11.5</td><td>1.6</td><td>-7.3</td><td>3.9E-13</td><td>1.6E-12</td><td>MB231/MB468</td><td>MP</td></tr>
<tr><td>>MI0001445|hsa-mir-423&WithFlank&17|+|30117073|30117178@23.45.23&[MIMAT0004748&hsa-miR-423-5p&offsets|0|0;m-141&17|+|30117095|30117117&offsets|0|0]|TGAGGGGCAGAGAGCGAGACTTT</td><td>>MI0001445|hsa-mir-423&WithFlank&17|+|30117073|30117178</td><td>23</td><td>45</td><td>23</td><td>TGAGGGGCAGAGAGCGAGACTTT</td><td>3035.5</td><td>5.0</td><td>0.7</td><td>7.2</td><td>4.6E-13</td><td>1.6E-12</td><td>MB231/MB468</td><td>NUC</td></tr>
<tr><td>>hg38_21_+_8433172_8446622_RNA45SN1_45S_50ntFlanks@11316.11334.19|CACCCCACGTCTCGTCGCG</td><td>>hg38_21_+_8433172_8446622_RNA45SN1_45S_50ntFlanks</td><td>11316</td><td>11334</td><td>19</td><td>CACCCCACGTCTCGTCGCG</td><td>1866.7</td><td>-9.5</td><td>1.3</td><td>-7.2</td><td>5.7E-13</td><td>1.6E-12</td><td>MB231/MB468</td><td>MITO</td></tr>
<tr><td>>hg38_21_+_8433172_8446622_RNA45SN1_45S_50ntFlanks@11708.11728.21|AGCGCGGGTAAACGGCGGGAG</td><td>>hg38_21_+_8433172_8446622_RNA45SN1_45S_50ntFlanks</td><td>11708</td><td>11728</td><td>21</td><td>AGCGCGGGTAAACGGCGGGAG</td><td>114.0</td><td>-10.7</td><td>1.5</td><td>-7.3</td><td>3.9E-13</td><td>1.6E-12</td><td>MB231/MB468</td><td>MP</td></tr>
<tr><td>>MI0000734|hsa-mir-106b&WithFlank&7|-|100093987|100094080@58.76.19(+1C)&[MIMAT0004672&hsa-miR-106b-3p&offsets|0|-3(+1C);m-87&7|-|100094003|100094023&offsets|0|-2(+1C)]|CCGCACTGTGGGTACTTGCC</td><td>>MI0000734|hsa-mir-106b&WithFlank&7|-|100093987|100094080</td><td>58</td><td>76</td><td>19(+1C)</td><td>CCGCACTGTGGGTACTTGCC</td><td>362.2</td><td>9.7</td><td>1.3</td><td>7.2</td><td>4.5E-13</td><td>1.6E-12</td><td>BT20/MB468</td><td>CYTO</td></tr>
<tr><td>>MI0003560|hsa-mir-92b&WithFlank&1|+|155195171|155195278@67.88.22&[MIMAT0003218&hsa-miR-92b-3p&offsets|0|0;m-115&1|+|155195237|155195258&offsets|0|0]|TATTGCACTCGTCCCGGCCTCC</td><td>>MI0003560|hsa-mir-92b&WithFlank&1|+|155195171|155195278</td><td>67</td><td>88</td><td>22</td><td>TATTGCACTCGTCCCGGCCTCC</td><td>2579.8</td><td>6.4</td><td>0.9</td><td>7.2</td><td>5.7E-13</td><td>1.7E-12</td><td>MB231/MB468</td><td>MITO</td></tr>
<tr><td>>hg38_21_+_8433172_8446622_RNA45SN1_45S_50ntFlanks@10131.10153.23|CGGCGGCGGCGGCGGCGGGGGTG</td><td>>hg38_21_+_8433172_8446622_RNA45SN1_45S_50ntFlanks</td><td>10131</td><td>10153</td><td>23</td><td>CGGCGGCGGCGGCGGCGGGGGTG</td><td>174.5</td><td>-11.3</td><td>1.6</td><td>-7.3</td><td>3.9E-13</td><td>1.7E-12</td><td>MB231/MB468</td><td>MP</td></tr>
<tr><td>>MI0000101|hsa-mir-99a&WithFlank&21|+|16539083|16539175@19.38.20(+1C)&[MIMAT0000097&hsa-miR-99a-5p&offsets|0|-2(+1C);m-64&21|+|16539101|16539121&offsets|0|-1(+1C)]|AACCCGTAGATCCGATCTTGC</td><td>>MI0000101|hsa-mir-99a&WithFlank&21|+|16539083|16539175</td><td>19</td><td>38</td><td>20(+1C)</td><td>AACCCGTAGATCCGATCTTGC</td><td>392.9</td><td>-11.7</td><td>1.5</td><td>-7.7</td><td>2.0E-14</td><td>1.7E-12</td><td>BT20/MB468</td><td>MP</td></tr>
<tr><td>>hg38_21_+_8433172_8446622_RNA45SN1_45S_50ntFlanks@7985.8003.19|GATCAGACGTGGCGACCCG</td><td>>hg38_21_+_8433172_8446622_RNA45SN1_45S_50ntFlanks</td><td>7985</td><td>8003</td><td>19</td><td>GATCAGACGTGGCGACCCG</td><td>451.7</td><td>-11.9</td><td>1.6</td><td>-7.7</td><td>2.0E-14</td><td>1.7E-12</td><td>BT20/MB468</td><td>MP</td></tr>
<tr><td>>trna14_LysTTT_11_-_59327808_59327880@1.33.33,trna2_LysTTT_17_+_8022473_8022545@1.33.33,trna54_LysTTT_1_+_204475655_204475727@1.33.33,trna5_LysTTT_11_+_59323902_59323974@1.33.33,trna62_LysTTT_1_-_204476158_204476230@1.33.33,trna76_LysTTT_6_+_28918806_28918878@1.33.33|GCCCGGATAGCTCAGTCGGTAGAGCATCAGACT</td><td>>trna14_LysTTT_11_-_59327808_59327880</td><td>1</td><td>33</td><td>33</td><td>GCCCGGATAGCTCAGTCGGTAGAGCATCAGACT</td><td>10480.4</td><td>5.9</td><td>0.8</td><td>7.3</td><td>3.9E-13</td><td>1.7E-12</td><td>MB231/MB468</td><td>MP</td></tr>
<tr><td>>hg38_21_+_8433172_8446622_RNA45SN1_45S_50ntFlanks@5446.5467.22|CGGATCGGCCCCGCCGGGGTCG</td><td>>hg38_21_+_8433172_8446622_RNA45SN1_45S_50ntFlanks</td><td>5446</td><td>5467</td><td>22</td><td>CGGATCGGCCCCGCCGGGGTCG</td><td>331.9</td><td>-10.5</td><td>1.4</td><td>-7.3</td><td>3.5E-13</td><td>1.7E-12</td><td>BT20/MB468</td><td>MITO</td></tr>
<tr><td>>hg38_21_+_8433172_8446622_RNA45SN1_45S_50ntFlanks@12615.12652.38|GCTGAGGAGCCAATGGGGCGAAGCTACCATCTGTGGGA</td><td>>hg38_21_+_8433172_8446622_RNA45SN1_45S_50ntFlanks</td><td>12615</td><td>12652</td><td>38</td><td>GCTGAGGAGCCAATGGGGCGAAGCTACCATCTGTGGGA</td><td>238.7</td><td>-11.7</td><td>1.6</td><td>-7.2</td><td>5.7E-13</td><td>1.7E-12</td><td>MB231/MB468</td><td>MITO</td></tr>
<tr><td>>MI0000087|hsa-mir-29a&WithFlank&7|-|130876741|130876816@47.69.23&[MIMAT0000086&hsa-miR-29a-3p&offsets|-1|0;m-26&7|-|130876749|130876769&offsets|-1|+1]|CTAGCACCATCTGAAATCGGTTA</td><td>>MI0000087|hsa-mir-29a&WithFlank&7|-|130876741|130876816</td><td>47</td><td>69</td><td>23</td><td>CTAGCACCATCTGAAATCGGTTA</td><td>471.6</td><td>10.6</td><td>1.5</td><td>7.2</td><td>4.6E-13</td><td>1.7E-12</td><td>MB231/MB468</td><td>CYTO</td></tr>
<tr><td>>MI0000285|hsa-mir-205&WithFlank&1|+|209432127|209432248@40.60.21(+1U)&[MIMAT0000266&hsa-miR-205-5p&offsets|0|-1(+1U);m-351&1|+|209432166|209432187&offsets|0|-1(+1U)]|TCCTTCATTCCACCGGAGTCTT</td><td>>MI0000285|hsa-mir-205&WithFlank&1|+|209432127|209432248</td><td>40</td><td>60</td><td>21(+1U)</td><td>TCCTTCATTCCACCGGAGTCTT</td><td>194.0</td><td>-11.5</td><td>1.6</td><td>-7.3</td><td>4.0E-13</td><td>1.7E-12</td><td>MB231/MB468</td><td>MP</td></tr>
<tr><td>>hg38_21_+_8433172_8446622_RNA45SN1_45S_50ntFlanks@11564.11606.43|CCTAGCAGCCGACTTAGAACTGGTGCGGACCAGGGGAATCCGA</td><td>>hg38_21_+_8433172_8446622_RNA45SN1_45S_50ntFlanks</td><td>11564</td><td>11606</td><td>43</td><td>CCTAGCAGCCGACTTAGAACTGGTGCGGACCAGGGGAATCCGA</td><td>275.5</td><td>-11.9</td><td>1.7</td><td>-7.2</td><td>5.8E-13</td><td>1.7E-12</td><td>MB231/MB468</td><td>MITO</td></tr>
<tr><td>>hg38_21_+_8433172_8446622_RNA45SN1_45S_50ntFlanks@12457.12476.20|CAAGCGTTGGATTGTTCACC</td><td>>hg38_21_+_8433172_8446622_RNA45SN1_45S_50ntFlanks</td><td>12457</td><td>12476</td><td>20</td><td>CAAGCGTTGGATTGTTCACC</td><td>4758.1</td><td>-5.5</td><td>0.8</td><td>-7.2</td><td>5.8E-13</td><td>1.7E-12</td><td>MB231/MB468</td><td>MITO</td></tr>
<tr><td>>hg38_21_+_8433172_8446622_RNA45SN1_45S_50ntFlanks@3776.3798.23|CCGGTACAGTGAAACTGCGAATG</td><td>>hg38_21_+_8433172_8446622_RNA45SN1_45S_50ntFlanks</td><td>3776</td><td>3798</td><td>23</td><td>CCGGTACAGTGAAACTGCGAATG</td><td>158.1</td><td>-11.2</td><td>1.5</td><td>-7.3</td><td>4.0E-13</td><td>1.7E-12</td><td>MB231/MB468</td><td>MP</td></tr>
<tr><td>>hg38_21_+_8433172_8446622_RNA45SN1_45S_50ntFlanks@10845.10867.23|ACTTCGGGATAAGGATTGGCTCT</td><td>>hg38_21_+_8433172_8446622_RNA45SN1_45S_50ntFlanks</td><td>10845</td><td>10867</td><td>23</td><td>ACTTCGGGATAAGGATTGGCTCT</td><td>589.4</td><td>-4.6</td><td>0.6</td><td>-7.2</td><td>4.7E-13</td><td>1.7E-12</td><td>MB231/MB468</td><td>NUC</td></tr>
<tr><td>>hg38_21_+_8433172_8446622_RNA45SN1_45S_50ntFlanks@11570.11600.31|AGCCGACTTAGAACTGGTGCGGACCAGGGGA</td><td>>hg38_21_+_8433172_8446622_RNA45SN1_45S_50ntFlanks</td><td>11570</td><td>11600</td><td>31</td><td>AGCCGACTTAGAACTGGTGCGGACCAGGGGA</td><td>1187.0</td><td>-4.2</td><td>0.6</td><td>-7.2</td><td>4.7E-13</td><td>1.7E-12</td><td>MB231/MB468</td><td>NUC</td></tr>
<tr><td>>hg38_21_+_8433172_8446622_RNA45SN1_45S_50ntFlanks@7975.8002.28|CGCGACCTCAGATCAGACGTGGCGACCC</td><td>>hg38_21_+_8433172_8446622_RNA45SN1_45S_50ntFlanks</td><td>7975</td><td>8002</td><td>28</td><td>CGCGACCTCAGATCAGACGTGGCGACCC</td><td>15193.0</td><td>6.4</td><td>0.9</td><td>7.3</td><td>3.5E-13</td><td>1.7E-12</td><td>BT20/MB468</td><td>MITO</td></tr>
<tr><td>>hg38_21_+_8433172_8446622_RNA45SN1_45S_50ntFlanks@10922.10941.20|CTGGACGAGGCGCCGCCGCC</td><td>>hg38_21_+_8433172_8446622_RNA45SN1_45S_50ntFlanks</td><td>10922</td><td>10941</td><td>20</td><td>CTGGACGAGGCGCCGCCGCC</td><td>287.2</td><td>-12.0</td><td>1.7</td><td>-7.3</td><td>4.0E-13</td><td>1.7E-12</td><td>MB231/MB468</td><td>MP</td></tr>
<tr><td>>hg38_21_+_8433172_8446622_RNA45SN1_45S_50ntFlanks@9740.9759.20|AACGATCTCAACCTATTCTC</td><td>>hg38_21_+_8433172_8446622_RNA45SN1_45S_50ntFlanks</td><td>9740</td><td>9759</td><td>20</td><td>AACGATCTCAACCTATTCTC</td><td>116.0</td><td>-10.7</td><td>1.5</td><td>-7.3</td><td>4.0E-13</td><td>1.7E-12</td><td>MB231/MB468</td><td>MP</td></tr>
<tr><td>>hg38_21_+_8433172_8446622_RNA45SN1_45S_50ntFlanks@10681.10710.30|TTCCGGCGGCGTCCGGTGAGCTCTCGCTGG</td><td>>hg38_21_+_8433172_8446622_RNA45SN1_45S_50ntFlanks</td><td>10681</td><td>10710</td><td>30</td><td>TTCCGGCGGCGTCCGGTGAGCTCTCGCTGG</td><td>1235.3</td><td>-9.4</td><td>1.3</td><td>-7.2</td><td>4.7E-13</td><td>1.7E-12</td><td>MB231/MB468</td><td>NUC</td></tr>
<tr><td>>hg38_21_+_8433172_8446622_RNA45SN1_45S_50ntFlanks@6780.6806.27|CGGGGCTACGCCTGTCTGAGCGTCGCT</td><td>>hg38_21_+_8433172_8446622_RNA45SN1_45S_50ntFlanks</td><td>6780</td><td>6806</td><td>27</td><td>CGGGGCTACGCCTGTCTGAGCGTCGCT</td><td>303.9</td><td>4.9</td><td>0.7</td><td>7.2</td><td>4.6E-13</td><td>1.7E-12</td><td>MB231/MB468</td><td>CYTO</td></tr>
<tr><td>>MI0000342|hsa-mir-200b&WithFlank&1|+|1167098|1167204@63.84.22&[MIMAT0000318&hsa-miR-200b-3p&offsets|0|0;m-93&1|+|1167160|1167182&offsets|0|-1]|TAATACTGCCTGGTAATGATGA</td><td>>MI0000342|hsa-mir-200b&WithFlank&1|+|1167098|1167204</td><td>63</td><td>84</td><td>22</td><td>TAATACTGCCTGGTAATGATGA</td><td>3830.3</td><td>-5.7</td><td>0.8</td><td>-7.3</td><td>4.0E-13</td><td>1.7E-12</td><td>MB231/MB468</td><td>MP</td></tr>
<tr><td>>hg38_21_+_8433172_8446622_RNA45SN1_45S_50ntFlanks@12675.12708.34|CAGAATCCCGCCCAGGCGGAACGATACGGCAGCG</td><td>>hg38_21_+_8433172_8446622_RNA45SN1_45S_50ntFlanks</td><td>12675</td><td>12708</td><td>34</td><td>CAGAATCCCGCCCAGGCGGAACGATACGGCAGCG</td><td>109.8</td><td>-10.6</td><td>1.5</td><td>-7.3</td><td>4.0E-13</td><td>1.7E-12</td><td>MB231/MB468</td><td>MP</td></tr>
<tr><td>>hg38_21_+_8433172_8446622_RNA45SN1_45S_50ntFlanks@3984.4005.22|GCGGCTTTGGTGACTCTAGATA</td><td>>hg38_21_+_8433172_8446622_RNA45SN1_45S_50ntFlanks</td><td>3984</td><td>4005</td><td>22</td><td>GCGGCTTTGGTGACTCTAGATA</td><td>714.9</td><td>-9.2</td><td>1.3</td><td>-7.2</td><td>4.7E-13</td><td>1.7E-12</td><td>MB231/MB468</td><td>NUC</td></tr>
<tr><td>>hg38_21_+_8433172_8446622_RNA45SN1_45S_50ntFlanks@12006.12033.28|GGCCGCCGGTGAAATACCACTACTCTGA</td><td>>hg38_21_+_8433172_8446622_RNA45SN1_45S_50ntFlanks</td><td>12006</td><td>12033</td><td>28</td><td>GGCCGCCGGTGAAATACCACTACTCTGA</td><td>845.3</td><td>6.1</td><td>0.8</td><td>7.3</td><td>3.5E-13</td><td>1.7E-12</td><td>BT20/MB468</td><td>MITO</td></tr>
<tr><td>>hg38_21_+_8433172_8446622_RNA45SN1_45S_50ntFlanks@4728.4759.32|ATCAAGAACGAAAGTCGGAGGTTCGAAGACGA</td><td>>hg38_21_+_8433172_8446622_RNA45SN1_45S_50ntFlanks</td><td>4728</td><td>4759</td><td>32</td><td>ATCAAGAACGAAAGTCGGAGGTTCGAAGACGA</td><td>192.1</td><td>-9.7</td><td>1.3</td><td>-7.3</td><td>3.5E-13</td><td>1.7E-12</td><td>BT20/MB468</td><td>MITO</td></tr>
<tr><td>>hg38_1_-_228634819_228635039_RNA5S12_5S_50ntFlanks@84.103.20|CTCGTCTGATCTCGGAAGCT</td><td>>hg38_1_-_228634819_228635039_RNA5S12_5S_50ntFlanks</td><td>84</td><td>103</td><td>20</td><td>CTCGTCTGATCTCGGAAGCT</td><td>2382.7</td><td>-10.4</td><td>1.4</td><td>-7.2</td><td>6.5E-13</td><td>1.7E-12</td><td>BT20/MB468</td><td>NUC</td></tr>
<tr><td>>hg38_21_+_8433172_8446622_RNA45SN1_45S_50ntFlanks@10729.10752.24|AGAGGGTGTAAATCTCGCGCCGGG</td><td>>hg38_21_+_8433172_8446622_RNA45SN1_45S_50ntFlanks</td><td>10729</td><td>10752</td><td>24</td><td>AGAGGGTGTAAATCTCGCGCCGGG</td><td>207.0</td><td>-6.7</td><td>0.9</td><td>-7.2</td><td>4.7E-13</td><td>1.7E-12</td><td>MB231/MB468</td><td>CYTO</td></tr>
<tr><td>>hg38_21_+_8433172_8446622_RNA45SN1_45S_50ntFlanks@12182.12211.30|CAAACGGTAACGCAGGTGTCCTAAGGCGAG</td><td>>hg38_21_+_8433172_8446622_RNA45SN1_45S_50ntFlanks</td><td>12182</td><td>12211</td><td>30</td><td>CAAACGGTAACGCAGGTGTCCTAAGGCGAG</td><td>152.9</td><td>-11.1</td><td>1.5</td><td>-7.3</td><td>4.0E-13</td><td>1.7E-12</td><td>MB231/MB468</td><td>MP</td></tr>
<tr><td>>hg38_21_+_8433172_8446622_RNA45SN1_45S_50ntFlanks@12476.12499.24|CCACTAATAGGGAACGTGAGCTGG</td><td>>hg38_21_+_8433172_8446622_RNA45SN1_45S_50ntFlanks</td><td>12476</td><td>12499</td><td>24</td><td>CCACTAATAGGGAACGTGAGCTGG</td><td>382.6</td><td>-8.4</td><td>1.2</td><td>-7.2</td><td>4.8E-13</td><td>1.7E-12</td><td>MB231/MB468</td><td>NUC</td></tr>
<tr><td>>am_trna128_GlyGCC_6_-_27870686_27870756@1.31.31,trna133_GlyCCC_1_-_16872434_16872504@1.31.31,trna18_GlyGCC_16_+_70822597_70822667@1.31.31,trna19_GlyGCC_16_+_70823410_70823480@1.31.31,trna19_GlyGCC_2_-_157257659_157257729@1.31.31,trna24_GlyGCC_16_-_70812942_70813012@1.31.31,trna25_GlyGCC_16_-_70812114_70812184@1.31.31,trna4_GlyCCC_1_+_17188416_17188486@1.31.31,trna5_GlyGCC_17_+_8029064_8029134@1.31.31,trna68_GlyGCC_1_-_161493637_161493707@1.31.31|GCATTGGTGGTTCAGTGGTAGAATTCTCGCC</td><td>>am_trna128_GlyGCC_6_-_27870686_27870756</td><td>1</td><td>31</td><td>31</td><td>GCATTGGTGGTTCAGTGGTAGAATTCTCGCC</td><td>95685.0</td><td>5.9</td><td>0.8</td><td>7.2</td><td>4.7E-13</td><td>1.7E-12</td><td>MB231/MB468</td><td>CYTO</td></tr>
<tr><td>>hg38_21_+_8433172_8446622_RNA45SN1_45S_50ntFlanks@8380.8401.22|CAAGAGGGCGTGAAACCGTTAA</td><td>>hg38_21_+_8433172_8446622_RNA45SN1_45S_50ntFlanks</td><td>8380</td><td>8401</td><td>22</td><td>CAAGAGGGCGTGAAACCGTTAA</td><td>335.7</td><td>-10.5</td><td>1.4</td><td>-7.3</td><td>3.6E-13</td><td>1.7E-12</td><td>BT20/MB468</td><td>MITO</td></tr>
<tr><td>>MI0000085|hsa-mir-27a&WithFlank&19|-|13836434|13836523@16.37.22&[MIMAT0004501&hsa-miR-27a-5p&offsets|0|0;m-270&19|-|13836487|13836508&offsets|0|0]|AGGGCTTAGCTGCTTGTGAGCA</td><td>>MI0000085|hsa-mir-27a&WithFlank&19|-|13836434|13836523</td><td>16</td><td>37</td><td>22</td><td>AGGGCTTAGCTGCTTGTGAGCA</td><td>446.2</td><td>6.2</td><td>0.9</td><td>7.2</td><td>4.8E-13</td><td>1.7E-12</td><td>MB231/MB468</td><td>NUC</td></tr>
<tr><td>>hg38_21_+_8433172_8446622_RNA45SN1_45S_50ntFlanks@9339.9374.36|CGCACGAAAGCCGCCGTGGCGCAATGAAGGTGAAGG</td><td>>hg38_21_+_8433172_8446622_RNA45SN1_45S_50ntFlanks</td><td>9339</td><td>9374</td><td>36</td><td>CGCACGAAAGCCGCCGTGGCGCAATGAAGGTGAAGG</td><td>183.5</td><td>-11.4</td><td>1.6</td><td>-7.3</td><td>4.1E-13</td><td>1.7E-12</td><td>MB231/MB468</td><td>MP</td></tr>
</table>

| | | | | | | | | | | | | | |
|---|---|---|---|---|---|---|---|---|---|---|---|---|---|
| >hg38_21_+_8433172_8446622_RNA45SN1_45S_50ntFlanks@10683.10708.26\|CCGGCGGCGTCCGGTGAGCTCTCGCT | >hg38_21_+_8433172_8446622_RNA45SN1_45S_50ntFlanks | 10683 | 10708 | 26 | CCGGCGGCGTCCGGTGAGCTCTCGCT | 196.3 | -5.9 | 0.8 | -7.2 | 4.7E-13 | 1.7E-12 | MB231/MB468 | CYTO |
| >hg38_21_+_8433172_8446622_RNA45SN1_45S_50ntFlanks@12203.12220.18\|TAAGGCGAGCTCAGGGAG | >hg38_21_+_8433172_8446622_RNA45SN1_45S_50ntFlanks | 12203 | 12220 | 18 | TAAGGCGAGCTCAGGGAG | 524.1 | -12.8 | 1.8 | -7.2 | 5.9E-13 | 1.7E-12 | MB231/MB468 | MITO |
| >hg38_21_+_8433172_8446622_RNA45SN1_45S_50ntFlanks@4734.4759.26\|AACGAAAGTCGGAGGTTCGAAGACGA | >hg38_21_+_8433172_8446622_RNA45SN1_45S_50ntFlanks | 4734 | 4759 | 26 | AACGAAAGTCGGAGGTTCGAAGACGA | 929.5 | -3.7 | 0.5 | -7.2 | 4.8E-13 | 1.7E-12 | MB231/MB468 | NUC |
| >hg38_21_+_8433172_8446622_RNA45SN1_45S_50ntFlanks@9738.9759.22\|GAAACGATCTCAACCTATTCTC | >hg38_21_+_8433172_8446622_RNA45SN1_45S_50ntFlanks | 9738 | 9759 | 22 | GAAACGATCTCAACCTATTCTC | 816.9 | -4.5 | 0.6 | -7.2 | 4.8E-13 | 1.7E-12 | MB231/MB468 | NUC |
| >hg38_21_+_8433172_8446622_RNA45SN1_45S_50ntFlanks@10402.10433.32\|AACATGGGTCAGTCGGTCCTGAGAGATGGGCG | >hg38_21_+_8433172_8446622_RNA45SN1_45S_50ntFlanks | 10402 | 10433 | 32 | AACATGGGTCAGTCGGTCCTGAGAGATGGGCG | 2240.2 | -10.7 | 1.5 | -7.2 | 6.6E-13 | 1.7E-12 | BT20/MB468 | NUC |
| >hg38_21_+_8433172_8446622_RNA45SN1_45S_50ntFlanks@12717.12753.37\|CCTCGGTTGGCCTCGGATAGCCGGTCCCCCGCCTGTC | >hg38_21_+_8433172_8446622_RNA45SN1_45S_50ntFlanks | 12717 | 12753 | 37 | CCTCGGTTGGCCTCGGATAGCCGGTCCCCCGCCTGTC | 346.5 | -8.4 | 1.2 | -7.2 | 6.0E-13 | 1.7E-12 | MB231/MB468 | MITO |
| >hg38_21_+_8433172_8446622_RNA45SN1_45S_50ntFlanks@8182.8223.42\|ATCGAGGCCCAGCCCGTGGACGGTGTGAGGCCGGTAGCGGCC | >hg38_21_+_8433172_8446622_RNA45SN1_45S_50ntFlanks | 8182 | 8223 | 42 | ATCGAGGCCCAGCCCGTGGACGGTGTGAGGCCGGTAGCGGCC | 312.5 | -10.6 | 1.5 | -7.2 | 4.7E-13 | 1.7E-12 | BT20/MB468 | CYTO |
| >hg38_21_+_8433172_8446622_RNA45SN1_45S_50ntFlanks@6721.6739.19\|AATTGCAGGACACATTGAT | >hg38_21_+_8433172_8446622_RNA45SN1_45S_50ntFlanks | 6721 | 6739 | 19 | AATTGCAGGACACATTGAT | 1452.9 | -9.2 | 1.3 | -7.2 | 4.7E-13 | 1.7E-12 | BT20/MB468 | CYTO |
| >hg38_21_+_8433172_8446622_RNA45SN1_45S_50ntFlanks@12437.12454.18\|CATTGTGAAGCAGAATTC | >hg38_21_+_8433172_8446622_RNA45SN1_45S_50ntFlanks | 12437 | 12454 | 18 | CATTGTGAAGCAGAATTC | 1012.0 | -7.2 | 1.0 | -7.2 | 6.0E-13 | 1.7E-12 | MB231/MB468 | MITO |
| >hg38_21_+_8433172_8446622_RNA45SN1_45S_50ntFlanks@12709.12743.35\|CCGCGGAGCCTCGGTTGGCCTCGGATAGCCGGTCC | >hg38_21_+_8433172_8446622_RNA45SN1_45S_50ntFlanks | 12709 | 12743 | 35 | CCGCGGAGCCTCGGTTGGCCTCGGATAGCCGGTCC | 116.5 | -10.7 | 1.5 | -7.3 | 4.1E-13 | 1.7E-12 | MB231/MB468 | MP |
| >hg38_21_+_8433172_8446622_RNA45SN1_45S_50ntFlanks@5487.5514.28\|CGCTGAGAAGACGGTCGAACTTGACTAT | >hg38_21_+_8433172_8446622_RNA45SN1_45S_50ntFlanks | 5487 | 5514 | 28 | CGCTGAGAAGACGGTCGAACTTGACTAT | 2341.0 | -8.7 | 1.2 | -7.2 | 6.7E-13 | 1.7E-12 | BT20/MB468 | NUC |
| >hg38_21_+_8433172_8446622_RNA45SN1_45S_50ntFlanks@12712.12742.31\|CGGAGCCTCGGTTGGCCTCGGATAGCCGGTC | >hg38_21_+_8433172_8446622_RNA45SN1_45S_50ntFlanks | 12712 | 12742 | 31 | CGGAGCCTCGGTTGGCCTCGGATAGCCGGTC | 357.4 | -10.6 | 1.5 | -7.3 | 3.6E-13 | 1.7E-12 | BT20/MB468 | MITO |
| >hg38_21_+_8433172_8446622_RNA45SN1_45S_50ntFlanks@9027.9068.42\|CGTCTCCTCGTGGGGGGGCCGGGCCACCCCTCCCACGGCGCG | >hg38_21_+_8433172_8446622_RNA45SN1_45S_50ntFlanks | 9027 | 9068 | 42 | CGTCTCCTCGTGGGGGGGCCGGGCCACCCCTCCCACGGCGCG | 222.2 | -11.6 | 1.6 | -7.2 | 6.0E-13 | 1.7E-12 | MB231/MB468 | MITO |
| >hg38_21_+_8433172_8446622_RNA45SN1_45S_50ntFlanks@11918.11952.35\|ACATGAGAGGTGTAGAATAAGTGGGAGGCCCCCGG | >hg38_21_+_8433172_8446622_RNA45SN1_45S_50ntFlanks | 11918 | 11952 | 35 | ACATGAGAGGTGTAGAATAAGTGGGAGGCCCCCGG | 176.2 | -11.3 | 1.6 | -7.3 | 4.1E-13 | 1.7E-12 | MB231/MB468 | MP |
| >hg38_21_+_8433172_8446622_RNA45SN1_45S_50ntFlanks@11576.11599.24\|CTTAGAACTGGTGCGGACCAGGGG | >hg38_21_+_8433172_8446622_RNA45SN1_45S_50ntFlanks | 11576 | 11599 | 24 | CTTAGAACTGGTGCGGACCAGGGG | 174.8 | -10.2 | 1.4 | -7.2 | 4.8E-13 | 1.7E-12 | MB231/MB468 | CYTO |
| >hg38_21_+_8433172_8446622_RNA45SN1_45S_50ntFlanks@10687.10709.23\|CGGCGTCCGGTGAGCTCTCGCTG | >hg38_21_+_8433172_8446622_RNA45SN1_45S_50ntFlanks | 10687 | 10709 | 23 | CGGCGTCCGGTGAGCTCTCGCTG | 5026.2 | -8.6 | 1.2 | -7.2 | 6.0E-13 | 1.7E-12 | MB231/MB468 | MITO |
| >hg38_21_+_8433172_8446622_RNA45SN1_45S_50ntFlanks@4739.4758.20\|AAGTCGGAGGTTCGAAGACG | >hg38_21_+_8433172_8446622_RNA45SN1_45S_50ntFlanks | 4739 | 4758 | 20 | AAGTCGGAGGTTCGAAGACG | 1148.1 | -4.4 | 0.6 | -7.3 | 4.2E-13 | 1.7E-12 | MB231/MB468 | MP |
| >MI0000446\|hsa-mir-125b-1&WithFlank&11\|-\|122099751\|122099850@22.42.21;MI0000470\|hsa-mir-125b-2&WithFlank&21\|+\|16590231\|16590331@24.44.21&[MIMAT0000423&hsa-miR-125b-5p&offsets\|+1\|0;m-67&11\|-\|122099809\|122099830&offsets\|+1\|0];[MIMAT0000423_1&hsa-miR-125b-5p&offsets\|+1\|0;m-68&21\|+\|16590253\|16590274&offsets\|+1\|0]\|CCCTGAGACCCTAACTTGTGA | >MI0000446\|hsa-mir-125b-1&WithFlank&11\|-\|122099751\|122099850 | 22 | 42 | 21 | CCCTGAGACCCTAACTTGTGA | 237.2 | -11.0 | 1.5 | -7.5 | 5.5E-14 | 1.7E-12 | BT20/MB231 | NUC |
| >hg38_21_+_8433172_8446622_RNA45SN1_45S_50ntFlanks@10876.10902.27\|GGTCGGTCGGGCTGGGGCGCGAAGCGG | >hg38_21_+_8433172_8446622_RNA45SN1_45S_50ntFlanks | 10876 | 10902 | 27 | GGTCGGTCGGGCTGGGGCGCGAAGCGG | 185.9 | -11.4 | 1.6 | -7.3 | 4.2E-13 | 1.7E-12 | MB231/MB468 | MP |
| >hg38_21_+_8433172_8446622_RNA45SN1_45S_50ntFlanks@12419.12458.40\|GATGTCGGCTCTTCCTATCATTGTGAAGCAGAATTCACCA | >hg38_21_+_8433172_8446622_RNA45SN1_45S_50ntFlanks | 12419 | 12458 | 40 | GATGTCGGCTCTTCCTATCATTGTGAAGCAGAATTCACCA | 1332.3 | 7.3 | 1.0 | 7.2 | 4.8E-13 | 1.7E-12 | MB231/MB468 | CYTO |
| >hg38_21_+_8433172_8446622_RNA45SN1_45S_50ntFlanks@4730.4765.36\|CAAGAACGAAAGTCGGAGGTTCGAAGACGATCAGAT | >hg38_21_+_8433172_8446622_RNA45SN1_45S_50ntFlanks | 4730 | 4765 | 36 | CAAGAACGAAAGTCGGAGGTTCGAAGACGATCAGAT | 1432.3 | 8.0 | 1.1 | 7.2 | 4.8E-13 | 1.7E-12 | BT20/MB468 | CYTO |
| >MI0003560\|hsa-mir-92b&WithFlank&1\|+\|155195171\|155195278@67.88.22&[MIMAT0003218&hsa-miR-92b-3p&offsets\|0\|0;m-115&1\|+\|155195237\|155195258&offsets\|0\|0]\|TATTGCACTCGTCCCGGCCTCC | >MI0003560\|hsa-mir-92b&WithFlank&1\|+\|155195171\|155195278 | 67 | 88 | 22 | TATTGCACTCGTCCCGGCCTCC | 2753.2 | 5.1 | 0.7 | 7.2 | 4.9E-13 | 1.7E-12 | MB231/MB468 | NUC |
| >MI0000078\|hsa-mir-22&WithFlank&17\|-\|1713897\|1713993@21.41.21&[MIMAT0004495&hsa-miR-22-5p&offsets\|0\|-1;m-217&17\|-\|1713953\|1713973&offsets\|0\|0]\|AGTTCTTCAGTGGCAAGCTTT | >MI0000078\|hsa-mir-22&WithFlank&17\|-\|1713897\|1713993 | 21 | 41 | 21 | AGTTCTTCAGTGGCAAGCTTT | 841.6 | 9.6 | 1.3 | 7.2 | 4.8E-13 | 1.7E-12 | BT20/MB468 | CYTO |
| >hg38_21_+_8433172_8446622_RNA45SN1_45S_50ntFlanks@6788.6806.19\|CGCCTGTCTGAGCGTCGCT | >hg38_21_+_8433172_8446622_RNA45SN1_45S_50ntFlanks | 6788 | 6806 | 19 | CGCCTGTCTGAGCGTCGCT | 11557.6 | 4.8 | 0.7 | 7.3 | 3.7E-13 | 1.8E-12 | BT20/MB468 | MITO |
| >MI0000255\|hsa-mir-30d&WithFlank&8\|-\|134804870\|134804951@12.32.21&[MIMAT0000245&hsa-miR-30d-5p&offsets\|0\|-1;m-21&8\|-\|134804919\|134804940&offsets\|0\|-1]\|TGTAAACATCCCCGACTGGAA | >MI0000255\|hsa-mir-30d&WithFlank&8\|-\|134804870\|134804951 | 12 | 32 | 21 | TGTAAACATCCCCGACTGGAA | 368.1 | 5.5 | 0.8 | 7.2 | 5.0E-13 | 1.8E-12 | MB231/MB468 | NUC |
| >hg38_21_+_8433172_8446622_RNA45SN1_45S_50ntFlanks@11729.11748.20\|TAACTATGACTCTCTTAAGG | >hg38_21_+_8433172_8446622_RNA45SN1_45S_50ntFlanks | 11729 | 11748 | 20 | TAACTATGACTCTCTTAAGG | 867.6 | -9.2 | 1.3 | -7.2 | 4.2E-13 | 1.8E-12 | MB231/MB468 | MP |
| >hg38_21_+_8433172_8446622_RNA45SN1_45S_50ntFlanks@11453.11477.25\|TCCTCGGCGCGCGGCGGCGGCGGCG | >hg38_21_+_8433172_8446622_RNA45SN1_45S_50ntFlanks | 11453 | 11477 | 25 | TCCTCGGCGCGCGGCGGCGGCGGCG | 230.9 | -7.9 | 1.1 | -7.2 | 4.8E-13 | 1.8E-12 | BT20/MB468 | CYTO |
| >MI0000483\|hsa-mir-186&WithFlank&1\|-\|71067625\|71067722@21.42.22&[MIMAT0000456&hsa-miR-186-5p&offsets\|0\|0;m-84&1\|-\|71067681\|71067702&offsets\|0\|0]\|CAAAGAATTCTCCTTTTGGGCT | >MI0000483\|hsa-mir-186&WithFlank&1\|-\|71067625\|71067722 | 21 | 42 | 22 | CAAAGAATTCTCCTTTTGGGCT | 1022.5 | 5.2 | 0.7 | 7.2 | 5.0E-13 | 1.8E-12 | MB231/MB468 | NUC |
| >hg38_21_+_8433172_8446622_RNA45SN1_45S_50ntFlanks@10688.10706.19\|GGCGTCCGGTGAGCTCTCG | >hg38_21_+_8433172_8446622_RNA45SN1_45S_50ntFlanks | 10688 | 10706 | 19 | GGCGTCCGGTGAGCTCTCG | 160.7 | -11.2 | 1.5 | -7.2 | 4.2E-13 | 1.8E-12 | MB231/MB468 | MP |
| >hg38_21_+_8433172_8446622_RNA45SN1_45S_50ntFlanks@9554.9574.21\|CGTAGCGGTCCTGACGTGCAA | >hg38_21_+_8433172_8446622_RNA45SN1_45S_50ntFlanks | 9554 | 9574 | 21 | CGTAGCGGTCCTGACGTGCAA | 1953.2 | -5.1 | 0.7 | -7.2 | 6.2E-13 | 1.8E-12 | MB231/MB468 | MITO |
| >am_trna128_GlyGCC_6_-_27870686_27870756@1.28.28,trna133_GlyCCC_1_-_16872434_16872504@1.28.28,trna18_GlyGCC_16_+_70822597_70822667@1.28.28,trna19_GlyGCC_16_+_70823410_70823480@1.28.28,trna19_GlyGCC_2_-_157257659_157257729@1.28.28,trna24_GlyGCC_16_-_70812942_70813012@1.28.28,trna25_GlyGCC_16_-_70812114_70812184@1.28.28,trna4_GlyCCC_1_+_17188416_17188486@1.28.28,trna5_GlyGCC_17_+_8029064_8029134@1.28.28,trna68_GlyGCC_1_-_161493637_161493707@1.28.28\|GCATTGGTGGTTCAGTGGTAGAATTCTC | >am_trna128_GlyGCC_6_-_27870686_27870756 | 1 | 28 | 28 | GCATTGGTGGTTCAGTGGTAGAATTCTC | 30097.0 | -5.3 | 0.7 | -7.2 | 4.2E-13 | 1.8E-12 | MB231/MB468 | MP |
| >hg38_21_+_8433172_8446622_RNA45SN1_45S_50ntFlanks@4027.4046.20\|CCCCGTGGCGGCGACGACCC | >hg38_21_+_8433172_8446622_RNA45SN1_45S_50ntFlanks | 4027 | 4046 | 20 | CCCCGTGGCGGCGACGACCC | 320.6 | -12.1 | 1.7 | -7.2 | 6.2E-13 | 1.8E-12 | MB231/MB468 | MITO |
| >hg38_21_+_8433172_8446622_RNA45SN1_45S_50ntFlanks@7985.8009.25\|GATCAGACGTGGCGACCCGCTGAAT | >hg38_21_+_8433172_8446622_RNA45SN1_45S_50ntFlanks | 7985 | 8009 | 25 | GATCAGACGTGGCGACCCGCTGAAT | 470.2 | -12.0 | 1.6 | -7.6 | 2.1E-14 | 1.8E-12 | BT20/MB468 | MP |
| >hg38_21_+_8433172_8446622_RNA45SN1_45S_50ntFlanks@11661.11681.21\|TGATTTCTGCCCAGTGCTCTG | >hg38_21_+_8433172_8446622_RNA45SN1_45S_50ntFlanks | 11661 | 11681 | 21 | TGATTTCTGCCCAGTGCTCTG | 15628.7 | -9.8 | 1.4 | -7.2 | 6.9E-13 | 1.8E-12 | BT20/MB468 | NUC |
| >hg38_21_+_8433172_8446622_RNA45SN1_45S_50ntFlanks@12303.12323.21\|CCTCACGATCCTTCTGACCTT | >hg38_21_+_8433172_8446622_RNA45SN1_45S_50ntFlanks | 12303 | 12323 | 21 | CCTCACGATCCTTCTGACCTT | 548.6 | -12.9 | 1.8 | -7.2 | 6.2E-13 | 1.8E-12 | MB231/MB468 | MITO |
| >hg38_21_+_8433172_8446622_RNA45SN1_45S_50ntFlanks@11200.11220.21\|CGGCGGCGGCGGCGACTCTGG | >hg38_21_+_8433172_8446622_RNA45SN1_45S_50ntFlanks | 11200 | 11220 | 21 | CGGCGGCGGCGGCGACTCTGG | 161.3 | -11.2 | 1.5 | -7.2 | 4.3E-13 | 1.8E-12 | MB231/MB468 | MP |
| >hg38_21_+_8433172_8446622_RNA45SN1_45S_50ntFlanks@3984.4002.19\|GCGGCTTTGGTGACTCTAG | >hg38_21_+_8433172_8446622_RNA45SN1_45S_50ntFlanks | 3984 | 4002 | 19 | GCGGCTTTGGTGACTCTAG | 4281.2 | -10.3 | 1.4 | -7.2 | 5.0E-13 | 1.8E-12 | MB231/MB468 | NUC |
| >hg38_21_+_8433172_8446622_RNA45SN1_45S_50ntFlanks@10116.10134.19\|TGCGTCGGAGGGCGGCGGC | >hg38_21_+_8433172_8446622_RNA45SN1_45S_50ntFlanks | 10116 | 10134 | 19 | TGCGTCGGAGGGCGGCGGC | 254.0 | -11.9 | 1.6 | -7.2 | 4.3E-13 | 1.8E-12 | MB231/MB468 | MP |
| >MI0000462\|hsa-mir-152&WithFlank&17\|-\|48037155\|48037253@60.80.21&[MIMAT0000438&hsa-miR-152-3p&offsets\|0\|0;m-97&17\|-\|48037174\|48037194&offsets\|0\|0]\|TCAGTGCATGACAGAACTTGG | >MI0000462\|hsa-mir-152&WithFlank&17\|-\|48037155\|48037253 | 60 | 80 | 21 | TCAGTGCATGACAGAACTTGG | 544.6 | 12.3 | 1.7 | 7.2 | 6.3E-13 | 1.8E-12 | MB231/MB468 | MITO |
| >trna111_HisGTG_1_-_147774845_147774916@-1G.32.33,trna118_HisGTG_1_-_145396881_145396952@-1G.32.33,trna16_HisGTG_1_+_146544773_146544844@-1G.32.33,trna1_HisGTG_15_+_45493349_45493420@-1G.32.33,trna21_HisGTG_1_+_147753471_147753542@-1G.32.33,trna33_HisGTG_6_+_27125906_27125977@-1G.32.33,trna7_HisGTG_9_-_14433938_14434009@-1G.32.33,trna8_HisGTG_15_-_45492611_45492682@-1G.32.33,trna9_HisGTG_15_-_45490804_45490875@-1G.32.33\|GGCCGTGATCGTATAGTGGTTAGTACTCTGCGT | >trna111_HisGTG_1_-_147774845_147774916 | -1G | 32 | 33 | GGCCGTGATCGTATAGTGGTTAGTACTCTGCGT | 3199.7 | 8.6 | 1.2 | 7.2 | 5.1E-13 | 1.8E-12 | MB231/MB468 | NUC |
| >hg38_21_+_8433172_8446622_RNA45SN1_45S_50ntFlanks@8839.8861.23\|CGGTGGCGGCGGCGGCGGCGGCG | >hg38_21_+_8433172_8446622_RNA45SN1_45S_50ntFlanks | 8839 | 8861 | 23 | CGGTGGCGGCGGCGGCGGCGGCG | 232.7 | -11.7 | 1.6 | -7.2 | 4.3E-13 | 1.8E-12 | MB231/MB468 | MP |
| >hg38_21_+_8433172_8446622_RNA45SN1_45S_50ntFlanks@11200.11233.34\|CGGCGGCGGCGGCGACTCTGGACGCGAGCCGGGC | >hg38_21_+_8433172_8446622_RNA45SN1_45S_50ntFlanks | 11200 | 11233 | 34 | CGGCGGCGGCGGCGACTCTGGACGCGAGCCGGGC | 182.4 | -11.4 | 1.6 | -7.2 | 4.3E-13 | 1.8E-12 | MB231/MB468 | MP |
| >hg38_21_+_8433172_8446622_RNA45SN1_45S_50ntFlanks@11602.11640.39\|TCCGACTGTTTAATTAAAACAAAGCATCGCGAAGGCCCG | >hg38_21_+_8433172_8446622_RNA45SN1_45S_50ntFlanks | 11602 | 11640 | 39 | TCCGACTGTTTAATTAAAACAAAGCATCGCGAAGGCCCG | 218.0 | -11.6 | 1.6 | -7.2 | 6.3E-13 | 1.8E-12 | MB231/MB468 | MITO |
| >hg38_21_+_8433172_8446622_RNA45SN1_45S_50ntFlanks@12658.12681.24\|ACTGAACGCCTCTAAGTCAGAATC | >hg38_21_+_8433172_8446622_RNA45SN1_45S_50ntFlanks | 12658 | 12681 | 24 | ACTGAACGCCTCTAAGTCAGAATC | 341.5 | -12.3 | 1.7 | -7.2 | 4.3E-13 | 1.8E-12 | MB231/MB468 | MP |
| >hg38_21_+_8433172_8446622_RNA45SN1_45S_50ntFlanks@11874.11904.31\|AGAAGACCCTGTTGAGCTTGACTCTAGTCTG | >hg38_21_+_8433172_8446622_RNA45SN1_45S_50ntFlanks | 11874 | 11904 | 31 | AGAAGACCCTGTTGAGCTTGACTCTAGTCTG | 221.0 | -10.1 | 1.4 | -7.2 | 5.0E-13 | 1.8E-12 | BT20/MB468 | CYTO |
| >MI0000342\|hsa-mir-200b&WithFlank&1\|+\|1167098\|1167204@63.84.22&[MIMAT0000318&hsa-miR-200b-3p&offsets\|0\|0;m-93&1\|+\|1167160\|1167182&offsets\|0\|-1]\|TAATACTGCCTGGTAATGATGA | >MI0000342\|hsa-mir-200b&WithFlank&1\|+\|1167098\|1167204 | 63 | 84 | 22 | TAATACTGCCTGGTAATGATGA | 913.1 | -4.7 | 0.6 | -7.2 | 5.1E-13 | 1.8E-12 | MB231/MB468 | NUC |

<table>
<tr><td>>hg38_21_+_8433172_8446622_RNA45SN1_45S_50ntFlanks@10351.10382.32|CGAGAACTTTGAAGGCCGAAGTGGAGAAGGGT</td><td>>hg38_21_+_8433172_8446622_RNA45SN1_45S_50ntFlanks</td><td>10351</td><td>10382</td><td>32</td><td>CGAGAACTTTGAAGGCCGAAGTGGAGAAGGGT</td><td>1861.5</td><td>-5.1</td><td>0.7</td><td>-7.2</td><td>6.3E-13</td><td>1.8E-12</td><td>MB231/MB468</td><td>MITO</td></tr>
<tr><td>>trna116_GluCTC_1_-_145399233_145399304@32.70.39,trna71_GluCTC_1_-_161439189_161439260@32.70.39,trna74_GluCTC_1_-_161431809_161431880@32.70.39,trna77_GluCTC_1_-_161424398_161424469@32.70.39,trna77_GluCTC_6_+_28949976_28950047@32.70.39,trna80_GluCTC_1_-_161417018_161417089@32.70.39,trna87_GluCTC_6_-_126101393_126101464@32.70.39|CTCTCACCGCCGCGGCCCGGGTTCGATTCCCGGTCAGGG</td><td>>trna116_GluCTC_1_-_145399233_145399304</td><td>32</td><td>70</td><td>39</td><td>CTCTCACCGCCGCGGCCCGGGTTCGATTCCCGGTCAGGG</td><td>238.8</td><td>-11.8</td><td>1.6</td><td>-7.2</td><td>4.3E-13</td><td>1.8E-12</td><td>MB231/MB468</td><td>MP</td></tr>
<tr><td>>hg38_21_+_8433172_8446622_RNA45SN1_45S_50ntFlanks@12141.12169.29|ACAGTGCCAGGTGGGGAGTTTGACTGGGG</td><td>>hg38_21_+_8433172_8446622_RNA45SN1_45S_50ntFlanks</td><td>12141</td><td>12169</td><td>29</td><td>ACAGTGCCAGGTGGGGAGTTTGACTGGGG</td><td>128.6</td><td>-10.9</td><td>1.5</td><td>-7.2</td><td>4.3E-13</td><td>1.8E-12</td><td>MB231/MB468</td><td>MP</td></tr>
<tr><td>>hg38_21_+_8433172_8446622_RNA45SN1_45S_50ntFlanks@6653.6679.27|ACTCTTAGCGGTGGATCACTCGGCTCG</td><td>>hg38_21_+_8433172_8446622_RNA45SN1_45S_50ntFlanks</td><td>6653</td><td>6679</td><td>27</td><td>ACTCTTAGCGGTGGATCACTCGGCTCG</td><td>527.2</td><td>-3.9</td><td>0.5</td><td>-7.2</td><td>7.0E-13</td><td>1.8E-12</td><td>BT20/MB468</td><td>NUC</td></tr>
<tr><td>>hg38_21_+_8433172_8446622_RNA45SN1_45S_50ntFlanks@8117.8136.20|CGGGACATGTGGCGTACGGA</td><td>>hg38_21_+_8433172_8446622_RNA45SN1_45S_50ntFlanks</td><td>8117</td><td>8136</td><td>20</td><td>CGGGACATGTGGCGTACGGA</td><td>184.1</td><td>-11.4</td><td>1.6</td><td>-7.2</td><td>4.4E-13</td><td>1.8E-12</td><td>MB231/MB468</td><td>MP</td></tr>
<tr><td>>hg38_21_+_8433172_8446622_RNA45SN1_45S_50ntFlanks@12306.12328.23|CACGATCCTTCTGACCTTTTGGG</td><td>>hg38_21_+_8433172_8446622_RNA45SN1_45S_50ntFlanks</td><td>12306</td><td>12328</td><td>23</td><td>CACGATCCTTCTGACCTTTTGGG</td><td>2940.8</td><td>-9.3</td><td>1.3</td><td>-7.2</td><td>6.4E-13</td><td>1.8E-12</td><td>MB231/MB468</td><td>MITO</td></tr>
<tr><td>>hg38_21_+_8433172_8446622_RNA45SN1_45S_50ntFlanks@12241.12269.29|CAGAAGGGCAAAAGCTCGCTTGATCTTGA</td><td>>hg38_21_+_8433172_8446622_RNA45SN1_45S_50ntFlanks</td><td>12241</td><td>12269</td><td>29</td><td>CAGAAGGGCAAAAGCTCGCTTGATCTTGA</td><td>767.7</td><td>-7.3</td><td>1.0</td><td>-7.2</td><td>6.4E-13</td><td>1.8E-12</td><td>MB231/MB468</td><td>MITO</td></tr>
<tr><td>>hg38_21_+_8433172_8446622_RNA45SN1_45S_50ntFlanks@5222.5244.23|CTGCACGCGCGCTACACTGACTG</td><td>>hg38_21_+_8433172_8446622_RNA45SN1_45S_50ntFlanks</td><td>5222</td><td>5244</td><td>23</td><td>CTGCACGCGCGCTACACTGACTG</td><td>1080.9</td><td>-13.9</td><td>1.9</td><td>-7.2</td><td>6.4E-13</td><td>1.8E-12</td><td>MB231/MB468</td><td>MITO</td></tr>
<tr><td>>trna10_ValCAC_5_-_180649395_180649467@35.54.20,trna12_ValAAC_5_-_180645270_180645342@35.54.20,trna132_ValAAC_6_-_27721179_27721251@35.54.20,trna13_ValCAC_19_-_4724647_4724719@35.54.20,trna152_ValCAC_6_-_27248049_27248121@35.54.20,trna157_ValCAC_6_-_27173867_27173939@35.54.20,trna15_ValAAC_5_-_180615416_180615488@35.54.20,trna18_ValCAC_5_-_180529253_180529325@35.54.20,trna2_ValAAC_3_+_169490018_169490090@35.54.20,trna2_ValCAC_5_+_180524070_180524142@35.54.20,trna37_ValAAC_6_+_27203288_27203360@35.54.20,trna4_ValAAC_5_+_180591154_180591226@35.54.20,trna5_ValAAC_5_+_180596610_180596682@35.54.20,trna6_ValCAC_5_+_180600650_180600722@35.54.20,trna85_ValCAC_1_-_161369490_161369562@35.54.20,trna98_ValCAC_1_-_149298555_149298627@35.54.20,trna9_ValCAC_6_+_26538282_26538354@35.54.20|ACACGCGAAAGGTCCCCGGT</td><td>>trna10_ValCAC_5_-_180649395_180649467</td><td>35</td><td>54</td><td>20</td><td>ACACGCGAAAGGTCCCCGGT</td><td>202.3</td><td>-7.9</td><td>1.1</td><td>-7.2</td><td>5.1E-13</td><td>1.8E-12</td><td>MB231/MB468</td><td>CYTO</td></tr>
<tr><td>>hg38_21_+_8433172_8446622_RNA45SN1_45S_50ntFlanks@4995.5023.29|ACAGATTGATAGCTCTTTCTCGATTCCGT</td><td>>hg38_21_+_8433172_8446622_RNA45SN1_45S_50ntFlanks</td><td>4995</td><td>5023</td><td>29</td><td>ACAGATTGATAGCTCTTTCTCGATTCCGT</td><td>394.8</td><td>-12.4</td><td>1.7</td><td>-7.2</td><td>6.4E-13</td><td>1.8E-12</td><td>MB231/MB468</td><td>MITO</td></tr>
<tr><td>>hg38_21_+_8433172_8446622_RNA45SN1_45S_50ntFlanks@6721.6752.32|AATTGCAGGACACATTGATCATCGACACTTCG</td><td>>hg38_21_+_8433172_8446622_RNA45SN1_45S_50ntFlanks</td><td>6721</td><td>6752</td><td>32</td><td>AATTGCAGGACACATTGATCATCGACACTTCG</td><td>673.5</td><td>-4.3</td><td>0.6</td><td>-7.2</td><td>5.2E-13</td><td>1.8E-12</td><td>MB231/MB468</td><td>NUC</td></tr>
<tr><td>>hg38_21_+_8433172_8446622_RNA45SN1_45S_50ntFlanks@6771.6797.27|GGTTCCTCCCGGGGCTACGCCTGTCTG</td><td>>hg38_21_+_8433172_8446622_RNA45SN1_45S_50ntFlanks</td><td>6771</td><td>6797</td><td>27</td><td>GGTTCCTCCCGGGGCTACGCCTGTCTG</td><td>220.9</td><td>-11.7</td><td>1.6</td><td>-7.2</td><td>4.4E-13</td><td>1.8E-12</td><td>MB231/MB468</td><td>MP</td></tr>
<tr><td>>hg38_21_+_8433172_8446622_RNA45SN1_45S_50ntFlanks@7968.7987.20|TCCGAGACGCGACCTCAGAT</td><td>>hg38_21_+_8433172_8446622_RNA45SN1_45S_50ntFlanks</td><td>7968</td><td>7987</td><td>20</td><td>TCCGAGACGCGACCTCAGAT</td><td>189.3</td><td>4.9</td><td>0.7</td><td>7.2</td><td>5.2E-13</td><td>1.8E-12</td><td>MB231/MB468</td><td>NUC</td></tr>
<tr><td>>hg38_21_+_8433172_8446622_RNA45SN1_45S_50ntFlanks@4069.4104.36|CTTTCGATGGTAGTCGCCGTGCCTACCATGGTGACC</td><td>>hg38_21_+_8433172_8446622_RNA45SN1_45S_50ntFlanks</td><td>4069</td><td>4104</td><td>36</td><td>CTTTCGATGGTAGTCGCCGTGCCTACCATGGTGACC</td><td>353.9</td><td>5.1</td><td>0.7</td><td>7.2</td><td>5.0E-13</td><td>1.8E-12</td><td>BT20/MB468</td><td>CYTO</td></tr>
<tr><td>>hg38_21_+_8433172_8446622_RNA45SN1_45S_50ntFlanks@4722.4753.32|TCATTAATCAAGAACGAAAGTCGGAGGTTCGA</td><td>>hg38_21_+_8433172_8446622_RNA45SN1_45S_50ntFlanks</td><td>4722</td><td>4753</td><td>32</td><td>TCATTAATCAAGAACGAAAGTCGGAGGTTCGA</td><td>161.5</td><td>-11.2</td><td>1.5</td><td>-7.2</td><td>4.4E-13</td><td>1.8E-12</td><td>MB231/MB468</td><td>MP</td></tr>
<tr><td>>MI0003513|hsa-mir-455&WithFlank&9|+|114209428|114209535@60.81.22&[MIMAT0004784&hsa-miR-455-3p&offsets|0|+1;m-135&9|+|114209487|114209508&offsets|0|0]|GCAGTCCATGGGCATATACACT</td><td>>MI0003513|hsa-mir-455&WithFlank&9|+|114209428|114209535</td><td>60</td><td>81</td><td>22</td><td>GCAGTCCATGGGCATATACACT</td><td>339.8</td><td>11.6</td><td>1.6</td><td>7.2</td><td>6.4E-13</td><td>1.8E-12</td><td>MB231/MB468</td><td>MITO</td></tr>
<tr><td>>MI0000272|hsa-mir-182&WithFlank&7|-|129770377|129770498@29.52.24(+1U)&[MIMAT0000259&hsa-miR-182-5p&offsets|0|0(+1U);m-22&7|-|129770449|129770470&offsets|0|+2(+1U)]|TTTGGCAATGGTAGAACTCACACTT</td><td>>MI0000272|hsa-mir-182&WithFlank&7|-|129770377|129770498</td><td>29</td><td>52</td><td>24(+1U)</td><td>TTTGGCAATGGTAGAACTCACACTT</td><td>699.1</td><td>9.2</td><td>1.3</td><td>7.2</td><td>5.0E-13</td><td>1.8E-12</td><td>BT20/MB468</td><td>CYTO</td></tr>
<tr><td>>hg38_21_+_8433172_8446622_RNA45SN1_45S_50ntFlanks@4445.4471.27|CTTGCCTCTCGGCGCCCCCTCGATGCT</td><td>>hg38_21_+_8433172_8446622_RNA45SN1_45S_50ntFlanks</td><td>4445</td><td>4471</td><td>27</td><td>CTTGCCTCTCGGCGCCCCCTCGATGCT</td><td>632.3</td><td>-4.0</td><td>0.6</td><td>-7.2</td><td>5.0E-13</td><td>1.8E-12</td><td>BT20/MB468</td><td>CYTO</td></tr>
<tr><td>>hg38_21_+_8433172_8446622_RNA45SN1_45S_50ntFlanks@10411.10435.25|CAGTCGGTCCTGAGAGATGGGCGAG</td><td>>hg38_21_+_8433172_8446622_RNA45SN1_45S_50ntFlanks</td><td>10411</td><td>10435</td><td>25</td><td>CAGTCGGTCCTGAGAGATGGGCGAG</td><td>333.2</td><td>-8.0</td><td>1.1</td><td>-7.2</td><td>5.1E-13</td><td>1.8E-12</td><td>BT20/MB468</td><td>CYTO</td></tr>
<tr><td>>hg38_21_+_8433172_8446622_RNA45SN1_45S_50ntFlanks@10458.10484.27|ATGGCCTCCGTTGCCCTCGGCCGATCG</td><td>>hg38_21_+_8433172_8446622_RNA45SN1_45S_50ntFlanks</td><td>10458</td><td>10484</td><td>27</td><td>ATGGCCTCCGTTGCCCTCGGCCGATCG</td><td>459.2</td><td>-8.8</td><td>1.2</td><td>-7.2</td><td>6.4E-13</td><td>1.8E-12</td><td>MB231/MB468</td><td>MITO</td></tr>
<tr><td>>hg38_21_+_8433172_8446622_RNA45SN1_45S_50ntFlanks@11581.11601.21|AACTGGTGCGGACCAGGGGAA</td><td>>hg38_21_+_8433172_8446622_RNA45SN1_45S_50ntFlanks</td><td>11581</td><td>11601</td><td>21</td><td>AACTGGTGCGGACCAGGGGAA</td><td>819.4</td><td>-8.8</td><td>1.2</td><td>-7.2</td><td>6.5E-13</td><td>1.8E-12</td><td>MB231/MB468</td><td>MITO</td></tr>
<tr><td>>MI0000108|hsa-mir-103a-2&WithFlank&20|+|3917488|3917577@54.76.23(+2U);MI0000109|hsa-mir-103a-1&WithFlank&5|-|168560890|168560979@54.76.23(+2U)&[MIMAT0000101&hsa-miR-103a-3p&offsets|0|0(+2U);m-12&20|+|3917541|3917563&offsets|0|0(+2U)];[MIMAT0000101_1&hsa-miR-103a-3p&offsets|0|0(+2U);m-13&5|-|168560904|168560926&offsets|0|0(+2U);m-44220&5|-|168560908|168560926&offsets|0|+4(+2U)]|AGCAGCATTGTACAGGGCTATGATT</td><td>>MI0000108|hsa-mir-103a-2&WithFlank&20|+|3917488|3917577</td><td>54</td><td>76</td><td>23(+2U)</td><td>AGCAGCATTGTACAGGGCTATGATT</td><td>1067.6</td><td>6.6</td><td>0.9</td><td>7.2</td><td>5.1E-13</td><td>1.8E-12</td><td>MB231/MB468</td><td>CYTO</td></tr>
<tr><td>>hg38_21_+_8433172_8446622_RNA45SN1_45S_50ntFlanks@5073.5093.21|AATTCCGATAACGAACGAGAC</td><td>>hg38_21_+_8433172_8446622_RNA45SN1_45S_50ntFlanks</td><td>5073</td><td>5093</td><td>21</td><td>AATTCCGATAACGAACGAGAC</td><td>1015.2</td><td>-13.8</td><td>1.9</td><td>-7.2</td><td>6.5E-13</td><td>1.8E-12</td><td>MB231/MB468</td><td>MITO</td></tr>
<tr><td>>hg38_21_+_8433172_8446622_RNA45SN1_45S_50ntFlanks@11325.11342.18|TCTCGTCGCGCGCGCGTC</td><td>>hg38_21_+_8433172_8446622_RNA45SN1_45S_50ntFlanks</td><td>11325</td><td>11342</td><td>18</td><td>TCTCGTCGCGCGCGCGTC</td><td>135.0</td><td>-10.9</td><td>1.5</td><td>-7.2</td><td>4.4E-13</td><td>1.9E-12</td><td>MB231/MB468</td><td>MP</td></tr>
<tr><td>>hg38_21_+_8433172_8446622_RNA45SN1_45S_50ntFlanks@12935.12961.27|TAAACCATTCGTAGACGACCTGCTTCT</td><td>>hg38_21_+_8433172_8446622_RNA45SN1_45S_50ntFlanks</td><td>12935</td><td>12961</td><td>27</td><td>TAAACCATTCGTAGACGACCTGCTTCT</td><td>864.8</td><td>-9.7</td><td>1.4</td><td>-7.2</td><td>7.2E-13</td><td>1.9E-12</td><td>BT20/MB468</td><td>NUC</td></tr>
<tr><td>>MI0000449|hsa-mir-132&WithFlank&17|-|2049902|2050014@65.86.22&[MIMAT0000426&hsa-miR-132-3p&offsets|0|0;m-138&17|-|2049929|2049950&offsets|0|0]|TAACAGTCTACAGCCATGGTCG</td><td>>MI0000449|hsa-mir-132&WithFlank&17|-|2049902|2050014</td><td>65</td><td>86</td><td>22</td><td>TAACAGTCTACAGCCATGGTCG</td><td>126.4</td><td>5.2</td><td>0.7</td><td>7.2</td><td>7.2E-13</td><td>1.9E-12</td><td>BT20/MB468</td><td>NUC</td></tr>
<tr><td>>hg38_21_+_8433172_8446622_RNA45SN1_45S_50ntFlanks@11918.11967.50|ACATGAGAGGTGTAGAATAAGTGGGAGGCCCCCGGCGCCCCCCCGGTGTC</td><td>>hg38_21_+_8433172_8446622_RNA45SN1_45S_50ntFlanks</td><td>11918</td><td>11967</td><td>50</td><td>ACATGAGAGGTGTAGAATAAGTGGGAGGCCCCCGGCGCCCCCCCGGTGTC</td><td>507.0</td><td>-11.3</td><td>1.6</td><td>-7.2</td><td>5.1E-13</td><td>1.9E-12</td><td>BT20/MB468</td><td>CYTO</td></tr>
<tr><td>>hg38_21_+_8433172_8446622_RNA45SN1_45S_50ntFlanks@13002.13030.29|CGATCTATTGAAAGTCAGCCCTCGACACA</td><td>>hg38_21_+_8433172_8446622_RNA45SN1_45S_50ntFlanks</td><td>13002</td><td>13030</td><td>29</td><td>CGATCTATTGAAAGTCAGCCCTCGACACA</td><td>206.8</td><td>-10.0</td><td>1.4</td><td>-7.2</td><td>5.1E-13</td><td>1.9E-12</td><td>BT20/MB468</td><td>CYTO</td></tr>
<tr><td>>hg38_21_+_8433172_8446622_RNA45SN1_45S_50ntFlanks@6745.6770.26|ACACTTCGAACGCACTTGCGGCCCCG</td><td>>hg38_21_+_8433172_8446622_RNA45SN1_45S_50ntFlanks</td><td>6745</td><td>6770</td><td>26</td><td>ACACTTCGAACGCACTTGCGGCCCCG</td><td>5062.8</td><td>-7.8</td><td>1.1</td><td>-7.2</td><td>5.1E-13</td><td>1.9E-12</td><td>BT20/MB468</td><td>CYTO</td></tr>
<tr><td>>hg38_1_-_228634819_228635039_RNA5S12_5S_50ntFlanks@51.85.35|GTCTACGGCCATACCACCCTGAACGCGCCCGATCT</td><td>>hg38_1_-_228634819_228635039_RNA5S12_5S_50ntFlanks</td><td>51</td><td>85</td><td>35</td><td>GTCTACGGCCATACCACCCTGAACGCGCCCGATCT</td><td>2591.9</td><td>5.9</td><td>0.8</td><td>7.2</td><td>5.1E-13</td><td>1.9E-12</td><td>BT20/MB468</td><td>CYTO</td></tr>
<tr><td>>hg38_21_+_8433172_8446622_RNA45SN1_45S_50ntFlanks@12008.12034.27|CCGCCGGTGAAATACCACTACTCTGAT</td><td>>hg38_21_+_8433172_8446622_RNA45SN1_45S_50ntFlanks</td><td>12008</td><td>12034</td><td>27</td><td>CCGCCGGTGAAATACCACTACTCTGAT</td><td>297.4</td><td>-3.4</td><td>0.5</td><td>-7.3</td><td>3.9E-13</td><td>1.9E-12</td><td>BT20/MB468</td><td>MITO</td></tr>
<tr><td>>hg38_21_+_8433172_8446622_RNA45SN1_45S_50ntFlanks@12957.12985.29|CTTCTGGGTCGGGGTTTCGTACGTAGCAG</td><td>>hg38_21_+_8433172_8446622_RNA45SN1_45S_50ntFlanks</td><td>12957</td><td>12985</td><td>29</td><td>CTTCTGGGTCGGGGTTTCGTACGTAGCAG</td><td>793.8</td><td>-9.8</td><td>1.4</td><td>-7.2</td><td>7.2E-13</td><td>1.9E-12</td><td>BT20/MB468</td><td>NUC</td></tr>
<tr><td>>hg38_21_+_8433172_8446622_RNA45SN1_45S_50ntFlanks@12087.12106.20|CTTCTGGCGCCAAGCGCCCG</td><td>>hg38_21_+_8433172_8446622_RNA45SN1_45S_50ntFlanks</td><td>12087</td><td>12106</td><td>20</td><td>CTTCTGGCGCCAAGCGCCCG</td><td>122.1</td><td>-10.8</td><td>1.5</td><td>-7.2</td><td>4.5E-13</td><td>1.9E-12</td><td>MB231/MB468</td><td>MP</td></tr>
<tr><td>>hg38_21_+_8433172_8446622_RNA45SN1_45S_50ntFlanks@5334.5352.19|ATGAACGAGGAATTCCCAG</td><td>>hg38_21_+_8433172_8446622_RNA45SN1_45S_50ntFlanks</td><td>5334</td><td>5352</td><td>19</td><td>ATGAACGAGGAATTCCCAG</td><td>123.5</td><td>-10.8</td><td>1.5</td><td>-7.2</td><td>4.5E-13</td><td>1.9E-12</td><td>MB231/MB468</td><td>MP</td></tr>
<tr><td>>Antisense_hg38_21_-_8433172_8446622_RNA45SN1_45S_50ntFlanks@7674.7691.18|CTCTCTCTCTCTCTCCCC</td><td>>Antisense_hg38_21_-_8433172_8446622_RNA45SN1_45S_50ntFlanks</td><td>7674</td><td>7691</td><td>18</td><td>CTCTCTCTCTCTCTCCCC</td><td>288.6</td><td>-11.1</td><td>1.5</td><td>-7.2</td><td>5.3E-13</td><td>1.9E-12</td><td>MB231/MB468</td><td>NUC</td></tr>
<tr><td>>hg38_21_+_8433172_8446622_RNA45SN1_45S_50ntFlanks@11027.11044.18|CTCTCTCTCTCTCTCCCC</td><td>>hg38_21_+_8433172_8446622_RNA45SN1_45S_50ntFlanks</td><td>11027</td><td>11044</td><td>18</td><td>CTCTCTCTCTCTCTCCCC</td><td>288.6</td><td>-11.1</td><td>1.5</td><td>-7.2</td><td>5.3E-13</td><td>1.9E-12</td><td>MB231/MB468</td><td>NUC</td></tr>
<tr><td>>MI0000077|hsa-mir-21&WithFlank&17|+|59841260|59841343@14.37.24&[MIMAT0000076&hsa-miR-21-5p&offsets|0|+2;m-1&17|+|59841273|59841294&offsets|0|+2]|TAGCTTATCAGACTGATGTTGACT</td><td>>MI0000077|hsa-mir-21&WithFlank&17|+|59841260|59841343</td><td>14</td><td>37</td><td>24</td><td>TAGCTTATCAGACTGATGTTGACT</td><td>38544.3</td><td>3.4</td><td>0.5</td><td>7.2</td><td>7.2E-13</td><td>1.9E-12</td><td>BT20/MB468</td><td>NUC</td></tr>
<tr><td>>hg38_21_+_8433172_8446622_RNA45SN1_45S_50ntFlanks@7985.8004.20|GATCAGACGTGGCGACCCGC</td><td>>hg38_21_+_8433172_8446622_RNA45SN1_45S_50ntFlanks</td><td>7985</td><td>8004</td><td>20</td><td>GATCAGACGTGGCGACCCGC</td><td>604.6</td><td>-12.3</td><td>1.6</td><td>-7.6</td><td>2.3E-14</td><td>1.9E-12</td><td>BT20/MB468</td><td>MP</td></tr>
<tr><td>>hg38_21_+_8433172_8446622_RNA45SN1_45S_50ntFlanks@7989.8009.21|AGACGTGGCGACCCGCTGAAT</td><td>>hg38_21_+_8433172_8446622_RNA45SN1_45S_50ntFlanks</td><td>7989</td><td>8009</td><td>21</td><td>AGACGTGGCGACCCGCTGAAT</td><td>1144.7</td><td>5.1</td><td>0.7</td><td>7.2</td><td>5.2E-13</td><td>1.9E-12</td><td>BT20/MB468</td><td>CYTO</td></tr>
<tr><td>>MI0000102|hsa-mir-100&WithFlank&11|-|122152223|122152314@19.37.19&[MIMAT0000098&hsa-miR-100-5p&offsets|0|-3;m-27&11|-|122152276|122152296&offsets|0|-2]|AACCCGTAGATCCGAACTT</td><td>>MI0000102|hsa-mir-100&WithFlank&11|-|122152223|122152314</td><td>19</td><td>37</td><td>19</td><td>AACCCGTAGATCCGAACTT</td><td>387.3</td><td>11.8</td><td>1.6</td><td>7.2</td><td>6.6E-13</td><td>1.9E-12</td><td>MB231/MB468</td><td>MITO</td></tr>
<tr><td>>hg38_21_+_8433172_8446622_RNA45SN1_45S_50ntFlanks@12942.12964.23|TTCGTAGACGACCTGCTTCTGGG</td><td>>hg38_21_+_8433172_8446622_RNA45SN1_45S_50ntFlanks</td><td>12942</td><td>12964</td><td>23</td><td>TTCGTAGACGACCTGCTTCTGGG</td><td>159.2</td><td>-7.3</td><td>1.0</td><td>-7.2</td><td>5.2E-13</td><td>1.9E-12</td><td>BT20/MB468</td><td>CYTO</td></tr>
<tr><td>>hg38_21_+_8433172_8446622_RNA45SN1_45S_50ntFlanks@3769.3792.24|CGCACGGCCGGTACAGTGAAACTG</td><td>>hg38_21_+_8433172_8446622_RNA45SN1_45S_50ntFlanks</td><td>3769</td><td>3792</td><td>24</td><td>CGCACGGCCGGTACAGTGAAACTG</td><td>178.1</td><td>-11.3</td><td>1.6</td><td>-7.2</td><td>4.5E-13</td><td>1.9E-12</td><td>MB231/MB468</td><td>MP</td></tr>
<tr><td>>MI0000087|hsa-mir-29a&WithFlank&7|-|130876741|130876816@48.68.21&[MIMAT0000086&hsa-miR-29a-3p&offsets|0|-1;m-26&7|-|130876749|130876769&offsets|0|0]|TAGCACCATCTGAAATCGGTT</td><td>>MI0000087|hsa-mir-29a&WithFlank&7|-|130876741|130876816</td><td>48</td><td>68</td><td>21</td><td>TAGCACCATCTGAAATCGGTT</td><td>29224.6</td><td>6.3</td><td>0.9</td><td>7.2</td><td>4.5E-13</td><td>1.9E-12</td><td>MB231/MB468</td><td>MP</td></tr>
</table>

<table>
<tr><td>>hg38_21_+_8433172_8446622_RNA45SN1_45S_50ntFlanks@6740.6769.30|CATCGACACTTCGAACGCACTTGCGGCCCC</td><td>>hg38_21_+_8433172_8446622_RNA45SN1_45S_50ntFlanks</td><td>6740</td><td>6769</td><td>30</td><td>CATCGACACTTCGAACGCACTTGCGGCCCC</td><td>112.2</td><td>-10.7</td><td>1.5</td><td>-7.2</td><td>4.5E-13</td><td>1.9E-12</td><td>MB231/MB468</td><td>MP</td></tr>
<tr><td>>hg38_21_+_8433172_8446622_RNA45SN1_45S_50ntFlanks@12676.12710.35|AGAATCCCGCCCAGGCGGAACGATACGGCAGCGCC</td><td>>hg38_21_+_8433172_8446622_RNA45SN1_45S_50ntFlanks</td><td>12676</td><td>12710</td><td>35</td><td>AGAATCCCGCCCAGGCGGAACGATACGGCAGCGCC</td><td>104.2</td><td>-10.6</td><td>1.5</td><td>-7.2</td><td>4.6E-13</td><td>1.9E-12</td><td>MB231/MB468</td><td>MP</td></tr>
<tr><td>>hg38_21_+_8433172_8446622_RNA45SN1_45S_50ntFlanks@4596.4616.21|TTGGTTTTCGGAACTGAGGCC</td><td>>hg38_21_+_8433172_8446622_RNA45SN1_45S_50ntFlanks</td><td>4596</td><td>4616</td><td>21</td><td>TTGGTTTTCGGAACTGAGGCC</td><td>126.6</td><td>-10.8</td><td>1.5</td><td>-7.2</td><td>4.6E-13</td><td>1.9E-12</td><td>MB231/MB468</td><td>MP</td></tr>
<tr><td>>hg38_21_+_8433172_8446622_RNA45SN1_45S_50ntFlanks@3910.3932.23|GGGATGCGTGCATTTATCAGATC</td><td>>hg38_21_+_8433172_8446622_RNA45SN1_45S_50ntFlanks</td><td>3910</td><td>3932</td><td>23</td><td>GGGATGCGTGCATTTATCAGATC</td><td>361.7</td><td>-12.4</td><td>1.7</td><td>-7.2</td><td>4.6E-13</td><td>1.9E-12</td><td>MB231/MB468</td><td>MP</td></tr>
<tr><td>>trna152_ValCAC_6_-_27248049_27248121@1.29.29|GCTTCTGTAGTGTAGTGGTTATCACGTTC</td><td>>trna152_ValCAC_6_-_27248049_27248121</td><td>1</td><td>29</td><td>29</td><td>GCTTCTGTAGTGTAGTGGTTATCACGTTC</td><td>23335.3</td><td>-6.1</td><td>0.8</td><td>-7.2</td><td>5.2E-13</td><td>1.9E-12</td><td>BT20/MB468</td><td>CYTO</td></tr>
<tr><td>>hg38_21_+_8433172_8446622_RNA45SN1_45S_50ntFlanks@10808.10826.19|CAATGTAGGTAAGGGAAGT</td><td>>hg38_21_+_8433172_8446622_RNA45SN1_45S_50ntFlanks</td><td>10808</td><td>10826</td><td>19</td><td>CAATGTAGGTAAGGGAAGT</td><td>1318.6</td><td>-8.0</td><td>1.1</td><td>-7.2</td><td>5.4E-13</td><td>1.9E-12</td><td>MB231/MB468</td><td>NUC</td></tr>
<tr><td>>hg38_21_+_8433172_8446622_RNA45SN1_45S_50ntFlanks@11578.11603.26|TAGAACTGGTGCGGACCAGGGGAATC</td><td>>hg38_21_+_8433172_8446622_RNA45SN1_45S_50ntFlanks</td><td>11578</td><td>11603</td><td>26</td><td>TAGAACTGGTGCGGACCAGGGGAATC</td><td>1119.7</td><td>-8.0</td><td>1.1</td><td>-7.2</td><td>4.6E-13</td><td>1.9E-12</td><td>MB231/MB468</td><td>MP</td></tr>
<tr><td>>hg38_21_+_8433172_8446622_RNA45SN1_45S_50ntFlanks@6787.6807.21|ACGCCTGTCTGAGCGTCGCTT</td><td>>hg38_21_+_8433172_8446622_RNA45SN1_45S_50ntFlanks</td><td>6787</td><td>6807</td><td>21</td><td>ACGCCTGTCTGAGCGTCGCTT</td><td>300.8</td><td>4.5</td><td>0.6</td><td>7.2</td><td>5.4E-13</td><td>1.9E-12</td><td>MB231/MB468</td><td>NUC</td></tr>
<tr><td>>hg38_21_+_8433172_8446622_RNA45SN1_45S_50ntFlanks@10781.10808.28|AGGTGAACAGCCTCTGGCATGTTGGAAC</td><td>>hg38_21_+_8433172_8446622_RNA45SN1_45S_50ntFlanks</td><td>10781</td><td>10808</td><td>28</td><td>AGGTGAACAGCCTCTGGCATGTTGGAAC</td><td>115.4</td><td>-10.7</td><td>1.5</td><td>-7.2</td><td>4.6E-13</td><td>1.9E-12</td><td>MB231/MB468</td><td>MP</td></tr>
<tr><td>>hg38_21_+_8433172_8446622_RNA45SN1_45S_50ntFlanks@12419.12448.30|GATGTCGGCTCTTCCTATCATTGTGAAGCA</td><td>>hg38_21_+_8433172_8446622_RNA45SN1_45S_50ntFlanks</td><td>12419</td><td>12448</td><td>30</td><td>GATGTCGGCTCTTCCTATCATTGTGAAGCA</td><td>1909.5</td><td>5.6</td><td>0.8</td><td>7.2</td><td>5.3E-13</td><td>1.9E-12</td><td>BT20/MB468</td><td>CYTO</td></tr>
<tr><td>>hg38_21_+_8433172_8446622_RNA45SN1_45S_50ntFlanks@12960.12982.23|CTGGGTCGGGGTTTCGTACGTAG</td><td>>hg38_21_+_8433172_8446622_RNA45SN1_45S_50ntFlanks</td><td>12960</td><td>12982</td><td>23</td><td>CTGGGTCGGGGTTTCGTACGTAG</td><td>629.5</td><td>-8.7</td><td>1.2</td><td>-7.2</td><td>5.4E-13</td><td>1.9E-12</td><td>MB231/MB468</td><td>NUC</td></tr>
<tr><td>>MI0003560|hsa-mir-92b&WithFlank&1|+|155195171|155195278@67.87.21&[MIMAT0003218&hsa-miR-92b-3p&offsets|0|-1;m-115&1|+|155195237|155195258&offsets|0|-1]|TATTGCACTCGTCCCGGCCTC</td><td>>MI0003560|hsa-mir-92b&WithFlank&1|+|155195171|155195278</td><td>67</td><td>87</td><td>21</td><td>TATTGCACTCGTCCCGGCCTC</td><td>1582.2</td><td>9.2</td><td>1.3</td><td>7.2</td><td>5.3E-13</td><td>1.9E-12</td><td>MB231/MB468</td><td>CYTO</td></tr>
<tr><td>>MI0003591|hsa-mir-584&WithFlank&5|-|149062307|149062415@22.41.20&[MIMAT0003249&hsa-miR-584-5p&offsets|0|-2;m-167&5|-|149062374|149062394&offsets|0|-1]|TTATGGTTTGCCTGGGACTG</td><td>>MI0003591|hsa-mir-584&WithFlank&5|-|149062307|149062415</td><td>22</td><td>41</td><td>20</td><td>TTATGGTTTGCCTGGGACTG</td><td>365.7</td><td>8.5</td><td>1.2</td><td>7.2</td><td>5.3E-13</td><td>1.9E-12</td><td>BT20/MB468</td><td>CYTO</td></tr>
<tr><td>>hg38_21_+_8433172_8446622_RNA45SN1_45S_50ntFlanks@12635.12667.33|AAGCTACCATCTGTGGGATTATGACTGAACGCC</td><td>>hg38_21_+_8433172_8446622_RNA45SN1_45S_50ntFlanks</td><td>12635</td><td>12667</td><td>33</td><td>AAGCTACCATCTGTGGGATTATGACTGAACGCC</td><td>329.3</td><td>6.8</td><td>0.9</td><td>7.2</td><td>5.3E-13</td><td>1.9E-12</td><td>BT20/MB468</td><td>CYTO</td></tr>
<tr><td>>MI0000683|hsa-mir-181b-2&WithFlank&9|+|124693704|124693804@22.46.25&[MIMAT0000257_1&hsa-miR-181b-5p&offsets|0|+2;m-72&9|+|124693725|124693747&offsets|0|+2]|AACATTCATTGCTGTCGGTGGGTTT</td><td>>MI0000683|hsa-mir-181b-2&WithFlank&9|+|124693704|124693804</td><td>22</td><td>46</td><td>25</td><td>AACATTCATTGCTGTCGGTGGGTTT</td><td>1493.6</td><td>7.2</td><td>1.0</td><td>7.2</td><td>6.7E-13</td><td>1.9E-12</td><td>MB231/MB468</td><td>MITO</td></tr>
<tr><td>>hg38_21_+_8433172_8446622_RNA45SN1_45S_50ntFlanks@11217.11246.30|CTGGACGCGAGCCGGGCCCTTCCCGTGGAT</td><td>>hg38_21_+_8433172_8446622_RNA45SN1_45S_50ntFlanks</td><td>11217</td><td>11246</td><td>30</td><td>CTGGACGCGAGCCGGGCCCTTCCCGTGGAT</td><td>239.7</td><td>7.3</td><td>1.0</td><td>7.2</td><td>5.3E-13</td><td>1.9E-12</td><td>MB231/MB468</td><td>CYTO</td></tr>
<tr><td>>hg38_21_+_8433172_8446622_RNA45SN1_45S_50ntFlanks@9548.9572.25|GAGGTCCGTAGCGGTCCTGACGTGC</td><td>>hg38_21_+_8433172_8446622_RNA45SN1_45S_50ntFlanks</td><td>9548</td><td>9572</td><td>25</td><td>GAGGTCCGTAGCGGTCCTGACGTGC</td><td>521.2</td><td>-4.8</td><td>0.7</td><td>-7.2</td><td>7.5E-13</td><td>1.9E-12</td><td>BT20/MB468</td><td>NUC</td></tr>
<tr><td>>hg38_21_+_8433172_8446622_RNA45SN1_45S_50ntFlanks@8814.8835.22|CGGGTCGGGGCGGCGGCGGCGG</td><td>>hg38_21_+_8433172_8446622_RNA45SN1_45S_50ntFlanks</td><td>8814</td><td>8835</td><td>22</td><td>CGGGTCGGGGCGGCGGCGGCGG</td><td>872.6</td><td>-12.9</td><td>1.7</td><td>-7.6</td><td>2.3E-14</td><td>1.9E-12</td><td>BT20/MB468</td><td>MP</td></tr>
<tr><td>>hg38_1_-_228634819_228635039_RNA5S12_5S_50ntFlanks@90.123.34|TGATCTCGGAAGCTAAGCAGGGTCGGGCCTGGTT</td><td>>hg38_1_-_228634819_228635039_RNA5S12_5S_50ntFlanks</td><td>90</td><td>123</td><td>34</td><td>TGATCTCGGAAGCTAAGCAGGGTCGGGCCTGGTT</td><td>646.1</td><td>8.7</td><td>1.2</td><td>7.2</td><td>5.4E-13</td><td>1.9E-12</td><td>MB231/MB468</td><td>CYTO</td></tr>
<tr><td>>hg38_21_+_8433172_8446622_RNA45SN1_45S_50ntFlanks@11769.11790.22|TAATTAGTGACGCGCATGAATG</td><td>>hg38_21_+_8433172_8446622_RNA45SN1_45S_50ntFlanks</td><td>11769</td><td>11790</td><td>22</td><td>TAATTAGTGACGCGCATGAATG</td><td>819.2</td><td>-3.8</td><td>0.5</td><td>-7.2</td><td>5.3E-13</td><td>1.9E-12</td><td>BT20/MB468</td><td>CYTO</td></tr>
<tr><td>>trnaMT_GluTTC_MT_-_14674_14742@53.72.20,trnalookalike8_GluTTC_5_-_93905172_93905240@53.72.20|TAGTCCGTGCGAGAATACCA</td><td>>trnaMT_GluTTC_MT_-_14674_14742</td><td>53</td><td>72</td><td>20</td><td>TAGTCCGTGCGAGAATACCA</td><td>497.7</td><td>-6.3</td><td>0.9</td><td>-7.2</td><td>7.5E-13</td><td>1.9E-12</td><td>BT20/MB468</td><td>NUC</td></tr>
<tr><td>>MI0000750|hsa-mir-26a-2&WithFlank&12|-|57824603|57824698@20.41.22(+2U);MI0000083|hsa-mir-26a-1&WithFlank&3|+|37969398|37969486@16.37.22(+2U)&[MIMAT0000082&hsa-miR-26a-5p&offsets|0|0(+2U);m-33&12|-|57824658|57824679&offsets|0|0(+2U)];[MIMAT0000082_1&hsa-miR-26a-5p&offsets|0|0(+2U);m-34&3|+|37969413|37969434&offsets|0|0(+2U)]|TTCAAGTAATCCAGGATAGGCTTT</td><td>>MI0000750|hsa-mir-26a-2&WithFlank&12|-|57824603|57824698</td><td>20</td><td>41</td><td>22(+2U)</td><td>TTCAAGTAATCCAGGATAGGCTTT</td><td>4758.7</td><td>7.4</td><td>1.0</td><td>7.2</td><td>6.8E-13</td><td>1.9E-12</td><td>MB231/MB468</td><td>MITO</td></tr>
<tr><td>>hg38_21_+_8433172_8446622_RNA45SN1_45S_50ntFlanks@12627.12653.27|ATGGGGCGAAGCTACCATCTGTGGGAT</td><td>>hg38_21_+_8433172_8446622_RNA45SN1_45S_50ntFlanks</td><td>12627</td><td>12653</td><td>27</td><td>ATGGGGCGAAGCTACCATCTGTGGGAT</td><td>2544.3</td><td>-7.0</td><td>1.0</td><td>-7.2</td><td>5.4E-13</td><td>1.9E-12</td><td>MB231/MB468</td><td>CYTO</td></tr>
<tr><td>>MI0001448|hsa-mir-425&WithFlank&3|-|49020142|49020240@21.44.24&[MIMAT0003393&hsa-miR-425-5p&offsets|+1|+2;m-95&3|-|49020200|49020221&offsets|+1|+3]|ATGACACGATCACTCCCGTTGAGT</td><td>>MI0001448|hsa-mir-425&WithFlank&3|-|49020142|49020240</td><td>21</td><td>44</td><td>24</td><td>ATGACACGATCACTCCCGTTGAGT</td><td>1207.9</td><td>8.2</td><td>1.1</td><td>7.2</td><td>5.4E-13</td><td>1.9E-12</td><td>BT20/MB468</td><td>CYTO</td></tr>
<tr><td>>hg38_21_+_8433172_8446622_RNA45SN1_45S_50ntFlanks@8381.8403.23|AAGAGGGCGTGAAACCGTTAAGA</td><td>>hg38_21_+_8433172_8446622_RNA45SN1_45S_50ntFlanks</td><td>8381</td><td>8403</td><td>23</td><td>AAGAGGGCGTGAAACCGTTAAGA</td><td>1375.4</td><td>-7.6</td><td>1.1</td><td>-7.2</td><td>4.7E-13</td><td>1.9E-12</td><td>MB231/MB468</td><td>MP</td></tr>
<tr><td>>MI0000299|hsa-mir-222&WithFlank&X|-|45747009|45747130@75.98.24(+1C)&[MIMAT0000279&hsa-miR-222-3p&offsets|0|+3(+1C);m-140&X|-|45747033|45747056&offsets|0|0(+1C)]|AGCTACATCTGGCTACTGGGTCTCC</td><td>>MI0000299|hsa-mir-222&WithFlank&X|-|45747009|45747130</td><td>75</td><td>98</td><td>24(+1C)</td><td>AGCTACATCTGGCTACTGGGTCTCC</td><td>6547.5</td><td>5.6</td><td>0.8</td><td>7.2</td><td>5.5E-13</td><td>1.9E-12</td><td>MB231/MB468</td><td>NUC</td></tr>
<tr><td>>hg38_21_+_8433172_8446622_RNA45SN1_45S_50ntFlanks@12589.12615.27|GTTCAGACATTTGGTGTATGTGCTTGG</td><td>>hg38_21_+_8433172_8446622_RNA45SN1_45S_50ntFlanks</td><td>12589</td><td>12615</td><td>27</td><td>GTTCAGACATTTGGTGTATGTGCTTGG</td><td>138.7</td><td>-11.0</td><td>1.5</td><td>-7.2</td><td>4.7E-13</td><td>1.9E-12</td><td>MB231/MB468</td><td>MP</td></tr>
<tr><td>>MI0000285|hsa-mir-205&WithFlank&1|+|209432127|209432248@40.60.21(+1A)&[MIMAT0000266&hsa-miR-205-5p&offsets|0|-1(+1A);m-351&1|+|209432166|209432187&offsets|0|-1(+1A)]|TCCTTCATTCCACCGGAGTCTA</td><td>>MI0000285|hsa-mir-205&WithFlank&1|+|209432127|209432248</td><td>40</td><td>60</td><td>21(+1A)</td><td>TCCTTCATTCCACCGGAGTCTA</td><td>184.8</td><td>-11.3</td><td>1.6</td><td>-7.2</td><td>6.8E-13</td><td>1.9E-12</td><td>MB231/MB468</td><td>MITO</td></tr>
<tr><td>>hg38_21_+_8433172_8446622_RNA45SN1_45S_50ntFlanks@10351.10377.27|CGAGAACTTTGAAGGCCGAAGTGGAGA</td><td>>hg38_21_+_8433172_8446622_RNA45SN1_45S_50ntFlanks</td><td>10351</td><td>10377</td><td>27</td><td>CGAGAACTTTGAAGGCCGAAGTGGAGA</td><td>3926.3</td><td>-4.0</td><td>0.6</td><td>-7.2</td><td>5.4E-13</td><td>1.9E-12</td><td>BT20/MB468</td><td>CYTO</td></tr>
<tr><td>>trnaMT_ValTAC_MT_+_1602_1670@48.71.24|CAACTTAACTTGACCGCTCTGACC</td><td>>trnaMT_ValTAC_MT_+_1602_1670</td><td>48</td><td>71</td><td>24</td><td>CAACTTAACTTGACCGCTCTGACC</td><td>1949.2</td><td>10.4</td><td>1.4</td><td>7.2</td><td>5.4E-13</td><td>1.9E-12</td><td>BT20/MB468</td><td>CYTO</td></tr>
<tr><td>>hg38_21_+_8433172_8446622_RNA45SN1_45S_50ntFlanks@8194.8215.22|CCCGTGGACGGTGTGAGGCCGG</td><td>>hg38_21_+_8433172_8446622_RNA45SN1_45S_50ntFlanks</td><td>8194</td><td>8215</td><td>22</td><td>CCCGTGGACGGTGTGAGGCCGG</td><td>1703.1</td><td>-9.4</td><td>1.3</td><td>-7.2</td><td>6.9E-13</td><td>2.0E-12</td><td>MB231/MB468</td><td>MITO</td></tr>
<tr><td>>hg38_21_+_8433172_8446622_RNA45SN1_45S_50ntFlanks@9580.9597.18|TCGTCCGACCTGGGTATA</td><td>>hg38_21_+_8433172_8446622_RNA45SN1_45S_50ntFlanks</td><td>9580</td><td>9597</td><td>18</td><td>TCGTCCGACCTGGGTATA</td><td>126.1</td><td>-10.8</td><td>1.5</td><td>-7.2</td><td>4.7E-13</td><td>2.0E-12</td><td>MB231/MB468</td><td>MP</td></tr>
<tr><td>>hg38_21_+_8433172_8446622_RNA45SN1_45S_50ntFlanks@4732.4766.35|AGAACGAAAGTCGGAGGTTCGAAGACGATCAGATA</td><td>>hg38_21_+_8433172_8446622_RNA45SN1_45S_50ntFlanks</td><td>4732</td><td>4766</td><td>35</td><td>AGAACGAAAGTCGGAGGTTCGAAGACGATCAGATA</td><td>794.7</td><td>7.0</td><td>1.0</td><td>7.2</td><td>5.4E-13</td><td>2.0E-12</td><td>MB231/MB468</td><td>CYTO</td></tr>
<tr><td>>hg38_21_+_8433172_8446622_RNA45SN1_45S_50ntFlanks@11768.11791.24|CTAATTAGTGACGCGCATGAATGG</td><td>>hg38_21_+_8433172_8446622_RNA45SN1_45S_50ntFlanks</td><td>11768</td><td>11791</td><td>24</td><td>CTAATTAGTGACGCGCATGAATGG</td><td>238.4</td><td>-10.0</td><td>1.4</td><td>-7.3</td><td>4.1E-13</td><td>2.0E-12</td><td>BT20/MB468</td><td>MITO</td></tr>
<tr><td>>MI0000483|hsa-mir-186&WithFlank&1|-|71067625|71067722@21.41.21&[MIMAT0000456&hsa-miR-186-5p&offsets|0|-1;m-84&1|-|71067681|71067702&offsets|0|-1]|CAAAGAATTCTCCTTTTGGGC</td><td>>MI0000483|hsa-mir-186&WithFlank&1|-|71067625|71067722</td><td>21</td><td>41</td><td>21</td><td>CAAAGAATTCTCCTTTTGGGC</td><td>106.0</td><td>2.7</td><td>0.4</td><td>7.2</td><td>7.6E-13</td><td>2.0E-12</td><td>BT20/MB468</td><td>NUC</td></tr>
<tr><td>>hg38_21_+_8433172_8446622_RNA45SN1_45S_50ntFlanks@5086.5120.35|AACGAGACTCTGGCATGCTAACTAGTTACGCGACC</td><td>>hg38_21_+_8433172_8446622_RNA45SN1_45S_50ntFlanks</td><td>5086</td><td>5120</td><td>35</td><td>AACGAGACTCTGGCATGCTAACTAGTTACGCGACC</td><td>307.5</td><td>5.0</td><td>0.7</td><td>7.2</td><td>5.4E-13</td><td>2.0E-12</td><td>MB231/MB468</td><td>CYTO</td></tr>
<tr><td>>hg38_21_+_8433172_8446622_RNA45SN1_45S_50ntFlanks@11208.11234.27|GCGGCGACTCTGGACGCGAGCCGGGCC</td><td>>hg38_21_+_8433172_8446622_RNA45SN1_45S_50ntFlanks</td><td>11208</td><td>11234</td><td>27</td><td>GCGGCGACTCTGGACGCGAGCCGGGCC</td><td>7870.3</td><td>-9.8</td><td>1.4</td><td>-7.2</td><td>5.4E-13</td><td>2.0E-12</td><td>MB231/MB468</td><td>CYTO</td></tr>
<tr><td>>hg38_21_+_8433172_8446622_RNA45SN1_45S_50ntFlanks@12940.12973.34|CATTCGTAGACGACCTGCTTCTGGGTCGGGGTTT</td><td>>hg38_21_+_8433172_8446622_RNA45SN1_45S_50ntFlanks</td><td>12940</td><td>12973</td><td>34</td><td>CATTCGTAGACGACCTGCTTCTGGGTCGGGGTTT</td><td>164.3</td><td>-11.2</td><td>1.6</td><td>-7.2</td><td>4.7E-13</td><td>2.0E-12</td><td>MB231/MB468</td><td>MP</td></tr>
<tr><td>>hg38_21_+_8433172_8446622_RNA45SN1_45S_50ntFlanks@4002.4024.23|GATAACCTCGGGCCGATCGCACG</td><td>>hg38_21_+_8433172_8446622_RNA45SN1_45S_50ntFlanks</td><td>4002</td><td>4024</td><td>23</td><td>GATAACCTCGGGCCGATCGCACG</td><td>5377.1</td><td>-11.4</td><td>1.6</td><td>-7.2</td><td>7.6E-13</td><td>2.0E-12</td><td>BT20/MB468</td><td>NUC</td></tr>
<tr><td>>hg38_21_+_8433172_8446622_RNA45SN1_45S_50ntFlanks@6772.6797.26|GTTCCTCCCGGGGCTACGCCTGTCTG</td><td>>hg38_21_+_8433172_8446622_RNA45SN1_45S_50ntFlanks</td><td>6772</td><td>6797</td><td>26</td><td>GTTCCTCCCGGGGCTACGCCTGTCTG</td><td>251.3</td><td>-8.3</td><td>1.1</td><td>-7.2</td><td>5.6E-13</td><td>2.0E-12</td><td>MB231/MB468</td><td>NUC</td></tr>
<tr><td>>trna119_LysCTT_1_-_145395522_145395594@1.33.33,trna11_LysCTT_5_-_180648979_180649051@1.33.33,trna13_LysCTT_6_+_26556774_26556846@1.33.33,trna7_LysCTT_16_+_3225692_3225764@1.33.33,trna9_LysCTT_5_+_180634755_180634827@1.33.33|GCCCGGCTAGCTCAGTCGGTAGAGCATGAGACT</td><td>>trna119_LysCTT_1_-_145395522_145395594</td><td>1</td><td>33</td><td>33</td><td>GCCCGGCTAGCTCAGTCGGTAGAGCATGAGACT</td><td>15971.2</td><td>8.4</td><td>1.2</td><td>7.2</td><td>5.6E-13</td><td>2.0E-12</td><td>MB231/MB468</td><td>NUC</td></tr>
<tr><td>>hg38_21_+_8433172_8446622_RNA45SN1_45S_50ntFlanks@12681.12703.23|CCCGCCCAGGCGGAACGATACGG</td><td>>hg38_21_+_8433172_8446622_RNA45SN1_45S_50ntFlanks</td><td>12681</td><td>12703</td><td>23</td><td>CCCGCCCAGGCGGAACGATACGG</td><td>208.7</td><td>-11.6</td><td>1.6</td><td>-7.2</td><td>4.8E-13</td><td>2.0E-12</td><td>MB231/MB468</td><td>MP</td></tr>
<tr><td>>hg38_21_+_8433172_8446622_RNA45SN1_45S_50ntFlanks@8527.8551.25|CCTTCCCCCGCCGCCCCTCCTCCTC</td><td>>hg38_21_+_8433172_8446622_RNA45SN1_45S_50ntFlanks</td><td>8527</td><td>8551</td><td>25</td><td>CCTTCCCCCGCCGCCCCTCCTCCTC</td><td>199.7</td><td>-10.6</td><td>1.5</td><td>-7.2</td><td>5.6E-13</td><td>2.0E-12</td><td>MB231/MB468</td><td>NUC</td></tr>
<tr><td>>hg38_21_+_8433172_8446622_RNA45SN1_45S_50ntFlanks@4289.4317.29|CGAGGATCCATTGGAGGGCAAGTCTGGTG</td><td>>hg38_21_+_8433172_8446622_RNA45SN1_45S_50ntFlanks</td><td>4289</td><td>4317</td><td>29</td><td>CGAGGATCCATTGGAGGGCAAGTCTGGTG</td><td>186.7</td><td>-10.5</td><td>1.5</td><td>-7.2</td><td>5.6E-13</td><td>2.0E-12</td><td>MB231/MB468</td><td>NUC</td></tr>
<tr><td>>hg38_21_+_8433172_8446622_RNA45SN1_45S_50ntFlanks@7977.8013.37|CGACCTCAGATCAGACGTGGCGACCCGCTGAATTTAA</td><td>>hg38_21_+_8433172_8446622_RNA45SN1_45S_50ntFlanks</td><td>7977</td><td>8013</td><td>37</td><td>CGACCTCAGATCAGACGTGGCGACCCGCTGAATTTAA</td><td>424.7</td><td>10.4</td><td>1.4</td><td>7.2</td><td>5.5E-13</td><td>2.0E-12</td><td>MB231/MB468</td><td>CYTO</td></tr>
<tr><td>>hg38_21_+_8433172_8446622_RNA45SN1_45S_50ntFlanks@12932.12972.41|CGCTAAACCATTCGTAGACGACCTGCTTCTGGGTCGGGGTT</td><td>>hg38_21_+_8433172_8446622_RNA45SN1_45S_50ntFlanks</td><td>12932</td><td>12972</td><td>41</td><td>CGCTAAACCATTCGTAGACGACCTGCTTCTGGGTCGGGGTT</td><td>377.9</td><td>-12.4</td><td>1.7</td><td>-7.2</td><td>4.8E-13</td><td>2.0E-12</td><td>MB231/MB468</td><td>MP</td></tr>
<tr><td>>hg38_21_+_8433172_8446622_RNA45SN1_45S_50ntFlanks@12433.12455.23|CTATCATTGTGAAGCAGAATTCA</td><td>>hg38_21_+_8433172_8446622_RNA45SN1_45S_50ntFlanks</td><td>12433</td><td>12455</td><td>23</td><td>CTATCATTGTGAAGCAGAATTCA</td><td>143.7</td><td>-10.1</td><td>1.4</td><td>-7.2</td><td>5.7E-13</td><td>2.0E-12</td><td>MB231/MB468</td><td>NUC</td></tr>
</table>

| | | | | | | | | | | | | | |
|---|---|---|---|---|---|---|---|---|---|---|---|---|---|
| >hg38_21_+_8433172_8446622_RNA45SN1_45S_50ntFlanks@12413.12448.36\|TCCTTCGATGTCGGCTCTTCCTATCATTGTGAAGCA | >hg38_21_+_8433172_8446622_RNA45SN1_45S_50ntFlanks | 12413 | 12448 | 36 | TCCTTCGATGTCGGCTCTTCCTATCATTGTGAAGCA | 250.9 | 6.2 | 0.9 | 7.2 | 5.5E-13 | 2.0E-12 | MB231/MB468 | CYTO |
| >MI0000476\|hsa-mir-138-1&WithFlank&3\|+\|44114206\|44114316@29.53.25&[MIMAT0000430_1&hsa-miR-138-5p&offsets\|0\|+2;m-497&3\|+\|44114234\|44114256&offsets\|0\|+2]\|AGCTGGTGTTGTGAATCAGGCCGTT | >MI0000476\|hsa-mir-138-1&WithFlank&3\|+\|44114206\|44114316 | 29 | 53 | 25 | AGCTGGTGTTGTGAATCAGGCCGTT | 160.0 | 10.5 | 1.5 | 7.2 | 4.8E-13 | 2.0E-12 | MB231/MB468 | MP |
| >hg38_21_+_8433172_8446622_RNA45SN1_45S_50ntFlanks@12488.12525.38\|AACGTGAGCTGGGTTTAGACCGTCGTGAGACAGGTTAG | >hg38_21_+_8433172_8446622_RNA45SN1_45S_50ntFlanks | 12488 | 12525 | 38 | AACGTGAGCTGGGTTTAGACCGTCGTGAGACAGGTTAG | 221.0 | -11.6 | 1.6 | -7.2 | 7.0E-13 | 2.0E-12 | MB231/MB468 | MITO |
| >hg38_21_+_8433172_8446622_RNA45SN1_45S_50ntFlanks@9204.9221.18\|TCGGGGCCACGCGCGCGT | >hg38_21_+_8433172_8446622_RNA45SN1_45S_50ntFlanks | 9204 | 9221 | 18 | TCGGGGCCACGCGCGCGT | 146.5 | -11.1 | 1.5 | -7.2 | 4.8E-13 | 2.0E-12 | MB231/MB468 | MP |
| >hg38_21_+_8433172_8446622_RNA45SN1_45S_50ntFlanks@11783.11810.28\|CATGAATGGATGAACGAGATTCCCACTG | >hg38_21_+_8433172_8446622_RNA45SN1_45S_50ntFlanks | 11783 | 11810 | 28 | CATGAATGGATGAACGAGATTCCCACTG | 295.7 | -12.1 | 1.7 | -7.2 | 4.8E-13 | 2.0E-12 | MB231/MB468 | MP |
| >trnaMT_PheGAA_MT_+_577_647@1.18.18\|GTTTATGTAGCTTACCTC | >trnaMT_PheGAA_MT_+_577_647 | 1 | 18 | 18 | GTTTATGTAGCTTACCTC | 374.3 | 7.1 | 1.0 | 7.2 | 5.7E-13 | 2.0E-12 | MB231/MB468 | NUC |
| >hg38_21_+_8433172_8446622_RNA45SN1_45S_50ntFlanks@11739.11768.30\|TCTCTTAAGGTAGCCAAATGCCTCGTCATC | >hg38_21_+_8433172_8446622_RNA45SN1_45S_50ntFlanks | 11739 | 11768 | 30 | TCTCTTAAGGTAGCCAAATGCCTCGTCATC | 150.7 | -5.1 | 0.7 | -7.2 | 5.6E-13 | 2.0E-12 | MB231/MB468 | CYTO |
| >trna5_AspGTC_12_+_98897281_98897352@37.73.37\|ACGCGGGAGACCGGGGTTCAATTCCCCGACGGGGAGC | >trna5_AspGTC_12_+_98897281_98897352 | 37 | 73 | 37 | ACGCGGGAGACCGGGGTTCAATTCCCCGACGGGGAGC | 4879.6 | 9.7 | 1.3 | 7.2 | 5.6E-13 | 2.0E-12 | MB231/MB468 | CYTO |
| >MI0000342\|hsa-mir-200b&WithFlank&1\|+\|1167098\|1167204@63.83.21&[MIMAT0000318&hsa-miR-200b-3p&offsets\|0\|-1;m-93&1\|+\|1167160\|1167182&offsets\|0\|-2]\|TAATACTGCCTGGTAATGATG | >MI0000342\|hsa-mir-200b&WithFlank&1\|+\|1167098\|1167204 | 63 | 83 | 21 | TAATACTGCCTGGTAATGATG | 246.0 | -5.3 | 0.7 | -7.2 | 5.6E-13 | 2.0E-12 | MB231/MB468 | CYTO |
| >hg38_21_+_8433172_8446622_RNA45SN1_45S_50ntFlanks@10127.10145.19\|GCGGCGGCGGCGGCGGCGG | >hg38_21_+_8433172_8446622_RNA45SN1_45S_50ntFlanks | 10127 | 10145 | 19 | GCGGCGGCGGCGGCGGCGG | 153.5 | -11.1 | 1.5 | -7.2 | 4.8E-13 | 2.0E-12 | MB231/MB468 | MP |
| >hg38_21_+_8433172_8446622_RNA45SN1_45S_50ntFlanks@10130.10148.19\|GCGGCGGCGGCGGCGGCGG | >hg38_21_+_8433172_8446622_RNA45SN1_45S_50ntFlanks | 10130 | 10148 | 19 | GCGGCGGCGGCGGCGGCGG | 153.5 | -11.1 | 1.5 | -7.2 | 4.8E-13 | 2.0E-12 | MB231/MB468 | MP |
| >hg38_21_+_8433172_8446622_RNA45SN1_45S_50ntFlanks@8823.8841.19\|GCGGCGGCGGCGGCGGCGG | >hg38_21_+_8433172_8446622_RNA45SN1_45S_50ntFlanks | 8823 | 8841 | 19 | GCGGCGGCGGCGGCGGCGG | 153.5 | -11.1 | 1.5 | -7.2 | 4.8E-13 | 2.0E-12 | MB231/MB468 | MP |
| >hg38_21_+_8433172_8446622_RNA45SN1_45S_50ntFlanks@8844.8862.19\|GCGGCGGCGGCGGCGGCGG | >hg38_21_+_8433172_8446622_RNA45SN1_45S_50ntFlanks | 8844 | 8862 | 19 | GCGGCGGCGGCGGCGGCGG | 153.5 | -11.1 | 1.5 | -7.2 | 4.8E-13 | 2.0E-12 | MB231/MB468 | MP |
| >hg38_21_+_8433172_8446622_RNA45SN1_45S_50ntFlanks@8847.8865.19\|GCGGCGGCGGCGGCGGCGG | >hg38_21_+_8433172_8446622_RNA45SN1_45S_50ntFlanks | 8847 | 8865 | 19 | GCGGCGGCGGCGGCGGCGG | 153.5 | -11.1 | 1.5 | -7.2 | 4.8E-13 | 2.0E-12 | MB231/MB468 | MP |
| >MI0000081\|hsa-mir-24-2&WithFlank&19\|-\|13836281\|13836365@56.75.20;MI0000080\|hsa-mir-24-1&WithFlank&9\|+\|95086015\|95086094@50.69.20&[MIMAT0000080&hsa-miR-24-3p&offsets\|0\|-2;m-36&19\|-\|13836288\|13836310&offsets\|0\|-3];[MIMAT0000080_1&hsa-miR-24-3p&offsets\|0\|-2;m-35&9\|+\|95086064\|95086086&offsets\|0\|-3]\|TGGCTCAGTTCAGCAGGAAC | >MI0000081\|hsa-mir-24-2&WithFlank&19\|-\|13836281\|13836365 | 56 | 75 | 20 | TGGCTCAGTTCAGCAGGAAC | 10624.0 | 4.7 | 0.6 | 7.2 | 5.7E-13 | 2.0E-12 | MB231/MB468 | NUC |
| >hg38_21_+_8433172_8446622_RNA45SN1_45S_50ntFlanks@5486.5526.41\|GCGCTGAGAAGACGGTCGAACTTGACTATCTAGAGGAAGTA | >hg38_21_+_8433172_8446622_RNA45SN1_45S_50ntFlanks | 5486 | 5526 | 41 | GCGCTGAGAAGACGGTCGAACTTGACTATCTAGAGGAAGTA | 350.8 | -12.3 | 1.7 | -7.2 | 4.9E-13 | 2.0E-12 | MB231/MB468 | MP |
| >hg38_1_-_228634819_228635039_RNA5S12_5S_50ntFlanks@84.103.20\|CTCGTCTGATCTCGGAAGCT | >hg38_1_-_228634819_228635039_RNA5S12_5S_50ntFlanks | 84 | 103 | 20 | CTCGTCTGATCTCGGAAGCT | 205.6 | -9.8 | 1.3 | -7.2 | 4.2E-13 | 2.0E-12 | BT20/MB468 | MITO |
| >MI0000088\|hsa-mir-30a&WithFlank&6\|-\|71403545\|71403627@12.35.24(+2U)&[MIMAT0000087&hsa-miR-30a-5p&offsets\|0\|+2(+2U);m-15&6\|-\|71403595\|71403616&offsets\|0\|+2(+2U)]\|TGTAAACATCCTCGACTGGAAGCTTT | >MI0000088\|hsa-mir-30a&WithFlank&6\|-\|71403545\|71403627 | 12 | 35 | 24(+2U) | TGTAAACATCCTCGACTGGAAGCTTT | 2703.3 | 5.3 | 0.7 | 7.2 | 4.9E-13 | 2.0E-12 | MB231/MB468 | MP |
| >hg38_21_+_8433172_8446622_RNA45SN1_45S_50ntFlanks@9562.9595.34\|TCCTGACGTGCAAATCGGTCGTCCGACCTGGGTA | >hg38_21_+_8433172_8446622_RNA45SN1_45S_50ntFlanks | 9562 | 9595 | 34 | TCCTGACGTGCAAATCGGTCGTCCGACCTGGGTA | 298.1 | -5.5 | 0.8 | -7.2 | 5.6E-13 | 2.0E-12 | MB231/MB468 | CYTO |
| >hg38_21_+_8433172_8446622_RNA45SN1_45S_50ntFlanks@10731.10755.25\|AGGGTGTAAATCTCGCGCCGGGCCG | >hg38_21_+_8433172_8446622_RNA45SN1_45S_50ntFlanks | 10731 | 10755 | 25 | AGGGTGTAAATCTCGCGCCGGGCCG | 981.0 | -9.0 | 1.3 | -7.2 | 5.8E-13 | 2.0E-12 | MB231/MB468 | NUC |
| >MI0000273\|hsa-mir-183&WithFlank&7\|-\|129774899\|129775020@33.55.23(+1C)&[MIMAT0000261&hsa-miR-183-5p&offsets\|0\|+1(+1C);m-41&7\|-\|129774966\|129774987&offsets\|-1\|0(+1C)]\|TATGGCACTGGTAGAATTCACTGC | >MI0000273\|hsa-mir-183&WithFlank&7\|-\|129774899\|129775020 | 33 | 55 | 23(+1C) | TATGGCACTGGTAGAATTCACTGC | 327.1 | 8.4 | 1.2 | 7.2 | 5.6E-13 | 2.0E-12 | BT20/MB468 | CYTO |
| >MI0000088\|hsa-mir-30a&WithFlank&6\|-\|71403545\|71403627@54.75.22&[MIMAT0000088&hsa-miR-30a-3p&offsets\|+1\|+1;m-38&6\|-\|71403554\|71403575&offsets\|+1\|+1]\|TTTCAGTCGGATGTTTGCAGCT | >MI0000088\|hsa-mir-30a&WithFlank&6\|-\|71403545\|71403627 | 54 | 75 | 22 | TTTCAGTCGGATGTTTGCAGCT | 990.3 | 7.8 | 1.1 | 7.2 | 5.6E-13 | 2.0E-12 | MB231/MB468 | CYTO |
| >MI0000253\|hsa-mir-148a&WithFlank&7\|-\|25949913\|25949992@50.71.22&[MIMAT0000243&hsa-miR-148a-3p&offsets\|0\|0;m-7&7\|-\|25949922\|25949943&offsets\|0\|0]\|TCAGTGCACTACAGAACTTTGT | >MI0000253\|hsa-mir-148a&WithFlank&7\|-\|25949913\|25949992 | 50 | 71 | 22 | TCAGTGCACTACAGAACTTTGT | 157153.5 | 4.2 | 0.6 | 7.5 | 6.5E-14 | 2.0E-12 | BT20/MB231 | NUC |
| >hg38_21_+_8433172_8446622_RNA45SN1_45S_50ntFlanks@6787.6807.21\|ACGCCTGTCTGAGCGTCGCTT | >hg38_21_+_8433172_8446622_RNA45SN1_45S_50ntFlanks | 6787 | 6807 | 21 | ACGCCTGTCTGAGCGTCGCTT | 283.1 | 4.7 | 0.7 | 7.2 | 5.6E-13 | 2.0E-12 | MB231/MB468 | CYTO |
| >hg38_21_+_8433172_8446622_RNA45SN1_45S_50ntFlanks@6651.6685.35\|CGACTCTTAGCGGTGGATCACTCGGCTCGTGCGTC | >hg38_21_+_8433172_8446622_RNA45SN1_45S_50ntFlanks | 6651 | 6685 | 35 | CGACTCTTAGCGGTGGATCACTCGGCTCGTGCGTC | 1751.9 | 5.2 | 0.7 | 7.2 | 5.6E-13 | 2.0E-12 | BT20/MB468 | CYTO |
| >hg38_21_+_8433172_8446622_RNA45SN1_45S_50ntFlanks@8240.8260.21\|CGGGTCTTCCCGGAGTCGGGT | >hg38_21_+_8433172_8446622_RNA45SN1_45S_50ntFlanks | 8240 | 8260 | 21 | CGGGTCTTCCCGGAGTCGGGT | 396.3 | 8.5 | 1.2 | 7.2 | 5.7E-13 | 2.0E-12 | MB231/MB468 | CYTO |
| >MI0000102\|hsa-mir-100&WithFlank&11\|-\|122152223\|122152314@19.40.22(+1U)&[MIMAT0000098&hsa-miR-100-5p&offsets\|0\|0(+1U);m-27&11\|-\|122152276\|122152296&offsets\|0\|+1(+1U)]\|AACCCGTAGATCCGAACTTGTGT | >MI0000102\|hsa-mir-100&WithFlank&11\|-\|122152223\|122152314 | 19 | 40 | 22(+1U) | AACCCGTAGATCCGAACTTGTGT | 67050.0 | -8.5 | 1.1 | -7.8 | 7.7E-15 | 2.0E-12 | BT20/MB231 | MP |
| >hg38_21_+_8433172_8446622_RNA45SN1_45S_50ntFlanks@11781.11804.24\|CGCATGAATGGATGAACGAGATTC | >hg38_21_+_8433172_8446622_RNA45SN1_45S_50ntFlanks | 11781 | 11804 | 24 | CGCATGAATGGATGAACGAGATTC | 2255.8 | -9.3 | 1.3 | -7.2 | 7.1E-13 | 2.0E-12 | MB231/MB468 | MITO |
| >hg38_21_+_8433172_8446622_RNA45SN1_45S_50ntFlanks@11671.11701.31\|CCAGTGCTCTGAATGTCAAAGTGAAGAAATT | >hg38_21_+_8433172_8446622_RNA45SN1_45S_50ntFlanks | 11671 | 11701 | 31 | CCAGTGCTCTGAATGTCAAAGTGAAGAAATT | 1205.7 | -4.1 | 0.6 | -7.2 | 5.8E-13 | 2.0E-12 | MB231/MB468 | NUC |
| >hg38_21_+_8433172_8446622_RNA45SN1_45S_50ntFlanks@11873.11920.48\|AAGAAGACCCTGTTGAGCTTGACTCTAGTCTGGCACGGTGAAGAGACA | >hg38_21_+_8433172_8446622_RNA45SN1_45S_50ntFlanks | 11873 | 11920 | 48 | AAGAAGACCCTGTTGAGCTTGACTCTAGTCTGGCACGGTGAAGAGACA | 297.2 | -12.0 | 1.7 | -7.2 | 7.2E-13 | 2.0E-12 | MB231/MB468 | MITO |
| >hg38_21_+_8433172_8446622_RNA45SN1_45S_50ntFlanks@11163.11188.26\|CGGTCCCCCGCGAGGGGGGCCCGGGC | >hg38_21_+_8433172_8446622_RNA45SN1_45S_50ntFlanks | 11163 | 11188 | 26 | CGGTCCCCCGCGAGGGGGGCCCGGGC | 474.8 | -12.8 | 1.8 | -7.2 | 4.9E-13 | 2.0E-12 | MB231/MB468 | MP |
| >hg38_21_+_8433172_8446622_RNA45SN1_45S_50ntFlanks@10841.10864.24\|CGTAACTTCGGGATAAGGATTGGC | >hg38_21_+_8433172_8446622_RNA45SN1_45S_50ntFlanks | 10841 | 10864 | 24 | CGTAACTTCGGGATAAGGATTGGC | 4164.3 | -10.6 | 1.5 | -7.2 | 4.9E-13 | 2.0E-12 | MB231/MB468 | MP |
| >hg38_21_+_8433172_8446622_RNA45SN1_45S_50ntFlanks@5467.5498.32\|GGCCCACGGCCCTGGCGGAGCGCTGAGAAGAC | >hg38_21_+_8433172_8446622_RNA45SN1_45S_50ntFlanks | 5467 | 5498 | 32 | GGCCCACGGCCCTGGCGGAGCGCTGAGAAGAC | 391.7 | -8.9 | 1.2 | -7.2 | 5.8E-13 | 2.0E-12 | MB231/MB468 | NUC |
| >hg38_21_+_8433172_8446622_RNA45SN1_45S_50ntFlanks@11561.11582.22\|GCGCCTAGCAGCCGACTTAGAA | >hg38_21_+_8433172_8446622_RNA45SN1_45S_50ntFlanks | 11561 | 11582 | 22 | GCGCCTAGCAGCCGACTTAGAA | 422.2 | -10.8 | 1.5 | -7.2 | 4.3E-13 | 2.0E-12 | BT20/MB468 | MITO |
| >hg38_21_+_8433172_8446622_RNA45SN1_45S_50ntFlanks@7988.8006.19\|CAGACGTGGCGACCCGCTG | >hg38_21_+_8433172_8446622_RNA45SN1_45S_50ntFlanks | 7988 | 8006 | 19 | CAGACGTGGCGACCCGCTG | 384.1 | -10.7 | 1.5 | -7.2 | 4.3E-13 | 2.0E-12 | BT20/MB468 | MITO |
| >MI0000082\|hsa-mir-25&WithFlank&7\|-\|100093554\|100093649@58.78.21&[MIMAT0000081&hsa-miR-25-3p&offsets\|0\|-1;m-23&7\|-\|100093571\|100093592&offsets\|0\|-1]\|CATTGCACTTGTCTCGGTCTG | >MI0000082\|hsa-mir-25&WithFlank&7\|-\|100093554\|100093649 | 58 | 78 | 21 | CATTGCACTTGTCTCGGTCTG | 4463.9 | 5.7 | 0.8 | 7.2 | 5.7E-13 | 2.0E-12 | MB231/MB468 | CYTO |
| >hg38_21_+_8433172_8446622_RNA45SN1_45S_50ntFlanks@5080.5097.18\|ATAACGAACGAGACTCTG | >hg38_21_+_8433172_8446622_RNA45SN1_45S_50ntFlanks | 5080 | 5097 | 18 | ATAACGAACGAGACTCTG | 413.0 | -12.6 | 1.7 | -7.2 | 5.0E-13 | 2.0E-12 | MB231/MB468 | MP |
| >hg38_21_+_8433172_8446622_RNA45SN1_45S_50ntFlanks@10391.10428.38\|AACAGCAGTTGAACATGGGTCAGTCGGTCCTGAGAGAT | >hg38_21_+_8433172_8446622_RNA45SN1_45S_50ntFlanks | 10391 | 10428 | 38 | AACAGCAGTTGAACATGGGTCAGTCGGTCCTGAGAGAT | 489.8 | -4.5 | 0.6 | -7.2 | 8.0E-13 | 2.0E-12 | BT20/MB468 | NUC |
| >MI0000750\|hsa-mir-26a-2&WithFlank&12\|-\|57824603\|57824698@20.40.21(+1G);MI0000083\|hsa-mir-26a-1&WithFlank&3\|+\|37969398\|37969486@16.36.21(+1G)&[MIMAT0000082&hsa-miR-26a-5p&offsets\|0\|-1(+1G);m-33&12\|-\|57824658\|57824679&offsets\|0\|-1(+1G)];[MIMAT0000082_1&hsa-miR-26a-5p&offsets\|0\|-1(+1G);m-34&3\|+\|37969413\|37969434&offsets\|0\|-1(+1G)]\|TTCAAGTAATCCAGGATAGGCG | >MI0000750\|hsa-mir-26a-2&WithFlank&12\|-\|57824603\|57824698 | 20 | 40 | 21(+1G) | TTCAAGTAATCCAGGATAGGCG | 2240.5 | 5.1 | 0.7 | 7.2 | 5.9E-13 | 2.1E-12 | MB231/MB468 | NUC |
| >hg38_21_+_8433172_8446622_RNA45SN1_45S_50ntFlanks@10912.10939.28\|CGCGCCGCGGCTGGACGAGGCGCCGCCG | >hg38_21_+_8433172_8446622_RNA45SN1_45S_50ntFlanks | 10912 | 10939 | 28 | CGCGCCGCGGCTGGACGAGGCGCCGCCG | 182.1 | -11.4 | 1.6 | -7.2 | 5.0E-13 | 2.1E-12 | MB231/MB468 | MP |
| >hg38_21_+_8433172_8446622_RNA45SN1_45S_50ntFlanks@12406.12431.26\|TTTTTGATCCTTCGATGTCGGCTCTT | >hg38_21_+_8433172_8446622_RNA45SN1_45S_50ntFlanks | 12406 | 12431 | 26 | TTTTTGATCCTTCGATGTCGGCTCTT | 167.1 | -11.2 | 1.6 | -7.2 | 5.0E-13 | 2.1E-12 | MB231/MB468 | MP |
| >hg38_21_+_8433172_8446622_RNA45SN1_45S_50ntFlanks@12476.12498.23\|CCACTAATAGGGAACGTGAGCTG | >hg38_21_+_8433172_8446622_RNA45SN1_45S_50ntFlanks | 12476 | 12498 | 23 | CCACTAATAGGGAACGTGAGCTG | 696.3 | -3.4 | 0.5 | -7.2 | 4.3E-13 | 2.1E-12 | BT20/MB468 | MITO |
| >hg38_21_+_8433172_8446622_RNA45SN1_45S_50ntFlanks@10534.10570.37\|CCGCGAGGCGTCCAGTGCGGTAACGCGACCGATCCCG | >hg38_21_+_8433172_8446622_RNA45SN1_45S_50ntFlanks | 10534 | 10570 | 37 | CCGCGAGGCGTCCAGTGCGGTAACGCGACCGATCCCG | 639.8 | -12.4 | 1.6 | -7.6 | 2.5E-14 | 2.1E-12 | BT20/MB468 | MP |
| >hg38_21_+_8433172_8446622_RNA45SN1_45S_50ntFlanks@8179.8211.33\|CTGATCGAGGCCCAGCCCGTGGACGGTGTGAGG | >hg38_21_+_8433172_8446622_RNA45SN1_45S_50ntFlanks | 8179 | 8211 | 33 | CTGATCGAGGCCCAGCCCGTGGACGGTGTGAGG | 908.6 | -12.9 | 1.7 | -7.6 | 2.5E-14 | 2.1E-12 | BT20/MB468 | MP |
| >hg38_21_+_8433172_8446622_RNA45SN1_45S_50ntFlanks@6745.6767.23\|ACACTTCGAACGCACTTGCGGCC | >hg38_21_+_8433172_8446622_RNA45SN1_45S_50ntFlanks | 6745 | 6767 | 23 | ACACTTCGAACGCACTTGCGGCC | 142.1 | -11.0 | 1.5 | -7.2 | 5.0E-13 | 2.1E-12 | MB231/MB468 | MP |
| >hg38_21_+_8433172_8446622_RNA45SN1_45S_50ntFlanks@8379.8400.22\|TCAAGAGGGCGTGAAACCGTTA | >hg38_21_+_8433172_8446622_RNA45SN1_45S_50ntFlanks | 8379 | 8400 | 22 | TCAAGAGGGCGTGAAACCGTTA | 153.7 | -6.7 | 0.9 | -7.2 | 5.8E-13 | 2.1E-12 | MB231/MB468 | CYTO |
| >trna10_LysCTT_16_+_3241501_3241573@1.36.36,trna13_LysCTT_14_-_58706613_58706685@1.36.36,trna2_LysCTT_15_+_79152904_79152976@1.36.36\|GCCCGGCTAGCTCAGTCGGTAGAGCATGGGACTCTT | >trna10_LysCTT_16_+_3241501_3241573 | 1 | 36 | 36 | GCCCGGCTAGCTCAGTCGGTAGAGCATGGGACTCTT | 526.6 | 7.0 | 1.0 | 7.2 | 5.9E-13 | 2.1E-12 | MB231/MB468 | NUC |
| >am_trna117_GlyTCC_1_-_145397864_145397935@1.30.30,trna45_GlyTCC_1_+_161500903_161500974@1.30.30,trna70_GlyTCC_1_-_161439547_161439618@1.30.30,trna73_GlyTCC_1_-_161432166_161432237@1.30.30,trna76_GlyTCC_1_-_161424756_161424827@1.30.30,trna79_GlyTCC_1_-_161417375_161417446@1.30.30\|GCGTTGGTGGTATAGTGGTGAGCATAGCTG | >am_trna117_GlyTCC_1_-_145397864_145397935 | 1 | 30 | 30 | GCGTTGGTGGTATAGTGGTGAGCATAGCTG | 1966.2 | -9.8 | 1.4 | -7.2 | 5.9E-13 | 2.1E-12 | MB231/MB468 | NUC |

<table>
<tr><td>>MI0000285|hsa-mir-205&WithFlank&1|+|209432127|209432248@40.60.21(+1C)&[MIMAT0000266&hsa-miR-205-5p&offsets|0|-1(+1C);m-351&1|+|209432166|209432187&offsets|0|-1(+1C)]|TCCTTCATTCCACCGGAGTCTC</td><td>>MI0000285|hsa-mir-205&WithFlank&1|+|209432127|209432248</td><td>40</td><td>60</td><td>21(+1C)</td><td>TCCTTCATTCCACCGGAGTCTC</td><td>439.0</td><td>6.4</td><td>0.9</td><td>7.2</td><td>5.7E-13</td><td>2.1E-12</td><td>BT20/MB468</td><td>CYTO</td></tr>
<tr><td>>hg38_21_+_8433172_8446622_RNA45SN1_45S_50ntFlanks@10793.10822.30|TCTGGCATGTTGGAACAATGTAGGTAAGGG</td><td>>hg38_21_+_8433172_8446622_RNA45SN1_45S_50ntFlanks</td><td>10793</td><td>10822</td><td>30</td><td>TCTGGCATGTTGGAACAATGTAGGTAAGGG</td><td>114.9</td><td>-10.7</td><td>1.5</td><td>-7.2</td><td>5.0E-13</td><td>2.1E-12</td><td>MB231/MB468</td><td>MP</td></tr>
<tr><td>>hg38_21_+_8433172_8446622_RNA45SN1_45S_50ntFlanks@10866.10893.28|CTAAGGGCTGGGTCGGTCGGGCTGGGGC</td><td>>hg38_21_+_8433172_8446622_RNA45SN1_45S_50ntFlanks</td><td>10866</td><td>10893</td><td>28</td><td>CTAAGGGCTGGGTCGGTCGGGCTGGGGC</td><td>144.9</td><td>-11.0</td><td>1.5</td><td>-7.2</td><td>5.0E-13</td><td>2.1E-12</td><td>MB231/MB468</td><td>MP</td></tr>
<tr><td>>hg38_21_+_8433172_8446622_RNA45SN1_45S_50ntFlanks@9032.9058.27|CCTCGTGGGGGGGCCGGGCCACCCCTC</td><td>>hg38_21_+_8433172_8446622_RNA45SN1_45S_50ntFlanks</td><td>9032</td><td>9058</td><td>27</td><td>CCTCGTGGGGGGGCCGGGCCACCCCTC</td><td>480.0</td><td>-12.0</td><td>1.6</td><td>-7.6</td><td>2.6E-14</td><td>2.1E-12</td><td>BT20/MB468</td><td>MP</td></tr>
<tr><td>>hg38_21_+_8433172_8446622_RNA45SN1_45S_50ntFlanks@12457.12479.23|CAAGCGTTGGATTGTTCACCCAC</td><td>>hg38_21_+_8433172_8446622_RNA45SN1_45S_50ntFlanks</td><td>12457</td><td>12479</td><td>23</td><td>CAAGCGTTGGATTGTTCACCCAC</td><td>128.6</td><td>-10.9</td><td>1.5</td><td>-7.2</td><td>5.0E-13</td><td>2.1E-12</td><td>MB231/MB468</td><td>MP</td></tr>
<tr><td>>trna10_AspGTC_12_-_125424193_125424264@36.74.39,trna12_AspGTC_12_-_125411891_125411962@36.74.39,trna144_AspGTC_6_-_27551236_27551307@36.74.39,trna38_AspGTC_17_-_8125556_8125627@36.74.39,trna45_AspGTC_6_+_27447453_27447524@36.74.39,trna48_AspGTC_6_+_27471523_27471594@36.74.39,trna4_AspGTC_12_+_96429799_96429870@36.74.39,trna69_AspGTC_1_-_161440205_161440276@36.74.39,trna72_AspGTC_1_-_161432824_161432895@36.74.39,trna75_AspGTC_1_-_161425414_161425485@36.74.39,trna78_AspGTC_1_-_161418033_161418104@36.74.39,trna81_AspGTC_1_-_161410615_161410686@36.74.39|CACGCGGGAGACCGGGGTTCGATTCCCCGACGGGGAGCC</td><td>>trna10_AspGTC_12_-_125424193_125424264</td><td>36</td><td>74</td><td>39</td><td>CACGCGGGAGACCGGGGTTCGATTCCCCGACGGGGAGCC</td><td>670.8</td><td>7.2</td><td>1.0</td><td>7.2</td><td>5.8E-13</td><td>2.1E-12</td><td>BT20/MB468</td><td>CYTO</td></tr>
<tr><td>>am_trna10_LysCTT_16_+_3241501_3241573@34.51.18,trna119_LysCTT_1_-_145395522_145395594@34.51.18,trna11_LysCTT_5_-_180648979_180649051@34.51.18,trna13_LysCTT_16_+_3246153_3246225@34.51.18,trna13_LysCTT_6_+_26556774_26556846@34.51.18,trna30_LysCTT_16_-_3230555_3230627@34.51.18,trna32_LysCTT_16_-_3207406_3207478@34.51.18,trna5_LysCTT_16_+_3214939_3215011@34.51.18,trna7_LysCTT_16_+_3225692_3225764@34.51.18,trna8_LysCTT_1_+_39970195_39970267@34.51.18,trna9_LysCTT_5_+_180634755_180634827@34.51.18|CTTAATCTCAGGGTCGTG</td><td>>am_trna10_LysCTT_16_+_3241501_3241573</td><td>34</td><td>51</td><td>18</td><td>CTTAATCTCAGGGTCGTG</td><td>319.1</td><td>-12.2</td><td>1.7</td><td>-7.2</td><td>5.1E-13</td><td>2.1E-12</td><td>MB231/MB468</td><td>MP</td></tr>
<tr><td>>MI0001448|hsa-mir-425&WithFlank&3|-|49020142|49020240@20.43.24(+1C)&[MIMAT0003393&hsa-miR-425-5p&offsets|0|+1(+1C);m-95&3|-|49020200|49020221&offsets|0|+2(+1C)]|AATGACACGATCACTCCCGTTGAGC</td><td>>MI0001448|hsa-mir-425&WithFlank&3|-|49020142|49020240</td><td>20</td><td>43</td><td>24(+1C)</td><td>AATGACACGATCACTCCCGTTGAGC</td><td>382.3</td><td>6.2</td><td>0.9</td><td>7.2</td><td>5.8E-13</td><td>2.1E-12</td><td>MB231/MB468</td><td>CYTO</td></tr>
<tr><td>>hg38_21_+_8433172_8446622_RNA45SN1_45S_50ntFlanks@12957.12980.24|CTTCTGGGTCGGGGTTTCGTACGT</td><td>>hg38_21_+_8433172_8446622_RNA45SN1_45S_50ntFlanks</td><td>12957</td><td>12980</td><td>24</td><td>CTTCTGGGTCGGGGTTTCGTACGT</td><td>139.6</td><td>-11.0</td><td>1.5</td><td>-7.2</td><td>5.1E-13</td><td>2.1E-12</td><td>MB231/MB468</td><td>MP</td></tr>
<tr><td>>hg38_21_+_8433172_8446622_RNA45SN1_45S_50ntFlanks@8378.8405.28|TTCAAGAGGGCGTGAAACCGTTAAGAGG</td><td>>hg38_21_+_8433172_8446622_RNA45SN1_45S_50ntFlanks</td><td>8378</td><td>8405</td><td>28</td><td>TTCAAGAGGGCGTGAAACCGTTAAGAGG</td><td>132.4</td><td>-10.9</td><td>1.5</td><td>-7.2</td><td>5.1E-13</td><td>2.1E-12</td><td>MB231/MB468</td><td>MP</td></tr>
<tr><td>>hg38_21_+_8433172_8446622_RNA45SN1_45S_50ntFlanks@11579.11600.22|AGAACTGGTGCGGACCAGGGGA</td><td>>hg38_21_+_8433172_8446622_RNA45SN1_45S_50ntFlanks</td><td>11579</td><td>11600</td><td>22</td><td>AGAACTGGTGCGGACCAGGGGA</td><td>283.4</td><td>-12.0</td><td>1.7</td><td>-7.2</td><td>5.1E-13</td><td>2.1E-12</td><td>MB231/MB468</td><td>MP</td></tr>
<tr><td>>MI0000760|hsa-mir-361&WithFlank&X|-|85903630|85903713@52.75.24&[MIMAT0004682&hsa-miR-361-3p&offsets|+1|+2;m-124&X|-|85903642|85903663&offsets|+1|+3]|CCCCCAGGTGTGATTCTGATTTGC</td><td>>MI0000760|hsa-mir-361&WithFlank&X|-|85903630|85903713</td><td>52</td><td>75</td><td>24</td><td>CCCCCAGGTGTGATTCTGATTTGC</td><td>626.4</td><td>5.8</td><td>0.8</td><td>7.2</td><td>5.9E-13</td><td>2.1E-12</td><td>MB231/MB468</td><td>CYTO</td></tr>
<tr><td>>MI0000264|hsa-mir-7-2&WithFlank&15|+|88611819|88611940@38.61.24(+1A);MI0000265|hsa-mir-7-3&WithFlank&19|+|4770664|4770785@37.60.24(+1A);MI0000263|hsa-mir-7-1&WithFlank&9|-|83969742|83969863@30.53.24(+1A)&[MIMAT0000252&hsa-miR-7-5p&offsets|0|0(+1A);m-400&15|+|88611856|88611879&offsets|0|0(+1A)];[MIMAT0000252_1&hsa-miR-7-5p&offsets|0|0(+1A);m-402&19|+|4770700|4770723&offsets|0|0(+1A)];[MIMAT0000252_2&hsa-miR-7-5p&offsets|0|0(+1A);m-401&9|-|83969811|83969834&offsets|0|0(+1A)]|TGGAAGACTAGTGATTTTGTTGTTA</td><td>>MI0000264|hsa-mir-7-2&WithFlank&15|+|88611819|88611940</td><td>38</td><td>61</td><td>24(+1A)</td><td>TGGAAGACTAGTGATTTTGTTGTTA</td><td>166.6</td><td>3.0</td><td>0.4</td><td>7.2</td><td>8.1E-13</td><td>2.1E-12</td><td>BT20/MB468</td><td>NUC</td></tr>
<tr><td>>hg38_21_+_8433172_8446622_RNA45SN1_45S_50ntFlanks@3985.4013.29|CGGCTTTGGTGACTCTAGATAACCTCGGG</td><td>>hg38_21_+_8433172_8446622_RNA45SN1_45S_50ntFlanks</td><td>3985</td><td>4013</td><td>29</td><td>CGGCTTTGGTGACTCTAGATAACCTCGGG</td><td>205.5</td><td>-11.5</td><td>1.6</td><td>-7.2</td><td>5.1E-13</td><td>2.1E-12</td><td>MB231/MB468</td><td>MP</td></tr>
<tr><td>>hg38_21_+_8433172_8446622_RNA45SN1_45S_50ntFlanks@5473.5493.21|CGGCCCTGGCGGAGCGCTGAG</td><td>>hg38_21_+_8433172_8446622_RNA45SN1_45S_50ntFlanks</td><td>5473</td><td>5493</td><td>21</td><td>CGGCCCTGGCGGAGCGCTGAG</td><td>121.7</td><td>-10.8</td><td>1.5</td><td>-7.2</td><td>5.1E-13</td><td>2.1E-12</td><td>MB231/MB468</td><td>MP</td></tr>
<tr><td>>hg38_21_+_8433172_8446622_RNA45SN1_45S_50ntFlanks@6718.6743.26|GTGAATTGCAGGACACATTGATCATC</td><td>>hg38_21_+_8433172_8446622_RNA45SN1_45S_50ntFlanks</td><td>6718</td><td>6743</td><td>26</td><td>GTGAATTGCAGGACACATTGATCATC</td><td>476.0</td><td>4.3</td><td>0.6</td><td>7.2</td><td>4.4E-13</td><td>2.1E-12</td><td>BT20/MB468</td><td>MITO</td></tr>
<tr><td>>hg38_21_+_8433172_8446622_RNA45SN1_45S_50ntFlanks@12360.12382.23|CAGGGATAACTGGCTTGTGGCGG</td><td>>hg38_21_+_8433172_8446622_RNA45SN1_45S_50ntFlanks</td><td>12360</td><td>12382</td><td>23</td><td>CAGGGATAACTGGCTTGTGGCGG</td><td>1057.3</td><td>-10.0</td><td>1.4</td><td>-7.2</td><td>8.2E-13</td><td>2.1E-12</td><td>BT20/MB468</td><td>NUC</td></tr>
<tr><td>>MI0000272|hsa-mir-182&WithFlank&7|-|129770377|129770498@29.53.25(+1U)&[MIMAT0000259&hsa-miR-182-5p&offsets|0|+1(+1U);m-22&7|-|129770449|129770470&offsets|0|+3(+1U)]|TTTGGCAATGGTAGAACTCACACTGT</td><td>>MI0000272|hsa-mir-182&WithFlank&7|-|129770377|129770498</td><td>29</td><td>53</td><td>25(+1U)</td><td>TTTGGCAATGGTAGAACTCACACTGT</td><td>452.7</td><td>2.9</td><td>0.4</td><td>7.2</td><td>8.2E-13</td><td>2.1E-12</td><td>BT20/MB468</td><td>NUC</td></tr>
<tr><td>>hg38_21_+_8433172_8446622_RNA45SN1_45S_50ntFlanks@12954.12972.19|CTGCTTCTGGGTCGGGGTT</td><td>>hg38_21_+_8433172_8446622_RNA45SN1_45S_50ntFlanks</td><td>12954</td><td>12972</td><td>19</td><td>CTGCTTCTGGGTCGGGGTT</td><td>1673.4</td><td>-10.5</td><td>1.5</td><td>-7.2</td><td>8.2E-13</td><td>2.1E-12</td><td>BT20/MB468</td><td>NUC</td></tr>
<tr><td>>hg38_21_+_8433172_8446622_RNA45SN1_45S_50ntFlanks@5417.5445.29|CTACCGATTGGATGGTTTAGTGAGGCCCT</td><td>>hg38_21_+_8433172_8446622_RNA45SN1_45S_50ntFlanks</td><td>5417</td><td>5445</td><td>29</td><td>CTACCGATTGGATGGTTTAGTGAGGCCCT</td><td>576.8</td><td>-8.9</td><td>1.2</td><td>-7.2</td><td>6.1E-13</td><td>2.1E-12</td><td>MB231/MB468</td><td>NUC</td></tr>
<tr><td>>hg38_21_+_8433172_8446622_RNA45SN1_45S_50ntFlanks@12712.12746.35|CGGAGCCTCGGTTGGCCTCGGATAGCCGGTCCCCC</td><td>>hg38_21_+_8433172_8446622_RNA45SN1_45S_50ntFlanks</td><td>12712</td><td>12746</td><td>35</td><td>CGGAGCCTCGGTTGGCCTCGGATAGCCGGTCCCCC</td><td>132.5</td><td>-10.9</td><td>1.5</td><td>-7.2</td><td>5.1E-13</td><td>2.1E-12</td><td>MB231/MB468</td><td>MP</td></tr>
<tr><td>>MI0000456|hsa-mir-140&WithFlank&16|+|69933075|69933186@69.90.22&[MIMAT0004597&hsa-miR-140-3p&offsets|+1|+2;m-45&16|+|69933143|69933164&offsets|0|0]|ACCACAGGGTAGAACCACGGAC</td><td>>MI0000456|hsa-mir-140&WithFlank&16|+|69933075|69933186</td><td>69</td><td>90</td><td>22</td><td>ACCACAGGGTAGAACCACGGAC</td><td>2024.2</td><td>10.2</td><td>1.4</td><td>7.2</td><td>5.9E-13</td><td>2.1E-12</td><td>MB231/MB468</td><td>CYTO</td></tr>
<tr><td>>hg38_21_+_8433172_8446622_RNA45SN1_45S_50ntFlanks@11661.11682.22|TGATTTCTGCCCAGTGCTCTGA</td><td>>hg38_21_+_8433172_8446622_RNA45SN1_45S_50ntFlanks</td><td>11661</td><td>11682</td><td>22</td><td>TGATTTCTGCCCAGTGCTCTGA</td><td>766.1</td><td>-6.2</td><td>0.9</td><td>-7.2</td><td>5.9E-13</td><td>2.1E-12</td><td>MB231/MB468</td><td>CYTO</td></tr>
<tr><td>>hg38_21_+_8433172_8446622_RNA45SN1_45S_50ntFlanks@4002.4020.19|GATAACCTCGGGCCGATCG</td><td>>hg38_21_+_8433172_8446622_RNA45SN1_45S_50ntFlanks</td><td>4002</td><td>4020</td><td>19</td><td>GATAACCTCGGGCCGATCG</td><td>1561.3</td><td>-9.7</td><td>1.3</td><td>-7.2</td><td>6.1E-13</td><td>2.1E-12</td><td>MB231/MB468</td><td>NUC</td></tr>
<tr><td>>hg38_21_+_8433172_8446622_RNA45SN1_45S_50ntFlanks@9538.9571.34|AACTCTGGTGGAGGTCCGTAGCGGTCCTGACGTG</td><td>>hg38_21_+_8433172_8446622_RNA45SN1_45S_50ntFlanks</td><td>9538</td><td>9571</td><td>34</td><td>AACTCTGGTGGAGGTCCGTAGCGGTCCTGACGTG</td><td>1848.9</td><td>-4.9</td><td>0.7</td><td>-7.2</td><td>6.1E-13</td><td>2.1E-12</td><td>MB231/MB468</td><td>NUC</td></tr>
<tr><td>>hg38_21_+_8433172_8446622_RNA45SN1_45S_50ntFlanks@5486.5512.27|GCGCTGAGAAGACGGTCGAACTTGACT</td><td>>hg38_21_+_8433172_8446622_RNA45SN1_45S_50ntFlanks</td><td>5486</td><td>5512</td><td>27</td><td>GCGCTGAGAAGACGGTCGAACTTGACT</td><td>1720.0</td><td>7.1</td><td>1.0</td><td>7.2</td><td>5.9E-13</td><td>2.1E-12</td><td>BT20/MB468</td><td>CYTO</td></tr>
<tr><td>>hg38_21_+_8433172_8446622_RNA45SN1_45S_50ntFlanks@11778.11804.27|ACGCGCATGAATGGATGAACGAGATTC</td><td>>hg38_21_+_8433172_8446622_RNA45SN1_45S_50ntFlanks</td><td>11778</td><td>11804</td><td>27</td><td>ACGCGCATGAATGGATGAACGAGATTC</td><td>1455.5</td><td>-7.8</td><td>1.1</td><td>-7.2</td><td>5.2E-13</td><td>2.1E-12</td><td>MB231/MB468</td><td>MP</td></tr>
<tr><td>>hg38_21_+_8433172_8446622_RNA45SN1_45S_50ntFlanks@8176.8193.18|CTTCTGATCGAGGCCCAG</td><td>>hg38_21_+_8433172_8446622_RNA45SN1_45S_50ntFlanks</td><td>8176</td><td>8193</td><td>18</td><td>CTTCTGATCGAGGCCCAG</td><td>448.6</td><td>-11.9</td><td>1.6</td><td>-7.6</td><td>2.7E-14</td><td>2.1E-12</td><td>BT20/MB468</td><td>MP</td></tr>
<tr><td>>hg38_21_+_8433172_8446622_RNA45SN1_45S_50ntFlanks@12156.12183.28|GAGTTTGACTGGGGCGGTACACCTGTCA</td><td>>hg38_21_+_8433172_8446622_RNA45SN1_45S_50ntFlanks</td><td>12156</td><td>12183</td><td>28</td><td>GAGTTTGACTGGGGCGGTACACCTGTCA</td><td>145.9</td><td>-11.1</td><td>1.5</td><td>-7.2</td><td>5.2E-13</td><td>2.1E-12</td><td>MB231/MB468</td><td>MP</td></tr>
<tr><td>>hg38_21_+_8433172_8446622_RNA45SN1_45S_50ntFlanks@4722.4747.26|TCATTAATCAAGAACGAAAGTCGGAG</td><td>>hg38_21_+_8433172_8446622_RNA45SN1_45S_50ntFlanks</td><td>4722</td><td>4747</td><td>26</td><td>TCATTAATCAAGAACGAAAGTCGGAG</td><td>135.9</td><td>-11.0</td><td>1.5</td><td>-7.2</td><td>5.2E-13</td><td>2.1E-12</td><td>MB231/MB468</td><td>MP</td></tr>
<tr><td>>hg38_21_+_8433172_8446622_RNA45SN1_45S_50ntFlanks@9457.9486.30|CGCCGGGGAGGTGGAGCACGAGCGCACGTG</td><td>>hg38_21_+_8433172_8446622_RNA45SN1_45S_50ntFlanks</td><td>9457</td><td>9486</td><td>30</td><td>CGCCGGGGAGGTGGAGCACGAGCGCACGTG</td><td>144.7</td><td>-11.0</td><td>1.5</td><td>-7.2</td><td>5.2E-13</td><td>2.1E-12</td><td>MB231/MB468</td><td>MP</td></tr>
<tr><td>>hg38_21_+_8433172_8446622_RNA45SN1_45S_50ntFlanks@9562.9592.31|TCCTGACGTGCAAATCGGTCGTCCGACCTGG</td><td>>hg38_21_+_8433172_8446622_RNA45SN1_45S_50ntFlanks</td><td>9562</td><td>9592</td><td>31</td><td>TCCTGACGTGCAAATCGGTCGTCCGACCTGG</td><td>256.1</td><td>-6.0</td><td>0.8</td><td>-7.2</td><td>6.0E-13</td><td>2.1E-12</td><td>MB231/MB468</td><td>CYTO</td></tr>
<tr><td>>hg38_21_+_8433172_8446622_RNA45SN1_45S_50ntFlanks@10571.10588.18|GAGAAGCCGGCGGGAGCC</td><td>>hg38_21_+_8433172_8446622_RNA45SN1_45S_50ntFlanks</td><td>10571</td><td>10588</td><td>18</td><td>GAGAAGCCGGCGGGAGCC</td><td>110.6</td><td>-10.7</td><td>1.5</td><td>-7.2</td><td>5.2E-13</td><td>2.1E-12</td><td>MB231/MB468</td><td>MP</td></tr>
<tr><td>>MI0000269|hsa-mir-181a-2&WithFlank&9|+|124692436|124692557@83.103.21&[MIMAT0004558&hsa-miR-181a-2-3p&offsets|0|-1;m-71&9|+|124692518|124692539&offsets|0|-1]|ACCACTGACCGTTGACTGTAC</td><td>>MI0000269|hsa-mir-181a-2&WithFlank&9|+|124692436|124692557</td><td>83</td><td>103</td><td>21</td><td>ACCACTGACCGTTGACTGTAC</td><td>341.0</td><td>9.8</td><td>1.4</td><td>7.2</td><td>6.0E-13</td><td>2.1E-12</td><td>MB231/MB468</td><td>CYTO</td></tr>
<tr><td>>hg38_21_+_8433172_8446622_RNA45SN1_45S_50ntFlanks@11233.11252.20|CCCTTCCCGTGGATCGCCCC</td><td>>hg38_21_+_8433172_8446622_RNA45SN1_45S_50ntFlanks</td><td>11233</td><td>11252</td><td>20</td><td>CCCTTCCCGTGGATCGCCCC</td><td>1433.9</td><td>-10.4</td><td>1.5</td><td>-7.2</td><td>8.3E-13</td><td>2.1E-12</td><td>BT20/MB468</td><td>NUC</td></tr>
<tr><td>>hg38_21_+_8433172_8446622_RNA45SN1_45S_50ntFlanks@11918.11967.50|ACATGAGAGGTGTAGAATAAGTGGGAGGCCCCCGGCGCCCCCCCGGTGTC</td><td>>hg38_21_+_8433172_8446622_RNA45SN1_45S_50ntFlanks</td><td>11918</td><td>11967</td><td>50</td><td>ACATGAGAGGTGTAGAATAAGTGGGAGGCCCCCGGCGCCCCCCCGGTGTC</td><td>957.4</td><td>-12.9</td><td>1.8</td><td>-7.2</td><td>6.2E-13</td><td>2.1E-12</td><td>MB231/MB468</td><td>NUC</td></tr>
<tr><td>>hg38_21_+_8433172_8446622_RNA45SN1_45S_50ntFlanks@12455.12478.24|ACCAAGCGTTGGATTGTTCACCCA</td><td>>hg38_21_+_8433172_8446622_RNA45SN1_45S_50ntFlanks</td><td>12455</td><td>12478</td><td>24</td><td>ACCAAGCGTTGGATTGTTCACCCA</td><td>136.1</td><td>-11.0</td><td>1.5</td><td>-7.2</td><td>5.2E-13</td><td>2.1E-12</td><td>MB231/MB468</td><td>MP</td></tr>
<tr><td>>am_trna13_LysCTT_14_-_58706613_58706685@31.51.21,trna2_LysCTT_15_+_79152904_79152976@31.51.21,trna4_LysCTT_18_-_43669271_43669343@31.51.21|ACTCTTAATCCCAGGGTCGTG</td><td>>am_trna13_LysCTT_14_-_58706613_58706685</td><td>31</td><td>51</td><td>21</td><td>ACTCTTAATCCCAGGGTCGTG</td><td>172.4</td><td>-10.4</td><td>1.4</td><td>-7.2</td><td>6.2E-13</td><td>2.2E-12</td><td>MB231/MB468</td><td>NUC</td></tr>
<tr><td>>hg38_21_+_8433172_8446622_RNA45SN1_45S_50ntFlanks@11729.11767.39|TAACTATGACTCTCTTAAGGTAGCCAAATGCCTCGTCAT</td><td>>hg38_21_+_8433172_8446622_RNA45SN1_45S_50ntFlanks</td><td>11729</td><td>11767</td><td>39</td><td>TAACTATGACTCTCTTAAGGTAGCCAAATGCCTCGTCAT</td><td>4519.1</td><td>-7.5</td><td>1.0</td><td>-7.2</td><td>6.1E-13</td><td>2.2E-12</td><td>MB231/MB468</td><td>CYTO</td></tr>
</table>

<table>
<tr><td>>MI0000272|hsa-mir-182&WithFlank&7|-|129770377|129770498@29.51.23(+1A)&[MIMAT0000259&hsa-miR-182-5p&offsets|0|-1(+1A);m-22&7|-|129770449|129770470&offsets|0|+1(+1A)]|TTTGGCAATGGTAGAACTCACACA</td><td>>MI0000272|hsa-mir-182&WithFlank&7|-|129770377|129770498</td><td>29</td><td>51</td><td>23(+1A)</td><td>TTTGGCAATGGTAGAACTCACACA</td><td>1990.4</td><td>8.2</td><td>1.1</td><td>7.2</td><td>4.6E-13</td><td>2.2E-12</td><td>BT20/MB468</td><td>MITO</td></tr>
<tr><td>>hg38_21_+_8433172_8446622_RNA45SN1_45S_50ntFlanks@11583.11603.21|CTGGTGCGGACCAGGGGAATC</td><td>>hg38_21_+_8433172_8446622_RNA45SN1_45S_50ntFlanks</td><td>11583</td><td>11603</td><td>21</td><td>CTGGTGCGGACCAGGGGAATC</td><td>1049.9</td><td>-3.4</td><td>0.5</td><td>-7.2</td><td>4.6E-13</td><td>2.2E-12</td><td>BT20/MB468</td><td>MITO</td></tr>
<tr><td>>hg38_21_+_8433172_8446622_RNA45SN1_45S_50ntFlanks@5152.5185.34|AGGGACAAGTGGCGTTCAGCCACCCGAGATTGAG</td><td>>hg38_21_+_8433172_8446622_RNA45SN1_45S_50ntFlanks</td><td>5152</td><td>5185</td><td>34</td><td>AGGGACAAGTGGCGTTCAGCCACCCGAGATTGAG</td><td>946.1</td><td>-14.3</td><td>2.0</td><td>-7.2</td><td>8.4E-13</td><td>2.2E-12</td><td>BT20/MB468</td><td>NUC</td></tr>
<tr><td>>trna111_HisGTG_1_-_147774845_147774916@2.30.29,trna118_HisGTG_1_-_145396881_145396952@2.30.29,trna16_HisGTG_1_+_146544773_146544844@2.30.29,trna1_HisGTG_15_+_45493349_45493420@2.30.29,trna21_HisGTG_1_+_147753471_147753542@2.30.29,trna33_HisGTG_6_+_27125906_27125977@2.30.29,trna7_HisGTG_9_-_14433938_14434009@2.30.29,trna8_HisGTG_15_-_45492611_45492682@2.30.29,trna9_HisGTG_15_-_45490804_45490875@2.30.29|CCGTGATCGTATAGTGGTTAGTACTCTGC</td><td>>trna111_HisGTG_1_-_147774845_147774916</td><td>2</td><td>30</td><td>29</td><td>CCGTGATCGTATAGTGGTTAGTACTCTGC</td><td>3844.9</td><td>5.9</td><td>0.8</td><td>7.2</td><td>6.0E-13</td><td>2.2E-12</td><td>BT20/MB468</td><td>CYTO</td></tr>
<tr><td>>hg38_21_+_8433172_8446622_RNA45SN1_45S_50ntFlanks@11768.11804.37|CTAATTAGTGACGCGCATGAATGGATGAACGAGATTC</td><td>>hg38_21_+_8433172_8446622_RNA45SN1_45S_50ntFlanks</td><td>11768</td><td>11804</td><td>37</td><td>CTAATTAGTGACGCGCATGAATGGATGAACGAGATTC</td><td>714.4</td><td>-9.0</td><td>1.3</td><td>-7.2</td><td>7.6E-13</td><td>2.2E-12</td><td>MB231/MB468</td><td>MITO</td></tr>
<tr><td>>hg38_21_+_8433172_8446622_RNA45SN1_45S_50ntFlanks@9572.9594.23|CAAATCGGTCGTCCGACCTGGGT</td><td>>hg38_21_+_8433172_8446622_RNA45SN1_45S_50ntFlanks</td><td>9572</td><td>9594</td><td>23</td><td>CAAATCGGTCGTCCGACCTGGGT</td><td>3821.9</td><td>-7.0</td><td>0.9</td><td>-7.6</td><td>2.7E-14</td><td>2.2E-12</td><td>BT20/MB468</td><td>MP</td></tr>
<tr><td>>hg38_21_+_8433172_8446622_RNA45SN1_45S_50ntFlanks@11781.11804.24|CGCATGAATGGATGAACGAGATTC</td><td>>hg38_21_+_8433172_8446622_RNA45SN1_45S_50ntFlanks</td><td>11781</td><td>11804</td><td>24</td><td>CGCATGAATGGATGAACGAGATTC</td><td>1993.8</td><td>-5.7</td><td>0.8</td><td>-7.2</td><td>6.1E-13</td><td>2.2E-12</td><td>MB231/MB468</td><td>CYTO</td></tr>
<tr><td>>hg38_21_+_8433172_8446622_RNA45SN1_45S_50ntFlanks@4979.5000.22|GACACGGACAGGATTGACAGAT</td><td>>hg38_21_+_8433172_8446622_RNA45SN1_45S_50ntFlanks</td><td>4979</td><td>5000</td><td>22</td><td>GACACGGACAGGATTGACAGAT</td><td>183.8</td><td>-11.4</td><td>1.6</td><td>-7.2</td><td>5.3E-13</td><td>2.2E-12</td><td>MB231/MB468</td><td>MP</td></tr>
<tr><td>>hg38_21_+_8433172_8446622_RNA45SN1_45S_50ntFlanks@11873.11900.28|AAGAAGACCCTGTTGAGCTTGACTCTAG</td><td>>hg38_21_+_8433172_8446622_RNA45SN1_45S_50ntFlanks</td><td>11873</td><td>11900</td><td>28</td><td>AAGAAGACCCTGTTGAGCTTGACTCTAG</td><td>1049.2</td><td>-6.5</td><td>0.9</td><td>-7.2</td><td>6.1E-13</td><td>2.2E-12</td><td>MB231/MB468</td><td>CYTO</td></tr>
<tr><td>>hg38_21_+_8433172_8446622_RNA45SN1_45S_50ntFlanks@4853.4875.23|AAGTCTTTGGGTTCCGGGGGGAG</td><td>>hg38_21_+_8433172_8446622_RNA45SN1_45S_50ntFlanks</td><td>4853</td><td>4875</td><td>23</td><td>AAGTCTTTGGGTTCCGGGGGGAG</td><td>118.6</td><td>-10.8</td><td>1.5</td><td>-7.2</td><td>5.3E-13</td><td>2.2E-12</td><td>MB231/MB468</td><td>MP</td></tr>
<tr><td>>hg38_21_+_8433172_8446622_RNA45SN1_45S_50ntFlanks@12616.12653.38|CTGAGGAGCCAATGGGGCGAAGCTACCATCTGTGGGAT</td><td>>hg38_21_+_8433172_8446622_RNA45SN1_45S_50ntFlanks</td><td>12616</td><td>12653</td><td>38</td><td>CTGAGGAGCCAATGGGGCGAAGCTACCATCTGTGGGAT</td><td>542.4</td><td>-8.9</td><td>1.2</td><td>-7.2</td><td>6.3E-13</td><td>2.2E-12</td><td>MB231/MB468</td><td>NUC</td></tr>
<tr><td>>MI0000082|hsa-mir-25&WithFlank&7|-|100093554|100093649@58.79.22(+1G)&[MIMAT0000081&hsa-miR-25-3p&offsets|0|0(+1G);m-23&7|-|100093571|100093592&offsets|0|0(+1G)]|CATTGCACTTGTCTCGGTCTGAG</td><td>>MI0000082|hsa-mir-25&WithFlank&7|-|100093554|100093649</td><td>58</td><td>79</td><td>22(+1G)</td><td>CATTGCACTTGTCTCGGTCTGAG</td><td>266.1</td><td>8.0</td><td>1.1</td><td>7.2</td><td>6.1E-13</td><td>2.2E-12</td><td>BT20/MB468</td><td>CYTO</td></tr>
<tr><td>>hg38_21_+_8433172_8446622_RNA45SN1_45S_50ntFlanks@12934.12970.37|CTAAACCATTCGTAGACGACCTGCTTCTGGGTCGGGG</td><td>>hg38_21_+_8433172_8446622_RNA45SN1_45S_50ntFlanks</td><td>12934</td><td>12970</td><td>37</td><td>CTAAACCATTCGTAGACGACCTGCTTCTGGGTCGGGG</td><td>582.5</td><td>-8.5</td><td>1.2</td><td>-7.2</td><td>6.1E-13</td><td>2.2E-12</td><td>BT20/MB468</td><td>CYTO</td></tr>
<tr><td>>hg38_21_+_8433172_8446622_RNA45SN1_45S_50ntFlanks@11771.11792.22|ATTAGTGACGCGCATGAATGGA</td><td>>hg38_21_+_8433172_8446622_RNA45SN1_45S_50ntFlanks</td><td>11771</td><td>11792</td><td>22</td><td>ATTAGTGACGCGCATGAATGGA</td><td>112.2</td><td>-10.7</td><td>1.5</td><td>-7.2</td><td>5.4E-13</td><td>2.2E-12</td><td>MB231/MB468</td><td>MP</td></tr>
<tr><td>>hg38_21_+_8433172_8446622_RNA45SN1_45S_50ntFlanks@4443.4470.28|CCCTTGCCTCTCGGCGCCCCCTCGATGC</td><td>>hg38_21_+_8433172_8446622_RNA45SN1_45S_50ntFlanks</td><td>4443</td><td>4470</td><td>28</td><td>CCCTTGCCTCTCGGCGCCCCCTCGATGC</td><td>504.6</td><td>-11.9</td><td>1.7</td><td>-7.2</td><td>6.4E-13</td><td>2.2E-12</td><td>MB231/MB468</td><td>NUC</td></tr>
<tr><td>>hg38_21_+_8433172_8446622_RNA45SN1_45S_50ntFlanks@5236.5267.32|CACTGACTGGCTCAGCGTGTGCCTACCCTACG</td><td>>hg38_21_+_8433172_8446622_RNA45SN1_45S_50ntFlanks</td><td>5236</td><td>5267</td><td>32</td><td>CACTGACTGGCTCAGCGTGTGCCTACCCTACG</td><td>1635.9</td><td>-10.0</td><td>1.4</td><td>-7.2</td><td>7.8E-13</td><td>2.2E-12</td><td>MB231/MB468</td><td>MITO</td></tr>
<tr><td>>MI0000063|hsa-let-7b&WithFlank&22|+|46113680|46113774@12.31.20(+1A)&[MIMAT0000063&hsa-let-7b-5p&offsets|0|-2(+1A);m-14&22|+|46113691|46113712&offsets|0|-2(+1A)]|TGAGGTAGTAGGTTGTGTGGA</td><td>>MI0000063|hsa-let-7b&WithFlank&22|+|46113680|46113774</td><td>12</td><td>31</td><td>20(+1A)</td><td>TGAGGTAGTAGGTTGTGTGGA</td><td>74.5</td><td>3.2</td><td>0.4</td><td>7.2</td><td>8.6E-13</td><td>2.2E-12</td><td>BT20/MB468</td><td>NUC</td></tr>
<tr><td>>hg38_21_+_8433172_8446622_RNA45SN1_45S_50ntFlanks@9790.9820.31|CTGGCGTGGAGCCGGGCGTGGAATGCGAGTG</td><td>>hg38_21_+_8433172_8446622_RNA45SN1_45S_50ntFlanks</td><td>9790</td><td>9820</td><td>31</td><td>CTGGCGTGGAGCCGGGCGTGGAATGCGAGTG</td><td>633.8</td><td>-9.1</td><td>1.3</td><td>-7.2</td><td>7.8E-13</td><td>2.2E-12</td><td>MB231/MB468</td><td>MITO</td></tr>
<tr><td>>hg38_21_+_8433172_8446622_RNA45SN1_45S_50ntFlanks@8191.8219.29|CAGCCCGTGGACGGTGTGAGGCCGGTAGC</td><td>>hg38_21_+_8433172_8446622_RNA45SN1_45S_50ntFlanks</td><td>8191</td><td>8219</td><td>29</td><td>CAGCCCGTGGACGGTGTGAGGCCGGTAGC</td><td>503.5</td><td>-9.2</td><td>1.3</td><td>-7.2</td><td>8.6E-13</td><td>2.2E-12</td><td>BT20/MB468</td><td>NUC</td></tr>
<tr><td>>trna14_LysTTT_11_-_59327808_59327880@1.33.33,trna2_LysTTT_17_+_8022473_8022545@1.33.33,trna54_LysTTT_1_+_204475655_204475727@1.33.33,trna5_LysTTT_11_+_59323902_59323974@1.33.33,trna62_LysTTT_1_-_204476158_204476230@1.33.33,trna76_LysTTT_6_+_28918806_28918878@1.33.33|GCCCGGATAGCTCAGTCGGTAGAGCATCAGACT</td><td>>trna14_LysTTT_11_-_59327808_59327880</td><td>1</td><td>33</td><td>33</td><td>GCCCGGATAGCTCAGTCGGTAGAGCATCAGACT</td><td>10756.1</td><td>7.3</td><td>1.0</td><td>7.2</td><td>7.8E-13</td><td>2.2E-12</td><td>MB231/MB468</td><td>MITO</td></tr>
<tr><td>>hg38_21_+_8433172_8446622_RNA45SN1_45S_50ntFlanks@8158.8181.24|TCGTGGGGGGCCCAAGTCCTTCTG</td><td>>hg38_21_+_8433172_8446622_RNA45SN1_45S_50ntFlanks</td><td>8158</td><td>8181</td><td>24</td><td>TCGTGGGGGGCCCAAGTCCTTCTG</td><td>489.1</td><td>-12.7</td><td>1.8</td><td>-7.2</td><td>7.8E-13</td><td>2.2E-12</td><td>MB231/MB468</td><td>MITO</td></tr>
<tr><td>>MI0000063|hsa-let-7b&WithFlank&22|+|46113680|46113774@12.33.22&[MIMAT0000063&hsa-let-7b-5p&offsets|0|0;m-14&22|+|46113691|46113712&offsets|0|0]|TGAGGTAGTAGGTTGTGTGGTT</td><td>>MI0000063|hsa-let-7b&WithFlank&22|+|46113680|46113774</td><td>12</td><td>33</td><td>22</td><td>TGAGGTAGTAGGTTGTGTGGTT</td><td>16535.5</td><td>2.8</td><td>0.4</td><td>7.1</td><td>8.7E-13</td><td>2.2E-12</td><td>BT20/MB468</td><td>NUC</td></tr>
<tr><td>>hg38_21_+_8433172_8446622_RNA45SN1_45S_50ntFlanks@10787.10815.29|ACAGCCTCTGGCATGTTGGAACAATGTAG</td><td>>hg38_21_+_8433172_8446622_RNA45SN1_45S_50ntFlanks</td><td>10787</td><td>10815</td><td>29</td><td>ACAGCCTCTGGCATGTTGGAACAATGTAG</td><td>653.4</td><td>-9.5</td><td>1.3</td><td>-7.1</td><td>8.7E-13</td><td>2.2E-12</td><td>BT20/MB468</td><td>NUC</td></tr>
<tr><td>>hg38_21_+_8433172_8446622_RNA45SN1_45S_50ntFlanks@7986.8014.29|ATCAGACGTGGCGACCCGCTGAATTTAAG</td><td>>hg38_21_+_8433172_8446622_RNA45SN1_45S_50ntFlanks</td><td>7986</td><td>8014</td><td>29</td><td>ATCAGACGTGGCGACCCGCTGAATTTAAG</td><td>377.4</td><td>-10.6</td><td>1.5</td><td>-7.2</td><td>4.7E-13</td><td>2.2E-12</td><td>BT20/MB468</td><td>MITO</td></tr>
<tr><td>>hg38_21_+_8433172_8446622_RNA45SN1_45S_50ntFlanks@13005.13025.21|TCTATTGAAAGTCAGCCCTCG</td><td>>hg38_21_+_8433172_8446622_RNA45SN1_45S_50ntFlanks</td><td>13005</td><td>13025</td><td>21</td><td>TCTATTGAAAGTCAGCCCTCG</td><td>209.6</td><td>-10.7</td><td>1.5</td><td>-7.2</td><td>6.5E-13</td><td>2.2E-12</td><td>MB231/MB468</td><td>NUC</td></tr>
<tr><td>>hg38_21_+_8433172_8446622_RNA45SN1_45S_50ntFlanks@10534.10562.29|CCGCGAGGCGTCCAGTGCGGTAACGCGAC</td><td>>hg38_21_+_8433172_8446622_RNA45SN1_45S_50ntFlanks</td><td>10534</td><td>10562</td><td>29</td><td>CCGCGAGGCGTCCAGTGCGGTAACGCGAC</td><td>116.9</td><td>-10.7</td><td>1.5</td><td>-7.2</td><td>5.5E-13</td><td>2.2E-12</td><td>MB231/MB468</td><td>MP</td></tr>
<tr><td>>hg38_21_+_8433172_8446622_RNA45SN1_45S_50ntFlanks@11820.11844.25|ACTATCCAGCGAAACCACAGCCAAG</td><td>>hg38_21_+_8433172_8446622_RNA45SN1_45S_50ntFlanks</td><td>11820</td><td>11844</td><td>25</td><td>ACTATCCAGCGAAACCACAGCCAAG</td><td>199.5</td><td>-11.5</td><td>1.6</td><td>-7.2</td><td>5.5E-13</td><td>2.2E-12</td><td>MB231/MB468</td><td>MP</td></tr>
<tr><td>>hg38_21_+_8433172_8446622_RNA45SN1_45S_50ntFlanks@11455.11477.23|CTCGGCGCGCGGCGGCGGCGGCG</td><td>>hg38_21_+_8433172_8446622_RNA45SN1_45S_50ntFlanks</td><td>11455</td><td>11477</td><td>23</td><td>CTCGGCGCGCGGCGGCGGCGGCG</td><td>232.8</td><td>-7.6</td><td>1.1</td><td>-7.2</td><td>6.3E-13</td><td>2.3E-12</td><td>MB231/MB468</td><td>CYTO</td></tr>
<tr><td>>hg38_21_+_8433172_8446622_RNA45SN1_45S_50ntFlanks@5484.5514.31|GAGCGCTGAGAAGACGGTCGAACTTGACTAT</td><td>>hg38_21_+_8433172_8446622_RNA45SN1_45S_50ntFlanks</td><td>5484</td><td>5514</td><td>31</td><td>GAGCGCTGAGAAGACGGTCGAACTTGACTAT</td><td>4803.7</td><td>-5.3</td><td>0.7</td><td>-7.2</td><td>6.3E-13</td><td>2.3E-12</td><td>MB231/MB468</td><td>CYTO</td></tr>
<tr><td>>hg38_21_+_8433172_8446622_RNA45SN1_45S_50ntFlanks@10926.10946.21|ACGAGGCGCCGCCGCCCCCCC</td><td>>hg38_21_+_8433172_8446622_RNA45SN1_45S_50ntFlanks</td><td>10926</td><td>10946</td><td>21</td><td>ACGAGGCGCCGCCGCCCCCCC</td><td>181.5</td><td>-11.3</td><td>1.6</td><td>-7.2</td><td>8.0E-13</td><td>2.3E-12</td><td>MB231/MB468</td><td>MITO</td></tr>
<tr><td>>hg38_21_+_8433172_8446622_RNA45SN1_45S_50ntFlanks@11930.11967.38|TAGAATAAGTGGGAGGCCCCCGGCGCCCCCCCGGTGTC</td><td>>hg38_21_+_8433172_8446622_RNA45SN1_45S_50ntFlanks</td><td>11930</td><td>11967</td><td>38</td><td>TAGAATAAGTGGGAGGCCCCCGGCGCCCCCCCGGTGTC</td><td>169.0</td><td>-11.3</td><td>1.6</td><td>-7.2</td><td>5.5E-13</td><td>2.3E-12</td><td>MB231/MB468</td><td>MP</td></tr>
<tr><td>>hg38_21_+_8433172_8446622_RNA45SN1_45S_50ntFlanks@4759.4780.22|ATCAGATACCGTCGTAGTTCCG</td><td>>hg38_21_+_8433172_8446622_RNA45SN1_45S_50ntFlanks</td><td>4759</td><td>4780</td><td>22</td><td>ATCAGATACCGTCGTAGTTCCG</td><td>9756.1</td><td>-10.4</td><td>1.5</td><td>-7.2</td><td>6.5E-13</td><td>2.3E-12</td><td>MB231/MB468</td><td>NUC</td></tr>
<tr><td>>hg38_21_+_8433172_8446622_RNA45SN1_45S_50ntFlanks@10515.10533.19|GGAGTGGCGGAGATGGGCG</td><td>>hg38_21_+_8433172_8446622_RNA45SN1_45S_50ntFlanks</td><td>10515</td><td>10533</td><td>19</td><td>GGAGTGGCGGAGATGGGCG</td><td>255.1</td><td>-11.9</td><td>1.6</td><td>-7.2</td><td>5.5E-13</td><td>2.3E-12</td><td>MB231/MB468</td><td>MP</td></tr>
<tr><td>>hg38_21_+_8433172_8446622_RNA45SN1_45S_50ntFlanks@4390.4423.34|TCTTGGGAGCGGGCGGGCGGTCCGCCGCGAGGCG</td><td>>hg38_21_+_8433172_8446622_RNA45SN1_45S_50ntFlanks</td><td>4390</td><td>4423</td><td>34</td><td>TCTTGGGAGCGGGCGGGCGGTCCGCCGCGAGGCG</td><td>132.8</td><td>-10.9</td><td>1.5</td><td>-7.2</td><td>5.5E-13</td><td>2.3E-12</td><td>MB231/MB468</td><td>MP</td></tr>
<tr><td>>hg38_21_+_8433172_8446622_RNA45SN1_45S_50ntFlanks@8196.8215.20|CGTGGACGGTGTGAGGCCGG</td><td>>hg38_21_+_8433172_8446622_RNA45SN1_45S_50ntFlanks</td><td>8196</td><td>8215</td><td>20</td><td>CGTGGACGGTGTGAGGCCGG</td><td>2653.9</td><td>-8.7</td><td>1.2</td><td>-7.1</td><td>8.8E-13</td><td>2.3E-12</td><td>BT20/MB468</td><td>NUC</td></tr>
<tr><td>>hg38_21_+_8433172_8446622_RNA45SN1_45S_50ntFlanks@4367.4394.28|CAGTTAAAAAGCTCGTAGTTGGATCTTG</td><td>>hg38_21_+_8433172_8446622_RNA45SN1_45S_50ntFlanks</td><td>4367</td><td>4394</td><td>28</td><td>CAGTTAAAAAGCTCGTAGTTGGATCTTG</td><td>264.5</td><td>-5.5</td><td>0.8</td><td>-7.2</td><td>6.4E-13</td><td>2.3E-12</td><td>MB231/MB468</td><td>CYTO</td></tr>
<tr><td>>hg38_21_+_8433172_8446622_RNA45SN1_45S_50ntFlanks@11783.11804.22|CATGAATGGATGAACGAGATTC</td><td>>hg38_21_+_8433172_8446622_RNA45SN1_45S_50ntFlanks</td><td>11783</td><td>11804</td><td>22</td><td>CATGAATGGATGAACGAGATTC</td><td>5551.3</td><td>-4.4</td><td>0.6</td><td>-7.2</td><td>4.8E-13</td><td>2.3E-12</td><td>BT20/MB468</td><td>MITO</td></tr>
<tr><td>>am_trnaMT_PheGAA_MT_+_577_647@40.57.18|TGTTTAGACGGGCTCACA</td><td>>am_trnaMT_PheGAA_MT_+_577_647</td><td>40</td><td>57</td><td>18</td><td>TGTTTAGACGGGCTCACA</td><td>420.0</td><td>7.2</td><td>1.0</td><td>7.2</td><td>6.6E-13</td><td>2.3E-12</td><td>MB231/MB468</td><td>NUC</td></tr>
<tr><td>>hg38_21_+_8433172_8446622_RNA45SN1_45S_50ntFlanks@6732.6770.39|ACATTGATCATCGACACTTCGAACGCACTTGCGGCCCCG</td><td>>hg38_21_+_8433172_8446622_RNA45SN1_45S_50ntFlanks</td><td>6732</td><td>6770</td><td>39</td><td>ACATTGATCATCGACACTTCGAACGCACTTGCGGCCCCG</td><td>1306.8</td><td>-8.6</td><td>1.2</td><td>-7.2</td><td>5.6E-13</td><td>2.3E-12</td><td>MB231/MB468</td><td>MP</td></tr>
<tr><td>>hg38_21_+_8433172_8446622_RNA45SN1_45S_50ntFlanks@11570.11601.32|AGCCGACTTAGAACTGGTGCGGACCAGGGGAA</td><td>>hg38_21_+_8433172_8446622_RNA45SN1_45S_50ntFlanks</td><td>11570</td><td>11601</td><td>32</td><td>AGCCGACTTAGAACTGGTGCGGACCAGGGGAA</td><td>2117.0</td><td>-7.9</td><td>1.1</td><td>-7.2</td><td>5.6E-13</td><td>2.3E-12</td><td>MB231/MB468</td><td>MP</td></tr>
<tr><td>>hg38_21_+_8433172_8446622_RNA45SN1_45S_50ntFlanks@6719.6747.29|TGAATTGCAGGACACATTGATCATCGACA</td><td>>hg38_21_+_8433172_8446622_RNA45SN1_45S_50ntFlanks</td><td>6719</td><td>6747</td><td>29</td><td>TGAATTGCAGGACACATTGATCATCGACA</td><td>2274.0</td><td>-7.5</td><td>1.0</td><td>-7.2</td><td>8.1E-13</td><td>2.3E-12</td><td>MB231/MB468</td><td>MITO</td></tr>
<tr><td>>trna10_AspGTC_12_-_125424193_125424264@31.72.42,trna12_AspGTC_12_-_125411891_125411962@31.72.42,trna38_AspGTC_17_-_8125556_8125627@31.72.42,trna45_AspGTC_6_+_27447453_27447524@31.72.42,trna48_AspGTC_6_+_27471523_27471594@31.72.42,trna4_AspGTC_12_+_96429799_96429870@31.72.42,trna69_AspGTC_1_-_161440205_161440276@31.72.42,trna72_AspGTC_1_-_161432824_161432895@31.72.42,trna75_AspGTC_1_-_161425414_161425485@31.72.42,trna78_AspGTC_1_-_161418033_161418104@31.72.42,trna81_AspGTC_1_-_161410615_161410686@31.72.42|CCTGTCACGCGGGAGACCGGGGTTCGATTCCCCGACGGGGAG</td><td>>trna10_AspGTC_12_-_125424193_125424264</td><td>31</td><td>72</td><td>42</td><td>CCTGTCACGCGGGAGACCGGGGTTCGATTCCCCGACGGGGAG</td><td>875.6</td><td>-13.6</td><td>1.9</td><td>-7.2</td><td>8.1E-13</td><td>2.3E-12</td><td>MB231/MB468</td><td>MITO</td></tr>
</table>

<table>
<tr><td>>hg38_21_+_8433172_8446622_RNA45SN1_45S_50ntFlanks@9622.9658.37|CTAGTAGCTGGTTCCCTCCGAAGTTTCCCTCAGGATA</td><td>>hg38_21_+_8433172_8446622_RNA45SN1_45S_50ntFlanks</td><td>9622</td><td>9658</td><td>37</td><td>CTAGTAGCTGGTTCCCTCCGAAGTTTCCCTCAGGATA</td><td>1151.0</td><td>-7.3</td><td>1.0</td><td>-7.2</td><td>8.1E-13</td><td>2.3E-12</td><td>MB231/MB468</td><td>MITO</td></tr>
<tr><td>>hg38_21_+_8433172_8446622_RNA45SN1_45S_50ntFlanks@3839.3862.24|TCTCCTACTTGGATAACTGTGGTA</td><td>>hg38_21_+_8433172_8446622_RNA45SN1_45S_50ntFlanks</td><td>3839</td><td>3862</td><td>24</td><td>TCTCCTACTTGGATAACTGTGGTA</td><td>1637.9</td><td>7.0</td><td>1.0</td><td>7.2</td><td>6.4E-13</td><td>2.3E-12</td><td>BT20/MB468</td><td>CYTO</td></tr>
<tr><td>>MI0000079|hsa-mir-23a&WithFlank&19|-|13836581|13836665@51.71.21&[MIMAT0000078&hsa-miR-23a-3p&offsets|0|0;m-31&19|-|13836595|13836615&offsets|0|0]|ATCACATTGCCAGGGATTTCC</td><td>>MI0000079|hsa-mir-23a&WithFlank&19|-|13836581|13836665</td><td>51</td><td>71</td><td>21</td><td>ATCACATTGCCAGGGATTTCC</td><td>126.1</td><td>-9.3</td><td>1.3</td><td>-7.2</td><td>6.4E-13</td><td>2.3E-12</td><td>BT20/MB468</td><td>CYTO</td></tr>
<tr><td>>hg38_21_+_8433172_8446622_RNA45SN1_45S_50ntFlanks@10391.10424.34|AACAGCAGTTGAACATGGGTCAGTCGGTCCTGAG</td><td>>hg38_21_+_8433172_8446622_RNA45SN1_45S_50ntFlanks</td><td>10391</td><td>10424</td><td>34</td><td>AACAGCAGTTGAACATGGGTCAGTCGGTCCTGAG</td><td>30570.1</td><td>4.7</td><td>0.7</td><td>7.2</td><td>4.9E-13</td><td>2.3E-12</td><td>BT20/MB468</td><td>MITO</td></tr>
<tr><td>>MI0000736|hsa-mir-30c-1&WithFlank&1|+|40757278|40757378@23.46.24;MI0000254|hsa-mir-30c-2&WithFlank&6|-|71376954|71377037@13.36.24&[MIMAT0000244&hsa-miR-30c-5p&offsets|0|+1;m-57&1|+|40757300|40757323&offsets|0|0];[MIMAT0000244_1&hsa-miR-30c-5p&offsets|0|+1;m-56&6|-|71377002|71377025&offsets|0|0]|TGTAAACATCCTACACTCTCAGCT</td><td>>MI0000736|hsa-mir-30c-1&WithFlank&1|+|40757278|40757378</td><td>23</td><td>46</td><td>24</td><td>TGTAAACATCCTACACTCTCAGCT</td><td>28793.3</td><td>4.1</td><td>0.6</td><td>7.2</td><td>6.7E-13</td><td>2.3E-12</td><td>MB231/MB468</td><td>NUC</td></tr>
<tr><td>>am_trna146_GlnCTG_6_-_27515531_27515602@1.29.29,trna1_GlnCTG_6_+_18836402_18836473@1.29.29,trna3_GlnCTG_17_+_8023070_8023141@1.29.29,trna42_GlnCTG_6_+_27263212_27263283@1.29.29,trna49_GlnCTG_6_+_27487308_27487379@1.29.29,trna7_GlnCTG_15_-_66161400_66161471@1.29.29,trna99_GlnCTG_6_-_28909378_28909449@1.29.29|GGTTCCATGGTGTAATGGTTAGCACTCTG</td><td>>am_trna146_GlnCTG_6_-_27515531_27515602</td><td>1</td><td>29</td><td>29</td><td>GGTTCCATGGTGTAATGGTTAGCACTCTG</td><td>8663.1</td><td>-6.8</td><td>0.9</td><td>-7.2</td><td>5.7E-13</td><td>2.3E-12</td><td>MB231/MB468</td><td>MP</td></tr>
<tr><td>>MI0000087|hsa-mir-29a&WithFlank&7|-|130876741|130876816@47.67.21(+1A)&[MIMAT0000086&hsa-miR-29a-3p&offsets|-1|-2(+1A);m-26&7|-|130876749|130876769&offsets|-1|-1(+1A)]|CTAGCACCATCTGAAATCGGTA</td><td>>MI0000087|hsa-mir-29a&WithFlank&7|-|130876741|130876816</td><td>47</td><td>67</td><td>21(+1A)</td><td>CTAGCACCATCTGAAATCGGTA</td><td>403.9</td><td>11.9</td><td>1.7</td><td>7.2</td><td>8.2E-13</td><td>2.3E-12</td><td>MB231/MB468</td><td>MITO</td></tr>
<tr><td>>hg38_21_+_8433172_8446622_RNA45SN1_45S_50ntFlanks@11641.11660.20|CGGCGGGTGTTGACGCGATG</td><td>>hg38_21_+_8433172_8446622_RNA45SN1_45S_50ntFlanks</td><td>11641</td><td>11660</td><td>20</td><td>CGGCGGGTGTTGACGCGATG</td><td>1777.0</td><td>-7.5</td><td>1.0</td><td>-7.2</td><td>6.5E-13</td><td>2.3E-12</td><td>MB231/MB468</td><td>CYTO</td></tr>
<tr><td>>hg38_21_+_8433172_8446622_RNA45SN1_45S_50ntFlanks@12408.12426.19|TTTGATCCTTCGATGTCGG</td><td>>hg38_21_+_8433172_8446622_RNA45SN1_45S_50ntFlanks</td><td>12408</td><td>12426</td><td>19</td><td>TTTGATCCTTCGATGTCGG</td><td>146.8</td><td>-6.5</td><td>0.9</td><td>-7.2</td><td>6.5E-13</td><td>2.3E-12</td><td>MB231/MB468</td><td>CYTO</td></tr>
<tr><td>>hg38_21_+_8433172_8446622_RNA45SN1_45S_50ntFlanks@12416.12441.26|TTCGATGTCGGCTCTTCCTATCATTG</td><td>>hg38_21_+_8433172_8446622_RNA45SN1_45S_50ntFlanks</td><td>12416</td><td>12441</td><td>26</td><td>TTCGATGTCGGCTCTTCCTATCATTG</td><td>823.6</td><td>-4.0</td><td>0.6</td><td>-7.1</td><td>9.0E-13</td><td>2.3E-12</td><td>BT20/MB468</td><td>NUC</td></tr>
<tr><td>>hg38_21_+_8433172_8446622_RNA45SN1_45S_50ntFlanks@10391.10430.40|AACAGCAGTTGAACATGGGTCAGTCGGTCCTGAGAGATGG</td><td>>hg38_21_+_8433172_8446622_RNA45SN1_45S_50ntFlanks</td><td>10391</td><td>10430</td><td>40</td><td>AACAGCAGTTGAACATGGGTCAGTCGGTCCTGAGAGATGG</td><td>1313.7</td><td>-9.7</td><td>1.4</td><td>-7.2</td><td>8.2E-13</td><td>2.3E-12</td><td>MB231/MB468</td><td>MITO</td></tr>
<tr><td>>hg38_21_+_8433172_8446622_RNA45SN1_45S_50ntFlanks@11999.12031.33|CCCTGCGGGCCGCCGGTGAAATACCACTACTCT</td><td>>hg38_21_+_8433172_8446622_RNA45SN1_45S_50ntFlanks</td><td>11999</td><td>12031</td><td>33</td><td>CCCTGCGGGCCGCCGGTGAAATACCACTACTCT</td><td>1433.1</td><td>-7.2</td><td>1.0</td><td>-7.2</td><td>6.6E-13</td><td>2.3E-12</td><td>MB231/MB468</td><td>CYTO</td></tr>
<tr><td>>hg38_21_+_8433172_8446622_RNA45SN1_45S_50ntFlanks@10393.10435.43|CAGCAGTTGAACATGGGTCAGTCGGTCCTGAGAGATGGGCGAG</td><td>>hg38_21_+_8433172_8446622_RNA45SN1_45S_50ntFlanks</td><td>10393</td><td>10435</td><td>43</td><td>CAGCAGTTGAACATGGGTCAGTCGGTCCTGAGAGATGGGCGAG</td><td>1016.3</td><td>-9.5</td><td>1.3</td><td>-7.2</td><td>8.2E-13</td><td>2.3E-12</td><td>MB231/MB468</td><td>MITO</td></tr>
<tr><td>>hg38_21_+_8433172_8446622_RNA45SN1_45S_50ntFlanks@476.501.26|TTCGTGATCGATGTGGTGACGTCGTG</td><td>>hg38_21_+_8433172_8446622_RNA45SN1_45S_50ntFlanks</td><td>476</td><td>501</td><td>26</td><td>TTCGTGATCGATGTGGTGACGTCGTG</td><td>3287.7</td><td>4.9</td><td>0.7</td><td>7.2</td><td>6.7E-13</td><td>2.3E-12</td><td>MB231/MB468</td><td>NUC</td></tr>
<tr><td>>hg38_21_+_8433172_8446622_RNA45SN1_45S_50ntFlanks@12539.12565.27|ATGATGTGTTGTTGCCATGGTAATCCT</td><td>>hg38_21_+_8433172_8446622_RNA45SN1_45S_50ntFlanks</td><td>12539</td><td>12565</td><td>27</td><td>ATGATGTGTTGTTGCCATGGTAATCCT</td><td>182.0</td><td>-7.5</td><td>1.0</td><td>-7.2</td><td>6.5E-13</td><td>2.3E-12</td><td>BT20/MB468</td><td>CYTO</td></tr>
<tr><td>>hg38_21_+_8433172_8446622_RNA45SN1_45S_50ntFlanks@11567.11602.36|AGCAGCCGACTTAGAACTGGTGCGGACCAGGGGAAT</td><td>>hg38_21_+_8433172_8446622_RNA45SN1_45S_50ntFlanks</td><td>11567</td><td>11602</td><td>36</td><td>AGCAGCCGACTTAGAACTGGTGCGGACCAGGGGAAT</td><td>1226.8</td><td>-6.1</td><td>0.9</td><td>-7.1</td><td>9.2E-13</td><td>2.4E-12</td><td>BT20/MB468</td><td>NUC</td></tr>
<tr><td>>hg38_21_+_8433172_8446622_RNA45SN1_45S_50ntFlanks@11233.11252.20|CCCTTCCCGTGGATCGCCCC</td><td>>hg38_21_+_8433172_8446622_RNA45SN1_45S_50ntFlanks</td><td>11233</td><td>11252</td><td>20</td><td>CCCTTCCCGTGGATCGCCCC</td><td>725.3</td><td>-9.3</td><td>1.3</td><td>-7.2</td><td>6.8E-13</td><td>2.4E-12</td><td>MB231/MB468</td><td>NUC</td></tr>
<tr><td>>hg38_21_+_8433172_8446622_RNA45SN1_45S_50ntFlanks@12605.12634.30|TATGTGCTTGGCTGAGGAGCCAATGGGGCG</td><td>>hg38_21_+_8433172_8446622_RNA45SN1_45S_50ntFlanks</td><td>12605</td><td>12634</td><td>30</td><td>TATGTGCTTGGCTGAGGAGCCAATGGGGCG</td><td>144.0</td><td>-11.0</td><td>1.5</td><td>-7.2</td><td>5.8E-13</td><td>2.4E-12</td><td>MB231/MB468</td><td>MP</td></tr>
<tr><td>>hg38_21_+_8433172_8446622_RNA45SN1_45S_50ntFlanks@6788.6807.20|CGCCTGTCTGAGCGTCGCTT</td><td>>hg38_21_+_8433172_8446622_RNA45SN1_45S_50ntFlanks</td><td>6788</td><td>6807</td><td>20</td><td>CGCCTGTCTGAGCGTCGCTT</td><td>1634.2</td><td>5.3</td><td>0.7</td><td>7.2</td><td>6.7E-13</td><td>2.4E-12</td><td>MB231/MB468</td><td>CYTO</td></tr>
<tr><td>>hg38_21_+_8433172_8446622_RNA45SN1_45S_50ntFlanks@12296.12323.28|AGCGGGGCCTCACGATCCTTCTGACCTT</td><td>>hg38_21_+_8433172_8446622_RNA45SN1_45S_50ntFlanks</td><td>12296</td><td>12323</td><td>28</td><td>AGCGGGGCCTCACGATCCTTCTGACCTT</td><td>336.2</td><td>-12.2</td><td>1.7</td><td>-7.2</td><td>8.4E-13</td><td>2.4E-12</td><td>MB231/MB468</td><td>MITO</td></tr>
<tr><td>>hg38_21_+_8433172_8446622_RNA45SN1_45S_50ntFlanks@12626.12652.27|AATGGGGCGAAGCTACCATCTGTGGGA</td><td>>hg38_21_+_8433172_8446622_RNA45SN1_45S_50ntFlanks</td><td>12626</td><td>12652</td><td>27</td><td>AATGGGGCGAAGCTACCATCTGTGGGA</td><td>651.7</td><td>-8.6</td><td>1.2</td><td>-7.2</td><td>8.4E-13</td><td>2.4E-12</td><td>MB231/MB468</td><td>MITO</td></tr>
<tr><td>>hg38_21_+_8433172_8446622_RNA45SN1_45S_50ntFlanks@4090.4121.32|CCTACCATGGTGACCACGGGTGACGGGGAATC</td><td>>hg38_21_+_8433172_8446622_RNA45SN1_45S_50ntFlanks</td><td>4090</td><td>4121</td><td>32</td><td>CCTACCATGGTGACCACGGGTGACGGGGAATC</td><td>114.8</td><td>-10.7</td><td>1.5</td><td>-7.2</td><td>5.8E-13</td><td>2.4E-12</td><td>MB231/MB468</td><td>MP</td></tr>
<tr><td>>MI0003560|hsa-mir-92b&WithFlank&1|+|155195171|155195278@67.87.21&[MIMAT0003218&hsa-miR-92b-3p&offsets|0|-1;m-115&1|+|155195237|155195258&offsets|0|-1]|TATTGCACTCGTCCCGGCCTC</td><td>>MI0003560|hsa-mir-92b&WithFlank&1|+|155195171|155195278</td><td>67</td><td>87</td><td>21</td><td>TATTGCACTCGTCCCGGCCTC</td><td>1285.8</td><td>10.0</td><td>1.4</td><td>7.2</td><td>5.0E-13</td><td>2.4E-12</td><td>BT20/MB468</td><td>MITO</td></tr>
<tr><td>>hg38_21_+_8433172_8446622_RNA45SN1_45S_50ntFlanks@12449.12480.32|GAATTCACCAAGCGTTGGATTGTTCACCCACT</td><td>>hg38_21_+_8433172_8446622_RNA45SN1_45S_50ntFlanks</td><td>12449</td><td>12480</td><td>32</td><td>GAATTCACCAAGCGTTGGATTGTTCACCCACT</td><td>855.9</td><td>7.3</td><td>1.0</td><td>7.2</td><td>6.7E-13</td><td>2.4E-12</td><td>MB231/MB468</td><td>CYTO</td></tr>
<tr><td>>hg38_21_+_8433172_8446622_RNA45SN1_45S_50ntFlanks@11874.11919.46|AGAAGACCCTGTTGAGCTTGACTCTAGTCTGGCACGGTGAAGAGAC</td><td>>hg38_21_+_8433172_8446622_RNA45SN1_45S_50ntFlanks</td><td>11874</td><td>11919</td><td>46</td><td>AGAAGACCCTGTTGAGCTTGACTCTAGTCTGGCACGGTGAAGAGAC</td><td>268.9</td><td>-11.9</td><td>1.7</td><td>-7.2</td><td>8.4E-13</td><td>2.4E-12</td><td>MB231/MB468</td><td>MITO</td></tr>
<tr><td>>hg38_21_+_8433172_8446622_RNA45SN1_45S_50ntFlanks@11881.11900.20|CCTGTTGAGCTTGACTCTAG</td><td>>hg38_21_+_8433172_8446622_RNA45SN1_45S_50ntFlanks</td><td>11881</td><td>11900</td><td>20</td><td>CCTGTTGAGCTTGACTCTAG</td><td>207.5</td><td>-5.6</td><td>0.8</td><td>-7.2</td><td>6.8E-13</td><td>2.4E-12</td><td>MB231/MB468</td><td>CYTO</td></tr>
<tr><td>>hg38_21_+_8433172_8446622_RNA45SN1_45S_50ntFlanks@4122.4144.23|AGGGTTCGATTCCGGAGAGGGAG</td><td>>hg38_21_+_8433172_8446622_RNA45SN1_45S_50ntFlanks</td><td>4122</td><td>4144</td><td>23</td><td>AGGGTTCGATTCCGGAGAGGGAG</td><td>266.1</td><td>-8.4</td><td>1.2</td><td>-7.2</td><td>6.9E-13</td><td>2.4E-12</td><td>MB231/MB468</td><td>NUC</td></tr>
<tr><td>>hg38_21_+_8433172_8446622_RNA45SN1_45S_50ntFlanks@4823.4852.30|ATGACCCGCCGGGCAGCTTCCGGGAAACCA</td><td>>hg38_21_+_8433172_8446622_RNA45SN1_45S_50ntFlanks</td><td>4823</td><td>4852</td><td>30</td><td>ATGACCCGCCGGGCAGCTTCCGGGAAACCA</td><td>175.4</td><td>4.6</td><td>0.6</td><td>7.2</td><td>6.8E-13</td><td>2.4E-12</td><td>MB231/MB468</td><td>CYTO</td></tr>
<tr><td>>hg38_21_+_8433172_8446622_RNA45SN1_45S_50ntFlanks@9029.9064.36|TCTCCTCGTGGGGGGGCCGGGCCACCCCTCCCACGG</td><td>>hg38_21_+_8433172_8446622_RNA45SN1_45S_50ntFlanks</td><td>9029</td><td>9064</td><td>36</td><td>TCTCCTCGTGGGGGGGCCGGGCCACCCCTCCCACGG</td><td>129.8</td><td>-10.9</td><td>1.5</td><td>-7.2</td><td>5.9E-13</td><td>2.4E-12</td><td>MB231/MB468</td><td>MP</td></tr>
<tr><td>>MI0000299|hsa-mir-222&WithFlank&X|-|45747009|45747130@75.97.23(+1A)&[MIMAT0000279&hsa-miR-222-3p&offsets|0|+2(+1A);m-140&X|-|45747033|45747056&offsets|0|-1(+1A)]|AGCTACATCTGGCTACTGGGTCTA</td><td>>MI0000299|hsa-mir-222&WithFlank&X|-|45747009|45747130</td><td>75</td><td>97</td><td>23(+1A)</td><td>AGCTACATCTGGCTACTGGGTCTA</td><td>1158.3</td><td>6.4</td><td>0.9</td><td>7.2</td><td>8.5E-13</td><td>2.4E-12</td><td>MB231/MB468</td><td>MITO</td></tr>
<tr><td>>hg38_21_+_8433172_8446622_RNA45SN1_45S_50ntFlanks@12412.12431.20|ATCCTTCGATGTCGGCTCTT</td><td>>hg38_21_+_8433172_8446622_RNA45SN1_45S_50ntFlanks</td><td>12412</td><td>12431</td><td>20</td><td>ATCCTTCGATGTCGGCTCTT</td><td>118.2</td><td>-10.7</td><td>1.5</td><td>-7.2</td><td>5.9E-13</td><td>2.4E-12</td><td>MB231/MB468</td><td>MP</td></tr>
<tr><td>>hg38_21_+_8433172_8446622_RNA45SN1_45S_50ntFlanks@12957.12990.34|CTTCTGGGTCGGGGTTTCGTACGTAGCAGAGCAG</td><td>>hg38_21_+_8433172_8446622_RNA45SN1_45S_50ntFlanks</td><td>12957</td><td>12990</td><td>34</td><td>CTTCTGGGTCGGGGTTTCGTACGTAGCAGAGCAG</td><td>199.7</td><td>-11.5</td><td>1.6</td><td>-7.2</td><td>5.9E-13</td><td>2.4E-12</td><td>MB231/MB468</td><td>MP</td></tr>
<tr><td>>hg38_21_+_8433172_8446622_RNA45SN1_45S_50ntFlanks@11669.11693.25|GCCCAGTGCTCTGAATGTCAAAGTG</td><td>>hg38_21_+_8433172_8446622_RNA45SN1_45S_50ntFlanks</td><td>11669</td><td>11693</td><td>25</td><td>GCCCAGTGCTCTGAATGTCAAAGTG</td><td>326.4</td><td>5.2</td><td>0.7</td><td>7.2</td><td>6.7E-13</td><td>2.4E-12</td><td>BT20/MB468</td><td>CYTO</td></tr>
<tr><td>>MI0000078|hsa-mir-22&WithFlank&17|-|1713897|1713993@21.41.21&[MIMAT0004495&hsa-miR-22-5p&offsets|0|-1;m-217&17|-|1713953|1713973&offsets|0|0]|AGTTCTTCAGTGGCAAGCTTT</td><td>>MI0000078|hsa-mir-22&WithFlank&17|-|1713897|1713993</td><td>21</td><td>41</td><td>21</td><td>AGTTCTTCAGTGGCAAGCTTT</td><td>2696.1</td><td>5.5</td><td>0.8</td><td>7.2</td><td>5.9E-13</td><td>2.4E-12</td><td>MB231/MB468</td><td>MP</td></tr>
<tr><td>>hg38_21_+_8433172_8446622_RNA45SN1_45S_50ntFlanks@10727.10747.21|GGAGAGGGTGTAAATCTCGCG</td><td>>hg38_21_+_8433172_8446622_RNA45SN1_45S_50ntFlanks</td><td>10727</td><td>10747</td><td>21</td><td>GGAGAGGGTGTAAATCTCGCG</td><td>175.6</td><td>-11.3</td><td>1.6</td><td>-7.2</td><td>5.9E-13</td><td>2.4E-12</td><td>MB231/MB468</td><td>MP</td></tr>
<tr><td>>hg38_21_+_8433172_8446622_RNA45SN1_45S_50ntFlanks@12044.12071.28|ACTGACCCGGTGAGGCGGGGGGGCGAGC</td><td>>hg38_21_+_8433172_8446622_RNA45SN1_45S_50ntFlanks</td><td>12044</td><td>12071</td><td>28</td><td>ACTGACCCGGTGAGGCGGGGGGGCGAGC</td><td>146.2</td><td>-7.4</td><td>1.0</td><td>-7.2</td><td>6.8E-13</td><td>2.4E-12</td><td>MB231/MB468</td><td>CYTO</td></tr>
<tr><td>>trna10_AspGTC_12_-_125424193_125424264@32.70.39,trna12_AspGTC_12_-_125411891_125411962@32.70.39,trna144_AspGTC_6_-_27551236_27551307@32.70.39,trna38_AspGTC_17_-_8125556_8125627@32.70.39,trna45_AspGTC_6_+_27447453_27447524@32.70.39,trna48_AspGTC_6_+_27471523_27471594@32.70.39,trna4_AspGTC_12_+_96429799_96429870@32.70.39,trna69_AspGTC_1_-_161440205_161440276@32.70.39,trna72_AspGTC_1_-_161432824_161432895@32.70.39,trna75_AspGTC_1_-_161425414_161425485@32.70.39,trna78_AspGTC_1_-_161418033_161418104@32.70.39,trna81_AspGTC_1_-_161410615_161410686@32.70.39|CTGTCACGCGGGAGACCGGGGTTCGATTCCCCGACGGGG</td><td>>trna10_AspGTC_12_-_125424193_125424264</td><td>32</td><td>70</td><td>39</td><td>CTGTCACGCGGGAGACCGGGGTTCGATTCCCCGACGGGG</td><td>1103.5</td><td>-10.3</td><td>1.4</td><td>-7.1</td><td>9.4E-13</td><td>2.4E-12</td><td>BT20/MB468</td><td>NUC</td></tr>
<tr><td>>hg38_21_+_8433172_8446622_RNA45SN1_45S_50ntFlanks@9860.9878.19|GGATGAACCGAACGCCGGG</td><td>>hg38_21_+_8433172_8446622_RNA45SN1_45S_50ntFlanks</td><td>9860</td><td>9878</td><td>19</td><td>GGATGAACCGAACGCCGGG</td><td>153.3</td><td>-11.1</td><td>1.5</td><td>-7.2</td><td>5.9E-13</td><td>2.4E-12</td><td>MB231/MB468</td><td>MP</td></tr>
<tr><td>>hg38_21_+_8433172_8446622_RNA45SN1_45S_50ntFlanks@10877.10902.26|GTCGGTCGGGCTGGGGCGCGAAGCGG</td><td>>hg38_21_+_8433172_8446622_RNA45SN1_45S_50ntFlanks</td><td>10877</td><td>10902</td><td>26</td><td>GTCGGTCGGGCTGGGGCGCGAAGCGG</td><td>194.1</td><td>-5.1</td><td>0.7</td><td>-7.2</td><td>6.8E-13</td><td>2.4E-12</td><td>MB231/MB468</td><td>CYTO</td></tr>
<tr><td>>hg38_21_+_8433172_8446622_RNA45SN1_45S_50ntFlanks@9572.9592.21|CAAATCGGTCGTCCGACCTGG</td><td>>hg38_21_+_8433172_8446622_RNA45SN1_45S_50ntFlanks</td><td>9572</td><td>9592</td><td>21</td><td>CAAATCGGTCGTCCGACCTGG</td><td>1146.1</td><td>-13.2</td><td>1.7</td><td>-7.6</td><td>3.1E-14</td><td>2.4E-12</td><td>BT20/MB468</td><td>MP</td></tr>
<tr><td>>hg38_21_+_8433172_8446622_RNA45SN1_45S_50ntFlanks@10461.10497.37|GCCTCCGTTGCCCTCGGCCGATCGAAAGGGAGTCGGG</td><td>>hg38_21_+_8433172_8446622_RNA45SN1_45S_50ntFlanks</td><td>10461</td><td>10497</td><td>37</td><td>GCCTCCGTTGCCCTCGGCCGATCGAAAGGGAGTCGGG</td><td>139.0</td><td>-11.0</td><td>1.5</td><td>-7.2</td><td>5.9E-13</td><td>2.4E-12</td><td>MB231/MB468</td><td>MP</td></tr>
<tr><td>>hg38_21_+_8433172_8446622_RNA45SN1_45S_50ntFlanks@11783.11806.24|CATGAATGGATGAACGAGATTCCC</td><td>>hg38_21_+_8433172_8446622_RNA45SN1_45S_50ntFlanks</td><td>11783</td><td>11806</td><td>24</td><td>CATGAATGGATGAACGAGATTCCC</td><td>3655.4</td><td>-7.8</td><td>1.1</td><td>-7.2</td><td>8.6E-13</td><td>2.4E-12</td><td>MB231/MB468</td><td>MITO</td></tr>
<tr><td>>hg38_21_+_8433172_8446622_RNA45SN1_45S_50ntFlanks@10384.10430.47|CCATGTGAACAGCAGTTGAACATGGGTCAGTCGGTCCTGAGAGATGG</td><td>>hg38_21_+_8433172_8446622_RNA45SN1_45S_50ntFlanks</td><td>10384</td><td>10430</td><td>47</td><td>CCATGTGAACAGCAGTTGAACATGGGTCAGTCGGTCCTGAGAGATGG</td><td>863.9</td><td>-14.2</td><td>2.0</td><td>-7.1</td><td>9.5E-13</td><td>2.4E-12</td><td>BT20/MB468</td><td>NUC</td></tr>
<tr><td>>am_MI0000809|hsa-mir-151a&WithFlank&8|-|140732558|140732659@53.73.21(+1C)&[MIMAT0000757&hsa-miR-151a-3p&offsets|0|0(+1C);m-30&8|-|140732587|140732607&offsets|0|0(+1C)]|CTAGACTGAAGCTCCTTGAGGC</td><td>>am_MI0000809|hsa-mir-151a&WithFlank&8|-|140732558|140732659</td><td>53</td><td>73</td><td>21(+1C)</td><td>CTAGACTGAAGCTCCTTGAGGC</td><td>987.7</td><td>6.9</td><td>1.0</td><td>7.2</td><td>8.6E-13</td><td>2.4E-12</td><td>MB231/MB468</td><td>MITO</td></tr>
</table>

| | | | | | | | | | | | | | |
|---|---|---|---|---|---|---|---|---|---|---|---|---|---|
| >hg38_21_+_8433172_8446622_RNA45SN1_45S_50ntFlanks@5487.5515.29\|CGCTGAGAAGACGGTCGAACTTGACTATC | >hg38_21_+_8433172_8446622_RNA45SN1_45S_50ntFlanks | 5487 | 5515 | 29 | CGCTGAGAAGACGGTCGAACTTGACTATC | 1007.3 | -7.5 | 1.0 | -7.2 | 8.6E-13 | 2.4E-12 | MB231/MB468 | MITO |
| >hg38_21_+_8433172_8446622_RNA45SN1_45S_50ntFlanks@10411.10442.32\|CAGTCGGTCCTGAGAGATGGGCGAGCGCCGTT | >hg38_21_+_8433172_8446622_RNA45SN1_45S_50ntFlanks | 10411 | 10442 | 32 | CAGTCGGTCCTGAGAGATGGGCGAGCGCCGTT | 637.2 | -4.1 | 0.6 | -7.2 | 6.8E-13 | 2.4E-12 | BT20/MB468 | CYTO |
| >hg38_21_+_8433172_8446622_RNA45SN1_45S_50ntFlanks@9454.9489.36\|CCGCGCCGGGGAGGTGGAGCACGAGCGCACGTGTTA | >hg38_21_+_8433172_8446622_RNA45SN1_45S_50ntFlanks | 9454 | 9489 | 36 | CCGCGCCGGGGAGGTGGAGCACGAGCGCACGTGTTA | 110.4 | -10.7 | 1.5 | -7.2 | 6.0E-13 | 2.4E-12 | MB231/MB468 | MP |
| >hg38_21_+_8433172_8446622_RNA45SN1_45S_50ntFlanks@3999.4024.26\|CTAGATAACCTCGGGCCGATCGCACG | >hg38_21_+_8433172_8446622_RNA45SN1_45S_50ntFlanks | 3999 | 4024 | 26 | CTAGATAACCTCGGGCCGATCGCACG | 139.3 | -11.0 | 1.5 | -7.2 | 6.0E-13 | 2.4E-12 | MB231/MB468 | MP |
| >MI0000746\|hsa-mir-99b&WithFlank&19\|+\|51692606\|51692687@13.32.20&[MIMAT0000689&hsa-miR-99b-5p&offsets\|0\|-2;m-10&19\|+\|51692618\|51692639&offsets\|0\|-2]\|CACCCGTAGAACCGACCTTG | >MI0000746\|hsa-mir-99b&WithFlank&19\|+\|51692606\|51692687 | 13 | 32 | 20 | CACCCGTAGAACCGACCTTG | 1184.7 | 6.9 | 1.0 | 7.2 | 6.9E-13 | 2.4E-12 | MB231/MB468 | CYTO |
| >hg38_21_+_8433172_8446622_RNA45SN1_45S_50ntFlanks@12838.12862.25\|GTGCGGAGTGCCCTTCGTCCTGGGA | >hg38_21_+_8433172_8446622_RNA45SN1_45S_50ntFlanks | 12838 | 12862 | 25 | GTGCGGAGTGCCCTTCGTCCTGGGA | 260.9 | -11.9 | 1.7 | -7.2 | 6.0E-13 | 2.4E-12 | MB231/MB468 | MP |
| >hg38_21_+_8433172_8446622_RNA45SN1_45S_50ntFlanks@4555.4572.18\|CGCAGCTAGGAATAATGG | >hg38_21_+_8433172_8446622_RNA45SN1_45S_50ntFlanks | 4555 | 4572 | 18 | CGCAGCTAGGAATAATGG | 1034.6 | -7.8 | 1.1 | -7.2 | 7.1E-13 | 2.4E-12 | MB231/MB468 | NUC |
| >hg38_21_+_8433172_8446622_RNA45SN1_45S_50ntFlanks@9563.9597.35\|CCTGACGTGCAAATCGGTCGTCCGACCTGGGTATA | >hg38_21_+_8433172_8446622_RNA45SN1_45S_50ntFlanks | 9563 | 9597 | 35 | CCTGACGTGCAAATCGGTCGTCCGACCTGGGTATA | 3613.2 | -7.7 | 1.1 | -7.2 | 8.7E-13 | 2.4E-12 | MB231/MB468 | MITO |
| >hg38_21_+_8433172_8446622_RNA45SN1_45S_50ntFlanks@12073.12110.38\|CCGAGGGGCTCTCGCTTCTGGCGCCAAGCGCCCGGCCG | >hg38_21_+_8433172_8446622_RNA45SN1_45S_50ntFlanks | 12073 | 12110 | 38 | CCGAGGGGCTCTCGCTTCTGGCGCCAAGCGCCCGGCCG | 226.2 | -8.1 | 1.1 | -7.2 | 6.9E-13 | 2.4E-12 | MB231/MB468 | CYTO |
| >hg38_21_+_8433172_8446622_RNA45SN1_45S_50ntFlanks@4367.4393.27\|CAGTTAAAAAGCTCGTAGTTGGATCTT | >hg38_21_+_8433172_8446622_RNA45SN1_45S_50ntFlanks | 4367 | 4393 | 27 | CAGTTAAAAAGCTCGTAGTTGGATCTT | 143.3 | -11.0 | 1.5 | -7.2 | 6.0E-13 | 2.5E-12 | MB231/MB468 | MP |
| >hg38_21_+_8433172_8446622_RNA45SN1_45S_50ntFlanks@12008.12035.28\|CCGCCGGTGAAATACCACTACTCTGATC | >hg38_21_+_8433172_8446622_RNA45SN1_45S_50ntFlanks | 12008 | 12035 | 28 | CCGCCGGTGAAATACCACTACTCTGATC | 711.0 | -8.7 | 1.2 | -7.1 | 8.7E-13 | 2.5E-12 | MB231/MB468 | MITO |
| >hg38_21_+_8433172_8446622_RNA45SN1_45S_50ntFlanks@12433.12455.23\|CTATCATTGTGAAGCAGAATTCA | >hg38_21_+_8433172_8446622_RNA45SN1_45S_50ntFlanks | 12433 | 12455 | 23 | CTATCATTGTGAAGCAGAATTCA | 158.0 | -10.0 | 1.4 | -7.2 | 7.0E-13 | 2.5E-12 | MB231/MB468 | CYTO |
| >MI0000342\|hsa-mir-200b&WithFlank&1\|+\|1167098\|1167204@63.86.24&[MIMAT0000318&hsa-miR-200b-3p&offsets\|0\|+2;m-93&1\|+\|1167160\|1167182&offsets\|0\|+1]\|TAATACTGCCTGGTAATGATGACG | >MI0000342\|hsa-mir-200b&WithFlank&1\|+\|1167098\|1167204 | 63 | 86 | 24 | TAATACTGCCTGGTAATGATGACG | 98.4 | -10.5 | 1.5 | -7.2 | 6.1E-13 | 2.5E-12 | MB231/MB468 | MP |
| >hg38_21_+_8433172_8446622_RNA45SN1_45S_50ntFlanks@10681.10712.32\|TTCCGGCGGCGTCCGGTGAGCTCTCGCTGGCC | >hg38_21_+_8433172_8446622_RNA45SN1_45S_50ntFlanks | 10681 | 10712 | 32 | TTCCGGCGGCGTCCGGTGAGCTCTCGCTGGCC | 444.2 | -8.5 | 1.2 | -7.2 | 6.1E-13 | 2.5E-12 | MB231/MB468 | MP |
| >MI0000095\|hsa-mir-93&WithFlank&7\|-\|100093762\|100093853@17.40.24&[MIMAT0000093&hsa-miR-93-5p&offsets\|0\|+1;m-29&7\|-\|100093815\|100093837&offsets\|0\|+1]\|CAAAGTGCTGTTCGTGCAGGTAGT | >MI0000095\|hsa-mir-93&WithFlank&7\|-\|100093762\|100093853 | 17 | 40 | 24 | CAAAGTGCTGTTCGTGCAGGTAGT | 2863.4 | 5.1 | 0.7 | 7.2 | 7.1E-13 | 2.5E-12 | MB231/MB468 | NUC |
| >hg38_21_+_8433172_8446622_RNA45SN1_45S_50ntFlanks@10906.10933.28\|TGGGCGCGCGCCGCGGCTGGACGAGGCG | >hg38_21_+_8433172_8446622_RNA45SN1_45S_50ntFlanks | 10906 | 10933 | 28 | TGGGCGCGCGCCGCGGCTGGACGAGGCG | 178.2 | -11.3 | 1.6 | -7.2 | 6.1E-13 | 2.5E-12 | MB231/MB468 | MP |
| >MI0000471\|hsa-mir-126&WithFlank&9\|+\|136670596\|136670692@58.79.22(+1U)&[MIMAT0000445&hsa-miR-126-3p&offsets\|0\|0(+1U);m-40&9\|+\|136670653\|136670674&offsets\|0\|0(+1U)]\|TCGTACCGTGAGTAATAATGCGT | >MI0000471\|hsa-mir-126&WithFlank&9\|+\|136670596\|136670692 | 58 | 79 | 22(+1U) | TCGTACCGTGAGTAATAATGCGT | 189.6 | 10.8 | 1.5 | 7.2 | 6.1E-13 | 2.5E-12 | MB231/MB468 | MP |
| >hg38_21_+_8433172_8446622_RNA45SN1_45S_50ntFlanks@8182.8204.23\|ATCGAGGCCCAGCCCGTGGACGG | >hg38_21_+_8433172_8446622_RNA45SN1_45S_50ntFlanks | 8182 | 8204 | 23 | ATCGAGGCCCAGCCCGTGGACGG | 208.1 | -9.8 | 1.4 | -7.2 | 5.2E-13 | 2.5E-12 | BT20/MB468 | MITO |
| >hg38_21_+_8433172_8446622_RNA45SN1_45S_50ntFlanks@12328.12346.19\|GTTTTAAGCAGGAGGTGTC | >hg38_21_+_8433172_8446622_RNA45SN1_45S_50ntFlanks | 12328 | 12346 | 19 | GTTTTAAGCAGGAGGTGTC | 177.4 | -11.3 | 1.6 | -7.2 | 6.1E-13 | 2.5E-12 | MB231/MB468 | MP |
| >hg38_21_+_8433172_8446622_RNA45SN1_45S_50ntFlanks@4615.4632.18\|CCATGATTAAGAGGGACG | >hg38_21_+_8433172_8446622_RNA45SN1_45S_50ntFlanks | 4615 | 4632 | 18 | CCATGATTAAGAGGGACG | 161.7 | -11.2 | 1.6 | -7.2 | 6.1E-13 | 2.5E-12 | MB231/MB468 | MP |
| >hg38_21_+_8433172_8446622_RNA45SN1_45S_50ntFlanks@10922.10945.24\|CTGGACGAGGCGCCGCCGCCCCCC | >hg38_21_+_8433172_8446622_RNA45SN1_45S_50ntFlanks | 10922 | 10945 | 24 | CTGGACGAGGCGCCGCCGCCCCCC | 1523.6 | -9.8 | 1.4 | -7.2 | 6.1E-13 | 2.5E-12 | MB231/MB468 | MP |
| >hg38_21_+_8433172_8446622_RNA45SN1_45S_50ntFlanks@12415.12432.18\|CTTCGATGTCGGCTCTTC | >hg38_21_+_8433172_8446622_RNA45SN1_45S_50ntFlanks | 12415 | 12432 | 18 | CTTCGATGTCGGCTCTTC | 2971.7 | -5.1 | 0.7 | -7.1 | 8.8E-13 | 2.5E-12 | MB231/MB468 | MITO |
| >MI0003643\|hsa-mir-629&WithFlank&15\|-\|70079366\|70079474@28.47.20(+1C)&[MIMAT0004810&hsa-miR-629-5p&offsets\|0\|-1(+1C);m-150&15\|-\|70079427\|70079447&offsets\|0\|-1(+1C)]\|TGGGTTTACGTTGGGAGAACC | >MI0003643\|hsa-mir-629&WithFlank&15\|-\|70079366\|70079474 | 28 | 47 | 20(+1C) | TGGGTTTACGTTGGGAGAACC | 881.1 | 13.5 | 1.8 | 7.6 | 3.2E-14 | 2.5E-12 | BT20/MB468 | MP |
| >hg38_21_+_8433172_8446622_RNA45SN1_45S_50ntFlanks@8168.8188.21\|CCCAAGTCCTTCTGATCGAGG | >hg38_21_+_8433172_8446622_RNA45SN1_45S_50ntFlanks | 8168 | 8188 | 21 | CCCAAGTCCTTCTGATCGAGG | 494.5 | -11.7 | 1.6 | -7.2 | 7.1E-13 | 2.5E-12 | MB231/MB468 | CYTO |
| >hg38_21_+_8433172_8446622_RNA45SN1_45S_50ntFlanks@8194.8211.18\|CCCGTGGACGGTGTGAGG | >hg38_21_+_8433172_8446622_RNA45SN1_45S_50ntFlanks | 8194 | 8211 | 18 | CCCGTGGACGGTGTGAGG | 5033.0 | -10.5 | 1.5 | -7.2 | 6.1E-13 | 2.5E-12 | MB231/MB468 | MP |
| >MI0000434\|hsa-let-7i&WithFlank&12\|+\|62603680\|62603775@12.30.19&[MIMAT0000415&hsa-let-7i-5p&offsets\|0\|-3;m-49&12\|+\|62603691\|62603712&offsets\|0\|-3]\|TGAGGTAGTAGTTTGTGCT | >MI0000434\|hsa-let-7i&WithFlank&12\|+\|62603680\|62603775 | 12 | 30 | 19 | TGAGGTAGTAGTTTGTGCT | 191.9 | 4.9 | 0.7 | 7.2 | 7.2E-13 | 2.5E-12 | MB231/MB468 | NUC |
| >hg38_21_+_8433172_8446622_RNA45SN1_45S_50ntFlanks@12395.12420.26\|AGCGACGTCGCTTTTTGATCCTTCGA | >hg38_21_+_8433172_8446622_RNA45SN1_45S_50ntFlanks | 12395 | 12420 | 26 | AGCGACGTCGCTTTTTGATCCTTCGA | 287.7 | 5.7 | 0.8 | 7.2 | 7.2E-13 | 2.5E-12 | MB231/MB468 | NUC |
| >hg38_21_+_8433172_8446622_RNA45SN1_45S_50ntFlanks@12154.12178.25\|GGGAGTTTGACTGGGGCGGTACACC | >hg38_21_+_8433172_8446622_RNA45SN1_45S_50ntFlanks | 12154 | 12178 | 25 | GGGAGTTTGACTGGGGCGGTACACC | 141.4 | -11.0 | 1.5 | -7.2 | 6.1E-13 | 2.5E-12 | MB231/MB468 | MP |
| >trna10_AspGTC_12_-_125424193_125424264@38.74.37,trna12_AspGTC_12_-_125411891_125411962@38.74.37,trna144_AspGTC_6_-_27551236_27551307@38.74.37,trna38_AspGTC_17_-_8125556_8125627@38.74.37,trna45_AspGTC_6_+_27447453_27447524@38.74.37,trna48_AspGTC_6_+_27471523_27471594@38.74.37,trna4_AspGTC_12_+_96429799_96429870@38.74.37,trna69_AspGTC_1_-_161440205_161440276@38.74.37,trna72_AspGTC_1_-_161432824_161432895@38.74.37,trna75_AspGTC_1_-_161425414_161425485@38.74.37,trna78_AspGTC_1_-_161418033_161418104@38.74.37,trna81_AspGTC_1_-_161410615_161410686@38.74.37\|CGCGGGAGACCGGGGTTCGATTCCCCGACGGGGAGCC | >trna10_AspGTC_12_-_125424193_125424264 | 38 | 74 | 37 | CGCGGGAGACCGGGGTTCGATTCCCCGACGGGGAGCC | 2221.6 | 7.1 | 1.0 | 7.2 | 7.1E-13 | 2.5E-12 | MB231/MB468 | CYTO |
| >hg38_21_+_8433172_8446622_RNA45SN1_45S_50ntFlanks@9171.9195.25\|CGGGTCGCGCCGTCGGGCCCGGGGG | >hg38_21_+_8433172_8446622_RNA45SN1_45S_50ntFlanks | 9171 | 9195 | 25 | CGGGTCGCGCCGTCGGGCCCGGGGG | 333.9 | -12.2 | 1.7 | -7.2 | 6.1E-13 | 2.5E-12 | MB231/MB468 | MP |
| >MI0000082\|hsa-mir-25&WithFlank&7\|-\|100093554\|100093649@58.77.20&[MIMAT0000081&hsa-miR-25-3p&offsets\|0\|-2;m-23&7\|-\|100093571\|100093592&offsets\|0\|-2]\|CATTGCACTTGTCTCGGTCT | >MI0000082\|hsa-mir-25&WithFlank&7\|-\|100093554\|100093649 | 58 | 77 | 20 | CATTGCACTTGTCTCGGTCT | 244.5 | 4.8 | 0.7 | 7.2 | 7.2E-13 | 2.5E-12 | MB231/MB468 | NUC |
| >trna22_MetCAT_16_-_87417628_87417700@15.42.28,trna75_MetCAT_6_+_28912352_28912424@15.42.28,trna97_MetCAT_6_-_28921042_28921114@15.42.28\|GTAGGCAGCGCGTCAGTCTCATAATCTG | >trna22_MetCAT_16_-_87417628_87417700 | 15 | 42 | 28 | GTAGGCAGCGCGTCAGTCTCATAATCTG | 189.3 | -10.5 | 1.5 | -7.2 | 7.3E-13 | 2.5E-12 | MB231/MB468 | NUC |
| >hg38_21_+_8433172_8446622_RNA45SN1_45S_50ntFlanks@10758.10785.28\|CCCATATCCGCAGCAGGTCTCCAAGGTG | >hg38_21_+_8433172_8446622_RNA45SN1_45S_50ntFlanks | 10758 | 10785 | 28 | CCCATATCCGCAGCAGGTCTCCAAGGTG | 189.0 | -11.4 | 1.6 | -7.2 | 6.2E-13 | 2.5E-12 | MB231/MB468 | MP |
| >hg38_21_+_8433172_8446622_RNA45SN1_45S_50ntFlanks@10238.10259.22\|CGCTACGCCGCGACGAGTAGGA | >hg38_21_+_8433172_8446622_RNA45SN1_45S_50ntFlanks | 10238 | 10259 | 22 | CGCTACGCCGCGACGAGTAGGA | 696.2 | 6.2 | 0.9 | 7.2 | 5.3E-13 | 2.5E-12 | BT20/MB468 | MITO |
| >MI0003628\|hsa-mir-615&WithFlank&12\|+\|54033944\|54034051@67.85.19&[MIMAT0003283&hsa-miR-615-3p&offsets\|0\|-3;m-495&12\|+\|54034010\|54034030&offsets\|0\|-2]\|TCCGAGCCTGGGTCTCCCT | >MI0003628\|hsa-mir-615&WithFlank&12\|+\|54033944\|54034051 | 67 | 85 | 19 | TCCGAGCCTGGGTCTCCCT | 394.9 | 5.2 | 0.7 | 7.5 | 7.2E-14 | 2.5E-12 | BT20/MB231 | MITO |
| >hg38_21_+_8433172_8446622_RNA45SN1_45S_50ntFlanks@12395.12421.27\|AGCGACGTCGCTTTTTGATCCTTCGAT | >hg38_21_+_8433172_8446622_RNA45SN1_45S_50ntFlanks | 12395 | 12421 | 27 | AGCGACGTCGCTTTTTGATCCTTCGAT | 209.1 | 5.1 | 0.7 | 7.2 | 7.3E-13 | 2.5E-12 | MB231/MB468 | NUC |
| >hg38_21_+_8433172_8446622_RNA45SN1_45S_50ntFlanks@10290.10319.30\|GCGCGGGCCCGGGTGGAGCCGCCGCAGGTG | >hg38_21_+_8433172_8446622_RNA45SN1_45S_50ntFlanks | 10290 | 10319 | 30 | GCGCGGGCCCGGGTGGAGCCGCCGCAGGTG | 313.6 | -12.2 | 1.7 | -7.2 | 6.2E-13 | 2.5E-12 | MB231/MB468 | MP |
| >hg38_21_+_8433172_8446622_RNA45SN1_45S_50ntFlanks@6644.6662.19\|CCTCGTACGACTCTTAGCG | >hg38_21_+_8433172_8446622_RNA45SN1_45S_50ntFlanks | 6644 | 6662 | 19 | CCTCGTACGACTCTTAGCG | 898.8 | 5.0 | 0.7 | 7.2 | 5.3E-13 | 2.5E-12 | BT20/MB468 | MITO |
| >hg38_21_+_8433172_8446622_RNA45SN1_45S_50ntFlanks@12678.12702.25\|AATCCCGCCCAGGCGGAACGATACG | >hg38_21_+_8433172_8446622_RNA45SN1_45S_50ntFlanks | 12678 | 12702 | 25 | AATCCCGCCCAGGCGGAACGATACG | 304.1 | -12.1 | 1.7 | -7.2 | 6.2E-13 | 2.5E-12 | MB231/MB468 | MP |
| >hg38_21_+_8433172_8446622_RNA45SN1_45S_50ntFlanks@11873.11890.18\|AAGAAGACCCTGTTGAGC | >hg38_21_+_8433172_8446622_RNA45SN1_45S_50ntFlanks | 11873 | 11890 | 18 | AAGAAGACCCTGTTGAGC | 793.1 | -8.8 | 1.2 | -7.1 | 9.0E-13 | 2.5E-12 | MB231/MB468 | MITO |
| >MI0000542\|hsa-mir-320a&WithFlank&8\|-\|22244960\|22245043@49.70.22(+1U)&[MIMAT0000510&hsa-miR-320a-3p&offsets\|+1\|+1(+1U);m-60&8\|-\|22244975\|22244996&offsets\|+1\|+1(+1U)]\|AAAGCTGGGTTGAGAGGGCGAAT | >MI0000542\|hsa-mir-320a&WithFlank&8\|-\|22244960\|22245043 | 49 | 70 | 22(+1U) | AAAGCTGGGTTGAGAGGGCGAAT | 515.8 | 10.2 | 1.4 | 7.2 | 5.4E-13 | 2.5E-12 | BT20/MB468 | MITO |
| >hg38_21_+_8433172_8446622_RNA45SN1_45S_50ntFlanks@12357.12376.20\|CCACAGGGATAACTGGCTTG | >hg38_21_+_8433172_8446622_RNA45SN1_45S_50ntFlanks | 12357 | 12376 | 20 | CCACAGGGATAACTGGCTTG | 437.0 | -10.9 | 1.5 | -7.2 | 5.4E-13 | 2.5E-12 | BT20/MB468 | MITO |
| >MI0000088\|hsa-mir-30a&WithFlank&6\|-\|71403545\|71403627@54.73.20&[MIMAT0000088&hsa-miR-30a-3p&offsets\|+1\|-1;m-38&6\|-\|71403554\|71403575&offsets\|+1\|-1]\|TTTCAGTCGGATGTTTGCAG | >MI0000088\|hsa-mir-30a&WithFlank&6\|-\|71403545\|71403627 | 54 | 73 | 20 | TTTCAGTCGGATGTTTGCAG | 256.3 | 8.9 | 1.2 | 7.2 | 7.2E-13 | 2.5E-12 | MB231/MB468 | CYTO |
| >hg38_21_+_8433172_8446622_RNA45SN1_45S_50ntFlanks@11027.11053.27\|CTCTCTCTCTCTCTCCCCCGCTCCCCG | >hg38_21_+_8433172_8446622_RNA45SN1_45S_50ntFlanks | 11027 | 11053 | 27 | CTCTCTCTCTCTCTCCCCCGCTCCCCG | 1350.5 | -8.4 | 1.2 | -7.2 | 7.2E-13 | 2.5E-12 | MB231/MB468 | CYTO |
| >hg38_21_+_8433172_8446622_RNA45SN1_45S_50ntFlanks@11707.11728.22\|AAGCGCGGGTAAACGGCGGGAG | >hg38_21_+_8433172_8446622_RNA45SN1_45S_50ntFlanks | 11707 | 11728 | 22 | AAGCGCGGGTAAACGGCGGGAG | 1194.5 | -10.0 | 1.4 | -7.1 | 9.9E-13 | 2.5E-12 | BT20/MB468 | NUC |

<table>
<tr><th></th><th></th><th></th><th></th><th></th><th></th><th></th><th></th><th></th><th></th><th></th><th></th><th></th><th></th><th></th></tr>
<tr><td>>am_trna116_GluCTC_1_-_145399233_145399304@7.29.23,trna59_GluCTC_1_+_249168447_249168518@7.29.23,trna71_GluCTC_1_-_161439189_161439260@7.29.23,trna74_GluCTC_1_-_161431809_161431880@7.29.23,trna77_GluCTC_1_-_161424398_161424469@7.29.23,trna77_GluCTC_6_+_28949976_28950047@7.29.23,trna80_GluCTC_1_-_161417018_161417089@7.29.23,trna87_GluCTC_6_-_126101393_126101464@7.29.23|GTGGTCTAGTGGTTAGGATTCGG</td><td>>am_trna116_GluCTC_1_-_145399233_145399304</td><td>7</td><td>29</td><td>23</td><td>GTGGTCTAGTGGTTAGGATTCGG</td><td>144.0</td><td>-11.0</td><td>1.5</td><td>-7.2</td><td>6.2E-13</td><td>2.5E-12</td><td>MB231/MB468</td><td>MP</td></tr>
<tr><td>>hg38_21_+_8433172_8446622_RNA45SN1_45S_50ntFlanks@12991.13017.27|CTCCCTCGCTGCGATCTATTGAAAGTC</td><td>>hg38_21_+_8433172_8446622_RNA45SN1_45S_50ntFlanks</td><td>12991</td><td>13017</td><td>27</td><td>CTCCCTCGCTGCGATCTATTGAAAGTC</td><td>1629.0</td><td>-4.6</td><td>0.6</td><td>-7.2</td><td>7.4E-13</td><td>2.5E-12</td><td>MB231/MB468</td><td>NUC</td></tr>
<tr><td>>hg38_21_+_8433172_8446622_RNA45SN1_45S_50ntFlanks@9504.9525.22|GTGAACTATGCCTGGGCAGGGC</td><td>>hg38_21_+_8433172_8446622_RNA45SN1_45S_50ntFlanks</td><td>9504</td><td>9525</td><td>22</td><td>GTGAACTATGCCTGGGCAGGGC</td><td>483.4</td><td>-6.9</td><td>1.0</td><td>-7.2</td><td>7.4E-13</td><td>2.5E-12</td><td>MB231/MB468</td><td>NUC</td></tr>
<tr><td>>hg38_21_+_8433172_8446622_RNA45SN1_45S_50ntFlanks@11580.11603.24|GAACTGGTGCGGACCAGGGGAATC</td><td>>hg38_21_+_8433172_8446622_RNA45SN1_45S_50ntFlanks</td><td>11580</td><td>11603</td><td>24</td><td>GAACTGGTGCGGACCAGGGGAATC</td><td>15030.8</td><td>-9.9</td><td>1.4</td><td>-7.2</td><td>6.2E-13</td><td>2.5E-12</td><td>MB231/MB468</td><td>MP</td></tr>
<tr><td>>hg38_21_+_8433172_8446622_RNA45SN1_45S_50ntFlanks@11613.11660.48|AATTAAAACAAAGCATCGCGAAGGCCCGCGGCGGGTGTTGACGCGATG</td><td>>hg38_21_+_8433172_8446622_RNA45SN1_45S_50ntFlanks</td><td>11613</td><td>11660</td><td>48</td><td>AATTAAAACAAAGCATCGCGAAGGCCCGCGGCGGGTGTTGACGCGATG</td><td>338.9</td><td>-12.2</td><td>1.7</td><td>-7.1</td><td>9.0E-13</td><td>2.5E-12</td><td>MB231/MB468</td><td>MITO</td></tr>
<tr><td>>hg38_21_+_8433172_8446622_RNA45SN1_45S_50ntFlanks@12998.13039.42|GCTGCGATCTATTGAAAGTCAGCCCTCGACACAAGGGTTTGT</td><td>>hg38_21_+_8433172_8446622_RNA45SN1_45S_50ntFlanks</td><td>12998</td><td>13039</td><td>42</td><td>GCTGCGATCTATTGAAAGTCAGCCCTCGACACAAGGGTTTGT</td><td>8452.2</td><td>6.9</td><td>1.0</td><td>7.2</td><td>7.2E-13</td><td>2.5E-12</td><td>MB231/MB468</td><td>CYTO</td></tr>
<tr><td>>hg38_21_+_8433172_8446622_RNA45SN1_45S_50ntFlanks@10391.10410.20|AACAGCAGTTGAACATGGGT</td><td>>hg38_21_+_8433172_8446622_RNA45SN1_45S_50ntFlanks</td><td>10391</td><td>10410</td><td>20</td><td>AACAGCAGTTGAACATGGGT</td><td>1514.4</td><td>-9.2</td><td>1.3</td><td>-7.1</td><td>9.0E-13</td><td>2.5E-12</td><td>MB231/MB468</td><td>MITO</td></tr>
<tr><td>>hg38_21_+_8433172_8446622_RNA45SN1_45S_50ntFlanks@12539.12566.28|ATGATGTGTTGTTGCCATGGTAATCCTG</td><td>>hg38_21_+_8433172_8446622_RNA45SN1_45S_50ntFlanks</td><td>12539</td><td>12566</td><td>28</td><td>ATGATGTGTTGTTGCCATGGTAATCCTG</td><td>1050.9</td><td>-7.0</td><td>1.0</td><td>-7.2</td><td>7.2E-13</td><td>2.5E-12</td><td>MB231/MB468</td><td>CYTO</td></tr>
<tr><td>>hg38_21_+_8433172_8446622_RNA45SN1_45S_50ntFlanks@11620.11640.21|ACAAAGCATCGCGAAGGCCCG</td><td>>hg38_21_+_8433172_8446622_RNA45SN1_45S_50ntFlanks</td><td>11620</td><td>11640</td><td>21</td><td>ACAAAGCATCGCGAAGGCCCG</td><td>185.8</td><td>-10.5</td><td>1.5</td><td>-7.2</td><td>7.4E-13</td><td>2.5E-12</td><td>MB231/MB468</td><td>NUC</td></tr>
<tr><td>>hg38_21_+_8433172_8446622_RNA45SN1_45S_50ntFlanks@11565.11601.37|CTAGCAGCCGACTTAGAACTGGTGCGGACCAGGGGAA</td><td>>hg38_21_+_8433172_8446622_RNA45SN1_45S_50ntFlanks</td><td>11565</td><td>11601</td><td>37</td><td>CTAGCAGCCGACTTAGAACTGGTGCGGACCAGGGGAA</td><td>482.7</td><td>-4.8</td><td>0.7</td><td>-7.2</td><td>7.4E-13</td><td>2.5E-12</td><td>MB231/MB468</td><td>NUC</td></tr>
<tr><td>>hg38_21_+_8433172_8446622_RNA45SN1_45S_50ntFlanks@4918.4939.22|ACCACCAGGAGTGGAGCCTGCG</td><td>>hg38_21_+_8433172_8446622_RNA45SN1_45S_50ntFlanks</td><td>4918</td><td>4939</td><td>22</td><td>ACCACCAGGAGTGGAGCCTGCG</td><td>118.0</td><td>-10.7</td><td>1.5</td><td>-7.2</td><td>6.3E-13</td><td>2.5E-12</td><td>MB231/MB468</td><td>MP</td></tr>
<tr><td>>hg38_21_+_8433172_8446622_RNA45SN1_45S_50ntFlanks@10786.10817.32|AACAGCCTCTGGCATGTTGGAACAATGTAGGT</td><td>>hg38_21_+_8433172_8446622_RNA45SN1_45S_50ntFlanks</td><td>10786</td><td>10817</td><td>32</td><td>AACAGCCTCTGGCATGTTGGAACAATGTAGGT</td><td>931.9</td><td>-4.5</td><td>0.6</td><td>-7.2</td><td>7.2E-13</td><td>2.5E-12</td><td>MB231/MB468</td><td>CYTO</td></tr>
<tr><td>>hg38_21_+_8433172_8446622_RNA45SN1_45S_50ntFlanks@5073.5110.38|AATTCCGATAACGAACGAGACTCTGGCATGCTAACTAG</td><td>>hg38_21_+_8433172_8446622_RNA45SN1_45S_50ntFlanks</td><td>5073</td><td>5110</td><td>38</td><td>AATTCCGATAACGAACGAGACTCTGGCATGCTAACTAG</td><td>1136.1</td><td>-8.4</td><td>1.2</td><td>-7.1</td><td>1.0E-12</td><td>2.5E-12</td><td>BT20/MB468</td><td>NUC</td></tr>
<tr><td>>MI0000087|hsa-mir-29a&WithFlank&7|-|130876741|130876816@48.67.20(+1C)&[MIMAT0000086&hsa-miR-29a-3p&offsets|0|-2(+1C);m-26&7|-|130876749|130876769&offsets|0|-1(+1C)]|TAGCACCATCTGAAATCGGTC</td><td>>MI0000087|hsa-mir-29a&WithFlank&7|-|130876741|130876816</td><td>48</td><td>67</td><td>20(+1C)</td><td>TAGCACCATCTGAAATCGGTC</td><td>678.8</td><td>10.1</td><td>1.4</td><td>7.2</td><td>7.4E-13</td><td>2.5E-12</td><td>MB231/MB468</td><td>NUC</td></tr>
<tr><td>>hg38_1_-_228634819_228635039_RNA5S12_5S_50ntFlanks@150.170.21|AATACCGGGTGCTGTAGGCTT</td><td>>hg38_1_-_228634819_228635039_RNA5S12_5S_50ntFlanks</td><td>150</td><td>170</td><td>21</td><td>AATACCGGGTGCTGTAGGCTT</td><td>5234.4</td><td>4.8</td><td>0.7</td><td>7.2</td><td>7.1E-13</td><td>2.6E-12</td><td>BT20/MB468</td><td>CYTO</td></tr>
<tr><td>>MI0000266|hsa-mir-10a&WithFlank&17|-|48579832|48579953@28.48.21&[MIMAT0000253&hsa-miR-10a-5p&offsets|0|-2;m-18&17|-|48579905|48579926&offsets|0|-1]|TACCCTGTAGATCCGAATTTG</td><td>>MI0000266|hsa-mir-10a&WithFlank&17|-|48579832|48579953</td><td>28</td><td>48</td><td>21</td><td>TACCCTGTAGATCCGAATTTG</td><td>233.1</td><td>11.1</td><td>1.5</td><td>7.2</td><td>6.3E-13</td><td>2.6E-12</td><td>MB231/MB468</td><td>MP</td></tr>
<tr><td>>hg38_21_+_8433172_8446622_RNA45SN1_45S_50ntFlanks@10731.10756.26|AGGGTGTAAATCTCGCGCCGGGCCGT</td><td>>hg38_21_+_8433172_8446622_RNA45SN1_45S_50ntFlanks</td><td>10731</td><td>10756</td><td>26</td><td>AGGGTGTAAATCTCGCGCCGGGCCGT</td><td>104.7</td><td>-10.6</td><td>1.5</td><td>-7.2</td><td>6.3E-13</td><td>2.6E-12</td><td>MB231/MB468</td><td>MP</td></tr>
<tr><td>>hg38_21_+_8433172_8446622_RNA45SN1_45S_50ntFlanks@11568.11602.35|GCAGCCGACTTAGAACTGGTGCGGACCAGGGGAAT</td><td>>hg38_21_+_8433172_8446622_RNA45SN1_45S_50ntFlanks</td><td>11568</td><td>11602</td><td>35</td><td>GCAGCCGACTTAGAACTGGTGCGGACCAGGGGAAT</td><td>1031.1</td><td>-3.9</td><td>0.5</td><td>-7.2</td><td>7.4E-13</td><td>2.6E-12</td><td>MB231/MB468</td><td>NUC</td></tr>
<tr><td>>MI0000102|hsa-mir-100&WithFlank&11|-|122152223|122152314@19.39.21(+1A)&[MIMAT0000098&hsa-miR-100-5p&offsets|0|-1(+1A);m-27&11|-|122152276|122152296&offsets|0|0(+1A)]|AACCCGTAGATCCGAACTTGTA</td><td>>MI0000102|hsa-mir-100&WithFlank&11|-|122152223|122152314</td><td>19</td><td>39</td><td>21(+1A)</td><td>AACCCGTAGATCCGAACTTGTA</td><td>22491.9</td><td>14.9</td><td>2.1</td><td>7.2</td><td>7.1E-13</td><td>2.6E-12</td><td>BT20/MB468</td><td>CYTO</td></tr>
<tr><td>>hg38_1_-_228634819_228635039_RNA5S12_5S_50ntFlanks@131.169.39|GGATGGGAGACCGCCTGGGAATACCGGGTGCTGTAGGCT</td><td>>hg38_1_-_228634819_228635039_RNA5S12_5S_50ntFlanks</td><td>131</td><td>169</td><td>39</td><td>GGATGGGAGACCGCCTGGGAATACCGGGTGCTGTAGGCT</td><td>576.0</td><td>7.8</td><td>1.1</td><td>7.2</td><td>5.5E-13</td><td>2.6E-12</td><td>BT20/MB468</td><td>MITO</td></tr>
<tr><td>>hg38_21_+_8433172_8446622_RNA45SN1_45S_50ntFlanks@8182.8224.43|ATCGAGGCCCAGCCCGTGGACGGTGTGAGGCCGGTAGCGGCCC</td><td>>hg38_21_+_8433172_8446622_RNA45SN1_45S_50ntFlanks</td><td>8182</td><td>8224</td><td>43</td><td>ATCGAGGCCCAGCCCGTGGACGGTGTGAGGCCGGTAGCGGCCC</td><td>220.2</td><td>-10.7</td><td>1.5</td><td>-7.2</td><td>7.5E-13</td><td>2.6E-12</td><td>MB231/MB468</td><td>NUC</td></tr>
<tr><td>>hg38_21_+_8433172_8446622_RNA45SN1_45S_50ntFlanks@13008.13028.21|ATTGAAAGTCAGCCCTCGACA</td><td>>hg38_21_+_8433172_8446622_RNA45SN1_45S_50ntFlanks</td><td>13008</td><td>13028</td><td>21</td><td>ATTGAAAGTCAGCCCTCGACA</td><td>539.2</td><td>-9.3</td><td>1.3</td><td>-7.1</td><td>1.0E-12</td><td>2.6E-12</td><td>BT20/MB468</td><td>NUC</td></tr>
<tr><td>>hg38_21_+_8433172_8446622_RNA45SN1_45S_50ntFlanks@4002.4022.21|GATAACCTCGGGCCGATCGCA</td><td>>hg38_21_+_8433172_8446622_RNA45SN1_45S_50ntFlanks</td><td>4002</td><td>4022</td><td>21</td><td>GATAACCTCGGGCCGATCGCA</td><td>697.0</td><td>-8.9</td><td>1.2</td><td>-7.1</td><td>9.1E-13</td><td>2.6E-12</td><td>MB231/MB468</td><td>MITO</td></tr>
<tr><td>>hg38_21_+_8433172_8446622_RNA45SN1_45S_50ntFlanks@10101.10125.25|CGCGCGCGTGTGGTGTGCGTCGGAG</td><td>>hg38_21_+_8433172_8446622_RNA45SN1_45S_50ntFlanks</td><td>10101</td><td>10125</td><td>25</td><td>CGCGCGCGTGTGGTGTGCGTCGGAG</td><td>692.8</td><td>-12.5</td><td>1.6</td><td>-7.6</td><td>3.3E-14</td><td>2.6E-12</td><td>BT20/MB468</td><td>MP</td></tr>
<tr><td>>hg38_21_+_8433172_8446622_RNA45SN1_45S_50ntFlanks@12633.12659.27|CGAAGCTACCATCTGTGGGATTATGAC</td><td>>hg38_21_+_8433172_8446622_RNA45SN1_45S_50ntFlanks</td><td>12633</td><td>12659</td><td>27</td><td>CGAAGCTACCATCTGTGGGATTATGAC</td><td>939.6</td><td>-5.3</td><td>0.7</td><td>-7.2</td><td>7.5E-13</td><td>2.6E-12</td><td>MB231/MB468</td><td>NUC</td></tr>
<tr><td>>hg38_21_+_8433172_8446622_RNA45SN1_45S_50ntFlanks@9577.9596.20|CGGTCGTCCGACCTGGGTAT</td><td>>hg38_21_+_8433172_8446622_RNA45SN1_45S_50ntFlanks</td><td>9577</td><td>9596</td><td>20</td><td>CGGTCGTCCGACCTGGGTAT</td><td>178.3</td><td>-11.3</td><td>1.6</td><td>-7.2</td><td>6.4E-13</td><td>2.6E-12</td><td>MB231/MB468</td><td>MP</td></tr>
<tr><td>>MI0000301|hsa-mir-224&WithFlank&X|-|151958572|151958664@14.36.23&[MIMAT0000281&hsa-miR-224-5p&offsets|+1|-1;m-238&X|-|151958630|151958651&offsets|0|+1]|CAAGTCACTAGTGGTTCCGTTTA</td><td>>MI0000301|hsa-mir-224&WithFlank&X|-|151958572|151958664</td><td>14</td><td>36</td><td>23</td><td>CAAGTCACTAGTGGTTCCGTTTA</td><td>322.7</td><td>11.6</td><td>1.6</td><td>7.1</td><td>9.2E-13</td><td>2.6E-12</td><td>MB231/MB468</td><td>MITO</td></tr>
<tr><td>>hg38_21_+_8433172_8446622_RNA45SN1_45S_50ntFlanks@11205.11234.30|GCGGCGGCGACTCTGGACGCGAGCCGGGCC</td><td>>hg38_21_+_8433172_8446622_RNA45SN1_45S_50ntFlanks</td><td>11205</td><td>11234</td><td>30</td><td>GCGGCGGCGACTCTGGACGCGAGCCGGGCC</td><td>261.0</td><td>-6.4</td><td>0.9</td><td>-7.2</td><td>7.4E-13</td><td>2.6E-12</td><td>MB231/MB468</td><td>CYTO</td></tr>
<tr><td>>hg38_21_+_8433172_8446622_RNA45SN1_45S_50ntFlanks@12456.12483.28|CCAAGCGTTGGATTGTTCACCCACTAAT</td><td>>hg38_21_+_8433172_8446622_RNA45SN1_45S_50ntFlanks</td><td>12456</td><td>12483</td><td>28</td><td>CCAAGCGTTGGATTGTTCACCCACTAAT</td><td>1336.0</td><td>-6.3</td><td>0.9</td><td>-7.2</td><td>7.4E-13</td><td>2.6E-12</td><td>MB231/MB468</td><td>CYTO</td></tr>
<tr><td>>hg38_21_+_8433172_8446622_RNA45SN1_45S_50ntFlanks@5472.5489.18|ACGGCCCTGGCGGAGCGC</td><td>>hg38_21_+_8433172_8446622_RNA45SN1_45S_50ntFlanks</td><td>5472</td><td>5489</td><td>18</td><td>ACGGCCCTGGCGGAGCGC</td><td>472.9</td><td>5.9</td><td>0.8</td><td>7.1</td><td>9.2E-13</td><td>2.6E-12</td><td>MB231/MB468</td><td>MITO</td></tr>
<tr><td>>hg38_21_+_8433172_8446622_RNA45SN1_45S_50ntFlanks@7986.8015.30|ATCAGACGTGGCGACCCGCTGAATTTAAGC</td><td>>hg38_21_+_8433172_8446622_RNA45SN1_45S_50ntFlanks</td><td>7986</td><td>8015</td><td>30</td><td>ATCAGACGTGGCGACCCGCTGAATTTAAGC</td><td>298.2</td><td>-10.3</td><td>1.4</td><td>-7.2</td><td>5.5E-13</td><td>2.6E-12</td><td>BT20/MB468</td><td>MITO</td></tr>
<tr><td>>MI0000299|hsa-mir-222&WithFlank&X|-|45747009|45747130@75.97.23(+2U)&[MIMAT0000279&hsa-miR-222-3p&offsets|0|+2(+2U);m-140&X|-|45747033|45747056&offsets|0|-1(+2U)]|AGCTACATCTGGCTACTGGGTCTTT</td><td>>MI0000299|hsa-mir-222&WithFlank&X|-|45747009|45747130</td><td>75</td><td>97</td><td>23(+2U)</td><td>AGCTACATCTGGCTACTGGGTCTTT</td><td>1088.4</td><td>6.5</td><td>0.9</td><td>7.2</td><td>7.5E-13</td><td>2.6E-12</td><td>MB231/MB468</td><td>NUC</td></tr>
<tr><td>>hg38_21_+_8433172_8446622_RNA45SN1_45S_50ntFlanks@10444.10461.18|CGAAGGGACGGGCGATGG</td><td>>hg38_21_+_8433172_8446622_RNA45SN1_45S_50ntFlanks</td><td>10444</td><td>10461</td><td>18</td><td>CGAAGGGACGGGCGATGG</td><td>122.7</td><td>-10.8</td><td>1.5</td><td>-7.2</td><td>6.4E-13</td><td>2.6E-12</td><td>MB231/MB468</td><td>MP</td></tr>
<tr><td>>hg38_21_+_8433172_8446622_RNA45SN1_45S_50ntFlanks@12510.12527.18|TCGTGAGACAGGTTAGTT</td><td>>hg38_21_+_8433172_8446622_RNA45SN1_45S_50ntFlanks</td><td>12510</td><td>12527</td><td>18</td><td>TCGTGAGACAGGTTAGTT</td><td>516.4</td><td>-8.7</td><td>1.2</td><td>-7.2</td><td>6.4E-13</td><td>2.6E-12</td><td>MB231/MB468</td><td>MP</td></tr>
<tr><td>>hg38_21_+_8433172_8446622_RNA45SN1_45S_50ntFlanks@11641.11669.29|CGGCGGGTGTTGACGCGATGTGATTTCTG</td><td>>hg38_21_+_8433172_8446622_RNA45SN1_45S_50ntFlanks</td><td>11641</td><td>11669</td><td>29</td><td>CGGCGGGTGTTGACGCGATGTGATTTCTG</td><td>910.5</td><td>-7.0</td><td>1.0</td><td>-7.2</td><td>7.4E-13</td><td>2.6E-12</td><td>MB231/MB468</td><td>CYTO</td></tr>
<tr><td>>hg38_21_+_8433172_8446622_RNA45SN1_45S_50ntFlanks@11766.11786.21|ATCTAATTAGTGACGCGCATG</td><td>>hg38_21_+_8433172_8446622_RNA45SN1_45S_50ntFlanks</td><td>11766</td><td>11786</td><td>21</td><td>ATCTAATTAGTGACGCGCATG</td><td>189.7</td><td>-10.5</td><td>1.5</td><td>-7.2</td><td>7.6E-13</td><td>2.6E-12</td><td>MB231/MB468</td><td>NUC</td></tr>
<tr><td>>hg38_21_+_8433172_8446622_RNA45SN1_45S_50ntFlanks@10922.10942.21|CTGGACGAGGCGCCGCCGCCC</td><td>>hg38_21_+_8433172_8446622_RNA45SN1_45S_50ntFlanks</td><td>10922</td><td>10942</td><td>21</td><td>CTGGACGAGGCGCCGCCGCCC</td><td>126.4</td><td>-9.7</td><td>1.4</td><td>-7.2</td><td>7.4E-13</td><td>2.6E-12</td><td>MB231/MB468</td><td>CYTO</td></tr>
<tr><td>>hg38_21_+_8433172_8446622_RNA45SN1_45S_50ntFlanks@3854.3881.28|ACTGTGGTAATTCTAGAGCTAATACATG</td><td>>hg38_21_+_8433172_8446622_RNA45SN1_45S_50ntFlanks</td><td>3854</td><td>3881</td><td>28</td><td>ACTGTGGTAATTCTAGAGCTAATACATG</td><td>1547.5</td><td>-9.8</td><td>1.4</td><td>-7.2</td><td>6.5E-13</td><td>2.6E-12</td><td>MB231/MB468</td><td>MP</td></tr>
<tr><td>>hg38_21_+_8433172_8446622_RNA45SN1_45S_50ntFlanks@8339.8373.35|TACCGTAAGGGAAAGTTGAAAAGAACTTTGAAGAG</td><td>>hg38_21_+_8433172_8446622_RNA45SN1_45S_50ntFlanks</td><td>8339</td><td>8373</td><td>35</td><td>TACCGTAAGGGAAAGTTGAAAAGAACTTTGAAGAG</td><td>525.9</td><td>12.8</td><td>1.7</td><td>7.6</td><td>3.3E-14</td><td>2.6E-12</td><td>BT20/MB468</td><td>MP</td></tr>
<tr><td>>hg38_21_+_8433172_8446622_RNA45SN1_45S_50ntFlanks@12409.12448.40|TTGATCCTTCGATGTCGGCTCTTCCTATCATTGTGAAGCA</td><td>>hg38_21_+_8433172_8446622_RNA45SN1_45S_50ntFlanks</td><td>12409</td><td>12448</td><td>40</td><td>TTGATCCTTCGATGTCGGCTCTTCCTATCATTGTGAAGCA</td><td>291.5</td><td>7.1</td><td>1.0</td><td>7.2</td><td>7.5E-13</td><td>2.6E-12</td><td>MB231/MB468</td><td>CYTO</td></tr>
<tr><td>>hg38_21_+_8433172_8446622_RNA45SN1_45S_50ntFlanks@5470.5516.47|CCACGGCCCTGGCGGAGCGCTGAGAAGACGGTCGAACTTGACTATCT</td><td>>hg38_21_+_8433172_8446622_RNA45SN1_45S_50ntFlanks</td><td>5470</td><td>5516</td><td>47</td><td>CCACGGCCCTGGCGGAGCGCTGAGAAGACGGTCGAACTTGACTATCT</td><td>1721.2</td><td>6.7</td><td>0.9</td><td>7.2</td><td>7.5E-13</td><td>2.6E-12</td><td>MB231/MB468</td><td>CYTO</td></tr>
<tr><td>>hg38_21_+_8433172_8446622_RNA45SN1_45S_50ntFlanks@5472.5509.38|ACGGCCCTGGCGGAGCGCTGAGAAGACGGTCGAACTTG</td><td>>hg38_21_+_8433172_8446622_RNA45SN1_45S_50ntFlanks</td><td>5472</td><td>5509</td><td>38</td><td>ACGGCCCTGGCGGAGCGCTGAGAAGACGGTCGAACTTG</td><td>5104.8</td><td>5.9</td><td>0.8</td><td>7.2</td><td>7.3E-13</td><td>2.6E-12</td><td>BT20/MB468</td><td>CYTO</td></tr>
<tr><td>>hg38_21_+_8433172_8446622_RNA45SN1_45S_50ntFlanks@8169.8197.29|CCAAGTCCTTCTGATCGAGGCCCAGCCCG</td><td>>hg38_21_+_8433172_8446622_RNA45SN1_45S_50ntFlanks</td><td>8169</td><td>8197</td><td>29</td><td>CCAAGTCCTTCTGATCGAGGCCCAGCCCG</td><td>926.3</td><td>-9.6</td><td>1.3</td><td>-7.1</td><td>9.3E-13</td><td>2.6E-12</td><td>MB231/MB468</td><td>MITO</td></tr>
<tr><td>>hg38_21_+_8433172_8446622_RNA45SN1_45S_50ntFlanks@12405.12430.26|CTTTTTGATCCTTCGATGTCGGCTCT</td><td>>hg38_21_+_8433172_8446622_RNA45SN1_45S_50ntFlanks</td><td>12405</td><td>12430</td><td>26</td><td>CTTTTTGATCCTTCGATGTCGGCTCT</td><td>483.2</td><td>-8.6</td><td>1.2</td><td>-7.1</td><td>9.3E-13</td><td>2.6E-12</td><td>MB231/MB468</td><td>MITO</td></tr>
<tr><td>>hg38_21_+_8433172_8446622_RNA45SN1_45S_50ntFlanks@10808.10836.29|CAATGTAGGTAAGGGAAGTCGGCAAGCCG</td><td>>hg38_21_+_8433172_8446622_RNA45SN1_45S_50ntFlanks</td><td>10808</td><td>10836</td><td>29</td><td>CAATGTAGGTAAGGGAAGTCGGCAAGCCG</td><td>103.0</td><td>-10.5</td><td>1.5</td><td>-7.2</td><td>6.5E-13</td><td>2.6E-12</td><td>MB231/MB468</td><td>MP</td></tr>
<tr><td>>hg38_21_+_8433172_8446622_RNA45SN1_45S_50ntFlanks@4069.4092.24|CTTTCGATGGTAGTCGCCGTGCCT</td><td>>hg38_21_+_8433172_8446622_RNA45SN1_45S_50ntFlanks</td><td>4069</td><td>4092</td><td>24</td><td>CTTTCGATGGTAGTCGCCGTGCCT</td><td>116.3</td><td>-10.7</td><td>1.5</td><td>-7.2</td><td>6.5E-13</td><td>2.6E-12</td><td>MB231/MB468</td><td>MP</td></tr>
<tr><td>>hg38_21_+_8433172_8446622_RNA45SN1_45S_50ntFlanks@10678.10712.35|CGGTTCCGGCGGCGTCCGGTGAGCTCTCGCTGGCC</td><td>>hg38_21_+_8433172_8446622_RNA45SN1_45S_50ntFlanks</td><td>10678</td><td>10712</td><td>35</td><td>CGGTTCCGGCGGCGTCCGGTGAGCTCTCGCTGGCC</td><td>8759.0</td><td>-7.6</td><td>1.1</td><td>-7.2</td><td>7.5E-13</td><td>2.6E-12</td><td>MB231/MB468</td><td>CYTO</td></tr>
</table>

<table>
<tr><td>>MI0000285|hsa-mir-205&WithFlank&1|+|209432127|209432248@40.62.23&[MIMAT0000266&hsa-miR-205-5p&offsets|0|+1;m-351&1|+|209432166|209432187&offsets|0|+1]|TCCTTCATTCCACCGGAGTCTGT</td><td>>MI0000285|hsa-mir-205&WithFlank&1|+|209432127|209432248</td><td>40</td><td>62</td><td>23</td><td>TCCTTCATTCCACCGGAGTCTGT</td><td>73187.4</td><td>7.5</td><td>1.0</td><td>7.2</td><td>5.6E-13</td><td>2.6E-12</td><td>BT20/MB468</td><td>MITO</td></tr>
<tr><td>>hg38_21_+_8433172_8446622_RNA45SN1_45S_50ntFlanks@12002.12032.31|TGCGGGCCGCCGGTGAAATACCACTACTCTG</td><td>>hg38_21_+_8433172_8446622_RNA45SN1_45S_50ntFlanks</td><td>12002</td><td>12032</td><td>31</td><td>TGCGGGCCGCCGGTGAAATACCACTACTCTG</td><td>5927.9</td><td>-6.3</td><td>0.9</td><td>-7.2</td><td>7.5E-13</td><td>2.6E-12</td><td>MB231/MB468</td><td>CYTO</td></tr>
<tr><td>>hg38_21_+_8433172_8446622_RNA45SN1_45S_50ntFlanks@10350.10376.27|ACGAGAACTTTGAAGGCCGAAGTGGAG</td><td>>hg38_21_+_8433172_8446622_RNA45SN1_45S_50ntFlanks</td><td>10350</td><td>10376</td><td>27</td><td>ACGAGAACTTTGAAGGCCGAAGTGGAG</td><td>1048.6</td><td>-3.6</td><td>0.5</td><td>-7.2</td><td>7.4E-13</td><td>2.6E-12</td><td>BT20/MB468</td><td>CYTO</td></tr>
<tr><td>>hg38_21_+_8433172_8446622_RNA45SN1_45S_50ntFlanks@11480.11502.23|CGGCGGAGGGGCCGCGGGCCGGT</td><td>>hg38_21_+_8433172_8446622_RNA45SN1_45S_50ntFlanks</td><td>11480</td><td>11502</td><td>23</td><td>CGGCGGAGGGGCCGCGGGCCGGT</td><td>229.7</td><td>-11.7</td><td>1.6</td><td>-7.2</td><td>6.5E-13</td><td>2.6E-12</td><td>MB231/MB468</td><td>MP</td></tr>
<tr><td>>hg38_21_+_8433172_8446622_RNA45SN1_45S_50ntFlanks@12007.12043.37|GCCGCCGGTGAAATACCACTACTCTGATCGTTTTTTC</td><td>>hg38_21_+_8433172_8446622_RNA45SN1_45S_50ntFlanks</td><td>12007</td><td>12043</td><td>37</td><td>GCCGCCGGTGAAATACCACTACTCTGATCGTTTTTTC</td><td>3419.3</td><td>-7.3</td><td>1.0</td><td>-7.2</td><td>7.7E-13</td><td>2.6E-12</td><td>MB231/MB468</td><td>NUC</td></tr>
<tr><td>>trna10_AspGTC_12_-_125424193_125424264@37.73.37,trna12_AspGTC_12_-_125411891_125411962@37.73.37,trna144_AspGTC_6_-_27551236_27551307@37.73.37,trna38_AspGTC_17_-_8125556_8125627@37.73.37,trna45_AspGTC_6_+_27447453_27447524@37.73.37,trna48_AspGTC_6_+_27471523_27471594@37.73.37,trna4_AspGTC_12_+_96429799_96429870@37.73.37,trna69_AspGTC_1_-_161440205_161440276@37.73.37,trna72_AspGTC_1_-_161432824_161432895@37.73.37,trna75_AspGTC_1_-_161425414_161425485@37.73.37,trna78_AspGTC_1_-_161418033_161418104@37.73.37,trna81_AspGTC_1_-_161410615_161410686@37.73.37|ACGCGGGAGACCGGGGTTCGATTCCCCGACGGGGAGC</td><td>>trna10_AspGTC_12_-_125424193_125424264</td><td>37</td><td>73</td><td>37</td><td>ACGCGGGAGACCGGGGTTCGATTCCCCGACGGGGAGC</td><td>39599.7</td><td>8.2</td><td>1.1</td><td>7.2</td><td>7.6E-13</td><td>2.6E-12</td><td>MB231/MB468</td><td>CYTO</td></tr>
<tr><td>>hg38_21_+_8433172_8446622_RNA45SN1_45S_50ntFlanks@11641.11685.45|CGGCGGGTGTTGACGCGATGTGATTTCTGCCCAGTGCTCTGAATG</td><td>>hg38_21_+_8433172_8446622_RNA45SN1_45S_50ntFlanks</td><td>11641</td><td>11685</td><td>45</td><td>CGGCGGGTGTTGACGCGATGTGATTTCTGCCCAGTGCTCTGAATG</td><td>887.7</td><td>-13.7</td><td>1.9</td><td>-7.2</td><td>6.6E-13</td><td>2.6E-12</td><td>MB231/MB468</td><td>MP</td></tr>
<tr><td>>trnaMT_GluTTC_MT_-_14674_14742@27.69.43,trnalookalike8_GluTTC_5_-_93905172_93905240@27.69.43|GTTTTTCATATCATTGGTCGTGGTTGTAGTCCGTGCGAGAATA</td><td>>trnaMT_GluTTC_MT_-_14674_14742</td><td>27</td><td>69</td><td>43</td><td>GTTTTTCATATCATTGGTCGTGGTTGTAGTCCGTGCGAGAATA</td><td>1166.4</td><td>-14.6</td><td>2.1</td><td>-7.1</td><td>1.0E-12</td><td>2.7E-12</td><td>BT20/MB468</td><td>NUC</td></tr>
<tr><td>>hg38_21_+_8433172_8446622_RNA45SN1_45S_50ntFlanks@4722.4752.31|TCATTAATCAAGAACGAAAGTCGGAGGTTCG</td><td>>hg38_21_+_8433172_8446622_RNA45SN1_45S_50ntFlanks</td><td>4722</td><td>4752</td><td>31</td><td>TCATTAATCAAGAACGAAAGTCGGAGGTTCG</td><td>730.8</td><td>-8.6</td><td>1.2</td><td>-7.2</td><td>7.8E-13</td><td>2.7E-12</td><td>MB231/MB468</td><td>NUC</td></tr>
<tr><td>>MI0000746|hsa-mir-99b&WithFlank&19|+|51692606|51692687@51.71.21(+1U)&[MIMAT0004678&hsa-miR-99b-3p&offsets|0|-1(+1U);m-151&19|+|51692656|51692678&offsets|0|-2(+1U)]|CAAGCTCGTGTCTGTGGGTCCT</td><td>>MI0000746|hsa-mir-99b&WithFlank&19|+|51692606|51692687</td><td>51</td><td>71</td><td>21(+1U)</td><td>CAAGCTCGTGTCTGTGGGTCCT</td><td>308.8</td><td>9.8</td><td>1.4</td><td>7.2</td><td>7.4E-13</td><td>2.7E-12</td><td>BT20/MB468</td><td>CYTO</td></tr>
<tr><td>>hg38_21_+_8433172_8446622_RNA45SN1_45S_50ntFlanks@10683.10702.20|CCGGCGGCGTCCGGTGAGCT</td><td>>hg38_21_+_8433172_8446622_RNA45SN1_45S_50ntFlanks</td><td>10683</td><td>10702</td><td>20</td><td>CCGGCGGCGTCCGGTGAGCT</td><td>108.0</td><td>-10.6</td><td>1.5</td><td>-7.2</td><td>6.6E-13</td><td>2.7E-12</td><td>MB231/MB468</td><td>MP</td></tr>
<tr><td>>hg38_21_+_8433172_8446622_RNA45SN1_45S_50ntFlanks@11881.11899.19|CCTGTTGAGCTTGACTCTA</td><td>>hg38_21_+_8433172_8446622_RNA45SN1_45S_50ntFlanks</td><td>11881</td><td>11899</td><td>19</td><td>CCTGTTGAGCTTGACTCTA</td><td>126.4</td><td>-10.8</td><td>1.5</td><td>-7.2</td><td>6.6E-13</td><td>2.7E-12</td><td>MB231/MB468</td><td>MP</td></tr>
<tr><td>>MI0003560|hsa-mir-92b&WithFlank&1|+|155195171|155195278@67.87.21(+1A)&[MIMAT0003218&hsa-miR-92b-3p&offsets|0|-1(+1A);m-115&1|+|155195237|155195258&offsets|0|-1(+1A)]|TATTGCACTCGTCCCGGCCTCA</td><td>>MI0003560|hsa-mir-92b&WithFlank&1|+|155195171|155195278</td><td>67</td><td>87</td><td>21(+1A)</td><td>TATTGCACTCGTCCCGGCCTCA</td><td>3662.1</td><td>11.8</td><td>1.6</td><td>7.2</td><td>7.5E-13</td><td>2.7E-12</td><td>BT20/MB468</td><td>CYTO</td></tr>
<tr><td>>hg38_21_+_8433172_8446622_RNA45SN1_45S_50ntFlanks@6740.6772.33|CATCGACACTTCGAACGCACTTGCGGCCCCGGG</td><td>>hg38_21_+_8433172_8446622_RNA45SN1_45S_50ntFlanks</td><td>6740</td><td>6772</td><td>33</td><td>CATCGACACTTCGAACGCACTTGCGGCCCCGGG</td><td>508.4</td><td>-8.8</td><td>1.2</td><td>-7.2</td><td>7.8E-13</td><td>2.7E-12</td><td>MB231/MB468</td><td>NUC</td></tr>
<tr><td>>hg38_1_-_228634819_228635039_RNA5S12_5S_50ntFlanks@51.77.27|GTCTACGGCCATACCACCCTGAACGCG</td><td>>hg38_1_-_228634819_228635039_RNA5S12_5S_50ntFlanks</td><td>51</td><td>77</td><td>27</td><td>GTCTACGGCCATACCACCCTGAACGCG</td><td>528.9</td><td>-3.5</td><td>0.5</td><td>-7.1</td><td>1.0E-12</td><td>2.7E-12</td><td>BT20/MB468</td><td>NUC</td></tr>
<tr><td>>hg38_21_+_8433172_8446622_RNA45SN1_45S_50ntFlanks@4003.4026.24|ATAACCTCGGGCCGATCGCACGCC</td><td>>hg38_21_+_8433172_8446622_RNA45SN1_45S_50ntFlanks</td><td>4003</td><td>4026</td><td>24</td><td>ATAACCTCGGGCCGATCGCACGCC</td><td>1043.4</td><td>-9.4</td><td>1.3</td><td>-7.2</td><td>7.8E-13</td><td>2.7E-12</td><td>MB231/MB468</td><td>NUC</td></tr>
<tr><td>>hg38_21_+_8433172_8446622_RNA45SN1_45S_50ntFlanks@11641.11665.25|CGGCGGGTGTTGACGCGATGTGATT</td><td>>hg38_21_+_8433172_8446622_RNA45SN1_45S_50ntFlanks</td><td>11641</td><td>11665</td><td>25</td><td>CGGCGGGTGTTGACGCGATGTGATT</td><td>1336.6</td><td>-6.8</td><td>0.9</td><td>-7.2</td><td>7.7E-13</td><td>2.7E-12</td><td>MB231/MB468</td><td>CYTO</td></tr>
<tr><td>>hg38_21_+_8433172_8446622_RNA45SN1_45S_50ntFlanks@10479.10512.34|CGATCGAAAGGGAGTCGGGTTCAGATCCCCGAAT</td><td>>hg38_21_+_8433172_8446622_RNA45SN1_45S_50ntFlanks</td><td>10479</td><td>10512</td><td>34</td><td>CGATCGAAAGGGAGTCGGGTTCAGATCCCCGAAT</td><td>147.7</td><td>-11.1</td><td>1.5</td><td>-7.2</td><td>6.6E-13</td><td>2.7E-12</td><td>MB231/MB468</td><td>MP</td></tr>
<tr><td>>am_trna106_PheGAA_6_-_28775610_28775682@31.52.22,trna109_PheGAA_6_-_28758499_28758571@31.52.22,trna11_PheGAA_12_-_125412389_125412461@31.52.22,trna13_PheGAA_11_-_59333853_59333925@31.52.22,trna14_PheGAA_19_-_1383361_1383433@31.52.22,trna15_PheGAA_11_-_59324970_59325042@31.52.22,trna1_PheGAA_13_-_95201904_95201976@31.52.22,trna72_PheGAA_6_+_28732164_28732236@31.52.22,trna96_PheGAA_6_-_28949449_28949521@31.52.22|ACTGAAGATCTAAAGGTCCCTG</td><td>>am_trna106_PheGAA_6_-_28775610_28775682</td><td>31</td><td>52</td><td>22</td><td>ACTGAAGATCTAAAGGTCCCTG</td><td>174.5</td><td>-6.9</td><td>1.0</td><td>-7.2</td><td>7.7E-13</td><td>2.7E-12</td><td>MB231/MB468</td><td>CYTO</td></tr>
<tr><td>>hg38_21_+_8433172_8446622_RNA45SN1_45S_50ntFlanks@5099.5136.38|CATGCTAACTAGTTACGCGACCCCCGAGCGGTCGGCGT</td><td>>hg38_21_+_8433172_8446622_RNA45SN1_45S_50ntFlanks</td><td>5099</td><td>5136</td><td>38</td><td>CATGCTAACTAGTTACGCGACCCCCGAGCGGTCGGCGT</td><td>92.3</td><td>-23.0</td><td>3.0</td><td>-7.6</td><td>3.5E-14</td><td>2.7E-12</td><td>BT20/MB468</td><td>MP</td></tr>
<tr><td>>MI0000736|hsa-mir-30c-1&WithFlank&1|+|40757278|40757378@23.45.23(+1C);MI0000254|hsa-mir-30c-2&WithFlank&6|-|71376954|71377037@13.35.23(+1C)&[MIMAT0000244&hsa-miR-30c-5p&offsets|0|0(+1C);m-57&1|+|40757300|40757323&offsets|0|-1(+1C)];[MIMAT0000244_1&hsa-miR-30c-5p&offsets|0|0(+1C);m-56&6|-|71377002|71377025&offsets|0|-1(+1C)]|TGTAAACATCCTACACTCTCAGCC</td><td>>MI0000736|hsa-mir-30c-1&WithFlank&1|+|40757278|40757378</td><td>23</td><td>45</td><td>23(+1C)</td><td>TGTAAACATCCTACACTCTCAGCC</td><td>3603.4</td><td>4.1</td><td>0.6</td><td>7.2</td><td>7.9E-13</td><td>2.7E-12</td><td>MB231/MB468</td><td>NUC</td></tr>
<tr><td>>trna116_GluCTC_1_-_145399233_145399304@32.73.42,trna71_GluCTC_1_-_161439189_161439260@32.73.42,trna74_GluCTC_1_-_161431809_161431880@32.73.42,trna77_GluCTC_1_-_161424398_161424469@32.73.42,trna77_GluCTC_6_+_28949976_28950047@32.73.42,trna80_GluCTC_1_-_161417018_161417089@32.73.42,trna87_GluCTC_6_-_126101393_126101464@32.73.42|CTCTCACCGCCGCGGCCCGGGTTCGATTCCCGGTCAGGGAAC</td><td>>trna116_GluCTC_1_-_145399233_145399304</td><td>32</td><td>73</td><td>42</td><td>CTCTCACCGCCGCGGCCCGGGTTCGATTCCCGGTCAGGGAAC</td><td>319.6</td><td>-12.1</td><td>1.7</td><td>-7.1</td><td>9.6E-13</td><td>2.7E-12</td><td>MB231/MB468</td><td>MITO</td></tr>
<tr><td>>trna111_HisGTG_1_-_147774845_147774916@1.32.32,trna118_HisGTG_1_-_145396881_145396952@1.32.32,trna16_HisGTG_1_+_146544773_146544844@1.32.32,trna1_HisGTG_15_+_45493349_45493420@1.32.32,trna21_HisGTG_1_+_147753471_147753542@1.32.32,trna33_HisGTG_6_+_27125906_27125977@1.32.32,trna7_HisGTG_9_-_14433938_14434009@1.32.32,trna8_HisGTG_15_-_45492611_45492682@1.32.32,trna9_HisGTG_15_-_45490804_45490875@1.32.32|GCCGTGATCGTATAGTGGTTAGTACTCTGCGT</td><td>>trna111_HisGTG_1_-_147774845_147774916</td><td>1</td><td>32</td><td>32</td><td>GCCGTGATCGTATAGTGGTTAGTACTCTGCGT</td><td>1342.1</td><td>7.7</td><td>1.1</td><td>7.2</td><td>7.8E-13</td><td>2.7E-12</td><td>MB231/MB468</td><td>CYTO</td></tr>
<tr><td>>hg38_21_+_8433172_8446622_RNA45SN1_45S_50ntFlanks@11027.11048.22|CTCTCTCTCTCTCTCCCCCGCT</td><td>>hg38_21_+_8433172_8446622_RNA45SN1_45S_50ntFlanks</td><td>11027</td><td>11048</td><td>22</td><td>CTCTCTCTCTCTCTCCCCCGCT</td><td>510.6</td><td>-12.0</td><td>1.7</td><td>-7.2</td><td>7.9E-13</td><td>2.7E-12</td><td>MB231/MB468</td><td>NUC</td></tr>
<tr><td>>hg38_21_+_8433172_8446622_RNA45SN1_45S_50ntFlanks@11314.11343.30|CCCACCCCACGTCTCGTCGCGCGCGCGTCC</td><td>>hg38_21_+_8433172_8446622_RNA45SN1_45S_50ntFlanks</td><td>11314</td><td>11343</td><td>30</td><td>CCCACCCCACGTCTCGTCGCGCGCGCGTCC</td><td>236.4</td><td>-10.8</td><td>1.5</td><td>-7.2</td><td>7.9E-13</td><td>2.7E-12</td><td>MB231/MB468</td><td>NUC</td></tr>
<tr><td>>MI0000086|hsa-mir-28&WithFlank&3|+|188688775|188688872@60.81.22(+1U)&[MIMAT0004502&hsa-miR-28-3p&offsets|0|0(+1U);m-28&3|+|188688834|188688855&offsets|0|0(+1U)]|CACTAGATTGTGAGCTCCTGGAT</td><td>>MI0000086|hsa-mir-28&WithFlank&3|+|188688775|188688872</td><td>60</td><td>81</td><td>22(+1U)</td><td>CACTAGATTGTGAGCTCCTGGAT</td><td>523.8</td><td>8.7</td><td>1.2</td><td>7.2</td><td>7.8E-13</td><td>2.7E-12</td><td>MB231/MB468</td><td>CYTO</td></tr>
<tr><td>>hg38_21_+_8433172_8446622_RNA45SN1_45S_50ntFlanks@10684.10712.29|CGGCGGCGTCCGGTGAGCTCTCGCTGGCC</td><td>>hg38_21_+_8433172_8446622_RNA45SN1_45S_50ntFlanks</td><td>10684</td><td>10712</td><td>29</td><td>CGGCGGCGTCCGGTGAGCTCTCGCTGGCC</td><td>859.0</td><td>-4.7</td><td>0.7</td><td>-7.1</td><td>9.7E-13</td><td>2.7E-12</td><td>MB231/MB468</td><td>MITO</td></tr>
<tr><td>>hg38_21_+_8433172_8446622_RNA45SN1_45S_50ntFlanks@8198.8237.40|TGGACGGTGTGAGGCCGGTAGCGGCCCCCGGCGCGCCGGG</td><td>>hg38_21_+_8433172_8446622_RNA45SN1_45S_50ntFlanks</td><td>8198</td><td>8237</td><td>40</td><td>TGGACGGTGTGAGGCCGGTAGCGGCCCCCGGCGCGCCGGG</td><td>131.4</td><td>-9.8</td><td>1.4</td><td>-7.2</td><td>7.8E-13</td><td>2.7E-12</td><td>MB231/MB468</td><td>CYTO</td></tr>
<tr><td>>hg38_21_+_8433172_8446622_RNA45SN1_45S_50ntFlanks@12073.12110.38|CCGAGGGGCTCTCGCTTCTGGCGCCAAGCGCCCGGCCG</td><td>>hg38_21_+_8433172_8446622_RNA45SN1_45S_50ntFlanks</td><td>12073</td><td>12110</td><td>38</td><td>CCGAGGGGCTCTCGCTTCTGGCGCCAAGCGCCCGGCCG</td><td>284.1</td><td>-11.1</td><td>1.6</td><td>-7.2</td><td>8.0E-13</td><td>2.7E-12</td><td>MB231/MB468</td><td>NUC</td></tr>
<tr><td>>hg38_21_+_8433172_8446622_RNA45SN1_45S_50ntFlanks@6761.6807.47|TGCGGCCCCGGGTTCCTCCCGGGGCTACGCCTGTCTGAGCGTCGCTT</td><td>>hg38_21_+_8433172_8446622_RNA45SN1_45S_50ntFlanks</td><td>6761</td><td>6807</td><td>47</td><td>TGCGGCCCCGGGTTCCTCCCGGGGCTACGCCTGTCTGAGCGTCGCTT</td><td>441.7</td><td>6.9</td><td>1.0</td><td>7.2</td><td>5.8E-13</td><td>2.7E-12</td><td>BT20/MB468</td><td>MITO</td></tr>
<tr><td>>hg38_21_+_8433172_8446622_RNA45SN1_45S_50ntFlanks@4729.4751.23|TCAAGAACGAAAGTCGGAGGTTC</td><td>>hg38_21_+_8433172_8446622_RNA45SN1_45S_50ntFlanks</td><td>4729</td><td>4751</td><td>23</td><td>TCAAGAACGAAAGTCGGAGGTTC</td><td>368.9</td><td>-3.9</td><td>0.6</td><td>-7.2</td><td>8.0E-13</td><td>2.7E-12</td><td>MB231/MB468</td><td>NUC</td></tr>
<tr><td>>MI0000084|hsa-mir-26b&WithFlank&2|+|218402640|218402728@18.38.21&[MIMAT0000083&hsa-miR-26b-5p&offsets|0|0;m-53&2|+|218402657|218402678&offsets|0|-1]|TTCAAGTAATTCAGGATAGGT</td><td>>MI0000084|hsa-mir-26b&WithFlank&2|+|218402640|218402728</td><td>18</td><td>38</td><td>21</td><td>TTCAAGTAATTCAGGATAGGT</td><td>370.6</td><td>2.0</td><td>0.3</td><td>7.1</td><td>1.1E-12</td><td>2.7E-12</td><td>BT20/MB468</td><td>NUC</td></tr>
<tr><td>>hg38_21_+_8433172_8446622_RNA45SN1_45S_50ntFlanks@10727.10763.37|GGAGAGGGTGTAAATCTCGCGCCGGGCCGTACCCATA</td><td>>hg38_21_+_8433172_8446622_RNA45SN1_45S_50ntFlanks</td><td>10727</td><td>10763</td><td>37</td><td>GGAGAGGGTGTAAATCTCGCGCCGGGCCGTACCCATA</td><td>162.4</td><td>-10.3</td><td>1.4</td><td>-7.2</td><td>8.0E-13</td><td>2.7E-12</td><td>MB231/MB468</td><td>NUC</td></tr>
<tr><td>>hg38_21_+_8433172_8446622_RNA45SN1_45S_50ntFlanks@9759.9786.28|CAAACTTTAAATGGGTAAGAAGCCCGGC</td><td>>hg38_21_+_8433172_8446622_RNA45SN1_45S_50ntFlanks</td><td>9759</td><td>9786</td><td>28</td><td>CAAACTTTAAATGGGTAAGAAGCCCGGC</td><td>544.3</td><td>-13.0</td><td>1.8</td><td>-7.2</td><td>6.8E-13</td><td>2.7E-12</td><td>MB231/MB468</td><td>MP</td></tr>
<tr><td>>hg38_21_+_8433172_8446622_RNA45SN1_45S_50ntFlanks@4064.4101.38|ATCAACTTTCGATGGTAGTCGCCGTGCCTACCATGGTG</td><td>>hg38_21_+_8433172_8446622_RNA45SN1_45S_50ntFlanks</td><td>4064</td><td>4101</td><td>38</td><td>ATCAACTTTCGATGGTAGTCGCCGTGCCTACCATGGTG</td><td>165.3</td><td>-11.2</td><td>1.6</td><td>-7.2</td><td>6.8E-13</td><td>2.7E-12</td><td>MB231/MB468</td><td>MP</td></tr>
<tr><td>>trna16_TyrGTA_14_-_21131351_21131444@59.76.18,trna17_TyrGTA_14_-_21128117_21128210@59.76.18,trna19_TyrGTA_14_-_21121258_21121351@59.76.18,trna2_TyrGTA_2_+_27273650_27273738@59.76.18,trna4_TyrGTA_8_+_67025602_67025694@59.76.18,trna5_TyrGTA_14_+_21151432_21151520@59.76.18,trna5_TyrGTA_8_+_67026223_67026311@59.76.18|TTCCGGCTCGAAGGACCA</td><td>>trna16_TyrGTA_14_-_21131351_21131444</td><td>59</td><td>76</td><td>18</td><td>TTCCGGCTCGAAGGACCA</td><td>2320.2</td><td>7.2</td><td>1.0</td><td>7.2</td><td>5.8E-13</td><td>2.7E-12</td><td>BT20/MB468</td><td>MITO</td></tr>
</table>

<table>
<tr><th></th><th></th><th></th><th></th><th></th><th></th><th></th><th></th><th></th><th></th><th></th><th></th><th></th><th></th></tr>
<tr><td>>hg38_21_+_8433172_8446622_RNA45SN1_45S_50ntFlanks@4723.4747.25|CATTAATCAAGAACGAAAGTCGGAG</td><td>>hg38_21_+_8433172_8446622_RNA45SN1_45S_50ntFlanks</td><td>4723</td><td>4747</td><td>25</td><td>CATTAATCAAGAACGAAAGTCGGAG</td><td>154.0</td><td>-11.1</td><td>1.5</td><td>-7.2</td><td>6.8E-13</td><td>2.7E-12</td><td>MB231/MB468</td><td>MP</td></tr>
<tr><td>>hg38_21_+_8433172_8446622_RNA45SN1_45S_50ntFlanks@10410.10428.19|TCAGTCGGTCCTGAGAGAT</td><td>>hg38_21_+_8433172_8446622_RNA45SN1_45S_50ntFlanks</td><td>10410</td><td>10428</td><td>19</td><td>TCAGTCGGTCCTGAGAGAT</td><td>523.2</td><td>-8.9</td><td>1.3</td><td>-7.1</td><td>9.8E-13</td><td>2.7E-12</td><td>MB231/MB468</td><td>MITO</td></tr>
<tr><td>>MI0003834|hsa-mir-769&WithFlank&19|+|46018926|46019055@36.57.22&[MIMAT0003886&hsa-miR-769-5p&offsets|0|0;m-216&19|+|46018961|46018982&offsets|0|0]|TGAGACCTCTGGGTTCTGAGCT</td><td>>MI0003834|hsa-mir-769&WithFlank&19|+|46018926|46019055</td><td>36</td><td>57</td><td>22</td><td>TGAGACCTCTGGGTTCTGAGCT</td><td>876.1</td><td>5.2</td><td>0.7</td><td>7.2</td><td>8.1E-13</td><td>2.7E-12</td><td>MB231/MB468</td><td>NUC</td></tr>
<tr><td>>hg38_21_+_8433172_8446622_RNA45SN1_45S_50ntFlanks@11206.11225.20|CGGCGGCGACTCTGGACGCG</td><td>>hg38_21_+_8433172_8446622_RNA45SN1_45S_50ntFlanks</td><td>11206</td><td>11225</td><td>20</td><td>CGGCGGCGACTCTGGACGCG</td><td>506.5</td><td>-8.7</td><td>1.2</td><td>-7.2</td><td>6.8E-13</td><td>2.7E-12</td><td>MB231/MB468</td><td>MP</td></tr>
<tr><td>>hg38_21_+_8433172_8446622_RNA45SN1_45S_50ntFlanks@13004.13025.22|ATCTATTGAAAGTCAGCCCTCG</td><td>>hg38_21_+_8433172_8446622_RNA45SN1_45S_50ntFlanks</td><td>13004</td><td>13025</td><td>22</td><td>ATCTATTGAAAGTCAGCCCTCG</td><td>203.8</td><td>-9.8</td><td>1.4</td><td>-7.2</td><td>5.9E-13</td><td>2.7E-12</td><td>BT20/MB468</td><td>MITO</td></tr>
<tr><td>>hg38_21_+_8433172_8446622_RNA45SN1_45S_50ntFlanks@10463.10481.19|CTCCGTTGCCCTCGGCCGA</td><td>>hg38_21_+_8433172_8446622_RNA45SN1_45S_50ntFlanks</td><td>10463</td><td>10481</td><td>19</td><td>CTCCGTTGCCCTCGGCCGA</td><td>129.3</td><td>-10.9</td><td>1.5</td><td>-7.2</td><td>6.8E-13</td><td>2.7E-12</td><td>MB231/MB468</td><td>MP</td></tr>
<tr><td>>hg38_21_+_8433172_8446622_RNA45SN1_45S_50ntFlanks@9572.9594.23|CAAATCGGTCGTCCGACCTGGGT</td><td>>hg38_21_+_8433172_8446622_RNA45SN1_45S_50ntFlanks</td><td>9572</td><td>9594</td><td>23</td><td>CAAATCGGTCGTCCGACCTGGGT</td><td>5526.3</td><td>-10.6</td><td>1.5</td><td>-7.2</td><td>6.8E-13</td><td>2.8E-12</td><td>MB231/MB468</td><td>MP</td></tr>
<tr><td>>hg38_21_+_8433172_8446622_RNA45SN1_45S_50ntFlanks@6719.6756.38|TGAATTGCAGGACACATTGATCATCGACACTTCGAACG</td><td>>hg38_21_+_8433172_8446622_RNA45SN1_45S_50ntFlanks</td><td>6719</td><td>6756</td><td>38</td><td>TGAATTGCAGGACACATTGATCATCGACACTTCGAACG</td><td>215.3</td><td>-11.6</td><td>1.6</td><td>-7.2</td><td>6.9E-13</td><td>2.8E-12</td><td>MB231/MB468</td><td>MP</td></tr>
<tr><td>>MI0000279|hsa-mir-196a-2&WithFlank&12|+|53991732|53991853@31.53.23&[MIMAT0000226&hsa-miR-196a-5p&offsets|0|+1;m-261&12|+|53991762|53991783&offsets|0|+1]|TAGGTAGTTTCATGTTGTTGGGA</td><td>>MI0000279|hsa-mir-196a-2&WithFlank&12|+|53991732|53991853</td><td>31</td><td>53</td><td>23</td><td>TAGGTAGTTTCATGTTGTTGGGA</td><td>886.6</td><td>13.1</td><td>1.7</td><td>7.7</td><td>1.1E-14</td><td>2.8E-12</td><td>BT20/MB231</td><td>MP</td></tr>
<tr><td>>hg38_MT_+_1621_3279_RNR2_16S_50ntFlanks@51.79.29|GCTAAACCTAGCCCCAAACCCACTCCACC</td><td>>hg38_MT_+_1621_3279_RNR2_16S_50ntFlanks</td><td>51</td><td>79</td><td>29</td><td>GCTAAACCTAGCCCCAAACCCACTCCACC</td><td>12986.2</td><td>7.0</td><td>1.0</td><td>7.2</td><td>5.9E-13</td><td>2.8E-12</td><td>BT20/MB468</td><td>MITO</td></tr>
<tr><td>>MI0000077|hsa-mir-21&WithFlank&17|+|59841260|59841343@15.36.22&[MIMAT0000076&hsa-miR-21-5p&offsets|+1|+1;m-1&17|+|59841273|59841294&offsets|+1|+1]|AGCTTATCAGACTGATGTTGAC</td><td>>MI0000077|hsa-mir-21&WithFlank&17|+|59841260|59841343</td><td>15</td><td>36</td><td>22</td><td>AGCTTATCAGACTGATGTTGAC</td><td>330.4</td><td>3.6</td><td>0.5</td><td>7.1</td><td>1.1E-12</td><td>2.8E-12</td><td>BT20/MB468</td><td>NUC</td></tr>
<tr><td>>hg38_21_+_8433172_8446622_RNA45SN1_45S_50ntFlanks@11561.11602.42|GCGCCTAGCAGCCGACTTAGAACTGGTGCGGACCAGGGGAAT</td><td>>hg38_21_+_8433172_8446622_RNA45SN1_45S_50ntFlanks</td><td>11561</td><td>11602</td><td>42</td><td>GCGCCTAGCAGCCGACTTAGAACTGGTGCGGACCAGGGGAAT</td><td>4568.6</td><td>-11.4</td><td>1.6</td><td>-7.1</td><td>1.1E-12</td><td>2.8E-12</td><td>BT20/MB468</td><td>NUC</td></tr>
<tr><td>>hg38_21_+_8433172_8446622_RNA45SN1_45S_50ntFlanks@12324.12347.24|TTGGGTTTTAAGCAGGAGGTGTCA</td><td>>hg38_21_+_8433172_8446622_RNA45SN1_45S_50ntFlanks</td><td>12324</td><td>12347</td><td>24</td><td>TTGGGTTTTAAGCAGGAGGTGTCA</td><td>176.1</td><td>-5.2</td><td>0.7</td><td>-7.2</td><td>8.0E-13</td><td>2.8E-12</td><td>MB231/MB468</td><td>CYTO</td></tr>
<tr><td>>MI0000298|hsa-mir-221&WithFlank&X|-|45746151|45746272@31.56.26&[MIMAT0004568&hsa-miR-221-5p&offsets|0|+4;m-405&X|-|45746220|45746242&offsets|0|+3]|ACCTGGCATACAATGTAGATTTCTGT</td><td>>MI0000298|hsa-mir-221&WithFlank&X|-|45746151|45746272</td><td>31</td><td>56</td><td>26</td><td>ACCTGGCATACAATGTAGATTTCTGT</td><td>1378.1</td><td>6.3</td><td>0.9</td><td>7.1</td><td>9.9E-13</td><td>2.8E-12</td><td>MB231/MB468</td><td>MITO</td></tr>
<tr><td>>hg38_21_+_8433172_8446622_RNA45SN1_45S_50ntFlanks@6651.6673.23|CGACTCTTAGCGGTGGATCACTC</td><td>>hg38_21_+_8433172_8446622_RNA45SN1_45S_50ntFlanks</td><td>6651</td><td>6673</td><td>23</td><td>CGACTCTTAGCGGTGGATCACTC</td><td>2003.3</td><td>5.0</td><td>0.7</td><td>7.2</td><td>7.8E-13</td><td>2.8E-12</td><td>BT20/MB468</td><td>CYTO</td></tr>
<tr><td>>hg38_21_+_8433172_8446622_RNA45SN1_45S_50ntFlanks@10781.10826.46|AGGTGAACAGCCTCTGGCATGTTGGAACAATGTAGGTAAGGGAAGT</td><td>>hg38_21_+_8433172_8446622_RNA45SN1_45S_50ntFlanks</td><td>10781</td><td>10826</td><td>46</td><td>AGGTGAACAGCCTCTGGCATGTTGGAACAATGTAGGTAAGGGAAGT</td><td>313.7</td><td>-12.1</td><td>1.7</td><td>-7.1</td><td>9.9E-13</td><td>2.8E-12</td><td>MB231/MB468</td><td>MITO</td></tr>
<tr><td>>hg38_21_+_8433172_8446622_RNA45SN1_45S_50ntFlanks@8017.8042.26|TATTAGTCAGCGGAGGAGAAGAAACT</td><td>>hg38_21_+_8433172_8446622_RNA45SN1_45S_50ntFlanks</td><td>8017</td><td>8042</td><td>26</td><td>TATTAGTCAGCGGAGGAGAAGAAACT</td><td>363.8</td><td>9.6</td><td>1.3</td><td>7.2</td><td>8.0E-13</td><td>2.8E-12</td><td>MB231/MB468</td><td>CYTO</td></tr>
<tr><td>>hg38_21_+_8433172_8446622_RNA45SN1_45S_50ntFlanks@10787.10828.42|ACAGCCTCTGGCATGTTGGAACAATGTAGGTAAGGGAAGTCG</td><td>>hg38_21_+_8433172_8446622_RNA45SN1_45S_50ntFlanks</td><td>10787</td><td>10828</td><td>42</td><td>ACAGCCTCTGGCATGTTGGAACAATGTAGGTAAGGGAAGTCG</td><td>232.4</td><td>-11.6</td><td>1.6</td><td>-7.1</td><td>1.0E-12</td><td>2.8E-12</td><td>MB231/MB468</td><td>MITO</td></tr>
<tr><td>>hg38_21_+_8433172_8446622_RNA45SN1_45S_50ntFlanks@12510.12546.37|TCGTGAGACAGGTTAGTTTTACCCTACTGATGATGTG</td><td>>hg38_21_+_8433172_8446622_RNA45SN1_45S_50ntFlanks</td><td>12510</td><td>12546</td><td>37</td><td>TCGTGAGACAGGTTAGTTTTACCCTACTGATGATGTG</td><td>189.1</td><td>-11.3</td><td>1.6</td><td>-7.1</td><td>1.0E-12</td><td>2.8E-12</td><td>MB231/MB468</td><td>MITO</td></tr>
<tr><td>>hg38_21_+_8433172_8446622_RNA45SN1_45S_50ntFlanks@12903.12930.28|CACGCACCGCACGTTCGTGGGGAACCTG</td><td>>hg38_21_+_8433172_8446622_RNA45SN1_45S_50ntFlanks</td><td>12903</td><td>12930</td><td>28</td><td>CACGCACCGCACGTTCGTGGGGAACCTG</td><td>101.6</td><td>-10.5</td><td>1.5</td><td>-7.2</td><td>6.9E-13</td><td>2.8E-12</td><td>MB231/MB468</td><td>MP</td></tr>
<tr><td>>hg38_21_+_8433172_8446622_RNA45SN1_45S_50ntFlanks@4731.4759.29|AAGAACGAAAGTCGGAGGTTCGAAGACGA</td><td>>hg38_21_+_8433172_8446622_RNA45SN1_45S_50ntFlanks</td><td>4731</td><td>4759</td><td>29</td><td>AAGAACGAAAGTCGGAGGTTCGAAGACGA</td><td>2494.7</td><td>-5.7</td><td>0.8</td><td>-7.1</td><td>1.0E-12</td><td>2.8E-12</td><td>MB231/MB468</td><td>MITO</td></tr>
<tr><td>>MI0003560|hsa-mir-92b&WithFlank&1|+|155195171|155195278@67.87.21(+1U)&[MIMAT0003218&hsa-miR-92b-3p&offsets|0|-1(+1U);m-115&1|+|155195237|155195258&offsets|0|-1(+1U)]|TATTGCACTCGTCCCGGCCTCT</td><td>>MI0003560|hsa-mir-92b&WithFlank&1|+|155195171|155195278</td><td>67</td><td>87</td><td>21(+1U)</td><td>TATTGCACTCGTCCCGGCCTCT</td><td>1719.6</td><td>7.1</td><td>1.0</td><td>7.1</td><td>1.0E-12</td><td>2.8E-12</td><td>MB231/MB468</td><td>MITO</td></tr>
<tr><td>>MI0000065|hsa-let-7d&WithFlank&9|+|94178828|94178926@14.33.20(+1C)&[MIMAT0000065&hsa-let-7d-5p&offsets|0|-2(+1C);m-90&9|+|94178841|94178862&offsets|0|-2(+1C)]|AGAGGTAGTAGGTTGCATAGC</td><td>>MI0000065|hsa-let-7d&WithFlank&9|+|94178828|94178926</td><td>14</td><td>33</td><td>20(+1C)</td><td>AGAGGTAGTAGGTTGCATAGC</td><td>424.3</td><td>9.1</td><td>1.3</td><td>7.2</td><td>7.9E-13</td><td>2.8E-12</td><td>BT20/MB468</td><td>CYTO</td></tr>
<tr><td>>MI0000108|hsa-mir-103a-2&WithFlank&20|+|3917488|3917577@54.77.24;MI0000109|hsa-mir-103a-1&WithFlank&5|-|168560890|168560979@54.77.24&[MIMAT0000101&hsa-miR-103a-3p&offsets|0|+1;m-12&20|+|3917541|3917563&offsets|0|+1];[MIMAT0000101_1&hsa-miR-103a-3p&offsets|0|+1;m-13&5|-|168560904|168560926&offsets|0|+1;m-44220&5|-|168560908|168560926&offsets|0|+5]|AGCAGCATTGTACAGGGCTATGAA</td><td>>MI0000108|hsa-mir-103a-2&WithFlank&20|+|3917488|3917577</td><td>54</td><td>77</td><td>24</td><td>AGCAGCATTGTACAGGGCTATGAA</td><td>888.0</td><td>5.0</td><td>0.7</td><td>7.2</td><td>8.3E-13</td><td>2.8E-12</td><td>MB231/MB468</td><td>NUC</td></tr>
<tr><td>>hg38_21_+_8433172_8446622_RNA45SN1_45S_50ntFlanks@10761.10793.33|ATATCCGCAGCAGGTCTCCAAGGTGAACAGCCT</td><td>>hg38_21_+_8433172_8446622_RNA45SN1_45S_50ntFlanks</td><td>10761</td><td>10793</td><td>33</td><td>ATATCCGCAGCAGGTCTCCAAGGTGAACAGCCT</td><td>195.9</td><td>-10.6</td><td>1.5</td><td>-7.2</td><td>8.3E-13</td><td>2.8E-12</td><td>MB231/MB468</td><td>NUC</td></tr>
<tr><td>>hg38_21_+_8433172_8446622_RNA45SN1_45S_50ntFlanks@5236.5260.25|CACTGACTGGCTCAGCGTGTGCCTA</td><td>>hg38_21_+_8433172_8446622_RNA45SN1_45S_50ntFlanks</td><td>5236</td><td>5260</td><td>25</td><td>CACTGACTGGCTCAGCGTGTGCCTA</td><td>877.2</td><td>-13.6</td><td>1.9</td><td>-7.1</td><td>1.0E-12</td><td>2.8E-12</td><td>MB231/MB468</td><td>MITO</td></tr>
<tr><td>>MI0000070|hsa-mir-16-1&WithFlank&13|-|50048967|50049067@20.39.20(+1U);MI0000115|hsa-mir-16-2&WithFlank&3|+|160404739|160404831@16.35.20(+1U)&[MIMAT0000069&hsa-miR-16-5p&offsets|0|-2(+1U);m-62&13|-|50049027|50049048&offsets|0|-2(+1U)];[MIMAT0000069_1&hsa-miR-16-5p&offsets|0|-2(+1U);m-61&3|+|160404754|160404775&offsets|0|-2(+1U)]|TAGCAGCACGTAAATATTGGT</td><td>>MI0000070|hsa-mir-16-1&WithFlank&13|-|50048967|50049067</td><td>20</td><td>39</td><td>20(+1U)</td><td>TAGCAGCACGTAAATATTGGT</td><td>1098.3</td><td>8.5</td><td>1.2</td><td>7.2</td><td>7.9E-13</td><td>2.8E-12</td><td>BT20/MB468</td><td>CYTO</td></tr>
<tr><td>>hg38_21_+_8433172_8446622_RNA45SN1_45S_50ntFlanks@13004.13024.21|ATCTATTGAAAGTCAGCCCTC</td><td>>hg38_21_+_8433172_8446622_RNA45SN1_45S_50ntFlanks</td><td>13004</td><td>13024</td><td>21</td><td>ATCTATTGAAAGTCAGCCCTC</td><td>141.3</td><td>-9.5</td><td>1.3</td><td>-7.2</td><td>7.9E-13</td><td>2.8E-12</td><td>BT20/MB468</td><td>CYTO</td></tr>
<tr><td>>hg38_21_+_8433172_8446622_RNA45SN1_45S_50ntFlanks@10849.10868.20|CGGGATAAGGATTGGCTCTA</td><td>>hg38_21_+_8433172_8446622_RNA45SN1_45S_50ntFlanks</td><td>10849</td><td>10868</td><td>20</td><td>CGGGATAAGGATTGGCTCTA</td><td>738.2</td><td>-8.9</td><td>1.2</td><td>-7.2</td><td>8.3E-13</td><td>2.8E-12</td><td>MB231/MB468</td><td>NUC</td></tr>
<tr><td>>hg38_21_+_8433172_8446622_RNA45SN1_45S_50ntFlanks@6648.6667.20|GTACGACTCTTAGCGGTGGA</td><td>>hg38_21_+_8433172_8446622_RNA45SN1_45S_50ntFlanks</td><td>6648</td><td>6667</td><td>20</td><td>GTACGACTCTTAGCGGTGGA</td><td>1972.9</td><td>6.0</td><td>0.8</td><td>7.2</td><td>6.0E-13</td><td>2.8E-12</td><td>BT20/MB468</td><td>MITO</td></tr>
<tr><td>>hg38_21_+_8433172_8446622_RNA45SN1_45S_50ntFlanks@9575.9601.27|ATCGGTCGTCCGACCTGGGTATAGGGG</td><td>>hg38_21_+_8433172_8446622_RNA45SN1_45S_50ntFlanks</td><td>9575</td><td>9601</td><td>27</td><td>ATCGGTCGTCCGACCTGGGTATAGGGG</td><td>174.3</td><td>-9.5</td><td>1.3</td><td>-7.2</td><td>6.1E-13</td><td>2.8E-12</td><td>BT20/MB468</td><td>MITO</td></tr>
<tr><td>>hg38_21_+_8433172_8446622_RNA45SN1_45S_50ntFlanks@11767.11792.26|TCTAATTAGTGACGCGCATGAATGGA</td><td>>hg38_21_+_8433172_8446622_RNA45SN1_45S_50ntFlanks</td><td>11767</td><td>11792</td><td>26</td><td>TCTAATTAGTGACGCGCATGAATGGA</td><td>170.4</td><td>-10.4</td><td>1.5</td><td>-7.2</td><td>8.3E-13</td><td>2.8E-12</td><td>MB231/MB468</td><td>NUC</td></tr>
<tr><td>>hg38_21_+_8433172_8446622_RNA45SN1_45S_50ntFlanks@10384.10406.23|CCATGTGAACAGCAGTTGAACAT</td><td>>hg38_21_+_8433172_8446622_RNA45SN1_45S_50ntFlanks</td><td>10384</td><td>10406</td><td>23</td><td>CCATGTGAACAGCAGTTGAACAT</td><td>317.4</td><td>-12.2</td><td>1.7</td><td>-7.2</td><td>7.0E-13</td><td>2.8E-12</td><td>MB231/MB468</td><td>MP</td></tr>
<tr><td>>hg38_21_+_8433172_8446622_RNA45SN1_45S_50ntFlanks@9565.9597.33|TGACGTGCAAATCGGTCGTCCGACCTGGGTATA</td><td>>hg38_21_+_8433172_8446622_RNA45SN1_45S_50ntFlanks</td><td>9565</td><td>9597</td><td>33</td><td>TGACGTGCAAATCGGTCGTCCGACCTGGGTATA</td><td>3983.2</td><td>-4.7</td><td>0.7</td><td>-7.2</td><td>8.3E-13</td><td>2.8E-12</td><td>MB231/MB468</td><td>NUC</td></tr>
<tr><td>>hg38_21_+_8433172_8446622_RNA45SN1_45S_50ntFlanks@12073.12091.19|CCGAGGGGCTCTCGCTTCT</td><td>>hg38_21_+_8433172_8446622_RNA45SN1_45S_50ntFlanks</td><td>12073</td><td>12091</td><td>19</td><td>CCGAGGGGCTCTCGCTTCT</td><td>215.0</td><td>-11.5</td><td>1.6</td><td>-7.1</td><td>1.0E-12</td><td>2.8E-12</td><td>MB231/MB468</td><td>MITO</td></tr>
<tr><td>>hg38_21_+_8433172_8446622_RNA45SN1_45S_50ntFlanks@3939.3958.20|AACCCGGTCAGCCCCTCTCC</td><td>>hg38_21_+_8433172_8446622_RNA45SN1_45S_50ntFlanks</td><td>3939</td><td>3958</td><td>20</td><td>AACCCGGTCAGCCCCTCTCC</td><td>382.2</td><td>-11.5</td><td>1.6</td><td>-7.2</td><td>8.4E-13</td><td>2.8E-12</td><td>MB231/MB468</td><td>NUC</td></tr>
<tr><td>>hg38_21_+_8433172_8446622_RNA45SN1_45S_50ntFlanks@4223.4243.21|AAAAATAACAATACAGGACTC</td><td>>hg38_21_+_8433172_8446622_RNA45SN1_45S_50ntFlanks</td><td>4223</td><td>4243</td><td>21</td><td>AAAAATAACAATACAGGACTC</td><td>138.1</td><td>-11.0</td><td>1.5</td><td>-7.2</td><td>7.1E-13</td><td>2.8E-12</td><td>MB231/MB468</td><td>MP</td></tr>
<tr><td>>hg38_21_+_8433172_8446622_RNA45SN1_45S_50ntFlanks@4367.4397.31|CAGTTAAAAAGCTCGTAGTTGGATCTTGGGA</td><td>>hg38_21_+_8433172_8446622_RNA45SN1_45S_50ntFlanks</td><td>4367</td><td>4397</td><td>31</td><td>CAGTTAAAAAGCTCGTAGTTGGATCTTGGGA</td><td>514.1</td><td>-9.2</td><td>1.3</td><td>-7.1</td><td>1.1E-12</td><td>2.8E-12</td><td>BT20/MB468</td><td>NUC</td></tr>
<tr><td>>hg38_21_+_8433172_8446622_RNA45SN1_45S_50ntFlanks@4983.5016.34|CGGACAGGATTGACAGATTGATAGCTCTTTCTCG</td><td>>hg38_21_+_8433172_8446622_RNA45SN1_45S_50ntFlanks</td><td>4983</td><td>5016</td><td>34</td><td>CGGACAGGATTGACAGATTGATAGCTCTTTCTCG</td><td>3378.3</td><td>-11.1</td><td>1.6</td><td>-7.1</td><td>1.0E-12</td><td>2.8E-12</td><td>MB231/MB468</td><td>MITO</td></tr>
<tr><td>>hg38_21_+_8433172_8446622_RNA45SN1_45S_50ntFlanks@12003.12043.41|GCGGGCCGCCGGTGAAATACCACTACTCTGATCGTTTTTTC</td><td>>hg38_21_+_8433172_8446622_RNA45SN1_45S_50ntFlanks</td><td>12003</td><td>12043</td><td>41</td><td>GCGGGCCGCCGGTGAAATACCACTACTCTGATCGTTTTTTC</td><td>1078.3</td><td>-10.3</td><td>1.4</td><td>-7.2</td><td>8.4E-13</td><td>2.8E-12</td><td>MB231/MB468</td><td>NUC</td></tr>
<tr><td>>hg38_21_+_8433172_8446622_RNA45SN1_45S_50ntFlanks@9575.9605.31|ATCGGTCGTCCGACCTGGGTATAGGGGCGAA</td><td>>hg38_21_+_8433172_8446622_RNA45SN1_45S_50ntFlanks</td><td>9575</td><td>9605</td><td>31</td><td>ATCGGTCGTCCGACCTGGGTATAGGGGCGAA</td><td>257.2</td><td>-10.1</td><td>1.4</td><td>-7.2</td><td>6.1E-13</td><td>2.9E-12</td><td>BT20/MB468</td><td>MITO</td></tr>
<tr><td>>hg38_21_+_8433172_8446622_RNA45SN1_45S_50ntFlanks@3988.4026.39|CTTTGGTGACTCTAGATAACCTCGGGCCGATCGCACGCC</td><td>>hg38_21_+_8433172_8446622_RNA45SN1_45S_50ntFlanks</td><td>3988</td><td>4026</td><td>39</td><td>CTTTGGTGACTCTAGATAACCTCGGGCCGATCGCACGCC</td><td>210.7</td><td>-11.6</td><td>1.6</td><td>-7.2</td><td>7.1E-13</td><td>2.9E-12</td><td>MB231/MB468</td><td>MP</td></tr>
<tr><td>>MI0000298|hsa-mir-221&WithFlank&X|-|45746151|45746272@31.53.23&[MIMAT0004568&hsa-miR-221-5p&offsets|0|+1;m-405&X|-|45746220|45746242&offsets|0|0]|ACCTGGCATACAATGTAGATTTC</td><td>>MI0000298|hsa-mir-221&WithFlank&X|-|45746151|45746272</td><td>31</td><td>53</td><td>23</td><td>ACCTGGCATACAATGTAGATTTC</td><td>614.7</td><td>6.2</td><td>0.9</td><td>7.2</td><td>8.2E-13</td><td>2.9E-12</td><td>MB231/MB468</td><td>CYTO</td></tr>
<tr><td>>hg38_21_+_8433172_8446622_RNA45SN1_45S_50ntFlanks@12892.12930.39|CCCTCGCCCGTCACGCACCGCACGTTCGTGGGGAACCTG</td><td>>hg38_21_+_8433172_8446622_RNA45SN1_45S_50ntFlanks</td><td>12892</td><td>12930</td><td>39</td><td>CCCTCGCCCGTCACGCACCGCACGTTCGTGGGGAACCTG</td><td>369.6</td><td>-12.4</td><td>1.7</td><td>-7.2</td><td>7.1E-13</td><td>2.9E-12</td><td>MB231/MB468</td><td>MP</td></tr>
<tr><td>>hg38_21_+_8433172_8446622_RNA45SN1_45S_50ntFlanks@12894.12931.38|CTCGCCCGTCACGCACCGCACGTTCGTGGGGAACCTGG</td><td>>hg38_21_+_8433172_8446622_RNA45SN1_45S_50ntFlanks</td><td>12894</td><td>12931</td><td>38</td><td>CTCGCCCGTCACGCACCGCACGTTCGTGGGGAACCTGG</td><td>482.7</td><td>-8.5</td><td>1.2</td><td>-7.1</td><td>1.0E-12</td><td>2.9E-12</td><td>MB231/MB468</td><td>MITO</td></tr>
<tr><td>>hg38_21_+_8433172_8446622_RNA45SN1_45S_50ntFlanks@11576.11594.19|CTTAGAACTGGTGCGGACC</td><td>>hg38_21_+_8433172_8446622_RNA45SN1_45S_50ntFlanks</td><td>11576</td><td>11594</td><td>19</td><td>CTTAGAACTGGTGCGGACC</td><td>1840.1</td><td>-4.3</td><td>0.6</td><td>-7.2</td><td>8.4E-13</td><td>2.9E-12</td><td>MB231/MB468</td><td>NUC</td></tr>
<tr><td>>hg38_21_+_8433172_8446622_RNA45SN1_45S_50ntFlanks@3782.3801.20|CAGTGAAACTGCGAATGGCT</td><td>>hg38_21_+_8433172_8446622_RNA45SN1_45S_50ntFlanks</td><td>3782</td><td>3801</td><td>20</td><td>CAGTGAAACTGCGAATGGCT</td><td>3224.0</td><td>-6.7</td><td>0.9</td><td>-7.2</td><td>7.1E-13</td><td>2.9E-12</td><td>MB231/MB468</td><td>MP</td></tr>
<tr><td>>MI0000749|hsa-mir-30e&WithFlank&1|+|40754349|40754452@23.46.24(+1U)&[MIMAT0000692&hsa-miR-30e-5p&offsets|0|+2(+1U);m-24&1|+|40754371|40754394&offsets|0|0(+1U)]|TGTAAACATCCTTGACTGGAAGCTT</td><td>>MI0000749|hsa-mir-30e&WithFlank&1|+|40754349|40754452</td><td>23</td><td>46</td><td>24(+1U)</td><td>TGTAAACATCCTTGACTGGAAGCTT</td><td>365.4</td><td>6.0</td><td>0.8</td><td>7.2</td><td>8.5E-13</td><td>2.9E-12</td><td>MB231/MB468</td><td>NUC</td></tr>
</table>

| | | | | | | | | | | | | | |
|---|---|---|---|---|---|---|---|---|---|---|---|---|---|
| >trna111_HisGTG_1_-_147774845_147774916@-1G.32.33,trna118_HisGTG_1_-_145396881_145396952@-1G.32.33,trna16_HisGTG_1_+_146544773_146544844@-1G.32.33,trna1_HisGTG_15_+_45493349_45493420@-1G.32.33,trna21_HisGTG_1_+_147753471_147753542@-1G.32.33,trna33_HisGTG_6_+_27125906_27125977@-1G.32.33,trna7_HisGTG_9_-_14433938_14434009@-1G.32.33,trna8_HisGTG_15_-_45492611_45492682@-1G.32.33,trna9_HisGTG_15_-_45490804_45490875@-1G.32.33\|GGCCGTGATCGTATAGTGGTTAGTACTCTGCGT | >trna111_HisGTG_1_-_147774845_147774916 | -1G | 32 | 33 | GGCCGTGATCGTATAGTGGTTAGTACTCTGCGT | 1972.8 | 8.4 | 1.2 | 7.1 | 1.0E-12 | 2.9E-12 | MB231/MB468 | MITO |
| >hg38_21_+_8433172_8446622_RNA45SN1_45S_50ntFlanks@6689.6720.32\|GAAGAACGCAGCTAGCTGCGAGAATTAATGTG | >hg38_21_+_8433172_8446622_RNA45SN1_45S_50ntFlanks | 6689 | 6720 | 32 | GAAGAACGCAGCTAGCTGCGAGAATTAATGTG | 110.4 | -10.7 | 1.5 | -7.2 | 7.2E-13 | 2.9E-12 | MB231/MB468 | MP |
| >hg38_21_+_8433172_8446622_RNA45SN1_45S_50ntFlanks@12073.12098.26\|CCGAGGGGCTCTCGCTTCTGGCGCCA | >hg38_21_+_8433172_8446622_RNA45SN1_45S_50ntFlanks | 12073 | 12098 | 26 | CCGAGGGGCTCTCGCTTCTGGCGCCA | 449.0 | -8.6 | 1.2 | -7.1 | 1.0E-12 | 2.9E-12 | MB231/MB468 | MITO |
| >MI0000542\|hsa-mir-320a&WithFlank&8\|-\|22244960\|22245043@48.68.21(+3U)&[MIMAT0000510&hsa-miR-320a-3p&offsets\|0\|-1(+3U);m-60&8\|-\|22244975\|22244996&offsets\|0\|-1(+3U)]\|AAAAGCTGGGTTGAGAGGGCGTTT | >MI0000542\|hsa-mir-320a&WithFlank&8\|-\|22244960\|22245043 | 48 | 68 | 21(+3U) | AAAAGCTGGGTTGAGAGGGCGTTT | 171.1 | 4.6 | 0.7 | 7.1 | 1.1E-12 | 2.9E-12 | BT20/MB468 | NUC |
| >MI0000255\|hsa-mir-30d&WithFlank&8\|-\|134804870\|134804951@12.35.24(+1U)&[MIMAT0000245&hsa-miR-30d-5p&offsets\|0\|+2(+1U);m-21&8\|-\|134804919\|134804940&offsets\|0\|+2(+1U)]\|TGTAAACATCCCCGACTGGAAGCTT | >MI0000255\|hsa-mir-30d&WithFlank&8\|-\|134804870\|134804951 | 12 | 35 | 24(+1U) | TGTAAACATCCCCGACTGGAAGCTT | 3057.7 | 7.2 | 1.0 | 7.1 | 1.0E-12 | 2.9E-12 | MB231/MB468 | MITO |
| >hg38_21_+_8433172_8446622_RNA45SN1_45S_50ntFlanks@8836.8858.23\|CGGCGGTGGCGGCGGCGGCGG | >hg38_21_+_8433172_8446622_RNA45SN1_45S_50ntFlanks | 8836 | 8858 | 23 | CGGCGGTGGCGGCGGCGGCGG | 153.1 | -11.1 | 1.5 | -7.2 | 7.2E-13 | 2.9E-12 | MB231/MB468 | MP |
| >hg38_21_+_8433172_8446622_RNA45SN1_45S_50ntFlanks@4102.4121.20\|ACCACGGGTGACGGGGAATC | >hg38_21_+_8433172_8446622_RNA45SN1_45S_50ntFlanks | 4102 | 4121 | 20 | ACCACGGGTGACGGGGAATC | 2039.2 | -6.5 | 0.9 | -7.2 | 8.5E-13 | 2.9E-12 | MB231/MB468 | NUC |
| >hg38_21_+_8433172_8446622_RNA45SN1_45S_50ntFlanks@4486.4511.26\|CCGCGGGGCCCGAAGCGTTTACTTTG | >hg38_21_+_8433172_8446622_RNA45SN1_45S_50ntFlanks | 4486 | 4511 | 26 | CCGCGGGGCCCGAAGCGTTTACTTTG | 564.7 | -12.1 | 1.7 | -7.2 | 8.5E-13 | 2.9E-12 | MB231/MB468 | NUC |
| >am_trna128_GlyGCC_6_-_27870686_27870756@1.28.28,trna133_GlyCCC_1_-_16872434_16872504@1.28.28,trna18_GlyGCC_16_+_70822597_70822667@1.28.28,trna19_GlyGCC_16_+_70823410_70823480@1.28.28,trna19_GlyGCC_2_-_157257659_157257729@1.28.28,trna24_GlyGCC_16_-_70812942_70813012@1.28.28,trna25_GlyGCC_16_-_70812114_70812184@1.28.28,trna4_GlyCCC_1_+_17188416_17188486@1.28.28,trna5_GlyGCC_17_+_8029064_8029134@1.28.28,trna68_GlyGCC_1_-_161493637_161493707@1.28.28\|GCATTGGTGGTTCAGTGGTAGAATTCTC | >am_trna128_GlyGCC_6_-_27870686_27870756 | 1 | 28 | 28 | GCATTGGTGGTTCAGTGGTAGAATTCTC | 56591.6 | -3.9 | 0.5 | -7.2 | 8.1E-13 | 2.9E-12 | BT20/MB468 | CYTO |
| >MI0000809\|hsa-mir-151a&WithFlank&8\|-\|140732558\|140732659@53.74.22(+1U)&[MIMAT0000757&hsa-miR-151a-3p&offsets\|0\|+1(+1U);m-30&8\|-\|140732587\|140732607&offsets\|0\|+1(+1U)]\|CTAGACTGAAGCTCCTTGAGGAT | >MI0000809\|hsa-mir-151a&WithFlank&8\|-\|140732558\|140732659 | 53 | 74 | 22(+1U) | CTAGACTGAAGCTCCTTGAGGAT | 1279.5 | 6.3 | 0.9 | 7.1 | 1.0E-12 | 2.9E-12 | MB231/MB468 | MITO |
| >hg38_21_+_8433172_8446622_RNA45SN1_45S_50ntFlanks@6738.6769.32\|ATCATCGACACTTCGAACGCACTTGCGGCCCC | >hg38_21_+_8433172_8446622_RNA45SN1_45S_50ntFlanks | 6738 | 6769 | 32 | ATCATCGACACTTCGAACGCACTTGCGGCCCC | 173.2 | -9.5 | 1.3 | -7.2 | 6.2E-13 | 2.9E-12 | BT20/MB468 | MITO |
| >hg38_21_+_8433172_8446622_RNA45SN1_45S_50ntFlanks@10349.10373.25\|AACGAGAACTTTGAAGGCCGAAGTG | >hg38_21_+_8433172_8446622_RNA45SN1_45S_50ntFlanks | 10349 | 10373 | 25 | AACGAGAACTTTGAAGGCCGAAGTG | 508.3 | -4.0 | 0.6 | -7.1 | 1.1E-12 | 2.9E-12 | BT20/MB468 | NUC |
| >MI0006444\|hsa-mir-1307&WithFlank&10\|-\|103394247\|103394407@87.109.23&[MIMAT0005951&hsa-miR-1307-3p&offsets\|+1\|+2;m-66&10\|-\|103394300\|103394322&offsets\|+1\|+1]\|CTCGGCGTGGCGTCGGTCGTGGT | >MI0006444\|hsa-mir-1307&WithFlank&10\|-\|103394247\|103394407 | 87 | 109 | 23 | CTCGGCGTGGCGTCGGTCGTGGT | 495.1 | 8.5 | 1.2 | 7.2 | 8.2E-13 | 2.9E-12 | BT20/MB468 | CYTO |
| >hg38_21_+_8433172_8446622_RNA45SN1_45S_50ntFlanks@12938.12956.19\|ACCATTCGTAGACGACCTG | >hg38_21_+_8433172_8446622_RNA45SN1_45S_50ntFlanks | 12938 | 12956 | 19 | ACCATTCGTAGACGACCTG | 2004.5 | -4.8 | 0.7 | -7.2 | 8.2E-13 | 2.9E-12 | BT20/MB468 | CYTO |
| >hg38_21_+_8433172_8446622_RNA45SN1_45S_50ntFlanks@10458.10493.36\|ATGGCCTCCGTTGCCCTCGGCCGATCGAAAGGGAGT | >hg38_21_+_8433172_8446622_RNA45SN1_45S_50ntFlanks | 10458 | 10493 | 36 | ATGGCCTCCGTTGCCCTCGGCCGATCGAAAGGGAGT | 157.4 | -11.2 | 1.6 | -7.2 | 7.2E-13 | 2.9E-12 | MB231/MB468 | MP |
| >hg38_21_+_8433172_8446622_RNA45SN1_45S_50ntFlanks@11572.11602.31\|CCGACTTAGAACTGGTGCGGACCAGGGGAAT | >hg38_21_+_8433172_8446622_RNA45SN1_45S_50ntFlanks | 11572 | 11602 | 31 | CCGACTTAGAACTGGTGCGGACCAGGGGAAT | 5311.2 | -4.3 | 0.6 | -7.2 | 8.2E-13 | 2.9E-12 | BT20/MB468 | CYTO |
| >trna108_AlaAGC_6_-_28763741_28763812@58.75.18\|TCCCCAGTACCTCCACCA | >trna108_AlaAGC_6_-_28763741_28763812 | 58 | 75 | 18 | TCCCCAGTACCTCCACCA | 2654.3 | 6.9 | 1.0 | 7.2 | 8.4E-13 | 2.9E-12 | MB231/MB468 | CYTO |
| >hg38_21_+_8433172_8446622_RNA45SN1_45S_50ntFlanks@7975.8012.38\|CGCGACCTCAGATCAGACGTGGCGACCCGCTGAATTTA | >hg38_21_+_8433172_8446622_RNA45SN1_45S_50ntFlanks | 7975 | 8012 | 38 | CGCGACCTCAGATCAGACGTGGCGACCCGCTGAATTTA | 19056.1 | 4.6 | 0.6 | 7.2 | 8.4E-13 | 2.9E-12 | MB231/MB468 | CYTO |
| >hg38_21_+_8433172_8446622_RNA45SN1_45S_50ntFlanks@12615.12653.39\|GCTGAGGAGCCAATGGGGCGAAGCTACCATCTGTGGGAT | >hg38_21_+_8433172_8446622_RNA45SN1_45S_50ntFlanks | 12615 | 12653 | 39 | GCTGAGGAGCCAATGGGGCGAAGCTACCATCTGTGGGAT | 4075.8 | -11.4 | 1.6 | -7.1 | 1.1E-12 | 2.9E-12 | BT20/MB468 | NUC |
| >hg38_21_+_8433172_8446622_RNA45SN1_45S_50ntFlanks@10237.10262.26\|ACGCTACGCCGCGACGAGTAGGAGGG | >hg38_21_+_8433172_8446622_RNA45SN1_45S_50ntFlanks | 10237 | 10262 | 26 | ACGCTACGCCGCGACGAGTAGGAGGG | 146.5 | -11.1 | 1.5 | -7.2 | 7.3E-13 | 2.9E-12 | MB231/MB468 | MP |
| >hg38_21_+_8433172_8446622_RNA45SN1_45S_50ntFlanks@13008.13025.18\|ATTGAAAGTCAGCCCTCG | >hg38_21_+_8433172_8446622_RNA45SN1_45S_50ntFlanks | 13008 | 13025 | 18 | ATTGAAAGTCAGCCCTCG | 221.2 | -9.9 | 1.4 | -7.2 | 6.3E-13 | 2.9E-12 | BT20/MB468 | MITO |
| >hg38_21_+_8433172_8446622_RNA45SN1_45S_50ntFlanks@10795.10825.31\|TGGCATGTTGGAACAATGTAGGTAAGGGAAG | >hg38_21_+_8433172_8446622_RNA45SN1_45S_50ntFlanks | 10795 | 10825 | 31 | TGGCATGTTGGAACAATGTAGGTAAGGGAAG | 110.8 | -10.7 | 1.5 | -7.2 | 7.3E-13 | 2.9E-12 | MB231/MB468 | MP |
| >hg38_21_+_8433172_8446622_RNA45SN1_45S_50ntFlanks@11999.12042.44\|CCCTGCGGGCCGCCGGTGAAATACCACTACTCTGATCGTTTTTT | >hg38_21_+_8433172_8446622_RNA45SN1_45S_50ntFlanks | 11999 | 12042 | 44 | CCCTGCGGGCCGCCGGTGAAATACCACTACTCTGATCGTTTTTT | 387.8 | -12.4 | 1.7 | -7.1 | 1.1E-12 | 2.9E-12 | MB231/MB468 | MITO |
| >hg38_21_+_8433172_8446622_RNA45SN1_45S_50ntFlanks@3993.4013.21\|GTGACTCTAGATAACCTCGGG | >hg38_21_+_8433172_8446622_RNA45SN1_45S_50ntFlanks | 3993 | 4013 | 21 | GTGACTCTAGATAACCTCGGG | 799.8 | -7.6 | 1.1 | -7.2 | 7.3E-13 | 2.9E-12 | MB231/MB468 | MP |
| >hg38_21_+_8433172_8446622_RNA45SN1_45S_50ntFlanks@12366.12384.19\|TAACTGGCTTGTGGCGGCC | >hg38_21_+_8433172_8446622_RNA45SN1_45S_50ntFlanks | 12366 | 12384 | 19 | TAACTGGCTTGTGGCGGCC | 419.6 | -10.8 | 1.5 | -7.2 | 6.3E-13 | 2.9E-12 | BT20/MB468 | MITO |
| >hg38_21_+_8433172_8446622_RNA45SN1_45S_50ntFlanks@5082.5104.23\|AACGAACGAGACTCTGGCATGCT | >hg38_21_+_8433172_8446622_RNA45SN1_45S_50ntFlanks | 5082 | 5104 | 23 | AACGAACGAGACTCTGGCATGCT | 464.9 | 5.3 | 0.7 | 7.2 | 8.5E-13 | 2.9E-12 | MB231/MB468 | CYTO |
| >hg38_21_+_8433172_8446622_RNA45SN1_45S_50ntFlanks@5227.5244.18\|CGCGCGCTACACTGACTG | >hg38_21_+_8433172_8446622_RNA45SN1_45S_50ntFlanks | 5227 | 5244 | 18 | CGCGCGCTACACTGACTG | 479.4 | -12.8 | 1.8 | -7.2 | 7.4E-13 | 3.0E-12 | MB231/MB468 | MP |
| >hg38_21_+_8433172_8446622_RNA45SN1_45S_50ntFlanks@6652.6676.25\|GACTCTTAGCGGTGGATCACTCGGC | >hg38_21_+_8433172_8446622_RNA45SN1_45S_50ntFlanks | 6652 | 6676 | 25 | GACTCTTAGCGGTGGATCACTCGGC | 7605.6 | 5.4 | 0.7 | 7.2 | 6.3E-13 | 3.0E-12 | BT20/MB468 | MITO |
| >hg38_21_+_8433172_8446622_RNA45SN1_45S_50ntFlanks@3802.3834.33\|CATTAAATCAGTTATGGTTCCTTTGGTCGCTCG | >hg38_21_+_8433172_8446622_RNA45SN1_45S_50ntFlanks | 3802 | 3834 | 33 | CATTAAATCAGTTATGGTTCCTTTGGTCGCTCG | 642.4 | -4.4 | 0.6 | -7.2 | 8.5E-13 | 3.0E-12 | MB231/MB468 | CYTO |
| >hg38_21_+_8433172_8446622_RNA45SN1_45S_50ntFlanks@3813.3834.22\|TTATGGTTCCTTTGGTCGCTCG | >hg38_21_+_8433172_8446622_RNA45SN1_45S_50ntFlanks | 3813 | 3834 | 22 | TTATGGTTCCTTTGGTCGCTCG | 362.7 | -4.0 | 0.6 | -7.2 | 8.5E-13 | 3.0E-12 | MB231/MB468 | CYTO |
| >MI0000078\|hsa-mir-22&WithFlank&17\|-\|1713897\|1713993@59.79.21(+1C)&[MIMAT0000077&hsa-miR-22-3p&offsets\|0\|-1(+1C);m-2&17\|-\|1713914\|1713935&offsets\|0\|-1(+1C)]\|AAGCTGCCAGTTGAAGAACTGC | >MI0000078\|hsa-mir-22&WithFlank&17\|-\|1713897\|1713993 | 59 | 79 | 21(+1C) | AAGCTGCCAGTTGAAGAACTGC | 1877.4 | 7.1 | 1.0 | 7.1 | 1.1E-12 | 3.0E-12 | MB231/MB468 | MITO |
| >hg38_21_+_8433172_8446622_RNA45SN1_45S_50ntFlanks@8105.8134.30\|CGCGGCGGGGCGCGGGACATGTGGCGTACG | >hg38_21_+_8433172_8446622_RNA45SN1_45S_50ntFlanks | 8105 | 8134 | 30 | CGCGGCGGGGCGCGGGACATGTGGCGTACG | 274.0 | -12.0 | 1.7 | -7.2 | 7.4E-13 | 3.0E-12 | MB231/MB468 | MP |
| >hg38_21_+_8433172_8446622_RNA45SN1_45S_50ntFlanks@12944.12972.29\|CGTAGACGACCTGCTTCTGGGTCGGGGTT | >hg38_21_+_8433172_8446622_RNA45SN1_45S_50ntFlanks | 12944 | 12972 | 29 | CGTAGACGACCTGCTTCTGGGTCGGGGTT | 363.3 | -7.9 | 1.1 | -7.2 | 8.5E-13 | 3.0E-12 | MB231/MB468 | CYTO |
| >hg38_21_+_8433172_8446622_RNA45SN1_45S_50ntFlanks@8439.8469.31\|GATTCAACCCGGCGGCGGGTCCGGCCGTGTC | >hg38_21_+_8433172_8446622_RNA45SN1_45S_50ntFlanks | 8439 | 8469 | 31 | GATTCAACCCGGCGGCGGGTCCGGCCGTGTC | 140.3 | -11.0 | 1.5 | -7.2 | 7.4E-13 | 3.0E-12 | MB231/MB468 | MP |
| >hg38_21_+_8433172_8446622_RNA45SN1_45S_50ntFlanks@12356.12375.20\|ACCACAGGGATAACTGGCTT | >hg38_21_+_8433172_8446622_RNA45SN1_45S_50ntFlanks | 12356 | 12375 | 20 | ACCACAGGGATAACTGGCTT | 112.4 | -10.7 | 1.5 | -7.2 | 7.4E-13 | 3.0E-12 | MB231/MB468 | MP |
| >hg38_21_+_8433172_8446622_RNA45SN1_45S_50ntFlanks@4980.4999.20\|ACACGGACAGGATTGACAGA | >hg38_21_+_8433172_8446622_RNA45SN1_45S_50ntFlanks | 4980 | 4999 | 20 | ACACGGACAGGATTGACAGA | 494.1 | -6.9 | 1.0 | -7.2 | 7.4E-13 | 3.0E-12 | MB231/MB468 | MP |
| >hg38_21_+_8433172_8446622_RNA45SN1_45S_50ntFlanks@10781.10805.25\|AGGTGAACAGCCTCTGGCATGTTGG | >hg38_21_+_8433172_8446622_RNA45SN1_45S_50ntFlanks | 10781 | 10805 | 25 | AGGTGAACAGCCTCTGGCATGTTGG | 524.9 | -8.3 | 1.2 | -7.2 | 8.6E-13 | 3.0E-12 | MB231/MB468 | CYTO |
| >MI0000086\|hsa-mir-28&WithFlank&3\|+\|188688775\|188688872@61.82.22&[MIMAT0004502&hsa-miR-28-3p&offsets\|+1\|+1;m-28&3\|+\|188688834\|188688855&offsets\|+1\|+1]\|ACTAGATTGTGAGCTCCTGGAG | >MI0000086\|hsa-mir-28&WithFlank&3\|+\|188688775\|188688872 | 61 | 82 | 22 | ACTAGATTGTGAGCTCCTGGAG | 301.0 | 8.4 | 1.2 | 7.2 | 8.6E-13 | 3.0E-12 | MB231/MB468 | CYTO |
| >trnaMT_GluTTC_MT_-_14674_14742@26.58.33,trnalookalike8_GluTTC_5_-_93905172_93905240@26.58.33\|GGTTTTTCATATCATTGGTCGTGGTTGTAGTCC | >trnaMT_GluTTC_MT_-_14674_14742 | 26 | 58 | 33 | GGTTTTTCATATCATTGGTCGTGGTTGTAGTCC | 725.4 | 11.0 | 1.5 | 7.2 | 6.4E-13 | 3.0E-12 | BT20/MB468 | MITO |
| >hg38_21_+_8433172_8446622_RNA45SN1_45S_50ntFlanks@8472.8494.23\|CGGCCCGGCGGATCTTTCCCGCC | >hg38_21_+_8433172_8446622_RNA45SN1_45S_50ntFlanks | 8472 | 8494 | 23 | CGGCCCGGCGGATCTTTCCCGCC | 1208.6 | -12.3 | 1.7 | -7.2 | 6.4E-13 | 3.0E-12 | BT20/MB468 | MITO |
| >hg38_21_+_8433172_8446622_RNA45SN1_45S_50ntFlanks@10410.10432.23\|TCAGTCGGTCCTGAGAGATGGGC | >hg38_21_+_8433172_8446622_RNA45SN1_45S_50ntFlanks | 10410 | 10432 | 23 | TCAGTCGGTCCTGAGAGATGGGC | 1449.8 | -10.1 | 1.4 | -7.1 | 1.2E-12 | 3.0E-12 | BT20/MB468 | NUC |
| >am_trnaMT_GluTTC_MT_-_14674_14742@27.49.23,trnalookalike8_GluTTC_5_-_93905172_93905240@27.49.23\|GTTTTTCATATCATTGGTCGTGG | >am_trnaMT_GluTTC_MT_-_14674_14742 | 27 | 49 | 23 | GTTTTTCATATCATTGGTCGTGG | 119.0 | -9.9 | 1.4 | -7.1 | 8.8E-13 | 3.0E-12 | MB231/MB468 | NUC |
| >hg38_21_+_8433172_8446622_RNA45SN1_45S_50ntFlanks@12437.12467.31\|CATTGTGAAGCAGAATTCACCAAGCGTTGGA | >hg38_21_+_8433172_8446622_RNA45SN1_45S_50ntFlanks | 12437 | 12467 | 31 | CATTGTGAAGCAGAATTCACCAAGCGTTGGA | 313.5 | -12.2 | 1.7 | -7.2 | 7.5E-13 | 3.0E-12 | MB231/MB468 | MP |
| >hg38_21_+_8433172_8446622_RNA45SN1_45S_50ntFlanks@12957.12987.31\|CTTCTGGGTCGGGGTTTCGTACGTAGCAGAG | >hg38_21_+_8433172_8446622_RNA45SN1_45S_50ntFlanks | 12957 | 12987 | 31 | CTTCTGGGTCGGGGTTTCGTACGTAGCAGAG | 2121.7 | -10.7 | 1.5 | -7.1 | 1.2E-12 | 3.0E-12 | BT20/MB468 | NUC |
| >hg38_21_+_8433172_8446622_RNA45SN1_45S_50ntFlanks@10845.10866.22\|ACTTCGGGATAAGGATTGGCTC | >hg38_21_+_8433172_8446622_RNA45SN1_45S_50ntFlanks | 10845 | 10866 | 22 | ACTTCGGGATAAGGATTGGCTC | 134.6 | -10.9 | 1.5 | -7.2 | 7.5E-13 | 3.0E-12 | MB231/MB468 | MP |

<table>
<tr><td></td><td></td><td></td><td></td><td></td><td></td><td></td><td></td><td></td><td></td><td></td><td></td><td></td><td></td></tr>
<tr><td>>hg38_21_+_8433172_8446622_RNA45SN1_45S_50ntFlanks@11876.11902.27|AAGACCCTGTTGAGCTTGACTCTAGTC</td><td>>hg38_21_+_8433172_8446622_RNA45SN1_45S_50ntFlanks</td><td>11876</td><td>11902</td><td>27</td><td>AAGACCCTGTTGAGCTTGACTCTAGTC</td><td>107.8</td><td>-10.6</td><td>1.5</td><td>-7.2</td><td>7.5E-13</td><td>3.0E-12</td><td>MB231/MB468</td><td>MP</td></tr>
<tr><td>>hg38_21_+_8433172_8446622_RNA45SN1_45S_50ntFlanks@4722.4755.34|TCATTAATCAAGAACGAAAGTCGGAGGTTCGAAG</td><td>>hg38_21_+_8433172_8446622_RNA45SN1_45S_50ntFlanks</td><td>4722</td><td>4755</td><td>34</td><td>TCATTAATCAAGAACGAAAGTCGGAGGTTCGAAG</td><td>172.2</td><td>-11.3</td><td>1.6</td><td>-7.2</td><td>7.5E-13</td><td>3.0E-12</td><td>MB231/MB468</td><td>MP</td></tr>
<tr><td>>MI0000650|hsa-mir-200c&WithFlank&12|+|6963693|6963772@50.72.23(+1U)&[MIMAT0000617&hsa-miR-200c-3p&offsets|0|0(+1U);m-54&12|+|6963742|6963764&offsets|0|0(+1U)]|TAATACTGCCGGGTAATGATGGAT</td><td>>MI0000650|hsa-mir-200c&WithFlank&12|+|6963693|6963772</td><td>50</td><td>72</td><td>23(+1U)</td><td>TAATACTGCCGGGTAATGATGGAT</td><td>2650.7</td><td>2.8</td><td>0.4</td><td>7.1</td><td>1.2E-12</td><td>3.0E-12</td><td>BT20/MB468</td><td>NUC</td></tr>
<tr><td>>hg38_21_+_8433172_8446622_RNA45SN1_45S_50ntFlanks@8170.8197.28|CAAGTCCTTCTGATCGAGGCCCAGCCCG</td><td>>hg38_21_+_8433172_8446622_RNA45SN1_45S_50ntFlanks</td><td>8170</td><td>8197</td><td>28</td><td>CAAGTCCTTCTGATCGAGGCCCAGCCCG</td><td>4066.9</td><td>-6.4</td><td>0.9</td><td>-7.2</td><td>6.5E-13</td><td>3.0E-12</td><td>BT20/MB468</td><td>MITO</td></tr>
<tr><td>>hg38_21_+_8433172_8446622_RNA45SN1_45S_50ntFlanks@11705.11728.24|TGAAGCGCGGGTAAACGGCGGGAG</td><td>>hg38_21_+_8433172_8446622_RNA45SN1_45S_50ntFlanks</td><td>11705</td><td>11728</td><td>24</td><td>TGAAGCGCGGGTAAACGGCGGGAG</td><td>1474.4</td><td>-9.2</td><td>1.3</td><td>-7.1</td><td>8.9E-13</td><td>3.0E-12</td><td>MB231/MB468</td><td>NUC</td></tr>
<tr><td>>hg38_21_+_8433172_8446622_RNA45SN1_45S_50ntFlanks@12412.12436.25|ATCCTTCGATGTCGGCTCTTCCTAT</td><td>>hg38_21_+_8433172_8446622_RNA45SN1_45S_50ntFlanks</td><td>12412</td><td>12436</td><td>25</td><td>ATCCTTCGATGTCGGCTCTTCCTAT</td><td>398.2</td><td>-10.9</td><td>1.5</td><td>-7.2</td><td>8.5E-13</td><td>3.0E-12</td><td>BT20/MB468</td><td>CYTO</td></tr>
<tr><td>>hg38_21_+_8433172_8446622_RNA45SN1_45S_50ntFlanks@12233.12258.26|CCGTGGAGCAGAAGGGCAAAAGCTCG</td><td>>hg38_21_+_8433172_8446622_RNA45SN1_45S_50ntFlanks</td><td>12233</td><td>12258</td><td>26</td><td>CCGTGGAGCAGAAGGGCAAAAGCTCG</td><td>300.6</td><td>-12.0</td><td>1.7</td><td>-7.1</td><td>1.1E-12</td><td>3.0E-12</td><td>MB231/MB468</td><td>MITO</td></tr>
<tr><td>>trna112_GlnCTG_1_-_147737382_147737453@30.52.23,trna131_GlnCTG_6_-_27759135_27759206@30.52.23,trna146_GlnCTG_6_-_27515531_27515602@30.52.23,trna15_GlnCTG_1_+_145963304_145963375@30.52.23,trna19_GlnCTG_1_+_147505038_147505109@30.52.23,trna1_GlnCTG_6_+_18836402_18836473@30.52.23,trna28_GlnCTG_1_+_149186125_149186196@30.52.23,trna3_GlnCTG_17_+_8023070_8023141@30.52.23,trna49_GlnCTG_6_+_27487308_27487379@30.52.23,trna7_GlnCTG_15_-_66161400_66161471@30.52.23,trna99_GlnCTG_6_-_28909378_28909449@30.52.23|GACTCTGAATCCAGCGATCCGAG</td><td>>trna112_GlnCTG_1_-_147737382_147737453</td><td>30</td><td>52</td><td>23</td><td>GACTCTGAATCCAGCGATCCGAG</td><td>293.4</td><td>-12.1</td><td>1.7</td><td>-7.2</td><td>7.6E-13</td><td>3.0E-12</td><td>MB231/MB468</td><td>MP</td></tr>
<tr><td>>hg38_21_+_8433172_8446622_RNA45SN1_45S_50ntFlanks@4472.4497.26|CTTAGCTGAGTGTCCCGCGGGGCCCG</td><td>>hg38_21_+_8433172_8446622_RNA45SN1_45S_50ntFlanks</td><td>4472</td><td>4497</td><td>26</td><td>CTTAGCTGAGTGTCCCGCGGGGCCCG</td><td>338.1</td><td>-10.5</td><td>1.5</td><td>-7.2</td><td>6.5E-13</td><td>3.0E-12</td><td>BT20/MB468</td><td>MITO</td></tr>
<tr><td>>hg38_21_+_8433172_8446622_RNA45SN1_45S_50ntFlanks@9545.9566.22|GTGGAGGTCCGTAGCGGTCCTG</td><td>>hg38_21_+_8433172_8446622_RNA45SN1_45S_50ntFlanks</td><td>9545</td><td>9566</td><td>22</td><td>GTGGAGGTCCGTAGCGGTCCTG</td><td>311.4</td><td>-10.4</td><td>1.4</td><td>-7.2</td><td>6.5E-13</td><td>3.0E-12</td><td>BT20/MB468</td><td>MITO</td></tr>
<tr><td>>MI0000284|hsa-mir-204&WithFlank&9|-|70809969|70810090@39.60.22&[MIMAT0000265&hsa-miR-204-5p&offsets|0|0;m-296&9|-|70810031|70810052&offsets|0|0]|TTCCCTTTGTCATCCTATGCCT</td><td>>MI0000284|hsa-mir-204&WithFlank&9|-|70809969|70810090</td><td>39</td><td>60</td><td>22</td><td>TTCCCTTTGTCATCCTATGCCT</td><td>161.1</td><td>-10.6</td><td>1.4</td><td>-7.5</td><td>8.9E-14</td><td>3.0E-12</td><td>BT20/MB231</td><td>MITO</td></tr>
<tr><td>>hg38_21_+_8433172_8446622_RNA45SN1_45S_50ntFlanks@4853.4870.18|AAGTCTTTGGGTTCCGGG</td><td>>hg38_21_+_8433172_8446622_RNA45SN1_45S_50ntFlanks</td><td>4853</td><td>4870</td><td>18</td><td>AAGTCTTTGGGTTCCGGG</td><td>134.8</td><td>-10.9</td><td>1.5</td><td>-7.2</td><td>7.6E-13</td><td>3.0E-12</td><td>MB231/MB468</td><td>MP</td></tr>
<tr><td>>hg38_21_+_8433172_8446622_RNA45SN1_45S_50ntFlanks@3987.4010.24|GCTTTGGTGACTCTAGATAACCTC</td><td>>hg38_21_+_8433172_8446622_RNA45SN1_45S_50ntFlanks</td><td>3987</td><td>4010</td><td>24</td><td>GCTTTGGTGACTCTAGATAACCTC</td><td>346.0</td><td>4.2</td><td>0.6</td><td>7.2</td><td>6.6E-13</td><td>3.0E-12</td><td>BT20/MB468</td><td>MITO</td></tr>
<tr><td>>hg38_21_+_8433172_8446622_RNA45SN1_45S_50ntFlanks@11212.11254.43|CGACTCTGGACGCGAGCCGGGCCCTTCCCGTGGATCGCCCCAG</td><td>>hg38_21_+_8433172_8446622_RNA45SN1_45S_50ntFlanks</td><td>11212</td><td>11254</td><td>43</td><td>CGACTCTGGACGCGAGCCGGGCCCTTCCCGTGGATCGCCCCAG</td><td>106.2</td><td>-10.6</td><td>1.5</td><td>-7.2</td><td>7.6E-13</td><td>3.0E-12</td><td>MB231/MB468</td><td>MP</td></tr>
<tr><td>>hg38_21_+_8433172_8446622_RNA45SN1_45S_50ntFlanks@6667.6686.20|ATCACTCGGCTCGTGCGTCG</td><td>>hg38_21_+_8433172_8446622_RNA45SN1_45S_50ntFlanks</td><td>6667</td><td>6686</td><td>20</td><td>ATCACTCGGCTCGTGCGTCG</td><td>379.7</td><td>-11.6</td><td>1.5</td><td>-7.6</td><td>3.9E-14</td><td>3.0E-12</td><td>BT20/MB468</td><td>MP</td></tr>
<tr><td>>hg38_21_+_8433172_8446622_RNA45SN1_45S_50ntFlanks@10839.10865.27|TCCGTAACTTCGGGATAAGGATTGGCT</td><td>>hg38_21_+_8433172_8446622_RNA45SN1_45S_50ntFlanks</td><td>10839</td><td>10865</td><td>27</td><td>TCCGTAACTTCGGGATAAGGATTGGCT</td><td>1544.7</td><td>-6.8</td><td>1.0</td><td>-7.1</td><td>8.8E-13</td><td>3.0E-12</td><td>MB231/MB468</td><td>CYTO</td></tr>
<tr><td>>hg38_21_+_8433172_8446622_RNA45SN1_45S_50ntFlanks@9459.9503.45|CCGGGGAGGTGGAGCACGAGCGCACGTGTTAGGACCCGAAAGATG</td><td>>hg38_21_+_8433172_8446622_RNA45SN1_45S_50ntFlanks</td><td>9459</td><td>9503</td><td>45</td><td>CCGGGGAGGTGGAGCACGAGCGCACGTGTTAGGACCCGAAAGATG</td><td>276.9</td><td>-11.1</td><td>1.6</td><td>-7.1</td><td>9.0E-13</td><td>3.1E-12</td><td>MB231/MB468</td><td>NUC</td></tr>
<tr><td>>am_trnaMT_SerGCT_MT_+_12207_12265@13.38.26|AAGAACTGCTAACTCATGCCCCCATG</td><td>>am_trnaMT_SerGCT_MT_+_12207_12265</td><td>13</td><td>38</td><td>26</td><td>AAGAACTGCTAACTCATGCCCCCATG</td><td>2242.2</td><td>-9.4</td><td>1.3</td><td>-7.1</td><td>1.2E-12</td><td>3.1E-12</td><td>BT20/MB468</td><td>NUC</td></tr>
<tr><td>>MI0000767|hsa-mir-365a&WithFlank&16|+|14309279|14309377@62.81.20;MI0000769|hsa-mir-365b&WithFlank&17|+|31575405|31575527@74.93.20&[MIMAT0000710&hsa-miR-365a-3p&offsets|0|-2;m-176&16|+|14309340|14309361&offsets|0|-2];[MIMAT0022834&hsa-miR-365b-3p&offsets|0|-2;m-175&17|+|31575478|31575499&offsets|0|-2]|TAATGCCCCTAAAAATCCTT</td><td>>MI0000767|hsa-mir-365a&WithFlank&16|+|14309279|14309377</td><td>62</td><td>81</td><td>20</td><td>TAATGCCCCTAAAAATCCTT</td><td>481.1</td><td>10.1</td><td>1.4</td><td>7.2</td><td>8.7E-13</td><td>3.1E-12</td><td>BT20/MB468</td><td>CYTO</td></tr>
<tr><td>>trnaMT_GluTTC_MT_-_14674_14742@40.71.32,trnalookalike8_GluTTC_5_-_93905172_93905240@40.71.32|TTGGTCGTGGTTGTAGTCCGTGCGAGAATACC</td><td>>trnaMT_GluTTC_MT_-_14674_14742</td><td>40</td><td>71</td><td>32</td><td>TTGGTCGTGGTTGTAGTCCGTGCGAGAATACC</td><td>1560.6</td><td>5.3</td><td>0.7</td><td>7.1</td><td>9.2E-13</td><td>3.1E-12</td><td>MB231/MB468</td><td>NUC</td></tr>
<tr><td>>MI0000087|hsa-mir-29a&WithFlank&7|-|130876741|130876816@48.66.19&[MIMAT0000086&hsa-miR-29a-3p&offsets|0|-3;m-26&7|-|130876749|130876769&offsets|0|-2]|TAGCACCATCTGAAATCGG</td><td>>MI0000087|hsa-mir-29a&WithFlank&7|-|130876741|130876816</td><td>48</td><td>66</td><td>19</td><td>TAGCACCATCTGAAATCGG</td><td>180.4</td><td>9.1</td><td>1.3</td><td>7.1</td><td>9.2E-13</td><td>3.1E-12</td><td>MB231/MB468</td><td>NUC</td></tr>
<tr><td>>hg38_21_+_8433172_8446622_RNA45SN1_45S_50ntFlanks@5152.5185.34|AGGGACAAGTGGCGTTCAGCCACCCGAGATTGAG</td><td>>hg38_21_+_8433172_8446622_RNA45SN1_45S_50ntFlanks</td><td>5152</td><td>5185</td><td>34</td><td>AGGGACAAGTGGCGTTCAGCCACCCGAGATTGAG</td><td>841.1</td><td>-13.6</td><td>1.9</td><td>-7.2</td><td>7.7E-13</td><td>3.1E-12</td><td>MB231/MB468</td><td>MP</td></tr>
<tr><td>>MI0000433|hsa-let-7g&WithFlank&3|-|52268272|52268367@11.31.21&[MIMAT0000414&hsa-let-7g-5p&offsets|0|-1;m-58&3|-|52268336|52268357&offsets|0|-1]|TGAGGTAGTAGTTTGTACAGT</td><td>>MI0000433|hsa-let-7g&WithFlank&3|-|52268272|52268367</td><td>11</td><td>31</td><td>21</td><td>TGAGGTAGTAGTTTGTACAGT</td><td>2374.6</td><td>5.5</td><td>0.8</td><td>7.1</td><td>9.0E-13</td><td>3.1E-12</td><td>MB231/MB468</td><td>CYTO</td></tr>
<tr><td>>hg38_21_+_8433172_8446622_RNA45SN1_45S_50ntFlanks@12654.12671.18|TATGACTGAACGCCTCTA</td><td>>hg38_21_+_8433172_8446622_RNA45SN1_45S_50ntFlanks</td><td>12654</td><td>12671</td><td>18</td><td>TATGACTGAACGCCTCTA</td><td>120.2</td><td>-10.8</td><td>1.5</td><td>-7.2</td><td>7.8E-13</td><td>3.1E-12</td><td>MB231/MB468</td><td>MP</td></tr>
<tr><td>>hg38_21_+_8433172_8446622_RNA45SN1_45S_50ntFlanks@6743.6771.29|CGACACTTCGAACGCACTTGCGGCCCCGG</td><td>>hg38_21_+_8433172_8446622_RNA45SN1_45S_50ntFlanks</td><td>6743</td><td>6771</td><td>29</td><td>CGACACTTCGAACGCACTTGCGGCCCCGG</td><td>1893.1</td><td>-8.3</td><td>1.2</td><td>-7.1</td><td>1.2E-12</td><td>3.1E-12</td><td>BT20/MB468</td><td>NUC</td></tr>
<tr><td>>MI0000088|hsa-mir-30a&WithFlank&6|-|71403545|71403627@12.35.24(+1C)&[MIMAT0000087&hsa-miR-30a-5p&offsets|0|+2(+1C);m-15&6|-|71403595|71403616&offsets|0|+2(+1C)]|TGTAAACATCCTCGACTGGAAGCTC</td><td>>MI0000088|hsa-mir-30a&WithFlank&6|-|71403545|71403627</td><td>12</td><td>35</td><td>24(+1C)</td><td>TGTAAACATCCTCGACTGGAAGCTC</td><td>2681.4</td><td>5.5</td><td>0.8</td><td>7.2</td><td>7.8E-13</td><td>3.1E-12</td><td>MB231/MB468</td><td>MP</td></tr>
<tr><td>>hg38_21_+_8433172_8446622_RNA45SN1_45S_50ntFlanks@4723.4753.31|CATTAATCAAGAACGAAAGTCGGAGGTTCGA</td><td>>hg38_21_+_8433172_8446622_RNA45SN1_45S_50ntFlanks</td><td>4723</td><td>4753</td><td>31</td><td>CATTAATCAAGAACGAAAGTCGGAGGTTCGA</td><td>149.3</td><td>-11.1</td><td>1.5</td><td>-7.2</td><td>7.8E-13</td><td>3.1E-12</td><td>MB231/MB468</td><td>MP</td></tr>
<tr><td>>hg38_21_+_8433172_8446622_RNA45SN1_45S_50ntFlanks@5472.5512.41|ACGGCCCTGGCGGAGCGCTGAGAAGACGGTCGAACTTGACT</td><td>>hg38_21_+_8433172_8446622_RNA45SN1_45S_50ntFlanks</td><td>5472</td><td>5512</td><td>41</td><td>ACGGCCCTGGCGGAGCGCTGAGAAGACGGTCGAACTTGACT</td><td>3053.1</td><td>6.3</td><td>0.9</td><td>7.1</td><td>9.2E-13</td><td>3.1E-12</td><td>MB231/MB468</td><td>NUC</td></tr>
<tr><td>>hg38_21_+_8433172_8446622_RNA45SN1_45S_50ntFlanks@8799.8820.22|CCTCGGGAGGGCGCGCGGGTCG</td><td>>hg38_21_+_8433172_8446622_RNA45SN1_45S_50ntFlanks</td><td>8799</td><td>8820</td><td>22</td><td>CCTCGGGAGGGCGCGCGGGTCG</td><td>253.9</td><td>-11.9</td><td>1.7</td><td>-7.2</td><td>7.8E-13</td><td>3.1E-12</td><td>MB231/MB468</td><td>MP</td></tr>
<tr><td>>hg38_21_+_8433172_8446622_RNA45SN1_45S_50ntFlanks@4066.4095.30|CAACTTTCGATGGTAGTCGCCGTGCCTACC</td><td>>hg38_21_+_8433172_8446622_RNA45SN1_45S_50ntFlanks</td><td>4066</td><td>4095</td><td>30</td><td>CAACTTTCGATGGTAGTCGCCGTGCCTACC</td><td>133.4</td><td>-10.9</td><td>1.5</td><td>-7.2</td><td>7.8E-13</td><td>3.1E-12</td><td>MB231/MB468</td><td>MP</td></tr>
<tr><td>>hg38_21_+_8433172_8446622_RNA45SN1_45S_50ntFlanks@8170.8193.24|CAAGTCCTTCTGATCGAGGCCCAG</td><td>>hg38_21_+_8433172_8446622_RNA45SN1_45S_50ntFlanks</td><td>8170</td><td>8193</td><td>24</td><td>CAAGTCCTTCTGATCGAGGCCCAG</td><td>168.2</td><td>-11.3</td><td>1.6</td><td>-7.2</td><td>7.8E-13</td><td>3.1E-12</td><td>MB231/MB468</td><td>MP</td></tr>
<tr><td>>hg38_21_+_8433172_8446622_RNA45SN1_45S_50ntFlanks@6671.6696.26|CTCGGCTCGTGCGTCGATGAAGAACG</td><td>>hg38_21_+_8433172_8446622_RNA45SN1_45S_50ntFlanks</td><td>6671</td><td>6696</td><td>26</td><td>CTCGGCTCGTGCGTCGATGAAGAACG</td><td>8891.8</td><td>-6.9</td><td>1.0</td><td>-7.1</td><td>1.1E-12</td><td>3.1E-12</td><td>MB231/MB468</td><td>MITO</td></tr>
<tr><td>>hg38_21_+_8433172_8446622_RNA45SN1_45S_50ntFlanks@12488.12509.22|AACGTGAGCTGGGTTTAGACCG</td><td>>hg38_21_+_8433172_8446622_RNA45SN1_45S_50ntFlanks</td><td>12488</td><td>12509</td><td>22</td><td>AACGTGAGCTGGGTTTAGACCG</td><td>5667.8</td><td>-8.3</td><td>1.2</td><td>-7.1</td><td>9.2E-13</td><td>3.1E-12</td><td>MB231/MB468</td><td>NUC</td></tr>
<tr><td>>hg38_21_+_8433172_8446622_RNA45SN1_45S_50ntFlanks@12151.12178.28|GTGGGGAGTTTGACTGGGGCGGTACACC</td><td>>hg38_21_+_8433172_8446622_RNA45SN1_45S_50ntFlanks</td><td>12151</td><td>12178</td><td>28</td><td>GTGGGGAGTTTGACTGGGGCGGTACACC</td><td>124.5</td><td>-10.8</td><td>1.5</td><td>-7.2</td><td>7.8E-13</td><td>3.1E-12</td><td>MB231/MB468</td><td>MP</td></tr>
<tr><td>>hg38_21_+_8433172_8446622_RNA45SN1_45S_50ntFlanks@9176.9197.22|CGCGCCGTCGGGCCCGGGGGAG</td><td>>hg38_21_+_8433172_8446622_RNA45SN1_45S_50ntFlanks</td><td>9176</td><td>9197</td><td>22</td><td>CGCGCCGTCGGGCCCGGGGGAG</td><td>156.2</td><td>-11.2</td><td>1.6</td><td>-7.2</td><td>7.8E-13</td><td>3.1E-12</td><td>MB231/MB468</td><td>MP</td></tr>
<tr><td>>hg38_21_+_8433172_8446622_RNA45SN1_45S_50ntFlanks@3851.3881.31|ATAACTGTGGTAATTCTAGAGCTAATACATG</td><td>>hg38_21_+_8433172_8446622_RNA45SN1_45S_50ntFlanks</td><td>3851</td><td>3881</td><td>31</td><td>ATAACTGTGGTAATTCTAGAGCTAATACATG</td><td>149.6</td><td>-11.1</td><td>1.5</td><td>-7.2</td><td>7.8E-13</td><td>3.1E-12</td><td>MB231/MB468</td><td>MP</td></tr>
<tr><td>>MI0015866|hsa-mir-4326&WithFlank&20|+|63286802|63286872@17.40.24&[MIMAT0016888&hsa-miR-4326&offsets|0|+4;m-339&20|+|63286818|63286841&offsets|0|0]|TGTTCCTCTGTCTCCCAGACTCTG</td><td>>MI0015866|hsa-mir-4326&WithFlank&20|+|63286802|63286872</td><td>17</td><td>40</td><td>24</td><td>TGTTCCTCTGTCTCCCAGACTCTG</td><td>357.8</td><td>9.7</td><td>1.3</td><td>7.2</td><td>6.8E-13</td><td>3.1E-12</td><td>BT20/MB468</td><td>MITO</td></tr>
<tr><td>>hg38_21_+_8433172_8446622_RNA45SN1_45S_50ntFlanks@10362.10391.30|AAGGCCGAAGTGGAGAAGGGTTCCATGTGA</td><td>>hg38_21_+_8433172_8446622_RNA45SN1_45S_50ntFlanks</td><td>10362</td><td>10391</td><td>30</td><td>AAGGCCGAAGTGGAGAAGGGTTCCATGTGA</td><td>271.7</td><td>-11.0</td><td>1.5</td><td>-7.1</td><td>9.3E-13</td><td>3.1E-12</td><td>MB231/MB468</td><td>NUC</td></tr>
<tr><td>>MI0016005|hsa-mir-3615&WithFlank&17|+|74748607|74748705@57.77.21(+1A)&[MIMAT0017994&hsa-miR-3615&offsets|0|0(+1A);m-271&17|+|74748663|74748684&offsets|0|-1(+1A)]|TCTCTCGGCTCCTCGCGGCTCA</td><td>>MI0016005|hsa-mir-3615&WithFlank&17|+|74748607|74748705</td><td>57</td><td>77</td><td>21(+1A)</td><td>TCTCTCGGCTCCTCGCGGCTCA</td><td>424.1</td><td>6.7</td><td>0.9</td><td>7.1</td><td>8.9E-13</td><td>3.1E-12</td><td>BT20/MB468</td><td>CYTO</td></tr>
<tr><td>>hg38_21_+_8433172_8446622_RNA45SN1_45S_50ntFlanks@11575.11602.28|ACTTAGAACTGGTGCGGACCAGGGGAAT</td><td>>hg38_21_+_8433172_8446622_RNA45SN1_45S_50ntFlanks</td><td>11575</td><td>11602</td><td>28</td><td>ACTTAGAACTGGTGCGGACCAGGGGAAT</td><td>1574.9</td><td>-4.2</td><td>0.6</td><td>-7.1</td><td>8.9E-13</td><td>3.2E-12</td><td>BT20/MB468</td><td>CYTO</td></tr>
<tr><td>>hg38_21_+_8433172_8446622_RNA45SN1_45S_50ntFlanks@12934.12958.25|CTAAACCATTCGTAGACGACCTGCT</td><td>>hg38_21_+_8433172_8446622_RNA45SN1_45S_50ntFlanks</td><td>12934</td><td>12958</td><td>25</td><td>CTAAACCATTCGTAGACGACCTGCT</td><td>1437.1</td><td>-10.1</td><td>1.4</td><td>-7.1</td><td>1.2E-12</td><td>3.2E-12</td><td>BT20/MB468</td><td>NUC</td></tr>
<tr><td>>am_MI0000809|hsa-mir-151a&WithFlank&8|-|140732558|140732659@53.72.20(+1C)&[MIMAT0000757&hsa-miR-151a-3p&offsets|0|-1(+1C);m-30&8|-|140732587|140732607&offsets|0|-1(+1C)]|CTAGACTGAAGCTCCTTGAGC</td><td>>am_MI0000809|hsa-mir-151a&WithFlank&8|-|140732558|140732659</td><td>53</td><td>72</td><td>20(+1C)</td><td>CTAGACTGAAGCTCCTTGAGC</td><td>993.8</td><td>9.3</td><td>1.3</td><td>7.2</td><td>6.9E-13</td><td>3.2E-12</td><td>BT20/MB468</td><td>MITO</td></tr>
<tr><td>>MI0000085|hsa-mir-27a&WithFlank&19|-|13836434|13836523@57.76.20&[MIMAT0000084&hsa-miR-27a-3p&offsets|0|-1;m-43&19|-|13836447|13836467&offsets|0|-1]|TTCACAGTGGCTAAGTTCCG</td><td>>MI0000085|hsa-mir-27a&WithFlank&19|-|13836434|13836523</td><td>57</td><td>76</td><td>20</td><td>TTCACAGTGGCTAAGTTCCG</td><td>14410.9</td><td>5.8</td><td>0.8</td><td>7.1</td><td>9.4E-13</td><td>3.2E-12</td><td>MB231/MB468</td><td>NUC</td></tr>
<tr><td>>hg38_21_+_8433172_8446622_RNA45SN1_45S_50ntFlanks@12459.12480.22|AGCGTTGGATTGTTCACCCACT</td><td>>hg38_21_+_8433172_8446622_RNA45SN1_45S_50ntFlanks</td><td>12459</td><td>12480</td><td>22</td><td>AGCGTTGGATTGTTCACCCACT</td><td>131.1</td><td>-10.9</td><td>1.5</td><td>-7.2</td><td>8.0E-13</td><td>3.2E-12</td><td>MB231/MB468</td><td>MP</td></tr>
<tr><td>>hg38_21_+_8433172_8446622_RNA45SN1_45S_50ntFlanks@12008.12042.35|CCGCCGGTGAAATACCACTACTCTGATCGTTTTTT</td><td>>hg38_21_+_8433172_8446622_RNA45SN1_45S_50ntFlanks</td><td>12008</td><td>12042</td><td>35</td><td>CCGCCGGTGAAATACCACTACTCTGATCGTTTTTT</td><td>377.0</td><td>-12.4</td><td>1.7</td><td>-7.2</td><td>8.0E-13</td><td>3.2E-12</td><td>MB231/MB468</td><td>MP</td></tr>
<tr><td>>hg38_21_+_8433172_8446622_RNA45SN1_45S_50ntFlanks@12394.12417.24|TAGCGACGTCGCTTTTTGATCCTT</td><td>>hg38_21_+_8433172_8446622_RNA45SN1_45S_50ntFlanks</td><td>12394</td><td>12417</td><td>24</td><td>TAGCGACGTCGCTTTTTGATCCTT</td><td>275.5</td><td>-12.0</td><td>1.7</td><td>-7.2</td><td>8.0E-13</td><td>3.2E-12</td><td>MB231/MB468</td><td>MP</td></tr>
</table>

<table>
<tr><th></th><th></th><th></th><th></th><th></th><th></th><th></th><th></th><th></th><th></th><th></th><th></th><th></th><th></th></tr>
<tr><td>>hg38_21_+_8433172_8446622_RNA45SN1_45S_50ntFlanks@12934.12966.33|CTAAACCATTCGTAGACGACCTGCTTCTGGGTC</td><td>>hg38_21_+_8433172_8446622_RNA45SN1_45S_50ntFlanks</td><td>12934</td><td>12966</td><td>33</td><td>CTAAACCATTCGTAGACGACCTGCTTCTGGGTC</td><td>305.1</td><td>-12.1</td><td>1.7</td><td>-7.2</td><td>8.0E-13</td><td>3.2E-12</td><td>MB231/MB468</td><td>MP</td></tr>
<tr><td>>hg38_21_+_8433172_8446622_RNA45SN1_45S_50ntFlanks@9605.9622.18|AAGACTAATCGAACCATC</td><td>>hg38_21_+_8433172_8446622_RNA45SN1_45S_50ntFlanks</td><td>9605</td><td>9622</td><td>18</td><td>AAGACTAATCGAACCATC</td><td>1652.6</td><td>-8.2</td><td>1.1</td><td>-7.2</td><td>8.0E-13</td><td>3.2E-12</td><td>MB231/MB468</td><td>MP</td></tr>
<tr><td>>hg38_21_+_8433172_8446622_RNA45SN1_45S_50ntFlanks@9339.9368.30|CGCACGAAAGCCGCCGTGGCGCAATGAAGG</td><td>>hg38_21_+_8433172_8446622_RNA45SN1_45S_50ntFlanks</td><td>9339</td><td>9368</td><td>30</td><td>CGCACGAAAGCCGCCGTGGCGCAATGAAGG</td><td>201.0</td><td>-11.4</td><td>1.6</td><td>-7.1</td><td>1.1E-12</td><td>3.2E-12</td><td>MB231/MB468</td><td>MITO</td></tr>
<tr><td>>hg38_21_+_8433172_8446622_RNA45SN1_45S_50ntFlanks@12957.12975.19|CTTCTGGGTCGGGGTTTCG</td><td>>hg38_21_+_8433172_8446622_RNA45SN1_45S_50ntFlanks</td><td>12957</td><td>12975</td><td>19</td><td>CTTCTGGGTCGGGGTTTCG</td><td>123.7</td><td>-10.8</td><td>1.5</td><td>-7.2</td><td>8.0E-13</td><td>3.2E-12</td><td>MB231/MB468</td><td>MP</td></tr>
<tr><td>>hg38_21_+_8433172_8446622_RNA45SN1_45S_50ntFlanks@10408.10437.30|GGTCAGTCGGTCCTGAGAGATGGGCGAGCG</td><td>>hg38_21_+_8433172_8446622_RNA45SN1_45S_50ntFlanks</td><td>10408</td><td>10437</td><td>30</td><td>GGTCAGTCGGTCCTGAGAGATGGGCGAGCG</td><td>156.0</td><td>-4.8</td><td>0.7</td><td>-7.1</td><td>9.2E-13</td><td>3.2E-12</td><td>MB231/MB468</td><td>CYTO</td></tr>
<tr><td>>hg38_21_+_8433172_8446622_RNA45SN1_45S_50ntFlanks@4002.4020.19|GATAACCTCGGGCCGATCG</td><td>>hg38_21_+_8433172_8446622_RNA45SN1_45S_50ntFlanks</td><td>4002</td><td>4020</td><td>19</td><td>GATAACCTCGGGCCGATCG</td><td>3080.8</td><td>-11.1</td><td>1.6</td><td>-7.1</td><td>1.3E-12</td><td>3.2E-12</td><td>BT20/MB468</td><td>NUC</td></tr>
<tr><td>>hg38_21_+_8433172_8446622_RNA45SN1_45S_50ntFlanks@7984.8008.25|AGATCAGACGTGGCGACCCGCTGAA</td><td>>hg38_21_+_8433172_8446622_RNA45SN1_45S_50ntFlanks</td><td>7984</td><td>8008</td><td>25</td><td>AGATCAGACGTGGCGACCCGCTGAA</td><td>541.4</td><td>-4.4</td><td>0.6</td><td>-7.1</td><td>9.5E-13</td><td>3.2E-12</td><td>MB231/MB468</td><td>NUC</td></tr>
<tr><td>>hg38_21_+_8433172_8446622_RNA45SN1_45S_50ntFlanks@8155.8175.21|CGCTCGTGGGGGGCCCAAGTC</td><td>>hg38_21_+_8433172_8446622_RNA45SN1_45S_50ntFlanks</td><td>8155</td><td>8175</td><td>21</td><td>CGCTCGTGGGGGGCCCAAGTC</td><td>133.2</td><td>-10.9</td><td>1.5</td><td>-7.2</td><td>8.1E-13</td><td>3.2E-12</td><td>MB231/MB468</td><td>MP</td></tr>
<tr><td>>hg38_21_+_8433172_8446622_RNA45SN1_45S_50ntFlanks@12994.13013.20|CCTCGCTGCGATCTATTGAA</td><td>>hg38_21_+_8433172_8446622_RNA45SN1_45S_50ntFlanks</td><td>12994</td><td>13013</td><td>20</td><td>CCTCGCTGCGATCTATTGAA</td><td>5680.6</td><td>-11.3</td><td>1.6</td><td>-7.1</td><td>1.3E-12</td><td>3.2E-12</td><td>BT20/MB468</td><td>NUC</td></tr>
<tr><td>>hg38_21_+_8433172_8446622_RNA45SN1_45S_50ntFlanks@3839.3861.23|TCTCCTACTTGGATAACTGTGGT</td><td>>hg38_21_+_8433172_8446622_RNA45SN1_45S_50ntFlanks</td><td>3839</td><td>3861</td><td>23</td><td>TCTCCTACTTGGATAACTGTGGT</td><td>702.2</td><td>5.2</td><td>0.7</td><td>7.1</td><td>9.1E-13</td><td>3.2E-12</td><td>BT20/MB468</td><td>CYTO</td></tr>
<tr><td>>hg38_21_+_8433172_8446622_RNA45SN1_45S_50ntFlanks@10041.10072.32|CCATACCCGGCCGTCGCCGGCAGTCGAGAGTG</td><td>>hg38_21_+_8433172_8446622_RNA45SN1_45S_50ntFlanks</td><td>10041</td><td>10072</td><td>32</td><td>CCATACCCGGCCGTCGCCGGCAGTCGAGAGTG</td><td>155.4</td><td>-11.1</td><td>1.6</td><td>-7.2</td><td>8.1E-13</td><td>3.2E-12</td><td>MB231/MB468</td><td>MP</td></tr>
<tr><td>>hg38_21_+_8433172_8446622_RNA45SN1_45S_50ntFlanks@4445.4472.28|CTTGCCTCTCGGCGCCCCCTCGATGCTC</td><td>>hg38_21_+_8433172_8446622_RNA45SN1_45S_50ntFlanks</td><td>4445</td><td>4472</td><td>28</td><td>CTTGCCTCTCGGCGCCCCCTCGATGCTC</td><td>273.9</td><td>-10.2</td><td>1.4</td><td>-7.2</td><td>7.0E-13</td><td>3.2E-12</td><td>BT20/MB468</td><td>MITO</td></tr>
<tr><td>>hg38_21_+_8433172_8446622_RNA45SN1_45S_50ntFlanks@12415.12434.20|CTTCGATGTCGGCTCTTCCT</td><td>>hg38_21_+_8433172_8446622_RNA45SN1_45S_50ntFlanks</td><td>12415</td><td>12434</td><td>20</td><td>CTTCGATGTCGGCTCTTCCT</td><td>985.3</td><td>-5.6</td><td>0.8</td><td>-7.2</td><td>8.1E-13</td><td>3.2E-12</td><td>MB231/MB468</td><td>MP</td></tr>
<tr><td>>hg38_21_+_8433172_8446622_RNA45SN1_45S_50ntFlanks@10350.10376.27|ACGAGAACTTTGAAGGCCGAAGTGGAG</td><td>>hg38_21_+_8433172_8446622_RNA45SN1_45S_50ntFlanks</td><td>10350</td><td>10376</td><td>27</td><td>ACGAGAACTTTGAAGGCCGAAGTGGAG</td><td>1269.0</td><td>-7.3</td><td>1.0</td><td>-7.1</td><td>1.2E-12</td><td>3.2E-12</td><td>MB231/MB468</td><td>MITO</td></tr>
<tr><td>>hg38_21_+_8433172_8446622_RNA45SN1_45S_50ntFlanks@8170.8199.30|CAAGTCCTTCTGATCGAGGCCCAGCCCGTG</td><td>>hg38_21_+_8433172_8446622_RNA45SN1_45S_50ntFlanks</td><td>8170</td><td>8199</td><td>30</td><td>CAAGTCCTTCTGATCGAGGCCCAGCCCGTG</td><td>625.7</td><td>-8.8</td><td>1.2</td><td>-7.1</td><td>1.2E-12</td><td>3.2E-12</td><td>MB231/MB468</td><td>MITO</td></tr>
<tr><td>>hg38_21_+_8433172_8446622_RNA45SN1_45S_50ntFlanks@3777.3810.34|CGGTACAGTGAAACTGCGAATGGCTCATTAAATC</td><td>>hg38_21_+_8433172_8446622_RNA45SN1_45S_50ntFlanks</td><td>3777</td><td>3810</td><td>34</td><td>CGGTACAGTGAAACTGCGAATGGCTCATTAAATC</td><td>595.2</td><td>-9.4</td><td>1.3</td><td>-7.1</td><td>1.3E-12</td><td>3.2E-12</td><td>BT20/MB468</td><td>NUC</td></tr>
<tr><td>>hg38_21_+_8433172_8446622_RNA45SN1_45S_50ntFlanks@12007.12031.25|GCCGCCGGTGAAATACCACTACTCT</td><td>>hg38_21_+_8433172_8446622_RNA45SN1_45S_50ntFlanks</td><td>12007</td><td>12031</td><td>25</td><td>GCCGCCGGTGAAATACCACTACTCT</td><td>7772.3</td><td>-5.3</td><td>0.7</td><td>-7.1</td><td>1.2E-12</td><td>3.2E-12</td><td>MB231/MB468</td><td>MITO</td></tr>
<tr><td>>hg38_21_+_8433172_8446622_RNA45SN1_45S_50ntFlanks@10498.10521.24|TTCAGATCCCCGAATCCGGAGTGG</td><td>>hg38_21_+_8433172_8446622_RNA45SN1_45S_50ntFlanks</td><td>10498</td><td>10521</td><td>24</td><td>TTCAGATCCCCGAATCCGGAGTGG</td><td>282.2</td><td>-12.0</td><td>1.7</td><td>-7.2</td><td>8.1E-13</td><td>3.2E-12</td><td>MB231/MB468</td><td>MP</td></tr>
<tr><td>>hg38_21_+_8433172_8446622_RNA45SN1_45S_50ntFlanks@10402.10429.28|AACATGGGTCAGTCGGTCCTGAGAGATG</td><td>>hg38_21_+_8433172_8446622_RNA45SN1_45S_50ntFlanks</td><td>10402</td><td>10429</td><td>28</td><td>AACATGGGTCAGTCGGTCCTGAGAGATG</td><td>3470.2</td><td>-4.1</td><td>0.6</td><td>-7.1</td><td>9.1E-13</td><td>3.2E-12</td><td>BT20/MB468</td><td>CYTO</td></tr>
<tr><td>>MI0000088|hsa-mir-30a&WithFlank&6|-|71403545|71403627@12.34.23(+1A)&[MIMAT0000087&hsa-miR-30a-5p&offsets|0|+1(+1A);m-15&6|-|71403595|71403616&offsets|0|+1(+1A)]|TGTAAACATCCTCGACTGGAAGCA</td><td>>MI0000088|hsa-mir-30a&WithFlank&6|-|71403545|71403627</td><td>12</td><td>34</td><td>23(+1A)</td><td>TGTAAACATCCTCGACTGGAAGCA</td><td>3856.1</td><td>7.0</td><td>1.0</td><td>7.1</td><td>9.1E-13</td><td>3.2E-12</td><td>BT20/MB468</td><td>CYTO</td></tr>
<tr><td>>hg38_21_+_8433172_8446622_RNA45SN1_45S_50ntFlanks@9739.9767.29|AAACGATCTCAACCTATTCTCAAACTTTA</td><td>>hg38_21_+_8433172_8446622_RNA45SN1_45S_50ntFlanks</td><td>9739</td><td>9767</td><td>29</td><td>AAACGATCTCAACCTATTCTCAAACTTTA</td><td>5563.8</td><td>-8.0</td><td>1.1</td><td>-7.1</td><td>1.2E-12</td><td>3.2E-12</td><td>MB231/MB468</td><td>MITO</td></tr>
<tr><td>>hg38_21_+_8433172_8446622_RNA45SN1_45S_50ntFlanks@6746.6769.24|CACTTCGAACGCACTTGCGGCCCC</td><td>>hg38_21_+_8433172_8446622_RNA45SN1_45S_50ntFlanks</td><td>6746</td><td>6769</td><td>24</td><td>CACTTCGAACGCACTTGCGGCCCC</td><td>118.0</td><td>-10.7</td><td>1.5</td><td>-7.2</td><td>8.2E-13</td><td>3.2E-12</td><td>MB231/MB468</td><td>MP</td></tr>
<tr><td>>hg38_21_+_8433172_8446622_RNA45SN1_45S_50ntFlanks@3913.3938.26|ATGCGTGCATTTATCAGATCAAAACC</td><td>>hg38_21_+_8433172_8446622_RNA45SN1_45S_50ntFlanks</td><td>3913</td><td>3938</td><td>26</td><td>ATGCGTGCATTTATCAGATCAAAACC</td><td>193.7</td><td>-11.5</td><td>1.6</td><td>-7.2</td><td>8.2E-13</td><td>3.3E-12</td><td>MB231/MB468</td><td>MP</td></tr>
<tr><td>>hg38_21_+_8433172_8446622_RNA45SN1_45S_50ntFlanks@12970.12990.21|GTTTCGTACGTAGCAGAGCAG</td><td>>hg38_21_+_8433172_8446622_RNA45SN1_45S_50ntFlanks</td><td>12970</td><td>12990</td><td>21</td><td>GTTTCGTACGTAGCAGAGCAG</td><td>167.3</td><td>-5.6</td><td>0.8</td><td>-7.1</td><td>9.4E-13</td><td>3.3E-12</td><td>MB231/MB468</td><td>CYTO</td></tr>
<tr><td>>hg38_21_+_8433172_8446622_RNA45SN1_45S_50ntFlanks@5467.5484.18|GGCCCACGGCCCTGGCGG</td><td>>hg38_21_+_8433172_8446622_RNA45SN1_45S_50ntFlanks</td><td>5467</td><td>5484</td><td>18</td><td>GGCCCACGGCCCTGGCGG</td><td>252.4</td><td>-7.7</td><td>1.1</td><td>-7.1</td><td>9.5E-13</td><td>3.3E-12</td><td>MB231/MB468</td><td>CYTO</td></tr>
<tr><td>>hg38_21_+_8433172_8446622_RNA45SN1_45S_50ntFlanks@6739.6770.32|TCATCGACACTTCGAACGCACTTGCGGCCCCG</td><td>>hg38_21_+_8433172_8446622_RNA45SN1_45S_50ntFlanks</td><td>6739</td><td>6770</td><td>32</td><td>TCATCGACACTTCGAACGCACTTGCGGCCCCG</td><td>283.5</td><td>-5.6</td><td>0.8</td><td>-7.1</td><td>9.5E-13</td><td>3.3E-12</td><td>MB231/MB468</td><td>CYTO</td></tr>
<tr><td>>hg38_21_+_8433172_8446622_RNA45SN1_45S_50ntFlanks@12033.12071.39|ATCGTTTTTTCACTGACCCGGTGAGGCGGGGGGGCGAGC</td><td>>hg38_21_+_8433172_8446622_RNA45SN1_45S_50ntFlanks</td><td>12033</td><td>12071</td><td>39</td><td>ATCGTTTTTTCACTGACCCGGTGAGGCGGGGGGGCGAGC</td><td>259.9</td><td>-11.8</td><td>1.7</td><td>-7.1</td><td>1.2E-12</td><td>3.3E-12</td><td>MB231/MB468</td><td>MITO</td></tr>
<tr><td>>hg38_21_+_8433172_8446622_RNA45SN1_45S_50ntFlanks@4287.4317.31|AACGAGGATCCATTGGAGGGCAAGTCTGGTG</td><td>>hg38_21_+_8433172_8446622_RNA45SN1_45S_50ntFlanks</td><td>4287</td><td>4317</td><td>31</td><td>AACGAGGATCCATTGGAGGGCAAGTCTGGTG</td><td>193.9</td><td>-10.6</td><td>1.5</td><td>-7.1</td><td>9.7E-13</td><td>3.3E-12</td><td>MB231/MB468</td><td>NUC</td></tr>
<tr><td>>hg38_21_+_8433172_8446622_RNA45SN1_45S_50ntFlanks@10838.10868.31|ATCCGTAACTTCGGGATAAGGATTGGCTCTA</td><td>>hg38_21_+_8433172_8446622_RNA45SN1_45S_50ntFlanks</td><td>10838</td><td>10868</td><td>31</td><td>ATCCGTAACTTCGGGATAAGGATTGGCTCTA</td><td>554.1</td><td>-11.4</td><td>1.6</td><td>-7.1</td><td>9.3E-13</td><td>3.3E-12</td><td>BT20/MB468</td><td>CYTO</td></tr>
<tr><td>>hg38_21_+_8433172_8446622_RNA45SN1_45S_50ntFlanks@12141.12182.42|ACAGTGCCAGGTGGGGAGTTTGACTGGGGCGGTACACCTGTC</td><td>>hg38_21_+_8433172_8446622_RNA45SN1_45S_50ntFlanks</td><td>12141</td><td>12182</td><td>42</td><td>ACAGTGCCAGGTGGGGAGTTTGACTGGGGCGGTACACCTGTC</td><td>252.4</td><td>-11.8</td><td>1.7</td><td>-7.1</td><td>1.2E-12</td><td>3.3E-12</td><td>MB231/MB468</td><td>MITO</td></tr>
<tr><td>>hg38_21_+_8433172_8446622_RNA45SN1_45S_50ntFlanks@6660.6681.22|GCGGTGGATCACTCGGCTCGTG</td><td>>hg38_21_+_8433172_8446622_RNA45SN1_45S_50ntFlanks</td><td>6660</td><td>6681</td><td>22</td><td>GCGGTGGATCACTCGGCTCGTG</td><td>1947.5</td><td>-8.4</td><td>1.2</td><td>-7.2</td><td>8.3E-13</td><td>3.3E-12</td><td>MB231/MB468</td><td>MP</td></tr>
<tr><td>>hg38_21_+_8433172_8446622_RNA45SN1_45S_50ntFlanks@9339.9373.35|CGCACGAAAGCCGCCGTGGCGCAATGAAGGTGAAG</td><td>>hg38_21_+_8433172_8446622_RNA45SN1_45S_50ntFlanks</td><td>9339</td><td>9373</td><td>35</td><td>CGCACGAAAGCCGCCGTGGCGCAATGAAGGTGAAG</td><td>165.8</td><td>-11.2</td><td>1.6</td><td>-7.2</td><td>8.3E-13</td><td>3.3E-12</td><td>MB231/MB468</td><td>MP</td></tr>
<tr><td>>hg38_21_+_8433172_8446622_RNA45SN1_45S_50ntFlanks@10411.10440.30|CAGTCGGTCCTGAGAGATGGGCGAGCGCCG</td><td>>hg38_21_+_8433172_8446622_RNA45SN1_45S_50ntFlanks</td><td>10411</td><td>10440</td><td>30</td><td>CAGTCGGTCCTGAGAGATGGGCGAGCGCCG</td><td>282.1</td><td>-5.5</td><td>0.8</td><td>-7.1</td><td>9.5E-13</td><td>3.3E-12</td><td>MB231/MB468</td><td>CYTO</td></tr>
<tr><td>>MI0000434|hsa-let-7i&WithFlank&12|+|62603680|62603775@12.32.21(+1C)&[MIMAT0000415&hsa-let-7i-5p&offsets|0|-1(+1C);m-49&12|+|62603691|62603712&offsets|0|-1(+1C)]|TGAGGTAGTAGTTTGTGCTGTC</td><td>>MI0000434|hsa-let-7i&WithFlank&12|+|62603680|62603775</td><td>12</td><td>32</td><td>21(+1C)</td><td>TGAGGTAGTAGTTTGTGCTGTC</td><td>7834.6</td><td>5.4</td><td>0.8</td><td>7.1</td><td>9.5E-13</td><td>3.3E-12</td><td>MB231/MB468</td><td>CYTO</td></tr>
<tr><td>>hg38_21_+_8433172_8446622_RNA45SN1_45S_50ntFlanks@5084.5106.23|CGAACGAGACTCTGGCATGCTAA</td><td>>hg38_21_+_8433172_8446622_RNA45SN1_45S_50ntFlanks</td><td>5084</td><td>5106</td><td>23</td><td>CGAACGAGACTCTGGCATGCTAA</td><td>645.3</td><td>-13.1</td><td>1.8</td><td>-7.1</td><td>1.2E-12</td><td>3.3E-12</td><td>MB231/MB468</td><td>MITO</td></tr>
<tr><td>>MI0000294|hsa-mir-218-1&WithFlank&4|+|20528269|20528390@31.51.21;MI0000295|hsa-mir-218-2&WithFlank&5|-|168768140|168768261@31.51.21&[MIMAT0000275&hsa-miR-218-5p&offsets|0|0;m-185&4|+|20528299|20528319&offsets|0|0];[MIMAT0000275_1&hsa-miR-218-5p&offsets|0|0;m-187&5|-|168768211|168768231&offsets|0|0]|TTGTGCTTGATCTAACCATGT</td><td>>MI0000294|hsa-mir-218-1&WithFlank&4|+|20528269|20528390</td><td>31</td><td>51</td><td>21</td><td>TTGTGCTTGATCTAACCATGT</td><td>392.8</td><td>11.8</td><td>1.7</td><td>7.1</td><td>1.2E-12</td><td>3.3E-12</td><td>MB231/MB468</td><td>MITO</td></tr>
<tr><td>>hg38_21_+_8433172_8446622_RNA45SN1_45S_50ntFlanks@10663.10710.48|CCTTGGAAAGCGTCGCGGTTCCGGCGGCGTCCGGTGAGCTCTCGCTGG</td><td>>hg38_21_+_8433172_8446622_RNA45SN1_45S_50ntFlanks</td><td>10663</td><td>10710</td><td>48</td><td>CCTTGGAAAGCGTCGCGGTTCCGGCGGCGTCCGGTGAGCTCTCGCTGG</td><td>392.2</td><td>-11.3</td><td>1.6</td><td>-7.1</td><td>9.6E-13</td><td>3.3E-12</td><td>MB231/MB468</td><td>CYTO</td></tr>
<tr><td>>hg38_21_+_8433172_8446622_RNA45SN1_45S_50ntFlanks@12073.12096.24|CCGAGGGGCTCTCGCTTCTGGCGC</td><td>>hg38_21_+_8433172_8446622_RNA45SN1_45S_50ntFlanks</td><td>12073</td><td>12096</td><td>24</td><td>CCGAGGGGCTCTCGCTTCTGGCGC</td><td>1208.8</td><td>-10.2</td><td>1.4</td><td>-7.1</td><td>1.3E-12</td><td>3.3E-12</td><td>BT20/MB468</td><td>NUC</td></tr>
<tr><td>>hg38_21_+_8433172_8446622_RNA45SN1_45S_50ntFlanks@10678.10708.31|CGGTTCCGGCGGCGTCCGGTGAGCTCTCGCT</td><td>>hg38_21_+_8433172_8446622_RNA45SN1_45S_50ntFlanks</td><td>10678</td><td>10708</td><td>31</td><td>CGGTTCCGGCGGCGTCCGGTGAGCTCTCGCT</td><td>10998.9</td><td>-5.9</td><td>0.8</td><td>-7.1</td><td>9.6E-13</td><td>3.3E-12</td><td>MB231/MB468</td><td>CYTO</td></tr>
<tr><td>>hg38_21_+_8433172_8446622_RNA45SN1_45S_50ntFlanks@12938.12961.24|ACCATTCGTAGACGACCTGCTTCT</td><td>>hg38_21_+_8433172_8446622_RNA45SN1_45S_50ntFlanks</td><td>12938</td><td>12961</td><td>24</td><td>ACCATTCGTAGACGACCTGCTTCT</td><td>268.6</td><td>-3.7</td><td>0.5</td><td>-7.1</td><td>9.9E-13</td><td>3.3E-12</td><td>MB231/MB468</td><td>NUC</td></tr>
<tr><td>>hg38_21_+_8433172_8446622_RNA45SN1_45S_50ntFlanks@4733.4764.32|GAACGAAAGTCGGAGGTTCGAAGACGATCAGA</td><td>>hg38_21_+_8433172_8446622_RNA45SN1_45S_50ntFlanks</td><td>4733</td><td>4764</td><td>32</td><td>GAACGAAAGTCGGAGGTTCGAAGACGATCAGA</td><td>439.9</td><td>9.2</td><td>1.3</td><td>7.1</td><td>9.4E-13</td><td>3.3E-12</td><td>BT20/MB468</td><td>CYTO</td></tr>
<tr><td>>hg38_21_+_8433172_8446622_RNA45SN1_45S_50ntFlanks@11208.11253.46|GCGGCGACTCTGGACGCGAGCCGGGCCCTTCCCGTGGATCGCCCCA</td><td>>hg38_21_+_8433172_8446622_RNA45SN1_45S_50ntFlanks</td><td>11208</td><td>11253</td><td>46</td><td>GCGGCGACTCTGGACGCGAGCCGGGCCCTTCCCGTGGATCGCCCCA</td><td>209.1</td><td>-11.5</td><td>1.6</td><td>-7.1</td><td>1.2E-12</td><td>3.3E-12</td><td>MB231/MB468</td><td>MITO</td></tr>
<tr><td>>MI0000456|hsa-mir-140&WithFlank&16|+|69933075|69933186@68.90.23&[MIMAT0004597&hsa-miR-140-3p&offsets|0|+2;m-45&16|+|69933143|69933164&offsets|-1|0]|TACCACAGGGTAGAACCACGGAC</td><td>>MI0000456|hsa-mir-140&WithFlank&16|+|69933075|69933186</td><td>68</td><td>90</td><td>23</td><td>TACCACAGGGTAGAACCACGGAC</td><td>411.1</td><td>11.9</td><td>1.7</td><td>7.1</td><td>1.2E-12</td><td>3.3E-12</td><td>MB231/MB468</td><td>MITO</td></tr>
<tr><td>>hg38_21_+_8433172_8446622_RNA45SN1_45S_50ntFlanks@4919.4940.22|CCACCAGGAGTGGAGCCTGCGG</td><td>>hg38_21_+_8433172_8446622_RNA45SN1_45S_50ntFlanks</td><td>4919</td><td>4940</td><td>22</td><td>CCACCAGGAGTGGAGCCTGCGG</td><td>609.0</td><td>-13.0</td><td>1.8</td><td>-7.1</td><td>1.2E-12</td><td>3.3E-12</td><td>MB231/MB468</td><td>MITO</td></tr>
<tr><td>>hg38_21_+_8433172_8446622_RNA45SN1_45S_50ntFlanks@12420.12442.23|ATGTCGGCTCTTCCTATCATTGT</td><td>>hg38_21_+_8433172_8446622_RNA45SN1_45S_50ntFlanks</td><td>12420</td><td>12442</td><td>23</td><td>ATGTCGGCTCTTCCTATCATTGT</td><td>1185.3</td><td>-5.5</td><td>0.8</td><td>-7.1</td><td>1.3E-12</td><td>3.3E-12</td><td>BT20/MB468</td><td>NUC</td></tr>
<tr><td>>MI0000272|hsa-mir-182&WithFlank&7|-|129770377|129770498@29.51.23(+1G)&[MIMAT0000259&hsa-miR-182-5p&offsets|0|-1(+1G);m-22&7|-|129770449|129770470&offsets|0|+1(+1G)]|TTTGGCAATGGTAGAACTCACACG</td><td>>MI0000272|hsa-mir-182&WithFlank&7|-|129770377|129770498</td><td>29</td><td>51</td><td>23(+1G)</td><td>TTTGGCAATGGTAGAACTCACACG</td><td>1578.7</td><td>4.2</td><td>0.6</td><td>7.4</td><td>1.1E-13</td><td>3.3E-12</td><td>BT20/MB231</td><td>NUC</td></tr>
<tr><td>>hg38_21_+_8433172_8446622_RNA45SN1_45S_50ntFlanks@4555.4587.33|CGCAGCTAGGAATAATGGAATAGGACCGCGGTT</td><td>>hg38_21_+_8433172_8446622_RNA45SN1_45S_50ntFlanks</td><td>4555</td><td>4587</td><td>33</td><td>CGCAGCTAGGAATAATGGAATAGGACCGCGGTT</td><td>163.3</td><td>-11.2</td><td>1.6</td><td>-7.2</td><td>8.4E-13</td><td>3.3E-12</td><td>MB231/MB468</td><td>MP</td></tr>
<tr><td>>hg38_21_+_8433172_8446622_RNA45SN1_45S_50ntFlanks@11873.11917.45|AAGAAGACCCTGTTGAGCTTGACTCTAGTCTGGCACGGTGAAGAG</td><td>>hg38_21_+_8433172_8446622_RNA45SN1_45S_50ntFlanks</td><td>11873</td><td>11917</td><td>45</td><td>AAGAAGACCCTGTTGAGCTTGACTCTAGTCTGGCACGGTGAAGAG</td><td>370.1</td><td>-12.3</td><td>1.7</td><td>-7.1</td><td>1.2E-12</td><td>3.3E-12</td><td>MB231/MB468</td><td>MITO</td></tr>
<tr><td>>hg38_21_+_8433172_8446622_RNA45SN1_45S_50ntFlanks@4763.4780.18|GATACCGTCGTAGTTCCG</td><td>>hg38_21_+_8433172_8446622_RNA45SN1_45S_50ntFlanks</td><td>4763</td><td>4780</td><td>18</td><td>GATACCGTCGTAGTTCCG</td><td>1003.8</td><td>-9.9</td><td>1.4</td><td>-7.1</td><td>1.3E-12</td><td>3.3E-12</td><td>BT20/MB468</td><td>NUC</td></tr>
<tr><td>>hg38_21_+_8433172_8446622_RNA45SN1_45S_50ntFlanks@4753.4784.32|AAGACGATCAGATACCGTCGTAGTTCCGACCA</td><td>>hg38_21_+_8433172_8446622_RNA45SN1_45S_50ntFlanks</td><td>4753</td><td>4784</td><td>32</td><td>AAGACGATCAGATACCGTCGTAGTTCCGACCA</td><td>338.9</td><td>-12.2</td><td>1.7</td><td>-7.1</td><td>1.2E-12</td><td>3.3E-12</td><td>MB231/MB468</td><td>MITO</td></tr>
</table>

<table>
<tr><td>>hg38_21_+_8433172_8446622_RNA45SN1_45S_50ntFlanks@6646.6671.26|TCGTACGACTCTTAGCGGTGGATCAC</td><td>>hg38_21_+_8433172_8446622_RNA45SN1_45S_50ntFlanks</td><td>6646</td><td>6671</td><td>26</td><td>TCGTACGACTCTTAGCGGTGGATCAC</td><td>2854.8</td><td>4.9</td><td>0.7</td><td>7.2</td><td>7.2E-13</td><td>3.3E-12</td><td>BT20/MB468</td><td>MITO</td></tr>
<tr><td>>MI0000082|hsa-mir-25&WithFlank&7|-|100093554|100093649@58.77.20&[MIMAT0000081&hsa-miR-25-3p&offsets|0|-2;m-23&7|-|100093571|100093592&offsets|0|-2]|CATTGCACTTGTCTCGGTCT</td><td>>MI0000082|hsa-mir-25&WithFlank&7|-|100093554|100093649</td><td>58</td><td>77</td><td>20</td><td>CATTGCACTTGTCTCGGTCT</td><td>748.3</td><td>9.2</td><td>1.3</td><td>7.1</td><td>9.4E-13</td><td>3.3E-12</td><td>BT20/MB468</td><td>CYTO</td></tr>
<tr><td>>MI0000063|hsa-let-7b&WithFlank&22|+|46113680|46113774@12.33.22(+1G)&[MIMAT0000063&hsa-let-7b-5p&offsets|0|0(+1G);m-14&22|+|46113691|46113712&offsets|0|0(+1G)]|TGAGGTAGTAGGTTGTGTGGTTG</td><td>>MI0000063|hsa-let-7b&WithFlank&22|+|46113680|46113774</td><td>12</td><td>33</td><td>22(+1G)</td><td>TGAGGTAGTAGGTTGTGTGGTTG</td><td>108.2</td><td>2.8</td><td>0.4</td><td>7.1</td><td>1.3E-12</td><td>3.3E-12</td><td>BT20/MB468</td><td>NUC</td></tr>
<tr><td>>hg38_21_+_8433172_8446622_RNA45SN1_45S_50ntFlanks@6661.6689.29|CGGTGGATCACTCGGCTCGTGCGTCGATG</td><td>>hg38_21_+_8433172_8446622_RNA45SN1_45S_50ntFlanks</td><td>6661</td><td>6689</td><td>29</td><td>CGGTGGATCACTCGGCTCGTGCGTCGATG</td><td>8859.4</td><td>-2.7</td><td>0.4</td><td>-7.1</td><td>1.3E-12</td><td>3.3E-12</td><td>BT20/MB468</td><td>NUC</td></tr>
<tr><td>>am_trna128_GlyGCC_6_-_27870686_27870756@1.34.34,trna18_GlyGCC_16_+_70822597_70822667@1.34.34,trna19_GlyGCC_16_+_70823410_70823480@1.34.34,trna19_GlyGCC_2_-_157257659_157257729@1.34.34,trna24_GlyGCC_16_-_70812942_70813012@1.34.34,trna25_GlyGCC_16_-_70812114_70812184@1.34.34,trna5_GlyGCC_17_+_8029064_8029134@1.34.34,trna68_GlyGCC_1_-_161493637_161493707@1.34.34|GCATTGGTGGTTCAGTGGTAGAATTCTCGCCTGC</td><td>>am_trna128_GlyGCC_6_-_27870686_27870756</td><td>1</td><td>34</td><td>34</td><td>GCATTGGTGGTTCAGTGGTAGAATTCTCGCCTGC</td><td>1568.0</td><td>9.2</td><td>1.3</td><td>7.1</td><td>1.0E-12</td><td>3.3E-12</td><td>MB231/MB468</td><td>NUC</td></tr>
<tr><td>>hg38_21_+_8433172_8446622_RNA45SN1_45S_50ntFlanks@10408.10427.20|GGTCAGTCGGTCCTGAGAGA</td><td>>hg38_21_+_8433172_8446622_RNA45SN1_45S_50ntFlanks</td><td>10408</td><td>10427</td><td>20</td><td>GGTCAGTCGGTCCTGAGAGA</td><td>160.2</td><td>-11.2</td><td>1.6</td><td>-7.2</td><td>8.4E-13</td><td>3.3E-12</td><td>MB231/MB468</td><td>MP</td></tr>
<tr><td>>hg38_21_+_8433172_8446622_RNA45SN1_45S_50ntFlanks@7992.8019.28|CGTGGCGACCCGCTGAATTTAAGCATAT</td><td>>hg38_21_+_8433172_8446622_RNA45SN1_45S_50ntFlanks</td><td>7992</td><td>8019</td><td>28</td><td>CGTGGCGACCCGCTGAATTTAAGCATAT</td><td>266.9</td><td>9.4</td><td>1.3</td><td>7.1</td><td>9.7E-13</td><td>3.3E-12</td><td>MB231/MB468</td><td>CYTO</td></tr>
<tr><td>>hg38_21_+_8433172_8446622_RNA45SN1_45S_50ntFlanks@5488.5512.25|GCTGAGAAGACGGTCGAACTTGACT</td><td>>hg38_21_+_8433172_8446622_RNA45SN1_45S_50ntFlanks</td><td>5488</td><td>5512</td><td>25</td><td>GCTGAGAAGACGGTCGAACTTGACT</td><td>1324.3</td><td>9.9</td><td>1.4</td><td>7.1</td><td>9.5E-13</td><td>3.4E-12</td><td>BT20/MB468</td><td>CYTO</td></tr>
<tr><td>>hg38_21_+_8433172_8446622_RNA45SN1_45S_50ntFlanks@9573.9595.23|AAATCGGTCGTCCGACCTGGGTA</td><td>>hg38_21_+_8433172_8446622_RNA45SN1_45S_50ntFlanks</td><td>9573</td><td>9595</td><td>23</td><td>AAATCGGTCGTCCGACCTGGGTA</td><td>1632.9</td><td>-3.9</td><td>0.5</td><td>-7.1</td><td>9.5E-13</td><td>3.4E-12</td><td>BT20/MB468</td><td>CYTO</td></tr>
<tr><td>>MI0000085|hsa-mir-27a&WithFlank&19|-|13836434|13836523@57.75.19(+1A)&[MIMAT0000084&hsa-miR-27a-3p&offsets|0|-2(+1A);m-43&19|-|13836447|13836467&offsets|0|-2(+1A)]|TTCACAGTGGCTAAGTTCCA</td><td>>MI0000085|hsa-mir-27a&WithFlank&19|-|13836434|13836523</td><td>57</td><td>75</td><td>19(+1A)</td><td>TTCACAGTGGCTAAGTTCCA</td><td>1885.4</td><td>6.0</td><td>0.8</td><td>7.1</td><td>1.0E-12</td><td>3.4E-12</td><td>MB231/MB468</td><td>NUC</td></tr>
<tr><td>>hg38_21_+_8433172_8446622_RNA45SN1_45S_50ntFlanks@11319.11338.20|CCCACGTCTCGTCGCGCGCG</td><td>>hg38_21_+_8433172_8446622_RNA45SN1_45S_50ntFlanks</td><td>11319</td><td>11338</td><td>20</td><td>CCCACGTCTCGTCGCGCGCG</td><td>150.0</td><td>-10.2</td><td>1.4</td><td>-7.1</td><td>1.0E-12</td><td>3.4E-12</td><td>MB231/MB468</td><td>NUC</td></tr>
<tr><td>>hg38_21_+_8433172_8446622_RNA45SN1_45S_50ntFlanks@3777.3801.25|CGGTACAGTGAAACTGCGAATGGCT</td><td>>hg38_21_+_8433172_8446622_RNA45SN1_45S_50ntFlanks</td><td>3777</td><td>3801</td><td>25</td><td>CGGTACAGTGAAACTGCGAATGGCT</td><td>1208.6</td><td>-7.8</td><td>1.1</td><td>-7.2</td><td>8.5E-13</td><td>3.4E-12</td><td>MB231/MB468</td><td>MP</td></tr>
<tr><td>>hg38_21_+_8433172_8446622_RNA45SN1_45S_50ntFlanks@10367.10384.18|CGAAGTGGAGAAGGGTTC</td><td>>hg38_21_+_8433172_8446622_RNA45SN1_45S_50ntFlanks</td><td>10367</td><td>10384</td><td>18</td><td>CGAAGTGGAGAAGGGTTC</td><td>163.1</td><td>-11.2</td><td>1.6</td><td>-7.2</td><td>8.5E-13</td><td>3.4E-12</td><td>MB231/MB468</td><td>MP</td></tr>
<tr><td>>MI0000482|hsa-mir-185&WithFlank&22|+|20033133|20033226@21.42.22(+1A)&[MIMAT0000455&hsa-miR-185-5p&offsets|0|0(+1A);m-145&22|+|20033153|20033174&offsets|0|0(+1A)]|TGGAGAGAAAGGCAGTTCCTGAA</td><td>>MI0000482|hsa-mir-185&WithFlank&22|+|20033133|20033226</td><td>21</td><td>42</td><td>22(+1A)</td><td>TGGAGAGAAAGGCAGTTCCTGAA</td><td>318.8</td><td>8.3</td><td>1.2</td><td>7.2</td><td>7.4E-13</td><td>3.4E-12</td><td>BT20/MB468</td><td>MITO</td></tr>
<tr><td>>hg38_21_+_8433172_8446622_RNA45SN1_45S_50ntFlanks@5421.5444.24|CGATTGGATGGTTTAGTGAGGCCC</td><td>>hg38_21_+_8433172_8446622_RNA45SN1_45S_50ntFlanks</td><td>5421</td><td>5444</td><td>24</td><td>CGATTGGATGGTTTAGTGAGGCCC</td><td>150.7</td><td>-4.6</td><td>0.7</td><td>-7.1</td><td>9.9E-13</td><td>3.4E-12</td><td>MB231/MB468</td><td>CYTO</td></tr>
<tr><td>>hg38_21_+_8433172_8446622_RNA45SN1_45S_50ntFlanks@9554.9584.31|CGTAGCGGTCCTGACGTGCAAATCGGTCGTC</td><td>>hg38_21_+_8433172_8446622_RNA45SN1_45S_50ntFlanks</td><td>9554</td><td>9584</td><td>31</td><td>CGTAGCGGTCCTGACGTGCAAATCGGTCGTC</td><td>8957.4</td><td>-9.6</td><td>1.4</td><td>-7.1</td><td>1.3E-12</td><td>3.4E-12</td><td>BT20/MB468</td><td>NUC</td></tr>
<tr><td>>hg38_21_+_8433172_8446622_RNA45SN1_45S_50ntFlanks@11920.11952.33|ATGAGAGGTGTAGAATAAGTGGGAGGCCCCCGG</td><td>>hg38_21_+_8433172_8446622_RNA45SN1_45S_50ntFlanks</td><td>11920</td><td>11952</td><td>33</td><td>ATGAGAGGTGTAGAATAAGTGGGAGGCCCCCGG</td><td>132.0</td><td>-10.9</td><td>1.5</td><td>-7.2</td><td>8.6E-13</td><td>3.4E-12</td><td>MB231/MB468</td><td>MP</td></tr>
<tr><td>>hg38_21_+_8433172_8446622_RNA45SN1_45S_50ntFlanks@12936.12959.24|AAACCATTCGTAGACGACCTGCTT</td><td>>hg38_21_+_8433172_8446622_RNA45SN1_45S_50ntFlanks</td><td>12936</td><td>12959</td><td>24</td><td>AAACCATTCGTAGACGACCTGCTT</td><td>107.6</td><td>-10.6</td><td>1.5</td><td>-7.2</td><td>8.6E-13</td><td>3.4E-12</td><td>MB231/MB468</td><td>MP</td></tr>
<tr><td>>hg38_21_+_8433172_8446622_RNA45SN1_45S_50ntFlanks@11933.11957.25|AATAAGTGGGAGGCCCCCGGCGCCC</td><td>>hg38_21_+_8433172_8446622_RNA45SN1_45S_50ntFlanks</td><td>11933</td><td>11957</td><td>25</td><td>AATAAGTGGGAGGCCCCCGGCGCCC</td><td>183.2</td><td>-11.4</td><td>1.6</td><td>-7.2</td><td>8.6E-13</td><td>3.4E-12</td><td>MB231/MB468</td><td>MP</td></tr>
<tr><td>>hg38_21_+_8433172_8446622_RNA45SN1_45S_50ntFlanks@12578.12601.24|AGGAACCGCAGGTTCAGACATTTG</td><td>>hg38_21_+_8433172_8446622_RNA45SN1_45S_50ntFlanks</td><td>12578</td><td>12601</td><td>24</td><td>AGGAACCGCAGGTTCAGACATTTG</td><td>102.2</td><td>-10.5</td><td>1.5</td><td>-7.2</td><td>8.6E-13</td><td>3.4E-12</td><td>MB231/MB468</td><td>MP</td></tr>
<tr><td>>hg38_21_+_8433172_8446622_RNA45SN1_45S_50ntFlanks@9241.9273.33|CGGAGCGAGCGCACGGGGTCGGCGGCGACGTCG</td><td>>hg38_21_+_8433172_8446622_RNA45SN1_45S_50ntFlanks</td><td>9241</td><td>9273</td><td>33</td><td>CGGAGCGAGCGCACGGGGTCGGCGGCGACGTCG</td><td>108.0</td><td>-10.6</td><td>1.5</td><td>-7.2</td><td>8.6E-13</td><td>3.4E-12</td><td>MB231/MB468</td><td>MP</td></tr>
<tr><td>>hg38_21_+_8433172_8446622_RNA45SN1_45S_50ntFlanks@3840.3864.25|CTCCTACTTGGATAACTGTGGTAAT</td><td>>hg38_21_+_8433172_8446622_RNA45SN1_45S_50ntFlanks</td><td>3840</td><td>3864</td><td>25</td><td>CTCCTACTTGGATAACTGTGGTAAT</td><td>501.9</td><td>6.7</td><td>0.9</td><td>7.1</td><td>9.7E-13</td><td>3.4E-12</td><td>BT20/MB468</td><td>CYTO</td></tr>
<tr><td>>hg38_21_+_8433172_8446622_RNA45SN1_45S_50ntFlanks@4161.4182.22|CACATCCAAGGAAGGCAGCAGG</td><td>>hg38_21_+_8433172_8446622_RNA45SN1_45S_50ntFlanks</td><td>4161</td><td>4182</td><td>22</td><td>CACATCCAAGGAAGGCAGCAGG</td><td>195.8</td><td>-11.5</td><td>1.6</td><td>-7.2</td><td>8.6E-13</td><td>3.4E-12</td><td>MB231/MB468</td><td>MP</td></tr>
<tr><td>>hg38_21_+_8433172_8446622_RNA45SN1_45S_50ntFlanks@3777.3801.25|CGGTACAGTGAAACTGCGAATGGCT</td><td>>hg38_21_+_8433172_8446622_RNA45SN1_45S_50ntFlanks</td><td>3777</td><td>3801</td><td>25</td><td>CGGTACAGTGAAACTGCGAATGGCT</td><td>2199.9</td><td>-8.7</td><td>1.2</td><td>-7.1</td><td>1.4E-12</td><td>3.4E-12</td><td>BT20/MB468</td><td>NUC</td></tr>
<tr><td>>hg38_21_+_8433172_8446622_RNA45SN1_45S_50ntFlanks@4145.4162.18|CCTGAGAAACGGCTACCA</td><td>>hg38_21_+_8433172_8446622_RNA45SN1_45S_50ntFlanks</td><td>4145</td><td>4162</td><td>18</td><td>CCTGAGAAACGGCTACCA</td><td>196.9</td><td>-11.5</td><td>1.6</td><td>-7.2</td><td>8.7E-13</td><td>3.4E-12</td><td>MB231/MB468</td><td>MP</td></tr>
<tr><td>>hg38_21_+_8433172_8446622_RNA45SN1_45S_50ntFlanks@3990.4013.24|TTGGTGACTCTAGATAACCTCGGG</td><td>>hg38_21_+_8433172_8446622_RNA45SN1_45S_50ntFlanks</td><td>3990</td><td>4013</td><td>24</td><td>TTGGTGACTCTAGATAACCTCGGG</td><td>272.3</td><td>-4.6</td><td>0.6</td><td>-7.1</td><td>1.0E-12</td><td>3.4E-12</td><td>MB231/MB468</td><td>NUC</td></tr>
<tr><td>>MI0000270|hsa-mir-181b-1&WithFlank&1|-|198858867|198858988@42.63.22;MI0000683|hsa-mir-181b-2&WithFlank&9|+|124693704|124693804@22.43.22&[MIMAT0000257&hsa-miR-181b-5p&offsets|0|-1;m-73&1|-|198858925|198858947&offsets|0|-1];[MIMAT0000257_1&hsa-miR-181b-5p&offsets|0|-1;m-72&9|+|124693725|124693747&offsets|0|-1]|AACATTCATTGCTGTCGGTGGG</td><td>>MI0000270|hsa-mir-181b-1&WithFlank&1|-|198858867|198858988</td><td>42</td><td>63</td><td>22</td><td>AACATTCATTGCTGTCGGTGGG</td><td>335.2</td><td>5.1</td><td>0.7</td><td>7.1</td><td>9.7E-13</td><td>3.4E-12</td><td>BT20/MB468</td><td>CYTO</td></tr>
<tr><td>>hg38_21_+_8433172_8446622_RNA45SN1_45S_50ntFlanks@4547.4564.18|CTGGATACCGCAGCTAGG</td><td>>hg38_21_+_8433172_8446622_RNA45SN1_45S_50ntFlanks</td><td>4547</td><td>4564</td><td>18</td><td>CTGGATACCGCAGCTAGG</td><td>155.6</td><td>-11.1</td><td>1.6</td><td>-7.1</td><td>8.7E-13</td><td>3.4E-12</td><td>MB231/MB468</td><td>MP</td></tr>
<tr><td>>hg38_21_+_8433172_8446622_RNA45SN1_45S_50ntFlanks@4657.4690.34|CTAGAGGTGAAATTCTTGGACCGGCGCAAGACGG</td><td>>hg38_21_+_8433172_8446622_RNA45SN1_45S_50ntFlanks</td><td>4657</td><td>4690</td><td>34</td><td>CTAGAGGTGAAATTCTTGGACCGGCGCAAGACGG</td><td>207.9</td><td>-11.6</td><td>1.6</td><td>-7.1</td><td>8.7E-13</td><td>3.4E-12</td><td>MB231/MB468</td><td>MP</td></tr>
<tr><td>>hg38_21_+_8433172_8446622_RNA45SN1_45S_50ntFlanks@9716.9737.22|AATGATTAGAGGTCTTGGGGCC</td><td>>hg38_21_+_8433172_8446622_RNA45SN1_45S_50ntFlanks</td><td>9716</td><td>9737</td><td>22</td><td>AATGATTAGAGGTCTTGGGGCC</td><td>1528.0</td><td>-9.8</td><td>1.4</td><td>-7.1</td><td>1.3E-12</td><td>3.5E-12</td><td>MB231/MB468</td><td>MITO</td></tr>
<tr><td>>hg38_21_+_8433172_8446622_RNA45SN1_45S_50ntFlanks@10393.10420.28|CAGCAGTTGAACATGGGTCAGTCGGTCC</td><td>>hg38_21_+_8433172_8446622_RNA45SN1_45S_50ntFlanks</td><td>10393</td><td>10420</td><td>28</td><td>CAGCAGTTGAACATGGGTCAGTCGGTCC</td><td>442.8</td><td>-8.6</td><td>1.2</td><td>-7.1</td><td>1.0E-12</td><td>3.5E-12</td><td>MB231/MB468</td><td>NUC</td></tr>
<tr><td>>MI0000255|hsa-mir-30d&WithFlank&8|-|134804870|134804951@12.34.23(+1A)&[MIMAT0000245&hsa-miR-30d-5p&offsets|0|+1(+1A);m-21&8|-|134804919|134804940&offsets|0|+1(+1A)]|TGTAAACATCCCCGACTGGAAGCA</td><td>>MI0000255|hsa-mir-30d&WithFlank&8|-|134804870|134804951</td><td>12</td><td>34</td><td>23(+1A)</td><td>TGTAAACATCCCCGACTGGAAGCA</td><td>2367.2</td><td>6.9</td><td>1.0</td><td>7.1</td><td>1.0E-12</td><td>3.5E-12</td><td>MB231/MB468</td><td>CYTO</td></tr>
<tr><td>>hg38_21_+_8433172_8446622_RNA45SN1_45S_50ntFlanks@9741.9758.18|ACGATCTCAACCTATTCT</td><td>>hg38_21_+_8433172_8446622_RNA45SN1_45S_50ntFlanks</td><td>9741</td><td>9758</td><td>18</td><td>ACGATCTCAACCTATTCT</td><td>104.7</td><td>-10.6</td><td>1.5</td><td>-7.1</td><td>8.8E-13</td><td>3.5E-12</td><td>MB231/MB468</td><td>MP</td></tr>
<tr><td>>hg38_21_+_8433172_8446622_RNA45SN1_45S_50ntFlanks@8686.8706.21|CCGGCTCCGGGACGGCTGGGA</td><td>>hg38_21_+_8433172_8446622_RNA45SN1_45S_50ntFlanks</td><td>8686</td><td>8706</td><td>21</td><td>CCGGCTCCGGGACGGCTGGGA</td><td>363.7</td><td>10.1</td><td>1.4</td><td>7.1</td><td>1.0E-12</td><td>3.5E-12</td><td>MB231/MB468</td><td>NUC</td></tr>
<tr><td>>hg38_21_+_8433172_8446622_RNA45SN1_45S_50ntFlanks@9312.9338.27|GTCTAACACGTGCGCGAGTCGGGGGCT</td><td>>hg38_21_+_8433172_8446622_RNA45SN1_45S_50ntFlanks</td><td>9312</td><td>9338</td><td>27</td><td>GTCTAACACGTGCGCGAGTCGGGGGCT</td><td>216.5</td><td>-11.6</td><td>1.6</td><td>-7.1</td><td>8.8E-13</td><td>3.5E-12</td><td>MB231/MB468</td><td>MP</td></tr>
<tr><td>>hg38_21_+_8433172_8446622_RNA45SN1_45S_50ntFlanks@11564.11590.27|CCTAGCAGCCGACTTAGAACTGGTGCG</td><td>>hg38_21_+_8433172_8446622_RNA45SN1_45S_50ntFlanks</td><td>11564</td><td>11590</td><td>27</td><td>CCTAGCAGCCGACTTAGAACTGGTGCG</td><td>529.6</td><td>-6.7</td><td>0.9</td><td>-7.1</td><td>1.3E-12</td><td>3.5E-12</td><td>MB231/MB468</td><td>MITO</td></tr>
<tr><td>>hg38_21_+_8433172_8446622_RNA45SN1_45S_50ntFlanks@9574.9595.22|AATCGGTCGTCCGACCTGGGTA</td><td>>hg38_21_+_8433172_8446622_RNA45SN1_45S_50ntFlanks</td><td>9574</td><td>9595</td><td>22</td><td>AATCGGTCGTCCGACCTGGGTA</td><td>4302.3</td><td>-6.5</td><td>0.9</td><td>-7.1</td><td>1.3E-12</td><td>3.5E-12</td><td>MB231/MB468</td><td>MITO</td></tr>
<tr><td>>hg38_21_+_8433172_8446622_RNA45SN1_45S_50ntFlanks@11643.11669.27|GCGGGTGTTGACGCGATGTGATTTCTG</td><td>>hg38_21_+_8433172_8446622_RNA45SN1_45S_50ntFlanks</td><td>11643</td><td>11669</td><td>27</td><td>GCGGGTGTTGACGCGATGTGATTTCTG</td><td>229.0</td><td>-11.7</td><td>1.6</td><td>-7.1</td><td>8.8E-13</td><td>3.5E-12</td><td>MB231/MB468</td><td>MP</td></tr>
<tr><td>>MI0000066|hsa-let-7e&WithFlank&19|+|51692780|51692870@14.35.22(+1U)&[MIMAT0000066&hsa-let-7e-5p&offsets|0|0(+1U);m-50&19|+|51692793|51692814&offsets|0|0(+1U)]|TGAGGTAGGAGGTTGTATAGTTT</td><td>>MI0000066|hsa-let-7e&WithFlank&19|+|51692780|51692870</td><td>14</td><td>35</td><td>22(+1U)</td><td>TGAGGTAGGAGGTTGTATAGTTT</td><td>1095.8</td><td>6.4</td><td>0.9</td><td>7.1</td><td>1.0E-12</td><td>3.5E-12</td><td>MB231/MB468</td><td>CYTO</td></tr>
<tr><td>>hg38_21_+_8433172_8446622_RNA45SN1_45S_50ntFlanks@9534.9565.32|AGGAAACTCTGGTGGAGGTCCGTAGCGGTCCT</td><td>>hg38_21_+_8433172_8446622_RNA45SN1_45S_50ntFlanks</td><td>9534</td><td>9565</td><td>32</td><td>AGGAAACTCTGGTGGAGGTCCGTAGCGGTCCT</td><td>1123.4</td><td>-3.1</td><td>0.4</td><td>-7.1</td><td>9.9E-13</td><td>3.5E-12</td><td>BT20/MB468</td><td>CYTO</td></tr>
<tr><td>>hg38_21_+_8433172_8446622_RNA45SN1_45S_50ntFlanks@12605.12625.21|TATGTGCTTGGCTGAGGAGCC</td><td>>hg38_21_+_8433172_8446622_RNA45SN1_45S_50ntFlanks</td><td>12605</td><td>12625</td><td>21</td><td>TATGTGCTTGGCTGAGGAGCC</td><td>19938.9</td><td>-12.3</td><td>1.7</td><td>-7.1</td><td>1.4E-12</td><td>3.5E-12</td><td>BT20/MB468</td><td>NUC</td></tr>
<tr><td>>hg38_21_+_8433172_8446622_RNA45SN1_45S_50ntFlanks@13000.13027.28|TGCGATCTATTGAAAGTCAGCCCTCGAC</td><td>>hg38_21_+_8433172_8446622_RNA45SN1_45S_50ntFlanks</td><td>13000</td><td>13027</td><td>28</td><td>TGCGATCTATTGAAAGTCAGCCCTCGAC</td><td>1671.9</td><td>-8.1</td><td>1.1</td><td>-7.1</td><td>1.4E-12</td><td>3.5E-12</td><td>BT20/MB468</td><td>NUC</td></tr>
<tr><td>>hg38_21_+_8433172_8446622_RNA45SN1_45S_50ntFlanks@3993.4026.34|GTGACTCTAGATAACCTCGGGCCGATCGCACGCC</td><td>>hg38_21_+_8433172_8446622_RNA45SN1_45S_50ntFlanks</td><td>3993</td><td>4026</td><td>34</td><td>GTGACTCTAGATAACCTCGGGCCGATCGCACGCC</td><td>1522.2</td><td>-7.6</td><td>1.1</td><td>-7.1</td><td>1.3E-12</td><td>3.5E-12</td><td>MB231/MB468</td><td>MITO</td></tr>
<tr><td>>hg38_21_+_8433172_8446622_RNA45SN1_45S_50ntFlanks@7980.8005.26|CCTCAGATCAGACGTGGCGACCCGCT</td><td>>hg38_21_+_8433172_8446622_RNA45SN1_45S_50ntFlanks</td><td>7980</td><td>8005</td><td>26</td><td>CCTCAGATCAGACGTGGCGACCCGCT</td><td>916.1</td><td>-7.0</td><td>1.0</td><td>-7.1</td><td>1.3E-12</td><td>3.5E-12</td><td>MB231/MB468</td><td>MITO</td></tr>
<tr><td>>hg38_21_+_8433172_8446622_RNA45SN1_45S_50ntFlanks@3789.3810.22|ACTGCGAATGGCTCATTAAATC</td><td>>hg38_21_+_8433172_8446622_RNA45SN1_45S_50ntFlanks</td><td>3789</td><td>3810</td><td>22</td><td>ACTGCGAATGGCTCATTAAATC</td><td>1268.8</td><td>-10.3</td><td>1.4</td><td>-7.1</td><td>1.4E-12</td><td>3.5E-12</td><td>BT20/MB468</td><td>NUC</td></tr>
<tr><td>>hg38_21_+_8433172_8446622_RNA45SN1_45S_50ntFlanks@4659.4678.20|AGAGGTGAAATTCTTGGACC</td><td>>hg38_21_+_8433172_8446622_RNA45SN1_45S_50ntFlanks</td><td>4659</td><td>4678</td><td>20</td><td>AGAGGTGAAATTCTTGGACC</td><td>172.5</td><td>11.3</td><td>1.6</td><td>7.1</td><td>1.1E-12</td><td>3.5E-12</td><td>MB231/MB468</td><td>NUC</td></tr>
<tr><td>>trnaMT_GluTTC_MT_-_14674_14742@31.72.42,trnalookalike8_GluTTC_5_-_93905172_93905240@31.72.42|TTCATATCATTGGTCGTGGTTGTAGTCCGTGCGAGAATACCA</td><td>>trnaMT_GluTTC_MT_-_14674_14742</td><td>31</td><td>72</td><td>42</td><td>TTCATATCATTGGTCGTGGTTGTAGTCCGTGCGAGAATACCA</td><td>930.7</td><td>5.5</td><td>0.8</td><td>7.2</td><td>7.6E-13</td><td>3.5E-12</td><td>BT20/MB468</td><td>MITO</td></tr>
</table>

| | | | | | | | | | | | | | |
|---|---|---|---|---|---|---|---|---|---|---|---|---|---|
| >am_trna117_GlyTCC_1_-_145397864_145397935@1.29.29,trna45_GlyTCC_1_+_161500903_161500974@1.29.29,trna70_GlyTCC_1_-_161439547_161439618@1.29.29,trna73_GlyTCC_1_-_161432166_161432237@1.29.29,trna76_GlyTCC_1_-_161424756_161424827@1.29.29,trna79_GlyTCC_1_-_161417375_161417446@1.29.29\|GCGTTGGTGGTATAGTGGTGAGCATAGCT | >am_trna117_GlyTCC_1_-_145397864_145397935 | 1 | 29 | 29 | GCGTTGGTGGTATAGTGGTGAGCATAGCT | 2219.1 | -10.2 | 1.4 | -7.1 | 1.1E-12 | 3.5E-12 | MB231/MB468 | NUC |
| >hg38_21_+_8433172_8446622_RNA45SN1_45S_50ntFlanks@11538.11560.23\|CGCGCGGCGCCTCGCCTCGGCCG | >hg38_21_+_8433172_8446622_RNA45SN1_45S_50ntFlanks | 11538 | 11560 | 23 | CGCGCGGCGCCTCGCCTCGGCCG | 142.7 | -6.4 | 0.9 | -7.1 | 1.0E-12 | 3.5E-12 | MB231/MB468 | CYTO |
| >hg38_21_+_8433172_8446622_RNA45SN1_45S_50ntFlanks@3802.3842.41\|CATTAAATCAGTTATGGTTCCTTTGGTCGCTCGCTCCTCTC | >hg38_21_+_8433172_8446622_RNA45SN1_45S_50ntFlanks | 3802 | 3842 | 41 | CATTAAATCAGTTATGGTTCCTTTGGTCGCTCGCTCCTCTC | 298.4 | -11.2 | 1.6 | -7.1 | 1.1E-12 | 3.5E-12 | MB231/MB468 | NUC |
| >hg38_21_+_8433172_8446622_RNA45SN1_45S_50ntFlanks@13002.13039.38\|CGATCTATTGAAAGTCAGCCCTCGACACAAGGGTTTGT | >hg38_21_+_8433172_8446622_RNA45SN1_45S_50ntFlanks | 13002 | 13039 | 38 | CGATCTATTGAAAGTCAGCCCTCGACACAAGGGTTTGT | 808.4 | -12.7 | 1.7 | -7.5 | 4.6E-14 | 3.5E-12 | BT20/MB468 | MP |
| >hg38_21_+_8433172_8446622_RNA45SN1_45S_50ntFlanks@7974.8001.28\|ACGCGACCTCAGATCAGACGTGGCGACC | >hg38_21_+_8433172_8446622_RNA45SN1_45S_50ntFlanks | 7974 | 8001 | 28 | ACGCGACCTCAGATCAGACGTGGCGACC | 153.8 | 4.4 | 0.6 | 7.1 | 1.4E-12 | 3.5E-12 | BT20/MB468 | NUC |
| >hg38_21_+_8433172_8446622_RNA45SN1_45S_50ntFlanks@10735.10757.23\|TGTAAATCTCGCGCCGGGCCGTA | >hg38_21_+_8433172_8446622_RNA45SN1_45S_50ntFlanks | 10735 | 10757 | 23 | TGTAAATCTCGCGCCGGGCCGTA | 95.1 | -10.4 | 1.5 | -7.1 | 8.9E-13 | 3.5E-12 | MB231/MB468 | MP |
| >hg38_21_+_8433172_8446622_RNA45SN1_45S_50ntFlanks@12033.12072.40\|ATCGTTTTTTCACTGACCCGGTGAGGCGGGGGGGCGAGCC | >hg38_21_+_8433172_8446622_RNA45SN1_45S_50ntFlanks | 12033 | 12072 | 40 | ATCGTTTTTTCACTGACCCGGTGAGGCGGGGGGGCGAGCC | 220.8 | -11.6 | 1.6 | -7.1 | 1.3E-12 | 3.5E-12 | MB231/MB468 | MITO |
| >hg38_21_+_8433172_8446622_RNA45SN1_45S_50ntFlanks@6735.6771.37\|TTGATCATCGACACTTCGAACGCACTTGCGGCCCCGG | >hg38_21_+_8433172_8446622_RNA45SN1_45S_50ntFlanks | 6735 | 6771 | 37 | TTGATCATCGACACTTCGAACGCACTTGCGGCCCCGG | 1332.1 | -7.2 | 1.0 | -7.1 | 1.3E-12 | 3.5E-12 | MB231/MB468 | MITO |
| >hg38_21_+_8433172_8446622_RNA45SN1_45S_50ntFlanks@12748.12769.22\|CCTGTCCCCGCCGGCGGGCCGC | >hg38_21_+_8433172_8446622_RNA45SN1_45S_50ntFlanks | 12748 | 12769 | 22 | CCTGTCCCCGCCGGCGGGCCGC | 445.0 | -8.8 | 1.2 | -7.1 | 1.3E-12 | 3.5E-12 | MB231/MB468 | MITO |
| >hg38_21_+_8433172_8446622_RNA45SN1_45S_50ntFlanks@10408.10428.21\|GGTCAGTCGGTCCTGAGAGAT | >hg38_21_+_8433172_8446622_RNA45SN1_45S_50ntFlanks | 10408 | 10428 | 21 | GGTCAGTCGGTCCTGAGAGAT | 784.6 | -7.5 | 1.1 | -7.1 | 1.1E-12 | 3.5E-12 | MB231/MB468 | NUC |
| >hg38_21_+_8433172_8446622_RNA45SN1_45S_50ntFlanks@12126.12146.21\|CGACCCGCTCCGGGGACAGTG | >hg38_21_+_8433172_8446622_RNA45SN1_45S_50ntFlanks | 12126 | 12146 | 21 | CGACCCGCTCCGGGGACAGTG | 738.3 | -11.6 | 1.6 | -7.2 | 7.7E-13 | 3.5E-12 | BT20/MB468 | MITO |
| >hg38_21_+_8433172_8446622_RNA45SN1_45S_50ntFlanks@9556.9596.41\|TAGCGGTCCTGACGTGCAAATCGGTCGTCCGACCTGGGTAT | >hg38_21_+_8433172_8446622_RNA45SN1_45S_50ntFlanks | 9556 | 9596 | 41 | TAGCGGTCCTGACGTGCAAATCGGTCGTCCGACCTGGGTAT | 412.9 | -12.5 | 1.8 | -7.1 | 1.3E-12 | 3.6E-12 | MB231/MB468 | MITO |
| >hg38_21_+_8433172_8446622_RNA45SN1_45S_50ntFlanks@11319.11342.24\|CCCACGTCTCGTCGCGCGCGCGTC | >hg38_21_+_8433172_8446622_RNA45SN1_45S_50ntFlanks | 11319 | 11342 | 24 | CCCACGTCTCGTCGCGCGCGCGTC | 138.5 | -11.0 | 1.5 | -7.1 | 9.0E-13 | 3.6E-12 | MB231/MB468 | MP |
| >hg38_21_+_8433172_8446622_RNA45SN1_45S_50ntFlanks@10408.10434.27\|GGTCAGTCGGTCCTGAGAGATGGGCGA | >hg38_21_+_8433172_8446622_RNA45SN1_45S_50ntFlanks | 10408 | 10434 | 27 | GGTCAGTCGGTCCTGAGAGATGGGCGA | 272.8 | -12.0 | 1.7 | -7.1 | 9.0E-13 | 3.6E-12 | MB231/MB468 | MP |
| >hg38_21_+_8433172_8446622_RNA45SN1_45S_50ntFlanks@11829.11872.44\|CGAAACCACAGCCAAGGGAACGGGCTTGGCGGAATCAGCGGGGA | >hg38_21_+_8433172_8446622_RNA45SN1_45S_50ntFlanks | 11829 | 11872 | 44 | CGAAACCACAGCCAAGGGAACGGGCTTGGCGGAATCAGCGGGGA | 212.4 | -11.5 | 1.6 | -7.1 | 1.3E-12 | 3.6E-12 | MB231/MB468 | MITO |
| >hg38_21_+_8433172_8446622_RNA45SN1_45S_50ntFlanks@11872.11890.19\|AAAGAAGACCCTGTTGAGC | >hg38_21_+_8433172_8446622_RNA45SN1_45S_50ntFlanks | 11872 | 11890 | 19 | AAAGAAGACCCTGTTGAGC | 136.1 | -11.0 | 1.5 | -7.1 | 9.0E-13 | 3.6E-12 | MB231/MB468 | MP |
| >hg38_21_+_8433172_8446622_RNA45SN1_45S_50ntFlanks@10411.10445.35\|CAGTCGGTCCTGAGAGATGGGCGAGCGCCGTTCCG | >hg38_21_+_8433172_8446622_RNA45SN1_45S_50ntFlanks | 10411 | 10445 | 35 | CAGTCGGTCCTGAGAGATGGGCGAGCGCCGTTCCG | 189.8 | -5.2 | 0.7 | -7.1 | 1.0E-12 | 3.6E-12 | MB231/MB468 | CYTO |
| >trna119_LysCTT_1_-_145395522_145395594@2.33.32,trna11_LysCTT_5_-_180648979_180649051@2.33.32,trna13_LysCTT_6_+_26556774_26556846@2.33.32,trna7_LysCTT_16_+_3225692_3225764@2.33.32,trna9_LysCTT_5_+_180634755_180634827@2.33.32\|CCCGGCTAGCTCAGTCGGTAGAGCATGAGACT | >trna119_LysCTT_1_-_145395522_145395594 | 2 | 33 | 32 | CCCGGCTAGCTCAGTCGGTAGAGCATGAGACT | 977.4 | 12.0 | 1.7 | 7.1 | 1.1E-12 | 3.6E-12 | MB231/MB468 | NUC |
| >hg38_21_+_8433172_8446622_RNA45SN1_45S_50ntFlanks@5074.5102.29\|ATTCCGATAACGAACGAGACTCTGGCATG | >hg38_21_+_8433172_8446622_RNA45SN1_45S_50ntFlanks | 5074 | 5102 | 29 | ATTCCGATAACGAACGAGACTCTGGCATG | 534.4 | -12.9 | 1.8 | -7.1 | 9.1E-13 | 3.6E-12 | MB231/MB468 | MP |
| >hg38_21_+_8433172_8446622_RNA45SN1_45S_50ntFlanks@12274.12317.44\|CAGTACGAATACAGACCGTGAAAGCGGGGCCTCACGATCCTTCT | >hg38_21_+_8433172_8446622_RNA45SN1_45S_50ntFlanks | 12274 | 12317 | 44 | CAGTACGAATACAGACCGTGAAAGCGGGGCCTCACGATCCTTCT | 229.2 | -11.6 | 1.6 | -7.1 | 1.3E-12 | 3.6E-12 | MB231/MB468 | MITO |
| >hg38_21_+_8433172_8446622_RNA45SN1_45S_50ntFlanks@12285.12319.35\|CAGACCGTGAAAGCGGGGCCTCACGATCCTTCTGA | >hg38_21_+_8433172_8446622_RNA45SN1_45S_50ntFlanks | 12285 | 12319 | 35 | CAGACCGTGAAAGCGGGGCCTCACGATCCTTCTGA | 1117.0 | -9.5 | 1.3 | -7.1 | 1.3E-12 | 3.6E-12 | MB231/MB468 | MITO |
| >hg38_21_+_8433172_8446622_RNA45SN1_45S_50ntFlanks@7982.8004.23\|TCAGATCAGACGTGGCGACCCGC | >hg38_21_+_8433172_8446622_RNA45SN1_45S_50ntFlanks | 7982 | 8004 | 23 | TCAGATCAGACGTGGCGACCCGC | 838.0 | -7.1 | 1.0 | -7.1 | 1.3E-12 | 3.6E-12 | MB231/MB468 | MITO |
| >hg38_21_+_8433172_8446622_RNA45SN1_45S_50ntFlanks@10389.10407.19\|TGAACAGCAGTTGAACATG | >hg38_21_+_8433172_8446622_RNA45SN1_45S_50ntFlanks | 10389 | 10407 | 19 | TGAACAGCAGTTGAACATG | 3025.7 | -8.3 | 1.2 | -7.1 | 1.0E-12 | 3.6E-12 | MB231/MB468 | CYTO |
| >hg38_21_+_8433172_8446622_RNA45SN1_45S_50ntFlanks@10498.10533.36\|TTCAGATCCCCGAATCCGGAGTGGCGGAGATGGGCG | >hg38_21_+_8433172_8446622_RNA45SN1_45S_50ntFlanks | 10498 | 10533 | 36 | TTCAGATCCCCGAATCCGGAGTGGCGGAGATGGGCG | 191.7 | -11.4 | 1.6 | -7.1 | 1.3E-12 | 3.6E-12 | MB231/MB468 | MITO |
| >MI0000088\|hsa-mir-30a&WithFlank&6\|-\|71403545\|71403627@12.35.24(+1U)&[MIMAT0000087&hsa-miR-30a-5p&offsets\|0\|+2(+1U);m-15&6\|-\|71403595\|71403616&offsets\|0\|+2(+1U)]\|TGTAAACATCCTCGACTGGAAGCTT | >MI0000088\|hsa-mir-30a&WithFlank&6\|-\|71403545\|71403627 | 12 | 35 | 24(+1U) | TGTAAACATCCTCGACTGGAAGCTT | 17979.4 | 5.3 | 0.7 | 7.1 | 9.2E-13 | 3.6E-12 | MB231/MB468 | MP |
| >hg38_21_+_8433172_8446622_RNA45SN1_45S_50ntFlanks@11880.11904.25\|CCCTGTTGAGCTTGACTCTAGTCTG | >hg38_21_+_8433172_8446622_RNA45SN1_45S_50ntFlanks | 11880 | 11904 | 25 | CCCTGTTGAGCTTGACTCTAGTCTG | 166.3 | -11.2 | 1.6 | -7.1 | 9.1E-13 | 3.6E-12 | MB231/MB468 | MP |
| >hg38_21_+_8433172_8446622_RNA45SN1_45S_50ntFlanks@5527.5553.27\|AAAGTCGTAACAAGGTTTCCGTAGGTG | >hg38_21_+_8433172_8446622_RNA45SN1_45S_50ntFlanks | 5527 | 5553 | 27 | AAAGTCGTAACAAGGTTTCCGTAGGTG | 103.0 | -10.5 | 1.5 | -7.1 | 9.2E-13 | 3.6E-12 | MB231/MB468 | MP |
| >hg38_21_+_8433172_8446622_RNA45SN1_45S_50ntFlanks@9539.9563.25\|ACTCTGGTGGAGGTCCGTAGCGGTC | >hg38_21_+_8433172_8446622_RNA45SN1_45S_50ntFlanks | 9539 | 9563 | 25 | ACTCTGGTGGAGGTCCGTAGCGGTC | 137.6 | -11.0 | 1.5 | -7.1 | 9.2E-13 | 3.6E-12 | MB231/MB468 | MP |
| >hg38_21_+_8433172_8446622_RNA45SN1_45S_50ntFlanks@12157.12183.27\|AGTTTGACTGGGGCGGTACACCTGTCA | >hg38_21_+_8433172_8446622_RNA45SN1_45S_50ntFlanks | 12157 | 12183 | 27 | AGTTTGACTGGGGCGGTACACCTGTCA | 568.8 | -12.2 | 1.6 | -7.5 | 4.7E-14 | 3.6E-12 | BT20/MB468 | MP |
| >hg38_21_+_8433172_8446622_RNA45SN1_45S_50ntFlanks@12957.12974.18\|CTTCTGGGTCGGGGTTTC | >hg38_21_+_8433172_8446622_RNA45SN1_45S_50ntFlanks | 12957 | 12974 | 18 | CTTCTGGGTCGGGGTTTC | 188.2 | -11.4 | 1.6 | -7.1 | 9.2E-13 | 3.6E-12 | MB231/MB468 | MP |
| >hg38_21_+_8433172_8446622_RNA45SN1_45S_50ntFlanks@6655.6689.35\|TCTTAGCGGTGGATCACTCGGCTCGTGCGTCGATG | >hg38_21_+_8433172_8446622_RNA45SN1_45S_50ntFlanks | 6655 | 6689 | 35 | TCTTAGCGGTGGATCACTCGGCTCGTGCGTCGATG | 3094.3 | -5.0 | 0.7 | -7.1 | 1.3E-12 | 3.6E-12 | MB231/MB468 | MITO |
| >MI0000298\|hsa-mir-221&WithFlank&X\|-\|45746151\|45746272@71.90.20&[MIMAT0000278&hsa-miR-221-3p&offsets\|0\|-3;m-91&X\|-\|45746181\|45746202&offsets\|0\|-2]\|AGCTACATTGTCTGCTGGGT | >MI0000298\|hsa-mir-221&WithFlank&X\|-\|45746151\|45746272 | 71 | 90 | 20 | AGCTACATTGTCTGCTGGGT | 646.0 | 6.4 | 0.9 | 7.1 | 1.3E-12 | 3.6E-12 | MB231/MB468 | MITO |
| >hg38_21_+_8433172_8446622_RNA45SN1_45S_50ntFlanks@11873.11898.26\|AAGAAGACCCTGTTGAGCTTGACTCT | >hg38_21_+_8433172_8446622_RNA45SN1_45S_50ntFlanks | 11873 | 11898 | 26 | AAGAAGACCCTGTTGAGCTTGACTCT | 1082.4 | -9.2 | 1.3 | -7.1 | 9.2E-13 | 3.6E-12 | MB231/MB468 | MP |
| >hg38_21_+_8433172_8446622_RNA45SN1_45S_50ntFlanks@11575.11603.29\|ACTTAGAACTGGTGCGGACCAGGGGAATC | >hg38_21_+_8433172_8446622_RNA45SN1_45S_50ntFlanks | 11575 | 11603 | 29 | ACTTAGAACTGGTGCGGACCAGGGGAATC | 6219.1 | -10.7 | 1.5 | -7.1 | 9.2E-13 | 3.6E-12 | MB231/MB468 | MP |
| >hg38_21_+_8433172_8446622_RNA45SN1_45S_50ntFlanks@10678.10704.27\|CGGTTCCGGCGGCGTCCGGTGAGCTCT | >hg38_21_+_8433172_8446622_RNA45SN1_45S_50ntFlanks | 10678 | 10704 | 27 | CGGTTCCGGCGGCGTCCGGTGAGCTCT | 149.7 | -11.1 | 1.6 | -7.1 | 9.2E-13 | 3.6E-12 | MB231/MB468 | MP |
| >hg38_21_+_8433172_8446622_RNA45SN1_45S_50ntFlanks@8194.8212.19\|CCCGTGGACGGTGTGAGGC | >hg38_21_+_8433172_8446622_RNA45SN1_45S_50ntFlanks | 8194 | 8212 | 19 | CCCGTGGACGGTGTGAGGC | 3448.8 | -8.5 | 1.2 | -7.1 | 9.3E-13 | 3.6E-12 | MB231/MB468 | MP |
| >hg38_21_+_8433172_8446622_RNA45SN1_45S_50ntFlanks@12417.12440.24\|TCGATGTCGGCTCTTCCTATCATT | >hg38_21_+_8433172_8446622_RNA45SN1_45S_50ntFlanks | 12417 | 12440 | 24 | TCGATGTCGGCTCTTCCTATCATT | 811.2 | -4.3 | 0.6 | -7.1 | 1.4E-12 | 3.6E-12 | BT20/MB468 | NUC |
| >hg38_21_+_8433172_8446622_RNA45SN1_45S_50ntFlanks@12991.13028.38\|CTCCCTCGCTGCGATCTATTGAAAGTCAGCCCTCGACA | >hg38_21_+_8433172_8446622_RNA45SN1_45S_50ntFlanks | 12991 | 13028 | 38 | CTCCCTCGCTGCGATCTATTGAAAGTCAGCCCTCGACA | 513.8 | -11.1 | 1.5 | -7.2 | 7.9E-13 | 3.6E-12 | BT20/MB468 | MITO |
| >hg38_21_+_8433172_8446622_RNA45SN1_45S_50ntFlanks@4731.4761.31\|AAGAACGAAAGTCGGAGGTTCGAAGACGATC | >hg38_21_+_8433172_8446622_RNA45SN1_45S_50ntFlanks | 4731 | 4761 | 31 | AAGAACGAAAGTCGGAGGTTCGAAGACGATC | 3351.8 | 4.8 | 0.7 | 7.1 | 1.1E-12 | 3.6E-12 | MB231/MB468 | CYTO |
| >hg38_21_+_8433172_8446622_RNA45SN1_45S_50ntFlanks@9564.9597.34\|CTGACGTGCAAATCGGTCGTCCGACCTGGGTATA | >hg38_21_+_8433172_8446622_RNA45SN1_45S_50ntFlanks | 9564 | 9597 | 34 | CTGACGTGCAAATCGGTCGTCCGACCTGGGTATA | 42338.4 | -8.9 | 1.3 | -7.1 | 1.3E-12 | 3.6E-12 | MB231/MB468 | MITO |
| >hg38_21_+_8433172_8446622_RNA45SN1_45S_50ntFlanks@10360.10388.29\|TGAAGGCCGAAGTGGAGAAGGGTTCCATG | >hg38_21_+_8433172_8446622_RNA45SN1_45S_50ntFlanks | 10360 | 10388 | 29 | TGAAGGCCGAAGTGGAGAAGGGTTCCATG | 206.6 | -11.6 | 1.6 | -7.1 | 9.3E-13 | 3.6E-12 | MB231/MB468 | MP |
| >hg38_21_+_8433172_8446622_RNA45SN1_45S_50ntFlanks@10794.10816.23\|CTGGCATGTTGGAACAATGTAGG | >hg38_21_+_8433172_8446622_RNA45SN1_45S_50ntFlanks | 10794 | 10816 | 23 | CTGGCATGTTGGAACAATGTAGG | 2153.2 | -9.5 | 1.3 | -7.1 | 9.3E-13 | 3.7E-12 | MB231/MB468 | MP |
| >hg38_21_+_8433172_8446622_RNA45SN1_45S_50ntFlanks@11872.11904.33\|AAAGAAGACCCTGTTGAGCTTGACTCTAGTCTG | >hg38_21_+_8433172_8446622_RNA45SN1_45S_50ntFlanks | 11872 | 11904 | 33 | AAAGAAGACCCTGTTGAGCTTGACTCTAGTCTG | 137.2 | -11.0 | 1.5 | -7.1 | 9.4E-13 | 3.7E-12 | MB231/MB468 | MP |
| >hg38_21_+_8433172_8446622_RNA45SN1_45S_50ntFlanks@4099.4116.18\|GTGACCACGGGTGACGGG | >hg38_21_+_8433172_8446622_RNA45SN1_45S_50ntFlanks | 4099 | 4116 | 18 | GTGACCACGGGTGACGGG | 207.8 | -11.6 | 1.6 | -7.1 | 9.3E-13 | 3.7E-12 | MB231/MB468 | MP |
| >hg38_21_+_8433172_8446622_RNA45SN1_45S_50ntFlanks@11200.11234.35\|CGGCGGCGGCGGCGACTCTGGACGCGAGCCGGGCC | >hg38_21_+_8433172_8446622_RNA45SN1_45S_50ntFlanks | 11200 | 11234 | 35 | CGGCGGCGGCGGCGACTCTGGACGCGAGCCGGGCC | 149.1 | -11.1 | 1.6 | -7.1 | 9.4E-13 | 3.7E-12 | MB231/MB468 | MP |
| >MI0000087\|hsa-mir-29a&WithFlank&7\|-\|130876741\|130876816@48.69.22&[MIMAT0000086&hsa-miR-29a-3p&offsets\|0\|0;m-26&7\|-\|130876749\|130876769&offsets\|0\|+1]\|TAGCACCATCTGAAATCGGTTA | >MI0000087\|hsa-mir-29a&WithFlank&7\|-\|130876741\|130876816 | 48 | 69 | 22 | TAGCACCATCTGAAATCGGTTA | 2140.4 | 6.9 | 1.0 | 7.1 | 1.5E-12 | 3.7E-12 | BT20/MB468 | NUC |
| >hg38_21_+_8433172_8446622_RNA45SN1_45S_50ntFlanks@8912.8936.25\|CGCCGAATCCCGGGGCCGAGGGAGC | >hg38_21_+_8433172_8446622_RNA45SN1_45S_50ntFlanks | 8912 | 8936 | 25 | CGCCGAATCCCGGGGCCGAGGGAGC | 928.6 | 7.4 | 1.0 | 7.1 | 1.1E-12 | 3.7E-12 | MB231/MB468 | NUC |
| >hg38_21_+_8433172_8446622_RNA45SN1_45S_50ntFlanks@11821.11844.24\|CTATCCAGCGAAACCACAGCCAAG | >hg38_21_+_8433172_8446622_RNA45SN1_45S_50ntFlanks | 11821 | 11844 | 24 | CTATCCAGCGAAACCACAGCCAAG | 316.4 | -8.1 | 1.1 | -7.1 | 1.1E-12 | 3.7E-12 | MB231/MB468 | NUC |

| | | | | | | | | | | | | | |
|---|---|---|---|---|---|---|---|---|---|---|---|---|---|
| >hg38_21_+_8433172_8446622_RNA45SN1_45S_50ntFlanks@4723.4751.29\|CATTAATCAAGAACGAAAGTCGGAGGTTC | >hg38_21_+_8433172_8446622_RNA45SN1_45S_50ntFlanks | 4723 | 4751 | 29 | CATTAATCAAGAACGAAAGTCGGAGGTTC | 520.5 | -6.6 | 0.9 | -7.1 | 1.1E-12 | 3.7E-12 | MB231/MB468 | NUC |
| >hg38_21_+_8433172_8446622_RNA45SN1_45S_50ntFlanks@4734.4765.32\|AACGAAAGTCGGAGGTTCGAAGACGATCAGAT | >hg38_21_+_8433172_8446622_RNA45SN1_45S_50ntFlanks | 4734 | 4765 | 32 | AACGAAAGTCGGAGGTTCGAAGACGATCAGAT | 817.9 | 7.5 | 1.1 | 7.1 | 1.0E-12 | 3.7E-12 | BT20/MB468 | CYTO |
| >hg38_21_+_8433172_8446622_RNA45SN1_45S_50ntFlanks@4760.4783.24\|TCAGATACCGTCGTAGTTCCGACC | >hg38_21_+_8433172_8446622_RNA45SN1_45S_50ntFlanks | 4760 | 4783 | 24 | TCAGATACCGTCGTAGTTCCGACC | 1599.9 | -8.4 | 1.2 | -7.1 | 1.3E-12 | 3.7E-12 | MB231/MB468 | MITO |
| >am_trna117_GlyTCC_1_-_145397864_145397935@1.28.28,trna45_GlyTCC_1_+_161500903_161500974@1.28.28,trna70_GlyTCC_1_-_161439547_161439618@1.28.28,trna73_GlyTCC_1_-_161432166_161432237@1.28.28,trna76_GlyTCC_1_-_161424756_161424827@1.28.28,trna79_GlyTCC_1_-_161417375_161417446@1.28.28\|GCGTTGGTGGTATAGTGGTGAGCATAGC | >am_trna117_GlyTCC_1_-_145397864_145397935 | 1 | 28 | 28 | GCGTTGGTGGTATAGTGGTGAGCATAGC | 319.2 | -11.3 | 1.6 | -7.1 | 1.1E-12 | 3.7E-12 | MB231/MB468 | NUC |
| >hg38_21_+_8433172_8446622_RNA45SN1_45S_50ntFlanks@8195.8213.19\|CCGTGGACGGTGTGAGGCC | >hg38_21_+_8433172_8446622_RNA45SN1_45S_50ntFlanks | 8195 | 8213 | 19 | CCGTGGACGGTGTGAGGCC | 2175.0 | 5.3 | 0.7 | 7.1 | 1.0E-12 | 3.7E-12 | BT20/MB468 | CYTO |
| >hg38_21_+_8433172_8446622_RNA45SN1_45S_50ntFlanks@12141.12181.41\|ACAGTGCCAGGTGGGGAGTTTGACTGGGGCGGTACACCTGT | >hg38_21_+_8433172_8446622_RNA45SN1_45S_50ntFlanks | 12141 | 12181 | 41 | ACAGTGCCAGGTGGGGAGTTTGACTGGGGCGGTACACCTGT | 186.9 | -11.3 | 1.6 | -7.1 | 1.3E-12 | 3.7E-12 | MB231/MB468 | MITO |
| >hg38_21_+_8433172_8446622_RNA45SN1_45S_50ntFlanks@12932.12972.41\|CGCTAAACCATTCGTAGACGACCTGCTTCTGGGTCGGGGTT | >hg38_21_+_8433172_8446622_RNA45SN1_45S_50ntFlanks | 12932 | 12972 | 41 | CGCTAAACCATTCGTAGACGACCTGCTTCTGGGTCGGGGTT | 1006.5 | -7.7 | 1.1 | -7.1 | 1.3E-12 | 3.7E-12 | MB231/MB468 | MITO |
| >hg38_21_+_8433172_8446622_RNA45SN1_45S_50ntFlanks@9033.9061.29\|CTCGTGGGGGGGCCGGGCCACCCCTCCCA | >hg38_21_+_8433172_8446622_RNA45SN1_45S_50ntFlanks | 9033 | 9061 | 29 | CTCGTGGGGGGGCCGGGCCACCCCTCCCA | 184.0 | -11.4 | 1.6 | -7.1 | 9.4E-13 | 3.7E-12 | MB231/MB468 | MP |
| >hg38_21_+_8433172_8446622_RNA45SN1_45S_50ntFlanks@9572.9594.23\|CAAATCGGTCGTCCGACCTGGGT | >hg38_21_+_8433172_8446622_RNA45SN1_45S_50ntFlanks | 9572 | 9594 | 23 | CAAATCGGTCGTCCGACCTGGGT | 679.4 | -4.5 | 0.6 | -7.1 | 1.1E-12 | 3.7E-12 | BT20/MB468 | CYTO |
| >hg38_21_+_8433172_8446622_RNA45SN1_45S_50ntFlanks@5059.5080.22\|CGATTTGTCTGGTTAATTCCGA | >hg38_21_+_8433172_8446622_RNA45SN1_45S_50ntFlanks | 5059 | 5080 | 22 | CGATTTGTCTGGTTAATTCCGA | 1323.4 | -14.2 | 2.0 | -7.1 | 1.4E-12 | 3.7E-12 | MB231/MB468 | MITO |
| >hg38_21_+_8433172_8446622_RNA45SN1_45S_50ntFlanks@11683.11706.24\|ATGTCAAAGTGAAGAAATTCAATG | >hg38_21_+_8433172_8446622_RNA45SN1_45S_50ntFlanks | 11683 | 11706 | 24 | ATGTCAAAGTGAAGAAATTCAATG | 139.9 | -11.0 | 1.5 | -7.1 | 9.5E-13 | 3.7E-12 | MB231/MB468 | MP |
| >hg38_21_+_8433172_8446622_RNA45SN1_45S_50ntFlanks@11792.11816.25\|ATGAACGAGATTCCCACTGTCCCTA | >hg38_21_+_8433172_8446622_RNA45SN1_45S_50ntFlanks | 11792 | 11816 | 25 | ATGAACGAGATTCCCACTGTCCCTA | 132.8 | -10.9 | 1.5 | -7.1 | 9.5E-13 | 3.7E-12 | MB231/MB468 | MP |
| >hg38_21_+_8433172_8446622_RNA45SN1_45S_50ntFlanks@9623.9649.27\|TAGTAGCTGGTTCCCTCCGAAGTTTCC | >hg38_21_+_8433172_8446622_RNA45SN1_45S_50ntFlanks | 9623 | 9649 | 27 | TAGTAGCTGGTTCCCTCCGAAGTTTCC | 97.2 | -10.5 | 1.5 | -7.1 | 9.5E-13 | 3.7E-12 | MB231/MB468 | MP |
| >hg38_21_+_8433172_8446622_RNA45SN1_45S_50ntFlanks@9534.9563.30\|AGGAAACTCTGGTGGAGGTCCGTAGCGGTC | >hg38_21_+_8433172_8446622_RNA45SN1_45S_50ntFlanks | 9534 | 9563 | 30 | AGGAAACTCTGGTGGAGGTCCGTAGCGGTC | 8388.6 | -10.2 | 1.4 | -7.1 | 1.4E-12 | 3.7E-12 | MB231/MB468 | MITO |
| >hg38_21_+_8433172_8446622_RNA45SN1_45S_50ntFlanks@4592.4614.23\|TTTGTTGGTTTTCGGAACTGAGG | >hg38_21_+_8433172_8446622_RNA45SN1_45S_50ntFlanks | 4592 | 4614 | 23 | TTTGTTGGTTTTCGGAACTGAGG | 111.6 | -10.7 | 1.5 | -7.1 | 9.5E-13 | 3.7E-12 | MB231/MB468 | MP |
| >hg38_21_+_8433172_8446622_RNA45SN1_45S_50ntFlanks@8182.8210.29\|ATCGAGGCCCAGCCCGTGGACGGTGTGAG | >hg38_21_+_8433172_8446622_RNA45SN1_45S_50ntFlanks | 8182 | 8210 | 29 | ATCGAGGCCCAGCCCGTGGACGGTGTGAG | 172.4 | -9.7 | 1.4 | -7.1 | 1.1E-12 | 3.7E-12 | BT20/MB468 | CYTO |
| >hg38_21_+_8433172_8446622_RNA45SN1_45S_50ntFlanks@10793.10826.34\|TCTGGCATGTTGGAACAATGTAGGTAAGGGAAGT | >hg38_21_+_8433172_8446622_RNA45SN1_45S_50ntFlanks | 10793 | 10826 | 34 | TCTGGCATGTTGGAACAATGTAGGTAAGGGAAGT | 133.0 | -10.9 | 1.5 | -7.1 | 9.5E-13 | 3.7E-12 | MB231/MB468 | MP |
| >hg38_21_+_8433172_8446622_RNA45SN1_45S_50ntFlanks@3813.3842.30\|TTATGGTTCCTTTGGTCGCTCGCTCCTCTC | >hg38_21_+_8433172_8446622_RNA45SN1_45S_50ntFlanks | 3813 | 3842 | 30 | TTATGGTTCCTTTGGTCGCTCGCTCCTCTC | 259.3 | -10.1 | 1.4 | -7.2 | 8.1E-13 | 3.7E-12 | BT20/MB468 | MITO |
| >MI0000462\|hsa-mir-152&WithFlank&17\|-\|48037155\|48037253@60.79.20(+1A)&[MIMAT0000438&hsa-miR-152-3p&offsets\|0\|-1(+1A);m-97&17\|-\|48037174\|48037194&offsets\|0\|-1(+1A)]\|TCAGTGCATGACAGAACTTGA | >MI0000462\|hsa-mir-152&WithFlank&17\|-\|48037155\|48037253 | 60 | 79 | 20(+1A) | TCAGTGCATGACAGAACTTGA | 173.1 | 10.7 | 1.5 | 7.1 | 9.6E-13 | 3.8E-12 | MB231/MB468 | MP |
| >hg38_21_+_8433172_8446622_RNA45SN1_45S_50ntFlanks@4225.4242.18\|AAATAACAATACAGGACT | >hg38_21_+_8433172_8446622_RNA45SN1_45S_50ntFlanks | 4225 | 4242 | 18 | AAATAACAATACAGGACT | 760.4 | -5.5 | 0.8 | -7.1 | 1.4E-12 | 3.8E-12 | MB231/MB468 | MITO |
| >hg38_21_+_8433172_8446622_RNA45SN1_45S_50ntFlanks@11602.11619.18\|TCCGACTGTTTAATTAAA | >hg38_21_+_8433172_8446622_RNA45SN1_45S_50ntFlanks | 11602 | 11619 | 18 | TCCGACTGTTTAATTAAA | 2651.3 | -10.8 | 1.5 | -7.1 | 1.5E-12 | 3.8E-12 | BT20/MB468 | NUC |
| >hg38_21_+_8433172_8446622_RNA45SN1_45S_50ntFlanks@7985.8007.23\|GATCAGACGTGGCGACCCGCTGA | >hg38_21_+_8433172_8446622_RNA45SN1_45S_50ntFlanks | 7985 | 8007 | 23 | GATCAGACGTGGCGACCCGCTGA | 296.2 | -10.3 | 1.4 | -7.2 | 8.2E-13 | 3.8E-12 | BT20/MB468 | MITO |
| >hg38_21_+_8433172_8446622_RNA45SN1_45S_50ntFlanks@10873.10893.21\|CTGGGTCGGTCGGGCTGGGGC | >hg38_21_+_8433172_8446622_RNA45SN1_45S_50ntFlanks | 10873 | 10893 | 21 | CTGGGTCGGTCGGGCTGGGGC | 159.0 | -11.2 | 1.6 | -7.1 | 9.7E-13 | 3.8E-12 | MB231/MB468 | MP |
| >hg38_21_+_8433172_8446622_RNA45SN1_45S_50ntFlanks@12456.12481.26\|CCAAGCGTTGGATTGTTCACCCACTA | >hg38_21_+_8433172_8446622_RNA45SN1_45S_50ntFlanks | 12456 | 12481 | 26 | CCAAGCGTTGGATTGTTCACCCACTA | 907.5 | -8.7 | 1.2 | -7.1 | 1.4E-12 | 3.8E-12 | MB231/MB468 | MITO |
| >hg38_21_+_8433172_8446622_RNA45SN1_45S_50ntFlanks@6738.6760.23\|ATCATCGACACTTCGAACGCACT | >hg38_21_+_8433172_8446622_RNA45SN1_45S_50ntFlanks | 6738 | 6760 | 23 | ATCATCGACACTTCGAACGCACT | 101.1 | -10.5 | 1.5 | -7.1 | 9.7E-13 | 3.8E-12 | MB231/MB468 | MP |
| >MI0000433\|hsa-let-7g&WithFlank&3\|-\|52268272\|52268367@11.32.22(+1C)&[MIMAT0000414&hsa-let-7g-5p&offsets\|0\|0(+1C);m-58&3\|-\|52268336\|52268357&offsets\|0\|0(+1C)]\|TGAGGTAGTAGTTTGTACAGTTC | >MI0000433\|hsa-let-7g&WithFlank&3\|-\|52268272\|52268367 | 11 | 32 | 22(+1C) | TGAGGTAGTAGTTTGTACAGTTC | 190.4 | 4.8 | 0.7 | 7.1 | 1.1E-12 | 3.8E-12 | MB231/MB468 | NUC |
| >hg38_21_+_8433172_8446622_RNA45SN1_45S_50ntFlanks@12710.12743.34\|CGCGGAGCCTCGGTTGGCCTCGGATAGCCGGTCC | >hg38_21_+_8433172_8446622_RNA45SN1_45S_50ntFlanks | 12710 | 12743 | 34 | CGCGGAGCCTCGGTTGGCCTCGGATAGCCGGTCC | 115.0 | -10.7 | 1.5 | -7.1 | 9.7E-13 | 3.8E-12 | MB231/MB468 | MP |
| >hg38_21_+_8433172_8446622_RNA45SN1_45S_50ntFlanks@3793.3812.20\|CGAATGGCTCATTAAATCAG | >hg38_21_+_8433172_8446622_RNA45SN1_45S_50ntFlanks | 3793 | 3812 | 20 | CGAATGGCTCATTAAATCAG | 116.8 | -10.7 | 1.5 | -7.1 | 9.7E-13 | 3.8E-12 | MB231/MB468 | MP |
| >hg38_21_+_8433172_8446622_RNA45SN1_45S_50ntFlanks@6649.6670.22\|TACGACTCTTAGCGGTGGATCA | >hg38_21_+_8433172_8446622_RNA45SN1_45S_50ntFlanks | 6649 | 6670 | 22 | TACGACTCTTAGCGGTGGATCA | 130.9 | -10.9 | 1.5 | -7.1 | 9.8E-13 | 3.8E-12 | MB231/MB468 | MP |
| >trna146_GlnCTG_6_-_27515531_27515602@17.53.37,trna1_GlnCTG_6_+_18836402_18836473@17.53.37,trna3_GlnCTG_17_+_8023070_8023141@17.53.37,trna49_GlnCTG_6_+_27487308_27487379@17.53.37,trna7_GlnCTG_15_-_66161400_66161471@17.53.37,trna99_GlnCTG_6_-_28909378_28909449@17.53.37\|GGTTAGCACTCTGGACTCTGAATCCAGCGATCCGAGT | >trna146_GlnCTG_6_-_27515531_27515602 | 17 | 53 | 37 | GGTTAGCACTCTGGACTCTGAATCCAGCGATCCGAGT | 567.6 | 10.6 | 1.5 | 7.1 | 1.1E-12 | 3.8E-12 | BT20/MB468 | CYTO |
| >MI0000266\|hsa-mir-10a&WithFlank&17\|-\|48579832\|48579953@28.49.22(+1A)&[MIMAT0000253&hsa-miR-10a-5p&offsets\|0\|-1(+1A);m-18&17\|-\|48579905\|48579926&offsets\|0\|0(+1A)]\|TACCCTGTAGATCCGAATTTGTA | >MI0000266\|hsa-mir-10a&WithFlank&17\|-\|48579832\|48579953 | 28 | 49 | 22(+1A) | TACCCTGTAGATCCGAATTTGTA | 739.2 | 13.3 | 1.8 | 7.5 | 5.1E-14 | 3.8E-12 | BT20/MB468 | MP |
| >trna10_AspGTC_12_-_125424193_125424264@1.34.34,trna12_AspGTC_12_-_125411891_125411962@1.34.34,trna38_AspGTC_17_-_8125556_8125627@1.34.34,trna45_AspGTC_6_+_27447453_27447524@1.34.34,trna48_AspGTC_6_+_27471523_27471594@1.34.34,trna4_AspGTC_12_+_96429799_96429870@1.34.34,trna69_AspGTC_1_-_161440205_161440276@1.34.34,trna72_AspGTC_1_-_161432824_161432895@1.34.34,trna75_AspGTC_1_-_161425414_161425485@1.34.34,trna78_AspGTC_1_-_161418033_161418104@1.34.34,trna81_AspGTC_1_-_161410615_161410686@1.34.34\|TCCTCGTTAGTATAGTGGTGAGTATCCCCGCCTG | >trna10_AspGTC_12_-_125424193_125424264 | 1 | 34 | 34 | TCCTCGTTAGTATAGTGGTGAGTATCCCCGCCTG | 1387.0 | 9.5 | 1.3 | 7.1 | 1.1E-12 | 3.8E-12 | MB231/MB468 | CYTO |
| >MI0000440\|hsa-mir-27b&WithFlank&9\|+\|95085439\|95085547@67.85.19&[MIMAT0000419&hsa-miR-27b-3p&offsets\|0\|-2;m-39&9\|+\|95085505\|95085525&offsets\|0\|-2]\|TTCACAGTGGCTAAGTTCT | >MI0000440\|hsa-mir-27b&WithFlank&9\|+\|95085439\|95085547 | 67 | 85 | 19 | TTCACAGTGGCTAAGTTCT | 1744.2 | 5.1 | 0.7 | 7.1 | 1.2E-12 | 3.8E-12 | MB231/MB468 | NUC |
| >hg38_21_+_8433172_8446622_RNA45SN1_45S_50ntFlanks@5500.5526.27\|GTCGAACTTGACTATCTAGAGGAAGTA | >hg38_21_+_8433172_8446622_RNA45SN1_45S_50ntFlanks | 5500 | 5526 | 27 | GTCGAACTTGACTATCTAGAGGAAGTA | 324.8 | -12.1 | 1.7 | -7.1 | 1.4E-12 | 3.9E-12 | MB231/MB468 | MITO |
| >trna10_LysCTT_16_+_3241501_3241573@1.34.34,trna13_LysCTT_14_-_58706613_58706685@1.34.34,trna2_LysCTT_15_+_79152904_79152976@1.34.34\|GCCCGGCTAGCTCAGTCGGTAGAGCATGGGACTC | >trna10_LysCTT_16_+_3241501_3241573 | 1 | 34 | 34 | GCCCGGCTAGCTCAGTCGGTAGAGCATGGGACTC | 5661.4 | 8.6 | 1.2 | 7.1 | 1.1E-12 | 3.9E-12 | MB231/MB468 | CYTO |
| >MI0003513\|hsa-mir-455&WithFlank&9\|+\|114209428\|114209535@22.43.22(+1C)&[MIMAT0003150&hsa-miR-455-5p&offsets\|0\|0(+1C);m-232&9\|+\|114209449\|114209470&offsets\|0\|0(+1C)]\|TATGTGCCTTTGGACTACATCGC | >MI0003513\|hsa-mir-455&WithFlank&9\|+\|114209428\|114209535 | 22 | 43 | 22(+1C) | TATGTGCCTTTGGACTACATCGC | 146.1 | 10.4 | 1.5 | 7.1 | 9.9E-13 | 3.9E-12 | MB231/MB468 | MP |
| >MI0001641\|hsa-mir-429&WithFlank&1\|+\|1168999\|1169093@57.78.22&[MIMAT0001536&hsa-miR-429&offsets\|0\|0;m-207&1\|+\|1169055\|1169076&offsets\|0\|0]\|TAATACTGTCTGGTAAAACCGT | >MI0001641\|hsa-mir-429&WithFlank&1\|+\|1168999\|1169093 | 57 | 78 | 22 | TAATACTGTCTGGTAAAACCGT | 324.7 | -6.1 | 0.9 | -7.1 | 1.1E-12 | 3.9E-12 | MB231/MB468 | CYTO |
| >hg38_21_+_8433172_8446622_RNA45SN1_45S_50ntFlanks@6722.6771.50\|ATTGCAGGACACATTGATCATCGACACTTCGAACGCACTTGCGGCCCCGG | >hg38_21_+_8433172_8446622_RNA45SN1_45S_50ntFlanks | 6722 | 6771 | 50 | ATTGCAGGACACATTGATCATCGACACTTCGAACGCACTTGCGGCCCCGG | 291.2 | -12.0 | 1.7 | -7.1 | 1.4E-12 | 3.9E-12 | MB231/MB468 | MITO |
| >hg38_21_+_8433172_8446622_RNA45SN1_45S_50ntFlanks@11874.11910.37\|AGAAGACCCTGTTGAGCTTGACTCTAGTCTGGCACGG | >hg38_21_+_8433172_8446622_RNA45SN1_45S_50ntFlanks | 11874 | 11910 | 37 | AGAAGACCCTGTTGAGCTTGACTCTAGTCTGGCACGG | 869.4 | -4.0 | 0.6 | -7.2 | 8.4E-13 | 3.9E-12 | BT20/MB468 | MITO |
| >hg38_21_+_8433172_8446622_RNA45SN1_45S_50ntFlanks@10845.10862.18\|ACTTCGGGATAAGGATTG | >hg38_21_+_8433172_8446622_RNA45SN1_45S_50ntFlanks | 10845 | 10862 | 18 | ACTTCGGGATAAGGATTG | 1337.0 | -6.7 | 0.9 | -7.1 | 1.1E-12 | 3.9E-12 | MB231/MB468 | CYTO |
| >hg38_21_+_8433172_8446622_RNA45SN1_45S_50ntFlanks@10830.10856.27\|CAAGCCGGATCCGTAACTTCGGGATAA | >hg38_21_+_8433172_8446622_RNA45SN1_45S_50ntFlanks | 10830 | 10856 | 27 | CAAGCCGGATCCGTAACTTCGGGATAA | 174.6 | -11.3 | 1.6 | -7.1 | 9.9E-13 | 3.9E-12 | MB231/MB468 | MP |
| >hg38_21_+_8433172_8446622_RNA45SN1_45S_50ntFlanks@12150.12182.33\|GGTGGGGAGTTTGACTGGGGCGGTACACCTGTC | >hg38_21_+_8433172_8446622_RNA45SN1_45S_50ntFlanks | 12150 | 12182 | 33 | GGTGGGGAGTTTGACTGGGGCGGTACACCTGTC | 115.7 | -10.7 | 1.5 | -7.1 | 9.9E-13 | 3.9E-12 | MB231/MB468 | MP |
| >hg38_21_+_8433172_8446622_RNA45SN1_45S_50ntFlanks@6719.6750.32\|TGAATTGCAGGACACATTGATCATCGACACTT | >hg38_21_+_8433172_8446622_RNA45SN1_45S_50ntFlanks | 6719 | 6750 | 32 | TGAATTGCAGGACACATTGATCATCGACACTT | 154.7 | -11.1 | 1.6 | -7.1 | 9.9E-13 | 3.9E-12 | MB231/MB468 | MP |

| | | | | | | | | | | | | | |
|---|---|---|---|---|---|---|---|---|---|---|---|---|---|
| >trna11_GluTTC_15_-_26327381_26327452@39.73.35,trna3_GluTTC_13_-_45492062_45492133@39.73.35\|CCAGGCGGCCCGGGTTCGACTCCCGGTGTGGGAAC | >trna11_GluTTC_15_-_26327381_26327452 | 39 | 73 | 35 | CCAGGCGGCCCGGGTTCGACTCCCGGTGTGGGAAC | 286.2 | 8.7 | 1.2 | 7.1 | 1.1E-12 | 3.9E-12 | MB231/MB468 | CYTO |
| >am_trna116_GluCTC_1_-_145399233_145399304@1.23.23,trna59_GluCTC_1_+_249168447_249168518@1.23.23,trna71_GluCTC_1_-_161439189_161439260@1.23.23,trna74_GluCTC_1_-_161431809_161431880@1.23.23,trna77_GluCTC_1_-_161424398_161424469@1.23.23,trna77_GluCTC_6_+_28949976_28950047@1.23.23,trna80_GluCTC_1_-_161417018_161417089@1.23.23,trna87_GluCTC_6_-_126101393_126101464@1.23.23\|TCCCTGGTGGTCTAGTGGTTAGG | >am_trna116_GluCTC_1_-_145399233_145399304 | 1 | 23 | 23 | TCCCTGGTGGTCTAGTGGTTAGG | 371.8 | 4.9 | 0.7 | 7.1 | 1.1E-12 | 3.9E-12 | BT20/MB468 | CYTO |
| >MI0000298\|hsa-mir-221&WithFlank&X\|-\|45746151\|45746272@31.54.24&[MIMAT0004568&hsa-miR-221-5p&offsets\|0\|+2;m-405&X\|-\|45746220\|45746242&offsets\|0\|+1]\|ACCTGGCATACAATGTAGATTTCT | >MI0000298\|hsa-mir-221&WithFlank&X\|-\|45746151\|45746272 | 31 | 54 | 24 | ACCTGGCATACAATGTAGATTTCT | 1883.6 | 6.7 | 0.9 | 7.1 | 1.4E-12 | 3.9E-12 | MB231/MB468 | MITO |
| >MI0000477\|hsa-mir-146a&WithFlank&5\|+\|160485346\|160485456@26.47.22&[MIMAT0000449&hsa-miR-146a-5p&offsets\|-1\|-1;m-129&5\|+\|160485372\|160485393&offsets\|-1\|-1]\|TTGAGAACTGAATTCCATGGGT | >MI0000477\|hsa-mir-146a&WithFlank&5\|+\|160485346\|160485456 | 26 | 47 | 22 | TTGAGAACTGAATTCCATGGGT | 248.7 | 11.2 | 1.6 | 7.1 | 1.0E-12 | 3.9E-12 | MB231/MB468 | MP |
| >MI0000809\|hsa-mir-151a&WithFlank&8\|-\|140732558\|140732659@51.72.22(+1U)&[MIMAT0000757&hsa-miR-151a-3p&offsets\|-2\|-1(+1U);m-30&8\|-\|140732587\|140732607&offsets\|-2\|-1(+1U)]\|TACTAGACTGAAGCTCCTTGAGT | >MI0000809\|hsa-mir-151a&WithFlank&8\|-\|140732558\|140732659 | 51 | 72 | 22(+1U) | TACTAGACTGAAGCTCCTTGAGT | 305.4 | 6.5 | 0.9 | 7.1 | 1.1E-12 | 3.9E-12 | MB231/MB468 | CYTO |
| >hg38_21_+_8433172_8446622_RNA45SN1_45S_50ntFlanks@8170.8196.27\|CAAGTCCTTCTGATCGAGGCCCAGCCC | >hg38_21_+_8433172_8446622_RNA45SN1_45S_50ntFlanks | 8170 | 8196 | 27 | CAAGTCCTTCTGATCGAGGCCCAGCCC | 693.8 | -9.0 | 1.3 | -7.1 | 1.4E-12 | 3.9E-12 | MB231/MB468 | MITO |
| >hg38_21_+_8433172_8446622_RNA45SN1_45S_50ntFlanks@11498.11516.19\|CCGGTCCCCCCCGCCGGGT | >hg38_21_+_8433172_8446622_RNA45SN1_45S_50ntFlanks | 11498 | 11516 | 19 | CCGGTCCCCCCCGCCGGGT | 203.9 | -11.5 | 1.6 | -7.1 | 1.0E-12 | 3.9E-12 | MB231/MB468 | MP |
| >hg38_21_+_8433172_8446622_RNA45SN1_45S_50ntFlanks@11575.11599.25\|ACTTAGAACTGGTGCGGACCAGGGG | >hg38_21_+_8433172_8446622_RNA45SN1_45S_50ntFlanks | 11575 | 11599 | 25 | ACTTAGAACTGGTGCGGACCAGGGG | 220.0 | -11.6 | 1.6 | -7.1 | 1.0E-12 | 3.9E-12 | MB231/MB468 | MP |
| >hg38_21_+_8433172_8446622_RNA45SN1_45S_50ntFlanks@9576.9596.21\|TCGGTCGTCCGACCTGGGTAT | >hg38_21_+_8433172_8446622_RNA45SN1_45S_50ntFlanks | 9576 | 9596 | 21 | TCGGTCGTCCGACCTGGGTAT | 546.4 | -12.2 | 1.6 | -7.5 | 5.2E-14 | 3.9E-12 | BT20/MB468 | MP |
| >MI0003776\|hsa-mir-320b-1&WithFlank&1\|+\|116671740\|116671823@48.68.21;MI0003839\|hsa-mir-320b-2&WithFlank&1\|-\|224257034\|224257116@47.67.21&[MIMAT0005792&hsa-miR-320b&offsets\|0\|-1;m-183&1\|+\|116671787\|116671806&offsets\|0\|+1];[MIMAT0005792_1&hsa-miR-320b&offsets\|0\|-1;m-183&1\|-\|224257051\|224257070&offsets\|0\|+1]\|AAAAGCTGGGTTGAGAGGGCA | >MI0003776\|hsa-mir-320b-1&WithFlank&1\|+\|116671740\|116671823 | 48 | 68 | 21 | AAAAGCTGGGTTGAGAGGGCA | 266.3 | 3.3 | 0.5 | 7.1 | 1.6E-12 | 3.9E-12 | BT20/MB468 | NUC |
| >hg38_21_+_8433172_8446622_RNA45SN1_45S_50ntFlanks@10409.10431.23\|GTCAGTCGGTCCTGAGAGATGGG | >hg38_21_+_8433172_8446622_RNA45SN1_45S_50ntFlanks | 10409 | 10431 | 23 | GTCAGTCGGTCCTGAGAGATGGG | 144.5 | -11.0 | 1.5 | -7.1 | 1.0E-12 | 3.9E-12 | MB231/MB468 | MP |
| >hg38_21_+_8433172_8446622_RNA45SN1_45S_50ntFlanks@6698.6719.22\|AGCTAGCTGCGAGAATTAATGT | >hg38_21_+_8433172_8446622_RNA45SN1_45S_50ntFlanks | 6698 | 6719 | 22 | AGCTAGCTGCGAGAATTAATGT | 3649.0 | -10.9 | 1.5 | -7.1 | 1.6E-12 | 3.9E-12 | BT20/MB468 | NUC |
| >hg38_21_+_8433172_8446622_RNA45SN1_45S_50ntFlanks@11572.11592.21\|CCGACTTAGAACTGGTGCGGA | >hg38_21_+_8433172_8446622_RNA45SN1_45S_50ntFlanks | 11572 | 11592 | 21 | CCGACTTAGAACTGGTGCGGA | 886.1 | -6.7 | 0.9 | -7.1 | 1.2E-12 | 3.9E-12 | MB231/MB468 | CYTO |
| >hg38_21_+_8433172_8446622_RNA45SN1_45S_50ntFlanks@11580.11602.23\|GAACTGGTGCGGACCAGGGGAAT | >hg38_21_+_8433172_8446622_RNA45SN1_45S_50ntFlanks | 11580 | 11602 | 23 | GAACTGGTGCGGACCAGGGGAAT | 615.7 | -6.9 | 1.0 | -7.1 | 1.4E-12 | 3.9E-12 | MB231/MB468 | MITO |
| >hg38_21_+_8433172_8446622_RNA45SN1_45S_50ntFlanks@10402.10420.19\|AACATGGGTCAGTCGGTCC | >hg38_21_+_8433172_8446622_RNA45SN1_45S_50ntFlanks | 10402 | 10420 | 19 | AACATGGGTCAGTCGGTCC | 2528.3 | -9.7 | 1.4 | -7.1 | 1.0E-12 | 3.9E-12 | MB231/MB468 | MP |
| >MI0000066\|hsa-let-7e&WithFlank&19\|+\|51692780\|51692870@59.80.22&[MIMAT0004485&hsa-let-7e-3p&offsets\|0\|0;m-260&19\|+\|51692838\|51692859&offsets\|0\|0]\|CTATACGGCCTCCTAGCTTTCC | >MI0000066\|hsa-let-7e&WithFlank&19\|+\|51692780\|51692870 | 59 | 80 | 22 | CTATACGGCCTCCTAGCTTTCC | 291.3 | 8.1 | 1.1 | 7.1 | 1.1E-12 | 3.9E-12 | BT20/MB468 | CYTO |
| >hg38_1_-_228634819_228635039_RNA5S12_5S_50ntFlanks@141.169.29\|CCGCCTGGGAATACCGGGTGCTGTAGGCT | >hg38_1_-_228634819_228635039_RNA5S12_5S_50ntFlanks | 141 | 169 | 29 | CCGCCTGGGAATACCGGGTGCTGTAGGCT | 2628.5 | 5.0 | 0.7 | 7.1 | 1.1E-12 | 3.9E-12 | BT20/MB468 | CYTO |
| >hg38_21_+_8433172_8446622_RNA45SN1_45S_50ntFlanks@12154.12182.29\|GGGAGTTTGACTGGGGCGGTACACCTGTC | >hg38_21_+_8433172_8446622_RNA45SN1_45S_50ntFlanks | 12154 | 12182 | 29 | GGGAGTTTGACTGGGGCGGTACACCTGTC | 1067.5 | -7.6 | 1.1 | -7.1 | 1.0E-12 | 4.0E-12 | MB231/MB468 | MP |
| >hg38_21_+_8433172_8446622_RNA45SN1_45S_50ntFlanks@10534.10564.31\|CCGCGAGGCGTCCAGTGCGGTAACGCGACCG | >hg38_21_+_8433172_8446622_RNA45SN1_45S_50ntFlanks | 10534 | 10564 | 31 | CCGCGAGGCGTCCAGTGCGGTAACGCGACCG | 423.9 | -11.8 | 1.6 | -7.5 | 5.3E-14 | 4.0E-12 | BT20/MB468 | MP |
| >hg38_21_+_8433172_8446622_RNA45SN1_45S_50ntFlanks@8191.8214.24\|CAGCCCGTGGACGGTGTGAGGCCG | >hg38_21_+_8433172_8446622_RNA45SN1_45S_50ntFlanks | 8191 | 8214 | 24 | CAGCCCGTGGACGGTGTGAGGCCG | 3377.7 | -11.1 | 1.6 | -7.1 | 1.6E-12 | 4.0E-12 | BT20/MB468 | NUC |
| >hg38_21_+_8433172_8446622_RNA45SN1_45S_50ntFlanks@10410.10443.34\|TCAGTCGGTCCTGAGAGATGGGCGAGCGCCGTTC | >hg38_21_+_8433172_8446622_RNA45SN1_45S_50ntFlanks | 10410 | 10443 | 34 | TCAGTCGGTCCTGAGAGATGGGCGAGCGCCGTTC | 701.8 | -9.4 | 1.3 | -7.1 | 1.5E-12 | 4.0E-12 | MB231/MB468 | MITO |
| >hg38_21_+_8433172_8446622_RNA45SN1_45S_50ntFlanks@6651.6681.31\|CGACTCTTAGCGGTGGATCACTCGGCTCGTG | >hg38_21_+_8433172_8446622_RNA45SN1_45S_50ntFlanks | 6651 | 6681 | 31 | CGACTCTTAGCGGTGGATCACTCGGCTCGTG | 65023.4 | -3.6 | 0.5 | -7.1 | 1.6E-12 | 4.0E-12 | BT20/MB468 | NUC |
| >hg38_21_+_8433172_8446622_RNA45SN1_45S_50ntFlanks@10808.10826.19\|CAATGTAGGTAAGGGAAGT | >hg38_21_+_8433172_8446622_RNA45SN1_45S_50ntFlanks | 10808 | 10826 | 19 | CAATGTAGGTAAGGGAAGT | 2584.4 | -10.9 | 1.5 | -7.1 | 1.6E-12 | 4.0E-12 | BT20/MB468 | NUC |
| >hg38_21_+_8433172_8446622_RNA45SN1_45S_50ntFlanks@482.530.49\|ATCGATGTGGTGACGTCGTGCTCTCCCGGGCCGGGTCCGAGCCGCGACG | >hg38_21_+_8433172_8446622_RNA45SN1_45S_50ntFlanks | 482 | 530 | 49 | ATCGATGTGGTGACGTCGTGCTCTCCCGGGCCGGGTCCGAGCCGCGACG | 1212.2 | -14.7 | 2.1 | -7.1 | 1.6E-12 | 4.0E-12 | BT20/MB468 | NUC |
| >hg38_21_+_8433172_8446622_RNA45SN1_45S_50ntFlanks@12126.12149.24\|CGACCCGCTCCGGGGACAGTGCCA | >hg38_21_+_8433172_8446622_RNA45SN1_45S_50ntFlanks | 12126 | 12149 | 24 | CGACCCGCTCCGGGGACAGTGCCA | 124.4 | -10.8 | 1.5 | -7.1 | 1.0E-12 | 4.0E-12 | MB231/MB468 | MP |
| >hg38_21_+_8433172_8446622_RNA45SN1_45S_50ntFlanks@11104.11129.26\|AAGGGTCGGGGCGGCAGGGGCCGGCG | >hg38_21_+_8433172_8446622_RNA45SN1_45S_50ntFlanks | 11104 | 11129 | 26 | AAGGGTCGGGGCGGCAGGGGCCGGCG | 170.0 | -11.3 | 1.6 | -7.1 | 1.0E-12 | 4.0E-12 | MB231/MB468 | MP |
| >hg38_21_+_8433172_8446622_RNA45SN1_45S_50ntFlanks@8115.8139.25\|CGCGGGACATGTGGCGTACGGAAGA | >hg38_21_+_8433172_8446622_RNA45SN1_45S_50ntFlanks | 8115 | 8139 | 25 | CGCGGGACATGTGGCGTACGGAAGA | 203.2 | 11.3 | 1.5 | 7.4 | 1.2E-13 | 4.0E-12 | BT20/MB231 | MITO |
| >MI0003599\|hsa-mir-589&WithFlank&7\|-\|5495813\|5495923@30.52.23&[MIMAT0004799&hsa-miR-589-5p&offsets\|0\|+1;m-162&7\|-\|5495874\|5495894&offsets\|0\|+2]\|TGAGAACCACGTCTGCTCTGAGC | >MI0003599\|hsa-mir-589&WithFlank&7\|-\|5495813\|5495923 | 30 | 52 | 23 | TGAGAACCACGTCTGCTCTGAGC | 1083.0 | 10.8 | 1.5 | 7.1 | 8.7E-13 | 4.0E-12 | BT20/MB468 | MITO |
| >MI0000253\|hsa-mir-148a&WithFlank&7\|-\|25949913\|25949992@12.33.22&[MIMAT0004549&hsa-miR-148a-5p&offsets\|0\|0;m-189&7\|-\|25949960\|25949981&offsets\|0\|0]\|AAAGTTCTGAGACACTCCGACT | >MI0000253\|hsa-mir-148a&WithFlank&7\|-\|25949913\|25949992 | 12 | 33 | 22 | AAAGTTCTGAGACACTCCGACT | 197.2 | 3.2 | 0.5 | 7.1 | 1.6E-12 | 4.0E-12 | BT20/MB468 | NUC |
| >hg38_21_+_8433172_8446622_RNA45SN1_45S_50ntFlanks@12638.12656.19\|CTACCATCTGTGGGATTAT | >hg38_21_+_8433172_8446622_RNA45SN1_45S_50ntFlanks | 12638 | 12656 | 19 | CTACCATCTGTGGGATTAT | 97.8 | -10.5 | 1.5 | -7.1 | 1.0E-12 | 4.0E-12 | MB231/MB468 | MP |
| >hg38_21_+_8433172_8446622_RNA45SN1_45S_50ntFlanks@10682.10711.30\|TCCGGCGGCGTCCGGTGAGCTCTCGCTGGC | >hg38_21_+_8433172_8446622_RNA45SN1_45S_50ntFlanks | 10682 | 10711 | 30 | TCCGGCGGCGTCCGGTGAGCTCTCGCTGGC | 101.2 | -10.5 | 1.5 | -7.1 | 1.0E-12 | 4.0E-12 | MB231/MB468 | MP |
| >hg38_21_+_8433172_8446622_RNA45SN1_45S_50ntFlanks@6719.6742.24\|TGAATTGCAGGACACATTGATCAT | >hg38_21_+_8433172_8446622_RNA45SN1_45S_50ntFlanks | 6719 | 6742 | 24 | TGAATTGCAGGACACATTGATCAT | 781.1 | -7.2 | 1.0 | -7.1 | 1.2E-12 | 4.0E-12 | MB231/MB468 | NUC |
| >hg38_1_-_228634819_228635039_RNA5S12_5S_50ntFlanks@138.169.32\|AGACCGCCTGGGAATACCGGGTGCTGTAGGCT | >hg38_1_-_228634819_228635039_RNA5S12_5S_50ntFlanks | 138 | 169 | 32 | AGACCGCCTGGGAATACCGGGTGCTGTAGGCT | 519.2 | 4.5 | 0.6 | 7.1 | 8.8E-13 | 4.0E-12 | BT20/MB468 | MITO |
| >hg38_21_+_8433172_8446622_RNA45SN1_45S_50ntFlanks@11036.11060.25\|TCTCTCCCCCGCTCCCCGTCCTCCC | >hg38_21_+_8433172_8446622_RNA45SN1_45S_50ntFlanks | 11036 | 11060 | 25 | TCTCTCCCCCGCTCCCCGTCCTCCC | 351.9 | -11.4 | 1.6 | -7.1 | 1.2E-12 | 4.0E-12 | MB231/MB468 | NUC |
| >hg38_21_+_8433172_8446622_RNA45SN1_45S_50ntFlanks@11027.11061.35\|CTCTCTCTCTCTCTCCCCCGCTCCCCGTCCTCCCC | >hg38_21_+_8433172_8446622_RNA45SN1_45S_50ntFlanks | 11027 | 11061 | 35 | CTCTCTCTCTCTCTCCCCCGCTCCCCGTCCTCCCC | 375.9 | -7.8 | 1.1 | -7.1 | 1.2E-12 | 4.0E-12 | MB231/MB468 | CYTO |
| >hg38_21_+_8433172_8446622_RNA45SN1_45S_50ntFlanks@10389.10435.47\|TGAACAGCAGTTGAACATGGGTCAGTCGGTCCTGAGAGATGGGCGAG | >hg38_21_+_8433172_8446622_RNA45SN1_45S_50ntFlanks | 10389 | 10435 | 47 | TGAACAGCAGTTGAACATGGGTCAGTCGGTCCTGAGAGATGGGCGAG | 424.6 | -13.2 | 1.9 | -7.1 | 1.6E-12 | 4.0E-12 | BT20/MB468 | NUC |
| >hg38_21_+_8433172_8446622_RNA45SN1_45S_50ntFlanks@6652.6689.38\|GACTCTTAGCGGTGGATCACTCGGCTCGTGCGTCGATG | >hg38_21_+_8433172_8446622_RNA45SN1_45S_50ntFlanks | 6652 | 6689 | 38 | GACTCTTAGCGGTGGATCACTCGGCTCGTGCGTCGATG | 78845.5 | -5.3 | 0.7 | -7.1 | 1.5E-12 | 4.0E-12 | MB231/MB468 | MITO |
| >hg38_21_+_8433172_8446622_RNA45SN1_45S_50ntFlanks@9534.9570.37\|AGGAAACTCTGGTGGAGGTCCGTAGCGGTCCTGACGT | >hg38_21_+_8433172_8446622_RNA45SN1_45S_50ntFlanks | 9534 | 9570 | 37 | AGGAAACTCTGGTGGAGGTCCGTAGCGGTCCTGACGT | 1411.5 | -5.8 | 0.8 | -7.1 | 1.5E-12 | 4.0E-12 | MB231/MB468 | MITO |
| >hg38_21_+_8433172_8446622_RNA45SN1_45S_50ntFlanks@11037.11060.24\|CTCTCCCCCGCTCCCCGTCCTCCC | >hg38_21_+_8433172_8446622_RNA45SN1_45S_50ntFlanks | 11037 | 11060 | 24 | CTCTCCCCCGCTCCCCGTCCTCCC | 1796.8 | -10.2 | 1.4 | -7.1 | 1.2E-12 | 4.0E-12 | MB231/MB468 | NUC |
| >hg38_21_+_8433172_8446622_RNA45SN1_45S_50ntFlanks@6666.6689.24\|GATCACTCGGCTCGTGCGTCGATG | >hg38_21_+_8433172_8446622_RNA45SN1_45S_50ntFlanks | 6666 | 6689 | 24 | GATCACTCGGCTCGTGCGTCGATG | 237.6 | -10.0 | 1.4 | -7.1 | 8.8E-13 | 4.0E-12 | BT20/MB468 | MITO |
| >hg38_21_+_8433172_8446622_RNA45SN1_45S_50ntFlanks@12072.12093.22\|CCCGAGGGGCTCTCGCTTCTGG | >hg38_21_+_8433172_8446622_RNA45SN1_45S_50ntFlanks | 12072 | 12093 | 22 | CCCGAGGGGCTCTCGCTTCTGG | 119.3 | -10.8 | 1.5 | -7.1 | 1.0E-12 | 4.0E-12 | MB231/MB468 | MP |
| >hg38_21_+_8433172_8446622_RNA45SN1_45S_50ntFlanks@5492.5514.23\|AGAAGACGGTCGAACTTGACTAT | >hg38_21_+_8433172_8446622_RNA45SN1_45S_50ntFlanks | 5492 | 5514 | 23 | AGAAGACGGTCGAACTTGACTAT | 2355.6 | -8.3 | 1.2 | -7.1 | 1.5E-12 | 4.0E-12 | MB231/MB468 | MITO |
| >hg38_21_+_8433172_8446622_RNA45SN1_45S_50ntFlanks@5449.5466.18\|ATCGGCCCCGCCGGGGTC | >hg38_21_+_8433172_8446622_RNA45SN1_45S_50ntFlanks | 5449 | 5466 | 18 | ATCGGCCCCGCCGGGGTC | 185.9 | -9.8 | 1.4 | -7.1 | 1.2E-12 | 4.0E-12 | BT20/MB468 | CYTO |
| >hg38_21_+_8433172_8446622_RNA45SN1_45S_50ntFlanks@10357.10383.27\|CTTTGAAGGCCGAAGTGGAGAAGGGTT | >hg38_21_+_8433172_8446622_RNA45SN1_45S_50ntFlanks | 10357 | 10383 | 27 | CTTTGAAGGCCGAAGTGGAGAAGGGTT | 795.3 | -7.5 | 1.1 | -7.1 | 1.0E-12 | 4.0E-12 | MB231/MB468 | MP |
| >hg38_21_+_8433172_8446622_RNA45SN1_45S_50ntFlanks@12936.12966.31\|AAACCATTCGTAGACGACCTGCTTCTGGGTC | >hg38_21_+_8433172_8446622_RNA45SN1_45S_50ntFlanks | 12936 | 12966 | 31 | AAACCATTCGTAGACGACCTGCTTCTGGGTC | 184.4 | -11.4 | 1.6 | -7.1 | 1.0E-12 | 4.0E-12 | MB231/MB468 | MP |
| >hg38_21_+_8433172_8446622_RNA45SN1_45S_50ntFlanks@10905.10939.35\|CTGGGCGCGCGCCGCGGCTGGACGAGGCGCCGCCG | >hg38_21_+_8433172_8446622_RNA45SN1_45S_50ntFlanks | 10905 | 10939 | 35 | CTGGGCGCGCGCCGCGGCTGGACGAGGCGCCGCCG | 249.9 | -10.7 | 1.5 | -7.1 | 1.2E-12 | 4.1E-12 | MB231/MB468 | CYTO |

<table>
<tr><td>>MI0000542|hsa-mir-320a&WithFlank&8|-|22244960|22245043@49.70.22&[MIMAT0000510&hsa-miR-320a-3p&offsets|+1|+1;m-60&8|-|22244975|22244996&offsets|+1|+1]|AAAGCTGGGTTGAGAGGGCGAA</td><td>>MI0000542|hsa-mir-320a&WithFlank&8|-|22244960|22245043</td><td>49</td><td>70</td><td>22</td><td>AAAGCTGGGTTGAGAGGGCGAA</td><td>402.7</td><td>9.9</td><td>1.4</td><td>7.1</td><td>8.9E-13</td><td>4.1E-12</td><td>BT20/MB468</td><td>MITO</td></tr>
<tr><td>>hg38_21_+_8433172_8446622_RNA45SN1_45S_50ntFlanks@12418.12443.26|CGATGTCGGCTCTTCCTATCATTGTG</td><td>>hg38_21_+_8433172_8446622_RNA45SN1_45S_50ntFlanks</td><td>12418</td><td>12443</td><td>26</td><td>CGATGTCGGCTCTTCCTATCATTGTG</td><td>6335.5</td><td>-4.3</td><td>0.6</td><td>-7.1</td><td>8.9E-13</td><td>4.1E-12</td><td>BT20/MB468</td><td>MITO</td></tr>
<tr><td>>hg38_21_+_8433172_8446622_RNA45SN1_45S_50ntFlanks@8191.8215.25|CAGCCCGTGGACGGTGTGAGGCCGG</td><td>>hg38_21_+_8433172_8446622_RNA45SN1_45S_50ntFlanks</td><td>8191</td><td>8215</td><td>25</td><td>CAGCCCGTGGACGGTGTGAGGCCGG</td><td>982.1</td><td>-5.5</td><td>0.7</td><td>-7.5</td><td>5.4E-14</td><td>4.1E-12</td><td>BT20/MB468</td><td>MP</td></tr>
<tr><td>>hg38_21_+_8433172_8446622_RNA45SN1_45S_50ntFlanks@11208.11229.22|GCGGCGACTCTGGACGCGAGCC</td><td>>hg38_21_+_8433172_8446622_RNA45SN1_45S_50ntFlanks</td><td>11208</td><td>11229</td><td>22</td><td>GCGGCGACTCTGGACGCGAGCC</td><td>806.1</td><td>-4.6</td><td>0.6</td><td>-7.1</td><td>1.2E-12</td><td>4.1E-12</td><td>MB231/MB468</td><td>NUC</td></tr>
<tr><td>>MI0000811|hsa-mir-148b&WithFlank&12|+|54337210|54337320@69.89.21&[MIMAT0000759&hsa-miR-148b-3p&offsets|0|-1;m-77&12|+|54337278|54337299&offsets|0|-1]|TCAGTGCATCACAGAACTTTG</td><td>>MI0000811|hsa-mir-148b&WithFlank&12|+|54337210|54337320</td><td>69</td><td>89</td><td>21</td><td>TCAGTGCATCACAGAACTTTG</td><td>481.4</td><td>6.0</td><td>0.8</td><td>7.1</td><td>1.2E-12</td><td>4.1E-12</td><td>MB231/MB468</td><td>NUC</td></tr>
<tr><td>>hg38_21_+_8433172_8446622_RNA45SN1_45S_50ntFlanks@10408.10432.25|GGTCAGTCGGTCCTGAGAGATGGGC</td><td>>hg38_21_+_8433172_8446622_RNA45SN1_45S_50ntFlanks</td><td>10408</td><td>10432</td><td>25</td><td>GGTCAGTCGGTCCTGAGAGATGGGC</td><td>1392.6</td><td>-9.5</td><td>1.3</td><td>-7.1</td><td>1.2E-12</td><td>4.1E-12</td><td>MB231/MB468</td><td>NUC</td></tr>
<tr><td>>hg38_21_+_8433172_8446622_RNA45SN1_45S_50ntFlanks@11925.11949.25|AGGTGTAGAATAAGTGGGAGGCCCC</td><td>>hg38_21_+_8433172_8446622_RNA45SN1_45S_50ntFlanks</td><td>11925</td><td>11949</td><td>25</td><td>AGGTGTAGAATAAGTGGGAGGCCCC</td><td>102.9</td><td>-10.5</td><td>1.5</td><td>-7.1</td><td>1.0E-12</td><td>4.1E-12</td><td>MB231/MB468</td><td>MP</td></tr>
<tr><td>>hg38_21_+_8433172_8446622_RNA45SN1_45S_50ntFlanks@4473.4493.21|TTAGCTGAGTGTCCCGCGGGG</td><td>>hg38_21_+_8433172_8446622_RNA45SN1_45S_50ntFlanks</td><td>4473</td><td>4493</td><td>21</td><td>TTAGCTGAGTGTCCCGCGGGG</td><td>161.0</td><td>-10.3</td><td>1.4</td><td>-7.1</td><td>1.2E-12</td><td>4.1E-12</td><td>MB231/MB468</td><td>NUC</td></tr>
<tr><td>>hg38_21_+_8433172_8446622_RNA45SN1_45S_50ntFlanks@9419.9452.34|TCCGCCGAGGGCGCACCACCGGCCCGTCTCGCCC</td><td>>hg38_21_+_8433172_8446622_RNA45SN1_45S_50ntFlanks</td><td>9419</td><td>9452</td><td>34</td><td>TCCGCCGAGGGCGCACCACCGGCCCGTCTCGCCC</td><td>106.7</td><td>-10.6</td><td>1.5</td><td>-7.1</td><td>1.0E-12</td><td>4.1E-12</td><td>MB231/MB468</td><td>MP</td></tr>
<tr><td>>hg38_21_+_8433172_8446622_RNA45SN1_45S_50ntFlanks@10408.10430.23|GGTCAGTCGGTCCTGAGAGATGG</td><td>>hg38_21_+_8433172_8446622_RNA45SN1_45S_50ntFlanks</td><td>10408</td><td>10430</td><td>23</td><td>GGTCAGTCGGTCCTGAGAGATGG</td><td>257.7</td><td>-10.1</td><td>1.4</td><td>-7.1</td><td>9.0E-13</td><td>4.1E-12</td><td>BT20/MB468</td><td>MITO</td></tr>
<tr><td>>MI0000289|hsa-mir-181a-1&WithFlank&1|-|198859038|198859159@70.90.21(+1A)&[MIMAT0000270&hsa-miR-181a-3p&offsets|0|-1(+1A);m-125&1|-|198859069|198859090&offsets|0|-1(+1A)]|ACCATCGACCGTTGATTGTACA</td><td>>MI0000289|hsa-mir-181a-1&WithFlank&1|-|198859038|198859159</td><td>70</td><td>90</td><td>21(+1A)</td><td>ACCATCGACCGTTGATTGTACA</td><td>408.7</td><td>8.8</td><td>1.2</td><td>7.1</td><td>1.2E-12</td><td>4.1E-12</td><td>MB231/MB468</td><td>CYTO</td></tr>
<tr><td>>MI0000085|hsa-mir-27a&WithFlank&19|-|13836434|13836523@57.76.20(+1A)&[MIMAT0000084&hsa-miR-27a-3p&offsets|0|-1(+1A);m-43&19|-|13836447|13836467&offsets|0|-1(+1A)]|TTCACAGTGGCTAAGTTCCGA</td><td>>MI0000085|hsa-mir-27a&WithFlank&19|-|13836434|13836523</td><td>57</td><td>76</td><td>20(+1A)</td><td>TTCACAGTGGCTAAGTTCCGA</td><td>2149.2</td><td>5.6</td><td>0.8</td><td>7.1</td><td>1.2E-12</td><td>4.1E-12</td><td>BT20/MB468</td><td>CYTO</td></tr>
<tr><td>>hg38_21_+_8433172_8446622_RNA45SN1_45S_50ntFlanks@12159.12183.25|TTTGACTGGGGCGGTACACCTGTCA</td><td>>hg38_21_+_8433172_8446622_RNA45SN1_45S_50ntFlanks</td><td>12159</td><td>12183</td><td>25</td><td>TTTGACTGGGGCGGTACACCTGTCA</td><td>643.1</td><td>-12.4</td><td>1.7</td><td>-7.5</td><td>5.6E-14</td><td>4.1E-12</td><td>BT20/MB468</td><td>MP</td></tr>
<tr><td>>hg38_21_+_8433172_8446622_RNA45SN1_45S_50ntFlanks@10534.10564.31|CCGCGAGGCGTCCAGTGCGGTAACGCGACCG</td><td>>hg38_21_+_8433172_8446622_RNA45SN1_45S_50ntFlanks</td><td>10534</td><td>10564</td><td>31</td><td>CCGCGAGGCGTCCAGTGCGGTAACGCGACCG</td><td>235.5</td><td>-11.7</td><td>1.6</td><td>-7.1</td><td>1.5E-12</td><td>4.1E-12</td><td>MB231/MB468</td><td>MITO</td></tr>
<tr><td>>hg38_21_+_8433172_8446622_RNA45SN1_45S_50ntFlanks@4760.4778.19|TCAGATACCGTCGTAGTTC</td><td>>hg38_21_+_8433172_8446622_RNA45SN1_45S_50ntFlanks</td><td>4760</td><td>4778</td><td>19</td><td>TCAGATACCGTCGTAGTTC</td><td>298.4</td><td>-10.3</td><td>1.4</td><td>-7.1</td><td>9.0E-13</td><td>4.1E-12</td><td>BT20/MB468</td><td>MITO</td></tr>
<tr><td>>hg38_21_+_8433172_8446622_RNA45SN1_45S_50ntFlanks@11570.11605.36|AGCCGACTTAGAACTGGTGCGGACCAGGGGAATCCG</td><td>>hg38_21_+_8433172_8446622_RNA45SN1_45S_50ntFlanks</td><td>11570</td><td>11605</td><td>36</td><td>AGCCGACTTAGAACTGGTGCGGACCAGGGGAATCCG</td><td>838.6</td><td>-9.3</td><td>1.3</td><td>-7.1</td><td>1.5E-12</td><td>4.1E-12</td><td>MB231/MB468</td><td>MITO</td></tr>
<tr><td>>hg38_21_+_8433172_8446622_RNA45SN1_45S_50ntFlanks@8335.8373.39|CAAGTACCGTAAGGGAAAGTTGAAAAGAACTTTGAAGAG</td><td>>hg38_21_+_8433172_8446622_RNA45SN1_45S_50ntFlanks</td><td>8335</td><td>8373</td><td>39</td><td>CAAGTACCGTAAGGGAAAGTTGAAAAGAACTTTGAAGAG</td><td>327.9</td><td>12.1</td><td>1.6</td><td>7.5</td><td>5.6E-14</td><td>4.1E-12</td><td>BT20/MB468</td><td>MP</td></tr>
<tr><td>>hg38_21_+_8433172_8446622_RNA45SN1_45S_50ntFlanks@12072.12106.35|CCCGAGGGGCTCTCGCTTCTGGCGCCAAGCGCCCG</td><td>>hg38_21_+_8433172_8446622_RNA45SN1_45S_50ntFlanks</td><td>12072</td><td>12106</td><td>35</td><td>CCCGAGGGGCTCTCGCTTCTGGCGCCAAGCGCCCG</td><td>199.5</td><td>-7.1</td><td>1.0</td><td>-7.1</td><td>1.2E-12</td><td>4.1E-12</td><td>MB231/MB468</td><td>CYTO</td></tr>
<tr><td>>hg38_21_+_8433172_8446622_RNA45SN1_45S_50ntFlanks@9533.9563.31|GAGGAAACTCTGGTGGAGGTCCGTAGCGGTC</td><td>>hg38_21_+_8433172_8446622_RNA45SN1_45S_50ntFlanks</td><td>9533</td><td>9563</td><td>31</td><td>GAGGAAACTCTGGTGGAGGTCCGTAGCGGTC</td><td>2086.9</td><td>-6.8</td><td>1.0</td><td>-7.1</td><td>1.1E-12</td><td>4.1E-12</td><td>MB231/MB468</td><td>MP</td></tr>
<tr><td>>hg38_21_+_8433172_8446622_RNA45SN1_45S_50ntFlanks@11853.11873.21|CTTGGCGGAATCAGCGGGGAA</td><td>>hg38_21_+_8433172_8446622_RNA45SN1_45S_50ntFlanks</td><td>11853</td><td>11873</td><td>21</td><td>CTTGGCGGAATCAGCGGGGAA</td><td>98.5</td><td>-10.5</td><td>1.5</td><td>-7.1</td><td>1.1E-12</td><td>4.1E-12</td><td>MB231/MB468</td><td>MP</td></tr>
<tr><td>>MI0000088|hsa-mir-30a&WithFlank&6|-|71403545|71403627@12.35.24&[MIMAT0000087&hsa-miR-30a-5p&offsets|0|+2;m-15&6|-|71403595|71403616&offsets|0|+2]|TGTAAACATCCTCGACTGGAAGCT</td><td>>MI0000088|hsa-mir-30a&WithFlank&6|-|71403545|71403627</td><td>12</td><td>35</td><td>24</td><td>TGTAAACATCCTCGACTGGAAGCT</td><td>142148.4</td><td>7.3</td><td>1.0</td><td>7.1</td><td>1.5E-12</td><td>4.1E-12</td><td>MB231/MB468</td><td>MITO</td></tr>
<tr><td>>hg38_21_+_8433172_8446622_RNA45SN1_45S_50ntFlanks@3875.3899.25|ATACATGCCGACGGGCGCTGACCCC</td><td>>hg38_21_+_8433172_8446622_RNA45SN1_45S_50ntFlanks</td><td>3875</td><td>3899</td><td>25</td><td>ATACATGCCGACGGGCGCTGACCCC</td><td>701.3</td><td>-4.2</td><td>0.6</td><td>-7.1</td><td>9.1E-13</td><td>4.1E-12</td><td>BT20/MB468</td><td>MITO</td></tr>
<tr><td>>trnaMT_PheGAA_MT_+_577_647@43.74.32|TTAGACGGGCTCACATCACCCCATAAACACCA</td><td>>trnaMT_PheGAA_MT_+_577_647</td><td>43</td><td>74</td><td>32</td><td>TTAGACGGGCTCACATCACCCCATAAACACCA</td><td>313.5</td><td>9.5</td><td>1.3</td><td>7.1</td><td>1.2E-12</td><td>4.1E-12</td><td>BT20/MB468</td><td>CYTO</td></tr>
<tr><td>>hg38_21_+_8433172_8446622_RNA45SN1_45S_50ntFlanks@5073.5098.26|AATTCCGATAACGAACGAGACTCTGG</td><td>>hg38_21_+_8433172_8446622_RNA45SN1_45S_50ntFlanks</td><td>5073</td><td>5098</td><td>26</td><td>AATTCCGATAACGAACGAGACTCTGG</td><td>4778.6</td><td>-11.0</td><td>1.6</td><td>-7.1</td><td>1.5E-12</td><td>4.1E-12</td><td>MB231/MB468</td><td>MITO</td></tr>
<tr><td>>MI0000073|hsa-mir-19a&WithFlank&13|+|91350885|91350978@55.78.24&[MIMAT0000073&hsa-miR-19a-3p&offsets|0|+1;m-201&13|+|91350939|91350961&offsets|0|+1]|TGTGCAAATCTATGCAAAACTGAT</td><td>>MI0000073|hsa-mir-19a&WithFlank&13|+|91350885|91350978</td><td>55</td><td>78</td><td>24</td><td>TGTGCAAATCTATGCAAAACTGAT</td><td>176.6</td><td>6.6</td><td>0.9</td><td>7.1</td><td>1.3E-12</td><td>4.1E-12</td><td>MB231/MB468</td><td>NUC</td></tr>
<tr><td>>MI0000088|hsa-mir-30a&WithFlank&6|-|71403545|71403627@54.73.20(+2U)&[MIMAT0000088&hsa-miR-30a-3p&offsets|+1|-1(+2U);m-38&6|-|71403554|71403575&offsets|+1|-1(+2U)]|TTTCAGTCGGATGTTTGCAGTT</td><td>>MI0000088|hsa-mir-30a&WithFlank&6|-|71403545|71403627</td><td>54</td><td>73</td><td>20(+2U)</td><td>TTTCAGTCGGATGTTTGCAGTT</td><td>281.6</td><td>11.4</td><td>1.6</td><td>7.1</td><td>1.5E-12</td><td>4.1E-12</td><td>MB231/MB468</td><td>MITO</td></tr>
<tr><td>>hg38_21_+_8433172_8446622_RNA45SN1_45S_50ntFlanks@4367.4396.30|CAGTTAAAAAGCTCGTAGTTGGATCTTGGG</td><td>>hg38_21_+_8433172_8446622_RNA45SN1_45S_50ntFlanks</td><td>4367</td><td>4396</td><td>30</td><td>CAGTTAAAAAGCTCGTAGTTGGATCTTGGG</td><td>174.9</td><td>-11.3</td><td>1.6</td><td>-7.1</td><td>1.1E-12</td><td>4.1E-12</td><td>MB231/MB468</td><td>MP</td></tr>
<tr><td>>hg38_21_+_8433172_8446622_RNA45SN1_45S_50ntFlanks@3822.3850.29|CTTTGGTCGCTCGCTCCTCTCCTACTTGG</td><td>>hg38_21_+_8433172_8446622_RNA45SN1_45S_50ntFlanks</td><td>3822</td><td>3850</td><td>29</td><td>CTTTGGTCGCTCGCTCCTCTCCTACTTGG</td><td>515.7</td><td>-9.2</td><td>1.3</td><td>-7.1</td><td>1.7E-12</td><td>4.2E-12</td><td>BT20/MB468</td><td>NUC</td></tr>
<tr><td>>hg38_21_+_8433172_8446622_RNA45SN1_45S_50ntFlanks@4378.4396.19|CTCGTAGTTGGATCTTGGG</td><td>>hg38_21_+_8433172_8446622_RNA45SN1_45S_50ntFlanks</td><td>4378</td><td>4396</td><td>19</td><td>CTCGTAGTTGGATCTTGGG</td><td>131.3</td><td>-10.9</td><td>1.5</td><td>-7.1</td><td>1.1E-12</td><td>4.2E-12</td><td>MB231/MB468</td><td>MP</td></tr>
<tr><td>>MI0000446|hsa-mir-125b-1&WithFlank&11|-|122099751|122099850@21.43.23(+1U)&[MIMAT0000423&hsa-miR-125b-5p&offsets|0|+1(+1U);m-67&11|-|122099809|122099830&offsets|0|+1(+1U)]|TCCCTGAGACCCTAACTTGTGATT</td><td>>MI0000446|hsa-mir-125b-1&WithFlank&11|-|122099751|122099850</td><td>21</td><td>43</td><td>23(+1U)</td><td>TCCCTGAGACCCTAACTTGTGATT</td><td>416.1</td><td>11.9</td><td>1.7</td><td>7.1</td><td>1.5E-12</td><td>4.2E-12</td><td>MB231/MB468</td><td>MITO</td></tr>
<tr><td>>hg38_21_+_8433172_8446622_RNA45SN1_45S_50ntFlanks@6790.6807.18|CCTGTCTGAGCGTCGCTT</td><td>>hg38_21_+_8433172_8446622_RNA45SN1_45S_50ntFlanks</td><td>6790</td><td>6807</td><td>18</td><td>CCTGTCTGAGCGTCGCTT</td><td>1673.4</td><td>4.2</td><td>0.6</td><td>7.1</td><td>1.2E-12</td><td>4.2E-12</td><td>MB231/MB468</td><td>CYTO</td></tr>
<tr><td>>hg38_21_+_8433172_8446622_RNA45SN1_45S_50ntFlanks@6744.6770.27|GACACTTCGAACGCACTTGCGGCCCCG</td><td>>hg38_21_+_8433172_8446622_RNA45SN1_45S_50ntFlanks</td><td>6744</td><td>6770</td><td>27</td><td>GACACTTCGAACGCACTTGCGGCCCCG</td><td>2845.2</td><td>-7.1</td><td>1.0</td><td>-7.1</td><td>1.2E-12</td><td>4.2E-12</td><td>MB231/MB468</td><td>CYTO</td></tr>
<tr><td>>MI0000102|hsa-mir-100&WithFlank&11|-|122152223|122152314@19.41.23&[MIMAT0000098&hsa-miR-100-5p&offsets|0|+1;m-27&11|-|122152276|122152296&offsets|0|+2]|AACCCGTAGATCCGAACTTGTGG</td><td>>MI0000102|hsa-mir-100&WithFlank&11|-|122152223|122152314</td><td>19</td><td>41</td><td>23</td><td>AACCCGTAGATCCGAACTTGTGG</td><td>2341.9</td><td>-6.4</td><td>0.9</td><td>-7.4</td><td>1.2E-13</td><td>4.2E-12</td><td>BT20/MB231</td><td>MITO</td></tr>
<tr><td>>hg38_21_+_8433172_8446622_RNA45SN1_45S_50ntFlanks@12008.12039.32|CCGCCGGTGAAATACCACTACTCTGATCGTTT</td><td>>hg38_21_+_8433172_8446622_RNA45SN1_45S_50ntFlanks</td><td>12008</td><td>12039</td><td>32</td><td>CCGCCGGTGAAATACCACTACTCTGATCGTTT</td><td>1088.3</td><td>-10.1</td><td>1.4</td><td>-7.1</td><td>1.7E-12</td><td>4.2E-12</td><td>BT20/MB468</td><td>NUC</td></tr>
<tr><td>>MI0000087|hsa-mir-29a&WithFlank&7|-|130876741|130876816@47.68.22(+2U)&[MIMAT0000086&hsa-miR-29a-3p&offsets|-1|-1(+2U);m-26&7|-|130876749|130876769&offsets|-1|0(+2U)]|CTAGCACCATCTGAAATCGGTTTT</td><td>>MI0000087|hsa-mir-29a&WithFlank&7|-|130876741|130876816</td><td>47</td><td>68</td><td>22(+2U)</td><td>CTAGCACCATCTGAAATCGGTTTT</td><td>186.3</td><td>10.8</td><td>1.5</td><td>7.1</td><td>1.1E-12</td><td>4.2E-12</td><td>MB231/MB468</td><td>MP</td></tr>
<tr><td>>hg38_21_+_8433172_8446622_RNA45SN1_45S_50ntFlanks@12957.12977.21|CTTCTGGGTCGGGGTTTCGTA</td><td>>hg38_21_+_8433172_8446622_RNA45SN1_45S_50ntFlanks</td><td>12957</td><td>12977</td><td>21</td><td>CTTCTGGGTCGGGGTTTCGTA</td><td>152.8</td><td>-11.1</td><td>1.6</td><td>-7.1</td><td>1.1E-12</td><td>4.2E-12</td><td>MB231/MB468</td><td>MP</td></tr>
<tr><td>>hg38_1_-_228634819_228635039_RNA5S12_5S_50ntFlanks@137.171.35|GAGACCGCCTGGGAATACCGGGTGCTGTAGGCTTT</td><td>>hg38_1_-_228634819_228635039_RNA5S12_5S_50ntFlanks</td><td>137</td><td>171</td><td>35</td><td>GAGACCGCCTGGGAATACCGGGTGCTGTAGGCTTT</td><td>1759.5</td><td>5.8</td><td>0.8</td><td>7.1</td><td>1.3E-12</td><td>4.2E-12</td><td>MB231/MB468</td><td>NUC</td></tr>
<tr><td>>hg38_21_+_8433172_8446622_RNA45SN1_45S_50ntFlanks@12973.13001.29|TCGTACGTAGCAGAGCAGCTCCCTCGCTG</td><td>>hg38_21_+_8433172_8446622_RNA45SN1_45S_50ntFlanks</td><td>12973</td><td>13001</td><td>29</td><td>TCGTACGTAGCAGAGCAGCTCCCTCGCTG</td><td>199.5</td><td>-9.7</td><td>1.4</td><td>-7.1</td><td>9.2E-13</td><td>4.2E-12</td><td>BT20/MB468</td><td>MITO</td></tr>
<tr><td>>hg38_21_+_8433172_8446622_RNA45SN1_45S_50ntFlanks@8191.8215.25|CAGCCCGTGGACGGTGTGAGGCCGG</td><td>>hg38_21_+_8433172_8446622_RNA45SN1_45S_50ntFlanks</td><td>8191</td><td>8215</td><td>25</td><td>CAGCCCGTGGACGGTGTGAGGCCGG</td><td>2322.4</td><td>-9.6</td><td>1.4</td><td>-7.1</td><td>1.3E-12</td><td>4.2E-12</td><td>MB231/MB468</td><td>NUC</td></tr>
<tr><td>>hg38_21_+_8433172_8446622_RNA45SN1_45S_50ntFlanks@3766.3798.33|GTACGCACGGCCGGTACAGTGAAACTGCGAATG</td><td>>hg38_21_+_8433172_8446622_RNA45SN1_45S_50ntFlanks</td><td>3766</td><td>3798</td><td>33</td><td>GTACGCACGGCCGGTACAGTGAAACTGCGAATG</td><td>404.0</td><td>7.1</td><td>1.0</td><td>7.1</td><td>1.2E-12</td><td>4.2E-12</td><td>MB231/MB468</td><td>CYTO</td></tr>
<tr><td>>hg38_21_+_8433172_8446622_RNA45SN1_45S_50ntFlanks@12129.12146.18|CCCGCTCCGGGGACAGTG</td><td>>hg38_21_+_8433172_8446622_RNA45SN1_45S_50ntFlanks</td><td>12129</td><td>12146</td><td>18</td><td>CCCGCTCCGGGGACAGTG</td><td>151.6</td><td>-9.6</td><td>1.3</td><td>-7.1</td><td>1.2E-12</td><td>4.2E-12</td><td>BT20/MB468</td><td>CYTO</td></tr>
<tr><td>>hg38_21_+_8433172_8446622_RNA45SN1_45S_50ntFlanks@11921.11954.34|TGAGAGGTGTAGAATAAGTGGGAGGCCCCCGGCG</td><td>>hg38_21_+_8433172_8446622_RNA45SN1_45S_50ntFlanks</td><td>11921</td><td>11954</td><td>34</td><td>TGAGAGGTGTAGAATAAGTGGGAGGCCCCCGGCG</td><td>188.5</td><td>-11.4</td><td>1.6</td><td>-7.1</td><td>1.1E-12</td><td>4.2E-12</td><td>MB231/MB468</td><td>MP</td></tr>
<tr><td>>hg38_21_+_8433172_8446622_RNA45SN1_45S_50ntFlanks@10479.10511.33|CGATCGAAAGGGAGTCGGGTTCAGATCCCCGAA</td><td>>hg38_21_+_8433172_8446622_RNA45SN1_45S_50ntFlanks</td><td>10479</td><td>10511</td><td>33</td><td>CGATCGAAAGGGAGTCGGGTTCAGATCCCCGAA</td><td>145.4</td><td>-9.5</td><td>1.3</td><td>-7.1</td><td>1.2E-12</td><td>4.2E-12</td><td>BT20/MB468</td><td>CYTO</td></tr>
<tr><td>>hg38_21_+_8433172_8446622_RNA45SN1_45S_50ntFlanks@12974.12991.18|CGTACGTAGCAGAGCAGC</td><td>>hg38_21_+_8433172_8446622_RNA45SN1_45S_50ntFlanks</td><td>12974</td><td>12991</td><td>18</td><td>CGTACGTAGCAGAGCAGC</td><td>3547.0</td><td>-4.3</td><td>0.6</td><td>-7.1</td><td>1.3E-12</td><td>4.3E-12</td><td>MB231/MB468</td><td>CYTO</td></tr>
<tr><td>>hg38_MT_+_1621_3279_RNR2_16S_50ntFlanks@1438.1477.40|TACGTGATCTGAGTTCAGACCGGAGTAATCCAGGTCGGTT</td><td>>hg38_MT_+_1621_3279_RNR2_16S_50ntFlanks</td><td>1438</td><td>1477</td><td>40</td><td>TACGTGATCTGAGTTCAGACCGGAGTAATCCAGGTCGGTT</td><td>230.4</td><td>8.2</td><td>1.2</td><td>7.1</td><td>1.3E-12</td><td>4.3E-12</td><td>MB231/MB468</td><td>CYTO</td></tr>
<tr><td>>hg38_21_+_8433172_8446622_RNA45SN1_45S_50ntFlanks@12646.12669.24|TGTGGGATTATGACTGAACGCCTC</td><td>>hg38_21_+_8433172_8446622_RNA45SN1_45S_50ntFlanks</td><td>12646</td><td>12669</td><td>24</td><td>TGTGGGATTATGACTGAACGCCTC</td><td>392.2</td><td>8.5</td><td>1.2</td><td>7.1</td><td>1.2E-12</td><td>4.3E-12</td><td>BT20/MB468</td><td>CYTO</td></tr>
<tr><td>>hg38_21_+_8433172_8446622_RNA45SN1_45S_50ntFlanks@4008.4042.35|CTCGGGCCGATCGCACGCCCCCCGTGGCGGCGACG</td><td>>hg38_21_+_8433172_8446622_RNA45SN1_45S_50ntFlanks</td><td>4008</td><td>4042</td><td>35</td><td>CTCGGGCCGATCGCACGCCCCCCGTGGCGGCGACG</td><td>117.8</td><td>-10.7</td><td>1.5</td><td>-7.1</td><td>1.1E-12</td><td>4.3E-12</td><td>MB231/MB468</td><td>MP</td></tr>
<tr><td>>hg38_21_+_8433172_8446622_RNA45SN1_45S_50ntFlanks@12626.12667.42|AATGGGGCGAAGCTACCATCTGTGGGATTATGACTGAACGCC</td><td>>hg38_21_+_8433172_8446622_RNA45SN1_45S_50ntFlanks</td><td>12626</td><td>12667</td><td>42</td><td>AATGGGGCGAAGCTACCATCTGTGGGATTATGACTGAACGCC</td><td>1162.5</td><td>7.7</td><td>1.1</td><td>7.1</td><td>1.3E-12</td><td>4.3E-12</td><td>MB231/MB468</td><td>CYTO</td></tr>
<tr><td>>hg38_21_+_8433172_8446622_RNA45SN1_45S_50ntFlanks@475.503.29|CTTCGTGATCGATGTGGTGACGTCGTGCT</td><td>>hg38_21_+_8433172_8446622_RNA45SN1_45S_50ntFlanks</td><td>475</td><td>503</td><td>29</td><td>CTTCGTGATCGATGTGGTGACGTCGTGCT</td><td>1327.3</td><td>9.6</td><td>1.4</td><td>7.1</td><td>1.2E-12</td><td>4.3E-12</td><td>BT20/MB468</td><td>CYTO</td></tr>
</table>

<table>
<tr><td></td><td></td><td></td><td></td><td></td><td></td><td></td><td></td><td></td><td></td><td></td><td></td><td></td><td></td></tr>
<tr><td>>hg38_21_+_8433172_8446622_RNA45SN1_45S_50ntFlanks@3875.3899.25|ATACATGCCGACGGGCGCTGACCCC</td><td>>hg38_21_+_8433172_8446622_RNA45SN1_45S_50ntFlanks</td><td>3875</td><td>3899</td><td>25</td><td>ATACATGCCGACGGGCGCTGACCCC</td><td>793.0</td><td>-3.1</td><td>0.4</td><td>-7.1</td><td>1.2E-12</td><td>4.3E-12</td><td>BT20/MB468</td><td>CYTO</td></tr>
<tr><td>>hg38_21_+_8433172_8446622_RNA45SN1_45S_50ntFlanks@6719.6751.33|TGAATTGCAGGACACATTGATCATCGACACTTC</td><td>>hg38_21_+_8433172_8446622_RNA45SN1_45S_50ntFlanks</td><td>6719</td><td>6751</td><td>33</td><td>TGAATTGCAGGACACATTGATCATCGACACTTC</td><td>1071.7</td><td>-10.0</td><td>1.4</td><td>-7.1</td><td>1.7E-12</td><td>4.3E-12</td><td>BT20/MB468</td><td>NUC</td></tr>
<tr><td>>hg38_21_+_8433172_8446622_RNA45SN1_45S_50ntFlanks@10357.10376.20|CTTTGAAGGCCGAAGTGGAG</td><td>>hg38_21_+_8433172_8446622_RNA45SN1_45S_50ntFlanks</td><td>10357</td><td>10376</td><td>20</td><td>CTTTGAAGGCCGAAGTGGAG</td><td>133.5</td><td>-10.9</td><td>1.5</td><td>-7.1</td><td>1.1E-12</td><td>4.3E-12</td><td>MB231/MB468</td><td>MP</td></tr>
<tr><td>>MI0000066|hsa-let-7e&WithFlank&19|+|51692780|51692870@14.35.22&[MIMAT0000066&hsa-let-7e-5p&offsets|0|0;m-50&19|+|51692793|51692814&offsets|0|0]|TGAGGTAGGAGGTTGTATAGTT</td><td>>MI0000066|hsa-let-7e&WithFlank&19|+|51692780|51692870</td><td>14</td><td>35</td><td>22</td><td>TGAGGTAGGAGGTTGTATAGTT</td><td>13144.7</td><td>6.3</td><td>0.9</td><td>7.1</td><td>1.3E-12</td><td>4.3E-12</td><td>MB231/MB468</td><td>CYTO</td></tr>
<tr><td>>hg38_21_+_8433172_8446622_RNA45SN1_45S_50ntFlanks@9929.9954.26|GATATAGACAGCAGGACGGTGGCCAT</td><td>>hg38_21_+_8433172_8446622_RNA45SN1_45S_50ntFlanks</td><td>9929</td><td>9954</td><td>26</td><td>GATATAGACAGCAGGACGGTGGCCAT</td><td>252.6</td><td>-11.8</td><td>1.7</td><td>-7.1</td><td>1.1E-12</td><td>4.3E-12</td><td>MB231/MB468</td><td>MP</td></tr>
<tr><td>>hg38_21_+_8433172_8446622_RNA45SN1_45S_50ntFlanks@11561.11580.20|GCGCCTAGCAGCCGACTTAG</td><td>>hg38_21_+_8433172_8446622_RNA45SN1_45S_50ntFlanks</td><td>11561</td><td>11580</td><td>20</td><td>GCGCCTAGCAGCCGACTTAG</td><td>167.5</td><td>-11.3</td><td>1.6</td><td>-7.1</td><td>1.1E-12</td><td>4.3E-12</td><td>MB231/MB468</td><td>MP</td></tr>
<tr><td>>hg38_21_+_8433172_8446622_RNA45SN1_45S_50ntFlanks@3920.3958.39|CATTTATCAGATCAAAACCAACCCGGTCAGCCCCTCTCC</td><td>>hg38_21_+_8433172_8446622_RNA45SN1_45S_50ntFlanks</td><td>3920</td><td>3958</td><td>39</td><td>CATTTATCAGATCAAAACCAACCCGGTCAGCCCCTCTCC</td><td>204.5</td><td>-11.5</td><td>1.6</td><td>-7.1</td><td>1.6E-12</td><td>4.3E-12</td><td>MB231/MB468</td><td>MITO</td></tr>
<tr><td>>hg38_21_+_8433172_8446622_RNA45SN1_45S_50ntFlanks@9252.9273.22|CACGGGGTCGGCGGCGACGTCG</td><td>>hg38_21_+_8433172_8446622_RNA45SN1_45S_50ntFlanks</td><td>9252</td><td>9273</td><td>22</td><td>CACGGGGTCGGCGGCGACGTCG</td><td>594.0</td><td>-9.3</td><td>1.3</td><td>-7.1</td><td>1.6E-12</td><td>4.3E-12</td><td>MB231/MB468</td><td>MITO</td></tr>
<tr><td>>hg38_21_+_8433172_8446622_RNA45SN1_45S_50ntFlanks@11671.11702.32|CCAGTGCTCTGAATGTCAAAGTGAAGAAATTC</td><td>>hg38_21_+_8433172_8446622_RNA45SN1_45S_50ntFlanks</td><td>11671</td><td>11702</td><td>32</td><td>CCAGTGCTCTGAATGTCAAAGTGAAGAAATTC</td><td>3625.0</td><td>-4.0</td><td>0.6</td><td>-7.1</td><td>1.3E-12</td><td>4.3E-12</td><td>MB231/MB468</td><td>NUC</td></tr>
<tr><td>>hg38_21_+_8433172_8446622_RNA45SN1_45S_50ntFlanks@9544.9571.28|GGTGGAGGTCCGTAGCGGTCCTGACGTG</td><td>>hg38_21_+_8433172_8446622_RNA45SN1_45S_50ntFlanks</td><td>9544</td><td>9571</td><td>28</td><td>GGTGGAGGTCCGTAGCGGTCCTGACGTG</td><td>6039.7</td><td>-4.6</td><td>0.7</td><td>-7.1</td><td>1.3E-12</td><td>4.3E-12</td><td>MB231/MB468</td><td>NUC</td></tr>
<tr><td>>MI0000093|hsa-mir-92a-1&WithFlank&13|+|91351308|91351397@54.75.22(+2A);MI0000094|hsa-mir-92a-2&WithFlank&X|-|134169532|134169618@54.75.22(+2A)&[MIMAT0000092&hsa-miR-92a-3p&offsets|0|0(+2A);m-9&13|+|91351361|91351382&offsets|0|0(+2A)];[MIMAT0000092_1&hsa-miR-92a-3p&offsets|0|0(+2A);m-11&X|-|134169544|134169565&offsets|0|0(+2A)]|TATTGCACTTGTCCCGGCCTGTAA</td><td>>MI0000093|hsa-mir-92a-1&WithFlank&13|+|91351308|91351397</td><td>54</td><td>75</td><td>22(+2A)</td><td>TATTGCACTTGTCCCGGCCTGTAA</td><td>582.4</td><td>5.5</td><td>0.8</td><td>7.1</td><td>1.3E-12</td><td>4.3E-12</td><td>MB231/MB468</td><td>CYTO</td></tr>
<tr><td>>hg38_21_+_8433172_8446622_RNA45SN1_45S_50ntFlanks@10391.10408.18|AACAGCAGTTGAACATGG</td><td>>hg38_21_+_8433172_8446622_RNA45SN1_45S_50ntFlanks</td><td>10391</td><td>10408</td><td>18</td><td>AACAGCAGTTGAACATGG</td><td>470.9</td><td>-8.2</td><td>1.2</td><td>-7.1</td><td>1.2E-12</td><td>4.3E-12</td><td>BT20/MB468</td><td>CYTO</td></tr>
<tr><td>>hg38_21_+_8433172_8446622_RNA45SN1_45S_50ntFlanks@11196.11225.30|GGGCCGGCGGCGGCGGCGACTCTGGACGCG</td><td>>hg38_21_+_8433172_8446622_RNA45SN1_45S_50ntFlanks</td><td>11196</td><td>11225</td><td>30</td><td>GGGCCGGCGGCGGCGGCGACTCTGGACGCG</td><td>169.7</td><td>-11.3</td><td>1.6</td><td>-7.1</td><td>1.1E-12</td><td>4.3E-12</td><td>MB231/MB468</td><td>MP</td></tr>
<tr><td>>hg38_21_+_8433172_8446622_RNA45SN1_45S_50ntFlanks@3711.3737.27|GTTGATCCTGCCAGTAGCATATGCTTG</td><td>>hg38_21_+_8433172_8446622_RNA45SN1_45S_50ntFlanks</td><td>3711</td><td>3737</td><td>27</td><td>GTTGATCCTGCCAGTAGCATATGCTTG</td><td>511.5</td><td>-3.8</td><td>0.5</td><td>-7.1</td><td>1.2E-12</td><td>4.4E-12</td><td>BT20/MB468</td><td>CYTO</td></tr>
<tr><td>>hg38_21_+_8433172_8446622_RNA45SN1_45S_50ntFlanks@8079.8102.24|AAGAGCCCAGCGCCGAATCCCCGC</td><td>>hg38_21_+_8433172_8446622_RNA45SN1_45S_50ntFlanks</td><td>8079</td><td>8102</td><td>24</td><td>AAGAGCCCAGCGCCGAATCCCCGC</td><td>347.3</td><td>-12.3</td><td>1.7</td><td>-7.1</td><td>1.1E-12</td><td>4.4E-12</td><td>MB231/MB468</td><td>MP</td></tr>
<tr><td>>am_trnaMT_ValTAC_MT_+_1602_1670@48.67.20|CAACTTAACTTGACCGCTCT</td><td>>am_trnaMT_ValTAC_MT_+_1602_1670</td><td>48</td><td>67</td><td>20</td><td>CAACTTAACTTGACCGCTCT</td><td>1242.0</td><td>5.3</td><td>0.7</td><td>7.1</td><td>1.3E-12</td><td>4.4E-12</td><td>MB231/MB468</td><td>NUC</td></tr>
<tr><td>>MI0000239|hsa-mir-197&WithFlank&1|+|109598887|109598973@54.72.19&[MIMAT0000227&hsa-miR-197-3p&offsets|0|-3;m-99&1|+|109598940|109598961&offsets|0|-3]|TTCACCACCTTCTCCACCC</td><td>>MI0000239|hsa-mir-197&WithFlank&1|+|109598887|109598973</td><td>54</td><td>72</td><td>19</td><td>TTCACCACCTTCTCCACCC</td><td>782.2</td><td>10.3</td><td>1.5</td><td>7.1</td><td>1.3E-12</td><td>4.4E-12</td><td>BT20/MB468</td><td>CYTO</td></tr>
<tr><td>>hg38_21_+_8433172_8446622_RNA45SN1_45S_50ntFlanks@12255.12272.18|CTCGCTTGATCTTGATTT</td><td>>hg38_21_+_8433172_8446622_RNA45SN1_45S_50ntFlanks</td><td>12255</td><td>12272</td><td>18</td><td>CTCGCTTGATCTTGATTT</td><td>111.1</td><td>-10.7</td><td>1.5</td><td>-7.1</td><td>1.1E-12</td><td>4.4E-12</td><td>MB231/MB468</td><td>MP</td></tr>
<tr><td>>hg38_21_+_8433172_8446622_RNA45SN1_45S_50ntFlanks@12676.12705.30|AGAATCCCGCCCAGGCGGAACGATACGGCA</td><td>>hg38_21_+_8433172_8446622_RNA45SN1_45S_50ntFlanks</td><td>12676</td><td>12705</td><td>30</td><td>AGAATCCCGCCCAGGCGGAACGATACGGCA</td><td>142.4</td><td>-11.0</td><td>1.5</td><td>-7.1</td><td>1.1E-12</td><td>4.4E-12</td><td>MB231/MB468</td><td>MP</td></tr>
<tr><td>>MI0000746|hsa-mir-99b&WithFlank&19|+|51692606|51692687@13.34.22(+1C)&[MIMAT0000689&hsa-miR-99b-5p&offsets|0|0(+1C);m-10&19|+|51692618|51692639&offsets|0|0(+1C)]|CACCCGTAGAACCGACCTTGCGC</td><td>>MI0000746|hsa-mir-99b&WithFlank&19|+|51692606|51692687</td><td>13</td><td>34</td><td>22(+1C)</td><td>CACCCGTAGAACCGACCTTGCGC</td><td>257.1</td><td>5.1</td><td>0.7</td><td>7.1</td><td>1.3E-12</td><td>4.4E-12</td><td>MB231/MB468</td><td>NUC</td></tr>
<tr><td>>hg38_21_+_8433172_8446622_RNA45SN1_45S_50ntFlanks@11208.11253.46|GCGGCGACTCTGGACGCGAGCCGGGCCCTTCCCGTGGATCGCCCCA</td><td>>hg38_21_+_8433172_8446622_RNA45SN1_45S_50ntFlanks</td><td>11208</td><td>11253</td><td>46</td><td>GCGGCGACTCTGGACGCGAGCCGGGCCCTTCCCGTGGATCGCCCCA</td><td>183.5</td><td>-10.2</td><td>1.4</td><td>-7.1</td><td>1.3E-12</td><td>4.4E-12</td><td>MB231/MB468</td><td>CYTO</td></tr>
<tr><td>>hg38_21_+_8433172_8446622_RNA45SN1_45S_50ntFlanks@9165.9183.19|CCCGGGCGGGTCGCGCCGT</td><td>>hg38_21_+_8433172_8446622_RNA45SN1_45S_50ntFlanks</td><td>9165</td><td>9183</td><td>19</td><td>CCCGGGCGGGTCGCGCCGT</td><td>113.3</td><td>-10.7</td><td>1.5</td><td>-7.1</td><td>1.1E-12</td><td>4.4E-12</td><td>MB231/MB468</td><td>MP</td></tr>
<tr><td>>hg38_21_+_8433172_8446622_RNA45SN1_45S_50ntFlanks@3921.3959.39|ATTTATCAGATCAAAACCAACCCGGTCAGCCCCTCTCCG</td><td>>hg38_21_+_8433172_8446622_RNA45SN1_45S_50ntFlanks</td><td>3921</td><td>3959</td><td>39</td><td>ATTTATCAGATCAAAACCAACCCGGTCAGCCCCTCTCCG</td><td>357.6</td><td>-11.4</td><td>1.6</td><td>-7.1</td><td>1.3E-12</td><td>4.4E-12</td><td>MB231/MB468</td><td>NUC</td></tr>
<tr><td>>MI0000065|hsa-let-7d&WithFlank&9|+|94178828|94178926@69.89.21&[MIMAT0004484&hsa-let-7d-3p&offsets|+1|0;m-98&9|+|94178895|94178916&offsets|+1|0]|TATACGACCTGCTGCCTTTCT</td><td>>MI0000065|hsa-let-7d&WithFlank&9|+|94178828|94178926</td><td>69</td><td>89</td><td>21</td><td>TATACGACCTGCTGCCTTTCT</td><td>287.3</td><td>8.1</td><td>1.1</td><td>7.1</td><td>1.3E-12</td><td>4.4E-12</td><td>BT20/MB468</td><td>CYTO</td></tr>
<tr><td>>hg38_21_+_8433172_8446622_RNA45SN1_45S_50ntFlanks@12416.12449.34|TTCGATGTCGGCTCTTCCTATCATTGTGAAGCAG</td><td>>hg38_21_+_8433172_8446622_RNA45SN1_45S_50ntFlanks</td><td>12416</td><td>12449</td><td>34</td><td>TTCGATGTCGGCTCTTCCTATCATTGTGAAGCAG</td><td>1174.0</td><td>5.3</td><td>0.7</td><td>7.1</td><td>1.3E-12</td><td>4.4E-12</td><td>MB231/MB468</td><td>CYTO</td></tr>
<tr><td>>hg38_21_+_8433172_8446622_RNA45SN1_45S_50ntFlanks@12619.12652.34|AGGAGCCAATGGGGCGAAGCTACCATCTGTGGGA</td><td>>hg38_21_+_8433172_8446622_RNA45SN1_45S_50ntFlanks</td><td>12619</td><td>12652</td><td>34</td><td>AGGAGCCAATGGGGCGAAGCTACCATCTGTGGGA</td><td>110.9</td><td>-10.7</td><td>1.5</td><td>-7.1</td><td>1.1E-12</td><td>4.4E-12</td><td>MB231/MB468</td><td>MP</td></tr>
<tr><td>>hg38_21_+_8433172_8446622_RNA45SN1_45S_50ntFlanks@12936.12955.20|AAACCATTCGTAGACGACCT</td><td>>hg38_21_+_8433172_8446622_RNA45SN1_45S_50ntFlanks</td><td>12936</td><td>12955</td><td>20</td><td>AAACCATTCGTAGACGACCT</td><td>344.2</td><td>-8.4</td><td>1.2</td><td>-7.1</td><td>1.6E-12</td><td>4.4E-12</td><td>MB231/MB468</td><td>MITO</td></tr>
<tr><td>>hg38_21_+_8433172_8446622_RNA45SN1_45S_50ntFlanks@4403.4423.21|CGGGCGGTCCGCCGCGAGGCG</td><td>>hg38_21_+_8433172_8446622_RNA45SN1_45S_50ntFlanks</td><td>4403</td><td>4423</td><td>21</td><td>CGGGCGGTCCGCCGCGAGGCG</td><td>606.5</td><td>-13.0</td><td>1.8</td><td>-7.1</td><td>1.6E-12</td><td>4.4E-12</td><td>MB231/MB468</td><td>MITO</td></tr>
<tr><td>>hg38_21_+_8433172_8446622_RNA45SN1_45S_50ntFlanks@5443.5466.24|CCTCGGATCGGCCCCGCCGGGGTC</td><td>>hg38_21_+_8433172_8446622_RNA45SN1_45S_50ntFlanks</td><td>5443</td><td>5466</td><td>24</td><td>CCTCGGATCGGCCCCGCCGGGGTC</td><td>1062.7</td><td>-4.7</td><td>0.7</td><td>-7.1</td><td>1.3E-12</td><td>4.4E-12</td><td>MB231/MB468</td><td>NUC</td></tr>
<tr><td>>hg38_21_+_8433172_8446622_RNA45SN1_45S_50ntFlanks@8114.8143.30|GCGCGGGACATGTGGCGTACGGAAGACCCG</td><td>>hg38_21_+_8433172_8446622_RNA45SN1_45S_50ntFlanks</td><td>8114</td><td>8143</td><td>30</td><td>GCGCGGGACATGTGGCGTACGGAAGACCCG</td><td>3282.4</td><td>-9.1</td><td>1.3</td><td>-7.1</td><td>9.7E-13</td><td>4.4E-12</td><td>BT20/MB468</td><td>MITO</td></tr>
<tr><td>>hg38_21_+_8433172_8446622_RNA45SN1_45S_50ntFlanks@4006.4024.19|ACCTCGGGCCGATCGCACG</td><td>>hg38_21_+_8433172_8446622_RNA45SN1_45S_50ntFlanks</td><td>4006</td><td>4024</td><td>19</td><td>ACCTCGGGCCGATCGCACG</td><td>1085.8</td><td>-7.2</td><td>1.0</td><td>-7.1</td><td>1.6E-12</td><td>4.4E-12</td><td>MB231/MB468</td><td>MITO</td></tr>
<tr><td>>trna152_ValCAC_6_-_27248049_27248121@1.32.32|GCTTCTGTAGTGTAGTGGTTATCACGTTCGCC</td><td>>trna152_ValCAC_6_-_27248049_27248121</td><td>1</td><td>32</td><td>32</td><td>GCTTCTGTAGTGTAGTGGTTATCACGTTCGCC</td><td>1547.1</td><td>6.3</td><td>0.9</td><td>7.1</td><td>1.1E-12</td><td>4.4E-12</td><td>MB231/MB468</td><td>MP</td></tr>
<tr><td>>hg38_21_+_8433172_8446622_RNA45SN1_45S_50ntFlanks@6719.6739.21|TGAATTGCAGGACACATTGAT</td><td>>hg38_21_+_8433172_8446622_RNA45SN1_45S_50ntFlanks</td><td>6719</td><td>6739</td><td>21</td><td>TGAATTGCAGGACACATTGAT</td><td>596.9</td><td>-5.7</td><td>0.8</td><td>-7.1</td><td>1.6E-12</td><td>4.4E-12</td><td>MB231/MB468</td><td>MITO</td></tr>
<tr><td>>MI0000102|hsa-mir-100&WithFlank&11|-|122152223|122152314@19.37.19(+1U)&[MIMAT0000098&hsa-miR-100-5p&offsets|0|-3(+1U);m-27&11|-|122152276|122152296&offsets|0|-2(+1U)]|AACCCGTAGATCCGAACTTT</td><td>>MI0000102|hsa-mir-100&WithFlank&11|-|122152223|122152314</td><td>19</td><td>37</td><td>19(+1U)</td><td>AACCCGTAGATCCGAACTTT</td><td>185.8</td><td>11.4</td><td>1.6</td><td>7.1</td><td>1.3E-12</td><td>4.4E-12</td><td>MB231/MB468</td><td>NUC</td></tr>
<tr><td>>MI0000456|hsa-mir-140&WithFlank&16|+|69933075|69933186@68.89.22&[MIMAT0004597&hsa-miR-140-3p&offsets|0|+1;m-45&16|+|69933143|69933164&offsets|-1|-1]|TACCACAGGGTAGAACCACGGA</td><td>>MI0000456|hsa-mir-140&WithFlank&16|+|69933075|69933186</td><td>68</td><td>89</td><td>22</td><td>TACCACAGGGTAGAACCACGGA</td><td>1359.1</td><td>10.1</td><td>1.4</td><td>7.1</td><td>1.3E-12</td><td>4.4E-12</td><td>MB231/MB468</td><td>CYTO</td></tr>
<tr><td>>hg38_21_+_8433172_8446622_RNA45SN1_45S_50ntFlanks@6721.6747.27|AATTGCAGGACACATTGATCATCGACA</td><td>>hg38_21_+_8433172_8446622_RNA45SN1_45S_50ntFlanks</td><td>6721</td><td>6747</td><td>27</td><td>AATTGCAGGACACATTGATCATCGACA</td><td>3666.3</td><td>-8.9</td><td>1.3</td><td>-7.1</td><td>1.8E-12</td><td>4.4E-12</td><td>BT20/MB468</td><td>NUC</td></tr>
<tr><td>>hg38_21_+_8433172_8446622_RNA45SN1_45S_50ntFlanks@5035.5060.26|CATGGCCGTTCTTAGTTGGTGGAGCG</td><td>>hg38_21_+_8433172_8446622_RNA45SN1_45S_50ntFlanks</td><td>5035</td><td>5060</td><td>26</td><td>CATGGCCGTTCTTAGTTGGTGGAGCG</td><td>962.6</td><td>-13.7</td><td>1.9</td><td>-7.1</td><td>1.6E-12</td><td>4.4E-12</td><td>MB231/MB468</td><td>MITO</td></tr>
<tr><td>>hg38_1_-_228634819_228635039_RNA5S12_5S_50ntFlanks@127.169.43|ACTTGGATGGGAGACCGCCTGGGAATACCGGGTGCTGTAGGCT</td><td>>hg38_1_-_228634819_228635039_RNA5S12_5S_50ntFlanks</td><td>127</td><td>169</td><td>43</td><td>ACTTGGATGGGAGACCGCCTGGGAATACCGGGTGCTGTAGGCT</td><td>868.2</td><td>10.0</td><td>1.4</td><td>7.1</td><td>1.3E-12</td><td>4.4E-12</td><td>MB231/MB468</td><td>CYTO</td></tr>
<tr><td>>trna112_GlnCTG_1_-_147737382_147737453@31.50.20,trna131_GlnCTG_6_-_27759135_27759206@31.50.20,trna146_GlnCTG_6_-_27515531_27515602@31.50.20,trna15_GlnCTG_1_+_145963304_145963375@31.50.20,trna19_GlnCTG_1_+_147505038_147505109@31.50.20,trna1_GlnCTG_6_+_18836402_18836473@31.50.20,trna28_GlnCTG_1_+_149186125_149186196@31.50.20,trna3_GlnCTG_17_+_8023070_8023141@31.50.20,trna49_GlnCTG_6_+_27487308_27487379@31.50.20,trna7_GlnCTG_15_-_66161400_66161471@31.50.20,trna99_GlnCTG_6_-_28909378_28909449@31.50.20|ACTCTGAATCCAGCGATCCG</td><td>>trna112_GlnCTG_1_-_147737382_147737453</td><td>31</td><td>50</td><td>20</td><td>ACTCTGAATCCAGCGATCCG</td><td>143.9</td><td>-11.0</td><td>1.6</td><td>-7.1</td><td>1.1E-12</td><td>4.4E-12</td><td>MB231/MB468</td><td>MP</td></tr>
<tr><td>>hg38_21_+_8433172_8446622_RNA45SN1_45S_50ntFlanks@12722.12742.21|GTTGGCCTCGGATAGCCGGTC</td><td>>hg38_21_+_8433172_8446622_RNA45SN1_45S_50ntFlanks</td><td>12722</td><td>12742</td><td>21</td><td>GTTGGCCTCGGATAGCCGGTC</td><td>143.7</td><td>-11.0</td><td>1.6</td><td>-7.1</td><td>1.1E-12</td><td>4.4E-12</td><td>MB231/MB468</td><td>MP</td></tr>
<tr><td>>hg38_21_+_8433172_8446622_RNA45SN1_45S_50ntFlanks@7992.8016.25|CGTGGCGACCCGCTGAATTTAAGCA</td><td>>hg38_21_+_8433172_8446622_RNA45SN1_45S_50ntFlanks</td><td>7992</td><td>8016</td><td>25</td><td>CGTGGCGACCCGCTGAATTTAAGCA</td><td>718.1</td><td>5.8</td><td>0.8</td><td>7.1</td><td>1.3E-12</td><td>4.4E-12</td><td>MB231/MB468</td><td>CYTO</td></tr>
<tr><td>>hg38_21_+_8433172_8446622_RNA45SN1_45S_50ntFlanks@9339.9365.27|CGCACGAAAGCCGCCGTGGCGCAATGA</td><td>>hg38_21_+_8433172_8446622_RNA45SN1_45S_50ntFlanks</td><td>9339</td><td>9365</td><td>27</td><td>CGCACGAAAGCCGCCGTGGCGCAATGA</td><td>208.9</td><td>-11.5</td><td>1.6</td><td>-7.1</td><td>1.6E-12</td><td>4.4E-12</td><td>MB231/MB468</td><td>MITO</td></tr>
<tr><td>>hg38_21_+_8433172_8446622_RNA45SN1_45S_50ntFlanks@12568.12587.20|TCAGTACGAGAGGAACCGCA</td><td>>hg38_21_+_8433172_8446622_RNA45SN1_45S_50ntFlanks</td><td>12568</td><td>12587</td><td>20</td><td>TCAGTACGAGAGGAACCGCA</td><td>108.8</td><td>-5.3</td><td>0.7</td><td>-7.1</td><td>1.3E-12</td><td>4.4E-12</td><td>MB231/MB468</td><td>CYTO</td></tr>
<tr><td>>hg38_21_+_8433172_8446622_RNA45SN1_45S_50ntFlanks@10729.10757.29|AGAGGGTGTAAATCTCGCGCCGGGCCGTA</td><td>>hg38_21_+_8433172_8446622_RNA45SN1_45S_50ntFlanks</td><td>10729</td><td>10757</td><td>29</td><td>AGAGGGTGTAAATCTCGCGCCGGGCCGTA</td><td>102.4</td><td>-10.5</td><td>1.5</td><td>-7.1</td><td>1.2E-12</td><td>4.4E-12</td><td>MB231/MB468</td><td>MP</td></tr>
<tr><td>>hg38_21_+_8433172_8446622_RNA45SN1_45S_50ntFlanks@3874.3898.25|AATACATGCCGACGGGCGCTGACCC</td><td>>hg38_21_+_8433172_8446622_RNA45SN1_45S_50ntFlanks</td><td>3874</td><td>3898</td><td>25</td><td>AATACATGCCGACGGGCGCTGACCC</td><td>4494.6</td><td>-5.3</td><td>0.8</td><td>-7.1</td><td>1.3E-12</td><td>4.5E-12</td><td>MB231/MB468</td><td>CYTO</td></tr>
<tr><td>>hg38_21_+_8433172_8446622_RNA45SN1_45S_50ntFlanks@5227.5245.19|CGCGCGCTACACTGACTGG</td><td>>hg38_21_+_8433172_8446622_RNA45SN1_45S_50ntFlanks</td><td>5227</td><td>5245</td><td>19</td><td>CGCGCGCTACACTGACTGG</td><td>1157.1</td><td>-14.0</td><td>2.0</td><td>-7.1</td><td>1.6E-12</td><td>4.5E-12</td><td>MB231/MB468</td><td>MITO</td></tr>
<tr><td>>hg38_MT_+_1621_3279_RNR2_16S_50ntFlanks@1185.1216.32|ATTTCGGTTGGGGCGACCTCGGAGCAGAACCC</td><td>>hg38_MT_+_1621_3279_RNR2_16S_50ntFlanks</td><td>1185</td><td>1216</td><td>32</td><td>ATTTCGGTTGGGGCGACCTCGGAGCAGAACCC</td><td>655.7</td><td>8.0</td><td>1.1</td><td>7.1</td><td>9.8E-13</td><td>4.5E-12</td><td>BT20/MB468</td><td>MITO</td></tr>
</table>

<table>
<tr><th></th><th></th><th></th><th></th><th></th><th></th><th></th><th></th><th></th><th></th><th></th><th></th><th></th><th></th></tr>
<tr><td>>MI0003560|hsa-mir-92b&WithFlank&1|+|155195171|155195278@67.88.22(+1U)&[MIMAT0003218&hsa-miR-92b-3p&offsets|0|0(+1U);m-115&1|+|155195237|155195258&offsets|0|0(+1U)]|TATTGCACTCGTCCCGGCCTCCT</td><td>>MI0003560|hsa-mir-92b&WithFlank&1|+|155195171|155195278</td><td>67</td><td>88</td><td>22(+1U)</td><td>TATTGCACTCGTCCCGGCCTCCT</td><td>348.1</td><td>5.6</td><td>0.8</td><td>7.1</td><td>1.4E-12</td><td>4.5E-12</td><td>MB231/MB468</td><td>NUC</td></tr>
<tr><td>>hg38_1_-_228634819_228635039_RNA5S12_5S_50ntFlanks@154.171.18|CCGGGTGCTGTAGGCTTT</td><td>>hg38_1_-_228634819_228635039_RNA5S12_5S_50ntFlanks</td><td>154</td><td>171</td><td>18</td><td>CCGGGTGCTGTAGGCTTT</td><td>5778.5</td><td>4.7</td><td>0.7</td><td>7.1</td><td>1.3E-12</td><td>4.5E-12</td><td>BT20/MB468</td><td>CYTO</td></tr>
<tr><td>>hg38_21_+_8433172_8446622_RNA45SN1_45S_50ntFlanks@10498.10533.36|TTCAGATCCCCGAATCCGGAGTGGCGGAGATGGGCG</td><td>>hg38_21_+_8433172_8446622_RNA45SN1_45S_50ntFlanks</td><td>10498</td><td>10533</td><td>36</td><td>TTCAGATCCCCGAATCCGGAGTGGCGGAGATGGGCG</td><td>121.9</td><td>-10.8</td><td>1.5</td><td>-7.1</td><td>1.2E-12</td><td>4.5E-12</td><td>MB231/MB468</td><td>MP</td></tr>
<tr><td>>hg38_21_+_8433172_8446622_RNA45SN1_45S_50ntFlanks@10737.10757.21|TAAATCTCGCGCCGGGCCGTA</td><td>>hg38_21_+_8433172_8446622_RNA45SN1_45S_50ntFlanks</td><td>10737</td><td>10757</td><td>21</td><td>TAAATCTCGCGCCGGGCCGTA</td><td>428.0</td><td>-8.7</td><td>1.2</td><td>-7.1</td><td>1.7E-12</td><td>4.5E-12</td><td>MB231/MB468</td><td>MITO</td></tr>
<tr><td>>hg38_21_+_8433172_8446622_RNA45SN1_45S_50ntFlanks@475.510.36|CTTCGTGATCGATGTGGTGACGTCGTGCTCTCCCGG</td><td>>hg38_21_+_8433172_8446622_RNA45SN1_45S_50ntFlanks</td><td>475</td><td>510</td><td>36</td><td>CTTCGTGATCGATGTGGTGACGTCGTGCTCTCCCGG</td><td>3520.0</td><td>7.2</td><td>1.0</td><td>7.1</td><td>9.9E-13</td><td>4.5E-12</td><td>BT20/MB468</td><td>MITO</td></tr>
<tr><td>>hg38_21_+_8433172_8446622_RNA45SN1_45S_50ntFlanks@9003.9026.24|GTCTCCCCCGCGGGGGCGCGCCGG</td><td>>hg38_21_+_8433172_8446622_RNA45SN1_45S_50ntFlanks</td><td>9003</td><td>9026</td><td>24</td><td>GTCTCCCCCGCGGGGGCGCGCCGG</td><td>297.7</td><td>-10.3</td><td>1.4</td><td>-7.1</td><td>9.9E-13</td><td>4.5E-12</td><td>BT20/MB468</td><td>MITO</td></tr>
<tr><td>>hg38_21_+_8433172_8446622_RNA45SN1_45S_50ntFlanks@9203.9221.19|CTCGGGGCCACGCGCGCGT</td><td>>hg38_21_+_8433172_8446622_RNA45SN1_45S_50ntFlanks</td><td>9203</td><td>9221</td><td>19</td><td>CTCGGGGCCACGCGCGCGT</td><td>134.0</td><td>-10.9</td><td>1.5</td><td>-7.1</td><td>1.2E-12</td><td>4.5E-12</td><td>MB231/MB468</td><td>MP</td></tr>
<tr><td>>hg38_21_+_8433172_8446622_RNA45SN1_45S_50ntFlanks@11641.11667.27|CGGCGGGTGTTGACGCGATGTGATTTC</td><td>>hg38_21_+_8433172_8446622_RNA45SN1_45S_50ntFlanks</td><td>11641</td><td>11667</td><td>27</td><td>CGGCGGGTGTTGACGCGATGTGATTTC</td><td>2417.5</td><td>-8.5</td><td>1.2</td><td>-7.1</td><td>1.7E-12</td><td>4.5E-12</td><td>MB231/MB468</td><td>MITO</td></tr>
<tr><td>>hg38_21_+_8433172_8446622_RNA45SN1_45S_50ntFlanks@7984.8005.22|AGATCAGACGTGGCGACCCGCT</td><td>>hg38_21_+_8433172_8446622_RNA45SN1_45S_50ntFlanks</td><td>7984</td><td>8005</td><td>22</td><td>AGATCAGACGTGGCGACCCGCT</td><td>486.6</td><td>-4.2</td><td>0.6</td><td>-7.0</td><td>1.8E-12</td><td>4.5E-12</td><td>BT20/MB468</td><td>NUC</td></tr>
<tr><td>>hg38_21_+_8433172_8446622_RNA45SN1_45S_50ntFlanks@12932.12952.21|CGCTAAACCATTCGTAGACGA</td><td>>hg38_21_+_8433172_8446622_RNA45SN1_45S_50ntFlanks</td><td>12932</td><td>12952</td><td>21</td><td>CGCTAAACCATTCGTAGACGA</td><td>96.5</td><td>-10.5</td><td>1.5</td><td>-7.1</td><td>1.2E-12</td><td>4.5E-12</td><td>MB231/MB468</td><td>MP</td></tr>
<tr><td>>hg38_21_+_8433172_8446622_RNA45SN1_45S_50ntFlanks@8850.8867.18|GCGGCGGCGGCGGCGGGA</td><td>>hg38_21_+_8433172_8446622_RNA45SN1_45S_50ntFlanks</td><td>8850</td><td>8867</td><td>18</td><td>GCGGCGGCGGCGGCGGGA</td><td>249.8</td><td>-11.8</td><td>1.7</td><td>-7.1</td><td>1.2E-12</td><td>4.5E-12</td><td>MB231/MB468</td><td>MP</td></tr>
<tr><td>>hg38_21_+_8433172_8446622_RNA45SN1_45S_50ntFlanks@3900.3937.38|CTTCGCGGGGGGGATGCGTGCATTTATCAGATCAAAAC</td><td>>hg38_21_+_8433172_8446622_RNA45SN1_45S_50ntFlanks</td><td>3900</td><td>3937</td><td>38</td><td>CTTCGCGGGGGGGATGCGTGCATTTATCAGATCAAAAC</td><td>498.0</td><td>-12.8</td><td>1.8</td><td>-7.1</td><td>1.2E-12</td><td>4.5E-12</td><td>MB231/MB468</td><td>MP</td></tr>
<tr><td>>hg38_21_+_8433172_8446622_RNA45SN1_45S_50ntFlanks@11561.11585.25|GCGCCTAGCAGCCGACTTAGAACTG</td><td>>hg38_21_+_8433172_8446622_RNA45SN1_45S_50ntFlanks</td><td>11561</td><td>11585</td><td>25</td><td>GCGCCTAGCAGCCGACTTAGAACTG</td><td>2008.4</td><td>-4.8</td><td>0.7</td><td>-7.1</td><td>1.0E-12</td><td>4.5E-12</td><td>BT20/MB468</td><td>MITO</td></tr>
<tr><td>>MI0005756|hsa-mir-934&WithFlank&X|+|136550872|136550966@21.42.22(+1C)&[MIMAT0004977&hsa-miR-934&offsets|0|0(+1C);m-943&X|+|136550892|136550913&offsets|0|0(+1C)]|TGTCTACTACTGGAGACACTGGC</td><td>>MI0005756|hsa-mir-934&WithFlank&X|+|136550872|136550966</td><td>21</td><td>42</td><td>22(+1C)</td><td>TGTCTACTACTGGAGACACTGGC</td><td>206.9</td><td>11.1</td><td>1.5</td><td>7.4</td><td>9.9E-14</td><td>4.5E-12</td><td>BT20/MB231</td><td>CYTO</td></tr>
<tr><td>>hg38_21_+_8433172_8446622_RNA45SN1_45S_50ntFlanks@11781.11804.24|CGCATGAATGGATGAACGAGATTC</td><td>>hg38_21_+_8433172_8446622_RNA45SN1_45S_50ntFlanks</td><td>11781</td><td>11804</td><td>24</td><td>CGCATGAATGGATGAACGAGATTC</td><td>1700.3</td><td>-4.9</td><td>0.7</td><td>-7.1</td><td>1.3E-12</td><td>4.5E-12</td><td>BT20/MB468</td><td>CYTO</td></tr>
<tr><td>>hg38_21_+_8433172_8446622_RNA45SN1_45S_50ntFlanks@4378.4402.25|CTCGTAGTTGGATCTTGGGAGCGGG</td><td>>hg38_21_+_8433172_8446622_RNA45SN1_45S_50ntFlanks</td><td>4378</td><td>4402</td><td>25</td><td>CTCGTAGTTGGATCTTGGGAGCGGG</td><td>158.1</td><td>-11.2</td><td>1.6</td><td>-7.1</td><td>1.2E-12</td><td>4.5E-12</td><td>MB231/MB468</td><td>MP</td></tr>
<tr><td>>hg38_21_+_8433172_8446622_RNA45SN1_45S_50ntFlanks@5047.5078.32|TAGTTGGTGGAGCGATTTGTCTGGTTAATTCC</td><td>>hg38_21_+_8433172_8446622_RNA45SN1_45S_50ntFlanks</td><td>5047</td><td>5078</td><td>32</td><td>TAGTTGGTGGAGCGATTTGTCTGGTTAATTCC</td><td>152.2</td><td>7.9</td><td>1.1</td><td>7.1</td><td>1.4E-12</td><td>4.6E-12</td><td>MB231/MB468</td><td>CYTO</td></tr>
<tr><td>>am_trna116_GluCTC_1_-_145399233_145399304@12.29.18,trna25_GluCTC_2_-_71273488_71273560@12.29.18,trna59_GluCTC_1_+_249168447_249168518@12.29.18,trna71_GluCTC_1_-_161439189_161439260@12.29.18,trna74_GluCTC_1_-_161431809_161431880@12.29.18,trna77_GluCTC_1_-_161424398_161424469@12.29.18,trna77_GluCTC_6_+_28949976_28950047@12.29.18,trna80_GluCTC_1_-_161417018_161417089@12.29.18,trna87_GluCTC_6_-_126101393_126101464@12.29.18|CTAGTGGTTAGGATTCGG</td><td>>am_trna116_GluCTC_1_-_145399233_145399304</td><td>12</td><td>29</td><td>18</td><td>CTAGTGGTTAGGATTCGG</td><td>161.1</td><td>-11.2</td><td>1.6</td><td>-7.1</td><td>1.2E-12</td><td>4.6E-12</td><td>MB231/MB468</td><td>MP</td></tr>
<tr><td>>hg38_21_+_8433172_8446622_RNA45SN1_45S_50ntFlanks@6743.6770.28|CGACACTTCGAACGCACTTGCGGCCCCG</td><td>>hg38_21_+_8433172_8446622_RNA45SN1_45S_50ntFlanks</td><td>6743</td><td>6770</td><td>28</td><td>CGACACTTCGAACGCACTTGCGGCCCCG</td><td>2608.1</td><td>-7.2</td><td>1.0</td><td>-7.1</td><td>1.4E-12</td><td>4.6E-12</td><td>MB231/MB468</td><td>CYTO</td></tr>
<tr><td>>MI0000082|hsa-mir-25&WithFlank&7|-|100093554|100093649@58.79.22(+2A)&[MIMAT0000081&hsa-miR-25-3p&offsets|0|0(+2A);m-23&7|-|100093571|100093592&offsets|0|0(+2A)]|CATTGCACTTGTCTCGGTCTGAAA</td><td>>MI0000082|hsa-mir-25&WithFlank&7|-|100093554|100093649</td><td>58</td><td>79</td><td>22(+2A)</td><td>CATTGCACTTGTCTCGGTCTGAAA</td><td>1119.8</td><td>7.3</td><td>1.0</td><td>7.1</td><td>1.3E-12</td><td>4.6E-12</td><td>BT20/MB468</td><td>CYTO</td></tr>
<tr><td>>hg38_21_+_8433172_8446622_RNA45SN1_45S_50ntFlanks@4370.4389.20|TTAAAAAGCTCGTAGTTGGA</td><td>>hg38_21_+_8433172_8446622_RNA45SN1_45S_50ntFlanks</td><td>4370</td><td>4389</td><td>20</td><td>TTAAAAAGCTCGTAGTTGGA</td><td>240.0</td><td>-11.8</td><td>1.7</td><td>-7.1</td><td>1.2E-12</td><td>4.6E-12</td><td>MB231/MB468</td><td>MP</td></tr>
<tr><td>>hg38_21_+_8433172_8446622_RNA45SN1_45S_50ntFlanks@7991.8012.22|ACGTGGCGACCCGCTGAATTTA</td><td>>hg38_21_+_8433172_8446622_RNA45SN1_45S_50ntFlanks</td><td>7991</td><td>8012</td><td>22</td><td>ACGTGGCGACCCGCTGAATTTA</td><td>10978.9</td><td>-5.0</td><td>0.7</td><td>-7.1</td><td>1.0E-12</td><td>4.6E-12</td><td>BT20/MB468</td><td>MITO</td></tr>
<tr><td>>hg38_21_+_8433172_8446622_RNA45SN1_45S_50ntFlanks@11666.11692.27|TCTGCCCAGTGCTCTGAATGTCAAAGT</td><td>>hg38_21_+_8433172_8446622_RNA45SN1_45S_50ntFlanks</td><td>11666</td><td>11692</td><td>27</td><td>TCTGCCCAGTGCTCTGAATGTCAAAGT</td><td>252.1</td><td>4.4</td><td>0.6</td><td>7.1</td><td>1.4E-12</td><td>4.6E-12</td><td>MB231/MB468</td><td>CYTO</td></tr>
<tr><td>>hg38_21_+_8433172_8446622_RNA45SN1_45S_50ntFlanks@9562.9597.36|TCCTGACGTGCAAATCGGTCGTCCGACCTGGGTATA</td><td>>hg38_21_+_8433172_8446622_RNA45SN1_45S_50ntFlanks</td><td>9562</td><td>9597</td><td>36</td><td>TCCTGACGTGCAAATCGGTCGTCCGACCTGGGTATA</td><td>232.1</td><td>-11.7</td><td>1.6</td><td>-7.1</td><td>1.2E-12</td><td>4.6E-12</td><td>MB231/MB468</td><td>MP</td></tr>
<tr><td>>MI0003137|hsa-mir-193b&WithFlank&16|+|14303961|14304055@57.77.21(+1C)&[MIMAT0002819&hsa-miR-193b-3p&offsets|0|-1(+1C);m-166&16|+|14304017|14304038&offsets|0|-1(+1C)]|AACTGGCCCTCAAAGTCCCGCC</td><td>>MI0003137|hsa-mir-193b&WithFlank&16|+|14303961|14304055</td><td>57</td><td>77</td><td>21(+1C)</td><td>AACTGGCCCTCAAAGTCCCGCC</td><td>669.2</td><td>10.2</td><td>1.4</td><td>7.1</td><td>1.3E-12</td><td>4.6E-12</td><td>BT20/MB468</td><td>CYTO</td></tr>
<tr><td>>hg38_21_+_8433172_8446622_RNA45SN1_45S_50ntFlanks@10845.10865.21|ACTTCGGGATAAGGATTGGCT</td><td>>hg38_21_+_8433172_8446622_RNA45SN1_45S_50ntFlanks</td><td>10845</td><td>10865</td><td>21</td><td>ACTTCGGGATAAGGATTGGCT</td><td>14804.2</td><td>-12.1</td><td>1.7</td><td>-7.0</td><td>1.8E-12</td><td>4.6E-12</td><td>BT20/MB468</td><td>NUC</td></tr>
<tr><td>>hg38_21_+_8433172_8446622_RNA45SN1_45S_50ntFlanks@12934.12965.32|CTAAACCATTCGTAGACGACCTGCTTCTGGGT</td><td>>hg38_21_+_8433172_8446622_RNA45SN1_45S_50ntFlanks</td><td>12934</td><td>12965</td><td>32</td><td>CTAAACCATTCGTAGACGACCTGCTTCTGGGT</td><td>1149.0</td><td>-5.5</td><td>0.8</td><td>-7.1</td><td>1.4E-12</td><td>4.6E-12</td><td>MB231/MB468</td><td>CYTO</td></tr>
<tr><td>>hg38_21_+_8433172_8446622_RNA45SN1_45S_50ntFlanks@6699.6719.21|GCTAGCTGCGAGAATTAATGT</td><td>>hg38_21_+_8433172_8446622_RNA45SN1_45S_50ntFlanks</td><td>6699</td><td>6719</td><td>21</td><td>GCTAGCTGCGAGAATTAATGT</td><td>1740.1</td><td>-7.7</td><td>1.1</td><td>-7.1</td><td>1.7E-12</td><td>4.6E-12</td><td>MB231/MB468</td><td>MITO</td></tr>
<tr><td>>hg38_21_+_8433172_8446622_RNA45SN1_45S_50ntFlanks@10876.10894.19|GGTCGGTCGGGCTGGGGCG</td><td>>hg38_21_+_8433172_8446622_RNA45SN1_45S_50ntFlanks</td><td>10876</td><td>10894</td><td>19</td><td>GGTCGGTCGGGCTGGGGCG</td><td>154.7</td><td>-11.1</td><td>1.6</td><td>-7.1</td><td>1.2E-12</td><td>4.6E-12</td><td>MB231/MB468</td><td>MP</td></tr>
<tr><td>>hg38_21_+_8433172_8446622_RNA45SN1_45S_50ntFlanks@4742.4761.20|TCGGAGGTTCGAAGACGATC</td><td>>hg38_21_+_8433172_8446622_RNA45SN1_45S_50ntFlanks</td><td>4742</td><td>4761</td><td>20</td><td>TCGGAGGTTCGAAGACGATC</td><td>303.8</td><td>5.1</td><td>0.7</td><td>7.1</td><td>1.4E-12</td><td>4.6E-12</td><td>MB231/MB468</td><td>CYTO</td></tr>
<tr><td>>trna116_GluCTC_1_-_145399233_145399304@13.33.21,trna25_GluCTC_2_-_71273488_71273560@13.33.21,trna59_GluCTC_1_+_249168447_249168518@13.33.21,trna71_GluCTC_1_-_161439189_161439260@13.33.21,trna74_GluCTC_1_-_161431809_161431880@13.33.21,trna77_GluCTC_1_-_161424398_161424469@13.33.21,trna77_GluCTC_6_+_28949976_28950047@13.33.21,trna80_GluCTC_1_-_161417018_161417089@13.33.21,trna87_GluCTC_6_-_126101393_126101464@13.33.21|TAGTGGTTAGGATTCGGCGCT</td><td>>trna116_GluCTC_1_-_145399233_145399304</td><td>13</td><td>33</td><td>21</td><td>TAGTGGTTAGGATTCGGCGCT</td><td>481.0</td><td>5.0</td><td>0.7</td><td>7.1</td><td>1.3E-12</td><td>4.6E-12</td><td>BT20/MB468</td><td>CYTO</td></tr>
<tr><td>>hg38_21_+_8433172_8446622_RNA45SN1_45S_50ntFlanks@12944.12964.21|CGTAGACGACCTGCTTCTGGG</td><td>>hg38_21_+_8433172_8446622_RNA45SN1_45S_50ntFlanks</td><td>12944</td><td>12964</td><td>21</td><td>CGTAGACGACCTGCTTCTGGG</td><td>1078.2</td><td>-10.0</td><td>1.4</td><td>-7.0</td><td>1.9E-12</td><td>4.6E-12</td><td>BT20/MB468</td><td>NUC</td></tr>
<tr><td>>hg38_21_+_8433172_8446622_RNA45SN1_45S_50ntFlanks@9562.9595.34|TCCTGACGTGCAAATCGGTCGTCCGACCTGGGTA</td><td>>hg38_21_+_8433172_8446622_RNA45SN1_45S_50ntFlanks</td><td>9562</td><td>9595</td><td>34</td><td>TCCTGACGTGCAAATCGGTCGTCCGACCTGGGTA</td><td>1022.7</td><td>-9.1</td><td>1.3</td><td>-7.1</td><td>1.7E-12</td><td>4.6E-12</td><td>MB231/MB468</td><td>MITO</td></tr>
<tr><td>>am_MI0003581|hsa-mir-574&WithFlank&4|+|38868026|38868133@31.53.23&[MIMAT0004795&hsa-miR-574-5p&offsets|0|0;m-213&4|+|38868056|38868078&offsets|0|0]|TGAGTGTGTGTGTGTGAGTGTGT</td><td>>am_MI0003581|hsa-mir-574&WithFlank&4|+|38868026|38868133</td><td>31</td><td>53</td><td>23</td><td>TGAGTGTGTGTGTGTGAGTGTGT</td><td>767.1</td><td>6.8</td><td>1.0</td><td>7.1</td><td>1.7E-12</td><td>4.6E-12</td><td>MB231/MB468</td><td>MITO</td></tr>
<tr><td>>hg38_21_+_8433172_8446622_RNA45SN1_45S_50ntFlanks@8319.8356.38|CGAGACCGATAGTCAACAAGTACCGTAAGGGAAAGTTG</td><td>>hg38_21_+_8433172_8446622_RNA45SN1_45S_50ntFlanks</td><td>8319</td><td>8356</td><td>38</td><td>CGAGACCGATAGTCAACAAGTACCGTAAGGGAAAGTTG</td><td>595.1</td><td>-6.7</td><td>0.9</td><td>-7.1</td><td>1.7E-12</td><td>4.6E-12</td><td>MB231/MB468</td><td>MITO</td></tr>
<tr><td>>hg38_21_+_8433172_8446622_RNA45SN1_45S_50ntFlanks@4588.4608.21|CTATTTTGTTGGTTTTCGGAA</td><td>>hg38_21_+_8433172_8446622_RNA45SN1_45S_50ntFlanks</td><td>4588</td><td>4608</td><td>21</td><td>CTATTTTGTTGGTTTTCGGAA</td><td>499.7</td><td>-7.1</td><td>1.0</td><td>-7.1</td><td>1.4E-12</td><td>4.7E-12</td><td>MB231/MB468</td><td>CYTO</td></tr>
<tr><td>>hg38_21_+_8433172_8446622_RNA45SN1_45S_50ntFlanks@12163.12181.19|ACTGGGGCGGTACACCTGT</td><td>>hg38_21_+_8433172_8446622_RNA45SN1_45S_50ntFlanks</td><td>12163</td><td>12181</td><td>19</td><td>ACTGGGGCGGTACACCTGT</td><td>547.4</td><td>-9.3</td><td>1.3</td><td>-7.0</td><td>1.9E-12</td><td>4.7E-12</td><td>BT20/MB468</td><td>NUC</td></tr>
<tr><td>>hg38_21_+_8433172_8446622_RNA45SN1_45S_50ntFlanks@3885.3915.31|ACGGGCGCTGACCCCCTTCGCGGGGGGGATG</td><td>>hg38_21_+_8433172_8446622_RNA45SN1_45S_50ntFlanks</td><td>3885</td><td>3915</td><td>31</td><td>ACGGGCGCTGACCCCCTTCGCGGGGGGGATG</td><td>313.0</td><td>-6.3</td><td>0.9</td><td>-7.1</td><td>1.4E-12</td><td>4.7E-12</td><td>MB231/MB468</td><td>NUC</td></tr>
<tr><td>>hg38_21_+_8433172_8446622_RNA45SN1_45S_50ntFlanks@10807.10840.34|ACAATGTAGGTAAGGGAAGTCGGCAAGCCGGATC</td><td>>hg38_21_+_8433172_8446622_RNA45SN1_45S_50ntFlanks</td><td>10807</td><td>10840</td><td>34</td><td>ACAATGTAGGTAAGGGAAGTCGGCAAGCCGGATC</td><td>317.2</td><td>-8.6</td><td>1.2</td><td>-7.1</td><td>1.4E-12</td><td>4.7E-12</td><td>MB231/MB468</td><td>NUC</td></tr>
<tr><td>>hg38_21_+_8433172_8446622_RNA45SN1_45S_50ntFlanks@11207.11233.27|GGCGGCGACTCTGGACGCGAGCCGGGC</td><td>>hg38_21_+_8433172_8446622_RNA45SN1_45S_50ntFlanks</td><td>11207</td><td>11233</td><td>27</td><td>GGCGGCGACTCTGGACGCGAGCCGGGC</td><td>140.6</td><td>-11.0</td><td>1.5</td><td>-7.1</td><td>1.2E-12</td><td>4.7E-12</td><td>MB231/MB468</td><td>MP</td></tr>
<tr><td>>hg38_21_+_8433172_8446622_RNA45SN1_45S_50ntFlanks@12163.12180.18|ACTGGGGCGGTACACCTG</td><td>>hg38_21_+_8433172_8446622_RNA45SN1_45S_50ntFlanks</td><td>12163</td><td>12180</td><td>18</td><td>ACTGGGGCGGTACACCTG</td><td>7192.5</td><td>-10.8</td><td>1.4</td><td>-7.5</td><td>6.4E-14</td><td>4.7E-12</td><td>BT20/MB468</td><td>MP</td></tr>
<tr><td>>am_trnaMT_ArgTCG_MT_+_10405_10469@28.48.21|ATTTCGACTCATTAAATTATG</td><td>>am_trnaMT_ArgTCG_MT_+_10405_10469</td><td>28</td><td>48</td><td>21</td><td>ATTTCGACTCATTAAATTATG</td><td>2278.2</td><td>-11.3</td><td>1.6</td><td>-7.1</td><td>1.4E-12</td><td>4.7E-12</td><td>MB231/MB468</td><td>NUC</td></tr>
<tr><td>>hg38_21_+_8433172_8446622_RNA45SN1_45S_50ntFlanks@12033.12068.36|ATCGTTTTTTCACTGACCCGGTGAGGCGGGGGGGCG</td><td>>hg38_21_+_8433172_8446622_RNA45SN1_45S_50ntFlanks</td><td>12033</td><td>12068</td><td>36</td><td>ATCGTTTTTTCACTGACCCGGTGAGGCGGGGGGGCG</td><td>151.1</td><td>-11.1</td><td>1.6</td><td>-7.1</td><td>1.2E-12</td><td>4.7E-12</td><td>MB231/MB468</td><td>MP</td></tr>
<tr><td>>hg38_21_+_8433172_8446622_RNA45SN1_45S_50ntFlanks@12939.12966.28|CCATTCGTAGACGACCTGCTTCTGGGTC</td><td>>hg38_21_+_8433172_8446622_RNA45SN1_45S_50ntFlanks</td><td>12939</td><td>12966</td><td>28</td><td>CCATTCGTAGACGACCTGCTTCTGGGTC</td><td>192.9</td><td>-11.5</td><td>1.6</td><td>-7.1</td><td>1.2E-12</td><td>4.7E-12</td><td>MB231/MB468</td><td>MP</td></tr>
<tr><td>>hg38_21_+_8433172_8446622_RNA45SN1_45S_50ntFlanks@4723.4748.26|CATTAATCAAGAACGAAAGTCGGAGG</td><td>>hg38_21_+_8433172_8446622_RNA45SN1_45S_50ntFlanks</td><td>4723</td><td>4748</td><td>26</td><td>CATTAATCAAGAACGAAAGTCGGAGG</td><td>720.8</td><td>-9.0</td><td>1.3</td><td>-7.1</td><td>1.2E-12</td><td>4.7E-12</td><td>MB231/MB468</td><td>MP</td></tr>
<tr><td>>hg38_21_+_8433172_8446622_RNA45SN1_45S_50ntFlanks@11876.11908.33|AAGACCCTGTTGAGCTTGACTCTAGTCTGGCAC</td><td>>hg38_21_+_8433172_8446622_RNA45SN1_45S_50ntFlanks</td><td>11876</td><td>11908</td><td>33</td><td>AAGACCCTGTTGAGCTTGACTCTAGTCTGGCAC</td><td>169.0</td><td>-5.0</td><td>0.7</td><td>-7.1</td><td>1.4E-12</td><td>4.7E-12</td><td>MB231/MB468</td><td>CYTO</td></tr>
</table>

<table>
<tr><th></th><th></th><th></th><th></th><th></th><th></th><th></th><th></th><th></th><th></th><th></th><th></th><th></th><th></th></tr>
<tr><td>>MI0000070|hsa-mir-16-1&WithFlank&13|-|50048967|50049067@20.39.20(+1U);MI0000115|hsa-mir-16-2&WithFlank&3|+|160404739|160404831@16.35.20(+1U)&[MIMAT0000069&hsa-miR-16-5p&offsets|0|-2(+1U);m-62&13|-|50049027|50049048&offsets|0|-2(+1U)];[MIMAT0000069_1&hsa-miR-16-5p&offsets|0|-2(+1U);m-61&3|+|160404754|160404775&offsets|0|-2(+1U)]|TAGCAGCACGTAAATATTGGT</td><td>>MI0000070|hsa-mir-16-1&WithFlank&13|-|50048967|50049067</td><td>20</td><td>39</td><td>20(+1U)</td><td>TAGCAGCACGTAAATATTGGT</td><td>187.6</td><td>4.6</td><td>0.7</td><td>7.1</td><td>1.4E-12</td><td>4.7E-12</td><td>MB231/MB468</td><td>NUC</td></tr>
<tr><td>>hg38_21_+_8433172_8446622_RNA45SN1_45S_50ntFlanks@10663.10709.47|CCTTGGAAAGCGTCGCGGTTCCGGCGGCGTCCGGTGAGCTCTCGCTG</td><td>>hg38_21_+_8433172_8446622_RNA45SN1_45S_50ntFlanks</td><td>10663</td><td>10709</td><td>47</td><td>CCTTGGAAAGCGTCGCGGTTCCGGCGGCGTCCGGTGAGCTCTCGCTG</td><td>400.8</td><td>-12.4</td><td>1.8</td><td>-7.1</td><td>1.7E-12</td><td>4.7E-12</td><td>MB231/MB468</td><td>MITO</td></tr>
<tr><td>>hg38_21_+_8433172_8446622_RNA45SN1_45S_50ntFlanks@476.505.30|TTCGTGATCGATGTGGTGACGTCGTGCTCT</td><td>>hg38_21_+_8433172_8446622_RNA45SN1_45S_50ntFlanks</td><td>476</td><td>505</td><td>30</td><td>TTCGTGATCGATGTGGTGACGTCGTGCTCT</td><td>26911.5</td><td>5.1</td><td>0.7</td><td>7.1</td><td>1.4E-12</td><td>4.7E-12</td><td>MB231/MB468</td><td>NUC</td></tr>
<tr><td>>hg38_21_+_8433172_8446622_RNA45SN1_45S_50ntFlanks@12405.12439.35|CTTTTTGATCCTTCGATGTCGGCTCTTCCTATCAT</td><td>>hg38_21_+_8433172_8446622_RNA45SN1_45S_50ntFlanks</td><td>12405</td><td>12439</td><td>35</td><td>CTTTTTGATCCTTCGATGTCGGCTCTTCCTATCAT</td><td>1210.7</td><td>-4.6</td><td>0.7</td><td>-7.1</td><td>1.4E-12</td><td>4.7E-12</td><td>MB231/MB468</td><td>NUC</td></tr>
<tr><td>>hg38_21_+_8433172_8446622_RNA45SN1_45S_50ntFlanks@10869.10889.21|AGGGCTGGGTCGGTCGGGCTG</td><td>>hg38_21_+_8433172_8446622_RNA45SN1_45S_50ntFlanks</td><td>10869</td><td>10889</td><td>21</td><td>AGGGCTGGGTCGGTCGGGCTG</td><td>878.4</td><td>-9.4</td><td>1.3</td><td>-7.1</td><td>1.2E-12</td><td>4.7E-12</td><td>MB231/MB468</td><td>MP</td></tr>
<tr><td>>am_trnaMT_SerGCT_MT_+_12207_12265@16.38.23|AACTGCTAACTCATGCCCCCATG</td><td>>am_trnaMT_SerGCT_MT_+_12207_12265</td><td>16</td><td>38</td><td>23</td><td>AACTGCTAACTCATGCCCCCATG</td><td>752.1</td><td>-7.9</td><td>1.1</td><td>-7.1</td><td>1.4E-12</td><td>4.7E-12</td><td>MB231/MB468</td><td>NUC</td></tr>
<tr><td>>hg38_21_+_8433172_8446622_RNA45SN1_45S_50ntFlanks@10463.10480.18|CTCCGTTGCCCTCGGCCG</td><td>>hg38_21_+_8433172_8446622_RNA45SN1_45S_50ntFlanks</td><td>10463</td><td>10480</td><td>18</td><td>CTCCGTTGCCCTCGGCCG</td><td>138.6</td><td>-11.0</td><td>1.5</td><td>-7.1</td><td>1.2E-12</td><td>4.7E-12</td><td>MB231/MB468</td><td>MP</td></tr>
<tr><td>>hg38_21_+_8433172_8446622_RNA45SN1_45S_50ntFlanks@11853.11871.19|CTTGGCGGAATCAGCGGGG</td><td>>hg38_21_+_8433172_8446622_RNA45SN1_45S_50ntFlanks</td><td>11853</td><td>11871</td><td>19</td><td>CTTGGCGGAATCAGCGGGG</td><td>119.3</td><td>-10.8</td><td>1.5</td><td>-7.1</td><td>1.2E-12</td><td>4.8E-12</td><td>MB231/MB468</td><td>MP</td></tr>
<tr><td>>hg38_21_+_8433172_8446622_RNA45SN1_45S_50ntFlanks@6661.6688.28|CGGTGGATCACTCGGCTCGTGCGTCGAT</td><td>>hg38_21_+_8433172_8446622_RNA45SN1_45S_50ntFlanks</td><td>6661</td><td>6688</td><td>28</td><td>CGGTGGATCACTCGGCTCGTGCGTCGAT</td><td>14574.5</td><td>4.3</td><td>0.6</td><td>7.1</td><td>1.0E-12</td><td>4.8E-12</td><td>BT20/MB468</td><td>MITO</td></tr>
<tr><td>>hg38_21_+_8433172_8446622_RNA45SN1_45S_50ntFlanks@10905.10936.32|CTGGGCGCGCGCCGCGGCTGGACGAGGCGCCG</td><td>>hg38_21_+_8433172_8446622_RNA45SN1_45S_50ntFlanks</td><td>10905</td><td>10936</td><td>32</td><td>CTGGGCGCGCGCCGCGGCTGGACGAGGCGCCG</td><td>257.3</td><td>-11.0</td><td>1.5</td><td>-7.1</td><td>1.5E-12</td><td>4.8E-12</td><td>MB231/MB468</td><td>NUC</td></tr>
<tr><td>>hg38_21_+_8433172_8446622_RNA45SN1_45S_50ntFlanks@9534.9551.18|AGGAAACTCTGGTGGAGG</td><td>>hg38_21_+_8433172_8446622_RNA45SN1_45S_50ntFlanks</td><td>9534</td><td>9551</td><td>18</td><td>AGGAAACTCTGGTGGAGG</td><td>6494.1</td><td>-7.3</td><td>1.0</td><td>-7.1</td><td>1.4E-12</td><td>4.8E-12</td><td>MB231/MB468</td><td>CYTO</td></tr>
<tr><td>>hg38_1_-_228634819_228635039_RNA5S12_5S_50ntFlanks@153.170.18|ACCGGGTGCTGTAGGCTT</td><td>>hg38_1_-_228634819_228635039_RNA5S12_5S_50ntFlanks</td><td>153</td><td>170</td><td>18</td><td>ACCGGGTGCTGTAGGCTT</td><td>32004.9</td><td>6.9</td><td>1.0</td><td>7.1</td><td>1.1E-12</td><td>4.8E-12</td><td>BT20/MB468</td><td>MITO</td></tr>
<tr><td>>MI0000077|hsa-mir-21&WithFlank&17|+|59841260|59841343@14.34.21(+2C)&[MIMAT0000076&hsa-miR-21-5p&offsets|0|-1(+2C);m-1&17|+|59841273|59841294&offsets|0|-1(+2C)]|TAGCTTATCAGACTGATGTTGCC</td><td>>MI0000077|hsa-mir-21&WithFlank&17|+|59841260|59841343</td><td>14</td><td>34</td><td>21(+2C)</td><td>TAGCTTATCAGACTGATGTTGCC</td><td>790.4</td><td>6.4</td><td>0.9</td><td>7.1</td><td>1.4E-12</td><td>4.8E-12</td><td>BT20/MB468</td><td>CYTO</td></tr>
<tr><td>>hg38_21_+_8433172_8446622_RNA45SN1_45S_50ntFlanks@11561.11590.30|GCGCCTAGCAGCCGACTTAGAACTGGTGCG</td><td>>hg38_21_+_8433172_8446622_RNA45SN1_45S_50ntFlanks</td><td>11561</td><td>11590</td><td>30</td><td>GCGCCTAGCAGCCGACTTAGAACTGGTGCG</td><td>273.2</td><td>-5.4</td><td>0.8</td><td>-7.1</td><td>1.4E-12</td><td>4.8E-12</td><td>MB231/MB468</td><td>CYTO</td></tr>
<tr><td>>hg38_21_+_8433172_8446622_RNA45SN1_45S_50ntFlanks@12405.12441.37|CTTTTTGATCCTTCGATGTCGGCTCTTCCTATCATTG</td><td>>hg38_21_+_8433172_8446622_RNA45SN1_45S_50ntFlanks</td><td>12405</td><td>12441</td><td>37</td><td>CTTTTTGATCCTTCGATGTCGGCTCTTCCTATCATTG</td><td>4967.3</td><td>-7.8</td><td>1.1</td><td>-7.1</td><td>1.8E-12</td><td>4.8E-12</td><td>MB231/MB468</td><td>MITO</td></tr>
<tr><td>>hg38_21_+_8433172_8446622_RNA45SN1_45S_50ntFlanks@10793.10818.26|TCTGGCATGTTGGAACAATGTAGGTA</td><td>>hg38_21_+_8433172_8446622_RNA45SN1_45S_50ntFlanks</td><td>10793</td><td>10818</td><td>26</td><td>TCTGGCATGTTGGAACAATGTAGGTA</td><td>471.7</td><td>-8.6</td><td>1.2</td><td>-7.1</td><td>1.2E-12</td><td>4.8E-12</td><td>MB231/MB468</td><td>MP</td></tr>
<tr><td>>MI0003763|hsa-mir-767&WithFlank&X|-|152393415|152393535@33.52.20&[MIMAT0003882&hsa-miR-767-5p&offsets|0|-3;m-788&X|-|152393481|152393503&offsets|0|-3]|TGCACCATGGTTGTCTGAGC</td><td>>MI0003763|hsa-mir-767&WithFlank&X|-|152393415|152393535</td><td>33</td><td>52</td><td>20</td><td>TGCACCATGGTTGTCTGAGC</td><td>153.9</td><td>10.6</td><td>1.4</td><td>7.4</td><td>1.1E-13</td><td>4.8E-12</td><td>BT20/MB231</td><td>CYTO</td></tr>
<tr><td>>MI0000088|hsa-mir-30a&WithFlank&6|-|71403545|71403627@12.34.23(+1C)&[MIMAT0000087&hsa-miR-30a-5p&offsets|0|+1(+1C);m-15&6|-|71403595|71403616&offsets|0|+1(+1C)]|TGTAAACATCCTCGACTGGAAGCC</td><td>>MI0000088|hsa-mir-30a&WithFlank&6|-|71403545|71403627</td><td>12</td><td>34</td><td>23(+1C)</td><td>TGTAAACATCCTCGACTGGAAGCC</td><td>25134.5</td><td>7.6</td><td>1.1</td><td>7.1</td><td>1.8E-12</td><td>4.8E-12</td><td>MB231/MB468</td><td>MITO</td></tr>
<tr><td>>hg38_21_+_8433172_8446622_RNA45SN1_45S_50ntFlanks@6719.6742.24|TGAATTGCAGGACACATTGATCAT</td><td>>hg38_21_+_8433172_8446622_RNA45SN1_45S_50ntFlanks</td><td>6719</td><td>6742</td><td>24</td><td>TGAATTGCAGGACACATTGATCAT</td><td>404.3</td><td>-6.4</td><td>0.9</td><td>-7.1</td><td>1.4E-12</td><td>4.8E-12</td><td>MB231/MB468</td><td>CYTO</td></tr>
<tr><td>>hg38_21_+_8433172_8446622_RNA45SN1_45S_50ntFlanks@9759.9789.31|CAAACTTTAAATGGGTAAGAAGCCCGGCTCG</td><td>>hg38_21_+_8433172_8446622_RNA45SN1_45S_50ntFlanks</td><td>9759</td><td>9789</td><td>31</td><td>CAAACTTTAAATGGGTAAGAAGCCCGGCTCG</td><td>170.3</td><td>-11.3</td><td>1.6</td><td>-7.1</td><td>1.2E-12</td><td>4.8E-12</td><td>MB231/MB468</td><td>MP</td></tr>
<tr><td>>trna27_GlyCCC_2_-_70476123_70476193@1.35.35,trna34_GlyCCC_16_-_686736_686806@1.35.35|GCGCCGCTGGTGTAGTGGTATCATGCAAGATTCCC</td><td>>trna27_GlyCCC_2_-_70476123_70476193</td><td>1</td><td>35</td><td>35</td><td>GCGCCGCTGGTGTAGTGGTATCATGCAAGATTCCC</td><td>335.1</td><td>10.1</td><td>1.4</td><td>7.1</td><td>1.4E-12</td><td>4.8E-12</td><td>MB231/MB468</td><td>CYTO</td></tr>
<tr><td>>hg38_21_+_8433172_8446622_RNA45SN1_45S_50ntFlanks@6719.6758.40|TGAATTGCAGGACACATTGATCATCGACACTTCGAACGCA</td><td>>hg38_21_+_8433172_8446622_RNA45SN1_45S_50ntFlanks</td><td>6719</td><td>6758</td><td>40</td><td>TGAATTGCAGGACACATTGATCATCGACACTTCGAACGCA</td><td>711.8</td><td>-7.5</td><td>1.1</td><td>-7.1</td><td>1.8E-12</td><td>4.8E-12</td><td>MB231/MB468</td><td>MITO</td></tr>
<tr><td>>hg38_21_+_8433172_8446622_RNA45SN1_45S_50ntFlanks@10846.10868.23|CTTCGGGATAAGGATTGGCTCTA</td><td>>hg38_21_+_8433172_8446622_RNA45SN1_45S_50ntFlanks</td><td>10846</td><td>10868</td><td>23</td><td>CTTCGGGATAAGGATTGGCTCTA</td><td>1340.1</td><td>-7.6</td><td>1.1</td><td>-7.1</td><td>1.5E-12</td><td>4.8E-12</td><td>MB231/MB468</td><td>NUC</td></tr>
<tr><td>>hg38_21_+_8433172_8446622_RNA45SN1_45S_50ntFlanks@10787.10820.34|ACAGCCTCTGGCATGTTGGAACAATGTAGGTAAG</td><td>>hg38_21_+_8433172_8446622_RNA45SN1_45S_50ntFlanks</td><td>10787</td><td>10820</td><td>34</td><td>ACAGCCTCTGGCATGTTGGAACAATGTAGGTAAG</td><td>100.2</td><td>-10.5</td><td>1.5</td><td>-7.1</td><td>1.2E-12</td><td>4.8E-12</td><td>MB231/MB468</td><td>MP</td></tr>
<tr><td>>hg38_21_+_8433172_8446622_RNA45SN1_45S_50ntFlanks@8472.8491.20|CGGCCCGGCGGATCTTTCCC</td><td>>hg38_21_+_8433172_8446622_RNA45SN1_45S_50ntFlanks</td><td>8472</td><td>8491</td><td>20</td><td>CGGCCCGGCGGATCTTTCCC</td><td>104.5</td><td>-10.6</td><td>1.5</td><td>-7.1</td><td>1.2E-12</td><td>4.8E-12</td><td>MB231/MB468</td><td>MP</td></tr>
<tr><td>>hg38_21_+_8433172_8446622_RNA45SN1_45S_50ntFlanks@11317.11340.24|ACCCCACGTCTCGTCGCGCGCGCG</td><td>>hg38_21_+_8433172_8446622_RNA45SN1_45S_50ntFlanks</td><td>11317</td><td>11340</td><td>24</td><td>ACCCCACGTCTCGTCGCGCGCGCG</td><td>205.1</td><td>-6.9</td><td>1.0</td><td>-7.1</td><td>1.4E-12</td><td>4.8E-12</td><td>MB231/MB468</td><td>CYTO</td></tr>
<tr><td>>MI0000087|hsa-mir-29a&WithFlank&7|-|130876741|130876816@47.69.23&[MIMAT0000086&hsa-miR-29a-3p&offsets|-1|0;m-26&7|-|130876749|130876769&offsets|-1|+1]|CTAGCACCATCTGAAATCGGTTA</td><td>>MI0000087|hsa-mir-29a&WithFlank&7|-|130876741|130876816</td><td>47</td><td>69</td><td>23</td><td>CTAGCACCATCTGAAATCGGTTA</td><td>2589.1</td><td>9.0</td><td>1.3</td><td>7.1</td><td>1.8E-12</td><td>4.8E-12</td><td>MB231/MB468</td><td>MITO</td></tr>
<tr><td>>MI0000070|hsa-mir-16-1&WithFlank&13|-|50048967|50049067@20.39.20;MI0000115|hsa-mir-16-2&WithFlank&3|+|160404739|160404831@16.35.20&[MIMAT0000069&hsa-miR-16-5p&offsets|0|-2;m-62&13|-|50049027|50049048&offsets|0|-2];[MIMAT0000069_1&hsa-miR-16-5p&offsets|0|-2;m-61&3|+|160404754|160404775&offsets|0|-2]|TAGCAGCACGTAAATATTGG</td><td>>MI0000070|hsa-mir-16-1&WithFlank&13|-|50048967|50049067</td><td>20</td><td>39</td><td>20</td><td>TAGCAGCACGTAAATATTGG</td><td>793.4</td><td>8.3</td><td>1.2</td><td>7.1</td><td>1.4E-12</td><td>4.8E-12</td><td>BT20/MB468</td><td>CYTO</td></tr>
<tr><td>>hg38_21_+_8433172_8446622_RNA45SN1_45S_50ntFlanks@4727.4766.40|AATCAAGAACGAAAGTCGGAGGTTCGAAGACGATCAGATA</td><td>>hg38_21_+_8433172_8446622_RNA45SN1_45S_50ntFlanks</td><td>4727</td><td>4766</td><td>40</td><td>AATCAAGAACGAAAGTCGGAGGTTCGAAGACGATCAGATA</td><td>1285.4</td><td>9.0</td><td>1.3</td><td>7.1</td><td>1.4E-12</td><td>4.8E-12</td><td>BT20/MB468</td><td>CYTO</td></tr>
<tr><td>>hg38_21_+_8433172_8446622_RNA45SN1_45S_50ntFlanks@4758.4780.23|GATCAGATACCGTCGTAGTTCCG</td><td>>hg38_21_+_8433172_8446622_RNA45SN1_45S_50ntFlanks</td><td>4758</td><td>4780</td><td>23</td><td>GATCAGATACCGTCGTAGTTCCG</td><td>178.9</td><td>-10.4</td><td>1.5</td><td>-7.1</td><td>1.5E-12</td><td>4.8E-12</td><td>MB231/MB468</td><td>NUC</td></tr>
<tr><td>>hg38_21_+_8433172_8446622_RNA45SN1_45S_50ntFlanks@11207.11225.19|GGCGGCGACTCTGGACGCG</td><td>>hg38_21_+_8433172_8446622_RNA45SN1_45S_50ntFlanks</td><td>11207</td><td>11225</td><td>19</td><td>GGCGGCGACTCTGGACGCG</td><td>361.7</td><td>-8.4</td><td>1.2</td><td>-7.0</td><td>1.8E-12</td><td>4.8E-12</td><td>MB231/MB468</td><td>MITO</td></tr>
<tr><td>>hg38_21_+_8433172_8446622_RNA45SN1_45S_50ntFlanks@5084.5113.30|CGAACGAGACTCTGGCATGCTAACTAGTTA</td><td>>hg38_21_+_8433172_8446622_RNA45SN1_45S_50ntFlanks</td><td>5084</td><td>5113</td><td>30</td><td>CGAACGAGACTCTGGCATGCTAACTAGTTA</td><td>393.1</td><td>-12.4</td><td>1.8</td><td>-7.0</td><td>1.8E-12</td><td>4.8E-12</td><td>MB231/MB468</td><td>MITO</td></tr>
<tr><td>>hg38_21_+_8433172_8446622_RNA45SN1_45S_50ntFlanks@4730.4759.30|CAAGAACGAAAGTCGGAGGTTCGAAGACGA</td><td>>hg38_21_+_8433172_8446622_RNA45SN1_45S_50ntFlanks</td><td>4730</td><td>4759</td><td>30</td><td>CAAGAACGAAAGTCGGAGGTTCGAAGACGA</td><td>1334.7</td><td>-8.0</td><td>1.1</td><td>-7.0</td><td>1.9E-12</td><td>4.8E-12</td><td>BT20/MB468</td><td>NUC</td></tr>
<tr><td>>hg38_21_+_8433172_8446622_RNA45SN1_45S_50ntFlanks@10392.10419.28|ACAGCAGTTGAACATGGGTCAGTCGGTC</td><td>>hg38_21_+_8433172_8446622_RNA45SN1_45S_50ntFlanks</td><td>10392</td><td>10419</td><td>28</td><td>ACAGCAGTTGAACATGGGTCAGTCGGTC</td><td>1342.3</td><td>-9.2</td><td>1.3</td><td>-7.1</td><td>1.5E-12</td><td>4.8E-12</td><td>MB231/MB468</td><td>NUC</td></tr>
<tr><td>>hg38_21_+_8433172_8446622_RNA45SN1_45S_50ntFlanks@11621.11640.20|CAAAGCATCGCGAAGGCCCG</td><td>>hg38_21_+_8433172_8446622_RNA45SN1_45S_50ntFlanks</td><td>11621</td><td>11640</td><td>20</td><td>CAAAGCATCGCGAAGGCCCG</td><td>175.4</td><td>-10.4</td><td>1.5</td><td>-7.1</td><td>1.5E-12</td><td>4.8E-12</td><td>MB231/MB468</td><td>NUC</td></tr>
<tr><td>>hg38_21_+_8433172_8446622_RNA45SN1_45S_50ntFlanks@11718.11751.34|AACGGCGGGAGTAACTATGACTCTCTTAAGGTAG</td><td>>hg38_21_+_8433172_8446622_RNA45SN1_45S_50ntFlanks</td><td>11718</td><td>11751</td><td>34</td><td>AACGGCGGGAGTAACTATGACTCTCTTAAGGTAG</td><td>183.7</td><td>-9.6</td><td>1.3</td><td>-7.1</td><td>1.1E-12</td><td>4.9E-12</td><td>BT20/MB468</td><td>MITO</td></tr>
<tr><td>>hg38_21_+_8433172_8446622_RNA45SN1_45S_50ntFlanks@9459.9491.33|CCGGGGAGGTGGAGCACGAGCGCACGTGTTAGG</td><td>>hg38_21_+_8433172_8446622_RNA45SN1_45S_50ntFlanks</td><td>9459</td><td>9491</td><td>33</td><td>CCGGGGAGGTGGAGCACGAGCGCACGTGTTAGG</td><td>102.0</td><td>-10.5</td><td>1.5</td><td>-7.1</td><td>1.3E-12</td><td>4.9E-12</td><td>MB231/MB468</td><td>MP</td></tr>
<tr><td>>trnaMT_GluTTC_MT_-_14674_14742@34.72.39,trnalookalike8_GluTTC_5_-_93905172_93905240@34.72.39|ATATCATTGGTCGTGGTTGTAGTCCGTGCGAGAATACCA</td><td>>trnaMT_GluTTC_MT_-_14674_14742</td><td>34</td><td>72</td><td>39</td><td>ATATCATTGGTCGTGGTTGTAGTCCGTGCGAGAATACCA</td><td>3065.4</td><td>5.7</td><td>0.8</td><td>7.1</td><td>1.5E-12</td><td>4.9E-12</td><td>MB231/MB468</td><td>NUC</td></tr>
<tr><td>>hg38_21_+_8433172_8446622_RNA45SN1_45S_50ntFlanks@6652.6684.33|GACTCTTAGCGGTGGATCACTCGGCTCGTGCGT</td><td>>hg38_21_+_8433172_8446622_RNA45SN1_45S_50ntFlanks</td><td>6652</td><td>6684</td><td>33</td><td>GACTCTTAGCGGTGGATCACTCGGCTCGTGCGT</td><td>407.4</td><td>4.5</td><td>0.6</td><td>7.1</td><td>1.5E-12</td><td>4.9E-12</td><td>MB231/MB468</td><td>CYTO</td></tr>
<tr><td>>hg38_21_+_8433172_8446622_RNA45SN1_45S_50ntFlanks@9504.9532.29|GTGAACTATGCCTGGGCAGGGCGAAGCCA</td><td>>hg38_21_+_8433172_8446622_RNA45SN1_45S_50ntFlanks</td><td>9504</td><td>9532</td><td>29</td><td>GTGAACTATGCCTGGGCAGGGCGAAGCCA</td><td>201.6</td><td>-9.7</td><td>1.4</td><td>-7.1</td><td>1.1E-12</td><td>4.9E-12</td><td>BT20/MB468</td><td>MITO</td></tr>
<tr><td>>hg38_21_+_8433172_8446622_RNA45SN1_45S_50ntFlanks@4760.4783.24|TCAGATACCGTCGTAGTTCCGACC</td><td>>hg38_21_+_8433172_8446622_RNA45SN1_45S_50ntFlanks</td><td>4760</td><td>4783</td><td>24</td><td>TCAGATACCGTCGTAGTTCCGACC</td><td>581.7</td><td>-3.7</td><td>0.5</td><td>-7.1</td><td>1.4E-12</td><td>4.9E-12</td><td>BT20/MB468</td><td>CYTO</td></tr>
<tr><td>>hg38_21_+_8433172_8446622_RNA45SN1_45S_50ntFlanks@10805.10840.36|GAACAATGTAGGTAAGGGAAGTCGGCAAGCCGGATC</td><td>>hg38_21_+_8433172_8446622_RNA45SN1_45S_50ntFlanks</td><td>10805</td><td>10840</td><td>36</td><td>GAACAATGTAGGTAAGGGAAGTCGGCAAGCCGGATC</td><td>695.3</td><td>-7.0</td><td>1.0</td><td>-7.0</td><td>1.8E-12</td><td>4.9E-12</td><td>MB231/MB468</td><td>MITO</td></tr>
<tr><td>>trnaMT_MetCAT_MT_+_4402_4469@51.71.21|TATACCCTTCCCGTACTACCA</td><td>>trnaMT_MetCAT_MT_+_4402_4469</td><td>51</td><td>71</td><td>21</td><td>TATACCCTTCCCGTACTACCA</td><td>650.5</td><td>9.9</td><td>1.4</td><td>7.1</td><td>1.5E-12</td><td>4.9E-12</td><td>MB231/MB468</td><td>CYTO</td></tr>
<tr><td>>am_trnaMT_LysTTT_MT_+_8295_8364@51.69.19,trnalookalike7_LysTTT_1_+_568844_568913@51.69.19|CCAACACCTCTTTACAGTG</td><td>>am_trnaMT_LysTTT_MT_+_8295_8364</td><td>51</td><td>69</td><td>19</td><td>CCAACACCTCTTTACAGTG</td><td>125.4</td><td>-10.8</td><td>1.5</td><td>-7.1</td><td>1.3E-12</td><td>4.9E-12</td><td>MB231/MB468</td><td>MP</td></tr>
<tr><td>>hg38_21_+_8433172_8446622_RNA45SN1_45S_50ntFlanks@10787.10822.36|ACAGCCTCTGGCATGTTGGAACAATGTAGGTAAGGG</td><td>>hg38_21_+_8433172_8446622_RNA45SN1_45S_50ntFlanks</td><td>10787</td><td>10822</td><td>36</td><td>ACAGCCTCTGGCATGTTGGAACAATGTAGGTAAGGG</td><td>449.7</td><td>-8.7</td><td>1.2</td><td>-7.0</td><td>1.8E-12</td><td>4.9E-12</td><td>MB231/MB468</td><td>MITO</td></tr>
<tr><td>>hg38_21_+_8433172_8446622_RNA45SN1_45S_50ntFlanks@10411.10437.27|CAGTCGGTCCTGAGAGATGGGCGAGCG</td><td>>hg38_21_+_8433172_8446622_RNA45SN1_45S_50ntFlanks</td><td>10411</td><td>10437</td><td>27</td><td>CAGTCGGTCCTGAGAGATGGGCGAGCG</td><td>282.7</td><td>-12.0</td><td>1.7</td><td>-7.1</td><td>1.3E-12</td><td>4.9E-12</td><td>MB231/MB468</td><td>MP</td></tr>
<tr><td>>hg38_21_+_8433172_8446622_RNA45SN1_45S_50ntFlanks@10572.10612.41|AGAAGCCGGCGGGAGCCCCGGGGAGAGTTCTCTTTTCTTTG</td><td>>hg38_21_+_8433172_8446622_RNA45SN1_45S_50ntFlanks</td><td>10572</td><td>10612</td><td>41</td><td>AGAAGCCGGCGGGAGCCCCGGGGAGAGTTCTCTTTTCTTTG</td><td>195.1</td><td>-10.6</td><td>1.5</td><td>-7.1</td><td>1.5E-12</td><td>4.9E-12</td><td>MB231/MB468</td><td>NUC</td></tr>
<tr><td>>hg38_21_+_8433172_8446622_RNA45SN1_45S_50ntFlanks@4898.4916.19|AAAGGAATTGACGGAAGGG</td><td>>hg38_21_+_8433172_8446622_RNA45SN1_45S_50ntFlanks</td><td>4898</td><td>4916</td><td>19</td><td>AAAGGAATTGACGGAAGGG</td><td>201.4</td><td>-11.5</td><td>1.6</td><td>-7.1</td><td>1.3E-12</td><td>4.9E-12</td><td>MB231/MB468</td><td>MP</td></tr>
<tr><td>>hg38_21_+_8433172_8446622_RNA45SN1_45S_50ntFlanks@11564.11582.19|CCTAGCAGCCGACTTAGAA</td><td>>hg38_21_+_8433172_8446622_RNA45SN1_45S_50ntFlanks</td><td>11564</td><td>11582</td><td>19</td><td>CCTAGCAGCCGACTTAGAA</td><td>2999.5</td><td>-8.0</td><td>1.1</td><td>-7.1</td><td>1.5E-12</td><td>5.0E-12</td><td>MB231/MB468</td><td>NUC</td></tr>
<tr><td>>MI0000253|hsa-mir-148a&WithFlank&7|-|25949913|25949992@12.33.22&[MIMAT0004549&hsa-miR-148a-5p&offsets|0|0;m-189&7|-|25949960|25949981&offsets|0|0]|AAAGTTCTGAGACACTCCGACT</td><td>>MI0000253|hsa-mir-148a&WithFlank&7|-|25949913|25949992</td><td>12</td><td>33</td><td>22</td><td>AAAGTTCTGAGACACTCCGACT</td><td>600.1</td><td>5.0</td><td>0.7</td><td>7.4</td><td>1.6E-13</td><td>5.0E-12</td><td>BT20/MB231</td><td>NUC</td></tr>
<tr><td>>hg38_21_+_8433172_8446622_RNA45SN1_45S_50ntFlanks@12893.12927.35|CCTCGCCCGTCACGCACCGCACGTTCGTGGGGAAC</td><td>>hg38_21_+_8433172_8446622_RNA45SN1_45S_50ntFlanks</td><td>12893</td><td>12927</td><td>35</td><td>CCTCGCCCGTCACGCACCGCACGTTCGTGGGGAAC</td><td>411.1</td><td>-11.8</td><td>1.6</td><td>-7.5</td><td>6.8E-14</td><td>5.0E-12</td><td>BT20/MB468</td><td>MP</td></tr>
</table>

| | | | | | | | | | | | | | |
|---|---|---|---|---|---|---|---|---|---|---|---|---|---|
| >hg38_21_+_8433172_8446622_RNA45SN1_45S_50ntFlanks@4641.4662.22\|CATTCGTATTGCGCCGCTAGAG | >hg38_21_+_8433172_8446622_RNA45SN1_45S_50ntFlanks | 4641 | 4662 | 22 | CATTCGTATTGCGCCGCTAGAG | 338.0 | -11.5 | 1.5 | -7.5 | 6.8E-14 | 5.0E-12 | BT20/MB468 | MP |
| >MI0000063\|hsa-let-7b&WithFlank&22\|+\|46113680\|46113774@12.34.23&[MIMAT0000063&hsa-let-7b-5p&offsets\|0\|+1;m-14&22\|+\|46113691\|46113712&offsets\|0\|+1]\|TGAGGTAGTAGGTTGTGTGGTTT | >MI0000063\|hsa-let-7b&WithFlank&22\|+\|46113680\|46113774 | 12 | 34 | 23 | TGAGGTAGTAGGTTGTGTGGTTT | 2056.5 | 2.8 | 0.4 | 7.0 | 2.0E-12 | 5.0E-12 | BT20/MB468 | NUC |
| >hg38_21_+_8433172_8446622_RNA45SN1_45S_50ntFlanks@10687.10706.20\|CGGCGTCCGGTGAGCTCTCG | >hg38_21_+_8433172_8446622_RNA45SN1_45S_50ntFlanks | 10687 | 10706 | 20 | CGGCGTCCGGTGAGCTCTCG | 2555.4 | -4.8 | 0.7 | -7.1 | 1.1E-12 | 5.0E-12 | BT20/MB468 | MITO |
| >hg38_21_+_8433172_8446622_RNA45SN1_45S_50ntFlanks@10846.10865.20\|CTTCGGGATAAGGATTGGCT | >hg38_21_+_8433172_8446622_RNA45SN1_45S_50ntFlanks | 10846 | 10865 | 20 | CTTCGGGATAAGGATTGGCT | 6592.5 | -11.4 | 1.6 | -7.0 | 2.0E-12 | 5.0E-12 | BT20/MB468 | NUC |
| >hg38_21_+_8433172_8446622_RNA45SN1_45S_50ntFlanks@3872.3899.28\|CTAATACATGCCGACGGGCGCTGACCCC | >hg38_21_+_8433172_8446622_RNA45SN1_45S_50ntFlanks | 3872 | 3899 | 28 | CTAATACATGCCGACGGGCGCTGACCCC | 215.4 | -9.8 | 1.4 | -7.1 | 1.1E-12 | 5.0E-12 | BT20/MB468 | MITO |
| >hg38_21_+_8433172_8446622_RNA45SN1_45S_50ntFlanks@4588.4614.27\|CTATTTTGTTGGTTTTCGGAACTGAGG | >hg38_21_+_8433172_8446622_RNA45SN1_45S_50ntFlanks | 4588 | 4614 | 27 | CTATTTTGTTGGTTTTCGGAACTGAGG | 481.2 | -8.4 | 1.2 | -7.0 | 1.8E-12 | 5.0E-12 | MB231/MB468 | MITO |
| >hg38_21_+_8433172_8446622_RNA45SN1_45S_50ntFlanks@11670.11702.33\|CCCAGTGCTCTGAATGTCAAAGTGAAGAAATTC | >hg38_21_+_8433172_8446622_RNA45SN1_45S_50ntFlanks | 11670 | 11702 | 33 | CCCAGTGCTCTGAATGTCAAAGTGAAGAAATTC | 2644.7 | -7.3 | 1.0 | -7.0 | 1.9E-12 | 5.0E-12 | MB231/MB468 | MITO |
| >hg38_21_+_8433172_8446622_RNA45SN1_45S_50ntFlanks@10808.10831.24\|CAATGTAGGTAAGGGAAGTCGGCA | >hg38_21_+_8433172_8446622_RNA45SN1_45S_50ntFlanks | 10808 | 10831 | 24 | CAATGTAGGTAAGGGAAGTCGGCA | 591.8 | -8.7 | 1.2 | -7.0 | 1.9E-12 | 5.0E-12 | MB231/MB468 | MITO |
| >hg38_21_+_8433172_8446622_RNA45SN1_45S_50ntFlanks@11629.11662.34\|CGCGAAGGCCCGCGGCGGGTGTTGACGCGATGTG | >hg38_21_+_8433172_8446622_RNA45SN1_45S_50ntFlanks | 11629 | 11662 | 34 | CGCGAAGGCCCGCGGCGGGTGTTGACGCGATGTG | 718.8 | -5.5 | 0.8 | -7.1 | 1.3E-12 | 5.0E-12 | MB231/MB468 | MP |
| >MI0000299\|hsa-mir-222&WithFlank&X\|-\|45747009\|45747130@75.96.22(+1G)&[MIMAT0000279&hsa-miR-222-3p&offsets\|0\|+1(+1G);m-140&X\|-\|45747033\|45747056&offsets\|0\|-2(+1G)]\|AGCTACATCTGGCTACTGGGTCG | >MI0000299\|hsa-mir-222&WithFlank&X\|-\|45747009\|45747130 | 75 | 96 | 22(+1G) | AGCTACATCTGGCTACTGGGTCG | 170.5 | 6.3 | 0.9 | 7.1 | 1.5E-12 | 5.0E-12 | MB231/MB468 | NUC |
| >hg38_21_+_8433172_8446622_RNA45SN1_45S_50ntFlanks@12163.12180.18\|ACTGGGGCGGTACACCTG | >hg38_21_+_8433172_8446622_RNA45SN1_45S_50ntFlanks | 12163 | 12180 | 18 | ACTGGGGCGGTACACCTG | 485.3 | -8.7 | 1.2 | -7.1 | 1.5E-12 | 5.0E-12 | MB231/MB468 | NUC |
| >hg38_21_+_8433172_8446622_RNA45SN1_45S_50ntFlanks@10420.10437.18\|CTGAGAGATGGGCGAGCG | >hg38_21_+_8433172_8446622_RNA45SN1_45S_50ntFlanks | 10420 | 10437 | 18 | CTGAGAGATGGGCGAGCG | 146.4 | -11.1 | 1.6 | -7.1 | 1.3E-12 | 5.0E-12 | MB231/MB468 | MP |
| >hg38_21_+_8433172_8446622_RNA45SN1_45S_50ntFlanks@12075.12095.21\|GAGGGGCTCTCGCTTCTGGCG | >hg38_21_+_8433172_8446622_RNA45SN1_45S_50ntFlanks | 12075 | 12095 | 21 | GAGGGGCTCTCGCTTCTGGCG | 178.9 | -11.3 | 1.6 | -7.1 | 1.3E-12 | 5.0E-12 | MB231/MB468 | MP |
| >MI0000809\|hsa-mir-151a&WithFlank&8\|-\|140732558\|140732659@53.74.22(+2U)&[MIMAT0000757&hsa-miR-151a-3p&offsets\|0\|+1(+2U);m-30&8\|-\|140732587\|140732607&offsets\|0\|+1(+2U)]\|CTAGACTGAAGCTCCTTGAGGATT | >MI0000809\|hsa-mir-151a&WithFlank&8\|-\|140732558\|140732659 | 53 | 74 | 22(+2U) | CTAGACTGAAGCTCCTTGAGGATT | 823.5 | 9.1 | 1.3 | 7.1 | 1.5E-12 | 5.0E-12 | BT20/MB468 | CYTO |
| >hg38_21_+_8433172_8446622_RNA45SN1_45S_50ntFlanks@11875.11898.24\|GAAGACCCTGTTGAGCTTGACTCT | >hg38_21_+_8433172_8446622_RNA45SN1_45S_50ntFlanks | 11875 | 11898 | 24 | GAAGACCCTGTTGAGCTTGACTCT | 120.5 | -10.8 | 1.5 | -7.1 | 1.3E-12 | 5.0E-12 | MB231/MB468 | MP |
| >hg38_21_+_8433172_8446622_RNA45SN1_45S_50ntFlanks@3843.3862.20\|CTACTTGGATAACTGTGGTA | >hg38_21_+_8433172_8446622_RNA45SN1_45S_50ntFlanks | 3843 | 3862 | 20 | CTACTTGGATAACTGTGGTA | 196.8 | -11.5 | 1.6 | -7.1 | 1.3E-12 | 5.0E-12 | MB231/MB468 | MP |
| >hg38_21_+_8433172_8446622_RNA45SN1_45S_50ntFlanks@8191.8211.21\|CAGCCCGTGGACGGTGTGAGG | >hg38_21_+_8433172_8446622_RNA45SN1_45S_50ntFlanks | 8191 | 8211 | 21 | CAGCCCGTGGACGGTGTGAGG | 888.8 | -12.9 | 1.7 | -7.5 | 7.0E-14 | 5.0E-12 | BT20/MB468 | MP |
| >hg38_21_+_8433172_8446622_RNA45SN1_45S_50ntFlanks@10482.10500.19\|TCGAAAGGGAGTCGGGTTC | >hg38_21_+_8433172_8446622_RNA45SN1_45S_50ntFlanks | 10482 | 10500 | 19 | TCGAAAGGGAGTCGGGTTC | 239.8 | -11.8 | 1.7 | -7.1 | 1.3E-12 | 5.0E-12 | MB231/MB468 | MP |
| >hg38_21_+_8433172_8446622_RNA45SN1_45S_50ntFlanks@4047.4081.35\|ATTCGAACGTCTGCCCTATCAACTTTCGATGGTAG | >hg38_21_+_8433172_8446622_RNA45SN1_45S_50ntFlanks | 4047 | 4081 | 35 | ATTCGAACGTCTGCCCTATCAACTTTCGATGGTAG | 242.4 | -11.8 | 1.7 | -7.1 | 1.3E-12 | 5.0E-12 | MB231/MB468 | MP |
| >hg38_21_+_8433172_8446622_RNA45SN1_45S_50ntFlanks@9572.9591.20\|CAAATCGGTCGTCCGACCTG | >hg38_21_+_8433172_8446622_RNA45SN1_45S_50ntFlanks | 9572 | 9591 | 20 | CAAATCGGTCGTCCGACCTG | 179.9 | -11.4 | 1.6 | -7.1 | 1.3E-12 | 5.0E-12 | MB231/MB468 | MP |
| >hg38_21_+_8433172_8446622_RNA45SN1_45S_50ntFlanks@10466.10488.23\|CGTTGCCCTCGGCCGATCGAAAG | >hg38_21_+_8433172_8446622_RNA45SN1_45S_50ntFlanks | 10466 | 10488 | 23 | CGTTGCCCTCGGCCGATCGAAAG | 131.8 | -10.9 | 1.5 | -7.1 | 1.3E-12 | 5.0E-12 | MB231/MB468 | MP |
| >hg38_21_+_8433172_8446622_RNA45SN1_45S_50ntFlanks@10392.10425.34\|ACAGCAGTTGAACATGGGTCAGTCGGTCCTGAGA | >hg38_21_+_8433172_8446622_RNA45SN1_45S_50ntFlanks | 10392 | 10425 | 34 | ACAGCAGTTGAACATGGGTCAGTCGGTCCTGAGA | 152.0 | -11.1 | 1.6 | -7.1 | 1.3E-12 | 5.1E-12 | MB231/MB468 | MP |
| >hg38_21_+_8433172_8446622_RNA45SN1_45S_50ntFlanks@12274.12307.34\|CAGTACGAATACAGACCGTGAAAGCGGGGCCTCA | >hg38_21_+_8433172_8446622_RNA45SN1_45S_50ntFlanks | 12274 | 12307 | 34 | CAGTACGAATACAGACCGTGAAAGCGGGGCCTCA | 195.1 | -11.5 | 1.6 | -7.1 | 1.3E-12 | 5.1E-12 | MB231/MB468 | MP |
| >hg38_21_+_8433172_8446622_RNA45SN1_45S_50ntFlanks@4759.4779.21\|ATCAGATACCGTCGTAGTTCC | >hg38_21_+_8433172_8446622_RNA45SN1_45S_50ntFlanks | 4759 | 4779 | 21 | ATCAGATACCGTCGTAGTTCC | 394.0 | -10.7 | 1.5 | -7.1 | 1.1E-12 | 5.1E-12 | BT20/MB468 | MITO |
| >hg38_21_+_8433172_8446622_RNA45SN1_45S_50ntFlanks@9858.9880.23\|CGGGATGAACCGAACGCCGGGTT | >hg38_21_+_8433172_8446622_RNA45SN1_45S_50ntFlanks | 9858 | 9880 | 23 | CGGGATGAACCGAACGCCGGGTT | 420.2 | -12.6 | 1.8 | -7.1 | 1.3E-12 | 5.1E-12 | MB231/MB468 | MP |
| >MI0000255\|hsa-mir-30d&WithFlank&8\|-\|134804870\|134804951@12.35.24(+2U)&[MIMAT0000245&hsa-miR-30d-5p&offsets\|0\|+2(+2U);m-21&8\|-\|134804919\|134804940&offsets\|0\|+2(+2U)]\|TGTAAACATCCCCGACTGGAAGCTTT | >MI0000255\|hsa-mir-30d&WithFlank&8\|-\|134804870\|134804951 | 12 | 35 | 24(+2U) | TGTAAACATCCCCGACTGGAAGCTTT | 424.4 | 8.3 | 1.2 | 7.1 | 1.5E-12 | 5.1E-12 | BT20/MB468 | CYTO |
| >hg38_21_+_8433172_8446622_RNA45SN1_45S_50ntFlanks@11881.11905.25\|CCTGTTGAGCTTGACTCTAGTCTGG | >hg38_21_+_8433172_8446622_RNA45SN1_45S_50ntFlanks | 11881 | 11905 | 25 | CCTGTTGAGCTTGACTCTAGTCTGG | 205.0 | -10.0 | 1.4 | -7.1 | 1.5E-12 | 5.1E-12 | BT20/MB468 | CYTO |
| >hg38_21_+_8433172_8446622_RNA45SN1_45S_50ntFlanks@9804.9831.28\|GGCGTGGAATGCGAGTGCCTAGTGGGCC | >hg38_21_+_8433172_8446622_RNA45SN1_45S_50ntFlanks | 9804 | 9831 | 28 | GGCGTGGAATGCGAGTGCCTAGTGGGCC | 151.7 | -11.1 | 1.6 | -7.1 | 1.3E-12 | 5.1E-12 | MB231/MB468 | MP |
| >hg38_21_+_8433172_8446622_RNA45SN1_45S_50ntFlanks@12934.12970.37\|CTAAACCATTCGTAGACGACCTGCTTCTGGGTCGGGG | >hg38_21_+_8433172_8446622_RNA45SN1_45S_50ntFlanks | 12934 | 12970 | 37 | CTAAACCATTCGTAGACGACCTGCTTCTGGGTCGGGG | 225.5 | -11.7 | 1.6 | -7.1 | 1.3E-12 | 5.1E-12 | MB231/MB468 | MP |
| >hg38_21_+_8433172_8446622_RNA45SN1_45S_50ntFlanks@3776.3794.19\|CCGGTACAGTGAAACTGCG | >hg38_21_+_8433172_8446622_RNA45SN1_45S_50ntFlanks | 3776 | 3794 | 19 | CCGGTACAGTGAAACTGCG | 127.1 | -10.9 | 1.5 | -7.1 | 1.3E-12 | 5.1E-12 | MB231/MB468 | MP |
| >hg38_21_+_8433172_8446622_RNA45SN1_45S_50ntFlanks@12893.12930.38\|CCTCGCCCGTCACGCACCGCACGTTCGTGGGGAACCTG | >hg38_21_+_8433172_8446622_RNA45SN1_45S_50ntFlanks | 12893 | 12930 | 38 | CCTCGCCCGTCACGCACCGCACGTTCGTGGGGAACCTG | 4130.2 | -8.5 | 1.2 | -7.0 | 1.9E-12 | 5.1E-12 | MB231/MB468 | MITO |
| >hg38_21_+_8433172_8446622_RNA45SN1_45S_50ntFlanks@4045.4066.22\|CCATTCGAACGTCTGCCCTATC | >hg38_21_+_8433172_8446622_RNA45SN1_45S_50ntFlanks | 4045 | 4066 | 22 | CCATTCGAACGTCTGCCCTATC | 2725.9 | -10.7 | 1.5 | -7.0 | 2.0E-12 | 5.1E-12 | BT20/MB468 | NUC |
| >hg38_21_+_8433172_8446622_RNA45SN1_45S_50ntFlanks@11577.11603.27\|TTAGAACTGGTGCGGACCAGGGGAATC | >hg38_21_+_8433172_8446622_RNA45SN1_45S_50ntFlanks | 11577 | 11603 | 27 | TTAGAACTGGTGCGGACCAGGGGAATC | 2424.3 | -10.7 | 1.5 | -7.0 | 2.0E-12 | 5.1E-12 | BT20/MB468 | NUC |
| >hg38_21_+_8433172_8446622_RNA45SN1_45S_50ntFlanks@10727.10745.19\|GGAGAGGGTGTAAATCTCG | >hg38_21_+_8433172_8446622_RNA45SN1_45S_50ntFlanks | 10727 | 10745 | 19 | GGAGAGGGTGTAAATCTCG | 209.7 | -11.6 | 1.6 | -7.1 | 1.3E-12 | 5.1E-12 | MB231/MB468 | MP |
| >hg38_21_+_8433172_8446622_RNA45SN1_45S_50ntFlanks@11453.11478.26\|TCCTCGGCGCGCGGCGGCGGCGGCGG | >hg38_21_+_8433172_8446622_RNA45SN1_45S_50ntFlanks | 11453 | 11478 | 26 | TCCTCGGCGCGCGGCGGCGGCGG | 131.1 | -10.9 | 1.5 | -7.1 | 1.3E-12 | 5.1E-12 | MB231/MB468 | MP |
| >trna111_HisGTG_1_-_147774845_147774916@-1G.35.36,trna118_HisGTG_1_-_145396881_145396952@-1G.35.36,trna16_HisGTG_1_+_146544773_146544844@-1G.35.36,trna1_HisGTG_15_+_45493349_45493420@-1G.35.36,trna21_HisGTG_1_+_147753471_147753542@-1G.35.36,trna33_HisGTG_6_+_27125906_27125977@-1G.35.36,trna7_HisGTG_9_-_14433938_14434009@-1G.35.36,trna8_HisGTG_15_-_45492611_45492682@-1G.35.36,trna9_HisGTG_15_-_45490804_45490875@-1G.35.36\|GGCCGTGATCGTATAGTGGTTAGTACTCTGCGTTGT | >trna111_HisGTG_1_-_147774845_147774916 | -1G | 35 | 36 | GGCCGTGATCGTATAGTGGTTAGTACTCTGCGTTGT | 2221.8 | 11.1 | 1.6 | 7.1 | 1.6E-12 | 5.1E-12 | MB231/MB468 | NUC |
| >hg38_21_+_8433172_8446622_RNA45SN1_45S_50ntFlanks@5125.5158.34\|AGCGGTCGGCGTCCCCCAACTTCTTAGAGGGACA | >hg38_21_+_8433172_8446622_RNA45SN1_45S_50ntFlanks | 5125 | 5158 | 34 | AGCGGTCGGCGTCCCCCAACTTCTTAGAGGGACA | 3034.8 | 8.9 | 1.3 | 7.1 | 1.5E-12 | 5.1E-12 | BT20/MB468 | CYTO |
| >MI0000471\|hsa-mir-126&WithFlank&9\|+\|136670596\|136670692@58.78.21(+1A)&[MIMAT0000445&hsa-miR-126-3p&offsets\|0\|-1(+1A);m-40&9\|+\|136670653\|136670674&offsets\|0\|-1(+1A)]\|TCGTACCGTGAGTAATAATGCA | >MI0000471\|hsa-mir-126&WithFlank&9\|+\|136670596\|136670692 | 58 | 78 | 21(+1A) | TCGTACCGTGAGTAATAATGCA | 921.3 | 10.7 | 1.5 | 7.1 | 1.1E-12 | 5.1E-12 | BT20/MB468 | MITO |
| >hg38_21_+_8433172_8446622_RNA45SN1_45S_50ntFlanks@4759.4781.23\|ATCAGATACCGTCGTAGTTCCGA | >hg38_21_+_8433172_8446622_RNA45SN1_45S_50ntFlanks | 4759 | 4781 | 23 | ATCAGATACCGTCGTAGTTCCGA | 519.1 | -8.1 | 1.1 | -7.1 | 1.5E-12 | 5.1E-12 | MB231/MB468 | CYTO |
| >hg38_21_+_8433172_8446622_RNA45SN1_45S_50ntFlanks@4090.4121.32\|CCTACCATGGTGACCACGGGTGACGGGGAATC | >hg38_21_+_8433172_8446622_RNA45SN1_45S_50ntFlanks | 4090 | 4121 | 32 | CCTACCATGGTGACCACGGGTGACGGGGAATC | 468.8 | -12.7 | 1.8 | -7.0 | 1.9E-12 | 5.1E-12 | MB231/MB468 | MITO |
| >hg38_1_-_228634819_228635039_RNA5S12_5S_50ntFlanks@141.170.30\|CCGCCTGGGAATACCGGGTGCTGTAGGCTT | >hg38_1_-_228634819_228635039_RNA5S12_5S_50ntFlanks | 141 | 170 | 30 | CCGCCTGGGAATACCGGGTGCTGTAGGCTT | 1854.5 | 4.3 | 0.6 | 7.1 | 1.6E-12 | 5.1E-12 | MB231/MB468 | NUC |
| >hg38_21_+_8433172_8446622_RNA45SN1_45S_50ntFlanks@4760.4782.23\|TCAGATACCGTCGTAGTTCCGAC | >hg38_21_+_8433172_8446622_RNA45SN1_45S_50ntFlanks | 4760 | 4782 | 23 | TCAGATACCGTCGTAGTTCCGAC | 1280.8 | -12.4 | 1.7 | -7.1 | 1.1E-12 | 5.1E-12 | BT20/MB468 | MITO |
| >MI0000088\|hsa-mir-30a&WithFlank&6\|-\|71403545\|71403627@12.35.24(+2U)&[MIMAT0000087&hsa-miR-30a-5p&offsets\|0\|+2(+2U);m-15&6\|-\|71403595\|71403616&offsets\|0\|+2(+2U)]\|TGTAAACATCCTCGACTGGAAGCTTT | >MI0000088\|hsa-mir-30a&WithFlank&6\|-\|71403545\|71403627 | 12 | 35 | 24(+2U) | TGTAAACATCCTCGACTGGAAGCTTT | 484.4 | 7.2 | 1.0 | 7.1 | 1.5E-12 | 5.1E-12 | MB231/MB468 | CYTO |
| >hg38_21_+_8433172_8446622_RNA45SN1_45S_50ntFlanks@4046.4067.22\|CATTCGAACGTCTGCCCTATCA | >hg38_21_+_8433172_8446622_RNA45SN1_45S_50ntFlanks | 4046 | 4067 | 22 | CATTCGAACGTCTGCCCTATCA | 95.5 | -10.4 | 1.5 | -7.1 | 1.3E-12 | 5.1E-12 | MB231/MB468 | MP |
| >hg38_21_+_8433172_8446622_RNA45SN1_45S_50ntFlanks@10877.10894.18\|GTCGGTCGGGCTGGGGCG | >hg38_21_+_8433172_8446622_RNA45SN1_45S_50ntFlanks | 10877 | 10894 | 18 | GTCGGTCGGGCTGGGGCG | 115.9 | -10.7 | 1.5 | -7.1 | 1.3E-12 | 5.1E-12 | MB231/MB468 | MP |
| >hg38_21_+_8433172_8446622_RNA45SN1_45S_50ntFlanks@11209.11229.21\|CGGCGACTCTGGACGCGAGCC | >hg38_21_+_8433172_8446622_RNA45SN1_45S_50ntFlanks | 11209 | 11229 | 21 | CGGCGACTCTGGACGCGAGCC | 745.5 | -8.9 | 1.3 | -7.1 | 1.3E-12 | 5.1E-12 | MB231/MB468 | MP |
| >hg38_21_+_8433172_8446622_RNA45SN1_45S_50ntFlanks@11885.11906.22\|TTGAGCTTGACTCTAGTCTGGC | >hg38_21_+_8433172_8446622_RNA45SN1_45S_50ntFlanks | 11885 | 11906 | 22 | TTGAGCTTGACTCTAGTCTGGC | 155.4 | -11.1 | 1.6 | -7.1 | 1.4E-12 | 5.2E-12 | MB231/MB468 | MP |

| | | | | | | | | | | | | | |
|---|---|---|---|---|---|---|---|---|---|---|---|---|---|
| >MI0000085\|hsa-mir-27a&WithFlank&19\|-\|13836434\|13836523@16.36.21&[MIMAT0004501&hsa-miR-27a-5p&offsets\|0\|-1;m-270&19\|-\|13836487\|13836508&offsets\|0\|-1]\|AGGGCTTAGCTGCTTGTGAGC | >MI0000085\|hsa-mir-27a&WithFlank&19\|-\|13836434\|13836523 | 16 | 36 | 21 | AGGGCTTAGCTGCTTGTGAGC | 667.9 | 5.8 | 0.8 | 7.1 | 1.5E-12 | 5.2E-12 | BT20/MB468 | CYTO |
| >MI0000266\|hsa-mir-10a&WithFlank&17\|-\|48579832\|48579953@29.50.22&[MIMAT0000253&hsa-miR-10a-5p&offsets\|+1\|0;m-18&17\|-\|48579905\|48579926&offsets\|+1\|+1]\|ACCCTGTAGATCCGAATTTGTG | >MI0000266\|hsa-mir-10a&WithFlank&17\|-\|48579832\|48579953 | 29 | 50 | 22 | ACCCTGTAGATCCGAATTTGTG | 1379.5 | 9.9 | 1.4 | 7.0 | 2.1E-12 | 5.2E-12 | BT20/MB468 | NUC |
| >hg38_21_+_8433172_8446622_RNA45SN1_45S_50ntFlanks@6738.6758.21\|ATCATCGACACTTCGAACGCA | >hg38_21_+_8433172_8446622_RNA45SN1_45S_50ntFlanks | 6738 | 6758 | 21 | ATCATCGACACTTCGAACGCA | 223.9 | -10.1 | 1.4 | -7.1 | 1.5E-12 | 5.2E-12 | BT20/MB468 | CYTO |
| >hg38_21_+_8433172_8446622_RNA45SN1_45S_50ntFlanks@10866.10886.21\|CTAAGGGCTGGGTCGGTCGGG | >hg38_21_+_8433172_8446622_RNA45SN1_45S_50ntFlanks | 10866 | 10886 | 21 | CTAAGGGCTGGGTCGGTCGGG | 544.0 | -7.6 | 1.1 | -7.1 | 1.5E-12 | 5.2E-12 | MB231/MB468 | CYTO |
| >hg38_21_+_8433172_8446622_RNA45SN1_45S_50ntFlanks@11768.11790.23\|CTAATTAGTGACGCGCATGAATG | >hg38_21_+_8433172_8446622_RNA45SN1_45S_50ntFlanks | 11768 | 11790 | 23 | CTAATTAGTGACGCGCATGAATG | 759.6 | -9.1 | 1.3 | -7.1 | 1.4E-12 | 5.2E-12 | MB231/MB468 | MP |
| >hg38_1_-_228634819_228635039_RNA5S12_5S_50ntFlanks@84.109.26\|CTCGTCTGATCTCGGAAGCTAAGCAG | >hg38_1_-_228634819_228635039_RNA5S12_5S_50ntFlanks | 84 | 109 | 26 | CTCGTCTGATCTCGGAAGCTAAGCAG | 97.9 | -10.5 | 1.5 | -7.1 | 1.4E-12 | 5.2E-12 | MB231/MB468 | MP |
| >MI0000111\|hsa-mir-105-1&WithFlank&X\|-\|152392213\|152392305@19.40.22;MI0000112\|hsa-mir-105-2&WithFlank&X\|-\|152394406\|152394498@19.40.22&[MIMAT0000102&hsa-miR-105-5p&offsets\|0\|-1;m-735&X\|-\|152392266\|152392287&offsets\|0\|0];[MIMAT0000102_1&hsa-miR-105-5p&offsets\|0\|-1;m-735&X\|-\|152394459\|152394480&offsets\|0\|0]\|TCAAATGCTCAGACTCCTGTGG | >MI0000111\|hsa-mir-105-1&WithFlank&X\|-\|152392213\|152392305 | 19 | 40 | 22 | TCAAATGCTCAGACTCCTGTGG | 219.1 | -11.6 | 1.6 | -7.1 | 1.4E-12 | 5.2E-12 | MB231/MB468 | MP |
| >MI0000102\|hsa-mir-100&WithFlank&11\|-\|122152223\|122152314@19.39.21(+1U)&[MIMAT0000098&hsa-miR-100-5p&offsets\|0\|-1(+1U);m-27&11\|-\|122152276\|122152296&offsets\|0\|0(+1U)]\|AACCCGTAGATCCGAACTTGTT | >MI0000102\|hsa-mir-100&WithFlank&11\|-\|122152223\|122152314 | 19 | 39 | 21(+1U) | AACCCGTAGATCCGAACTTGTT | 2036.7 | 15.2 | 2.2 | 7.1 | 1.5E-12 | 5.2E-12 | BT20/MB468 | CYTO |
| >am_trna106_HisGTG_1_-_149155828_149155899@7.29.23,trna111_HisGTG_1_-_147774845_147774916@7.29.23,trna118_HisGTG_1_-_145396881_145396952@7.29.23,trna16_HisGTG_1_+_146544773_146544844@7.29.23,trna1_HisGTG_15_+_45493349_45493420@7.29.23,trna21_HisGTG_1_+_147753471_147753542@7.29.23,trna33_HisGTG_6_+_27125906_27125977@7.29.23,trna7_HisGTG_9_-_14433938_14434009@7.29.23,trna8_HisGTG_15_-_45492611_45492682@7.29.23,trna9_HisGTG_15_-_45490804_45490875@7.29.23\|ATCGTATAGTGGTTAGTACTCTG | >am_trna106_HisGTG_1_-_149155828_149155899 | 7 | 29 | 23 | ATCGTATAGTGGTTAGTACTCTG | 621.7 | -8.6 | 1.2 | -7.1 | 1.6E-12 | 5.2E-12 | MB231/MB468 | CYTO |
| >hg38_21_+_8433172_8446622_RNA45SN1_45S_50ntFlanks@3854.3889.36\|ACTGTGGTAATTCTAGAGCTAATACATGCCGACGGG | >hg38_21_+_8433172_8446622_RNA45SN1_45S_50ntFlanks | 3854 | 3889 | 36 | ACTGTGGTAATTCTAGAGCTAATACATGCCGACGGG | 1201.8 | -10.0 | 1.4 | -7.0 | 2.1E-12 | 5.2E-12 | BT20/MB468 | NUC |
| >hg38_21_+_8433172_8446622_RNA45SN1_45S_50ntFlanks@9623.9650.28\|TAGTAGCTGGTTCCCTCCGAAGTTTCCC | >hg38_21_+_8433172_8446622_RNA45SN1_45S_50ntFlanks | 9623 | 9650 | 28 | TAGTAGCTGGTTCCCTCCGAAGTTTCCC | 432.0 | -9.0 | 1.3 | -7.0 | 2.1E-12 | 5.2E-12 | BT20/MB468 | NUC |
| >hg38_21_+_8433172_8446622_RNA45SN1_45S_50ntFlanks@5449.5466.18\|ATCGGCCCCGCCGGGGTC | >hg38_21_+_8433172_8446622_RNA45SN1_45S_50ntFlanks | 5449 | 5466 | 18 | ATCGGCCCCGCCGGGGTC | 208.9 | -9.8 | 1.4 | -7.1 | 1.2E-12 | 5.2E-12 | BT20/MB468 | MITO |
| >hg38_21_+_8433172_8446622_RNA45SN1_45S_50ntFlanks@6730.6770.41\|ACACATTGATCATCGACACTTCGAACGCACTTGCGGCCCCG | >hg38_21_+_8433172_8446622_RNA45SN1_45S_50ntFlanks | 6730 | 6770 | 41 | ACACATTGATCATCGACACTTCGAACGCACTTGCGGCCCCG | 1219.2 | -10.4 | 1.5 | -7.0 | 2.1E-12 | 5.2E-12 | BT20/MB468 | NUC |
| >hg38_21_+_8433172_8446622_RNA45SN1_45S_50ntFlanks@6733.6757.25\|CATTGATCATCGACACTTCGAACGC | >hg38_21_+_8433172_8446622_RNA45SN1_45S_50ntFlanks | 6733 | 6757 | 25 | CATTGATCATCGACACTTCGAACGC | 167.2 | -11.2 | 1.6 | -7.1 | 1.4E-12 | 5.2E-12 | MB231/MB468 | MP |
| >hg38_21_+_8433172_8446622_RNA45SN1_45S_50ntFlanks@10839.10868.30\|TCCGTAACTTCGGGATAAGGATTGGCTCTA | >hg38_21_+_8433172_8446622_RNA45SN1_45S_50ntFlanks | 10839 | 10868 | 30 | TCCGTAACTTCGGGATAAGGATTGGCTCTA | 142.3 | -11.0 | 1.6 | -7.1 | 1.4E-12 | 5.2E-12 | MB231/MB468 | MP |
| >am_trna128_GlyGCC_6_-_27870686_27870756@1.29.29,trna133_GlyCCC_1_-_16872434_16872504@1.29.29,trna18_GlyGCC_16_+_70822597_70822667@1.29.29,trna19_GlyGCC_16_+_70823410_70823480@1.29.29,trna19_GlyGCC_2_-_157257659_157257729@1.29.29,trna24_GlyGCC_16_-_70812942_70813012@1.29.29,trna25_GlyGCC_16_-_70812114_70812184@1.29.29,trna4_GlyCCC_1_+_17188416_17188486@1.29.29,trna5_GlyGCC_17_+_8029064_8029134@1.29.29,trna68_GlyGCC_1_-_161493637_161493707@1.29.29\|GCATTGGTGGTTCAGTGGTAGAATTCTCG | >am_trna128_GlyGCC_6_-_27870686_27870756 | 1 | 29 | 29 | GCATTGGTGGTTCAGTGGTAGAATTCTCG | 20187.9 | -5.8 | 0.8 | -7.1 | 1.6E-12 | 5.2E-12 | MB231/MB468 | NUC |
| >hg38_21_+_8433172_8446622_RNA45SN1_45S_50ntFlanks@11569.11596.28\|CAGCCGACTTAGAACTGGTGCGGACCAG | >hg38_21_+_8433172_8446622_RNA45SN1_45S_50ntFlanks | 11569 | 11596 | 28 | CAGCCGACTTAGAACTGGTGCGGACCAG | 116.1 | -10.7 | 1.5 | -7.1 | 1.4E-12 | 5.2E-12 | MB231/MB468 | MP |
| >hg38_21_+_8433172_8446622_RNA45SN1_45S_50ntFlanks@4560.4582.23\|CTAGGAATAATGGAATAGGACCG | >hg38_21_+_8433172_8446622_RNA45SN1_45S_50ntFlanks | 4560 | 4582 | 23 | CTAGGAATAATGGAATAGGACCG | 124.7 | -4.5 | 0.6 | -7.1 | 1.6E-12 | 5.2E-12 | MB231/MB468 | CYTO |
| >hg38_21_+_8433172_8446622_RNA45SN1_45S_50ntFlanks@10389.10419.31\|TGAACAGCAGTTGAACATGGGTCAGTCGGTC | >hg38_21_+_8433172_8446622_RNA45SN1_45S_50ntFlanks | 10389 | 10419 | 31 | TGAACAGCAGTTGAACATGGGTCAGTCGGTC | 118.4 | -10.8 | 1.5 | -7.1 | 1.4E-12 | 5.2E-12 | MB231/MB468 | MP |
| >hg38_21_+_8433172_8446622_RNA45SN1_45S_50ntFlanks@10463.10497.35\|CTCCGTTGCCCTCGGCCGATCGAAAGGGAGTCGGG | >hg38_21_+_8433172_8446622_RNA45SN1_45S_50ntFlanks | 10463 | 10497 | 35 | CTCCGTTGCCCTCGGCCGATCGAAAGGGAGTCGGG | 215.7 | -9.8 | 1.4 | -7.1 | 1.2E-12 | 5.3E-12 | BT20/MB468 | MITO |
| >hg38_21_+_8433172_8446622_RNA45SN1_45S_50ntFlanks@12420.12446.27\|ATGTCGGCTCTTCCTATCATTGTGAAG | >hg38_21_+_8433172_8446622_RNA45SN1_45S_50ntFlanks | 12420 | 12446 | 27 | ATGTCGGCTCTTCCTATCATTGTGAAG | 1489.3 | -4.0 | 0.6 | -7.1 | 1.2E-12 | 5.3E-12 | BT20/MB468 | MITO |
| >MI0000477\|hsa-mir-146a&WithFlank&5\|+\|160485346\|160485456@27.46.20&[MIMAT0000449&hsa-miR-146a-5p&offsets\|0\|-2;m-129&5\|+\|160485372\|160485393&offsets\|0\|-2]\|TGAGAACTGAATTCCATGGG | >MI0000477\|hsa-mir-146a&WithFlank&5\|+\|160485346\|160485456 | 27 | 46 | 20 | TGAGAACTGAATTCCATGGG | 886.5 | -13.2 | 1.7 | -7.6 | 2.1E-14 | 5.3E-12 | BT20/MB231 | MP |
| >hg38_21_+_8433172_8446622_RNA45SN1_45S_50ntFlanks@3985.4013.29\|CGGCTTTGGTGACTCTAGATAACCTCGGG | >hg38_21_+_8433172_8446622_RNA45SN1_45S_50ntFlanks | 3985 | 4013 | 29 | CGGCTTTGGTGACTCTAGATAACCTCGGG | 564.1 | -8.5 | 1.2 | -7.1 | 1.6E-12 | 5.3E-12 | MB231/MB468 | NUC |
| >hg38_21_+_8433172_8446622_RNA45SN1_45S_50ntFlanks@11233.11254.22\|CCCTTCCCGTGGATCGCCCCAG | >hg38_21_+_8433172_8446622_RNA45SN1_45S_50ntFlanks | 11233 | 11254 | 22 | CCCTTCCCGTGGATCGCCCCAG | 685.8 | -12.5 | 1.7 | -7.5 | 7.4E-14 | 5.3E-12 | BT20/MB468 | MP |
| >hg38_21_+_8433172_8446622_RNA45SN1_45S_50ntFlanks@10873.10901.29\|CTGGGTCGGTCGGGCTGGGGCGCGAAGCG | >hg38_21_+_8433172_8446622_RNA45SN1_45S_50ntFlanks | 10873 | 10901 | 29 | CTGGGTCGGTCGGGCTGGGGCGCGAAGCG | 264.7 | -7.1 | 1.0 | -7.1 | 1.6E-12 | 5.3E-12 | MB231/MB468 | CYTO |
| >hg38_21_+_8433172_8446622_RNA45SN1_45S_50ntFlanks@10808.10825.18\|CAATGTAGGTAAGGGAAG | >hg38_21_+_8433172_8446622_RNA45SN1_45S_50ntFlanks | 10808 | 10825 | 18 | CAATGTAGGTAAGGGAAG | 1341.0 | -8.6 | 1.2 | -7.1 | 1.2E-12 | 5.3E-12 | BT20/MB468 | MITO |
| >hg38_21_+_8433172_8446622_RNA45SN1_45S_50ntFlanks@10683.10706.24\|CCGGCGGCGTCCGGTGAGCTCTCG | >hg38_21_+_8433172_8446622_RNA45SN1_45S_50ntFlanks | 10683 | 10706 | 24 | CCGGCGGCGTCCGGTGAGCTCTCG | 8908.5 | -6.0 | 0.8 | -7.1 | 1.6E-12 | 5.3E-12 | MB231/MB468 | CYTO |
| >MI0000285\|hsa-mir-205&WithFlank&1\|+\|209432127\|209432248@40.61.22(+2A)&[MIMAT0000266&hsa-miR-205-5p&offsets\|0\|0(+2A);m-351&1\|+\|209432166\|209432187&offsets\|0\|0(+2A)]\|TCCTTCATTCCACCGGAGTCTGAA | >MI0000285\|hsa-mir-205&WithFlank&1\|+\|209432127\|209432248 | 40 | 61 | 22(+2A) | TCCTTCATTCCACCGGAGTCTGAA | 1776.9 | 6.4 | 0.9 | 7.1 | 1.2E-12 | 5.3E-12 | BT20/MB468 | MITO |
| >hg38_21_+_8433172_8446622_RNA45SN1_45S_50ntFlanks@11729.11764.36\|TAACTATGACTCTCTTAAGGTAGCCAAATGCCTCGT | >hg38_21_+_8433172_8446622_RNA45SN1_45S_50ntFlanks | 11729 | 11764 | 36 | TAACTATGACTCTCTTAAGGTAGCCAAATGCCTCGT | 1936.0 | -7.2 | 1.0 | -7.1 | 1.6E-12 | 5.3E-12 | MB231/MB468 | CYTO |
| >hg38_21_+_8433172_8446622_RNA45SN1_45S_50ntFlanks@3990.4026.37\|TTGGTGACTCTAGATAACCTCGGGCCGATCGCACGCC | >hg38_21_+_8433172_8446622_RNA45SN1_45S_50ntFlanks | 3990 | 4026 | 37 | TTGGTGACTCTAGATAACCTCGGGCCGATCGCACGCC | 192.9 | -11.5 | 1.6 | -7.1 | 1.4E-12 | 5.3E-12 | MB231/MB468 | MP |
| >hg38_21_+_8433172_8446622_RNA45SN1_45S_50ntFlanks@10478.10512.35\|CCGATCGAAAGGGAGTCGGGTTCAGATCCCCGAAT | >hg38_21_+_8433172_8446622_RNA45SN1_45S_50ntFlanks | 10478 | 10512 | 35 | CCGATCGAAAGGGAGTCGGGTTCAGATCCCCGAAT | 314.9 | -5.4 | 0.8 | -7.1 | 1.6E-12 | 5.3E-12 | MB231/MB468 | CYTO |
| >hg38_21_+_8433172_8446622_RNA45SN1_45S_50ntFlanks@9738.9758.21\|GAAACGATCTCAACCTATTCT | >hg38_21_+_8433172_8446622_RNA45SN1_45S_50ntFlanks | 9738 | 9758 | 21 | GAAACGATCTCAACCTATTCT | 4297.8 | -5.9 | 0.8 | -7.1 | 1.6E-12 | 5.3E-12 | MB231/MB468 | CYTO |
| >hg38_21_+_8433172_8446622_RNA45SN1_45S_50ntFlanks@7989.8010.22\|AGACGTGGCGACCCGCTGAATT | >hg38_21_+_8433172_8446622_RNA45SN1_45S_50ntFlanks | 7989 | 8010 | 22 | AGACGTGGCGACCCGCTGAATT | 964.3 | 5.8 | 0.8 | 7.1 | 1.6E-12 | 5.3E-12 | BT20/MB468 | CYTO |
| >hg38_21_+_8433172_8446622_RNA45SN1_45S_50ntFlanks@12605.12623.19\|TATGTGCTTGGCTGAGGAG | >hg38_21_+_8433172_8446622_RNA45SN1_45S_50ntFlanks | 12605 | 12623 | 19 | TATGTGCTTGGCTGAGGAG | 2190.4 | -10.4 | 1.5 | -7.0 | 2.1E-12 | 5.3E-12 | BT20/MB468 | NUC |
| >hg38_21_+_8433172_8446622_RNA45SN1_45S_50ntFlanks@5484.5514.31\|GAGCGCTGAGAAGACGGTCGAACTTGACTAT | >hg38_21_+_8433172_8446622_RNA45SN1_45S_50ntFlanks | 5484 | 5514 | 31 | GAGCGCTGAGAAGACGGTCGAACTTGACTAT | 4162.3 | -4.2 | 0.6 | -7.1 | 1.6E-12 | 5.4E-12 | BT20/MB468 | CYTO |
| >am_trna2_GlyGCC_21_-_18827107_18827177@1.22.22,trna35_GlyGCC_1_+_161413094_161413164@1.22.22,trna37_GlyGCC_1_+_161420467_161420537@1.22.22,trna39_GlyGCC_1_+_161427898_161427968@1.22.22,trna41_GlyGCC_1_+_161435258_161435328@1.22.22\|GCATGGGTGGTTCAGTGGTAGA | >am_trna2_GlyGCC_21_-_18827107_18827177 | 1 | 22 | 22 | GCATGGGTGGTTCAGTGGTAGA | 705.4 | 5.2 | 0.7 | 7.1 | 1.6E-12 | 5.4E-12 | BT20/MB468 | CYTO |
| >hg38_21_+_8433172_8446622_RNA45SN1_45S_50ntFlanks@9472.9503.32\|GCACGAGCGCACGTGTTAGGACCCGAAAGATG | >hg38_21_+_8433172_8446622_RNA45SN1_45S_50ntFlanks | 9472 | 9503 | 32 | GCACGAGCGCACGTGTTAGGACCCGAAAGATG | 163.6 | -11.2 | 1.6 | -7.1 | 1.4E-12 | 5.4E-12 | MB231/MB468 | MP |
| >hg38_21_+_8433172_8446622_RNA45SN1_45S_50ntFlanks@9556.9582.27\|TAGCGGTCCTGACGTGCAAATCGGTCG | >hg38_21_+_8433172_8446622_RNA45SN1_45S_50ntFlanks | 9556 | 9582 | 27 | TAGCGGTCCTGACGTGCAAATCGGTCG | 155.2 | -11.1 | 1.6 | -7.1 | 1.4E-12 | 5.4E-12 | MB231/MB468 | MP |
| >hg38_21_+_8433172_8446622_RNA45SN1_45S_50ntFlanks@6707.6736.30\|CGAGAATTAATGTGAATTGCAGGACACATT | >hg38_21_+_8433172_8446622_RNA45SN1_45S_50ntFlanks | 6707 | 6736 | 30 | CGAGAATTAATGTGAATTGCAGGACACATT | 120.0 | -10.8 | 1.5 | -7.1 | 1.4E-12 | 5.4E-12 | MB231/MB468 | MP |
| >hg38_21_+_8433172_8446622_RNA45SN1_45S_50ntFlanks@10482.10499.18\|TCGAAAGGGAGTCGGGTT | >hg38_21_+_8433172_8446622_RNA45SN1_45S_50ntFlanks | 10482 | 10499 | 18 | TCGAAAGGGAGTCGGGTT | 154.9 | -11.1 | 1.6 | -7.1 | 1.4E-12 | 5.4E-12 | MB231/MB468 | MP |
| >hg38_21_+_8433172_8446622_RNA45SN1_45S_50ntFlanks@12946.12964.19\|TAGACGACCTGCTTCTGGG | >hg38_21_+_8433172_8446622_RNA45SN1_45S_50ntFlanks | 12946 | 12964 | 19 | TAGACGACCTGCTTCTGGG | 160.4 | -11.2 | 1.6 | -7.1 | 1.4E-12 | 5.4E-12 | MB231/MB468 | MP |
| >hg38_21_+_8433172_8446622_RNA45SN1_45S_50ntFlanks@10411.10428.18\|CAGTCGGTCCTGAGAGAT | >hg38_21_+_8433172_8446622_RNA45SN1_45S_50ntFlanks | 10411 | 10428 | 18 | CAGTCGGTCCTGAGAGAT | 4397.4 | -9.1 | 1.3 | -7.0 | 2.0E-12 | 5.4E-12 | MB231/MB468 | MITO |
| >hg38_21_+_8433172_8446622_RNA45SN1_45S_50ntFlanks@8972.8990.19\|CCACCCCCGCGGGGAATCC | >hg38_21_+_8433172_8446622_RNA45SN1_45S_50ntFlanks | 8972 | 8990 | 19 | CCACCCCCGCGGGGAATCC | 354.9 | -12.3 | 1.7 | -7.0 | 2.0E-12 | 5.4E-12 | MB231/MB468 | MITO |
| >hg38_21_+_8433172_8446622_RNA45SN1_45S_50ntFlanks@9790.9816.27\|CTGGCGTGGAGCCGGGCGTGGAATGCG | >hg38_21_+_8433172_8446622_RNA45SN1_45S_50ntFlanks | 9790 | 9816 | 27 | CTGGCGTGGAGCCGGGCGTGGAATGCG | 131.0 | -10.9 | 1.5 | -7.1 | 1.4E-12 | 5.4E-12 | MB231/MB468 | MP |
| >hg38_21_+_8433172_8446622_RNA45SN1_45S_50ntFlanks@12539.12564.26\|ATGATGTGTTGTTGCCATGGTAATCC | >hg38_21_+_8433172_8446622_RNA45SN1_45S_50ntFlanks | 12539 | 12564 | 26 | ATGATGTGTTGTTGCCATGGTAATCC | 118.6 | -10.8 | 1.5 | -7.1 | 1.4E-12 | 5.4E-12 | MB231/MB468 | MP |

<table>
<tr><td>>hg38_21_+_8433172_8446622_RNA45SN1_45S_50ntFlanks@11641.11663.23|CGGCGGGTGTTGACGCGATGTGA</td><td>>hg38_21_+_8433172_8446622_RNA45SN1_45S_50ntFlanks</td><td>11641</td><td>11663</td><td>23</td><td>CGGCGGGTGTTGACGCGATGTGA</td><td>843.2</td><td>-9.9</td><td>1.4</td><td>-7.0</td><td>2.2E-12</td><td>5.4E-12</td><td>BT20/MB468</td><td>NUC</td></tr>
<tr><td>>hg38_21_+_8433172_8446622_RNA45SN1_45S_50ntFlanks@3900.3921.22|CTTCGCGGGGGGGATGCGTGCA</td><td>>hg38_21_+_8433172_8446622_RNA45SN1_45S_50ntFlanks</td><td>3900</td><td>3921</td><td>22</td><td>CTTCGCGGGGGGGATGCGTGCA</td><td>139.3</td><td>-11.0</td><td>1.6</td><td>-7.1</td><td>1.4E-12</td><td>5.4E-12</td><td>MB231/MB468</td><td>MP</td></tr>
<tr><td>>hg38_21_+_8433172_8446622_RNA45SN1_45S_50ntFlanks@11568.11587.20|GCAGCCGACTTAGAACTGGT</td><td>>hg38_21_+_8433172_8446622_RNA45SN1_45S_50ntFlanks</td><td>11568</td><td>11587</td><td>20</td><td>GCAGCCGACTTAGAACTGGT</td><td>385.7</td><td>3.7</td><td>0.5</td><td>7.1</td><td>1.2E-12</td><td>5.4E-12</td><td>BT20/MB468</td><td>MITO</td></tr>
<tr><td>>hg38_21_+_8433172_8446622_RNA45SN1_45S_50ntFlanks@3987.4024.38|GCTTTGGTGACTCTAGATAACCTCGGGCCGATCGCACG</td><td>>hg38_21_+_8433172_8446622_RNA45SN1_45S_50ntFlanks</td><td>3987</td><td>4024</td><td>38</td><td>GCTTTGGTGACTCTAGATAACCTCGGGCCGATCGCACG</td><td>2612.9</td><td>-11.0</td><td>1.5</td><td>-7.1</td><td>1.4E-12</td><td>5.4E-12</td><td>MB231/MB468</td><td>MP</td></tr>
<tr><td>>hg38_21_+_8433172_8446622_RNA45SN1_45S_50ntFlanks@11564.11592.29|CCTAGCAGCCGACTTAGAACTGGTGCGGA</td><td>>hg38_21_+_8433172_8446622_RNA45SN1_45S_50ntFlanks</td><td>11564</td><td>11592</td><td>29</td><td>CCTAGCAGCCGACTTAGAACTGGTGCGGA</td><td>709.5</td><td>-8.8</td><td>1.2</td><td>-7.0</td><td>2.0E-12</td><td>5.4E-12</td><td>MB231/MB468</td><td>MITO</td></tr>
<tr><td>>hg38_21_+_8433172_8446622_RNA45SN1_45S_50ntFlanks@12324.12343.20|TTGGGTTTTAAGCAGGAGGT</td><td>>hg38_21_+_8433172_8446622_RNA45SN1_45S_50ntFlanks</td><td>12324</td><td>12343</td><td>20</td><td>TTGGGTTTTAAGCAGGAGGT</td><td>243.7</td><td>-7.0</td><td>1.0</td><td>-7.1</td><td>1.6E-12</td><td>5.4E-12</td><td>BT20/MB468</td><td>CYTO</td></tr>
<tr><td>>hg38_21_+_8433172_8446622_RNA45SN1_45S_50ntFlanks@11208.11231.24|GCGGCGACTCTGGACGCGAGCCGG</td><td>>hg38_21_+_8433172_8446622_RNA45SN1_45S_50ntFlanks</td><td>11208</td><td>11231</td><td>24</td><td>GCGGCGACTCTGGACGCGAGCCGG</td><td>2360.1</td><td>-6.2</td><td>0.9</td><td>-7.1</td><td>1.6E-12</td><td>5.4E-12</td><td>MB231/MB468</td><td>CYTO</td></tr>
<tr><td>>MI0000285|hsa-mir-205&WithFlank&1|+|209432127|209432248@40.61.22(+1C)&[MIMAT0000266&hsa-miR-205-5p&offsets|0|0(+1C);m-351&1|+|209432166|209432187&offsets|0|0(+1C)]|TCCTTCATTCCACCGGAGTCTGC</td><td>>MI0000285|hsa-mir-205&WithFlank&1|+|209432127|209432248</td><td>40</td><td>61</td><td>22(+1C)</td><td>TCCTTCATTCCACCGGAGTCTGC</td><td>10842.1</td><td>7.6</td><td>1.1</td><td>7.1</td><td>1.2E-12</td><td>5.4E-12</td><td>BT20/MB468</td><td>MITO</td></tr>
<tr><td>>hg38_21_+_8433172_8446622_RNA45SN1_45S_50ntFlanks@4265.4282.18|ATGAGTCCACTTTAAATC</td><td>>hg38_21_+_8433172_8446622_RNA45SN1_45S_50ntFlanks</td><td>4265</td><td>4282</td><td>18</td><td>ATGAGTCCACTTTAAATC</td><td>95.4</td><td>-10.4</td><td>1.5</td><td>-7.1</td><td>1.4E-12</td><td>5.4E-12</td><td>MB231/MB468</td><td>MP</td></tr>
<tr><td>>hg38_21_+_8433172_8446622_RNA45SN1_45S_50ntFlanks@8847.8867.21|GCGGCGGCGGCGGCGGCGGGA</td><td>>hg38_21_+_8433172_8446622_RNA45SN1_45S_50ntFlanks</td><td>8847</td><td>8867</td><td>21</td><td>GCGGCGGCGGCGGCGGCGGGA</td><td>144.0</td><td>-11.0</td><td>1.6</td><td>-7.1</td><td>1.4E-12</td><td>5.4E-12</td><td>MB231/MB468</td><td>MP</td></tr>
<tr><td>>hg38_21_+_8433172_8446622_RNA45SN1_45S_50ntFlanks@6699.6717.19|GCTAGCTGCGAGAATTAAT</td><td>>hg38_21_+_8433172_8446622_RNA45SN1_45S_50ntFlanks</td><td>6699</td><td>6717</td><td>19</td><td>GCTAGCTGCGAGAATTAAT</td><td>416.0</td><td>-5.7</td><td>0.8</td><td>-7.1</td><td>1.6E-12</td><td>5.5E-12</td><td>BT20/MB468</td><td>CYTO</td></tr>
<tr><td>>hg38_21_+_8433172_8446622_RNA45SN1_45S_50ntFlanks@12437.12454.18|CATTGTGAAGCAGAATTC</td><td>>hg38_21_+_8433172_8446622_RNA45SN1_45S_50ntFlanks</td><td>12437</td><td>12454</td><td>18</td><td>CATTGTGAAGCAGAATTC</td><td>2570.9</td><td>-5.5</td><td>0.8</td><td>-7.1</td><td>1.6E-12</td><td>5.5E-12</td><td>MB231/MB468</td><td>CYTO</td></tr>
<tr><td>>hg38_21_+_8433172_8446622_RNA45SN1_45S_50ntFlanks@10793.10823.31|TCTGGCATGTTGGAACAATGTAGGTAAGGGA</td><td>>hg38_21_+_8433172_8446622_RNA45SN1_45S_50ntFlanks</td><td>10793</td><td>10823</td><td>31</td><td>TCTGGCATGTTGGAACAATGTAGGTAAGGGA</td><td>103.7</td><td>-10.6</td><td>1.5</td><td>-7.1</td><td>1.4E-12</td><td>5.5E-12</td><td>MB231/MB468</td><td>MP</td></tr>
<tr><td>>hg38_21_+_8433172_8446622_RNA45SN1_45S_50ntFlanks@5074.5097.24|ATTCCGATAACGAACGAGACTCTG</td><td>>hg38_21_+_8433172_8446622_RNA45SN1_45S_50ntFlanks</td><td>5074</td><td>5097</td><td>24</td><td>ATTCCGATAACGAACGAGACTCTG</td><td>727.4</td><td>-13.4</td><td>1.9</td><td>-7.1</td><td>1.4E-12</td><td>5.5E-12</td><td>MB231/MB468</td><td>MP</td></tr>
<tr><td>>trnaMT_GluTTC_MT_-_14674_14742@38.71.34,trnalookalike8_GluTTC_5_-_93905172_93905240@38.71.34|CATTGGTCGTGGTTGTAGTCCGTGCGAGAATACC</td><td>>trnaMT_GluTTC_MT_-_14674_14742</td><td>38</td><td>71</td><td>34</td><td>CATTGGTCGTGGTTGTAGTCCGTGCGAGAATACC</td><td>462.5</td><td>5.4</td><td>0.8</td><td>7.1</td><td>1.7E-12</td><td>5.5E-12</td><td>MB231/MB468</td><td>NUC</td></tr>
<tr><td>>hg38_21_+_8433172_8446622_RNA45SN1_45S_50ntFlanks@10362.10384.23|AAGGCCGAAGTGGAGAAGGGTTC</td><td>>hg38_21_+_8433172_8446622_RNA45SN1_45S_50ntFlanks</td><td>10362</td><td>10384</td><td>23</td><td>AAGGCCGAAGTGGAGAAGGGTTC</td><td>641.9</td><td>-8.6</td><td>1.2</td><td>-7.0</td><td>2.0E-12</td><td>5.5E-12</td><td>MB231/MB468</td><td>MITO</td></tr>
<tr><td>>hg38_21_+_8433172_8446622_RNA45SN1_45S_50ntFlanks@11317.11340.24|ACCCCACGTCTCGTCGCGCGCGCG</td><td>>hg38_21_+_8433172_8446622_RNA45SN1_45S_50ntFlanks</td><td>11317</td><td>11340</td><td>24</td><td>ACCCCACGTCTCGTCGCGCGCGCG</td><td>578.1</td><td>-8.6</td><td>1.2</td><td>-7.1</td><td>1.7E-12</td><td>5.5E-12</td><td>MB231/MB468</td><td>NUC</td></tr>
<tr><td>>hg38_21_+_8433172_8446622_RNA45SN1_45S_50ntFlanks@12685.12708.24|CCCAGGCGGAACGATACGGCAGCG</td><td>>hg38_21_+_8433172_8446622_RNA45SN1_45S_50ntFlanks</td><td>12685</td><td>12708</td><td>24</td><td>CCCAGGCGGAACGATACGGCAGCG</td><td>103.3</td><td>-10.6</td><td>1.5</td><td>-7.1</td><td>1.5E-12</td><td>5.5E-12</td><td>MB231/MB468</td><td>MP</td></tr>
<tr><td>>hg38_21_+_8433172_8446622_RNA45SN1_45S_50ntFlanks@10410.10431.22|TCAGTCGGTCCTGAGAGATGGG</td><td>>hg38_21_+_8433172_8446622_RNA45SN1_45S_50ntFlanks</td><td>10410</td><td>10431</td><td>22</td><td>TCAGTCGGTCCTGAGAGATGGG</td><td>2751.1</td><td>-6.0</td><td>0.8</td><td>-7.1</td><td>1.7E-12</td><td>5.5E-12</td><td>MB231/MB468</td><td>CYTO</td></tr>
<tr><td>>hg38_21_+_8433172_8446622_RNA45SN1_45S_50ntFlanks@5186.5207.22|CAATAACAGGTCTGTGATGCCC</td><td>>hg38_21_+_8433172_8446622_RNA45SN1_45S_50ntFlanks</td><td>5186</td><td>5207</td><td>22</td><td>CAATAACAGGTCTGTGATGCCC</td><td>891.4</td><td>-13.6</td><td>1.9</td><td>-7.0</td><td>2.1E-12</td><td>5.5E-12</td><td>MB231/MB468</td><td>MITO</td></tr>
<tr><td>>am_trna128_GlyGCC_6_-_27870686_27870756@6.28.23,trna133_GlyCCC_1_-_16872434_16872504@6.28.23,trna17_GluTTC_2_-_204229131_204229201@6.28.23,trna18_GlyGCC_16_+_70822597_70822667@6.28.23,trna19_GlyGCC_16_+_70823410_70823480@6.28.23,trna19_GlyGCC_2_-_157257659_157257729@6.28.23,trna24_GlyGCC_16_-_70812942_70813012@6.28.23,trna25_GlyGCC_16_-_70812114_70812184@6.28.23,trna2_GlyGCC_21_-_18827107_18827177@6.28.23,trna35_GlyGCC_1_+_161413094_161413164@6.28.23,trna37_GlyGCC_1_+_161420467_161420537@6.28.23,trna39_GlyGCC_1_+_161427898_161427968@6.28.23,trna41_GlyGCC_1_+_161435258_161435328@6.28.23,trna4_GlyCCC_1_+_17188416_17188486@6.28.23,trna5_GlyGCC_17_+_8029064_8029134@6.28.23,trna68_GlyGCC_1_-_161493637_161493707@6.28.23,trna91_GlyCCC_1_-_149680210_149680280@6.28.23,trna99_ValCAC_1_-_149294666_149294736@6.28.23|GGTGGTTCAGTGGTAGAATTCTC</td><td>>am_trna128_GlyGCC_6_-_27870686_27870756</td><td>6</td><td>28</td><td>23</td><td>GGTGGTTCAGTGGTAGAATTCTC</td><td>193.4</td><td>-11.5</td><td>1.6</td><td>-7.1</td><td>1.5E-12</td><td>5.5E-12</td><td>MB231/MB468</td><td>MP</td></tr>
<tr><td>>hg38_21_+_8433172_8446622_RNA45SN1_45S_50ntFlanks@12074.12095.22|CGAGGGGCTCTCGCTTCTGGCG</td><td>>hg38_21_+_8433172_8446622_RNA45SN1_45S_50ntFlanks</td><td>12074</td><td>12095</td><td>22</td><td>CGAGGGGCTCTCGCTTCTGGCG</td><td>928.1</td><td>-9.1</td><td>1.3</td><td>-7.1</td><td>1.5E-12</td><td>5.5E-12</td><td>MB231/MB468</td><td>MP</td></tr>
<tr><td>>MI0000447|hsa-mir-128-1&WithFlank&2|+|135665391|135665484@56.75.20(+1A);MI0000727|hsa-mir-128-2&WithFlank&3|+|35744470|35744565@58.77.20(+1A)&[MIMAT0000424&hsa-miR-128-3p&offsets|0|-1(+1A);m-108&2|+|135665446|135665466&offsets|0|-1(+1A)];[MIMAT0000424_1&hsa-miR-128-3p&offsets|0|-1(+1A);m-117&3|+|35744527|35744547&offsets|0|-1(+1A)]|TCACAGTGAACCGGTCTCTTA</td><td>>MI0000447|hsa-mir-128-1&WithFlank&2|+|135665391|135665484</td><td>56</td><td>75</td><td>20(+1A)</td><td>TCACAGTGAACCGGTCTCTTA</td><td>540.8</td><td>6.9</td><td>1.0</td><td>7.1</td><td>1.6E-12</td><td>5.5E-12</td><td>BT20/MB468</td><td>CYTO</td></tr>
<tr><td>>hg38_21_+_8433172_8446622_RNA45SN1_45S_50ntFlanks@9552.9571.20|TCCGTAGCGGTCCTGACGTG</td><td>>hg38_21_+_8433172_8446622_RNA45SN1_45S_50ntFlanks</td><td>9552</td><td>9571</td><td>20</td><td>TCCGTAGCGGTCCTGACGTG</td><td>924.8</td><td>-3.5</td><td>0.5</td><td>-7.1</td><td>1.7E-12</td><td>5.5E-12</td><td>MB231/MB468</td><td>CYTO</td></tr>
<tr><td>>hg38_21_+_8433172_8446622_RNA45SN1_45S_50ntFlanks@4240.4258.19|ACTCTTTCGAGGCCCTGTA</td><td>>hg38_21_+_8433172_8446622_RNA45SN1_45S_50ntFlanks</td><td>4240</td><td>4258</td><td>19</td><td>ACTCTTTCGAGGCCCTGTA</td><td>680.9</td><td>-9.0</td><td>1.3</td><td>-7.0</td><td>2.1E-12</td><td>5.5E-12</td><td>MB231/MB468</td><td>MITO</td></tr>
<tr><td>>hg38_21_+_8433172_8446622_RNA45SN1_45S_50ntFlanks@10351.10376.26|CGAGAACTTTGAAGGCCGAAGTGGAG</td><td>>hg38_21_+_8433172_8446622_RNA45SN1_45S_50ntFlanks</td><td>10351</td><td>10376</td><td>26</td><td>CGAGAACTTTGAAGGCCGAAGTGGAG</td><td>1407.5</td><td>-6.3</td><td>0.9</td><td>-7.1</td><td>1.7E-12</td><td>5.6E-12</td><td>MB231/MB468</td><td>CYTO</td></tr>
<tr><td>>hg38_21_+_8433172_8446622_RNA45SN1_45S_50ntFlanks@11835.11852.18|CACAGCCAAGGGAACGGG</td><td>>hg38_21_+_8433172_8446622_RNA45SN1_45S_50ntFlanks</td><td>11835</td><td>11852</td><td>18</td><td>CACAGCCAAGGGAACGGG</td><td>116.8</td><td>-10.7</td><td>1.5</td><td>-7.1</td><td>1.5E-12</td><td>5.6E-12</td><td>MB231/MB468</td><td>MP</td></tr>
<tr><td>>hg38_21_+_8433172_8446622_RNA45SN1_45S_50ntFlanks@12436.12454.19|TCATTGTGAAGCAGAATTC</td><td>>hg38_21_+_8433172_8446622_RNA45SN1_45S_50ntFlanks</td><td>12436</td><td>12454</td><td>19</td><td>TCATTGTGAAGCAGAATTC</td><td>2787.8</td><td>-8.3</td><td>1.2</td><td>-7.1</td><td>1.5E-12</td><td>5.6E-12</td><td>MB231/MB468</td><td>MP</td></tr>
<tr><td>>hg38_21_+_8433172_8446622_RNA45SN1_45S_50ntFlanks@10348.10368.21|AAACGAGAACTTTGAAGGCCG</td><td>>hg38_21_+_8433172_8446622_RNA45SN1_45S_50ntFlanks</td><td>10348</td><td>10368</td><td>21</td><td>AAACGAGAACTTTGAAGGCCG</td><td>1116.8</td><td>-5.3</td><td>0.8</td><td>-7.0</td><td>2.1E-12</td><td>5.6E-12</td><td>MB231/MB468</td><td>MITO</td></tr>
<tr><td>>MI0000299|hsa-mir-222&WithFlank&X|-|45747009|45747130@75.96.22&[MIMAT0000279&hsa-miR-222-3p&offsets|0|+1;m-140&X|-|45747033|45747056&offsets|0|-2]|AGCTACATCTGGCTACTGGGTC</td><td>>MI0000299|hsa-mir-222&WithFlank&X|-|45747009|45747130</td><td>75</td><td>96</td><td>22</td><td>AGCTACATCTGGCTACTGGGTC</td><td>1730.3</td><td>5.8</td><td>0.8</td><td>7.0</td><td>2.1E-12</td><td>5.6E-12</td><td>MB231/MB468</td><td>MITO</td></tr>
<tr><td>>hg38_21_+_8433172_8446622_RNA45SN1_45S_50ntFlanks@11583.11605.23|CTGGTGCGGACCAGGGGAATCCG</td><td>>hg38_21_+_8433172_8446622_RNA45SN1_45S_50ntFlanks</td><td>11583</td><td>11605</td><td>23</td><td>CTGGTGCGGACCAGGGGAATCCG</td><td>714.8</td><td>-6.9</td><td>1.0</td><td>-7.1</td><td>1.7E-12</td><td>5.6E-12</td><td>MB231/MB468</td><td>NUC</td></tr>
<tr><td>>hg38_21_+_8433172_8446622_RNA45SN1_45S_50ntFlanks@9790.9821.32|CTGGCGTGGAGCCGGGCGTGGAATGCGAGTGC</td><td>>hg38_21_+_8433172_8446622_RNA45SN1_45S_50ntFlanks</td><td>9790</td><td>9821</td><td>32</td><td>CTGGCGTGGAGCCGGGCGTGGAATGCGAGTGC</td><td>99.0</td><td>-10.5</td><td>1.5</td><td>-7.1</td><td>1.5E-12</td><td>5.6E-12</td><td>MB231/MB468</td><td>MP</td></tr>
<tr><td>>MI0000063|hsa-let-7b&WithFlank&22|+|46113680|46113774@12.33.22(+1C)&[MIMAT0000063&hsa-let-7b-5p&offsets|0|0(+1C);m-14&22|+|46113691|46113712&offsets|0|0(+1C)]|TGAGGTAGTAGGTTGTGTGGTTC</td><td>>MI0000063|hsa-let-7b&WithFlank&22|+|46113680|46113774</td><td>12</td><td>33</td><td>22(+1C)</td><td>TGAGGTAGTAGGTTGTGTGGTTC</td><td>293.5</td><td>5.7</td><td>0.8</td><td>7.1</td><td>1.7E-12</td><td>5.6E-12</td><td>MB231/MB468</td><td>CYTO</td></tr>
<tr><td>>hg38_21_+_8433172_8446622_RNA45SN1_45S_50ntFlanks@5084.5110.27|CGAACGAGACTCTGGCATGCTAACTAG</td><td>>hg38_21_+_8433172_8446622_RNA45SN1_45S_50ntFlanks</td><td>5084</td><td>5110</td><td>27</td><td>CGAACGAGACTCTGGCATGCTAACTAG</td><td>2486.6</td><td>-9.1</td><td>1.3</td><td>-7.0</td><td>2.2E-12</td><td>5.6E-12</td><td>BT20/MB468</td><td>NUC</td></tr>
<tr><td>>MI0000809|hsa-mir-151a&WithFlank&8|-|140732558|140732659@53.74.22(+1A)&[MIMAT0000757&hsa-miR-151a-3p&offsets|0|+1(+1A);m-30&8|-|140732587|140732607&offsets|0|+1(+1A)]|CTAGACTGAAGCTCCTTGAGGAA</td><td>>MI0000809|hsa-mir-151a&WithFlank&8|-|140732558|140732659</td><td>53</td><td>74</td><td>22(+1A)</td><td>CTAGACTGAAGCTCCTTGAGGAA</td><td>4740.8</td><td>5.5</td><td>0.7</td><td>7.5</td><td>7.8E-14</td><td>5.6E-12</td><td>BT20/MB468</td><td>MP</td></tr>
<tr><td>>hg38_21_+_8433172_8446622_RNA45SN1_45S_50ntFlanks@9339.9364.26|CGCACGAAAGCCGCCGTGGCGCAATG</td><td>>hg38_21_+_8433172_8446622_RNA45SN1_45S_50ntFlanks</td><td>9339</td><td>9364</td><td>26</td><td>CGCACGAAAGCCGCCGTGGCGCAATG</td><td>336.2</td><td>-12.3</td><td>1.7</td><td>-7.1</td><td>1.5E-12</td><td>5.6E-12</td><td>MB231/MB468</td><td>MP</td></tr>
<tr><td>>MI0000065|hsa-let-7d&WithFlank&9|+|94178828|94178926@68.88.21(+1A)&[MIMAT0004484&hsa-let-7d-3p&offsets|0|-1(+1A);m-98&9|+|94178895|94178916&offsets|0|-1(+1A)]|CTATACGACCTGCTGCCTTTCA</td><td>>MI0000065|hsa-let-7d&WithFlank&9|+|94178828|94178926</td><td>68</td><td>88</td><td>21(+1A)</td><td>CTATACGACCTGCTGCCTTTCA</td><td>600.3</td><td>10.1</td><td>1.4</td><td>7.1</td><td>1.6E-12</td><td>5.6E-12</td><td>BT20/MB468</td><td>CYTO</td></tr>
<tr><td>>hg38_21_+_8433172_8446622_RNA45SN1_45S_50ntFlanks@12941.12972.32|ATTCGTAGACGACCTGCTTCTGGGTCGGGGTT</td><td>>hg38_21_+_8433172_8446622_RNA45SN1_45S_50ntFlanks</td><td>12941</td><td>12972</td><td>32</td><td>ATTCGTAGACGACCTGCTTCTGGGTCGGGGTT</td><td>295.4</td><td>-7.9</td><td>1.1</td><td>-7.1</td><td>1.7E-12</td><td>5.6E-12</td><td>MB231/MB468</td><td>CYTO</td></tr>
<tr><td>>hg38_21_+_8433172_8446622_RNA45SN1_45S_50ntFlanks@3985.4027.43|CGGCTTTGGTGACTCTAGATAACCTCGGGCCGATCGCACGCCC</td><td>>hg38_21_+_8433172_8446622_RNA45SN1_45S_50ntFlanks</td><td>3985</td><td>4027</td><td>43</td><td>CGGCTTTGGTGACTCTAGATAACCTCGGGCCGATCGCACGCCC</td><td>226.9</td><td>-11.6</td><td>1.7</td><td>-7.0</td><td>2.1E-12</td><td>5.6E-12</td><td>MB231/MB468</td><td>MITO</td></tr>
<tr><td>>am_MI0000809|hsa-mir-151a&WithFlank&8|-|140732558|140732659@53.73.21(+1G)&[MIMAT0000757&hsa-miR-151a-3p&offsets|0|0(+1G);m-30&8|-|140732587|140732607&offsets|0|0(+1G)]|CTAGACTGAAGCTCCTTGAGGG</td><td>>am_MI0000809|hsa-mir-151a&WithFlank&8|-|140732558|140732659</td><td>53</td><td>73</td><td>21(+1G)</td><td>CTAGACTGAAGCTCCTTGAGGG</td><td>431.8</td><td>8.7</td><td>1.2</td><td>7.1</td><td>1.3E-12</td><td>5.6E-12</td><td>BT20/MB468</td><td>MITO</td></tr>
<tr><td>>hg38_21_+_8433172_8446622_RNA45SN1_45S_50ntFlanks@9225.9249.25|CCGAAGAGGGGGACGGCGGAGCGAG</td><td>>hg38_21_+_8433172_8446622_RNA45SN1_45S_50ntFlanks</td><td>9225</td><td>9249</td><td>25</td><td>CCGAAGAGGGGGACGGCGGAGCGAG</td><td>104.2</td><td>-10.6</td><td>1.5</td><td>-7.1</td><td>1.5E-12</td><td>5.6E-12</td><td>MB231/MB468</td><td>MP</td></tr>
<tr><td>>MI0000285|hsa-mir-205&WithFlank&1|+|209432127|209432248@40.61.22(+1G)&[MIMAT0000266&hsa-miR-205-5p&offsets|0|0(+1G);m-351&1|+|209432166|209432187&offsets|0|0(+1G)]|TCCTTCATTCCACCGGAGTCTGG</td><td>>MI0000285|hsa-mir-205&WithFlank&1|+|209432127|209432248</td><td>40</td><td>61</td><td>22(+1G)</td><td>TCCTTCATTCCACCGGAGTCTGG</td><td>190.9</td><td>-11.4</td><td>1.6</td><td>-7.1</td><td>1.5E-12</td><td>5.6E-12</td><td>MB231/MB468</td><td>MP</td></tr>
<tr><td>>hg38_21_+_8433172_8446622_RNA45SN1_45S_50ntFlanks@11585.11623.39|GGTGCGGACCAGGGGAATCCGACTGTTTAATTAAAACAA</td><td>>hg38_21_+_8433172_8446622_RNA45SN1_45S_50ntFlanks</td><td>11585</td><td>11623</td><td>39</td><td>GGTGCGGACCAGGGGAATCCGACTGTTTAATTAAAACAA</td><td>660.2</td><td>13.1</td><td>1.8</td><td>7.5</td><td>7.9E-14</td><td>5.6E-12</td><td>BT20/MB468</td><td>MP</td></tr>
<tr><td>>hg38_21_+_8433172_8446622_RNA45SN1_45S_50ntFlanks@12415.12455.41|CTTCGATGTCGGCTCTTCCTATCATTGTGAAGCAGAATTCA</td><td>>hg38_21_+_8433172_8446622_RNA45SN1_45S_50ntFlanks</td><td>12415</td><td>12455</td><td>41</td><td>CTTCGATGTCGGCTCTTCCTATCATTGTGAAGCAGAATTCA</td><td>3278.6</td><td>-5.6</td><td>0.8</td><td>-7.0</td><td>2.1E-12</td><td>5.7E-12</td><td>MB231/MB468</td><td>MITO</td></tr>
</table>

<table>
<tr><td>>hg38_21_+_8433172_8446622_RNA45SN1_45S_50ntFlanks@9858.9902.45|CGGGATGAACCGAACGCCGGGTTAAGGCGCCCGATGCCGACGCTC</td><td>>hg38_21_+_8433172_8446622_RNA45SN1_45S_50ntFlanks</td><td>9858</td><td>9902</td><td>45</td><td>CGGGATGAACCGAACGCCGGGTTAAGGCGCCCGATGCCGACGCTC</td><td>440.5</td><td>-12.6</td><td>1.8</td><td>-7.0</td><td>2.1E-12</td><td>5.7E-12</td><td>MB231/MB468</td><td>MITO</td></tr>
<tr><td>>MI0005756|hsa-mir-934&WithFlank&X|+|136550872|136550966@21.42.22(+1A)&[MIMAT0004977&hsa-miR-934&offsets|0|0(+1A);m-943&X|+|136550892|136550913&offsets|0|0(+1A)]|TGTCTACTACTGGAGACACTGGA</td><td>>MI0005756|hsa-mir-934&WithFlank&X|+|136550872|136550966</td><td>21</td><td>42</td><td>22(+1A)</td><td>TGTCTACTACTGGAGACACTGGA</td><td>552.4</td><td>4.6</td><td>0.7</td><td>7.0</td><td>2.3E-12</td><td>5.7E-12</td><td>BT20/MB468</td><td>NUC</td></tr>
<tr><td>>hg38_21_+_8433172_8446622_RNA45SN1_45S_50ntFlanks@12971.12990.20|TTTCGTACGTAGCAGAGCAG</td><td>>hg38_21_+_8433172_8446622_RNA45SN1_45S_50ntFlanks</td><td>12971</td><td>12990</td><td>20</td><td>TTTCGTACGTAGCAGAGCAG</td><td>355.8</td><td>-8.5</td><td>1.2</td><td>-7.0</td><td>2.1E-12</td><td>5.7E-12</td><td>MB231/MB468</td><td>MITO</td></tr>
<tr><td>>hg38_21_+_8433172_8446622_RNA45SN1_45S_50ntFlanks@12685.12711.27|CCCAGGCGGAACGATACGGCAGCGCCG</td><td>>hg38_21_+_8433172_8446622_RNA45SN1_45S_50ntFlanks</td><td>12685</td><td>12711</td><td>27</td><td>CCCAGGCGGAACGATACGGCAGCGCCG</td><td>225.2</td><td>-11.7</td><td>1.7</td><td>-7.1</td><td>1.5E-12</td><td>5.7E-12</td><td>MB231/MB468</td><td>MP</td></tr>
<tr><td>>hg38_21_+_8433172_8446622_RNA45SN1_45S_50ntFlanks@4047.4065.19|ATTCGAACGTCTGCCCTAT</td><td>>hg38_21_+_8433172_8446622_RNA45SN1_45S_50ntFlanks</td><td>4047</td><td>4065</td><td>19</td><td>ATTCGAACGTCTGCCCTAT</td><td>429.7</td><td>-3.5</td><td>0.5</td><td>-7.1</td><td>1.7E-12</td><td>5.7E-12</td><td>BT20/MB468</td><td>CYTO</td></tr>
<tr><td>>hg38_21_+_8433172_8446622_RNA45SN1_45S_50ntFlanks@11161.11189.29|CACGGTCCCCCGCGAGGGGGGCCCGGGCA</td><td>>hg38_21_+_8433172_8446622_RNA45SN1_45S_50ntFlanks</td><td>11161</td><td>11189</td><td>29</td><td>CACGGTCCCCCGCGAGGGGGGCCCGGGCA</td><td>310.0</td><td>-11.2</td><td>1.6</td><td>-7.1</td><td>1.8E-12</td><td>5.7E-12</td><td>MB231/MB468</td><td>NUC</td></tr>
<tr><td>>MI0000299|hsa-mir-222&WithFlank&X|-|45747009|45747130@75.96.22(+1C)&[MIMAT0000279&hsa-miR-222-3p&offsets|0|+1(+1C);m-140&X|-|45747033|45747056&offsets|0|-2(+1C)]|AGCTACATCTGGCTACTGGGTCC</td><td>>MI0000299|hsa-mir-222&WithFlank&X|-|45747009|45747130</td><td>75</td><td>96</td><td>22(+1C)</td><td>AGCTACATCTGGCTACTGGGTCC</td><td>1549.1</td><td>6.9</td><td>1.0</td><td>7.0</td><td>2.1E-12</td><td>5.7E-12</td><td>MB231/MB468</td><td>MITO</td></tr>
<tr><td>>hg38_21_+_8433172_8446622_RNA45SN1_45S_50ntFlanks@4761.4782.22|CAGATACCGTCGTAGTTCCGAC</td><td>>hg38_21_+_8433172_8446622_RNA45SN1_45S_50ntFlanks</td><td>4761</td><td>4782</td><td>22</td><td>CAGATACCGTCGTAGTTCCGAC</td><td>97.1</td><td>-10.5</td><td>1.5</td><td>-7.1</td><td>1.5E-12</td><td>5.7E-12</td><td>MB231/MB468</td><td>MP</td></tr>
<tr><td>>hg38_21_+_8433172_8446622_RNA45SN1_45S_50ntFlanks@8202.8223.22|CGGTGTGAGGCCGGTAGCGGCC</td><td>>hg38_21_+_8433172_8446622_RNA45SN1_45S_50ntFlanks</td><td>8202</td><td>8223</td><td>22</td><td>CGGTGTGAGGCCGGTAGCGGCC</td><td>103.6</td><td>-10.6</td><td>1.5</td><td>-7.1</td><td>1.5E-12</td><td>5.7E-12</td><td>MB231/MB468</td><td>MP</td></tr>
<tr><td>>hg38_21_+_8433172_8446622_RNA45SN1_45S_50ntFlanks@12709.12747.39|CCGCGGAGCCTCGGTTGGCCTCGGATAGCCGGTCCCCCG</td><td>>hg38_21_+_8433172_8446622_RNA45SN1_45S_50ntFlanks</td><td>12709</td><td>12747</td><td>39</td><td>CCGCGGAGCCTCGGTTGGCCTCGGATAGCCGGTCCCCCG</td><td>173.4</td><td>-11.3</td><td>1.6</td><td>-7.1</td><td>1.5E-12</td><td>5.7E-12</td><td>MB231/MB468</td><td>MP</td></tr>
<tr><td>>MI0000477|hsa-mir-146a&WithFlank&5|+|160485346|160485456@27.44.18&[MIMAT0000449&hsa-miR-146a-5p&offsets|0|-4;m-129&5|+|160485372|160485393&offsets|0|-4]|TGAGAACTGAATTCCATG</td><td>>MI0000477|hsa-mir-146a&WithFlank&5|+|160485346|160485456</td><td>27</td><td>44</td><td>18</td><td>TGAGAACTGAATTCCATG</td><td>474.1</td><td>12.1</td><td>1.7</td><td>7.1</td><td>1.5E-12</td><td>5.7E-12</td><td>MB231/MB468</td><td>MP</td></tr>
<tr><td>>hg38_21_+_8433172_8446622_RNA45SN1_45S_50ntFlanks@4297.4320.24|CATTGGAGGGCAAGTCTGGTGCCA</td><td>>hg38_21_+_8433172_8446622_RNA45SN1_45S_50ntFlanks</td><td>4297</td><td>4320</td><td>24</td><td>CATTGGAGGGCAAGTCTGGTGCCA</td><td>194.6</td><td>-5.1</td><td>0.7</td><td>-7.1</td><td>1.7E-12</td><td>5.7E-12</td><td>MB231/MB468</td><td>CYTO</td></tr>
<tr><td>>hg38_21_+_8433172_8446622_RNA45SN1_45S_50ntFlanks@10465.10495.31|CCGTTGCCCTCGGCCGATCGAAAGGGAGTCG</td><td>>hg38_21_+_8433172_8446622_RNA45SN1_45S_50ntFlanks</td><td>10465</td><td>10495</td><td>31</td><td>CCGTTGCCCTCGGCCGATCGAAAGGGAGTCG</td><td>110.4</td><td>-10.6</td><td>1.5</td><td>-7.1</td><td>1.5E-12</td><td>5.7E-12</td><td>MB231/MB468</td><td>MP</td></tr>
<tr><td>>hg38_21_+_8433172_8446622_RNA45SN1_45S_50ntFlanks@7979.8007.29|ACCTCAGATCAGACGTGGCGACCCGCTGA</td><td>>hg38_21_+_8433172_8446622_RNA45SN1_45S_50ntFlanks</td><td>7979</td><td>8007</td><td>29</td><td>ACCTCAGATCAGACGTGGCGACCCGCTGA</td><td>412.4</td><td>-8.4</td><td>1.2</td><td>-7.1</td><td>1.5E-12</td><td>5.7E-12</td><td>MB231/MB468</td><td>MP</td></tr>
<tr><td>>hg38_21_+_8433172_8446622_RNA45SN1_45S_50ntFlanks@6774.6802.29|TCCTCCCGGGGCTACGCCTGTCTGAGCGT</td><td>>hg38_21_+_8433172_8446622_RNA45SN1_45S_50ntFlanks</td><td>6774</td><td>6802</td><td>29</td><td>TCCTCCCGGGGCTACGCCTGTCTGAGCGT</td><td>701.0</td><td>6.7</td><td>0.9</td><td>7.1</td><td>1.3E-12</td><td>5.8E-12</td><td>BT20/MB468</td><td>MITO</td></tr>
<tr><td>>hg38_21_+_8433172_8446622_RNA45SN1_45S_50ntFlanks@3593.3610.18|CCCGTCCGTCCGTCCGTC</td><td>>hg38_21_+_8433172_8446622_RNA45SN1_45S_50ntFlanks</td><td>3593</td><td>3610</td><td>18</td><td>CCCGTCCGTCCGTCCGTC</td><td>364.8</td><td>-7.6</td><td>1.1</td><td>-7.1</td><td>1.7E-12</td><td>5.8E-12</td><td>BT20/MB468</td><td>CYTO</td></tr>
<tr><td>>hg38_21_+_8433172_8446622_RNA45SN1_45S_50ntFlanks@8737.8754.18|CCCGTCCGTCCGTCCGTC</td><td>>hg38_21_+_8433172_8446622_RNA45SN1_45S_50ntFlanks</td><td>8737</td><td>8754</td><td>18</td><td>CCCGTCCGTCCGTCCGTC</td><td>364.8</td><td>-7.6</td><td>1.1</td><td>-7.1</td><td>1.7E-12</td><td>5.8E-12</td><td>BT20/MB468</td><td>CYTO</td></tr>
<tr><td>>hg38_21_+_8433172_8446622_RNA45SN1_45S_50ntFlanks@8836.8859.24|CGGCGGTGGCGGCGGCGGCGGCGG</td><td>>hg38_21_+_8433172_8446622_RNA45SN1_45S_50ntFlanks</td><td>8836</td><td>8859</td><td>24</td><td>CGGCGGTGGCGGCGGCGGCGGCGG</td><td>168.2</td><td>-11.3</td><td>1.6</td><td>-7.1</td><td>1.5E-12</td><td>5.8E-12</td><td>MB231/MB468</td><td>MP</td></tr>
<tr><td>>hg38_21_+_8433172_8446622_RNA45SN1_45S_50ntFlanks@4817.4835.19|ATTCCCATGACCCGCCGGG</td><td>>hg38_21_+_8433172_8446622_RNA45SN1_45S_50ntFlanks</td><td>4817</td><td>4835</td><td>19</td><td>ATTCCCATGACCCGCCGGG</td><td>110.2</td><td>-10.6</td><td>1.5</td><td>-7.1</td><td>1.5E-12</td><td>5.8E-12</td><td>MB231/MB468</td><td>MP</td></tr>
<tr><td>>hg38_21_+_8433172_8446622_RNA45SN1_45S_50ntFlanks@10869.10894.26|AGGGCTGGGTCGGTCGGGCTGGGGCG</td><td>>hg38_21_+_8433172_8446622_RNA45SN1_45S_50ntFlanks</td><td>10869</td><td>10894</td><td>26</td><td>AGGGCTGGGTCGGTCGGGCTGGGGCG</td><td>666.0</td><td>-9.3</td><td>1.3</td><td>-7.0</td><td>2.2E-12</td><td>5.8E-12</td><td>MB231/MB468</td><td>MITO</td></tr>
<tr><td>>trna10_MetCAT_8_-_124169470_124169542@31.51.21,trna22_MetCAT_16_-_87417628_87417700@31.51.21|TCTCATAATCTGAAGGTCGTG</td><td>>trna10_MetCAT_8_-_124169470_124169542</td><td>31</td><td>51</td><td>21</td><td>TCTCATAATCTGAAGGTCGTG</td><td>206.4</td><td>-9.8</td><td>1.4</td><td>-7.1</td><td>1.3E-12</td><td>5.8E-12</td><td>BT20/MB468</td><td>MITO</td></tr>
<tr><td>>hg38_21_+_8433172_8446622_RNA45SN1_45S_50ntFlanks@4224.4242.19|AAAATAACAATACAGGACT</td><td>>hg38_21_+_8433172_8446622_RNA45SN1_45S_50ntFlanks</td><td>4224</td><td>4242</td><td>19</td><td>AAAATAACAATACAGGACT</td><td>2107.3</td><td>-8.2</td><td>1.2</td><td>-7.1</td><td>1.5E-12</td><td>5.8E-12</td><td>MB231/MB468</td><td>MP</td></tr>
<tr><td>>hg38_21_+_8433172_8446622_RNA45SN1_45S_50ntFlanks@10664.10709.46|CTTGGAAAGCGTCGCGGTTCCGGCGGCGTCCGGTGAGCTCTCGCTG</td><td>>hg38_21_+_8433172_8446622_RNA45SN1_45S_50ntFlanks</td><td>10664</td><td>10709</td><td>46</td><td>CTTGGAAAGCGTCGCGGTTCCGGCGGCGTCCGGTGAGCTCTCGCTG</td><td>302.1</td><td>-12.0</td><td>1.7</td><td>-7.0</td><td>2.2E-12</td><td>5.8E-12</td><td>MB231/MB468</td><td>MITO</td></tr>
<tr><td>>hg38_21_+_8433172_8446622_RNA45SN1_45S_50ntFlanks@13004.13023.20|ATCTATTGAAAGTCAGCCCT</td><td>>hg38_21_+_8433172_8446622_RNA45SN1_45S_50ntFlanks</td><td>13004</td><td>13023</td><td>20</td><td>ATCTATTGAAAGTCAGCCCT</td><td>302.7</td><td>-7.1</td><td>1.0</td><td>-7.1</td><td>1.8E-12</td><td>5.8E-12</td><td>MB231/MB468</td><td>CYTO</td></tr>
<tr><td>>hg38_21_+_8433172_8446622_RNA45SN1_45S_50ntFlanks@11686.11717.32|TCAAAGTGAAGAAATTCAATGAAGCGCGGGTA</td><td>>hg38_21_+_8433172_8446622_RNA45SN1_45S_50ntFlanks</td><td>11686</td><td>11717</td><td>32</td><td>TCAAAGTGAAGAAATTCAATGAAGCGCGGGTA</td><td>200.8</td><td>-10.6</td><td>1.5</td><td>-7.0</td><td>1.8E-12</td><td>5.8E-12</td><td>MB231/MB468</td><td>NUC</td></tr>
<tr><td>>hg38_21_+_8433172_8446622_RNA45SN1_45S_50ntFlanks@9468.9503.36|TGGAGCACGAGCGCACGTGTTAGGACCCGAAAGATG</td><td>>hg38_21_+_8433172_8446622_RNA45SN1_45S_50ntFlanks</td><td>9468</td><td>9503</td><td>36</td><td>TGGAGCACGAGCGCACGTGTTAGGACCCGAAAGATG</td><td>200.6</td><td>-11.5</td><td>1.6</td><td>-7.1</td><td>1.5E-12</td><td>5.8E-12</td><td>MB231/MB468</td><td>MP</td></tr>
<tr><td>>hg38_21_+_8433172_8446622_RNA45SN1_45S_50ntFlanks@6651.6676.26|CGACTCTTAGCGGTGGATCACTCGGC</td><td>>hg38_21_+_8433172_8446622_RNA45SN1_45S_50ntFlanks</td><td>6651</td><td>6676</td><td>26</td><td>CGACTCTTAGCGGTGGATCACTCGGC</td><td>61390.1</td><td>9.2</td><td>1.3</td><td>7.1</td><td>1.7E-12</td><td>5.9E-12</td><td>BT20/MB468</td><td>CYTO</td></tr>
<tr><td>>hg38_21_+_8433172_8446622_RNA45SN1_45S_50ntFlanks@9202.9219.18|TCTCGGGGCCACGCGCGC</td><td>>hg38_21_+_8433172_8446622_RNA45SN1_45S_50ntFlanks</td><td>9202</td><td>9219</td><td>18</td><td>TCTCGGGGCCACGCGCGC</td><td>123.0</td><td>-10.8</td><td>1.5</td><td>-7.1</td><td>1.6E-12</td><td>5.9E-12</td><td>MB231/MB468</td><td>MP</td></tr>
<tr><td>>hg38_21_+_8433172_8446622_RNA45SN1_45S_50ntFlanks@5471.5514.44|CACGGCCCTGGCGGAGCGCTGAGAAGACGGTCGAACTTGACTAT</td><td>>hg38_21_+_8433172_8446622_RNA45SN1_45S_50ntFlanks</td><td>5471</td><td>5514</td><td>44</td><td>CACGGCCCTGGCGGAGCGCTGAGAAGACGGTCGAACTTGACTAT</td><td>833.6</td><td>-13.6</td><td>1.9</td><td>-7.1</td><td>1.6E-12</td><td>5.9E-12</td><td>MB231/MB468</td><td>MP</td></tr>
<tr><td>>hg38_21_+_8433172_8446622_RNA45SN1_45S_50ntFlanks@10841.10862.22|CGTAACTTCGGGATAAGGATTG</td><td>>hg38_21_+_8433172_8446622_RNA45SN1_45S_50ntFlanks</td><td>10841</td><td>10862</td><td>22</td><td>CGTAACTTCGGGATAAGGATTG</td><td>1357.8</td><td>-9.1</td><td>1.3</td><td>-7.0</td><td>2.2E-12</td><td>5.9E-12</td><td>MB231/MB468</td><td>MITO</td></tr>
<tr><td>>hg38_21_+_8433172_8446622_RNA45SN1_45S_50ntFlanks@5449.5466.18|ATCGGCCCCGCCGGGGTC</td><td>>hg38_21_+_8433172_8446622_RNA45SN1_45S_50ntFlanks</td><td>5449</td><td>5466</td><td>18</td><td>ATCGGCCCCGCCGGGGTC</td><td>222.8</td><td>-7.0</td><td>1.0</td><td>-7.1</td><td>1.8E-12</td><td>5.9E-12</td><td>MB231/MB468</td><td>CYTO</td></tr>
<tr><td>>hg38_21_+_8433172_8446622_RNA45SN1_45S_50ntFlanks@7992.8020.29|CGTGGCGACCCGCTGAATTTAAGCATATT</td><td>>hg38_21_+_8433172_8446622_RNA45SN1_45S_50ntFlanks</td><td>7992</td><td>8020</td><td>29</td><td>CGTGGCGACCCGCTGAATTTAAGCATATT</td><td>142.6</td><td>7.0</td><td>1.0</td><td>7.1</td><td>1.8E-12</td><td>5.9E-12</td><td>MB231/MB468</td><td>CYTO</td></tr>
<tr><td>>hg38_21_+_8433172_8446622_RNA45SN1_45S_50ntFlanks@6748.6771.24|CTTCGAACGCACTTGCGGCCCCGG</td><td>>hg38_21_+_8433172_8446622_RNA45SN1_45S_50ntFlanks</td><td>6748</td><td>6771</td><td>24</td><td>CTTCGAACGCACTTGCGGCCCCGG</td><td>196.8</td><td>-9.7</td><td>1.4</td><td>-7.1</td><td>1.3E-12</td><td>5.9E-12</td><td>BT20/MB468</td><td>MITO</td></tr>
<tr><td>>hg38_21_+_8433172_8446622_RNA45SN1_45S_50ntFlanks@10098.10117.20|GCGCGCGCGCGTGTGGTGTG</td><td>>hg38_21_+_8433172_8446622_RNA45SN1_45S_50ntFlanks</td><td>10098</td><td>10117</td><td>20</td><td>GCGCGCGCGCGTGTGGTGTG</td><td>130.1</td><td>-10.9</td><td>1.5</td><td>-7.1</td><td>1.6E-12</td><td>5.9E-12</td><td>MB231/MB468</td><td>MP</td></tr>
<tr><td>>hg38_21_+_8433172_8446622_RNA45SN1_45S_50ntFlanks@5493.5513.21|GAAGACGGTCGAACTTGACTA</td><td>>hg38_21_+_8433172_8446622_RNA45SN1_45S_50ntFlanks</td><td>5493</td><td>5513</td><td>21</td><td>GAAGACGGTCGAACTTGACTA</td><td>123.7</td><td>-10.8</td><td>1.5</td><td>-7.1</td><td>1.6E-12</td><td>5.9E-12</td><td>MB231/MB468</td><td>MP</td></tr>
<tr><td>>hg38_21_+_8433172_8446622_RNA45SN1_45S_50ntFlanks@9568.9592.25|CGTGCAAATCGGTCGTCCGACCTGG</td><td>>hg38_21_+_8433172_8446622_RNA45SN1_45S_50ntFlanks</td><td>9568</td><td>9592</td><td>25</td><td>CGTGCAAATCGGTCGTCCGACCTGG</td><td>435.3</td><td>-9.0</td><td>1.3</td><td>-7.0</td><td>2.4E-12</td><td>5.9E-12</td><td>BT20/MB468</td><td>NUC</td></tr>
<tr><td>>hg38_21_+_8433172_8446622_RNA45SN1_45S_50ntFlanks@12623.12653.31|GCCAATGGGGCGAAGCTACCATCTGTGGGAT</td><td>>hg38_21_+_8433172_8446622_RNA45SN1_45S_50ntFlanks</td><td>12623</td><td>12653</td><td>31</td><td>GCCAATGGGGCGAAGCTACCATCTGTGGGAT</td><td>100.8</td><td>-10.5</td><td>1.5</td><td>-7.1</td><td>1.6E-12</td><td>5.9E-12</td><td>MB231/MB468</td><td>MP</td></tr>
<tr><td>>hg38_21_+_8433172_8446622_RNA45SN1_45S_50ntFlanks@4729.4755.27|TCAAGAACGAAAGTCGGAGGTTCGAAG</td><td>>hg38_21_+_8433172_8446622_RNA45SN1_45S_50ntFlanks</td><td>4729</td><td>4755</td><td>27</td><td>TCAAGAACGAAAGTCGGAGGTTCGAAG</td><td>104.6</td><td>-10.6</td><td>1.5</td><td>-7.1</td><td>1.6E-12</td><td>5.9E-12</td><td>MB231/MB468</td><td>MP</td></tr>
<tr><td>>trna16_ValTAC_11_-_59318460_59318532@37.75.39,trna17_ValTAC_11_-_59318102_59318174@37.75.39,trna4_ValTAC_X_-_18693029_18693101@37.75.39|ACGCAGAAGGTCCTGGGTTCGAGCCCCAGTGGAACCACC</td><td>>trna16_ValTAC_11_-_59318460_59318532</td><td>37</td><td>75</td><td>39</td><td>ACGCAGAAGGTCCTGGGTTCGAGCCCCAGTGGAACCACC</td><td>1064.3</td><td>10.6</td><td>1.5</td><td>7.0</td><td>1.8E-12</td><td>5.9E-12</td><td>MB231/MB468</td><td>CYTO</td></tr>
<tr><td>>MI0000098|hsa-mir-96&WithFlank&7|-|129774686|129774775@15.36.22(+1A)&[MIMAT0000095&hsa-miR-96-5p&offsets|0|-1(+1A);m-248&7|-|129774739|129774761&offsets|0|-1(+1A)]|TTTGGCACTAGCACATTTTTGCA</td><td>>MI0000098|hsa-mir-96&WithFlank&7|-|129774686|129774775</td><td>15</td><td>36</td><td>22(+1A)</td><td>TTTGGCACTAGCACATTTTTGCA</td><td>198.6</td><td>4.8</td><td>0.7</td><td>7.0</td><td>2.4E-12</td><td>5.9E-12</td><td>BT20/MB468</td><td>NUC</td></tr>
<tr><td>>hg38_21_+_8433172_8446622_RNA45SN1_45S_50ntFlanks@5420.5444.25|CCGATTGGATGGTTTAGTGAGGCCC</td><td>>hg38_21_+_8433172_8446622_RNA45SN1_45S_50ntFlanks</td><td>5420</td><td>5444</td><td>25</td><td>CCGATTGGATGGTTTAGTGAGGCCC</td><td>843.3</td><td>-6.4</td><td>0.9</td><td>-7.0</td><td>1.8E-12</td><td>6.0E-12</td><td>MB231/MB468</td><td>CYTO</td></tr>
<tr><td>>hg38_21_+_8433172_8446622_RNA45SN1_45S_50ntFlanks@8170.8213.44|CAAGTCCTTCTGATCGAGGCCCAGCCCGTGGACGGTGTGAGGCC</td><td>>hg38_21_+_8433172_8446622_RNA45SN1_45S_50ntFlanks</td><td>8170</td><td>8213</td><td>44</td><td>CAAGTCCTTCTGATCGAGGCCCAGCCCGTGGACGGTGTGAGGCC</td><td>1598.9</td><td>-8.3</td><td>1.2</td><td>-7.0</td><td>2.4E-12</td><td>6.0E-12</td><td>BT20/MB468</td><td>NUC</td></tr>
<tr><td>>hg38_1_-_228634819_228635039_RNA5S12_5S_50ntFlanks@142.171.30|CGCCTGGGAATACCGGGTGCTGTAGGCTTT</td><td>>hg38_1_-_228634819_228635039_RNA5S12_5S_50ntFlanks</td><td>142</td><td>171</td><td>30</td><td>CGCCTGGGAATACCGGGTGCTGTAGGCTTT</td><td>3373.1</td><td>5.5</td><td>0.8</td><td>7.0</td><td>1.8E-12</td><td>6.0E-12</td><td>MB231/MB468</td><td>CYTO</td></tr>
<tr><td>>am_trna10_ValCAC_5_-_180649395_180649467@1.32.32,trna12_ValAAC_5_-_180645270_180645342@1.32.32,trna132_ValAAC_6_-_27721179_27721251@1.32.32,trna136_ValAAC_6_-_27648885_27648957@1.32.32,trna139_ValAAC_6_-_27618707_27618779@1.32.32,trna18_ValCAC_5_-_180529253_180529325@1.32.32,trna2_ValAAC_3_+_169490018_169490090@1.32.32,trna2_ValCAC_5_+_180524070_180524142@1.32.32,trna4_ValAAC_5_+_180591154_180591226@1.32.32,trna5_ValAAC_5_+_180596610_180596682@1.32.32,trna6_ValCAC_5_+_180600650_180600722@1.32.32,trna85_ValCAC_1_-_161369490_161369562@1.32.32,trna90_ValCAC_1_-_149684088_149684161@1.32.32,trna98_ValCAC_1_-_149298555_149298627@1.32.32,trna9_ValCAC_6_+_26538282_26538354@1.32.32|GTTTCCGTAGTGTAGTGGTTATCACGTTCGCC</td><td>>am_trna10_ValCAC_5_-_180649395_180649467</td><td>1</td><td>32</td><td>32</td><td>GTTTCCGTAGTGTAGTGGTTATCACGTTCGCC</td><td>6143.0</td><td>8.2</td><td>1.2</td><td>7.1</td><td>1.3E-12</td><td>6.0E-12</td><td>BT20/MB468</td><td>MITO</td></tr>
<tr><td>>hg38_21_+_8433172_8446622_RNA45SN1_45S_50ntFlanks@3782.3800.19|CAGTGAAACTGCGAATGGC</td><td>>hg38_21_+_8433172_8446622_RNA45SN1_45S_50ntFlanks</td><td>3782</td><td>3800</td><td>19</td><td>CAGTGAAACTGCGAATGGC</td><td>1485.6</td><td>-10.5</td><td>1.5</td><td>-7.0</td><td>2.4E-12</td><td>6.0E-12</td><td>BT20/MB468</td><td>NUC</td></tr>
<tr><td>>MI0000234|hsa-mir-192&WithFlank&11|-|64891131|64891252@30.51.22(+1U)&[MIMAT0000222&hsa-miR-192-5p&offsets|0|+1(+1U);m-59&11|-|64891203|64891223&offsets|0|+1(+1U)]|CTGACCTATGAATTGACAGCCAT</td><td>>MI0000234|hsa-mir-192&WithFlank&11|-|64891131|64891252</td><td>30</td><td>51</td><td>22(+1U)</td><td>CTGACCTATGAATTGACAGCCAT</td><td>954.8</td><td>5.1</td><td>0.7</td><td>7.0</td><td>1.8E-12</td><td>6.0E-12</td><td>MB231/MB468</td><td>NUC</td></tr>
<tr><td>>hg38_21_+_8433172_8446622_RNA45SN1_45S_50ntFlanks@12436.12455.20|TCATTGTGAAGCAGAATTCA</td><td>>hg38_21_+_8433172_8446622_RNA45SN1_45S_50ntFlanks</td><td>12436</td><td>12455</td><td>20</td><td>TCATTGTGAAGCAGAATTCA</td><td>3349.7</td><td>-10.3</td><td>1.5</td><td>-7.1</td><td>1.7E-12</td><td>6.0E-12</td><td>BT20/MB468</td><td>CYTO</td></tr>
</table>

<table>
<tr><th></th><th></th><th></th><th></th><th></th><th></th><th></th><th></th><th></th><th></th><th></th><th></th><th></th><th></th></tr>
<tr><td>>hg38_21_+_8433172_8446622_RNA45SN1_45S_50ntFlanks@3738.3756.19|TCTCAAAGATTAAGCCATG</td><td>>hg38_21_+_8433172_8446622_RNA45SN1_45S_50ntFlanks</td><td>3738</td><td>3756</td><td>19</td><td>TCTCAAAGATTAAGCCATG</td><td>146.8</td><td>-9.5</td><td>1.3</td><td>-7.1</td><td>1.7E-12</td><td>6.0E-12</td><td>BT20/MB468</td><td>CYTO</td></tr>
<tr><td>>MI0000263|hsa-mir-7-1&WithFlank&9|-|83969742|83969863@30.54.25&[MIMAT0000252_2&hsa-miR-7-5p&offsets|0|+1;m-401&9|-|83969811|83969834&offsets|0|+1]|TGGAAGACTAGTGATTTTGTTGTTT</td><td>>MI0000263|hsa-mir-7-1&WithFlank&9|-|83969742|83969863</td><td>30</td><td>54</td><td>25</td><td>TGGAAGACTAGTGATTTTGTTGTTT</td><td>1743.8</td><td>6.7</td><td>0.9</td><td>7.1</td><td>1.8E-12</td><td>6.0E-12</td><td>BT20/MB468</td><td>CYTO</td></tr>
<tr><td>>hg38_21_+_8433172_8446622_RNA45SN1_45S_50ntFlanks@9548.9576.29|GAGGTCCGTAGCGGTCCTGACGTGCAAAT</td><td>>hg38_21_+_8433172_8446622_RNA45SN1_45S_50ntFlanks</td><td>9548</td><td>9576</td><td>29</td><td>GAGGTCCGTAGCGGTCCTGACGTGCAAAT</td><td>202.5</td><td>5.7</td><td>0.8</td><td>7.0</td><td>1.8E-12</td><td>6.0E-12</td><td>MB231/MB468</td><td>CYTO</td></tr>
<tr><td>>hg38_21_+_8433172_8446622_RNA45SN1_45S_50ntFlanks@4051.4071.21|GAACGTCTGCCCTATCAACTT</td><td>>hg38_21_+_8433172_8446622_RNA45SN1_45S_50ntFlanks</td><td>4051</td><td>4071</td><td>21</td><td>GAACGTCTGCCCTATCAACTT</td><td>141.9</td><td>6.9</td><td>1.0</td><td>7.0</td><td>1.8E-12</td><td>6.0E-12</td><td>MB231/MB468</td><td>CYTO</td></tr>
<tr><td>>MI0000750|hsa-mir-26a-2&WithFlank&12|-|57824603|57824698@20.40.21(+1C);MI0000083|hsa-mir-26a-1&WithFlank&3|+|37969398|37969486@16.36.21(+1C)&[MIMAT0000082&hsa-miR-26a-5p&offsets|0|-1(+1C);m-33&12|-|57824658|57824679&offsets|0|-1(+1C)];[MIMAT0000082_1&hsa-miR-26a-5p&offsets|0|-1(+1C);m-34&3|+|37969413|37969434&offsets|0|-1(+1C)]|TTCAAGTAATCCAGGATAGGCC</td><td>>MI0000750|hsa-mir-26a-2&WithFlank&12|-|57824603|57824698</td><td>20</td><td>40</td><td>21(+1C)</td><td>TTCAAGTAATCCAGGATAGGCC</td><td>9731.5</td><td>4.9</td><td>0.7</td><td>7.0</td><td>1.9E-12</td><td>6.0E-12</td><td>MB231/MB468</td><td>NUC</td></tr>
<tr><td>>hg38_21_+_8433172_8446622_RNA45SN1_45S_50ntFlanks@11933.11958.26|AATAAGTGGGAGGCCCCCGGCGCCCC</td><td>>hg38_21_+_8433172_8446622_RNA45SN1_45S_50ntFlanks</td><td>11933</td><td>11958</td><td>26</td><td>AATAAGTGGGAGGCCCCCGGCGCCCC</td><td>139.8</td><td>-11.0</td><td>1.6</td><td>-7.1</td><td>1.6E-12</td><td>6.0E-12</td><td>MB231/MB468</td><td>MP</td></tr>
<tr><td>>hg38_21_+_8433172_8446622_RNA45SN1_45S_50ntFlanks@3843.3874.32|CTACTTGGATAACTGTGGTAATTCTAGAGCTA</td><td>>hg38_21_+_8433172_8446622_RNA45SN1_45S_50ntFlanks</td><td>3843</td><td>3874</td><td>32</td><td>CTACTTGGATAACTGTGGTAATTCTAGAGCTA</td><td>666.4</td><td>-7.3</td><td>1.0</td><td>-7.0</td><td>2.3E-12</td><td>6.0E-12</td><td>MB231/MB468</td><td>MITO</td></tr>
<tr><td>>hg38_21_+_8433172_8446622_RNA45SN1_45S_50ntFlanks@4006.4024.19|ACCTCGGGCCGATCGCACG</td><td>>hg38_21_+_8433172_8446622_RNA45SN1_45S_50ntFlanks</td><td>4006</td><td>4024</td><td>19</td><td>ACCTCGGGCCGATCGCACG</td><td>1763.2</td><td>-10.4</td><td>1.5</td><td>-7.0</td><td>2.4E-12</td><td>6.0E-12</td><td>BT20/MB468</td><td>NUC</td></tr>
<tr><td>>am_trna116_GluCTC_1_-_145399233_145399304@12.29.18,trna25_GluCTC_2_-_71273488_71273560@12.29.18,trna59_GluCTC_1_+_249168447_249168518@12.29.18,trna71_GluCTC_1_-_161439189_161439260@12.29.18,trna74_GluCTC_1_-_161431809_161431880@12.29.18,trna77_GluCTC_1_-_161424398_161424469@12.29.18,trna77_GluCTC_6_+_28949976_28950047@12.29.18,trna80_GluCTC_1_-_161417018_161417089@12.29.18,trna87_GluCTC_6_-_126101393_126101464@12.29.18|CTAGTGGTTAGGATTCGG</td><td>>am_trna116_GluCTC_1_-_145399233_145399304</td><td>12</td><td>29</td><td>18</td><td>CTAGTGGTTAGGATTCGG</td><td>174.4</td><td>-6.9</td><td>1.0</td><td>-7.0</td><td>1.8E-12</td><td>6.0E-12</td><td>MB231/MB468</td><td>CYTO</td></tr>
<tr><td>>hg38_21_+_8433172_8446622_RNA45SN1_45S_50ntFlanks@9534.9566.33|AGGAAACTCTGGTGGAGGTCCGTAGCGGTCCTG</td><td>>hg38_21_+_8433172_8446622_RNA45SN1_45S_50ntFlanks</td><td>9534</td><td>9566</td><td>33</td><td>AGGAAACTCTGGTGGAGGTCCGTAGCGGTCCTG</td><td>2562.3</td><td>-6.7</td><td>1.0</td><td>-7.0</td><td>1.8E-12</td><td>6.0E-12</td><td>MB231/MB468</td><td>CYTO</td></tr>
<tr><td>>MI0000270|hsa-mir-181b-1&WithFlank&1|-|198858867|198858988@42.64.23(+1C);MI0000683|hsa-mir-181b-2&WithFlank&9|+|124693704|124693804@22.44.23(+1C)&[MIMAT0000257&hsa-miR-181b-5p&offsets|0|0(+1C);m-73&1|-|198858925|198858947&offsets|0|0(+1C)];[MIMAT0000257_1&hsa-miR-181b-5p&offsets|0|0(+1C);m-72&9|+|124693725|124693747&offsets|0|0(+1C)]|AACATTCATTGCTGTCGGTGGGTC</td><td>>MI0000270|hsa-mir-181b-1&WithFlank&1|-|198858867|198858988</td><td>42</td><td>64</td><td>23(+1C)</td><td>AACATTCATTGCTGTCGGTGGGTC</td><td>222.9</td><td>7.0</td><td>1.0</td><td>7.0</td><td>1.9E-12</td><td>6.0E-12</td><td>MB231/MB468</td><td>NUC</td></tr>
<tr><td>>hg38_21_+_8433172_8446622_RNA45SN1_45S_50ntFlanks@6663.6685.23|GTGGATCACTCGGCTCGTGCGTC</td><td>>hg38_21_+_8433172_8446622_RNA45SN1_45S_50ntFlanks</td><td>6663</td><td>6685</td><td>23</td><td>GTGGATCACTCGGCTCGTGCGTC</td><td>119.8</td><td>-10.8</td><td>1.5</td><td>-7.1</td><td>1.6E-12</td><td>6.0E-12</td><td>MB231/MB468</td><td>MP</td></tr>
<tr><td>>hg38_21_+_8433172_8446622_RNA45SN1_45S_50ntFlanks@4225.4242.18|AAATAACAATACAGGACT</td><td>>hg38_21_+_8433172_8446622_RNA45SN1_45S_50ntFlanks</td><td>4225</td><td>4242</td><td>18</td><td>AAATAACAATACAGGACT</td><td>1691.2</td><td>-5.7</td><td>0.8</td><td>-7.1</td><td>1.6E-12</td><td>6.1E-12</td><td>MB231/MB468</td><td>MP</td></tr>
<tr><td>>hg38_21_+_8433172_8446622_RNA45SN1_45S_50ntFlanks@12655.12673.19|ATGACTGAACGCCTCTAAG</td><td>>hg38_21_+_8433172_8446622_RNA45SN1_45S_50ntFlanks</td><td>12655</td><td>12673</td><td>19</td><td>ATGACTGAACGCCTCTAAG</td><td>2572.1</td><td>-10.8</td><td>1.5</td><td>-7.0</td><td>2.5E-12</td><td>6.1E-12</td><td>BT20/MB468</td><td>NUC</td></tr>
<tr><td>>hg38_21_+_8433172_8446622_RNA45SN1_45S_50ntFlanks@12902.12933.32|TCACGCACCGCACGTTCGTGGGGAACCTGGCG</td><td>>hg38_21_+_8433172_8446622_RNA45SN1_45S_50ntFlanks</td><td>12902</td><td>12933</td><td>32</td><td>TCACGCACCGCACGTTCGTGGGGAACCTGGCG</td><td>127.1</td><td>-10.9</td><td>1.5</td><td>-7.1</td><td>1.6E-12</td><td>6.1E-12</td><td>MB231/MB468</td><td>MP</td></tr>
<tr><td>>hg38_21_+_8433172_8446622_RNA45SN1_45S_50ntFlanks@11821.11852.32|CTATCCAGCGAAACCACAGCCAAGGGAACGGG</td><td>>hg38_21_+_8433172_8446622_RNA45SN1_45S_50ntFlanks</td><td>11821</td><td>11852</td><td>32</td><td>CTATCCAGCGAAACCACAGCCAAGGGAACGGG</td><td>95.0</td><td>-10.4</td><td>1.5</td><td>-7.1</td><td>1.6E-12</td><td>6.1E-12</td><td>MB231/MB468</td><td>MP</td></tr>
<tr><td>>hg38_21_+_8433172_8446622_RNA45SN1_45S_50ntFlanks@12324.12348.25|TTGGGTTTTAAGCAGGAGGTGTCAG</td><td>>hg38_21_+_8433172_8446622_RNA45SN1_45S_50ntFlanks</td><td>12324</td><td>12348</td><td>25</td><td>TTGGGTTTTAAGCAGGAGGTGTCAG</td><td>225.9</td><td>-9.9</td><td>1.4</td><td>-7.1</td><td>1.4E-12</td><td>6.1E-12</td><td>BT20/MB468</td><td>MITO</td></tr>
<tr><td>>MI0000088|hsa-mir-30a&WithFlank&6|-|71403545|71403627@54.73.20&[MIMAT0000088&hsa-miR-30a-3p&offsets|+1|-1;m-38&6|-|71403554|71403575&offsets|+1|-1]|TTTCAGTCGGATGTTTGCAG</td><td>>MI0000088|hsa-mir-30a&WithFlank&6|-|71403545|71403627</td><td>54</td><td>73</td><td>20</td><td>TTTCAGTCGGATGTTTGCAG</td><td>358.4</td><td>7.1</td><td>1.0</td><td>7.0</td><td>1.9E-12</td><td>6.1E-12</td><td>MB231/MB468</td><td>NUC</td></tr>
<tr><td>>MI0003599|hsa-mir-589&WithFlank&7|-|5495813|5495923@30.52.23&[MIMAT0004799&hsa-miR-589-5p&offsets|0|+1;m-162&7|-|5495874|5495894&offsets|0|+2]|TGAGAACCACGTCTGCTCTGAGC</td><td>>MI0003599|hsa-mir-589&WithFlank&7|-|5495813|5495923</td><td>30</td><td>52</td><td>23</td><td>TGAGAACCACGTCTGCTCTGAGC</td><td>1116.1</td><td>9.1</td><td>1.3</td><td>7.0</td><td>1.9E-12</td><td>6.1E-12</td><td>MB231/MB468</td><td>CYTO</td></tr>
<tr><td>>hg38_1_-_228634819_228635039_RNA5S12_5S_50ntFlanks@142.170.29|CGCCTGGGAATACCGGGTGCTGTAGGCTT</td><td>>hg38_1_-_228634819_228635039_RNA5S12_5S_50ntFlanks</td><td>142</td><td>170</td><td>29</td><td>CGCCTGGGAATACCGGGTGCTGTAGGCTT</td><td>1739.7</td><td>5.2</td><td>0.7</td><td>7.0</td><td>1.9E-12</td><td>6.1E-12</td><td>MB231/MB468</td><td>CYTO</td></tr>
<tr><td>>hg38_21_+_8433172_8446622_RNA45SN1_45S_50ntFlanks@5071.5098.28|TTAATTCCGATAACGAACGAGACTCTGG</td><td>>hg38_21_+_8433172_8446622_RNA45SN1_45S_50ntFlanks</td><td>5071</td><td>5098</td><td>28</td><td>TTAATTCCGATAACGAACGAGACTCTGG</td><td>1761.6</td><td>-10.5</td><td>1.5</td><td>-7.0</td><td>2.3E-12</td><td>6.2E-12</td><td>MB231/MB468</td><td>MITO</td></tr>
<tr><td>>hg38_21_+_8433172_8446622_RNA45SN1_45S_50ntFlanks@10392.10430.39|ACAGCAGTTGAACATGGGTCAGTCGGTCCTGAGAGATGG</td><td>>hg38_21_+_8433172_8446622_RNA45SN1_45S_50ntFlanks</td><td>10392</td><td>10430</td><td>39</td><td>ACAGCAGTTGAACATGGGTCAGTCGGTCCTGAGAGATGG</td><td>1317.2</td><td>-7.6</td><td>1.1</td><td>-7.0</td><td>2.3E-12</td><td>6.2E-12</td><td>MB231/MB468</td><td>MITO</td></tr>
<tr><td>>hg38_21_+_8433172_8446622_RNA45SN1_45S_50ntFlanks@10903.10935.33|GGCTGGGCGCGCGCCGCGGCTGGACGAGGCGCC</td><td>>hg38_21_+_8433172_8446622_RNA45SN1_45S_50ntFlanks</td><td>10903</td><td>10935</td><td>33</td><td>GGCTGGGCGCGCGCCGCGGCTGGACGAGGCGCC</td><td>305.9</td><td>-11.2</td><td>1.6</td><td>-7.0</td><td>1.9E-12</td><td>6.2E-12</td><td>MB231/MB468</td><td>NUC</td></tr>
<tr><td>>trna116_GluCTC_1_-_145399233_145399304@35.74.40,trna134_GluTTC_1_-_16861774_16861845@35.74.40,trna71_GluCTC_1_-_161439189_161439260@35.74.40,trna74_GluCTC_1_-_161431809_161431880@35.74.40,trna77_GluCTC_1_-_161424398_161424469@35.74.40,trna77_GluCTC_6_+_28949976_28950047@35.74.40,trna80_GluCTC_1_-_161417018_161417089@35.74.40,trna84_GluTTC_1_-_161391883_161391954@35.74.40,trna87_GluCTC_6_-_126101393_126101464@35.74.40|TCACCGCCGCGGCCCGGGTTCGATTCCCGGTCAGGGAACC</td><td>>trna116_GluCTC_1_-_145399233_145399304</td><td>35</td><td>74</td><td>40</td><td>TCACCGCCGCGGCCCGGGTTCGATTCCCGGTCAGGGAACC</td><td>3034.1</td><td>8.6</td><td>1.2</td><td>7.0</td><td>1.9E-12</td><td>6.2E-12</td><td>MB231/MB468</td><td>CYTO</td></tr>
<tr><td>>hg38_21_+_8433172_8446622_RNA45SN1_45S_50ntFlanks@6782.6803.22|GGGCTACGCCTGTCTGAGCGTC</td><td>>hg38_21_+_8433172_8446622_RNA45SN1_45S_50ntFlanks</td><td>6782</td><td>6803</td><td>22</td><td>GGGCTACGCCTGTCTGAGCGTC</td><td>440.3</td><td>-11.9</td><td>1.6</td><td>-7.5</td><td>8.7E-14</td><td>6.2E-12</td><td>BT20/MB468</td><td>MP</td></tr>
<tr><td>>hg38_21_+_8433172_8446622_RNA45SN1_45S_50ntFlanks@12073.12106.34|CCGAGGGGCTCTCGCTTCTGGCGCCAAGCGCCCG</td><td>>hg38_21_+_8433172_8446622_RNA45SN1_45S_50ntFlanks</td><td>12073</td><td>12106</td><td>34</td><td>CCGAGGGGCTCTCGCTTCTGGCGCCAAGCGCCCG</td><td>264.7</td><td>-7.6</td><td>1.1</td><td>-7.0</td><td>1.9E-12</td><td>6.2E-12</td><td>MB231/MB468</td><td>CYTO</td></tr>
<tr><td>>am_trnaMT_LysTTT_MT_+_8295_8364@30.52.23,trnalookalike7_LysTTT_1_+_568844_568913@30.52.23|TTAAGTTAAAGATTAAGAGAACC</td><td>>am_trnaMT_LysTTT_MT_+_8295_8364</td><td>30</td><td>52</td><td>23</td><td>TTAAGTTAAAGATTAAGAGAACC</td><td>831.7</td><td>9.1</td><td>1.3</td><td>7.0</td><td>1.8E-12</td><td>6.2E-12</td><td>BT20/MB468</td><td>CYTO</td></tr>
<tr><td>>MI0000065|hsa-let-7d&WithFlank&9|+|94178828|94178926@14.34.21(+1A)&[MIMAT0000065&hsa-let-7d-5p&offsets|0|-1(+1A);m-90&9|+|94178841|94178862&offsets|0|-1(+1A)]|AGAGGTAGTAGGTTGCATAGTA</td><td>>MI0000065|hsa-let-7d&WithFlank&9|+|94178828|94178926</td><td>14</td><td>34</td><td>21(+1A)</td><td>AGAGGTAGTAGGTTGCATAGTA</td><td>89.7</td><td>3.2</td><td>0.5</td><td>7.0</td><td>2.5E-12</td><td>6.2E-12</td><td>BT20/MB468</td><td>NUC</td></tr>
<tr><td>>hg38_21_+_8433172_8446622_RNA45SN1_45S_50ntFlanks@12605.12624.20|TATGTGCTTGGCTGAGGAGC</td><td>>hg38_21_+_8433172_8446622_RNA45SN1_45S_50ntFlanks</td><td>12605</td><td>12624</td><td>20</td><td>TATGTGCTTGGCTGAGGAGC</td><td>12667.8</td><td>-9.8</td><td>1.4</td><td>-7.0</td><td>2.5E-12</td><td>6.2E-12</td><td>BT20/MB468</td><td>NUC</td></tr>
<tr><td>>hg38_21_+_8433172_8446622_RNA45SN1_45S_50ntFlanks@7979.8003.25|ACCTCAGATCAGACGTGGCGACCCG</td><td>>hg38_21_+_8433172_8446622_RNA45SN1_45S_50ntFlanks</td><td>7979</td><td>8003</td><td>25</td><td>ACCTCAGATCAGACGTGGCGACCCG</td><td>2699.7</td><td>-8.4</td><td>1.2</td><td>-7.0</td><td>2.5E-12</td><td>6.2E-12</td><td>BT20/MB468</td><td>NUC</td></tr>
<tr><td>>am_trnaMT_SerGCT_MT_+_12207_12265@18.38.21|CTGCTAACTCATGCCCCCATG</td><td>>am_trnaMT_SerGCT_MT_+_12207_12265</td><td>18</td><td>38</td><td>21</td><td>CTGCTAACTCATGCCCCCATG</td><td>236.5</td><td>-10.8</td><td>1.5</td><td>-7.0</td><td>1.9E-12</td><td>6.2E-12</td><td>MB231/MB468</td><td>NUC</td></tr>
<tr><td>>hg38_21_+_8433172_8446622_RNA45SN1_45S_50ntFlanks@12332.12349.18|TAAGCAGGAGGTGTCAGA</td><td>>hg38_21_+_8433172_8446622_RNA45SN1_45S_50ntFlanks</td><td>12332</td><td>12349</td><td>18</td><td>TAAGCAGGAGGTGTCAGA</td><td>107.4</td><td>-10.6</td><td>1.5</td><td>-7.1</td><td>1.6E-12</td><td>6.2E-12</td><td>MB231/MB468</td><td>MP</td></tr>
<tr><td>>MI0003628|hsa-mir-615&WithFlank&12|+|54033944|54034051@67.85.19&[MIMAT0003283&hsa-miR-615-3p&offsets|0|-3;m-495&12|+|54034010|54034030&offsets|0|-2]|TCCGAGCCTGGGTCTCCCT</td><td>>MI0003628|hsa-mir-615&WithFlank&12|+|54033944|54034051</td><td>67</td><td>85</td><td>19</td><td>TCCGAGCCTGGGTCTCCCT</td><td>707.7</td><td>7.7</td><td>1.1</td><td>7.1</td><td>1.4E-12</td><td>6.2E-12</td><td>BT20/MB468</td><td>MITO</td></tr>
<tr><td>>am_MI0003639|hsa-mir-625&WithFlank&14|+|65471096|65471192@58.80.23&[MIMAT0004808&hsa-miR-625-3p&offsets|0|+1;m-137&14|+|65471153|65471174&offsets|0|+1]|GACTATAGAACTTTCCCCCTCAT</td><td>>am_MI0003639|hsa-mir-625&WithFlank&14|+|65471096|65471192</td><td>58</td><td>80</td><td>23</td><td>GACTATAGAACTTTCCCCCTCAT</td><td>413.9</td><td>-4.3</td><td>0.6</td><td>-7.3</td><td>2.1E-13</td><td>6.2E-12</td><td>BT20/MB231</td><td>NUC</td></tr>
<tr><td>>hg38_21_+_8433172_8446622_RNA45SN1_45S_50ntFlanks@12991.13019.29|CTCCCTCGCTGCGATCTATTGAAAGTCAG</td><td>>hg38_21_+_8433172_8446622_RNA45SN1_45S_50ntFlanks</td><td>12991</td><td>13019</td><td>29</td><td>CTCCCTCGCTGCGATCTATTGAAAGTCAG</td><td>241.9</td><td>-10.0</td><td>1.4</td><td>-7.1</td><td>1.4E-12</td><td>6.2E-12</td><td>BT20/MB468</td><td>MITO</td></tr>
<tr><td>>hg38_21_+_8433172_8446622_RNA45SN1_45S_50ntFlanks@6785.6806.22|CTACGCCTGTCTGAGCGTCGCT</td><td>>hg38_21_+_8433172_8446622_RNA45SN1_45S_50ntFlanks</td><td>6785</td><td>6806</td><td>22</td><td>CTACGCCTGTCTGAGCGTCGCT</td><td>2047.4</td><td>4.7</td><td>0.7</td><td>7.0</td><td>1.8E-12</td><td>6.2E-12</td><td>BT20/MB468</td><td>CYTO</td></tr>
<tr><td>>hg38_21_+_8433172_8446622_RNA45SN1_45S_50ntFlanks@9741.9765.25|ACGATCTCAACCTATTCTCAAACTT</td><td>>hg38_21_+_8433172_8446622_RNA45SN1_45S_50ntFlanks</td><td>9741</td><td>9765</td><td>25</td><td>ACGATCTCAACCTATTCTCAAACTT</td><td>519.1</td><td>-11.1</td><td>1.6</td><td>-7.1</td><td>1.4E-12</td><td>6.2E-12</td><td>BT20/MB468</td><td>MITO</td></tr>
<tr><td>>hg38_21_+_8433172_8446622_RNA45SN1_45S_50ntFlanks@5485.5524.40|AGCGCTGAGAAGACGGTCGAACTTGACTATCTAGAGGAAG</td><td>>hg38_21_+_8433172_8446622_RNA45SN1_45S_50ntFlanks</td><td>5485</td><td>5524</td><td>40</td><td>AGCGCTGAGAAGACGGTCGAACTTGACTATCTAGAGGAAG</td><td>153.8</td><td>-11.1</td><td>1.6</td><td>-7.1</td><td>1.7E-12</td><td>6.2E-12</td><td>MB231/MB468</td><td>MP</td></tr>
<tr><td>>hg38_21_+_8433172_8446622_RNA45SN1_45S_50ntFlanks@8191.8215.25|CAGCCCGTGGACGGTGTGAGGCCGG</td><td>>hg38_21_+_8433172_8446622_RNA45SN1_45S_50ntFlanks</td><td>8191</td><td>8215</td><td>25</td><td>CAGCCCGTGGACGGTGTGAGGCCGG</td><td>4657.7</td><td>-9.0</td><td>1.3</td><td>-7.0</td><td>2.5E-12</td><td>6.2E-12</td><td>BT20/MB468</td><td>NUC</td></tr>
<tr><td>>hg38_21_+_8433172_8446622_RNA45SN1_45S_50ntFlanks@10869.10902.34|AGGGCTGGGTCGGTCGGGCTGGGGCGCGAAGCGG</td><td>>hg38_21_+_8433172_8446622_RNA45SN1_45S_50ntFlanks</td><td>10869</td><td>10902</td><td>34</td><td>AGGGCTGGGTCGGTCGGGCTGGGGCGCGAAGCGG</td><td>3454.2</td><td>-9.9</td><td>1.4</td><td>-7.0</td><td>1.9E-12</td><td>6.2E-12</td><td>MB231/MB468</td><td>NUC</td></tr>
<tr><td>>hg38_21_+_8433172_8446622_RNA45SN1_45S_50ntFlanks@12300.12318.19|GGGCCTCACGATCCTTCTG</td><td>>hg38_21_+_8433172_8446622_RNA45SN1_45S_50ntFlanks</td><td>12300</td><td>12318</td><td>19</td><td>GGGCCTCACGATCCTTCTG</td><td>212.5</td><td>-11.6</td><td>1.6</td><td>-7.1</td><td>1.7E-12</td><td>6.2E-12</td><td>MB231/MB468</td><td>MP</td></tr>
<tr><td>>hg38_21_+_8433172_8446622_RNA45SN1_45S_50ntFlanks@476.510.35|TTCGTGATCGATGTGGTGACGTCGTGCTCTCCCGG</td><td>>hg38_21_+_8433172_8446622_RNA45SN1_45S_50ntFlanks</td><td>476</td><td>510</td><td>35</td><td>TTCGTGATCGATGTGGTGACGTCGTGCTCTCCCGG</td><td>509.2</td><td>8.3</td><td>1.2</td><td>7.0</td><td>1.9E-12</td><td>6.2E-12</td><td>MB231/MB468</td><td>CYTO</td></tr>
<tr><td>>hg38_21_+_8433172_8446622_RNA45SN1_45S_50ntFlanks@12457.12477.21|CAAGCGTTGGATTGTTCACCC</td><td>>hg38_21_+_8433172_8446622_RNA45SN1_45S_50ntFlanks</td><td>12457</td><td>12477</td><td>21</td><td>CAAGCGTTGGATTGTTCACCC</td><td>2255.9</td><td>-5.6</td><td>0.8</td><td>-7.0</td><td>2.3E-12</td><td>6.2E-12</td><td>MB231/MB468</td><td>MITO</td></tr>
<tr><td>>MI0000465|hsa-mir-191&WithFlank&3|-|49020612|49020715@22.45.24&[MIMAT0000440&hsa-miR-191-5p&offsets|0|+1;m-48&3|-|49020672|49020694&offsets|0|+1]|CAACGGAATCCCAAAAGCAGCTGT</td><td>>MI0000465|hsa-mir-191&WithFlank&3|-|49020612|49020715</td><td>22</td><td>45</td><td>24</td><td>CAACGGAATCCCAAAAGCAGCTGT</td><td>5713.9</td><td>7.2</td><td>1.0</td><td>7.0</td><td>2.3E-12</td><td>6.3E-12</td><td>MB231/MB468</td><td>MITO</td></tr>
<tr><td>>hg38_21_+_8433172_8446622_RNA45SN1_45S_50ntFlanks@12627.12667.41|ATGGGGCGAAGCTACCATCTGTGGGATTATGACTGAACGCC</td><td>>hg38_21_+_8433172_8446622_RNA45SN1_45S_50ntFlanks</td><td>12627</td><td>12667</td><td>41</td><td>ATGGGGCGAAGCTACCATCTGTGGGATTATGACTGAACGCC</td><td>341.4</td><td>6.9</td><td>1.0</td><td>7.0</td><td>1.9E-12</td><td>6.3E-12</td><td>MB231/MB468</td><td>CYTO</td></tr>
</table>

<table>
<tr><td>>hg38_21_+_8433172_8446622_RNA45SN1_45S_50ntFlanks@6641.6681.41|CGACCTCGTACGACTCTTAGCGGTGGATCACTCGGCTCGTG</td><td>>hg38_21_+_8433172_8446622_RNA45SN1_45S_50ntFlanks</td><td>6641</td><td>6681</td><td>41</td><td>CGACCTCGTACGACTCTTAGCGGTGGATCACTCGGCTCGTG</td><td>205.8</td><td>9.7</td><td>1.4</td><td>7.0</td><td>1.9E-12</td><td>6.3E-12</td><td>MB231/MB468</td><td>NUC</td></tr>
<tr><td>>hg38_21_+_8433172_8446622_RNA45SN1_45S_50ntFlanks@4047.4066.20|ATTCGAACGTCTGCCCTATC</td><td>>hg38_21_+_8433172_8446622_RNA45SN1_45S_50ntFlanks</td><td>4047</td><td>4066</td><td>20</td><td>ATTCGAACGTCTGCCCTATC</td><td>3884.4</td><td>-4.4</td><td>0.6</td><td>-7.0</td><td>1.9E-12</td><td>6.3E-12</td><td>MB231/MB468</td><td>NUC</td></tr>
<tr><td>>hg38_21_+_8433172_8446622_RNA45SN1_45S_50ntFlanks@10367.10388.22|CGAAGTGGAGAAGGGTTCCATG</td><td>>hg38_21_+_8433172_8446622_RNA45SN1_45S_50ntFlanks</td><td>10367</td><td>10388</td><td>22</td><td>CGAAGTGGAGAAGGGTTCCATG</td><td>115.6</td><td>-10.7</td><td>1.5</td><td>-7.1</td><td>1.7E-12</td><td>6.3E-12</td><td>MB231/MB468</td><td>MP</td></tr>
<tr><td>>hg38_21_+_8433172_8446622_RNA45SN1_45S_50ntFlanks@12635.12657.23|AAGCTACCATCTGTGGGATTATG</td><td>>hg38_21_+_8433172_8446622_RNA45SN1_45S_50ntFlanks</td><td>12635</td><td>12657</td><td>23</td><td>AAGCTACCATCTGTGGGATTATG</td><td>2798.1</td><td>-9.7</td><td>1.4</td><td>-7.0</td><td>2.4E-12</td><td>6.3E-12</td><td>MB231/MB468</td><td>MITO</td></tr>
<tr><td>>hg38_21_+_8433172_8446622_RNA45SN1_45S_50ntFlanks@12397.12418.22|CGACGTCGCTTTTTGATCCTTC</td><td>>hg38_21_+_8433172_8446622_RNA45SN1_45S_50ntFlanks</td><td>12397</td><td>12418</td><td>22</td><td>CGACGTCGCTTTTTGATCCTTC</td><td>716.5</td><td>-4.0</td><td>0.6</td><td>-7.0</td><td>2.5E-12</td><td>6.3E-12</td><td>BT20/MB468</td><td>NUC</td></tr>
<tr><td>>hg38_21_+_8433172_8446622_RNA45SN1_45S_50ntFlanks@10806.10828.23|AACAATGTAGGTAAGGGAAGTCG</td><td>>hg38_21_+_8433172_8446622_RNA45SN1_45S_50ntFlanks</td><td>10806</td><td>10828</td><td>23</td><td>AACAATGTAGGTAAGGGAAGTCG</td><td>361.8</td><td>-5.7</td><td>0.8</td><td>-7.0</td><td>1.9E-12</td><td>6.3E-12</td><td>MB231/MB468</td><td>CYTO</td></tr>
<tr><td>>hg38_21_+_8433172_8446622_RNA45SN1_45S_50ntFlanks@9578.9602.25|GGTCGTCCGACCTGGGTATAGGGGC</td><td>>hg38_21_+_8433172_8446622_RNA45SN1_45S_50ntFlanks</td><td>9578</td><td>9602</td><td>25</td><td>GGTCGTCCGACCTGGGTATAGGGGC</td><td>270.3</td><td>11.8</td><td>1.6</td><td>7.5</td><td>8.9E-14</td><td>6.3E-12</td><td>BT20/MB468</td><td>MP</td></tr>
<tr><td>>hg38_21_+_8433172_8446622_RNA45SN1_45S_50ntFlanks@12479.12509.31|CTAATAGGGAACGTGAGCTGGGTTTAGACCG</td><td>>hg38_21_+_8433172_8446622_RNA45SN1_45S_50ntFlanks</td><td>12479</td><td>12509</td><td>31</td><td>CTAATAGGGAACGTGAGCTGGGTTTAGACCG</td><td>276.4</td><td>-11.1</td><td>1.6</td><td>-7.0</td><td>1.9E-12</td><td>6.3E-12</td><td>MB231/MB468</td><td>NUC</td></tr>
<tr><td>>hg38_21_+_8433172_8446622_RNA45SN1_45S_50ntFlanks@10910.10928.19|CGCGCGCCGCGGCTGGACG</td><td>>hg38_21_+_8433172_8446622_RNA45SN1_45S_50ntFlanks</td><td>10910</td><td>10928</td><td>19</td><td>CGCGCGCCGCGGCTGGACG</td><td>195.9</td><td>-11.5</td><td>1.6</td><td>-7.1</td><td>1.7E-12</td><td>6.3E-12</td><td>MB231/MB468</td><td>MP</td></tr>
<tr><td>>hg38_21_+_8433172_8446622_RNA45SN1_45S_50ntFlanks@10846.10865.20|CTTCGGGATAAGGATTGGCT</td><td>>hg38_21_+_8433172_8446622_RNA45SN1_45S_50ntFlanks</td><td>10846</td><td>10865</td><td>20</td><td>CTTCGGGATAAGGATTGGCT</td><td>5316.8</td><td>-6.0</td><td>0.8</td><td>-7.0</td><td>1.8E-12</td><td>6.3E-12</td><td>BT20/MB468</td><td>CYTO</td></tr>
<tr><td>>am_trna128_GlyGCC_6_-_27870686_27870756@7.35.29,trna18_GlyGCC_16_+_70822597_70822667@7.35.29,trna19_GlyGCC_16_+_70823410_70823480@7.35.29,trna19_GlyGCC_2_-_157257659_157257729@7.35.29,trna24_GlyGCC_16_-_70812942_70813012@7.35.29,trna25_GlyGCC_16_-_70812114_70812184@7.35.29,trna2_GlyGCC_21_-_18827107_18827177@7.35.29,trna35_GlyGCC_1_+_161413094_161413164@7.35.29,trna37_GlyGCC_1_+_161420467_161420537@7.35.29,trna39_GlyGCC_1_+_161427898_161427968@7.35.29,trna41_GlyGCC_1_+_161435258_161435328@7.35.29,trna5_GlyGCC_17_+_8029064_8029134@7.35.29,trna68_GlyGCC_1_-_161493637_161493707@7.35.29|GTGGTTCAGTGGTAGAATTCTCGCCTGCC</td><td>>am_trna128_GlyGCC_6_-_27870686_27870756</td><td>7</td><td>35</td><td>29</td><td>GTGGTTCAGTGGTAGAATTCTCGCCTGCC</td><td>202.1</td><td>5.2</td><td>0.7</td><td>7.0</td><td>2.0E-12</td><td>6.3E-12</td><td>MB231/MB468</td><td>NUC</td></tr>
<tr><td>>hg38_21_+_8433172_8446622_RNA45SN1_45S_50ntFlanks@11211.11254.44|GCGACTCTGGACGCGAGCCGGGCCCTTCCCGTGGATCGCCCCAG</td><td>>hg38_21_+_8433172_8446622_RNA45SN1_45S_50ntFlanks</td><td>11211</td><td>11254</td><td>44</td><td>GCGACTCTGGACGCGAGCCGGGCCCTTCCCGTGGATCGCCCCAG</td><td>331.2</td><td>-5.8</td><td>0.8</td><td>-7.0</td><td>1.9E-12</td><td>6.3E-12</td><td>MB231/MB468</td><td>CYTO</td></tr>
<tr><td>>hg38_21_+_8433172_8446622_RNA45SN1_45S_50ntFlanks@10349.10372.24|AACGAGAACTTTGAAGGCCGAAGT</td><td>>hg38_21_+_8433172_8446622_RNA45SN1_45S_50ntFlanks</td><td>10349</td><td>10372</td><td>24</td><td>AACGAGAACTTTGAAGGCCGAAGT</td><td>1012.9</td><td>3.5</td><td>0.5</td><td>7.1</td><td>1.4E-12</td><td>6.3E-12</td><td>BT20/MB468</td><td>MITO</td></tr>
<tr><td>>hg38_21_+_8433172_8446622_RNA45SN1_45S_50ntFlanks@12482.12500.19|ATAGGGAACGTGAGCTGGG</td><td>>hg38_21_+_8433172_8446622_RNA45SN1_45S_50ntFlanks</td><td>12482</td><td>12500</td><td>19</td><td>ATAGGGAACGTGAGCTGGG</td><td>274.7</td><td>-4.1</td><td>0.6</td><td>-7.0</td><td>2.0E-12</td><td>6.3E-12</td><td>MB231/MB468</td><td>NUC</td></tr>
<tr><td>>hg38_21_+_8433172_8446622_RNA45SN1_45S_50ntFlanks@8351.8375.25|AAGTTGAAAAGAACTTTGAAGAGAG</td><td>>hg38_21_+_8433172_8446622_RNA45SN1_45S_50ntFlanks</td><td>8351</td><td>8375</td><td>25</td><td>AAGTTGAAAAGAACTTTGAAGAGAG</td><td>3975.9</td><td>9.3</td><td>1.2</td><td>7.5</td><td>9.0E-14</td><td>6.3E-12</td><td>BT20/MB468</td><td>MP</td></tr>
<tr><td>>hg38_21_+_8433172_8446622_RNA45SN1_45S_50ntFlanks@12640.12657.18|ACCATCTGTGGGATTATG</td><td>>hg38_21_+_8433172_8446622_RNA45SN1_45S_50ntFlanks</td><td>12640</td><td>12657</td><td>18</td><td>ACCATCTGTGGGATTATG</td><td>319.6</td><td>-11.4</td><td>1.5</td><td>-7.5</td><td>9.1E-14</td><td>6.3E-12</td><td>BT20/MB468</td><td>MP</td></tr>
<tr><td>>hg38_21_+_8433172_8446622_RNA45SN1_45S_50ntFlanks@11918.11952.35|ACATGAGAGGTGTAGAATAAGTGGGAGGCCCCCGG</td><td>>hg38_21_+_8433172_8446622_RNA45SN1_45S_50ntFlanks</td><td>11918</td><td>11952</td><td>35</td><td>ACATGAGAGGTGTAGAATAAGTGGGAGGCCCCCGG</td><td>361.2</td><td>-7.8</td><td>1.1</td><td>-7.0</td><td>1.9E-12</td><td>6.3E-12</td><td>BT20/MB468</td><td>CYTO</td></tr>
<tr><td>>hg38_21_+_8433172_8446622_RNA45SN1_45S_50ntFlanks@485.529.45|GATGTGGTGACGTCGTGCTCTCCCGGGCCGGGTCCGAGCCGCGAC</td><td>>hg38_21_+_8433172_8446622_RNA45SN1_45S_50ntFlanks</td><td>485</td><td>529</td><td>45</td><td>GATGTGGTGACGTCGTGCTCTCCCGGGCCGGGTCCGAGCCGCGAC</td><td>354.1</td><td>11.3</td><td>1.6</td><td>7.0</td><td>2.6E-12</td><td>6.3E-12</td><td>BT20/MB468</td><td>NUC</td></tr>
<tr><td>>hg38_21_+_8433172_8446622_RNA45SN1_45S_50ntFlanks@12940.12972.33|CATTCGTAGACGACCTGCTTCTGGGTCGGGGTT</td><td>>hg38_21_+_8433172_8446622_RNA45SN1_45S_50ntFlanks</td><td>12940</td><td>12972</td><td>33</td><td>CATTCGTAGACGACCTGCTTCTGGGTCGGGGTT</td><td>406.5</td><td>-11.7</td><td>1.6</td><td>-7.5</td><td>9.1E-14</td><td>6.3E-12</td><td>BT20/MB468</td><td>MP</td></tr>
<tr><td>>hg38_21_+_8433172_8446622_RNA45SN1_45S_50ntFlanks@10391.10429.39|AACAGCAGTTGAACATGGGTCAGTCGGTCCTGAGAGATG</td><td>>hg38_21_+_8433172_8446622_RNA45SN1_45S_50ntFlanks</td><td>10391</td><td>10429</td><td>39</td><td>AACAGCAGTTGAACATGGGTCAGTCGGTCCTGAGAGATG</td><td>2504.4</td><td>-5.4</td><td>0.8</td><td>-7.0</td><td>2.0E-12</td><td>6.3E-12</td><td>MB231/MB468</td><td>NUC</td></tr>
<tr><td>>trna10_AspGTC_12_-_125424193_125424264@31.75.45,trna12_AspGTC_12_-_125411891_125411962@31.75.45,trna38_AspGTC_17_-_8125556_8125627@31.75.45,trna45_AspGTC_6_+_27447453_27447524@31.75.45,trna48_AspGTC_6_+_27471523_27471594@31.75.45,trna4_AspGTC_12_+_96429799_96429870@31.75.45,trna69_AspGTC_1_-_161440205_161440276@31.75.45,trna72_AspGTC_1_-_161432824_161432895@31.75.45,trna75_AspGTC_1_-_161425414_161425485@31.75.45,trna78_AspGTC_1_-_161418033_161418104@31.75.45,trna81_AspGTC_1_-_161410615_161410686@31.75.45|CCTGTCACGCGGGAGACCGGGGTTCGATTCCCCGACGGGGAGCCA</td><td>>trna10_AspGTC_12_-_125424193_125424264</td><td>31</td><td>75</td><td>45</td><td>CCTGTCACGCGGGAGACCGGGGTTCGATTCCCCGACGGGGAGCCA</td><td>461.1</td><td>-11.8</td><td>1.7</td><td>-7.0</td><td>2.0E-12</td><td>6.3E-12</td><td>MB231/MB468</td><td>NUC</td></tr>
<tr><td>>hg38_21_+_8433172_8446622_RNA45SN1_45S_50ntFlanks@12415.12436.22|CTTCGATGTCGGCTCTTCCTAT</td><td>>hg38_21_+_8433172_8446622_RNA45SN1_45S_50ntFlanks</td><td>12415</td><td>12436</td><td>22</td><td>CTTCGATGTCGGCTCTTCCTAT</td><td>6217.1</td><td>-5.1</td><td>0.7</td><td>-7.1</td><td>1.4E-12</td><td>6.3E-12</td><td>BT20/MB468</td><td>MITO</td></tr>
<tr><td>>hg38_21_+_8433172_8446622_RNA45SN1_45S_50ntFlanks@4823.4851.29|ATGACCCGCCGGGCAGCTTCCGGGAAACC</td><td>>hg38_21_+_8433172_8446622_RNA45SN1_45S_50ntFlanks</td><td>4823</td><td>4851</td><td>29</td><td>ATGACCCGCCGGGCAGCTTCCGGGAAACC</td><td>331.4</td><td>7.9</td><td>1.1</td><td>7.0</td><td>1.9E-12</td><td>6.3E-12</td><td>MB231/MB468</td><td>CYTO</td></tr>
<tr><td>>hg38_21_+_8433172_8446622_RNA45SN1_45S_50ntFlanks@11783.11800.18|CATGAATGGATGAACGAG</td><td>>hg38_21_+_8433172_8446622_RNA45SN1_45S_50ntFlanks</td><td>11783</td><td>11800</td><td>18</td><td>CATGAATGGATGAACGAG</td><td>1021.0</td><td>-5.1</td><td>0.7</td><td>-7.0</td><td>2.0E-12</td><td>6.4E-12</td><td>MB231/MB468</td><td>NUC</td></tr>
<tr><td>>trna10_AspGTC_12_-_125424193_125424264@32.71.40,trna12_AspGTC_12_-_125411891_125411962@32.71.40,trna144_AspGTC_6_-_27551236_27551307@32.71.40,trna38_AspGTC_17_-_8125556_8125627@32.71.40,trna45_AspGTC_6_+_27447453_27447524@32.71.40,trna48_AspGTC_6_+_27471523_27471594@32.71.40,trna4_AspGTC_12_+_96429799_96429870@32.71.40,trna69_AspGTC_1_-_161440205_161440276@32.71.40,trna72_AspGTC_1_-_161432824_161432895@32.71.40,trna75_AspGTC_1_-_161425414_161425485@32.71.40,trna78_AspGTC_1_-_161418033_161418104@32.71.40,trna81_AspGTC_1_-_161410615_161410686@32.71.40|CTGTCACGCGGGAGACCGGGGTTCGATTCCCCGACGGGGA</td><td>>trna10_AspGTC_12_-_125424193_125424264</td><td>32</td><td>71</td><td>40</td><td>CTGTCACGCGGGAGACCGGGGTTCGATTCCCCGACGGGGA</td><td>230.4</td><td>-10.8</td><td>1.5</td><td>-7.0</td><td>2.0E-12</td><td>6.4E-12</td><td>MB231/MB468</td><td>NUC</td></tr>
<tr><td>>hg38_21_+_8433172_8446622_RNA45SN1_45S_50ntFlanks@4917.4938.22|CACCACCAGGAGTGGAGCCTGC</td><td>>hg38_21_+_8433172_8446622_RNA45SN1_45S_50ntFlanks</td><td>4917</td><td>4938</td><td>22</td><td>CACCACCAGGAGTGGAGCCTGC</td><td>115.2</td><td>-10.7</td><td>1.5</td><td>-7.1</td><td>1.7E-12</td><td>6.4E-12</td><td>MB231/MB468</td><td>MP</td></tr>
<tr><td>>hg38_21_+_8433172_8446622_RNA45SN1_45S_50ntFlanks@11857.11875.19|GCGGAATCAGCGGGGAAAG</td><td>>hg38_21_+_8433172_8446622_RNA45SN1_45S_50ntFlanks</td><td>11857</td><td>11875</td><td>19</td><td>GCGGAATCAGCGGGGAAAG</td><td>138.6</td><td>-11.0</td><td>1.6</td><td>-7.1</td><td>1.7E-12</td><td>6.4E-12</td><td>MB231/MB468</td><td>MP</td></tr>
<tr><td>>trna111_HisGTG_1_-_147774845_147774916@1.32.32,trna118_HisGTG_1_-_145396881_145396952@1.32.32,trna16_HisGTG_1_+_146544773_146544844@1.32.32,trna1_HisGTG_15_+_45493349_45493420@1.32.32,trna21_HisGTG_1_+_147753471_147753542@1.32.32,trna33_HisGTG_6_+_27125906_27125977@1.32.32,trna7_HisGTG_9_-_14433938_14434009@1.32.32,trna8_HisGTG_15_-_45492611_45492682@1.32.32,trna9_HisGTG_15_-_45490804_45490875@1.32.32|GCCGTGATCGTATAGTGGTTAGTACTCTGCGT</td><td>>trna111_HisGTG_1_-_147774845_147774916</td><td>1</td><td>32</td><td>32</td><td>GCCGTGATCGTATAGTGGTTAGTACTCTGCGT</td><td>829.7</td><td>7.3</td><td>1.0</td><td>7.0</td><td>1.9E-12</td><td>6.4E-12</td><td>BT20/MB468</td><td>CYTO</td></tr>
<tr><td>>hg38_21_+_8433172_8446622_RNA45SN1_45S_50ntFlanks@10397.10424.28|AGTTGAACATGGGTCAGTCGGTCCTGAG</td><td>>hg38_21_+_8433172_8446622_RNA45SN1_45S_50ntFlanks</td><td>10397</td><td>10424</td><td>28</td><td>AGTTGAACATGGGTCAGTCGGTCCTGAG</td><td>315.3</td><td>7.7</td><td>1.1</td><td>7.0</td><td>1.9E-12</td><td>6.4E-12</td><td>MB231/MB468</td><td>CYTO</td></tr>
<tr><td>>hg38_21_+_8433172_8446622_RNA45SN1_45S_50ntFlanks@11667.11692.26|CTGCCCAGTGCTCTGAATGTCAAAGT</td><td>>hg38_21_+_8433172_8446622_RNA45SN1_45S_50ntFlanks</td><td>11667</td><td>11692</td><td>26</td><td>CTGCCCAGTGCTCTGAATGTCAAAGT</td><td>429.5</td><td>8.0</td><td>1.1</td><td>7.0</td><td>1.9E-12</td><td>6.4E-12</td><td>MB231/MB468</td><td>CYTO</td></tr>
<tr><td>>am_trnaMT_GluTTC_MT_-_14674_14742@41.58.18,trnalookalike8_GluTTC_5_-_93905172_93905240@41.58.18|TGGTCGTGGTTGTAGTCC</td><td>>am_trnaMT_GluTTC_MT_-_14674_14742</td><td>41</td><td>58</td><td>18</td><td>TGGTCGTGGTTGTAGTCC</td><td>448.8</td><td>7.4</td><td>1.0</td><td>7.1</td><td>1.4E-12</td><td>6.4E-12</td><td>BT20/MB468</td><td>MITO</td></tr>
<tr><td>>hg38_21_+_8433172_8446622_RNA45SN1_45S_50ntFlanks@12004.12030.27|CGGGCCGCCGGTGAAATACCACTACTC</td><td>>hg38_21_+_8433172_8446622_RNA45SN1_45S_50ntFlanks</td><td>12004</td><td>12030</td><td>27</td><td>CGGGCCGCCGGTGAAATACCACTACTC</td><td>129.6</td><td>-10.9</td><td>1.5</td><td>-7.1</td><td>1.7E-12</td><td>6.4E-12</td><td>MB231/MB468</td><td>MP</td></tr>
<tr><td>>hg38_21_+_8433172_8446622_RNA45SN1_45S_50ntFlanks@12912.12935.24|CACGTTCGTGGGGAACCTGGCGCT</td><td>>hg38_21_+_8433172_8446622_RNA45SN1_45S_50ntFlanks</td><td>12912</td><td>12935</td><td>24</td><td>CACGTTCGTGGGGAACCTGGCGCT</td><td>612.9</td><td>-8.9</td><td>1.3</td><td>-7.0</td><td>2.4E-12</td><td>6.4E-12</td><td>MB231/MB468</td><td>MITO</td></tr>
<tr><td>>hg38_21_+_8433172_8446622_RNA45SN1_45S_50ntFlanks@12935.12952.18|TAAACCATTCGTAGACGA</td><td>>hg38_21_+_8433172_8446622_RNA45SN1_45S_50ntFlanks</td><td>12935</td><td>12952</td><td>18</td><td>TAAACCATTCGTAGACGA</td><td>103.3</td><td>-10.6</td><td>1.5</td><td>-7.1</td><td>1.7E-12</td><td>6.4E-12</td><td>MB231/MB468</td><td>MP</td></tr>
<tr><td>>hg38_21_+_8433172_8446622_RNA45SN1_45S_50ntFlanks@10392.10410.19|ACAGCAGTTGAACATGGGT</td><td>>hg38_21_+_8433172_8446622_RNA45SN1_45S_50ntFlanks</td><td>10392</td><td>10410</td><td>19</td><td>ACAGCAGTTGAACATGGGT</td><td>2691.8</td><td>-6.1</td><td>0.8</td><td>-7.5</td><td>9.3E-14</td><td>6.4E-12</td><td>BT20/MB468</td><td>MP</td></tr>
<tr><td>>hg38_21_+_8433172_8446622_RNA45SN1_45S_50ntFlanks@10401.10435.35|GAACATGGGTCAGTCGGTCCTGAGAGATGGGCGAG</td><td>>hg38_21_+_8433172_8446622_RNA45SN1_45S_50ntFlanks</td><td>10401</td><td>10435</td><td>35</td><td>GAACATGGGTCAGTCGGTCCTGAGAGATGGGCGAG</td><td>214.0</td><td>-11.6</td><td>1.6</td><td>-7.1</td><td>1.7E-12</td><td>6.4E-12</td><td>MB231/MB468</td><td>MP</td></tr>
<tr><td>>MI0000102|hsa-mir-100&WithFlank&11|-|122152223|122152314@19.37.19(+1A)&[MIMAT0000098&hsa-miR-100-5p&offsets|0|-3(+1A);m-27&11|-|122152276|122152296&offsets|0|-2(+1A)]|AACCCGTAGATCCGAACTTA</td><td>>MI0000102|hsa-mir-100&WithFlank&11|-|122152223|122152314</td><td>19</td><td>37</td><td>19(+1A)</td><td>AACCCGTAGATCCGAACTTA</td><td>5250.2</td><td>12.0</td><td>1.7</td><td>7.1</td><td>1.7E-12</td><td>6.4E-12</td><td>MB231/MB468</td><td>MP</td></tr>
<tr><td>>hg38_21_+_8433172_8446622_RNA45SN1_45S_50ntFlanks@12332.12365.34|TAAGCAGGAGGTGTCAGAAAAGTTACCACAGGGA</td><td>>hg38_21_+_8433172_8446622_RNA45SN1_45S_50ntFlanks</td><td>12332</td><td>12365</td><td>34</td><td>TAAGCAGGAGGTGTCAGAAAAGTTACCACAGGGA</td><td>153.1</td><td>-11.1</td><td>1.6</td><td>-7.1</td><td>1.7E-12</td><td>6.4E-12</td><td>MB231/MB468</td><td>MP</td></tr>
<tr><td>>hg38_21_+_8433172_8446622_RNA45SN1_45S_50ntFlanks@9622.9656.35|CTAGTAGCTGGTTCCCTCCGAAGTTTCCCTCAGGA</td><td>>hg38_21_+_8433172_8446622_RNA45SN1_45S_50ntFlanks</td><td>9622</td><td>9656</td><td>35</td><td>CTAGTAGCTGGTTCCCTCCGAAGTTTCCCTCAGGA</td><td>338.8</td><td>-8.4</td><td>1.2</td><td>-7.0</td><td>2.4E-12</td><td>6.4E-12</td><td>MB231/MB468</td><td>MITO</td></tr>
</table>

| | | | | | | | | | | | | | |
|---|---|---|---|---|---|---|---|---|---|---|---|---|---|
| >hg38_21_+_8433172_8446622_RNA45SN1_45S_50ntFlanks@475.496.22\|CTTCGTGATCGATGTGGTGACG | >hg38_21_+_8433172_8446622_RNA45SN1_45S_50ntFlanks | 475 | 496 | 22 | CTTCGTGATCGATGTGGTGACG | 24899.8 | 5.6 | 0.8 | 7.0 | 2.0E-12 | 6.4E-12 | MB231/MB468 | NUC |
| >hg38_21_+_8433172_8446622_RNA45SN1_45S_50ntFlanks@12971.12992.22\|TTTCGTACGTAGCAGAGCAGCT | >hg38_21_+_8433172_8446622_RNA45SN1_45S_50ntFlanks | 12971 | 12992 | 22 | TTTCGTACGTAGCAGAGCAGCT | 289.5 | -3.3 | 0.5 | -7.0 | 2.0E-12 | 6.4E-12 | MB231/MB468 | NUC |
| >hg38_21_+_8433172_8446622_RNA45SN1_45S_50ntFlanks@8194.8214.21\|CCCGTGGACGGTGTGAGGCCG | >hg38_21_+_8433172_8446622_RNA45SN1_45S_50ntFlanks | 8194 | 8214 | 21 | CCCGTGGACGGTGTGAGGCCG | 3354.2 | -10.8 | 1.5 | -7.0 | 2.6E-12 | 6.4E-12 | BT20/MB468 | NUC |
| >hg38_21_+_8433172_8446622_RNA45SN1_45S_50ntFlanks@10463.10488.26\|CTCCGTTGCCCTCGGCCGATCGAAAG | >hg38_21_+_8433172_8446622_RNA45SN1_45S_50ntFlanks | 10463 | 10488 | 26 | CTCCGTTGCCCTCGGCCGATCGAAAG | 98.5 | -10.5 | 1.5 | -7.1 | 1.7E-12 | 6.5E-12 | MB231/MB468 | MP |
| >MI0000299\|hsa-mir-222&WithFlank&X\|-\|45747009\|45747130@75.97.23&[MIMAT0000279&hsa-miR-222-3p&offsets\|0\|+2;m-140&X\|-\|45747033\|45747056&offsets\|0\|-1]\|AGCTACATCTGGCTACTGGGTCT | >MI0000299\|hsa-mir-222&WithFlank&X\|-\|45747009\|45747130 | 75 | 97 | 23 | AGCTACATCTGGCTACTGGGTCT | 8574.1 | 6.2 | 0.9 | 7.0 | 2.4E-12 | 6.5E-12 | MB231/MB468 | MITO |
| >hg38_21_+_8433172_8446622_RNA45SN1_45S_50ntFlanks@9504.9522.19\|GTGAACTATGCCTGGGCAG | >hg38_21_+_8433172_8446622_RNA45SN1_45S_50ntFlanks | 9504 | 9522 | 19 | GTGAACTATGCCTGGGCAG | 2110.3 | -10.5 | 1.5 | -7.0 | 2.6E-12 | 6.5E-12 | BT20/MB468 | NUC |
| >hg38_21_+_8433172_8446622_RNA45SN1_45S_50ntFlanks@11999.12039.41\|CCCTGCGGGCCGCCGGTGAAATACCACTACTCTGATCGTTT | >hg38_21_+_8433172_8446622_RNA45SN1_45S_50ntFlanks | 11999 | 12039 | 41 | CCCTGCGGGCCGCCGGTGAAATACCACTACTCTGATCGTTT | 178.7 | -10.2 | 1.4 | -7.0 | 2.0E-12 | 6.5E-12 | MB231/MB468 | CYTO |
| >hg38_21_+_8433172_8446622_RNA45SN1_45S_50ntFlanks@6650.6677.28\|ACGACTCTTAGCGGTGGATCACTCGGCT | >hg38_21_+_8433172_8446622_RNA45SN1_45S_50ntFlanks | 6650 | 6677 | 28 | ACGACTCTTAGCGGTGGATCACTCGGCT | 5882.9 | 4.6 | 0.6 | 7.1 | 1.5E-12 | 6.5E-12 | BT20/MB468 | MITO |
| >MI0000433\|hsa-let-7g&WithFlank&3\|-\|52268272\|52268367@10.32.23&[MIMAT0000414&hsa-let-7g-5p&offsets\|-1\|0;m-58&3\|-\|52268336\|52268357&offsets\|-1\|0]\|CTGAGGTAGTAGTTTGTACAGTT | >MI0000433\|hsa-let-7g&WithFlank&3\|-\|52268272\|52268367 | 10 | 32 | 23 | CTGAGGTAGTAGTTTGTACAGTT | 440.5 | 7.3 | 1.0 | 7.1 | 1.5E-12 | 6.5E-12 | BT20/MB468 | MITO |
| >am_MI0006405\|hsa-mir-1268a&WithFlank&15\|-\|22225272\|22225335@11.28.18;MI0016748\|hsa-mir-1268b&WithFlank&17\|+\|80098822\|80098883@10.27.18&[MIMAT0005922&hsa-miR-1268a&offsets\|0\|0;m-386&15\|-\|22225308\|22225325&offsets\|0\|0];[MIMAT0018925&hsa-miR-1268b&offsets\|0\|-2;m-350&17\|+\|80098831\|80098848&offsets\|0\|0]\|CGGGCGTGGTGGTGGGGG | >am_MI0006405\|hsa-mir-1268a&WithFlank&15\|-\|22225272\|22225335 | 11 | 28 | 18 | CGGGCGTGGTGGTGGGGG | 306.3 | 9.1 | 1.3 | 7.0 | 2.0E-12 | 6.5E-12 | MB231/MB468 | CYTO |
| >hg38_21_+_8433172_8446622_RNA45SN1_45S_50ntFlanks@13148.13165.18\|CGGGCGTGGTGGTGGGGG | >hg38_21_+_8433172_8446622_RNA45SN1_45S_50ntFlanks | 13148 | 13165 | 18 | CGGGCGTGGTGGTGGGGG | 306.3 | 9.1 | 1.3 | 7.0 | 2.0E-12 | 6.5E-12 | MB231/MB468 | CYTO |
| >hg38_21_+_8433172_8446622_RNA45SN1_45S_50ntFlanks@11566.11583.18\|TAGCAGCCGACTTAGAAC | >hg38_21_+_8433172_8446622_RNA45SN1_45S_50ntFlanks | 11566 | 11583 | 18 | TAGCAGCCGACTTAGAAC | 234.2 | 4.9 | 0.7 | 7.0 | 2.0E-12 | 6.6E-12 | MB231/MB468 | NUC |
| >hg38_21_+_8433172_8446622_RNA45SN1_45S_50ntFlanks@12539.12587.49\|ATGATGTGTTGTTGCCATGGTAATCCTGCTCAGTACGAGAGGAACCGCA | >hg38_21_+_8433172_8446622_RNA45SN1_45S_50ntFlanks | 12539 | 12587 | 49 | ATGATGTGTTGTTGCCATGGTAATCCTGCTCAGTACGAGAGGAACCGCA | 739.1 | -12.5 | 1.8 | -7.0 | 2.0E-12 | 6.6E-12 | MB231/MB468 | NUC |
| >hg38_21_+_8433172_8446622_RNA45SN1_45S_50ntFlanks@5487.5518.32\|CGCTGAGAAGACGGTCGAACTTGACTATCTAG | >hg38_21_+_8433172_8446622_RNA45SN1_45S_50ntFlanks | 5487 | 5518 | 32 | CGCTGAGAAGACGGTCGAACTTGACTATCTAG | 735.7 | -12.6 | 1.7 | -7.4 | 9.6E-14 | 6.6E-12 | BT20/MB468 | MP |
| >hg38_21_+_8433172_8446622_RNA45SN1_45S_50ntFlanks@12405.12424.20\|CTTTTTGATCCTTCGATGTC | >hg38_21_+_8433172_8446622_RNA45SN1_45S_50ntFlanks | 12405 | 12424 | 20 | CTTTTTGATCCTTCGATGTC | 433.0 | -10.8 | 1.5 | -7.1 | 1.5E-12 | 6.6E-12 | BT20/MB468 | MITO |
| >hg38_21_+_8433172_8446622_RNA45SN1_45S_50ntFlanks@3746.3763.18\|ATTAAGCCATGCATGTCT | >hg38_21_+_8433172_8446622_RNA45SN1_45S_50ntFlanks | 3746 | 3763 | 18 | ATTAAGCCATGCATGTCT | 118.1 | -10.7 | 1.5 | -7.1 | 1.8E-12 | 6.6E-12 | MB231/MB468 | MP |
| >hg38_21_+_8433172_8446622_RNA45SN1_45S_50ntFlanks@12083.12107.25\|CTCGCTTCTGGCGCCAAGCGCCCGG | >hg38_21_+_8433172_8446622_RNA45SN1_45S_50ntFlanks | 12083 | 12107 | 25 | CTCGCTTCTGGCGCCAAGCGCCCGG | 333.4 | -11.5 | 1.5 | -7.4 | 9.7E-14 | 6.6E-12 | BT20/MB468 | MP |
| >hg38_21_+_8433172_8446622_RNA45SN1_45S_50ntFlanks@6698.6720.23\|AGCTAGCTGCGAGAATTAATGTG | >hg38_21_+_8433172_8446622_RNA45SN1_45S_50ntFlanks | 6698 | 6720 | 23 | AGCTAGCTGCGAGAATTAATGTG | 33353.9 | -6.8 | 1.0 | -7.1 | 1.8E-12 | 6.6E-12 | MB231/MB468 | MP |
| >hg38_21_+_8433172_8446622_RNA45SN1_45S_50ntFlanks@12686.12711.26\|CCAGGCGGAACGATACGGCAGCGCCG | >hg38_21_+_8433172_8446622_RNA45SN1_45S_50ntFlanks | 12686 | 12711 | 26 | CCAGGCGGAACGATACGGCAGCGCCG | 736.7 | -9.2 | 1.3 | -7.0 | 2.5E-12 | 6.6E-12 | MB231/MB468 | MITO |
| >hg38_21_+_8433172_8446622_RNA45SN1_45S_50ntFlanks@12008.12037.30\|CCGCCGGTGAAATACCACTACTCTGATCGT | >hg38_21_+_8433172_8446622_RNA45SN1_45S_50ntFlanks | 12008 | 12037 | 30 | CCGCCGGTGAAATACCACTACTCTGATCGT | 671.0 | -8.6 | 1.2 | -7.0 | 2.5E-12 | 6.6E-12 | MB231/MB468 | MITO |
| >MI0000477\|hsa-mir-146a&WithFlank&5\|+\|160485346\|160485456@27.47.21(+1A)&[MIMAT0000449&hsa-miR-146a-5p&offsets\|0\|-1(+1A);m-129&5\|+\|160485372\|160485393&offsets\|0\|-1(+1A)]\|TGAGAACTGAATTCCATGGGTA | >MI0000477\|hsa-mir-146a&WithFlank&5\|+\|160485346\|160485456 | 27 | 47 | 21(+1A) | TGAGAACTGAATTCCATGGGTA | 541.5 | -12.2 | 1.7 | -7.3 | 2.2E-13 | 6.6E-12 | BT20/MB231 | NUC |
| >am_trna128_GlyGCC_6_-_27870686_27870756@1.25.25,trna133_GlyCCC_1_-_16872434_16872504@1.25.25,trna18_GlyGCC_16_+_70822597_70822667@1.25.25,trna19_GlyGCC_16_+_70823410_70823480@1.25.25,trna19_GlyGCC_2_-_157257659_157257729@1.25.25,trna24_GlyGCC_16_-_70812942_70813012@1.25.25,trna25_GlyGCC_16_-_70812114_70812184@1.25.25,trna4_GlyCCC_1_+_17188416_17188486@1.25.25,trna5_GlyGCC_17_+_8029064_8029134@1.25.25,trna68_GlyGCC_1_-_161493637_161493707@1.25.25\|GCATTGGTGGTTCAGTGGTAGAATT | >am_trna128_GlyGCC_6_-_27870686_27870756 | 1 | 25 | 25 | GCATTGGTGGTTCAGTGGTAGAATT | 1376.0 | -6.1 | 0.9 | -7.0 | 2.7E-12 | 6.6E-12 | BT20/MB468 | NUC |
| >hg38_21_+_8433172_8446622_RNA45SN1_45S_50ntFlanks@9462.9503.42\|GGGAGGTGGAGCACGAGCGCACGTGTTAGGACCCGAAAGATG | >hg38_21_+_8433172_8446622_RNA45SN1_45S_50ntFlanks | 9462 | 9503 | 42 | GGGAGGTGGAGCACGAGCGCACGTGTTAGGACCCGAAAGATG | 312.2 | -12.2 | 1.7 | -7.1 | 1.8E-12 | 6.6E-12 | MB231/MB468 | MP |
| >hg38_21_+_8433172_8446622_RNA45SN1_45S_50ntFlanks@3841.3879.39\|TCCTACTTGGATAACTGTGGTAATTCTAGAGCTAATACA | >hg38_21_+_8433172_8446622_RNA45SN1_45S_50ntFlanks | 3841 | 3879 | 39 | TCCTACTTGGATAACTGTGGTAATTCTAGAGCTAATACA | 998.9 | 7.2 | 1.0 | 7.0 | 2.0E-12 | 6.7E-12 | BT20/MB468 | CYTO |
| >MI0000734\|hsa-mir-106b&WithFlank&7\|-\|100093987\|100094080@56.77.22&[MIMAT0004672&hsa-miR-106b-3p&offsets\|-2\|-2;m-87&7\|-\|100094003\|100094023&offsets\|-2\|-1]\|TACCGCACTGTGGGTACTTGCT | >MI0000734\|hsa-mir-106b&WithFlank&7\|-\|100093987\|100094080 | 56 | 77 | 22 | TACCGCACTGTGGGTACTTGCT | 982.6 | 8.0 | 1.1 | 7.0 | 2.0E-12 | 6.7E-12 | MB231/MB468 | CYTO |
| >hg38_21_+_8433172_8446622_RNA45SN1_45S_50ntFlanks@4728.4752.25\|ATCAAGAACGAAAGTCGGAGGTTCG | >hg38_21_+_8433172_8446622_RNA45SN1_45S_50ntFlanks | 4728 | 4752 | 25 | ATCAAGAACGAAAGTCGGAGGTTCG | 166.7 | -9.5 | 1.3 | -7.1 | 1.5E-12 | 6.7E-12 | BT20/MB468 | MITO |
| >hg38_21_+_8433172_8446622_RNA45SN1_45S_50ntFlanks@7989.8011.23\|AGACGTGGCGACCCGCTGAATTT | >hg38_21_+_8433172_8446622_RNA45SN1_45S_50ntFlanks | 7989 | 8011 | 23 | AGACGTGGCGACCCGCTGAATTT | 5702.2 | 6.8 | 1.0 | 7.0 | 2.0E-12 | 6.7E-12 | MB231/MB468 | CYTO |
| >hg38_21_+_8433172_8446622_RNA45SN1_45S_50ntFlanks@4678.4707.30\|CGGCGCAAGACGGACCAGAGCGAAAGCATT | >hg38_21_+_8433172_8446622_RNA45SN1_45S_50ntFlanks | 4678 | 4707 | 30 | CGGCGCAAGACGGACCAGAGCGAAAGCATT | 183.5 | 7.4 | 1.1 | 7.0 | 2.0E-12 | 6.7E-12 | MB231/MB468 | CYTO |
| >MI0000462\|hsa-mir-152&WithFlank&17\|-\|48037155\|48037253@60.81.22&[MIMAT0000438&hsa-miR-152-3p&offsets\|0\|+1;m-97&17\|-\|48037174\|48037194&offsets\|0\|+1]\|TCAGTGCATGACAGAACTTGGG | >MI0000462\|hsa-mir-152&WithFlank&17\|-\|48037155\|48037253 | 60 | 81 | 22 | TCAGTGCATGACAGAACTTGGG | 277.3 | 11.3 | 1.6 | 7.0 | 2.5E-12 | 6.7E-12 | MB231/MB468 | MITO |
| >MI0000076\|hsa-mir-20a&WithFlank&13\|+\|91351059\|91351141@14.37.24&[MIMAT0000075&hsa-miR-20a-5p&offsets\|0\|+1;m-96&13\|+\|91351072\|91351094&offsets\|0\|+1]\|TAAAGTGCTTATAGTGCAGGTAGT | >MI0000076\|hsa-mir-20a&WithFlank&13\|+\|91351059\|91351141 | 14 | 37 | 24 | TAAAGTGCTTATAGTGCAGGTAGT | 1205.7 | 6.1 | 0.9 | 7.0 | 2.1E-12 | 6.7E-12 | MB231/MB468 | NUC |
| >hg38_21_+_8433172_8446622_RNA45SN1_45S_50ntFlanks@11592.11613.22\|ACCAGGGGAATCCGACTGTTTA | >hg38_21_+_8433172_8446622_RNA45SN1_45S_50ntFlanks | 11592 | 11613 | 22 | ACCAGGGGAATCCGACTGTTTA | 656.7 | -8.6 | 1.2 | -7.0 | 2.5E-12 | 6.7E-12 | MB231/MB468 | MITO |
| >hg38_21_+_8433172_8446622_RNA45SN1_45S_50ntFlanks@4773.4793.21\|TAGTTCCGACCATAAACGATG | >hg38_21_+_8433172_8446622_RNA45SN1_45S_50ntFlanks | 4773 | 4793 | 21 | TAGTTCCGACCATAAACGATG | 5006.6 | -11.6 | 1.7 | -7.0 | 2.7E-12 | 6.7E-12 | BT20/MB468 | NUC |
| >hg38_21_+_8433172_8446622_RNA45SN1_45S_50ntFlanks@10431.10456.26\|GCGAGCGCCGTTCCGAAGGGACGGGC | >hg38_21_+_8433172_8446622_RNA45SN1_45S_50ntFlanks | 10431 | 10456 | 26 | GCGAGCGCCGTTCCGAAGGGACGGGC | 167.4 | -9.7 | 1.4 | -7.0 | 2.0E-12 | 6.7E-12 | BT20/MB468 | CYTO |
| >hg38_21_+_8433172_8446622_RNA45SN1_45S_50ntFlanks@11873.11908.36\|AAGAAGACCCTGTTGAGCTTGACTCTAGTCTGGCAC | >hg38_21_+_8433172_8446622_RNA45SN1_45S_50ntFlanks | 11873 | 11908 | 36 | AAGAAGACCCTGTTGAGCTTGACTCTAGTCTGGCAC | 175.3 | -11.3 | 1.6 | -7.0 | 1.8E-12 | 6.7E-12 | MB231/MB468 | MP |
| >hg38_21_+_8433172_8446622_RNA45SN1_45S_50ntFlanks@10731.10748.18\|AGGGTGTAAATCTCGCGC | >hg38_21_+_8433172_8446622_RNA45SN1_45S_50ntFlanks | 10731 | 10748 | 18 | AGGGTGTAAATCTCGCGC | 110.5 | -10.7 | 1.5 | -7.0 | 1.8E-12 | 6.7E-12 | MB231/MB468 | MP |
| >hg38_21_+_8433172_8446622_RNA45SN1_45S_50ntFlanks@3882.3915.34\|CCGACGGGCGCTGACCCCCTTCGCGGGGGGGATG | >hg38_21_+_8433172_8446622_RNA45SN1_45S_50ntFlanks | 3882 | 3915 | 34 | CCGACGGGCGCTGACCCCCTTCGCGGGGGGGATG | 127.5 | -10.9 | 1.5 | -7.0 | 1.8E-12 | 6.7E-12 | MB231/MB468 | MP |
| >hg38_21_+_8433172_8446622_RNA45SN1_45S_50ntFlanks@9534.9553.20\|AGGAAACTCTGGTGGAGGTC | >hg38_21_+_8433172_8446622_RNA45SN1_45S_50ntFlanks | 9534 | 9553 | 20 | AGGAAACTCTGGTGGAGGTC | 106236.2 | -7.0 | 1.0 | -7.0 | 2.0E-12 | 6.7E-12 | BT20/MB468 | CYTO |
| >MI0000279\|hsa-mir-196a-2&WithFlank&12\|+\|53991732\|53991853@31.50.20(+1A);MI0000238\|hsa-mir-196a-1&WithFlank&17\|-\|48632484\|48632565@13.32.20(+1A)&[MIMAT0000226&hsa-miR-196a-5p&offsets\|0\|-2(+1A);m-261&12\|+\|53991762\|53991783&offsets\|0\|-2(+1A)];[MIMAT0000226_1&hsa-miR-196a-5p&offsets\|0\|-2(+1A);m-268&17\|-\|48632532\|48632553&offsets\|0\|-2(+1A)]\|TAGGTAGTTTCATGTTGTTGA | >MI0000279\|hsa-mir-196a-2&WithFlank&12\|+\|53991732\|53991853 | 31 | 50 | 20(+1A) | TAGGTAGTTTCATGTTGTTGA | 593.7 | 8.4 | 1.2 | 7.0 | 2.0E-12 | 6.7E-12 | BT20/MB468 | CYTO |
| >hg38_21_+_8433172_8446622_RNA45SN1_45S_50ntFlanks@12714.12743.30\|GAGCCTCGGTTGGCCTCGGATAGCCGGTCC | >hg38_21_+_8433172_8446622_RNA45SN1_45S_50ntFlanks | 12714 | 12743 | 30 | GAGCCTCGGTTGGCCTCGGATAGCCGGTCC | 88.7 | -10.3 | 1.5 | -7.0 | 1.8E-12 | 6.7E-12 | MB231/MB468 | MP |
| >hg38_21_+_8433172_8446622_RNA45SN1_45S_50ntFlanks@11874.11912.39\|AGAAGACCCTGTTGAGCTTGACTCTAGTCTGGCACGGTG | >hg38_21_+_8433172_8446622_RNA45SN1_45S_50ntFlanks | 11874 | 11912 | 39 | AGAAGACCCTGTTGAGCTTGACTCTAGTCTGGCACGGTG | 360.3 | -10.6 | 1.5 | -7.1 | 1.5E-12 | 6.7E-12 | BT20/MB468 | MITO |
| >hg38_21_+_8433172_8446622_RNA45SN1_45S_50ntFlanks@12141.12179.39\|ACAGTGCCAGGTGGGGAGTTTGACTGGGGCGGTACACCT | >hg38_21_+_8433172_8446622_RNA45SN1_45S_50ntFlanks | 12141 | 12179 | 39 | ACAGTGCCAGGTGGGGAGTTTGACTGGGGCGGTACACCT | 269.2 | -11.9 | 1.7 | -7.0 | 1.8E-12 | 6.7E-12 | MB231/MB468 | MP |
| >hg38_21_+_8433172_8446622_RNA45SN1_45S_50ntFlanks@12934.12957.24\|CTAAACCATTCGTAGACGACCTGC | >hg38_21_+_8433172_8446622_RNA45SN1_45S_50ntFlanks | 12934 | 12957 | 24 | CTAAACCATTCGTAGACGACCTGC | 93.4 | -10.4 | 1.5 | -7.0 | 1.8E-12 | 6.8E-12 | MB231/MB468 | MP |
| >hg38_21_+_8433172_8446622_RNA45SN1_45S_50ntFlanks@12623.12654.32\|GCCAATGGGGCGAAGCTACCATCTGTGGGATT | >hg38_21_+_8433172_8446622_RNA45SN1_45S_50ntFlanks | 12623 | 12654 | 32 | GCCAATGGGGCGAAGCTACCATCTGTGGGATT | 305.5 | -11.3 | 1.5 | -7.4 | 1.0E-13 | 6.8E-12 | BT20/MB468 | MP |
| >hg38_21_+_8433172_8446622_RNA45SN1_45S_50ntFlanks@9033.9076.44\|CTCGTGGGGGGGCCGGGCCACCCCTCCCACGGCGCGACCGCTCT | >hg38_21_+_8433172_8446622_RNA45SN1_45S_50ntFlanks | 9033 | 9076 | 44 | CTCGTGGGGGGGCCGGGCCACCCCTCCCACGGCGCGACCGCTCT | 81.0 | -22.8 | 3.1 | -7.4 | 1.0E-13 | 6.8E-12 | BT20/MB468 | MP |
| >hg38_21_+_8433172_8446622_RNA45SN1_45S_50ntFlanks@8401.8422.22\|AGAGGTAAACGGGTGGGGTCCG | >hg38_21_+_8433172_8446622_RNA45SN1_45S_50ntFlanks | 8401 | 8422 | 22 | AGAGGTAAACGGGTGGGGTCCG | 127.5 | -10.9 | 1.5 | -7.0 | 1.8E-12 | 6.8E-12 | MB231/MB468 | MP |
| >hg38_21_+_8433172_8446622_RNA45SN1_45S_50ntFlanks@12674.12704.31\|TCAGAATCCCGCCCAGGCGGAACGATACGGC | >hg38_21_+_8433172_8446622_RNA45SN1_45S_50ntFlanks | 12674 | 12704 | 31 | TCAGAATCCCGCCCAGGCGGAACGATACGGC | 114.2 | -10.7 | 1.5 | -7.0 | 1.8E-12 | 6.8E-12 | MB231/MB468 | MP |
| >hg38_21_+_8433172_8446622_RNA45SN1_45S_50ntFlanks@12274.12306.33\|CAGTACGAATACAGACCGTGAAAGCGGGGCCTC | >hg38_21_+_8433172_8446622_RNA45SN1_45S_50ntFlanks | 12274 | 12306 | 33 | CAGTACGAATACAGACCGTGAAAGCGGGGCCTC | 181.2 | -11.3 | 1.6 | -7.0 | 2.6E-12 | 6.8E-12 | MB231/MB468 | MITO |

| | | | | | | | | | | | | | |
|---|---|---|---|---|---|---|---|---|---|---|---|---|---|
| >hg38_21_+_8433172_8446622_RNA45SN1_45S_50ntFlanks@6721.6744.24\|AATTGCAGGACACATTGATCATCG | >hg38_21_+_8433172_8446622_RNA45SN1_45S_50ntFlanks | 6721 | 6744 | 24 | AATTGCAGGACACATTGATCATCG | 27828.6 | -7.0 | 1.0 | -7.0 | 2.0E-12 | 6.8E-12 | BT20/MB468 | CYTO |
| >hg38_21_+_8433172_8446622_RNA45SN1_45S_50ntFlanks@6647.6667.21\|CGTACGACTCTTAGCGGTGGA | >hg38_21_+_8433172_8446622_RNA45SN1_45S_50ntFlanks | 6647 | 6667 | 21 | CGTACGACTCTTAGCGGTGGA | 641.7 | 3.8 | 0.5 | 7.0 | 2.1E-12 | 6.8E-12 | MB231/MB468 | NUC |
| >MI0000102\|hsa-mir-100&WithFlank&11\|-\|122152223\|122152314@19.39.21(+1A)&[MIMAT0000098&hsa-miR-100-5p&offsets\|0\|-1(+1A);m-27&11\|-\|122152276\|122152296&offsets\|0\|0(+1A)]\|AACCCGTAGATCCGAACTTGTA | >MI0000102\|hsa-mir-100&WithFlank&11\|-\|122152223\|122152314 | 19 | 39 | 21(+1A) | AACCCGTAGATCCGAACTTGTA | 126984.9 | -8.3 | 1.1 | -7.6 | 2.8E-14 | 6.8E-12 | BT20/MB231 | MP |
| >hg38_21_+_8433172_8446622_RNA45SN1_45S_50ntFlanks@11210.11254.45\|GGCGACTCTGGACGCGAGCCGGGCCCTTCCCGTGGATCGCCCCAG | >hg38_21_+_8433172_8446622_RNA45SN1_45S_50ntFlanks | 11210 | 11254 | 45 | GGCGACTCTGGACGCGAGCCGGGCCCTTCCCGTGGATCGCCCCAG | 4712.3 | -9.7 | 1.4 | -7.0 | 2.8E-12 | 6.8E-12 | BT20/MB468 | NUC |
| >MI0003591\|hsa-mir-584&WithFlank&5\|-\|149062307\|149062415@22.40.19&[MIMAT0003249&hsa-miR-584-5p&offsets\|0\|-3;m-167&5\|-\|149062374\|149062394&offsets\|0\|-2]\|TTATGGTTTGCCTGGGACT | >MI0003591\|hsa-mir-584&WithFlank&5\|-\|149062307\|149062415 | 22 | 40 | 19 | TTATGGTTTGCCTGGGACT | 333.3 | 7.9 | 1.1 | 7.0 | 2.1E-12 | 6.8E-12 | MB231/MB468 | CYTO |
| >MI0000298\|hsa-mir-221&WithFlank&X\|-\|45746151\|45746272@71.91.21(+1G)&[MIMAT0000278&hsa-miR-221-3p&offsets\|0\|-2(+1G);m-91&X\|-\|45746181\|45746202&offsets\|0\|-1(+1G)]\|AGCTACATTGTCTGCTGGGTTG | >MI0000298\|hsa-mir-221&WithFlank&X\|-\|45746151\|45746272 | 71 | 91 | 21(+1G) | AGCTACATTGTCTGCTGGGTTG | 187.1 | 7.8 | 1.1 | 7.0 | 2.1E-12 | 6.8E-12 | MB231/MB468 | NUC |
| >hg38_21_+_8433172_8446622_RNA45SN1_45S_50ntFlanks@6664.6687.24\|TGGATCACTCGGCTCGTGCGTCGA | >hg38_21_+_8433172_8446622_RNA45SN1_45S_50ntFlanks | 6664 | 6687 | 24 | TGGATCACTCGGCTCGTGCGTCGA | 109.4 | -10.6 | 1.5 | -7.0 | 1.8E-12 | 6.8E-12 | MB231/MB468 | MP |
| >hg38_21_+_8433172_8446622_RNA45SN1_45S_50ntFlanks@9122.9142.21\|CCGCGCGCGGGTCGGGGGGCG | >hg38_21_+_8433172_8446622_RNA45SN1_45S_50ntFlanks | 9122 | 9142 | 21 | CCGCGCGCGGGTCGGGGGGCG | 184.1 | -11.4 | 1.6 | -7.0 | 1.8E-12 | 6.9E-12 | MB231/MB468 | MP |
| >hg38_21_+_8433172_8446622_RNA45SN1_45S_50ntFlanks@4589.4616.28\|TATTTTGTTGGTTTTCGGAACTGAGGCC | >hg38_21_+_8433172_8446622_RNA45SN1_45S_50ntFlanks | 4589 | 4616 | 28 | TATTTTGTTGGTTTTCGGAACTGAGGCC | 194.4 | -11.5 | 1.6 | -7.0 | 1.8E-12 | 6.9E-12 | MB231/MB468 | MP |
| >hg38_21_+_8433172_8446622_RNA45SN1_45S_50ntFlanks@3874.3899.26\|AATACATGCCGACGGGCGCTGACCCC | >hg38_21_+_8433172_8446622_RNA45SN1_45S_50ntFlanks | 3874 | 3899 | 26 | AATACATGCCGACGGGCGCTGACCCC | 177.8 | -11.3 | 1.6 | -7.0 | 1.8E-12 | 6.9E-12 | MB231/MB468 | MP |
| >hg38_21_+_8433172_8446622_RNA45SN1_45S_50ntFlanks@5485.5511.27\|AGCGCTGAGAAGACGGTCGAACTTGAC | >hg38_21_+_8433172_8446622_RNA45SN1_45S_50ntFlanks | 5485 | 5511 | 27 | AGCGCTGAGAAGACGGTCGAACTTGAC | 128.0 | -10.9 | 1.5 | -7.0 | 1.8E-12 | 6.9E-12 | MB231/MB468 | MP |
| >hg38_21_+_8433172_8446622_RNA45SN1_45S_50ntFlanks@11907.11929.23\|ACGGTGAAGAGACATGAGAGGTG | >hg38_21_+_8433172_8446622_RNA45SN1_45S_50ntFlanks | 11907 | 11929 | 23 | ACGGTGAAGAGACATGAGAGGTG | 165.1 | -11.2 | 1.6 | -7.0 | 1.9E-12 | 6.9E-12 | MB231/MB468 | MP |
| >hg38_21_+_8433172_8446622_RNA45SN1_45S_50ntFlanks@12461.12487.27\|CGTTGGATTGTTCACCCACTAATAGGG | >hg38_21_+_8433172_8446622_RNA45SN1_45S_50ntFlanks | 12461 | 12487 | 27 | CGTTGGATTGTTCACCCACTAATAGGG | 146.5 | -11.1 | 1.6 | -7.0 | 1.9E-12 | 6.9E-12 | MB231/MB468 | MP |
| >hg38_21_+_8433172_8446622_RNA45SN1_45S_50ntFlanks@9554.9575.22\|CGTAGCGGTCCTGACGTGCAAA | >hg38_21_+_8433172_8446622_RNA45SN1_45S_50ntFlanks | 9554 | 9575 | 22 | CGTAGCGGTCCTGACGTGCAAA | 1258.2 | -4.9 | 0.7 | -7.0 | 2.6E-12 | 6.9E-12 | MB231/MB468 | MITO |
| >hg38_21_+_8433172_8446622_RNA45SN1_45S_50ntFlanks@11766.11803.38\|ATCTAATTAGTGACGCGCATGAATGGATGAACGAGATT | >hg38_21_+_8433172_8446622_RNA45SN1_45S_50ntFlanks | 11766 | 11803 | 38 | ATCTAATTAGTGACGCGCATGAATGGATGAACGAGATT | 244.8 | -10.2 | 1.5 | -7.0 | 2.0E-12 | 6.9E-12 | BT20/MB468 | CYTO |
| >am_trna10_ValCAC_5_-_180649395_180649467@1.31.31,trna12_ValAAC_5_-_180645270_180645342@1.31.31,trna132_ValAAC_6_-_27721179_27721251@1.31.31,trna136_ValAAC_6_-_27648885_27648957@1.31.31,trna139_ValAAC_6_-_27618707_27618779@1.31.31,trna18_ValCAC_5_-_180529253_180529325@1.31.31,trna2_ValAAC_3_+_169490018_169490090@1.31.31,trna2_ValCAC_5_+_180524070_180524142@1.31.31,trna4_ValAAC_5_+_180591154_180591226@1.31.31,trna5_ValAAC_5_+_180596610_180596682@1.31.31,trna6_ValCAC_5_+_180600650_180600722@1.31.31,trna85_ValCAC_1_-_161369490_161369562@1.31.31,trna90_ValCAC_1_-_149684088_149684161@1.31.31,trna98_ValCAC_1_-_149298555_149298627@1.31.31,trna9_ValCAC_6_+_26538282_26538354@1.31.31\|GTTTCCGTAGTGTAGTGGTTATCACGTTCGC | >am_trna10_ValCAC_5_-_180649395_180649467 | 1 | 31 | 31 | GTTTCCGTAGTGTAGTGGTTATCACGTTCGC | 872.1 | -9.9 | 1.4 | -7.0 | 2.8E-12 | 6.9E-12 | BT20/MB468 | NUC |
| >MI0000077\|hsa-mir-21&WithFlank&17\|+\|59841260\|59841343@14.36.23(+1C)&[MIMAT0000076&hsa-miR-21-5p&offsets\|0\|+1(+1C);m-1&17\|+\|59841273\|59841294&offsets\|0\|+1(+1C)]\|TAGCTTATCAGACTGATGTTGACC | >MI0000077\|hsa-mir-21&WithFlank&17\|+\|59841260\|59841343 | 14 | 36 | 23(+1C) | TAGCTTATCAGACTGATGTTGACC | 7877.2 | 3.7 | 0.5 | 7.0 | 2.8E-12 | 6.9E-12 | BT20/MB468 | NUC |
| >hg38_21_+_8433172_8446622_RNA45SN1_45S_50ntFlanks@483.524.42\|TCGATGTGGTGACGTCGTGCTCTCCCGGGCCGGGTCCGAGCC | >hg38_21_+_8433172_8446622_RNA45SN1_45S_50ntFlanks | 483 | 524 | 42 | TCGATGTGGTGACGTCGTGCTCTCCCGGGCCGGGTCCGAGCC | 1347.1 | 5.9 | 0.8 | 7.0 | 2.2E-12 | 6.9E-12 | MB231/MB468 | NUC |
| >hg38_MT_+_598_1651_RNR1_12S_50ntFlanks@428.468.41\|AAAGTGGCTTTAACATATCTGAACACACAATAGCTAAGACC | >hg38_MT_+_598_1651_RNR1_12S_50ntFlanks | 428 | 468 | 41 | AAAGTGGCTTTAACATATCTGAACACACAATAGCTAAGACC | 1266.7 | 7.9 | 1.1 | 7.1 | 1.6E-12 | 6.9E-12 | BT20/MB468 | MITO |
| >hg38_21_+_8433172_8446622_RNA45SN1_45S_50ntFlanks@11233.11251.19\|CCCTTCCCGTGGATCGCCC | >hg38_21_+_8433172_8446622_RNA45SN1_45S_50ntFlanks | 11233 | 11251 | 19 | CCCTTCCCGTGGATCGCCC | 234.2 | -10.6 | 1.5 | -7.0 | 2.1E-12 | 6.9E-12 | MB231/MB468 | CYTO |
| >trna10_LysCTT_16_+_3241501_3241573@4.33.30,trna13_LysCTT_14_-_58706613_58706685@4.33.30,trna2_LysCTT_15_+_79152904_79152976@4.33.30\|CGGCTAGCTCAGTCGGTAGAGCATGGGACT | >trna10_LysCTT_16_+_3241501_3241573 | 4 | 33 | 30 | CGGCTAGCTCAGTCGGTAGAGCATGGGACT | 2061.1 | 6.6 | 0.9 | 7.0 | 2.6E-12 | 6.9E-12 | MB231/MB468 | MITO |
| >hg38_21_+_8433172_8446622_RNA45SN1_45S_50ntFlanks@7974.7997.24\|ACGCGACCTCAGATCAGACGTGGC | >hg38_21_+_8433172_8446622_RNA45SN1_45S_50ntFlanks | 7974 | 7997 | 24 | ACGCGACCTCAGATCAGACGTGGC | 1573.0 | 3.7 | 0.5 | 7.0 | 2.2E-12 | 7.0E-12 | MB231/MB468 | NUC |
| >hg38_21_+_8433172_8446622_RNA45SN1_45S_50ntFlanks@11208.11254.47\|GCGGCGACTCTGGACGCGAGCCGGGCCCTTCCCGTGGATCGCCCCAG | >hg38_21_+_8433172_8446622_RNA45SN1_45S_50ntFlanks | 11208 | 11254 | 47 | GCGGCGACTCTGGACGCGAGCCGGGCCCTTCCCGTGGATCGCCCCAG | 31517.4 | -9.9 | 1.4 | -7.0 | 2.2E-12 | 7.0E-12 | MB231/MB468 | NUC |
| >MI0000088\|hsa-mir-30a&WithFlank&6\|-\|71403545\|71403627@53.75.23&[MIMAT0000088&hsa-miR-30a-3p&offsets\|0\|+1;m-38&6\|-\|71403554\|71403575&offsets\|0\|+1]\|CTTTCAGTCGGATGTTTGCAGCT | >MI0000088\|hsa-mir-30a&WithFlank&6\|-\|71403545\|71403627 | 53 | 75 | 23 | CTTTCAGTCGGATGTTTGCAGCT | 531.1 | 8.6 | 1.2 | 7.0 | 2.1E-12 | 7.0E-12 | MB231/MB468 | CYTO |
| >hg38_21_+_8433172_8446622_RNA45SN1_45S_50ntFlanks@11188.11208.21\|CACCCGGGGGGCCGGCGGCGG | >hg38_21_+_8433172_8446622_RNA45SN1_45S_50ntFlanks | 11188 | 11208 | 21 | CACCCGGGGGGCCGGCGGCGG | 185.7 | -11.4 | 1.6 | -7.0 | 1.9E-12 | 7.0E-12 | MB231/MB468 | MP |
| >hg38_21_+_8433172_8446622_RNA45SN1_45S_50ntFlanks@3920.3938.19\|CATTTATCAGATCAAAACC | >hg38_21_+_8433172_8446622_RNA45SN1_45S_50ntFlanks | 3920 | 3938 | 19 | CATTTATCAGATCAAAACC | 122.0 | -10.8 | 1.5 | -7.0 | 1.9E-12 | 7.0E-12 | MB231/MB468 | MP |
| >hg38_21_+_8433172_8446622_RNA45SN1_45S_50ntFlanks@6653.6681.29\|ACTCTTAGCGGTGGATCACTCGGCTCGTG | >hg38_21_+_8433172_8446622_RNA45SN1_45S_50ntFlanks | 6653 | 6681 | 29 | ACTCTTAGCGGTGGATCACTCGGCTCGTG | 3505.8 | -6.9 | 1.0 | -7.0 | 1.9E-12 | 7.0E-12 | MB231/MB468 | MP |
| >hg38_21_+_8433172_8446622_RNA45SN1_45S_50ntFlanks@11787.11804.18\|AATGGATGAACGAGATTC | >hg38_21_+_8433172_8446622_RNA45SN1_45S_50ntFlanks | 11787 | 11804 | 18 | AATGGATGAACGAGATTC | 9708.2 | -6.1 | 0.8 | -7.4 | 1.0E-13 | 7.0E-12 | BT20/MB468 | MP |
| >hg38_21_+_8433172_8446622_RNA45SN1_45S_50ntFlanks@7975.8010.36\|CGCGACCTCAGATCAGACGTGGCGACCCGCTGAATT | >hg38_21_+_8433172_8446622_RNA45SN1_45S_50ntFlanks | 7975 | 8010 | 36 | CGCGACCTCAGATCAGACGTGGCGACCCGCTGAATT | 67446.7 | 4.3 | 0.6 | 7.0 | 2.2E-12 | 7.1E-12 | MB231/MB468 | NUC |
| >hg38_21_+_8433172_8446622_RNA45SN1_45S_50ntFlanks@10772.10805.34\|AGGTCTCCAAGGTGAACAGCCTCTGGCATGTTGG | >hg38_21_+_8433172_8446622_RNA45SN1_45S_50ntFlanks | 10772 | 10805 | 34 | AGGTCTCCAAGGTGAACAGCCTCTGGCATGTTGG | 103.1 | -10.6 | 1.5 | -7.0 | 1.9E-12 | 7.1E-12 | MB231/MB468 | MP |
| >hg38_21_+_8433172_8446622_RNA45SN1_45S_50ntFlanks@11209.11234.26\|CGGCGACTCTGGACGCGAGCCGGGCC | >hg38_21_+_8433172_8446622_RNA45SN1_45S_50ntFlanks | 11209 | 11234 | 26 | CGGCGACTCTGGACGCGAGCCGGGCC | 4490.6 | -9.3 | 1.3 | -7.0 | 2.2E-12 | 7.1E-12 | MB231/MB468 | CYTO |
| >hg38_21_+_8433172_8446622_RNA45SN1_45S_50ntFlanks@8528.8551.24\|CTTCCCCCGCCGCCCCTCCTCCTC | >hg38_21_+_8433172_8446622_RNA45SN1_45S_50ntFlanks | 8528 | 8551 | 24 | CTTCCCCCGCCGCCCCTCCTCCTC | 199.8 | -10.6 | 1.5 | -7.0 | 2.2E-12 | 7.1E-12 | MB231/MB468 | NUC |
| >hg38_21_+_8433172_8446622_RNA45SN1_45S_50ntFlanks@9806.9829.24\|CGTGGAATGCGAGTGCCTAGTGGG | >hg38_21_+_8433172_8446622_RNA45SN1_45S_50ntFlanks | 9806 | 9829 | 24 | CGTGGAATGCGAGTGCCTAGTGGG | 182.9 | -11.4 | 1.6 | -7.0 | 1.9E-12 | 7.1E-12 | MB231/MB468 | MP |
| >hg38_21_+_8433172_8446622_RNA45SN1_45S_50ntFlanks@9533.9571.39\|GAGGAAACTCTGGTGGAGGTCCGTAGCGGTCCTGACGTG | >hg38_21_+_8433172_8446622_RNA45SN1_45S_50ntFlanks | 9533 | 9571 | 39 | GAGGAAACTCTGGTGGAGGTCCGTAGCGGTCCTGACGTG | 11092.3 | -5.0 | 0.7 | -7.0 | 2.2E-12 | 7.1E-12 | MB231/MB468 | NUC |
| >hg38_21_+_8433172_8446622_RNA45SN1_45S_50ntFlanks@12450.12480.31\|AATTCACCAAGCGTTGGATTGTTCACCCACT | >hg38_21_+_8433172_8446622_RNA45SN1_45S_50ntFlanks | 12450 | 12480 | 31 | AATTCACCAAGCGTTGGATTGTTCACCCACT | 965.5 | 6.4 | 0.9 | 7.0 | 2.1E-12 | 7.1E-12 | BT20/MB468 | CYTO |
| >hg38_21_+_8433172_8446622_RNA45SN1_45S_50ntFlanks@7974.7997.24\|ACGCGACCTCAGATCAGACGTGGC | >hg38_21_+_8433172_8446622_RNA45SN1_45S_50ntFlanks | 7974 | 7997 | 24 | ACGCGACCTCAGATCAGACGTGGC | 1660.9 | 5.2 | 0.7 | 7.0 | 2.1E-12 | 7.1E-12 | BT20/MB468 | CYTO |
| >hg38_21_+_8433172_8446622_RNA45SN1_45S_50ntFlanks@12357.12375.19\|CCACAGGGATAACTGGCTT | >hg38_21_+_8433172_8446622_RNA45SN1_45S_50ntFlanks | 12357 | 12375 | 19 | CCACAGGGATAACTGGCTT | 327.8 | -7.1 | 1.0 | -7.0 | 2.2E-12 | 7.2E-12 | MB231/MB468 | CYTO |
| >hg38_21_+_8433172_8446622_RNA45SN1_45S_50ntFlanks@10781.10806.26\|AGGTGAACAGCCTCTGGCATGTTGGA | >hg38_21_+_8433172_8446622_RNA45SN1_45S_50ntFlanks | 10781 | 10806 | 26 | AGGTGAACAGCCTCTGGCATGTTGGA | 223.4 | -8.1 | 1.2 | -7.0 | 2.2E-12 | 7.2E-12 | MB231/MB468 | NUC |
| >hg38_21_+_8433172_8446622_RNA45SN1_45S_50ntFlanks@12259.12281.23\|CTTGATCTTGATTTTCAGTACGA | >hg38_21_+_8433172_8446622_RNA45SN1_45S_50ntFlanks | 12259 | 12281 | 23 | CTTGATCTTGATTTTCAGTACGA | 147.3 | -11.1 | 1.6 | -7.0 | 1.9E-12 | 7.2E-12 | MB231/MB468 | MP |
| >hg38_21_+_8433172_8446622_RNA45SN1_45S_50ntFlanks@10388.10409.22\|GTGAACAGCAGTTGAACATGGG | >hg38_21_+_8433172_8446622_RNA45SN1_45S_50ntFlanks | 10388 | 10409 | 22 | GTGAACAGCAGTTGAACATGGG | 168.7 | -11.3 | 1.6 | -7.0 | 1.9E-12 | 7.2E-12 | MB231/MB468 | MP |
| >hg38_21_+_8433172_8446622_RNA45SN1_45S_50ntFlanks@4730.4761.32\|CAAGAACGAAAGTCGGAGGTTCGAAGACGATC | >hg38_21_+_8433172_8446622_RNA45SN1_45S_50ntFlanks | 4730 | 4761 | 32 | CAAGAACGAAAGTCGGAGGTTCGAAGACGATC | 561.9 | 4.6 | 0.7 | 7.0 | 2.2E-12 | 7.2E-12 | MB231/MB468 | CYTO |
| >hg38_21_+_8433172_8446622_RNA45SN1_45S_50ntFlanks@9696.9722.27\|CAGTTTTATCCGGTAAAGCGAATGATT | >hg38_21_+_8433172_8446622_RNA45SN1_45S_50ntFlanks | 9696 | 9722 | 27 | CAGTTTTATCCGGTAAAGCGAATGATT | 347.3 | -12.2 | 1.7 | -7.0 | 2.7E-12 | 7.2E-12 | MB231/MB468 | MITO |
| >hg38_21_+_8433172_8446622_RNA45SN1_45S_50ntFlanks@10806.10828.23\|AACAATGTAGGTAAGGGAAGTCG | >hg38_21_+_8433172_8446622_RNA45SN1_45S_50ntFlanks | 10806 | 10828 | 23 | AACAATGTAGGTAAGGGAAGTCG | 820.3 | -8.6 | 1.2 | -7.0 | 2.7E-12 | 7.2E-12 | MB231/MB468 | MITO |
| >hg38_21_+_8433172_8446622_RNA45SN1_45S_50ntFlanks@7985.8007.23\|GATCAGACGTGGCGACCCGCTGA | >hg38_21_+_8433172_8446622_RNA45SN1_45S_50ntFlanks | 7985 | 8007 | 23 | GATCAGACGTGGCGACCCGCTGA | 410.0 | -11.8 | 1.6 | -7.4 | 1.1E-13 | 7.2E-12 | BT20/MB468 | MP |
| >hg38_21_+_8433172_8446622_RNA45SN1_45S_50ntFlanks@11205.11223.19\|GCGGCGGCGACTCTGGACG | >hg38_21_+_8433172_8446622_RNA45SN1_45S_50ntFlanks | 11205 | 11223 | 19 | GCGGCGGCGACTCTGGACG | 137.3 | -11.0 | 1.6 | -7.0 | 1.9E-12 | 7.2E-12 | MB231/MB468 | MP |
| >hg38_21_+_8433172_8446622_RNA45SN1_45S_50ntFlanks@8105.8140.36\|CGCGGCGGGGCGCGGGACATGTGGCGTACGGAAGAC | >hg38_21_+_8433172_8446622_RNA45SN1_45S_50ntFlanks | 8105 | 8140 | 36 | CGCGGCGGGGCGCGGGACATGTGGCGTACGGAAGAC | 15712.3 | -9.8 | 1.4 | -7.0 | 1.9E-12 | 7.2E-12 | MB231/MB468 | MP |

| | | | | | | | | | | | | | |
|---|---|---|---|---|---|---|---|---|---|---|---|---|---|
| >trnaMT_GluTTC_MT_-_14674_14742@41.71.31,trnalookalike8_GluTTC_5_-_93905172_93905240@41.71.31\|TGGTCGTGGTTGTAGTCCGTGCGAGAATACC | >trnaMT_GluTTC_MT_-_14674_14742 | 41 | 71 | 31 | TGGTCGTGGTTGTAGTCCGTGCGAGAATACC | 835.2 | 8.8 | 1.2 | 7.0 | 2.2E-12 | 7.2E-12 | MB231/MB468 | CYTO |
| >hg38_21_+_8433172_8446622_RNA45SN1_45S_50ntFlanks@3900.3919.20\|CTTCGCGGGGGGGATGCGTG | >hg38_21_+_8433172_8446622_RNA45SN1_45S_50ntFlanks | 3900 | 3919 | 20 | CTTCGCGGGGGGGATGCGTG | 529.6 | -8.8 | 1.3 | -7.0 | 2.3E-12 | 7.2E-12 | MB231/MB468 | NUC |
| >hg38_21_+_8433172_8446622_RNA45SN1_45S_50ntFlanks@11847.11869.23\|AACGGGCTTGGCGGAATCAGCGG | >hg38_21_+_8433172_8446622_RNA45SN1_45S_50ntFlanks | 11847 | 11869 | 23 | AACGGGCTTGGCGGAATCAGCGG | 121.7 | -10.8 | 1.5 | -7.0 | 1.9E-12 | 7.2E-12 | MB231/MB468 | MP |
| >hg38_21_+_8433172_8446622_RNA45SN1_45S_50ntFlanks@8079.8103.25\|AAGAGCCCAGCGCCGAATCCCCGCC | >hg38_21_+_8433172_8446622_RNA45SN1_45S_50ntFlanks | 8079 | 8103 | 25 | AAGAGCCCAGCGCCGAATCCCCGCC | 2457.8 | -10.2 | 1.5 | -7.0 | 1.9E-12 | 7.2E-12 | MB231/MB468 | MP |
| >hg38_21_+_8433172_8446622_RNA45SN1_45S_50ntFlanks@11315.11334.20\|CCACCCCACGTCTCGTCGCG | >hg38_21_+_8433172_8446622_RNA45SN1_45S_50ntFlanks | 11315 | 11334 | 20 | CCACCCCACGTCTCGTCGCG | 535.6 | -8.8 | 1.3 | -7.0 | 2.7E-12 | 7.2E-12 | MB231/MB468 | MITO |
| >hg38_21_+_8433172_8446622_RNA45SN1_45S_50ntFlanks@10407.10435.29\|GGGTCAGTCGGTCCTGAGAGATGGGCGAG | >hg38_21_+_8433172_8446622_RNA45SN1_45S_50ntFlanks | 10407 | 10435 | 29 | GGGTCAGTCGGTCCTGAGAGATGGGCGAG | 120.5 | -10.8 | 1.5 | -7.0 | 1.9E-12 | 7.2E-12 | MB231/MB468 | MP |
| >hg38_21_+_8433172_8446622_RNA45SN1_45S_50ntFlanks@12242.12269.28\|AGAAGGGCAAAAGCTCGCTTGATCTTGA | >hg38_21_+_8433172_8446622_RNA45SN1_45S_50ntFlanks | 12242 | 12269 | 28 | AGAAGGGCAAAAGCTCGCTTGATCTTGA | 124.3 | -10.8 | 1.5 | -7.0 | 1.9E-12 | 7.2E-12 | MB231/MB468 | MP |
| >hg38_21_+_8433172_8446622_RNA45SN1_45S_50ntFlanks@7981.8005.25\|CTCAGATCAGACGTGGCGACCCGCT | >hg38_21_+_8433172_8446622_RNA45SN1_45S_50ntFlanks | 7981 | 8005 | 25 | CTCAGATCAGACGTGGCGACCCGCT | 753.7 | -6.8 | 1.0 | -7.0 | 2.7E-12 | 7.2E-12 | MB231/MB468 | MITO |
| >hg38_21_+_8433172_8446622_RNA45SN1_45S_50ntFlanks@9250.9273.24\|CGCACGGGGTCGGCGGCGACGTCG | >hg38_21_+_8433172_8446622_RNA45SN1_45S_50ntFlanks | 9250 | 9273 | 24 | CGCACGGGGTCGGCGGCGACGTCG | 148.1 | -11.1 | 1.6 | -7.0 | 2.0E-12 | 7.2E-12 | MB231/MB468 | MP |
| >hg38_21_+_8433172_8446622_RNA45SN1_45S_50ntFlanks@10916.10936.21\|CCGCGGCTGGACGAGGCGCCG | >hg38_21_+_8433172_8446622_RNA45SN1_45S_50ntFlanks | 10916 | 10936 | 21 | CCGCGGCTGGACGAGGCGCCG | 244.0 | -10.9 | 1.6 | -7.0 | 2.3E-12 | 7.3E-12 | MB231/MB468 | NUC |
| >MI0005559\|hsa-mir-744&WithFlank&17\|+\|12081893\|12082002@17.36.20(+1U)&[MIMAT0004945&hsa-miR-744-5p&offsets\|0\|-2(+1U);m-172&17\|+\|12081909\|12081929&offsets\|0\|-1(+1U)]\|TGCGGGGCTAGGGCTAACAGT | >MI0005559\|hsa-mir-744&WithFlank&17\|+\|12081893\|12082002 | 17 | 36 | 20(+1U) | TGCGGGGCTAGGGCTAACAGT | 100.7 | 3.5 | 0.5 | 7.0 | 3.0E-12 | 7.3E-12 | BT20/MB468 | NUC |
| >hg38_21_+_8433172_8446622_RNA45SN1_45S_50ntFlanks@3988.4023.36\|CTTTGGTGACTCTAGATAACCTCGGGCCGATCGCAC | >hg38_21_+_8433172_8446622_RNA45SN1_45S_50ntFlanks | 3988 | 4023 | 36 | CTTTGGTGACTCTAGATAACCTCGGGCCGATCGCAC | 1366.1 | 5.4 | 0.8 | 7.0 | 2.2E-12 | 7.3E-12 | MB231/MB468 | CYTO |
| >hg38_21_+_8433172_8446622_RNA45SN1_45S_50ntFlanks@10686.10710.25\|GCGGCGTCCGGTGAGCTCTCGCTGG | >hg38_21_+_8433172_8446622_RNA45SN1_45S_50ntFlanks | 10686 | 10710 | 25 | GCGGCGTCCGGTGAGCTCTCGCTGG | 4954.1 | -5.4 | 0.8 | -7.0 | 2.8E-12 | 7.3E-12 | MB231/MB468 | MITO |
| >hg38_21_+_8433172_8446622_RNA45SN1_45S_50ntFlanks@12539.12563.25\|ATGATGTGTTGTTGCCATGGTAATC | >hg38_21_+_8433172_8446622_RNA45SN1_45S_50ntFlanks | 12539 | 12563 | 25 | ATGATGTGTTGTTGCCATGGTAATC | 710.3 | -8.6 | 1.2 | -7.0 | 2.8E-12 | 7.3E-12 | MB231/MB468 | MITO |
| >hg38_21_+_8433172_8446622_RNA45SN1_45S_50ntFlanks@10392.10431.40\|ACAGCAGTTGAACATGGGTCAGTCGGTCCTGAGAGATGGG | >hg38_21_+_8433172_8446622_RNA45SN1_45S_50ntFlanks | 10392 | 10431 | 40 | ACAGCAGTTGAACATGGGTCAGTCGGTCCTGAGAGATGGG | 248.8 | -11.8 | 1.7 | -7.0 | 2.0E-12 | 7.3E-12 | MB231/MB468 | MP |
| >hg38_21_+_8433172_8446622_RNA45SN1_45S_50ntFlanks@4047.4067.21\|ATTCGAACGTCTGCCCTATCA | >hg38_21_+_8433172_8446622_RNA45SN1_45S_50ntFlanks | 4047 | 4067 | 21 | ATTCGAACGTCTGCCCTATCA | 745.2 | -5.5 | 0.8 | -7.0 | 2.3E-12 | 7.3E-12 | MB231/MB468 | NUC |
| >MI0000289\|hsa-mir-181a-1&WithFlank&1\|-\|198859038\|198859159@30.53.24;MI0000269\|hsa-mir-181a-2&WithFlank&9\|+\|124692436\|124692557@45.68.24&[MIMAT0000256&hsa-miR-181a-5p&offsets\|0\|+1;m-32&1\|-\|198859108\|198859130&offsets\|0\|+1];[MIMAT0000256_1&hsa-miR-181a-5p&offsets\|0\|+1;m-32&9\|+\|124692480\|124692502&offsets\|0\|+1]\|AACATTCAACGCTGTCGGTGAGTT | >MI0000289\|hsa-mir-181a-1&WithFlank&1\|-\|198859038\|198859159 | 30 | 53 | 24 | AACATTCAACGCTGTCGGTGAGTT | 3297.9 | 8.0 | 1.1 | 7.0 | 2.8E-12 | 7.3E-12 | MB231/MB468 | MITO |
| >hg38_21_+_8433172_8446622_RNA45SN1_45S_50ntFlanks@12841.12871.31\|CGGAGTGCCCTTCGTCCTGGGAAACGGGGCG | >hg38_21_+_8433172_8446622_RNA45SN1_45S_50ntFlanks | 12841 | 12871 | 31 | CGGAGTGCCCTTCGTCCTGGGAAACGGGGCG | 148.5 | -11.1 | 1.6 | -7.0 | 2.0E-12 | 7.3E-12 | MB231/MB468 | MP |
| >hg38_21_+_8433172_8446622_RNA45SN1_45S_50ntFlanks@12332.12356.25\|TAAGCAGGAGGTGTCAGAAAAGTTA | >hg38_21_+_8433172_8446622_RNA45SN1_45S_50ntFlanks | 12332 | 12356 | 25 | TAAGCAGGAGGTGTCAGAAAAGTTA | 93.9 | -10.4 | 1.5 | -7.0 | 2.0E-12 | 7.3E-12 | MB231/MB468 | MP |
| >hg38_21_+_8433172_8446622_RNA45SN1_45S_50ntFlanks@4591.4614.24\|TTTTGTTGGTTTTCGGAACTGAGG | >hg38_21_+_8433172_8446622_RNA45SN1_45S_50ntFlanks | 4591 | 4614 | 24 | TTTTGTTGGTTTTCGGAACTGAGG | 117.9 | -10.7 | 1.5 | -7.0 | 2.0E-12 | 7.3E-12 | MB231/MB468 | MP |
| >hg38_21_+_8433172_8446622_RNA45SN1_45S_50ntFlanks@7975.8004.30\|CGCGACCTCAGATCAGACGTGGCGACCCGC | >hg38_21_+_8433172_8446622_RNA45SN1_45S_50ntFlanks | 7975 | 8004 | 30 | CGCGACCTCAGATCAGACGTGGCGACCCGC | 80500.5 | 5.9 | 0.8 | 7.1 | 1.7E-12 | 7.3E-12 | BT20/MB468 | MITO |
| >hg38_21_+_8433172_8446622_RNA45SN1_45S_50ntFlanks@9536.9571.36\|GAAACTCTGGTGGAGGTCCGTAGCGGTCCTGACGTG | >hg38_21_+_8433172_8446622_RNA45SN1_45S_50ntFlanks | 9536 | 9571 | 36 | GAAACTCTGGTGGAGGTCCGTAGCGGTCCTGACGTG | 1310.6 | -3.7 | 0.5 | -7.0 | 2.3E-12 | 7.3E-12 | MB231/MB468 | NUC |
| >hg38_21_+_8433172_8446622_RNA45SN1_45S_50ntFlanks@9651.9679.29\|TCAGGATAGCTGGCGCTCTCGCAGACCCG | >hg38_21_+_8433172_8446622_RNA45SN1_45S_50ntFlanks | 9651 | 9679 | 29 | TCAGGATAGCTGGCGCTCTCGCAGACCCG | 88.3 | -10.3 | 1.5 | -7.0 | 2.0E-12 | 7.4E-12 | MB231/MB468 | MP |
| >hg38_21_+_8433172_8446622_RNA45SN1_45S_50ntFlanks@4730.4765.36\|CAAGAACGAAAGTCGGAGGTTCGAAGACGATCAGAT | >hg38_21_+_8433172_8446622_RNA45SN1_45S_50ntFlanks | 4730 | 4765 | 36 | CAAGAACGAAAGTCGGAGGTTCGAAGACGATCAGAT | 660.1 | 6.6 | 0.9 | 7.0 | 2.3E-12 | 7.4E-12 | MB231/MB468 | CYTO |
| >MI0000100\|hsa-mir-98&WithFlank&X\|-\|53556217\|53556347@28.48.21&[MIMAT0000096&hsa-miR-98-5p&offsets\|0\|-1;m-143&X\|-\|53556299\|53556320&offsets\|0\|-1]\|TGAGGTAGTAAGTTGTATTGT | >MI0000100\|hsa-mir-98&WithFlank&X\|-\|53556217\|53556347 | 28 | 48 | 21 | TGAGGTAGTAAGTTGTATTGT | 335.5 | 3.6 | 0.5 | 7.0 | 2.3E-12 | 7.4E-12 | MB231/MB468 | NUC |
| >hg38_21_+_8433172_8446622_RNA45SN1_45S_50ntFlanks@12937.12962.26\|AACCATTCGTAGACGACCTGCTTCTG | >hg38_21_+_8433172_8446622_RNA45SN1_45S_50ntFlanks | 12937 | 12962 | 26 | AACCATTCGTAGACGACCTGCTTCTG | 97.0 | -10.5 | 1.5 | -7.0 | 2.0E-12 | 7.4E-12 | MB231/MB468 | MP |
| >hg38_21_+_8433172_8446622_RNA45SN1_45S_50ntFlanks@8229.8253.25\|CGCGCCGGGCCCGGGTCTTCCCGGA | >hg38_21_+_8433172_8446622_RNA45SN1_45S_50ntFlanks | 8229 | 8253 | 25 | CGCGCCGGGCCCGGGTCTTCCCGGA | 331.2 | 9.7 | 1.4 | 7.0 | 2.3E-12 | 7.4E-12 | MB231/MB468 | CYTO |
| >MI0000457\|hsa-mir-141&WithFlank&12\|+\|6964091\|6964197@65.85.21&[MIMAT0000432&hsa-miR-141-3p&offsets\|0\|-1;m-134&12\|+\|6964155\|6964175&offsets\|0\|0]\|TAACACTGTCTGGTAAAGATG | >MI0000457\|hsa-mir-141&WithFlank&12\|+\|6964091\|6964197 | 65 | 85 | 21 | TAACACTGTCTGGTAAAGATG | 141.6 | -11.0 | 1.6 | -7.0 | 2.0E-12 | 7.4E-12 | MB231/MB468 | MP |
| >hg38_21_+_8433172_8446622_RNA45SN1_45S_50ntFlanks@12073.12096.24\|CCGAGGGGCTCTCGCTTCTGGCGC | >hg38_21_+_8433172_8446622_RNA45SN1_45S_50ntFlanks | 12073 | 12096 | 24 | CCGAGGGGCTCTCGCTTCTGGCGC | 587.1 | -7.1 | 1.0 | -7.0 | 2.3E-12 | 7.4E-12 | MB231/MB468 | NUC |
| >hg38_21_+_8433172_8446622_RNA45SN1_45S_50ntFlanks@10905.10933.29\|CTGGGCGCGCGCCGCGGCTGGACGAGGCG | >hg38_21_+_8433172_8446622_RNA45SN1_45S_50ntFlanks | 10905 | 10933 | 29 | CTGGGCGCGCGCCGCGGCTGGACGAGGCG | 287.0 | -7.4 | 1.1 | -7.0 | 2.3E-12 | 7.4E-12 | MB231/MB468 | CYTO |
| >hg38_21_+_8433172_8446622_RNA45SN1_45S_50ntFlanks@11729.11750.22\|TAACTATGACTCTCTTAAGGTA | >hg38_21_+_8433172_8446622_RNA45SN1_45S_50ntFlanks | 11729 | 11750 | 22 | TAACTATGACTCTCTTAAGGTA | 523.9 | -8.5 | 1.2 | -7.0 | 2.0E-12 | 7.4E-12 | MB231/MB468 | MP |
| >hg38_21_+_8433172_8446622_RNA45SN1_45S_50ntFlanks@12615.12632.18\|GCTGAGGAGCCAATGGGG | >hg38_21_+_8433172_8446622_RNA45SN1_45S_50ntFlanks | 12615 | 12632 | 18 | GCTGAGGAGCCAATGGGG | 106.7 | -10.6 | 1.5 | -7.0 | 2.0E-12 | 7.5E-12 | MB231/MB468 | MP |
| >MI0001445\|hsa-mir-423&WithFlank&17\|+\|30117073\|30117178@59.80.22(+1G)&[MIMAT0001340&hsa-miR-423-3p&offsets\|0\|-1(+1G);m-100&17\|+\|30117131\|30117153&offsets\|0\|-1(+1G)]\|AGCTCGGTCTGAGGCCCCTCAGG | >MI0001445\|hsa-mir-423&WithFlank&17\|+\|30117073\|30117178 | 59 | 80 | 22(+1G) | AGCTCGGTCTGAGGCCCCTCAGG | 223.0 | 5.5 | 0.8 | 7.0 | 2.3E-12 | 7.5E-12 | MB231/MB468 | NUC |
| >hg38_21_+_8433172_8446622_RNA45SN1_45S_50ntFlanks@4985.5002.18\|GACAGGATTGACAGATTG | >hg38_21_+_8433172_8446622_RNA45SN1_45S_50ntFlanks | 4985 | 5002 | 18 | GACAGGATTGACAGATTG | 192.4 | -11.5 | 1.6 | -7.0 | 2.0E-12 | 7.5E-12 | MB231/MB468 | MP |
| >hg38_21_+_8433172_8446622_RNA45SN1_45S_50ntFlanks@10389.10407.19\|TGAACAGCAGTTGAACATG | >hg38_21_+_8433172_8446622_RNA45SN1_45S_50ntFlanks | 10389 | 10407 | 19 | TGAACAGCAGTTGAACATG | 790.9 | -8.8 | 1.3 | -7.0 | 2.8E-12 | 7.5E-12 | MB231/MB468 | MITO |
| >hg38_21_+_8433172_8446622_RNA45SN1_45S_50ntFlanks@9759.9788.30\|CAAACTTTAAATGGGTAAGAAGCCCGGCTC | >hg38_21_+_8433172_8446622_RNA45SN1_45S_50ntFlanks | 9759 | 9788 | 30 | CAAACTTTAAATGGGTAAGAAGCCCGGCTC | 169.6 | -11.3 | 1.6 | -7.0 | 2.0E-12 | 7.5E-12 | MB231/MB468 | MP |
| >hg38_21_+_8433172_8446622_RNA45SN1_45S_50ntFlanks@12936.12969.34\|AAACCATTCGTAGACGACCTGCTTCTGGGTCGGG | >hg38_21_+_8433172_8446622_RNA45SN1_45S_50ntFlanks | 12936 | 12969 | 34 | AAACCATTCGTAGACGACCTGCTTCTGGGTCGGG | 212.6 | -6.5 | 0.9 | -7.0 | 2.3E-12 | 7.5E-12 | MB231/MB468 | CYTO |
| >hg38_21_+_8433172_8446622_RNA45SN1_45S_50ntFlanks@11208.11238.31\|GCGGCGACTCTGGACGCGAGCCGGGCCCTTC | >hg38_21_+_8433172_8446622_RNA45SN1_45S_50ntFlanks | 11208 | 11238 | 31 | GCGGCGACTCTGGACGCGAGCCGGGCCCTTC | 137.4 | -11.0 | 1.6 | -7.0 | 2.0E-12 | 7.5E-12 | MB231/MB468 | MP |
| >hg38_21_+_8433172_8446622_RNA45SN1_45S_50ntFlanks@10869.10890.22\|AGGGCTGGGTCGGTCGGGCTGG | >hg38_21_+_8433172_8446622_RNA45SN1_45S_50ntFlanks | 10869 | 10890 | 22 | AGGGCTGGGTCGGTCGGGCTGG | 1118.6 | -7.6 | 1.1 | -7.0 | 2.0E-12 | 7.5E-12 | MB231/MB468 | MP |
| >trnaMT_PheGAA_MT_+_577_647@35.62.28\|GAAAATGTTTAGACGGGCTCACATCACC | >trnaMT_PheGAA_MT_+_577_647 | 35 | 62 | 28 | GAAAATGTTTAGACGGGCTCACATCACC | 232.4 | 7.9 | 1.1 | 7.0 | 2.4E-12 | 7.5E-12 | MB231/MB468 | NUC |
| >hg38_21_+_8433172_8446622_RNA45SN1_45S_50ntFlanks@5078.5105.28\|CGATAACGAACGAGACTCTGGCATGCTA | >hg38_21_+_8433172_8446622_RNA45SN1_45S_50ntFlanks | 5078 | 5105 | 28 | CGATAACGAACGAGACTCTGGCATGCTA | 370.9 | -12.4 | 1.8 | -7.0 | 2.0E-12 | 7.5E-12 | MB231/MB468 | MP |
| >hg38_21_+_8433172_8446622_RNA45SN1_45S_50ntFlanks@12126.12147.22\|CGACCCGCTCCGGGGACAGTGC | >hg38_21_+_8433172_8446622_RNA45SN1_45S_50ntFlanks | 12126 | 12147 | 22 | CGACCCGCTCCGGGGACAGTGC | 128.2 | -10.9 | 1.5 | -7.0 | 2.0E-12 | 7.5E-12 | MB231/MB468 | MP |
| >am_trna134_GluTTC_1_-_16861774_16861845@1.28.28,trna5_GluTTC_1_+_17199078_17199149@1.28.28,trna84_GluTTC_1_-_161391883_161391954@1.28.28,trna94_GluTTC_1_-_149664355_149664427@1.28.28\|TCCCTGGTGGTCTAGTGGCTAGGATTCG | >am_trna134_GluTTC_1_-_16861774_16861845 | 1 | 28 | 28 | TCCCTGGTGGTCTAGTGGCTAGGATTCG | 843.5 | -9.1 | 1.3 | -7.0 | 2.4E-12 | 7.5E-12 | MB231/MB468 | NUC |
| >hg38_21_+_8433172_8446622_RNA45SN1_45S_50ntFlanks@12399.12417.19\|ACGTCGCTTTTTGATCCTT | >hg38_21_+_8433172_8446622_RNA45SN1_45S_50ntFlanks | 12399 | 12417 | 19 | ACGTCGCTTTTTGATCCTT | 113.3 | -10.7 | 1.5 | -7.0 | 2.0E-12 | 7.5E-12 | MB231/MB468 | MP |
| >hg38_21_+_8433172_8446622_RNA45SN1_45S_50ntFlanks@4102.4121.20\|ACCACGGGTGACGGGGAATC | >hg38_21_+_8433172_8446622_RNA45SN1_45S_50ntFlanks | 4102 | 4121 | 20 | ACCACGGGTGACGGGGAATC | 3917.6 | -11.1 | 1.6 | -7.0 | 3.1E-12 | 7.6E-12 | BT20/MB468 | NUC |
| >hg38_21_+_8433172_8446622_RNA45SN1_45S_50ntFlanks@11594.11611.18\|CAGGGGAATCCGACTGTT | >hg38_21_+_8433172_8446622_RNA45SN1_45S_50ntFlanks | 11594 | 11611 | 18 | CAGGGGAATCCGACTGTT | 2266.4 | -3.8 | 0.5 | -7.0 | 2.3E-12 | 7.6E-12 | MB231/MB468 | CYTO |
| >trna119_LysCTT_1_-_145395522_145395594@5.36.32,trna11_LysCTT_5_-_180648979_180649051@5.36.32,trna13_LysCTT_6_+_26556774_26556846@5.36.32,trna7_LysCTT_16_+_3225692_3225764@5.36.32,trna9_LysCTT_5_+_180634755_180634827@5.36.32\|GGCTAGCTCAGTCGGTAGAGCATGAGACTCTT | >trna119_LysCTT_1_-_145395522_145395594 | 5 | 36 | 32 | GGCTAGCTCAGTCGGTAGAGCATGAGACTCTT | 364.0 | 7.6 | 1.1 | 7.0 | 2.4E-12 | 7.6E-12 | MB231/MB468 | NUC |
| >hg38_21_+_8433172_8446622_RNA45SN1_45S_50ntFlanks@12626.12657.32\|AATGGGGCGAAGCTACCATCTGTGGGATTATG | >hg38_21_+_8433172_8446622_RNA45SN1_45S_50ntFlanks | 12626 | 12657 | 32 | AATGGGGCGAAGCTACCATCTGTGGGATTATG | 1142.9 | -9.2 | 1.3 | -7.0 | 2.1E-12 | 7.6E-12 | MB231/MB468 | MP |
| >hg38_21_+_8433172_8446622_RNA45SN1_45S_50ntFlanks@9034.9056.23\|TCGTGGGGGGGCCGGGCCACCCC | >hg38_21_+_8433172_8446622_RNA45SN1_45S_50ntFlanks | 9034 | 9056 | 23 | TCGTGGGGGGGCCGGGCCACCCC | 163.4 | -11.2 | 1.6 | -7.0 | 2.1E-12 | 7.6E-12 | MB231/MB468 | MP |

<table>
<tr><th></th><th></th><th></th><th></th><th></th><th></th><th></th><th></th><th></th><th></th><th></th><th></th><th></th><th></th></tr>
<tr><td>>hg38_21_+_8433172_8446622_RNA45SN1_45S_50ntFlanks@10391.10429.39|AACAGCAGTTGAACATGGGTCAGTCGGTCCTGAGAGATG</td><td>>hg38_21_+_8433172_8446622_RNA45SN1_45S_50ntFlanks</td><td>10391</td><td>10429</td><td>39</td><td>AACAGCAGTTGAACATGGGTCAGTCGGTCCTGAGAGATG</td><td>2118.5</td><td>-5.7</td><td>0.8</td><td>-7.0</td><td>2.9E-12</td><td>7.6E-12</td><td>MB231/MB468</td><td>MITO</td></tr>
<tr><td>>trnaMT_HisGTG_MT_+_12138_12206@28.71.44|ATTGTGAATCTGACAACAGAGGCTTACGACCCCTTATTTACCCC</td><td>>trnaMT_HisGTG_MT_+_12138_12206</td><td>28</td><td>71</td><td>44</td><td>ATTGTGAATCTGACAACAGAGGCTTACGACCCCTTATTTACCCC</td><td>365.0</td><td>-12.9</td><td>1.9</td><td>-7.0</td><td>3.1E-12</td><td>7.6E-12</td><td>BT20/MB468</td><td>NUC</td></tr>
<tr><td>>hg38_21_+_8433172_8446622_RNA45SN1_45S_50ntFlanks@9564.9595.32|CTGACGTGCAAATCGGTCGTCCGACCTGGGTA</td><td>>hg38_21_+_8433172_8446622_RNA45SN1_45S_50ntFlanks</td><td>9564</td><td>9595</td><td>32</td><td>CTGACGTGCAAATCGGTCGTCCGACCTGGGTA</td><td>2173.0</td><td>-4.3</td><td>0.6</td><td>-7.0</td><td>2.3E-12</td><td>7.6E-12</td><td>MB231/MB468</td><td>CYTO</td></tr>
<tr><td>>trna11_GluTTC_15_-_26327381_26327452@1.34.34,trna3_GluTTC_13_-_45492062_45492133@1.34.34|TCCCACATGGTCTAGCGGTTAGGATTCCTGGTTT</td><td>>trna11_GluTTC_15_-_26327381_26327452</td><td>1</td><td>34</td><td>34</td><td>TCCCACATGGTCTAGCGGTTAGGATTCCTGGTTT</td><td>608.4</td><td>10.4</td><td>1.5</td><td>7.0</td><td>2.4E-12</td><td>7.6E-12</td><td>MB231/MB468</td><td>CYTO</td></tr>
<tr><td>>hg38_21_+_8433172_8446622_RNA45SN1_45S_50ntFlanks@6731.6756.26|CACATTGATCATCGACACTTCGAACG</td><td>>hg38_21_+_8433172_8446622_RNA45SN1_45S_50ntFlanks</td><td>6731</td><td>6756</td><td>26</td><td>CACATTGATCATCGACACTTCGAACG</td><td>716.1</td><td>-3.9</td><td>0.6</td><td>-7.0</td><td>2.4E-12</td><td>7.6E-12</td><td>MB231/MB468</td><td>NUC</td></tr>
<tr><td>>hg38_21_+_8433172_8446622_RNA45SN1_45S_50ntFlanks@9167.9197.31|CGGGCGGGTCGCGCCGTCGGGCCCGGGGGAG</td><td>>hg38_21_+_8433172_8446622_RNA45SN1_45S_50ntFlanks</td><td>9167</td><td>9197</td><td>31</td><td>CGGGCGGGTCGCGCCGTCGGGCCCGGGGGAG</td><td>216.7</td><td>-10.7</td><td>1.5</td><td>-7.0</td><td>2.4E-12</td><td>7.6E-12</td><td>MB231/MB468</td><td>NUC</td></tr>
<tr><td>>hg38_21_+_8433172_8446622_RNA45SN1_45S_50ntFlanks@5472.5514.43|ACGGCCCTGGCGGAGCGCTGAGAAGACGGTCGAACTTGACTAT</td><td>>hg38_21_+_8433172_8446622_RNA45SN1_45S_50ntFlanks</td><td>5472</td><td>5514</td><td>43</td><td>ACGGCCCTGGCGGAGCGCTGAGAAGACGGTCGAACTTGACTAT</td><td>2198.8</td><td>-6.1</td><td>0.9</td><td>-7.0</td><td>3.1E-12</td><td>7.6E-12</td><td>BT20/MB468</td><td>NUC</td></tr>
<tr><td>>MI0000737|hsa-mir-200a&WithFlank&1|+|1167857|1167958@60.81.22&[MIMAT0000682&hsa-miR-200a-3p&offsets|0|0;m-142&1|+|1167916|1167938&offsets|0|-1]|TAACACTGTCTGGTAACGATGT</td><td>>MI0000737|hsa-mir-200a&WithFlank&1|+|1167857|1167958</td><td>60</td><td>81</td><td>22</td><td>TAACACTGTCTGGTAACGATGT</td><td>1767.4</td><td>-5.9</td><td>0.8</td><td>-7.0</td><td>2.1E-12</td><td>7.7E-12</td><td>MB231/MB468</td><td>MP</td></tr>
<tr><td>>hg38_21_+_8433172_8446622_RNA45SN1_45S_50ntFlanks@9535.9553.19|GGAAACTCTGGTGGAGGTC</td><td>>hg38_21_+_8433172_8446622_RNA45SN1_45S_50ntFlanks</td><td>9535</td><td>9553</td><td>19</td><td>GGAAACTCTGGTGGAGGTC</td><td>240.1</td><td>-11.8</td><td>1.7</td><td>-7.0</td><td>2.1E-12</td><td>7.7E-12</td><td>MB231/MB468</td><td>MP</td></tr>
<tr><td>>hg38_21_+_8433172_8446622_RNA45SN1_45S_50ntFlanks@11037.11075.39|CTCTCCCCCGCTCCCCGTCCTCCCCCCTCCCCGGGGGAG</td><td>>hg38_21_+_8433172_8446622_RNA45SN1_45S_50ntFlanks</td><td>11037</td><td>11075</td><td>39</td><td>CTCTCCCCCGCTCCCCGTCCTCCCCCCTCCCCGGGGGAG</td><td>415.2</td><td>-8.9</td><td>1.3</td><td>-7.0</td><td>2.4E-12</td><td>7.7E-12</td><td>MB231/MB468</td><td>NUC</td></tr>
<tr><td>>hg38_21_+_8433172_8446622_RNA45SN1_45S_50ntFlanks@10389.10424.36|TGAACAGCAGTTGAACATGGGTCAGTCGGTCCTGAG</td><td>>hg38_21_+_8433172_8446622_RNA45SN1_45S_50ntFlanks</td><td>10389</td><td>10424</td><td>36</td><td>TGAACAGCAGTTGAACATGGGTCAGTCGGTCCTGAG</td><td>24675.4</td><td>5.1</td><td>0.7</td><td>7.1</td><td>1.7E-12</td><td>7.7E-12</td><td>BT20/MB468</td><td>MITO</td></tr>
<tr><td>>hg38_21_+_8433172_8446622_RNA45SN1_45S_50ntFlanks@4664.4695.32|TGAAATTCTTGGACCGGCGCAAGACGGACCAG</td><td>>hg38_21_+_8433172_8446622_RNA45SN1_45S_50ntFlanks</td><td>4664</td><td>4695</td><td>32</td><td>TGAAATTCTTGGACCGGCGCAAGACGGACCAG</td><td>106.4</td><td>-10.6</td><td>1.5</td><td>-7.0</td><td>2.1E-12</td><td>7.7E-12</td><td>MB231/MB468</td><td>MP</td></tr>
<tr><td>>hg38_21_+_8433172_8446622_RNA45SN1_45S_50ntFlanks@11710.11728.19|CGCGGGTAAACGGCGGGAG</td><td>>hg38_21_+_8433172_8446622_RNA45SN1_45S_50ntFlanks</td><td>11710</td><td>11728</td><td>19</td><td>CGCGGGTAAACGGCGGGAG</td><td>677.4</td><td>-6.9</td><td>1.0</td><td>-7.0</td><td>2.4E-12</td><td>7.7E-12</td><td>MB231/MB468</td><td>NUC</td></tr>
<tr><td>>hg38_21_+_8433172_8446622_RNA45SN1_45S_50ntFlanks@6779.6806.28|CCGGGGCTACGCCTGTCTGAGCGTCGCT</td><td>>hg38_21_+_8433172_8446622_RNA45SN1_45S_50ntFlanks</td><td>6779</td><td>6806</td><td>28</td><td>CCGGGGCTACGCCTGTCTGAGCGTCGCT</td><td>421.5</td><td>3.7</td><td>0.5</td><td>7.0</td><td>2.4E-12</td><td>7.7E-12</td><td>MB231/MB468</td><td>NUC</td></tr>
<tr><td>>hg38_21_+_8433172_8446622_RNA45SN1_45S_50ntFlanks@10800.10818.19|TGTTGGAACAATGTAGGTA</td><td>>hg38_21_+_8433172_8446622_RNA45SN1_45S_50ntFlanks</td><td>10800</td><td>10818</td><td>19</td><td>TGTTGGAACAATGTAGGTA</td><td>129.1</td><td>-10.9</td><td>1.5</td><td>-7.0</td><td>2.1E-12</td><td>7.7E-12</td><td>MB231/MB468</td><td>MP</td></tr>
<tr><td>>hg38_21_+_8433172_8446622_RNA45SN1_45S_50ntFlanks@4240.4258.19|ACTCTTTCGAGGCCCTGTA</td><td>>hg38_21_+_8433172_8446622_RNA45SN1_45S_50ntFlanks</td><td>4240</td><td>4258</td><td>19</td><td>ACTCTTTCGAGGCCCTGTA</td><td>637.7</td><td>-5.0</td><td>0.7</td><td>-7.0</td><td>2.4E-12</td><td>7.7E-12</td><td>MB231/MB468</td><td>CYTO</td></tr>
<tr><td>>hg38_21_+_8433172_8446622_RNA45SN1_45S_50ntFlanks@12081.12109.29|CTCTCGCTTCTGGCGCCAAGCGCCCGGCC</td><td>>hg38_21_+_8433172_8446622_RNA45SN1_45S_50ntFlanks</td><td>12081</td><td>12109</td><td>29</td><td>CTCTCGCTTCTGGCGCCAAGCGCCCGGCC</td><td>115.5</td><td>-10.7</td><td>1.5</td><td>-7.0</td><td>2.1E-12</td><td>7.7E-12</td><td>MB231/MB468</td><td>MP</td></tr>
<tr><td>>hg38_21_+_8433172_8446622_RNA45SN1_45S_50ntFlanks@10355.10383.29|AACTTTGAAGGCCGAAGTGGAGAAGGGTT</td><td>>hg38_21_+_8433172_8446622_RNA45SN1_45S_50ntFlanks</td><td>10355</td><td>10383</td><td>29</td><td>AACTTTGAAGGCCGAAGTGGAGAAGGGTT</td><td>553.3</td><td>-6.7</td><td>1.0</td><td>-7.0</td><td>2.5E-12</td><td>7.8E-12</td><td>MB231/MB468</td><td>NUC</td></tr>
<tr><td>>hg38_21_+_8433172_8446622_RNA45SN1_45S_50ntFlanks@10820.10840.21|GGGAAGTCGGCAAGCCGGATC</td><td>>hg38_21_+_8433172_8446622_RNA45SN1_45S_50ntFlanks</td><td>10820</td><td>10840</td><td>21</td><td>GGGAAGTCGGCAAGCCGGATC</td><td>1094.5</td><td>-7.0</td><td>1.0</td><td>-7.0</td><td>2.9E-12</td><td>7.8E-12</td><td>MB231/MB468</td><td>MITO</td></tr>
<tr><td>>hg38_21_+_8433172_8446622_RNA45SN1_45S_50ntFlanks@11205.11232.28|GCGGCGGCGACTCTGGACGCGAGCCGGG</td><td>>hg38_21_+_8433172_8446622_RNA45SN1_45S_50ntFlanks</td><td>11205</td><td>11232</td><td>28</td><td>GCGGCGGCGACTCTGGACGCGAGCCGGG</td><td>1152.2</td><td>-6.6</td><td>0.9</td><td>-7.0</td><td>2.4E-12</td><td>7.8E-12</td><td>MB231/MB468</td><td>CYTO</td></tr>
<tr><td>>hg38_21_+_8433172_8446622_RNA45SN1_45S_50ntFlanks@3985.4009.25|CGGCTTTGGTGACTCTAGATAACCT</td><td>>hg38_21_+_8433172_8446622_RNA45SN1_45S_50ntFlanks</td><td>3985</td><td>4009</td><td>25</td><td>CGGCTTTGGTGACTCTAGATAACCT</td><td>180.3</td><td>-11.4</td><td>1.6</td><td>-7.0</td><td>2.1E-12</td><td>7.8E-12</td><td>MB231/MB468</td><td>MP</td></tr>
<tr><td>>hg38_21_+_8433172_8446622_RNA45SN1_45S_50ntFlanks@8170.8195.26|CAAGTCCTTCTGATCGAGGCCCAGCC</td><td>>hg38_21_+_8433172_8446622_RNA45SN1_45S_50ntFlanks</td><td>8170</td><td>8195</td><td>26</td><td>CAAGTCCTTCTGATCGAGGCCCAGCC</td><td>2895.8</td><td>-10.0</td><td>1.4</td><td>-7.0</td><td>2.1E-12</td><td>7.8E-12</td><td>MB231/MB468</td><td>MP</td></tr>
<tr><td>>hg38_21_+_8433172_8446622_RNA45SN1_45S_50ntFlanks@11212.11234.23|CGACTCTGGACGCGAGCCGGGCC</td><td>>hg38_21_+_8433172_8446622_RNA45SN1_45S_50ntFlanks</td><td>11212</td><td>11234</td><td>23</td><td>CGACTCTGGACGCGAGCCGGGCC</td><td>127.0</td><td>-10.9</td><td>1.5</td><td>-7.0</td><td>2.1E-12</td><td>7.8E-12</td><td>MB231/MB468</td><td>MP</td></tr>
<tr><td>>hg38_21_+_8433172_8446622_RNA45SN1_45S_50ntFlanks@12399.12436.38|ACGTCGCTTTTTGATCCTTCGATGTCGGCTCTTCCTAT</td><td>>hg38_21_+_8433172_8446622_RNA45SN1_45S_50ntFlanks</td><td>12399</td><td>12436</td><td>38</td><td>ACGTCGCTTTTTGATCCTTCGATGTCGGCTCTTCCTAT</td><td>855.9</td><td>-7.2</td><td>1.0</td><td>-7.0</td><td>3.0E-12</td><td>7.8E-12</td><td>MB231/MB468</td><td>MITO</td></tr>
<tr><td>>hg38_21_+_8433172_8446622_RNA45SN1_45S_50ntFlanks@6671.6696.26|CTCGGCTCGTGCGTCGATGAAGAACG</td><td>>hg38_21_+_8433172_8446622_RNA45SN1_45S_50ntFlanks</td><td>6671</td><td>6696</td><td>26</td><td>CTCGGCTCGTGCGTCGATGAAGAACG</td><td>3777.5</td><td>-8.0</td><td>1.1</td><td>-7.0</td><td>2.1E-12</td><td>7.8E-12</td><td>MB231/MB468</td><td>MP</td></tr>
<tr><td>>hg38_21_+_8433172_8446622_RNA45SN1_45S_50ntFlanks@12448.12480.33|AGAATTCACCAAGCGTTGGATTGTTCACCCACT</td><td>>hg38_21_+_8433172_8446622_RNA45SN1_45S_50ntFlanks</td><td>12448</td><td>12480</td><td>33</td><td>AGAATTCACCAAGCGTTGGATTGTTCACCCACT</td><td>257.2</td><td>6.7</td><td>1.0</td><td>7.0</td><td>2.5E-12</td><td>7.8E-12</td><td>MB231/MB468</td><td>NUC</td></tr>
<tr><td>>hg38_21_+_8433172_8446622_RNA45SN1_45S_50ntFlanks@12201.12221.21|CCTAAGGCGAGCTCAGGGAGG</td><td>>hg38_21_+_8433172_8446622_RNA45SN1_45S_50ntFlanks</td><td>12201</td><td>12221</td><td>21</td><td>CCTAAGGCGAGCTCAGGGAGG</td><td>120.7</td><td>-10.8</td><td>1.5</td><td>-7.0</td><td>2.1E-12</td><td>7.8E-12</td><td>MB231/MB468</td><td>MP</td></tr>
<tr><td>>hg38_21_+_8433172_8446622_RNA45SN1_45S_50ntFlanks@12941.12964.24|ATTCGTAGACGACCTGCTTCTGGG</td><td>>hg38_21_+_8433172_8446622_RNA45SN1_45S_50ntFlanks</td><td>12941</td><td>12964</td><td>24</td><td>ATTCGTAGACGACCTGCTTCTGGG</td><td>121.2</td><td>-10.8</td><td>1.5</td><td>-7.0</td><td>2.1E-12</td><td>7.8E-12</td><td>MB231/MB468</td><td>MP</td></tr>
<tr><td>>hg38_21_+_8433172_8446622_RNA45SN1_45S_50ntFlanks@7984.8006.23|AGATCAGACGTGGCGACCCGCTG</td><td>>hg38_21_+_8433172_8446622_RNA45SN1_45S_50ntFlanks</td><td>7984</td><td>8006</td><td>23</td><td>AGATCAGACGTGGCGACCCGCTG</td><td>1689.1</td><td>-7.8</td><td>1.1</td><td>-7.0</td><td>2.5E-12</td><td>7.8E-12</td><td>MB231/MB468</td><td>NUC</td></tr>
<tr><td>>hg38_21_+_8433172_8446622_RNA45SN1_45S_50ntFlanks@9029.9065.37|TCTCCTCGTGGGGGGGCCGGGCCACCCCTCCCACGGC</td><td>>hg38_21_+_8433172_8446622_RNA45SN1_45S_50ntFlanks</td><td>9029</td><td>9065</td><td>37</td><td>TCTCCTCGTGGGGGGGCCGGGCCACCCCTCCCACGGC</td><td>105.0</td><td>-10.6</td><td>1.5</td><td>-7.0</td><td>2.1E-12</td><td>7.8E-12</td><td>MB231/MB468</td><td>MP</td></tr>
<tr><td>>MI0000077|hsa-mir-21&WithFlank&17|+|59841260|59841343@14.34.21(+3C)&[MIMAT0000076&hsa-miR-21-5p&offsets|0|-1(+3C);m-1&17|+|59841273|59841294&offsets|0|-1(+3C)]|TAGCTTATCAGACTGATGTTGCCC</td><td>>MI0000077|hsa-mir-21&WithFlank&17|+|59841260|59841343</td><td>14</td><td>34</td><td>21(+3C)</td><td>TAGCTTATCAGACTGATGTTGCCC</td><td>281.7</td><td>8.2</td><td>1.2</td><td>7.0</td><td>2.5E-12</td><td>7.9E-12</td><td>MB231/MB468</td><td>NUC</td></tr>
<tr><td>>hg38_21_+_8433172_8446622_RNA45SN1_45S_50ntFlanks@10385.10405.21|CATGTGAACAGCAGTTGAACA</td><td>>hg38_21_+_8433172_8446622_RNA45SN1_45S_50ntFlanks</td><td>10385</td><td>10405</td><td>21</td><td>CATGTGAACAGCAGTTGAACA</td><td>366.7</td><td>-12.4</td><td>1.8</td><td>-7.0</td><td>2.1E-12</td><td>7.9E-12</td><td>MB231/MB468</td><td>MP</td></tr>
<tr><td>>hg38_21_+_8433172_8446622_RNA45SN1_45S_50ntFlanks@4889.4917.29|CTGAAACTTAAAGGAATTGACGGAAGGGC</td><td>>hg38_21_+_8433172_8446622_RNA45SN1_45S_50ntFlanks</td><td>4889</td><td>4917</td><td>29</td><td>CTGAAACTTAAAGGAATTGACGGAAGGGC</td><td>107.5</td><td>-10.6</td><td>1.5</td><td>-7.0</td><td>2.1E-12</td><td>7.9E-12</td><td>MB231/MB468</td><td>MP</td></tr>
<tr><td>>hg38_21_+_8433172_8446622_RNA45SN1_45S_50ntFlanks@7983.8005.23|CAGATCAGACGTGGCGACCCGCT</td><td>>hg38_21_+_8433172_8446622_RNA45SN1_45S_50ntFlanks</td><td>7983</td><td>8005</td><td>23</td><td>CAGATCAGACGTGGCGACCCGCT</td><td>1477.1</td><td>-9.2</td><td>1.3</td><td>-7.0</td><td>2.1E-12</td><td>7.9E-12</td><td>MB231/MB468</td><td>MP</td></tr>
<tr><td>>hg38_21_+_8433172_8446622_RNA45SN1_45S_50ntFlanks@10410.10431.22|TCAGTCGGTCCTGAGAGATGGG</td><td>>hg38_21_+_8433172_8446622_RNA45SN1_45S_50ntFlanks</td><td>10410</td><td>10431</td><td>22</td><td>TCAGTCGGTCCTGAGAGATGGG</td><td>2295.4</td><td>-6.9</td><td>1.0</td><td>-7.0</td><td>2.3E-12</td><td>7.9E-12</td><td>BT20/MB468</td><td>CYTO</td></tr>
<tr><td>>hg38_21_+_8433172_8446622_RNA45SN1_45S_50ntFlanks@485.504.20|GATGTGGTGACGTCGTGCTC</td><td>>hg38_21_+_8433172_8446622_RNA45SN1_45S_50ntFlanks</td><td>485</td><td>504</td><td>20</td><td>GATGTGGTGACGTCGTGCTC</td><td>635.9</td><td>13.1</td><td>1.8</td><td>7.4</td><td>1.2E-13</td><td>7.9E-12</td><td>BT20/MB468</td><td>MP</td></tr>
<tr><td>>hg38_21_+_8433172_8446622_RNA45SN1_45S_50ntFlanks@8836.8861.26|CGGCGGTGGCGGCGGCGGCGGCGGCG</td><td>>hg38_21_+_8433172_8446622_RNA45SN1_45S_50ntFlanks</td><td>8836</td><td>8861</td><td>26</td><td>CGGCGGTGGCGGCGGCGGCGGCGGCG</td><td>156.5</td><td>-11.2</td><td>1.6</td><td>-7.0</td><td>2.2E-12</td><td>7.9E-12</td><td>MB231/MB468</td><td>MP</td></tr>
<tr><td>>hg38_21_+_8433172_8446622_RNA45SN1_45S_50ntFlanks@3984.4001.18|GCGGCTTTGGTGACTCTA</td><td>>hg38_21_+_8433172_8446622_RNA45SN1_45S_50ntFlanks</td><td>3984</td><td>4001</td><td>18</td><td>GCGGCTTTGGTGACTCTA</td><td>1479.3</td><td>-5.3</td><td>0.8</td><td>-7.0</td><td>1.8E-12</td><td>7.9E-12</td><td>BT20/MB468</td><td>MITO</td></tr>
<tr><td>>hg38_21_+_8433172_8446622_RNA45SN1_45S_50ntFlanks@10806.10825.20|AACAATGTAGGTAAGGGAAG</td><td>>hg38_21_+_8433172_8446622_RNA45SN1_45S_50ntFlanks</td><td>10806</td><td>10825</td><td>20</td><td>AACAATGTAGGTAAGGGAAG</td><td>632.2</td><td>-9.5</td><td>1.4</td><td>-7.0</td><td>2.5E-12</td><td>7.9E-12</td><td>MB231/MB468</td><td>NUC</td></tr>
<tr><td>>hg38_21_+_8433172_8446622_RNA45SN1_45S_50ntFlanks@4006.4024.19|ACCTCGGGCCGATCGCACG</td><td>>hg38_21_+_8433172_8446622_RNA45SN1_45S_50ntFlanks</td><td>4006</td><td>4024</td><td>19</td><td>ACCTCGGGCCGATCGCACG</td><td>877.8</td><td>-4.8</td><td>0.7</td><td>-7.0</td><td>2.5E-12</td><td>7.9E-12</td><td>MB231/MB468</td><td>NUC</td></tr>
<tr><td>>hg38_21_+_8433172_8446622_RNA45SN1_45S_50ntFlanks@9120.9138.19|TGCCGCGCGCGGGTCGGGG</td><td>>hg38_21_+_8433172_8446622_RNA45SN1_45S_50ntFlanks</td><td>9120</td><td>9138</td><td>19</td><td>TGCCGCGCGCGGGTCGGGG</td><td>155.6</td><td>-11.1</td><td>1.6</td><td>-7.0</td><td>2.2E-12</td><td>7.9E-12</td><td>MB231/MB468</td><td>MP</td></tr>
<tr><td>>hg38_21_+_8433172_8446622_RNA45SN1_45S_50ntFlanks@11873.11901.29|AAGAAGACCCTGTTGAGCTTGACTCTAGT</td><td>>hg38_21_+_8433172_8446622_RNA45SN1_45S_50ntFlanks</td><td>11873</td><td>11901</td><td>29</td><td>AAGAAGACCCTGTTGAGCTTGACTCTAGT</td><td>91.5</td><td>-10.4</td><td>1.5</td><td>-7.0</td><td>2.2E-12</td><td>7.9E-12</td><td>MB231/MB468</td><td>MP</td></tr>
<tr><td>>hg38_21_+_8433172_8446622_RNA45SN1_45S_50ntFlanks@10905.10931.27|CTGGGCGCGCGCCGCGGCTGGACGAGG</td><td>>hg38_21_+_8433172_8446622_RNA45SN1_45S_50ntFlanks</td><td>10905</td><td>10931</td><td>27</td><td>CTGGGCGCGCGCCGCGGCTGGACGAGG</td><td>111.6</td><td>-10.7</td><td>1.5</td><td>-7.0</td><td>2.2E-12</td><td>7.9E-12</td><td>MB231/MB468</td><td>MP</td></tr>
<tr><td>>hg38_21_+_8433172_8446622_RNA45SN1_45S_50ntFlanks@9739.9758.20|AAACGATCTCAACCTATTCT</td><td>>hg38_21_+_8433172_8446622_RNA45SN1_45S_50ntFlanks</td><td>9739</td><td>9758</td><td>20</td><td>AAACGATCTCAACCTATTCT</td><td>4625.1</td><td>-5.2</td><td>0.7</td><td>-7.0</td><td>1.8E-12</td><td>8.0E-12</td><td>BT20/MB468</td><td>MITO</td></tr>
<tr><td>>trna32_LysCTT_16_-_3207406_3207478@4.33.30|CGGCTAGCTCAGTCGGTAGAGCATGAGACC</td><td>>trna32_LysCTT_16_-_3207406_3207478</td><td>4</td><td>33</td><td>30</td><td>CGGCTAGCTCAGTCGGTAGAGCATGAGACC</td><td>434.7</td><td>12.0</td><td>1.7</td><td>7.0</td><td>3.0E-12</td><td>8.0E-12</td><td>MB231/MB468</td><td>MITO</td></tr>
<tr><td>>hg38_21_+_8433172_8446622_RNA45SN1_45S_50ntFlanks@12405.12430.26|CTTTTTGATCCTTCGATGTCGGCTCT</td><td>>hg38_21_+_8433172_8446622_RNA45SN1_45S_50ntFlanks</td><td>12405</td><td>12430</td><td>26</td><td>CTTTTTGATCCTTCGATGTCGGCTCT</td><td>2162.6</td><td>-4.7</td><td>0.7</td><td>-7.0</td><td>2.5E-12</td><td>8.0E-12</td><td>MB231/MB468</td><td>CYTO</td></tr>
<tr><td>>hg38_21_+_8433172_8446622_RNA45SN1_45S_50ntFlanks@10391.10420.30|AACAGCAGTTGAACATGGGTCAGTCGGTCC</td><td>>hg38_21_+_8433172_8446622_RNA45SN1_45S_50ntFlanks</td><td>10391</td><td>10420</td><td>30</td><td>AACAGCAGTTGAACATGGGTCAGTCGGTCC</td><td>853.8</td><td>-8.8</td><td>1.3</td><td>-7.0</td><td>2.5E-12</td><td>8.0E-12</td><td>MB231/MB468</td><td>NUC</td></tr>
<tr><td>>am_MI0000786|hsa-mir-378a&WithFlank&5|+|149732819|149732896@49.70.22(+1U)&[MIMAT0000732&hsa-miR-378a-3p&offsets|0|0(+1U);m-44&5|+|149732867|149732888&offsets|0|0(+1U)]|ACTGGACTTGGAGTCAGAAGGCT</td><td>>am_MI0000786|hsa-mir-378a&WithFlank&5|+|149732819|149732896</td><td>49</td><td>70</td><td>22(+1U)</td><td>ACTGGACTTGGAGTCAGAAGGCT</td><td>1428.2</td><td>4.9</td><td>0.7</td><td>7.0</td><td>1.8E-12</td><td>8.0E-12</td><td>BT20/MB468</td><td>MITO</td></tr>
<tr><td>>hg38_21_+_8433172_8446622_RNA45SN1_45S_50ntFlanks@10845.10865.21|ACTTCGGGATAAGGATTGGCT</td><td>>hg38_21_+_8433172_8446622_RNA45SN1_45S_50ntFlanks</td><td>10845</td><td>10865</td><td>21</td><td>ACTTCGGGATAAGGATTGGCT</td><td>10682.8</td><td>-6.0</td><td>0.9</td><td>-7.0</td><td>2.5E-12</td><td>8.0E-12</td><td>MB231/MB468</td><td>CYTO</td></tr>
<tr><td>>hg38_21_+_8433172_8446622_RNA45SN1_45S_50ntFlanks@11919.11950.32|CATGAGAGGTGTAGAATAAGTGGGAGGCCCCC</td><td>>hg38_21_+_8433172_8446622_RNA45SN1_45S_50ntFlanks</td><td>11919</td><td>11950</td><td>32</td><td>CATGAGAGGTGTAGAATAAGTGGGAGGCCCCC</td><td>312.9</td><td>-5.8</td><td>0.8</td><td>-7.0</td><td>2.5E-12</td><td>8.0E-12</td><td>MB231/MB468</td><td>CYTO</td></tr>
<tr><td>>hg38_21_+_8433172_8446622_RNA45SN1_45S_50ntFlanks@3988.4024.37|CTTTGGTGACTCTAGATAACCTCGGGCCGATCGCACG</td><td>>hg38_21_+_8433172_8446622_RNA45SN1_45S_50ntFlanks</td><td>3988</td><td>4024</td><td>37</td><td>CTTTGGTGACTCTAGATAACCTCGGGCCGATCGCACG</td><td>890.1</td><td>-6.9</td><td>1.0</td><td>-7.0</td><td>3.0E-12</td><td>8.0E-12</td><td>MB231/MB468</td><td>MITO</td></tr>
<tr><td>>hg38_21_+_8433172_8446622_RNA45SN1_45S_50ntFlanks@10874.10892.19|TGGGTCGGTCGGGCTGGGG</td><td>>hg38_21_+_8433172_8446622_RNA45SN1_45S_50ntFlanks</td><td>10874</td><td>10892</td><td>19</td><td>TGGGTCGGTCGGGCTGGGG</td><td>150.7</td><td>-11.1</td><td>1.6</td><td>-7.0</td><td>2.2E-12</td><td>8.0E-12</td><td>MB231/MB468</td><td>MP</td></tr>
<tr><td>>hg38_21_+_8433172_8446622_RNA45SN1_45S_50ntFlanks@9344.9370.27|GAAAGCCGCCGTGGCGCAATGAAGGTG</td><td>>hg38_21_+_8433172_8446622_RNA45SN1_45S_50ntFlanks</td><td>9344</td><td>9370</td><td>27</td><td>GAAAGCCGCCGTGGCGCAATGAAGGTG</td><td>118.6</td><td>-10.8</td><td>1.5</td><td>-7.0</td><td>2.2E-12</td><td>8.0E-12</td><td>MB231/MB468</td><td>MP</td></tr>
</table>

<table>
<tr><td>>hg38_21_+_8433172_8446622_RNA45SN1_45S_50ntFlanks@12690.12708.19|GCGGAACGATACGGCAGCG</td><td>>hg38_21_+_8433172_8446622_RNA45SN1_45S_50ntFlanks</td><td>12690</td><td>12708</td><td>19</td><td>GCGGAACGATACGGCAGCG</td><td>249.7</td><td>-11.8</td><td>1.7</td><td>-7.0</td><td>2.2E-12</td><td>8.0E-12</td><td>MB231/MB468</td><td>MP</td></tr>
<tr><td>>MI0000077|hsa-mir-21&WithFlank&17|+|59841260|59841343@16.36.21&[MIMAT0000076&hsa-miR-21-5p&offsets|+2|+1;m-1&17|+|59841273|59841294&offsets|+2|+1]|GCTTATCAGACTGATGTTGAC</td><td>>MI0000077|hsa-mir-21&WithFlank&17|+|59841260|59841343</td><td>16</td><td>36</td><td>21</td><td>GCTTATCAGACTGATGTTGAC</td><td>103.6</td><td>7.6</td><td>1.1</td><td>7.0</td><td>3.3E-12</td><td>8.1E-12</td><td>BT20/MB468</td><td>NUC</td></tr>
<tr><td>>hg38_21_+_8433172_8446622_RNA45SN1_45S_50ntFlanks@11970.11998.29|CGCGAGGGGCCCGGGGCGGGGTCCGCCGG</td><td>>hg38_21_+_8433172_8446622_RNA45SN1_45S_50ntFlanks</td><td>11970</td><td>11998</td><td>29</td><td>CGCGAGGGGCCCGGGGCGGGGTCCGCCGG</td><td>234.3</td><td>-11.7</td><td>1.7</td><td>-7.0</td><td>2.2E-12</td><td>8.1E-12</td><td>MB231/MB468</td><td>MP</td></tr>
<tr><td>>hg38_21_+_8433172_8446622_RNA45SN1_45S_50ntFlanks@4163.4182.20|CATCCAAGGAAGGCAGCAGG</td><td>>hg38_21_+_8433172_8446622_RNA45SN1_45S_50ntFlanks</td><td>4163</td><td>4182</td><td>20</td><td>CATCCAAGGAAGGCAGCAGG</td><td>211.1</td><td>-9.8</td><td>1.4</td><td>-7.0</td><td>1.8E-12</td><td>8.1E-12</td><td>BT20/MB468</td><td>MITO</td></tr>
<tr><td>>MI0000272|hsa-mir-182&WithFlank&7|-|129770377|129770498@29.49.21&[MIMAT0000259&hsa-miR-182-5p&offsets|0|-3;m-22&7|-|129770449|129770470&offsets|0|-1]|TTTGGCAATGGTAGAACTCAC</td><td>>MI0000272|hsa-mir-182&WithFlank&7|-|129770377|129770498</td><td>29</td><td>49</td><td>21</td><td>TTTGGCAATGGTAGAACTCAC</td><td>1646.9</td><td>9.8</td><td>1.4</td><td>7.0</td><td>2.4E-12</td><td>8.1E-12</td><td>BT20/MB468</td><td>CYTO</td></tr>
<tr><td>>hg38_21_+_8433172_8446622_RNA45SN1_45S_50ntFlanks@11668.11692.25|TGCCCAGTGCTCTGAATGTCAAAGT</td><td>>hg38_21_+_8433172_8446622_RNA45SN1_45S_50ntFlanks</td><td>11668</td><td>11692</td><td>25</td><td>TGCCCAGTGCTCTGAATGTCAAAGT</td><td>249.6</td><td>3.5</td><td>0.5</td><td>7.0</td><td>2.6E-12</td><td>8.1E-12</td><td>MB231/MB468</td><td>NUC</td></tr>
<tr><td>>hg38_21_+_8433172_8446622_RNA45SN1_45S_50ntFlanks@10411.10436.26|CAGTCGGTCCTGAGAGATGGGCGAGC</td><td>>hg38_21_+_8433172_8446622_RNA45SN1_45S_50ntFlanks</td><td>10411</td><td>10436</td><td>26</td><td>CAGTCGGTCCTGAGAGATGGGCGAGC</td><td>149.4</td><td>-11.1</td><td>1.6</td><td>-7.0</td><td>2.2E-12</td><td>8.1E-12</td><td>MB231/MB468</td><td>MP</td></tr>
<tr><td>>hg38_21_+_8433172_8446622_RNA45SN1_45S_50ntFlanks@7975.8017.43|CGCGACCTCAGATCAGACGTGGCGACCCGCTGAATTTAAGCAT</td><td>>hg38_21_+_8433172_8446622_RNA45SN1_45S_50ntFlanks</td><td>7975</td><td>8017</td><td>43</td><td>CGCGACCTCAGATCAGACGTGGCGACCCGCTGAATTTAAGCAT</td><td>20489.5</td><td>5.4</td><td>0.8</td><td>7.0</td><td>1.8E-12</td><td>8.1E-12</td><td>BT20/MB468</td><td>MITO</td></tr>
<tr><td>>am_trnaMT_ProTGG_MT_-_15956_16023@52.71.20|AAAGACTTTTTCTCTGACCA</td><td>>am_trnaMT_ProTGG_MT_-_15956_16023</td><td>52</td><td>71</td><td>20</td><td>AAAGACTTTTTCTCTGACCA</td><td>1591.3</td><td>6.9</td><td>1.0</td><td>7.0</td><td>2.6E-12</td><td>8.1E-12</td><td>MB231/MB468</td><td>NUC</td></tr>
<tr><td>>MI0000465|hsa-mir-191&WithFlank&3|-|49020612|49020715@22.44.23(+1C)&[MIMAT0000440&hsa-miR-191-5p&offsets|0|0(+1C);m-48&3|-|49020672|49020694&offsets|0|0(+1C)]|CAACGGAATCCCAAAAGCAGCTGC</td><td>>MI0000465|hsa-mir-191&WithFlank&3|-|49020612|49020715</td><td>22</td><td>44</td><td>23(+1C)</td><td>CAACGGAATCCCAAAAGCAGCTGC</td><td>510.5</td><td>6.1</td><td>0.9</td><td>7.0</td><td>2.6E-12</td><td>8.2E-12</td><td>MB231/MB468</td><td>NUC</td></tr>
<tr><td>>hg38_21_+_8433172_8446622_RNA45SN1_45S_50ntFlanks@10137.10157.21|CGGCGGCGGCGGGGGTGTGGG</td><td>>hg38_21_+_8433172_8446622_RNA45SN1_45S_50ntFlanks</td><td>10137</td><td>10157</td><td>21</td><td>CGGCGGCGGCGGGGGTGTGGG</td><td>170.0</td><td>-11.3</td><td>1.6</td><td>-7.0</td><td>2.2E-12</td><td>8.2E-12</td><td>MB231/MB468</td><td>MP</td></tr>
<tr><td>>trna20_GluTTC_2_-_131094701_131094772@1.33.33,trna5_GluTTC_13_-_41634874_41634945@1.33.33|TCCCATATGGTCTAGCGGTTAGGATTCCTGGTT</td><td>>trna20_GluTTC_2_-_131094701_131094772</td><td>1</td><td>33</td><td>33</td><td>TCCCATATGGTCTAGCGGTTAGGATTCCTGGTT</td><td>1352.1</td><td>10.2</td><td>1.5</td><td>7.0</td><td>2.2E-12</td><td>8.2E-12</td><td>MB231/MB468</td><td>MP</td></tr>
<tr><td>>MI0000456|hsa-mir-140&WithFlank&16|+|69933075|69933186@68.90.23&[MIMAT0004597&hsa-miR-140-3p&offsets|0|+2;m-45&16|+|69933143|69933164&offsets|-1|0]|TACCACAGGGTAGAACCACGGAC</td><td>>MI0000456|hsa-mir-140&WithFlank&16|+|69933075|69933186</td><td>68</td><td>90</td><td>23</td><td>TACCACAGGGTAGAACCACGGAC</td><td>1266.4</td><td>10.0</td><td>1.4</td><td>7.0</td><td>2.5E-12</td><td>8.2E-12</td><td>MB231/MB468</td><td>CYTO</td></tr>
<tr><td>>hg38_21_+_8433172_8446622_RNA45SN1_45S_50ntFlanks@12714.12751.38|GAGCCTCGGTTGGCCTCGGATAGCCGGTCCCCCGCCTG</td><td>>hg38_21_+_8433172_8446622_RNA45SN1_45S_50ntFlanks</td><td>12714</td><td>12751</td><td>38</td><td>GAGCCTCGGTTGGCCTCGGATAGCCGGTCCCCCGCCTG</td><td>430.4</td><td>-12.6</td><td>1.8</td><td>-7.0</td><td>2.2E-12</td><td>8.2E-12</td><td>MB231/MB468</td><td>MP</td></tr>
<tr><td>>MI0000440|hsa-mir-27b&WithFlank&9|+|95085439|95085547@67.87.21&[MIMAT0000419&hsa-miR-27b-3p&offsets|0|0;m-39&9|+|95085505|95085525&offsets|0|0]|TTCACAGTGGCTAAGTTCTGC</td><td>>MI0000440|hsa-mir-27b&WithFlank&9|+|95085439|95085547</td><td>67</td><td>87</td><td>21</td><td>TTCACAGTGGCTAAGTTCTGC</td><td>46231.1</td><td>6.2</td><td>0.9</td><td>7.0</td><td>3.1E-12</td><td>8.2E-12</td><td>MB231/MB468</td><td>MITO</td></tr>
<tr><td>>hg38_21_+_8433172_8446622_RNA45SN1_45S_50ntFlanks@9738.9756.19|GAAACGATCTCAACCTATT</td><td>>hg38_21_+_8433172_8446622_RNA45SN1_45S_50ntFlanks</td><td>9738</td><td>9756</td><td>19</td><td>GAAACGATCTCAACCTATT</td><td>132.1</td><td>-10.9</td><td>1.6</td><td>-7.0</td><td>2.2E-12</td><td>8.2E-12</td><td>MB231/MB468</td><td>MP</td></tr>
<tr><td>>hg38_21_+_8433172_8446622_RNA45SN1_45S_50ntFlanks@10882.10901.20|TCGGGCTGGGGCGCGAAGCG</td><td>>hg38_21_+_8433172_8446622_RNA45SN1_45S_50ntFlanks</td><td>10882</td><td>10901</td><td>20</td><td>TCGGGCTGGGGCGCGAAGCG</td><td>135.6</td><td>-10.9</td><td>1.6</td><td>-7.0</td><td>2.2E-12</td><td>8.2E-12</td><td>MB231/MB468</td><td>MP</td></tr>
<tr><td>>hg38_21_+_8433172_8446622_RNA45SN1_45S_50ntFlanks@11933.11956.24|AATAAGTGGGAGGCCCCCGGCGCC</td><td>>hg38_21_+_8433172_8446622_RNA45SN1_45S_50ntFlanks</td><td>11933</td><td>11956</td><td>24</td><td>AATAAGTGGGAGGCCCCCGGCGCC</td><td>158.8</td><td>-11.2</td><td>1.6</td><td>-7.0</td><td>2.2E-12</td><td>8.2E-12</td><td>MB231/MB468</td><td>MP</td></tr>
<tr><td>>hg38_21_+_8433172_8446622_RNA45SN1_45S_50ntFlanks@6772.6801.30|GTTCCTCCCGGGGCTACGCCTGTCTGAGCG</td><td>>hg38_21_+_8433172_8446622_RNA45SN1_45S_50ntFlanks</td><td>6772</td><td>6801</td><td>30</td><td>GTTCCTCCCGGGGCTACGCCTGTCTGAGCG</td><td>96.9</td><td>-10.5</td><td>1.5</td><td>-7.0</td><td>2.2E-12</td><td>8.2E-12</td><td>MB231/MB468</td><td>MP</td></tr>
<tr><td>>MI0000446|hsa-mir-125b-1&WithFlank&11|-|122099751|122099850@21.42.22(+1C);MI0000470|hsa-mir-125b-2&WithFlank&21|+|16590231|16590331@23.44.22(+1C)&[MIMAT0000423&hsa-miR-125b-5p&offsets|0|0(+1C);m-67&11|-|122099809|122099830&offsets|0|0(+1C)];[MIMAT0000423_1&hsa-miR-125b-5p&offsets|0|0(+1C);m-68&21|+|16590253|16590274&offsets|0|0(+1C)]|TCCCTGAGACCCTAACTTGTGAC</td><td>>MI0000446|hsa-mir-125b-1&WithFlank&11|-|122099751|122099850</td><td>21</td><td>42</td><td>22(+1C)</td><td>TCCCTGAGACCCTAACTTGTGAC</td><td>444.0</td><td>12.0</td><td>1.7</td><td>7.0</td><td>3.1E-12</td><td>8.2E-12</td><td>MB231/MB468</td><td>MITO</td></tr>
<tr><td>>MI0000087|hsa-mir-29a&WithFlank&7|-|130876741|130876816@47.68.22(+1U)&[MIMAT0000086&hsa-miR-29a-3p&offsets|-1|-1(+1U);m-26&7|-|130876749|130876769&offsets|-1|0(+1U)]|CTAGCACCATCTGAAATCGGTTT</td><td>>MI0000087|hsa-mir-29a&WithFlank&7|-|130876741|130876816</td><td>47</td><td>68</td><td>22(+1U)</td><td>CTAGCACCATCTGAAATCGGTTT</td><td>409.4</td><td>10.0</td><td>1.4</td><td>7.0</td><td>2.5E-12</td><td>8.2E-12</td><td>MB231/MB468</td><td>CYTO</td></tr>
<tr><td>>hg38_21_+_8433172_8446622_RNA45SN1_45S_50ntFlanks@10916.10935.20|CCGCGGCTGGACGAGGCGCC</td><td>>hg38_21_+_8433172_8446622_RNA45SN1_45S_50ntFlanks</td><td>10916</td><td>10935</td><td>20</td><td>CCGCGGCTGGACGAGGCGCC</td><td>290.3</td><td>-6.4</td><td>0.9</td><td>-7.0</td><td>2.5E-12</td><td>8.2E-12</td><td>MB231/MB468</td><td>CYTO</td></tr>
<tr><td>>hg38_21_+_8433172_8446622_RNA45SN1_45S_50ntFlanks@10458.10494.37|ATGGCCTCCGTTGCCCTCGGCCGATCGAAAGGGAGTC</td><td>>hg38_21_+_8433172_8446622_RNA45SN1_45S_50ntFlanks</td><td>10458</td><td>10494</td><td>37</td><td>ATGGCCTCCGTTGCCCTCGGCCGATCGAAAGGGAGTC</td><td>120.4</td><td>-10.8</td><td>1.5</td><td>-7.0</td><td>2.3E-12</td><td>8.2E-12</td><td>MB231/MB468</td><td>MP</td></tr>
<tr><td>>hg38_21_+_8433172_8446622_RNA45SN1_45S_50ntFlanks@10349.10368.20|AACGAGAACTTTGAAGGCCG</td><td>>hg38_21_+_8433172_8446622_RNA45SN1_45S_50ntFlanks</td><td>10349</td><td>10368</td><td>20</td><td>AACGAGAACTTTGAAGGCCG</td><td>1771.5</td><td>-5.0</td><td>0.7</td><td>-7.0</td><td>3.1E-12</td><td>8.3E-12</td><td>MB231/MB468</td><td>MITO</td></tr>
<tr><td>>hg38_21_+_8433172_8446622_RNA45SN1_45S_50ntFlanks@11845.11875.31|GGAACGGGCTTGGCGGAATCAGCGGGGAAAG</td><td>>hg38_21_+_8433172_8446622_RNA45SN1_45S_50ntFlanks</td><td>11845</td><td>11875</td><td>31</td><td>GGAACGGGCTTGGCGGAATCAGCGGGGAAAG</td><td>171.4</td><td>-11.3</td><td>1.6</td><td>-7.0</td><td>2.3E-12</td><td>8.3E-12</td><td>MB231/MB468</td><td>MP</td></tr>
<tr><td>>MI0000081|hsa-mir-24-2&WithFlank&19|-|13836281|13836365@56.74.19(+1U);MI0000080|hsa-mir-24-1&WithFlank&9|+|95086015|95086094@50.68.19(+1U)&[MIMAT0000080&hsa-miR-24-3p&offsets|0|-3(+1U);m-36&19|-|13836288|13836310&offsets|0|-4(+1U)];[MIMAT0000080_1&hsa-miR-24-3p&offsets|0|-3(+1U);m-35&9|+|95086064|95086086&offsets|0|-4(+1U)]|TGGCTCAGTTCAGCAGGAAT</td><td>>MI0000081|hsa-mir-24-2&WithFlank&19|-|13836281|13836365</td><td>56</td><td>74</td><td>19(+1U)</td><td>TGGCTCAGTTCAGCAGGAAT</td><td>511.2</td><td>5.0</td><td>0.7</td><td>7.0</td><td>2.6E-12</td><td>8.3E-12</td><td>MB231/MB468</td><td>NUC</td></tr>
<tr><td>>hg38_21_+_8433172_8446622_RNA45SN1_45S_50ntFlanks@10403.10424.22|ACATGGGTCAGTCGGTCCTGAG</td><td>>hg38_21_+_8433172_8446622_RNA45SN1_45S_50ntFlanks</td><td>10403</td><td>10424</td><td>22</td><td>ACATGGGTCAGTCGGTCCTGAG</td><td>7145.7</td><td>5.8</td><td>0.8</td><td>7.4</td><td>1.3E-13</td><td>8.3E-12</td><td>BT20/MB468</td><td>MP</td></tr>
<tr><td>>hg38_21_+_8433172_8446622_RNA45SN1_45S_50ntFlanks@6734.6774.41|ATTGATCATCGACACTTCGAACGCACTTGCGGCCCCGGGTT</td><td>>hg38_21_+_8433172_8446622_RNA45SN1_45S_50ntFlanks</td><td>6734</td><td>6774</td><td>41</td><td>ATTGATCATCGACACTTCGAACGCACTTGCGGCCCCGGGTT</td><td>676.1</td><td>6.2</td><td>0.9</td><td>7.0</td><td>2.6E-12</td><td>8.3E-12</td><td>MB231/MB468</td><td>NUC</td></tr>
<tr><td>>hg38_21_+_8433172_8446622_RNA45SN1_45S_50ntFlanks@12203.12220.18|TAAGGCGAGCTCAGGGAG</td><td>>hg38_21_+_8433172_8446622_RNA45SN1_45S_50ntFlanks</td><td>12203</td><td>12220</td><td>18</td><td>TAAGGCGAGCTCAGGGAG</td><td>195.7</td><td>-11.5</td><td>1.6</td><td>-7.0</td><td>2.3E-12</td><td>8.3E-12</td><td>MB231/MB468</td><td>MP</td></tr>
<tr><td>>hg38_21_+_8433172_8446622_RNA45SN1_45S_50ntFlanks@4657.4680.24|CTAGAGGTGAAATTCTTGGACCGG</td><td>>hg38_21_+_8433172_8446622_RNA45SN1_45S_50ntFlanks</td><td>4657</td><td>4680</td><td>24</td><td>CTAGAGGTGAAATTCTTGGACCGG</td><td>146.7</td><td>-11.1</td><td>1.6</td><td>-7.0</td><td>2.3E-12</td><td>8.3E-12</td><td>MB231/MB468</td><td>MP</td></tr>
<tr><td>>hg38_21_+_8433172_8446622_RNA45SN1_45S_50ntFlanks@6652.6682.31|GACTCTTAGCGGTGGATCACTCGGCTCGTGC</td><td>>hg38_21_+_8433172_8446622_RNA45SN1_45S_50ntFlanks</td><td>6652</td><td>6682</td><td>31</td><td>GACTCTTAGCGGTGGATCACTCGGCTCGTGC</td><td>3422.6</td><td>5.3</td><td>0.8</td><td>7.0</td><td>1.9E-12</td><td>8.3E-12</td><td>BT20/MB468</td><td>MITO</td></tr>
<tr><td>>hg38_21_+_8433172_8446622_RNA45SN1_45S_50ntFlanks@10734.10755.22|GTGTAAATCTCGCGCCGGGCCG</td><td>>hg38_21_+_8433172_8446622_RNA45SN1_45S_50ntFlanks</td><td>10734</td><td>10755</td><td>22</td><td>GTGTAAATCTCGCGCCGGGCCG</td><td>100.2</td><td>-10.5</td><td>1.5</td><td>-7.0</td><td>2.3E-12</td><td>8.3E-12</td><td>MB231/MB468</td><td>MP</td></tr>
<tr><td>>hg38_21_+_8433172_8446622_RNA45SN1_45S_50ntFlanks@12336.12365.30|CAGGAGGTGTCAGAAAAGTTACCACAGGGA</td><td>>hg38_21_+_8433172_8446622_RNA45SN1_45S_50ntFlanks</td><td>12336</td><td>12365</td><td>30</td><td>CAGGAGGTGTCAGAAAAGTTACCACAGGGA</td><td>99.6</td><td>-10.5</td><td>1.5</td><td>-7.0</td><td>2.3E-12</td><td>8.3E-12</td><td>MB231/MB468</td><td>MP</td></tr>
<tr><td>>am_MI0005763|hsa-mir-941-1&WithFlank&20|+|63919443|63919526@53.74.22;MI0005764|hsa-mir-941-2&WithFlank&20|+|63919499|63919582@53.74.22;MI0005765|hsa-mir-941-3&WithFlank&20|+|63919555|63919638@53.74.22;MI0005766|hsa-mir-941-4&WithFlank&20|+|63919750|63919833@53.74.22;MI0031520|hsa-mir-941-5&WithFlank&20|+|63919862|63919945@53.74.22&[MIMAT0004984&hsa-miR-941&offsets|0|-1;m-263&20|+|63919495|63919517&offsets|0|-1];[MIMAT0004984_1&hsa-miR-941&offsets|0|-1;m-263&20|+|63919551|63919573&offsets|0|-1];[MIMAT0004984_2&hsa-miR-941&offsets|0|-1;m-263&20|+|63919607|63919629&offsets|0|-1];[MIMAT0004984_3&hsa-miR-941&offsets|0|-1;m-263&20|+|63919802|63919824&offsets|0|-1];[MIMAT0004984_4&hsa-miR-941&offsets|0|-1;m-263&20|+|63919914|63919936&offsets|0|-1]|CACCCGGCTGTGTGCACATGTG</td><td>>am_MI0005763|hsa-mir-941-1&WithFlank&20|+|63919443|63919526</td><td>53</td><td>74</td><td>22</td><td>CACCCGGCTGTGTGCACATGTG</td><td>846.5</td><td>6.6</td><td>0.9</td><td>7.0</td><td>2.6E-12</td><td>8.3E-12</td><td>MB231/MB468</td><td>CYTO</td></tr>
<tr><td>>hg38_1_-_228634819_228635039_RNA5S12_5S_50ntFlanks@150.169.20|AATACCGGGTGCTGTAGGCT</td><td>>hg38_1_-_228634819_228635039_RNA5S12_5S_50ntFlanks</td><td>150</td><td>169</td><td>20</td><td>AATACCGGGTGCTGTAGGCT</td><td>4573.8</td><td>-5.3</td><td>0.8</td><td>-7.0</td><td>2.3E-12</td><td>8.3E-12</td><td>MB231/MB468</td><td>MP</td></tr>
<tr><td>>hg38_21_+_8433172_8446622_RNA45SN1_45S_50ntFlanks@5473.5496.24|CGGCCCTGGCGGAGCGCTGAGAAG</td><td>>hg38_21_+_8433172_8446622_RNA45SN1_45S_50ntFlanks</td><td>5473</td><td>5496</td><td>24</td><td>CGGCCCTGGCGGAGCGCTGAGAAG</td><td>158.2</td><td>6.8</td><td>1.0</td><td>7.0</td><td>2.6E-12</td><td>8.4E-12</td><td>MB231/MB468</td><td>CYTO</td></tr>
<tr><td>>hg38_21_+_8433172_8446622_RNA45SN1_45S_50ntFlanks@7973.8012.40|GACGCGACCTCAGATCAGACGTGGCGACCCGCTGAATTTA</td><td>>hg38_21_+_8433172_8446622_RNA45SN1_45S_50ntFlanks</td><td>7973</td><td>8012</td><td>40</td><td>GACGCGACCTCAGATCAGACGTGGCGACCCGCTGAATTTA</td><td>427.2</td><td>4.7</td><td>0.7</td><td>7.0</td><td>2.6E-12</td><td>8.4E-12</td><td>MB231/MB468</td><td>CYTO</td></tr>
<tr><td>>hg38_21_+_8433172_8446622_RNA45SN1_45S_50ntFlanks@11718.11751.34|AACGGCGGGAGTAACTATGACTCTCTTAAGGTAG</td><td>>hg38_21_+_8433172_8446622_RNA45SN1_45S_50ntFlanks</td><td>11718</td><td>11751</td><td>34</td><td>AACGGCGGGAGTAACTATGACTCTCTTAAGGTAG</td><td>1120.7</td><td>-9.9</td><td>1.4</td><td>-7.0</td><td>3.4E-12</td><td>8.4E-12</td><td>BT20/MB468</td><td>NUC</td></tr>
<tr><td>>hg38_21_+_8433172_8446622_RNA45SN1_45S_50ntFlanks@12973.12992.20|TCGTACGTAGCAGAGCAGCT</td><td>>hg38_21_+_8433172_8446622_RNA45SN1_45S_50ntFlanks</td><td>12973</td><td>12992</td><td>20</td><td>TCGTACGTAGCAGAGCAGCT</td><td>4093.1</td><td>-10.9</td><td>1.6</td><td>-7.0</td><td>3.4E-12</td><td>8.4E-12</td><td>BT20/MB468</td><td>NUC</td></tr>
<tr><td>>am_MI0005763|hsa-mir-941-1&WithFlank&20|+|63919443|63919526@53.75.23;MI0005764|hsa-mir-941-2&WithFlank&20|+|63919499|63919582@53.75.23;MI0005765|hsa-mir-941-3&WithFlank&20|+|63919555|63919638@53.75.23;MI0005766|hsa-mir-941-4&WithFlank&20|+|63919750|63919833@53.75.23;MI0031520|hsa-mir-941-5&WithFlank&20|+|63919862|63919945@53.75.23&[MIMAT0004984&hsa-miR-941&offsets|0|0;m-263&20|+|63919495|63919517&offsets|0|0];[MIMAT0004984_1&hsa-miR-941&offsets|0|0;m-263&20|+|63919551|63919573&offsets|0|0];[MIMAT0004984_2&hsa-miR-941&offsets|0|0;m-263&20|+|63919607|63919629&offsets|0|0];[MIMAT0004984_3&hsa-miR-941&offsets|0|0;m-263&20|+|63919802|63919824&offsets|0|0];[MIMAT0004984_4&hsa-miR-941&offsets|0|0;m-263&20|+|63919914|63919936&offsets|0|0]|CACCCGGCTGTGTGCACATGTGC</td><td>>am_MI0005763|hsa-mir-941-1&WithFlank&20|+|63919443|63919526</td><td>53</td><td>75</td><td>23</td><td>CACCCGGCTGTGTGCACATGTGC</td><td>12540.9</td><td>5.1</td><td>0.7</td><td>7.0</td><td>2.7E-12</td><td>8.4E-12</td><td>MB231/MB468</td><td>NUC</td></tr>
<tr><td>>hg38_21_+_8433172_8446622_RNA45SN1_45S_50ntFlanks@10436.10453.18|CGCCGTTCCGAAGGGACG</td><td>>hg38_21_+_8433172_8446622_RNA45SN1_45S_50ntFlanks</td><td>10436</td><td>10453</td><td>18</td><td>CGCCGTTCCGAAGGGACG</td><td>389.8</td><td>-11.7</td><td>1.6</td><td>-7.4</td><td>1.3E-13</td><td>8.4E-12</td><td>BT20/MB468</td><td>MP</td></tr>
</table>

<table>
<tr><td>>trna10_AspGTC_12_-_125424193_125424264@37.74.38,trna12_AspGTC_12_-_125411891_125411962@37.74.38,trna144_AspGTC_6_-_27551236_27551307@37.74.38,trna38_AspGTC_17_-_8125556_8125627@37.74.38,trna45_AspGTC_6_+_27447453_27447524@37.74.38,trna48_AspGTC_6_+_27471523_27471594@37.74.38,trna4_AspGTC_12_+_96429799_96429870@37.74.38,trna69_AspGTC_1_-_161440205_161440276@37.74.38,trna72_AspGTC_1_-_161432824_161432895@37.74.38,trna75_AspGTC_1_-_161425414_161425485@37.74.38,trna78_AspGTC_1_-_161418033_161418104@37.74.38,trna81_AspGTC_1_-_161410615_161410686@37.74.38|ACGCGGGAGACCGGGGTTCGATTCCCCGACGGGGAGCC</td><td>>trna10_AspGTC_12_-_125424193_125424264</td><td>37</td><td>74</td><td>38</td><td>ACGCGGGAGACCGGGGTTCGATTCCCCGACGGGGAGCC</td><td>442.4</td><td>4.9</td><td>0.7</td><td>7.0</td><td>2.5E-12</td><td>8.4E-12</td><td>BT20/MB468</td><td>CYTO</td></tr>
<tr><td>>hg38_21_+_8433172_8446622_RNA45SN1_45S_50ntFlanks@11879.11904.26|ACCCTGTTGAGCTTGACTCTAGTCTG</td><td>>hg38_21_+_8433172_8446622_RNA45SN1_45S_50ntFlanks</td><td>11879</td><td>11904</td><td>26</td><td>ACCCTGTTGAGCTTGACTCTAGTCTG</td><td>161.6</td><td>-11.2</td><td>1.6</td><td>-7.0</td><td>2.3E-12</td><td>8.5E-12</td><td>MB231/MB468</td><td>MP</td></tr>
<tr><td>>hg38_21_+_8433172_8446622_RNA45SN1_45S_50ntFlanks@12151.12179.29|GTGGGGAGTTTGACTGGGGCGGTACACCT</td><td>>hg38_21_+_8433172_8446622_RNA45SN1_45S_50ntFlanks</td><td>12151</td><td>12179</td><td>29</td><td>GTGGGGAGTTTGACTGGGGCGGTACACCT</td><td>194.2</td><td>-5.3</td><td>0.8</td><td>-7.0</td><td>2.6E-12</td><td>8.5E-12</td><td>MB231/MB468</td><td>CYTO</td></tr>
<tr><td>>hg38_21_+_8433172_8446622_RNA45SN1_45S_50ntFlanks@6721.6746.26|AATTGCAGGACACATTGATCATCGAC</td><td>>hg38_21_+_8433172_8446622_RNA45SN1_45S_50ntFlanks</td><td>6721</td><td>6746</td><td>26</td><td>AATTGCAGGACACATTGATCATCGAC</td><td>3666.2</td><td>-10.8</td><td>1.6</td><td>-7.0</td><td>3.4E-12</td><td>8.5E-12</td><td>BT20/MB468</td><td>NUC</td></tr>
<tr><td>>trna116_GluCTC_1_-_145399233_145399304@32.69.38,trna59_GluCTC_1_+_249168447_249168518@32.69.38,trna71_GluCTC_1_-_161439189_161439260@32.69.38,trna74_GluCTC_1_-_161431809_161431880@32.69.38,trna77_GluCTC_1_-_161424398_161424469@32.69.38,trna77_GluCTC_6_+_28949976_28950047@32.69.38,trna80_GluCTC_1_-_161417018_161417089@32.69.38,trna87_GluCTC_6_-_126101393_126101464@32.69.38|CTCTCACCGCCGCGGCCCGGGTTCGATTCCCGGTCAGG</td><td>>trna116_GluCTC_1_-_145399233_145399304</td><td>32</td><td>69</td><td>38</td><td>CTCTCACCGCCGCGGCCCGGGTTCGATTCCCGGTCAGG</td><td>235.6</td><td>-11.7</td><td>1.7</td><td>-7.0</td><td>3.2E-12</td><td>8.5E-12</td><td>MB231/MB468</td><td>MITO</td></tr>
<tr><td>>hg38_21_+_8433172_8446622_RNA45SN1_45S_50ntFlanks@11705.11740.36|TGAAGCGCGGGTAAACGGCGGGAGTAACTATGACTC</td><td>>hg38_21_+_8433172_8446622_RNA45SN1_45S_50ntFlanks</td><td>11705</td><td>11740</td><td>36</td><td>TGAAGCGCGGGTAAACGGCGGGAGTAACTATGACTC</td><td>672.4</td><td>-9.6</td><td>1.4</td><td>-7.0</td><td>3.5E-12</td><td>8.5E-12</td><td>BT20/MB468</td><td>NUC</td></tr>
<tr><td>>hg38_21_+_8433172_8446622_RNA45SN1_45S_50ntFlanks@9562.9584.23|TCCTGACGTGCAAATCGGTCGTC</td><td>>hg38_21_+_8433172_8446622_RNA45SN1_45S_50ntFlanks</td><td>9562</td><td>9584</td><td>23</td><td>TCCTGACGTGCAAATCGGTCGTC</td><td>115.0</td><td>-10.7</td><td>1.5</td><td>-7.0</td><td>2.3E-12</td><td>8.5E-12</td><td>MB231/MB468</td><td>MP</td></tr>
<tr><td>>hg38_1_-_228634819_228635039_RNA5S12_5S_50ntFlanks@144.171.28|CCTGGGAATACCGGGTGCTGTAGGCTTT</td><td>>hg38_1_-_228634819_228635039_RNA5S12_5S_50ntFlanks</td><td>144</td><td>171</td><td>28</td><td>CCTGGGAATACCGGGTGCTGTAGGCTTT</td><td>7839.6</td><td>6.1</td><td>0.9</td><td>7.0</td><td>2.6E-12</td><td>8.5E-12</td><td>MB231/MB468</td><td>CYTO</td></tr>
<tr><td>>hg38_21_+_8433172_8446622_RNA45SN1_45S_50ntFlanks@8144.8164.21|CTCCCCGGCGCCGCTCGTGGG</td><td>>hg38_21_+_8433172_8446622_RNA45SN1_45S_50ntFlanks</td><td>8144</td><td>8164</td><td>21</td><td>CTCCCCGGCGCCGCTCGTGGG</td><td>308.6</td><td>-12.1</td><td>1.7</td><td>-7.0</td><td>2.3E-12</td><td>8.5E-12</td><td>MB231/MB468</td><td>MP</td></tr>
<tr><td>>hg38_21_+_8433172_8446622_RNA45SN1_45S_50ntFlanks@3870.3898.29|AGCTAATACATGCCGACGGGCGCTGACCC</td><td>>hg38_21_+_8433172_8446622_RNA45SN1_45S_50ntFlanks</td><td>3870</td><td>3898</td><td>29</td><td>AGCTAATACATGCCGACGGGCGCTGACCC</td><td>384.7</td><td>-11.6</td><td>1.7</td><td>-7.0</td><td>2.7E-12</td><td>8.5E-12</td><td>MB231/MB468</td><td>NUC</td></tr>
<tr><td>>hg38_21_+_8433172_8446622_RNA45SN1_45S_50ntFlanks@11671.11703.33|CCAGTGCTCTGAATGTCAAAGTGAAGAAATTCA</td><td>>hg38_21_+_8433172_8446622_RNA45SN1_45S_50ntFlanks</td><td>11671</td><td>11703</td><td>33</td><td>CCAGTGCTCTGAATGTCAAAGTGAAGAAATTCA</td><td>1402.8</td><td>-3.8</td><td>0.5</td><td>-7.0</td><td>2.7E-12</td><td>8.6E-12</td><td>MB231/MB468</td><td>NUC</td></tr>
<tr><td>>hg38_21_+_8433172_8446622_RNA45SN1_45S_50ntFlanks@11565.11591.27|CTAGCAGCCGACTTAGAACTGGTGCGG</td><td>>hg38_21_+_8433172_8446622_RNA45SN1_45S_50ntFlanks</td><td>11565</td><td>11591</td><td>27</td><td>CTAGCAGCCGACTTAGAACTGGTGCGG</td><td>168.4</td><td>-11.3</td><td>1.6</td><td>-7.0</td><td>2.3E-12</td><td>8.6E-12</td><td>MB231/MB468</td><td>MP</td></tr>
<tr><td>>hg38_21_+_8433172_8446622_RNA45SN1_45S_50ntFlanks@8839.8856.18|CGGTGGCGGCGGCGGCGG</td><td>>hg38_21_+_8433172_8446622_RNA45SN1_45S_50ntFlanks</td><td>8839</td><td>8856</td><td>18</td><td>CGGTGGCGGCGGCGGCGG</td><td>126.3</td><td>-10.8</td><td>1.5</td><td>-7.0</td><td>2.4E-12</td><td>8.6E-12</td><td>MB231/MB468</td><td>MP</td></tr>
<tr><td>>MI0000061|hsa-let-7a-2&WithFlank&11|-|122146516|122146599@11.30.20(+1C);MI0000062|hsa-let-7a-3&WithFlank&22|+|46112743|46112828@10.29.20(+1C);MI0000060|hsa-let-7a-1&WithFlank&9|+|94175951|94176042@12.31.20(+1C)&[MIMAT0000062&hsa-let-7a-5p&offsets|0|-2(+1C);m-6&11|-|122146568|122146589&offsets|0|-2(+1C)];[MIMAT0000062_1&hsa-let-7a-5p&offsets|0|-2(+1C);m-5&22|+|46112752|46112773&offsets|0|-2(+1C)];[MIMAT0000062_2&hsa-let-7a-5p&offsets|0|-2(+1C);m-4&9|+|94175962|94175983&offsets|0|-2(+1C)]|TGAGGTAGTAGGTTGTATAGC</td><td>>MI0000061|hsa-let-7a-2&WithFlank&11|-|122146516|122146599</td><td>11</td><td>30</td><td>20(+1C)</td><td>TGAGGTAGTAGGTTGTATAGC</td><td>340.9</td><td>2.7</td><td>0.4</td><td>7.0</td><td>3.5E-12</td><td>8.6E-12</td><td>BT20/MB468</td><td>NUC</td></tr>
<tr><td>>hg38_21_+_8433172_8446622_RNA45SN1_45S_50ntFlanks@12399.12419.21|ACGTCGCTTTTTGATCCTTCG</td><td>>hg38_21_+_8433172_8446622_RNA45SN1_45S_50ntFlanks</td><td>12399</td><td>12419</td><td>21</td><td>ACGTCGCTTTTTGATCCTTCG</td><td>109.0</td><td>-10.6</td><td>1.5</td><td>-7.0</td><td>2.4E-12</td><td>8.6E-12</td><td>MB231/MB468</td><td>MP</td></tr>
<tr><td>>hg38_21_+_8433172_8446622_RNA45SN1_45S_50ntFlanks@4002.4023.22|GATAACCTCGGGCCGATCGCAC</td><td>>hg38_21_+_8433172_8446622_RNA45SN1_45S_50ntFlanks</td><td>4002</td><td>4023</td><td>22</td><td>GATAACCTCGGGCCGATCGCAC</td><td>3464.4</td><td>-6.4</td><td>0.9</td><td>-7.0</td><td>3.5E-12</td><td>8.6E-12</td><td>BT20/MB468</td><td>NUC</td></tr>
<tr><td>>hg38_21_+_8433172_8446622_RNA45SN1_45S_50ntFlanks@12459.12476.18|AGCGTTGGATTGTTCACC</td><td>>hg38_21_+_8433172_8446622_RNA45SN1_45S_50ntFlanks</td><td>12459</td><td>12476</td><td>18</td><td>AGCGTTGGATTGTTCACC</td><td>2106.4</td><td>-5.0</td><td>0.7</td><td>-7.0</td><td>2.7E-12</td><td>8.6E-12</td><td>MB231/MB468</td><td>NUC</td></tr>
<tr><td>>am_trnaMT_ArgTCG_MT_+_10405_10469@28.48.21|ATTTCGACTCATTAAATTATG</td><td>>am_trnaMT_ArgTCG_MT_+_10405_10469</td><td>28</td><td>48</td><td>21</td><td>ATTTCGACTCATTAAATTATG</td><td>452.4</td><td>-11.9</td><td>1.6</td><td>-7.4</td><td>1.3E-13</td><td>8.7E-12</td><td>BT20/MB468</td><td>MP</td></tr>
<tr><td>>hg38_21_+_8433172_8446622_RNA45SN1_45S_50ntFlanks@12676.12702.27|AGAATCCCGCCCAGGCGGAACGATACG</td><td>>hg38_21_+_8433172_8446622_RNA45SN1_45S_50ntFlanks</td><td>12676</td><td>12702</td><td>27</td><td>AGAATCCCGCCCAGGCGGAACGATACG</td><td>386.8</td><td>-12.5</td><td>1.8</td><td>-7.0</td><td>2.4E-12</td><td>8.7E-12</td><td>MB231/MB468</td><td>MP</td></tr>
<tr><td>>MI0000102|hsa-mir-100&WithFlank&11|-|122152223|122152314@19.40.22(+3A)&[MIMAT0000098&hsa-miR-100-5p&offsets|0|0(+3A);m-27&11|-|122152276|122152296&offsets|0|+1(+3A)]|AACCCGTAGATCCGAACTTGTGAAA</td><td>>MI0000102|hsa-mir-100&WithFlank&11|-|122152223|122152314</td><td>19</td><td>40</td><td>22(+3A)</td><td>AACCCGTAGATCCGAACTTGTGAAA</td><td>224.7</td><td>11.0</td><td>1.6</td><td>7.0</td><td>3.3E-12</td><td>8.7E-12</td><td>MB231/MB468</td><td>MITO</td></tr>
<tr><td>>hg38_21_+_8433172_8446622_RNA45SN1_45S_50ntFlanks@5515.5533.19|CTAGAGGAAGTAAAAGTCG</td><td>>hg38_21_+_8433172_8446622_RNA45SN1_45S_50ntFlanks</td><td>5515</td><td>5533</td><td>19</td><td>CTAGAGGAAGTAAAAGTCG</td><td>294.2</td><td>-11.3</td><td>1.5</td><td>-7.4</td><td>1.3E-13</td><td>8.7E-12</td><td>BT20/MB468</td><td>MP</td></tr>
<tr><td>>trnaMT_PheGAA_MT_+_577_647@39.62.24|ATGTTTAGACGGGCTCACATCACC</td><td>>trnaMT_PheGAA_MT_+_577_647</td><td>39</td><td>62</td><td>24</td><td>ATGTTTAGACGGGCTCACATCACC</td><td>303.5</td><td>5.2</td><td>0.7</td><td>7.0</td><td>2.8E-12</td><td>8.7E-12</td><td>MB231/MB468</td><td>NUC</td></tr>
<tr><td>>hg38_21_+_8433172_8446622_RNA45SN1_45S_50ntFlanks@4760.4778.19|TCAGATACCGTCGTAGTTC</td><td>>hg38_21_+_8433172_8446622_RNA45SN1_45S_50ntFlanks</td><td>4760</td><td>4778</td><td>19</td><td>TCAGATACCGTCGTAGTTC</td><td>653.3</td><td>-5.7</td><td>0.8</td><td>-7.0</td><td>2.7E-12</td><td>8.7E-12</td><td>MB231/MB468</td><td>CYTO</td></tr>
<tr><td>>MI0001445|hsa-mir-423&WithFlank&17|+|30117073|30117178@58.79.22&[MIMAT0001340&hsa-miR-423-3p&offsets|-1|-2;m-100&17|+|30117131|30117153&offsets|-1|-2]|AAGCTCGGTCTGAGGCCCCTCA</td><td>>MI0001445|hsa-mir-423&WithFlank&17|+|30117073|30117178</td><td>58</td><td>79</td><td>22</td><td>AAGCTCGGTCTGAGGCCCCTCA</td><td>698.9</td><td>7.8</td><td>1.1</td><td>7.0</td><td>2.0E-12</td><td>8.7E-12</td><td>BT20/MB468</td><td>MITO</td></tr>
<tr><td>>MI0000750|hsa-mir-26a-2&WithFlank&12|-|57824603|57824698@20.41.22;MI0000083|hsa-mir-26a-1&WithFlank&3|+|37969398|37969486@16.37.22&[MIMAT0000082&hsa-miR-26a-5p&offsets|0|0;m-33&12|-|57824658|57824679&offsets|0|0];[MIMAT0000082_1&hsa-miR-26a-5p&offsets|0|0;m-34&3|+|37969413|37969434&offsets|0|0]|TTCAAGTAATCCAGGATAGGCT</td><td>>MI0000750|hsa-mir-26a-2&WithFlank&12|-|57824603|57824698</td><td>20</td><td>41</td><td>22</td><td>TTCAAGTAATCCAGGATAGGCT</td><td>82812.5</td><td>4.9</td><td>0.7</td><td>7.0</td><td>2.8E-12</td><td>8.7E-12</td><td>MB231/MB468</td><td>NUC</td></tr>
<tr><td>>hg38_21_+_8433172_8446622_RNA45SN1_45S_50ntFlanks@3939.3959.21|AACCCGGTCAGCCCCTCTCCG</td><td>>hg38_21_+_8433172_8446622_RNA45SN1_45S_50ntFlanks</td><td>3939</td><td>3959</td><td>21</td><td>AACCCGGTCAGCCCCTCTCCG</td><td>900.8</td><td>-9.5</td><td>1.4</td><td>-7.0</td><td>2.8E-12</td><td>8.7E-12</td><td>MB231/MB468</td><td>NUC</td></tr>
<tr><td>>hg38_21_+_8433172_8446622_RNA45SN1_45S_50ntFlanks@8799.8823.25|CCTCGGGAGGGCGCGCGGGTCGGGG</td><td>>hg38_21_+_8433172_8446622_RNA45SN1_45S_50ntFlanks</td><td>8799</td><td>8823</td><td>25</td><td>CCTCGGGAGGGCGCGCGGGTCGGGG</td><td>425.0</td><td>-11.8</td><td>1.6</td><td>-7.4</td><td>1.3E-13</td><td>8.8E-12</td><td>BT20/MB468</td><td>MP</td></tr>
<tr><td>>am_MI0000786|hsa-mir-378a&WithFlank&5|+|149732819|149732896@50.70.21(+1U)&[MIMAT0000732&hsa-miR-378a-3p&offsets|+1|0(+1U);m-44&5|+|149732867|149732888&offsets|+1|0(+1U)]|CTGGACTTGGAGTCAGAAGGCT</td><td>>am_MI0000786|hsa-mir-378a&WithFlank&5|+|149732819|149732896</td><td>50</td><td>70</td><td>21(+1U)</td><td>CTGGACTTGGAGTCAGAAGGCT</td><td>639.5</td><td>5.2</td><td>0.7</td><td>7.0</td><td>2.0E-12</td><td>8.8E-12</td><td>BT20/MB468</td><td>MITO</td></tr>
<tr><td>>hg38_21_+_8433172_8446622_RNA45SN1_45S_50ntFlanks@6740.6772.33|CATCGACACTTCGAACGCACTTGCGGCCCCGGG</td><td>>hg38_21_+_8433172_8446622_RNA45SN1_45S_50ntFlanks</td><td>6740</td><td>6772</td><td>33</td><td>CATCGACACTTCGAACGCACTTGCGGCCCCGGG</td><td>1213.0</td><td>-4.3</td><td>0.6</td><td>-7.0</td><td>2.0E-12</td><td>8.8E-12</td><td>BT20/MB468</td><td>MITO</td></tr>
<tr><td>>hg38_21_+_8433172_8446622_RNA45SN1_45S_50ntFlanks@10420.10453.34|CTGAGAGATGGGCGAGCGCCGTTCCGAAGGGACG</td><td>>hg38_21_+_8433172_8446622_RNA45SN1_45S_50ntFlanks</td><td>10420</td><td>10453</td><td>34</td><td>CTGAGAGATGGGCGAGCGCCGTTCCGAAGGGACG</td><td>1402.8</td><td>-4.4</td><td>0.6</td><td>-7.0</td><td>2.6E-12</td><td>8.8E-12</td><td>BT20/MB468</td><td>CYTO</td></tr>
<tr><td>>trna104_AlaTGC_6_-_28785012_28785083@58.75.18,trna107_AlaTGC_6_-_28770577_28770647@57.74.18|TCCCCGACACCTCCACCA</td><td>>trna104_AlaTGC_6_-_28785012_28785083</td><td>58</td><td>75</td><td>18</td><td>TCCCCGACACCTCCACCA</td><td>346.5</td><td>9.8</td><td>1.4</td><td>7.0</td><td>2.6E-12</td><td>8.8E-12</td><td>BT20/MB468</td><td>CYTO</td></tr>
<tr><td>>hg38_21_+_8433172_8446622_RNA45SN1_45S_50ntFlanks@9623.9656.34|TAGTAGCTGGTTCCCTCCGAAGTTTCCCTCAGGA</td><td>>hg38_21_+_8433172_8446622_RNA45SN1_45S_50ntFlanks</td><td>9623</td><td>9656</td><td>34</td><td>TAGTAGCTGGTTCCCTCCGAAGTTTCCCTCAGGA</td><td>606.0</td><td>-8.5</td><td>1.2</td><td>-7.0</td><td>3.4E-12</td><td>8.8E-12</td><td>MB231/MB468</td><td>MITO</td></tr>
<tr><td>>hg38_21_+_8433172_8446622_RNA45SN1_45S_50ntFlanks@12285.12318.34|CAGACCGTGAAAGCGGGGCCTCACGATCCTTCTG</td><td>>hg38_21_+_8433172_8446622_RNA45SN1_45S_50ntFlanks</td><td>12285</td><td>12318</td><td>34</td><td>CAGACCGTGAAAGCGGGGCCTCACGATCCTTCTG</td><td>368.0</td><td>-10.6</td><td>1.5</td><td>-7.0</td><td>2.0E-12</td><td>8.8E-12</td><td>BT20/MB468</td><td>MITO</td></tr>
<tr><td>>hg38_21_+_8433172_8446622_RNA45SN1_45S_50ntFlanks@11207.11224.18|GGCGGCGACTCTGGACGC</td><td>>hg38_21_+_8433172_8446622_RNA45SN1_45S_50ntFlanks</td><td>11207</td><td>11224</td><td>18</td><td>GGCGGCGACTCTGGACGC</td><td>110.7</td><td>-10.7</td><td>1.5</td><td>-7.0</td><td>2.4E-12</td><td>8.8E-12</td><td>MB231/MB468</td><td>MP</td></tr>
<tr><td>>hg38_21_+_8433172_8446622_RNA45SN1_45S_50ntFlanks@12183.12211.29|AAACGGTAACGCAGGTGTCCTAAGGCGAG</td><td>>hg38_21_+_8433172_8446622_RNA45SN1_45S_50ntFlanks</td><td>12183</td><td>12211</td><td>29</td><td>AAACGGTAACGCAGGTGTCCTAAGGCGAG</td><td>124.0</td><td>-10.8</td><td>1.5</td><td>-7.0</td><td>2.4E-12</td><td>8.8E-12</td><td>MB231/MB468</td><td>MP</td></tr>
<tr><td>>hg38_21_+_8433172_8446622_RNA45SN1_45S_50ntFlanks@4899.4918.20|AAGGAATTGACGGAAGGGCA</td><td>>hg38_21_+_8433172_8446622_RNA45SN1_45S_50ntFlanks</td><td>4899</td><td>4918</td><td>20</td><td>AAGGAATTGACGGAAGGGCA</td><td>220.7</td><td>-11.6</td><td>1.7</td><td>-7.0</td><td>2.4E-12</td><td>8.8E-12</td><td>MB231/MB468</td><td>MP</td></tr>
<tr><td>>trna111_HisGTG_1_-_147774845_147774916@-1A.29.30,trna118_HisGTG_1_-_145396881_145396952@-1A.29.30,trna16_HisGTG_1_+_146544773_146544844@-1A.29.30,trna1_HisGTG_15_+_45493349_45493420@-1A.29.30,trna21_HisGTG_1_+_147753471_147753542@-1A.29.30,trna33_HisGTG_6_+_27125906_27125977@-1A.29.30,trna7_HisGTG_9_-_14433938_14434009@-1A.29.30,trna8_HisGTG_15_-_45492611_45492682@-1A.29.30,trna9_HisGTG_15_-_45490804_45490875@-1A.29.30|AGCCGTGATCGTATAGTGGTTAGTACTCTG</td><td>>trna111_HisGTG_1_-_147774845_147774916</td><td>-1A</td><td>29</td><td>30</td><td>AGCCGTGATCGTATAGTGGTTAGTACTCTG</td><td>1183.3</td><td>-5.5</td><td>0.8</td><td>-7.0</td><td>2.7E-12</td><td>8.8E-12</td><td>MB231/MB468</td><td>CYTO</td></tr>
<tr><td>>hg38_21_+_8433172_8446622_RNA45SN1_45S_50ntFlanks@3850.3873.24|GATAACTGTGGTAATTCTAGAGCT</td><td>>hg38_21_+_8433172_8446622_RNA45SN1_45S_50ntFlanks</td><td>3850</td><td>3873</td><td>24</td><td>GATAACTGTGGTAATTCTAGAGCT</td><td>636.5</td><td>-5.0</td><td>0.7</td><td>-7.0</td><td>2.4E-12</td><td>8.9E-12</td><td>MB231/MB468</td><td>MP</td></tr>
<tr><td>>hg38_21_+_8433172_8446622_RNA45SN1_45S_50ntFlanks@3728.3748.21|CATATGCTTGTCTCAAAGATT</td><td>>hg38_21_+_8433172_8446622_RNA45SN1_45S_50ntFlanks</td><td>3728</td><td>3748</td><td>21</td><td>CATATGCTTGTCTCAAAGATT</td><td>91.0</td><td>-10.4</td><td>1.5</td><td>-7.0</td><td>2.4E-12</td><td>8.9E-12</td><td>MB231/MB468</td><td>MP</td></tr>
<tr><td>>hg38_21_+_8433172_8446622_RNA45SN1_45S_50ntFlanks@4895.4917.23|CTTAAAGGAATTGACGGAAGGGC</td><td>>hg38_21_+_8433172_8446622_RNA45SN1_45S_50ntFlanks</td><td>4895</td><td>4917</td><td>23</td><td>CTTAAAGGAATTGACGGAAGGGC</td><td>188.1</td><td>-11.4</td><td>1.6</td><td>-7.0</td><td>2.4E-12</td><td>8.9E-12</td><td>MB231/MB468</td><td>MP</td></tr>
<tr><td>>hg38_21_+_8433172_8446622_RNA45SN1_45S_50ntFlanks@11324.11344.21|GTCTCGTCGCGCGCGCGTCCG</td><td>>hg38_21_+_8433172_8446622_RNA45SN1_45S_50ntFlanks</td><td>11324</td><td>11344</td><td>21</td><td>GTCTCGTCGCGCGCGCGTCCG</td><td>432.2</td><td>-8.7</td><td>1.2</td><td>-7.0</td><td>3.4E-12</td><td>8.9E-12</td><td>MB231/MB468</td><td>MITO</td></tr>
</table>

| | | | | | | | | | | | | | |
|---|---|---|---|---|---|---|---|---|---|---|---|---|---|
| >hg38_21_+_8433172_8446622_RNA45SN1_45S_50ntFlanks@8357.8378.22\|AAAAGAACTTTGAAGAGAGAGT | >hg38_21_+_8433172_8446622_RNA45SN1_45S_50ntFlanks | 8357 | 8378 | 22 | AAAAGAACTTTGAAGAGAGAGT | 31720.4 | -8.1 | 1.1 | -7.4 | 1.4E-13 | 8.9E-12 | BT20/MB468 | MP |
| >hg38_21_+_8433172_8446622_RNA45SN1_45S_50ntFlanks@9304.9323.20\|ACCAAGGAGTCTAACACGTG | >hg38_21_+_8433172_8446622_RNA45SN1_45S_50ntFlanks | 9304 | 9323 | 20 | ACCAAGGAGTCTAACACGTG | 210.8 | -11.6 | 1.7 | -7.0 | 2.4E-12 | 8.9E-12 | MB231/MB468 | MP |
| >hg38_21_+_8433172_8446622_RNA45SN1_45S_50ntFlanks@12044.12064.21\|ACTGACCCGGTGAGGCGGGGG | >hg38_21_+_8433172_8446622_RNA45SN1_45S_50ntFlanks | 12044 | 12064 | 21 | ACTGACCCGGTGAGGCGGGGG | 237.0 | -11.8 | 1.7 | -7.0 | 2.5E-12 | 8.9E-12 | MB231/MB468 | MP |
| >hg38_21_+_8433172_8446622_RNA45SN1_45S_50ntFlanks@9352.9373.22\|CCGTGGCGCAATGAAGGTGAAG | >hg38_21_+_8433172_8446622_RNA45SN1_45S_50ntFlanks | 9352 | 9373 | 22 | CCGTGGCGCAATGAAGGTGAAG | 96.9 | -10.5 | 1.5 | -7.0 | 2.5E-12 | 8.9E-12 | MB231/MB468 | MP |
| >hg38_21_+_8433172_8446622_RNA45SN1_45S_50ntFlanks@10458.10488.31\|ATGGCCTCCGTTGCCCTCGGCCGATCGAAAG | >hg38_21_+_8433172_8446622_RNA45SN1_45S_50ntFlanks | 10458 | 10488 | 31 | ATGGCCTCCGTTGCCCTCGGCCGATCGAAAG | 680.7 | -4.1 | 0.6 | -7.0 | 2.0E-12 | 8.9E-12 | BT20/MB468 | MITO |
| >hg38_21_+_8433172_8446622_RNA45SN1_45S_50ntFlanks@12686.12708.23\|CCAGGCGGAACGATACGGCAGCG | >hg38_21_+_8433172_8446622_RNA45SN1_45S_50ntFlanks | 12686 | 12708 | 23 | CCAGGCGGAACGATACGGCAGCG | 153.4 | -7.6 | 1.1 | -7.0 | 2.8E-12 | 8.9E-12 | MB231/MB468 | NUC |
| >hg38_21_+_8433172_8446622_RNA45SN1_45S_50ntFlanks@10427.10453.27\|ATGGGCGAGCGCCGTTCCGAAGGGACG | >hg38_21_+_8433172_8446622_RNA45SN1_45S_50ntFlanks | 10427 | 10453 | 27 | ATGGGCGAGCGCCGTTCCGAAGGGACG | 993.1 | -5.7 | 0.8 | -7.0 | 2.9E-12 | 8.9E-12 | MB231/MB468 | NUC |
| >MI0000298\|hsa-mir-221&WithFlank&X\|-\|45746151\|45746272@71.90.20(+1C)&[MIMAT0000278&hsa-miR-221-3p&offsets\|0\|-3(+1C);m-91&X\|-\|45746181\|45746202&offsets\|0\|-2(+1C)]\|AGCTACATTGTCTGCTGGGTC | >MI0000298\|hsa-mir-221&WithFlank&X\|-\|45746151\|45746272 | 71 | 90 | 20(+1C) | AGCTACATTGTCTGCTGGGTC | 184.2 | 7.8 | 1.1 | 7.0 | 2.9E-12 | 8.9E-12 | MB231/MB468 | NUC |
| >hg38_21_+_8433172_8446622_RNA45SN1_45S_50ntFlanks@11235.11268.34\|CTTCCCGTGGATCGCCCCAGCTGCGGCGGGCGTC | >hg38_21_+_8433172_8446622_RNA45SN1_45S_50ntFlanks | 11235 | 11268 | 34 | CTTCCCGTGGATCGCCCCAGCTGCGGCGGGCGTC | 90.9 | -10.4 | 1.5 | -7.0 | 2.5E-12 | 9.0E-12 | MB231/MB468 | MP |
| >hg38_21_+_8433172_8446622_RNA45SN1_45S_50ntFlanks@6664.6686.23\|TGGATCACTCGGCTCGTGCGTCG | >hg38_21_+_8433172_8446622_RNA45SN1_45S_50ntFlanks | 6664 | 6686 | 23 | TGGATCACTCGGCTCGTGCGTCG | 1140.9 | -7.0 | 1.0 | -7.0 | 3.4E-12 | 9.0E-12 | MB231/MB468 | MITO |
| >hg38_21_+_8433172_8446622_RNA45SN1_45S_50ntFlanks@12717.12751.35\|CCTCGGTTGGCCTCGGATAGCCGGTCCCCCGCCTG | >hg38_21_+_8433172_8446622_RNA45SN1_45S_50ntFlanks | 12717 | 12751 | 35 | CCTCGGTTGGCCTCGGATAGCCGGTCCCCCGCCTG | 342.3 | -12.3 | 1.8 | -7.0 | 2.5E-12 | 9.0E-12 | MB231/MB468 | MP |
| >hg38_21_+_8433172_8446622_RNA45SN1_45S_50ntFlanks@7973.8004.32\|GACGCGACCTCAGATCAGACGTGGCGACCCGC | >hg38_21_+_8433172_8446622_RNA45SN1_45S_50ntFlanks | 7973 | 8004 | 32 | GACGCGACCTCAGATCAGACGTGGCGACCCGC | 937.4 | 5.6 | 0.8 | 7.0 | 2.1E-12 | 9.0E-12 | BT20/MB468 | MITO |
| >hg38_21_+_8433172_8446622_RNA45SN1_45S_50ntFlanks@3835.3861.27\|CTCCTCTCCTACTTGGATAACTGTGGT | >hg38_21_+_8433172_8446622_RNA45SN1_45S_50ntFlanks | 3835 | 3861 | 27 | CTCCTCTCCTACTTGGATAACTGTGGT | 592.1 | -4.4 | 0.6 | -7.0 | 2.9E-12 | 9.0E-12 | MB231/MB468 | NUC |
| >hg38_21_+_8433172_8446622_RNA45SN1_45S_50ntFlanks@4163.4186.24\|CATCCAAGGAAGGCAGCAGGCGCG | >hg38_21_+_8433172_8446622_RNA45SN1_45S_50ntFlanks | 4163 | 4186 | 24 | CATCCAAGGAAGGCAGCAGGCGCG | 749.3 | -7.0 | 1.0 | -7.0 | 2.9E-12 | 9.0E-12 | MB231/MB468 | NUC |
| >hg38_21_+_8433172_8446622_RNA45SN1_45S_50ntFlanks@10513.10546.34\|CCGGAGTGGCGGAGATGGGCGCCGCGAGGCGTCC | >hg38_21_+_8433172_8446622_RNA45SN1_45S_50ntFlanks | 10513 | 10546 | 34 | CCGGAGTGGCGGAGATGGGCGCCGCGAGGCGTCC | 105.1 | -10.6 | 1.5 | -7.0 | 2.5E-12 | 9.0E-12 | MB231/MB468 | MP |
| >hg38_21_+_8433172_8446622_RNA45SN1_45S_50ntFlanks@10411.10453.43\|CAGTCGGTCCTGAGAGATGGGCGAGCGCCGTTCCGAAGGGACG | >hg38_21_+_8433172_8446622_RNA45SN1_45S_50ntFlanks | 10411 | 10453 | 43 | CAGTCGGTCCTGAGAGATGGGCGAGCGCCGTTCCGAAGGGACG | 3469.2 | -8.1 | 1.2 | -7.0 | 3.4E-12 | 9.0E-12 | MB231/MB468 | MITO |
| >hg38_21_+_8433172_8446622_RNA45SN1_45S_50ntFlanks@12158.12183.26\|GTTTGACTGGGGCGGTACACCTGTCA | >hg38_21_+_8433172_8446622_RNA45SN1_45S_50ntFlanks | 12158 | 12183 | 26 | GTTTGACTGGGGCGGTACACCTGTCA | 116.4 | -10.7 | 1.5 | -7.0 | 2.5E-12 | 9.0E-12 | MB231/MB468 | MP |
| >hg38_21_+_8433172_8446622_RNA45SN1_45S_50ntFlanks@12654.12673.20\|TATGACTGAACGCCTCTAAG | >hg38_21_+_8433172_8446622_RNA45SN1_45S_50ntFlanks | 12654 | 12673 | 20 | TATGACTGAACGCCTCTAAG | 619.0 | -9.5 | 1.4 | -6.9 | 3.7E-12 | 9.0E-12 | BT20/MB468 | NUC |
| >MI0000809\|hsa-mir-151a&WithFlank&8\|-\|140732558\|140732659@53.74.22&[MIMAT0000757&hsa-miR-151a-3p&offsets\|0\|+1;m-30&8\|-\|140732587\|140732607&offsets\|0\|+1]\|CTAGACTGAAGCTCCTTGAGGA | >MI0000809\|hsa-mir-151a&WithFlank&8\|-\|140732558\|140732659 | 53 | 74 | 22 | CTAGACTGAAGCTCCTTGAGGA | 6678.2 | 5.6 | 0.8 | 7.0 | 3.4E-12 | 9.1E-12 | MB231/MB468 | MITO |
| >hg38_21_+_8433172_8446622_RNA45SN1_45S_50ntFlanks@11255.11281.27\|CTGCGGCGGGCGTCGCGGCCGCCCCCG | >hg38_21_+_8433172_8446622_RNA45SN1_45S_50ntFlanks | 11255 | 11281 | 27 | CTGCGGCGGGCGTCGCGGCCGCCCCCG | 125.7 | -10.8 | 1.5 | -7.0 | 2.5E-12 | 9.1E-12 | MB231/MB468 | MP |
| >MI0000298\|hsa-mir-221&WithFlank&X\|-\|45746151\|45746272@71.94.24&[MIMAT0000278&hsa-miR-221-3p&offsets\|0\|+1;m-91&X\|-\|45746181\|45746202&offsets\|0\|+2]\|AGCTACATTGTCTGCTGGGTTTCA | >MI0000298\|hsa-mir-221&WithFlank&X\|-\|45746151\|45746272 | 71 | 94 | 24 | AGCTACATTGTCTGCTGGGTTTCA | 2772.5 | 5.9 | 0.8 | 7.0 | 2.8E-12 | 9.1E-12 | MB231/MB468 | CYTO |
| >hg38_21_+_8433172_8446622_RNA45SN1_45S_50ntFlanks@5086.5122.37\|AACGAGACTCTGGCATGCTAACTAGTTACGCGACCCC | >hg38_21_+_8433172_8446622_RNA45SN1_45S_50ntFlanks | 5086 | 5122 | 37 | AACGAGACTCTGGCATGCTAACTAGTTACGCGACCCC | 304.5 | 8.8 | 1.3 | 7.0 | 2.8E-12 | 9.1E-12 | MB231/MB468 | CYTO |
| >hg38_21_+_8433172_8446622_RNA45SN1_45S_50ntFlanks@13006.13025.20\|CTATTGAAAGTCAGCCCTCG | >hg38_21_+_8433172_8446622_RNA45SN1_45S_50ntFlanks | 13006 | 13025 | 20 | CTATTGAAAGTCAGCCCTCG | 164.2 | -9.4 | 1.3 | -7.0 | 2.1E-12 | 9.1E-12 | BT20/MB468 | MITO |
| >MI0000298\|hsa-mir-221&WithFlank&X\|-\|45746151\|45746272@71.94.24&[MIMAT0000278&hsa-miR-221-3p&offsets\|0\|+1;m-91&X\|-\|45746181\|45746202&offsets\|0\|+2]\|AGCTACATTGTCTGCTGGGTTTCA | >MI0000298\|hsa-mir-221&WithFlank&X\|-\|45746151\|45746272 | 71 | 94 | 24 | AGCTACATTGTCTGCTGGGTTTCA | 2118.2 | 5.1 | 0.7 | 7.0 | 2.9E-12 | 9.1E-12 | MB231/MB468 | NUC |
| >hg38_21_+_8433172_8446622_RNA45SN1_45S_50ntFlanks@7980.8012.33\|CCTCAGATCAGACGTGGCGACCCGCTGAATTTA | >hg38_21_+_8433172_8446622_RNA45SN1_45S_50ntFlanks | 7980 | 8012 | 33 | CCTCAGATCAGACGTGGCGACCCGCTGAATTTA | 1548.7 | -7.2 | 1.0 | -7.0 | 3.5E-12 | 9.2E-12 | MB231/MB468 | MITO |
| >MI0000433\|hsa-let-7g&WithFlank&3\|-\|52268272\|52268367@11.33.23&[MIMAT0000414&hsa-let-7g-5p&offsets\|0\|+1;m-58&3\|-\|52268336\|52268357&offsets\|0\|+1]\|TGAGGTAGTAGTTTGTACAGTTT | >MI0000433\|hsa-let-7g&WithFlank&3\|-\|52268272\|52268367 | 11 | 33 | 23 | TGAGGTAGTAGTTTGTACAGTTT | 831.8 | 5.5 | 0.8 | 7.0 | 2.8E-12 | 9.2E-12 | MB231/MB468 | CYTO |
| >hg38_21_+_8433172_8446622_RNA45SN1_45S_50ntFlanks@12960.12993.34\|CTGGGTCGGGGTTTCGTACGTAGCAGAGCAGCTC | >hg38_21_+_8433172_8446622_RNA45SN1_45S_50ntFlanks | 12960 | 12993 | 34 | CTGGGTCGGGGTTTCGTACGTAGCAGAGCAGCTC | 220.7 | 8.2 | 1.2 | 7.0 | 2.9E-12 | 9.2E-12 | MB231/MB468 | CYTO |
| >hg38_21_+_8433172_8446622_RNA45SN1_45S_50ntFlanks@10771.10798.28\|CAGGTCTCCAAGGTGAACAGCCTCTGGC | >hg38_21_+_8433172_8446622_RNA45SN1_45S_50ntFlanks | 10771 | 10798 | 28 | CAGGTCTCCAAGGTGAACAGCCTCTGGC | 88.1 | -10.3 | 1.5 | -7.0 | 2.5E-12 | 9.2E-12 | MB231/MB468 | MP |
| >hg38_21_+_8433172_8446622_RNA45SN1_45S_50ntFlanks@5467.5491.25\|GGCCCACGGCCCTGGCGGAGCGCTG | >hg38_21_+_8433172_8446622_RNA45SN1_45S_50ntFlanks | 5467 | 5491 | 25 | GGCCCACGGCCCTGGCGGAGCGCTG | 172.2 | -11.3 | 1.6 | -7.0 | 2.5E-12 | 9.2E-12 | MB231/MB468 | MP |
| >hg38_1_-_228634819_228635039_RNA5S12_5S_50ntFlanks@51.76.26\|GTCTACGGCCATACCACCCTGAACGC | >hg38_1_-_228634819_228635039_RNA5S12_5S_50ntFlanks | 51 | 76 | 26 | GTCTACGGCCATACCACCCTGAACGC | 1100.1 | 6.2 | 0.9 | 7.0 | 2.7E-12 | 9.2E-12 | BT20/MB468 | CYTO |
| >hg38_21_+_8433172_8446622_RNA45SN1_45S_50ntFlanks@11921.11967.47\|TGAGAGGTGTAGAATAAGTGGGAGGCCCCCGGCGCCCCCCCGGTGTC | >hg38_21_+_8433172_8446622_RNA45SN1_45S_50ntFlanks | 11921 | 11967 | 47 | TGAGAGGTGTAGAATAAGTGGGAGGCCCCCGGCGCCCCCCCGGTGTC | 241.3 | -10.2 | 1.5 | -7.0 | 2.7E-12 | 9.2E-12 | BT20/MB468 | CYTO |
| >am_MI0005763\|hsa-mir-941-1&WithFlank&20\|+\|63919443\|63919526@53.75.23(+1U);MI0005764\|hsa-mir-941-2&WithFlank&20\|+\|63919499\|63919582@53.75.23(+1U);MI0005765\|hsa-mir-941-3&WithFlank&20\|+\|63919555\|63919638@53.75.23(+1U);MI0005766\|hsa-mir-941-4&WithFlank&20\|+\|63919750\|63919833@53.75.23(+1U);MI0031520\|hsa-mir-941-5&WithFlank&20\|+\|63919862\|63919945@53.75.23(+1U)&[MIMAT0004984&hsa-miR-941&offsets\|0\|0(+1U);m-263&20\|+\|63919495\|63919517&offsets\|0\|0(+1U)];[MIMAT0004984_1&hsa-miR-941&offsets\|0\|0(+1U);m-263&20\|+\|63919551\|63919573&offsets\|0\|0(+1U)];[MIMAT0004984_2&hsa-miR-941&offsets\|0\|0(+1U);m-263&20\|+\|63919607\|63919629&offsets\|0\|0(+1U)];[MIMAT0004984_3&hsa-miR-941&offsets\|0\|0(+1U);m-263&20\|+\|63919802\|63919824&offsets\|0\|0(+1U)];[MIMAT0004984_4&hsa-miR-941&offsets\|0\|0(+1U);m-263&20\|+\|63919914\|63919936&offsets\|0\|0(+1U)]\|CACCCGGCTGTGTGCACATGTGCT | >am_MI0005763\|hsa-mir-941-1&WithFlank&20\|+\|63919443\|63919526 | 53 | 75 | 23(+1U) | CACCCGGCTGTGTGCACATGTGCT | 611.2 | 9.2 | 1.3 | 7.0 | 2.1E-12 | 9.2E-12 | BT20/MB468 | MITO |
| >hg38_21_+_8433172_8446622_RNA45SN1_45S_50ntFlanks@12589.12614.26\|GTTCAGACATTTGGTGTATGTGCTTG | >hg38_21_+_8433172_8446622_RNA45SN1_45S_50ntFlanks | 12589 | 12614 | 26 | GTTCAGACATTTGGTGTATGTGCTTG | 219.5 | -9.9 | 1.4 | -7.0 | 2.1E-12 | 9.2E-12 | BT20/MB468 | MITO |
| >hg38_21_+_8433172_8446622_RNA45SN1_45S_50ntFlanks@5558.5580.23\|TGCGGAAGGATCATTAACGGAGC | >hg38_21_+_8433172_8446622_RNA45SN1_45S_50ntFlanks | 5558 | 5580 | 23 | TGCGGAAGGATCATTAACGGAGC | 189.7 | 6.9 | 1.0 | 7.0 | 2.9E-12 | 9.3E-12 | MB231/MB468 | CYTO |
| >hg38_21_+_8433172_8446622_RNA45SN1_45S_50ntFlanks@6722.6746.25\|ATTGCAGGACACATTGATCATCGAC | >hg38_21_+_8433172_8446622_RNA45SN1_45S_50ntFlanks | 6722 | 6746 | 25 | ATTGCAGGACACATTGATCATCGAC | 395.8 | -8.8 | 1.3 | -6.9 | 3.8E-12 | 9.3E-12 | BT20/MB468 | NUC |
| >hg38_21_+_8433172_8446622_RNA45SN1_45S_50ntFlanks@3960.3984.25\|GCCCCGGCCGGGGGGCGGGCGCCGG | >hg38_21_+_8433172_8446622_RNA45SN1_45S_50ntFlanks | 3960 | 3984 | 25 | GCCCCGGCCGGGGGGCGGGCGCCGG | 112.2 | -9.5 | 1.4 | -7.0 | 2.9E-12 | 9.3E-12 | MB231/MB468 | CYTO |
| >hg38_21_+_8433172_8446622_RNA45SN1_45S_50ntFlanks@5485.5520.36\|AGCGCTGAGAAGACGGTCGAACTTGACTATCTAGAG | >hg38_21_+_8433172_8446622_RNA45SN1_45S_50ntFlanks | 5485 | 5520 | 36 | AGCGCTGAGAAGACGGTCGAACTTGACTATCTAGAG | 130.0 | -10.9 | 1.6 | -7.0 | 2.6E-12 | 9.3E-12 | MB231/MB468 | MP |
| >hg38_21_+_8433172_8446622_RNA45SN1_45S_50ntFlanks@4367.4406.40\|CAGTTAAAAAGCTCGTAGTTGGATCTTGGGAGCGGGCGGG | >hg38_21_+_8433172_8446622_RNA45SN1_45S_50ntFlanks | 4367 | 4406 | 40 | CAGTTAAAAAGCTCGTAGTTGGATCTTGGGAGCGGGCGGG | 256.7 | -11.0 | 1.6 | -7.0 | 3.0E-12 | 9.3E-12 | MB231/MB468 | NUC |
| >hg38_21_+_8433172_8446622_RNA45SN1_45S_50ntFlanks@6667.6689.23\|ATCACTCGGCTCGTGCGTCGATG | >hg38_21_+_8433172_8446622_RNA45SN1_45S_50ntFlanks | 6667 | 6689 | 23 | ATCACTCGGCTCGTGCGTCGATG | 210.0 | -9.8 | 1.4 | -7.0 | 2.1E-12 | 9.3E-12 | BT20/MB468 | MITO |
| >MI0000487\|hsa-mir-193a&WithFlank&17\|+\|31559990\|31560089@27.47.21&[MIMAT0004614&hsa-miR-193a-5p&offsets\|0\|-1;m-123&17\|+\|31560016\|31560037&offsets\|0\|-1]\|TGGGTCTTTGCGGGCGAGATG | >MI0000487\|hsa-mir-193a&WithFlank&17\|+\|31559990\|31560089 | 27 | 47 | 21 | TGGGTCTTTGCGGGCGAGATG | 676.1 | 8.9 | 1.3 | 7.0 | 2.8E-12 | 9.3E-12 | BT20/MB468 | CYTO |
| >hg38_21_+_8433172_8446622_RNA45SN1_45S_50ntFlanks@9534.9556.23\|AGGAAACTCTGGTGGAGGTCCGT | >hg38_21_+_8433172_8446622_RNA45SN1_45S_50ntFlanks | 9534 | 9556 | 23 | AGGAAACTCTGGTGGAGGTCCGT | 292.8 | -12.1 | 1.7 | -7.0 | 2.6E-12 | 9.3E-12 | MB231/MB468 | MP |
| >hg38_21_+_8433172_8446622_RNA45SN1_45S_50ntFlanks@7983.8012.30\|CAGATCAGACGTGGCGACCCGCTGAATTTA | >hg38_21_+_8433172_8446622_RNA45SN1_45S_50ntFlanks | 7983 | 8012 | 30 | CAGATCAGACGTGGCGACCCGCTGAATTTA | 1668.5 | -7.5 | 1.1 | -7.0 | 3.6E-12 | 9.3E-12 | MB231/MB468 | MITO |
| >hg38_21_+_8433172_8446622_RNA45SN1_45S_50ntFlanks@475.498.24\|CTTCGTGATCGATGTGGTGACGTC | >hg38_21_+_8433172_8446622_RNA45SN1_45S_50ntFlanks | 475 | 498 | 24 | CTTCGTGATCGATGTGGTGACGTC | 16582.9 | 6.9 | 1.0 | 7.0 | 3.6E-12 | 9.3E-12 | MB231/MB468 | MITO |
| >hg38_21_+_8433172_8446622_RNA45SN1_45S_50ntFlanks@12001.12032.32\|CTGCGGGCCGCCGGTGAAATACCACTACTCTG | >hg38_21_+_8433172_8446622_RNA45SN1_45S_50ntFlanks | 12001 | 12032 | 32 | CTGCGGGCCGCCGGTGAAATACCACTACTCTG | 5440.4 | -8.3 | 1.2 | -7.0 | 3.6E-12 | 9.3E-12 | MB231/MB468 | MITO |

<table>
<tr><th></th><th></th><th></th><th></th><th></th><th></th><th></th><th></th><th></th><th></th><th></th><th></th><th></th><th></th></tr>
<tr><td>>hg38_21_+_8433172_8446622_RNA45SN1_45S_50ntFlanks@11857.11880.24|GCGGAATCAGCGGGGAAAGAAGAC</td><td>>hg38_21_+_8433172_8446622_RNA45SN1_45S_50ntFlanks</td><td>11857</td><td>11880</td><td>24</td><td>GCGGAATCAGCGGGGAAAGAAGAC</td><td>97.6</td><td>-10.5</td><td>1.5</td><td>-7.0</td><td>2.6E-12</td><td>9.3E-12</td><td>MB231/MB468</td><td>MP</td></tr>
<tr><td>>hg38_21_+_8433172_8446622_RNA45SN1_45S_50ntFlanks@12630.12661.32|GGGCGAAGCTACCATCTGTGGGATTATGACTG</td><td>>hg38_21_+_8433172_8446622_RNA45SN1_45S_50ntFlanks</td><td>12630</td><td>12661</td><td>32</td><td>GGGCGAAGCTACCATCTGTGGGATTATGACTG</td><td>411.5</td><td>-12.5</td><td>1.8</td><td>-7.0</td><td>2.6E-12</td><td>9.4E-12</td><td>MB231/MB468</td><td>MP</td></tr>
<tr><td>>hg38_21_+_8433172_8446622_RNA45SN1_45S_50ntFlanks@9391.9412.22|CCGAGGTGGGATCCCGAGGCCT</td><td>>hg38_21_+_8433172_8446622_RNA45SN1_45S_50ntFlanks</td><td>9391</td><td>9412</td><td>22</td><td>CCGAGGTGGGATCCCGAGGCCT</td><td>130.3</td><td>-10.9</td><td>1.6</td><td>-7.0</td><td>2.6E-12</td><td>9.4E-12</td><td>MB231/MB468</td><td>MP</td></tr>
<tr><td>>hg38_21_+_8433172_8446622_RNA45SN1_45S_50ntFlanks@12084.12106.23|TCGCTTCTGGCGCCAAGCGCCCG</td><td>>hg38_21_+_8433172_8446622_RNA45SN1_45S_50ntFlanks</td><td>12084</td><td>12106</td><td>23</td><td>TCGCTTCTGGCGCCAAGCGCCCG</td><td>108.4</td><td>-10.6</td><td>1.5</td><td>-7.0</td><td>2.6E-12</td><td>9.4E-12</td><td>MB231/MB468</td><td>MP</td></tr>
<tr><td>>hg38_21_+_8433172_8446622_RNA45SN1_45S_50ntFlanks@10411.10443.33|CAGTCGGTCCTGAGAGATGGGCGAGCGCCGTTC</td><td>>hg38_21_+_8433172_8446622_RNA45SN1_45S_50ntFlanks</td><td>10411</td><td>10443</td><td>33</td><td>CAGTCGGTCCTGAGAGATGGGCGAGCGCCGTTC</td><td>1896.6</td><td>-10.5</td><td>1.5</td><td>-6.9</td><td>3.8E-12</td><td>9.4E-12</td><td>BT20/MB468</td><td>NUC</td></tr>
<tr><td>>trna17_ValTAC_11_-_59318102_59318174@1.30.30,trna4_ValTAC_X_-_18693029_18693101@1.30.30|GGTTCCATAGTGTAGTGGTTATCACGTCTG</td><td>>trna17_ValTAC_11_-_59318102_59318174</td><td>1</td><td>30</td><td>30</td><td>GGTTCCATAGTGTAGTGGTTATCACGTCTG</td><td>124.1</td><td>-10.8</td><td>1.5</td><td>-7.0</td><td>2.6E-12</td><td>9.4E-12</td><td>MB231/MB468</td><td>MP</td></tr>
<tr><td>>trna119_LysCTT_1_-_145395522_145395594@4.33.30,trna11_LysCTT_5_-_180648979_180649051@4.33.30,trna13_LysCTT_6_+_26556774_26556846@4.33.30,trna7_LysCTT_16_+_3225692_3225764@4.33.30,trna9_LysCTT_5_+_180634755_180634827@4.33.30|CGGCTAGCTCAGTCGGTAGAGCATGAGACT</td><td>>trna119_LysCTT_1_-_145395522_145395594</td><td>4</td><td>33</td><td>30</td><td>CGGCTAGCTCAGTCGGTAGAGCATGAGACT</td><td>2823.8</td><td>6.8</td><td>1.0</td><td>7.0</td><td>3.6E-12</td><td>9.4E-12</td><td>MB231/MB468</td><td>MITO</td></tr>
<tr><td>>hg38_21_+_8433172_8446622_RNA45SN1_45S_50ntFlanks@3831.3850.20|CTCGCTCCTCTCCTACTTGG</td><td>>hg38_21_+_8433172_8446622_RNA45SN1_45S_50ntFlanks</td><td>3831</td><td>3850</td><td>20</td><td>CTCGCTCCTCTCCTACTTGG</td><td>176.1</td><td>-7.8</td><td>1.1</td><td>-7.0</td><td>3.0E-12</td><td>9.4E-12</td><td>MB231/MB468</td><td>NUC</td></tr>
<tr><td>>MI0000082|hsa-mir-25&WithFlank&7|-|100093554|100093649@58.77.20(+1A)&[MIMAT0000081&hsa-miR-25-3p&offsets|0|-2(+1A);m-23&7|-|100093571|100093592&offsets|0|-2(+1A)]|CATTGCACTTGTCTCGGTCTA</td><td>>MI0000082|hsa-mir-25&WithFlank&7|-|100093554|100093649</td><td>58</td><td>77</td><td>20(+1A)</td><td>CATTGCACTTGTCTCGGTCTA</td><td>544.1</td><td>6.2</td><td>0.9</td><td>7.0</td><td>2.9E-12</td><td>9.4E-12</td><td>MB231/MB468</td><td>CYTO</td></tr>
<tr><td>>hg38_21_+_8433172_8446622_RNA45SN1_45S_50ntFlanks@11573.11592.20|CGACTTAGAACTGGTGCGGA</td><td>>hg38_21_+_8433172_8446622_RNA45SN1_45S_50ntFlanks</td><td>11573</td><td>11592</td><td>20</td><td>CGACTTAGAACTGGTGCGGA</td><td>1711.0</td><td>-8.4</td><td>1.2</td><td>-7.0</td><td>2.6E-12</td><td>9.5E-12</td><td>MB231/MB468</td><td>MP</td></tr>
<tr><td>>hg38_21_+_8433172_8446622_RNA45SN1_45S_50ntFlanks@4547.4573.27|CTGGATACCGCAGCTAGGAATAATGGA</td><td>>hg38_21_+_8433172_8446622_RNA45SN1_45S_50ntFlanks</td><td>4547</td><td>4573</td><td>27</td><td>CTGGATACCGCAGCTAGGAATAATGGA</td><td>172.3</td><td>-11.3</td><td>1.6</td><td>-7.0</td><td>2.6E-12</td><td>9.5E-12</td><td>MB231/MB468</td><td>MP</td></tr>
<tr><td>>hg38_21_+_8433172_8446622_RNA45SN1_45S_50ntFlanks@5482.5509.28|CGGAGCGCTGAGAAGACGGTCGAACTTG</td><td>>hg38_21_+_8433172_8446622_RNA45SN1_45S_50ntFlanks</td><td>5482</td><td>5509</td><td>28</td><td>CGGAGCGCTGAGAAGACGGTCGAACTTG</td><td>115.5</td><td>-10.7</td><td>1.5</td><td>-7.0</td><td>2.6E-12</td><td>9.5E-12</td><td>MB231/MB468</td><td>MP</td></tr>
<tr><td>>am_trna128_GlyGCC_6_-_27870686_27870756@1.33.33,trna18_GlyGCC_16_+_70822597_70822667@1.33.33,trna19_GlyGCC_16_+_70823410_70823480@1.33.33,trna19_GlyGCC_2_-_157257659_157257729@1.33.33,trna24_GlyGCC_16_-_70812942_70813012@1.33.33,trna25_GlyGCC_16_-_70812114_70812184@1.33.33,trna5_GlyGCC_17_+_8029064_8029134@1.33.33,trna68_GlyGCC_1_-_161493637_161493707@1.33.33|GCATTGGTGGTTCAGTGGTAGAATTCTCGCCTG</td><td>>am_trna128_GlyGCC_6_-_27870686_27870756</td><td>1</td><td>33</td><td>33</td><td>GCATTGGTGGTTCAGTGGTAGAATTCTCGCCTG</td><td>5847.8</td><td>5.8</td><td>0.8</td><td>7.0</td><td>3.0E-12</td><td>9.5E-12</td><td>MB231/MB468</td><td>CYTO</td></tr>
<tr><td>>hg38_21_+_8433172_8446622_RNA45SN1_45S_50ntFlanks@7982.7999.18|TCAGATCAGACGTGGCGA</td><td>>hg38_21_+_8433172_8446622_RNA45SN1_45S_50ntFlanks</td><td>7982</td><td>7999</td><td>18</td><td>TCAGATCAGACGTGGCGA</td><td>102.2</td><td>-10.5</td><td>1.5</td><td>-7.0</td><td>2.6E-12</td><td>9.5E-12</td><td>MB231/MB468</td><td>MP</td></tr>
<tr><td>>hg38_21_+_8433172_8446622_RNA45SN1_45S_50ntFlanks@7988.8019.32|CAGACGTGGCGACCCGCTGAATTTAAGCATAT</td><td>>hg38_21_+_8433172_8446622_RNA45SN1_45S_50ntFlanks</td><td>7988</td><td>8019</td><td>32</td><td>CAGACGTGGCGACCCGCTGAATTTAAGCATAT</td><td>363.2</td><td>8.1</td><td>1.2</td><td>7.0</td><td>3.0E-12</td><td>9.5E-12</td><td>MB231/MB468</td><td>CYTO</td></tr>
<tr><td>>hg38_21_+_8433172_8446622_RNA45SN1_45S_50ntFlanks@5125.5145.21|AGCGGTCGGCGTCCCCCAACT</td><td>>hg38_21_+_8433172_8446622_RNA45SN1_45S_50ntFlanks</td><td>5125</td><td>5145</td><td>21</td><td>AGCGGTCGGCGTCCCCCAACT</td><td>822.2</td><td>9.0</td><td>1.3</td><td>7.0</td><td>2.8E-12</td><td>9.5E-12</td><td>BT20/MB468</td><td>CYTO</td></tr>
<tr><td>>hg38_21_+_8433172_8446622_RNA45SN1_45S_50ntFlanks@11829.11855.27|CGAAACCACAGCCAAGGGAACGGGCTT</td><td>>hg38_21_+_8433172_8446622_RNA45SN1_45S_50ntFlanks</td><td>11829</td><td>11855</td><td>27</td><td>CGAAACCACAGCCAAGGGAACGGGCTT</td><td>126.2</td><td>-10.8</td><td>1.5</td><td>-7.0</td><td>2.6E-12</td><td>9.5E-12</td><td>MB231/MB468</td><td>MP</td></tr>
<tr><td>>MI0006444|hsa-mir-1307&WithFlank&10|-|103394247|103394407@87.109.23&[MIMAT0005951&hsa-miR-1307-3p&offsets|+1|+2;m-66&10|-|103394300|103394322&offsets|+1|+1]|CTCGGCGTGGCGTCGGTCGTGGT</td><td>>MI0006444|hsa-mir-1307&WithFlank&10|-|103394247|103394407</td><td>87</td><td>109</td><td>23</td><td>CTCGGCGTGGCGTCGGTCGTGGT</td><td>351.2</td><td>7.8</td><td>1.1</td><td>7.0</td><td>3.0E-12</td><td>9.5E-12</td><td>MB231/MB468</td><td>CYTO</td></tr>
<tr><td>>hg38_21_+_8433172_8446622_RNA45SN1_45S_50ntFlanks@10794.10816.23|CTGGCATGTTGGAACAATGTAGG</td><td>>hg38_21_+_8433172_8446622_RNA45SN1_45S_50ntFlanks</td><td>10794</td><td>10816</td><td>23</td><td>CTGGCATGTTGGAACAATGTAGG</td><td>1404.8</td><td>-8.1</td><td>1.2</td><td>-6.9</td><td>3.9E-12</td><td>9.5E-12</td><td>BT20/MB468</td><td>NUC</td></tr>
<tr><td>>hg38_21_+_8433172_8446622_RNA45SN1_45S_50ntFlanks@6641.6683.43|CGACCTCGTACGACTCTTAGCGGTGGATCACTCGGCTCGTGCG</td><td>>hg38_21_+_8433172_8446622_RNA45SN1_45S_50ntFlanks</td><td>6641</td><td>6683</td><td>43</td><td>CGACCTCGTACGACTCTTAGCGGTGGATCACTCGGCTCGTGCG</td><td>587.4</td><td>7.9</td><td>1.1</td><td>7.0</td><td>3.1E-12</td><td>9.5E-12</td><td>MB231/MB468</td><td>NUC</td></tr>
<tr><td>>trnaMT_GluTTC_MT_-_14674_14742@46.71.26,trnalookalike8_GluTTC_5_-_93905172_93905240@46.71.26|GTGGTTGTAGTCCGTGCGAGAATACC</td><td>>trnaMT_GluTTC_MT_-_14674_14742</td><td>46</td><td>71</td><td>26</td><td>GTGGTTGTAGTCCGTGCGAGAATACC</td><td>202.0</td><td>6.7</td><td>1.0</td><td>7.0</td><td>3.1E-12</td><td>9.5E-12</td><td>MB231/MB468</td><td>NUC</td></tr>
<tr><td>>hg38_21_+_8433172_8446622_RNA45SN1_45S_50ntFlanks@6738.6762.25|ATCATCGACACTTCGAACGCACTTG</td><td>>hg38_21_+_8433172_8446622_RNA45SN1_45S_50ntFlanks</td><td>6738</td><td>6762</td><td>25</td><td>ATCATCGACACTTCGAACGCACTTG</td><td>93.7</td><td>-10.4</td><td>1.5</td><td>-7.0</td><td>2.6E-12</td><td>9.6E-12</td><td>MB231/MB468</td><td>MP</td></tr>
<tr><td>>hg38_21_+_8433172_8446622_RNA45SN1_45S_50ntFlanks@12456.12488.33|CCAAGCGTTGGATTGTTCACCCACTAATAGGGA</td><td>>hg38_21_+_8433172_8446622_RNA45SN1_45S_50ntFlanks</td><td>12456</td><td>12488</td><td>33</td><td>CCAAGCGTTGGATTGTTCACCCACTAATAGGGA</td><td>6557.3</td><td>-7.5</td><td>1.1</td><td>-7.0</td><td>3.0E-12</td><td>9.6E-12</td><td>MB231/MB468</td><td>CYTO</td></tr>
<tr><td>>hg38_21_+_8433172_8446622_RNA45SN1_45S_50ntFlanks@4919.4939.21|CCACCAGGAGTGGAGCCTGCG</td><td>>hg38_21_+_8433172_8446622_RNA45SN1_45S_50ntFlanks</td><td>4919</td><td>4939</td><td>21</td><td>CCACCAGGAGTGGAGCCTGCG</td><td>549.9</td><td>-12.9</td><td>1.9</td><td>-6.9</td><td>3.7E-12</td><td>9.6E-12</td><td>MB231/MB468</td><td>MITO</td></tr>
<tr><td>>MI0000087|hsa-mir-29a&WithFlank&7|-|130876741|130876816@48.67.20(+1C)&[MIMAT0000086&hsa-miR-29a-3p&offsets|0|-2(+1C);m-26&7|-|130876749|130876769&offsets|0|-1(+1C)]|TAGCACCATCTGAAATCGGTC</td><td>>MI0000087|hsa-mir-29a&WithFlank&7|-|130876741|130876816</td><td>48</td><td>67</td><td>20(+1C)</td><td>TAGCACCATCTGAAATCGGTC</td><td>436.7</td><td>10.5</td><td>1.5</td><td>7.0</td><td>3.0E-12</td><td>9.6E-12</td><td>MB231/MB468</td><td>CYTO</td></tr>
<tr><td>>hg38_21_+_8433172_8446622_RNA45SN1_45S_50ntFlanks@11316.11342.27|CACCCCACGTCTCGTCGCGCGCGCGTC</td><td>>hg38_21_+_8433172_8446622_RNA45SN1_45S_50ntFlanks</td><td>11316</td><td>11342</td><td>27</td><td>CACCCCACGTCTCGTCGCGCGCGCGTC</td><td>1015.6</td><td>-13.1</td><td>1.8</td><td>-7.4</td><td>1.5E-13</td><td>9.6E-12</td><td>BT20/MB468</td><td>MP</td></tr>
<tr><td>>MI0005756|hsa-mir-934&WithFlank&X|+|136550872|136550966@21.43.23&[MIMAT0004977&hsa-miR-934&offsets|0|+1;m-943&X|+|136550892|136550913&offsets|0|+1]|TGTCTACTACTGGAGACACTGGT</td><td>>MI0005756|hsa-mir-934&WithFlank&X|+|136550872|136550966</td><td>21</td><td>43</td><td>23</td><td>TGTCTACTACTGGAGACACTGGT</td><td>670.3</td><td>12.7</td><td>1.7</td><td>7.6</td><td>4.1E-14</td><td>9.6E-12</td><td>BT20/MB231</td><td>MP</td></tr>
<tr><td>>hg38_21_+_8433172_8446622_RNA45SN1_45S_50ntFlanks@9739.9759.21|AAACGATCTCAACCTATTCTC</td><td>>hg38_21_+_8433172_8446622_RNA45SN1_45S_50ntFlanks</td><td>9739</td><td>9759</td><td>21</td><td>AAACGATCTCAACCTATTCTC</td><td>979.8</td><td>-3.2</td><td>0.5</td><td>-7.0</td><td>3.1E-12</td><td>9.6E-12</td><td>MB231/MB468</td><td>NUC</td></tr>
<tr><td>>hg38_21_+_8433172_8446622_RNA45SN1_45S_50ntFlanks@12625.12657.33|CAATGGGGCGAAGCTACCATCTGTGGGATTATG</td><td>>hg38_21_+_8433172_8446622_RNA45SN1_45S_50ntFlanks</td><td>12625</td><td>12657</td><td>33</td><td>CAATGGGGCGAAGCTACCATCTGTGGGATTATG</td><td>268.4</td><td>-4.4</td><td>0.6</td><td>-7.0</td><td>3.1E-12</td><td>9.6E-12</td><td>MB231/MB468</td><td>NUC</td></tr>
<tr><td>>hg38_21_+_8433172_8446622_RNA45SN1_45S_50ntFlanks@10781.10807.27|AGGTGAACAGCCTCTGGCATGTTGGAA</td><td>>hg38_21_+_8433172_8446622_RNA45SN1_45S_50ntFlanks</td><td>10781</td><td>10807</td><td>27</td><td>AGGTGAACAGCCTCTGGCATGTTGGAA</td><td>449.7</td><td>-8.4</td><td>1.2</td><td>-7.0</td><td>3.0E-12</td><td>9.6E-12</td><td>MB231/MB468</td><td>CYTO</td></tr>
<tr><td>>MI0005756|hsa-mir-934&WithFlank&X|+|136550872|136550966@21.42.22(+1A)&[MIMAT0004977&hsa-miR-934&offsets|0|0(+1A);m-943&X|+|136550892|136550913&offsets|0|0(+1A)]|TGTCTACTACTGGAGACACTGGA</td><td>>MI0005756|hsa-mir-934&WithFlank&X|+|136550872|136550966</td><td>21</td><td>42</td><td>22(+1A)</td><td>TGTCTACTACTGGAGACACTGGA</td><td>431.8</td><td>-12.6</td><td>1.8</td><td>-7.0</td><td>2.7E-12</td><td>9.6E-12</td><td>MB231/MB468</td><td>MP</td></tr>
<tr><td>>hg38_21_+_8433172_8446622_RNA45SN1_45S_50ntFlanks@10234.10263.30|CGGACGCTACGCCGCGACGAGTAGGAGGGC</td><td>>hg38_21_+_8433172_8446622_RNA45SN1_45S_50ntFlanks</td><td>10234</td><td>10263</td><td>30</td><td>CGGACGCTACGCCGCGACGAGTAGGAGGGC</td><td>93.5</td><td>-10.4</td><td>1.5</td><td>-7.0</td><td>2.7E-12</td><td>9.6E-12</td><td>MB231/MB468</td><td>MP</td></tr>
<tr><td>>MI0000750|hsa-mir-26a-2&WithFlank&12|-|57824603|57824698@20.41.22(+2U);MI0000083|hsa-mir-26a-1&WithFlank&3|+|37969398|37969486@16.37.22(+2U)&[MIMAT0000082&hsa-miR-26a-5p&offsets|0|0(+2U);m-33&12|-|57824658|57824679&offsets|0|0(+2U)];[MIMAT0000082_1&hsa-miR-26a-5p&offsets|0|0(+2U);m-34&3|+|37969413|37969434&offsets|0|0(+2U)]|TTCAAGTAATCCAGGATAGGCTTT</td><td>>MI0000750|hsa-mir-26a-2&WithFlank&12|-|57824603|57824698</td><td>20</td><td>41</td><td>22(+2U)</td><td>TTCAAGTAATCCAGGATAGGCTTT</td><td>1547.2</td><td>5.9</td><td>0.8</td><td>7.0</td><td>3.0E-12</td><td>9.6E-12</td><td>MB231/MB468</td><td>CYTO</td></tr>
<tr><td>>hg38_21_+_8433172_8446622_RNA45SN1_45S_50ntFlanks@11703.11736.34|AATGAAGCGCGGGTAAACGGCGGGAGTAACTATG</td><td>>hg38_21_+_8433172_8446622_RNA45SN1_45S_50ntFlanks</td><td>11703</td><td>11736</td><td>34</td><td>AATGAAGCGCGGGTAAACGGCGGGAGTAACTATG</td><td>719.4</td><td>-5.5</td><td>0.8</td><td>-6.9</td><td>3.9E-12</td><td>9.6E-12</td><td>BT20/MB468</td><td>NUC</td></tr>
<tr><td>>hg38_21_+_8433172_8446622_RNA45SN1_45S_50ntFlanks@11209.11239.31|CGGCGACTCTGGACGCGAGCCGGGCCCTTCC</td><td>>hg38_21_+_8433172_8446622_RNA45SN1_45S_50ntFlanks</td><td>11209</td><td>11239</td><td>31</td><td>CGGCGACTCTGGACGCGAGCCGGGCCCTTCC</td><td>106.3</td><td>-10.6</td><td>1.5</td><td>-7.0</td><td>2.7E-12</td><td>9.7E-12</td><td>MB231/MB468</td><td>MP</td></tr>
<tr><td>>hg38_21_+_8433172_8446622_RNA45SN1_45S_50ntFlanks@3813.3834.22|TTATGGTTCCTTTGGTCGCTCG</td><td>>hg38_21_+_8433172_8446622_RNA45SN1_45S_50ntFlanks</td><td>3813</td><td>3834</td><td>22</td><td>TTATGGTTCCTTTGGTCGCTCG</td><td>728.7</td><td>-9.3</td><td>1.3</td><td>-6.9</td><td>3.7E-12</td><td>9.7E-12</td><td>MB231/MB468</td><td>MITO</td></tr>
<tr><td>>hg38_21_+_8433172_8446622_RNA45SN1_45S_50ntFlanks@3593.3610.18|CCCGTCCGTCCGTCCGTC</td><td>>hg38_21_+_8433172_8446622_RNA45SN1_45S_50ntFlanks</td><td>3593</td><td>3610</td><td>18</td><td>CCCGTCCGTCCGTCCGTC</td><td>207.5</td><td>-9.8</td><td>1.4</td><td>-7.0</td><td>2.2E-12</td><td>9.7E-12</td><td>BT20/MB468</td><td>MITO</td></tr>
<tr><td>>hg38_21_+_8433172_8446622_RNA45SN1_45S_50ntFlanks@8737.8754.18|CCCGTCCGTCCGTCCGTC</td><td>>hg38_21_+_8433172_8446622_RNA45SN1_45S_50ntFlanks</td><td>8737</td><td>8754</td><td>18</td><td>CCCGTCCGTCCGTCCGTC</td><td>207.5</td><td>-9.8</td><td>1.4</td><td>-7.0</td><td>2.2E-12</td><td>9.7E-12</td><td>BT20/MB468</td><td>MITO</td></tr>
<tr><td>>hg38_1_-_228634819_228635039_RNA5S12_5S_50ntFlanks@148.170.23|GGAATACCGGGTGCTGTAGGCTT</td><td>>hg38_1_-_228634819_228635039_RNA5S12_5S_50ntFlanks</td><td>148</td><td>170</td><td>23</td><td>GGAATACCGGGTGCTGTAGGCTT</td><td>6194.2</td><td>3.7</td><td>0.5</td><td>7.0</td><td>2.9E-12</td><td>9.7E-12</td><td>BT20/MB468</td><td>CYTO</td></tr>
<tr><td>>hg38_21_+_8433172_8446622_RNA45SN1_45S_50ntFlanks@9564.9596.33|CTGACGTGCAAATCGGTCGTCCGACCTGGGTAT</td><td>>hg38_21_+_8433172_8446622_RNA45SN1_45S_50ntFlanks</td><td>9564</td><td>9596</td><td>33</td><td>CTGACGTGCAAATCGGTCGTCCGACCTGGGTAT</td><td>17461.5</td><td>-5.9</td><td>0.8</td><td>-7.0</td><td>3.0E-12</td><td>9.7E-12</td><td>MB231/MB468</td><td>CYTO</td></tr>
<tr><td>>hg38_21_+_8433172_8446622_RNA45SN1_45S_50ntFlanks@4202.4219.18|CGACCCGGGGAGGTAGTG</td><td>>hg38_21_+_8433172_8446622_RNA45SN1_45S_50ntFlanks</td><td>4202</td><td>4219</td><td>18</td><td>CGACCCGGGGAGGTAGTG</td><td>787.1</td><td>-8.8</td><td>1.3</td><td>-7.0</td><td>2.7E-12</td><td>9.7E-12</td><td>MB231/MB468</td><td>MP</td></tr>
<tr><td>>hg38_21_+_8433172_8446622_RNA45SN1_45S_50ntFlanks@10427.10457.31|ATGGGCGAGCGCCGTTCCGAAGGGACGGGCG</td><td>>hg38_21_+_8433172_8446622_RNA45SN1_45S_50ntFlanks</td><td>10427</td><td>10457</td><td>31</td><td>ATGGGCGAGCGCCGTTCCGAAGGGACGGGCG</td><td>298.4</td><td>-11.2</td><td>1.6</td><td>-7.0</td><td>3.1E-12</td><td>9.7E-12</td><td>MB231/MB468</td><td>NUC</td></tr>
<tr><td>>hg38_21_+_8433172_8446622_RNA45SN1_45S_50ntFlanks@12458.12496.39|AAGCGTTGGATTGTTCACCCACTAATAGGGAACGTGAGC</td><td>>hg38_21_+_8433172_8446622_RNA45SN1_45S_50ntFlanks</td><td>12458</td><td>12496</td><td>39</td><td>AAGCGTTGGATTGTTCACCCACTAATAGGGAACGTGAGC</td><td>380.2</td><td>4.7</td><td>0.7</td><td>7.0</td><td>2.9E-12</td><td>9.8E-12</td><td>BT20/MB468</td><td>CYTO</td></tr>
<tr><td>>hg38_21_+_8433172_8446622_RNA45SN1_45S_50ntFlanks@8111.8143.33|GGGGCGCGGGACATGTGGCGTACGGAAGACCCG</td><td>>hg38_21_+_8433172_8446622_RNA45SN1_45S_50ntFlanks</td><td>8111</td><td>8143</td><td>33</td><td>GGGGCGCGGGACATGTGGCGTACGGAAGACCCG</td><td>148.0</td><td>-11.1</td><td>1.6</td><td>-7.0</td><td>2.7E-12</td><td>9.8E-12</td><td>MB231/MB468</td><td>MP</td></tr>
<tr><td>>hg38_21_+_8433172_8446622_RNA45SN1_45S_50ntFlanks@7986.8015.30|ATCAGACGTGGCGACCCGCTGAATTTAAGC</td><td>>hg38_21_+_8433172_8446622_RNA45SN1_45S_50ntFlanks</td><td>7986</td><td>8015</td><td>30</td><td>ATCAGACGTGGCGACCCGCTGAATTTAAGC</td><td>310.1</td><td>-11.4</td><td>1.5</td><td>-7.4</td><td>1.5E-13</td><td>9.8E-12</td><td>BT20/MB468</td><td>MP</td></tr>
<tr><td>>hg38_21_+_8433172_8446622_RNA45SN1_45S_50ntFlanks@3843.3881.39|CTACTTGGATAACTGTGGTAATTCTAGAGCTAATACATG</td><td>>hg38_21_+_8433172_8446622_RNA45SN1_45S_50ntFlanks</td><td>3843</td><td>3881</td><td>39</td><td>CTACTTGGATAACTGTGGTAATTCTAGAGCTAATACATG</td><td>381.8</td><td>-8.5</td><td>1.2</td><td>-6.9</td><td>3.7E-12</td><td>9.8E-12</td><td>MB231/MB468</td><td>MITO</td></tr>
<tr><td>>hg38_21_+_8433172_8446622_RNA45SN1_45S_50ntFlanks@3996.4026.31|ACTCTAGATAACCTCGGGCCGATCGCACGCC</td><td>>hg38_21_+_8433172_8446622_RNA45SN1_45S_50ntFlanks</td><td>3996</td><td>4026</td><td>31</td><td>ACTCTAGATAACCTCGGGCCGATCGCACGCC</td><td>98.0</td><td>-10.5</td><td>1.5</td><td>-7.0</td><td>2.7E-12</td><td>9.8E-12</td><td>MB231/MB468</td><td>MP</td></tr>
</table>

<table>
<tr><td>>hg38_21_+_8433172_8446622_RNA45SN1_45S_50ntFlanks@9790.9832.43|CTGGCGTGGAGCCGGGCGTGGAATGCGAGTGCCTAGTGGGCCA</td><td>>hg38_21_+_8433172_8446622_RNA45SN1_45S_50ntFlanks</td><td>9790</td><td>9832</td><td>43</td><td>CTGGCGTGGAGCCGGGCGTGGAATGCGAGTGCCTAGTGGGCCA</td><td>244.6</td><td>-11.7</td><td>1.7</td><td>-6.9</td><td>3.7E-12</td><td>9.8E-12</td><td>MB231/MB468</td><td>MITO</td></tr>
<tr><td>>hg38_21_+_8433172_8446622_RNA45SN1_45S_50ntFlanks@10794.10818.25|CTGGCATGTTGGAACAATGTAGGTA</td><td>>hg38_21_+_8433172_8446622_RNA45SN1_45S_50ntFlanks</td><td>10794</td><td>10818</td><td>25</td><td>CTGGCATGTTGGAACAATGTAGGTA</td><td>2177.3</td><td>-7.0</td><td>1.0</td><td>-7.0</td><td>3.1E-12</td><td>9.8E-12</td><td>MB231/MB468</td><td>CYTO</td></tr>
<tr><td>>hg38_21_+_8433172_8446622_RNA45SN1_45S_50ntFlanks@10841.10860.20|CGTAACTTCGGGATAAGGAT</td><td>>hg38_21_+_8433172_8446622_RNA45SN1_45S_50ntFlanks</td><td>10841</td><td>10860</td><td>20</td><td>CGTAACTTCGGGATAAGGAT</td><td>114.3</td><td>-10.7</td><td>1.5</td><td>-7.0</td><td>2.7E-12</td><td>9.8E-12</td><td>MB231/MB468</td><td>MP</td></tr>
<tr><td>>hg38_21_+_8433172_8446622_RNA45SN1_45S_50ntFlanks@10685.10707.23|GGCGGCGTCCGGTGAGCTCTCGC</td><td>>hg38_21_+_8433172_8446622_RNA45SN1_45S_50ntFlanks</td><td>10685</td><td>10707</td><td>23</td><td>GGCGGCGTCCGGTGAGCTCTCGC</td><td>983.0</td><td>7.3</td><td>1.0</td><td>7.0</td><td>2.3E-12</td><td>9.8E-12</td><td>BT20/MB468</td><td>MITO</td></tr>
<tr><td>>trna111_HisGTG_1_-_147774845_147774916@-1T.32.33,trna118_HisGTG_1_-_145396881_145396952@-1T.32.33,trna16_HisGTG_1_+_146544773_146544844@-1T.32.33,trna1_HisGTG_15_+_45493349_45493420@-1T.32.33,trna21_HisGTG_1_+_147753471_147753542@-1T.32.33,trna33_HisGTG_6_+_27125906_27125977@-1T.32.33,trna7_HisGTG_9_-_14433938_14434009@-1T.32.33,trna8_HisGTG_15_-_45492611_45492682@-1T.32.33,trna9_HisGTG_15_-_45490804_45490875@-1T.32.33|TGCCGTGATCGTATAGTGGTTAGTACTCTGCGT</td><td>>trna111_HisGTG_1_-_147774845_147774916</td><td>-1T</td><td>32</td><td>33</td><td>TGCCGTGATCGTATAGTGGTTAGTACTCTGCGT</td><td>231.6</td><td>11.1</td><td>1.6</td><td>6.9</td><td>3.8E-12</td><td>9.8E-12</td><td>MB231/MB468</td><td>MITO</td></tr>
<tr><td>>hg38_21_+_8433172_8446622_RNA45SN1_45S_50ntFlanks@10389.10422.34|TGAACAGCAGTTGAACATGGGTCAGTCGGTCCTG</td><td>>hg38_21_+_8433172_8446622_RNA45SN1_45S_50ntFlanks</td><td>10389</td><td>10422</td><td>34</td><td>TGAACAGCAGTTGAACATGGGTCAGTCGGTCCTG</td><td>149.9</td><td>-11.1</td><td>1.6</td><td>-7.0</td><td>2.7E-12</td><td>9.8E-12</td><td>MB231/MB468</td><td>MP</td></tr>
<tr><td>>hg38_21_+_8433172_8446622_RNA45SN1_45S_50ntFlanks@11930.11950.21|TAGAATAAGTGGGAGGCCCCC</td><td>>hg38_21_+_8433172_8446622_RNA45SN1_45S_50ntFlanks</td><td>11930</td><td>11950</td><td>21</td><td>TAGAATAAGTGGGAGGCCCCC</td><td>367.2</td><td>-11.6</td><td>1.6</td><td>-7.4</td><td>1.5E-13</td><td>9.9E-12</td><td>BT20/MB468</td><td>MP</td></tr>
<tr><td>>hg38_21_+_8433172_8446622_RNA45SN1_45S_50ntFlanks@7973.8009.37|GACGCGACCTCAGATCAGACGTGGCGACCCGCTGAAT</td><td>>hg38_21_+_8433172_8446622_RNA45SN1_45S_50ntFlanks</td><td>7973</td><td>8009</td><td>37</td><td>GACGCGACCTCAGATCAGACGTGGCGACCCGCTGAAT</td><td>1944.7</td><td>3.9</td><td>0.6</td><td>7.0</td><td>3.2E-12</td><td>9.9E-12</td><td>MB231/MB468</td><td>NUC</td></tr>
<tr><td>>hg38_21_+_8433172_8446622_RNA45SN1_45S_50ntFlanks@6708.6731.24|GAGAATTAATGTGAATTGCAGGAC</td><td>>hg38_21_+_8433172_8446622_RNA45SN1_45S_50ntFlanks</td><td>6708</td><td>6731</td><td>24</td><td>GAGAATTAATGTGAATTGCAGGAC</td><td>325.2</td><td>5.2</td><td>0.8</td><td>7.0</td><td>3.1E-12</td><td>9.9E-12</td><td>MB231/MB468</td><td>CYTO</td></tr>
<tr><td>>hg38_21_+_8433172_8446622_RNA45SN1_45S_50ntFlanks@12991.13009.19|CTCCCTCGCTGCGATCTAT</td><td>>hg38_21_+_8433172_8446622_RNA45SN1_45S_50ntFlanks</td><td>12991</td><td>13009</td><td>19</td><td>CTCCCTCGCTGCGATCTAT</td><td>215.0</td><td>-9.8</td><td>1.4</td><td>-7.0</td><td>2.3E-12</td><td>9.9E-12</td><td>BT20/MB468</td><td>MITO</td></tr>
<tr><td>>hg38_21_+_8433172_8446622_RNA45SN1_45S_50ntFlanks@9903.9928.26|ATCAGACCCCAGAAAAGGTGTTGGTT</td><td>>hg38_21_+_8433172_8446622_RNA45SN1_45S_50ntFlanks</td><td>9903</td><td>9928</td><td>26</td><td>ATCAGACCCCAGAAAAGGTGTTGGTT</td><td>211.9</td><td>-10.7</td><td>1.5</td><td>-7.0</td><td>3.2E-12</td><td>9.9E-12</td><td>MB231/MB468</td><td>NUC</td></tr>
<tr><td>>hg38_1_-_228634819_228635039_RNA5S12_5S_50ntFlanks@137.170.34|GAGACCGCCTGGGAATACCGGGTGCTGTAGGCTT</td><td>>hg38_1_-_228634819_228635039_RNA5S12_5S_50ntFlanks</td><td>137</td><td>170</td><td>34</td><td>GAGACCGCCTGGGAATACCGGGTGCTGTAGGCTT</td><td>335.6</td><td>5.2</td><td>0.7</td><td>7.0</td><td>2.3E-12</td><td>9.9E-12</td><td>BT20/MB468</td><td>MITO</td></tr>
<tr><td>>hg38_21_+_8433172_8446622_RNA45SN1_45S_50ntFlanks@12436.12455.20|TCATTGTGAAGCAGAATTCA</td><td>>hg38_21_+_8433172_8446622_RNA45SN1_45S_50ntFlanks</td><td>12436</td><td>12455</td><td>20</td><td>TCATTGTGAAGCAGAATTCA</td><td>3190.7</td><td>-8.6</td><td>1.2</td><td>-6.9</td><td>3.8E-12</td><td>9.9E-12</td><td>MB231/MB468</td><td>MITO</td></tr>
<tr><td>>hg38_21_+_8433172_8446622_RNA45SN1_45S_50ntFlanks@5492.5513.22|AGAAGACGGTCGAACTTGACTA</td><td>>hg38_21_+_8433172_8446622_RNA45SN1_45S_50ntFlanks</td><td>5492</td><td>5513</td><td>22</td><td>AGAAGACGGTCGAACTTGACTA</td><td>1367.6</td><td>-9.5</td><td>1.4</td><td>-7.0</td><td>2.8E-12</td><td>9.9E-12</td><td>MB231/MB468</td><td>MP</td></tr>
<tr><td>>trna116_GluCTC_1_-_145399233_145399304@32.68.37,trna59_GluCTC_1_+_249168447_249168518@32.68.37,trna71_GluCTC_1_-_161439189_161439260@32.68.37,trna74_GluCTC_1_-_161431809_161431880@32.68.37,trna77_GluCTC_1_-_161424398_161424469@32.68.37,trna77_GluCTC_6_+_28949976_28950047@32.68.37,trna80_GluCTC_1_-_161417018_161417089@32.68.37,trna87_GluCTC_6_-_126101393_126101464@32.68.37|CTCTCACCGCCGCGGCCCGGGTTCGATTCCCGGTCAG</td><td>>trna116_GluCTC_1_-_145399233_145399304</td><td>32</td><td>68</td><td>37</td><td>CTCTCACCGCCGCGGCCCGGGTTCGATTCCCGGTCAG</td><td>129.4</td><td>-9.7</td><td>1.4</td><td>-7.0</td><td>3.1E-12</td><td>1.0E-11</td><td>MB231/MB468</td><td>CYTO</td></tr>
<tr><td>>hg38_21_+_8433172_8446622_RNA45SN1_45S_50ntFlanks@10186.10209.24|TCCTCCCCTCCTCCCGCCCACGCC</td><td>>hg38_21_+_8433172_8446622_RNA45SN1_45S_50ntFlanks</td><td>10186</td><td>10209</td><td>24</td><td>TCCTCCCCTCCTCCCGCCCACGCC</td><td>225.1</td><td>-7.1</td><td>1.0</td><td>-7.0</td><td>3.1E-12</td><td>1.0E-11</td><td>MB231/MB468</td><td>CYTO</td></tr>
<tr><td>>hg38_21_+_8433172_8446622_RNA45SN1_45S_50ntFlanks@11671.11700.30|CCAGTGCTCTGAATGTCAAAGTGAAGAAAT</td><td>>hg38_21_+_8433172_8446622_RNA45SN1_45S_50ntFlanks</td><td>11671</td><td>11700</td><td>30</td><td>CCAGTGCTCTGAATGTCAAAGTGAAGAAAT</td><td>165.7</td><td>-11.2</td><td>1.6</td><td>-7.0</td><td>2.8E-12</td><td>1.0E-11</td><td>MB231/MB468</td><td>MP</td></tr>
<tr><td>>hg38_21_+_8433172_8446622_RNA45SN1_45S_50ntFlanks@11321.11343.23|CACGTCTCGTCGCGCGCGCGTCC</td><td>>hg38_21_+_8433172_8446622_RNA45SN1_45S_50ntFlanks</td><td>11321</td><td>11343</td><td>23</td><td>CACGTCTCGTCGCGCGCGCGTCC</td><td>1322.5</td><td>-5.8</td><td>0.8</td><td>-6.9</td><td>3.8E-12</td><td>1.0E-11</td><td>MB231/MB468</td><td>MITO</td></tr>
<tr><td>>hg38_21_+_8433172_8446622_RNA45SN1_45S_50ntFlanks@12912.12931.20|CACGTTCGTGGGGAACCTGG</td><td>>hg38_21_+_8433172_8446622_RNA45SN1_45S_50ntFlanks</td><td>12912</td><td>12931</td><td>20</td><td>CACGTTCGTGGGGAACCTGG</td><td>129.6</td><td>-10.9</td><td>1.6</td><td>-7.0</td><td>2.8E-12</td><td>1.0E-11</td><td>MB231/MB468</td><td>MP</td></tr>
<tr><td>>hg38_21_+_8433172_8446622_RNA45SN1_45S_50ntFlanks@12347.12364.18|AGAAAAGTTACCACAGGG</td><td>>hg38_21_+_8433172_8446622_RNA45SN1_45S_50ntFlanks</td><td>12347</td><td>12364</td><td>18</td><td>AGAAAAGTTACCACAGGG</td><td>153.3</td><td>-9.6</td><td>1.4</td><td>-7.0</td><td>3.0E-12</td><td>1.0E-11</td><td>BT20/MB468</td><td>CYTO</td></tr>
<tr><td>>hg38_21_+_8433172_8446622_RNA45SN1_45S_50ntFlanks@11212.11233.22|CGACTCTGGACGCGAGCCGGGC</td><td>>hg38_21_+_8433172_8446622_RNA45SN1_45S_50ntFlanks</td><td>11212</td><td>11233</td><td>22</td><td>CGACTCTGGACGCGAGCCGGGC</td><td>91.0</td><td>-10.4</td><td>1.5</td><td>-7.0</td><td>2.8E-12</td><td>1.0E-11</td><td>MB231/MB468</td><td>MP</td></tr>
<tr><td>>hg38_21_+_8433172_8446622_RNA45SN1_45S_50ntFlanks@11363.11381.19|CGGGCGGCGGCGGTCGGCG</td><td>>hg38_21_+_8433172_8446622_RNA45SN1_45S_50ntFlanks</td><td>11363</td><td>11381</td><td>19</td><td>CGGGCGGCGGCGGTCGGCG</td><td>171.2</td><td>-11.3</td><td>1.6</td><td>-7.0</td><td>2.8E-12</td><td>1.0E-11</td><td>MB231/MB468</td><td>MP</td></tr>
<tr><td>>hg38_21_+_8433172_8446622_RNA45SN1_45S_50ntFlanks@10792.10823.32|CTCTGGCATGTTGGAACAATGTAGGTAAGGGA</td><td>>hg38_21_+_8433172_8446622_RNA45SN1_45S_50ntFlanks</td><td>10792</td><td>10823</td><td>32</td><td>CTCTGGCATGTTGGAACAATGTAGGTAAGGGA</td><td>106.4</td><td>-10.6</td><td>1.5</td><td>-7.0</td><td>2.8E-12</td><td>1.0E-11</td><td>MB231/MB468</td><td>MP</td></tr>
<tr><td>>hg38_21_+_8433172_8446622_RNA45SN1_45S_50ntFlanks@10805.10826.22|GAACAATGTAGGTAAGGGAAGT</td><td>>hg38_21_+_8433172_8446622_RNA45SN1_45S_50ntFlanks</td><td>10805</td><td>10826</td><td>22</td><td>GAACAATGTAGGTAAGGGAAGT</td><td>163.2</td><td>-7.3</td><td>1.1</td><td>-7.0</td><td>3.0E-12</td><td>1.0E-11</td><td>BT20/MB468</td><td>CYTO</td></tr>
<tr><td>>hg38_21_+_8433172_8446622_RNA45SN1_45S_50ntFlanks@11927.11948.22|GTGTAGAATAAGTGGGAGGCCC</td><td>>hg38_21_+_8433172_8446622_RNA45SN1_45S_50ntFlanks</td><td>11927</td><td>11948</td><td>22</td><td>GTGTAGAATAAGTGGGAGGCCC</td><td>107.5</td><td>-10.6</td><td>1.5</td><td>-7.0</td><td>2.8E-12</td><td>1.0E-11</td><td>MB231/MB468</td><td>MP</td></tr>
<tr><td>>hg38_21_+_8433172_8446622_RNA45SN1_45S_50ntFlanks@12934.12962.29|CTAAACCATTCGTAGACGACCTGCTTCTG</td><td>>hg38_21_+_8433172_8446622_RNA45SN1_45S_50ntFlanks</td><td>12934</td><td>12962</td><td>29</td><td>CTAAACCATTCGTAGACGACCTGCTTCTG</td><td>313.9</td><td>-8.3</td><td>1.2</td><td>-6.9</td><td>3.8E-12</td><td>1.0E-11</td><td>MB231/MB468</td><td>MITO</td></tr>
<tr><td>>hg38_21_+_8433172_8446622_RNA45SN1_45S_50ntFlanks@5421.5444.24|CGATTGGATGGTTTAGTGAGGCCC</td><td>>hg38_21_+_8433172_8446622_RNA45SN1_45S_50ntFlanks</td><td>5421</td><td>5444</td><td>24</td><td>CGATTGGATGGTTTAGTGAGGCCC</td><td>203.0</td><td>-11.5</td><td>1.6</td><td>-7.0</td><td>2.8E-12</td><td>1.0E-11</td><td>MB231/MB468</td><td>MP</td></tr>
<tr><td>>hg38_21_+_8433172_8446622_RNA45SN1_45S_50ntFlanks@11890.11907.18|CTTGACTCTAGTCTGGCA</td><td>>hg38_21_+_8433172_8446622_RNA45SN1_45S_50ntFlanks</td><td>11890</td><td>11907</td><td>18</td><td>CTTGACTCTAGTCTGGCA</td><td>1227.2</td><td>-6.8</td><td>1.0</td><td>-7.0</td><td>3.2E-12</td><td>1.0E-11</td><td>MB231/MB468</td><td>CYTO</td></tr>
<tr><td>>hg38_21_+_8433172_8446622_RNA45SN1_45S_50ntFlanks@11920.11967.48|ATGAGAGGTGTAGAATAAGTGGGAGGCCCCCGGCGCCCCCCCGGTGTC</td><td>>hg38_21_+_8433172_8446622_RNA45SN1_45S_50ntFlanks</td><td>11920</td><td>11967</td><td>48</td><td>ATGAGAGGTGTAGAATAAGTGGGAGGCCCCCGGCGCCCCCCCGGTGTC</td><td>265.7</td><td>-10.4</td><td>1.5</td><td>-7.0</td><td>3.0E-12</td><td>1.0E-11</td><td>BT20/MB468</td><td>CYTO</td></tr>
<tr><td>>hg38_21_+_8433172_8446622_RNA45SN1_45S_50ntFlanks@3841.3868.28|TCCTACTTGGATAACTGTGGTAATTCTA</td><td>>hg38_21_+_8433172_8446622_RNA45SN1_45S_50ntFlanks</td><td>3841</td><td>3868</td><td>28</td><td>TCCTACTTGGATAACTGTGGTAATTCTA</td><td>995.6</td><td>2.8</td><td>0.4</td><td>7.0</td><td>3.0E-12</td><td>1.0E-11</td><td>BT20/MB468</td><td>CYTO</td></tr>
<tr><td>>hg38_21_+_8433172_8446622_RNA45SN1_45S_50ntFlanks@4443.4470.28|CCCTTGCCTCTCGGCGCCCCCTCGATGC</td><td>>hg38_21_+_8433172_8446622_RNA45SN1_45S_50ntFlanks</td><td>4443</td><td>4470</td><td>28</td><td>CCCTTGCCTCTCGGCGCCCCCTCGATGC</td><td>172.7</td><td>-5.8</td><td>0.8</td><td>-7.0</td><td>3.2E-12</td><td>1.0E-11</td><td>MB231/MB468</td><td>CYTO</td></tr>
<tr><td>>hg38_21_+_8433172_8446622_RNA45SN1_45S_50ntFlanks@12433.12455.23|CTATCATTGTGAAGCAGAATTCA</td><td>>hg38_21_+_8433172_8446622_RNA45SN1_45S_50ntFlanks</td><td>12433</td><td>12455</td><td>23</td><td>CTATCATTGTGAAGCAGAATTCA</td><td>472.7</td><td>-12.0</td><td>1.6</td><td>-7.4</td><td>1.6E-13</td><td>1.0E-11</td><td>BT20/MB468</td><td>MP</td></tr>
<tr><td>>hg38_21_+_8433172_8446622_RNA45SN1_45S_50ntFlanks@6671.6699.29|CTCGGCTCGTGCGTCGATGAAGAACGCAG</td><td>>hg38_21_+_8433172_8446622_RNA45SN1_45S_50ntFlanks</td><td>6671</td><td>6699</td><td>29</td><td>CTCGGCTCGTGCGTCGATGAAGAACGCAG</td><td>1858.9</td><td>-6.8</td><td>1.0</td><td>-7.0</td><td>3.0E-12</td><td>1.0E-11</td><td>BT20/MB468</td><td>CYTO</td></tr>
<tr><td>>hg38_21_+_8433172_8446622_RNA45SN1_45S_50ntFlanks@11570.11602.33|AGCCGACTTAGAACTGGTGCGGACCAGGGGAAT</td><td>>hg38_21_+_8433172_8446622_RNA45SN1_45S_50ntFlanks</td><td>11570</td><td>11602</td><td>33</td><td>AGCCGACTTAGAACTGGTGCGGACCAGGGGAAT</td><td>1561.3</td><td>-4.2</td><td>0.6</td><td>-7.0</td><td>3.3E-12</td><td>1.0E-11</td><td>MB231/MB468</td><td>NUC</td></tr>
<tr><td>>hg38_21_+_8433172_8446622_RNA45SN1_45S_50ntFlanks@12678.12708.31|AATCCCGCCCAGGCGGAACGATACGGCAGCG</td><td>>hg38_21_+_8433172_8446622_RNA45SN1_45S_50ntFlanks</td><td>12678</td><td>12708</td><td>31</td><td>AATCCCGCCCAGGCGGAACGATACGGCAGCG</td><td>334.1</td><td>-11.5</td><td>1.6</td><td>-7.4</td><td>1.6E-13</td><td>1.0E-11</td><td>BT20/MB468</td><td>MP</td></tr>
<tr><td>>hg38_21_+_8433172_8446622_RNA45SN1_45S_50ntFlanks@7986.8007.22|ATCAGACGTGGCGACCCGCTGA</td><td>>hg38_21_+_8433172_8446622_RNA45SN1_45S_50ntFlanks</td><td>7986</td><td>8007</td><td>22</td><td>ATCAGACGTGGCGACCCGCTGA</td><td>686.9</td><td>-12.5</td><td>1.7</td><td>-7.4</td><td>1.6E-13</td><td>1.0E-11</td><td>BT20/MB468</td><td>MP</td></tr>
<tr><td>>hg38_21_+_8433172_8446622_RNA45SN1_45S_50ntFlanks@9575.9601.27|ATCGGTCGTCCGACCTGGGTATAGGGG</td><td>>hg38_21_+_8433172_8446622_RNA45SN1_45S_50ntFlanks</td><td>9575</td><td>9601</td><td>27</td><td>ATCGGTCGTCCGACCTGGGTATAGGGG</td><td>310.5</td><td>-11.4</td><td>1.5</td><td>-7.4</td><td>1.6E-13</td><td>1.0E-11</td><td>BT20/MB468</td><td>MP</td></tr>
<tr><td>>hg38_21_+_8433172_8446622_RNA45SN1_45S_50ntFlanks@6741.6772.32|ATCGACACTTCGAACGCACTTGCGGCCCCGGG</td><td>>hg38_21_+_8433172_8446622_RNA45SN1_45S_50ntFlanks</td><td>6741</td><td>6772</td><td>32</td><td>ATCGACACTTCGAACGCACTTGCGGCCCCGGG</td><td>978.8</td><td>-4.1</td><td>0.6</td><td>-7.0</td><td>2.3E-12</td><td>1.0E-11</td><td>BT20/MB468</td><td>MITO</td></tr>
<tr><td>>hg38_21_+_8433172_8446622_RNA45SN1_45S_50ntFlanks@3996.4024.29|ACTCTAGATAACCTCGGGCCGATCGCACG</td><td>>hg38_21_+_8433172_8446622_RNA45SN1_45S_50ntFlanks</td><td>3996</td><td>4024</td><td>29</td><td>ACTCTAGATAACCTCGGGCCGATCGCACG</td><td>2028.2</td><td>-10.6</td><td>1.5</td><td>-6.9</td><td>4.2E-12</td><td>1.0E-11</td><td>BT20/MB468</td><td>NUC</td></tr>
<tr><td>>hg38_21_+_8433172_8446622_RNA45SN1_45S_50ntFlanks@12478.12499.22|ACTAATAGGGAACGTGAGCTGG</td><td>>hg38_21_+_8433172_8446622_RNA45SN1_45S_50ntFlanks</td><td>12478</td><td>12499</td><td>22</td><td>ACTAATAGGGAACGTGAGCTGG</td><td>433.6</td><td>-4.5</td><td>0.6</td><td>-7.0</td><td>3.3E-12</td><td>1.0E-11</td><td>MB231/MB468</td><td>NUC</td></tr>
<tr><td>>hg38_21_+_8433172_8446622_RNA45SN1_45S_50ntFlanks@5477.5516.40|CCTGGCGGAGCGCTGAGAAGACGGTCGAACTTGACTATCT</td><td>>hg38_21_+_8433172_8446622_RNA45SN1_45S_50ntFlanks</td><td>5477</td><td>5516</td><td>40</td><td>CCTGGCGGAGCGCTGAGAAGACGGTCGAACTTGACTATCT</td><td>269.0</td><td>11.8</td><td>1.6</td><td>7.4</td><td>1.6E-13</td><td>1.0E-11</td><td>BT20/MB468</td><td>MP</td></tr>
<tr><td>>hg38_21_+_8433172_8446622_RNA45SN1_45S_50ntFlanks@11663.11685.23|ATTTCTGCCCAGTGCTCTGAATG</td><td>>hg38_21_+_8433172_8446622_RNA45SN1_45S_50ntFlanks</td><td>11663</td><td>11685</td><td>23</td><td>ATTTCTGCCCAGTGCTCTGAATG</td><td>684.4</td><td>-4.2</td><td>0.6</td><td>-7.0</td><td>3.2E-12</td><td>1.0E-11</td><td>MB231/MB468</td><td>CYTO</td></tr>
<tr><td>>hg38_21_+_8433172_8446622_RNA45SN1_45S_50ntFlanks@5168.5185.18|CAGCCACCCGAGATTGAG</td><td>>hg38_21_+_8433172_8446622_RNA45SN1_45S_50ntFlanks</td><td>5168</td><td>5185</td><td>18</td><td>CAGCCACCCGAGATTGAG</td><td>589.1</td><td>-13.0</td><td>1.9</td><td>-6.9</td><td>3.9E-12</td><td>1.0E-11</td><td>MB231/MB468</td><td>MITO</td></tr>
<tr><td>>hg38_21_+_8433172_8446622_RNA45SN1_45S_50ntFlanks@7975.8005.31|CGCGACCTCAGATCAGACGTGGCGACCCGCT</td><td>>hg38_21_+_8433172_8446622_RNA45SN1_45S_50ntFlanks</td><td>7975</td><td>8005</td><td>31</td><td>CGCGACCTCAGATCAGACGTGGCGACCCGCT</td><td>73122.9</td><td>3.9</td><td>0.6</td><td>6.9</td><td>4.2E-12</td><td>1.0E-11</td><td>BT20/MB468</td><td>NUC</td></tr>
<tr><td>>hg38_21_+_8433172_8446622_RNA45SN1_45S_50ntFlanks@12539.12560.22|ATGATGTGTTGTTGCCATGGTA</td><td>>hg38_21_+_8433172_8446622_RNA45SN1_45S_50ntFlanks</td><td>12539</td><td>12560</td><td>22</td><td>ATGATGTGTTGTTGCCATGGTA</td><td>116.1</td><td>-9.2</td><td>1.3</td><td>-7.0</td><td>3.1E-12</td><td>1.0E-11</td><td>BT20/MB468</td><td>CYTO</td></tr>
<tr><td>>hg38_21_+_8433172_8446622_RNA45SN1_45S_50ntFlanks@12046.12070.25|TGACCCGGTGAGGCGGGGGGGCGAG</td><td>>hg38_21_+_8433172_8446622_RNA45SN1_45S_50ntFlanks</td><td>12046</td><td>12070</td><td>25</td><td>TGACCCGGTGAGGCGGGGGGGCGAG</td><td>129.6</td><td>-10.9</td><td>1.6</td><td>-7.0</td><td>2.9E-12</td><td>1.0E-11</td><td>MB231/MB468</td><td>MP</td></tr>
<tr><td>>MI0000089|hsa-mir-31&WithFlank&9|-|21512109|21512191@14.36.23&[MIMAT0000089&hsa-miR-31-5p&offsets|0|+2;m-360&9|-|21512157|21512178&offsets|0|+1]|AGGCAAGATGCTGGCATAGCTGT</td><td>>MI0000089|hsa-mir-31&WithFlank&9|-|21512109|21512191</td><td>14</td><td>36</td><td>23</td><td>AGGCAAGATGCTGGCATAGCTGT</td><td>240.6</td><td>-11.8</td><td>1.7</td><td>-7.0</td><td>2.9E-12</td><td>1.0E-11</td><td>MB231/MB468</td><td>MP</td></tr>
</table>

<table>
<tr><th></th><th></th><th></th><th></th><th></th><th></th><th></th><th></th><th></th><th></th><th></th><th></th><th></th><th></th></tr>
<tr><td>>hg38_21_+_8433172_8446622_RNA45SN1_45S_50ntFlanks@5491.5525.35|GAGAAGACGGTCGAACTTGACTATCTAGAGGAAGT</td><td>>hg38_21_+_8433172_8446622_RNA45SN1_45S_50ntFlanks</td><td>5491</td><td>5525</td><td>35</td><td>GAGAAGACGGTCGAACTTGACTATCTAGAGGAAGT</td><td>1745.8</td><td>10.6</td><td>1.5</td><td>7.0</td><td>3.1E-12</td><td>1.0E-11</td><td>BT20/MB468</td><td>CYTO</td></tr>
<tr><td>>hg38_21_+_8433172_8446622_RNA45SN1_45S_50ntFlanks@12936.12962.27|AAACCATTCGTAGACGACCTGCTTCTG</td><td>>hg38_21_+_8433172_8446622_RNA45SN1_45S_50ntFlanks</td><td>12936</td><td>12962</td><td>27</td><td>AAACCATTCGTAGACGACCTGCTTCTG</td><td>101.0</td><td>-10.5</td><td>1.5</td><td>-7.0</td><td>2.9E-12</td><td>1.0E-11</td><td>MB231/MB468</td><td>MP</td></tr>
<tr><td>>hg38_21_+_8433172_8446622_RNA45SN1_45S_50ntFlanks@11766.11794.29|ATCTAATTAGTGACGCGCATGAATGGATG</td><td>>hg38_21_+_8433172_8446622_RNA45SN1_45S_50ntFlanks</td><td>11766</td><td>11794</td><td>29</td><td>ATCTAATTAGTGACGCGCATGAATGGATG</td><td>209.3</td><td>-10.0</td><td>1.4</td><td>-7.0</td><td>3.1E-12</td><td>1.0E-11</td><td>BT20/MB468</td><td>CYTO</td></tr>
<tr><td>>hg38_21_+_8433172_8446622_RNA45SN1_45S_50ntFlanks@10725.10752.28|GGGGAGAGGGTGTAAATCTCGCGCCGGG</td><td>>hg38_21_+_8433172_8446622_RNA45SN1_45S_50ntFlanks</td><td>10725</td><td>10752</td><td>28</td><td>GGGGAGAGGGTGTAAATCTCGCGCCGGG</td><td>127.8</td><td>-6.7</td><td>1.0</td><td>-7.0</td><td>3.2E-12</td><td>1.0E-11</td><td>MB231/MB468</td><td>CYTO</td></tr>
<tr><td>>hg38_21_+_8433172_8446622_RNA45SN1_45S_50ntFlanks@12285.12323.39|CAGACCGTGAAAGCGGGGCCTCACGATCCTTCTGACCTT</td><td>>hg38_21_+_8433172_8446622_RNA45SN1_45S_50ntFlanks</td><td>12285</td><td>12323</td><td>39</td><td>CAGACCGTGAAAGCGGGGCCTCACGATCCTTCTGACCTT</td><td>1021.0</td><td>-5.2</td><td>0.7</td><td>-7.0</td><td>2.4E-12</td><td>1.0E-11</td><td>BT20/MB468</td><td>MITO</td></tr>
<tr><td>>hg38_21_+_8433172_8446622_RNA45SN1_45S_50ntFlanks@12419.12457.39|GATGTCGGCTCTTCCTATCATTGTGAAGCAGAATTCACC</td><td>>hg38_21_+_8433172_8446622_RNA45SN1_45S_50ntFlanks</td><td>12419</td><td>12457</td><td>39</td><td>GATGTCGGCTCTTCCTATCATTGTGAAGCAGAATTCACC</td><td>2556.0</td><td>7.7</td><td>1.1</td><td>7.0</td><td>3.3E-12</td><td>1.0E-11</td><td>MB231/MB468</td><td>CYTO</td></tr>
<tr><td>>hg38_21_+_8433172_8446622_RNA45SN1_45S_50ntFlanks@11619.11660.42|AACAAAGCATCGCGAAGGCCCGCGGCGGGTGTTGACGCGATG</td><td>>hg38_21_+_8433172_8446622_RNA45SN1_45S_50ntFlanks</td><td>11619</td><td>11660</td><td>42</td><td>AACAAAGCATCGCGAAGGCCCGCGGCGGGTGTTGACGCGATG</td><td>211.2</td><td>-11.5</td><td>1.7</td><td>-6.9</td><td>4.0E-12</td><td>1.0E-11</td><td>MB231/MB468</td><td>MITO</td></tr>
<tr><td>>hg38_21_+_8433172_8446622_RNA45SN1_45S_50ntFlanks@4445.4470.26|CTTGCCTCTCGGCGCCCCCTCGATGC</td><td>>hg38_21_+_8433172_8446622_RNA45SN1_45S_50ntFlanks</td><td>4445</td><td>4470</td><td>26</td><td>CTTGCCTCTCGGCGCCCCCTCGATGC</td><td>153.1</td><td>-4.5</td><td>0.6</td><td>-7.0</td><td>3.3E-12</td><td>1.0E-11</td><td>MB231/MB468</td><td>CYTO</td></tr>
<tr><td>>hg38_21_+_8433172_8446622_RNA45SN1_45S_50ntFlanks@10389.10411.23|TGAACAGCAGTTGAACATGGGTC</td><td>>hg38_21_+_8433172_8446622_RNA45SN1_45S_50ntFlanks</td><td>10389</td><td>10411</td><td>23</td><td>TGAACAGCAGTTGAACATGGGTC</td><td>244.8</td><td>-3.9</td><td>0.6</td><td>-7.0</td><td>3.3E-12</td><td>1.0E-11</td><td>MB231/MB468</td><td>NUC</td></tr>
<tr><td>>hg38_21_+_8433172_8446622_RNA45SN1_45S_50ntFlanks@6697.6720.24|CAGCTAGCTGCGAGAATTAATGTG</td><td>>hg38_21_+_8433172_8446622_RNA45SN1_45S_50ntFlanks</td><td>6697</td><td>6720</td><td>24</td><td>CAGCTAGCTGCGAGAATTAATGTG</td><td>2697.2</td><td>-6.0</td><td>0.9</td><td>-6.9</td><td>4.2E-12</td><td>1.0E-11</td><td>BT20/MB468</td><td>NUC</td></tr>
<tr><td>>MI0000542|hsa-mir-320a&WithFlank&8|-|22244960|22245043@48.71.24(+1U)&[MIMAT0000510&hsa-miR-320a-3p&offsets|0|+2(+1U);m-60&8|-|22244975|22244996&offsets|0|+2(+1U)]|AAAAGCTGGGTTGAGAGGGCGAAAT</td><td>>MI0000542|hsa-mir-320a&WithFlank&8|-|22244960|22245043</td><td>48</td><td>71</td><td>24(+1U)</td><td>AAAAGCTGGGTTGAGAGGGCGAAAT</td><td>510.5</td><td>10.2</td><td>1.5</td><td>7.0</td><td>2.4E-12</td><td>1.0E-11</td><td>BT20/MB468</td><td>MITO</td></tr>
<tr><td>>hg38_21_+_8433172_8446622_RNA45SN1_45S_50ntFlanks@9226.9247.22|CGAAGAGGGGGACGGCGGAGCG</td><td>>hg38_21_+_8433172_8446622_RNA45SN1_45S_50ntFlanks</td><td>9226</td><td>9247</td><td>22</td><td>CGAAGAGGGGGACGGCGGAGCG</td><td>104.9</td><td>-10.6</td><td>1.5</td><td>-7.0</td><td>2.9E-12</td><td>1.0E-11</td><td>MB231/MB468</td><td>MP</td></tr>
<tr><td>>hg38_21_+_8433172_8446622_RNA45SN1_45S_50ntFlanks@12405.12433.29|CTTTTTGATCCTTCGATGTCGGCTCTTCC</td><td>>hg38_21_+_8433172_8446622_RNA45SN1_45S_50ntFlanks</td><td>12405</td><td>12433</td><td>29</td><td>CTTTTTGATCCTTCGATGTCGGCTCTTCC</td><td>1609.8</td><td>-2.7</td><td>0.4</td><td>-7.0</td><td>2.4E-12</td><td>1.0E-11</td><td>BT20/MB468</td><td>MITO</td></tr>
<tr><td>>hg38_21_+_8433172_8446622_RNA45SN1_45S_50ntFlanks@11561.11601.41|GCGCCTAGCAGCCGACTTAGAACTGGTGCGGACCAGGGGAA</td><td>>hg38_21_+_8433172_8446622_RNA45SN1_45S_50ntFlanks</td><td>11561</td><td>11601</td><td>41</td><td>GCGCCTAGCAGCCGACTTAGAACTGGTGCGGACCAGGGGAA</td><td>3788.9</td><td>-8.6</td><td>1.2</td><td>-7.0</td><td>3.4E-12</td><td>1.0E-11</td><td>MB231/MB468</td><td>NUC</td></tr>
<tr><td>>hg38_21_+_8433172_8446622_RNA45SN1_45S_50ntFlanks@11675.11707.33|TGCTCTGAATGTCAAAGTGAAGAAATTCAATGA</td><td>>hg38_21_+_8433172_8446622_RNA45SN1_45S_50ntFlanks</td><td>11675</td><td>11707</td><td>33</td><td>TGCTCTGAATGTCAAAGTGAAGAAATTCAATGA</td><td>443.6</td><td>4.0</td><td>0.6</td><td>7.0</td><td>3.4E-12</td><td>1.0E-11</td><td>MB231/MB468</td><td>NUC</td></tr>
<tr><td>>hg38_21_+_8433172_8446622_RNA45SN1_45S_50ntFlanks@11577.11600.24|TTAGAACTGGTGCGGACCAGGGGA</td><td>>hg38_21_+_8433172_8446622_RNA45SN1_45S_50ntFlanks</td><td>11577</td><td>11600</td><td>24</td><td>TTAGAACTGGTGCGGACCAGGGGA</td><td>196.2</td><td>-7.6</td><td>1.1</td><td>-7.0</td><td>3.3E-12</td><td>1.0E-11</td><td>MB231/MB468</td><td>CYTO</td></tr>
<tr><td>>MI0000477|hsa-mir-146a&WithFlank&5|+|160485346|160485456@27.50.24&[MIMAT0000449&hsa-miR-146a-5p&offsets|0|+2;m-129&5|+|160485372|160485393&offsets|0|+2]|TGAGAACTGAATTCCATGGGTTGT</td><td>>MI0000477|hsa-mir-146a&WithFlank&5|+|160485346|160485456</td><td>27</td><td>50</td><td>24</td><td>TGAGAACTGAATTCCATGGGTTGT</td><td>385.7</td><td>-11.7</td><td>1.6</td><td>-7.3</td><td>3.6E-13</td><td>1.0E-11</td><td>BT20/MB231</td><td>NUC</td></tr>
<tr><td>>am_MI0000809|hsa-mir-151a&WithFlank&8|-|140732558|140732659@53.73.21(+1U)&[MIMAT0000757&hsa-miR-151a-3p&offsets|0|0(+1U);m-30&8|-|140732587|140732607&offsets|0|0(+1U)]|CTAGACTGAAGCTCCTTGAGGT</td><td>>am_MI0000809|hsa-mir-151a&WithFlank&8|-|140732558|140732659</td><td>53</td><td>73</td><td>21(+1U)</td><td>CTAGACTGAAGCTCCTTGAGGT</td><td>5200.8</td><td>5.0</td><td>0.7</td><td>7.0</td><td>2.9E-12</td><td>1.0E-11</td><td>MB231/MB468</td><td>MP</td></tr>
<tr><td>>hg38_21_+_8433172_8446622_RNA45SN1_45S_50ntFlanks@11718.11740.23|AACGGCGGGAGTAACTATGACTC</td><td>>hg38_21_+_8433172_8446622_RNA45SN1_45S_50ntFlanks</td><td>11718</td><td>11740</td><td>23</td><td>AACGGCGGGAGTAACTATGACTC</td><td>13880.7</td><td>-12.1</td><td>1.7</td><td>-6.9</td><td>4.3E-12</td><td>1.0E-11</td><td>BT20/MB468</td><td>NUC</td></tr>
<tr><td>>MI0000077|hsa-mir-21&WithFlank&17|+|59841260|59841343@14.32.19&[MIMAT0000076&hsa-miR-21-5p&offsets|0|-3;m-1&17|+|59841273|59841294&offsets|0|-3]|TAGCTTATCAGACTGATGT</td><td>>MI0000077|hsa-mir-21&WithFlank&17|+|59841260|59841343</td><td>14</td><td>32</td><td>19</td><td>TAGCTTATCAGACTGATGT</td><td>410.1</td><td>4.7</td><td>0.7</td><td>6.9</td><td>4.3E-12</td><td>1.1E-11</td><td>BT20/MB468</td><td>NUC</td></tr>
<tr><td>>hg38_21_+_8433172_8446622_RNA45SN1_45S_50ntFlanks@12567.12587.21|CTCAGTACGAGAGGAACCGCA</td><td>>hg38_21_+_8433172_8446622_RNA45SN1_45S_50ntFlanks</td><td>12567</td><td>12587</td><td>21</td><td>CTCAGTACGAGAGGAACCGCA</td><td>407.0</td><td>-8.1</td><td>1.2</td><td>-7.0</td><td>3.4E-12</td><td>1.1E-11</td><td>MB231/MB468</td><td>NUC</td></tr>
<tr><td>>hg38_21_+_8433172_8446622_RNA45SN1_45S_50ntFlanks@10678.10710.33|CGGTTCCGGCGGCGTCCGGTGAGCTCTCGCTGG</td><td>>hg38_21_+_8433172_8446622_RNA45SN1_45S_50ntFlanks</td><td>10678</td><td>10710</td><td>33</td><td>CGGTTCCGGCGGCGTCCGGTGAGCTCTCGCTGG</td><td>172660.3</td><td>-7.4</td><td>1.1</td><td>-7.0</td><td>3.3E-12</td><td>1.1E-11</td><td>MB231/MB468</td><td>CYTO</td></tr>
<tr><td>>hg38_21_+_8433172_8446622_RNA45SN1_45S_50ntFlanks@9903.9928.26|ATCAGACCCCAGAAAAGGTGTTGGTT</td><td>>hg38_21_+_8433172_8446622_RNA45SN1_45S_50ntFlanks</td><td>9903</td><td>9928</td><td>26</td><td>ATCAGACCCCAGAAAAGGTGTTGGTT</td><td>134.3</td><td>-10.9</td><td>1.6</td><td>-7.0</td><td>2.9E-12</td><td>1.1E-11</td><td>MB231/MB468</td><td>MP</td></tr>
<tr><td>>hg38_21_+_8433172_8446622_RNA45SN1_45S_50ntFlanks@10407.10430.24|GGGTCAGTCGGTCCTGAGAGATGG</td><td>>hg38_21_+_8433172_8446622_RNA45SN1_45S_50ntFlanks</td><td>10407</td><td>10430</td><td>24</td><td>GGGTCAGTCGGTCCTGAGAGATGG</td><td>148.6</td><td>-11.1</td><td>1.6</td><td>-7.0</td><td>2.9E-12</td><td>1.1E-11</td><td>MB231/MB468</td><td>MP</td></tr>
<tr><td>>hg38_21_+_8433172_8446622_RNA45SN1_45S_50ntFlanks@4983.5003.21|CGGACAGGATTGACAGATTGA</td><td>>hg38_21_+_8433172_8446622_RNA45SN1_45S_50ntFlanks</td><td>4983</td><td>5003</td><td>21</td><td>CGGACAGGATTGACAGATTGA</td><td>1055.8</td><td>-13.8</td><td>2.0</td><td>-6.9</td><td>4.1E-12</td><td>1.1E-11</td><td>MB231/MB468</td><td>MITO</td></tr>
<tr><td>>MI0000294|hsa-mir-218-1&WithFlank&4|+|20528269|20528390@31.52.22(+1U);MI0000295|hsa-mir-218-2&WithFlank&5|-|168768140|168768261@31.52.22(+1U)&[MIMAT0000275&hsa-miR-218-5p&offsets|0|+1(+1U);m-185&4|+|20528299|20528319&offsets|0|+1(+1U)];[MIMAT0000275_1&hsa-miR-218-5p&offsets|0|+1(+1U);m-187&5|-|168768211|168768231&offsets|0|+1(+1U)]|TTGTGCTTGATCTAACCATGTGT</td><td>>MI0000294|hsa-mir-218-1&WithFlank&4|+|20528269|20528390</td><td>31</td><td>52</td><td>22(+1U)</td><td>TTGTGCTTGATCTAACCATGTGT</td><td>199.8</td><td>10.9</td><td>1.6</td><td>7.0</td><td>3.0E-12</td><td>1.1E-11</td><td>MB231/MB468</td><td>MP</td></tr>
<tr><td>>hg38_21_+_8433172_8446622_RNA45SN1_45S_50ntFlanks@11661.11678.18|TGATTTCTGCCCAGTGCT</td><td>>hg38_21_+_8433172_8446622_RNA45SN1_45S_50ntFlanks</td><td>11661</td><td>11678</td><td>18</td><td>TGATTTCTGCCCAGTGCT</td><td>133.3</td><td>-10.9</td><td>1.6</td><td>-7.0</td><td>3.0E-12</td><td>1.1E-11</td><td>MB231/MB468</td><td>MP</td></tr>
<tr><td>>hg38_21_+_8433172_8446622_RNA45SN1_45S_50ntFlanks@8171.8197.27|AAGTCCTTCTGATCGAGGCCCAGCCCG</td><td>>hg38_21_+_8433172_8446622_RNA45SN1_45S_50ntFlanks</td><td>8171</td><td>8197</td><td>27</td><td>AAGTCCTTCTGATCGAGGCCCAGCCCG</td><td>1612.2</td><td>-9.9</td><td>1.4</td><td>-6.9</td><td>4.1E-12</td><td>1.1E-11</td><td>MB231/MB468</td><td>MITO</td></tr>
<tr><td>>hg38_21_+_8433172_8446622_RNA45SN1_45S_50ntFlanks@10392.10410.19|ACAGCAGTTGAACATGGGT</td><td>>hg38_21_+_8433172_8446622_RNA45SN1_45S_50ntFlanks</td><td>10392</td><td>10410</td><td>19</td><td>ACAGCAGTTGAACATGGGT</td><td>756.9</td><td>-3.9</td><td>0.6</td><td>-7.0</td><td>2.5E-12</td><td>1.1E-11</td><td>BT20/MB468</td><td>MITO</td></tr>
<tr><td>>hg38_21_+_8433172_8446622_RNA45SN1_45S_50ntFlanks@4223.4242.20|AAAAATAACAATACAGGACT</td><td>>hg38_21_+_8433172_8446622_RNA45SN1_45S_50ntFlanks</td><td>4223</td><td>4242</td><td>20</td><td>AAAAATAACAATACAGGACT</td><td>794.9</td><td>-9.0</td><td>1.3</td><td>-7.0</td><td>3.0E-12</td><td>1.1E-11</td><td>MB231/MB468</td><td>MP</td></tr>
<tr><td>>hg38_21_+_8433172_8446622_RNA45SN1_45S_50ntFlanks@3866.3898.33|CTAGAGCTAATACATGCCGACGGGCGCTGACCC</td><td>>hg38_21_+_8433172_8446622_RNA45SN1_45S_50ntFlanks</td><td>3866</td><td>3898</td><td>33</td><td>CTAGAGCTAATACATGCCGACGGGCGCTGACCC</td><td>310.6</td><td>-11.2</td><td>1.6</td><td>-7.0</td><td>3.4E-12</td><td>1.1E-11</td><td>MB231/MB468</td><td>NUC</td></tr>
<tr><td>>hg38_21_+_8433172_8446622_RNA45SN1_45S_50ntFlanks@11705.11726.22|TGAAGCGCGGGTAAACGGCGGG</td><td>>hg38_21_+_8433172_8446622_RNA45SN1_45S_50ntFlanks</td><td>11705</td><td>11726</td><td>22</td><td>TGAAGCGCGGGTAAACGGCGGG</td><td>131.4</td><td>-10.9</td><td>1.6</td><td>-7.0</td><td>3.0E-12</td><td>1.1E-11</td><td>MB231/MB468</td><td>MP</td></tr>
<tr><td>>hg38_21_+_8433172_8446622_RNA45SN1_45S_50ntFlanks@11778.11800.23|ACGCGCATGAATGGATGAACGAG</td><td>>hg38_21_+_8433172_8446622_RNA45SN1_45S_50ntFlanks</td><td>11778</td><td>11800</td><td>23</td><td>ACGCGCATGAATGGATGAACGAG</td><td>132.5</td><td>-10.9</td><td>1.6</td><td>-7.0</td><td>3.0E-12</td><td>1.1E-11</td><td>MB231/MB468</td><td>MP</td></tr>
<tr><td>>hg38_21_+_8433172_8446622_RNA45SN1_45S_50ntFlanks@12249.12266.18|CAAAAGCTCGCTTGATCT</td><td>>hg38_21_+_8433172_8446622_RNA45SN1_45S_50ntFlanks</td><td>12249</td><td>12266</td><td>18</td><td>CAAAAGCTCGCTTGATCT</td><td>107.5</td><td>-10.6</td><td>1.5</td><td>-7.0</td><td>3.0E-12</td><td>1.1E-11</td><td>MB231/MB468</td><td>MP</td></tr>
<tr><td>>MI0000446|hsa-mir-125b-1&WithFlank&11|-|122099751|122099850@21.41.21(+1G);MI0000470|hsa-mir-125b-2&WithFlank&21|+|16590231|16590331@23.43.21(+1G)&[MIMAT0000423&hsa-miR-125b-5p&offsets|0|-1(+1G);m-67&11|-|122099809|122099830&offsets|0|-1(+1G)];[MIMAT0000423_1&hsa-miR-125b-5p&offsets|0|-1(+1G);m-68&21|+|16590253|16590274&offsets|0|-1(+1G)]|TCCCTGAGACCCTAACTTGTGG</td><td>>MI0000446|hsa-mir-125b-1&WithFlank&11|-|122099751|122099850</td><td>21</td><td>41</td><td>21(+1G)</td><td>TCCCTGAGACCCTAACTTGTGG</td><td>400.9</td><td>8.5</td><td>1.2</td><td>7.0</td><td>3.5E-12</td><td>1.1E-11</td><td>MB231/MB468</td><td>NUC</td></tr>
<tr><td>>am_trna116_GluCTC_1_-_145399233_145399304@7.28.22,trna59_GluCTC_1_+_249168447_249168518@7.28.22,trna71_GluCTC_1_-_161439189_161439260@7.28.22,trna74_GluCTC_1_-_161431809_161431880@7.28.22,trna77_GluCTC_1_-_161424398_161424469@7.28.22,trna77_GluCTC_6_+_28949976_28950047@7.28.22,trna80_GluCTC_1_-_161417018_161417089@7.28.22,trna87_GluCTC_6_-_126101393_126101464@7.28.22|GTGGTCTAGTGGTTAGGATTCG</td><td>>am_trna116_GluCTC_1_-_145399233_145399304</td><td>7</td><td>28</td><td>22</td><td>GTGGTCTAGTGGTTAGGATTCG</td><td>209.6</td><td>-11.6</td><td>1.7</td><td>-7.0</td><td>3.0E-12</td><td>1.1E-11</td><td>MB231/MB468</td><td>MP</td></tr>
<tr><td>>hg38_21_+_8433172_8446622_RNA45SN1_45S_50ntFlanks@7977.8010.34|CGACCTCAGATCAGACGTGGCGACCCGCTGAATT</td><td>>hg38_21_+_8433172_8446622_RNA45SN1_45S_50ntFlanks</td><td>7977</td><td>8010</td><td>34</td><td>CGACCTCAGATCAGACGTGGCGACCCGCTGAATT</td><td>1300.1</td><td>9.0</td><td>1.3</td><td>7.0</td><td>3.4E-12</td><td>1.1E-11</td><td>MB231/MB468</td><td>CYTO</td></tr>
<tr><td>>MI0000061|hsa-let-7a-2&WithFlank&11|-|122146516|122146599@11.30.20;MI0000062|hsa-let-7a-3&WithFlank&22|+|46112743|46112828@10.29.20;MI0000060|hsa-let-7a-1&WithFlank&9|+|94175951|94176042@12.31.20&[MIMAT0000062&hsa-let-7a-5p&offsets|0|-2;m-6&11|-|122146568|122146589&offsets|0|-2];[MIMAT0000062_1&hsa-let-7a-5p&offsets|0|-2;m-5&22|+|46112752|46112773&offsets|0|-2];[MIMAT0000062_2&hsa-let-7a-5p&offsets|0|-2;m-4&9|+|94175962|94175983&offsets|0|-2]|TGAGGTAGTAGGTTGTATAG</td><td>>MI0000061|hsa-let-7a-2&WithFlank&11|-|122146516|122146599</td><td>11</td><td>30</td><td>20</td><td>TGAGGTAGTAGGTTGTATAG</td><td>493.5</td><td>8.8</td><td>1.3</td><td>7.0</td><td>3.2E-12</td><td>1.1E-11</td><td>BT20/MB468</td><td>CYTO</td></tr>
<tr><td>>hg38_21_+_8433172_8446622_RNA45SN1_45S_50ntFlanks@11199.11225.27|CCGGCGGCGGCGGCGACTCTGGACGCG</td><td>>hg38_21_+_8433172_8446622_RNA45SN1_45S_50ntFlanks</td><td>11199</td><td>11225</td><td>27</td><td>CCGGCGGCGGCGGCGACTCTGGACGCG</td><td>5749.4</td><td>-11.2</td><td>1.6</td><td>-6.9</td><td>4.4E-12</td><td>1.1E-11</td><td>BT20/MB468</td><td>NUC</td></tr>
<tr><td>>hg38_21_+_8433172_8446622_RNA45SN1_45S_50ntFlanks@6647.6674.28|CGTACGACTCTTAGCGGTGGATCACTCG</td><td>>hg38_21_+_8433172_8446622_RNA45SN1_45S_50ntFlanks</td><td>6647</td><td>6674</td><td>28</td><td>CGTACGACTCTTAGCGGTGGATCACTCG</td><td>1553.0</td><td>4.7</td><td>0.7</td><td>7.0</td><td>2.5E-12</td><td>1.1E-11</td><td>BT20/MB468</td><td>MITO</td></tr>
<tr><td>>hg38_21_+_8433172_8446622_RNA45SN1_45S_50ntFlanks@3851.3873.23|ATAACTGTGGTAATTCTAGAGCT</td><td>>hg38_21_+_8433172_8446622_RNA45SN1_45S_50ntFlanks</td><td>3851</td><td>3873</td><td>23</td><td>ATAACTGTGGTAATTCTAGAGCT</td><td>332.8</td><td>-6.6</td><td>0.9</td><td>-7.0</td><td>3.0E-12</td><td>1.1E-11</td><td>MB231/MB468</td><td>MP</td></tr>
<tr><td>>hg38_21_+_8433172_8446622_RNA45SN1_45S_50ntFlanks@10727.10757.31|GGAGAGGGTGTAAATCTCGCGCCGGGCCGTA</td><td>>hg38_21_+_8433172_8446622_RNA45SN1_45S_50ntFlanks</td><td>10727</td><td>10757</td><td>31</td><td>GGAGAGGGTGTAAATCTCGCGCCGGGCCGTA</td><td>352.9</td><td>-6.1</td><td>0.9</td><td>-7.0</td><td>3.4E-12</td><td>1.1E-11</td><td>MB231/MB468</td><td>CYTO</td></tr>
<tr><td>>MI0001448|hsa-mir-425&WithFlank&3|-|49020142|49020240@20.42.23(+1A)&[MIMAT0003393&hsa-miR-425-5p&offsets|0|0(+1A);m-95&3|-|49020200|49020221&offsets|0|+1(+1A)]|AATGACACGATCACTCCCGTTGAA</td><td>>MI0001448|hsa-mir-425&WithFlank&3|-|49020142|49020240</td><td>20</td><td>42</td><td>23(+1A)</td><td>AATGACACGATCACTCCCGTTGAA</td><td>450.5</td><td>10.0</td><td>1.4</td><td>7.0</td><td>3.2E-12</td><td>1.1E-11</td><td>BT20/MB468</td><td>CYTO</td></tr>
</table>

| | | | | | | | | | | | | | |
|---|---|---|---|---|---|---|---|---|---|---|---|---|---|
| >hg38_21_+_8433172_8446622_RNA45SN1_45S_50ntFlanks@11906.11929.24\|CACGGTGAAGAGACATGAGAGGTG | >hg38_21_+_8433172_8446622_RNA45SN1_45S_50ntFlanks | 11906 | 11929 | 24 | CACGGTGAAGAGACATGAGAGGTG | 426.3 | -8.9 | 1.3 | -6.9 | 4.4E-12 | 1.1E-11 | BT20/MB468 | NUC |
| >hg38_21_+_8433172_8446622_RNA45SN1_45S_50ntFlanks@12539.12580.42\|ATGATGTGTTGTTGCCATGGTAATCCTGCTCAGTACGAGAGG | >hg38_21_+_8433172_8446622_RNA45SN1_45S_50ntFlanks | 12539 | 12580 | 42 | ATGATGTGTTGTTGCCATGGTAATCCTGCTCAGTACGAGAGG | 369.7 | -12.3 | 1.8 | -6.9 | 4.2E-12 | 1.1E-11 | MB231/MB468 | MITO |
| >hg38_21_+_8433172_8446622_RNA45SN1_45S_50ntFlanks@11572.11605.34\|CCGACTTAGAACTGGTGCGGACCAGGGGAATCCG | >hg38_21_+_8433172_8446622_RNA45SN1_45S_50ntFlanks | 11572 | 11605 | 34 | CCGACTTAGAACTGGTGCGGACCAGGGGAATCCG | 11213.5 | -11.7 | 1.7 | -6.9 | 4.5E-12 | 1.1E-11 | BT20/MB468 | NUC |
| >hg38_21_+_8433172_8446622_RNA45SN1_45S_50ntFlanks@8168.8193.26\|CCCAAGTCCTTCTGATCGAGGCCCAG | >hg38_21_+_8433172_8446622_RNA45SN1_45S_50ntFlanks | 8168 | 8193 | 26 | CCCAAGTCCTTCTGATCGAGGCCCAG | 118.2 | -10.7 | 1.5 | -7.0 | 3.0E-12 | 1.1E-11 | MB231/MB468 | MP |
| >hg38_21_+_8433172_8446622_RNA45SN1_45S_50ntFlanks@8196.8224.29\|CGTGGACGGTGTGAGGCCGGTAGCGGCCC | >hg38_21_+_8433172_8446622_RNA45SN1_45S_50ntFlanks | 8196 | 8224 | 29 | CGTGGACGGTGTGAGGCCGGTAGCGGCCC | 910.4 | -7.6 | 1.1 | -7.0 | 3.0E-12 | 1.1E-11 | MB231/MB468 | MP |
| >hg38_21_+_8433172_8446622_RNA45SN1_45S_50ntFlanks@5143.5162.20\|ACTTCTTAGAGGGACAAGTG | >hg38_21_+_8433172_8446622_RNA45SN1_45S_50ntFlanks | 5143 | 5162 | 20 | ACTTCTTAGAGGGACAAGTG | 704.4 | -13.2 | 1.9 | -6.9 | 4.2E-12 | 1.1E-11 | MB231/MB468 | MITO |
| >hg38_21_+_8433172_8446622_RNA45SN1_45S_50ntFlanks@6667.6689.23\|ATCACTCGGCTCGTGCGTCGATG | >hg38_21_+_8433172_8446622_RNA45SN1_45S_50ntFlanks | 6667 | 6689 | 23 | ATCACTCGGCTCGTGCGTCGATG | 590.2 | -12.3 | 1.7 | -7.4 | 1.8E-13 | 1.1E-11 | BT20/MB468 | MP |
| >MI0000471\|hsa-mir-126&WithFlank&9\|+\|136670596\|136670692@57.79.23&[MIMAT0000445&hsa-miR-126-3p&offsets\|-1\|0;m-40&9\|+\|136670653\|136670674&offsets\|-1\|0]\|CTCGTACCGTGAGTAATAATGCG | >MI0000471\|hsa-mir-126&WithFlank&9\|+\|136670596\|136670692 | 57 | 79 | 23 | CTCGTACCGTGAGTAATAATGCG | 125.6 | 7.5 | 1.1 | 6.9 | 4.5E-12 | 1.1E-11 | BT20/MB468 | NUC |
| >MI0000542\|hsa-mir-320a&WithFlank&8\|-\|22244960\|22245043@48.68.21(+1U)&[MIMAT0000510&hsa-miR-320a-3p&offsets\|0\|-1(+1U);m-60&8\|-\|22244975\|22244996&offsets\|0\|-1(+1U)]\|AAAAGCTGGGTTGAGAGGGCGT | >MI0000542\|hsa-mir-320a&WithFlank&8\|-\|22244960\|22245043 | 48 | 68 | 21(+1U) | AAAAGCTGGGTTGAGAGGGCGT | 1564.8 | 5.2 | 0.7 | 7.0 | 3.5E-12 | 1.1E-11 | MB231/MB468 | NUC |
| >hg38_21_+_8433172_8446622_RNA45SN1_45S_50ntFlanks@10771.10796.26\|CAGGTCTCCAAGGTGAACAGCCTCTG | >hg38_21_+_8433172_8446622_RNA45SN1_45S_50ntFlanks | 10771 | 10796 | 26 | CAGGTCTCCAAGGTGAACAGCCTCTG | 546.5 | -4.5 | 0.6 | -7.0 | 3.5E-12 | 1.1E-11 | MB231/MB468 | NUC |
| >hg38_21_+_8433172_8446622_RNA45SN1_45S_50ntFlanks@9805.9831.27\|GCGTGGAATGCGAGTGCCTAGTGGGCC | >hg38_21_+_8433172_8446622_RNA45SN1_45S_50ntFlanks | 9805 | 9831 | 27 | GCGTGGAATGCGAGTGCCTAGTGGGCC | 142.0 | -11.0 | 1.6 | -7.0 | 3.1E-12 | 1.1E-11 | MB231/MB468 | MP |
| >trna152_ValCAC_6_-_27248049_27248121@1.42.42\|GCTTCTGTAGTGTAGTGGTTATCACGTTCGCCTCACACGCGA | >trna152_ValCAC_6_-_27248049_27248121 | 1 | 42 | 42 | GCTTCTGTAGTGTAGTGGTTATCACGTTCGCCTCACACGCGA | 1133.6 | 7.9 | 1.1 | 7.0 | 3.5E-12 | 1.1E-11 | MB231/MB468 | CYTO |
| >hg38_21_+_8433172_8446622_RNA45SN1_45S_50ntFlanks@6772.6793.22\|GTTCCTCCCGGGGCTACGCCTG | >hg38_21_+_8433172_8446622_RNA45SN1_45S_50ntFlanks | 6772 | 6793 | 22 | GTTCCTCCCGGGGCTACGCCTG | 310.3 | -10.4 | 1.5 | -7.0 | 2.5E-12 | 1.1E-11 | BT20/MB468 | MITO |
| >hg38_21_+_8433172_8446622_RNA45SN1_45S_50ntFlanks@9738.9755.18\|GAAACGATCTCAACCTAT | >hg38_21_+_8433172_8446622_RNA45SN1_45S_50ntFlanks | 9738 | 9755 | 18 | GAAACGATCTCAACCTAT | 741.3 | -9.2 | 1.3 | -7.0 | 3.3E-12 | 1.1E-11 | BT20/MB468 | CYTO |
| >hg38_21_+_8433172_8446622_RNA45SN1_45S_50ntFlanks@12539.12562.24\|ATGATGTGTTGTTGCCATGGTAAT | >hg38_21_+_8433172_8446622_RNA45SN1_45S_50ntFlanks | 12539 | 12562 | 24 | ATGATGTGTTGTTGCCATGGTAAT | 94.4 | -10.4 | 1.5 | -7.0 | 3.1E-12 | 1.1E-11 | MB231/MB468 | MP |
| >hg38_21_+_8433172_8446622_RNA45SN1_45S_50ntFlanks@4099.4128.30\|GTGACCACGGGTGACGGGGAATCAGGGTTC | >hg38_21_+_8433172_8446622_RNA45SN1_45S_50ntFlanks | 4099 | 4128 | 30 | GTGACCACGGGTGACGGGGAATCAGGGTTC | 95.5 | -10.4 | 1.5 | -7.0 | 3.1E-12 | 1.1E-11 | MB231/MB468 | MP |
| >hg38_21_+_8433172_8446622_RNA45SN1_45S_50ntFlanks@7974.7997.24\|ACGCGACCTCAGATCAGACGTGGC | >hg38_21_+_8433172_8446622_RNA45SN1_45S_50ntFlanks | 7974 | 7997 | 24 | ACGCGACCTCAGATCAGACGTGGC | 1567.9 | 4.8 | 0.7 | 7.0 | 3.5E-12 | 1.1E-11 | MB231/MB468 | CYTO |
| >hg38_21_+_8433172_8446622_RNA45SN1_45S_50ntFlanks@4367.4408.42\|CAGTTAAAAAGCTCGTAGTTGGATCTTGGGAGCGGGCGGGCG | >hg38_21_+_8433172_8446622_RNA45SN1_45S_50ntFlanks | 4367 | 4408 | 42 | CAGTTAAAAAGCTCGTAGTTGGATCTTGGGAGCGGGCGGGCG | 251.7 | -10.9 | 1.6 | -7.0 | 3.6E-12 | 1.1E-11 | MB231/MB468 | NUC |
| >hg38_21_+_8433172_8446622_RNA45SN1_45S_50ntFlanks@8170.8213.44\|CAAGTCCTTCTGATCGAGGCCCAGCCCGTGGACGGTGTGAGGCC | >hg38_21_+_8433172_8446622_RNA45SN1_45S_50ntFlanks | 8170 | 8213 | 44 | CAAGTCCTTCTGATCGAGGCCCAGCCCGTGGACGGTGTGAGGCC | 360.5 | -12.4 | 1.8 | -7.0 | 3.1E-12 | 1.1E-11 | MB231/MB468 | MP |
| >hg38_21_+_8433172_8446622_RNA45SN1_45S_50ntFlanks@10404.10435.32\|CATGGGTCAGTCGGTCCTGAGAGATGGGCGAG | >hg38_21_+_8433172_8446622_RNA45SN1_45S_50ntFlanks | 10404 | 10435 | 32 | CATGGGTCAGTCGGTCCTGAGAGATGGGCGAG | 848.9 | -9.7 | 1.4 | -6.9 | 4.5E-12 | 1.1E-11 | BT20/MB468 | NUC |
| >hg38_21_+_8433172_8446622_RNA45SN1_45S_50ntFlanks@11233.11269.37\|CCCTTCCCGTGGATCGCCCCAGCTGCGGCGGGCGTCG | >hg38_21_+_8433172_8446622_RNA45SN1_45S_50ntFlanks | 11233 | 11269 | 37 | CCCTTCCCGTGGATCGCCCCAGCTGCGGCGGGCGTCG | 226.3 | -10.8 | 1.6 | -7.0 | 3.6E-12 | 1.1E-11 | MB231/MB468 | NUC |
| >hg38_21_+_8433172_8446622_RNA45SN1_45S_50ntFlanks@11593.11611.19\|CCAGGGGAATCCGACTGTT | >hg38_21_+_8433172_8446622_RNA45SN1_45S_50ntFlanks | 11593 | 11611 | 19 | CCAGGGGAATCCGACTGTT | 2422.6 | -6.0 | 0.8 | -7.4 | 1.8E-13 | 1.1E-11 | BT20/MB468 | MP |
| >hg38_21_+_8433172_8446622_RNA45SN1_45S_50ntFlanks@12436.12454.19\|TCATTGTGAAGCAGAATTC | >hg38_21_+_8433172_8446622_RNA45SN1_45S_50ntFlanks | 12436 | 12454 | 19 | TCATTGTGAAGCAGAATTC | 1896.0 | -10.1 | 1.4 | -7.4 | 1.8E-13 | 1.1E-11 | BT20/MB468 | MP |
| >hg38_21_+_8433172_8446622_RNA45SN1_45S_50ntFlanks@10868.10894.27\|AAGGGCTGGGTCGGTCGGGCTGGGGCG | >hg38_21_+_8433172_8446622_RNA45SN1_45S_50ntFlanks | 10868 | 10894 | 27 | AAGGGCTGGGTCGGTCGGGCTGGGGCG | 147.0 | -11.1 | 1.6 | -7.0 | 3.1E-12 | 1.1E-11 | MB231/MB468 | MP |
| >MI0000077\|hsa-mir-21&WithFlank&17\|+\|59841260\|59841343@52.72.21&[MIMAT0004494&hsa-miR-21-3p&offsets\|0\|0;m-52&17\|+\|59841311\|59841332&offsets\|0\|-1]\|CAACACCAGTCGATGGGCTGT | >MI0000077\|hsa-mir-21&WithFlank&17\|+\|59841260\|59841343 | 52 | 72 | 21 | CAACACCAGTCGATGGGCTGT | 630.4 | 12.5 | 1.8 | 6.9 | 4.3E-12 | 1.1E-11 | MB231/MB468 | MITO |
| >hg38_21_+_8433172_8446622_RNA45SN1_45S_50ntFlanks@11729.11758.30\|TAACTATGACTCTCTTAAGGTAGCCAAATG | >hg38_21_+_8433172_8446622_RNA45SN1_45S_50ntFlanks | 11729 | 11758 | 30 | TAACTATGACTCTCTTAAGGTAGCCAAATG | 204.2 | -11.5 | 1.7 | -7.0 | 3.1E-12 | 1.1E-11 | MB231/MB468 | MP |
| >hg38_21_+_8433172_8446622_RNA45SN1_45S_50ntFlanks@12932.12956.25\|CGCTAAACCATTCGTAGACGACCTG | >hg38_21_+_8433172_8446622_RNA45SN1_45S_50ntFlanks | 12932 | 12956 | 25 | CGCTAAACCATTCGTAGACGACCTG | 624.2 | -3.0 | 0.4 | -7.0 | 2.6E-12 | 1.1E-11 | BT20/MB468 | MITO |
| >hg38_21_+_8433172_8446622_RNA45SN1_45S_50ntFlanks@7988.8007.20\|CAGACGTGGCGACCCGCTGA | >hg38_21_+_8433172_8446622_RNA45SN1_45S_50ntFlanks | 7988 | 8007 | 20 | CAGACGTGGCGACCCGCTGA | 2745.7 | -9.7 | 1.4 | -7.0 | 3.1E-12 | 1.1E-11 | MB231/MB468 | MP |
| >hg38_1_-_228634819_228635039_RNA5S12_5S_50ntFlanks@134.168.35\|TGGGAGACCGCCTGGGAATACCGGGTGCTGTAGGC | >hg38_1_-_228634819_228635039_RNA5S12_5S_50ntFlanks | 134 | 168 | 35 | TGGGAGACCGCCTGGGAATACCGGGTGCTGTAGGC | 523.6 | 6.7 | 1.0 | 7.0 | 3.3E-12 | 1.1E-11 | BT20/MB468 | CYTO |
| >hg38_21_+_8433172_8446622_RNA45SN1_45S_50ntFlanks@11592.11611.20\|ACCAGGGGAATCCGACTGTT | >hg38_21_+_8433172_8446622_RNA45SN1_45S_50ntFlanks | 11592 | 11611 | 20 | ACCAGGGGAATCCGACTGTT | 573.6 | -7.3 | 1.0 | -7.0 | 3.5E-12 | 1.1E-11 | MB231/MB468 | CYTO |
| >MI0006444\|hsa-mir-1307&WithFlank&10\|-\|103394247\|103394407@87.107.21(+1A)&[MIMAT0005951&hsa-miR-1307-3p&offsets\|+1\|0(+1A);m-66&10\|-\|103394300\|103394322&offsets\|+1\|-1(+1A)]\|CTCGGCGTGGCGTCGGTCGTGA | >MI0006444\|hsa-mir-1307&WithFlank&10\|-\|103394247\|103394407 | 87 | 107 | 21(+1A) | CTCGGCGTGGCGTCGGTCGTGA | 544.5 | 8.6 | 1.2 | 7.0 | 3.3E-12 | 1.1E-11 | BT20/MB468 | CYTO |
| >hg38_21_+_8433172_8446622_RNA45SN1_45S_50ntFlanks@12512.12538.27\|GTGAGACAGGTTAGTTTTACCCTACTG | >hg38_21_+_8433172_8446622_RNA45SN1_45S_50ntFlanks | 12512 | 12538 | 27 | GTGAGACAGGTTAGTTTTACCCTACTG | 100.3 | -10.5 | 1.5 | -7.0 | 3.1E-12 | 1.1E-11 | MB231/MB468 | MP |
| >hg38_21_+_8433172_8446622_RNA45SN1_45S_50ntFlanks@10498.10524.27\|TTCAGATCCCCGAATCCGGAGTGGCGG | >hg38_21_+_8433172_8446622_RNA45SN1_45S_50ntFlanks | 10498 | 10524 | 27 | TTCAGATCCCCGAATCCGGAGTGGCGG | 120.3 | -10.8 | 1.5 | -7.0 | 3.2E-12 | 1.1E-11 | MB231/MB468 | MP |
| >MI0000266\|hsa-mir-10a&WithFlank&17\|-\|48579832\|48579953@69.89.21&[MIMAT0004555&hsa-miR-10a-3p&offsets\|0\|-1;m-468&17\|-\|48579865\|48579885&offsets\|0\|0]\|CAAATTCGTATCTAGGGGAAT | >MI0000266\|hsa-mir-10a&WithFlank&17\|-\|48579832\|48579953 | 69 | 89 | 21 | CAAATTCGTATCTAGGGGAAT | 198.8 | 10.9 | 1.6 | 7.0 | 3.2E-12 | 1.1E-11 | MB231/MB468 | MP |
| >hg38_21_+_8433172_8446622_RNA45SN1_45S_50ntFlanks@10500.10524.25\|CAGATCCCCGAATCCGGAGTGGCGG | >hg38_21_+_8433172_8446622_RNA45SN1_45S_50ntFlanks | 10500 | 10524 | 25 | CAGATCCCCGAATCCGGAGTGGCGG | 100.6 | -10.5 | 1.5 | -7.0 | 3.2E-12 | 1.1E-11 | MB231/MB468 | MP |
| >MI0000433\|hsa-let-7g&WithFlank&3\|-\|52268272\|52268367@11.30.20&[MIMAT0000414&hsa-let-7g-5p&offsets\|0\|-2;m-58&3\|-\|52268336\|52268357&offsets\|0\|-2]\|TGAGGTAGTAGTTTGTACAG | >MI0000433\|hsa-let-7g&WithFlank&3\|-\|52268272\|52268367 | 11 | 30 | 20 | TGAGGTAGTAGTTTGTACAG | 200.0 | 5.6 | 0.8 | 6.9 | 3.7E-12 | 1.1E-11 | MB231/MB468 | NUC |
| >hg38_21_+_8433172_8446622_RNA45SN1_45S_50ntFlanks@4121.4145.25\|CAGGGTTCGATTCCGGAGAGGGAGC | >hg38_21_+_8433172_8446622_RNA45SN1_45S_50ntFlanks | 4121 | 4145 | 25 | CAGGGTTCGATTCCGGAGAGGGAGC | 100.4 | -10.5 | 1.5 | -7.0 | 3.2E-12 | 1.1E-11 | MB231/MB468 | MP |
| >MI0000102\|hsa-mir-100&WithFlank&11\|-\|122152223\|122152314@19.38.20(+1A)&[MIMAT0000098&hsa-miR-100-5p&offsets\|0\|-2(+1A);m-27&11\|-\|122152276\|122152296&offsets\|0\|-1(+1A)]\|AACCCGTAGATCCGAACTTGA | >MI0000102\|hsa-mir-100&WithFlank&11\|-\|122152223\|122152314 | 19 | 38 | 20(+1A) | AACCCGTAGATCCGAACTTGA | 1222.7 | 14.5 | 2.1 | 7.0 | 3.4E-12 | 1.1E-11 | BT20/MB468 | CYTO |
| >hg38_21_+_8433172_8446622_RNA45SN1_45S_50ntFlanks@9738.9758.21\|GAAACGATCTCAACCTATTCT | >hg38_21_+_8433172_8446622_RNA45SN1_45S_50ntFlanks | 9738 | 9758 | 21 | GAAACGATCTCAACCTATTCT | 3662.2 | -6.2 | 0.9 | -7.0 | 3.4E-12 | 1.1E-11 | BT20/MB468 | CYTO |
| >hg38_21_+_8433172_8446622_RNA45SN1_45S_50ntFlanks@11669.11702.34\|GCCCAGTGCTCTGAATGTCAAAGTGAAGAAATTC | >hg38_21_+_8433172_8446622_RNA45SN1_45S_50ntFlanks | 11669 | 11702 | 34 | GCCCAGTGCTCTGAATGTCAAAGTGAAGAAATTC | 1229.9 | 4.5 | 0.6 | 7.0 | 3.4E-12 | 1.1E-11 | BT20/MB468 | CYTO |
| >hg38_21_+_8433172_8446622_RNA45SN1_45S_50ntFlanks@10481.10512.32\|ATCGAAAGGGAGTCGGGTTCAGATCCCCGAAT | >hg38_21_+_8433172_8446622_RNA45SN1_45S_50ntFlanks | 10481 | 10512 | 32 | ATCGAAAGGGAGTCGGGTTCAGATCCCCGAAT | 297.4 | -10.5 | 1.5 | -7.0 | 3.4E-12 | 1.1E-11 | BT20/MB468 | CYTO |
| >hg38_21_+_8433172_8446622_RNA45SN1_45S_50ntFlanks@12151.12181.31\|GTGGGGAGTTTGACTGGGGCGGTACACCTGT | >hg38_21_+_8433172_8446622_RNA45SN1_45S_50ntFlanks | 12151 | 12181 | 31 | GTGGGGAGTTTGACTGGGGCGGTACACCTGT | 164.1 | -9.4 | 1.4 | -7.0 | 2.6E-12 | 1.1E-11 | BT20/MB468 | MITO |
| >hg38_21_+_8433172_8446622_RNA45SN1_45S_50ntFlanks@12082.12110.29\|TCTCGCTTCTGGCGCCAAGCGCCCGGCCG | >hg38_21_+_8433172_8446622_RNA45SN1_45S_50ntFlanks | 12082 | 12110 | 29 | TCTCGCTTCTGGCGCCAAGCGCCCGGCCG | 94.6 | -10.4 | 1.5 | -7.0 | 3.2E-12 | 1.1E-11 | MB231/MB468 | MP |
| >hg38_21_+_8433172_8446622_RNA45SN1_45S_50ntFlanks@8357.8374.18\|AAAAGAACTTTGAAGAGA | >hg38_21_+_8433172_8446622_RNA45SN1_45S_50ntFlanks | 8357 | 8374 | 18 | AAAAGAACTTTGAAGAGA | 105.7 | -10.6 | 1.5 | -7.0 | 3.2E-12 | 1.1E-11 | MB231/MB468 | MP |
| >hg38_21_+_8433172_8446622_RNA45SN1_45S_50ntFlanks@12419.12452.34\|GATGTCGGCTCTTCCTATCATTGTGAAGCAGAAT | >hg38_21_+_8433172_8446622_RNA45SN1_45S_50ntFlanks | 12419 | 12452 | 34 | GATGTCGGCTCTTCCTATCATTGTGAAGCAGAAT | 1435.2 | 7.1 | 1.0 | 7.0 | 3.6E-12 | 1.1E-11 | MB231/MB468 | CYTO |
| >hg38_21_+_8433172_8446622_RNA45SN1_45S_50ntFlanks@7983.8004.22\|CAGATCAGACGTGGCGACCCGC | >hg38_21_+_8433172_8446622_RNA45SN1_45S_50ntFlanks | 7983 | 8004 | 22 | CAGATCAGACGTGGCGACCCGC | 1891.1 | -4.4 | 0.6 | -7.0 | 2.6E-12 | 1.1E-11 | BT20/MB468 | MITO |
| >MI0000449\|hsa-mir-132&WithFlank&17\|-\|2049902\|2050014@65.86.22&[MIMAT0000426&hsa-miR-132-3p&offsets\|0\|0;m-138&17\|-\|2049929\|2049950&offsets\|0\|0]\|TAACAGTCTACAGCCATGGTCG | >MI0000449\|hsa-mir-132&WithFlank&17\|-\|2049902\|2050014 | 65 | 86 | 22 | TAACAGTCTACAGCCATGGTCG | 188.6 | 10.8 | 1.5 | 7.0 | 3.2E-12 | 1.1E-11 | MB231/MB468 | MP |
| >hg38_21_+_8433172_8446622_RNA45SN1_45S_50ntFlanks@12129.12146.18\|CCCGCTCCGGGGACAGTG | >hg38_21_+_8433172_8446622_RNA45SN1_45S_50ntFlanks | 12129 | 12146 | 18 | CCCGCTCCGGGGACAGTG | 94.9 | -10.4 | 1.5 | -7.0 | 3.2E-12 | 1.1E-11 | MB231/MB468 | MP |
| >hg38_21_+_8433172_8446622_RNA45SN1_45S_50ntFlanks@10827.10848.22\|CGGCAAGCCGGATCCGTAACTT | >hg38_21_+_8433172_8446622_RNA45SN1_45S_50ntFlanks | 10827 | 10848 | 22 | CGGCAAGCCGGATCCGTAACTT | 95.5 | -10.4 | 1.5 | -7.0 | 3.2E-12 | 1.1E-11 | MB231/MB468 | MP |
| >hg38_21_+_8433172_8446622_RNA45SN1_45S_50ntFlanks@6646.6676.31\|TCGTACGACTCTTAGCGGTGGATCACTCGGC | >hg38_21_+_8433172_8446622_RNA45SN1_45S_50ntFlanks | 6646 | 6676 | 31 | TCGTACGACTCTTAGCGGTGGATCACTCGGC | 3906.8 | 6.5 | 0.9 | 7.0 | 3.4E-12 | 1.1E-11 | BT20/MB468 | CYTO |
| >hg38_21_+_8433172_8446622_RNA45SN1_45S_50ntFlanks@10819.10839.21\|AGGGAAGTCGGCAAGCCGGAT | >hg38_21_+_8433172_8446622_RNA45SN1_45S_50ntFlanks | 10819 | 10839 | 21 | AGGGAAGTCGGCAAGCCGGAT | 231.2 | -8.1 | 1.2 | -6.9 | 3.7E-12 | 1.1E-11 | MB231/MB468 | NUC |
| >hg38_21_+_8433172_8446622_RNA45SN1_45S_50ntFlanks@11233.11250.18\|CCCTTCCCGTGGATCGCC | >hg38_21_+_8433172_8446622_RNA45SN1_45S_50ntFlanks | 11233 | 11250 | 18 | CCCTTCCCGTGGATCGCC | 570.8 | -8.9 | 1.3 | -6.9 | 3.7E-12 | 1.1E-11 | MB231/MB468 | NUC |

| | | | | | | | | | | | | | |
|---|---|---|---|---|---|---|---|---|---|---|---|---|---|
| >hg38_21_+_8433172_8446622_RNA45SN1_45S_50ntFlanks@9565.9597.33\|TGACGTGCAAATCGGTCGTCCGACCTGGGTATA | >hg38_21_+_8433172_8446622_RNA45SN1_45S_50ntFlanks | 9565 | 9597 | 33 | TGACGTGCAAATCGGTCGTCCGACCTGGGTATA | 4175.5 | -4.3 | 0.6 | -7.0 | 2.7E-12 | 1.1E-11 | BT20/MB468 | MITO |
| >hg38_21_+_8433172_8446622_RNA45SN1_45S_50ntFlanks@10393.10433.41\|CAGCAGTTGAACATGGGTCAGTCGGTCCTGAGAGATGGGCG | >hg38_21_+_8433172_8446622_RNA45SN1_45S_50ntFlanks | 10393 | 10433 | 41 | CAGCAGTTGAACATGGGTCAGTCGGTCCTGAGAGATGGGCG | 228.8 | -11.7 | 1.7 | -7.0 | 3.2E-12 | 1.1E-11 | MB231/MB468 | MP |
| >trnaMT_SerGCT_MT_+_12207_12265@8.30.23\|CTCACAAGAACTGCTAACTCATG | >trnaMT_SerGCT_MT_+_12207_12265 | 8 | 30 | 23 | CTCACAAGAACTGCTAACTCATG | 145.1 | -11.0 | 1.6 | -7.0 | 3.2E-12 | 1.1E-11 | MB231/MB468 | MP |
| >hg38_21_+_8433172_8446622_RNA45SN1_45S_50ntFlanks@12082.12107.26\|TCTCGCTTCTGGCGCCAAGCGCCCGG | >hg38_21_+_8433172_8446622_RNA45SN1_45S_50ntFlanks | 12082 | 12107 | 26 | TCTCGCTTCTGGCGCCAAGCGCCCGG | 93.0 | -10.4 | 1.5 | -7.0 | 3.2E-12 | 1.1E-11 | MB231/MB468 | MP |
| >hg38_21_+_8433172_8446622_RNA45SN1_45S_50ntFlanks@9167.9195.29\|CGGGCGGGTCGCGCCGTCGGGCCCGGGGG | >hg38_21_+_8433172_8446622_RNA45SN1_45S_50ntFlanks | 9167 | 9195 | 29 | CGGGCGGGTCGCGCCGTCGGGCCCGGGGG | 257.4 | -11.9 | 1.7 | -7.0 | 3.2E-12 | 1.1E-11 | MB231/MB468 | MP |
| >hg38_21_+_8433172_8446622_RNA45SN1_45S_50ntFlanks@9564.9591.28\|CTGACGTGCAAATCGGTCGTCCGACCTG | >hg38_21_+_8433172_8446622_RNA45SN1_45S_50ntFlanks | 9564 | 9591 | 28 | CTGACGTGCAAATCGGTCGTCCGACCTG | 86.4 | -10.3 | 1.5 | -7.0 | 3.2E-12 | 1.1E-11 | MB231/MB468 | MP |
| >hg38_21_+_8433172_8446622_RNA45SN1_45S_50ntFlanks@4566.4587.22\|ATAATGGAATAGGACCGCGGTT | >hg38_21_+_8433172_8446622_RNA45SN1_45S_50ntFlanks | 4566 | 4587 | 22 | ATAATGGAATAGGACCGCGGTT | 1123.7 | -9.9 | 1.4 | -6.9 | 4.7E-12 | 1.1E-11 | BT20/MB468 | NUC |
| >hg38_21_+_8433172_8446622_RNA45SN1_45S_50ntFlanks@10458.10495.38\|ATGGCCTCCGTTGCCCTCGGCCGATCGAAAGGGAGTCG | >hg38_21_+_8433172_8446622_RNA45SN1_45S_50ntFlanks | 10458 | 10495 | 38 | ATGGCCTCCGTTGCCCTCGGCCGATCGAAAGGGAGTCG | 540.1 | -8.4 | 1.2 | -6.9 | 4.4E-12 | 1.2E-11 | MB231/MB468 | MITO |
| >hg38_21_+_8433172_8446622_RNA45SN1_45S_50ntFlanks@4283.4314.32\|CTTTAACGAGGATCCATTGGAGGGCAAGTCTG | >hg38_21_+_8433172_8446622_RNA45SN1_45S_50ntFlanks | 4283 | 4314 | 32 | CTTTAACGAGGATCCATTGGAGGGCAAGTCTG | 906.6 | -7.5 | 1.1 | -6.9 | 3.7E-12 | 1.2E-11 | MB231/MB468 | NUC |
| >hg38_21_+_8433172_8446622_RNA45SN1_45S_50ntFlanks@10866.10902.37\|CTAAGGGCTGGGTCGGTCGGGCTGGGGCGCGAAGCGG | >hg38_21_+_8433172_8446622_RNA45SN1_45S_50ntFlanks | 10866 | 10902 | 37 | CTAAGGGCTGGGTCGGTCGGGCTGGGGCGCGAAGCGG | 6804.0 | -11.5 | 1.7 | -6.9 | 4.7E-12 | 1.2E-11 | BT20/MB468 | NUC |
| >hg38_21_+_8433172_8446622_RNA45SN1_45S_50ntFlanks@3705.3746.42\|TACCTGGTTGATCCTGCCAGTAGCATATGCTTGTCTCAAAGA | >hg38_21_+_8433172_8446622_RNA45SN1_45S_50ntFlanks | 3705 | 3746 | 42 | TACCTGGTTGATCCTGCCAGTAGCATATGCTTGTCTCAAAGA | 14737.0 | 11.1 | 1.6 | 7.0 | 3.4E-12 | 1.2E-11 | BT20/MB468 | CYTO |
| >trnaMT_GluTTC_MT_-_14674_14742@39.72.34,trnalookalike8_GluTTC_5_-_93905172_93905240@39.72.34\|ATTGGTCGTGGTTGTAGTCCGTGCGAGAATACCA | >trnaMT_GluTTC_MT_-_14674_14742 | 39 | 72 | 34 | ATTGGTCGTGGTTGTAGTCCGTGCGAGAATACCA | 27872.4 | 5.9 | 0.9 | 6.9 | 3.7E-12 | 1.2E-11 | MB231/MB468 | NUC |
| >hg38_21_+_8433172_8446622_RNA45SN1_45S_50ntFlanks@9573.9603.31\|AAATCGGTCGTCCGACCTGGGTATAGGGGCG | >hg38_21_+_8433172_8446622_RNA45SN1_45S_50ntFlanks | 9573 | 9603 | 31 | AAATCGGTCGTCCGACCTGGGTATAGGGGCG | 1683.4 | -5.1 | 0.7 | -6.9 | 4.7E-12 | 1.2E-11 | BT20/MB468 | NUC |
| >hg38_21_+_8433172_8446622_RNA45SN1_45S_50ntFlanks@9534.9563.30\|AGGAAACTCTGGTGGAGGTCCGTAGCGGTC | >hg38_21_+_8433172_8446622_RNA45SN1_45S_50ntFlanks | 9534 | 9563 | 30 | AGGAAACTCTGGTGGAGGTCCGTAGCGGTC | 64463.1 | -13.0 | 1.9 | -6.9 | 4.7E-12 | 1.2E-11 | BT20/MB468 | NUC |
| >hg38_21_+_8433172_8446622_RNA45SN1_45S_50ntFlanks@9004.9026.23\|TCTCCCCCGCGGGGGCGCGCCGG | >hg38_21_+_8433172_8446622_RNA45SN1_45S_50ntFlanks | 9004 | 9026 | 23 | TCTCCCCCGCGGGGGCGCGCCGG | 198.0 | -11.5 | 1.6 | -7.0 | 3.2E-12 | 1.2E-11 | MB231/MB468 | MP |
| >hg38_21_+_8433172_8446622_RNA45SN1_45S_50ntFlanks@10829.10855.27\|GCAAGCCGGATCCGTAACTTCGGGATA | >hg38_21_+_8433172_8446622_RNA45SN1_45S_50ntFlanks | 10829 | 10855 | 27 | GCAAGCCGGATCCGTAACTTCGGGATA | 120.3 | -10.8 | 1.5 | -7.0 | 3.2E-12 | 1.2E-11 | MB231/MB468 | MP |
| >hg38_21_+_8433172_8446622_RNA45SN1_45S_50ntFlanks@8179.8199.21\|CTGATCGAGGCCCAGCCCGTG | >hg38_21_+_8433172_8446622_RNA45SN1_45S_50ntFlanks | 8179 | 8199 | 21 | CTGATCGAGGCCCAGCCCGTG | 155.2 | -11.1 | 1.6 | -7.0 | 3.2E-12 | 1.2E-11 | MB231/MB468 | MP |
| >trna134_GluTTC_1_-_16861774_16861845@1.34.34,trna5_GluTTC_1_+_17199078_17199149@1.34.34,trna84_GluTTC_1_-_161391883_161391954@1.34.34,trna94_GluTTC_1_-_149664355_149664427@1.34.34\|TCCCTGGTGGTCTAGTGGCTAGGATTCGGCGCTT | >trna134_GluTTC_1_-_16861774_16861845 | 1 | 34 | 34 | TCCCTGGTGGTCTAGTGGCTAGGATTCGGCGCTT | 927.6 | 10.6 | 1.5 | 7.0 | 3.5E-12 | 1.2E-11 | BT20/MB468 | CYTO |
| >hg38_21_+_8433172_8446622_RNA45SN1_45S_50ntFlanks@4977.5000.24\|CGGACACGGACAGGATTGACAGAT | >hg38_21_+_8433172_8446622_RNA45SN1_45S_50ntFlanks | 4977 | 5000 | 24 | CGGACACGGACAGGATTGACAGAT | 101.1 | -10.5 | 1.5 | -7.0 | 3.3E-12 | 1.2E-11 | MB231/MB468 | MP |
| >hg38_21_+_8433172_8446622_RNA45SN1_45S_50ntFlanks@12241.12273.33\|CAGAAGGGCAAAAGCTCGCTTGATCTTGATTTT | >hg38_21_+_8433172_8446622_RNA45SN1_45S_50ntFlanks | 12241 | 12273 | 33 | CAGAAGGGCAAAAGCTCGCTTGATCTTGATTTT | 1874.7 | -7.4 | 1.1 | -6.9 | 4.4E-12 | 1.2E-11 | MB231/MB468 | MITO |
| >MI0000065\|hsa-let-7d&WithFlank&9\|+\|94178828\|94178926@14.36.23&[MIMAT0000065&hsa-let-7d-5p&offsets\|0\|+1;m-90&9\|+\|94178841\|94178862&offsets\|0\|+1]\|AGAGGTAGTAGGTTGCATAGTTT | >MI0000065\|hsa-let-7d&WithFlank&9\|+\|94178828\|94178926 | 14 | 36 | 23 | AGAGGTAGTAGGTTGCATAGTTT | 261.1 | 5.7 | 0.8 | 6.9 | 3.7E-12 | 1.2E-11 | MB231/MB468 | CYTO |
| >hg38_21_+_8433172_8446622_RNA45SN1_45S_50ntFlanks@12405.12423.19\|CTTTTTGATCCTTCGATGT | >hg38_21_+_8433172_8446622_RNA45SN1_45S_50ntFlanks | 12405 | 12423 | 19 | CTTTTTGATCCTTCGATGT | 549.4 | -7.6 | 1.1 | -6.9 | 3.7E-12 | 1.2E-11 | MB231/MB468 | CYTO |
| >hg38_21_+_8433172_8446622_RNA45SN1_45S_50ntFlanks@9736.9755.20\|CCGAAACGATCTCAACCTAT | >hg38_21_+_8433172_8446622_RNA45SN1_45S_50ntFlanks | 9736 | 9755 | 20 | CCGAAACGATCTCAACCTAT | 286.8 | -7.0 | 1.0 | -7.0 | 3.5E-12 | 1.2E-11 | BT20/MB468 | CYTO |
| >hg38_21_+_8433172_8446622_RNA45SN1_45S_50ntFlanks@12204.12223.20\|AAGGCGAGCTCAGGGAGGAC | >hg38_21_+_8433172_8446622_RNA45SN1_45S_50ntFlanks | 12204 | 12223 | 20 | AAGGCGAGCTCAGGGAGGAC | 703.8 | 9.0 | 1.3 | 6.9 | 3.7E-12 | 1.2E-11 | MB231/MB468 | CYTO |
| >MI0000270\|hsa-mir-181b-1&WithFlank&1\|-\|198858867\|198858988@42.63.22(+1C);MI0000683\|hsa-mir-181b-2&WithFlank&9\|+\|124693704\|124693804@22.43.22(+1C)&[MIMAT0000257&hsa-miR-181b-5p&offsets\|0\|-1(+1C);m-73&1\|-\|198858925\|198858947&offsets\|0\|-1(+1C)];[MIMAT0000257_1&hsa-miR-181b-5p&offsets\|0\|-1(+1C);m-72&9\|+\|124693725\|124693747&offsets\|0\|-1(+1C)]\|AACATTCATTGCTGTCGGTGGGC | >MI0000270\|hsa-mir-181b-1&WithFlank&1\|-\|198858867\|198858988 | 42 | 63 | 22(+1C) | AACATTCATTGCTGTCGGTGGGC | 349.5 | 5.8 | 0.8 | 7.0 | 3.5E-12 | 1.2E-11 | BT20/MB468 | CYTO |
| >hg38_21_+_8433172_8446622_RNA45SN1_45S_50ntFlanks@10384.10403.20\|CCATGTGAACAGCAGTTGAA | >hg38_21_+_8433172_8446622_RNA45SN1_45S_50ntFlanks | 10384 | 10403 | 20 | CCATGTGAACAGCAGTTGAA | 178.0 | -11.3 | 1.6 | -7.0 | 3.3E-12 | 1.2E-11 | MB231/MB468 | MP |
| >hg38_21_+_8433172_8446622_RNA45SN1_45S_50ntFlanks@9473.9503.31\|CACGAGCGCACGTGTTAGGACCCGAAAGATG | >hg38_21_+_8433172_8446622_RNA45SN1_45S_50ntFlanks | 9473 | 9503 | 31 | CACGAGCGCACGTGTTAGGACCCGAAAGATG | 1502.5 | -9.9 | 1.4 | -6.9 | 4.5E-12 | 1.2E-11 | MB231/MB468 | MITO |
| >hg38_21_+_8433172_8446622_RNA45SN1_45S_50ntFlanks@6670.6689.20\|ACTCGGCTCGTGCGTCGATG | >hg38_21_+_8433172_8446622_RNA45SN1_45S_50ntFlanks | 6670 | 6689 | 20 | ACTCGGCTCGTGCGTCGATG | 333.0 | -12.2 | 1.8 | -7.0 | 3.3E-12 | 1.2E-11 | MB231/MB468 | MP |
| >hg38_21_+_8433172_8446622_RNA45SN1_45S_50ntFlanks@8196.8225.30\|CGTGGACGGTGTGAGGCCGGTAGCGGCCCC | >hg38_21_+_8433172_8446622_RNA45SN1_45S_50ntFlanks | 8196 | 8225 | 30 | CGTGGACGGTGTGAGGCCGGTAGCGGCCCC | 994.8 | -4.6 | 0.7 | -6.9 | 3.8E-12 | 1.2E-11 | MB231/MB468 | NUC |
| >hg38_MT_+_1621_3279_RNR2_16S_50ntFlanks@51.70.20\|GCTAAACCTAGCCCCAAACC | >hg38_MT_+_1621_3279_RNR2_16S_50ntFlanks | 51 | 70 | 20 | GCTAAACCTAGCCCCAAACC | 173.3 | 7.6 | 1.1 | 6.9 | 3.8E-12 | 1.2E-11 | MB231/MB468 | NUC |
| >hg38_21_+_8433172_8446622_RNA45SN1_45S_50ntFlanks@9951.9968.18\|CCATGGAAGTCGGAATCC | >hg38_21_+_8433172_8446622_RNA45SN1_45S_50ntFlanks | 9951 | 9968 | 18 | CCATGGAAGTCGGAATCC | 134.4 | -10.9 | 1.6 | -7.0 | 3.3E-12 | 1.2E-11 | MB231/MB468 | MP |
| >hg38_21_+_8433172_8446622_RNA45SN1_45S_50ntFlanks@12874.12891.18\|GCTGGAAAGGCGGCCGCC | >hg38_21_+_8433172_8446622_RNA45SN1_45S_50ntFlanks | 12874 | 12891 | 18 | GCTGGAAAGGCGGCCGCC | 173.1 | -10.4 | 1.5 | -6.9 | 3.8E-12 | 1.2E-11 | MB231/MB468 | NUC |
| >hg38_21_+_8433172_8446622_RNA45SN1_45S_50ntFlanks@12940.12965.26\|CATTCGTAGACGACCTGCTTCTGGGT | >hg38_21_+_8433172_8446622_RNA45SN1_45S_50ntFlanks | 12940 | 12965 | 26 | CATTCGTAGACGACCTGCTTCTGGGT | 346.6 | -8.6 | 1.3 | -6.9 | 4.8E-12 | 1.2E-11 | BT20/MB468 | NUC |
| >hg38_21_+_8433172_8446622_RNA45SN1_45S_50ntFlanks@7990.8009.20\|GACGTGGCGACCCGCTGAAT | >hg38_21_+_8433172_8446622_RNA45SN1_45S_50ntFlanks | 7990 | 8009 | 20 | GACGTGGCGACCCGCTGAAT | 493.0 | 5.1 | 0.7 | 6.9 | 3.7E-12 | 1.2E-11 | MB231/MB468 | CYTO |
| >hg38_21_+_8433172_8446622_RNA45SN1_45S_50ntFlanks@11876.11909.34\|AAGACCCTGTTGAGCTTGACTCTAGTCTGGCACG | >hg38_21_+_8433172_8446622_RNA45SN1_45S_50ntFlanks | 11876 | 11909 | 34 | AAGACCCTGTTGAGCTTGACTCTAGTCTGGCACG | 112.6 | -10.7 | 1.5 | -7.0 | 3.3E-12 | 1.2E-11 | MB231/MB468 | MP |
| >hg38_21_+_8433172_8446622_RNA45SN1_45S_50ntFlanks@7980.8005.26\|CCTCAGATCAGACGTGGCGACCCGCT | >hg38_21_+_8433172_8446622_RNA45SN1_45S_50ntFlanks | 7980 | 8005 | 26 | CCTCAGATCAGACGTGGCGACCCGCT | 385.1 | -4.0 | 0.6 | -6.9 | 3.8E-12 | 1.2E-11 | MB231/MB468 | NUC |
| >hg38_21_+_8433172_8446622_RNA45SN1_45S_50ntFlanks@12618.12653.36\|GAGGAGCCAATGGGGCGAAGCTACCATCTGTGGGAT | >hg38_21_+_8433172_8446622_RNA45SN1_45S_50ntFlanks | 12618 | 12653 | 36 | GAGGAGCCAATGGGGCGAAGCTACCATCTGTGGGAT | 103.9 | -10.6 | 1.5 | -7.0 | 3.3E-12 | 1.2E-11 | MB231/MB468 | MP |
| >hg38_21_+_8433172_8446622_RNA45SN1_45S_50ntFlanks@12836.12866.31\|CGGTGCGGAGTGCCCTTCGTCCTGGGAAACG | >hg38_21_+_8433172_8446622_RNA45SN1_45S_50ntFlanks | 12836 | 12866 | 31 | CGGTGCGGAGTGCCCTTCGTCCTGGGAAACG | 84.4 | -10.3 | 1.5 | -7.0 | 3.3E-12 | 1.2E-11 | MB231/MB468 | MP |
| >hg38_21_+_8433172_8446622_RNA45SN1_45S_50ntFlanks@4730.4752.23\|CAAGAACGAAAGTCGGAGGTTCG | >hg38_21_+_8433172_8446622_RNA45SN1_45S_50ntFlanks | 4730 | 4752 | 23 | CAAGAACGAAAGTCGGAGGTTCG | 441.3 | -6.7 | 1.0 | -7.0 | 3.3E-12 | 1.2E-11 | MB231/MB468 | MP |
| >hg38_21_+_8433172_8446622_RNA45SN1_45S_50ntFlanks@11579.11605.27\|AGAACTGGTGCGGACCAGGGGAATCCG | >hg38_21_+_8433172_8446622_RNA45SN1_45S_50ntFlanks | 11579 | 11605 | 27 | AGAACTGGTGCGGACCAGGGGAATCCG | 3480.8 | -10.8 | 1.6 | -6.9 | 4.9E-12 | 1.2E-11 | BT20/MB468 | NUC |
| >hg38_21_+_8433172_8446622_RNA45SN1_45S_50ntFlanks@476.499.24\|TTCGTGATCGATGTGGTGACGTCG | >hg38_21_+_8433172_8446622_RNA45SN1_45S_50ntFlanks | 476 | 499 | 24 | TTCGTGATCGATGTGGTGACGTCG | 706.6 | 7.4 | 1.1 | 6.9 | 3.8E-12 | 1.2E-11 | MB231/MB468 | CYTO |
| >trna152_ValCAC_6_-_27248049_27248121@1.34.34\|GCTTCTGTAGTGTAGTGGTTATCACGTTCGCCTC | >trna152_ValCAC_6_-_27248049_27248121 | 1 | 34 | 34 | GCTTCTGTAGTGTAGTGGTTATCACGTTCGCCTC | 1194.3 | 7.2 | 1.0 | 7.0 | 3.3E-12 | 1.2E-11 | MB231/MB468 | MP |
| >hg38_21_+_8433172_8446622_RNA45SN1_45S_50ntFlanks@10916.10946.31\|CCGCGGCTGGACGAGGCGCCGCCGCCCCCCC | >hg38_21_+_8433172_8446622_RNA45SN1_45S_50ntFlanks | 10916 | 10946 | 31 | CCGCGGCTGGACGAGGCGCCGCCGCCCCCCC | 133.1 | -10.9 | 1.6 | -7.0 | 3.3E-12 | 1.2E-11 | MB231/MB468 | MP |
| >MI0000465\|hsa-mir-191&WithFlank&3\|-\|49020612\|49020715@22.43.22(+1U)&[MIMAT0000440&hsa-miR-191-5p&offsets\|0\|-1(+1U);m-48&3\|-\|49020672\|49020694&offsets\|0\|-1(+1U)]\|CAACGGAATCCCAAAAGCAGCTT | >MI0000465\|hsa-mir-191&WithFlank&3\|-\|49020612\|49020715 | 22 | 43 | 22(+1U) | CAACGGAATCCCAAAAGCAGCTT | 620.5 | 8.1 | 1.2 | 7.0 | 2.8E-12 | 1.2E-11 | BT20/MB468 | MITO |
| >hg38_21_+_8433172_8446622_RNA45SN1_45S_50ntFlanks@5471.5498.28\|CACGGCCCTGGCGGAGCGCTGAGAAGAC | >hg38_21_+_8433172_8446622_RNA45SN1_45S_50ntFlanks | 5471 | 5498 | 28 | CACGGCCCTGGCGGAGCGCTGAGAAGAC | 150.5 | -11.1 | 1.6 | -7.0 | 3.4E-12 | 1.2E-11 | MB231/MB468 | MP |
| >hg38_21_+_8433172_8446622_RNA45SN1_45S_50ntFlanks@5482.5503.22\|CGGAGCGCTGAGAAGACGGTCG | >hg38_21_+_8433172_8446622_RNA45SN1_45S_50ntFlanks | 5482 | 5503 | 22 | CGGAGCGCTGAGAAGACGGTCG | 122.1 | -10.8 | 1.6 | -7.0 | 3.4E-12 | 1.2E-11 | MB231/MB468 | MP |
| >hg38_21_+_8433172_8446622_RNA45SN1_45S_50ntFlanks@10350.10377.28\|ACGAGAACTTTGAAGGCCGAAGTGGAGA | >hg38_21_+_8433172_8446622_RNA45SN1_45S_50ntFlanks | 10350 | 10377 | 28 | ACGAGAACTTTGAAGGCCGAAGTGGAGA | 8898.4 | -9.1 | 1.3 | -7.0 | 3.4E-12 | 1.2E-11 | MB231/MB468 | MP |
| >hg38_21_+_8433172_8446622_RNA45SN1_45S_50ntFlanks@10425.10453.29\|AGATGGGCGAGCGCCGTTCCGAAGGGACG | >hg38_21_+_8433172_8446622_RNA45SN1_45S_50ntFlanks | 10425 | 10453 | 29 | AGATGGGCGAGCGCCGTTCCGAAGGGACG | 118.7 | -10.8 | 1.5 | -7.0 | 3.4E-12 | 1.2E-11 | MB231/MB468 | MP |
| >hg38_21_+_8433172_8446622_RNA45SN1_45S_50ntFlanks@8195.8213.19\|CCGTGGACGGTGTGAGGCC | >hg38_21_+_8433172_8446622_RNA45SN1_45S_50ntFlanks | 8195 | 8213 | 19 | CCGTGGACGGTGTGAGGCC | 393.4 | 3.3 | 0.5 | 7.0 | 2.8E-12 | 1.2E-11 | BT20/MB468 | MITO |
| >hg38_21_+_8433172_8446622_RNA45SN1_45S_50ntFlanks@6730.6770.41\|ACACATTGATCATCGACACTTCGAACGCACTTGCGGCCCCG | >hg38_21_+_8433172_8446622_RNA45SN1_45S_50ntFlanks | 6730 | 6770 | 41 | ACACATTGATCATCGACACTTCGAACGCACTTGCGGCCCCG | 310.3 | -12.1 | 1.7 | -7.0 | 3.4E-12 | 1.2E-11 | MB231/MB468 | MP |
| >hg38_21_+_8433172_8446622_RNA45SN1_45S_50ntFlanks@11729.11761.33\|TAACTATGACTCTCTTAAGGTAGCCAAATGCCT | >hg38_21_+_8433172_8446622_RNA45SN1_45S_50ntFlanks | 11729 | 11761 | 33 | TAACTATGACTCTCTTAAGGTAGCCAAATGCCT | 173.7 | -11.3 | 1.6 | -7.0 | 3.4E-12 | 1.2E-11 | MB231/MB468 | MP |
| >hg38_21_+_8433172_8446622_RNA45SN1_45S_50ntFlanks@9545.9573.29\|GTGGAGGTCCGTAGCGGTCCTGACGTGCA | >hg38_21_+_8433172_8446622_RNA45SN1_45S_50ntFlanks | 9545 | 9573 | 29 | GTGGAGGTCCGTAGCGGTCCTGACGTGCA | 4430.5 | -8.9 | 1.3 | -6.9 | 4.9E-12 | 1.2E-11 | BT20/MB468 | NUC |

| | | | | | | | | | | | | | |
|---|---|---|---|---|---|---|---|---|---|---|---|---|---|
| >hg38_21_+_8433172_8446622_RNA45SN1_45S_50ntFlanks@5494.5526.33\|AAGACGGTCGAACTTGACTATCTAGAGGAAGTA | >hg38_21_+_8433172_8446622_RNA45SN1_45S_50ntFlanks | 5494 | 5526 | 33 | AAGACGGTCGAACTTGACTATCTAGAGGAAGTA | 1216.9 | -9.0 | 1.3 | -6.9 | 4.6E-12 | 1.2E-11 | MB231/MB468 | MITO |
| >hg38_21_+_8433172_8446622_RNA45SN1_45S_50ntFlanks@12822.12847.26\|CCGGGACCGGGGTCCGGTGCGGAGTG | >hg38_21_+_8433172_8446622_RNA45SN1_45S_50ntFlanks | 12822 | 12847 | 26 | CCGGGACCGGGGTCCGGTGCGGAGTG | 427.2 | -12.6 | 1.8 | -7.0 | 3.4E-12 | 1.2E-11 | MB231/MB468 | MP |
| >trnaMT_PheGAA_MT_+_577_647@35.57.23\|GAAAATGTTTAGACGGGCTCACA | >trnaMT_PheGAA_MT_+_577_647 | 35 | 57 | 23 | GAAAATGTTTAGACGGGCTCACA | 579.7 | 7.0 | 1.0 | 6.9 | 5.0E-12 | 1.2E-11 | BT20/MB468 | NUC |
| >hg38_21_+_8433172_8446622_RNA45SN1_45S_50ntFlanks@9554.9576.23\|CGTAGCGGTCCTGACGTGCAAAT | >hg38_21_+_8433172_8446622_RNA45SN1_45S_50ntFlanks | 9554 | 9576 | 23 | CGTAGCGGTCCTGACGTGCAAAT | 568.4 | -2.9 | 0.4 | -6.9 | 5.0E-12 | 1.2E-11 | BT20/MB468 | NUC |
| >hg38_21_+_8433172_8446622_RNA45SN1_45S_50ntFlanks@6649.6668.20\|TACGACTCTTAGCGGTGGAT | >hg38_21_+_8433172_8446622_RNA45SN1_45S_50ntFlanks | 6649 | 6668 | 20 | TACGACTCTTAGCGGTGGAT | 5179.0 | 6.1 | 0.9 | 7.0 | 2.8E-12 | 1.2E-11 | BT20/MB468 | MITO |
| >hg38_21_+_8433172_8446622_RNA45SN1_45S_50ntFlanks@5071.5097.27\|TTAATTCCGATAACGAACGAGACTCTG | >hg38_21_+_8433172_8446622_RNA45SN1_45S_50ntFlanks | 5071 | 5097 | 27 | TTAATTCCGATAACGAACGAGACTCTG | 722.8 | -13.4 | 1.9 | -7.0 | 3.4E-12 | 1.2E-11 | MB231/MB468 | MP |
| >MI0000298\|hsa-mir-221&WithFlank&X\|-\|45746151\|45746272@71.93.23&[MIMAT0000278&hsa-miR-221-3p&offsets\|0\|0;m-91&X\|-\|45746181\|45746202&offsets\|0\|+1]\|AGCTACATTGTCTGCTGGGTTTC | >MI0000298\|hsa-mir-221&WithFlank&X\|-\|45746151\|45746272 | 71 | 93 | 23 | AGCTACATTGTCTGCTGGGTTTC | 31365.1 | 5.6 | 0.8 | 6.9 | 4.6E-12 | 1.2E-11 | MB231/MB468 | MITO |
| >hg38_21_+_8433172_8446622_RNA45SN1_45S_50ntFlanks@5083.5117.35\|ACGAACGAGACTCTGGCATGCTAACTAGTTACGCG | >hg38_21_+_8433172_8446622_RNA45SN1_45S_50ntFlanks | 5083 | 5117 | 35 | ACGAACGAGACTCTGGCATGCTAACTAGTTACGCG | 315.2 | -12.2 | 1.7 | -7.0 | 3.4E-12 | 1.2E-11 | MB231/MB468 | MP |
| >hg38_21_+_8433172_8446622_RNA45SN1_45S_50ntFlanks@9796.9831.36\|TGGAGCCGGGCGTGGAATGCGAGTGCCTAGTGGGCC | >hg38_21_+_8433172_8446622_RNA45SN1_45S_50ntFlanks | 9796 | 9831 | 36 | TGGAGCCGGGCGTGGAATGCGAGTGCCTAGTGGGCC | 108.5 | -10.6 | 1.5 | -7.0 | 3.4E-12 | 1.2E-11 | MB231/MB468 | MP |
| >hg38_21_+_8433172_8446622_RNA45SN1_45S_50ntFlanks@10841.10858.18\|CGTAACTTCGGGATAAGG | >hg38_21_+_8433172_8446622_RNA45SN1_45S_50ntFlanks | 10841 | 10858 | 18 | CGTAACTTCGGGATAAGG | 298.0 | -8.5 | 1.2 | -6.9 | 3.9E-12 | 1.2E-11 | MB231/MB468 | NUC |
| >trna116_GluCTC_1_-_145399233_145399304@1.39.39,trna59_GluCTC_1_+_249168447_249168518@1.39.39,trna71_GluCTC_1_-_161439189_161439260@1.39.39,trna74_GluCTC_1_-_161431809_161431880@1.39.39,trna77_GluCTC_1_-_161424398_161424469@1.39.39,trna77_GluCTC_6_+_28949976_28950047@1.39.39,trna80_GluCTC_1_-_161417018_161417089@1.39.39,trna87_GluCTC_6_-_126101393_126101464@1.39.39\|TCCCTGGTGGTCTAGTGGTTAGGATTCGGCGCTCTCACC | >trna116_GluCTC_1_-_145399233_145399304 | 1 | 39 | 39 | TCCCTGGTGGTCTAGTGGTTAGGATTCGGCGCTCTCACC | 12151.9 | 6.0 | 0.9 | 7.0 | 3.6E-12 | 1.2E-11 | BT20/MB468 | CYTO |
| >MI0000650\|hsa-mir-200c&WithFlank&12\|+\|6963693\|6963772@50.70.21&[MIMAT0000617&hsa-miR-200c-3p&offsets\|0\|-2;m-54&12\|+\|6963742\|6963764&offsets\|0\|-2]\|TAATACTGCCGGGTAATGATG | >MI0000650\|hsa-mir-200c&WithFlank&12\|+\|6963693\|6963772 | 50 | 70 | 21 | TAATACTGCCGGGTAATGATG | 103.7 | -10.6 | 1.5 | -7.0 | 3.4E-12 | 1.2E-11 | MB231/MB468 | MP |
| >MI0000085\|hsa-mir-27a&WithFlank&19\|-\|13836434\|13836523@57.77.21&[MIMAT0000084&hsa-miR-27a-3p&offsets\|0\|0;m-43&19\|-\|13836447\|13836467&offsets\|0\|0]\|TTCACAGTGGCTAAGTTCCGC | >MI0000085\|hsa-mir-27a&WithFlank&19\|-\|13836434\|13836523 | 57 | 77 | 21 | TTCACAGTGGCTAAGTTCCGC | 988.6 | 5.2 | 0.7 | 7.0 | 3.6E-12 | 1.2E-11 | BT20/MB468 | CYTO |
| >MI0000273\|hsa-mir-183&WithFlank&7\|-\|129774899\|129775020@34.53.20(+1A)&[MIMAT0000261&hsa-miR-183-5p&offsets\|+1\|-1(+1A);m-41&7\|-\|129774966\|129774987&offsets\|0\|-2(+1A)]\|ATGGCACTGGTAGAATTCACA | >MI0000273\|hsa-mir-183&WithFlank&7\|-\|129774899\|129775020 | 34 | 53 | 20(+1A) | ATGGCACTGGTAGAATTCACA | 617.1 | 8.8 | 1.3 | 7.0 | 3.6E-12 | 1.2E-11 | BT20/MB468 | CYTO |
| >hg38_21_+_8433172_8446622_RNA45SN1_45S_50ntFlanks@12414.12444.31\|CCTTCGATGTCGGCTCTTCCTATCATTGTGA | >hg38_21_+_8433172_8446622_RNA45SN1_45S_50ntFlanks | 12414 | 12444 | 31 | CCTTCGATGTCGGCTCTTCCTATCATTGTGA | 391.7 | 4.5 | 0.6 | 6.9 | 3.9E-12 | 1.2E-11 | MB231/MB468 | CYTO |
| >hg38_21_+_8433172_8446622_RNA45SN1_45S_50ntFlanks@11877.11905.29\|AGACCCTGTTGAGCTTGACTCTAGTCTGG | >hg38_21_+_8433172_8446622_RNA45SN1_45S_50ntFlanks | 11877 | 11905 | 29 | AGACCCTGTTGAGCTTGACTCTAGTCTGG | 99.6 | -10.5 | 1.5 | -7.0 | 3.4E-12 | 1.2E-11 | MB231/MB468 | MP |
| >hg38_21_+_8433172_8446622_RNA45SN1_45S_50ntFlanks@13006.13023.18\|CTATTGAAAGTCAGCCCT | >hg38_21_+_8433172_8446622_RNA45SN1_45S_50ntFlanks | 13006 | 13023 | 18 | CTATTGAAAGTCAGCCCT | 182.2 | -7.5 | 1.1 | -6.9 | 3.7E-12 | 1.2E-11 | BT20/MB468 | CYTO |
| >hg38_21_+_8433172_8446622_RNA45SN1_45S_50ntFlanks@10771.10806.36\|CAGGTCTCCAAGGTGAACAGCCTCTGGCATGTTGGA | >hg38_21_+_8433172_8446622_RNA45SN1_45S_50ntFlanks | 10771 | 10806 | 36 | CAGGTCTCCAAGGTGAACAGCCTCTGGCATGTTGGA | 550.3 | -12.2 | 1.7 | -7.3 | 2.0E-13 | 1.2E-11 | BT20/MB468 | MP |
| >hg38_21_+_8433172_8446622_RNA45SN1_45S_50ntFlanks@12357.12375.19\|CCACAGGGATAACTGGCTT | >hg38_21_+_8433172_8446622_RNA45SN1_45S_50ntFlanks | 12357 | 12375 | 19 | CCACAGGGATAACTGGCTT | 329.9 | -11.4 | 1.6 | -7.3 | 2.0E-13 | 1.2E-11 | BT20/MB468 | MP |
| >MI0000269\|hsa-mir-181a-2&WithFlank&9\|+\|124692436\|124692557@83.104.22&[MIMAT0004558&hsa-miR-181a-2-3p&offsets\|0\|0;m-71&9\|+\|124692518\|124692539&offsets\|0\|0]\|ACCACTGACCGTTGACTGTACC | >MI0000269\|hsa-mir-181a-2&WithFlank&9\|+\|124692436\|124692557 | 83 | 104 | 22 | ACCACTGACCGTTGACTGTACC | 1492.8 | 5.2 | 0.7 | 6.9 | 4.0E-12 | 1.2E-11 | MB231/MB468 | NUC |
| >trnaMT_GluTTC_MT_-_14674_14742@24.58.35,trnalookalike8_GluTTC_5_-_93905172_93905240@24.58.35\|ATGGTTTTTCATATCATTGGTCGTGGTTGTAGTCC | >trnaMT_GluTTC_MT_-_14674_14742 | 24 | 58 | 35 | ATGGTTTTTCATATCATTGGTCGTGGTTGTAGTCC | 929.0 | 9.2 | 1.3 | 7.0 | 2.8E-12 | 1.2E-11 | BT20/MB468 | MITO |
| >hg38_21_+_8433172_8446622_RNA45SN1_45S_50ntFlanks@10869.10892.24\|AGGGCTGGGTCGGTCGGGCTGGGG | >hg38_21_+_8433172_8446622_RNA45SN1_45S_50ntFlanks | 10869 | 10892 | 24 | AGGGCTGGGTCGGTCGGGCTGGGG | 596.3 | -9.0 | 1.3 | -6.9 | 4.0E-12 | 1.2E-11 | MB231/MB468 | NUC |
| >hg38_21_+_8433172_8446622_RNA45SN1_45S_50ntFlanks@8352.8375.24\|AGTTGAAAAGAACTTTGAAGAGAG | >hg38_21_+_8433172_8446622_RNA45SN1_45S_50ntFlanks | 8352 | 8375 | 24 | AGTTGAAAAGAACTTTGAAGAGAG | 4655.9 | 9.2 | 1.2 | 7.3 | 2.0E-13 | 1.2E-11 | BT20/MB468 | MP |
| >hg38_21_+_8433172_8446622_RNA45SN1_45S_50ntFlanks@3782.3801.20\|CAGTGAAACTGCGAATGGCT | >hg38_21_+_8433172_8446622_RNA45SN1_45S_50ntFlanks | 3782 | 3801 | 20 | CAGTGAAACTGCGAATGGCT | 1053.2 | -3.1 | 0.4 | -6.9 | 3.7E-12 | 1.2E-11 | BT20/MB468 | CYTO |
| >hg38_21_+_8433172_8446622_RNA45SN1_45S_50ntFlanks@11316.11343.28\|CACCCCACGTCTCGTCGCGCGCGCGTCC | >hg38_21_+_8433172_8446622_RNA45SN1_45S_50ntFlanks | 11316 | 11343 | 28 | CACCCCACGTCTCGTCGCGCGCGCGTCC | 134.9 | -10.9 | 1.6 | -7.0 | 3.5E-12 | 1.2E-11 | MB231/MB468 | MP |
| >MI0000433\|hsa-let-7g&WithFlank&3\|-\|52268272\|52268367@11.28.18&[MIMAT0000414&hsa-let-7g-5p&offsets\|0\|-4;m-58&3\|-\|52268336\|52268357&offsets\|0\|-4]\|TGAGGTAGTAGTTTGTAC | >MI0000433\|hsa-let-7g&WithFlank&3\|-\|52268272\|52268367 | 11 | 28 | 18 | TGAGGTAGTAGTTTGTAC | 4121.4 | 5.4 | 0.8 | 6.9 | 4.0E-12 | 1.2E-11 | MB231/MB468 | NUC |
| >hg38_21_+_8433172_8446622_RNA45SN1_45S_50ntFlanks@4241.4258.18\|CTCTTTCGAGGCCCTGTA | >hg38_21_+_8433172_8446622_RNA45SN1_45S_50ntFlanks | 4241 | 4258 | 18 | CTCTTTCGAGGCCCTGTA | 268.0 | -10.2 | 1.5 | -7.0 | 2.9E-12 | 1.2E-11 | BT20/MB468 | MITO |
| >hg38_21_+_8433172_8446622_RNA45SN1_45S_50ntFlanks@11933.11967.35\|AATAAGTGGGAGGCCCCCGGCGCCCCCCCGGTGTC | >hg38_21_+_8433172_8446622_RNA45SN1_45S_50ntFlanks | 11933 | 11967 | 35 | AATAAGTGGGAGGCCCCCGGCGCCCCCCCGGTGTC | 349.1 | -11.5 | 1.6 | -7.3 | 2.1E-13 | 1.2E-11 | BT20/MB468 | MP |
| >hg38_21_+_8433172_8446622_RNA45SN1_45S_50ntFlanks@13002.13029.28\|CGATCTATTGAAAGTCAGCCCTCGACAC | >hg38_21_+_8433172_8446622_RNA45SN1_45S_50ntFlanks | 13002 | 13029 | 28 | CGATCTATTGAAAGTCAGCCCTCGACAC | 279.2 | -11.2 | 1.5 | -7.3 | 2.1E-13 | 1.2E-11 | BT20/MB468 | MP |
| >MI0000650\|hsa-mir-200c&WithFlank&12\|+\|6963693\|6963772@49.71.23&[MIMAT0000617&hsa-miR-200c-3p&offsets\|-1\|-1;m-54&12\|+\|6963742\|6963764&offsets\|-1\|-1]\|CTAATACTGCCGGGTAATGATGG | >MI0000650\|hsa-mir-200c&WithFlank&12\|+\|6963693\|6963772 | 49 | 71 | 23 | CTAATACTGCCGGGTAATGATGG | 162.9 | 3.1 | 0.4 | 6.9 | 5.1E-12 | 1.2E-11 | BT20/MB468 | NUC |
| >hg38_21_+_8433172_8446622_RNA45SN1_45S_50ntFlanks@7986.8007.22\|ATCAGACGTGGCGACCCGCTGA | >hg38_21_+_8433172_8446622_RNA45SN1_45S_50ntFlanks | 7986 | 8007 | 22 | ATCAGACGTGGCGACCCGCTGA | 297.6 | -10.3 | 1.5 | -7.0 | 2.9E-12 | 1.2E-11 | BT20/MB468 | MITO |
| >hg38_21_+_8433172_8446622_RNA45SN1_45S_50ntFlanks@10408.10425.18\|GGTCAGTCGGTCCTGAGA | >hg38_21_+_8433172_8446622_RNA45SN1_45S_50ntFlanks | 10408 | 10425 | 18 | GGTCAGTCGGTCCTGAGA | 365.4 | -5.3 | 0.8 | -6.9 | 3.9E-12 | 1.2E-11 | MB231/MB468 | CYTO |
| >hg38_21_+_8433172_8446622_RNA45SN1_45S_50ntFlanks@11581.11600.20\|AACTGGTGCGGACCAGGGGA | >hg38_21_+_8433172_8446622_RNA45SN1_45S_50ntFlanks | 11581 | 11600 | 20 | AACTGGTGCGGACCAGGGGA | 951.3 | -5.3 | 0.8 | -6.9 | 3.9E-12 | 1.2E-11 | MB231/MB468 | CYTO |
| >hg38_21_+_8433172_8446622_RNA45SN1_45S_50ntFlanks@9548.9577.30\|GAGGTCCGTAGCGGTCCTGACGTGCAAATC | >hg38_21_+_8433172_8446622_RNA45SN1_45S_50ntFlanks | 9548 | 9577 | 30 | GAGGTCCGTAGCGGTCCTGACGTGCAAATC | 161.5 | 5.7 | 0.8 | 6.9 | 3.9E-12 | 1.2E-11 | MB231/MB468 | CYTO |
| >hg38_21_+_8433172_8446622_RNA45SN1_45S_50ntFlanks@10874.10902.29\|TGGGTCGGTCGGGCTGGGGCGCGAAGCGG | >hg38_21_+_8433172_8446622_RNA45SN1_45S_50ntFlanks | 10874 | 10902 | 29 | TGGGTCGGTCGGGCTGGGGCGCGAAGCGG | 118.8 | -10.8 | 1.5 | -7.0 | 3.5E-12 | 1.2E-11 | MB231/MB468 | MP |
| >MI0000542\|hsa-mir-320a&WithFlank&8\|-\|22244960\|22245043@48.68.21(+1C)&[MIMAT0000510&hsa-miR-320a-3p&offsets\|0\|-1(+1C);m-60&8\|-\|22244975\|22244996&offsets\|0\|-1(+1C)]\|AAAAGCTGGGTTGAGAGGGCGC | >MI0000542\|hsa-mir-320a&WithFlank&8\|-\|22244960\|22245043 | 48 | 68 | 21(+1C) | AAAAGCTGGGTTGAGAGGGCGC | 165.2 | 3.6 | 0.5 | 6.9 | 5.1E-12 | 1.2E-11 | BT20/MB468 | NUC |
| >hg38_21_+_8433172_8446622_RNA45SN1_45S_50ntFlanks@5485.5525.41\|AGCGCTGAGAAGACGGTCGAACTTGACTATCTAGAGGAAGT | >hg38_21_+_8433172_8446622_RNA45SN1_45S_50ntFlanks | 5485 | 5525 | 41 | AGCGCTGAGAAGACGGTCGAACTTGACTATCTAGAGGAAGT | 160.3 | -11.2 | 1.6 | -7.0 | 3.5E-12 | 1.2E-11 | MB231/MB468 | MP |
| >hg38_21_+_8433172_8446622_RNA45SN1_45S_50ntFlanks@4064.4081.18\|ATCAACTTTCGATGGTAG | >hg38_21_+_8433172_8446622_RNA45SN1_45S_50ntFlanks | 4064 | 4081 | 18 | ATCAACTTTCGATGGTAG | 561.1 | -11.4 | 1.6 | -6.9 | 3.7E-12 | 1.2E-11 | BT20/MB468 | CYTO |
| >hg38_21_+_8433172_8446622_RNA45SN1_45S_50ntFlanks@6670.6698.29\|ACTCGGCTCGTGCGTCGATGAAGAACGCA | >hg38_21_+_8433172_8446622_RNA45SN1_45S_50ntFlanks | 6670 | 6698 | 29 | ACTCGGCTCGTGCGTCGATGAAGAACGCA | 2923.7 | -6.5 | 0.9 | -6.9 | 4.8E-12 | 1.2E-11 | MB231/MB468 | MITO |
| >hg38_21_+_8433172_8446622_RNA45SN1_45S_50ntFlanks@8659.8699.41\|CATTTCCACCGCGGCGGTGCGCCGCGACCGGCTCCGGGACG | >hg38_21_+_8433172_8446622_RNA45SN1_45S_50ntFlanks | 8659 | 8699 | 41 | CATTTCCACCGCGGCGGTGCGCCGCGACCGGCTCCGGGACG | 258.1 | -11.8 | 1.7 | -6.9 | 4.8E-12 | 1.2E-11 | MB231/MB468 | MITO |
| >hg38_21_+_8433172_8446622_RNA45SN1_45S_50ntFlanks@8170.8212.43\|CAAGTCCTTCTGATCGAGGCCCAGCCCGTGGACGGTGTGAGGC | >hg38_21_+_8433172_8446622_RNA45SN1_45S_50ntFlanks | 8170 | 8212 | 43 | CAAGTCCTTCTGATCGAGGCCCAGCCCGTGGACGGTGTGAGGC | 2532.0 | -11.0 | 1.6 | -7.0 | 3.5E-12 | 1.2E-11 | MB231/MB468 | MP |
| >hg38_21_+_8433172_8446622_RNA45SN1_45S_50ntFlanks@10071.10098.28\|TGGACGGGAGCGGCGGGGGCGGCGCG | >hg38_21_+_8433172_8446622_RNA45SN1_45S_50ntFlanks | 10071 | 10098 | 28 | TGGACGGGAGCGGCGGGGGCGGCGCG | 143.2 | -11.0 | 1.6 | -7.0 | 3.5E-12 | 1.2E-11 | MB231/MB468 | MP |
| >hg38_21_+_8433172_8446622_RNA45SN1_45S_50ntFlanks@11561.11605.45\|GCGCCTAGCAGCCGACTTAGAACTGGTGCGGACCAGGGGAATCCG | >hg38_21_+_8433172_8446622_RNA45SN1_45S_50ntFlanks | 11561 | 11605 | 45 | GCGCCTAGCAGCCGACTTAGAACTGGTGCGGACCAGGGGAATCCG | 294.6 | -11.2 | 1.6 | -6.9 | 4.1E-12 | 1.2E-11 | MB231/MB468 | NUC |
| >trna111_HisGTG_1_-_147774845_147774916@-1T.33.34,trna118_HisGTG_1_-_145396881_145396952@-1T.33.34,trna16_HisGTG_1_+_146544773_146544844@-1T.33.34,trna1_HisGTG_15_+_45493349_45493420@-1T.33.34,trna21_HisGTG_1_+_147753471_147753542@-1T.33.34,trna33_HisGTG_6_+_27125906_27125977@-1T.33.34,trna7_HisGTG_9_-_14433938_14434009@-1T.33.34,trna8_HisGTG_15_-_45492611_45492682@-1T.33.34,trna9_HisGTG_15_-_45490804_45490875@-1T.33.34\|TGCCGTGATCGTATAGTGGTTAGTACTCTGCGTT | >trna111_HisGTG_1_-_147774845_147774916 | -1T | 33 | 34 | TGCCGTGATCGTATAGTGGTTAGTACTCTGCGTT | 2508.9 | 7.8 | 1.1 | 6.9 | 4.1E-12 | 1.3E-11 | MB231/MB468 | NUC |
| >hg38_21_+_8433172_8446622_RNA45SN1_45S_50ntFlanks@11076.11103.28\|CGCCGCGTGGGGGCGGCGGCGGGGGGAG | >hg38_21_+_8433172_8446622_RNA45SN1_45S_50ntFlanks | 11076 | 11103 | 28 | CGCCGCGTGGGGGCGGCGGCGGGGGGAG | 231.2 | -11.7 | 1.7 | -7.0 | 3.5E-12 | 1.3E-11 | MB231/MB468 | MP |
| >hg38_21_+_8433172_8446622_RNA45SN1_45S_50ntFlanks@6735.6771.37\|TTGATCATCGACACTTCGAACGCACTTGCGGCCCCGG | >hg38_21_+_8433172_8446622_RNA45SN1_45S_50ntFlanks | 6735 | 6771 | 37 | TTGATCATCGACACTTCGAACGCACTTGCGGCCCCGG | 224.8 | -11.7 | 1.7 | -7.0 | 3.5E-12 | 1.3E-11 | MB231/MB468 | MP |
| >hg38_21_+_8433172_8446622_RNA45SN1_45S_50ntFlanks@12458.12496.39\|AAGCGTTGGATTGTTCACCCACTAATAGGGAACGTGAGC | >hg38_21_+_8433172_8446622_RNA45SN1_45S_50ntFlanks | 12458 | 12496 | 39 | AAGCGTTGGATTGTTCACCCACTAATAGGGAACGTGAGC | 240.4 | 4.0 | 0.6 | 6.9 | 4.1E-12 | 1.3E-11 | MB231/MB468 | NUC |
| >hg38_21_+_8433172_8446622_RNA45SN1_45S_50ntFlanks@10351.10389.39\|CGAGAACTTTGAAGGCCGAAGTGGAGAAGGGTTCCATGT | >hg38_21_+_8433172_8446622_RNA45SN1_45S_50ntFlanks | 10351 | 10389 | 39 | CGAGAACTTTGAAGGCCGAAGTGGAGAAGGGTTCCATGT | 180.8 | -11.4 | 1.6 | -7.0 | 3.6E-12 | 1.3E-11 | MB231/MB468 | MP |

| | | | | | | | | | | | | | |
|---|---|---|---|---|---|---|---|---|---|---|---|---|---|
| >hg38_1_-_228634819_228635039_RNA5S12_5S_50ntFlanks@144.171.28\|CCTGGGAATACCGGGTGCTGTAGGCTTT | >hg38_1_-_228634819_228635039_RNA5S12_5S_50ntFlanks | 144 | 171 | 28 | CCTGGGAATACCGGGTGCTGTAGGCTTT | 2829.9 | 4.9 | 0.7 | 6.9 | 3.8E-12 | 1.3E-11 | BT20/MB468 | CYTO |
| >hg38_21_+_8433172_8446622_RNA45SN1_45S_50ntFlanks@12621.12653.33\|GAGCCAATGGGGCGAAGCTACCATCTGTGGGAT | >hg38_21_+_8433172_8446622_RNA45SN1_45S_50ntFlanks | 12621 | 12653 | 33 | GAGCCAATGGGGCGAAGCTACCATCTGTGGGAT | 101.8 | -10.5 | 1.5 | -7.0 | 3.6E-12 | 1.3E-11 | MB231/MB468 | MP |
| >hg38_1_-_228634819_228635039_RNA5S12_5S_50ntFlanks@134.172.39\|TGGGAGACCGCCTGGGAATACCGGGTGCTGTAGGCTTTT | >hg38_1_-_228634819_228635039_RNA5S12_5S_50ntFlanks | 134 | 172 | 39 | TGGGAGACCGCCTGGGAATACCGGGTGCTGTAGGCTTTT | 2451.2 | 9.1 | 1.3 | 6.9 | 4.0E-12 | 1.3E-11 | MB231/MB468 | CYTO |
| >MI0000750\|hsa-mir-26a-2&WithFlank&12\|-\|57824603\|57824698@20.41.22(+1A);MI0000083\|hsa-mir-26a-1&WithFlank&3\|+\|37969398\|37969486@16.37.22(+1A)&[MIMAT0000082&hsa-miR-26a-5p&offsets\|0\|0(+1A);m-33&12\|-\|57824658\|57824679&offsets\|0\|0(+1A)];[MIMAT0000082_1&hsa-miR-26a-5p&offsets\|0\|0(+1A);m-34&3\|+\|37969413\|37969434&offsets\|0\|0(+1A)]\|TTCAAGTAATCCAGGATAGGCTA | >MI0000750\|hsa-mir-26a-2&WithFlank&12\|-\|57824603\|57824698 | 20 | 41 | 22(+1A) | TTCAAGTAATCCAGGATAGGCTA | 692.8 | 4.5 | 0.7 | 6.9 | 4.1E-12 | 1.3E-11 | MB231/MB468 | NUC |
| >hg38_21_+_8433172_8446622_RNA45SN1_45S_50ntFlanks@11163.11188.26\|CGGTCCCCCGCGAGGGGGGCCCGGGC | >hg38_21_+_8433172_8446622_RNA45SN1_45S_50ntFlanks | 11163 | 11188 | 26 | CGGTCCCCCGCGAGGGGGGCCCGGGC | 184.5 | -10.5 | 1.5 | -6.9 | 4.1E-12 | 1.3E-11 | MB231/MB468 | NUC |
| >hg38_21_+_8433172_8446622_RNA45SN1_45S_50ntFlanks@9737.9765.29\|CGAAACGATCTCAACCTATTCTCAAACTT | >hg38_21_+_8433172_8446622_RNA45SN1_45S_50ntFlanks | 9737 | 9765 | 29 | CGAAACGATCTCAACCTATTCTCAAACTT | 447.8 | -4.8 | 0.7 | -6.9 | 5.2E-12 | 1.3E-11 | BT20/MB468 | NUC |
| >hg38_21_+_8433172_8446622_RNA45SN1_45S_50ntFlanks@12456.12491.36\|CCAAGCGTTGGATTGTTCACCCACTAATAGGGAACG | >hg38_21_+_8433172_8446622_RNA45SN1_45S_50ntFlanks | 12456 | 12491 | 36 | CCAAGCGTTGGATTGTTCACCCACTAATAGGGAACG | 174.8 | -9.5 | 1.4 | -7.0 | 2.9E-12 | 1.3E-11 | BT20/MB468 | MITO |
| >hg38_21_+_8433172_8446622_RNA45SN1_45S_50ntFlanks@4041.4059.19\|CGACCCATTCGAACGTCTG | >hg38_21_+_8433172_8446622_RNA45SN1_45S_50ntFlanks | 4041 | 4059 | 19 | CGACCCATTCGAACGTCTG | 106.9 | -10.6 | 1.5 | -7.0 | 3.6E-12 | 1.3E-11 | MB231/MB468 | MP |
| >hg38_21_+_8433172_8446622_RNA45SN1_45S_50ntFlanks@11586.11607.22\|GTGCGGACCAGGGGAATCCGAC | >hg38_21_+_8433172_8446622_RNA45SN1_45S_50ntFlanks | 11586 | 11607 | 22 | GTGCGGACCAGGGGAATCCGAC | 228.3 | -11.7 | 1.7 | -7.0 | 3.6E-12 | 1.3E-11 | MB231/MB468 | MP |
| >hg38_21_+_8433172_8446622_RNA45SN1_45S_50ntFlanks@3782.3812.31\|CAGTGAAACTGCGAATGGCTCATTAAATCAG | >hg38_21_+_8433172_8446622_RNA45SN1_45S_50ntFlanks | 3782 | 3812 | 31 | CAGTGAAACTGCGAATGGCTCATTAAATCAG | 384.8 | -12.5 | 1.8 | -7.0 | 3.6E-12 | 1.3E-11 | MB231/MB468 | MP |
| >hg38_21_+_8433172_8446622_RNA45SN1_45S_50ntFlanks@10786.10816.31\|AACAGCCTCTGGCATGTTGGAACAATGTAGG | >hg38_21_+_8433172_8446622_RNA45SN1_45S_50ntFlanks | 10786 | 10816 | 31 | AACAGCCTCTGGCATGTTGGAACAATGTAGG | 2539.8 | -10.5 | 1.5 | -6.9 | 5.2E-12 | 1.3E-11 | BT20/MB468 | NUC |
| >hg38_21_+_8433172_8446622_RNA45SN1_45S_50ntFlanks@6644.6661.18\|CCTCGTACGACTCTTAGC | >hg38_21_+_8433172_8446622_RNA45SN1_45S_50ntFlanks | 6644 | 6661 | 18 | CCTCGTACGACTCTTAGC | 668.3 | 6.2 | 0.9 | 7.0 | 3.0E-12 | 1.3E-11 | BT20/MB468 | MITO |
| >hg38_21_+_8433172_8446622_RNA45SN1_45S_50ntFlanks@12456.12481.26\|CCAAGCGTTGGATTGTTCACCCACTA | >hg38_21_+_8433172_8446622_RNA45SN1_45S_50ntFlanks | 12456 | 12481 | 26 | CCAAGCGTTGGATTGTTCACCCACTA | 732.1 | -7.8 | 1.1 | -6.9 | 4.1E-12 | 1.3E-11 | MB231/MB468 | NUC |
| >MI0016887\|hsa-mir-4521&WithFlank&17\|+\|8186939\|8187010@13.34.22&[MIMAT0019058&hsa-miR-4521&offsets\|+3\|+3;m-765&17\|+\|8186948\|8186969&offsets\|+3\|+3]\|AAGGAAGTCCTGTGCTCAGTTT | >MI0016887\|hsa-mir-4521&WithFlank&17\|+\|8186939\|8187010 | 13 | 34 | 22 | AAGGAAGTCCTGTGCTCAGTTT | 834.2 | 10.7 | 1.5 | 6.9 | 3.8E-12 | 1.3E-11 | BT20/MB468 | CYTO |
| >hg38_21_+_8433172_8446622_RNA45SN1_45S_50ntFlanks@7983.8008.26\|CAGATCAGACGTGGCGACCCGCTGAA | >hg38_21_+_8433172_8446622_RNA45SN1_45S_50ntFlanks | 7983 | 8008 | 26 | CAGATCAGACGTGGCGACCCGCTGAA | 4724.6 | -4.6 | 0.7 | -7.0 | 3.0E-12 | 1.3E-11 | BT20/MB468 | MITO |
| >hg38_21_+_8433172_8446622_RNA45SN1_45S_50ntFlanks@9711.9737.27\|AAGCGAATGATTAGAGGTCTTGGGGCC | >hg38_21_+_8433172_8446622_RNA45SN1_45S_50ntFlanks | 9711 | 9737 | 27 | AAGCGAATGATTAGAGGTCTTGGGGCC | 179.0 | -11.3 | 1.6 | -7.0 | 3.6E-12 | 1.3E-11 | MB231/MB468 | MP |
| >hg38_21_+_8433172_8446622_RNA45SN1_45S_50ntFlanks@11933.11967.35\|AATAAGTGGGAGGCCCCCGGCGCCCCCCCGGTGTC | >hg38_21_+_8433172_8446622_RNA45SN1_45S_50ntFlanks | 11933 | 11967 | 35 | AATAAGTGGGAGGCCCCCGGCGCCCCCCCGGTGTC | 390.7 | -12.4 | 1.8 | -6.9 | 4.9E-12 | 1.3E-11 | MB231/MB468 | MITO |
| >hg38_21_+_8433172_8446622_RNA45SN1_45S_50ntFlanks@8104.8138.35\|CCGCGGCGGGGCGCGGGACATGTGGCGTACGGAAG | >hg38_21_+_8433172_8446622_RNA45SN1_45S_50ntFlanks | 8104 | 8138 | 35 | CCGCGGCGGGGCGCGGGACATGTGGCGTACGGAAG | 1809.0 | -8.2 | 1.2 | -6.9 | 4.9E-12 | 1.3E-11 | MB231/MB468 | MITO |
| >hg38_21_+_8433172_8446622_RNA45SN1_45S_50ntFlanks@4657.4683.27\|CTAGAGGTGAAATTCTTGGACCGGCGC | >hg38_21_+_8433172_8446622_RNA45SN1_45S_50ntFlanks | 4657 | 4683 | 27 | CTAGAGGTGAAATTCTTGGACCGGCGC | 126.3 | -10.8 | 1.6 | -7.0 | 3.6E-12 | 1.3E-11 | MB231/MB468 | MP |
| >hg38_21_+_8433172_8446622_RNA45SN1_45S_50ntFlanks@8687.8706.20\|CGGCTCCGGGACGGCTGGGA | >hg38_21_+_8433172_8446622_RNA45SN1_45S_50ntFlanks | 8687 | 8706 | 20 | CGGCTCCGGGACGGCTGGGA | 318.3 | 9.5 | 1.4 | 6.9 | 3.9E-12 | 1.3E-11 | BT20/MB468 | CYTO |
| >hg38_21_+_8433172_8446622_RNA45SN1_45S_50ntFlanks@5473.5507.35\|CGGCCCTGGCGGAGCGCTGAGAAGACGGTCGAACT | >hg38_21_+_8433172_8446622_RNA45SN1_45S_50ntFlanks | 5473 | 5507 | 35 | CGGCCCTGGCGGAGCGCTGAGAAGACGGTCGAACT | 1647.3 | 8.7 | 1.3 | 6.9 | 3.9E-12 | 1.3E-11 | BT20/MB468 | CYTO |
| >hg38_21_+_8433172_8446622_RNA45SN1_45S_50ntFlanks@8170.8215.46\|CAAGTCCTTCTGATCGAGGCCCAGCCCGTGGACGGTGTGAGGCCGG | >hg38_21_+_8433172_8446622_RNA45SN1_45S_50ntFlanks | 8170 | 8215 | 46 | CAAGTCCTTCTGATCGAGGCCCAGCCCGTGGACGGTGTGAGGCCGG | 478.5 | -12.8 | 1.8 | -7.0 | 3.6E-12 | 1.3E-11 | MB231/MB468 | MP |
| >trna10_ValCAC_5_-_180649395_180649467@36.54.19,trna12_ValAAC_5_-_180645270_180645342@36.54.19,trna132_ValAAC_6_-_27721179_27721251@36.54.19,trna13_ValCAC_19_-_4724647_4724719@36.54.19,trna152_ValCAC_6_-_27248049_27248121@36.54.19,trna157_ValCAC_6_-_27173867_27173939@36.54.19,trna15_ValAAC_5_-_180615416_180615488@36.54.19,trna18_ValCAC_5_-_180529253_180529325@36.54.19,trna2_ValAAC_3_+_169490018_169490090@36.54.19,trna2_ValCAC_5_+_180524070_180524142@36.54.19,trna37_ValAAC_6_+_27203288_27203360@36.54.19,trna4_ValAAC_5_+_180591154_180591226@36.54.19,trna5_ValAAC_5_+_180596610_180596682@36.54.19,trna6_ValCAC_5_+_180600650_180600722@36.54.19,trna85_ValCAC_1_-_161369490_161369562@36.54.19,trna98_ValCAC_1_-_149298555_149298627@36.54.19,trna9_ValCAC_6_+_26538282_26538354@36.54.19\|CACGCGAAAGGTCCCCGGT | >trna10_ValCAC_5_-_180649395_180649467 | 36 | 54 | 19 | CACGCGAAAGGTCCCCGGT | 368.6 | -12.4 | 1.8 | -7.0 | 3.6E-12 | 1.3E-11 | MB231/MB468 | MP |
| >hg38_21_+_8433172_8446622_RNA45SN1_45S_50ntFlanks@8195.8223.29\|CCGTGGACGGTGTGAGGCCGGTAGCGGCC | >hg38_21_+_8433172_8446622_RNA45SN1_45S_50ntFlanks | 8195 | 8223 | 29 | CCGTGGACGGTGTGAGGCCGGTAGCGGCC | 100.9 | -10.5 | 1.5 | -7.0 | 3.6E-12 | 1.3E-11 | MB231/MB468 | MP |
| >hg38_21_+_8433172_8446622_RNA45SN1_45S_50ntFlanks@8154.8184.31\|CCGCTCGTGGGGGGCCCAAGTCCTTCTGATC | >hg38_21_+_8433172_8446622_RNA45SN1_45S_50ntFlanks | 8154 | 8184 | 31 | CCGCTCGTGGGGGGCCCAAGTCCTTCTGATC | 186.6 | -11.4 | 1.6 | -7.0 | 3.6E-12 | 1.3E-11 | MB231/MB468 | MP |
| >hg38_21_+_8433172_8446622_RNA45SN1_45S_50ntFlanks@11453.11478.26\|TCCTCGGCGCGCGGCGGCGGCGG | >hg38_21_+_8433172_8446622_RNA45SN1_45S_50ntFlanks | 11453 | 11478 | 26 | TCCTCGGCGCGCGGCGGCGGCGG | 141.9 | -9.5 | 1.4 | -6.9 | 3.9E-12 | 1.3E-11 | BT20/MB468 | CYTO |
| >MI0000087\|hsa-mir-29a&WithFlank&7\|-\|130876741\|130876816@48.67.20(+1C)&[MIMAT0000086&hsa-miR-29a-3p&offsets\|0\|-2(+1C);m-26&7\|-\|130876749\|130876769&offsets\|0\|-1(+1C)]\|TAGCACCATCTGAAATCGGTC | >MI0000087\|hsa-mir-29a&WithFlank&7\|-\|130876741\|130876816 | 48 | 67 | 20(+1C) | TAGCACCATCTGAAATCGGTC | 1995.7 | 9.2 | 1.3 | 6.9 | 5.0E-12 | 1.3E-11 | MB231/MB468 | MITO |
| >hg38_21_+_8433172_8446622_RNA45SN1_45S_50ntFlanks@12352.12376.25\|AGTTACCACAGGGATAACTGGCTTG | >hg38_21_+_8433172_8446622_RNA45SN1_45S_50ntFlanks | 12352 | 12376 | 25 | AGTTACCACAGGGATAACTGGCTTG | 91.9 | -10.4 | 1.5 | -6.9 | 3.7E-12 | 1.3E-11 | MB231/MB468 | MP |
| >MI0000478\|hsa-mir-149&WithFlank&2\|+\|240455995\|240456095@21.43.23&[MIMAT0000450&hsa-miR-149-5p&offsets\|0\|0;m-195&2\|+\|240456015\|240456037&offsets\|0\|0]\|TCTGGCTCCGTGTCTTCACTCCC | >MI0000478\|hsa-mir-149&WithFlank&2\|+\|240455995\|240456095 | 21 | 43 | 23 | TCTGGCTCCGTGTCTTCACTCCC | 201.0 | 5.1 | 0.7 | 6.9 | 4.2E-12 | 1.3E-11 | MB231/MB468 | NUC |
| >hg38_21_+_8433172_8446622_RNA45SN1_45S_50ntFlanks@7984.8004.21\|AGATCAGACGTGGCGACCCGC | >hg38_21_+_8433172_8446622_RNA45SN1_45S_50ntFlanks | 7984 | 8004 | 21 | AGATCAGACGTGGCGACCCGC | 1781.6 | -9.3 | 1.3 | -6.9 | 5.0E-12 | 1.3E-11 | MB231/MB468 | MITO |
| >hg38_21_+_8433172_8446622_RNA45SN1_45S_50ntFlanks@12605.12626.22\|TATGTGCTTGGCTGAGGAGCCA | >hg38_21_+_8433172_8446622_RNA45SN1_45S_50ntFlanks | 12605 | 12626 | 22 | TATGTGCTTGGCTGAGGAGCCA | 4146.7 | -9.9 | 1.4 | -6.9 | 4.2E-12 | 1.3E-11 | MB231/MB468 | NUC |
| >hg38_21_+_8433172_8446622_RNA45SN1_45S_50ntFlanks@10877.10901.25\|GTCGGTCGGGCTGGGGCGCGAAGCG | >hg38_21_+_8433172_8446622_RNA45SN1_45S_50ntFlanks | 10877 | 10901 | 25 | GTCGGTCGGGCTGGGGCGCGAAGCG | 122.2 | -10.8 | 1.6 | -6.9 | 3.7E-12 | 1.3E-11 | MB231/MB468 | MP |
| >MI0000342\|hsa-mir-200b&WithFlank&1\|+\|1167098\|1167204@64.85.22(+1A)&[MIMAT0000318&hsa-miR-200b-3p&offsets\|+1\|+1(+1A);m-93&1\|+\|1167160\|1167182&offsets\|+1\|0(+1A)]\|AATACTGCCTGGTAATGATGACA | >MI0000342\|hsa-mir-200b&WithFlank&1\|+\|1167098\|1167204 | 64 | 85 | 22(+1A) | AATACTGCCTGGTAATGATGACA | 92.5 | -10.4 | 1.5 | -6.9 | 3.7E-12 | 1.3E-11 | MB231/MB468 | MP |
| >hg38_21_+_8433172_8446622_RNA45SN1_45S_50ntFlanks@8111.8140.30\|GGGGCGCGGGACATGTGGCGTACGGAAGAC | >hg38_21_+_8433172_8446622_RNA45SN1_45S_50ntFlanks | 8111 | 8140 | 30 | GGGGCGCGGGACATGTGGCGTACGGAAGAC | 189.3 | -11.4 | 1.6 | -6.9 | 3.7E-12 | 1.3E-11 | MB231/MB468 | MP |
| >hg38_21_+_8433172_8446622_RNA45SN1_45S_50ntFlanks@6731.6771.41\|CACATTGATCATCGACACTTCGAACGCACTTGCGGCCCCGG | >hg38_21_+_8433172_8446622_RNA45SN1_45S_50ntFlanks | 6731 | 6771 | 41 | CACATTGATCATCGACACTTCGAACGCACTTGCGGCCCCGG | 449.8 | -12.7 | 1.8 | -6.9 | 3.7E-12 | 1.3E-11 | MB231/MB468 | MP |
| >hg38_21_+_8433172_8446622_RNA45SN1_45S_50ntFlanks@11233.11251.19\|CCCTTCCCGTGGATCGCCC | >hg38_21_+_8433172_8446622_RNA45SN1_45S_50ntFlanks | 11233 | 11251 | 19 | CCCTTCCCGTGGATCGCCC | 208.8 | -10.7 | 1.5 | -6.9 | 4.3E-12 | 1.3E-11 | MB231/MB468 | NUC |
| >MI0000471\|hsa-mir-126&WithFlank&9\|+\|136670596\|136670692@59.77.19(+1U)&[MIMAT0000445&hsa-miR-126-3p&offsets\|+1\|-2(+1U);m-40&9\|+\|136670653\|136670674&offsets\|+1\|-2(+1U)]\|CGTACCGTGAGTAATAATGT | >MI0000471\|hsa-mir-126&WithFlank&9\|+\|136670596\|136670692 | 59 | 77 | 19(+1U) | CGTACCGTGAGTAATAATGT | 296.3 | 11.4 | 1.6 | 6.9 | 3.7E-12 | 1.3E-11 | MB231/MB468 | MP |
| >hg38_21_+_8433172_8446622_RNA45SN1_45S_50ntFlanks@10469.10497.29\|TGCCCTCGGCCGATCGAAAGGGAGTCGGG | >hg38_21_+_8433172_8446622_RNA45SN1_45S_50ntFlanks | 10469 | 10497 | 29 | TGCCCTCGGCCGATCGAAAGGGAGTCGGG | 96.3 | -10.5 | 1.5 | -6.9 | 3.7E-12 | 1.3E-11 | MB231/MB468 | MP |
| >hg38_21_+_8433172_8446622_RNA45SN1_45S_50ntFlanks@9808.9830.23\|TGGAATGCGAGTGCCTAGTGGGC | >hg38_21_+_8433172_8446622_RNA45SN1_45S_50ntFlanks | 9808 | 9830 | 23 | TGGAATGCGAGTGCCTAGTGGGC | 140.5 | -11.0 | 1.6 | -6.9 | 3.7E-12 | 1.3E-11 | MB231/MB468 | MP |
| >trna37_ValAAC_6_+_27203288_27203360@1.29.29\|GTTTCCGTAGTGTAGTGGTTATCACGTTT | >trna37_ValAAC_6_+_27203288_27203360 | 1 | 29 | 29 | GTTTCCGTAGTGTAGTGGTTATCACGTTT | 1419.7 | -6.3 | 0.9 | -6.9 | 4.2E-12 | 1.3E-11 | MB231/MB468 | CYTO |
| >MI0000342\|hsa-mir-200b&WithFlank&1\|+\|1167098\|1167204@64.85.22(+1U)&[MIMAT0000318&hsa-miR-200b-3p&offsets\|+1\|+1(+1U);m-93&1\|+\|1167160\|1167182&offsets\|+1\|0(+1U)]\|AATACTGCCTGGTAATGATGACT | >MI0000342\|hsa-mir-200b&WithFlank&1\|+\|1167098\|1167204 | 64 | 85 | 22(+1U) | AATACTGCCTGGTAATGATGACT | 244.8 | -5.2 | 0.8 | -6.9 | 4.2E-12 | 1.3E-11 | MB231/MB468 | CYTO |
| >hg38_21_+_8433172_8446622_RNA45SN1_45S_50ntFlanks@11729.11765.37\|TAACTATGACTCTCTTAAGGTAGCCAAATGCCTCGTC | >hg38_21_+_8433172_8446622_RNA45SN1_45S_50ntFlanks | 11729 | 11765 | 37 | TAACTATGACTCTCTTAAGGTAGCCAAATGCCTCGTC | 1463.2 | -5.0 | 0.7 | -7.0 | 3.1E-12 | 1.3E-11 | BT20/MB468 | MITO |
| >hg38_21_+_8433172_8446622_RNA45SN1_45S_50ntFlanks@10481.10512.32\|ATCGAAAGGGAGTCGGGTTCAGATCCCCGAAT | >hg38_21_+_8433172_8446622_RNA45SN1_45S_50ntFlanks | 10481 | 10512 | 32 | ATCGAAAGGGAGTCGGGTTCAGATCCCCGAAT | 351.4 | -7.9 | 1.1 | -6.9 | 4.2E-12 | 1.3E-11 | MB231/MB468 | CYTO |

<table>
<tr><td>>hg38_21_+_8433172_8446622_RNA45SN1_45S_50ntFlanks@11572.11609.38|CCGACTTAGAACTGGTGCGGACCAGGGGAATCCGACTG</td><td>>hg38_21_+_8433172_8446622_RNA45SN1_45S_50ntFlanks</td><td>11572</td><td>11609</td><td>38</td><td>CCGACTTAGAACTGGTGCGGACCAGGGGAATCCGACTG</td><td>1804.6</td><td>-5.8</td><td>0.8</td><td>-6.9</td><td>5.1E-12</td><td>1.3E-11</td><td>MB231/MB468</td><td>MITO</td></tr>
<tr><td>>hg38_21_+_8433172_8446622_RNA45SN1_45S_50ntFlanks@3815.3842.28|ATGGTTCCTTTGGTCGCTCGCTCCTCTC</td><td>>hg38_21_+_8433172_8446622_RNA45SN1_45S_50ntFlanks</td><td>3815</td><td>3842</td><td>28</td><td>ATGGTTCCTTTGGTCGCTCGCTCCTCTC</td><td>540.8</td><td>-9.3</td><td>1.3</td><td>-6.9</td><td>5.5E-12</td><td>1.3E-11</td><td>BT20/MB468</td><td>NUC</td></tr>
<tr><td>>hg38_21_+_8433172_8446622_RNA45SN1_45S_50ntFlanks@8381.8400.20|AAGAGGGCGTGAAACCGTTA</td><td>>hg38_21_+_8433172_8446622_RNA45SN1_45S_50ntFlanks</td><td>8381</td><td>8400</td><td>20</td><td>AAGAGGGCGTGAAACCGTTA</td><td>3224.4</td><td>-10.2</td><td>1.5</td><td>-6.9</td><td>3.8E-12</td><td>1.3E-11</td><td>MB231/MB468</td><td>MP</td></tr>
<tr><td>>hg38_21_+_8433172_8446622_RNA45SN1_45S_50ntFlanks@9574.9592.19|AATCGGTCGTCCGACCTGG</td><td>>hg38_21_+_8433172_8446622_RNA45SN1_45S_50ntFlanks</td><td>9574</td><td>9592</td><td>19</td><td>AATCGGTCGTCCGACCTGG</td><td>160.2</td><td>-9.6</td><td>1.4</td><td>-6.9</td><td>4.0E-12</td><td>1.3E-11</td><td>BT20/MB468</td><td>CYTO</td></tr>
<tr><td>>hg38_21_+_8433172_8446622_RNA45SN1_45S_50ntFlanks@4685.4726.42|AGACGGACCAGAGCGAAAGCATTTGCCAAGAATGTTTTCATT</td><td>>hg38_21_+_8433172_8446622_RNA45SN1_45S_50ntFlanks</td><td>4685</td><td>4726</td><td>42</td><td>AGACGGACCAGAGCGAAAGCATTTGCCAAGAATGTTTTCATT</td><td>466.8</td><td>8.4</td><td>1.2</td><td>6.9</td><td>4.2E-12</td><td>1.3E-11</td><td>MB231/MB468</td><td>CYTO</td></tr>
<tr><td>>hg38_21_+_8433172_8446622_RNA45SN1_45S_50ntFlanks@10392.10433.42|ACAGCAGTTGAACATGGGTCAGTCGGTCCTGAGAGATGGGCG</td><td>>hg38_21_+_8433172_8446622_RNA45SN1_45S_50ntFlanks</td><td>10392</td><td>10433</td><td>42</td><td>ACAGCAGTTGAACATGGGTCAGTCGGTCCTGAGAGATGGGCG</td><td>2027.2</td><td>-10.9</td><td>1.6</td><td>-6.9</td><td>5.5E-12</td><td>1.3E-11</td><td>BT20/MB468</td><td>NUC</td></tr>
<tr><td>>trnaMT_GlnTTG_MT_-_4329_4400@1.33.33,trnalookalike2_GlnTTG_1_-_564879_564950@1.33.33|TAGGATGGGGTGTGATAGGTGGCACGGAGAATT</td><td>>trnaMT_GlnTTG_MT_-_4329_4400</td><td>1</td><td>33</td><td>33</td><td>TAGGATGGGGTGTGATAGGTGGCACGGAGAATT</td><td>75.6</td><td>5.3</td><td>0.8</td><td>6.9</td><td>5.5E-12</td><td>1.3E-11</td><td>BT20/MB468</td><td>NUC</td></tr>
<tr><td>>hg38_21_+_8433172_8446622_RNA45SN1_45S_50ntFlanks@9465.9488.24|AGGTGGAGCACGAGCGCACGTGTT</td><td>>hg38_21_+_8433172_8446622_RNA45SN1_45S_50ntFlanks</td><td>9465</td><td>9488</td><td>24</td><td>AGGTGGAGCACGAGCGCACGTGTT</td><td>333.0</td><td>-12.2</td><td>1.8</td><td>-6.9</td><td>3.8E-12</td><td>1.3E-11</td><td>MB231/MB468</td><td>MP</td></tr>
<tr><td>>hg38_21_+_8433172_8446622_RNA45SN1_45S_50ntFlanks@11890.11907.18|CTTGACTCTAGTCTGGCA</td><td>>hg38_21_+_8433172_8446622_RNA45SN1_45S_50ntFlanks</td><td>11890</td><td>11907</td><td>18</td><td>CTTGACTCTAGTCTGGCA</td><td>1014.6</td><td>-9.2</td><td>1.3</td><td>-6.9</td><td>5.1E-12</td><td>1.3E-11</td><td>MB231/MB468</td><td>MITO</td></tr>
<tr><td>>hg38_21_+_8433172_8446622_RNA45SN1_45S_50ntFlanks@13001.13027.27|GCGATCTATTGAAAGTCAGCCCTCGAC</td><td>>hg38_21_+_8433172_8446622_RNA45SN1_45S_50ntFlanks</td><td>13001</td><td>13027</td><td>27</td><td>GCGATCTATTGAAAGTCAGCCCTCGAC</td><td>160.8</td><td>-9.4</td><td>1.4</td><td>-7.0</td><td>3.1E-12</td><td>1.3E-11</td><td>BT20/MB468</td><td>MITO</td></tr>
<tr><td>>MI0000433|hsa-let-7g&WithFlank&3|-|52268272|52268367@11.32.22(+1A)&[MIMAT0000414&hsa-let-7g-5p&offsets|0|0(+1A);m-58&3|-|52268336|52268357&offsets|0|0(+1A)]|TGAGGTAGTAGTTTGTACAGTTA</td><td>>MI0000433|hsa-let-7g&WithFlank&3|-|52268272|52268367</td><td>11</td><td>32</td><td>22(+1A)</td><td>TGAGGTAGTAGTTTGTACAGTTA</td><td>1225.5</td><td>4.4</td><td>0.6</td><td>6.9</td><td>4.4E-12</td><td>1.3E-11</td><td>MB231/MB468</td><td>NUC</td></tr>
<tr><td>>MI0000456|hsa-mir-140&WithFlank&16|+|69933075|69933186@69.89.21(+1U)&[MIMAT0004597&hsa-miR-140-3p&offsets|+1|+1(+1U);m-45&16|+|69933143|69933164&offsets|0|-1(+1U)]|ACCACAGGGTAGAACCACGGAT</td><td>>MI0000456|hsa-mir-140&WithFlank&16|+|69933075|69933186</td><td>69</td><td>89</td><td>21(+1U)</td><td>ACCACAGGGTAGAACCACGGAT</td><td>50.5</td><td>8.5</td><td>1.2</td><td>6.9</td><td>5.5E-12</td><td>1.3E-11</td><td>BT20/MB468</td><td>NUC</td></tr>
<tr><td>>hg38_21_+_8433172_8446622_RNA45SN1_45S_50ntFlanks@10808.10833.26|CAATGTAGGTAAGGGAAGTCGGCAAG</td><td>>hg38_21_+_8433172_8446622_RNA45SN1_45S_50ntFlanks</td><td>10808</td><td>10833</td><td>26</td><td>CAATGTAGGTAAGGGAAGTCGGCAAG</td><td>97.8</td><td>-10.5</td><td>1.5</td><td>-6.9</td><td>3.8E-12</td><td>1.3E-11</td><td>MB231/MB468</td><td>MP</td></tr>
<tr><td>>hg38_21_+_8433172_8446622_RNA45SN1_45S_50ntFlanks@4100.4117.18|TGACCACGGGTGACGGGG</td><td>>hg38_21_+_8433172_8446622_RNA45SN1_45S_50ntFlanks</td><td>4100</td><td>4117</td><td>18</td><td>TGACCACGGGTGACGGGG</td><td>131.9</td><td>-10.9</td><td>1.6</td><td>-6.9</td><td>3.8E-12</td><td>1.3E-11</td><td>MB231/MB468</td><td>MP</td></tr>
<tr><td>>hg38_21_+_8433172_8446622_RNA45SN1_45S_50ntFlanks@6690.6719.30|AAGAACGCAGCTAGCTGCGAGAATTAATGT</td><td>>hg38_21_+_8433172_8446622_RNA45SN1_45S_50ntFlanks</td><td>6690</td><td>6719</td><td>30</td><td>AAGAACGCAGCTAGCTGCGAGAATTAATGT</td><td>140.1</td><td>-9.4</td><td>1.4</td><td>-6.9</td><td>4.0E-12</td><td>1.3E-11</td><td>BT20/MB468</td><td>CYTO</td></tr>
<tr><td>>hg38_21_+_8433172_8446622_RNA45SN1_45S_50ntFlanks@11623.11660.38|AAGCATCGCGAAGGCCCGCGGCGGGTGTTGACGCGATG</td><td>>hg38_21_+_8433172_8446622_RNA45SN1_45S_50ntFlanks</td><td>11623</td><td>11660</td><td>38</td><td>AAGCATCGCGAAGGCCCGCGGCGGGTGTTGACGCGATG</td><td>101.3</td><td>-10.5</td><td>1.5</td><td>-6.9</td><td>3.8E-12</td><td>1.3E-11</td><td>MB231/MB468</td><td>MP</td></tr>
<tr><td>>trna16_GlnTTG_17_+_47269890_47269961@1.29.29,trna64_GlnTTG_6_+_28557156_28557227@1.29.29,trna84_GlnTTG_6_+_145503859_145503930@1.29.29|GGTCCCATGGTGTAATGGTTAGCACTCTG</td><td>>trna16_GlnTTG_17_+_47269890_47269961</td><td>1</td><td>29</td><td>29</td><td>GGTCCCATGGTGTAATGGTTAGCACTCTG</td><td>962.6</td><td>-9.4</td><td>1.4</td><td>-6.9</td><td>5.1E-12</td><td>1.3E-11</td><td>MB231/MB468</td><td>MITO</td></tr>
<tr><td>>hg38_21_+_8433172_8446622_RNA45SN1_45S_50ntFlanks@10687.10711.25|CGGCGTCCGGTGAGCTCTCGCTGGC</td><td>>hg38_21_+_8433172_8446622_RNA45SN1_45S_50ntFlanks</td><td>10687</td><td>10711</td><td>25</td><td>CGGCGTCCGGTGAGCTCTCGCTGGC</td><td>2909.3</td><td>-3.9</td><td>0.6</td><td>-7.0</td><td>3.1E-12</td><td>1.3E-11</td><td>BT20/MB468</td><td>MITO</td></tr>
<tr><td>>hg38_21_+_8433172_8446622_RNA45SN1_45S_50ntFlanks@11604.11622.19|CGACTGTTTAATTAAAACA</td><td>>hg38_21_+_8433172_8446622_RNA45SN1_45S_50ntFlanks</td><td>11604</td><td>11622</td><td>19</td><td>CGACTGTTTAATTAAAACA</td><td>106.8</td><td>-10.6</td><td>1.5</td><td>-6.9</td><td>3.8E-12</td><td>1.3E-11</td><td>MB231/MB468</td><td>MP</td></tr>
<tr><td>>hg38_21_+_8433172_8446622_RNA45SN1_45S_50ntFlanks@11146.11166.21|CCCCGGCGGCGGGGGCACGGT</td><td>>hg38_21_+_8433172_8446622_RNA45SN1_45S_50ntFlanks</td><td>11146</td><td>11166</td><td>21</td><td>CCCCGGCGGCGGGGGCACGGT</td><td>164.9</td><td>-10.3</td><td>1.5</td><td>-6.9</td><td>4.4E-12</td><td>1.3E-11</td><td>MB231/MB468</td><td>NUC</td></tr>
<tr><td>>hg38_21_+_8433172_8446622_RNA45SN1_45S_50ntFlanks@9421.9453.33|CGCCGAGGGCGCACCACCGGCCCGTCTCGCCCG</td><td>>hg38_21_+_8433172_8446622_RNA45SN1_45S_50ntFlanks</td><td>9421</td><td>9453</td><td>33</td><td>CGCCGAGGGCGCACCACCGGCCCGTCTCGCCCG</td><td>115.4</td><td>-10.7</td><td>1.5</td><td>-6.9</td><td>3.8E-12</td><td>1.3E-11</td><td>MB231/MB468</td><td>MP</td></tr>
<tr><td>>hg38_21_+_8433172_8446622_RNA45SN1_45S_50ntFlanks@4917.4940.24|CACCACCAGGAGTGGAGCCTGCGG</td><td>>hg38_21_+_8433172_8446622_RNA45SN1_45S_50ntFlanks</td><td>4917</td><td>4940</td><td>24</td><td>CACCACCAGGAGTGGAGCCTGCGG</td><td>705.7</td><td>-9.0</td><td>1.3</td><td>-6.9</td><td>3.8E-12</td><td>1.3E-11</td><td>MB231/MB468</td><td>MP</td></tr>
<tr><td>>hg38_21_+_8433172_8446622_RNA45SN1_45S_50ntFlanks@11314.11344.31|CCCACCCCACGTCTCGTCGCGCGCGCGTCCG</td><td>>hg38_21_+_8433172_8446622_RNA45SN1_45S_50ntFlanks</td><td>11314</td><td>11344</td><td>31</td><td>CCCACCCCACGTCTCGTCGCGCGCGCGTCCG</td><td>93.1</td><td>-10.4</td><td>1.5</td><td>-6.9</td><td>3.8E-12</td><td>1.3E-11</td><td>MB231/MB468</td><td>MP</td></tr>
<tr><td>>hg38_21_+_8433172_8446622_RNA45SN1_45S_50ntFlanks@8170.8188.19|CAAGTCCTTCTGATCGAGG</td><td>>hg38_21_+_8433172_8446622_RNA45SN1_45S_50ntFlanks</td><td>8170</td><td>8188</td><td>19</td><td>CAAGTCCTTCTGATCGAGG</td><td>544.7</td><td>-11.4</td><td>1.6</td><td>-6.9</td><td>4.1E-12</td><td>1.3E-11</td><td>BT20/MB468</td><td>CYTO</td></tr>
<tr><td>>hg38_21_+_8433172_8446622_RNA45SN1_45S_50ntFlanks@12245.12268.24|AGGGCAAAAGCTCGCTTGATCTTG</td><td>>hg38_21_+_8433172_8446622_RNA45SN1_45S_50ntFlanks</td><td>12245</td><td>12268</td><td>24</td><td>AGGGCAAAAGCTCGCTTGATCTTG</td><td>125.1</td><td>-10.8</td><td>1.6</td><td>-6.9</td><td>3.8E-12</td><td>1.3E-11</td><td>MB231/MB468</td><td>MP</td></tr>
<tr><td>>hg38_21_+_8433172_8446622_RNA45SN1_45S_50ntFlanks@8552.8579.28|CTCCCCGGAGGGGGCGGGCTCCGGCGGG</td><td>>hg38_21_+_8433172_8446622_RNA45SN1_45S_50ntFlanks</td><td>8552</td><td>8579</td><td>28</td><td>CTCCCCGGAGGGGGCGGGCTCCGGCGGG</td><td>190.0</td><td>-10.5</td><td>1.5</td><td>-6.9</td><td>4.4E-12</td><td>1.3E-11</td><td>MB231/MB468</td><td>NUC</td></tr>
<tr><td>>hg38_21_+_8433172_8446622_RNA45SN1_45S_50ntFlanks@3787.3812.26|AAACTGCGAATGGCTCATTAAATCAG</td><td>>hg38_21_+_8433172_8446622_RNA45SN1_45S_50ntFlanks</td><td>3787</td><td>3812</td><td>26</td><td>AAACTGCGAATGGCTCATTAAATCAG</td><td>317.8</td><td>-12.2</td><td>1.8</td><td>-6.9</td><td>3.8E-12</td><td>1.3E-11</td><td>MB231/MB468</td><td>MP</td></tr>
<tr><td>>hg38_21_+_8433172_8446622_RNA45SN1_45S_50ntFlanks@11768.11798.31|CTAATTAGTGACGCGCATGAATGGATGAACG</td><td>>hg38_21_+_8433172_8446622_RNA45SN1_45S_50ntFlanks</td><td>11768</td><td>11798</td><td>31</td><td>CTAATTAGTGACGCGCATGAATGGATGAACG</td><td>103.4</td><td>-10.6</td><td>1.5</td><td>-6.9</td><td>3.9E-12</td><td>1.3E-11</td><td>MB231/MB468</td><td>MP</td></tr>
<tr><td>>hg38_21_+_8433172_8446622_RNA45SN1_45S_50ntFlanks@11583.11602.20|CTGGTGCGGACCAGGGGAAT</td><td>>hg38_21_+_8433172_8446622_RNA45SN1_45S_50ntFlanks</td><td>11583</td><td>11602</td><td>20</td><td>CTGGTGCGGACCAGGGGAAT</td><td>914.6</td><td>-5.3</td><td>0.7</td><td>-7.3</td><td>2.3E-13</td><td>1.3E-11</td><td>BT20/MB468</td><td>MP</td></tr>
<tr><td>>hg38_21_+_8433172_8446622_RNA45SN1_45S_50ntFlanks@11821.11844.24|CTATCCAGCGAAACCACAGCCAAG</td><td>>hg38_21_+_8433172_8446622_RNA45SN1_45S_50ntFlanks</td><td>11821</td><td>11844</td><td>24</td><td>CTATCCAGCGAAACCACAGCCAAG</td><td>623.5</td><td>-12.4</td><td>1.7</td><td>-7.3</td><td>2.3E-13</td><td>1.3E-11</td><td>BT20/MB468</td><td>MP</td></tr>
<tr><td>>hg38_21_+_8433172_8446622_RNA45SN1_45S_50ntFlanks@12588.12610.23|GGTTCAGACATTTGGTGTATGTG</td><td>>hg38_21_+_8433172_8446622_RNA45SN1_45S_50ntFlanks</td><td>12588</td><td>12610</td><td>23</td><td>GGTTCAGACATTTGGTGTATGTG</td><td>409.0</td><td>-11.8</td><td>1.6</td><td>-7.3</td><td>2.3E-13</td><td>1.3E-11</td><td>BT20/MB468</td><td>MP</td></tr>
<tr><td>>hg38_21_+_8433172_8446622_RNA45SN1_45S_50ntFlanks@8182.8204.23|ATCGAGGCCCAGCCCGTGGACGG</td><td>>hg38_21_+_8433172_8446622_RNA45SN1_45S_50ntFlanks</td><td>8182</td><td>8204</td><td>23</td><td>ATCGAGGCCCAGCCCGTGGACGG</td><td>370.0</td><td>-11.6</td><td>1.6</td><td>-7.3</td><td>2.3E-13</td><td>1.3E-11</td><td>BT20/MB468</td><td>MP</td></tr>
<tr><td>>hg38_21_+_8433172_8446622_RNA45SN1_45S_50ntFlanks@8155.8178.24|CGCTCGTGGGGGGCCCAAGTCCTT</td><td>>hg38_21_+_8433172_8446622_RNA45SN1_45S_50ntFlanks</td><td>8155</td><td>8178</td><td>24</td><td>CGCTCGTGGGGGGCCCAAGTCCTT</td><td>274.6</td><td>-10.2</td><td>1.5</td><td>-7.0</td><td>3.2E-12</td><td>1.3E-11</td><td>BT20/MB468</td><td>MITO</td></tr>
<tr><td>>hg38_21_+_8433172_8446622_RNA45SN1_45S_50ntFlanks@10895.10913.19|CGAAGCGGGGCTGGGCGCG</td><td>>hg38_21_+_8433172_8446622_RNA45SN1_45S_50ntFlanks</td><td>10895</td><td>10913</td><td>19</td><td>CGAAGCGGGGCTGGGCGCG</td><td>105.8</td><td>-10.6</td><td>1.5</td><td>-6.9</td><td>3.9E-12</td><td>1.4E-11</td><td>MB231/MB468</td><td>MP</td></tr>
<tr><td>>hg38_21_+_8433172_8446622_RNA45SN1_45S_50ntFlanks@11784.11804.21|ATGAATGGATGAACGAGATTC</td><td>>hg38_21_+_8433172_8446622_RNA45SN1_45S_50ntFlanks</td><td>11784</td><td>11804</td><td>21</td><td>ATGAATGGATGAACGAGATTC</td><td>5736.7</td><td>-5.6</td><td>0.8</td><td>-6.9</td><td>4.3E-12</td><td>1.4E-11</td><td>MB231/MB468</td><td>CYTO</td></tr>
<tr><td>>hg38_21_+_8433172_8446622_RNA45SN1_45S_50ntFlanks@10350.10368.19|ACGAGAACTTTGAAGGCCG</td><td>>hg38_21_+_8433172_8446622_RNA45SN1_45S_50ntFlanks</td><td>10350</td><td>10368</td><td>19</td><td>ACGAGAACTTTGAAGGCCG</td><td>8157.3</td><td>-7.0</td><td>1.0</td><td>-7.3</td><td>2.3E-13</td><td>1.4E-11</td><td>BT20/MB468</td><td>MP</td></tr>
<tr><td>>hg38_21_+_8433172_8446622_RNA45SN1_45S_50ntFlanks@12991.13011.21|CTCCCTCGCTGCGATCTATTG</td><td>>hg38_21_+_8433172_8446622_RNA45SN1_45S_50ntFlanks</td><td>12991</td><td>13011</td><td>21</td><td>CTCCCTCGCTGCGATCTATTG</td><td>89.8</td><td>-10.4</td><td>1.5</td><td>-6.9</td><td>3.9E-12</td><td>1.4E-11</td><td>MB231/MB468</td><td>MP</td></tr>
<tr><td>>hg38_21_+_8433172_8446622_RNA45SN1_45S_50ntFlanks@10389.10406.18|TGAACAGCAGTTGAACAT</td><td>>hg38_21_+_8433172_8446622_RNA45SN1_45S_50ntFlanks</td><td>10389</td><td>10406</td><td>18</td><td>TGAACAGCAGTTGAACAT</td><td>106.0</td><td>-10.6</td><td>1.5</td><td>-6.9</td><td>3.9E-12</td><td>1.4E-11</td><td>MB231/MB468</td><td>MP</td></tr>
<tr><td>>hg38_21_+_8433172_8446622_RNA45SN1_45S_50ntFlanks@9801.9820.20|CCGGGCGTGGAATGCGAGTG</td><td>>hg38_21_+_8433172_8446622_RNA45SN1_45S_50ntFlanks</td><td>9801</td><td>9820</td><td>20</td><td>CCGGGCGTGGAATGCGAGTG</td><td>117.1</td><td>-10.7</td><td>1.5</td><td>-6.9</td><td>3.9E-12</td><td>1.4E-11</td><td>MB231/MB468</td><td>MP</td></tr>
<tr><td>>hg38_21_+_8433172_8446622_RNA45SN1_45S_50ntFlanks@3920.3959.40|CATTTATCAGATCAAAACCAACCCGGTCAGCCCCTCTCCG</td><td>>hg38_21_+_8433172_8446622_RNA45SN1_45S_50ntFlanks</td><td>3920</td><td>3959</td><td>40</td><td>CATTTATCAGATCAAAACCAACCCGGTCAGCCCCTCTCCG</td><td>480.1</td><td>-11.9</td><td>1.7</td><td>-6.9</td><td>4.5E-12</td><td>1.4E-11</td><td>MB231/MB468</td><td>NUC</td></tr>
<tr><td>>hg38_21_+_8433172_8446622_RNA45SN1_45S_50ntFlanks@10912.10932.21|CGCGCCGCGGCTGGACGAGGC</td><td>>hg38_21_+_8433172_8446622_RNA45SN1_45S_50ntFlanks</td><td>10912</td><td>10932</td><td>21</td><td>CGCGCCGCGGCTGGACGAGGC</td><td>509.6</td><td>-8.7</td><td>1.3</td><td>-6.9</td><td>4.3E-12</td><td>1.4E-11</td><td>MB231/MB468</td><td>CYTO</td></tr>
<tr><td>>hg38_21_+_8433172_8446622_RNA45SN1_45S_50ntFlanks@10841.10867.27|CGTAACTTCGGGATAAGGATTGGCTCT</td><td>>hg38_21_+_8433172_8446622_RNA45SN1_45S_50ntFlanks</td><td>10841</td><td>10867</td><td>27</td><td>CGTAACTTCGGGATAAGGATTGGCTCT</td><td>8169.7</td><td>-4.8</td><td>0.7</td><td>-6.9</td><td>5.6E-12</td><td>1.4E-11</td><td>BT20/MB468</td><td>NUC</td></tr>
<tr><td>>hg38_21_+_8433172_8446622_RNA45SN1_45S_50ntFlanks@4390.4410.21|TCTTGGGAGCGGGCGGGCGGT</td><td>>hg38_21_+_8433172_8446622_RNA45SN1_45S_50ntFlanks</td><td>4390</td><td>4410</td><td>21</td><td>TCTTGGGAGCGGGCGGGCGGT</td><td>119.9</td><td>-7.1</td><td>1.0</td><td>-6.9</td><td>4.4E-12</td><td>1.4E-11</td><td>MB231/MB468</td><td>CYTO</td></tr>
<tr><td>>hg38_21_+_8433172_8446622_RNA45SN1_45S_50ntFlanks@11890.11908.19|CTTGACTCTAGTCTGGCAC</td><td>>hg38_21_+_8433172_8446622_RNA45SN1_45S_50ntFlanks</td><td>11890</td><td>11908</td><td>19</td><td>CTTGACTCTAGTCTGGCAC</td><td>102.4</td><td>-10.5</td><td>1.5</td><td>-6.9</td><td>3.9E-12</td><td>1.4E-11</td><td>MB231/MB468</td><td>MP</td></tr>
<tr><td>>hg38_21_+_8433172_8446622_RNA45SN1_45S_50ntFlanks@12937.12959.23|AACCATTCGTAGACGACCTGCTT</td><td>>hg38_21_+_8433172_8446622_RNA45SN1_45S_50ntFlanks</td><td>12937</td><td>12959</td><td>23</td><td>AACCATTCGTAGACGACCTGCTT</td><td>88.6</td><td>-10.3</td><td>1.5</td><td>-6.9</td><td>3.9E-12</td><td>1.4E-11</td><td>MB231/MB468</td><td>MP</td></tr>
<tr><td>>hg38_21_+_8433172_8446622_RNA45SN1_45S_50ntFlanks@4102.4127.26|ACCACGGGTGACGGGGAATCAGGGTT</td><td>>hg38_21_+_8433172_8446622_RNA45SN1_45S_50ntFlanks</td><td>4102</td><td>4127</td><td>26</td><td>ACCACGGGTGACGGGGAATCAGGGTT</td><td>1371.5</td><td>-10.0</td><td>1.5</td><td>-6.9</td><td>5.7E-12</td><td>1.4E-11</td><td>BT20/MB468</td><td>NUC</td></tr>
<tr><td>>hg38_21_+_8433172_8446622_RNA45SN1_45S_50ntFlanks@9534.9558.25|AGGAAACTCTGGTGGAGGTCCGTAG</td><td>>hg38_21_+_8433172_8446622_RNA45SN1_45S_50ntFlanks</td><td>9534</td><td>9558</td><td>25</td><td>AGGAAACTCTGGTGGAGGTCCGTAG</td><td>181.9</td><td>-9.6</td><td>1.4</td><td>-7.0</td><td>3.2E-12</td><td>1.4E-11</td><td>BT20/MB468</td><td>MITO</td></tr>
<tr><td>>MI0005756|hsa-mir-934&WithFlank&X|+|136550872|136550966@21.40.20(+1A)&[MIMAT0004977&hsa-miR-934&offsets|0|-2(+1A);m-943&X|+|136550892|136550913&offsets|0|-2(+1A)]|TGTCTACTACTGGAGACACTA</td><td>>MI0005756|hsa-mir-934&WithFlank&X|+|136550872|136550966</td><td>21</td><td>40</td><td>20(+1A)</td><td>TGTCTACTACTGGAGACACTA</td><td>165.2</td><td>10.7</td><td>1.5</td><td>7.3</td><td>3.1E-13</td><td>1.4E-11</td><td>BT20/MB231</td><td>CYTO</td></tr>
<tr><td>>hg38_21_+_8433172_8446622_RNA45SN1_45S_50ntFlanks@10346.10388.43|TCAAACGAGAACTTTGAAGGCCGAAGTGGAGAAGGGTTCCATG</td><td>>hg38_21_+_8433172_8446622_RNA45SN1_45S_50ntFlanks</td><td>10346</td><td>10388</td><td>43</td><td>TCAAACGAGAACTTTGAAGGCCGAAGTGGAGAAGGGTTCCATG</td><td>1240.0</td><td>-5.8</td><td>0.8</td><td>-6.9</td><td>5.3E-12</td><td>1.4E-11</td><td>MB231/MB468</td><td>MITO</td></tr>
<tr><td>>hg38_21_+_8433172_8446622_RNA45SN1_45S_50ntFlanks@4069.4089.21|CTTTCGATGGTAGTCGCCGTG</td><td>>hg38_21_+_8433172_8446622_RNA45SN1_45S_50ntFlanks</td><td>4069</td><td>4089</td><td>21</td><td>CTTTCGATGGTAGTCGCCGTG</td><td>637.5</td><td>-5.1</td><td>0.7</td><td>-6.9</td><td>3.9E-12</td><td>1.4E-11</td><td>MB231/MB468</td><td>MP</td></tr>
<tr><td>>hg38_21_+_8433172_8446622_RNA45SN1_45S_50ntFlanks@6671.6693.23|CTCGGCTCGTGCGTCGATGAAGA</td><td>>hg38_21_+_8433172_8446622_RNA45SN1_45S_50ntFlanks</td><td>6671</td><td>6693</td><td>23</td><td>CTCGGCTCGTGCGTCGATGAAGA</td><td>114.8</td><td>-10.7</td><td>1.5</td><td>-6.9</td><td>3.9E-12</td><td>1.4E-11</td><td>MB231/MB468</td><td>MP</td></tr>
<tr><td>>hg38_21_+_8433172_8446622_RNA45SN1_45S_50ntFlanks@12451.12480.30|ATTCACCAAGCGTTGGATTGTTCACCCACT</td><td>>hg38_21_+_8433172_8446622_RNA45SN1_45S_50ntFlanks</td><td>12451</td><td>12480</td><td>30</td><td>ATTCACCAAGCGTTGGATTGTTCACCCACT</td><td>215.5</td><td>6.2</td><td>0.9</td><td>6.9</td><td>4.4E-12</td><td>1.4E-11</td><td>MB231/MB468</td><td>CYTO</td></tr>
</table>

<table>
<tr><th></th><th></th><th></th><th></th><th></th><th></th><th></th><th></th><th></th><th></th><th></th><th></th><th></th><th></th></tr>
<tr><td>>MI0000736|hsa-mir-30c-1&WithFlank&1|+|40757278|40757378@23.48.26;MI0000254|hsa-mir-30c-2&WithFlank&6|-|71376954|71377037@13.38.26&[MIMAT0000244&hsa-miR-30c-5p&offsets|0|+3;m-57&1|+|40757300|40757323&offsets|0|+2];[MIMAT0000244_1&hsa-miR-30c-5p&offsets|0|+3;m-56&6|-|71377002|71377025&offsets|0|+2]|TGTAAACATCCTACACTCTCAGCTGT</td><td>>MI0000736|hsa-mir-30c-1&WithFlank&1|+|40757278|40757378</td><td>23</td><td>48</td><td>26</td><td>TGTAAACATCCTACACTCTCAGCTGT</td><td>350.8</td><td>5.0</td><td>0.7</td><td>6.9</td><td>4.5E-12</td><td>1.4E-11</td><td>MB231/MB468</td><td>NUC</td></tr>
<tr><td>>hg38_21_+_8433172_8446622_RNA45SN1_45S_50ntFlanks@10107.10127.21|CGTGTGGTGTGCGTCGGAGGG</td><td>>hg38_21_+_8433172_8446622_RNA45SN1_45S_50ntFlanks</td><td>10107</td><td>10127</td><td>21</td><td>CGTGTGGTGTGCGTCGGAGGG</td><td>481.7</td><td>-7.5</td><td>1.1</td><td>-6.9</td><td>4.4E-12</td><td>1.4E-11</td><td>MB231/MB468</td><td>CYTO</td></tr>
<tr><td>>hg38_21_+_8433172_8446622_RNA45SN1_45S_50ntFlanks@8168.8197.30|CCCAAGTCCTTCTGATCGAGGCCCAGCCCG</td><td>>hg38_21_+_8433172_8446622_RNA45SN1_45S_50ntFlanks</td><td>8168</td><td>8197</td><td>30</td><td>CCCAAGTCCTTCTGATCGAGGCCCAGCCCG</td><td>118.4</td><td>-10.8</td><td>1.5</td><td>-6.9</td><td>3.9E-12</td><td>1.4E-11</td><td>MB231/MB468</td><td>MP</td></tr>
<tr><td>>hg38_21_+_8433172_8446622_RNA45SN1_45S_50ntFlanks@9004.9026.23|TCTCCCCCGCGGGGGCGCGCCGG</td><td>>hg38_21_+_8433172_8446622_RNA45SN1_45S_50ntFlanks</td><td>9004</td><td>9026</td><td>23</td><td>TCTCCCCCGCGGGGGCGCGCCGG</td><td>2275.6</td><td>-10.5</td><td>1.5</td><td>-6.9</td><td>5.7E-12</td><td>1.4E-11</td><td>BT20/MB468</td><td>NUC</td></tr>
<tr><td>>hg38_21_+_8433172_8446622_RNA45SN1_45S_50ntFlanks@7977.8008.32|CGACCTCAGATCAGACGTGGCGACCCGCTGAA</td><td>>hg38_21_+_8433172_8446622_RNA45SN1_45S_50ntFlanks</td><td>7977</td><td>8008</td><td>32</td><td>CGACCTCAGATCAGACGTGGCGACCCGCTGAA</td><td>696.8</td><td>-5.2</td><td>0.7</td><td>-6.9</td><td>4.0E-12</td><td>1.4E-11</td><td>MB231/MB468</td><td>MP</td></tr>
<tr><td>>hg38_21_+_8433172_8446622_RNA45SN1_45S_50ntFlanks@10341.10373.33|AATATTCAAACGAGAACTTTGAAGGCCGAAGTG</td><td>>hg38_21_+_8433172_8446622_RNA45SN1_45S_50ntFlanks</td><td>10341</td><td>10373</td><td>33</td><td>AATATTCAAACGAGAACTTTGAAGGCCGAAGTG</td><td>371.5</td><td>5.5</td><td>0.8</td><td>7.0</td><td>3.2E-12</td><td>1.4E-11</td><td>BT20/MB468</td><td>MITO</td></tr>
<tr><td>>MI0000102|hsa-mir-100&WithFlank&11|-|122152223|122152314@19.40.22(+1C)&[MIMAT0000098&hsa-miR-100-5p&offsets|0|0(+1C);m-27&11|-|122152276|122152296&offsets|0|+1(+1C)]|AACCCGTAGATCCGAACTTGTGC</td><td>>MI0000102|hsa-mir-100&WithFlank&11|-|122152223|122152314</td><td>19</td><td>40</td><td>22(+1C)</td><td>AACCCGTAGATCCGAACTTGTGC</td><td>997.0</td><td>14.2</td><td>2.1</td><td>6.9</td><td>4.2E-12</td><td>1.4E-11</td><td>BT20/MB468</td><td>CYTO</td></tr>
<tr><td>>MI0000746|hsa-mir-99b&WithFlank&19|+|51692606|51692687@13.34.22&[MIMAT0000689&hsa-miR-99b-5p&offsets|0|0;m-10&19|+|51692618|51692639&offsets|0|0]|CACCCGTAGAACCGACCTTGCG</td><td>>MI0000746|hsa-mir-99b&WithFlank&19|+|51692606|51692687</td><td>13</td><td>34</td><td>22</td><td>CACCCGTAGAACCGACCTTGCG</td><td>37273.0</td><td>5.6</td><td>0.8</td><td>6.9</td><td>4.4E-12</td><td>1.4E-11</td><td>MB231/MB468</td><td>CYTO</td></tr>
<tr><td>>hg38_21_+_8433172_8446622_RNA45SN1_45S_50ntFlanks@12405.12432.28|CTTTTTGATCCTTCGATGTCGGCTCTTC</td><td>>hg38_21_+_8433172_8446622_RNA45SN1_45S_50ntFlanks</td><td>12405</td><td>12432</td><td>28</td><td>CTTTTTGATCCTTCGATGTCGGCTCTTC</td><td>15254.6</td><td>-6.5</td><td>0.9</td><td>-6.9</td><td>4.4E-12</td><td>1.4E-11</td><td>MB231/MB468</td><td>CYTO</td></tr>
<tr><td>>hg38_21_+_8433172_8446622_RNA45SN1_45S_50ntFlanks@10761.10785.25|ATATCCGCAGCAGGTCTCCAAGGTG</td><td>>hg38_21_+_8433172_8446622_RNA45SN1_45S_50ntFlanks</td><td>10761</td><td>10785</td><td>25</td><td>ATATCCGCAGCAGGTCTCCAAGGTG</td><td>298.9</td><td>-8.2</td><td>1.2</td><td>-6.9</td><td>5.4E-12</td><td>1.4E-11</td><td>MB231/MB468</td><td>MITO</td></tr>
<tr><td>>hg38_21_+_8433172_8446622_RNA45SN1_45S_50ntFlanks@4730.4758.29|CAAGAACGAAAGTCGGAGGTTCGAAGACG</td><td>>hg38_21_+_8433172_8446622_RNA45SN1_45S_50ntFlanks</td><td>4730</td><td>4758</td><td>29</td><td>CAAGAACGAAAGTCGGAGGTTCGAAGACG</td><td>5698.0</td><td>-5.9</td><td>0.9</td><td>-6.9</td><td>5.4E-12</td><td>1.4E-11</td><td>MB231/MB468</td><td>MITO</td></tr>
<tr><td>>hg38_21_+_8433172_8446622_RNA45SN1_45S_50ntFlanks@10572.10589.18|AGAAGCCGGCGGGAGCCC</td><td>>hg38_21_+_8433172_8446622_RNA45SN1_45S_50ntFlanks</td><td>10572</td><td>10589</td><td>18</td><td>AGAAGCCGGCGGGAGCCC</td><td>119.8</td><td>-10.8</td><td>1.6</td><td>-6.9</td><td>4.0E-12</td><td>1.4E-11</td><td>MB231/MB468</td><td>MP</td></tr>
<tr><td>>hg38_21_+_8433172_8446622_RNA45SN1_45S_50ntFlanks@12941.12973.33|ATTCGTAGACGACCTGCTTCTGGGTCGGGGTTT</td><td>>hg38_21_+_8433172_8446622_RNA45SN1_45S_50ntFlanks</td><td>12941</td><td>12973</td><td>33</td><td>ATTCGTAGACGACCTGCTTCTGGGTCGGGGTTT</td><td>110.0</td><td>-10.6</td><td>1.5</td><td>-6.9</td><td>4.0E-12</td><td>1.4E-11</td><td>MB231/MB468</td><td>MP</td></tr>
<tr><td>>hg38_21_+_8433172_8446622_RNA45SN1_45S_50ntFlanks@10680.10705.26|GTTCCGGCGGCGTCCGGTGAGCTCTC</td><td>>hg38_21_+_8433172_8446622_RNA45SN1_45S_50ntFlanks</td><td>10680</td><td>10705</td><td>26</td><td>GTTCCGGCGGCGTCCGGTGAGCTCTC</td><td>952.7</td><td>-4.5</td><td>0.6</td><td>-6.9</td><td>4.5E-12</td><td>1.4E-11</td><td>MB231/MB468</td><td>CYTO</td></tr>
<tr><td>>hg38_21_+_8433172_8446622_RNA45SN1_45S_50ntFlanks@10805.10829.25|GAACAATGTAGGTAAGGGAAGTCGG</td><td>>hg38_21_+_8433172_8446622_RNA45SN1_45S_50ntFlanks</td><td>10805</td><td>10829</td><td>25</td><td>GAACAATGTAGGTAAGGGAAGTCGG</td><td>112.8</td><td>-10.7</td><td>1.5</td><td>-6.9</td><td>4.0E-12</td><td>1.4E-11</td><td>MB231/MB468</td><td>MP</td></tr>
<tr><td>>hg38_21_+_8433172_8446622_RNA45SN1_45S_50ntFlanks@4882.4903.22|TGCAAAGCTGAAACTTAAAGGA</td><td>>hg38_21_+_8433172_8446622_RNA45SN1_45S_50ntFlanks</td><td>4882</td><td>4903</td><td>22</td><td>TGCAAAGCTGAAACTTAAAGGA</td><td>421.9</td><td>9.8</td><td>1.4</td><td>7.0</td><td>3.3E-12</td><td>1.4E-11</td><td>BT20/MB468</td><td>MITO</td></tr>
<tr><td>>trna112_GlnCTG_1_-_147737382_147737453@30.50.21,trna131_GlnCTG_6_-_27759135_27759206@30.50.21,trna146_GlnCTG_6_-_27515531_27515602@30.50.21,trna15_GlnCTG_1_+_145963304_145963375@30.50.21,trna19_GlnCTG_1_+_147505038_147505109@30.50.21,trna1_GlnCTG_6_+_18836402_18836473@30.50.21,trna28_GlnCTG_1_+_149186125_149186196@30.50.21,trna3_GlnCTG_17_+_8023070_8023141@30.50.21,trna49_GlnCTG_6_+_27487308_27487379@30.50.21,trna7_GlnCTG_15_-_66161400_66161471@30.50.21,trna99_GlnCTG_6_-_28909378_28909449@30.50.21|GACTCTGAATCCAGCGATCCG</td><td>>trna112_GlnCTG_1_-_147737382_147737453</td><td>30</td><td>50</td><td>21</td><td>GACTCTGAATCCAGCGATCCG</td><td>404.1</td><td>-8.9</td><td>1.3</td><td>-6.9</td><td>4.6E-12</td><td>1.4E-11</td><td>MB231/MB468</td><td>NUC</td></tr>
<tr><td>>MI0000734|hsa-mir-106b&WithFlank&7|-|100093987|100094080@58.77.20&[MIMAT0004672&hsa-miR-106b-3p&offsets|0|-2;m-87&7|-|100094003|100094023&offsets|0|-1]|CCGCACTGTGGGTACTTGCT</td><td>>MI0000734|hsa-mir-106b&WithFlank&7|-|100093987|100094080</td><td>58</td><td>77</td><td>20</td><td>CCGCACTGTGGGTACTTGCT</td><td>501.6</td><td>6.1</td><td>0.9</td><td>6.9</td><td>5.4E-12</td><td>1.4E-11</td><td>MB231/MB468</td><td>MITO</td></tr>
<tr><td>>hg38_21_+_8433172_8446622_RNA45SN1_45S_50ntFlanks@12438.12455.18|ATTGTGAAGCAGAATTCA</td><td>>hg38_21_+_8433172_8446622_RNA45SN1_45S_50ntFlanks</td><td>12438</td><td>12455</td><td>18</td><td>ATTGTGAAGCAGAATTCA</td><td>1100.1</td><td>-5.5</td><td>0.8</td><td>-7.0</td><td>3.3E-12</td><td>1.4E-11</td><td>BT20/MB468</td><td>MITO</td></tr>
<tr><td>>MI0003643|hsa-mir-629&WithFlank&15|-|70079366|70079474@28.47.20(+1C)&[MIMAT0004810&hsa-miR-629-5p&offsets|0|-1(+1C);m-150&15|-|70079427|70079447&offsets|0|-1(+1C)]|TGGGTTTACGTTGGGAGAACC</td><td>>MI0003643|hsa-mir-629&WithFlank&15|-|70079366|70079474</td><td>28</td><td>47</td><td>20(+1C)</td><td>TGGGTTTACGTTGGGAGAACC</td><td>195.3</td><td>10.8</td><td>1.6</td><td>6.9</td><td>4.0E-12</td><td>1.4E-11</td><td>MB231/MB468</td><td>MP</td></tr>
<tr><td>>hg38_21_+_8433172_8446622_RNA45SN1_45S_50ntFlanks@9299.9338.40|CACGGACCAAGGAGTCTAACACGTGCGCGAGTCGGGGGCT</td><td>>hg38_21_+_8433172_8446622_RNA45SN1_45S_50ntFlanks</td><td>9299</td><td>9338</td><td>40</td><td>CACGGACCAAGGAGTCTAACACGTGCGCGAGTCGGGGGCT</td><td>126.7</td><td>-10.9</td><td>1.6</td><td>-6.9</td><td>4.0E-12</td><td>1.4E-11</td><td>MB231/MB468</td><td>MP</td></tr>
<tr><td>>hg38_21_+_8433172_8446622_RNA45SN1_45S_50ntFlanks@12714.12742.29|GAGCCTCGGTTGGCCTCGGATAGCCGGTC</td><td>>hg38_21_+_8433172_8446622_RNA45SN1_45S_50ntFlanks</td><td>12714</td><td>12742</td><td>29</td><td>GAGCCTCGGTTGGCCTCGGATAGCCGGTC</td><td>861.5</td><td>-7.2</td><td>1.0</td><td>-6.9</td><td>4.0E-12</td><td>1.4E-11</td><td>MB231/MB468</td><td>MP</td></tr>
<tr><td>>hg38_21_+_8433172_8446622_RNA45SN1_45S_50ntFlanks@11210.11229.20|GGCGACTCTGGACGCGAGCC</td><td>>hg38_21_+_8433172_8446622_RNA45SN1_45S_50ntFlanks</td><td>11210</td><td>11229</td><td>20</td><td>GGCGACTCTGGACGCGAGCC</td><td>147.5</td><td>-11.1</td><td>1.6</td><td>-6.9</td><td>4.0E-12</td><td>1.4E-11</td><td>MB231/MB468</td><td>MP</td></tr>
<tr><td>>hg38_21_+_8433172_8446622_RNA45SN1_45S_50ntFlanks@9457.9503.47|CGCCGGGGAGGTGGAGCACGAGCGCACGTGTTAGGACCCGAAAGATG</td><td>>hg38_21_+_8433172_8446622_RNA45SN1_45S_50ntFlanks</td><td>9457</td><td>9503</td><td>47</td><td>CGCCGGGGAGGTGGAGCACGAGCGCACGTGTTAGGACCCGAAAGATG</td><td>337.5</td><td>-11.4</td><td>1.6</td><td>-6.9</td><td>4.6E-12</td><td>1.4E-11</td><td>MB231/MB468</td><td>NUC</td></tr>
<tr><td>>MI0000299|hsa-mir-222&WithFlank&X|-|45747009|45747130@75.99.25(+1A)&[MIMAT0000279&hsa-miR-222-3p&offsets|0|+4(+1A);m-140&X|-|45747033|45747056&offsets|0|+1(+1A)]|AGCTACATCTGGCTACTGGGTCTCTA</td><td>>MI0000299|hsa-mir-222&WithFlank&X|-|45747009|45747130</td><td>75</td><td>99</td><td>25(+1A)</td><td>AGCTACATCTGGCTACTGGGTCTCTA</td><td>295.7</td><td>11.4</td><td>1.7</td><td>6.9</td><td>5.4E-12</td><td>1.4E-11</td><td>MB231/MB468</td><td>MITO</td></tr>
<tr><td>>hg38_21_+_8433172_8446622_RNA45SN1_45S_50ntFlanks@12583.12601.19|CCGCAGGTTCAGACATTTG</td><td>>hg38_21_+_8433172_8446622_RNA45SN1_45S_50ntFlanks</td><td>12583</td><td>12601</td><td>19</td><td>CCGCAGGTTCAGACATTTG</td><td>110.6</td><td>-10.7</td><td>1.5</td><td>-6.9</td><td>4.0E-12</td><td>1.4E-11</td><td>MB231/MB468</td><td>MP</td></tr>
<tr><td>>MI0000440|hsa-mir-27b&WithFlank&9|+|95085439|95085547@67.86.20&[MIMAT0000419&hsa-miR-27b-3p&offsets|0|-1;m-39&9|+|95085505|95085525&offsets|0|-1]|TTCACAGTGGCTAAGTTCTG</td><td>>MI0000440|hsa-mir-27b&WithFlank&9|+|95085439|95085547</td><td>67</td><td>86</td><td>20</td><td>TTCACAGTGGCTAAGTTCTG</td><td>4676.4</td><td>6.0</td><td>0.9</td><td>6.9</td><td>5.5E-12</td><td>1.4E-11</td><td>MB231/MB468</td><td>MITO</td></tr>
<tr><td>>hg38_21_+_8433172_8446622_RNA45SN1_45S_50ntFlanks@9573.9595.23|AAATCGGTCGTCCGACCTGGGTA</td><td>>hg38_21_+_8433172_8446622_RNA45SN1_45S_50ntFlanks</td><td>9573</td><td>9595</td><td>23</td><td>AAATCGGTCGTCCGACCTGGGTA</td><td>2760.9</td><td>-6.8</td><td>1.0</td><td>-6.9</td><td>5.5E-12</td><td>1.4E-11</td><td>MB231/MB468</td><td>MITO</td></tr>
<tr><td>>hg38_21_+_8433172_8446622_RNA45SN1_45S_50ntFlanks@6651.6678.28|CGACTCTTAGCGGTGGATCACTCGGCTC</td><td>>hg38_21_+_8433172_8446622_RNA45SN1_45S_50ntFlanks</td><td>6651</td><td>6678</td><td>28</td><td>CGACTCTTAGCGGTGGATCACTCGGCTC</td><td>6768.7</td><td>4.3</td><td>0.6</td><td>6.9</td><td>4.3E-12</td><td>1.4E-11</td><td>BT20/MB468</td><td>CYTO</td></tr>
<tr><td>>MI0000440|hsa-mir-27b&WithFlank&9|+|95085439|95085547@67.88.22(+2U)&[MIMAT0000419&hsa-miR-27b-3p&offsets|0|+1(+2U);m-39&9|+|95085505|95085525&offsets|0|+1(+2U)]|TTCACAGTGGCTAAGTTCTGCATT</td><td>>MI0000440|hsa-mir-27b&WithFlank&9|+|95085439|95085547</td><td>67</td><td>88</td><td>22(+2U)</td><td>TTCACAGTGGCTAAGTTCTGCATT</td><td>187.5</td><td>5.1</td><td>0.7</td><td>6.9</td><td>4.6E-12</td><td>1.4E-11</td><td>MB231/MB468</td><td>NUC</td></tr>
<tr><td>>hg38_21_+_8433172_8446622_RNA45SN1_45S_50ntFlanks@12728.12747.20|CTCGGATAGCCGGTCCCCCG</td><td>>hg38_21_+_8433172_8446622_RNA45SN1_45S_50ntFlanks</td><td>12728</td><td>12747</td><td>20</td><td>CTCGGATAGCCGGTCCCCCG</td><td>213.2</td><td>-11.6</td><td>1.7</td><td>-6.9</td><td>4.1E-12</td><td>1.4E-11</td><td>MB231/MB468</td><td>MP</td></tr>
<tr><td>>hg38_21_+_8433172_8446622_RNA45SN1_45S_50ntFlanks@10482.10512.31|TCGAAAGGGAGTCGGGTTCAGATCCCCGAAT</td><td>>hg38_21_+_8433172_8446622_RNA45SN1_45S_50ntFlanks</td><td>10482</td><td>10512</td><td>31</td><td>TCGAAAGGGAGTCGGGTTCAGATCCCCGAAT</td><td>106.2</td><td>-10.6</td><td>1.5</td><td>-6.9</td><td>4.1E-12</td><td>1.4E-11</td><td>MB231/MB468</td><td>MP</td></tr>
<tr><td>>hg38_21_+_8433172_8446622_RNA45SN1_45S_50ntFlanks@11579.11601.23|AGAACTGGTGCGGACCAGGGGAA</td><td>>hg38_21_+_8433172_8446622_RNA45SN1_45S_50ntFlanks</td><td>11579</td><td>11601</td><td>23</td><td>AGAACTGGTGCGGACCAGGGGAA</td><td>715.2</td><td>-12.6</td><td>1.7</td><td>-7.3</td><td>2.4E-13</td><td>1.4E-11</td><td>BT20/MB468</td><td>MP</td></tr>
<tr><td>>hg38_21_+_8433172_8446622_RNA45SN1_45S_50ntFlanks@3900.3935.36|CTTCGCGGGGGGGATGCGTGCATTTATCAGATCAAA</td><td>>hg38_21_+_8433172_8446622_RNA45SN1_45S_50ntFlanks</td><td>3900</td><td>3935</td><td>36</td><td>CTTCGCGGGGGGGATGCGTGCATTTATCAGATCAAA</td><td>223.9</td><td>-11.7</td><td>1.7</td><td>-6.9</td><td>4.1E-12</td><td>1.4E-11</td><td>MB231/MB468</td><td>MP</td></tr>
<tr><td>>trnaMT_ValTAC_MT_+_1602_1670@53.72.20|TAACTTGACCGCTCTGACCA</td><td>>trnaMT_ValTAC_MT_+_1602_1670</td><td>53</td><td>72</td><td>20</td><td>TAACTTGACCGCTCTGACCA</td><td>7043.1</td><td>6.2</td><td>0.9</td><td>7.0</td><td>3.3E-12</td><td>1.4E-11</td><td>BT20/MB468</td><td>MITO</td></tr>
<tr><td>>hg38_21_+_8433172_8446622_RNA45SN1_45S_50ntFlanks@11564.11587.24|CCTAGCAGCCGACTTAGAACTGGT</td><td>>hg38_21_+_8433172_8446622_RNA45SN1_45S_50ntFlanks</td><td>11564</td><td>11587</td><td>24</td><td>CCTAGCAGCCGACTTAGAACTGGT</td><td>398.4</td><td>-6.6</td><td>1.0</td><td>-6.9</td><td>4.1E-12</td><td>1.4E-11</td><td>MB231/MB468</td><td>MP</td></tr>
<tr><td>>hg38_21_+_8433172_8446622_RNA45SN1_45S_50ntFlanks@10512.10534.23|TCCGGAGTGGCGGAGATGGGCGC</td><td>>hg38_21_+_8433172_8446622_RNA45SN1_45S_50ntFlanks</td><td>10512</td><td>10534</td><td>23</td><td>TCCGGAGTGGCGGAGATGGGCGC</td><td>238.2</td><td>-10.0</td><td>1.4</td><td>-7.0</td><td>3.4E-12</td><td>1.4E-11</td><td>BT20/MB468</td><td>MITO</td></tr>
<tr><td>>hg38_21_+_8433172_8446622_RNA45SN1_45S_50ntFlanks@5084.5109.26|CGAACGAGACTCTGGCATGCTAACTA</td><td>>hg38_21_+_8433172_8446622_RNA45SN1_45S_50ntFlanks</td><td>5084</td><td>5109</td><td>26</td><td>CGAACGAGACTCTGGCATGCTAACTA</td><td>507.1</td><td>-12.8</td><td>1.9</td><td>-6.9</td><td>5.5E-12</td><td>1.4E-11</td><td>MB231/MB468</td><td>MITO</td></tr>
<tr><td>>hg38_21_+_8433172_8446622_RNA45SN1_45S_50ntFlanks@10735.10752.18|TGTAAATCTCGCGCCGGG</td><td>>hg38_21_+_8433172_8446622_RNA45SN1_45S_50ntFlanks</td><td>10735</td><td>10752</td><td>18</td><td>TGTAAATCTCGCGCCGGG</td><td>232.4</td><td>-8.2</td><td>1.2</td><td>-6.9</td><td>4.7E-12</td><td>1.4E-11</td><td>MB231/MB468</td><td>NUC</td></tr>
<tr><td>>hg38_21_+_8433172_8446622_RNA45SN1_45S_50ntFlanks@4728.4748.21|ATCAAGAACGAAAGTCGGAGG</td><td>>hg38_21_+_8433172_8446622_RNA45SN1_45S_50ntFlanks</td><td>4728</td><td>4748</td><td>21</td><td>ATCAAGAACGAAAGTCGGAGG</td><td>194.4</td><td>-9.9</td><td>1.4</td><td>-6.9</td><td>4.3E-12</td><td>1.4E-11</td><td>BT20/MB468</td><td>CYTO</td></tr>
<tr><td>>hg38_21_+_8433172_8446622_RNA45SN1_45S_50ntFlanks@8552.8581.30|CTCCCCGGAGGGGGCGGGCTCCGGCGGGTG</td><td>>hg38_21_+_8433172_8446622_RNA45SN1_45S_50ntFlanks</td><td>8552</td><td>8581</td><td>30</td><td>CTCCCCGGAGGGGGCGGGCTCCGGCGGGTG</td><td>226.7</td><td>-9.9</td><td>1.4</td><td>-7.0</td><td>3.4E-12</td><td>1.4E-11</td><td>BT20/MB468</td><td>MITO</td></tr>
<tr><td>>hg38_21_+_8433172_8446622_RNA45SN1_45S_50ntFlanks@5470.5499.30|CCACGGCCCTGGCGGAGCGCTGAGAAGACG</td><td>>hg38_21_+_8433172_8446622_RNA45SN1_45S_50ntFlanks</td><td>5470</td><td>5499</td><td>30</td><td>CCACGGCCCTGGCGGAGCGCTGAGAAGACG</td><td>819.6</td><td>-8.6</td><td>1.3</td><td>-6.9</td><td>5.6E-12</td><td>1.4E-11</td><td>MB231/MB468</td><td>MITO</td></tr>
<tr><td>>hg38_21_+_8433172_8446622_RNA45SN1_45S_50ntFlanks@9534.9558.25|AGGAAACTCTGGTGGAGGTCCGTAG</td><td>>hg38_21_+_8433172_8446622_RNA45SN1_45S_50ntFlanks</td><td>9534</td><td>9558</td><td>25</td><td>AGGAAACTCTGGTGGAGGTCCGTAG</td><td>425.2</td><td>-8.3</td><td>1.2</td><td>-6.9</td><td>5.6E-12</td><td>1.4E-11</td><td>MB231/MB468</td><td>MITO</td></tr>
<tr><td>>hg38_21_+_8433172_8446622_RNA45SN1_45S_50ntFlanks@10764.10785.22|TCCGCAGCAGGTCTCCAAGGTG</td><td>>hg38_21_+_8433172_8446622_RNA45SN1_45S_50ntFlanks</td><td>10764</td><td>10785</td><td>22</td><td>TCCGCAGCAGGTCTCCAAGGTG</td><td>101.3</td><td>-10.5</td><td>1.5</td><td>-6.9</td><td>4.1E-12</td><td>1.4E-11</td><td>MB231/MB468</td><td>MP</td></tr>
<tr><td>>hg38_21_+_8433172_8446622_RNA45SN1_45S_50ntFlanks@8168.8199.32|CCCAAGTCCTTCTGATCGAGGCCCAGCCCGTG</td><td>>hg38_21_+_8433172_8446622_RNA45SN1_45S_50ntFlanks</td><td>8168</td><td>8199</td><td>32</td><td>CCCAAGTCCTTCTGATCGAGGCCCAGCCCGTG</td><td>105.5</td><td>-10.6</td><td>1.5</td><td>-6.9</td><td>4.1E-12</td><td>1.4E-11</td><td>MB231/MB468</td><td>MP</td></tr>
<tr><td>>hg38_21_+_8433172_8446622_RNA45SN1_45S_50ntFlanks@6738.6756.19|ATCATCGACACTTCGAACG</td><td>>hg38_21_+_8433172_8446622_RNA45SN1_45S_50ntFlanks</td><td>6738</td><td>6756</td><td>19</td><td>ATCATCGACACTTCGAACG</td><td>130.3</td><td>-10.0</td><td>1.4</td><td>-6.9</td><td>4.7E-12</td><td>1.4E-11</td><td>MB231/MB468</td><td>NUC</td></tr>
<tr><td>>hg38_21_+_8433172_8446622_RNA45SN1_45S_50ntFlanks@12935.12966.32|TAAACCATTCGTAGACGACCTGCTTCTGGGTC</td><td>>hg38_21_+_8433172_8446622_RNA45SN1_45S_50ntFlanks</td><td>12935</td><td>12966</td><td>32</td><td>TAAACCATTCGTAGACGACCTGCTTCTGGGTC</td><td>410.0</td><td>-5.9</td><td>0.9</td><td>-6.9</td><td>4.6E-12</td><td>1.4E-11</td><td>MB231/MB468</td><td>CYTO</td></tr>
</table>

<table>
<tr><td>>MI0000342|hsa-mir-200b&WithFlank&1|+|1167098|1167204@63.85.23&[MIMAT0000318&hsa-miR-200b-3p&offsets|0|+1;m-93&1|+|1167160|1167182&offsets|0|0]|TAATACTGCCTGGTAATGATGAC</td><td>>MI0000342|hsa-mir-200b&WithFlank&1|+|1167098|1167204</td><td>63</td><td>85</td><td>23</td><td>TAATACTGCCTGGTAATGATGAC</td><td>3125.0</td><td>-5.3</td><td>0.8</td><td>-6.9</td><td>4.2E-12</td><td>1.4E-11</td><td>MB231/MB468</td><td>MP</td></tr>
<tr><td>>hg38_21_+_8433172_8446622_RNA45SN1_45S_50ntFlanks@11578.11602.25|TAGAACTGGTGCGGACCAGGGGAAT</td><td>>hg38_21_+_8433172_8446622_RNA45SN1_45S_50ntFlanks</td><td>11578</td><td>11602</td><td>25</td><td>TAGAACTGGTGCGGACCAGGGGAAT</td><td>103.4</td><td>-10.6</td><td>1.5</td><td>-6.9</td><td>4.2E-12</td><td>1.4E-11</td><td>MB231/MB468</td><td>MP</td></tr>
<tr><td>>MI0000088|hsa-mir-30a&WithFlank&6|-|71403545|71403627@13.35.23&[MIMAT0000087&hsa-miR-30a-5p&offsets|+1|+2;m-15&6|-|71403595|71403616&offsets|+1|+2]|GTAAACATCCTCGACTGGAAGCT</td><td>>MI0000088|hsa-mir-30a&WithFlank&6|-|71403545|71403627</td><td>13</td><td>35</td><td>23</td><td>GTAAACATCCTCGACTGGAAGCT</td><td>582.1</td><td>7.1</td><td>1.0</td><td>6.9</td><td>4.4E-12</td><td>1.4E-11</td><td>BT20/MB468</td><td>CYTO</td></tr>
<tr><td>>hg38_21_+_8433172_8446622_RNA45SN1_45S_50ntFlanks@4731.4763.33|AAGAACGAAAGTCGGAGGTTCGAAGACGATCAG</td><td>>hg38_21_+_8433172_8446622_RNA45SN1_45S_50ntFlanks</td><td>4731</td><td>4763</td><td>33</td><td>AAGAACGAAAGTCGGAGGTTCGAAGACGATCAG</td><td>590.3</td><td>5.3</td><td>0.8</td><td>6.9</td><td>4.4E-12</td><td>1.4E-11</td><td>BT20/MB468</td><td>CYTO</td></tr>
<tr><td>>hg38_21_+_8433172_8446622_RNA45SN1_45S_50ntFlanks@7990.8013.24|GACGTGGCGACCCGCTGAATTTAA</td><td>>hg38_21_+_8433172_8446622_RNA45SN1_45S_50ntFlanks</td><td>7990</td><td>8013</td><td>24</td><td>GACGTGGCGACCCGCTGAATTTAA</td><td>180.0</td><td>6.0</td><td>0.9</td><td>6.9</td><td>4.6E-12</td><td>1.4E-11</td><td>MB231/MB468</td><td>CYTO</td></tr>
<tr><td>>hg38_21_+_8433172_8446622_RNA45SN1_45S_50ntFlanks@9621.9658.38|TCTAGTAGCTGGTTCCCTCCGAAGTTTCCCTCAGGATA</td><td>>hg38_21_+_8433172_8446622_RNA45SN1_45S_50ntFlanks</td><td>9621</td><td>9658</td><td>38</td><td>TCTAGTAGCTGGTTCCCTCCGAAGTTTCCCTCAGGATA</td><td>189.2</td><td>-11.4</td><td>1.6</td><td>-6.9</td><td>4.2E-12</td><td>1.4E-11</td><td>MB231/MB468</td><td>MP</td></tr>
<tr><td>>MI0001445|hsa-mir-423&WithFlank&17|+|30117073|30117178@23.44.22(+1C)&[MIMAT0004748&hsa-miR-423-5p&offsets|0|-1(+1C);m-141&17|+|30117095|30117117&offsets|0|-1(+1C)]|TGAGGGGCAGAGAGCGAGACTTC</td><td>>MI0001445|hsa-mir-423&WithFlank&17|+|30117073|30117178</td><td>23</td><td>44</td><td>22(+1C)</td><td>TGAGGGGCAGAGAGCGAGACTTC</td><td>8583.4</td><td>7.7</td><td>1.0</td><td>7.3</td><td>2.5E-13</td><td>1.4E-11</td><td>BT20/MB468</td><td>MP</td></tr>
<tr><td>>hg38_21_+_8433172_8446622_RNA45SN1_45S_50ntFlanks@6721.6756.36|AATTGCAGGACACATTGATCATCGACACTTCGAACG</td><td>>hg38_21_+_8433172_8446622_RNA45SN1_45S_50ntFlanks</td><td>6721</td><td>6756</td><td>36</td><td>AATTGCAGGACACATTGATCATCGACACTTCGAACG</td><td>999.7</td><td>-7.7</td><td>1.1</td><td>-6.9</td><td>4.2E-12</td><td>1.4E-11</td><td>MB231/MB468</td><td>MP</td></tr>
<tr><td>>hg38_21_+_8433172_8446622_RNA45SN1_45S_50ntFlanks@6778.6802.25|CCCGGGGCTACGCCTGTCTGAGCGT</td><td>>hg38_21_+_8433172_8446622_RNA45SN1_45S_50ntFlanks</td><td>6778</td><td>6802</td><td>25</td><td>CCCGGGGCTACGCCTGTCTGAGCGT</td><td>2828.5</td><td>6.1</td><td>0.9</td><td>6.9</td><td>4.6E-12</td><td>1.4E-11</td><td>MB231/MB468</td><td>CYTO</td></tr>
<tr><td>>hg38_21_+_8433172_8446622_RNA45SN1_45S_50ntFlanks@11572.11612.41|CCGACTTAGAACTGGTGCGGACCAGGGGAATCCGACTGTTT</td><td>>hg38_21_+_8433172_8446622_RNA45SN1_45S_50ntFlanks</td><td>11572</td><td>11612</td><td>41</td><td>CCGACTTAGAACTGGTGCGGACCAGGGGAATCCGACTGTTT</td><td>10269.9</td><td>-6.2</td><td>0.9</td><td>-6.9</td><td>5.6E-12</td><td>1.5E-11</td><td>MB231/MB468</td><td>MITO</td></tr>
<tr><td>>hg38_21_+_8433172_8446622_RNA45SN1_45S_50ntFlanks@7973.7997.25|GACGCGACCTCAGATCAGACGTGGC</td><td>>hg38_21_+_8433172_8446622_RNA45SN1_45S_50ntFlanks</td><td>7973</td><td>7997</td><td>25</td><td>GACGCGACCTCAGATCAGACGTGGC</td><td>1421.8</td><td>6.9</td><td>1.0</td><td>7.0</td><td>3.4E-12</td><td>1.5E-11</td><td>BT20/MB468</td><td>MITO</td></tr>
<tr><td>>MI0000298|hsa-mir-221&WithFlank&X|-|45746151|45746272@71.91.21(+1A)&[MIMAT0000278&hsa-miR-221-3p&offsets|0|-2(+1A);m-91&X|-|45746181|45746202&offsets|0|-1(+1A)]|AGCTACATTGTCTGCTGGGTTA</td><td>>MI0000298|hsa-mir-221&WithFlank&X|-|45746151|45746272</td><td>71</td><td>91</td><td>21(+1A)</td><td>AGCTACATTGTCTGCTGGGTTA</td><td>1049.3</td><td>7.0</td><td>1.0</td><td>6.9</td><td>5.6E-12</td><td>1.5E-11</td><td>MB231/MB468</td><td>MITO</td></tr>
<tr><td>>hg38_21_+_8433172_8446622_RNA45SN1_45S_50ntFlanks@11104.11122.19|AAGGGTCGGGGCGGCAGGG</td><td>>hg38_21_+_8433172_8446622_RNA45SN1_45S_50ntFlanks</td><td>11104</td><td>11122</td><td>19</td><td>AAGGGTCGGGGCGGCAGGG</td><td>149.0</td><td>-11.1</td><td>1.6</td><td>-6.9</td><td>4.2E-12</td><td>1.5E-11</td><td>MB231/MB468</td><td>MP</td></tr>
<tr><td>>trna116_GluCTC_1_-_145399233_145399304@3.35.33,trna59_GluCTC_1_+_249168447_249168518@3.35.33,trna71_GluCTC_1_-_161439189_161439260@3.35.33,trna74_GluCTC_1_-_161431809_161431880@3.35.33,trna77_GluCTC_1_-_161424398_161424469@3.35.33,trna77_GluCTC_6_+_28949976_28950047@3.35.33,trna80_GluCTC_1_-_161417018_161417089@3.35.33,trna87_GluCTC_6_-_126101393_126101464@3.35.33|CCTGGTGGTCTAGTGGTTAGGATTCGGCGCTCT</td><td>>trna116_GluCTC_1_-_145399233_145399304</td><td>3</td><td>35</td><td>33</td><td>CCTGGTGGTCTAGTGGTTAGGATTCGGCGCTCT</td><td>251.5</td><td>5.2</td><td>0.7</td><td>6.9</td><td>4.8E-12</td><td>1.5E-11</td><td>MB231/MB468</td><td>NUC</td></tr>
<tr><td>>hg38_21_+_8433172_8446622_RNA45SN1_45S_50ntFlanks@4895.4916.22|CTTAAAGGAATTGACGGAAGGG</td><td>>hg38_21_+_8433172_8446622_RNA45SN1_45S_50ntFlanks</td><td>4895</td><td>4916</td><td>22</td><td>CTTAAAGGAATTGACGGAAGGG</td><td>124.0</td><td>-10.8</td><td>1.6</td><td>-6.9</td><td>4.2E-12</td><td>1.5E-11</td><td>MB231/MB468</td><td>MP</td></tr>
<tr><td>>hg38_21_+_8433172_8446622_RNA45SN1_45S_50ntFlanks@6752.6770.19|GAACGCACTTGCGGCCCCG</td><td>>hg38_21_+_8433172_8446622_RNA45SN1_45S_50ntFlanks</td><td>6752</td><td>6770</td><td>19</td><td>GAACGCACTTGCGGCCCCG</td><td>105.3</td><td>-10.6</td><td>1.5</td><td>-6.9</td><td>4.2E-12</td><td>1.5E-11</td><td>MB231/MB468</td><td>MP</td></tr>
<tr><td>>hg38_21_+_8433172_8446622_RNA45SN1_45S_50ntFlanks@4980.5000.21|ACACGGACAGGATTGACAGAT</td><td>>hg38_21_+_8433172_8446622_RNA45SN1_45S_50ntFlanks</td><td>4980</td><td>5000</td><td>21</td><td>ACACGGACAGGATTGACAGAT</td><td>106.2</td><td>-10.6</td><td>1.5</td><td>-6.9</td><td>4.2E-12</td><td>1.5E-11</td><td>MB231/MB468</td><td>MP</td></tr>
<tr><td>>hg38_21_+_8433172_8446622_RNA45SN1_45S_50ntFlanks@10726.10755.30|GGGAGAGGGTGTAAATCTCGCGCCGGGCCG</td><td>>hg38_21_+_8433172_8446622_RNA45SN1_45S_50ntFlanks</td><td>10726</td><td>10755</td><td>30</td><td>GGGAGAGGGTGTAAATCTCGCGCCGGGCCG</td><td>296.1</td><td>-11.3</td><td>1.5</td><td>-7.3</td><td>2.5E-13</td><td>1.5E-11</td><td>BT20/MB468</td><td>MP</td></tr>
<tr><td>>hg38_21_+_8433172_8446622_RNA45SN1_45S_50ntFlanks@10922.10941.20|CTGGACGAGGCGCCGCCGCC</td><td>>hg38_21_+_8433172_8446622_RNA45SN1_45S_50ntFlanks</td><td>10922</td><td>10941</td><td>20</td><td>CTGGACGAGGCGCCGCCGCC</td><td>890.0</td><td>-9.0</td><td>1.3</td><td>-6.9</td><td>5.7E-12</td><td>1.5E-11</td><td>MB231/MB468</td><td>MITO</td></tr>
<tr><td>>hg38_21_+_8433172_8446622_RNA45SN1_45S_50ntFlanks@9177.9197.21|GCGCCGTCGGGCCCGGGGGAG</td><td>>hg38_21_+_8433172_8446622_RNA45SN1_45S_50ntFlanks</td><td>9177</td><td>9197</td><td>21</td><td>GCGCCGTCGGGCCCGGGGGAG</td><td>232.3</td><td>-11.7</td><td>1.7</td><td>-6.9</td><td>4.2E-12</td><td>1.5E-11</td><td>MB231/MB468</td><td>MP</td></tr>
<tr><td>>hg38_21_+_8433172_8446622_RNA45SN1_45S_50ntFlanks@5379.5419.41|ATTAAGTCCCTGCCCTTTGTACACACCGCCCGTCGCTACTA</td><td>>hg38_21_+_8433172_8446622_RNA45SN1_45S_50ntFlanks</td><td>5379</td><td>5419</td><td>41</td><td>ATTAAGTCCCTGCCCTTTGTACACACCGCCCGTCGCTACTA</td><td>303.4</td><td>-10.3</td><td>1.5</td><td>-7.0</td><td>3.4E-12</td><td>1.5E-11</td><td>BT20/MB468</td><td>MITO</td></tr>
<tr><td>>hg38_21_+_8433172_8446622_RNA45SN1_45S_50ntFlanks@9736.9758.23|CCGAAACGATCTCAACCTATTCT</td><td>>hg38_21_+_8433172_8446622_RNA45SN1_45S_50ntFlanks</td><td>9736</td><td>9758</td><td>23</td><td>CCGAAACGATCTCAACCTATTCT</td><td>176.1</td><td>-9.8</td><td>1.4</td><td>-6.9</td><td>4.4E-12</td><td>1.5E-11</td><td>BT20/MB468</td><td>CYTO</td></tr>
<tr><td>>hg38_21_+_8433172_8446622_RNA45SN1_45S_50ntFlanks@9129.9148.20|CGGGTCGGGGGGCGGGGCGG</td><td>>hg38_21_+_8433172_8446622_RNA45SN1_45S_50ntFlanks</td><td>9129</td><td>9148</td><td>20</td><td>CGGGTCGGGGGGCGGGGCGG</td><td>223.8</td><td>-11.7</td><td>1.7</td><td>-6.9</td><td>4.2E-12</td><td>1.5E-11</td><td>MB231/MB468</td><td>MP</td></tr>
<tr><td>>hg38_21_+_8433172_8446622_RNA45SN1_45S_50ntFlanks@12640.12669.30|ACCATCTGTGGGATTATGACTGAACGCCTC</td><td>>hg38_21_+_8433172_8446622_RNA45SN1_45S_50ntFlanks</td><td>12640</td><td>12669</td><td>30</td><td>ACCATCTGTGGGATTATGACTGAACGCCTC</td><td>675.2</td><td>3.8</td><td>0.5</td><td>7.0</td><td>3.5E-12</td><td>1.5E-11</td><td>BT20/MB468</td><td>MITO</td></tr>
<tr><td>>trnaMT_GluTTC_MT_-_14674_14742@26.67.42,trnalookalike8_GluTTC_5_-_93905172_93905240@26.67.42|GGTTTTTCATATCATTGGTCGTGGTTGTAGTCCGTGCGAGAA</td><td>>trnaMT_GluTTC_MT_-_14674_14742</td><td>26</td><td>67</td><td>42</td><td>GGTTTTTCATATCATTGGTCGTGGTTGTAGTCCGTGCGAGAA</td><td>360.7</td><td>8.4</td><td>1.2</td><td>7.0</td><td>3.5E-12</td><td>1.5E-11</td><td>BT20/MB468</td><td>MITO</td></tr>
<tr><td>>MI0000078|hsa-mir-22&WithFlank&17|-|1713897|1713993@59.80.22(+1A)&[MIMAT0000077&hsa-miR-22-3p&offsets|0|0(+1A);m-2&17|-|1713914|1713935&offsets|0|0(+1A)]|AAGCTGCCAGTTGAAGAACTGTA</td><td>>MI0000078|hsa-mir-22&WithFlank&17|-|1713897|1713993</td><td>59</td><td>80</td><td>22(+1A)</td><td>AAGCTGCCAGTTGAAGAACTGTA</td><td>343.1</td><td>8.5</td><td>1.2</td><td>7.0</td><td>3.5E-12</td><td>1.5E-11</td><td>BT20/MB468</td><td>MITO</td></tr>
<tr><td>>MI0000743|hsa-mir-34c&WithFlank&11|+|111513433|111513521@19.40.22(+1U)&[MIMAT0000686&hsa-miR-34c-5p&offsets|0|-1(+1U);m-336&11|+|111513451|111513473&offsets|0|-1(+1U)]|AGGCAGTGTAGTTAGCTGATTGT</td><td>>MI0000743|hsa-mir-34c&WithFlank&11|+|111513433|111513521</td><td>19</td><td>40</td><td>22(+1U)</td><td>AGGCAGTGTAGTTAGCTGATTGT</td><td>537.9</td><td>-12.5</td><td>1.7</td><td>-7.5</td><td>6.4E-14</td><td>1.5E-11</td><td>BT20/MB231</td><td>MP</td></tr>
<tr><td>>hg38_21_+_8433172_8446622_RNA45SN1_45S_50ntFlanks@10349.10368.20|AACGAGAACTTTGAAGGCCG</td><td>>hg38_21_+_8433172_8446622_RNA45SN1_45S_50ntFlanks</td><td>10349</td><td>10368</td><td>20</td><td>AACGAGAACTTTGAAGGCCG</td><td>1980.3</td><td>-3.8</td><td>0.5</td><td>-6.9</td><td>4.4E-12</td><td>1.5E-11</td><td>BT20/MB468</td><td>CYTO</td></tr>
<tr><td>>hg38_21_+_8433172_8446622_RNA45SN1_45S_50ntFlanks@6719.6736.18|TGAATTGCAGGACACATT</td><td>>hg38_21_+_8433172_8446622_RNA45SN1_45S_50ntFlanks</td><td>6719</td><td>6736</td><td>18</td><td>TGAATTGCAGGACACATT</td><td>1823.6</td><td>-5.4</td><td>0.8</td><td>-6.9</td><td>4.4E-12</td><td>1.5E-11</td><td>BT20/MB468</td><td>CYTO</td></tr>
<tr><td>>hg38_21_+_8433172_8446622_RNA45SN1_45S_50ntFlanks@11874.11898.25|AGAAGACCCTGTTGAGCTTGACTCT</td><td>>hg38_21_+_8433172_8446622_RNA45SN1_45S_50ntFlanks</td><td>11874</td><td>11898</td><td>25</td><td>AGAAGACCCTGTTGAGCTTGACTCT</td><td>2778.8</td><td>-6.0</td><td>0.9</td><td>-6.9</td><td>5.7E-12</td><td>1.5E-11</td><td>MB231/MB468</td><td>MITO</td></tr>
<tr><td>>hg38_21_+_8433172_8446622_RNA45SN1_45S_50ntFlanks@8196.8232.37|CGTGGACGGTGTGAGGCCGGTAGCGGCCCCCGGCGCG</td><td>>hg38_21_+_8433172_8446622_RNA45SN1_45S_50ntFlanks</td><td>8196</td><td>8232</td><td>37</td><td>CGTGGACGGTGTGAGGCCGGTAGCGGCCCCCGGCGCG</td><td>103.0</td><td>-10.6</td><td>1.5</td><td>-6.9</td><td>4.2E-12</td><td>1.5E-11</td><td>MB231/MB468</td><td>MP</td></tr>
<tr><td>>hg38_21_+_8433172_8446622_RNA45SN1_45S_50ntFlanks@7990.8019.30|GACGTGGCGACCCGCTGAATTTAAGCATAT</td><td>>hg38_21_+_8433172_8446622_RNA45SN1_45S_50ntFlanks</td><td>7990</td><td>8019</td><td>30</td><td>GACGTGGCGACCCGCTGAATTTAAGCATAT</td><td>813.8</td><td>10.1</td><td>1.5</td><td>6.9</td><td>4.7E-12</td><td>1.5E-11</td><td>MB231/MB468</td><td>CYTO</td></tr>
<tr><td>>hg38_21_+_8433172_8446622_RNA45SN1_45S_50ntFlanks@10805.10828.24|GAACAATGTAGGTAAGGGAAGTCG</td><td>>hg38_21_+_8433172_8446622_RNA45SN1_45S_50ntFlanks</td><td>10805</td><td>10828</td><td>24</td><td>GAACAATGTAGGTAAGGGAAGTCG</td><td>143.5</td><td>-9.3</td><td>1.3</td><td>-7.0</td><td>3.5E-12</td><td>1.5E-11</td><td>BT20/MB468</td><td>MITO</td></tr>
<tr><td>>hg38_21_+_8433172_8446622_RNA45SN1_45S_50ntFlanks@10786.10826.41|AACAGCCTCTGGCATGTTGGAACAATGTAGGTAAGGGAAGT</td><td>>hg38_21_+_8433172_8446622_RNA45SN1_45S_50ntFlanks</td><td>10786</td><td>10826</td><td>41</td><td>AACAGCCTCTGGCATGTTGGAACAATGTAGGTAAGGGAAGT</td><td>828.9</td><td>-5.6</td><td>0.8</td><td>-6.9</td><td>4.8E-12</td><td>1.5E-11</td><td>MB231/MB468</td><td>NUC</td></tr>
<tr><td>>hg38_21_+_8433172_8446622_RNA45SN1_45S_50ntFlanks@5425.5450.26|TGGATGGTTTAGTGAGGCCCTCGGAT</td><td>>hg38_21_+_8433172_8446622_RNA45SN1_45S_50ntFlanks</td><td>5425</td><td>5450</td><td>26</td><td>TGGATGGTTTAGTGAGGCCCTCGGAT</td><td>600.1</td><td>10.4</td><td>1.5</td><td>6.9</td><td>4.5E-12</td><td>1.5E-11</td><td>BT20/MB468</td><td>CYTO</td></tr>
<tr><td>>MI0016887|hsa-mir-4521&WithFlank&17|+|8186939|8187010@13.35.23&[MIMAT0019058&hsa-miR-4521&offsets|+3|+4;m-765&17|+|8186948|8186969&offsets|+3|+4]|AAGGAAGTCCTGTGCTCAGTTTT</td><td>>MI0016887|hsa-mir-4521&WithFlank&17|+|8186939|8187010</td><td>13</td><td>35</td><td>23</td><td>AAGGAAGTCCTGTGCTCAGTTTT</td><td>799.5</td><td>10.9</td><td>1.6</td><td>6.9</td><td>4.5E-12</td><td>1.5E-11</td><td>BT20/MB468</td><td>CYTO</td></tr>
<tr><td>>hg38_21_+_8433172_8446622_RNA45SN1_45S_50ntFlanks@11575.11605.31|ACTTAGAACTGGTGCGGACCAGGGGAATCCG</td><td>>hg38_21_+_8433172_8446622_RNA45SN1_45S_50ntFlanks</td><td>11575</td><td>11605</td><td>31</td><td>ACTTAGAACTGGTGCGGACCAGGGGAATCCG</td><td>4140.7</td><td>-10.9</td><td>1.6</td><td>-6.9</td><td>6.1E-12</td><td>1.5E-11</td><td>BT20/MB468</td><td>NUC</td></tr>
<tr><td>>hg38_21_+_8433172_8446622_RNA45SN1_45S_50ntFlanks@12033.12068.36|ATCGTTTTTTCACTGACCCGGTGAGGCGGGGGGGCG</td><td>>hg38_21_+_8433172_8446622_RNA45SN1_45S_50ntFlanks</td><td>12033</td><td>12068</td><td>36</td><td>ATCGTTTTTTCACTGACCCGGTGAGGCGGGGGGGCG</td><td>253.7</td><td>-10.1</td><td>1.4</td><td>-7.0</td><td>3.5E-12</td><td>1.5E-11</td><td>BT20/MB468</td><td>MITO</td></tr>
<tr><td>>hg38_21_+_8433172_8446622_RNA45SN1_45S_50ntFlanks@10808.10825.18|CAATGTAGGTAAGGGAAG</td><td>>hg38_21_+_8433172_8446622_RNA45SN1_45S_50ntFlanks</td><td>10808</td><td>10825</td><td>18</td><td>CAATGTAGGTAAGGGAAG</td><td>1695.6</td><td>-8.5</td><td>1.2</td><td>-6.9</td><td>4.7E-12</td><td>1.5E-11</td><td>MB231/MB468</td><td>CYTO</td></tr>
<tr><td>>hg38_21_+_8433172_8446622_RNA45SN1_45S_50ntFlanks@6789.6807.19|GCCTGTCTGAGCGTCGCTT</td><td>>hg38_21_+_8433172_8446622_RNA45SN1_45S_50ntFlanks</td><td>6789</td><td>6807</td><td>19</td><td>GCCTGTCTGAGCGTCGCTT</td><td>625.4</td><td>5.2</td><td>0.7</td><td>6.9</td><td>4.8E-12</td><td>1.5E-11</td><td>MB231/MB468</td><td>CYTO</td></tr>
<tr><td>>hg38_21_+_8433172_8446622_RNA45SN1_45S_50ntFlanks@10362.10384.23|AAGGCCGAAGTGGAGAAGGGTTC</td><td>>hg38_21_+_8433172_8446622_RNA45SN1_45S_50ntFlanks</td><td>10362</td><td>10384</td><td>23</td><td>AAGGCCGAAGTGGAGAAGGGTTC</td><td>1504.0</td><td>-7.8</td><td>1.1</td><td>-6.9</td><td>4.3E-12</td><td>1.5E-11</td><td>MB231/MB468</td><td>MP</td></tr>
<tr><td>>hg38_21_+_8433172_8446622_RNA45SN1_45S_50ntFlanks@11778.11803.26|ACGCGCATGAATGGATGAACGAGATT</td><td>>hg38_21_+_8433172_8446622_RNA45SN1_45S_50ntFlanks</td><td>11778</td><td>11803</td><td>26</td><td>ACGCGCATGAATGGATGAACGAGATT</td><td>1164.9</td><td>-7.4</td><td>1.1</td><td>-6.9</td><td>4.3E-12</td><td>1.5E-11</td><td>MB231/MB468</td><td>MP</td></tr>
<tr><td>>hg38_21_+_8433172_8446622_RNA45SN1_45S_50ntFlanks@10687.10708.22|CGGCGTCCGGTGAGCTCTCGCT</td><td>>hg38_21_+_8433172_8446622_RNA45SN1_45S_50ntFlanks</td><td>10687</td><td>10708</td><td>22</td><td>CGGCGTCCGGTGAGCTCTCGCT</td><td>2579.8</td><td>-4.4</td><td>0.6</td><td>-6.9</td><td>4.9E-12</td><td>1.5E-11</td><td>MB231/MB468</td><td>NUC</td></tr>
<tr><td>>hg38_21_+_8433172_8446622_RNA45SN1_45S_50ntFlanks@10927.10945.19|CGAGGCGCCGCCGCCCCCC</td><td>>hg38_21_+_8433172_8446622_RNA45SN1_45S_50ntFlanks</td><td>10927</td><td>10945</td><td>19</td><td>CGAGGCGCCGCCGCCCCCC</td><td>517.3</td><td>-12.9</td><td>1.9</td><td>-6.9</td><td>4.3E-12</td><td>1.5E-11</td><td>MB231/MB468</td><td>MP</td></tr>
<tr><td>>hg38_21_+_8433172_8446622_RNA45SN1_45S_50ntFlanks@12043.12070.28|CACTGACCCGGTGAGGCGGGGGGGGCGAG</td><td>>hg38_21_+_8433172_8446622_RNA45SN1_45S_50ntFlanks</td><td>12043</td><td>12070</td><td>28</td><td>CACTGACCCGGTGAGGCGGGGGGGCGAG</td><td>124.8</td><td>-10.8</td><td>1.6</td><td>-6.9</td><td>4.3E-12</td><td>1.5E-11</td><td>MB231/MB468</td><td>MP</td></tr>
<tr><td>>hg38_21_+_8433172_8446622_RNA45SN1_45S_50ntFlanks@6733.6770.38|CATTGATCATCGACACTTCGAACGCACTTGCGGCCCCG</td><td>>hg38_21_+_8433172_8446622_RNA45SN1_45S_50ntFlanks</td><td>6733</td><td>6770</td><td>38</td><td>CATTGATCATCGACACTTCGAACGCACTTGCGGCCCCG</td><td>1980.9</td><td>-10.1</td><td>1.5</td><td>-6.9</td><td>4.3E-12</td><td>1.5E-11</td><td>MB231/MB468</td><td>MP</td></tr>
<tr><td>>hg38_21_+_8433172_8446622_RNA45SN1_45S_50ntFlanks@9033.9057.25|CTCGTGGGGGGGCCGGGCCACCCCT</td><td>>hg38_21_+_8433172_8446622_RNA45SN1_45S_50ntFlanks</td><td>9033</td><td>9057</td><td>25</td><td>CTCGTGGGGGGGCCGGGCCACCCCT</td><td>495.4</td><td>-12.0</td><td>1.6</td><td>-7.3</td><td>2.6E-13</td><td>1.5E-11</td><td>BT20/MB468</td><td>MP</td></tr>
<tr><td>>hg38_21_+_8433172_8446622_RNA45SN1_45S_50ntFlanks@9929.9948.20|GATATAGACAGCAGGACGGT</td><td>>hg38_21_+_8433172_8446622_RNA45SN1_45S_50ntFlanks</td><td>9929</td><td>9948</td><td>20</td><td>GATATAGACAGCAGGACGGT</td><td>1515.0</td><td>-5.9</td><td>0.9</td><td>-6.9</td><td>4.3E-12</td><td>1.5E-11</td><td>MB231/MB468</td><td>MP</td></tr>
</table>

<table>
<tr><td></td><td></td><td></td><td></td><td></td><td></td><td></td><td></td><td></td><td></td><td></td><td></td><td></td><td></td></tr>
<tr><td>>hg38_21_+_8433172_8446622_RNA45SN1_45S_50ntFlanks@11321.11343.23|CACGTCTCGTCGCGCGCGCGTCC</td><td>>hg38_21_+_8433172_8446622_RNA45SN1_45S_50ntFlanks</td><td>11321</td><td>11343</td><td>23</td><td>CACGTCTCGTCGCGCGCGCGTCC</td><td>97.7</td><td>-10.5</td><td>1.5</td><td>-6.9</td><td>4.3E-12</td><td>1.5E-11</td><td>MB231/MB468</td><td>MP</td></tr>
<tr><td>>hg38_21_+_8433172_8446622_RNA45SN1_45S_50ntFlanks@12825.12847.23|GGACCGGGGTCCGGTGCGGAGTG</td><td>>hg38_21_+_8433172_8446622_RNA45SN1_45S_50ntFlanks</td><td>12825</td><td>12847</td><td>23</td><td>GGACCGGGGTCCGGTGCGGAGTG</td><td>191.8</td><td>-11.4</td><td>1.7</td><td>-6.9</td><td>4.3E-12</td><td>1.5E-11</td><td>MB231/MB468</td><td>MP</td></tr>
<tr><td>>hg38_21_+_8433172_8446622_RNA45SN1_45S_50ntFlanks@6651.6693.43|CGACTCTTAGCGGTGGATCACTCGGCTCGTGCGTCGATGAAGA</td><td>>hg38_21_+_8433172_8446622_RNA45SN1_45S_50ntFlanks</td><td>6651</td><td>6693</td><td>43</td><td>CGACTCTTAGCGGTGGATCACTCGGCTCGTGCGTCGATGAAGA</td><td>1370.7</td><td>4.9</td><td>0.7</td><td>6.9</td><td>4.8E-12</td><td>1.5E-11</td><td>MB231/MB468</td><td>CYTO</td></tr>
<tr><td>>MI0000264|hsa-mir-7-2&WithFlank&15|+|88611819|88611940@38.60.23(+1G);MI0000265|hsa-mir-7-3&WithFlank&19|+|4770664|4770785@37.59.23(+1G);MI0000263|hsa-mir-7-1&WithFlank&9|-|83969742|83969863@30.52.23(+1G)&[MIMAT0000252&hsa-miR-7-5p&offsets|0|-1(+1G);m-400&15|+|88611856|88611879&offsets|0|-1(+1G)];[MIMAT0000252_1&hsa-miR-7-5p&offsets|0|-1(+1G);m-402&19|+|4770700|4770723&offsets|0|-1(+1G)];[MIMAT0000252_2&hsa-miR-7-5p&offsets|0|-1(+1G);m-401&9|-|83969811|83969834&offsets|0|-1(+1G)]|TGGAAGACTAGTGATTTTGTTGTG</td><td>>MI0000264|hsa-mir-7-2&WithFlank&15|+|88611819|88611940</td><td>38</td><td>60</td><td>23(+1G)</td><td>TGGAAGACTAGTGATTTTGTTGTG</td><td>537.1</td><td>7.0</td><td>1.0</td><td>7.0</td><td>3.6E-12</td><td>1.5E-11</td><td>BT20/MB468</td><td>MITO</td></tr>
<tr><td>>MI0000078|hsa-mir-22&WithFlank&17|-|1713897|1713993@21.41.21&[MIMAT0004495&hsa-miR-22-5p&offsets|0|-1;m-217&17|-|1713953|1713973&offsets|0|0]|AGTTCTTCAGTGGCAAGCTTT</td><td>>MI0000078|hsa-mir-22&WithFlank&17|-|1713897|1713993</td><td>21</td><td>41</td><td>21</td><td>AGTTCTTCAGTGGCAAGCTTT</td><td>322.6</td><td>7.9</td><td>1.1</td><td>6.9</td><td>4.8E-12</td><td>1.5E-11</td><td>MB231/MB468</td><td>CYTO</td></tr>
<tr><td>>hg38_21_+_8433172_8446622_RNA45SN1_45S_50ntFlanks@4640.4665.26|GCATTCGTATTGCGCCGCTAGAGGTG</td><td>>hg38_21_+_8433172_8446622_RNA45SN1_45S_50ntFlanks</td><td>4640</td><td>4665</td><td>26</td><td>GCATTCGTATTGCGCCGCTAGAGGTG</td><td>85.8</td><td>-10.3</td><td>1.5</td><td>-6.9</td><td>4.3E-12</td><td>1.5E-11</td><td>MB231/MB468</td><td>MP</td></tr>
<tr><td>>MI0003513|hsa-mir-455&WithFlank&9|+|114209428|114209535@60.80.21(+1C)&[MIMAT0004784&hsa-miR-455-3p&offsets|0|0(+1C);m-135&9|+|114209487|114209508&offsets|0|-1(+1C)]|GCAGTCCATGGGCATATACACC</td><td>>MI0003513|hsa-mir-455&WithFlank&9|+|114209428|114209535</td><td>60</td><td>80</td><td>21(+1C)</td><td>GCAGTCCATGGGCATATACACC</td><td>215.0</td><td>11.0</td><td>1.6</td><td>6.9</td><td>5.8E-12</td><td>1.5E-11</td><td>MB231/MB468</td><td>MITO</td></tr>
<tr><td>>hg38_21_+_8433172_8446622_RNA45SN1_45S_50ntFlanks@10402.10431.30|AACATGGGTCAGTCGGTCCTGAGAGATGGG</td><td>>hg38_21_+_8433172_8446622_RNA45SN1_45S_50ntFlanks</td><td>10402</td><td>10431</td><td>30</td><td>AACATGGGTCAGTCGGTCCTGAGAGATGGG</td><td>2545.8</td><td>-7.8</td><td>1.1</td><td>-6.9</td><td>5.0E-12</td><td>1.5E-11</td><td>MB231/MB468</td><td>NUC</td></tr>
<tr><td>>hg38_21_+_8433172_8446622_RNA45SN1_45S_50ntFlanks@12933.12953.21|GCTAAACCATTCGTAGACGAC</td><td>>hg38_21_+_8433172_8446622_RNA45SN1_45S_50ntFlanks</td><td>12933</td><td>12953</td><td>21</td><td>GCTAAACCATTCGTAGACGAC</td><td>139.5</td><td>-11.0</td><td>1.6</td><td>-6.9</td><td>4.4E-12</td><td>1.5E-11</td><td>MB231/MB468</td><td>MP</td></tr>
<tr><td>>hg38_21_+_8433172_8446622_RNA45SN1_45S_50ntFlanks@10771.10792.22|CAGGTCTCCAAGGTGAACAGCC</td><td>>hg38_21_+_8433172_8446622_RNA45SN1_45S_50ntFlanks</td><td>10771</td><td>10792</td><td>22</td><td>CAGGTCTCCAAGGTGAACAGCC</td><td>88.7</td><td>-10.3</td><td>1.5</td><td>-6.9</td><td>4.4E-12</td><td>1.5E-11</td><td>MB231/MB468</td><td>MP</td></tr>
<tr><td>>hg38_21_+_8433172_8446622_RNA45SN1_45S_50ntFlanks@5017.5034.18|ATTCCGTGGGTGGTGGTG</td><td>>hg38_21_+_8433172_8446622_RNA45SN1_45S_50ntFlanks</td><td>5017</td><td>5034</td><td>18</td><td>ATTCCGTGGGTGGTGGTG</td><td>178.9</td><td>-11.3</td><td>1.6</td><td>-6.9</td><td>4.4E-12</td><td>1.5E-11</td><td>MB231/MB468</td><td>MP</td></tr>
<tr><td>>hg38_21_+_8433172_8446622_RNA45SN1_45S_50ntFlanks@11205.11232.28|GCGGCGGCGACTCTGGACGCGAGCCGGG</td><td>>hg38_21_+_8433172_8446622_RNA45SN1_45S_50ntFlanks</td><td>11205</td><td>11232</td><td>28</td><td>GCGGCGGCGACTCTGGACGCGAGCCGGG</td><td>441.6</td><td>-8.3</td><td>1.2</td><td>-6.9</td><td>5.9E-12</td><td>1.5E-11</td><td>MB231/MB468</td><td>MITO</td></tr>
<tr><td>>hg38_21_+_8433172_8446622_RNA45SN1_45S_50ntFlanks@10781.10801.21|AGGTGAACAGCCTCTGGCATG</td><td>>hg38_21_+_8433172_8446622_RNA45SN1_45S_50ntFlanks</td><td>10781</td><td>10801</td><td>21</td><td>AGGTGAACAGCCTCTGGCATG</td><td>836.2</td><td>-6.1</td><td>0.9</td><td>-6.9</td><td>4.9E-12</td><td>1.5E-11</td><td>MB231/MB468</td><td>CYTO</td></tr>
<tr><td>>hg38_21_+_8433172_8446622_RNA45SN1_45S_50ntFlanks@10107.10130.24|CGTGTGGTGTGCGTCGGAGGGCGG</td><td>>hg38_21_+_8433172_8446622_RNA45SN1_45S_50ntFlanks</td><td>10107</td><td>10130</td><td>24</td><td>CGTGTGGTGTGCGTCGGAGGGCGG</td><td>108.5</td><td>-10.6</td><td>1.5</td><td>-6.9</td><td>4.4E-12</td><td>1.5E-11</td><td>MB231/MB468</td><td>MP</td></tr>
<tr><td>>hg38_21_+_8433172_8446622_RNA45SN1_45S_50ntFlanks@11565.11603.39|CTAGCAGCCGACTTAGAACTGGTGCGGACCAGGGGAATC</td><td>>hg38_21_+_8433172_8446622_RNA45SN1_45S_50ntFlanks</td><td>11565</td><td>11603</td><td>39</td><td>CTAGCAGCCGACTTAGAACTGGTGCGGACCAGGGGAATC</td><td>185.3</td><td>-11.4</td><td>1.6</td><td>-6.9</td><td>4.4E-12</td><td>1.5E-11</td><td>MB231/MB468</td><td>MP</td></tr>
<tr><td>>hg38_21_+_8433172_8446622_RNA45SN1_45S_50ntFlanks@11705.11728.24|TGAAGCGCGGGTAAACGGCGGGAG</td><td>>hg38_21_+_8433172_8446622_RNA45SN1_45S_50ntFlanks</td><td>11705</td><td>11728</td><td>24</td><td>TGAAGCGCGGGTAAACGGCGGGAG</td><td>615.3</td><td>-7.4</td><td>1.1</td><td>-6.9</td><td>4.9E-12</td><td>1.5E-11</td><td>MB231/MB468</td><td>CYTO</td></tr>
<tr><td>>MI0000469|hsa-mir-125a&WithFlank&19|+|51693248|51693345@21.43.23(+1U)&[MIMAT0000443&hsa-miR-125a-5p&offsets|0|-1(+1U);m-63&19|+|51693268|51693289&offsets|0|+1(+1U)]|TCCCTGAGACCCTTTAACCTGTGT</td><td>>MI0000469|hsa-mir-125a&WithFlank&19|+|51693248|51693345</td><td>21</td><td>43</td><td>23(+1U)</td><td>TCCCTGAGACCCTTTAACCTGTGT</td><td>776.6</td><td>5.7</td><td>0.8</td><td>6.9</td><td>4.9E-12</td><td>1.5E-11</td><td>MB231/MB468</td><td>CYTO</td></tr>
<tr><td>>hg38_21_+_8433172_8446622_RNA45SN1_45S_50ntFlanks@8439.8467.29|GATTCAACCCGGCGGCGGGTCCGGCCGTG</td><td>>hg38_21_+_8433172_8446622_RNA45SN1_45S_50ntFlanks</td><td>8439</td><td>8467</td><td>29</td><td>GATTCAACCCGGCGGCGGGTCCGGCCGTG</td><td>100.7</td><td>-10.5</td><td>1.5</td><td>-6.9</td><td>4.4E-12</td><td>1.5E-11</td><td>MB231/MB468</td><td>MP</td></tr>
<tr><td>>am_trnaMT_ValTAC_MT_+_1602_1670@48.65.18|CAACTTAACTTGACCGCT</td><td>>am_trnaMT_ValTAC_MT_+_1602_1670</td><td>48</td><td>65</td><td>18</td><td>CAACTTAACTTGACCGCT</td><td>583.0</td><td>10.4</td><td>1.5</td><td>6.9</td><td>4.6E-12</td><td>1.5E-11</td><td>BT20/MB468</td><td>CYTO</td></tr>
<tr><td>>MI0000748|hsa-mir-130b&WithFlank&22|+|21653298|21653391@19.40.22&[MIMAT0004680&hsa-miR-130b-5p&offsets|0|+1;m-234&22|+|21653316|21653337&offsets|0|0]|ACTCTTTCCCTGTTGCACTACT</td><td>>MI0000748|hsa-mir-130b&WithFlank&22|+|21653298|21653391</td><td>19</td><td>40</td><td>22</td><td>ACTCTTTCCCTGTTGCACTACT</td><td>817.1</td><td>6.2</td><td>0.9</td><td>6.9</td><td>5.9E-12</td><td>1.5E-11</td><td>MB231/MB468</td><td>MITO</td></tr>
<tr><td>>MI0000750|hsa-mir-26a-2&WithFlank&12|-|57824603|57824698@20.40.21(+1G);MI0000083|hsa-mir-26a-1&WithFlank&3|+|37969398|37969486@16.36.21(+1G)&[MIMAT0000082&hsa-miR-26a-5p&offsets|0|-1(+1G);m-33&12|-|57824658|57824679&offsets|0|-1(+1G)];[MIMAT0000082_1&hsa-miR-26a-5p&offsets|0|-1(+1G);m-34&3|+|37969413|37969434&offsets|0|-1(+1G)]|TTCAAGTAATCCAGGATAGGCG</td><td>>MI0000750|hsa-mir-26a-2&WithFlank&12|-|57824603|57824698</td><td>20</td><td>40</td><td>21(+1G)</td><td>TTCAAGTAATCCAGGATAGGCG</td><td>1194.8</td><td>4.7</td><td>0.7</td><td>7.0</td><td>3.6E-12</td><td>1.5E-11</td><td>BT20/MB468</td><td>MITO</td></tr>
<tr><td>>hg38_21_+_8433172_8446622_RNA45SN1_45S_50ntFlanks@10137.10158.22|CGGCGGCGGCGGGGGTGTGGGG</td><td>>hg38_21_+_8433172_8446622_RNA45SN1_45S_50ntFlanks</td><td>10137</td><td>10158</td><td>22</td><td>CGGCGGCGGCGGGGGTGTGGGG</td><td>142.1</td><td>-11.0</td><td>1.6</td><td>-6.9</td><td>4.4E-12</td><td>1.5E-11</td><td>MB231/MB468</td><td>MP</td></tr>
<tr><td>>MI0000542|hsa-mir-320a&WithFlank&8|-|22244960|22245043@48.69.22(+2U)&[MIMAT0000510&hsa-miR-320a-3p&offsets|0|0(+2U);m-60&8|-|22244975|22244996&offsets|0|0(+2U)]|AAAAGCTGGGTTGAGAGGGCGATT</td><td>>MI0000542|hsa-mir-320a&WithFlank&8|-|22244960|22245043</td><td>48</td><td>69</td><td>22(+2U)</td><td>AAAAGCTGGGTTGAGAGGGCGATT</td><td>1054.3</td><td>9.6</td><td>1.4</td><td>7.0</td><td>3.6E-12</td><td>1.5E-11</td><td>BT20/MB468</td><td>MITO</td></tr>
<tr><td>>hg38_21_+_8433172_8446622_RNA45SN1_45S_50ntFlanks@12944.12988.45|CGTAGACGACCTGCTTCTGGGTCGGGGTTTCGTACGTAGCAGAGC</td><td>>hg38_21_+_8433172_8446622_RNA45SN1_45S_50ntFlanks</td><td>12944</td><td>12988</td><td>45</td><td>CGTAGACGACCTGCTTCTGGGTCGGGGTTTCGTACGTAGCAGAGC</td><td>170.3</td><td>-10.9</td><td>1.5</td><td>-7.3</td><td>3.5E-13</td><td>1.5E-11</td><td>BT20/MB231</td><td>CYTO</td></tr>
<tr><td>>hg38_21_+_8433172_8446622_RNA45SN1_45S_50ntFlanks@3854.3874.21|ACTGTGGTAATTCTAGAGCTA</td><td>>hg38_21_+_8433172_8446622_RNA45SN1_45S_50ntFlanks</td><td>3854</td><td>3874</td><td>21</td><td>ACTGTGGTAATTCTAGAGCTA</td><td>719.5</td><td>-8.7</td><td>1.3</td><td>-6.9</td><td>4.4E-12</td><td>1.5E-11</td><td>MB231/MB468</td><td>MP</td></tr>
<tr><td>>hg38_21_+_8433172_8446622_RNA45SN1_45S_50ntFlanks@9507.9526.20|AACTATGCCTGGGCAGGGCG</td><td>>hg38_21_+_8433172_8446622_RNA45SN1_45S_50ntFlanks</td><td>9507</td><td>9526</td><td>20</td><td>AACTATGCCTGGGCAGGGCG</td><td>149.6</td><td>-9.5</td><td>1.4</td><td>-6.9</td><td>4.7E-12</td><td>1.5E-11</td><td>BT20/MB468</td><td>CYTO</td></tr>
<tr><td>>hg38_21_+_8433172_8446622_RNA45SN1_45S_50ntFlanks@12674.12703.30|TCAGAATCCCGCCCAGGCGGAACGATACGG</td><td>>hg38_21_+_8433172_8446622_RNA45SN1_45S_50ntFlanks</td><td>12674</td><td>12703</td><td>30</td><td>TCAGAATCCCGCCCAGGCGGAACGATACGG</td><td>199.1</td><td>-11.5</td><td>1.7</td><td>-6.9</td><td>4.4E-12</td><td>1.5E-11</td><td>MB231/MB468</td><td>MP</td></tr>
<tr><td>>hg38_21_+_8433172_8446622_RNA45SN1_45S_50ntFlanks@11671.11699.29|CCAGTGCTCTGAATGTCAAAGTGAAGAAA</td><td>>hg38_21_+_8433172_8446622_RNA45SN1_45S_50ntFlanks</td><td>11671</td><td>11699</td><td>29</td><td>CCAGTGCTCTGAATGTCAAAGTGAAGAAA</td><td>145.2</td><td>-11.0</td><td>1.6</td><td>-6.9</td><td>4.4E-12</td><td>1.5E-11</td><td>MB231/MB468</td><td>MP</td></tr>
<tr><td>>hg38_21_+_8433172_8446622_RNA45SN1_45S_50ntFlanks@11586.11611.26|GTGCGGACCAGGGGAATCCGACTGTT</td><td>>hg38_21_+_8433172_8446622_RNA45SN1_45S_50ntFlanks</td><td>11586</td><td>11611</td><td>26</td><td>GTGCGGACCAGGGGAATCCGACTGTT</td><td>2099.6</td><td>-3.7</td><td>0.5</td><td>-6.9</td><td>5.1E-12</td><td>1.5E-11</td><td>MB231/MB468</td><td>NUC</td></tr>
<tr><td>>hg38_21_+_8433172_8446622_RNA45SN1_45S_50ntFlanks@12971.13003.33|TTTCGTACGTAGCAGAGCAGCTCCCTCGCTGCG</td><td>>hg38_21_+_8433172_8446622_RNA45SN1_45S_50ntFlanks</td><td>12971</td><td>13003</td><td>33</td><td>TTTCGTACGTAGCAGAGCAGCTCCCTCGCTGCG</td><td>199.9</td><td>-5.8</td><td>0.8</td><td>-6.9</td><td>4.9E-12</td><td>1.5E-11</td><td>MB231/MB468</td><td>CYTO</td></tr>
<tr><td>>hg38_21_+_8433172_8446622_RNA45SN1_45S_50ntFlanks@3985.4005.21|CGGCTTTGGTGACTCTAGATA</td><td>>hg38_21_+_8433172_8446622_RNA45SN1_45S_50ntFlanks</td><td>3985</td><td>4005</td><td>21</td><td>CGGCTTTGGTGACTCTAGATA</td><td>2579.5</td><td>-10.5</td><td>1.5</td><td>-6.9</td><td>6.4E-12</td><td>1.5E-11</td><td>BT20/MB468</td><td>NUC</td></tr>
<tr><td>>MI0000542|hsa-mir-320a&WithFlank&8|-|22244960|22245043@48.71.24&[MIMAT0000510&hsa-miR-320a-3p&offsets|0|+2;m-60&8|-|22244975|22244996&offsets|0|+2]|AAAAGCTGGGTTGAGAGGGCGAAA</td><td>>MI0000542|hsa-mir-320a&WithFlank&8|-|22244960|22245043</td><td>48</td><td>71</td><td>24</td><td>AAAAGCTGGGTTGAGAGGGCGAAA</td><td>742.9</td><td>7.5</td><td>1.1</td><td>6.9</td><td>4.9E-12</td><td>1.5E-11</td><td>MB231/MB468</td><td>CYTO</td></tr>
<tr><td>>hg38_21_+_8433172_8446622_RNA45SN1_45S_50ntFlanks@11564.11581.18|CCTAGCAGCCGACTTAGA</td><td>>hg38_21_+_8433172_8446622_RNA45SN1_45S_50ntFlanks</td><td>11564</td><td>11581</td><td>18</td><td>CCTAGCAGCCGACTTAGA</td><td>1175.7</td><td>-3.8</td><td>0.5</td><td>-7.0</td><td>3.6E-12</td><td>1.5E-11</td><td>BT20/MB468</td><td>MITO</td></tr>
<tr><td>>hg38_21_+_8433172_8446622_RNA45SN1_45S_50ntFlanks@9526.9573.48|GAAGCCAGAGGAAACTCTGGTGGAGGTCCGTAGCGGTCCTGACGTGCA</td><td>>hg38_21_+_8433172_8446622_RNA45SN1_45S_50ntFlanks</td><td>9526</td><td>9573</td><td>48</td><td>GAAGCCAGAGGAAACTCTGGTGGAGGTCCGTAGCGGTCCTGACGTGCA</td><td>324.9</td><td>8.7</td><td>1.3</td><td>6.9</td><td>4.9E-12</td><td>1.5E-11</td><td>MB231/MB468</td><td>CYTO</td></tr>
<tr><td>>trna10_LysCTT_16_+_3241501_3241573@1.30.30,trna13_LysCTT_14_-_58706613_58706685@1.30.30,trna2_LysCTT_15_+_79152904_79152976@1.30.30|GCCCGGCTAGCTCAGTCGGTAGAGCATGGG</td><td>>trna10_LysCTT_16_+_3241501_3241573</td><td>1</td><td>30</td><td>30</td><td>GCCCGGCTAGCTCAGTCGGTAGAGCATGGG</td><td>514.9</td><td>-8.1</td><td>1.2</td><td>-6.9</td><td>4.7E-12</td><td>1.5E-11</td><td>BT20/MB468</td><td>CYTO</td></tr>
<tr><td>>hg38_21_+_8433172_8446622_RNA45SN1_45S_50ntFlanks@6731.6756.26|CACATTGATCATCGACACTTCGAACG</td><td>>hg38_21_+_8433172_8446622_RNA45SN1_45S_50ntFlanks</td><td>6731</td><td>6756</td><td>26</td><td>CACATTGATCATCGACACTTCGAACG</td><td>1484.2</td><td>-8.2</td><td>1.2</td><td>-6.9</td><td>6.4E-12</td><td>1.5E-11</td><td>BT20/MB468</td><td>NUC</td></tr>
<tr><td>>hg38_21_+_8433172_8446622_RNA45SN1_45S_50ntFlanks@10234.10258.25|CGGACGCTACGCCGCGACGAGTAGG</td><td>>hg38_21_+_8433172_8446622_RNA45SN1_45S_50ntFlanks</td><td>10234</td><td>10258</td><td>25</td><td>CGGACGCTACGCCGCGACGAGTAGG</td><td>96.4</td><td>-10.5</td><td>1.5</td><td>-6.9</td><td>4.5E-12</td><td>1.5E-11</td><td>MB231/MB468</td><td>MP</td></tr>
<tr><td>>hg38_21_+_8433172_8446622_RNA45SN1_45S_50ntFlanks@5431.5466.36|GTTTAGTGAGGCCCTCGGATCGGCCCCGCCGGGGTC</td><td>>hg38_21_+_8433172_8446622_RNA45SN1_45S_50ntFlanks</td><td>5431</td><td>5466</td><td>36</td><td>GTTTAGTGAGGCCCTCGGATCGGCCCCGCCGGGGTC</td><td>248.1</td><td>-5.7</td><td>0.8</td><td>-6.9</td><td>5.0E-12</td><td>1.5E-11</td><td>MB231/MB468</td><td>CYTO</td></tr>
<tr><td>>hg38_21_+_8433172_8446622_RNA45SN1_45S_50ntFlanks@11586.11609.24|GTGCGGACCAGGGGAATCCGACTG</td><td>>hg38_21_+_8433172_8446622_RNA45SN1_45S_50ntFlanks</td><td>11586</td><td>11609</td><td>24</td><td>GTGCGGACCAGGGGAATCCGACTG</td><td>533.2</td><td>-4.6</td><td>0.7</td><td>-6.9</td><td>6.0E-12</td><td>1.5E-11</td><td>MB231/MB468</td><td>MITO</td></tr>
<tr><td>>hg38_21_+_8433172_8446622_RNA45SN1_45S_50ntFlanks@12973.13000.28|TCGTACGTAGCAGAGCAGCTCCCTCGCT</td><td>>hg38_21_+_8433172_8446622_RNA45SN1_45S_50ntFlanks</td><td>12973</td><td>13000</td><td>28</td><td>TCGTACGTAGCAGAGCAGCTCCCTCGCT</td><td>278.4</td><td>-4.3</td><td>0.6</td><td>-6.9</td><td>5.1E-12</td><td>1.6E-11</td><td>MB231/MB468</td><td>NUC</td></tr>
<tr><td>>hg38_21_+_8433172_8446622_RNA45SN1_45S_50ntFlanks@12655.12673.19|ATGACTGAACGCCTCTAAG</td><td>>hg38_21_+_8433172_8446622_RNA45SN1_45S_50ntFlanks</td><td>12655</td><td>12673</td><td>19</td><td>ATGACTGAACGCCTCTAAG</td><td>442.3</td><td>-8.3</td><td>1.2</td><td>-6.9</td><td>6.0E-12</td><td>1.6E-11</td><td>MB231/MB468</td><td>MITO</td></tr>
<tr><td>>MI0000065|hsa-let-7d&WithFlank&9|+|94178828|94178926@68.89.22&[MIMAT0004484&hsa-let-7d-3p&offsets|0|0;m-98&9|+|94178895|94178916&offsets|0|0]|CTATACGACCTGCTGCCTTTCT</td><td>>MI0000065|hsa-let-7d&WithFlank&9|+|94178828|94178926</td><td>68</td><td>89</td><td>22</td><td>CTATACGACCTGCTGCCTTTCT</td><td>2805.6</td><td>4.9</td><td>0.7</td><td>6.9</td><td>5.1E-12</td><td>1.6E-11</td><td>MB231/MB468</td><td>NUC</td></tr>
<tr><td>>MI0000298|hsa-mir-221&WithFlank&X|-|45746151|45746272@71.92.22&[MIMAT0000278&hsa-miR-221-3p&offsets|0|-1;m-91&X|-|45746181|45746202&offsets|0|0]|AGCTACATTGTCTGCTGGGTTT</td><td>>MI0000298|hsa-mir-221&WithFlank&X|-|45746151|45746272</td><td>71</td><td>92</td><td>22</td><td>AGCTACATTGTCTGCTGGGTTT</td><td>12873.1</td><td>6.4</td><td>0.9</td><td>6.9</td><td>5.0E-12</td><td>1.6E-11</td><td>MB231/MB468</td><td>CYTO</td></tr>
<tr><td>>hg38_21_+_8433172_8446622_RNA45SN1_45S_50ntFlanks@11581.11609.29|AACTGGTGCGGACCAGGGGAATCCGACTG</td><td>>hg38_21_+_8433172_8446622_RNA45SN1_45S_50ntFlanks</td><td>11581</td><td>11609</td><td>29</td><td>AACTGGTGCGGACCAGGGGAATCCGACTG</td><td>2223.0</td><td>-5.1</td><td>0.7</td><td>-6.9</td><td>6.5E-12</td><td>1.6E-11</td><td>BT20/MB468</td><td>NUC</td></tr>
<tr><td>>hg38_21_+_8433172_8446622_RNA45SN1_45S_50ntFlanks@10344.10362.19|ATTCAAACGAGAACTTTGA</td><td>>hg38_21_+_8433172_8446622_RNA45SN1_45S_50ntFlanks</td><td>10344</td><td>10362</td><td>19</td><td>ATTCAAACGAGAACTTTGA</td><td>425.8</td><td>7.5</td><td>1.1</td><td>6.9</td><td>3.7E-12</td><td>1.6E-11</td><td>BT20/MB468</td><td>MITO</td></tr>
<tr><td>>hg38_21_+_8433172_8446622_RNA45SN1_45S_50ntFlanks@11768.11790.23|CTAATTAGTGACGCGCATGAATG</td><td>>hg38_21_+_8433172_8446622_RNA45SN1_45S_50ntFlanks</td><td>11768</td><td>11790</td><td>23</td><td>CTAATTAGTGACGCGCATGAATG</td><td>640.8</td><td>-8.5</td><td>1.2</td><td>-6.9</td><td>6.1E-12</td><td>1.6E-11</td><td>MB231/MB468</td><td>MITO</td></tr>
</table>

| | | | | | | | | | | | | | |
|---|---|---|---|---|---|---|---|---|---|---|---|---|---|
| >hg38_21_+_8433172_8446622_RNA45SN1_45S_50ntFlanks@4892.4916.25\|AAACTTAAAGGAATTGACGGAAGGG | >hg38_21_+_8433172_8446622_RNA45SN1_45S_50ntFlanks | 4892 | 4916 | 25 | AAACTTAAAGGAATTGACGGAAGGG | 142.0 | -11.0 | 1.6 | -6.9 | 4.5E-12 | 1.6E-11 | MB231/MB468 | MP |
| >hg38_21_+_8433172_8446622_RNA45SN1_45S_50ntFlanks@5484.5518.35\|GAGCGCTGAGAAGACGGTCGAACTTGACTATCTAG | >hg38_21_+_8433172_8446622_RNA45SN1_45S_50ntFlanks | 5484 | 5518 | 35 | GAGCGCTGAGAAGACGGTCGAACTTGACTATCTAG | 194.8 | -9.7 | 1.4 | -6.9 | 3.7E-12 | 1.6E-11 | BT20/MB468 | MITO |
| >hg38_21_+_8433172_8446622_RNA45SN1_45S_50ntFlanks@12444.12483.40\|AAGCAGAATTCACCAAGCGTTGGATTGTTCACCCACTAAT | >hg38_21_+_8433172_8446622_RNA45SN1_45S_50ntFlanks | 12444 | 12483 | 40 | AAGCAGAATTCACCAAGCGTTGGATTGTTCACCCACTAAT | 732.3 | -7.8 | 1.1 | -6.9 | 6.5E-12 | 1.6E-11 | BT20/MB468 | NUC |
| >hg38_21_+_8433172_8446622_RNA45SN1_45S_50ntFlanks@11671.11701.31\|CCAGTGCTCTGAATGTCAAAGTGAAGAAATT | >hg38_21_+_8433172_8446622_RNA45SN1_45S_50ntFlanks | 11671 | 11701 | 31 | CCAGTGCTCTGAATGTCAAAGTGAAGAAATT | 2440.0 | -8.3 | 1.2 | -6.9 | 6.5E-12 | 1.6E-11 | BT20/MB468 | NUC |
| >trna116_GluCTC_1_-_145399233_145399304@30.71.42,trna71_GluCTC_1_-_161439189_161439260@30.71.42,trna74_GluCTC_1_-_161431809_161431880@30.71.42,trna77_GluCTC_1_-_161424398_161424469@30.71.42,trna77_GluCTC_6_+_28949976_28950047@30.71.42,trna80_GluCTC_1_-_161417018_161417089@30.71.42,trna87_GluCTC_6_-_126101393_126101464@30.71.42\|CGCTCTCACCGCCGCGGCCCGGGTTCGATTCCCGGTCAGGGA | >trna116_GluCTC_1_-_145399233_145399304 | 30 | 71 | 42 | CGCTCTCACCGCCGCGGCCCGGGTTCGATTCCCGGTCAGGGA | 258.7 | -7.1 | 1.0 | -6.9 | 4.8E-12 | 1.6E-11 | BT20/MB468 | CYTO |
| >hg38_21_+_8433172_8446622_RNA45SN1_45S_50ntFlanks@11918.11956.39\|ACATGAGAGGTGTAGAATAAGTGGGAGGCCCCCGGCGCC | >hg38_21_+_8433172_8446622_RNA45SN1_45S_50ntFlanks | 11918 | 11956 | 39 | ACATGAGAGGTGTAGAATAAGTGGGAGGCCCCCGGCGCC | 242.4 | -8.2 | 1.2 | -6.9 | 5.2E-12 | 1.6E-11 | MB231/MB468 | NUC |
| >hg38_21_+_8433172_8446622_RNA45SN1_45S_50ntFlanks@8423.8471.49\|CGCAGTCCGCCCGGAGGATTCAACCCGGCGGCGGGTCCGGCCGTGTCGG | >hg38_21_+_8433172_8446622_RNA45SN1_45S_50ntFlanks | 8423 | 8471 | 49 | CGCAGTCCGCCCGGAGGATTCAACCCGGCGGCGGGTCCGGCCGTGTCGG | 480.7 | -12.8 | 1.8 | -6.9 | 4.6E-12 | 1.6E-11 | MB231/MB468 | MP |
| >hg38_21_+_8433172_8446622_RNA45SN1_45S_50ntFlanks@8170.8195.26\|CAAGTCCTTCTGATCGAGGCCCAGCC | >hg38_21_+_8433172_8446622_RNA45SN1_45S_50ntFlanks | 8170 | 8195 | 26 | CAAGTCCTTCTGATCGAGGCCCAGCC | 687.6 | -5.6 | 0.8 | -6.9 | 5.2E-12 | 1.6E-11 | MB231/MB468 | NUC |
| >hg38_21_+_8433172_8446622_RNA45SN1_45S_50ntFlanks@10756.10785.30\|TACCCATATCCGCAGCAGGTCTCCAAGGTG | >hg38_21_+_8433172_8446622_RNA45SN1_45S_50ntFlanks | 10756 | 10785 | 30 | TACCCATATCCGCAGCAGGTCTCCAAGGTG | 111.7 | -10.7 | 1.5 | -6.9 | 4.6E-12 | 1.6E-11 | MB231/MB468 | MP |
| >hg38_21_+_8433172_8446622_RNA45SN1_45S_50ntFlanks@10905.10941.37\|CTGGGCGCGCGCCGCGGCTGGACGAGGCGCCGCCGCC | >hg38_21_+_8433172_8446622_RNA45SN1_45S_50ntFlanks | 10905 | 10941 | 37 | CTGGGCGCGCGCCGCGGCTGGACGAGGCGCCGCCGCC | 118.3 | -10.8 | 1.6 | -6.9 | 4.6E-12 | 1.6E-11 | MB231/MB468 | MP |
| >hg38_21_+_8433172_8446622_RNA45SN1_45S_50ntFlanks@8198.8226.29\|TGGACGGTGTGAGGCCGGTAGCGGCCCCC | >hg38_21_+_8433172_8446622_RNA45SN1_45S_50ntFlanks | 8198 | 8226 | 29 | TGGACGGTGTGAGGCCGGTAGCGGCCCCC | 85.9 | -10.3 | 1.5 | -6.9 | 4.6E-12 | 1.6E-11 | MB231/MB468 | MP |
| >hg38_21_+_8433172_8446622_RNA45SN1_45S_50ntFlanks@12054.12072.19\|TGAGGCGGGGGGGCGAGCC | >hg38_21_+_8433172_8446622_RNA45SN1_45S_50ntFlanks | 12054 | 12072 | 19 | TGAGGCGGGGGGGCGAGCC | 138.0 | -11.0 | 1.6 | -6.9 | 4.6E-12 | 1.6E-11 | MB231/MB468 | MP |
| >hg38_21_+_8433172_8446622_RNA45SN1_45S_50ntFlanks@8168.8189.22\|CCCAAGTCCTTCTGATCGAGGC | >hg38_21_+_8433172_8446622_RNA45SN1_45S_50ntFlanks | 8168 | 8189 | 22 | CCCAAGTCCTTCTGATCGAGGC | 104.1 | -10.6 | 1.5 | -6.9 | 4.6E-12 | 1.6E-11 | MB231/MB468 | MP |
| >MI0000076\|hsa-mir-20a&WithFlank&13\|+\|91351059\|91351141@14.36.23(+1A)&[MIMAT0000075&hsa-miR-20a-5p&offsets\|0\|0(+1A);m-96&13\|+\|91351072\|91351094&offsets\|0\|0(+1A)]\|TAAAGTGCTTATAGTGCAGGTAGA | >MI0000076\|hsa-mir-20a&WithFlank&13\|+\|91351059\|91351141 | 14 | 36 | 23(+1A) | TAAAGTGCTTATAGTGCAGGTAGA | 1359.9 | 5.0 | 0.7 | 6.9 | 4.8E-12 | 1.6E-11 | BT20/MB468 | CYTO |
| >MI0000650\|hsa-mir-200c&WithFlank&12\|+\|6963693\|6963772@50.72.23&[MIMAT0000617&hsa-miR-200c-3p&offsets\|0\|0;m-54&12\|+\|6963742\|6963764&offsets\|0\|0]\|TAATACTGCCGGGTAATGATGGA | >MI0000650\|hsa-mir-200c&WithFlank&12\|+\|6963693\|6963772 | 50 | 72 | 23 | TAATACTGCCGGGTAATGATGGA | 2562.7 | -4.3 | 0.6 | -6.9 | 5.3E-12 | 1.6E-11 | MB231/MB468 | NUC |
| >am_trna111_HisGTG_1_-_147774845_147774916@-1G.29.30,trna118_HisGTG_1_-_145396881_145396952@-1G.29.30,trna16_HisGTG_1_+_146544773_146544844@-1G.29.30,trna1_HisGTG_15_+_45493349_45493420@-1G.29.30,trna21_HisGTG_1_+_147753471_147753542@-1G.29.30,trna33_HisGTG_6_+_27125906_27125977@-1G.29.30,trna7_HisGTG_9_-_14433938_14434009@-1G.29.30,trna8_HisGTG_15_-_45492611_45492682@-1G.29.30,trna9_HisGTG_15_-_45490804_45490875@-1G.29.30\|GGCCGTGATCGTATAGTGGTTAGTACTCTG | >am_trna111_HisGTG_1_-_147774845_147774916 | -1G | 29 | 30 | GGCCGTGATCGTATAGTGGTTAGTACTCTG | 14966.7 | -5.3 | 0.8 | -6.9 | 5.3E-12 | 1.6E-11 | MB231/MB468 | NUC |
| >hg38_21_+_8433172_8446622_RNA45SN1_45S_50ntFlanks@9739.9766.28\|AAACGATCTCAACCTATTCTCAAACTTT | >hg38_21_+_8433172_8446622_RNA45SN1_45S_50ntFlanks | 9739 | 9766 | 28 | AAACGATCTCAACCTATTCTCAAACTTT | 114.9 | -10.7 | 1.5 | -6.9 | 4.6E-12 | 1.6E-11 | MB231/MB468 | MP |
| >hg38_21_+_8433172_8446622_RNA45SN1_45S_50ntFlanks@476.504.29\|TTCGTGATCGATGTGGTGACGTCGTGCTC | >hg38_21_+_8433172_8446622_RNA45SN1_45S_50ntFlanks | 476 | 504 | 29 | TTCGTGATCGATGTGGTGACGTCGTGCTC | 10927.3 | 5.1 | 0.7 | 6.9 | 5.3E-12 | 1.6E-11 | MB231/MB468 | NUC |
| >hg38_MT_+_1621_3279_RNR2_16S_50ntFlanks@493.517.25\|GCTCTTTGGACACTAGGAAAAAACC | >hg38_MT_+_1621_3279_RNR2_16S_50ntFlanks | 493 | 517 | 25 | GCTCTTTGGACACTAGGAAAAAACC | 1002.9 | 5.9 | 0.8 | 6.9 | 3.8E-12 | 1.6E-11 | BT20/MB468 | MITO |
| >hg38_21_+_8433172_8446622_RNA45SN1_45S_50ntFlanks@11215.11234.20\|CTCTGGACGCGAGCCGGGCC | >hg38_21_+_8433172_8446622_RNA45SN1_45S_50ntFlanks | 11215 | 11234 | 20 | CTCTGGACGCGAGCCGGGCC | 197.6 | -11.5 | 1.7 | -6.9 | 4.6E-12 | 1.6E-11 | MB231/MB468 | MP |
| >hg38_21_+_8433172_8446622_RNA45SN1_45S_50ntFlanks@5471.5488.18\|CACGGCCCTGGCGGAGCG | >hg38_21_+_8433172_8446622_RNA45SN1_45S_50ntFlanks | 5471 | 5488 | 18 | CACGGCCCTGGCGGAGCG | 137.1 | -11.0 | 1.6 | -6.9 | 4.6E-12 | 1.6E-11 | MB231/MB468 | MP |
| >hg38_21_+_8433172_8446622_RNA45SN1_45S_50ntFlanks@9714.9732.19\|CGAATGATTAGAGGTCTTG | >hg38_21_+_8433172_8446622_RNA45SN1_45S_50ntFlanks | 9714 | 9732 | 19 | CGAATGATTAGAGGTCTTG | 117.9 | -10.7 | 1.6 | -6.9 | 4.6E-12 | 1.6E-11 | MB231/MB468 | MP |
| >MI0000285\|hsa-mir-205&WithFlank&1\|+\|209432127\|209432248@40.59.20(+1A)&[MIMAT0000266&hsa-miR-205-5p&offsets\|0\|-2(+1A);m-351&1\|+\|209432166\|209432187&offsets\|0\|-2(+1A)]\|TCCTTCATTCCACCGGAGTCA | >MI0000285\|hsa-mir-205&WithFlank&1\|+\|209432127\|209432248 | 40 | 59 | 20(+1A) | TCCTTCATTCCACCGGAGTCA | 160.6 | -11.2 | 1.6 | -6.9 | 4.6E-12 | 1.6E-11 | MB231/MB468 | MP |
| >hg38_21_+_8433172_8446622_RNA45SN1_45S_50ntFlanks@11569.11594.26\|CAGCCGACTTAGAACTGGTGCGGACC | >hg38_21_+_8433172_8446622_RNA45SN1_45S_50ntFlanks | 11569 | 11594 | 26 | CAGCCGACTTAGAACTGGTGCGGACC | 560.8 | -4.3 | 0.6 | -6.9 | 6.2E-12 | 1.6E-11 | MB231/MB468 | MITO |
| >MI0000077\|hsa-mir-21&WithFlank&17\|+\|59841260\|59841343@14.36.23(+1G)&[MIMAT0000076&hsa-miR-21-5p&offsets\|0\|+1(+1G);m-1&17\|+\|59841273\|59841294&offsets\|0\|+1(+1G)]\|TAGCTTATCAGACTGATGTTGACG | >MI0000077\|hsa-mir-21&WithFlank&17\|+\|59841260\|59841343 | 14 | 36 | 23(+1G) | TAGCTTATCAGACTGATGTTGACG | 25924.4 | 6.7 | 1.0 | 6.9 | 4.9E-12 | 1.6E-11 | BT20/MB468 | CYTO |
| >hg38_21_+_8433172_8446622_RNA45SN1_45S_50ntFlanks@12941.12972.32\|ATTCGTAGACGACCTGCTTCTGGGTCGGGGTT | >hg38_21_+_8433172_8446622_RNA45SN1_45S_50ntFlanks | 12941 | 12972 | 32 | ATTCGTAGACGACCTGCTTCTGGGTCGGGGTT | 392.4 | -11.7 | 1.6 | -7.3 | 2.8E-13 | 1.6E-11 | BT20/MB468 | MP |
| >hg38_21_+_8433172_8446622_RNA45SN1_45S_50ntFlanks@5156.5185.30\|ACAAGTGGCGTTCAGCCACCCGAGATTGAG | >hg38_21_+_8433172_8446622_RNA45SN1_45S_50ntFlanks | 5156 | 5185 | 30 | ACAAGTGGCGTTCAGCCACCCGAGATTGAG | 563.7 | -12.9 | 1.9 | -6.9 | 6.2E-12 | 1.6E-11 | MB231/MB468 | MITO |
| >hg38_21_+_8433172_8446622_RNA45SN1_45S_50ntFlanks@11656.11685.30\|CGATGTGATTTCTGCCCAGTGCTCTGAATG | >hg38_21_+_8433172_8446622_RNA45SN1_45S_50ntFlanks | 11656 | 11685 | 30 | CGATGTGATTTCTGCCCAGTGCTCTGAATG | 98.2 | -10.5 | 1.5 | -6.9 | 4.7E-12 | 1.6E-11 | MB231/MB468 | MP |
| >hg38_21_+_8433172_8446622_RNA45SN1_45S_50ntFlanks@4922.4944.23\|CCAGGAGTGGAGCCTGCGGCTTA | >hg38_21_+_8433172_8446622_RNA45SN1_45S_50ntFlanks | 4922 | 4944 | 23 | CCAGGAGTGGAGCCTGCGGCTTA | 438.8 | -12.6 | 1.8 | -6.9 | 6.3E-12 | 1.6E-11 | MB231/MB468 | MITO |
| >hg38_21_+_8433172_8446622_RNA45SN1_45S_50ntFlanks@475.495.21\|CTTCGTGATCGATGTGGTGAC | >hg38_21_+_8433172_8446622_RNA45SN1_45S_50ntFlanks | 475 | 495 | 21 | CTTCGTGATCGATGTGGTGAC | 6811.6 | 4.6 | 0.7 | 6.9 | 5.3E-12 | 1.6E-11 | MB231/MB468 | NUC |
| >hg38_21_+_8433172_8446622_RNA45SN1_45S_50ntFlanks@11768.11806.39\|CTAATTAGTGACGCGCATGAATGGATGAACGAGATTCCC | >hg38_21_+_8433172_8446622_RNA45SN1_45S_50ntFlanks | 11768 | 11806 | 39 | CTAATTAGTGACGCGCATGAATGGATGAACGAGATTCCC | 161.9 | -11.2 | 1.6 | -6.9 | 4.7E-12 | 1.6E-11 | MB231/MB468 | MP |
| >MI0000462\|hsa-mir-152&WithFlank&17\|-\|48037155\|48037253@60.81.22(+1U)&[MIMAT0000438&hsa-miR-152-3p&offsets\|0\|+1(+1U);m-97&17\|-\|48037174\|48037194&offsets\|0\|+1(+1U)]\|TCAGTGCATGACAGAACTTGGGT | >MI0000462\|hsa-mir-152&WithFlank&17\|-\|48037155\|48037253 | 60 | 81 | 22(+1U) | TCAGTGCATGACAGAACTTGGGT | 343.5 | 11.6 | 1.7 | 6.9 | 6.3E-12 | 1.6E-11 | MB231/MB468 | MITO |
| >MI0000088\|hsa-mir-30a&WithFlank&6\|-\|71403545\|71403627@53.73.21&[MIMAT0000088&hsa-miR-30a-3p&offsets\|0\|-1;m-38&6\|-\|71403554\|71403575&offsets\|0\|-1]\|CTTTCAGTCGGATGTTTGCAG | >MI0000088\|hsa-mir-30a&WithFlank&6\|-\|71403545\|71403627 | 53 | 73 | 21 | CTTTCAGTCGGATGTTTGCAG | 602.3 | 5.8 | 0.8 | 6.9 | 6.3E-12 | 1.6E-11 | MB231/MB468 | MITO |
| >MI0000085\|hsa-mir-27a&WithFlank&19\|-\|13836434\|13836523@57.75.19&[MIMAT0000084&hsa-miR-27a-3p&offsets\|0\|-2;m-43&19\|-\|13836447\|13836467&offsets\|0\|-2]\|TTCACAGTGGCTAAGTTCC | >MI0000085\|hsa-mir-27a&WithFlank&19\|-\|13836434\|13836523 | 57 | 75 | 19 | TTCACAGTGGCTAAGTTCC | 674.2 | 5.2 | 0.7 | 6.9 | 5.4E-12 | 1.6E-11 | MB231/MB468 | NUC |
| >MI0000734\|hsa-mir-106b&WithFlank&7\|-\|100093987\|100094080@58.79.22&[MIMAT0004672&hsa-miR-106b-3p&offsets\|0\|0;m-87&7\|-\|100094003\|100094023&offsets\|0\|+1]\|CCGCACTGTGGGTACTTGCTGC | >MI0000734\|hsa-mir-106b&WithFlank&7\|-\|100093987\|100094080 | 58 | 79 | 22 | CCGCACTGTGGGTACTTGCTGC | 6975.1 | 4.4 | 0.6 | 6.9 | 5.4E-12 | 1.6E-11 | MB231/MB468 | NUC |
| >hg38_21_+_8433172_8446622_RNA45SN1_45S_50ntFlanks@10411.10443.33\|CAGTCGGTCCTGAGAGATGGGCGAGCGCCGTTC | >hg38_21_+_8433172_8446622_RNA45SN1_45S_50ntFlanks | 10411 | 10443 | 33 | CAGTCGGTCCTGAGAGATGGGCGAGCGCCGTTC | 1339.0 | -6.3 | 0.9 | -6.9 | 5.2E-12 | 1.6E-11 | MB231/MB468 | CYTO |
| >hg38_21_+_8433172_8446622_RNA45SN1_45S_50ntFlanks@10781.10804.24\|AGGTGAACAGCCTCTGGCATGTTG | >hg38_21_+_8433172_8446622_RNA45SN1_45S_50ntFlanks | 10781 | 10804 | 24 | AGGTGAACAGCCTCTGGCATGTTG | 355.9 | -11.6 | 1.6 | -7.3 | 2.8E-13 | 1.6E-11 | BT20/MB468 | MP |
| >hg38_21_+_8433172_8446622_RNA45SN1_45S_50ntFlanks@8182.8208.27\|ATCGAGGCCCAGCCCGTGGACGGTGTG | >hg38_21_+_8433172_8446622_RNA45SN1_45S_50ntFlanks | 8182 | 8208 | 27 | ATCGAGGCCCAGCCCGTGGACGGTGTG | 428.4 | -11.8 | 1.6 | -7.3 | 2.8E-13 | 1.6E-11 | BT20/MB468 | MP |
| >MI0000298\|hsa-mir-221&WithFlank&X\|-\|45746151\|45746272@71.93.23(+2U)&[MIMAT0000278&hsa-miR-221-3p&offsets\|0\|0(+2U);m-91&X\|-\|45746181\|45746202&offsets\|0\|+1(+2U)]\|AGCTACATTGTCTGCTGGGTTTCTT | >MI0000298\|hsa-mir-221&WithFlank&X\|-\|45746151\|45746272 | 71 | 93 | 23(+2U) | AGCTACATTGTCTGCTGGGTTTCTT | 202.5 | 5.6 | 0.8 | 6.9 | 5.4E-12 | 1.6E-11 | MB231/MB468 | NUC |
| >hg38_21_+_8433172_8446622_RNA45SN1_45S_50ntFlanks@11661.11682.22\|TGATTTCTGCCCAGTGCTCTGA | >hg38_21_+_8433172_8446622_RNA45SN1_45S_50ntFlanks | 11661 | 11682 | 22 | TGATTTCTGCCCAGTGCTCTGA | 1769.1 | -8.0 | 1.2 | -6.9 | 6.7E-12 | 1.6E-11 | BT20/MB468 | NUC |
| >hg38_21_+_8433172_8446622_RNA45SN1_45S_50ntFlanks@12449.12480.32\|GAATTCACCAAGCGTTGGATTGTTCACCCACT | >hg38_21_+_8433172_8446622_RNA45SN1_45S_50ntFlanks | 12449 | 12480 | 32 | GAATTCACCAAGCGTTGGATTGTTCACCCACT | 1128.2 | 8.0 | 1.2 | 6.9 | 5.0E-12 | 1.6E-11 | BT20/MB468 | CYTO |
| >hg38_21_+_8433172_8446622_RNA45SN1_45S_50ntFlanks@12786.12805.20\|CCCGCGCGCGCGGGAGGGCG | >hg38_21_+_8433172_8446622_RNA45SN1_45S_50ntFlanks | 12786 | 12805 | 20 | CCCGCGCGCGCGGGAGGGCG | 131.3 | -10.9 | 1.6 | -6.9 | 4.7E-12 | 1.6E-11 | MB231/MB468 | MP |
| >hg38_21_+_8433172_8446622_RNA45SN1_45S_50ntFlanks@12149.12182.34\|AGGTGGGGAGTTTGACTGGGGCGGTACACCTGTC | >hg38_21_+_8433172_8446622_RNA45SN1_45S_50ntFlanks | 12149 | 12182 | 34 | AGGTGGGGAGTTTGACTGGGGCGGTACACCTGTC | 100.0 | -10.5 | 1.5 | -6.9 | 4.7E-12 | 1.6E-11 | MB231/MB468 | MP |
| >hg38_21_+_8433172_8446622_RNA45SN1_45S_50ntFlanks@11927.11951.25\|GTGTAGAATAAGTGGGAGGCCCCCG | >hg38_21_+_8433172_8446622_RNA45SN1_45S_50ntFlanks | 11927 | 11951 | 25 | GTGTAGAATAAGTGGGAGGCCCCCG | 113.2 | -10.7 | 1.5 | -6.9 | 4.8E-12 | 1.6E-11 | MB231/MB468 | MP |
| >hg38_21_+_8433172_8446622_RNA45SN1_45S_50ntFlanks@8357.8379.23\|AAAAGAACTTTGAAGAGAGAGTT | >hg38_21_+_8433172_8446622_RNA45SN1_45S_50ntFlanks | 8357 | 8379 | 23 | AAAAGAACTTTGAAGAGAGAGTT | 20894.9 | -5.9 | 0.9 | -6.9 | 3.9E-12 | 1.6E-11 | BT20/MB468 | MITO |

| | | | | | | | | | | | | | |
|---|---|---|---|---|---|---|---|---|---|---|---|---|---|
| >hg38_21_+_8433172_8446622_RNA45SN1_45S_50ntFlanks@6648.6681.34\|GTACGACTCTTAGCGGTGGATCACTCGGCTCGTG | >hg38_21_+_8433172_8446622_RNA45SN1_45S_50ntFlanks | 6648 | 6681 | 34 | GTACGACTCTTAGCGGTGGATCACTCGGCTCGTG | 5489.7 | -6.1 | 0.9 | -6.9 | 4.8E-12 | 1.6E-11 | MB231/MB468 | MP |
| >am_MI0000815\|hsa-mir-339&WithFlank&7\|-\|1022927\|1023032@21.42.22&[MIMAT0000764&hsa-miR-339-5p&offsets\|0\|-1;m-163&7\|-\|1022990\|1023012&offsets\|0\|-1]\|TCCCTGTCCTCCAGGAGCTCAC | >am_MI0000815\|hsa-mir-339&WithFlank&7\|-\|1022927\|1023032 | 21 | 42 | 22 | TCCCTGTCCTCCAGGAGCTCAC | 470.0 | 8.6 | 1.2 | 6.9 | 5.0E-12 | 1.6E-11 | BT20/MB468 | CYTO |
| >hg38_21_+_8433172_8446622_RNA45SN1_45S_50ntFlanks@11209.11230.22\|CGGCGACTCTGGACGCGAGCCG | >hg38_21_+_8433172_8446622_RNA45SN1_45S_50ntFlanks | 11209 | 11230 | 22 | CGGCGACTCTGGACGCGAGCCG | 1045.6 | -7.1 | 1.0 | -6.9 | 6.4E-12 | 1.6E-11 | MB231/MB468 | MITO |
| >hg38_21_+_8433172_8446622_RNA45SN1_45S_50ntFlanks@10686.10710.25\|GCGGCGTCCGGTGAGCTCTCGCTGG | >hg38_21_+_8433172_8446622_RNA45SN1_45S_50ntFlanks | 10686 | 10710 | 25 | GCGGCGTCCGGTGAGCTCTCGCTGG | 3606.1 | -4.9 | 0.7 | -6.9 | 5.4E-12 | 1.6E-11 | MB231/MB468 | NUC |
| >hg38_1_-_228634819_228635039_RNA5S12_5S_50ntFlanks@128.169.42\|CTTGGATGGGAGACCGCCTGGGAATACCGGGTGCTGTAGGCT | >hg38_1_-_228634819_228635039_RNA5S12_5S_50ntFlanks | 128 | 169 | 42 | CTTGGATGGGAGACCGCCTGGGAATACCGGGTGCTGTAGGCT | 658.2 | 7.3 | 1.1 | 6.9 | 5.3E-12 | 1.6E-11 | MB231/MB468 | CYTO |
| >MI0000462\|hsa-mir-152&WithFlank&17\|-\|48037155\|48037253@60.80.21(+1A)&[MIMAT0000438&hsa-miR-152-3p&offsets\|0\|0(+1A);m-97&17\|-\|48037174\|48037194&offsets\|0\|0(+1A)]\|TCAGTGCATGACAGAACTTGGA | >MI0000462\|hsa-mir-152&WithFlank&17\|-\|48037155\|48037253 | 60 | 80 | 21(+1A) | TCAGTGCATGACAGAACTTGGA | 176.6 | 10.7 | 1.5 | 6.9 | 4.8E-12 | 1.6E-11 | MB231/MB468 | MP |
| >hg38_21_+_8433172_8446622_RNA45SN1_45S_50ntFlanks@6647.6672.26\|CGTACGACTCTTAGCGGTGGATCACT | >hg38_21_+_8433172_8446622_RNA45SN1_45S_50ntFlanks | 6647 | 6672 | 26 | CGTACGACTCTTAGCGGTGGATCACT | 368.4 | 4.7 | 0.7 | 6.9 | 5.3E-12 | 1.6E-11 | MB231/MB468 | CYTO |
| >hg38_21_+_8433172_8446622_RNA45SN1_45S_50ntFlanks@9475.9503.29\|CGAGCGCACGTGTTAGGACCCGAAAGATG | >hg38_21_+_8433172_8446622_RNA45SN1_45S_50ntFlanks | 9475 | 9503 | 29 | CGAGCGCACGTGTTAGGACCCGAAAGATG | 196.1 | -11.5 | 1.7 | -6.9 | 4.8E-12 | 1.7E-11 | MB231/MB468 | MP |
| >am_trna128_GlyGCC_6_-_27870686_27870756@1.29.29,trna133_GlyCCC_1_-_16872434_16872504@1.29.29,trna18_GlyGCC_16_+_70822597_70822667@1.29.29,trna19_GlyGCC_16_+_70823410_70823480@1.29.29,trna19_GlyGCC_2_-_157257659_157257729@1.29.29,trna24_GlyGCC_16_-_70812942_70813012@1.29.29,trna25_GlyGCC_16_-_70812114_70812184@1.29.29,trna4_GlyCCC_1_+_17188416_17188486@1.29.29,trna5_GlyGCC_17_+_8029064_8029134@1.29.29,trna68_GlyGCC_1_-_161493637_161493707@1.29.29\|GCATTGGTGGTTCAGTGGTAGAATTCTCG | >am_trna128_GlyGCC_6_-_27870686_27870756 | 1 | 29 | 29 | GCATTGGTGGTTCAGTGGTAGAATTCTCG | 38828.1 | -5.2 | 0.7 | -6.9 | 5.0E-12 | 1.7E-11 | BT20/MB468 | CYTO |
| >hg38_21_+_8433172_8446622_RNA45SN1_45S_50ntFlanks@12347.12366.20\|AGAAAAGTTACCACAGGGAT | >hg38_21_+_8433172_8446622_RNA45SN1_45S_50ntFlanks | 12347 | 12366 | 20 | AGAAAAGTTACCACAGGGAT | 150.6 | -9.5 | 1.4 | -6.9 | 5.1E-12 | 1.7E-11 | BT20/MB468 | CYTO |
| >hg38_21_+_8433172_8446622_RNA45SN1_45S_50ntFlanks@13002.13031.30\|CGATCTATTGAAAGTCAGCCCTCGACACAA | >hg38_21_+_8433172_8446622_RNA45SN1_45S_50ntFlanks | 13002 | 13031 | 30 | CGATCTATTGAAAGTCAGCCCTCGACACAA | 119.1 | -9.2 | 1.3 | -6.9 | 5.0E-12 | 1.7E-11 | BT20/MB468 | CYTO |
| >hg38_21_+_8433172_8446622_RNA45SN1_45S_50ntFlanks@4002.4026.25\|GATAACCTCGGGCCGATCGCACGCC | >hg38_21_+_8433172_8446622_RNA45SN1_45S_50ntFlanks | 4002 | 4026 | 25 | GATAACCTCGGGCCGATCGCACGCC | 238.1 | -10.0 | 1.4 | -6.9 | 3.9E-12 | 1.7E-11 | BT20/MB468 | MITO |
| >hg38_21_+_8433172_8446622_RNA45SN1_45S_50ntFlanks@10134.10158.25\|CGGCGGCGGCGGCGGGGGTGTGGGG | >hg38_21_+_8433172_8446622_RNA45SN1_45S_50ntFlanks | 10134 | 10158 | 25 | CGGCGGCGGCGGCGGGGGTGTGGGG | 128.1 | -10.9 | 1.6 | -6.9 | 4.8E-12 | 1.7E-11 | MB231/MB468 | MP |
| >hg38_21_+_8433172_8446622_RNA45SN1_45S_50ntFlanks@8200.8223.24\|GACGGTGTGAGGCCGGTAGCGGCC | >hg38_21_+_8433172_8446622_RNA45SN1_45S_50ntFlanks | 8200 | 8223 | 24 | GACGGTGTGAGGCCGGTAGCGGCC | 130.1 | -10.9 | 1.6 | -6.9 | 4.8E-12 | 1.7E-11 | MB231/MB468 | MP |
| >hg38_21_+_8433172_8446622_RNA45SN1_45S_50ntFlanks@12488.12512.25\|AACGTGAGCTGGGTTTAGACCGTCG | >hg38_21_+_8433172_8446622_RNA45SN1_45S_50ntFlanks | 12488 | 12512 | 25 | AACGTGAGCTGGGTTTAGACCGTCG | 153.6 | -11.1 | 1.6 | -6.9 | 4.8E-12 | 1.7E-11 | MB231/MB468 | MP |
| >hg38_21_+_8433172_8446622_RNA45SN1_45S_50ntFlanks@12432.12455.24\|CCTATCATTGTGAAGCAGAATTCA | >hg38_21_+_8433172_8446622_RNA45SN1_45S_50ntFlanks | 12432 | 12455 | 24 | CCTATCATTGTGAAGCAGAATTCA | 480.7 | -12.0 | 1.6 | -7.3 | 2.9E-13 | 1.7E-11 | BT20/MB468 | MP |
| >MI0000294\|hsa-mir-218-1&WithFlank&4\|+\|20528269\|20528390@31.52.22;MI0000295\|hsa-mir-218-2&WithFlank&5\|-\|168768140\|168768261@31.52.22&[MIMAT0000275&hsa-miR-218-5p&offsets\|0\|+1;m-185&4\|+\|20528299\|20528319&offsets\|0\|+1];[MIMAT0000275_1&hsa-miR-218-5p&offsets\|0\|+1;m-187&5\|-\|168768211\|168768231&offsets\|0\|+1]\|TTGTGCTTGATCTAACCATGTG | >MI0000294\|hsa-mir-218-1&WithFlank&4\|+\|20528269\|20528390 | 31 | 52 | 22 | TTGTGCTTGATCTAACCATGTG | 195.0 | 10.8 | 1.6 | 6.9 | 4.9E-12 | 1.7E-11 | MB231/MB468 | MP |
| >hg38_1_-_228634819_228635039_RNA5S12_5S_50ntFlanks@141.171.31\|CCGCCTGGGAATACCGGGTGCTGTAGGCTTT | >hg38_1_-_228634819_228635039_RNA5S12_5S_50ntFlanks | 141 | 171 | 31 | CCGCCTGGGAATACCGGGTGCTGTAGGCTTT | 5504.7 | 5.8 | 0.8 | 6.9 | 5.4E-12 | 1.7E-11 | MB231/MB468 | CYTO |
| >hg38_21_+_8433172_8446622_RNA45SN1_45S_50ntFlanks@10787.10815.29\|ACAGCCTCTGGCATGTTGGAACAATGTAG | >hg38_21_+_8433172_8446622_RNA45SN1_45S_50ntFlanks | 10787 | 10815 | 29 | ACAGCCTCTGGCATGTTGGAACAATGTAG | 121.1 | -10.8 | 1.6 | -6.9 | 4.9E-12 | 1.7E-11 | MB231/MB468 | MP |
| >hg38_21_+_8433172_8446622_RNA45SN1_45S_50ntFlanks@11570.11601.32\|AGCCGACTTAGAACTGGTGCGGACCAGGGGAA | >hg38_21_+_8433172_8446622_RNA45SN1_45S_50ntFlanks | 11570 | 11601 | 32 | AGCCGACTTAGAACTGGTGCGGACCAGGGGAA | 1961.7 | -7.6 | 1.1 | -6.9 | 6.5E-12 | 1.7E-11 | MB231/MB468 | MITO |
| >hg38_21_+_8433172_8446622_RNA45SN1_45S_50ntFlanks@4385.4406.22\|TTGGATCTTGGGAGCGGGCGGG | >hg38_21_+_8433172_8446622_RNA45SN1_45S_50ntFlanks | 4385 | 4406 | 22 | TTGGATCTTGGGAGCGGGCGGG | 151.9 | -11.1 | 1.6 | -6.9 | 4.9E-12 | 1.7E-11 | MB231/MB468 | MP |
| >trna10_GlyTCC_17_+_8124866_8124937@31.70.40,trna117_GlyTCC_1_-_145397864_145397935@31.70.40,trna2_GlyTCC_19_+_4724082_4724153@31.70.40,trna45_GlyTCC_1_+_161500903_161500974@31.70.40,trna70_GlyTCC_1_-_161439547_161439618@31.70.40,trna73_GlyTCC_1_-_161432166_161432237@31.70.40,trna76_GlyTCC_1_-_161424756_161424827@31.70.40,trna79_GlyTCC_1_-_161417375_161417446@31.70.40\|CCTTCCAAGCAGTTGACCCGGGTTCGATTCCCGGCCAACG | >trna10_GlyTCC_17_+_8124866_8124937 | 31 | 70 | 40 | CCTTCCAAGCAGTTGACCCGGGTTCGATTCCCGGCCAACG | 193.2 | -10.3 | 1.5 | -6.9 | 5.4E-12 | 1.7E-11 | MB231/MB468 | CYTO |
| >hg38_21_+_8433172_8446622_RNA45SN1_45S_50ntFlanks@10677.10711.35\|GCGGTTCCGGCGGCGTCCGGTGAGCTCTCGCTGGC | >hg38_21_+_8433172_8446622_RNA45SN1_45S_50ntFlanks | 10677 | 10711 | 35 | GCGGTTCCGGCGGCGTCCGGTGAGCTCTCGCTGGC | 132.1 | -10.9 | 1.6 | -6.9 | 4.9E-12 | 1.7E-11 | MB231/MB468 | MP |
| >hg38_21_+_8433172_8446622_RNA45SN1_45S_50ntFlanks@8356.8379.24\|GAAAAGAACTTTGAAGAGAGAGTT | >hg38_21_+_8433172_8446622_RNA45SN1_45S_50ntFlanks | 8356 | 8379 | 24 | GAAAAGAACTTTGAAGAGAGAGTT | 5243.1 | -6.1 | 0.9 | -6.9 | 5.1E-12 | 1.7E-11 | BT20/MB468 | CYTO |
| >MI0000446\|hsa-mir-125b-1&WithFlank&11\|-\|122099751\|122099850@21.43.23(+1U)&[MIMAT0000423&hsa-miR-125b-5p&offsets\|0\|+1(+1U);m-67&11\|-\|122099809\|122099830&offsets\|0\|+1(+1U)]\|TCCCTGAGACCCTAACTTGTGATT | >MI0000446\|hsa-mir-125b-1&WithFlank&11\|-\|122099751\|122099850 | 21 | 43 | 23(+1U) | TCCCTGAGACCCTAACTTGTGATT | 205.8 | -10.8 | 1.5 | -7.2 | 5.8E-13 | 1.7E-11 | BT20/MB231 | NUC |
| >MI0006444\|hsa-mir-1307&WithFlank&10\|-\|103394247\|103394407@86.107.22&[MIMAT0005951&hsa-miR-1307-3p&offsets\|0\|0;m-66&10\|-\|103394300\|103394322&offsets\|0\|-1]\|ACTCGGCGTGGCGTCGGTCGTG | >MI0006444\|hsa-mir-1307&WithFlank&10\|-\|103394247\|103394407 | 86 | 107 | 22 | ACTCGGCGTGGCGTCGGTCGTG | 801.9 | 5.3 | 0.8 | 6.9 | 5.6E-12 | 1.7E-11 | MB231/MB468 | NUC |
| >hg38_21_+_8433172_8446622_RNA45SN1_45S_50ntFlanks@10366.10383.18\|CCGAAGTGGAGAAGGGTT | >hg38_21_+_8433172_8446622_RNA45SN1_45S_50ntFlanks | 10366 | 10383 | 18 | CCGAAGTGGAGAAGGGTT | 146.2 | -7.5 | 1.1 | -6.9 | 5.6E-12 | 1.7E-11 | MB231/MB468 | NUC |
| >hg38_21_+_8433172_8446622_RNA45SN1_45S_50ntFlanks@10458.10478.21\|ATGGCCTCCGTTGCCCTCGGC | >hg38_21_+_8433172_8446622_RNA45SN1_45S_50ntFlanks | 10458 | 10478 | 21 | ATGGCCTCCGTTGCCCTCGGC | 2696.1 | -9.2 | 1.3 | -6.9 | 6.6E-12 | 1.7E-11 | MB231/MB468 | MITO |
| >hg38_21_+_8433172_8446622_RNA45SN1_45S_50ntFlanks@4049.4069.21\|TCGAACGTCTGCCCTATCAAC | >hg38_21_+_8433172_8446622_RNA45SN1_45S_50ntFlanks | 4049 | 4069 | 21 | TCGAACGTCTGCCCTATCAAC | 306.9 | 6.7 | 1.0 | 6.9 | 5.4E-12 | 1.7E-11 | MB231/MB468 | CYTO |
| >hg38_21_+_8433172_8446622_RNA45SN1_45S_50ntFlanks@4923.4944.22\|CAGGAGTGGAGCCTGCGGCTTA | >hg38_21_+_8433172_8446622_RNA45SN1_45S_50ntFlanks | 4923 | 4944 | 22 | CAGGAGTGGAGCCTGCGGCTTA | 576.8 | -13.0 | 1.9 | -6.9 | 6.6E-12 | 1.7E-11 | MB231/MB468 | MITO |
| >hg38_21_+_8433172_8446622_RNA45SN1_45S_50ntFlanks@10410.10429.20\|TCAGTCGGTCCTGAGAGATG | >hg38_21_+_8433172_8446622_RNA45SN1_45S_50ntFlanks | 10410 | 10429 | 20 | TCAGTCGGTCCTGAGAGATG | 7690.8 | -5.5 | 0.8 | -6.9 | 5.4E-12 | 1.7E-11 | MB231/MB468 | CYTO |
| >hg38_21_+_8433172_8446622_RNA45SN1_45S_50ntFlanks@4588.4621.34\|CTATTTTGTTGGTTTTCGGAACTGAGGCCATGAT | >hg38_21_+_8433172_8446622_RNA45SN1_45S_50ntFlanks | 4588 | 4621 | 34 | CTATTTTGTTGGTTTTCGGAACTGAGGCCATGAT | 571.5 | -8.5 | 1.2 | -6.9 | 6.6E-12 | 1.7E-11 | MB231/MB468 | MITO |
| >hg38_21_+_8433172_8446622_RNA45SN1_45S_50ntFlanks@10788.10805.18\|CAGCCTCTGGCATGTTGG | >hg38_21_+_8433172_8446622_RNA45SN1_45S_50ntFlanks | 10788 | 10805 | 18 | CAGCCTCTGGCATGTTGG | 104.0 | -10.6 | 1.5 | -6.9 | 4.9E-12 | 1.7E-11 | MB231/MB468 | MP |
| >hg38_21_+_8433172_8446622_RNA45SN1_45S_50ntFlanks@9559.9597.39\|CGGTCCTGACGTGCAAATCGGTCGTCCGACCTGGGTATA | >hg38_21_+_8433172_8446622_RNA45SN1_45S_50ntFlanks | 9559 | 9597 | 39 | CGGTCCTGACGTGCAAATCGGTCGTCCGACCTGGGTATA | 259.7 | -11.8 | 1.7 | -6.9 | 6.6E-12 | 1.7E-11 | MB231/MB468 | MITO |
| >hg38_21_+_8433172_8446622_RNA45SN1_45S_50ntFlanks@9572.9595.24\|CAAATCGGTCGTCCGACCTGGGTA | >hg38_21_+_8433172_8446622_RNA45SN1_45S_50ntFlanks | 9572 | 9595 | 24 | CAAATCGGTCGTCCGACCTGGGTA | 1237.1 | -10.4 | 1.5 | -6.9 | 7.0E-12 | 1.7E-11 | BT20/MB468 | NUC |
| >hg38_21_+_8433172_8446622_RNA45SN1_45S_50ntFlanks@10683.10709.27\|CCGGCGGCGTCCGGTGAGCTCTCGCTG | >hg38_21_+_8433172_8446622_RNA45SN1_45S_50ntFlanks | 10683 | 10709 | 27 | CCGGCGGCGTCCGGTGAGCTCTCGCTG | 3639.5 | -8.0 | 1.2 | -6.9 | 6.6E-12 | 1.7E-11 | MB231/MB468 | MITO |
| >hg38_21_+_8433172_8446622_RNA45SN1_45S_50ntFlanks@12417.12434.18\|TCGATGTCGGCTCTTCCT | >hg38_21_+_8433172_8446622_RNA45SN1_45S_50ntFlanks | 12417 | 12434 | 18 | TCGATGTCGGCTCTTCCT | 5207.4 | -3.5 | 0.5 | -6.9 | 5.7E-12 | 1.7E-11 | MB231/MB468 | NUC |
| >MI0000441\|hsa-mir-30b&WithFlank&8\|-\|134800514\|134800613@23.43.21(+1A)&[MIMAT0000420&hsa-miR-30b-5p&offsets\|0\|-1(+1A);m-76&8\|-\|134800570\|134800591&offsets\|0\|-1(+1A)]\|TGTAAACATCCTACACTCAGCA | >MI0000441\|hsa-mir-30b&WithFlank&8\|-\|134800514\|134800613 | 23 | 43 | 21(+1A) | TGTAAACATCCTACACTCAGCA | 223.4 | 6.9 | 1.0 | 6.9 | 5.7E-12 | 1.7E-11 | MB231/MB468 | NUC |
| >hg38_21_+_8433172_8446622_RNA45SN1_45S_50ntFlanks@6745.6771.27\|ACACTTCGAACGCACTTGCGGCCCCGG | >hg38_21_+_8433172_8446622_RNA45SN1_45S_50ntFlanks | 6745 | 6771 | 27 | ACACTTCGAACGCACTTGCGGCCCCGG | 2591.1 | -6.8 | 1.0 | -6.9 | 5.5E-12 | 1.7E-11 | MB231/MB468 | CYTO |
| >MI0000063\|hsa-let-7b&WithFlank&22\|+\|46113680\|46113774@12.29.18&[MIMAT0000063&hsa-let-7b-5p&offsets\|0\|-4;m-14&22\|+\|46113691\|46113712&offsets\|0\|-4]\|TGAGGTAGTAGGTTGTGT | >MI0000063\|hsa-let-7b&WithFlank&22\|+\|46113680\|46113774 | 12 | 29 | 18 | TGAGGTAGTAGGTTGTGT | 83.7 | 3.2 | 0.5 | 6.9 | 7.1E-12 | 1.7E-11 | BT20/MB468 | NUC |
| >hg38_21_+_8433172_8446622_RNA45SN1_45S_50ntFlanks@12635.12654.20\|AAGCTACCATCTGTGGGATT | >hg38_21_+_8433172_8446622_RNA45SN1_45S_50ntFlanks | 12635 | 12654 | 20 | AAGCTACCATCTGTGGGATT | 1565.5 | -10.1 | 1.5 | -6.9 | 7.1E-12 | 1.7E-11 | BT20/MB468 | NUC |
| >trnaMT_GluTTC_MT_-_14674_14742@33.72.40,trnalookalike8_GluTTC_5_-_93905172_93905240@33.72.40\|CATATCATTGGTCGTGGTTGTAGTCCGTGCGAGAATACCA | >trnaMT_GluTTC_MT_-_14674_14742 | 33 | 72 | 40 | CATATCATTGGTCGTGGTTGTAGTCCGTGCGAGAATACCA | 1517.4 | 5.4 | 0.8 | 6.9 | 4.1E-12 | 1.7E-11 | BT20/MB468 | MITO |
| >MI0000734\|hsa-mir-106b&WithFlank&7\|-\|100093987\|100094080@58.77.20(+1A)&[MIMAT0004672&hsa-miR-106b-3p&offsets\|0\|-2(+1A);m-87&7\|-\|100094003\|100094023&offsets\|0\|-1(+1A)]\|CCGCACTGTGGGTACTTGCTA | >MI0000734\|hsa-mir-106b&WithFlank&7\|-\|100093987\|100094080 | 58 | 77 | 20(+1A) | CCGCACTGTGGGTACTTGCTA | 425.0 | 8.5 | 1.2 | 6.9 | 5.6E-12 | 1.7E-11 | MB231/MB468 | CYTO |

| | | | | | | | | | | | | | |
|---|---|---|---|---|---|---|---|---|---|---|---|---|---|
| >hg38_21_+_8433172_8446622_RNA45SN1_45S_50ntFlanks@10471.10488.18\|CCCTCGGCCGATCGAAAG | >hg38_21_+_8433172_8446622_RNA45SN1_45S_50ntFlanks | 10471 | 10488 | 18 | CCCTCGGCCGATCGAAAG | 1520.4 | -7.8 | 1.1 | -6.9 | 6.7E-12 | 1.7E-11 | MB231/MB468 | MITO |
| >hg38_21_+_8433172_8446622_RNA45SN1_45S_50ntFlanks@3925.3959.35\|ATCAGATCAAAACCAACCCGGTCAGCCCCTCTCCG | >hg38_21_+_8433172_8446622_RNA45SN1_45S_50ntFlanks | 3925 | 3959 | 35 | ATCAGATCAAAACCAACCCGGTCAGCCCCTCTCCG | 106.3 | -9.0 | 1.3 | -6.9 | 5.3E-12 | 1.7E-11 | BT20/MB468 | CYTO |
| >hg38_21_+_8433172_8446622_RNA45SN1_45S_50ntFlanks@12345.12365.21\|TCAGAAAAGTTACCACAGGGA | >hg38_21_+_8433172_8446622_RNA45SN1_45S_50ntFlanks | 12345 | 12365 | 21 | TCAGAAAAGTTACCACAGGGA | 175.1 | -7.5 | 1.1 | -6.9 | 5.3E-12 | 1.7E-11 | BT20/MB468 | CYTO |
| >hg38_21_+_8433172_8446622_RNA45SN1_45S_50ntFlanks@10794.10816.23\|CTGGCATGTTGGAACAATGTAGG | >hg38_21_+_8433172_8446622_RNA45SN1_45S_50ntFlanks | 10794 | 10816 | 23 | CTGGCATGTTGGAACAATGTAGG | 1062.3 | -7.3 | 1.1 | -6.9 | 6.7E-12 | 1.7E-11 | MB231/MB468 | MITO |
| >hg38_21_+_8433172_8446622_RNA45SN1_45S_50ntFlanks@12936.12969.34\|AAACCATTCGTAGACGACCTGCTTCTGGGTCGGG | >hg38_21_+_8433172_8446622_RNA45SN1_45S_50ntFlanks | 12936 | 12969 | 34 | AAACCATTCGTAGACGACCTGCTTCTGGGTCGGG | 138.0 | -11.0 | 1.6 | -6.9 | 5.0E-12 | 1.7E-11 | MB231/MB468 | MP |
| >hg38_21_+_8433172_8446622_RNA45SN1_45S_50ntFlanks@8179.8209.31\|CTGATCGAGGCCCAGCCCGTGGACGGTGTGA | >hg38_21_+_8433172_8446622_RNA45SN1_45S_50ntFlanks | 8179 | 8209 | 31 | CTGATCGAGGCCCAGCCCGTGGACGGTGTGA | 101.3 | -10.5 | 1.5 | -6.9 | 5.0E-12 | 1.7E-11 | MB231/MB468 | MP |
| >MI0000449\|hsa-mir-132&WithFlank&17\|-\|2049902\|2050014@29.50.22&[MIMAT0004594&hsa-miR-132-5p&offsets\|0\|0;m-414&17\|-\|2049965\|2049986&offsets\|0\|0]\|ACCGTGGCTTTCGATTGTTACT | >MI0000449\|hsa-mir-132&WithFlank&17\|-\|2049902\|2050014 | 29 | 50 | 22 | ACCGTGGCTTTCGATTGTTACT | 52.2 | 6.6 | 1.0 | 6.9 | 7.2E-12 | 1.7E-11 | BT20/MB468 | NUC |
| >MI0000102\|hsa-mir-100&WithFlank&11\|-\|122152223\|122152314@19.38.20&[MIMAT0000098&hsa-miR-100-5p&offsets\|0\|-2;m-27&11\|-\|122152276\|122152296&offsets\|0\|-1]\|AACCCGTAGATCCGAACTTG | >MI0000102\|hsa-mir-100&WithFlank&11\|-\|122152223\|122152314 | 19 | 38 | 20 | AACCCGTAGATCCGAACTTG | 23746.4 | -6.5 | 0.9 | -7.2 | 5.2E-13 | 1.7E-11 | BT20/MB231 | MITO |
| >hg38_21_+_8433172_8446622_RNA45SN1_45S_50ntFlanks@5186.5207.22\|CAATAACAGGTCTGTGATGCCC | >hg38_21_+_8433172_8446622_RNA45SN1_45S_50ntFlanks | 5186 | 5207 | 22 | CAATAACAGGTCTGTGATGCCC | 221.9 | -11.7 | 1.7 | -6.9 | 5.1E-12 | 1.7E-11 | MB231/MB468 | MP |
| >hg38_21_+_8433172_8446622_RNA45SN1_45S_50ntFlanks@6721.6738.18\|AATTGCAGGACACATTGA | >hg38_21_+_8433172_8446622_RNA45SN1_45S_50ntFlanks | 6721 | 6738 | 18 | AATTGCAGGACACATTGA | 206.0 | -7.4 | 1.1 | -6.9 | 5.6E-12 | 1.7E-11 | MB231/MB468 | CYTO |
| >hg38_21_+_8433172_8446622_RNA45SN1_45S_50ntFlanks@4923.4941.19\|CAGGAGTGGAGCCTGCGGC | >hg38_21_+_8433172_8446622_RNA45SN1_45S_50ntFlanks | 4923 | 4941 | 19 | CAGGAGTGGAGCCTGCGGC | 112.8 | -10.7 | 1.5 | -6.9 | 5.1E-12 | 1.7E-11 | MB231/MB468 | MP |
| >hg38_21_+_8433172_8446622_RNA45SN1_45S_50ntFlanks@5470.5516.47\|CCACGGCCCTGGCGGAGCGCTGAGAAGACGGTCGAACTTGACTATCT | >hg38_21_+_8433172_8446622_RNA45SN1_45S_50ntFlanks | 5470 | 5516 | 47 | CCACGGCCCTGGCGGAGCGCTGAGAAGACGGTCGAACTTGACTATCT | 1218.4 | -5.7 | 0.8 | -7.3 | 4.0E-13 | 1.7E-11 | BT20/MB231 | CYTO |
| >hg38_1_-_228634819_228635039_RNA5S12_5S_50ntFlanks@130.170.41\|TGGATGGGAGACCGCCTGGGAATACCGGGTGCTGTAGGCTT | >hg38_1_-_228634819_228635039_RNA5S12_5S_50ntFlanks | 130 | 170 | 41 | TGGATGGGAGACCGCCTGGGAATACCGGGTGCTGTAGGCTT | 3711.3 | 8.5 | 1.2 | 6.9 | 4.1E-12 | 1.7E-11 | BT20/MB468 | MITO |
| >hg38_21_+_8433172_8446622_RNA45SN1_45S_50ntFlanks@10794.10822.29\|CTGGCATGTTGGAACAATGTAGGTAAGGG | >hg38_21_+_8433172_8446622_RNA45SN1_45S_50ntFlanks | 10794 | 10822 | 29 | CTGGCATGTTGGAACAATGTAGGTAAGGG | 336.2 | -5.8 | 0.8 | -6.9 | 5.6E-12 | 1.7E-11 | MB231/MB468 | CYTO |
| >MI0000061\|hsa-let-7a-2&WithFlank&11\|-\|122146516\|122146599@11.31.21(+1A);MI0000062\|hsa-let-7a-3&WithFlank&22\|+\|46112743\|46112828@10.30.21(+1A);MI0000060\|hsa-let-7a-1&WithFlank&9\|+\|94175951\|94176042@12.32.21(+1A)&[MIMAT0000062&hsa-let-7a-5p&offsets\|0\|-1(+1A);m-6&11\|-\|122146568\|122146589&offsets\|0\|-1(+1A)];[MIMAT0000062_1&hsa-let-7a-5p&offsets\|0\|-1(+1A);m-5&22\|+\|46112752\|46112773&offsets\|0\|-1(+1A)];[MIMAT0000062_2&hsa-let-7a-5p&offsets\|0\|-1(+1A);m-4&9\|+\|94175962\|94175983&offsets\|0\|-1(+1A)]\|TGAGGTAGTAGGTTGTATAGTA | >MI0000061\|hsa-let-7a-2&WithFlank&11\|-\|122146516\|122146599 | 11 | 31 | 21(+1A) | TGAGGTAGTAGGTTGTATAGTA | 2182.2 | 6.4 | 0.9 | 6.9 | 5.6E-12 | 1.7E-11 | MB231/MB468 | CYTO |
| >hg38_21_+_8433172_8446622_RNA45SN1_45S_50ntFlanks@4723.4757.35\|CATTAATCAAGAACGAAAGTCGGAGGTTCGAAGAC | >hg38_21_+_8433172_8446622_RNA45SN1_45S_50ntFlanks | 4723 | 4757 | 35 | CATTAATCAAGAACGAAAGTCGGAGGTTCGAAGAC | 174.4 | -11.3 | 1.6 | -6.9 | 5.1E-12 | 1.7E-11 | MB231/MB468 | MP |
| >hg38_21_+_8433172_8446622_RNA45SN1_45S_50ntFlanks@10362.10390.29\|AAGGCCGAAGTGGAGAAGGGTTCCATGTG | >hg38_21_+_8433172_8446622_RNA45SN1_45S_50ntFlanks | 10362 | 10390 | 29 | AAGGCCGAAGTGGAGAAGGGTTCCATGTG | 2818.2 | -7.1 | 1.0 | -6.9 | 5.7E-12 | 1.7E-11 | MB231/MB468 | CYTO |
| >hg38_21_+_8433172_8446622_RNA45SN1_45S_50ntFlanks@10834.10853.20\|CCGGATCCGTAACTTCGGGA | >hg38_21_+_8433172_8446622_RNA45SN1_45S_50ntFlanks | 10834 | 10853 | 20 | CCGGATCCGTAACTTCGGGA | 91.4 | -10.4 | 1.5 | -6.9 | 5.1E-12 | 1.7E-11 | MB231/MB468 | MP |
| >hg38_21_+_8433172_8446622_RNA45SN1_45S_50ntFlanks@9352.9371.20\|CCGTGGCGCAATGAAGGTGA | >hg38_21_+_8433172_8446622_RNA45SN1_45S_50ntFlanks | 9352 | 9371 | 20 | CCGTGGCGCAATGAAGGTGA | 228.2 | -9.9 | 1.4 | -6.9 | 4.2E-12 | 1.8E-11 | BT20/MB468 | MITO |
| >hg38_21_+_8433172_8446622_RNA45SN1_45S_50ntFlanks@9862.9902.41\|ATGAACCGAACGCCGGGTTAAGGCGCCCGATGCCGACGCTC | >hg38_21_+_8433172_8446622_RNA45SN1_45S_50ntFlanks | 9862 | 9902 | 41 | ATGAACCGAACGCCGGGTTAAGGCGCCCGATGCCGACGCTC | 133.6 | -10.9 | 1.6 | -6.9 | 5.1E-12 | 1.8E-11 | MB231/MB468 | MP |
| >hg38_21_+_8433172_8446622_RNA45SN1_45S_50ntFlanks@4983.5002.20\|CGGACAGGATTGACAGATTG | >hg38_21_+_8433172_8446622_RNA45SN1_45S_50ntFlanks | 4983 | 5002 | 20 | CGGACAGGATTGACAGATTG | 947.7 | -9.2 | 1.3 | -6.9 | 5.1E-12 | 1.8E-11 | MB231/MB468 | MP |
| >hg38_21_+_8433172_8446622_RNA45SN1_45S_50ntFlanks@4854.4871.18\|AGTCTTTGGGTTCCGGGG | >hg38_21_+_8433172_8446622_RNA45SN1_45S_50ntFlanks | 4854 | 4871 | 18 | AGTCTTTGGGTTCCGGGG | 116.0 | -10.7 | 1.6 | -6.9 | 5.1E-12 | 1.8E-11 | MB231/MB468 | MP |
| >hg38_21_+_8433172_8446622_RNA45SN1_45S_50ntFlanks@10342.10360.19\|ATATTCAAACGAGAACTTT | >hg38_21_+_8433172_8446622_RNA45SN1_45S_50ntFlanks | 10342 | 10360 | 19 | ATATTCAAACGAGAACTTT | 108.9 | -10.6 | 1.5 | -6.9 | 5.1E-12 | 1.8E-11 | MB231/MB468 | MP |
| >am_trnaMT_LysTTT_MT_+_8295_8364@34.52.19,trnalookalike7_LysTTT_1_+_568844_568913@34.52.19\|GTTAAAGATTAAGAGAACC | >am_trnaMT_LysTTT_MT_+_8295_8364 | 34 | 52 | 19 | GTTAAAGATTAAGAGAACC | 1059.3 | 9.2 | 1.3 | 6.9 | 5.4E-12 | 1.8E-11 | BT20/MB468 | CYTO |
| >trna111_HisGTG_1_-_147774845_147774916@-1A.33.34,trna118_HisGTG_1_-_145396881_145396952@-1A.33.34,trna16_HisGTG_1_+_146544773_146544844@-1A.33.34,trna1_HisGTG_15_+_45493349_45493420@-1A.33.34,trna21_HisGTG_1_+_147753471_147753542@-1A.33.34,trna33_HisGTG_6_+_27125906_27125977@-1A.33.34,trna7_HisGTG_9_-_14433938_14434009@-1A.33.34,trna8_HisGTG_15_-_45492611_45492682@-1A.33.34,trna9_HisGTG_15_-_45490804_45490875@-1A.33.34\|AGCCGTGATCGTATAGTGGTTAGTACTCTGCGTT | >trna111_HisGTG_1_-_147774845_147774916 | -1A | 33 | 34 | AGCCGTGATCGTATAGTGGTTAGTACTCTGCGTT | 551.6 | 8.9 | 1.3 | 6.9 | 5.4E-12 | 1.8E-11 | BT20/MB468 | CYTO |
| >MI0000261\|hsa-mir-139&WithFlank&11\|-\|72615057\|72615136@13.34.22(+1C)&[MIMAT0000250&hsa-miR-139-5p&offsets\|0\|-1(+1C);m-186&11\|-\|72615102\|72615124&offsets\|0\|-1(+1C)]\|TCTACAGTGCACGTGTCTCCAGC | >MI0000261\|hsa-mir-139&WithFlank&11\|-\|72615057\|72615136 | 13 | 34 | 22(+1C) | TCTACAGTGCACGTGTCTCCAGC | 175.1 | 10.7 | 1.5 | 6.9 | 5.2E-12 | 1.8E-11 | MB231/MB468 | MP |
| >hg38_21_+_8433172_8446622_RNA45SN1_45S_50ntFlanks@6698.6718.21\|AGCTAGCTGCGAGAATTAATG | >hg38_21_+_8433172_8446622_RNA45SN1_45S_50ntFlanks | 6698 | 6718 | 21 | AGCTAGCTGCGAGAATTAATG | 11474.3 | -7.1 | 1.0 | -7.3 | 3.1E-13 | 1.8E-11 | BT20/MB468 | MP |
| >hg38_21_+_8433172_8446622_RNA45SN1_45S_50ntFlanks@11875.11900.26\|GAAGACCCTGTTGAGCTTGACTCTAG | >hg38_21_+_8433172_8446622_RNA45SN1_45S_50ntFlanks | 11875 | 11900 | 26 | GAAGACCCTGTTGAGCTTGACTCTAG | 97.6 | -10.5 | 1.5 | -6.9 | 5.2E-12 | 1.8E-11 | MB231/MB468 | MP |
| >hg38_21_+_8433172_8446622_RNA45SN1_45S_50ntFlanks@3993.4023.31\|GTGACTCTAGATAACCTCGGGCCGATCGCAC | >hg38_21_+_8433172_8446622_RNA45SN1_45S_50ntFlanks | 3993 | 4023 | 31 | GTGACTCTAGATAACCTCGGGCCGATCGCAC | 1059.7 | 4.7 | 0.7 | 6.9 | 5.7E-12 | 1.8E-11 | MB231/MB468 | CYTO |
| >hg38_21_+_8433172_8446622_RNA45SN1_45S_50ntFlanks@6784.6801.18\|GCTACGCCTGTCTGAGCG | >hg38_21_+_8433172_8446622_RNA45SN1_45S_50ntFlanks | 6784 | 6801 | 18 | GCTACGCCTGTCTGAGCG | 269.0 | -11.2 | 1.5 | -7.3 | 3.1E-13 | 1.8E-11 | BT20/MB468 | MP |
| >hg38_21_+_8433172_8446622_RNA45SN1_45S_50ntFlanks@11233.11252.20\|CCCTTCCCGTGGATCGCCCC | >hg38_21_+_8433172_8446622_RNA45SN1_45S_50ntFlanks | 11233 | 11252 | 20 | CCCTTCCCGTGGATCGCCCC | 510.2 | -7.9 | 1.1 | -6.9 | 5.7E-12 | 1.8E-11 | MB231/MB468 | CYTO |
| >hg38_21_+_8433172_8446622_RNA45SN1_45S_50ntFlanks@5447.5466.20\|GGATCGGCCCCGCCGGGGTC | >hg38_21_+_8433172_8446622_RNA45SN1_45S_50ntFlanks | 5447 | 5466 | 20 | GGATCGGCCCCGCCGGGGTC | 193.2 | -11.5 | 1.7 | -6.9 | 5.2E-12 | 1.8E-11 | MB231/MB468 | MP |
| >am_trna134_GluTTC_1_-_16861774_16861845@1.29.29,trna5_GluTTC_1_+_17199078_17199149@1.29.29,trna84_GluTTC_1_-_161391883_161391954@1.29.29,trna94_GluTTC_1_-_149664355_149664427@1.29.29\|TCCCTGGTGGTCTAGTGGCTAGGATTCGG | >am_trna134_GluTTC_1_-_16861774_16861845 | 1 | 29 | 29 | TCCCTGGTGGTCTAGTGGCTAGGATTCGG | 3733.4 | -4.4 | 0.6 | -6.9 | 5.4E-12 | 1.8E-11 | BT20/MB468 | CYTO |
| >hg38_21_+_8433172_8446622_RNA45SN1_45S_50ntFlanks@5487.5525.39\|CGCTGAGAAGACGGTCGAACTTGACTATCTAGAGGAAGT | >hg38_21_+_8433172_8446622_RNA45SN1_45S_50ntFlanks | 5487 | 5525 | 39 | CGCTGAGAAGACGGTCGAACTTGACTATCTAGAGGAAGT | 146.7 | -11.1 | 1.6 | -6.9 | 5.2E-12 | 1.8E-11 | MB231/MB468 | MP |
| >hg38_21_+_8433172_8446622_RNA45SN1_45S_50ntFlanks@10793.10825.33\|TCTGGCATGTTGGAACAATGTAGGTAAGGGAAG | >hg38_21_+_8433172_8446622_RNA45SN1_45S_50ntFlanks | 10793 | 10825 | 33 | TCTGGCATGTTGGAACAATGTAGGTAAGGGAAG | 91.7 | -10.4 | 1.5 | -6.9 | 5.2E-12 | 1.8E-11 | MB231/MB468 | MP |
| >hg38_21_+_8433172_8446622_RNA45SN1_45S_50ntFlanks@12893.12931.39\|CCTCGCCCGTCACGCACCGCACGTTCGTGGGGAACCTGG | >hg38_21_+_8433172_8446622_RNA45SN1_45S_50ntFlanks | 12893 | 12931 | 39 | CCTCGCCCGTCACGCACCGCACGTTCGTGGGGAACCTGG | 123.9 | -10.8 | 1.6 | -6.9 | 5.2E-12 | 1.8E-11 | MB231/MB468 | MP |
| >hg38_21_+_8433172_8446622_RNA45SN1_45S_50ntFlanks@10713.10738.26\|CTTGAAAATCCGGGGGAGAGGGTGTA | >hg38_21_+_8433172_8446622_RNA45SN1_45S_50ntFlanks | 10713 | 10738 | 26 | CTTGAAAATCCGGGGGAGAGGGTGTA | 98.3 | -10.5 | 1.5 | -6.9 | 5.2E-12 | 1.8E-11 | MB231/MB468 | MP |
| >hg38_21_+_8433172_8446622_RNA45SN1_45S_50ntFlanks@10392.10433.42\|ACAGCAGTTGAACATGGGTCAGTCGGTCCTGAGAGATGGGCG | >hg38_21_+_8433172_8446622_RNA45SN1_45S_50ntFlanks | 10392 | 10433 | 42 | ACAGCAGTTGAACATGGGTCAGTCGGTCCTGAGAGATGGGCG | 1005.1 | -6.3 | 0.9 | -6.9 | 5.9E-12 | 1.8E-11 | MB231/MB468 | NUC |
| >MI0000077\|hsa-mir-21&WithFlank&17\|+\|59841260\|59841343@52.72.21&[MIMAT0004494&hsa-miR-21-3p&offsets\|0\|0;m-52&17\|+\|59841311\|59841332&offsets\|0\|-1]\|CAACACCAGTCGATGGGCTGT | >MI0000077\|hsa-mir-21&WithFlank&17\|+\|59841260\|59841343 | 52 | 72 | 21 | CAACACCAGTCGATGGGCTGT | 209.5 | 8.3 | 1.2 | 6.9 | 5.9E-12 | 1.8E-11 | MB231/MB468 | NUC |
| >MI0005566\|hsa-mir-374b&WithFlank&X\|-\|74218541\|74218624@17.37.21&[MIMAT0004955&hsa-miR-374b-5p&offsets\|0\|-1;m-153&X\|-\|74218587\|74218608&offsets\|0\|-1]\|ATATAATACAACCTGCTAAGT | >MI0005566\|hsa-mir-374b&WithFlank&X\|-\|74218541\|74218624 | 17 | 37 | 21 | ATATAATACAACCTGCTAAGT | 822.7 | 5.2 | 0.8 | 6.9 | 5.9E-12 | 1.8E-11 | MB231/MB468 | NUC |
| >hg38_21_+_8433172_8446622_RNA45SN1_45S_50ntFlanks@4892.4917.26\|AAACTTAAAGGAATTGACGGAAGGGC | >hg38_21_+_8433172_8446622_RNA45SN1_45S_50ntFlanks | 4892 | 4917 | 26 | AAACTTAAAGGAATTGACGGAAGGGC | 246.3 | -11.8 | 1.7 | -6.9 | 5.2E-12 | 1.8E-11 | MB231/MB468 | MP |
| >hg38_21_+_8433172_8446622_RNA45SN1_45S_50ntFlanks@11785.11804.20\|TGAATGGATGAACGAGATTC | >hg38_21_+_8433172_8446622_RNA45SN1_45S_50ntFlanks | 11785 | 11804 | 20 | TGAATGGATGAACGAGATTC | 8663.3 | -6.4 | 0.9 | -6.9 | 5.8E-12 | 1.8E-11 | MB231/MB468 | CYTO |
| >hg38_21_+_8433172_8446622_RNA45SN1_45S_50ntFlanks@12658.12675.18\|ACTGAACGCCTCTAAGTC | >hg38_21_+_8433172_8446622_RNA45SN1_45S_50ntFlanks | 12658 | 12675 | 18 | ACTGAACGCCTCTAAGTC | 3067.8 | -10.7 | 1.6 | -6.8 | 7.4E-12 | 1.8E-11 | BT20/MB468 | NUC |
| >hg38_21_+_8433172_8446622_RNA45SN1_45S_50ntFlanks@5431.5451.21\|GTTTAGTGAGGCCCTCGGATC | >hg38_21_+_8433172_8446622_RNA45SN1_45S_50ntFlanks | 5431 | 5451 | 21 | GTTTAGTGAGGCCCTCGGATC | 240.3 | -8.2 | 1.2 | -6.9 | 6.0E-12 | 1.8E-11 | MB231/MB468 | NUC |

<table>
<tr><td>>am_trna11_ProAGG_16_+_3241989_3242060@1.29.29,trna12_ProAGG_6_+_26555498_26555569@1.29.29,trna12_ProTGG_11_-_75946869_75946940@1.29.29,trna14_ProTGG_5_-_180615854_180615925@1.29.29,trna22_ProAGG_14_-_21081560_21081631@1.29.29,trna23_ProAGG_14_-_21077495_21077566@1.29.29,trna28_ProTGG_16_-_3234133_3234204@1.29.29,trna29_ProAGG_16_-_3232635_3232706@1.29.29,trna2_ProAGG_7_+_128423504_128423575@1.29.29,trna30_ProCGG_6_+_27059521_27059592@1.29.29,trna37_ProCGG_17_-_8126151_8126222@1.29.29,trna3_ProTGG_16_+_3208923_3208994@1.29.29,trna52_ProCGG_1_+_167683962_167684033@1.29.29,trna65_ProAGG_1_-_167684725_167684796@1.29.29,trna6_ProCGG_16_+_3222049_3222120@1.29.29,trna6_ProTGG_14_+_21152175_21152246@1.29.29,trna8_ProTGG_16_+_3238094_3238165@1.29.29,trna9_ProAGG_11_+_75946557_75946628@1.29.29,trna9_ProAGG_16_+_3239634_3239705@1.29.29|GGCTCGTTGGTCTAGGGGTATGATTCTCG</td><td>>am_trna11_ProAGG_16_+_3241989_3242060</td><td>1</td><td>29</td><td>29</td><td>GGCTCGTTGGTCTAGGGGTATGATTCTCG</td><td>5806.0</td><td>-11.1</td><td>1.6</td><td>-6.8</td><td>7.4E-12</td><td>1.8E-11</td><td>BT20/MB468</td><td>NUC</td></tr>
<tr><td>>hg38_21_+_8433172_8446622_RNA45SN1_45S_50ntFlanks@12712.12745.34|CGGAGCCTCGGTTGGCCTCGGATAGCCGGTCCCC</td><td>>hg38_21_+_8433172_8446622_RNA45SN1_45S_50ntFlanks</td><td>12712</td><td>12745</td><td>34</td><td>CGGAGCCTCGGTTGGCCTCGGATAGCCGGTCCCC</td><td>331.2</td><td>-10.5</td><td>1.5</td><td>-6.9</td><td>4.3E-12</td><td>1.8E-11</td><td>BT20/MB468</td><td>MITO</td></tr>
<tr><td>>am_trna10_ValCAC_5_-_180649395_180649467@1.33.33,trna12_ValAAC_5_-_180645270_180645342@1.33.33,trna132_ValAAC_6_-_27721179_27721251@1.33.33,trna136_ValAAC_6_-_27648885_27648957@1.33.33,trna139_ValAAC_6_-_27618707_27618779@1.33.33,trna18_ValCAC_5_-_180529253_180529325@1.33.33,trna2_ValAAC_3_+_169490018_169490090@1.33.33,trna2_ValCAC_5_+_180524070_180524142@1.33.33,trna4_ValAAC_5_+_180591154_180591226@1.33.33,trna5_ValAAC_5_+_180596610_180596682@1.33.33,trna6_ValCAC_5_+_180600650_180600722@1.33.33,trna85_ValCAC_1_-_161369490_161369562@1.33.33,trna90_ValCAC_1_-_149684088_149684161@1.33.33,trna98_ValCAC_1_-_149298555_149298627@1.33.33,trna9_ValCAC_6_+_26538282_26538354@1.33.33|GTTTCCGTAGTGTAGTGGTTATCACGTTCGCCT</td><td>>am_trna10_ValCAC_5_-_180649395_180649467</td><td>1</td><td>33</td><td>33</td><td>GTTTCCGTAGTGTAGTGGTTATCACGTTCGCCT</td><td>2010.2</td><td>6.9</td><td>1.0</td><td>6.9</td><td>7.0E-12</td><td>1.8E-11</td><td>MB231/MB468</td><td>MITO</td></tr>
<tr><td>>MI0000095|hsa-mir-93&WithFlank&7|-|100093762|100093853@17.38.22(+1A)&[MIMAT0000093&hsa-miR-93-5p&offsets|0|-1(+1A);m-29&7|-|100093815|100093837&offsets|0|-1(+1A)]|CAAAGTGCTGTTCGTGCAGGTAA</td><td>>MI0000095|hsa-mir-93&WithFlank&7|-|100093762|100093853</td><td>17</td><td>38</td><td>22(+1A)</td><td>CAAAGTGCTGTTCGTGCAGGTAA</td><td>628.7</td><td>5.9</td><td>0.9</td><td>6.9</td><td>4.3E-12</td><td>1.8E-11</td><td>BT20/MB468</td><td>MITO</td></tr>
<tr><td>>hg38_21_+_8433172_8446622_RNA45SN1_45S_50ntFlanks@10690.10709.20|CGTCCGGTGAGCTCTCGCTG</td><td>>hg38_21_+_8433172_8446622_RNA45SN1_45S_50ntFlanks</td><td>10690</td><td>10709</td><td>20</td><td>CGTCCGGTGAGCTCTCGCTG</td><td>1846.0</td><td>-9.3</td><td>1.4</td><td>-6.9</td><td>6.0E-12</td><td>1.8E-11</td><td>MB231/MB468</td><td>NUC</td></tr>
<tr><td>>hg38_21_+_8433172_8446622_RNA45SN1_45S_50ntFlanks@12539.12567.29|ATGATGTGTTGTTGCCATGGTAATCCTGC</td><td>>hg38_21_+_8433172_8446622_RNA45SN1_45S_50ntFlanks</td><td>12539</td><td>12567</td><td>29</td><td>ATGATGTGTTGTTGCCATGGTAATCCTGC</td><td>2290.5</td><td>-10.4</td><td>1.5</td><td>-6.8</td><td>7.5E-12</td><td>1.8E-11</td><td>BT20/MB468</td><td>NUC</td></tr>
<tr><td>>MI0000471|hsa-mir-126&WithFlank&9|+|136670596|136670692@59.79.21&[MIMAT0000445&hsa-miR-126-3p&offsets|+1|0;m-40&9|+|136670653|136670674&offsets|+1|0]|CGTACCGTGAGTAATAATGCG</td><td>>MI0000471|hsa-mir-126&WithFlank&9|+|136670596|136670692</td><td>59</td><td>79</td><td>21</td><td>CGTACCGTGAGTAATAATGCG</td><td>77.4</td><td>7.1</td><td>1.0</td><td>6.8</td><td>7.5E-12</td><td>1.8E-11</td><td>BT20/MB468</td><td>NUC</td></tr>
<tr><td>>hg38_21_+_8433172_8446622_RNA45SN1_45S_50ntFlanks@10794.10823.30|CTGGCATGTTGGAACAATGTAGGTAAGGGA</td><td>>hg38_21_+_8433172_8446622_RNA45SN1_45S_50ntFlanks</td><td>10794</td><td>10823</td><td>30</td><td>CTGGCATGTTGGAACAATGTAGGTAAGGGA</td><td>501.8</td><td>-9.2</td><td>1.3</td><td>-6.8</td><td>7.5E-12</td><td>1.8E-11</td><td>BT20/MB468</td><td>NUC</td></tr>
<tr><td>>hg38_21_+_8433172_8446622_RNA45SN1_45S_50ntFlanks@4442.4471.30|CCCCTTGCCTCTCGGCGCCCCCTCGATGCT</td><td>>hg38_21_+_8433172_8446622_RNA45SN1_45S_50ntFlanks</td><td>4442</td><td>4471</td><td>30</td><td>CCCCTTGCCTCTCGGCGCCCCCTCGATGCT</td><td>139.5</td><td>-9.4</td><td>1.4</td><td>-6.9</td><td>5.6E-12</td><td>1.8E-11</td><td>BT20/MB468</td><td>CYTO</td></tr>
<tr><td>>hg38_21_+_8433172_8446622_RNA45SN1_45S_50ntFlanks@12615.12659.45|GCTGAGGAGCCAATGGGGCGAAGCTACCATCTGTGGGATTATGAC</td><td>>hg38_21_+_8433172_8446622_RNA45SN1_45S_50ntFlanks</td><td>12615</td><td>12659</td><td>45</td><td>GCTGAGGAGCCAATGGGGCGAAGCTACCATCTGTGGGATTATGAC</td><td>301.3</td><td>-11.2</td><td>1.6</td><td>-6.9</td><td>6.0E-12</td><td>1.8E-11</td><td>MB231/MB468</td><td>NUC</td></tr>
<tr><td>>hg38_21_+_8433172_8446622_RNA45SN1_45S_50ntFlanks@4547.4566.20|CTGGATACCGCAGCTAGGAA</td><td>>hg38_21_+_8433172_8446622_RNA45SN1_45S_50ntFlanks</td><td>4547</td><td>4566</td><td>20</td><td>CTGGATACCGCAGCTAGGAA</td><td>251.6</td><td>-11.8</td><td>1.7</td><td>-6.9</td><td>5.3E-12</td><td>1.8E-11</td><td>MB231/MB468</td><td>MP</td></tr>
<tr><td>>hg38_21_+_8433172_8446622_RNA45SN1_45S_50ntFlanks@4583.4601.19|CGGTTCTATTTTGTTGGTT</td><td>>hg38_21_+_8433172_8446622_RNA45SN1_45S_50ntFlanks</td><td>4583</td><td>4601</td><td>19</td><td>CGGTTCTATTTTGTTGGTT</td><td>110.1</td><td>-10.6</td><td>1.5</td><td>-6.9</td><td>5.3E-12</td><td>1.8E-11</td><td>MB231/MB468</td><td>MP</td></tr>
<tr><td>>hg38_21_+_8433172_8446622_RNA45SN1_45S_50ntFlanks@7974.8010.37|ACGCGACCTCAGATCAGACGTGGCGACCCGCTGAATT</td><td>>hg38_21_+_8433172_8446622_RNA45SN1_45S_50ntFlanks</td><td>7974</td><td>8010</td><td>37</td><td>ACGCGACCTCAGATCAGACGTGGCGACCCGCTGAATT</td><td>18612.7</td><td>4.3</td><td>0.6</td><td>6.8</td><td>7.6E-12</td><td>1.8E-11</td><td>BT20/MB468</td><td>NUC</td></tr>
<tr><td>>hg38_21_+_8433172_8446622_RNA45SN1_45S_50ntFlanks@10791.10817.27|CCTCTGGCATGTTGGAACAATGTAGGT</td><td>>hg38_21_+_8433172_8446622_RNA45SN1_45S_50ntFlanks</td><td>10791</td><td>10817</td><td>27</td><td>CCTCTGGCATGTTGGAACAATGTAGGT</td><td>88.4</td><td>-10.3</td><td>1.5</td><td>-6.9</td><td>5.3E-12</td><td>1.8E-11</td><td>MB231/MB468</td><td>MP</td></tr>
<tr><td>>hg38_21_+_8433172_8446622_RNA45SN1_45S_50ntFlanks@9575.9595.21|ATCGGTCGTCCGACCTGGGTA</td><td>>hg38_21_+_8433172_8446622_RNA45SN1_45S_50ntFlanks</td><td>9575</td><td>9595</td><td>21</td><td>ATCGGTCGTCCGACCTGGGTA</td><td>4804.1</td><td>-10.7</td><td>1.6</td><td>-6.9</td><td>5.3E-12</td><td>1.8E-11</td><td>MB231/MB468</td><td>MP</td></tr>
<tr><td>>hg38_21_+_8433172_8446622_RNA45SN1_45S_50ntFlanks@11875.11906.32|GAAGACCCTGTTGAGCTTGACTCTAGTCTGGC</td><td>>hg38_21_+_8433172_8446622_RNA45SN1_45S_50ntFlanks</td><td>11875</td><td>11906</td><td>32</td><td>GAAGACCCTGTTGAGCTTGACTCTAGTCTGGC</td><td>360.5</td><td>-8.4</td><td>1.2</td><td>-6.9</td><td>7.1E-12</td><td>1.8E-11</td><td>MB231/MB468</td><td>MITO</td></tr>
<tr><td>>hg38_21_+_8433172_8446622_RNA45SN1_45S_50ntFlanks@10069.10092.24|AGTGGACGGGAGCGGCGGGGGCGG</td><td>>hg38_21_+_8433172_8446622_RNA45SN1_45S_50ntFlanks</td><td>10069</td><td>10092</td><td>24</td><td>AGTGGACGGGAGCGGCGGGGGCGG</td><td>176.0</td><td>-11.3</td><td>1.6</td><td>-6.9</td><td>5.3E-12</td><td>1.8E-11</td><td>MB231/MB468</td><td>MP</td></tr>
<tr><td>>hg38_21_+_8433172_8446622_RNA45SN1_45S_50ntFlanks@9738.9764.27|GAAACGATCTCAACCTATTCTCAAACT</td><td>>hg38_21_+_8433172_8446622_RNA45SN1_45S_50ntFlanks</td><td>9738</td><td>9764</td><td>27</td><td>GAAACGATCTCAACCTATTCTCAAACT</td><td>103.2</td><td>-10.6</td><td>1.5</td><td>-6.9</td><td>5.3E-12</td><td>1.8E-11</td><td>MB231/MB468</td><td>MP</td></tr>
<tr><td>>hg38_21_+_8433172_8446622_RNA45SN1_45S_50ntFlanks@9339.9378.40|CGCACGAAAGCCGCCGTGGCGCAATGAAGGTGAAGGCCGG</td><td>>hg38_21_+_8433172_8446622_RNA45SN1_45S_50ntFlanks</td><td>9339</td><td>9378</td><td>40</td><td>CGCACGAAAGCCGCCGTGGCGCAATGAAGGTGAAGGCCGG</td><td>225.8</td><td>-11.6</td><td>1.7</td><td>-6.9</td><td>7.1E-12</td><td>1.8E-11</td><td>MB231/MB468</td><td>MITO</td></tr>
<tr><td>>hg38_21_+_8433172_8446622_RNA45SN1_45S_50ntFlanks@11314.11336.23|CCCACCCCACGTCTCGTCGCGCG</td><td>>hg38_21_+_8433172_8446622_RNA45SN1_45S_50ntFlanks</td><td>11314</td><td>11336</td><td>23</td><td>CCCACCCCACGTCTCGTCGCGCG</td><td>219.6</td><td>-10.7</td><td>1.6</td><td>-6.9</td><td>6.1E-12</td><td>1.8E-11</td><td>MB231/MB468</td><td>NUC</td></tr>
<tr><td>>hg38_21_+_8433172_8446622_RNA45SN1_45S_50ntFlanks@12931.12960.30|GCGCTAAACCATTCGTAGACGACCTGCTTC</td><td>>hg38_21_+_8433172_8446622_RNA45SN1_45S_50ntFlanks</td><td>12931</td><td>12960</td><td>30</td><td>GCGCTAAACCATTCGTAGACGACCTGCTTC</td><td>105.9</td><td>-10.6</td><td>1.5</td><td>-6.9</td><td>5.4E-12</td><td>1.8E-11</td><td>MB231/MB468</td><td>MP</td></tr>
<tr><td>>hg38_1_-_228634819_228635039_RNA5S12_5S_50ntFlanks@138.169.32|AGACCGCCTGGGAATACCGGGTGCTGTAGGCT</td><td>>hg38_1_-_228634819_228635039_RNA5S12_5S_50ntFlanks</td><td>138</td><td>169</td><td>32</td><td>AGACCGCCTGGGAATACCGGGTGCTGTAGGCT</td><td>521.4</td><td>5.7</td><td>0.8</td><td>6.9</td><td>5.9E-12</td><td>1.8E-11</td><td>MB231/MB468</td><td>CYTO</td></tr>
<tr><td>>hg38_21_+_8433172_8446622_RNA45SN1_45S_50ntFlanks@5334.5362.29|ATGAACGAGGAATTCCCAGTAAGTGCGGG</td><td>>hg38_21_+_8433172_8446622_RNA45SN1_45S_50ntFlanks</td><td>5334</td><td>5362</td><td>29</td><td>ATGAACGAGGAATTCCCAGTAAGTGCGGG</td><td>432.9</td><td>-12.5</td><td>1.8</td><td>-6.9</td><td>7.2E-12</td><td>1.8E-11</td><td>MB231/MB468</td><td>MITO</td></tr>
<tr><td>>hg38_21_+_8433172_8446622_RNA45SN1_45S_50ntFlanks@4444.4472.29|CCTTGCCTCTCGGCGCCCCCTCGATGCTC</td><td>>hg38_21_+_8433172_8446622_RNA45SN1_45S_50ntFlanks</td><td>4444</td><td>4472</td><td>29</td><td>CCTTGCCTCTCGGCGCCCCCTCGATGCTC</td><td>307.7</td><td>-11.2</td><td>1.6</td><td>-6.9</td><td>6.1E-12</td><td>1.8E-11</td><td>MB231/MB468</td><td>NUC</td></tr>
<tr><td>>MI0000255|hsa-mir-30d&WithFlank&8|-|134804870|134804951@12.35.24&[MIMAT0000245&hsa-miR-30d-5p&offsets|0|+2;m-21&8|-|134804919|134804940&offsets|0|+2]|TGTAAACATCCCCGACTGGAAGCT</td><td>>MI0000255|hsa-mir-30d&WithFlank&8|-|134804870|134804951</td><td>12</td><td>35</td><td>24</td><td>TGTAAACATCCCCGACTGGAAGCT</td><td>44640.4</td><td>6.5</td><td>0.9</td><td>6.9</td><td>7.2E-12</td><td>1.8E-11</td><td>MB231/MB468</td><td>MITO</td></tr>
<tr><td>>hg38_21_+_8433172_8446622_RNA45SN1_45S_50ntFlanks@6671.6698.28|CTCGGCTCGTGCGTCGATGAAGAACGCA</td><td>>hg38_21_+_8433172_8446622_RNA45SN1_45S_50ntFlanks</td><td>6671</td><td>6698</td><td>28</td><td>CTCGGCTCGTGCGTCGATGAAGAACGCA</td><td>771.5</td><td>-5.2</td><td>0.8</td><td>-6.9</td><td>6.1E-12</td><td>1.8E-11</td><td>MB231/MB468</td><td>NUC</td></tr>
<tr><td>>hg38_21_+_8433172_8446622_RNA45SN1_45S_50ntFlanks@6772.6798.27|GTTCCTCCCGGGGCTACGCCTGTCTGA</td><td>>hg38_21_+_8433172_8446622_RNA45SN1_45S_50ntFlanks</td><td>6772</td><td>6798</td><td>27</td><td>GTTCCTCCCGGGGCTACGCCTGTCTGA</td><td>104.1</td><td>-10.6</td><td>1.5</td><td>-6.9</td><td>5.4E-12</td><td>1.8E-11</td><td>MB231/MB468</td><td>MP</td></tr>
<tr><td>>hg38_21_+_8433172_8446622_RNA45SN1_45S_50ntFlanks@10770.10805.36|GCAGGTCTCCAAGGTGAACAGCCTCTGGCATGTTGG</td><td>>hg38_21_+_8433172_8446622_RNA45SN1_45S_50ntFlanks</td><td>10770</td><td>10805</td><td>36</td><td>GCAGGTCTCCAAGGTGAACAGCCTCTGGCATGTTGG</td><td>97.3</td><td>-10.5</td><td>1.5</td><td>-6.9</td><td>5.4E-12</td><td>1.8E-11</td><td>MB231/MB468</td><td>MP</td></tr>
<tr><td>>trnaMT_ValTAC_MT_+_1602_1670@53.72.20|TAACTTGACCGCTCTGACCA</td><td>>trnaMT_ValTAC_MT_+_1602_1670</td><td>53</td><td>72</td><td>20</td><td>TAACTTGACCGCTCTGACCA</td><td>2238.4</td><td>11.5</td><td>1.7</td><td>6.9</td><td>5.7E-12</td><td>1.8E-11</td><td>BT20/MB468</td><td>CYTO</td></tr>
<tr><td>>hg38_21_+_8433172_8446622_RNA45SN1_45S_50ntFlanks@12475.12496.22|CCCACTAATAGGGAACGTGAGC</td><td>>hg38_21_+_8433172_8446622_RNA45SN1_45S_50ntFlanks</td><td>12475</td><td>12496</td><td>22</td><td>CCCACTAATAGGGAACGTGAGC</td><td>698.9</td><td>-4.9</td><td>0.7</td><td>-6.8</td><td>7.7E-12</td><td>1.8E-11</td><td>BT20/MB468</td><td>NUC</td></tr>
<tr><td>>hg38_1_-_228634819_228635039_RNA5S12_5S_50ntFlanks@147.170.24|GGGAATACCGGGTGCTGTAGGCTT</td><td>>hg38_1_-_228634819_228635039_RNA5S12_5S_50ntFlanks</td><td>147</td><td>170</td><td>24</td><td>GGGAATACCGGGTGCTGTAGGCTT</td><td>4321.9</td><td>3.9</td><td>0.6</td><td>6.9</td><td>4.4E-12</td><td>1.8E-11</td><td>BT20/MB468</td><td>MITO</td></tr>
<tr><td>>hg38_21_+_8433172_8446622_RNA45SN1_45S_50ntFlanks@4486.4511.26|CCGCGGGGCCCGAAGCGTTTACTTTG</td><td>>hg38_21_+_8433172_8446622_RNA45SN1_45S_50ntFlanks</td><td>4486</td><td>4511</td><td>26</td><td>CCGCGGGGCCCGAAGCGTTTACTTTG</td><td>428.1</td><td>-10.8</td><td>1.6</td><td>-6.9</td><td>4.4E-12</td><td>1.8E-11</td><td>BT20/MB468</td><td>MITO</td></tr>
<tr><td>>hg38_21_+_8433172_8446622_RNA45SN1_45S_50ntFlanks@9226.9251.26|CGAAGAGGGGGACGGCGGAGCGAGCG</td><td>>hg38_21_+_8433172_8446622_RNA45SN1_45S_50ntFlanks</td><td>9226</td><td>9251</td><td>26</td><td>CGAAGAGGGGGACGGCGGAGCGAGCG</td><td>101.2</td><td>-10.5</td><td>1.5</td><td>-6.9</td><td>5.5E-12</td><td>1.9E-11</td><td>MB231/MB468</td><td>MP</td></tr>
<tr><td>>hg38_21_+_8433172_8446622_RNA45SN1_45S_50ntFlanks@10798.10826.29|CATGTTGGAACAATGTAGGTAAGGGAAGT</td><td>>hg38_21_+_8433172_8446622_RNA45SN1_45S_50ntFlanks</td><td>10798</td><td>10826</td><td>29</td><td>CATGTTGGAACAATGTAGGTAAGGGAAGT</td><td>572.7</td><td>-6.6</td><td>1.0</td><td>-6.9</td><td>7.3E-12</td><td>1.9E-11</td><td>MB231/MB468</td><td>MITO</td></tr>
<tr><td>>hg38_21_+_8433172_8446622_RNA45SN1_45S_50ntFlanks@12033.12055.23|ATCGTTTTTTCACTGACCCGGTG</td><td>>hg38_21_+_8433172_8446622_RNA45SN1_45S_50ntFlanks</td><td>12033</td><td>12055</td><td>23</td><td>ATCGTTTTTTCACTGACCCGGTG</td><td>349.8</td><td>-11.5</td><td>1.6</td><td>-7.3</td><td>3.3E-13</td><td>1.9E-11</td><td>BT20/MB468</td><td>MP</td></tr>
<tr><td>>hg38_21_+_8433172_8446622_RNA45SN1_45S_50ntFlanks@11200.11224.25|CGGCGGCGGCGGCGACTCTGGACGC</td><td>>hg38_21_+_8433172_8446622_RNA45SN1_45S_50ntFlanks</td><td>11200</td><td>11224</td><td>25</td><td>CGGCGGCGGCGGCGACTCTGGACGC</td><td>106.0</td><td>-10.6</td><td>1.5</td><td>-6.9</td><td>5.5E-12</td><td>1.9E-11</td><td>MB231/MB468</td><td>MP</td></tr>
<tr><td>>hg38_21_+_8433172_8446622_RNA45SN1_45S_50ntFlanks@12628.12647.20|TGGGGCGAAGCTACCATCTG</td><td>>hg38_21_+_8433172_8446622_RNA45SN1_45S_50ntFlanks</td><td>12628</td><td>12647</td><td>20</td><td>TGGGGCGAAGCTACCATCTG</td><td>124.2</td><td>-10.8</td><td>1.6</td><td>-6.9</td><td>5.5E-12</td><td>1.9E-11</td><td>MB231/MB468</td><td>MP</td></tr>
<tr><td>>MI0000065|hsa-let-7d&WithFlank&9|+|94178828|94178926@14.34.21(+1C)&[MIMAT0000065&hsa-let-7d-5p&offsets|0|-1(+1C);m-90&9|+|94178841|94178862&offsets|0|-1(+1C)]|AGAGGTAGTAGGTTGCATAGTC</td><td>>MI0000065|hsa-let-7d&WithFlank&9|+|94178828|94178926</td><td>14</td><td>34</td><td>21(+1C)</td><td>AGAGGTAGTAGGTTGCATAGTC</td><td>772.0</td><td>4.6</td><td>0.7</td><td>6.9</td><td>6.2E-12</td><td>1.9E-11</td><td>MB231/MB468</td><td>NUC</td></tr>
<tr><td>>MI0000088|hsa-mir-30a&WithFlank&6|-|71403545|71403627@53.71.19&[MIMAT0000088&hsa-miR-30a-3p&offsets|0|-3;m-38&6|-|71403554|71403575&offsets|0|-3]|CTTTCAGTCGGATGTTTGC</td><td>>MI0000088|hsa-mir-30a&WithFlank&6|-|71403545|71403627</td><td>53</td><td>71</td><td>19</td><td>CTTTCAGTCGGATGTTTGC</td><td>3156.4</td><td>4.8</td><td>0.7</td><td>6.9</td><td>6.2E-12</td><td>1.9E-11</td><td>MB231/MB468</td><td>NUC</td></tr>
<tr><td>>hg38_21_+_8433172_8446622_RNA45SN1_45S_50ntFlanks@10866.10894.29|CTAAGGGCTGGGTCGGTCGGGCTGGGGCG</td><td>>hg38_21_+_8433172_8446622_RNA45SN1_45S_50ntFlanks</td><td>10866</td><td>10894</td><td>29</td><td>CTAAGGGCTGGGTCGGTCGGGCTGGGGCG</td><td>1717.1</td><td>-10.2</td><td>1.5</td><td>-6.8</td><td>7.8E-12</td><td>1.9E-11</td><td>BT20/MB468</td><td>NUC</td></tr>
</table>

| | | | | | | | | | | | | | |
|---|---|---|---|---|---|---|---|---|---|---|---|---|---|
| >hg38_MT_+_598_1651_RNR1_12S_50ntFlanks@372.404.33\|AAAGCTAAAACTCACCTGAGTTGTAAAAAACTC | >hg38_MT_+_598_1651_RNR1_12S_50ntFlanks | 372 | 404 | 33 | AAAGCTAAAACTCACCTGAGTTGTAAAAAACTC | 680.1 | 5.2 | 0.8 | 6.9 | 4.5E-12 | 1.9E-11 | BT20/MB468 | MITO |
| >trna16_TyrGTA_14_-_21131351_21131444@59.76.18,trna17_TyrGTA_14_-_21128117_21128210@59.76.18,trna19_TyrGTA_14_-_21121258_21121351@59.76.18,trna2_TyrGTA_2_+_27273650_27273738@59.76.18,trna4_TyrGTA_8_+_67025602_67025694@59.76.18,trna5_TyrGTA_14_+_21151432_21151520@59.76.18,trna5_TyrGTA_8_+_67026223_67026311@59.76.18\|TTCCGGCTCGAAGGACCA | >trna16_TyrGTA_14_-_21131351_21131444 | 59 | 76 | 18 | TTCCGGCTCGAAGGACCA | 918.1 | 5.3 | 0.8 | 6.9 | 6.1E-12 | 1.9E-11 | MB231/MB468 | CYTO |
| >hg38_21_+_8433172_8446622_RNA45SN1_45S_50ntFlanks@4047.4067.21\|ATTCGAACGTCTGCCCTATCA | >hg38_21_+_8433172_8446622_RNA45SN1_45S_50ntFlanks | 4047 | 4067 | 21 | ATTCGAACGTCTGCCCTATCA | 483.7 | -11.0 | 1.6 | -6.9 | 4.5E-12 | 1.9E-11 | BT20/MB468 | MITO |
| >am_trna106_HisGTG_1_-_149155828_149155899@7.28.22,trna111_HisGTG_1_-_147774845_147774916@7.28.22,trna118_HisGTG_1_-_145396881_145396952@7.28.22,trna16_HisGTG_1_+_146544773_146544844@7.28.22,trna1_HisGTG_15_+_45493349_45493420@7.28.22,trna21_HisGTG_1_+_147753471_147753542@7.28.22,trna33_HisGTG_6_+_27125906_27125977@7.28.22,trna7_HisGTG_9_-_14433938_14434009@7.28.22,trna8_HisGTG_15_-_45492611_45492682@7.28.22,trna9_HisGTG_15_-_45490804_45490875@7.28.22\|ATCGTATAGTGGTTAGTACTCT | >am_trna106_HisGTG_1_-_149155828_149155899 | 7 | 28 | 22 | ATCGTATAGTGGTTAGTACTCT | 264.8 | -11.1 | 1.5 | -7.3 | 3.3E-13 | 1.9E-11 | BT20/MB468 | MP |
| >MI0000342\|hsa-mir-200b&WithFlank&1\|+\|1167098\|1167204@64.85.22(+1C)&[MIMAT0000318&hsa-miR-200b-3p&offsets\|+1\|+1(+1C);m-93&1\|+\|1167160\|1167182&offsets\|+1\|0(+1C)]\|AATACTGCCTGGTAATGATGACC | >MI0000342\|hsa-mir-200b&WithFlank&1\|+\|1167098\|1167204 | 64 | 85 | 22(+1C) | AATACTGCCTGGTAATGATGACC | 94.8 | -10.4 | 1.5 | -6.9 | 5.5E-12 | 1.9E-11 | MB231/MB468 | MP |
| >hg38_21_+_8433172_8446622_RNA45SN1_45S_50ntFlanks@8192.8214.23\|AGCCCGTGGACGGTGTGAGGCCG | >hg38_21_+_8433172_8446622_RNA45SN1_45S_50ntFlanks | 8192 | 8214 | 23 | AGCCCGTGGACGGTGTGAGGCCG | 159.7 | -11.2 | 1.6 | -6.9 | 5.5E-12 | 1.9E-11 | MB231/MB468 | MP |
| >hg38_21_+_8433172_8446622_RNA45SN1_45S_50ntFlanks@10792.10816.25\|CTCTGGCATGTTGGAACAATGTAGG | >hg38_21_+_8433172_8446622_RNA45SN1_45S_50ntFlanks | 10792 | 10816 | 25 | CTCTGGCATGTTGGAACAATGTAGG | 425.6 | -8.4 | 1.2 | -6.9 | 5.5E-12 | 1.9E-11 | MB231/MB468 | MP |
| >hg38_21_+_8433172_8446622_RNA45SN1_45S_50ntFlanks@4367.4407.41\|CAGTTAAAAAGCTCGTAGTTGGATCTTGGGAGCGGGCGGGC | >hg38_21_+_8433172_8446622_RNA45SN1_45S_50ntFlanks | 4367 | 4407 | 41 | CAGTTAAAAAGCTCGTAGTTGGATCTTGGGAGCGGGCGGGC | 197.9 | -10.6 | 1.5 | -6.9 | 6.3E-12 | 1.9E-11 | MB231/MB468 | NUC |
| >hg38_21_+_8433172_8446622_RNA45SN1_45S_50ntFlanks@9504.9531.28\|GTGAACTATGCCTGGGCAGGGCGAAGCC | >hg38_21_+_8433172_8446622_RNA45SN1_45S_50ntFlanks | 9504 | 9531 | 28 | GTGAACTATGCCTGGGCAGGGCGAAGCC | 504.9 | -7.0 | 1.0 | -6.9 | 7.3E-12 | 1.9E-11 | MB231/MB468 | MITO |
| >hg38_21_+_8433172_8446622_RNA45SN1_45S_50ntFlanks@12709.12739.31\|CCGCGGAGCCTCGGTTGGCCTCGGATAGCCG | >hg38_21_+_8433172_8446622_RNA45SN1_45S_50ntFlanks | 12709 | 12739 | 31 | CCGCGGAGCCTCGGTTGGCCTCGGATAGCCG | 86.8 | -10.3 | 1.5 | -6.9 | 5.5E-12 | 1.9E-11 | MB231/MB468 | MP |
| >hg38_21_+_8433172_8446622_RNA45SN1_45S_50ntFlanks@10234.10268.35\|CGGACGCTACGCCGCGACGAGTAGGAGGGCCGCTG | >hg38_21_+_8433172_8446622_RNA45SN1_45S_50ntFlanks | 10234 | 10268 | 35 | CGGACGCTACGCCGCGACGAGTAGGAGGGCCGCTG | 85.9 | -10.3 | 1.5 | -6.9 | 5.5E-12 | 1.9E-11 | MB231/MB468 | MP |
| >hg38_21_+_8433172_8446622_RNA45SN1_45S_50ntFlanks@4591.4613.23\|TTTTGTTGGTTTTCGGAACTGAG | >hg38_21_+_8433172_8446622_RNA45SN1_45S_50ntFlanks | 4591 | 4613 | 23 | TTTTGTTGGTTTTCGGAACTGAG | 117.0 | -10.7 | 1.6 | -6.9 | 5.5E-12 | 1.9E-11 | MB231/MB468 | MP |
| >hg38_21_+_8433172_8446622_RNA45SN1_45S_50ntFlanks@12934.12966.33\|CTAAACCATTCGTAGACGACCTGCTTCTGGGTC | >hg38_21_+_8433172_8446622_RNA45SN1_45S_50ntFlanks | 12934 | 12966 | 33 | CTAAACCATTCGTAGACGACCTGCTTCTGGGTC | 371.9 | -8.3 | 1.2 | -6.9 | 7.4E-12 | 1.9E-11 | MB231/MB468 | MITO |
| >hg38_21_+_8433172_8446622_RNA45SN1_45S_50ntFlanks@10366.10384.19\|CCGAAGTGGAGAAGGGTTC | >hg38_21_+_8433172_8446622_RNA45SN1_45S_50ntFlanks | 10366 | 10384 | 19 | CCGAAGTGGAGAAGGGTTC | 125.9 | -10.8 | 1.6 | -6.9 | 5.5E-12 | 1.9E-11 | MB231/MB468 | MP |
| >hg38_21_+_8433172_8446622_RNA45SN1_45S_50ntFlanks@9720.9748.29\|ATTAGAGGTCTTGGGGCCGAAACGATCTC | >hg38_21_+_8433172_8446622_RNA45SN1_45S_50ntFlanks | 9720 | 9748 | 29 | ATTAGAGGTCTTGGGGCCGAAACGATCTC | 120.8 | -10.8 | 1.6 | -6.9 | 5.6E-12 | 1.9E-11 | MB231/MB468 | MP |
| >hg38_21_+_8433172_8446622_RNA45SN1_45S_50ntFlanks@3852.3881.30\|TAACTGTGGTAATTCTAGAGCTAATACATG | >hg38_21_+_8433172_8446622_RNA45SN1_45S_50ntFlanks | 3852 | 3881 | 30 | TAACTGTGGTAATTCTAGAGCTAATACATG | 446.2 | -8.6 | 1.3 | -6.8 | 7.4E-12 | 1.9E-11 | MB231/MB468 | MITO |
| >hg38_21_+_8433172_8446622_RNA45SN1_45S_50ntFlanks@12539.12573.35\|ATGATGTGTTGTTGCCATGGTAATCCTGCTCAGTA | >hg38_21_+_8433172_8446622_RNA45SN1_45S_50ntFlanks | 12539 | 12573 | 35 | ATGATGTGTTGTTGCCATGGTAATCCTGCTCAGTA | 725.0 | -9.7 | 1.4 | -6.8 | 7.9E-12 | 1.9E-11 | BT20/MB468 | NUC |
| >hg38_21_+_8433172_8446622_RNA45SN1_45S_50ntFlanks@4919.4938.20\|CCACCAGGAGTGGAGCCTGC | >hg38_21_+_8433172_8446622_RNA45SN1_45S_50ntFlanks | 4919 | 4938 | 20 | CCACCAGGAGTGGAGCCTGC | 561.2 | -12.9 | 1.9 | -6.8 | 7.4E-12 | 1.9E-11 | MB231/MB468 | MITO |
| >MI0000298\|hsa-mir-221&WithFlank&X\|-\|45746151\|45746272@71.93.23(+1U)&[MIMAT0000278&hsa-miR-221-3p&offsets\|0\|0(+1U);m-91&X\|-\|45746181\|45746202&offsets\|0\|+1(+1U)]\|AGCTACATTGTCTGCTGGGTTTCT | >MI0000298\|hsa-mir-221&WithFlank&X\|-\|45746151\|45746272 | 71 | 93 | 23(+1U) | AGCTACATTGTCTGCTGGGTTTCT | 10416.3 | 6.0 | 0.9 | 6.8 | 7.4E-12 | 1.9E-11 | MB231/MB468 | MITO |
| >hg38_21_+_8433172_8446622_RNA45SN1_45S_50ntFlanks@12934.12969.36\|CTAAACCATTCGTAGACGACCTGCTTCTGGGTCGGG | >hg38_21_+_8433172_8446622_RNA45SN1_45S_50ntFlanks | 12934 | 12969 | 36 | CTAAACCATTCGTAGACGACCTGCTTCTGGGTCGGG | 399.7 | -8.4 | 1.2 | -6.8 | 7.4E-12 | 1.9E-11 | MB231/MB468 | MITO |
| >hg38_21_+_8433172_8446622_RNA45SN1_45S_50ntFlanks@6646.6673.28\|TCGTACGACTCTTAGCGGTGGATCACTC | >hg38_21_+_8433172_8446622_RNA45SN1_45S_50ntFlanks | 6646 | 6673 | 28 | TCGTACGACTCTTAGCGGTGGATCACTC | 1653.2 | 5.6 | 0.8 | 6.9 | 5.8E-12 | 1.9E-11 | BT20/MB468 | CYTO |
| >MI0000088\|hsa-mir-30a&WithFlank&6\|-\|71403545\|71403627@53.74.22(+1A)&[MIMAT0000088&hsa-miR-30a-3p&offsets\|0\|0(+1A);m-38&6\|-\|71403554\|71403575&offsets\|0\|0(+1A)]\|CTTTCAGTCGGATGTTTGCAGCA | >MI0000088\|hsa-mir-30a&WithFlank&6\|-\|71403545\|71403627 | 53 | 74 | 22(+1A) | CTTTCAGTCGGATGTTTGCAGCA | 363.1 | 4.7 | 0.7 | 6.9 | 6.3E-12 | 1.9E-11 | MB231/MB468 | NUC |
| >hg38_21_+_8433172_8446622_RNA45SN1_45S_50ntFlanks@10832.10852.21\|AGCCGGATCCGTAACTTCGGG | >hg38_21_+_8433172_8446622_RNA45SN1_45S_50ntFlanks | 10832 | 10852 | 21 | AGCCGGATCCGTAACTTCGGG | 105.2 | -10.6 | 1.5 | -6.9 | 5.6E-12 | 1.9E-11 | MB231/MB468 | MP |
| >MI0000266\|hsa-mir-10a&WithFlank&17\|-\|48579832\|48579953@29.50.22(+1A)&[MIMAT0000253&hsa-miR-10a-5p&offsets\|+1\|0(+1A);m-18&17\|-\|48579905\|48579926&offsets\|+1\|+1(+1A)]\|ACCCTGTAGATCCGAATTTGTGA | >MI0000266\|hsa-mir-10a&WithFlank&17\|-\|48579832\|48579953 | 29 | 50 | 22(+1A) | ACCCTGTAGATCCGAATTTGTGA | 260.7 | 11.8 | 1.6 | 7.3 | 3.4E-13 | 1.9E-11 | BT20/MB468 | MP |
| >hg38_21_+_8433172_8446622_RNA45SN1_45S_50ntFlanks@8109.8143.35\|GCGGGGCGCGGGACATGTGGCGTACGGAAGACCCG | >hg38_21_+_8433172_8446622_RNA45SN1_45S_50ntFlanks | 8109 | 8143 | 35 | GCGGGGCGCGGGACATGTGGCGTACGGAAGACCCG | 309.0 | -11.4 | 1.6 | -7.3 | 3.4E-13 | 1.9E-11 | BT20/MB468 | MP |
| >hg38_21_+_8433172_8446622_RNA45SN1_45S_50ntFlanks@10421.10453.33\|TGAGAGATGGGCGAGCGCCGTTCCGAAGGGACG | >hg38_21_+_8433172_8446622_RNA45SN1_45S_50ntFlanks | 10421 | 10453 | 33 | TGAGAGATGGGCGAGCGCCGTTCCGAAGGGACG | 104.6 | -10.6 | 1.5 | -6.9 | 5.6E-12 | 1.9E-11 | MB231/MB468 | MP |
| >hg38_21_+_8433172_8446622_RNA45SN1_45S_50ntFlanks@10794.10829.36\|CTGGCATGTTGGAACAATGTAGGTAAGGGAAGTCGG | >hg38_21_+_8433172_8446622_RNA45SN1_45S_50ntFlanks | 10794 | 10829 | 36 | CTGGCATGTTGGAACAATGTAGGTAAGGGAAGTCGG | 122.2 | -10.8 | 1.6 | -6.9 | 5.6E-12 | 1.9E-11 | MB231/MB468 | MP |
| >hg38_21_+_8433172_8446622_RNA45SN1_45S_50ntFlanks@5484.5506.23\|GAGCGCTGAGAAGACGGTCGAAC | >hg38_21_+_8433172_8446622_RNA45SN1_45S_50ntFlanks | 5484 | 5506 | 23 | GAGCGCTGAGAAGACGGTCGAAC | 129.4 | -10.9 | 1.6 | -6.9 | 5.6E-12 | 1.9E-11 | MB231/MB468 | MP |
| >hg38_21_+_8433172_8446622_RNA45SN1_45S_50ntFlanks@11792.11820.29\|ATGAACGAGATTCCCACTGTCCCTACCTA | >hg38_21_+_8433172_8446622_RNA45SN1_45S_50ntFlanks | 11792 | 11820 | 29 | ATGAACGAGATTCCCACTGTCCCTACCTA | 192.8 | -9.7 | 1.4 | -6.9 | 4.6E-12 | 1.9E-11 | BT20/MB468 | MITO |
| >hg38_21_+_8433172_8446622_RNA45SN1_45S_50ntFlanks@12083.12110.28\|CTCGCTTCTGGCGCCAAGCGCCCGGCCG | >hg38_21_+_8433172_8446622_RNA45SN1_45S_50ntFlanks | 12083 | 12110 | 28 | CTCGCTTCTGGCGCCAAGCGCCCGGCCG | 164.7 | -9.5 | 1.4 | -6.9 | 4.6E-12 | 1.9E-11 | BT20/MB468 | MITO |
| >hg38_21_+_8433172_8446622_RNA45SN1_45S_50ntFlanks@10403.10426.24\|ACATGGGTCAGTCGGTCCTGAGAG | >hg38_21_+_8433172_8446622_RNA45SN1_45S_50ntFlanks | 10403 | 10426 | 24 | ACATGGGTCAGTCGGTCCTGAGAG | 91.6 | -10.4 | 1.5 | -6.9 | 5.6E-12 | 1.9E-11 | MB231/MB468 | MP |
| >hg38_21_+_8433172_8446622_RNA45SN1_45S_50ntFlanks@12044.12072.29\|ACTGACCCGGTGAGGCGGGGGGGCGAGCC | >hg38_21_+_8433172_8446622_RNA45SN1_45S_50ntFlanks | 12044 | 12072 | 29 | ACTGACCCGGTGAGGCGGGGGGGCGAGCC | 1396.0 | -8.7 | 1.3 | -6.9 | 4.6E-12 | 1.9E-11 | BT20/MB468 | MITO |
| >hg38_21_+_8433172_8446622_RNA45SN1_45S_50ntFlanks@11821.11854.34\|CTATCCAGCGAAACCACAGCCAAGGGAACGGGCT | >hg38_21_+_8433172_8446622_RNA45SN1_45S_50ntFlanks | 11821 | 11854 | 34 | CTATCCAGCGAAACCACAGCCAAGGGAACGGGCT | 100.8 | -10.5 | 1.5 | -6.9 | 5.7E-12 | 1.9E-11 | MB231/MB468 | MP |
| >am_trna128_GlyGCC_6_-_27870686_27870756@10.28.19,trna133_GlyCCC_1_-_16872434_16872504@10.28.19,trna17_GluTTC_2_-_204229131_204229201@10.28.19,trna18_GlyGCC_16_+_70822597_70822667@10.28.19,trna19_GlyGCC_16_+_70823410_70823480@10.28.19,trna19_GlyGCC_2_-_157257659_157257729@10.28.19,trna24_GlyGCC_16_-_70812942_70813012@10.28.19,trna25_GlyGCC_16_-_70812114_70812184@10.28.19,trna2_GlyGCC_21_-_18827107_18827177@10.28.19,trna35_GlyGCC_1_+_161413094_161413164@10.28.19,trna37_GlyGCC_1_+_161420467_161420537@10.28.19,trna39_GlyGCC_1_+_161427898_161427968@10.28.19,trna41_GlyGCC_1_+_161435258_161435328@10.28.19,trna4_GlyCCC_1_+_17188416_17188486@10.28.19,trna5_GlyGCC_17_+_8029064_8029134@10.28.19,trna68_GlyGCC_1_-_161493637_161493707@10.28.19,trna91_GlyCCC_1_-_149680210_149680280@10.28.19,trna99_ValCAC_1_-_149294666_149294736@10.28.19\|GTTCAGTGGTAGAATTCTC | >am_trna128_GlyGCC_6_-_27870686_27870756 | 10 | 28 | 19 | GTTCAGTGGTAGAATTCTC | 238.7 | -11.8 | 1.7 | -6.9 | 5.7E-12 | 1.9E-11 | MB231/MB468 | MP |
| >hg38_21_+_8433172_8446622_RNA45SN1_45S_50ntFlanks@11589.11611.23\|CGGACCAGGGGAATCCGACTGTT | >hg38_21_+_8433172_8446622_RNA45SN1_45S_50ntFlanks | 11589 | 11611 | 23 | CGGACCAGGGGAATCCGACTGTT | 951.6 | -4.1 | 0.6 | -6.8 | 7.6E-12 | 1.9E-11 | MB231/MB468 | MITO |
| >trna111_HisGTG_1_-_147774845_147774916@-1T.29.30,trna118_HisGTG_1_-_145396881_145396952@-1T.29.30,trna16_HisGTG_1_+_146544773_146544844@-1T.29.30,trna1_HisGTG_15_+_45493349_45493420@-1T.29.30,trna21_HisGTG_1_+_147753471_147753542@-1T.29.30,trna33_HisGTG_6_+_27125906_27125977@-1T.29.30,trna7_HisGTG_9_-_14433938_14434009@-1T.29.30,trna8_HisGTG_15_-_45492611_45492682@-1T.29.30,trna9_HisGTG_15_-_45490804_45490875@-1T.29.30\|TGCCGTGATCGTATAGTGGTTAGTACTCTG | >trna111_HisGTG_1_-_147774845_147774916 | -1T | 29 | 30 | TGCCGTGATCGTATAGTGGTTAGTACTCTG | 9280.1 | -6.3 | 0.9 | -6.9 | 5.9E-12 | 1.9E-11 | BT20/MB468 | CYTO |
| >hg38_21_+_8433172_8446622_RNA45SN1_45S_50ntFlanks@11848.11868.21\|ACGGGCTTGGCGGAATCAGCG | >hg38_21_+_8433172_8446622_RNA45SN1_45S_50ntFlanks | 11848 | 11868 | 21 | ACGGGCTTGGCGGAATCAGCG | 127.8 | -10.9 | 1.6 | -6.9 | 5.7E-12 | 1.9E-11 | MB231/MB468 | MP |
| >hg38_21_+_8433172_8446622_RNA45SN1_45S_50ntFlanks@6744.6771.28\|GACACTTCGAACGCACTTGCGGCCCCGG | >hg38_21_+_8433172_8446622_RNA45SN1_45S_50ntFlanks | 6744 | 6771 | 28 | GACACTTCGAACGCACTTGCGGCCCCGG | 848.6 | -6.2 | 0.9 | -6.9 | 6.3E-12 | 1.9E-11 | MB231/MB468 | CYTO |

<table>
<tr><td>>hg38_21_+_8433172_8446622_RNA45SN1_45S_50ntFlanks@4975.4994.20|CCCGGACACGGACAGGATTG</td><td>>hg38_21_+_8433172_8446622_RNA45SN1_45S_50ntFlanks</td><td>4975</td><td>4994</td><td>20</td><td>CCCGGACACGGACAGGATTG</td><td>1261.3</td><td>-10.4</td><td>1.5</td><td>-6.8</td><td>7.6E-12</td><td>1.9E-11</td><td>MB231/MB468</td><td>MITO</td></tr>
<tr><td>>hg38_21_+_8433172_8446622_RNA45SN1_45S_50ntFlanks@6738.6756.19|ATCATCGACACTTCGAACG</td><td>>hg38_21_+_8433172_8446622_RNA45SN1_45S_50ntFlanks</td><td>6738</td><td>6756</td><td>19</td><td>ATCATCGACACTTCGAACG</td><td>176.9</td><td>-9.6</td><td>1.4</td><td>-6.9</td><td>4.7E-12</td><td>1.9E-11</td><td>BT20/MB468</td><td>MITO</td></tr>
<tr><td>>hg38_21_+_8433172_8446622_RNA45SN1_45S_50ntFlanks@9736.9765.30|CCGAAACGATCTCAACCTATTCTCAAACTT</td><td>>hg38_21_+_8433172_8446622_RNA45SN1_45S_50ntFlanks</td><td>9736</td><td>9765</td><td>30</td><td>CCGAAACGATCTCAACCTATTCTCAAACTT</td><td>2328.2</td><td>-5.9</td><td>0.9</td><td>-6.9</td><td>4.7E-12</td><td>1.9E-11</td><td>BT20/MB468</td><td>MITO</td></tr>
<tr><td>>MI0000342|hsa-mir-200b&WithFlank&1|+|1167098|1167204@64.85.22(+1U)&[MIMAT0000318&hsa-miR-200b-3p&offsets|+1|+1(+1U);m-93&1|+|1167160|1167182&offsets|+1|0(+1U)]|AATACTGCCTGGTAATGATGACT</td><td>>MI0000342|hsa-mir-200b&WithFlank&1|+|1167098|1167204</td><td>64</td><td>85</td><td>22(+1U)</td><td>AATACTGCCTGGTAATGATGACT</td><td>289.8</td><td>-11.3</td><td>1.5</td><td>-7.3</td><td>3.5E-13</td><td>1.9E-11</td><td>BT20/MB468</td><td>MP</td></tr>
<tr><td>>hg38_21_+_8433172_8446622_RNA45SN1_45S_50ntFlanks@9341.9367.27|CACGAAAGCCGCCGTGGCGCAATGAAG</td><td>>hg38_21_+_8433172_8446622_RNA45SN1_45S_50ntFlanks</td><td>9341</td><td>9367</td><td>27</td><td>CACGAAAGCCGCCGTGGCGCAATGAAG</td><td>102.5</td><td>-10.5</td><td>1.5</td><td>-6.9</td><td>5.7E-12</td><td>1.9E-11</td><td>MB231/MB468</td><td>MP</td></tr>
<tr><td>>hg38_21_+_8433172_8446622_RNA45SN1_45S_50ntFlanks@12076.12110.35|AGGGGCTCTCGCTTCTGGCGCCAAGCGCCCGGCCG</td><td>>hg38_21_+_8433172_8446622_RNA45SN1_45S_50ntFlanks</td><td>12076</td><td>12110</td><td>35</td><td>AGGGGCTCTCGCTTCTGGCGCCAAGCGCCCGGCCG</td><td>591.1</td><td>-6.8</td><td>1.0</td><td>-6.8</td><td>7.7E-12</td><td>1.9E-11</td><td>MB231/MB468</td><td>MITO</td></tr>
<tr><td>>hg38_21_+_8433172_8446622_RNA45SN1_45S_50ntFlanks@11844.11864.21|GGGAACGGGCTTGGCGGAATC</td><td>>hg38_21_+_8433172_8446622_RNA45SN1_45S_50ntFlanks</td><td>11844</td><td>11864</td><td>21</td><td>GGGAACGGGCTTGGCGGAATC</td><td>131.2</td><td>-10.9</td><td>1.6</td><td>-6.9</td><td>5.8E-12</td><td>1.9E-11</td><td>MB231/MB468</td><td>MP</td></tr>
<tr><td>>hg38_21_+_8433172_8446622_RNA45SN1_45S_50ntFlanks@11452.11471.20|CTCCTCGGCGCGCGGCGGCG</td><td>>hg38_21_+_8433172_8446622_RNA45SN1_45S_50ntFlanks</td><td>11452</td><td>11471</td><td>20</td><td>CTCCTCGGCGCGCGGCGGCG</td><td>271.2</td><td>-5.3</td><td>0.8</td><td>-6.9</td><td>6.0E-12</td><td>1.9E-11</td><td>BT20/MB468</td><td>CYTO</td></tr>
<tr><td>>hg38_21_+_8433172_8446622_RNA45SN1_45S_50ntFlanks@10679.10711.33|GGTTCCGGCGGCGTCCGGTGAGCTCTCGCTGGC</td><td>>hg38_21_+_8433172_8446622_RNA45SN1_45S_50ntFlanks</td><td>10679</td><td>10711</td><td>33</td><td>GGTTCCGGCGGCGTCCGGTGAGCTCTCGCTGGC</td><td>18616.2</td><td>-10.3</td><td>1.5</td><td>-6.8</td><td>8.1E-12</td><td>1.9E-11</td><td>BT20/MB468</td><td>NUC</td></tr>
<tr><td>>hg38_21_+_8433172_8446622_RNA45SN1_45S_50ntFlanks@4995.5024.30|ACAGATTGATAGCTCTTTCTCGATTCCGTG</td><td>>hg38_21_+_8433172_8446622_RNA45SN1_45S_50ntFlanks</td><td>4995</td><td>5024</td><td>30</td><td>ACAGATTGATAGCTCTTTCTCGATTCCGTG</td><td>2162.4</td><td>-9.8</td><td>1.4</td><td>-6.8</td><td>7.7E-12</td><td>2.0E-11</td><td>MB231/MB468</td><td>MITO</td></tr>
<tr><td>>hg38_21_+_8433172_8446622_RNA45SN1_45S_50ntFlanks@6651.6700.50|CGACTCTTAGCGGTGGATCACTCGGCTCGTGCGTCGATGAAGAACGCAGC</td><td>>hg38_21_+_8433172_8446622_RNA45SN1_45S_50ntFlanks</td><td>6651</td><td>6700</td><td>50</td><td>CGACTCTTAGCGGTGGATCACTCGGCTCGTGCGTCGATGAAGAACGCAGC</td><td>29245.0</td><td>7.3</td><td>1.1</td><td>6.9</td><td>4.7E-12</td><td>2.0E-11</td><td>BT20/MB468</td><td>MITO</td></tr>
<tr><td>>MI0000342|hsa-mir-200b&WithFlank&1|+|1167098|1167204@63.83.21(+1U)&[MIMAT0000318&hsa-miR-200b-3p&offsets|0|-1(+1U);m-93&1|+|1167160|1167182&offsets|0|-2(+1U)]|TAATACTGCCTGGTAATGATGT</td><td>>MI0000342|hsa-mir-200b&WithFlank&1|+|1167098|1167204</td><td>63</td><td>83</td><td>21(+1U)</td><td>TAATACTGCCTGGTAATGATGT</td><td>102.6</td><td>-10.5</td><td>1.5</td><td>-6.9</td><td>5.8E-12</td><td>2.0E-11</td><td>MB231/MB468</td><td>MP</td></tr>
<tr><td>>hg38_21_+_8433172_8446622_RNA45SN1_45S_50ntFlanks@4006.4039.34|ACCTCGGGCCGATCGCACGCCCCCCGTGGCGGCG</td><td>>hg38_21_+_8433172_8446622_RNA45SN1_45S_50ntFlanks</td><td>4006</td><td>4039</td><td>34</td><td>ACCTCGGGCCGATCGCACGCCCCCCGTGGCGGCG</td><td>590.6</td><td>-9.2</td><td>1.3</td><td>-6.8</td><td>8.2E-12</td><td>2.0E-11</td><td>BT20/MB468</td><td>NUC</td></tr>
<tr><td>>hg38_21_+_8433172_8446622_RNA45SN1_45S_50ntFlanks@6649.6683.35|TACGACTCTTAGCGGTGGATCACTCGGCTCGTGCG</td><td>>hg38_21_+_8433172_8446622_RNA45SN1_45S_50ntFlanks</td><td>6649</td><td>6683</td><td>35</td><td>TACGACTCTTAGCGGTGGATCACTCGGCTCGTGCG</td><td>6393.0</td><td>6.0</td><td>0.9</td><td>6.9</td><td>6.6E-12</td><td>2.0E-11</td><td>MB231/MB468</td><td>NUC</td></tr>
<tr><td>>trna128_GlyGCC_6_-_27870686_27870756@16.52.37,trna19_GlyGCC_16_+_70823410_70823480@16.52.37,trna19_GlyGCC_2_-_157257659_157257729@16.52.37,trna24_GlyGCC_16_-_70812942_70813012@16.52.37,trna25_GlyGCC_16_-_70812114_70812184@16.52.37,trna2_GlyGCC_21_-_18827107_18827177@16.52.37,trna35_GlyGCC_1_+_161413094_161413164@16.52.37,trna37_GlyGCC_1_+_161420467_161420537@16.52.37,trna39_GlyGCC_1_+_161427898_161427968@16.52.37,trna41_GlyGCC_1_+_161435258_161435328@16.52.37,trna5_GlyGCC_17_+_8029064_8029134@16.52.37,trna68_GlyGCC_1_-_161493637_161493707@16.52.37|TGGTAGAATTCTCGCCTGCCACGCGGGAGGCCCGGGT</td><td>>trna128_GlyGCC_6_-_27870686_27870756</td><td>16</td><td>52</td><td>37</td><td>TGGTAGAATTCTCGCCTGCCACGCGGGAGGCCCGGGT</td><td>981.4</td><td>7.4</td><td>1.1</td><td>6.9</td><td>6.1E-12</td><td>2.0E-11</td><td>BT20/MB468</td><td>CYTO</td></tr>
<tr><td>>hg38_21_+_8433172_8446622_RNA45SN1_45S_50ntFlanks@4725.4748.24|TTAATCAAGAACGAAAGTCGGAGG</td><td>>hg38_21_+_8433172_8446622_RNA45SN1_45S_50ntFlanks</td><td>4725</td><td>4748</td><td>24</td><td>TTAATCAAGAACGAAAGTCGGAGG</td><td>102.1</td><td>-10.5</td><td>1.5</td><td>-6.9</td><td>5.8E-12</td><td>2.0E-11</td><td>MB231/MB468</td><td>MP</td></tr>
<tr><td>>hg38_21_+_8433172_8446622_RNA45SN1_45S_50ntFlanks@10805.10827.23|GAACAATGTAGGTAAGGGAAGTC</td><td>>hg38_21_+_8433172_8446622_RNA45SN1_45S_50ntFlanks</td><td>10805</td><td>10827</td><td>23</td><td>GAACAATGTAGGTAAGGGAAGTC</td><td>105.2</td><td>-10.6</td><td>1.5</td><td>-6.9</td><td>5.8E-12</td><td>2.0E-11</td><td>MB231/MB468</td><td>MP</td></tr>
<tr><td>>hg38_21_+_8433172_8446622_RNA45SN1_45S_50ntFlanks@11583.11603.21|CTGGTGCGGACCAGGGGAATC</td><td>>hg38_21_+_8433172_8446622_RNA45SN1_45S_50ntFlanks</td><td>11583</td><td>11603</td><td>21</td><td>CTGGTGCGGACCAGGGGAATC</td><td>4622.9</td><td>-6.0</td><td>0.8</td><td>-7.3</td><td>3.6E-13</td><td>2.0E-11</td><td>BT20/MB468</td><td>MP</td></tr>
<tr><td>>hg38_21_+_8433172_8446622_RNA45SN1_45S_50ntFlanks@10362.10388.27|AAGGCCGAAGTGGAGAAGGGTTCCATG</td><td>>hg38_21_+_8433172_8446622_RNA45SN1_45S_50ntFlanks</td><td>10362</td><td>10388</td><td>27</td><td>AAGGCCGAAGTGGAGAAGGGTTCCATG</td><td>2686.0</td><td>-6.7</td><td>0.9</td><td>-7.3</td><td>3.6E-13</td><td>2.0E-11</td><td>BT20/MB468</td><td>MP</td></tr>
<tr><td>>hg38_21_+_8433172_8446622_RNA45SN1_45S_50ntFlanks@8168.8190.23|CCCAAGTCCTTCTGATCGAGGCC</td><td>>hg38_21_+_8433172_8446622_RNA45SN1_45S_50ntFlanks</td><td>8168</td><td>8190</td><td>23</td><td>CCCAAGTCCTTCTGATCGAGGCC</td><td>278.7</td><td>-11.2</td><td>1.5</td><td>-7.3</td><td>3.6E-13</td><td>2.0E-11</td><td>BT20/MB468</td><td>MP</td></tr>
<tr><td>>hg38_21_+_8433172_8446622_RNA45SN1_45S_50ntFlanks@9545.9563.19|GTGGAGGTCCGTAGCGGTC</td><td>>hg38_21_+_8433172_8446622_RNA45SN1_45S_50ntFlanks</td><td>9545</td><td>9563</td><td>19</td><td>GTGGAGGTCCGTAGCGGTC</td><td>1483.8</td><td>-9.1</td><td>1.3</td><td>-6.8</td><td>7.8E-12</td><td>2.0E-11</td><td>MB231/MB468</td><td>MITO</td></tr>
<tr><td>>hg38_21_+_8433172_8446622_RNA45SN1_45S_50ntFlanks@10234.10267.34|CGGACGCTACGCCGCGACGAGTAGGAGGGCCGCT</td><td>>hg38_21_+_8433172_8446622_RNA45SN1_45S_50ntFlanks</td><td>10234</td><td>10267</td><td>34</td><td>CGGACGCTACGCCGCGACGAGTAGGAGGGCCGCT</td><td>92.9</td><td>-10.4</td><td>1.5</td><td>-6.9</td><td>5.8E-12</td><td>2.0E-11</td><td>MB231/MB468</td><td>MP</td></tr>
<tr><td>>hg38_21_+_8433172_8446622_RNA45SN1_45S_50ntFlanks@12408.12426.19|TTTGATCCTTCGATGTCGG</td><td>>hg38_21_+_8433172_8446622_RNA45SN1_45S_50ntFlanks</td><td>12408</td><td>12426</td><td>19</td><td>TTTGATCCTTCGATGTCGG</td><td>125.9</td><td>-10.8</td><td>1.6</td><td>-6.9</td><td>5.8E-12</td><td>2.0E-11</td><td>MB231/MB468</td><td>MP</td></tr>
<tr><td>>hg38_21_+_8433172_8446622_RNA45SN1_45S_50ntFlanks@10414.10443.30|TCGGTCCTGAGAGATGGGCGAGCGCCGTTC</td><td>>hg38_21_+_8433172_8446622_RNA45SN1_45S_50ntFlanks</td><td>10414</td><td>10443</td><td>30</td><td>TCGGTCCTGAGAGATGGGCGAGCGCCGTTC</td><td>129.4</td><td>-10.9</td><td>1.6</td><td>-6.9</td><td>5.9E-12</td><td>2.0E-11</td><td>MB231/MB468</td><td>MP</td></tr>
<tr><td>>hg38_21_+_8433172_8446622_RNA45SN1_45S_50ntFlanks@4050.4081.32|CGAACGTCTGCCCTATCAACTTTCGATGGTAG</td><td>>hg38_21_+_8433172_8446622_RNA45SN1_45S_50ntFlanks</td><td>4050</td><td>4081</td><td>32</td><td>CGAACGTCTGCCCTATCAACTTTCGATGGTAG</td><td>180.2</td><td>-11.4</td><td>1.7</td><td>-6.9</td><td>5.9E-12</td><td>2.0E-11</td><td>MB231/MB468</td><td>MP</td></tr>
<tr><td>>hg38_21_+_8433172_8446622_RNA45SN1_45S_50ntFlanks@9339.9378.40|CGCACGAAAGCCGCCGTGGCGCAATGAAGGTGAAGGCCGG</td><td>>hg38_21_+_8433172_8446622_RNA45SN1_45S_50ntFlanks</td><td>9339</td><td>9378</td><td>40</td><td>CGCACGAAAGCCGCCGTGGCGCAATGAAGGTGAAGGCCGG</td><td>140.9</td><td>-11.0</td><td>1.6</td><td>-6.9</td><td>5.9E-12</td><td>2.0E-11</td><td>MB231/MB468</td><td>MP</td></tr>
<tr><td>>hg38_21_+_8433172_8446622_RNA45SN1_45S_50ntFlanks@10770.10797.28|GCAGGTCTCCAAGGTGAACAGCCTCTGG</td><td>>hg38_21_+_8433172_8446622_RNA45SN1_45S_50ntFlanks</td><td>10770</td><td>10797</td><td>28</td><td>GCAGGTCTCCAAGGTGAACAGCCTCTGG</td><td>89.0</td><td>-10.3</td><td>1.5</td><td>-6.9</td><td>5.9E-12</td><td>2.0E-11</td><td>MB231/MB468</td><td>MP</td></tr>
<tr><td>>hg38_21_+_8433172_8446622_RNA45SN1_45S_50ntFlanks@3885.3915.31|ACGGGCGCTGACCCCCTTCGCGGGGGGGATG</td><td>>hg38_21_+_8433172_8446622_RNA45SN1_45S_50ntFlanks</td><td>3885</td><td>3915</td><td>31</td><td>ACGGGCGCTGACCCCCTTCGCGGGGGGGATG</td><td>656.5</td><td>-9.3</td><td>1.4</td><td>-6.8</td><td>8.3E-12</td><td>2.0E-11</td><td>BT20/MB468</td><td>NUC</td></tr>
<tr><td>>hg38_21_+_8433172_8446622_RNA45SN1_45S_50ntFlanks@12163.12183.21|ACTGGGGCGGTACACCTGTCA</td><td>>hg38_21_+_8433172_8446622_RNA45SN1_45S_50ntFlanks</td><td>12163</td><td>12183</td><td>21</td><td>ACTGGGGCGGTACACCTGTCA</td><td>835.3</td><td>-5.5</td><td>0.8</td><td>-7.3</td><td>3.7E-13</td><td>2.0E-11</td><td>BT20/MB468</td><td>MP</td></tr>
<tr><td>>hg38_21_+_8433172_8446622_RNA45SN1_45S_50ntFlanks@12367.12384.18|AACTGGCTTGTGGCGGCC</td><td>>hg38_21_+_8433172_8446622_RNA45SN1_45S_50ntFlanks</td><td>12367</td><td>12384</td><td>18</td><td>AACTGGCTTGTGGCGGCC</td><td>588.0</td><td>-12.3</td><td>1.7</td><td>-7.3</td><td>3.7E-13</td><td>2.0E-11</td><td>BT20/MB468</td><td>MP</td></tr>
<tr><td>>hg38_21_+_8433172_8446622_RNA45SN1_45S_50ntFlanks@12420.12446.27|ATGTCGGCTCTTCCTATCATTGTGAAG</td><td>>hg38_21_+_8433172_8446622_RNA45SN1_45S_50ntFlanks</td><td>12420</td><td>12446</td><td>27</td><td>ATGTCGGCTCTTCCTATCATTGTGAAG</td><td>1747.5</td><td>-5.6</td><td>0.8</td><td>-7.3</td><td>3.7E-13</td><td>2.0E-11</td><td>BT20/MB468</td><td>MP</td></tr>
<tr><td>>hg38_21_+_8433172_8446622_RNA45SN1_45S_50ntFlanks@6772.6806.35|GTTCCTCCCGGGGCTACGCCTGTCTGAGCGTCGCT</td><td>>hg38_21_+_8433172_8446622_RNA45SN1_45S_50ntFlanks</td><td>6772</td><td>6806</td><td>35</td><td>GTTCCTCCCGGGGCTACGCCTGTCTGAGCGTCGCT</td><td>2337.5</td><td>-4.6</td><td>0.7</td><td>-6.9</td><td>5.9E-12</td><td>2.0E-11</td><td>MB231/MB468</td><td>MP</td></tr>
<tr><td>>hg38_MT_+_1621_3279_RNR2_16S_50ntFlanks@459.492.34|CTTGTAAATTTAACTGTTAGTCCAAAGAGGAACA</td><td>>hg38_MT_+_1621_3279_RNR2_16S_50ntFlanks</td><td>459</td><td>492</td><td>34</td><td>CTTGTAAATTTAACTGTTAGTCCAAAGAGGAACA</td><td>831.4</td><td>5.9</td><td>0.9</td><td>6.9</td><td>4.8E-12</td><td>2.0E-11</td><td>BT20/MB468</td><td>MITO</td></tr>
<tr><td>>hg38_21_+_8433172_8446622_RNA45SN1_45S_50ntFlanks@11208.11230.23|GCGGCGACTCTGGACGCGAGCCG</td><td>>hg38_21_+_8433172_8446622_RNA45SN1_45S_50ntFlanks</td><td>11208</td><td>11230</td><td>23</td><td>GCGGCGACTCTGGACGCGAGCCG</td><td>1581.3</td><td>-7.0</td><td>1.0</td><td>-6.9</td><td>6.5E-12</td><td>2.0E-11</td><td>MB231/MB468</td><td>CYTO</td></tr>
<tr><td>>hg38_21_+_8433172_8446622_RNA45SN1_45S_50ntFlanks@7974.8002.29|ACGCGACCTCAGATCAGACGTGGCGACCC</td><td>>hg38_21_+_8433172_8446622_RNA45SN1_45S_50ntFlanks</td><td>7974</td><td>8002</td><td>29</td><td>ACGCGACCTCAGATCAGACGTGGCGACCC</td><td>1947.3</td><td>6.2</td><td>0.9</td><td>6.9</td><td>4.8E-12</td><td>2.0E-11</td><td>BT20/MB468</td><td>MITO</td></tr>
<tr><td>>MI0000483|hsa-mir-186&WithFlank&1|-|71067625|71067722@21.41.21(+1C)&[MIMAT0000456&hsa-miR-186-5p&offsets|0|-1(+1C);m-84&1|-|71067681|71067702&offsets|0|-1(+1C)]|CAAAGAATTCTCCTTTTGGGCC</td><td>>MI0000483|hsa-mir-186&WithFlank&1|-|71067625|71067722</td><td>21</td><td>41</td><td>21(+1C)</td><td>CAAAGAATTCTCCTTTTGGGCC</td><td>389.6</td><td>7.6</td><td>1.1</td><td>6.9</td><td>6.1E-12</td><td>2.0E-11</td><td>BT20/MB468</td><td>CYTO</td></tr>
<tr><td>>hg38_21_+_8433172_8446622_RNA45SN1_45S_50ntFlanks@7975.8004.30|CGCGACCTCAGATCAGACGTGGCGACCCGC</td><td>>hg38_21_+_8433172_8446622_RNA45SN1_45S_50ntFlanks</td><td>7975</td><td>8004</td><td>30</td><td>CGCGACCTCAGATCAGACGTGGCGACCCGC</td><td>32485.1</td><td>7.7</td><td>1.1</td><td>6.9</td><td>6.1E-12</td><td>2.0E-11</td><td>BT20/MB468</td><td>CYTO</td></tr>
<tr><td>>hg38_21_+_8433172_8446622_RNA45SN1_45S_50ntFlanks@4687.4707.21|ACGGACCAGAGCGAAAGCATT</td><td>>hg38_21_+_8433172_8446622_RNA45SN1_45S_50ntFlanks</td><td>4687</td><td>4707</td><td>21</td><td>ACGGACCAGAGCGAAAGCATT</td><td>952.2</td><td>-9.4</td><td>1.4</td><td>-6.8</td><td>7.8E-12</td><td>2.0E-11</td><td>MB231/MB468</td><td>MITO</td></tr>
<tr><td>>hg38_21_+_8433172_8446622_RNA45SN1_45S_50ntFlanks@12456.12480.25|CCAAGCGTTGGATTGTTCACCCACT</td><td>>hg38_21_+_8433172_8446622_RNA45SN1_45S_50ntFlanks</td><td>12456</td><td>12480</td><td>25</td><td>CCAAGCGTTGGATTGTTCACCCACT</td><td>2683.3</td><td>-6.8</td><td>1.0</td><td>-6.8</td><td>8.3E-12</td><td>2.0E-11</td><td>BT20/MB468</td><td>NUC</td></tr>
<tr><td>>hg38_21_+_8433172_8446622_RNA45SN1_45S_50ntFlanks@11999.12039.41|CCCTGCGGGCCGCCGGTGAAATACCACTACTCTGATCGTTT</td><td>>hg38_21_+_8433172_8446622_RNA45SN1_45S_50ntFlanks</td><td>11999</td><td>12039</td><td>41</td><td>CCCTGCGGGCCGCCGGTGAAATACCACTACTCTGATCGTTT</td><td>281.9</td><td>-11.1</td><td>1.6</td><td>-6.9</td><td>6.7E-12</td><td>2.0E-11</td><td>MB231/MB468</td><td>NUC</td></tr>
<tr><td>>hg38_21_+_8433172_8446622_RNA45SN1_45S_50ntFlanks@12615.12654.40|GCTGAGGAGCCAATGGGGCGAAGCTACCATCTGTGGGATT</td><td>>hg38_21_+_8433172_8446622_RNA45SN1_45S_50ntFlanks</td><td>12615</td><td>12654</td><td>40</td><td>GCTGAGGAGCCAATGGGGCGAAGCTACCATCTGTGGGATT</td><td>22341.5</td><td>-12.7</td><td>1.9</td><td>-6.8</td><td>8.4E-12</td><td>2.0E-11</td><td>BT20/MB468</td><td>NUC</td></tr>
<tr><td>>hg38_21_+_8433172_8446622_RNA45SN1_45S_50ntFlanks@4603.4622.20|TCGGAACTGAGGCCATGATT</td><td>>hg38_21_+_8433172_8446622_RNA45SN1_45S_50ntFlanks</td><td>4603</td><td>4622</td><td>20</td><td>TCGGAACTGAGGCCATGATT</td><td>86.0</td><td>-10.3</td><td>1.5</td><td>-6.9</td><td>5.9E-12</td><td>2.0E-11</td><td>MB231/MB468</td><td>MP</td></tr>
<tr><td>>hg38_21_+_8433172_8446622_RNA45SN1_45S_50ntFlanks@4554.4574.21|CCGCAGCTAGGAATAATGGAA</td><td>>hg38_21_+_8433172_8446622_RNA45SN1_45S_50ntFlanks</td><td>4554</td><td>4574</td><td>21</td><td>CCGCAGCTAGGAATAATGGAA</td><td>133.6</td><td>-10.9</td><td>1.6</td><td>-6.9</td><td>5.9E-12</td><td>2.0E-11</td><td>MB231/MB468</td><td>MP</td></tr>
<tr><td>>hg38_21_+_8433172_8446622_RNA45SN1_45S_50ntFlanks@6733.6771.39|CATTGATCATCGACACTTCGAACGCACTTGCGGCCCCGG</td><td>>hg38_21_+_8433172_8446622_RNA45SN1_45S_50ntFlanks</td><td>6733</td><td>6771</td><td>39</td><td>CATTGATCATCGACACTTCGAACGCACTTGCGGCCCCGG</td><td>2948.5</td><td>-5.4</td><td>0.8</td><td>-6.9</td><td>6.7E-12</td><td>2.0E-11</td><td>MB231/MB468</td><td>NUC</td></tr>
<tr><td>>hg38_21_+_8433172_8446622_RNA45SN1_45S_50ntFlanks@9562.9595.34|TCCTGACGTGCAAATCGGTCGTCCGACCTGGGTA</td><td>>hg38_21_+_8433172_8446622_RNA45SN1_45S_50ntFlanks</td><td>9562</td><td>9595</td><td>34</td><td>TCCTGACGTGCAAATCGGTCGTCCGACCTGGGTA</td><td>119.1</td><td>-10.8</td><td>1.6</td><td>-6.9</td><td>6.0E-12</td><td>2.0E-11</td><td>MB231/MB468</td><td>MP</td></tr>
<tr><td>>hg38_21_+_8433172_8446622_RNA45SN1_45S_50ntFlanks@12893.12926.34|CCTCGCCCGTCACGCACCGCACGTTCGTGGGGAA</td><td>>hg38_21_+_8433172_8446622_RNA45SN1_45S_50ntFlanks</td><td>12893</td><td>12926</td><td>34</td><td>CCTCGCCCGTCACGCACCGCACGTTCGTGGGGAA</td><td>387.8</td><td>-11.7</td><td>1.6</td><td>-7.3</td><td>3.7E-13</td><td>2.0E-11</td><td>BT20/MB468</td><td>MP</td></tr>
<tr><td>>hg38_21_+_8433172_8446622_RNA45SN1_45S_50ntFlanks@10458.10486.29|ATGGCCTCCGTTGCCCTCGGCCGATCGAA</td><td>>hg38_21_+_8433172_8446622_RNA45SN1_45S_50ntFlanks</td><td>10458</td><td>10486</td><td>29</td><td>ATGGCCTCCGTTGCCCTCGGCCGATCGAA</td><td>514.9</td><td>-6.9</td><td>1.0</td><td>-6.8</td><td>7.9E-12</td><td>2.0E-11</td><td>MB231/MB468</td><td>MITO</td></tr>
<tr><td>>trna119_LysCTT_1_-_145395522_145395594@4.35.32,trna11_LysCTT_5_-_180648979_180649051@4.35.32,trna13_LysCTT_6_+_26556774_26556846@4.35.32,trna7_LysCTT_16_+_3225692_3225764@4.35.32,trna9_LysCTT_5_+_180634755_180634827@4.35.32|CGGCTAGCTCAGTCGGTAGAGCATGAGACTCT</td><td>>trna119_LysCTT_1_-_145395522_145395594</td><td>4</td><td>35</td><td>32</td><td>CGGCTAGCTCAGTCGGTAGAGCATGAGACTCT</td><td>634.6</td><td>5.0</td><td>0.7</td><td>6.9</td><td>6.0E-12</td><td>2.0E-11</td><td>MB231/MB468</td><td>MP</td></tr>
</table>

<table>
<tr><th></th><th></th><th></th><th></th><th></th><th></th><th></th><th></th><th></th><th></th><th></th><th></th><th></th><th></th></tr>
<tr><td>>am_MI0000809|hsa-mir-151a&WithFlank&8|-|140732558|140732659@53.73.21(+2U)&[MIMAT0000757&hsa-miR-151a-3p&offsets|0|0(+2U);m-30&8|-|140732587|140732607&offsets|0|0(+2U)]|CTAGACTGAAGCTCCTTGAGGTT</td><td>>am_MI0000809|hsa-mir-151a&WithFlank&8|-|140732558|140732659</td><td>53</td><td>73</td><td>21(+2U)</td><td>CTAGACTGAAGCTCCTTGAGGTT</td><td>626.8</td><td>8.6</td><td>1.2</td><td>6.9</td><td>6.6E-12</td><td>2.0E-11</td><td>MB231/MB468</td><td>CYTO</td></tr>
<tr><td>>hg38_21_+_8433172_8446622_RNA45SN1_45S_50ntFlanks@11768.11786.19|CTAATTAGTGACGCGCATG</td><td>>hg38_21_+_8433172_8446622_RNA45SN1_45S_50ntFlanks</td><td>11768</td><td>11786</td><td>19</td><td>CTAATTAGTGACGCGCATG</td><td>1680.9</td><td>-10.3</td><td>1.5</td><td>-6.8</td><td>8.4E-12</td><td>2.0E-11</td><td>BT20/MB468</td><td>NUC</td></tr>
<tr><td>>hg38_21_+_8433172_8446622_RNA45SN1_45S_50ntFlanks@5225.5244.20|CACGCGCGCTACACTGACTG</td><td>>hg38_21_+_8433172_8446622_RNA45SN1_45S_50ntFlanks</td><td>5225</td><td>5244</td><td>20</td><td>CACGCGCGCTACACTGACTG</td><td>244.5</td><td>-11.8</td><td>1.7</td><td>-6.9</td><td>6.0E-12</td><td>2.0E-11</td><td>MB231/MB468</td><td>MP</td></tr>
<tr><td>>MI0000272|hsa-mir-182&WithFlank&7|-|129770377|129770498@29.50.22(+2U)&[MIMAT0000259&hsa-miR-182-5p&offsets|0|-2(+2U);m-22&7|-|129770449|129770470&offsets|0|0(+2U)]|TTTGGCAATGGTAGAACTCACATT</td><td>>MI0000272|hsa-mir-182&WithFlank&7|-|129770377|129770498</td><td>29</td><td>50</td><td>22(+2U)</td><td>TTTGGCAATGGTAGAACTCACATT</td><td>562.1</td><td>10.7</td><td>1.6</td><td>6.9</td><td>6.2E-12</td><td>2.0E-11</td><td>BT20/MB468</td><td>CYTO</td></tr>
<tr><td>>hg38_21_+_8433172_8446622_RNA45SN1_45S_50ntFlanks@10768.10785.18|CAGCAGGTCTCCAAGGTG</td><td>>hg38_21_+_8433172_8446622_RNA45SN1_45S_50ntFlanks</td><td>10768</td><td>10785</td><td>18</td><td>CAGCAGGTCTCCAAGGTG</td><td>506.1</td><td>-3.9</td><td>0.6</td><td>-6.9</td><td>6.2E-12</td><td>2.0E-11</td><td>BT20/MB468</td><td>CYTO</td></tr>
<tr><td>>hg38_21_+_8433172_8446622_RNA45SN1_45S_50ntFlanks@9740.9767.28|AACGATCTCAACCTATTCTCAAACTTTA</td><td>>hg38_21_+_8433172_8446622_RNA45SN1_45S_50ntFlanks</td><td>9740</td><td>9767</td><td>28</td><td>AACGATCTCAACCTATTCTCAAACTTTA</td><td>88.5</td><td>-10.3</td><td>1.5</td><td>-6.9</td><td>6.0E-12</td><td>2.0E-11</td><td>MB231/MB468</td><td>MP</td></tr>
<tr><td>>hg38_21_+_8433172_8446622_RNA45SN1_45S_50ntFlanks@9413.9453.41|CTCCAGTCCGCCGAGGGCGCACCACCGGCCCGTCTCGCCCG</td><td>>hg38_21_+_8433172_8446622_RNA45SN1_45S_50ntFlanks</td><td>9413</td><td>9453</td><td>41</td><td>CTCCAGTCCGCCGAGGGCGCACCACCGGCCCGTCTCGCCCG</td><td>157.8</td><td>-9.4</td><td>1.4</td><td>-6.9</td><td>4.9E-12</td><td>2.0E-11</td><td>BT20/MB468</td><td>MITO</td></tr>
<tr><td>>MI0000082|hsa-mir-25&WithFlank&7|-|100093554|100093649@58.77.20&[MIMAT0000081&hsa-miR-25-3p&offsets|0|-2;m-23&7|-|100093571|100093592&offsets|0|-2]|CATTGCACTTGTCTCGGTCT</td><td>>MI0000082|hsa-mir-25&WithFlank&7|-|100093554|100093649</td><td>58</td><td>77</td><td>20</td><td>CATTGCACTTGTCTCGGTCT</td><td>92.9</td><td>2.0</td><td>0.3</td><td>6.8</td><td>8.5E-12</td><td>2.0E-11</td><td>BT20/MB468</td><td>NUC</td></tr>
<tr><td>>hg38_21_+_8433172_8446622_RNA45SN1_45S_50ntFlanks@11881.11903.23|CCTGTTGAGCTTGACTCTAGTCT</td><td>>hg38_21_+_8433172_8446622_RNA45SN1_45S_50ntFlanks</td><td>11881</td><td>11903</td><td>23</td><td>CCTGTTGAGCTTGACTCTAGTCT</td><td>444.7</td><td>-9.0</td><td>1.3</td><td>-6.8</td><td>8.5E-12</td><td>2.0E-11</td><td>BT20/MB468</td><td>NUC</td></tr>
<tr><td>>hg38_21_+_8433172_8446622_RNA45SN1_45S_50ntFlanks@12944.12964.21|CGTAGACGACCTGCTTCTGGG</td><td>>hg38_21_+_8433172_8446622_RNA45SN1_45S_50ntFlanks</td><td>12944</td><td>12964</td><td>21</td><td>CGTAGACGACCTGCTTCTGGG</td><td>229.4</td><td>-5.0</td><td>0.7</td><td>-6.9</td><td>6.6E-12</td><td>2.0E-11</td><td>MB231/MB468</td><td>CYTO</td></tr>
<tr><td>>hg38_21_+_8433172_8446622_RNA45SN1_45S_50ntFlanks@12686.12704.19|CCAGGCGGAACGATACGGC</td><td>>hg38_21_+_8433172_8446622_RNA45SN1_45S_50ntFlanks</td><td>12686</td><td>12704</td><td>19</td><td>CCAGGCGGAACGATACGGC</td><td>87.9</td><td>-10.3</td><td>1.5</td><td>-6.9</td><td>6.0E-12</td><td>2.0E-11</td><td>MB231/MB468</td><td>MP</td></tr>
<tr><td>>hg38_21_+_8433172_8446622_RNA45SN1_45S_50ntFlanks@4753.4782.30|AAGACGATCAGATACCGTCGTAGTTCCGAC</td><td>>hg38_21_+_8433172_8446622_RNA45SN1_45S_50ntFlanks</td><td>4753</td><td>4782</td><td>30</td><td>AAGACGATCAGATACCGTCGTAGTTCCGAC</td><td>1875.9</td><td>-10.1</td><td>1.5</td><td>-6.8</td><td>8.0E-12</td><td>2.0E-11</td><td>MB231/MB468</td><td>MITO</td></tr>
<tr><td>>hg38_21_+_8433172_8446622_RNA45SN1_45S_50ntFlanks@6661.6695.35|CGGTGGATCACTCGGCTCGTGCGTCGATGAAGAAC</td><td>>hg38_21_+_8433172_8446622_RNA45SN1_45S_50ntFlanks</td><td>6661</td><td>6695</td><td>35</td><td>CGGTGGATCACTCGGCTCGTGCGTCGATGAAGAAC</td><td>259.9</td><td>5.8</td><td>0.8</td><td>6.9</td><td>6.6E-12</td><td>2.0E-11</td><td>MB231/MB468</td><td>CYTO</td></tr>
<tr><td>>MI0000086|hsa-mir-28&WithFlank&3|+|188688775|188688872@60.80.21&[MIMAT0004502&hsa-miR-28-3p&offsets|0|-1;m-28&3|+|188688834|188688855&offsets|0|-1]|CACTAGATTGTGAGCTCCTGG</td><td>>MI0000086|hsa-mir-28&WithFlank&3|+|188688775|188688872</td><td>60</td><td>80</td><td>21</td><td>CACTAGATTGTGAGCTCCTGG</td><td>975.6</td><td>10.3</td><td>1.5</td><td>6.9</td><td>4.9E-12</td><td>2.0E-11</td><td>BT20/MB468</td><td>MITO</td></tr>
<tr><td>>hg38_21_+_8433172_8446622_RNA45SN1_45S_50ntFlanks@10781.10805.25|AGGTGAACAGCCTCTGGCATGTTGG</td><td>>hg38_21_+_8433172_8446622_RNA45SN1_45S_50ntFlanks</td><td>10781</td><td>10805</td><td>25</td><td>AGGTGAACAGCCTCTGGCATGTTGG</td><td>155.3</td><td>-7.6</td><td>1.1</td><td>-6.9</td><td>6.8E-12</td><td>2.0E-11</td><td>MB231/MB468</td><td>NUC</td></tr>
<tr><td>>hg38_21_+_8433172_8446622_RNA45SN1_45S_50ntFlanks@9537.9563.27|AAACTCTGGTGGAGGTCCGTAGCGGTC</td><td>>hg38_21_+_8433172_8446622_RNA45SN1_45S_50ntFlanks</td><td>9537</td><td>9563</td><td>27</td><td>AAACTCTGGTGGAGGTCCGTAGCGGTC</td><td>110.5</td><td>-10.7</td><td>1.5</td><td>-6.9</td><td>6.0E-12</td><td>2.0E-11</td><td>MB231/MB468</td><td>MP</td></tr>
<tr><td>>hg38_21_+_8433172_8446622_RNA45SN1_45S_50ntFlanks@5238.5266.29|CTGACTGGCTCAGCGTGTGCCTACCCTAC</td><td>>hg38_21_+_8433172_8446622_RNA45SN1_45S_50ntFlanks</td><td>5238</td><td>5266</td><td>29</td><td>CTGACTGGCTCAGCGTGTGCCTACCCTAC</td><td>187.8</td><td>4.4</td><td>0.6</td><td>6.9</td><td>6.8E-12</td><td>2.0E-11</td><td>MB231/MB468</td><td>NUC</td></tr>
<tr><td>>hg38_21_+_8433172_8446622_RNA45SN1_45S_50ntFlanks@8106.8142.37|GCGGCGGGGCGCGGGACATGTGGCGTACGGAAGACCC</td><td>>hg38_21_+_8433172_8446622_RNA45SN1_45S_50ntFlanks</td><td>8106</td><td>8142</td><td>37</td><td>GCGGCGGGGCGCGGGACATGTGGCGTACGGAAGACCC</td><td>156.3</td><td>-11.2</td><td>1.6</td><td>-6.9</td><td>6.1E-12</td><td>2.0E-11</td><td>MB231/MB468</td><td>MP</td></tr>
<tr><td>>MI0000542|hsa-mir-320a&WithFlank&8|-|22244960|22245043@48.70.23(+1U)&[MIMAT0000510&hsa-miR-320a-3p&offsets|0|+1(+1U);m-60&8|-|22244975|22244996&offsets|0|+1(+1U)]|AAAAGCTGGGTTGAGAGGGCGAAT</td><td>>MI0000542|hsa-mir-320a&WithFlank&8|-|22244960|22245043</td><td>48</td><td>70</td><td>23(+1U)</td><td>AAAAGCTGGGTTGAGAGGGCGAAT</td><td>8922.7</td><td>6.8</td><td>0.9</td><td>7.3</td><td>3.9E-13</td><td>2.0E-11</td><td>BT20/MB468</td><td>MP</td></tr>
<tr><td>>hg38_21_+_8433172_8446622_RNA45SN1_45S_50ntFlanks@4995.5012.18|ACAGATTGATAGCTCTTT</td><td>>hg38_21_+_8433172_8446622_RNA45SN1_45S_50ntFlanks</td><td>4995</td><td>5012</td><td>18</td><td>ACAGATTGATAGCTCTTT</td><td>533.7</td><td>-12.1</td><td>1.7</td><td>-7.3</td><td>3.9E-13</td><td>2.0E-11</td><td>BT20/MB468</td><td>MP</td></tr>
<tr><td>>hg38_21_+_8433172_8446622_RNA45SN1_45S_50ntFlanks@6721.6740.20|AATTGCAGGACACATTGATC</td><td>>hg38_21_+_8433172_8446622_RNA45SN1_45S_50ntFlanks</td><td>6721</td><td>6740</td><td>20</td><td>AATTGCAGGACACATTGATC</td><td>4837.9</td><td>-6.3</td><td>0.9</td><td>-7.3</td><td>3.8E-13</td><td>2.0E-11</td><td>BT20/MB468</td><td>MP</td></tr>
<tr><td>>MI0000456|hsa-mir-140&WithFlank&16|+|69933075|69933186@69.91.23&[MIMAT0004597&hsa-miR-140-3p&offsets|+1|+3;m-45&16|+|69933143|69933164&offsets|0|+1]|ACCACAGGGTAGAACCACGGACA</td><td>>MI0000456|hsa-mir-140&WithFlank&16|+|69933075|69933186</td><td>69</td><td>91</td><td>23</td><td>ACCACAGGGTAGAACCACGGACA</td><td>622.1</td><td>10.2</td><td>1.5</td><td>6.9</td><td>6.7E-12</td><td>2.0E-11</td><td>MB231/MB468</td><td>CYTO</td></tr>
<tr><td>>MI0001445|hsa-mir-423&WithFlank&17|+|30117073|30117178@59.81.23(+1U)&[MIMAT0001340&hsa-miR-423-3p&offsets|0|0(+1U);m-100&17|+|30117131|30117153&offsets|0|0(+1U)]|AGCTCGGTCTGAGGCCCCTCAGTT</td><td>>MI0001445|hsa-mir-423&WithFlank&17|+|30117073|30117178</td><td>59</td><td>81</td><td>23(+1U)</td><td>AGCTCGGTCTGAGGCCCCTCAGTT</td><td>1405.0</td><td>5.5</td><td>0.8</td><td>6.9</td><td>6.9E-12</td><td>2.1E-11</td><td>MB231/MB468</td><td>NUC</td></tr>
<tr><td>>hg38_21_+_8433172_8446622_RNA45SN1_45S_50ntFlanks@11203.11223.21|CGGCGGCGGCGACTCTGGACG</td><td>>hg38_21_+_8433172_8446622_RNA45SN1_45S_50ntFlanks</td><td>11203</td><td>11223</td><td>21</td><td>CGGCGGCGGCGACTCTGGACG</td><td>106.8</td><td>-10.6</td><td>1.5</td><td>-6.9</td><td>6.1E-12</td><td>2.1E-11</td><td>MB231/MB468</td><td>MP</td></tr>
<tr><td>>hg38_21_+_8433172_8446622_RNA45SN1_45S_50ntFlanks@10786.10828.43|AACAGCCTCTGGCATGTTGGAACAATGTAGGTAAGGGAAGTCG</td><td>>hg38_21_+_8433172_8446622_RNA45SN1_45S_50ntFlanks</td><td>10786</td><td>10828</td><td>43</td><td>AACAGCCTCTGGCATGTTGGAACAATGTAGGTAAGGGAAGTCG</td><td>196.8</td><td>-11.5</td><td>1.7</td><td>-6.9</td><td>6.1E-12</td><td>2.1E-11</td><td>MB231/MB468</td><td>MP</td></tr>
<tr><td>>hg38_21_+_8433172_8446622_RNA45SN1_45S_50ntFlanks@10411.10433.23|CAGTCGGTCCTGAGAGATGGGCG</td><td>>hg38_21_+_8433172_8446622_RNA45SN1_45S_50ntFlanks</td><td>10411</td><td>10433</td><td>23</td><td>CAGTCGGTCCTGAGAGATGGGCG</td><td>262.9</td><td>-8.3</td><td>1.2</td><td>-6.9</td><td>6.9E-12</td><td>2.1E-11</td><td>MB231/MB468</td><td>NUC</td></tr>
<tr><td>>hg38_21_+_8433172_8446622_RNA45SN1_45S_50ntFlanks@10420.10455.36|CTGAGAGATGGGCGAGCGCCGTTCCGAAGGGACGGG</td><td>>hg38_21_+_8433172_8446622_RNA45SN1_45S_50ntFlanks</td><td>10420</td><td>10455</td><td>36</td><td>CTGAGAGATGGGCGAGCGCCGTTCCGAAGGGACGGG</td><td>439.4</td><td>-9.0</td><td>1.3</td><td>-6.9</td><td>6.9E-12</td><td>2.1E-11</td><td>MB231/MB468</td><td>NUC</td></tr>
<tr><td>>hg38_21_+_8433172_8446622_RNA45SN1_45S_50ntFlanks@11821.11843.23|CTATCCAGCGAAACCACAGCCAA</td><td>>hg38_21_+_8433172_8446622_RNA45SN1_45S_50ntFlanks</td><td>11821</td><td>11843</td><td>23</td><td>CTATCCAGCGAAACCACAGCCAA</td><td>118.0</td><td>-10.7</td><td>1.6</td><td>-6.9</td><td>6.1E-12</td><td>2.1E-11</td><td>MB231/MB468</td><td>MP</td></tr>
<tr><td>>hg38_21_+_8433172_8446622_RNA45SN1_45S_50ntFlanks@10421.10456.36|TGAGAGATGGGCGAGCGCCGTTCCGAAGGGACGGGC</td><td>>hg38_21_+_8433172_8446622_RNA45SN1_45S_50ntFlanks</td><td>10421</td><td>10456</td><td>36</td><td>TGAGAGATGGGCGAGCGCCGTTCCGAAGGGACGGGC</td><td>1997.8</td><td>-4.4</td><td>0.6</td><td>-6.9</td><td>6.4E-12</td><td>2.1E-11</td><td>BT20/MB468</td><td>CYTO</td></tr>
<tr><td>>hg38_21_+_8433172_8446622_RNA45SN1_45S_50ntFlanks@6726.6744.19|CAGGACACATTGATCATCG</td><td>>hg38_21_+_8433172_8446622_RNA45SN1_45S_50ntFlanks</td><td>6726</td><td>6744</td><td>19</td><td>CAGGACACATTGATCATCG</td><td>209.4</td><td>-11.6</td><td>1.7</td><td>-6.9</td><td>6.1E-12</td><td>2.1E-11</td><td>MB231/MB468</td><td>MP</td></tr>
<tr><td>>hg38_21_+_8433172_8446622_RNA45SN1_45S_50ntFlanks@10350.10390.41|ACGAGAACTTTGAAGGCCGAAGTGGAGAAGGGTTCCATGTG</td><td>>hg38_21_+_8433172_8446622_RNA45SN1_45S_50ntFlanks</td><td>10350</td><td>10390</td><td>41</td><td>ACGAGAACTTTGAAGGCCGAAGTGGAGAAGGGTTCCATGTG</td><td>9236.5</td><td>-9.6</td><td>1.4</td><td>-6.8</td><td>8.6E-12</td><td>2.1E-11</td><td>BT20/MB468</td><td>NUC</td></tr>
<tr><td>>hg38_21_+_8433172_8446622_RNA45SN1_45S_50ntFlanks@11572.11607.36|CCGACTTAGAACTGGTGCGGACCAGGGGAATCCGAC</td><td>>hg38_21_+_8433172_8446622_RNA45SN1_45S_50ntFlanks</td><td>11572</td><td>11607</td><td>36</td><td>CCGACTTAGAACTGGTGCGGACCAGGGGAATCCGAC</td><td>560.6</td><td>-5.2</td><td>0.8</td><td>-6.8</td><td>8.6E-12</td><td>2.1E-11</td><td>BT20/MB468</td><td>NUC</td></tr>
<tr><td>>trna10_AlaCGC_6_+_26553731_26553802@1.19.19,trna119_AlaCGC_6_-_28641613_28641684@1.19.19,trna120_AlaAGC_6_-_28626014_28626085@1.19.19,trna13_AlaCGC_2_+_157257281_157257352@1.19.19,trna13_AlaTGC_12_-_125406301_125406372@1.19.19,trna66_AlaTGC_6_+_28611222_28611293@1.19.19,trna8_AlaTGC_12_+_125424512_125424583@1.19.19,trna8_AlaTGC_5_+_180633868_180633939@1.19.19|GGGGATGTAGCTCAGTGGT</td><td>>trna10_AlaCGC_6_+_26553731_26553802</td><td>1</td><td>19</td><td>19</td><td>GGGGATGTAGCTCAGTGGT</td><td>392.3</td><td>4.1</td><td>0.6</td><td>6.9</td><td>6.4E-12</td><td>2.1E-11</td><td>BT20/MB468</td><td>CYTO</td></tr>
<tr><td>>hg38_21_+_8433172_8446622_RNA45SN1_45S_50ntFlanks@4892.4918.27|AAACTTAAAGGAATTGACGGAAGGGCA</td><td>>hg38_21_+_8433172_8446622_RNA45SN1_45S_50ntFlanks</td><td>4892</td><td>4918</td><td>27</td><td>AAACTTAAAGGAATTGACGGAAGGGCA</td><td>147.5</td><td>-11.1</td><td>1.6</td><td>-6.9</td><td>6.1E-12</td><td>2.1E-11</td><td>MB231/MB468</td><td>MP</td></tr>
<tr><td>>MI0000477|hsa-mir-146a&WithFlank&5|+|160485346|160485456@27.49.23(+1A)&[MIMAT0000449&hsa-miR-146a-5p&offsets|0|+1(+1A);m-129&5|+|160485372|160485393&offsets|0|+1(+1A)]|TGAGAACTGAATTCCATGGGTTGA</td><td>>MI0000477|hsa-mir-146a&WithFlank&5|+|160485346|160485456</td><td>27</td><td>49</td><td>23(+1A)</td><td>TGAGAACTGAATTCCATGGGTTGA</td><td>168.9</td><td>10.6</td><td>1.5</td><td>6.9</td><td>6.2E-12</td><td>2.1E-11</td><td>MB231/MB468</td><td>MP</td></tr>
<tr><td>>hg38_21_+_8433172_8446622_RNA45SN1_45S_50ntFlanks@3738.3763.26|TCTCAAAGATTAAGCCATGCATGTCT</td><td>>hg38_21_+_8433172_8446622_RNA45SN1_45S_50ntFlanks</td><td>3738</td><td>3763</td><td>26</td><td>TCTCAAAGATTAAGCCATGCATGTCT</td><td>96.5</td><td>-10.5</td><td>1.5</td><td>-6.9</td><td>6.2E-12</td><td>2.1E-11</td><td>MB231/MB468</td><td>MP</td></tr>
<tr><td>>hg38_21_+_8433172_8446622_RNA45SN1_45S_50ntFlanks@4240.4258.19|ACTCTTTCGAGGCCCTGTA</td><td>>hg38_21_+_8433172_8446622_RNA45SN1_45S_50ntFlanks</td><td>4240</td><td>4258</td><td>19</td><td>ACTCTTTCGAGGCCCTGTA</td><td>179.0</td><td>-7.8</td><td>1.1</td><td>-6.9</td><td>7.0E-12</td><td>2.1E-11</td><td>MB231/MB468</td><td>NUC</td></tr>
<tr><td>>hg38_21_+_8433172_8446622_RNA45SN1_45S_50ntFlanks@10463.10483.21|CTCCGTTGCCCTCGGCCGATC</td><td>>hg38_21_+_8433172_8446622_RNA45SN1_45S_50ntFlanks</td><td>10463</td><td>10483</td><td>21</td><td>CTCCGTTGCCCTCGGCCGATC</td><td>120.1</td><td>-10.8</td><td>1.6</td><td>-6.9</td><td>6.2E-12</td><td>2.1E-11</td><td>MB231/MB468</td><td>MP</td></tr>
<tr><td>>hg38_21_+_8433172_8446622_RNA45SN1_45S_50ntFlanks@9534.9574.41|AGGAAACTCTGGTGGAGGTCCGTAGCGGTCCTGACGTGCAA</td><td>>hg38_21_+_8433172_8446622_RNA45SN1_45S_50ntFlanks</td><td>9534</td><td>9574</td><td>41</td><td>AGGAAACTCTGGTGGAGGTCCGTAGCGGTCCTGACGTGCAA</td><td>669.4</td><td>-8.9</td><td>1.3</td><td>-6.8</td><td>8.2E-12</td><td>2.1E-11</td><td>MB231/MB468</td><td>MITO</td></tr>
<tr><td>>MI0016005|hsa-mir-3615&WithFlank&17|+|74748607|74748705@57.78.22&[MIMAT0017994&hsa-miR-3615&offsets|0|+1;m-271&17|+|74748663|74748684&offsets|0|0]|TCTCTCGGCTCCTCGCGGCTCG</td><td>>MI0016005|hsa-mir-3615&WithFlank&17|+|74748607|74748705</td><td>57</td><td>78</td><td>22</td><td>TCTCTCGGCTCCTCGCGGCTCG</td><td>1670.0</td><td>5.4</td><td>0.8</td><td>6.9</td><td>6.8E-12</td><td>2.1E-11</td><td>MB231/MB468</td><td>CYTO</td></tr>
<tr><td>>hg38_21_+_8433172_8446622_RNA45SN1_45S_50ntFlanks@5222.5244.23|CTGCACGCGCGCTACACTGACTG</td><td>>hg38_21_+_8433172_8446622_RNA45SN1_45S_50ntFlanks</td><td>5222</td><td>5244</td><td>23</td><td>CTGCACGCGCGCTACACTGACTG</td><td>163.7</td><td>-11.2</td><td>1.6</td><td>-6.9</td><td>6.2E-12</td><td>2.1E-11</td><td>MB231/MB468</td><td>MP</td></tr>
<tr><td>>hg38_21_+_8433172_8446622_RNA45SN1_45S_50ntFlanks@4898.4917.20|AAAGGAATTGACGGAAGGGC</td><td>>hg38_21_+_8433172_8446622_RNA45SN1_45S_50ntFlanks</td><td>4898</td><td>4917</td><td>20</td><td>AAAGGAATTGACGGAAGGGC</td><td>343.3</td><td>-12.3</td><td>1.8</td><td>-6.9</td><td>6.2E-12</td><td>2.1E-11</td><td>MB231/MB468</td><td>MP</td></tr>
<tr><td>>hg38_21_+_8433172_8446622_RNA45SN1_45S_50ntFlanks@9568.9586.19|CGTGCAAATCGGTCGTCCG</td><td>>hg38_21_+_8433172_8446622_RNA45SN1_45S_50ntFlanks</td><td>9568</td><td>9586</td><td>19</td><td>CGTGCAAATCGGTCGTCCG</td><td>421.4</td><td>-5.9</td><td>0.9</td><td>-6.9</td><td>6.8E-12</td><td>2.1E-11</td><td>MB231/MB468</td><td>CYTO</td></tr>
<tr><td>>hg38_21_+_8433172_8446622_RNA45SN1_45S_50ntFlanks@11615.11640.26|TTAAAACAAAGCATCGCGAAGGCCCG</td><td>>hg38_21_+_8433172_8446622_RNA45SN1_45S_50ntFlanks</td><td>11615</td><td>11640</td><td>26</td><td>TTAAAACAAAGCATCGCGAAGGCCCG</td><td>257.9</td><td>-11.1</td><td>1.5</td><td>-7.3</td><td>3.9E-13</td><td>2.1E-11</td><td>BT20/MB468</td><td>MP</td></tr>
<tr><td>>hg38_21_+_8433172_8446622_RNA45SN1_45S_50ntFlanks@4728.4748.21|ATCAAGAACGAAAGTCGGAGG</td><td>>hg38_21_+_8433172_8446622_RNA45SN1_45S_50ntFlanks</td><td>4728</td><td>4748</td><td>21</td><td>ATCAAGAACGAAAGTCGGAGG</td><td>428.0</td><td>-11.8</td><td>1.6</td><td>-7.3</td><td>3.9E-13</td><td>2.1E-11</td><td>BT20/MB468</td><td>MP</td></tr>
<tr><td>>hg38_21_+_8433172_8446622_RNA45SN1_45S_50ntFlanks@12201.12222.22|CCTAAGGCGAGCTCAGGGAGGA</td><td>>hg38_21_+_8433172_8446622_RNA45SN1_45S_50ntFlanks</td><td>12201</td><td>12222</td><td>22</td><td>CCTAAGGCGAGCTCAGGGAGGA</td><td>94.9</td><td>-10.4</td><td>1.5</td><td>-6.9</td><td>6.2E-12</td><td>2.1E-11</td><td>MB231/MB468</td><td>MP</td></tr>
<tr><td>>MI0016887|hsa-mir-4521&WithFlank&17|+|8186939|8187010@10.34.25&[MIMAT0019058&hsa-miR-4521&offsets|0|+3;m-765&17|+|8186948|8186969&offsets|0|+3]|GCTAAGGAAGTCCTGTGCTCAGTTT</td><td>>MI0016887|hsa-mir-4521&WithFlank&17|+|8186939|8187010</td><td>10</td><td>34</td><td>25</td><td>GCTAAGGAAGTCCTGTGCTCAGTTT</td><td>3779.3</td><td>6.9</td><td>1.0</td><td>6.9</td><td>6.4E-12</td><td>2.1E-11</td><td>BT20/MB468</td><td>CYTO</td></tr>
<tr><td>>hg38_21_+_8433172_8446622_RNA45SN1_45S_50ntFlanks@3979.4001.23|CGCCGGCGGCTTTGGTGACTCTA</td><td>>hg38_21_+_8433172_8446622_RNA45SN1_45S_50ntFlanks</td><td>3979</td><td>4001</td><td>23</td><td>CGCCGGCGGCTTTGGTGACTCTA</td><td>762.6</td><td>-9.7</td><td>1.4</td><td>-6.8</td><td>8.7E-12</td><td>2.1E-11</td><td>BT20/MB468</td><td>NUC</td></tr>
<tr><td>>hg38_21_+_8433172_8446622_RNA45SN1_45S_50ntFlanks@8168.8215.48|CCCAAGTCCTTCTGATCGAGGCCCAGCCCGTGGACGGTGTGAGGCCGG</td><td>>hg38_21_+_8433172_8446622_RNA45SN1_45S_50ntFlanks</td><td>8168</td><td>8215</td><td>48</td><td>CCCAAGTCCTTCTGATCGAGGCCCAGCCCGTGGACGGTGTGAGGCCGG</td><td>1131.6</td><td>-10.3</td><td>1.5</td><td>-6.9</td><td>7.0E-12</td><td>2.1E-11</td><td>MB231/MB468</td><td>NUC</td></tr>
</table>

<table>
<tr><td>>MI0000088|hsa-mir-30a&WithFlank&6|-|71403545|71403627@53.74.22&[MIMAT0000088&hsa-miR-30a-3p&offsets|0|0;m-38&6|-|71403554|71403575&offsets|0|0]|CTTTCAGTCGGATGTTTGCAGC</td><td>>MI0000088|hsa-mir-30a&WithFlank&6|-|71403545|71403627</td><td>53</td><td>74</td><td>22</td><td>CTTTCAGTCGGATGTTTGCAGC</td><td>7758.3</td><td>5.9</td><td>0.9</td><td>6.8</td><td>8.2E-12</td><td>2.1E-11</td><td>MB231/MB468</td><td>MITO</td></tr>
<tr><td>>MI0000088|hsa-mir-30a&WithFlank&6|-|71403545|71403627@12.35.24&[MIMAT0000087&hsa-miR-30a-5p&offsets|0|+2;m-15&6|-|71403595|71403616&offsets|0|+2]|TGTAAACATCCTCGACTGGAAGCT</td><td>>MI0000088|hsa-mir-30a&WithFlank&6|-|71403545|71403627</td><td>12</td><td>35</td><td>24</td><td>TGTAAACATCCTCGACTGGAAGCT</td><td>25642.8</td><td>6.1</td><td>0.9</td><td>6.9</td><td>6.8E-12</td><td>2.1E-11</td><td>MB231/MB468</td><td>CYTO</td></tr>
<tr><td>>MI0000108|hsa-mir-103a-2&WithFlank&20|+|3917488|3917577@54.76.23(+1U);MI0000109|hsa-mir-103a-1&WithFlank&5|-|168560890|168560979@54.76.23(+1U)&[MIMAT0000101&hsa-miR-103a-3p&offsets|0|0(+1U);m-12&20|+|3917541|3917563&offsets|0|0(+1U)];[MIMAT0000101_1&hsa-miR-103a-3p&offsets|0|0(+1U);m-13&5|-|168560904|168560926&offsets|0|0(+1U);m-44220&5|-|168560908|168560926&offsets|0|+4(+1U)]|AGCAGCATTGTACAGGGCTATGAT</td><td>>MI0000108|hsa-mir-103a-2&WithFlank&20|+|3917488|3917577</td><td>54</td><td>76</td><td>23(+1U)</td><td>AGCAGCATTGTACAGGGCTATGAT</td><td>2093.0</td><td>6.2</td><td>0.9</td><td>6.9</td><td>6.8E-12</td><td>2.1E-11</td><td>MB231/MB468</td><td>CYTO</td></tr>
<tr><td>>hg38_21_+_8433172_8446622_RNA45SN1_45S_50ntFlanks@10389.10411.23|TGAACAGCAGTTGAACATGGGTC</td><td>>hg38_21_+_8433172_8446622_RNA45SN1_45S_50ntFlanks</td><td>10389</td><td>10411</td><td>23</td><td>TGAACAGCAGTTGAACATGGGTC</td><td>116.9</td><td>-10.7</td><td>1.6</td><td>-6.9</td><td>6.2E-12</td><td>2.1E-11</td><td>MB231/MB468</td><td>MP</td></tr>
<tr><td>>trnaMT_PheGAA_MT_+_577_647@38.56.19|AATGTTTAGACGGGCTCAC</td><td>>trnaMT_PheGAA_MT_+_577_647</td><td>38</td><td>56</td><td>19</td><td>AATGTTTAGACGGGCTCAC</td><td>351.5</td><td>5.7</td><td>0.8</td><td>6.8</td><td>8.3E-12</td><td>2.1E-11</td><td>MB231/MB468</td><td>MITO</td></tr>
<tr><td>>hg38_21_+_8433172_8446622_RNA45SN1_45S_50ntFlanks@12580.12604.25|GAACCGCAGGTTCAGACATTTGGTG</td><td>>hg38_21_+_8433172_8446622_RNA45SN1_45S_50ntFlanks</td><td>12580</td><td>12604</td><td>25</td><td>GAACCGCAGGTTCAGACATTTGGTG</td><td>88.0</td><td>-10.3</td><td>1.5</td><td>-6.9</td><td>6.3E-12</td><td>2.1E-11</td><td>MB231/MB468</td><td>MP</td></tr>
<tr><td>>hg38_21_+_8433172_8446622_RNA45SN1_45S_50ntFlanks@12893.12929.37|CCTCGCCCGTCACGCACCGCACGTTCGTGGGGAACCT</td><td>>hg38_21_+_8433172_8446622_RNA45SN1_45S_50ntFlanks</td><td>12893</td><td>12929</td><td>37</td><td>CCTCGCCCGTCACGCACCGCACGTTCGTGGGGAACCT</td><td>137.8</td><td>-11.0</td><td>1.6</td><td>-6.9</td><td>6.3E-12</td><td>2.1E-11</td><td>MB231/MB468</td><td>MP</td></tr>
<tr><td>>MI0000070|hsa-mir-16-1&WithFlank&13|-|50048967|50049067@20.41.22;MI0000115|hsa-mir-16-2&WithFlank&3|+|160404739|160404831@16.37.22&[MIMAT0000069&hsa-miR-16-5p&offsets|0|0;m-62&13|-|50049027|50049048&offsets|0|0];[MIMAT0000069_1&hsa-miR-16-5p&offsets|0|0;m-61&3|+|160404754|160404775&offsets|0|0]|TAGCAGCACGTAAATATTGGCG</td><td>>MI0000070|hsa-mir-16-1&WithFlank&13|-|50048967|50049067</td><td>20</td><td>41</td><td>22</td><td>TAGCAGCACGTAAATATTGGCG</td><td>910.3</td><td>5.3</td><td>0.8</td><td>6.9</td><td>5.1E-12</td><td>2.1E-11</td><td>BT20/MB468</td><td>MITO</td></tr>
<tr><td>>hg38_21_+_8433172_8446622_RNA45SN1_45S_50ntFlanks@9566.9596.31|GACGTGCAAATCGGTCGTCCGACCTGGGTAT</td><td>>hg38_21_+_8433172_8446622_RNA45SN1_45S_50ntFlanks</td><td>9566</td><td>9596</td><td>31</td><td>GACGTGCAAATCGGTCGTCCGACCTGGGTAT</td><td>2854.7</td><td>-4.2</td><td>0.6</td><td>-6.9</td><td>6.9E-12</td><td>2.1E-11</td><td>MB231/MB468</td><td>CYTO</td></tr>
<tr><td>>MI0000294|hsa-mir-218-1&WithFlank&4|+|20528269|20528390@31.52.22;MI0000295|hsa-mir-218-2&WithFlank&5|-|168768140|168768261@31.52.22&[MIMAT0000275&hsa-miR-218-5p&offsets|0|+1;m-185&4|+|20528299|20528319&offsets|0|+1];[MIMAT0000275_1&hsa-miR-218-5p&offsets|0|+1;m-187&5|-|168768211|168768231&offsets|0|+1]|TTGTGCTTGATCTAACCATGTG</td><td>>MI0000294|hsa-mir-218-1&WithFlank&4|+|20528269|20528390</td><td>31</td><td>52</td><td>22</td><td>TTGTGCTTGATCTAACCATGTG</td><td>150.9</td><td>-10.5</td><td>1.5</td><td>-7.2</td><td>6.4E-13</td><td>2.1E-11</td><td>BT20/MB231</td><td>MITO</td></tr>
<tr><td>>MI0000746|hsa-mir-99b&WithFlank&19|+|51692606|51692687@13.34.22(+1A)&[MIMAT0000689&hsa-miR-99b-5p&offsets|0|0(+1A);m-10&19|+|51692618|51692639&offsets|0|0(+1A)]|CACCCGTAGAACCGACCTTGCGA</td><td>>MI0000746|hsa-mir-99b&WithFlank&19|+|51692606|51692687</td><td>13</td><td>34</td><td>22(+1A)</td><td>CACCCGTAGAACCGACCTTGCGA</td><td>1213.0</td><td>5.8</td><td>0.8</td><td>6.9</td><td>6.9E-12</td><td>2.1E-11</td><td>MB231/MB468</td><td>CYTO</td></tr>
<tr><td>>MI0003628|hsa-mir-615&WithFlank&12|+|54033944|54034051@67.87.21(+1A)&[MIMAT0003283&hsa-miR-615-3p&offsets|0|-1(+1A);m-495&12|+|54034010|54034030&offsets|0|0(+1A)]|TCCGAGCCTGGGTCTCCCTCTA</td><td>>MI0003628|hsa-mir-615&WithFlank&12|+|54033944|54034051</td><td>67</td><td>87</td><td>21(+1A)</td><td>TCCGAGCCTGGGTCTCCCTCTA</td><td>190.4</td><td>7.8</td><td>1.1</td><td>7.2</td><td>4.9E-13</td><td>2.1E-11</td><td>BT20/MB231</td><td>CYTO</td></tr>
<tr><td>>hg38_21_+_8433172_8446622_RNA45SN1_45S_50ntFlanks@11781.11807.27|CGCATGAATGGATGAACGAGATTCCCA</td><td>>hg38_21_+_8433172_8446622_RNA45SN1_45S_50ntFlanks</td><td>11781</td><td>11807</td><td>27</td><td>CGCATGAATGGATGAACGAGATTCCCA</td><td>232.3</td><td>-11.7</td><td>1.7</td><td>-6.9</td><td>6.3E-12</td><td>2.1E-11</td><td>MB231/MB468</td><td>MP</td></tr>
<tr><td>>hg38_21_+_8433172_8446622_RNA45SN1_45S_50ntFlanks@8423.8469.47|CGCAGTCCGCCCGGAGGATTCAACCCGGCGGCGGGTCCGGCCGTGTC</td><td>>hg38_21_+_8433172_8446622_RNA45SN1_45S_50ntFlanks</td><td>8423</td><td>8469</td><td>47</td><td>CGCAGTCCGCCCGGAGGATTCAACCCGGCGGCGGGTCCGGCCGTGTC</td><td>653.1</td><td>-13.8</td><td>2.0</td><td>-6.8</td><td>8.9E-12</td><td>2.1E-11</td><td>BT20/MB468</td><td>NUC</td></tr>
<tr><td>>MI0000088|hsa-mir-30a&WithFlank&6|-|71403545|71403627@12.35.24(+3U)&[MIMAT0000087&hsa-miR-30a-5p&offsets|0|+2(+3U);m-15&6|-|71403595|71403616&offsets|0|+2(+3U)]|TGTAAACATCCTCGACTGGAAGCTTTT</td><td>>MI0000088|hsa-mir-30a&WithFlank&6|-|71403545|71403627</td><td>12</td><td>35</td><td>24(+3U)</td><td>TGTAAACATCCTCGACTGGAAGCTTTT</td><td>461.2</td><td>12.1</td><td>1.8</td><td>6.8</td><td>8.4E-12</td><td>2.1E-11</td><td>MB231/MB468</td><td>MITO</td></tr>
<tr><td>>am_MI0000786|hsa-mir-378a&WithFlank&5|+|149732819|149732896@49.69.21&[MIMAT0000732&hsa-miR-378a-3p&offsets|0|-1;m-44&5|+|149732867|149732888&offsets|0|-1]|ACTGGACTTGGAGTCAGAAGG</td><td>>am_MI0000786|hsa-mir-378a&WithFlank&5|+|149732819|149732896</td><td>49</td><td>69</td><td>21</td><td>ACTGGACTTGGAGTCAGAAGG</td><td>2200.8</td><td>3.8</td><td>0.6</td><td>6.9</td><td>7.2E-12</td><td>2.1E-11</td><td>MB231/MB468</td><td>NUC</td></tr>
<tr><td>>MI0016746|hsa-mir-548o-2&WithFlank&20|+|38516557|38516638@51.72.22;MI0006402|hsa-mir-548o&WithFlank&7|-|102405736|102405861@93.114.22&[MIMAT0005919&hsa-miR-548o-3p&offsets|0|0;m-431&20|+|38516607|38516628&offsets|0|0];[MIMAT0005919_1&hsa-miR-548o-3p&offsets|0|0;m-430&7|-|102405748|102405769&offsets|0|0]|CCAAAACTGCAGTTACTTTTGC</td><td>>MI0016746|hsa-mir-548o-2&WithFlank&20|+|38516557|38516638</td><td>51</td><td>72</td><td>22</td><td>CCAAAACTGCAGTTACTTTTGC</td><td>349.0</td><td>5.0</td><td>0.7</td><td>6.9</td><td>7.2E-12</td><td>2.1E-11</td><td>MB231/MB468</td><td>NUC</td></tr>
<tr><td>>hg38_21_+_8433172_8446622_RNA45SN1_45S_50ntFlanks@11564.11600.37|CCTAGCAGCCGACTTAGAACTGGTGCGGACCAGGGGA</td><td>>hg38_21_+_8433172_8446622_RNA45SN1_45S_50ntFlanks</td><td>11564</td><td>11600</td><td>37</td><td>CCTAGCAGCCGACTTAGAACTGGTGCGGACCAGGGGA</td><td>3093.8</td><td>-8.6</td><td>1.3</td><td>-6.8</td><td>8.9E-12</td><td>2.1E-11</td><td>BT20/MB468</td><td>NUC</td></tr>
<tr><td>>hg38_21_+_8433172_8446622_RNA45SN1_45S_50ntFlanks@10535.10568.34|CGCGAGGCGTCCAGTGCGGTAACGCGACCGATCC</td><td>>hg38_21_+_8433172_8446622_RNA45SN1_45S_50ntFlanks</td><td>10535</td><td>10568</td><td>34</td><td>CGCGAGGCGTCCAGTGCGGTAACGCGACCGATCC</td><td>87.4</td><td>-10.3</td><td>1.5</td><td>-6.9</td><td>6.4E-12</td><td>2.1E-11</td><td>MB231/MB468</td><td>MP</td></tr>
<tr><td>>hg38_21_+_8433172_8446622_RNA45SN1_45S_50ntFlanks@11643.11665.23|GCGGGTGTTGACGCGATGTGATT</td><td>>hg38_21_+_8433172_8446622_RNA45SN1_45S_50ntFlanks</td><td>11643</td><td>11665</td><td>23</td><td>GCGGGTGTTGACGCGATGTGATT</td><td>108.2</td><td>-10.6</td><td>1.5</td><td>-6.9</td><td>6.4E-12</td><td>2.1E-11</td><td>MB231/MB468</td><td>MP</td></tr>
<tr><td>>hg38_21_+_8433172_8446622_RNA45SN1_45S_50ntFlanks@9104.9128.25|CCGGCGACGGGGGGGGTGCCGCGCG</td><td>>hg38_21_+_8433172_8446622_RNA45SN1_45S_50ntFlanks</td><td>9104</td><td>9128</td><td>25</td><td>CCGGCGACGGGGGGGGTGCCGCGCG</td><td>119.1</td><td>-10.8</td><td>1.6</td><td>-6.9</td><td>6.4E-12</td><td>2.1E-11</td><td>MB231/MB468</td><td>MP</td></tr>
<tr><td>>hg38_MT_+_1621_3279_RNR2_16S_50ntFlanks@51.81.31|GCTAAACCTAGCCCCAAACCCACTCCACCTT</td><td>>hg38_MT_+_1621_3279_RNR2_16S_50ntFlanks</td><td>51</td><td>81</td><td>31</td><td>GCTAAACCTAGCCCCAAACCCACTCCACCTT</td><td>5486.8</td><td>6.7</td><td>1.0</td><td>6.9</td><td>5.2E-12</td><td>2.1E-11</td><td>BT20/MB468</td><td>MITO</td></tr>
<tr><td>>MI0000078|hsa-mir-22&WithFlank&17|-|1713897|1713993@59.78.20&[MIMAT0000077&hsa-miR-22-3p&offsets|0|-2;m-2&17|-|1713914|1713935&offsets|0|-2]|AAGCTGCCAGTTGAAGAACT</td><td>>MI0000078|hsa-mir-22&WithFlank&17|-|1713897|1713993</td><td>59</td><td>78</td><td>20</td><td>AAGCTGCCAGTTGAAGAACT</td><td>365.7</td><td>8.2</td><td>1.2</td><td>6.9</td><td>7.0E-12</td><td>2.1E-11</td><td>MB231/MB468</td><td>CYTO</td></tr>
<tr><td>>hg38_21_+_8433172_8446622_RNA45SN1_45S_50ntFlanks@12717.12755.39|CCTCGGTTGGCCTCGGATAGCCGGTCCCCCGCCTGTCCC</td><td>>hg38_21_+_8433172_8446622_RNA45SN1_45S_50ntFlanks</td><td>12717</td><td>12755</td><td>39</td><td>CCTCGGTTGGCCTCGGATAGCCGGTCCCCCGCCTGTCCC</td><td>412.7</td><td>-8.9</td><td>1.3</td><td>-6.8</td><td>9.0E-12</td><td>2.1E-11</td><td>BT20/MB468</td><td>NUC</td></tr>
<tr><td>>hg38_21_+_8433172_8446622_RNA45SN1_45S_50ntFlanks@12716.12747.32|GCCTCGGTTGGCCTCGGATAGCCGGTCCCCCG</td><td>>hg38_21_+_8433172_8446622_RNA45SN1_45S_50ntFlanks</td><td>12716</td><td>12747</td><td>32</td><td>GCCTCGGTTGGCCTCGGATAGCCGGTCCCCCG</td><td>182.5</td><td>-11.4</td><td>1.7</td><td>-6.9</td><td>6.4E-12</td><td>2.1E-11</td><td>MB231/MB468</td><td>MP</td></tr>
<tr><td>>hg38_21_+_8433172_8446622_RNA45SN1_45S_50ntFlanks@10727.10748.22|GGAGAGGGTGTAAATCTCGCGC</td><td>>hg38_21_+_8433172_8446622_RNA45SN1_45S_50ntFlanks</td><td>10727</td><td>10748</td><td>22</td><td>GGAGAGGGTGTAAATCTCGCGC</td><td>175.2</td><td>-11.3</td><td>1.6</td><td>-6.9</td><td>6.4E-12</td><td>2.1E-11</td><td>MB231/MB468</td><td>MP</td></tr>
<tr><td>>MI0000264|hsa-mir-7-2&WithFlank&15|+|88611819|88611940@38.59.22(+1A);MI0000265|hsa-mir-7-3&WithFlank&19|+|4770664|4770785@37.58.22(+1A);MI0000263|hsa-mir-7-1&WithFlank&9|-|83969742|83969863@30.51.22(+1A)&[MIMAT0000252&hsa-miR-7-5p&offsets|0|-2(+1A);m-400&15|+|88611856|88611879&offsets|0|-2(+1A)];[MIMAT0000252_1&hsa-miR-7-5p&offsets|0|-2(+1A);m-402&19|+|4770700|4770723&offsets|0|-2(+1A)];[MIMAT0000252_2&hsa-miR-7-5p&offsets|0|-2(+1A);m-401&9|-|83969811|83969834&offsets|0|-2(+1A)]|TGGAAGACTAGTGATTTTGTTGA</td><td>>MI0000264|hsa-mir-7-2&WithFlank&15|+|88611819|88611940</td><td>38</td><td>59</td><td>22(+1A)</td><td>TGGAAGACTAGTGATTTTGTTGA</td><td>79.5</td><td>5.7</td><td>0.8</td><td>6.8</td><td>9.0E-12</td><td>2.1E-11</td><td>BT20/MB468</td><td>NUC</td></tr>
<tr><td>>hg38_21_+_8433172_8446622_RNA45SN1_45S_50ntFlanks@4007.4026.20|CCTCGGGCCGATCGCACGCC</td><td>>hg38_21_+_8433172_8446622_RNA45SN1_45S_50ntFlanks</td><td>4007</td><td>4026</td><td>20</td><td>CCTCGGGCCGATCGCACGCC</td><td>543.6</td><td>-9.0</td><td>1.3</td><td>-6.8</td><td>8.5E-12</td><td>2.2E-11</td><td>MB231/MB468</td><td>MITO</td></tr>
<tr><td>>hg38_21_+_8433172_8446622_RNA45SN1_45S_50ntFlanks@11584.11623.40|TGGTGCGGACCAGGGGAATCCGACTGTTTAATTAAAACAA</td><td>>hg38_21_+_8433172_8446622_RNA45SN1_45S_50ntFlanks</td><td>11584</td><td>11623</td><td>40</td><td>TGGTGCGGACCAGGGGAATCCGACTGTTTAATTAAAACAA</td><td>4468.9</td><td>10.5</td><td>1.5</td><td>6.9</td><td>5.2E-12</td><td>2.2E-11</td><td>BT20/MB468</td><td>MITO</td></tr>
<tr><td>>trnaMT_LeuTAA_MT_+_3230_3304@16.45.30|CCCGGTAATCGCATAAAACTTAAAACTTTA</td><td>>trnaMT_LeuTAA_MT_+_3230_3304</td><td>16</td><td>45</td><td>30</td><td>CCCGGTAATCGCATAAAACTTAAAACTTTA</td><td>138.5</td><td>-9.8</td><td>1.4</td><td>-6.9</td><td>7.1E-12</td><td>2.2E-11</td><td>MB231/MB468</td><td>CYTO</td></tr>
<tr><td>>MI0000542|hsa-mir-320a&WithFlank&8|-|22244960|22245043@48.68.21&[MIMAT0000510&hsa-miR-320a-3p&offsets|0|-1;m-60&8|-|22244975|22244996&offsets|0|-1]|AAAAGCTGGGTTGAGAGGGCG</td><td>>MI0000542|hsa-mir-320a&WithFlank&8|-|22244960|22245043</td><td>48</td><td>68</td><td>21</td><td>AAAAGCTGGGTTGAGAGGGCG</td><td>1437.2</td><td>6.0</td><td>0.9</td><td>6.9</td><td>7.1E-12</td><td>2.2E-11</td><td>MB231/MB468</td><td>CYTO</td></tr>
<tr><td>>hg38_21_+_8433172_8446622_RNA45SN1_45S_50ntFlanks@10771.10795.25|CAGGTCTCCAAGGTGAACAGCCTCT</td><td>>hg38_21_+_8433172_8446622_RNA45SN1_45S_50ntFlanks</td><td>10771</td><td>10795</td><td>25</td><td>CAGGTCTCCAAGGTGAACAGCCTCT</td><td>80.6</td><td>-10.2</td><td>1.5</td><td>-6.9</td><td>6.5E-12</td><td>2.2E-11</td><td>MB231/MB468</td><td>MP</td></tr>
<tr><td>>hg38_21_+_8433172_8446622_RNA45SN1_45S_50ntFlanks@8168.8206.39|CCCAAGTCCTTCTGATCGAGGCCCAGCCCGTGGACGGTG</td><td>>hg38_21_+_8433172_8446622_RNA45SN1_45S_50ntFlanks</td><td>8168</td><td>8206</td><td>39</td><td>CCCAAGTCCTTCTGATCGAGGCCCAGCCCGTGGACGGTG</td><td>192.6</td><td>-11.5</td><td>1.7</td><td>-6.9</td><td>6.5E-12</td><td>2.2E-11</td><td>MB231/MB468</td><td>MP</td></tr>
<tr><td>>MI0001445|hsa-mir-423&WithFlank&17|+|30117073|30117178@23.45.23&[MIMAT0004748&hsa-miR-423-5p&offsets|0|0;m-141&17|+|30117095|30117117&offsets|0|0]|TGAGGGGCAGAGAGCGAGACTTT</td><td>>MI0001445|hsa-mir-423&WithFlank&17|+|30117073|30117178</td><td>23</td><td>45</td><td>23</td><td>TGAGGGGCAGAGAGCGAGACTTT</td><td>76503.4</td><td>7.6</td><td>1.0</td><td>7.3</td><td>4.1E-13</td><td>2.2E-11</td><td>BT20/MB468</td><td>MP</td></tr>
<tr><td>>hg38_21_+_8433172_8446622_RNA45SN1_45S_50ntFlanks@12892.12931.40|CCCTCGCCCGTCACGCACCGCACGTTCGTGGGGAACCTGG</td><td>>hg38_21_+_8433172_8446622_RNA45SN1_45S_50ntFlanks</td><td>12892</td><td>12931</td><td>40</td><td>CCCTCGCCCGTCACGCACCGCACGTTCGTGGGGAACCTGG</td><td>186.6</td><td>-9.6</td><td>1.4</td><td>-6.9</td><td>5.2E-12</td><td>2.2E-11</td><td>BT20/MB468</td><td>MITO</td></tr>
<tr><td>>hg38_21_+_8433172_8446622_RNA45SN1_45S_50ntFlanks@12456.12483.28|CCAAGCGTTGGATTGTTCACCCACTAAT</td><td>>hg38_21_+_8433172_8446622_RNA45SN1_45S_50ntFlanks</td><td>12456</td><td>12483</td><td>28</td><td>CCAAGCGTTGGATTGTTCACCCACTAAT</td><td>2628.4</td><td>-8.0</td><td>1.2</td><td>-6.9</td><td>7.3E-12</td><td>2.2E-11</td><td>MB231/MB468</td><td>NUC</td></tr>
<tr><td>>hg38_21_+_8433172_8446622_RNA45SN1_45S_50ntFlanks@7990.8018.29|GACGTGGCGACCCGCTGAATTTAAGCATA</td><td>>hg38_21_+_8433172_8446622_RNA45SN1_45S_50ntFlanks</td><td>7990</td><td>8018</td><td>29</td><td>GACGTGGCGACCCGCTGAATTTAAGCATA</td><td>284.9</td><td>9.3</td><td>1.4</td><td>6.9</td><td>7.1E-12</td><td>2.2E-11</td><td>MB231/MB468</td><td>CYTO</td></tr>
<tr><td>>hg38_21_+_8433172_8446622_RNA45SN1_45S_50ntFlanks@11933.11953.21|AATAAGTGGGAGGCCCCCGGC</td><td>>hg38_21_+_8433172_8446622_RNA45SN1_45S_50ntFlanks</td><td>11933</td><td>11953</td><td>21</td><td>AATAAGTGGGAGGCCCCCGGC</td><td>97.3</td><td>-10.5</td><td>1.5</td><td>-6.9</td><td>6.5E-12</td><td>2.2E-11</td><td>MB231/MB468</td><td>MP</td></tr>
<tr><td>>hg38_21_+_8433172_8446622_RNA45SN1_45S_50ntFlanks@6651.6683.33|CGACTCTTAGCGGTGGATCACTCGGCTCGTGCG</td><td>>hg38_21_+_8433172_8446622_RNA45SN1_45S_50ntFlanks</td><td>6651</td><td>6683</td><td>33</td><td>CGACTCTTAGCGGTGGATCACTCGGCTCGTGCG</td><td>31658.8</td><td>5.9</td><td>0.9</td><td>6.9</td><td>7.3E-12</td><td>2.2E-11</td><td>MB231/MB468</td><td>NUC</td></tr>
<tr><td>>hg38_21_+_8433172_8446622_RNA45SN1_45S_50ntFlanks@11911.11929.19|TGAAGAGACATGAGAGGTG</td><td>>hg38_21_+_8433172_8446622_RNA45SN1_45S_50ntFlanks</td><td>11911</td><td>11929</td><td>19</td><td>TGAAGAGACATGAGAGGTG</td><td>151.3</td><td>-11.1</td><td>1.6</td><td>-6.9</td><td>6.5E-12</td><td>2.2E-11</td><td>MB231/MB468</td><td>MP</td></tr>
<tr><td>>hg38_21_+_8433172_8446622_RNA45SN1_45S_50ntFlanks@5421.5443.23|CGATTGGATGGTTTAGTGAGGCC</td><td>>hg38_21_+_8433172_8446622_RNA45SN1_45S_50ntFlanks</td><td>5421</td><td>5443</td><td>23</td><td>CGATTGGATGGTTTAGTGAGGCC</td><td>1431.5</td><td>-5.2</td><td>0.8</td><td>-6.9</td><td>7.1E-12</td><td>2.2E-11</td><td>MB231/MB468</td><td>CYTO</td></tr>
<tr><td>>MI0000737|hsa-mir-200a&WithFlank&1|+|1167857|1167958@60.82.23&[MIMAT0000682&hsa-miR-200a-3p&offsets|0|+1;m-142&1|+|1167916|1167938&offsets|0|0]|TAACACTGTCTGGTAACGATGTT</td><td>>MI0000737|hsa-mir-200a&WithFlank&1|+|1167857|1167958</td><td>60</td><td>82</td><td>23</td><td>TAACACTGTCTGGTAACGATGTT</td><td>307.0</td><td>-7.3</td><td>1.1</td><td>-6.9</td><td>7.1E-12</td><td>2.2E-11</td><td>MB231/MB468</td><td>CYTO</td></tr>
<tr><td>>hg38_21_+_8433172_8446622_RNA45SN1_45S_50ntFlanks@12456.12490.35|CCAAGCGTTGGATTGTTCACCCACTAATAGGGAAC</td><td>>hg38_21_+_8433172_8446622_RNA45SN1_45S_50ntFlanks</td><td>12456</td><td>12490</td><td>35</td><td>CCAAGCGTTGGATTGTTCACCCACTAATAGGGAAC</td><td>1419.7</td><td>-9.3</td><td>1.4</td><td>-6.9</td><td>6.5E-12</td><td>2.2E-11</td><td>MB231/MB468</td><td>MP</td></tr>
<tr><td>>hg38_21_+_8433172_8446622_RNA45SN1_45S_50ntFlanks@12347.12364.18|AGAAAAGTTACCACAGGG</td><td>>hg38_21_+_8433172_8446622_RNA45SN1_45S_50ntFlanks</td><td>12347</td><td>12364</td><td>18</td><td>AGAAAAGTTACCACAGGG</td><td>164.2</td><td>-11.2</td><td>1.6</td><td>-6.9</td><td>6.5E-12</td><td>2.2E-11</td><td>MB231/MB468</td><td>MP</td></tr>
</table>

<table>
<tr><td>>trna119_LysCTT_1_-_145395522_145395594@3.33.31,trna11_LysCTT_5_-_180648979_180649051@3.33.31,trna13_LysCTT_6_+_26556774_26556846@3.33.31,trna7_LysCTT_16_+_3225692_3225764@3.33.31,trna9_LysCTT_5_+_180634755_180634827@3.33.31|CCGGCTAGCTCAGTCGGTAGAGCATGAGACT</td><td>>trna119_LysCTT_1_-_145395522_145395594</td><td>3</td><td>33</td><td>31</td><td>CCGGCTAGCTCAGTCGGTAGAGCATGAGACT</td><td>1731.0</td><td>10.3</td><td>1.5</td><td>6.9</td><td>7.3E-12</td><td>2.2E-11</td><td>MB231/MB468</td><td>NUC</td></tr>
<tr><td>>hg38_21_+_8433172_8446622_RNA45SN1_45S_50ntFlanks@11874.11908.35|AGAAGACCCTGTTGAGCTTGACTCTAGTCTGGCAC</td><td>>hg38_21_+_8433172_8446622_RNA45SN1_45S_50ntFlanks</td><td>11874</td><td>11908</td><td>35</td><td>AGAAGACCCTGTTGAGCTTGACTCTAGTCTGGCAC</td><td>203.9</td><td>-11.5</td><td>1.7</td><td>-6.9</td><td>6.5E-12</td><td>2.2E-11</td><td>MB231/MB468</td><td>MP</td></tr>
<tr><td>>hg38_21_+_8433172_8446622_RNA45SN1_45S_50ntFlanks@3984.4024.41|GCGGCTTTGGTGACTCTAGATAACCTCGGGCCGATCGCACG</td><td>>hg38_21_+_8433172_8446622_RNA45SN1_45S_50ntFlanks</td><td>3984</td><td>4024</td><td>41</td><td>GCGGCTTTGGTGACTCTAGATAACCTCGGGCCGATCGCACG</td><td>973.2</td><td>-8.6</td><td>1.2</td><td>-6.9</td><td>7.1E-12</td><td>2.2E-11</td><td>MB231/MB468</td><td>CYTO</td></tr>
<tr><td>>MI0000469|hsa-mir-125a&WithFlank&19|+|51693248|51693345@21.42.22&[MIMAT0000443&hsa-miR-125a-5p&offsets|0|-2;m-63&19|+|51693268|51693289&offsets|0|0]|TCCCTGAGACCCTTTAACCTGT</td><td>>MI0000469|hsa-mir-125a&WithFlank&19|+|51693248|51693345</td><td>21</td><td>42</td><td>22</td><td>TCCCTGAGACCCTTTAACCTGT</td><td>2709.4</td><td>6.0</td><td>0.9</td><td>6.9</td><td>7.2E-12</td><td>2.2E-11</td><td>MB231/MB468</td><td>CYTO</td></tr>
<tr><td>>hg38_21_+_8433172_8446622_RNA45SN1_45S_50ntFlanks@6740.6770.31|CATCGACACTTCGAACGCACTTGCGGCCCCG</td><td>>hg38_21_+_8433172_8446622_RNA45SN1_45S_50ntFlanks</td><td>6740</td><td>6770</td><td>31</td><td>CATCGACACTTCGAACGCACTTGCGGCCCCG</td><td>2519.2</td><td>-10.5</td><td>1.5</td><td>-6.8</td><td>9.2E-12</td><td>2.2E-11</td><td>BT20/MB468</td><td>NUC</td></tr>
<tr><td>>hg38_21_+_8433172_8446622_RNA45SN1_45S_50ntFlanks@12259.12278.20|CTTGATCTTGATTTTCAGTA</td><td>>hg38_21_+_8433172_8446622_RNA45SN1_45S_50ntFlanks</td><td>12259</td><td>12278</td><td>20</td><td>CTTGATCTTGATTTTCAGTA</td><td>96.8</td><td>-10.5</td><td>1.5</td><td>-6.9</td><td>6.6E-12</td><td>2.2E-11</td><td>MB231/MB468</td><td>MP</td></tr>
<tr><td>>hg38_21_+_8433172_8446622_RNA45SN1_45S_50ntFlanks@11718.11742.25|AACGGCGGGAGTAACTATGACTCTC</td><td>>hg38_21_+_8433172_8446622_RNA45SN1_45S_50ntFlanks</td><td>11718</td><td>11742</td><td>25</td><td>AACGGCGGGAGTAACTATGACTCTC</td><td>4368.1</td><td>-5.6</td><td>0.8</td><td>-6.9</td><td>7.2E-12</td><td>2.2E-11</td><td>MB231/MB468</td><td>CYTO</td></tr>
<tr><td>>hg38_21_+_8433172_8446622_RNA45SN1_45S_50ntFlanks@5219.5244.26|GGGCTGCACGCGCGCTACACTGACTG</td><td>>hg38_21_+_8433172_8446622_RNA45SN1_45S_50ntFlanks</td><td>5219</td><td>5244</td><td>26</td><td>GGGCTGCACGCGCGCTACACTGACTG</td><td>652.8</td><td>-13.1</td><td>1.9</td><td>-6.8</td><td>8.7E-12</td><td>2.2E-11</td><td>MB231/MB468</td><td>MITO</td></tr>
<tr><td>>trnaMT_ValTAC_MT_+_1602_1670@44.71.28|ATTTCAACTTAACTTGACCGCTCTGACC</td><td>>trnaMT_ValTAC_MT_+_1602_1670</td><td>44</td><td>71</td><td>28</td><td>ATTTCAACTTAACTTGACCGCTCTGACC</td><td>323.7</td><td>4.7</td><td>0.7</td><td>6.8</td><td>7.4E-12</td><td>2.2E-11</td><td>MB231/MB468</td><td>NUC</td></tr>
<tr><td>>trna9_ArgTCT_1_+_94313129_94313213@1.29.29|GGCTCCGTGGCGCAATGGATAGCGCATTG</td><td>>trna9_ArgTCT_1_+_94313129_94313213</td><td>1</td><td>29</td><td>29</td><td>GGCTCCGTGGCGCAATGGATAGCGCATTG</td><td>92.0</td><td>-10.4</td><td>1.5</td><td>-6.9</td><td>6.6E-12</td><td>2.2E-11</td><td>MB231/MB468</td><td>MP</td></tr>
<tr><td>>hg38_21_+_8433172_8446622_RNA45SN1_45S_50ntFlanks@4445.4462.18|CTTGCCTCTCGGCGCCCC</td><td>>hg38_21_+_8433172_8446622_RNA45SN1_45S_50ntFlanks</td><td>4445</td><td>4462</td><td>18</td><td>CTTGCCTCTCGGCGCCCC</td><td>142.6</td><td>-11.0</td><td>1.6</td><td>-6.9</td><td>6.6E-12</td><td>2.2E-11</td><td>MB231/MB468</td><td>MP</td></tr>
<tr><td>>MI0000088|hsa-mir-30a&WithFlank&6|-|71403545|71403627@13.35.23&[MIMAT0000087&hsa-miR-30a-5p&offsets|+1|+2;m-15&6|-|71403595|71403616&offsets|+1|+2]|GTAAACATCCTCGACTGGAAGCT</td><td>>MI0000088|hsa-mir-30a&WithFlank&6|-|71403545|71403627</td><td>13</td><td>35</td><td>23</td><td>GTAAACATCCTCGACTGGAAGCT</td><td>1604.2</td><td>7.1</td><td>1.0</td><td>6.8</td><td>8.7E-12</td><td>2.2E-11</td><td>MB231/MB468</td><td>MITO</td></tr>
<tr><td>>hg38_21_+_8433172_8446622_RNA45SN1_45S_50ntFlanks@11873.11893.21|AAGAAGACCCTGTTGAGCTTG</td><td>>hg38_21_+_8433172_8446622_RNA45SN1_45S_50ntFlanks</td><td>11873</td><td>11893</td><td>21</td><td>AAGAAGACCCTGTTGAGCTTG</td><td>3781.7</td><td>-8.2</td><td>1.2</td><td>-6.8</td><td>7.4E-12</td><td>2.2E-11</td><td>MB231/MB468</td><td>NUC</td></tr>
<tr><td>>hg38_1_-_228634819_228635039_RNA5S12_5S_50ntFlanks@135.170.36|GGGAGACCGCCTGGGAATACCGGGTGCTGTAGGCTT</td><td>>hg38_1_-_228634819_228635039_RNA5S12_5S_50ntFlanks</td><td>135</td><td>170</td><td>36</td><td>GGGAGACCGCCTGGGAATACCGGGTGCTGTAGGCTT</td><td>26710.6</td><td>5.7</td><td>0.8</td><td>6.9</td><td>7.3E-12</td><td>2.2E-11</td><td>MB231/MB468</td><td>CYTO</td></tr>
<tr><td>>hg38_21_+_8433172_8446622_RNA45SN1_45S_50ntFlanks@10590.10607.18|CGGGGAGAGTTCTCTTTT</td><td>>hg38_21_+_8433172_8446622_RNA45SN1_45S_50ntFlanks</td><td>10590</td><td>10607</td><td>18</td><td>CGGGGAGAGTTCTCTTTT</td><td>153.3</td><td>-11.1</td><td>1.6</td><td>-6.9</td><td>6.6E-12</td><td>2.2E-11</td><td>MB231/MB468</td><td>MP</td></tr>
<tr><td>>hg38_21_+_8433172_8446622_RNA45SN1_45S_50ntFlanks@3802.3829.28|CATTAAATCAGTTATGGTTCCTTTGGTC</td><td>>hg38_21_+_8433172_8446622_RNA45SN1_45S_50ntFlanks</td><td>3802</td><td>3829</td><td>28</td><td>CATTAAATCAGTTATGGTTCCTTTGGTC</td><td>146.0</td><td>-11.1</td><td>1.6</td><td>-6.9</td><td>6.6E-12</td><td>2.2E-11</td><td>MB231/MB468</td><td>MP</td></tr>
<tr><td>>hg38_21_+_8433172_8446622_RNA45SN1_45S_50ntFlanks@8182.8211.30|ATCGAGGCCCAGCCCGTGGACGGTGTGAGG</td><td>>hg38_21_+_8433172_8446622_RNA45SN1_45S_50ntFlanks</td><td>8182</td><td>8211</td><td>30</td><td>ATCGAGGCCCAGCCCGTGGACGGTGTGAGG</td><td>383.1</td><td>-11.7</td><td>1.6</td><td>-7.2</td><td>4.2E-13</td><td>2.2E-11</td><td>BT20/MB468</td><td>MP</td></tr>
<tr><td>>MI0000071|hsa-mir-17&WithFlank&13|+|91350599|91350694@20.41.22&[MIMAT0000070&hsa-miR-17-5p&offsets|0|-1;m-88&13|+|91350618|91350640&offsets|0|-1]|CAAAGTGCTTACAGTGCAGGTA</td><td>>MI0000071|hsa-mir-17&WithFlank&13|+|91350599|91350694</td><td>20</td><td>41</td><td>22</td><td>CAAAGTGCTTACAGTGCAGGTA</td><td>589.7</td><td>4.6</td><td>0.7</td><td>6.8</td><td>7.5E-12</td><td>2.2E-11</td><td>MB231/MB468</td><td>NUC</td></tr>
<tr><td>>hg38_21_+_8433172_8446622_RNA45SN1_45S_50ntFlanks@11561.11588.28|GCGCCTAGCAGCCGACTTAGAACTGGTG</td><td>>hg38_21_+_8433172_8446622_RNA45SN1_45S_50ntFlanks</td><td>11561</td><td>11588</td><td>28</td><td>GCGCCTAGCAGCCGACTTAGAACTGGTG</td><td>1138.0</td><td>-6.6</td><td>1.0</td><td>-6.9</td><td>7.3E-12</td><td>2.2E-11</td><td>MB231/MB468</td><td>CYTO</td></tr>
<tr><td>>hg38_21_+_8433172_8446622_RNA45SN1_45S_50ntFlanks@5073.5102.30|AATTCCGATAACGAACGAGACTCTGGCATG</td><td>>hg38_21_+_8433172_8446622_RNA45SN1_45S_50ntFlanks</td><td>5073</td><td>5102</td><td>30</td><td>AATTCCGATAACGAACGAGACTCTGGCATG</td><td>1025.0</td><td>-9.8</td><td>1.4</td><td>-6.8</td><td>8.8E-12</td><td>2.2E-11</td><td>MB231/MB468</td><td>MITO</td></tr>
<tr><td>>hg38_21_+_8433172_8446622_RNA45SN1_45S_50ntFlanks@8194.8225.32|CCCGTGGACGGTGTGAGGCCGGTAGCGGCCCC</td><td>>hg38_21_+_8433172_8446622_RNA45SN1_45S_50ntFlanks</td><td>8194</td><td>8225</td><td>32</td><td>CCCGTGGACGGTGTGAGGCCGGTAGCGGCCCC</td><td>479.0</td><td>-5.6</td><td>0.8</td><td>-6.9</td><td>7.3E-12</td><td>2.2E-11</td><td>MB231/MB468</td><td>CYTO</td></tr>
<tr><td>>hg38_21_+_8433172_8446622_RNA45SN1_45S_50ntFlanks@12489.12518.30|ACGTGAGCTGGGTTTAGACCGTCGTGAGAC</td><td>>hg38_21_+_8433172_8446622_RNA45SN1_45S_50ntFlanks</td><td>12489</td><td>12518</td><td>30</td><td>ACGTGAGCTGGGTTTAGACCGTCGTGAGAC</td><td>111.8</td><td>-10.7</td><td>1.6</td><td>-6.9</td><td>6.7E-12</td><td>2.2E-11</td><td>MB231/MB468</td><td>MP</td></tr>
<tr><td>>hg38_21_+_8433172_8446622_RNA45SN1_45S_50ntFlanks@8168.8191.24|CCCAAGTCCTTCTGATCGAGGCCC</td><td>>hg38_21_+_8433172_8446622_RNA45SN1_45S_50ntFlanks</td><td>8168</td><td>8191</td><td>24</td><td>CCCAAGTCCTTCTGATCGAGGCCC</td><td>112.2</td><td>-10.7</td><td>1.6</td><td>-6.9</td><td>6.7E-12</td><td>2.2E-11</td><td>MB231/MB468</td><td>MP</td></tr>
<tr><td>>hg38_21_+_8433172_8446622_RNA45SN1_45S_50ntFlanks@6669.6689.21|CACTCGGCTCGTGCGTCGATG</td><td>>hg38_21_+_8433172_8446622_RNA45SN1_45S_50ntFlanks</td><td>6669</td><td>6689</td><td>21</td><td>CACTCGGCTCGTGCGTCGATG</td><td>234.4</td><td>-11.7</td><td>1.7</td><td>-6.9</td><td>6.7E-12</td><td>2.2E-11</td><td>MB231/MB468</td><td>MP</td></tr>
<tr><td>>MI0000477|hsa-mir-146a&WithFlank&5|+|160485346|160485456@27.48.22(+1A)&[MIMAT0000449&hsa-miR-146a-5p&offsets|0|0(+1A);m-129&5|+|160485372|160485393&offsets|0|0(+1A)]|TGAGAACTGAATTCCATGGGTTA</td><td>>MI0000477|hsa-mir-146a&WithFlank&5|+|160485346|160485456</td><td>27</td><td>48</td><td>22(+1A)</td><td>TGAGAACTGAATTCCATGGGTTA</td><td>494.9</td><td>-12.0</td><td>1.7</td><td>-7.2</td><td>7.8E-13</td><td>2.2E-11</td><td>BT20/MB231</td><td>NUC</td></tr>
<tr><td>>MI0000809|hsa-mir-151a&WithFlank&8|-|140732558|140732659@53.74.22(+1U)&[MIMAT0000757&hsa-miR-151a-3p&offsets|0|+1(+1U);m-30&8|-|140732587|140732607&offsets|0|+1(+1U)]|CTAGACTGAAGCTCCTTGAGGAT</td><td>>MI0000809|hsa-mir-151a&WithFlank&8|-|140732558|140732659</td><td>53</td><td>74</td><td>22(+1U)</td><td>CTAGACTGAAGCTCCTTGAGGAT</td><td>702.7</td><td>6.7</td><td>1.0</td><td>6.9</td><td>7.3E-12</td><td>2.2E-11</td><td>MB231/MB468</td><td>CYTO</td></tr>
<tr><td>>hg38_21_+_8433172_8446622_RNA45SN1_45S_50ntFlanks@12588.12605.18|GGTTCAGACATTTGGTGT</td><td>>hg38_21_+_8433172_8446622_RNA45SN1_45S_50ntFlanks</td><td>12588</td><td>12605</td><td>18</td><td>GGTTCAGACATTTGGTGT</td><td>91.1</td><td>-10.4</td><td>1.5</td><td>-6.9</td><td>6.7E-12</td><td>2.2E-11</td><td>MB231/MB468</td><td>MP</td></tr>
<tr><td>>hg38_21_+_8433172_8446622_RNA45SN1_45S_50ntFlanks@483.505.23|TCGATGTGGTGACGTCGTGCTCT</td><td>>hg38_21_+_8433172_8446622_RNA45SN1_45S_50ntFlanks</td><td>483</td><td>505</td><td>23</td><td>TCGATGTGGTGACGTCGTGCTCT</td><td>1294.0</td><td>5.0</td><td>0.7</td><td>6.8</td><td>7.6E-12</td><td>2.2E-11</td><td>MB231/MB468</td><td>NUC</td></tr>
<tr><td>>trna16_GlnTTG_17_+_47269890_47269961@1.29.29,trna64_GlnTTG_6_+_28557156_28557227@1.29.29,trna84_GlnTTG_6_+_145503859_145503930@1.29.29|GGTCCCATGGTGTAATGGTTAGCACTCTG</td><td>>trna16_GlnTTG_17_+_47269890_47269961</td><td>1</td><td>29</td><td>29</td><td>GGTCCCATGGTGTAATGGTTAGCACTCTG</td><td>1010.8</td><td>-8.8</td><td>1.3</td><td>-6.8</td><td>7.6E-12</td><td>2.2E-11</td><td>MB231/MB468</td><td>NUC</td></tr>
<tr><td>>hg38_21_+_8433172_8446622_RNA45SN1_45S_50ntFlanks@10458.10477.20|ATGGCCTCCGTTGCCCTCGG</td><td>>hg38_21_+_8433172_8446622_RNA45SN1_45S_50ntFlanks</td><td>10458</td><td>10477</td><td>20</td><td>ATGGCCTCCGTTGCCCTCGG</td><td>5743.6</td><td>-8.1</td><td>1.2</td><td>-6.8</td><td>7.4E-12</td><td>2.2E-11</td><td>MB231/MB468</td><td>CYTO</td></tr>
<tr><td>>hg38_21_+_8433172_8446622_RNA45SN1_45S_50ntFlanks@7983.8004.22|CAGATCAGACGTGGCGACCCGC</td><td>>hg38_21_+_8433172_8446622_RNA45SN1_45S_50ntFlanks</td><td>7983</td><td>8004</td><td>22</td><td>CAGATCAGACGTGGCGACCCGC</td><td>1609.5</td><td>-9.3</td><td>1.4</td><td>-6.9</td><td>6.7E-12</td><td>2.2E-11</td><td>MB231/MB468</td><td>MP</td></tr>
<tr><td>>hg38_21_+_8433172_8446622_RNA45SN1_45S_50ntFlanks@10830.10858.29|CAAGCCGGATCCGTAACTTCGGGATAAGG</td><td>>hg38_21_+_8433172_8446622_RNA45SN1_45S_50ntFlanks</td><td>10830</td><td>10858</td><td>29</td><td>CAAGCCGGATCCGTAACTTCGGGATAAGG</td><td>551.0</td><td>-6.8</td><td>1.0</td><td>-6.8</td><td>7.4E-12</td><td>2.3E-11</td><td>MB231/MB468</td><td>CYTO</td></tr>
<tr><td>>hg38_21_+_8433172_8446622_RNA45SN1_45S_50ntFlanks@3854.3884.31|ACTGTGGTAATTCTAGAGCTAATACATGCCG</td><td>>hg38_21_+_8433172_8446622_RNA45SN1_45S_50ntFlanks</td><td>3854</td><td>3884</td><td>31</td><td>ACTGTGGTAATTCTAGAGCTAATACATGCCG</td><td>177.4</td><td>-11.3</td><td>1.7</td><td>-6.9</td><td>6.8E-12</td><td>2.3E-11</td><td>MB231/MB468</td><td>MP</td></tr>
<tr><td>>hg38_21_+_8433172_8446622_RNA45SN1_45S_50ntFlanks@12444.12475.32|AAGCAGAATTCACCAAGCGTTGGATTGTTCAC</td><td>>hg38_21_+_8433172_8446622_RNA45SN1_45S_50ntFlanks</td><td>12444</td><td>12475</td><td>32</td><td>AAGCAGAATTCACCAAGCGTTGGATTGTTCAC</td><td>193.1</td><td>-11.5</td><td>1.7</td><td>-6.9</td><td>6.8E-12</td><td>2.3E-11</td><td>MB231/MB468</td><td>MP</td></tr>
<tr><td>>MI0000239|hsa-mir-197&WithFlank&1|+|109598887|109598973@54.74.21(+1A)&[MIMAT0000227&hsa-miR-197-3p&offsets|0|-1(+1A);m-99&1|+|109598940|109598961&offsets|0|-1(+1A)]|TTCACCACCTTCTCCACCCAGA</td><td>>MI0000239|hsa-mir-197&WithFlank&1|+|109598887|109598973</td><td>54</td><td>74</td><td>21(+1A)</td><td>TTCACCACCTTCTCCACCCAGA</td><td>205.9</td><td>3.2</td><td>0.5</td><td>6.8</td><td>9.5E-12</td><td>2.3E-11</td><td>BT20/MB468</td><td>NUC</td></tr>
<tr><td>>hg38_21_+_8433172_8446622_RNA45SN1_45S_50ntFlanks@12640.12667.28|ACCATCTGTGGGATTATGACTGAACGCC</td><td>>hg38_21_+_8433172_8446622_RNA45SN1_45S_50ntFlanks</td><td>12640</td><td>12667</td><td>28</td><td>ACCATCTGTGGGATTATGACTGAACGCC</td><td>291.1</td><td>7.1</td><td>1.0</td><td>6.9</td><td>7.0E-12</td><td>2.3E-11</td><td>BT20/MB468</td><td>CYTO</td></tr>
<tr><td>>trnaMT_SerGCT_MT_+_12207_12265@8.38.31|CTCACAAGAACTGCTAACTCATGCCCCCATG</td><td>>trnaMT_SerGCT_MT_+_12207_12265</td><td>8</td><td>38</td><td>31</td><td>CTCACAAGAACTGCTAACTCATGCCCCCATG</td><td>167.8</td><td>-11.3</td><td>1.6</td><td>-6.9</td><td>6.8E-12</td><td>2.3E-11</td><td>MB231/MB468</td><td>MP</td></tr>
<tr><td>>hg38_21_+_8433172_8446622_RNA45SN1_45S_50ntFlanks@10478.10512.35|CCGATCGAAAGGGAGTCGGGTTCAGATCCCCGAAT</td><td>>hg38_21_+_8433172_8446622_RNA45SN1_45S_50ntFlanks</td><td>10478</td><td>10512</td><td>35</td><td>CCGATCGAAAGGGAGTCGGGTTCAGATCCCCGAAT</td><td>115.5</td><td>-10.7</td><td>1.6</td><td>-6.9</td><td>6.8E-12</td><td>2.3E-11</td><td>MB231/MB468</td><td>MP</td></tr>
<tr><td>>hg38_21_+_8433172_8446622_RNA45SN1_45S_50ntFlanks@4602.4623.22|TTCGGAACTGAGGCCATGATTA</td><td>>hg38_21_+_8433172_8446622_RNA45SN1_45S_50ntFlanks</td><td>4602</td><td>4623</td><td>22</td><td>TTCGGAACTGAGGCCATGATTA</td><td>95.3</td><td>-10.4</td><td>1.5</td><td>-6.9</td><td>6.8E-12</td><td>2.3E-11</td><td>MB231/MB468</td><td>MP</td></tr>
<tr><td>>hg38_21_+_8433172_8446622_RNA45SN1_45S_50ntFlanks@10689.10710.22|GCGTCCGGTGAGCTCTCGCTGG</td><td>>hg38_21_+_8433172_8446622_RNA45SN1_45S_50ntFlanks</td><td>10689</td><td>10710</td><td>22</td><td>GCGTCCGGTGAGCTCTCGCTGG</td><td>207.2</td><td>-10.0</td><td>1.5</td><td>-6.9</td><td>7.0E-12</td><td>2.3E-11</td><td>BT20/MB468</td><td>CYTO</td></tr>
<tr><td>>trna129_MetCAT_6_-_27870271_27870342@29.51.23,trna142_MetCAT_6_-_27560600_27560671@29.51.23,trna150_MetCAT_6_-_27300764_27300835@29.51.23,trna169_MetCAT_6_-_26330529_26330600@29.51.23,trna171_MetCAT_6_-_26313352_26313423@29.51.23,trna20_MetCAT_17_-_80452597_80452668@29.51.23,trna2_MetCAT_6_+_26286754_26286825@29.51.23,trna32_MetCAT_1_+_153643726_153643797@29.51.23,trna61_MetCAT_6_+_27745664_27745735@29.51.23|GGCCCATAACCCAGAGGTCGATG</td><td>>trna129_MetCAT_6_-_27870271_27870342</td><td>29</td><td>51</td><td>23</td><td>GGCCCATAACCCAGAGGTCGATG</td><td>158.6</td><td>-11.2</td><td>1.6</td><td>-6.9</td><td>6.8E-12</td><td>2.3E-11</td><td>MB231/MB468</td><td>MP</td></tr>
<tr><td>>hg38_21_+_8433172_8446622_RNA45SN1_45S_50ntFlanks@4047.4065.19|ATTCGAACGTCTGCCCTAT</td><td>>hg38_21_+_8433172_8446622_RNA45SN1_45S_50ntFlanks</td><td>4047</td><td>4065</td><td>19</td><td>ATTCGAACGTCTGCCCTAT</td><td>702.2</td><td>-9.1</td><td>1.3</td><td>-6.8</td><td>9.0E-12</td><td>2.3E-11</td><td>MB231/MB468</td><td>MITO</td></tr>
<tr><td>>hg38_21_+_8433172_8446622_RNA45SN1_45S_50ntFlanks@10361.10383.23|GAAGGCCGAAGTGGAGAAGGGTT</td><td>>hg38_21_+_8433172_8446622_RNA45SN1_45S_50ntFlanks</td><td>10361</td><td>10383</td><td>23</td><td>GAAGGCCGAAGTGGAGAAGGGTT</td><td>894.3</td><td>-7.4</td><td>1.1</td><td>-6.9</td><td>6.8E-12</td><td>2.3E-11</td><td>MB231/MB468</td><td>MP</td></tr>
<tr><td>>hg38_21_+_8433172_8446622_RNA45SN1_45S_50ntFlanks@12525.12546.22|GTTTTACCCTACTGATGATGTG</td><td>>hg38_21_+_8433172_8446622_RNA45SN1_45S_50ntFlanks</td><td>12525</td><td>12546</td><td>22</td><td>GTTTTACCCTACTGATGATGTG</td><td>157.6</td><td>-11.2</td><td>1.6</td><td>-6.9</td><td>6.9E-12</td><td>2.3E-11</td><td>MB231/MB468</td><td>MP</td></tr>
<tr><td>>hg38_21_+_8433172_8446622_RNA45SN1_45S_50ntFlanks@12959.12990.32|TCTGGGTCGGGGTTTCGTACGTAGCAGAGCAG</td><td>>hg38_21_+_8433172_8446622_RNA45SN1_45S_50ntFlanks</td><td>12959</td><td>12990</td><td>32</td><td>TCTGGGTCGGGGTTTCGTACGTAGCAGAGCAG</td><td>580.0</td><td>-9.2</td><td>1.3</td><td>-6.8</td><td>9.6E-12</td><td>2.3E-11</td><td>BT20/MB468</td><td>NUC</td></tr>
<tr><td>>hg38_21_+_8433172_8446622_RNA45SN1_45S_50ntFlanks@12306.12326.21|CACGATCCTTCTGACCTTTTG</td><td>>hg38_21_+_8433172_8446622_RNA45SN1_45S_50ntFlanks</td><td>12306</td><td>12326</td><td>21</td><td>CACGATCCTTCTGACCTTTTG</td><td>1834.7</td><td>-8.2</td><td>1.2</td><td>-6.8</td><td>9.0E-12</td><td>2.3E-11</td><td>MB231/MB468</td><td>MITO</td></tr>
<tr><td>>hg38_21_+_8433172_8446622_RNA45SN1_45S_50ntFlanks@7984.8010.27|AGATCAGACGTGGCGACCCGCTGAATT</td><td>>hg38_21_+_8433172_8446622_RNA45SN1_45S_50ntFlanks</td><td>7984</td><td>8010</td><td>27</td><td>AGATCAGACGTGGCGACCCGCTGAATT</td><td>154.0</td><td>-11.1</td><td>1.6</td><td>-6.9</td><td>6.9E-12</td><td>2.3E-11</td><td>MB231/MB468</td><td>MP</td></tr>
</table>

| | | | | | | | | | | | | | |
|---|---|---|---|---|---|---|---|---|---|---|---|---|---|
| >MI0000266\|hsa-mir-10a&WithFlank&17\|-\|48579832\|48579953@29.50.22&[MIMAT0000253&hsa-miR-10a-5p&offsets\|+1\|0;m-18&17\|-\|48579905\|48579926&offsets\|+1\|+1]\|ACCCTGTAGATCCGAATTTGTG | >MI0000266\|hsa-mir-10a&WithFlank&17\|-\|48579832\|48579953 | 29 | 50 | 22 | ACCCTGTAGATCCGAATTTGTG | 1744.7 | 9.2 | 1.3 | 6.9 | 6.9E-12 | 2.3E-11 | MB231/MB468 | MP |
| >hg38_21_+_8433172_8446622_RNA45SN1_45S_50ntFlanks@6668.6699.32\|TCACTCGGCTCGTGCGTCGATGAAGAACGCAG | >hg38_21_+_8433172_8446622_RNA45SN1_45S_50ntFlanks | 6668 | 6699 | 32 | TCACTCGGCTCGTGCGTCGATGAAGAACGCAG | 117.8 | -10.7 | 1.6 | -6.9 | 6.9E-12 | 2.3E-11 | MB231/MB468 | MP |
| >hg38_21_+_8433172_8446622_RNA45SN1_45S_50ntFlanks@3996.4024.29\|ACTCTAGATAACCTCGGGCCGATCGCACG | >hg38_21_+_8433172_8446622_RNA45SN1_45S_50ntFlanks | 3996 | 4024 | 29 | ACTCTAGATAACCTCGGGCCGATCGCACG | 1051.7 | -5.0 | 0.7 | -6.8 | 7.8E-12 | 2.3E-11 | MB231/MB468 | NUC |
| >hg38_21_+_8433172_8446622_RNA45SN1_45S_50ntFlanks@8324.8356.33\|CCGATAGTCAACAAGTACCGTAAGGGAAAGTTG | >hg38_21_+_8433172_8446622_RNA45SN1_45S_50ntFlanks | 8324 | 8356 | 33 | CCGATAGTCAACAAGTACCGTAAGGGAAAGTTG | 297.2 | -11.3 | 1.6 | -7.2 | 4.4E-13 | 2.3E-11 | BT20/MB468 | MP |
| >hg38_21_+_8433172_8446622_RNA45SN1_45S_50ntFlanks@11874.11897.24\|AGAAGACCCTGTTGAGCTTGACTC | >hg38_21_+_8433172_8446622_RNA45SN1_45S_50ntFlanks | 11874 | 11897 | 24 | AGAAGACCCTGTTGAGCTTGACTC | 658.3 | -8.6 | 1.3 | -6.9 | 6.9E-12 | 2.3E-11 | MB231/MB468 | MP |
| >hg38_21_+_8433172_8446622_RNA45SN1_45S_50ntFlanks@10907.10930.24\|GGGCGCGCGCCGCGGCTGGACGAG | >hg38_21_+_8433172_8446622_RNA45SN1_45S_50ntFlanks | 10907 | 10930 | 24 | GGGCGCGCGCCGCGGCTGGACGAG | 4479.5 | 12.2 | 1.8 | 6.9 | 5.6E-12 | 2.3E-11 | BT20/MB468 | MITO |
| >trnaMT_GluTTC_MT_-_14674_14742@39.57.19,trnalookalike8_GluTTC_5_-_93905172_93905240@39.57.19\|ATTGGTCGTGGTTGTAGTC | >trnaMT_GluTTC_MT_-_14674_14742 | 39 | 57 | 19 | ATTGGTCGTGGTTGTAGTC | 782.4 | 8.0 | 1.2 | 6.9 | 5.6E-12 | 2.3E-11 | BT20/MB468 | MITO |
| >hg38_21_+_8433172_8446622_RNA45SN1_45S_50ntFlanks@5468.5499.32\|GCCCACGGCCCTGGCGGAGCGCTGAGAAGACG | >hg38_21_+_8433172_8446622_RNA45SN1_45S_50ntFlanks | 5468 | 5499 | 32 | GCCCACGGCCCTGGCGGAGCGCTGAGAAGACG | 209.1 | -11.6 | 1.7 | -6.9 | 7.0E-12 | 2.3E-11 | MB231/MB468 | MP |
| >hg38_21_+_8433172_8446622_RNA45SN1_45S_50ntFlanks@12412.12441.30\|ATCCTTCGATGTCGGCTCTTCCTATCATTG | >hg38_21_+_8433172_8446622_RNA45SN1_45S_50ntFlanks | 12412 | 12441 | 30 | ATCCTTCGATGTCGGCTCTTCCTATCATTG | 199.3 | -9.7 | 1.4 | -6.9 | 5.6E-12 | 2.3E-11 | BT20/MB468 | MITO |
| >hg38_21_+_8433172_8446622_RNA45SN1_45S_50ntFlanks@6723.6744.22\|TTGCAGGACACATTGATCATCG | >hg38_21_+_8433172_8446622_RNA45SN1_45S_50ntFlanks | 6723 | 6744 | 22 | TTGCAGGACACATTGATCATCG | 674.9 | -4.7 | 0.7 | -6.8 | 7.8E-12 | 2.3E-11 | MB231/MB468 | NUC |
| >hg38_21_+_8433172_8446622_RNA45SN1_45S_50ntFlanks@8357.8378.22\|AAAAGAACTTTGAAGAGAGAGT | >hg38_21_+_8433172_8446622_RNA45SN1_45S_50ntFlanks | 8357 | 8378 | 22 | AAAAGAACTTTGAAGAGAGAGT | 4375.1 | -5.1 | 0.7 | -6.9 | 7.2E-12 | 2.3E-11 | BT20/MB468 | CYTO |
| >hg38_21_+_8433172_8446622_RNA45SN1_45S_50ntFlanks@11170.11188.19\|CCGCGAGGGGGGCCCGGGC | >hg38_21_+_8433172_8446622_RNA45SN1_45S_50ntFlanks | 11170 | 11188 | 19 | CCGCGAGGGGGGCCCGGGC | 379.3 | 8.6 | 1.3 | 6.9 | 7.2E-12 | 2.3E-11 | BT20/MB468 | CYTO |
| >hg38_21_+_8433172_8446622_RNA45SN1_45S_50ntFlanks@12407.12425.19\|TTTTGATCCTTCGATGTCG | >hg38_21_+_8433172_8446622_RNA45SN1_45S_50ntFlanks | 12407 | 12425 | 19 | TTTTGATCCTTCGATGTCG | 108.0 | -10.6 | 1.5 | -6.9 | 7.0E-12 | 2.3E-11 | MB231/MB468 | MP |
| >trna10_AspGTC_12_-_125424193_125424264@32.70.39,trna12_AspGTC_12_-_125411891_125411962@32.70.39,trna144_AspGTC_6_-_27551236_27551307@32.70.39,trna38_AspGTC_17_-_8125556_8125627@32.70.39,trna45_AspGTC_6_+_27447453_27447524@32.70.39,trna48_AspGTC_6_+_27471523_27471594@32.70.39,trna4_AspGTC_12_+_96429799_96429870@32.70.39,trna69_AspGTC_1_-_161440205_161440276@32.70.39,trna72_AspGTC_1_-_161432824_161432895@32.70.39,trna75_AspGTC_1_-_161425414_161425485@32.70.39,trna78_AspGTC_1_-_161418033_161418104@32.70.39,trna81_AspGTC_1_-_161410615_161410686@32.70.39\|CTGTCACGCGGGAGACCGGGGTTCGATTCCCCGACGGGG | >trna10_AspGTC_12_-_125424193_125424264 | 32 | 70 | 39 | CTGTCACGCGGGAGACCGGGGTTCGATTCCCCGACGGGG | 237.5 | -11.7 | 1.7 | -6.8 | 9.2E-12 | 2.3E-11 | MB231/MB468 | MITO |
| >MI0000542\|hsa-mir-320a&WithFlank&8\|-\|22244960\|22245043@48.68.21(+2U)&[MIMAT0000510&hsa-miR-320a-3p&offsets\|0\|-1(+2U);m-60&8\|-\|22244975\|22244996&offsets\|0\|-1(+2U)]\|AAAAGCTGGGTTGAGAGGGCGTT | >MI0000542\|hsa-mir-320a&WithFlank&8\|-\|22244960\|22245043 | 48 | 68 | 21(+2U) | AAAAGCTGGGTTGAGAGGGCGTT | 520.5 | 5.3 | 0.8 | 6.8 | 7.9E-12 | 2.3E-11 | MB231/MB468 | NUC |
| >hg38_21_+_8433172_8446622_RNA45SN1_45S_50ntFlanks@10905.10936.32\|CTGGGCGCGCGCCGCGGCTGGACGAGGCGCCG | >hg38_21_+_8433172_8446622_RNA45SN1_45S_50ntFlanks | 10905 | 10936 | 32 | CTGGGCGCGCGCCGCGGCTGGACGAGGCGCCG | 273.5 | -11.2 | 1.5 | -7.2 | 4.5E-13 | 2.3E-11 | BT20/MB468 | MP |
| >hg38_21_+_8433172_8446622_RNA45SN1_45S_50ntFlanks@6690.6720.31\|AAGAACGCAGCTAGCTGCGAGAATTAATGTG | >hg38_21_+_8433172_8446622_RNA45SN1_45S_50ntFlanks | 6690 | 6720 | 31 | AAGAACGCAGCTAGCTGCGAGAATTAATGTG | 1688.8 | -7.2 | 1.0 | -6.9 | 5.7E-12 | 2.3E-11 | BT20/MB468 | MITO |
| >hg38_21_+_8433172_8446622_RNA45SN1_45S_50ntFlanks@10391.10419.29\|AACAGCAGTTGAACATGGGTCAGTCGGTC | >hg38_21_+_8433172_8446622_RNA45SN1_45S_50ntFlanks | 10391 | 10419 | 29 | AACAGCAGTTGAACATGGGTCAGTCGGTC | 809.7 | -8.8 | 1.3 | -6.8 | 7.9E-12 | 2.3E-11 | MB231/MB468 | NUC |
| >hg38_21_+_8433172_8446622_RNA45SN1_45S_50ntFlanks@9625.9658.34\|GTAGCTGGTTCCCTCCGAAGTTTCCCTCAGGATA | >hg38_21_+_8433172_8446622_RNA45SN1_45S_50ntFlanks | 9625 | 9658 | 34 | GTAGCTGGTTCCCTCCGAAGTTTCCCTCAGGATA | 372.1 | -8.2 | 1.2 | -6.8 | 9.3E-12 | 2.3E-11 | MB231/MB468 | MITO |
| >trna116_GluCTC_1_-_145399233_145399304@30.71.42,trna71_GluCTC_1_-_161439189_161439260@30.71.42,trna74_GluCTC_1_-_161431809_161431880@30.71.42,trna77_GluCTC_1_-_161424398_161424469@30.71.42,trna77_GluCTC_6_+_28949976_28950047@30.71.42,trna80_GluCTC_1_-_161417018_161417089@30.71.42,trna87_GluCTC_6_-_126101393_126101464@30.71.42\|CGCTCTCACCGCCGCGGCCCGGGTTCGATTCCCGGTCAGGGA | >trna116_GluCTC_1_-_145399233_145399304 | 30 | 71 | 42 | CGCTCTCACCGCCGCGGCCCGGGTTCGATTCCCGGTCAGGGA | 173.1 | -10.4 | 1.5 | -6.8 | 8.0E-12 | 2.3E-11 | MB231/MB468 | NUC |
| >MI0000446\|hsa-mir-125b-1&WithFlank&11\|-\|122099751\|122099850@21.42.22(+1C);MI0000470\|hsa-mir-125b-2&WithFlank&21\|+\|16590231\|16590331@23.44.22(+1C)&[MIMAT0000423&hsa-miR-125b-5p&offsets\|0\|0(+1C);m-67&11\|-\|122099809\|122099830&offsets\|0\|0(+1C)];[MIMAT0000423_1&hsa-miR-125b-5p&offsets\|0\|0(+1C);m-68&21\|+\|16590253\|16590274&offsets\|0\|0(+1C)]\|TCCCTGAGACCCTAACTTGTGAC | >MI0000446\|hsa-mir-125b-1&WithFlank&11\|-\|122099751\|122099850 | 21 | 42 | 22(+1C) | TCCCTGAGACCCTAACTTGTGAC | 187.4 | -10.6 | 1.5 | -7.2 | 8.3E-13 | 2.4E-11 | BT20/MB231 | NUC |
| >hg38_21_+_8433172_8446622_RNA45SN1_45S_50ntFlanks@11729.11750.22\|TAACTATGACTCTCTTAAGGTA | >hg38_21_+_8433172_8446622_RNA45SN1_45S_50ntFlanks | 11729 | 11750 | 22 | TAACTATGACTCTCTTAAGGTA | 1923.1 | -4.1 | 0.6 | -6.8 | 7.8E-12 | 2.4E-11 | MB231/MB468 | CYTO |
| >hg38_21_+_8433172_8446622_RNA45SN1_45S_50ntFlanks@11874.11893.20\|AGAAGACCCTGTTGAGCTTG | >hg38_21_+_8433172_8446622_RNA45SN1_45S_50ntFlanks | 11874 | 11893 | 20 | AGAAGACCCTGTTGAGCTTG | 5756.2 | -6.1 | 0.9 | -6.8 | 7.8E-12 | 2.4E-11 | MB231/MB468 | CYTO |
| >hg38_21_+_8433172_8446622_RNA45SN1_45S_50ntFlanks@12938.12955.18\|ACCATTCGTAGACGACCT | >hg38_21_+_8433172_8446622_RNA45SN1_45S_50ntFlanks | 12938 | 12955 | 18 | ACCATTCGTAGACGACCT | 327.6 | -8.6 | 1.3 | -6.8 | 9.9E-12 | 2.4E-11 | BT20/MB468 | NUC |
| >trnaMT_ValTAC_MT_+_1602_1670@46.72.27\|TTCAACTTAACTTGACCGCTCTGACCA | >trnaMT_ValTAC_MT_+_1602_1670 | 46 | 72 | 27 | TTCAACTTAACTTGACCGCTCTGACCA | 936.0 | 5.2 | 0.8 | 6.8 | 8.0E-12 | 2.4E-11 | MB231/MB468 | NUC |
| >hg38_21_+_8433172_8446622_RNA45SN1_45S_50ntFlanks@10392.10435.44\|ACAGCAGTTGAACATGGGTCAGTCGGTCCTGAGAGATGGGCGAG | >hg38_21_+_8433172_8446622_RNA45SN1_45S_50ntFlanks | 10392 | 10435 | 44 | ACAGCAGTTGAACATGGGTCAGTCGGTCCTGAGAGATGGGCGAG | 335.2 | -12.3 | 1.8 | -6.9 | 7.1E-12 | 2.4E-11 | MB231/MB468 | MP |
| >hg38_21_+_8433172_8446622_RNA45SN1_45S_50ntFlanks@10922.10946.25\|CTGGACGAGGCGCCGCCGCCCCCCC | >hg38_21_+_8433172_8446622_RNA45SN1_45S_50ntFlanks | 10922 | 10946 | 25 | CTGGACGAGGCGCCGCCGCCCCCCC | 1014.9 | -8.8 | 1.3 | -6.8 | 8.1E-12 | 2.4E-11 | MB231/MB468 | NUC |
| >hg38_21_+_8433172_8446622_RNA45SN1_45S_50ntFlanks@3866.3899.34\|CTAGAGCTAATACATGCCGACGGGCGCTGACCCC | >hg38_21_+_8433172_8446622_RNA45SN1_45S_50ntFlanks | 3866 | 3899 | 34 | CTAGAGCTAATACATGCCGACGGGCGCTGACCCC | 141.3 | -9.2 | 1.3 | -6.9 | 5.8E-12 | 2.4E-11 | BT20/MB468 | MITO |
| >hg38_21_+_8433172_8446622_RNA45SN1_45S_50ntFlanks@11583.11623.41\|CTGGTGCGGACCAGGGGAATCCGACTGTTTAATTAAAACAA | >hg38_21_+_8433172_8446622_RNA45SN1_45S_50ntFlanks | 11583 | 11623 | 41 | CTGGTGCGGACCAGGGGAATCCGACTGTTTAATTAAAACAA | 1994.4 | 7.1 | 1.0 | 6.9 | 5.8E-12 | 2.4E-11 | BT20/MB468 | MITO |
| >MI0001448\|hsa-mir-425&WithFlank&3\|-\|49020142\|49020240@20.44.25&[MIMAT0003393&hsa-miR-425-5p&offsets\|0\|+2;m-95&3\|-\|49020200\|49020221&offsets\|0\|+3]\|AATGACACGATCACTCCCGTTGAGT | >MI0001448\|hsa-mir-425&WithFlank&3\|-\|49020142\|49020240 | 20 | 44 | 25 | AATGACACGATCACTCCCGTTGAGT | 5577.3 | 6.6 | 1.0 | 6.8 | 9.5E-12 | 2.4E-11 | MB231/MB468 | MITO |
| >hg38_21_+_8433172_8446622_RNA45SN1_45S_50ntFlanks@5086.5108.23\|AACGAGACTCTGGCATGCTAACT | >hg38_21_+_8433172_8446622_RNA45SN1_45S_50ntFlanks | 5086 | 5108 | 23 | AACGAGACTCTGGCATGCTAACT | 157.5 | 6.5 | 1.0 | 6.8 | 7.9E-12 | 2.4E-11 | MB231/MB468 | CYTO |
| >hg38_21_+_8433172_8446622_RNA45SN1_45S_50ntFlanks@13004.13028.25\|ATCTATTGAAAGTCAGCCCTCGACA | >hg38_21_+_8433172_8446622_RNA45SN1_45S_50ntFlanks | 13004 | 13028 | 25 | ATCTATTGAAAGTCAGCCCTCGACA | 243.2 | -10.0 | 1.5 | -6.9 | 5.8E-12 | 2.4E-11 | BT20/MB468 | MITO |
| >MI0002470\|hsa-mir-486-1&WithFlank&8\|-\|41660435\|41660514@10.31.22(+1U);MI0023622\|hsa-mir-486-2&WithFlank&8\|+\|41660438\|41660513@7.28.22(+1U)&[MIMAT0002177&hsa-miR-486-5p&offsets\|0\|0(+1U);m-107&8\|-\|41660484\|41660505&offsets\|0\|0(+1U)];[MIMAT0002177_1&hsa-miR-486-5p&offsets\|0\|0(+1U);TJU_CMC_MD2.ID03138.5p-miR&offsets\|0\|-2(+1U);m-106&8\|+\|41660444\|41660465&offsets\|0\|0(+1U)]\|TCCTGTACTGAGCTGCCCCGAGT | >MI0002470\|hsa-mir-486-1&WithFlank&8\|-\|41660435\|41660514 | 10 | 31 | 22(+1U) | TCCTGTACTGAGCTGCCCCGAGT | 187.5 | -10.6 | 1.5 | -7.2 | 8.6E-13 | 2.4E-11 | BT20/MB231 | NUC |
| >hg38_21_+_8433172_8446622_RNA45SN1_45S_50ntFlanks@5079.5104.26\|GATAACGAACGAGACTCTGGCATGCT | >hg38_21_+_8433172_8446622_RNA45SN1_45S_50ntFlanks | 5079 | 5104 | 26 | GATAACGAACGAGACTCTGGCATGCT | 280.6 | 7.3 | 1.1 | 6.8 | 8.0E-12 | 2.4E-11 | MB231/MB468 | CYTO |
| >hg38_MT_+_598_1651_RNR1_12S_50ntFlanks@914.946.33\|CAAAGGACATTTAACTAAAACCCCTACGCATTT | >hg38_MT_+_598_1651_RNR1_12S_50ntFlanks | 914 | 946 | 33 | CAAAGGACATTTAACTAAAACCCCTACGCATTT | 564.0 | 5.6 | 0.8 | 6.9 | 5.9E-12 | 2.4E-11 | BT20/MB468 | MITO |
| >hg38_21_+_8433172_8446622_RNA45SN1_45S_50ntFlanks@7987.8006.20\|TCAGACGTGGCGACCCGCTG | >hg38_21_+_8433172_8446622_RNA45SN1_45S_50ntFlanks | 7987 | 8006 | 20 | TCAGACGTGGCGACCCGCTG | 300.0 | -10.3 | 1.5 | -6.9 | 5.9E-12 | 2.4E-11 | BT20/MB468 | MITO |
| >hg38_21_+_8433172_8446622_RNA45SN1_45S_50ntFlanks@9534.9567.34\|AGGAAACTCTGGTGGAGGTCCGTAGCGGTCCTGA | >hg38_21_+_8433172_8446622_RNA45SN1_45S_50ntFlanks | 9534 | 9567 | 34 | AGGAAACTCTGGTGGAGGTCCGTAGCGGTCCTGA | 1362.1 | -7.0 | 1.0 | -6.8 | 8.0E-12 | 2.4E-11 | MB231/MB468 | CYTO |
| >hg38_21_+_8433172_8446622_RNA45SN1_45S_50ntFlanks@10236.10260.25\|GACGCTACGCCGCGACGAGTAGGAG | >hg38_21_+_8433172_8446622_RNA45SN1_45S_50ntFlanks | 10236 | 10260 | 25 | GACGCTACGCCGCGACGAGTAGGAG | 114.1 | -10.7 | 1.6 | -6.9 | 7.3E-12 | 2.4E-11 | MB231/MB468 | MP |
| >hg38_21_+_8433172_8446622_RNA45SN1_45S_50ntFlanks@12588.12606.19\|GGTTCAGACATTTGGTGTA | >hg38_21_+_8433172_8446622_RNA45SN1_45S_50ntFlanks | 12588 | 12606 | 19 | GGTTCAGACATTTGGTGTA | 87.2 | -10.3 | 1.5 | -6.9 | 7.3E-12 | 2.4E-11 | MB231/MB468 | MP |
| >hg38_21_+_8433172_8446622_RNA45SN1_45S_50ntFlanks@8212.8230.19\|CCGGTAGCGGCCCCCGGCG | >hg38_21_+_8433172_8446622_RNA45SN1_45S_50ntFlanks | 8212 | 8230 | 19 | CCGGTAGCGGCCCCCGGCG | 129.5 | -9.3 | 1.4 | -6.8 | 7.5E-12 | 2.4E-11 | BT20/MB468 | CYTO |
| >hg38_21_+_8433172_8446622_RNA45SN1_45S_50ntFlanks@4002.4025.24\|GATAACCTCGGGCCGATCGCACGC | >hg38_21_+_8433172_8446622_RNA45SN1_45S_50ntFlanks | 4002 | 4025 | 24 | GATAACCTCGGGCCGATCGCACGC | 102.8 | -10.5 | 1.5 | -6.9 | 7.3E-12 | 2.4E-11 | MB231/MB468 | MP |
| >MI0003591\|hsa-mir-584&WithFlank&5\|-\|149062307\|149062415@22.41.20(+1U)&[MIMAT0003249&hsa-miR-584-5p&offsets\|0\|-2(+1U);m-167&5\|-\|149062374\|149062394&offsets\|0\|-1(+1U)]\|TTATGGTTTGCCTGGGACTGT | >MI0003591\|hsa-mir-584&WithFlank&5\|-\|149062307\|149062415 | 22 | 41 | 20(+1U) | TTATGGTTTGCCTGGGACTGT | 408.8 | 8.5 | 1.2 | 6.8 | 8.0E-12 | 2.4E-11 | MB231/MB468 | CYTO |
| >hg38_21_+_8433172_8446622_RNA45SN1_45S_50ntFlanks@12476.12493.18\|CCACTAATAGGGAACGTG | >hg38_21_+_8433172_8446622_RNA45SN1_45S_50ntFlanks | 12476 | 12493 | 18 | CCACTAATAGGGAACGTG | 354.4 | -8.5 | 1.2 | -6.8 | 9.6E-12 | 2.4E-11 | MB231/MB468 | MITO |
| >hg38_21_+_8433172_8446622_RNA45SN1_45S_50ntFlanks@12942.12969.28\|TTCGTAGACGACCTGCTTCTGGGTCGGG | >hg38_21_+_8433172_8446622_RNA45SN1_45S_50ntFlanks | 12942 | 12969 | 28 | TTCGTAGACGACCTGCTTCTGGGTCGGG | 89.0 | -10.3 | 1.5 | -6.9 | 7.3E-12 | 2.4E-11 | MB231/MB468 | MP |

| | | | | | | | | | | | | | |
|---|---|---|---|---|---|---|---|---|---|---|---|---|---|
| >hg38_21_+_8433172_8446622_RNA45SN1_45S_50ntFlanks@12163.12181.19\|ACTGGGGCGGTACACCTGT | >hg38_21_+_8433172_8446622_RNA45SN1_45S_50ntFlanks | 12163 | 12181 | 19 | ACTGGGGCGGTACACCTGT | 261.9 | -7.8 | 1.1 | -6.8 | 8.2E-12 | 2.4E-11 | MB231/MB468 | NUC |
| >MI0003760\|hsa-mir-671&WithFlank&7\|+\|151238415\|151238544@35.58.24&[MIMAT0003880&hsa-miR-671-5p&offsets\|0\|+1;m-316&7\|+\|151238449\|151238472&offsets\|0\|0]\|AGGAAGCCCTGGAGGGGCTGGAGG | >MI0003760\|hsa-mir-671&WithFlank&7\|+\|151238415\|151238544 | 35 | 58 | 24 | AGGAAGCCCTGGAGGGGCTGGAGG | 422.7 | 8.4 | 1.2 | 6.8 | 7.5E-12 | 2.4E-11 | BT20/MB468 | CYTO |
| >hg38_21_+_8433172_8446622_RNA45SN1_45S_50ntFlanks@10355.10388.34\|AACTTTGAAGGCCGAAGTGGAGAAGGGTTCCATG | >hg38_21_+_8433172_8446622_RNA45SN1_45S_50ntFlanks | 10355 | 10388 | 34 | AACTTTGAAGGCCGAAGTGGAGAAGGGTTCCATG | 554.8 | -8.9 | 1.3 | -6.8 | 9.6E-12 | 2.4E-11 | MB231/MB468 | MITO |
| >hg38_21_+_8433172_8446622_RNA45SN1_45S_50ntFlanks@9567.9592.26\|ACGTGCAAATCGGTCGTCCGACCTGG | >hg38_21_+_8433172_8446622_RNA45SN1_45S_50ntFlanks | 9567 | 9592 | 26 | ACGTGCAAATCGGTCGTCCGACCTGG | 1632.2 | -6.6 | 1.0 | -6.8 | 8.1E-12 | 2.4E-11 | MB231/MB468 | CYTO |
| >hg38_21_+_8433172_8446622_RNA45SN1_45S_50ntFlanks@11730.11751.22\|AACTATGACTCTCTTAAGGTAG | >hg38_21_+_8433172_8446622_RNA45SN1_45S_50ntFlanks | 11730 | 11751 | 22 | AACTATGACTCTCTTAAGGTAG | 309.9 | -4.0 | 0.6 | -6.8 | 8.3E-12 | 2.4E-11 | MB231/MB468 | NUC |
| >hg38_21_+_8433172_8446622_RNA45SN1_45S_50ntFlanks@4160.4181.22\|CCACATCCAAGGAAGGCAGCAG | >hg38_21_+_8433172_8446622_RNA45SN1_45S_50ntFlanks | 4160 | 4181 | 22 | CCACATCCAAGGAAGGCAGCAG | 144.2 | -11.0 | 1.6 | -6.9 | 7.4E-12 | 2.4E-11 | MB231/MB468 | MP |
| >hg38_21_+_8433172_8446622_RNA45SN1_45S_50ntFlanks@3851.3881.31\|ATAACTGTGGTAATTCTAGAGCTAATACATG | >hg38_21_+_8433172_8446622_RNA45SN1_45S_50ntFlanks | 3851 | 3881 | 31 | ATAACTGTGGTAATTCTAGAGCTAATACATG | 374.2 | -8.6 | 1.3 | -6.8 | 9.7E-12 | 2.4E-11 | MB231/MB468 | MITO |
| >hg38_21_+_8433172_8446622_RNA45SN1_45S_50ntFlanks@5515.5535.21\|CTAGAGGAAGTAAAAGTCGTA | >hg38_21_+_8433172_8446622_RNA45SN1_45S_50ntFlanks | 5515 | 5535 | 21 | CTAGAGGAAGTAAAAGTCGTA | 658.3 | -8.9 | 1.3 | -6.8 | 9.7E-12 | 2.4E-11 | MB231/MB468 | MITO |
| >hg38_21_+_8433172_8446622_RNA45SN1_45S_50ntFlanks@4641.4663.23\|CATTCGTATTGCGCCGCTAGAGG | >hg38_21_+_8433172_8446622_RNA45SN1_45S_50ntFlanks | 4641 | 4663 | 23 | CATTCGTATTGCGCCGCTAGAGG | 366.5 | -5.4 | 0.8 | -6.8 | 9.7E-12 | 2.5E-11 | MB231/MB468 | MITO |
| >hg38_21_+_8433172_8446622_RNA45SN1_45S_50ntFlanks@5472.5516.45\|ACGGCCCTGGCGGAGCGCTGAGAAGACGGTCGAACTTGACTATCT | >hg38_21_+_8433172_8446622_RNA45SN1_45S_50ntFlanks | 5472 | 5516 | 45 | ACGGCCCTGGCGGAGCGCTGAGAAGACGGTCGAACTTGACTATCT | 2289.7 | 6.2 | 0.9 | 6.8 | 8.3E-12 | 2.5E-11 | MB231/MB468 | NUC |
| >hg38_21_+_8433172_8446622_RNA45SN1_45S_50ntFlanks@10348.10379.32\|AAACGAGAACTTTGAAGGCCGAAGTGGAGAAG | >hg38_21_+_8433172_8446622_RNA45SN1_45S_50ntFlanks | 10348 | 10379 | 32 | AAACGAGAACTTTGAAGGCCGAAGTGGAGAAG | 304.1 | 3.0 | 0.4 | 6.8 | 7.6E-12 | 2.5E-11 | BT20/MB468 | CYTO |
| >hg38_21_+_8433172_8446622_RNA45SN1_45S_50ntFlanks@9720.9737.18\|ATTAGAGGTCTTGGGGCC | >hg38_21_+_8433172_8446622_RNA45SN1_45S_50ntFlanks | 9720 | 9737 | 18 | ATTAGAGGTCTTGGGGCC | 501.6 | -11.1 | 1.6 | -6.9 | 6.0E-12 | 2.5E-11 | BT20/MB468 | MITO |
| >hg38_21_+_8433172_8446622_RNA45SN1_45S_50ntFlanks@4729.4765.37\|TCAAGAACGAAAGTCGGAGGTTCGAAGACGATCAGAT | >hg38_21_+_8433172_8446622_RNA45SN1_45S_50ntFlanks | 4729 | 4765 | 37 | TCAAGAACGAAAGTCGGAGGTTCGAAGACGATCAGAT | 309.7 | 5.9 | 0.9 | 6.8 | 8.2E-12 | 2.5E-11 | MB231/MB468 | CYTO |
| >trna5_AspGTC_12_+_98897281_98897352@35.73.39\|TCACGCGGGAGACCGGGGTTCAATTCCCCGACGGGGAGC | >trna5_AspGTC_12_+_98897281_98897352 | 35 | 73 | 39 | TCACGCGGGAGACCGGGGTTCAATTCCCCGACGGGGAGC | 1133.4 | 9.6 | 1.4 | 6.8 | 8.2E-12 | 2.5E-11 | MB231/MB468 | CYTO |
| >hg38_21_+_8433172_8446622_RNA45SN1_45S_50ntFlanks@12490.12509.20\|CGTGAGCTGGGTTTAGACCG | >hg38_21_+_8433172_8446622_RNA45SN1_45S_50ntFlanks | 12490 | 12509 | 20 | CGTGAGCTGGGTTTAGACCG | 129.6 | -10.9 | 1.6 | -6.8 | 7.4E-12 | 2.5E-11 | MB231/MB468 | MP |
| >MI0000093\|hsa-mir-92a-1&WithFlank&13\|+\|91351308\|91351397@54.76.23(+1U)&[MIMAT0000092&hsa-miR-92a-3p&offsets\|0\|+1(+1U);m-9&13\|+\|91351361\|91351382&offsets\|0\|+1(+1U)]\|TATTGCACTTGTCCCGGCCTGTTT | >MI0000093\|hsa-mir-92a-1&WithFlank&13\|+\|91351308\|91351397 | 54 | 76 | 23(+1U) | TATTGCACTTGTCCCGGCCTGTTT | 536.2 | 6.6 | 1.0 | 6.8 | 8.2E-12 | 2.5E-11 | MB231/MB468 | CYTO |
| >hg38_21_+_8433172_8446622_RNA45SN1_45S_50ntFlanks@10408.10436.29\|GGTCAGTCGGTCCTGAGAGATGGGCGAGC | >hg38_21_+_8433172_8446622_RNA45SN1_45S_50ntFlanks | 10408 | 10436 | 29 | GGTCAGTCGGTCCTGAGAGATGGGCGAGC | 118.5 | -10.8 | 1.6 | -6.8 | 7.5E-12 | 2.5E-11 | MB231/MB468 | MP |
| >hg38_21_+_8433172_8446622_RNA45SN1_45S_50ntFlanks@12071.12095.25\|CCCCGAGGGGCTCTCGCTTCTGGCG | >hg38_21_+_8433172_8446622_RNA45SN1_45S_50ntFlanks | 12071 | 12095 | 25 | CCCCGAGGGGCTCTCGCTTCTGGCG | 873.7 | -7.0 | 1.0 | -6.8 | 8.4E-12 | 2.5E-11 | MB231/MB468 | NUC |
| >hg38_21_+_8433172_8446622_RNA45SN1_45S_50ntFlanks@6661.6680.20\|CGGTGGATCACTCGGCTCGT | >hg38_21_+_8433172_8446622_RNA45SN1_45S_50ntFlanks | 6661 | 6680 | 20 | CGGTGGATCACTCGGCTCGT | 508.8 | 2.8 | 0.4 | 6.8 | 7.7E-12 | 2.5E-11 | BT20/MB468 | CYTO |
| >hg38_21_+_8433172_8446622_RNA45SN1_45S_50ntFlanks@10514.10534.21\|CGGAGTGGCGGAGATGGGCGC | >hg38_21_+_8433172_8446622_RNA45SN1_45S_50ntFlanks | 10514 | 10534 | 21 | CGGAGTGGCGGAGATGGGCGC | 762.5 | -9.0 | 1.3 | -6.8 | 7.5E-12 | 2.5E-11 | MB231/MB468 | MP |
| >hg38_21_+_8433172_8446622_RNA45SN1_45S_50ntFlanks@11542.11584.43\|CGGCGCCTCGCCTCGGCCGGCGCCTAGCAGCCGACTTAGAACT | >hg38_21_+_8433172_8446622_RNA45SN1_45S_50ntFlanks | 11542 | 11584 | 43 | CGGCGCCTCGCCTCGGCCGGCGCCTAGCAGCCGACTTAGAACT | 361.5 | 7.5 | 1.1 | 6.8 | 8.2E-12 | 2.5E-11 | MB231/MB468 | CYTO |
| >hg38_21_+_8433172_8446622_RNA45SN1_45S_50ntFlanks@9526.9570.45\|GAAGCCAGAGGAAACTCTGGTGGAGGTCCGTAGCGGTCCTGACGT | >hg38_21_+_8433172_8446622_RNA45SN1_45S_50ntFlanks | 9526 | 9570 | 45 | GAAGCCAGAGGAAACTCTGGTGGAGGTCCGTAGCGGTCCTGACGT | 800.9 | 6.9 | 1.0 | 6.8 | 8.2E-12 | 2.5E-11 | MB231/MB468 | CYTO |
| >hg38_21_+_8433172_8446622_RNA45SN1_45S_50ntFlanks@12478.12500.23\|ACTAATAGGGAACGTGAGCTGGG | >hg38_21_+_8433172_8446622_RNA45SN1_45S_50ntFlanks | 12478 | 12500 | 23 | ACTAATAGGGAACGTGAGCTGGG | 300.1 | -3.5 | 0.5 | -6.8 | 8.5E-12 | 2.5E-11 | MB231/MB468 | NUC |
| >hg38_21_+_8433172_8446622_RNA45SN1_45S_50ntFlanks@8472.8489.18\|CGGCCCGGCGGATCTTTC | >hg38_21_+_8433172_8446622_RNA45SN1_45S_50ntFlanks | 8472 | 8489 | 18 | CGGCCCGGCGGATCTTTC | 99.3 | -10.5 | 1.5 | -6.8 | 7.5E-12 | 2.5E-11 | MB231/MB468 | MP |
| >hg38_21_+_8433172_8446622_RNA45SN1_45S_50ntFlanks@10571.10589.19\|GAGAAGCCGGCGGGAGCCC | >hg38_21_+_8433172_8446622_RNA45SN1_45S_50ntFlanks | 10571 | 10589 | 19 | GAGAAGCCGGCGGGAGCCC | 118.7 | -10.8 | 1.6 | -6.8 | 7.5E-12 | 2.5E-11 | MB231/MB468 | MP |
| >hg38_21_+_8433172_8446622_RNA45SN1_45S_50ntFlanks@12850.12871.22\|CTTCGTCCTGGGAAACGGGGCG | >hg38_21_+_8433172_8446622_RNA45SN1_45S_50ntFlanks | 12850 | 12871 | 22 | CTTCGTCCTGGGAAACGGGGCG | 222.6 | -11.7 | 1.7 | -6.8 | 7.5E-12 | 2.5E-11 | MB231/MB468 | MP |
| >hg38_21_+_8433172_8446622_RNA45SN1_45S_50ntFlanks@6650.6678.29\|ACGACTCTTAGCGGTGGATCACTCGGCTC | >hg38_21_+_8433172_8446622_RNA45SN1_45S_50ntFlanks | 6650 | 6678 | 29 | ACGACTCTTAGCGGTGGATCACTCGGCTC | 571.1 | 3.9 | 0.6 | 6.8 | 7.7E-12 | 2.5E-11 | BT20/MB468 | CYTO |
| >hg38_21_+_8433172_8446622_RNA45SN1_45S_50ntFlanks@4732.4758.27\|AGAACGAAAGTCGGAGGTTCGAAGACG | >hg38_21_+_8433172_8446622_RNA45SN1_45S_50ntFlanks | 4732 | 4758 | 27 | AGAACGAAAGTCGGAGGTTCGAAGACG | 4707.2 | -5.1 | 0.8 | -6.8 | 7.5E-12 | 2.5E-11 | MB231/MB468 | MP |
| >hg38_21_+_8433172_8446622_RNA45SN1_45S_50ntFlanks@475.509.35\|CTTCGTGATCGATGTGGTGACGTCGTGCTCTCCCG | >hg38_21_+_8433172_8446622_RNA45SN1_45S_50ntFlanks | 475 | 509 | 35 | CTTCGTGATCGATGTGGTGACGTCGTGCTCTCCCG | 2514.1 | 7.3 | 1.1 | 6.9 | 6.1E-12 | 2.5E-11 | BT20/MB468 | MITO |
| >am_trna10_ValCAC_5_-_180649395_180649467@2.32.31,trna12_ValAAC_5_-_180645270_180645342@2.32.31,trna132_ValAAC_6_-_27721179_27721251@2.32.31,trna136_ValAAC_6_-_27648885_27648957@2.32.31,trna139_ValAAC_6_-_27618707_27618779@2.32.31,trna18_ValCAC_5_-_180529253_180529325@2.32.31,trna2_ValAAC_3_+_169490018_169490090@2.32.31,trna2_ValCAC_5_+_180524070_180524142@2.32.31,trna4_ValAAC_5_+_180591154_180591226@2.32.31,trna5_ValAAC_5_+_180596610_180596682@2.32.31,trna6_ValCAC_5_+_180600650_180600722@2.32.31,trna85_ValCAC_1_-_161369490_161369562@2.32.31,trna90_ValCAC_1_-_149684088_149684161@2.32.31,trna98_ValCAC_1_-_149298555_149298627@2.32.31,trna9_ValCAC_6_+_26538282_26538354@2.32.31\|TTTCCGTAGTGTAGTGGTTATCACGTTCGCC | >am_trna10_ValCAC_5_-_180649395_180649467 | 2 | 32 | 31 | TTTCCGTAGTGTAGTGGTTATCACGTTCGCC | 276.0 | 11.3 | 1.7 | 6.8 | 9.9E-12 | 2.5E-11 | MB231/MB468 | MITO |
| >hg38_1_-_228634819_228635039_RNA5S12_5S_50ntFlanks@139.169.31\|GACCGCCTGGGAATACCGGGTGCTGTAGGCT | >hg38_1_-_228634819_228635039_RNA5S12_5S_50ntFlanks | 139 | 169 | 31 | GACCGCCTGGGAATACCGGGTGCTGTAGGCT | 816.1 | 5.3 | 0.8 | 6.8 | 8.3E-12 | 2.5E-11 | MB231/MB468 | CYTO |
| >hg38_21_+_8433172_8446622_RNA45SN1_45S_50ntFlanks@4981.4999.19\|CACGGACAGGATTGACAGA | >hg38_21_+_8433172_8446622_RNA45SN1_45S_50ntFlanks | 4981 | 4999 | 19 | CACGGACAGGATTGACAGA | 418.2 | -4.7 | 0.7 | -6.8 | 7.6E-12 | 2.5E-11 | MB231/MB468 | MP |
| >hg38_21_+_8433172_8446622_RNA45SN1_45S_50ntFlanks@12625.12653.29\|CAATGGGGCGAAGCTACCATCTGTGGGAT | >hg38_21_+_8433172_8446622_RNA45SN1_45S_50ntFlanks | 12625 | 12653 | 29 | CAATGGGGCGAAGCTACCATCTGTGGGAT | 2137.5 | -10.3 | 1.5 | -6.8 | 1.1E-11 | 2.5E-11 | BT20/MB468 | NUC |
| >hg38_21_+_8433172_8446622_RNA45SN1_45S_50ntFlanks@4588.4610.23\|CTATTTTGTTGGTTTTCGGAACT | >hg38_21_+_8433172_8446622_RNA45SN1_45S_50ntFlanks | 4588 | 4610 | 23 | CTATTTTGTTGGTTTTCGGAACT | 123.5 | -10.8 | 1.6 | -6.8 | 7.6E-12 | 2.5E-11 | MB231/MB468 | MP |
| >trnaMT_GlyTCC_MT_+_9991_10058@28.68.41\|ACTTCCAATTAACTAGTTTTGACAACATTCAAAAAAGAGTA | >trnaMT_GlyTCC_MT_+_9991_10058 | 28 | 68 | 41 | ACTTCCAATTAACTAGTTTTGACAACATTCAAAAAAGAGTA | 881.6 | -14.2 | 2.1 | -6.8 | 1.1E-11 | 2.5E-11 | BT20/MB468 | NUC |
| >hg38_21_+_8433172_8446622_RNA45SN1_45S_50ntFlanks@12457.12476.20\|CAAGCGTTGGATTGTTCACC | >hg38_21_+_8433172_8446622_RNA45SN1_45S_50ntFlanks | 12457 | 12476 | 20 | CAAGCGTTGGATTGTTCACC | 2065.0 | -3.3 | 0.5 | -6.9 | 6.1E-12 | 2.5E-11 | BT20/MB468 | MITO |
| >hg38_21_+_8433172_8446622_RNA45SN1_45S_50ntFlanks@6746.6771.26\|CACTTCGAACGCACTTGCGGCCCCGG | >hg38_21_+_8433172_8446622_RNA45SN1_45S_50ntFlanks | 6746 | 6771 | 26 | CACTTCGAACGCACTTGCGGCCCCGG | 364.2 | -7.9 | 1.2 | -6.8 | 8.6E-12 | 2.5E-11 | MB231/MB468 | NUC |
| >hg38_21_+_8433172_8446622_RNA45SN1_45S_50ntFlanks@12259.12284.26\|CTTGATCTTGATTTTCAGTACGAATA | >hg38_21_+_8433172_8446622_RNA45SN1_45S_50ntFlanks | 12259 | 12284 | 26 | CTTGATCTTGATTTTCAGTACGAATA | 3467.1 | -8.5 | 1.2 | -6.8 | 1.0E-11 | 2.5E-11 | MB231/MB468 | MITO |
| >MI0000767\|hsa-mir-365a&WithFlank&16\|+\|14309279\|14309377@22.44.23&[MIMAT0009199&hsa-miR-365a-5p&offsets\|0\|0;m-595&16\|+\|14309300\|14309322&offsets\|0\|0]\|AGGGACTTTTGGGGGCAGATGTG | >MI0000767\|hsa-mir-365a&WithFlank&16\|+\|14309279\|14309377 | 22 | 44 | 23 | AGGGACTTTTGGGGGCAGATGTG | 444.9 | 8.5 | 1.2 | 6.8 | 7.8E-12 | 2.5E-11 | BT20/MB468 | CYTO |
| >hg38_21_+_8433172_8446622_RNA45SN1_45S_50ntFlanks@10393.10424.32\|CAGCAGTTGAACATGGGTCAGTCGGTCCTGAG | >hg38_21_+_8433172_8446622_RNA45SN1_45S_50ntFlanks | 10393 | 10424 | 32 | CAGCAGTTGAACATGGGTCAGTCGGTCCTGAG | 4349.6 | -3.8 | 0.6 | -6.8 | 1.1E-11 | 2.5E-11 | BT20/MB468 | NUC |
| >trnaMT_GlyTCC_MT_+_9991_10058@27.68.42\|AACTTCCAATTAACTAGTTTTGACAACATTCAAAAAAGAGTA | >trnaMT_GlyTCC_MT_+_9991_10058 | 27 | 68 | 42 | AACTTCCAATTAACTAGTTTTGACAACATTCAAAAAAGAGTA | 306.1 | -12.7 | 1.9 | -6.8 | 1.1E-11 | 2.5E-11 | BT20/MB468 | NUC |
| >MI0000455\|hsa-mir-138-2&WithFlank&16\|+\|56858512\|56858607@16.38.23;MI0000476\|hsa-mir-138-1&WithFlank&3\|+\|44114206\|44114316@29.51.23&[MIMAT0000430&hsa-miR-138-5p&offsets\|0\|0;m-509&16\|+\|56858527\|56858549&offsets\|0\|0];[MIMAT0000430_1&hsa-miR-138-5p&offsets\|0\|0;m-497&3\|+\|44114234\|44114256&offsets\|0\|0]\|AGCTGGTGTTGTGAATCAGGCCG | >MI0000455\|hsa-mir-138-2&WithFlank&16\|+\|56858512\|56858607 | 16 | 38 | 23 | AGCTGGTGTTGTGAATCAGGCCG | 535.6 | -12.5 | 1.7 | -7.4 | 1.1E-13 | 2.5E-11 | BT20/MB231 | MP |
| >MI0000102\|hsa-mir-100&WithFlank&11\|-\|122152223\|122152314@22.40.19&[MIMAT0000098&hsa-miR-100-5p&offsets\|+3\|0;m-27&11\|-\|122152276\|122152296&offsets\|+3\|+1]\|CCGTAGATCCGAACTTGTG | >MI0000102\|hsa-mir-100&WithFlank&11\|-\|122152223\|122152314 | 22 | 40 | 19 | CCGTAGATCCGAACTTGTG | 562.7 | 12.4 | 1.8 | 6.8 | 1.0E-11 | 2.5E-11 | MB231/MB468 | MITO |
| >hg38_21_+_8433172_8446622_RNA45SN1_45S_50ntFlanks@10427.10456.30\|ATGGGCGAGCGCCGTTCCGAAGGGACGGGC | >hg38_21_+_8433172_8446622_RNA45SN1_45S_50ntFlanks | 10427 | 10456 | 30 | ATGGGCGAGCGCCGTTCCGAAGGGACGGGC | 1003.8 | -4.7 | 0.7 | -6.8 | 8.6E-12 | 2.5E-11 | MB231/MB468 | NUC |
| >hg38_21_+_8433172_8446622_RNA45SN1_45S_50ntFlanks@4918.4937.20\|ACCACCAGGAGTGGAGCCTG | >hg38_21_+_8433172_8446622_RNA45SN1_45S_50ntFlanks | 4918 | 4937 | 20 | ACCACCAGGAGTGGAGCCTG | 106.5 | -10.6 | 1.5 | -6.8 | 7.7E-12 | 2.5E-11 | MB231/MB468 | MP |
| >hg38_21_+_8433172_8446622_RNA45SN1_45S_50ntFlanks@12396.12417.22\|GCGACGTCGCTTTTTGATCCTT | >hg38_21_+_8433172_8446622_RNA45SN1_45S_50ntFlanks | 12396 | 12417 | 22 | GCGACGTCGCTTTTTGATCCTT | 1561.2 | -4.6 | 0.7 | -6.8 | 8.6E-12 | 2.5E-11 | MB231/MB468 | NUC |

<table>
<tr><th></th><th></th><th></th><th></th><th></th><th></th><th></th><th></th><th></th><th></th><th></th><th></th><th></th><th></th></tr>
<tr><td>>MI0000298|hsa-mir-221&WithFlank&X|-|45746151|45746272@31.56.26(+1U)&[MIMAT0004568&hsa-miR-221-5p&offsets|0|+4(+1U);m-405&X|-|45746220|45746242&offsets|0|+3(+1U)]|ACCTGGCATACAATGTAGATTTCTGTT</td><td>>MI0000298|hsa-mir-221&WithFlank&X|-|45746151|45746272</td><td>31</td><td>56</td><td>26(+1U)</td><td>ACCTGGCATACAATGTAGATTTCTGTT</td><td>323.9</td><td>7.9</td><td>1.2</td><td>6.8</td><td>8.4E-12</td><td>2.5E-11</td><td>MB231/MB468</td><td>CYTO</td></tr>
<tr><td>>trnaMT_GluTTC_MT_-_14674_14742@46.72.27,trnalookalike8_GluTTC_5_-_93905172_93905240@46.72.27|GTGGTTGTAGTCCGTGCGAGAATACCA</td><td>>trnaMT_GluTTC_MT_-_14674_14742</td><td>46</td><td>72</td><td>27</td><td>GTGGTTGTAGTCCGTGCGAGAATACCA</td><td>803.2</td><td>8.9</td><td>1.3</td><td>6.8</td><td>8.4E-12</td><td>2.5E-11</td><td>MB231/MB468</td><td>CYTO</td></tr>
<tr><td>>hg38_21_+_8433172_8446622_RNA45SN1_45S_50ntFlanks@12892.12932.41|CCCTCGCCCGTCACGCACCGCACGTTCGTGGGGAACCTGGC</td><td>>hg38_21_+_8433172_8446622_RNA45SN1_45S_50ntFlanks</td><td>12892</td><td>12932</td><td>41</td><td>CCCTCGCCCGTCACGCACCGCACGTTCGTGGGGAACCTGGC</td><td>478.1</td><td>-12.8</td><td>1.9</td><td>-6.8</td><td>7.7E-12</td><td>2.5E-11</td><td>MB231/MB468</td><td>MP</td></tr>
<tr><td>>hg38_21_+_8433172_8446622_RNA45SN1_45S_50ntFlanks@12366.12404.39|TAACTGGCTTGTGGCGGCCAAGCGTTCATAGCGACGTCG</td><td>>hg38_21_+_8433172_8446622_RNA45SN1_45S_50ntFlanks</td><td>12366</td><td>12404</td><td>39</td><td>TAACTGGCTTGTGGCGGCCAAGCGTTCATAGCGACGTCG</td><td>345.2</td><td>-11.1</td><td>1.6</td><td>-6.8</td><td>8.4E-12</td><td>2.5E-11</td><td>MB231/MB468</td><td>CYTO</td></tr>
<tr><td>>hg38_21_+_8433172_8446622_RNA45SN1_45S_50ntFlanks@11217.11254.38|CTGGACGCGAGCCGGGCCCTTCCCGTGGATCGCCCCAG</td><td>>hg38_21_+_8433172_8446622_RNA45SN1_45S_50ntFlanks</td><td>11217</td><td>11254</td><td>38</td><td>CTGGACGCGAGCCGGGCCCTTCCCGTGGATCGCCCCAG</td><td>94.6</td><td>-10.4</td><td>1.5</td><td>-6.8</td><td>7.7E-12</td><td>2.5E-11</td><td>MB231/MB468</td><td>MP</td></tr>
<tr><td>>hg38_21_+_8433172_8446622_RNA45SN1_45S_50ntFlanks@8191.8214.24|CAGCCCGTGGACGGTGTGAGGCCG</td><td>>hg38_21_+_8433172_8446622_RNA45SN1_45S_50ntFlanks</td><td>8191</td><td>8214</td><td>24</td><td>CAGCCCGTGGACGGTGTGAGGCCG</td><td>1140.5</td><td>-7.1</td><td>1.0</td><td>-6.8</td><td>1.0E-11</td><td>2.5E-11</td><td>MB231/MB468</td><td>MITO</td></tr>
<tr><td>>hg38_21_+_8433172_8446622_RNA45SN1_45S_50ntFlanks@12044.12071.28|ACTGACCCGGTGAGGCGGGGGGCGAGC</td><td>>hg38_21_+_8433172_8446622_RNA45SN1_45S_50ntFlanks</td><td>12044</td><td>12071</td><td>28</td><td>ACTGACCCGGTGAGGCGGGGGGGCGAGC</td><td>1046.4</td><td>-8.4</td><td>1.2</td><td>-6.9</td><td>6.2E-12</td><td>2.5E-11</td><td>BT20/MB468</td><td>MITO</td></tr>
<tr><td>>hg38_21_+_8433172_8446622_RNA45SN1_45S_50ntFlanks@12008.12038.31|CCGCCGGTGAAATACCACTACTCTGATCGTT</td><td>>hg38_21_+_8433172_8446622_RNA45SN1_45S_50ntFlanks</td><td>12008</td><td>12038</td><td>31</td><td>CCGCCGGTGAAATACCACTACTCTGATCGTT</td><td>163.3</td><td>-5.9</td><td>0.9</td><td>-6.8</td><td>8.5E-12</td><td>2.5E-11</td><td>MB231/MB468</td><td>CYTO</td></tr>
<tr><td>>hg38_21_+_8433172_8446622_RNA45SN1_45S_50ntFlanks@8170.8191.22|CAAGTCCTTCTGATCGAGGCCC</td><td>>hg38_21_+_8433172_8446622_RNA45SN1_45S_50ntFlanks</td><td>8170</td><td>8191</td><td>22</td><td>CAAGTCCTTCTGATCGAGGCCC</td><td>124.4</td><td>-10.8</td><td>1.6</td><td>-6.8</td><td>7.7E-12</td><td>2.5E-11</td><td>MB231/MB468</td><td>MP</td></tr>
<tr><td>>MI0016887|hsa-mir-4521&WithFlank&17|+|8186939|8187010@13.35.23&[MIMAT0019058&hsa-miR-4521&offsets|+3|+4;m-765&17|+|8186948|8186969&offsets|+3|+4]|AAGGAAGTCCTGTGCTCAGTTTT</td><td>>MI0016887|hsa-mir-4521&WithFlank&17|+|8186939|8187010</td><td>13</td><td>35</td><td>23</td><td>AAGGAAGTCCTGTGCTCAGTTTT</td><td>95.2</td><td>9.4</td><td>1.4</td><td>6.8</td><td>1.1E-11</td><td>2.5E-11</td><td>BT20/MB468</td><td>NUC</td></tr>
<tr><td>>hg38_21_+_8433172_8446622_RNA45SN1_45S_50ntFlanks@12417.12436.20|TCGATGTCGGCTCTTCCTAT</td><td>>hg38_21_+_8433172_8446622_RNA45SN1_45S_50ntFlanks</td><td>12417</td><td>12436</td><td>20</td><td>TCGATGTCGGCTCTTCCTAT</td><td>7650.0</td><td>-5.1</td><td>0.7</td><td>-6.8</td><td>7.9E-12</td><td>2.6E-11</td><td>BT20/MB468</td><td>CYTO</td></tr>
<tr><td>>MI0000102|hsa-mir-100&WithFlank&11|-|122152223|122152314@19.40.22&[MIMAT0000098&hsa-miR-100-5p&offsets|0|0;m-27&11|-|122152276|122152296&offsets|0|+1]|AACCCGTAGATCCGAACTTGTG</td><td>>MI0000102|hsa-mir-100&WithFlank&11|-|122152223|122152314</td><td>19</td><td>40</td><td>22</td><td>AACCCGTAGATCCGAACTTGTG</td><td>463896.6</td><td>10.7</td><td>1.6</td><td>6.8</td><td>7.8E-12</td><td>2.6E-11</td><td>MB231/MB468</td><td>MP</td></tr>
<tr><td>>hg38_21_+_8433172_8446622_RNA45SN1_45S_50ntFlanks@10664.10706.43|CTTGGAAAGCGTCGCGGTTCCGGCGGCGTCCGGTGAGCTCTCG</td><td>>hg38_21_+_8433172_8446622_RNA45SN1_45S_50ntFlanks</td><td>10664</td><td>10706</td><td>43</td><td>CTTGGAAAGCGTCGCGGTTCCGGCGGCGTCCGGTGAGCTCTCG</td><td>328.2</td><td>-12.2</td><td>1.8</td><td>-6.8</td><td>7.8E-12</td><td>2.6E-11</td><td>MB231/MB468</td><td>MP</td></tr>
<tr><td>>MI0000542|hsa-mir-320a&WithFlank&8|-|22244960|22245043@48.68.21&[MIMAT0000510&hsa-miR-320a-3p&offsets|0|-1;m-60&8|-|22244975|22244996&offsets|0|-1]|AAAAGCTGGGTTGAGAGGGCG</td><td>>MI0000542|hsa-mir-320a&WithFlank&8|-|22244960|22245043</td><td>48</td><td>68</td><td>21</td><td>AAAAGCTGGGTTGAGAGGGCG</td><td>22876.8</td><td>7.5</td><td>1.0</td><td>7.2</td><td>5.0E-13</td><td>2.6E-11</td><td>BT20/MB468</td><td>MP</td></tr>
<tr><td>>hg38_21_+_8433172_8446622_RNA45SN1_45S_50ntFlanks@11669.11688.20|GCCCAGTGCTCTGAATGTCA</td><td>>hg38_21_+_8433172_8446622_RNA45SN1_45S_50ntFlanks</td><td>11669</td><td>11688</td><td>20</td><td>GCCCAGTGCTCTGAATGTCA</td><td>231.8</td><td>4.7</td><td>0.7</td><td>6.8</td><td>8.0E-12</td><td>2.6E-11</td><td>BT20/MB468</td><td>CYTO</td></tr>
<tr><td>>hg38_1_-_228634819_228635039_RNA5S12_5S_50ntFlanks@148.169.22|GGAATACCGGGTGCTGTAGGCT</td><td>>hg38_1_-_228634819_228635039_RNA5S12_5S_50ntFlanks</td><td>148</td><td>169</td><td>22</td><td>GGAATACCGGGTGCTGTAGGCT</td><td>1456.7</td><td>-4.8</td><td>0.7</td><td>-6.8</td><td>1.0E-11</td><td>2.6E-11</td><td>MB231/MB468</td><td>MITO</td></tr>
<tr><td>>hg38_21_+_8433172_8446622_RNA45SN1_45S_50ntFlanks@6719.6740.22|TGAATTGCAGGACACATTGATC</td><td>>hg38_21_+_8433172_8446622_RNA45SN1_45S_50ntFlanks</td><td>6719</td><td>6740</td><td>22</td><td>TGAATTGCAGGACACATTGATC</td><td>2085.9</td><td>-7.4</td><td>1.1</td><td>-6.8</td><td>1.0E-11</td><td>2.6E-11</td><td>MB231/MB468</td><td>MITO</td></tr>
<tr><td>>hg38_21_+_8433172_8446622_RNA45SN1_45S_50ntFlanks@8171.8212.42|AAGTCCTTCTGATCGAGGCCCAGCCCGTGGACGGTGTGAGGC</td><td>>hg38_21_+_8433172_8446622_RNA45SN1_45S_50ntFlanks</td><td>8171</td><td>8212</td><td>42</td><td>AAGTCCTTCTGATCGAGGCCCAGCCCGTGGACGGTGTGAGGC</td><td>1308.8</td><td>-7.2</td><td>1.1</td><td>-6.8</td><td>1.0E-11</td><td>2.6E-11</td><td>MB231/MB468</td><td>MITO</td></tr>
<tr><td>>hg38_21_+_8433172_8446622_RNA45SN1_45S_50ntFlanks@9243.9273.31|GAGCGAGCGCACGGGGTCGGCGGCGACGTCG</td><td>>hg38_21_+_8433172_8446622_RNA45SN1_45S_50ntFlanks</td><td>9243</td><td>9273</td><td>31</td><td>GAGCGAGCGCACGGGGTCGGCGGCGACGTCG</td><td>151.4</td><td>-11.1</td><td>1.6</td><td>-6.8</td><td>7.8E-12</td><td>2.6E-11</td><td>MB231/MB468</td><td>MP</td></tr>
<tr><td>>hg38_21_+_8433172_8446622_RNA45SN1_45S_50ntFlanks@10420.10443.24|CTGAGAGATGGGCGAGCGCCGTTC</td><td>>hg38_21_+_8433172_8446622_RNA45SN1_45S_50ntFlanks</td><td>10420</td><td>10443</td><td>24</td><td>CTGAGAGATGGGCGAGCGCCGTTC</td><td>658.9</td><td>-6.1</td><td>0.9</td><td>-6.8</td><td>8.1E-12</td><td>2.6E-11</td><td>BT20/MB468</td><td>CYTO</td></tr>
<tr><td>>hg38_21_+_8433172_8446622_RNA45SN1_45S_50ntFlanks@6646.6678.33|TCGTACGACTCTTAGCGGTGGATCACTCGGCTC</td><td>>hg38_21_+_8433172_8446622_RNA45SN1_45S_50ntFlanks</td><td>6646</td><td>6678</td><td>33</td><td>TCGTACGACTCTTAGCGGTGGATCACTCGGCTC</td><td>16750.0</td><td>4.9</td><td>0.7</td><td>6.9</td><td>6.3E-12</td><td>2.6E-11</td><td>BT20/MB468</td><td>MITO</td></tr>
<tr><td>>MI0000102|hsa-mir-100&WithFlank&11|-|122152223|122152314@19.40.22(+1C)&[MIMAT0000098&hsa-miR-100-5p&offsets|0|0(+1C);m-27&11|-|122152276|122152296&offsets|0|+1(+1C)]|AACCCGTAGATCCGAACTTGTGC</td><td>>MI0000102|hsa-mir-100&WithFlank&11|-|122152223|122152314</td><td>19</td><td>40</td><td>22(+1C)</td><td>AACCCGTAGATCCGAACTTGTGC</td><td>8342.9</td><td>-8.1</td><td>1.1</td><td>-7.2</td><td>8.0E-13</td><td>2.6E-11</td><td>BT20/MB231</td><td>MITO</td></tr>
<tr><td>>hg38_21_+_8433172_8446622_RNA45SN1_45S_50ntFlanks@10781.10817.37|AGGTGAACAGCCTCTGGCATGTTGGAACAATGTAGGT</td><td>>hg38_21_+_8433172_8446622_RNA45SN1_45S_50ntFlanks</td><td>10781</td><td>10817</td><td>37</td><td>AGGTGAACAGCCTCTGGCATGTTGGAACAATGTAGGT</td><td>501.0</td><td>-6.6</td><td>1.0</td><td>-6.8</td><td>1.0E-11</td><td>2.6E-11</td><td>MB231/MB468</td><td>MITO</td></tr>
<tr><td>>hg38_21_+_8433172_8446622_RNA45SN1_45S_50ntFlanks@9504.9523.20|GTGAACTATGCCTGGGCAGG</td><td>>hg38_21_+_8433172_8446622_RNA45SN1_45S_50ntFlanks</td><td>9504</td><td>9523</td><td>20</td><td>GTGAACTATGCCTGGGCAGG</td><td>2340.0</td><td>-7.1</td><td>1.0</td><td>-6.8</td><td>8.1E-12</td><td>2.6E-11</td><td>BT20/MB468</td><td>CYTO</td></tr>
<tr><td>>hg38_21_+_8433172_8446622_RNA45SN1_45S_50ntFlanks@9164.9183.20|CCCCGGGCGGGTCGCGCCGT</td><td>>hg38_21_+_8433172_8446622_RNA45SN1_45S_50ntFlanks</td><td>9164</td><td>9183</td><td>20</td><td>CCCCGGGCGGGTCGCGCCGT</td><td>93.8</td><td>-10.4</td><td>1.5</td><td>-6.8</td><td>7.9E-12</td><td>2.6E-11</td><td>MB231/MB468</td><td>MP</td></tr>
<tr><td>>hg38_21_+_8433172_8446622_RNA45SN1_45S_50ntFlanks@8190.8211.22|CCAGCCCGTGGACGGTGTGAGG</td><td>>hg38_21_+_8433172_8446622_RNA45SN1_45S_50ntFlanks</td><td>8190</td><td>8211</td><td>22</td><td>CCAGCCCGTGGACGGTGTGAGG</td><td>182.2</td><td>-11.4</td><td>1.7</td><td>-6.8</td><td>7.9E-12</td><td>2.6E-11</td><td>MB231/MB468</td><td>MP</td></tr>
<tr><td>>MI0003643|hsa-mir-629&WithFlank&15|-|70079366|70079474@28.50.23&[MIMAT0004810&hsa-miR-629-5p&offsets|0|+2;m-150&15|-|70079427|70079447&offsets|0|+2]|TGGGTTTACGTTGGGAGAACTTT</td><td>>MI0003643|hsa-mir-629&WithFlank&15|-|70079366|70079474</td><td>28</td><td>50</td><td>23</td><td>TGGGTTTACGTTGGGAGAACTTT</td><td>127.2</td><td>10.2</td><td>1.5</td><td>6.8</td><td>7.9E-12</td><td>2.6E-11</td><td>MB231/MB468</td><td>MP</td></tr>
<tr><td>>hg38_21_+_8433172_8446622_RNA45SN1_45S_50ntFlanks@11843.11864.22|AGGGAACGGGCTTGGCGGAATC</td><td>>hg38_21_+_8433172_8446622_RNA45SN1_45S_50ntFlanks</td><td>11843</td><td>11864</td><td>22</td><td>AGGGAACGGGCTTGGCGGAATC</td><td>150.2</td><td>-11.1</td><td>1.6</td><td>-6.8</td><td>7.9E-12</td><td>2.6E-11</td><td>MB231/MB468</td><td>MP</td></tr>
<tr><td>>hg38_21_+_8433172_8446622_RNA45SN1_45S_50ntFlanks@11658.11685.28|ATGTGATTTCTGCCCAGTGCTCTGAATG</td><td>>hg38_21_+_8433172_8446622_RNA45SN1_45S_50ntFlanks</td><td>11658</td><td>11685</td><td>28</td><td>ATGTGATTTCTGCCCAGTGCTCTGAATG</td><td>1086.0</td><td>-7.4</td><td>1.1</td><td>-6.8</td><td>1.0E-11</td><td>2.6E-11</td><td>MB231/MB468</td><td>MITO</td></tr>
<tr><td>>hg38_21_+_8433172_8446622_RNA45SN1_45S_50ntFlanks@12619.12653.35|AGGAGCCAATGGGGCGAAGCTACCATCTGTGGGAT</td><td>>hg38_21_+_8433172_8446622_RNA45SN1_45S_50ntFlanks</td><td>12619</td><td>12653</td><td>35</td><td>AGGAGCCAATGGGGCGAAGCTACCATCTGTGGGAT</td><td>778.4</td><td>-8.5</td><td>1.2</td><td>-6.8</td><td>9.0E-12</td><td>2.6E-11</td><td>MB231/MB468</td><td>NUC</td></tr>
<tr><td>>hg38_21_+_8433172_8446622_RNA45SN1_45S_50ntFlanks@9929.9953.25|GATATAGACAGCAGGACGGTGGCCA</td><td>>hg38_21_+_8433172_8446622_RNA45SN1_45S_50ntFlanks</td><td>9929</td><td>9953</td><td>25</td><td>GATATAGACAGCAGGACGGTGGCCA</td><td>133.6</td><td>-10.9</td><td>1.6</td><td>-6.8</td><td>8.0E-12</td><td>2.6E-11</td><td>MB231/MB468</td><td>MP</td></tr>
<tr><td>>MI0000469|hsa-mir-125a&WithFlank&19|+|51693248|51693345@21.41.21&[MIMAT0000443&hsa-miR-125a-5p&offsets|0|-3;m-63&19|+|51693268|51693289&offsets|0|-1]|TCCCTGAGACCCTTTAACCTG</td><td>>MI0000469|hsa-mir-125a&WithFlank&19|+|51693248|51693345</td><td>21</td><td>41</td><td>21</td><td>TCCCTGAGACCCTTTAACCTG</td><td>974.8</td><td>9.6</td><td>1.4</td><td>6.8</td><td>8.2E-12</td><td>2.6E-11</td><td>BT20/MB468</td><td>CYTO</td></tr>
<tr><td>>hg38_1_-_228634819_228635039_RNA5S12_5S_50ntFlanks@149.169.21|GAATACCGGGTGCTGTAGGCT</td><td>>hg38_1_-_228634819_228635039_RNA5S12_5S_50ntFlanks</td><td>149</td><td>169</td><td>21</td><td>GAATACCGGGTGCTGTAGGCT</td><td>950.8</td><td>5.3</td><td>0.8</td><td>6.8</td><td>8.8E-12</td><td>2.6E-11</td><td>MB231/MB468</td><td>CYTO</td></tr>
<tr><td>>hg38_21_+_8433172_8446622_RNA45SN1_45S_50ntFlanks@9553.9577.25|CCGTAGCGGTCCTGACGTGCAAATC</td><td>>hg38_21_+_8433172_8446622_RNA45SN1_45S_50ntFlanks</td><td>9553</td><td>9577</td><td>25</td><td>CCGTAGCGGTCCTGACGTGCAAATC</td><td>258.9</td><td>5.7</td><td>0.8</td><td>6.8</td><td>8.8E-12</td><td>2.6E-11</td><td>MB231/MB468</td><td>CYTO</td></tr>
<tr><td>>hg38_21_+_8433172_8446622_RNA45SN1_45S_50ntFlanks@12956.12982.27|GCTTCTGGGTCGGGGTTTCGTACGTAG</td><td>>hg38_21_+_8433172_8446622_RNA45SN1_45S_50ntFlanks</td><td>12956</td><td>12982</td><td>27</td><td>GCTTCTGGGTCGGGGTTTCGTACGTAG</td><td>850.9</td><td>-9.7</td><td>1.4</td><td>-6.8</td><td>1.1E-11</td><td>2.6E-11</td><td>BT20/MB468</td><td>NUC</td></tr>
<tr><td>>hg38_21_+_8433172_8446622_RNA45SN1_45S_50ntFlanks@11575.11607.33|ACTTAGAACTGGTGCGGACCAGGGGAATCCGAC</td><td>>hg38_21_+_8433172_8446622_RNA45SN1_45S_50ntFlanks</td><td>11575</td><td>11607</td><td>33</td><td>ACTTAGAACTGGTGCGGACCAGGGGAATCCGAC</td><td>431.6</td><td>-3.9</td><td>0.6</td><td>-6.8</td><td>9.0E-12</td><td>2.6E-11</td><td>MB231/MB468</td><td>NUC</td></tr>
<tr><td>>hg38_21_+_8433172_8446622_RNA45SN1_45S_50ntFlanks@11593.11613.21|CCAGGGGAATCCGACTGTTTA</td><td>>hg38_21_+_8433172_8446622_RNA45SN1_45S_50ntFlanks</td><td>11593</td><td>11613</td><td>21</td><td>CCAGGGGAATCCGACTGTTTA</td><td>511.7</td><td>-8.6</td><td>1.3</td><td>-6.8</td><td>8.0E-12</td><td>2.6E-11</td><td>MB231/MB468</td><td>MP</td></tr>
<tr><td>>MI0000266|hsa-mir-10a&WithFlank&17|-|48579832|48579953@28.49.22&[MIMAT0000253&hsa-miR-10a-5p&offsets|0|-1;m-18&17|-|48579905|48579926&offsets|0|0]|TACCCTGTAGATCCGAATTTGT</td><td>>MI0000266|hsa-mir-10a&WithFlank&17|-|48579832|48579953</td><td>28</td><td>49</td><td>22</td><td>TACCCTGTAGATCCGAATTTGT</td><td>6745.5</td><td>11.1</td><td>1.6</td><td>6.8</td><td>1.1E-11</td><td>2.6E-11</td><td>MB231/MB468</td><td>MITO</td></tr>
<tr><td>>hg38_21_+_8433172_8446622_RNA45SN1_45S_50ntFlanks@12405.12440.36|CTTTTTGATCCTTCGATGTCGGCTCTTCCTATCATT</td><td>>hg38_21_+_8433172_8446622_RNA45SN1_45S_50ntFlanks</td><td>12405</td><td>12440</td><td>36</td><td>CTTTTTGATCCTTCGATGTCGGCTCTTCCTATCATT</td><td>595.9</td><td>-4.3</td><td>0.6</td><td>-6.8</td><td>1.1E-11</td><td>2.6E-11</td><td>MB231/MB468</td><td>MITO</td></tr>
<tr><td>>MI0000285|hsa-mir-205&WithFlank&1|+|209432127|209432248@40.62.23(+1A)&[MIMAT0000266&hsa-miR-205-5p&offsets|0|+1(+1A);m-351&1|+|209432166|209432187&offsets|0|+1(+1A)]|TCCTTCATTCCACCGGAGTCTGTA</td><td>>MI0000285|hsa-mir-205&WithFlank&1|+|209432127|209432248</td><td>40</td><td>62</td><td>23(+1A)</td><td>TCCTTCATTCCACCGGAGTCTGTA</td><td>2774.1</td><td>6.9</td><td>1.0</td><td>6.9</td><td>6.5E-12</td><td>2.7E-11</td><td>BT20/MB468</td><td>MITO</td></tr>
<tr><td>>hg38_21_+_8433172_8446622_RNA45SN1_45S_50ntFlanks@5157.5185.29|CAAGTGGCGTTCAGCCACCCGAGATTGAG</td><td>>hg38_21_+_8433172_8446622_RNA45SN1_45S_50ntFlanks</td><td>5157</td><td>5185</td><td>29</td><td>CAAGTGGCGTTCAGCCACCCGAGATTGAG</td><td>730.4</td><td>-13.3</td><td>2.0</td><td>-6.8</td><td>1.1E-11</td><td>2.7E-11</td><td>MB231/MB468</td><td>MITO</td></tr>
<tr><td>>hg38_21_+_8433172_8446622_RNA45SN1_45S_50ntFlanks@11755.11777.23|AATGCCTCGTCATCTAATTAGTG</td><td>>hg38_21_+_8433172_8446622_RNA45SN1_45S_50ntFlanks</td><td>11755</td><td>11777</td><td>23</td><td>AATGCCTCGTCATCTAATTAGTG</td><td>85.0</td><td>-10.3</td><td>1.5</td><td>-6.8</td><td>8.1E-12</td><td>2.7E-11</td><td>MB231/MB468</td><td>MP</td></tr>
<tr><td>>hg38_21_+_8433172_8446622_RNA45SN1_45S_50ntFlanks@5045.5072.28|CTTAGTTGGTGGAGCGATTTGTCTGGTT</td><td>>hg38_21_+_8433172_8446622_RNA45SN1_45S_50ntFlanks</td><td>5045</td><td>5072</td><td>28</td><td>CTTAGTTGGTGGAGCGATTTGTCTGGTT</td><td>453.8</td><td>-12.6</td><td>1.9</td><td>-6.8</td><td>1.1E-11</td><td>2.7E-11</td><td>MB231/MB468</td><td>MITO</td></tr>
<tr><td>>trna134_GluTTC_1_-_16861774_16861845@1.31.31,trna5_GluTTC_1_+_17199078_17199149@1.31.31,trna84_GluTTC_1_-_161391883_161391954@1.31.31,trna94_GluTTC_1_-_149664355_149664427@1.31.31|TCCCTGGTGGTCTAGTGGCTAGGATTCGGCG</td><td>>trna134_GluTTC_1_-_16861774_16861845</td><td>1</td><td>31</td><td>31</td><td>TCCCTGGTGGTCTAGTGGCTAGGATTCGGCG</td><td>12241.6</td><td>-9.8</td><td>1.4</td><td>-6.8</td><td>1.1E-11</td><td>2.7E-11</td><td>BT20/MB468</td><td>NUC</td></tr>
<tr><td>>hg38_21_+_8433172_8446622_RNA45SN1_45S_50ntFlanks@476.516.41|TTCGTGATCGATGTGGTGACGTCGTGCTCTCCCGGGCCGGG</td><td>>hg38_21_+_8433172_8446622_RNA45SN1_45S_50ntFlanks</td><td>476</td><td>516</td><td>41</td><td>TTCGTGATCGATGTGGTGACGTCGTGCTCTCCCGGGCCGGG</td><td>1480.3</td><td>7.8</td><td>1.1</td><td>6.9</td><td>6.5E-12</td><td>2.7E-11</td><td>BT20/MB468</td><td>MITO</td></tr>
<tr><td>>trna10_LysCTT_16_+_3241501_3241573@8.35.28,trna13_LysCTT_14_-_58706613_58706685@8.35.28,trna2_LysCTT_15_+_79152904_79152976@8.35.28,trna4_LysCTT_18_-_43669271_43669343@8.35.28|TAGCTCAGTCGGTAGAGCATGGGACTCT</td><td>>trna10_LysCTT_16_+_3241501_3241573</td><td>8</td><td>35</td><td>28</td><td>TAGCTCAGTCGGTAGAGCATGGGACTCT</td><td>240.9</td><td>11.1</td><td>1.6</td><td>6.8</td><td>1.1E-11</td><td>2.7E-11</td><td>MB231/MB468</td><td>MITO</td></tr>
<tr><td>>hg38_21_+_8433172_8446622_RNA45SN1_45S_50ntFlanks@12606.12626.21|ATGTGCTTGGCTGAGGAGCCA</td><td>>hg38_21_+_8433172_8446622_RNA45SN1_45S_50ntFlanks</td><td>12606</td><td>12626</td><td>21</td><td>ATGTGCTTGGCTGAGGAGCCA</td><td>711.4</td><td>-9.6</td><td>1.4</td><td>-6.8</td><td>1.1E-11</td><td>2.7E-11</td><td>BT20/MB468</td><td>NUC</td></tr>
<tr><td>>hg38_21_+_8433172_8446622_RNA45SN1_45S_50ntFlanks@10099.10126.28|CGCGCGCGCGTGTGGTGTGCGTCGGAGG</td><td>>hg38_21_+_8433172_8446622_RNA45SN1_45S_50ntFlanks</td><td>10099</td><td>10126</td><td>28</td><td>CGCGCGCGCGTGTGGTGTGCGTCGGAGG</td><td>156.2</td><td>-11.1</td><td>1.6</td><td>-6.8</td><td>8.1E-12</td><td>2.7E-11</td><td>MB231/MB468</td><td>MP</td></tr>
</table>

| | | | | | | | | | | | | | |
|---|---|---|---|---|---|---|---|---|---|---|---|---|---|
| >hg38_21_+_8433172_8446622_RNA45SN1_45S_50ntFlanks@8170.8200.31\|CAAGTCCTTCTGATCGAGGCCCAGCCCGTGG | >hg38_21_+_8433172_8446622_RNA45SN1_45S_50ntFlanks | 8170 | 8200 | 31 | CAAGTCCTTCTGATCGAGGCCCAGCCCGTGG | 111.0 | -10.7 | 1.6 | -6.8 | 8.2E-12 | 2.7E-11 | MB231/MB468 | MP |
| >hg38_21_+_8433172_8446622_RNA45SN1_45S_50ntFlanks@12646.12670.25\|TGTGGGATTATGACTGAACGCCTCT | >hg38_21_+_8433172_8446622_RNA45SN1_45S_50ntFlanks | 12646 | 12670 | 25 | TGTGGGATTATGACTGAACGCCTCT | 399.8 | 9.8 | 1.4 | 6.8 | 9.0E-12 | 2.7E-11 | MB231/MB468 | CYTO |
| >hg38_21_+_8433172_8446622_RNA45SN1_45S_50ntFlanks@6771.6803.33\|GGTTCCTCCCGGGGCTACGCCTGTCTGAGCGTC | >hg38_21_+_8433172_8446622_RNA45SN1_45S_50ntFlanks | 6771 | 6803 | 33 | GGTTCCTCCCGGGGCTACGCCTGTCTGAGCGTC | 707.3 | -9.0 | 1.3 | -6.8 | 8.2E-12 | 2.7E-11 | MB231/MB468 | MP |
| >hg38_21_+_8433172_8446622_RNA45SN1_45S_50ntFlanks@10335.10361.27\|GTAGCAAATATTCAAACGAGAACTTTG | >hg38_21_+_8433172_8446622_RNA45SN1_45S_50ntFlanks | 10335 | 10361 | 27 | GTAGCAAATATTCAAACGAGAACTTTG | 80.4 | -10.2 | 1.5 | -6.8 | 8.2E-12 | 2.7E-11 | MB231/MB468 | MP |
| >hg38_21_+_8433172_8446622_RNA45SN1_45S_50ntFlanks@4641.4662.22\|CATTCGTATTGCGCCGCTAGAG | >hg38_21_+_8433172_8446622_RNA45SN1_45S_50ntFlanks | 4641 | 4662 | 22 | CATTCGTATTGCGCCGCTAGAG | 511.4 | -8.3 | 1.2 | -6.8 | 1.1E-11 | 2.7E-11 | MB231/MB468 | MITO |
| >hg38_21_+_8433172_8446622_RNA45SN1_45S_50ntFlanks@10367.10390.24\|CGAAGTGGAGAAGGGTTCCATGTG | >hg38_21_+_8433172_8446622_RNA45SN1_45S_50ntFlanks | 10367 | 10390 | 24 | CGAAGTGGAGAAGGGTTCCATGTG | 111.9 | -10.7 | 1.6 | -6.8 | 8.2E-12 | 2.7E-11 | MB231/MB468 | MP |
| >hg38_21_+_8433172_8446622_RNA45SN1_45S_50ntFlanks@6745.6769.25\|ACACTTCGAACGCACTTGCGGCCCC | >hg38_21_+_8433172_8446622_RNA45SN1_45S_50ntFlanks | 6745 | 6769 | 25 | ACACTTCGAACGCACTTGCGGCCCC | 582.5 | -6.7 | 1.0 | -6.8 | 1.1E-11 | 2.7E-11 | MB231/MB468 | MITO |
| >hg38_21_+_8433172_8446622_RNA45SN1_45S_50ntFlanks@10768.10797.30\|CAGCAGGTCTCCAAGGTGAACAGCCTCTGG | >hg38_21_+_8433172_8446622_RNA45SN1_45S_50ntFlanks | 10768 | 10797 | 30 | CAGCAGGTCTCCAAGGTGAACAGCCTCTGG | 168.5 | -6.7 | 1.0 | -6.8 | 8.4E-12 | 2.7E-11 | BT20/MB468 | CYTO |
| >hg38_21_+_8433172_8446622_RNA45SN1_45S_50ntFlanks@12008.12037.30\|CCGCCGGTGAAATACCACTACTCTGATCGT | >hg38_21_+_8433172_8446622_RNA45SN1_45S_50ntFlanks | 12008 | 12037 | 30 | CCGCCGGTGAAATACCACTACTCTGATCGT | 666.8 | -9.4 | 1.4 | -6.8 | 1.1E-11 | 2.7E-11 | BT20/MB468 | NUC |
| >MI0000809\|hsa-mir-151a&WithFlank&8\|-\|140732558\|140732659@53.74.22&[MIMAT0000757&hsa-miR-151a-3p&offsets\|0\|+1;m-30&8\|-\|140732587\|140732607&offsets\|0\|+1]\|CTAGACTGAAGCTCCTTGAGGA | >MI0000809\|hsa-mir-151a&WithFlank&8\|-\|140732558\|140732659 | 53 | 74 | 22 | CTAGACTGAAGCTCCTTGAGGA | 3899.5 | 2.5 | 0.4 | 6.8 | 1.1E-11 | 2.7E-11 | BT20/MB468 | NUC |
| >MI0003667\|hsa-mir-652&WithFlank&X\|+\|110055323\|110055432@67.87.21&[MIMAT0003322&hsa-miR-652-3p&offsets\|0\|0;m-148&X\|+\|110055389\|110055410&offsets\|0\|-1]\|AATGGCGCCACTAGGGTTGTG | >MI0003667\|hsa-mir-652&WithFlank&X\|+\|110055323\|110055432 | 67 | 87 | 21 | AATGGCGCCACTAGGGTTGTG | 599.0 | 8.7 | 1.3 | 6.8 | 8.4E-12 | 2.7E-11 | BT20/MB468 | CYTO |
| >MI0000466\|hsa-mir-9-1&WithFlank&1\|-\|156420335\|156420435@22.43.22;MI0000468\|hsa-mir-9-3&WithFlank&15\|+\|89368011\|89368112@22.43.22;MI0000467\|hsa-mir-9-2&WithFlank&5\|-\|88666847\|88666945@22.43.22&[MIMAT0000441&hsa-miR-9-5p&offsets\|0\|-1;m-104&1\|-\|156420392\|156420414&offsets\|0\|-1];[MIMAT0000441_1&hsa-miR-9-5p&offsets\|0\|-1;m-105&15\|+\|89368032\|89368054&offsets\|0\|-1];[MIMAT0000441_2&hsa-miR-9-5p&offsets\|0\|-1;m-104&5\|-\|88666902\|88666924&offsets\|0\|-1]\|TCTTTGGTTATCTAGCTGTATG | >MI0000466\|hsa-mir-9-1&WithFlank&1\|-\|156420335\|156420435 | 22 | 43 | 22 | TCTTTGGTTATCTAGCTGTATG | 1145.8 | 9.4 | 1.4 | 6.9 | 6.6E-12 | 2.7E-11 | BT20/MB468 | MITO |
| >hg38_21_+_8433172_8446622_RNA45SN1_45S_50ntFlanks@12934.12953.20\|CTAAACCATTCGTAGACGAC | >hg38_21_+_8433172_8446622_RNA45SN1_45S_50ntFlanks | 12934 | 12953 | 20 | CTAAACCATTCGTAGACGAC | 1671.1 | -6.4 | 0.9 | -6.8 | 9.0E-12 | 2.7E-11 | MB231/MB468 | CYTO |
| >trna59_GluCTC_1_+_249168447_249168518@32.70.39\|CTCTCACCGCCGCGGCCCGGGTTCGATTCCCGGTCAGGA | >trna59_GluCTC_1_+_249168447_249168518 | 32 | 70 | 39 | CTCTCACCGCCGCGGCCCGGGTTCGATTCCCGGTCAGGA | 226.0 | -11.6 | 1.7 | -6.8 | 1.1E-11 | 2.7E-11 | MB231/MB468 | MITO |
| >hg38_21_+_8433172_8446622_RNA45SN1_45S_50ntFlanks@4444.4462.19\|CCTTGCCTCTCGGCGCCCC | >hg38_21_+_8433172_8446622_RNA45SN1_45S_50ntFlanks | 4444 | 4462 | 19 | CCTTGCCTCTCGGCGCCCC | 131.8 | -10.9 | 1.6 | -6.8 | 8.2E-12 | 2.7E-11 | MB231/MB468 | MP |
| >MI0000746\|hsa-mir-99b&WithFlank&19\|+\|51692606\|51692687@13.33.21(+1U)&[MIMAT0000689&hsa-miR-99b-5p&offsets\|0\|-1(+1U);m-10&19\|+\|51692618\|51692639&offsets\|0\|-1(+1U)]\|CACCCGTAGAACCGACCTTGCT | >MI0000746\|hsa-mir-99b&WithFlank&19\|+\|51692606\|51692687 | 13 | 33 | 21(+1U) | CACCCGTAGAACCGACCTTGCT | 378.0 | 4.4 | 0.6 | 6.8 | 9.2E-12 | 2.7E-11 | MB231/MB468 | NUC |
| >hg38_21_+_8433172_8446622_RNA45SN1_45S_50ntFlanks@12639.12657.19\|TACCATCTGTGGGATTATG | >hg38_21_+_8433172_8446622_RNA45SN1_45S_50ntFlanks | 12639 | 12657 | 19 | TACCATCTGTGGGATTATG | 94.7 | -10.4 | 1.5 | -6.8 | 8.2E-12 | 2.7E-11 | MB231/MB468 | MP |
| >hg38_21_+_8433172_8446622_RNA45SN1_45S_50ntFlanks@11759.11780.22\|CCTCGTCATCTAATTAGTGACG | >hg38_21_+_8433172_8446622_RNA45SN1_45S_50ntFlanks | 11759 | 11780 | 22 | CCTCGTCATCTAATTAGTGACG | 156.1 | -11.1 | 1.6 | -6.8 | 8.3E-12 | 2.7E-11 | MB231/MB468 | MP |
| >hg38_21_+_8433172_8446622_RNA45SN1_45S_50ntFlanks@10869.10900.32\|AGGGCTGGGTCGGTCGGGCTGGGGCGCGAAGC | >hg38_21_+_8433172_8446622_RNA45SN1_45S_50ntFlanks | 10869 | 10900 | 32 | AGGGCTGGGTCGGTCGGGCTGGGGCGCGAAGC | 171.0 | -11.3 | 1.7 | -6.8 | 8.3E-12 | 2.7E-11 | MB231/MB468 | MP |
| >hg38_21_+_8433172_8446622_RNA45SN1_45S_50ntFlanks@11316.11336.21\|CACCCCACGTCTCGTCGCGCG | >hg38_21_+_8433172_8446622_RNA45SN1_45S_50ntFlanks | 11316 | 11336 | 21 | CACCCCACGTCTCGTCGCGCG | 3277.2 | -9.9 | 1.5 | -6.8 | 9.3E-12 | 2.7E-11 | MB231/MB468 | NUC |
| >hg38_21_+_8433172_8446622_RNA45SN1_45S_50ntFlanks@12822.12849.28\|CCGGGACCGGGGTCCGGTGCGGAGTGCC | >hg38_21_+_8433172_8446622_RNA45SN1_45S_50ntFlanks | 12822 | 12849 | 28 | CCGGGACCGGGGTCCGGTGCGGAGTGCC | 307.6 | -12.1 | 1.8 | -6.8 | 8.3E-12 | 2.7E-11 | MB231/MB468 | MP |
| >hg38_21_+_8433172_8446622_RNA45SN1_45S_50ntFlanks@11670.11696.27\|CCCAGTGCTCTGAATGTCAAAGTGAAG | >hg38_21_+_8433172_8446622_RNA45SN1_45S_50ntFlanks | 11670 | 11696 | 27 | CCCAGTGCTCTGAATGTCAAAGTGAAG | 86.3 | -10.3 | 1.5 | -6.8 | 8.3E-12 | 2.7E-11 | MB231/MB468 | MP |
| >hg38_1_-_228634819_228635039_RNA5S12_5S_50ntFlanks@134.169.36\|TGGGAGACCGCCTGGGAATACCGGGTGCTGTAGGCT | >hg38_1_-_228634819_228635039_RNA5S12_5S_50ntFlanks | 134 | 169 | 36 | TGGGAGACCGCCTGGGAATACCGGGTGCTGTAGGCT | 8761.2 | 7.2 | 1.0 | 6.9 | 6.7E-12 | 2.7E-11 | BT20/MB468 | MITO |
| >trna16_ValTAC_11_-_59318460_59318532@1.34.34\|GGTTCCATAGTGTAGCGGTTATCACGTCTGCTTT | >trna16_ValTAC_11_-_59318460_59318532 | 1 | 34 | 34 | GGTTCCATAGTGTAGCGGTTATCACGTCTGCTTT | 379.2 | 11.8 | 1.7 | 6.8 | 1.1E-11 | 2.7E-11 | MB231/MB468 | MITO |
| >hg38_21_+_8433172_8446622_RNA45SN1_45S_50ntFlanks@476.507.32\|TTCGTGATCGATGTGGTGACGTCGTGCTCTCC | >hg38_21_+_8433172_8446622_RNA45SN1_45S_50ntFlanks | 476 | 507 | 32 | TTCGTGATCGATGTGGTGACGTCGTGCTCTCC | 566.6 | 8.5 | 1.2 | 6.8 | 8.5E-12 | 2.7E-11 | BT20/MB468 | CYTO |
| >hg38_21_+_8433172_8446622_RNA45SN1_45S_50ntFlanks@11577.11602.26\|TTAGAACTGGTGCGGACCAGGGGAAT | >hg38_21_+_8433172_8446622_RNA45SN1_45S_50ntFlanks | 11577 | 11602 | 26 | TTAGAACTGGTGCGGACCAGGGGAAT | 133.5 | -10.9 | 1.6 | -6.8 | 8.3E-12 | 2.7E-11 | MB231/MB468 | MP |
| >hg38_21_+_8433172_8446622_RNA45SN1_45S_50ntFlanks@5084.5120.37\|CGAACGAGACTCTGGCATGCTAACTAGTTACGCGACC | >hg38_21_+_8433172_8446622_RNA45SN1_45S_50ntFlanks | 5084 | 5120 | 37 | CGAACGAGACTCTGGCATGCTAACTAGTTACGCGACC | 985.8 | -6.4 | 0.9 | -6.8 | 1.1E-11 | 2.7E-11 | MB231/MB468 | MITO |
| >hg38_21_+_8433172_8446622_RNA45SN1_45S_50ntFlanks@12662.12703.42\|AACGCCTCTAAGTCAGAATCCCGCCCAGGCGGAACGATACGG | >hg38_21_+_8433172_8446622_RNA45SN1_45S_50ntFlanks | 12662 | 12703 | 42 | AACGCCTCTAAGTCAGAATCCCGCCCAGGCGGAACGATACGG | 251.5 | -11.8 | 1.7 | -6.8 | 1.1E-11 | 2.7E-11 | MB231/MB468 | MITO |
| >hg38_21_+_8433172_8446622_RNA45SN1_45S_50ntFlanks@6734.6774.41\|ATTGATCATCGACACTTCGAACGCACTTGCGGCCCCGGGTT | >hg38_21_+_8433172_8446622_RNA45SN1_45S_50ntFlanks | 6734 | 6774 | 41 | ATTGATCATCGACACTTCGAACGCACTTGCGGCCCCGGGTT | 583.7 | 9.1 | 1.3 | 6.8 | 8.6E-12 | 2.7E-11 | BT20/MB468 | CYTO |
| >hg38_21_+_8433172_8446622_RNA45SN1_45S_50ntFlanks@3738.3756.19\|TCTCAAAGATTAAGCCATG | >hg38_21_+_8433172_8446622_RNA45SN1_45S_50ntFlanks | 3738 | 3756 | 19 | TCTCAAAGATTAAGCCATG | 177.4 | -6.5 | 1.0 | -6.8 | 9.2E-12 | 2.8E-11 | MB231/MB468 | CYTO |
| >hg38_21_+_8433172_8446622_RNA45SN1_45S_50ntFlanks@12676.12703.28\|AGAATCCCGCCCAGGCGGAACGATACGG | >hg38_21_+_8433172_8446622_RNA45SN1_45S_50ntFlanks | 12676 | 12703 | 28 | AGAATCCCGCCCAGGCGGAACGATACGG | 364.7 | -8.1 | 1.2 | -6.8 | 8.6E-12 | 2.8E-11 | BT20/MB468 | CYTO |
| >hg38_21_+_8433172_8446622_RNA45SN1_45S_50ntFlanks@10410.10437.28\|TCAGTCGGTCCTGAGAGATGGGCGAGCG | >hg38_21_+_8433172_8446622_RNA45SN1_45S_50ntFlanks | 10410 | 10437 | 28 | TCAGTCGGTCCTGAGAGATGGGCGAGCG | 165.4 | -11.2 | 1.6 | -6.8 | 8.4E-12 | 2.8E-11 | MB231/MB468 | MP |
| >hg38_21_+_8433172_8446622_RNA45SN1_45S_50ntFlanks@4240.4258.19\|ACTCTTTCGAGGCCCTGTA | >hg38_21_+_8433172_8446622_RNA45SN1_45S_50ntFlanks | 4240 | 4258 | 19 | ACTCTTTCGAGGCCCTGTA | 298.5 | -10.3 | 1.5 | -6.9 | 6.8E-12 | 2.8E-11 | BT20/MB468 | MITO |
| >hg38_21_+_8433172_8446622_RNA45SN1_45S_50ntFlanks@8381.8404.24\|AAGAGGGCGTGAAACCGTTAAGAG | >hg38_21_+_8433172_8446622_RNA45SN1_45S_50ntFlanks | 8381 | 8404 | 24 | AAGAGGGCGTGAAACCGTTAAGAG | 570.1 | -8.7 | 1.3 | -6.8 | 8.4E-12 | 2.8E-11 | MB231/MB468 | MP |
| >hg38_21_+_8433172_8446622_RNA45SN1_45S_50ntFlanks@485.528.44\|GATGTGGTGACGTCGTGCTCTCCCGGGCCGGGTCCGAGCCGCGA | >hg38_21_+_8433172_8446622_RNA45SN1_45S_50ntFlanks | 485 | 528 | 44 | GATGTGGTGACGTCGTGCTCTCCCGGGCCGGGTCCGAGCCGCGA | 309.5 | 11.1 | 1.6 | 6.8 | 1.2E-11 | 2.8E-11 | BT20/MB468 | NUC |
| >hg38_21_+_8433172_8446622_RNA45SN1_45S_50ntFlanks@11891.11910.20\|TTGACTCTAGTCTGGCACGG | >hg38_21_+_8433172_8446622_RNA45SN1_45S_50ntFlanks | 11891 | 11910 | 20 | TTGACTCTAGTCTGGCACGG | 532.4 | -9.0 | 1.3 | -6.8 | 1.2E-11 | 2.8E-11 | BT20/MB468 | NUC |
| >MI0000085\|hsa-mir-27a&WithFlank&19\|-\|13836434\|13836523@57.76.20(+1G)&[MIMAT0000084&hsa-miR-27a-3p&offsets\|0\|-1(+1G);m-43&19\|-\|13836447\|13836467&offsets\|0\|-1(+1G)]\|TTCACAGTGGCTAAGTTCCGG | >MI0000085\|hsa-mir-27a&WithFlank&19\|-\|13836434\|13836523 | 57 | 76 | 20(+1G) | TTCACAGTGGCTAAGTTCCGG | 170.9 | 5.8 | 0.8 | 6.8 | 9.5E-12 | 2.8E-11 | MB231/MB468 | NUC |
| >MI0000438\|hsa-mir-15b&WithFlank&3\|+\|160404582\|160404691@64.84.21&[MIMAT0004586&hsa-miR-15b-3p&offsets\|0\|-1;m-204&3\|+\|160404645\|160404665&offsets\|0\|0]\|CGAATCATTATTTGCTGCTCT | >MI0000438\|hsa-mir-15b&WithFlank&3\|+\|160404582\|160404691 | 64 | 84 | 21 | CGAATCATTATTTGCTGCTCT | 283.8 | 4.1 | 0.6 | 6.8 | 9.5E-12 | 2.8E-11 | MB231/MB468 | NUC |
| >MI0000065\|hsa-let-7d&WithFlank&9\|+\|94178828\|94178926@14.34.21(+1G)&[MIMAT0000065&hsa-let-7d-5p&offsets\|0\|-1(+1G);m-90&9\|+\|94178841\|94178862&offsets\|0\|-1(+1G)]\|AGAGGTAGTAGGTTGCATAGTG | >MI0000065\|hsa-let-7d&WithFlank&9\|+\|94178828\|94178926 | 14 | 34 | 21(+1G) | AGAGGTAGTAGGTTGCATAGTG | 356.9 | 7.5 | 1.1 | 6.8 | 8.7E-12 | 2.8E-11 | BT20/MB468 | CYTO |
| >hg38_21_+_8433172_8446622_RNA45SN1_45S_50ntFlanks@3787.3812.26\|AAACTGCGAATGGCTCATTAAATCAG | >hg38_21_+_8433172_8446622_RNA45SN1_45S_50ntFlanks | 3787 | 3812 | 26 | AAACTGCGAATGGCTCATTAAATCAG | 1620.4 | -10.2 | 1.5 | -6.8 | 1.1E-11 | 2.8E-11 | MB231/MB468 | MITO |
| >hg38_21_+_8433172_8446622_RNA45SN1_45S_50ntFlanks@12416.12442.27\|TTCGATGTCGGCTCTTCCTATCATTGT | >hg38_21_+_8433172_8446622_RNA45SN1_45S_50ntFlanks | 12416 | 12442 | 27 | TTCGATGTCGGCTCTTCCTATCATTGT | 392.8 | 5.2 | 0.8 | 6.9 | 6.8E-12 | 2.8E-11 | BT20/MB468 | MITO |
| >hg38_21_+_8433172_8446622_RNA45SN1_45S_50ntFlanks@9244.9274.31\|AGCGAGCGCACGGGGTCGGCGGCGACGTCGG | >hg38_21_+_8433172_8446622_RNA45SN1_45S_50ntFlanks | 9244 | 9274 | 31 | AGCGAGCGCACGGGGTCGGCGGCGACGTCGG | 113.6 | -10.7 | 1.6 | -6.8 | 8.5E-12 | 2.8E-11 | MB231/MB468 | MP |
| >MI0000441\|hsa-mir-30b&WithFlank&8\|-\|134800514\|134800613@23.44.22&[MIMAT0000420&hsa-miR-30b-5p&offsets\|0\|0;m-76&8\|-\|134800570\|134800591&offsets\|0\|0]\|TGTAAACATCCTACACTCAGCT | >MI0000441\|hsa-mir-30b&WithFlank&8\|-\|134800514\|134800613 | 23 | 44 | 22 | TGTAAACATCCTACACTCAGCT | 1808.7 | 4.6 | 0.7 | 6.8 | 1.2E-11 | 2.8E-11 | BT20/MB468 | NUC |
| >hg38_21_+_8433172_8446622_RNA45SN1_45S_50ntFlanks@8168.8195.28\|CCCAAGTCCTTCTGATCGAGGCCCAGCC | >hg38_21_+_8433172_8446622_RNA45SN1_45S_50ntFlanks | 8168 | 8195 | 28 | CCCAAGTCCTTCTGATCGAGGCCCAGCC | 665.5 | -9.6 | 1.4 | -6.8 | 1.2E-11 | 2.8E-11 | BT20/MB468 | NUC |
| >hg38_21_+_8433172_8446622_RNA45SN1_45S_50ntFlanks@11584.11611.28\|TGGTGCGGACCAGGGGAATCCGACTGTT | >hg38_21_+_8433172_8446622_RNA45SN1_45S_50ntFlanks | 11584 | 11611 | 28 | TGGTGCGGACCAGGGGAATCCGACTGTT | 270.3 | 3.7 | 0.5 | 6.8 | 8.7E-12 | 2.8E-11 | BT20/MB468 | CYTO |
| >hg38_21_+_8433172_8446622_RNA45SN1_45S_50ntFlanks@11998.12032.35\|GCCCTGCGGGCCGCCGGTGAAATACCACTACTCTG | >hg38_21_+_8433172_8446622_RNA45SN1_45S_50ntFlanks | 11998 | 12032 | 35 | GCCCTGCGGGCCGCCGGTGAAATACCACTACTCTG | 259.7 | -10.1 | 1.5 | -6.9 | 6.9E-12 | 2.8E-11 | BT20/MB468 | MITO |
| >hg38_21_+_8433172_8446622_RNA45SN1_45S_50ntFlanks@4657.4686.30\|CTAGAGGTGAAATTCTTGGACCGGCGCAAG | >hg38_21_+_8433172_8446622_RNA45SN1_45S_50ntFlanks | 4657 | 4686 | 30 | CTAGAGGTGAAATTCTTGGACCGGCGCAAG | 371.4 | -10.6 | 1.5 | -6.9 | 6.9E-12 | 2.8E-11 | BT20/MB468 | MITO |
| >hg38_21_+_8433172_8446622_RNA45SN1_45S_50ntFlanks@4735.4765.31\|ACGAAAGTCGGAGGTTCGAAGACGATCAGAT | >hg38_21_+_8433172_8446622_RNA45SN1_45S_50ntFlanks | 4735 | 4765 | 31 | ACGAAAGTCGGAGGTTCGAAGACGATCAGAT | 564.3 | 8.7 | 1.3 | 6.8 | 8.8E-12 | 2.8E-11 | BT20/MB468 | CYTO |

| | | | | | | | | | | | | | |
|---|---|---|---|---|---|---|---|---|---|---|---|---|---|
| >hg38_21_+_8433172_8446622_RNA45SN1_45S_50ntFlanks@11569.11600.32\|CAGCCGACTTAGAACTGGTGCGGACCAGGGGA | >hg38_21_+_8433172_8446622_RNA45SN1_45S_50ntFlanks | 11569 | 11600 | 32 | CAGCCGACTTAGAACTGGTGCGGACCAGGGGA | 13840.1 | -9.7 | 1.4 | -6.8 | 1.2E-11 | 2.8E-11 | BT20/MB468 | NUC |
| >hg38_21_+_8433172_8446622_RNA45SN1_45S_50ntFlanks@12719.12742.24\|TCGGTTGGCCTCGGATAGCCGGTC | >hg38_21_+_8433172_8446622_RNA45SN1_45S_50ntFlanks | 12719 | 12742 | 24 | TCGGTTGGCCTCGGATAGCCGGTC | 136.6 | -11.0 | 1.6 | -6.8 | 8.6E-12 | 2.8E-11 | MB231/MB468 | MP |
| >hg38_21_+_8433172_8446622_RNA45SN1_45S_50ntFlanks@12892.12927.36\|CCCTCGCCCGTCACGCACCGCACGTTCGTGGGGAAC | >hg38_21_+_8433172_8446622_RNA45SN1_45S_50ntFlanks | 12892 | 12927 | 36 | CCCTCGCCCGTCACGCACCGCACGTTCGTGGGGAAC | 90.7 | -10.4 | 1.5 | -6.8 | 8.6E-12 | 2.8E-11 | MB231/MB468 | MP |
| >hg38_21_+_8433172_8446622_RNA45SN1_45S_50ntFlanks@4574.4611.38\|ATAGGACCGCGGTTCTATTTTGTTGGTTTTCGGAACTG | >hg38_21_+_8433172_8446622_RNA45SN1_45S_50ntFlanks | 4574 | 4611 | 38 | ATAGGACCGCGGTTCTATTTTGTTGGTTTTCGGAACTG | 292.1 | -12.0 | 1.8 | -6.8 | 1.1E-11 | 2.8E-11 | MB231/MB468 | MITO |
| >hg38_21_+_8433172_8446622_RNA45SN1_45S_50ntFlanks@10802.10829.28\|TTGGAACAATGTAGGTAAGGGAAGTCGG | >hg38_21_+_8433172_8446622_RNA45SN1_45S_50ntFlanks | 10802 | 10829 | 28 | TTGGAACAATGTAGGTAAGGGAAGTCGG | 106.6 | -10.6 | 1.6 | -6.8 | 8.6E-12 | 2.8E-11 | MB231/MB468 | MP |
| >hg38_21_+_8433172_8446622_RNA45SN1_45S_50ntFlanks@9505.9525.21\|TGAACTATGCCTGGGCAGGGC | >hg38_21_+_8433172_8446622_RNA45SN1_45S_50ntFlanks | 9505 | 9525 | 21 | TGAACTATGCCTGGGCAGGGC | 200.5 | -11.5 | 1.7 | -6.8 | 8.6E-12 | 2.8E-11 | MB231/MB468 | MP |
| >hg38_21_+_8433172_8446622_RNA45SN1_45S_50ntFlanks@9576.9597.22\|TCGGTCGTCCGACCTGGGTATA | >hg38_21_+_8433172_8446622_RNA45SN1_45S_50ntFlanks | 9576 | 9597 | 22 | TCGGTCGTCCGACCTGGGTATA | 263.8 | -10.1 | 1.5 | -6.9 | 6.9E-12 | 2.8E-11 | BT20/MB468 | MITO |
| >hg38_21_+_8433172_8446622_RNA45SN1_45S_50ntFlanks@6719.6750.32\|TGAATTGCAGGACACATTGATCATCGACACTT | >hg38_21_+_8433172_8446622_RNA45SN1_45S_50ntFlanks | 6719 | 6750 | 32 | TGAATTGCAGGACACATTGATCATCGACACTT | 433.5 | -9.0 | 1.3 | -6.8 | 1.2E-11 | 2.8E-11 | BT20/MB468 | NUC |
| >hg38_21_+_8433172_8446622_RNA45SN1_45S_50ntFlanks@10922.10944.23\|CTGGACGAGGCGCCGCCGCCCCC | >hg38_21_+_8433172_8446622_RNA45SN1_45S_50ntFlanks | 10922 | 10944 | 23 | CTGGACGAGGCGCCGCCGCCCCC | 230.3 | -6.9 | 1.0 | -6.8 | 9.4E-12 | 2.8E-11 | MB231/MB468 | CYTO |
| >hg38_21_+_8433172_8446622_RNA45SN1_45S_50ntFlanks@6662.6689.28\|GGTGGATCACTCGGCTCGTGCGTCGATG | >hg38_21_+_8433172_8446622_RNA45SN1_45S_50ntFlanks | 6662 | 6689 | 28 | GGTGGATCACTCGGCTCGTGCGTCGATG | 409.9 | -4.9 | 0.7 | -6.8 | 1.1E-11 | 2.8E-11 | MB231/MB468 | MITO |
| >hg38_21_+_8433172_8446622_RNA45SN1_45S_50ntFlanks@10806.10829.24\|AACAATGTAGGTAAGGGAAGTCGG | >hg38_21_+_8433172_8446622_RNA45SN1_45S_50ntFlanks | 10806 | 10829 | 24 | AACAATGTAGGTAAGGGAAGTCGG | 182.1 | -6.2 | 0.9 | -6.8 | 9.5E-12 | 2.8E-11 | MB231/MB468 | CYTO |
| >hg38_21_+_8433172_8446622_RNA45SN1_45S_50ntFlanks@6753.6770.18\|AACGCACTTGCGGCCCCG | >hg38_21_+_8433172_8446622_RNA45SN1_45S_50ntFlanks | 6753 | 6770 | 18 | AACGCACTTGCGGCCCCG | 173.1 | -5.1 | 0.7 | -6.8 | 9.5E-12 | 2.8E-11 | MB231/MB468 | CYTO |
| >hg38_21_+_8433172_8446622_RNA45SN1_45S_50ntFlanks@11881.11904.24\|CCTGTTGAGCTTGACTCTAGTCTG | >hg38_21_+_8433172_8446622_RNA45SN1_45S_50ntFlanks | 11881 | 11904 | 24 | CCTGTTGAGCTTGACTCTAGTCTG | 1350.3 | -8.3 | 1.2 | -6.8 | 8.9E-12 | 2.8E-11 | BT20/MB468 | CYTO |
| >hg38_21_+_8433172_8446622_RNA45SN1_45S_50ntFlanks@10402.10422.21\|AACATGGGTCAGTCGGTCCTG | >hg38_21_+_8433172_8446622_RNA45SN1_45S_50ntFlanks | 10402 | 10422 | 21 | AACATGGGTCAGTCGGTCCTG | 1319.2 | -7.5 | 1.1 | -6.8 | 8.7E-12 | 2.8E-11 | MB231/MB468 | MP |
| >hg38_21_+_8433172_8446622_RNA45SN1_45S_50ntFlanks@4555.4575.21\|CGCAGCTAGGAATAATGGAAT | >hg38_21_+_8433172_8446622_RNA45SN1_45S_50ntFlanks | 4555 | 4575 | 21 | CGCAGCTAGGAATAATGGAAT | 549.1 | -7.5 | 1.1 | -6.8 | 1.2E-11 | 2.8E-11 | BT20/MB468 | NUC |
| >hg38_21_+_8433172_8446622_RNA45SN1_45S_50ntFlanks@12212.12232.21\|CTCAGGGAGGACAGAAACCTC | >hg38_21_+_8433172_8446622_RNA45SN1_45S_50ntFlanks | 12212 | 12232 | 21 | CTCAGGGAGGACAGAAACCTC | 112.1 | -10.7 | 1.6 | -6.8 | 8.7E-12 | 2.8E-11 | MB231/MB468 | MP |
| >MI0000299\|hsa-mir-222&WithFlank&X\|-\|45747009\|45747130@75.95.21&[MIMAT0000279&hsa-miR-222-3p&offsets\|0\|0;m-140&X\|-\|45747033\|45747056&offsets\|0\|-3]\|AGCTACATCTGGCTACTGGGT | >MI0000299\|hsa-mir-222&WithFlank&X\|-\|45747009\|45747130 | 75 | 95 | 21 | AGCTACATCTGGCTACTGGGT | 2157.7 | 5.1 | 0.8 | 6.8 | 9.7E-12 | 2.8E-11 | MB231/MB468 | NUC |
| >hg38_21_+_8433172_8446622_RNA45SN1_45S_50ntFlanks@3802.3830.29\|CATTAAATCAGTTATGGTTCCTTTGGTCG | >hg38_21_+_8433172_8446622_RNA45SN1_45S_50ntFlanks | 3802 | 3830 | 29 | CATTAAATCAGTTATGGTTCCTTTGGTCG | 101.5 | -10.5 | 1.5 | -6.8 | 8.7E-12 | 2.8E-11 | MB231/MB468 | MP |
| >MI0000289\|hsa-mir-181a-1&WithFlank&1\|-\|198859038\|198859159@30.51.22;MI0000269\|hsa-mir-181a-2&WithFlank&9\|+\|124692436\|124692557@45.66.22&[MIMAT0000256&hsa-miR-181a-5p&offsets\|0\|-1;m-32&1\|-\|198859108\|198859130&offsets\|0\|-1];[MIMAT0000256_1&hsa-miR-181a-5p&offsets\|0\|-1;m-32&9\|+\|124692480\|124692502&offsets\|0\|-1]\|AACATTCAACGCTGTCGGTGAG | >MI0000289\|hsa-mir-181a-1&WithFlank&1\|-\|198859038\|198859159 | 30 | 51 | 22 | AACATTCAACGCTGTCGGTGAG | 568.2 | 5.2 | 0.8 | 6.8 | 9.7E-12 | 2.8E-11 | MB231/MB468 | NUC |
| >hg38_21_+_8433172_8446622_RNA45SN1_45S_50ntFlanks@10798.10818.21\|CATGTTGGAACAATGTAGGTA | >hg38_21_+_8433172_8446622_RNA45SN1_45S_50ntFlanks | 10798 | 10818 | 21 | CATGTTGGAACAATGTAGGTA | 3556.3 | -4.8 | 0.7 | -6.8 | 8.9E-12 | 2.8E-11 | BT20/MB468 | CYTO |
| >MI0000088\|hsa-mir-30a&WithFlank&6\|-\|71403545\|71403627@12.33.22(+2U)&[MIMAT0000087&hsa-miR-30a-5p&offsets\|0\|0(+2U);m-15&6\|-\|71403595\|71403616&offsets\|0\|0(+2U)]\|TGTAAACATCCTCGACTGGAAGTT | >MI0000088\|hsa-mir-30a&WithFlank&6\|-\|71403545\|71403627 | 12 | 33 | 22(+2U) | TGTAAACATCCTCGACTGGAAGTT | 221.1 | 6.9 | 1.0 | 6.8 | 9.7E-12 | 2.8E-11 | MB231/MB468 | NUC |
| >hg38_21_+_8433172_8446622_RNA45SN1_45S_50ntFlanks@7987.8008.22\|TCAGACGTGGCGACCCGCTGAA | >hg38_21_+_8433172_8446622_RNA45SN1_45S_50ntFlanks | 7987 | 8008 | 22 | TCAGACGTGGCGACCCGCTGAA | 436.0 | -11.8 | 1.6 | -7.2 | 5.5E-13 | 2.8E-11 | BT20/MB468 | MP |
| >hg38_21_+_8433172_8446622_RNA45SN1_45S_50ntFlanks@9736.9767.32\|CCGAAACGATCTCAACCTATTCTCAAACTTTA | >hg38_21_+_8433172_8446622_RNA45SN1_45S_50ntFlanks | 9736 | 9767 | 32 | CCGAAACGATCTCAACCTATTCTCAAACTTTA | 720.6 | -9.5 | 1.4 | -6.8 | 1.2E-11 | 2.8E-11 | BT20/MB468 | NUC |
| >hg38_21_+_8433172_8446622_RNA45SN1_45S_50ntFlanks@3853.3871.19\|AACTGTGGTAATTCTAGAG | >hg38_21_+_8433172_8446622_RNA45SN1_45S_50ntFlanks | 3853 | 3871 | 19 | AACTGTGGTAATTCTAGAG | 367.4 | -11.6 | 1.6 | -7.2 | 5.6E-13 | 2.9E-11 | BT20/MB468 | MP |
| >hg38_21_+_8433172_8446622_RNA45SN1_45S_50ntFlanks@12942.12972.31\|TTCGTAGACGACCTGCTTCTGGGTCGGGGTT | >hg38_21_+_8433172_8446622_RNA45SN1_45S_50ntFlanks | 12942 | 12972 | 31 | TTCGTAGACGACCTGCTTCTGGGTCGGGGTT | 183.7 | -9.6 | 1.4 | -6.9 | 7.0E-12 | 2.9E-11 | BT20/MB468 | MITO |
| >hg38_21_+_8433172_8446622_RNA45SN1_45S_50ntFlanks@4557.4579.23\|CAGCTAGGAATAATGGAATAGGA | >hg38_21_+_8433172_8446622_RNA45SN1_45S_50ntFlanks | 4557 | 4579 | 23 | CAGCTAGGAATAATGGAATAGGA | 98.7 | -10.5 | 1.5 | -6.8 | 8.7E-12 | 2.9E-11 | MB231/MB468 | MP |
| >hg38_21_+_8433172_8446622_RNA45SN1_45S_50ntFlanks@12274.12318.45\|CAGTACGAATACAGACCGTGAAAGCGGGGCCTCACGATCCTTCTG | >hg38_21_+_8433172_8446622_RNA45SN1_45S_50ntFlanks | 12274 | 12318 | 45 | CAGTACGAATACAGACCGTGAAAGCGGGGCCTCACGATCCTTCTG | 247.8 | -11.7 | 1.7 | -6.8 | 1.1E-11 | 2.9E-11 | MB231/MB468 | MITO |
| >hg38_21_+_8433172_8446622_RNA45SN1_45S_50ntFlanks@8799.8827.29\|CCTCGGGAGGGCGCGCGGGTCGGGGCGGC | >hg38_21_+_8433172_8446622_RNA45SN1_45S_50ntFlanks | 8799 | 8827 | 29 | CCTCGGGAGGGCGCGCGGGTCGGGGCGGC | 150.7 | -11.1 | 1.6 | -6.8 | 8.7E-12 | 2.9E-11 | MB231/MB468 | MP |
| >MI0001445\|hsa-mir-423&WithFlank&17\|+\|30117073\|30117178@59.77.19(+1U)&[MIMAT0001340&hsa-miR-423-3p&offsets\|0\|-4(+1U);m-100&17\|+\|30117131\|30117153&offsets\|0\|-4(+1U)]\|AGCTCGGTCTGAGGCCCCTT | >MI0001445\|hsa-mir-423&WithFlank&17\|+\|30117073\|30117178 | 59 | 77 | 19(+1U) | AGCTCGGTCTGAGGCCCCTT | 307.4 | 5.6 | 0.8 | 6.8 | 9.8E-12 | 2.9E-11 | MB231/MB468 | NUC |
| >MI0000095\|hsa-mir-93&WithFlank&7\|-\|100093762\|100093853@56.78.23&[MIMAT0004509&hsa-miR-93-3p&offsets\|0\|+1;m-250&7\|-\|100093776\|100093798&offsets\|0\|0]\|ACTGCTGAGCTAGCACTTCCCGA | >MI0000095\|hsa-mir-93&WithFlank&7\|-\|100093762\|100093853 | 56 | 78 | 23 | ACTGCTGAGCTAGCACTTCCCGA | 330.0 | 8.0 | 1.2 | 6.8 | 9.0E-12 | 2.9E-11 | BT20/MB468 | CYTO |
| >hg38_21_+_8433172_8446622_RNA45SN1_45S_50ntFlanks@12420.12457.38\|ATGTCGGCTCTTCCTATCATTGTGAAGCAGAATTCACC | >hg38_21_+_8433172_8446622_RNA45SN1_45S_50ntFlanks | 12420 | 12457 | 38 | ATGTCGGCTCTTCCTATCATTGTGAAGCAGAATTCACC | 545.9 | 2.9 | 0.4 | 6.8 | 9.0E-12 | 2.9E-11 | BT20/MB468 | CYTO |
| >hg38_21_+_8433172_8446622_RNA45SN1_45S_50ntFlanks@11206.11223.18\|CGGCGGCGACTCTGGACG | >hg38_21_+_8433172_8446622_RNA45SN1_45S_50ntFlanks | 11206 | 11223 | 18 | CGGCGGCGACTCTGGACG | 107.7 | -10.6 | 1.6 | -6.8 | 8.8E-12 | 2.9E-11 | MB231/MB468 | MP |
| >hg38_21_+_8433172_8446622_RNA45SN1_45S_50ntFlanks@4045.4062.18\|CCATTCGAACGTCTGCCC | >hg38_21_+_8433172_8446622_RNA45SN1_45S_50ntFlanks | 4045 | 4062 | 18 | CCATTCGAACGTCTGCCC | 263.2 | -10.1 | 1.5 | -6.9 | 7.1E-12 | 2.9E-11 | BT20/MB468 | MITO |
| >hg38_21_+_8433172_8446622_RNA45SN1_45S_50ntFlanks@10805.10825.21\|GAACAATGTAGGTAAGGGAAG | >hg38_21_+_8433172_8446622_RNA45SN1_45S_50ntFlanks | 10805 | 10825 | 21 | GAACAATGTAGGTAAGGGAAG | 166.7 | -9.5 | 1.4 | -6.9 | 7.1E-12 | 2.9E-11 | BT20/MB468 | MITO |
| >hg38_21_+_8433172_8446622_RNA45SN1_45S_50ntFlanks@10425.10453.29\|AGATGGGCGAGCGCCGTTCCGAAGGGACG | >hg38_21_+_8433172_8446622_RNA45SN1_45S_50ntFlanks | 10425 | 10453 | 29 | AGATGGGCGAGCGCCGTTCCGAAGGGACG | 1047.6 | -9.7 | 1.4 | -6.8 | 1.2E-11 | 2.9E-11 | BT20/MB468 | NUC |
| >hg38_21_+_8433172_8446622_RNA45SN1_45S_50ntFlanks@8194.8213.20\|CCCGTGGACGGTGTGAGGCC | >hg38_21_+_8433172_8446622_RNA45SN1_45S_50ntFlanks | 8194 | 8213 | 20 | CCCGTGGACGGTGTGAGGCC | 2474.2 | -4.4 | 0.6 | -6.8 | 9.9E-12 | 2.9E-11 | MB231/MB468 | NUC |
| >MI0000255\|hsa-mir-30d&WithFlank&8\|-\|134804870\|134804951@12.33.22(+1U)&[MIMAT0000245&hsa-miR-30d-5p&offsets\|0\|0(+1U);m-21&8\|-\|134804919\|134804940&offsets\|0\|0(+1U)]\|TGTAAACATCCCCGACTGGAAGT | >MI0000255\|hsa-mir-30d&WithFlank&8\|-\|134804870\|134804951 | 12 | 33 | 22(+1U) | TGTAAACATCCCCGACTGGAAGT | 1814.5 | 6.2 | 0.9 | 6.8 | 9.7E-12 | 2.9E-11 | MB231/MB468 | CYTO |
| >hg38_MT_+_1621_3279_RNR2_16S_50ntFlanks@51.80.30\|GCTAAACCTAGCCCCAAACCCACTCCACCT | >hg38_MT_+_1621_3279_RNR2_16S_50ntFlanks | 51 | 80 | 30 | GCTAAACCTAGCCCCAAACCCACTCCACCT | 3580.7 | 6.5 | 1.0 | 6.9 | 7.1E-12 | 2.9E-11 | BT20/MB468 | MITO |
| >hg38_21_+_8433172_8446622_RNA45SN1_45S_50ntFlanks@5470.5499.30\|CCACGGCCCTGGCGGAGCGCTGAGAAGACG | >hg38_21_+_8433172_8446622_RNA45SN1_45S_50ntFlanks | 5470 | 5499 | 30 | CCACGGCCCTGGCGGAGCGCTGAGAAGACG | 1007.4 | -7.1 | 1.0 | -6.8 | 1.0E-11 | 2.9E-11 | MB231/MB468 | NUC |
| >hg38_21_+_8433172_8446622_RNA45SN1_45S_50ntFlanks@8342.8376.35\|CGTAAGGGAAAGTTGAAAAGAACTTTGAAGAGAGA | >hg38_21_+_8433172_8446622_RNA45SN1_45S_50ntFlanks | 8342 | 8376 | 35 | CGTAAGGGAAAGTTGAAAAGAACTTTGAAGAGAGA | 580.6 | 8.5 | 1.2 | 6.8 | 9.7E-12 | 2.9E-11 | MB231/MB468 | CYTO |
| >hg38_21_+_8433172_8446622_RNA45SN1_45S_50ntFlanks@9943.9968.26\|GACGGTGGCCATGGAAGTCGGAATCC | >hg38_21_+_8433172_8446622_RNA45SN1_45S_50ntFlanks | 9943 | 9968 | 26 | GACGGTGGCCATGGAAGTCGGAATCC | 169.7 | -11.3 | 1.7 | -6.8 | 8.9E-12 | 2.9E-11 | MB231/MB468 | MP |
| >hg38_21_+_8433172_8446622_RNA45SN1_45S_50ntFlanks@11569.11597.29\|CAGCCGACTTAGAACTGGTGCGGACCAGG | >hg38_21_+_8433172_8446622_RNA45SN1_45S_50ntFlanks | 11569 | 11597 | 29 | CAGCCGACTTAGAACTGGTGCGGACCAGG | 432.4 | -4.5 | 0.7 | -6.8 | 1.2E-11 | 2.9E-11 | MB231/MB468 | MITO |
| >hg38_21_+_8433172_8446622_RNA45SN1_45S_50ntFlanks@5423.5443.21\|ATTGGATGGTTTAGTGAGGCC | >hg38_21_+_8433172_8446622_RNA45SN1_45S_50ntFlanks | 5423 | 5443 | 21 | ATTGGATGGTTTAGTGAGGCC | 211.2 | -5.4 | 0.8 | -6.8 | 9.7E-12 | 2.9E-11 | MB231/MB468 | CYTO |
| >hg38_21_+_8433172_8446622_RNA45SN1_45S_50ntFlanks@8171.8214.44\|AAGTCCTTCTGATCGAGGCCCAGCCCGTGGACGGTGTGAGGCCG | >hg38_21_+_8433172_8446622_RNA45SN1_45S_50ntFlanks | 8171 | 8214 | 44 | AAGTCCTTCTGATCGAGGCCCAGCCCGTGGACGGTGTGAGGCCG | 2342.4 | -8.4 | 1.2 | -6.8 | 1.2E-11 | 2.9E-11 | BT20/MB468 | NUC |
| >hg38_21_+_8433172_8446622_RNA45SN1_45S_50ntFlanks@11568.11588.21\|GCAGCCGACTTAGAACTGGTG | >hg38_21_+_8433172_8446622_RNA45SN1_45S_50ntFlanks | 11568 | 11588 | 21 | GCAGCCGACTTAGAACTGGTG | 670.2 | -5.4 | 0.8 | -6.8 | 1.2E-11 | 2.9E-11 | BT20/MB468 | NUC |
| >hg38_21_+_8433172_8446622_RNA45SN1_45S_50ntFlanks@10782.10801.20\|GGTGAACAGCCTCTGGCATG | >hg38_21_+_8433172_8446622_RNA45SN1_45S_50ntFlanks | 10782 | 10801 | 20 | GGTGAACAGCCTCTGGCATG | 685.6 | -5.6 | 0.8 | -6.8 | 9.8E-12 | 2.9E-11 | MB231/MB468 | CYTO |
| >hg38_21_+_8433172_8446622_RNA45SN1_45S_50ntFlanks@10291.10322.32\|CGCGGGCCCGGGTGGAGCCGCCGCAGGTGCAG | >hg38_21_+_8433172_8446622_RNA45SN1_45S_50ntFlanks | 10291 | 10322 | 32 | CGCGGGCCCGGGTGGAGCCGCCGCAGGTGCAG | 163.9 | -11.2 | 1.6 | -6.8 | 8.9E-12 | 2.9E-11 | MB231/MB468 | MP |
| >hg38_21_+_8433172_8446622_RNA45SN1_45S_50ntFlanks@476.495.20\|TTCGTGATCGATGTGGTGAC | >hg38_21_+_8433172_8446622_RNA45SN1_45S_50ntFlanks | 476 | 495 | 20 | TTCGTGATCGATGTGGTGAC | 6710.4 | 4.9 | 0.7 | 6.8 | 1.0E-11 | 2.9E-11 | MB231/MB468 | NUC |
| >hg38_21_+_8433172_8446622_RNA45SN1_45S_50ntFlanks@5467.5494.28\|GGCCCACGGCCCTGGCGGAGCGCTGAGA | >hg38_21_+_8433172_8446622_RNA45SN1_45S_50ntFlanks | 5467 | 5494 | 28 | GGCCCACGGCCCTGGCGGAGCGCTGAGA | 185.8 | -7.8 | 1.2 | -6.8 | 1.0E-11 | 2.9E-11 | MB231/MB468 | NUC |
| >MI0000465\|hsa-mir-191&WithFlank&3\|-\|49020612\|49020715@23.44.22&[MIMAT0000440&hsa-miR-191-5p&offsets\|+1\|0;m-48&3\|-\|49020672\|49020694&offsets\|+1\|0]\|AACGGAATCCCAAAAGCAGCTG | >MI0000465\|hsa-mir-191&WithFlank&3\|-\|49020612\|49020715 | 23 | 44 | 22 | AACGGAATCCCAAAAGCAGCTG | 288.6 | 5.6 | 0.8 | 6.8 | 9.8E-12 | 2.9E-11 | MB231/MB468 | CYTO |
| >MI0000089\|hsa-mir-31&WithFlank&9\|-\|21512109\|21512191@14.36.23&[MIMAT0000089&hsa-miR-31-5p&offsets\|0\|+2;m-360&9\|-\|21512157\|21512178&offsets\|0\|+1]\|AGGCAAGATGCTGGCATAGCTGT | >MI0000089\|hsa-mir-31&WithFlank&9\|-\|21512109\|21512191 | 14 | 36 | 23 | AGGCAAGATGCTGGCATAGCTGT | 1020.0 | 13.3 | 1.8 | 7.4 | 1.4E-13 | 2.9E-11 | BT20/MB231 | MP |

| | | | | | | | | | | | | | |
|---|---|---|---|---|---|---|---|---|---|---|---|---|---|
| >hg38_21_+_8433172_8446622_RNA45SN1_45S_50ntFlanks@3985.4010.26\|CGGCTTTGGTGACTCTAGATAACCTC | >hg38_21_+_8433172_8446622_RNA45SN1_45S_50ntFlanks | 3985 | 4010 | 26 | CGGCTTTGGTGACTCTAGATAACCTC | 171.1 | -11.3 | 1.7 | -6.8 | 9.0E-12 | 2.9E-11 | MB231/MB468 | MP |
| >hg38_21_+_8433172_8446622_RNA45SN1_45S_50ntFlanks@12082.12108.27\|TCTCGCTTCTGGCGCCAAGCGCCCGGC | >hg38_21_+_8433172_8446622_RNA45SN1_45S_50ntFlanks | 12082 | 12108 | 27 | TCTCGCTTCTGGCGCCAAGCGCCCGGC | 92.8 | -10.4 | 1.5 | -6.8 | 9.0E-12 | 2.9E-11 | MB231/MB468 | MP |
| >hg38_21_+_8433172_8446622_RNA45SN1_45S_50ntFlanks@3776.3801.26\|CCGGTACAGTGAAACTGCGAATGGCT | >hg38_21_+_8433172_8446622_RNA45SN1_45S_50ntFlanks | 3776 | 3801 | 26 | CCGGTACAGTGAAACTGCGAATGGCT | 830.7 | -9.5 | 1.4 | -6.8 | 1.2E-11 | 2.9E-11 | BT20/MB468 | NUC |
| >hg38_21_+_8433172_8446622_RNA45SN1_45S_50ntFlanks@12625.12657.33\|CAATGGGGCGAAGCTACCATCTGTGGGATTATG | >hg38_21_+_8433172_8446622_RNA45SN1_45S_50ntFlanks | 12625 | 12657 | 33 | CAATGGGGCGAAGCTACCATCTGTGGGATTATG | 1319.0 | -7.5 | 1.1 | -6.8 | 1.2E-11 | 2.9E-11 | MB231/MB468 | MITO |
| >hg38_21_+_8433172_8446622_RNA45SN1_45S_50ntFlanks@5239.5256.18\|TGACTGGCTCAGCGTGTG | >hg38_21_+_8433172_8446622_RNA45SN1_45S_50ntFlanks | 5239 | 5256 | 18 | TGACTGGCTCAGCGTGTG | 169.6 | -11.3 | 1.7 | -6.8 | 9.0E-12 | 2.9E-11 | MB231/MB468 | MP |
| >hg38_21_+_8433172_8446622_RNA45SN1_45S_50ntFlanks@9157.9187.31\|CAGTGCGCCCCGGGCGGGTCGCGCCGTCGGG | >hg38_21_+_8433172_8446622_RNA45SN1_45S_50ntFlanks | 9157 | 9187 | 31 | CAGTGCGCCCCGGGCGGGTCGCGCCGTCGGG | 624.1 | -8.4 | 1.2 | -6.8 | 1.2E-11 | 2.9E-11 | MB231/MB468 | MITO |
| >am_MI0000786\|hsa-mir-378a&WithFlank&5\|+\|149732819\|149732896@49.70.22(+2A)&[MIMAT0000732&hsa-miR-378a-3p&offsets\|0\|0(+2A);m-44&5\|+\|149732867\|149732888&offsets\|0\|0(+2A)]\|ACTGGACTTGGAGTCAGAAGGCAA | >am_MI0000786\|hsa-mir-378a&WithFlank&5\|+\|149732819\|149732896 | 49 | 70 | 22(+2A) | ACTGGACTTGGAGTCAGAAGGCAA | 265.3 | 6.1 | 0.9 | 6.8 | 9.2E-12 | 2.9E-11 | BT20/MB468 | CYTO |
| >hg38_21_+_8433172_8446622_RNA45SN1_45S_50ntFlanks@5158.5185.28\|AAGTGGCGTTCAGCCACCCGAGATTGAG | >hg38_21_+_8433172_8446622_RNA45SN1_45S_50ntFlanks | 5158 | 5185 | 28 | AAGTGGCGTTCAGCCACCCGAGATTGAG | 978.2 | -13.7 | 2.0 | -6.8 | 1.2E-11 | 2.9E-11 | MB231/MB468 | MITO |
| >hg38_21_+_8433172_8446622_RNA45SN1_45S_50ntFlanks@12992.13039.48\|TCCCTCGCTGCGATCTATTGAAAGTCAGCCCTCGACACAAGGGTTTGT | >hg38_21_+_8433172_8446622_RNA45SN1_45S_50ntFlanks | 12992 | 13039 | 48 | TCCCTCGCTGCGATCTATTGAAAGTCAGCCCTCGACACAAGGGTTTGT | 5948.4 | -9.3 | 1.4 | -6.8 | 1.3E-11 | 2.9E-11 | BT20/MB468 | NUC |
| >hg38_21_+_8433172_8446622_RNA45SN1_45S_50ntFlanks@10384.10401.18\|CCATGTGAACAGCAGTTG | >hg38_21_+_8433172_8446622_RNA45SN1_45S_50ntFlanks | 10384 | 10401 | 18 | CCATGTGAACAGCAGTTG | 225.6 | -11.7 | 1.7 | -6.8 | 9.0E-12 | 3.0E-11 | MB231/MB468 | MP |
| >trnaMT_GluTTC_MT_-_14674_14742@35.71.37,trnalookalike8_GluTTC_5_-_93905172_93905240@35.71.37\|TATCATTGGTCGTGGTTGTAGTCCGTGCGAGAATACC | >trnaMT_GluTTC_MT_-_14674_14742 | 35 | 71 | 37 | TATCATTGGTCGTGGTTGTAGTCCGTGCGAGAATACC | 346.3 | 4.9 | 0.7 | 6.8 | 1.2E-11 | 3.0E-11 | MB231/MB468 | MITO |
| >am_trna10_ValCAC_5_-_180649395_180649467@1.34.34,trna18_ValCAC_5_-_180529253_180529325@1.34.34,trna2_ValCAC_5_+_180524070_180524142@1.34.34,trna6_ValCAC_5_+_180600650_180600722@1.34.34,trna85_ValCAC_1_-_161369490_161369562@1.34.34,trna90_ValCAC_1_-_149684088_149684161@1.34.34,trna98_ValCAC_1_-_149298555_149298627@1.34.34,trna9_ValCAC_6_+_26538282_26538354@1.34.34\|GTTTCCGTAGTGTAGTGGTTATCACGTTCGCCTC | >am_trna10_ValCAC_5_-_180649395_180649467 | 1 | 34 | 34 | GTTTCCGTAGTGTAGTGGTTATCACGTTCGCCTC | 9971.2 | 11.7 | 1.7 | 6.8 | 9.9E-12 | 3.0E-11 | MB231/MB468 | CYTO |
| >hg38_21_+_8433172_8446622_RNA45SN1_45S_50ntFlanks@10391.10420.30\|AACAGCAGTTGAACATGGGTCAGTCGGTCC | >hg38_21_+_8433172_8446622_RNA45SN1_45S_50ntFlanks | 10391 | 10420 | 30 | AACAGCAGTTGAACATGGGTCAGTCGGTCC | 589.0 | -5.5 | 0.8 | -6.8 | 9.9E-12 | 3.0E-11 | MB231/MB468 | CYTO |
| >hg38_21_+_8433172_8446622_RNA45SN1_45S_50ntFlanks@11189.11207.19\|ACCCGGGGGGCCGGCGGCG | >hg38_21_+_8433172_8446622_RNA45SN1_45S_50ntFlanks | 11189 | 11207 | 19 | ACCCGGGGGGCCGGCGGCG | 134.8 | -10.9 | 1.6 | -6.8 | 9.0E-12 | 3.0E-11 | MB231/MB468 | MP |
| >hg38_21_+_8433172_8446622_RNA45SN1_45S_50ntFlanks@8170.8196.27\|CAAGTCCTTCTGATCGAGGCCCAGCCC | >hg38_21_+_8433172_8446622_RNA45SN1_45S_50ntFlanks | 8170 | 8196 | 27 | CAAGTCCTTCTGATCGAGGCCCAGCCC | 164.1 | -11.2 | 1.6 | -6.8 | 9.1E-12 | 3.0E-11 | MB231/MB468 | MP |
| >hg38_21_+_8433172_8446622_RNA45SN1_45S_50ntFlanks@8168.8204.37\|CCCAAGTCCTTCTGATCGAGGCCCAGCCCGTGGACGG | >hg38_21_+_8433172_8446622_RNA45SN1_45S_50ntFlanks | 8168 | 8204 | 37 | CCCAAGTCCTTCTGATCGAGGCCCAGCCCGTGGACGG | 210.0 | -11.6 | 1.7 | -6.8 | 9.1E-12 | 3.0E-11 | MB231/MB468 | MP |
| >hg38_21_+_8433172_8446622_RNA45SN1_45S_50ntFlanks@4005.4022.18\|AACCTCGGGCCGATCGCA | >hg38_21_+_8433172_8446622_RNA45SN1_45S_50ntFlanks | 4005 | 4022 | 18 | AACCTCGGGCCGATCGCA | 97.6 | -10.5 | 1.5 | -6.8 | 9.1E-12 | 3.0E-11 | MB231/MB468 | MP |
| >hg38_21_+_8433172_8446622_RNA45SN1_45S_50ntFlanks@4725.4759.35\|TTAATCAAGAACGAAAGTCGGAGGTTCGAAGACGA | >hg38_21_+_8433172_8446622_RNA45SN1_45S_50ntFlanks | 4725 | 4759 | 35 | TTAATCAAGAACGAAAGTCGGAGGTTCGAAGACGA | 1161.2 | -9.8 | 1.4 | -6.8 | 1.3E-11 | 3.0E-11 | BT20/MB468 | NUC |
| >hg38_21_+_8433172_8446622_RNA45SN1_45S_50ntFlanks@4147.4175.29\|TGAGAAACGGCTACCACATCCAAGGAAGG | >hg38_21_+_8433172_8446622_RNA45SN1_45S_50ntFlanks | 4147 | 4175 | 29 | TGAGAAACGGCTACCACATCCAAGGAAGG | 126.7 | -10.8 | 1.6 | -6.8 | 9.1E-12 | 3.0E-11 | MB231/MB468 | MP |
| >hg38_21_+_8433172_8446622_RNA45SN1_45S_50ntFlanks@3993.4026.34\|GTGACTCTAGATAACCTCGGGCCGATCGCACGCC | >hg38_21_+_8433172_8446622_RNA45SN1_45S_50ntFlanks | 3993 | 4026 | 34 | GTGACTCTAGATAACCTCGGGCCGATCGCACGCC | 1330.9 | -8.3 | 1.2 | -6.8 | 1.3E-11 | 3.0E-11 | BT20/MB468 | NUC |
| >hg38_21_+_8433172_8446622_RNA45SN1_45S_50ntFlanks@3878.3898.21\|CATGCCGACGGGCGCTGACCC | >hg38_21_+_8433172_8446622_RNA45SN1_45S_50ntFlanks | 3878 | 3898 | 21 | CATGCCGACGGGCGCTGACCC | 123.2 | -10.8 | 1.6 | -6.8 | 9.1E-12 | 3.0E-11 | MB231/MB468 | MP |
| >hg38_21_+_8433172_8446622_RNA45SN1_45S_50ntFlanks@9564.9593.30\|CTGACGTGCAAATCGGTCGTCCGACCTGGG | >hg38_21_+_8433172_8446622_RNA45SN1_45S_50ntFlanks | 9564 | 9593 | 30 | CTGACGTGCAAATCGGTCGTCCGACCTGGG | 80.0 | -10.2 | 1.5 | -6.8 | 9.1E-12 | 3.0E-11 | MB231/MB468 | MP |
| >hg38_21_+_8433172_8446622_RNA45SN1_45S_50ntFlanks@11197.11232.36\|GGCCGGCGGCGGCGGCGACTCTGGACGCGAGCCGGG | >hg38_21_+_8433172_8446622_RNA45SN1_45S_50ntFlanks | 11197 | 11232 | 36 | GGCCGGCGGCGGCGGCGACTCTGGACGCGAGCCGGG | 98.3 | -10.5 | 1.5 | -6.8 | 9.2E-12 | 3.0E-11 | MB231/MB468 | MP |
| >hg38_21_+_8433172_8446622_RNA45SN1_45S_50ntFlanks@11755.11777.23\|AATGCCTCGTCATCTAATTAGTG | >hg38_21_+_8433172_8446622_RNA45SN1_45S_50ntFlanks | 11755 | 11777 | 23 | AATGCCTCGTCATCTAATTAGTG | 519.0 | -9.2 | 1.4 | -6.8 | 1.3E-11 | 3.0E-11 | BT20/MB468 | NUC |
| >hg38_21_+_8433172_8446622_RNA45SN1_45S_50ntFlanks@4681.4720.40\|CGCAAGACGGACCAGAGCGAAAGCATTTGCCAAGAATGTT | >hg38_21_+_8433172_8446622_RNA45SN1_45S_50ntFlanks | 4681 | 4720 | 40 | CGCAAGACGGACCAGAGCGAAAGCATTTGCCAAGAATGTT | 381.0 | 8.0 | 1.2 | 6.8 | 1.0E-11 | 3.0E-11 | MB231/MB468 | CYTO |
| >hg38_21_+_8433172_8446622_RNA45SN1_45S_50ntFlanks@10406.10424.19\|TGGGTCAGTCGGTCCTGAG | >hg38_21_+_8433172_8446622_RNA45SN1_45S_50ntFlanks | 10406 | 10424 | 19 | TGGGTCAGTCGGTCCTGAG | 365.4 | 4.4 | 0.6 | 6.8 | 1.0E-11 | 3.0E-11 | MB231/MB468 | CYTO |
| >am_trna128_GlyGCC_6_-_27870686_27870756@1.31.31,trna133_GlyCCC_1_-_16872434_16872504@1.31.31,trna18_GlyGCC_16_+_70822597_70822667@1.31.31,trna19_GlyGCC_16_+_70823410_70823480@1.31.31,trna19_GlyGCC_2_-_157257659_157257729@1.31.31,trna24_GlyGCC_16_-_70812942_70813012@1.31.31,trna25_GlyGCC_16_-_70812114_70812184@1.31.31,trna4_GlyCCC_1_+_17188416_17188486@1.31.31,trna5_GlyGCC_17_+_8029064_8029134@1.31.31,trna68_GlyGCC_1_-_161493637_161493707@1.31.31\|GCATTGGTGGTTCAGTGGTAGAATTCTCGCC | >am_trna128_GlyGCC_6_-_27870686_27870756 | 1 | 31 | 31 | GCATTGGTGGTTCAGTGGTAGAATTCTCGCC | 23818.5 | 5.9 | 0.9 | 6.8 | 1.2E-11 | 3.0E-11 | MB231/MB468 | MITO |
| >hg38_21_+_8433172_8446622_RNA45SN1_45S_50ntFlanks@11571.11613.43\|GCCGACTTAGAACTGGTGCGGACCAGGGGAATCCGACTGTTTA | >hg38_21_+_8433172_8446622_RNA45SN1_45S_50ntFlanks | 11571 | 11613 | 43 | GCCGACTTAGAACTGGTGCGGACCAGGGGAATCCGACTGTTTA | 251.2 | 8.9 | 1.3 | 6.8 | 1.0E-11 | 3.0E-11 | MB231/MB468 | CYTO |
| >hg38_21_+_8433172_8446622_RNA45SN1_45S_50ntFlanks@12539.12567.29\|ATGATGTGTTGTTGCCATGGTAATCCTGC | >hg38_21_+_8433172_8446622_RNA45SN1_45S_50ntFlanks | 12539 | 12567 | 29 | ATGATGTGTTGTTGCCATGGTAATCCTGC | 1124.1 | -8.8 | 1.3 | -6.8 | 1.0E-11 | 3.0E-11 | MB231/MB468 | NUC |
| >hg38_21_+_8433172_8446622_RNA45SN1_45S_50ntFlanks@10411.10441.31\|CAGTCGGTCCTGAGAGATGGGCGAGCGCCGT | >hg38_21_+_8433172_8446622_RNA45SN1_45S_50ntFlanks | 10411 | 10441 | 31 | CAGTCGGTCCTGAGAGATGGGCGAGCGCCGT | 177.0 | -4.4 | 0.6 | -6.8 | 1.0E-11 | 3.0E-11 | MB231/MB468 | CYTO |
| >hg38_21_+_8433172_8446622_RNA45SN1_45S_50ntFlanks@6648.6665.18\|GTACGACTCTTAGCGGTG | >hg38_21_+_8433172_8446622_RNA45SN1_45S_50ntFlanks | 6648 | 6665 | 18 | GTACGACTCTTAGCGGTG | 818.7 | 5.2 | 0.8 | 6.8 | 9.4E-12 | 3.0E-11 | BT20/MB468 | CYTO |
| >hg38_21_+_8433172_8446622_RNA45SN1_45S_50ntFlanks@8381.8415.35\|AAGAGGGCGTGAAACCGTTAAGAGGTAAACGGGTG | >hg38_21_+_8433172_8446622_RNA45SN1_45S_50ntFlanks | 8381 | 8415 | 35 | AAGAGGGCGTGAAACCGTTAAGAGGTAAACGGGTG | 114.5 | -10.7 | 1.6 | -6.8 | 9.2E-12 | 3.0E-11 | MB231/MB468 | MP |
| >hg38_21_+_8433172_8446622_RNA45SN1_45S_50ntFlanks@12071.12108.38\|CCCCGAGGGGCTCTCGCTTCTGGCGCCAAGCGCCCGGC | >hg38_21_+_8433172_8446622_RNA45SN1_45S_50ntFlanks | 12071 | 12108 | 38 | CCCCGAGGGGCTCTCGCTTCTGGCGCCAAGCGCCCGGC | 391.0 | -6.5 | 1.0 | -6.8 | 1.2E-11 | 3.0E-11 | MB231/MB468 | MITO |
| >MI0000446\|hsa-mir-125b-1&WithFlank&11\|-\|122099751\|122099850@21.39.19(+1C);MI0000470\|hsa-mir-125b-2&WithFlank&21\|+\|16590231\|16590331@23.41.19(+1C)&[MIMAT0000423&hsa-miR-125b-5p&offsets\|0\|-3(+1C);m-67&11\|-\|122099809\|122099830&offsets\|0\|-3(+1C)];[MIMAT0000423_1&hsa-miR-125b-5p&offsets\|0\|-3(+1C);m-68&21\|+\|16590253\|16590274&offsets\|0\|-3(+1C)]\|TCCCTGAGACCCTAACTTGC | >MI0000446\|hsa-mir-125b-1&WithFlank&11\|-\|122099751\|122099850 | 21 | 39 | 19(+1C) | TCCCTGAGACCCTAACTTGC | 300.5 | 11.5 | 1.7 | 6.8 | 1.2E-11 | 3.0E-11 | MB231/MB468 | MITO |
| >hg38_21_+_8433172_8446622_RNA45SN1_45S_50ntFlanks@10781.10801.21\|AGGTGAACAGCCTCTGGCATG | >hg38_21_+_8433172_8446622_RNA45SN1_45S_50ntFlanks | 10781 | 10801 | 21 | AGGTGAACAGCCTCTGGCATG | 633.3 | -8.5 | 1.3 | -6.8 | 1.2E-11 | 3.0E-11 | MB231/MB468 | MITO |
| >hg38_21_+_8433172_8446622_RNA45SN1_45S_50ntFlanks@10846.10866.21\|CTTCGGGATAAGGATTGGCTC | >hg38_21_+_8433172_8446622_RNA45SN1_45S_50ntFlanks | 10846 | 10866 | 21 | CTTCGGGATAAGGATTGGCTC | 97.0 | -10.5 | 1.5 | -6.8 | 9.2E-12 | 3.0E-11 | MB231/MB468 | MP |
| >hg38_21_+_8433172_8446622_RNA45SN1_45S_50ntFlanks@4322.4346.25\|CAGCCGCGGTAATTCCAGCTCCAAT | >hg38_21_+_8433172_8446622_RNA45SN1_45S_50ntFlanks | 4322 | 4346 | 25 | CAGCCGCGGTAATTCCAGCTCCAAT | 90.1 | -10.4 | 1.5 | -6.8 | 9.2E-12 | 3.0E-11 | MB231/MB468 | MP |
| >trna10_AspGTC_12_-_125424193_125424264@32.72.41,trna12_AspGTC_12_-_125411891_125411962@32.72.41,trna144_AspGTC_6_-_27551236_27551307@32.72.41,trna38_AspGTC_17_-_8125556_8125627@32.72.41,trna45_AspGTC_6_+_27447453_27447524@32.72.41,trna48_AspGTC_6_+_27471523_27471594@32.72.41,trna4_AspGTC_12_+_96429799_96429870@32.72.41,trna69_AspGTC_1_-_161440205_161440276@32.72.41,trna72_AspGTC_1_-_161432824_161432895@32.72.41,trna75_AspGTC_1_-_161425414_161425485@32.72.41,trna78_AspGTC_1_-_161418033_161418104@32.72.41,trna81_AspGTC_1_-_161410615_161410686@32.72.41\|CTGTCACGCGGGAGACCGGGGTTCGATTCCCCGACGGGGAG | >trna10_AspGTC_12_-_125424193_125424264 | 32 | 72 | 41 | CTGTCACGCGGGAGACCGGGGTTCGATTCCCCGACGGGGAG | 3319.1 | -11.2 | 1.6 | -6.8 | 1.3E-11 | 3.0E-11 | BT20/MB468 | NUC |

<table>
<tr><td>>trna10_AspGTC_12_-_125424193_125424264@1.33.33,trna12_AspGTC_12_-_125411891_125411962@1.33.33,trna38_AspGTC_17_-_8125556_8125627@1.33.33,trna45_AspGTC_6_+_27447453_27447524@1.33.33,trna48_AspGTC_6_+_27471523_27471594@1.33.33,trna4_AspGTC_12_+_96429799_96429870@1.33.33,trna69_AspGTC_1_-_161440205_161440276@1.33.33,trna72_AspGTC_1_-_161432824_161432895@1.33.33,trna75_AspGTC_1_-_161425414_161425485@1.33.33,trna78_AspGTC_1_-_161418033_161418104@1.33.33,trna81_AspGTC_1_-_161410615_161410686@1.33.33|TCCTCGTTAGTATAGTGGTGAGTATCCCCGCCT</td><td>>trna10_AspGTC_12_-_125424193_125424264</td><td>1</td><td>33</td><td>33</td><td>TCCTCGTTAGTATAGTGGTGAGTATCCCCGCCT</td><td>667.5</td><td>10.7</td><td>1.6</td><td>6.8</td><td>1.0E-11</td><td>3.0E-11</td><td>MB231/MB468</td><td>CYTO</td></tr>
<tr><td>>hg38_21_+_8433172_8446622_RNA45SN1_45S_50ntFlanks@10393.10430.38|CAGCAGTTGAACATGGGTCAGTCGGTCCTGAGAGATGG</td><td>>hg38_21_+_8433172_8446622_RNA45SN1_45S_50ntFlanks</td><td>10393</td><td>10430</td><td>38</td><td>CAGCAGTTGAACATGGGTCAGTCGGTCCTGAGAGATGG</td><td>960.3</td><td>-7.2</td><td>1.1</td><td>-6.8</td><td>1.2E-11</td><td>3.0E-11</td><td>MB231/MB468</td><td>MITO</td></tr>
<tr><td>>hg38_21_+_8433172_8446622_RNA45SN1_45S_50ntFlanks@4759.4787.29|ATCAGATACCGTCGTAGTTCCGACCATAA</td><td>>hg38_21_+_8433172_8446622_RNA45SN1_45S_50ntFlanks</td><td>4759</td><td>4787</td><td>29</td><td>ATCAGATACCGTCGTAGTTCCGACCATAA</td><td>115.4</td><td>-10.7</td><td>1.6</td><td>-6.8</td><td>9.3E-12</td><td>3.0E-11</td><td>MB231/MB468</td><td>MP</td></tr>
<tr><td>>hg38_21_+_8433172_8446622_RNA45SN1_45S_50ntFlanks@7986.8009.24|ATCAGACGTGGCGACCCGCTGAAT</td><td>>hg38_21_+_8433172_8446622_RNA45SN1_45S_50ntFlanks</td><td>7986</td><td>8009</td><td>24</td><td>ATCAGACGTGGCGACCCGCTGAAT</td><td>220.9</td><td>-9.9</td><td>1.4</td><td>-6.8</td><td>7.5E-12</td><td>3.0E-11</td><td>BT20/MB468</td><td>MITO</td></tr>
<tr><td>>hg38_21_+_8433172_8446622_RNA45SN1_45S_50ntFlanks@9027.9059.33|CGTCTCCTCGTGGGGGGGCCGGGCCACCCCTCC</td><td>>hg38_21_+_8433172_8446622_RNA45SN1_45S_50ntFlanks</td><td>9027</td><td>9059</td><td>33</td><td>CGTCTCCTCGTGGGGGGGCCGGGCCACCCCTCC</td><td>786.1</td><td>-7.8</td><td>1.1</td><td>-6.8</td><td>7.5E-12</td><td>3.0E-11</td><td>BT20/MB468</td><td>MITO</td></tr>
<tr><td>>hg38_21_+_8433172_8446622_RNA45SN1_45S_50ntFlanks@11552.11592.41|CCTCGGCCGGCGCCTAGCAGCCGACTTAGAACTGGTGCGGA</td><td>>hg38_21_+_8433172_8446622_RNA45SN1_45S_50ntFlanks</td><td>11552</td><td>11592</td><td>41</td><td>CCTCGGCCGGCGCCTAGCAGCCGACTTAGAACTGGTGCGGA</td><td>700.0</td><td>-5.3</td><td>0.8</td><td>-6.8</td><td>1.2E-11</td><td>3.0E-11</td><td>MB231/MB468</td><td>MITO</td></tr>
<tr><td>>hg38_21_+_8433172_8446622_RNA45SN1_45S_50ntFlanks@8383.8400.18|GAGGGCGTGAAACCGTTA</td><td>>hg38_21_+_8433172_8446622_RNA45SN1_45S_50ntFlanks</td><td>8383</td><td>8400</td><td>18</td><td>GAGGGCGTGAAACCGTTA</td><td>142.7</td><td>-11.0</td><td>1.6</td><td>-6.8</td><td>9.3E-12</td><td>3.0E-11</td><td>MB231/MB468</td><td>MP</td></tr>
<tr><td>>hg38_21_+_8433172_8446622_RNA45SN1_45S_50ntFlanks@11212.11253.42|CGACTCTGGACGCGAGCCGGGCCCTTCCCGTGGATCGCCCCA</td><td>>hg38_21_+_8433172_8446622_RNA45SN1_45S_50ntFlanks</td><td>11212</td><td>11253</td><td>42</td><td>CGACTCTGGACGCGAGCCGGGCCCTTCCCGTGGATCGCCCCA</td><td>607.1</td><td>-9.0</td><td>1.3</td><td>-6.8</td><td>1.2E-11</td><td>3.0E-11</td><td>MB231/MB468</td><td>MITO</td></tr>
<tr><td>>hg38_21_+_8433172_8446622_RNA45SN1_45S_50ntFlanks@12323.12344.22|TTTGGGTTTTAAGCAGGAGGTG</td><td>>hg38_21_+_8433172_8446622_RNA45SN1_45S_50ntFlanks</td><td>12323</td><td>12344</td><td>22</td><td>TTTGGGTTTTAAGCAGGAGGTG</td><td>128.9</td><td>-10.9</td><td>1.6</td><td>-6.8</td><td>9.3E-12</td><td>3.0E-11</td><td>MB231/MB468</td><td>MP</td></tr>
<tr><td>>MI0000542|hsa-mir-320a&WithFlank&8|-|22244960|22245043@48.69.22(+1U)&[MIMAT0000510&hsa-miR-320a-3p&offsets|0|0(+1U);m-60&8|-|22244975|22244996&offsets|0|0(+1U)]|AAAAGCTGGGTTGAGAGGGCGAT</td><td>>MI0000542|hsa-mir-320a&WithFlank&8|-|22244960|22245043</td><td>48</td><td>69</td><td>22(+1U)</td><td>AAAAGCTGGGTTGAGAGGGCGAT</td><td>2194.4</td><td>3.8</td><td>0.6</td><td>6.8</td><td>1.3E-11</td><td>3.0E-11</td><td>BT20/MB468</td><td>NUC</td></tr>
<tr><td>>hg38_21_+_8433172_8446622_RNA45SN1_45S_50ntFlanks@11568.11601.34|GCAGCCGACTTAGAACTGGTGCGGACCAGGGGAA</td><td>>hg38_21_+_8433172_8446622_RNA45SN1_45S_50ntFlanks</td><td>11568</td><td>11601</td><td>34</td><td>GCAGCCGACTTAGAACTGGTGCGGACCAGGGGAA</td><td>4215.1</td><td>-6.9</td><td>1.0</td><td>-6.8</td><td>1.3E-11</td><td>3.0E-11</td><td>BT20/MB468</td><td>NUC</td></tr>
<tr><td>>MI0000469|hsa-mir-125a&WithFlank&19|+|51693248|51693345@59.79.21&[MIMAT0004602&hsa-miR-125a-3p&offsets|0|-1;m-333&19|+|51693306|51693326&offsets|0|0]|ACAGGTGAGGTTCTTGGGAGC</td><td>>MI0000469|hsa-mir-125a&WithFlank&19|+|51693248|51693345</td><td>59</td><td>79</td><td>21</td><td>ACAGGTGAGGTTCTTGGGAGC</td><td>1185.3</td><td>9.1</td><td>1.3</td><td>6.8</td><td>9.6E-12</td><td>3.1E-11</td><td>BT20/MB468</td><td>CYTO</td></tr>
<tr><td>>hg38_21_+_8433172_8446622_RNA45SN1_45S_50ntFlanks@11564.11581.18|CCTAGCAGCCGACTTAGA</td><td>>hg38_21_+_8433172_8446622_RNA45SN1_45S_50ntFlanks</td><td>11564</td><td>11581</td><td>18</td><td>CCTAGCAGCCGACTTAGA</td><td>937.6</td><td>-6.0</td><td>0.9</td><td>-6.8</td><td>1.0E-11</td><td>3.1E-11</td><td>MB231/MB468</td><td>CYTO</td></tr>
<tr><td>>hg38_MT_+_1621_3279_RNR2_16S_50ntFlanks@493.518.26|GCTCTTTGGACACTAGGAAAAAACCT</td><td>>hg38_MT_+_1621_3279_RNR2_16S_50ntFlanks</td><td>493</td><td>518</td><td>26</td><td>GCTCTTTGGACACTAGGAAAAAACCT</td><td>1061.7</td><td>6.3</td><td>0.9</td><td>6.8</td><td>7.5E-12</td><td>3.1E-11</td><td>BT20/MB468</td><td>MITO</td></tr>
<tr><td>>hg38_21_+_8433172_8446622_RNA45SN1_45S_50ntFlanks@8839.8862.24|CGGTGGCGGCGGCGGCGGCGGCGG</td><td>>hg38_21_+_8433172_8446622_RNA45SN1_45S_50ntFlanks</td><td>8839</td><td>8862</td><td>24</td><td>CGGTGGCGGCGGCGGCGGCGGCGG</td><td>160.2</td><td>-11.2</td><td>1.6</td><td>-6.8</td><td>9.4E-12</td><td>3.1E-11</td><td>MB231/MB468</td><td>MP</td></tr>
<tr><td>>hg38_21_+_8433172_8446622_RNA45SN1_45S_50ntFlanks@10689.10706.18|GCGTCCGGTGAGCTCTCG</td><td>>hg38_21_+_8433172_8446622_RNA45SN1_45S_50ntFlanks</td><td>10689</td><td>10706</td><td>18</td><td>GCGTCCGGTGAGCTCTCG</td><td>799.2</td><td>-7.7</td><td>1.1</td><td>-6.8</td><td>7.6E-12</td><td>3.1E-11</td><td>BT20/MB468</td><td>MITO</td></tr>
<tr><td>>am_trna11_ProAGG_16_+_3241989_3242060@1.28.28,trna12_ProAGG_6_+_26555498_26555569@1.28.28,trna12_ProTGG_11_-_75946869_75946940@1.28.28,trna14_ProTGG_5_-_180615854_180615925@1.28.28,trna22_ProAGG_14_-_21081560_21081631@1.28.28,trna23_ProAGG_14_-_21077495_21077566@1.28.28,trna28_ProTGG_16_-_3234133_3234204@1.28.28,trna29_ProAGG_16_-_3232635_3232706@1.28.28,trna2_ProAGG_7_+_128423504_128423575@1.28.28,trna30_ProCGG_6_+_27059521_27059592@1.28.28,trna37_ProCGG_17_-_8126151_8126222@1.28.28,trna3_ProTGG_16_+_3208923_3208994@1.28.28,trna52_ProCGG_1_+_167683962_167684033@1.28.28,trna65_ProAGG_1_-_167684725_167684796@1.28.28,trna6_ProCGG_16_+_3222049_3222120@1.28.28,trna6_ProTGG_14_+_21152175_21152246@1.28.28,trna8_ProTGG_16_+_3238094_3238165@1.28.28,trna9_ProAGG_11_+_75946557_75946628@1.28.28,trna9_ProAGG_16_+_3239634_3239705@1.28.28|GGCTCGTTGGTCTAGGGGTATGATTCTC</td><td>>am_trna11_ProAGG_16_+_3241989_3242060</td><td>1</td><td>28</td><td>28</td><td>GGCTCGTTGGTCTAGGGGTATGATTCTC</td><td>204.3</td><td>-11.5</td><td>1.7</td><td>-6.8</td><td>9.4E-12</td><td>3.1E-11</td><td>MB231/MB468</td><td>MP</td></tr>
<tr><td>>hg38_21_+_8433172_8446622_RNA45SN1_45S_50ntFlanks@12627.12657.31|ATGGGGCGAAGCTACCATCTGTGGGATTATG</td><td>>hg38_21_+_8433172_8446622_RNA45SN1_45S_50ntFlanks</td><td>12627</td><td>12657</td><td>31</td><td>ATGGGGCGAAGCTACCATCTGTGGGATTATG</td><td>1951.1</td><td>-8.8</td><td>1.3</td><td>-6.8</td><td>7.6E-12</td><td>3.1E-11</td><td>BT20/MB468</td><td>MITO</td></tr>
<tr><td>>hg38_21_+_8433172_8446622_RNA45SN1_45S_50ntFlanks@8687.8707.21|CGGCTCCGGGACGGCTGGGAA</td><td>>hg38_21_+_8433172_8446622_RNA45SN1_45S_50ntFlanks</td><td>8687</td><td>8707</td><td>21</td><td>CGGCTCCGGGACGGCTGGGAA</td><td>224.9</td><td>9.8</td><td>1.4</td><td>6.8</td><td>1.1E-11</td><td>3.1E-11</td><td>MB231/MB468</td><td>NUC</td></tr>
<tr><td>>hg38_21_+_8433172_8446622_RNA45SN1_45S_50ntFlanks@8191.8212.22|CAGCCCGTGGACGGTGTGAGGC</td><td>>hg38_21_+_8433172_8446622_RNA45SN1_45S_50ntFlanks</td><td>8191</td><td>8212</td><td>22</td><td>CAGCCCGTGGACGGTGTGAGGC</td><td>2413.5</td><td>-8.0</td><td>1.2</td><td>-6.8</td><td>9.4E-12</td><td>3.1E-11</td><td>MB231/MB468</td><td>MP</td></tr>
<tr><td>>hg38_21_+_8433172_8446622_RNA45SN1_45S_50ntFlanks@11587.11611.25|TGCGGACCAGGGGAATCCGACTGTT</td><td>>hg38_21_+_8433172_8446622_RNA45SN1_45S_50ntFlanks</td><td>11587</td><td>11611</td><td>25</td><td>TGCGGACCAGGGGAATCCGACTGTT</td><td>876.4</td><td>-4.2</td><td>0.6</td><td>-6.8</td><td>1.3E-11</td><td>3.1E-11</td><td>BT20/MB468</td><td>NUC</td></tr>
<tr><td>>hg38_21_+_8433172_8446622_RNA45SN1_45S_50ntFlanks@6741.6769.29|ATCGACACTTCGAACGCACTTGCGGCCCC</td><td>>hg38_21_+_8433172_8446622_RNA45SN1_45S_50ntFlanks</td><td>6741</td><td>6769</td><td>29</td><td>ATCGACACTTCGAACGCACTTGCGGCCCC</td><td>99.4</td><td>-10.5</td><td>1.5</td><td>-6.8</td><td>9.5E-12</td><td>3.1E-11</td><td>MB231/MB468</td><td>MP</td></tr>
<tr><td>>hg38_21_+_8433172_8446622_RNA45SN1_45S_50ntFlanks@12617.12652.36|TGAGGAGCCAATGGGGCGAAGCTACCATCTGTGGGA</td><td>>hg38_21_+_8433172_8446622_RNA45SN1_45S_50ntFlanks</td><td>12617</td><td>12652</td><td>36</td><td>TGAGGAGCCAATGGGGCGAAGCTACCATCTGTGGGA</td><td>96.6</td><td>-10.5</td><td>1.5</td><td>-6.8</td><td>9.5E-12</td><td>3.1E-11</td><td>MB231/MB468</td><td>MP</td></tr>
<tr><td>>MI0000289|hsa-mir-181a-1&WithFlank&1|-|198859038|198859159@70.90.21(+1U)&[MIMAT0000270&hsa-miR-181a-3p&offsets|0|-1(+1U);m-125&1|-|198859069|198859090&offsets|0|-1(+1U)]|ACCATCGACCGTTGATTGTACT</td><td>>MI0000289|hsa-mir-181a-1&WithFlank&1|-|198859038|198859159</td><td>70</td><td>90</td><td>21(+1U)</td><td>ACCATCGACCGTTGATTGTACT</td><td>165.8</td><td>-10.5</td><td>1.5</td><td>-7.1</td><td>1.1E-12</td><td>3.1E-11</td><td>BT20/MB231</td><td>NUC</td></tr>
<tr><td>>hg38_21_+_8433172_8446622_RNA45SN1_45S_50ntFlanks@12936.12956.21|AAACCATTCGTAGACGACCTG</td><td>>hg38_21_+_8433172_8446622_RNA45SN1_45S_50ntFlanks</td><td>12936</td><td>12956</td><td>21</td><td>AAACCATTCGTAGACGACCTG</td><td>5067.7</td><td>-5.2</td><td>0.8</td><td>-6.8</td><td>9.7E-12</td><td>3.1E-11</td><td>BT20/MB468</td><td>CYTO</td></tr>
<tr><td>>hg38_21_+_8433172_8446622_RNA45SN1_45S_50ntFlanks@7988.8018.31|CAGACGTGGCGACCCGCTGAATTTAAGCATA</td><td>>hg38_21_+_8433172_8446622_RNA45SN1_45S_50ntFlanks</td><td>7988</td><td>8018</td><td>31</td><td>CAGACGTGGCGACCCGCTGAATTTAAGCATA</td><td>609.7</td><td>-8.9</td><td>1.3</td><td>-6.8</td><td>9.5E-12</td><td>3.1E-11</td><td>MB231/MB468</td><td>MP</td></tr>
<tr><td>>hg38_21_+_8433172_8446622_RNA45SN1_45S_50ntFlanks@8336.8373.38|AAGTACCGTAAGGGAAAGTTGAAAAGAACTTTGAAGAG</td><td>>hg38_21_+_8433172_8446622_RNA45SN1_45S_50ntFlanks</td><td>8336</td><td>8373</td><td>38</td><td>AAGTACCGTAAGGGAAAGTTGAAAAGAACTTTGAAGAG</td><td>330.1</td><td>12.1</td><td>1.7</td><td>7.2</td><td>6.1E-13</td><td>3.1E-11</td><td>BT20/MB468</td><td>MP</td></tr>
<tr><td>>hg38_21_+_8433172_8446622_RNA45SN1_45S_50ntFlanks@12935.12952.18|TAAACCATTCGTAGACGA</td><td>>hg38_21_+_8433172_8446622_RNA45SN1_45S_50ntFlanks</td><td>12935</td><td>12952</td><td>18</td><td>TAAACCATTCGTAGACGA</td><td>265.2</td><td>-7.1</td><td>1.0</td><td>-6.8</td><td>1.0E-11</td><td>3.1E-11</td><td>MB231/MB468</td><td>CYTO</td></tr>
<tr><td>>hg38_21_+_8433172_8446622_RNA45SN1_45S_50ntFlanks@10572.10591.20|AGAAGCCGGCGGGAGCCCCG</td><td>>hg38_21_+_8433172_8446622_RNA45SN1_45S_50ntFlanks</td><td>10572</td><td>10591</td><td>20</td><td>AGAAGCCGGCGGGAGCCCCG</td><td>984.4</td><td>-9.0</td><td>1.3</td><td>-6.8</td><td>1.1E-11</td><td>3.1E-11</td><td>MB231/MB468</td><td>NUC</td></tr>
<tr><td>>hg38_21_+_8433172_8446622_RNA45SN1_45S_50ntFlanks@10458.10491.34|ATGGCCTCCGTTGCCCTCGGCCGATCGAAAGGGA</td><td>>hg38_21_+_8433172_8446622_RNA45SN1_45S_50ntFlanks</td><td>10458</td><td>10491</td><td>34</td><td>ATGGCCTCCGTTGCCCTCGGCCGATCGAAAGGGA</td><td>112.0</td><td>-10.7</td><td>1.6</td><td>-6.8</td><td>9.5E-12</td><td>3.1E-11</td><td>MB231/MB468</td><td>MP</td></tr>
<tr><td>>hg38_21_+_8433172_8446622_RNA45SN1_45S_50ntFlanks@11881.11900.20|CCTGTTGAGCTTGACTCTAG</td><td>>hg38_21_+_8433172_8446622_RNA45SN1_45S_50ntFlanks</td><td>11881</td><td>11900</td><td>20</td><td>CCTGTTGAGCTTGACTCTAG</td><td>173.0</td><td>-9.7</td><td>1.4</td><td>-6.8</td><td>9.7E-12</td><td>3.1E-11</td><td>BT20/MB468</td><td>CYTO</td></tr>
<tr><td>>hg38_21_+_8433172_8446622_RNA45SN1_45S_50ntFlanks@10794.10828.35|CTGGCATGTTGGAACAATGTAGGTAAGGGAAGTCG</td><td>>hg38_21_+_8433172_8446622_RNA45SN1_45S_50ntFlanks</td><td>10794</td><td>10828</td><td>35</td><td>CTGGCATGTTGGAACAATGTAGGTAAGGGAAGTCG</td><td>448.1</td><td>-8.4</td><td>1.2</td><td>-6.8</td><td>1.2E-11</td><td>3.1E-11</td><td>MB231/MB468</td><td>MITO</td></tr>
<tr><td>>hg38_21_+_8433172_8446622_RNA45SN1_45S_50ntFlanks@12627.12656.30|ATGGGGCGAAGCTACCATCTGTGGGATTAT</td><td>>hg38_21_+_8433172_8446622_RNA45SN1_45S_50ntFlanks</td><td>12627</td><td>12656</td><td>30</td><td>ATGGGGCGAAGCTACCATCTGTGGGATTAT</td><td>555.0</td><td>-9.2</td><td>1.4</td><td>-6.8</td><td>1.2E-11</td><td>3.1E-11</td><td>MB231/MB468</td><td>MITO</td></tr>
<tr><td>>MI0000264|hsa-mir-7-2&WithFlank&15|+|88611819|88611940@38.59.22;MI0000265|hsa-mir-7-3&WithFlank&19|+|4770664|4770785@37.58.22;MI0000263|hsa-mir-7-1&WithFlank&9|-|83969742|83969863@30.51.22&[MIMAT0000252&hsa-miR-7-5p&offsets|0|-2;m-400&15|+|88611856|88611879&offsets|0|-2];[MIMAT0000252_1&hsa-miR-7-5p&offsets|0|-2;m-402&19|+|4770700|4770723&offsets|0|-2];[MIMAT0000252_2&hsa-miR-7-5p&offsets|0|-2;m-401&9|-|83969811|83969834&offsets|0|-2]|TGGAAGACTAGTGATTTTGTTG</td><td>>MI0000264|hsa-mir-7-2&WithFlank&15|+|88611819|88611940</td><td>38</td><td>59</td><td>22</td><td>TGGAAGACTAGTGATTTTGTTG</td><td>767.8</td><td>7.6</td><td>1.1</td><td>6.8</td><td>7.7E-12</td><td>3.1E-11</td><td>BT20/MB468</td><td>MITO</td></tr>
<tr><td>>hg38_21_+_8433172_8446622_RNA45SN1_45S_50ntFlanks@11583.11609.27|CTGGTGCGGACCAGGGGAATCCGACTG</td><td>>hg38_21_+_8433172_8446622_RNA45SN1_45S_50ntFlanks</td><td>11583</td><td>11609</td><td>27</td><td>CTGGTGCGGACCAGGGGAATCCGACTG</td><td>1490.3</td><td>-5.3</td><td>0.8</td><td>-6.8</td><td>1.3E-11</td><td>3.1E-11</td><td>MB231/MB468</td><td>MITO</td></tr>
<tr><td>>hg38_21_+_8433172_8446622_RNA45SN1_45S_50ntFlanks@8178.8211.34|TCTGATCGAGGCCCAGCCCGTGGACGGTGTGAGG</td><td>>hg38_21_+_8433172_8446622_RNA45SN1_45S_50ntFlanks</td><td>8178</td><td>8211</td><td>34</td><td>TCTGATCGAGGCCCAGCCCGTGGACGGTGTGAGG</td><td>92.1</td><td>-10.4</td><td>1.5</td><td>-6.8</td><td>9.6E-12</td><td>3.1E-11</td><td>MB231/MB468</td><td>MP</td></tr>
<tr><td>>hg38_21_+_8433172_8446622_RNA45SN1_45S_50ntFlanks@10404.10422.19|CATGGGTCAGTCGGTCCTG</td><td>>hg38_21_+_8433172_8446622_RNA45SN1_45S_50ntFlanks</td><td>10404</td><td>10422</td><td>19</td><td>CATGGGTCAGTCGGTCCTG</td><td>645.1</td><td>-5.3</td><td>0.8</td><td>-6.8</td><td>9.6E-12</td><td>3.1E-11</td><td>MB231/MB468</td><td>MP</td></tr>
<tr><td>>trna146_GlnCTG_6_-_27515531_27515602@15.43.29,trna1_GlnCTG_6_+_18836402_18836473@15.43.29,trna3_GlnCTG_17_+_8023070_8023141@15.43.29,trna49_GlnCTG_6_+_27487308_27487379@15.43.29,trna7_GlnCTG_15_-_66161400_66161471@15.43.29,trna99_GlnCTG_6_-_28909378_28909449@15.43.29|ATGGTTAGCACTCTGGACTCTGAATCCAG</td><td>>trna146_GlnCTG_6_-_27515531_27515602</td><td>15</td><td>43</td><td>29</td><td>ATGGTTAGCACTCTGGACTCTGAATCCAG</td><td>94.3</td><td>-10.4</td><td>1.5</td><td>-6.8</td><td>9.6E-12</td><td>3.1E-11</td><td>MB231/MB468</td><td>MP</td></tr>
<tr><td>>MI0000239|hsa-mir-197&WithFlank&1|+|109598887|109598973@54.75.22&[MIMAT0000227&hsa-miR-197-3p&offsets|0|0;m-99&1|+|109598940|109598961&offsets|0|0]|TTCACCACCTTCTCCACCCAGC</td><td>>MI0000239|hsa-mir-197&WithFlank&1|+|109598887|109598973</td><td>54</td><td>75</td><td>22</td><td>TTCACCACCTTCTCCACCCAGC</td><td>2740.7</td><td>6.0</td><td>0.9</td><td>6.8</td><td>1.1E-11</td><td>3.1E-11</td><td>MB231/MB468</td><td>CYTO</td></tr>
</table>

| | | | | | | | | | | | | | |
|---|---|---|---|---|---|---|---|---|---|---|---|---|---|
| >hg38_21_+_8433172_8446622_RNA45SN1_45S_50ntFlanks@10883.10905.23\|CGGGCTGGGGCGCGAAGCGGGGC | >hg38_21_+_8433172_8446622_RNA45SN1_45S_50ntFlanks | 10883 | 10905 | 23 | CGGGCTGGGGCGCGAAGCGGGGC | 348.0 | 5.4 | 0.8 | 6.8 | 1.1E-11 | 3.1E-11 | MB231/MB468 | NUC |
| >hg38_21_+_8433172_8446622_RNA45SN1_45S_50ntFlanks@5489.5512.24\|CTGAGAAGACGGTCGAACTTGACT | >hg38_21_+_8433172_8446622_RNA45SN1_45S_50ntFlanks | 5489 | 5512 | 24 | CTGAGAAGACGGTCGAACTTGACT | 411.8 | 3.9 | 0.6 | 6.8 | 1.1E-11 | 3.1E-11 | MB231/MB468 | CYTO |
| >hg38_21_+_8433172_8446622_RNA45SN1_45S_50ntFlanks@12453.12476.24\|TCACCAAGCGTTGGATTGTTCACC | >hg38_21_+_8433172_8446622_RNA45SN1_45S_50ntFlanks | 12453 | 12476 | 24 | TCACCAAGCGTTGGATTGTTCACC | 630.6 | 4.4 | 0.6 | 6.8 | 1.1E-11 | 3.1E-11 | MB231/MB468 | CYTO |
| >hg38_21_+_8433172_8446622_RNA45SN1_45S_50ntFlanks@8115.8143.29\|CGCGGGACATGTGGCGTACGGAAGACCCG | >hg38_21_+_8433172_8446622_RNA45SN1_45S_50ntFlanks | 8115 | 8143 | 29 | CGCGGGACATGTGGCGTACGGAAGACCCG | 215.1 | -6.6 | 1.0 | -6.8 | 1.1E-11 | 3.1E-11 | MB231/MB468 | CYTO |
| >am_trna111_HisGTG_1_-_147774845_147774916@1.29.29,trna118_HisGTG_1_-_145396881_145396952@1.29.29,trna16_HisGTG_1_+_146544773_146544844@1.29.29,trna1_HisGTG_15_+_45493349_45493420@1.29.29,trna21_HisGTG_1_+_147753471_147753542@1.29.29,trna33_HisGTG_6_+_27125906_27125977@1.29.29,trna7_HisGTG_9_-_14433938_14434009@1.29.29,trna8_HisGTG_15_-_45492611_45492682@1.29.29,trna9_HisGTG_15_-_45490804_45490875@1.29.29\|GCCGTGATCGTATAGTGGTTAGTACTCTG | >am_trna111_HisGTG_1_-_147774845_147774916 | 1 | 29 | 29 | GCCGTGATCGTATAGTGGTTAGTACTCTG | 9409.3 | -5.9 | 0.9 | -6.8 | 1.1E-11 | 3.1E-11 | MB231/MB468 | NUC |
| >am_trnaMT_ValTAC_MT_+_1602_1670@51.69.19\|CTTAACTTGACCGCTCTGA | >am_trnaMT_ValTAC_MT_+_1602_1670 | 51 | 69 | 19 | CTTAACTTGACCGCTCTGA | 160.2 | -11.2 | 1.6 | -6.8 | 9.7E-12 | 3.1E-11 | MB231/MB468 | MP |
| >hg38_21_+_8433172_8446622_RNA45SN1_45S_50ntFlanks@10788.10825.38\|CAGCCTCTGGCATGTTGGAACAATGTAGGTAAGGGAAG | >hg38_21_+_8433172_8446622_RNA45SN1_45S_50ntFlanks | 10788 | 10825 | 38 | CAGCCTCTGGCATGTTGGAACAATGTAGGTAAGGGAAG | 156.9 | -11.2 | 1.6 | -6.8 | 9.7E-12 | 3.1E-11 | MB231/MB468 | MP |
| >hg38_21_+_8433172_8446622_RNA45SN1_45S_50ntFlanks@3793.3810.18\|CGAATGGCTCATTAAATC | >hg38_21_+_8433172_8446622_RNA45SN1_45S_50ntFlanks | 3793 | 3810 | 18 | CGAATGGCTCATTAAATC | 298.2 | -8.4 | 1.2 | -6.8 | 1.3E-11 | 3.1E-11 | BT20/MB468 | NUC |
| >MI0001445\|hsa-mir-423&WithFlank&17\|+\|30117073\|30117178@59.81.23(+1A)&[MIMAT0001340&hsa-miR-423-3p&offsets\|0\|0(+1A);m-100&17\|+\|30117131\|30117153&offsets\|0\|0(+1A)]\|AGCTCGGTCTGAGGCCCCTCAGTA | >MI0001445\|hsa-mir-423&WithFlank&17\|+\|30117073\|30117178 | 59 | 81 | 23(+1A) | AGCTCGGTCTGAGGCCCCTCAGTA | 467.2 | 5.8 | 0.9 | 6.8 | 1.1E-11 | 3.1E-11 | MB231/MB468 | CYTO |
| >hg38_21_+_8433172_8446622_RNA45SN1_45S_50ntFlanks@11874.11910.37\|AGAAGACCCTGTTGAGCTTGACTCTAGTCTGGCACGG | >hg38_21_+_8433172_8446622_RNA45SN1_45S_50ntFlanks | 11874 | 11910 | 37 | AGAAGACCCTGTTGAGCTTGACTCTAGTCTGGCACGG | 236.8 | -5.7 | 0.8 | -6.8 | 1.1E-11 | 3.1E-11 | MB231/MB468 | CYTO |
| >hg38_21_+_8433172_8446622_RNA45SN1_45S_50ntFlanks@9576.9596.21\|TCGGTCGTCCGACCTGGGTAT | >hg38_21_+_8433172_8446622_RNA45SN1_45S_50ntFlanks | 9576 | 9596 | 21 | TCGGTCGTCCGACCTGGGTAT | 227.9 | -9.9 | 1.5 | -6.8 | 7.8E-12 | 3.1E-11 | BT20/MB468 | MITO |
| >MI0000446\|hsa-mir-125b-1&WithFlank&11\|-\|122099751\|122099850@21.42.22(+1C);MI0000470\|hsa-mir-125b-2&WithFlank&21\|+\|16590231\|16590331@23.44.22(+1C)&[MIMAT0000423&hsa-miR-125b-5p&offsets\|0\|0(+1C);m-67&11\|-\|122099809\|122099830&offsets\|0\|0(+1C)];[MIMAT0000423_1&hsa-miR-125b-5p&offsets\|0\|0(+1C);m-68&21\|+\|16590253\|16590274&offsets\|0\|0(+1C)]\|TCCCTGAGACCCTAACTTGTGAC | >MI0000446\|hsa-mir-125b-1&WithFlank&11\|-\|122099751\|122099850 | 21 | 42 | 22(+1C) | TCCCTGAGACCCTAACTTGTGAC | 281.2 | 9.3 | 1.4 | 6.8 | 1.1E-11 | 3.2E-11 | MB231/MB468 | NUC |
| >am_trna117_GlyTCC_1_-_145397864_145397935@1.29.29,trna45_GlyTCC_1_+_161500903_161500974@1.29.29,trna70_GlyTCC_1_-_161439547_161439618@1.29.29,trna73_GlyTCC_1_-_161432166_161432237@1.29.29,trna76_GlyTCC_1_-_161424756_161424827@1.29.29,trna79_GlyTCC_1_-_161417375_161417446@1.29.29\|GCGTTGGTGGTATAGTGGTGAGCATAGCT | >am_trna117_GlyTCC_1_-_145397864_145397935 | 1 | 29 | 29 | GCGTTGGTGGTATAGTGGTGAGCATAGCT | 4308.6 | -11.3 | 1.7 | -6.8 | 1.3E-11 | 3.2E-11 | BT20/MB468 | NUC |
| >hg38_21_+_8433172_8446622_RNA45SN1_45S_50ntFlanks@10357.10388.32\|CTTTGAAGGCCGAAGTGGAGAAGGGTTCCATG | >hg38_21_+_8433172_8446622_RNA45SN1_45S_50ntFlanks | 10357 | 10388 | 32 | CTTTGAAGGCCGAAGTGGAGAAGGGTTCCATG | 637.1 | -8.9 | 1.3 | -6.8 | 9.7E-12 | 3.2E-11 | MB231/MB468 | MP |
| >hg38_21_+_8433172_8446622_RNA45SN1_45S_50ntFlanks@10420.10453.34\|CTGAGAGATGGGCGAGCGCCGTTCCGAAGGGACG | >hg38_21_+_8433172_8446622_RNA45SN1_45S_50ntFlanks | 10420 | 10453 | 34 | CTGAGAGATGGGCGAGCGCCGTTCCGAAGGGACG | 2028.6 | -7.9 | 1.2 | -6.8 | 1.1E-11 | 3.2E-11 | MB231/MB468 | NUC |
| >hg38_21_+_8433172_8446622_RNA45SN1_45S_50ntFlanks@11569.11599.31\|CAGCCGACTTAGAACTGGTGCGGACCAGGGG | >hg38_21_+_8433172_8446622_RNA45SN1_45S_50ntFlanks | 11569 | 11599 | 31 | CAGCCGACTTAGAACTGGTGCGGACCAGGGG | 644.3 | -4.3 | 0.6 | -6.8 | 1.1E-11 | 3.2E-11 | MB231/MB468 | NUC |
| >MI0000455\|hsa-mir-138-2&WithFlank&16\|+\|56858512\|56858607@16.38.23(+1C);MI0000476\|hsa-mir-138-1&WithFlank&3\|+\|44114206\|44114316@29.51.23(+1C)&[MIMAT0000430&hsa-miR-138-5p&offsets\|0\|0(+1C);m-509&16\|+\|56858527\|56858549&offsets\|0\|0(+1C)];[MIMAT0000430_1&hsa-miR-138-5p&offsets\|0\|0(+1C);m-497&3\|+\|44114234\|44114256&offsets\|0\|0(+1C)]\|AGCTGGTGTTGTGAATCAGGCCGC | >MI0000455\|hsa-mir-138-2&WithFlank&16\|+\|56858512\|56858607 | 16 | 38 | 23(+1C) | AGCTGGTGTTGTGAATCAGGCCGC | 122.8 | 10.2 | 1.5 | 6.8 | 9.7E-12 | 3.2E-11 | MB231/MB468 | MP |
| >hg38_21_+_8433172_8446622_RNA45SN1_45S_50ntFlanks@3875.3897.23\|ATACATGCCGACGGGCGCTGACC | >hg38_21_+_8433172_8446622_RNA45SN1_45S_50ntFlanks | 3875 | 3897 | 23 | ATACATGCCGACGGGCGCTGACC | 105.3 | -10.6 | 1.6 | -6.8 | 9.7E-12 | 3.2E-11 | MB231/MB468 | MP |
| >hg38_21_+_8433172_8446622_RNA45SN1_45S_50ntFlanks@6743.6769.27\|CGACACTTCGAACGCACTTGCGGCCCC | >hg38_21_+_8433172_8446622_RNA45SN1_45S_50ntFlanks | 6743 | 6769 | 27 | CGACACTTCGAACGCACTTGCGGCCCC | 90.3 | -10.4 | 1.5 | -6.8 | 9.7E-12 | 3.2E-11 | MB231/MB468 | MP |
| >hg38_21_+_8433172_8446622_RNA45SN1_45S_50ntFlanks@4072.4089.18\|TCGATGGTAGTCGCCGTG | >hg38_21_+_8433172_8446622_RNA45SN1_45S_50ntFlanks | 4072 | 4089 | 18 | TCGATGGTAGTCGCCGTG | 88.9 | -10.3 | 1.5 | -6.8 | 9.8E-12 | 3.2E-11 | MB231/MB468 | MP |
| >hg38_21_+_8433172_8446622_RNA45SN1_45S_50ntFlanks@4787.4812.26\|AACGATGCCGACCGGCGATGCGGCGG | >hg38_21_+_8433172_8446622_RNA45SN1_45S_50ntFlanks | 4787 | 4812 | 26 | AACGATGCCGACCGGCGATGCGGCGG | 173.5 | -11.3 | 1.7 | -6.8 | 9.8E-12 | 3.2E-11 | MB231/MB468 | MP |
| >hg38_MT_+_598_1651_RNR1_12S_50ntFlanks@371.404.34\|TAAAGCTAAAACTCACCTGAGTTGTAAAAAACTC | >hg38_MT_+_598_1651_RNR1_12S_50ntFlanks | 371 | 404 | 34 | TAAAGCTAAAACTCACCTGAGTTGTAAAAAACTC | 737.3 | 5.4 | 0.8 | 6.8 | 7.8E-12 | 3.2E-11 | BT20/MB468 | MITO |
| >hg38_21_+_8433172_8446622_RNA45SN1_45S_50ntFlanks@11665.11685.21\|TTCTGCCCAGTGCTCTGAATG | >hg38_21_+_8433172_8446622_RNA45SN1_45S_50ntFlanks | 11665 | 11685 | 21 | TTCTGCCCAGTGCTCTGAATG | 519.2 | -4.4 | 0.6 | -6.8 | 9.8E-12 | 3.2E-11 | MB231/MB468 | MP |
| >hg38_21_+_8433172_8446622_RNA45SN1_45S_50ntFlanks@8111.8138.28\|GGGGCGCGGGACATGTGGCGTACGGAAG | >hg38_21_+_8433172_8446622_RNA45SN1_45S_50ntFlanks | 8111 | 8138 | 28 | GGGGCGCGGGACATGTGGCGTACGGAAG | 183.5 | -11.4 | 1.7 | -6.8 | 9.8E-12 | 3.2E-11 | MB231/MB468 | MP |
| >hg38_21_+_8433172_8446622_RNA45SN1_45S_50ntFlanks@12654.12675.22\|TATGACTGAACGCCTCTAAGTC | >hg38_21_+_8433172_8446622_RNA45SN1_45S_50ntFlanks | 12654 | 12675 | 22 | TATGACTGAACGCCTCTAAGTC | 7228.9 | -7.9 | 1.2 | -6.8 | 1.1E-11 | 3.2E-11 | MB231/MB468 | CYTO |
| >hg38_21_+_8433172_8446622_RNA45SN1_45S_50ntFlanks@8168.8187.20\|CCCAAGTCCTTCTGATCGAG | >hg38_21_+_8433172_8446622_RNA45SN1_45S_50ntFlanks | 8168 | 8187 | 20 | CCCAAGTCCTTCTGATCGAG | 1107.3 | -7.4 | 1.1 | -6.8 | 1.1E-11 | 3.2E-11 | MB231/MB468 | NUC |
| >hg38_21_+_8433172_8446622_RNA45SN1_45S_50ntFlanks@9738.9765.28\|GAAACGATCTCAACCTATTCTCAAACTT | >hg38_21_+_8433172_8446622_RNA45SN1_45S_50ntFlanks | 9738 | 9765 | 28 | GAAACGATCTCAACCTATTCTCAAACTT | 6906.9 | -9.1 | 1.3 | -6.8 | 1.4E-11 | 3.2E-11 | BT20/MB468 | NUC |
| >hg38_21_+_8433172_8446622_RNA45SN1_45S_50ntFlanks@10421.10443.23\|TGAGAGATGGGCGAGCGCCGTTC | >hg38_21_+_8433172_8446622_RNA45SN1_45S_50ntFlanks | 10421 | 10443 | 23 | TGAGAGATGGGCGAGCGCCGTTC | 353.1 | -12.3 | 1.8 | -6.8 | 9.9E-12 | 3.2E-11 | MB231/MB468 | MP |
| >MI0000750\|hsa-mir-26a-2&WithFlank&12\|-\|57824603\|57824698@20.41.22(+1U);MI0000083\|hsa-mir-26a-1&WithFlank&3\|+\|37969398\|37969486@16.37.22(+1U)&[MIMAT0000082&hsa-miR-26a-5p&offsets\|0\|0(+1U);m-33&12\|-\|57824658\|57824679&offsets\|0\|0(+1U)];[MIMAT0000082_1&hsa-miR-26a-5p&offsets\|0\|0(+1U);m-34&3\|+\|37969413\|37969434&offsets\|0\|0(+1U)]\|TTCAAGTAATCCAGGATAGGCTT | >MI0000750\|hsa-mir-26a-2&WithFlank&12\|-\|57824603\|57824698 | 20 | 41 | 22(+1U) | TTCAAGTAATCCAGGATAGGCTT | 5292.1 | 6.6 | 1.0 | 6.8 | 1.3E-11 | 3.2E-11 | MB231/MB468 | MITO |
| >MI0000102\|hsa-mir-100&WithFlank&11\|-\|122152223\|122152314@19.37.19(+1U)&[MIMAT0000098&hsa-miR-100-5p&offsets\|0\|-3(+1U);m-27&11\|-\|122152276\|122152296&offsets\|0\|-2(+1U)]\|AACCCGTAGATCCGAACTTT | >MI0000102\|hsa-mir-100&WithFlank&11\|-\|122152223\|122152314 | 19 | 37 | 19(+1U) | AACCCGTAGATCCGAACTTT | 157.0 | -10.5 | 1.5 | -7.1 | 1.0E-12 | 3.2E-11 | BT20/MB231 | MITO |
| >hg38_21_+_8433172_8446622_RNA45SN1_45S_50ntFlanks@11895.11912.18\|CTCTAGTCTGGCACGGTG | >hg38_21_+_8433172_8446622_RNA45SN1_45S_50ntFlanks | 11895 | 11912 | 18 | CTCTAGTCTGGCACGGTG | 103.5 | -10.6 | 1.6 | -6.8 | 9.9E-12 | 3.2E-11 | MB231/MB468 | MP |
| >hg38_21_+_8433172_8446622_RNA45SN1_45S_50ntFlanks@475.508.34\|CTTCGTGATCGATGTGGTGACGTCGTGCTCTCCC | >hg38_21_+_8433172_8446622_RNA45SN1_45S_50ntFlanks | 475 | 508 | 34 | CTTCGTGATCGATGTGGTGACGTCGTGCTCTCCC | 551.0 | 8.5 | 1.3 | 6.8 | 1.0E-11 | 3.2E-11 | BT20/MB468 | CYTO |
| >trna119_LysCTT_1_-_145395522_145395594@3.33.31,trna11_LysCTT_5_-_180648979_180649051@3.33.31,trna13_LysCTT_6_+_26556774_26556846@3.33.31,trna7_LysCTT_16_+_3225692_3225764@3.33.31,trna9_LysCTT_5_+_180634755_180634827@3.33.31\|CCGGCTAGCTCAGTCGGTAGAGCATGAGACT | >trna119_LysCTT_1_-_145395522_145395594 | 3 | 33 | 31 | CCGGCTAGCTCAGTCGGTAGAGCATGAGACT | 1151.1 | 8.4 | 1.2 | 6.8 | 1.1E-11 | 3.2E-11 | MB231/MB468 | CYTO |
| >hg38_21_+_8433172_8446622_RNA45SN1_45S_50ntFlanks@12626.12661.36\|AATGGGGCGAAGCTACCATCTGTGGGATTATGACTG | >hg38_21_+_8433172_8446622_RNA45SN1_45S_50ntFlanks | 12626 | 12661 | 36 | AATGGGGCGAAGCTACCATCTGTGGGATTATGACTG | 9419.5 | -8.4 | 1.2 | -6.8 | 1.3E-11 | 3.2E-11 | MB231/MB468 | MITO |
| >hg38_21_+_8433172_8446622_RNA45SN1_45S_50ntFlanks@12945.12988.44\|GTAGACGACCTGCTTCTGGGTCGGGGTTTCGTACGTAGCAGAGC | >hg38_21_+_8433172_8446622_RNA45SN1_45S_50ntFlanks | 12945 | 12988 | 44 | GTAGACGACCTGCTTCTGGGTCGGGGTTTCGTACGTAGCAGAGC | 893.3 | 5.6 | 0.8 | 6.8 | 1.1E-11 | 3.2E-11 | MB231/MB468 | CYTO |
| >hg38_MT_+_1621_3279_RNR2_16S_50ntFlanks@1184.1216.33\|AATTTCGGTTGGGGCGACCTCGGAGCAGAACCC | >hg38_MT_+_1621_3279_RNR2_16S_50ntFlanks | 1184 | 1216 | 33 | AATTTCGGTTGGGGCGACCTCGGAGCAGAACCC | 898.6 | 8.2 | 1.2 | 6.8 | 8.0E-12 | 3.2E-11 | BT20/MB468 | MITO |
| >trna3_ProTGG_14_+_21101165_21101236@1.29.29\|GGCTCGTTGGTCTAGTGGTATGATTCTCG | >trna3_ProTGG_14_+_21101165_21101236 | 1 | 29 | 29 | GGCTCGTTGGTCTAGTGGTATGATTCTCG | 175.7 | -6.5 | 1.0 | -6.8 | 1.1E-11 | 3.2E-11 | MB231/MB468 | CYTO |
| >MI0000736\|hsa-mir-30c-1&WithFlank&1\|+\|40757278\|40757378@23.45.23;MI0000254\|hsa-mir-30c-2&WithFlank&6\|-\|71376954\|71377037@13.35.23&[MIMAT0000244&hsa-miR-30c-5p&offsets\|0\|0;m-57&1\|+\|40757300\|40757323&offsets\|0\|-1];[MIMAT0000244_1&hsa-miR-30c-5p&offsets\|0\|0;m-56&6\|-\|71377002\|71377025&offsets\|0\|-1]\|TGTAAACATCCTACACTCTCAGC | >MI0000736\|hsa-mir-30c-1&WithFlank&1\|+\|40757278\|40757378 | 23 | 45 | 23 | TGTAAACATCCTACACTCTCAGC | 11635.9 | 5.5 | 0.8 | 6.8 | 1.1E-11 | 3.2E-11 | MB231/MB468 | CYTO |
| >hg38_21_+_8433172_8446622_RNA45SN1_45S_50ntFlanks@8182.8210.29\|ATCGAGGCCCAGCCCGTGGACGGTGTGAG | >hg38_21_+_8433172_8446622_RNA45SN1_45S_50ntFlanks | 8182 | 8210 | 29 | ATCGAGGCCCAGCCCGTGGACGGTGTGAG | 400.9 | -11.7 | 1.6 | -7.2 | 6.4E-13 | 3.2E-11 | BT20/MB468 | MP |
| >hg38_MT_+_1621_3279_RNR2_16S_50ntFlanks@51.84.34\|GCTAAACCTAGCCCCAAACCCACTCCACCTTACT | >hg38_MT_+_1621_3279_RNR2_16S_50ntFlanks | 51 | 84 | 34 | GCTAAACCTAGCCCCAAACCCACTCCACCTTACT | 16435.2 | 7.0 | 1.0 | 6.8 | 8.1E-12 | 3.3E-11 | BT20/MB468 | MITO |
| >hg38_21_+_8433172_8446622_RNA45SN1_45S_50ntFlanks@11318.11343.26\|CCCCACGTCTCGTCGCGCGCGCGTCC | >hg38_21_+_8433172_8446622_RNA45SN1_45S_50ntFlanks | 11318 | 11343 | 26 | CCCCACGTCTCGTCGCGCGCGCGTCC | 2303.1 | -9.6 | 1.4 | -6.8 | 1.1E-11 | 3.3E-11 | MB231/MB468 | NUC |
| >hg38_21_+_8433172_8446622_RNA45SN1_45S_50ntFlanks@4760.4783.24\|TCAGATACCGTCGTAGTTCCGACC | >hg38_21_+_8433172_8446622_RNA45SN1_45S_50ntFlanks | 4760 | 4783 | 24 | TCAGATACCGTCGTAGTTCCGACC | 930.5 | -9.0 | 1.3 | -6.8 | 1.0E-11 | 3.3E-11 | MB231/MB468 | MP |
| >hg38_21_+_8433172_8446622_RNA45SN1_45S_50ntFlanks@6698.6718.21\|AGCTAGCTGCGAGAATTAATG | >hg38_21_+_8433172_8446622_RNA45SN1_45S_50ntFlanks | 6698 | 6718 | 21 | AGCTAGCTGCGAGAATTAATG | 18354.3 | -9.1 | 1.3 | -6.8 | 1.0E-11 | 3.3E-11 | MB231/MB468 | MP |

<table>
<tr><td>>hg38_21_+_8433172_8446622_RNA45SN1_45S_50ntFlanks@9027.9063.37|CGTCTCCTCGTGGGGGGGCCGGGCCACCCCTCCCACG</td><td>>hg38_21_+_8433172_8446622_RNA45SN1_45S_50ntFlanks</td><td>9027</td><td>9063</td><td>37</td><td>CGTCTCCTCGTGGGGGGGCCGGGCCACCCCTCCCACG</td><td>99.4</td><td>-10.5</td><td>1.5</td><td>-6.8</td><td>1.0E-11</td><td>3.3E-11</td><td>MB231/MB468</td><td>MP</td></tr>
<tr><td>>hg38_21_+_8433172_8446622_RNA45SN1_45S_50ntFlanks@10410.10441.32|TCAGTCGGTCCTGAGAGATGGGCGAGCGCCGT</td><td>>hg38_21_+_8433172_8446622_RNA45SN1_45S_50ntFlanks</td><td>10410</td><td>10441</td><td>32</td><td>TCAGTCGGTCCTGAGAGATGGGCGAGCGCCGT</td><td>99.5</td><td>-10.5</td><td>1.5</td><td>-6.8</td><td>1.0E-11</td><td>3.3E-11</td><td>MB231/MB468</td><td>MP</td></tr>
<tr><td>>hg38_21_+_8433172_8446622_RNA45SN1_45S_50ntFlanks@12347.12366.20|AGAAAAGTTACCACAGGGAT</td><td>>hg38_21_+_8433172_8446622_RNA45SN1_45S_50ntFlanks</td><td>12347</td><td>12366</td><td>20</td><td>AGAAAAGTTACCACAGGGAT</td><td>128.2</td><td>-10.9</td><td>1.6</td><td>-6.8</td><td>1.0E-11</td><td>3.3E-11</td><td>MB231/MB468</td><td>MP</td></tr>
<tr><td>>trnaMT_HisGTG_MT_+_12138_12206@32.70.39|TGAATCTGACAACAGAGGCTTACGACCCCTTATTTACCC</td><td>>trnaMT_HisGTG_MT_+_12138_12206</td><td>32</td><td>70</td><td>39</td><td>TGAATCTGACAACAGAGGCTTACGACCCCTTATTTACCC</td><td>862.8</td><td>8.1</td><td>1.2</td><td>6.8</td><td>1.1E-11</td><td>3.3E-11</td><td>MB231/MB468</td><td>CYTO</td></tr>
<tr><td>>hg38_21_+_8433172_8446622_RNA45SN1_45S_50ntFlanks@9243.9272.30|GAGCGAGCGCACGGGGTCGGCGGCGACGTC</td><td>>hg38_21_+_8433172_8446622_RNA45SN1_45S_50ntFlanks</td><td>9243</td><td>9272</td><td>30</td><td>GAGCGAGCGCACGGGGTCGGCGGCGACGTC</td><td>91.8</td><td>-10.4</td><td>1.5</td><td>-6.8</td><td>1.0E-11</td><td>3.3E-11</td><td>MB231/MB468</td><td>MP</td></tr>
<tr><td>>hg38_21_+_8433172_8446622_RNA45SN1_45S_50ntFlanks@4473.4494.22|TTAGCTGAGTGTCCCGCGGGGC</td><td>>hg38_21_+_8433172_8446622_RNA45SN1_45S_50ntFlanks</td><td>4473</td><td>4494</td><td>22</td><td>TTAGCTGAGTGTCCCGCGGGGC</td><td>219.6</td><td>-7.8</td><td>1.1</td><td>-6.8</td><td>1.0E-11</td><td>3.3E-11</td><td>BT20/MB468</td><td>CYTO</td></tr>
<tr><td>>MI0001445|hsa-mir-423&WithFlank&17|+|30117073|30117178@59.80.22(+1C)&[MIMAT0001340&hsa-miR-423-3p&offsets|0|-1(+1C);m-100&17|+|30117131|30117153&offsets|0|-1(+1C)]|AGCTCGGTCTGAGGCCCCTCAGC</td><td>>MI0001445|hsa-mir-423&WithFlank&17|+|30117073|30117178</td><td>59</td><td>80</td><td>22(+1C)</td><td>AGCTCGGTCTGAGGCCCCTCAGC</td><td>733.8</td><td>3.6</td><td>0.5</td><td>6.8</td><td>1.4E-11</td><td>3.3E-11</td><td>BT20/MB468</td><td>NUC</td></tr>
<tr><td>>trnaMT_LeuTAA_MT_+_3230_3304@16.44.29|CCCGGTAATCGCATAAAACTTAAAACTTT</td><td>>trnaMT_LeuTAA_MT_+_3230_3304</td><td>16</td><td>44</td><td>29</td><td>CCCGGTAATCGCATAAAACTTAAAACTTT</td><td>2449.1</td><td>-7.7</td><td>1.1</td><td>-6.8</td><td>1.1E-11</td><td>3.3E-11</td><td>MB231/MB468</td><td>NUC</td></tr>
<tr><td>>hg38_21_+_8433172_8446622_RNA45SN1_45S_50ntFlanks@10866.10894.29|CTAAGGGCTGGGTCGGTCGGGCTGGGGCG</td><td>>hg38_21_+_8433172_8446622_RNA45SN1_45S_50ntFlanks</td><td>10866</td><td>10894</td><td>29</td><td>CTAAGGGCTGGGTCGGTCGGGCTGGGGCG</td><td>413.3</td><td>-6.0</td><td>0.9</td><td>-6.8</td><td>1.1E-11</td><td>3.3E-11</td><td>MB231/MB468</td><td>CYTO</td></tr>
<tr><td>>MI0000433|hsa-let-7g&WithFlank&3|-|52268272|52268367@11.28.18&[MIMAT0000414&hsa-let-7g-5p&offsets|0|-4;m-58&3|-|52268336|52268357&offsets|0|-4]|TGAGGTAGTAGTTTGTAC</td><td>>MI0000433|hsa-let-7g&WithFlank&3|-|52268272|52268367</td><td>11</td><td>28</td><td>18</td><td>TGAGGTAGTAGTTTGTAC</td><td>1416.6</td><td>5.9</td><td>0.9</td><td>6.8</td><td>1.1E-11</td><td>3.3E-11</td><td>MB231/MB468</td><td>CYTO</td></tr>
<tr><td>>hg38_21_+_8433172_8446622_RNA45SN1_45S_50ntFlanks@11579.11607.29|AGAACTGGTGCGGACCAGGGGAATCCGAC</td><td>>hg38_21_+_8433172_8446622_RNA45SN1_45S_50ntFlanks</td><td>11579</td><td>11607</td><td>29</td><td>AGAACTGGTGCGGACCAGGGGAATCCGAC</td><td>778.5</td><td>-4.5</td><td>0.7</td><td>-6.8</td><td>1.3E-11</td><td>3.3E-11</td><td>MB231/MB468</td><td>MITO</td></tr>
<tr><td>>hg38_21_+_8433172_8446622_RNA45SN1_45S_50ntFlanks@12636.12656.21|AGCTACCATCTGTGGGATTAT</td><td>>hg38_21_+_8433172_8446622_RNA45SN1_45S_50ntFlanks</td><td>12636</td><td>12656</td><td>21</td><td>AGCTACCATCTGTGGGATTAT</td><td>100.6</td><td>-10.5</td><td>1.5</td><td>-6.8</td><td>1.0E-11</td><td>3.3E-11</td><td>MB231/MB468</td><td>MP</td></tr>
<tr><td>>hg38_21_+_8433172_8446622_RNA45SN1_45S_50ntFlanks@6787.6806.20|ACGCCTGTCTGAGCGTCGCT</td><td>>hg38_21_+_8433172_8446622_RNA45SN1_45S_50ntFlanks</td><td>6787</td><td>6806</td><td>20</td><td>ACGCCTGTCTGAGCGTCGCT</td><td>4433.0</td><td>5.6</td><td>0.8</td><td>6.8</td><td>8.2E-12</td><td>3.3E-11</td><td>BT20/MB468</td><td>MITO</td></tr>
<tr><td>>MI0000070|hsa-mir-16-1&WithFlank&13|-|50048967|50049067@20.42.23;MI0000115|hsa-mir-16-2&WithFlank&3|+|160404739|160404831@16.38.23&[MIMAT0000069&hsa-miR-16-5p&offsets|0|+1;m-62&13|-|50049027|50049048&offsets|0|+1];[MIMAT0000069_1&hsa-miR-16-5p&offsets|0|+1;m-61&3|+|160404754|160404775&offsets|0|+1]|TAGCAGCACGTAAATATTGGCGT</td><td>>MI0000070|hsa-mir-16-1&WithFlank&13|-|50048967|50049067</td><td>20</td><td>42</td><td>23</td><td>TAGCAGCACGTAAATATTGGCGT</td><td>733.9</td><td>5.7</td><td>0.8</td><td>6.8</td><td>8.2E-12</td><td>3.3E-11</td><td>BT20/MB468</td><td>MITO</td></tr>
<tr><td>>hg38_21_+_8433172_8446622_RNA45SN1_45S_50ntFlanks@11718.11736.19|AACGGCGGGAGTAACTATG</td><td>>hg38_21_+_8433172_8446622_RNA45SN1_45S_50ntFlanks</td><td>11718</td><td>11736</td><td>19</td><td>AACGGCGGGAGTAACTATG</td><td>822.3</td><td>-6.2</td><td>0.9</td><td>-6.8</td><td>1.3E-11</td><td>3.3E-11</td><td>MB231/MB468</td><td>MITO</td></tr>
<tr><td>>MI0000803|hsa-mir-330&WithFlank&19|-|45638988|45639093@63.86.24(+1U)&[MIMAT0000751&hsa-miR-330-3p&offsets|0|+1(+1U);m-347&19|-|45639009|45639030&offsets|-1|+1(+1U)]|GCAAAGCACACGGCCTGCAGAGAGT</td><td>>MI0000803|hsa-mir-330&WithFlank&19|-|45638988|45639093</td><td>63</td><td>86</td><td>24(+1U)</td><td>GCAAAGCACACGGCCTGCAGAGAGT</td><td>801.6</td><td>7.1</td><td>1.0</td><td>6.8</td><td>1.0E-11</td><td>3.3E-11</td><td>BT20/MB468</td><td>CYTO</td></tr>
<tr><td>>hg38_21_+_8433172_8446622_RNA45SN1_45S_50ntFlanks@9419.9453.35|TCCGCCGAGGGCGCACCACCGGCCCGTCTCGCCCG</td><td>>hg38_21_+_8433172_8446622_RNA45SN1_45S_50ntFlanks</td><td>9419</td><td>9453</td><td>35</td><td>TCCGCCGAGGGCGCACCACCGGCCCGTCTCGCCCG</td><td>190.8</td><td>-9.7</td><td>1.4</td><td>-6.8</td><td>8.2E-12</td><td>3.3E-11</td><td>BT20/MB468</td><td>MITO</td></tr>
<tr><td>>MI0025242|hsa-mir-7706&WithFlank&15|+|85380590|85380668@50.71.22&[MIMAT0030021&hsa-miR-7706&offsets|0|-2;m-390&15|+|85380639|85380662&offsets|0|-2]|TGAAGCGCCTGTGCTCTGCCGA</td><td>>MI0025242|hsa-mir-7706&WithFlank&15|+|85380590|85380668</td><td>50</td><td>71</td><td>22</td><td>TGAAGCGCCTGTGCTCTGCCGA</td><td>350.2</td><td>8.2</td><td>1.2</td><td>6.8</td><td>8.2E-12</td><td>3.3E-11</td><td>BT20/MB468</td><td>MITO</td></tr>
<tr><td>>MI0000299|hsa-mir-222&WithFlank&X|-|45747009|45747130@75.95.21&[MIMAT0000279&hsa-miR-222-3p&offsets|0|0;m-140&X|-|45747033|45747056&offsets|0|-3]|AGCTACATCTGGCTACTGGGT</td><td>>MI0000299|hsa-mir-222&WithFlank&X|-|45747009|45747130</td><td>75</td><td>95</td><td>21</td><td>AGCTACATCTGGCTACTGGGT</td><td>2934.4</td><td>6.0</td><td>0.9</td><td>6.8</td><td>1.3E-11</td><td>3.3E-11</td><td>MB231/MB468</td><td>MITO</td></tr>
<tr><td>>hg38_21_+_8433172_8446622_RNA45SN1_45S_50ntFlanks@13004.13027.24|ATCTATTGAAAGTCAGCCCTCGAC</td><td>>hg38_21_+_8433172_8446622_RNA45SN1_45S_50ntFlanks</td><td>13004</td><td>13027</td><td>24</td><td>ATCTATTGAAAGTCAGCCCTCGAC</td><td>3003.6</td><td>-8.7</td><td>1.3</td><td>-6.8</td><td>1.1E-11</td><td>3.3E-11</td><td>MB231/MB468</td><td>CYTO</td></tr>
<tr><td>>hg38_21_+_8433172_8446622_RNA45SN1_45S_50ntFlanks@4045.4066.22|CCATTCGAACGTCTGCCCTATC</td><td>>hg38_21_+_8433172_8446622_RNA45SN1_45S_50ntFlanks</td><td>4045</td><td>4066</td><td>22</td><td>CCATTCGAACGTCTGCCCTATC</td><td>208.7</td><td>-6.2</td><td>0.9</td><td>-6.8</td><td>1.0E-11</td><td>3.3E-11</td><td>MB231/MB468</td><td>MP</td></tr>
<tr><td>>MI0000285|hsa-mir-205&WithFlank&1|+|209432127|209432248@40.61.22(+1C)&[MIMAT0000266&hsa-miR-205-5p&offsets|0|0(+1C);m-351&1|+|209432166|209432187&offsets|0|0(+1C)]|TCCTTCATTCCACCGGAGTCTGC</td><td>>MI0000285|hsa-mir-205&WithFlank&1|+|209432127|209432248</td><td>40</td><td>61</td><td>22(+1C)</td><td>TCCTTCATTCCACCGGAGTCTGC</td><td>490.0</td><td>12.3</td><td>1.7</td><td>7.4</td><td>1.6E-13</td><td>3.3E-11</td><td>BT20/MB231</td><td>MP</td></tr>
<tr><td>>hg38_21_+_8433172_8446622_RNA45SN1_45S_50ntFlanks@9860.9893.34|GGATGAACCGAACGCCGGGTTAAGGCGCCCGATG</td><td>>hg38_21_+_8433172_8446622_RNA45SN1_45S_50ntFlanks</td><td>9860</td><td>9893</td><td>34</td><td>GGATGAACCGAACGCCGGGTTAAGGCGCCCGATG</td><td>165.9</td><td>-11.2</td><td>1.7</td><td>-6.8</td><td>1.0E-11</td><td>3.3E-11</td><td>MB231/MB468</td><td>MP</td></tr>
<tr><td>>hg38_1_-_228634819_228635039_RNA5S12_5S_50ntFlanks@137.170.34|GAGACCGCCTGGGAATACCGGGTGCTGTAGGCTT</td><td>>hg38_1_-_228634819_228635039_RNA5S12_5S_50ntFlanks</td><td>137</td><td>170</td><td>34</td><td>GAGACCGCCTGGGAATACCGGGTGCTGTAGGCTT</td><td>468.1</td><td>5.1</td><td>0.7</td><td>6.8</td><td>1.2E-11</td><td>3.3E-11</td><td>MB231/MB468</td><td>NUC</td></tr>
<tr><td>>hg38_21_+_8433172_8446622_RNA45SN1_45S_50ntFlanks@9563.9595.33|CCTGACGTGCAAATCGGTCGTCCGACCTGGGTA</td><td>>hg38_21_+_8433172_8446622_RNA45SN1_45S_50ntFlanks</td><td>9563</td><td>9595</td><td>33</td><td>CCTGACGTGCAAATCGGTCGTCCGACCTGGGTA</td><td>91.6</td><td>-10.4</td><td>1.5</td><td>-6.8</td><td>1.0E-11</td><td>3.3E-11</td><td>MB231/MB468</td><td>MP</td></tr>
<tr><td>>hg38_21_+_8433172_8446622_RNA45SN1_45S_50ntFlanks@4559.4587.29|GCTAGGAATAATGGAATAGGACCGCGGTT</td><td>>hg38_21_+_8433172_8446622_RNA45SN1_45S_50ntFlanks</td><td>4559</td><td>4587</td><td>29</td><td>GCTAGGAATAATGGAATAGGACCGCGGTT</td><td>203.5</td><td>-8.0</td><td>1.2</td><td>-6.8</td><td>1.2E-11</td><td>3.3E-11</td><td>MB231/MB468</td><td>NUC</td></tr>
<tr><td>>hg38_21_+_8433172_8446622_RNA45SN1_45S_50ntFlanks@6732.6754.23|ACATTGATCATCGACACTTCGAA</td><td>>hg38_21_+_8433172_8446622_RNA45SN1_45S_50ntFlanks</td><td>6732</td><td>6754</td><td>23</td><td>ACATTGATCATCGACACTTCGAA</td><td>86.8</td><td>-10.3</td><td>1.5</td><td>-6.8</td><td>1.0E-11</td><td>3.3E-11</td><td>MB231/MB468</td><td>MP</td></tr>
<tr><td>>hg38_21_+_8433172_8446622_RNA45SN1_45S_50ntFlanks@6651.6673.23|CGACTCTTAGCGGTGGATCACTC</td><td>>hg38_21_+_8433172_8446622_RNA45SN1_45S_50ntFlanks</td><td>6651</td><td>6673</td><td>23</td><td>CGACTCTTAGCGGTGGATCACTC</td><td>2059.2</td><td>4.8</td><td>0.7</td><td>6.8</td><td>1.1E-11</td><td>3.4E-11</td><td>MB231/MB468</td><td>CYTO</td></tr>
<tr><td>>hg38_21_+_8433172_8446622_RNA45SN1_45S_50ntFlanks@12542.12573.32|ATGTGTTGTTGCCATGGTAATCCTGCTCAGTA</td><td>>hg38_21_+_8433172_8446622_RNA45SN1_45S_50ntFlanks</td><td>12542</td><td>12573</td><td>32</td><td>ATGTGTTGTTGCCATGGTAATCCTGCTCAGTA</td><td>92.4</td><td>-10.4</td><td>1.5</td><td>-6.8</td><td>1.0E-11</td><td>3.4E-11</td><td>MB231/MB468</td><td>MP</td></tr>
<tr><td>>hg38_21_+_8433172_8446622_RNA45SN1_45S_50ntFlanks@7977.7995.19|CGACCTCAGATCAGACGTG</td><td>>hg38_21_+_8433172_8446622_RNA45SN1_45S_50ntFlanks</td><td>7977</td><td>7995</td><td>19</td><td>CGACCTCAGATCAGACGTG</td><td>430.9</td><td>-8.2</td><td>1.2</td><td>-6.8</td><td>1.0E-11</td><td>3.4E-11</td><td>MB231/MB468</td><td>MP</td></tr>
<tr><td>>hg38_21_+_8433172_8446622_RNA45SN1_45S_50ntFlanks@11873.11897.25|AAGAAGACCCTGTTGAGCTTGACTC</td><td>>hg38_21_+_8433172_8446622_RNA45SN1_45S_50ntFlanks</td><td>11873</td><td>11897</td><td>25</td><td>AAGAAGACCCTGTTGAGCTTGACTC</td><td>593.3</td><td>-3.4</td><td>0.5</td><td>-6.8</td><td>1.1E-11</td><td>3.4E-11</td><td>BT20/MB468</td><td>CYTO</td></tr>
<tr><td>>MI0002470|hsa-mir-486-1&WithFlank&8|-|41660435|41660514@10.31.22(+1U);MI0023622|hsa-mir-486-2&WithFlank&8|+|41660438|41660513@7.28.22(+1U)&[MIMAT0002177&hsa-miR-486-5p&offsets|0|0(+1U);m-107&8|-|41660484|41660505&offsets|0|0(+1U)];[MIMAT0002177_1&hsa-miR-486-5p&offsets|0|0(+1U);TJU_CMC_MD2.ID03138.5p-miR&offsets|0|-2(+1U);m-106&8|+|41660444|41660465&offsets|0|0(+1U)]|TCCTGTACTGAGCTGCCCCGAGT</td><td>>MI0002470|hsa-mir-486-1&WithFlank&8|-|41660435|41660514</td><td>10</td><td>31</td><td>22(+1U)</td><td>TCCTGTACTGAGCTGCCCCGAGT</td><td>235.7</td><td>11.1</td><td>1.6</td><td>6.8</td><td>1.4E-11</td><td>3.4E-11</td><td>MB231/MB468</td><td>MITO</td></tr>
<tr><td>>hg38_21_+_8433172_8446622_RNA45SN1_45S_50ntFlanks@4722.4752.31|TCATTAATCAAGAACGAAAGTCGGAGGTTCG</td><td>>hg38_21_+_8433172_8446622_RNA45SN1_45S_50ntFlanks</td><td>4722</td><td>4752</td><td>31</td><td>TCATTAATCAAGAACGAAAGTCGGAGGTTCG</td><td>1152.2</td><td>-7.0</td><td>1.0</td><td>-6.8</td><td>1.1E-11</td><td>3.4E-11</td><td>MB231/MB468</td><td>CYTO</td></tr>
<tr><td>>hg38_21_+_8433172_8446622_RNA45SN1_45S_50ntFlanks@10683.10709.27|CCGGCGGCGTCCGGTGAGCTCTCGCTG</td><td>>hg38_21_+_8433172_8446622_RNA45SN1_45S_50ntFlanks</td><td>10683</td><td>10709</td><td>27</td><td>CCGGCGGCGTCCGGTGAGCTCTCGCTG</td><td>1597.9</td><td>-4.5</td><td>0.7</td><td>-6.8</td><td>8.4E-12</td><td>3.4E-11</td><td>BT20/MB468</td><td>MITO</td></tr>
<tr><td>>hg38_21_+_8433172_8446622_RNA45SN1_45S_50ntFlanks@9766.9789.24|TAAATGGGTAAGAAGCCCGGCTCG</td><td>>hg38_21_+_8433172_8446622_RNA45SN1_45S_50ntFlanks</td><td>9766</td><td>9789</td><td>24</td><td>TAAATGGGTAAGAAGCCCGGCTCG</td><td>130.4</td><td>-10.9</td><td>1.6</td><td>-6.8</td><td>1.0E-11</td><td>3.4E-11</td><td>MB231/MB468</td><td>MP</td></tr>
<tr><td>>hg38_21_+_8433172_8446622_RNA45SN1_45S_50ntFlanks@3929.3959.31|GATCAAAACCAACCCGGTCAGCCCCTCTCCG</td><td>>hg38_21_+_8433172_8446622_RNA45SN1_45S_50ntFlanks</td><td>3929</td><td>3959</td><td>31</td><td>GATCAAAACCAACCCGGTCAGCCCCTCTCCG</td><td>97.4</td><td>-8.9</td><td>1.3</td><td>-6.8</td><td>1.1E-11</td><td>3.4E-11</td><td>BT20/MB468</td><td>CYTO</td></tr>
<tr><td>>hg38_21_+_8433172_8446622_RNA45SN1_45S_50ntFlanks@8439.8471.33|GATTCAACCCGGCGGCGGGTCCGGCCGTGTCGG</td><td>>hg38_21_+_8433172_8446622_RNA45SN1_45S_50ntFlanks</td><td>8439</td><td>8471</td><td>33</td><td>GATTCAACCCGGCGGCGGGTCCGGCCGTGTCGG</td><td>271.0</td><td>-11.2</td><td>1.6</td><td>-7.2</td><td>6.7E-13</td><td>3.4E-11</td><td>BT20/MB468</td><td>MP</td></tr>
<tr><td>>hg38_21_+_8433172_8446622_RNA45SN1_45S_50ntFlanks@11572.11591.20|CCGACTTAGAACTGGTGCGG</td><td>>hg38_21_+_8433172_8446622_RNA45SN1_45S_50ntFlanks</td><td>11572</td><td>11591</td><td>20</td><td>CCGACTTAGAACTGGTGCGG</td><td>3701.3</td><td>-10.7</td><td>1.6</td><td>-6.8</td><td>1.4E-11</td><td>3.4E-11</td><td>BT20/MB468</td><td>NUC</td></tr>
<tr><td>>hg38_21_+_8433172_8446622_RNA45SN1_45S_50ntFlanks@5188.5218.31|ATAACAGGTCTGTGATGCCCTTAGATGTCCG</td><td>>hg38_21_+_8433172_8446622_RNA45SN1_45S_50ntFlanks</td><td>5188</td><td>5218</td><td>31</td><td>ATAACAGGTCTGTGATGCCCTTAGATGTCCG</td><td>534.4</td><td>-12.8</td><td>1.9</td><td>-6.8</td><td>1.4E-11</td><td>3.4E-11</td><td>MB231/MB468</td><td>MITO</td></tr>
<tr><td>>MI0000070|hsa-mir-16-1&WithFlank&13|-|50048967|50049067@20.40.21(+1A);MI0000115|hsa-mir-16-2&WithFlank&3|+|160404739|160404831@16.36.21(+1A)&[MIMAT0000069&hsa-miR-16-5p&offsets|0|-1(+1A);m-62&13|-|50049027|50049048&offsets|0|-1(+1A)];[MIMAT0000069_1&hsa-miR-16-5p&offsets|0|-1(+1A);m-61&3|+|160404754|160404775&offsets|0|-1(+1A)]|TAGCAGCACGTAAATATTGGCA</td><td>>MI0000070|hsa-mir-16-1&WithFlank&13|-|50048967|50049067</td><td>20</td><td>40</td><td>21(+1A)</td><td>TAGCAGCACGTAAATATTGGCA</td><td>385.1</td><td>5.1</td><td>0.8</td><td>6.8</td><td>1.2E-11</td><td>3.4E-11</td><td>MB231/MB468</td><td>NUC</td></tr>
<tr><td>>hg38_MT_+_1621_3279_RNR2_16S_50ntFlanks@714.740.27|CATAAGCCTGCGTCAGATTAAAACACT</td><td>>hg38_MT_+_1621_3279_RNR2_16S_50ntFlanks</td><td>714</td><td>740</td><td>27</td><td>CATAAGCCTGCGTCAGATTAAAACACT</td><td>140.1</td><td>10.4</td><td>1.5</td><td>6.8</td><td>1.4E-11</td><td>3.4E-11</td><td>MB231/MB468</td><td>MITO</td></tr>
<tr><td>>MI0000342|hsa-mir-200b&WithFlank&1|+|1167098|1167204@63.83.21(+1G)&[MIMAT0000318&hsa-miR-200b-3p&offsets|0|-1(+1G);m-93&1|+|1167160|1167182&offsets|0|-2(+1G)]|TAATACTGCCTGGTAATGATGG</td><td>>MI0000342|hsa-mir-200b&WithFlank&1|+|1167098|1167204</td><td>63</td><td>83</td><td>21(+1G)</td><td>TAATACTGCCTGGTAATGATGG</td><td>111.1</td><td>-10.7</td><td>1.6</td><td>-6.8</td><td>1.0E-11</td><td>3.4E-11</td><td>MB231/MB468</td><td>MP</td></tr>
<tr><td>>hg38_21_+_8433172_8446622_RNA45SN1_45S_50ntFlanks@4283.4310.28|CTTTAACGAGGATCCATTGGAGGGCAAG</td><td>>hg38_21_+_8433172_8446622_RNA45SN1_45S_50ntFlanks</td><td>4283</td><td>4310</td><td>28</td><td>CTTTAACGAGGATCCATTGGAGGGCAAG</td><td>109.2</td><td>-10.6</td><td>1.6</td><td>-6.8</td><td>1.0E-11</td><td>3.4E-11</td><td>MB231/MB468</td><td>MP</td></tr>
<tr><td>>hg38_21_+_8433172_8446622_RNA45SN1_45S_50ntFlanks@12626.12653.28|AATGGGGCGAAGCTACCATCTGTGGGAT</td><td>>hg38_21_+_8433172_8446622_RNA45SN1_45S_50ntFlanks</td><td>12626</td><td>12653</td><td>28</td><td>AATGGGGCGAAGCTACCATCTGTGGGAT</td><td>6157.1</td><td>-11.2</td><td>1.7</td><td>-6.8</td><td>1.5E-11</td><td>3.4E-11</td><td>BT20/MB468</td><td>NUC</td></tr>
<tr><td>>MI0000254|hsa-mir-30c-2&WithFlank&6|-|71376954|71377037@53.74.22&[MIMAT0004550&hsa-miR-30c-2-3p&offsets|0|0;m-206&6|-|71376964|71376985&offsets|0|0]|CTGGGAGAAGGCTGTTTACTCT</td><td>>MI0000254|hsa-mir-30c-2&WithFlank&6|-|71376954|71377037</td><td>53</td><td>74</td><td>22</td><td>CTGGGAGAAGGCTGTTTACTCT</td><td>801.9</td><td>6.3</td><td>0.9</td><td>6.8</td><td>1.4E-11</td><td>3.4E-11</td><td>MB231/MB468</td><td>MITO</td></tr>
<tr><td>>hg38_21_+_8433172_8446622_RNA45SN1_45S_50ntFlanks@9862.9902.41|ATGAACCGAACGCCGGGTTAAGGCGCCCGATGCCGACGCTC</td><td>>hg38_21_+_8433172_8446622_RNA45SN1_45S_50ntFlanks</td><td>9862</td><td>9902</td><td>41</td><td>ATGAACCGAACGCCGGGTTAAGGCGCCCGATGCCGACGCTC</td><td>279.0</td><td>-11.9</td><td>1.8</td><td>-6.8</td><td>1.4E-11</td><td>3.4E-11</td><td>MB231/MB468</td><td>MITO</td></tr>
</table>

| | | | | | | | | | | | | | |
|---|---|---|---|---|---|---|---|---|---|---|---|---|---|
| >hg38_21_+_8433172_8446622_RNA45SN1_45S_50ntFlanks@12578.12604.27\|AGGAACCGCAGGTTCAGACATTTGGTG | >hg38_21_+_8433172_8446622_RNA45SN1_45S_50ntFlanks | 12578 | 12604 | 27 | AGGAACCGCAGGTTCAGACATTTGGTG | 167.5 | -9.5 | 1.4 | -6.8 | 8.5E-12 | 3.4E-11 | BT20/MB468 | MITO |
| >trna10_LysCTT_16_+_3241501_3241573@9.35.27,trna13_LysCTT_14_-_58706613_58706685@9.35.27,trna2_LysCTT_15_+_79152904_79152976@9.35.27,trna4_LysCTT_18_-_43669271_43669343@9.35.27\|AGCTCAGTCGGTAGAGCATGGGACTCT | >trna10_LysCTT_16_+_3241501_3241573 | 9 | 35 | 27 | AGCTCAGTCGGTAGAGCATGGGACTCT | 257.6 | 11.2 | 1.7 | 6.8 | 1.4E-11 | 3.4E-11 | MB231/MB468 | MITO |
| >hg38_21_+_8433172_8446622_RNA45SN1_45S_50ntFlanks@10826.10844.19\|TCGGCAAGCCGGATCCGTA | >hg38_21_+_8433172_8446622_RNA45SN1_45S_50ntFlanks | 10826 | 10844 | 19 | TCGGCAAGCCGGATCCGTA | 705.0 | -8.9 | 1.3 | -6.8 | 1.2E-11 | 3.4E-11 | MB231/MB468 | CYTO |
| >hg38_21_+_8433172_8446622_RNA45SN1_45S_50ntFlanks@12456.12486.31\|CCAAGCGTTGGATTGTTCACCCACTAATAGG | >hg38_21_+_8433172_8446622_RNA45SN1_45S_50ntFlanks | 12456 | 12486 | 31 | CCAAGCGTTGGATTGTTCACCCACTAATAGG | 6315.7 | -7.1 | 1.0 | -6.8 | 1.2E-11 | 3.4E-11 | MB231/MB468 | CYTO |
| >hg38_21_+_8433172_8446622_RNA45SN1_45S_50ntFlanks@10919.10946.28\|CGGCTGGACGAGGCGCCGCCGCCCCCCC | >hg38_21_+_8433172_8446622_RNA45SN1_45S_50ntFlanks | 10919 | 10946 | 28 | CGGCTGGACGAGGCGCCGCCGCCCCCCC | 157.6 | -11.2 | 1.6 | -6.8 | 1.1E-11 | 3.4E-11 | MB231/MB468 | MP |
| >MI0000462\|hsa-mir-152&WithFlank&17\|-\|48037155\|48037253@60.78.19&[MIMAT0000438&hsa-miR-152-3p&offsets\|0\|-2;m-97&17\|-\|48037174\|48037194&offsets\|0\|-2]\|TCAGTGCATGACAGAACTT | >MI0000462\|hsa-mir-152&WithFlank&17\|-\|48037155\|48037253 | 60 | 78 | 19 | TCAGTGCATGACAGAACTT | 159.9 | 10.5 | 1.6 | 6.8 | 1.1E-11 | 3.4E-11 | MB231/MB468 | MP |
| >trnaMT_GluTTC_MT_-_14674_14742@45.72.28,trnalookalike8_GluTTC_5_-_93905172_93905240@45.72.28\|CGTGGTTGTAGTCCGTGCGAGAATACCA | >trnaMT_GluTTC_MT_-_14674_14742 | 45 | 72 | 28 | CGTGGTTGTAGTCCGTGCGAGAATACCA | 1395.0 | 5.4 | 0.8 | 6.8 | 8.5E-12 | 3.4E-11 | BT20/MB468 | MITO |
| >hg38_21_+_8433172_8446622_RNA45SN1_45S_50ntFlanks@8170.8208.39\|CAAGTCCTTCTGATCGAGGCCCAGCCCGTGGACGGTGTG | >hg38_21_+_8433172_8446622_RNA45SN1_45S_50ntFlanks | 8170 | 8208 | 39 | CAAGTCCTTCTGATCGAGGCCCAGCCCGTGGACGGTGTG | 204.1 | -11.5 | 1.7 | -6.8 | 1.1E-11 | 3.4E-11 | MB231/MB468 | MP |
| >hg38_21_+_8433172_8446622_RNA45SN1_45S_50ntFlanks@12628.12661.34\|TGGGGCGAAGCTACCATCTGTGGGATTATGACTG | >hg38_21_+_8433172_8446622_RNA45SN1_45S_50ntFlanks | 12628 | 12661 | 34 | TGGGGCGAAGCTACCATCTGTGGGATTATGACTG | 2259.8 | -10.5 | 1.6 | -6.8 | 1.5E-11 | 3.4E-11 | BT20/MB468 | NUC |
| >MI0000102\|hsa-mir-100&WithFlank&11\|-\|122152223\|122152314@19.37.19(+1U)&[MIMAT0000098&hsa-miR-100-5p&offsets\|0\|-3(+1U);m-27&11\|-\|122152276\|122152296&offsets\|0\|-2(+1U)]\|AACCCGTAGATCCGAACTTT | >MI0000102\|hsa-mir-100&WithFlank&11\|-\|122152223\|122152314 | 19 | 37 | 19(+1U) | AACCCGTAGATCCGAACTTT | 409.9 | 11.9 | 1.8 | 6.8 | 1.1E-11 | 3.4E-11 | MB231/MB468 | MP |
| >hg38_21_+_8433172_8446622_RNA45SN1_45S_50ntFlanks@12435.12473.39\|ATCATTGTGAAGCAGAATTCACCAAGCGTTGGATTGTTC | >hg38_21_+_8433172_8446622_RNA45SN1_45S_50ntFlanks | 12435 | 12473 | 39 | ATCATTGTGAAGCAGAATTCACCAAGCGTTGGATTGTTC | 233.9 | -10.2 | 1.5 | -6.8 | 1.1E-11 | 3.4E-11 | BT20/MB468 | CYTO |
| >hg38_21_+_8433172_8446622_RNA45SN1_45S_50ntFlanks@9381.9415.35\|CGCTCGCCGGCCGAGGTGGGATCCCGAGGCCTCTC | >hg38_21_+_8433172_8446622_RNA45SN1_45S_50ntFlanks | 9381 | 9415 | 35 | CGCTCGCCGGCCGAGGTGGGATCCCGAGGCCTCTC | 93.4 | -10.4 | 1.5 | -6.8 | 1.1E-11 | 3.5E-11 | MB231/MB468 | MP |
| >MI0000088\|hsa-mir-30a&WithFlank&6\|-\|71403545\|71403627@12.31.20&[MIMAT0000087&hsa-miR-30a-5p&offsets\|0\|-2;m-15&6\|-\|71403595\|71403616&offsets\|0\|-2]\|TGTAAACATCCTCGACTGGA | >MI0000088\|hsa-mir-30a&WithFlank&6\|-\|71403545\|71403627 | 12 | 31 | 20 | TGTAAACATCCTCGACTGGA | 10299.6 | 4.7 | 0.7 | 6.8 | 1.1E-11 | 3.5E-11 | MB231/MB468 | MP |
| >MI0000737\|hsa-mir-200a&WithFlank&1\|+\|1167857\|1167958@60.81.22(+1G)&[MIMAT0000682&hsa-miR-200a-3p&offsets\|0\|0(+1G);m-142&1\|+\|1167916\|1167938&offsets\|0\|-1(+1G)]\|TAACACTGTCTGGTAACGATGTG | >MI0000737\|hsa-mir-200a&WithFlank&1\|+\|1167857\|1167958 | 60 | 81 | 22(+1G) | TAACACTGTCTGGTAACGATGTG | 180.6 | -11.4 | 1.7 | -6.8 | 1.1E-11 | 3.5E-11 | MB231/MB468 | MP |
| >hg38_21_+_8433172_8446622_RNA45SN1_45S_50ntFlanks@12630.12659.30\|GGGCGAAGCTACCATCTGTGGGATTATGAC | >hg38_21_+_8433172_8446622_RNA45SN1_45S_50ntFlanks | 12630 | 12659 | 30 | GGGCGAAGCTACCATCTGTGGGATTATGAC | 261.5 | -11.9 | 1.8 | -6.8 | 1.1E-11 | 3.5E-11 | MB231/MB468 | MP |
| >hg38_21_+_8433172_8446622_RNA45SN1_45S_50ntFlanks@4380.4405.26\|CGTAGTTGGATCTTGGGAGCGGGCGG | >hg38_21_+_8433172_8446622_RNA45SN1_45S_50ntFlanks | 4380 | 4405 | 26 | CGTAGTTGGATCTTGGGAGCGGGCGG | 115.4 | -10.7 | 1.6 | -6.8 | 1.1E-11 | 3.5E-11 | MB231/MB468 | MP |
| >MI0000078\|hsa-mir-22&WithFlank&17\|-\|1713897\|1713993@59.80.22&[MIMAT0000077&hsa-miR-22-3p&offsets\|0\|0;m-2&17\|-\|1713914\|1713935&offsets\|0\|0]\|AAGCTGCCAGTTGAAGAACTGT | >MI0000078\|hsa-mir-22&WithFlank&17\|-\|1713897\|1713993 | 59 | 80 | 22 | AAGCTGCCAGTTGAAGAACTGT | 3626.5 | 6.3 | 0.9 | 6.8 | 1.2E-11 | 3.5E-11 | MB231/MB468 | CYTO |
| >hg38_21_+_8433172_8446622_RNA45SN1_45S_50ntFlanks@12654.12674.21\|TATGACTGAACGCCTCTAAGT | >hg38_21_+_8433172_8446622_RNA45SN1_45S_50ntFlanks | 12654 | 12674 | 21 | TATGACTGAACGCCTCTAAGT | 2320.7 | -7.5 | 1.1 | -6.8 | 1.2E-11 | 3.5E-11 | MB231/MB468 | NUC |
| >hg38_21_+_8433172_8446622_RNA45SN1_45S_50ntFlanks@4726.4755.30\|TAATCAAGAACGAAAGTCGGAGGTTCGAAG | >hg38_21_+_8433172_8446622_RNA45SN1_45S_50ntFlanks | 4726 | 4755 | 30 | TAATCAAGAACGAAAGTCGGAGGTTCGAAG | 105.9 | -10.6 | 1.6 | -6.8 | 1.1E-11 | 3.5E-11 | MB231/MB468 | MP |
| >hg38_21_+_8433172_8446622_RNA45SN1_45S_50ntFlanks@8552.8571.20\|CTCCCCGGAGGGGGCGGGCT | >hg38_21_+_8433172_8446622_RNA45SN1_45S_50ntFlanks | 8552 | 8571 | 20 | CTCCCCGGAGGGGGCGGGCT | 85.4 | -10.3 | 1.5 | -6.8 | 1.1E-11 | 3.5E-11 | MB231/MB468 | MP |
| >hg38_21_+_8433172_8446622_RNA45SN1_45S_50ntFlanks@9602.9621.20\|CGAAAGACTAATCGAACCAT | >hg38_21_+_8433172_8446622_RNA45SN1_45S_50ntFlanks | 9602 | 9621 | 20 | CGAAAGACTAATCGAACCAT | 106.2 | -10.6 | 1.6 | -6.8 | 1.1E-11 | 3.5E-11 | MB231/MB468 | MP |
| >hg38_21_+_8433172_8446622_RNA45SN1_45S_50ntFlanks@8170.8218.49\|CAAGTCCTTCTGATCGAGGCCCAGCCCGTGGACGGTGTGAGGCCGGTAG | >hg38_21_+_8433172_8446622_RNA45SN1_45S_50ntFlanks | 8170 | 8218 | 49 | CAAGTCCTTCTGATCGAGGCCCAGCCCGTGGACGGTGTGAGGCCGGTAG | 229.8 | -11.6 | 1.7 | -6.8 | 1.4E-11 | 3.5E-11 | MB231/MB468 | MITO |
| >hg38_21_+_8433172_8446622_RNA45SN1_45S_50ntFlanks@4006.4042.37\|ACCTCGGGCCGATCGCACGCCCCCCGTGGCGGCGACG | >hg38_21_+_8433172_8446622_RNA45SN1_45S_50ntFlanks | 4006 | 4042 | 37 | ACCTCGGGCCGATCGCACGCCCCCCGTGGCGGCGACG | 1443.3 | -10.0 | 1.5 | -6.7 | 1.5E-11 | 3.5E-11 | BT20/MB468 | NUC |
| >hg38_21_+_8433172_8446622_RNA45SN1_45S_50ntFlanks@10343.10373.31\|TATTCAAACGAGAACTTTGAAGGCCGAAGTG | >hg38_21_+_8433172_8446622_RNA45SN1_45S_50ntFlanks | 10343 | 10373 | 31 | TATTCAAACGAGAACTTTGAAGGCCGAAGTG | 420.3 | 5.3 | 0.8 | 6.8 | 1.2E-11 | 3.5E-11 | MB231/MB468 | CYTO |
| >hg38_21_+_8433172_8446622_RNA45SN1_45S_50ntFlanks@12934.12968.35\|CTAAACCATTCGTAGACGACCTGCTTCTGGGTCGG | >hg38_21_+_8433172_8446622_RNA45SN1_45S_50ntFlanks | 12934 | 12968 | 35 | CTAAACCATTCGTAGACGACCTGCTTCTGGGTCGG | 139.6 | -9.4 | 1.4 | -6.8 | 1.1E-11 | 3.5E-11 | BT20/MB468 | CYTO |
| >hg38_21_+_8433172_8446622_RNA45SN1_45S_50ntFlanks@476.496.21\|TTCGTGATCGATGTGGTGACG | >hg38_21_+_8433172_8446622_RNA45SN1_45S_50ntFlanks | 476 | 496 | 21 | TTCGTGATCGATGTGGTGACG | 5294.6 | 6.2 | 0.9 | 6.8 | 1.2E-11 | 3.5E-11 | MB231/MB468 | NUC |
| >am_MI0003581\|hsa-mir-574&WithFlank&4\|+\|38868026\|38868133@31.53.23&[MIMAT0004795&hsa-miR-574-5p&offsets\|0\|0;m-213&4\|+\|38868056\|38868078&offsets\|0\|0]\|TGAGTGTGTGTGTGTGAGTGTGT | >am_MI0003581\|hsa-mir-574&WithFlank&4\|+\|38868026\|38868133 | 31 | 53 | 23 | TGAGTGTGTGTGTGTGAGTGTGT | 1343.8 | 9.5 | 1.4 | 6.8 | 1.1E-11 | 3.5E-11 | BT20/MB468 | CYTO |
| >hg38_21_+_8433172_8446622_RNA45SN1_45S_50ntFlanks@6736.6770.35\|TGATCATCGACACTTCGAACGCACTTGCGGCCCCG | >hg38_21_+_8433172_8446622_RNA45SN1_45S_50ntFlanks | 6736 | 6770 | 35 | TGATCATCGACACTTCGAACGCACTTGCGGCCCCG | 337.2 | -8.2 | 1.2 | -6.8 | 1.4E-11 | 3.5E-11 | MB231/MB468 | MITO |
| >hg38_21_+_8433172_8446622_RNA45SN1_45S_50ntFlanks@3831.3860.30\|CTCGCTCCTCTCCTACTTGGATAACTGTGG | >hg38_21_+_8433172_8446622_RNA45SN1_45S_50ntFlanks | 3831 | 3860 | 30 | CTCGCTCCTCTCCTACTTGGATAACTGTGG | 726.4 | -7.0 | 1.0 | -6.8 | 1.4E-11 | 3.5E-11 | MB231/MB468 | MITO |
| >hg38_21_+_8433172_8446622_RNA45SN1_45S_50ntFlanks@10686.10711.26\|GCGGCGTCCGGTGAGCTCTCGCTGGC | >hg38_21_+_8433172_8446622_RNA45SN1_45S_50ntFlanks | 10686 | 10711 | 26 | GCGGCGTCCGGTGAGCTCTCGCTGGC | 680.8 | -4.4 | 0.6 | -6.8 | 1.2E-11 | 3.5E-11 | MB231/MB468 | CYTO |
| >hg38_21_+_8433172_8446622_RNA45SN1_45S_50ntFlanks@12991.13008.18\|CTCCCTCGCTGCGATCTA | >hg38_21_+_8433172_8446622_RNA45SN1_45S_50ntFlanks | 12991 | 13008 | 18 | CTCCCTCGCTGCGATCTA | 171.0 | -9.5 | 1.4 | -6.8 | 8.8E-12 | 3.5E-11 | BT20/MB468 | MITO |
| >hg38_21_+_8433172_8446622_RNA45SN1_45S_50ntFlanks@7973.7993.21\|GACGCGACCTCAGATCAGACG | >hg38_21_+_8433172_8446622_RNA45SN1_45S_50ntFlanks | 7973 | 7993 | 21 | GACGCGACCTCAGATCAGACG | 5055.5 | 5.3 | 0.8 | 6.8 | 8.8E-12 | 3.5E-11 | BT20/MB468 | MITO |
| >hg38_21_+_8433172_8446622_RNA45SN1_45S_50ntFlanks@7974.7993.20\|ACGCGACCTCAGATCAGACG | >hg38_21_+_8433172_8446622_RNA45SN1_45S_50ntFlanks | 7974 | 7993 | 20 | ACGCGACCTCAGATCAGACG | 27257.5 | 4.9 | 0.7 | 6.8 | 8.8E-12 | 3.5E-11 | BT20/MB468 | MITO |
| >hg38_1_-_228634819_228635039_RNA5S12_5S_50ntFlanks@84.101.18\|CTCGTCTGATCTCGGAAG | >hg38_1_-_228634819_228635039_RNA5S12_5S_50ntFlanks | 84 | 101 | 18 | CTCGTCTGATCTCGGAAG | 1385.3 | -5.6 | 0.8 | -6.8 | 1.2E-11 | 3.5E-11 | MB231/MB468 | NUC |
| >am_trna2_LeuTAA_X_-_55207755_55207829@1.18.18,trna4_LeuTAA_20_-_55933428_55933502@1.18.18,trna60_LeuTAA_1_-_238106956_238107030@1.18.18,trna9_LeuTAA_2_+_117781417_117781491@1.18.18,trnaMT_LeuTAA_MT_+_3230_3304@1.18.18\|GTTAAGATGGCAGAGCCC | >am_trna2_LeuTAA_X_-_55207755_55207829 | 1 | 18 | 18 | GTTAAGATGGCAGAGCCC | 71.7 | 4.4 | 0.7 | 6.7 | 1.5E-11 | 3.5E-11 | BT20/MB468 | NUC |
| >hg38_21_+_8433172_8446622_RNA45SN1_45S_50ntFlanks@10921.10939.19\|GCTGGACGAGGCGCCGCCG | >hg38_21_+_8433172_8446622_RNA45SN1_45S_50ntFlanks | 10921 | 10939 | 19 | GCTGGACGAGGCGCCGCCG | 231.6 | -7.3 | 1.1 | -6.8 | 1.2E-11 | 3.5E-11 | MB231/MB468 | CYTO |
| >hg38_21_+_8433172_8446622_RNA45SN1_45S_50ntFlanks@9381.9406.26\|CGCTCGCCGGCCGAGGTGGGATCCCG | >hg38_21_+_8433172_8446622_RNA45SN1_45S_50ntFlanks | 9381 | 9406 | 26 | CGCTCGCCGGCCGAGGTGGGATCCCG | 367.4 | -12.3 | 1.8 | -6.8 | 1.4E-11 | 3.5E-11 | MB231/MB468 | MITO |
| >hg38_21_+_8433172_8446622_RNA45SN1_45S_50ntFlanks@10716.10733.18\|GAAAATCCGGGGGAGAGG | >hg38_21_+_8433172_8446622_RNA45SN1_45S_50ntFlanks | 10716 | 10733 | 18 | GAAAATCCGGGGGAGAGG | 145.1 | -11.0 | 1.6 | -6.8 | 1.1E-11 | 3.5E-11 | MB231/MB468 | MP |
| >hg38_21_+_8433172_8446622_RNA45SN1_45S_50ntFlanks@12499.12521.23\|GGTTTAGACCGTCGTGAGACAGG | >hg38_21_+_8433172_8446622_RNA45SN1_45S_50ntFlanks | 12499 | 12521 | 23 | GGTTTAGACCGTCGTGAGACAGG | 208.9 | -11.6 | 1.7 | -6.8 | 1.1E-11 | 3.5E-11 | MB231/MB468 | MP |
| >hg38_21_+_8433172_8446622_RNA45SN1_45S_50ntFlanks@10352.10382.31\|GAGAACTTTGAAGGCCGAAGTGGAGAAGGGT | >hg38_21_+_8433172_8446622_RNA45SN1_45S_50ntFlanks | 10352 | 10382 | 31 | GAGAACTTTGAAGGCCGAAGTGGAGAAGGGT | 210.3 | 4.0 | 0.6 | 6.8 | 1.2E-11 | 3.5E-11 | MB231/MB468 | CYTO |
| >hg38_21_+_8433172_8446622_RNA45SN1_45S_50ntFlanks@10350.10377.28\|ACGAGAACTTTGAAGGCCGAAGTGGAGA | >hg38_21_+_8433172_8446622_RNA45SN1_45S_50ntFlanks | 10350 | 10377 | 28 | ACGAGAACTTTGAAGGCCGAAGTGGAGA | 1508.2 | -3.6 | 0.5 | -6.8 | 1.2E-11 | 3.5E-11 | MB231/MB468 | CYTO |
| >trnaMT_SerGCT_MT_+_12207_12265@23.62.40\|AACTCATGCCCCCATGTCTAACAACATGGCTTTCTCACCA | >trnaMT_SerGCT_MT_+_12207_12265 | 23 | 62 | 40 | AACTCATGCCCCCATGTCTAACAACATGGCTTTCTCACCA | 940.7 | 6.9 | 1.0 | 6.8 | 8.9E-12 | 3.5E-11 | BT20/MB468 | MITO |
| >MI0000446\|hsa-mir-125b-1&WithFlank&11\|-\|122099751\|122099850@61.82.22&[MIMAT0004592&hsa-miR-125b-1-3p&offsets\|0\|0;m-485&11\|-\|122099769\|122099790&offsets\|0\|0]\|ACGGGTTAGGCTCTTGGGAGCT | >MI0000446\|hsa-mir-125b-1&WithFlank&11\|-\|122099751\|122099850 | 61 | 82 | 22 | ACGGGTTAGGCTCTTGGGAGCT | 921.7 | 10.0 | 1.5 | 6.8 | 1.1E-11 | 3.5E-11 | MB231/MB468 | MP |
| >hg38_21_+_8433172_8446622_RNA45SN1_45S_50ntFlanks@11921.11952.32\|TGAGAGGTGTAGAATAAGTGGGAGGCCCCCGG | >hg38_21_+_8433172_8446622_RNA45SN1_45S_50ntFlanks | 11921 | 11952 | 32 | TGAGAGGTGTAGAATAAGTGGGAGGCCCCCGG | 96.8 | -10.5 | 1.5 | -6.8 | 1.1E-11 | 3.6E-11 | MB231/MB468 | MP |
| >hg38_21_+_8433172_8446622_RNA45SN1_45S_50ntFlanks@10534.10569.36\|CCGCGAGGCGTCCAGTGCGGTAACGCGACCGATCCC | >hg38_21_+_8433172_8446622_RNA45SN1_45S_50ntFlanks | 10534 | 10569 | 36 | CCGCGAGGCGTCCAGTGCGGTAACGCGACCGATCCC | 190.0 | -9.7 | 1.4 | -6.8 | 8.9E-12 | 3.6E-11 | BT20/MB468 | MITO |
| >hg38_21_+_8433172_8446622_RNA45SN1_45S_50ntFlanks@10468.10497.30\|TTGCCCTCGGCCGATCGAAAGGGAGTCGGG | >hg38_21_+_8433172_8446622_RNA45SN1_45S_50ntFlanks | 10468 | 10497 | 30 | TTGCCCTCGGCCGATCGAAAGGGAGTCGGG | 530.2 | -9.0 | 1.3 | -6.7 | 1.5E-11 | 3.6E-11 | BT20/MB468 | NUC |
| >hg38_21_+_8433172_8446622_RNA45SN1_45S_50ntFlanks@11784.11805.22\|ATGAATGGATGAACGAGATTCC | >hg38_21_+_8433172_8446622_RNA45SN1_45S_50ntFlanks | 11784 | 11805 | 22 | ATGAATGGATGAACGAGATTCC | 880.0 | -7.1 | 1.1 | -6.8 | 1.1E-11 | 3.6E-11 | MB231/MB468 | MP |
| >hg38_21_+_8433172_8446622_RNA45SN1_45S_50ntFlanks@10765.10785.21\|CCGCAGCAGGTCTCCAAGGTG | >hg38_21_+_8433172_8446622_RNA45SN1_45S_50ntFlanks | 10765 | 10785 | 21 | CCGCAGCAGGTCTCCAAGGTG | 108.7 | -10.6 | 1.6 | -6.8 | 1.1E-11 | 3.6E-11 | MB231/MB468 | MP |
| >hg38_21_+_8433172_8446622_RNA45SN1_45S_50ntFlanks@6751.6772.22\|CGAACGCACTTGCGGCCCCGGG | >hg38_21_+_8433172_8446622_RNA45SN1_45S_50ntFlanks | 6751 | 6772 | 22 | CGAACGCACTTGCGGCCCCGGG | 113.6 | -10.7 | 1.6 | -6.8 | 1.1E-11 | 3.6E-11 | MB231/MB468 | MP |
| >hg38_21_+_8433172_8446622_RNA45SN1_45S_50ntFlanks@10798.10822.25\|CATGTTGGAACAATGTAGGTAAGGG | >hg38_21_+_8433172_8446622_RNA45SN1_45S_50ntFlanks | 10798 | 10822 | 25 | CATGTTGGAACAATGTAGGTAAGGG | 762.7 | -8.7 | 1.3 | -6.8 | 1.1E-11 | 3.6E-11 | MB231/MB468 | MP |

| | | | | | | | | | | | | | |
|---|---|---|---|---|---|---|---|---|---|---|---|---|---|
| >hg38_21_+_8433172_8446622_RNA45SN1_45S_50ntFlanks@11190.11207.18\|CCCGGGGGGCCGGCGGCG | >hg38_21_+_8433172_8446622_RNA45SN1_45S_50ntFlanks | 11190 | 11207 | 18 | CCCGGGGGGCCGGCGGCG | 524.7 | -7.6 | 1.1 | -6.8 | 1.2E-11 | 3.6E-11 | MB231/MB468 | CYTO |
| >hg38_21_+_8433172_8446622_RNA45SN1_45S_50ntFlanks@4439.4471.33\|CCGCCCCTTGCCTCTCGGCGCCCCCTCGATGCT | >hg38_21_+_8433172_8446622_RNA45SN1_45S_50ntFlanks | 4439 | 4471 | 33 | CCGCCCCTTGCCTCTCGGCGCCCCCTCGATGCT | 153.0 | -6.3 | 0.9 | -6.8 | 1.2E-11 | 3.6E-11 | MB231/MB468 | CYTO |
| >trnaMT_PheGAA_MT_+_577_647@35.56.22\|GAAAATGTTTAGACGGGCTCAC | >trnaMT_PheGAA_MT_+_577_647 | 35 | 56 | 22 | GAAAATGTTTAGACGGGCTCAC | 255.7 | 9.3 | 1.4 | 6.8 | 1.2E-11 | 3.6E-11 | MB231/MB468 | NUC |
| >hg38_21_+_8433172_8446622_RNA45SN1_45S_50ntFlanks@6731.6754.24\|CACATTGATCATCGACACTTCGAA | >hg38_21_+_8433172_8446622_RNA45SN1_45S_50ntFlanks | 6731 | 6754 | 24 | CACATTGATCATCGACACTTCGAA | 99.8 | -10.5 | 1.5 | -6.8 | 1.1E-11 | 3.6E-11 | MB231/MB468 | MP |
| >hg38_21_+_8433172_8446622_RNA45SN1_45S_50ntFlanks@12053.12071.19\|GTGAGGCGGGGGGGCGAGC | >hg38_21_+_8433172_8446622_RNA45SN1_45S_50ntFlanks | 12053 | 12071 | 19 | GTGAGGCGGGGGGGCGAGC | 116.2 | -10.7 | 1.6 | -6.8 | 1.1E-11 | 3.6E-11 | MB231/MB468 | MP |
| >hg38_21_+_8433172_8446622_RNA45SN1_45S_50ntFlanks@10782.10801.20\|GGTGAACAGCCTCTGGCATG | >hg38_21_+_8433172_8446622_RNA45SN1_45S_50ntFlanks | 10782 | 10801 | 20 | GGTGAACAGCCTCTGGCATG | 1092.5 | -9.3 | 1.4 | -6.7 | 1.5E-11 | 3.6E-11 | BT20/MB468 | NUC |
| >hg38_21_+_8433172_8446622_RNA45SN1_45S_50ntFlanks@10921.10941.21\|GCTGGACGAGGCGCCGCCGCC | >hg38_21_+_8433172_8446622_RNA45SN1_45S_50ntFlanks | 10921 | 10941 | 21 | GCTGGACGAGGCGCCGCCGCC | 141.7 | -7.4 | 1.1 | -6.8 | 1.2E-11 | 3.6E-11 | MB231/MB468 | CYTO |
| >hg38_21_+_8433172_8446622_RNA45SN1_45S_50ntFlanks@9574.9592.19\|AATCGGTCGTCCGACCTGG | >hg38_21_+_8433172_8446622_RNA45SN1_45S_50ntFlanks | 9574 | 9592 | 19 | AATCGGTCGTCCGACCTGG | 1492.7 | -10.0 | 1.5 | -6.8 | 1.1E-11 | 3.6E-11 | MB231/MB468 | MP |
| >MI0000067\|hsa-let-7f-1&WithFlank&9\|+\|94176341\|94176439@13.30.18;MI0000068\|hsa-let-7f-2&WithFlank&X\|-\|53557186\|53557280@14.31.18&[MIMAT0000067&hsa-let-7f-5p&offsets\|0\|-4;m-20&9\|+\|94176353\|94176374&offsets\|0\|-4];[MIMAT0000067_1&hsa-let-7f-5p&offsets\|0\|-4;m-19&X\|-\|53557246\|53557267&offsets\|0\|-4]\|TGAGGTAGTAGATTGTAT | >MI0000067\|hsa-let-7f-1&WithFlank&9\|+\|94176341\|94176439 | 13 | 30 | 18 | TGAGGTAGTAGATTGTAT | 1997.6 | 5.1 | 0.8 | 6.8 | 1.2E-11 | 3.6E-11 | MB231/MB468 | CYTO |
| >hg38_21_+_8433172_8446622_RNA45SN1_45S_50ntFlanks@3843.3871.29\|CTACTTGGATAACTGTGGTAATTCTAGAG | >hg38_21_+_8433172_8446622_RNA45SN1_45S_50ntFlanks | 3843 | 3871 | 29 | CTACTTGGATAACTGTGGTAATTCTAGAG | 594.1 | -8.6 | 1.3 | -6.8 | 1.1E-11 | 3.6E-11 | MB231/MB468 | MP |
| >hg38_21_+_8433172_8446622_RNA45SN1_45S_50ntFlanks@5485.5502.18\|AGCGCTGAGAAGACGGTC | >hg38_21_+_8433172_8446622_RNA45SN1_45S_50ntFlanks | 5485 | 5502 | 18 | AGCGCTGAGAAGACGGTC | 163.1 | -11.2 | 1.7 | -6.8 | 1.1E-11 | 3.6E-11 | MB231/MB468 | MP |
| >hg38_21_+_8433172_8446622_RNA45SN1_45S_50ntFlanks@10355.10388.34\|AACTTTGAAGGCCGAAGTGGAGAAGGGTTCCATG | >hg38_21_+_8433172_8446622_RNA45SN1_45S_50ntFlanks | 10355 | 10388 | 34 | AACTTTGAAGGCCGAAGTGGAGAAGGGTTCCATG | 755.6 | -7.5 | 1.1 | -6.8 | 1.2E-11 | 3.6E-11 | MB231/MB468 | CYTO |
| >hg38_21_+_8433172_8446622_RNA45SN1_45S_50ntFlanks@4722.4755.34\|TCATTAATCAAGAACGAAAGTCGGAGGTTCGAAG | >hg38_21_+_8433172_8446622_RNA45SN1_45S_50ntFlanks | 4722 | 4755 | 34 | TCATTAATCAAGAACGAAAGTCGGAGGTTCGAAG | 180.5 | -9.6 | 1.4 | -6.8 | 9.0E-12 | 3.6E-11 | BT20/MB468 | MITO |
| >hg38_21_+_8433172_8446622_RNA45SN1_45S_50ntFlanks@10095.10115.21\|CGCGCGCGCGCGCGTGTGGTG | >hg38_21_+_8433172_8446622_RNA45SN1_45S_50ntFlanks | 10095 | 10115 | 21 | CGCGCGCGCGCGCGTGTGGTG | 112.8 | -10.7 | 1.6 | -6.8 | 1.1E-11 | 3.6E-11 | MB231/MB468 | MP |
| >hg38_21_+_8433172_8446622_RNA45SN1_45S_50ntFlanks@4046.4067.22\|CATTCGAACGTCTGCCCTATCA | >hg38_21_+_8433172_8446622_RNA45SN1_45S_50ntFlanks | 4046 | 4067 | 22 | CATTCGAACGTCTGCCCTATCA | 614.8 | -7.5 | 1.1 | -6.8 | 1.5E-11 | 3.6E-11 | MB231/MB468 | MITO |
| >hg38_21_+_8433172_8446622_RNA45SN1_45S_50ntFlanks@12249.12269.21\|CAAAAGCTCGCTTGATCTTGA | >hg38_21_+_8433172_8446622_RNA45SN1_45S_50ntFlanks | 12249 | 12269 | 21 | CAAAAGCTCGCTTGATCTTGA | 121.5 | -10.8 | 1.6 | -6.8 | 1.1E-11 | 3.6E-11 | MB231/MB468 | MP |
| >hg38_21_+_8433172_8446622_RNA45SN1_45S_50ntFlanks@11272.11295.24\|GCCGCCCCCGGGGAGCCCGGCGGG | >hg38_21_+_8433172_8446622_RNA45SN1_45S_50ntFlanks | 11272 | 11295 | 24 | GCCGCCCCCGGGGAGCCCGGCGGG | 159.6 | -11.2 | 1.6 | -6.8 | 1.1E-11 | 3.6E-11 | MB231/MB468 | MP |
| >hg38_21_+_8433172_8446622_RNA45SN1_45S_50ntFlanks@476.499.24\|TTCGTGATCGATGTGGTGACGTCG | >hg38_21_+_8433172_8446622_RNA45SN1_45S_50ntFlanks | 476 | 499 | 24 | TTCGTGATCGATGTGGTGACGTCG | 2891.9 | 7.1 | 1.0 | 6.8 | 9.0E-12 | 3.6E-11 | BT20/MB468 | MITO |
| >hg38_21_+_8433172_8446622_RNA45SN1_45S_50ntFlanks@4679.4707.29\|GGCGCAAGACGGACCAGAGCGAAAGCATT | >hg38_21_+_8433172_8446622_RNA45SN1_45S_50ntFlanks | 4679 | 4707 | 29 | GGCGCAAGACGGACCAGAGCGAAAGCATT | 368.1 | 8.3 | 1.2 | 6.8 | 1.2E-11 | 3.6E-11 | MB231/MB468 | CYTO |
| >hg38_21_+_8433172_8446622_RNA45SN1_45S_50ntFlanks@11235.11252.18\|CTTCCCGTGGATCGCCCC | >hg38_21_+_8433172_8446622_RNA45SN1_45S_50ntFlanks | 11235 | 11252 | 18 | CTTCCCGTGGATCGCCCC | 2329.4 | -7.1 | 1.0 | -6.8 | 1.2E-11 | 3.6E-11 | MB231/MB468 | CYTO |
| >MI0000456\|hsa-mir-140&WithFlank&16\|+\|69933075\|69933186@68.90.23&[MIMAT0004597&hsa-miR-140-3p&offsets\|0\|+2;m-45&16\|+\|69933143\|69933164&offsets\|-1\|0]\|TACCACAGGGTAGAACCACGGAC | >MI0000456\|hsa-mir-140&WithFlank&16\|+\|69933075\|69933186 | 68 | 90 | 23 | TACCACAGGGTAGAACCACGGAC | 99.9 | 5.5 | 0.8 | 6.7 | 1.6E-11 | 3.6E-11 | BT20/MB468 | NUC |
| >hg38_21_+_8433172_8446622_RNA45SN1_45S_50ntFlanks@6722.6743.22\|ATTGCAGGACACATTGATCATC | >hg38_21_+_8433172_8446622_RNA45SN1_45S_50ntFlanks | 6722 | 6743 | 22 | ATTGCAGGACACATTGATCATC | 2806.4 | -8.6 | 1.3 | -6.7 | 1.6E-11 | 3.6E-11 | BT20/MB468 | NUC |
| >hg38_21_+_8433172_8446622_RNA45SN1_45S_50ntFlanks@12605.12640.36\|TATGTGCTTGGCTGAGGAGCCAATGGGGCGAAGCTA | >hg38_21_+_8433172_8446622_RNA45SN1_45S_50ntFlanks | 12605 | 12640 | 36 | TATGTGCTTGGCTGAGGAGCCAATGGGGCGAAGCTA | 148.4 | -10.2 | 1.5 | -6.8 | 1.3E-11 | 3.6E-11 | MB231/MB468 | NUC |
| >hg38_21_+_8433172_8446622_RNA45SN1_45S_50ntFlanks@12412.12433.22\|ATCCTTCGATGTCGGCTCTTCC | >hg38_21_+_8433172_8446622_RNA45SN1_45S_50ntFlanks | 12412 | 12433 | 22 | ATCCTTCGATGTCGGCTCTTCC | 239.0 | -10.2 | 1.5 | -6.8 | 1.1E-11 | 3.6E-11 | BT20/MB468 | CYTO |
| >MI0000825\|hsa-mir-345&WithFlank&14\|+\|100307853\|100307962@24.45.22(+1U)&[MIMAT0000772&hsa-miR-345-5p&offsets\|0\|0(+1U);m-193&14\|+\|100307876\|100307896&offsets\|0\|+1(+1U)]\|GCTGACTCCTAGTCCAGGGCTCT | >MI0000825\|hsa-mir-345&WithFlank&14\|+\|100307853\|100307962 | 24 | 45 | 22(+1U) | GCTGACTCCTAGTCCAGGGCTCT | 328.1 | 11.6 | 1.7 | 6.8 | 1.5E-11 | 3.6E-11 | MB231/MB468 | MITO |
| >hg38_21_+_8433172_8446622_RNA45SN1_45S_50ntFlanks@6664.6681.18\|TGGATCACTCGGCTCGTG | >hg38_21_+_8433172_8446622_RNA45SN1_45S_50ntFlanks | 6664 | 6681 | 18 | TGGATCACTCGGCTCGTG | 3340.8 | -7.5 | 1.1 | -6.8 | 1.5E-11 | 3.6E-11 | MB231/MB468 | MITO |
| >hg38_21_+_8433172_8446622_RNA45SN1_45S_50ntFlanks@4661.4684.24\|AGGTGAAATTCTTGGACCGGCGCA | >hg38_21_+_8433172_8446622_RNA45SN1_45S_50ntFlanks | 4661 | 4684 | 24 | AGGTGAAATTCTTGGACCGGCGCA | 140.4 | -11.0 | 1.6 | -6.8 | 1.1E-11 | 3.6E-11 | MB231/MB468 | MP |
| >hg38_21_+_8433172_8446622_RNA45SN1_45S_50ntFlanks@10387.10410.24\|TGTGAACAGCAGTTGAACATGGGT | >hg38_21_+_8433172_8446622_RNA45SN1_45S_50ntFlanks | 10387 | 10410 | 24 | TGTGAACAGCAGTTGAACATGGGT | 537.2 | -6.5 | 1.0 | -6.8 | 1.3E-11 | 3.6E-11 | MB231/MB468 | NUC |
| >hg38_21_+_8433172_8446622_RNA45SN1_45S_50ntFlanks@11561.11592.32\|GCGCCTAGCAGCCGACTTAGAACTGGTGCGGA | >hg38_21_+_8433172_8446622_RNA45SN1_45S_50ntFlanks | 11561 | 11592 | 32 | GCGCCTAGCAGCCGACTTAGAACTGGTGCGGA | 1271.0 | -4.4 | 0.6 | -6.8 | 9.1E-12 | 3.6E-11 | BT20/MB468 | MITO |
| >hg38_21_+_8433172_8446622_RNA45SN1_45S_50ntFlanks@9419.9451.33\|TCCGCCGAGGGCGCACCACCGGCCCGTCTCGCC | >hg38_21_+_8433172_8446622_RNA45SN1_45S_50ntFlanks | 9419 | 9451 | 33 | TCCGCCGAGGGCGCACCACCGGCCCGTCTCGCC | 92.8 | -10.4 | 1.5 | -6.8 | 1.1E-11 | 3.6E-11 | MB231/MB468 | MP |
| >hg38_21_+_8433172_8446622_RNA45SN1_45S_50ntFlanks@8196.8213.18\|CGTGGACGGTGTGAGGCC | >hg38_21_+_8433172_8446622_RNA45SN1_45S_50ntFlanks | 8196 | 8213 | 18 | CGTGGACGGTGTGAGGCC | 3913.1 | -5.1 | 0.8 | -6.7 | 1.6E-11 | 3.6E-11 | BT20/MB468 | NUC |
| >hg38_21_+_8433172_8446622_RNA45SN1_45S_50ntFlanks@10537.10571.35\|CGAGGCGTCCAGTGCGGTAACGCGACCGATCCCGG | >hg38_21_+_8433172_8446622_RNA45SN1_45S_50ntFlanks | 10537 | 10571 | 35 | CGAGGCGTCCAGTGCGGTAACGCGACCGATCCCGG | 81.7 | -10.2 | 1.5 | -6.8 | 1.1E-11 | 3.6E-11 | MB231/MB468 | MP |
| >hg38_21_+_8433172_8446622_RNA45SN1_45S_50ntFlanks@11873.11897.25\|AAGAAGACCCTGTTGAGCTTGACTC | >hg38_21_+_8433172_8446622_RNA45SN1_45S_50ntFlanks | 11873 | 11897 | 25 | AAGAAGACCCTGTTGAGCTTGACTC | 683.4 | -6.7 | 1.0 | -6.7 | 1.5E-11 | 3.7E-11 | MB231/MB468 | MITO |
| >hg38_21_+_8433172_8446622_RNA45SN1_45S_50ntFlanks@6731.6756.26\|CACATTGATCATCGACACTTCGAACG | >hg38_21_+_8433172_8446622_RNA45SN1_45S_50ntFlanks | 6731 | 6756 | 26 | CACATTGATCATCGACACTTCGAACG | 639.4 | -3.5 | 0.5 | -6.8 | 1.2E-11 | 3.7E-11 | BT20/MB468 | CYTO |
| >hg38_21_+_8433172_8446622_RNA45SN1_45S_50ntFlanks@8176.8204.29\|CTTCTGATCGAGGCCCAGCCCGTGGACGG | >hg38_21_+_8433172_8446622_RNA45SN1_45S_50ntFlanks | 8176 | 8204 | 29 | CTTCTGATCGAGGCCCAGCCCGTGGACGG | 3342.6 | -8.5 | 1.3 | -6.7 | 1.6E-11 | 3.7E-11 | BT20/MB468 | NUC |
| >trnaMT_LeuTAG_MT_+_12266_12336@38.74.37\|CCCAAAAAATTTTGGTGCAACTCCAAATAAAAGTACCA | >trnaMT_LeuTAG_MT_+_12266_12336 | 38 | 74 | 37 | CCCAAAAAATTTTGGTGCAACTCCAAATAAAAGTACCA | 237.3 | -11.1 | 1.6 | -7.1 | 1.2E-12 | 3.7E-11 | BT20/MB231 | MITO |
| >MI0000301\|hsa-mir-224&WithFlank&X\|-\|151958572\|151958664@14.34.21&[MIMAT0000281&hsa-miR-224-5p&offsets\|+1\|-3;m-238&X\|-\|151958630\|151958651&offsets\|0\|-1]\|CAAGTCACTAGTGGTTCCGTT | >MI0000301\|hsa-mir-224&WithFlank&X\|-\|151958572\|151958664 | 14 | 34 | 21 | CAAGTCACTAGTGGTTCCGTT | 204.1 | 10.9 | 1.6 | 6.7 | 1.5E-11 | 3.7E-11 | MB231/MB468 | MITO |
| >hg38_21_+_8433172_8446622_RNA45SN1_45S_50ntFlanks@12838.12871.34\|GTGCGGAGTGCCCTTCGTCCTGGGAAACGGGGCG | >hg38_21_+_8433172_8446622_RNA45SN1_45S_50ntFlanks | 12838 | 12871 | 34 | GTGCGGAGTGCCCTTCGTCCTGGGAAACGGGGCG | 90.4 | -10.4 | 1.5 | -6.8 | 1.1E-11 | 3.7E-11 | MB231/MB468 | MP |
| >MI0000078\|hsa-mir-22&WithFlank&17\|-\|1713897\|1713993@59.80.22(+1A)&[MIMAT0000077&hsa-miR-22-3p&offsets\|0\|0(+1A);m-2&17\|-\|1713914\|1713935&offsets\|0\|0(+1A)]\|AAGCTGCCAGTTGAAGAACTGTA | >MI0000078\|hsa-mir-22&WithFlank&17\|-\|1713897\|1713993 | 59 | 80 | 22(+1A) | AAGCTGCCAGTTGAAGAACTGTA | 100.1 | 4.2 | 0.6 | 6.7 | 1.6E-11 | 3.7E-11 | BT20/MB468 | NUC |
| >hg38_21_+_8433172_8446622_RNA45SN1_45S_50ntFlanks@11581.11603.23\|AACTGGTGCGGACCAGGGGAATC | >hg38_21_+_8433172_8446622_RNA45SN1_45S_50ntFlanks | 11581 | 11603 | 23 | AACTGGTGCGGACCAGGGGAATC | 13273.8 | -7.8 | 1.1 | -7.2 | 7.3E-13 | 3.7E-11 | BT20/MB468 | MP |
| >hg38_21_+_8433172_8446622_RNA45SN1_45S_50ntFlanks@4759.4777.19\|ATCAGATACCGTCGTAGTT | >hg38_21_+_8433172_8446622_RNA45SN1_45S_50ntFlanks | 4759 | 4777 | 19 | ATCAGATACCGTCGTAGTT | 309.4 | -11.4 | 1.6 | -7.2 | 7.3E-13 | 3.7E-11 | BT20/MB468 | MP |
| >hg38_21_+_8433172_8446622_RNA45SN1_45S_50ntFlanks@12456.12480.25\|CCAAGCGTTGGATTGTTCACCCACT | >hg38_21_+_8433172_8446622_RNA45SN1_45S_50ntFlanks | 12456 | 12480 | 25 | CCAAGCGTTGGATTGTTCACCCACT | 2373.9 | -4.7 | 0.7 | -6.8 | 1.2E-11 | 3.7E-11 | BT20/MB468 | CYTO |
| >MI0000783\|hsa-mir-375&WithFlank&2\|-\|219001639\|219001714@46.66.21&[MIMAT0000728&hsa-miR-375-3p&offsets\|0\|-1;m-92&2\|-\|219001648\|219001669&offsets\|0\|-1]\|TTTGTTCGTTCGGCTCGCGTG | >MI0000783\|hsa-mir-375&WithFlank&2\|-\|219001639\|219001714 | 46 | 66 | 21 | TTTGTTCGTTCGGCTCGCGTG | 116.4 | -10.7 | 1.6 | -6.8 | 1.1E-11 | 3.7E-11 | MB231/MB468 | MP |
| >am_MI0005763\|hsa-mir-941-1&WithFlank&20\|+\|63919443\|63919526@53.74.22;MI0005764\|hsa-mir-941-2&WithFlank&20\|+\|63919499\|63919582@53.74.22;MI0005765\|hsa-mir-941-3&WithFlank&20\|+\|63919555\|63919638@53.74.22;MI0005766\|hsa-mir-941-4&WithFlank&20\|+\|63919750\|63919833@53.74.22;MI0031520\|hsa-mir-941-5&WithFlank&20\|+\|63919862\|63919945@53.74.22&[MIMAT0004984&hsa-miR-941&offsets\|0\|-1;m-263&20\|+\|63919495\|63919517&offsets\|0\|-1];[MIMAT0004984_1&hsa-miR-941&offsets\|0\|-1;m-263&20\|+\|63919551\|63919573&offsets\|0\|-1];[MIMAT0004984_2&hsa-miR-941&offsets\|0\|-1;m-263&20\|+\|63919607\|63919629&offsets\|0\|-1];[MIMAT0004984_3&hsa-miR-941&offsets\|0\|-1;m-263&20\|+\|63919802\|63919824&offsets\|0\|-1];[MIMAT0004984_4&hsa-miR-941&offsets\|0\|-1;m-263&20\|+\|63919914\|63919936&offsets\|0\|-1]\|CACCCGGCTGTGTGCACATGTG | >am_MI0005763\|hsa-mir-941-1&WithFlank&20\|+\|63919443\|63919526 | 53 | 74 | 22 | CACCCGGCTGTGTGCACATGTG | 515.6 | 8.6 | 1.3 | 6.8 | 9.2E-12 | 3.7E-11 | BT20/MB468 | MITO |
| >hg38_21_+_8433172_8446622_RNA45SN1_45S_50ntFlanks@10421.10456.36\|TGAGAGATGGGCGAGCGCCGTTCCGAAGGGACGGGC | >hg38_21_+_8433172_8446622_RNA45SN1_45S_50ntFlanks | 10421 | 10456 | 36 | TGAGAGATGGGCGAGCGCCGTTCCGAAGGGACGGGC | 376.1 | -8.6 | 1.3 | -6.7 | 1.5E-11 | 3.7E-11 | MB231/MB468 | MITO |
| >hg38_21_+_8433172_8446622_RNA45SN1_45S_50ntFlanks@3765.3792.28\|AGTACGCACGGCCGGTACAGTGAAACTG | >hg38_21_+_8433172_8446622_RNA45SN1_45S_50ntFlanks | 3765 | 3792 | 28 | AGTACGCACGGCCGGTACAGTGAAACTG | 628.5 | -9.3 | 1.4 | -6.7 | 1.6E-11 | 3.7E-11 | BT20/MB468 | NUC |
| >hg38_1_-_228634819_228635039_RNA5S12_5S_50ntFlanks@140.171.32\|ACCGCCTGGGAATACCGGGTGCTGTAGGCTTT | >hg38_1_-_228634819_228635039_RNA5S12_5S_50ntFlanks | 140 | 171 | 32 | ACCGCCTGGGAATACCGGGTGCTGTAGGCTTT | 4802.7 | 5.3 | 0.8 | 6.8 | 1.3E-11 | 3.7E-11 | MB231/MB468 | NUC |
| >trna152_ValCAC_6_-_27248049_27248121@1.32.32\|GCTTCTGTAGTGTAGTGGTTATCACGTTCGCC | >trna152_ValCAC_6_-_27248049_27248121 | 1 | 32 | 32 | GCTTCTGTAGTGTAGTGGTTATCACGTTCGCC | 1851.7 | 7.4 | 1.1 | 6.8 | 1.3E-11 | 3.7E-11 | MB231/MB468 | NUC |

| | | | | | | | | | | | | | |
|---|---|---|---|---|---|---|---|---|---|---|---|---|---|
| >hg38_21_+_8433172_8446622_RNA45SN1_45S_50ntFlanks@8170.8214.45\|CAAGTCCTTCTGATCGAGGCCCAGCCCGTGGACGGTGTGAGGCCG | >hg38_21_+_8433172_8446622_RNA45SN1_45S_50ntFlanks | 8170 | 8214 | 45 | CAAGTCCTTCTGATCGAGGCCCAGCCCGTGGACGGTGTGAGGCCG | 3124.5 | -5.8 | 0.9 | -6.8 | 1.3E-11 | 3.7E-11 | MB231/MB468 | NUC |
| >trnaMT_ValTAC_MT_+_1602_1670@51.72.22\|CTTAACTTGACCGCTCTGACCA | >trnaMT_ValTAC_MT_+_1602_1670 | 51 | 72 | 22 | CTTAACTTGACCGCTCTGACCA | 1838.3 | -5.1 | 0.7 | -6.8 | 1.2E-11 | 3.7E-11 | MB231/MB468 | MP |
| >hg38_21_+_8433172_8446622_RNA45SN1_45S_50ntFlanks@6650.6695.46\|ACGACTCTTAGCGGTGGATCACTCGGCTCGTGCGTCGATGAAGAAC | >hg38_21_+_8433172_8446622_RNA45SN1_45S_50ntFlanks | 6650 | 6695 | 46 | ACGACTCTTAGCGGTGGATCACTCGGCTCGTGCGTCGATGAAGAAC | 444.7 | 7.9 | 1.2 | 6.8 | 1.3E-11 | 3.7E-11 | MB231/MB468 | CYTO |
| >hg38_21_+_8433172_8446622_RNA45SN1_45S_50ntFlanks@11729.11747.19\|TAACTATGACTCTCTTAAG | >hg38_21_+_8433172_8446622_RNA45SN1_45S_50ntFlanks | 11729 | 11747 | 19 | TAACTATGACTCTCTTAAG | 7827.9 | -4.9 | 0.7 | -6.8 | 1.3E-11 | 3.7E-11 | MB231/MB468 | NUC |
| >hg38_21_+_8433172_8446622_RNA45SN1_45S_50ntFlanks@12455.12487.33\|ACCAAGCGTTGGATTGTTCACCCACTAATAGGG | >hg38_21_+_8433172_8446622_RNA45SN1_45S_50ntFlanks | 12455 | 12487 | 33 | ACCAAGCGTTGGATTGTTCACCCACTAATAGGG | 148.9 | -11.1 | 1.6 | -6.8 | 1.2E-11 | 3.7E-11 | MB231/MB468 | MP |
| >hg38_21_+_8433172_8446622_RNA45SN1_45S_50ntFlanks@10841.10863.23\|CGTAACTTCGGGATAAGGATTGG | >hg38_21_+_8433172_8446622_RNA45SN1_45S_50ntFlanks | 10841 | 10863 | 23 | CGTAACTTCGGGATAAGGATTGG | 4600.3 | -10.1 | 1.5 | -6.8 | 1.2E-11 | 3.7E-11 | MB231/MB468 | MP |
| >hg38_21_+_8433172_8446622_RNA45SN1_45S_50ntFlanks@4129.4150.22\|GATTCCGGAGAGGGAGCCTGAG | >hg38_21_+_8433172_8446622_RNA45SN1_45S_50ntFlanks | 4129 | 4150 | 22 | GATTCCGGAGAGGGAGCCTGAG | 75.7 | -10.1 | 1.5 | -6.8 | 1.2E-11 | 3.7E-11 | MB231/MB468 | MP |
| >trnaMT_GlnTTG_MT_-_4329_4400@1.36.36,trnalookalike2_GlnTTG_1_-_564879_564950@1.36.36\|TAGGATGGGGTGTGATAGGTGGCACGGAGAATTTTG | >trnaMT_GlnTTG_MT_-_4329_4400 | 1 | 36 | 36 | TAGGATGGGGTGTGATAGGTGGCACGGAGAATTTTG | 1041.5 | 10.1 | 1.5 | 6.8 | 9.3E-12 | 3.7E-11 | BT20/MB468 | MITO |
| >hg38_21_+_8433172_8446622_RNA45SN1_45S_50ntFlanks@11874.11898.25\|AGAAGACCCTGTTGAGCTTGACTCT | >hg38_21_+_8433172_8446622_RNA45SN1_45S_50ntFlanks | 11874 | 11898 | 25 | AGAAGACCCTGTTGAGCTTGACTCT | 4979.1 | -8.6 | 1.3 | -6.7 | 1.6E-11 | 3.7E-11 | BT20/MB468 | NUC |
| >hg38_21_+_8433172_8446622_RNA45SN1_45S_50ntFlanks@10571.10598.28\|GAGAAGCCGGCGGGAGCCCCGGGGAGAG | >hg38_21_+_8433172_8446622_RNA45SN1_45S_50ntFlanks | 10571 | 10598 | 28 | GAGAAGCCGGCGGGAGCCCCGGGGAGAG | 132.7 | -10.9 | 1.6 | -6.8 | 1.2E-11 | 3.7E-11 | MB231/MB468 | MP |
| >hg38_1_-_228634819_228635039_RNA5S12_5S_50ntFlanks@128.170.43\|CTTGGATGGGAGACCGCCTGGGAATACCGGGTGCTGTAGGCTT | >hg38_1_-_228634819_228635039_RNA5S12_5S_50ntFlanks | 128 | 170 | 43 | CTTGGATGGGAGACCGCCTGGGAATACCGGGTGCTGTAGGCTT | 1696.7 | 6.9 | 1.0 | 6.8 | 9.3E-12 | 3.7E-11 | BT20/MB468 | MITO |
| >hg38_21_+_8433172_8446622_RNA45SN1_45S_50ntFlanks@10906.10925.20\|TGGGCGCGCGCCGCGGCTGG | >hg38_21_+_8433172_8446622_RNA45SN1_45S_50ntFlanks | 10906 | 10925 | 20 | TGGGCGCGCGCCGCGGCTGG | 117.3 | -10.7 | 1.6 | -6.8 | 1.2E-11 | 3.7E-11 | MB231/MB468 | MP |
| >hg38_21_+_8433172_8446622_RNA45SN1_45S_50ntFlanks@11913.11951.39\|AAGAGACATGAGAGGTGTAGAATAAGTGGGAGGCCCCCG | >hg38_21_+_8433172_8446622_RNA45SN1_45S_50ntFlanks | 11913 | 11951 | 39 | AAGAGACATGAGAGGTGTAGAATAAGTGGGAGGCCCCCG | 160.3 | -11.2 | 1.6 | -6.8 | 1.2E-11 | 3.7E-11 | MB231/MB468 | MP |
| >hg38_21_+_8433172_8446622_RNA45SN1_45S_50ntFlanks@11729.11765.37\|TAACTATGACTCTCTTAAGGTAGCCAAATGCCTCGTC | >hg38_21_+_8433172_8446622_RNA45SN1_45S_50ntFlanks | 11729 | 11765 | 37 | TAACTATGACTCTCTTAAGGTAGCCAAATGCCTCGTC | 884.1 | -9.4 | 1.4 | -6.8 | 1.2E-11 | 3.7E-11 | MB231/MB468 | MP |
| >hg38_21_+_8433172_8446622_RNA45SN1_45S_50ntFlanks@11601.11619.19\|ATCCGACTGTTTAATTAAA | >hg38_21_+_8433172_8446622_RNA45SN1_45S_50ntFlanks | 11601 | 11619 | 19 | ATCCGACTGTTTAATTAAA | 209.2 | -10.0 | 1.5 | -6.8 | 1.2E-11 | 3.7E-11 | BT20/MB468 | CYTO |
| >trna111_HisGTG_1_-_147774845_147774916@-1T.22.23,trna118_HisGTG_1_-_145396881_145396952@-1T.22.23,trna16_HisGTG_1_+_146544773_146544844@-1T.22.23,trna1_HisGTG_15_+_45493349_45493420@-1T.22.23,trna21_HisGTG_1_+_147753471_147753542@-1T.22.23,trna33_HisGTG_6_+_27125906_27125977@-1T.22.23,trna7_HisGTG_9_-_14433938_14434009@-1T.22.23,trna8_HisGTG_15_-_45492611_45492682@-1T.22.23,trna9_HisGTG_15_-_45490804_45490875@-1T.22.23\|TGCCGTGATCGTATAGTGGTTAG | >trna111_HisGTG_1_-_147774845_147774916 | -1T | 22 | 23 | TGCCGTGATCGTATAGTGGTTAG | 185.8 | 10.8 | 1.6 | 6.7 | 1.5E-11 | 3.7E-11 | MB231/MB468 | MITO |
| >hg38_21_+_8433172_8446622_RNA45SN1_45S_50ntFlanks@11576.11593.18\|CTTAGAACTGGTGCGGAC | >hg38_21_+_8433172_8446622_RNA45SN1_45S_50ntFlanks | 11576 | 11593 | 18 | CTTAGAACTGGTGCGGAC | 1286.2 | -7.4 | 1.1 | -6.8 | 1.3E-11 | 3.7E-11 | MB231/MB468 | NUC |
| >hg38_21_+_8433172_8446622_RNA45SN1_45S_50ntFlanks@11728.11751.24\|GTAACTATGACTCTCTTAAGGTAG | >hg38_21_+_8433172_8446622_RNA45SN1_45S_50ntFlanks | 11728 | 11751 | 24 | GTAACTATGACTCTCTTAAGGTAG | 2435.6 | -5.9 | 0.9 | -6.8 | 1.3E-11 | 3.7E-11 | MB231/MB468 | NUC |
| >hg38_21_+_8433172_8446622_RNA45SN1_45S_50ntFlanks@4557.4575.19\|CAGCTAGGAATAATGGAAT | >hg38_21_+_8433172_8446622_RNA45SN1_45S_50ntFlanks | 4557 | 4575 | 19 | CAGCTAGGAATAATGGAAT | 103.2 | -10.6 | 1.6 | -6.8 | 1.2E-11 | 3.7E-11 | MB231/MB468 | MP |
| >hg38_21_+_8433172_8446622_RNA45SN1_45S_50ntFlanks@11768.11791.24\|CTAATTAGTGACGCGCATGAATGG | >hg38_21_+_8433172_8446622_RNA45SN1_45S_50ntFlanks | 11768 | 11791 | 24 | CTAATTAGTGACGCGCATGAATGG | 258.3 | -11.1 | 1.5 | -7.2 | 7.5E-13 | 3.7E-11 | BT20/MB468 | MP |
| >hg38_21_+_8433172_8446622_RNA45SN1_45S_50ntFlanks@3775.3799.25\|GCCGGTACAGTGAAACTGCGAATGG | >hg38_21_+_8433172_8446622_RNA45SN1_45S_50ntFlanks | 3775 | 3799 | 25 | GCCGGTACAGTGAAACTGCGAATGG | 255.9 | -11.1 | 1.5 | -7.2 | 7.5E-13 | 3.7E-11 | BT20/MB468 | MP |
| >hg38_21_+_8433172_8446622_RNA45SN1_45S_50ntFlanks@4385.4402.18\|TTGGATCTTGGGAGCGGG | >hg38_21_+_8433172_8446622_RNA45SN1_45S_50ntFlanks | 4385 | 4402 | 18 | TTGGATCTTGGGAGCGGG | 120.5 | -10.8 | 1.6 | -6.8 | 1.2E-11 | 3.7E-11 | MB231/MB468 | MP |
| >hg38_21_+_8433172_8446622_RNA45SN1_45S_50ntFlanks@8556.8577.22\|CCGGAGGGGGCGGGCTCCGGCG | >hg38_21_+_8433172_8446622_RNA45SN1_45S_50ntFlanks | 8556 | 8577 | 22 | CCGGAGGGGGCGGGCTCCGGCG | 130.4 | -10.9 | 1.6 | -6.8 | 1.2E-11 | 3.7E-11 | MB231/MB468 | MP |
| >hg38_21_+_8433172_8446622_RNA45SN1_45S_50ntFlanks@10788.10818.31\|CAGCCTCTGGCATGTTGGAACAATGTAGGTA | >hg38_21_+_8433172_8446622_RNA45SN1_45S_50ntFlanks | 10788 | 10818 | 31 | CAGCCTCTGGCATGTTGGAACAATGTAGGTA | 1393.1 | -4.8 | 0.7 | -6.8 | 1.3E-11 | 3.7E-11 | MB231/MB468 | CYTO |
| >MI0000299\|hsa-mir-222&WithFlank&X\|-\|45747009\|45747130@75.96.22(+1A)&[MIMAT0000279&hsa-miR-222-3p&offsets\|0\|+1(+1A);m-140&X\|-\|45747033\|45747056&offsets\|0\|-2(+1A)]\|AGCTACATCTGGCTACTGGGTCA | >MI0000299\|hsa-mir-222&WithFlank&X\|-\|45747009\|45747130 | 75 | 96 | 22(+1A) | AGCTACATCTGGCTACTGGGTCA | 1138.1 | 6.6 | 1.0 | 6.7 | 1.5E-11 | 3.7E-11 | MB231/MB468 | MITO |
| >MI0000270\|hsa-mir-181b-1&WithFlank&1\|-\|198858867\|198858988@42.64.23;MI0000683\|hsa-mir-181b-2&WithFlank&9\|+\|124693704\|124693804@22.44.23&[MIMAT0000257&hsa-miR-181b-5p&offsets\|0\|0;m-73&1\|-\|198858925\|198858947&offsets\|0\|0];[MIMAT0000257_1&hsa-miR-181b-5p&offsets\|0\|0;m-72&9\|+\|124693725\|124693747&offsets\|0\|0]\|AACATTCATTGCTGTCGGTGGGT | >MI0000270\|hsa-mir-181b-1&WithFlank&1\|-\|198858867\|198858988 | 42 | 64 | 23 | AACATTCATTGCTGTCGGTGGGT | 3057.6 | 5.4 | 0.8 | 6.8 | 1.3E-11 | 3.8E-11 | MB231/MB468 | NUC |
| >hg38_21_+_8433172_8446622_RNA45SN1_45S_50ntFlanks@6651.6668.18\|CGACTCTTAGCGGTGGAT | >hg38_21_+_8433172_8446622_RNA45SN1_45S_50ntFlanks | 6651 | 6668 | 18 | CGACTCTTAGCGGTGGAT | 3587.4 | 3.6 | 0.5 | 6.8 | 1.3E-11 | 3.8E-11 | MB231/MB468 | NUC |
| >hg38_21_+_8433172_8446622_RNA45SN1_45S_50ntFlanks@4047.4073.27\|ATTCGAACGTCTGCCCTATCAACTTTC | >hg38_21_+_8433172_8446622_RNA45SN1_45S_50ntFlanks | 4047 | 4073 | 27 | ATTCGAACGTCTGCCCTATCAACTTTC | 435.1 | -7.0 | 1.0 | -6.7 | 1.6E-11 | 3.8E-11 | BT20/MB468 | NUC |
| >hg38_21_+_8433172_8446622_RNA45SN1_45S_50ntFlanks@9711.9729.19\|AAGCGAATGATTAGAGGTC | >hg38_21_+_8433172_8446622_RNA45SN1_45S_50ntFlanks | 9711 | 9729 | 19 | AAGCGAATGATTAGAGGTC | 148.7 | -11.1 | 1.6 | -6.8 | 1.2E-11 | 3.8E-11 | MB231/MB468 | MP |
| >hg38_21_+_8433172_8446622_RNA45SN1_45S_50ntFlanks@11115.11133.19\|CGGCAGGGGCCGGCGGCGG | >hg38_21_+_8433172_8446622_RNA45SN1_45S_50ntFlanks | 11115 | 11133 | 19 | CGGCAGGGGCCGGCGGCGG | 385.0 | -7.2 | 1.1 | -6.8 | 1.3E-11 | 3.8E-11 | MB231/MB468 | CYTO |
| >MI0000749\|hsa-mir-30e&WithFlank&1\|+\|40754349\|40754452@23.46.24(+2U)&[MIMAT0000692&hsa-miR-30e-5p&offsets\|0\|+2(+2U);m-24&1\|+\|40754371\|40754394&offsets\|0\|0(+2U)]\|TGTAAACATCCTTGACTGGAAGCTTT | >MI0000749\|hsa-mir-30e&WithFlank&1\|+\|40754349\|40754452 | 23 | 46 | 24(+2U) | TGTAAACATCCTTGACTGGAAGCTTT | 194.2 | 6.1 | 0.9 | 6.8 | 1.3E-11 | 3.8E-11 | MB231/MB468 | NUC |
| >hg38_21_+_8433172_8446622_RNA45SN1_45S_50ntFlanks@3993.4025.33\|GTGACTCTAGATAACCTCGGGCCGATCGCACGC | >hg38_21_+_8433172_8446622_RNA45SN1_45S_50ntFlanks | 3993 | 4025 | 33 | GTGACTCTAGATAACCTCGGGCCGATCGCACGC | 99.3 | -10.5 | 1.5 | -6.8 | 1.2E-11 | 3.8E-11 | MB231/MB468 | MP |
| >hg38_21_+_8433172_8446622_RNA45SN1_45S_50ntFlanks@8191.8212.22\|CAGCCCGTGGACGGTGTGAGGC | >hg38_21_+_8433172_8446622_RNA45SN1_45S_50ntFlanks | 8191 | 8212 | 22 | CAGCCCGTGGACGGTGTGAGGC | 493.6 | -3.2 | 0.5 | -6.8 | 1.2E-11 | 3.8E-11 | BT20/MB468 | CYTO |
| >MI0000446\|hsa-mir-125b-1&WithFlank&11\|-\|122099751\|122099850@22.42.21;MI0000470\|hsa-mir-125b-2&WithFlank&21\|+\|16590231\|16590331@24.44.21&[MIMAT0000423&hsa-miR-125b-5p&offsets\|+1\|0;m-67&11\|-\|122099809\|122099830&offsets\|+1\|0];[MIMAT0000423_1&hsa-miR-125b-5p&offsets\|+1\|0;m-68&21\|+\|16590253\|16590274&offsets\|+1\|0]\|CCCTGAGACCCTAACTTGTGA | >MI0000446\|hsa-mir-125b-1&WithFlank&11\|-\|122099751\|122099850 | 22 | 42 | 21 | CCCTGAGACCCTAACTTGTGA | 276.7 | 11.3 | 1.7 | 6.7 | 1.5E-11 | 3.8E-11 | MB231/MB468 | MITO |
| >hg38_21_+_8433172_8446622_RNA45SN1_45S_50ntFlanks@4683.4723.41\|CAAGACGGACCAGAGCGAAAGCATTTGCCAAGAATGTTTTC | >hg38_21_+_8433172_8446622_RNA45SN1_45S_50ntFlanks | 4683 | 4723 | 41 | CAAGACGGACCAGAGCGAAAGCATTTGCCAAGAATGTTTTC | 1176.0 | -7.7 | 1.1 | -6.7 | 1.5E-11 | 3.8E-11 | MB231/MB468 | MITO |
| >hg38_21_+_8433172_8446622_RNA45SN1_45S_50ntFlanks@8242.8270.29\|GGTCTTCCCGGAGTCGGGTTGCTTGGGAA | >hg38_21_+_8433172_8446622_RNA45SN1_45S_50ntFlanks | 8242 | 8270 | 29 | GGTCTTCCCGGAGTCGGGTTGCTTGGGAA | 105.8 | -10.6 | 1.6 | -6.8 | 1.2E-11 | 3.8E-11 | MB231/MB468 | MP |
| >hg38_21_+_8433172_8446622_RNA45SN1_45S_50ntFlanks@3771.3799.29\|CACGGCCGGTACAGTGAAACTGCGAATGG | >hg38_21_+_8433172_8446622_RNA45SN1_45S_50ntFlanks | 3771 | 3799 | 29 | CACGGCCGGTACAGTGAAACTGCGAATGG | 355.0 | -6.2 | 0.9 | -6.8 | 1.3E-11 | 3.8E-11 | MB231/MB468 | NUC |
| >hg38_21_+_8433172_8446622_RNA45SN1_45S_50ntFlanks@10402.10430.29\|AACATGGGTCAGTCGGTCCTGAGAGATGG | >hg38_21_+_8433172_8446622_RNA45SN1_45S_50ntFlanks | 10402 | 10430 | 29 | AACATGGGTCAGTCGGTCCTGAGAGATGG | 1515.3 | -4.9 | 0.7 | -7.2 | 7.6E-13 | 3.8E-11 | BT20/MB468 | MP |
| >MI0003760\|hsa-mir-671&WithFlank&7\|+\|151238415\|151238544@35.58.24&[MIMAT0003880&hsa-miR-671-5p&offsets\|0\|+1;m-316&7\|+\|151238449\|151238472&offsets\|0\|0]\|AGGAAGCCCTGGAGGGGCTGGAGG | >MI0003760\|hsa-mir-671&WithFlank&7\|+\|151238415\|151238544 | 35 | 58 | 24 | AGGAAGCCCTGGAGGGGCTGGAGG | 669.6 | 9.0 | 1.3 | 6.8 | 9.5E-12 | 3.8E-11 | BT20/MB468 | MITO |
| >hg38_21_+_8433172_8446622_RNA45SN1_45S_50ntFlanks@10681.10711.31\|TTCCGGCGGCGTCCGGTGAGCTCTCGCTGGC | >hg38_21_+_8433172_8446622_RNA45SN1_45S_50ntFlanks | 10681 | 10711 | 31 | TTCCGGCGGCGTCCGGTGAGCTCTCGCTGGC | 1115.2 | -7.5 | 1.1 | -6.8 | 1.3E-11 | 3.8E-11 | MB231/MB468 | NUC |
| >hg38_21_+_8433172_8446622_RNA45SN1_45S_50ntFlanks@11884.11904.21\|GTTGAGCTTGACTCTAGTCTG | >hg38_21_+_8433172_8446622_RNA45SN1_45S_50ntFlanks | 11884 | 11904 | 21 | GTTGAGCTTGACTCTAGTCTG | 219.8 | -7.7 | 1.1 | -6.8 | 1.3E-11 | 3.8E-11 | MB231/MB468 | CYTO |
| >hg38_1_-_228634819_228635039_RNA5S12_5S_50ntFlanks@127.169.43\|ACTTGGATGGGAGACCGCCTGGGAATACCGGGTGCTGTAGGCT | >hg38_1_-_228634819_228635039_RNA5S12_5S_50ntFlanks | 127 | 169 | 43 | ACTTGGATGGGAGACCGCCTGGGAATACCGGGTGCTGTAGGCT | 223.8 | 11.6 | 1.6 | 7.2 | 7.7E-13 | 3.8E-11 | BT20/MB468 | MP |
| >hg38_21_+_8433172_8446622_RNA45SN1_45S_50ntFlanks@12419.12447.29\|GATGTCGGCTCTTCCTATCATTGTGAAGC | >hg38_21_+_8433172_8446622_RNA45SN1_45S_50ntFlanks | 12419 | 12447 | 29 | GATGTCGGCTCTTCCTATCATTGTGAAGC | 658.3 | 5.0 | 0.7 | 6.8 | 1.2E-11 | 3.8E-11 | BT20/MB468 | CYTO |
| >hg38_21_+_8433172_8446622_RNA45SN1_45S_50ntFlanks@10408.10453.46\|GGTCAGTCGGTCCTGAGAGATGGGCGAGCGCCGTTCCGAAGGGACG | >hg38_21_+_8433172_8446622_RNA45SN1_45S_50ntFlanks | 10408 | 10453 | 46 | GGTCAGTCGGTCCTGAGAGATGGGCGAGCGCCGTTCCGAAGGGACG | 198.9 | -9.9 | 1.5 | -6.8 | 1.2E-11 | 3.8E-11 | BT20/MB468 | CYTO |
| >hg38_21_+_8433172_8446622_RNA45SN1_45S_50ntFlanks@12405.12438.34\|CTTTTTGATCCTTCGATGTCGGCTCTTCCTATCA | >hg38_21_+_8433172_8446622_RNA45SN1_45S_50ntFlanks | 12405 | 12438 | 34 | CTTTTTGATCCTTCGATGTCGGCTCTTCCTATCA | 2017.8 | -5.6 | 0.8 | -6.7 | 1.5E-11 | 3.8E-11 | MB231/MB468 | MITO |
| >MI0000067\|hsa-let-7f-1&WithFlank&9\|+\|94176341\|94176439@13.31.19;MI0000068\|hsa-let-7f-2&WithFlank&X\|-\|53557186\|53557280@14.32.19&[MIMAT0000067&hsa-let-7f-5p&offsets\|0\|-3;m-20&9\|+\|94176353\|94176374&offsets\|0\|-3];[MIMAT0000067_1&hsa-let-7f-5p&offsets\|0\|-3;m-19&X\|-\|53557246\|53557267&offsets\|0\|-3]\|TGAGGTAGTAGATTGTATA | >MI0000067\|hsa-let-7f-1&WithFlank&9\|+\|94176341\|94176439 | 13 | 31 | 19 | TGAGGTAGTAGATTGTATA | 396.9 | 9.7 | 1.4 | 6.8 | 9.6E-12 | 3.8E-11 | BT20/MB468 | MITO |
| >hg38_21_+_8433172_8446622_RNA45SN1_45S_50ntFlanks@6661.6695.35\|CGGTGGATCACTCGGCTCGTGCGTCGATGAAGAAC | >hg38_21_+_8433172_8446622_RNA45SN1_45S_50ntFlanks | 6661 | 6695 | 35 | CGGTGGATCACTCGGCTCGTGCGTCGATGAAGAAC | 362.8 | 3.7 | 0.5 | 6.8 | 1.3E-11 | 3.8E-11 | MB231/MB468 | NUC |

| | | | | | | | | | | | | | |
|---|---|---|---|---|---|---|---|---|---|---|---|---|---|
| >hg38_21_+_8433172_8446622_RNA45SN1_45S_50ntFlanks@12420.12459.40\|ATGTCGGCTCTTCCTATCATTGTGAAGCAGAATTCACCAA | >hg38_21_+_8433172_8446622_RNA45SN1_45S_50ntFlanks | 12420 | 12459 | 40 | ATGTCGGCTCTTCCTATCATTGTGAAGCAGAATTCACCAA | 312.1 | 4.5 | 0.7 | 6.8 | 1.3E-11 | 3.8E-11 | MB231/MB468 | CYTO |
| >hg38_21_+_8433172_8446622_RNA45SN1_45S_50ntFlanks@4064.4089.26\|ATCAACTTTCGATGGTAGTCGCCGTG | >hg38_21_+_8433172_8446622_RNA45SN1_45S_50ntFlanks | 4064 | 4089 | 26 | ATCAACTTTCGATGGTAGTCGCCGTG | 85.8 | -10.3 | 1.5 | -6.8 | 1.2E-11 | 3.8E-11 | MB231/MB468 | MP |
| >MI0000462\|hsa-mir-152&WithFlank&17\|-\|48037155\|48037253@60.81.22(+1A)&[MIMAT0000438&hsa-miR-152-3p&offsets\|0\|+1(+1A);m-97&17\|-\|48037174\|48037194&offsets\|0\|+1(+1A)]\|TCAGTGCATGACAGAACTTGGGA | >MI0000462\|hsa-mir-152&WithFlank&17\|-\|48037155\|48037253 | 60 | 81 | 22(+1A) | TCAGTGCATGACAGAACTTGGGA | 138.0 | 10.3 | 1.5 | 6.8 | 1.2E-11 | 3.8E-11 | MB231/MB468 | MP |
| >hg38_21_+_8433172_8446622_RNA45SN1_45S_50ntFlanks@10789.10816.28\|AGCCTCTGGCATGTTGGAACAATGTAGG | >hg38_21_+_8433172_8446622_RNA45SN1_45S_50ntFlanks | 10789 | 10816 | 28 | AGCCTCTGGCATGTTGGAACAATGTAGG | 118.2 | -10.7 | 1.6 | -6.8 | 1.2E-11 | 3.8E-11 | MB231/MB468 | MP |
| >hg38_21_+_8433172_8446622_RNA45SN1_45S_50ntFlanks@4283.4306.24\|CTTTAACGAGGATCCATTGGAGGG | >hg38_21_+_8433172_8446622_RNA45SN1_45S_50ntFlanks | 4283 | 4306 | 24 | CTTTAACGAGGATCCATTGGAGGG | 125.5 | -10.8 | 1.6 | -6.8 | 1.2E-11 | 3.9E-11 | MB231/MB468 | MP |
| >trna11_GluTTC_15_-_26327381_26327452@1.31.31,trna3_GluTTC_13_-_45492062_45492133@1.31.31\|TCCCACATGGTCTAGCGGTTAGGATTCCTGG | >trna11_GluTTC_15_-_26327381_26327452 | 1 | 31 | 31 | TCCCACATGGTCTAGCGGTTAGGATTCCTGG | 1144.7 | -7.6 | 1.1 | -6.7 | 1.7E-11 | 3.9E-11 | BT20/MB468 | NUC |
| >MI0000065\|hsa-let-7d&WithFlank&9\|+\|94178828\|94178926@14.35.22(+1A)&[MIMAT0000065&hsa-let-7d-5p&offsets\|0\|0(+1A);m-90&9\|+\|94178841\|94178862&offsets\|0\|0(+1A)]\|AGAGGTAGTAGGTTGCATAGTTA | >MI0000065\|hsa-let-7d&WithFlank&9\|+\|94178828\|94178926 | 14 | 35 | 22(+1A) | AGAGGTAGTAGGTTGCATAGTTA | 498.6 | 9.0 | 1.3 | 6.8 | 1.2E-11 | 3.9E-11 | BT20/MB468 | CYTO |
| >MI0000283\|hsa-mir-203a&WithFlank&14\|+\|104117399\|104117520@72.92.21&[MIMAT0000264&hsa-miR-203a-3p&offsets\|+1\|0;m-79&14\|+\|104117469\|104117490&offsets\|+1\|0]\|TGAAATGTTTAGGACCACTAG | >MI0000283\|hsa-mir-203a&WithFlank&14\|+\|104117399\|104117520 | 72 | 92 | 21 | TGAAATGTTTAGGACCACTAG | 567.4 | 12.5 | 1.7 | 7.4 | 1.9E-13 | 3.9E-11 | BT20/MB231 | MP |
| >hg38_21_+_8433172_8446622_RNA45SN1_45S_50ntFlanks@3881.3898.18\|GCCGACGGGCGCTGACCC | >hg38_21_+_8433172_8446622_RNA45SN1_45S_50ntFlanks | 3881 | 3898 | 18 | GCCGACGGGCGCTGACCC | 144.7 | -11.0 | 1.6 | -6.8 | 1.2E-11 | 3.9E-11 | MB231/MB468 | MP |
| >hg38_21_+_8433172_8446622_RNA45SN1_45S_50ntFlanks@3785.3810.26\|TGAAACTGCGAATGGCTCATTAAATC | >hg38_21_+_8433172_8446622_RNA45SN1_45S_50ntFlanks | 3785 | 3810 | 26 | TGAAACTGCGAATGGCTCATTAAATC | 246.1 | -11.8 | 1.7 | -6.8 | 1.2E-11 | 3.9E-11 | MB231/MB468 | MP |
| >MI0000802\|hsa-mir-340&WithFlank&5\|-\|180015297\|180015403@22.43.22&[MIMAT0004692&hsa-miR-340-5p&offsets\|0\|0;m-220&5\|-\|180015361\|180015382&offsets\|0\|0]\|TTATAAAGCAATGAGACTGATT | >MI0000802\|hsa-mir-340&WithFlank&5\|-\|180015297\|180015403 | 22 | 43 | 22 | TTATAAAGCAATGAGACTGATT | 2034.9 | 5.9 | 0.9 | 6.8 | 1.2E-11 | 3.9E-11 | BT20/MB468 | CYTO |
| >hg38_21_+_8433172_8446622_RNA45SN1_45S_50ntFlanks@9564.9594.31\|CTGACGTGCAAATCGGTCGTCCGACCTGGGT | >hg38_21_+_8433172_8446622_RNA45SN1_45S_50ntFlanks | 9564 | 9594 | 31 | CTGACGTGCAAATCGGTCGTCCGACCTGGGT | 2098.6 | -8.2 | 1.2 | -6.7 | 1.7E-11 | 3.9E-11 | BT20/MB468 | NUC |
| >hg38_21_+_8433172_8446622_RNA45SN1_45S_50ntFlanks@9554.9597.44\|CGTAGCGGTCCTGACGTGCAAATCGGTCGTCCGACCTGGGTATA | >hg38_21_+_8433172_8446622_RNA45SN1_45S_50ntFlanks | 9554 | 9597 | 44 | CGTAGCGGTCCTGACGTGCAAATCGGTCGTCCGACCTGGGTATA | 6312.5 | -8.3 | 1.2 | -6.7 | 1.6E-11 | 3.9E-11 | MB231/MB468 | MITO |
| >hg38_21_+_8433172_8446622_RNA45SN1_45S_50ntFlanks@10572.10591.20\|AGAAGCCGGCGGGAGCCCCG | >hg38_21_+_8433172_8446622_RNA45SN1_45S_50ntFlanks | 10572 | 10591 | 20 | AGAAGCCGGCGGGAGCCCCG | 1977.1 | -10.4 | 1.5 | -6.7 | 1.7E-11 | 3.9E-11 | BT20/MB468 | NUC |
| >hg38_21_+_8433172_8446622_RNA45SN1_45S_50ntFlanks@10420.10457.38\|CTGAGAGATGGGCGAGCGCCGTTCCGAAGGGACGGGCG | >hg38_21_+_8433172_8446622_RNA45SN1_45S_50ntFlanks | 10420 | 10457 | 38 | CTGAGAGATGGGCGAGCGCCGTTCCGAAGGGACGGGCG | 124.1 | -10.8 | 1.6 | -6.8 | 1.2E-11 | 3.9E-11 | MB231/MB468 | MP |
| >hg38_21_+_8433172_8446622_RNA45SN1_45S_50ntFlanks@9534.9551.18\|AGGAAACTCTGGTGGAGG | >hg38_21_+_8433172_8446622_RNA45SN1_45S_50ntFlanks | 9534 | 9551 | 18 | AGGAAACTCTGGTGGAGG | 9581.4 | -11.5 | 1.7 | -6.7 | 1.7E-11 | 3.9E-11 | BT20/MB468 | NUC |
| >hg38_21_+_8433172_8446622_RNA45SN1_45S_50ntFlanks@5485.5525.41\|AGCGCTGAGAAGACGGTCGAACTTGACTATCTAGAGGAAGT | >hg38_21_+_8433172_8446622_RNA45SN1_45S_50ntFlanks | 5485 | 5525 | 41 | AGCGCTGAGAAGACGGTCGAACTTGACTATCTAGAGGAAGT | 474.0 | -8.4 | 1.3 | -6.7 | 1.6E-11 | 3.9E-11 | MB231/MB468 | MITO |
| >hg38_21_+_8433172_8446622_RNA45SN1_45S_50ntFlanks@10351.10391.41\|CGAGAACTTTGAAGGCCGAAGTGGAGAAGGGTTCCATGTGA | >hg38_21_+_8433172_8446622_RNA45SN1_45S_50ntFlanks | 10351 | 10391 | 41 | CGAGAACTTTGAAGGCCGAAGTGGAGAAGGGTTCCATGTGA | 1347.6 | -5.6 | 0.8 | -6.8 | 1.4E-11 | 3.9E-11 | MB231/MB468 | NUC |
| >hg38_21_+_8433172_8446622_RNA45SN1_45S_50ntFlanks@10781.10813.33\|AGGTGAACAGCCTCTGGCATGTTGGAACAATGT | >hg38_21_+_8433172_8446622_RNA45SN1_45S_50ntFlanks | 10781 | 10813 | 33 | AGGTGAACAGCCTCTGGCATGTTGGAACAATGT | 129.0 | -10.9 | 1.6 | -6.8 | 1.2E-11 | 3.9E-11 | MB231/MB468 | MP |
| >trna119_LysCTT_1_-_145395522_145395594@5.35.31,trna11_LysCTT_5_-_180648979_180649051@5.35.31,trna13_LysCTT_6_+_26556774_26556846@5.35.31,trna7_LysCTT_16_+_3225692_3225764@5.35.31,trna9_LysCTT_5_+_180634755_180634827@5.35.31\|GGCTAGCTCAGTCGGTAGAGCATGAGACTCT | >trna119_LysCTT_1_-_145395522_145395594 | 5 | 35 | 31 | GGCTAGCTCAGTCGGTAGAGCATGAGACTCT | 280.5 | 8.0 | 1.2 | 6.8 | 1.4E-11 | 3.9E-11 | MB231/MB468 | NUC |
| >hg38_21_+_8433172_8446622_RNA45SN1_45S_50ntFlanks@10668.10706.39\|GAAAGCGTCGCGGTTCCGGCGGCGTCCGGTGAGCTCTCG | >hg38_21_+_8433172_8446622_RNA45SN1_45S_50ntFlanks | 10668 | 10706 | 39 | GAAAGCGTCGCGGTTCCGGCGGCGTCCGGTGAGCTCTCG | 758.3 | -9.2 | 1.4 | -6.7 | 1.6E-11 | 3.9E-11 | MB231/MB468 | MITO |
| >hg38_21_+_8433172_8446622_RNA45SN1_45S_50ntFlanks@10684.10711.28\|CGGCGGCGTCCGGTGAGCTCTCGCTGGC | >hg38_21_+_8433172_8446622_RNA45SN1_45S_50ntFlanks | 10684 | 10711 | 28 | CGGCGGCGTCCGGTGAGCTCTCGCTGGC | 4427.2 | -4.5 | 0.7 | -6.8 | 1.2E-11 | 3.9E-11 | BT20/MB468 | CYTO |
| >trna10_AspGTC_12_-_125424193_125424264@35.73.39,trna12_AspGTC_12_-_125411891_125411962@35.73.39,trna144_AspGTC_6_-_27551236_27551307@35.73.39,trna38_AspGTC_17_-_8125556_8125627@35.73.39,trna45_AspGTC_6_+_27447453_27447524@35.73.39,trna48_AspGTC_6_+_27471523_27471594@35.73.39,trna4_AspGTC_12_+_96429799_96429870@35.73.39,trna69_AspGTC_1_-_161440205_161440276@35.73.39,trna72_AspGTC_1_-_161432824_161432895@35.73.39,trna75_AspGTC_1_-_161425414_161425485@35.73.39,trna78_AspGTC_1_-_161418033_161418104@35.73.39,trna81_AspGTC_1_-_161410615_161410686@35.73.39\|TCACGCGGGAGACCGGGGTTCGATTCCCCGACGGGGAGC | >trna10_AspGTC_12_-_125424193_125424264 | 35 | 73 | 39 | TCACGCGGGAGACCGGGGTTCGATTCCCCGACGGGGAGC | 6245.7 | 7.0 | 1.0 | 6.8 | 1.3E-11 | 3.9E-11 | MB231/MB468 | CYTO |
| >MI0000811\|hsa-mir-148b&WithFlank&12\|+\|54337210\|54337320@69.90.22(+1A)&[MIMAT0000759&hsa-miR-148b-3p&offsets\|0\|0(+1A);m-77&12\|+\|54337278\|54337299&offsets\|0\|0(+1A)]\|TCAGTGCATCACAGAACTTTGTA | >MI0000811\|hsa-mir-148b&WithFlank&12\|+\|54337210\|54337320 | 69 | 90 | 22(+1A) | TCAGTGCATCACAGAACTTTGTA | 467.4 | 7.1 | 1.1 | 6.8 | 1.2E-11 | 3.9E-11 | BT20/MB468 | CYTO |
| >hg38_21_+_8433172_8446622_RNA45SN1_45S_50ntFlanks@10678.10712.35\|CGGTTCCGGCGGCGTCCGGTGAGCTCTCGCTGGCC | >hg38_21_+_8433172_8446622_RNA45SN1_45S_50ntFlanks | 10678 | 10712 | 35 | CGGTTCCGGCGGCGTCCGGTGAGCTCTCGCTGGCC | 13614.1 | -6.2 | 0.9 | -7.2 | 7.9E-13 | 3.9E-11 | BT20/MB468 | MP |
| >trna11_GluTTC_15_-_26327381_26327452@1.35.35,trna3_GluTTC_13_-_45492062_45492133@1.35.35\|TCCCACATGGTCTAGCGGTTAGGATTCCTGGTTTT | >trna11_GluTTC_15_-_26327381_26327452 | 1 | 35 | 35 | TCCCACATGGTCTAGCGGTTAGGATTCCTGGTTTT | 423.1 | 10.1 | 1.5 | 6.8 | 1.3E-11 | 3.9E-11 | MB231/MB468 | CYTO |
| >hg38_21_+_8433172_8446622_RNA45SN1_45S_50ntFlanks@12626.12651.26\|AATGGGGCGAAGCTACCATCTGTGGG | >hg38_21_+_8433172_8446622_RNA45SN1_45S_50ntFlanks | 12626 | 12651 | 26 | AATGGGGCGAAGCTACCATCTGTGGG | 350.6 | -11.5 | 1.6 | -7.2 | 8.0E-13 | 3.9E-11 | BT20/MB468 | MP |
| >hg38_21_+_8433172_8446622_RNA45SN1_45S_50ntFlanks@12637.12659.23\|GCTACCATCTGTGGGATTATGAC | >hg38_21_+_8433172_8446622_RNA45SN1_45S_50ntFlanks | 12637 | 12659 | 23 | GCTACCATCTGTGGGATTATGAC | 3929.0 | -8.6 | 1.3 | -6.7 | 1.6E-11 | 3.9E-11 | MB231/MB468 | MITO |
| >hg38_21_+_8433172_8446622_RNA45SN1_45S_50ntFlanks@5486.5518.33\|GCGCTGAGAAGACGGTCGAACTTGACTATCTAG | >hg38_21_+_8433172_8446622_RNA45SN1_45S_50ntFlanks | 5486 | 5518 | 33 | GCGCTGAGAAGACGGTCGAACTTGACTATCTAG | 314.0 | -8.2 | 1.2 | -6.7 | 1.6E-11 | 3.9E-11 | MB231/MB468 | MITO |
| >hg38_21_+_8433172_8446622_RNA45SN1_45S_50ntFlanks@7982.8007.26\|TCAGATCAGACGTGGCGACCCGCTGA | >hg38_21_+_8433172_8446622_RNA45SN1_45S_50ntFlanks | 7982 | 8007 | 26 | TCAGATCAGACGTGGCGACCCGCTGA | 698.1 | -6.9 | 1.0 | -6.7 | 1.6E-11 | 3.9E-11 | MB231/MB468 | MITO |
| >hg38_21_+_8433172_8446622_RNA45SN1_45S_50ntFlanks@4753.4782.30\|AAGACGATCAGATACCGTCGTAGTTCCGAC | >hg38_21_+_8433172_8446622_RNA45SN1_45S_50ntFlanks | 4753 | 4782 | 30 | AAGACGATCAGATACCGTCGTAGTTCCGAC | 780.6 | -11.7 | 1.7 | -6.8 | 9.9E-12 | 4.0E-11 | BT20/MB468 | MITO |
| >hg38_21_+_8433172_8446622_RNA45SN1_45S_50ntFlanks@10681.10705.25\|TTCCGGCGGCGTCCGGTGAGCTCTC | >hg38_21_+_8433172_8446622_RNA45SN1_45S_50ntFlanks | 10681 | 10705 | 25 | TTCCGGCGGCGTCCGGTGAGCTCTC | 87.3 | -10.3 | 1.5 | -6.8 | 1.2E-11 | 4.0E-11 | MB231/MB468 | MP |
| >MI0003820\|hsa-mir-454&WithFlank&17\|-\|59137752\|59137878@70.94.25&[MIMAT0003885&hsa-miR-454-3p&offsets\|0\|+2;m-272&17\|-\|59137787\|59137809&offsets\|0\|+2]\|TAGTGCAATATTGCTTATAGGGTTT | >MI0003820\|hsa-mir-454&WithFlank&17\|-\|59137752\|59137878 | 70 | 94 | 25 | TAGTGCAATATTGCTTATAGGGTTT | 530.2 | 6.9 | 1.0 | 6.8 | 1.4E-11 | 4.0E-11 | MB231/MB468 | NUC |
| >hg38_21_+_8433172_8446622_RNA45SN1_45S_50ntFlanks@11538.11560.23\|CGCGCGGCGCCTCGCCTCGGCCG | >hg38_21_+_8433172_8446622_RNA45SN1_45S_50ntFlanks | 11538 | 11560 | 23 | CGCGCGGCGCCTCGCCTCGGCCG | 119.0 | -10.8 | 1.6 | -6.8 | 1.2E-11 | 4.0E-11 | MB231/MB468 | MP |
| >hg38_21_+_8433172_8446622_RNA45SN1_45S_50ntFlanks@12457.12474.18\|CAAGCGTTGGATTGTTCA | >hg38_21_+_8433172_8446622_RNA45SN1_45S_50ntFlanks | 12457 | 12474 | 18 | CAAGCGTTGGATTGTTCA | 455.3 | -7.1 | 1.1 | -6.7 | 1.7E-11 | 4.0E-11 | BT20/MB468 | NUC |
| >hg38_21_+_8433172_8446622_RNA45SN1_45S_50ntFlanks@4000.4026.27\|TAGATAACCTCGGGCCGATCGCACGCC | >hg38_21_+_8433172_8446622_RNA45SN1_45S_50ntFlanks | 4000 | 4026 | 27 | TAGATAACCTCGGGCCGATCGCACGCC | 103.2 | -10.6 | 1.6 | -6.8 | 1.2E-11 | 4.0E-11 | MB231/MB468 | MP |
| >hg38_21_+_8433172_8446622_RNA45SN1_45S_50ntFlanks@6736.6770.35\|TGATCATCGACACTTCGAACGCACTTGCGGCCCCG | >hg38_21_+_8433172_8446622_RNA45SN1_45S_50ntFlanks | 6736 | 6770 | 35 | TGATCATCGACACTTCGAACGCACTTGCGGCCCCG | 123.4 | -10.8 | 1.6 | -6.8 | 1.3E-11 | 4.0E-11 | MB231/MB468 | MP |
| >hg38_21_+_8433172_8446622_RNA45SN1_45S_50ntFlanks@12511.12538.28\|CGTGAGACAGGTTAGTTTTACCCTACTG | >hg38_21_+_8433172_8446622_RNA45SN1_45S_50ntFlanks | 12511 | 12538 | 28 | CGTGAGACAGGTTAGTTTTACCCTACTG | 193.0 | -9.7 | 1.4 | -6.8 | 1.0E-11 | 4.0E-11 | BT20/MB468 | MITO |
| >hg38_21_+_8433172_8446622_RNA45SN1_45S_50ntFlanks@9125.9142.18\|CGCGCGGGTCGGGGGGCG | >hg38_21_+_8433172_8446622_RNA45SN1_45S_50ntFlanks | 9125 | 9142 | 18 | CGCGCGGGTCGGGGGGCG | 138.0 | -11.0 | 1.6 | -6.8 | 1.3E-11 | 4.0E-11 | MB231/MB468 | MP |
| >hg38_21_+_8433172_8446622_RNA45SN1_45S_50ntFlanks@9556.9597.42\|TAGCGGTCCTGACGTGCAAATCGGTCGTCCGACCTGGGTATA | >hg38_21_+_8433172_8446622_RNA45SN1_45S_50ntFlanks | 9556 | 9597 | 42 | TAGCGGTCCTGACGTGCAAATCGGTCGTCCGACCTGGGTATA | 287.7 | -12.0 | 1.8 | -6.7 | 1.6E-11 | 4.0E-11 | MB231/MB468 | MITO |
| >hg38_21_+_8433172_8446622_RNA45SN1_45S_50ntFlanks@9573.9594.22\|AAATCGGTCGTCCGACCTGGGT | >hg38_21_+_8433172_8446622_RNA45SN1_45S_50ntFlanks | 9573 | 9594 | 22 | AAATCGGTCGTCCGACCTGGGT | 2673.4 | -5.7 | 0.8 | -6.8 | 1.3E-11 | 4.0E-11 | MB231/MB468 | MP |
| >hg38_21_+_8433172_8446622_RNA45SN1_45S_50ntFlanks@5423.5466.44\|ATTGGATGGTTTAGTGAGGCCCTCGGATCGGCCCCGCCGGGGTC | >hg38_21_+_8433172_8446622_RNA45SN1_45S_50ntFlanks | 5423 | 5466 | 44 | ATTGGATGGTTTAGTGAGGCCCTCGGATCGGCCCCGCCGGGGTC | 530.1 | -12.9 | 1.9 | -6.8 | 1.3E-11 | 4.0E-11 | MB231/MB468 | MP |
| >hg38_21_+_8433172_8446622_RNA45SN1_45S_50ntFlanks@5471.5514.44\|CACGGCCCTGGCGGAGCGCTGAGAAGACGGTCGAACTTGACTAT | >hg38_21_+_8433172_8446622_RNA45SN1_45S_50ntFlanks | 5471 | 5514 | 44 | CACGGCCCTGGCGGAGCGCTGAGAAGACGGTCGAACTTGACTAT | 3269.2 | -11.1 | 1.7 | -6.7 | 1.7E-11 | 4.0E-11 | BT20/MB468 | NUC |
| >hg38_21_+_8433172_8446622_RNA45SN1_45S_50ntFlanks@6653.6676.24\|ACTCTTAGCGGTGGATCACTCGGC | >hg38_21_+_8433172_8446622_RNA45SN1_45S_50ntFlanks | 6653 | 6676 | 24 | ACTCTTAGCGGTGGATCACTCGGC | 629.2 | 5.9 | 0.9 | 6.8 | 1.4E-11 | 4.0E-11 | MB231/MB468 | CYTO |
| >hg38_21_+_8433172_8446622_RNA45SN1_45S_50ntFlanks@10349.10388.40\|AACGAGAACTTTGAAGGCCGAAGTGGAGAAGGGTTCCATG | >hg38_21_+_8433172_8446622_RNA45SN1_45S_50ntFlanks | 10349 | 10388 | 40 | AACGAGAACTTTGAAGGCCGAAGTGGAGAAGGGTTCCATG | 974.8 | -5.6 | 0.8 | -6.8 | 1.3E-11 | 4.0E-11 | MB231/MB468 | MP |

| | | | | | | | | | | | | | |
|---|---|---|---|---|---|---|---|---|---|---|---|---|---|
| >hg38_21_+_8433172_8446622_RNA45SN1_45S_50ntFlanks@12420.12456.37\|ATGTCGGCTCTTCCTATCATTGTGAAGCAGAATTCAC | >hg38_21_+_8433172_8446622_RNA45SN1_45S_50ntFlanks | 12420 | 12456 | 37 | ATGTCGGCTCTTCCTATCATTGTGAAGCAGAATTCAC | 4362.1 | -9.0 | 1.3 | -6.7 | 1.7E-11 | 4.0E-11 | BT20/MB468 | NUC |
| >hg38_21_+_8433172_8446622_RNA45SN1_45S_50ntFlanks@11575.11600.26\|ACTTAGAACTGGTGCGGACCAGGGGA | >hg38_21_+_8433172_8446622_RNA45SN1_45S_50ntFlanks | 11575 | 11600 | 26 | ACTTAGAACTGGTGCGGACCAGGGGA | 336.2 | -8.2 | 1.2 | -6.7 | 1.6E-11 | 4.0E-11 | MB231/MB468 | MITO |
| >hg38_21_+_8433172_8446622_RNA45SN1_45S_50ntFlanks@6699.6720.22\|GCTAGCTGCGAGAATTAATGTG | >hg38_21_+_8433172_8446622_RNA45SN1_45S_50ntFlanks | 6699 | 6720 | 22 | GCTAGCTGCGAGAATTAATGTG | 14715.4 | -4.6 | 0.7 | -6.8 | 1.0E-11 | 4.0E-11 | BT20/MB468 | MITO |
| >hg38_21_+_8433172_8446622_RNA45SN1_45S_50ntFlanks@11190.11207.18\|CCCGGGGGGCCGGCGGCG | >hg38_21_+_8433172_8446622_RNA45SN1_45S_50ntFlanks | 11190 | 11207 | 18 | CCCGGGGGGCCGGCGGCG | 536.1 | -12.1 | 1.7 | -7.2 | 8.2E-13 | 4.0E-11 | BT20/MB468 | MP |
| >hg38_21_+_8433172_8446622_RNA45SN1_45S_50ntFlanks@10351.10371.21\|CGAGAACTTTGAAGGCCGAAG | >hg38_21_+_8433172_8446622_RNA45SN1_45S_50ntFlanks | 10351 | 10371 | 21 | CGAGAACTTTGAAGGCCGAAG | 713.1 | -3.2 | 0.5 | -6.8 | 1.4E-11 | 4.0E-11 | MB231/MB468 | CYTO |
| >hg38_21_+_8433172_8446622_RNA45SN1_45S_50ntFlanks@12991.13028.38\|CTCCCTCGCTGCGATCTATTGAAAGTCAGCCCTCGACA | >hg38_21_+_8433172_8446622_RNA45SN1_45S_50ntFlanks | 12991 | 13028 | 38 | CTCCCTCGCTGCGATCTATTGAAAGTCAGCCCTCGACA | 615.1 | -9.5 | 1.4 | -6.8 | 1.4E-11 | 4.0E-11 | MB231/MB468 | NUC |
| >trna10_GlyTCC_17_+_8124866_8124937@58.75.18,trna117_GlyTCC_1_-_145397864_145397935@58.75.18,trna2_GlyTCC_19_+_4724082_4724153@58.75.18,trna45_GlyTCC_1_+_161500903_161500974@58.75.18,trna70_GlyTCC_1_-_161439547_161439618@58.75.18,trna73_GlyTCC_1_-_161432166_161432237@58.75.18,trna76_GlyTCC_1_-_161424756_161424827@58.75.18,trna79_GlyTCC_1_-_161417375_161417446@58.75.18\|TTCCCGGCCAACGCACCA | >trna10_GlyTCC_17_+_8124866_8124937 | 58 | 75 | 18 | TTCCCGGCCAACGCACCA | 417.9 | 8.4 | 1.2 | 6.8 | 1.3E-11 | 4.0E-11 | BT20/MB468 | CYTO |
| >hg38_21_+_8433172_8446622_RNA45SN1_45S_50ntFlanks@6722.6739.18\|ATTGCAGGACACATTGAT | >hg38_21_+_8433172_8446622_RNA45SN1_45S_50ntFlanks | 6722 | 6739 | 18 | ATTGCAGGACACATTGAT | 96.6 | -10.5 | 1.5 | -6.8 | 1.3E-11 | 4.1E-11 | MB231/MB468 | MP |
| >hg38_21_+_8433172_8446622_RNA45SN1_45S_50ntFlanks@8343.8373.31\|GTAAGGGAAAGTTGAAAAGAACTTTGAAGAG | >hg38_21_+_8433172_8446622_RNA45SN1_45S_50ntFlanks | 8343 | 8373 | 31 | GTAAGGGAAAGTTGAAAAGAACTTTGAAGAG | 419.0 | 12.5 | 1.7 | 7.2 | 8.3E-13 | 4.1E-11 | BT20/MB468 | MP |
| >hg38_21_+_8433172_8446622_RNA45SN1_45S_50ntFlanks@11729.11753.25\|TAACTATGACTCTCTTAAGGTAGCC | >hg38_21_+_8433172_8446622_RNA45SN1_45S_50ntFlanks | 11729 | 11753 | 25 | TAACTATGACTCTCTTAAGGTAGCC | 274.5 | -10.2 | 1.5 | -6.8 | 1.0E-11 | 4.1E-11 | BT20/MB468 | MITO |
| >trna10_AspGTC_12_-_125424193_125424264@31.72.42,trna12_AspGTC_12_-_125411891_125411962@31.72.42,trna38_AspGTC_17_-_8125556_8125627@31.72.42,trna45_AspGTC_6_+_27447453_27447524@31.72.42,trna48_AspGTC_6_+_27471523_27471594@31.72.42,trna4_AspGTC_12_+_96429799_96429870@31.72.42,trna69_AspGTC_1_-_161440205_161440276@31.72.42,trna72_AspGTC_1_-_161432824_161432895@31.72.42,trna75_AspGTC_1_-_161425414_161425485@31.72.42,trna78_AspGTC_1_-_161418033_161418104@31.72.42,trna81_AspGTC_1_-_161410615_161410686@31.72.42\|CCTGTCACGCGGGAGACCGGGGTTCGATTCCCCGACGGGGAG | >trna10_AspGTC_12_-_125424193_125424264 | 31 | 72 | 42 | CCTGTCACGCGGGAGACCGGGGTTCGATTCCCCGACGGGGAG | 1581.0 | -10.6 | 1.6 | -6.7 | 1.8E-11 | 4.1E-11 | BT20/MB468 | NUC |
| >MI0006273\|hsa-mir-1180&WithFlank&17\|-\|19344500\|19344580@47.66.20&[MIMAT0005825&hsa-miR-1180-3p&offsets\|0\|-2;m-214&17\|-\|19344513\|19344534&offsets\|0\|-2]\|TTTCCGGCTCGCGTGGGTGT | >MI0006273\|hsa-mir-1180&WithFlank&17\|-\|19344500\|19344580 | 47 | 66 | 20 | TTTCCGGCTCGCGTGGGTGT | 454.4 | 8.4 | 1.2 | 6.8 | 1.0E-11 | 4.1E-11 | BT20/MB468 | MITO |
| >am_trna111_HisGTG_1_-_147774845_147774916@3.29.27,trna118_HisGTG_1_-_145396881_145396952@3.29.27,trna16_HisGTG_1_+_146544773_146544844@3.29.27,trna1_HisGTG_15_+_45493349_45493420@3.29.27,trna21_HisGTG_1_+_147753471_147753542@3.29.27,trna33_HisGTG_6_+_27125906_27125977@3.29.27,trna7_HisGTG_9_-_14433938_14434009@3.29.27,trna8_HisGTG_15_-_45492611_45492682@3.29.27,trna9_HisGTG_15_-_45490804_45490875@3.29.27\|CGTGATCGTATAGTGGTTAGTACTCTG | >am_trna111_HisGTG_1_-_147774845_147774916 | 3 | 29 | 27 | CGTGATCGTATAGTGGTTAGTACTCTG | 398.3 | -8.3 | 1.2 | -6.7 | 1.7E-11 | 4.1E-11 | MB231/MB468 | MITO |
| >hg38_21_+_8433172_8446622_RNA45SN1_45S_50ntFlanks@11564.11591.28\|CCTAGCAGCCGACTTAGAACTGGTGCGG | >hg38_21_+_8433172_8446622_RNA45SN1_45S_50ntFlanks | 11564 | 11591 | 28 | CCTAGCAGCCGACTTAGAACTGGTGCGG | 761.9 | -6.7 | 1.0 | -6.8 | 1.4E-11 | 4.1E-11 | MB231/MB468 | NUC |
| >hg38_21_+_8433172_8446622_RNA45SN1_45S_50ntFlanks@9534.9561.28\|AGGAAACTCTGGTGGAGGTCCGTAGCGG | >hg38_21_+_8433172_8446622_RNA45SN1_45S_50ntFlanks | 9534 | 9561 | 28 | AGGAAACTCTGGTGGAGGTCCGTAGCGG | 3074.9 | -6.9 | 1.0 | -6.8 | 1.4E-11 | 4.1E-11 | MB231/MB468 | CYTO |
| >hg38_21_+_8433172_8446622_RNA45SN1_45S_50ntFlanks@11208.11250.43\|GCGGCGACTCTGGACGCGAGCCGGGCCCTTCCCGTGGATCGCC | >hg38_21_+_8433172_8446622_RNA45SN1_45S_50ntFlanks | 11208 | 11250 | 43 | GCGGCGACTCTGGACGCGAGCCGGGCCCTTCCCGTGGATCGCC | 604.3 | -6.7 | 1.0 | -6.8 | 1.4E-11 | 4.1E-11 | MB231/MB468 | CYTO |
| >hg38_21_+_8433172_8446622_RNA45SN1_45S_50ntFlanks@12404.12431.28\|GCTTTTTGATCCTTCGATGTCGGCTCTT | >hg38_21_+_8433172_8446622_RNA45SN1_45S_50ntFlanks | 12404 | 12431 | 28 | GCTTTTTGATCCTTCGATGTCGGCTCTT | 103.0 | -10.5 | 1.6 | -6.8 | 1.3E-11 | 4.1E-11 | MB231/MB468 | MP |
| >hg38_21_+_8433172_8446622_RNA45SN1_45S_50ntFlanks@10412.10432.21\|AGTCGGTCCTGAGAGATGGGC | >hg38_21_+_8433172_8446622_RNA45SN1_45S_50ntFlanks | 10412 | 10432 | 21 | AGTCGGTCCTGAGAGATGGGC | 100.0 | -10.5 | 1.6 | -6.8 | 1.3E-11 | 4.1E-11 | MB231/MB468 | MP |
| >hg38_21_+_8433172_8446622_RNA45SN1_45S_50ntFlanks@11579.11603.25\|AGAACTGGTGCGGACCAGGGGAATC | >hg38_21_+_8433172_8446622_RNA45SN1_45S_50ntFlanks | 11579 | 11603 | 25 | AGAACTGGTGCGGACCAGGGGAATC | 9903.2 | -7.9 | 1.1 | -7.2 | 8.4E-13 | 4.1E-11 | BT20/MB468 | MP |
| >hg38_21_+_8433172_8446622_RNA45SN1_45S_50ntFlanks@5515.5543.29\|CTAGAGGAAGTAAAAGTCGTAACAAGGTT | >hg38_21_+_8433172_8446622_RNA45SN1_45S_50ntFlanks | 5515 | 5543 | 29 | CTAGAGGAAGTAAAAGTCGTAACAAGGTT | 113.4 | -10.7 | 1.6 | -6.8 | 1.3E-11 | 4.1E-11 | MB231/MB468 | MP |
| >hg38_21_+_8433172_8446622_RNA45SN1_45S_50ntFlanks@12836.12867.32\|CGGTGCGGAGTGCCCTTCGTCCTGGGAAACGG | >hg38_21_+_8433172_8446622_RNA45SN1_45S_50ntFlanks | 12836 | 12867 | 32 | CGGTGCGGAGTGCCCTTCGTCCTGGGAAACGG | 89.8 | -10.4 | 1.5 | -6.8 | 1.3E-11 | 4.1E-11 | MB231/MB468 | MP |
| >MI0000457\|hsa-mir-141&WithFlank&12\|+\|6964091\|6964197@65.85.21&[MIMAT0000432&hsa-miR-141-3p&offsets\|0\|-1;m-134&12\|+\|6964155\|6964175&offsets\|0\|0]\|TAACACTGTCTGGTAAAGATG | >MI0000457\|hsa-mir-141&WithFlank&12\|+\|6964091\|6964197 | 65 | 85 | 21 | TAACACTGTCTGGTAAAGATG | 870.7 | 13.1 | 1.8 | 7.3 | 2.1E-13 | 4.1E-11 | BT20/MB231 | MP |
| >hg38_21_+_8433172_8446622_RNA45SN1_45S_50ntFlanks@9651.9668.18\|TCAGGATAGCTGGCGCTC | >hg38_21_+_8433172_8446622_RNA45SN1_45S_50ntFlanks | 9651 | 9668 | 18 | TCAGGATAGCTGGCGCTC | 120.6 | -10.8 | 1.6 | -6.8 | 1.3E-11 | 4.1E-11 | MB231/MB468 | MP |
| >MI0000434\|hsa-let-7i&WithFlank&12\|+\|62603680\|62603775@12.33.22(+1C)&[MIMAT0000415&hsa-let-7i-5p&offsets\|0\|0(+1C);m-49&12\|+\|62603691\|62603712&offsets\|0\|0(+1C)]\|TGAGGTAGTAGTTTGTGCTGTTC | >MI0000434\|hsa-let-7i&WithFlank&12\|+\|62603680\|62603775 | 12 | 33 | 22(+1C) | TGAGGTAGTAGTTTGTGCTGTTC | 389.7 | 5.9 | 0.9 | 6.8 | 1.4E-11 | 4.1E-11 | MB231/MB468 | CYTO |
| >hg38_21_+_8433172_8446622_RNA45SN1_45S_50ntFlanks@5442.5466.25\|CCCTCGGATCGGCCCCGCCGGGGTC | >hg38_21_+_8433172_8446622_RNA45SN1_45S_50ntFlanks | 5442 | 5466 | 25 | CCCTCGGATCGGCCCCGCCGGGGTC | 115.7 | -10.7 | 1.6 | -6.8 | 1.3E-11 | 4.1E-11 | MB231/MB468 | MP |
| >hg38_21_+_8433172_8446622_RNA45SN1_45S_50ntFlanks@10432.10456.25\|CGAGCGCCGTTCCGAAGGGACGGGC | >hg38_21_+_8433172_8446622_RNA45SN1_45S_50ntFlanks | 10432 | 10456 | 25 | CGAGCGCCGTTCCGAAGGGACGGGC | 88.3 | -10.3 | 1.5 | -6.8 | 1.3E-11 | 4.1E-11 | MB231/MB468 | MP |
| >hg38_21_+_8433172_8446622_RNA45SN1_45S_50ntFlanks@10458.10488.31\|ATGGCCTCCGTTGCCCTCGGCCGATCGAAAG | >hg38_21_+_8433172_8446622_RNA45SN1_45S_50ntFlanks | 10458 | 10488 | 31 | ATGGCCTCCGTTGCCCTCGGCCGATCGAAAG | 1588.9 | -7.2 | 1.1 | -6.7 | 1.7E-11 | 4.1E-11 | MB231/MB468 | MITO |
| >hg38_21_+_8433172_8446622_RNA45SN1_45S_50ntFlanks@12642.12661.20\|CATCTGTGGGATTATGACTG | >hg38_21_+_8433172_8446622_RNA45SN1_45S_50ntFlanks | 12642 | 12661 | 20 | CATCTGTGGGATTATGACTG | 1762.2 | -7.9 | 1.2 | -6.7 | 1.7E-11 | 4.1E-11 | MB231/MB468 | MITO |
| >hg38_21_+_8433172_8446622_RNA45SN1_45S_50ntFlanks@4159.4184.26\|ACCACATCCAAGGAAGGCAGCAGGCG | >hg38_21_+_8433172_8446622_RNA45SN1_45S_50ntFlanks | 4159 | 4184 | 26 | ACCACATCCAAGGAAGGCAGCAGGCG | 172.5 | -11.3 | 1.7 | -6.8 | 1.3E-11 | 4.1E-11 | MB231/MB468 | MP |
| >hg38_21_+_8433172_8446622_RNA45SN1_45S_50ntFlanks@12893.12924.32\|CCTCGCCCGTCACGCACCGCACGTTCGTGGGG | >hg38_21_+_8433172_8446622_RNA45SN1_45S_50ntFlanks | 12893 | 12924 | 32 | CCTCGCCCGTCACGCACCGCACGTTCGTGGGG | 114.5 | -10.7 | 1.6 | -6.8 | 1.3E-11 | 4.1E-11 | MB231/MB468 | MP |
| >trna134_GluTTC_1_-_16861774_16861845@1.33.33,trna5_GluTTC_1_+_17199078_17199149@1.33.33,trna84_GluTTC_1_-_161391883_161391954@1.33.33,trna94_GluTTC_1_-_149664355_149664427@1.33.33\|TCCCTGGTGGTCTAGTGGCTAGGATTCGGCGCT | >trna134_GluTTC_1_-_16861774_16861845 | 1 | 33 | 33 | TCCCTGGTGGTCTAGTGGCTAGGATTCGGCGCT | 1010.3 | 6.0 | 0.9 | 6.8 | 1.4E-11 | 4.1E-11 | MB231/MB468 | CYTO |
| >hg38_21_+_8433172_8446622_RNA45SN1_45S_50ntFlanks@3837.3872.36\|CCTCTCCTACTTGGATAACTGTGGTAATTCTAGAGC | >hg38_21_+_8433172_8446622_RNA45SN1_45S_50ntFlanks | 3837 | 3872 | 36 | CCTCTCCTACTTGGATAACTGTGGTAATTCTAGAGC | 466.2 | -9.1 | 1.3 | -6.7 | 1.8E-11 | 4.2E-11 | BT20/MB468 | NUC |
| >hg38_21_+_8433172_8446622_RNA45SN1_45S_50ntFlanks@10421.10443.23\|TGAGAGATGGGCGAGCGCCGTTC | >hg38_21_+_8433172_8446622_RNA45SN1_45S_50ntFlanks | 10421 | 10443 | 23 | TGAGAGATGGGCGAGCGCCGTTC | 192.4 | -7.6 | 1.1 | -6.8 | 1.3E-11 | 4.2E-11 | BT20/MB468 | CYTO |
| >hg38_21_+_8433172_8446622_RNA45SN1_45S_50ntFlanks@6669.6686.18\|CACTCGGCTCGTGCGTCG | >hg38_21_+_8433172_8446622_RNA45SN1_45S_50ntFlanks | 6669 | 6686 | 18 | CACTCGGCTCGTGCGTCG | 591.4 | -8.4 | 1.2 | -6.8 | 1.3E-11 | 4.2E-11 | MB231/MB468 | MP |
| >am_trna2_GlyGCC_21_-_18827107_18827177@1.26.26,trna35_GlyGCC_1_+_161413094_161413164@1.26.26,trna37_GlyGCC_1_+_161420467_161420537@1.26.26,trna39_GlyGCC_1_+_161427898_161427968@1.26.26,trna41_GlyGCC_1_+_161435258_161435328@1.26.26\|GCATGGGTGGTTCAGTGGTAGAATTC | >am_trna2_GlyGCC_21_-_18827107_18827177 | 1 | 26 | 26 | GCATGGGTGGTTCAGTGGTAGAATTC | 887.6 | 3.5 | 0.5 | 6.8 | 1.3E-11 | 4.2E-11 | BT20/MB468 | CYTO |
| >hg38_21_+_8433172_8446622_RNA45SN1_45S_50ntFlanks@12081.12106.26\|CTCTCGCTTCTGGCGCCAAGCGCCCG | >hg38_21_+_8433172_8446622_RNA45SN1_45S_50ntFlanks | 12081 | 12106 | 26 | CTCTCGCTTCTGGCGCCAAGCGCCCG | 113.0 | -10.7 | 1.6 | -6.8 | 1.3E-11 | 4.2E-11 | MB231/MB468 | MP |
| >hg38_21_+_8433172_8446622_RNA45SN1_45S_50ntFlanks@12800.12817.18\|AGGGCGCGTGCCCCGCCG | >hg38_21_+_8433172_8446622_RNA45SN1_45S_50ntFlanks | 12800 | 12817 | 18 | AGGGCGCGTGCCCCGCCG | 164.4 | -11.2 | 1.7 | -6.8 | 1.3E-11 | 4.2E-11 | MB231/MB468 | MP |
| >MI0000272\|hsa-mir-182&WithFlank&7\|-\|129770377\|129770498@29.52.24&[MIMAT0000259&hsa-miR-182-5p&offsets\|0\|0;m-22&7\|-\|129770449\|129770470&offsets\|0\|+2]\|TTTGGCAATGGTAGAACTCACACT | >MI0000272\|hsa-mir-182&WithFlank&7\|-\|129770377\|129770498 | 29 | 52 | 24 | TTTGGCAATGGTAGAACTCACACT | 51338.5 | 3.5 | 0.5 | 7.1 | 1.5E-12 | 4.2E-11 | BT20/MB231 | NUC |
| >MI0000449\|hsa-mir-132&WithFlank&17\|-\|2049902\|2050014@65.86.22&[MIMAT0000426&hsa-miR-132-3p&offsets\|0\|0;m-138&17\|-\|2049929\|2049950&offsets\|0\|0]\|TAACAGTCTACAGCCATGGTCG | >MI0000449\|hsa-mir-132&WithFlank&17\|-\|2049902\|2050014 | 65 | 86 | 22 | TAACAGTCTACAGCCATGGTCG | 385.8 | 9.0 | 1.3 | 6.8 | 1.4E-11 | 4.2E-11 | MB231/MB468 | CYTO |
| >hg38_21_+_8433172_8446622_RNA45SN1_45S_50ntFlanks@12243.12280.38\|GAAGGGCAAAAGCTCGCTTGATCTTGATTTTCAGTACG | >hg38_21_+_8433172_8446622_RNA45SN1_45S_50ntFlanks | 12243 | 12280 | 38 | GAAGGGCAAAAGCTCGCTTGATCTTGATTTTCAGTACG | 176.1 | 8.6 | 1.3 | 6.8 | 1.4E-11 | 4.2E-11 | MB231/MB468 | CYTO |
| >hg38_21_+_8433172_8446622_RNA45SN1_45S_50ntFlanks@7989.8019.31\|AGACGTGGCGACCCGCTGAATTTAAGCATAT | >hg38_21_+_8433172_8446622_RNA45SN1_45S_50ntFlanks | 7989 | 8019 | 31 | AGACGTGGCGACCCGCTGAATTTAAGCATAT | 874.6 | 9.7 | 1.4 | 6.8 | 1.4E-11 | 4.2E-11 | MB231/MB468 | CYTO |
| >hg38_21_+_8433172_8446622_RNA45SN1_45S_50ntFlanks@8839.8867.29\|CGGTGGCGGCGGCGGCGGCGGCGGCGGGA | >hg38_21_+_8433172_8446622_RNA45SN1_45S_50ntFlanks | 8839 | 8867 | 29 | CGGTGGCGGCGGCGGCGGCGGCGGCGGGA | 170.8 | -11.3 | 1.7 | -6.8 | 1.3E-11 | 4.2E-11 | MB231/MB468 | MP |

<table>
<tr><td>>MI0000264|hsa-mir-7-2&WithFlank&15|+|88611819|88611940@38.60.23(+1G);MI0000265|hsa-mir-7-3&WithFlank&19|+|4770664|4770785@37.59.23(+1G);MI0000263|hsa-mir-7-1&WithFlank&9|-|83969742|83969863@30.52.23(+1G)&[MIMAT0000252&hsa-miR-7-5p&offsets|0|-1(+1G);m-400&15|+|88611856|88611879&offsets|0|-1(+1G)];[MIMAT0000252_1&hsa-miR-7-5p&offsets|0|-1(+1G);m-402&19|+|4770700|4770723&offsets|0|-1(+1G)];[MIMAT0000252_2&hsa-miR-7-5p&offsets|0|-1(+1G);m-401&9|-|83969811|83969834&offsets|0|-1(+1G)]|TGGAAGACTAGTGATTTTGTTGTG</td><td>>MI0000264|hsa-mir-7-2&WithFlank&15|+|88611819|88611940</td><td>38</td><td>60</td><td>23(+1G)</td><td>TGGAAGACTAGTGATTTTGTTGTG</td><td>4029.6</td><td>6.9</td><td>1.0</td><td>6.8</td><td>1.3E-11</td><td>4.2E-11</td><td>BT20/MB468</td><td>CYTO</td></tr>
<tr><td>>MI0000477|hsa-mir-146a&WithFlank&5|+|160485346|160485456@27.47.21(+1G)&[MIMAT0000449&hsa-miR-146a-5p&offsets|0|-1(+1G);m-129&5|+|160485372|160485393&offsets|0|-1(+1G)]|TGAGAACTGAATTCCATGGGTG</td><td>>MI0000477|hsa-mir-146a&WithFlank&5|+|160485346|160485456</td><td>27</td><td>47</td><td>21(+1G)</td><td>TGAGAACTGAATTCCATGGGTG</td><td>206.6</td><td>-10.9</td><td>1.5</td><td>-7.1</td><td>1.3E-12</td><td>4.2E-11</td><td>BT20/MB231</td><td>MITO</td></tr>
<tr><td>>MI0000446|hsa-mir-125b-1&WithFlank&11|-|122099751|122099850@61.81.21(+1C)&[MIMAT0004592&hsa-miR-125b-1-3p&offsets|0|-1(+1C);m-485&11|-|122099769|122099790&offsets|0|-1(+1C)]|ACGGGTTAGGCTCTTGGGAGCC</td><td>>MI0000446|hsa-mir-125b-1&WithFlank&11|-|122099751|122099850</td><td>61</td><td>81</td><td>21(+1C)</td><td>ACGGGTTAGGCTCTTGGGAGCC</td><td>191.6</td><td>10.8</td><td>1.6</td><td>6.8</td><td>1.3E-11</td><td>4.2E-11</td><td>MB231/MB468</td><td>MP</td></tr>
<tr><td>>hg38_21_+_8433172_8446622_RNA45SN1_45S_50ntFlanks@13001.13027.27|GCGATCTATTGAAAGTCAGCCCTCGAC</td><td>>hg38_21_+_8433172_8446622_RNA45SN1_45S_50ntFlanks</td><td>13001</td><td>13027</td><td>27</td><td>GCGATCTATTGAAAGTCAGCCCTCGAC</td><td>80.1</td><td>-10.2</td><td>1.5</td><td>-6.8</td><td>1.3E-11</td><td>4.2E-11</td><td>MB231/MB468</td><td>MP</td></tr>
<tr><td>>hg38_21_+_8433172_8446622_RNA45SN1_45S_50ntFlanks@6721.6758.38|AATTGCAGGACACATTGATCATCGACACTTCGAACGCA</td><td>>hg38_21_+_8433172_8446622_RNA45SN1_45S_50ntFlanks</td><td>6721</td><td>6758</td><td>38</td><td>AATTGCAGGACACATTGATCATCGACACTTCGAACGCA</td><td>3504.8</td><td>-7.6</td><td>1.1</td><td>-6.7</td><td>1.7E-11</td><td>4.2E-11</td><td>MB231/MB468</td><td>MITO</td></tr>
<tr><td>>hg38_21_+_8433172_8446622_RNA45SN1_45S_50ntFlanks@6722.6745.24|ATTGCAGGACACATTGATCATCGA</td><td>>hg38_21_+_8433172_8446622_RNA45SN1_45S_50ntFlanks</td><td>6722</td><td>6745</td><td>24</td><td>ATTGCAGGACACATTGATCATCGA</td><td>295.9</td><td>-5.8</td><td>0.9</td><td>-6.8</td><td>1.5E-11</td><td>4.2E-11</td><td>MB231/MB468</td><td>CYTO</td></tr>
<tr><td>>hg38_21_+_8433172_8446622_RNA45SN1_45S_50ntFlanks@11908.11929.22|CGGTGAAGAGACATGAGAGGTG</td><td>>hg38_21_+_8433172_8446622_RNA45SN1_45S_50ntFlanks</td><td>11908</td><td>11929</td><td>22</td><td>CGGTGAAGAGACATGAGAGGTG</td><td>159.9</td><td>-11.2</td><td>1.7</td><td>-6.8</td><td>1.3E-11</td><td>4.2E-11</td><td>MB231/MB468</td><td>MP</td></tr>
<tr><td>>hg38_21_+_8433172_8446622_RNA45SN1_45S_50ntFlanks@4590.4615.26|ATTTTGTTGGTTTTCGGAACTGAGGC</td><td>>hg38_21_+_8433172_8446622_RNA45SN1_45S_50ntFlanks</td><td>4590</td><td>4615</td><td>26</td><td>ATTTTGTTGGTTTTCGGAACTGAGGC</td><td>98.1</td><td>-10.5</td><td>1.5</td><td>-6.8</td><td>1.3E-11</td><td>4.2E-11</td><td>MB231/MB468</td><td>MP</td></tr>
<tr><td>>hg38_21_+_8433172_8446622_RNA45SN1_45S_50ntFlanks@3985.4002.18|CGGCTTTGGTGACTCTAG</td><td>>hg38_21_+_8433172_8446622_RNA45SN1_45S_50ntFlanks</td><td>3985</td><td>4002</td><td>18</td><td>CGGCTTTGGTGACTCTAG</td><td>2093.0</td><td>-4.8</td><td>0.7</td><td>-6.8</td><td>1.4E-11</td><td>4.2E-11</td><td>BT20/MB468</td><td>CYTO</td></tr>
<tr><td>>hg38_21_+_8433172_8446622_RNA45SN1_45S_50ntFlanks@12934.12952.19|CTAAACCATTCGTAGACGA</td><td>>hg38_21_+_8433172_8446622_RNA45SN1_45S_50ntFlanks</td><td>12934</td><td>12952</td><td>19</td><td>CTAAACCATTCGTAGACGA</td><td>782.1</td><td>-8.0</td><td>1.2</td><td>-6.8</td><td>1.4E-11</td><td>4.3E-11</td><td>BT20/MB468</td><td>CYTO</td></tr>
<tr><td>>hg38_21_+_8433172_8446622_RNA45SN1_45S_50ntFlanks@6669.6699.31|CACTCGGCTCGTGCGTCGATGAAGAACGCAG</td><td>>hg38_21_+_8433172_8446622_RNA45SN1_45S_50ntFlanks</td><td>6669</td><td>6699</td><td>31</td><td>CACTCGGCTCGTGCGTCGATGAAGAACGCAG</td><td>433.5</td><td>-8.5</td><td>1.3</td><td>-6.8</td><td>1.3E-11</td><td>4.3E-11</td><td>MB231/MB468</td><td>MP</td></tr>
<tr><td>>hg38_21_+_8433172_8446622_RNA45SN1_45S_50ntFlanks@4003.4039.37|ATAACCTCGGGCCGATCGCACGCCCCCCGTGGCGGCG</td><td>>hg38_21_+_8433172_8446622_RNA45SN1_45S_50ntFlanks</td><td>4003</td><td>4039</td><td>37</td><td>ATAACCTCGGGCCGATCGCACGCCCCCCGTGGCGGCG</td><td>813.0</td><td>-9.5</td><td>1.4</td><td>-6.7</td><td>1.8E-11</td><td>4.3E-11</td><td>BT20/MB468</td><td>NUC</td></tr>
<tr><td>>hg38_21_+_8433172_8446622_RNA45SN1_45S_50ntFlanks@9504.9525.22|GTGAACTATGCCTGGGCAGGGC</td><td>>hg38_21_+_8433172_8446622_RNA45SN1_45S_50ntFlanks</td><td>9504</td><td>9525</td><td>22</td><td>GTGAACTATGCCTGGGCAGGGC</td><td>1353.7</td><td>-8.9</td><td>1.3</td><td>-6.7</td><td>1.7E-11</td><td>4.3E-11</td><td>MB231/MB468</td><td>MITO</td></tr>
<tr><td>>hg38_21_+_8433172_8446622_RNA45SN1_45S_50ntFlanks@13009.13027.19|TTGAAAGTCAGCCCTCGAC</td><td>>hg38_21_+_8433172_8446622_RNA45SN1_45S_50ntFlanks</td><td>13009</td><td>13027</td><td>19</td><td>TTGAAAGTCAGCCCTCGAC</td><td>107.1</td><td>-10.6</td><td>1.6</td><td>-6.8</td><td>1.4E-11</td><td>4.3E-11</td><td>MB231/MB468</td><td>MP</td></tr>
<tr><td>>hg38_21_+_8433172_8446622_RNA45SN1_45S_50ntFlanks@5421.5445.25|CGATTGGATGGTTTAGTGAGGCCCT</td><td>>hg38_21_+_8433172_8446622_RNA45SN1_45S_50ntFlanks</td><td>5421</td><td>5445</td><td>25</td><td>CGATTGGATGGTTTAGTGAGGCCCT</td><td>690.4</td><td>-6.2</td><td>0.9</td><td>-6.8</td><td>1.5E-11</td><td>4.3E-11</td><td>MB231/MB468</td><td>CYTO</td></tr>
<tr><td>>hg38_21_+_8433172_8446622_RNA45SN1_45S_50ntFlanks@4895.4918.24|CTTAAAGGAATTGACGGAAGGGCA</td><td>>hg38_21_+_8433172_8446622_RNA45SN1_45S_50ntFlanks</td><td>4895</td><td>4918</td><td>24</td><td>CTTAAAGGAATTGACGGAAGGGCA</td><td>125.7</td><td>-10.8</td><td>1.6</td><td>-6.8</td><td>1.4E-11</td><td>4.3E-11</td><td>MB231/MB468</td><td>MP</td></tr>
<tr><td>>trnaMT_MetCAT_MT_+_4402_4469@38.71.34|CGAAAATGTTGGTTATACCCTTCCCGTACTACCA</td><td>>trnaMT_MetCAT_MT_+_4402_4469</td><td>38</td><td>71</td><td>34</td><td>CGAAAATGTTGGTTATACCCTTCCCGTACTACCA</td><td>269.2</td><td>9.3</td><td>1.4</td><td>6.7</td><td>1.5E-11</td><td>4.3E-11</td><td>MB231/MB468</td><td>NUC</td></tr>
<tr><td>>hg38_21_+_8433172_8446622_RNA45SN1_45S_50ntFlanks@7973.8011.39|GACGCGACCTCAGATCAGACGTGGCGACCCGCTGAATTT</td><td>>hg38_21_+_8433172_8446622_RNA45SN1_45S_50ntFlanks</td><td>7973</td><td>8011</td><td>39</td><td>GACGCGACCTCAGATCAGACGTGGCGACCCGCTGAATTT</td><td>3244.1</td><td>4.7</td><td>0.7</td><td>6.7</td><td>1.5E-11</td><td>4.3E-11</td><td>MB231/MB468</td><td>NUC</td></tr>
<tr><td>>am_trnaMT_SerGCT_MT_+_12207_12265@12.38.27|CAAGAACTGCTAACTCATGCCCCCATG</td><td>>am_trnaMT_SerGCT_MT_+_12207_12265</td><td>12</td><td>38</td><td>27</td><td>CAAGAACTGCTAACTCATGCCCCCATG</td><td>1331.9</td><td>-6.1</td><td>0.9</td><td>-6.7</td><td>1.5E-11</td><td>4.3E-11</td><td>MB231/MB468</td><td>NUC</td></tr>
<tr><td>>trnaMT_LeuTAG_MT_+_12266_12336@37.74.38|CCCCAAAAATTTTGGTGCAACTCCAAATAAAAGTACCA</td><td>>trnaMT_LeuTAG_MT_+_12266_12336</td><td>37</td><td>74</td><td>38</td><td>CCCCAAAAATTTTGGTGCAACTCCAAATAAAAGTACCA</td><td>207.6</td><td>-10.9</td><td>1.5</td><td>-7.1</td><td>1.4E-12</td><td>4.3E-11</td><td>BT20/MB231</td><td>MITO</td></tr>
<tr><td>>hg38_21_+_8433172_8446622_RNA45SN1_45S_50ntFlanks@9507.9525.19|AACTATGCCTGGGCAGGGC</td><td>>hg38_21_+_8433172_8446622_RNA45SN1_45S_50ntFlanks</td><td>9507</td><td>9525</td><td>19</td><td>AACTATGCCTGGGCAGGGC</td><td>208.8</td><td>-11.6</td><td>1.7</td><td>-6.8</td><td>1.4E-11</td><td>4.3E-11</td><td>MB231/MB468</td><td>MP</td></tr>
<tr><td>>hg38_21_+_8433172_8446622_RNA45SN1_45S_50ntFlanks@12431.12455.25|TCCTATCATTGTGAAGCAGAATTCA</td><td>>hg38_21_+_8433172_8446622_RNA45SN1_45S_50ntFlanks</td><td>12431</td><td>12455</td><td>25</td><td>TCCTATCATTGTGAAGCAGAATTCA</td><td>103.0</td><td>-10.5</td><td>1.6</td><td>-6.8</td><td>1.4E-11</td><td>4.3E-11</td><td>MB231/MB468</td><td>MP</td></tr>
<tr><td>>hg38_21_+_8433172_8446622_RNA45SN1_45S_50ntFlanks@12940.12964.25|CATTCGTAGACGACCTGCTTCTGGG</td><td>>hg38_21_+_8433172_8446622_RNA45SN1_45S_50ntFlanks</td><td>12940</td><td>12964</td><td>25</td><td>CATTCGTAGACGACCTGCTTCTGGG</td><td>77.8</td><td>-10.1</td><td>1.5</td><td>-6.8</td><td>1.4E-11</td><td>4.3E-11</td><td>MB231/MB468</td><td>MP</td></tr>
<tr><td>>hg38_21_+_8433172_8446622_RNA45SN1_45S_50ntFlanks@10838.10867.30|ATCCGTAACTTCGGGATAAGGATTGGCTCT</td><td>>hg38_21_+_8433172_8446622_RNA45SN1_45S_50ntFlanks</td><td>10838</td><td>10867</td><td>30</td><td>ATCCGTAACTTCGGGATAAGGATTGGCTCT</td><td>171.5</td><td>-9.7</td><td>1.4</td><td>-6.8</td><td>1.4E-11</td><td>4.3E-11</td><td>BT20/MB468</td><td>CYTO</td></tr>
<tr><td>>hg38_21_+_8433172_8446622_RNA45SN1_45S_50ntFlanks@11562.11590.29|CGCCTAGCAGCCGACTTAGAACTGGTGCG</td><td>>hg38_21_+_8433172_8446622_RNA45SN1_45S_50ntFlanks</td><td>11562</td><td>11590</td><td>29</td><td>CGCCTAGCAGCCGACTTAGAACTGGTGCG</td><td>112.2</td><td>-10.7</td><td>1.6</td><td>-6.8</td><td>1.4E-11</td><td>4.3E-11</td><td>MB231/MB468</td><td>MP</td></tr>
<tr><td>>hg38_21_+_8433172_8446622_RNA45SN1_45S_50ntFlanks@11569.11612.44|CAGCCGACTTAGAACTGGTGCGGACCAGGGGAATCCGACTGTTT</td><td>>hg38_21_+_8433172_8446622_RNA45SN1_45S_50ntFlanks</td><td>11569</td><td>11612</td><td>44</td><td>CAGCCGACTTAGAACTGGTGCGGACCAGGGGAATCCGACTGTTT</td><td>7055.7</td><td>-6.5</td><td>1.0</td><td>-6.7</td><td>1.8E-11</td><td>4.3E-11</td><td>MB231/MB468</td><td>MITO</td></tr>
<tr><td>>trnaMT_PheGAA_MT_+_577_647@36.62.27|AAAATGTTTAGACGGGCTCACATCACC</td><td>>trnaMT_PheGAA_MT_+_577_647</td><td>36</td><td>62</td><td>27</td><td>AAAATGTTTAGACGGGCTCACATCACC</td><td>208.1</td><td>6.8</td><td>1.0</td><td>6.7</td><td>1.5E-11</td><td>4.3E-11</td><td>MB231/MB468</td><td>NUC</td></tr>
<tr><td>>hg38_21_+_8433172_8446622_RNA45SN1_45S_50ntFlanks@12954.12972.19|CTGCTTCTGGGTCGGGGTT</td><td>>hg38_21_+_8433172_8446622_RNA45SN1_45S_50ntFlanks</td><td>12954</td><td>12972</td><td>19</td><td>CTGCTTCTGGGTCGGGGTT</td><td>839.9</td><td>-8.8</td><td>1.3</td><td>-6.7</td><td>1.5E-11</td><td>4.3E-11</td><td>MB231/MB468</td><td>NUC</td></tr>
<tr><td>>hg38_21_+_8433172_8446622_RNA45SN1_45S_50ntFlanks@11759.11777.19|CCTCGTCATCTAATTAGTG</td><td>>hg38_21_+_8433172_8446622_RNA45SN1_45S_50ntFlanks</td><td>11759</td><td>11777</td><td>19</td><td>CCTCGTCATCTAATTAGTG</td><td>504.4</td><td>-7.8</td><td>1.2</td><td>-6.7</td><td>1.5E-11</td><td>4.3E-11</td><td>MB231/MB468</td><td>CYTO</td></tr>
<tr><td>>hg38_1_-_228634819_228635039_RNA5S12_5S_50ntFlanks@146.171.26|TGGGAATACCGGGTGCTGTAGGCTTT</td><td>>hg38_1_-_228634819_228635039_RNA5S12_5S_50ntFlanks</td><td>146</td><td>171</td><td>26</td><td>TGGGAATACCGGGTGCTGTAGGCTTT</td><td>1162.6</td><td>4.5</td><td>0.7</td><td>6.8</td><td>1.1E-11</td><td>4.3E-11</td><td>BT20/MB468</td><td>MITO</td></tr>
<tr><td>>hg38_21_+_8433172_8446622_RNA45SN1_45S_50ntFlanks@12008.12043.36|CCGCCGGTGAAATACCACTACTCTGATCGTTTTTTC</td><td>>hg38_21_+_8433172_8446622_RNA45SN1_45S_50ntFlanks</td><td>12008</td><td>12043</td><td>36</td><td>CCGCCGGTGAAATACCACTACTCTGATCGTTTTTTC</td><td>11256.3</td><td>-8.2</td><td>1.2</td><td>-6.8</td><td>1.1E-11</td><td>4.3E-11</td><td>BT20/MB468</td><td>MITO</td></tr>
<tr><td>>MI0000102|hsa-mir-100&WithFlank&11|-|122152223|122152314@22.40.19&[MIMAT0000098&hsa-miR-100-5p&offsets|+3|0;m-27&11|-|122152276|122152296&offsets|+3|+1]|CCGTAGATCCGAACTTGTG</td><td>>MI0000102|hsa-mir-100&WithFlank&11|-|122152223|122152314</td><td>22</td><td>40</td><td>19</td><td>CCGTAGATCCGAACTTGTG</td><td>389.4</td><td>11.8</td><td>1.7</td><td>6.8</td><td>1.4E-11</td><td>4.3E-11</td><td>MB231/MB468</td><td>MP</td></tr>
<tr><td>>hg38_21_+_8433172_8446622_RNA45SN1_45S_50ntFlanks@12321.12344.24|CTTTTGGGTTTTAAGCAGGAGGTG</td><td>>hg38_21_+_8433172_8446622_RNA45SN1_45S_50ntFlanks</td><td>12321</td><td>12344</td><td>24</td><td>CTTTTGGGTTTTAAGCAGGAGGTG</td><td>81.8</td><td>-10.2</td><td>1.5</td><td>-6.8</td><td>1.4E-11</td><td>4.4E-11</td><td>MB231/MB468</td><td>MP</td></tr>
<tr><td>>MI0000065|hsa-let-7d&WithFlank&9|+|94178828|94178926@14.34.21&[MIMAT0000065&hsa-let-7d-5p&offsets|0|-1;m-90&9|+|94178841|94178862&offsets|0|-1]|AGAGGTAGTAGGTTGCATAGT</td><td>>MI0000065|hsa-let-7d&WithFlank&9|+|94178828|94178926</td><td>14</td><td>34</td><td>21</td><td>AGAGGTAGTAGGTTGCATAGT</td><td>596.9</td><td>2.7</td><td>0.4</td><td>6.7</td><td>1.9E-11</td><td>4.4E-11</td><td>BT20/MB468</td><td>NUC</td></tr>
<tr><td>>hg38_21_+_8433172_8446622_RNA45SN1_45S_50ntFlanks@3874.3898.25|AATACATGCCGACGGGCGCTGACCC</td><td>>hg38_21_+_8433172_8446622_RNA45SN1_45S_50ntFlanks</td><td>3874</td><td>3898</td><td>25</td><td>AATACATGCCGACGGGCGCTGACCC</td><td>8237.9</td><td>-9.6</td><td>1.4</td><td>-6.7</td><td>1.9E-11</td><td>4.4E-11</td><td>BT20/MB468</td><td>NUC</td></tr>
<tr><td>>trna10_LysCTT_16_+_3241501_3241573@1.34.34,trna13_LysCTT_14_-_58706613_58706685@1.34.34,trna2_LysCTT_15_+_79152904_79152976@1.34.34|GCCCGGCTAGCTCAGTCGGTAGAGCATGGGACTC</td><td>>trna10_LysCTT_16_+_3241501_3241573</td><td>1</td><td>34</td><td>34</td><td>GCCCGGCTAGCTCAGTCGGTAGAGCATGGGACTC</td><td>748.0</td><td>8.1</td><td>1.2</td><td>6.7</td><td>1.5E-11</td><td>4.4E-11</td><td>MB231/MB468</td><td>NUC</td></tr>
<tr><td>>trna152_ValCAC_6_-_27248049_27248121@1.34.34|GCTTCTGTAGTGTAGTGGTTATCACGTTCGCCTC</td><td>>trna152_ValCAC_6_-_27248049_27248121</td><td>1</td><td>34</td><td>34</td><td>GCTTCTGTAGTGTAGTGGTTATCACGTTCGCCTC</td><td>6202.0</td><td>10.3</td><td>1.5</td><td>6.7</td><td>1.5E-11</td><td>4.4E-11</td><td>MB231/MB468</td><td>NUC</td></tr>
<tr><td>>hg38_21_+_8433172_8446622_RNA45SN1_45S_50ntFlanks@10843.10863.21|TAACTTCGGGATAAGGATTGG</td><td>>hg38_21_+_8433172_8446622_RNA45SN1_45S_50ntFlanks</td><td>10843</td><td>10863</td><td>21</td><td>TAACTTCGGGATAAGGATTGG</td><td>724.7</td><td>-5.9</td><td>0.9</td><td>-6.7</td><td>1.5E-11</td><td>4.4E-11</td><td>MB231/MB468</td><td>CYTO</td></tr>
<tr><td>>hg38_21_+_8433172_8446622_RNA45SN1_45S_50ntFlanks@11188.11225.38|CACCCGGGGGGCCGGCGGCGGCGGCGACTCTGGACGCG</td><td>>hg38_21_+_8433172_8446622_RNA45SN1_45S_50ntFlanks</td><td>11188</td><td>11225</td><td>38</td><td>CACCCGGGGGGCCGGCGGCGGCGGCGACTCTGGACGCG</td><td>319.3</td><td>-7.1</td><td>1.0</td><td>-6.7</td><td>1.5E-11</td><td>4.4E-11</td><td>MB231/MB468</td><td>CYTO</td></tr>
<tr><td>>hg38_21_+_8433172_8446622_RNA45SN1_45S_50ntFlanks@12589.12606.18|GTTCAGACATTTGGTGTA</td><td>>hg38_21_+_8433172_8446622_RNA45SN1_45S_50ntFlanks</td><td>12589</td><td>12606</td><td>18</td><td>GTTCAGACATTTGGTGTA</td><td>80.2</td><td>-10.2</td><td>1.5</td><td>-6.8</td><td>1.4E-11</td><td>4.4E-11</td><td>MB231/MB468</td><td>MP</td></tr>
<tr><td>>hg38_21_+_8433172_8446622_RNA45SN1_45S_50ntFlanks@12457.12487.31|CAAGCGTTGGATTGTTCACCCACTAATAGGG</td><td>>hg38_21_+_8433172_8446622_RNA45SN1_45S_50ntFlanks</td><td>12457</td><td>12487</td><td>31</td><td>CAAGCGTTGGATTGTTCACCCACTAATAGGG</td><td>1875.8</td><td>-9.4</td><td>1.4</td><td>-6.7</td><td>1.8E-11</td><td>4.4E-11</td><td>MB231/MB468</td><td>MITO</td></tr>
<tr><td>>hg38_21_+_8433172_8446622_RNA45SN1_45S_50ntFlanks@10678.10711.34|CGGTTCCGGCGGCGTCCGGTGAGCTCTCGCTGGC</td><td>>hg38_21_+_8433172_8446622_RNA45SN1_45S_50ntFlanks</td><td>10678</td><td>10711</td><td>34</td><td>CGGTTCCGGCGGCGTCCGGTGAGCTCTCGCTGGC</td><td>80477.5</td><td>-6.3</td><td>0.9</td><td>-6.8</td><td>1.1E-11</td><td>4.4E-11</td><td>BT20/MB468</td><td>MITO</td></tr>
<tr><td>>hg38_21_+_8433172_8446622_RNA45SN1_45S_50ntFlanks@12628.12653.26|TGGGGCGAAGCTACCATCTGTGGGAT</td><td>>hg38_21_+_8433172_8446622_RNA45SN1_45S_50ntFlanks</td><td>12628</td><td>12653</td><td>26</td><td>TGGGGCGAAGCTACCATCTGTGGGAT</td><td>1398.7</td><td>-8.2</td><td>1.2</td><td>-6.8</td><td>1.1E-11</td><td>4.4E-11</td><td>BT20/MB468</td><td>MITO</td></tr>
<tr><td>>hg38_21_+_8433172_8446622_RNA45SN1_45S_50ntFlanks@12635.12661.27|AAGCTACCATCTGTGGGATTATGACTG</td><td>>hg38_21_+_8433172_8446622_RNA45SN1_45S_50ntFlanks</td><td>12635</td><td>12661</td><td>27</td><td>AAGCTACCATCTGTGGGATTATGACTG</td><td>9925.9</td><td>-4.7</td><td>0.7</td><td>-6.8</td><td>1.1E-11</td><td>4.4E-11</td><td>BT20/MB468</td><td>MITO</td></tr>
<tr><td>>hg38_21_+_8433172_8446622_RNA45SN1_45S_50ntFlanks@12909.12933.25|CCGCACGTTCGTGGGGAACCTGGCG</td><td>>hg38_21_+_8433172_8446622_RNA45SN1_45S_50ntFlanks</td><td>12909</td><td>12933</td><td>25</td><td>CCGCACGTTCGTGGGGAACCTGGCG</td><td>558.5</td><td>-8.2</td><td>1.2</td><td>-6.7</td><td>1.8E-11</td><td>4.4E-11</td><td>MB231/MB468</td><td>MITO</td></tr>
<tr><td>>hg38_21_+_8433172_8446622_RNA45SN1_45S_50ntFlanks@9563.9597.35|CCTGACGTGCAAATCGGTCGTCCGACCTGGGTATA</td><td>>hg38_21_+_8433172_8446622_RNA45SN1_45S_50ntFlanks</td><td>9563</td><td>9597</td><td>35</td><td>CCTGACGTGCAAATCGGTCGTCCGACCTGGGTATA</td><td>672.5</td><td>-4.3</td><td>0.6</td><td>-6.7</td><td>1.5E-11</td><td>4.4E-11</td><td>MB231/MB468</td><td>NUC</td></tr>
<tr><td>>MI0000266|hsa-mir-10a&WithFlank&17|-|48579832|48579953@29.48.20(+1C)&[MIMAT0000253&hsa-miR-10a-5p&offsets|+1|-2(+1C);m-18&17|-|48579905|48579926&offsets|+1|-1(+1C)]|ACCCTGTAGATCCGAATTTGC</td><td>>MI0000266|hsa-mir-10a&WithFlank&17|-|48579832|48579953</td><td>29</td><td>48</td><td>20(+1C)</td><td>ACCCTGTAGATCCGAATTTGC</td><td>130.0</td><td>10.2</td><td>1.5</td><td>6.8</td><td>1.4E-11</td><td>4.4E-11</td><td>MB231/MB468</td><td>MP</td></tr>
<tr><td>>MI0000478|hsa-mir-149&WithFlank&2|+|240455995|240456095@21.41.21(+1U)&[MIMAT0000450&hsa-miR-149-5p&offsets|0|-2(+1U);m-195&2|+|240456015|240456037&offsets|0|-2(+1U)]|TCTGGCTCCGTGTCTTCACTCT</td><td>>MI0000478|hsa-mir-149&WithFlank&2|+|240455995|240456095</td><td>21</td><td>41</td><td>21(+1U)</td><td>TCTGGCTCCGTGTCTTCACTCT</td><td>153.7</td><td>7.5</td><td>1.1</td><td>6.7</td><td>1.9E-11</td><td>4.4E-11</td><td>BT20/MB468</td><td>NUC</td></tr>
<tr><td>>hg38_21_+_8433172_8446622_RNA45SN1_45S_50ntFlanks@4163.4183.21|CATCCAAGGAAGGCAGCAGGC</td><td>>hg38_21_+_8433172_8446622_RNA45SN1_45S_50ntFlanks</td><td>4163</td><td>4183</td><td>21</td><td>CATCCAAGGAAGGCAGCAGGC</td><td>120.9</td><td>-10.8</td><td>1.6</td><td>-6.8</td><td>1.4E-11</td><td>4.4E-11</td><td>MB231/MB468</td><td>MP</td></tr>
</table>

| | | | | | | | | | | | | | |
|---|---|---|---|---|---|---|---|---|---|---|---|---|---|
| >trna119_LysCTT_1_-_145395522_145395594@5.35.31,trna11_LysCTT_5_-_180648979_180649051@5.35.31,trna13_LysCTT_6_+_26556774_26556846@5.35.31,trna7_LysCTT_16_+_3225692_3225764@5.35.31,trna9_LysCTT_5_+_180634755_180634827@5.35.31\|GGCTAGCTCAGTCGGTAGAGCATGAGACTCT | >trna119_LysCTT_1_-_145395522_145395594 | 5 | 35 | 31 | GGCTAGCTCAGTCGGTAGAGCATGAGACTCT | 1799.7 | 5.9 | 0.9 | 6.7 | 1.8E-11 | 4.4E-11 | MB231/MB468 | MITO |
| >hg38_21_+_8433172_8446622_RNA45SN1_45S_50ntFlanks@5003.5024.22\|ATAGCTCTTTCTCGATTCCGTG | >hg38_21_+_8433172_8446622_RNA45SN1_45S_50ntFlanks | 5003 | 5024 | 22 | ATAGCTCTTTCTCGATTCCGTG | 99.3 | -10.5 | 1.6 | -6.8 | 1.4E-11 | 4.4E-11 | MB231/MB468 | MP |
| >hg38_21_+_8433172_8446622_RNA45SN1_45S_50ntFlanks@4005.4042.38\|AACCTCGGGCCGATCGCACGCCCCCCGTGGCGGCGACG | >hg38_21_+_8433172_8446622_RNA45SN1_45S_50ntFlanks | 4005 | 4042 | 38 | AACCTCGGGCCGATCGCACGCCCCCCGTGGCGGCGACG | 1459.0 | -8.1 | 1.2 | -6.7 | 1.9E-11 | 4.4E-11 | BT20/MB468 | NUC |
| >hg38_21_+_8433172_8446622_RNA45SN1_45S_50ntFlanks@9554.9579.26\|CGTAGCGGTCCTGACGTGCAAATCGG | >hg38_21_+_8433172_8446622_RNA45SN1_45S_50ntFlanks | 9554 | 9579 | 26 | CGTAGCGGTCCTGACGTGCAAATCGG | 91.1 | -10.4 | 1.5 | -6.8 | 1.4E-11 | 4.4E-11 | MB231/MB468 | MP |
| >hg38_21_+_8433172_8446622_RNA45SN1_45S_50ntFlanks@12941.12965.25\|ATTCGTAGACGACCTGCTTCTGGGT | >hg38_21_+_8433172_8446622_RNA45SN1_45S_50ntFlanks | 12941 | 12965 | 25 | ATTCGTAGACGACCTGCTTCTGGGT | 366.3 | -3.6 | 0.5 | -6.7 | 1.5E-11 | 4.4E-11 | MB231/MB468 | CYTO |
| >hg38_21_+_8433172_8446622_RNA45SN1_45S_50ntFlanks@9534.9564.31\|AGGAAACTCTGGTGGAGGTCCGTAGCGGTCC | >hg38_21_+_8433172_8446622_RNA45SN1_45S_50ntFlanks | 9534 | 9564 | 31 | AGGAAACTCTGGTGGAGGTCCGTAGCGGTCC | 1753.7 | -3.9 | 0.6 | -6.7 | 1.5E-11 | 4.4E-11 | MB231/MB468 | CYTO |
| >MI0000093\|hsa-mir-92a-1&WithFlank&13\|+\|91351308\|91351397@54.75.22(+1A);MI0000094\|hsa-mir-92a-2&WithFlank&X\|-\|134169532\|134169618@54.75.22(+1A)&[MIMAT0000092&hsa-miR-92a-3p&offsets\|0\|0(+1A);m-9&13\|+\|91351361\|91351382&offsets\|0\|0(+1A)];[MIMAT0000092_1&hsa-miR-92a-3p&offsets\|0\|0(+1A);m-11&X\|-\|134169544\|134169565&offsets\|0\|0(+1A)]\|TATTGCACTTGTCCCGGCCTGTA | >MI0000093\|hsa-mir-92a-1&WithFlank&13\|+\|91351308\|91351397 | 54 | 75 | 22(+1A) | TATTGCACTTGTCCCGGCCTGTA | 10288.6 | 5.5 | 0.8 | 6.7 | 1.5E-11 | 4.4E-11 | MB231/MB468 | CYTO |
| >hg38_21_+_8433172_8446622_RNA45SN1_45S_50ntFlanks@4003.4025.23\|ATAACCTCGGGCCGATCGCACGC | >hg38_21_+_8433172_8446622_RNA45SN1_45S_50ntFlanks | 4003 | 4025 | 23 | ATAACCTCGGGCCGATCGCACGC | 933.3 | -4.8 | 0.7 | -6.7 | 1.5E-11 | 4.4E-11 | MB231/MB468 | CYTO |
| >hg38_21_+_8433172_8446622_RNA45SN1_45S_50ntFlanks@13006.13028.23\|CTATTGAAAGTCAGCCCTCGACA | >hg38_21_+_8433172_8446622_RNA45SN1_45S_50ntFlanks | 13006 | 13028 | 23 | CTATTGAAAGTCAGCCCTCGACA | 233.6 | -8.1 | 1.2 | -6.7 | 1.6E-11 | 4.4E-11 | MB231/MB468 | NUC |
| >hg38_21_+_8433172_8446622_RNA45SN1_45S_50ntFlanks@11586.11603.18\|GTGCGGACCAGGGGAATC | >hg38_21_+_8433172_8446622_RNA45SN1_45S_50ntFlanks | 11586 | 11603 | 18 | GTGCGGACCAGGGGAATC | 6590.1 | -11.1 | 1.7 | -6.7 | 1.9E-11 | 4.4E-11 | BT20/MB468 | NUC |
| >hg38_21_+_8433172_8446622_RNA45SN1_45S_50ntFlanks@4560.4582.23\|CTAGGAATAATGGAATAGGACCG | >hg38_21_+_8433172_8446622_RNA45SN1_45S_50ntFlanks | 4560 | 4582 | 23 | CTAGGAATAATGGAATAGGACCG | 93.7 | -10.4 | 1.5 | -6.8 | 1.4E-11 | 4.5E-11 | MB231/MB468 | MP |
| >hg38_21_+_8433172_8446622_RNA45SN1_45S_50ntFlanks@12988.13012.25\|CAGCTCCCTCGCTGCGATCTATTGA | >hg38_21_+_8433172_8446622_RNA45SN1_45S_50ntFlanks | 12988 | 13012 | 25 | CAGCTCCCTCGCTGCGATCTATTGA | 201.6 | 3.8 | 0.6 | 6.7 | 1.5E-11 | 4.5E-11 | MB231/MB468 | CYTO |
| >MI0000239\|hsa-mir-197&WithFlank&1\|+\|109598887\|109598973@54.74.21(+1U)&[MIMAT0000227&hsa-miR-197-3p&offsets\|0\|-1(+1U);m-99&1\|+\|109598940\|109598961&offsets\|0\|-1(+1U)]\|TTCACCACCTTCTCCACCCAGT | >MI0000239\|hsa-mir-197&WithFlank&1\|+\|109598887\|109598973 | 54 | 74 | 21(+1U) | TTCACCACCTTCTCCACCCAGT | 2736.1 | 3.9 | 0.6 | 7.1 | 1.6E-12 | 4.5E-11 | BT20/MB231 | NUC |
| >hg38_21_+_8433172_8446622_RNA45SN1_45S_50ntFlanks@12450.12476.27\|AATTCACCAAGCGTTGGATTGTTCACC | >hg38_21_+_8433172_8446622_RNA45SN1_45S_50ntFlanks | 12450 | 12476 | 27 | AATTCACCAAGCGTTGGATTGTTCACC | 211.1 | 5.2 | 0.8 | 6.7 | 1.5E-11 | 4.5E-11 | MB231/MB468 | CYTO |
| >hg38_21_+_8433172_8446622_RNA45SN1_45S_50ntFlanks@11199.11229.31\|CCGGCGGCGGCGGCGACTCTGGACGCGAGCC | >hg38_21_+_8433172_8446622_RNA45SN1_45S_50ntFlanks | 11199 | 11229 | 31 | CCGGCGGCGGCGGCGACTCTGGACGCGAGCC | 265.9 | -5.3 | 0.8 | -6.7 | 1.5E-11 | 4.5E-11 | MB231/MB468 | CYTO |
| >hg38_21_+_8433172_8446622_RNA45SN1_45S_50ntFlanks@10689.10711.23\|GCGTCCGGTGAGCTCTCGCTGGC | >hg38_21_+_8433172_8446622_RNA45SN1_45S_50ntFlanks | 10689 | 10711 | 23 | GCGTCCGGTGAGCTCTCGCTGGC | 227.5 | -10.1 | 1.5 | -6.8 | 1.4E-11 | 4.5E-11 | BT20/MB468 | CYTO |
| >hg38_21_+_8433172_8446622_RNA45SN1_45S_50ntFlanks@10731.10754.24\|AGGGTGTAAATCTCGCGCCGGGCC | >hg38_21_+_8433172_8446622_RNA45SN1_45S_50ntFlanks | 10731 | 10754 | 24 | AGGGTGTAAATCTCGCGCCGGGCC | 79.4 | -10.2 | 1.5 | -6.8 | 1.4E-11 | 4.5E-11 | MB231/MB468 | MP |
| >am_trna119_LysCTT_1_-_145395522_145395594@14.32.19,trna11_LysCTT_5_-_180648979_180649051@14.32.19,trna128_LysCTT_1_-_55423542_55423614@14.32.19,trna13_LysCTT_6_+_26556774_26556846@14.32.19,trna32_LysCTT_16_-_3207406_3207478@14.32.19,trna6_LysCTT_19_-_52425396_52425468@14.32.19,trna7_LysCTT_16_+_3225692_3225764@14.32.19,trna9_LysCTT_5_+_180634755_180634827@14.32.19\|AGTCGGTAGAGCATGAGAC | >am_trna119_LysCTT_1_-_145395522_145395594 | 14 | 32 | 19 | AGTCGGTAGAGCATGAGAC | 895.0 | 5.1 | 0.8 | 6.8 | 1.4E-11 | 4.5E-11 | BT20/MB468 | CYTO |
| >hg38_21_+_8433172_8446622_RNA45SN1_45S_50ntFlanks@12072.12094.23\|CCCGAGGGGCTCTCGCTTCTGGC | >hg38_21_+_8433172_8446622_RNA45SN1_45S_50ntFlanks | 12072 | 12094 | 23 | CCCGAGGGGCTCTCGCTTCTGGC | 94.2 | -10.4 | 1.5 | -6.8 | 1.4E-11 | 4.5E-11 | MB231/MB468 | MP |
| >hg38_21_+_8433172_8446622_RNA45SN1_45S_50ntFlanks@12804.12821.18\|CGCGTGCCCCGCCGCGCG | >hg38_21_+_8433172_8446622_RNA45SN1_45S_50ntFlanks | 12804 | 12821 | 18 | CGCGTGCCCCGCCGCGCG | 191.4 | -9.7 | 1.4 | -6.8 | 1.1E-11 | 4.5E-11 | BT20/MB468 | MITO |
| >MI0000298\|hsa-mir-221&WithFlank&X\|-\|45746151\|45746272@71.91.21(+1C)&[MIMAT0000278&hsa-miR-221-3p&offsets\|0\|-2(+1C);m-91&X\|-\|45746181\|45746202&offsets\|0\|-1(+1C)]\|AGCTACATTGTCTGCTGGGTTC | >MI0000298\|hsa-mir-221&WithFlank&X\|-\|45746151\|45746272 | 71 | 91 | 21(+1C) | AGCTACATTGTCTGCTGGGTTC | 1560.3 | 6.3 | 0.9 | 6.7 | 1.6E-11 | 4.5E-11 | MB231/MB468 | CYTO |
| >hg38_21_+_8433172_8446622_RNA45SN1_45S_50ntFlanks@11579.11602.24\|AGAACTGGTGCGGACCAGGGGAAT | >hg38_21_+_8433172_8446622_RNA45SN1_45S_50ntFlanks | 11579 | 11602 | 24 | AGAACTGGTGCGGACCAGGGGAAT | 887.5 | -9.2 | 1.4 | -6.8 | 1.4E-11 | 4.5E-11 | MB231/MB468 | MP |
| >hg38_21_+_8433172_8446622_RNA45SN1_45S_50ntFlanks@11575.11602.28\|ACTTAGAACTGGTGCGGACCAGGGGAAT | >hg38_21_+_8433172_8446622_RNA45SN1_45S_50ntFlanks | 11575 | 11602 | 28 | ACTTAGAACTGGTGCGGACCAGGGGAAT | 1213.1 | -7.1 | 1.0 | -6.7 | 1.6E-11 | 4.5E-11 | MB231/MB468 | NUC |
| >hg38_21_+_8433172_8446622_RNA45SN1_45S_50ntFlanks@12709.12747.39\|CCGCGGAGCCTCGGTTGGCCTCGGATAGCCGGTCCCCCG | >hg38_21_+_8433172_8446622_RNA45SN1_45S_50ntFlanks | 12709 | 12747 | 39 | CCGCGGAGCCTCGGTTGGCCTCGGATAGCCGGTCCCCCG | 245.8 | -10.0 | 1.5 | -6.8 | 1.1E-11 | 4.5E-11 | BT20/MB468 | MITO |
| >hg38_21_+_8433172_8446622_RNA45SN1_45S_50ntFlanks@10512.10533.22\|TCCGGAGTGGCGGAGATGGGCG | >hg38_21_+_8433172_8446622_RNA45SN1_45S_50ntFlanks | 10512 | 10533 | 22 | TCCGGAGTGGCGGAGATGGGCG | 5908.1 | -11.6 | 1.7 | -6.7 | 2.0E-11 | 4.5E-11 | BT20/MB468 | NUC |
| >hg38_21_+_8433172_8446622_RNA45SN1_45S_50ntFlanks@11561.11601.41\|GCGCCTAGCAGCCGACTTAGAACTGGTGCGGACCAGGGGAA | >hg38_21_+_8433172_8446622_RNA45SN1_45S_50ntFlanks | 11561 | 11601 | 41 | GCGCCTAGCAGCCGACTTAGAACTGGTGCGGACCAGGGGAA | 7502.8 | -11.4 | 1.7 | -6.7 | 2.0E-11 | 4.5E-11 | BT20/MB468 | NUC |
| >hg38_21_+_8433172_8446622_RNA45SN1_45S_50ntFlanks@9739.9757.19\|AAACGATCTCAACCTATTC | >hg38_21_+_8433172_8446622_RNA45SN1_45S_50ntFlanks | 9739 | 9757 | 19 | AAACGATCTCAACCTATTC | 2730.3 | -6.3 | 0.9 | -6.8 | 1.1E-11 | 4.5E-11 | BT20/MB468 | MITO |
| >hg38_21_+_8433172_8446622_RNA45SN1_45S_50ntFlanks@11769.11790.22\|TAATTAGTGACGCGCATGAATG | >hg38_21_+_8433172_8446622_RNA45SN1_45S_50ntFlanks | 11769 | 11790 | 22 | TAATTAGTGACGCGCATGAATG | 1420.5 | -7.7 | 1.1 | -6.7 | 2.0E-11 | 4.5E-11 | BT20/MB468 | NUC |
| >hg38_21_+_8433172_8446622_RNA45SN1_45S_50ntFlanks@3876.3898.23\|TACATGCCGACGGGCGCTGACCC | >hg38_21_+_8433172_8446622_RNA45SN1_45S_50ntFlanks | 3876 | 3898 | 23 | TACATGCCGACGGGCGCTGACCC | 2151.5 | -8.6 | 1.3 | -6.7 | 2.0E-11 | 4.5E-11 | BT20/MB468 | NUC |
| >hg38_21_+_8433172_8446622_RNA45SN1_45S_50ntFlanks@12993.13013.21\|CCCTCGCTGCGATCTATTGAA | >hg38_21_+_8433172_8446622_RNA45SN1_45S_50ntFlanks | 12993 | 13013 | 21 | CCCTCGCTGCGATCTATTGAA | 1840.8 | -4.7 | 0.7 | -6.7 | 1.6E-11 | 4.5E-11 | MB231/MB468 | NUC |
| >hg38_21_+_8433172_8446622_RNA45SN1_45S_50ntFlanks@4378.4408.31\|CTCGTAGTTGGATCTTGGGAGCGGGCGGGCG | >hg38_21_+_8433172_8446622_RNA45SN1_45S_50ntFlanks | 4378 | 4408 | 31 | CTCGTAGTTGGATCTTGGGAGCGGGCGGGCG | 107.8 | -10.6 | 1.6 | -6.8 | 1.4E-11 | 4.5E-11 | MB231/MB468 | MP |
| >hg38_21_+_8433172_8446622_RNA45SN1_45S_50ntFlanks@5420.5442.23\|CCGATTGGATGGTTTAGTGAGGC | >hg38_21_+_8433172_8446622_RNA45SN1_45S_50ntFlanks | 5420 | 5442 | 23 | CCGATTGGATGGTTTAGTGAGGC | 111.0 | -10.7 | 1.6 | -6.8 | 1.4E-11 | 4.5E-11 | MB231/MB468 | MP |
| >hg38_21_+_8433172_8446622_RNA45SN1_45S_50ntFlanks@12416.12447.32\|TTCGATGTCGGCTCTTCCTATCATTGTGAAGC | >hg38_21_+_8433172_8446622_RNA45SN1_45S_50ntFlanks | 12416 | 12447 | 32 | TTCGATGTCGGCTCTTCCTATCATTGTGAAGC | 2325.3 | 7.2 | 1.1 | 6.7 | 1.6E-11 | 4.5E-11 | MB231/MB468 | CYTO |
| >MI0000456\|hsa-mir-140&WithFlank&16\|+\|69933075\|69933186@68.89.22&[MIMAT0004597&hsa-miR-140-3p&offsets\|0\|+1;m-45&16\|+\|69933143\|69933164&offsets\|-1\|-1]\|TACCACAGGGTAGAACCACGGA | >MI0000456\|hsa-mir-140&WithFlank&16\|+\|69933075\|69933186 | 68 | 89 | 22 | TACCACAGGGTAGAACCACGGA | 378.2 | 8.3 | 1.2 | 6.7 | 1.6E-11 | 4.6E-11 | MB231/MB468 | NUC |
| >hg38_21_+_8433172_8446622_RNA45SN1_45S_50ntFlanks@11573.11603.31\|CGACTTAGAACTGGTGCGGACCAGGGGAATC | >hg38_21_+_8433172_8446622_RNA45SN1_45S_50ntFlanks | 11573 | 11603 | 31 | CGACTTAGAACTGGTGCGGACCAGGGGAATC | 2379.3 | -8.1 | 1.2 | -6.7 | 2.0E-11 | 4.6E-11 | BT20/MB468 | NUC |
| >hg38_21_+_8433172_8446622_RNA45SN1_45S_50ntFlanks@6747.6770.24\|ACTTCGAACGCACTTGCGGCCCCG | >hg38_21_+_8433172_8446622_RNA45SN1_45S_50ntFlanks | 6747 | 6770 | 24 | ACTTCGAACGCACTTGCGGCCCCG | 617.1 | -9.2 | 1.4 | -6.7 | 2.0E-11 | 4.6E-11 | BT20/MB468 | NUC |
| >hg38_21_+_8433172_8446622_RNA45SN1_45S_50ntFlanks@11570.11601.32\|AGCCGACTTAGAACTGGTGCGGACCAGGGGAA | >hg38_21_+_8433172_8446622_RNA45SN1_45S_50ntFlanks | 11570 | 11601 | 32 | AGCCGACTTAGAACTGGTGCGGACCAGGGGAA | 6538.2 | -9.0 | 1.3 | -6.7 | 2.0E-11 | 4.6E-11 | BT20/MB468 | NUC |
| >hg38_21_+_8433172_8446622_RNA45SN1_45S_50ntFlanks@9910.9928.19\|CCCAGAAAAGGTGTTGGTT | >hg38_21_+_8433172_8446622_RNA45SN1_45S_50ntFlanks | 9910 | 9928 | 19 | CCCAGAAAAGGTGTTGGTT | 139.7 | -11.0 | 1.6 | -6.8 | 1.4E-11 | 4.6E-11 | MB231/MB468 | MP |
| >hg38_21_+_8433172_8446622_RNA45SN1_45S_50ntFlanks@9547.9570.24\|GGAGGTCCGTAGCGGTCCTGACGT | >hg38_21_+_8433172_8446622_RNA45SN1_45S_50ntFlanks | 9547 | 9570 | 24 | GGAGGTCCGTAGCGGTCCTGACGT | 317.2 | 4.0 | 0.6 | 6.7 | 1.6E-11 | 4.6E-11 | MB231/MB468 | CYTO |
| >hg38_21_+_8433172_8446622_RNA45SN1_45S_50ntFlanks@11565.11592.28\|CTAGCAGCCGACTTAGAACTGGTGCGGA | >hg38_21_+_8433172_8446622_RNA45SN1_45S_50ntFlanks | 11565 | 11592 | 28 | CTAGCAGCCGACTTAGAACTGGTGCGGA | 234.4 | -11.7 | 1.7 | -6.8 | 1.4E-11 | 4.6E-11 | MB231/MB468 | MP |
| >hg38_21_+_8433172_8446622_RNA45SN1_45S_50ntFlanks@12353.12376.24\|GTTACCACAGGGATAACTGGCTTG | >hg38_21_+_8433172_8446622_RNA45SN1_45S_50ntFlanks | 12353 | 12376 | 24 | GTTACCACAGGGATAACTGGCTTG | 95.4 | -10.4 | 1.5 | -6.8 | 1.4E-11 | 4.6E-11 | MB231/MB468 | MP |
| >hg38_21_+_8433172_8446622_RNA45SN1_45S_50ntFlanks@11885.11905.21\|TTGAGCTTGACTCTAGTCTGG | >hg38_21_+_8433172_8446622_RNA45SN1_45S_50ntFlanks | 11885 | 11905 | 21 | TTGAGCTTGACTCTAGTCTGG | 1154.3 | -8.2 | 1.2 | -6.7 | 1.6E-11 | 4.6E-11 | MB231/MB468 | CYTO |
| >hg38_21_+_8433172_8446622_RNA45SN1_45S_50ntFlanks@12436.12454.19\|TCATTGTGAAGCAGAATTC | >hg38_21_+_8433172_8446622_RNA45SN1_45S_50ntFlanks | 12436 | 12454 | 19 | TCATTGTGAAGCAGAATTC | 1318.9 | -7.5 | 1.1 | -6.7 | 1.6E-11 | 4.6E-11 | MB231/MB468 | CYTO |
| >hg38_21_+_8433172_8446622_RNA45SN1_45S_50ntFlanks@11317.11344.28\|ACCCCACGTCTCGTCGCGCGCGCGTCCG | >hg38_21_+_8433172_8446622_RNA45SN1_45S_50ntFlanks | 11317 | 11344 | 28 | ACCCCACGTCTCGTCGCGCGCGCGTCCG | 337.3 | -11.5 | 1.6 | -7.1 | 9.4E-13 | 4.6E-11 | BT20/MB468 | MP |
| >hg38_21_+_8433172_8446622_RNA45SN1_45S_50ntFlanks@12907.12935.29\|CACCGCACGTTCGTGGGGAACCTGGCGCT | >hg38_21_+_8433172_8446622_RNA45SN1_45S_50ntFlanks | 12907 | 12935 | 29 | CACCGCACGTTCGTGGGGAACCTGGCGCT | 998.8 | -7.0 | 1.0 | -6.7 | 1.9E-11 | 4.6E-11 | MB231/MB468 | MITO |
| >hg38_21_+_8433172_8446622_RNA45SN1_45S_50ntFlanks@11875.11905.31\|GAAGACCCTGTTGAGCTTGACTCTAGTCTGG | >hg38_21_+_8433172_8446622_RNA45SN1_45S_50ntFlanks | 11875 | 11905 | 31 | GAAGACCCTGTTGAGCTTGACTCTAGTCTGG | 117.8 | -10.7 | 1.6 | -6.8 | 1.5E-11 | 4.6E-11 | MB231/MB468 | MP |
| >hg38_21_+_8433172_8446622_RNA45SN1_45S_50ntFlanks@11273.11293.21\|CCGCCCCCGGGGAGCCCGGCG | >hg38_21_+_8433172_8446622_RNA45SN1_45S_50ntFlanks | 11273 | 11293 | 21 | CCGCCCCCGGGGAGCCCGGCG | 124.6 | -10.8 | 1.6 | -6.8 | 1.5E-11 | 4.6E-11 | MB231/MB468 | MP |

| | | | | | | | | | | | | | |
|---|---|---|---|---|---|---|---|---|---|---|---|---|---|
| >hg38_21_+_8433172_8446622_RNA45SN1_45S_50ntFlanks@12182.12200.19\|CAAACGGTAACGCAGGTGT | >hg38_21_+_8433172_8446622_RNA45SN1_45S_50ntFlanks | 12182 | 12200 | 19 | CAAACGGTAACGCAGGTGT | 148.9 | -11.1 | 1.6 | -6.8 | 1.5E-11 | 4.6E-11 | MB231/MB468 | MP |
| >hg38_21_+_8433172_8446622_RNA45SN1_45S_50ntFlanks@8403.8442.40\|AGGTAAACGGGTGGGGTCCGCGCAGTCCGCCCGGAGGATT | >hg38_21_+_8433172_8446622_RNA45SN1_45S_50ntFlanks | 8403 | 8442 | 40 | AGGTAAACGGGTGGGGTCCGCGCAGTCCGCCCGGAGGATT | 119.1 | -10.8 | 1.6 | -6.8 | 1.5E-11 | 4.6E-11 | MB231/MB468 | MP |
| >MI0000825\|hsa-mir-345&WithFlank&14\|+\|100307853\|100307962@24.45.22&[MIMAT0000772&hsa-miR-345-5p&offsets\|0\|0;m-193&14\|+\|100307876\|100307896&offsets\|0\|+1]\|GCTGACTCCTAGTCCAGGGCTC | >MI0000825\|hsa-mir-345&WithFlank&14\|+\|100307853\|100307962 | 24 | 45 | 22 | GCTGACTCCTAGTCCAGGGCTC | 315.7 | 5.2 | 0.8 | 6.7 | 1.6E-11 | 4.6E-11 | MB231/MB468 | NUC |
| >trnaMT_PheGAA_MT_+_577_647@1.19.19\|GTTTATGTAGCTTACCTCC | >trnaMT_PheGAA_MT_+_577_647 | 1 | 19 | 19 | GTTTATGTAGCTTACCTCC | 406.9 | 9.1 | 1.3 | 6.7 | 1.6E-11 | 4.6E-11 | MB231/MB468 | CYTO |
| >hg38_21_+_8433172_8446622_RNA45SN1_45S_50ntFlanks@11905.11929.25\|GCACGGTGAAGAGACATGAGAGGTG | >hg38_21_+_8433172_8446622_RNA45SN1_45S_50ntFlanks | 11905 | 11929 | 25 | GCACGGTGAAGAGACATGAGAGGTG | 184.4 | -11.4 | 1.7 | -6.8 | 1.5E-11 | 4.6E-11 | MB231/MB468 | MP |
| >hg38_21_+_8433172_8446622_RNA45SN1_45S_50ntFlanks@11703.11722.20\|AATGAAGCGCGGGTAAACGG | >hg38_21_+_8433172_8446622_RNA45SN1_45S_50ntFlanks | 11703 | 11722 | 20 | AATGAAGCGCGGGTAAACGG | 107.8 | -10.6 | 1.6 | -6.8 | 1.5E-11 | 4.6E-11 | MB231/MB468 | MP |
| >hg38_21_+_8433172_8446622_RNA45SN1_45S_50ntFlanks@10694.10712.19\|CGGTGAGCTCTCGCTGGCC | >hg38_21_+_8433172_8446622_RNA45SN1_45S_50ntFlanks | 10694 | 10712 | 19 | CGGTGAGCTCTCGCTGGCC | 175.8 | -11.3 | 1.7 | -6.7 | 1.5E-11 | 4.6E-11 | MB231/MB468 | MP |
| >hg38_21_+_8433172_8446622_RNA45SN1_45S_50ntFlanks@10392.10420.29\|ACAGCAGTTGAACATGGGTCAGTCGGTCC | >hg38_21_+_8433172_8446622_RNA45SN1_45S_50ntFlanks | 10392 | 10420 | 29 | ACAGCAGTTGAACATGGGTCAGTCGGTCC | 977.8 | -9.0 | 1.3 | -6.7 | 1.6E-11 | 4.7E-11 | MB231/MB468 | NUC |
| >hg38_21_+_8433172_8446622_RNA45SN1_45S_50ntFlanks@12457.12477.21\|CAAGCGTTGGATTGTTCACCC | >hg38_21_+_8433172_8446622_RNA45SN1_45S_50ntFlanks | 12457 | 12477 | 21 | CAAGCGTTGGATTGTTCACCC | 3191.1 | -4.7 | 0.7 | -6.7 | 1.6E-11 | 4.7E-11 | MB231/MB468 | NUC |
| >hg38_21_+_8433172_8446622_RNA45SN1_45S_50ntFlanks@9387.9409.23\|CCGGCCGAGGTGGGATCCCGAGG | >hg38_21_+_8433172_8446622_RNA45SN1_45S_50ntFlanks | 9387 | 9409 | 23 | CCGGCCGAGGTGGGATCCCGAGG | 146.2 | -11.1 | 1.6 | -6.7 | 1.5E-11 | 4.7E-11 | MB231/MB468 | MP |
| >hg38_21_+_8433172_8446622_RNA45SN1_45S_50ntFlanks@4760.4793.34\|TCAGATACCGTCGTAGTTCCGACCATAAACGATG | >hg38_21_+_8433172_8446622_RNA45SN1_45S_50ntFlanks | 4760 | 4793 | 34 | TCAGATACCGTCGTAGTTCCGACCATAAACGATG | 297.7 | -6.5 | 1.0 | -6.7 | 1.9E-11 | 4.7E-11 | MB231/MB468 | MITO |
| >hg38_21_+_8433172_8446622_RNA45SN1_45S_50ntFlanks@9568.9595.28\|CGTGCAAATCGGTCGTCCGACCTGGGTA | >hg38_21_+_8433172_8446622_RNA45SN1_45S_50ntFlanks | 9568 | 9595 | 28 | CGTGCAAATCGGTCGTCCGACCTGGGTA | 3451.7 | -6.8 | 1.0 | -6.7 | 1.6E-11 | 4.7E-11 | MB231/MB468 | CYTO |
| >hg38_21_+_8433172_8446622_RNA45SN1_45S_50ntFlanks@6774.6806.33\|TCCTCCCGGGGCTACGCCTGTCTGAGCGTCGCT | >hg38_21_+_8433172_8446622_RNA45SN1_45S_50ntFlanks | 6774 | 6806 | 33 | TCCTCCCGGGGCTACGCCTGTCTGAGCGTCGCT | 633.9 | 4.6 | 0.7 | 6.7 | 1.6E-11 | 4.7E-11 | MB231/MB468 | NUC |
| >hg38_21_+_8433172_8446622_RNA45SN1_45S_50ntFlanks@11602.11619.18\|TCCGACTGTTTAATTAAA | >hg38_21_+_8433172_8446622_RNA45SN1_45S_50ntFlanks | 11602 | 11619 | 18 | TCCGACTGTTTAATTAAA | 628.9 | -6.7 | 1.0 | -6.7 | 1.5E-11 | 4.7E-11 | MB231/MB468 | MP |
| >hg38_21_+_8433172_8446622_RNA45SN1_45S_50ntFlanks@3838.3860.23\|CTCTCCTACTTGGATAACTGTGG | >hg38_21_+_8433172_8446622_RNA45SN1_45S_50ntFlanks | 3838 | 3860 | 23 | CTCTCCTACTTGGATAACTGTGG | 3551.7 | -8.5 | 1.3 | -6.7 | 2.0E-11 | 4.7E-11 | BT20/MB468 | NUC |
| >hg38_21_+_8433172_8446622_RNA45SN1_45S_50ntFlanks@9469.9503.35\|GGAGCACGAGCGCACGTGTTAGGACCCGAAAGATG | >hg38_21_+_8433172_8446622_RNA45SN1_45S_50ntFlanks | 9469 | 9503 | 35 | GGAGCACGAGCGCACGTGTTAGGACCCGAAAGATG | 181.5 | -11.4 | 1.7 | -6.7 | 1.5E-11 | 4.7E-11 | MB231/MB468 | MP |
| >hg38_1_-_228634819_228635039_RNA5S12_5S_50ntFlanks@150.171.22\|AATACCGGGTGCTGTAGGCTTT | >hg38_1_-_228634819_228635039_RNA5S12_5S_50ntFlanks | 150 | 171 | 22 | AATACCGGGTGCTGTAGGCTTT | 23469.5 | -5.6 | 0.8 | -6.7 | 1.5E-11 | 4.7E-11 | MB231/MB468 | MP |
| >hg38_21_+_8433172_8446622_RNA45SN1_45S_50ntFlanks@10786.10816.31\|AACAGCCTCTGGCATGTTGGAACAATGTAGG | >hg38_21_+_8433172_8446622_RNA45SN1_45S_50ntFlanks | 10786 | 10816 | 31 | AACAGCCTCTGGCATGTTGGAACAATGTAGG | 2208.8 | -4.9 | 0.7 | -6.7 | 1.5E-11 | 4.7E-11 | BT20/MB468 | CYTO |
| >hg38_21_+_8433172_8446622_RNA45SN1_45S_50ntFlanks@5078.5115.38\|CGATAACGAACGAGACTCTGGCATGCTAACTAGTTACG | >hg38_21_+_8433172_8446622_RNA45SN1_45S_50ntFlanks | 5078 | 5115 | 38 | CGATAACGAACGAGACTCTGGCATGCTAACTAGTTACG | 488.1 | 4.0 | 0.6 | 6.7 | 1.6E-11 | 4.7E-11 | MB231/MB468 | CYTO |
| >hg38_21_+_8433172_8446622_RNA45SN1_45S_50ntFlanks@8154.8188.35\|CCGCTCGTGGGGGGCCCAAGTCCTTCTGATCGAGG | >hg38_21_+_8433172_8446622_RNA45SN1_45S_50ntFlanks | 8154 | 8188 | 35 | CCGCTCGTGGGGGGCCCAAGTCCTTCTGATCGAGG | 88.0 | -10.3 | 1.5 | -6.7 | 1.5E-11 | 4.7E-11 | MB231/MB468 | MP |
| >hg38_21_+_8433172_8446622_RNA45SN1_45S_50ntFlanks@10470.10497.28\|GCCCTCGGCCGATCGAAAGGGAGTCGGG | >hg38_21_+_8433172_8446622_RNA45SN1_45S_50ntFlanks | 10470 | 10497 | 28 | GCCCTCGGCCGATCGAAAGGGAGTCGGG | 76.5 | -10.1 | 1.5 | -6.7 | 1.5E-11 | 4.7E-11 | MB231/MB468 | MP |
| >hg38_21_+_8433172_8446622_RNA45SN1_45S_50ntFlanks@8356.8378.23\|GAAAAGAACTTTGAAGAGAGAGT | >hg38_21_+_8433172_8446622_RNA45SN1_45S_50ntFlanks | 8356 | 8378 | 23 | GAAAAGAACTTTGAAGAGAGAGT | 2532.6 | -5.2 | 0.8 | -6.7 | 1.7E-11 | 4.7E-11 | MB231/MB468 | NUC |
| >hg38_21_+_8433172_8446622_RNA45SN1_45S_50ntFlanks@4555.4576.22\|CGCAGCTAGGAATAATGGAATA | >hg38_21_+_8433172_8446622_RNA45SN1_45S_50ntFlanks | 4555 | 4576 | 22 | CGCAGCTAGGAATAATGGAATA | 100.8 | -10.5 | 1.6 | -6.7 | 1.5E-11 | 4.7E-11 | MB231/MB468 | MP |
| >hg38_21_+_8433172_8446622_RNA45SN1_45S_50ntFlanks@8839.8866.28\|CGGTGGCGGCGGCGGCGGCGGCGGCGGG | >hg38_21_+_8433172_8446622_RNA45SN1_45S_50ntFlanks | 8839 | 8866 | 28 | CGGTGGCGGCGGCGGCGGCGGCGGCGGG | 170.6 | -11.3 | 1.7 | -6.7 | 1.5E-11 | 4.7E-11 | MB231/MB468 | MP |
| >hg38_21_+_8433172_8446622_RNA45SN1_45S_50ntFlanks@11675.11706.32\|TGCTCTGAATGTCAAAGTGAAGAAATTCAATG | >hg38_21_+_8433172_8446622_RNA45SN1_45S_50ntFlanks | 11675 | 11706 | 32 | TGCTCTGAATGTCAAAGTGAAGAAATTCAATG | 311.1 | 4.3 | 0.6 | 6.8 | 1.2E-11 | 4.7E-11 | BT20/MB468 | MITO |
| >hg38_21_+_8433172_8446622_RNA45SN1_45S_50ntFlanks@12658.12685.28\|ACTGAACGCCTCTAAGTCAGAATCCCGC | >hg38_21_+_8433172_8446622_RNA45SN1_45S_50ntFlanks | 12658 | 12685 | 28 | ACTGAACGCCTCTAAGTCAGAATCCCGC | 99.8 | -10.5 | 1.6 | -6.7 | 1.5E-11 | 4.7E-11 | MB231/MB468 | MP |
| >hg38_21_+_8433172_8446622_RNA45SN1_45S_50ntFlanks@10380.10424.45\|GGTTCCATGTGAACAGCAGTTGAACATGGGTCAGTCGGTCCTGAG | >hg38_21_+_8433172_8446622_RNA45SN1_45S_50ntFlanks | 10380 | 10424 | 45 | GGTTCCATGTGAACAGCAGTTGAACATGGGTCAGTCGGTCCTGAG | 268.5 | 11.8 | 1.7 | 7.1 | 9.8E-13 | 4.7E-11 | BT20/MB468 | MP |
| >MI0005561\|hsa-mir-877&WithFlank&6\|+\|30584326\|30584423@72.92.21(+1U)&[MIMAT0004950&hsa-miR-877-3p&offsets\|0\|0(+1U);m-607&6\|+\|30584397\|30584418&offsets\|0\|-1(+1U)]\|TCCTCTTCTCCCTCCTCCCAGT | >MI0005561\|hsa-mir-877&WithFlank&6\|+\|30584326\|30584423 | 72 | 92 | 21(+1U) | TCCTCTTCTCCCTCCTCCCAGT | 1002.2 | 9.5 | 1.4 | 6.7 | 1.5E-11 | 4.7E-11 | BT20/MB468 | CYTO |
| >hg38_21_+_8433172_8446622_RNA45SN1_45S_50ntFlanks@10687.10712.26\|CGGCGTCCGGTGAGCTCTCGCTGGCC | >hg38_21_+_8433172_8446622_RNA45SN1_45S_50ntFlanks | 10687 | 10712 | 26 | CGGCGTCCGGTGAGCTCTCGCTGGCC | 993.5 | -8.8 | 1.3 | -6.7 | 1.5E-11 | 4.7E-11 | MB231/MB468 | MP |
| >hg38_21_+_8433172_8446622_RNA45SN1_45S_50ntFlanks@10439.10460.22\|CGTTCCGAAGGGACGGGCGATG | >hg38_21_+_8433172_8446622_RNA45SN1_45S_50ntFlanks | 10439 | 10460 | 22 | CGTTCCGAAGGGACGGGCGATG | 93.2 | -10.4 | 1.5 | -6.7 | 1.5E-11 | 4.7E-11 | MB231/MB468 | MP |
| >hg38_21_+_8433172_8446622_RNA45SN1_45S_50ntFlanks@8402.8422.21\|GAGGTAAACGGGTGGGGTCCG | >hg38_21_+_8433172_8446622_RNA45SN1_45S_50ntFlanks | 8402 | 8422 | 21 | GAGGTAAACGGGTGGGGTCCG | 98.5 | -10.5 | 1.6 | -6.7 | 1.5E-11 | 4.7E-11 | MB231/MB468 | MP |
| >hg38_21_+_8433172_8446622_RNA45SN1_45S_50ntFlanks@8839.8858.20\|CGGTGGCGGCGGCGGCGGCG | >hg38_21_+_8433172_8446622_RNA45SN1_45S_50ntFlanks | 8839 | 8858 | 20 | CGGTGGCGGCGGCGGCGGCG | 134.0 | -10.9 | 1.6 | -6.7 | 1.5E-11 | 4.7E-11 | MB231/MB468 | MP |
| >hg38_21_+_8433172_8446622_RNA45SN1_45S_50ntFlanks@8012.8042.31\|AAGCATATTAGTCAGCGGAGGAGAAGAAACT | >hg38_21_+_8433172_8446622_RNA45SN1_45S_50ntFlanks | 8012 | 8042 | 31 | AAGCATATTAGTCAGCGGAGGAGAAGAAACT | 260.0 | 8.7 | 1.3 | 6.7 | 1.7E-11 | 4.8E-11 | MB231/MB468 | CYTO |
| >hg38_21_+_8433172_8446622_RNA45SN1_45S_50ntFlanks@9202.9225.24\|TCTCGGGGCCACGCGCGCGTCCCC | >hg38_21_+_8433172_8446622_RNA45SN1_45S_50ntFlanks | 9202 | 9225 | 24 | TCTCGGGGCCACGCGCGCGTCCCC | 390.7 | -11.7 | 1.6 | -7.1 | 9.9E-13 | 4.8E-11 | BT20/MB468 | MP |
| >hg38_21_+_8433172_8446622_RNA45SN1_45S_50ntFlanks@3705.3741.37\|TACCTGGTTGATCCTGCCAGTAGCATATGCTTGTCTC | >hg38_21_+_8433172_8446622_RNA45SN1_45S_50ntFlanks | 3705 | 3741 | 37 | TACCTGGTTGATCCTGCCAGTAGCATATGCTTGTCTC | 2624.7 | 4.5 | 0.7 | 6.7 | 1.7E-11 | 4.8E-11 | MB231/MB468 | CYTO |
| >hg38_1_-_228634819_228635039_RNA5S12_5S_50ntFlanks@129.170.42\|TTGGATGGGAGACCGCCTGGGAATACCGGGTGCTGTAGGCTT | >hg38_1_-_228634819_228635039_RNA5S12_5S_50ntFlanks | 129 | 170 | 42 | TTGGATGGGAGACCGCCTGGGAATACCGGGTGCTGTAGGCTT | 2745.3 | 8.3 | 1.2 | 6.8 | 1.2E-11 | 4.8E-11 | BT20/MB468 | MITO |
| >hg38_21_+_8433172_8446622_RNA45SN1_45S_50ntFlanks@9507.9526.20\|AACTATGCCTGGGCAGGGCG | >hg38_21_+_8433172_8446622_RNA45SN1_45S_50ntFlanks | 9507 | 9526 | 20 | AACTATGCCTGGGCAGGGCG | 1047.1 | -13.1 | 1.8 | -7.1 | 9.9E-13 | 4.8E-11 | BT20/MB468 | MP |
| >hg38_21_+_8433172_8446622_RNA45SN1_45S_50ntFlanks@12405.12426.22\|CTTTTTGATCCTTCGATGTCGG | >hg38_21_+_8433172_8446622_RNA45SN1_45S_50ntFlanks | 12405 | 12426 | 22 | CTTTTTGATCCTTCGATGTCGG | 1181.5 | -4.2 | 0.6 | -6.8 | 1.2E-11 | 4.8E-11 | BT20/MB468 | MITO |
| >hg38_21_+_8433172_8446622_RNA45SN1_45S_50ntFlanks@11786.11806.21\|GAATGGATGAACGAGATTCCC | >hg38_21_+_8433172_8446622_RNA45SN1_45S_50ntFlanks | 11786 | 11806 | 21 | GAATGGATGAACGAGATTCCC | 143.7 | -11.0 | 1.6 | -6.7 | 1.5E-11 | 4.8E-11 | MB231/MB468 | MP |
| >hg38_21_+_8433172_8446622_RNA45SN1_45S_50ntFlanks@3705.3740.36\|TACCTGGTTGATCCTGCCAGTAGCATATGCTTGTCT | >hg38_21_+_8433172_8446622_RNA45SN1_45S_50ntFlanks | 3705 | 3740 | 36 | TACCTGGTTGATCCTGCCAGTAGCATATGCTTGTCT | 1136.4 | 9.5 | 1.4 | 6.7 | 1.5E-11 | 4.8E-11 | BT20/MB468 | CYTO |
| >hg38_21_+_8433172_8446622_RNA45SN1_45S_50ntFlanks@4006.4025.20\|ACCTCGGGCCGATCGCACGC | >hg38_21_+_8433172_8446622_RNA45SN1_45S_50ntFlanks | 4006 | 4025 | 20 | ACCTCGGGCCGATCGCACGC | 96.5 | -10.5 | 1.6 | -6.7 | 1.5E-11 | 4.8E-11 | MB231/MB468 | MP |
| >hg38_21_+_8433172_8446622_RNA45SN1_45S_50ntFlanks@7986.8018.33\|ATCAGACGTGGCGACCCGCTGAATTTAAGCATA | >hg38_21_+_8433172_8446622_RNA45SN1_45S_50ntFlanks | 7986 | 8018 | 33 | ATCAGACGTGGCGACCCGCTGAATTTAAGCATA | 416.6 | -11.8 | 1.7 | -7.1 | 1.0E-12 | 4.8E-11 | BT20/MB468 | MP |
| >hg38_21_+_8433172_8446622_RNA45SN1_45S_50ntFlanks@8179.8208.30\|CTGATCGAGGCCCAGCCCGTGGACGGTGTG | >hg38_21_+_8433172_8446622_RNA45SN1_45S_50ntFlanks | 8179 | 8208 | 30 | CTGATCGAGGCCCAGCCCGTGGACGGTGTG | 417.5 | -8.4 | 1.2 | -6.7 | 1.5E-11 | 4.8E-11 | MB231/MB468 | MP |
| >hg38_21_+_8433172_8446622_RNA45SN1_45S_50ntFlanks@11874.11892.19\|AGAAGACCCTGTTGAGCTT | >hg38_21_+_8433172_8446622_RNA45SN1_45S_50ntFlanks | 11874 | 11892 | 19 | AGAAGACCCTGTTGAGCTT | 503.7 | -4.6 | 0.7 | -6.7 | 1.7E-11 | 4.8E-11 | MB231/MB468 | CYTO |
| >hg38_21_+_8433172_8446622_RNA45SN1_45S_50ntFlanks@12627.12653.27\|ATGGGGCGAAGCTACCATCTGTGGGAT | >hg38_21_+_8433172_8446622_RNA45SN1_45S_50ntFlanks | 12627 | 12653 | 27 | ATGGGGCGAAGCTACCATCTGTGGGAT | 7421.6 | -10.6 | 1.5 | -7.1 | 1.0E-12 | 4.8E-11 | BT20/MB468 | MP |
| >MI0000086\|hsa-mir-28&WithFlank&3\|+\|188688775\|188688872@60.81.22&[MIMAT0004502&hsa-miR-28-3p&offsets\|0\|0;m-28&3\|+\|188688834\|188688855&offsets\|0\|0]\|CACTAGATTGTGAGCTCCTGGA | >MI0000086\|hsa-mir-28&WithFlank&3\|+\|188688775\|188688872 | 60 | 81 | 22 | CACTAGATTGTGAGCTCCTGGA | 6368.5 | 6.5 | 0.9 | 7.1 | 1.0E-12 | 4.8E-11 | BT20/MB468 | MP |
| >trnaMT_GluTTC_MT_-_14674_14742@40.59.20,trnalookalike8_GluTTC_5_-_93905172_93905240@40.59.20\|TTGGTCGTGGTTGTAGTCCG | >trnaMT_GluTTC_MT_-_14674_14742 | 40 | 59 | 20 | TTGGTCGTGGTTGTAGTCCG | 347.2 | 8.4 | 1.2 | 6.8 | 1.2E-11 | 4.8E-11 | BT20/MB468 | MITO |
| >hg38_21_+_8433172_8446622_RNA45SN1_45S_50ntFlanks@11576.11609.34\|CTTAGAACTGGTGCGGACCAGGGGAATCCGACTG | >hg38_21_+_8433172_8446622_RNA45SN1_45S_50ntFlanks | 11576 | 11609 | 34 | CTTAGAACTGGTGCGGACCAGGGGAATCCGACTG | 978.9 | -7.1 | 1.1 | -6.7 | 2.0E-11 | 4.8E-11 | MB231/MB468 | MITO |
| >hg38_21_+_8433172_8446622_RNA45SN1_45S_50ntFlanks@4732.4757.26\|AGAACGAAAGTCGGAGGTTCGAAGAC | >hg38_21_+_8433172_8446622_RNA45SN1_45S_50ntFlanks | 4732 | 4757 | 26 | AGAACGAAAGTCGGAGGTTCGAAGAC | 416.9 | 5.0 | 0.7 | 6.7 | 1.5E-11 | 4.8E-11 | BT20/MB468 | CYTO |
| >MI0000289\|hsa-mir-181a-1&WithFlank&1\|-\|198859038\|198859159@30.54.25;MI0000269\|hsa-mir-181a-2&WithFlank&9\|+\|124692436\|124692557@45.69.25&[MIMAT0000256&hsa-miR-181a-5p&offsets\|0\|+2;m-32&1\|-\|198859108\|198859130&offsets\|0\|+2];[MIMAT0000256_1&hsa-miR-181a-5p&offsets\|0\|+2;m-32&9\|+\|124692480\|124692502&offsets\|0\|+2]\|AACATTCAACGCTGTCGGTGAGTTT | >MI0000289\|hsa-mir-181a-1&WithFlank&1\|-\|198859038\|198859159 | 30 | 54 | 25 | AACATTCAACGCTGTCGGTGAGTTT | 3860.6 | 7.5 | 1.1 | 6.7 | 2.0E-11 | 4.8E-11 | MB231/MB468 | MITO |
| >hg38_21_+_8433172_8446622_RNA45SN1_45S_50ntFlanks@10574.10596.23\|AAGCCGGCGGGAGCCCCGGGGAG | >hg38_21_+_8433172_8446622_RNA45SN1_45S_50ntFlanks | 10574 | 10596 | 23 | AAGCCGGCGGGAGCCCCGGGGAG | 522.7 | -9.0 | 1.3 | -6.7 | 2.1E-11 | 4.8E-11 | BT20/MB468 | NUC |

| | | | | | | | | | | | | | |
|---|---|---|---|---|---|---|---|---|---|---|---|---|---|
| >hg38_21_+_8433172_8446622_RNA45SN1_45S_50ntFlanks@10781.10804.24\|AGGTGAACAGCCTCTGGCATGTTG | >hg38_21_+_8433172_8446622_RNA45SN1_45S_50ntFlanks | 10781 | 10804 | 24 | AGGTGAACAGCCTCTGGCATGTTG | 467.0 | -8.4 | 1.3 | -6.7 | 1.7E-11 | 4.8E-11 | MB231/MB468 | CYTO |
| >hg38_21_+_8433172_8446622_RNA45SN1_45S_50ntFlanks@12160.12178.19\|TTGACTGGGGCGGTACACC | >hg38_21_+_8433172_8446622_RNA45SN1_45S_50ntFlanks | 12160 | 12178 | 19 | TTGACTGGGGCGGTACACC | 94.0 | -10.4 | 1.5 | -6.7 | 1.5E-11 | 4.8E-11 | MB231/MB468 | MP |
| >hg38_21_+_8433172_8446622_RNA45SN1_45S_50ntFlanks@3793.3810.18\|CGAATGGCTCATTAAATC | >hg38_21_+_8433172_8446622_RNA45SN1_45S_50ntFlanks | 3793 | 3810 | 18 | CGAATGGCTCATTAAATC | 594.3 | -9.2 | 1.4 | -6.7 | 2.0E-11 | 4.8E-11 | MB231/MB468 | MITO |
| >hg38_21_+_8433172_8446622_RNA45SN1_45S_50ntFlanks@5334.5362.29\|ATGAACGAGGAATTCCCAGTAAGTGCGGG | >hg38_21_+_8433172_8446622_RNA45SN1_45S_50ntFlanks | 5334 | 5362 | 29 | ATGAACGAGGAATTCCCAGTAAGTGCGGG | 291.9 | -12.1 | 1.8 | -6.7 | 1.5E-11 | 4.8E-11 | MB231/MB468 | MP |
| >hg38_21_+_8433172_8446622_RNA45SN1_45S_50ntFlanks@11579.11609.31\|AGAACTGGTGCGGACCAGGGGAATCCGACTG | >hg38_21_+_8433172_8446622_RNA45SN1_45S_50ntFlanks | 11579 | 11609 | 31 | AGAACTGGTGCGGACCAGGGGAATCCGACTG | 629.8 | -4.5 | 0.7 | -6.7 | 2.1E-11 | 4.9E-11 | BT20/MB468 | NUC |
| >hg38_21_+_8433172_8446622_RNA45SN1_45S_50ntFlanks@12149.12180.32\|AGGTGGGGAGTTTGACTGGGGCGGTACACCTG | >hg38_21_+_8433172_8446622_RNA45SN1_45S_50ntFlanks | 12149 | 12180 | 32 | AGGTGGGGAGTTTGACTGGGGCGGTACACCTG | 82.1 | -10.2 | 1.5 | -6.7 | 1.6E-11 | 4.9E-11 | MB231/MB468 | MP |
| >hg38_21_+_8433172_8446622_RNA45SN1_45S_50ntFlanks@5475.5516.42\|GCCCTGGCGGAGCGCTGAGAAGACGGTCGAACTTGACTATCT | >hg38_21_+_8433172_8446622_RNA45SN1_45S_50ntFlanks | 5475 | 5516 | 42 | GCCCTGGCGGAGCGCTGAGAAGACGGTCGAACTTGACTATCT | 559.2 | 10.0 | 1.5 | 6.7 | 1.6E-11 | 4.9E-11 | BT20/MB468 | CYTO |
| >MI0000650\|hsa-mir-200c&WithFlank&12\|+\|6963693\|6963772@50.70.21(+1U)&[MIMAT0000617&hsa-miR-200c-3p&offsets\|0\|-2(+1U);m-54&12\|+\|6963742\|6963764&offsets\|0\|-2(+1U)]\|TAATACTGCCGGGTAATGATGT | >MI0000650\|hsa-mir-200c&WithFlank&12\|+\|6963693\|6963772 | 50 | 70 | 21(+1U) | TAATACTGCCGGGTAATGATGT | 175.6 | 4.3 | 0.6 | 6.7 | 2.1E-11 | 4.9E-11 | BT20/MB468 | NUC |
| >MI0001445\|hsa-mir-423&WithFlank&17\|+\|30117073\|30117178@23.46.24&[MIMAT0004748&hsa-miR-423-5p&offsets\|0\|+1;m-141&17\|+\|30117095\|30117117&offsets\|0\|+1]\|TGAGGGGCAGAGAGCGAGACTTTT | >MI0001445\|hsa-mir-423&WithFlank&17\|+\|30117073\|30117178 | 23 | 46 | 24 | TGAGGGGCAGAGAGCGAGACTTTT | 223.8 | 2.7 | 0.4 | 6.7 | 2.1E-11 | 4.9E-11 | BT20/MB468 | NUC |
| >am_trna116_GluCTC_1_-_145399233_145399304@1.30.30,trna59_GluCTC_1_+_249168447_249168518@1.30.30,trna71_GluCTC_1_-_161439189_161439260@1.30.30,trna74_GluCTC_1_-_161431809_161431880@1.30.30,trna77_GluCTC_1_-_161424398_161424469@1.30.30,trna77_GluCTC_6_+_28949976_28950047@1.30.30,trna80_GluCTC_1_-_161417018_161417089@1.30.30,trna87_GluCTC_6_-_126101393_126101464@1.30.30\|TCCCTGGTGGTCTAGTGGTTAGGATTCGGC | >am_trna116_GluCTC_1_-_145399233_145399304 | 1 | 30 | 30 | TCCCTGGTGGTCTAGTGGTTAGGATTCGGC | 36723.2 | -6.5 | 1.0 | -6.7 | 2.1E-11 | 4.9E-11 | BT20/MB468 | NUC |
| >MI0000284\|hsa-mir-204&WithFlank&9\|-\|70809969\|70810090@39.60.22&[MIMAT0000265&hsa-miR-204-5p&offsets\|0\|0;m-296&9\|-\|70810031\|70810052&offsets\|0\|0]\|TTCCCTTTGTCATCCTATGCCT | >MI0000284\|hsa-mir-204&WithFlank&9\|-\|70809969\|70810090 | 39 | 60 | 22 | TTCCCTTTGTCATCCTATGCCT | 160.7 | -10.4 | 1.5 | -7.0 | 1.8E-12 | 4.9E-11 | BT20/MB231 | NUC |
| >hg38_21_+_8433172_8446622_RNA45SN1_45S_50ntFlanks@5523.5550.28\|AGTAAAAGTCGTAACAAGGTTTCCGTAG | >hg38_21_+_8433172_8446622_RNA45SN1_45S_50ntFlanks | 5523 | 5550 | 28 | AGTAAAAGTCGTAACAAGGTTTCCGTAG | 109.8 | -10.6 | 1.6 | -6.7 | 1.6E-11 | 4.9E-11 | MB231/MB468 | MP |
| >am_trna116_GluCTC_1_-_145399233_145399304@1.28.28,trna59_GluCTC_1_+_249168447_249168518@1.28.28,trna71_GluCTC_1_-_161439189_161439260@1.28.28,trna74_GluCTC_1_-_161431809_161431880@1.28.28,trna77_GluCTC_1_-_161424398_161424469@1.28.28,trna77_GluCTC_6_+_28949976_28950047@1.28.28,trna80_GluCTC_1_-_161417018_161417089@1.28.28,trna87_GluCTC_6_-_126101393_126101464@1.28.28\|TCCCTGGTGGTCTAGTGGTTAGGATTCG | >am_trna116_GluCTC_1_-_145399233_145399304 | 1 | 28 | 28 | TCCCTGGTGGTCTAGTGGTTAGGATTCG | 6371.5 | -5.3 | 0.8 | -6.7 | 1.6E-11 | 4.9E-11 | MB231/MB468 | MP |
| >hg38_21_+_8433172_8446622_RNA45SN1_45S_50ntFlanks@6719.6736.18\|TGAATTGCAGGACACATT | >hg38_21_+_8433172_8446622_RNA45SN1_45S_50ntFlanks | 6719 | 6736 | 18 | TGAATTGCAGGACACATT | 1478.4 | -7.0 | 1.0 | -6.7 | 2.0E-11 | 4.9E-11 | MB231/MB468 | MITO |
| >hg38_21_+_8433172_8446622_RNA45SN1_45S_50ntFlanks@4046.4063.18\|CATTCGAACGTCTGCCCT | >hg38_21_+_8433172_8446622_RNA45SN1_45S_50ntFlanks | 4046 | 4063 | 18 | CATTCGAACGTCTGCCCT | 1757.1 | -5.1 | 0.8 | -6.8 | 1.2E-11 | 4.9E-11 | BT20/MB468 | MITO |
| >hg38_21_+_8433172_8446622_RNA45SN1_45S_50ntFlanks@6774.6806.33\|TCCTCCCGGGGCTACGCCTGTCTGAGCGTCGCT | >hg38_21_+_8433172_8446622_RNA45SN1_45S_50ntFlanks | 6774 | 6806 | 33 | TCCTCCCGGGGCTACGCCTGTCTGAGCGTCGCT | 933.8 | 5.6 | 0.8 | 6.7 | 1.7E-11 | 4.9E-11 | MB231/MB468 | CYTO |
| >hg38_21_+_8433172_8446622_RNA45SN1_45S_50ntFlanks@8170.8195.26\|CAAGTCCTTCTGATCGAGGCCCAGCC | >hg38_21_+_8433172_8446622_RNA45SN1_45S_50ntFlanks | 8170 | 8195 | 26 | CAAGTCCTTCTGATCGAGGCCCAGCC | 2035.0 | -9.1 | 1.4 | -6.7 | 2.0E-11 | 4.9E-11 | MB231/MB468 | MITO |
| >hg38_21_+_8433172_8446622_RNA45SN1_45S_50ntFlanks@7968.7994.27\|TCCGAGACGCGACCTCAGATCAGACGT | >hg38_21_+_8433172_8446622_RNA45SN1_45S_50ntFlanks | 7968 | 7994 | 27 | TCCGAGACGCGACCTCAGATCAGACGT | 303.4 | 7.2 | 1.1 | 6.8 | 1.2E-11 | 4.9E-11 | BT20/MB468 | MITO |
| >hg38_21_+_8433172_8446622_RNA45SN1_45S_50ntFlanks@11573.11592.20\|CGACTTAGAACTGGTGCGGA | >hg38_21_+_8433172_8446622_RNA45SN1_45S_50ntFlanks | 11573 | 11592 | 20 | CGACTTAGAACTGGTGCGGA | 461.2 | -4.1 | 0.6 | -6.7 | 1.7E-11 | 4.9E-11 | MB231/MB468 | CYTO |
| >hg38_21_+_8433172_8446622_RNA45SN1_45S_50ntFlanks@12419.12445.27\|GATGTCGGCTCTTCCTATCATTGTGAA | >hg38_21_+_8433172_8446622_RNA45SN1_45S_50ntFlanks | 12419 | 12445 | 27 | GATGTCGGCTCTTCCTATCATTGTGAA | 420.1 | -4.0 | 0.6 | -6.7 | 2.1E-11 | 4.9E-11 | BT20/MB468 | NUC |
| >hg38_1_-_228634819_228635039_RNA5S12_5S_50ntFlanks@136.171.36\|GGAGACCGCCTGGGAATACCGGGTGCTGTAGGCTTT | >hg38_1_-_228634819_228635039_RNA5S12_5S_50ntFlanks | 136 | 171 | 36 | GGAGACCGCCTGGGAATACCGGGTGCTGTAGGCTTT | 2806.5 | 5.2 | 0.8 | 6.7 | 1.7E-11 | 4.9E-11 | MB231/MB468 | NUC |
| >hg38_21_+_8433172_8446622_RNA45SN1_45S_50ntFlanks@6656.6689.34\|CTTAGCGGTGGATCACTCGGCTCGTGCGTCGATG | >hg38_21_+_8433172_8446622_RNA45SN1_45S_50ntFlanks | 6656 | 6689 | 34 | CTTAGCGGTGGATCACTCGGCTCGTGCGTCGATG | 1755.3 | -7.0 | 1.1 | -6.7 | 2.0E-11 | 4.9E-11 | MB231/MB468 | MITO |
| >hg38_21_+_8433172_8446622_RNA45SN1_45S_50ntFlanks@4734.4751.18\|AACGAAAGTCGGAGGTTC | >hg38_21_+_8433172_8446622_RNA45SN1_45S_50ntFlanks | 4734 | 4751 | 18 | AACGAAAGTCGGAGGTTC | 1347.1 | -5.1 | 0.8 | -6.7 | 1.6E-11 | 4.9E-11 | MB231/MB468 | MP |
| >hg38_21_+_8433172_8446622_RNA45SN1_45S_50ntFlanks@10819.10865.47\|AGGGAAGTCGGCAAGCCGGATCCGTAACTTCGGGATAAGGATTGGCT | >hg38_21_+_8433172_8446622_RNA45SN1_45S_50ntFlanks | 10819 | 10865 | 47 | AGGGAAGTCGGCAAGCCGGATCCGTAACTTCGGGATAAGGATTGGCT | 537.6 | -12.0 | 1.8 | -6.7 | 1.7E-11 | 4.9E-11 | MB231/MB468 | NUC |
| >hg38_21_+_8433172_8446622_RNA45SN1_45S_50ntFlanks@3813.3837.25\|TTATGGTTCCTTTGGTCGCTCGCTC | >hg38_21_+_8433172_8446622_RNA45SN1_45S_50ntFlanks | 3813 | 3837 | 25 | TTATGGTTCCTTTGGTCGCTCGCTC | 677.5 | -8.2 | 1.2 | -6.7 | 1.7E-11 | 4.9E-11 | MB231/MB468 | NUC |
| >hg38_21_+_8433172_8446622_RNA45SN1_45S_50ntFlanks@11932.11951.20\|GAATAAGTGGGAGGCCCCCG | >hg38_21_+_8433172_8446622_RNA45SN1_45S_50ntFlanks | 11932 | 11951 | 20 | GAATAAGTGGGAGGCCCCCG | 128.5 | -10.9 | 1.6 | -6.7 | 1.6E-11 | 4.9E-11 | MB231/MB468 | MP |
| >trna116_GluCTC_1_-_145399233_145399304@35.75.41,trna134_GluTTC_1_-_16861774_16861845@35.75.41,trna71_GluCTC_1_-_161439189_161439260@35.75.41,trna74_GluCTC_1_-_161431809_161431880@35.75.41,trna77_GluCTC_1_-_161424398_161424469@35.75.41,trna77_GluCTC_6_+_28949976_28950047@35.75.41,trna80_GluCTC_1_-_161417018_161417089@35.75.41,trna84_GluTTC_1_-_161391883_161391954@35.75.41,trna87_GluCTC_6_-_126101393_126101464@35.75.41\|TCACCGCCGCGGCCCGGGTTCGATTCCCGGTCAGGGAACCA | >trna116_GluCTC_1_-_145399233_145399304 | 35 | 75 | 41 | TCACCGCCGCGGCCCGGGTTCGATTCCCGGTCAGGGAACCA | 5797.7 | 7.4 | 1.1 | 6.7 | 1.7E-11 | 4.9E-11 | MB231/MB468 | CYTO |
| >am_MI0000809\|hsa-mir-151a&WithFlank&8\|-\|140732558\|140732659@53.70.18&[MIMAT0000757&hsa-miR-151a-3p&offsets\|0\|-3;m-30&8\|-\|140732587\|140732607&offsets\|0\|-3]\|CTAGACTGAAGCTCCTTG | >am_MI0000809\|hsa-mir-151a&WithFlank&8\|-\|140732558\|140732659 | 53 | 70 | 18 | CTAGACTGAAGCTCCTTG | 594.2 | 8.2 | 1.2 | 6.7 | 1.7E-11 | 4.9E-11 | MB231/MB468 | CYTO |
| >hg38_21_+_8433172_8446622_RNA45SN1_45S_50ntFlanks@9575.9605.31\|ATCGGTCGTCCGACCTGGGTATAGGGGCGAA | >hg38_21_+_8433172_8446622_RNA45SN1_45S_50ntFlanks | 9575 | 9605 | 31 | ATCGGTCGTCCGACCTGGGTATAGGGGCGAA | 384.8 | -11.7 | 1.6 | -7.1 | 1.0E-12 | 5.0E-11 | BT20/MB468 | MP |
| >hg38_21_+_8433172_8446622_RNA45SN1_45S_50ntFlanks@12416.12437.22\|TTCGATGTCGGCTCTTCCTATC | >hg38_21_+_8433172_8446622_RNA45SN1_45S_50ntFlanks | 12416 | 12437 | 22 | TTCGATGTCGGCTCTTCCTATC | 6848.2 | 4.2 | 0.6 | 6.7 | 1.7E-11 | 5.0E-11 | MB231/MB468 | CYTO |
| >hg38_21_+_8433172_8446622_RNA45SN1_45S_50ntFlanks@8171.8215.45\|AAGTCCTTCTGATCGAGGCCCAGCCCGTGGACGGTGTGAGGCCGG | >hg38_21_+_8433172_8446622_RNA45SN1_45S_50ntFlanks | 8171 | 8215 | 45 | AAGTCCTTCTGATCGAGGCCCAGCCCGTGGACGGTGTGAGGCCGG | 1495.8 | -10.5 | 1.6 | -6.7 | 2.2E-11 | 5.0E-11 | BT20/MB468 | NUC |
| >hg38_21_+_8433172_8446622_RNA45SN1_45S_50ntFlanks@10821.10840.20\|GGAAGTCGGCAAGCCGGATC | >hg38_21_+_8433172_8446622_RNA45SN1_45S_50ntFlanks | 10821 | 10840 | 20 | GGAAGTCGGCAAGCCGGATC | 3805.3 | -4.9 | 0.7 | -6.7 | 1.7E-11 | 5.0E-11 | MB231/MB468 | CYTO |
| >hg38_21_+_8433172_8446622_RNA45SN1_45S_50ntFlanks@4589.4616.28\|TATTTTGTTGGTTTTCGGAACTGAGGCC | >hg38_21_+_8433172_8446622_RNA45SN1_45S_50ntFlanks | 4589 | 4616 | 28 | TATTTTGTTGGTTTTCGGAACTGAGGCC | 303.8 | -8.5 | 1.3 | -6.7 | 2.2E-11 | 5.0E-11 | BT20/MB468 | NUC |
| >hg38_21_+_8433172_8446622_RNA45SN1_45S_50ntFlanks@12002.12033.32\|TGCGGGCCGCCGGTGAAATACCACTACTCTGA | >hg38_21_+_8433172_8446622_RNA45SN1_45S_50ntFlanks | 12002 | 12033 | 32 | TGCGGGCCGCCGGTGAAATACCACTACTCTGA | 92.8 | -10.4 | 1.5 | -6.7 | 1.6E-11 | 5.0E-11 | MB231/MB468 | MP |
| >hg38_21_+_8433172_8446622_RNA45SN1_45S_50ntFlanks@6772.6806.35\|GTTCCTCCCGGGGCTACGCCTGTCTGAGCGTCGCT | >hg38_21_+_8433172_8446622_RNA45SN1_45S_50ntFlanks | 6772 | 6806 | 35 | GTTCCTCCCGGGGCTACGCCTGTCTGAGCGTCGCT | 1522.5 | -3.9 | 0.6 | -6.7 | 2.2E-11 | 5.0E-11 | BT20/MB468 | NUC |
| >trnaMT_GlyTCC_MT_+_9991_10058@28.66.39\|ACTTCCAATTAACTAGTTTTGACAACATTCAAAAAAGAG | >trnaMT_GlyTCC_MT_+_9991_10058 | 28 | 66 | 39 | ACTTCCAATTAACTAGTTTTGACAACATTCAAAAAAGAG | 466.2 | -13.3 | 2.0 | -6.7 | 2.2E-11 | 5.0E-11 | BT20/MB468 | NUC |
| >hg38_21_+_8433172_8446622_RNA45SN1_45S_50ntFlanks@11667.11706.40\|CTGCCCAGTGCTCTGAATGTCAAAGTGAAGAAATTCAATG | >hg38_21_+_8433172_8446622_RNA45SN1_45S_50ntFlanks | 11667 | 11706 | 40 | CTGCCCAGTGCTCTGAATGTCAAAGTGAAGAAATTCAATG | 546.7 | 10.1 | 1.5 | 6.8 | 1.3E-11 | 5.0E-11 | BT20/MB468 | MITO |
| >hg38_21_+_8433172_8446622_RNA45SN1_45S_50ntFlanks@8443.8471.29\|CAACCCGGCGGCGGGTCCGGCCGTGTCGG | >hg38_21_+_8433172_8446622_RNA45SN1_45S_50ntFlanks | 8443 | 8471 | 29 | CAACCCGGCGGCGGGTCCGGCCGTGTCGG | 110.7 | -10.7 | 1.6 | -6.7 | 1.6E-11 | 5.0E-11 | MB231/MB468 | MP |
| >hg38_21_+_8433172_8446622_RNA45SN1_45S_50ntFlanks@12905.12933.29\|CGCACCGCACGTTCGTGGGGAACCTGGCG | >hg38_21_+_8433172_8446622_RNA45SN1_45S_50ntFlanks | 12905 | 12933 | 29 | CGCACCGCACGTTCGTGGGGAACCTGGCG | 209.4 | -11.6 | 1.7 | -6.7 | 1.6E-11 | 5.0E-11 | MB231/MB468 | MP |
| >hg38_21_+_8433172_8446622_RNA45SN1_45S_50ntFlanks@10843.10862.20\|TAACTTCGGGATAAGGATTG | >hg38_21_+_8433172_8446622_RNA45SN1_45S_50ntFlanks | 10843 | 10862 | 20 | TAACTTCGGGATAAGGATTG | 980.9 | -6.2 | 0.9 | -6.7 | 1.8E-11 | 5.0E-11 | MB231/MB468 | CYTO |
| >hg38_21_+_8433172_8446622_RNA45SN1_45S_50ntFlanks@9534.9569.36\|AGGAAACTCTGGTGGAGGTCCGTAGCGGTCCTGACG | >hg38_21_+_8433172_8446622_RNA45SN1_45S_50ntFlanks | 9534 | 9569 | 36 | AGGAAACTCTGGTGGAGGTCCGTAGCGGTCCTGACG | 2670.2 | -7.7 | 1.2 | -6.7 | 2.1E-11 | 5.0E-11 | MB231/MB468 | MITO |
| >hg38_21_+_8433172_8446622_RNA45SN1_45S_50ntFlanks@11454.11478.25\|CCTCGGCGCGCGGCGGCGGCGGCGG | >hg38_21_+_8433172_8446622_RNA45SN1_45S_50ntFlanks | 11454 | 11478 | 25 | CCTCGGCGCGCGGCGGCGGCGGCGG | 114.7 | -10.7 | 1.6 | -6.7 | 1.6E-11 | 5.0E-11 | MB231/MB468 | MP |
| >hg38_21_+_8433172_8446622_RNA45SN1_45S_50ntFlanks@11886.11905.20\|TGAGCTTGACTCTAGTCTGG | >hg38_21_+_8433172_8446622_RNA45SN1_45S_50ntFlanks | 11886 | 11905 | 20 | TGAGCTTGACTCTAGTCTGG | 305.6 | -11.3 | 1.6 | -7.1 | 1.1E-12 | 5.0E-11 | BT20/MB468 | MP |
| >hg38_21_+_8433172_8446622_RNA45SN1_45S_50ntFlanks@6649.6668.20\|TACGACTCTTAGCGGTGGAT | >hg38_21_+_8433172_8446622_RNA45SN1_45S_50ntFlanks | 6649 | 6668 | 20 | TACGACTCTTAGCGGTGGAT | 523.8 | 3.7 | 0.5 | 6.7 | 1.6E-11 | 5.0E-11 | BT20/MB468 | CYTO |

<table>
<tr><th></th><th></th><th></th><th></th><th></th><th></th><th></th><th></th><th></th><th></th><th></th><th></th><th></th><th></th></tr>
<tr><td>>hg38_21_+_8433172_8446622_RNA45SN1_45S_50ntFlanks@10738.10757.20|AAATCTCGCGCCGGGCCGTA</td><td>>hg38_21_+_8433172_8446622_RNA45SN1_45S_50ntFlanks</td><td>10738</td><td>10757</td><td>20</td><td>AAATCTCGCGCCGGGCCGTA</td><td>633.4</td><td>-4.4</td><td>0.7</td><td>-6.7</td><td>1.8E-11</td><td>5.0E-11</td><td>MB231/MB468</td><td>NUC</td></tr>
<tr><td>>hg38_21_+_8433172_8446622_RNA45SN1_45S_50ntFlanks@7976.8010.35|GCGACCTCAGATCAGACGTGGCGACCCGCTGAATT</td><td>>hg38_21_+_8433172_8446622_RNA45SN1_45S_50ntFlanks</td><td>7976</td><td>8010</td><td>35</td><td>GCGACCTCAGATCAGACGTGGCGACCCGCTGAATT</td><td>404.8</td><td>8.3</td><td>1.2</td><td>6.7</td><td>1.6E-11</td><td>5.0E-11</td><td>BT20/MB468</td><td>CYTO</td></tr>
<tr><td>>MI0000441|hsa-mir-30b&WithFlank&8|-|134800514|134800613@23.44.22&[MIMAT0000420&hsa-miR-30b-5p&offsets|0|0;m-76&8|-|134800570|134800591&offsets|0|0]|TGTAAACATCCTACACTCAGCT</td><td>>MI0000441|hsa-mir-30b&WithFlank&8|-|134800514|134800613</td><td>23</td><td>44</td><td>22</td><td>TGTAAACATCCTACACTCAGCT</td><td>1573.3</td><td>6.7</td><td>1.0</td><td>6.8</td><td>1.3E-11</td><td>5.0E-11</td><td>BT20/MB468</td><td>MITO</td></tr>
<tr><td>>hg38_21_+_8433172_8446622_RNA45SN1_45S_50ntFlanks@6668.6689.22|TCACTCGGCTCGTGCGTCGATG</td><td>>hg38_21_+_8433172_8446622_RNA45SN1_45S_50ntFlanks</td><td>6668</td><td>6689</td><td>22</td><td>TCACTCGGCTCGTGCGTCGATG</td><td>123.1</td><td>-10.8</td><td>1.6</td><td>-6.7</td><td>1.6E-11</td><td>5.0E-11</td><td>MB231/MB468</td><td>MP</td></tr>
<tr><td>>MI0000061|hsa-let-7a-2&WithFlank&11|-|122146516|122146599@11.32.22(+1C);MI0000062|hsa-let-7a-3&WithFlank&22|+|46112743|46112828@10.31.22(+1C);MI0000060|hsa-let-7a-1&WithFlank&9|+|94175951|94176042@12.33.22(+1C)&[MIMAT0000062&hsa-let-7a-5p&offsets|0|0(+1C);m-6&11|-|122146568|122146589&offsets|0|0(+1C)];[MIMAT0000062_1&hsa-let-7a-5p&offsets|0|0(+1C);m-5&22|+|46112752|46112773&offsets|0|0(+1C)];[MIMAT0000062_2&hsa-let-7a-5p&offsets|0|0(+1C);m-4&9|+|94175962|94175983&offsets|0|0(+1C)]|TGAGGTAGTAGGTTGTATAGTTC</td><td>>MI0000061|hsa-let-7a-2&WithFlank&11|-|122146516|122146599</td><td>11</td><td>32</td><td>22(+1C)</td><td>TGAGGTAGTAGGTTGTATAGTTC</td><td>292.6</td><td>6.4</td><td>1.0</td><td>6.7</td><td>1.8E-11</td><td>5.0E-11</td><td>MB231/MB468</td><td>CYTO</td></tr>
<tr><td>>hg38_21_+_8433172_8446622_RNA45SN1_45S_50ntFlanks@10838.10865.28|ATCCGTAACTTCGGGATAAGGATTGGCT</td><td>>hg38_21_+_8433172_8446622_RNA45SN1_45S_50ntFlanks</td><td>10838</td><td>10865</td><td>28</td><td>ATCCGTAACTTCGGGATAAGGATTGGCT</td><td>232.0</td><td>-10.9</td><td>1.5</td><td>-7.1</td><td>1.1E-12</td><td>5.1E-11</td><td>BT20/MB468</td><td>MP</td></tr>
<tr><td>>hg38_21_+_8433172_8446622_RNA45SN1_45S_50ntFlanks@5486.5506.21|GCGCTGAGAAGACGGTCGAAC</td><td>>hg38_21_+_8433172_8446622_RNA45SN1_45S_50ntFlanks</td><td>5486</td><td>5506</td><td>21</td><td>GCGCTGAGAAGACGGTCGAAC</td><td>376.4</td><td>5.3</td><td>0.8</td><td>6.7</td><td>1.6E-11</td><td>5.1E-11</td><td>BT20/MB468</td><td>CYTO</td></tr>
<tr><td>>hg38_21_+_8433172_8446622_RNA45SN1_45S_50ntFlanks@10771.10797.27|CAGGTCTCCAAGGTGAACAGCCTCTGG</td><td>>hg38_21_+_8433172_8446622_RNA45SN1_45S_50ntFlanks</td><td>10771</td><td>10797</td><td>27</td><td>CAGGTCTCCAAGGTGAACAGCCTCTGG</td><td>597.9</td><td>-7.9</td><td>1.2</td><td>-6.7</td><td>1.6E-11</td><td>5.1E-11</td><td>BT20/MB468</td><td>CYTO</td></tr>
<tr><td>>trna152_ValCAC_6_-_27248049_27248121@1.29.29|GCTTCTGTAGTGTAGTGGTTATCACGTTC</td><td>>trna152_ValCAC_6_-_27248049_27248121</td><td>1</td><td>29</td><td>29</td><td>GCTTCTGTAGTGTAGTGGTTATCACGTTC</td><td>28073.6</td><td>-6.2</td><td>0.9</td><td>-6.7</td><td>1.8E-11</td><td>5.1E-11</td><td>MB231/MB468</td><td>CYTO</td></tr>
<tr><td>>hg38_21_+_8433172_8446622_RNA45SN1_45S_50ntFlanks@11199.11224.26|CCGGCGGCGGCGGCGACTCTGGACGC</td><td>>hg38_21_+_8433172_8446622_RNA45SN1_45S_50ntFlanks</td><td>11199</td><td>11224</td><td>26</td><td>CCGGCGGCGGCGGCGACTCTGGACGC</td><td>150.1</td><td>-11.1</td><td>1.6</td><td>-6.7</td><td>1.6E-11</td><td>5.1E-11</td><td>MB231/MB468</td><td>MP</td></tr>
<tr><td>>MI0000082|hsa-mir-25&WithFlank&7|-|100093554|100093649@58.78.21(+1U)&[MIMAT0000081&hsa-miR-25-3p&offsets|0|-1(+1U);m-23&7|-|100093571|100093592&offsets|0|-1(+1U)]|CATTGCACTTGTCTCGGTCTGT</td><td>>MI0000082|hsa-mir-25&WithFlank&7|-|100093554|100093649</td><td>58</td><td>78</td><td>21(+1U)</td><td>CATTGCACTTGTCTCGGTCTGT</td><td>1825.5</td><td>6.2</td><td>0.9</td><td>6.7</td><td>2.1E-11</td><td>5.1E-11</td><td>MB231/MB468</td><td>MITO</td></tr>
<tr><td>>hg38_21_+_8433172_8446622_RNA45SN1_45S_50ntFlanks@11576.11611.36|CTTAGAACTGGTGCGGACCAGGGGAATCCGACTGTT</td><td>>hg38_21_+_8433172_8446622_RNA45SN1_45S_50ntFlanks</td><td>11576</td><td>11611</td><td>36</td><td>CTTAGAACTGGTGCGGACCAGGGGAATCCGACTGTT</td><td>343.7</td><td>-4.0</td><td>0.6</td><td>-6.7</td><td>1.8E-11</td><td>5.1E-11</td><td>MB231/MB468</td><td>NUC</td></tr>
<tr><td>>hg38_1_-_228634819_228635039_RNA5S12_5S_50ntFlanks@135.169.35|GGGAGACCGCCTGGGAATACCGGGTGCTGTAGGCT</td><td>>hg38_1_-_228634819_228635039_RNA5S12_5S_50ntFlanks</td><td>135</td><td>169</td><td>35</td><td>GGGAGACCGCCTGGGAATACCGGGTGCTGTAGGCT</td><td>1727.6</td><td>4.9</td><td>0.7</td><td>6.7</td><td>1.8E-11</td><td>5.1E-11</td><td>MB231/MB468</td><td>CYTO</td></tr>
<tr><td>>trna10_LysCTT_16_+_3241501_3241573@11.35.25,trna13_LysCTT_14_-_58706613_58706685@11.35.25,trna2_LysCTT_15_+_79152904_79152976@11.35.25,trna4_LysCTT_18_-_43669271_43669343@11.35.25|CTCAGTCGGTAGAGCATGGGACTCT</td><td>>trna10_LysCTT_16_+_3241501_3241573</td><td>11</td><td>35</td><td>25</td><td>CTCAGTCGGTAGAGCATGGGACTCT</td><td>268.8</td><td>11.3</td><td>1.7</td><td>6.7</td><td>2.1E-11</td><td>5.1E-11</td><td>MB231/MB468</td><td>MITO</td></tr>
<tr><td>>hg38_21_+_8433172_8446622_RNA45SN1_45S_50ntFlanks@11727.11744.18|AGTAACTATGACTCTCTT</td><td>>hg38_21_+_8433172_8446622_RNA45SN1_45S_50ntFlanks</td><td>11727</td><td>11744</td><td>18</td><td>AGTAACTATGACTCTCTT</td><td>1579.6</td><td>-5.3</td><td>0.8</td><td>-6.7</td><td>2.2E-11</td><td>5.1E-11</td><td>BT20/MB468</td><td>NUC</td></tr>
<tr><td>>trna111_HisGTG_1_-_147774845_147774916@1.35.35,trna118_HisGTG_1_-_145396881_145396952@1.35.35,trna16_HisGTG_1_+_146544773_146544844@1.35.35,trna1_HisGTG_15_+_45493349_45493420@1.35.35,trna21_HisGTG_1_+_147753471_147753542@1.35.35,trna33_HisGTG_6_+_27125906_27125977@1.35.35,trna7_HisGTG_9_-_14433938_14434009@1.35.35,trna8_HisGTG_15_-_45492611_45492682@1.35.35,trna9_HisGTG_15_-_45490804_45490875@1.35.35|GCCGTGATCGTATAGTGGTTAGTACTCTGCGTTGT</td><td>>trna111_HisGTG_1_-_147774845_147774916</td><td>1</td><td>35</td><td>35</td><td>GCCGTGATCGTATAGTGGTTAGTACTCTGCGTTGT</td><td>571.2</td><td>8.8</td><td>1.3</td><td>6.7</td><td>1.8E-11</td><td>5.1E-11</td><td>MB231/MB468</td><td>CYTO</td></tr>
<tr><td>>hg38_21_+_8433172_8446622_RNA45SN1_45S_50ntFlanks@12637.12657.21|GCTACCATCTGTGGGATTATG</td><td>>hg38_21_+_8433172_8446622_RNA45SN1_45S_50ntFlanks</td><td>12637</td><td>12657</td><td>21</td><td>GCTACCATCTGTGGGATTATG</td><td>628.9</td><td>-9.0</td><td>1.3</td><td>-6.7</td><td>2.1E-11</td><td>5.1E-11</td><td>MB231/MB468</td><td>MITO</td></tr>
<tr><td>>MI0000477|hsa-mir-146a&WithFlank&5|+|160485346|160485456@27.48.22(+2U)&[MIMAT0000449&hsa-miR-146a-5p&offsets|0|0(+2U);m-129&5|+|160485372|160485393&offsets|0|0(+2U)]|TGAGAACTGAATTCCATGGGTTTT</td><td>>MI0000477|hsa-mir-146a&WithFlank&5|+|160485346|160485456</td><td>27</td><td>48</td><td>22(+2U)</td><td>TGAGAACTGAATTCCATGGGTTTT</td><td>175.8</td><td>-10.7</td><td>1.5</td><td>-7.1</td><td>1.7E-12</td><td>5.1E-11</td><td>BT20/MB231</td><td>MITO</td></tr>
<tr><td>>hg38_21_+_8433172_8446622_RNA45SN1_45S_50ntFlanks@4367.4400.34|CAGTTAAAAAGCTCGTAGTTGGATCTTGGGAGCG</td><td>>hg38_21_+_8433172_8446622_RNA45SN1_45S_50ntFlanks</td><td>4367</td><td>4400</td><td>34</td><td>CAGTTAAAAAGCTCGTAGTTGGATCTTGGGAGCG</td><td>159.6</td><td>-11.2</td><td>1.7</td><td>-6.7</td><td>1.6E-11</td><td>5.1E-11</td><td>MB231/MB468</td><td>MP</td></tr>
<tr><td>>hg38_21_+_8433172_8446622_RNA45SN1_45S_50ntFlanks@5420.5466.47|CCGATTGGATGGTTTAGTGAGGCCCTCGGATCGGCCCCGCCGGGGTC</td><td>>hg38_21_+_8433172_8446622_RNA45SN1_45S_50ntFlanks</td><td>5420</td><td>5466</td><td>47</td><td>CCGATTGGATGGTTTAGTGAGGCCCTCGGATCGGCCCCGCCGGGGTC</td><td>154.4</td><td>-10.0</td><td>1.5</td><td>-6.7</td><td>1.8E-11</td><td>5.1E-11</td><td>MB231/MB468</td><td>CYTO</td></tr>
<tr><td>>hg38_21_+_8433172_8446622_RNA45SN1_45S_50ntFlanks@11564.11588.25|CCTAGCAGCCGACTTAGAACTGGTG</td><td>>hg38_21_+_8433172_8446622_RNA45SN1_45S_50ntFlanks</td><td>11564</td><td>11588</td><td>25</td><td>CCTAGCAGCCGACTTAGAACTGGTG</td><td>2104.3</td><td>-9.3</td><td>1.4</td><td>-6.7</td><td>1.6E-11</td><td>5.1E-11</td><td>MB231/MB468</td><td>MP</td></tr>
<tr><td>>hg38_21_+_8433172_8446622_RNA45SN1_45S_50ntFlanks@10903.10935.33|GGCTGGGCGCGCGCCGCGGCTGGACGAGGCGCC</td><td>>hg38_21_+_8433172_8446622_RNA45SN1_45S_50ntFlanks</td><td>10903</td><td>10935</td><td>33</td><td>GGCTGGGCGCGCGCCGCGGCTGGACGAGGCGCC</td><td>157.0</td><td>-10.0</td><td>1.5</td><td>-6.7</td><td>1.8E-11</td><td>5.1E-11</td><td>MB231/MB468</td><td>CYTO</td></tr>
<tr><td>>MI0000266|hsa-mir-10a&WithFlank&17|-|48579832|48579953@28.49.22&[MIMAT0000253&hsa-miR-10a-5p&offsets|0|-1;m-18&17|-|48579905|48579926&offsets|0|0]|TACCCTGTAGATCCGAATTTGT</td><td>>MI0000266|hsa-mir-10a&WithFlank&17|-|48579832|48579953</td><td>28</td><td>49</td><td>22</td><td>TACCCTGTAGATCCGAATTTGT</td><td>14454.7</td><td>10.2</td><td>1.4</td><td>7.1</td><td>1.1E-12</td><td>5.1E-11</td><td>BT20/MB468</td><td>MP</td></tr>
<tr><td>>hg38_21_+_8433172_8446622_RNA45SN1_45S_50ntFlanks@11623.11640.18|AAGCATCGCGAAGGCCCG</td><td>>hg38_21_+_8433172_8446622_RNA45SN1_45S_50ntFlanks</td><td>11623</td><td>11640</td><td>18</td><td>AAGCATCGCGAAGGCCCG</td><td>328.0</td><td>-11.4</td><td>1.6</td><td>-7.1</td><td>1.1E-12</td><td>5.1E-11</td><td>BT20/MB468</td><td>MP</td></tr>
<tr><td>>hg38_21_+_8433172_8446622_RNA45SN1_45S_50ntFlanks@476.504.29|TTCGTGATCGATGTGGTGACGTCGTGCTC</td><td>>hg38_21_+_8433172_8446622_RNA45SN1_45S_50ntFlanks</td><td>476</td><td>504</td><td>29</td><td>TTCGTGATCGATGTGGTGACGTCGTGCTC</td><td>1031.6</td><td>9.2</td><td>1.4</td><td>6.7</td><td>1.6E-11</td><td>5.1E-11</td><td>BT20/MB468</td><td>CYTO</td></tr>
<tr><td>>hg38_21_+_8433172_8446622_RNA45SN1_45S_50ntFlanks@8176.8204.29|CTTCTGATCGAGGCCCAGCCCGTGGACGG</td><td>>hg38_21_+_8433172_8446622_RNA45SN1_45S_50ntFlanks</td><td>8176</td><td>8204</td><td>29</td><td>CTTCTGATCGAGGCCCAGCCCGTGGACGG</td><td>927.6</td><td>-2.9</td><td>0.4</td><td>-6.7</td><td>1.6E-11</td><td>5.1E-11</td><td>BT20/MB468</td><td>CYTO</td></tr>
<tr><td>>hg38_21_+_8433172_8446622_RNA45SN1_45S_50ntFlanks@3789.3812.24|ACTGCGAATGGCTCATTAAATCAG</td><td>>hg38_21_+_8433172_8446622_RNA45SN1_45S_50ntFlanks</td><td>3789</td><td>3812</td><td>24</td><td>ACTGCGAATGGCTCATTAAATCAG</td><td>159.8</td><td>-11.2</td><td>1.7</td><td>-6.7</td><td>1.6E-11</td><td>5.1E-11</td><td>MB231/MB468</td><td>MP</td></tr>
<tr><td>>am_trnaMT_PheGAA_MT_+_577_647@40.62.23|TGTTTAGACGGGCTCACATCACC</td><td>>am_trnaMT_PheGAA_MT_+_577_647</td><td>40</td><td>62</td><td>23</td><td>TGTTTAGACGGGCTCACATCACC</td><td>280.5</td><td>5.3</td><td>0.8</td><td>6.7</td><td>2.1E-11</td><td>5.1E-11</td><td>MB231/MB468</td><td>MITO</td></tr>
<tr><td>>hg38_21_+_8433172_8446622_RNA45SN1_45S_50ntFlanks@5419.5443.25|ACCGATTGGATGGTTTAGTGAGGCC</td><td>>hg38_21_+_8433172_8446622_RNA45SN1_45S_50ntFlanks</td><td>5419</td><td>5443</td><td>25</td><td>ACCGATTGGATGGTTTAGTGAGGCC</td><td>126.7</td><td>-10.8</td><td>1.6</td><td>-6.7</td><td>1.6E-11</td><td>5.1E-11</td><td>MB231/MB468</td><td>MP</td></tr>
<tr><td>>hg38_21_+_8433172_8446622_RNA45SN1_45S_50ntFlanks@9535.9571.37|GGAAACTCTGGTGGAGGTCCGTAGCGGTCCTGACGTG</td><td>>hg38_21_+_8433172_8446622_RNA45SN1_45S_50ntFlanks</td><td>9535</td><td>9571</td><td>37</td><td>GGAAACTCTGGTGGAGGTCCGTAGCGGTCCTGACGTG</td><td>2957.7</td><td>-8.4</td><td>1.3</td><td>-6.7</td><td>2.2E-11</td><td>5.1E-11</td><td>BT20/MB468</td><td>NUC</td></tr>
<tr><td>>hg38_21_+_8433172_8446622_RNA45SN1_45S_50ntFlanks@12627.12656.30|ATGGGGCGAAGCTACCATCTGTGGGATTAT</td><td>>hg38_21_+_8433172_8446622_RNA45SN1_45S_50ntFlanks</td><td>12627</td><td>12656</td><td>30</td><td>ATGGGGCGAAGCTACCATCTGTGGGATTAT</td><td>127.6</td><td>-10.9</td><td>1.6</td><td>-6.7</td><td>1.7E-11</td><td>5.1E-11</td><td>MB231/MB468</td><td>MP</td></tr>
<tr><td>>hg38_21_+_8433172_8446622_RNA45SN1_45S_50ntFlanks@12994.13038.45|CCTCGCTGCGATCTATTGAAAGTCAGCCCTCGACACAAGGGTTTG</td><td>>hg38_21_+_8433172_8446622_RNA45SN1_45S_50ntFlanks</td><td>12994</td><td>13038</td><td>45</td><td>CCTCGCTGCGATCTATTGAAAGTCAGCCCTCGACACAAGGGTTTG</td><td>601.9</td><td>9.2</td><td>1.4</td><td>6.7</td><td>1.8E-11</td><td>5.1E-11</td><td>MB231/MB468</td><td>CYTO</td></tr>
<tr><td>>am_trnaMT_PheGAA_MT_+_577_647@41.58.18|GTTTAGACGGGCTCACAT</td><td>>am_trnaMT_PheGAA_MT_+_577_647</td><td>41</td><td>58</td><td>18</td><td>GTTTAGACGGGCTCACAT</td><td>354.5</td><td>9.5</td><td>1.4</td><td>6.7</td><td>1.7E-11</td><td>5.1E-11</td><td>BT20/MB468</td><td>CYTO</td></tr>
<tr><td>>hg38_21_+_8433172_8446622_RNA45SN1_45S_50ntFlanks@5492.5523.32|AGAAGACGGTCGAACTTGACTATCTAGAGGAA</td><td>>hg38_21_+_8433172_8446622_RNA45SN1_45S_50ntFlanks</td><td>5492</td><td>5523</td><td>32</td><td>AGAAGACGGTCGAACTTGACTATCTAGAGGAA</td><td>152.2</td><td>-11.1</td><td>1.7</td><td>-6.7</td><td>1.7E-11</td><td>5.2E-11</td><td>MB231/MB468</td><td>MP</td></tr>
<tr><td>>hg38_21_+_8433172_8446622_RNA45SN1_45S_50ntFlanks@9505.9526.22|TGAACTATGCCTGGGCAGGGCG</td><td>>hg38_21_+_8433172_8446622_RNA45SN1_45S_50ntFlanks</td><td>9505</td><td>9526</td><td>22</td><td>TGAACTATGCCTGGGCAGGGCG</td><td>213.2</td><td>-9.8</td><td>1.5</td><td>-6.8</td><td>1.3E-11</td><td>5.2E-11</td><td>BT20/MB468</td><td>MITO</td></tr>
<tr><td>>hg38_21_+_8433172_8446622_RNA45SN1_45S_50ntFlanks@6776.6797.22|CTCCCGGGGCTACGCCTGTCTG</td><td>>hg38_21_+_8433172_8446622_RNA45SN1_45S_50ntFlanks</td><td>6776</td><td>6797</td><td>22</td><td>CTCCCGGGGCTACGCCTGTCTG</td><td>87.4</td><td>-10.3</td><td>1.5</td><td>-6.7</td><td>1.7E-11</td><td>5.2E-11</td><td>MB231/MB468</td><td>MP</td></tr>
<tr><td>>hg38_21_+_8433172_8446622_RNA45SN1_45S_50ntFlanks@11666.11692.27|TCTGCCCAGTGCTCTGAATGTCAAAGT</td><td>>hg38_21_+_8433172_8446622_RNA45SN1_45S_50ntFlanks</td><td>11666</td><td>11692</td><td>27</td><td>TCTGCCCAGTGCTCTGAATGTCAAAGT</td><td>392.5</td><td>7.0</td><td>1.0</td><td>6.8</td><td>1.3E-11</td><td>5.2E-11</td><td>BT20/MB468</td><td>MITO</td></tr>
<tr><td>>hg38_21_+_8433172_8446622_RNA45SN1_45S_50ntFlanks@12717.12741.25|CCTCGGTTGGCCTCGGATAGCCGGT</td><td>>hg38_21_+_8433172_8446622_RNA45SN1_45S_50ntFlanks</td><td>12717</td><td>12741</td><td>25</td><td>CCTCGGTTGGCCTCGGATAGCCGGT</td><td>81.9</td><td>-10.2</td><td>1.5</td><td>-6.7</td><td>1.7E-11</td><td>5.2E-11</td><td>MB231/MB468</td><td>MP</td></tr>
<tr><td>>hg38_21_+_8433172_8446622_RNA45SN1_45S_50ntFlanks@4099.4120.22|GTGACCACGGGTGACGGGGAAT</td><td>>hg38_21_+_8433172_8446622_RNA45SN1_45S_50ntFlanks</td><td>4099</td><td>4120</td><td>22</td><td>GTGACCACGGGTGACGGGGAAT</td><td>95.4</td><td>-10.4</td><td>1.6</td><td>-6.7</td><td>1.7E-11</td><td>5.2E-11</td><td>MB231/MB468</td><td>MP</td></tr>
<tr><td>>hg38_21_+_8433172_8446622_RNA45SN1_45S_50ntFlanks@8153.8181.29|GCCGCTCGTGGGGGGCCCAAGTCCTTCTG</td><td>>hg38_21_+_8433172_8446622_RNA45SN1_45S_50ntFlanks</td><td>8153</td><td>8181</td><td>29</td><td>GCCGCTCGTGGGGGGCCCAAGTCCTTCTG</td><td>199.8</td><td>-11.5</td><td>1.7</td><td>-6.7</td><td>1.7E-11</td><td>5.2E-11</td><td>MB231/MB468</td><td>MP</td></tr>
<tr><td>>hg38_1_-_228634819_228635039_RNA5S12_5S_50ntFlanks@151.169.19|ATACCGGGTGCTGTAGGCT</td><td>>hg38_1_-_228634819_228635039_RNA5S12_5S_50ntFlanks</td><td>151</td><td>169</td><td>19</td><td>ATACCGGGTGCTGTAGGCT</td><td>817.7</td><td>4.8</td><td>0.7</td><td>6.7</td><td>1.8E-11</td><td>5.2E-11</td><td>MB231/MB468</td><td>CYTO</td></tr>
<tr><td>>MI0000085|hsa-mir-27a&WithFlank&19|-|13836434|13836523@16.37.22&[MIMAT0004501&hsa-miR-27a-5p&offsets|0|0;m-270&19|-|13836487|13836508&offsets|0|0]|AGGGCTTAGCTGCTTGTGAGCA</td><td>>MI0000085|hsa-mir-27a&WithFlank&19|-|13836434|13836523</td><td>16</td><td>37</td><td>22</td><td>AGGGCTTAGCTGCTTGTGAGCA</td><td>1799.5</td><td>6.3</td><td>0.9</td><td>6.7</td><td>1.7E-11</td><td>5.2E-11</td><td>BT20/MB468</td><td>CYTO</td></tr>
<tr><td>>hg38_21_+_8433172_8446622_RNA45SN1_45S_50ntFlanks@10408.10435.28|GGTCAGTCGGTCCTGAGAGATGGGCGAG</td><td>>hg38_21_+_8433172_8446622_RNA45SN1_45S_50ntFlanks</td><td>10408</td><td>10435</td><td>28</td><td>GGTCAGTCGGTCCTGAGAGATGGGCGAG</td><td>1862.4</td><td>-8.5</td><td>1.3</td><td>-6.7</td><td>1.7E-11</td><td>5.2E-11</td><td>BT20/MB468</td><td>CYTO</td></tr>
<tr><td>>hg38_21_+_8433172_8446622_RNA45SN1_45S_50ntFlanks@12616.12634.19|CTGAGGAGCCAATGGGGCG</td><td>>hg38_21_+_8433172_8446622_RNA45SN1_45S_50ntFlanks</td><td>12616</td><td>12634</td><td>19</td><td>CTGAGGAGCCAATGGGGCG</td><td>116.6</td><td>-10.7</td><td>1.6</td><td>-6.7</td><td>1.7E-11</td><td>5.2E-11</td><td>MB231/MB468</td><td>MP</td></tr>
<tr><td>>hg38_21_+_8433172_8446622_RNA45SN1_45S_50ntFlanks@4731.4766.36|AAGAACGAAAGTCGGAGGTTCGAAGACGATCAGATA</td><td>>hg38_21_+_8433172_8446622_RNA45SN1_45S_50ntFlanks</td><td>4731</td><td>4766</td><td>36</td><td>AAGAACGAAAGTCGGAGGTTCGAAGACGATCAGATA</td><td>4291.2</td><td>8.6</td><td>1.3</td><td>6.7</td><td>1.7E-11</td><td>5.2E-11</td><td>BT20/MB468</td><td>CYTO</td></tr>
<tr><td>>MI0001733|hsa-mir-452&WithFlank&X|-|151959622|151959718@20.41.22&[MIMAT0001635&hsa-miR-452-5p&offsets|0|0;m-188&X|-|151959677|151959699&offsets|0|-1]|AACTGTTTGCAGAGGAAACTGA</td><td>>MI0001733|hsa-mir-452&WithFlank&X|-|151959622|151959718</td><td>20</td><td>41</td><td>22</td><td>AACTGTTTGCAGAGGAAACTGA</td><td>201.6</td><td>7.4</td><td>1.0</td><td>7.1</td><td>1.2E-12</td><td>5.2E-11</td><td>BT20/MB231</td><td>CYTO</td></tr>
</table>

| | | | | | | | | | | | | | |
|---|---|---|---|---|---|---|---|---|---|---|---|---|---|
| >hg38_21_+_8433172_8446622_RNA45SN1_45S_50ntFlanks@9568.9595.28\|CGTGCAAATCGGTCGTCCGACCTGGGTA | >hg38_21_+_8433172_8446622_RNA45SN1_45S_50ntFlanks | 9568 | 9595 | 28 | CGTGCAAATCGGTCGTCCGACCTGGGTA | 1480.9 | -7.4 | 1.1 | -6.7 | 1.8E-11 | 5.2E-11 | MB231/MB468 | NUC |
| >hg38_21_+_8433172_8446622_RNA45SN1_45S_50ntFlanks@5143.5162.20\|ACTTCTTAGAGGGACAAGTG | >hg38_21_+_8433172_8446622_RNA45SN1_45S_50ntFlanks | 5143 | 5162 | 20 | ACTTCTTAGAGGGACAAGTG | 534.6 | -12.9 | 1.9 | -6.7 | 1.7E-11 | 5.2E-11 | MB231/MB468 | MP |
| >hg38_21_+_8433172_8446622_RNA45SN1_45S_50ntFlanks@11602.11623.22\|TCCGACTGTTTAATTAAAACAA | >hg38_21_+_8433172_8446622_RNA45SN1_45S_50ntFlanks | 11602 | 11623 | 22 | TCCGACTGTTTAATTAAAACAA | 1849.7 | 5.0 | 0.7 | 6.8 | 1.3E-11 | 5.2E-11 | BT20/MB468 | MITO |
| >trna15_ValAAC_5_-_180615416_180615488@1.23.23\|GTTTCCGTAGTGTAGTGGTCATC | >trna15_ValAAC_5_-_180615416_180615488 | 1 | 23 | 23 | GTTTCCGTAGTGTAGTGGTCATC | 440.4 | -3.3 | 0.5 | -6.7 | 2.3E-11 | 5.2E-11 | BT20/MB468 | NUC |
| >MI0003776\|hsa-mir-320b-1&WithFlank&1\|+\|116671740\|116671823@48.66.19(+1A);MI0003839\|hsa-mir-320b-2&WithFlank&1\|-\|224257034\|224257116@47.65.19(+1A);MI0003778\|hsa-mir-320c-1&WithFlank&18\|+\|21683512\|21683595@48.66.19(+1A);MI0008191\|hsa-mir-320c-2&WithFlank&18\|+\|24321669\|24321752@48.66.19(+1A);MI0000542\|hsa-mir-320a&WithFlank&8\|-\|22244960\|22245043@48.66.19(+1A)&[MIMAT0005792&hsa-miR-320b&offsets\|0\|-3(+1A);m-183&1\|+\|116671787\|116671806&offsets\|0\|-1(+1A)];[MIMAT0005792_1&hsa-miR-320b&offsets\|0\|-3(+1A);m-183&1\|-\|224257051\|224257070&offsets\|0\|-1(+1A)];[MIMAT0005793&hsa-miR-320c&offsets\|0\|-1(+1A);m-222&18\|+\|21683559\|21683578&offsets\|0\|-1(+1A)];[MIMAT0005793_1&hsa-miR-320c&offsets\|0\|-1(+1A);m-219&18\|+\|24321716\|24321735&offsets\|0\|-1(+1A)];[MIMAT0000510&hsa-miR-320a-3p&offsets\|0\|-3(+1A);m-60&8\|-\|22244975\|22244996&offsets\|0\|-3(+1A)]\|AAAAGCTGGGTTGAGAGGGA | >MI0003776\|hsa-mir-320b-1&WithFlank&1\|+\|116671740\|116671823 | 48 | 66 | 19(+1A) | AAAAGCTGGGTTGAGAGGGA | 77.3 | 4.2 | 0.6 | 6.7 | 2.3E-11 | 5.2E-11 | BT20/MB468 | NUC |
| >hg38_21_+_8433172_8446622_RNA45SN1_45S_50ntFlanks@10463.10489.27\|CTCCGTTGCCCTCGGCCGATCGAAAGG | >hg38_21_+_8433172_8446622_RNA45SN1_45S_50ntFlanks | 10463 | 10489 | 27 | CTCCGTTGCCCTCGGCCGATCGAAAGG | 448.0 | -9.0 | 1.3 | -6.7 | 2.3E-11 | 5.2E-11 | BT20/MB468 | NUC |
| >hg38_21_+_8433172_8446622_RNA45SN1_45S_50ntFlanks@9200.9218.19\|TCTCTCGGGGCCACGCGCG | >hg38_21_+_8433172_8446622_RNA45SN1_45S_50ntFlanks | 9200 | 9218 | 19 | TCTCTCGGGGCCACGCGCG | 106.0 | -10.6 | 1.6 | -6.7 | 1.7E-11 | 5.2E-11 | MB231/MB468 | MP |
| >hg38_21_+_8433172_8446622_RNA45SN1_45S_50ntFlanks@11729.11753.25\|TAACTATGACTCTCTTAAGGTAGCC | >hg38_21_+_8433172_8446622_RNA45SN1_45S_50ntFlanks | 11729 | 11753 | 25 | TAACTATGACTCTCTTAAGGTAGCC | 321.2 | -6.6 | 1.0 | -6.7 | 1.8E-11 | 5.2E-11 | MB231/MB468 | NUC |
| >hg38_21_+_8433172_8446622_RNA45SN1_45S_50ntFlanks@12436.12455.20\|TCATTGTGAAGCAGAATTCA | >hg38_21_+_8433172_8446622_RNA45SN1_45S_50ntFlanks | 12436 | 12455 | 20 | TCATTGTGAAGCAGAATTCA | 1668.2 | -7.3 | 1.1 | -6.7 | 1.9E-11 | 5.2E-11 | MB231/MB468 | NUC |
| >hg38_21_+_8433172_8446622_RNA45SN1_45S_50ntFlanks@10679.10710.32\|GGTTCCGGCGGCGTCCGGTGAGCTCTCGCTGG | >hg38_21_+_8433172_8446622_RNA45SN1_45S_50ntFlanks | 10679 | 10710 | 32 | GGTTCCGGCGGCGTCCGGTGAGCTCTCGCTGG | 41137.5 | -6.2 | 0.9 | -6.8 | 1.3E-11 | 5.2E-11 | BT20/MB468 | MITO |
| >hg38_21_+_8433172_8446622_RNA45SN1_45S_50ntFlanks@9623.9648.26\|TAGTAGCTGGTTCCCTCCGAAGTTTC | >hg38_21_+_8433172_8446622_RNA45SN1_45S_50ntFlanks | 9623 | 9648 | 26 | TAGTAGCTGGTTCCCTCCGAAGTTTC | 2120.1 | -9.1 | 1.4 | -6.7 | 2.2E-11 | 5.2E-11 | MB231/MB468 | MITO |
| >hg38_21_+_8433172_8446622_RNA45SN1_45S_50ntFlanks@11709.11728.20\|GCGCGGGTAAACGGCGGGAG | >hg38_21_+_8433172_8446622_RNA45SN1_45S_50ntFlanks | 11709 | 11728 | 20 | GCGCGGGTAAACGGCGGGAG | 87.0 | -10.3 | 1.5 | -6.7 | 1.7E-11 | 5.2E-11 | MB231/MB468 | MP |
| >hg38_21_+_8433172_8446622_RNA45SN1_45S_50ntFlanks@3775.3792.18\|GCCGGTACAGTGAAACTG | >hg38_21_+_8433172_8446622_RNA45SN1_45S_50ntFlanks | 3775 | 3792 | 18 | GCCGGTACAGTGAAACTG | 2437.9 | -5.0 | 0.7 | -6.7 | 1.8E-11 | 5.2E-11 | MB231/MB468 | CYTO |
| >hg38_21_+_8433172_8446622_RNA45SN1_45S_50ntFlanks@11454.11477.24\|CCTCGGCGCGCGGCGGCGGCGGCG | >hg38_21_+_8433172_8446622_RNA45SN1_45S_50ntFlanks | 11454 | 11477 | 24 | CCTCGGCGCGCGGCGGCGGCGGCG | 93.9 | -10.4 | 1.5 | -6.7 | 1.7E-11 | 5.3E-11 | MB231/MB468 | MP |
| >hg38_21_+_8433172_8446622_RNA45SN1_45S_50ntFlanks@4473.4497.25\|TTAGCTGAGTGTCCCGCGGGGCCCG | >hg38_21_+_8433172_8446622_RNA45SN1_45S_50ntFlanks | 4473 | 4497 | 25 | TTAGCTGAGTGTCCCGCGGGGCCCG | 101.3 | -10.5 | 1.6 | -6.7 | 1.7E-11 | 5.3E-11 | MB231/MB468 | MP |
| >hg38_21_+_8433172_8446622_RNA45SN1_45S_50ntFlanks@11199.11223.25\|CCGGCGGCGGCGGCGACTCTGGACG | >hg38_21_+_8433172_8446622_RNA45SN1_45S_50ntFlanks | 11199 | 11223 | 25 | CCGGCGGCGGCGGCGACTCTGGACG | 1683.3 | -10.5 | 1.6 | -6.7 | 2.3E-11 | 5.3E-11 | BT20/MB468 | NUC |
| >hg38_21_+_8433172_8446622_RNA45SN1_45S_50ntFlanks@12640.12661.22\|ACCATCTGTGGGATTATGACTG | >hg38_21_+_8433172_8446622_RNA45SN1_45S_50ntFlanks | 12640 | 12661 | 22 | ACCATCTGTGGGATTATGACTG | 2492.5 | -6.1 | 0.9 | -6.7 | 2.2E-11 | 5.3E-11 | MB231/MB468 | MITO |
| >hg38_21_+_8433172_8446622_RNA45SN1_45S_50ntFlanks@12283.12318.36\|TACAGACCGTGAAAGCGGGGCCTCACGATCCTTCTG | >hg38_21_+_8433172_8446622_RNA45SN1_45S_50ntFlanks | 12283 | 12318 | 36 | TACAGACCGTGAAAGCGGGGCCTCACGATCCTTCTG | 107.1 | -10.6 | 1.6 | -6.7 | 1.7E-11 | 5.3E-11 | MB231/MB468 | MP |
| >hg38_21_+_8433172_8446622_RNA45SN1_45S_50ntFlanks@12477.12498.22\|CACTAATAGGGAACGTGAGCTG | >hg38_21_+_8433172_8446622_RNA45SN1_45S_50ntFlanks | 12477 | 12498 | 22 | CACTAATAGGGAACGTGAGCTG | 2607.5 | -8.3 | 1.2 | -6.7 | 2.3E-11 | 5.3E-11 | BT20/MB468 | NUC |
| >MI0000253\|hsa-mir-148a&WithFlank&7\|-\|25949913\|25949992@50.70.21(+1A)&[MIMAT0000243&hsa-miR-148a-3p&offsets\|0\|-1(+1A);m-7&7\|-\|25949922\|25949943&offsets\|0\|-1(+1A)]\|TCAGTGCACTACAGAACTTTGA | >MI0000253\|hsa-mir-148a&WithFlank&7\|-\|25949913\|25949992 | 50 | 70 | 21(+1A) | TCAGTGCACTACAGAACTTTGA | 5543.1 | 5.9 | 0.8 | 7.1 | 1.3E-12 | 5.3E-11 | BT20/MB231 | CYTO |
| >hg38_21_+_8433172_8446622_RNA45SN1_45S_50ntFlanks@12033.12053.21\|ATCGTTTTTTCACTGACCCGG | >hg38_21_+_8433172_8446622_RNA45SN1_45S_50ntFlanks | 12033 | 12053 | 21 | ATCGTTTTTTCACTGACCCGG | 438.6 | -7.5 | 1.1 | -6.7 | 1.9E-11 | 5.3E-11 | MB231/MB468 | CYTO |
| >hg38_21_+_8433172_8446622_RNA45SN1_45S_50ntFlanks@10389.10410.22\|TGAACAGCAGTTGAACATGGGT | >hg38_21_+_8433172_8446622_RNA45SN1_45S_50ntFlanks | 10389 | 10410 | 22 | TGAACAGCAGTTGAACATGGGT | 1803.9 | -5.1 | 0.8 | -6.7 | 1.7E-11 | 5.3E-11 | BT20/MB468 | CYTO |
| >hg38_21_+_8433172_8446622_RNA45SN1_45S_50ntFlanks@11622.11640.19\|AAAGCATCGCGAAGGCCCG | >hg38_21_+_8433172_8446622_RNA45SN1_45S_50ntFlanks | 11622 | 11640 | 19 | AAAGCATCGCGAAGGCCCG | 264.3 | -10.1 | 1.5 | -6.8 | 1.4E-11 | 5.3E-11 | BT20/MB468 | MITO |
| >hg38_21_+_8433172_8446622_RNA45SN1_45S_50ntFlanks@10783.10815.33\|GTGAACAGCCTCTGGCATGTTGGAACAATGTAG | >hg38_21_+_8433172_8446622_RNA45SN1_45S_50ntFlanks | 10783 | 10815 | 33 | GTGAACAGCCTCTGGCATGTTGGAACAATGTAG | 102.6 | -10.5 | 1.6 | -6.7 | 1.7E-11 | 5.3E-11 | MB231/MB468 | MP |
| >MI0000286\|hsa-mir-210&WithFlank&11\|-\|568083\|568204@72.94.23&[MIMAT0000267&hsa-miR-210-3p&offsets\|0\|+1;m-102&11\|-\|568112\|568133&offsets\|0\|+1]\|CTGTGCGTGTGACAGCGGCTGAT | >MI0000286\|hsa-mir-210&WithFlank&11\|-\|568083\|568204 | 72 | 94 | 23 | CTGTGCGTGTGACAGCGGCTGAT | 426.9 | 12.0 | 1.8 | 6.7 | 2.2E-11 | 5.3E-11 | MB231/MB468 | MITO |
| >hg38_21_+_8433172_8446622_RNA45SN1_45S_50ntFlanks@10572.10607.36\|AGAAGCCGGCGGGAGCCCCGGGGAGAGTTCTCTTTT | >hg38_21_+_8433172_8446622_RNA45SN1_45S_50ntFlanks | 10572 | 10607 | 36 | AGAAGCCGGCGGGAGCCCCGGGGAGAGTTCTCTTTT | 100.2 | -10.5 | 1.6 | -6.7 | 1.7E-11 | 5.3E-11 | MB231/MB468 | MP |
| >hg38_21_+_8433172_8446622_RNA45SN1_45S_50ntFlanks@11933.11968.36\|AATAAGTGGGAGGCCCCCGGCGCCCCCCCGGTGTCC | >hg38_21_+_8433172_8446622_RNA45SN1_45S_50ntFlanks | 11933 | 11968 | 36 | AATAAGTGGGAGGCCCCCGGCGCCCCCCCGGTGTCC | 249.0 | -11.7 | 1.8 | -6.7 | 2.2E-11 | 5.3E-11 | MB231/MB468 | MITO |
| >MI0002470\|hsa-mir-486-1&WithFlank&8\|-\|41660435\|41660514@10.31.22(+1U);MI0023622\|hsa-mir-486-2&WithFlank&8\|+\|41660438\|41660513@7.28.22(+1U)&[MIMAT0002177&hsa-miR-486-5p&offsets\|0\|0(+1U);m-107&8\|-\|41660484\|41660505&offsets\|0\|0(+1U)];[MIMAT0002177_1&hsa-miR-486-5p&offsets\|0\|0(+1U);TJU_CMC_MD2.ID03138.5p-miR&offsets\|0\|-2(+1U);m-106&8\|+\|41660444\|41660465&offsets\|0\|0(+1U)]\|TCCTGTACTGAGCTGCCCCGAGT | >MI0002470\|hsa-mir-486-1&WithFlank&8\|-\|41660435\|41660514 | 10 | 31 | 22(+1U) | TCCTGTACTGAGCTGCCCCGAGT | 561.8 | -12.5 | 1.7 | -7.3 | 2.7E-13 | 5.3E-11 | BT20/MB231 | MP |
| >MI0000102\|hsa-mir-100&WithFlank&11\|-\|122152223\|122152314@20.39.20(+1A)&[MIMAT0000098&hsa-miR-100-5p&offsets\|+1\|-1(+1A);m-27&11\|-\|122152276\|122152296&offsets\|+1\|0(+1A)]\|ACCCGTAGATCCGAACTTGTA | >MI0000102\|hsa-mir-100&WithFlank&11\|-\|122152223\|122152314 | 20 | 39 | 20(+1A) | ACCCGTAGATCCGAACTTGTA | 160.1 | 10.5 | 1.6 | 6.7 | 1.7E-11 | 5.3E-11 | MB231/MB468 | MP |
| >hg38_21_+_8433172_8446622_RNA45SN1_45S_50ntFlanks@5421.5466.46\|CGATTGGATGGTTTAGTGAGGCCCTCGGATCGGCCCCGCCGGGGTC | >hg38_21_+_8433172_8446622_RNA45SN1_45S_50ntFlanks | 5421 | 5466 | 46 | CGATTGGATGGTTTAGTGAGGCCCTCGGATCGGCCCCGCCGGGGTC | 178.1 | -11.3 | 1.7 | -6.7 | 2.2E-11 | 5.3E-11 | MB231/MB468 | MITO |
| >trna10_LysCTT_16_+_3241501_3241573@3.35.33,trna13_LysCTT_14_-_58706613_58706685@3.35.33,trna2_LysCTT_15_+_79152904_79152976@3.35.33\|CCGGCTAGCTCAGTCGGTAGAGCATGGGACTCT | >trna10_LysCTT_16_+_3241501_3241573 | 3 | 35 | 33 | CCGGCTAGCTCAGTCGGTAGAGCATGGGACTCT | 2038.1 | 6.9 | 1.0 | 6.7 | 2.2E-11 | 5.3E-11 | MB231/MB468 | MITO |
| >hg38_21_+_8433172_8446622_RNA45SN1_45S_50ntFlanks@11718.11742.25\|AACGGCGGGAGTAACTATGACTCTC | >hg38_21_+_8433172_8446622_RNA45SN1_45S_50ntFlanks | 11718 | 11742 | 25 | AACGGCGGGAGTAACTATGACTCTC | 1401.2 | -9.1 | 1.4 | -6.7 | 1.7E-11 | 5.3E-11 | MB231/MB468 | MP |
| >hg38_21_+_8433172_8446622_RNA45SN1_45S_50ntFlanks@10808.10830.23\|CAATGTAGGTAAGGGAAGTCGGC | >hg38_21_+_8433172_8446622_RNA45SN1_45S_50ntFlanks | 10808 | 10830 | 23 | CAATGTAGGTAAGGGAAGTCGGC | 517.3 | -7.2 | 1.1 | -6.7 | 1.9E-11 | 5.3E-11 | MB231/MB468 | CYTO |
| >hg38_21_+_8433172_8446622_RNA45SN1_45S_50ntFlanks@9545.9570.26\|GTGGAGGTCCGTAGCGGTCCTGACGT | >hg38_21_+_8433172_8446622_RNA45SN1_45S_50ntFlanks | 9545 | 9570 | 26 | GTGGAGGTCCGTAGCGGTCCTGACGT | 728.3 | -5.0 | 0.8 | -6.7 | 2.2E-11 | 5.3E-11 | MB231/MB468 | MITO |
| >hg38_21_+_8433172_8446622_RNA45SN1_45S_50ntFlanks@11581.11599.19\|AACTGGTGCGGACCAGGGG | >hg38_21_+_8433172_8446622_RNA45SN1_45S_50ntFlanks | 11581 | 11599 | 19 | AACTGGTGCGGACCAGGGG | 184.3 | -11.4 | 1.7 | -6.7 | 1.7E-11 | 5.4E-11 | MB231/MB468 | MP |
| >hg38_21_+_8433172_8446622_RNA45SN1_45S_50ntFlanks@5419.5445.27\|ACCGATTGGATGGTTTAGTGAGGCCCT | >hg38_21_+_8433172_8446622_RNA45SN1_45S_50ntFlanks | 5419 | 5445 | 27 | ACCGATTGGATGGTTTAGTGAGGCCCT | 214.5 | -3.6 | 0.5 | -6.7 | 1.7E-11 | 5.4E-11 | BT20/MB468 | CYTO |
| >hg38_21_+_8433172_8446622_RNA45SN1_45S_50ntFlanks@4297.4321.25\|CATTGGAGGGCAAGTCTGGTGCCAG | >hg38_21_+_8433172_8446622_RNA45SN1_45S_50ntFlanks | 4297 | 4321 | 25 | CATTGGAGGGCAAGTCTGGTGCCAG | 442.5 | -5.4 | 0.8 | -6.7 | 1.9E-11 | 5.4E-11 | MB231/MB468 | CYTO |
| >hg38_21_+_8433172_8446622_RNA45SN1_45S_50ntFlanks@10681.10706.26\|TTCCGGCGGCGTCCGGTGAGCTCTCG | >hg38_21_+_8433172_8446622_RNA45SN1_45S_50ntFlanks | 10681 | 10706 | 26 | TTCCGGCGGCGTCCGGTGAGCTCTCG | 1869.2 | -8.9 | 1.3 | -6.7 | 2.2E-11 | 5.4E-11 | MB231/MB468 | MITO |
| >hg38_21_+_8433172_8446622_RNA45SN1_45S_50ntFlanks@6739.6772.34\|TCATCGACACTTCGAACGCACTTGCGGCCCCGGG | >hg38_21_+_8433172_8446622_RNA45SN1_45S_50ntFlanks | 6739 | 6772 | 34 | TCATCGACACTTCGAACGCACTTGCGGCCCCGGG | 754.6 | -8.6 | 1.3 | -6.7 | 2.2E-11 | 5.4E-11 | MB231/MB468 | MITO |
| >hg38_21_+_8433172_8446622_RNA45SN1_45S_50ntFlanks@11711.11728.18\|GCGGGTAAACGGCGGGAG | >hg38_21_+_8433172_8446622_RNA45SN1_45S_50ntFlanks | 11711 | 11728 | 18 | GCGGGTAAACGGCGGGAG | 90.3 | -10.4 | 1.5 | -6.7 | 1.7E-11 | 5.4E-11 | MB231/MB468 | MP |
| >hg38_21_+_8433172_8446622_RNA45SN1_45S_50ntFlanks@12605.12625.21\|TATGTGCTTGGCTGAGGAGCC | >hg38_21_+_8433172_8446622_RNA45SN1_45S_50ntFlanks | 12605 | 12625 | 21 | TATGTGCTTGGCTGAGGAGCC | 3835.4 | -7.9 | 1.2 | -6.7 | 1.7E-11 | 5.4E-11 | MB231/MB468 | MP |
| >MI0000088\|hsa-mir-30a&WithFlank&6\|-\|71403545\|71403627@13.35.23&[MIMAT0000087&hsa-miR-30a-5p&offsets\|+1\|+2;m-15&6\|-\|71403595\|71403616&offsets\|+1\|+2]\|GTAAACATCCTCGACTGGAAGCT | >MI0000088\|hsa-mir-30a&WithFlank&6\|-\|71403545\|71403627 | 13 | 35 | 23 | GTAAACATCCTCGACTGGAAGCT | 3346.1 | 4.3 | 0.6 | 6.7 | 1.7E-11 | 5.4E-11 | MB231/MB468 | MP |
| >hg38_21_+_8433172_8446622_RNA45SN1_45S_50ntFlanks@4370.4398.29\|TTAAAAAGCTCGTAGTTGGATCTTGGGAG | >hg38_21_+_8433172_8446622_RNA45SN1_45S_50ntFlanks | 4370 | 4398 | 29 | TTAAAAAGCTCGTAGTTGGATCTTGGGAG | 177.9 | -11.3 | 1.7 | -6.7 | 1.7E-11 | 5.4E-11 | MB231/MB468 | MP |
| >hg38_21_+_8433172_8446622_RNA45SN1_45S_50ntFlanks@4549.4573.25\|GGATACCGCAGCTAGGAATAATGGA | >hg38_21_+_8433172_8446622_RNA45SN1_45S_50ntFlanks | 4549 | 4573 | 25 | GGATACCGCAGCTAGGAATAATGGA | 94.3 | -10.4 | 1.5 | -6.7 | 1.7E-11 | 5.4E-11 | MB231/MB468 | MP |
| >hg38_21_+_8433172_8446622_RNA45SN1_45S_50ntFlanks@9786.9803.18\|CTCGCTGGCGTGGAGCCG | >hg38_21_+_8433172_8446622_RNA45SN1_45S_50ntFlanks | 9786 | 9803 | 18 | CTCGCTGGCGTGGAGCCG | 110.3 | -10.6 | 1.6 | -6.7 | 1.7E-11 | 5.4E-11 | MB231/MB468 | MP |

| | | | | | | | | | | | | | |
|---|---|---|---|---|---|---|---|---|---|---|---|---|---|
| >hg38_21_+_8433172_8446622_RNA45SN1_45S_50ntFlanks@11587.11611.25\|TGCGGACCAGGGGAATCCGACTGTT | >hg38_21_+_8433172_8446622_RNA45SN1_45S_50ntFlanks | 11587 | 11611 | 25 | TGCGGACCAGGGGAATCCGACTGTT | 905.7 | -4.9 | 0.7 | -6.7 | 1.7E-11 | 5.4E-11 | MB231/MB468 | MP |
| >trna116_GluCTC_1_-_145399233_145399304@30.71.42,trna71_GluCTC_1_-_161439189_161439260@30.71.42,trna74_GluCTC_1_-_161431809_161431880@30.71.42,trna77_GluCTC_1_-_161424398_161424469@30.71.42,trna77_GluCTC_6_+_28949976_28950047@30.71.42,trna80_GluCTC_1_-_161417018_161417089@30.71.42,trna87_GluCTC_6_-_126101393_126101464@30.71.42\|CGCTCTCACCGCCGCGGCCCGGGTTCGATTCCCGGTCAGGGA | >trna116_GluCTC_1_-_145399233_145399304 | 30 | 71 | 42 | CGCTCTCACCGCCGCGGCCCGGGTTCGATTCCCGGTCAGGGA | 308.1 | -6.9 | 1.0 | -6.7 | 1.9E-11 | 5.4E-11 | MB231/MB468 | CYTO |
| >hg38_21_+_8433172_8446622_RNA45SN1_45S_50ntFlanks@5485.5515.31\|AGCGCTGAGAAGACGGTCGAACTTGACTATC | >hg38_21_+_8433172_8446622_RNA45SN1_45S_50ntFlanks | 5485 | 5515 | 31 | AGCGCTGAGAAGACGGTCGAACTTGACTATC | 987.1 | -8.7 | 1.3 | -6.7 | 2.2E-11 | 5.4E-11 | MB231/MB468 | MITO |
| >hg38_21_+_8433172_8446622_RNA45SN1_45S_50ntFlanks@6648.6687.40\|GTACGACTCTTAGCGGTGGATCACTCGGCTCGTGCGTCGA | >hg38_21_+_8433172_8446622_RNA45SN1_45S_50ntFlanks | 6648 | 6687 | 40 | GTACGACTCTTAGCGGTGGATCACTCGGCTCGTGCGTCGA | 347.9 | 6.0 | 0.9 | 6.7 | 1.9E-11 | 5.4E-11 | MB231/MB468 | NUC |
| >hg38_21_+_8433172_8446622_RNA45SN1_45S_50ntFlanks@11805.11830.26\|CCACTGTCCCTACCTACTATCCAGCG | >hg38_21_+_8433172_8446622_RNA45SN1_45S_50ntFlanks | 11805 | 11830 | 26 | CCACTGTCCCTACCTACTATCCAGCG | 108.9 | -10.6 | 1.6 | -6.7 | 1.8E-11 | 5.4E-11 | MB231/MB468 | MP |
| >hg38_21_+_8433172_8446622_RNA45SN1_45S_50ntFlanks@9166.9183.18\|CCGGGCGGGTCGCGCCGT | >hg38_21_+_8433172_8446622_RNA45SN1_45S_50ntFlanks | 9166 | 9183 | 18 | CCGGGCGGGTCGCGCCGT | 252.1 | -10.1 | 1.5 | -6.8 | 1.4E-11 | 5.4E-11 | BT20/MB468 | MITO |
| >hg38_21_+_8433172_8446622_RNA45SN1_45S_50ntFlanks@9604.9622.19\|AAAGACTAATCGAACCATC | >hg38_21_+_8433172_8446622_RNA45SN1_45S_50ntFlanks | 9604 | 9622 | 19 | AAAGACTAATCGAACCATC | 1417.2 | -3.3 | 0.5 | -6.7 | 1.9E-11 | 5.4E-11 | MB231/MB468 | CYTO |
| >hg38_21_+_8433172_8446622_RNA45SN1_45S_50ntFlanks@8824.8844.21\|CGGCGGCGGCGGCGGCGGTGG | >hg38_21_+_8433172_8446622_RNA45SN1_45S_50ntFlanks | 8824 | 8844 | 21 | CGGCGGCGGCGGCGGCGGTGG | 137.9 | -11.0 | 1.6 | -6.7 | 1.8E-11 | 5.4E-11 | MB231/MB468 | MP |
| >hg38_21_+_8433172_8446622_RNA45SN1_45S_50ntFlanks@4449.4471.23\|CCTCTCGGCGCCCCCTCGATGCT | >hg38_21_+_8433172_8446622_RNA45SN1_45S_50ntFlanks | 4449 | 4471 | 23 | CCTCTCGGCGCCCCCTCGATGCT | 1646.9 | -7.1 | 1.0 | -6.8 | 1.4E-11 | 5.4E-11 | BT20/MB468 | MITO |
| >hg38_21_+_8433172_8446622_RNA45SN1_45S_50ntFlanks@11189.11225.37\|ACCCGGGGGGCCGGCGGCGGCGGCGACTCTGGACGCG | >hg38_21_+_8433172_8446622_RNA45SN1_45S_50ntFlanks | 11189 | 11225 | 37 | ACCCGGGGGGCCGGCGGCGGCGGCGACTCTGGACGCG | 222.7 | -11.7 | 1.7 | -6.7 | 1.8E-11 | 5.5E-11 | MB231/MB468 | MP |
| >hg38_21_+_8433172_8446622_RNA45SN1_45S_50ntFlanks@11641.11666.26\|CGGCGGGTGTTGACGCGATGTGATTT | >hg38_21_+_8433172_8446622_RNA45SN1_45S_50ntFlanks | 11641 | 11666 | 26 | CGGCGGGTGTTGACGCGATGTGATTT | 263.4 | -11.1 | 1.6 | -7.1 | 1.2E-12 | 5.5E-11 | BT20/MB468 | MP |
| >hg38_21_+_8433172_8446622_RNA45SN1_45S_50ntFlanks@7979.8005.27\|ACCTCAGATCAGACGTGGCGACCCGCT | >hg38_21_+_8433172_8446622_RNA45SN1_45S_50ntFlanks | 7979 | 8005 | 27 | ACCTCAGATCAGACGTGGCGACCCGCT | 742.3 | -4.5 | 0.7 | -6.7 | 2.3E-11 | 5.5E-11 | MB231/MB468 | MITO |
| >hg38_21_+_8433172_8446622_RNA45SN1_45S_50ntFlanks@3854.3882.29\|ACTGTGGTAATTCTAGAGCTAATACATGC | >hg38_21_+_8433172_8446622_RNA45SN1_45S_50ntFlanks | 3854 | 3882 | 29 | ACTGTGGTAATTCTAGAGCTAATACATGC | 168.0 | -11.3 | 1.7 | -6.7 | 1.8E-11 | 5.5E-11 | MB231/MB468 | MP |
| >hg38_21_+_8433172_8446622_RNA45SN1_45S_50ntFlanks@10687.10710.24\|CGGCGTCCGGTGAGCTCTCGCTGG | >hg38_21_+_8433172_8446622_RNA45SN1_45S_50ntFlanks | 10687 | 10710 | 24 | CGGCGTCCGGTGAGCTCTCGCTGG | 3691.8 | -4.8 | 0.7 | -6.8 | 1.4E-11 | 5.5E-11 | BT20/MB468 | MITO |
| >hg38_21_+_8433172_8446622_RNA45SN1_45S_50ntFlanks@12412.12434.23\|ATCCTTCGATGTCGGCTCTTCCT | >hg38_21_+_8433172_8446622_RNA45SN1_45S_50ntFlanks | 12412 | 12434 | 23 | ATCCTTCGATGTCGGCTCTTCCT | 90.8 | -10.4 | 1.5 | -6.7 | 1.8E-11 | 5.5E-11 | MB231/MB468 | MP |
| >trnaMT_PheGAA_MT_+_577_647@35.55.21\|GAAAATGTTTAGACGGGCTCA | >trnaMT_PheGAA_MT_+_577_647 | 35 | 55 | 21 | GAAAATGTTTAGACGGGCTCA | 2000.8 | 7.0 | 1.0 | 7.1 | 1.2E-12 | 5.5E-11 | BT20/MB468 | MP |
| >hg38_21_+_8433172_8446622_RNA45SN1_45S_50ntFlanks@6782.6799.18\|GGGCTACGCCTGTCTGAG | >hg38_21_+_8433172_8446622_RNA45SN1_45S_50ntFlanks | 6782 | 6799 | 18 | GGGCTACGCCTGTCTGAG | 327.1 | -11.4 | 1.6 | -7.1 | 1.2E-12 | 5.5E-11 | BT20/MB468 | MP |
| >hg38_21_+_8433172_8446622_RNA45SN1_45S_50ntFlanks@6648.6678.31\|GTACGACTCTTAGCGGTGGATCACTCGGCTC | >hg38_21_+_8433172_8446622_RNA45SN1_45S_50ntFlanks | 6648 | 6678 | 31 | GTACGACTCTTAGCGGTGGATCACTCGGCTC | 5824.3 | 4.8 | 0.7 | 6.8 | 1.4E-11 | 5.5E-11 | BT20/MB468 | MITO |
| >hg38_21_+_8433172_8446622_RNA45SN1_45S_50ntFlanks@10458.10484.27\|ATGGCCTCCGTTGCCCTCGGCCGATCG | >hg38_21_+_8433172_8446622_RNA45SN1_45S_50ntFlanks | 10458 | 10484 | 27 | ATGGCCTCCGTTGCCCTCGGCCGATCG | 199.3 | -9.7 | 1.4 | -6.8 | 1.4E-11 | 5.5E-11 | BT20/MB468 | MITO |
| >hg38_21_+_8433172_8446622_RNA45SN1_45S_50ntFlanks@10864.10891.28\|CTCTAAGGGCTGGGTCGGTCGGGCTGGG | >hg38_21_+_8433172_8446622_RNA45SN1_45S_50ntFlanks | 10864 | 10891 | 28 | CTCTAAGGGCTGGGTCGGTCGGGCTGGG | 102.3 | -10.5 | 1.6 | -6.7 | 1.8E-11 | 5.5E-11 | MB231/MB468 | MP |
| >hg38_21_+_8433172_8446622_RNA45SN1_45S_50ntFlanks@9196.9218.23\|AGGTTCTCTCGGGGCCACGCGCG | >hg38_21_+_8433172_8446622_RNA45SN1_45S_50ntFlanks | 9196 | 9218 | 23 | AGGTTCTCTCGGGGCCACGCGCG | 131.0 | -10.9 | 1.6 | -6.7 | 1.8E-11 | 5.5E-11 | MB231/MB468 | MP |
| >hg38_21_+_8433172_8446622_RNA45SN1_45S_50ntFlanks@12539.12569.31\|ATGATGTGTTGTTGCCATGGTAATCCTGCTC | >hg38_21_+_8433172_8446622_RNA45SN1_45S_50ntFlanks | 12539 | 12569 | 31 | ATGATGTGTTGTTGCCATGGTAATCCTGCTC | 1981.6 | -6.6 | 1.0 | -6.7 | 1.9E-11 | 5.5E-11 | MB231/MB468 | CYTO |
| >hg38_21_+_8433172_8446622_RNA45SN1_45S_50ntFlanks@5445.5464.20\|TCGGATCGGCCCCGCCGGGG | >hg38_21_+_8433172_8446622_RNA45SN1_45S_50ntFlanks | 5445 | 5464 | 20 | TCGGATCGGCCCCGCCGGGG | 86.4 | -10.3 | 1.5 | -6.7 | 1.8E-11 | 5.5E-11 | MB231/MB468 | MP |
| >hg38_21_+_8433172_8446622_RNA45SN1_45S_50ntFlanks@12488.12505.18\|AACGTGAGCTGGGTTTAG | >hg38_21_+_8433172_8446622_RNA45SN1_45S_50ntFlanks | 12488 | 12505 | 18 | AACGTGAGCTGGGTTTAG | 160.8 | -11.2 | 1.7 | -6.7 | 1.8E-11 | 5.5E-11 | MB231/MB468 | MP |
| >hg38_21_+_8433172_8446622_RNA45SN1_45S_50ntFlanks@12448.12480.33\|AGAATTCACCAAGCGTTGGATTGTTCACCCACT | >hg38_21_+_8433172_8446622_RNA45SN1_45S_50ntFlanks | 12448 | 12480 | 33 | AGAATTCACCAAGCGTTGGATTGTTCACCCACT | 300.2 | 7.0 | 1.0 | 6.7 | 1.8E-11 | 5.5E-11 | BT20/MB468 | CYTO |
| >hg38_21_+_8433172_8446622_RNA45SN1_45S_50ntFlanks@7986.8004.19\|ATCAGACGTGGCGACCCGC | >hg38_21_+_8433172_8446622_RNA45SN1_45S_50ntFlanks | 7986 | 8004 | 19 | ATCAGACGTGGCGACCCGC | 151.0 | -9.5 | 1.4 | -6.7 | 1.8E-11 | 5.5E-11 | BT20/MB468 | CYTO |
| >hg38_1_-_228634819_228635039_RNA5S12_5S_50ntFlanks@69.107.39\|CTGAACGCGCCCGATCTCGTCTGATCTCGGAAGCTAAGC | >hg38_1_-_228634819_228635039_RNA5S12_5S_50ntFlanks | 69 | 107 | 39 | CTGAACGCGCCCGATCTCGTCTGATCTCGGAAGCTAAGC | 116.7 | -10.7 | 1.6 | -6.7 | 1.8E-11 | 5.5E-11 | MB231/MB468 | MP |
| >hg38_21_+_8433172_8446622_RNA45SN1_45S_50ntFlanks@5414.5443.30\|CTACTACCGATTGGATGGTTTAGTGAGGCC | >hg38_21_+_8433172_8446622_RNA45SN1_45S_50ntFlanks | 5414 | 5443 | 30 | CTACTACCGATTGGATGGTTTAGTGAGGCC | 95.0 | -10.4 | 1.6 | -6.7 | 1.8E-11 | 5.5E-11 | MB231/MB468 | MP |
| >hg38_21_+_8433172_8446622_RNA45SN1_45S_50ntFlanks@10792.10826.35\|CTCTGGCATGTTGGAACAATGTAGGTAAGGGAAGT | >hg38_21_+_8433172_8446622_RNA45SN1_45S_50ntFlanks | 10792 | 10826 | 35 | CTCTGGCATGTTGGAACAATGTAGGTAAGGGAAGT | 108.9 | -10.6 | 1.6 | -6.7 | 1.8E-11 | 5.5E-11 | MB231/MB468 | MP |
| >hg38_21_+_8433172_8446622_RNA45SN1_45S_50ntFlanks@8179.8212.34\|CTGATCGAGGCCCAGCCCGTGGACGGTGTGAGGC | >hg38_21_+_8433172_8446622_RNA45SN1_45S_50ntFlanks | 8179 | 8212 | 34 | CTGATCGAGGCCCAGCCCGTGGACGGTGTGAGGC | 985.0 | -9.2 | 1.4 | -6.7 | 1.8E-11 | 5.5E-11 | MB231/MB468 | MP |
| >hg38_21_+_8433172_8446622_RNA45SN1_45S_50ntFlanks@3854.3877.24\|ACTGTGGTAATTCTAGAGCTAATA | >hg38_21_+_8433172_8446622_RNA45SN1_45S_50ntFlanks | 3854 | 3877 | 24 | ACTGTGGTAATTCTAGAGCTAATA | 125.4 | -10.8 | 1.6 | -6.7 | 1.8E-11 | 5.5E-11 | MB231/MB468 | MP |
| >hg38_21_+_8433172_8446622_RNA45SN1_45S_50ntFlanks@10853.10873.21\|ATAAGGATTGGCTCTAAGGGC | >hg38_21_+_8433172_8446622_RNA45SN1_45S_50ntFlanks | 10853 | 10873 | 21 | ATAAGGATTGGCTCTAAGGGC | 430.9 | -8.9 | 1.3 | -6.7 | 2.4E-11 | 5.5E-11 | BT20/MB468 | NUC |
| >hg38_21_+_8433172_8446622_RNA45SN1_45S_50ntFlanks@10350.10374.25\|ACGAGAACTTTGAAGGCCGAAGTGG | >hg38_21_+_8433172_8446622_RNA45SN1_45S_50ntFlanks | 10350 | 10374 | 25 | ACGAGAACTTTGAAGGCCGAAGTGG | 1618.2 | -4.3 | 0.6 | -6.7 | 1.8E-11 | 5.5E-11 | BT20/MB468 | CYTO |
| >hg38_21_+_8433172_8446622_RNA45SN1_45S_50ntFlanks@10926.10946.21\|ACGAGGCGCCGCCGCCCCCCC | >hg38_21_+_8433172_8446622_RNA45SN1_45S_50ntFlanks | 10926 | 10946 | 21 | ACGAGGCGCCGCCGCCCCCCC | 148.3 | -10.2 | 1.5 | -6.7 | 2.0E-11 | 5.5E-11 | MB231/MB468 | NUC |
| >hg38_21_+_8433172_8446622_RNA45SN1_45S_50ntFlanks@12083.12108.26\|CTCGCTTCTGGCGCCAAGCGCCCGGC | >hg38_21_+_8433172_8446622_RNA45SN1_45S_50ntFlanks | 12083 | 12108 | 26 | CTCGCTTCTGGCGCCAAGCGCCCGGC | 756.6 | -8.6 | 1.3 | -6.7 | 2.3E-11 | 5.6E-11 | MB231/MB468 | MITO |
| >hg38_21_+_8433172_8446622_RNA45SN1_45S_50ntFlanks@12944.12966.23\|CGTAGACGACCTGCTTCTGGGTC | >hg38_21_+_8433172_8446622_RNA45SN1_45S_50ntFlanks | 12944 | 12966 | 23 | CGTAGACGACCTGCTTCTGGGTC | 125.0 | -10.8 | 1.6 | -6.7 | 1.8E-11 | 5.6E-11 | MB231/MB468 | MP |
| >hg38_21_+_8433172_8446622_RNA45SN1_45S_50ntFlanks@12525.12549.25\|GTTTTACCCTACTGATGATGTGTTG | >hg38_21_+_8433172_8446622_RNA45SN1_45S_50ntFlanks | 12525 | 12549 | 25 | GTTTTACCCTACTGATGATGTGTTG | 106.5 | -10.6 | 1.6 | -6.7 | 1.8E-11 | 5.6E-11 | MB231/MB468 | MP |
| >hg38_21_+_8433172_8446622_RNA45SN1_45S_50ntFlanks@3984.4010.27\|GCGGCTTTGGTGACTCTAGATAACCTC | >hg38_21_+_8433172_8446622_RNA45SN1_45S_50ntFlanks | 3984 | 4010 | 27 | GCGGCTTTGGTGACTCTAGATAACCTC | 179.8 | -11.4 | 1.7 | -6.7 | 1.8E-11 | 5.6E-11 | MB231/MB468 | MP |
| >hg38_21_+_8433172_8446622_RNA45SN1_45S_50ntFlanks@12522.12546.25\|TTAGTTTTACCCTACTGATGATGTG | >hg38_21_+_8433172_8446622_RNA45SN1_45S_50ntFlanks | 12522 | 12546 | 25 | TTAGTTTTACCCTACTGATGATGTG | 101.5 | -10.5 | 1.6 | -6.7 | 1.8E-11 | 5.6E-11 | MB231/MB468 | MP |
| >am_trnaMT_ValTAC_MT_+_1602_1670@30.68.39\|CTTACACTTAGGAGATTTCAACTTAACTTGACCGCTCTG | >am_trnaMT_ValTAC_MT_+_1602_1670 | 30 | 68 | 39 | CTTACACTTAGGAGATTTCAACTTAACTTGACCGCTCTG | 191.0 | -11.4 | 1.7 | -6.7 | 2.3E-11 | 5.6E-11 | MB231/MB468 | MITO |
| >hg38_21_+_8433172_8446622_RNA45SN1_45S_50ntFlanks@13003.13027.25\|GATCTATTGAAAGTCAGCCCTCGAC | >hg38_21_+_8433172_8446622_RNA45SN1_45S_50ntFlanks | 13003 | 13027 | 25 | GATCTATTGAAAGTCAGCCCTCGAC | 170.6 | -9.7 | 1.4 | -6.7 | 1.8E-11 | 5.6E-11 | BT20/MB468 | CYTO |
| >hg38_21_+_8433172_8446622_RNA45SN1_45S_50ntFlanks@10458.10483.26\|ATGGCCTCCGTTGCCCTCGGCCGATC | >hg38_21_+_8433172_8446622_RNA45SN1_45S_50ntFlanks | 10458 | 10483 | 26 | ATGGCCTCCGTTGCCCTCGGCCGATC | 197.3 | -11.5 | 1.7 | -6.7 | 1.8E-11 | 5.6E-11 | MB231/MB468 | MP |
| >hg38_21_+_8433172_8446622_RNA45SN1_45S_50ntFlanks@4758.4780.23\|GATCAGATACCGTCGTAGTTCCG | >hg38_21_+_8433172_8446622_RNA45SN1_45S_50ntFlanks | 4758 | 4780 | 23 | GATCAGATACCGTCGTAGTTCCG | 151.4 | -11.1 | 1.7 | -6.7 | 1.8E-11 | 5.6E-11 | MB231/MB468 | MP |
| >MI0000299\|hsa-mir-222&WithFlank&X\|-\|45747009\|45747130@75.96.22(+1C)&[MIMAT0000279&hsa-miR-222-3p&offsets\|0\|+1(+1C);m-140&X\|-\|45747033\|45747056&offsets\|0\|-2(+1C)]\|AGCTACATCTGGCTACTGGGTCC | >MI0000299\|hsa-mir-222&WithFlank&X\|-\|45747009\|45747130 | 75 | 96 | 22(+1C) | AGCTACATCTGGCTACTGGGTCC | 775.5 | 5.3 | 0.8 | 6.7 | 2.0E-11 | 5.6E-11 | MB231/MB468 | NUC |
| >hg38_21_+_8433172_8446622_RNA45SN1_45S_50ntFlanks@12129.12146.18\|CCCGCTCCGGGGACAGTG | >hg38_21_+_8433172_8446622_RNA45SN1_45S_50ntFlanks | 12129 | 12146 | 18 | CCCGCTCCGGGGACAGTG | 180.8 | -6.4 | 1.0 | -6.7 | 2.0E-11 | 5.6E-11 | MB231/MB468 | CYTO |
| >hg38_21_+_8433172_8446622_RNA45SN1_45S_50ntFlanks@12359.12382.24\|ACAGGGATAACTGGCTTGTGGCGG | >hg38_21_+_8433172_8446622_RNA45SN1_45S_50ntFlanks | 12359 | 12382 | 24 | ACAGGGATAACTGGCTTGTGGCGG | 106.5 | -10.6 | 1.6 | -6.7 | 1.8E-11 | 5.6E-11 | MB231/MB468 | MP |
| >hg38_21_+_8433172_8446622_RNA45SN1_45S_50ntFlanks@9858.9878.21\|CGGGATGAACCGAACGCCGGG | >hg38_21_+_8433172_8446622_RNA45SN1_45S_50ntFlanks | 9858 | 9878 | 21 | CGGGATGAACCGAACGCCGGG | 161.8 | -11.2 | 1.7 | -6.7 | 1.8E-11 | 5.6E-11 | MB231/MB468 | MP |
| >hg38_21_+_8433172_8446622_RNA45SN1_45S_50ntFlanks@12072.12095.24\|CCCGAGGGGCTCTCGCTTCTGGCG | >hg38_21_+_8433172_8446622_RNA45SN1_45S_50ntFlanks | 12072 | 12095 | 24 | CCCGAGGGGCTCTCGCTTCTGGCG | 1385.7 | -9.9 | 1.5 | -6.7 | 2.5E-11 | 5.6E-11 | BT20/MB468 | NUC |
| >hg38_21_+_8433172_8446622_RNA45SN1_45S_50ntFlanks@10347.10374.28\|CAAACGAGAACTTTGAAGGCCGAAGTGG | >hg38_21_+_8433172_8446622_RNA45SN1_45S_50ntFlanks | 10347 | 10374 | 28 | CAAACGAGAACTTTGAAGGCCGAAGTGG | 1240.6 | -6.4 | 1.0 | -6.7 | 1.8E-11 | 5.6E-11 | MB231/MB468 | MP |
| >hg38_21_+_8433172_8446622_RNA45SN1_45S_50ntFlanks@4001.4023.23\|AGATAACCTCGGGCCGATCGCAC | >hg38_21_+_8433172_8446622_RNA45SN1_45S_50ntFlanks | 4001 | 4023 | 23 | AGATAACCTCGGGCCGATCGCAC | 750.6 | 6.2 | 0.9 | 6.7 | 1.8E-11 | 5.6E-11 | BT20/MB468 | CYTO |
| >hg38_21_+_8433172_8446622_RNA45SN1_45S_50ntFlanks@12971.12990.20\|TTTCGTACGTAGCAGAGCAG | >hg38_21_+_8433172_8446622_RNA45SN1_45S_50ntFlanks | 12971 | 12990 | 20 | TTTCGTACGTAGCAGAGCAG | 80.2 | -10.2 | 1.5 | -6.7 | 1.8E-11 | 5.7E-11 | MB231/MB468 | MP |

| | | | | | | | | | | | | | |
|---|---|---|---|---|---|---|---|---|---|---|---|---|---|
| >hg38_1_-_228634819_228635039_RNA5S12_5S_50ntFlanks@51.92.42\|GTCTACGGCCATACCACCCTGAACGCGCCCGATCTCGTCTGA | >hg38_1_-_228634819_228635039_RNA5S12_5S_50ntFlanks | 51 | 92 | 42 | GTCTACGGCCATACCACCCTGAACGCGCCCGATCTCGTCTGA | 988.1 | 7.0 | 1.0 | 6.7 | 2.0E-11 | 5.7E-11 | MB231/MB468 | CYTO |
| >hg38_21_+_8433172_8446622_RNA45SN1_45S_50ntFlanks@9027.9057.31\|CGTCTCCTCGTGGGGGGGCCGGGCCACCCCT | >hg38_21_+_8433172_8446622_RNA45SN1_45S_50ntFlanks | 9027 | 9057 | 31 | CGTCTCCTCGTGGGGGGGCCGGGCCACCCCT | 104.7 | -10.6 | 1.6 | -6.7 | 1.8E-11 | 5.7E-11 | MB231/MB468 | MP |
| >hg38_21_+_8433172_8446622_RNA45SN1_45S_50ntFlanks@11785.11805.21\|TGAATGGATGAACGAGATTCC | >hg38_21_+_8433172_8446622_RNA45SN1_45S_50ntFlanks | 11785 | 11805 | 21 | TGAATGGATGAACGAGATTCC | 1027.9 | -9.6 | 1.4 | -6.7 | 2.5E-11 | 5.7E-11 | BT20/MB468 | NUC |
| >hg38_21_+_8433172_8446622_RNA45SN1_45S_50ntFlanks@7986.8015.30\|ATCAGACGTGGCGACCCGCTGAATTTAAGC | >hg38_21_+_8433172_8446622_RNA45SN1_45S_50ntFlanks | 7986 | 8015 | 30 | ATCAGACGTGGCGACCCGCTGAATTTAAGC | 446.1 | -8.3 | 1.2 | -6.7 | 1.8E-11 | 5.7E-11 | MB231/MB468 | MP |
| >hg38_21_+_8433172_8446622_RNA45SN1_45S_50ntFlanks@9572.9600.29\|CAAATCGGTCGTCCGACCTGGGTATAGGG | >hg38_21_+_8433172_8446622_RNA45SN1_45S_50ntFlanks | 9572 | 9600 | 29 | CAAATCGGTCGTCCGACCTGGGTATAGGG | 112.5 | -10.7 | 1.6 | -6.7 | 1.8E-11 | 5.7E-11 | MB231/MB468 | MP |
| >MI0003137\|hsa-mir-193b&WithFlank&16\|+\|14303961\|14304055@57.77.21(+1A)&[MIMAT0002819&hsa-miR-193b-3p&offsets\|0\|-1(+1A);m-166&16\|+\|14304017\|14304038&offsets\|0\|-1(+1A)]\|AACTGGCCCTCAAAGTCCCGCA | >MI0003137\|hsa-mir-193b&WithFlank&16\|+\|14303961\|14304055 | 57 | 77 | 21(+1A) | AACTGGCCCTCAAAGTCCCGCA | 145.1 | 7.4 | 1.1 | 6.7 | 2.5E-11 | 5.7E-11 | BT20/MB468 | NUC |
| >hg38_21_+_8433172_8446622_RNA45SN1_45S_50ntFlanks@12488.12512.25\|AACGTGAGCTGGGTTTAGACCGTCG | >hg38_21_+_8433172_8446622_RNA45SN1_45S_50ntFlanks | 12488 | 12512 | 25 | AACGTGAGCTGGGTTTAGACCGTCG | 249.2 | -7.0 | 1.0 | -6.7 | 2.0E-11 | 5.7E-11 | MB231/MB468 | CYTO |
| >hg38_21_+_8433172_8446622_RNA45SN1_45S_50ntFlanks@12182.12201.20\|CAAACGGTAACGCAGGTGTC | >hg38_21_+_8433172_8446622_RNA45SN1_45S_50ntFlanks | 12182 | 12201 | 20 | CAAACGGTAACGCAGGTGTC | 147.8 | -11.1 | 1.6 | -6.7 | 1.8E-11 | 5.7E-11 | MB231/MB468 | MP |
| >MI0000746\|hsa-mir-99b&WithFlank&19\|+\|51692606\|51692687@13.33.21(+1A)&[MIMAT0000689&hsa-miR-99b-5p&offsets\|0\|-1(+1A);m-10&19\|+\|51692618\|51692639&offsets\|0\|-1(+1A)]\|CACCCGTAGAACCGACCTTGCA | >MI0000746\|hsa-mir-99b&WithFlank&19\|+\|51692606\|51692687 | 13 | 33 | 21(+1A) | CACCCGTAGAACCGACCTTGCA | 6820.0 | 5.9 | 0.9 | 6.7 | 2.0E-11 | 5.7E-11 | MB231/MB468 | CYTO |
| >MI0000270\|hsa-mir-181b-1&WithFlank&1\|-\|198858867\|198858988@42.62.21;MI0000683\|hsa-mir-181b-2&WithFlank&9\|+\|124693704\|124693804@22.42.21&[MIMAT0000257&hsa-miR-181b-5p&offsets\|0\|-2;m-73&1\|-\|198858925\|198858947&offsets\|0\|-2];[MIMAT0000257_1&hsa-miR-181b-5p&offsets\|0\|-2;m-72&9\|+\|124693725\|124693747&offsets\|0\|-2]\|AACATTCATTGCTGTCGGTGG | >MI0000270\|hsa-mir-181b-1&WithFlank&1\|-\|198858867\|198858988 | 42 | 62 | 21 | AACATTCATTGCTGTCGGTGG | 519.1 | 5.7 | 0.9 | 6.7 | 2.0E-11 | 5.7E-11 | MB231/MB468 | NUC |
| >hg38_21_+_8433172_8446622_RNA45SN1_45S_50ntFlanks@9240.9274.35\|GCGGAGCGAGCGCACGGGGTCGGCGGCGACGTCGG | >hg38_21_+_8433172_8446622_RNA45SN1_45S_50ntFlanks | 9240 | 9274 | 35 | GCGGAGCGAGCGCACGGGGTCGGCGGCGACGTCGG | 322.7 | -11.4 | 1.6 | -7.1 | 1.2E-12 | 5.7E-11 | BT20/MB468 | MP |
| >MI0000440\|hsa-mir-27b&WithFlank&9\|+\|95085439\|95085547@66.87.22&[MIMAT0000419&hsa-miR-27b-3p&offsets\|-1\|0;m-39&9\|+\|95085505\|95085525&offsets\|-1\|0]\|GTTCACAGTGGCTAAGTTCTGC | >MI0000440\|hsa-mir-27b&WithFlank&9\|+\|95085439\|95085547 | 66 | 87 | 22 | GTTCACAGTGGCTAAGTTCTGC | 477.0 | 8.6 | 1.3 | 6.8 | 1.5E-11 | 5.7E-11 | BT20/MB468 | MITO |
| >hg38_21_+_8433172_8446622_RNA45SN1_45S_50ntFlanks@11212.11232.21\|CGACTCTGGACGCGAGCCGGG | >hg38_21_+_8433172_8446622_RNA45SN1_45S_50ntFlanks | 11212 | 11232 | 21 | CGACTCTGGACGCGAGCCGGG | 930.9 | -9.5 | 1.4 | -6.7 | 2.5E-11 | 5.7E-11 | BT20/MB468 | NUC |
| >hg38_21_+_8433172_8446622_RNA45SN1_45S_50ntFlanks@11209.11226.18\|CGGCGACTCTGGACGCGA | >hg38_21_+_8433172_8446622_RNA45SN1_45S_50ntFlanks | 11209 | 11226 | 18 | CGGCGACTCTGGACGCGA | 127.9 | -10.9 | 1.6 | -6.7 | 1.8E-11 | 5.7E-11 | MB231/MB468 | MP |
| >hg38_21_+_8433172_8446622_RNA45SN1_45S_50ntFlanks@8527.8551.25\|CCTTCCCCCGCCGCCCCTCCTCCTC | >hg38_21_+_8433172_8446622_RNA45SN1_45S_50ntFlanks | 8527 | 8551 | 25 | CCTTCCCCCGCCGCCCCTCCTCCTC | 142.5 | -11.0 | 1.6 | -6.7 | 1.9E-11 | 5.7E-11 | MB231/MB468 | MP |
| >hg38_21_+_8433172_8446622_RNA45SN1_45S_50ntFlanks@11704.11728.25\|ATGAAGCGCGGGTAAACGGCGGGAG | >hg38_21_+_8433172_8446622_RNA45SN1_45S_50ntFlanks | 11704 | 11728 | 25 | ATGAAGCGCGGGTAAACGGCGGGAG | 4056.0 | -10.7 | 1.6 | -6.7 | 2.5E-11 | 5.7E-11 | BT20/MB468 | NUC |
| >trna116_GluCTC_1_-_145399233_145399304@1.42.42,trna59_GluCTC_1_+_249168447_249168518@1.42.42,trna71_GluCTC_1_-_161439189_161439260@1.42.42,trna74_GluCTC_1_-_161431809_161431880@1.42.42,trna77_GluCTC_1_-_161424398_161424469@1.42.42,trna77_GluCTC_6_+_28949976_28950047@1.42.42,trna80_GluCTC_1_-_161417018_161417089@1.42.42,trna87_GluCTC_6_-_126101393_126101464@1.42.42\|TCCCTGGTGGTCTAGTGGTTAGGATTCGGCGCTCTCACCGCC | >trna116_GluCTC_1_-_145399233_145399304 | 1 | 42 | 42 | TCCCTGGTGGTCTAGTGGTTAGGATTCGGCGCTCTCACCGCC | 338.6 | 7.4 | 1.1 | 6.8 | 1.5E-11 | 5.7E-11 | BT20/MB468 | MITO |
| >hg38_21_+_8433172_8446622_RNA45SN1_45S_50ntFlanks@9034.9055.22\|TCGTGGGGGGGCCGGGCCACCC | >hg38_21_+_8433172_8446622_RNA45SN1_45S_50ntFlanks | 9034 | 9055 | 22 | TCGTGGGGGGGCCGGGCCACCC | 118.2 | -10.7 | 1.6 | -6.7 | 1.9E-11 | 5.7E-11 | MB231/MB468 | MP |
| >hg38_21_+_8433172_8446622_RNA45SN1_45S_50ntFlanks@6779.6803.25\|CCGGGGCTACGCCTGTCTGAGCGTC | >hg38_21_+_8433172_8446622_RNA45SN1_45S_50ntFlanks | 6779 | 6803 | 25 | CCGGGGCTACGCCTGTCTGAGCGTC | 1172.2 | -7.4 | 1.1 | -6.7 | 1.9E-11 | 5.7E-11 | MB231/MB468 | MP |
| >hg38_21_+_8433172_8446622_RNA45SN1_45S_50ntFlanks@10781.10822.42\|AGGTGAACAGCCTCTGGCATGTTGGAACAATGTAGGTAAGGG | >hg38_21_+_8433172_8446622_RNA45SN1_45S_50ntFlanks | 10781 | 10822 | 42 | AGGTGAACAGCCTCTGGCATGTTGGAACAATGTAGGTAAGGG | 171.0 | -11.3 | 1.7 | -6.7 | 1.9E-11 | 5.7E-11 | MB231/MB468 | MP |
| >hg38_21_+_8433172_8446622_RNA45SN1_45S_50ntFlanks@10369.10393.25\|AAGTGGAGAAGGGTTCCATGTGAAC | >hg38_21_+_8433172_8446622_RNA45SN1_45S_50ntFlanks | 10369 | 10393 | 25 | AAGTGGAGAAGGGTTCCATGTGAAC | 81.5 | -10.2 | 1.5 | -6.7 | 1.9E-11 | 5.7E-11 | MB231/MB468 | MP |
| >hg38_21_+_8433172_8446622_RNA45SN1_45S_50ntFlanks@9572.9597.26\|CAAATCGGTCGTCCGACCTGGGTATA | >hg38_21_+_8433172_8446622_RNA45SN1_45S_50ntFlanks | 9572 | 9597 | 26 | CAAATCGGTCGTCCGACCTGGGTATA | 12305.4 | -10.2 | 1.5 | -6.8 | 1.5E-11 | 5.7E-11 | BT20/MB468 | MITO |
| >trna116_GluCTC_1_-_145399233_145399304@1.36.36,trna59_GluCTC_1_+_249168447_249168518@1.36.36,trna71_GluCTC_1_-_161439189_161439260@1.36.36,trna74_GluCTC_1_-_161431809_161431880@1.36.36,trna77_GluCTC_1_-_161424398_161424469@1.36.36,trna77_GluCTC_6_+_28949976_28950047@1.36.36,trna80_GluCTC_1_-_161417018_161417089@1.36.36,trna87_GluCTC_6_-_126101393_126101464@1.36.36\|TCCCTGGTGGTCTAGTGGTTAGGATTCGGCGCTCTC | >trna116_GluCTC_1_-_145399233_145399304 | 1 | 36 | 36 | TCCCTGGTGGTCTAGTGGTTAGGATTCGGCGCTCTC | 3784.4 | 7.3 | 1.1 | 6.7 | 2.0E-11 | 5.7E-11 | MB231/MB468 | NUC |
| >hg38_21_+_8433172_8446622_RNA45SN1_45S_50ntFlanks@9537.9571.35\|AAACTCTGGTGGAGGTCCGTAGCGGTCCTGACGTG | >hg38_21_+_8433172_8446622_RNA45SN1_45S_50ntFlanks | 9537 | 9571 | 35 | AAACTCTGGTGGAGGTCCGTAGCGGTCCTGACGTG | 1095.8 | -3.8 | 0.6 | -6.7 | 2.0E-11 | 5.7E-11 | MB231/MB468 | NUC |
| >hg38_21_+_8433172_8446622_RNA45SN1_45S_50ntFlanks@4934.4951.18\|CCTGCGGCTTAATTTGAC | >hg38_21_+_8433172_8446622_RNA45SN1_45S_50ntFlanks | 4934 | 4951 | 18 | CCTGCGGCTTAATTTGAC | 412.2 | -12.5 | 1.9 | -6.7 | 2.4E-11 | 5.7E-11 | MB231/MB468 | MITO |
| >hg38_21_+_8433172_8446622_RNA45SN1_45S_50ntFlanks@10830.10859.30\|CAAGCCGGATCCGTAACTTCGGGATAAGGA | >hg38_21_+_8433172_8446622_RNA45SN1_45S_50ntFlanks | 10830 | 10859 | 30 | CAAGCCGGATCCGTAACTTCGGGATAAGGA | 331.3 | -12.2 | 1.8 | -6.7 | 1.9E-11 | 5.7E-11 | MB231/MB468 | MP |
| >hg38_21_+_8433172_8446622_RNA45SN1_45S_50ntFlanks@4724.4757.34\|ATTAATCAAGAACGAAAGTCGGAGGTTCGAAGAC | >hg38_21_+_8433172_8446622_RNA45SN1_45S_50ntFlanks | 4724 | 4757 | 34 | ATTAATCAAGAACGAAAGTCGGAGGTTCGAAGAC | 253.3 | -10.1 | 1.5 | -6.8 | 1.5E-11 | 5.7E-11 | BT20/MB468 | MITO |
| >hg38_21_+_8433172_8446622_RNA45SN1_45S_50ntFlanks@10396.10420.25\|CAGTTGAACATGGGTCAGTCGGTCC | >hg38_21_+_8433172_8446622_RNA45SN1_45S_50ntFlanks | 10396 | 10420 | 25 | CAGTTGAACATGGGTCAGTCGGTCC | 123.9 | -10.8 | 1.6 | -6.7 | 1.9E-11 | 5.7E-11 | MB231/MB468 | MP |
| >hg38_21_+_8433172_8446622_RNA45SN1_45S_50ntFlanks@9459.9495.37\|CCGGGGAGGTGGAGCACGAGCGCACGTGTTAGGACCC | >hg38_21_+_8433172_8446622_RNA45SN1_45S_50ntFlanks | 9459 | 9495 | 37 | CCGGGGAGGTGGAGCACGAGCGCACGTGTTAGGACCC | 100.2 | -10.5 | 1.6 | -6.7 | 1.9E-11 | 5.7E-11 | MB231/MB468 | MP |
| >hg38_21_+_8433172_8446622_RNA45SN1_45S_50ntFlanks@9559.9596.38\|CGGTCCTGACGTGCAAATCGGTCGTCCGACCTGGGTAT | >hg38_21_+_8433172_8446622_RNA45SN1_45S_50ntFlanks | 9559 | 9596 | 38 | CGGTCCTGACGTGCAAATCGGTCGTCCGACCTGGGTAT | 257.9 | -11.8 | 1.8 | -6.7 | 2.4E-11 | 5.7E-11 | MB231/MB468 | MITO |
| >hg38_21_+_8433172_8446622_RNA45SN1_45S_50ntFlanks@11580.11603.24\|GAACTGGTGCGGACCAGGGGAATC | >hg38_21_+_8433172_8446622_RNA45SN1_45S_50ntFlanks | 11580 | 11603 | 24 | GAACTGGTGCGGACCAGGGGAATC | 1676.7 | -4.1 | 0.6 | -6.8 | 1.5E-11 | 5.7E-11 | BT20/MB468 | MITO |
| >hg38_21_+_8433172_8446622_RNA45SN1_45S_50ntFlanks@13009.13027.19\|TTGAAAGTCAGCCCTCGAC | >hg38_21_+_8433172_8446622_RNA45SN1_45S_50ntFlanks | 13009 | 13027 | 19 | TTGAAAGTCAGCCCTCGAC | 684.2 | -7.7 | 1.1 | -6.7 | 2.0E-11 | 5.8E-11 | MB231/MB468 | CYTO |
| >hg38_21_+_8433172_8446622_RNA45SN1_45S_50ntFlanks@10421.10453.33\|TGAGAGATGGGCGAGCGCCGTTCCGAAGGGACG | >hg38_21_+_8433172_8446622_RNA45SN1_45S_50ntFlanks | 10421 | 10453 | 33 | TGAGAGATGGGCGAGCGCCGTTCCGAAGGGACG | 1696.5 | -10.1 | 1.5 | -6.7 | 2.5E-11 | 5.8E-11 | BT20/MB468 | NUC |
| >hg38_21_+_8433172_8446622_RNA45SN1_45S_50ntFlanks@10511.10533.23\|ATCCGGAGTGGCGGAGATGGGCG | >hg38_21_+_8433172_8446622_RNA45SN1_45S_50ntFlanks | 10511 | 10533 | 23 | ATCCGGAGTGGCGGAGATGGGCG | 270.2 | -10.2 | 1.5 | -6.7 | 1.5E-11 | 5.8E-11 | BT20/MB468 | MITO |
| >hg38_21_+_8433172_8446622_RNA45SN1_45S_50ntFlanks@11739.11768.30\|TCTCTTAAGGTAGCCAAATGCCTCGTCATC | >hg38_21_+_8433172_8446622_RNA45SN1_45S_50ntFlanks | 11739 | 11768 | 30 | TCTCTTAAGGTAGCCAAATGCCTCGTCATC | 247.9 | -6.1 | 0.9 | -6.7 | 2.0E-11 | 5.8E-11 | MB231/MB468 | NUC |
| >hg38_21_+_8433172_8446622_RNA45SN1_45S_50ntFlanks@4364.4389.26\|CTGCAGTTAAAAAGCTCGTAGTTGGA | >hg38_21_+_8433172_8446622_RNA45SN1_45S_50ntFlanks | 4364 | 4389 | 26 | CTGCAGTTAAAAAGCTCGTAGTTGGA | 79.7 | -10.2 | 1.5 | -6.7 | 1.9E-11 | 5.8E-11 | MB231/MB468 | MP |
| >hg38_21_+_8433172_8446622_RNA45SN1_45S_50ntFlanks@3851.3869.19\|ATAACTGTGGTAATTCTAG | >hg38_21_+_8433172_8446622_RNA45SN1_45S_50ntFlanks | 3851 | 3869 | 19 | ATAACTGTGGTAATTCTAG | 93.3 | -10.4 | 1.6 | -6.7 | 1.9E-11 | 5.8E-11 | MB231/MB468 | MP |
| >hg38_21_+_8433172_8446622_RNA45SN1_45S_50ntFlanks@11575.11609.35\|ACTTAGAACTGGTGCGGACCAGGGGAATCCGACTG | >hg38_21_+_8433172_8446622_RNA45SN1_45S_50ntFlanks | 11575 | 11609 | 35 | ACTTAGAACTGGTGCGGACCAGGGGAATCCGACTG | 927.0 | -7.5 | 1.1 | -6.7 | 1.9E-11 | 5.8E-11 | MB231/MB468 | MP |
| >hg38_21_+_8433172_8446622_RNA45SN1_45S_50ntFlanks@9536.9553.18\|GAAACTCTGGTGGAGGTC | >hg38_21_+_8433172_8446622_RNA45SN1_45S_50ntFlanks | 9536 | 9553 | 18 | GAAACTCTGGTGGAGGTC | 206.0 | -11.6 | 1.7 | -6.7 | 1.9E-11 | 5.8E-11 | MB231/MB468 | MP |
| >hg38_21_+_8433172_8446622_RNA45SN1_45S_50ntFlanks@10027.10072.46\|CTGGAGCGTCGGGCCCATACCCGGCCGTCGCCGGCAGTCGAGAGTG | >hg38_21_+_8433172_8446622_RNA45SN1_45S_50ntFlanks | 10027 | 10072 | 46 | CTGGAGCGTCGGGCCCATACCCGGCCGTCGCCGGCAGTCGAGAGTG | 371.8 | -12.3 | 1.8 | -6.7 | 2.4E-11 | 5.8E-11 | MB231/MB468 | MITO |
| >hg38_21_+_8433172_8446622_RNA45SN1_45S_50ntFlanks@7985.8009.25\|GATCAGACGTGGCGACCCGCTGAAT | >hg38_21_+_8433172_8446622_RNA45SN1_45S_50ntFlanks | 7985 | 8009 | 25 | GATCAGACGTGGCGACCCGCTGAAT | 199.8 | -9.7 | 1.4 | -6.7 | 1.5E-11 | 5.8E-11 | BT20/MB468 | MITO |
| >hg38_21_+_8433172_8446622_RNA45SN1_45S_50ntFlanks@11323.11344.22\|CGTCTCGTCGCGCGCGCGTCCG | >hg38_21_+_8433172_8446622_RNA45SN1_45S_50ntFlanks | 11323 | 11344 | 22 | CGTCTCGTCGCGCGCGCGTCCG | 17899.5 | -8.7 | 1.3 | -6.7 | 2.0E-11 | 5.8E-11 | MB231/MB468 | CYTO |
| >trna10_LysCTT_16_+_3241501_3241573@13.35.23,trna13_LysCTT_14_-_58706613_58706685@13.35.23,trna2_LysCTT_15_+_79152904_79152976@13.35.23,trna4_LysCTT_18_-_43669271_43669343@13.35.23\|CAGTCGGTAGAGCATGGGACTCT | >trna10_LysCTT_16_+_3241501_3241573 | 13 | 35 | 23 | CAGTCGGTAGAGCATGGGACTCT | 270.2 | 11.3 | 1.7 | 6.7 | 2.4E-11 | 5.8E-11 | MB231/MB468 | MITO |
| >hg38_21_+_8433172_8446622_RNA45SN1_45S_50ntFlanks@485.530.46\|GATGTGGTGACGTCGTGCTCTCCCGGGCCGGGTCCGAGCCGCGACG | >hg38_21_+_8433172_8446622_RNA45SN1_45S_50ntFlanks | 485 | 530 | 46 | GATGTGGTGACGTCGTGCTCTCCCGGGCCGGGTCCGAGCCGCGACG | 259.9 | 10.9 | 1.6 | 6.7 | 2.6E-11 | 5.8E-11 | BT20/MB468 | NUC |
| >hg38_21_+_8433172_8446622_RNA45SN1_45S_50ntFlanks@9035.9056.22\|CGTGGGGGGGCCGGGCCACCCC | >hg38_21_+_8433172_8446622_RNA45SN1_45S_50ntFlanks | 9035 | 9056 | 22 | CGTGGGGGGGCCGGGCCACCCC | 400.9 | -11.7 | 1.7 | -7.1 | 1.3E-12 | 5.8E-11 | BT20/MB468 | MP |

| | | | | | | | | | | | | | |
|---|---|---|---|---|---|---|---|---|---|---|---|---|---|
| >hg38_21_+_8433172_8446622_RNA45SN1_45S_50ntFlanks@10351.10380.30\|CGAGAACTTTGAAGGCCGAAGTGGAGAAGG | >hg38_21_+_8433172_8446622_RNA45SN1_45S_50ntFlanks | 10351 | 10380 | 30 | CGAGAACTTTGAAGGCCGAAGTGGAGAAGG | 1940.0 | -8.0 | 1.2 | -6.7 | 1.9E-11 | 5.8E-11 | MB231/MB468 | MP |
| >hg38_21_+_8433172_8446622_RNA45SN1_45S_50ntFlanks@9360.9380.21\|CAATGAAGGTGAAGGCCGGCG | >hg38_21_+_8433172_8446622_RNA45SN1_45S_50ntFlanks | 9360 | 9380 | 21 | CAATGAAGGTGAAGGCCGGCG | 194.5 | -11.5 | 1.7 | -6.7 | 1.9E-11 | 5.8E-11 | MB231/MB468 | MP |
| >hg38_21_+_8433172_8446622_RNA45SN1_45S_50ntFlanks@11845.11862.18\|GGAACGGGCTTGGCGGAA | >hg38_21_+_8433172_8446622_RNA45SN1_45S_50ntFlanks | 11845 | 11862 | 18 | GGAACGGGCTTGGCGGAA | 115.9 | -10.7 | 1.6 | -6.7 | 1.9E-11 | 5.8E-11 | MB231/MB468 | MP |
| >hg38_21_+_8433172_8446622_RNA45SN1_45S_50ntFlanks@12539.12579.41\|ATGATGTGTTGTTGCCATGGTAATCCTGCTCAGTACGAGAG | >hg38_21_+_8433172_8446622_RNA45SN1_45S_50ntFlanks | 12539 | 12579 | 41 | ATGATGTGTTGTTGCCATGGTAATCCTGCTCAGTACGAGAG | 193.4 | -11.4 | 1.7 | -6.7 | 2.4E-11 | 5.9E-11 | MB231/MB468 | MITO |
| >hg38_21_+_8433172_8446622_RNA45SN1_45S_50ntFlanks@12072.12092.21\|CCCGAGGGGCTCTCGCTTCTG | >hg38_21_+_8433172_8446622_RNA45SN1_45S_50ntFlanks | 12072 | 12092 | 21 | CCCGAGGGGCTCTCGCTTCTG | 99.3 | -10.5 | 1.6 | -6.7 | 1.9E-11 | 5.9E-11 | MB231/MB468 | MP |
| >hg38_21_+_8433172_8446622_RNA45SN1_45S_50ntFlanks@12072.12090.19\|CCCGAGGGGCTCTCGCTTC | >hg38_21_+_8433172_8446622_RNA45SN1_45S_50ntFlanks | 12072 | 12090 | 19 | CCCGAGGGGCTCTCGCTTC | 331.8 | -5.2 | 0.8 | -6.7 | 2.1E-11 | 5.9E-11 | MB231/MB468 | CYTO |
| >MI0001445\|hsa-mir-423&WithFlank&17\|+\|30117073\|30117178@23.43.21&[MIMAT0004748&hsa-miR-423-5p&offsets\|0\|-2;m-141&17\|+\|30117095\|30117117&offsets\|0\|-2]\|TGAGGGGCAGAGAGCGAGACT | >MI0001445\|hsa-mir-423&WithFlank&17\|+\|30117073\|30117178 | 23 | 43 | 21 | TGAGGGGCAGAGAGCGAGACT | 527.1 | 9.8 | 1.5 | 6.7 | 1.5E-11 | 5.9E-11 | BT20/MB468 | MITO |
| >MI0000079\|hsa-mir-23a&WithFlank&19\|-\|13836581\|13836665@51.71.21&[MIMAT0000078&hsa-miR-23a-3p&offsets\|0\|0;m-31&19\|-\|13836595\|13836615&offsets\|0\|0]\|ATCACATTGCCAGGGATTTCC | >MI0000079\|hsa-mir-23a&WithFlank&19\|-\|13836581\|13836665 | 51 | 71 | 21 | ATCACATTGCCAGGGATTTCC | 471.1 | -12.0 | 1.7 | -7.1 | 1.3E-12 | 5.9E-11 | BT20/MB468 | MP |
| >hg38_21_+_8433172_8446622_RNA45SN1_45S_50ntFlanks@10914.10941.28\|CGCCGCGGCTGGACGAGGCGCCGCCGCC | >hg38_21_+_8433172_8446622_RNA45SN1_45S_50ntFlanks | 10914 | 10941 | 28 | CGCCGCGGCTGGACGAGGCGCCGCCGCC | 130.0 | -10.9 | 1.6 | -6.7 | 1.9E-11 | 5.9E-11 | MB231/MB468 | MP |
| >hg38_21_+_8433172_8446622_RNA45SN1_45S_50ntFlanks@9031.9059.29\|TCCTCGTGGGGGGGCCGGGCCACCCCTCC | >hg38_21_+_8433172_8446622_RNA45SN1_45S_50ntFlanks | 9031 | 9059 | 29 | TCCTCGTGGGGGGGCCGGGCCACCCCTCC | 224.2 | -9.9 | 1.5 | -6.7 | 1.5E-11 | 5.9E-11 | BT20/MB468 | MITO |
| >hg38_21_+_8433172_8446622_RNA45SN1_45S_50ntFlanks@5239.5267.29\|TGACTGGCTCAGCGTGTGCCTACCCTACG | >hg38_21_+_8433172_8446622_RNA45SN1_45S_50ntFlanks | 5239 | 5267 | 29 | TGACTGGCTCAGCGTGTGCCTACCCTACG | 134.9 | -10.9 | 1.6 | -6.7 | 1.9E-11 | 5.9E-11 | MB231/MB468 | MP |
| >hg38_21_+_8433172_8446622_RNA45SN1_45S_50ntFlanks@11845.11863.19\|GGAACGGGCTTGGCGGAAT | >hg38_21_+_8433172_8446622_RNA45SN1_45S_50ntFlanks | 11845 | 11863 | 19 | GGAACGGGCTTGGCGGAAT | 87.7 | -10.3 | 1.5 | -6.7 | 1.9E-11 | 5.9E-11 | MB231/MB468 | MP |
| >hg38_21_+_8433172_8446622_RNA45SN1_45S_50ntFlanks@10389.10420.32\|TGAACAGCAGTTGAACATGGGTCAGTCGGTCC | >hg38_21_+_8433172_8446622_RNA45SN1_45S_50ntFlanks | 10389 | 10420 | 32 | TGAACAGCAGTTGAACATGGGTCAGTCGGTCC | 102.8 | -10.5 | 1.6 | -6.7 | 1.9E-11 | 5.9E-11 | MB231/MB468 | MP |
| >hg38_21_+_8433172_8446622_RNA45SN1_45S_50ntFlanks@12078.12095.18\|GGGCTCTCGCTTCTGGCG | >hg38_21_+_8433172_8446622_RNA45SN1_45S_50ntFlanks | 12078 | 12095 | 18 | GGGCTCTCGCTTCTGGCG | 128.4 | -10.9 | 1.6 | -6.7 | 1.9E-11 | 5.9E-11 | MB231/MB468 | MP |
| >MI0005756\|hsa-mir-934&WithFlank&X\|+\|136550872\|136550966@21.42.22(+2A)&[MIMAT0004977&hsa-miR-934&offsets\|0\|0(+2A);m-943&X\|+\|136550892\|136550913&offsets\|0\|0(+2A)]\|TGTCTACTACTGGAGACACTGGAA | >MI0005756\|hsa-mir-934&WithFlank&X\|+\|136550872\|136550966 | 21 | 42 | 22(+2A) | TGTCTACTACTGGAGACACTGGAA | 186.4 | 4.2 | 0.6 | 6.7 | 2.6E-11 | 5.9E-11 | BT20/MB468 | NUC |
| >hg38_21_+_8433172_8446622_RNA45SN1_45S_50ntFlanks@10866.10889.24\|CTAAGGGCTGGGTCGGTCGGGCTG | >hg38_21_+_8433172_8446622_RNA45SN1_45S_50ntFlanks | 10866 | 10889 | 24 | CTAAGGGCTGGGTCGGTCGGGCTG | 1047.4 | -8.9 | 1.3 | -6.7 | 2.1E-11 | 5.9E-11 | MB231/MB468 | NUC |
| >hg38_21_+_8433172_8446622_RNA45SN1_45S_50ntFlanks@8236.8266.31\|GGCCCGGGTCTTCCCGGAGTCGGGTTGCTTG | >hg38_21_+_8433172_8446622_RNA45SN1_45S_50ntFlanks | 8236 | 8266 | 31 | GGCCCGGGTCTTCCCGGAGTCGGGTTGCTTG | 106.4 | -10.6 | 1.6 | -6.7 | 1.9E-11 | 5.9E-11 | MB231/MB468 | MP |
| >hg38_21_+_8433172_8446622_RNA45SN1_45S_50ntFlanks@12289.12323.35\|CCGTGAAAGCGGGGCCTCACGATCCTTCTGACCTT | >hg38_21_+_8433172_8446622_RNA45SN1_45S_50ntFlanks | 12289 | 12323 | 35 | CCGTGAAAGCGGGGCCTCACGATCCTTCTGACCTT | 2251.9 | -6.6 | 1.0 | -6.7 | 1.5E-11 | 5.9E-11 | BT20/MB468 | MITO |
| >hg38_21_+_8433172_8446622_RNA45SN1_45S_50ntFlanks@8776.8796.21\|CCGCCCCCCGGCCCCGCGTCC | >hg38_21_+_8433172_8446622_RNA45SN1_45S_50ntFlanks | 8776 | 8796 | 21 | CCGCCCCCCGGCCCCGCGTCC | 473.8 | -12.8 | 1.9 | -6.7 | 1.9E-11 | 5.9E-11 | MB231/MB468 | MP |
| >hg38_21_+_8433172_8446622_RNA45SN1_45S_50ntFlanks@10786.10825.40\|AACAGCCTCTGGCATGTTGGAACAATGTAGGTAAGGGAAG | >hg38_21_+_8433172_8446622_RNA45SN1_45S_50ntFlanks | 10786 | 10825 | 40 | AACAGCCTCTGGCATGTTGGAACAATGTAGGTAAGGGAAG | 456.2 | -6.1 | 0.9 | -6.7 | 2.1E-11 | 5.9E-11 | MB231/MB468 | CYTO |
| >hg38_21_+_8433172_8446622_RNA45SN1_45S_50ntFlanks@4007.4025.19\|CCTCGGGCCGATCGCACGC | >hg38_21_+_8433172_8446622_RNA45SN1_45S_50ntFlanks | 4007 | 4025 | 19 | CCTCGGGCCGATCGCACGC | 122.4 | -10.8 | 1.6 | -6.7 | 1.9E-11 | 5.9E-11 | MB231/MB468 | MP |
| >MI0001448\|hsa-mir-425&WithFlank&3\|-\|49020142\|49020240@21.43.23&[MIMAT0003393&hsa-miR-425-5p&offsets\|+1\|+1;m-95&3\|-\|49020200\|49020221&offsets\|+1\|+2]\|ATGACACGATCACTCCCGTTGAG | >MI0001448\|hsa-mir-425&WithFlank&3\|-\|49020142\|49020240 | 21 | 43 | 23 | ATGACACGATCACTCCCGTTGAG | 349.7 | 8.3 | 1.2 | 6.7 | 1.9E-11 | 5.9E-11 | BT20/MB468 | CYTO |
| >hg38_21_+_8433172_8446622_RNA45SN1_45S_50ntFlanks@10873.10902.30\|CTGGGTCGGTCGGGCTGGGGCGCGAAGCGG | >hg38_21_+_8433172_8446622_RNA45SN1_45S_50ntFlanks | 10873 | 10902 | 30 | CTGGGTCGGTCGGGCTGGGGCGCGAAGCGG | 1277.3 | -8.9 | 1.3 | -6.7 | 2.1E-11 | 6.0E-11 | MB231/MB468 | NUC |
| >hg38_21_+_8433172_8446622_RNA45SN1_45S_50ntFlanks@11576.11602.27\|CTTAGAACTGGTGCGGACCAGGGGAAT | >hg38_21_+_8433172_8446622_RNA45SN1_45S_50ntFlanks | 11576 | 11602 | 27 | CTTAGAACTGGTGCGGACCAGGGGAAT | 1918.8 | -10.1 | 1.5 | -6.7 | 2.6E-11 | 6.0E-11 | BT20/MB468 | NUC |
| >hg38_21_+_8433172_8446622_RNA45SN1_45S_50ntFlanks@5086.5112.27\|AACGAGACTCTGGCATGCTAACTAGTT | >hg38_21_+_8433172_8446622_RNA45SN1_45S_50ntFlanks | 5086 | 5112 | 27 | AACGAGACTCTGGCATGCTAACTAGTT | 291.8 | 5.7 | 0.9 | 6.7 | 2.1E-11 | 6.0E-11 | MB231/MB468 | CYTO |
| >hg38_21_+_8433172_8446622_RNA45SN1_45S_50ntFlanks@9590.9610.21\|TGGGTATAGGGGCGAAAGACT | >hg38_21_+_8433172_8446622_RNA45SN1_45S_50ntFlanks | 9590 | 9610 | 21 | TGGGTATAGGGGCGAAAGACT | 309.3 | 7.4 | 1.1 | 6.7 | 2.1E-11 | 6.0E-11 | MB231/MB468 | CYTO |
| >trnaMT_SerGCT_MT_+_12207_12265@1.18.18\|GAGAAAGCTCACAAGAAC | >trnaMT_SerGCT_MT_+_12207_12265 | 1 | 18 | 18 | GAGAAAGCTCACAAGAAC | 3913.5 | 9.5 | 1.4 | 6.7 | 1.9E-11 | 6.0E-11 | BT20/MB468 | CYTO |
| >hg38_21_+_8433172_8446622_RNA45SN1_45S_50ntFlanks@11882.11906.25\|CTGTTGAGCTTGACTCTAGTCTGGC | >hg38_21_+_8433172_8446622_RNA45SN1_45S_50ntFlanks | 11882 | 11906 | 25 | CTGTTGAGCTTGACTCTAGTCTGGC | 82.0 | -10.2 | 1.5 | -6.7 | 2.0E-11 | 6.0E-11 | MB231/MB468 | MP |
| >hg38_21_+_8433172_8446622_RNA45SN1_45S_50ntFlanks@10883.10901.19\|CGGGCTGGGGCGCGAAGCG | >hg38_21_+_8433172_8446622_RNA45SN1_45S_50ntFlanks | 10883 | 10901 | 19 | CGGGCTGGGGCGCGAAGCG | 116.8 | -10.7 | 1.6 | -6.7 | 2.0E-11 | 6.0E-11 | MB231/MB468 | MP |
| >hg38_21_+_8433172_8446622_RNA45SN1_45S_50ntFlanks@5484.5521.38\|GAGCGCTGAGAAGACGGTCGAACTTGACTATCTAGAGG | >hg38_21_+_8433172_8446622_RNA45SN1_45S_50ntFlanks | 5484 | 5521 | 38 | GAGCGCTGAGAAGACGGTCGAACTTGACTATCTAGAGG | 132.7 | -10.9 | 1.6 | -6.7 | 2.0E-11 | 6.0E-11 | MB231/MB468 | MP |
| >hg38_21_+_8433172_8446622_RNA45SN1_45S_50ntFlanks@10408.10430.23\|GGTCAGTCGGTCCTGAGAGATGG | >hg38_21_+_8433172_8446622_RNA45SN1_45S_50ntFlanks | 10408 | 10430 | 23 | GGTCAGTCGGTCCTGAGAGATGG | 1999.7 | -7.0 | 1.0 | -6.7 | 2.1E-11 | 6.0E-11 | MB231/MB468 | CYTO |
| >MI0000465\|hsa-mir-191&WithFlank&3\|-\|49020612\|49020715@22.44.23(+1A)&[MIMAT0000440&hsa-miR-191-5p&offsets\|0\|0(+1A);m-48&3\|-\|49020672\|49020694&offsets\|0\|0(+1A)]\|CAACGGAATCCCAAAAGCAGCTGA | >MI0000465\|hsa-mir-191&WithFlank&3\|-\|49020612\|49020715 | 22 | 44 | 23(+1A) | CAACGGAATCCCAAAAGCAGCTGA | 481.8 | 4.2 | 0.6 | 6.7 | 2.1E-11 | 6.0E-11 | MB231/MB468 | NUC |
| >hg38_21_+_8433172_8446622_RNA45SN1_45S_50ntFlanks@12008.12039.32\|CCGCCGGTGAAATACCACTACTCTGATCGTTT | >hg38_21_+_8433172_8446622_RNA45SN1_45S_50ntFlanks | 12008 | 12039 | 32 | CCGCCGGTGAAATACCACTACTCTGATCGTTT | 550.3 | -6.9 | 1.0 | -6.7 | 2.1E-11 | 6.0E-11 | MB231/MB468 | NUC |
| >hg38_21_+_8433172_8446622_RNA45SN1_45S_50ntFlanks@11718.11751.34\|AACGGCGGGAGTAACTATGACTCTCTTAAGGTAG | >hg38_21_+_8433172_8446622_RNA45SN1_45S_50ntFlanks | 11718 | 11751 | 34 | AACGGCGGGAGTAACTATGACTCTCTTAAGGTAG | 337.1 | -11.5 | 1.6 | -7.1 | 1.3E-12 | 6.0E-11 | BT20/MB468 | MP |
| >hg38_21_+_8433172_8446622_RNA45SN1_45S_50ntFlanks@11578.11605.28\|TAGAACTGGTGCGGACCAGGGGAATCCG | >hg38_21_+_8433172_8446622_RNA45SN1_45S_50ntFlanks | 11578 | 11605 | 28 | TAGAACTGGTGCGGACCAGGGGAATCCG | 645.8 | -5.5 | 0.8 | -6.7 | 2.1E-11 | 6.0E-11 | MB231/MB468 | CYTO |
| >hg38_21_+_8433172_8446622_RNA45SN1_45S_50ntFlanks@7993.8010.18\|GTGGCGACCCGCTGAATT | >hg38_21_+_8433172_8446622_RNA45SN1_45S_50ntFlanks | 7993 | 8010 | 18 | GTGGCGACCCGCTGAATT | 84.1 | 3.7 | 0.6 | 6.7 | 2.6E-11 | 6.0E-11 | BT20/MB468 | NUC |
| >hg38_21_+_8433172_8446622_RNA45SN1_45S_50ntFlanks@10069.10089.21\|AGTGGACGGGAGCGGCGGGGG | >hg38_21_+_8433172_8446622_RNA45SN1_45S_50ntFlanks | 10069 | 10089 | 21 | AGTGGACGGGAGCGGCGGGGG | 144.2 | -11.0 | 1.6 | -6.7 | 2.0E-11 | 6.0E-11 | MB231/MB468 | MP |
| >hg38_21_+_8433172_8446622_RNA45SN1_45S_50ntFlanks@11999.12030.32\|CCCTGCGGGCCGCCGGTGAAATACCACTACTC | >hg38_21_+_8433172_8446622_RNA45SN1_45S_50ntFlanks | 11999 | 12030 | 32 | CCCTGCGGGCCGCCGGTGAAATACCACTACTC | 78.7 | -10.2 | 1.5 | -6.7 | 2.0E-11 | 6.0E-11 | MB231/MB468 | MP |
| >hg38_21_+_8433172_8446622_RNA45SN1_45S_50ntFlanks@12072.12096.25\|CCCGAGGGGCTCTCGCTTCTGGCGC | >hg38_21_+_8433172_8446622_RNA45SN1_45S_50ntFlanks | 12072 | 12096 | 25 | CCCGAGGGGCTCTCGCTTCTGGCGC | 200.9 | -11.5 | 1.7 | -6.7 | 2.0E-11 | 6.0E-11 | MB231/MB468 | MP |
| >hg38_21_+_8433172_8446622_RNA45SN1_45S_50ntFlanks@10795.10818.24\|TGGCATGTTGGAACAATGTAGGTA | >hg38_21_+_8433172_8446622_RNA45SN1_45S_50ntFlanks | 10795 | 10818 | 24 | TGGCATGTTGGAACAATGTAGGTA | 929.6 | -8.6 | 1.3 | -6.7 | 2.5E-11 | 6.0E-11 | MB231/MB468 | MITO |
| >hg38_21_+_8433172_8446622_RNA45SN1_45S_50ntFlanks@12289.12325.37\|CCGTGAAAGCGGGGCCTCACGATCCTTCTGACCTTTT | >hg38_21_+_8433172_8446622_RNA45SN1_45S_50ntFlanks | 12289 | 12325 | 37 | CCGTGAAAGCGGGGCCTCACGATCCTTCTGACCTTTT | 414.6 | -8.7 | 1.3 | -6.7 | 2.5E-11 | 6.0E-11 | MB231/MB468 | MITO |
| >MI0000063\|hsa-let-7b&WithFlank&22\|+\|46113680\|46113774@12.34.23&[MIMAT0000063&hsa-let-7b-5p&offsets\|0\|+1;m-14&22\|+\|46113691\|46113712&offsets\|0\|+1]\|TGAGGTAGTAGGTTGTGTGGTTT | >MI0000063\|hsa-let-7b&WithFlank&22\|+\|46113680\|46113774 | 12 | 34 | 23 | TGAGGTAGTAGGTTGTGTGGTTT | 2038.7 | 3.9 | 0.6 | 6.7 | 2.1E-11 | 6.0E-11 | MB231/MB468 | NUC |
| >hg38_21_+_8433172_8446622_RNA45SN1_45S_50ntFlanks@12628.12659.32\|TGGGGCGAAGCTACCATCTGTGGGATTATGAC | >hg38_21_+_8433172_8446622_RNA45SN1_45S_50ntFlanks | 12628 | 12659 | 32 | TGGGGCGAAGCTACCATCTGTGGGATTATGAC | 541.2 | -8.5 | 1.3 | -6.7 | 2.1E-11 | 6.0E-11 | MB231/MB468 | NUC |
| >hg38_21_+_8433172_8446622_RNA45SN1_45S_50ntFlanks@7986.8009.24\|ATCAGACGTGGCGACCCGCTGAAT | >hg38_21_+_8433172_8446622_RNA45SN1_45S_50ntFlanks | 7986 | 8009 | 24 | ATCAGACGTGGCGACCCGCTGAAT | 1299.3 | -9.3 | 1.4 | -6.7 | 2.0E-11 | 6.0E-11 | MB231/MB468 | MP |
| >hg38_21_+_8433172_8446622_RNA45SN1_45S_50ntFlanks@6730.6770.41\|ACACATTGATCATCGACACTTCGAACGCACTTGCGGCCCCG | >hg38_21_+_8433172_8446622_RNA45SN1_45S_50ntFlanks | 6730 | 6770 | 41 | ACACATTGATCATCGACACTTCGAACGCACTTGCGGCCCCG | 582.3 | -4.0 | 0.6 | -6.7 | 1.6E-11 | 6.1E-11 | BT20/MB468 | MITO |
| >hg38_21_+_8433172_8446622_RNA45SN1_45S_50ntFlanks@10362.10381.20\|AAGGCCGAAGTGGAGAAGGG | >hg38_21_+_8433172_8446622_RNA45SN1_45S_50ntFlanks | 10362 | 10381 | 20 | AAGGCCGAAGTGGAGAAGGG | 193.6 | -7.9 | 1.2 | -6.7 | 2.2E-11 | 6.1E-11 | MB231/MB468 | NUC |
| >hg38_21_+_8433172_8446622_RNA45SN1_45S_50ntFlanks@3831.3860.30\|CTCGCTCCTCTCCTACTTGGATAACTGTGG | >hg38_21_+_8433172_8446622_RNA45SN1_45S_50ntFlanks | 3831 | 3860 | 30 | CTCGCTCCTCTCCTACTTGGATAACTGTGG | 3837.7 | -10.6 | 1.6 | -6.7 | 2.7E-11 | 6.1E-11 | BT20/MB468 | NUC |
| >hg38_21_+_8433172_8446622_RNA45SN1_45S_50ntFlanks@12942.12973.32\|TTCGTAGACGACCTGCTTCTGGGTCGGGGTTT | >hg38_21_+_8433172_8446622_RNA45SN1_45S_50ntFlanks | 12942 | 12973 | 32 | TTCGTAGACGACCTGCTTCTGGGTCGGGGTTT | 265.0 | -11.1 | 1.6 | -7.1 | 1.3E-12 | 6.1E-11 | BT20/MB468 | MP |
| >hg38_21_+_8433172_8446622_RNA45SN1_45S_50ntFlanks@8437.8471.35\|AGGATTCAACCCGGCGGCGGGTCCGGCCGTGTCGG | >hg38_21_+_8433172_8446622_RNA45SN1_45S_50ntFlanks | 8437 | 8471 | 35 | AGGATTCAACCCGGCGGCGGGTCCGGCCGTGTCGG | 346.0 | -11.5 | 1.6 | -7.1 | 1.3E-12 | 6.1E-11 | BT20/MB468 | MP |
| >hg38_21_+_8433172_8446622_RNA45SN1_45S_50ntFlanks@10681.10709.29\|TTCCGGCGGCGTCCGGTGAGCTCTCGCTG | >hg38_21_+_8433172_8446622_RNA45SN1_45S_50ntFlanks | 10681 | 10709 | 29 | TTCCGGCGGCGTCCGGTGAGCTCTCGCTG | 673.5 | -8.4 | 1.3 | -6.7 | 2.5E-11 | 6.1E-11 | MB231/MB468 | MITO |
| >hg38_21_+_8433172_8446622_RNA45SN1_45S_50ntFlanks@12419.12454.36\|GATGTCGGCTCTTCCTATCATTGTGAAGCAGAATTC | >hg38_21_+_8433172_8446622_RNA45SN1_45S_50ntFlanks | 12419 | 12454 | 36 | GATGTCGGCTCTTCCTATCATTGTGAAGCAGAATTC | 885.5 | 4.9 | 0.7 | 6.7 | 2.1E-11 | 6.1E-11 | MB231/MB468 | CYTO |

<table>
<tr><td>>trna111_HisGTG_1_-_147774845_147774916@-1T.32.33,trna118_HisGTG_1_-_145396881_145396952@-1T.32.33,trna16_HisGTG_1_+_146544773_146544844@-1T.32.33,trna1_HisGTG_15_+_45493349_45493420@-1T.32.33,trna21_HisGTG_1_+_147753471_147753542@-1T.32.33,trna33_HisGTG_6_+_27125906_27125977@-1T.32.33,trna7_HisGTG_9_-_14433938_14434009@-1T.32.33,trna8_HisGTG_15_-_45492611_45492682@-1T.32.33,trna9_HisGTG_15_-_45490804_45490875@-1T.32.33|TGCCGTGATCGTATAGTGGTTAGTACTCTGCGT</td><td>>trna111_HisGTG_1_-_147774845_147774916</td><td>-1T</td><td>32</td><td>33</td><td>TGCCGTGATCGTATAGTGGTTAGTACTCTGCGT</td><td>184.6</td><td>10.7</td><td>1.6</td><td>6.7</td><td>2.0E-11</td><td>6.1E-11</td><td>MB231/MB468</td><td>MP</td></tr>
<tr><td>>hg38_21_+_8433172_8446622_RNA45SN1_45S_50ntFlanks@4103.4121.19|CCACGGGTGACGGGGAATC</td><td>>hg38_21_+_8433172_8446622_RNA45SN1_45S_50ntFlanks</td><td>4103</td><td>4121</td><td>19</td><td>CCACGGGTGACGGGGAATC</td><td>711.5</td><td>-5.6</td><td>0.8</td><td>-6.7</td><td>2.2E-11</td><td>6.1E-11</td><td>MB231/MB468</td><td>CYTO</td></tr>
<tr><td>>hg38_21_+_8433172_8446622_RNA45SN1_45S_50ntFlanks@12712.12743.32|CGGAGCCTCGGTTGGCCTCGGATAGCCGGTCC</td><td>>hg38_21_+_8433172_8446622_RNA45SN1_45S_50ntFlanks</td><td>12712</td><td>12743</td><td>32</td><td>CGGAGCCTCGGTTGGCCTCGGATAGCCGGTCC</td><td>320.7</td><td>-11.4</td><td>1.6</td><td>-7.1</td><td>1.3E-12</td><td>6.1E-11</td><td>BT20/MB468</td><td>MP</td></tr>
<tr><td>>hg38_21_+_8433172_8446622_RNA45SN1_45S_50ntFlanks@3705.3745.41|TACCTGGTTGATCCTGCCAGTAGCATATGCTTGTCTCAAAG</td><td>>hg38_21_+_8433172_8446622_RNA45SN1_45S_50ntFlanks</td><td>3705</td><td>3745</td><td>41</td><td>TACCTGGTTGATCCTGCCAGTAGCATATGCTTGTCTCAAAG</td><td>2604.2</td><td>7.2</td><td>1.1</td><td>6.7</td><td>2.0E-11</td><td>6.1E-11</td><td>BT20/MB468</td><td>CYTO</td></tr>
<tr><td>>MI0000088|hsa-mir-30a&WithFlank&6|-|71403545|71403627@13.35.23(+1C)&[MIMAT0000087&hsa-miR-30a-5p&offsets|+1|+2(+1C);m-15&6|-|71403595|71403616&offsets|+1|+2(+1C)]|GTAAACATCCTCGACTGGAAGCTC</td><td>>MI0000088|hsa-mir-30a&WithFlank&6|-|71403545|71403627</td><td>13</td><td>35</td><td>23(+1C)</td><td>GTAAACATCCTCGACTGGAAGCTC</td><td>114.5</td><td>10.1</td><td>1.5</td><td>6.7</td><td>2.0E-11</td><td>6.1E-11</td><td>MB231/MB468</td><td>MP</td></tr>
<tr><td>>hg38_21_+_8433172_8446622_RNA45SN1_45S_50ntFlanks@9200.9222.23|TCTCTCGGGGCCACGCGCGCGTC</td><td>>hg38_21_+_8433172_8446622_RNA45SN1_45S_50ntFlanks</td><td>9200</td><td>9222</td><td>23</td><td>TCTCTCGGGGCCACGCGCGCGTC</td><td>111.0</td><td>-10.7</td><td>1.6</td><td>-6.7</td><td>2.0E-11</td><td>6.1E-11</td><td>MB231/MB468</td><td>MP</td></tr>
<tr><td>>hg38_21_+_8433172_8446622_RNA45SN1_45S_50ntFlanks@12414.12447.34|CCTTCGATGTCGGCTCTTCCTATCATTGTGAAGC</td><td>>hg38_21_+_8433172_8446622_RNA45SN1_45S_50ntFlanks</td><td>12414</td><td>12447</td><td>34</td><td>CCTTCGATGTCGGCTCTTCCTATCATTGTGAAGC</td><td>347.4</td><td>9.5</td><td>1.4</td><td>6.7</td><td>1.6E-11</td><td>6.1E-11</td><td>BT20/MB468</td><td>MITO</td></tr>
<tr><td>>hg38_21_+_8433172_8446622_RNA45SN1_45S_50ntFlanks@9029.9059.31|TCTCCTCGTGGGGGGGCCGGGCCACCCCTCC</td><td>>hg38_21_+_8433172_8446622_RNA45SN1_45S_50ntFlanks</td><td>9029</td><td>9059</td><td>31</td><td>TCTCCTCGTGGGGGGGCCGGGCCACCCCTCC</td><td>258.8</td><td>-10.1</td><td>1.5</td><td>-6.7</td><td>1.6E-11</td><td>6.2E-11</td><td>BT20/MB468</td><td>MITO</td></tr>
<tr><td>>hg38_21_+_8433172_8446622_RNA45SN1_45S_50ntFlanks@11564.11586.23|CCTAGCAGCCGACTTAGAACTGG</td><td>>hg38_21_+_8433172_8446622_RNA45SN1_45S_50ntFlanks</td><td>11564</td><td>11586</td><td>23</td><td>CCTAGCAGCCGACTTAGAACTGG</td><td>1498.5</td><td>-7.2</td><td>1.1</td><td>-6.7</td><td>2.0E-11</td><td>6.2E-11</td><td>MB231/MB468</td><td>MP</td></tr>
<tr><td>>hg38_21_+_8433172_8446622_RNA45SN1_45S_50ntFlanks@12399.12419.21|ACGTCGCTTTTTGATCCTTCG</td><td>>hg38_21_+_8433172_8446622_RNA45SN1_45S_50ntFlanks</td><td>12399</td><td>12419</td><td>21</td><td>ACGTCGCTTTTTGATCCTTCG</td><td>227.5</td><td>-9.9</td><td>1.5</td><td>-6.7</td><td>1.6E-11</td><td>6.2E-11</td><td>BT20/MB468</td><td>MITO</td></tr>
<tr><td>>hg38_21_+_8433172_8446622_RNA45SN1_45S_50ntFlanks@12456.12487.32|CCAAGCGTTGGATTGTTCACCCACTAATAGGG</td><td>>hg38_21_+_8433172_8446622_RNA45SN1_45S_50ntFlanks</td><td>12456</td><td>12487</td><td>32</td><td>CCAAGCGTTGGATTGTTCACCCACTAATAGGG</td><td>40904.1</td><td>-8.5</td><td>1.3</td><td>-6.7</td><td>2.2E-11</td><td>6.2E-11</td><td>MB231/MB468</td><td>CYTO</td></tr>
<tr><td>>MI0000289|hsa-mir-181a-1&WithFlank&1|-|198859038|198859159@70.90.21(+1U)&[MIMAT0000270&hsa-miR-181a-3p&offsets|0|-1(+1U);m-125&1|-|198859069|198859090&offsets|0|-1(+1U)]|ACCATCGACCGTTGATTGTACT</td><td>>MI0000289|hsa-mir-181a-1&WithFlank&1|-|198859038|198859159</td><td>70</td><td>90</td><td>21(+1U)</td><td>ACCATCGACCGTTGATTGTACT</td><td>504.2</td><td>-12.4</td><td>1.7</td><td>-7.3</td><td>3.2E-13</td><td>6.2E-11</td><td>BT20/MB231</td><td>MP</td></tr>
<tr><td>>MI0000255|hsa-mir-30d&WithFlank&8|-|134804870|134804951@12.34.23(+1C)&[MIMAT0000245&hsa-miR-30d-5p&offsets|0|+1(+1C);m-21&8|-|134804919|134804940&offsets|0|+1(+1C)]|TGTAAACATCCCCGACTGGAAGCC</td><td>>MI0000255|hsa-mir-30d&WithFlank&8|-|134804870|134804951</td><td>12</td><td>34</td><td>23(+1C)</td><td>TGTAAACATCCCCGACTGGAAGCC</td><td>8051.4</td><td>6.7</td><td>1.0</td><td>6.7</td><td>2.6E-11</td><td>6.2E-11</td><td>MB231/MB468</td><td>MITO</td></tr>
<tr><td>>hg38_21_+_8433172_8446622_RNA45SN1_45S_50ntFlanks@4725.4766.42|TTAATCAAGAACGAAAGTCGGAGGTTCGAAGACGATCAGATA</td><td>>hg38_21_+_8433172_8446622_RNA45SN1_45S_50ntFlanks</td><td>4725</td><td>4766</td><td>42</td><td>TTAATCAAGAACGAAAGTCGGAGGTTCGAAGACGATCAGATA</td><td>291.6</td><td>7.2</td><td>1.1</td><td>6.7</td><td>2.2E-11</td><td>6.2E-11</td><td>MB231/MB468</td><td>CYTO</td></tr>
<tr><td>>hg38_21_+_8433172_8446622_RNA45SN1_45S_50ntFlanks@11579.11611.33|AGAACTGGTGCGGACCAGGGGAATCCGACTGTT</td><td>>hg38_21_+_8433172_8446622_RNA45SN1_45S_50ntFlanks</td><td>11579</td><td>11611</td><td>33</td><td>AGAACTGGTGCGGACCAGGGGAATCCGACTGTT</td><td>1684.9</td><td>-5.4</td><td>0.8</td><td>-6.7</td><td>2.0E-11</td><td>6.2E-11</td><td>MB231/MB468</td><td>MP</td></tr>
<tr><td>>hg38_21_+_8433172_8446622_RNA45SN1_45S_50ntFlanks@11821.11853.33|CTATCCAGCGAAACCACAGCCAAGGGAACGGGC</td><td>>hg38_21_+_8433172_8446622_RNA45SN1_45S_50ntFlanks</td><td>11821</td><td>11853</td><td>33</td><td>CTATCCAGCGAAACCACAGCCAAGGGAACGGGC</td><td>76.2</td><td>-10.1</td><td>1.5</td><td>-6.7</td><td>2.0E-11</td><td>6.2E-11</td><td>MB231/MB468</td><td>MP</td></tr>
<tr><td>>hg38_21_+_8433172_8446622_RNA45SN1_45S_50ntFlanks@6730.6752.23|ACACATTGATCATCGACACTTCG</td><td>>hg38_21_+_8433172_8446622_RNA45SN1_45S_50ntFlanks</td><td>6730</td><td>6752</td><td>23</td><td>ACACATTGATCATCGACACTTCG</td><td>95.1</td><td>-10.4</td><td>1.6</td><td>-6.7</td><td>2.0E-11</td><td>6.2E-11</td><td>MB231/MB468</td><td>MP</td></tr>
<tr><td>>hg38_21_+_8433172_8446622_RNA45SN1_45S_50ntFlanks@12407.12431.25|TTTTGATCCTTCGATGTCGGCTCTT</td><td>>hg38_21_+_8433172_8446622_RNA45SN1_45S_50ntFlanks</td><td>12407</td><td>12431</td><td>25</td><td>TTTTGATCCTTCGATGTCGGCTCTT</td><td>726.0</td><td>-7.2</td><td>1.1</td><td>-6.7</td><td>2.0E-11</td><td>6.2E-11</td><td>MB231/MB468</td><td>MP</td></tr>
<tr><td>>MI0000063|hsa-let-7b&WithFlank&22|+|46113680|46113774@12.33.22&[MIMAT0000063&hsa-let-7b-5p&offsets|0|0;m-14&22|+|46113691|46113712&offsets|0|0]|TGAGGTAGTAGGTTGTGTGGTT</td><td>>MI0000063|hsa-let-7b&WithFlank&22|+|46113680|46113774</td><td>12</td><td>33</td><td>22</td><td>TGAGGTAGTAGGTTGTGTGGTT</td><td>12156.5</td><td>5.3</td><td>0.8</td><td>6.7</td><td>2.2E-11</td><td>6.2E-11</td><td>MB231/MB468</td><td>CYTO</td></tr>
<tr><td>>hg38_21_+_8433172_8446622_RNA45SN1_45S_50ntFlanks@11576.11607.32|CTTAGAACTGGTGCGGACCAGGGGAATCCGAC</td><td>>hg38_21_+_8433172_8446622_RNA45SN1_45S_50ntFlanks</td><td>11576</td><td>11607</td><td>32</td><td>CTTAGAACTGGTGCGGACCAGGGGAATCCGAC</td><td>348.0</td><td>-4.8</td><td>0.7</td><td>-6.7</td><td>2.6E-11</td><td>6.2E-11</td><td>MB231/MB468</td><td>MITO</td></tr>
<tr><td>>hg38_21_+_8433172_8446622_RNA45SN1_45S_50ntFlanks@12365.12382.18|ATAACTGGCTTGTGGCGG</td><td>>hg38_21_+_8433172_8446622_RNA45SN1_45S_50ntFlanks</td><td>12365</td><td>12382</td><td>18</td><td>ATAACTGGCTTGTGGCGG</td><td>1258.2</td><td>-7.9</td><td>1.2</td><td>-6.7</td><td>2.2E-11</td><td>6.2E-11</td><td>MB231/MB468</td><td>CYTO</td></tr>
<tr><td>>hg38_21_+_8433172_8446622_RNA45SN1_45S_50ntFlanks@4783.4800.18|CATAAACGATGCCGACCG</td><td>>hg38_21_+_8433172_8446622_RNA45SN1_45S_50ntFlanks</td><td>4783</td><td>4800</td><td>18</td><td>CATAAACGATGCCGACCG</td><td>124.6</td><td>-5.9</td><td>0.9</td><td>-6.7</td><td>2.2E-11</td><td>6.2E-11</td><td>MB231/MB468</td><td>CYTO</td></tr>
<tr><td>>hg38_21_+_8433172_8446622_RNA45SN1_45S_50ntFlanks@12938.12956.19|ACCATTCGTAGACGACCTG</td><td>>hg38_21_+_8433172_8446622_RNA45SN1_45S_50ntFlanks</td><td>12938</td><td>12956</td><td>19</td><td>ACCATTCGTAGACGACCTG</td><td>2459.1</td><td>-3.7</td><td>0.5</td><td>-6.7</td><td>2.2E-11</td><td>6.2E-11</td><td>MB231/MB468</td><td>CYTO</td></tr>
<tr><td>>MI0000077|hsa-mir-21&WithFlank&17|+|59841260|59841343@14.31.18&[MIMAT0000076&hsa-miR-21-5p&offsets|0|-4;m-1&17|+|59841273|59841294&offsets|0|-4]|TAGCTTATCAGACTGATG</td><td>>MI0000077|hsa-mir-21&WithFlank&17|+|59841260|59841343</td><td>14</td><td>31</td><td>18</td><td>TAGCTTATCAGACTGATG</td><td>366.6</td><td>4.1</td><td>0.6</td><td>6.7</td><td>2.7E-11</td><td>6.2E-11</td><td>BT20/MB468</td><td>NUC</td></tr>
<tr><td>>hg38_21_+_8433172_8446622_RNA45SN1_45S_50ntFlanks@10914.10939.26|CGCCGCGGCTGGACGAGGCGCCGCCG</td><td>>hg38_21_+_8433172_8446622_RNA45SN1_45S_50ntFlanks</td><td>10914</td><td>10939</td><td>26</td><td>CGCCGCGGCTGGACGAGGCGCCGCCG</td><td>318.8</td><td>-11.4</td><td>1.6</td><td>-7.1</td><td>1.4E-12</td><td>6.2E-11</td><td>BT20/MB468</td><td>MP</td></tr>
<tr><td>>hg38_21_+_8433172_8446622_RNA45SN1_45S_50ntFlanks@13006.13024.19|CTATTGAAAGTCAGCCCTC</td><td>>hg38_21_+_8433172_8446622_RNA45SN1_45S_50ntFlanks</td><td>13006</td><td>13024</td><td>19</td><td>CTATTGAAAGTCAGCCCTC</td><td>175.8</td><td>-6.2</td><td>0.9</td><td>-6.7</td><td>2.2E-11</td><td>6.2E-11</td><td>MB231/MB468</td><td>CYTO</td></tr>
<tr><td>>hg38_MT_+_1621_3279_RNR2_16S_50ntFlanks@1185.1215.31|ATTTCGGTTGGGGCGACCTCGGAGCAGAACC</td><td>>hg38_MT_+_1621_3279_RNR2_16S_50ntFlanks</td><td>1185</td><td>1215</td><td>31</td><td>ATTTCGGTTGGGGCGACCTCGGAGCAGAACC</td><td>694.1</td><td>9.4</td><td>1.4</td><td>6.7</td><td>1.6E-11</td><td>6.2E-11</td><td>BT20/MB468</td><td>MITO</td></tr>
<tr><td>>hg38_21_+_8433172_8446622_RNA45SN1_45S_50ntFlanks@9391.9416.26|CCGAGGTGGGATCCCGAGGCCTCTCC</td><td>>hg38_21_+_8433172_8446622_RNA45SN1_45S_50ntFlanks</td><td>9391</td><td>9416</td><td>26</td><td>CCGAGGTGGGATCCCGAGGCCTCTCC</td><td>98.3</td><td>-10.5</td><td>1.6</td><td>-6.7</td><td>2.0E-11</td><td>6.2E-11</td><td>MB231/MB468</td><td>MP</td></tr>
<tr><td>>hg38_21_+_8433172_8446622_RNA45SN1_45S_50ntFlanks@474.496.23|TCTTCGTGATCGATGTGGTGACG</td><td>>hg38_21_+_8433172_8446622_RNA45SN1_45S_50ntFlanks</td><td>474</td><td>496</td><td>23</td><td>TCTTCGTGATCGATGTGGTGACG</td><td>774.1</td><td>5.9</td><td>0.9</td><td>6.7</td><td>2.2E-11</td><td>6.2E-11</td><td>MB231/MB468</td><td>NUC</td></tr>
<tr><td>>hg38_21_+_8433172_8446622_RNA45SN1_45S_50ntFlanks@11567.11603.37|AGCAGCCGACTTAGAACTGGTGCGGACCAGGGGAATC</td><td>>hg38_21_+_8433172_8446622_RNA45SN1_45S_50ntFlanks</td><td>11567</td><td>11603</td><td>37</td><td>AGCAGCCGACTTAGAACTGGTGCGGACCAGGGGAATC</td><td>968.1</td><td>-4.1</td><td>0.6</td><td>-6.7</td><td>2.2E-11</td><td>6.2E-11</td><td>MB231/MB468</td><td>NUC</td></tr>
<tr><td>>hg38_21_+_8433172_8446622_RNA45SN1_45S_50ntFlanks@10462.10497.36|CCTCCGTTGCCCTCGGCCGATCGAAAGGGAGTCGGG</td><td>>hg38_21_+_8433172_8446622_RNA45SN1_45S_50ntFlanks</td><td>10462</td><td>10497</td><td>36</td><td>CCTCCGTTGCCCTCGGCCGATCGAAAGGGAGTCGGG</td><td>99.4</td><td>-10.5</td><td>1.6</td><td>-6.7</td><td>2.0E-11</td><td>6.3E-11</td><td>MB231/MB468</td><td>MP</td></tr>
<tr><td>>hg38_21_+_8433172_8446622_RNA45SN1_45S_50ntFlanks@11576.11605.30|CTTAGAACTGGTGCGGACCAGGGGAATCCG</td><td>>hg38_21_+_8433172_8446622_RNA45SN1_45S_50ntFlanks</td><td>11576</td><td>11605</td><td>30</td><td>CTTAGAACTGGTGCGGACCAGGGGAATCCG</td><td>1784.5</td><td>-6.4</td><td>1.0</td><td>-6.7</td><td>2.2E-11</td><td>6.3E-11</td><td>MB231/MB468</td><td>CYTO</td></tr>
<tr><td>>hg38_21_+_8433172_8446622_RNA45SN1_45S_50ntFlanks@9653.9679.27|AGGATAGCTGGCGCTCTCGCAGACCCG</td><td>>hg38_21_+_8433172_8446622_RNA45SN1_45S_50ntFlanks</td><td>9653</td><td>9679</td><td>27</td><td>AGGATAGCTGGCGCTCTCGCAGACCCG</td><td>812.6</td><td>-7.2</td><td>1.1</td><td>-6.7</td><td>2.6E-11</td><td>6.3E-11</td><td>MB231/MB468</td><td>MITO</td></tr>
<tr><td>>hg38_21_+_8433172_8446622_RNA45SN1_45S_50ntFlanks@7983.8002.20|CAGATCAGACGTGGCGACCC</td><td>>hg38_21_+_8433172_8446622_RNA45SN1_45S_50ntFlanks</td><td>7983</td><td>8002</td><td>20</td><td>CAGATCAGACGTGGCGACCC</td><td>142.1</td><td>-11.0</td><td>1.6</td><td>-6.7</td><td>2.1E-11</td><td>6.3E-11</td><td>MB231/MB468</td><td>MP</td></tr>
<tr><td>>hg38_21_+_8433172_8446622_RNA45SN1_45S_50ntFlanks@3899.3920.22|CCTTCGCGGGGGGGATGCGTGC</td><td>>hg38_21_+_8433172_8446622_RNA45SN1_45S_50ntFlanks</td><td>3899</td><td>3920</td><td>22</td><td>CCTTCGCGGGGGGGATGCGTGC</td><td>695.2</td><td>-4.1</td><td>0.6</td><td>-6.7</td><td>2.0E-11</td><td>6.3E-11</td><td>BT20/MB468</td><td>CYTO</td></tr>
<tr><td>>hg38_21_+_8433172_8446622_RNA45SN1_45S_50ntFlanks@12936.12959.24|AAACCATTCGTAGACGACCTGCTT</td><td>>hg38_21_+_8433172_8446622_RNA45SN1_45S_50ntFlanks</td><td>12936</td><td>12959</td><td>24</td><td>AAACCATTCGTAGACGACCTGCTT</td><td>347.4</td><td>-4.8</td><td>0.7</td><td>-6.7</td><td>2.6E-11</td><td>6.3E-11</td><td>MB231/MB468</td><td>MITO</td></tr>
<tr><td>>hg38_21_+_8433172_8446622_RNA45SN1_45S_50ntFlanks@10883.10905.23|CGGGCTGGGGCGCGAAGCGGGGC</td><td>>hg38_21_+_8433172_8446622_RNA45SN1_45S_50ntFlanks</td><td>10883</td><td>10905</td><td>23</td><td>CGGGCTGGGGCGCGAAGCGGGGC</td><td>479.6</td><td>7.1</td><td>1.1</td><td>6.7</td><td>2.2E-11</td><td>6.3E-11</td><td>MB231/MB468</td><td>CYTO</td></tr>
<tr><td>>hg38_21_+_8433172_8446622_RNA45SN1_45S_50ntFlanks@11786.11804.19|GAATGGATGAACGAGATTC</td><td>>hg38_21_+_8433172_8446622_RNA45SN1_45S_50ntFlanks</td><td>11786</td><td>11804</td><td>19</td><td>GAATGGATGAACGAGATTC</td><td>2562.7</td><td>-10.2</td><td>1.5</td><td>-6.7</td><td>2.8E-11</td><td>6.3E-11</td><td>BT20/MB468</td><td>NUC</td></tr>
<tr><td>>hg38_21_+_8433172_8446622_RNA45SN1_45S_50ntFlanks@12971.12990.20|TTTCGTACGTAGCAGAGCAG</td><td>>hg38_21_+_8433172_8446622_RNA45SN1_45S_50ntFlanks</td><td>12971</td><td>12990</td><td>20</td><td>TTTCGTACGTAGCAGAGCAG</td><td>148.3</td><td>-9.3</td><td>1.4</td><td>-6.7</td><td>1.6E-11</td><td>6.3E-11</td><td>BT20/MB468</td><td>MITO</td></tr>
<tr><td>>MI0000750|hsa-mir-26a-2&WithFlank&12|-|57824603|57824698@20.39.20(+2U);MI0000083|hsa-mir-26a-1&WithFlank&3|+|37969398|37969486@16.35.20(+2U)&[MIMAT0000082&hsa-miR-26a-5p&offsets|0|-2(+2U);m-33&12|-|57824658|57824679&offsets|0|-2(+2U)];[MIMAT0000082_1&hsa-miR-26a-5p&offsets|0|-2(+2U);m-34&3|+|37969413|37969434&offsets|0|-2(+2U)]|TTCAAGTAATCCAGGATAGGTT</td><td>>MI0000750|hsa-mir-26a-2&WithFlank&12|-|57824603|57824698</td><td>20</td><td>39</td><td>20(+2U)</td><td>TTCAAGTAATCCAGGATAGGTT</td><td>265.2</td><td>5.1</td><td>0.8</td><td>6.7</td><td>2.3E-11</td><td>6.3E-11</td><td>MB231/MB468</td><td>NUC</td></tr>
<tr><td>>hg38_21_+_8433172_8446622_RNA45SN1_45S_50ntFlanks@10874.10894.21|TGGGTCGGTCGGGCTGGGGCG</td><td>>hg38_21_+_8433172_8446622_RNA45SN1_45S_50ntFlanks</td><td>10874</td><td>10894</td><td>21</td><td>TGGGTCGGTCGGGCTGGGGCG</td><td>101.6</td><td>-10.5</td><td>1.6</td><td>-6.7</td><td>2.1E-11</td><td>6.3E-11</td><td>MB231/MB468</td><td>MP</td></tr>
<tr><td>>hg38_21_+_8433172_8446622_RNA45SN1_45S_50ntFlanks@12126.12144.19|CGACCCGCTCCGGGGACAG</td><td>>hg38_21_+_8433172_8446622_RNA45SN1_45S_50ntFlanks</td><td>12126</td><td>12144</td><td>19</td><td>CGACCCGCTCCGGGGACAG</td><td>128.8</td><td>-10.9</td><td>1.6</td><td>-6.7</td><td>2.1E-11</td><td>6.3E-11</td><td>MB231/MB468</td><td>MP</td></tr>
<tr><td>>hg38_21_+_8433172_8446622_RNA45SN1_45S_50ntFlanks@10128.10157.30|CGGCGGCGGCGGCGGCGGCGGGGGTGTGGG</td><td>>hg38_21_+_8433172_8446622_RNA45SN1_45S_50ntFlanks</td><td>10128</td><td>10157</td><td>30</td><td>CGGCGGCGGCGGCGGCGGCGGGGGTGTGGG</td><td>143.3</td><td>-11.0</td><td>1.6</td><td>-6.7</td><td>2.1E-11</td><td>6.3E-11</td><td>MB231/MB468</td><td>MP</td></tr>
<tr><td>>hg38_21_+_8433172_8446622_RNA45SN1_45S_50ntFlanks@4129.4158.30|GATTCCGGAGAGGGAGCCTGAGAAACGGCT</td><td>>hg38_21_+_8433172_8446622_RNA45SN1_45S_50ntFlanks</td><td>4129</td><td>4158</td><td>30</td><td>GATTCCGGAGAGGGAGCCTGAGAAACGGCT</td><td>109.0</td><td>-10.6</td><td>1.6</td><td>-6.7</td><td>2.1E-11</td><td>6.3E-11</td><td>MB231/MB468</td><td>MP</td></tr>
<tr><td>>trna17_ValTAC_11_-_59318102_59318174@1.34.34,trna4_ValTAC_X_-_18693029_18693101@1.34.34|GGTTCCATAGTGTAGTGGTTATCACGTCTGCTTT</td><td>>trna17_ValTAC_11_-_59318102_59318174</td><td>1</td><td>34</td><td>34</td><td>GGTTCCATAGTGTAGTGGTTATCACGTCTGCTTT</td><td>247.5</td><td>11.8</td><td>1.8</td><td>6.7</td><td>2.3E-11</td><td>6.3E-11</td><td>MB231/MB468</td><td>NUC</td></tr>
<tr><td>>hg38_21_+_8433172_8446622_RNA45SN1_45S_50ntFlanks@9352.9377.26|CCGTGGCGCAATGAAGGTGAAGGCCG</td><td>>hg38_21_+_8433172_8446622_RNA45SN1_45S_50ntFlanks</td><td>9352</td><td>9377</td><td>26</td><td>CCGTGGCGCAATGAAGGTGAAGGCCG</td><td>256.4</td><td>-10.1</td><td>1.5</td><td>-6.7</td><td>1.6E-11</td><td>6.3E-11</td><td>BT20/MB468</td><td>MITO</td></tr>
<tr><td>>hg38_21_+_8433172_8446622_RNA45SN1_45S_50ntFlanks@6650.6681.32|ACGACTCTTAGCGGTGGATCACTCGGCTCGTG</td><td>>hg38_21_+_8433172_8446622_RNA45SN1_45S_50ntFlanks</td><td>6650</td><td>6681</td><td>32</td><td>ACGACTCTTAGCGGTGGATCACTCGGCTCGTG</td><td>4469.6</td><td>-3.0</td><td>0.5</td><td>-6.7</td><td>2.8E-11</td><td>6.3E-11</td><td>BT20/MB468</td><td>NUC</td></tr>
<tr><td>>hg38_21_+_8433172_8446622_RNA45SN1_45S_50ntFlanks@4444.4471.28|CCTTGCCTCTCGGCGCCCCCTCGATGCT</td><td>>hg38_21_+_8433172_8446622_RNA45SN1_45S_50ntFlanks</td><td>4444</td><td>4471</td><td>28</td><td>CCTTGCCTCTCGGCGCCCCCTCGATGCT</td><td>339.3</td><td>-5.3</td><td>0.8</td><td>-6.7</td><td>2.3E-11</td><td>6.4E-11</td><td>MB231/MB468</td><td>CYTO</td></tr>
</table>

| | | | | | | | | | | | | | |
|---|---|---|---|---|---|---|---|---|---|---|---|---|---|
| >hg38_21_+_8433172_8446622_RNA45SN1_45S_50ntFlanks@9556.9573.18\|TAGCGGTCCTGACGTGCA | >hg38_21_+_8433172_8446622_RNA45SN1_45S_50ntFlanks | 9556 | 9573 | 18 | TAGCGGTCCTGACGTGCA | 316.0 | -11.4 | 1.6 | -7.1 | 1.4E-12 | 6.4E-11 | BT20/MB468 | MP |
| >hg38_21_+_8433172_8446622_RNA45SN1_45S_50ntFlanks@3705.3729.25\|TACCTGGTTGATCCTGCCAGTAGCA | >hg38_21_+_8433172_8446622_RNA45SN1_45S_50ntFlanks | 3705 | 3729 | 25 | TACCTGGTTGATCCTGCCAGTAGCA | 593.8 | 8.9 | 1.3 | 6.7 | 2.1E-11 | 6.4E-11 | BT20/MB468 | CYTO |
| >MI0000298\|hsa-mir-221&WithFlank&X\|-\|45746151\|45746272@31.55.25(+1C)&[MIMAT0004568&hsa-miR-221-5p&offsets\|0\|+3(+1C);m-405&X\|-\|45746220\|45746242&offsets\|0\|+2(+1C)]\|ACCTGGCATACAATGTAGATTTCTGC | >MI0000298\|hsa-mir-221&WithFlank&X\|-\|45746151\|45746272 | 31 | 55 | 25(+1C) | ACCTGGCATACAATGTAGATTTCTGC | 376.3 | 7.0 | 1.1 | 6.7 | 2.3E-11 | 6.4E-11 | MB231/MB468 | CYTO |
| >hg38_21_+_8433172_8446622_RNA45SN1_45S_50ntFlanks@12974.13001.28\|CGTACGTAGCAGAGCAGCTCCCTCGCTG | >hg38_21_+_8433172_8446622_RNA45SN1_45S_50ntFlanks | 12974 | 13001 | 28 | CGTACGTAGCAGAGCAGCTCCCTCGCTG | 139.8 | -9.2 | 1.4 | -6.7 | 1.7E-11 | 6.4E-11 | BT20/MB468 | MITO |
| >hg38_21_+_8433172_8446622_RNA45SN1_45S_50ntFlanks@10681.10706.26\|TTCCGGCGGCGTCCGGTGAGCTCTCG | >hg38_21_+_8433172_8446622_RNA45SN1_45S_50ntFlanks | 10681 | 10706 | 26 | TTCCGGCGGCGTCCGGTGAGCTCTCG | 2703.5 | -5.3 | 0.8 | -6.7 | 2.1E-11 | 6.4E-11 | BT20/MB468 | CYTO |
| >MI0000078\|hsa-mir-22&WithFlank&17\|-\|1713897\|1713993@59.78.20&[MIMAT0000077&hsa-miR-22-3p&offsets\|0\|-2;m-2&17\|-\|1713914\|1713935&offsets\|0\|-2]\|AAGCTGCCAGTTGAAGAACT | >MI0000078\|hsa-mir-22&WithFlank&17\|-\|1713897\|1713993 | 59 | 78 | 20 | AAGCTGCCAGTTGAAGAACT | 167.5 | 3.7 | 0.6 | 6.7 | 2.8E-11 | 6.4E-11 | BT20/MB468 | NUC |
| >hg38_21_+_8433172_8446622_RNA45SN1_45S_50ntFlanks@10841.10873.33\|CGTAACTTCGGGATAAGGATTGGCTCTAAGGGC | >hg38_21_+_8433172_8446622_RNA45SN1_45S_50ntFlanks | 10841 | 10873 | 33 | CGTAACTTCGGGATAAGGATTGGCTCTAAGGGC | 2311.1 | -8.2 | 1.2 | -6.7 | 2.8E-11 | 6.4E-11 | BT20/MB468 | NUC |
| >hg38_21_+_8433172_8446622_RNA45SN1_45S_50ntFlanks@4382.4408.27\|TAGTTGGATCTTGGGAGCGGGCGGGCG | >hg38_21_+_8433172_8446622_RNA45SN1_45S_50ntFlanks | 4382 | 4408 | 27 | TAGTTGGATCTTGGGAGCGGGCGGGCG | 89.4 | -10.3 | 1.5 | -6.7 | 2.1E-11 | 6.4E-11 | MB231/MB468 | MP |
| >hg38_21_+_8433172_8446622_RNA45SN1_45S_50ntFlanks@10821.10840.20\|GGAAGTCGGCAAGCCGGATC | >hg38_21_+_8433172_8446622_RNA45SN1_45S_50ntFlanks | 10821 | 10840 | 20 | GGAAGTCGGCAAGCCGGATC | 1806.7 | -7.2 | 1.1 | -6.7 | 2.7E-11 | 6.4E-11 | MB231/MB468 | MITO |
| >hg38_21_+_8433172_8446622_RNA45SN1_45S_50ntFlanks@12191.12221.31\|ACGCAGGTGTCCTAAGGCGAGCTCAGGGAGG | >hg38_21_+_8433172_8446622_RNA45SN1_45S_50ntFlanks | 12191 | 12221 | 31 | ACGCAGGTGTCCTAAGGCGAGCTCAGGGAGG | 94.1 | -10.4 | 1.6 | -6.7 | 2.1E-11 | 6.4E-11 | MB231/MB468 | MP |
| >hg38_21_+_8433172_8446622_RNA45SN1_45S_50ntFlanks@10375.10406.32\|AGAAGGGTTCCATGTGAACAGCAGTTGAACAT | >hg38_21_+_8433172_8446622_RNA45SN1_45S_50ntFlanks | 10375 | 10406 | 32 | AGAAGGGTTCCATGTGAACAGCAGTTGAACAT | 522.7 | -7.8 | 1.2 | -6.7 | 2.3E-11 | 6.4E-11 | MB231/MB468 | NUC |
| >hg38_21_+_8433172_8446622_RNA45SN1_45S_50ntFlanks@4917.4941.25\|CACCACCAGGAGTGGAGCCTGCGGC | >hg38_21_+_8433172_8446622_RNA45SN1_45S_50ntFlanks | 4917 | 4941 | 25 | CACCACCAGGAGTGGAGCCTGCGGC | 662.3 | -9.3 | 1.4 | -6.7 | 2.7E-11 | 6.4E-11 | MB231/MB468 | MITO |
| >hg38_21_+_8433172_8446622_RNA45SN1_45S_50ntFlanks@4657.4686.30\|CTAGAGGTGAAATTCTTGGACCGGCGCAAG | >hg38_21_+_8433172_8446622_RNA45SN1_45S_50ntFlanks | 4657 | 4686 | 30 | CTAGAGGTGAAATTCTTGGACCGGCGCAAG | 278.6 | -7.4 | 1.1 | -6.7 | 2.3E-11 | 6.5E-11 | MB231/MB468 | CYTO |
| >hg38_21_+_8433172_8446622_RNA45SN1_45S_50ntFlanks@4758.4780.23\|GATCAGATACCGTCGTAGTTCCG | >hg38_21_+_8433172_8446622_RNA45SN1_45S_50ntFlanks | 4758 | 4780 | 23 | GATCAGATACCGTCGTAGTTCCG | 103.8 | -9.0 | 1.3 | -6.7 | 2.1E-11 | 6.5E-11 | BT20/MB468 | CYTO |
| >hg38_21_+_8433172_8446622_RNA45SN1_45S_50ntFlanks@10664.10710.47\|CTTGGAAAGCGTCGCGGTTCCGGCGGCGTCCGGTGAGCTCTCGCTGG | >hg38_21_+_8433172_8446622_RNA45SN1_45S_50ntFlanks | 10664 | 10710 | 47 | CTTGGAAAGCGTCGCGGTTCCGGCGGCGTCCGGTGAGCTCTCGCTGG | 421.0 | -10.8 | 1.6 | -6.7 | 1.7E-11 | 6.5E-11 | BT20/MB468 | MITO |
| >hg38_21_+_8433172_8446622_RNA45SN1_45S_50ntFlanks@9381.9410.30\|CGCTCGCCGGCCGAGGTGGGATCCCGAGGC | >hg38_21_+_8433172_8446622_RNA45SN1_45S_50ntFlanks | 9381 | 9410 | 30 | CGCTCGCCGGCCGAGGTGGGATCCCGAGGC | 115.3 | -10.7 | 1.6 | -6.7 | 2.1E-11 | 6.5E-11 | MB231/MB468 | MP |
| >trna152_ValCAC_6_-_27248049_27248121@1.33.33\|GCTTCTGTAGTGTAGTGGTTATCACGTTCGCCT | >trna152_ValCAC_6_-_27248049_27248121 | 1 | 33 | 33 | GCTTCTGTAGTGTAGTGGTTATCACGTTCGCCT | 9478.7 | 11.5 | 1.7 | 6.7 | 2.3E-11 | 6.5E-11 | MB231/MB468 | CYTO |
| >hg38_21_+_8433172_8446622_RNA45SN1_45S_50ntFlanks@6661.6700.40\|CGGTGGATCACTCGGCTCGTGCGTCGATGAAGAACGCAGC | >hg38_21_+_8433172_8446622_RNA45SN1_45S_50ntFlanks | 6661 | 6700 | 40 | CGGTGGATCACTCGGCTCGTGCGTCGATGAAGAACGCAGC | 7099.4 | 7.2 | 1.0 | 7.1 | 1.5E-12 | 6.5E-11 | BT20/MB468 | MP |
| >hg38_21_+_8433172_8446622_RNA45SN1_45S_50ntFlanks@11161.11188.28\|CACGGTCCCCCGCGAGGGGGGCCCGGGC | >hg38_21_+_8433172_8446622_RNA45SN1_45S_50ntFlanks | 11161 | 11188 | 28 | CACGGTCCCCCGCGAGGGGGGCCCGGGC | 643.9 | -12.4 | 1.8 | -7.1 | 1.5E-12 | 6.5E-11 | BT20/MB468 | MP |
| >hg38_21_+_8433172_8446622_RNA45SN1_45S_50ntFlanks@6689.6706.18\|GAAGAACGCAGCTAGCTG | >hg38_21_+_8433172_8446622_RNA45SN1_45S_50ntFlanks | 6689 | 6706 | 18 | GAAGAACGCAGCTAGCTG | 117.0 | -10.7 | 1.6 | -6.7 | 2.1E-11 | 6.5E-11 | MB231/MB468 | MP |
| >hg38_21_+_8433172_8446622_RNA45SN1_45S_50ntFlanks@8358.8379.22\|AAAGAACTTTGAAGAGAGAGTT | >hg38_21_+_8433172_8446622_RNA45SN1_45S_50ntFlanks | 8358 | 8379 | 22 | AAAGAACTTTGAAGAGAGAGTT | 1467.2 | -9.5 | 1.4 | -6.7 | 2.1E-11 | 6.5E-11 | MB231/MB468 | MP |
| >hg38_21_+_8433172_8446622_RNA45SN1_45S_50ntFlanks@9347.9370.24\|AGCCGCCGTGGCGCAATGAAGGTG | >hg38_21_+_8433172_8446622_RNA45SN1_45S_50ntFlanks | 9347 | 9370 | 24 | AGCCGCCGTGGCGCAATGAAGGTG | 99.2 | -10.5 | 1.6 | -6.7 | 2.1E-11 | 6.5E-11 | MB231/MB468 | MP |
| >hg38_21_+_8433172_8446622_RNA45SN1_45S_50ntFlanks@10350.10374.25\|ACGAGAACTTTGAAGGCCGAAGTGG | >hg38_21_+_8433172_8446622_RNA45SN1_45S_50ntFlanks | 10350 | 10374 | 25 | ACGAGAACTTTGAAGGCCGAAGTGG | 1933.0 | -4.1 | 0.6 | -6.7 | 2.3E-11 | 6.5E-11 | MB231/MB468 | CYTO |
| >hg38_21_+_8433172_8446622_RNA45SN1_45S_50ntFlanks@8552.8581.30\|CTCCCCGGAGGGGGCGGGCTCCGGCGGGTG | >hg38_21_+_8433172_8446622_RNA45SN1_45S_50ntFlanks | 8552 | 8581 | 30 | CTCCCCGGAGGGGGCGGGCTCCGGCGGGTG | 202.8 | -11.5 | 1.7 | -6.7 | 2.1E-11 | 6.5E-11 | MB231/MB468 | MP |
| >hg38_21_+_8433172_8446622_RNA45SN1_45S_50ntFlanks@11612.11660.49\|TAATTAAAACAAAGCATCGCGAAGGCCCGCGGCGGGTGTTGACGCGATG | >hg38_21_+_8433172_8446622_RNA45SN1_45S_50ntFlanks | 11612 | 11660 | 49 | TAATTAAAACAAAGCATCGCGAAGGCCCGCGGCGGGTGTTGACGCGATG | 244.7 | -11.7 | 1.8 | -6.7 | 2.7E-11 | 6.5E-11 | MB231/MB468 | MITO |
| >hg38_21_+_8433172_8446622_RNA45SN1_45S_50ntFlanks@10683.10712.30\|CCGGCGGCGTCCGGTGAGCTCTCGCTGGCC | >hg38_21_+_8433172_8446622_RNA45SN1_45S_50ntFlanks | 10683 | 10712 | 30 | CCGGCGGCGTCCGGTGAGCTCTCGCTGGCC | 4749.5 | -8.9 | 1.3 | -6.7 | 2.9E-11 | 6.5E-11 | BT20/MB468 | NUC |
| >trna16_TyrGTA_14_-_21131351_21131444@59.76.18,trna17_TyrGTA_14_-_21128117_21128210@59.76.18,trna19_TyrGTA_14_-_21121258_21121351@59.76.18,trna2_TyrGTA_2_+_27273650_27273738@59.76.18,trna4_TyrGTA_8_+_67025602_67025694@59.76.18,trna5_TyrGTA_14_+_21151432_21151520@59.76.18,trna5_TyrGTA_8_+_67026223_67026311@59.76.18\|TTCCGGCTCGAAGGACCA | >trna16_TyrGTA_14_-_21131351_21131444 | 59 | 76 | 18 | TTCCGGCTCGAAGGACCA | 464.9 | 4.6 | 0.7 | 6.7 | 2.3E-11 | 6.5E-11 | MB231/MB468 | NUC |
| >MI0000734\|hsa-mir-106b&WithFlank&7\|-\|100093987\|100094080@56.77.22&[MIMAT0004672&hsa-miR-106b-3p&offsets\|-2\|-2;m-87&7\|-\|100094003\|100094023&offsets\|-2\|-1]\|TACCGCACTGTGGGTACTTGCT | >MI0000734\|hsa-mir-106b&WithFlank&7\|-\|100093987\|100094080 | 56 | 77 | 22 | TACCGCACTGTGGGTACTTGCT | 243.4 | 4.3 | 0.7 | 6.7 | 2.9E-11 | 6.5E-11 | BT20/MB468 | NUC |
| >hg38_21_+_8433172_8446622_RNA45SN1_45S_50ntFlanks@10817.10840.24\|TAAGGGAAGTCGGCAAGCCGGATC | >hg38_21_+_8433172_8446622_RNA45SN1_45S_50ntFlanks | 10817 | 10840 | 24 | TAAGGGAAGTCGGCAAGCCGGATC | 314.4 | -3.2 | 0.5 | -6.7 | 2.3E-11 | 6.5E-11 | MB231/MB468 | CYTO |
| >hg38_21_+_8433172_8446622_RNA45SN1_45S_50ntFlanks@6739.6771.33\|TCATCGACACTTCGAACGCACTTGCGGCCCCGG | >hg38_21_+_8433172_8446622_RNA45SN1_45S_50ntFlanks | 6739 | 6771 | 33 | TCATCGACACTTCGAACGCACTTGCGGCCCCGG | 158.1 | -5.2 | 0.8 | -6.7 | 2.3E-11 | 6.5E-11 | MB231/MB468 | CYTO |
| >trna10_AspGTC_12_-_125424193_125424264@32.71.40,trna12_AspGTC_12_-_125411891_125411962@32.71.40,trna144_AspGTC_6_-_27551236_27551307@32.71.40,trna38_AspGTC_17_-_8125556_8125627@32.71.40,trna45_AspGTC_6_+_27447453_27447524@32.71.40,trna48_AspGTC_6_+_27471523_27471594@32.71.40,trna4_AspGTC_12_+_96429799_96429870@32.71.40,trna69_AspGTC_1_-_161440205_161440276@32.71.40,trna72_AspGTC_1_-_161432824_161432895@32.71.40,trna75_AspGTC_1_-_161425414_161425485@32.71.40,trna78_AspGTC_1_-_161418033_161418104@32.71.40,trna81_AspGTC_1_-_161410615_161410686@32.71.40\|CTGTCACGCGGGAGACCGGGGTTCGATTCCCCGACGGGGA | >trna10_AspGTC_12_-_125424193_125424264 | 32 | 71 | 40 | CTGTCACGCGGGAGACCGGGGTTCGATTCCCCGACGGGGA | 140.3 | -11.0 | 1.6 | -6.7 | 2.1E-11 | 6.5E-11 | MB231/MB468 | MP |
| >hg38_MT_+_1621_3279_RNR2_16S_50ntFlanks@51.87.37\|GCTAAACCTAGCCCCAAACCCACTCCACCTTACTACC | >hg38_MT_+_1621_3279_RNR2_16S_50ntFlanks | 51 | 87 | 37 | GCTAAACCTAGCCCCAAACCCACTCCACCTTACTACC | 975.4 | 5.9 | 0.9 | 6.7 | 2.3E-11 | 6.5E-11 | MB231/MB468 | NUC |
| >am_trnaMT_PheGAA_MT_+_577_647@41.61.21\|GTTTAGACGGGCTCACATCAC | >am_trnaMT_PheGAA_MT_+_577_647 | 41 | 61 | 21 | GTTTAGACGGGCTCACATCAC | 239.9 | 9.6 | 1.4 | 6.7 | 2.1E-11 | 6.5E-11 | BT20/MB468 | CYTO |
| >hg38_21_+_8433172_8446622_RNA45SN1_45S_50ntFlanks@10408.10434.27\|GGTCAGTCGGTCCTGAGAGATGGGCGA | >hg38_21_+_8433172_8446622_RNA45SN1_45S_50ntFlanks | 10408 | 10434 | 27 | GGTCAGTCGGTCCTGAGAGATGGGCGA | 732.6 | -9.7 | 1.5 | -6.7 | 2.9E-11 | 6.6E-11 | BT20/MB468 | NUC |
| >hg38_21_+_8433172_8446622_RNA45SN1_45S_50ntFlanks@12957.12986.30\|CTTCTGGGTCGGGGTTTCGTACGTAGCAGA | >hg38_21_+_8433172_8446622_RNA45SN1_45S_50ntFlanks | 12957 | 12986 | 30 | CTTCTGGGTCGGGGTTTCGTACGTAGCAGA | 147.9 | -11.1 | 1.7 | -6.7 | 2.2E-11 | 6.6E-11 | MB231/MB468 | MP |
| >hg38_1_-_228634819_228635039_RNA5S12_5S_50ntFlanks@126.169.44\|TACTTGGATGGGAGACCGCCTGGGAATACCGGGTGCTGTAGGCT | >hg38_1_-_228634819_228635039_RNA5S12_5S_50ntFlanks | 126 | 169 | 44 | TACTTGGATGGGAGACCGCCTGGGAATACCGGGTGCTGTAGGCT | 904.7 | 7.3 | 1.1 | 6.7 | 2.3E-11 | 6.6E-11 | MB231/MB468 | CYTO |
| >hg38_21_+_8433172_8446622_RNA45SN1_45S_50ntFlanks@5421.5443.23\|CGATTGGATGGTTTAGTGAGGCC | >hg38_21_+_8433172_8446622_RNA45SN1_45S_50ntFlanks | 5421 | 5443 | 23 | CGATTGGATGGTTTAGTGAGGCC | 1698.4 | -10.2 | 1.5 | -6.7 | 2.9E-11 | 6.6E-11 | BT20/MB468 | NUC |
| >hg38_21_+_8433172_8446622_RNA45SN1_45S_50ntFlanks@7974.8004.31\|ACGCGACCTCAGATCAGACGTGGCGACCCGC | >hg38_21_+_8433172_8446622_RNA45SN1_45S_50ntFlanks | 7974 | 8004 | 31 | ACGCGACCTCAGATCAGACGTGGCGACCCGC | 5067.2 | 8.0 | 1.2 | 6.7 | 2.1E-11 | 6.6E-11 | BT20/MB468 | CYTO |
| >hg38_21_+_8433172_8446622_RNA45SN1_45S_50ntFlanks@10393.10429.37\|CAGCAGTTGAACATGGGTCAGTCGGTCCTGAGAGATG | >hg38_21_+_8433172_8446622_RNA45SN1_45S_50ntFlanks | 10393 | 10429 | 37 | CAGCAGTTGAACATGGGTCAGTCGGTCCTGAGAGATG | 1540.7 | -6.3 | 0.9 | -6.7 | 2.2E-11 | 6.6E-11 | MB231/MB468 | MP |
| >am_trnaMT_ValTAC_MT_+_1602_1670@48.67.20\|CAACTTAACTTGACCGCTCT | >am_trnaMT_ValTAC_MT_+_1602_1670 | 48 | 67 | 20 | CAACTTAACTTGACCGCTCT | 4297.6 | 7.4 | 1.1 | 6.7 | 1.7E-11 | 6.6E-11 | BT20/MB468 | MITO |
| >hg38_21_+_8433172_8446622_RNA45SN1_45S_50ntFlanks@9032.9060.29\|CCTCGTGGGGGGGCCGGGCCACCCCTCCC | >hg38_21_+_8433172_8446622_RNA45SN1_45S_50ntFlanks | 9032 | 9060 | 29 | CCTCGTGGGGGGGCCGGGCCACCCCTCCC | 92.8 | -10.4 | 1.6 | -6.7 | 2.2E-11 | 6.6E-11 | MB231/MB468 | MP |
| >MI0000750\|hsa-mir-26a-2&WithFlank&12\|-\|57824603\|57824698@20.41.22(+2U);MI0000083\|hsa-mir-26a-1&WithFlank&3\|+\|37969398\|37969486@16.37.22(+2U)&[MIMAT0000082&hsa-miR-26a-5p&offsets\|0\|0(+2U);m-33&12\|-\|57824658\|57824679&offsets\|0\|0(+2U)];[MIMAT0000082_1&hsa-miR-26a-5p&offsets\|0\|0(+2U);m-34&3\|+\|37969413\|37969434&offsets\|0\|0(+2U)]\|TTCAAGTAATCCAGGATAGGCTTT | >MI0000750\|hsa-mir-26a-2&WithFlank&12\|-\|57824603\|57824698 | 20 | 41 | 22(+2U) | TTCAAGTAATCCAGGATAGGCTTT | 1106.5 | 6.5 | 1.0 | 6.7 | 1.7E-11 | 6.6E-11 | BT20/MB468 | MITO |
| >hg38_21_+_8433172_8446622_RNA45SN1_45S_50ntFlanks@11874.11903.30\|AGAAGACCCTGTTGAGCTTGACTCTAGTCT | >hg38_21_+_8433172_8446622_RNA45SN1_45S_50ntFlanks | 11874 | 11903 | 30 | AGAAGACCCTGTTGAGCTTGACTCTAGTCT | 205.3 | -6.5 | 1.0 | -6.7 | 2.4E-11 | 6.6E-11 | MB231/MB468 | CYTO |
| >hg38_21_+_8433172_8446622_RNA45SN1_45S_50ntFlanks@6652.6670.19\|GACTCTTAGCGGTGGATCA | >hg38_21_+_8433172_8446622_RNA45SN1_45S_50ntFlanks | 6652 | 6670 | 19 | GACTCTTAGCGGTGGATCA | 107.6 | -10.6 | 1.6 | -6.7 | 2.2E-11 | 6.6E-11 | MB231/MB468 | MP |
| >hg38_21_+_8433172_8446622_RNA45SN1_45S_50ntFlanks@10291.10317.27\|CGCGGGCCCGGGTGGAGCCGCCGCAGG | >hg38_21_+_8433172_8446622_RNA45SN1_45S_50ntFlanks | 10291 | 10317 | 27 | CGCGGGCCCGGGTGGAGCCGCCGCAGG | 260.5 | -11.9 | 1.8 | -6.7 | 2.2E-11 | 6.6E-11 | MB231/MB468 | MP |

<table>
<tr><td>>hg38_21_+_8433172_8446622_RNA45SN1_45S_50ntFlanks@4679.4707.29|GGCGCAAGACGGACCAGAGCGAAAGCATT</td><td>>hg38_21_+_8433172_8446622_RNA45SN1_45S_50ntFlanks</td><td>4679</td><td>4707</td><td>29</td><td>GGCGCAAGACGGACCAGAGCGAAAGCATT</td><td>252.2</td><td>8.1</td><td>1.2</td><td>6.7</td><td>2.2E-11</td><td>6.6E-11</td><td>BT20/MB468</td><td>CYTO</td></tr>
<tr><td>>MI0002468|hsa-mir-484&WithFlank&16|+|15643288|15643378@14.35.22&[MIMAT0002174&hsa-miR-484&offsets|0|0;m-127&16|+|15643301|15643321&offsets|0|+1]|TCAGGCTCAGTCCCCTCCCGAT</td><td>>MI0002468|hsa-mir-484&WithFlank&16|+|15643288|15643378</td><td>14</td><td>35</td><td>22</td><td>TCAGGCTCAGTCCCCTCCCGAT</td><td>952.3</td><td>5.8</td><td>0.9</td><td>6.7</td><td>2.4E-11</td><td>6.6E-11</td><td>MB231/MB468</td><td>CYTO</td></tr>
<tr><td>>hg38_21_+_8433172_8446622_RNA45SN1_45S_50ntFlanks@8354.8391.38|TTGAAAAGAACTTTGAAGAGAGAGTTCAAGAGGGCGTG</td><td>>hg38_21_+_8433172_8446622_RNA45SN1_45S_50ntFlanks</td><td>8354</td><td>8391</td><td>38</td><td>TTGAAAAGAACTTTGAAGAGAGAGTTCAAGAGGGCGTG</td><td>193.0</td><td>11.3</td><td>1.6</td><td>7.1</td><td>1.5E-12</td><td>6.6E-11</td><td>BT20/MB468</td><td>MP</td></tr>
<tr><td>>hg38_21_+_8433172_8446622_RNA45SN1_45S_50ntFlanks@3782.3812.31|CAGTGAAACTGCGAATGGCTCATTAAATCAG</td><td>>hg38_21_+_8433172_8446622_RNA45SN1_45S_50ntFlanks</td><td>3782</td><td>3812</td><td>31</td><td>CAGTGAAACTGCGAATGGCTCATTAAATCAG</td><td>397.5</td><td>-10.7</td><td>1.6</td><td>-6.7</td><td>1.7E-11</td><td>6.7E-11</td><td>BT20/MB468</td><td>MITO</td></tr>
<tr><td>>hg38_21_+_8433172_8446622_RNA45SN1_45S_50ntFlanks@12615.12652.38|GCTGAGGAGCCAATGGGGCGAAGCTACCATCTGTGGGA</td><td>>hg38_21_+_8433172_8446622_RNA45SN1_45S_50ntFlanks</td><td>12615</td><td>12652</td><td>38</td><td>GCTGAGGAGCCAATGGGGCGAAGCTACCATCTGTGGGA</td><td>102.2</td><td>-10.5</td><td>1.6</td><td>-6.7</td><td>2.2E-11</td><td>6.7E-11</td><td>MB231/MB468</td><td>MP</td></tr>
<tr><td>>hg38_21_+_8433172_8446622_RNA45SN1_45S_50ntFlanks@4884.4914.31|CAAAGCTGAAACTTAAAGGAATTGACGGAAG</td><td>>hg38_21_+_8433172_8446622_RNA45SN1_45S_50ntFlanks</td><td>4884</td><td>4914</td><td>31</td><td>CAAAGCTGAAACTTAAAGGAATTGACGGAAG</td><td>98.6</td><td>-10.5</td><td>1.6</td><td>-6.7</td><td>2.2E-11</td><td>6.7E-11</td><td>MB231/MB468</td><td>MP</td></tr>
<tr><td>>hg38_21_+_8433172_8446622_RNA45SN1_45S_50ntFlanks@10726.10757.32|GGGAGAGGGTGTAAATCTCGCGCCGGGCCGTA</td><td>>hg38_21_+_8433172_8446622_RNA45SN1_45S_50ntFlanks</td><td>10726</td><td>10757</td><td>32</td><td>GGGAGAGGGTGTAAATCTCGCGCCGGGCCGTA</td><td>81.3</td><td>-10.2</td><td>1.5</td><td>-6.7</td><td>2.2E-11</td><td>6.7E-11</td><td>MB231/MB468</td><td>MP</td></tr>
<tr><td>>MI0000088|hsa-mir-30a&WithFlank&6|-|71403545|71403627@53.73.21(+1A)&[MIMAT0000088&hsa-miR-30a-3p&offsets|0|-1(+1A);m-38&6|-|71403554|71403575&offsets|0|-1(+1A)]|CTTTCAGTCGGATGTTTGCAGA</td><td>>MI0000088|hsa-mir-30a&WithFlank&6|-|71403545|71403627</td><td>53</td><td>73</td><td>21(+1A)</td><td>CTTTCAGTCGGATGTTTGCAGA</td><td>1038.7</td><td>7.7</td><td>1.2</td><td>6.7</td><td>2.4E-11</td><td>6.7E-11</td><td>MB231/MB468</td><td>CYTO</td></tr>
<tr><td>>hg38_21_+_8433172_8446622_RNA45SN1_45S_50ntFlanks@6653.6689.37|ACTCTTAGCGGTGGATCACTCGGCTCGTGCGTCGATG</td><td>>hg38_21_+_8433172_8446622_RNA45SN1_45S_50ntFlanks</td><td>6653</td><td>6689</td><td>37</td><td>ACTCTTAGCGGTGGATCACTCGGCTCGTGCGTCGATG</td><td>6952.1</td><td>-4.6</td><td>0.7</td><td>-6.7</td><td>2.8E-11</td><td>6.7E-11</td><td>MB231/MB468</td><td>MITO</td></tr>
<tr><td>>hg38_21_+_8433172_8446622_RNA45SN1_45S_50ntFlanks@12456.12473.18|CCAAGCGTTGGATTGTTC</td><td>>hg38_21_+_8433172_8446622_RNA45SN1_45S_50ntFlanks</td><td>12456</td><td>12473</td><td>18</td><td>CCAAGCGTTGGATTGTTC</td><td>2218.0</td><td>-6.2</td><td>0.9</td><td>-6.7</td><td>2.8E-11</td><td>6.7E-11</td><td>MB231/MB468</td><td>MITO</td></tr>
<tr><td>>hg38_21_+_8433172_8446622_RNA45SN1_45S_50ntFlanks@9032.9055.24|CCTCGTGGGGGGGCCGGGCCACCC</td><td>>hg38_21_+_8433172_8446622_RNA45SN1_45S_50ntFlanks</td><td>9032</td><td>9055</td><td>24</td><td>CCTCGTGGGGGGGCCGGGCCACCC</td><td>126.2</td><td>-10.8</td><td>1.6</td><td>-6.7</td><td>2.2E-11</td><td>6.7E-11</td><td>MB231/MB468</td><td>MP</td></tr>
<tr><td>>hg38_21_+_8433172_8446622_RNA45SN1_45S_50ntFlanks@9467.9488.22|GTGGAGCACGAGCGCACGTGTT</td><td>>hg38_21_+_8433172_8446622_RNA45SN1_45S_50ntFlanks</td><td>9467</td><td>9488</td><td>22</td><td>GTGGAGCACGAGCGCACGTGTT</td><td>267.9</td><td>-11.9</td><td>1.8</td><td>-6.7</td><td>2.2E-11</td><td>6.7E-11</td><td>MB231/MB468</td><td>MP</td></tr>
<tr><td>>hg38_21_+_8433172_8446622_RNA45SN1_45S_50ntFlanks@12259.12282.24|CTTGATCTTGATTTTCAGTACGAA</td><td>>hg38_21_+_8433172_8446622_RNA45SN1_45S_50ntFlanks</td><td>12259</td><td>12282</td><td>24</td><td>CTTGATCTTGATTTTCAGTACGAA</td><td>1213.0</td><td>-7.9</td><td>1.2</td><td>-6.7</td><td>2.8E-11</td><td>6.7E-11</td><td>MB231/MB468</td><td>MITO</td></tr>
<tr><td>>trna14_ThrCGT_17_+_29877093_29877164@58.75.18,trna15_ThrCGT_16_+_14379750_14379821@58.75.18|ACCCCGTCCGTGCCTCCA</td><td>>trna14_ThrCGT_17_+_29877093_29877164</td><td>58</td><td>75</td><td>18</td><td>ACCCCGTCCGTGCCTCCA</td><td>450.4</td><td>8.6</td><td>1.3</td><td>6.7</td><td>2.2E-11</td><td>6.7E-11</td><td>BT20/MB468</td><td>CYTO</td></tr>
<tr><td>>hg38_21_+_8433172_8446622_RNA45SN1_45S_50ntFlanks@4243.4273.31|CTTTCGAGGCCCTGTAATTGGAATGAGTCCA</td><td>>hg38_21_+_8433172_8446622_RNA45SN1_45S_50ntFlanks</td><td>4243</td><td>4273</td><td>31</td><td>CTTTCGAGGCCCTGTAATTGGAATGAGTCCA</td><td>121.2</td><td>-10.8</td><td>1.6</td><td>-6.7</td><td>2.2E-11</td><td>6.7E-11</td><td>MB231/MB468</td><td>MP</td></tr>
<tr><td>>hg38_21_+_8433172_8446622_RNA45SN1_45S_50ntFlanks@4882.4903.22|TGCAAAGCTGAAACTTAAAGGA</td><td>>hg38_21_+_8433172_8446622_RNA45SN1_45S_50ntFlanks</td><td>4882</td><td>4903</td><td>22</td><td>TGCAAAGCTGAAACTTAAAGGA</td><td>709.2</td><td>9.0</td><td>1.4</td><td>6.7</td><td>2.4E-11</td><td>6.7E-11</td><td>MB231/MB468</td><td>CYTO</td></tr>
<tr><td>>hg38_21_+_8433172_8446622_RNA45SN1_45S_50ntFlanks@9343.9368.26|CGAAAGCCGCCGTGGCGCAATGAAGG</td><td>>hg38_21_+_8433172_8446622_RNA45SN1_45S_50ntFlanks</td><td>9343</td><td>9368</td><td>26</td><td>CGAAAGCCGCCGTGGCGCAATGAAGG</td><td>102.5</td><td>-10.5</td><td>1.6</td><td>-6.7</td><td>2.2E-11</td><td>6.7E-11</td><td>MB231/MB468</td><td>MP</td></tr>
<tr><td>>MI0000440|hsa-mir-27b&WithFlank&9|+|95085439|95085547@67.86.20&[MIMAT0000419&hsa-miR-27b-3p&offsets|0|-1;m-39&9|+|95085505|95085525&offsets|0|-1]|TTCACAGTGGCTAAGTTCTG</td><td>>MI0000440|hsa-mir-27b&WithFlank&9|+|95085439|95085547</td><td>67</td><td>86</td><td>20</td><td>TTCACAGTGGCTAAGTTCTG</td><td>4391.3</td><td>5.9</td><td>0.9</td><td>6.7</td><td>2.4E-11</td><td>6.7E-11</td><td>MB231/MB468</td><td>CYTO</td></tr>
<tr><td>>hg38_21_+_8433172_8446622_RNA45SN1_45S_50ntFlanks@10410.10435.26|TCAGTCGGTCCTGAGAGATGGGCGAG</td><td>>hg38_21_+_8433172_8446622_RNA45SN1_45S_50ntFlanks</td><td>10410</td><td>10435</td><td>26</td><td>TCAGTCGGTCCTGAGAGATGGGCGAG</td><td>259.5</td><td>-10.1</td><td>1.5</td><td>-6.7</td><td>1.8E-11</td><td>6.8E-11</td><td>BT20/MB468</td><td>MITO</td></tr>
<tr><td>>hg38_21_+_8433172_8446622_RNA45SN1_45S_50ntFlanks@9306.9323.18|CAAGGAGTCTAACACGTG</td><td>>hg38_21_+_8433172_8446622_RNA45SN1_45S_50ntFlanks</td><td>9306</td><td>9323</td><td>18</td><td>CAAGGAGTCTAACACGTG</td><td>143.3</td><td>-6.4</td><td>1.0</td><td>-6.7</td><td>2.4E-11</td><td>6.8E-11</td><td>MB231/MB468</td><td>CYTO</td></tr>
<tr><td>>trnaMT_PheGAA_MT_+_577_647@39.59.21|ATGTTTAGACGGGCTCACATC</td><td>>trnaMT_PheGAA_MT_+_577_647</td><td>39</td><td>59</td><td>21</td><td>ATGTTTAGACGGGCTCACATC</td><td>187.0</td><td>5.5</td><td>0.8</td><td>6.7</td><td>2.4E-11</td><td>6.8E-11</td><td>MB231/MB468</td><td>NUC</td></tr>
<tr><td>>hg38_21_+_8433172_8446622_RNA45SN1_45S_50ntFlanks@11768.11803.36|CTAATTAGTGACGCGCATGAATGGATGAACGAGATT</td><td>>hg38_21_+_8433172_8446622_RNA45SN1_45S_50ntFlanks</td><td>11768</td><td>11803</td><td>36</td><td>CTAATTAGTGACGCGCATGAATGGATGAACGAGATT</td><td>2203.9</td><td>-7.3</td><td>1.1</td><td>-6.7</td><td>2.4E-11</td><td>6.8E-11</td><td>MB231/MB468</td><td>NUC</td></tr>
<tr><td>>hg38_21_+_8433172_8446622_RNA45SN1_45S_50ntFlanks@10795.10816.22|TGGCATGTTGGAACAATGTAGG</td><td>>hg38_21_+_8433172_8446622_RNA45SN1_45S_50ntFlanks</td><td>10795</td><td>10816</td><td>22</td><td>TGGCATGTTGGAACAATGTAGG</td><td>78.8</td><td>-10.2</td><td>1.5</td><td>-6.7</td><td>2.2E-11</td><td>6.8E-11</td><td>MB231/MB468</td><td>MP</td></tr>
<tr><td>>hg38_21_+_8433172_8446622_RNA45SN1_45S_50ntFlanks@4243.4275.33|CTTTCGAGGCCCTGTAATTGGAATGAGTCCACT</td><td>>hg38_21_+_8433172_8446622_RNA45SN1_45S_50ntFlanks</td><td>4243</td><td>4275</td><td>33</td><td>CTTTCGAGGCCCTGTAATTGGAATGAGTCCACT</td><td>157.8</td><td>-11.2</td><td>1.7</td><td>-6.7</td><td>2.2E-11</td><td>6.8E-11</td><td>MB231/MB468</td><td>MP</td></tr>
<tr><td>>hg38_21_+_8433172_8446622_RNA45SN1_45S_50ntFlanks@6682.6718.37|CGTCGATGAAGAACGCAGCTAGCTGCGAGAATTAATG</td><td>>hg38_21_+_8433172_8446622_RNA45SN1_45S_50ntFlanks</td><td>6682</td><td>6718</td><td>37</td><td>CGTCGATGAAGAACGCAGCTAGCTGCGAGAATTAATG</td><td>130.0</td><td>-7.0</td><td>1.0</td><td>-6.7</td><td>2.2E-11</td><td>6.8E-11</td><td>BT20/MB468</td><td>CYTO</td></tr>
<tr><td>>hg38_21_+_8433172_8446622_RNA45SN1_45S_50ntFlanks@11200.11223.24|CGGCGGCGGCGGCGACTCTGGACG</td><td>>hg38_21_+_8433172_8446622_RNA45SN1_45S_50ntFlanks</td><td>11200</td><td>11223</td><td>24</td><td>CGGCGGCGGCGGCGACTCTGGACG</td><td>368.1</td><td>-8.1</td><td>1.2</td><td>-6.7</td><td>2.8E-11</td><td>6.8E-11</td><td>MB231/MB468</td><td>MITO</td></tr>
<tr><td>>MI0001445|hsa-mir-423&WithFlank&17|+|30117073|30117178@23.44.22(+1C)&[MIMAT0004748&hsa-miR-423-5p&offsets|0|-1(+1C);m-141&17|+|30117095|30117117&offsets|0|-1(+1C)]|TGAGGGGCAGAGAGCGAGACTTC</td><td>>MI0001445|hsa-mir-423&WithFlank&17|+|30117073|30117178</td><td>23</td><td>44</td><td>22(+1C)</td><td>TGAGGGGCAGAGAGCGAGACTTC</td><td>618.3</td><td>8.3</td><td>1.2</td><td>6.7</td><td>2.4E-11</td><td>6.8E-11</td><td>MB231/MB468</td><td>CYTO</td></tr>
<tr><td>>hg38_21_+_8433172_8446622_RNA45SN1_45S_50ntFlanks@10686.10706.21|GCGGCGTCCGGTGAGCTCTCG</td><td>>hg38_21_+_8433172_8446622_RNA45SN1_45S_50ntFlanks</td><td>10686</td><td>10706</td><td>21</td><td>GCGGCGTCCGGTGAGCTCTCG</td><td>1366.4</td><td>-8.7</td><td>1.3</td><td>-6.7</td><td>2.8E-11</td><td>6.8E-11</td><td>MB231/MB468</td><td>MITO</td></tr>
<tr><td>>hg38_21_+_8433172_8446622_RNA45SN1_45S_50ntFlanks@10574.10591.18|AAGCCGGCGGGAGCCCCG</td><td>>hg38_21_+_8433172_8446622_RNA45SN1_45S_50ntFlanks</td><td>10574</td><td>10591</td><td>18</td><td>AAGCCGGCGGGAGCCCCG</td><td>187.9</td><td>-9.6</td><td>1.4</td><td>-6.7</td><td>1.8E-11</td><td>6.8E-11</td><td>BT20/MB468</td><td>MITO</td></tr>
<tr><td>>hg38_21_+_8433172_8446622_RNA45SN1_45S_50ntFlanks@9343.9369.27|CGAAAGCCGCCGTGGCGCAATGAAGGT</td><td>>hg38_21_+_8433172_8446622_RNA45SN1_45S_50ntFlanks</td><td>9343</td><td>9369</td><td>27</td><td>CGAAAGCCGCCGTGGCGCAATGAAGGT</td><td>94.3</td><td>-10.4</td><td>1.6</td><td>-6.7</td><td>2.3E-11</td><td>6.8E-11</td><td>MB231/MB468</td><td>MP</td></tr>
<tr><td>>hg38_21_+_8433172_8446622_RNA45SN1_45S_50ntFlanks@6789.6807.19|GCCTGTCTGAGCGTCGCTT</td><td>>hg38_21_+_8433172_8446622_RNA45SN1_45S_50ntFlanks</td><td>6789</td><td>6807</td><td>19</td><td>GCCTGTCTGAGCGTCGCTT</td><td>1907.6</td><td>4.7</td><td>0.7</td><td>6.7</td><td>1.8E-11</td><td>6.8E-11</td><td>BT20/MB468</td><td>MITO</td></tr>
<tr><td>>hg38_21_+_8433172_8446622_RNA45SN1_45S_50ntFlanks@12939.12965.27|CCATTCGTAGACGACCTGCTTCTGGGT</td><td>>hg38_21_+_8433172_8446622_RNA45SN1_45S_50ntFlanks</td><td>12939</td><td>12965</td><td>27</td><td>CCATTCGTAGACGACCTGCTTCTGGGT</td><td>96.5</td><td>-10.5</td><td>1.6</td><td>-6.7</td><td>2.3E-11</td><td>6.9E-11</td><td>MB231/MB468</td><td>MP</td></tr>
<tr><td>>MI0000065|hsa-let-7d&WithFlank&9|+|94178828|94178926@14.34.21(+1C)&[MIMAT0000065&hsa-let-7d-5p&offsets|0|-1(+1C);m-90&9|+|94178841|94178862&offsets|0|-1(+1C)]|AGAGGTAGTAGGTTGCATAGTC</td><td>>MI0000065|hsa-let-7d&WithFlank&9|+|94178828|94178926</td><td>14</td><td>34</td><td>21(+1C)</td><td>AGAGGTAGTAGGTTGCATAGTC</td><td>569.0</td><td>5.5</td><td>0.8</td><td>6.7</td><td>2.5E-11</td><td>6.9E-11</td><td>MB231/MB468</td><td>CYTO</td></tr>
<tr><td>>am_trna10_ValCAC_5_-_180649395_180649467@1.33.33,trna12_ValAAC_5_-_180645270_180645342@1.33.33,trna132_ValAAC_6_-_27721179_27721251@1.33.33,trna136_ValAAC_6_-_27648885_27648957@1.33.33,trna139_ValAAC_6_-_27618707_27618779@1.33.33,trna18_ValCAC_5_-_180529253_180529325@1.33.33,trna2_ValAAC_3_+_169490018_169490090@1.33.33,trna2_ValCAC_5_+_180524070_180524142@1.33.33,trna4_ValAAC_5_+_180591154_180591226@1.33.33,trna5_ValAAC_5_+_180596610_180596682@1.33.33,trna6_ValCAC_5_+_180600650_180600722@1.33.33,trna85_ValCAC_1_-_161369490_161369562@1.33.33,trna90_ValCAC_1_-_149684088_149684161@1.33.33,trna98_ValCAC_1_-_149298555_149298627@1.33.33,trna9_ValCAC_6_+_26538282_26538354@1.33.33|GTTTCCGTAGTGTAGTGGTTATCACGTTCGCCT</td><td>>am_trna10_ValCAC_5_-_180649395_180649467</td><td>1</td><td>33</td><td>33</td><td>GTTTCCGTAGTGTAGTGGTTATCACGTTCGCCT</td><td>11942.5</td><td>10.4</td><td>1.6</td><td>6.7</td><td>2.5E-11</td><td>6.9E-11</td><td>MB231/MB468</td><td>CYTO</td></tr>
<tr><td>>hg38_21_+_8433172_8446622_RNA45SN1_45S_50ntFlanks@10713.10730.18|CTTGAAAATCCGGGGGAG</td><td>>hg38_21_+_8433172_8446622_RNA45SN1_45S_50ntFlanks</td><td>10713</td><td>10730</td><td>18</td><td>CTTGAAAATCCGGGGGAG</td><td>130.6</td><td>-10.9</td><td>1.6</td><td>-6.7</td><td>2.3E-11</td><td>6.9E-11</td><td>MB231/MB468</td><td>MP</td></tr>
<tr><td>>hg38_21_+_8433172_8446622_RNA45SN1_45S_50ntFlanks@9505.9533.29|TGAACTATGCCTGGGCAGGGCGAAGCCAG</td><td>>hg38_21_+_8433172_8446622_RNA45SN1_45S_50ntFlanks</td><td>9505</td><td>9533</td><td>29</td><td>TGAACTATGCCTGGGCAGGGCGAAGCCAG</td><td>157.9</td><td>-11.2</td><td>1.7</td><td>-6.7</td><td>2.3E-11</td><td>6.9E-11</td><td>MB231/MB468</td><td>MP</td></tr>
<tr><td>>MI0000088|hsa-mir-30a&WithFlank&6|-|71403545|71403627@12.34.23(+1A)&[MIMAT0000087&hsa-miR-30a-5p&offsets|0|+1(+1A);m-15&6|-|71403595|71403616&offsets|0|+1(+1A)]|TGTAAACATCCTCGACTGGAAGCA</td><td>>MI0000088|hsa-mir-30a&WithFlank&6|-|71403545|71403627</td><td>12</td><td>34</td><td>23(+1A)</td><td>TGTAAACATCCTCGACTGGAAGCA</td><td>14739.0</td><td>7.2</td><td>1.1</td><td>6.7</td><td>2.9E-11</td><td>6.9E-11</td><td>MB231/MB468</td><td>MITO</td></tr>
<tr><td>>hg38_21_+_8433172_8446622_RNA45SN1_45S_50ntFlanks@11575.11609.35|ACTTAGAACTGGTGCGGACCAGGGGAATCCGACTG</td><td>>hg38_21_+_8433172_8446622_RNA45SN1_45S_50ntFlanks</td><td>11575</td><td>11609</td><td>35</td><td>ACTTAGAACTGGTGCGGACCAGGGGAATCCGACTG</td><td>395.9</td><td>-3.9</td><td>0.6</td><td>-6.7</td><td>2.5E-11</td><td>6.9E-11</td><td>MB231/MB468</td><td>NUC</td></tr>
<tr><td>>MI0000102|hsa-mir-100&WithFlank&11|-|122152223|122152314@19.37.19&[MIMAT0000098&hsa-miR-100-5p&offsets|0|-3;m-27&11|-|122152276|122152296&offsets|0|-2]|AACCCGTAGATCCGAACTT</td><td>>MI0000102|hsa-mir-100&WithFlank&11|-|122152223|122152314</td><td>19</td><td>37</td><td>19</td><td>AACCCGTAGATCCGAACTT</td><td>420.2</td><td>-7.6</td><td>1.1</td><td>-7.0</td><td>2.3E-12</td><td>6.9E-11</td><td>BT20/MB231</td><td>MITO</td></tr>
<tr><td>>hg38_21_+_8433172_8446622_RNA45SN1_45S_50ntFlanks@9562.9592.31|TCCTGACGTGCAAATCGGTCGTCCGACCTGG</td><td>>hg38_21_+_8433172_8446622_RNA45SN1_45S_50ntFlanks</td><td>9562</td><td>9592</td><td>31</td><td>TCCTGACGTGCAAATCGGTCGTCCGACCTGG</td><td>99.9</td><td>-10.5</td><td>1.6</td><td>-6.7</td><td>2.3E-11</td><td>6.9E-11</td><td>MB231/MB468</td><td>MP</td></tr>
<tr><td>>hg38_21_+_8433172_8446622_RNA45SN1_45S_50ntFlanks@6645.6663.19|CTCGTACGACTCTTAGCGG</td><td>>hg38_21_+_8433172_8446622_RNA45SN1_45S_50ntFlanks</td><td>6645</td><td>6663</td><td>19</td><td>CTCGTACGACTCTTAGCGG</td><td>9205.0</td><td>4.1</td><td>0.6</td><td>6.7</td><td>1.8E-11</td><td>6.9E-11</td><td>BT20/MB468</td><td>MITO</td></tr>
<tr><td>>hg38_21_+_8433172_8446622_RNA45SN1_45S_50ntFlanks@4283.4301.19|CTTTAACGAGGATCCATTG</td><td>>hg38_21_+_8433172_8446622_RNA45SN1_45S_50ntFlanks</td><td>4283</td><td>4301</td><td>19</td><td>CTTTAACGAGGATCCATTG</td><td>78.4</td><td>-10.2</td><td>1.5</td><td>-6.7</td><td>2.3E-11</td><td>6.9E-11</td><td>MB231/MB468</td><td>MP</td></tr>
<tr><td>>hg38_21_+_8433172_8446622_RNA45SN1_45S_50ntFlanks@3863.3898.36|ATTCTAGAGCTAATACATGCCGACGGGCGCTGACCC</td><td>>hg38_21_+_8433172_8446622_RNA45SN1_45S_50ntFlanks</td><td>3863</td><td>3898</td><td>36</td><td>ATTCTAGAGCTAATACATGCCGACGGGCGCTGACCC</td><td>225.7</td><td>-11.7</td><td>1.7</td><td>-6.7</td><td>2.3E-11</td><td>6.9E-11</td><td>MB231/MB468</td><td>MP</td></tr>
<tr><td>>hg38_21_+_8433172_8446622_RNA45SN1_45S_50ntFlanks@6648.6677.30|GTACGACTCTTAGCGGTGGATCACTCGGCT</td><td>>hg38_21_+_8433172_8446622_RNA45SN1_45S_50ntFlanks</td><td>6648</td><td>6677</td><td>30</td><td>GTACGACTCTTAGCGGTGGATCACTCGGCT</td><td>4003.0</td><td>4.5</td><td>0.7</td><td>6.7</td><td>1.8E-11</td><td>6.9E-11</td><td>BT20/MB468</td><td>MITO</td></tr>
<tr><td>>hg38_21_+_8433172_8446622_RNA45SN1_45S_50ntFlanks@12420.12443.24|ATGTCGGCTCTTCCTATCATTGTG</td><td>>hg38_21_+_8433172_8446622_RNA45SN1_45S_50ntFlanks</td><td>12420</td><td>12443</td><td>24</td><td>ATGTCGGCTCTTCCTATCATTGTG</td><td>12032.8</td><td>-4.8</td><td>0.7</td><td>-6.7</td><td>2.5E-11</td><td>6.9E-11</td><td>MB231/MB468</td><td>NUC</td></tr>
<tr><td>>trnaMT_PheGAA_MT_+_577_647@39.56.18|ATGTTTAGACGGGCTCAC</td><td>>trnaMT_PheGAA_MT_+_577_647</td><td>39</td><td>56</td><td>18</td><td>ATGTTTAGACGGGCTCAC</td><td>4113.8</td><td>4.6</td><td>0.7</td><td>6.7</td><td>2.9E-11</td><td>6.9E-11</td><td>MB231/MB468</td><td>MITO</td></tr>
<tr><td>>hg38_21_+_8433172_8446622_RNA45SN1_45S_50ntFlanks@11683.11702.20|ATGTCAAAGTGAAGAAATTC</td><td>>hg38_21_+_8433172_8446622_RNA45SN1_45S_50ntFlanks</td><td>11683</td><td>11702</td><td>20</td><td>ATGTCAAAGTGAAGAAATTC</td><td>726.7</td><td>-12.6</td><td>1.8</td><td>-7.1</td><td>1.6E-12</td><td>6.9E-11</td><td>BT20/MB468</td><td>MP</td></tr>
<tr><td>>hg38_21_+_8433172_8446622_RNA45SN1_45S_50ntFlanks@11857.11881.25|GCGGAATCAGCGGGGAAAGAAGACC</td><td>>hg38_21_+_8433172_8446622_RNA45SN1_45S_50ntFlanks</td><td>11857</td><td>11881</td><td>25</td><td>GCGGAATCAGCGGGGAAAGAAGACC</td><td>92.1</td><td>-10.4</td><td>1.6</td><td>-6.7</td><td>2.3E-11</td><td>6.9E-11</td><td>MB231/MB468</td><td>MP</td></tr>
<tr><td>>trnaMT_PheGAA_MT_+_577_647@38.62.25|AATGTTTAGACGGGCTCACATCACC</td><td>>trnaMT_PheGAA_MT_+_577_647</td><td>38</td><td>62</td><td>25</td><td>AATGTTTAGACGGGCTCACATCACC</td><td>724.4</td><td>8.1</td><td>1.2</td><td>6.7</td><td>1.8E-11</td><td>7.0E-11</td><td>BT20/MB468</td><td>MITO</td></tr>
</table>

| | | | | | | | | | | | | | |
|---|---|---|---|---|---|---|---|---|---|---|---|---|---|
| >hg38_21_+_8433172_8446622_RNA45SN1_45S_50ntFlanks@11569.11591.23\|CAGCCGACTTAGAACTGGTGCGG | >hg38_21_+_8433172_8446622_RNA45SN1_45S_50ntFlanks | 11569 | 11591 | 23 | CAGCCGACTTAGAACTGGTGCGG | 300.4 | -6.5 | 1.0 | -6.7 | 2.5E-11 | 7.0E-11 | MB231/MB468 | CYTO |
| >hg38_21_+_8433172_8446622_RNA45SN1_45S_50ntFlanks@11316.11336.21\|CACCCCACGTCTCGTCGCGCG | >hg38_21_+_8433172_8446622_RNA45SN1_45S_50ntFlanks | 11316 | 11336 | 21 | CACCCCACGTCTCGTCGCGCG | 2372.9 | -8.3 | 1.2 | -6.7 | 2.5E-11 | 7.0E-11 | MB231/MB468 | CYTO |
| >hg38_21_+_8433172_8446622_RNA45SN1_45S_50ntFlanks@10378.10406.29\|AGGGTTCCATGTGAACAGCAGTTGAACAT | >hg38_21_+_8433172_8446622_RNA45SN1_45S_50ntFlanks | 10378 | 10406 | 29 | AGGGTTCCATGTGAACAGCAGTTGAACAT | 83.1 | -10.2 | 1.5 | -6.7 | 2.3E-11 | 7.0E-11 | MB231/MB468 | MP |
| >hg38_21_+_8433172_8446622_RNA45SN1_45S_50ntFlanks@12457.12474.18\|CAAGCGTTGGATTGTTCA | >hg38_21_+_8433172_8446622_RNA45SN1_45S_50ntFlanks | 12457 | 12474 | 18 | CAAGCGTTGGATTGTTCA | 2057.0 | -6.1 | 0.9 | -6.7 | 2.5E-11 | 7.0E-11 | MB231/MB468 | CYTO |
| >hg38_21_+_8433172_8446622_RNA45SN1_45S_50ntFlanks@12445.12472.28\|AGCAGAATTCACCAAGCGTTGGATTGTT | >hg38_21_+_8433172_8446622_RNA45SN1_45S_50ntFlanks | 12445 | 12472 | 28 | AGCAGAATTCACCAAGCGTTGGATTGTT | 81.5 | -10.2 | 1.5 | -6.7 | 2.3E-11 | 7.0E-11 | MB231/MB468 | MP |
| >hg38_21_+_8433172_8446622_RNA45SN1_45S_50ntFlanks@10905.10930.26\|CTGGGCGCGCGCCGCGGCTGGACGAG | >hg38_21_+_8433172_8446622_RNA45SN1_45S_50ntFlanks | 10905 | 10930 | 26 | CTGGGCGCGCGCCGCGGCTGGACGAG | 87.8 | -10.3 | 1.5 | -6.7 | 2.3E-11 | 7.0E-11 | MB231/MB468 | MP |
| >hg38_21_+_8433172_8446622_RNA45SN1_45S_50ntFlanks@8379.8402.24\|TCAAGAGGGCGTGAAACCGTTAAG | >hg38_21_+_8433172_8446622_RNA45SN1_45S_50ntFlanks | 8379 | 8402 | 24 | TCAAGAGGGCGTGAAACCGTTAAG | 449.4 | -8.3 | 1.3 | -6.7 | 2.9E-11 | 7.0E-11 | MB231/MB468 | MITO |
| >hg38_21_+_8433172_8446622_RNA45SN1_45S_50ntFlanks@10768.10797.30\|CAGCAGGTCTCCAAGGTGAACAGCCTCTGG | >hg38_21_+_8433172_8446622_RNA45SN1_45S_50ntFlanks | 10768 | 10797 | 30 | CAGCAGGTCTCCAAGGTGAACAGCCTCTGG | 251.6 | -11.1 | 1.6 | -7.1 | 1.6E-12 | 7.0E-11 | BT20/MB468 | MP |
| >hg38_21_+_8433172_8446622_RNA45SN1_45S_50ntFlanks@10071.10100.30\|TGGACGGGAGCGGCGGGGGCGGCGCGCG | >hg38_21_+_8433172_8446622_RNA45SN1_45S_50ntFlanks | 10071 | 10100 | 30 | TGGACGGGAGCGGCGGGGGCGGCGCGCGCG | 120.3 | -10.8 | 1.6 | -6.7 | 2.3E-11 | 7.0E-11 | MB231/MB468 | MP |
| >trna2_SerTGA_10_+_69524261_69524342@68.85.18\|ACCCTGCTCGCTGCGCCA | >trna2_SerTGA_10_+_69524261_69524342 | 68 | 85 | 18 | ACCCTGCTCGCTGCGCCA | 316.8 | 8.2 | 1.2 | 6.7 | 2.3E-11 | 7.0E-11 | BT20/MB468 | CYTO |
| >hg38_21_+_8433172_8446622_RNA45SN1_45S_50ntFlanks@9624.9652.29\|AGTAGCTGGTTCCCTCCGAAGTTTCCCTC | >hg38_21_+_8433172_8446622_RNA45SN1_45S_50ntFlanks | 9624 | 9652 | 29 | AGTAGCTGGTTCCCTCCGAAGTTTCCCTC | 86.9 | -10.3 | 1.5 | -6.7 | 2.3E-11 | 7.0E-11 | MB231/MB468 | MP |
| >hg38_1_-_228634819_228635039_RNA5S12_5S_50ntFlanks@138.171.34\|AGACCGCCTGGGAATACCGGGTGCTGTAGGCTTT | >hg38_1_-_228634819_228635039_RNA5S12_5S_50ntFlanks | 138 | 171 | 34 | AGACCGCCTGGGAATACCGGGTGCTGTAGGCTTT | 1295.3 | 5.7 | 0.9 | 6.7 | 2.5E-11 | 7.0E-11 | MB231/MB468 | NUC |
| >hg38_21_+_8433172_8446622_RNA45SN1_45S_50ntFlanks@11878.11904.27\|GACCCTGTTGAGCTTGACTCTAGTCTG | >hg38_21_+_8433172_8446622_RNA45SN1_45S_50ntFlanks | 11878 | 11904 | 27 | GACCCTGTTGAGCTTGACTCTAGTCTG | 84.6 | -10.3 | 1.5 | -6.7 | 2.3E-11 | 7.0E-11 | MB231/MB468 | MP |
| >hg38_21_+_8433172_8446622_RNA45SN1_45S_50ntFlanks@12003.12034.32\|GCGGGCCGCCGGTGAAATACCACTACTCTGAT | >hg38_21_+_8433172_8446622_RNA45SN1_45S_50ntFlanks | 12003 | 12034 | 32 | GCGGGCCGCCGGTGAAATACCACTACTCTGAT | 2492.3 | 6.4 | 1.0 | 6.7 | 2.5E-11 | 7.0E-11 | MB231/MB468 | CYTO |
| >hg38_21_+_8433172_8446622_RNA45SN1_45S_50ntFlanks@10726.10752.27\|GGGAGAGGGTGTAAATCTCGCGCCGGG | >hg38_21_+_8433172_8446622_RNA45SN1_45S_50ntFlanks | 10726 | 10752 | 27 | GGGAGAGGGTGTAAATCTCGCGCCGGG | 265.7 | -11.1 | 1.6 | -7.1 | 1.6E-12 | 7.0E-11 | BT20/MB468 | MP |
| >hg38_21_+_8433172_8446622_RNA45SN1_45S_50ntFlanks@8437.8469.33\|AGGATTCAACCCGGCGGCGGGTCCGGCCGTGTC | >hg38_21_+_8433172_8446622_RNA45SN1_45S_50ntFlanks | 8437 | 8469 | 33 | AGGATTCAACCCGGCGGCGGGTCCGGCCGTGTC | 279.3 | -11.2 | 1.6 | -7.1 | 1.6E-12 | 7.0E-11 | BT20/MB468 | MP |
| >hg38_21_+_8433172_8446622_RNA45SN1_45S_50ntFlanks@4922.4943.22\|CCAGGAGTGGAGCCTGCGGCTT | >hg38_21_+_8433172_8446622_RNA45SN1_45S_50ntFlanks | 4922 | 4943 | 22 | CCAGGAGTGGAGCCTGCGGCTT | 94.3 | -10.4 | 1.6 | -6.7 | 2.3E-11 | 7.1E-11 | MB231/MB468 | MP |
| >hg38_21_+_8433172_8446622_RNA45SN1_45S_50ntFlanks@10350.10381.32\|ACGAGAACTTTGAAGGCCGAAGTGGAGAAGGG | >hg38_21_+_8433172_8446622_RNA45SN1_45S_50ntFlanks | 10350 | 10381 | 32 | ACGAGAACTTTGAAGGCCGAAGTGGAGAAGGG | 1450.6 | -9.8 | 1.5 | -6.6 | 3.1E-11 | 7.1E-11 | BT20/MB468 | NUC |
| >trnaMT_SerGCT_MT_+_12207_12265@1.21.21\|GAGAAAGCTCACAAGAACTGC | >trnaMT_SerGCT_MT_+_12207_12265 | 1 | 21 | 21 | GAGAAAGCTCACAAGAACTGC | 494.1 | 7.5 | 1.1 | 6.7 | 2.3E-11 | 7.1E-11 | BT20/MB468 | CYTO |
| >hg38_21_+_8433172_8446622_RNA45SN1_45S_50ntFlanks@8687.8706.20\|CGGCTCCGGGACGGCTGGGA | >hg38_21_+_8433172_8446622_RNA45SN1_45S_50ntFlanks | 8687 | 8706 | 20 | CGGCTCCGGGACGGCTGGGA | 653.4 | 9.0 | 1.4 | 6.7 | 2.5E-11 | 7.1E-11 | MB231/MB468 | NUC |
| >hg38_21_+_8433172_8446622_RNA45SN1_45S_50ntFlanks@6773.6803.31\|TTCCTCCCGGGGCTACGCCTGTCTGAGCGTC | >hg38_21_+_8433172_8446622_RNA45SN1_45S_50ntFlanks | 6773 | 6803 | 31 | TTCCTCCCGGGGCTACGCCTGTCTGAGCGTC | 871.9 | -7.0 | 1.1 | -6.7 | 2.5E-11 | 7.1E-11 | MB231/MB468 | NUC |
| >hg38_21_+_8433172_8446622_RNA45SN1_45S_50ntFlanks@4555.4579.25\|CGCAGCTAGGAATAATGGAATAGGA | >hg38_21_+_8433172_8446622_RNA45SN1_45S_50ntFlanks | 4555 | 4579 | 25 | CGCAGCTAGGAATAATGGAATAGGA | 106.8 | -10.6 | 1.6 | -6.7 | 2.4E-11 | 7.1E-11 | MB231/MB468 | MP |
| >MI0000077\|hsa-mir-21&WithFlank&17\|+\|59841260\|59841343@14.39.26(+1U)&[MIMAT0000076&hsa-miR-21-5p&offsets\|0\|+4(+1U);m-1&17\|+\|59841273\|59841294&offsets\|0\|+4(+1U)]\|TAGCTTATCAGACTGATGTTGACTGTT | >MI0000077\|hsa-mir-21&WithFlank&17\|+\|59841260\|59841343 | 14 | 39 | 26(+1U) | TAGCTTATCAGACTGATGTTGACTGTT | 431.2 | 5.2 | 0.8 | 6.7 | 2.3E-11 | 7.1E-11 | BT20/MB468 | CYTO |
| >hg38_21_+_8433172_8446622_RNA45SN1_45S_50ntFlanks@8552.8579.28\|CTCCCCGGAGGGGGCGGGCTCCGGCGGG | >hg38_21_+_8433172_8446622_RNA45SN1_45S_50ntFlanks | 8552 | 8579 | 28 | CTCCCCGGAGGGGGCGGGCTCCGGCGGG | 147.8 | -9.3 | 1.4 | -6.7 | 1.9E-11 | 7.1E-11 | BT20/MB468 | MITO |
| >hg38_21_+_8433172_8446622_RNA45SN1_45S_50ntFlanks@12259.12283.25\|CTTGATCTTGATTTTCAGTACGAAT | >hg38_21_+_8433172_8446622_RNA45SN1_45S_50ntFlanks | 12259 | 12283 | 25 | CTTGATCTTGATTTTCAGTACGAAT | 117.2 | -10.7 | 1.6 | -6.7 | 2.4E-11 | 7.1E-11 | MB231/MB468 | MP |
| >trna111_HisGTG_1_-_147774845_147774916@-1A.26.27,trna118_HisGTG_1_-_145396881_145396952@-1A.26.27,trna16_HisGTG_1_+_146544773_146544844@-1A.26.27,trna1_HisGTG_15_+_45493349_45493420@-1A.26.27,trna21_HisGTG_1_+_147753471_147753542@-1A.26.27,trna33_HisGTG_6_+_27125906_27125977@-1A.26.27,trna7_HisGTG_9_-_14433938_14434009@-1A.26.27,trna8_HisGTG_15_-_45492611_45492682@-1A.26.27,trna9_HisGTG_15_-_45490804_45490875@-1A.26.27\|AGCCGTGATCGTATAGTGGTTAGTACT | >trna111_HisGTG_1_-_147774845_147774916 | -1A | 26 | 27 | AGCCGTGATCGTATAGTGGTTAGTACT | 553.1 | 5.4 | 0.8 | 6.7 | 2.6E-11 | 7.1E-11 | MB231/MB468 | CYTO |
| >hg38_21_+_8433172_8446622_RNA45SN1_45S_50ntFlanks@12946.12965.20\|TAGACGACCTGCTTCTGGGT | >hg38_21_+_8433172_8446622_RNA45SN1_45S_50ntFlanks | 12946 | 12965 | 20 | TAGACGACCTGCTTCTGGGT | 89.6 | -10.4 | 1.5 | -6.7 | 2.4E-11 | 7.2E-11 | MB231/MB468 | MP |
| >hg38_21_+_8433172_8446622_RNA45SN1_45S_50ntFlanks@12242.12273.32\|AGAAGGGCAAAAGCTCGCTTGATCTTGATTTT | >hg38_21_+_8433172_8446622_RNA45SN1_45S_50ntFlanks | 12242 | 12273 | 32 | AGAAGGGCAAAAGCTCGCTTGATCTTGATTTT | 137.6 | -11.0 | 1.6 | -6.7 | 2.4E-11 | 7.2E-11 | MB231/MB468 | MP |
| >hg38_21_+_8433172_8446622_RNA45SN1_45S_50ntFlanks@4588.4613.26\|CTATTTTGTTGGTTTTCGGAACTGAG | >hg38_21_+_8433172_8446622_RNA45SN1_45S_50ntFlanks | 4588 | 4613 | 26 | CTATTTTGTTGGTTTTCGGAACTGAG | 128.0 | -9.1 | 1.4 | -6.7 | 1.9E-11 | 7.2E-11 | BT20/MB468 | MITO |
| >trna129_MetCAT_6_-_27870271_27870342@31.50.20,trna142_MetCAT_6_-_27560600_27560671@31.50.20,trna150_MetCAT_6_-_27300764_27300835@31.50.20,trna169_MetCAT_6_-_26330529_26330600@31.50.20,trna171_MetCAT_6_-_26313352_26313423@31.50.20,trna20_MetCAT_17_-_80452597_80452668@31.50.20,trna2_MetCAT_6_+_26286754_26286825@31.50.20,trna32_MetCAT_1_+_153643726_153643797@31.50.20,trna61_MetCAT_6_+_27745664_27745735@31.50.20\|CCCATAACCCAGAGGTCGAT | >trna129_MetCAT_6_-_27870271_27870342 | 31 | 50 | 20 | CCCATAACCCAGAGGTCGAT | 92.7 | -10.4 | 1.6 | -6.7 | 2.4E-11 | 7.2E-11 | MB231/MB468 | MP |
| >hg38_21_+_8433172_8446622_RNA45SN1_45S_50ntFlanks@12141.12162.22\|ACAGTGCCAGGTGGGGAGTTTG | >hg38_21_+_8433172_8446622_RNA45SN1_45S_50ntFlanks | 12141 | 12162 | 22 | ACAGTGCCAGGTGGGGAGTTTG | 102.6 | -9.0 | 1.3 | -6.7 | 2.3E-11 | 7.2E-11 | BT20/MB468 | CYTO |
| >hg38_21_+_8433172_8446622_RNA45SN1_45S_50ntFlanks@7983.8012.30\|CAGATCAGACGTGGCGACCCGCTGAATTTA | >hg38_21_+_8433172_8446622_RNA45SN1_45S_50ntFlanks | 7983 | 8012 | 30 | CAGATCAGACGTGGCGACCCGCTGAATTTA | 690.4 | -8.6 | 1.3 | -6.7 | 2.4E-11 | 7.2E-11 | MB231/MB468 | MP |
| >hg38_21_+_8433172_8446622_RNA45SN1_45S_50ntFlanks@10791.10818.28\|CCTCTGGCATGTTGGAACAATGTAGGTA | >hg38_21_+_8433172_8446622_RNA45SN1_45S_50ntFlanks | 10791 | 10818 | 28 | CCTCTGGCATGTTGGAACAATGTAGGTA | 330.2 | -6.3 | 0.9 | -6.6 | 3.0E-11 | 7.2E-11 | MB231/MB468 | MITO |
| >hg38_21_+_8433172_8446622_RNA45SN1_45S_50ntFlanks@11576.11600.25\|CTTAGAACTGGTGCGGACCAGGGGA | >hg38_21_+_8433172_8446622_RNA45SN1_45S_50ntFlanks | 11576 | 11600 | 25 | CTTAGAACTGGTGCGGACCAGGGGA | 894.6 | -8.1 | 1.2 | -6.7 | 2.6E-11 | 7.2E-11 | MB231/MB468 | CYTO |
| >MI0000750\|hsa-mir-26a-2&WithFlank&12\|-\|57824603\|57824698@20.41.22(+1A);MI0000083\|hsa-mir-26a-1&WithFlank&3\|+\|37969398\|37969486@16.37.22(+1A)&[MIMAT0000082&hsa-miR-26a-5p&offsets\|0\|0(+1A);m-33&12\|-\|57824658\|57824679&offsets\|0\|0(+1A)];[MIMAT0000082_1&hsa-miR-26a-5p&offsets\|0\|0(+1A);m-34&3\|+\|37969413\|37969434&offsets\|0\|0(+1A)]\|TTCAAGTAATCCAGGATAGGCTA | >MI0000750\|hsa-mir-26a-2&WithFlank&12\|-\|57824603\|57824698 | 20 | 41 | 22(+1A) | TTCAAGTAATCCAGGATAGGCTA | 1011.6 | 6.0 | 0.9 | 6.7 | 1.9E-11 | 7.2E-11 | BT20/MB468 | MITO |
| >hg38_21_+_8433172_8446622_RNA45SN1_45S_50ntFlanks@11320.11342.23\|CCACGTCTCGTCGCGCGCGCGTC | >hg38_21_+_8433172_8446622_RNA45SN1_45S_50ntFlanks | 11320 | 11342 | 23 | CCACGTCTCGTCGCGCGCGCGTC | 1744.2 | -9.0 | 1.4 | -6.7 | 2.6E-11 | 7.2E-11 | MB231/MB468 | NUC |
| >hg38_21_+_8433172_8446622_RNA45SN1_45S_50ntFlanks@5492.5526.35\|AGAAGACGGTCGAACTTGACTATCTAGAGGAAGTA | >hg38_21_+_8433172_8446622_RNA45SN1_45S_50ntFlanks | 5492 | 5526 | 35 | AGAAGACGGTCGAACTTGACTATCTAGAGGAAGTA | 1321.1 | -3.9 | 0.6 | -6.7 | 1.9E-11 | 7.2E-11 | BT20/MB468 | MITO |
| >hg38_21_+_8433172_8446622_RNA45SN1_45S_50ntFlanks@12274.12319.46\|CAGTACGAATACAGACCGTGAAAGCGGGGCCTCACGATCCTTCTGA | >hg38_21_+_8433172_8446622_RNA45SN1_45S_50ntFlanks | 12274 | 12319 | 46 | CAGTACGAATACAGACCGTGAAAGCGGGGCCTCACGATCCTTCTGA | 204.8 | -11.5 | 1.7 | -6.6 | 3.0E-11 | 7.2E-11 | MB231/MB468 | MITO |
| >hg38_21_+_8433172_8446622_RNA45SN1_45S_50ntFlanks@10807.10828.22\|ACAATGTAGGTAAGGGAAGTCG | >hg38_21_+_8433172_8446622_RNA45SN1_45S_50ntFlanks | 10807 | 10828 | 22 | ACAATGTAGGTAAGGGAAGTCG | 629.6 | -8.4 | 1.3 | -6.6 | 3.0E-11 | 7.2E-11 | MB231/MB468 | MITO |
| >MI0000088\|hsa-mir-30a&WithFlank&6\|-\|71403545\|71403627@12.35.24(+2U)&[MIMAT0000087&hsa-miR-30a-5p&offsets\|0\|+2(+2U);m-15&6\|-\|71403595\|71403616&offsets\|0\|+2(+2U)]\|TGTAAACATCCTCGACTGGAAGCTTT | >MI0000088\|hsa-mir-30a&WithFlank&6\|-\|71403545\|71403627 | 12 | 35 | 24(+2U) | TGTAAACATCCTCGACTGGAAGCTTT | 5090.1 | -6.4 | 0.9 | -7.3 | 3.9E-13 | 7.2E-11 | BT20/MB231 | MP |
| >hg38_21_+_8433172_8446622_RNA45SN1_45S_50ntFlanks@6735.6770.36\|TTGATCATCGACACTTCGAACGCACTTGCGGCCCCG | >hg38_21_+_8433172_8446622_RNA45SN1_45S_50ntFlanks | 6735 | 6770 | 36 | TTGATCATCGACACTTCGAACGCACTTGCGGCCCCG | 484.4 | -4.7 | 0.7 | -6.7 | 2.6E-11 | 7.2E-11 | MB231/MB468 | NUC |
| >hg38_21_+_8433172_8446622_RNA45SN1_45S_50ntFlanks@12624.12654.31\|CCAATGGGGCGAAGCTACCATCTGTGGGATT | >hg38_21_+_8433172_8446622_RNA45SN1_45S_50ntFlanks | 12624 | 12654 | 31 | CCAATGGGGCGAAGCTACCATCTGTGGGATT | 7112.5 | -8.7 | 1.2 | -7.1 | 1.7E-12 | 7.3E-11 | BT20/MB468 | MP |
| >hg38_21_+_8433172_8446622_RNA45SN1_45S_50ntFlanks@11921.11968.48\|TGAGAGGTGTAGAATAAGTGGGAGGCCCCCGGCGCCCCCCCGGTGTCC | >hg38_21_+_8433172_8446622_RNA45SN1_45S_50ntFlanks | 11921 | 11968 | 48 | TGAGAGGTGTAGAATAAGTGGGAGGCCCCCGGCGCCCCCCCGGTGTCC | 302.6 | -11.2 | 1.7 | -6.7 | 2.6E-11 | 7.3E-11 | MB231/MB468 | NUC |

<table>
<tr><td>>hg38_21_+_8433172_8446622_RNA45SN1_45S_50ntFlanks@12633.12651.19|CGAAGCTACCATCTGTGGG</td><td>>hg38_21_+_8433172_8446622_RNA45SN1_45S_50ntFlanks</td><td>12633</td><td>12651</td><td>19</td><td>CGAAGCTACCATCTGTGGG</td><td>747.1</td><td>-7.4</td><td>1.1</td><td>-6.6</td><td>3.2E-11</td><td>7.3E-11</td><td>BT20/MB468</td><td>NUC</td></tr>
<tr><td>>hg38_21_+_8433172_8446622_RNA45SN1_45S_50ntFlanks@12874.12892.19|GCTGGAAAGGCGGCCGCCC</td><td>>hg38_21_+_8433172_8446622_RNA45SN1_45S_50ntFlanks</td><td>12874</td><td>12892</td><td>19</td><td>GCTGGAAAGGCGGCCGCCC</td><td>108.4</td><td>-10.6</td><td>1.6</td><td>-6.7</td><td>2.4E-11</td><td>7.3E-11</td><td>MB231/MB468</td><td>MP</td></tr>
<tr><td>>hg38_21_+_8433172_8446622_RNA45SN1_45S_50ntFlanks@10802.10822.21|TTGGAACAATGTAGGTAAGGG</td><td>>hg38_21_+_8433172_8446622_RNA45SN1_45S_50ntFlanks</td><td>10802</td><td>10822</td><td>21</td><td>TTGGAACAATGTAGGTAAGGG</td><td>230.6</td><td>-6.4</td><td>1.0</td><td>-6.7</td><td>2.6E-11</td><td>7.3E-11</td><td>MB231/MB468</td><td>CYTO</td></tr>
<tr><td>>hg38_21_+_8433172_8446622_RNA45SN1_45S_50ntFlanks@9477.9501.25|AGCGCACGTGTTAGGACCCGAAAGA</td><td>>hg38_21_+_8433172_8446622_RNA45SN1_45S_50ntFlanks</td><td>9477</td><td>9501</td><td>25</td><td>AGCGCACGTGTTAGGACCCGAAAGA</td><td>221.5</td><td>7.0</td><td>1.0</td><td>6.7</td><td>2.6E-11</td><td>7.3E-11</td><td>MB231/MB468</td><td>NUC</td></tr>
<tr><td>>hg38_21_+_8433172_8446622_RNA45SN1_45S_50ntFlanks@11641.11658.18|CGGCGGGTGTTGACGCGA</td><td>>hg38_21_+_8433172_8446622_RNA45SN1_45S_50ntFlanks</td><td>11641</td><td>11658</td><td>18</td><td>CGGCGGGTGTTGACGCGA</td><td>319.0</td><td>-10.4</td><td>1.6</td><td>-6.7</td><td>1.9E-11</td><td>7.3E-11</td><td>BT20/MB468</td><td>MITO</td></tr>
<tr><td>>hg38_21_+_8433172_8446622_RNA45SN1_45S_50ntFlanks@5446.5466.21|CGGATCGGCCCCGCCGGGGTC</td><td>>hg38_21_+_8433172_8446622_RNA45SN1_45S_50ntFlanks</td><td>5446</td><td>5466</td><td>21</td><td>CGGATCGGCCCCGCCGGGGTC</td><td>9142.4</td><td>-8.1</td><td>1.2</td><td>-6.7</td><td>2.6E-11</td><td>7.3E-11</td><td>MB231/MB468</td><td>CYTO</td></tr>
<tr><td>>hg38_21_+_8433172_8446622_RNA45SN1_45S_50ntFlanks@11208.11254.47|GCGGCGACTCTGGACGCGAGCCGGGCCCTTCCCGTGGATCGCCCCAG</td><td>>hg38_21_+_8433172_8446622_RNA45SN1_45S_50ntFlanks</td><td>11208</td><td>11254</td><td>47</td><td>GCGGCGACTCTGGACGCGAGCCGGGCCCTTCCCGTGGATCGCCCCAG</td><td>8749.6</td><td>-6.4</td><td>1.0</td><td>-6.7</td><td>2.4E-11</td><td>7.3E-11</td><td>BT20/MB468</td><td>CYTO</td></tr>
<tr><td>>hg38_21_+_8433172_8446622_RNA45SN1_45S_50ntFlanks@12045.12068.24|CTGACCCGGTGAGGCGGGGGGGCG</td><td>>hg38_21_+_8433172_8446622_RNA45SN1_45S_50ntFlanks</td><td>12045</td><td>12068</td><td>24</td><td>CTGACCCGGTGAGGCGGGGGGGCG</td><td>277.4</td><td>-11.2</td><td>1.6</td><td>-7.1</td><td>1.7E-12</td><td>7.3E-11</td><td>BT20/MB468</td><td>MP</td></tr>
<tr><td>>hg38_21_+_8433172_8446622_RNA45SN1_45S_50ntFlanks@12455.12475.21|ACCAAGCGTTGGATTGTTCAC</td><td>>hg38_21_+_8433172_8446622_RNA45SN1_45S_50ntFlanks</td><td>12455</td><td>12475</td><td>21</td><td>ACCAAGCGTTGGATTGTTCAC</td><td>110.2</td><td>-10.6</td><td>1.6</td><td>-6.7</td><td>2.4E-11</td><td>7.3E-11</td><td>MB231/MB468</td><td>MP</td></tr>
<tr><td>>hg38_21_+_8433172_8446622_RNA45SN1_45S_50ntFlanks@11583.11605.23|CTGGTGCGGACCAGGGGAATCCG</td><td>>hg38_21_+_8433172_8446622_RNA45SN1_45S_50ntFlanks</td><td>11583</td><td>11605</td><td>23</td><td>CTGGTGCGGACCAGGGGAATCCG</td><td>865.4</td><td>-4.9</td><td>0.7</td><td>-7.1</td><td>1.7E-12</td><td>7.3E-11</td><td>BT20/MB468</td><td>MP</td></tr>
<tr><td>>hg38_21_+_8433172_8446622_RNA45SN1_45S_50ntFlanks@12476.12509.34|CCACTAATAGGGAACGTGAGCTGGGTTTAGACCG</td><td>>hg38_21_+_8433172_8446622_RNA45SN1_45S_50ntFlanks</td><td>12476</td><td>12509</td><td>34</td><td>CCACTAATAGGGAACGTGAGCTGGGTTTAGACCG</td><td>115.9</td><td>-7.1</td><td>1.1</td><td>-6.7</td><td>2.6E-11</td><td>7.3E-11</td><td>MB231/MB468</td><td>CYTO</td></tr>
<tr><td>>hg38_21_+_8433172_8446622_RNA45SN1_45S_50ntFlanks@12418.12455.38|CGATGTCGGCTCTTCCTATCATTGTGAAGCAGAATTCA</td><td>>hg38_21_+_8433172_8446622_RNA45SN1_45S_50ntFlanks</td><td>12418</td><td>12455</td><td>38</td><td>CGATGTCGGCTCTTCCTATCATTGTGAAGCAGAATTCA</td><td>3276.1</td><td>-5.3</td><td>0.8</td><td>-6.7</td><td>2.4E-11</td><td>7.3E-11</td><td>BT20/MB468</td><td>CYTO</td></tr>
<tr><td>>hg38_21_+_8433172_8446622_RNA45SN1_45S_50ntFlanks@12605.12626.22|TATGTGCTTGGCTGAGGAGCCA</td><td>>hg38_21_+_8433172_8446622_RNA45SN1_45S_50ntFlanks</td><td>12605</td><td>12626</td><td>22</td><td>TATGTGCTTGGCTGAGGAGCCA</td><td>4554.6</td><td>-7.1</td><td>1.1</td><td>-6.7</td><td>2.4E-11</td><td>7.3E-11</td><td>BT20/MB468</td><td>CYTO</td></tr>
<tr><td>>hg38_21_+_8433172_8446622_RNA45SN1_45S_50ntFlanks@4723.4757.35|CATTAATCAAGAACGAAAGTCGGAGGTTCGAAGAC</td><td>>hg38_21_+_8433172_8446622_RNA45SN1_45S_50ntFlanks</td><td>4723</td><td>4757</td><td>35</td><td>CATTAATCAAGAACGAAAGTCGGAGGTTCGAAGAC</td><td>427.9</td><td>-8.7</td><td>1.3</td><td>-6.6</td><td>3.1E-11</td><td>7.3E-11</td><td>MB231/MB468</td><td>MITO</td></tr>
<tr><td>>hg38_21_+_8433172_8446622_RNA45SN1_45S_50ntFlanks@10571.10593.23|GAGAAGCCGGCGGGAGCCCCGGG</td><td>>hg38_21_+_8433172_8446622_RNA45SN1_45S_50ntFlanks</td><td>10571</td><td>10593</td><td>23</td><td>GAGAAGCCGGCGGGAGCCCCGGG</td><td>115.6</td><td>-10.7</td><td>1.6</td><td>-6.7</td><td>2.4E-11</td><td>7.4E-11</td><td>MB231/MB468</td><td>MP</td></tr>
<tr><td>>hg38_21_+_8433172_8446622_RNA45SN1_45S_50ntFlanks@11569.11595.27|CAGCCGACTTAGAACTGGTGCGGACCA</td><td>>hg38_21_+_8433172_8446622_RNA45SN1_45S_50ntFlanks</td><td>11569</td><td>11595</td><td>27</td><td>CAGCCGACTTAGAACTGGTGCGGACCA</td><td>126.0</td><td>-10.8</td><td>1.6</td><td>-6.7</td><td>2.4E-11</td><td>7.4E-11</td><td>MB231/MB468</td><td>MP</td></tr>
<tr><td>>trna12_ArgCCT_16_+_3243918_3243990@35.52.18,trna18_ArgCCT_17_+_73030001_73030073@35.52.18,trna21_ArgCCT_17_-_73030526_73030598@35.52.18,trna2_ArgCCT_16_+_3202901_3202973@35.52.18,trna3_ArgCCT_7_+_139025446_139025518@35.52.18|CTAAGCCAGGGATTGTGG</td><td>>trna12_ArgCCT_16_+_3243918_3243990</td><td>35</td><td>52</td><td>18</td><td>CTAAGCCAGGGATTGTGG</td><td>96.7</td><td>-10.5</td><td>1.6</td><td>-6.7</td><td>2.4E-11</td><td>7.4E-11</td><td>MB231/MB468</td><td>MP</td></tr>
<tr><td>>hg38_21_+_8433172_8446622_RNA45SN1_45S_50ntFlanks@12850.12868.19|CTTCGTCCTGGGAAACGGG</td><td>>hg38_21_+_8433172_8446622_RNA45SN1_45S_50ntFlanks</td><td>12850</td><td>12868</td><td>19</td><td>CTTCGTCCTGGGAAACGGG</td><td>189.5</td><td>-11.4</td><td>1.7</td><td>-6.7</td><td>2.4E-11</td><td>7.4E-11</td><td>MB231/MB468</td><td>MP</td></tr>
<tr><td>>hg38_21_+_8433172_8446622_RNA45SN1_45S_50ntFlanks@12935.12962.28|TAAACCATTCGTAGACGACCTGCTTCTG</td><td>>hg38_21_+_8433172_8446622_RNA45SN1_45S_50ntFlanks</td><td>12935</td><td>12962</td><td>28</td><td>TAAACCATTCGTAGACGACCTGCTTCTG</td><td>419.8</td><td>-8.7</td><td>1.3</td><td>-6.6</td><td>3.3E-11</td><td>7.4E-11</td><td>BT20/MB468</td><td>NUC</td></tr>
<tr><td>>hg38_21_+_8433172_8446622_RNA45SN1_45S_50ntFlanks@10375.10407.33|AGAAGGGTTCCATGTGAACAGCAGTTGAACATG</td><td>>hg38_21_+_8433172_8446622_RNA45SN1_45S_50ntFlanks</td><td>10375</td><td>10407</td><td>33</td><td>AGAAGGGTTCCATGTGAACAGCAGTTGAACATG</td><td>147.7</td><td>-9.3</td><td>1.4</td><td>-6.7</td><td>1.9E-11</td><td>7.4E-11</td><td>BT20/MB468</td><td>MITO</td></tr>
<tr><td>>hg38_21_+_8433172_8446622_RNA45SN1_45S_50ntFlanks@4892.4911.20|AAACTTAAAGGAATTGACGG</td><td>>hg38_21_+_8433172_8446622_RNA45SN1_45S_50ntFlanks</td><td>4892</td><td>4911</td><td>20</td><td>AAACTTAAAGGAATTGACGG</td><td>294.8</td><td>-4.8</td><td>0.7</td><td>-6.7</td><td>2.4E-11</td><td>7.4E-11</td><td>MB231/MB468</td><td>MP</td></tr>
<tr><td>>hg38_21_+_8433172_8446622_RNA45SN1_45S_50ntFlanks@12150.12179.30|GGTGGGGAGTTTGACTGGGGCGGTACACCT</td><td>>hg38_21_+_8433172_8446622_RNA45SN1_45S_50ntFlanks</td><td>12150</td><td>12179</td><td>30</td><td>GGTGGGGAGTTTGACTGGGGCGGTACACCT</td><td>76.4</td><td>-10.1</td><td>1.5</td><td>-6.7</td><td>2.5E-11</td><td>7.4E-11</td><td>MB231/MB468</td><td>MP</td></tr>
<tr><td>>am_trna162_MetCAT_6_-_26758550_26758622@37.54.18,trna164_MetCAT_6_-_26735574_26735646@37.54.18,trna20_MetCAT_16_+_71460396_71460468@37.54.18,trna21_MetCAT_6_+_26701712_26701784@37.54.18,trna27_MetCAT_6_+_26766444_26766516@37.54.18,trna75_MetCAT_6_+_28912352_28912424@37.54.18,trna97_MetCAT_6_-_28921042_28921114@37.54.18|AATCTGAAGGTCCTGAGT</td><td>>am_trna162_MetCAT_6_-_26758550_26758622</td><td>37</td><td>54</td><td>18</td><td>AATCTGAAGGTCCTGAGT</td><td>108.9</td><td>-10.6</td><td>1.6</td><td>-6.7</td><td>2.5E-11</td><td>7.4E-11</td><td>MB231/MB468</td><td>MP</td></tr>
<tr><td>>hg38_21_+_8433172_8446622_RNA45SN1_45S_50ntFlanks@3782.3799.18|CAGTGAAACTGCGAATGG</td><td>>hg38_21_+_8433172_8446622_RNA45SN1_45S_50ntFlanks</td><td>3782</td><td>3799</td><td>18</td><td>CAGTGAAACTGCGAATGG</td><td>216.5</td><td>-10.8</td><td>1.5</td><td>-7.1</td><td>1.7E-12</td><td>7.4E-11</td><td>BT20/MB468</td><td>MP</td></tr>
<tr><td>>hg38_21_+_8433172_8446622_RNA45SN1_45S_50ntFlanks@10806.10826.21|AACAATGTAGGTAAGGGAAGT</td><td>>hg38_21_+_8433172_8446622_RNA45SN1_45S_50ntFlanks</td><td>10806</td><td>10826</td><td>21</td><td>AACAATGTAGGTAAGGGAAGT</td><td>568.7</td><td>-8.5</td><td>1.3</td><td>-6.7</td><td>2.7E-11</td><td>7.4E-11</td><td>MB231/MB468</td><td>NUC</td></tr>
<tr><td>>hg38_21_+_8433172_8446622_RNA45SN1_45S_50ntFlanks@12416.12450.35|TTCGATGTCGGCTCTTCCTATCATTGTGAAGCAGA</td><td>>hg38_21_+_8433172_8446622_RNA45SN1_45S_50ntFlanks</td><td>12416</td><td>12450</td><td>35</td><td>TTCGATGTCGGCTCTTCCTATCATTGTGAAGCAGA</td><td>204.7</td><td>4.8</td><td>0.7</td><td>6.7</td><td>2.7E-11</td><td>7.4E-11</td><td>MB231/MB468</td><td>NUC</td></tr>
<tr><td>>hg38_21_+_8433172_8446622_RNA45SN1_45S_50ntFlanks@11604.11640.37|CGACTGTTTAATTAAAACAAAGCATCGCGAAGGCCCG</td><td>>hg38_21_+_8433172_8446622_RNA45SN1_45S_50ntFlanks</td><td>11604</td><td>11640</td><td>37</td><td>CGACTGTTTAATTAAAACAAAGCATCGCGAAGGCCCG</td><td>152.3</td><td>-11.1</td><td>1.7</td><td>-6.7</td><td>2.5E-11</td><td>7.4E-11</td><td>MB231/MB468</td><td>MP</td></tr>
<tr><td>>hg38_21_+_8433172_8446622_RNA45SN1_45S_50ntFlanks@8347.8375.29|GGGAAAGTTGAAAAGAACTTTGAAGAGAG</td><td>>hg38_21_+_8433172_8446622_RNA45SN1_45S_50ntFlanks</td><td>8347</td><td>8375</td><td>29</td><td>GGGAAAGTTGAAAAGAACTTTGAAGAGAG</td><td>1940.8</td><td>10.1</td><td>1.4</td><td>7.1</td><td>1.7E-12</td><td>7.4E-11</td><td>BT20/MB468</td><td>MP</td></tr>
<tr><td>>hg38_1_-_228634819_228635039_RNA5S12_5S_50ntFlanks@84.112.29|CTCGTCTGATCTCGGAAGCTAAGCAGGGT</td><td>>hg38_1_-_228634819_228635039_RNA5S12_5S_50ntFlanks</td><td>84</td><td>112</td><td>29</td><td>CTCGTCTGATCTCGGAAGCTAAGCAGGGT</td><td>897.9</td><td>-8.4</td><td>1.3</td><td>-6.7</td><td>2.7E-11</td><td>7.4E-11</td><td>MB231/MB468</td><td>NUC</td></tr>
<tr><td>>hg38_21_+_8433172_8446622_RNA45SN1_45S_50ntFlanks@9526.9571.46|GAAGCCAGAGGAAACTCTGGTGGAGGTCCGTAGCGGTCCTGACGTG</td><td>>hg38_21_+_8433172_8446622_RNA45SN1_45S_50ntFlanks</td><td>9526</td><td>9571</td><td>46</td><td>GAAGCCAGAGGAAACTCTGGTGGAGGTCCGTAGCGGTCCTGACGTG</td><td>1399.7</td><td>5.2</td><td>0.8</td><td>6.7</td><td>2.7E-11</td><td>7.4E-11</td><td>MB231/MB468</td><td>CYTO</td></tr>
<tr><td>>hg38_21_+_8433172_8446622_RNA45SN1_45S_50ntFlanks@11759.11777.19|CCTCGTCATCTAATTAGTG</td><td>>hg38_21_+_8433172_8446622_RNA45SN1_45S_50ntFlanks</td><td>11759</td><td>11777</td><td>19</td><td>CCTCGTCATCTAATTAGTG</td><td>428.8</td><td>-8.8</td><td>1.3</td><td>-6.7</td><td>2.4E-11</td><td>7.4E-11</td><td>BT20/MB468</td><td>CYTO</td></tr>
<tr><td>>hg38_21_+_8433172_8446622_RNA45SN1_45S_50ntFlanks@11581.11607.27|AACTGGTGCGGACCAGGGGAATCCGAC</td><td>>hg38_21_+_8433172_8446622_RNA45SN1_45S_50ntFlanks</td><td>11581</td><td>11607</td><td>27</td><td>AACTGGTGCGGACCAGGGGAATCCGAC</td><td>619.5</td><td>-4.6</td><td>0.7</td><td>-6.6</td><td>3.1E-11</td><td>7.4E-11</td><td>MB231/MB468</td><td>MITO</td></tr>
<tr><td>>hg38_21_+_8433172_8446622_RNA45SN1_45S_50ntFlanks@12788.12805.18|CGCGCGCGCGGGAGGGCG</td><td>>hg38_21_+_8433172_8446622_RNA45SN1_45S_50ntFlanks</td><td>12788</td><td>12805</td><td>18</td><td>CGCGCGCGCGGGAGGGCG</td><td>115.1</td><td>-10.7</td><td>1.6</td><td>-6.7</td><td>2.5E-11</td><td>7.4E-11</td><td>MB231/MB468</td><td>MP</td></tr>
<tr><td>>trna11_GluTTC_15_-_26327381_26327452@1.26.26,trna3_GluTTC_13_-_45492062_45492133@1.26.26|TCCCACATGGTCTAGCGGTTAGGATT</td><td>>trna11_GluTTC_15_-_26327381_26327452</td><td>1</td><td>26</td><td>26</td><td>TCCCACATGGTCTAGCGGTTAGGATT</td><td>925.6</td><td>5.6</td><td>0.8</td><td>6.7</td><td>2.4E-11</td><td>7.4E-11</td><td>BT20/MB468</td><td>CYTO</td></tr>
<tr><td>>hg38_21_+_8433172_8446622_RNA45SN1_45S_50ntFlanks@12960.12982.23|CTGGGTCGGGGTTTCGTACGTAG</td><td>>hg38_21_+_8433172_8446622_RNA45SN1_45S_50ntFlanks</td><td>12960</td><td>12982</td><td>23</td><td>CTGGGTCGGGGTTTCGTACGTAG</td><td>1284.8</td><td>-9.8</td><td>1.5</td><td>-6.6</td><td>3.3E-11</td><td>7.4E-11</td><td>BT20/MB468</td><td>NUC</td></tr>
<tr><td>>trna20_GluTTC_2_-_131094701_131094772@1.32.32,trna5_GluTTC_13_-_41634874_41634945@1.32.32|TCCCATATGGTCTAGCGGTTAGGATTCCTGGT</td><td>>trna20_GluTTC_2_-_131094701_131094772</td><td>1</td><td>32</td><td>32</td><td>TCCCATATGGTCTAGCGGTTAGGATTCCTGGT</td><td>280.7</td><td>11.4</td><td>1.7</td><td>6.6</td><td>3.1E-11</td><td>7.5E-11</td><td>MB231/MB468</td><td>MITO</td></tr>
<tr><td>>am_trnaMT_ValTAC_MT_+_1602_1670@42.69.28|AGATTTCAACTTAACTTGACCGCTCTGA</td><td>>am_trnaMT_ValTAC_MT_+_1602_1670</td><td>42</td><td>69</td><td>28</td><td>AGATTTCAACTTAACTTGACCGCTCTGA</td><td>441.9</td><td>5.7</td><td>0.9</td><td>6.7</td><td>2.0E-11</td><td>7.5E-11</td><td>BT20/MB468</td><td>MITO</td></tr>
<tr><td>>hg38_21_+_8433172_8446622_RNA45SN1_45S_50ntFlanks@10845.10862.18|ACTTCGGGATAAGGATTG</td><td>>hg38_21_+_8433172_8446622_RNA45SN1_45S_50ntFlanks</td><td>10845</td><td>10862</td><td>18</td><td>ACTTCGGGATAAGGATTG</td><td>1135.4</td><td>-6.6</td><td>1.0</td><td>-6.7</td><td>2.5E-11</td><td>7.5E-11</td><td>BT20/MB468</td><td>CYTO</td></tr>
<tr><td>>hg38_21_+_8433172_8446622_RNA45SN1_45S_50ntFlanks@12822.12859.38|CCGGGACCGGGGTCCGGTGCGGAGTGCCCTTCGTCCTG</td><td>>hg38_21_+_8433172_8446622_RNA45SN1_45S_50ntFlanks</td><td>12822</td><td>12859</td><td>38</td><td>CCGGGACCGGGGTCCGGTGCGGAGTGCCCTTCGTCCTG</td><td>79.9</td><td>-10.2</td><td>1.5</td><td>-6.7</td><td>2.5E-11</td><td>7.5E-11</td><td>MB231/MB468</td><td>MP</td></tr>
<tr><td>>hg38_21_+_8433172_8446622_RNA45SN1_45S_50ntFlanks@12938.12966.29|ACCATTCGTAGACGACCTGCTTCTGGGTC</td><td>>hg38_21_+_8433172_8446622_RNA45SN1_45S_50ntFlanks</td><td>12938</td><td>12966</td><td>29</td><td>ACCATTCGTAGACGACCTGCTTCTGGGTC</td><td>135.5</td><td>-10.9</td><td>1.6</td><td>-6.7</td><td>2.5E-11</td><td>7.6E-11</td><td>MB231/MB468</td><td>MP</td></tr>
<tr><td>>hg38_21_+_8433172_8446622_RNA45SN1_45S_50ntFlanks@6775.6806.32|CCTCCCGGGGCTACGCCTGTCTGAGCGTCGCT</td><td>>hg38_21_+_8433172_8446622_RNA45SN1_45S_50ntFlanks</td><td>6775</td><td>6806</td><td>32</td><td>CCTCCCGGGGCTACGCCTGTCTGAGCGTCGCT</td><td>321.5</td><td>4.6</td><td>0.7</td><td>6.7</td><td>2.7E-11</td><td>7.6E-11</td><td>MB231/MB468</td><td>CYTO</td></tr>
<tr><td>>hg38_21_+_8433172_8446622_RNA45SN1_45S_50ntFlanks@4289.4307.19|CGAGGATCCATTGGAGGGC</td><td>>hg38_21_+_8433172_8446622_RNA45SN1_45S_50ntFlanks</td><td>4289</td><td>4307</td><td>19</td><td>CGAGGATCCATTGGAGGGC</td><td>98.9</td><td>-10.5</td><td>1.6</td><td>-6.7</td><td>2.5E-11</td><td>7.6E-11</td><td>MB231/MB468</td><td>MP</td></tr>
<tr><td>>hg38_21_+_8433172_8446622_RNA45SN1_45S_50ntFlanks@11273.11295.23|CCGCCCCCGGGGAGCCCGGCGGG</td><td>>hg38_21_+_8433172_8446622_RNA45SN1_45S_50ntFlanks</td><td>11273</td><td>11295</td><td>23</td><td>CCGCCCCCGGGGAGCCCGGCGGG</td><td>164.4</td><td>-11.2</td><td>1.7</td><td>-6.7</td><td>2.5E-11</td><td>7.6E-11</td><td>MB231/MB468</td><td>MP</td></tr>
<tr><td>>hg38_21_+_8433172_8446622_RNA45SN1_45S_50ntFlanks@12894.12928.35|CTCGCCCGTCACGCACCGCACGTTCGTGGGGAACC</td><td>>hg38_21_+_8433172_8446622_RNA45SN1_45S_50ntFlanks</td><td>12894</td><td>12928</td><td>35</td><td>CTCGCCCGTCACGCACCGCACGTTCGTGGGGAACC</td><td>72.9</td><td>-10.1</td><td>1.5</td><td>-6.7</td><td>2.5E-11</td><td>7.6E-11</td><td>MB231/MB468</td><td>MP</td></tr>
<tr><td>>hg38_21_+_8433172_8446622_RNA45SN1_45S_50ntFlanks@6774.6806.33|TCCTCCCGGGGCTACGCCTGTCTGAGCGTCGCT</td><td>>hg38_21_+_8433172_8446622_RNA45SN1_45S_50ntFlanks</td><td>6774</td><td>6806</td><td>33</td><td>TCCTCCCGGGGCTACGCCTGTCTGAGCGTCGCT</td><td>13782.6</td><td>7.5</td><td>1.1</td><td>6.7</td><td>2.0E-11</td><td>7.6E-11</td><td>BT20/MB468</td><td>MITO</td></tr>
<tr><td>>hg38_21_+_8433172_8446622_RNA45SN1_45S_50ntFlanks@12357.12375.19|CCACAGGGATAACTGGCTT</td><td>>hg38_21_+_8433172_8446622_RNA45SN1_45S_50ntFlanks</td><td>12357</td><td>12375</td><td>19</td><td>CCACAGGGATAACTGGCTT</td><td>219.3</td><td>-9.9</td><td>1.5</td><td>-6.7</td><td>2.0E-11</td><td>7.6E-11</td><td>BT20/MB468</td><td>MITO</td></tr>
<tr><td>>hg38_21_+_8433172_8446622_RNA45SN1_45S_50ntFlanks@9554.9596.43|CGTAGCGGTCCTGACGTGCAAATCGGTCGTCCGACCTGGGTAT</td><td>>hg38_21_+_8433172_8446622_RNA45SN1_45S_50ntFlanks</td><td>9554</td><td>9596</td><td>43</td><td>CGTAGCGGTCCTGACGTGCAAATCGGTCGTCCGACCTGGGTAT</td><td>693.0</td><td>-12.4</td><td>1.9</td><td>-6.7</td><td>2.7E-11</td><td>7.6E-11</td><td>MB231/MB468</td><td>NUC</td></tr>
<tr><td>>trna10_LysCTT_16_+_3241501_3241573@4.35.32,trna13_LysCTT_14_-_58706613_58706685@4.35.32,trna2_LysCTT_15_+_79152904_79152976@4.35.32|CGGCTAGCTCAGTCGGTAGAGCATGGGACTCT</td><td>>trna10_LysCTT_16_+_3241501_3241573</td><td>4</td><td>35</td><td>32</td><td>CGGCTAGCTCAGTCGGTAGAGCATGGGACTCT</td><td>386.8</td><td>7.2</td><td>1.1</td><td>6.7</td><td>2.7E-11</td><td>7.6E-11</td><td>MB231/MB468</td><td>NUC</td></tr>
<tr><td>>am_trnaMT_PheGAA_MT_+_577_647@40.57.18|TGTTTAGACGGGCTCACA</td><td>>am_trnaMT_PheGAA_MT_+_577_647</td><td>40</td><td>57</td><td>18</td><td>TGTTTAGACGGGCTCACA</td><td>299.5</td><td>8.5</td><td>1.3</td><td>6.7</td><td>2.7E-11</td><td>7.6E-11</td><td>MB231/MB468</td><td>CYTO</td></tr>
<tr><td>>hg38_21_+_8433172_8446622_RNA45SN1_45S_50ntFlanks@9713.9735.23|GCGAATGATTAGAGGTCTTGGGG</td><td>>hg38_21_+_8433172_8446622_RNA45SN1_45S_50ntFlanks</td><td>9713</td><td>9735</td><td>23</td><td>GCGAATGATTAGAGGTCTTGGGG</td><td>106.6</td><td>-10.6</td><td>1.6</td><td>-6.7</td><td>2.5E-11</td><td>7.6E-11</td><td>MB231/MB468</td><td>MP</td></tr>
<tr><td>>MI0000477|hsa-mir-146a&WithFlank&5|+|160485346|160485456@27.49.23(+1C)&[MIMAT0000449&hsa-miR-146a-5p&offsets|0|+1(+1C);m-129&5|+|160485372|160485393&offsets|0|+1(+1C)]|TGAGAACTGAATTCCATGGGTTGC</td><td>>MI0000477|hsa-mir-146a&WithFlank&5|+|160485346|160485456</td><td>27</td><td>49</td><td>23(+1C)</td><td>TGAGAACTGAATTCCATGGGTTGC</td><td>219.3</td><td>11.0</td><td>1.6</td><td>6.7</td><td>2.5E-11</td><td>7.6E-11</td><td>MB231/MB468</td><td>MP</td></tr>
</table>

| | | | | | | | | | | | | | |
|---|---|---|---|---|---|---|---|---|---|---|---|---|---|
| >hg38_21_+_8433172_8446622_RNA45SN1_45S_50ntFlanks@8109.8141.33\|GCGGGGCGCGGGACATGTGGCGTACGGAAGACC | >hg38_21_+_8433172_8446622_RNA45SN1_45S_50ntFlanks | 8109 | 8141 | 33 | GCGGGGCGCGGGACATGTGGCGTACGGAAGACC | 191.1 | -11.4 | 1.7 | -6.7 | 2.5E-11 | 7.6E-11 | MB231/MB468 | MP |
| >hg38_21_+_8433172_8446622_RNA45SN1_45S_50ntFlanks@8194.8215.22\|CCCGTGGACGGTGTGAGGCCGG | >hg38_21_+_8433172_8446622_RNA45SN1_45S_50ntFlanks | 8194 | 8215 | 22 | CCCGTGGACGGTGTGAGGCCGG | 2749.8 | -5.9 | 0.9 | -6.7 | 2.8E-11 | 7.6E-11 | MB231/MB468 | CYTO |
| >hg38_21_+_8433172_8446622_RNA45SN1_45S_50ntFlanks@11205.11225.21\|GCGGCGGCGACTCTGGACGCG | >hg38_21_+_8433172_8446622_RNA45SN1_45S_50ntFlanks | 11205 | 11225 | 21 | GCGGCGGCGACTCTGGACGCG | 359.7 | -8.2 | 1.2 | -6.6 | 3.2E-11 | 7.6E-11 | MB231/MB468 | MITO |
| >MI0001641\|hsa-mir-429&WithFlank&1\|+\|1168999\|1169093@57.77.21(+1C)&[MIMAT0001536&hsa-miR-429&offsets\|0\|-1(+1C);m-207&1\|+\|1169055\|1169076&offsets\|0\|-1(+1C)]\|TAATACTGTCTGGTAAAACCGC | >MI0001641\|hsa-mir-429&WithFlank&1\|+\|1168999\|1169093 | 57 | 77 | 21(+1C) | TAATACTGTCTGGTAAAACCGC | 212.7 | -10.8 | 1.5 | -7.1 | 1.8E-12 | 7.7E-11 | BT20/MB468 | MP |
| >hg38_21_+_8433172_8446622_RNA45SN1_45S_50ntFlanks@4372.4398.27\|AAAAAGCTCGTAGTTGGATCTTGGGAG | >hg38_21_+_8433172_8446622_RNA45SN1_45S_50ntFlanks | 4372 | 4398 | 27 | AAAAAGCTCGTAGTTGGATCTTGGGAG | 187.0 | -11.4 | 1.7 | -6.7 | 2.6E-11 | 7.7E-11 | MB231/MB468 | MP |
| >hg38_21_+_8433172_8446622_RNA45SN1_45S_50ntFlanks@10345.10372.28\|TTCAAACGAGAACTTTGAAGGCCGAAGT | >hg38_21_+_8433172_8446622_RNA45SN1_45S_50ntFlanks | 10345 | 10372 | 28 | TTCAAACGAGAACTTTGAAGGCCGAAGT | 191.8 | 4.3 | 0.6 | 6.7 | 2.8E-11 | 7.7E-11 | MB231/MB468 | CYTO |
| >hg38_21_+_8433172_8446622_RNA45SN1_45S_50ntFlanks@11363.11383.21\|CGGGCGGCGGCGGTCGGCGGG | >hg38_21_+_8433172_8446622_RNA45SN1_45S_50ntFlanks | 11363 | 11383 | 21 | CGGGCGGCGGCGGTCGGCGGG | 470.3 | -12.0 | 1.7 | -7.0 | 1.8E-12 | 7.7E-11 | BT20/MB468 | MP |
| >hg38_21_+_8433172_8446622_RNA45SN1_45S_50ntFlanks@12444.12473.30\|AAGCAGAATTCACCAAGCGTTGGATTGTTC | >hg38_21_+_8433172_8446622_RNA45SN1_45S_50ntFlanks | 12444 | 12473 | 30 | AAGCAGAATTCACCAAGCGTTGGATTGTTC | 385.0 | -11.7 | 1.7 | -7.0 | 1.8E-12 | 7.7E-11 | BT20/MB468 | MP |
| >hg38_21_+_8433172_8446622_RNA45SN1_45S_50ntFlanks@5448.5466.19\|GATCGGCCCCGCCGGGGTC | >hg38_21_+_8433172_8446622_RNA45SN1_45S_50ntFlanks | 5448 | 5466 | 19 | GATCGGCCCCGCCGGGGTC | 540.4 | -12.2 | 1.7 | -7.0 | 1.8E-12 | 7.7E-11 | BT20/MB468 | MP |
| >hg38_21_+_8433172_8446622_RNA45SN1_45S_50ntFlanks@9567.9592.26\|ACGTGCAAATCGGTCGTCCGACCTGG | >hg38_21_+_8433172_8446622_RNA45SN1_45S_50ntFlanks | 9567 | 9592 | 26 | ACGTGCAAATCGGTCGTCCGACCTGG | 665.3 | -8.3 | 1.2 | -6.6 | 3.2E-11 | 7.7E-11 | MB231/MB468 | MITO |
| >hg38_21_+_8433172_8446622_RNA45SN1_45S_50ntFlanks@12974.13001.28\|CGTACGTAGCAGAGCAGCTCCCTCGCTG | >hg38_21_+_8433172_8446622_RNA45SN1_45S_50ntFlanks | 12974 | 13001 | 28 | CGTACGTAGCAGAGCAGCTCCCTCGCTG | 136.2 | -4.6 | 0.7 | -6.7 | 2.8E-11 | 7.7E-11 | MB231/MB468 | CYTO |
| >hg38_21_+_8433172_8446622_RNA45SN1_45S_50ntFlanks@6650.6667.18\|ACGACTCTTAGCGGTGGA | >hg38_21_+_8433172_8446622_RNA45SN1_45S_50ntFlanks | 6650 | 6667 | 18 | ACGACTCTTAGCGGTGGA | 1745.8 | 6.1 | 0.9 | 6.7 | 2.0E-11 | 7.7E-11 | BT20/MB468 | MITO |
| >hg38_21_+_8433172_8446622_RNA45SN1_45S_50ntFlanks@11569.11605.37\|CAGCCGACTTAGAACTGGTGCGGACCAGGGGAATCCG | >hg38_21_+_8433172_8446622_RNA45SN1_45S_50ntFlanks | 11569 | 11605 | 37 | CAGCCGACTTAGAACTGGTGCGGACCAGGGGAATCCG | 1462.8 | -9.0 | 1.3 | -6.7 | 2.8E-11 | 7.7E-11 | MB231/MB468 | NUC |
| >hg38_21_+_8433172_8446622_RNA45SN1_45S_50ntFlanks@3912.3938.27\|GATGCGTGCATTTATCAGATCAAAACC | >hg38_21_+_8433172_8446622_RNA45SN1_45S_50ntFlanks | 3912 | 3938 | 27 | GATGCGTGCATTTATCAGATCAAAACC | 148.6 | -11.1 | 1.7 | -6.7 | 2.6E-11 | 7.7E-11 | MB231/MB468 | MP |
| >hg38_21_+_8433172_8446622_RNA45SN1_45S_50ntFlanks@10663.10710.48\|CCTTGGAAAGCGTCGCGGTTCCGGCGGCGTCCGGTGAGCTCTCGCTGG | >hg38_21_+_8433172_8446622_RNA45SN1_45S_50ntFlanks | 10663 | 10710 | 48 | CCTTGGAAAGCGTCGCGGTTCCGGCGGCGTCCGGTGAGCTCTCGCTGG | 720.3 | -12.5 | 1.9 | -6.7 | 2.8E-11 | 7.7E-11 | MB231/MB468 | NUC |
| >MI0000102\|hsa-mir-100&WithFlank&11\|-\|122152223\|122152314@20.39.20&[MIMAT0000098&hsa-miR-100-5p&offsets\|+1\|-1;m-27&11\|-\|122152276\|122152296&offsets\|+1\|0]\|ACCCGTAGATCCGAACTTGT | >MI0000102\|hsa-mir-100&WithFlank&11\|-\|122152223\|122152314 | 20 | 39 | 20 | ACCCGTAGATCCGAACTTGT | 223.6 | 11.0 | 1.7 | 6.6 | 3.2E-11 | 7.7E-11 | MB231/MB468 | MITO |
| >hg38_21_+_8433172_8446622_RNA45SN1_45S_50ntFlanks@11092.11114.23\|CGGCGGGGGGAGAAGGGTCGGGG | >hg38_21_+_8433172_8446622_RNA45SN1_45S_50ntFlanks | 11092 | 11114 | 23 | CGGCGGGGGGAGAAGGGTCGGGG | 125.3 | -10.8 | 1.6 | -6.7 | 2.6E-11 | 7.8E-11 | MB231/MB468 | MP |
| >hg38_21_+_8433172_8446622_RNA45SN1_45S_50ntFlanks@7988.8017.30\|CAGACGTGGCGACCCGCTGAATTTAAGCAT | >hg38_21_+_8433172_8446622_RNA45SN1_45S_50ntFlanks | 7988 | 8017 | 30 | CAGACGTGGCGACCCGCTGAATTTAAGCAT | 1644.8 | -4.3 | 0.6 | -6.6 | 3.2E-11 | 7.8E-11 | MB231/MB468 | MITO |
| >hg38_21_+_8433172_8446622_RNA45SN1_45S_50ntFlanks@5035.5061.27\|CATGGCCGTTCTTAGTTGGTGGAGCGA | >hg38_21_+_8433172_8446622_RNA45SN1_45S_50ntFlanks | 5035 | 5061 | 27 | CATGGCCGTTCTTAGTTGGTGGAGCGA | 449.4 | -12.6 | 1.9 | -6.6 | 3.2E-11 | 7.8E-11 | MB231/MB468 | MITO |
| >hg38_21_+_8433172_8446622_RNA45SN1_45S_50ntFlanks@11569.11586.18\|CAGCCGACTTAGAACTGG | >hg38_21_+_8433172_8446622_RNA45SN1_45S_50ntFlanks | 11569 | 11586 | 18 | CAGCCGACTTAGAACTGG | 2282.1 | -5.3 | 0.8 | -6.7 | 2.6E-11 | 7.8E-11 | MB231/MB468 | MP |
| >hg38_21_+_8433172_8446622_RNA45SN1_45S_50ntFlanks@5000.5024.25\|TTGATAGCTCTTTCTCGATTCCGTG | >hg38_21_+_8433172_8446622_RNA45SN1_45S_50ntFlanks | 5000 | 5024 | 25 | TTGATAGCTCTTTCTCGATTCCGTG | 79.6 | -10.2 | 1.5 | -6.7 | 2.6E-11 | 7.8E-11 | MB231/MB468 | MP |
| >hg38_21_+_8433172_8446622_RNA45SN1_45S_50ntFlanks@10544.10563.20\|TCCAGTGCGGTAACGCGACC | >hg38_21_+_8433172_8446622_RNA45SN1_45S_50ntFlanks | 10544 | 10563 | 20 | TCCAGTGCGGTAACGCGACC | 80.5 | -10.2 | 1.5 | -6.7 | 2.6E-11 | 7.8E-11 | MB231/MB468 | MP |
| >hg38_21_+_8433172_8446622_RNA45SN1_45S_50ntFlanks@10787.10823.37\|ACAGCCTCTGGCATGTTGGAACAATGTAGGTAAGGGA | >hg38_21_+_8433172_8446622_RNA45SN1_45S_50ntFlanks | 10787 | 10823 | 37 | ACAGCCTCTGGCATGTTGGAACAATGTAGGTAAGGGA | 377.5 | -8.3 | 1.2 | -6.6 | 3.3E-11 | 7.8E-11 | MB231/MB468 | MITO |
| >hg38_21_+_8433172_8446622_RNA45SN1_45S_50ntFlanks@9225.9247.23\|CCGAAGAGGGGGACGGCGGAGCG | >hg38_21_+_8433172_8446622_RNA45SN1_45S_50ntFlanks | 9225 | 9247 | 23 | CCGAAGAGGGGGACGGCGGAGCG | 85.3 | -10.3 | 1.5 | -6.7 | 2.6E-11 | 7.8E-11 | MB231/MB468 | MP |
| >hg38_21_+_8433172_8446622_RNA45SN1_45S_50ntFlanks@4128.4162.35\|CGATTCCGGAGAGGGAGCCTGAGAAACGGCTACCA | >hg38_21_+_8433172_8446622_RNA45SN1_45S_50ntFlanks | 4128 | 4162 | 35 | CGATTCCGGAGAGGGAGCCTGAGAAACGGCTACCA | 197.3 | -11.5 | 1.7 | -6.7 | 2.6E-11 | 7.8E-11 | MB231/MB468 | MP |
| >hg38_21_+_8433172_8446622_RNA45SN1_45S_50ntFlanks@9494.9526.33\|CCGAAAGATGGTGAACTATGCCTGGGCAGGGCG | >hg38_21_+_8433172_8446622_RNA45SN1_45S_50ntFlanks | 9494 | 9526 | 33 | CCGAAAGATGGTGAACTATGCCTGGGCAGGGCG | 102.0 | -10.5 | 1.6 | -6.7 | 2.6E-11 | 7.8E-11 | MB231/MB468 | MP |
| >MI0000289\|hsa-mir-181a-1&WithFlank&1\|-\|198859038\|198859159@30.54.25(+1U);MI0000269\|hsa-mir-181a-2&WithFlank&9\|+\|124692436\|124692557@45.69.25(+1U)&[MIMAT0000256&hsa-miR-181a-5p&offsets\|0\|+2(+1U);m-32&1\|-\|198859108\|198859130&offsets\|0\|+2(+1U)];[MIMAT0000256_1&hsa-miR-181a-5p&offsets\|0\|+2(+1U);m-32&9\|+\|124692480\|124692502&offsets\|0\|+2(+1U)]\|AACATTCAACGCTGTCGGTGAGTTTT | >MI0000289\|hsa-mir-181a-1&WithFlank&1\|-\|198859038\|198859159 | 30 | 54 | 25(+1U) | AACATTCAACGCTGTCGGTGAGTTTT | 465.5 | 7.7 | 1.2 | 6.7 | 2.8E-11 | 7.8E-11 | MB231/MB468 | NUC |
| >hg38_21_+_8433172_8446622_RNA45SN1_45S_50ntFlanks@10783.10816.34\|GTGAACAGCCTCTGGCATGTTGGAACAATGTAGG | >hg38_21_+_8433172_8446622_RNA45SN1_45S_50ntFlanks | 10783 | 10816 | 34 | GTGAACAGCCTCTGGCATGTTGGAACAATGTAGG | 91.3 | -10.4 | 1.6 | -6.7 | 2.6E-11 | 7.8E-11 | MB231/MB468 | MP |
| >hg38_21_+_8433172_8446622_RNA45SN1_45S_50ntFlanks@11854.11872.19\|TTGGCGGAATCAGCGGGGA | >hg38_21_+_8433172_8446622_RNA45SN1_45S_50ntFlanks | 11854 | 11872 | 19 | TTGGCGGAATCAGCGGGGA | 82.0 | -10.2 | 1.5 | -6.7 | 2.6E-11 | 7.8E-11 | MB231/MB468 | MP |
| >MI0000283\|hsa-mir-203a&WithFlank&14\|+\|104117399\|104117520@71.92.22&[MIMAT0000264&hsa-miR-203a-3p&offsets\|0\|0;m-79&14\|+\|104117469\|104117490&offsets\|0\|0]\|GTGAAATGTTTAGGACCACTAG | >MI0000283\|hsa-mir-203a&WithFlank&14\|+\|104117399\|104117520 | 71 | 92 | 22 | GTGAAATGTTTAGGACCACTAG | 545.6 | 12.4 | 1.7 | 7.2 | 4.3E-13 | 7.8E-11 | BT20/MB231 | MP |
| >MI0000446\|hsa-mir-125b-1&WithFlank&11\|-\|122099751\|122099850@21.42.22;MI0000470\|hsa-mir-125b-2&WithFlank&21\|+\|16590231\|16590331@23.44.22&[MIMAT0000423&hsa-miR-125b-5p&offsets\|0\|0;m-67&11\|-\|122099809\|122099830&offsets\|0\|0];[MIMAT0000423_1&hsa-miR-125b-5p&offsets\|0\|0;m-68&21\|+\|16590253\|16590274&offsets\|0\|0]\|TCCCTGAGACCCTAACTTGTGA | >MI0000446\|hsa-mir-125b-1&WithFlank&11\|-\|122099751\|122099850 | 21 | 42 | 22 | TCCCTGAGACCCTAACTTGTGA | 23822.0 | -7.1 | 1.0 | -7.0 | 2.6E-12 | 7.9E-11 | BT20/MB231 | MITO |
| >trna119_LysCTT_1_-_145395522_145395594@3.33.31,trna11_LysCTT_5_-_180648979_180649051@3.33.31,trna13_LysCTT_6_+_26556774_26556846@3.33.31,trna7_LysCTT_16_+_3225692_3225764@3.33.31,trna9_LysCTT_5_+_180634755_180634827@3.33.31\|CCGGCTAGCTCAGTCGGTAGAGCATGAGACT | >trna119_LysCTT_1_-_145395522_145395594 | 3 | 33 | 31 | CCGGCTAGCTCAGTCGGTAGAGCATGAGACT | 1168.9 | 7.7 | 1.2 | 6.6 | 3.3E-11 | 7.9E-11 | MB231/MB468 | MITO |
| >hg38_21_+_8433172_8446622_RNA45SN1_45S_50ntFlanks@9568.9594.27\|CGTGCAAATCGGTCGTCCGACCTGGGT | >hg38_21_+_8433172_8446622_RNA45SN1_45S_50ntFlanks | 9568 | 9594 | 27 | CGTGCAAATCGGTCGTCCGACCTGGGT | 469.3 | -3.8 | 0.6 | -6.7 | 2.6E-11 | 7.9E-11 | BT20/MB468 | CYTO |
| >hg38_21_+_8433172_8446622_RNA45SN1_45S_50ntFlanks@11362.11383.22\|TCGGGCGGCGGCGGTCGGCGGG | >hg38_21_+_8433172_8446622_RNA45SN1_45S_50ntFlanks | 11362 | 11383 | 22 | TCGGGCGGCGGCGGTCGGCGGG | 134.4 | -10.9 | 1.6 | -6.7 | 2.6E-11 | 7.9E-11 | MB231/MB468 | MP |
| >hg38_21_+_8433172_8446622_RNA45SN1_45S_50ntFlanks@9178.9195.18\|CGCCGTCGGGCCCGGGGG | >hg38_21_+_8433172_8446622_RNA45SN1_45S_50ntFlanks | 9178 | 9195 | 18 | CGCCGTCGGGCCCGGGGG | 163.9 | -11.2 | 1.7 | -6.7 | 2.6E-11 | 7.9E-11 | MB231/MB468 | MP |
| >hg38_21_+_8433172_8446622_RNA45SN1_45S_50ntFlanks@8182.8208.27\|ATCGAGGCCCAGCCCGTGGACGGTGTG | >hg38_21_+_8433172_8446622_RNA45SN1_45S_50ntFlanks | 8182 | 8208 | 27 | ATCGAGGCCCAGCCCGTGGACGGTGTG | 122.3 | -9.0 | 1.3 | -6.7 | 2.1E-11 | 7.9E-11 | BT20/MB468 | MITO |
| >hg38_21_+_8433172_8446622_RNA45SN1_45S_50ntFlanks@9305.9338.34\|CCAAGGAGTCTAACACGTGCGCGAGTCGGGGGCT | >hg38_21_+_8433172_8446622_RNA45SN1_45S_50ntFlanks | 9305 | 9338 | 34 | CCAAGGAGTCTAACACGTGCGCGAGTCGGGGGCT | 1610.5 | -6.9 | 1.0 | -6.7 | 2.1E-11 | 7.9E-11 | BT20/MB468 | MITO |
| >MI0000440\|hsa-mir-27b&WithFlank&9\|+\|95085439\|95085547@25.47.23&[MIMAT0004588&hsa-miR-27b-5p&offsets\|0\|+1;m-240&9\|+\|95085463\|95085484&offsets\|0\|+1]\|AGAGCTTAGCTGATTGGTGAACA | >MI0000440\|hsa-mir-27b&WithFlank&9\|+\|95085439\|95085547 | 25 | 47 | 23 | AGAGCTTAGCTGATTGGTGAACA | 522.9 | 8.8 | 1.3 | 6.7 | 2.6E-11 | 8.0E-11 | BT20/MB468 | CYTO |
| >hg38_21_+_8433172_8446622_RNA45SN1_45S_50ntFlanks@4774.4793.20\|AGTTCCGACCATAAACGATG | >hg38_21_+_8433172_8446622_RNA45SN1_45S_50ntFlanks | 4774 | 4793 | 20 | AGTTCCGACCATAAACGATG | 87.7 | -10.3 | 1.5 | -6.7 | 2.7E-11 | 8.0E-11 | MB231/MB468 | MP |
| >MI0000108\|hsa-mir-103a-2&WithFlank&20\|+\|3917488\|3917577@54.76.23(+1U);MI0000109\|hsa-mir-103a-1&WithFlank&5\|-\|168560890\|168560979@54.76.23(+1U)&[MIMAT0000101&hsa-miR-103a-3p&offsets\|0\|0(+1U);m-12&20\|+\|3917541\|3917563&offsets\|0\|0(+1U)];[MIMAT0000101_1&hsa-miR-103a-3p&offsets\|0\|0(+1U);m-13&5\|-\|168560904\|168560926&offsets\|0\|0(+1U);m-44220&5\|-\|168560908\|168560926&offsets\|0\|+4(+1U)]\|AGCAGCATTGTACAGGGCTATGAT | >MI0000108\|hsa-mir-103a-2&WithFlank&20\|+\|3917488\|3917577 | 54 | 76 | 23(+1U) | AGCAGCATTGTACAGGGCTATGAT | 5008.6 | 6.7 | 1.0 | 6.6 | 3.3E-11 | 8.0E-11 | MB231/MB468 | MITO |
| >MI0003188\|hsa-mir-503&WithFlank&X\|-\|134546322\|134546404@12.31.20&[MIMAT0002874&hsa-miR-503-5p&offsets\|0\|-3;m-297&X\|-\|134546371\|134546393&offsets\|0\|-3]\|TAGCAGCGGGAACAGTTCTG | >MI0003188\|hsa-mir-503&WithFlank&X\|-\|134546322\|134546404 | 12 | 31 | 20 | TAGCAGCGGGAACAGTTCTG | 188.2 | 10.8 | 1.6 | 6.6 | 3.3E-11 | 8.0E-11 | MB231/MB468 | MITO |
| >hg38_21_+_8433172_8446622_RNA45SN1_45S_50ntFlanks@10397.10424.28\|AGTTGAACATGGGTCAGTCGGTCCTGAG | >hg38_21_+_8433172_8446622_RNA45SN1_45S_50ntFlanks | 10397 | 10424 | 28 | AGTTGAACATGGGTCAGTCGGTCCTGAG | 341.7 | 8.1 | 1.2 | 6.7 | 2.6E-11 | 8.0E-11 | BT20/MB468 | CYTO |
| >hg38_21_+_8433172_8446622_RNA45SN1_45S_50ntFlanks@11759.11777.19\|CCTCGTCATCTAATTAGTG | >hg38_21_+_8433172_8446622_RNA45SN1_45S_50ntFlanks | 11759 | 11777 | 19 | CCTCGTCATCTAATTAGTG | 155.5 | -9.4 | 1.4 | -6.7 | 2.1E-11 | 8.0E-11 | BT20/MB468 | MITO |
| >MI0000650\|hsa-mir-200c&WithFlank&12\|+\|6963693\|6963772@50.71.22(+1G)&[MIMAT0000617&hsa-miR-200c-3p&offsets\|0\|-1(+1G);m-54&12\|+\|6963742\|6963764&offsets\|0\|-1(+1G)]\|TAATACTGCCGGGTAATGATGGG | >MI0000650\|hsa-mir-200c&WithFlank&12\|+\|6963693\|6963772 | 50 | 71 | 22(+1G) | TAATACTGCCGGGTAATGATGGG | 94.7 | -10.4 | 1.6 | -6.7 | 2.7E-11 | 8.0E-11 | MB231/MB468 | MP |
| >MI0003560\|hsa-mir-92b&WithFlank&1\|+\|155195171\|155195278@67.86.20(+1U)&[MIMAT0003218&hsa-miR-92b-3p&offsets\|0\|-2(+1U);m-115&1\|+\|155195237\|155195258&offsets\|0\|-2(+1U)]\|TATTGCACTCGTCCCGGCCTT | >MI0003560\|hsa-mir-92b&WithFlank&1\|+\|155195171\|155195278 | 67 | 86 | 20(+1U) | TATTGCACTCGTCCCGGCCTT | 260.2 | 11.2 | 1.7 | 6.6 | 3.4E-11 | 8.0E-11 | MB231/MB468 | MITO |
| >hg38_21_+_8433172_8446622_RNA45SN1_45S_50ntFlanks@12274.12299.26\|CAGTACGAATACAGACCGTGAAAGCG | >hg38_21_+_8433172_8446622_RNA45SN1_45S_50ntFlanks | 12274 | 12299 | 26 | CAGTACGAATACAGACCGTGAAAGCG | 1362.5 | -10.1 | 1.5 | -6.6 | 3.4E-11 | 8.0E-11 | MB231/MB468 | MITO |

<table>
<tr><td>>hg38_1_-_228634819_228635039_RNA5S12_5S_50ntFlanks@130.169.40|TGGATGGGAGACCGCCTGGGAATACCGGGTGCTGTAGGCT</td><td>>hg38_1_-_228634819_228635039_RNA5S12_5S_50ntFlanks</td><td>130</td><td>169</td><td>40</td><td>TGGATGGGAGACCGCCTGGGAATACCGGGTGCTGTAGGCT</td><td>854.4</td><td>6.3</td><td>0.9</td><td>7.0</td><td>1.9E-12</td><td>8.0E-11</td><td>BT20/MB468</td><td>MP</td></tr>
<tr><td>>trna10_GluCTC_3_-_103879524_103879595@16.33.18,trna116_GluCTC_1_-_145399233_145399304@16.33.18,trna25_GluCTC_2_-_71273488_71273560@16.33.18,trna59_GluCTC_1_+_249168447_249168518@16.33.18,trna71_GluCTC_1_-_161439189_161439260@16.33.18,trna74_GluCTC_1_-_161431809_161431880@16.33.18,trna77_GluCTC_1_-_161424398_161424469@16.33.18,trna77_GluCTC_6_+_28949976_28950047@16.33.18,trna80_GluCTC_1_-_161417018_161417089@16.33.18,trna87_GluCTC_6_-_126101393_126101464@16.33.18|TGGTTAGGATTCGGCGCT</td><td>>trna10_GluCTC_3_-_103879524_103879595</td><td>16</td><td>33</td><td>18</td><td>TGGTTAGGATTCGGCGCT</td><td>1420.5</td><td>5.8</td><td>0.9</td><td>6.7</td><td>2.6E-11</td><td>8.0E-11</td><td>BT20/MB468</td><td>CYTO</td></tr>
<tr><td>>hg38_21_+_8433172_8446622_RNA45SN1_45S_50ntFlanks@6719.6737.19|TGAATTGCAGGACACATTG</td><td>>hg38_21_+_8433172_8446622_RNA45SN1_45S_50ntFlanks</td><td>6719</td><td>6737</td><td>19</td><td>TGAATTGCAGGACACATTG</td><td>2331.0</td><td>-6.7</td><td>1.0</td><td>-6.7</td><td>2.9E-11</td><td>8.0E-11</td><td>MB231/MB468</td><td>CYTO</td></tr>
<tr><td>>hg38_21_+_8433172_8446622_RNA45SN1_45S_50ntFlanks@10393.10431.39|CAGCAGTTGAACATGGGTCAGTCGGTCCTGAGAGATGGG</td><td>>hg38_21_+_8433172_8446622_RNA45SN1_45S_50ntFlanks</td><td>10393</td><td>10431</td><td>39</td><td>CAGCAGTTGAACATGGGTCAGTCGGTCCTGAGAGATGGG</td><td>169.4</td><td>-11.3</td><td>1.7</td><td>-6.7</td><td>2.7E-11</td><td>8.0E-11</td><td>MB231/MB468</td><td>MP</td></tr>
<tr><td>>hg38_21_+_8433172_8446622_RNA45SN1_45S_50ntFlanks@4385.4410.26|TTGGATCTTGGGAGCGGGCGGGCGGT</td><td>>hg38_21_+_8433172_8446622_RNA45SN1_45S_50ntFlanks</td><td>4385</td><td>4410</td><td>26</td><td>TTGGATCTTGGGAGCGGGCGGGCGGT</td><td>138.2</td><td>-11.0</td><td>1.6</td><td>-6.7</td><td>2.7E-11</td><td>8.1E-11</td><td>MB231/MB468</td><td>MP</td></tr>
<tr><td>>am_trna128_GlyGCC_6_-_27870686_27870756@1.35.35,trna18_GlyGCC_16_+_70822597_70822667@1.35.35,trna19_GlyGCC_16_+_70823410_70823480@1.35.35,trna19_GlyGCC_2_-_157257659_157257729@1.35.35,trna24_GlyGCC_16_-_70812942_70813012@1.35.35,trna25_GlyGCC_16_-_70812114_70812184@1.35.35,trna5_GlyGCC_17_+_8029064_8029134@1.35.35,trna68_GlyGCC_1_-_161493637_161493707@1.35.35|GCATTGGTGGTTCAGTGGTAGAATTCTCGCCTGCC</td><td>>am_trna128_GlyGCC_6_-_27870686_27870756</td><td>1</td><td>35</td><td>35</td><td>GCATTGGTGGTTCAGTGGTAGAATTCTCGCCTGCC</td><td>2071.2</td><td>5.1</td><td>0.8</td><td>6.7</td><td>2.6E-11</td><td>8.1E-11</td><td>BT20/MB468</td><td>CYTO</td></tr>
<tr><td>>hg38_21_+_8433172_8446622_RNA45SN1_45S_50ntFlanks@6646.6681.36|TCGTACGACTCTTAGCGGTGGATCACTCGGCTCGTG</td><td>>hg38_21_+_8433172_8446622_RNA45SN1_45S_50ntFlanks</td><td>6646</td><td>6681</td><td>36</td><td>TCGTACGACTCTTAGCGGTGGATCACTCGGCTCGTG</td><td>33238.2</td><td>-6.0</td><td>0.9</td><td>-6.7</td><td>2.7E-11</td><td>8.1E-11</td><td>MB231/MB468</td><td>MP</td></tr>
<tr><td>>hg38_21_+_8433172_8446622_RNA45SN1_45S_50ntFlanks@12418.12446.29|CGATGTCGGCTCTTCCTATCATTGTGAAG</td><td>>hg38_21_+_8433172_8446622_RNA45SN1_45S_50ntFlanks</td><td>12418</td><td>12446</td><td>29</td><td>CGATGTCGGCTCTTCCTATCATTGTGAAG</td><td>3156.1</td><td>-4.9</td><td>0.7</td><td>-6.6</td><td>3.4E-11</td><td>8.1E-11</td><td>MB231/MB468</td><td>MITO</td></tr>
<tr><td>>hg38_21_+_8433172_8446622_RNA45SN1_45S_50ntFlanks@4380.4400.21|CGTAGTTGGATCTTGGGAGCG</td><td>>hg38_21_+_8433172_8446622_RNA45SN1_45S_50ntFlanks</td><td>4380</td><td>4400</td><td>21</td><td>CGTAGTTGGATCTTGGGAGCG</td><td>103.0</td><td>-10.5</td><td>1.6</td><td>-6.7</td><td>2.7E-11</td><td>8.1E-11</td><td>MB231/MB468</td><td>MP</td></tr>
<tr><td>>hg38_21_+_8433172_8446622_RNA45SN1_45S_50ntFlanks@3873.3898.26|TAATACATGCCGACGGGCGCTGACCC</td><td>>hg38_21_+_8433172_8446622_RNA45SN1_45S_50ntFlanks</td><td>3873</td><td>3898</td><td>26</td><td>TAATACATGCCGACGGGCGCTGACCC</td><td>2309.8</td><td>-8.1</td><td>1.2</td><td>-6.7</td><td>2.9E-11</td><td>8.1E-11</td><td>MB231/MB468</td><td>NUC</td></tr>
<tr><td>>MI0003560|hsa-mir-92b&WithFlank&1|+|155195171|155195278@67.88.22(+3U)&[MIMAT0003218&hsa-miR-92b-3p&offsets|0|0(+3U);m-115&1|+|155195237|155195258&offsets|0|0(+3U)]|TATTGCACTCGTCCCGGCCTCCTTT</td><td>>MI0003560|hsa-mir-92b&WithFlank&1|+|155195171|155195278</td><td>67</td><td>88</td><td>22(+3U)</td><td>TATTGCACTCGTCCCGGCCTCCTTT</td><td>251.7</td><td>11.2</td><td>1.7</td><td>6.6</td><td>3.4E-11</td><td>8.1E-11</td><td>MB231/MB468</td><td>MITO</td></tr>
<tr><td>>MI0000809|hsa-mir-151a&WithFlank&8|-|140732558|140732659@51.73.23(+1U)&[MIMAT0000757&hsa-miR-151a-3p&offsets|-2|0(+1U);m-30&8|-|140732587|140732607&offsets|-2|0(+1U)]|TACTAGACTGAAGCTCCTTGAGGT</td><td>>MI0000809|hsa-mir-151a&WithFlank&8|-|140732558|140732659</td><td>51</td><td>73</td><td>23(+1U)</td><td>TACTAGACTGAAGCTCCTTGAGGT</td><td>364.3</td><td>6.2</td><td>0.9</td><td>6.7</td><td>2.9E-11</td><td>8.1E-11</td><td>MB231/MB468</td><td>CYTO</td></tr>
<tr><td>>hg38_21_+_8433172_8446622_RNA45SN1_45S_50ntFlanks@5431.5466.36|GTTTAGTGAGGCCCTCGGATCGGCCCCGCCGGGGTC</td><td>>hg38_21_+_8433172_8446622_RNA45SN1_45S_50ntFlanks</td><td>5431</td><td>5466</td><td>36</td><td>GTTTAGTGAGGCCCTCGGATCGGCCCCGCCGGGGTC</td><td>87.0</td><td>-10.3</td><td>1.5</td><td>-6.7</td><td>2.7E-11</td><td>8.1E-11</td><td>MB231/MB468</td><td>MP</td></tr>
<tr><td>>hg38_21_+_8433172_8446622_RNA45SN1_45S_50ntFlanks@12932.12970.39|CGCTAAACCATTCGTAGACGACCTGCTTCTGGGTCGGGG</td><td>>hg38_21_+_8433172_8446622_RNA45SN1_45S_50ntFlanks</td><td>12932</td><td>12970</td><td>39</td><td>CGCTAAACCATTCGTAGACGACCTGCTTCTGGGTCGGGG</td><td>229.2</td><td>-5.7</td><td>0.9</td><td>-6.6</td><td>2.9E-11</td><td>8.1E-11</td><td>MB231/MB468</td><td>CYTO</td></tr>
<tr><td>>hg38_21_+_8433172_8446622_RNA45SN1_45S_50ntFlanks@12992.13011.20|TCCCTCGCTGCGATCTATTG</td><td>>hg38_21_+_8433172_8446622_RNA45SN1_45S_50ntFlanks</td><td>12992</td><td>13011</td><td>20</td><td>TCCCTCGCTGCGATCTATTG</td><td>654.6</td><td>-5.2</td><td>0.8</td><td>-6.6</td><td>2.9E-11</td><td>8.1E-11</td><td>MB231/MB468</td><td>NUC</td></tr>
<tr><td>>hg38_21_+_8433172_8446622_RNA45SN1_45S_50ntFlanks@12632.12653.22|GCGAAGCTACCATCTGTGGGAT</td><td>>hg38_21_+_8433172_8446622_RNA45SN1_45S_50ntFlanks</td><td>12632</td><td>12653</td><td>22</td><td>GCGAAGCTACCATCTGTGGGAT</td><td>288.0</td><td>-8.4</td><td>1.3</td><td>-6.6</td><td>2.9E-11</td><td>8.2E-11</td><td>MB231/MB468</td><td>NUC</td></tr>
<tr><td>>hg38_21_+_8433172_8446622_RNA45SN1_45S_50ntFlanks@8179.8214.36|CTGATCGAGGCCCAGCCCGTGGACGGTGTGAGGCCG</td><td>>hg38_21_+_8433172_8446622_RNA45SN1_45S_50ntFlanks</td><td>8179</td><td>8214</td><td>36</td><td>CTGATCGAGGCCCAGCCCGTGGACGGTGTGAGGCCG</td><td>123.7</td><td>-10.8</td><td>1.6</td><td>-6.7</td><td>2.7E-11</td><td>8.2E-11</td><td>MB231/MB468</td><td>MP</td></tr>
<tr><td>>hg38_21_+_8433172_8446622_RNA45SN1_45S_50ntFlanks@10713.10736.24|CTTGAAAATCCGGGGGAGAGGGTG</td><td>>hg38_21_+_8433172_8446622_RNA45SN1_45S_50ntFlanks</td><td>10713</td><td>10736</td><td>24</td><td>CTTGAAAATCCGGGGGAGAGGGTG</td><td>293.2</td><td>-6.6</td><td>1.0</td><td>-6.6</td><td>3.4E-11</td><td>8.2E-11</td><td>MB231/MB468</td><td>MITO</td></tr>
<tr><td>>hg38_21_+_8433172_8446622_RNA45SN1_45S_50ntFlanks@10784.10818.35|TGAACAGCCTCTGGCATGTTGGAACAATGTAGGTA</td><td>>hg38_21_+_8433172_8446622_RNA45SN1_45S_50ntFlanks</td><td>10784</td><td>10818</td><td>35</td><td>TGAACAGCCTCTGGCATGTTGGAACAATGTAGGTA</td><td>769.7</td><td>-6.6</td><td>1.0</td><td>-6.6</td><td>3.4E-11</td><td>8.2E-11</td><td>MB231/MB468</td><td>MITO</td></tr>
<tr><td>>hg38_21_+_8433172_8446622_RNA45SN1_45S_50ntFlanks@8358.8378.21|AAAGAACTTTGAAGAGAGAGT</td><td>>hg38_21_+_8433172_8446622_RNA45SN1_45S_50ntFlanks</td><td>8358</td><td>8378</td><td>21</td><td>AAAGAACTTTGAAGAGAGAGT</td><td>820.2</td><td>-5.8</td><td>0.9</td><td>-6.6</td><td>3.6E-11</td><td>8.2E-11</td><td>BT20/MB468</td><td>NUC</td></tr>
<tr><td>>hg38_21_+_8433172_8446622_RNA45SN1_45S_50ntFlanks@3900.3918.19|CTTCGCGGGGGGGATGCGT</td><td>>hg38_21_+_8433172_8446622_RNA45SN1_45S_50ntFlanks</td><td>3900</td><td>3918</td><td>19</td><td>CTTCGCGGGGGGGATGCGT</td><td>125.6</td><td>-10.8</td><td>1.6</td><td>-6.7</td><td>2.7E-11</td><td>8.2E-11</td><td>MB231/MB468</td><td>MP</td></tr>
<tr><td>>hg38_21_+_8433172_8446622_RNA45SN1_45S_50ntFlanks@6650.6669.20|ACGACTCTTAGCGGTGGATC</td><td>>hg38_21_+_8433172_8446622_RNA45SN1_45S_50ntFlanks</td><td>6650</td><td>6669</td><td>20</td><td>ACGACTCTTAGCGGTGGATC</td><td>13636.9</td><td>4.6</td><td>0.7</td><td>6.7</td><td>2.2E-11</td><td>8.2E-11</td><td>BT20/MB468</td><td>MITO</td></tr>
<tr><td>>hg38_21_+_8433172_8446622_RNA45SN1_45S_50ntFlanks@6734.6773.40|ATTGATCATCGACACTTCGAACGCACTTGCGGCCCCGGGT</td><td>>hg38_21_+_8433172_8446622_RNA45SN1_45S_50ntFlanks</td><td>6734</td><td>6773</td><td>40</td><td>ATTGATCATCGACACTTCGAACGCACTTGCGGCCCCGGGT</td><td>465.3</td><td>7.8</td><td>1.2</td><td>6.6</td><td>3.0E-11</td><td>8.2E-11</td><td>MB231/MB468</td><td>CYTO</td></tr>
<tr><td>>hg38_21_+_8433172_8446622_RNA45SN1_45S_50ntFlanks@11579.11601.23|AGAACTGGTGCGGACCAGGGGAA</td><td>>hg38_21_+_8433172_8446622_RNA45SN1_45S_50ntFlanks</td><td>11579</td><td>11601</td><td>23</td><td>AGAACTGGTGCGGACCAGGGGAA</td><td>1094.0</td><td>-6.0</td><td>0.9</td><td>-6.6</td><td>3.0E-11</td><td>8.2E-11</td><td>MB231/MB468</td><td>CYTO</td></tr>
<tr><td>>am_trnaMT_PheGAA_MT_+_577_647@41.62.22|GTTTAGACGGGCTCACATCACC</td><td>>am_trnaMT_PheGAA_MT_+_577_647</td><td>41</td><td>62</td><td>22</td><td>GTTTAGACGGGCTCACATCACC</td><td>404.7</td><td>5.1</td><td>0.8</td><td>6.6</td><td>3.0E-11</td><td>8.2E-11</td><td>MB231/MB468</td><td>NUC</td></tr>
<tr><td>>hg38_21_+_8433172_8446622_RNA45SN1_45S_50ntFlanks@8421.8442.22|CGCGCAGTCCGCCCGGAGGATT</td><td>>hg38_21_+_8433172_8446622_RNA45SN1_45S_50ntFlanks</td><td>8421</td><td>8442</td><td>22</td><td>CGCGCAGTCCGCCCGGAGGATT</td><td>88.8</td><td>-10.3</td><td>1.6</td><td>-6.7</td><td>2.7E-11</td><td>8.2E-11</td><td>MB231/MB468</td><td>MP</td></tr>
<tr><td>>hg38_21_+_8433172_8446622_RNA45SN1_45S_50ntFlanks@12008.12041.34|CCGCCGGTGAAATACCACTACTCTGATCGTTTTT</td><td>>hg38_21_+_8433172_8446622_RNA45SN1_45S_50ntFlanks</td><td>12008</td><td>12041</td><td>34</td><td>CCGCCGGTGAAATACCACTACTCTGATCGTTTTT</td><td>172.7</td><td>-11.3</td><td>1.7</td><td>-6.7</td><td>2.8E-11</td><td>8.2E-11</td><td>MB231/MB468</td><td>MP</td></tr>
<tr><td>>hg38_21_+_8433172_8446622_RNA45SN1_45S_50ntFlanks@12488.12509.22|AACGTGAGCTGGGTTTAGACCG</td><td>>hg38_21_+_8433172_8446622_RNA45SN1_45S_50ntFlanks</td><td>12488</td><td>12509</td><td>22</td><td>AACGTGAGCTGGGTTTAGACCG</td><td>1084.2</td><td>-8.5</td><td>1.3</td><td>-6.6</td><td>3.5E-11</td><td>8.2E-11</td><td>MB231/MB468</td><td>MITO</td></tr>
<tr><td>>hg38_21_+_8433172_8446622_RNA45SN1_45S_50ntFlanks@10869.10889.21|AGGGCTGGGTCGGTCGGGCTG</td><td>>hg38_21_+_8433172_8446622_RNA45SN1_45S_50ntFlanks</td><td>10869</td><td>10889</td><td>21</td><td>AGGGCTGGGTCGGTCGGGCTG</td><td>817.3</td><td>-7.9</td><td>1.2</td><td>-6.6</td><td>3.0E-11</td><td>8.2E-11</td><td>MB231/MB468</td><td>CYTO</td></tr>
<tr><td>>hg38_21_+_8433172_8446622_RNA45SN1_45S_50ntFlanks@11872.11893.22|AAAGAAGACCCTGTTGAGCTTG</td><td>>hg38_21_+_8433172_8446622_RNA45SN1_45S_50ntFlanks</td><td>11872</td><td>11893</td><td>22</td><td>AAAGAAGACCCTGTTGAGCTTG</td><td>504.4</td><td>-8.5</td><td>1.3</td><td>-6.7</td><td>2.8E-11</td><td>8.2E-11</td><td>MB231/MB468</td><td>MP</td></tr>
<tr><td>>hg38_21_+_8433172_8446622_RNA45SN1_45S_50ntFlanks@8191.8210.20|CAGCCCGTGGACGGTGTGAG</td><td>>hg38_21_+_8433172_8446622_RNA45SN1_45S_50ntFlanks</td><td>8191</td><td>8210</td><td>20</td><td>CAGCCCGTGGACGGTGTGAG</td><td>117.4</td><td>-10.7</td><td>1.6</td><td>-6.7</td><td>2.8E-11</td><td>8.2E-11</td><td>MB231/MB468</td><td>MP</td></tr>
<tr><td>>hg38_21_+_8433172_8446622_RNA45SN1_45S_50ntFlanks@4444.4471.28|CCTTGCCTCTCGGCGCCCCCTCGATGCT</td><td>>hg38_21_+_8433172_8446622_RNA45SN1_45S_50ntFlanks</td><td>4444</td><td>4471</td><td>28</td><td>CCTTGCCTCTCGGCGCCCCCTCGATGCT</td><td>822.7</td><td>-8.0</td><td>1.2</td><td>-6.6</td><td>3.6E-11</td><td>8.2E-11</td><td>BT20/MB468</td><td>NUC</td></tr>
<tr><td>>hg38_21_+_8433172_8446622_RNA45SN1_45S_50ntFlanks@9351.9370.20|GCCGTGGCGCAATGAAGGTG</td><td>>hg38_21_+_8433172_8446622_RNA45SN1_45S_50ntFlanks</td><td>9351</td><td>9370</td><td>20</td><td>GCCGTGGCGCAATGAAGGTG</td><td>110.7</td><td>-10.7</td><td>1.6</td><td>-6.7</td><td>2.8E-11</td><td>8.3E-11</td><td>MB231/MB468</td><td>MP</td></tr>
<tr><td>>MI0000471|hsa-mir-126&WithFlank&9|+|136670596|136670692@59.79.21(+1U)&[MIMAT0000445&hsa-miR-126-3p&offsets|+1|0(+1U);m-40&9|+|136670653|136670674&offsets|+1|0(+1U)]|CGTACCGTGAGTAATAATGCGT</td><td>>MI0000471|hsa-mir-126&WithFlank&9|+|136670596|136670692</td><td>59</td><td>79</td><td>21(+1U)</td><td>CGTACCGTGAGTAATAATGCGT</td><td>129.7</td><td>10.2</td><td>1.5</td><td>6.7</td><td>2.8E-11</td><td>8.3E-11</td><td>MB231/MB468</td><td>MP</td></tr>
<tr><td>>hg38_21_+_8433172_8446622_RNA45SN1_45S_50ntFlanks@4283.4317.35|CTTTAACGAGGATCCATTGGAGGGCAAGTCTGGTG</td><td>>hg38_21_+_8433172_8446622_RNA45SN1_45S_50ntFlanks</td><td>4283</td><td>4317</td><td>35</td><td>CTTTAACGAGGATCCATTGGAGGGCAAGTCTGGTG</td><td>1626.8</td><td>-4.6</td><td>0.7</td><td>-6.6</td><td>3.0E-11</td><td>8.3E-11</td><td>MB231/MB468</td><td>CYTO</td></tr>
<tr><td>>hg38_21_+_8433172_8446622_RNA45SN1_45S_50ntFlanks@11784.11806.23|ATGAATGGATGAACGAGATTCCC</td><td>>hg38_21_+_8433172_8446622_RNA45SN1_45S_50ntFlanks</td><td>11784</td><td>11806</td><td>23</td><td>ATGAATGGATGAACGAGATTCCC</td><td>1646.2</td><td>-7.3</td><td>1.1</td><td>-6.7</td><td>2.8E-11</td><td>8.3E-11</td><td>MB231/MB468</td><td>MP</td></tr>
<tr><td>>hg38_21_+_8433172_8446622_RNA45SN1_45S_50ntFlanks@12526.12546.21|TTTTACCCTACTGATGATGTG</td><td>>hg38_21_+_8433172_8446622_RNA45SN1_45S_50ntFlanks</td><td>12526</td><td>12546</td><td>21</td><td>TTTTACCCTACTGATGATGTG</td><td>186.1</td><td>-7.8</td><td>1.2</td><td>-6.6</td><td>3.0E-11</td><td>8.3E-11</td><td>MB231/MB468</td><td>CYTO</td></tr>
<tr><td>>hg38_21_+_8433172_8446622_RNA45SN1_45S_50ntFlanks@12447.12467.21|CAGAATTCACCAAGCGTTGGA</td><td>>hg38_21_+_8433172_8446622_RNA45SN1_45S_50ntFlanks</td><td>12447</td><td>12467</td><td>21</td><td>CAGAATTCACCAAGCGTTGGA</td><td>119.5</td><td>-10.8</td><td>1.6</td><td>-6.7</td><td>2.8E-11</td><td>8.3E-11</td><td>MB231/MB468</td><td>MP</td></tr>
<tr><td>>hg38_21_+_8433172_8446622_RNA45SN1_45S_50ntFlanks@4588.4614.27|CTATTTTGTTGGTTTTCGGAACTGAGG</td><td>>hg38_21_+_8433172_8446622_RNA45SN1_45S_50ntFlanks</td><td>4588</td><td>4614</td><td>27</td><td>CTATTTTGTTGGTTTTCGGAACTGAGG</td><td>637.6</td><td>-4.9</td><td>0.7</td><td>-6.6</td><td>3.0E-11</td><td>8.3E-11</td><td>MB231/MB468</td><td>CYTO</td></tr>
<tr><td>>hg38_21_+_8433172_8446622_RNA45SN1_45S_50ntFlanks@12606.12625.20|ATGTGCTTGGCTGAGGAGCC</td><td>>hg38_21_+_8433172_8446622_RNA45SN1_45S_50ntFlanks</td><td>12606</td><td>12625</td><td>20</td><td>ATGTGCTTGGCTGAGGAGCC</td><td>482.2</td><td>-6.6</td><td>1.0</td><td>-6.7</td><td>2.8E-11</td><td>8.3E-11</td><td>MB231/MB468</td><td>MP</td></tr>
<tr><td>>hg38_21_+_8433172_8446622_RNA45SN1_45S_50ntFlanks@9346.9370.25|AAGCCGCCGTGGCGCAATGAAGGTG</td><td>>hg38_21_+_8433172_8446622_RNA45SN1_45S_50ntFlanks</td><td>9346</td><td>9370</td><td>25</td><td>AAGCCGCCGTGGCGCAATGAAGGTG</td><td>118.7</td><td>-10.8</td><td>1.6</td><td>-6.7</td><td>2.8E-11</td><td>8.3E-11</td><td>MB231/MB468</td><td>MP</td></tr>
<tr><td>>MI0000809|hsa-mir-151a&WithFlank&8|-|140732558|140732659@51.72.22(+1A)&[MIMAT0000757&hsa-miR-151a-3p&offsets|-2|-1(+1A);m-30&8|-|140732587|140732607&offsets|-2|-1(+1A)]|TACTAGACTGAAGCTCCTTGAGA</td><td>>MI0000809|hsa-mir-151a&WithFlank&8|-|140732558|140732659</td><td>51</td><td>72</td><td>22(+1A)</td><td>TACTAGACTGAAGCTCCTTGAGA</td><td>367.5</td><td>6.3</td><td>0.9</td><td>6.6</td><td>3.0E-11</td><td>8.3E-11</td><td>MB231/MB468</td><td>CYTO</td></tr>
<tr><td>>hg38_21_+_8433172_8446622_RNA45SN1_45S_50ntFlanks@9339.9367.29|CGCACGAAAGCCGCCGTGGCGCAATGAAG</td><td>>hg38_21_+_8433172_8446622_RNA45SN1_45S_50ntFlanks</td><td>9339</td><td>9367</td><td>29</td><td>CGCACGAAAGCCGCCGTGGCGCAATGAAG</td><td>511.9</td><td>-12.1</td><td>1.7</td><td>-7.0</td><td>2.0E-12</td><td>8.3E-11</td><td>BT20/MB468</td><td>MP</td></tr>
<tr><td>>hg38_21_+_8433172_8446622_RNA45SN1_45S_50ntFlanks@9787.9818.32|TCGCTGGCGTGGAGCCGGGCGTGGAATGCGAG</td><td>>hg38_21_+_8433172_8446622_RNA45SN1_45S_50ntFlanks</td><td>9787</td><td>9818</td><td>32</td><td>TCGCTGGCGTGGAGCCGGGCGTGGAATGCGAG</td><td>89.9</td><td>-10.4</td><td>1.6</td><td>-6.7</td><td>2.8E-11</td><td>8.3E-11</td><td>MB231/MB468</td><td>MP</td></tr>
<tr><td>>hg38_21_+_8433172_8446622_RNA45SN1_45S_50ntFlanks@4738.4757.20|AAAGTCGGAGGTTCGAAGAC</td><td>>hg38_21_+_8433172_8446622_RNA45SN1_45S_50ntFlanks</td><td>4738</td><td>4757</td><td>20</td><td>AAAGTCGGAGGTTCGAAGAC</td><td>543.2</td><td>4.3</td><td>0.6</td><td>6.7</td><td>2.7E-11</td><td>8.4E-11</td><td>BT20/MB468</td><td>CYTO</td></tr>
</table>

| | | | | | | | | | | | | | |
|---|---|---|---|---|---|---|---|---|---|---|---|---|---|
| >hg38_21_+_8433172_8446622_RNA45SN1_45S_50ntFlanks@6721.6749.29\|AATTGCAGGACACATTGATCATCGACACT | >hg38_21_+_8433172_8446622_RNA45SN1_45S_50ntFlanks | 6721 | 6749 | 29 | AATTGCAGGACACATTGATCATCGACACT | 743.3 | -7.5 | 1.1 | -6.6 | 3.7E-11 | 8.4E-11 | BT20/MB468 | NUC |
| >hg38_21_+_8433172_8446622_RNA45SN1_45S_50ntFlanks@8836.8855.20\|CGGCGGTGGCGGCGGCGGCG | >hg38_21_+_8433172_8446622_RNA45SN1_45S_50ntFlanks | 8836 | 8855 | 20 | CGGCGGTGGCGGCGGCGGCG | 102.0 | -10.5 | 1.6 | -6.7 | 2.8E-11 | 8.4E-11 | MB231/MB468 | MP |
| >hg38_21_+_8433172_8446622_RNA45SN1_45S_50ntFlanks@10794.10825.32\|CTGGCATGTTGGAACAATGTAGGTAAGGGAAG | >hg38_21_+_8433172_8446622_RNA45SN1_45S_50ntFlanks | 10794 | 10825 | 32 | CTGGCATGTTGGAACAATGTAGGTAAGGGAAG | 413.4 | -8.4 | 1.3 | -6.6 | 3.0E-11 | 8.4E-11 | MB231/MB468 | NUC |
| >MI0000087\|hsa-mir-29a&WithFlank&7\|-\|130876741\|130876816@48.66.19&[MIMAT0000086&hsa-miR-29a-3p&offsets\|0\|-3;m-26&7\|-\|130876749\|130876769&offsets\|0\|-2]\|TAGCACCATCTGAAATCGG | >MI0000087\|hsa-mir-29a&WithFlank&7\|-\|130876741\|130876816 | 48 | 66 | 19 | TAGCACCATCTGAAATCGG | 207.6 | 10.9 | 1.6 | 6.7 | 2.8E-11 | 8.4E-11 | MB231/MB468 | MP |
| >hg38_21_+_8433172_8446622_RNA45SN1_45S_50ntFlanks@11849.11875.27\|CGGGCTTGGCGGAATCAGCGGGGAAAG | >hg38_21_+_8433172_8446622_RNA45SN1_45S_50ntFlanks | 11849 | 11875 | 27 | CGGGCTTGGCGGAATCAGCGGGGAAAG | 80.2 | -10.2 | 1.5 | -6.7 | 2.8E-11 | 8.4E-11 | MB231/MB468 | MP |
| >hg38_21_+_8433172_8446622_RNA45SN1_45S_50ntFlanks@5144.5162.19\|CTTCTTAGAGGGACAAGTG | >hg38_21_+_8433172_8446622_RNA45SN1_45S_50ntFlanks | 5144 | 5162 | 19 | CTTCTTAGAGGGACAAGTG | 309.6 | -12.1 | 1.8 | -6.7 | 2.8E-11 | 8.4E-11 | MB231/MB468 | MP |
| >hg38_21_+_8433172_8446622_RNA45SN1_45S_50ntFlanks@10420.10456.37\|CTGAGAGATGGGCGAGCGCCGTTCCGAAGGGACGGGC | >hg38_21_+_8433172_8446622_RNA45SN1_45S_50ntFlanks | 10420 | 10456 | 37 | CTGAGAGATGGGCGAGCGCCGTTCCGAAGGGACGGGC | 696.7 | -6.6 | 1.0 | -6.6 | 3.5E-11 | 8.4E-11 | MB231/MB468 | MITO |
| >hg38_21_+_8433172_8446622_RNA45SN1_45S_50ntFlanks@9563.9597.35\|CCTGACGTGCAAATCGGTCGTCCGACCTGGGTATA | >hg38_21_+_8433172_8446622_RNA45SN1_45S_50ntFlanks | 9563 | 9597 | 35 | CCTGACGTGCAAATCGGTCGTCCGACCTGGGTATA | 1326.7 | -9.8 | 1.5 | -6.6 | 3.7E-11 | 8.4E-11 | BT20/MB468 | NUC |
| >am_trna11_ProAGG_16_+_3241989_3242060@1.29.29,trna12_ProAGG_6_+_26555498_26555569@1.29.29,trna12_ProTGG_11_-_75946869_75946940@1.29.29,trna14_ProTGG_5_-_180615854_180615925@1.29.29,trna22_ProAGG_14_-_21081560_21081631@1.29.29,trna23_ProAGG_14_-_21077495_21077566@1.29.29,trna28_ProTGG_16_-_3234133_3234204@1.29.29,trna29_ProAGG_16_-_3232635_3232706@1.29.29,trna2_ProAGG_7_+_128423504_128423575@1.29.29,trna30_ProCGG_6_+_27059521_27059592@1.29.29,trna37_ProCGG_17_-_8126151_8126222@1.29.29,trna3_ProTGG_16_+_3208923_3208994@1.29.29,trna52_ProCGG_1_+_167683962_167684033@1.29.29,trna65_ProAGG_1_-_167684725_167684796@1.29.29,trna6_ProCGG_16_+_3222049_3222120@1.29.29,trna6_ProTGG_14_+_21152175_21152246@1.29.29,trna8_ProTGG_16_+_3238094_3238165@1.29.29,trna9_ProAGG_11_+_75946557_75946628@1.29.29,trna9_ProAGG_16_+_3239634_3239705@1.29.29\|GGCTCGTTGGTCTAGGGGTATGATTCTCG | >am_trna11_ProAGG_16_+_3241989_3242060 | 1 | 29 | 29 | GGCTCGTTGGTCTAGGGGTATGATTCTCG | 6245.6 | -5.9 | 0.9 | -6.7 | 2.8E-11 | 8.4E-11 | BT20/MB468 | CYTO |
| >hg38_21_+_8433172_8446622_RNA45SN1_45S_50ntFlanks@12456.12475.20\|CCAAGCGTTGGATTGTTCAC | >hg38_21_+_8433172_8446622_RNA45SN1_45S_50ntFlanks | 12456 | 12475 | 20 | CCAAGCGTTGGATTGTTCAC | 22464.1 | -9.0 | 1.4 | -6.7 | 2.8E-11 | 8.4E-11 | BT20/MB468 | CYTO |
| >hg38_21_+_8433172_8446622_RNA45SN1_45S_50ntFlanks@10794.10822.29\|CTGGCATGTTGGAACAATGTAGGTAAGGG | >hg38_21_+_8433172_8446622_RNA45SN1_45S_50ntFlanks | 10794 | 10822 | 29 | CTGGCATGTTGGAACAATGTAGGTAAGGG | 600.8 | -9.2 | 1.4 | -6.6 | 3.7E-11 | 8.4E-11 | BT20/MB468 | NUC |
| >hg38_21_+_8433172_8446622_RNA45SN1_45S_50ntFlanks@10817.10840.24\|TAAGGGAAGTCGGCAAGCCGGATC | >hg38_21_+_8433172_8446622_RNA45SN1_45S_50ntFlanks | 10817 | 10840 | 24 | TAAGGGAAGTCGGCAAGCCGGATC | 346.9 | -8.4 | 1.3 | -6.6 | 3.5E-11 | 8.4E-11 | MB231/MB468 | MITO |
| >hg38_21_+_8433172_8446622_RNA45SN1_45S_50ntFlanks@4983.5002.20\|CGGACAGGATTGACAGATTG | >hg38_21_+_8433172_8446622_RNA45SN1_45S_50ntFlanks | 4983 | 5002 | 20 | CGGACAGGATTGACAGATTG | 2526.9 | -10.7 | 1.6 | -6.6 | 3.6E-11 | 8.4E-11 | MB231/MB468 | MITO |
| >hg38_21_+_8433172_8446622_RNA45SN1_45S_50ntFlanks@11564.11601.38\|CCTAGCAGCCGACTTAGAACTGGTGCGGACCAGGGGAA | >hg38_21_+_8433172_8446622_RNA45SN1_45S_50ntFlanks | 11564 | 11601 | 38 | CCTAGCAGCCGACTTAGAACTGGTGCGGACCAGGGGAA | 4502.9 | -10.4 | 1.6 | -6.7 | 2.8E-11 | 8.5E-11 | MB231/MB468 | MP |
| >trna16_LeuAAG_16_+_22308461_22308542@1.20.20,trna16_LeuAAG_5_-_180601044_180601125@1.20.20,trna19_LeuAAG_5_-_180524474_180524555@1.20.20,trna1_LeuAAG_14_+_21078291_21078372@1.20.20,trna3_LeuAAG_5_+_180528840_180528921@1.20.20,trna42_LeuTAG_17_-_8023632_8023713@1.20.20,trna78_LeuAAG_6_+_28956779_28956860@1.20.20,trna7_LeuAAG_5_+_180614701_180614782@1.20.20,trna98_LeuAAG_6_-_28911399_28911480@1.20.20\|GGTAGCGTGGCCGAGCGGTC | >trna16_LeuAAG_16_+_22308461_22308542 | 1 | 20 | 20 | GGTAGCGTGGCCGAGCGGTC | 363.2 | 7.5 | 1.1 | 6.7 | 2.8E-11 | 8.5E-11 | BT20/MB468 | CYTO |
| >hg38_21_+_8433172_8446622_RNA45SN1_45S_50ntFlanks@10388.10424.37\|GTGAACAGCAGTTGAACATGGGTCAGTCGGTCCTGAG | >hg38_21_+_8433172_8446622_RNA45SN1_45S_50ntFlanks | 10388 | 10424 | 37 | GTGAACAGCAGTTGAACATGGGTCAGTCGGTCCTGAG | 2004.1 | 4.6 | 0.7 | 6.6 | 3.1E-11 | 8.5E-11 | MB231/MB468 | CYTO |
| >hg38_21_+_8433172_8446622_RNA45SN1_45S_50ntFlanks@5423.5441.19\|ATTGGATGGTTTAGTGAGG | >hg38_21_+_8433172_8446622_RNA45SN1_45S_50ntFlanks | 5423 | 5441 | 19 | ATTGGATGGTTTAGTGAGG | 136.7 | -11.0 | 1.6 | -6.7 | 2.9E-11 | 8.5E-11 | MB231/MB468 | MP |
| >hg38_21_+_8433172_8446622_RNA45SN1_45S_50ntFlanks@10069.10096.28\|AGTGGACGGGAGCGGCGGGGGCGGCGCG | >hg38_21_+_8433172_8446622_RNA45SN1_45S_50ntFlanks | 10069 | 10096 | 28 | AGTGGACGGGAGCGGCGGGGGCGGCGCG | 128.2 | -10.9 | 1.6 | -6.7 | 2.9E-11 | 8.5E-11 | MB231/MB468 | MP |
| >hg38_21_+_8433172_8446622_RNA45SN1_45S_50ntFlanks@12437.12473.37\|CATTGTGAAGCAGAATTCACCAAGCGTTGGATTGTTC | >hg38_21_+_8433172_8446622_RNA45SN1_45S_50ntFlanks | 12437 | 12473 | 37 | CATTGTGAAGCAGAATTCACCAAGCGTTGGATTGTTC | 6286.4 | -4.9 | 0.7 | -6.6 | 3.1E-11 | 8.5E-11 | MB231/MB468 | CYTO |
| >hg38_21_+_8433172_8446622_RNA45SN1_45S_50ntFlanks@9504.9526.23\|GTGAACTATGCCTGGGCAGGGCG | >hg38_21_+_8433172_8446622_RNA45SN1_45S_50ntFlanks | 9504 | 9526 | 23 | GTGAACTATGCCTGGGCAGGGCG | 3236.3 | -7.2 | 1.1 | -6.7 | 2.3E-11 | 8.5E-11 | BT20/MB468 | MITO |
| >MI0000736\|hsa-mir-30c-1&WithFlank&1\|+\|40757278\|40757378@23.45.23(+1G);MI0000254\|hsa-mir-30c-2&WithFlank&6\|-\|71376954\|71377037@13.35.23(+1G)&[MIMAT0000244&hsa-miR-30c-5p&offsets\|0\|0(+1G);m-57&1\|+\|40757300\|40757323&offsets\|0\|-1(+1G)];[MIMAT0000244_1&hsa-miR-30c-5p&offsets\|0\|0(+1G);m-56&6\|-\|71377002\|71377025&offsets\|0\|-1(+1G)]\|TGTAAACATCCTACACTCTCAGCG | >MI0000736\|hsa-mir-30c-1&WithFlank&1\|+\|40757278\|40757378 | 23 | 45 | 23(+1G) | TGTAAACATCCTACACTCTCAGCG | 558.4 | 4.2 | 0.6 | 6.7 | 2.8E-11 | 8.5E-11 | BT20/MB468 | CYTO |
| >hg38_21_+_8433172_8446622_RNA45SN1_45S_50ntFlanks@10664.10710.47\|CTTGGAAAGCGTCGCGGTTCCGGCGGCGTCCGGTGAGCTCTCGCTGG | >hg38_21_+_8433172_8446622_RNA45SN1_45S_50ntFlanks | 10664 | 10710 | 47 | CTTGGAAAGCGTCGCGGTTCCGGCGGCGTCCGGTGAGCTCTCGCTGG | 588.7 | -12.2 | 1.8 | -6.6 | 3.1E-11 | 8.6E-11 | MB231/MB468 | NUC |
| >hg38_21_+_8433172_8446622_RNA45SN1_45S_50ntFlanks@6719.6743.25\|TGAATTGCAGGACACATTGATCATC | >hg38_21_+_8433172_8446622_RNA45SN1_45S_50ntFlanks | 6719 | 6743 | 25 | TGAATTGCAGGACACATTGATCATC | 3652.0 | -7.3 | 1.1 | -6.6 | 3.6E-11 | 8.6E-11 | MB231/MB468 | MITO |
| >hg38_21_+_8433172_8446622_RNA45SN1_45S_50ntFlanks@12456.12493.38\|CCAAGCGTTGGATTGTTCACCCACTAATAGGGAACGTG | >hg38_21_+_8433172_8446622_RNA45SN1_45S_50ntFlanks | 12456 | 12493 | 38 | CCAAGCGTTGGATTGTTCACCCACTAATAGGGAACGTG | 1763.5 | -8.7 | 1.3 | -6.6 | 3.8E-11 | 8.6E-11 | BT20/MB468 | NUC |
| >hg38_21_+_8433172_8446622_RNA45SN1_45S_50ntFlanks@12413.12431.19\|TCCTTCGATGTCGGCTCTT | >hg38_21_+_8433172_8446622_RNA45SN1_45S_50ntFlanks | 12413 | 12431 | 19 | TCCTTCGATGTCGGCTCTT | 89.1 | -10.3 | 1.6 | -6.7 | 2.9E-11 | 8.6E-11 | MB231/MB468 | MP |
| >hg38_21_+_8433172_8446622_RNA45SN1_45S_50ntFlanks@11718.11744.27\|AACGGCGGGAGTAACTATGACTCTCTT | >hg38_21_+_8433172_8446622_RNA45SN1_45S_50ntFlanks | 11718 | 11744 | 27 | AACGGCGGGAGTAACTATGACTCTCTT | 893.8 | -7.2 | 1.1 | -6.7 | 2.9E-11 | 8.6E-11 | MB231/MB468 | MP |
| >hg38_21_+_8433172_8446622_RNA45SN1_45S_50ntFlanks@4403.4423.21\|CGGGCGGTCCGCCGCGAGGCG | >hg38_21_+_8433172_8446622_RNA45SN1_45S_50ntFlanks | 4403 | 4423 | 21 | CGGGCGGTCCGCCGCGAGGCG | 162.7 | -11.2 | 1.7 | -6.7 | 2.9E-11 | 8.6E-11 | MB231/MB468 | MP |
| >hg38_21_+_8433172_8446622_RNA45SN1_45S_50ntFlanks@6671.6698.28\|CTCGGCTCGTGCGTCGATGAAGAACGCA | >hg38_21_+_8433172_8446622_RNA45SN1_45S_50ntFlanks | 6671 | 6698 | 28 | CTCGGCTCGTGCGTCGATGAAGAACGCA | 679.7 | -5.6 | 0.8 | -6.7 | 2.8E-11 | 8.6E-11 | BT20/MB468 | CYTO |
| >hg38_21_+_8433172_8446622_RNA45SN1_45S_50ntFlanks@4670.4689.20\|TCTTGGACCGGCGCAAGACG | >hg38_21_+_8433172_8446622_RNA45SN1_45S_50ntFlanks | 4670 | 4689 | 20 | TCTTGGACCGGCGCAAGACG | 99.1 | -10.5 | 1.6 | -6.7 | 2.9E-11 | 8.6E-11 | MB231/MB468 | MP |
| >trnaMT_GluTTC_MT_-_14674_14742@40.57.18,trnalookalike8_GluTTC_5_-_93905172_93905240@40.57.18\|TTGGTCGTGGTTGTAGTC | >trnaMT_GluTTC_MT_-_14674_14742 | 40 | 57 | 18 | TTGGTCGTGGTTGTAGTC | 579.0 | 7.5 | 1.1 | 6.7 | 2.3E-11 | 8.6E-11 | BT20/MB468 | MITO |
| >hg38_21_+_8433172_8446622_RNA45SN1_45S_50ntFlanks@6652.6673.22\|GACTCTTAGCGGTGGATCACTC | >hg38_21_+_8433172_8446622_RNA45SN1_45S_50ntFlanks | 6652 | 6673 | 22 | GACTCTTAGCGGTGGATCACTC | 1625.2 | 5.5 | 0.8 | 6.7 | 2.3E-11 | 8.6E-11 | BT20/MB468 | MITO |
| >hg38_21_+_8433172_8446622_RNA45SN1_45S_50ntFlanks@12412.12435.24\|ATCCTTCGATGTCGGCTCTTCCTA | >hg38_21_+_8433172_8446622_RNA45SN1_45S_50ntFlanks | 12412 | 12435 | 24 | ATCCTTCGATGTCGGCTCTTCCTA | 215.0 | -9.8 | 1.5 | -6.7 | 2.3E-11 | 8.6E-11 | BT20/MB468 | MITO |
| >hg38_21_+_8433172_8446622_RNA45SN1_45S_50ntFlanks@6733.6758.26\|CATTGATCATCGACACTTCGAACGCA | >hg38_21_+_8433172_8446622_RNA45SN1_45S_50ntFlanks | 6733 | 6758 | 26 | CATTGATCATCGACACTTCGAACGCA | 718.4 | -7.0 | 1.1 | -6.7 | 2.9E-11 | 8.6E-11 | MB231/MB468 | MP |
| >hg38_21_+_8433172_8446622_RNA45SN1_45S_50ntFlanks@9105.9128.24\|CGGCGACGGGGGGGGTGCCGCGCG | >hg38_21_+_8433172_8446622_RNA45SN1_45S_50ntFlanks | 9105 | 9128 | 24 | CGGCGACGGGGGGGGTGCCGCGCG | 128.4 | -10.9 | 1.6 | -6.7 | 2.9E-11 | 8.6E-11 | MB231/MB468 | MP |
| >hg38_21_+_8433172_8446622_RNA45SN1_45S_50ntFlanks@9821.9838.18\|CCTAGTGGGCCACTTTTG | >hg38_21_+_8433172_8446622_RNA45SN1_45S_50ntFlanks | 9821 | 9838 | 18 | CCTAGTGGGCCACTTTTG | 275.3 | -10.2 | 1.5 | -6.7 | 2.3E-11 | 8.7E-11 | BT20/MB468 | MITO |
| >hg38_21_+_8433172_8446622_RNA45SN1_45S_50ntFlanks@12159.12180.22\|TTTGACTGGGGCGGTACACCTG | >hg38_21_+_8433172_8446622_RNA45SN1_45S_50ntFlanks | 12159 | 12180 | 22 | TTTGACTGGGGCGGTACACCTG | 685.3 | -8.5 | 1.3 | -6.6 | 3.1E-11 | 8.7E-11 | MB231/MB468 | NUC |
| >hg38_21_+_8433172_8446622_RNA45SN1_45S_50ntFlanks@11576.11599.24\|CTTAGAACTGGTGCGGACCAGGGG | >hg38_21_+_8433172_8446622_RNA45SN1_45S_50ntFlanks | 11576 | 11599 | 24 | CTTAGAACTGGTGCGGACCAGGGG | 120.4 | -10.8 | 1.6 | -6.7 | 2.9E-11 | 8.7E-11 | MB231/MB468 | MP |
| >trna116_GluCTC_1_-_145399233_145399304@32.72.41,trna71_GluCTC_1_-_161439189_161439260@32.72.41,trna74_GluCTC_1_-_161431809_161431880@32.72.41,trna77_GluCTC_1_-_161424398_161424469@32.72.41,trna77_GluCTC_6_+_28949976_28950047@32.72.41,trna80_GluCTC_1_-_161417018_161417089@32.72.41,trna87_GluCTC_6_-_126101393_126101464@32.72.41\|CTCTCACCGCCGCGGCCCGGGTTCGATTCCCGGTCAGGGAA | >trna116_GluCTC_1_-_145399233_145399304 | 32 | 72 | 41 | CTCTCACCGCCGCGGCCCGGGTTCGATTCCCGGTCAGGGAA | 243.4 | -11.7 | 1.8 | -6.6 | 3.7E-11 | 8.7E-11 | MB231/MB468 | MITO |
| >hg38_21_+_8433172_8446622_RNA45SN1_45S_50ntFlanks@4663.4695.33\|GTGAAATTCTTGGACCGGCGCAAGACGGACCAG | >hg38_21_+_8433172_8446622_RNA45SN1_45S_50ntFlanks | 4663 | 4695 | 33 | GTGAAATTCTTGGACCGGCGCAAGACGGACCAG | 79.6 | -10.2 | 1.5 | -6.7 | 2.9E-11 | 8.7E-11 | MB231/MB468 | MP |
| >MI0000434\|hsa-let-7i&WithFlank&12\|+\|62603680\|62603775@12.32.21(+1G)&[MIMAT0000415&hsa-let-7i-5p&offsets\|0\|-1(+1G);m-49&12\|+\|62603691\|62603712&offsets\|0\|-1(+1G)]\|TGAGGTAGTAGTTTGTGCTGTG | >MI0000434\|hsa-let-7i&WithFlank&12\|+\|62603680\|62603775 | 12 | 32 | 21(+1G) | TGAGGTAGTAGTTTGTGCTGTG | 2333.7 | 5.3 | 0.8 | 6.7 | 2.8E-11 | 8.7E-11 | BT20/MB468 | CYTO |
| >hg38_21_+_8433172_8446622_RNA45SN1_45S_50ntFlanks@10679.10704.26\|GGTTCCGGCGGCGTCCGGTGAGCTCT | >hg38_21_+_8433172_8446622_RNA45SN1_45S_50ntFlanks | 10679 | 10704 | 26 | GGTTCCGGCGGCGTCCGGTGAGCTCT | 93.7 | -10.4 | 1.6 | -6.7 | 2.9E-11 | 8.7E-11 | MB231/MB468 | MP |

| | | | | | | | | | | | | | |
|---|---|---|---|---|---|---|---|---|---|---|---|---|---|
| >hg38_21_+_8433172_8446622_RNA45SN1_45S_50ntFlanks@9572.9595.24\|CAAATCGGTCGTCCGACCTGGGTA | >hg38_21_+_8433172_8446622_RNA45SN1_45S_50ntFlanks | 9572 | 9595 | 24 | CAAATCGGTCGTCCGACCTGGGTA | 3410.9 | -6.3 | 0.9 | -6.7 | 2.3E-11 | 8.7E-11 | BT20/MB468 | MITO |
| >hg38_21_+_8433172_8446622_RNA45SN1_45S_50ntFlanks@6655.6681.27\|TCTTAGCGGTGGATCACTCGGCTCGTG | >hg38_21_+_8433172_8446622_RNA45SN1_45S_50ntFlanks | 6655 | 6681 | 27 | TCTTAGCGGTGGATCACTCGGCTCGTG | 2414.8 | -8.1 | 1.2 | -6.6 | 2.9E-11 | 8.7E-11 | MB231/MB468 | MP |
| >MI0000736\|hsa-mir-30c-1&WithFlank&1\|+\|40757278\|40757378@23.44.22(+1U);MI0000254\|hsa-mir-30c-2&WithFlank&6\|-\|71376954\|71377037@13.34.22(+1U)&[MIMAT0000244&hsa-miR-30c-5p&offsets\|0\|-1(+1U);m-57&1\|+\|40757300\|40757323&offsets\|0\|-2(+1U)];[MIMAT0000244_1&hsa-miR-30c-5p&offsets\|0\|-1(+1U);m-56&6\|-\|71377002\|71377025&offsets\|0\|-2(+1U)]\|TGTAAACATCCTACACTCTCAGT | >MI0000736\|hsa-mir-30c-1&WithFlank&1\|+\|40757278\|40757378 | 23 | 44 | 22(+1U) | TGTAAACATCCTACACTCTCAGT | 2369.4 | 5.7 | 0.9 | 6.6 | 3.2E-11 | 8.7E-11 | MB231/MB468 | CYTO |
| >hg38_21_+_8433172_8446622_RNA45SN1_45S_50ntFlanks@12635.12654.20\|AAGCTACCATCTGTGGGATT | >hg38_21_+_8433172_8446622_RNA45SN1_45S_50ntFlanks | 12635 | 12654 | 20 | AAGCTACCATCTGTGGGATT | 1476.1 | -6.9 | 1.0 | -6.6 | 3.2E-11 | 8.7E-11 | MB231/MB468 | CYTO |
| >hg38_21_+_8433172_8446622_RNA45SN1_45S_50ntFlanks@10806.10831.26\|AACAATGTAGGTAAGGGAAGTCGGCA | >hg38_21_+_8433172_8446622_RNA45SN1_45S_50ntFlanks | 10806 | 10831 | 26 | AACAATGTAGGTAAGGGAAGTCGGCA | 88.0 | -10.3 | 1.6 | -6.6 | 3.0E-11 | 8.8E-11 | MB231/MB468 | MP |
| >hg38_21_+_8433172_8446622_RNA45SN1_45S_50ntFlanks@11535.11553.19\|TTCCGCGCGGCGCCTCGCC | >hg38_21_+_8433172_8446622_RNA45SN1_45S_50ntFlanks | 11535 | 11553 | 19 | TTCCGCGCGGCGCCTCGCC | 248.6 | -8.2 | 1.2 | -6.6 | 3.2E-11 | 8.8E-11 | MB231/MB468 | NUC |
| >hg38_21_+_8433172_8446622_RNA45SN1_45S_50ntFlanks@12407.12431.25\|TTTTGATCCTTCGATGTCGGCTCTT | >hg38_21_+_8433172_8446622_RNA45SN1_45S_50ntFlanks | 12407 | 12431 | 25 | TTTTGATCCTTCGATGTCGGCTCTT | 696.7 | -4.0 | 0.6 | -6.6 | 3.2E-11 | 8.8E-11 | MB231/MB468 | NUC |
| >hg38_21_+_8433172_8446622_RNA45SN1_45S_50ntFlanks@6650.6696.47\|ACGACTCTTAGCGGTGGATCACTCGGCTCGTGCGTCGATGAAGAACG | >hg38_21_+_8433172_8446622_RNA45SN1_45S_50ntFlanks | 6650 | 6696 | 47 | ACGACTCTTAGCGGTGGATCACTCGGCTCGTGCGTCGATGAAGAACG | 440.5 | 5.5 | 0.8 | 6.6 | 3.2E-11 | 8.8E-11 | MB231/MB468 | CYTO |
| >hg38_21_+_8433172_8446622_RNA45SN1_45S_50ntFlanks@9929.9949.21\|GATATAGACAGCAGGACGGTG | >hg38_21_+_8433172_8446622_RNA45SN1_45S_50ntFlanks | 9929 | 9949 | 21 | GATATAGACAGCAGGACGGTG | 711.0 | -5.8 | 0.9 | -6.6 | 3.0E-11 | 8.8E-11 | MB231/MB468 | MP |
| >hg38_21_+_8433172_8446622_RNA45SN1_45S_50ntFlanks@11921.11969.49\|TGAGAGGTGTAGAATAAGTGGGAGGCCCCCGGCGCCCCCCCGGTGTCCC | >hg38_21_+_8433172_8446622_RNA45SN1_45S_50ntFlanks | 11921 | 11969 | 49 | TGAGAGGTGTAGAATAAGTGGGAGGCCCCCGGCGCCCCCCCGGTGTCCC | 243.2 | -10.9 | 1.6 | -6.6 | 3.2E-11 | 8.8E-11 | MB231/MB468 | NUC |
| >hg38_21_+_8433172_8446622_RNA45SN1_45S_50ntFlanks@3831.3853.23\|CTCGCTCCTCTCCTACTTGGATA | >hg38_21_+_8433172_8446622_RNA45SN1_45S_50ntFlanks | 3831 | 3853 | 23 | CTCGCTCCTCTCCTACTTGGATA | 339.4 | -8.6 | 1.3 | -6.6 | 3.9E-11 | 8.8E-11 | BT20/MB468 | NUC |
| >MI0000811\|hsa-mir-148b&WithFlank&12\|+\|54337210\|54337320@69.89.21(+1C)&[MIMAT0000759&hsa-miR-148b-3p&offsets\|0\|-1(+1C);m-77&12\|+\|54337278\|54337299&offsets\|0\|-1(+1C)]\|TCAGTGCATCACAGAACTTTGC | >MI0000811\|hsa-mir-148b&WithFlank&12\|+\|54337210\|54337320 | 69 | 89 | 21(+1C) | TCAGTGCATCACAGAACTTTGC | 209.3 | 3.4 | 0.5 | 6.6 | 3.9E-11 | 8.8E-11 | BT20/MB468 | NUC |
| >hg38_21_+_8433172_8446622_RNA45SN1_45S_50ntFlanks@10906.10936.31\|TGGGCGCGCGCCGCGGCTGGACGAGGCGCCG | >hg38_21_+_8433172_8446622_RNA45SN1_45S_50ntFlanks | 10906 | 10936 | 31 | TGGGCGCGCGCCGCGGCTGGACGAGGCGCCG | 87.9 | -10.3 | 1.6 | -6.6 | 3.0E-11 | 8.8E-11 | MB231/MB468 | MP |
| >hg38_21_+_8433172_8446622_RNA45SN1_45S_50ntFlanks@9343.9367.25\|CGAAAGCCGCCGTGGCGCAATGAAG | >hg38_21_+_8433172_8446622_RNA45SN1_45S_50ntFlanks | 9343 | 9367 | 25 | CGAAAGCCGCCGTGGCGCAATGAAG | 101.4 | -10.5 | 1.6 | -6.6 | 3.0E-11 | 8.8E-11 | MB231/MB468 | MP |
| >hg38_21_+_8433172_8446622_RNA45SN1_45S_50ntFlanks@11576.11606.31\|CTTAGAACTGGTGCGGACCAGGGGAATCCGA | >hg38_21_+_8433172_8446622_RNA45SN1_45S_50ntFlanks | 11576 | 11606 | 31 | CTTAGAACTGGTGCGGACCAGGGGAATCCGA | 121.0 | -10.8 | 1.6 | -6.6 | 3.0E-11 | 8.8E-11 | MB231/MB468 | MP |
| >hg38_21_+_8433172_8446622_RNA45SN1_45S_50ntFlanks@11190.11227.38\|CCCGGGGGGCCGGCGGCGGCGGCGACTCTGGACGCGAG | >hg38_21_+_8433172_8446622_RNA45SN1_45S_50ntFlanks | 11190 | 11227 | 38 | CCCGGGGGGCCGGCGGCGGCGGCGACTCTGGACGCGAG | 211.8 | -11.6 | 1.7 | -6.6 | 3.0E-11 | 8.8E-11 | MB231/MB468 | MP |
| >hg38_21_+_8433172_8446622_RNA45SN1_45S_50ntFlanks@11316.11344.29\|CACCCCACGTCTCGTCGCGCGCGCGTCCG | >hg38_21_+_8433172_8446622_RNA45SN1_45S_50ntFlanks | 11316 | 11344 | 29 | CACCCCACGTCTCGTCGCGCGCGCGTCCG | 1116.3 | -9.4 | 1.3 | -7.0 | 2.1E-12 | 8.9E-11 | BT20/MB468 | MP |
| >hg38_21_+_8433172_8446622_RNA45SN1_45S_50ntFlanks@13008.13028.21\|ATTGAAAGTCAGCCCTCGACA | >hg38_21_+_8433172_8446622_RNA45SN1_45S_50ntFlanks | 13008 | 13028 | 21 | ATTGAAAGTCAGCCCTCGACA | 264.7 | -6.2 | 0.9 | -6.6 | 3.2E-11 | 8.9E-11 | MB231/MB468 | NUC |
| >hg38_21_+_8433172_8446622_RNA45SN1_45S_50ntFlanks@11873.11898.26\|AAGAAGACCCTGTTGAGCTTGACTCT | >hg38_21_+_8433172_8446622_RNA45SN1_45S_50ntFlanks | 11873 | 11898 | 26 | AAGAAGACCCTGTTGAGCTTGACTCT | 3467.6 | -8.3 | 1.3 | -6.6 | 3.9E-11 | 8.9E-11 | BT20/MB468 | NUC |
| >hg38_21_+_8433172_8446622_RNA45SN1_45S_50ntFlanks@3767.3791.25\|TACGCACGGCCGGTACAGTGAAACT | >hg38_21_+_8433172_8446622_RNA45SN1_45S_50ntFlanks | 3767 | 3791 | 25 | TACGCACGGCCGGTACAGTGAAACT | 298.6 | 6.6 | 1.0 | 6.6 | 3.2E-11 | 8.9E-11 | MB231/MB468 | CYTO |
| >hg38_21_+_8433172_8446622_RNA45SN1_45S_50ntFlanks@10357.10384.28\|CTTTGAAGGCCGAAGTGGAGAAGGGTTC | >hg38_21_+_8433172_8446622_RNA45SN1_45S_50ntFlanks | 10357 | 10384 | 28 | CTTTGAAGGCCGAAGTGGAGAAGGGTTC | 440.5 | -8.8 | 1.3 | -6.6 | 3.9E-11 | 8.9E-11 | BT20/MB468 | NUC |
| >hg38_21_+_8433172_8446622_RNA45SN1_45S_50ntFlanks@5471.5514.44\|CACGGCCCTGGCGGAGCGCTGAGAAGACGGTCGAACTTGACTAT | >hg38_21_+_8433172_8446622_RNA45SN1_45S_50ntFlanks | 5471 | 5514 | 44 | CACGGCCCTGGCGGAGCGCTGAGAAGACGGTCGAACTTGACTAT | 2301.0 | -6.0 | 0.9 | -6.7 | 2.9E-11 | 8.9E-11 | BT20/MB468 | CYTO |
| >hg38_21_+_8433172_8446622_RNA45SN1_45S_50ntFlanks@4005.4026.22\|AACCTCGGGCCGATCGCACGCC | >hg38_21_+_8433172_8446622_RNA45SN1_45S_50ntFlanks | 4005 | 4026 | 22 | AACCTCGGGCCGATCGCACGCC | 298.5 | -10.3 | 1.5 | -6.7 | 2.4E-11 | 8.9E-11 | BT20/MB468 | MITO |
| >hg38_21_+_8433172_8446622_RNA45SN1_45S_50ntFlanks@11581.11613.33\|AACTGGTGCGGACCAGGGGAATCCGACTGTTTA | >hg38_21_+_8433172_8446622_RNA45SN1_45S_50ntFlanks | 11581 | 11613 | 33 | AACTGGTGCGGACCAGGGGAATCCGACTGTTTA | 1383.3 | -4.7 | 0.7 | -6.6 | 3.8E-11 | 8.9E-11 | MB231/MB468 | MITO |
| >hg38_21_+_8433172_8446622_RNA45SN1_45S_50ntFlanks@12588.12615.28\|GGTTCAGACATTTGGTGTATGTGCTTGG | >hg38_21_+_8433172_8446622_RNA45SN1_45S_50ntFlanks | 12588 | 12615 | 28 | GGTTCAGACATTTGGTGTATGTGCTTGG | 102.0 | -10.5 | 1.6 | -6.6 | 3.0E-11 | 8.9E-11 | MB231/MB468 | MP |
| >hg38_21_+_8433172_8446622_RNA45SN1_45S_50ntFlanks@8423.8469.47\|CGCAGTCCGCCCGGAGGATTCAACCCGGCGGCGGGTCCGGCCGTGTC | >hg38_21_+_8433172_8446622_RNA45SN1_45S_50ntFlanks | 8423 | 8469 | 47 | CGCAGTCCGCCCGGAGGATTCAACCCGGCGGCGGGTCCGGCCGTGTC | 310.6 | -12.1 | 1.8 | -6.6 | 3.0E-11 | 8.9E-11 | MB231/MB468 | MP |
| >hg38_21_+_8433172_8446622_RNA45SN1_45S_50ntFlanks@10351.10381.31\|CGAGAACTTTGAAGGCCGAAGTGGAGAAGGG | >hg38_21_+_8433172_8446622_RNA45SN1_45S_50ntFlanks | 10351 | 10381 | 31 | CGAGAACTTTGAAGGCCGAAGTGGAGAAGGG | 949.8 | -6.9 | 1.0 | -6.6 | 3.8E-11 | 8.9E-11 | MB231/MB468 | MITO |
| >hg38_21_+_8433172_8446622_RNA45SN1_45S_50ntFlanks@9204.9225.22\|TCGGGGCCACGCGCGCGTCCCC | >hg38_21_+_8433172_8446622_RNA45SN1_45S_50ntFlanks | 9204 | 9225 | 22 | TCGGGGCCACGCGCGCGTCCCC | 209.4 | -11.6 | 1.7 | -6.6 | 3.0E-11 | 8.9E-11 | MB231/MB468 | MP |
| >hg38_21_+_8433172_8446622_RNA45SN1_45S_50ntFlanks@11051.11075.25\|CCGTCCTCCCCCCTCCCCGGGGGAG | >hg38_21_+_8433172_8446622_RNA45SN1_45S_50ntFlanks | 11051 | 11075 | 25 | CCGTCCTCCCCCCTCCCCGGGGGAG | 161.5 | -11.2 | 1.7 | -6.6 | 3.0E-11 | 8.9E-11 | MB231/MB468 | MP |
| >am_trna116_GluCTC_1_-_145399233_145399304@1.29.29,trna59_GluCTC_1_+_249168447_249168518@1.29.29,trna71_GluCTC_1_-_161439189_161439260@1.29.29,trna74_GluCTC_1_-_161431809_161431880@1.29.29,trna77_GluCTC_1_-_161424398_161424469@1.29.29,trna77_GluCTC_6_+_28949976_28950047@1.29.29,trna80_GluCTC_1_-_161417018_161417089@1.29.29,trna87_GluCTC_6_-_126101393_126101464@1.29.29\|TCCCTGGTGGTCTAGTGGTTAGGATTCGG | >am_trna116_GluCTC_1_-_145399233_145399304 | 1 | 29 | 29 | TCCCTGGTGGTCTAGTGGTTAGGATTCGG | 65111.8 | -7.7 | 1.2 | -6.6 | 4.0E-11 | 8.9E-11 | BT20/MB468 | NUC |
| >hg38_21_+_8433172_8446622_RNA45SN1_45S_50ntFlanks@10535.10570.36\|CGCGAGGCGTCCAGTGCGGTAACGCGACCGATCCCG | >hg38_21_+_8433172_8446622_RNA45SN1_45S_50ntFlanks | 10535 | 10570 | 36 | CGCGAGGCGTCCAGTGCGGTAACGCGACCGATCCCG | 260.7 | -11.1 | 1.6 | -7.0 | 2.1E-12 | 9.0E-11 | BT20/MB468 | MP |
| >hg38_21_+_8433172_8446622_RNA45SN1_45S_50ntFlanks@10805.10830.26\|GAACAATGTAGGTAAGGGAAGTCGGC | >hg38_21_+_8433172_8446622_RNA45SN1_45S_50ntFlanks | 10805 | 10830 | 26 | GAACAATGTAGGTAAGGGAAGTCGGC | 86.2 | -10.3 | 1.5 | -6.6 | 3.0E-11 | 9.0E-11 | MB231/MB468 | MP |
| >hg38_21_+_8433172_8446622_RNA45SN1_45S_50ntFlanks@11874.11906.33\|AGAAGACCCTGTTGAGCTTGACTCTAGTCTGGC | >hg38_21_+_8433172_8446622_RNA45SN1_45S_50ntFlanks | 11874 | 11906 | 33 | AGAAGACCCTGTTGAGCTTGACTCTAGTCTGGC | 1957.4 | -4.5 | 0.7 | -6.7 | 2.4E-11 | 9.0E-11 | BT20/MB468 | MITO |
| >hg38_21_+_8433172_8446622_RNA45SN1_45S_50ntFlanks@6648.6674.27\|GTACGACTCTTAGCGGTGGATCACTCG | >hg38_21_+_8433172_8446622_RNA45SN1_45S_50ntFlanks | 6648 | 6674 | 27 | GTACGACTCTTAGCGGTGGATCACTCG | 1798.7 | 5.0 | 0.8 | 6.7 | 2.4E-11 | 9.0E-11 | BT20/MB468 | MITO |
| >hg38_21_+_8433172_8446622_RNA45SN1_45S_50ntFlanks@9169.9197.29\|GGCGGGTCGCGCCGTCGGGCCCGGGGGAG | >hg38_21_+_8433172_8446622_RNA45SN1_45S_50ntFlanks | 9169 | 9197 | 29 | GGCGGGTCGCGCCGTCGGGCCCGGGGGAG | 238.4 | -11.8 | 1.8 | -6.6 | 3.0E-11 | 9.0E-11 | MB231/MB468 | MP |
| >hg38_21_+_8433172_8446622_RNA45SN1_45S_50ntFlanks@5525.5550.26\|TAAAAGTCGTAACAAGGTTTCCGTAG | >hg38_21_+_8433172_8446622_RNA45SN1_45S_50ntFlanks | 5525 | 5550 | 26 | TAAAAGTCGTAACAAGGTTTCCGTAG | 288.7 | -5.0 | 0.8 | -6.6 | 3.3E-11 | 9.0E-11 | MB231/MB468 | CYTO |
| >MI0000746\|hsa-mir-99b&WithFlank&19\|+\|51692606\|51692687@13.33.21(+1U)&[MIMAT0000689&hsa-miR-99b-5p&offsets\|0\|-1(+1U);m-10&19\|+\|51692618\|51692639&offsets\|0\|-1(+1U)]\|CACCCGTAGAACCGACCTTGCT | >MI0000746\|hsa-mir-99b&WithFlank&19\|+\|51692606\|51692687 | 13 | 33 | 21(+1U) | CACCCGTAGAACCGACCTTGCT | 623.2 | 5.9 | 0.9 | 6.6 | 3.3E-11 | 9.0E-11 | MB231/MB468 | CYTO |
| >hg38_21_+_8433172_8446622_RNA45SN1_45S_50ntFlanks@3843.3881.39\|CTACTTGGATAACTGTGGTAATTCTAGAGCTAATACATG | >hg38_21_+_8433172_8446622_RNA45SN1_45S_50ntFlanks | 3843 | 3881 | 39 | CTACTTGGATAACTGTGGTAATTCTAGAGCTAATACATG | 161.5 | -9.4 | 1.4 | -6.7 | 2.4E-11 | 9.0E-11 | BT20/MB468 | MITO |
| >MI0000095\|hsa-mir-93&WithFlank&7\|-\|100093762\|100093853@17.37.21&[MIMAT0000093&hsa-miR-93-5p&offsets\|0\|-2;m-29&7\|-\|100093815\|100093837&offsets\|0\|-2]\|CAAAGTGCTGTTCGTGCAGGT | >MI0000095\|hsa-mir-93&WithFlank&7\|-\|100093762\|100093853 | 17 | 37 | 21 | CAAAGTGCTGTTCGTGCAGGT | 1026.7 | 5.1 | 0.8 | 6.6 | 3.3E-11 | 9.0E-11 | MB231/MB468 | NUC |
| >MI0000088\|hsa-mir-30a&WithFlank&6\|-\|71403545\|71403627@12.34.23&[MIMAT0000087&hsa-miR-30a-5p&offsets\|0\|+1;m-15&6\|-\|71403595\|71403616&offsets\|0\|+1]\|TGTAAACATCCTCGACTGGAAGC | >MI0000088\|hsa-mir-30a&WithFlank&6\|-\|71403545\|71403627 | 12 | 34 | 23 | TGTAAACATCCTCGACTGGAAGC | 26787.1 | 6.4 | 1.0 | 6.6 | 3.8E-11 | 9.0E-11 | MB231/MB468 | MITO |
| >hg38_21_+_8433172_8446622_RNA45SN1_45S_50ntFlanks@11582.11609.28\|ACTGGTGCGGACCAGGGGAATCCGACTG | >hg38_21_+_8433172_8446622_RNA45SN1_45S_50ntFlanks | 11582 | 11609 | 28 | ACTGGTGCGGACCAGGGGAATCCGACTG | 439.5 | -4.5 | 0.7 | -6.6 | 3.0E-11 | 9.0E-11 | MB231/MB468 | MP |
| >hg38_21_+_8433172_8446622_RNA45SN1_45S_50ntFlanks@12602.12626.25\|GTGTATGTGCTTGGCTGAGGAGCCA | >hg38_21_+_8433172_8446622_RNA45SN1_45S_50ntFlanks | 12602 | 12626 | 25 | GTGTATGTGCTTGGCTGAGGAGCCA | 171.5 | -7.7 | 1.2 | -6.6 | 3.3E-11 | 9.0E-11 | MB231/MB468 | NUC |
| >hg38_21_+_8433172_8446622_RNA45SN1_45S_50ntFlanks@10798.10830.33\|CATGTTGGAACAATGTAGGTAAGGGAAGTCGGC | >hg38_21_+_8433172_8446622_RNA45SN1_45S_50ntFlanks | 10798 | 10830 | 33 | CATGTTGGAACAATGTAGGTAAGGGAAGTCGGC | 78.4 | -10.2 | 1.5 | -6.6 | 3.1E-11 | 9.0E-11 | MB231/MB468 | MP |
| >hg38_21_+_8433172_8446622_RNA45SN1_45S_50ntFlanks@4999.5016.18\|ATTGATAGCTCTTTCTCG | >hg38_21_+_8433172_8446622_RNA45SN1_45S_50ntFlanks | 4999 | 5016 | 18 | ATTGATAGCTCTTTCTCG | 698.0 | -13.2 | 2.0 | -6.6 | 3.8E-11 | 9.0E-11 | MB231/MB468 | MITO |
| >hg38_21_+_8433172_8446622_RNA45SN1_45S_50ntFlanks@10830.10852.23\|CAAGCCGGATCCGTAACTTCGGG | >hg38_21_+_8433172_8446622_RNA45SN1_45S_50ntFlanks | 10830 | 10852 | 23 | CAAGCCGGATCCGTAACTTCGGG | 332.5 | -6.3 | 0.9 | -6.6 | 3.3E-11 | 9.0E-11 | MB231/MB468 | CYTO |
| >hg38_21_+_8433172_8446622_RNA45SN1_45S_50ntFlanks@11683.11702.20\|ATGTCAAAGTGAAGAAATTC | >hg38_21_+_8433172_8446622_RNA45SN1_45S_50ntFlanks | 11683 | 11702 | 20 | ATGTCAAAGTGAAGAAATTC | 1041.4 | -8.1 | 1.2 | -6.6 | 3.1E-11 | 9.0E-11 | MB231/MB468 | MP |

<table>
<tr><td>>hg38_21_+_8433172_8446622_RNA45SN1_45S_50ntFlanks@10805.10828.24|GAACAATGTAGGTAAGGGAAGTCG</td><td>>hg38_21_+_8433172_8446622_RNA45SN1_45S_50ntFlanks</td><td>10805</td><td>10828</td><td>24</td><td>GAACAATGTAGGTAAGGGAAGTCG</td><td>343.4</td><td>-8.0</td><td>1.2</td><td>-6.6</td><td>3.8E-11</td><td>9.1E-11</td><td>MB231/MB468</td><td>MITO</td></tr>
<tr><td>>hg38_MT_+_598_1651_RNR1_12S_50ntFlanks@368.404.37|CAATAAAGCTAAAACTCACCTGAGTTGTAAAAAACTC</td><td>>hg38_MT_+_598_1651_RNR1_12S_50ntFlanks</td><td>368</td><td>404</td><td>37</td><td>CAATAAAGCTAAAACTCACCTGAGTTGTAAAAAACTC</td><td>1616.1</td><td>5.3</td><td>0.8</td><td>6.7</td><td>2.4E-11</td><td>9.1E-11</td><td>BT20/MB468</td><td>MITO</td></tr>
<tr><td>>MI0000077|hsa-mir-21&WithFlank&17|+|59841260|59841343@14.36.23(+1C)&[MIMAT0000076&hsa-miR-21-5p&offsets|0|+1(+1C);m-1&17|+|59841273|59841294&offsets|0|+1(+1C)]|TAGCTTATCAGACTGATGTTGACC</td><td>>MI0000077|hsa-mir-21&WithFlank&17|+|59841260|59841343</td><td>14</td><td>36</td><td>23(+1C)</td><td>TAGCTTATCAGACTGATGTTGACC</td><td>19138.0</td><td>6.1</td><td>0.9</td><td>6.6</td><td>3.3E-11</td><td>9.1E-11</td><td>MB231/MB468</td><td>CYTO</td></tr>
<tr><td>>hg38_21_+_8433172_8446622_RNA45SN1_45S_50ntFlanks@9391.9410.20|CCGAGGTGGGATCCCGAGGC</td><td>>hg38_21_+_8433172_8446622_RNA45SN1_45S_50ntFlanks</td><td>9391</td><td>9410</td><td>20</td><td>CCGAGGTGGGATCCCGAGGC</td><td>139.5</td><td>-11.0</td><td>1.7</td><td>-6.6</td><td>3.1E-11</td><td>9.1E-11</td><td>MB231/MB468</td><td>MP</td></tr>
<tr><td>>hg38_21_+_8433172_8446622_RNA45SN1_45S_50ntFlanks@10786.10826.41|AACAGCCTCTGGCATGTTGGAACAATGTAGGTAAGGGAAGT</td><td>>hg38_21_+_8433172_8446622_RNA45SN1_45S_50ntFlanks</td><td>10786</td><td>10826</td><td>41</td><td>AACAGCCTCTGGCATGTTGGAACAATGTAGGTAAGGGAAGT</td><td>1689.1</td><td>-10.3</td><td>1.6</td><td>-6.6</td><td>4.0E-11</td><td>9.1E-11</td><td>BT20/MB468</td><td>NUC</td></tr>
<tr><td>>hg38_21_+_8433172_8446622_RNA45SN1_45S_50ntFlanks@11581.11609.29|AACTGGTGCGGACCAGGGGAATCCGACTG</td><td>>hg38_21_+_8433172_8446622_RNA45SN1_45S_50ntFlanks</td><td>11581</td><td>11609</td><td>29</td><td>AACTGGTGCGGACCAGGGGAATCCGACTG</td><td>1795.9</td><td>-3.0</td><td>0.5</td><td>-6.6</td><td>3.3E-11</td><td>9.1E-11</td><td>MB231/MB468</td><td>CYTO</td></tr>
<tr><td>>hg38_21_+_8433172_8446622_RNA45SN1_45S_50ntFlanks@11874.11899.26|AGAAGACCCTGTTGAGCTTGACTCTA</td><td>>hg38_21_+_8433172_8446622_RNA45SN1_45S_50ntFlanks</td><td>11874</td><td>11899</td><td>26</td><td>AGAAGACCCTGTTGAGCTTGACTCTA</td><td>797.2</td><td>-8.4</td><td>1.3</td><td>-6.6</td><td>3.9E-11</td><td>9.1E-11</td><td>MB231/MB468</td><td>MITO</td></tr>
<tr><td>>trna15_ValAAC_5_-_180615416_180615488@1.32.32|GTTTCCGTAGTGTAGTGGTCATCACGTTCGCC</td><td>>trna15_ValAAC_5_-_180615416_180615488</td><td>1</td><td>32</td><td>32</td><td>GTTTCCGTAGTGTAGTGGTCATCACGTTCGCC</td><td>586.2</td><td>6.5</td><td>1.0</td><td>6.6</td><td>3.9E-11</td><td>9.2E-11</td><td>MB231/MB468</td><td>MITO</td></tr>
<tr><td>>hg38_21_+_8433172_8446622_RNA45SN1_45S_50ntFlanks@12511.12538.28|CGTGAGACAGGTTAGTTTTACCCTACTG</td><td>>hg38_21_+_8433172_8446622_RNA45SN1_45S_50ntFlanks</td><td>12511</td><td>12538</td><td>28</td><td>CGTGAGACAGGTTAGTTTTACCCTACTG</td><td>227.8</td><td>-10.9</td><td>1.6</td><td>-7.0</td><td>2.2E-12</td><td>9.2E-11</td><td>BT20/MB468</td><td>MP</td></tr>
<tr><td>>hg38_21_+_8433172_8446622_RNA45SN1_45S_50ntFlanks@10713.10736.24|CTTGAAAATCCGGGGGAGAGGGTG</td><td>>hg38_21_+_8433172_8446622_RNA45SN1_45S_50ntFlanks</td><td>10713</td><td>10736</td><td>24</td><td>CTTGAAAATCCGGGGGAGAGGGTG</td><td>550.0</td><td>-9.1</td><td>1.4</td><td>-6.6</td><td>4.1E-11</td><td>9.2E-11</td><td>BT20/MB468</td><td>NUC</td></tr>
<tr><td>>MI0000294|hsa-mir-218-1&WithFlank&4|+|20528269|20528390@31.52.22;MI0000295|hsa-mir-218-2&WithFlank&5|-|168768140|168768261@31.52.22&[MIMAT0000275&hsa-miR-218-5p&offsets|0|+1;m-185&4|+|20528299|20528319&offsets|0|+1];[MIMAT0000275_1&hsa-miR-218-5p&offsets|0|+1;m-187&5|-|168768211|168768231&offsets|0|+1]|TTGTGCTTGATCTAACCATGTG</td><td>>MI0000294|hsa-mir-218-1&WithFlank&4|+|20528269|20528390</td><td>31</td><td>52</td><td>22</td><td>TTGTGCTTGATCTAACCATGTG</td><td>172.1</td><td>-10.5</td><td>1.5</td><td>-7.0</td><td>3.4E-12</td><td>9.2E-11</td><td>BT20/MB231</td><td>NUC</td></tr>
<tr><td>>hg38_21_+_8433172_8446622_RNA45SN1_45S_50ntFlanks@11316.11336.21|CACCCCACGTCTCGTCGCGCG</td><td>>hg38_21_+_8433172_8446622_RNA45SN1_45S_50ntFlanks</td><td>11316</td><td>11336</td><td>21</td><td>CACCCCACGTCTCGTCGCGCG</td><td>305.2</td><td>-11.3</td><td>1.6</td><td>-7.0</td><td>2.2E-12</td><td>9.2E-11</td><td>BT20/MB468</td><td>MP</td></tr>
<tr><td>>MI0000440|hsa-mir-27b&WithFlank&9|+|95085439|95085547@67.86.20(+1U)&[MIMAT0000419&hsa-miR-27b-3p&offsets|0|-1(+1U);m-39&9|+|95085505|95085525&offsets|0|-1(+1U)]|TTCACAGTGGCTAAGTTCTGT</td><td>>MI0000440|hsa-mir-27b&WithFlank&9|+|95085439|95085547</td><td>67</td><td>86</td><td>20(+1U)</td><td>TTCACAGTGGCTAAGTTCTGT</td><td>12442.3</td><td>6.8</td><td>1.0</td><td>6.6</td><td>3.9E-11</td><td>9.2E-11</td><td>MB231/MB468</td><td>MITO</td></tr>
<tr><td>>am_trnaMT_LeuTAG_MT_+_12266_12336@43.61.19|AAATTTTGGTGCAACTCCA</td><td>>am_trnaMT_LeuTAG_MT_+_12266_12336</td><td>43</td><td>61</td><td>19</td><td>AAATTTTGGTGCAACTCCA</td><td>96.6</td><td>-10.5</td><td>1.6</td><td>-6.6</td><td>3.1E-11</td><td>9.2E-11</td><td>MB231/MB468</td><td>MP</td></tr>
<tr><td>>hg38_21_+_8433172_8446622_RNA45SN1_45S_50ntFlanks@12488.12522.35|AACGTGAGCTGGGTTTAGACCGTCGTGAGACAGGT</td><td>>hg38_21_+_8433172_8446622_RNA45SN1_45S_50ntFlanks</td><td>12488</td><td>12522</td><td>35</td><td>AACGTGAGCTGGGTTTAGACCGTCGTGAGACAGGT</td><td>429.2</td><td>-6.9</td><td>1.0</td><td>-6.6</td><td>3.4E-11</td><td>9.2E-11</td><td>MB231/MB468</td><td>NUC</td></tr>
<tr><td>>hg38_21_+_8433172_8446622_RNA45SN1_45S_50ntFlanks@11671.11704.34|CCAGTGCTCTGAATGTCAAAGTGAAGAAATTCAA</td><td>>hg38_21_+_8433172_8446622_RNA45SN1_45S_50ntFlanks</td><td>11671</td><td>11704</td><td>34</td><td>CCAGTGCTCTGAATGTCAAAGTGAAGAAATTCAA</td><td>1689.5</td><td>-10.1</td><td>1.5</td><td>-6.6</td><td>4.1E-11</td><td>9.3E-11</td><td>BT20/MB468</td><td>NUC</td></tr>
<tr><td>>hg38_21_+_8433172_8446622_RNA45SN1_45S_50ntFlanks@11581.11607.27|AACTGGTGCGGACCAGGGGAATCCGAC</td><td>>hg38_21_+_8433172_8446622_RNA45SN1_45S_50ntFlanks</td><td>11581</td><td>11607</td><td>27</td><td>AACTGGTGCGGACCAGGGGAATCCGAC</td><td>626.9</td><td>-5.5</td><td>0.8</td><td>-6.6</td><td>3.1E-11</td><td>9.3E-11</td><td>MB231/MB468</td><td>MP</td></tr>
<tr><td>>MI0000734|hsa-mir-106b&WithFlank&7|-|100093987|100094080@18.39.22&[MIMAT0000680&hsa-miR-106b-5p&offsets|0|+1;m-103&7|-|100094043|100094063&offsets|0|+1]|TAAAGTGCTGACAGTGCAGATA</td><td>>MI0000734|hsa-mir-106b&WithFlank&7|-|100093987|100094080</td><td>18</td><td>39</td><td>22</td><td>TAAAGTGCTGACAGTGCAGATA</td><td>806.8</td><td>5.7</td><td>0.9</td><td>6.6</td><td>3.0E-11</td><td>9.3E-11</td><td>BT20/MB468</td><td>CYTO</td></tr>
<tr><td>>hg38_21_+_8433172_8446622_RNA45SN1_45S_50ntFlanks@6776.6803.28|CTCCCGGGGCTACGCCTGTCTGAGCGTC</td><td>>hg38_21_+_8433172_8446622_RNA45SN1_45S_50ntFlanks</td><td>6776</td><td>6803</td><td>28</td><td>CTCCCGGGGCTACGCCTGTCTGAGCGTC</td><td>4499.4</td><td>-5.9</td><td>0.9</td><td>-6.6</td><td>3.9E-11</td><td>9.3E-11</td><td>MB231/MB468</td><td>MITO</td></tr>
<tr><td>>hg38_MT_+_598_1651_RNR1_12S_50ntFlanks@373.404.32|AAGCTAAAACTCACCTGAGTTGTAAAAAACTC</td><td>>hg38_MT_+_598_1651_RNR1_12S_50ntFlanks</td><td>373</td><td>404</td><td>32</td><td>AAGCTAAAACTCACCTGAGTTGTAAAAAACTC</td><td>875.0</td><td>5.4</td><td>0.8</td><td>6.7</td><td>2.5E-11</td><td>9.3E-11</td><td>BT20/MB468</td><td>MITO</td></tr>
<tr><td>>MI0000803|hsa-mir-330&WithFlank&19|-|45638988|45639093@63.85.23&[MIMAT0000751&hsa-miR-330-3p&offsets|0|0;m-347&19|-|45639009|45639030&offsets|-1|0]|GCAAAGCACACGGCCTGCAGAGA</td><td>>MI0000803|hsa-mir-330&WithFlank&19|-|45638988|45639093</td><td>63</td><td>85</td><td>23</td><td>GCAAAGCACACGGCCTGCAGAGA</td><td>520.8</td><td>9.7</td><td>1.5</td><td>6.6</td><td>3.1E-11</td><td>9.3E-11</td><td>BT20/MB468</td><td>CYTO</td></tr>
<tr><td>>MI0003628|hsa-mir-615&WithFlank&12|+|54033944|54034051@67.87.21(+1C)&[MIMAT0003283&hsa-miR-615-3p&offsets|0|-1(+1C);m-495&12|+|54034010|54034030&offsets|0|0(+1C)]|TCCGAGCCTGGGTCTCCCTCTC</td><td>>MI0003628|hsa-mir-615&WithFlank&12|+|54033944|54034051</td><td>67</td><td>87</td><td>21(+1C)</td><td>TCCGAGCCTGGGTCTCCCTCTC</td><td>327.5</td><td>9.4</td><td>1.4</td><td>6.6</td><td>3.1E-11</td><td>9.3E-11</td><td>BT20/MB468</td><td>CYTO</td></tr>
<tr><td>>hg38_21_+_8433172_8446622_RNA45SN1_45S_50ntFlanks@10409.10429.21|GTCAGTCGGTCCTGAGAGATG</td><td>>hg38_21_+_8433172_8446622_RNA45SN1_45S_50ntFlanks</td><td>10409</td><td>10429</td><td>21</td><td>GTCAGTCGGTCCTGAGAGATG</td><td>1120.1</td><td>-2.5</td><td>0.4</td><td>-6.6</td><td>3.1E-11</td><td>9.3E-11</td><td>BT20/MB468</td><td>CYTO</td></tr>
<tr><td>>hg38_21_+_8433172_8446622_RNA45SN1_45S_50ntFlanks@11453.11477.25|TCCTCGGCGCGCGGCGGCGGCGGCG</td><td>>hg38_21_+_8433172_8446622_RNA45SN1_45S_50ntFlanks</td><td>11453</td><td>11477</td><td>25</td><td>TCCTCGGCGCGCGGCGGCGGCGGCG</td><td>92.6</td><td>-10.4</td><td>1.6</td><td>-6.6</td><td>3.1E-11</td><td>9.3E-11</td><td>MB231/MB468</td><td>MP</td></tr>
<tr><td>>hg38_21_+_8433172_8446622_RNA45SN1_45S_50ntFlanks@7984.8006.23|AGATCAGACGTGGCGACCCGCTG</td><td>>hg38_21_+_8433172_8446622_RNA45SN1_45S_50ntFlanks</td><td>7984</td><td>8006</td><td>23</td><td>AGATCAGACGTGGCGACCCGCTG</td><td>5691.7</td><td>-7.8</td><td>1.2</td><td>-6.7</td><td>2.5E-11</td><td>9.3E-11</td><td>BT20/MB468</td><td>MITO</td></tr>
<tr><td>>hg38_21_+_8433172_8446622_RNA45SN1_45S_50ntFlanks@9805.9838.34|GCGTGGAATGCGAGTGCCTAGTGGGCCACTTTTG</td><td>>hg38_21_+_8433172_8446622_RNA45SN1_45S_50ntFlanks</td><td>9805</td><td>9838</td><td>34</td><td>GCGTGGAATGCGAGTGCCTAGTGGGCCACTTTTG</td><td>169.6</td><td>-9.5</td><td>1.4</td><td>-6.7</td><td>2.5E-11</td><td>9.3E-11</td><td>BT20/MB468</td><td>MITO</td></tr>
<tr><td>>hg38_21_+_8433172_8446622_RNA45SN1_45S_50ntFlanks@3921.3959.39|ATTTATCAGATCAAAACCAACCCGGTCAGCCCCTCTCCG</td><td>>hg38_21_+_8433172_8446622_RNA45SN1_45S_50ntFlanks</td><td>3921</td><td>3959</td><td>39</td><td>ATTTATCAGATCAAAACCAACCCGGTCAGCCCCTCTCCG</td><td>205.3</td><td>-9.8</td><td>1.5</td><td>-6.7</td><td>2.5E-11</td><td>9.3E-11</td><td>BT20/MB468</td><td>MITO</td></tr>
<tr><td>>hg38_21_+_8433172_8446622_RNA45SN1_45S_50ntFlanks@11573.11602.30|CGACTTAGAACTGGTGCGGACCAGGGGAAT</td><td>>hg38_21_+_8433172_8446622_RNA45SN1_45S_50ntFlanks</td><td>11573</td><td>11602</td><td>30</td><td>CGACTTAGAACTGGTGCGGACCAGGGGAAT</td><td>507.0</td><td>-4.8</td><td>0.7</td><td>-6.6</td><td>3.9E-11</td><td>9.3E-11</td><td>MB231/MB468</td><td>MITO</td></tr>
<tr><td>>MI0000067|hsa-let-7f-1&WithFlank&9|+|94176341|94176439@13.34.22;MI0000068|hsa-let-7f-2&WithFlank&X|-|53557186|53557280@14.35.22&[MIMAT0000067&hsa-let-7f-5p&offsets|0|0;m-20&9|+|94176353|94176374&offsets|0|0];[MIMAT0000067_1&hsa-let-7f-5p&offsets|0|0;m-19&X|-|53557246|53557267&offsets|0|0]|TGAGGTAGTAGATTGTATAGTT</td><td>>MI0000067|hsa-let-7f-1&WithFlank&9|+|94176341|94176439</td><td>13</td><td>34</td><td>22</td><td>TGAGGTAGTAGATTGTATAGTT</td><td>53625.1</td><td>4.4</td><td>0.7</td><td>6.6</td><td>3.4E-11</td><td>9.3E-11</td><td>MB231/MB468</td><td>CYTO</td></tr>
<tr><td>>hg38_21_+_8433172_8446622_RNA45SN1_45S_50ntFlanks@6651.6684.34|CGACTCTTAGCGGTGGATCACTCGGCTCGTGCGT</td><td>>hg38_21_+_8433172_8446622_RNA45SN1_45S_50ntFlanks</td><td>6651</td><td>6684</td><td>34</td><td>CGACTCTTAGCGGTGGATCACTCGGCTCGTGCGT</td><td>1280.9</td><td>3.9</td><td>0.6</td><td>6.6</td><td>3.4E-11</td><td>9.3E-11</td><td>MB231/MB468</td><td>CYTO</td></tr>
<tr><td>>hg38_21_+_8433172_8446622_RNA45SN1_45S_50ntFlanks@8158.8181.24|TCGTGGGGGGCCCAAGTCCTTCTG</td><td>>hg38_21_+_8433172_8446622_RNA45SN1_45S_50ntFlanks</td><td>8158</td><td>8181</td><td>24</td><td>TCGTGGGGGGCCCAAGTCCTTCTG</td><td>75.6</td><td>-10.1</td><td>1.5</td><td>-6.6</td><td>3.2E-11</td><td>9.3E-11</td><td>MB231/MB468</td><td>MP</td></tr>
<tr><td>>am_trna119_LysCTT_1_-_145395522_145395594@13.35.23,trna11_LysCTT_5_-_180648979_180649051@13.35.23,trna128_LysCTT_1_-_55423542_55423614@13.35.23,trna13_LysCTT_6_+_26556774_26556846@13.35.23,trna6_LysCTT_19_-_52425396_52425468@13.35.23,trna7_LysCTT_16_+_3225692_3225764@13.35.23,trna9_LysCTT_5_+_180634755_180634827@13.35.23|CAGTCGGTAGAGCATGAGACTCT</td><td>>am_trna119_LysCTT_1_-_145395522_145395594</td><td>13</td><td>35</td><td>23</td><td>CAGTCGGTAGAGCATGAGACTCT</td><td>687.6</td><td>-7.0</td><td>1.0</td><td>-7.0</td><td>3.1E-12</td><td>9.3E-11</td><td>BT20/MB231</td><td>MITO</td></tr>
<tr><td>>MI0000066|hsa-let-7e&WithFlank&19|+|51692780|51692870@14.33.20&[MIMAT0000066&hsa-let-7e-5p&offsets|0|-2;m-50&19|+|51692793|51692814&offsets|0|-2]|TGAGGTAGGAGGTTGTATAG</td><td>>MI0000066|hsa-let-7e&WithFlank&19|+|51692780|51692870</td><td>14</td><td>33</td><td>20</td><td>TGAGGTAGGAGGTTGTATAG</td><td>348.4</td><td>8.2</td><td>1.2</td><td>6.6</td><td>3.1E-11</td><td>9.3E-11</td><td>BT20/MB468</td><td>CYTO</td></tr>
<tr><td>>hg38_21_+_8433172_8446622_RNA45SN1_45S_50ntFlanks@12526.12548.23|TTTTACCCTACTGATGATGTGTT</td><td>>hg38_21_+_8433172_8446622_RNA45SN1_45S_50ntFlanks</td><td>12526</td><td>12548</td><td>23</td><td>TTTTACCCTACTGATGATGTGTT</td><td>85.7</td><td>-10.3</td><td>1.5</td><td>-6.6</td><td>3.2E-11</td><td>9.3E-11</td><td>MB231/MB468</td><td>MP</td></tr>
<tr><td>>hg38_21_+_8433172_8446622_RNA45SN1_45S_50ntFlanks@11273.11293.21|CCGCCCCCGGGGAGCCCGGCG</td><td>>hg38_21_+_8433172_8446622_RNA45SN1_45S_50ntFlanks</td><td>11273</td><td>11293</td><td>21</td><td>CCGCCCCCGGGGAGCCCGGCG</td><td>274.2</td><td>-8.4</td><td>1.3</td><td>-6.6</td><td>3.4E-11</td><td>9.4E-11</td><td>MB231/MB468</td><td>NUC</td></tr>
<tr><td>>MI0000266|hsa-mir-10a&WithFlank&17|-|48579832|48579953@29.49.21(+1A)&[MIMAT0000253&hsa-miR-10a-5p&offsets|+1|-1(+1A);m-18&17|-|48579905|48579926&offsets|+1|0(+1A)]|ACCCTGTAGATCCGAATTTGTA</td><td>>MI0000266|hsa-mir-10a&WithFlank&17|-|48579832|48579953</td><td>29</td><td>49</td><td>21(+1A)</td><td>ACCCTGTAGATCCGAATTTGTA</td><td>285.4</td><td>11.9</td><td>1.7</td><td>7.0</td><td>2.2E-12</td><td>9.4E-11</td><td>BT20/MB468</td><td>MP</td></tr>
<tr><td>>hg38_21_+_8433172_8446622_RNA45SN1_45S_50ntFlanks@10116.10138.23|TGCGTCGGAGGGCGGCGGCGGCG</td><td>>hg38_21_+_8433172_8446622_RNA45SN1_45S_50ntFlanks</td><td>10116</td><td>10138</td><td>23</td><td>TGCGTCGGAGGGCGGCGGCGGCG</td><td>164.0</td><td>-9.7</td><td>1.5</td><td>-6.6</td><td>3.1E-11</td><td>9.4E-11</td><td>BT20/MB468</td><td>CYTO</td></tr>
<tr><td>>hg38_21_+_8433172_8446622_RNA45SN1_45S_50ntFlanks@12640.12657.18|ACCATCTGTGGGATTATG</td><td>>hg38_21_+_8433172_8446622_RNA45SN1_45S_50ntFlanks</td><td>12640</td><td>12657</td><td>18</td><td>ACCATCTGTGGGATTATG</td><td>470.0</td><td>-8.3</td><td>1.2</td><td>-6.6</td><td>3.2E-11</td><td>9.4E-11</td><td>MB231/MB468</td><td>MP</td></tr>
<tr><td>>hg38_21_+_8433172_8446622_RNA45SN1_45S_50ntFlanks@6666.6686.21|GATCACTCGGCTCGTGCGTCG</td><td>>hg38_21_+_8433172_8446622_RNA45SN1_45S_50ntFlanks</td><td>6666</td><td>6686</td><td>21</td><td>GATCACTCGGCTCGTGCGTCG</td><td>238.7</td><td>-11.0</td><td>1.6</td><td>-7.0</td><td>2.3E-12</td><td>9.4E-11</td><td>BT20/MB468</td><td>MP</td></tr>
<tr><td>>hg38_21_+_8433172_8446622_RNA45SN1_45S_50ntFlanks@11670.11694.25|CCCAGTGCTCTGAATGTCAAAGTGA</td><td>>hg38_21_+_8433172_8446622_RNA45SN1_45S_50ntFlanks</td><td>11670</td><td>11694</td><td>25</td><td>CCCAGTGCTCTGAATGTCAAAGTGA</td><td>74.7</td><td>-10.1</td><td>1.5</td><td>-6.6</td><td>3.2E-11</td><td>9.4E-11</td><td>MB231/MB468</td><td>MP</td></tr>
<tr><td>>hg38_21_+_8433172_8446622_RNA45SN1_45S_50ntFlanks@3984.4014.31|GCGGCTTTGGTGACTCTAGATAACCTCGGGC</td><td>>hg38_21_+_8433172_8446622_RNA45SN1_45S_50ntFlanks</td><td>3984</td><td>4014</td><td>31</td><td>GCGGCTTTGGTGACTCTAGATAACCTCGGGC</td><td>112.5</td><td>-10.7</td><td>1.6</td><td>-6.6</td><td>3.2E-11</td><td>9.4E-11</td><td>MB231/MB468</td><td>MP</td></tr>
<tr><td>>hg38_21_+_8433172_8446622_RNA45SN1_45S_50ntFlanks@3851.3871.21|ATAACTGTGGTAATTCTAGAG</td><td>>hg38_21_+_8433172_8446622_RNA45SN1_45S_50ntFlanks</td><td>3851</td><td>3871</td><td>21</td><td>ATAACTGTGGTAATTCTAGAG</td><td>104.8</td><td>-10.6</td><td>1.6</td><td>-6.6</td><td>3.2E-11</td><td>9.4E-11</td><td>MB231/MB468</td><td>MP</td></tr>
<tr><td>>hg38_21_+_8433172_8446622_RNA45SN1_45S_50ntFlanks@11188.11225.38|CACCCGGGGGGCCGGCGGCGGCGGCGACTCTGGACGCG</td><td>>hg38_21_+_8433172_8446622_RNA45SN1_45S_50ntFlanks</td><td>11188</td><td>11225</td><td>38</td><td>CACCCGGGGGGCCGGCGGCGGCGGCGACTCTGGACGCG</td><td>173.8</td><td>-11.3</td><td>1.7</td><td>-6.6</td><td>3.2E-11</td><td>9.4E-11</td><td>MB231/MB468</td><td>MP</td></tr>
<tr><td>>hg38_1_-_228634819_228635039_RNA5S12_5S_50ntFlanks@154.171.18|CCGGGTGCTGTAGGCTTT</td><td>>hg38_1_-_228634819_228635039_RNA5S12_5S_50ntFlanks</td><td>154</td><td>171</td><td>18</td><td>CCGGGTGCTGTAGGCTTT</td><td>7598.3</td><td>4.9</td><td>0.7</td><td>6.6</td><td>3.4E-11</td><td>9.4E-11</td><td>MB231/MB468</td><td>CYTO</td></tr>
<tr><td>>hg38_21_+_8433172_8446622_RNA45SN1_45S_50ntFlanks@7991.8018.28|ACGTGGCGACCCGCTGAATTTAAGCATA</td><td>>hg38_21_+_8433172_8446622_RNA45SN1_45S_50ntFlanks</td><td>7991</td><td>8018</td><td>28</td><td>ACGTGGCGACCCGCTGAATTTAAGCATA</td><td>258.3</td><td>-11.1</td><td>1.6</td><td>-7.0</td><td>2.3E-12</td><td>9.4E-11</td><td>BT20/MB468</td><td>MP</td></tr>
<tr><td>>hg38_1_-_228634819_228635039_RNA5S12_5S_50ntFlanks@138.170.33|AGACCGCCTGGGAATACCGGGTGCTGTAGGCTT</td><td>>hg38_1_-_228634819_228635039_RNA5S12_5S_50ntFlanks</td><td>138</td><td>170</td><td>33</td><td>AGACCGCCTGGGAATACCGGGTGCTGTAGGCTT</td><td>908.9</td><td>4.3</td><td>0.6</td><td>6.7</td><td>2.5E-11</td><td>9.4E-11</td><td>BT20/MB468</td><td>MITO</td></tr>
<tr><td>>hg38_21_+_8433172_8446622_RNA45SN1_45S_50ntFlanks@11564.11612.49|CCTAGCAGCCGACTTAGAACTGGTGCGGACCAGGGGAATCCGACTGTTT</td><td>>hg38_21_+_8433172_8446622_RNA45SN1_45S_50ntFlanks</td><td>11564</td><td>11612</td><td>49</td><td>CCTAGCAGCCGACTTAGAACTGGTGCGGACCAGGGGAATCCGACTGTTT</td><td>220.0</td><td>-11.6</td><td>1.8</td><td>-6.6</td><td>4.0E-11</td><td>9.5E-11</td><td>MB231/MB468</td><td>MITO</td></tr>
</table>

<table>
<tr><td>>MI0000088|hsa-mir-30a&WithFlank&6|-|71403545|71403627@53.73.21(+2U)&[MIMAT0000088&hsa-miR-30a-3p&offsets|0|-1(+2U);m-38&6|-|71403554|71403575&offsets|0|-1(+2U)]|CTTTCAGTCGGATGTTTGCAGTT</td><td>>MI0000088|hsa-mir-30a&WithFlank&6|-|71403545|71403627</td><td>53</td><td>73</td><td>21(+2U)</td><td>CTTTCAGTCGGATGTTTGCAGTT</td><td>512.5</td><td>8.5</td><td>1.3</td><td>6.6</td><td>3.1E-11</td><td>9.5E-11</td><td>BT20/MB468</td><td>CYTO</td></tr>
<tr><td>>hg38_1_-_228634819_228635039_RNA5S12_5S_50ntFlanks@146.171.26|TGGGAATACCGGGTGCTGTAGGCTTT</td><td>>hg38_1_-_228634819_228635039_RNA5S12_5S_50ntFlanks</td><td>146</td><td>171</td><td>26</td><td>TGGGAATACCGGGTGCTGTAGGCTTT</td><td>1097.6</td><td>4.8</td><td>0.7</td><td>6.6</td><td>3.4E-11</td><td>9.5E-11</td><td>MB231/MB468</td><td>NUC</td></tr>
<tr><td>>trna10_MetCAT_8_-_124169470_124169542@32.51.20,trna22_MetCAT_16_-_87417628_87417700@32.51.20|CTCATAATCTGAAGGTCGTG</td><td>>trna10_MetCAT_8_-_124169470_124169542</td><td>32</td><td>51</td><td>20</td><td>CTCATAATCTGAAGGTCGTG</td><td>105.3</td><td>-10.6</td><td>1.6</td><td>-6.6</td><td>3.2E-11</td><td>9.5E-11</td><td>MB231/MB468</td><td>MP</td></tr>
<tr><td>>hg38_21_+_8433172_8446622_RNA45SN1_45S_50ntFlanks@9470.9488.19|GAGCACGAGCGCACGTGTT</td><td>>hg38_21_+_8433172_8446622_RNA45SN1_45S_50ntFlanks</td><td>9470</td><td>9488</td><td>19</td><td>GAGCACGAGCGCACGTGTT</td><td>257.2</td><td>-11.9</td><td>1.8</td><td>-6.6</td><td>3.2E-11</td><td>9.5E-11</td><td>MB231/MB468</td><td>MP</td></tr>
<tr><td>>hg38_21_+_8433172_8446622_RNA45SN1_45S_50ntFlanks@11877.11906.30|AGACCCTGTTGAGCTTGACTCTAGTCTGGC</td><td>>hg38_21_+_8433172_8446622_RNA45SN1_45S_50ntFlanks</td><td>11877</td><td>11906</td><td>30</td><td>AGACCCTGTTGAGCTTGACTCTAGTCTGGC</td><td>73.0</td><td>-10.1</td><td>1.5</td><td>-6.6</td><td>3.2E-11</td><td>9.5E-11</td><td>MB231/MB468</td><td>MP</td></tr>
<tr><td>>hg38_21_+_8433172_8446622_RNA45SN1_45S_50ntFlanks@8638.8658.21|GCGACCGGCCGCCGCCGGGCG</td><td>>hg38_21_+_8433172_8446622_RNA45SN1_45S_50ntFlanks</td><td>8638</td><td>8658</td><td>21</td><td>GCGACCGGCCGCCGCCGGGCG</td><td>278.0</td><td>-12.0</td><td>1.8</td><td>-6.6</td><td>3.2E-11</td><td>9.5E-11</td><td>MB231/MB468</td><td>MP</td></tr>
<tr><td>>hg38_21_+_8433172_8446622_RNA45SN1_45S_50ntFlanks@10458.10478.21|ATGGCCTCCGTTGCCCTCGGC</td><td>>hg38_21_+_8433172_8446622_RNA45SN1_45S_50ntFlanks</td><td>10458</td><td>10478</td><td>21</td><td>ATGGCCTCCGTTGCCCTCGGC</td><td>9169.9</td><td>-11.4</td><td>1.7</td><td>-6.6</td><td>4.2E-11</td><td>9.5E-11</td><td>BT20/MB468</td><td>NUC</td></tr>
<tr><td>>hg38_21_+_8433172_8446622_RNA45SN1_45S_50ntFlanks@8687.8705.19|CGGCTCCGGGACGGCTGGG</td><td>>hg38_21_+_8433172_8446622_RNA45SN1_45S_50ntFlanks</td><td>8687</td><td>8705</td><td>19</td><td>CGGCTCCGGGACGGCTGGG</td><td>416.7</td><td>9.0</td><td>1.4</td><td>6.6</td><td>3.5E-11</td><td>9.5E-11</td><td>MB231/MB468</td><td>CYTO</td></tr>
<tr><td>>hg38_1_-_228634819_228635039_RNA5S12_5S_50ntFlanks@128.169.42|CTTGGATGGGAGACCGCCTGGGAATACCGGGTGCTGTAGGCT</td><td>>hg38_1_-_228634819_228635039_RNA5S12_5S_50ntFlanks</td><td>128</td><td>169</td><td>42</td><td>CTTGGATGGGAGACCGCCTGGGAATACCGGGTGCTGTAGGCT</td><td>1916.0</td><td>9.1</td><td>1.4</td><td>6.6</td><td>3.1E-11</td><td>9.5E-11</td><td>BT20/MB468</td><td>CYTO</td></tr>
<tr><td>>hg38_21_+_8433172_8446622_RNA45SN1_45S_50ntFlanks@4102.4121.20|ACCACGGGTGACGGGGAATC</td><td>>hg38_21_+_8433172_8446622_RNA45SN1_45S_50ntFlanks</td><td>4102</td><td>4121</td><td>20</td><td>ACCACGGGTGACGGGGAATC</td><td>2864.9</td><td>-5.7</td><td>0.9</td><td>-6.6</td><td>3.2E-11</td><td>9.5E-11</td><td>MB231/MB468</td><td>MP</td></tr>
<tr><td>>hg38_21_+_8433172_8446622_RNA45SN1_45S_50ntFlanks@11768.11794.27|CTAATTAGTGACGCGCATGAATGGATG</td><td>>hg38_21_+_8433172_8446622_RNA45SN1_45S_50ntFlanks</td><td>11768</td><td>11794</td><td>27</td><td>CTAATTAGTGACGCGCATGAATGGATG</td><td>14471.3</td><td>-11.6</td><td>1.8</td><td>-6.6</td><td>4.2E-11</td><td>9.5E-11</td><td>BT20/MB468</td><td>NUC</td></tr>
<tr><td>>MI0000439|hsa-mir-23b&WithFlank&9|+|95085202|95085310@67.84.18&[MIMAT0000418&hsa-miR-23b-3p&offsets|+3|-2;m-42&9|+|95085265|95085285&offsets|+3|0]|ACATTGCCAGGGATTACC</td><td>>MI0000439|hsa-mir-23b&WithFlank&9|+|95085202|95085310</td><td>67</td><td>84</td><td>18</td><td>ACATTGCCAGGGATTACC</td><td>259.0</td><td>11.8</td><td>1.7</td><td>7.0</td><td>2.3E-12</td><td>9.5E-11</td><td>BT20/MB468</td><td>MP</td></tr>
<tr><td>>MI0000062|hsa-let-7a-3&WithFlank&22|+|46112743|46112828@58.78.21(+1U);MI0000060|hsa-let-7a-1&WithFlank&9|+|94175951|94176042@63.83.21(+1U)&[MIMAT0004481&hsa-let-7a-3p&offsets|0|0(+1U);m-180&22|+|46112800|46112821&offsets|0|-1(+1U)];[MIMAT0004481_1&hsa-let-7a-3p&offsets|0|0(+1U);m-181&9|+|94176013|94176034&offsets|0|-1(+1U)]|CTATACAATCTACTGTCTTTCT</td><td>>MI0000062|hsa-let-7a-3&WithFlank&22|+|46112743|46112828</td><td>58</td><td>78</td><td>21(+1U)</td><td>CTATACAATCTACTGTCTTTCT</td><td>506.2</td><td>4.6</td><td>0.7</td><td>6.6</td><td>3.5E-11</td><td>9.6E-11</td><td>MB231/MB468</td><td>CYTO</td></tr>
<tr><td>>hg38_21_+_8433172_8446622_RNA45SN1_45S_50ntFlanks@9602.9622.21|CGAAAGACTAATCGAACCATC</td><td>>hg38_21_+_8433172_8446622_RNA45SN1_45S_50ntFlanks</td><td>9602</td><td>9622</td><td>21</td><td>CGAAAGACTAATCGAACCATC</td><td>498.0</td><td>-3.7</td><td>0.6</td><td>-6.6</td><td>3.1E-11</td><td>9.6E-11</td><td>BT20/MB468</td><td>CYTO</td></tr>
<tr><td>>MI0000809|hsa-mir-151a&WithFlank&8|-|140732558|140732659@53.74.22(+2U)&[MIMAT0000757&hsa-miR-151a-3p&offsets|0|+1(+2U);m-30&8|-|140732587|140732607&offsets|0|+1(+2U)]|CTAGACTGAAGCTCCTTGAGGATT</td><td>>MI0000809|hsa-mir-151a&WithFlank&8|-|140732558|140732659</td><td>53</td><td>74</td><td>22(+2U)</td><td>CTAGACTGAAGCTCCTTGAGGATT</td><td>418.5</td><td>7.9</td><td>1.2</td><td>6.6</td><td>3.5E-11</td><td>9.6E-11</td><td>MB231/MB468</td><td>CYTO</td></tr>
<tr><td>>MI0000433|hsa-let-7g&WithFlank&3|-|52268272|52268367@10.32.23&[MIMAT0000414&hsa-let-7g-5p&offsets|-1|0;m-58&3|-|52268336|52268357&offsets|-1|0]|CTGAGGTAGTAGTTTGTACAGTT</td><td>>MI0000433|hsa-let-7g&WithFlank&3|-|52268272|52268367</td><td>10</td><td>32</td><td>23</td><td>CTGAGGTAGTAGTTTGTACAGTT</td><td>459.5</td><td>4.6</td><td>0.7</td><td>6.6</td><td>3.5E-11</td><td>9.6E-11</td><td>MB231/MB468</td><td>NUC</td></tr>
<tr><td>>hg38_21_+_8433172_8446622_RNA45SN1_45S_50ntFlanks@9562.9595.34|TCCTGACGTGCAAATCGGTCGTCCGACCTGGGTA</td><td>>hg38_21_+_8433172_8446622_RNA45SN1_45S_50ntFlanks</td><td>9562</td><td>9595</td><td>34</td><td>TCCTGACGTGCAAATCGGTCGTCCGACCTGGGTA</td><td>610.4</td><td>-9.2</td><td>1.4</td><td>-6.6</td><td>4.3E-11</td><td>9.6E-11</td><td>BT20/MB468</td><td>NUC</td></tr>
<tr><td>>MI0003591|hsa-mir-584&WithFlank&5|-|149062307|149062415@22.41.20(+1U)&[MIMAT0003249&hsa-miR-584-5p&offsets|0|-2(+1U);m-167&5|-|149062374|149062394&offsets|0|-1(+1U)]|TTATGGTTTGCCTGGGACTGT</td><td>>MI0003591|hsa-mir-584&WithFlank&5|-|149062307|149062415</td><td>22</td><td>41</td><td>20(+1U)</td><td>TTATGGTTTGCCTGGGACTGT</td><td>867.3</td><td>5.5</td><td>0.8</td><td>6.6</td><td>3.3E-11</td><td>9.6E-11</td><td>MB231/MB468</td><td>MP</td></tr>
<tr><td>>hg38_21_+_8433172_8446622_RNA45SN1_45S_50ntFlanks@12476.12500.25|CCACTAATAGGGAACGTGAGCTGGG</td><td>>hg38_21_+_8433172_8446622_RNA45SN1_45S_50ntFlanks</td><td>12476</td><td>12500</td><td>25</td><td>CCACTAATAGGGAACGTGAGCTGGG</td><td>590.0</td><td>-8.5</td><td>1.3</td><td>-6.6</td><td>3.3E-11</td><td>9.6E-11</td><td>MB231/MB468</td><td>MP</td></tr>
<tr><td>>hg38_21_+_8433172_8446622_RNA45SN1_45S_50ntFlanks@7974.8015.42|ACGCGACCTCAGATCAGACGTGGCGACCCGCTGAATTTAAGC</td><td>>hg38_21_+_8433172_8446622_RNA45SN1_45S_50ntFlanks</td><td>7974</td><td>8015</td><td>42</td><td>ACGCGACCTCAGATCAGACGTGGCGACCCGCTGAATTTAAGC</td><td>8711.6</td><td>6.5</td><td>1.0</td><td>6.7</td><td>2.6E-11</td><td>9.6E-11</td><td>BT20/MB468</td><td>MITO</td></tr>
<tr><td>>hg38_21_+_8433172_8446622_RNA45SN1_45S_50ntFlanks@11323.11341.19|CGTCTCGTCGCGCGCGCGT</td><td>>hg38_21_+_8433172_8446622_RNA45SN1_45S_50ntFlanks</td><td>11323</td><td>11341</td><td>19</td><td>CGTCTCGTCGCGCGCGCGT</td><td>88.5</td><td>-10.3</td><td>1.6</td><td>-6.6</td><td>3.3E-11</td><td>9.6E-11</td><td>MB231/MB468</td><td>MP</td></tr>
<tr><td>>hg38_21_+_8433172_8446622_RNA45SN1_45S_50ntFlanks@12997.13040.44|CGCTGCGATCTATTGAAAGTCAGCCCTCGACACAAGGGTTTGTC</td><td>>hg38_21_+_8433172_8446622_RNA45SN1_45S_50ntFlanks</td><td>12997</td><td>13040</td><td>44</td><td>CGCTGCGATCTATTGAAAGTCAGCCCTCGACACAAGGGTTTGTC</td><td>219.6</td><td>8.9</td><td>1.3</td><td>6.6</td><td>3.5E-11</td><td>9.6E-11</td><td>MB231/MB468</td><td>NUC</td></tr>
<tr><td>>hg38_21_+_8433172_8446622_RNA45SN1_45S_50ntFlanks@12635.12659.25|AAGCTACCATCTGTGGGATTATGAC</td><td>>hg38_21_+_8433172_8446622_RNA45SN1_45S_50ntFlanks</td><td>12635</td><td>12659</td><td>25</td><td>AAGCTACCATCTGTGGGATTATGAC</td><td>2317.8</td><td>-10.5</td><td>1.6</td><td>-6.6</td><td>4.3E-11</td><td>9.6E-11</td><td>BT20/MB468</td><td>NUC</td></tr>
<tr><td>>hg38_21_+_8433172_8446622_RNA45SN1_45S_50ntFlanks@10343.10361.19|TATTCAAACGAGAACTTTG</td><td>>hg38_21_+_8433172_8446622_RNA45SN1_45S_50ntFlanks</td><td>10343</td><td>10361</td><td>19</td><td>TATTCAAACGAGAACTTTG</td><td>358.0</td><td>-6.5</td><td>1.0</td><td>-6.6</td><td>3.3E-11</td><td>9.7E-11</td><td>MB231/MB468</td><td>MP</td></tr>
<tr><td>>hg38_21_+_8433172_8446622_RNA45SN1_45S_50ntFlanks@5491.5532.42|GAGAAGACGGTCGAACTTGACTATCTAGAGGAAGTAAAAGTC</td><td>>hg38_21_+_8433172_8446622_RNA45SN1_45S_50ntFlanks</td><td>5491</td><td>5532</td><td>42</td><td>GAGAAGACGGTCGAACTTGACTATCTAGAGGAAGTAAAAGTC</td><td>656.4</td><td>10.5</td><td>1.6</td><td>6.6</td><td>3.2E-11</td><td>9.7E-11</td><td>BT20/MB468</td><td>CYTO</td></tr>
<tr><td>>am_MI0000809|hsa-mir-151a&WithFlank&8|-|140732558|140732659@53.72.20&[MIMAT0000757&hsa-miR-151a-3p&offsets|0|-1;m-30&8|-|140732587|140732607&offsets|0|-1]|CTAGACTGAAGCTCCTTGAG</td><td>>am_MI0000809|hsa-mir-151a&WithFlank&8|-|140732558|140732659</td><td>53</td><td>72</td><td>20</td><td>CTAGACTGAAGCTCCTTGAG</td><td>1088.0</td><td>5.7</td><td>0.9</td><td>6.6</td><td>4.1E-11</td><td>9.7E-11</td><td>MB231/MB468</td><td>MITO</td></tr>
<tr><td>>hg38_21_+_8433172_8446622_RNA45SN1_45S_50ntFlanks@11768.11786.19|CTAATTAGTGACGCGCATG</td><td>>hg38_21_+_8433172_8446622_RNA45SN1_45S_50ntFlanks</td><td>11768</td><td>11786</td><td>19</td><td>CTAATTAGTGACGCGCATG</td><td>236.6</td><td>-11.0</td><td>1.6</td><td>-7.0</td><td>2.3E-12</td><td>9.7E-11</td><td>BT20/MB468</td><td>MP</td></tr>
<tr><td>>hg38_21_+_8433172_8446622_RNA45SN1_45S_50ntFlanks@3899.3935.37|CCTTCGCGGGGGGGATGCGTGCATTTATCAGATCAAA</td><td>>hg38_21_+_8433172_8446622_RNA45SN1_45S_50ntFlanks</td><td>3899</td><td>3935</td><td>37</td><td>CCTTCGCGGGGGGGATGCGTGCATTTATCAGATCAAA</td><td>317.9</td><td>-12.2</td><td>1.8</td><td>-6.6</td><td>3.3E-11</td><td>9.7E-11</td><td>MB231/MB468</td><td>MP</td></tr>
<tr><td>>hg38_21_+_8433172_8446622_RNA45SN1_45S_50ntFlanks@10475.10494.20|CGGCCGATCGAAAGGGAGTC</td><td>>hg38_21_+_8433172_8446622_RNA45SN1_45S_50ntFlanks</td><td>10475</td><td>10494</td><td>20</td><td>CGGCCGATCGAAAGGGAGTC</td><td>137.4</td><td>-11.0</td><td>1.7</td><td>-6.6</td><td>3.3E-11</td><td>9.7E-11</td><td>MB231/MB468</td><td>MP</td></tr>
<tr><td>>hg38_21_+_8433172_8446622_RNA45SN1_45S_50ntFlanks@10905.10939.35|CTGGGCGCGCGCCGCGGCTGGACGAGGCGCCGCCG</td><td>>hg38_21_+_8433172_8446622_RNA45SN1_45S_50ntFlanks</td><td>10905</td><td>10939</td><td>35</td><td>CTGGGCGCGCGCCGCGGCTGGACGAGGCGCCGCCG</td><td>173.3</td><td>-11.3</td><td>1.7</td><td>-6.6</td><td>3.3E-11</td><td>9.7E-11</td><td>MB231/MB468</td><td>MP</td></tr>
<tr><td>>hg38_21_+_8433172_8446622_RNA45SN1_45S_50ntFlanks@11210.11232.23|GGCGACTCTGGACGCGAGCCGGG</td><td>>hg38_21_+_8433172_8446622_RNA45SN1_45S_50ntFlanks</td><td>11210</td><td>11232</td><td>23</td><td>GGCGACTCTGGACGCGAGCCGGG</td><td>1251.8</td><td>-8.3</td><td>1.3</td><td>-6.6</td><td>3.5E-11</td><td>9.7E-11</td><td>MB231/MB468</td><td>CYTO</td></tr>
<tr><td>>hg38_21_+_8433172_8446622_RNA45SN1_45S_50ntFlanks@11317.11338.22|ACCCCACGTCTCGTCGCGCGCG</td><td>>hg38_21_+_8433172_8446622_RNA45SN1_45S_50ntFlanks</td><td>11317</td><td>11338</td><td>22</td><td>ACCCCACGTCTCGTCGCGCGCG</td><td>77.6</td><td>-10.1</td><td>1.5</td><td>-6.6</td><td>3.3E-11</td><td>9.7E-11</td><td>MB231/MB468</td><td>MP</td></tr>
<tr><td>>hg38_21_+_8433172_8446622_RNA45SN1_45S_50ntFlanks@11212.11238.27|CGACTCTGGACGCGAGCCGGGCCCTTC</td><td>>hg38_21_+_8433172_8446622_RNA45SN1_45S_50ntFlanks</td><td>11212</td><td>11238</td><td>27</td><td>CGACTCTGGACGCGAGCCGGGCCCTTC</td><td>146.4</td><td>-11.1</td><td>1.7</td><td>-6.6</td><td>3.3E-11</td><td>9.7E-11</td><td>MB231/MB468</td><td>MP</td></tr>
<tr><td>>hg38_MT_+_1621_3279_RNR2_16S_50ntFlanks@729.759.31|GATTAAAACACTGAACTGACAATTAACAGCC</td><td>>hg38_MT_+_1621_3279_RNR2_16S_50ntFlanks</td><td>729</td><td>759</td><td>31</td><td>GATTAAAACACTGAACTGACAATTAACAGCC</td><td>176.7</td><td>10.7</td><td>1.6</td><td>6.6</td><td>3.3E-11</td><td>9.7E-11</td><td>MB231/MB468</td><td>MP</td></tr>
<tr><td>>hg38_21_+_8433172_8446622_RNA45SN1_45S_50ntFlanks@11932.11950.19|GAATAAGTGGGAGGCCCCC</td><td>>hg38_21_+_8433172_8446622_RNA45SN1_45S_50ntFlanks</td><td>11932</td><td>11950</td><td>19</td><td>GAATAAGTGGGAGGCCCCC</td><td>127.0</td><td>-10.9</td><td>1.6</td><td>-6.6</td><td>3.3E-11</td><td>9.7E-11</td><td>MB231/MB468</td><td>MP</td></tr>
<tr><td>>hg38_21_+_8433172_8446622_RNA45SN1_45S_50ntFlanks@12165.12182.18|TGGGGCGGTACACCTGTC</td><td>>hg38_21_+_8433172_8446622_RNA45SN1_45S_50ntFlanks</td><td>12165</td><td>12182</td><td>18</td><td>TGGGGCGGTACACCTGTC</td><td>472.7</td><td>-8.4</td><td>1.3</td><td>-6.6</td><td>3.3E-11</td><td>9.7E-11</td><td>MB231/MB468</td><td>MP</td></tr>
<tr><td>>hg38_21_+_8433172_8446622_RNA45SN1_45S_50ntFlanks@4680.4697.18|GCGCAAGACGGACCAGAG</td><td>>hg38_21_+_8433172_8446622_RNA45SN1_45S_50ntFlanks</td><td>4680</td><td>4697</td><td>18</td><td>GCGCAAGACGGACCAGAG</td><td>80.6</td><td>-10.2</td><td>1.5</td><td>-6.6</td><td>3.3E-11</td><td>9.8E-11</td><td>MB231/MB468</td><td>MP</td></tr>
<tr><td>>hg38_21_+_8433172_8446622_RNA45SN1_45S_50ntFlanks@5080.5105.26|ATAACGAACGAGACTCTGGCATGCTA</td><td>>hg38_21_+_8433172_8446622_RNA45SN1_45S_50ntFlanks</td><td>5080</td><td>5105</td><td>26</td><td>ATAACGAACGAGACTCTGGCATGCTA</td><td>180.8</td><td>-11.4</td><td>1.7</td><td>-6.6</td><td>3.3E-11</td><td>9.8E-11</td><td>MB231/MB468</td><td>MP</td></tr>
<tr><td>>hg38_21_+_8433172_8446622_RNA45SN1_45S_50ntFlanks@11315.11342.28|CCACCCCACGTCTCGTCGCGCGCGCGTC</td><td>>hg38_21_+_8433172_8446622_RNA45SN1_45S_50ntFlanks</td><td>11315</td><td>11342</td><td>28</td><td>CCACCCCACGTCTCGTCGCGCGCGCGTC</td><td>654.0</td><td>-7.6</td><td>1.1</td><td>-6.6</td><td>3.6E-11</td><td>9.8E-11</td><td>MB231/MB468</td><td>CYTO</td></tr>
<tr><td>>hg38_21_+_8433172_8446622_RNA45SN1_45S_50ntFlanks@8194.8214.21|CCCGTGGACGGTGTGAGGCCG</td><td>>hg38_21_+_8433172_8446622_RNA45SN1_45S_50ntFlanks</td><td>8194</td><td>8214</td><td>21</td><td>CCCGTGGACGGTGTGAGGCCG</td><td>1663.2</td><td>-7.3</td><td>1.1</td><td>-6.6</td><td>3.6E-11</td><td>9.8E-11</td><td>MB231/MB468</td><td>NUC</td></tr>
<tr><td>>hg38_21_+_8433172_8446622_RNA45SN1_45S_50ntFlanks@12513.12546.34|TGAGACAGGTTAGTTTTACCCTACTGATGATGTG</td><td>>hg38_21_+_8433172_8446622_RNA45SN1_45S_50ntFlanks</td><td>12513</td><td>12546</td><td>34</td><td>TGAGACAGGTTAGTTTTACCCTACTGATGATGTG</td><td>101.9</td><td>-10.5</td><td>1.6</td><td>-6.6</td><td>3.3E-11</td><td>9.8E-11</td><td>MB231/MB468</td><td>MP</td></tr>
<tr><td>>hg38_21_+_8433172_8446622_RNA45SN1_45S_50ntFlanks@12000.12032.33|CCTGCGGGCCGCCGGTGAAATACCACTACTCTG</td><td>>hg38_21_+_8433172_8446622_RNA45SN1_45S_50ntFlanks</td><td>12000</td><td>12032</td><td>33</td><td>CCTGCGGGCCGCCGGTGAAATACCACTACTCTG</td><td>1271.2</td><td>-8.8</td><td>1.3</td><td>-6.6</td><td>3.6E-11</td><td>9.8E-11</td><td>MB231/MB468</td><td>NUC</td></tr>
<tr><td>>hg38_21_+_8433172_8446622_RNA45SN1_45S_50ntFlanks@10349.10374.26|AACGAGAACTTTGAAGGCCGAAGTGG</td><td>>hg38_21_+_8433172_8446622_RNA45SN1_45S_50ntFlanks</td><td>10349</td><td>10374</td><td>26</td><td>AACGAGAACTTTGAAGGCCGAAGTGG</td><td>1216.6</td><td>-4.2</td><td>0.6</td><td>-6.6</td><td>3.6E-11</td><td>9.8E-11</td><td>MB231/MB468</td><td>NUC</td></tr>
<tr><td>>hg38_21_+_8433172_8446622_RNA45SN1_45S_50ntFlanks@11849.11872.24|CGGGCTTGGCGGAATCAGCGGGGA</td><td>>hg38_21_+_8433172_8446622_RNA45SN1_45S_50ntFlanks</td><td>11849</td><td>11872</td><td>24</td><td>CGGGCTTGGCGGAATCAGCGGGGA</td><td>647.0</td><td>-8.2</td><td>1.2</td><td>-6.6</td><td>3.6E-11</td><td>9.8E-11</td><td>MB231/MB468</td><td>NUC</td></tr>
<tr><td>>hg38_21_+_8433172_8446622_RNA45SN1_45S_50ntFlanks@12090.12110.21|CTGGCGCCAAGCGCCCGGCCG</td><td>>hg38_21_+_8433172_8446622_RNA45SN1_45S_50ntFlanks</td><td>12090</td><td>12110</td><td>21</td><td>CTGGCGCCAAGCGCCCGGCCG</td><td>171.0</td><td>-11.3</td><td>1.7</td><td>-6.6</td><td>3.3E-11</td><td>9.8E-11</td><td>MB231/MB468</td><td>MP</td></tr>
<tr><td>>am_trna2_GlyGCC_21_-_18827107_18827177@1.19.19,trna35_GlyGCC_1_+_161413094_161413164@1.19.19,trna37_GlyGCC_1_+_161420467_161420537@1.19.19,trna39_GlyGCC_1_+_161427898_161427968@1.19.19,trna41_GlyGCC_1_+_161435258_161435328@1.19.19|GCATGGGTGGTTCAGTGGT</td><td>>am_trna2_GlyGCC_21_-_18827107_18827177</td><td>1</td><td>19</td><td>19</td><td>GCATGGGTGGTTCAGTGGT</td><td>93.4</td><td>-10.4</td><td>1.6</td><td>-6.6</td><td>3.3E-11</td><td>9.8E-11</td><td>MB231/MB468</td><td>MP</td></tr>
<tr><td>>hg38_21_+_8433172_8446622_RNA45SN1_45S_50ntFlanks@10849.10866.18|CGGGATAAGGATTGGCTC</td><td>>hg38_21_+_8433172_8446622_RNA45SN1_45S_50ntFlanks</td><td>10849</td><td>10866</td><td>18</td><td>CGGGATAAGGATTGGCTC</td><td>87.3</td><td>-10.3</td><td>1.6</td><td>-6.6</td><td>3.3E-11</td><td>9.8E-11</td><td>MB231/MB468</td><td>MP</td></tr>
<tr><td>>trnaMT_ValTAC_MT_+_1602_1670@45.71.27|TTTCAACTTAACTTGACCGCTCTGACC</td><td>>trnaMT_ValTAC_MT_+_1602_1670</td><td>45</td><td>71</td><td>27</td><td>TTTCAACTTAACTTGACCGCTCTGACC</td><td>253.7</td><td>6.6</td><td>1.0</td><td>6.6</td><td>3.6E-11</td><td>9.8E-11</td><td>MB231/MB468</td><td>NUC</td></tr>
<tr><td>>hg38_21_+_8433172_8446622_RNA45SN1_45S_50ntFlanks@6773.6799.27|TTCCTCCCGGGGCTACGCCTGTCTGAG</td><td>>hg38_21_+_8433172_8446622_RNA45SN1_45S_50ntFlanks</td><td>6773</td><td>6799</td><td>27</td><td>TTCCTCCCGGGGCTACGCCTGTCTGAG</td><td>279.8</td><td>-6.0</td><td>0.9</td><td>-6.6</td><td>3.6E-11</td><td>9.9E-11</td><td>MB231/MB468</td><td>NUC</td></tr>
<tr><td>>hg38_21_+_8433172_8446622_RNA45SN1_45S_50ntFlanks@12418.12441.24|CGATGTCGGCTCTTCCTATCATTG</td><td>>hg38_21_+_8433172_8446622_RNA45SN1_45S_50ntFlanks</td><td>12418</td><td>12441</td><td>24</td><td>CGATGTCGGCTCTTCCTATCATTG</td><td>1516.7</td><td>-4.3</td><td>0.6</td><td>-6.6</td><td>3.6E-11</td><td>9.9E-11</td><td>MB231/MB468</td><td>NUC</td></tr>
<tr><td>>hg38_21_+_8433172_8446622_RNA45SN1_45S_50ntFlanks@10379.10424.46|GGGTTCCATGTGAACAGCAGTTGAACATGGGTCAGTCGGTCCTGAG</td><td>>hg38_21_+_8433172_8446622_RNA45SN1_45S_50ntFlanks</td><td>10379</td><td>10424</td><td>46</td><td>GGGTTCCATGTGAACAGCAGTTGAACATGGGTCAGTCGGTCCTGAG</td><td>391.2</td><td>6.5</td><td>1.0</td><td>6.6</td><td>3.6E-11</td><td>9.9E-11</td><td>MB231/MB468</td><td>CYTO</td></tr>
<tr><td>>hg38_21_+_8433172_8446622_RNA45SN1_45S_50ntFlanks@11883.11905.23|TGTTGAGCTTGACTCTAGTCTGG</td><td>>hg38_21_+_8433172_8446622_RNA45SN1_45S_50ntFlanks</td><td>11883</td><td>11905</td><td>23</td><td>TGTTGAGCTTGACTCTAGTCTGG</td><td>94.7</td><td>-10.4</td><td>1.6</td><td>-6.6</td><td>3.4E-11</td><td>9.9E-11</td><td>MB231/MB468</td><td>MP</td></tr>
</table>

<table>
<tr><td>>hg38_21_+_8433172_8446622_RNA45SN1_45S_50ntFlanks@11603.11623.21|CCGACTGTTTAATTAAAACAA</td><td>>hg38_21_+_8433172_8446622_RNA45SN1_45S_50ntFlanks</td><td>11603</td><td>11623</td><td>21</td><td>CCGACTGTTTAATTAAAACAA</td><td>628.1</td><td>7.4</td><td>1.1</td><td>6.6</td><td>3.3E-11</td><td>9.9E-11</td><td>BT20/MB468</td><td>CYTO</td></tr>
<tr><td>>hg38_21_+_8433172_8446622_RNA45SN1_45S_50ntFlanks@4003.4023.21|ATAACCTCGGGCCGATCGCAC</td><td>>hg38_21_+_8433172_8446622_RNA45SN1_45S_50ntFlanks</td><td>4003</td><td>4023</td><td>21</td><td>ATAACCTCGGGCCGATCGCAC</td><td>2280.3</td><td>-4.6</td><td>0.7</td><td>-6.6</td><td>3.6E-11</td><td>1.0E-10</td><td>MB231/MB468</td><td>NUC</td></tr>
<tr><td>>hg38_21_+_8433172_8446622_RNA45SN1_45S_50ntFlanks@4995.5023.29|ACAGATTGATAGCTCTTTCTCGATTCCGT</td><td>>hg38_21_+_8433172_8446622_RNA45SN1_45S_50ntFlanks</td><td>4995</td><td>5023</td><td>29</td><td>ACAGATTGATAGCTCTTTCTCGATTCCGT</td><td>260.5</td><td>-11.9</td><td>1.8</td><td>-6.6</td><td>3.4E-11</td><td>1.0E-10</td><td>MB231/MB468</td><td>MP</td></tr>
<tr><td>>hg38_21_+_8433172_8446622_RNA45SN1_45S_50ntFlanks@8154.8179.26|CCGCTCGTGGGGGGCCCAAGTCCTTC</td><td>>hg38_21_+_8433172_8446622_RNA45SN1_45S_50ntFlanks</td><td>8154</td><td>8179</td><td>26</td><td>CCGCTCGTGGGGGGCCCAAGTCCTTC</td><td>89.7</td><td>-10.3</td><td>1.6</td><td>-6.6</td><td>3.4E-11</td><td>1.0E-10</td><td>MB231/MB468</td><td>MP</td></tr>
<tr><td>>hg38_21_+_8433172_8446622_RNA45SN1_45S_50ntFlanks@4299.4317.19|TTGGAGGGCAAGTCTGGTG</td><td>>hg38_21_+_8433172_8446622_RNA45SN1_45S_50ntFlanks</td><td>4299</td><td>4317</td><td>19</td><td>TTGGAGGGCAAGTCTGGTG</td><td>91.6</td><td>-10.4</td><td>1.6</td><td>-6.6</td><td>3.4E-11</td><td>1.0E-10</td><td>MB231/MB468</td><td>MP</td></tr>
<tr><td>>hg38_21_+_8433172_8446622_RNA45SN1_45S_50ntFlanks@12363.12382.20|GGATAACTGGCTTGTGGCGG</td><td>>hg38_21_+_8433172_8446622_RNA45SN1_45S_50ntFlanks</td><td>12363</td><td>12382</td><td>20</td><td>GGATAACTGGCTTGTGGCGG</td><td>140.5</td><td>-11.0</td><td>1.7</td><td>-6.6</td><td>3.4E-11</td><td>1.0E-10</td><td>MB231/MB468</td><td>MP</td></tr>
<tr><td>>hg38_21_+_8433172_8446622_RNA45SN1_45S_50ntFlanks@11312.11344.33|CCCCCACCCCACGTCTCGTCGCGCGCGCGTCCG</td><td>>hg38_21_+_8433172_8446622_RNA45SN1_45S_50ntFlanks</td><td>11312</td><td>11344</td><td>33</td><td>CCCCCACCCCACGTCTCGTCGCGCGCGCGTCCG</td><td>157.7</td><td>-11.2</td><td>1.7</td><td>-6.6</td><td>3.4E-11</td><td>1.0E-10</td><td>MB231/MB468</td><td>MP</td></tr>
<tr><td>>am_MI0000086|hsa-mir-28&WithFlank&3|+|188688775|188688872@20.39.20&[MIMAT0000085&hsa-miR-28-5p&offsets|0|-2;m-133&3|+|188688794|188688815&offsets|0|-2]|AAGGAGCTCACAGTCTATTG</td><td>>am_MI0000086|hsa-mir-28&WithFlank&3|+|188688775|188688872</td><td>20</td><td>39</td><td>20</td><td>AAGGAGCTCACAGTCTATTG</td><td>329.0</td><td>8.5</td><td>1.3</td><td>6.6</td><td>3.3E-11</td><td>1.0E-10</td><td>BT20/MB468</td><td>CYTO</td></tr>
<tr><td>>hg38_21_+_8433172_8446622_RNA45SN1_45S_50ntFlanks@8342.8378.37|CGTAAGGGAAAGTTGAAAAGAACTTTGAAGAGAGAGT</td><td>>hg38_21_+_8433172_8446622_RNA45SN1_45S_50ntFlanks</td><td>8342</td><td>8378</td><td>37</td><td>CGTAAGGGAAAGTTGAAAAGAACTTTGAAGAGAGAGT</td><td>835.3</td><td>7.5</td><td>1.1</td><td>6.6</td><td>3.3E-11</td><td>1.0E-10</td><td>BT20/MB468</td><td>CYTO</td></tr>
<tr><td>>MI0000289|hsa-mir-181a-1&WithFlank&1|-|198859038|198859159@30.52.23;MI0000269|hsa-mir-181a-2&WithFlank&9|+|124692436|124692557@45.67.23&[MIMAT0000256&hsa-miR-181a-5p&offsets|0|0;m-32&1|-|198859108|198859130&offsets|0|0];[MIMAT0000256_1&hsa-miR-181a-5p&offsets|0|0;m-32&9|+|124692480|124692502&offsets|0|0]|AACATTCAACGCTGTCGGTGAGT</td><td>>MI0000289|hsa-mir-181a-1&WithFlank&1|-|198859038|198859159</td><td>30</td><td>52</td><td>23</td><td>AACATTCAACGCTGTCGGTGAGT</td><td>5354.1</td><td>6.1</td><td>0.9</td><td>6.6</td><td>3.7E-11</td><td>1.0E-10</td><td>MB231/MB468</td><td>NUC</td></tr>
<tr><td>>hg38_21_+_8433172_8446622_RNA45SN1_45S_50ntFlanks@10690.10710.21|CGTCCGGTGAGCTCTCGCTGG</td><td>>hg38_21_+_8433172_8446622_RNA45SN1_45S_50ntFlanks</td><td>10690</td><td>10710</td><td>21</td><td>CGTCCGGTGAGCTCTCGCTGG</td><td>1155.0</td><td>-8.4</td><td>1.3</td><td>-6.6</td><td>3.3E-11</td><td>1.0E-10</td><td>BT20/MB468</td><td>CYTO</td></tr>
<tr><td>>trna11_GluTTC_15_-_26327381_26327452@3.36.34,trna3_GluTTC_13_-_45492062_45492133@3.36.34|CCACATGGTCTAGCGGTTAGGATTCCTGGTTTTC</td><td>>trna11_GluTTC_15_-_26327381_26327452</td><td>3</td><td>36</td><td>34</td><td>CCACATGGTCTAGCGGTTAGGATTCCTGGTTTTC</td><td>226.5</td><td>11.0</td><td>1.7</td><td>6.6</td><td>4.3E-11</td><td>1.0E-10</td><td>MB231/MB468</td><td>MITO</td></tr>
<tr><td>>hg38_21_+_8433172_8446622_RNA45SN1_45S_50ntFlanks@4588.4616.29|CTATTTTGTTGGTTTTCGGAACTGAGGCC</td><td>>hg38_21_+_8433172_8446622_RNA45SN1_45S_50ntFlanks</td><td>4588</td><td>4616</td><td>29</td><td>CTATTTTGTTGGTTTTCGGAACTGAGGCC</td><td>2557.0</td><td>-4.7</td><td>0.7</td><td>-6.6</td><td>3.7E-11</td><td>1.0E-10</td><td>MB231/MB468</td><td>CYTO</td></tr>
<tr><td>>hg38_21_+_8433172_8446622_RNA45SN1_45S_50ntFlanks@11918.11967.50|ACATGAGAGGTGTAGAATAAGTGGGAGGCCCCCGGCGCCCCCCCGGTGTC</td><td>>hg38_21_+_8433172_8446622_RNA45SN1_45S_50ntFlanks</td><td>11918</td><td>11967</td><td>50</td><td>ACATGAGAGGTGTAGAATAAGTGGGAGGCCCCCGGCGCCCCCCCGGTGTC</td><td>391.2</td><td>-12.4</td><td>1.9</td><td>-6.6</td><td>4.3E-11</td><td>1.0E-10</td><td>MB231/MB468</td><td>MITO</td></tr>
<tr><td>>hg38_21_+_8433172_8446622_RNA45SN1_45S_50ntFlanks@12273.12299.27|TCAGTACGAATACAGACCGTGAAAGCG</td><td>>hg38_21_+_8433172_8446622_RNA45SN1_45S_50ntFlanks</td><td>12273</td><td>12299</td><td>27</td><td>TCAGTACGAATACAGACCGTGAAAGCG</td><td>89.9</td><td>-10.4</td><td>1.6</td><td>-6.6</td><td>3.4E-11</td><td>1.0E-10</td><td>MB231/MB468</td><td>MP</td></tr>
<tr><td>>hg38_21_+_8433172_8446622_RNA45SN1_45S_50ntFlanks@9533.9551.19|GAGGAAACTCTGGTGGAGG</td><td>>hg38_21_+_8433172_8446622_RNA45SN1_45S_50ntFlanks</td><td>9533</td><td>9551</td><td>19</td><td>GAGGAAACTCTGGTGGAGG</td><td>447.6</td><td>-4.4</td><td>0.7</td><td>-6.6</td><td>3.7E-11</td><td>1.0E-10</td><td>MB231/MB468</td><td>NUC</td></tr>
<tr><td>>hg38_21_+_8433172_8446622_RNA45SN1_45S_50ntFlanks@12324.12353.30|TTGGGTTTTAAGCAGGAGGTGTCAGAAAAG</td><td>>hg38_21_+_8433172_8446622_RNA45SN1_45S_50ntFlanks</td><td>12324</td><td>12353</td><td>30</td><td>TTGGGTTTTAAGCAGGAGGTGTCAGAAAAG</td><td>81.0</td><td>-10.2</td><td>1.5</td><td>-6.6</td><td>3.5E-11</td><td>1.0E-10</td><td>MB231/MB468</td><td>MP</td></tr>
<tr><td>>MI0000301|hsa-mir-224&WithFlank&X|-|151958572|151958664@14.37.24&[MIMAT0000281&hsa-miR-224-5p&offsets|+1|0;m-238&X|-|151958630|151958651&offsets|0|+2]|CAAGTCACTAGTGGTTCCGTTTAG</td><td>>MI0000301|hsa-mir-224&WithFlank&X|-|151958572|151958664</td><td>14</td><td>37</td><td>24</td><td>CAAGTCACTAGTGGTTCCGTTTAG</td><td>694.8</td><td>6.1</td><td>0.9</td><td>6.6</td><td>4.5E-11</td><td>1.0E-10</td><td>BT20/MB468</td><td>NUC</td></tr>
<tr><td>>hg38_21_+_8433172_8446622_RNA45SN1_45S_50ntFlanks@9540.9571.32|CTCTGGTGGAGGTCCGTAGCGGTCCTGACGTG</td><td>>hg38_21_+_8433172_8446622_RNA45SN1_45S_50ntFlanks</td><td>9540</td><td>9571</td><td>32</td><td>CTCTGGTGGAGGTCCGTAGCGGTCCTGACGTG</td><td>443.0</td><td>-4.9</td><td>0.7</td><td>-6.6</td><td>4.3E-11</td><td>1.0E-10</td><td>MB231/MB468</td><td>MITO</td></tr>
<tr><td>>hg38_21_+_8433172_8446622_RNA45SN1_45S_50ntFlanks@10393.10419.27|CAGCAGTTGAACATGGGTCAGTCGGTC</td><td>>hg38_21_+_8433172_8446622_RNA45SN1_45S_50ntFlanks</td><td>10393</td><td>10419</td><td>27</td><td>CAGCAGTTGAACATGGGTCAGTCGGTC</td><td>675.2</td><td>-6.8</td><td>1.0</td><td>-6.6</td><td>3.7E-11</td><td>1.0E-10</td><td>MB231/MB468</td><td>NUC</td></tr>
<tr><td>>hg38_21_+_8433172_8446622_RNA45SN1_45S_50ntFlanks@10403.10435.33|ACATGGGTCAGTCGGTCCTGAGAGATGGGCGAG</td><td>>hg38_21_+_8433172_8446622_RNA45SN1_45S_50ntFlanks</td><td>10403</td><td>10435</td><td>33</td><td>ACATGGGTCAGTCGGTCCTGAGAGATGGGCGAG</td><td>105.9</td><td>-10.6</td><td>1.6</td><td>-6.6</td><td>3.5E-11</td><td>1.0E-10</td><td>MB231/MB468</td><td>MP</td></tr>
<tr><td>>trna10_LysCTT_16_+_3241501_3241573@14.31.18,trna13_LysCTT_14_-_58706613_58706685@14.31.18,trna2_LysCTT_15_+_79152904_79152976@14.31.18,trna4_LysCTT_18_-_43669271_43669343@14.31.18|AGTCGGTAGAGCATGGGA</td><td>>trna10_LysCTT_16_+_3241501_3241573</td><td>14</td><td>31</td><td>18</td><td>AGTCGGTAGAGCATGGGA</td><td>1325.0</td><td>-9.0</td><td>1.4</td><td>-6.6</td><td>3.7E-11</td><td>1.0E-10</td><td>MB231/MB468</td><td>NUC</td></tr>
<tr><td>>hg38_21_+_8433172_8446622_RNA45SN1_45S_50ntFlanks@12660.12681.22|TGAACGCCTCTAAGTCAGAATC</td><td>>hg38_21_+_8433172_8446622_RNA45SN1_45S_50ntFlanks</td><td>12660</td><td>12681</td><td>22</td><td>TGAACGCCTCTAAGTCAGAATC</td><td>136.6</td><td>-11.0</td><td>1.7</td><td>-6.6</td><td>3.5E-11</td><td>1.0E-10</td><td>MB231/MB468</td><td>MP</td></tr>
<tr><td>>hg38_21_+_8433172_8446622_RNA45SN1_45S_50ntFlanks@4995.5022.28|ACAGATTGATAGCTCTTTCTCGATTCCG</td><td>>hg38_21_+_8433172_8446622_RNA45SN1_45S_50ntFlanks</td><td>4995</td><td>5022</td><td>28</td><td>ACAGATTGATAGCTCTTTCTCGATTCCG</td><td>698.6</td><td>-11.5</td><td>1.7</td><td>-6.7</td><td>2.7E-11</td><td>1.0E-10</td><td>BT20/MB468</td><td>MITO</td></tr>
<tr><td>>hg38_21_+_8433172_8446622_RNA45SN1_45S_50ntFlanks@9538.9563.26|AACTCTGGTGGAGGTCCGTAGCGGTC</td><td>>hg38_21_+_8433172_8446622_RNA45SN1_45S_50ntFlanks</td><td>9538</td><td>9563</td><td>26</td><td>AACTCTGGTGGAGGTCCGTAGCGGTC</td><td>100.8</td><td>-10.5</td><td>1.6</td><td>-6.6</td><td>3.5E-11</td><td>1.0E-10</td><td>MB231/MB468</td><td>MP</td></tr>
<tr><td>>hg38_21_+_8433172_8446622_RNA45SN1_45S_50ntFlanks@11562.11601.40|CGCCTAGCAGCCGACTTAGAACTGGTGCGGACCAGGGGAA</td><td>>hg38_21_+_8433172_8446622_RNA45SN1_45S_50ntFlanks</td><td>11562</td><td>11601</td><td>40</td><td>CGCCTAGCAGCCGACTTAGAACTGGTGCGGACCAGGGGAA</td><td>271.4</td><td>-11.9</td><td>1.8</td><td>-6.6</td><td>3.5E-11</td><td>1.0E-10</td><td>MB231/MB468</td><td>MP</td></tr>
<tr><td>>hg38_21_+_8433172_8446622_RNA45SN1_45S_50ntFlanks@11874.11909.36|AGAAGACCCTGTTGAGCTTGACTCTAGTCTGGCACG</td><td>>hg38_21_+_8433172_8446622_RNA45SN1_45S_50ntFlanks</td><td>11874</td><td>11909</td><td>36</td><td>AGAAGACCCTGTTGAGCTTGACTCTAGTCTGGCACG</td><td>296.8</td><td>-4.2</td><td>0.6</td><td>-6.6</td><td>3.7E-11</td><td>1.0E-10</td><td>MB231/MB468</td><td>CYTO</td></tr>
<tr><td>>hg38_21_+_8433172_8446622_RNA45SN1_45S_50ntFlanks@12004.12035.32|CGGGCCGCCGGTGAAATACCACTACTCTGATC</td><td>>hg38_21_+_8433172_8446622_RNA45SN1_45S_50ntFlanks</td><td>12004</td><td>12035</td><td>32</td><td>CGGGCCGCCGGTGAAATACCACTACTCTGATC</td><td>2542.3</td><td>5.8</td><td>0.8</td><td>7.0</td><td>3.5E-12</td><td>1.0E-10</td><td>BT20/MB231</td><td>MITO</td></tr>
<tr><td>>MI0000477|hsa-mir-146a&WithFlank&5|+|160485346|160485456@27.48.22(+1C)&[MIMAT0000449&hsa-miR-146a-5p&offsets|0|0(+1C);m-129&5|+|160485372|160485393&offsets|0|0(+1C)]|TGAGAACTGAATTCCATGGGTTC</td><td>>MI0000477|hsa-mir-146a&WithFlank&5|+|160485346|160485456</td><td>27</td><td>48</td><td>22(+1C)</td><td>TGAGAACTGAATTCCATGGGTTC</td><td>189.9</td><td>-10.8</td><td>1.6</td><td>-7.0</td><td>3.5E-12</td><td>1.0E-10</td><td>BT20/MB231</td><td>MITO</td></tr>
<tr><td>>MI0003205|hsa-mir-532&WithFlank&X|+|50003142|50003244@26.48.23&[MIMAT0002888&hsa-miR-532-5p&offsets|0|+1;m-65&X|+|50003167|50003188&offsets|0|+1]|CATGCCTTGAGTGTAGGACCGTT</td><td>>MI0003205|hsa-mir-532&WithFlank&X|+|50003142|50003244</td><td>26</td><td>48</td><td>23</td><td>CATGCCTTGAGTGTAGGACCGTT</td><td>255.9</td><td>6.2</td><td>0.9</td><td>6.6</td><td>3.4E-11</td><td>1.0E-10</td><td>BT20/MB468</td><td>CYTO</td></tr>
<tr><td>>hg38_21_+_8433172_8446622_RNA45SN1_45S_50ntFlanks@6732.6773.42|ACATTGATCATCGACACTTCGAACGCACTTGCGGCCCCGGGT</td><td>>hg38_21_+_8433172_8446622_RNA45SN1_45S_50ntFlanks</td><td>6732</td><td>6773</td><td>42</td><td>ACATTGATCATCGACACTTCGAACGCACTTGCGGCCCCGGGT</td><td>786.6</td><td>7.9</td><td>1.2</td><td>6.6</td><td>3.4E-11</td><td>1.0E-10</td><td>BT20/MB468</td><td>CYTO</td></tr>
<tr><td>>hg38_21_+_8433172_8446622_RNA45SN1_45S_50ntFlanks@9738.9759.22|GAAACGATCTCAACCTATTCTC</td><td>>hg38_21_+_8433172_8446622_RNA45SN1_45S_50ntFlanks</td><td>9738</td><td>9759</td><td>22</td><td>GAAACGATCTCAACCTATTCTC</td><td>757.6</td><td>-4.7</td><td>0.7</td><td>-6.6</td><td>3.4E-11</td><td>1.0E-10</td><td>BT20/MB468</td><td>CYTO</td></tr>
<tr><td>>hg38_21_+_8433172_8446622_RNA45SN1_45S_50ntFlanks@10402.10422.21|AACATGGGTCAGTCGGTCCTG</td><td>>hg38_21_+_8433172_8446622_RNA45SN1_45S_50ntFlanks</td><td>10402</td><td>10422</td><td>21</td><td>AACATGGGTCAGTCGGTCCTG</td><td>324.7</td><td>-5.1</td><td>0.8</td><td>-6.6</td><td>3.7E-11</td><td>1.0E-10</td><td>MB231/MB468</td><td>CYTO</td></tr>
<tr><td>>hg38_21_+_8433172_8446622_RNA45SN1_45S_50ntFlanks@11208.11229.22|GCGGCGACTCTGGACGCGAGCC</td><td>>hg38_21_+_8433172_8446622_RNA45SN1_45S_50ntFlanks</td><td>11208</td><td>11229</td><td>22</td><td>GCGGCGACTCTGGACGCGAGCC</td><td>500.1</td><td>-4.8</td><td>0.7</td><td>-6.6</td><td>4.3E-11</td><td>1.0E-10</td><td>MB231/MB468</td><td>MITO</td></tr>
<tr><td>>hg38_21_+_8433172_8446622_RNA45SN1_45S_50ntFlanks@12637.12661.25|GCTACCATCTGTGGGATTATGACTG</td><td>>hg38_21_+_8433172_8446622_RNA45SN1_45S_50ntFlanks</td><td>12637</td><td>12661</td><td>25</td><td>GCTACCATCTGTGGGATTATGACTG</td><td>4417.8</td><td>-8.1</td><td>1.2</td><td>-6.6</td><td>4.3E-11</td><td>1.0E-10</td><td>MB231/MB468</td><td>MITO</td></tr>
<tr><td>>hg38_21_+_8433172_8446622_RNA45SN1_45S_50ntFlanks@11104.11130.27|AAGGGTCGGGGCGGCAGGGGCCGGCGG</td><td>>hg38_21_+_8433172_8446622_RNA45SN1_45S_50ntFlanks</td><td>11104</td><td>11130</td><td>27</td><td>AAGGGTCGGGGCGGCAGGGGCCGGCGG</td><td>107.6</td><td>-10.6</td><td>1.6</td><td>-6.6</td><td>3.5E-11</td><td>1.0E-10</td><td>MB231/MB468</td><td>MP</td></tr>
<tr><td>>hg38_21_+_8433172_8446622_RNA45SN1_45S_50ntFlanks@10434.10453.20|AGCGCCGTTCCGAAGGGACG</td><td>>hg38_21_+_8433172_8446622_RNA45SN1_45S_50ntFlanks</td><td>10434</td><td>10453</td><td>20</td><td>AGCGCCGTTCCGAAGGGACG</td><td>115.3</td><td>-10.7</td><td>1.6</td><td>-6.6</td><td>3.5E-11</td><td>1.0E-10</td><td>MB231/MB468</td><td>MP</td></tr>
<tr><td>>hg38_21_+_8433172_8446622_RNA45SN1_45S_50ntFlanks@11930.11954.25|TAGAATAAGTGGGAGGCCCCCGGCG</td><td>>hg38_21_+_8433172_8446622_RNA45SN1_45S_50ntFlanks</td><td>11930</td><td>11954</td><td>25</td><td>TAGAATAAGTGGGAGGCCCCCGGCG</td><td>235.7</td><td>-11.0</td><td>1.6</td><td>-7.0</td><td>2.5E-12</td><td>1.0E-10</td><td>BT20/MB468</td><td>MP</td></tr>
<tr><td>>hg38_21_+_8433172_8446622_RNA45SN1_45S_50ntFlanks@4260.4282.23|TTGGAATGAGTCCACTTTAAATC</td><td>>hg38_21_+_8433172_8446622_RNA45SN1_45S_50ntFlanks</td><td>4260</td><td>4282</td><td>23</td><td>TTGGAATGAGTCCACTTTAAATC</td><td>212.4</td><td>-10.8</td><td>1.5</td><td>-7.0</td><td>2.5E-12</td><td>1.0E-10</td><td>BT20/MB468</td><td>MP</td></tr>
<tr><td>>hg38_21_+_8433172_8446622_RNA45SN1_45S_50ntFlanks@12605.12624.20|TATGTGCTTGGCTGAGGAGC</td><td>>hg38_21_+_8433172_8446622_RNA45SN1_45S_50ntFlanks</td><td>12605</td><td>12624</td><td>20</td><td>TATGTGCTTGGCTGAGGAGC</td><td>678.5</td><td>-4.5</td><td>0.7</td><td>-6.6</td><td>4.4E-11</td><td>1.0E-10</td><td>MB231/MB468</td><td>MITO</td></tr>
<tr><td>>hg38_21_+_8433172_8446622_RNA45SN1_45S_50ntFlanks@10922.10945.24|CTGGACGAGGCGCCGCCGCCCCCC</td><td>>hg38_21_+_8433172_8446622_RNA45SN1_45S_50ntFlanks</td><td>10922</td><td>10945</td><td>24</td><td>CTGGACGAGGCGCCGCCGCCCCCC</td><td>616.1</td><td>-8.1</td><td>1.2</td><td>-6.6</td><td>3.8E-11</td><td>1.0E-10</td><td>MB231/MB468</td><td>NUC</td></tr>
<tr><td>>hg38_21_+_8433172_8446622_RNA45SN1_45S_50ntFlanks@4006.4026.21|ACCTCGGGCCGATCGCACGCC</td><td>>hg38_21_+_8433172_8446622_RNA45SN1_45S_50ntFlanks</td><td>4006</td><td>4026</td><td>21</td><td>ACCTCGGGCCGATCGCACGCC</td><td>240.7</td><td>-10.0</td><td>1.5</td><td>-6.7</td><td>2.8E-11</td><td>1.0E-10</td><td>BT20/MB468</td><td>MITO</td></tr>
<tr><td>>hg38_21_+_8433172_8446622_RNA45SN1_45S_50ntFlanks@11785.11820.36|TGAATGGATGAACGAGATTCCCACTGTCCCTACCTA</td><td>>hg38_21_+_8433172_8446622_RNA45SN1_45S_50ntFlanks</td><td>11785</td><td>11820</td><td>36</td><td>TGAATGGATGAACGAGATTCCCACTGTCCCTACCTA</td><td>103.6</td><td>-10.6</td><td>1.6</td><td>-6.6</td><td>3.6E-11</td><td>1.0E-10</td><td>MB231/MB468</td><td>MP</td></tr>
<tr><td>>hg38_21_+_8433172_8446622_RNA45SN1_45S_50ntFlanks@9573.9597.25|AAATCGGTCGTCCGACCTGGGTATA</td><td>>hg38_21_+_8433172_8446622_RNA45SN1_45S_50ntFlanks</td><td>9573</td><td>9597</td><td>25</td><td>AAATCGGTCGTCCGACCTGGGTATA</td><td>519.3</td><td>-4.6</td><td>0.7</td><td>-6.6</td><td>3.8E-11</td><td>1.0E-10</td><td>MB231/MB468</td><td>NUC</td></tr>
<tr><td>>hg38_21_+_8433172_8446622_RNA45SN1_45S_50ntFlanks@10388.10410.23|GTGAACAGCAGTTGAACATGGGT</td><td>>hg38_21_+_8433172_8446622_RNA45SN1_45S_50ntFlanks</td><td>10388</td><td>10410</td><td>23</td><td>GTGAACAGCAGTTGAACATGGGT</td><td>423.1</td><td>-3.5</td><td>0.5</td><td>-6.6</td><td>3.5E-11</td><td>1.0E-10</td><td>BT20/MB468</td><td>CYTO</td></tr>
<tr><td>>hg38_21_+_8433172_8446622_RNA45SN1_45S_50ntFlanks@11228.11254.27|CCGGGCCCTTCCCGTGGATCGCCCCAG</td><td>>hg38_21_+_8433172_8446622_RNA45SN1_45S_50ntFlanks</td><td>11228</td><td>11254</td><td>27</td><td>CCGGGCCCTTCCCGTGGATCGCCCCAG</td><td>149.5</td><td>-11.1</td><td>1.7</td><td>-6.6</td><td>3.6E-11</td><td>1.0E-10</td><td>MB231/MB468</td><td>MP</td></tr>
<tr><td>>hg38_21_+_8433172_8446622_RNA45SN1_45S_50ntFlanks@9534.9554.21|AGGAAACTCTGGTGGAGGTCC</td><td>>hg38_21_+_8433172_8446622_RNA45SN1_45S_50ntFlanks</td><td>9534</td><td>9554</td><td>21</td><td>AGGAAACTCTGGTGGAGGTCC</td><td>583.8</td><td>-8.4</td><td>1.3</td><td>-6.6</td><td>3.8E-11</td><td>1.0E-10</td><td>MB231/MB468</td><td>CYTO</td></tr>
<tr><td>>hg38_21_+_8433172_8446622_RNA45SN1_45S_50ntFlanks@9738.9763.26|GAAACGATCTCAACCTATTCTCAAAC</td><td>>hg38_21_+_8433172_8446622_RNA45SN1_45S_50ntFlanks</td><td>9738</td><td>9763</td><td>26</td><td>GAAACGATCTCAACCTATTCTCAAAC</td><td>1070.3</td><td>-7.5</td><td>1.1</td><td>-6.6</td><td>4.7E-11</td><td>1.0E-10</td><td>BT20/MB468</td><td>NUC</td></tr>
<tr><td>>hg38_21_+_8433172_8446622_RNA45SN1_45S_50ntFlanks@10733.10752.20|GGTGTAAATCTCGCGCCGGG</td><td>>hg38_21_+_8433172_8446622_RNA45SN1_45S_50ntFlanks</td><td>10733</td><td>10752</td><td>20</td><td>GGTGTAAATCTCGCGCCGGG</td><td>78.8</td><td>-10.2</td><td>1.5</td><td>-6.6</td><td>3.6E-11</td><td>1.0E-10</td><td>MB231/MB468</td><td>MP</td></tr>
<tr><td>>hg38_21_+_8433172_8446622_RNA45SN1_45S_50ntFlanks@12008.12026.19|CCGCCGGTGAAATACCACT</td><td>>hg38_21_+_8433172_8446622_RNA45SN1_45S_50ntFlanks</td><td>12008</td><td>12026</td><td>19</td><td>CCGCCGGTGAAATACCACT</td><td>117.9</td><td>-10.7</td><td>1.6</td><td>-6.6</td><td>3.6E-11</td><td>1.0E-10</td><td>MB231/MB468</td><td>MP</td></tr>
</table>

<table>
<tr><td>>hg38_21_+_8433172_8446622_RNA45SN1_45S_50ntFlanks@12893.12933.41|CCTCGCCCGTCACGCACCGCACGTTCGTGGGGAACCTGGCG</td><td>>hg38_21_+_8433172_8446622_RNA45SN1_45S_50ntFlanks</td><td>12893</td><td>12933</td><td>41</td><td>CCTCGCCCGTCACGCACCGCACGTTCGTGGGGAACCTGGCG</td><td>1100.9</td><td>-9.7</td><td>1.5</td><td>-6.6</td><td>4.7E-11</td><td>1.0E-10</td><td>BT20/MB468</td><td>NUC</td></tr>
<tr><td>>hg38_21_+_8433172_8446622_RNA45SN1_45S_50ntFlanks@11822.11844.23|TATCCAGCGAAACCACAGCCAAG</td><td>>hg38_21_+_8433172_8446622_RNA45SN1_45S_50ntFlanks</td><td>11822</td><td>11844</td><td>23</td><td>TATCCAGCGAAACCACAGCCAAG</td><td>111.0</td><td>-10.7</td><td>1.6</td><td>-6.6</td><td>3.6E-11</td><td>1.0E-10</td><td>MB231/MB468</td><td>MP</td></tr>
<tr><td>>hg38_21_+_8433172_8446622_RNA45SN1_45S_50ntFlanks@10375.10407.33|AGAAGGGTTCCATGTGAACAGCAGTTGAACATG</td><td>>hg38_21_+_8433172_8446622_RNA45SN1_45S_50ntFlanks</td><td>10375</td><td>10407</td><td>33</td><td>AGAAGGGTTCCATGTGAACAGCAGTTGAACATG</td><td>479.9</td><td>-7.6</td><td>1.1</td><td>-6.6</td><td>3.5E-11</td><td>1.0E-10</td><td>BT20/MB468</td><td>CYTO</td></tr>
<tr><td>>hg38_21_+_8433172_8446622_RNA45SN1_45S_50ntFlanks@6722.6744.23|ATTGCAGGACACATTGATCATCG</td><td>>hg38_21_+_8433172_8446622_RNA45SN1_45S_50ntFlanks</td><td>6722</td><td>6744</td><td>23</td><td>ATTGCAGGACACATTGATCATCG</td><td>3062.1</td><td>-5.4</td><td>0.8</td><td>-6.6</td><td>3.5E-11</td><td>1.1E-10</td><td>BT20/MB468</td><td>CYTO</td></tr>
<tr><td>>trna5_AspGTC_12_+_98897281_98897352@37.75.39|ACGCGGGAGACCGGGGTTCAATTCCCCGACGGGGAGCCA</td><td>>trna5_AspGTC_12_+_98897281_98897352</td><td>37</td><td>75</td><td>39</td><td>ACGCGGGAGACCGGGGTTCAATTCCCCGACGGGGAGCCA</td><td>4469.8</td><td>9.7</td><td>1.5</td><td>6.6</td><td>3.9E-11</td><td>1.1E-10</td><td>MB231/MB468</td><td>CYTO</td></tr>
<tr><td>>hg38_21_+_8433172_8446622_RNA45SN1_45S_50ntFlanks@3997.4024.28|CTCTAGATAACCTCGGGCCGATCGCACG</td><td>>hg38_21_+_8433172_8446622_RNA45SN1_45S_50ntFlanks</td><td>3997</td><td>4024</td><td>28</td><td>CTCTAGATAACCTCGGGCCGATCGCACG</td><td>102.8</td><td>-10.5</td><td>1.6</td><td>-6.6</td><td>3.6E-11</td><td>1.1E-10</td><td>MB231/MB468</td><td>MP</td></tr>
<tr><td>>hg38_21_+_8433172_8446622_RNA45SN1_45S_50ntFlanks@10412.10443.32|AGTCGGTCCTGAGAGATGGGCGAGCGCCGTTC</td><td>>hg38_21_+_8433172_8446622_RNA45SN1_45S_50ntFlanks</td><td>10412</td><td>10443</td><td>32</td><td>AGTCGGTCCTGAGAGATGGGCGAGCGCCGTTC</td><td>108.5</td><td>-10.6</td><td>1.6</td><td>-6.6</td><td>3.6E-11</td><td>1.1E-10</td><td>MB231/MB468</td><td>MP</td></tr>
<tr><td>>hg38_21_+_8433172_8446622_RNA45SN1_45S_50ntFlanks@10393.10411.19|CAGCAGTTGAACATGGGTC</td><td>>hg38_21_+_8433172_8446622_RNA45SN1_45S_50ntFlanks</td><td>10393</td><td>10411</td><td>19</td><td>CAGCAGTTGAACATGGGTC</td><td>369.4</td><td>-6.7</td><td>1.0</td><td>-6.6</td><td>4.7E-11</td><td>1.1E-10</td><td>BT20/MB468</td><td>NUC</td></tr>
<tr><td>>hg38_21_+_8433172_8446622_RNA45SN1_45S_50ntFlanks@12894.12932.39|CTCGCCCGTCACGCACCGCACGTTCGTGGGGAACCTGGC</td><td>>hg38_21_+_8433172_8446622_RNA45SN1_45S_50ntFlanks</td><td>12894</td><td>12932</td><td>39</td><td>CTCGCCCGTCACGCACCGCACGTTCGTGGGGAACCTGGC</td><td>202.5</td><td>-11.5</td><td>1.7</td><td>-6.6</td><td>3.6E-11</td><td>1.1E-10</td><td>MB231/MB468</td><td>MP</td></tr>
<tr><td>>hg38_21_+_8433172_8446622_RNA45SN1_45S_50ntFlanks@6737.6772.36|GATCATCGACACTTCGAACGCACTTGCGGCCCCGGG</td><td>>hg38_21_+_8433172_8446622_RNA45SN1_45S_50ntFlanks</td><td>6737</td><td>6772</td><td>36</td><td>GATCATCGACACTTCGAACGCACTTGCGGCCCCGGG</td><td>221.6</td><td>-9.9</td><td>1.5</td><td>-6.7</td><td>2.8E-11</td><td>1.1E-10</td><td>BT20/MB468</td><td>MITO</td></tr>
<tr><td>>MI0001733|hsa-mir-452&WithFlank&X|-|151959622|151959718@20.42.23&[MIMAT0001635&hsa-miR-452-5p&offsets|0|+1;m-188&X|-|151959677|151959699&offsets|0|0]|AACTGTTTGCAGAGGAAACTGAG</td><td>>MI0001733|hsa-mir-452&WithFlank&X|-|151959622|151959718</td><td>20</td><td>42</td><td>23</td><td>AACTGTTTGCAGAGGAAACTGAG</td><td>1214.9</td><td>3.8</td><td>0.5</td><td>6.9</td><td>4.0E-12</td><td>1.1E-10</td><td>BT20/MB231</td><td>NUC</td></tr>
<tr><td>>hg38_21_+_8433172_8446622_RNA45SN1_45S_50ntFlanks@5074.5110.37|ATTCCGATAACGAACGAGACTCTGGCATGCTAACTAG</td><td>>hg38_21_+_8433172_8446622_RNA45SN1_45S_50ntFlanks</td><td>5074</td><td>5110</td><td>37</td><td>ATTCCGATAACGAACGAGACTCTGGCATGCTAACTAG</td><td>504.0</td><td>-9.0</td><td>1.4</td><td>-6.6</td><td>4.5E-11</td><td>1.1E-10</td><td>MB231/MB468</td><td>MITO</td></tr>
<tr><td>>hg38_21_+_8433172_8446622_RNA45SN1_45S_50ntFlanks@10846.10863.18|CTTCGGGATAAGGATTGG</td><td>>hg38_21_+_8433172_8446622_RNA45SN1_45S_50ntFlanks</td><td>10846</td><td>10863</td><td>18</td><td>CTTCGGGATAAGGATTGG</td><td>1039.6</td><td>-4.6</td><td>0.7</td><td>-6.6</td><td>3.9E-11</td><td>1.1E-10</td><td>MB231/MB468</td><td>CYTO</td></tr>
<tr><td>>hg38_21_+_8433172_8446622_RNA45SN1_45S_50ntFlanks@10482.10512.31|TCGAAAGGGAGTCGGGTTCAGATCCCCGAAT</td><td>>hg38_21_+_8433172_8446622_RNA45SN1_45S_50ntFlanks</td><td>10482</td><td>10512</td><td>31</td><td>TCGAAAGGGAGTCGGGTTCAGATCCCCGAAT</td><td>824.1</td><td>-6.5</td><td>1.0</td><td>-6.6</td><td>3.9E-11</td><td>1.1E-10</td><td>MB231/MB468</td><td>CYTO</td></tr>
<tr><td>>hg38_21_+_8433172_8446622_RNA45SN1_45S_50ntFlanks@11316.11340.25|CACCCCACGTCTCGTCGCGCGCGCG</td><td>>hg38_21_+_8433172_8446622_RNA45SN1_45S_50ntFlanks</td><td>11316</td><td>11340</td><td>25</td><td>CACCCCACGTCTCGTCGCGCGCGCG</td><td>479.0</td><td>-7.3</td><td>1.1</td><td>-6.6</td><td>3.9E-11</td><td>1.1E-10</td><td>MB231/MB468</td><td>CYTO</td></tr>
<tr><td>>hg38_21_+_8433172_8446622_RNA45SN1_45S_50ntFlanks@5469.5514.46|CCCACGGCCCTGGCGGAGCGCTGAGAAGACGGTCGAACTTGACTAT</td><td>>hg38_21_+_8433172_8446622_RNA45SN1_45S_50ntFlanks</td><td>5469</td><td>5514</td><td>46</td><td>CCCACGGCCCTGGCGGAGCGCTGAGAAGACGGTCGAACTTGACTAT</td><td>225.0</td><td>-10.8</td><td>1.6</td><td>-6.6</td><td>3.9E-11</td><td>1.1E-10</td><td>MB231/MB468</td><td>NUC</td></tr>
<tr><td>>hg38_21_+_8433172_8446622_RNA45SN1_45S_50ntFlanks@9736.9754.19|CCGAAACGATCTCAACCTA</td><td>>hg38_21_+_8433172_8446622_RNA45SN1_45S_50ntFlanks</td><td>9736</td><td>9754</td><td>19</td><td>CCGAAACGATCTCAACCTA</td><td>758.1</td><td>-8.1</td><td>1.2</td><td>-6.6</td><td>3.5E-11</td><td>1.1E-10</td><td>BT20/MB468</td><td>CYTO</td></tr>
<tr><td>>hg38_21_+_8433172_8446622_RNA45SN1_45S_50ntFlanks@4261.4282.22|TGGAATGAGTCCACTTTAAATC</td><td>>hg38_21_+_8433172_8446622_RNA45SN1_45S_50ntFlanks</td><td>4261</td><td>4282</td><td>22</td><td>TGGAATGAGTCCACTTTAAATC</td><td>83.5</td><td>-10.2</td><td>1.5</td><td>-6.6</td><td>3.6E-11</td><td>1.1E-10</td><td>MB231/MB468</td><td>MP</td></tr>
<tr><td>>hg38_21_+_8433172_8446622_RNA45SN1_45S_50ntFlanks@4378.4406.29|CTCGTAGTTGGATCTTGGGAGCGGGCGGG</td><td>>hg38_21_+_8433172_8446622_RNA45SN1_45S_50ntFlanks</td><td>4378</td><td>4406</td><td>29</td><td>CTCGTAGTTGGATCTTGGGAGCGGGCGGG</td><td>138.3</td><td>-11.0</td><td>1.7</td><td>-6.6</td><td>3.6E-11</td><td>1.1E-10</td><td>MB231/MB468</td><td>MP</td></tr>
<tr><td>>hg38_21_+_8433172_8446622_RNA45SN1_45S_50ntFlanks@12002.12043.42|TGCGGGCCGCCGGTGAAATACCACTACTCTGATCGTTTTTTC</td><td>>hg38_21_+_8433172_8446622_RNA45SN1_45S_50ntFlanks</td><td>12002</td><td>12043</td><td>42</td><td>TGCGGGCCGCCGGTGAAATACCACTACTCTGATCGTTTTTTC</td><td>394.8</td><td>-12.5</td><td>1.9</td><td>-6.6</td><td>3.6E-11</td><td>1.1E-10</td><td>MB231/MB468</td><td>MP</td></tr>
<tr><td>>hg38_21_+_8433172_8446622_RNA45SN1_45S_50ntFlanks@13006.13027.22|CTATTGAAAGTCAGCCCTCGAC</td><td>>hg38_21_+_8433172_8446622_RNA45SN1_45S_50ntFlanks</td><td>13006</td><td>13027</td><td>22</td><td>CTATTGAAAGTCAGCCCTCGAC</td><td>156.7</td><td>-9.4</td><td>1.4</td><td>-6.7</td><td>2.8E-11</td><td>1.1E-10</td><td>BT20/MB468</td><td>MITO</td></tr>
<tr><td>>MI0000111|hsa-mir-105-1&WithFlank&X|-|152392213|152392305@19.40.22;MI0000112|hsa-mir-105-2&WithFlank&X|-|152394406|152394498@19.40.22&[MIMAT0000102&hsa-miR-105-5p&offsets|0|-1;m-735&X|-|152392266|152392287&offsets|0|0];[MIMAT0000102_1&hsa-miR-105-5p&offsets|0|-1;m-735&X|-|152394459|152394480&offsets|0|0]|TCAAATGCTCAGACTCCTGTGG</td><td>>MI0000111|hsa-mir-105-1&WithFlank&X|-|152392213|152392305</td><td>19</td><td>40</td><td>22</td><td>TCAAATGCTCAGACTCCTGTGG</td><td>509.8</td><td>4.4</td><td>0.7</td><td>6.7</td><td>2.8E-11</td><td>1.1E-10</td><td>BT20/MB468</td><td>MITO</td></tr>
<tr><td>>MI0003513|hsa-mir-455&WithFlank&9|+|114209428|114209535@22.45.24&[MIMAT0003150&hsa-miR-455-5p&offsets|0|+2;m-232&9|+|114209449|114209470&offsets|0|+2]|TATGTGCCTTTGGACTACATCGTG</td><td>>MI0003513|hsa-mir-455&WithFlank&9|+|114209428|114209535</td><td>22</td><td>45</td><td>24</td><td>TATGTGCCTTTGGACTACATCGTG</td><td>206.8</td><td>10.9</td><td>1.7</td><td>6.6</td><td>4.5E-11</td><td>1.1E-10</td><td>MB231/MB468</td><td>MITO</td></tr>
<tr><td>>MI0005566|hsa-mir-374b&WithFlank&X|-|74218541|74218624@17.39.23&[MIMAT0004955&hsa-miR-374b-5p&offsets|0|+1;m-153&X|-|74218587|74218608&offsets|0|+1]|ATATAATACAACCTGCTAAGTGT</td><td>>MI0005566|hsa-mir-374b&WithFlank&X|-|74218541|74218624</td><td>17</td><td>39</td><td>23</td><td>ATATAATACAACCTGCTAAGTGT</td><td>292.6</td><td>6.9</td><td>1.0</td><td>6.6</td><td>3.9E-11</td><td>1.1E-10</td><td>MB231/MB468</td><td>NUC</td></tr>
<tr><td>>hg38_21_+_8433172_8446622_RNA45SN1_45S_50ntFlanks@8115.8135.21|CGCGGGACATGTGGCGTACGG</td><td>>hg38_21_+_8433172_8446622_RNA45SN1_45S_50ntFlanks</td><td>8115</td><td>8135</td><td>21</td><td>CGCGGGACATGTGGCGTACGG</td><td>88.5</td><td>-10.3</td><td>1.6</td><td>-6.6</td><td>3.7E-11</td><td>1.1E-10</td><td>MB231/MB468</td><td>MP</td></tr>
<tr><td>>hg38_21_+_8433172_8446622_RNA45SN1_45S_50ntFlanks@12365.12384.20|ATAACTGGCTTGTGGCGGCC</td><td>>hg38_21_+_8433172_8446622_RNA45SN1_45S_50ntFlanks</td><td>12365</td><td>12384</td><td>20</td><td>ATAACTGGCTTGTGGCGGCC</td><td>118.4</td><td>-9.2</td><td>1.4</td><td>-6.6</td><td>3.5E-11</td><td>1.1E-10</td><td>BT20/MB468</td><td>CYTO</td></tr>
<tr><td>>MI0003560|hsa-mir-92b&WithFlank&1|+|155195171|155195278@67.87.21(+1A)&[MIMAT0003218&hsa-miR-92b-3p&offsets|0|-1(+1A);m-115&1|+|155195237|155195258&offsets|0|-1(+1A)]|TATTGCACTCGTCCCGGCCTCA</td><td>>MI0003560|hsa-mir-92b&WithFlank&1|+|155195171|155195278</td><td>67</td><td>87</td><td>21(+1A)</td><td>TATTGCACTCGTCCCGGCCTCA</td><td>1120.1</td><td>9.8</td><td>1.5</td><td>6.6</td><td>3.9E-11</td><td>1.1E-10</td><td>MB231/MB468</td><td>CYTO</td></tr>
<tr><td>>MI0000084|hsa-mir-26b&WithFlank&2|+|218402640|218402728@18.38.21&[MIMAT0000083&hsa-miR-26b-5p&offsets|0|0;m-53&2|+|218402657|218402678&offsets|0|-1]|TTCAAGTAATTCAGGATAGGT</td><td>>MI0000084|hsa-mir-26b&WithFlank&2|+|218402640|218402728</td><td>18</td><td>38</td><td>21</td><td>TTCAAGTAATTCAGGATAGGT</td><td>653.2</td><td>4.2</td><td>0.6</td><td>6.6</td><td>3.9E-11</td><td>1.1E-10</td><td>MB231/MB468</td><td>NUC</td></tr>
<tr><td>>hg38_21_+_8433172_8446622_RNA45SN1_45S_50ntFlanks@9578.9601.24|GGTCGTCCGACCTGGGTATAGGGG</td><td>>hg38_21_+_8433172_8446622_RNA45SN1_45S_50ntFlanks</td><td>9578</td><td>9601</td><td>24</td><td>GGTCGTCCGACCTGGGTATAGGGG</td><td>218.8</td><td>11.5</td><td>1.6</td><td>7.0</td><td>2.6E-12</td><td>1.1E-10</td><td>BT20/MB468</td><td>MP</td></tr>
<tr><td>>hg38_21_+_8433172_8446622_RNA45SN1_45S_50ntFlanks@12875.12892.18|CTGGAAAGGCGGCCGCCC</td><td>>hg38_21_+_8433172_8446622_RNA45SN1_45S_50ntFlanks</td><td>12875</td><td>12892</td><td>18</td><td>CTGGAAAGGCGGCCGCCC</td><td>331.0</td><td>-11.5</td><td>1.6</td><td>-7.0</td><td>2.6E-12</td><td>1.1E-10</td><td>BT20/MB468</td><td>MP</td></tr>
<tr><td>>MI0000086|hsa-mir-28&WithFlank&3|+|188688775|188688872@60.80.21&[MIMAT0004502&hsa-miR-28-3p&offsets|0|-1;m-28&3|+|188688834|188688855&offsets|0|-1]|CACTAGATTGTGAGCTCCTGG</td><td>>MI0000086|hsa-mir-28&WithFlank&3|+|188688775|188688872</td><td>60</td><td>80</td><td>21</td><td>CACTAGATTGTGAGCTCCTGG</td><td>1035.4</td><td>4.8</td><td>0.7</td><td>6.6</td><td>3.7E-11</td><td>1.1E-10</td><td>MB231/MB468</td><td>MP</td></tr>
<tr><td>>MI0002468|hsa-mir-484&WithFlank&16|+|15643288|15643378@14.35.22&[MIMAT0002174&hsa-miR-484&offsets|0|0;m-127&16|+|15643301|15643321&offsets|0|+1]|TCAGGCTCAGTCCCCTCCCGAT</td><td>>MI0002468|hsa-mir-484&WithFlank&16|+|15643288|15643378</td><td>14</td><td>35</td><td>22</td><td>TCAGGCTCAGTCCCCTCCCGAT</td><td>2052.9</td><td>5.7</td><td>0.9</td><td>6.7</td><td>2.9E-11</td><td>1.1E-10</td><td>BT20/MB468</td><td>MITO</td></tr>
<tr><td>>hg38_21_+_8433172_8446622_RNA45SN1_45S_50ntFlanks@12974.12997.24|CGTACGTAGCAGAGCAGCTCCCTC</td><td>>hg38_21_+_8433172_8446622_RNA45SN1_45S_50ntFlanks</td><td>12974</td><td>12997</td><td>24</td><td>CGTACGTAGCAGAGCAGCTCCCTC</td><td>279.1</td><td>3.4</td><td>0.5</td><td>6.6</td><td>3.9E-11</td><td>1.1E-10</td><td>MB231/MB468</td><td>CYTO</td></tr>
<tr><td>>hg38_21_+_8433172_8446622_RNA45SN1_45S_50ntFlanks@10314.10331.18|CAGGTGCAGATCTTGGTG</td><td>>hg38_21_+_8433172_8446622_RNA45SN1_45S_50ntFlanks</td><td>10314</td><td>10331</td><td>18</td><td>CAGGTGCAGATCTTGGTG</td><td>108.8</td><td>-10.6</td><td>1.6</td><td>-6.6</td><td>3.7E-11</td><td>1.1E-10</td><td>MB231/MB468</td><td>MP</td></tr>
<tr><td>>am_trna2_GlyGCC_21_-_18827107_18827177@57.74.18,trna35_GlyGCC_1_+_161413094_161413164@57.74.18,trna37_GlyGCC_1_+_161420467_161420537@57.74.18,trna39_GlyGCC_1_+_161427898_161427968@57.74.18,trna41_GlyGCC_1_+_161435258_161435328@57.74.18|TTCCCGGCCCATGCACCA</td><td>>am_trna2_GlyGCC_21_-_18827107_18827177</td><td>57</td><td>74</td><td>18</td><td>TTCCCGGCCCATGCACCA</td><td>63.6</td><td>4.7</td><td>0.7</td><td>6.6</td><td>4.8E-11</td><td>1.1E-10</td><td>BT20/MB468</td><td>NUC</td></tr>
<tr><td>>hg38_21_+_8433172_8446622_RNA45SN1_45S_50ntFlanks@4003.4042.40|ATAACCTCGGGCCGATCGCACGCCCCCCGTGGCGGCGACG</td><td>>hg38_21_+_8433172_8446622_RNA45SN1_45S_50ntFlanks</td><td>4003</td><td>4042</td><td>40</td><td>ATAACCTCGGGCCGATCGCACGCCCCCCGTGGCGGCGACG</td><td>2207.0</td><td>-7.6</td><td>1.2</td><td>-6.6</td><td>3.9E-11</td><td>1.1E-10</td><td>MB231/MB468</td><td>NUC</td></tr>
<tr><td>>hg38_21_+_8433172_8446622_RNA45SN1_45S_50ntFlanks@12934.12951.18|CTAAACCATTCGTAGACG</td><td>>hg38_21_+_8433172_8446622_RNA45SN1_45S_50ntFlanks</td><td>12934</td><td>12951</td><td>18</td><td>CTAAACCATTCGTAGACG</td><td>536.0</td><td>-4.3</td><td>0.6</td><td>-6.6</td><td>3.9E-11</td><td>1.1E-10</td><td>MB231/MB468</td><td>NUC</td></tr>
<tr><td>>hg38_21_+_8433172_8446622_RNA45SN1_45S_50ntFlanks@12526.12554.29|TTTTACCCTACTGATGATGTGTTGTTGCC</td><td>>hg38_21_+_8433172_8446622_RNA45SN1_45S_50ntFlanks</td><td>12526</td><td>12554</td><td>29</td><td>TTTTACCCTACTGATGATGTGTTGTTGCC</td><td>82.1</td><td>-10.2</td><td>1.5</td><td>-6.6</td><td>3.7E-11</td><td>1.1E-10</td><td>MB231/MB468</td><td>MP</td></tr>
<tr><td>>hg38_21_+_8433172_8446622_RNA45SN1_45S_50ntFlanks@6698.6717.20|AGCTAGCTGCGAGAATTAAT</td><td>>hg38_21_+_8433172_8446622_RNA45SN1_45S_50ntFlanks</td><td>6698</td><td>6717</td><td>20</td><td>AGCTAGCTGCGAGAATTAAT</td><td>366.1</td><td>-6.4</td><td>1.0</td><td>-6.6</td><td>4.6E-11</td><td>1.1E-10</td><td>MB231/MB468</td><td>MITO</td></tr>
<tr><td>>hg38_21_+_8433172_8446622_RNA45SN1_45S_50ntFlanks@3771.3788.18|CACGGCCGGTACAGTGAA</td><td>>hg38_21_+_8433172_8446622_RNA45SN1_45S_50ntFlanks</td><td>3771</td><td>3788</td><td>18</td><td>CACGGCCGGTACAGTGAA</td><td>84.1</td><td>-10.3</td><td>1.6</td><td>-6.6</td><td>3.7E-11</td><td>1.1E-10</td><td>MB231/MB468</td><td>MP</td></tr>
<tr><td>>hg38_21_+_8433172_8446622_RNA45SN1_45S_50ntFlanks@11104.11133.30|AAGGGTCGGGGCGGCAGGGGCCGGCGGCGG</td><td>>hg38_21_+_8433172_8446622_RNA45SN1_45S_50ntFlanks</td><td>11104</td><td>11133</td><td>30</td><td>AAGGGTCGGGGCGGCAGGGGCCGGCGGCGG</td><td>1473.5</td><td>-9.9</td><td>1.5</td><td>-6.6</td><td>4.8E-11</td><td>1.1E-10</td><td>BT20/MB468</td><td>NUC</td></tr>
<tr><td>>hg38_21_+_8433172_8446622_RNA45SN1_45S_50ntFlanks@6773.6793.21|TTCCTCCCGGGGCTACGCCTG</td><td>>hg38_21_+_8433172_8446622_RNA45SN1_45S_50ntFlanks</td><td>6773</td><td>6793</td><td>21</td><td>TTCCTCCCGGGGCTACGCCTG</td><td>996.6</td><td>-8.6</td><td>1.3</td><td>-6.6</td><td>4.6E-11</td><td>1.1E-10</td><td>MB231/MB468</td><td>MITO</td></tr>
<tr><td>>hg38_21_+_8433172_8446622_RNA45SN1_45S_50ntFlanks@6707.6734.28|CGAGAATTAATGTGAATTGCAGGACACA</td><td>>hg38_21_+_8433172_8446622_RNA45SN1_45S_50ntFlanks</td><td>6707</td><td>6734</td><td>28</td><td>CGAGAATTAATGTGAATTGCAGGACACA</td><td>93.1</td><td>-10.4</td><td>1.6</td><td>-6.6</td><td>3.7E-11</td><td>1.1E-10</td><td>MB231/MB468</td><td>MP</td></tr>
<tr><td>>hg38_21_+_8433172_8446622_RNA45SN1_45S_50ntFlanks@11881.11903.23|CCTGTTGAGCTTGACTCTAGTCT</td><td>>hg38_21_+_8433172_8446622_RNA45SN1_45S_50ntFlanks</td><td>11881</td><td>11903</td><td>23</td><td>CCTGTTGAGCTTGACTCTAGTCT</td><td>493.4</td><td>-4.4</td><td>0.7</td><td>-6.6</td><td>4.0E-11</td><td>1.1E-10</td><td>MB231/MB468</td><td>CYTO</td></tr>
<tr><td>>trna119_LysCTT_1_-_145395522_145395594@4.33.30,trna11_LysCTT_5_-_180648979_180649051@4.33.30,trna13_LysCTT_6_+_26556774_26556846@4.33.30,trna7_LysCTT_16_+_3225692_3225764@4.33.30,trna9_LysCTT_5_+_180634755_180634827@4.33.30|CGGCTAGCTCAGTCGGTAGAGCATGAGACT</td><td>>trna119_LysCTT_1_-_145395522_145395594</td><td>4</td><td>33</td><td>30</td><td>CGGCTAGCTCAGTCGGTAGAGCATGAGACT</td><td>2933.6</td><td>-5.8</td><td>0.8</td><td>-6.9</td><td>3.7E-12</td><td>1.1E-10</td><td>BT20/MB231</td><td>MITO</td></tr>
<tr><td>>hg38_21_+_8433172_8446622_RNA45SN1_45S_50ntFlanks@12488.12524.37|AACGTGAGCTGGGTTTAGACCGTCGTGAGACAGGTTA</td><td>>hg38_21_+_8433172_8446622_RNA45SN1_45S_50ntFlanks</td><td>12488</td><td>12524</td><td>37</td><td>AACGTGAGCTGGGTTTAGACCGTCGTGAGACAGGTTA</td><td>82.3</td><td>-10.2</td><td>1.5</td><td>-6.6</td><td>3.7E-11</td><td>1.1E-10</td><td>MB231/MB468</td><td>MP</td></tr>
<tr><td>>hg38_21_+_8433172_8446622_RNA45SN1_45S_50ntFlanks@11875.11904.30|GAAGACCCTGTTGAGCTTGACTCTAGTCTG</td><td>>hg38_21_+_8433172_8446622_RNA45SN1_45S_50ntFlanks</td><td>11875</td><td>11904</td><td>30</td><td>GAAGACCCTGTTGAGCTTGACTCTAGTCTG</td><td>167.4</td><td>-11.3</td><td>1.7</td><td>-6.6</td><td>3.7E-11</td><td>1.1E-10</td><td>MB231/MB468</td><td>MP</td></tr>
<tr><td>>MI0000088|hsa-mir-30a&WithFlank&6|-|71403545|71403627@53.75.23&[MIMAT0000088&hsa-miR-30a-3p&offsets|0|+1;m-38&6|-|71403554|71403575&offsets|0|+1]|CTTTCAGTCGGATGTTTGCAGCT</td><td>>MI0000088|hsa-mir-30a&WithFlank&6|-|71403545|71403627</td><td>53</td><td>75</td><td>23</td><td>CTTTCAGTCGGATGTTTGCAGCT</td><td>206.0</td><td>10.9</td><td>1.7</td><td>6.6</td><td>4.6E-11</td><td>1.1E-10</td><td>MB231/MB468</td><td>MITO</td></tr>
<tr><td>>hg38_21_+_8433172_8446622_RNA45SN1_45S_50ntFlanks@4006.4042.37|ACCTCGGGCCGATCGCACGCCCCCCGTGGCGGCGACG</td><td>>hg38_21_+_8433172_8446622_RNA45SN1_45S_50ntFlanks</td><td>4006</td><td>4042</td><td>37</td><td>ACCTCGGGCCGATCGCACGCCCCCCGTGGCGGCGACG</td><td>442.8</td><td>-4.5</td><td>0.7</td><td>-6.6</td><td>4.6E-11</td><td>1.1E-10</td><td>MB231/MB468</td><td>MITO</td></tr>
<tr><td>>MI0000650|hsa-mir-200c&WithFlank&12|+|6963693|6963772@50.70.21&[MIMAT0000617&hsa-miR-200c-3p&offsets|0|-2;m-54&12|+|6963742|6963764&offsets|0|-2]|TAATACTGCCGGGTAATGATG</td><td>>MI0000650|hsa-mir-200c&WithFlank&12|+|6963693|6963772</td><td>50</td><td>70</td><td>21</td><td>TAATACTGCCGGGTAATGATG</td><td>4594.5</td><td>6.4</td><td>0.9</td><td>7.0</td><td>2.7E-12</td><td>1.1E-10</td><td>BT20/MB468</td><td>MP</td></tr>
</table>

| | | | | | | | | | | | | | |
|---|---|---|---|---|---|---|---|---|---|---|---|---|---|
| >hg38_21_+_8433172_8446622_RNA45SN1_45S_50ntFlanks@12894.12925.32\|CTCGCCCGTCACGCACCGCACGTTCGTGGGGA | >hg38_21_+_8433172_8446622_RNA45SN1_45S_50ntFlanks | 12894 | 12925 | 32 | CTCGCCCGTCACGCACCGCACGTTCGTGGGGA | 131.2 | -10.9 | 1.6 | -6.6 | 3.8E-11 | 1.1E-10 | MB231/MB468 | MP |
| >hg38_21_+_8433172_8446622_RNA45SN1_45S_50ntFlanks@9457.9503.47\|CGCCGGGGAGGTGGAGCACGAGCGCACGTGTTAGGACCCGAAAGATG | >hg38_21_+_8433172_8446622_RNA45SN1_45S_50ntFlanks | 9457 | 9503 | 47 | CGCCGGGGAGGTGGAGCACGAGCGCACGTGTTAGGACCCGAAAGATG | 608.3 | -13.1 | 2.0 | -6.6 | 3.8E-11 | 1.1E-10 | MB231/MB468 | MP |
| >hg38_21_+_8433172_8446622_RNA45SN1_45S_50ntFlanks@10796.10826.31\|GGCATGTTGGAACAATGTAGGTAAGGGAAGT | >hg38_21_+_8433172_8446622_RNA45SN1_45S_50ntFlanks | 10796 | 10826 | 31 | GGCATGTTGGAACAATGTAGGTAAGGGAAGT | 84.3 | -10.3 | 1.6 | -6.6 | 3.8E-11 | 1.1E-10 | MB231/MB468 | MP |
| >hg38_21_+_8433172_8446622_RNA45SN1_45S_50ntFlanks@11669.11707.39\|GCCCAGTGCTCTGAATGTCAAAGTGAAGAAATTCAATGA | >hg38_21_+_8433172_8446622_RNA45SN1_45S_50ntFlanks | 11669 | 11707 | 39 | GCCCAGTGCTCTGAATGTCAAAGTGAAGAAATTCAATGA | 456.3 | 8.5 | 1.3 | 6.6 | 3.6E-11 | 1.1E-10 | BT20/MB468 | CYTO |
| >hg38_21_+_8433172_8446622_RNA45SN1_45S_50ntFlanks@12071.12095.25\|CCCCGAGGGGCTCTCGCTTCTGGCG | >hg38_21_+_8433172_8446622_RNA45SN1_45S_50ntFlanks | 12071 | 12095 | 25 | CCCCGAGGGGCTCTCGCTTCTGGCG | 1796.8 | -10.0 | 1.5 | -6.6 | 4.9E-11 | 1.1E-10 | BT20/MB468 | NUC |
| >hg38_21_+_8433172_8446622_RNA45SN1_45S_50ntFlanks@3788.3810.23\|AACTGCGAATGGCTCATTAAATC | >hg38_21_+_8433172_8446622_RNA45SN1_45S_50ntFlanks | 3788 | 3810 | 23 | AACTGCGAATGGCTCATTAAATC | 754.1 | -7.3 | 1.1 | -6.6 | 4.9E-11 | 1.1E-10 | BT20/MB468 | NUC |
| >hg38_21_+_8433172_8446622_RNA45SN1_45S_50ntFlanks@11769.11804.36\|TAATTAGTGACGCGCATGAATGGATGAACGAGATTC | >hg38_21_+_8433172_8446622_RNA45SN1_45S_50ntFlanks | 11769 | 11804 | 36 | TAATTAGTGACGCGCATGAATGGATGAACGAGATTC | 2082.8 | -10.1 | 1.5 | -6.6 | 4.9E-11 | 1.1E-10 | BT20/MB468 | NUC |
| >hg38_21_+_8433172_8446622_RNA45SN1_45S_50ntFlanks@11575.11594.20\|ACTTAGAACTGGTGCGGACC | >hg38_21_+_8433172_8446622_RNA45SN1_45S_50ntFlanks | 11575 | 11594 | 20 | ACTTAGAACTGGTGCGGACC | 2885.1 | -4.1 | 0.6 | -6.6 | 4.0E-11 | 1.1E-10 | MB231/MB468 | NUC |
| >hg38_21_+_8433172_8446622_RNA45SN1_45S_50ntFlanks@9568.9595.28\|CGTGCAAATCGGTCGTCCGACCTGGGTA | >hg38_21_+_8433172_8446622_RNA45SN1_45S_50ntFlanks | 9568 | 9595 | 28 | CGTGCAAATCGGTCGTCCGACCTGGGTA | 2080.5 | -4.4 | 0.7 | -6.6 | 3.0E-11 | 1.1E-10 | BT20/MB468 | MITO |
| >hg38_21_+_8433172_8446622_RNA45SN1_45S_50ntFlanks@8321.8356.36\|AGACCGATAGTCAACAAGTACCGTAAGGGAAAGTTG | >hg38_21_+_8433172_8446622_RNA45SN1_45S_50ntFlanks | 8321 | 8356 | 36 | AGACCGATAGTCAACAAGTACCGTAAGGGAAAGTTG | 213.5 | -6.5 | 1.0 | -6.6 | 4.1E-11 | 1.1E-10 | MB231/MB468 | CYTO |
| >hg38_21_+_8433172_8446622_RNA45SN1_45S_50ntFlanks@10910.10930.21\|CGCGCGCCGCGGCTGGACGAG | >hg38_21_+_8433172_8446622_RNA45SN1_45S_50ntFlanks | 10910 | 10930 | 21 | CGCGCGCCGCGGCTGGACGAG | 106.1 | -10.6 | 1.6 | -6.6 | 3.8E-11 | 1.1E-10 | MB231/MB468 | MP |
| >hg38_21_+_8433172_8446622_RNA45SN1_45S_50ntFlanks@12616.12661.46\|CTGAGGAGCCAATGGGGCGAAGCTACCATCTGTGGGATTATGACTG | >hg38_21_+_8433172_8446622_RNA45SN1_45S_50ntFlanks | 12616 | 12661 | 46 | CTGAGGAGCCAATGGGGCGAAGCTACCATCTGTGGGATTATGACTG | 1245.8 | -7.9 | 1.2 | -6.6 | 4.7E-11 | 1.1E-10 | MB231/MB468 | MITO |
| >hg38_21_+_8433172_8446622_RNA45SN1_45S_50ntFlanks@10351.10369.19\|CGAGAACTTTGAAGGCCGA | >hg38_21_+_8433172_8446622_RNA45SN1_45S_50ntFlanks | 10351 | 10369 | 19 | CGAGAACTTTGAAGGCCGA | 1978.2 | -4.8 | 0.7 | -6.6 | 3.8E-11 | 1.1E-10 | MB231/MB468 | MP |
| >hg38_21_+_8433172_8446622_RNA45SN1_45S_50ntFlanks@10826.10848.23\|TCGGCAAGCCGGATCCGTAACTT | >hg38_21_+_8433172_8446622_RNA45SN1_45S_50ntFlanks | 10826 | 10848 | 23 | TCGGCAAGCCGGATCCGTAACTT | 113.2 | -10.7 | 1.6 | -6.6 | 3.8E-11 | 1.1E-10 | MB231/MB468 | MP |
| >hg38_21_+_8433172_8446622_RNA45SN1_45S_50ntFlanks@5444.5466.23\|CTCGGATCGGCCCCGCCGGGGTC | >hg38_21_+_8433172_8446622_RNA45SN1_45S_50ntFlanks | 5444 | 5466 | 23 | CTCGGATCGGCCCCGCCGGGGTC | 28294.7 | -7.2 | 1.0 | -7.0 | 2.7E-12 | 1.1E-10 | BT20/MB468 | MP |
| >hg38_21_+_8433172_8446622_RNA45SN1_45S_50ntFlanks@9307.9338.32\|AAGGAGTCTAACACGTGCGCGAGTCGGGGGCT | >hg38_21_+_8433172_8446622_RNA45SN1_45S_50ntFlanks | 9307 | 9338 | 32 | AAGGAGTCTAACACGTGCGCGAGTCGGGGGCT | 667.9 | -5.6 | 0.9 | -6.6 | 4.7E-11 | 1.1E-10 | MB231/MB468 | MITO |
| >hg38_21_+_8433172_8446622_RNA45SN1_45S_50ntFlanks@9579.9596.18\|GTCGTCCGACCTGGGTAT | >hg38_21_+_8433172_8446622_RNA45SN1_45S_50ntFlanks | 9579 | 9596 | 18 | GTCGTCCGACCTGGGTAT | 92.1 | -10.4 | 1.6 | -6.6 | 3.8E-11 | 1.1E-10 | MB231/MB468 | MP |
| >hg38_21_+_8433172_8446622_RNA45SN1_45S_50ntFlanks@3984.4002.19\|GCGGCTTTGGTGACTCTAG | >hg38_21_+_8433172_8446622_RNA45SN1_45S_50ntFlanks | 3984 | 4002 | 19 | GCGGCTTTGGTGACTCTAG | 2118.3 | -9.3 | 1.4 | -6.6 | 3.0E-11 | 1.1E-10 | BT20/MB468 | MITO |
| >hg38_21_+_8433172_8446622_RNA45SN1_45S_50ntFlanks@11209.11231.23\|CGGCGACTCTGGACGCGAGCCGG | >hg38_21_+_8433172_8446622_RNA45SN1_45S_50ntFlanks | 11209 | 11231 | 23 | CGGCGACTCTGGACGCGAGCCGG | 1601.3 | -4.3 | 0.7 | -6.6 | 3.7E-11 | 1.1E-10 | BT20/MB468 | CYTO |
| >hg38_21_+_8433172_8446622_RNA45SN1_45S_50ntFlanks@5236.5267.32\|CACTGACTGGCTCAGCGTGTGCCTACCCTACG | >hg38_21_+_8433172_8446622_RNA45SN1_45S_50ntFlanks | 5236 | 5267 | 32 | CACTGACTGGCTCAGCGTGTGCCTACCCTACG | 484.1 | -8.9 | 1.4 | -6.6 | 5.0E-11 | 1.1E-10 | BT20/MB468 | NUC |
| >hg38_21_+_8433172_8446622_RNA45SN1_45S_50ntFlanks@11615.11640.26\|TTAAAACAAAGCATCGCGAAGGCCCG | >hg38_21_+_8433172_8446622_RNA45SN1_45S_50ntFlanks | 11615 | 11640 | 26 | TTAAAACAAAGCATCGCGAAGGCCCG | 193.9 | -9.7 | 1.5 | -6.6 | 3.0E-11 | 1.1E-10 | BT20/MB468 | MITO |
| >hg38_21_+_8433172_8446622_RNA45SN1_45S_50ntFlanks@4738.4762.25\|AAAGTCGGAGGTTCGAAGACGATCA | >hg38_21_+_8433172_8446622_RNA45SN1_45S_50ntFlanks | 4738 | 4762 | 25 | AAAGTCGGAGGTTCGAAGACGATCA | 487.8 | 7.1 | 1.1 | 6.6 | 3.7E-11 | 1.1E-10 | BT20/MB468 | CYTO |
| >hg38_21_+_8433172_8446622_RNA45SN1_45S_50ntFlanks@11576.11612.37\|CTTAGAACTGGTGCGGACCAGGGGAATCCGACTGTTT | >hg38_21_+_8433172_8446622_RNA45SN1_45S_50ntFlanks | 11576 | 11612 | 37 | CTTAGAACTGGTGCGGACCAGGGGAATCCGACTGTTT | 575.3 | -5.5 | 0.8 | -6.6 | 4.8E-11 | 1.1E-10 | MB231/MB468 | MITO |
| >hg38_21_+_8433172_8446622_RNA45SN1_45S_50ntFlanks@9540.9563.24\|CTCTGGTGGAGGTCCGTAGCGGTC | >hg38_21_+_8433172_8446622_RNA45SN1_45S_50ntFlanks | 9540 | 9563 | 24 | CTCTGGTGGAGGTCCGTAGCGGTC | 95.0 | -10.4 | 1.6 | -6.6 | 3.9E-11 | 1.1E-10 | MB231/MB468 | MP |
| >hg38_21_+_8433172_8446622_RNA45SN1_45S_50ntFlanks@11930.11949.20\|TAGAATAAGTGGGAGGCCCC | >hg38_21_+_8433172_8446622_RNA45SN1_45S_50ntFlanks | 11930 | 11949 | 20 | TAGAATAAGTGGGAGGCCCC | 231.9 | -10.9 | 1.6 | -7.0 | 2.8E-12 | 1.1E-10 | BT20/MB468 | MP |
| >hg38_21_+_8433172_8446622_RNA45SN1_45S_50ntFlanks@13002.13032.31\|CGATCTATTGAAAGTCAGCCCTCGACACAAG | >hg38_21_+_8433172_8446622_RNA45SN1_45S_50ntFlanks | 13002 | 13032 | 31 | CGATCTATTGAAAGTCAGCCCTCGACACAAG | 95.4 | -10.4 | 1.6 | -6.6 | 3.9E-11 | 1.1E-10 | MB231/MB468 | MP |
| >hg38_21_+_8433172_8446622_RNA45SN1_45S_50ntFlanks@10788.10818.31\|CAGCCTCTGGCATGTTGGAACAATGTAGGTA | >hg38_21_+_8433172_8446622_RNA45SN1_45S_50ntFlanks | 10788 | 10818 | 31 | CAGCCTCTGGCATGTTGGAACAATGTAGGTA | 614.6 | -7.2 | 1.1 | -6.6 | 5.0E-11 | 1.1E-10 | BT20/MB468 | NUC |
| >hg38_21_+_8433172_8446622_RNA45SN1_45S_50ntFlanks@6739.6756.18\|TCATCGACACTTCGAACG | >hg38_21_+_8433172_8446622_RNA45SN1_45S_50ntFlanks | 6739 | 6756 | 18 | TCATCGACACTTCGAACG | 118.0 | -10.7 | 1.6 | -6.6 | 3.9E-11 | 1.1E-10 | MB231/MB468 | MP |
| >hg38_21_+_8433172_8446622_RNA45SN1_45S_50ntFlanks@4257.4275.19\|TAATTGGAATGAGTCCACT | >hg38_21_+_8433172_8446622_RNA45SN1_45S_50ntFlanks | 4257 | 4275 | 19 | TAATTGGAATGAGTCCACT | 229.4 | -10.9 | 1.6 | -7.0 | 2.8E-12 | 1.1E-10 | BT20/MB468 | MP |
| >hg38_21_+_8433172_8446622_RNA45SN1_45S_50ntFlanks@4724.4759.36\|ATTAATCAAGAACGAAAGTCGGAGGTTCGAAGACGA | >hg38_21_+_8433172_8446622_RNA45SN1_45S_50ntFlanks | 4724 | 4759 | 36 | ATTAATCAAGAACGAAAGTCGGAGGTTCGAAGACGA | 3555.6 | -9.9 | 1.5 | -6.6 | 3.9E-11 | 1.1E-10 | MB231/MB468 | MP |
| >hg38_21_+_8433172_8446622_RNA45SN1_45S_50ntFlanks@6775.6799.25\|CCTCCCGGGGCTACGCCTGTCTGAG | >hg38_21_+_8433172_8446622_RNA45SN1_45S_50ntFlanks | 6775 | 6799 | 25 | CCTCCCGGGGCTACGCCTGTCTGAG | 75.2 | -10.1 | 1.5 | -6.6 | 3.9E-11 | 1.1E-10 | MB231/MB468 | MP |
| >hg38_21_+_8433172_8446622_RNA45SN1_45S_50ntFlanks@10400.10424.25\|TGAACATGGGTCAGTCGGTCCTGAG | >hg38_21_+_8433172_8446622_RNA45SN1_45S_50ntFlanks | 10400 | 10424 | 25 | TGAACATGGGTCAGTCGGTCCTGAG | 1623.8 | 7.7 | 1.2 | 6.6 | 3.8E-11 | 1.1E-10 | BT20/MB468 | CYTO |
| >hg38_21_+_8433172_8446622_RNA45SN1_45S_50ntFlanks@7984.8009.26\|AGATCAGACGTGGCGACCCGCTGAAT | >hg38_21_+_8433172_8446622_RNA45SN1_45S_50ntFlanks | 7984 | 8009 | 26 | AGATCAGACGTGGCGACCCGCTGAAT | 1239.9 | -4.4 | 0.7 | -6.6 | 3.9E-11 | 1.1E-10 | MB231/MB468 | MP |
| >hg38_21_+_8433172_8446622_RNA45SN1_45S_50ntFlanks@9454.9489.36\|CCGCGCCGGGGAGGTGGAGCACGAGCGCACGTGTTA | >hg38_21_+_8433172_8446622_RNA45SN1_45S_50ntFlanks | 9454 | 9489 | 36 | CCGCGCCGGGGAGGTGGAGCACGAGCGCACGTGTTA | 115.2 | -8.9 | 1.3 | -6.6 | 3.0E-11 | 1.1E-10 | BT20/MB468 | MITO |
| >hg38_21_+_8433172_8446622_RNA45SN1_45S_50ntFlanks@6772.6799.28\|GTTCCTCCCGGGGCTACGCCTGTCTGAG | >hg38_21_+_8433172_8446622_RNA45SN1_45S_50ntFlanks | 6772 | 6799 | 28 | GTTCCTCCCGGGGCTACGCCTGTCTGAG | 1609.7 | -9.9 | 1.5 | -6.6 | 5.1E-11 | 1.1E-10 | BT20/MB468 | NUC |
| >hg38_21_+_8433172_8446622_RNA45SN1_45S_50ntFlanks@9738.9767.30\|GAAACGATCTCAACCTATTCTCAAACTTTA | >hg38_21_+_8433172_8446622_RNA45SN1_45S_50ntFlanks | 9738 | 9767 | 30 | GAAACGATCTCAACCTATTCTCAAACTTTA | 2368.6 | -8.2 | 1.2 | -6.6 | 5.1E-11 | 1.1E-10 | BT20/MB468 | NUC |
| >MI0000067\|hsa-let-7f-1&WithFlank&9\|+\|94176341\|94176439@13.32.20(+1G);MI0000068\|hsa-let-7f-2&WithFlank&X\|-\|53557186\|53557280@14.33.20(+1G)&[MIMAT0000067&hsa-let-7f-5p&offsets\|0\|-2(+1G);m-20&9\|+\|94176353\|94176374&offsets\|0\|-2(+1G)];[MIMAT0000067_1&hsa-let-7f-5p&offsets\|0\|-2(+1G);m-19&X\|-\|53557246\|53557267&offsets\|0\|-2(+1G)]\|TGAGGTAGTAGATTGTATAGG | >MI0000067\|hsa-let-7f-1&WithFlank&9\|+\|94176341\|94176439 | 13 | 32 | 20(+1G) | TGAGGTAGTAGATTGTATAGG | 343.5 | 5.5 | 0.8 | 6.6 | 3.8E-11 | 1.1E-10 | BT20/MB468 | CYTO |
| >hg38_21_+_8433172_8446622_RNA45SN1_45S_50ntFlanks@475.508.34\|CTTCGTGATCGATGTGGTGACGTCGTGCTCTCCC | >hg38_21_+_8433172_8446622_RNA45SN1_45S_50ntFlanks | 475 | 508 | 34 | CTTCGTGATCGATGTGGTGACGTCGTGCTCTCCC | 544.6 | 8.2 | 1.2 | 6.6 | 4.2E-11 | 1.1E-10 | MB231/MB468 | CYTO |
| >hg38_21_+_8433172_8446622_RNA45SN1_45S_50ntFlanks@12308.12328.21\|CGATCCTTCTGACCTTTTGGG | >hg38_21_+_8433172_8446622_RNA45SN1_45S_50ntFlanks | 12308 | 12328 | 21 | CGATCCTTCTGACCTTTTGGG | 68.7 | -10.0 | 1.5 | -6.6 | 3.9E-11 | 1.1E-10 | MB231/MB468 | MP |
| >hg38_21_+_8433172_8446622_RNA45SN1_45S_50ntFlanks@9504.9534.31\|GTGAACTATGCCTGGGCAGGGCGAAGCCAGA | >hg38_21_+_8433172_8446622_RNA45SN1_45S_50ntFlanks | 9504 | 9534 | 31 | GTGAACTATGCCTGGGCAGGGCGAAGCCAGA | 149.0 | -11.1 | 1.7 | -6.6 | 3.9E-11 | 1.1E-10 | MB231/MB468 | MP |
| >hg38_21_+_8433172_8446622_RNA45SN1_45S_50ntFlanks@12181.12201.21\|TCAAACGGTAACGCAGGTGTC | >hg38_21_+_8433172_8446622_RNA45SN1_45S_50ntFlanks | 12181 | 12201 | 21 | TCAAACGGTAACGCAGGTGTC | 124.2 | -10.8 | 1.6 | -6.6 | 3.9E-11 | 1.1E-10 | MB231/MB468 | MP |
| >MI0000465\|hsa-mir-191&WithFlank&3\|-\|49020612\|49020715@22.44.23(+1A)&[MIMAT0000440&hsa-miR-191-5p&offsets\|0\|0(+1A);m-48&3\|-\|49020672\|49020694&offsets\|0\|0(+1A)]\|CAACGGAATCCCAAAAGCAGCTGA | >MI0000465\|hsa-mir-191&WithFlank&3\|-\|49020612\|49020715 | 22 | 44 | 23(+1A) | CAACGGAATCCCAAAAGCAGCTGA | 418.3 | 6.0 | 0.9 | 6.6 | 4.2E-11 | 1.1E-10 | MB231/MB468 | CYTO |
| >hg38_21_+_8433172_8446622_RNA45SN1_45S_50ntFlanks@11576.11593.18\|CTTAGAACTGGTGCGGAC | >hg38_21_+_8433172_8446622_RNA45SN1_45S_50ntFlanks | 11576 | 11593 | 18 | CTTAGAACTGGTGCGGAC | 636.3 | -8.3 | 1.3 | -6.6 | 4.8E-11 | 1.1E-10 | MB231/MB468 | MITO |
| >hg38_21_+_8433172_8446622_RNA45SN1_45S_50ntFlanks@10829.10848.20\|GCAAGCCGGATCCGTAACTT | >hg38_21_+_8433172_8446622_RNA45SN1_45S_50ntFlanks | 10829 | 10848 | 20 | GCAAGCCGGATCCGTAACTT | 84.0 | -10.3 | 1.6 | -6.6 | 3.9E-11 | 1.1E-10 | MB231/MB468 | MP |
| >hg38_21_+_8433172_8446622_RNA45SN1_45S_50ntFlanks@12404.12433.30\|GCTTTTTGATCCTTCGATGTCGGCTCTTCC | >hg38_21_+_8433172_8446622_RNA45SN1_45S_50ntFlanks | 12404 | 12433 | 30 | GCTTTTTGATCCTTCGATGTCGGCTCTTCC | 696.4 | -4.6 | 0.7 | -6.6 | 5.1E-11 | 1.1E-10 | BT20/MB468 | NUC |
| >MI0000279\|hsa-mir-196a-2&WithFlank&12\|+\|53991732\|53991853@31.52.22;MI0000238\|hsa-mir-196a-1&WithFlank&17\|-\|48632484\|48632565@13.34.22&[MIMAT0000226&hsa-miR-196a-5p&offsets\|0\|0;m-261&12\|+\|53991762\|53991783&offsets\|0\|0];[MIMAT0000226_1&hsa-miR-196a-5p&offsets\|0\|0;m-268&17\|-\|48632532\|48632553&offsets\|0\|0]\|TAGGTAGTTTCATGTTGTTGGG | >MI0000279\|hsa-mir-196a-2&WithFlank&12\|+\|53991732\|53991853 | 31 | 52 | 22 | TAGGTAGTTTCATGTTGTTGGG | 1296.7 | 3.1 | 0.5 | 6.6 | 5.1E-11 | 1.1E-10 | BT20/MB468 | NUC |
| >hg38_21_+_8433172_8446622_RNA45SN1_45S_50ntFlanks@5184.5201.18\|AGCAATAACAGGTCTGTG | >hg38_21_+_8433172_8446622_RNA45SN1_45S_50ntFlanks | 5184 | 5201 | 18 | AGCAATAACAGGTCTGTG | 148.2 | -11.1 | 1.7 | -6.6 | 3.9E-11 | 1.1E-10 | MB231/MB468 | MP |
| >hg38_21_+_8433172_8446622_RNA45SN1_45S_50ntFlanks@6721.6770.50\|AATTGCAGGACACATTGATCATCGACACTTCGAACGCACTTGCGGCCCCG | >hg38_21_+_8433172_8446622_RNA45SN1_45S_50ntFlanks | 6721 | 6770 | 50 | AATTGCAGGACACATTGATCATCGACACTTCGAACGCACTTGCGGCCCCG | 293.3 | -10.5 | 1.6 | -6.6 | 3.8E-11 | 1.1E-10 | BT20/MB468 | CYTO |
| >hg38_21_+_8433172_8446622_RNA45SN1_45S_50ntFlanks@12259.12284.26\|CTTGATCTTGATTTTCAGTACGAATA | >hg38_21_+_8433172_8446622_RNA45SN1_45S_50ntFlanks | 12259 | 12284 | 26 | CTTGATCTTGATTTTCAGTACGAATA | 2495.5 | -9.8 | 1.5 | -6.6 | 3.9E-11 | 1.1E-10 | MB231/MB468 | MP |
| >hg38_21_+_8433172_8446622_RNA45SN1_45S_50ntFlanks@10826.10864.39\|TCGGCAAGCCGGATCCGTAACTTCGGGATAAGGATTGGC | >hg38_21_+_8433172_8446622_RNA45SN1_45S_50ntFlanks | 10826 | 10864 | 39 | TCGGCAAGCCGGATCCGTAACTTCGGGATAAGGATTGGC | 227.9 | -10.1 | 1.5 | -6.6 | 3.8E-11 | 1.1E-10 | BT20/MB468 | CYTO |

<table>
<tr><td>>hg38_21_+_8433172_8446622_RNA45SN1_45S_50ntFlanks@11768.11806.39|CTAATTAGTGACGCGCATGAATGGATGAACGAGATTCCC</td><td>>hg38_21_+_8433172_8446622_RNA45SN1_45S_50ntFlanks</td><td>11768</td><td>11806</td><td>39</td><td>CTAATTAGTGACGCGCATGAATGGATGAACGAGATTCCC</td><td>472.5</td><td>-8.3</td><td>1.3</td><td>-6.6</td><td>4.8E-11</td><td>1.1E-10</td><td>MB231/MB468</td><td>MITO</td></tr>
<tr><td>>hg38_21_+_8433172_8446622_RNA45SN1_45S_50ntFlanks@4163.4180.18|CATCCAAGGAAGGCAGCA</td><td>>hg38_21_+_8433172_8446622_RNA45SN1_45S_50ntFlanks</td><td>4163</td><td>4180</td><td>18</td><td>CATCCAAGGAAGGCAGCA</td><td>134.7</td><td>-10.9</td><td>1.7</td><td>-6.6</td><td>3.9E-11</td><td>1.1E-10</td><td>MB231/MB468</td><td>MP</td></tr>
<tr><td>>hg38_21_+_8433172_8446622_RNA45SN1_45S_50ntFlanks@12636.12654.19|AGCTACCATCTGTGGGATT</td><td>>hg38_21_+_8433172_8446622_RNA45SN1_45S_50ntFlanks</td><td>12636</td><td>12654</td><td>19</td><td>AGCTACCATCTGTGGGATT</td><td>2665.8</td><td>-6.0</td><td>0.9</td><td>-6.6</td><td>3.1E-11</td><td>1.1E-10</td><td>BT20/MB468</td><td>MITO</td></tr>
<tr><td>>hg38_21_+_8433172_8446622_RNA45SN1_45S_50ntFlanks@10869.10901.33|AGGGCTGGGTCGGTCGGGCTGGGGCGCGAAGCG</td><td>>hg38_21_+_8433172_8446622_RNA45SN1_45S_50ntFlanks</td><td>10869</td><td>10901</td><td>33</td><td>AGGGCTGGGTCGGTCGGGCTGGGGCGCGAAGCG</td><td>1518.1</td><td>-8.6</td><td>1.3</td><td>-6.6</td><td>4.2E-11</td><td>1.1E-10</td><td>MB231/MB468</td><td>NUC</td></tr>
<tr><td>>hg38_21_+_8433172_8446622_RNA45SN1_45S_50ntFlanks@12046.12070.25|TGACCCGGTGAGGCGGGGGGGCGAG</td><td>>hg38_21_+_8433172_8446622_RNA45SN1_45S_50ntFlanks</td><td>12046</td><td>12070</td><td>25</td><td>TGACCCGGTGAGGCGGGGGGGCGAG</td><td>796.8</td><td>-9.8</td><td>1.5</td><td>-6.6</td><td>5.1E-11</td><td>1.1E-10</td><td>BT20/MB468</td><td>NUC</td></tr>
<tr><td>>hg38_21_+_8433172_8446622_RNA45SN1_45S_50ntFlanks@475.496.22|CTTCGTGATCGATGTGGTGACG</td><td>>hg38_21_+_8433172_8446622_RNA45SN1_45S_50ntFlanks</td><td>475</td><td>496</td><td>22</td><td>CTTCGTGATCGATGTGGTGACG</td><td>3023.9</td><td>7.2</td><td>1.1</td><td>6.6</td><td>3.1E-11</td><td>1.1E-10</td><td>BT20/MB468</td><td>MITO</td></tr>
<tr><td>>hg38_21_+_8433172_8446622_RNA45SN1_45S_50ntFlanks@4728.4751.24|ATCAAGAACGAAAGTCGGAGGTTC</td><td>>hg38_21_+_8433172_8446622_RNA45SN1_45S_50ntFlanks</td><td>4728</td><td>4751</td><td>24</td><td>ATCAAGAACGAAAGTCGGAGGTTC</td><td>138.0</td><td>-9.2</td><td>1.4</td><td>-6.6</td><td>3.1E-11</td><td>1.1E-10</td><td>BT20/MB468</td><td>MITO</td></tr>
<tr><td>>hg38_21_+_8433172_8446622_RNA45SN1_45S_50ntFlanks@11233.11255.23|CCCTTCCCGTGGATCGCCCCAGC</td><td>>hg38_21_+_8433172_8446622_RNA45SN1_45S_50ntFlanks</td><td>11233</td><td>11255</td><td>23</td><td>CCCTTCCCGTGGATCGCCCCAGC</td><td>205.9</td><td>-11.5</td><td>1.7</td><td>-6.6</td><td>3.9E-11</td><td>1.1E-10</td><td>MB231/MB468</td><td>MP</td></tr>
<tr><td>>MI0000088|hsa-mir-30a&WithFlank&6|-|71403545|71403627@13.35.23(+1U)&[MIMAT0000087&hsa-miR-30a-5p&offsets|+1|+2(+1U);m-15&6|-|71403595|71403616&offsets|+1|+2(+1U)]|GTAAACATCCTCGACTGGAAGCTT</td><td>>MI0000088|hsa-mir-30a&WithFlank&6|-|71403545|71403627</td><td>13</td><td>35</td><td>23(+1U)</td><td>GTAAACATCCTCGACTGGAAGCTT</td><td>270.1</td><td>11.3</td><td>1.7</td><td>6.6</td><td>4.9E-11</td><td>1.1E-10</td><td>MB231/MB468</td><td>MITO</td></tr>
<tr><td>>hg38_21_+_8433172_8446622_RNA45SN1_45S_50ntFlanks@12676.12704.29|AGAATCCCGCCCAGGCGGAACGATACGGC</td><td>>hg38_21_+_8433172_8446622_RNA45SN1_45S_50ntFlanks</td><td>12676</td><td>12704</td><td>29</td><td>AGAATCCCGCCCAGGCGGAACGATACGGC</td><td>114.4</td><td>-9.1</td><td>1.4</td><td>-6.6</td><td>3.8E-11</td><td>1.1E-10</td><td>BT20/MB468</td><td>CYTO</td></tr>
<tr><td>>hg38_21_+_8433172_8446622_RNA45SN1_45S_50ntFlanks@6732.6772.41|ACATTGATCATCGACACTTCGAACGCACTTGCGGCCCCGGG</td><td>>hg38_21_+_8433172_8446622_RNA45SN1_45S_50ntFlanks</td><td>6732</td><td>6772</td><td>41</td><td>ACATTGATCATCGACACTTCGAACGCACTTGCGGCCCCGGG</td><td>609.4</td><td>-4.9</td><td>0.8</td><td>-6.6</td><td>5.1E-11</td><td>1.1E-10</td><td>BT20/MB468</td><td>NUC</td></tr>
<tr><td>>hg38_21_+_8433172_8446622_RNA45SN1_45S_50ntFlanks@4046.4067.22|CATTCGAACGTCTGCCCTATCA</td><td>>hg38_21_+_8433172_8446622_RNA45SN1_45S_50ntFlanks</td><td>4046</td><td>4067</td><td>22</td><td>CATTCGAACGTCTGCCCTATCA</td><td>850.4</td><td>-9.4</td><td>1.4</td><td>-6.6</td><td>5.2E-11</td><td>1.1E-10</td><td>BT20/MB468</td><td>NUC</td></tr>
<tr><td>>hg38_21_+_8433172_8446622_RNA45SN1_45S_50ntFlanks@6730.6770.41|ACACATTGATCATCGACACTTCGAACGCACTTGCGGCCCCG</td><td>>hg38_21_+_8433172_8446622_RNA45SN1_45S_50ntFlanks</td><td>6730</td><td>6770</td><td>41</td><td>ACACATTGATCATCGACACTTCGAACGCACTTGCGGCCCCG</td><td>605.8</td><td>-5.7</td><td>0.9</td><td>-6.6</td><td>4.2E-11</td><td>1.2E-10</td><td>MB231/MB468</td><td>NUC</td></tr>
<tr><td>>hg38_21_+_8433172_8446622_RNA45SN1_45S_50ntFlanks@10808.10844.37|CAATGTAGGTAAGGGAAGTCGGCAAGCCGGATCCGTA</td><td>>hg38_21_+_8433172_8446622_RNA45SN1_45S_50ntFlanks</td><td>10808</td><td>10844</td><td>37</td><td>CAATGTAGGTAAGGGAAGTCGGCAAGCCGGATCCGTA</td><td>115.5</td><td>-10.7</td><td>1.6</td><td>-6.6</td><td>4.0E-11</td><td>1.2E-10</td><td>MB231/MB468</td><td>MP</td></tr>
<tr><td>>hg38_21_+_8433172_8446622_RNA45SN1_45S_50ntFlanks@8169.8197.29|CCAAGTCCTTCTGATCGAGGCCCAGCCCG</td><td>>hg38_21_+_8433172_8446622_RNA45SN1_45S_50ntFlanks</td><td>8169</td><td>8197</td><td>29</td><td>CCAAGTCCTTCTGATCGAGGCCCAGCCCG</td><td>103.7</td><td>-10.6</td><td>1.6</td><td>-6.6</td><td>4.0E-11</td><td>1.2E-10</td><td>MB231/MB468</td><td>MP</td></tr>
<tr><td>>trnaMT_PheGAA_MT_+_577_647@36.57.22|AAAATGTTTAGACGGGCTCACA</td><td>>trnaMT_PheGAA_MT_+_577_647</td><td>36</td><td>57</td><td>22</td><td>AAAATGTTTAGACGGGCTCACA</td><td>873.8</td><td>6.1</td><td>0.9</td><td>7.0</td><td>2.9E-12</td><td>1.2E-10</td><td>BT20/MB468</td><td>MP</td></tr>
<tr><td>>hg38_21_+_8433172_8446622_RNA45SN1_45S_50ntFlanks@11641.11676.36|CGGCGGGTGTTGACGCGATGTGATTTCTGCCCAGTG</td><td>>hg38_21_+_8433172_8446622_RNA45SN1_45S_50ntFlanks</td><td>11641</td><td>11676</td><td>36</td><td>CGGCGGGTGTTGACGCGATGTGATTTCTGCCCAGTG</td><td>286.4</td><td>-11.2</td><td>1.6</td><td>-7.0</td><td>2.9E-12</td><td>1.2E-10</td><td>BT20/MB468</td><td>MP</td></tr>
<tr><td>>hg38_21_+_8433172_8446622_RNA45SN1_45S_50ntFlanks@12356.12376.21|ACCACAGGGATAACTGGCTTG</td><td>>hg38_21_+_8433172_8446622_RNA45SN1_45S_50ntFlanks</td><td>12356</td><td>12376</td><td>21</td><td>ACCACAGGGATAACTGGCTTG</td><td>223.8</td><td>-10.9</td><td>1.6</td><td>-7.0</td><td>2.9E-12</td><td>1.2E-10</td><td>BT20/MB468</td><td>MP</td></tr>
<tr><td>>hg38_21_+_8433172_8446622_RNA45SN1_45S_50ntFlanks@8179.8203.25|CTGATCGAGGCCCAGCCCGTGGACG</td><td>>hg38_21_+_8433172_8446622_RNA45SN1_45S_50ntFlanks</td><td>8179</td><td>8203</td><td>25</td><td>CTGATCGAGGCCCAGCCCGTGGACG</td><td>600.8</td><td>-8.5</td><td>1.3</td><td>-6.6</td><td>4.0E-11</td><td>1.2E-10</td><td>MB231/MB468</td><td>MP</td></tr>
<tr><td>>MI0000471|hsa-mir-126&WithFlank&9|+|136670596|136670692@58.77.20&[MIMAT0000445&hsa-miR-126-3p&offsets|0|-2;m-40&9|+|136670653|136670674&offsets|0|-2]|TCGTACCGTGAGTAATAATG</td><td>>MI0000471|hsa-mir-126&WithFlank&9|+|136670596|136670692</td><td>58</td><td>77</td><td>20</td><td>TCGTACCGTGAGTAATAATG</td><td>200.1</td><td>10.9</td><td>1.7</td><td>6.6</td><td>4.9E-11</td><td>1.2E-10</td><td>MB231/MB468</td><td>MITO</td></tr>
<tr><td>>MI0001445|hsa-mir-423&WithFlank&17|+|30117073|30117178@23.44.22&[MIMAT0004748&hsa-miR-423-5p&offsets|0|-1;m-141&17|+|30117095|30117117&offsets|0|-1]|TGAGGGGCAGAGAGCGAGACTT</td><td>>MI0001445|hsa-mir-423&WithFlank&17|+|30117073|30117178</td><td>23</td><td>44</td><td>22</td><td>TGAGGGGCAGAGAGCGAGACTT</td><td>1280.9</td><td>8.8</td><td>1.3</td><td>6.6</td><td>4.3E-11</td><td>1.2E-10</td><td>MB231/MB468</td><td>CYTO</td></tr>
<tr><td>>hg38_21_+_8433172_8446622_RNA45SN1_45S_50ntFlanks@10788.10825.38|CAGCCTCTGGCATGTTGGAACAATGTAGGTAAGGGAAG</td><td>>hg38_21_+_8433172_8446622_RNA45SN1_45S_50ntFlanks</td><td>10788</td><td>10825</td><td>38</td><td>CAGCCTCTGGCATGTTGGAACAATGTAGGTAAGGGAAG</td><td>435.6</td><td>-8.2</td><td>1.3</td><td>-6.6</td><td>4.9E-11</td><td>1.2E-10</td><td>MB231/MB468</td><td>MITO</td></tr>
<tr><td>>hg38_21_+_8433172_8446622_RNA45SN1_45S_50ntFlanks@8212.8235.24|CCGGTAGCGGCCCCCGGCGCGCCG</td><td>>hg38_21_+_8433172_8446622_RNA45SN1_45S_50ntFlanks</td><td>8212</td><td>8235</td><td>24</td><td>CCGGTAGCGGCCCCCGGCGCGCCG</td><td>78.4</td><td>-10.2</td><td>1.5</td><td>-6.6</td><td>4.0E-11</td><td>1.2E-10</td><td>MB231/MB468</td><td>MP</td></tr>
<tr><td>>MI0000650|hsa-mir-200c&WithFlank&12|+|6963693|6963772@50.70.21(+1U)&[MIMAT0000617&hsa-miR-200c-3p&offsets|0|-2(+1U);m-54&12|+|6963742|6963764&offsets|0|-2(+1U)]|TAATACTGCCGGGTAATGATGT</td><td>>MI0000650|hsa-mir-200c&WithFlank&12|+|6963693|6963772</td><td>50</td><td>70</td><td>21(+1U)</td><td>TAATACTGCCGGGTAATGATGT</td><td>480.0</td><td>10.0</td><td>1.5</td><td>6.6</td><td>3.1E-11</td><td>1.2E-10</td><td>BT20/MB468</td><td>MITO</td></tr>
<tr><td>>hg38_21_+_8433172_8446622_RNA45SN1_45S_50ntFlanks@12044.12062.19|ACTGACCCGGTGAGGCGGG</td><td>>hg38_21_+_8433172_8446622_RNA45SN1_45S_50ntFlanks</td><td>12044</td><td>12062</td><td>19</td><td>ACTGACCCGGTGAGGCGGG</td><td>167.6</td><td>-11.3</td><td>1.7</td><td>-6.6</td><td>4.0E-11</td><td>1.2E-10</td><td>MB231/MB468</td><td>MP</td></tr>
<tr><td>>hg38_21_+_8433172_8446622_RNA45SN1_45S_50ntFlanks@5485.5523.39|AGCGCTGAGAAGACGGTCGAACTTGACTATCTAGAGGAA</td><td>>hg38_21_+_8433172_8446622_RNA45SN1_45S_50ntFlanks</td><td>5485</td><td>5523</td><td>39</td><td>AGCGCTGAGAAGACGGTCGAACTTGACTATCTAGAGGAA</td><td>117.2</td><td>-10.7</td><td>1.6</td><td>-6.6</td><td>4.0E-11</td><td>1.2E-10</td><td>MB231/MB468</td><td>MP</td></tr>
<tr><td>>hg38_21_+_8433172_8446622_RNA45SN1_45S_50ntFlanks@11205.11254.50|GCGGCGGCGACTCTGGACGCGAGCCGGGCCCTTCCCGTGGATCGCCCCAG</td><td>>hg38_21_+_8433172_8446622_RNA45SN1_45S_50ntFlanks</td><td>11205</td><td>11254</td><td>50</td><td>GCGGCGGCGACTCTGGACGCGAGCCGGGCCCTTCCCGTGGATCGCCCCAG</td><td>311.7</td><td>-11.2</td><td>1.7</td><td>-6.6</td><td>4.3E-11</td><td>1.2E-10</td><td>MB231/MB468</td><td>NUC</td></tr>
<tr><td>>MI0000746|hsa-mir-99b&WithFlank&19|+|51692606|51692687@13.34.22(+1C)&[MIMAT0000689&hsa-miR-99b-5p&offsets|0|0(+1C);m-10&19|+|51692618|51692639&offsets|0|0(+1C)]|CACCCGTAGAACCGACCTTGCGC</td><td>>MI0000746|hsa-mir-99b&WithFlank&19|+|51692606|51692687</td><td>13</td><td>34</td><td>22(+1C)</td><td>CACCCGTAGAACCGACCTTGCGC</td><td>898.9</td><td>8.9</td><td>1.3</td><td>6.6</td><td>3.9E-11</td><td>1.2E-10</td><td>BT20/MB468</td><td>CYTO</td></tr>
<tr><td>>hg38_21_+_8433172_8446622_RNA45SN1_45S_50ntFlanks@12934.12972.39|CTAAACCATTCGTAGACGACCTGCTTCTGGGTCGGGGTT</td><td>>hg38_21_+_8433172_8446622_RNA45SN1_45S_50ntFlanks</td><td>12934</td><td>12972</td><td>39</td><td>CTAAACCATTCGTAGACGACCTGCTTCTGGGTCGGGGTT</td><td>6752.4</td><td>-8.5</td><td>1.3</td><td>-6.6</td><td>3.9E-11</td><td>1.2E-10</td><td>BT20/MB468</td><td>CYTO</td></tr>
<tr><td>>hg38_21_+_8433172_8446622_RNA45SN1_45S_50ntFlanks@10392.10426.35|ACAGCAGTTGAACATGGGTCAGTCGGTCCTGAGAG</td><td>>hg38_21_+_8433172_8446622_RNA45SN1_45S_50ntFlanks</td><td>10392</td><td>10426</td><td>35</td><td>ACAGCAGTTGAACATGGGTCAGTCGGTCCTGAGAG</td><td>1090.7</td><td>-7.5</td><td>1.1</td><td>-6.6</td><td>4.3E-11</td><td>1.2E-10</td><td>MB231/MB468</td><td>NUC</td></tr>
<tr><td>>hg38_21_+_8433172_8446622_RNA45SN1_45S_50ntFlanks@10794.10823.30|CTGGCATGTTGGAACAATGTAGGTAAGGGA</td><td>>hg38_21_+_8433172_8446622_RNA45SN1_45S_50ntFlanks</td><td>10794</td><td>10823</td><td>30</td><td>CTGGCATGTTGGAACAATGTAGGTAAGGGA</td><td>191.2</td><td>-5.2</td><td>0.8</td><td>-6.6</td><td>4.3E-11</td><td>1.2E-10</td><td>MB231/MB468</td><td>CYTO</td></tr>
<tr><td>>MI0005757|hsa-mir-935&WithFlank&19|+|53982301|53982403@63.84.22&[MIMAT0004978&hsa-miR-935&offsets|+1|0;m-549&19|+|53982363|53982384&offsets|0|0]|CAGTTACCGCTTCCGCTACCGC</td><td>>MI0005757|hsa-mir-935&WithFlank&19|+|53982301|53982403</td><td>63</td><td>84</td><td>22</td><td>CAGTTACCGCTTCCGCTACCGC</td><td>185.3</td><td>8.4</td><td>1.2</td><td>6.9</td><td>4.5E-12</td><td>1.2E-10</td><td>BT20/MB231</td><td>NUC</td></tr>
<tr><td>>hg38_21_+_8433172_8446622_RNA45SN1_45S_50ntFlanks@6663.6681.19|GTGGATCACTCGGCTCGTG</td><td>>hg38_21_+_8433172_8446622_RNA45SN1_45S_50ntFlanks</td><td>6663</td><td>6681</td><td>19</td><td>GTGGATCACTCGGCTCGTG</td><td>1662.4</td><td>-2.4</td><td>0.4</td><td>-6.6</td><td>3.2E-11</td><td>1.2E-10</td><td>BT20/MB468</td><td>MITO</td></tr>
<tr><td>>hg38_21_+_8433172_8446622_RNA45SN1_45S_50ntFlanks@10134.10153.20|CGGCGGCGGCGGCGGGGGTG</td><td>>hg38_21_+_8433172_8446622_RNA45SN1_45S_50ntFlanks</td><td>10134</td><td>10153</td><td>20</td><td>CGGCGGCGGCGGCGGGGGTG</td><td>121.3</td><td>-10.8</td><td>1.6</td><td>-6.6</td><td>4.1E-11</td><td>1.2E-10</td><td>MB231/MB468</td><td>MP</td></tr>
<tr><td>>hg38_1_-_228634819_228635039_RNA5S12_5S_50ntFlanks@142.171.30|CGCCTGGGAATACCGGGTGCTGTAGGCTTT</td><td>>hg38_1_-_228634819_228635039_RNA5S12_5S_50ntFlanks</td><td>142</td><td>171</td><td>30</td><td>CGCCTGGGAATACCGGGTGCTGTAGGCTTT</td><td>1329.7</td><td>4.5</td><td>0.7</td><td>6.6</td><td>4.3E-11</td><td>1.2E-10</td><td>MB231/MB468</td><td>NUC</td></tr>
<tr><td>>hg38_21_+_8433172_8446622_RNA45SN1_45S_50ntFlanks@10411.10453.43|CAGTCGGTCCTGAGAGATGGGCGAGCGCCGTTCCGAAGGGACG</td><td>>hg38_21_+_8433172_8446622_RNA45SN1_45S_50ntFlanks</td><td>10411</td><td>10453</td><td>43</td><td>CAGTCGGTCCTGAGAGATGGGCGAGCGCCGTTCCGAAGGGACG</td><td>2695.9</td><td>-11.0</td><td>1.7</td><td>-6.6</td><td>5.3E-11</td><td>1.2E-10</td><td>BT20/MB468</td><td>NUC</td></tr>
<tr><td>>hg38_21_+_8433172_8446622_RNA45SN1_45S_50ntFlanks@4995.5024.30|ACAGATTGATAGCTCTTTCTCGATTCCGTG</td><td>>hg38_21_+_8433172_8446622_RNA45SN1_45S_50ntFlanks</td><td>4995</td><td>5024</td><td>30</td><td>ACAGATTGATAGCTCTTTCTCGATTCCGTG</td><td>5728.0</td><td>-11.9</td><td>1.8</td><td>-6.6</td><td>5.3E-11</td><td>1.2E-10</td><td>BT20/MB468</td><td>NUC</td></tr>
<tr><td>>hg38_21_+_8433172_8446622_RNA45SN1_45S_50ntFlanks@9504.9521.18|GTGAACTATGCCTGGGCA</td><td>>hg38_21_+_8433172_8446622_RNA45SN1_45S_50ntFlanks</td><td>9504</td><td>9521</td><td>18</td><td>GTGAACTATGCCTGGGCA</td><td>3442.5</td><td>-10.6</td><td>1.6</td><td>-6.6</td><td>5.3E-11</td><td>1.2E-10</td><td>BT20/MB468</td><td>NUC</td></tr>
<tr><td>>hg38_21_+_8433172_8446622_RNA45SN1_45S_50ntFlanks@11577.11603.27|TTAGAACTGGTGCGGACCAGGGGAATC</td><td>>hg38_21_+_8433172_8446622_RNA45SN1_45S_50ntFlanks</td><td>11577</td><td>11603</td><td>27</td><td>TTAGAACTGGTGCGGACCAGGGGAATC</td><td>1133.2</td><td>-7.1</td><td>1.1</td><td>-6.6</td><td>4.4E-11</td><td>1.2E-10</td><td>MB231/MB468</td><td>NUC</td></tr>
<tr><td>>hg38_21_+_8433172_8446622_RNA45SN1_45S_50ntFlanks@9143.9162.20|GGGCGGACTGTCCCCAGTGC</td><td>>hg38_21_+_8433172_8446622_RNA45SN1_45S_50ntFlanks</td><td>9143</td><td>9162</td><td>20</td><td>GGGCGGACTGTCCCCAGTGC</td><td>108.6</td><td>-10.6</td><td>1.6</td><td>-6.6</td><td>4.1E-11</td><td>1.2E-10</td><td>MB231/MB468</td><td>MP</td></tr>
<tr><td>>hg38_21_+_8433172_8446622_RNA45SN1_45S_50ntFlanks@6648.6695.48|GTACGACTCTTAGCGGTGGATCACTCGGCTCGTGCGTCGATGAAGAAC</td><td>>hg38_21_+_8433172_8446622_RNA45SN1_45S_50ntFlanks</td><td>6648</td><td>6695</td><td>48</td><td>GTACGACTCTTAGCGGTGGATCACTCGGCTCGTGCGTCGATGAAGAAC</td><td>4152.5</td><td>6.9</td><td>1.0</td><td>6.6</td><td>3.2E-11</td><td>1.2E-10</td><td>BT20/MB468</td><td>MITO</td></tr>
<tr><td>>hg38_21_+_8433172_8446622_RNA45SN1_45S_50ntFlanks@6722.6746.25|ATTGCAGGACACATTGATCATCGAC</td><td>>hg38_21_+_8433172_8446622_RNA45SN1_45S_50ntFlanks</td><td>6722</td><td>6746</td><td>25</td><td>ATTGCAGGACACATTGATCATCGAC</td><td>337.6</td><td>-8.1</td><td>1.2</td><td>-6.6</td><td>4.1E-11</td><td>1.2E-10</td><td>MB231/MB468</td><td>MP</td></tr>
<tr><td>>hg38_21_+_8433172_8446622_RNA45SN1_45S_50ntFlanks@12616.12652.37|CTGAGGAGCCAATGGGGCGAAGCTACCATCTGTGGGA</td><td>>hg38_21_+_8433172_8446622_RNA45SN1_45S_50ntFlanks</td><td>12616</td><td>12652</td><td>37</td><td>CTGAGGAGCCAATGGGGCGAAGCTACCATCTGTGGGA</td><td>88.2</td><td>-10.3</td><td>1.6</td><td>-6.6</td><td>4.1E-11</td><td>1.2E-10</td><td>MB231/MB468</td><td>MP</td></tr>
<tr><td>>hg38_21_+_8433172_8446622_RNA45SN1_45S_50ntFlanks@12632.12661.30|GCGAAGCTACCATCTGTGGGATTATGACTG</td><td>>hg38_21_+_8433172_8446622_RNA45SN1_45S_50ntFlanks</td><td>12632</td><td>12661</td><td>30</td><td>GCGAAGCTACCATCTGTGGGATTATGACTG</td><td>2720.3</td><td>-10.1</td><td>1.5</td><td>-6.6</td><td>4.1E-11</td><td>1.2E-10</td><td>MB231/MB468</td><td>MP</td></tr>
<tr><td>>hg38_21_+_8433172_8446622_RNA45SN1_45S_50ntFlanks@5084.5102.19|CGAACGAGACTCTGGCATG</td><td>>hg38_21_+_8433172_8446622_RNA45SN1_45S_50ntFlanks</td><td>5084</td><td>5102</td><td>19</td><td>CGAACGAGACTCTGGCATG</td><td>2174.7</td><td>-8.4</td><td>1.3</td><td>-6.6</td><td>4.1E-11</td><td>1.2E-10</td><td>MB231/MB468</td><td>MP</td></tr>
<tr><td>>hg38_21_+_8433172_8446622_RNA45SN1_45S_50ntFlanks@5420.5446.27|CCGATTGGATGGTTTAGTGAGGCCCTC</td><td>>hg38_21_+_8433172_8446622_RNA45SN1_45S_50ntFlanks</td><td>5420</td><td>5446</td><td>27</td><td>CCGATTGGATGGTTTAGTGAGGCCCTC</td><td>186.9</td><td>-11.4</td><td>1.7</td><td>-6.6</td><td>4.1E-11</td><td>1.2E-10</td><td>MB231/MB468</td><td>MP</td></tr>
<tr><td>>MI0000294|hsa-mir-218-1&WithFlank&4|+|20528269|20528390@31.52.22(+1U);MI0000295|hsa-mir-218-2&WithFlank&5|-|168768140|168768261@31.52.22(+1U)&[MIMAT0000275&hsa-miR-218-5p&offsets|0|+1(+1U);m-185&4|+|20528299|20528319&offsets|0|+1(+1U)];[MIMAT0000275_1&hsa-miR-218-5p&offsets|0|+1(+1U);m-187&5|-|168768211|168768231&offsets|0|+1(+1U)]|TTGTGCTTGATCTAACCATGTGT</td><td>>MI0000294|hsa-mir-218-1&WithFlank&4|+|20528269|20528390</td><td>31</td><td>52</td><td>22(+1U)</td><td>TTGTGCTTGATCTAACCATGTGT</td><td>155.5</td><td>-10.4</td><td>1.5</td><td>-6.9</td><td>4.6E-12</td><td>1.2E-10</td><td>BT20/MB231</td><td>NUC</td></tr>
<tr><td>>hg38_21_+_8433172_8446622_RNA45SN1_45S_50ntFlanks@4759.4782.24|ATCAGATACCGTCGTAGTTCCGAC</td><td>>hg38_21_+_8433172_8446622_RNA45SN1_45S_50ntFlanks</td><td>4759</td><td>4782</td><td>24</td><td>ATCAGATACCGTCGTAGTTCCGAC</td><td>5102.0</td><td>-9.8</td><td>1.5</td><td>-6.6</td><td>4.1E-11</td><td>1.2E-10</td><td>MB231/MB468</td><td>MP</td></tr>
<tr><td>>hg38_21_+_8433172_8446622_RNA45SN1_45S_50ntFlanks@11583.11605.23|CTGGTGCGGACCAGGGGAATCCG</td><td>>hg38_21_+_8433172_8446622_RNA45SN1_45S_50ntFlanks</td><td>11583</td><td>11605</td><td>23</td><td>CTGGTGCGGACCAGGGGAATCCG</td><td>808.0</td><td>-4.1</td><td>0.6</td><td>-6.6</td><td>3.2E-11</td><td>1.2E-10</td><td>BT20/MB468</td><td>MITO</td></tr>
<tr><td>>hg38_21_+_8433172_8446622_RNA45SN1_45S_50ntFlanks@6699.6717.19|GCTAGCTGCGAGAATTAAT</td><td>>hg38_21_+_8433172_8446622_RNA45SN1_45S_50ntFlanks</td><td>6699</td><td>6717</td><td>19</td><td>GCTAGCTGCGAGAATTAAT</td><td>994.3</td><td>-5.0</td><td>0.8</td><td>-6.6</td><td>4.4E-11</td><td>1.2E-10</td><td>MB231/MB468</td><td>NUC</td></tr>
<tr><td>>am_trnaMT_PheGAA_MT_+_577_647@41.59.19|GTTTAGACGGGCTCACATC</td><td>>am_trnaMT_PheGAA_MT_+_577_647</td><td>41</td><td>59</td><td>19</td><td>GTTTAGACGGGCTCACATC</td><td>344.5</td><td>8.5</td><td>1.3</td><td>6.6</td><td>4.4E-11</td><td>1.2E-10</td><td>MB231/MB468</td><td>CYTO</td></tr>
<tr><td>>hg38_21_+_8433172_8446622_RNA45SN1_45S_50ntFlanks@9569.9597.29|GTGCAAATCGGTCGTCCGACCTGGGTATA</td><td>>hg38_21_+_8433172_8446622_RNA45SN1_45S_50ntFlanks</td><td>9569</td><td>9597</td><td>29</td><td>GTGCAAATCGGTCGTCCGACCTGGGTATA</td><td>932.2</td><td>-9.5</td><td>1.4</td><td>-6.6</td><td>5.4E-11</td><td>1.2E-10</td><td>BT20/MB468</td><td>NUC</td></tr>
</table>

<table>
<tr><th></th><th></th><th></th><th></th><th></th><th></th><th></th><th></th><th></th><th></th><th></th><th></th><th></th><th></th><th></th></tr>
<tr><td>>MI0000434|hsa-let-7i&WithFlank&12|+|62603680|62603775@12.32.21(+1A)&[MIMAT0000415&hsa-let-7i-5p&offsets|0|-1(+1A);m-49&12|+|62603691|62603712&offsets|0|-1(+1A)]|TGAGGTAGTAGTTTGTGCTGTA</td><td>>MI0000434|hsa-let-7i&WithFlank&12|+|62603680|62603775</td><td>12</td><td>32</td><td>21(+1A)</td><td>TGAGGTAGTAGTTTGTGCTGTA</td><td>2809.9</td><td>5.7</td><td>0.9</td><td>6.6</td><td>4.4E-11</td><td>1.2E-10</td><td>MB231/MB468</td><td>CYTO</td></tr>
<tr><td>>hg38_21_+_8433172_8446622_RNA45SN1_45S_50ntFlanks@5083.5100.18|ACGAACGAGACTCTGGCA</td><td>>hg38_21_+_8433172_8446622_RNA45SN1_45S_50ntFlanks</td><td>5083</td><td>5100</td><td>18</td><td>ACGAACGAGACTCTGGCA</td><td>406.0</td><td>6.5</td><td>1.0</td><td>6.6</td><td>4.4E-11</td><td>1.2E-10</td><td>MB231/MB468</td><td>CYTO</td></tr>
<tr><td>>hg38_21_+_8433172_8446622_RNA45SN1_45S_50ntFlanks@12450.12480.31|AATTCACCAAGCGTTGGATTGTTCACCCACT</td><td>>hg38_21_+_8433172_8446622_RNA45SN1_45S_50ntFlanks</td><td>12450</td><td>12480</td><td>31</td><td>AATTCACCAAGCGTTGGATTGTTCACCCACT</td><td>673.0</td><td>5.6</td><td>0.9</td><td>6.6</td><td>4.4E-11</td><td>1.2E-10</td><td>MB231/MB468</td><td>CYTO</td></tr>
<tr><td>>hg38_21_+_8433172_8446622_RNA45SN1_45S_50ntFlanks@4723.4748.26|CATTAATCAAGAACGAAAGTCGGAGG</td><td>>hg38_21_+_8433172_8446622_RNA45SN1_45S_50ntFlanks</td><td>4723</td><td>4748</td><td>26</td><td>CATTAATCAAGAACGAAAGTCGGAGG</td><td>906.3</td><td>-8.7</td><td>1.3</td><td>-6.6</td><td>5.1E-11</td><td>1.2E-10</td><td>MB231/MB468</td><td>MITO</td></tr>
<tr><td>>MI0000749|hsa-mir-30e&WithFlank&1|+|40754349|40754452@65.83.19&[MIMAT0000693&hsa-miR-30e-3p&offsets|0|-3;m-25&1|+|40754413|40754434&offsets|0|-3]|CTTTCAGTCGGATGTTTAC</td><td>>MI0000749|hsa-mir-30e&WithFlank&1|+|40754349|40754452</td><td>65</td><td>83</td><td>19</td><td>CTTTCAGTCGGATGTTTAC</td><td>1019.9</td><td>4.6</td><td>0.7</td><td>6.6</td><td>4.0E-11</td><td>1.2E-10</td><td>BT20/MB468</td><td>CYTO</td></tr>
<tr><td>>hg38_21_+_8433172_8446622_RNA45SN1_45S_50ntFlanks@12948.12990.43|GACGACCTGCTTCTGGGTCGGGGTTTCGTACGTAGCAGAGCAG</td><td>>hg38_21_+_8433172_8446622_RNA45SN1_45S_50ntFlanks</td><td>12948</td><td>12990</td><td>43</td><td>GACGACCTGCTTCTGGGTCGGGGTTTCGTACGTAGCAGAGCAG</td><td>1463.3</td><td>-10.7</td><td>1.6</td><td>-6.6</td><td>5.4E-11</td><td>1.2E-10</td><td>BT20/MB468</td><td>NUC</td></tr>
<tr><td>>hg38_21_+_8433172_8446622_RNA45SN1_45S_50ntFlanks@7974.8016.43|ACGCGACCTCAGATCAGACGTGGCGACCCGCTGAATTTAAGCA</td><td>>hg38_21_+_8433172_8446622_RNA45SN1_45S_50ntFlanks</td><td>7974</td><td>8016</td><td>43</td><td>ACGCGACCTCAGATCAGACGTGGCGACCCGCTGAATTTAAGCA</td><td>8884.3</td><td>5.2</td><td>0.8</td><td>6.6</td><td>3.3E-11</td><td>1.2E-10</td><td>BT20/MB468</td><td>MITO</td></tr>
<tr><td>>hg38_21_+_8433172_8446622_RNA45SN1_45S_50ntFlanks@3875.3894.20|ATACATGCCGACGGGCGCTG</td><td>>hg38_21_+_8433172_8446622_RNA45SN1_45S_50ntFlanks</td><td>3875</td><td>3894</td><td>20</td><td>ATACATGCCGACGGGCGCTG</td><td>85.9</td><td>-10.3</td><td>1.6</td><td>-6.6</td><td>4.2E-11</td><td>1.2E-10</td><td>MB231/MB468</td><td>MP</td></tr>
<tr><td>>hg38_21_+_8433172_8446622_RNA45SN1_45S_50ntFlanks@9621.9655.35|TCTAGTAGCTGGTTCCCTCCGAAGTTTCCCTCAGG</td><td>>hg38_21_+_8433172_8446622_RNA45SN1_45S_50ntFlanks</td><td>9621</td><td>9655</td><td>35</td><td>TCTAGTAGCTGGTTCCCTCCGAAGTTTCCCTCAGG</td><td>104.5</td><td>-10.6</td><td>1.6</td><td>-6.6</td><td>4.2E-11</td><td>1.2E-10</td><td>MB231/MB468</td><td>MP</td></tr>
<tr><td>>hg38_21_+_8433172_8446622_RNA45SN1_45S_50ntFlanks@12676.12703.28|AGAATCCCGCCCAGGCGGAACGATACGG</td><td>>hg38_21_+_8433172_8446622_RNA45SN1_45S_50ntFlanks</td><td>12676</td><td>12703</td><td>28</td><td>AGAATCCCGCCCAGGCGGAACGATACGG</td><td>2533.8</td><td>-10.2</td><td>1.5</td><td>-7.0</td><td>3.1E-12</td><td>1.2E-10</td><td>BT20/MB468</td><td>MP</td></tr>
<tr><td>>hg38_1_-_228634819_228635039_RNA5S12_5S_50ntFlanks@84.103.20|CTCGTCTGATCTCGGAAGCT</td><td>>hg38_1_-_228634819_228635039_RNA5S12_5S_50ntFlanks</td><td>84</td><td>103</td><td>20</td><td>CTCGTCTGATCTCGGAAGCT</td><td>1129.6</td><td>-6.6</td><td>1.0</td><td>-6.6</td><td>4.5E-11</td><td>1.2E-10</td><td>MB231/MB468</td><td>CYTO</td></tr>
<tr><td>>hg38_21_+_8433172_8446622_RNA45SN1_45S_50ntFlanks@12053.12072.20|GTGAGGCGGGGGGGCGAGCC</td><td>>hg38_21_+_8433172_8446622_RNA45SN1_45S_50ntFlanks</td><td>12053</td><td>12072</td><td>20</td><td>GTGAGGCGGGGGGGCGAGCC</td><td>109.4</td><td>-10.6</td><td>1.6</td><td>-6.6</td><td>4.2E-11</td><td>1.2E-10</td><td>MB231/MB468</td><td>MP</td></tr>
<tr><td>>MI0000542|hsa-mir-320a&WithFlank&8|-|22244960|22245043@48.70.23(+1U)&[MIMAT0000510&hsa-miR-320a-3p&offsets|0|+1(+1U);m-60&8|-|22244975|22244996&offsets|0|+1(+1U)]|AAAAGCTGGGTTGAGAGGGCGAAT</td><td>>MI0000542|hsa-mir-320a&WithFlank&8|-|22244960|22245043</td><td>48</td><td>70</td><td>23(+1U)</td><td>AAAAGCTGGGTTGAGAGGGCGAAT</td><td>1286.5</td><td>3.1</td><td>0.5</td><td>6.6</td><td>5.5E-11</td><td>1.2E-10</td><td>BT20/MB468</td><td>NUC</td></tr>
<tr><td>>hg38_21_+_8433172_8446622_RNA45SN1_45S_50ntFlanks@6721.6743.23|AATTGCAGGACACATTGATCATC</td><td>>hg38_21_+_8433172_8446622_RNA45SN1_45S_50ntFlanks</td><td>6721</td><td>6743</td><td>23</td><td>AATTGCAGGACACATTGATCATC</td><td>5662.5</td><td>-5.5</td><td>0.8</td><td>-6.6</td><td>4.1E-11</td><td>1.2E-10</td><td>BT20/MB468</td><td>CYTO</td></tr>
<tr><td>>hg38_21_+_8433172_8446622_RNA45SN1_45S_50ntFlanks@4569.4587.19|ATGGAATAGGACCGCGGTT</td><td>>hg38_21_+_8433172_8446622_RNA45SN1_45S_50ntFlanks</td><td>4569</td><td>4587</td><td>19</td><td>ATGGAATAGGACCGCGGTT</td><td>566.4</td><td>-4.8</td><td>0.7</td><td>-6.6</td><td>4.5E-11</td><td>1.2E-10</td><td>MB231/MB468</td><td>NUC</td></tr>
<tr><td>>MI0000093|hsa-mir-92a-1&WithFlank&13|+|91351308|91351397@54.75.22(+1C);MI0000094|hsa-mir-92a-2&WithFlank&X|-|134169532|134169618@54.75.22(+1C)&[MIMAT0000092&hsa-miR-92a-3p&offsets|0|0(+1C);m-9&13|+|91351361|91351382&offsets|0|0(+1C)];[MIMAT0000092_1&hsa-miR-92a-3p&offsets|0|0(+1C);m-11&X|-|134169544|134169565&offsets|0|0(+1C)]|TATTGCACTTGTCCCGGCCTGTC</td><td>>MI0000093|hsa-mir-92a-1&WithFlank&13|+|91351308|91351397</td><td>54</td><td>75</td><td>22(+1C)</td><td>TATTGCACTTGTCCCGGCCTGTC</td><td>635.8</td><td>4.1</td><td>0.6</td><td>6.6</td><td>4.5E-11</td><td>1.2E-10</td><td>MB231/MB468</td><td>NUC</td></tr>
<tr><td>>hg38_21_+_8433172_8446622_RNA45SN1_45S_50ntFlanks@6721.6770.50|AATTGCAGGACACATTGATCATCGACACTTCGAACGCACTTGCGGCCCCG</td><td>>hg38_21_+_8433172_8446622_RNA45SN1_45S_50ntFlanks</td><td>6721</td><td>6770</td><td>50</td><td>AATTGCAGGACACATTGATCATCGACACTTCGAACGCACTTGCGGCCCCG</td><td>458.2</td><td>-12.7</td><td>1.9</td><td>-6.6</td><td>4.2E-11</td><td>1.2E-10</td><td>MB231/MB468</td><td>MP</td></tr>
<tr><td>>hg38_21_+_8433172_8446622_RNA45SN1_45S_50ntFlanks@10834.10862.29|CCGGATCCGTAACTTCGGGATAAGGATTG</td><td>>hg38_21_+_8433172_8446622_RNA45SN1_45S_50ntFlanks</td><td>10834</td><td>10862</td><td>29</td><td>CCGGATCCGTAACTTCGGGATAAGGATTG</td><td>121.1</td><td>-10.8</td><td>1.6</td><td>-6.6</td><td>4.2E-11</td><td>1.2E-10</td><td>MB231/MB468</td><td>MP</td></tr>
<tr><td>>hg38_21_+_8433172_8446622_RNA45SN1_45S_50ntFlanks@12457.12485.29|CAAGCGTTGGATTGTTCACCCACTAATAG</td><td>>hg38_21_+_8433172_8446622_RNA45SN1_45S_50ntFlanks</td><td>12457</td><td>12485</td><td>29</td><td>CAAGCGTTGGATTGTTCACCCACTAATAG</td><td>97.7</td><td>-10.5</td><td>1.6</td><td>-6.6</td><td>4.2E-11</td><td>1.2E-10</td><td>MB231/MB468</td><td>MP</td></tr>
<tr><td>>hg38_MT_+_1621_3279_RNR2_16S_50ntFlanks@51.76.26|GCTAAACCTAGCCCCAAACCCACTCC</td><td>>hg38_MT_+_1621_3279_RNR2_16S_50ntFlanks</td><td>51</td><td>76</td><td>26</td><td>GCTAAACCTAGCCCCAAACCCACTCC</td><td>2067.4</td><td>6.4</td><td>1.0</td><td>6.6</td><td>3.3E-11</td><td>1.2E-10</td><td>BT20/MB468</td><td>MITO</td></tr>
<tr><td>>hg38_21_+_8433172_8446622_RNA45SN1_45S_50ntFlanks@4369.4389.21|GTTAAAAAGCTCGTAGTTGGA</td><td>>hg38_21_+_8433172_8446622_RNA45SN1_45S_50ntFlanks</td><td>4369</td><td>4389</td><td>21</td><td>GTTAAAAAGCTCGTAGTTGGA</td><td>133.0</td><td>-10.9</td><td>1.7</td><td>-6.6</td><td>4.2E-11</td><td>1.2E-10</td><td>MB231/MB468</td><td>MP</td></tr>
<tr><td>>hg38_21_+_8433172_8446622_RNA45SN1_45S_50ntFlanks@10682.10709.28|TCCGGCGGCGTCCGGTGAGCTCTCGCTG</td><td>>hg38_21_+_8433172_8446622_RNA45SN1_45S_50ntFlanks</td><td>10682</td><td>10709</td><td>28</td><td>TCCGGCGGCGTCCGGTGAGCTCTCGCTG</td><td>68.3</td><td>-10.0</td><td>1.5</td><td>-6.6</td><td>4.3E-11</td><td>1.2E-10</td><td>MB231/MB468</td><td>MP</td></tr>
<tr><td>>hg38_21_+_8433172_8446622_RNA45SN1_45S_50ntFlanks@10796.10823.28|GGCATGTTGGAACAATGTAGGTAAGGGA</td><td>>hg38_21_+_8433172_8446622_RNA45SN1_45S_50ntFlanks</td><td>10796</td><td>10823</td><td>28</td><td>GGCATGTTGGAACAATGTAGGTAAGGGA</td><td>84.2</td><td>-10.3</td><td>1.6</td><td>-6.6</td><td>4.2E-11</td><td>1.2E-10</td><td>MB231/MB468</td><td>MP</td></tr>
<tr><td>>MI0000089|hsa-mir-31&WithFlank&9|-|21512109|21512191@14.35.22&[MIMAT0000089&hsa-miR-31-5p&offsets|0|+1;m-360&9|-|21512157|21512178&offsets|0|0]|AGGCAAGATGCTGGCATAGCTG</td><td>>MI0000089|hsa-mir-31&WithFlank&9|-|21512109|21512191</td><td>14</td><td>35</td><td>22</td><td>AGGCAAGATGCTGGCATAGCTG</td><td>333.3</td><td>4.2</td><td>0.6</td><td>6.6</td><td>5.5E-11</td><td>1.2E-10</td><td>BT20/MB468</td><td>NUC</td></tr>
<tr><td>>hg38_21_+_8433172_8446622_RNA45SN1_45S_50ntFlanks@6646.6670.25|TCGTACGACTCTTAGCGGTGGATCA</td><td>>hg38_21_+_8433172_8446622_RNA45SN1_45S_50ntFlanks</td><td>6646</td><td>6670</td><td>25</td><td>TCGTACGACTCTTAGCGGTGGATCA</td><td>3515.2</td><td>-2.7</td><td>0.4</td><td>-6.6</td><td>5.5E-11</td><td>1.2E-10</td><td>BT20/MB468</td><td>NUC</td></tr>
<tr><td>>MI0000088|hsa-mir-30a&WithFlank&6|-|71403545|71403627@53.74.22(+1A)&[MIMAT0000088&hsa-miR-30a-3p&offsets|0|0(+1A);m-38&6|-|71403554|71403575&offsets|0|0(+1A)]|CTTTCAGTCGGATGTTTGCAGCA</td><td>>MI0000088|hsa-mir-30a&WithFlank&6|-|71403545|71403627</td><td>53</td><td>74</td><td>22(+1A)</td><td>CTTTCAGTCGGATGTTTGCAGCA</td><td>562.7</td><td>6.8</td><td>1.0</td><td>6.6</td><td>4.1E-11</td><td>1.2E-10</td><td>BT20/MB468</td><td>CYTO</td></tr>
<tr><td>>MI0000064|hsa-let-7c&WithFlank&21|+|16539822|16539917@17.38.22&[MIMAT0000064&hsa-let-7c-5p&offsets|0|0;m-37&21|+|16539838|16539859&offsets|0|0]|TGAGGTAGTAGGTTGTATGGTT</td><td>>MI0000064|hsa-let-7c&WithFlank&21|+|16539822|16539917</td><td>17</td><td>38</td><td>22</td><td>TGAGGTAGTAGGTTGTATGGTT</td><td>1099.8</td><td>2.2</td><td>0.3</td><td>6.6</td><td>4.1E-11</td><td>1.2E-10</td><td>BT20/MB468</td><td>CYTO</td></tr>
<tr><td>>trna10_LysCTT_16_+_3241501_3241573@6.35.30,trna13_LysCTT_14_-_58706613_58706685@6.35.30,trna2_LysCTT_15_+_79152904_79152976@6.35.30,trna4_LysCTT_18_-_43669271_43669343@6.35.30|GCTAGCTCAGTCGGTAGAGCATGGGACTCT</td><td>>trna10_LysCTT_16_+_3241501_3241573</td><td>6</td><td>35</td><td>30</td><td>GCTAGCTCAGTCGGTAGAGCATGGGACTCT</td><td>245.6</td><td>11.2</td><td>1.7</td><td>6.6</td><td>5.3E-11</td><td>1.2E-10</td><td>MB231/MB468</td><td>MITO</td></tr>
<tr><td>>hg38_21_+_8433172_8446622_RNA45SN1_45S_50ntFlanks@4564.4587.24|GAATAATGGAATAGGACCGCGGTT</td><td>>hg38_21_+_8433172_8446622_RNA45SN1_45S_50ntFlanks</td><td>4564</td><td>4587</td><td>24</td><td>GAATAATGGAATAGGACCGCGGTT</td><td>405.2</td><td>-8.4</td><td>1.3</td><td>-6.6</td><td>4.5E-11</td><td>1.2E-10</td><td>MB231/MB468</td><td>NUC</td></tr>
<tr><td>>hg38_21_+_8433172_8446622_RNA45SN1_45S_50ntFlanks@4472.4493.22|CTTAGCTGAGTGTCCCGCGGGG</td><td>>hg38_21_+_8433172_8446622_RNA45SN1_45S_50ntFlanks</td><td>4472</td><td>4493</td><td>22</td><td>CTTAGCTGAGTGTCCCGCGGGG</td><td>374.9</td><td>-8.4</td><td>1.3</td><td>-6.6</td><td>5.3E-11</td><td>1.2E-10</td><td>MB231/MB468</td><td>MITO</td></tr>
<tr><td>>hg38_21_+_8433172_8446622_RNA45SN1_45S_50ntFlanks@6663.6689.27|GTGGATCACTCGGCTCGTGCGTCGATG</td><td>>hg38_21_+_8433172_8446622_RNA45SN1_45S_50ntFlanks</td><td>6663</td><td>6689</td><td>27</td><td>GTGGATCACTCGGCTCGTGCGTCGATG</td><td>781.3</td><td>-4.3</td><td>0.7</td><td>-6.6</td><td>4.6E-11</td><td>1.2E-10</td><td>MB231/MB468</td><td>CYTO</td></tr>
<tr><td>>hg38_21_+_8433172_8446622_RNA45SN1_45S_50ntFlanks@10535.10562.28|CGCGAGGCGTCCAGTGCGGTAACGCGAC</td><td>>hg38_21_+_8433172_8446622_RNA45SN1_45S_50ntFlanks</td><td>10535</td><td>10562</td><td>28</td><td>CGCGAGGCGTCCAGTGCGGTAACGCGAC</td><td>122.9</td><td>-10.8</td><td>1.6</td><td>-6.6</td><td>4.3E-11</td><td>1.2E-10</td><td>MB231/MB468</td><td>MP</td></tr>
<tr><td>>hg38_21_+_8433172_8446622_RNA45SN1_45S_50ntFlanks@11586.11623.38|GTGCGGACCAGGGGAATCCGACTGTTTAATTAAAACAA</td><td>>hg38_21_+_8433172_8446622_RNA45SN1_45S_50ntFlanks</td><td>11586</td><td>11623</td><td>38</td><td>GTGCGGACCAGGGGAATCCGACTGTTTAATTAAAACAA</td><td>730.7</td><td>6.4</td><td>1.0</td><td>6.6</td><td>3.3E-11</td><td>1.2E-10</td><td>BT20/MB468</td><td>MITO</td></tr>
<tr><td>>MI0000100|hsa-mir-98&WithFlank&X|-|53556217|53556347@28.48.21(+1C)&[MIMAT0000096&hsa-miR-98-5p&offsets|0|-1(+1C);m-143&X|-|53556299|53556320&offsets|0|-1(+1C)]|TGAGGTAGTAAGTTGTATTGTC</td><td>>MI0000100|hsa-mir-98&WithFlank&X|-|53556217|53556347</td><td>28</td><td>48</td><td>21(+1C)</td><td>TGAGGTAGTAAGTTGTATTGTC</td><td>422.2</td><td>3.1</td><td>0.5</td><td>6.6</td><td>4.6E-11</td><td>1.2E-10</td><td>MB231/MB468</td><td>NUC</td></tr>
<tr><td>>hg38_21_+_8433172_8446622_RNA45SN1_45S_50ntFlanks@11805.11823.19|CCACTGTCCCTACCTACTA</td><td>>hg38_21_+_8433172_8446622_RNA45SN1_45S_50ntFlanks</td><td>11805</td><td>11823</td><td>19</td><td>CCACTGTCCCTACCTACTA</td><td>301.6</td><td>-8.0</td><td>1.2</td><td>-6.6</td><td>4.6E-11</td><td>1.2E-10</td><td>MB231/MB468</td><td>NUC</td></tr>
<tr><td>>hg38_21_+_8433172_8446622_RNA45SN1_45S_50ntFlanks@11808.11828.21|CTGTCCCTACCTACTATCCAG</td><td>>hg38_21_+_8433172_8446622_RNA45SN1_45S_50ntFlanks</td><td>11808</td><td>11828</td><td>21</td><td>CTGTCCCTACCTACTATCCAG</td><td>131.4</td><td>-10.9</td><td>1.7</td><td>-6.6</td><td>4.3E-11</td><td>1.2E-10</td><td>MB231/MB468</td><td>MP</td></tr>
<tr><td>>MI0000061|hsa-let-7a-2&WithFlank&11|-|122146516|122146599@11.30.20;MI0000062|hsa-let-7a-3&WithFlank&22|+|46112743|46112828@10.29.20;MI0000060|hsa-let-7a-1&WithFlank&9|+|94175951|94176042@12.31.20&[MIMAT0000062&hsa-let-7a-5p&offsets|0|-2;m-6&11|-|122146568|122146589&offsets|0|-2];[MIMAT0000062_1&hsa-let-7a-5p&offsets|0|-2;m-5&22|+|46112752|46112773&offsets|0|-2];[MIMAT0000062_2&hsa-let-7a-5p&offsets|0|-2;m-4&9|+|94175962|94175983&offsets|0|-2]|TGAGGTAGTAGGTTGTATAG</td><td>>MI0000061|hsa-let-7a-2&WithFlank&11|-|122146516|122146599</td><td>11</td><td>30</td><td>20</td><td>TGAGGTAGTAGGTTGTATAG</td><td>293.8</td><td>7.7</td><td>1.2</td><td>6.6</td><td>4.6E-11</td><td>1.2E-10</td><td>MB231/MB468</td><td>CYTO</td></tr>
<tr><td>>trna10_AspGTC_12_-_125424193_125424264@37.75.39,trna12_AspGTC_12_-_125411891_125411962@37.75.39,trna144_AspGTC_6_-_27551236_27551307@37.75.39,trna38_AspGTC_17_-_8125556_8125627@37.75.39,trna45_AspGTC_6_+_27447453_27447524@37.75.39,trna48_AspGTC_6_+_27471523_27471594@37.75.39,trna4_AspGTC_12_+_96429799_96429870@37.75.39,trna69_AspGTC_1_-_161440205_161440276@37.75.39,trna72_AspGTC_1_-_161432824_161432895@37.75.39,trna75_AspGTC_1_-_161425414_161425485@37.75.39,trna78_AspGTC_1_-_161418033_161418104@37.75.39,trna81_AspGTC_1_-_161410615_161410686@37.75.39|ACGCGGGAGACCGGGGTTCGATTCCCCGACGGGGAGCCA</td><td>>trna10_AspGTC_12_-_125424193_125424264</td><td>37</td><td>75</td><td>39</td><td>ACGCGGGAGACCGGGGTTCGATTCCCCGACGGGGAGCCA</td><td>36795.4</td><td>8.9</td><td>1.4</td><td>6.6</td><td>4.6E-11</td><td>1.2E-10</td><td>MB231/MB468</td><td>CYTO</td></tr>
<tr><td>>hg38_MT_+_1621_3279_RNR2_16S_50ntFlanks@51.71.21|GCTAAACCTAGCCCCAAACCC</td><td>>hg38_MT_+_1621_3279_RNR2_16S_50ntFlanks</td><td>51</td><td>71</td><td>21</td><td>GCTAAACCTAGCCCCAAACCC</td><td>430.8</td><td>5.3</td><td>0.8</td><td>6.6</td><td>4.6E-11</td><td>1.2E-10</td><td>MB231/MB468</td><td>NUC</td></tr>
<tr><td>>hg38_21_+_8433172_8446622_RNA45SN1_45S_50ntFlanks@12927.12952.26|CCTGGCGCTAAACCATTCGTAGACGA</td><td>>hg38_21_+_8433172_8446622_RNA45SN1_45S_50ntFlanks</td><td>12927</td><td>12952</td><td>26</td><td>CCTGGCGCTAAACCATTCGTAGACGA</td><td>87.7</td><td>-10.3</td><td>1.6</td><td>-6.6</td><td>4.3E-11</td><td>1.2E-10</td><td>MB231/MB468</td><td>MP</td></tr>
<tr><td>>hg38_21_+_8433172_8446622_RNA45SN1_45S_50ntFlanks@10389.10410.22|TGAACAGCAGTTGAACATGGGT</td><td>>hg38_21_+_8433172_8446622_RNA45SN1_45S_50ntFlanks</td><td>10389</td><td>10410</td><td>22</td><td>TGAACAGCAGTTGAACATGGGT</td><td>7921.5</td><td>-11.1</td><td>1.7</td><td>-6.6</td><td>5.6E-11</td><td>1.2E-10</td><td>BT20/MB468</td><td>NUC</td></tr>
<tr><td>>hg38_21_+_8433172_8446622_RNA45SN1_45S_50ntFlanks@10404.10433.30|CATGGGTCAGTCGGTCCTGAGAGATGGGCG</td><td>>hg38_21_+_8433172_8446622_RNA45SN1_45S_50ntFlanks</td><td>10404</td><td>10433</td><td>30</td><td>CATGGGTCAGTCGGTCCTGAGAGATGGGCG</td><td>281.0</td><td>-8.4</td><td>1.3</td><td>-6.6</td><td>4.6E-11</td><td>1.2E-10</td><td>MB231/MB468</td><td>NUC</td></tr>
<tr><td>>hg38_21_+_8433172_8446622_RNA45SN1_45S_50ntFlanks@3771.3794.24|CACGGCCGGTACAGTGAAACTGCG</td><td>>hg38_21_+_8433172_8446622_RNA45SN1_45S_50ntFlanks</td><td>3771</td><td>3794</td><td>24</td><td>CACGGCCGGTACAGTGAAACTGCG</td><td>492.4</td><td>-4.2</td><td>0.6</td><td>-6.6</td><td>4.6E-11</td><td>1.2E-10</td><td>MB231/MB468</td><td>NUC</td></tr>
<tr><td>>hg38_21_+_8433172_8446622_RNA45SN1_45S_50ntFlanks@11655.11687.33|GCGATGTGATTTCTGCCCAGTGCTCTGAATGTC</td><td>>hg38_21_+_8433172_8446622_RNA45SN1_45S_50ntFlanks</td><td>11655</td><td>11687</td><td>33</td><td>GCGATGTGATTTCTGCCCAGTGCTCTGAATGTC</td><td>227.0</td><td>8.1</td><td>1.2</td><td>6.6</td><td>4.6E-11</td><td>1.2E-10</td><td>MB231/MB468</td><td>CYTO</td></tr>
<tr><td>>hg38_21_+_8433172_8446622_RNA45SN1_45S_50ntFlanks@4046.4066.21|CATTCGAACGTCTGCCCTATC</td><td>>hg38_21_+_8433172_8446622_RNA45SN1_45S_50ntFlanks</td><td>4046</td><td>4066</td><td>21</td><td>CATTCGAACGTCTGCCCTATC</td><td>599.3</td><td>-6.6</td><td>1.0</td><td>-6.6</td><td>4.3E-11</td><td>1.2E-10</td><td>MB231/MB468</td><td>MP</td></tr>
</table>

<table>
<tr><td></td><td></td><td></td><td></td><td></td><td></td><td></td><td></td><td></td><td></td><td></td><td></td><td></td><td></td></tr>
<tr><td>>trna10_AspGTC_12_-_125424193_125424264@37.64.28,trna12_AspGTC_12_-_125411891_125411962@37.64.28,trna144_AspGTC_6_-_27551236_27551307@37.64.28,trna38_AspGTC_17_-_8125556_8125627@37.64.28,trna45_AspGTC_6_+_27447453_27447524@37.64.28,trna48_AspGTC_6_+_27471523_27471594@37.64.28,trna4_AspGTC_12_+_96429799_96429870@37.64.28,trna69_AspGTC_1_-_161440205_161440276@37.64.28,trna72_AspGTC_1_-_161432824_161432895@37.64.28,trna75_AspGTC_1_-_161425414_161425485@37.64.28,trna78_AspGTC_1_-_161418033_161418104@37.64.28,trna81_AspGTC_1_-_161410615_161410686@37.64.28|ACGCGGGAGACCGGGGTTCGATTCCCCG</td><td>>trna10_AspGTC_12_-_125424193_125424264</td><td>37</td><td>64</td><td>28</td><td>ACGCGGGAGACCGGGGTTCGATTCCCCG</td><td>159.7</td><td>-11.2</td><td>1.7</td><td>-6.6</td><td>4.3E-11</td><td>1.3E-10</td><td>MB231/MB468</td><td>MP</td></tr>
<tr><td>>hg38_21_+_8433172_8446622_RNA45SN1_45S_50ntFlanks@4981.5000.20|CACGGACAGGATTGACAGAT</td><td>>hg38_21_+_8433172_8446622_RNA45SN1_45S_50ntFlanks</td><td>4981</td><td>5000</td><td>20</td><td>CACGGACAGGATTGACAGAT</td><td>79.5</td><td>-10.2</td><td>1.5</td><td>-6.6</td><td>4.3E-11</td><td>1.3E-10</td><td>MB231/MB468</td><td>MP</td></tr>
<tr><td>>hg38_21_+_8433172_8446622_RNA45SN1_45S_50ntFlanks@11671.11699.29|CCAGTGCTCTGAATGTCAAAGTGAAGAAA</td><td>>hg38_21_+_8433172_8446622_RNA45SN1_45S_50ntFlanks</td><td>11671</td><td>11699</td><td>29</td><td>CCAGTGCTCTGAATGTCAAAGTGAAGAAA</td><td>719.6</td><td>-9.3</td><td>1.4</td><td>-6.6</td><td>5.6E-11</td><td>1.3E-10</td><td>BT20/MB468</td><td>NUC</td></tr>
<tr><td>>MI0000298|hsa-mir-221&WithFlank&X|-|45746151|45746272@71.94.24&[MIMAT0000278&hsa-miR-221-3p&offsets|0|+1;m-91&X|-|45746181|45746202&offsets|0|+2]|AGCTACATTGTCTGCTGGGTTTCA</td><td>>MI0000298|hsa-mir-221&WithFlank&X|-|45746151|45746272</td><td>71</td><td>94</td><td>24</td><td>AGCTACATTGTCTGCTGGGTTTCA</td><td>921.7</td><td>4.7</td><td>0.7</td><td>6.6</td><td>3.4E-11</td><td>1.3E-10</td><td>BT20/MB468</td><td>MITO</td></tr>
<tr><td>>hg38_21_+_8433172_8446622_RNA45SN1_45S_50ntFlanks@12914.12933.20|CGTTCGTGGGGAACCTGGCG</td><td>>hg38_21_+_8433172_8446622_RNA45SN1_45S_50ntFlanks</td><td>12914</td><td>12933</td><td>20</td><td>CGTTCGTGGGGAACCTGGCG</td><td>1554.1</td><td>-9.4</td><td>1.4</td><td>-6.6</td><td>4.4E-11</td><td>1.3E-10</td><td>MB231/MB468</td><td>MP</td></tr>
<tr><td>>hg38_21_+_8433172_8446622_RNA45SN1_45S_50ntFlanks@482.505.24|ATCGATGTGGTGACGTCGTGCTCT</td><td>>hg38_21_+_8433172_8446622_RNA45SN1_45S_50ntFlanks</td><td>482</td><td>505</td><td>24</td><td>ATCGATGTGGTGACGTCGTGCTCT</td><td>404.7</td><td>-11.7</td><td>1.7</td><td>-7.0</td><td>3.2E-12</td><td>1.3E-10</td><td>BT20/MB468</td><td>MP</td></tr>
<tr><td>>hg38_21_+_8433172_8446622_RNA45SN1_45S_50ntFlanks@9534.9554.21|AGGAAACTCTGGTGGAGGTCC</td><td>>hg38_21_+_8433172_8446622_RNA45SN1_45S_50ntFlanks</td><td>9534</td><td>9554</td><td>21</td><td>AGGAAACTCTGGTGGAGGTCC</td><td>489.1</td><td>-12.0</td><td>1.7</td><td>-7.0</td><td>3.2E-12</td><td>1.3E-10</td><td>BT20/MB468</td><td>MP</td></tr>
<tr><td>>hg38_21_+_8433172_8446622_RNA45SN1_45S_50ntFlanks@8381.8406.26|AAGAGGGCGTGAAACCGTTAAGAGGT</td><td>>hg38_21_+_8433172_8446622_RNA45SN1_45S_50ntFlanks</td><td>8381</td><td>8406</td><td>26</td><td>AAGAGGGCGTGAAACCGTTAAGAGGT</td><td>480.3</td><td>-3.8</td><td>0.6</td><td>-6.6</td><td>5.7E-11</td><td>1.3E-10</td><td>BT20/MB468</td><td>NUC</td></tr>
<tr><td>>hg38_21_+_8433172_8446622_RNA45SN1_45S_50ntFlanks@476.494.19|TTCGTGATCGATGTGGTGA</td><td>>hg38_21_+_8433172_8446622_RNA45SN1_45S_50ntFlanks</td><td>476</td><td>494</td><td>19</td><td>TTCGTGATCGATGTGGTGA</td><td>1026.3</td><td>5.3</td><td>0.8</td><td>6.6</td><td>4.7E-11</td><td>1.3E-10</td><td>MB231/MB468</td><td>NUC</td></tr>
<tr><td>>hg38_21_+_8433172_8446622_RNA45SN1_45S_50ntFlanks@9759.9789.31|CAAACTTTAAATGGGTAAGAAGCCCGGCTCG</td><td>>hg38_21_+_8433172_8446622_RNA45SN1_45S_50ntFlanks</td><td>9759</td><td>9789</td><td>31</td><td>CAAACTTTAAATGGGTAAGAAGCCCGGCTCG</td><td>120.1</td><td>-9.0</td><td>1.4</td><td>-6.6</td><td>3.4E-11</td><td>1.3E-10</td><td>BT20/MB468</td><td>MITO</td></tr>
<tr><td>>hg38_21_+_8433172_8446622_RNA45SN1_45S_50ntFlanks@13007.13025.19|TATTGAAAGTCAGCCCTCG</td><td>>hg38_21_+_8433172_8446622_RNA45SN1_45S_50ntFlanks</td><td>13007</td><td>13025</td><td>19</td><td>TATTGAAAGTCAGCCCTCG</td><td>816.2</td><td>-6.9</td><td>1.0</td><td>-6.6</td><td>4.7E-11</td><td>1.3E-10</td><td>MB231/MB468</td><td>NUC</td></tr>
<tr><td>>hg38_21_+_8433172_8446622_RNA45SN1_45S_50ntFlanks@10350.10390.41|ACGAGAACTTTGAAGGCCGAAGTGGAGAAGGGTTCCATGTG</td><td>>hg38_21_+_8433172_8446622_RNA45SN1_45S_50ntFlanks</td><td>10350</td><td>10390</td><td>41</td><td>ACGAGAACTTTGAAGGCCGAAGTGGAGAAGGGTTCCATGTG</td><td>4632.1</td><td>-5.3</td><td>0.8</td><td>-6.6</td><td>4.7E-11</td><td>1.3E-10</td><td>MB231/MB468</td><td>NUC</td></tr>
<tr><td>>MI0000272|hsa-mir-182&WithFlank&7|-|129770377|129770498@29.51.23(+1G)&[MIMAT0000259&hsa-miR-182-5p&offsets|0|-1(+1G);m-22&7|-|129770449|129770470&offsets|0|+1(+1G)]|TTTGGCAATGGTAGAACTCACACG</td><td>>MI0000272|hsa-mir-182&WithFlank&7|-|129770377|129770498</td><td>29</td><td>51</td><td>23(+1G)</td><td>TTTGGCAATGGTAGAACTCACACG</td><td>494.5</td><td>3.7</td><td>0.6</td><td>6.6</td><td>5.7E-11</td><td>1.3E-10</td><td>BT20/MB468</td><td>NUC</td></tr>
<tr><td>>hg38_21_+_8433172_8446622_RNA45SN1_45S_50ntFlanks@9736.9755.20|CCGAAACGATCTCAACCTAT</td><td>>hg38_21_+_8433172_8446622_RNA45SN1_45S_50ntFlanks</td><td>9736</td><td>9755</td><td>20</td><td>CCGAAACGATCTCAACCTAT</td><td>346.2</td><td>-5.4</td><td>0.8</td><td>-6.6</td><td>4.7E-11</td><td>1.3E-10</td><td>MB231/MB468</td><td>CYTO</td></tr>
<tr><td>>hg38_21_+_8433172_8446622_RNA45SN1_45S_50ntFlanks@12151.12179.29|GTGGGGAGTTTGACTGGGGCGGTACACCT</td><td>>hg38_21_+_8433172_8446622_RNA45SN1_45S_50ntFlanks</td><td>12151</td><td>12179</td><td>29</td><td>GTGGGGAGTTTGACTGGGGCGGTACACCT</td><td>151.8</td><td>-9.3</td><td>1.4</td><td>-6.6</td><td>3.4E-11</td><td>1.3E-10</td><td>BT20/MB468</td><td>MITO</td></tr>
<tr><td>>hg38_21_+_8433172_8446622_RNA45SN1_45S_50ntFlanks@12159.12182.24|TTTGACTGGGGCGGTACACCTGTC</td><td>>hg38_21_+_8433172_8446622_RNA45SN1_45S_50ntFlanks</td><td>12159</td><td>12182</td><td>24</td><td>TTTGACTGGGGCGGTACACCTGTC</td><td>5019.3</td><td>-8.9</td><td>1.3</td><td>-7.0</td><td>3.2E-12</td><td>1.3E-10</td><td>BT20/MB468</td><td>MP</td></tr>
<tr><td>>MI0000255|hsa-mir-30d&WithFlank&8|-|134804870|134804951@12.34.23&[MIMAT0000245&hsa-miR-30d-5p&offsets|0|+1;m-21&8|-|134804919|134804940&offsets|0|+1]|TGTAAACATCCCCGACTGGAAGC</td><td>>MI0000255|hsa-mir-30d&WithFlank&8|-|134804870|134804951</td><td>12</td><td>34</td><td>23</td><td>TGTAAACATCCCCGACTGGAAGC</td><td>9162.3</td><td>5.9</td><td>0.9</td><td>6.6</td><td>4.7E-11</td><td>1.3E-10</td><td>MB231/MB468</td><td>CYTO</td></tr>
<tr><td>>MI0000102|hsa-mir-100&WithFlank&11|-|122152223|122152314@22.40.19&[MIMAT0000098&hsa-miR-100-5p&offsets|+3|0;m-27&11|-|122152276|122152296&offsets|+3|+1]|CCGTAGATCCGAACTTGTG</td><td>>MI0000102|hsa-mir-100&WithFlank&11|-|122152223|122152314</td><td>22</td><td>40</td><td>19</td><td>CCGTAGATCCGAACTTGTG</td><td>509.4</td><td>-12.1</td><td>1.7</td><td>-6.9</td><td>5.0E-12</td><td>1.3E-10</td><td>BT20/MB231</td><td>NUC</td></tr>
<tr><td>>hg38_21_+_8433172_8446622_RNA45SN1_45S_50ntFlanks@10796.10822.27|GGCATGTTGGAACAATGTAGGTAAGGG</td><td>>hg38_21_+_8433172_8446622_RNA45SN1_45S_50ntFlanks</td><td>10796</td><td>10822</td><td>27</td><td>GGCATGTTGGAACAATGTAGGTAAGGG</td><td>88.8</td><td>-10.3</td><td>1.6</td><td>-6.6</td><td>4.5E-11</td><td>1.3E-10</td><td>MB231/MB468</td><td>MP</td></tr>
<tr><td>>hg38_21_+_8433172_8446622_RNA45SN1_45S_50ntFlanks@4723.4752.30|CATTAATCAAGAACGAAAGTCGGAGGTTCG</td><td>>hg38_21_+_8433172_8446622_RNA45SN1_45S_50ntFlanks</td><td>4723</td><td>4752</td><td>30</td><td>CATTAATCAAGAACGAAAGTCGGAGGTTCG</td><td>974.7</td><td>-5.0</td><td>0.8</td><td>-6.6</td><td>4.8E-11</td><td>1.3E-10</td><td>MB231/MB468</td><td>CYTO</td></tr>
<tr><td>>hg38_21_+_8433172_8446622_RNA45SN1_45S_50ntFlanks@9534.9573.40|AGGAAACTCTGGTGGAGGTCCGTAGCGGTCCTGACGTGCA</td><td>>hg38_21_+_8433172_8446622_RNA45SN1_45S_50ntFlanks</td><td>9534</td><td>9573</td><td>40</td><td>AGGAAACTCTGGTGGAGGTCCGTAGCGGTCCTGACGTGCA</td><td>7346.4</td><td>-9.3</td><td>1.4</td><td>-6.5</td><td>5.8E-11</td><td>1.3E-10</td><td>BT20/MB468</td><td>NUC</td></tr>
<tr><td>>hg38_21_+_8433172_8446622_RNA45SN1_45S_50ntFlanks@12994.13012.19|CCTCGCTGCGATCTATTGA</td><td>>hg38_21_+_8433172_8446622_RNA45SN1_45S_50ntFlanks</td><td>12994</td><td>13012</td><td>19</td><td>CCTCGCTGCGATCTATTGA</td><td>1294.8</td><td>-9.7</td><td>1.5</td><td>-6.5</td><td>5.8E-11</td><td>1.3E-10</td><td>BT20/MB468</td><td>NUC</td></tr>
<tr><td>>hg38_21_+_8433172_8446622_RNA45SN1_45S_50ntFlanks@4738.4768.31|AAAGTCGGAGGTTCGAAGACGATCAGATACC</td><td>>hg38_21_+_8433172_8446622_RNA45SN1_45S_50ntFlanks</td><td>4738</td><td>4768</td><td>31</td><td>AAAGTCGGAGGTTCGAAGACGATCAGATACC</td><td>1950.7</td><td>9.5</td><td>1.4</td><td>6.6</td><td>4.3E-11</td><td>1.3E-10</td><td>BT20/MB468</td><td>CYTO</td></tr>
<tr><td>>hg38_21_+_8433172_8446622_RNA45SN1_45S_50ntFlanks@8354.8375.22|TTGAAAAGAACTTTGAAGAGAG</td><td>>hg38_21_+_8433172_8446622_RNA45SN1_45S_50ntFlanks</td><td>8354</td><td>8375</td><td>22</td><td>TTGAAAAGAACTTTGAAGAGAG</td><td>788.8</td><td>10.7</td><td>1.6</td><td>6.6</td><td>3.5E-11</td><td>1.3E-10</td><td>BT20/MB468</td><td>MITO</td></tr>
<tr><td>>MI0000477|hsa-mir-146a&WithFlank&5|+|160485346|160485456@26.48.23&[MIMAT0000449&hsa-miR-146a-5p&offsets|-1|0;m-129&5|+|160485372|160485393&offsets|-1|0]|TTGAGAACTGAATTCCATGGGTT</td><td>>MI0000477|hsa-mir-146a&WithFlank&5|+|160485346|160485456</td><td>26</td><td>48</td><td>23</td><td>TTGAGAACTGAATTCCATGGGTT</td><td>567.1</td><td>-12.5</td><td>1.7</td><td>-7.2</td><td>7.2E-13</td><td>1.3E-10</td><td>BT20/MB231</td><td>MP</td></tr>
<tr><td>>MI0000298|hsa-mir-221&WithFlank&X|-|45746151|45746272@71.92.22(+1U)&[MIMAT0000278&hsa-miR-221-3p&offsets|0|-1(+1U);m-91&X|-|45746181|45746202&offsets|0|0(+1U)]|AGCTACATTGTCTGCTGGGTTTT</td><td>>MI0000298|hsa-mir-221&WithFlank&X|-|45746151|45746272</td><td>71</td><td>92</td><td>22(+1U)</td><td>AGCTACATTGTCTGCTGGGTTTT</td><td>3165.5</td><td>6.2</td><td>0.9</td><td>6.6</td><td>4.8E-11</td><td>1.3E-10</td><td>MB231/MB468</td><td>CYTO</td></tr>
<tr><td>>hg38_21_+_8433172_8446622_RNA45SN1_45S_50ntFlanks@12157.12182.26|AGTTTGACTGGGGCGGTACACCTGTC</td><td>>hg38_21_+_8433172_8446622_RNA45SN1_45S_50ntFlanks</td><td>12157</td><td>12182</td><td>26</td><td>AGTTTGACTGGGGCGGTACACCTGTC</td><td>147.0</td><td>-6.9</td><td>1.1</td><td>-6.6</td><td>4.8E-11</td><td>1.3E-10</td><td>MB231/MB468</td><td>CYTO</td></tr>
<tr><td>>MI0000085|hsa-mir-27a&WithFlank&19|-|13836434|13836523@57.76.20&[MIMAT0000084&hsa-miR-27a-3p&offsets|0|-1;m-43&19|-|13836447|13836467&offsets|0|-1]|TTCACAGTGGCTAAGTTCCG</td><td>>MI0000085|hsa-mir-27a&WithFlank&19|-|13836434|13836523</td><td>57</td><td>76</td><td>20</td><td>TTCACAGTGGCTAAGTTCCG</td><td>6827.1</td><td>5.3</td><td>0.8</td><td>6.6</td><td>4.3E-11</td><td>1.3E-10</td><td>BT20/MB468</td><td>CYTO</td></tr>
<tr><td>>hg38_21_+_8433172_8446622_RNA45SN1_45S_50ntFlanks@6650.6683.34|ACGACTCTTAGCGGTGGATCACTCGGCTCGTGCG</td><td>>hg38_21_+_8433172_8446622_RNA45SN1_45S_50ntFlanks</td><td>6650</td><td>6683</td><td>34</td><td>ACGACTCTTAGCGGTGGATCACTCGGCTCGTGCG</td><td>8064.7</td><td>5.9</td><td>0.9</td><td>6.6</td><td>4.8E-11</td><td>1.3E-10</td><td>MB231/MB468</td><td>NUC</td></tr>
<tr><td>>hg38_21_+_8433172_8446622_RNA45SN1_45S_50ntFlanks@10690.10711.22|CGTCCGGTGAGCTCTCGCTGGC</td><td>>hg38_21_+_8433172_8446622_RNA45SN1_45S_50ntFlanks</td><td>10690</td><td>10711</td><td>22</td><td>CGTCCGGTGAGCTCTCGCTGGC</td><td>225.8</td><td>-10.1</td><td>1.5</td><td>-6.6</td><td>4.3E-11</td><td>1.3E-10</td><td>BT20/MB468</td><td>CYTO</td></tr>
<tr><td>>hg38_21_+_8433172_8446622_RNA45SN1_45S_50ntFlanks@3711.3739.29|GTTGATCCTGCCAGTAGCATATGCTTGTC</td><td>>hg38_21_+_8433172_8446622_RNA45SN1_45S_50ntFlanks</td><td>3711</td><td>3739</td><td>29</td><td>GTTGATCCTGCCAGTAGCATATGCTTGTC</td><td>642.3</td><td>-3.6</td><td>0.5</td><td>-6.5</td><td>5.9E-11</td><td>1.3E-10</td><td>BT20/MB468</td><td>NUC</td></tr>
<tr><td>>hg38_21_+_8433172_8446622_RNA45SN1_45S_50ntFlanks@12083.12106.24|CTCGCTTCTGGCGCCAAGCGCCCG</td><td>>hg38_21_+_8433172_8446622_RNA45SN1_45S_50ntFlanks</td><td>12083</td><td>12106</td><td>24</td><td>CTCGCTTCTGGCGCCAAGCGCCCG</td><td>1125.3</td><td>-7.3</td><td>1.1</td><td>-6.6</td><td>5.5E-11</td><td>1.3E-10</td><td>MB231/MB468</td><td>MITO</td></tr>
<tr><td>>trna14_LysTTT_11_-_59327808_59327880@1.30.30,trna2_LysTTT_17_+_8022473_8022545@1.30.30,trna54_LysTTT_1_+_204475655_204475727@1.30.30,trna5_LysTTT_11_+_59323902_59323974@1.30.30,trna62_LysTTT_1_-_204476158_204476230@1.30.30,trna76_LysTTT_6_+_28918806_28918878@1.30.30|GCCCGGATAGCTCAGTCGGTAGAGCATCAG</td><td>>trna14_LysTTT_11_-_59327808_59327880</td><td>1</td><td>30</td><td>30</td><td>GCCCGGATAGCTCAGTCGGTAGAGCATCAG</td><td>1119.3</td><td>-5.5</td><td>0.8</td><td>-6.6</td><td>4.8E-11</td><td>1.3E-10</td><td>MB231/MB468</td><td>CYTO</td></tr>
<tr><td>>trnaMT_ValTAC_MT_+_1602_1670@53.71.19|TAACTTGACCGCTCTGACC</td><td>>trnaMT_ValTAC_MT_+_1602_1670</td><td>53</td><td>71</td><td>19</td><td>TAACTTGACCGCTCTGACC</td><td>1136.5</td><td>5.4</td><td>0.8</td><td>6.6</td><td>3.5E-11</td><td>1.3E-10</td><td>BT20/MB468</td><td>MITO</td></tr>
<tr><td>>hg38_21_+_8433172_8446622_RNA45SN1_45S_50ntFlanks@11570.11591.22|AGCCGACTTAGAACTGGTGCGG</td><td>>hg38_21_+_8433172_8446622_RNA45SN1_45S_50ntFlanks</td><td>11570</td><td>11591</td><td>22</td><td>AGCCGACTTAGAACTGGTGCGG</td><td>582.5</td><td>-6.6</td><td>1.0</td><td>-6.6</td><td>5.6E-11</td><td>1.3E-10</td><td>MB231/MB468</td><td>MITO</td></tr>
<tr><td>>hg38_21_+_8433172_8446622_RNA45SN1_45S_50ntFlanks@8183.8223.41|TCGAGGCCCAGCCCGTGGACGGTGTGAGGCCGGTAGCGGCC</td><td>>hg38_21_+_8433172_8446622_RNA45SN1_45S_50ntFlanks</td><td>8183</td><td>8223</td><td>41</td><td>TCGAGGCCCAGCCCGTGGACGGTGTGAGGCCGGTAGCGGCC</td><td>788.0</td><td>-7.1</td><td>1.1</td><td>-6.6</td><td>4.8E-11</td><td>1.3E-10</td><td>MB231/MB468</td><td>NUC</td></tr>
<tr><td>>hg38_21_+_8433172_8446622_RNA45SN1_45S_50ntFlanks@8382.8399.18|AGAGGGCGTGAAACCGTT</td><td>>hg38_21_+_8433172_8446622_RNA45SN1_45S_50ntFlanks</td><td>8382</td><td>8399</td><td>18</td><td>AGAGGGCGTGAAACCGTT</td><td>125.2</td><td>-10.8</td><td>1.6</td><td>-6.6</td><td>4.5E-11</td><td>1.3E-10</td><td>MB231/MB468</td><td>MP</td></tr>
<tr><td>>hg38_21_+_8433172_8446622_RNA45SN1_45S_50ntFlanks@4444.4472.29|CCTTGCCTCTCGGCGCCCCCTCGATGCTC</td><td>>hg38_21_+_8433172_8446622_RNA45SN1_45S_50ntFlanks</td><td>4444</td><td>4472</td><td>29</td><td>CCTTGCCTCTCGGCGCCCCCTCGATGCTC</td><td>165.9</td><td>-9.5</td><td>1.4</td><td>-6.6</td><td>3.5E-11</td><td>1.3E-10</td><td>BT20/MB468</td><td>MITO</td></tr>
<tr><td>>MI0000093|hsa-mir-92a-1&WithFlank&13|+|91351308|91351397@54.76.23&[MIMAT0000092&hsa-miR-92a-3p&offsets|0|+1;m-9&13|+|91351361|91351382&offsets|0|+1]|TATTGCACTTGTCCCGGCCTGTT</td><td>>MI0000093|hsa-mir-92a-1&WithFlank&13|+|91351308|91351397</td><td>54</td><td>76</td><td>23</td><td>TATTGCACTTGTCCCGGCCTGTT</td><td>6042.0</td><td>4.2</td><td>0.6</td><td>6.6</td><td>4.8E-11</td><td>1.3E-10</td><td>MB231/MB468</td><td>NUC</td></tr>
<tr><td>>MI0000749|hsa-mir-30e&WithFlank&1|+|40754349|40754452@23.46.24&[MIMAT0000692&hsa-miR-30e-5p&offsets|0|+2;m-24&1|+|40754371|40754394&offsets|0|0]|TGTAAACATCCTTGACTGGAAGCT</td><td>>MI0000749|hsa-mir-30e&WithFlank&1|+|40754349|40754452</td><td>23</td><td>46</td><td>24</td><td>TGTAAACATCCTTGACTGGAAGCT</td><td>3965.6</td><td>5.0</td><td>0.8</td><td>6.6</td><td>4.8E-11</td><td>1.3E-10</td><td>MB231/MB468</td><td>NUC</td></tr>
<tr><td>>hg38_21_+_8433172_8446622_RNA45SN1_45S_50ntFlanks@4224.4242.19|AAAATAACAATACAGGACT</td><td>>hg38_21_+_8433172_8446622_RNA45SN1_45S_50ntFlanks</td><td>4224</td><td>4242</td><td>19</td><td>AAAATAACAATACAGGACT</td><td>434.7</td><td>-4.2</td><td>0.6</td><td>-6.6</td><td>3.5E-11</td><td>1.3E-10</td><td>BT20/MB468</td><td>MITO</td></tr>
<tr><td>>hg38_21_+_8433172_8446622_RNA45SN1_45S_50ntFlanks@3785.3812.28|TGAAACTGCGAATGGCTCATTAAATCAG</td><td>>hg38_21_+_8433172_8446622_RNA45SN1_45S_50ntFlanks</td><td>3785</td><td>3812</td><td>28</td><td>TGAAACTGCGAATGGCTCATTAAATCAG</td><td>370.2</td><td>-12.4</td><td>1.9</td><td>-6.6</td><td>4.5E-11</td><td>1.3E-10</td><td>MB231/MB468</td><td>MP</td></tr>
<tr><td>>MI0001445|hsa-mir-423&WithFlank&17|+|30117073|30117178@59.81.23&[MIMAT0001340&hsa-miR-423-3p&offsets|0|0;m-100&17|+|30117131|30117153&offsets|0|0]|AGCTCGGTCTGAGGCCCCTCAGT</td><td>>MI0001445|hsa-mir-423&WithFlank&17|+|30117073|30117178</td><td>59</td><td>81</td><td>23</td><td>AGCTCGGTCTGAGGCCCCTCAGT</td><td>13839.1</td><td>5.5</td><td>0.8</td><td>6.6</td><td>4.9E-11</td><td>1.3E-10</td><td>MB231/MB468</td><td>NUC</td></tr>
<tr><td>>MI0000469|hsa-mir-125a&WithFlank&19|+|51693248|51693345@21.44.24(+1A)&[MIMAT0000443&hsa-miR-125a-5p&offsets|0|0(+1A);m-63&19|+|51693268|51693289&offsets|0|+2(+1A)]|TCCCTGAGACCCTTTAACCTGTGAA</td><td>>MI0000469|hsa-mir-125a&WithFlank&19|+|51693248|51693345</td><td>21</td><td>44</td><td>24(+1A)</td><td>TCCCTGAGACCCTTTAACCTGTGAA</td><td>357.6</td><td>8.2</td><td>1.2</td><td>6.6</td><td>4.4E-11</td><td>1.3E-10</td><td>BT20/MB468</td><td>CYTO</td></tr>
<tr><td>>hg38_21_+_8433172_8446622_RNA45SN1_45S_50ntFlanks@12709.12730.22|CCGCGGAGCCTCGGTTGGCCTC</td><td>>hg38_21_+_8433172_8446622_RNA45SN1_45S_50ntFlanks</td><td>12709</td><td>12730</td><td>22</td><td>CCGCGGAGCCTCGGTTGGCCTC</td><td>175.7</td><td>-11.3</td><td>1.7</td><td>-6.6</td><td>4.6E-11</td><td>1.3E-10</td><td>MB231/MB468</td><td>MP</td></tr>
<tr><td>>hg38_21_+_8433172_8446622_RNA45SN1_45S_50ntFlanks@12456.12480.25|CCAAGCGTTGGATTGTTCACCCACT</td><td>>hg38_21_+_8433172_8446622_RNA45SN1_45S_50ntFlanks</td><td>12456</td><td>12480</td><td>25</td><td>CCAAGCGTTGGATTGTTCACCCACT</td><td>761.0</td><td>-3.7</td><td>0.6</td><td>-6.6</td><td>3.6E-11</td><td>1.3E-10</td><td>BT20/MB468</td><td>MITO</td></tr>
</table>

<table>
<tr><th></th><th></th><th></th><th></th><th></th><th></th><th></th><th></th><th></th><th></th><th></th><th></th><th></th><th></th></tr>
<tr><td>>MI0000088|hsa-mir-30a&WithFlank&6|-|71403545|71403627@12.34.23(+1C)&[MIMAT0000087&hsa-miR-30a-5p&offsets|0|+1(+1C);m-15&6|-|71403595|71403616&offsets|0|+1(+1C)]|TGTAAACATCCTCGACTGGAAGCC</td><td>>MI0000088|hsa-mir-30a&WithFlank&6|-|71403545|71403627</td><td>12</td><td>34</td><td>23(+1C)</td><td>TGTAAACATCCTCGACTGGAAGCC</td><td>3316.7</td><td>5.9</td><td>0.9</td><td>6.6</td><td>4.9E-11</td><td>1.3E-10</td><td>MB231/MB468</td><td>CYTO</td></tr>
<tr><td>>hg38_21_+_8433172_8446622_RNA45SN1_45S_50ntFlanks@4753.4780.28|AAGACGATCAGATACCGTCGTAGTTCCG</td><td>>hg38_21_+_8433172_8446622_RNA45SN1_45S_50ntFlanks</td><td>4753</td><td>4780</td><td>28</td><td>AAGACGATCAGATACCGTCGTAGTTCCG</td><td>93.2</td><td>-10.4</td><td>1.6</td><td>-6.6</td><td>4.6E-11</td><td>1.3E-10</td><td>MB231/MB468</td><td>MP</td></tr>
<tr><td>>hg38_21_+_8433172_8446622_RNA45SN1_45S_50ntFlanks@4760.4786.27|TCAGATACCGTCGTAGTTCCGACCATA</td><td>>hg38_21_+_8433172_8446622_RNA45SN1_45S_50ntFlanks</td><td>4760</td><td>4786</td><td>27</td><td>TCAGATACCGTCGTAGTTCCGACCATA</td><td>736.3</td><td>-8.6</td><td>1.3</td><td>-6.6</td><td>4.6E-11</td><td>1.3E-10</td><td>MB231/MB468</td><td>MP</td></tr>
<tr><td>>MI0000816|hsa-mir-335&WithFlank&7|+|130496105|130496210@58.78.21(+1A)&[MIMAT0004703&hsa-miR-335-3p&offsets|0|-1(+1A);m-159&7|+|130496162|130496183&offsets|0|-1(+1A)]|TTTTTCATTATTGCTCCTGACA</td><td>>MI0000816|hsa-mir-335&WithFlank&7|+|130496105|130496210</td><td>58</td><td>78</td><td>21(+1A)</td><td>TTTTTCATTATTGCTCCTGACA</td><td>620.5</td><td>7.4</td><td>1.1</td><td>6.6</td><td>4.9E-11</td><td>1.3E-10</td><td>MB231/MB468</td><td>CYTO</td></tr>
<tr><td>>hg38_21_+_8433172_8446622_RNA45SN1_45S_50ntFlanks@11536.11553.18|TCCGCGCGGCGCCTCGCC</td><td>>hg38_21_+_8433172_8446622_RNA45SN1_45S_50ntFlanks</td><td>11536</td><td>11553</td><td>18</td><td>TCCGCGCGGCGCCTCGCC</td><td>194.1</td><td>-5.4</td><td>0.8</td><td>-6.6</td><td>4.9E-11</td><td>1.3E-10</td><td>MB231/MB468</td><td>CYTO</td></tr>
<tr><td>>trnaMT_LeuTAG_MT_+_12266_12336@43.74.32|AAATTTTGGTGCAACTCCAAATAAAAGTACCA</td><td>>trnaMT_LeuTAG_MT_+_12266_12336</td><td>43</td><td>74</td><td>32</td><td>AAATTTTGGTGCAACTCCAAATAAAAGTACCA</td><td>238.6</td><td>-10.0</td><td>1.5</td><td>-6.6</td><td>3.6E-11</td><td>1.3E-10</td><td>BT20/MB468</td><td>MITO</td></tr>
<tr><td>>hg38_21_+_8433172_8446622_RNA45SN1_45S_50ntFlanks@4661.4686.26|AGGTGAAATTCTTGGACCGGCGCAAG</td><td>>hg38_21_+_8433172_8446622_RNA45SN1_45S_50ntFlanks</td><td>4661</td><td>4686</td><td>26</td><td>AGGTGAAATTCTTGGACCGGCGCAAG</td><td>140.1</td><td>-11.0</td><td>1.7</td><td>-6.6</td><td>4.6E-11</td><td>1.3E-10</td><td>MB231/MB468</td><td>MP</td></tr>
<tr><td>>hg38_21_+_8433172_8446622_RNA45SN1_45S_50ntFlanks@10761.10785.25|ATATCCGCAGCAGGTCTCCAAGGTG</td><td>>hg38_21_+_8433172_8446622_RNA45SN1_45S_50ntFlanks</td><td>10761</td><td>10785</td><td>25</td><td>ATATCCGCAGCAGGTCTCCAAGGTG</td><td>128.1</td><td>-9.1</td><td>1.4</td><td>-6.6</td><td>3.6E-11</td><td>1.3E-10</td><td>BT20/MB468</td><td>MITO</td></tr>
<tr><td>>hg38_21_+_8433172_8446622_RNA45SN1_45S_50ntFlanks@12274.12307.34|CAGTACGAATACAGACCGTGAAAGCGGGGCCTCA</td><td>>hg38_21_+_8433172_8446622_RNA45SN1_45S_50ntFlanks</td><td>12274</td><td>12307</td><td>34</td><td>CAGTACGAATACAGACCGTGAAAGCGGGGCCTCA</td><td>196.4</td><td>-11.4</td><td>1.7</td><td>-6.6</td><td>5.6E-11</td><td>1.3E-10</td><td>MB231/MB468</td><td>MITO</td></tr>
<tr><td>>hg38_21_+_8433172_8446622_RNA45SN1_45S_50ntFlanks@10393.10435.43|CAGCAGTTGAACATGGGTCAGTCGGTCCTGAGAGATGGGCGAG</td><td>>hg38_21_+_8433172_8446622_RNA45SN1_45S_50ntFlanks</td><td>10393</td><td>10435</td><td>43</td><td>CAGCAGTTGAACATGGGTCAGTCGGTCCTGAGAGATGGGCGAG</td><td>718.4</td><td>-9.7</td><td>1.5</td><td>-6.6</td><td>4.9E-11</td><td>1.3E-10</td><td>MB231/MB468</td><td>NUC</td></tr>
<tr><td>>hg38_21_+_8433172_8446622_RNA45SN1_45S_50ntFlanks@5311.5330.20|ATCGGGGATTGCAATTATTC</td><td>>hg38_21_+_8433172_8446622_RNA45SN1_45S_50ntFlanks</td><td>5311</td><td>5330</td><td>20</td><td>ATCGGGGATTGCAATTATTC</td><td>154.1</td><td>-11.1</td><td>1.7</td><td>-6.6</td><td>4.6E-11</td><td>1.3E-10</td><td>MB231/MB468</td><td>MP</td></tr>
<tr><td>>hg38_21_+_8433172_8446622_RNA45SN1_45S_50ntFlanks@10248.10273.26|CGACGAGTAGGAGGGCCGCTGCGGTG</td><td>>hg38_21_+_8433172_8446622_RNA45SN1_45S_50ntFlanks</td><td>10248</td><td>10273</td><td>26</td><td>CGACGAGTAGGAGGGCCGCTGCGGTG</td><td>128.3</td><td>-10.9</td><td>1.7</td><td>-6.6</td><td>4.6E-11</td><td>1.3E-10</td><td>MB231/MB468</td><td>MP</td></tr>
<tr><td>>MI0000283|hsa-mir-203a&WithFlank&14|+|104117399|104117520@72.92.21(+1U)&[MIMAT0000264&hsa-miR-203a-3p&offsets|+1|0(+1U);m-79&14|+|104117469|104117490&offsets|+1|0(+1U)]|TGAAATGTTTAGGACCACTAGT</td><td>>MI0000283|hsa-mir-203a&WithFlank&14|+|104117399|104117520</td><td>72</td><td>92</td><td>21(+1U)</td><td>TGAAATGTTTAGGACCACTAGT</td><td>769.8</td><td>8.8</td><td>1.3</td><td>6.9</td><td>5.2E-12</td><td>1.3E-10</td><td>BT20/MB231</td><td>NUC</td></tr>
<tr><td>>MI0000446|hsa-mir-125b-1&WithFlank&11|-|122099751|122099850@61.81.21&[MIMAT0004592&hsa-miR-125b-1-3p&offsets|0|-1;m-485&11|-|122099769|122099790&offsets|0|-1]|ACGGGTTAGGCTCTTGGGAGC</td><td>>MI0000446|hsa-mir-125b-1&WithFlank&11|-|122099751|122099850</td><td>61</td><td>81</td><td>21</td><td>ACGGGTTAGGCTCTTGGGAGC</td><td>155.3</td><td>-10.4</td><td>1.5</td><td>-6.9</td><td>5.3E-12</td><td>1.3E-10</td><td>BT20/MB231</td><td>NUC</td></tr>
<tr><td>>hg38_21_+_8433172_8446622_RNA45SN1_45S_50ntFlanks@9624.9650.27|AGTAGCTGGTTCCCTCCGAAGTTTCCC</td><td>>hg38_21_+_8433172_8446622_RNA45SN1_45S_50ntFlanks</td><td>9624</td><td>9650</td><td>27</td><td>AGTAGCTGGTTCCCTCCGAAGTTTCCC</td><td>93.0</td><td>-10.4</td><td>1.6</td><td>-6.6</td><td>4.6E-11</td><td>1.3E-10</td><td>MB231/MB468</td><td>MP</td></tr>
<tr><td>>hg38_21_+_8433172_8446622_RNA45SN1_45S_50ntFlanks@5035.5058.24|CATGGCCGTTCTTAGTTGGTGGAG</td><td>>hg38_21_+_8433172_8446622_RNA45SN1_45S_50ntFlanks</td><td>5035</td><td>5058</td><td>24</td><td>CATGGCCGTTCTTAGTTGGTGGAG</td><td>490.5</td><td>-12.7</td><td>1.9</td><td>-6.6</td><td>5.6E-11</td><td>1.3E-10</td><td>MB231/MB468</td><td>MITO</td></tr>
<tr><td>>hg38_21_+_8433172_8446622_RNA45SN1_45S_50ntFlanks@8198.8223.26|TGGACGGTGTGAGGCCGGTAGCGGCC</td><td>>hg38_21_+_8433172_8446622_RNA45SN1_45S_50ntFlanks</td><td>8198</td><td>8223</td><td>26</td><td>TGGACGGTGTGAGGCCGGTAGCGGCC</td><td>1632.3</td><td>-9.9</td><td>1.5</td><td>-6.5</td><td>6.0E-11</td><td>1.3E-10</td><td>BT20/MB468</td><td>NUC</td></tr>
<tr><td>>hg38_21_+_8433172_8446622_RNA45SN1_45S_50ntFlanks@12633.12652.20|CGAAGCTACCATCTGTGGGA</td><td>>hg38_21_+_8433172_8446622_RNA45SN1_45S_50ntFlanks</td><td>12633</td><td>12652</td><td>20</td><td>CGAAGCTACCATCTGTGGGA</td><td>433.8</td><td>-6.0</td><td>0.9</td><td>-6.6</td><td>4.9E-11</td><td>1.3E-10</td><td>MB231/MB468</td><td>CYTO</td></tr>
<tr><td>>hg38_21_+_8433172_8446622_RNA45SN1_45S_50ntFlanks@12941.12960.20|ATTCGTAGACGACCTGCTTC</td><td>>hg38_21_+_8433172_8446622_RNA45SN1_45S_50ntFlanks</td><td>12941</td><td>12960</td><td>20</td><td>ATTCGTAGACGACCTGCTTC</td><td>815.0</td><td>3.5</td><td>0.5</td><td>6.6</td><td>4.5E-11</td><td>1.3E-10</td><td>BT20/MB468</td><td>CYTO</td></tr>
<tr><td>>hg38_21_+_8433172_8446622_RNA45SN1_45S_50ntFlanks@10346.10372.27|TCAAACGAGAACTTTGAAGGCCGAAGT</td><td>>hg38_21_+_8433172_8446622_RNA45SN1_45S_50ntFlanks</td><td>10346</td><td>10372</td><td>27</td><td>TCAAACGAGAACTTTGAAGGCCGAAGT</td><td>775.7</td><td>-5.3</td><td>0.8</td><td>-6.6</td><td>4.6E-11</td><td>1.3E-10</td><td>MB231/MB468</td><td>MP</td></tr>
<tr><td>>hg38_21_+_8433172_8446622_RNA45SN1_45S_50ntFlanks@12927.12973.47|CCTGGCGCTAAACCATTCGTAGACGACCTGCTTCTGGGTCGGGGTTT</td><td>>hg38_21_+_8433172_8446622_RNA45SN1_45S_50ntFlanks</td><td>12927</td><td>12973</td><td>47</td><td>CCTGGCGCTAAACCATTCGTAGACGACCTGCTTCTGGGTCGGGGTTT</td><td>274.5</td><td>-11.1</td><td>1.7</td><td>-6.6</td><td>5.0E-11</td><td>1.3E-10</td><td>MB231/MB468</td><td>NUC</td></tr>
<tr><td>>hg38_21_+_8433172_8446622_RNA45SN1_45S_50ntFlanks@6721.6754.34|AATTGCAGGACACATTGATCATCGACACTTCGAA</td><td>>hg38_21_+_8433172_8446622_RNA45SN1_45S_50ntFlanks</td><td>6721</td><td>6754</td><td>34</td><td>AATTGCAGGACACATTGATCATCGACACTTCGAA</td><td>146.4</td><td>-11.1</td><td>1.7</td><td>-6.6</td><td>4.7E-11</td><td>1.3E-10</td><td>MB231/MB468</td><td>MP</td></tr>
<tr><td>>hg38_21_+_8433172_8446622_RNA45SN1_45S_50ntFlanks@6746.6770.25|CACTTCGAACGCACTTGCGGCCCCG</td><td>>hg38_21_+_8433172_8446622_RNA45SN1_45S_50ntFlanks</td><td>6746</td><td>6770</td><td>25</td><td>CACTTCGAACGCACTTGCGGCCCCG</td><td>1364.5</td><td>-6.7</td><td>1.0</td><td>-6.6</td><td>3.6E-11</td><td>1.3E-10</td><td>BT20/MB468</td><td>MITO</td></tr>
<tr><td>>hg38_21_+_8433172_8446622_RNA45SN1_45S_50ntFlanks@9352.9380.29|CCGTGGCGCAATGAAGGTGAAGGCCGGCG</td><td>>hg38_21_+_8433172_8446622_RNA45SN1_45S_50ntFlanks</td><td>9352</td><td>9380</td><td>29</td><td>CCGTGGCGCAATGAAGGTGAAGGCCGGCG</td><td>180.3</td><td>-9.6</td><td>1.4</td><td>-6.6</td><td>3.6E-11</td><td>1.3E-10</td><td>BT20/MB468</td><td>MITO</td></tr>
<tr><td>>hg38_21_+_8433172_8446622_RNA45SN1_45S_50ntFlanks@9309.9336.28|GGAGTCTAACACGTGCGCGAGTCGGGGG</td><td>>hg38_21_+_8433172_8446622_RNA45SN1_45S_50ntFlanks</td><td>9309</td><td>9336</td><td>28</td><td>GGAGTCTAACACGTGCGCGAGTCGGGGG</td><td>117.6</td><td>-10.7</td><td>1.6</td><td>-6.6</td><td>4.7E-11</td><td>1.3E-10</td><td>MB231/MB468</td><td>MP</td></tr>
<tr><td>>hg38_21_+_8433172_8446622_RNA45SN1_45S_50ntFlanks@5035.5072.38|CATGGCCGTTCTTAGTTGGTGGAGCGATTTGTCTGGTT</td><td>>hg38_21_+_8433172_8446622_RNA45SN1_45S_50ntFlanks</td><td>5035</td><td>5072</td><td>38</td><td>CATGGCCGTTCTTAGTTGGTGGAGCGATTTGTCTGGTT</td><td>418.7</td><td>-10.8</td><td>1.6</td><td>-6.6</td><td>3.7E-11</td><td>1.3E-10</td><td>BT20/MB468</td><td>MITO</td></tr>
<tr><td>>trna119_LysCTT_1_-_145395522_145395594@1.34.34,trna11_LysCTT_5_-_180648979_180649051@1.34.34,trna13_LysCTT_6_+_26556774_26556846@1.34.34,trna7_LysCTT_16_+_3225692_3225764@1.34.34,trna9_LysCTT_5_+_180634755_180634827@1.34.34|GCCCGGCTAGCTCAGTCGGTAGAGCATGAGACTC</td><td>>trna119_LysCTT_1_-_145395522_145395594</td><td>1</td><td>34</td><td>34</td><td>GCCCGGCTAGCTCAGTCGGTAGAGCATGAGACTC</td><td>6558.8</td><td>7.6</td><td>1.2</td><td>6.6</td><td>5.0E-11</td><td>1.3E-10</td><td>MB231/MB468</td><td>NUC</td></tr>
<tr><td>>trnaMT_ValTAC_MT_+_1602_1670@55.72.18|ACTTGACCGCTCTGACCA</td><td>>trnaMT_ValTAC_MT_+_1602_1670</td><td>55</td><td>72</td><td>18</td><td>ACTTGACCGCTCTGACCA</td><td>531.0</td><td>5.5</td><td>0.8</td><td>6.6</td><td>5.0E-11</td><td>1.3E-10</td><td>MB231/MB468</td><td>CYTO</td></tr>
<tr><td>>hg38_21_+_8433172_8446622_RNA45SN1_45S_50ntFlanks@4731.4762.32|AAGAACGAAAGTCGGAGGTTCGAAGACGATCA</td><td>>hg38_21_+_8433172_8446622_RNA45SN1_45S_50ntFlanks</td><td>4731</td><td>4762</td><td>32</td><td>AAGAACGAAAGTCGGAGGTTCGAAGACGATCA</td><td>1240.9</td><td>9.5</td><td>1.4</td><td>6.6</td><td>4.5E-11</td><td>1.3E-10</td><td>BT20/MB468</td><td>CYTO</td></tr>
<tr><td>>hg38_1_-_228634819_228635039_RNA5S12_5S_50ntFlanks@147.169.23|GGGAATACCGGGTGCTGTAGGCT</td><td>>hg38_1_-_228634819_228635039_RNA5S12_5S_50ntFlanks</td><td>147</td><td>169</td><td>23</td><td>GGGAATACCGGGTGCTGTAGGCT</td><td>1010.9</td><td>4.1</td><td>0.6</td><td>6.6</td><td>3.7E-11</td><td>1.3E-10</td><td>BT20/MB468</td><td>MITO</td></tr>
<tr><td>>hg38_21_+_8433172_8446622_RNA45SN1_45S_50ntFlanks@8185.8210.26|GAGGCCCAGCCCGTGGACGGTGTGAG</td><td>>hg38_21_+_8433172_8446622_RNA45SN1_45S_50ntFlanks</td><td>8185</td><td>8210</td><td>26</td><td>GAGGCCCAGCCCGTGGACGGTGTGAG</td><td>336.0</td><td>7.0</td><td>1.1</td><td>6.6</td><td>4.5E-11</td><td>1.3E-10</td><td>BT20/MB468</td><td>CYTO</td></tr>
<tr><td>>am_trnaMT_PheGAA_MT_+_577_647@40.58.19|TGTTTAGACGGGCTCACAT</td><td>>am_trnaMT_PheGAA_MT_+_577_647</td><td>40</td><td>58</td><td>19</td><td>TGTTTAGACGGGCTCACAT</td><td>230.4</td><td>6.6</td><td>1.0</td><td>6.6</td><td>5.0E-11</td><td>1.4E-10</td><td>MB231/MB468</td><td>NUC</td></tr>
<tr><td>>hg38_21_+_8433172_8446622_RNA45SN1_45S_50ntFlanks@10471.10488.18|CCCTCGGCCGATCGAAAG</td><td>>hg38_21_+_8433172_8446622_RNA45SN1_45S_50ntFlanks</td><td>10471</td><td>10488</td><td>18</td><td>CCCTCGGCCGATCGAAAG</td><td>168.1</td><td>-11.3</td><td>1.7</td><td>-6.6</td><td>4.7E-11</td><td>1.4E-10</td><td>MB231/MB468</td><td>MP</td></tr>
<tr><td>>hg38_21_+_8433172_8446622_RNA45SN1_45S_50ntFlanks@10393.10422.30|CAGCAGTTGAACATGGGTCAGTCGGTCCTG</td><td>>hg38_21_+_8433172_8446622_RNA45SN1_45S_50ntFlanks</td><td>10393</td><td>10422</td><td>30</td><td>CAGCAGTTGAACATGGGTCAGTCGGTCCTG</td><td>697.7</td><td>-4.3</td><td>0.7</td><td>-6.6</td><td>5.0E-11</td><td>1.4E-10</td><td>MB231/MB468</td><td>NUC</td></tr>
<tr><td>>hg38_21_+_8433172_8446622_RNA45SN1_45S_50ntFlanks@11881.11906.26|CCTGTTGAGCTTGACTCTAGTCTGGC</td><td>>hg38_21_+_8433172_8446622_RNA45SN1_45S_50ntFlanks</td><td>11881</td><td>11906</td><td>26</td><td>CCTGTTGAGCTTGACTCTAGTCTGGC</td><td>420.3</td><td>-7.4</td><td>1.1</td><td>-6.6</td><td>4.6E-11</td><td>1.4E-10</td><td>BT20/MB468</td><td>CYTO</td></tr>
<tr><td>>MI0000809|hsa-mir-151a&WithFlank&8|-|140732558|140732659@51.72.22(+1A)&[MIMAT0000757&hsa-miR-151a-3p&offsets|-2|-1(+1A);m-30&8|-|140732587|140732607&offsets|-2|-1(+1A)]|TACTAGACTGAAGCTCCTTGAGA</td><td>>MI0000809|hsa-mir-151a&WithFlank&8|-|140732558|140732659</td><td>51</td><td>72</td><td>22(+1A)</td><td>TACTAGACTGAAGCTCCTTGAGA</td><td>758.6</td><td>8.8</td><td>1.3</td><td>6.6</td><td>3.7E-11</td><td>1.4E-10</td><td>BT20/MB468</td><td>MITO</td></tr>
<tr><td>>hg38_21_+_8433172_8446622_RNA45SN1_45S_50ntFlanks@3705.3748.44|TACCTGGTTGATCCTGCCAGTAGCATATGCTTGTCTCAAAGATT</td><td>>hg38_21_+_8433172_8446622_RNA45SN1_45S_50ntFlanks</td><td>3705</td><td>3748</td><td>44</td><td>TACCTGGTTGATCCTGCCAGTAGCATATGCTTGTCTCAAAGATT</td><td>13151.8</td><td>8.7</td><td>1.3</td><td>6.6</td><td>4.6E-11</td><td>1.4E-10</td><td>BT20/MB468</td><td>CYTO</td></tr>
<tr><td>>hg38_21_+_8433172_8446622_RNA45SN1_45S_50ntFlanks@7974.8015.42|ACGCGACCTCAGATCAGACGTGGCGACCCGCTGAATTTAAGC</td><td>>hg38_21_+_8433172_8446622_RNA45SN1_45S_50ntFlanks</td><td>7974</td><td>8015</td><td>42</td><td>ACGCGACCTCAGATCAGACGTGGCGACCCGCTGAATTTAAGC</td><td>860.5</td><td>7.3</td><td>1.1</td><td>6.6</td><td>4.6E-11</td><td>1.4E-10</td><td>BT20/MB468</td><td>CYTO</td></tr>
<tr><td>>hg38_21_+_8433172_8446622_RNA45SN1_45S_50ntFlanks@12456.12484.29|CCAAGCGTTGGATTGTTCACCCACTAATA</td><td>>hg38_21_+_8433172_8446622_RNA45SN1_45S_50ntFlanks</td><td>12456</td><td>12484</td><td>29</td><td>CCAAGCGTTGGATTGTTCACCCACTAATA</td><td>7876.0</td><td>-7.7</td><td>1.2</td><td>-6.6</td><td>5.1E-11</td><td>1.4E-10</td><td>MB231/MB468</td><td>CYTO</td></tr>
<tr><td>>hg38_21_+_8433172_8446622_RNA45SN1_45S_50ntFlanks@9622.9639.18|CTAGTAGCTGGTTCCCTC</td><td>>hg38_21_+_8433172_8446622_RNA45SN1_45S_50ntFlanks</td><td>9622</td><td>9639</td><td>18</td><td>CTAGTAGCTGGTTCCCTC</td><td>99.3</td><td>-10.5</td><td>1.6</td><td>-6.6</td><td>4.8E-11</td><td>1.4E-10</td><td>MB231/MB468</td><td>MP</td></tr>
<tr><td>>hg38_21_+_8433172_8446622_RNA45SN1_45S_50ntFlanks@4759.4781.23|ATCAGATACCGTCGTAGTTCCGA</td><td>>hg38_21_+_8433172_8446622_RNA45SN1_45S_50ntFlanks</td><td>4759</td><td>4781</td><td>23</td><td>ATCAGATACCGTCGTAGTTCCGA</td><td>259.2</td><td>-11.1</td><td>1.6</td><td>-7.0</td><td>3.5E-12</td><td>1.4E-10</td><td>BT20/MB468</td><td>MP</td></tr>
<tr><td>>MI0000342|hsa-mir-200b&WithFlank&1|+|1167098|1167204@63.84.22(+1U)&[MIMAT0000318&hsa-miR-200b-3p&offsets|0|0(+1U);m-93&1|+|1167160|1167182&offsets|0|-1(+1U)]|TAATACTGCCTGGTAATGATGAT</td><td>>MI0000342|hsa-mir-200b&WithFlank&1|+|1167098|1167204</td><td>63</td><td>84</td><td>22(+1U)</td><td>TAATACTGCCTGGTAATGATGAT</td><td>244.2</td><td>-11.0</td><td>1.6</td><td>-7.0</td><td>3.5E-12</td><td>1.4E-10</td><td>BT20/MB468</td><td>MP</td></tr>
<tr><td>>trna129_MetCAT_6_-_27870271_27870342@32.51.20,trna142_MetCAT_6_-_27560600_27560671@32.51.20,trna150_MetCAT_6_-_27300764_27300835@32.51.20,trna169_MetCAT_6_-_26330529_26330600@32.51.20,trna171_MetCAT_6_-_26313352_26313423@32.51.20,trna20_MetCAT_17_-_80452597_80452668@32.51.20,trna2_MetCAT_6_+_26286754_26286825@32.51.20,trna32_MetCAT_1_+_153643726_153643797@32.51.20,trna61_MetCAT_6_+_27745664_27745735@32.51.20|CCATAACCCAGAGGTCGATG</td><td>>trna129_MetCAT_6_-_27870271_27870342</td><td>32</td><td>51</td><td>20</td><td>CCATAACCCAGAGGTCGATG</td><td>436.7</td><td>-11.9</td><td>1.7</td><td>-7.0</td><td>3.5E-12</td><td>1.4E-10</td><td>BT20/MB468</td><td>MP</td></tr>
<tr><td>>hg38_21_+_8433172_8446622_RNA45SN1_45S_50ntFlanks@9167.9187.21|CGGGCGGGTCGCGCCGTCGGG</td><td>>hg38_21_+_8433172_8446622_RNA45SN1_45S_50ntFlanks</td><td>9167</td><td>9187</td><td>21</td><td>CGGGCGGGTCGCGCCGTCGGG</td><td>338.3</td><td>-11.5</td><td>1.7</td><td>-7.0</td><td>3.5E-12</td><td>1.4E-10</td><td>BT20/MB468</td><td>MP</td></tr>
</table>

<table>
<tr><td>>trna116_GluCTC_1_-_145399233_145399304@1.37.37,trna59_GluCTC_1_+_249168447_249168518@1.37.37,trna71_GluCTC_1_-_161439189_161439260@1.37.37,trna74_GluCTC_1_-_161431809_161431880@1.37.37,trna77_GluCTC_1_-_161424398_161424469@1.37.37,trna77_GluCTC_6_+_28949976_28950047@1.37.37,trna80_GluCTC_1_-_161417018_161417089@1.37.37,trna87_GluCTC_6_-_126101393_126101464@1.37.37|TCCCTGGTGGTCTAGTGGTTAGGATTCGGCGCTCTCA</td><td>>trna116_GluCTC_1_-_145399233_145399304</td><td>1</td><td>37</td><td>37</td><td>TCCCTGGTGGTCTAGTGGTTAGGATTCGGCGCTCTCA</td><td>312.4</td><td>7.5</td><td>1.1</td><td>6.6</td><td>5.1E-11</td><td>1.4E-10</td><td>MB231/MB468</td><td>CYTO</td></tr>
<tr><td>>hg38_21_+_8433172_8446622_RNA45SN1_45S_50ntFlanks@10906.10926.21|TGGGCGCGCGCCGCGGCTGGA</td><td>>hg38_21_+_8433172_8446622_RNA45SN1_45S_50ntFlanks</td><td>10906</td><td>10926</td><td>21</td><td>TGGGCGCGCGCCGCGGCTGGA</td><td>82.1</td><td>-10.2</td><td>1.6</td><td>-6.6</td><td>4.8E-11</td><td>1.4E-10</td><td>MB231/MB468</td><td>MP</td></tr>
<tr><td>>hg38_21_+_8433172_8446622_RNA45SN1_45S_50ntFlanks@10690.10708.19|CGTCCGGTGAGCTCTCGCT</td><td>>hg38_21_+_8433172_8446622_RNA45SN1_45S_50ntFlanks</td><td>10690</td><td>10708</td><td>19</td><td>CGTCCGGTGAGCTCTCGCT</td><td>12971.7</td><td>-9.3</td><td>1.4</td><td>-6.5</td><td>6.2E-11</td><td>1.4E-10</td><td>BT20/MB468</td><td>NUC</td></tr>
<tr><td>>hg38_21_+_8433172_8446622_RNA45SN1_45S_50ntFlanks@12251.12271.21|AAAGCTCGCTTGATCTTGATT</td><td>>hg38_21_+_8433172_8446622_RNA45SN1_45S_50ntFlanks</td><td>12251</td><td>12271</td><td>21</td><td>AAAGCTCGCTTGATCTTGATT</td><td>245.7</td><td>4.0</td><td>0.6</td><td>6.6</td><td>4.6E-11</td><td>1.4E-10</td><td>BT20/MB468</td><td>CYTO</td></tr>
<tr><td>>hg38_21_+_8433172_8446622_RNA45SN1_45S_50ntFlanks@10349.10383.35|AACGAGAACTTTGAAGGCCGAAGTGGAGAAGGGTT</td><td>>hg38_21_+_8433172_8446622_RNA45SN1_45S_50ntFlanks</td><td>10349</td><td>10383</td><td>35</td><td>AACGAGAACTTTGAAGGCCGAAGTGGAGAAGGGTT</td><td>6758.6</td><td>-5.8</td><td>0.9</td><td>-6.6</td><td>4.8E-11</td><td>1.4E-10</td><td>MB231/MB468</td><td>MP</td></tr>
<tr><td>>trna1_GlnCTG_6_+_18836402_18836473@30.70.41,trna3_GlnCTG_17_+_8023070_8023141@30.70.41,trna49_GlnCTG_6_+_27487308_27487379@30.70.41,trna7_GlnCTG_15_-_66161400_66161471@30.70.41,trna99_GlnCTG_6_-_28909378_28909449@30.70.41|GACTCTGAATCCAGCGATCCGAGTTCAAATCTCGGTGGAAC</td><td>>trna1_GlnCTG_6_+_18836402_18836473</td><td>30</td><td>70</td><td>41</td><td>GACTCTGAATCCAGCGATCCGAGTTCAAATCTCGGTGGAAC</td><td>229.8</td><td>-10.2</td><td>1.5</td><td>-6.6</td><td>4.6E-11</td><td>1.4E-10</td><td>BT20/MB468</td><td>CYTO</td></tr>
<tr><td>>hg38_21_+_8433172_8446622_RNA45SN1_45S_50ntFlanks@11792.11814.23|ATGAACGAGATTCCCACTGTCCC</td><td>>hg38_21_+_8433172_8446622_RNA45SN1_45S_50ntFlanks</td><td>11792</td><td>11814</td><td>23</td><td>ATGAACGAGATTCCCACTGTCCC</td><td>93.6</td><td>-10.4</td><td>1.6</td><td>-6.6</td><td>4.8E-11</td><td>1.4E-10</td><td>MB231/MB468</td><td>MP</td></tr>
<tr><td>>hg38_21_+_8433172_8446622_RNA45SN1_45S_50ntFlanks@12457.12478.22|CAAGCGTTGGATTGTTCACCCA</td><td>>hg38_21_+_8433172_8446622_RNA45SN1_45S_50ntFlanks</td><td>12457</td><td>12478</td><td>22</td><td>CAAGCGTTGGATTGTTCACCCA</td><td>3962.4</td><td>-5.3</td><td>0.8</td><td>-6.6</td><td>4.6E-11</td><td>1.4E-10</td><td>BT20/MB468</td><td>CYTO</td></tr>
<tr><td>>hg38_21_+_8433172_8446622_RNA45SN1_45S_50ntFlanks@6651.6691.41|CGACTCTTAGCGGTGGATCACTCGGCTCGTGCGTCGATGAA</td><td>>hg38_21_+_8433172_8446622_RNA45SN1_45S_50ntFlanks</td><td>6651</td><td>6691</td><td>41</td><td>CGACTCTTAGCGGTGGATCACTCGGCTCGTGCGTCGATGAA</td><td>313.6</td><td>5.7</td><td>0.9</td><td>6.6</td><td>4.6E-11</td><td>1.4E-10</td><td>BT20/MB468</td><td>CYTO</td></tr>
<tr><td>>hg38_21_+_8433172_8446622_RNA45SN1_45S_50ntFlanks@11569.11600.32|CAGCCGACTTAGAACTGGTGCGGACCAGGGGA</td><td>>hg38_21_+_8433172_8446622_RNA45SN1_45S_50ntFlanks</td><td>11569</td><td>11600</td><td>32</td><td>CAGCCGACTTAGAACTGGTGCGGACCAGGGGA</td><td>2216.1</td><td>-7.1</td><td>1.1</td><td>-6.5</td><td>5.9E-11</td><td>1.4E-10</td><td>MB231/MB468</td><td>MITO</td></tr>
<tr><td>>MI0000469|hsa-mir-125a&WithFlank&19|+|51693248|51693345@21.41.21(+1C)&[MIMAT0000443&hsa-miR-125a-5p&offsets|0|-3(+1C);m-63&19|+|51693268|51693289&offsets|0|-1(+1C)]|TCCCTGAGACCCTTTAACCTGC</td><td>>MI0000469|hsa-mir-125a&WithFlank&19|+|51693248|51693345</td><td>21</td><td>41</td><td>21(+1C)</td><td>TCCCTGAGACCCTTTAACCTGC</td><td>276.5</td><td>5.9</td><td>0.9</td><td>6.6</td><td>5.2E-11</td><td>1.4E-10</td><td>MB231/MB468</td><td>CYTO</td></tr>
<tr><td>>hg38_21_+_8433172_8446622_RNA45SN1_45S_50ntFlanks@10572.10596.25|AGAAGCCGGCGGGAGCCCCGGGGAG</td><td>>hg38_21_+_8433172_8446622_RNA45SN1_45S_50ntFlanks</td><td>10572</td><td>10596</td><td>25</td><td>AGAAGCCGGCGGGAGCCCCGGGGAG</td><td>168.2</td><td>-6.3</td><td>1.0</td><td>-6.6</td><td>5.2E-11</td><td>1.4E-10</td><td>MB231/MB468</td><td>CYTO</td></tr>
<tr><td>>hg38_21_+_8433172_8446622_RNA45SN1_45S_50ntFlanks@7990.8010.21|GACGTGGCGACCCGCTGAATT</td><td>>hg38_21_+_8433172_8446622_RNA45SN1_45S_50ntFlanks</td><td>7990</td><td>8010</td><td>21</td><td>GACGTGGCGACCCGCTGAATT</td><td>383.0</td><td>6.1</td><td>0.9</td><td>6.6</td><td>5.2E-11</td><td>1.4E-10</td><td>MB231/MB468</td><td>CYTO</td></tr>
<tr><td>>hg38_21_+_8433172_8446622_RNA45SN1_45S_50ntFlanks@4730.4758.29|CAAGAACGAAAGTCGGAGGTTCGAAGACG</td><td>>hg38_21_+_8433172_8446622_RNA45SN1_45S_50ntFlanks</td><td>4730</td><td>4758</td><td>29</td><td>CAAGAACGAAAGTCGGAGGTTCGAAGACG</td><td>6567.9</td><td>-4.6</td><td>0.7</td><td>-6.6</td><td>4.8E-11</td><td>1.4E-10</td><td>MB231/MB468</td><td>MP</td></tr>
<tr><td>>hg38_21_+_8433172_8446622_RNA45SN1_45S_50ntFlanks@10382.10424.43|TTCCATGTGAACAGCAGTTGAACATGGGTCAGTCGGTCCTGAG</td><td>>hg38_21_+_8433172_8446622_RNA45SN1_45S_50ntFlanks</td><td>10382</td><td>10424</td><td>43</td><td>TTCCATGTGAACAGCAGTTGAACATGGGTCAGTCGGTCCTGAG</td><td>1228.8</td><td>9.0</td><td>1.4</td><td>6.6</td><td>4.7E-11</td><td>1.4E-10</td><td>BT20/MB468</td><td>CYTO</td></tr>
<tr><td>>hg38_21_+_8433172_8446622_RNA45SN1_45S_50ntFlanks@6719.6750.32|TGAATTGCAGGACACATTGATCATCGACACTT</td><td>>hg38_21_+_8433172_8446622_RNA45SN1_45S_50ntFlanks</td><td>6719</td><td>6750</td><td>32</td><td>TGAATTGCAGGACACATTGATCATCGACACTT</td><td>279.4</td><td>-6.3</td><td>1.0</td><td>-6.5</td><td>6.0E-11</td><td>1.4E-10</td><td>MB231/MB468</td><td>MITO</td></tr>
<tr><td>>hg38_21_+_8433172_8446622_RNA45SN1_45S_50ntFlanks@9533.9577.45|GAGGAAACTCTGGTGGAGGTCCGTAGCGGTCCTGACGTGCAAATC</td><td>>hg38_21_+_8433172_8446622_RNA45SN1_45S_50ntFlanks</td><td>9533</td><td>9577</td><td>45</td><td>GAGGAAACTCTGGTGGAGGTCCGTAGCGGTCCTGACGTGCAAATC</td><td>292.3</td><td>8.7</td><td>1.3</td><td>6.6</td><td>5.2E-11</td><td>1.4E-10</td><td>MB231/MB468</td><td>CYTO</td></tr>
<tr><td>>hg38_21_+_8433172_8446622_RNA45SN1_45S_50ntFlanks@10786.10823.38|AACAGCCTCTGGCATGTTGGAACAATGTAGGTAAGGGA</td><td>>hg38_21_+_8433172_8446622_RNA45SN1_45S_50ntFlanks</td><td>10786</td><td>10823</td><td>38</td><td>AACAGCCTCTGGCATGTTGGAACAATGTAGGTAAGGGA</td><td>370.9</td><td>-5.6</td><td>0.9</td><td>-6.6</td><td>5.2E-11</td><td>1.4E-10</td><td>MB231/MB468</td><td>CYTO</td></tr>
<tr><td>>hg38_21_+_8433172_8446622_RNA45SN1_45S_50ntFlanks@11933.11969.37|AATAAGTGGGAGGCCCCCGGCGCCCCCCCGGTGTCCC</td><td>>hg38_21_+_8433172_8446622_RNA45SN1_45S_50ntFlanks</td><td>11933</td><td>11969</td><td>37</td><td>AATAAGTGGGAGGCCCCCGGCGCCCCCCCGGTGTCCC</td><td>119.5</td><td>-10.8</td><td>1.6</td><td>-6.6</td><td>4.9E-11</td><td>1.4E-10</td><td>MB231/MB468</td><td>MP</td></tr>
<tr><td>>hg38_21_+_8433172_8446622_RNA45SN1_45S_50ntFlanks@4512.4532.21|AAAAAATTAGAGTGTTCAAAG</td><td>>hg38_21_+_8433172_8446622_RNA45SN1_45S_50ntFlanks</td><td>4512</td><td>4532</td><td>21</td><td>AAAAAATTAGAGTGTTCAAAG</td><td>95.2</td><td>-10.4</td><td>1.6</td><td>-6.6</td><td>4.9E-11</td><td>1.4E-10</td><td>MB231/MB468</td><td>MP</td></tr>
<tr><td>>hg38_21_+_8433172_8446622_RNA45SN1_45S_50ntFlanks@3769.3792.24|CGCACGGCCGGTACAGTGAAACTG</td><td>>hg38_21_+_8433172_8446622_RNA45SN1_45S_50ntFlanks</td><td>3769</td><td>3792</td><td>24</td><td>CGCACGGCCGGTACAGTGAAACTG</td><td>443.3</td><td>-10.9</td><td>1.6</td><td>-6.6</td><td>3.8E-11</td><td>1.4E-10</td><td>BT20/MB468</td><td>MITO</td></tr>
<tr><td>>hg38_21_+_8433172_8446622_RNA45SN1_45S_50ntFlanks@11568.11605.38|GCAGCCGACTTAGAACTGGTGCGGACCAGGGGAATCCG</td><td>>hg38_21_+_8433172_8446622_RNA45SN1_45S_50ntFlanks</td><td>11568</td><td>11605</td><td>38</td><td>GCAGCCGACTTAGAACTGGTGCGGACCAGGGGAATCCG</td><td>843.5</td><td>-7.0</td><td>1.1</td><td>-6.5</td><td>6.0E-11</td><td>1.4E-10</td><td>MB231/MB468</td><td>MITO</td></tr>
<tr><td>>hg38_21_+_8433172_8446622_RNA45SN1_45S_50ntFlanks@10348.10368.21|AAACGAGAACTTTGAAGGCCG</td><td>>hg38_21_+_8433172_8446622_RNA45SN1_45S_50ntFlanks</td><td>10348</td><td>10368</td><td>21</td><td>AAACGAGAACTTTGAAGGCCG</td><td>1147.5</td><td>-3.4</td><td>0.5</td><td>-6.6</td><td>4.7E-11</td><td>1.4E-10</td><td>BT20/MB468</td><td>CYTO</td></tr>
<tr><td>>hg38_21_+_8433172_8446622_RNA45SN1_45S_50ntFlanks@11315.11334.20|CCACCCCACGTCTCGTCGCG</td><td>>hg38_21_+_8433172_8446622_RNA45SN1_45S_50ntFlanks</td><td>11315</td><td>11334</td><td>20</td><td>CCACCCCACGTCTCGTCGCG</td><td>485.3</td><td>-6.8</td><td>1.0</td><td>-6.6</td><td>5.2E-11</td><td>1.4E-10</td><td>MB231/MB468</td><td>CYTO</td></tr>
<tr><td>>am_trna11_ProAGG_16_+_3241989_3242060@1.28.28,trna12_ProAGG_6_+_26555498_26555569@1.28.28,trna12_ProTGG_11_-_75946869_75946940@1.28.28,trna14_ProTGG_5_-_180615854_180615925@1.28.28,trna22_ProAGG_14_-_21081560_21081631@1.28.28,trna23_ProAGG_14_-_21077495_21077566@1.28.28,trna28_ProTGG_16_-_3234133_3234204@1.28.28,trna29_ProAGG_16_-_3232635_3232706@1.28.28,trna2_ProAGG_7_+_128423504_128423575@1.28.28,trna30_ProCGG_6_+_27059521_27059592@1.28.28,trna37_ProCGG_17_-_8126151_8126222@1.28.28,trna3_ProTGG_16_+_3208923_3208994@1.28.28,trna52_ProCGG_1_+_167683962_167684033@1.28.28,trna65_ProAGG_1_-_167684725_167684796@1.28.28,trna6_ProCGG_16_+_3222049_3222120@1.28.28,trna6_ProTGG_14_+_21152175_21152246@1.28.28,trna8_ProTGG_16_+_3238094_3238165@1.28.28,trna9_ProAGG_11_+_75946557_75946628@1.28.28,trna9_ProAGG_16_+_3239634_3239705@1.28.28|GGCTCGTTGGTCTAGGGGTATGATTCTC</td><td>>am_trna11_ProAGG_16_+_3241989_3242060</td><td>1</td><td>28</td><td>28</td><td>GGCTCGTTGGTCTAGGGGTATGATTCTC</td><td>210.3</td><td>-9.8</td><td>1.5</td><td>-6.6</td><td>3.8E-11</td><td>1.4E-10</td><td>BT20/MB468</td><td>MITO</td></tr>
<tr><td>>hg38_1_-_228634819_228635039_RNA5S12_5S_50ntFlanks@151.168.18|ATACCGGGTGCTGTAGGC</td><td>>hg38_1_-_228634819_228635039_RNA5S12_5S_50ntFlanks</td><td>151</td><td>168</td><td>18</td><td>ATACCGGGTGCTGTAGGC</td><td>92.8</td><td>-10.4</td><td>1.6</td><td>-6.6</td><td>4.9E-11</td><td>1.4E-10</td><td>MB231/MB468</td><td>MP</td></tr>
<tr><td>>hg38_21_+_8433172_8446622_RNA45SN1_45S_50ntFlanks@4241.4263.23|CTCTTTCGAGGCCCTGTAATTGG</td><td>>hg38_21_+_8433172_8446622_RNA45SN1_45S_50ntFlanks</td><td>4241</td><td>4263</td><td>23</td><td>CTCTTTCGAGGCCCTGTAATTGG</td><td>89.5</td><td>-10.3</td><td>1.6</td><td>-6.6</td><td>4.9E-11</td><td>1.4E-10</td><td>MB231/MB468</td><td>MP</td></tr>
<tr><td>>hg38_21_+_8433172_8446622_RNA45SN1_45S_50ntFlanks@5417.5438.22|CTACCGATTGGATGGTTTAGTG</td><td>>hg38_21_+_8433172_8446622_RNA45SN1_45S_50ntFlanks</td><td>5417</td><td>5438</td><td>22</td><td>CTACCGATTGGATGGTTTAGTG</td><td>80.4</td><td>-10.2</td><td>1.6</td><td>-6.6</td><td>4.9E-11</td><td>1.4E-10</td><td>MB231/MB468</td><td>MP</td></tr>
<tr><td>>hg38_21_+_8433172_8446622_RNA45SN1_45S_50ntFlanks@3985.4024.40|CGGCTTTGGTGACTCTAGATAACCTCGGGCCGATCGCACG</td><td>>hg38_21_+_8433172_8446622_RNA45SN1_45S_50ntFlanks</td><td>3985</td><td>4024</td><td>40</td><td>CGGCTTTGGTGACTCTAGATAACCTCGGGCCGATCGCACG</td><td>266.5</td><td>-7.8</td><td>1.2</td><td>-6.6</td><td>5.2E-11</td><td>1.4E-10</td><td>MB231/MB468</td><td>NUC</td></tr>
<tr><td>>hg38_21_+_8433172_8446622_RNA45SN1_45S_50ntFlanks@4771.4793.23|CGTAGTTCCGACCATAAACGATG</td><td>>hg38_21_+_8433172_8446622_RNA45SN1_45S_50ntFlanks</td><td>4771</td><td>4793</td><td>23</td><td>CGTAGTTCCGACCATAAACGATG</td><td>77.9</td><td>-10.1</td><td>1.5</td><td>-6.6</td><td>4.9E-11</td><td>1.4E-10</td><td>MB231/MB468</td><td>MP</td></tr>
<tr><td>>hg38_21_+_8433172_8446622_RNA45SN1_45S_50ntFlanks@11882.11899.18|CTGTTGAGCTTGACTCTA</td><td>>hg38_21_+_8433172_8446622_RNA45SN1_45S_50ntFlanks</td><td>11882</td><td>11899</td><td>18</td><td>CTGTTGAGCTTGACTCTA</td><td>91.9</td><td>-10.4</td><td>1.6</td><td>-6.6</td><td>4.9E-11</td><td>1.4E-10</td><td>MB231/MB468</td><td>MP</td></tr>
<tr><td>>MI0000803|hsa-mir-330&WithFlank&19|-|45638988|45639093@63.86.24(+1U)&[MIMAT0000751&hsa-miR-330-3p&offsets|0|+1(+1U);m-347&19|-|45639009|45639030&offsets|-1|+1(+1U)]|GCAAAGCACACGGCCTGCAGAGAGT</td><td>>MI0000803|hsa-mir-330&WithFlank&19|-|45638988|45639093</td><td>63</td><td>86</td><td>24(+1U)</td><td>GCAAAGCACACGGCCTGCAGAGAGT</td><td>958.9</td><td>7.0</td><td>1.1</td><td>6.6</td><td>5.3E-11</td><td>1.4E-10</td><td>MB231/MB468</td><td>CYTO</td></tr>
<tr><td>>hg38_21_+_8433172_8446622_RNA45SN1_45S_50ntFlanks@10864.10890.27|CTCTAAGGGCTGGGTCGGTCGGGCTGG</td><td>>hg38_21_+_8433172_8446622_RNA45SN1_45S_50ntFlanks</td><td>10864</td><td>10890</td><td>27</td><td>CTCTAAGGGCTGGGTCGGTCGGGCTGG</td><td>89.2</td><td>-10.3</td><td>1.6</td><td>-6.6</td><td>4.9E-11</td><td>1.4E-10</td><td>MB231/MB468</td><td>MP</td></tr>
<tr><td>>hg38_21_+_8433172_8446622_RNA45SN1_45S_50ntFlanks@9576.9597.22|TCGGTCGTCCGACCTGGGTATA</td><td>>hg38_21_+_8433172_8446622_RNA45SN1_45S_50ntFlanks</td><td>9576</td><td>9597</td><td>22</td><td>TCGGTCGTCCGACCTGGGTATA</td><td>308.0</td><td>-11.3</td><td>1.6</td><td>-7.0</td><td>3.6E-12</td><td>1.4E-10</td><td>BT20/MB468</td><td>MP</td></tr>
<tr><td>>hg38_21_+_8433172_8446622_RNA45SN1_45S_50ntFlanks@9568.9603.36|CGTGCAAATCGGTCGTCCGACCTGGGTATAGGGGCG</td><td>>hg38_21_+_8433172_8446622_RNA45SN1_45S_50ntFlanks</td><td>9568</td><td>9603</td><td>36</td><td>CGTGCAAATCGGTCGTCCGACCTGGGTATAGGGGCG</td><td>685.0</td><td>-3.6</td><td>0.6</td><td>-6.6</td><td>5.2E-11</td><td>1.4E-10</td><td>MB231/MB468</td><td>NUC</td></tr>
<tr><td>>hg38_21_+_8433172_8446622_RNA45SN1_45S_50ntFlanks@10392.10431.40|ACAGCAGTTGAACATGGGTCAGTCGGTCCTGAGAGATGGG</td><td>>hg38_21_+_8433172_8446622_RNA45SN1_45S_50ntFlanks</td><td>10392</td><td>10431</td><td>40</td><td>ACAGCAGTTGAACATGGGTCAGTCGGTCCTGAGAGATGGG</td><td>558.0</td><td>-7.5</td><td>1.1</td><td>-6.6</td><td>5.3E-11</td><td>1.4E-10</td><td>MB231/MB468</td><td>NUC</td></tr>
<tr><td>>hg38_21_+_8433172_8446622_RNA45SN1_45S_50ntFlanks@10534.10570.37|CCGCGAGGCGTCCAGTGCGGTAACGCGACCGATCCCG</td><td>>hg38_21_+_8433172_8446622_RNA45SN1_45S_50ntFlanks</td><td>10534</td><td>10570</td><td>37</td><td>CCGCGAGGCGTCCAGTGCGGTAACGCGACCGATCCCG</td><td>764.9</td><td>-8.0</td><td>1.2</td><td>-6.6</td><td>3.9E-11</td><td>1.4E-10</td><td>BT20/MB468</td><td>MITO</td></tr>
<tr><td>>hg38_21_+_8433172_8446622_RNA45SN1_45S_50ntFlanks@12971.12990.20|TTTCGTACGTAGCAGAGCAG</td><td>>hg38_21_+_8433172_8446622_RNA45SN1_45S_50ntFlanks</td><td>12971</td><td>12990</td><td>20</td><td>TTTCGTACGTAGCAGAGCAG</td><td>598.4</td><td>-5.8</td><td>0.9</td><td>-6.6</td><td>5.3E-11</td><td>1.4E-10</td><td>MB231/MB468</td><td>CYTO</td></tr>
<tr><td>>hg38_21_+_8433172_8446622_RNA45SN1_45S_50ntFlanks@10350.10378.29|ACGAGAACTTTGAAGGCCGAAGTGGAGAA</td><td>>hg38_21_+_8433172_8446622_RNA45SN1_45S_50ntFlanks</td><td>10350</td><td>10378</td><td>29</td><td>ACGAGAACTTTGAAGGCCGAAGTGGAGAA</td><td>2581.0</td><td>-10.2</td><td>1.6</td><td>-6.5</td><td>6.4E-11</td><td>1.4E-10</td><td>BT20/MB468</td><td>NUC</td></tr>
<tr><td>>hg38_21_+_8433172_8446622_RNA45SN1_45S_50ntFlanks@10788.10826.39|CAGCCTCTGGCATGTTGGAACAATGTAGGTAAGGGAAGT</td><td>>hg38_21_+_8433172_8446622_RNA45SN1_45S_50ntFlanks</td><td>10788</td><td>10826</td><td>39</td><td>CAGCCTCTGGCATGTTGGAACAATGTAGGTAAGGGAAGT</td><td>163.4</td><td>-11.2</td><td>1.7</td><td>-6.6</td><td>5.0E-11</td><td>1.4E-10</td><td>MB231/MB468</td><td>MP</td></tr>
<tr><td>>hg38_21_+_8433172_8446622_RNA45SN1_45S_50ntFlanks@10463.10484.22|CTCCGTTGCCCTCGGCCGATCG</td><td>>hg38_21_+_8433172_8446622_RNA45SN1_45S_50ntFlanks</td><td>10463</td><td>10484</td><td>22</td><td>CTCCGTTGCCCTCGGCCGATCG</td><td>84.8</td><td>-10.3</td><td>1.6</td><td>-6.6</td><td>5.0E-11</td><td>1.4E-10</td><td>MB231/MB468</td><td>MP</td></tr>
</table>

| | | | | | | | | | | | | | |
|---|---|---|---|---|---|---|---|---|---|---|---|---|---|
| >hg38_21_+_8433172_8446622_RNA45SN1_45S_50ntFlanks@9758.9786.29\|TCAAACTTTAAATGGGTAAGAAGCCCGGC | >hg38_21_+_8433172_8446622_RNA45SN1_45S_50ntFlanks | 9758 | 9786 | 29 | TCAAACTTTAAATGGGTAAGAAGCCCGGC | 311.1 | -12.1 | 1.8 | -6.6 | 5.0E-11 | 1.4E-10 | MB231/MB468 | MP |
| >hg38_21_+_8433172_8446622_RNA45SN1_45S_50ntFlanks@12638.12662.25\|CTACCATCTGTGGGATTATGACTGA | >hg38_21_+_8433172_8446622_RNA45SN1_45S_50ntFlanks | 12638 | 12662 | 25 | CTACCATCTGTGGGATTATGACTGA | 86.2 | -10.3 | 1.6 | -6.6 | 5.0E-11 | 1.4E-10 | MB231/MB468 | MP |
| >hg38_21_+_8433172_8446622_RNA45SN1_45S_50ntFlanks@10233.10262.30\|GCGGACGCTACGCCGCGACGAGTAGGAGGG | >hg38_21_+_8433172_8446622_RNA45SN1_45S_50ntFlanks | 10233 | 10262 | 30 | GCGGACGCTACGCCGCGACGAGTAGGAGGG | 76.7 | -10.1 | 1.5 | -6.6 | 5.0E-11 | 1.4E-10 | MB231/MB468 | MP |
| >hg38_21_+_8433172_8446622_RNA45SN1_45S_50ntFlanks@12540.12567.28\|TGATGTGTTGTTGCCATGGTAATCCTGC | >hg38_21_+_8433172_8446622_RNA45SN1_45S_50ntFlanks | 12540 | 12567 | 28 | TGATGTGTTGTTGCCATGGTAATCCTGC | 83.9 | -10.3 | 1.6 | -6.6 | 5.0E-11 | 1.4E-10 | MB231/MB468 | MP |
| >hg38_21_+_8433172_8446622_RNA45SN1_45S_50ntFlanks@11891.11909.19\|TTGACTCTAGTCTGGCACG | >hg38_21_+_8433172_8446622_RNA45SN1_45S_50ntFlanks | 11891 | 11909 | 19 | TTGACTCTAGTCTGGCACG | 311.7 | -3.0 | 0.5 | -6.6 | 4.8E-11 | 1.4E-10 | BT20/MB468 | CYTO |
| >hg38_21_+_8433172_8446622_RNA45SN1_45S_50ntFlanks@10411.10432.22\|CAGTCGGTCCTGAGAGATGGGC | >hg38_21_+_8433172_8446622_RNA45SN1_45S_50ntFlanks | 10411 | 10432 | 22 | CAGTCGGTCCTGAGAGATGGGC | 1044.6 | -9.5 | 1.5 | -6.5 | 6.5E-11 | 1.4E-10 | BT20/MB468 | NUC |
| >hg38_21_+_8433172_8446622_RNA45SN1_45S_50ntFlanks@11163.11189.27\|CGGTCCCCCGCGAGGGGGGCCCGGGCA | >hg38_21_+_8433172_8446622_RNA45SN1_45S_50ntFlanks | 11163 | 11189 | 27 | CGGTCCCCCGCGAGGGGGGCCCGGGCA | 250.7 | -11.8 | 1.8 | -6.6 | 5.0E-11 | 1.4E-10 | MB231/MB468 | MP |
| >hg38_21_+_8433172_8446622_RNA45SN1_45S_50ntFlanks@7983.8006.24\|CAGATCAGACGTGGCGACCCGCTG | >hg38_21_+_8433172_8446622_RNA45SN1_45S_50ntFlanks | 7983 | 8006 | 24 | CAGATCAGACGTGGCGACCCGCTG | 2834.4 | -5.3 | 0.8 | -6.6 | 5.3E-11 | 1.4E-10 | MB231/MB468 | CYTO |
| >hg38_21_+_8433172_8446622_RNA45SN1_45S_50ntFlanks@475.507.33\|CTTCGTGATCGATGTGGTGACGTCGTGCTCTCC | >hg38_21_+_8433172_8446622_RNA45SN1_45S_50ntFlanks | 475 | 507 | 33 | CTTCGTGATCGATGTGGTGACGTCGTGCTCTCC | 1957.6 | 8.9 | 1.4 | 6.6 | 5.3E-11 | 1.4E-10 | MB231/MB468 | CYTO |
| >hg38_21_+_8433172_8446622_RNA45SN1_45S_50ntFlanks@12124.12150.27\|CGCGACCCGCTCCGGGGACAGTGCCAG | >hg38_21_+_8433172_8446622_RNA45SN1_45S_50ntFlanks | 12124 | 12150 | 27 | CGCGACCCGCTCCGGGGACAGTGCCAG | 107.2 | -10.6 | 1.6 | -6.6 | 5.0E-11 | 1.4E-10 | MB231/MB468 | MP |
| >hg38_21_+_8433172_8446622_RNA45SN1_45S_50ntFlanks@9534.9557.24\|AGGAAACTCTGGTGGAGGTCCGTA | >hg38_21_+_8433172_8446622_RNA45SN1_45S_50ntFlanks | 9534 | 9557 | 24 | AGGAAACTCTGGTGGAGGTCCGTA | 278.8 | -12.0 | 1.8 | -6.6 | 5.0E-11 | 1.4E-10 | MB231/MB468 | MP |
| >hg38_21_+_8433172_8446622_RNA45SN1_45S_50ntFlanks@8423.8468.46\|CGCAGTCCGCCCGGAGGATTCAACCCGGCGGCGGGTCCGGCCGTGT | >hg38_21_+_8433172_8446622_RNA45SN1_45S_50ntFlanks | 8423 | 8468 | 46 | CGCAGTCCGCCCGGAGGATTCAACCCGGCGGCGGGTCCGGCCGTGT | 182.0 | -11.3 | 1.7 | -6.5 | 6.1E-11 | 1.4E-10 | MB231/MB468 | MITO |
| >hg38_21_+_8433172_8446622_RNA45SN1_45S_50ntFlanks@11581.11611.31\|AACTGGTGCGGACCAGGGGAATCCGACTGTT | >hg38_21_+_8433172_8446622_RNA45SN1_45S_50ntFlanks | 11581 | 11611 | 31 | AACTGGTGCGGACCAGGGGAATCCGACTGTT | 626.5 | -3.4 | 0.5 | -6.6 | 5.4E-11 | 1.4E-10 | MB231/MB468 | NUC |
| >hg38_21_+_8433172_8446622_RNA45SN1_45S_50ntFlanks@10351.10368.18\|CGAGAACTTTGAAGGCCG | >hg38_21_+_8433172_8446622_RNA45SN1_45S_50ntFlanks | 10351 | 10368 | 18 | CGAGAACTTTGAAGGCCG | 15924.4 | -6.1 | 0.9 | -6.6 | 5.4E-11 | 1.4E-10 | MB231/MB468 | CYTO |
| >hg38_21_+_8433172_8446622_RNA45SN1_45S_50ntFlanks@11781.11803.23\|CGCATGAATGGATGAACGAGATT | >hg38_21_+_8433172_8446622_RNA45SN1_45S_50ntFlanks | 11781 | 11803 | 23 | CGCATGAATGGATGAACGAGATT | 921.0 | -5.2 | 0.8 | -6.6 | 5.4E-11 | 1.4E-10 | MB231/MB468 | CYTO |
| >hg38_21_+_8433172_8446622_RNA45SN1_45S_50ntFlanks@7989.8009.21\|AGACGTGGCGACCCGCTGAAT | >hg38_21_+_8433172_8446622_RNA45SN1_45S_50ntFlanks | 7989 | 8009 | 21 | AGACGTGGCGACCCGCTGAAT | 4828.4 | 7.0 | 1.1 | 6.6 | 5.4E-11 | 1.4E-10 | MB231/MB468 | CYTO |
| >hg38_21_+_8433172_8446622_RNA45SN1_45S_50ntFlanks@12624.12654.31\|CCAATGGGGCGAAGCTACCATCTGTGGGATT | >hg38_21_+_8433172_8446622_RNA45SN1_45S_50ntFlanks | 12624 | 12654 | 31 | CCAATGGGGCGAAGCTACCATCTGTGGGATT | 1577.1 | -7.4 | 1.1 | -6.6 | 5.4E-11 | 1.4E-10 | MB231/MB468 | NUC |
| >hg38_1_-_228634819_228635039_RNA5S12_5S_50ntFlanks@129.169.41\|TTGGATGGGAGACCGCCTGGGAATACCGGGTGCTGTAGGCT | >hg38_1_-_228634819_228635039_RNA5S12_5S_50ntFlanks | 129 | 169 | 41 | TTGGATGGGAGACCGCCTGGGAATACCGGGTGCTGTAGGCT | 2091.5 | 10.2 | 1.6 | 6.6 | 4.9E-11 | 1.4E-10 | BT20/MB468 | CYTO |
| >hg38_21_+_8433172_8446622_RNA45SN1_45S_50ntFlanks@4453.4471.19\|TCGGCGCCCCCTCGATGCT | >hg38_21_+_8433172_8446622_RNA45SN1_45S_50ntFlanks | 4453 | 4471 | 19 | TCGGCGCCCCCTCGATGCT | 102.6 | -10.5 | 1.6 | -6.6 | 5.0E-11 | 1.4E-10 | MB231/MB468 | MP |
| >am_trnaMT_ProTGG_MT_-_15956_16023@52.70.19\|AAAGACTTTTTCTCTGACC | >am_trnaMT_ProTGG_MT_-_15956_16023 | 52 | 70 | 19 | AAAGACTTTTTCTCTGACC | 1379.6 | 6.4 | 1.0 | 6.6 | 3.9E-11 | 1.4E-10 | BT20/MB468 | MITO |
| >hg38_21_+_8433172_8446622_RNA45SN1_45S_50ntFlanks@12324.12351.28\|TTGGGTTTTAAGCAGGAGGTGTCAGAAA | >hg38_21_+_8433172_8446622_RNA45SN1_45S_50ntFlanks | 12324 | 12351 | 28 | TTGGGTTTTAAGCAGGAGGTGTCAGAAA | 84.4 | -10.3 | 1.6 | -6.6 | 5.0E-11 | 1.4E-10 | MB231/MB468 | MP |
| >am_trna119_LysCTT_1_-_145395522_145395594@16.33.18,trna11_LysCTT_5_-_180648979_180649051@16.33.18,trna128_LysCTT_1_-_55423542_55423614@16.33.18,trna13_LysCTT_6_+_26556774_26556846@16.33.18,trna6_LysCTT_19_-_52425396_52425468@16.33.18,trna7_LysCTT_16_+_3225692_3225764@16.33.18,trna9_LysCTT_5_+_180634755_180634827@16.33.18\|TCGGTAGAGCATGAGACT | >am_trna119_LysCTT_1_-_145395522_145395594 | 16 | 33 | 18 | TCGGTAGAGCATGAGACT | 526.4 | 7.4 | 1.1 | 6.6 | 5.4E-11 | 1.4E-10 | MB231/MB468 | NUC |
| >MI0000081\|hsa-mir-24-2&WithFlank&19\|-\|13836281\|13836365@56.77.22;MI0000080\|hsa-mir-24-1&WithFlank&9\|+\|95086015\|95086094@50.71.22&[MIMAT0000080&hsa-miR-24-3p&offsets\|0\|0;m-36&19\|-\|13836288\|13836310&offsets\|0\|-1];[MIMAT0000080_1&hsa-miR-24-3p&offsets\|0\|0;m-35&9\|+\|95086064\|95086086&offsets\|0\|-1]\|TGGCTCAGTTCAGCAGGAACAG | >MI0000081\|hsa-mir-24-2&WithFlank&19\|-\|13836281\|13836365 | 56 | 77 | 22 | TGGCTCAGTTCAGCAGGAACAG | 12012.7 | 4.3 | 0.7 | 6.6 | 5.4E-11 | 1.4E-10 | MB231/MB468 | NUC |
| >hg38_21_+_8433172_8446622_RNA45SN1_45S_50ntFlanks@9198.9224.27\|GTTCTCTCGGGGCCACGCGCGCGTCCC | >hg38_21_+_8433172_8446622_RNA45SN1_45S_50ntFlanks | 9198 | 9224 | 27 | GTTCTCTCGGGGCCACGCGCGCGTCCC | 299.7 | -11.3 | 1.6 | -6.9 | 3.7E-12 | 1.4E-10 | BT20/MB468 | MP |
| >hg38_21_+_8433172_8446622_RNA45SN1_45S_50ntFlanks@4368.4389.22\|AGTTAAAAAGCTCGTAGTTGGA | >hg38_21_+_8433172_8446622_RNA45SN1_45S_50ntFlanks | 4368 | 4389 | 22 | AGTTAAAAAGCTCGTAGTTGGA | 106.6 | -10.6 | 1.6 | -6.6 | 5.1E-11 | 1.4E-10 | MB231/MB468 | MP |
| >hg38_21_+_8433172_8446622_RNA45SN1_45S_50ntFlanks@3705.3745.41\|TACCTGGTTGATCCTGCCAGTAGCATATGCTTGTCTCAAAG | >hg38_21_+_8433172_8446622_RNA45SN1_45S_50ntFlanks | 3705 | 3745 | 41 | TACCTGGTTGATCCTGCCAGTAGCATATGCTTGTCTCAAAG | 1144.2 | 5.8 | 0.9 | 6.6 | 5.4E-11 | 1.4E-10 | MB231/MB468 | CYTO |
| >hg38_21_+_8433172_8446622_RNA45SN1_45S_50ntFlanks@6738.6757.20\|ATCATCGACACTTCGAACGC | >hg38_21_+_8433172_8446622_RNA45SN1_45S_50ntFlanks | 6738 | 6757 | 20 | ATCATCGACACTTCGAACGC | 88.9 | -10.3 | 1.6 | -6.6 | 5.1E-11 | 1.4E-10 | MB231/MB468 | MP |
| >hg38_21_+_8433172_8446622_RNA45SN1_45S_50ntFlanks@10245.10262.18\|CCGCGACGAGTAGGAGGG | >hg38_21_+_8433172_8446622_RNA45SN1_45S_50ntFlanks | 10245 | 10262 | 18 | CCGCGACGAGTAGGAGGG | 97.8 | -10.5 | 1.6 | -6.6 | 5.1E-11 | 1.4E-10 | MB231/MB468 | MP |
| >hg38_21_+_8433172_8446622_RNA45SN1_45S_50ntFlanks@10409.10428.20\|GTCAGTCGGTCCTGAGAGAT | >hg38_21_+_8433172_8446622_RNA45SN1_45S_50ntFlanks | 10409 | 10428 | 20 | GTCAGTCGGTCCTGAGAGAT | 556.0 | -7.1 | 1.1 | -6.5 | 6.6E-11 | 1.5E-10 | BT20/MB468 | NUC |
| >hg38_21_+_8433172_8446622_RNA45SN1_45S_50ntFlanks@11576.11600.25\|CTTAGAACTGGTGCGGACCAGGGGA | >hg38_21_+_8433172_8446622_RNA45SN1_45S_50ntFlanks | 11576 | 11600 | 25 | CTTAGAACTGGTGCGGACCAGGGGA | 1003.9 | -9.4 | 1.4 | -6.5 | 6.6E-11 | 1.5E-10 | BT20/MB468 | NUC |
| >hg38_21_+_8433172_8446622_RNA45SN1_45S_50ntFlanks@10871.10889.19\|GGCTGGGTCGGTCGGGCTG | >hg38_21_+_8433172_8446622_RNA45SN1_45S_50ntFlanks | 10871 | 10889 | 19 | GGCTGGGTCGGTCGGGCTG | 95.7 | -10.4 | 1.6 | -6.6 | 5.1E-11 | 1.5E-10 | MB231/MB468 | MP |
| >hg38_21_+_8433172_8446622_RNA45SN1_45S_50ntFlanks@3771.3794.24\|CACGGCCGGTACAGTGAAACTGCG | >hg38_21_+_8433172_8446622_RNA45SN1_45S_50ntFlanks | 3771 | 3794 | 24 | CACGGCCGGTACAGTGAAACTGCG | 958.6 | -9.5 | 1.5 | -6.5 | 6.6E-11 | 1.5E-10 | BT20/MB468 | NUC |
| >MI0000459\|hsa-mir-143&WithFlank&5\|+\|149428912\|149429029@67.86.20&[MIMAT0000435&hsa-miR-143-3p&offsets\|0\|-1;m-3&5\|+\|149428978\|149428997&offsets\|0\|0]\|TGAGATGAAGCACTGTAGCT | >MI0000459\|hsa-mir-143&WithFlank&5\|+\|149428912\|149429029 | 67 | 86 | 20 | TGAGATGAAGCACTGTAGCT | 105.5 | -10.6 | 1.6 | -6.6 | 5.1E-11 | 1.5E-10 | MB231/MB468 | MP |
| >hg38_21_+_8433172_8446622_RNA45SN1_45S_50ntFlanks@8335.8375.41\|CAAGTACCGTAAGGGAAAGTTGAAAAGAACTTTGAAGAGAG | >hg38_21_+_8433172_8446622_RNA45SN1_45S_50ntFlanks | 8335 | 8375 | 41 | CAAGTACCGTAAGGGAAAGTTGAAAAGAACTTTGAAGAGAG | 547.5 | 10.3 | 1.6 | 6.6 | 4.0E-11 | 1.5E-10 | BT20/MB468 | MITO |
| >hg38_21_+_8433172_8446622_RNA45SN1_45S_50ntFlanks@10768.10785.18\|CAGCAGGTCTCCAAGGTG | >hg38_21_+_8433172_8446622_RNA45SN1_45S_50ntFlanks | 10768 | 10785 | 18 | CAGCAGGTCTCCAAGGTG | 1088.9 | -8.9 | 1.4 | -6.6 | 5.1E-11 | 1.5E-10 | MB231/MB468 | MP |
| >hg38_21_+_8433172_8446622_RNA45SN1_45S_50ntFlanks@12627.12662.36\|ATGGGGCGAAGCTACCATCTGTGGGATTATGACTGA | >hg38_21_+_8433172_8446622_RNA45SN1_45S_50ntFlanks | 12627 | 12662 | 36 | ATGGGGCGAAGCTACCATCTGTGGGATTATGACTGA | 109.2 | -10.6 | 1.6 | -6.6 | 5.1E-11 | 1.5E-10 | MB231/MB468 | MP |
| >hg38_21_+_8433172_8446622_RNA45SN1_45S_50ntFlanks@11212.11233.22\|CGACTCTGGACGCGAGCCGGGC | >hg38_21_+_8433172_8446622_RNA45SN1_45S_50ntFlanks | 11212 | 11233 | 22 | CGACTCTGGACGCGAGCCGGGC | 214.4 | -7.1 | 1.1 | -6.6 | 4.9E-11 | 1.5E-10 | BT20/MB468 | CYTO |
| >MI0000298\|hsa-mir-221&WithFlank&X\|-\|45746151\|45746272@31.55.25&[MIMAT0004568&hsa-miR-221-5p&offsets\|0\|+3;m-405&X\|-\|45746220\|45746242&offsets\|0\|+2]\|ACCTGGCATACAATGTAGATTTCTG | >MI0000298\|hsa-mir-221&WithFlank&X\|-\|45746151\|45746272 | 31 | 55 | 25 | ACCTGGCATACAATGTAGATTTCTG | 536.8 | 6.9 | 1.1 | 6.6 | 5.5E-11 | 1.5E-10 | MB231/MB468 | CYTO |
| >hg38_21_+_8433172_8446622_RNA45SN1_45S_50ntFlanks@5059.5098.40\|CGATTTGTCTGGTTAATTCCGATAACGAACGAGACTCTGG | >hg38_21_+_8433172_8446622_RNA45SN1_45S_50ntFlanks | 5059 | 5098 | 40 | CGATTTGTCTGGTTAATTCCGATAACGAACGAGACTCTGG | 1141.1 | -14.0 | 2.1 | -6.6 | 5.1E-11 | 1.5E-10 | MB231/MB468 | MP |
| >hg38_21_+_8433172_8446622_RNA45SN1_45S_50ntFlanks@13002.13020.19\|CGATCTATTGAAAGTCAGC | >hg38_21_+_8433172_8446622_RNA45SN1_45S_50ntFlanks | 13002 | 13020 | 19 | CGATCTATTGAAAGTCAGC | 242.8 | -11.0 | 1.6 | -6.9 | 3.8E-12 | 1.5E-10 | BT20/MB468 | MP |
| >hg38_21_+_8433172_8446622_RNA45SN1_45S_50ntFlanks@8342.8375.34\|CGTAAGGGAAAGTTGAAAAGAACTTTGAAGAGAG | >hg38_21_+_8433172_8446622_RNA45SN1_45S_50ntFlanks | 8342 | 8375 | 34 | CGTAAGGGAAAGTTGAAAAGAACTTTGAAGAGAG | 315.2 | 8.6 | 1.3 | 6.6 | 5.5E-11 | 1.5E-10 | MB231/MB468 | CYTO |
| >hg38_21_+_8433172_8446622_RNA45SN1_45S_50ntFlanks@12638.12660.23\|CTACCATCTGTGGGATTATGACT | >hg38_21_+_8433172_8446622_RNA45SN1_45S_50ntFlanks | 12638 | 12660 | 23 | CTACCATCTGTGGGATTATGACT | 100.2 | -10.5 | 1.6 | -6.6 | 5.1E-11 | 1.5E-10 | MB231/MB468 | MP |
| >trna10_AspGTC_12_-_125424193_125424264@35.75.41,trna12_AspGTC_12_-_125411891_125411962@35.75.41,trna144_AspGTC_6_-_27551236_27551307@35.75.41,trna38_AspGTC_17_-_8125556_8125627@35.75.41,trna45_AspGTC_6_+_27447453_27447524@35.75.41,trna48_AspGTC_6_+_27471523_27471594@35.75.41,trna4_AspGTC_12_+_96429799_96429870@35.75.41,trna69_AspGTC_1_-_161440205_161440276@35.75.41,trna72_AspGTC_1_-_161432824_161432895@35.75.41,trna75_AspGTC_1_-_161425414_161425485@35.75.41,trna78_AspGTC_1_-_161418033_161418104@35.75.41,trna81_AspGTC_1_-_161410615_161410686@35.75.41\|TCACGCGGGAGACCGGGGTTCGATTCCCCGACGGGGAGCCA | >trna10_AspGTC_12_-_125424193_125424264 | 35 | 75 | 41 | TCACGCGGGAGACCGGGGTTCGATTCCCCGACGGGGAGCCA | 1148.0 | 5.7 | 0.9 | 6.6 | 4.9E-11 | 1.5E-10 | BT20/MB468 | CYTO |
| >MI0000088\|hsa-mir-30a&WithFlank&6\|-\|71403545\|71403627@54.74.21(+1A)&[MIMAT0000088&hsa-miR-30a-3p&offsets\|+1\|0(+1A);m-38&6\|-\|71403554\|71403575&offsets\|+1\|0(+1A)]\|TTTCAGTCGGATGTTTGCAGCA | >MI0000088\|hsa-mir-30a&WithFlank&6\|-\|71403545\|71403627 | 54 | 74 | 21(+1A) | TTTCAGTCGGATGTTTGCAGCA | 222.8 | 6.8 | 1.0 | 6.6 | 5.5E-11 | 1.5E-10 | MB231/MB468 | NUC |
| >hg38_21_+_8433172_8446622_RNA45SN1_45S_50ntFlanks@12634.12652.19\|GAAGCTACCATCTGTGGGA | >hg38_21_+_8433172_8446622_RNA45SN1_45S_50ntFlanks | 12634 | 12652 | 19 | GAAGCTACCATCTGTGGGA | 98.3 | -10.5 | 1.6 | -6.6 | 5.1E-11 | 1.5E-10 | MB231/MB468 | MP |

<table>
<tr><td>>MI0000285|hsa-mir-205&WithFlank&1|+|209432127|209432248@40.60.21(+1U)&[MIMAT0000266&hsa-miR-205-5p&offsets|0|-1(+1U);m-351&1|+|209432166|209432187&offsets|0|-1(+1U)]|TCCTTCATTCCACCGGAGTCTT</td><td>>MI0000285|hsa-mir-205&WithFlank&1|+|209432127|209432248</td><td>40</td><td>60</td><td>21(+1U)</td><td>TCCTTCATTCCACCGGAGTCTT</td><td>641.3</td><td>12.7</td><td>1.8</td><td>7.2</td><td>8.6E-13</td><td>1.5E-10</td><td>BT20/MB231</td><td>MP</td></tr>
<tr><td>>MI0000102|hsa-mir-100&WithFlank&11|-|122152223|122152314@20.39.20&[MIMAT0000098&hsa-miR-100-5p&offsets|+1|-1;m-27&11|-|122152276|122152296&offsets|+1|0]|ACCCGTAGATCCGAACTTGT</td><td>>MI0000102|hsa-mir-100&WithFlank&11|-|122152223|122152314</td><td>20</td><td>39</td><td>20</td><td>ACCCGTAGATCCGAACTTGT</td><td>475.2</td><td>-12.3</td><td>1.7</td><td>-7.2</td><td>8.5E-13</td><td>1.5E-10</td><td>BT20/MB231</td><td>MP</td></tr>
<tr><td>>hg38_21_+_8433172_8446622_RNA45SN1_45S_50ntFlanks@11920.11951.32|ATGAGAGGTGTAGAATAAGTGGGAGGCCCCCG</td><td>>hg38_21_+_8433172_8446622_RNA45SN1_45S_50ntFlanks</td><td>11920</td><td>11951</td><td>32</td><td>ATGAGAGGTGTAGAATAAGTGGGAGGCCCCCG</td><td>426.4</td><td>-5.6</td><td>0.9</td><td>-6.6</td><td>5.0E-11</td><td>1.5E-10</td><td>BT20/MB468</td><td>CYTO</td></tr>
<tr><td>>MI0000809|hsa-mir-151a&WithFlank&8|-|140732558|140732659@53.74.22(+1U)&[MIMAT0000757&hsa-miR-151a-3p&offsets|0|+1(+1U);m-30&8|-|140732587|140732607&offsets|0|+1(+1U)]|CTAGACTGAAGCTCCTTGAGGAT</td><td>>MI0000809|hsa-mir-151a&WithFlank&8|-|140732558|140732659</td><td>53</td><td>74</td><td>22(+1U)</td><td>CTAGACTGAAGCTCCTTGAGGAT</td><td>1149.1</td><td>4.5</td><td>0.7</td><td>6.6</td><td>5.2E-11</td><td>1.5E-10</td><td>MB231/MB468</td><td>MP</td></tr>
<tr><td>>hg38_21_+_8433172_8446622_RNA45SN1_45S_50ntFlanks@9736.9757.22|CCGAAACGATCTCAACCTATTC</td><td>>hg38_21_+_8433172_8446622_RNA45SN1_45S_50ntFlanks</td><td>9736</td><td>9757</td><td>22</td><td>CCGAAACGATCTCAACCTATTC</td><td>370.8</td><td>-11.6</td><td>1.7</td><td>-6.9</td><td>3.8E-12</td><td>1.5E-10</td><td>BT20/MB468</td><td>MP</td></tr>
<tr><td>>hg38_21_+_8433172_8446622_RNA45SN1_45S_50ntFlanks@7977.8011.35|CGACCTCAGATCAGACGTGGCGACCCGCTGAATTT</td><td>>hg38_21_+_8433172_8446622_RNA45SN1_45S_50ntFlanks</td><td>7977</td><td>8011</td><td>35</td><td>CGACCTCAGATCAGACGTGGCGACCCGCTGAATTT</td><td>1613.9</td><td>3.9</td><td>0.6</td><td>6.6</td><td>4.0E-11</td><td>1.5E-10</td><td>BT20/MB468</td><td>MITO</td></tr>
<tr><td>>hg38_21_+_8433172_8446622_RNA45SN1_45S_50ntFlanks@9027.9065.39|CGTCTCCTCGTGGGGGGGCCGGGCCACCCCTCCCACGGC</td><td>>hg38_21_+_8433172_8446622_RNA45SN1_45S_50ntFlanks</td><td>9027</td><td>9065</td><td>39</td><td>CGTCTCCTCGTGGGGGGGCCGGGCCACCCCTCCCACGGC</td><td>1013.6</td><td>-4.8</td><td>0.7</td><td>-6.6</td><td>4.0E-11</td><td>1.5E-10</td><td>BT20/MB468</td><td>MITO</td></tr>
<tr><td>>hg38_21_+_8433172_8446622_RNA45SN1_45S_50ntFlanks@10351.10380.30|CGAGAACTTTGAAGGCCGAAGTGGAGAAGG</td><td>>hg38_21_+_8433172_8446622_RNA45SN1_45S_50ntFlanks</td><td>10351</td><td>10380</td><td>30</td><td>CGAGAACTTTGAAGGCCGAAGTGGAGAAGG</td><td>1628.9</td><td>-7.8</td><td>1.2</td><td>-6.5</td><td>6.7E-11</td><td>1.5E-10</td><td>BT20/MB468</td><td>NUC</td></tr>
<tr><td>>MI0000737|hsa-mir-200a&WithFlank&1|+|1167857|1167958@22.43.22&[MIMAT0001620&hsa-miR-200a-5p&offsets|0|0;m-154&1|+|1167878|1167899&offsets|0|0]|CATCTTACCGGACAGTGCTGGA</td><td>>MI0000737|hsa-mir-200a&WithFlank&1|+|1167857|1167958</td><td>22</td><td>43</td><td>22</td><td>CATCTTACCGGACAGTGCTGGA</td><td>143.3</td><td>-4.7</td><td>0.7</td><td>-6.6</td><td>5.5E-11</td><td>1.5E-10</td><td>MB231/MB468</td><td>CYTO</td></tr>
<tr><td>>MI0000081|hsa-mir-24-2&WithFlank&19|-|13836281|13836365@56.76.21;MI0000080|hsa-mir-24-1&WithFlank&9|+|95086015|95086094@50.70.21&[MIMAT0000080&hsa-miR-24-3p&offsets|0|-1;m-36&19|-|13836288|13836310&offsets|0|-2];[MIMAT0000080_1&hsa-miR-24-3p&offsets|0|-1;m-35&9|+|95086064|95086086&offsets|0|-2]|TGGCTCAGTTCAGCAGGAACA</td><td>>MI0000081|hsa-mir-24-2&WithFlank&19|-|13836281|13836365</td><td>56</td><td>76</td><td>21</td><td>TGGCTCAGTTCAGCAGGAACA</td><td>2404.5</td><td>3.9</td><td>0.6</td><td>6.6</td><td>5.5E-11</td><td>1.5E-10</td><td>MB231/MB468</td><td>NUC</td></tr>
<tr><td>>am_trna116_GluCTC_1_-_145399233_145399304@1.29.29,trna59_GluCTC_1_+_249168447_249168518@1.29.29,trna71_GluCTC_1_-_161439189_161439260@1.29.29,trna74_GluCTC_1_-_161431809_161431880@1.29.29,trna77_GluCTC_1_-_161424398_161424469@1.29.29,trna77_GluCTC_6_+_28949976_28950047@1.29.29,trna80_GluCTC_1_-_161417018_161417089@1.29.29,trna87_GluCTC_6_-_126101393_126101464@1.29.29|TCCCTGGTGGTCTAGTGGTTAGGATTCGG</td><td>>am_trna116_GluCTC_1_-_145399233_145399304</td><td>1</td><td>29</td><td>29</td><td>TCCCTGGTGGTCTAGTGGTTAGGATTCGG</td><td>62813.2</td><td>-3.7</td><td>0.6</td><td>-6.6</td><td>5.0E-11</td><td>1.5E-10</td><td>BT20/MB468</td><td>CYTO</td></tr>
<tr><td>>hg38_21_+_8433172_8446622_RNA45SN1_45S_50ntFlanks@4759.4776.18|ATCAGATACCGTCGTAGT</td><td>>hg38_21_+_8433172_8446622_RNA45SN1_45S_50ntFlanks</td><td>4759</td><td>4776</td><td>18</td><td>ATCAGATACCGTCGTAGT</td><td>221.4</td><td>-10.1</td><td>1.5</td><td>-6.6</td><td>5.0E-11</td><td>1.5E-10</td><td>BT20/MB468</td><td>CYTO</td></tr>
<tr><td>>hg38_21_+_8433172_8446622_RNA45SN1_45S_50ntFlanks@9806.9830.25|CGTGGAATGCGAGTGCCTAGTGGGC</td><td>>hg38_21_+_8433172_8446622_RNA45SN1_45S_50ntFlanks</td><td>9806</td><td>9830</td><td>25</td><td>CGTGGAATGCGAGTGCCTAGTGGGC</td><td>491.0</td><td>-8.2</td><td>1.3</td><td>-6.5</td><td>6.3E-11</td><td>1.5E-10</td><td>MB231/MB468</td><td>MITO</td></tr>
<tr><td>>hg38_21_+_8433172_8446622_RNA45SN1_45S_50ntFlanks@10782.10801.20|GGTGAACAGCCTCTGGCATG</td><td>>hg38_21_+_8433172_8446622_RNA45SN1_45S_50ntFlanks</td><td>10782</td><td>10801</td><td>20</td><td>GGTGAACAGCCTCTGGCATG</td><td>530.2</td><td>-6.5</td><td>1.0</td><td>-6.6</td><td>5.5E-11</td><td>1.5E-10</td><td>MB231/MB468</td><td>NUC</td></tr>
<tr><td>>MI0000298|hsa-mir-221&WithFlank&X|-|45746151|45746272@71.93.23(+1C)&[MIMAT0000278&hsa-miR-221-3p&offsets|0|0(+1C);m-91&X|-|45746181|45746202&offsets|0|+1(+1C)]|AGCTACATTGTCTGCTGGGTTTCC</td><td>>MI0000298|hsa-mir-221&WithFlank&X|-|45746151|45746272</td><td>71</td><td>93</td><td>23(+1C)</td><td>AGCTACATTGTCTGCTGGGTTTCC</td><td>1689.9</td><td>6.2</td><td>0.9</td><td>6.5</td><td>6.3E-11</td><td>1.5E-10</td><td>MB231/MB468</td><td>MITO</td></tr>
<tr><td>>MI0000737|hsa-mir-200a&WithFlank&1|+|1167857|1167958@60.80.21&[MIMAT0000682&hsa-miR-200a-3p&offsets|0|-1;m-142&1|+|1167916|1167938&offsets|0|-2]|TAACACTGTCTGGTAACGATG</td><td>>MI0000737|hsa-mir-200a&WithFlank&1|+|1167857|1167958</td><td>60</td><td>80</td><td>21</td><td>TAACACTGTCTGGTAACGATG</td><td>536.5</td><td>-6.7</td><td>1.0</td><td>-6.6</td><td>5.2E-11</td><td>1.5E-10</td><td>MB231/MB468</td><td>MP</td></tr>
<tr><td>>hg38_21_+_8433172_8446622_RNA45SN1_45S_50ntFlanks@6646.6679.34|TCGTACGACTCTTAGCGGTGGATCACTCGGCTCG</td><td>>hg38_21_+_8433172_8446622_RNA45SN1_45S_50ntFlanks</td><td>6646</td><td>6679</td><td>34</td><td>TCGTACGACTCTTAGCGGTGGATCACTCGGCTCG</td><td>7401.8</td><td>-3.1</td><td>0.5</td><td>-6.5</td><td>6.7E-11</td><td>1.5E-10</td><td>BT20/MB468</td><td>NUC</td></tr>
<tr><td>>hg38_21_+_8433172_8446622_RNA45SN1_45S_50ntFlanks@12461.12478.18|CGTTGGATTGTTCACCCA</td><td>>hg38_21_+_8433172_8446622_RNA45SN1_45S_50ntFlanks</td><td>12461</td><td>12478</td><td>18</td><td>CGTTGGATTGTTCACCCA</td><td>95.5</td><td>-10.4</td><td>1.6</td><td>-6.6</td><td>5.2E-11</td><td>1.5E-10</td><td>MB231/MB468</td><td>MP</td></tr>
<tr><td>>hg38_21_+_8433172_8446622_RNA45SN1_45S_50ntFlanks@8359.8379.21|AAGAACTTTGAAGAGAGAGTT</td><td>>hg38_21_+_8433172_8446622_RNA45SN1_45S_50ntFlanks</td><td>8359</td><td>8379</td><td>21</td><td>AAGAACTTTGAAGAGAGAGTT</td><td>1018.2</td><td>-9.0</td><td>1.4</td><td>-6.6</td><td>5.2E-11</td><td>1.5E-10</td><td>MB231/MB468</td><td>MP</td></tr>
<tr><td>>hg38_21_+_8433172_8446622_RNA45SN1_45S_50ntFlanks@11718.11742.25|AACGGCGGGAGTAACTATGACTCTC</td><td>>hg38_21_+_8433172_8446622_RNA45SN1_45S_50ntFlanks</td><td>11718</td><td>11742</td><td>25</td><td>AACGGCGGGAGTAACTATGACTCTC</td><td>3731.1</td><td>-5.8</td><td>0.9</td><td>-6.6</td><td>5.0E-11</td><td>1.5E-10</td><td>BT20/MB468</td><td>CYTO</td></tr>
<tr><td>>hg38_21_+_8433172_8446622_RNA45SN1_45S_50ntFlanks@12006.12043.38|GGCCGCCGGTGAAATACCACTACTCTGATCGTTTTTTC</td><td>>hg38_21_+_8433172_8446622_RNA45SN1_45S_50ntFlanks</td><td>12006</td><td>12043</td><td>38</td><td>GGCCGCCGGTGAAATACCACTACTCTGATCGTTTTTTC</td><td>86.6</td><td>-21.2</td><td>3.0</td><td>-6.9</td><td>3.9E-12</td><td>1.5E-10</td><td>BT20/MB468</td><td>MP</td></tr>
<tr><td>>MI0000088|hsa-mir-30a&WithFlank&6|-|71403545|71403627@53.71.19&[MIMAT0000088&hsa-miR-30a-3p&offsets|0|-3;m-38&6|-|71403554|71403575&offsets|0|-3]|CTTTCAGTCGGATGTTTGC</td><td>>MI0000088|hsa-mir-30a&WithFlank&6|-|71403545|71403627</td><td>53</td><td>71</td><td>19</td><td>CTTTCAGTCGGATGTTTGC</td><td>665.4</td><td>5.6</td><td>0.8</td><td>6.6</td><td>4.1E-11</td><td>1.5E-10</td><td>BT20/MB468</td><td>MITO</td></tr>
<tr><td>>hg38_21_+_8433172_8446622_RNA45SN1_45S_50ntFlanks@12004.12026.23|CGGGCCGCCGGTGAAATACCACT</td><td>>hg38_21_+_8433172_8446622_RNA45SN1_45S_50ntFlanks</td><td>12004</td><td>12026</td><td>23</td><td>CGGGCCGCCGGTGAAATACCACT</td><td>641.8</td><td>7.5</td><td>1.1</td><td>6.6</td><td>4.1E-11</td><td>1.5E-10</td><td>BT20/MB468</td><td>MITO</td></tr>
<tr><td>>hg38_21_+_8433172_8446622_RNA45SN1_45S_50ntFlanks@8406.8442.37|TAAACGGGTGGGGTCCGCGCAGTCCGCCCGGAGGATT</td><td>>hg38_21_+_8433172_8446622_RNA45SN1_45S_50ntFlanks</td><td>8406</td><td>8442</td><td>37</td><td>TAAACGGGTGGGGTCCGCGCAGTCCGCCCGGAGGATT</td><td>93.2</td><td>-10.4</td><td>1.6</td><td>-6.6</td><td>5.2E-11</td><td>1.5E-10</td><td>MB231/MB468</td><td>MP</td></tr>
<tr><td>>hg38_1_-_228634819_228635039_RNA5S12_5S_50ntFlanks@144.169.26|CCTGGGAATACCGGGTGCTGTAGGCT</td><td>>hg38_1_-_228634819_228635039_RNA5S12_5S_50ntFlanks</td><td>144</td><td>169</td><td>26</td><td>CCTGGGAATACCGGGTGCTGTAGGCT</td><td>467.6</td><td>6.6</td><td>1.0</td><td>6.6</td><td>5.6E-11</td><td>1.5E-10</td><td>MB231/MB468</td><td>CYTO</td></tr>
<tr><td>>hg38_21_+_8433172_8446622_RNA45SN1_45S_50ntFlanks@10845.10865.21|ACTTCGGGATAAGGATTGGCT</td><td>>hg38_21_+_8433172_8446622_RNA45SN1_45S_50ntFlanks</td><td>10845</td><td>10865</td><td>21</td><td>ACTTCGGGATAAGGATTGGCT</td><td>8996.2</td><td>-6.1</td><td>0.9</td><td>-6.6</td><td>5.1E-11</td><td>1.5E-10</td><td>BT20/MB468</td><td>CYTO</td></tr>
<tr><td>>MI0000542|hsa-mir-320a&WithFlank&8|-|22244960|22245043@48.69.22(+2U)&[MIMAT0000510&hsa-miR-320a-3p&offsets|0|0(+2U);m-60&8|-|22244975|22244996&offsets|0|0(+2U)]|AAAAGCTGGGTTGAGAGGGCGATT</td><td>>MI0000542|hsa-mir-320a&WithFlank&8|-|22244960|22245043</td><td>48</td><td>69</td><td>22(+2U)</td><td>AAAAGCTGGGTTGAGAGGGCGATT</td><td>1131.4</td><td>5.2</td><td>0.8</td><td>6.6</td><td>5.6E-11</td><td>1.5E-10</td><td>MB231/MB468</td><td>NUC</td></tr>
<tr><td>>hg38_21_+_8433172_8446622_RNA45SN1_45S_50ntFlanks@9738.9762.25|GAAACGATCTCAACCTATTCTCAAA</td><td>>hg38_21_+_8433172_8446622_RNA45SN1_45S_50ntFlanks</td><td>9738</td><td>9762</td><td>25</td><td>GAAACGATCTCAACCTATTCTCAAA</td><td>92.9</td><td>-10.4</td><td>1.6</td><td>-6.6</td><td>5.3E-11</td><td>1.5E-10</td><td>MB231/MB468</td><td>MP</td></tr>
<tr><td>>hg38_21_+_8433172_8446622_RNA45SN1_45S_50ntFlanks@11879.11904.26|ACCCTGTTGAGCTTGACTCTAGTCTG</td><td>>hg38_21_+_8433172_8446622_RNA45SN1_45S_50ntFlanks</td><td>11879</td><td>11904</td><td>26</td><td>ACCCTGTTGAGCTTGACTCTAGTCTG</td><td>147.1</td><td>-4.3</td><td>0.7</td><td>-6.6</td><td>5.6E-11</td><td>1.5E-10</td><td>MB231/MB468</td><td>CYTO</td></tr>
<tr><td>>hg38_21_+_8433172_8446622_RNA45SN1_45S_50ntFlanks@6647.6677.31|CGTACGACTCTTAGCGGTGGATCACTCGGCT</td><td>>hg38_21_+_8433172_8446622_RNA45SN1_45S_50ntFlanks</td><td>6647</td><td>6677</td><td>31</td><td>CGTACGACTCTTAGCGGTGGATCACTCGGCT</td><td>3399.6</td><td>4.4</td><td>0.7</td><td>6.6</td><td>4.1E-11</td><td>1.5E-10</td><td>BT20/MB468</td><td>MITO</td></tr>
<tr><td>>trna116_GluCTC_1_-_145399233_145399304@32.51.20,trna59_GluCTC_1_+_249168447_249168518@32.51.20,trna71_GluCTC_1_-_161439189_161439260@32.51.20,trna74_GluCTC_1_-_161431809_161431880@32.51.20,trna77_GluCTC_1_-_161424398_161424469@32.51.20,trna77_GluCTC_6_+_28949976_28950047@32.51.20,trna80_GluCTC_1_-_161417018_161417089@32.51.20,trna87_GluCTC_6_-_126101393_126101464@32.51.20|CTCTCACCGCCGCGGCCCGG</td><td>>trna116_GluCTC_1_-_145399233_145399304</td><td>32</td><td>51</td><td>20</td><td>CTCTCACCGCCGCGGCCCGG</td><td>459.7</td><td>-11.8</td><td>1.8</td><td>-6.6</td><td>5.6E-11</td><td>1.5E-10</td><td>MB231/MB468</td><td>NUC</td></tr>
<tr><td>>hg38_21_+_8433172_8446622_RNA45SN1_45S_50ntFlanks@12539.12567.29|ATGATGTGTTGTTGCCATGGTAATCCTGC</td><td>>hg38_21_+_8433172_8446622_RNA45SN1_45S_50ntFlanks</td><td>12539</td><td>12567</td><td>29</td><td>ATGATGTGTTGTTGCCATGGTAATCCTGC</td><td>645.0</td><td>-6.1</td><td>0.9</td><td>-6.6</td><td>5.6E-11</td><td>1.5E-10</td><td>MB231/MB468</td><td>CYTO</td></tr>
<tr><td>>hg38_21_+_8433172_8446622_RNA45SN1_45S_50ntFlanks@6736.6771.36|TGATCATCGACACTTCGAACGCACTTGCGGCCCCGG</td><td>>hg38_21_+_8433172_8446622_RNA45SN1_45S_50ntFlanks</td><td>6736</td><td>6771</td><td>36</td><td>TGATCATCGACACTTCGAACGCACTTGCGGCCCCGG</td><td>429.8</td><td>-8.1</td><td>1.2</td><td>-6.5</td><td>6.4E-11</td><td>1.5E-10</td><td>MB231/MB468</td><td>MITO</td></tr>
<tr><td>>hg38_21_+_8433172_8446622_RNA45SN1_45S_50ntFlanks@10687.10711.25|CGGCGTCCGGTGAGCTCTCGCTGGC</td><td>>hg38_21_+_8433172_8446622_RNA45SN1_45S_50ntFlanks</td><td>10687</td><td>10711</td><td>25</td><td>CGGCGTCCGGTGAGCTCTCGCTGGC</td><td>10807.6</td><td>-9.4</td><td>1.4</td><td>-6.5</td><td>6.8E-11</td><td>1.5E-10</td><td>BT20/MB468</td><td>NUC</td></tr>
<tr><td>>hg38_21_+_8433172_8446622_RNA45SN1_45S_50ntFlanks@9240.9271.32|GCGGAGCGAGCGCACGGGGTCGGCGGCGACGT</td><td>>hg38_21_+_8433172_8446622_RNA45SN1_45S_50ntFlanks</td><td>9240</td><td>9271</td><td>32</td><td>GCGGAGCGAGCGCACGGGGTCGGCGGCGACGT</td><td>83.6</td><td>-10.2</td><td>1.6</td><td>-6.6</td><td>5.3E-11</td><td>1.5E-10</td><td>MB231/MB468</td><td>MP</td></tr>
<tr><td>>hg38_21_+_8433172_8446622_RNA45SN1_45S_50ntFlanks@12994.13014.21|CCTCGCTGCGATCTATTGAAA</td><td>>hg38_21_+_8433172_8446622_RNA45SN1_45S_50ntFlanks</td><td>12994</td><td>13014</td><td>21</td><td>CCTCGCTGCGATCTATTGAAA</td><td>1941.2</td><td>-10.0</td><td>1.5</td><td>-6.5</td><td>6.8E-11</td><td>1.5E-10</td><td>BT20/MB468</td><td>NUC</td></tr>
<tr><td>>hg38_21_+_8433172_8446622_RNA45SN1_45S_50ntFlanks@10404.10428.25|CATGGGTCAGTCGGTCCTGAGAGAT</td><td>>hg38_21_+_8433172_8446622_RNA45SN1_45S_50ntFlanks</td><td>10404</td><td>10428</td><td>25</td><td>CATGGGTCAGTCGGTCCTGAGAGAT</td><td>81.2</td><td>-10.2</td><td>1.6</td><td>-6.6</td><td>5.3E-11</td><td>1.5E-10</td><td>MB231/MB468</td><td>MP</td></tr>
<tr><td>>hg38_21_+_8433172_8446622_RNA45SN1_45S_50ntFlanks@3787.3810.24|AAACTGCGAATGGCTCATTAAATC</td><td>>hg38_21_+_8433172_8446622_RNA45SN1_45S_50ntFlanks</td><td>3787</td><td>3810</td><td>24</td><td>AAACTGCGAATGGCTCATTAAATC</td><td>1225.7</td><td>-5.6</td><td>0.9</td><td>-6.5</td><td>6.5E-11</td><td>1.5E-10</td><td>MB231/MB468</td><td>MITO</td></tr>
<tr><td>>hg38_21_+_8433172_8446622_RNA45SN1_45S_50ntFlanks@10663.10711.49|CCTTGGAAAGCGTCGCGGTTCCGGCGGCGTCCGGTGAGCTCTCGCTGGC</td><td>>hg38_21_+_8433172_8446622_RNA45SN1_45S_50ntFlanks</td><td>10663</td><td>10711</td><td>49</td><td>CCTTGGAAAGCGTCGCGGTTCCGGCGGCGTCCGGTGAGCTCTCGCTGGC</td><td>790.7</td><td>-12.6</td><td>1.9</td><td>-6.6</td><td>5.7E-11</td><td>1.5E-10</td><td>MB231/MB468</td><td>NUC</td></tr>
<tr><td>>MI0000471|hsa-mir-126&WithFlank&9|+|136670596|136670692@59.78.20&[MIMAT0000445&hsa-miR-126-3p&offsets|+1|-1;m-40&9|+|136670653|136670674&offsets|+1|-1]|CGTACCGTGAGTAATAATGC</td><td>>MI0000471|hsa-mir-126&WithFlank&9|+|136670596|136670692</td><td>59</td><td>78</td><td>20</td><td>CGTACCGTGAGTAATAATGC</td><td>828.8</td><td>8.8</td><td>1.3</td><td>6.6</td><td>5.3E-11</td><td>1.5E-10</td><td>MB231/MB468</td><td>MP</td></tr>
<tr><td>>hg38_21_+_8433172_8446622_RNA45SN1_45S_50ntFlanks@9266.9285.20|CGACGTCGGCTACCCACCCG</td><td>>hg38_21_+_8433172_8446622_RNA45SN1_45S_50ntFlanks</td><td>9266</td><td>9285</td><td>20</td><td>CGACGTCGGCTACCCACCCG</td><td>86.2</td><td>-10.3</td><td>1.6</td><td>-6.6</td><td>5.3E-11</td><td>1.5E-10</td><td>MB231/MB468</td><td>MP</td></tr>
<tr><td>>MI0003628|hsa-mir-615&WithFlank&12|+|54033944|54034051@67.88.22(+1U)&[MIMAT0003283&hsa-miR-615-3p&offsets|0|0(+1U);m-495&12|+|54034010|54034030&offsets|0|+1(+1U)]|TCCGAGCCTGGGTCTCCCTCTTT</td><td>>MI0003628|hsa-mir-615&WithFlank&12|+|54033944|54034051</td><td>67</td><td>88</td><td>22(+1U)</td><td>TCCGAGCCTGGGTCTCCCTCTTT</td><td>151.1</td><td>7.9</td><td>1.1</td><td>7.0</td><td>3.7E-12</td><td>1.5E-10</td><td>BT20/MB231</td><td>CYTO</td></tr>
<tr><td>>hg38_21_+_8433172_8446622_RNA45SN1_45S_50ntFlanks@12255.12273.19|CTCGCTTGATCTTGATTTT</td><td>>hg38_21_+_8433172_8446622_RNA45SN1_45S_50ntFlanks</td><td>12255</td><td>12273</td><td>19</td><td>CTCGCTTGATCTTGATTTT</td><td>72.0</td><td>-10.0</td><td>1.5</td><td>-6.6</td><td>5.3E-11</td><td>1.5E-10</td><td>MB231/MB468</td><td>MP</td></tr>
</table>

<table>
<tr><td>>am_MI0000809|hsa-mir-151a&WithFlank&8|-|140732558|140732659@53.70.18&[MIMAT0000757&hsa-miR-151a-3p&offsets|0|-3;m-30&8|-|140732587|140732607&offsets|0|-3]|CTAGACTGAAGCTCCTTG</td><td>>am_MI0000809|hsa-mir-151a&WithFlank&8|-|140732558|140732659</td><td>53</td><td>70</td><td>18</td><td>CTAGACTGAAGCTCCTTG</td><td>2487.6</td><td>9.7</td><td>1.5</td><td>6.6</td><td>4.2E-11</td><td>1.5E-10</td><td>BT20/MB468</td><td>MITO</td></tr>
<tr><td>>hg38_21_+_8433172_8446622_RNA45SN1_45S_50ntFlanks@11580.11603.24|GAACTGGTGCGGACCAGGGGAATC</td><td>>hg38_21_+_8433172_8446622_RNA45SN1_45S_50ntFlanks</td><td>11580</td><td>11603</td><td>24</td><td>GAACTGGTGCGGACCAGGGGAATC</td><td>10545.1</td><td>-7.9</td><td>1.1</td><td>-6.9</td><td>4.0E-12</td><td>1.5E-10</td><td>BT20/MB468</td><td>MP</td></tr>
<tr><td>>hg38_21_+_8433172_8446622_RNA45SN1_45S_50ntFlanks@4590.4616.27|ATTTTGTTGGTTTTCGGAACTGAGGCC</td><td>>hg38_21_+_8433172_8446622_RNA45SN1_45S_50ntFlanks</td><td>4590</td><td>4616</td><td>27</td><td>ATTTTGTTGGTTTTCGGAACTGAGGCC</td><td>457.1</td><td>-8.2</td><td>1.3</td><td>-6.5</td><td>6.5E-11</td><td>1.5E-10</td><td>MB231/MB468</td><td>MITO</td></tr>
<tr><td>>trna119_LysCTT_1_-_145395522_145395594@3.34.32,trna11_LysCTT_5_-_180648979_180649051@3.34.32,trna13_LysCTT_6_+_26556774_26556846@3.34.32,trna7_LysCTT_16_+_3225692_3225764@3.34.32,trna9_LysCTT_5_+_180634755_180634827@3.34.32|CCGGCTAGCTCAGTCGGTAGAGCATGAGACTC</td><td>>trna119_LysCTT_1_-_145395522_145395594</td><td>3</td><td>34</td><td>32</td><td>CCGGCTAGCTCAGTCGGTAGAGCATGAGACTC</td><td>351.6</td><td>9.4</td><td>1.4</td><td>6.6</td><td>5.7E-11</td><td>1.5E-10</td><td>MB231/MB468</td><td>NUC</td></tr>
<tr><td>>trna10_AspGTC_12_-_125424193_125424264@52.70.19,trna12_AspGTC_12_-_125411891_125411962@52.70.19,trna144_AspGTC_6_-_27551236_27551307@52.70.19,trna38_AspGTC_17_-_8125556_8125627@52.70.19,trna45_AspGTC_6_+_27447453_27447524@52.70.19,trna48_AspGTC_6_+_27471523_27471594@52.70.19,trna4_AspGTC_12_+_96429799_96429870@52.70.19,trna69_AspGTC_1_-_161440205_161440276@52.70.19,trna72_AspGTC_1_-_161432824_161432895@52.70.19,trna75_AspGTC_1_-_161425414_161425485@52.70.19,trna78_AspGTC_1_-_161418033_161418104@52.70.19,trna81_AspGTC_1_-_161410615_161410686@52.70.19|GTTCGATTCCCCGACGGGG</td><td>>trna10_AspGTC_12_-_125424193_125424264</td><td>52</td><td>70</td><td>19</td><td>GTTCGATTCCCCGACGGGG</td><td>120.3</td><td>-10.8</td><td>1.6</td><td>-6.6</td><td>5.4E-11</td><td>1.5E-10</td><td>MB231/MB468</td><td>MP</td></tr>
<tr><td>>hg38_21_+_8433172_8446622_RNA45SN1_45S_50ntFlanks@9862.9894.33|ATGAACCGAACGCCGGGTTAAGGCGCCCGATGC</td><td>>hg38_21_+_8433172_8446622_RNA45SN1_45S_50ntFlanks</td><td>9862</td><td>9894</td><td>33</td><td>ATGAACCGAACGCCGGGTTAAGGCGCCCGATGC</td><td>164.8</td><td>-11.2</td><td>1.7</td><td>-6.6</td><td>5.4E-11</td><td>1.5E-10</td><td>MB231/MB468</td><td>MP</td></tr>
<tr><td>>trna103_AsnGTT_1_-_149230570_149230643@33.53.21,trna107_AsnGTT_1_-_148760356_148760429@33.53.21,trna108_AsnGTT_1_-_148598314_148598387@33.53.21,trna1_AsnGTT_19_+_1383562_1383635@33.53.21,trna26_AsnGTT_1_+_148248115_148248188@33.53.21,trna31_AsnGTT_17_-_36908034_36908107@33.53.21,trna47_AsnGTT_1_+_161510031_161510104@33.53.21,trna4_AsnGTT_10_-_22518438_22518511@33.53.21,trna7_AsnGTT_13_-_31248101_31248174@33.53.21,trna83_AsnGTT_1_-_161397867_161397940@33.53.21|CTGTTAACCGAAAGGTTGGTG</td><td>>trna103_AsnGTT_1_-_149230570_149230643</td><td>33</td><td>53</td><td>21</td><td>CTGTTAACCGAAAGGTTGGTG</td><td>144.5</td><td>-11.0</td><td>1.7</td><td>-6.6</td><td>5.4E-11</td><td>1.5E-10</td><td>MB231/MB468</td><td>MP</td></tr>
<tr><td>>MI0000272|hsa-mir-182&WithFlank&7|-|129770377|129770498@29.52.24(+1A)&[MIMAT0000259&hsa-miR-182-5p&offsets|0|0(+1A);m-22&7|-|129770449|129770470&offsets|0|+2(+1A)]|TTTGGCAATGGTAGAACTCACACTA</td><td>>MI0000272|hsa-mir-182&WithFlank&7|-|129770377|129770498</td><td>29</td><td>52</td><td>24(+1A)</td><td>TTTGGCAATGGTAGAACTCACACTA</td><td>1296.2</td><td>3.5</td><td>0.5</td><td>6.9</td><td>6.2E-12</td><td>1.5E-10</td><td>BT20/MB231</td><td>NUC</td></tr>
<tr><td>>trna10_LysCTT_16_+_3241501_3241573@1.33.33,trna13_LysCTT_14_-_58706613_58706685@1.33.33,trna2_LysCTT_15_+_79152904_79152976@1.33.33|GCCCGGCTAGCTCAGTCGGTAGAGCATGGGACT</td><td>>trna10_LysCTT_16_+_3241501_3241573</td><td>1</td><td>33</td><td>33</td><td>GCCCGGCTAGCTCAGTCGGTAGAGCATGGGACT</td><td>1453.7</td><td>7.5</td><td>1.1</td><td>6.6</td><td>5.8E-11</td><td>1.5E-10</td><td>MB231/MB468</td><td>CYTO</td></tr>
<tr><td>>MI0003188|hsa-mir-503&WithFlank&X|-|134546322|134546404@12.30.19&[MIMAT0002874&hsa-miR-503-5p&offsets|0|-4;m-297&X|-|134546371|134546393&offsets|0|-4]|TAGCAGCGGGAACAGTTCT</td><td>>MI0003188|hsa-mir-503&WithFlank&X|-|134546322|134546404</td><td>12</td><td>30</td><td>19</td><td>TAGCAGCGGGAACAGTTCT</td><td>153.3</td><td>10.5</td><td>1.6</td><td>6.6</td><td>5.4E-11</td><td>1.5E-10</td><td>MB231/MB468</td><td>MP</td></tr>
<tr><td>>hg38_21_+_8433172_8446622_RNA45SN1_45S_50ntFlanks@4731.4758.28|AAGAACGAAAGTCGGAGGTTCGAAGACG</td><td>>hg38_21_+_8433172_8446622_RNA45SN1_45S_50ntFlanks</td><td>4731</td><td>4758</td><td>28</td><td>AAGAACGAAAGTCGGAGGTTCGAAGACG</td><td>10036.5</td><td>-5.8</td><td>0.9</td><td>-6.5</td><td>6.6E-11</td><td>1.5E-10</td><td>MB231/MB468</td><td>MITO</td></tr>
<tr><td>>hg38_21_+_8433172_8446622_RNA45SN1_45S_50ntFlanks@8105.8138.34|CGCGGCGGGGCGCGGGACATGTGGCGTACGGAAG</td><td>>hg38_21_+_8433172_8446622_RNA45SN1_45S_50ntFlanks</td><td>8105</td><td>8138</td><td>34</td><td>CGCGGCGGGGCGCGGGACATGTGGCGTACGGAAG</td><td>11937.0</td><td>-9.1</td><td>1.4</td><td>-6.6</td><td>5.4E-11</td><td>1.5E-10</td><td>MB231/MB468</td><td>MP</td></tr>
<tr><td>>hg38_21_+_8433172_8446622_RNA45SN1_45S_50ntFlanks@10348.10373.26|AAACGAGAACTTTGAAGGCCGAAGTG</td><td>>hg38_21_+_8433172_8446622_RNA45SN1_45S_50ntFlanks</td><td>10348</td><td>10373</td><td>26</td><td>AAACGAGAACTTTGAAGGCCGAAGTG</td><td>380.4</td><td>-3.9</td><td>0.6</td><td>-6.5</td><td>7.0E-11</td><td>1.5E-10</td><td>BT20/MB468</td><td>NUC</td></tr>
<tr><td>>hg38_21_+_8433172_8446622_RNA45SN1_45S_50ntFlanks@10572.10593.22|AGAAGCCGGCGGGAGCCCCGGG</td><td>>hg38_21_+_8433172_8446622_RNA45SN1_45S_50ntFlanks</td><td>10572</td><td>10593</td><td>22</td><td>AGAAGCCGGCGGGAGCCCCGGG</td><td>87.6</td><td>-10.3</td><td>1.6</td><td>-6.6</td><td>5.4E-11</td><td>1.5E-10</td><td>MB231/MB468</td><td>MP</td></tr>
<tr><td>>hg38_21_+_8433172_8446622_RNA45SN1_45S_50ntFlanks@10399.10424.26|TTGAACATGGGTCAGTCGGTCCTGAG</td><td>>hg38_21_+_8433172_8446622_RNA45SN1_45S_50ntFlanks</td><td>10399</td><td>10424</td><td>26</td><td>TTGAACATGGGTCAGTCGGTCCTGAG</td><td>338.8</td><td>4.7</td><td>0.7</td><td>6.6</td><td>5.2E-11</td><td>1.5E-10</td><td>BT20/MB468</td><td>CYTO</td></tr>
<tr><td>>hg38_21_+_8433172_8446622_RNA45SN1_45S_50ntFlanks@8177.8202.26|TTCTGATCGAGGCCCAGCCCGTGGAC</td><td>>hg38_21_+_8433172_8446622_RNA45SN1_45S_50ntFlanks</td><td>8177</td><td>8202</td><td>26</td><td>TTCTGATCGAGGCCCAGCCCGTGGAC</td><td>247.7</td><td>4.3</td><td>0.7</td><td>6.5</td><td>5.8E-11</td><td>1.5E-10</td><td>MB231/MB468</td><td>CYTO</td></tr>
<tr><td>>hg38_21_+_8433172_8446622_RNA45SN1_45S_50ntFlanks@8406.8442.37|TAAACGGGTGGGGTCCGCGCAGTCCGCCCGGAGGATT</td><td>>hg38_21_+_8433172_8446622_RNA45SN1_45S_50ntFlanks</td><td>8406</td><td>8442</td><td>37</td><td>TAAACGGGTGGGGTCCGCGCAGTCCGCCCGGAGGATT</td><td>127.3</td><td>-9.1</td><td>1.4</td><td>-6.6</td><td>4.2E-11</td><td>1.5E-10</td><td>BT20/MB468</td><td>MITO</td></tr>
<tr><td>>hg38_21_+_8433172_8446622_RNA45SN1_45S_50ntFlanks@8472.8496.25|CGGCCCGGCGGATCTTTCCCGCCCC</td><td>>hg38_21_+_8433172_8446622_RNA45SN1_45S_50ntFlanks</td><td>8472</td><td>8496</td><td>25</td><td>CGGCCCGGCGGATCTTTCCCGCCCC</td><td>146.5</td><td>-11.1</td><td>1.7</td><td>-6.6</td><td>5.4E-11</td><td>1.5E-10</td><td>MB231/MB468</td><td>MP</td></tr>
<tr><td>>hg38_21_+_8433172_8446622_RNA45SN1_45S_50ntFlanks@8196.8215.20|CGTGGACGGTGTGAGGCCGG</td><td>>hg38_21_+_8433172_8446622_RNA45SN1_45S_50ntFlanks</td><td>8196</td><td>8215</td><td>20</td><td>CGTGGACGGTGTGAGGCCGG</td><td>4250.2</td><td>-4.9</td><td>0.8</td><td>-6.5</td><td>5.8E-11</td><td>1.5E-10</td><td>MB231/MB468</td><td>CYTO</td></tr>
<tr><td>>hg38_21_+_8433172_8446622_RNA45SN1_45S_50ntFlanks@11572.11607.36|CCGACTTAGAACTGGTGCGGACCAGGGGAATCCGAC</td><td>>hg38_21_+_8433172_8446622_RNA45SN1_45S_50ntFlanks</td><td>11572</td><td>11607</td><td>36</td><td>CCGACTTAGAACTGGTGCGGACCAGGGGAATCCGAC</td><td>597.1</td><td>-4.9</td><td>0.8</td><td>-6.5</td><td>6.6E-11</td><td>1.5E-10</td><td>MB231/MB468</td><td>MITO</td></tr>
<tr><td>>hg38_21_+_8433172_8446622_RNA45SN1_45S_50ntFlanks@5217.5240.24|CGGGGCTGCACGCGCGCTACACTG</td><td>>hg38_21_+_8433172_8446622_RNA45SN1_45S_50ntFlanks</td><td>5217</td><td>5240</td><td>24</td><td>CGGGGCTGCACGCGCGCTACACTG</td><td>474.8</td><td>-12.7</td><td>1.9</td><td>-6.5</td><td>6.6E-11</td><td>1.5E-10</td><td>MB231/MB468</td><td>MITO</td></tr>
<tr><td>>trna116_GluCTC_1_-_145399233_145399304@32.70.39,trna71_GluCTC_1_-_161439189_161439260@32.70.39,trna74_GluCTC_1_-_161431809_161431880@32.70.39,trna77_GluCTC_1_-_161424398_161424469@32.70.39,trna77_GluCTC_6_+_28949976_28950047@32.70.39,trna80_GluCTC_1_-_161417018_161417089@32.70.39,trna87_GluCTC_6_-_126101393_126101464@32.70.39|CTCTCACCGCCGCGGCCCGGGTTCGATTCCCGGTCAGGG</td><td>>trna116_GluCTC_1_-_145399233_145399304</td><td>32</td><td>70</td><td>39</td><td>CTCTCACCGCCGCGGCCCGGGTTCGATTCCCGGTCAGGG</td><td>6656.0</td><td>-8.9</td><td>1.4</td><td>-6.5</td><td>5.8E-11</td><td>1.6E-10</td><td>MB231/MB468</td><td>CYTO</td></tr>
<tr><td>>hg38_21_+_8433172_8446622_RNA45SN1_45S_50ntFlanks@10458.10498.41|ATGGCCTCCGTTGCCCTCGGCCGATCGAAAGGGAGTCGGGT</td><td>>hg38_21_+_8433172_8446622_RNA45SN1_45S_50ntFlanks</td><td>10458</td><td>10498</td><td>41</td><td>ATGGCCTCCGTTGCCCTCGGCCGATCGAAAGGGAGTCGGGT</td><td>108.6</td><td>-10.6</td><td>1.6</td><td>-6.6</td><td>5.5E-11</td><td>1.6E-10</td><td>MB231/MB468</td><td>MP</td></tr>
<tr><td>>hg38_21_+_8433172_8446622_RNA45SN1_45S_50ntFlanks@5071.5102.32|TTAATTCCGATAACGAACGAGACTCTGGCATG</td><td>>hg38_21_+_8433172_8446622_RNA45SN1_45S_50ntFlanks</td><td>5071</td><td>5102</td><td>32</td><td>TTAATTCCGATAACGAACGAGACTCTGGCATG</td><td>267.8</td><td>-11.9</td><td>1.8</td><td>-6.6</td><td>5.5E-11</td><td>1.6E-10</td><td>MB231/MB468</td><td>MP</td></tr>
<tr><td>>hg38_21_+_8433172_8446622_RNA45SN1_45S_50ntFlanks@6682.6718.37|CGTCGATGAAGAACGCAGCTAGCTGCGAGAATTAATG</td><td>>hg38_21_+_8433172_8446622_RNA45SN1_45S_50ntFlanks</td><td>6682</td><td>6718</td><td>37</td><td>CGTCGATGAAGAACGCAGCTAGCTGCGAGAATTAATG</td><td>322.5</td><td>-10.4</td><td>1.6</td><td>-6.6</td><td>4.3E-11</td><td>1.6E-10</td><td>BT20/MB468</td><td>MITO</td></tr>
<tr><td>>hg38_21_+_8433172_8446622_RNA45SN1_45S_50ntFlanks@12605.12622.18|TATGTGCTTGGCTGAGGA</td><td>>hg38_21_+_8433172_8446622_RNA45SN1_45S_50ntFlanks</td><td>12605</td><td>12622</td><td>18</td><td>TATGTGCTTGGCTGAGGA</td><td>82.2</td><td>-10.2</td><td>1.6</td><td>-6.6</td><td>5.5E-11</td><td>1.6E-10</td><td>MB231/MB468</td><td>MP</td></tr>
<tr><td>>hg38_21_+_8433172_8446622_RNA45SN1_45S_50ntFlanks@12212.12231.20|CTCAGGGAGGACAGAAACCT</td><td>>hg38_21_+_8433172_8446622_RNA45SN1_45S_50ntFlanks</td><td>12212</td><td>12231</td><td>20</td><td>CTCAGGGAGGACAGAAACCT</td><td>71.8</td><td>-10.0</td><td>1.5</td><td>-6.6</td><td>5.5E-11</td><td>1.6E-10</td><td>MB231/MB468</td><td>MP</td></tr>
<tr><td>>hg38_21_+_8433172_8446622_RNA45SN1_45S_50ntFlanks@10362.10407.46|AAGGCCGAAGTGGAGAAGGGTTCCATGTGAACAGCAGTTGAACATG</td><td>>hg38_21_+_8433172_8446622_RNA45SN1_45S_50ntFlanks</td><td>10362</td><td>10407</td><td>46</td><td>AAGGCCGAAGTGGAGAAGGGTTCCATGTGAACAGCAGTTGAACATG</td><td>307.7</td><td>-11.2</td><td>1.7</td><td>-6.5</td><td>5.9E-11</td><td>1.6E-10</td><td>MB231/MB468</td><td>NUC</td></tr>
<tr><td>>trna10_AspGTC_12_-_125424193_125424264@31.72.42,trna12_AspGTC_12_-_125411891_125411962@31.72.42,trna38_AspGTC_17_-_8125556_8125627@31.72.42,trna45_AspGTC_6_+_27447453_27447524@31.72.42,trna48_AspGTC_6_+_27471523_27471594@31.72.42,trna4_AspGTC_12_+_96429799_96429870@31.72.42,trna69_AspGTC_1_-_161440205_161440276@31.72.42,trna72_AspGTC_1_-_161432824_161432895@31.72.42,trna75_AspGTC_1_-_161425414_161425485@31.72.42,trna78_AspGTC_1_-_161418033_161418104@31.72.42,trna81_AspGTC_1_-_161410615_161410686@31.72.42|CCTGTCACGCGGGAGACCGGGGTTCGATTCCCCGACGGGGAG</td><td>>trna10_AspGTC_12_-_125424193_125424264</td><td>31</td><td>72</td><td>42</td><td>CCTGTCACGCGGGAGACCGGGGTTCGATTCCCCGACGGGGAG</td><td>123.4</td><td>-10.8</td><td>1.6</td><td>-6.6</td><td>5.5E-11</td><td>1.6E-10</td><td>MB231/MB468</td><td>MP</td></tr>
<tr><td>>hg38_21_+_8433172_8446622_RNA45SN1_45S_50ntFlanks@11572.11589.18|CCGACTTAGAACTGGTGC</td><td>>hg38_21_+_8433172_8446622_RNA45SN1_45S_50ntFlanks</td><td>11572</td><td>11589</td><td>18</td><td>CCGACTTAGAACTGGTGC</td><td>509.0</td><td>-7.0</td><td>1.1</td><td>-6.5</td><td>7.1E-11</td><td>1.6E-10</td><td>BT20/MB468</td><td>NUC</td></tr>
<tr><td>>hg38_21_+_8433172_8446622_RNA45SN1_45S_50ntFlanks@9483.9503.21|CGTGTTAGGACCCGAAAGATG</td><td>>hg38_21_+_8433172_8446622_RNA45SN1_45S_50ntFlanks</td><td>9483</td><td>9503</td><td>21</td><td>CGTGTTAGGACCCGAAAGATG</td><td>84.4</td><td>-10.3</td><td>1.6</td><td>-6.6</td><td>5.5E-11</td><td>1.6E-10</td><td>MB231/MB468</td><td>MP</td></tr>
<tr><td>>hg38_21_+_8433172_8446622_RNA45SN1_45S_50ntFlanks@4764.4793.30|ATACCGTCGTAGTTCCGACCATAAACGATG</td><td>>hg38_21_+_8433172_8446622_RNA45SN1_45S_50ntFlanks</td><td>4764</td><td>4793</td><td>30</td><td>ATACCGTCGTAGTTCCGACCATAAACGATG</td><td>80.2</td><td>-10.2</td><td>1.6</td><td>-6.6</td><td>5.5E-11</td><td>1.6E-10</td><td>MB231/MB468</td><td>MP</td></tr>
<tr><td>>am_trna2_GlyGCC_21_-_18827107_18827177@1.27.27,trna35_GlyGCC_1_+_161413094_161413164@1.27.27,trna37_GlyGCC_1_+_161420467_161420537@1.27.27,trna39_GlyGCC_1_+_161427898_161427968@1.27.27,trna41_GlyGCC_1_+_161435258_161435328@1.27.27|GCATGGGTGGTTCAGTGGTAGAATTCT</td><td>>am_trna2_GlyGCC_21_-_18827107_18827177</td><td>1</td><td>27</td><td>27</td><td>GCATGGGTGGTTCAGTGGTAGAATTCT</td><td>950.0</td><td>-4.5</td><td>0.7</td><td>-6.5</td><td>5.9E-11</td><td>1.6E-10</td><td>MB231/MB468</td><td>CYTO</td></tr>
<tr><td>>hg38_21_+_8433172_8446622_RNA45SN1_45S_50ntFlanks@11885.11912.28|TTGAGCTTGACTCTAGTCTGGCACGGTG</td><td>>hg38_21_+_8433172_8446622_RNA45SN1_45S_50ntFlanks</td><td>11885</td><td>11912</td><td>28</td><td>TTGAGCTTGACTCTAGTCTGGCACGGTG</td><td>150.1</td><td>-9.3</td><td>1.4</td><td>-6.6</td><td>4.3E-11</td><td>1.6E-10</td><td>BT20/MB468</td><td>MITO</td></tr>
<tr><td>>hg38_21_+_8433172_8446622_RNA45SN1_45S_50ntFlanks@10827.10865.39|CGGCAAGCCGGATCCGTAACTTCGGGATAAGGATTGGCT</td><td>>hg38_21_+_8433172_8446622_RNA45SN1_45S_50ntFlanks</td><td>10827</td><td>10865</td><td>39</td><td>CGGCAAGCCGGATCCGTAACTTCGGGATAAGGATTGGCT</td><td>414.9</td><td>-8.3</td><td>1.3</td><td>-6.6</td><td>5.3E-11</td><td>1.6E-10</td><td>BT20/MB468</td><td>CYTO</td></tr>
<tr><td>>hg38_21_+_8433172_8446622_RNA45SN1_45S_50ntFlanks@11569.11593.25|CAGCCGACTTAGAACTGGTGCGGAC</td><td>>hg38_21_+_8433172_8446622_RNA45SN1_45S_50ntFlanks</td><td>11569</td><td>11593</td><td>25</td><td>CAGCCGACTTAGAACTGGTGCGGAC</td><td>722.9</td><td>-12.6</td><td>1.8</td><td>-6.9</td><td>4.1E-12</td><td>1.6E-10</td><td>BT20/MB468</td><td>MP</td></tr>
</table>

<table>
<tr><th></th><th></th><th></th><th></th><th></th><th></th><th></th><th></th><th></th><th></th><th></th><th></th><th></th><th></th></tr>
<tr><td>>hg38_21_+_8433172_8446622_RNA45SN1_45S_50ntFlanks@7989.8007.19|AGACGTGGCGACCCGCTGA</td><td>>hg38_21_+_8433172_8446622_RNA45SN1_45S_50ntFlanks</td><td>7989</td><td>8007</td><td>19</td><td>AGACGTGGCGACCCGCTGA</td><td>4117.1</td><td>-5.1</td><td>0.8</td><td>-6.5</td><td>6.8E-11</td><td>1.6E-10</td><td>MB231/MB468</td><td>MITO</td></tr>
<tr><td>>hg38_21_+_8433172_8446622_RNA45SN1_45S_50ntFlanks@11534.11552.19|GTTCCGCGCGGCGCCTCGC</td><td>>hg38_21_+_8433172_8446622_RNA45SN1_45S_50ntFlanks</td><td>11534</td><td>11552</td><td>19</td><td>GTTCCGCGCGGCGCCTCGC</td><td>92.3</td><td>-10.4</td><td>1.6</td><td>-6.6</td><td>5.6E-11</td><td>1.6E-10</td><td>MB231/MB468</td><td>MP</td></tr>
<tr><td>>MI0000734|hsa-mir-106b&WithFlank&7|-|100093987|100094080@58.76.19&[MIMAT0004672&hsa-miR-106b-3p&offsets|0|-3;m-87&7|-|100094003|100094023&offsets|0|-2]|CCGCACTGTGGGTACTTGC</td><td>>MI0000734|hsa-mir-106b&WithFlank&7|-|100093987|100094080</td><td>58</td><td>76</td><td>19</td><td>CCGCACTGTGGGTACTTGC</td><td>274.5</td><td>7.6</td><td>1.2</td><td>6.5</td><td>5.9E-11</td><td>1.6E-10</td><td>MB231/MB468</td><td>CYTO</td></tr>
<tr><td>>MI0000272|hsa-mir-182&WithFlank&7|-|129770377|129770498@29.53.25(+1U)&[MIMAT0000259&hsa-miR-182-5p&offsets|0|+1(+1U);m-22&7|-|129770449|129770470&offsets|0|+3(+1U)]|TTTGGCAATGGTAGAACTCACACTGT</td><td>>MI0000272|hsa-mir-182&WithFlank&7|-|129770377|129770498</td><td>29</td><td>53</td><td>25(+1U)</td><td>TTTGGCAATGGTAGAACTCACACTGT</td><td>595.6</td><td>4.5</td><td>0.7</td><td>6.5</td><td>5.9E-11</td><td>1.6E-10</td><td>MB231/MB468</td><td>NUC</td></tr>
<tr><td>>hg38_21_+_8433172_8446622_RNA45SN1_45S_50ntFlanks@10421.10443.23|TGAGAGATGGGCGAGCGCCGTTC</td><td>>hg38_21_+_8433172_8446622_RNA45SN1_45S_50ntFlanks</td><td>10421</td><td>10443</td><td>23</td><td>TGAGAGATGGGCGAGCGCCGTTC</td><td>235.9</td><td>-5.1</td><td>0.8</td><td>-6.5</td><td>6.0E-11</td><td>1.6E-10</td><td>MB231/MB468</td><td>CYTO</td></tr>
<tr><td>>hg38_21_+_8433172_8446622_RNA45SN1_45S_50ntFlanks@10824.10865.42|AGTCGGCAAGCCGGATCCGTAACTTCGGGATAAGGATTGGCT</td><td>>hg38_21_+_8433172_8446622_RNA45SN1_45S_50ntFlanks</td><td>10824</td><td>10865</td><td>42</td><td>AGTCGGCAAGCCGGATCCGTAACTTCGGGATAAGGATTGGCT</td><td>126.3</td><td>-9.7</td><td>1.5</td><td>-6.5</td><td>6.0E-11</td><td>1.6E-10</td><td>MB231/MB468</td><td>CYTO</td></tr>
<tr><td>>hg38_21_+_8433172_8446622_RNA45SN1_45S_50ntFlanks@7985.8003.19|GATCAGACGTGGCGACCCG</td><td>>hg38_21_+_8433172_8446622_RNA45SN1_45S_50ntFlanks</td><td>7985</td><td>8003</td><td>19</td><td>GATCAGACGTGGCGACCCG</td><td>141.8</td><td>-9.2</td><td>1.4</td><td>-6.6</td><td>4.4E-11</td><td>1.6E-10</td><td>BT20/MB468</td><td>MITO</td></tr>
<tr><td>>hg38_21_+_8433172_8446622_RNA45SN1_45S_50ntFlanks@4753.4783.31|AAGACGATCAGATACCGTCGTAGTTCCGACC</td><td>>hg38_21_+_8433172_8446622_RNA45SN1_45S_50ntFlanks</td><td>4753</td><td>4783</td><td>31</td><td>AAGACGATCAGATACCGTCGTAGTTCCGACC</td><td>1316.6</td><td>-7.8</td><td>1.2</td><td>-6.5</td><td>7.2E-11</td><td>1.6E-10</td><td>BT20/MB468</td><td>NUC</td></tr>
<tr><td>>hg38_21_+_8433172_8446622_RNA45SN1_45S_50ntFlanks@3850.3874.25|GATAACTGTGGTAATTCTAGAGCTA</td><td>>hg38_21_+_8433172_8446622_RNA45SN1_45S_50ntFlanks</td><td>3850</td><td>3874</td><td>25</td><td>GATAACTGTGGTAATTCTAGAGCTA</td><td>345.6</td><td>-6.4</td><td>1.0</td><td>-6.6</td><td>5.6E-11</td><td>1.6E-10</td><td>MB231/MB468</td><td>MP</td></tr>
<tr><td>>hg38_21_+_8433172_8446622_RNA45SN1_45S_50ntFlanks@6776.6803.28|CTCCCGGGGCTACGCCTGTCTGAGCGTC</td><td>>hg38_21_+_8433172_8446622_RNA45SN1_45S_50ntFlanks</td><td>6776</td><td>6803</td><td>28</td><td>CTCCCGGGGCTACGCCTGTCTGAGCGTC</td><td>651.1</td><td>-8.7</td><td>1.3</td><td>-6.6</td><td>5.6E-11</td><td>1.6E-10</td><td>MB231/MB468</td><td>MP</td></tr>
<tr><td>>hg38_21_+_8433172_8446622_RNA45SN1_45S_50ntFlanks@5227.5246.20|CGCGCGCTACACTGACTGGC</td><td>>hg38_21_+_8433172_8446622_RNA45SN1_45S_50ntFlanks</td><td>5227</td><td>5246</td><td>20</td><td>CGCGCGCTACACTGACTGGC</td><td>506.6</td><td>-12.8</td><td>2.0</td><td>-6.5</td><td>6.8E-11</td><td>1.6E-10</td><td>MB231/MB468</td><td>MITO</td></tr>
<tr><td>>hg38_21_+_8433172_8446622_RNA45SN1_45S_50ntFlanks@11568.11602.35|GCAGCCGACTTAGAACTGGTGCGGACCAGGGGAAT</td><td>>hg38_21_+_8433172_8446622_RNA45SN1_45S_50ntFlanks</td><td>11568</td><td>11602</td><td>35</td><td>GCAGCCGACTTAGAACTGGTGCGGACCAGGGGAAT</td><td>702.6</td><td>-4.8</td><td>0.7</td><td>-6.6</td><td>5.6E-11</td><td>1.6E-10</td><td>MB231/MB468</td><td>MP</td></tr>
<tr><td>>hg38_21_+_8433172_8446622_RNA45SN1_45S_50ntFlanks@10025.10072.48|CGCTGGAGCGTCGGGCCCATACCCGGCCGTCGCCGGCAGTCGAGAGTG</td><td>>hg38_21_+_8433172_8446622_RNA45SN1_45S_50ntFlanks</td><td>10025</td><td>10072</td><td>48</td><td>CGCTGGAGCGTCGGGCCCATACCCGGCCGTCGCCGGCAGTCGAGAGTG</td><td>279.0</td><td>-11.9</td><td>1.8</td><td>-6.5</td><td>6.8E-11</td><td>1.6E-10</td><td>MB231/MB468</td><td>MITO</td></tr>
<tr><td>>MI0003820|hsa-mir-454&WithFlank&17|-|59137752|59137878@70.95.26&[MIMAT0003885&hsa-miR-454-3p&offsets|0|+3;m-272&17|-|59137787|59137809&offsets|0|+3]|TAGTGCAATATTGCTTATAGGGTTTT</td><td>>MI0003820|hsa-mir-454&WithFlank&17|-|59137752|59137878</td><td>70</td><td>95</td><td>26</td><td>TAGTGCAATATTGCTTATAGGGTTTT</td><td>260.6</td><td>11.2</td><td>1.7</td><td>6.5</td><td>6.8E-11</td><td>1.6E-10</td><td>MB231/MB468</td><td>MITO</td></tr>
<tr><td>>hg38_21_+_8433172_8446622_RNA45SN1_45S_50ntFlanks@11576.11594.19|CTTAGAACTGGTGCGGACC</td><td>>hg38_21_+_8433172_8446622_RNA45SN1_45S_50ntFlanks</td><td>11576</td><td>11594</td><td>19</td><td>CTTAGAACTGGTGCGGACC</td><td>478.8</td><td>-4.2</td><td>0.7</td><td>-6.5</td><td>6.9E-11</td><td>1.6E-10</td><td>MB231/MB468</td><td>MITO</td></tr>
<tr><td>>hg38_21_+_8433172_8446622_RNA45SN1_45S_50ntFlanks@10731.10759.29|AGGGTGTAAATCTCGCGCCGGGCCGTACC</td><td>>hg38_21_+_8433172_8446622_RNA45SN1_45S_50ntFlanks</td><td>10731</td><td>10759</td><td>29</td><td>AGGGTGTAAATCTCGCGCCGGGCCGTACC</td><td>89.4</td><td>-10.3</td><td>1.6</td><td>-6.6</td><td>5.6E-11</td><td>1.6E-10</td><td>MB231/MB468</td><td>MP</td></tr>
<tr><td>>MI0000066|hsa-let-7e&WithFlank&19|+|51692780|51692870@14.34.21(+1C)&[MIMAT0000066&hsa-let-7e-5p&offsets|0|-1(+1C);m-50&19|+|51692793|51692814&offsets|0|-1(+1C)]|TGAGGTAGGAGGTTGTATAGTC</td><td>>MI0000066|hsa-let-7e&WithFlank&19|+|51692780|51692870</td><td>14</td><td>34</td><td>21(+1C)</td><td>TGAGGTAGGAGGTTGTATAGTC</td><td>1435.3</td><td>6.1</td><td>0.9</td><td>6.5</td><td>6.0E-11</td><td>1.6E-10</td><td>MB231/MB468</td><td>CYTO</td></tr>
<tr><td>>hg38_21_+_8433172_8446622_RNA45SN1_45S_50ntFlanks@10786.10819.34|AACAGCCTCTGGCATGTTGGAACAATGTAGGTAA</td><td>>hg38_21_+_8433172_8446622_RNA45SN1_45S_50ntFlanks</td><td>10786</td><td>10819</td><td>34</td><td>AACAGCCTCTGGCATGTTGGAACAATGTAGGTAA</td><td>117.9</td><td>-9.0</td><td>1.4</td><td>-6.6</td><td>4.4E-11</td><td>1.6E-10</td><td>BT20/MB468</td><td>MITO</td></tr>
<tr><td>>hg38_21_+_8433172_8446622_RNA45SN1_45S_50ntFlanks@9533.9554.22|GAGGAAACTCTGGTGGAGGTCC</td><td>>hg38_21_+_8433172_8446622_RNA45SN1_45S_50ntFlanks</td><td>9533</td><td>9554</td><td>22</td><td>GAGGAAACTCTGGTGGAGGTCC</td><td>406.4</td><td>-8.9</td><td>1.4</td><td>-6.5</td><td>7.3E-11</td><td>1.6E-10</td><td>BT20/MB468</td><td>NUC</td></tr>
<tr><td>>hg38_21_+_8433172_8446622_RNA45SN1_45S_50ntFlanks@12631.12657.27|GGCGAAGCTACCATCTGTGGGATTATG</td><td>>hg38_21_+_8433172_8446622_RNA45SN1_45S_50ntFlanks</td><td>12631</td><td>12657</td><td>27</td><td>GGCGAAGCTACCATCTGTGGGATTATG</td><td>171.4</td><td>-11.3</td><td>1.7</td><td>-6.6</td><td>5.6E-11</td><td>1.6E-10</td><td>MB231/MB468</td><td>MP</td></tr>
<tr><td>>hg38_21_+_8433172_8446622_RNA45SN1_45S_50ntFlanks@10764.10785.22|TCCGCAGCAGGTCTCCAAGGTG</td><td>>hg38_21_+_8433172_8446622_RNA45SN1_45S_50ntFlanks</td><td>10764</td><td>10785</td><td>22</td><td>TCCGCAGCAGGTCTCCAAGGTG</td><td>119.6</td><td>-9.0</td><td>1.4</td><td>-6.6</td><td>4.4E-11</td><td>1.6E-10</td><td>BT20/MB468</td><td>MITO</td></tr>
<tr><td>>hg38_21_+_8433172_8446622_RNA45SN1_45S_50ntFlanks@11927.11950.24|GTGTAGAATAAGTGGGAGGCCCCC</td><td>>hg38_21_+_8433172_8446622_RNA45SN1_45S_50ntFlanks</td><td>11927</td><td>11950</td><td>24</td><td>GTGTAGAATAAGTGGGAGGCCCCC</td><td>108.7</td><td>-10.6</td><td>1.6</td><td>-6.6</td><td>5.7E-11</td><td>1.6E-10</td><td>MB231/MB468</td><td>MP</td></tr>
<tr><td>>hg38_21_+_8433172_8446622_RNA45SN1_45S_50ntFlanks@12359.12376.18|ACAGGGATAACTGGCTTG</td><td>>hg38_21_+_8433172_8446622_RNA45SN1_45S_50ntFlanks</td><td>12359</td><td>12376</td><td>18</td><td>ACAGGGATAACTGGCTTG</td><td>75.7</td><td>-10.1</td><td>1.5</td><td>-6.6</td><td>5.7E-11</td><td>1.6E-10</td><td>MB231/MB468</td><td>MP</td></tr>
<tr><td>>hg38_21_+_8433172_8446622_RNA45SN1_45S_50ntFlanks@12241.12268.28|CAGAAGGGCAAAAGCTCGCTTGATCTTG</td><td>>hg38_21_+_8433172_8446622_RNA45SN1_45S_50ntFlanks</td><td>12241</td><td>12268</td><td>28</td><td>CAGAAGGGCAAAAGCTCGCTTGATCTTG</td><td>691.2</td><td>-8.8</td><td>1.3</td><td>-6.6</td><td>5.7E-11</td><td>1.6E-10</td><td>MB231/MB468</td><td>MP</td></tr>
<tr><td>>hg38_21_+_8433172_8446622_RNA45SN1_45S_50ntFlanks@7986.8011.26|ATCAGACGTGGCGACCCGCTGAATTT</td><td>>hg38_21_+_8433172_8446622_RNA45SN1_45S_50ntFlanks</td><td>7986</td><td>8011</td><td>26</td><td>ATCAGACGTGGCGACCCGCTGAATTT</td><td>970.8</td><td>-8.9</td><td>1.4</td><td>-6.6</td><td>5.7E-11</td><td>1.6E-10</td><td>MB231/MB468</td><td>MP</td></tr>
<tr><td>>hg38_21_+_8433172_8446622_RNA45SN1_45S_50ntFlanks@6776.6806.31|CTCCCGGGGCTACGCCTGTCTGAGCGTCGCT</td><td>>hg38_21_+_8433172_8446622_RNA45SN1_45S_50ntFlanks</td><td>6776</td><td>6806</td><td>31</td><td>CTCCCGGGGCTACGCCTGTCTGAGCGTCGCT</td><td>1438.2</td><td>3.8</td><td>0.6</td><td>6.5</td><td>6.0E-11</td><td>1.6E-10</td><td>MB231/MB468</td><td>CYTO</td></tr>
<tr><td>>trna116_GluCTC_1_-_145399233_145399304@3.33.31,trna59_GluCTC_1_+_249168447_249168518@3.33.31,trna71_GluCTC_1_-_161439189_161439260@3.33.31,trna74_GluCTC_1_-_161431809_161431880@3.33.31,trna77_GluCTC_1_-_161424398_161424469@3.33.31,trna77_GluCTC_6_+_28949976_28950047@3.33.31,trna80_GluCTC_1_-_161417018_161417089@3.33.31,trna87_GluCTC_6_-_126101393_126101464@3.33.31|CCTGGTGGTCTAGTGGTTAGGATTCGGCGCT</td><td>>trna116_GluCTC_1_-_145399233_145399304</td><td>3</td><td>33</td><td>31</td><td>CCTGGTGGTCTAGTGGTTAGGATTCGGCGCT</td><td>451.4</td><td>5.0</td><td>0.8</td><td>6.5</td><td>6.0E-11</td><td>1.6E-10</td><td>MB231/MB468</td><td>NUC</td></tr>
<tr><td>>hg38_21_+_8433172_8446622_RNA45SN1_45S_50ntFlanks@8528.8549.22|CTTCCCCCGCCGCCCCTCCTCC</td><td>>hg38_21_+_8433172_8446622_RNA45SN1_45S_50ntFlanks</td><td>8528</td><td>8549</td><td>22</td><td>CTTCCCCCGCCGCCCCTCCTCC</td><td>156.7</td><td>-10.3</td><td>1.6</td><td>-6.5</td><td>6.0E-11</td><td>1.6E-10</td><td>MB231/MB468</td><td>NUC</td></tr>
<tr><td>>hg38_21_+_8433172_8446622_RNA45SN1_45S_50ntFlanks@9858.9900.43|CGGGATGAACCGAACGCCGGGTTAAGGCGCCCGATGCCGACGC</td><td>>hg38_21_+_8433172_8446622_RNA45SN1_45S_50ntFlanks</td><td>9858</td><td>9900</td><td>43</td><td>CGGGATGAACCGAACGCCGGGTTAAGGCGCCCGATGCCGACGC</td><td>289.0</td><td>-12.0</td><td>1.8</td><td>-6.5</td><td>6.9E-11</td><td>1.6E-10</td><td>MB231/MB468</td><td>MITO</td></tr>
<tr><td>>hg38_21_+_8433172_8446622_RNA45SN1_45S_50ntFlanks@11834.11851.18|CCACAGCCAAGGGAACGG</td><td>>hg38_21_+_8433172_8446622_RNA45SN1_45S_50ntFlanks</td><td>11834</td><td>11851</td><td>18</td><td>CCACAGCCAAGGGAACGG</td><td>81.0</td><td>-10.2</td><td>1.6</td><td>-6.6</td><td>5.7E-11</td><td>1.6E-10</td><td>MB231/MB468</td><td>MP</td></tr>
<tr><td>>hg38_21_+_8433172_8446622_RNA45SN1_45S_50ntFlanks@12141.12180.40|ACAGTGCCAGGTGGGGAGTTTGACTGGGGCGGTACACCTG</td><td>>hg38_21_+_8433172_8446622_RNA45SN1_45S_50ntFlanks</td><td>12141</td><td>12180</td><td>40</td><td>ACAGTGCCAGGTGGGGAGTTTGACTGGGGCGGTACACCTG</td><td>149.7</td><td>-11.1</td><td>1.7</td><td>-6.6</td><td>5.7E-11</td><td>1.6E-10</td><td>MB231/MB468</td><td>MP</td></tr>
<tr><td>>hg38_21_+_8433172_8446622_RNA45SN1_45S_50ntFlanks@11729.11754.26|TAACTATGACTCTCTTAAGGTAGCCA</td><td>>hg38_21_+_8433172_8446622_RNA45SN1_45S_50ntFlanks</td><td>11729</td><td>11754</td><td>26</td><td>TAACTATGACTCTCTTAAGGTAGCCA</td><td>235.4</td><td>-8.1</td><td>1.2</td><td>-6.5</td><td>6.1E-11</td><td>1.6E-10</td><td>MB231/MB468</td><td>CYTO</td></tr>
<tr><td>>hg38_21_+_8433172_8446622_RNA45SN1_45S_50ntFlanks@12407.12455.49|TTTTGATCCTTCGATGTCGGCTCTTCCTATCATTGTGAAGCAGAATTCA</td><td>>hg38_21_+_8433172_8446622_RNA45SN1_45S_50ntFlanks</td><td>12407</td><td>12455</td><td>49</td><td>TTTTGATCCTTCGATGTCGGCTCTTCCTATCATTGTGAAGCAGAATTCA</td><td>210.3</td><td>-11.5</td><td>1.8</td><td>-6.5</td><td>6.9E-11</td><td>1.6E-10</td><td>MB231/MB468</td><td>MITO</td></tr>
<tr><td>>hg38_21_+_8433172_8446622_RNA45SN1_45S_50ntFlanks@12407.12434.28|TTTTGATCCTTCGATGTCGGCTCTTCCT</td><td>>hg38_21_+_8433172_8446622_RNA45SN1_45S_50ntFlanks</td><td>12407</td><td>12434</td><td>28</td><td>TTTTGATCCTTCGATGTCGGCTCTTCCT</td><td>2117.7</td><td>-4.2</td><td>0.6</td><td>-6.5</td><td>6.9E-11</td><td>1.6E-10</td><td>MB231/MB468</td><td>MITO</td></tr>
<tr><td>>MI0000440|hsa-mir-27b&WithFlank&9|+|95085439|95085547@67.87.21(+1C)&[MIMAT0000419&hsa-miR-27b-3p&offsets|0|0(+1C);m-39&9|+|95085505|95085525&offsets|0|0(+1C)]|TTCACAGTGGCTAAGTTCTGCC</td><td>>MI0000440|hsa-mir-27b&WithFlank&9|+|95085439|95085547</td><td>67</td><td>87</td><td>21(+1C)</td><td>TTCACAGTGGCTAAGTTCTGCC</td><td>346.3</td><td>11.7</td><td>1.8</td><td>6.5</td><td>6.9E-11</td><td>1.6E-10</td><td>MB231/MB468</td><td>MITO</td></tr>
<tr><td>>hg38_21_+_8433172_8446622_RNA45SN1_45S_50ntFlanks@4047.4065.19|ATTCGAACGTCTGCCCTAT</td><td>>hg38_21_+_8433172_8446622_RNA45SN1_45S_50ntFlanks</td><td>4047</td><td>4065</td><td>19</td><td>ATTCGAACGTCTGCCCTAT</td><td>295.2</td><td>-10.3</td><td>1.6</td><td>-6.6</td><td>4.4E-11</td><td>1.6E-10</td><td>BT20/MB468</td><td>MITO</td></tr>
<tr><td>>hg38_21_+_8433172_8446622_RNA45SN1_45S_50ntFlanks@4555.4587.33|CGCAGCTAGGAATAATGGAATAGGACCGCGGTT</td><td>>hg38_21_+_8433172_8446622_RNA45SN1_45S_50ntFlanks</td><td>4555</td><td>4587</td><td>33</td><td>CGCAGCTAGGAATAATGGAATAGGACCGCGGTT</td><td>201.4</td><td>-4.7</td><td>0.7</td><td>-6.5</td><td>6.1E-11</td><td>1.6E-10</td><td>MB231/MB468</td><td>CYTO</td></tr>
<tr><td>>trna111_HisGTG_1_-_147774845_147774916@-1T.29.30,trna118_HisGTG_1_-_145396881_145396952@-1T.29.30,trna16_HisGTG_1_+_146544773_146544844@-1T.29.30,trna1_HisGTG_15_+_45493349_45493420@-1T.29.30,trna21_HisGTG_1_+_147753471_147753542@-1T.29.30,trna33_HisGTG_6_+_27125906_27125977@-1T.29.30,trna7_HisGTG_9_-_14433938_14434009@-1T.29.30,trna8_HisGTG_15_-_45492611_45492682@-1T.29.30,trna9_HisGTG_15_-_45490804_45490875@-1T.29.30|TGCCGTGATCGTATAGTGGTTAGTACTCTG</td><td>>trna111_HisGTG_1_-_147774845_147774916</td><td>-1T</td><td>29</td><td>30</td><td>TGCCGTGATCGTATAGTGGTTAGTACTCTG</td><td>12949.1</td><td>-11.5</td><td>1.8</td><td>-6.5</td><td>7.3E-11</td><td>1.6E-10</td><td>BT20/MB468</td><td>NUC</td></tr>
<tr><td>>hg38_21_+_8433172_8446622_RNA45SN1_45S_50ntFlanks@10420.10456.37|CTGAGAGATGGGCGAGCGCCGTTCCGAAGGGACGGGC</td><td>>hg38_21_+_8433172_8446622_RNA45SN1_45S_50ntFlanks</td><td>10420</td><td>10456</td><td>37</td><td>CTGAGAGATGGGCGAGCGCCGTTCCGAAGGGACGGGC</td><td>9415.1</td><td>-11.6</td><td>1.8</td><td>-6.5</td><td>7.3E-11</td><td>1.6E-10</td><td>BT20/MB468</td><td>NUC</td></tr>
<tr><td>>hg38_21_+_8433172_8446622_RNA45SN1_45S_50ntFlanks@6719.6758.40|TGAATTGCAGGACACATTGATCATCGACACTTCGAACGCA</td><td>>hg38_21_+_8433172_8446622_RNA45SN1_45S_50ntFlanks</td><td>6719</td><td>6758</td><td>40</td><td>TGAATTGCAGGACACATTGATCATCGACACTTCGAACGCA</td><td>154.0</td><td>-11.1</td><td>1.7</td><td>-6.6</td><td>5.7E-11</td><td>1.6E-10</td><td>MB231/MB468</td><td>MP</td></tr>
<tr><td>>hg38_21_+_8433172_8446622_RNA45SN1_45S_50ntFlanks@10677.10707.31|GCGGTTCCGGCGGCGTCCGGTGAGCTCTCGC</td><td>>hg38_21_+_8433172_8446622_RNA45SN1_45S_50ntFlanks</td><td>10677</td><td>10707</td><td>31</td><td>GCGGTTCCGGCGGCGTCCGGTGAGCTCTCGC</td><td>1101.6</td><td>7.6</td><td>1.1</td><td>6.6</td><td>4.5E-11</td><td>1.6E-10</td><td>BT20/MB468</td><td>MITO</td></tr>
<tr><td>>MI0000471|hsa-mir-126&WithFlank&9|+|136670596|136670692@59.79.21&[MIMAT0000445&hsa-miR-126-3p&offsets|+1|0;m-40&9|+|136670653|136670674&offsets|+1|0]|CGTACCGTGAGTAATAATGCG</td><td>>MI0000471|hsa-mir-126&WithFlank&9|+|136670596|136670692</td><td>59</td><td>79</td><td>21</td><td>CGTACCGTGAGTAATAATGCG</td><td>454.6</td><td>12.6</td><td>1.8</td><td>6.9</td><td>4.2E-12</td><td>1.6E-10</td><td>BT20/MB468</td><td>MP</td></tr>
<tr><td>>hg38_21_+_8433172_8446622_RNA45SN1_45S_50ntFlanks@10350.10379.30|ACGAGAACTTTGAAGGCCGAAGTGGAGAAG</td><td>>hg38_21_+_8433172_8446622_RNA45SN1_45S_50ntFlanks</td><td>10350</td><td>10379</td><td>30</td><td>ACGAGAACTTTGAAGGCCGAAGTGGAGAAG</td><td>736.7</td><td>-3.7</td><td>0.6</td><td>-6.5</td><td>6.1E-11</td><td>1.6E-10</td><td>MB231/MB468</td><td>NUC</td></tr>
<tr><td>>hg38_21_+_8433172_8446622_RNA45SN1_45S_50ntFlanks@11233.11269.37|CCCTTCCCGTGGATCGCCCCAGCTGCGGCGGGCGTCG</td><td>>hg38_21_+_8433172_8446622_RNA45SN1_45S_50ntFlanks</td><td>11233</td><td>11269</td><td>37</td><td>CCCTTCCCGTGGATCGCCCCAGCTGCGGCGGGCGTCG</td><td>92.9</td><td>-10.4</td><td>1.6</td><td>-6.6</td><td>5.7E-11</td><td>1.6E-10</td><td>MB231/MB468</td><td>MP</td></tr>
<tr><td>>hg38_21_+_8433172_8446622_RNA45SN1_45S_50ntFlanks@6663.6681.19|GTGGATCACTCGGCTCGTG</td><td>>hg38_21_+_8433172_8446622_RNA45SN1_45S_50ntFlanks</td><td>6663</td><td>6681</td><td>19</td><td>GTGGATCACTCGGCTCGTG</td><td>3582.0</td><td>-8.2</td><td>1.3</td><td>-6.5</td><td>7.4E-11</td><td>1.6E-10</td><td>BT20/MB468</td><td>NUC</td></tr>
<tr><td>>MI0000285|hsa-mir-205&WithFlank&1|+|209432127|209432248@40.61.22&[MIMAT0000266&hsa-miR-205-5p&offsets|0|0;m-351&1|+|209432166|209432187&offsets|0|0]|TCCTTCATTCCACCGGAGTCTG</td><td>>MI0000285|hsa-mir-205&WithFlank&1|+|209432127|209432248</td><td>40</td><td>61</td><td>22</td><td>TCCTTCATTCCACCGGAGTCTG</td><td>50442.8</td><td>6.6</td><td>1.0</td><td>6.6</td><td>4.5E-11</td><td>1.6E-10</td><td>BT20/MB468</td><td>MITO</td></tr>
<tr><td>>hg38_21_+_8433172_8446622_RNA45SN1_45S_50ntFlanks@8079.8106.28|AAGAGCCCAGCGCCGAATCCCCGCCCCG</td><td>>hg38_21_+_8433172_8446622_RNA45SN1_45S_50ntFlanks</td><td>8079</td><td>8106</td><td>28</td><td>AAGAGCCCAGCGCCGAATCCCCGCCCCG</td><td>146.0</td><td>-11.1</td><td>1.7</td><td>-6.5</td><td>5.8E-11</td><td>1.6E-10</td><td>MB231/MB468</td><td>MP</td></tr>
<tr><td>>hg38_21_+_8433172_8446622_RNA45SN1_45S_50ntFlanks@4724.4755.32|ATTAATCAAGAACGAAAGTCGGAGGTTCGAAG</td><td>>hg38_21_+_8433172_8446622_RNA45SN1_45S_50ntFlanks</td><td>4724</td><td>4755</td><td>32</td><td>ATTAATCAAGAACGAAAGTCGGAGGTTCGAAG</td><td>159.2</td><td>-9.4</td><td>1.4</td><td>-6.6</td><td>4.5E-11</td><td>1.6E-10</td><td>BT20/MB468</td><td>MITO</td></tr>
</table>

| | | | | | | | | | | | | | |
|---|---|---|---|---|---|---|---|---|---|---|---|---|---|
| >hg38_21_+_8433172_8446622_RNA45SN1_45S_50ntFlanks@4917.4937.21\|CACCACCAGGAGTGGAGCCTG | >hg38_21_+_8433172_8446622_RNA45SN1_45S_50ntFlanks | 4917 | 4937 | 21 | CACCACCAGGAGTGGAGCCTG | 73.0 | -10.1 | 1.5 | -6.5 | 5.8E-11 | 1.6E-10 | MB231/MB468 | MP |
| >hg38_21_+_8433172_8446622_RNA45SN1_45S_50ntFlanks@10512.10533.22\|TCCGGAGTGGCGGAGATGGGCG | >hg38_21_+_8433172_8446622_RNA45SN1_45S_50ntFlanks | 10512 | 10533 | 22 | TCCGGAGTGGCGGAGATGGGCG | 1151.9 | -6.9 | 1.1 | -6.5 | 6.2E-11 | 1.6E-10 | MB231/MB468 | CYTO |
| >hg38_21_+_8433172_8446622_RNA45SN1_45S_50ntFlanks@12456.12473.18\|CCAAGCGTTGGATTGTTC | >hg38_21_+_8433172_8446622_RNA45SN1_45S_50ntFlanks | 12456 | 12473 | 18 | CCAAGCGTTGGATTGTTC | 3896.7 | -10.9 | 1.7 | -6.5 | 7.5E-11 | 1.6E-10 | BT20/MB468 | NUC |
| >hg38_21_+_8433172_8446622_RNA45SN1_45S_50ntFlanks@6651.6672.22\|CGACTCTTAGCGGTGGATCACT | >hg38_21_+_8433172_8446622_RNA45SN1_45S_50ntFlanks | 6651 | 6672 | 22 | CGACTCTTAGCGGTGGATCACT | 2551.4 | 4.7 | 0.7 | 6.5 | 6.2E-11 | 1.6E-10 | MB231/MB468 | CYTO |
| >MI0000255\|hsa-mir-30d&WithFlank&8\|-\|134804870\|134804951@12.35.24(+2U)&[MIMAT0000245&hsa-miR-30d-5p&offsets\|0\|+2(+2U);m-21&8\|-\|134804919\|134804940&offsets\|0\|+2(+2U)]\|TGTAAACATCCCCGACTGGAAGCTTT | >MI0000255\|hsa-mir-30d&WithFlank&8\|-\|134804870\|134804951 | 12 | 35 | 24(+2U) | TGTAAACATCCCCGACTGGAAGCTTT | 477.0 | 8.2 | 1.3 | 6.5 | 6.2E-11 | 1.6E-10 | MB231/MB468 | CYTO |
| >hg38_21_+_8433172_8446622_RNA45SN1_45S_50ntFlanks@12539.12570.32\|ATGATGTGTTGTTGCCATGGTAATCCTGCTCA | >hg38_21_+_8433172_8446622_RNA45SN1_45S_50ntFlanks | 12539 | 12570 | 32 | ATGATGTGTTGTTGCCATGGTAATCCTGCTCA | 79.6 | -10.2 | 1.6 | -6.5 | 5.8E-11 | 1.6E-10 | MB231/MB468 | MP |
| >hg38_1_-_228634819_228635039_RNA5S12_5S_50ntFlanks@51.84.34\|GTCTACGGCCATACCACCCTGAACGCGCCCGATC | >hg38_1_-_228634819_228635039_RNA5S12_5S_50ntFlanks | 51 | 84 | 34 | GTCTACGGCCATACCACCCTGAACGCGCCCGATC | 17930.2 | 6.0 | 0.9 | 6.5 | 6.2E-11 | 1.6E-10 | MB231/MB468 | CYTO |
| >am_trnaMT_SerGCT_MT_+_12207_12265@12.38.27\|CAAGAACTGCTAACTCATGCCCCCATG | >am_trnaMT_SerGCT_MT_+_12207_12265 | 12 | 38 | 27 | CAAGAACTGCTAACTCATGCCCCCATG | 2995.1 | -11.1 | 1.7 | -6.5 | 7.5E-11 | 1.6E-10 | BT20/MB468 | NUC |
| >hg38_21_+_8433172_8446622_RNA45SN1_45S_50ntFlanks@6721.6743.23\|AATTGCAGGACACATTGATCATC | >hg38_21_+_8433172_8446622_RNA45SN1_45S_50ntFlanks | 6721 | 6743 | 23 | AATTGCAGGACACATTGATCATC | 4639.6 | -7.8 | 1.2 | -6.5 | 5.8E-11 | 1.6E-10 | MB231/MB468 | MP |
| >MI0003628\|hsa-mir-615&WithFlank&12\|+\|54033944\|54034051@67.87.21(+1C)&[MIMAT0003283&hsa-miR-615-3p&offsets\|0\|-1(+1C);m-495&12\|+\|54034010\|54034030&offsets\|0\|0(+1C)]\|TCCGAGCCTGGGTCTCCCTCTC | >MI0003628\|hsa-mir-615&WithFlank&12\|+\|54033944\|54034051 | 67 | 87 | 21(+1C) | TCCGAGCCTGGGTCTCCCTCTC | 418.6 | 8.2 | 1.3 | 6.6 | 4.5E-11 | 1.6E-10 | BT20/MB468 | MITO |
| >hg38_21_+_8433172_8446622_RNA45SN1_45S_50ntFlanks@9504.9529.26\|GTGAACTATGCCTGGGCAGGGCGAAG | >hg38_21_+_8433172_8446622_RNA45SN1_45S_50ntFlanks | 9504 | 9529 | 26 | GTGAACTATGCCTGGGCAGGGCGAAG | 198.7 | -9.7 | 1.5 | -6.6 | 4.6E-11 | 1.7E-10 | BT20/MB468 | MITO |
| >hg38_21_+_8433172_8446622_RNA45SN1_45S_50ntFlanks@11573.11605.33\|CGACTTAGAACTGGTGCGGACCAGGGGAATCCG | >hg38_21_+_8433172_8446622_RNA45SN1_45S_50ntFlanks | 11573 | 11605 | 33 | CGACTTAGAACTGGTGCGGACCAGGGGAATCCG | 538.8 | -8.7 | 1.3 | -6.5 | 5.8E-11 | 1.7E-10 | MB231/MB468 | MP |
| >hg38_21_+_8433172_8446622_RNA45SN1_45S_50ntFlanks@13002.13030.29\|CGATCTATTGAAAGTCAGCCCTCGACACA | >hg38_21_+_8433172_8446622_RNA45SN1_45S_50ntFlanks | 13002 | 13030 | 29 | CGATCTATTGAAAGTCAGCCCTCGACACA | 83.1 | -10.2 | 1.6 | -6.5 | 5.9E-11 | 1.7E-10 | MB231/MB468 | MP |
| >hg38_21_+_8433172_8446622_RNA45SN1_45S_50ntFlanks@3845.3868.24\|ACTTGGATAACTGTGGTAATTCTA | >hg38_21_+_8433172_8446622_RNA45SN1_45S_50ntFlanks | 3845 | 3868 | 24 | ACTTGGATAACTGTGGTAATTCTA | 573.2 | -3.9 | 0.6 | -6.5 | 7.5E-11 | 1.7E-10 | BT20/MB468 | NUC |
| >hg38_21_+_8433172_8446622_RNA45SN1_45S_50ntFlanks@10393.10435.43\|CAGCAGTTGAACATGGGTCAGTCGGTCCTGAGAGATGGGCGAG | >hg38_21_+_8433172_8446622_RNA45SN1_45S_50ntFlanks | 10393 | 10435 | 43 | CAGCAGTTGAACATGGGTCAGTCGGTCCTGAGAGATGGGCGAG | 1478.3 | -10.7 | 1.6 | -6.5 | 7.5E-11 | 1.7E-10 | BT20/MB468 | NUC |
| >hg38_21_+_8433172_8446622_RNA45SN1_45S_50ntFlanks@12936.12953.18\|AAACCATTCGTAGACGAC | >hg38_21_+_8433172_8446622_RNA45SN1_45S_50ntFlanks | 12936 | 12953 | 18 | AAACCATTCGTAGACGAC | 1208.6 | -9.7 | 1.5 | -6.5 | 7.6E-11 | 1.7E-10 | BT20/MB468 | NUC |
| >hg38_21_+_8433172_8446622_RNA45SN1_45S_50ntFlanks@12640.12660.21\|ACCATCTGTGGGATTATGACT | >hg38_21_+_8433172_8446622_RNA45SN1_45S_50ntFlanks | 12640 | 12660 | 21 | ACCATCTGTGGGATTATGACT | 145.0 | -11.0 | 1.7 | -6.5 | 5.9E-11 | 1.7E-10 | MB231/MB468 | MP |
| >hg38_21_+_8433172_8446622_RNA45SN1_45S_50ntFlanks@12416.12449.34\|TTCGATGTCGGCTCTTCCTATCATTGTGAAGCAG | >hg38_21_+_8433172_8446622_RNA45SN1_45S_50ntFlanks | 12416 | 12449 | 34 | TTCGATGTCGGCTCTTCCTATCATTGTGAAGCAG | 485.5 | 4.2 | 0.6 | 6.6 | 5.6E-11 | 1.7E-10 | BT20/MB468 | CYTO |
| >hg38_21_+_8433172_8446622_RNA45SN1_45S_50ntFlanks@11661.11685.25\|TGATTTCTGCCCAGTGCTCTGAATG | >hg38_21_+_8433172_8446622_RNA45SN1_45S_50ntFlanks | 11661 | 11685 | 25 | TGATTTCTGCCCAGTGCTCTGAATG | 7522.1 | -7.1 | 1.1 | -6.5 | 6.3E-11 | 1.7E-10 | MB231/MB468 | CYTO |
| >hg38_21_+_8433172_8446622_RNA45SN1_45S_50ntFlanks@3782.3802.21\|CAGTGAAACTGCGAATGGCTC | >hg38_21_+_8433172_8446622_RNA45SN1_45S_50ntFlanks | 3782 | 3802 | 21 | CAGTGAAACTGCGAATGGCTC | 661.2 | -8.8 | 1.3 | -6.5 | 7.2E-11 | 1.7E-10 | MB231/MB468 | MITO |
| >hg38_21_+_8433172_8446622_RNA45SN1_45S_50ntFlanks@12678.12711.34\|AATCCCGCCCAGGCGGAACGATACGGCAGCGCCG | >hg38_21_+_8433172_8446622_RNA45SN1_45S_50ntFlanks | 12678 | 12711 | 34 | AATCCCGCCCAGGCGGAACGATACGGCAGCGCCG | 94.7 | -10.4 | 1.6 | -6.5 | 5.9E-11 | 1.7E-10 | MB231/MB468 | MP |
| >hg38_21_+_8433172_8446622_RNA45SN1_45S_50ntFlanks@12957.12986.30\|CTTCTGGGTCGGGGTTTCGTACGTAGCAGA | >hg38_21_+_8433172_8446622_RNA45SN1_45S_50ntFlanks | 12957 | 12986 | 30 | CTTCTGGGTCGGGGTTTCGTACGTAGCAGA | 377.0 | -8.1 | 1.2 | -6.5 | 7.2E-11 | 1.7E-10 | MB231/MB468 | MITO |
| >hg38_21_+_8433172_8446622_RNA45SN1_45S_50ntFlanks@3835.3871.37\|CTCCTCTCCTACTTGGATAACTGTGGTAATTCTAGAG | >hg38_21_+_8433172_8446622_RNA45SN1_45S_50ntFlanks | 3835 | 3871 | 37 | CTCCTCTCCTACTTGGATAACTGTGGTAATTCTAGAG | 1026.6 | -5.3 | 0.8 | -6.5 | 6.3E-11 | 1.7E-10 | MB231/MB468 | NUC |
| >hg38_21_+_8433172_8446622_RNA45SN1_45S_50ntFlanks@12588.12614.27\|GGTTCAGACATTTGGTGTATGTGCTTG | >hg38_21_+_8433172_8446622_RNA45SN1_45S_50ntFlanks | 12588 | 12614 | 27 | GGTTCAGACATTTGGTGTATGTGCTTG | 378.5 | -11.6 | 1.7 | -6.9 | 4.4E-12 | 1.7E-10 | BT20/MB468 | MP |
| >hg38_21_+_8433172_8446622_RNA45SN1_45S_50ntFlanks@11170.11189.20\|CCGCGAGGGGGGCCCGGGCA | >hg38_21_+_8433172_8446622_RNA45SN1_45S_50ntFlanks | 11170 | 11189 | 20 | CCGCGAGGGGGGCCCGGGCA | 502.6 | 9.4 | 1.4 | 6.5 | 6.3E-11 | 1.7E-10 | MB231/MB468 | CYTO |
| >trna116_GluCTC_1_-_145399233_145399304@32.71.40,trna71_GluCTC_1_-_161439189_161439260@32.71.40,trna74_GluCTC_1_-_161431809_161431880@32.71.40,trna77_GluCTC_1_-_161424398_161424469@32.71.40,trna77_GluCTC_6_+_28949976_28950047@32.71.40,trna80_GluCTC_1_-_161417018_161417089@32.71.40,trna87_GluCTC_6_-_126101393_126101464@32.71.40\|CTCTCACCGCCGCGGCCCGGGTTCGATTCCCGGTCAGGGA | >trna116_GluCTC_1_-_145399233_145399304 | 32 | 71 | 40 | CTCTCACCGCCGCGGCCCGGGTTCGATTCCCGGTCAGGGA | 102.5 | -10.5 | 1.6 | -6.5 | 5.9E-11 | 1.7E-10 | MB231/MB468 | MP |
| >hg38_21_+_8433172_8446622_RNA45SN1_45S_50ntFlanks@11314.11344.31\|CCCACCCCACGTCTCGTCGCGCGCGCGTCCG | >hg38_21_+_8433172_8446622_RNA45SN1_45S_50ntFlanks | 11314 | 11344 | 31 | CCCACCCCACGTCTCGTCGCGCGCGCGTCCG | 682.7 | -7.5 | 1.2 | -6.5 | 6.3E-11 | 1.7E-10 | MB231/MB468 | CYTO |
| >hg38_21_+_8433172_8446622_RNA45SN1_45S_50ntFlanks@5061.5082.22\|ATTTGTCTGGTTAATTCCGATA | >hg38_21_+_8433172_8446622_RNA45SN1_45S_50ntFlanks | 5061 | 5082 | 22 | ATTTGTCTGGTTAATTCCGATA | 145.5 | -11.0 | 1.7 | -6.5 | 6.0E-11 | 1.7E-10 | MB231/MB468 | MP |
| >hg38_21_+_8433172_8446622_RNA45SN1_45S_50ntFlanks@11568.11603.36\|GCAGCCGACTTAGAACTGGTGCGGACCAGGGGAATC | >hg38_21_+_8433172_8446622_RNA45SN1_45S_50ntFlanks | 11568 | 11603 | 36 | GCAGCCGACTTAGAACTGGTGCGGACCAGGGGAATC | 3888.7 | -6.7 | 1.0 | -6.5 | 7.7E-11 | 1.7E-10 | BT20/MB468 | NUC |
| >trna10_AspGTC_12_-_125424193_125424264@1.29.29,trna12_AspGTC_12_-_125411891_125411962@1.29.29,trna38_AspGTC_17_-_8125556_8125627@1.29.29,trna45_AspGTC_6_+_27447453_27447524@1.29.29,trna48_AspGTC_6_+_27471523_27471594@1.29.29,trna4_AspGTC_12_+_96429799_96429870@1.29.29,trna69_AspGTC_1_-_161440205_161440276@1.29.29,trna72_AspGTC_1_-_161432824_161432895@1.29.29,trna75_AspGTC_1_-_161425414_161425485@1.29.29,trna78_AspGTC_1_-_161418033_161418104@1.29.29,trna81_AspGTC_1_-_161410615_161410686@1.29.29\|TCCTCGTTAGTATAGTGGTGAGTATCCCC | >trna10_AspGTC_12_-_125424193_125424264 | 1 | 29 | 29 | TCCTCGTTAGTATAGTGGTGAGTATCCCC | 601.5 | -4.7 | 0.7 | -6.5 | 6.4E-11 | 1.7E-10 | MB231/MB468 | CYTO |
| >hg38_21_+_8433172_8446622_RNA45SN1_45S_50ntFlanks@10823.10865.43\|AAGTCGGCAAGCCGGATCCGTAACTTCGGGATAAGGATTGGCT | >hg38_21_+_8433172_8446622_RNA45SN1_45S_50ntFlanks | 10823 | 10865 | 43 | AAGTCGGCAAGCCGGATCCGTAACTTCGGGATAAGGATTGGCT | 798.8 | -6.9 | 1.1 | -6.5 | 6.4E-11 | 1.7E-10 | MB231/MB468 | CYTO |
| >trna16_ValTAC_11_-_59318460_59318532@1.34.34\|GGTTCCATAGTGTAGCGGTTATCACGTCTGCTTT | >trna16_ValTAC_11_-_59318460_59318532 | 1 | 34 | 34 | GGTTCCATAGTGTAGCGGTTATCACGTCTGCTTT | 501.1 | 10.3 | 1.6 | 6.5 | 6.4E-11 | 1.7E-10 | MB231/MB468 | CYTO |
| >MI0000746\|hsa-mir-99b&WithFlank&19\|+\|51692606\|51692687@13.33.21(+1A)&[MIMAT0000689&hsa-miR-99b-5p&offsets\|0\|-1(+1A);m-10&19\|+\|51692618\|51692639&offsets\|0\|-1(+1A)]\|CACCCGTAGAACCGACCTTGCA | >MI0000746\|hsa-mir-99b&WithFlank&19\|+\|51692606\|51692687 | 13 | 33 | 21(+1A) | CACCCGTAGAACCGACCTTGCA | 3408.6 | 4.1 | 0.6 | 6.5 | 6.3E-11 | 1.7E-10 | MB231/MB468 | NUC |
| >hg38_21_+_8433172_8446622_RNA45SN1_45S_50ntFlanks@10729.10751.23\|AGAGGGTGTAAATCTCGCGCCGG | >hg38_21_+_8433172_8446622_RNA45SN1_45S_50ntFlanks | 10729 | 10751 | 23 | AGAGGGTGTAAATCTCGCGCCGG | 100.8 | -10.5 | 1.6 | -6.5 | 6.0E-11 | 1.7E-10 | MB231/MB468 | MP |
| >hg38_21_+_8433172_8446622_RNA45SN1_45S_50ntFlanks@9349.9370.22\|CCGCCGTGGCGCAATGAAGGTG | >hg38_21_+_8433172_8446622_RNA45SN1_45S_50ntFlanks | 9349 | 9370 | 22 | CCGCCGTGGCGCAATGAAGGTG | 292.7 | -11.3 | 1.6 | -6.9 | 4.5E-12 | 1.7E-10 | BT20/MB468 | MP |
| >MI0000063\|hsa-let-7b&WithFlank&22\|+\|46113680\|46113774@12.32.21(+1C)&[MIMAT0000063&hsa-let-7b-5p&offsets\|0\|-1(+1C);m-14&22\|+\|46113691\|46113712&offsets\|0\|-1(+1C)]\|TGAGGTAGTAGGTTGTGTGGTC | >MI0000063\|hsa-let-7b&WithFlank&22\|+\|46113680\|46113774 | 12 | 32 | 21(+1C) | TGAGGTAGTAGGTTGTGTGGTC | 1336.4 | 5.3 | 0.8 | 6.5 | 6.4E-11 | 1.7E-10 | MB231/MB468 | CYTO |
| >MI0000111\|hsa-mir-105-1&WithFlank&X\|-\|152392213\|152392305@21.42.22;MI0000112\|hsa-mir-105-2&WithFlank&X\|-\|152394406\|152394498@21.42.22&[MIMAT0000102&hsa-miR-105-5p&offsets\|+2\|+1;m-735&X\|-\|152392266\|152392287&offsets\|+2\|+2];[MIMAT0000102_1&hsa-miR-105-5p&offsets\|+2\|+1;m-735&X\|-\|152394459\|152394480&offsets\|+2\|+2]\|AAATGCTCAGACTCCTGTGGTG | >MI0000111\|hsa-mir-105-1&WithFlank&X\|-\|152392213\|152392305 | 21 | 42 | 22 | AAATGCTCAGACTCCTGTGGTG | 98.8 | -10.5 | 1.6 | -6.5 | 6.0E-11 | 1.7E-10 | MB231/MB468 | MP |
| >MI0003591\|hsa-mir-584&WithFlank&5\|-\|149062307\|149062415@22.40.19&[MIMAT0003249&hsa-miR-584-5p&offsets\|0\|-3;m-167&5\|-\|149062374\|149062394&offsets\|0\|-2]\|TTATGGTTTGCCTGGGACT | >MI0003591\|hsa-mir-584&WithFlank&5\|-\|149062307\|149062415 | 22 | 40 | 19 | TTATGGTTTGCCTGGGACT | 665.7 | 4.9 | 0.8 | 6.5 | 6.0E-11 | 1.7E-10 | MB231/MB468 | MP |
| >hg38_21_+_8433172_8446622_RNA45SN1_45S_50ntFlanks@12627.12651.25\|ATGGGGCGAAGCTACCATCTGTGGG | >hg38_21_+_8433172_8446622_RNA45SN1_45S_50ntFlanks | 12627 | 12651 | 25 | ATGGGGCGAAGCTACCATCTGTGGG | 291.4 | -11.3 | 1.6 | -6.9 | 4.5E-12 | 1.7E-10 | BT20/MB468 | MP |
| >hg38_21_+_8433172_8446622_RNA45SN1_45S_50ntFlanks@12202.12222.21\|CTAAGGCGAGCTCAGGGAGGA | >hg38_21_+_8433172_8446622_RNA45SN1_45S_50ntFlanks | 12202 | 12222 | 21 | CTAAGGCGAGCTCAGGGAGGA | 563.5 | -9.1 | 1.4 | -6.5 | 7.8E-11 | 1.7E-10 | BT20/MB468 | NUC |
| >hg38_21_+_8433172_8446622_RNA45SN1_45S_50ntFlanks@9250.9272.23\|CGCACGGGGTCGGCGGCGACGTC | >hg38_21_+_8433172_8446622_RNA45SN1_45S_50ntFlanks | 9250 | 9272 | 23 | CGCACGGGGTCGGCGGCGACGTC | 88.7 | -10.3 | 1.6 | -6.5 | 6.0E-11 | 1.7E-10 | MB231/MB468 | MP |
| >hg38_21_+_8433172_8446622_RNA45SN1_45S_50ntFlanks@4442.4471.30\|CCCCTTGCCTCTCGGCGCCCCCTCGATGCT | >hg38_21_+_8433172_8446622_RNA45SN1_45S_50ntFlanks | 4442 | 4471 | 30 | CCCCTTGCCTCTCGGCGCCCCCTCGATGCT | 99.0 | -10.5 | 1.6 | -6.5 | 6.1E-11 | 1.7E-10 | MB231/MB468 | MP |
| >hg38_21_+_8433172_8446622_RNA45SN1_45S_50ntFlanks@9454.9492.39\|CCGCGCCGGGGAGGTGGAGCACGAGCGCACGTGTTAGGA | >hg38_21_+_8433172_8446622_RNA45SN1_45S_50ntFlanks | 9454 | 9492 | 39 | CCGCGCCGGGGAGGTGGAGCACGAGCGCACGTGTTAGGA | 75.9 | -10.1 | 1.5 | -6.5 | 6.1E-11 | 1.7E-10 | MB231/MB468 | MP |
| >hg38_21_+_8433172_8446622_RNA45SN1_45S_50ntFlanks@9825.9850.26\|GTGGGCCACTTTTGGTAAGCAGAACT | >hg38_21_+_8433172_8446622_RNA45SN1_45S_50ntFlanks | 9825 | 9850 | 26 | GTGGGCCACTTTTGGTAAGCAGAACT | 83.2 | -10.2 | 1.6 | -6.5 | 6.1E-11 | 1.7E-10 | MB231/MB468 | MP |

| | | | | | | | | | | | | | |
|---|---|---|---|---|---|---|---|---|---|---|---|---|---|
| >hg38_21_+_8433172_8446622_RNA45SN1_45S_50ntFlanks@10365.10390.26\|GCCGAAGTGGAGAAGGGTTCCATGTG | >hg38_21_+_8433172_8446622_RNA45SN1_45S_50ntFlanks | 10365 | 10390 | 26 | GCCGAAGTGGAGAAGGGTTCCATGTG | 70.9 | -10.0 | 1.5 | -6.5 | 6.1E-11 | 1.7E-10 | MB231/MB468 | MP |
| >hg38_21_+_8433172_8446622_RNA45SN1_45S_50ntFlanks@4666.4686.21\|AAATTCTTGGACCGGCGCAAG | >hg38_21_+_8433172_8446622_RNA45SN1_45S_50ntFlanks | 4666 | 4686 | 21 | AAATTCTTGGACCGGCGCAAG | 111.1 | -10.7 | 1.6 | -6.5 | 6.1E-11 | 1.7E-10 | MB231/MB468 | MP |
| >hg38_21_+_8433172_8446622_RNA45SN1_45S_50ntFlanks@9416.9453.38\|CAGTCCGCCGAGGGCGCACCACCGGCCCGTCTCGCCCG | >hg38_21_+_8433172_8446622_RNA45SN1_45S_50ntFlanks | 9416 | 9453 | 38 | CAGTCCGCCGAGGGCGCACCACCGGCCCGTCTCGCCCG | 74.5 | -10.1 | 1.5 | -6.5 | 6.1E-11 | 1.7E-10 | MB231/MB468 | MP |
| >hg38_21_+_8433172_8446622_RNA45SN1_45S_50ntFlanks@9604.9622.19\|AAAGACTAATCGAACCATC | >hg38_21_+_8433172_8446622_RNA45SN1_45S_50ntFlanks | 9604 | 9622 | 19 | AAAGACTAATCGAACCATC | 3638.8 | -8.1 | 1.2 | -6.5 | 6.1E-11 | 1.7E-10 | MB231/MB468 | MP |
| >hg38_21_+_8433172_8446622_RNA45SN1_45S_50ntFlanks@11562.11603.42\|CGCCTAGCAGCCGACTTAGAACTGGTGCGGACCAGGGGAATC | >hg38_21_+_8433172_8446622_RNA45SN1_45S_50ntFlanks | 11562 | 11603 | 42 | CGCCTAGCAGCCGACTTAGAACTGGTGCGGACCAGGGGAATC | 1244.6 | -9.0 | 1.4 | -6.5 | 7.4E-11 | 1.7E-10 | MB231/MB468 | MITO |
| >hg38_21_+_8433172_8446622_RNA45SN1_45S_50ntFlanks@9760.9787.28\|AAACTTTAAATGGGTAAGAAGCCCGGCT | >hg38_21_+_8433172_8446622_RNA45SN1_45S_50ntFlanks | 9760 | 9787 | 28 | AAACTTTAAATGGGTAAGAAGCCCGGCT | 278.1 | -12.0 | 1.8 | -6.5 | 6.1E-11 | 1.7E-10 | MB231/MB468 | MP |
| >hg38_21_+_8433172_8446622_RNA45SN1_45S_50ntFlanks@12008.12042.35\|CCGCCGGTGAAATACCACTACTCTGATCGTTTTTT | >hg38_21_+_8433172_8446622_RNA45SN1_45S_50ntFlanks | 12008 | 12042 | 35 | CCGCCGGTGAAATACCACTACTCTGATCGTTTTTT | 1049.1 | -9.4 | 1.4 | -6.5 | 6.5E-11 | 1.7E-10 | MB231/MB468 | NUC |
| >hg38_21_+_8433172_8446622_RNA45SN1_45S_50ntFlanks@9240.9275.36\|GCGGAGCGAGCGCACGGGGTCGGCGGCGACGTCGGC | >hg38_21_+_8433172_8446622_RNA45SN1_45S_50ntFlanks | 9240 | 9275 | 36 | GCGGAGCGAGCGCACGGGGTCGGCGGCGACGTCGGC | 95.5 | -10.4 | 1.6 | -6.5 | 6.1E-11 | 1.7E-10 | MB231/MB468 | MP |
| >hg38_21_+_8433172_8446622_RNA45SN1_45S_50ntFlanks@8836.8862.27\|CGGCGGTGGCGGCGGCGGCGGCGGCGG | >hg38_21_+_8433172_8446622_RNA45SN1_45S_50ntFlanks | 8836 | 8862 | 27 | CGGCGGTGGCGGCGGCGGCGGCGGCGG | 107.1 | -10.6 | 1.6 | -6.5 | 6.2E-11 | 1.7E-10 | MB231/MB468 | MP |
| >hg38_21_+_8433172_8446622_RNA45SN1_45S_50ntFlanks@9862.9880.19\|ATGAACCGAACGCCGGGTT | >hg38_21_+_8433172_8446622_RNA45SN1_45S_50ntFlanks | 9862 | 9880 | 19 | ATGAACCGAACGCCGGGTT | 448.3 | -12.7 | 1.9 | -6.5 | 6.2E-11 | 1.7E-10 | MB231/MB468 | MP |
| >hg38_21_+_8433172_8446622_RNA45SN1_45S_50ntFlanks@7988.8016.29\|CAGACGTGGCGACCCGCTGAATTTAAGCA | >hg38_21_+_8433172_8446622_RNA45SN1_45S_50ntFlanks | 7988 | 8016 | 29 | CAGACGTGGCGACCCGCTGAATTTAAGCA | 1086.7 | -6.9 | 1.1 | -6.5 | 6.2E-11 | 1.7E-10 | MB231/MB468 | MP |
| >MI0000434\|hsa-let-7i&WithFlank&12\|+\|62603680\|62603775@12.32.21&[MIMAT0000415&hsa-let-7i-5p&offsets\|0\|-1;m-49&12\|+\|62603691\|62603712&offsets\|0\|-1]\|TGAGGTAGTAGTTTGTGCTGT | >MI0000434\|hsa-let-7i&WithFlank&12\|+\|62603680\|62603775 | 12 | 32 | 21 | TGAGGTAGTAGTTTGTGCTGT | 1078.9 | 2.2 | 0.3 | 6.5 | 7.9E-11 | 1.7E-10 | BT20/MB468 | NUC |
| >hg38_21_+_8433172_8446622_RNA45SN1_45S_50ntFlanks@12008.12034.27\|CCGCCGGTGAAATACCACTACTCTGAT | >hg38_21_+_8433172_8446622_RNA45SN1_45S_50ntFlanks | 12008 | 12034 | 27 | CCGCCGGTGAAATACCACTACTCTGAT | 516.6 | -5.8 | 0.9 | -6.5 | 5.9E-11 | 1.7E-10 | BT20/MB468 | CYTO |
| >hg38_21_+_8433172_8446622_RNA45SN1_45S_50ntFlanks@5444.5466.23\|CTCGGATCGGCCCCGCCGGGGTC | >hg38_21_+_8433172_8446622_RNA45SN1_45S_50ntFlanks | 5444 | 5466 | 23 | CTCGGATCGGCCCCGCCGGGGTC | 24462.5 | -5.9 | 0.9 | -6.5 | 6.6E-11 | 1.7E-10 | MB231/MB468 | CYTO |
| >hg38_21_+_8433172_8446622_RNA45SN1_45S_50ntFlanks@12934.12953.20\|CTAAACCATTCGTAGACGAC | >hg38_21_+_8433172_8446622_RNA45SN1_45S_50ntFlanks | 12934 | 12953 | 20 | CTAAACCATTCGTAGACGAC | 1412.1 | -8.4 | 1.3 | -6.5 | 5.9E-11 | 1.7E-10 | BT20/MB468 | CYTO |
| >hg38_21_+_8433172_8446622_RNA45SN1_45S_50ntFlanks@11586.11603.18\|GTGCGGACCAGGGGAATC | >hg38_21_+_8433172_8446622_RNA45SN1_45S_50ntFlanks | 11586 | 11603 | 18 | GTGCGGACCAGGGGAATC | 1689.8 | -6.0 | 0.9 | -6.5 | 6.6E-11 | 1.7E-10 | MB231/MB468 | CYTO |
| >hg38_21_+_8433172_8446622_RNA45SN1_45S_50ntFlanks@11534.11555.22\|GTTCCGCGCGGCGCCTCGCCTC | >hg38_21_+_8433172_8446622_RNA45SN1_45S_50ntFlanks | 11534 | 11555 | 22 | GTTCCGCGCGGCGCCTCGCCTC | 723.5 | -9.5 | 1.5 | -6.5 | 8.0E-11 | 1.7E-10 | BT20/MB468 | NUC |
| >hg38_21_+_8433172_8446622_RNA45SN1_45S_50ntFlanks@10866.10886.21\|CTAAGGGCTGGGTCGGTCGGG | >hg38_21_+_8433172_8446622_RNA45SN1_45S_50ntFlanks | 10866 | 10886 | 21 | CTAAGGGCTGGGTCGGTCGGG | 536.4 | -8.4 | 1.3 | -6.5 | 6.6E-11 | 1.8E-10 | MB231/MB468 | NUC |
| >hg38_21_+_8433172_8446622_RNA45SN1_45S_50ntFlanks@10869.10886.18\|AGGGCTGGGTCGGTCGGG | >hg38_21_+_8433172_8446622_RNA45SN1_45S_50ntFlanks | 10869 | 10886 | 18 | AGGGCTGGGTCGGTCGGG | 615.2 | -9.0 | 1.4 | -6.5 | 6.6E-11 | 1.8E-10 | MB231/MB468 | NUC |
| >hg38_21_+_8433172_8446622_RNA45SN1_45S_50ntFlanks@6734.6773.40\|ATTGATCATCGACACTTCGAACGCACTTGCGGCCCCGGGT | >hg38_21_+_8433172_8446622_RNA45SN1_45S_50ntFlanks | 6734 | 6773 | 40 | ATTGATCATCGACACTTCGAACGCACTTGCGGCCCCGGGT | 1185.2 | 5.8 | 0.9 | 6.5 | 6.6E-11 | 1.8E-10 | MB231/MB468 | NUC |
| >hg38_21_+_8433172_8446622_RNA45SN1_45S_50ntFlanks@9552.9572.21\|TCCGTAGCGGTCCTGACGTGC | >hg38_21_+_8433172_8446622_RNA45SN1_45S_50ntFlanks | 9552 | 9572 | 21 | TCCGTAGCGGTCCTGACGTGC | 254.8 | -2.9 | 0.4 | -6.5 | 6.6E-11 | 1.8E-10 | MB231/MB468 | NUC |
| >MI0000477\|hsa-mir-146a&WithFlank&5\|+\|160485346\|160485456@27.46.20(+1A)&[MIMAT0000449&hsa-miR-146a-5p&offsets\|0\|-2(+1A);m-129&5\|+\|160485372\|160485393&offsets\|0\|-2(+1A)]\|TGAGAACTGAATTCCATGGGA | >MI0000477\|hsa-mir-146a&WithFlank&5\|+\|160485346\|160485456 | 27 | 46 | 20(+1A) | TGAGAACTGAATTCCATGGGA | 737.6 | -12.9 | 1.8 | -7.1 | 1.0E-12 | 1.8E-10 | BT20/MB231 | MP |
| >hg38_21_+_8433172_8446622_RNA45SN1_45S_50ntFlanks@9534.9561.28\|AGGAAACTCTGGTGGAGGTCCGTAGCGG | >hg38_21_+_8433172_8446622_RNA45SN1_45S_50ntFlanks | 9534 | 9561 | 28 | AGGAAACTCTGGTGGAGGTCCGTAGCGG | 12431.6 | -11.7 | 1.8 | -6.5 | 8.0E-11 | 1.8E-10 | BT20/MB468 | NUC |
| >MI0000449\|hsa-mir-132&WithFlank&17\|-\|2049902\|2050014@65.86.22&[MIMAT0000426&hsa-miR-132-3p&offsets\|0\|0;m-138&17\|-\|2049929\|2049950&offsets\|0\|0]\|TAACAGTCTACAGCCATGGTCG | >MI0000449\|hsa-mir-132&WithFlank&17\|-\|2049902\|2050014 | 65 | 86 | 22 | TAACAGTCTACAGCCATGGTCG | 795.6 | 13.4 | 1.9 | 6.9 | 4.7E-12 | 1.8E-10 | BT20/MB468 | MP |
| >hg38_21_+_8433172_8446622_RNA45SN1_45S_50ntFlanks@5422.5450.29\|GATTGGATGGTTTAGTGAGGCCCTCGGAT | >hg38_21_+_8433172_8446622_RNA45SN1_45S_50ntFlanks | 5422 | 5450 | 29 | GATTGGATGGTTTAGTGAGGCCCTCGGAT | 673.7 | 13.1 | 1.9 | 6.9 | 4.7E-12 | 1.8E-10 | BT20/MB468 | MP |
| >hg38_21_+_8433172_8446622_RNA45SN1_45S_50ntFlanks@12676.12704.29\|AGAATCCCGCCCAGGCGGAACGATACGGC | >hg38_21_+_8433172_8446622_RNA45SN1_45S_50ntFlanks | 12676 | 12704 | 29 | AGAATCCCGCCCAGGCGGAACGATACGGC | 427.0 | -11.8 | 1.7 | -6.9 | 4.7E-12 | 1.8E-10 | BT20/MB468 | MP |
| >trnaMT_GluTTC_MT_-_14674_14742@46.72.27,trnalookalike8_GluTTC_5_-_93905172_93905240@46.72.27\|GTGGTTGTAGTCCGTGCGAGAATACCA | >trnaMT_GluTTC_MT_-_14674_14742 | 46 | 72 | 27 | GTGGTTGTAGTCCGTGCGAGAATACCA | 2368.6 | 6.7 | 1.0 | 6.6 | 4.9E-11 | 1.8E-10 | BT20/MB468 | MITO |
| >hg38_21_+_8433172_8446622_RNA45SN1_45S_50ntFlanks@6745.6763.19\|ACACTTCGAACGCACTTGC | >hg38_21_+_8433172_8446622_RNA45SN1_45S_50ntFlanks | 6745 | 6763 | 19 | ACACTTCGAACGCACTTGC | 98.9 | -10.5 | 1.6 | -6.5 | 6.3E-11 | 1.8E-10 | MB231/MB468 | MP |
| >hg38_21_+_8433172_8446622_RNA45SN1_45S_50ntFlanks@3777.3802.26\|CGGTACAGTGAAACTGCGAATGGCTC | >hg38_21_+_8433172_8446622_RNA45SN1_45S_50ntFlanks | 3777 | 3802 | 26 | CGGTACAGTGAAACTGCGAATGGCTC | 103.3 | -10.6 | 1.6 | -6.5 | 6.3E-11 | 1.8E-10 | MB231/MB468 | MP |
| >hg38_21_+_8433172_8446622_RNA45SN1_45S_50ntFlanks@9930.9948.19\|ATATAGACAGCAGGACGGT | >hg38_21_+_8433172_8446622_RNA45SN1_45S_50ntFlanks | 9930 | 9948 | 19 | ATATAGACAGCAGGACGGT | 375.1 | -4.8 | 0.7 | -6.5 | 6.3E-11 | 1.8E-10 | MB231/MB468 | MP |
| >hg38_21_+_8433172_8446622_RNA45SN1_45S_50ntFlanks@11321.11338.18\|CACGTCTCGTCGCGCGCG | >hg38_21_+_8433172_8446622_RNA45SN1_45S_50ntFlanks | 11321 | 11338 | 18 | CACGTCTCGTCGCGCGCG | 665.4 | -8.2 | 1.3 | -6.5 | 6.6E-11 | 1.8E-10 | MB231/MB468 | NUC |
| >MI0000439\|hsa-mir-23b&WithFlank&9\|+\|95085202\|95085310@67.86.20&[MIMAT0000418&hsa-miR-23b-3p&offsets\|+3\|0;m-42&9\|+\|95085265\|95085285&offsets\|+3\|+2]\|ACATTGCCAGGGATTACCAC | >MI0000439\|hsa-mir-23b&WithFlank&9\|+\|95085202\|95085310 | 67 | 86 | 20 | ACATTGCCAGGGATTACCAC | 190.8 | 11.3 | 1.6 | 6.9 | 4.7E-12 | 1.8E-10 | BT20/MB468 | MP |
| >hg38_21_+_8433172_8446622_RNA45SN1_45S_50ntFlanks@3777.3794.18\|CGGTACAGTGAAACTGCG | >hg38_21_+_8433172_8446622_RNA45SN1_45S_50ntFlanks | 3777 | 3794 | 18 | CGGTACAGTGAAACTGCG | 204.1 | -10.8 | 1.6 | -6.9 | 4.7E-12 | 1.8E-10 | BT20/MB468 | MP |
| >trna10_LysCTT_16_+_3241501_3241573@4.33.30,trna13_LysCTT_14_-_58706613_58706685@4.33.30,trna2_LysCTT_15_+_79152904_79152976@4.33.30\|CGGCTAGCTCAGTCGGTAGAGCATGGGACT | >trna10_LysCTT_16_+_3241501_3241573 | 4 | 33 | 30 | CGGCTAGCTCAGTCGGTAGAGCATGGGACT | 1181.9 | 6.3 | 1.0 | 6.5 | 6.6E-11 | 1.8E-10 | MB231/MB468 | NUC |
| >hg38_21_+_8433172_8446622_RNA45SN1_45S_50ntFlanks@11208.11229.22\|GCGGCGACTCTGGACGCGAGCC | >hg38_21_+_8433172_8446622_RNA45SN1_45S_50ntFlanks | 11208 | 11229 | 22 | GCGGCGACTCTGGACGCGAGCC | 772.5 | -8.7 | 1.3 | -6.5 | 6.3E-11 | 1.8E-10 | MB231/MB468 | MP |
| >MI0000095\|hsa-mir-93&WithFlank&7\|-\|100093762\|100093853@17.39.23(+1C)&[MIMAT0000093&hsa-miR-93-5p&offsets\|0\|0(+1C);m-29&7\|-\|100093815\|100093837&offsets\|0\|0(+1C)]\|CAAAGTGCTGTTCGTGCAGGTAGC | >MI0000095\|hsa-mir-93&WithFlank&7\|-\|100093762\|100093853 | 17 | 39 | 23(+1C) | CAAAGTGCTGTTCGTGCAGGTAGC | 308.1 | 4.8 | 0.7 | 6.5 | 6.6E-11 | 1.8E-10 | MB231/MB468 | NUC |
| >hg38_21_+_8433172_8446622_RNA45SN1_45S_50ntFlanks@3782.3802.21\|CAGTGAAACTGCGAATGGCTC | >hg38_21_+_8433172_8446622_RNA45SN1_45S_50ntFlanks | 3782 | 3802 | 21 | CAGTGAAACTGCGAATGGCTC | 608.0 | -9.0 | 1.4 | -6.5 | 6.3E-11 | 1.8E-10 | MB231/MB468 | MP |
| >MI0000077\|hsa-mir-21&WithFlank&17\|+\|59841260\|59841343@15.36.22(+1A)&[MIMAT0000076&hsa-miR-21-5p&offsets\|+1\|+1(+1A);m-1&17\|+\|59841273\|59841294&offsets\|+1\|+1(+1A)]\|AGCTTATCAGACTGATGTTGACA | >MI0000077\|hsa-mir-21&WithFlank&17\|+\|59841260\|59841343 | 15 | 36 | 22(+1A) | AGCTTATCAGACTGATGTTGACA | 101.6 | 3.2 | 0.5 | 6.5 | 8.1E-11 | 1.8E-10 | BT20/MB468 | NUC |
| >hg38_21_+_8433172_8446622_RNA45SN1_45S_50ntFlanks@12625.12659.35\|CAATGGGGCGAAGCTACCATCTGTGGGATTATGAC | >hg38_21_+_8433172_8446622_RNA45SN1_45S_50ntFlanks | 12625 | 12659 | 35 | CAATGGGGCGAAGCTACCATCTGTGGGATTATGAC | 1802.1 | -8.0 | 1.2 | -6.5 | 7.7E-11 | 1.8E-10 | MB231/MB468 | MITO |
| >MI0001445\|hsa-mir-423&WithFlank&17\|+\|30117073\|30117178@58.81.24&[MIMAT0001340&hsa-miR-423-3p&offsets\|-1\|0;m-100&17\|+\|30117131\|30117153&offsets\|-1\|0]\|AAGCTCGGTCTGAGGCCCCTCAGT | >MI0001445\|hsa-mir-423&WithFlank&17\|+\|30117073\|30117178 | 58 | 81 | 24 | AAGCTCGGTCTGAGGCCCCTCAGT | 3345.4 | 6.1 | 0.9 | 6.5 | 7.7E-11 | 1.8E-10 | MB231/MB468 | MITO |
| >hg38_21_+_8433172_8446622_RNA45SN1_45S_50ntFlanks@6666.6688.23\|GATCACTCGGCTCGTGCGTCGAT | >hg38_21_+_8433172_8446622_RNA45SN1_45S_50ntFlanks | 6666 | 6688 | 23 | GATCACTCGGCTCGTGCGTCGAT | 139.1 | -9.2 | 1.4 | -6.6 | 4.9E-11 | 1.8E-10 | BT20/MB468 | MITO |
| >hg38_21_+_8433172_8446622_RNA45SN1_45S_50ntFlanks@11568.11592.25\|GCAGCCGACTTAGAACTGGTGCGGA | >hg38_21_+_8433172_8446622_RNA45SN1_45S_50ntFlanks | 11568 | 11592 | 25 | GCAGCCGACTTAGAACTGGTGCGGA | 2212.2 | -8.7 | 1.3 | -6.5 | 6.4E-11 | 1.8E-10 | MB231/MB468 | MP |
| >hg38_21_+_8433172_8446622_RNA45SN1_45S_50ntFlanks@11670.11693.24\|CCCAGTGCTCTGAATGTCAAAGTG | >hg38_21_+_8433172_8446622_RNA45SN1_45S_50ntFlanks | 11670 | 11693 | 24 | CCCAGTGCTCTGAATGTCAAAGTG | 526.1 | -5.2 | 0.8 | -6.5 | 6.4E-11 | 1.8E-10 | MB231/MB468 | MP |
| >trna116_GluCTC_1_-_145399233_145399304@30.70.41,trna71_GluCTC_1_-_161439189_161439260@30.70.41,trna74_GluCTC_1_-_161431809_161431880@30.70.41,trna77_GluCTC_1_-_161424398_161424469@30.70.41,trna77_GluCTC_6_+_28949976_28950047@30.70.41,trna80_GluCTC_1_-_161417018_161417089@30.70.41,trna87_GluCTC_6_-_126101393_126101464@30.70.41\|CGCTCTCACCGCCGCGGCCCGGGTTCGATTCCCGGTCAGGG | >trna116_GluCTC_1_-_145399233_145399304 | 30 | 70 | 41 | CGCTCTCACCGCCGCGGCCCGGGTTCGATTCCCGGTCAGGG | 212.3 | -11.5 | 1.8 | -6.5 | 7.7E-11 | 1.8E-10 | MB231/MB468 | MITO |
| >hg38_21_+_8433172_8446622_RNA45SN1_45S_50ntFlanks@9598.9622.25\|GGGGCGAAAGACTAATCGAACCATC | >hg38_21_+_8433172_8446622_RNA45SN1_45S_50ntFlanks | 9598 | 9622 | 25 | GGGGCGAAAGACTAATCGAACCATC | 210.7 | -11.6 | 1.8 | -6.5 | 6.4E-11 | 1.8E-10 | MB231/MB468 | MP |
| >hg38_21_+_8433172_8446622_RNA45SN1_45S_50ntFlanks@3773.3794.22\|CGGCCGGTACAGTGAAACTGCG | >hg38_21_+_8433172_8446622_RNA45SN1_45S_50ntFlanks | 3773 | 3794 | 22 | CGGCCGGTACAGTGAAACTGCG | 142.9 | -11.0 | 1.7 | -6.5 | 6.4E-11 | 1.8E-10 | MB231/MB468 | MP |
| >hg38_21_+_8433172_8446622_RNA45SN1_45S_50ntFlanks@12006.12043.38\|GGCCGCCGGTGAAATACCACTACTCTGATCGTTTTTTC | >hg38_21_+_8433172_8446622_RNA45SN1_45S_50ntFlanks | 12006 | 12043 | 38 | GGCCGCCGGTGAAATACCACTACTCTGATCGTTTTTTC | 281.3 | -11.1 | 1.7 | -6.5 | 6.8E-11 | 1.8E-10 | MB231/MB468 | NUC |
| >hg38_21_+_8433172_8446622_RNA45SN1_45S_50ntFlanks@11821.11856.36\|CTATCCAGCGAAACCACAGCCAAGGGAACGGGCTTG | >hg38_21_+_8433172_8446622_RNA45SN1_45S_50ntFlanks | 11821 | 11856 | 36 | CTATCCAGCGAAACCACAGCCAAGGGAACGGGCTTG | 131.0 | -9.1 | 1.4 | -6.6 | 5.0E-11 | 1.8E-10 | BT20/MB468 | MITO |
| >hg38_21_+_8433172_8446622_RNA45SN1_45S_50ntFlanks@12626.12657.32\|AATGGGGCGAAGCTACCATCTGTGGGATTATG | >hg38_21_+_8433172_8446622_RNA45SN1_45S_50ntFlanks | 12626 | 12657 | 32 | AATGGGGCGAAGCTACCATCTGTGGGATTATG | 1113.1 | -4.9 | 0.7 | -6.6 | 5.0E-11 | 1.8E-10 | BT20/MB468 | MITO |

| | | | | | | | | | | | | | |
|---|---|---|---|---|---|---|---|---|---|---|---|---|---|
| >am_trna116_GluCTC_1_-_145399233_145399304@1.29.29,trna59_GluCTC_1_+_249168447_249168518@1.29.29,trna71_GluCTC_1_-_161439189_161439260@1.29.29,trna74_GluCTC_1_-_161431809_161431880@1.29.29,trna77_GluCTC_1_-_161424398_161424469@1.29.29,trna77_GluCTC_6_+_28949976_28950047@1.29.29,trna80_GluCTC_1_-_161417018_161417089@1.29.29,trna87_GluCTC_6_-_126101393_126101464@1.29.29\|TCCCTGGTGGTCTAGTGGTTAGGATTCGG | >am_trna116_GluCTC_1_-_145399233_145399304 | 1 | 29 | 29 | TCCCTGGTGGTCTAGTGGTTAGGATTCGG | 18622.2 | -6.1 | 0.9 | -6.5 | 6.4E-11 | 1.8E-10 | MB231/MB468 | MP |
| >hg38_21_+_8433172_8446622_RNA45SN1_45S_50ntFlanks@3882.3899.18\|CCGACGGGCGCTGACCCC | >hg38_21_+_8433172_8446622_RNA45SN1_45S_50ntFlanks | 3882 | 3899 | 18 | CCGACGGGCGCTGACCCC | 2250.3 | -5.8 | 0.9 | -6.6 | 5.0E-11 | 1.8E-10 | BT20/MB468 | MITO |
| >hg38_21_+_8433172_8446622_RNA45SN1_45S_50ntFlanks@11583.11607.25\|CTGGTGCGGACCAGGGGAATCCGAC | >hg38_21_+_8433172_8446622_RNA45SN1_45S_50ntFlanks | 11583 | 11607 | 25 | CTGGTGCGGACCAGGGGAATCCGAC | 435.5 | -4.3 | 0.7 | -6.5 | 7.8E-11 | 1.8E-10 | MB231/MB468 | MITO |
| >hg38_21_+_8433172_8446622_RNA45SN1_45S_50ntFlanks@8423.8469.47\|CGCAGTCCGCCCGGAGGATTCAACCCGGCGGCGGGTCCGGCCGTGTC | >hg38_21_+_8433172_8446622_RNA45SN1_45S_50ntFlanks | 8423 | 8469 | 47 | CGCAGTCCGCCCGGAGGATTCAACCCGGCGGCGGGTCCGGCCGTGTC | 1344.7 | -8.6 | 1.3 | -6.6 | 5.0E-11 | 1.8E-10 | BT20/MB468 | MITO |
| >hg38_21_+_8433172_8446622_RNA45SN1_45S_50ntFlanks@6722.6744.23\|ATTGCAGGACACATTGATCATCG | >hg38_21_+_8433172_8446622_RNA45SN1_45S_50ntFlanks | 6722 | 6744 | 23 | ATTGCAGGACACATTGATCATCG | 3608.4 | -5.2 | 0.8 | -6.5 | 6.8E-11 | 1.8E-10 | MB231/MB468 | CYTO |
| >hg38_21_+_8433172_8446622_RNA45SN1_45S_50ntFlanks@4283.4315.33\|CTTTAACGAGGATCCATTGGAGGGCAAGTCTGG | >hg38_21_+_8433172_8446622_RNA45SN1_45S_50ntFlanks | 4283 | 4315 | 33 | CTTTAACGAGGATCCATTGGAGGGCAAGTCTGG | 154.2 | -11.1 | 1.7 | -6.5 | 6.4E-11 | 1.8E-10 | MB231/MB468 | MP |
| >hg38_21_+_8433172_8446622_RNA45SN1_45S_50ntFlanks@4685.4707.23\|AGACGGACCAGAGCGAAAGCATT | >hg38_21_+_8433172_8446622_RNA45SN1_45S_50ntFlanks | 4685 | 4707 | 23 | AGACGGACCAGAGCGAAAGCATT | 410.2 | -6.7 | 1.0 | -6.5 | 6.4E-11 | 1.8E-10 | MB231/MB468 | MP |
| >hg38_21_+_8433172_8446622_RNA45SN1_45S_50ntFlanks@8439.8470.32\|GATTCAACCCGGCGGCGGGTCCGGCCGTGTCG | >hg38_21_+_8433172_8446622_RNA45SN1_45S_50ntFlanks | 8439 | 8470 | 32 | GATTCAACCCGGCGGCGGGTCCGGCCGTGTCG | 75.9 | -10.1 | 1.5 | -6.5 | 6.5E-11 | 1.8E-10 | MB231/MB468 | MP |
| >hg38_21_+_8433172_8446622_RNA45SN1_45S_50ntFlanks@12073.12098.26\|CCGAGGGGCTCTCGCTTCTGGCGCCA | >hg38_21_+_8433172_8446622_RNA45SN1_45S_50ntFlanks | 12073 | 12098 | 26 | CCGAGGGGCTCTCGCTTCTGGCGCCA | 80.5 | -10.2 | 1.6 | -6.5 | 6.5E-11 | 1.8E-10 | MB231/MB468 | MP |
| >hg38_21_+_8433172_8446622_RNA45SN1_45S_50ntFlanks@13002.13030.29\|CGATCTATTGAAAGTCAGCCCTCGACACA | >hg38_21_+_8433172_8446622_RNA45SN1_45S_50ntFlanks | 13002 | 13030 | 29 | CGATCTATTGAAAGTCAGCCCTCGACACA | 156.7 | -9.4 | 1.4 | -6.6 | 5.0E-11 | 1.8E-10 | BT20/MB468 | MITO |
| >am_MI0000809\|hsa-mir-151a&WithFlank&8\|-\|140732558\|140732659@53.72.20(+1U)&[MIMAT0000757&hsa-miR-151a-3p&offsets\|0\|-1(+1U);m-30&8\|-\|140732587\|140732607&offsets\|0\|-1(+1U)]\|CTAGACTGAAGCTCCTTGAGT | >am_MI0000809\|hsa-mir-151a&WithFlank&8\|-\|140732558\|140732659 | 53 | 72 | 20(+1U) | CTAGACTGAAGCTCCTTGAGT | 4521.2 | 6.1 | 0.9 | 6.9 | 4.9E-12 | 1.8E-10 | BT20/MB468 | MP |
| >hg38_21_+_8433172_8446622_RNA45SN1_45S_50ntFlanks@11677.11702.26\|CTCTGAATGTCAAAGTGAAGAAATTC | >hg38_21_+_8433172_8446622_RNA45SN1_45S_50ntFlanks | 11677 | 11702 | 26 | CTCTGAATGTCAAAGTGAAGAAATTC | 774.5 | -7.6 | 1.2 | -6.5 | 8.3E-11 | 1.8E-10 | BT20/MB468 | NUC |
| >hg38_21_+_8433172_8446622_RNA45SN1_45S_50ntFlanks@6644.6669.26\|CCTCGTACGACTCTTAGCGGTGGATC | >hg38_21_+_8433172_8446622_RNA45SN1_45S_50ntFlanks | 6644 | 6669 | 26 | CCTCGTACGACTCTTAGCGGTGGATC | 1678.6 | 4.6 | 0.7 | 6.6 | 5.0E-11 | 1.8E-10 | BT20/MB468 | MITO |
| >hg38_21_+_8433172_8446622_RNA45SN1_45S_50ntFlanks@3900.3919.20\|CTTCGCGGGGGGGATGCGTG | >hg38_21_+_8433172_8446622_RNA45SN1_45S_50ntFlanks | 3900 | 3919 | 20 | CTTCGCGGGGGGGATGCGTG | 1059.3 | -7.7 | 1.2 | -6.5 | 8.3E-11 | 1.8E-10 | BT20/MB468 | NUC |
| >hg38_21_+_8433172_8446622_RNA45SN1_45S_50ntFlanks@8465.8482.18\|GTGTCGGCGGCCCGGCGG | >hg38_21_+_8433172_8446622_RNA45SN1_45S_50ntFlanks | 8465 | 8482 | 18 | GTGTCGGCGGCCCGGCGG | 106.3 | -10.6 | 1.6 | -6.5 | 6.5E-11 | 1.8E-10 | MB231/MB468 | MP |
| >hg38_21_+_8433172_8446622_RNA45SN1_45S_50ntFlanks@9032.9058.27\|CCTCGTGGGGGGGCCGGGCCACCCCTC | >hg38_21_+_8433172_8446622_RNA45SN1_45S_50ntFlanks | 9032 | 9058 | 27 | CCTCGTGGGGGGGCCGGGCCACCCCTC | 180.8 | -9.6 | 1.5 | -6.6 | 5.1E-11 | 1.8E-10 | BT20/MB468 | MITO |
| >hg38_21_+_8433172_8446622_RNA45SN1_45S_50ntFlanks@8380.8405.26\|CAAGAGGGCGTGAAACCGTTAAGAGG | >hg38_21_+_8433172_8446622_RNA45SN1_45S_50ntFlanks | 8380 | 8405 | 26 | CAAGAGGGCGTGAAACCGTTAAGAGG | 1221.9 | -7.2 | 1.1 | -6.5 | 6.9E-11 | 1.8E-10 | MB231/MB468 | NUC |
| >hg38_21_+_8433172_8446622_RNA45SN1_45S_50ntFlanks@10840.10866.27\|CCGTAACTTCGGGATAAGGATTGGCTC | >hg38_21_+_8433172_8446622_RNA45SN1_45S_50ntFlanks | 10840 | 10866 | 27 | CCGTAACTTCGGGATAAGGATTGGCTC | 79.6 | -10.2 | 1.6 | -6.5 | 6.5E-11 | 1.8E-10 | MB231/MB468 | MP |
| >hg38_1_-_228634819_228635039_RNA5S12_5S_50ntFlanks@131.169.39\|GGATGGGAGACCGCCTGGGAATACCGGGTGCTGTAGGCT | >hg38_1_-_228634819_228635039_RNA5S12_5S_50ntFlanks | 131 | 169 | 39 | GGATGGGAGACCGCCTGGGAATACCGGGTGCTGTAGGCT | 658.7 | 8.0 | 1.2 | 6.5 | 6.9E-11 | 1.8E-10 | MB231/MB468 | CYTO |
| >hg38_21_+_8433172_8446622_RNA45SN1_45S_50ntFlanks@5473.5490.18\|CGGCCCTGGCGGAGCGCT | >hg38_21_+_8433172_8446622_RNA45SN1_45S_50ntFlanks | 5473 | 5490 | 18 | CGGCCCTGGCGGAGCGCT | 371.0 | 6.5 | 1.0 | 6.5 | 7.0E-11 | 1.8E-10 | MB231/MB468 | CYTO |
| >MI0006444\|hsa-mir-1307&WithFlank&10\|-\|103394247\|103394407@86.108.23&[MIMAT0005951&hsa-miR-1307-3p&offsets\|0\|+1;m-66&10\|-\|103394300\|103394322&offsets\|0\|0]\|ACTCGGCGTGGCGTCGGTCGTGG | >MI0006444\|hsa-mir-1307&WithFlank&10\|-\|103394247\|103394407 | 86 | 108 | 23 | ACTCGGCGTGGCGTCGGTCGTGG | 2936.1 | 5.1 | 0.8 | 6.5 | 6.9E-11 | 1.8E-10 | MB231/MB468 | NUC |
| >hg38_21_+_8433172_8446622_RNA45SN1_45S_50ntFlanks@12072.12095.24\|CCCGAGGGGCTCTCGCTTCTGGCG | >hg38_21_+_8433172_8446622_RNA45SN1_45S_50ntFlanks | 12072 | 12095 | 24 | CCCGAGGGGCTCTCGCTTCTGGCG | 666.7 | -6.7 | 1.0 | -6.5 | 6.9E-11 | 1.8E-10 | MB231/MB468 | NUC |
| >hg38_21_+_8433172_8446622_RNA45SN1_45S_50ntFlanks@11792.11810.19\|ATGAACGAGATTCCCACTG | >hg38_21_+_8433172_8446622_RNA45SN1_45S_50ntFlanks | 11792 | 11810 | 19 | ATGAACGAGATTCCCACTG | 320.8 | -8.1 | 1.2 | -6.5 | 7.0E-11 | 1.8E-10 | MB231/MB468 | CYTO |
| >hg38_21_+_8433172_8446622_RNA45SN1_45S_50ntFlanks@12076.12097.22\|AGGGGCTCTCGCTTCTGGCGCC | >hg38_21_+_8433172_8446622_RNA45SN1_45S_50ntFlanks | 12076 | 12097 | 22 | AGGGGCTCTCGCTTCTGGCGCC | 111.0 | -10.7 | 1.6 | -6.5 | 6.6E-11 | 1.8E-10 | MB231/MB468 | MP |
| >hg38_21_+_8433172_8446622_RNA45SN1_45S_50ntFlanks@12642.12659.18\|CATCTGTGGGATTATGAC | >hg38_21_+_8433172_8446622_RNA45SN1_45S_50ntFlanks | 12642 | 12659 | 18 | CATCTGTGGGATTATGAC | 2135.0 | -9.7 | 1.5 | -6.5 | 7.9E-11 | 1.8E-10 | MB231/MB468 | MITO |
| >hg38_21_+_8433172_8446622_RNA45SN1_45S_50ntFlanks@3773.3792.20\|CGGCCGGTACAGTGAAACTG | >hg38_21_+_8433172_8446622_RNA45SN1_45S_50ntFlanks | 3773 | 3792 | 20 | CGGCCGGTACAGTGAAACTG | 482.4 | -6.5 | 1.0 | -6.5 | 7.0E-11 | 1.8E-10 | MB231/MB468 | NUC |
| >hg38_21_+_8433172_8446622_RNA45SN1_45S_50ntFlanks@5527.5549.23\|AAAGTCGTAACAAGGTTTCCGTA | >hg38_21_+_8433172_8446622_RNA45SN1_45S_50ntFlanks | 5527 | 5549 | 23 | AAAGTCGTAACAAGGTTTCCGTA | 191.1 | -7.4 | 1.1 | -6.5 | 7.0E-11 | 1.8E-10 | MB231/MB468 | NUC |
| >MI0000289\|hsa-mir-181a-1&WithFlank&1\|-\|198859038\|198859159@70.91.22&[MIMAT0000270&hsa-miR-181a-3p&offsets\|0\|0;m-125&1\|-\|198859069\|198859090&offsets\|0\|0]\|ACCATCGACCGTTGATTGTACC | >MI0000289\|hsa-mir-181a-1&WithFlank&1\|-\|198859038\|198859159 | 70 | 91 | 22 | ACCATCGACCGTTGATTGTACC | 1047.1 | 6.3 | 1.0 | 6.5 | 8.0E-11 | 1.8E-10 | MB231/MB468 | MITO |
| >hg38_21_+_8433172_8446622_RNA45SN1_45S_50ntFlanks@12072.12097.26\|CCCGAGGGGCTCTCGCTTCTGGCGCC | >hg38_21_+_8433172_8446622_RNA45SN1_45S_50ntFlanks | 12072 | 12097 | 26 | CCCGAGGGGCTCTCGCTTCTGGCGCC | 134.2 | -10.9 | 1.7 | -6.5 | 6.6E-11 | 1.8E-10 | MB231/MB468 | MP |
| >hg38_21_+_8433172_8446622_RNA45SN1_45S_50ntFlanks@8191.8214.24\|CAGCCCGTGGACGGTGTGAGGCCG | >hg38_21_+_8433172_8446622_RNA45SN1_45S_50ntFlanks | 8191 | 8214 | 24 | CAGCCCGTGGACGGTGTGAGGCCG | 1713.1 | -7.5 | 1.1 | -6.5 | 7.0E-11 | 1.8E-10 | MB231/MB468 | NUC |
| >hg38_21_+_8433172_8446622_RNA45SN1_45S_50ntFlanks@475.497.23\|CTTCGTGATCGATGTGGTGACGT | >hg38_21_+_8433172_8446622_RNA45SN1_45S_50ntFlanks | 475 | 497 | 23 | CTTCGTGATCGATGTGGTGACGT | 7084.6 | 7.9 | 1.2 | 6.6 | 5.1E-11 | 1.8E-10 | BT20/MB468 | MITO |
| >hg38_21_+_8433172_8446622_RNA45SN1_45S_50ntFlanks@3771.3798.28\|CACGGCCGGTACAGTGAAACTGCGAATG | >hg38_21_+_8433172_8446622_RNA45SN1_45S_50ntFlanks | 3771 | 3798 | 28 | CACGGCCGGTACAGTGAAACTGCGAATG | 397.6 | -6.7 | 1.0 | -6.5 | 8.4E-11 | 1.8E-10 | BT20/MB468 | NUC |
| >hg38_21_+_8433172_8446622_RNA45SN1_45S_50ntFlanks@10535.10569.35\|CGCGAGGCGTCCAGTGCGGTAACGCGACCGATCCC | >hg38_21_+_8433172_8446622_RNA45SN1_45S_50ntFlanks | 10535 | 10569 | 35 | CGCGAGGCGTCCAGTGCGGTAACGCGACCGATCCC | 74.7 | -10.1 | 1.5 | -6.5 | 6.6E-11 | 1.8E-10 | MB231/MB468 | MP |
| >hg38_21_+_8433172_8446622_RNA45SN1_45S_50ntFlanks@11704.11727.24\|ATGAAGCGCGGGTAAACGGCGGGA | >hg38_21_+_8433172_8446622_RNA45SN1_45S_50ntFlanks | 11704 | 11727 | 24 | ATGAAGCGCGGGTAAACGGCGGGA | 117.6 | -10.7 | 1.6 | -6.5 | 6.6E-11 | 1.8E-10 | MB231/MB468 | MP |
| >hg38_21_+_8433172_8446622_RNA45SN1_45S_50ntFlanks@12931.12951.21\|GCGCTAAACCATTCGTAGACG | >hg38_21_+_8433172_8446622_RNA45SN1_45S_50ntFlanks | 12931 | 12951 | 21 | GCGCTAAACCATTCGTAGACG | 75.0 | -10.1 | 1.5 | -6.5 | 6.6E-11 | 1.9E-10 | MB231/MB468 | MP |
| >trna10_AspGTC_12_-_125424193_125424264@1.29.29,trna12_AspGTC_12_-_125411891_125411962@1.29.29,trna38_AspGTC_17_-_8125556_8125627@1.29.29,trna45_AspGTC_6_+_27447453_27447524@1.29.29,trna48_AspGTC_6_+_27471523_27471594@1.29.29,trna4_AspGTC_12_+_96429799_96429870@1.29.29,trna69_AspGTC_1_-_161440205_161440276@1.29.29,trna72_AspGTC_1_-_161432824_161432895@1.29.29,trna75_AspGTC_1_-_161425414_161425485@1.29.29,trna78_AspGTC_1_-_161418033_161418104@1.29.29,trna81_AspGTC_1_-_161410615_161410686@1.29.29\|TCCTCGTTAGTATAGTGGTGAGTATCCCC | >trna10_AspGTC_12_-_125424193_125424264 | 1 | 29 | 29 | TCCTCGTTAGTATAGTGGTGAGTATCCCC | 225.2 | -10.9 | 1.6 | -6.9 | 5.0E-12 | 1.9E-10 | BT20/MB468 | MP |
| >hg38_21_+_8433172_8446622_RNA45SN1_45S_50ntFlanks@11707.11726.20\|AAGCGCGGGTAAACGGCGGG | >hg38_21_+_8433172_8446622_RNA45SN1_45S_50ntFlanks | 11707 | 11726 | 20 | AAGCGCGGGTAAACGGCGGG | 83.7 | -10.3 | 1.6 | -6.5 | 6.6E-11 | 1.9E-10 | MB231/MB468 | MP |
| >am_trnaMT_PheGAA_MT_+_577_647@40.57.18\|TGTTTAGACGGGCTCACA | >am_trnaMT_PheGAA_MT_+_577_647 | 40 | 57 | 18 | TGTTTAGACGGGCTCACA | 698.3 | 10.0 | 1.5 | 6.5 | 6.3E-11 | 1.9E-10 | BT20/MB468 | CYTO |
| >hg38_21_+_8433172_8446622_RNA45SN1_45S_50ntFlanks@12990.13013.24\|GCTCCCTCGCTGCGATCTATTGAA | >hg38_21_+_8433172_8446622_RNA45SN1_45S_50ntFlanks | 12990 | 13013 | 24 | GCTCCCTCGCTGCGATCTATTGAA | 86.2 | -10.3 | 1.6 | -6.5 | 6.6E-11 | 1.9E-10 | MB231/MB468 | MP |
| >hg38_21_+_8433172_8446622_RNA45SN1_45S_50ntFlanks@9129.9149.21\|CGGGTCGGGGGGCGGGGCGGA | >hg38_21_+_8433172_8446622_RNA45SN1_45S_50ntFlanks | 9129 | 9149 | 21 | CGGGTCGGGGGGCGGGGCGGA | 462.4 | -11.9 | 1.7 | -6.9 | 5.0E-12 | 1.9E-10 | BT20/MB468 | MP |
| >hg38_21_+_8433172_8446622_RNA45SN1_45S_50ntFlanks@12446.12480.35\|GCAGAATTCACCAAGCGTTGGATTGTTCACCCACT | >hg38_21_+_8433172_8446622_RNA45SN1_45S_50ntFlanks | 12446 | 12480 | 35 | GCAGAATTCACCAAGCGTTGGATTGTTCACCCACT | 472.4 | 4.2 | 0.6 | 6.5 | 7.1E-11 | 1.9E-10 | MB231/MB468 | CYTO |
| >hg38_21_+_8433172_8446622_RNA45SN1_45S_50ntFlanks@12366.12404.39\|TAACTGGCTTGTGGCGGCCAAGCGTTCATAGCGACGTCG | >hg38_21_+_8433172_8446622_RNA45SN1_45S_50ntFlanks | 12366 | 12404 | 39 | TAACTGGCTTGTGGCGGCCAAGCGTTCATAGCGACGTCG | 147.9 | -9.3 | 1.4 | -6.6 | 5.2E-11 | 1.9E-10 | BT20/MB468 | MITO |
| >hg38_21_+_8433172_8446622_RNA45SN1_45S_50ntFlanks@11586.11606.21\|GTGCGGACCAGGGGAATCCGA | >hg38_21_+_8433172_8446622_RNA45SN1_45S_50ntFlanks | 11586 | 11606 | 21 | GTGCGGACCAGGGGAATCCGA | 123.4 | -10.8 | 1.7 | -6.5 | 6.7E-11 | 1.9E-10 | MB231/MB468 | MP |
| >MI0000102\|hsa-mir-100&WithFlank&11\|-\|122152223\|122152314@19.40.22(+3U)&[MIMAT0000098&hsa-miR-100-5p&offsets\|0\|0(+3U);m-27&11\|-\|122152276\|122152296&offsets\|0\|+1(+3U)]\|AACCCGTAGATCCGAACTTGTGTTT | >MI0000102\|hsa-mir-100&WithFlank&11\|-\|122152223\|122152314 | 19 | 40 | 22(+3U) | AACCCGTAGATCCGAACTTGTGTTT | 185.9 | 10.8 | 1.7 | 6.5 | 8.0E-11 | 1.9E-10 | MB231/MB468 | MITO |
| >MI0000289\|hsa-mir-181a-1&WithFlank&1\|-\|198859038\|198859159@70.90.21(+1U)&[MIMAT0000270&hsa-miR-181a-3p&offsets\|0\|-1(+1U);m-125&1\|-\|198859069\|198859090&offsets\|0\|-1(+1U)]\|ACCATCGACCGTTGATTGTACT | >MI0000289\|hsa-mir-181a-1&WithFlank&1\|-\|198859038\|198859159 | 70 | 90 | 21(+1U) | ACCATCGACCGTTGATTGTACT | 231.0 | 6.3 | 1.0 | 6.5 | 7.0E-11 | 1.9E-10 | MB231/MB468 | NUC |

<table>
<tr><td>>hg38_21_+_8433172_8446622_RNA45SN1_45S_50ntFlanks@8555.8581.27|CCCGGAGGGGGCGGGCTCCGGCGGGTG</td><td>>hg38_21_+_8433172_8446622_RNA45SN1_45S_50ntFlanks</td><td>8555</td><td>8581</td><td>27</td><td>CCCGGAGGGGGCGGGCTCCGGCGGGTG</td><td>110.8</td><td>-10.7</td><td>1.6</td><td>-6.5</td><td>6.7E-11</td><td>1.9E-10</td><td>MB231/MB468</td><td>MP</td></tr>
<tr><td>>hg38_21_+_8433172_8446622_RNA45SN1_45S_50ntFlanks@9570.9593.24|TGCAAATCGGTCGTCCGACCTGGG</td><td>>hg38_21_+_8433172_8446622_RNA45SN1_45S_50ntFlanks</td><td>9570</td><td>9593</td><td>24</td><td>TGCAAATCGGTCGTCCGACCTGGG</td><td>84.1</td><td>-10.3</td><td>1.6</td><td>-6.5</td><td>6.7E-11</td><td>1.9E-10</td><td>MB231/MB468</td><td>MP</td></tr>
<tr><td>>hg38_21_+_8433172_8446622_RNA45SN1_45S_50ntFlanks@9243.9275.33|GAGCGAGCGCACGGGGTCGGCGGCGACGTCGGC</td><td>>hg38_21_+_8433172_8446622_RNA45SN1_45S_50ntFlanks</td><td>9243</td><td>9275</td><td>33</td><td>GAGCGAGCGCACGGGGTCGGCGGCGACGTCGGC</td><td>91.6</td><td>-10.4</td><td>1.6</td><td>-6.5</td><td>6.7E-11</td><td>1.9E-10</td><td>MB231/MB468</td><td>MP</td></tr>
<tr><td>>hg38_21_+_8433172_8446622_RNA45SN1_45S_50ntFlanks@9620.9648.29|ATCTAGTAGCTGGTTCCCTCCGAAGTTTC</td><td>>hg38_21_+_8433172_8446622_RNA45SN1_45S_50ntFlanks</td><td>9620</td><td>9648</td><td>29</td><td>ATCTAGTAGCTGGTTCCCTCCGAAGTTTC</td><td>77.6</td><td>-10.1</td><td>1.6</td><td>-6.5</td><td>6.7E-11</td><td>1.9E-10</td><td>MB231/MB468</td><td>MP</td></tr>
<tr><td>>MI0000273|hsa-mir-183&WithFlank&7|-|129774899|129775020@34.54.21(+1A)&[MIMAT0000261&hsa-miR-183-5p&offsets|+1|0(+1A);m-41&7|-|129774966|129774987&offsets|0|-1(+1A)]|ATGGCACTGGTAGAATTCACTA</td><td>>MI0000273|hsa-mir-183&WithFlank&7|-|129774899|129775020</td><td>34</td><td>54</td><td>21(+1A)</td><td>ATGGCACTGGTAGAATTCACTA</td><td>440.6</td><td>4.0</td><td>0.6</td><td>6.8</td><td>7.6E-12</td><td>1.9E-10</td><td>BT20/MB231</td><td>NUC</td></tr>
<tr><td>>hg38_21_+_8433172_8446622_RNA45SN1_45S_50ntFlanks@11785.11806.22|TGAATGGATGAACGAGATTCCC</td><td>>hg38_21_+_8433172_8446622_RNA45SN1_45S_50ntFlanks</td><td>11785</td><td>11806</td><td>22</td><td>TGAATGGATGAACGAGATTCCC</td><td>1291.7</td><td>-7.1</td><td>1.1</td><td>-6.5</td><td>6.7E-11</td><td>1.9E-10</td><td>MB231/MB468</td><td>MP</td></tr>
<tr><td>>hg38_21_+_8433172_8446622_RNA45SN1_45S_50ntFlanks@5083.5113.31|ACGAACGAGACTCTGGCATGCTAACTAGTTA</td><td>>hg38_21_+_8433172_8446622_RNA45SN1_45S_50ntFlanks</td><td>5083</td><td>5113</td><td>31</td><td>ACGAACGAGACTCTGGCATGCTAACTAGTTA</td><td>161.8</td><td>-11.2</td><td>1.7</td><td>-6.5</td><td>6.7E-11</td><td>1.9E-10</td><td>MB231/MB468</td><td>MP</td></tr>
<tr><td>>hg38_21_+_8433172_8446622_RNA45SN1_45S_50ntFlanks@7985.8017.33|GATCAGACGTGGCGACCCGCTGAATTTAAGCAT</td><td>>hg38_21_+_8433172_8446622_RNA45SN1_45S_50ntFlanks</td><td>7985</td><td>8017</td><td>33</td><td>GATCAGACGTGGCGACCCGCTGAATTTAAGCAT</td><td>201.4</td><td>-9.7</td><td>1.5</td><td>-6.6</td><td>5.2E-11</td><td>1.9E-10</td><td>BT20/MB468</td><td>MITO</td></tr>
<tr><td>>MI0001445|hsa-mir-423&WithFlank&17|+|30117073|30117178@58.81.24&[MIMAT0001340&hsa-miR-423-3p&offsets|-1|0;m-100&17|+|30117131|30117153&offsets|-1|0]|AAGCTCGGTCTGAGGCCCCTCAGT</td><td>>MI0001445|hsa-mir-423&WithFlank&17|+|30117073|30117178</td><td>58</td><td>81</td><td>24</td><td>AAGCTCGGTCTGAGGCCCCTCAGT</td><td>1861.0</td><td>5.8</td><td>0.8</td><td>6.9</td><td>5.1E-12</td><td>1.9E-10</td><td>BT20/MB468</td><td>MP</td></tr>
<tr><td>>hg38_21_+_8433172_8446622_RNA45SN1_45S_50ntFlanks@11792.11810.19|ATGAACGAGATTCCCACTG</td><td>>hg38_21_+_8433172_8446622_RNA45SN1_45S_50ntFlanks</td><td>11792</td><td>11810</td><td>19</td><td>ATGAACGAGATTCCCACTG</td><td>414.2</td><td>-11.8</td><td>1.7</td><td>-6.9</td><td>5.1E-12</td><td>1.9E-10</td><td>BT20/MB468</td><td>MP</td></tr>
<tr><td>>hg38_21_+_8433172_8446622_RNA45SN1_45S_50ntFlanks@11999.12031.33|CCCTGCGGGCCGCCGGTGAAATACCACTACTCT</td><td>>hg38_21_+_8433172_8446622_RNA45SN1_45S_50ntFlanks</td><td>11999</td><td>12031</td><td>33</td><td>CCCTGCGGGCCGCCGGTGAAATACCACTACTCT</td><td>1154.9</td><td>-6.5</td><td>1.0</td><td>-6.6</td><td>5.2E-11</td><td>1.9E-10</td><td>BT20/MB468</td><td>MITO</td></tr>
<tr><td>>hg38_1_-_228634819_228635039_RNA5S12_5S_50ntFlanks@141.170.30|CCGCCTGGGAATACCGGGTGCTGTAGGCTT</td><td>>hg38_1_-_228634819_228635039_RNA5S12_5S_50ntFlanks</td><td>141</td><td>170</td><td>30</td><td>CCGCCTGGGAATACCGGGTGCTGTAGGCTT</td><td>3709.1</td><td>5.2</td><td>0.8</td><td>6.5</td><td>7.2E-11</td><td>1.9E-10</td><td>MB231/MB468</td><td>CYTO</td></tr>
<tr><td>>hg38_21_+_8433172_8446622_RNA45SN1_45S_50ntFlanks@4728.4755.28|ATCAAGAACGAAAGTCGGAGGTTCGAAG</td><td>>hg38_21_+_8433172_8446622_RNA45SN1_45S_50ntFlanks</td><td>4728</td><td>4755</td><td>28</td><td>ATCAAGAACGAAAGTCGGAGGTTCGAAG</td><td>213.6</td><td>-10.8</td><td>1.6</td><td>-6.9</td><td>5.1E-12</td><td>1.9E-10</td><td>BT20/MB468</td><td>MP</td></tr>
<tr><td>>hg38_21_+_8433172_8446622_RNA45SN1_45S_50ntFlanks@12991.13027.37|CTCCCTCGCTGCGATCTATTGAAAGTCAGCCCTCGAC</td><td>>hg38_21_+_8433172_8446622_RNA45SN1_45S_50ntFlanks</td><td>12991</td><td>13027</td><td>37</td><td>CTCCCTCGCTGCGATCTATTGAAAGTCAGCCCTCGAC</td><td>964.3</td><td>-8.5</td><td>1.3</td><td>-6.5</td><td>7.1E-11</td><td>1.9E-10</td><td>MB231/MB468</td><td>NUC</td></tr>
<tr><td>>hg38_21_+_8433172_8446622_RNA45SN1_45S_50ntFlanks@9941.9978.38|AGGACGGTGGCCATGGAAGTCGGAATCCGCTAAGGAGT</td><td>>hg38_21_+_8433172_8446622_RNA45SN1_45S_50ntFlanks</td><td>9941</td><td>9978</td><td>38</td><td>AGGACGGTGGCCATGGAAGTCGGAATCCGCTAAGGAGT</td><td>210.2</td><td>11.5</td><td>1.7</td><td>6.9</td><td>5.2E-12</td><td>1.9E-10</td><td>BT20/MB468</td><td>MP</td></tr>
<tr><td>>hg38_21_+_8433172_8446622_RNA45SN1_45S_50ntFlanks@8369.8402.34|AAGAGAGAGTTCAAGAGGGCGTGAAACCGTTAAG</td><td>>hg38_21_+_8433172_8446622_RNA45SN1_45S_50ntFlanks</td><td>8369</td><td>8402</td><td>34</td><td>AAGAGAGAGTTCAAGAGGGCGTGAAACCGTTAAG</td><td>79.2</td><td>-10.2</td><td>1.6</td><td>-6.5</td><td>6.7E-11</td><td>1.9E-10</td><td>MB231/MB468</td><td>MP</td></tr>
<tr><td>>hg38_21_+_8433172_8446622_RNA45SN1_45S_50ntFlanks@6700.6718.19|CTAGCTGCGAGAATTAATG</td><td>>hg38_21_+_8433172_8446622_RNA45SN1_45S_50ntFlanks</td><td>6700</td><td>6718</td><td>19</td><td>CTAGCTGCGAGAATTAATG</td><td>11588.2</td><td>-10.7</td><td>1.6</td><td>-6.5</td><td>6.8E-11</td><td>1.9E-10</td><td>MB231/MB468</td><td>MP</td></tr>
<tr><td>>hg38_21_+_8433172_8446622_RNA45SN1_45S_50ntFlanks@8198.8223.26|TGGACGGTGTGAGGCCGGTAGCGGCC</td><td>>hg38_21_+_8433172_8446622_RNA45SN1_45S_50ntFlanks</td><td>8198</td><td>8223</td><td>26</td><td>TGGACGGTGTGAGGCCGGTAGCGGCC</td><td>844.3</td><td>-9.0</td><td>1.4</td><td>-6.5</td><td>6.8E-11</td><td>1.9E-10</td><td>MB231/MB468</td><td>MP</td></tr>
<tr><td>>trna15_ValAAC_5_-_180615416_180615488@1.32.32|GTTTCCGTAGTGTAGTGGTCATCACGTTCGCC</td><td>>trna15_ValAAC_5_-_180615416_180615488</td><td>1</td><td>32</td><td>32</td><td>GTTTCCGTAGTGTAGTGGTCATCACGTTCGCC</td><td>2074.0</td><td>8.6</td><td>1.3</td><td>6.5</td><td>7.2E-11</td><td>1.9E-10</td><td>MB231/MB468</td><td>CYTO</td></tr>
<tr><td>>hg38_21_+_8433172_8446622_RNA45SN1_45S_50ntFlanks@11604.11621.18|CGACTGTTTAATTAAAAC</td><td>>hg38_21_+_8433172_8446622_RNA45SN1_45S_50ntFlanks</td><td>11604</td><td>11621</td><td>18</td><td>CGACTGTTTAATTAAAAC</td><td>886.7</td><td>-5.5</td><td>0.8</td><td>-6.5</td><td>8.6E-11</td><td>1.9E-10</td><td>BT20/MB468</td><td>NUC</td></tr>
<tr><td>>trna10_AspGTC_12_-_125424193_125424264@40.73.34,trna12_AspGTC_12_-_125411891_125411962@40.73.34,trna144_AspGTC_6_-_27551236_27551307@40.73.34,trna38_AspGTC_17_-_8125556_8125627@40.73.34,trna45_AspGTC_6_+_27447453_27447524@40.73.34,trna48_AspGTC_6_+_27471523_27471594@40.73.34,trna4_AspGTC_12_+_96429799_96429870@40.73.34,trna69_AspGTC_1_-_161440205_161440276@40.73.34,trna72_AspGTC_1_-_161432824_161432895@40.73.34,trna75_AspGTC_1_-_161425414_161425485@40.73.34,trna78_AspGTC_1_-_161418033_161418104@40.73.34,trna81_AspGTC_1_-_161410615_161410686@40.73.34|CGGGAGACCGGGGTTCGATTCCCCGACGGGGAGC</td><td>>trna10_AspGTC_12_-_125424193_125424264</td><td>40</td><td>73</td><td>34</td><td>CGGGAGACCGGGGTTCGATTCCCCGACGGGGAGC</td><td>3102.0</td><td>7.2</td><td>1.1</td><td>6.5</td><td>7.2E-11</td><td>1.9E-10</td><td>MB231/MB468</td><td>CYTO</td></tr>
<tr><td>>hg38_21_+_8433172_8446622_RNA45SN1_45S_50ntFlanks@8171.8197.27|AAGTCCTTCTGATCGAGGCCCAGCCCG</td><td>>hg38_21_+_8433172_8446622_RNA45SN1_45S_50ntFlanks</td><td>8171</td><td>8197</td><td>27</td><td>AAGTCCTTCTGATCGAGGCCCAGCCCG</td><td>178.7</td><td>-11.3</td><td>1.7</td><td>-6.5</td><td>6.8E-11</td><td>1.9E-10</td><td>MB231/MB468</td><td>MP</td></tr>
<tr><td>>hg38_21_+_8433172_8446622_RNA45SN1_45S_50ntFlanks@6657.6681.25|TTAGCGGTGGATCACTCGGCTCGTG</td><td>>hg38_21_+_8433172_8446622_RNA45SN1_45S_50ntFlanks</td><td>6657</td><td>6681</td><td>25</td><td>TTAGCGGTGGATCACTCGGCTCGTG</td><td>494.0</td><td>-4.4</td><td>0.7</td><td>-6.5</td><td>7.2E-11</td><td>1.9E-10</td><td>MB231/MB468</td><td>CYTO</td></tr>
<tr><td>>hg38_21_+_8433172_8446622_RNA45SN1_45S_50ntFlanks@12997.13014.18|CGCTGCGATCTATTGAAA</td><td>>hg38_21_+_8433172_8446622_RNA45SN1_45S_50ntFlanks</td><td>12997</td><td>13014</td><td>18</td><td>CGCTGCGATCTATTGAAA</td><td>5362.5</td><td>4.3</td><td>0.7</td><td>6.5</td><td>7.2E-11</td><td>1.9E-10</td><td>MB231/MB468</td><td>CYTO</td></tr>
<tr><td>>hg38_21_+_8433172_8446622_RNA45SN1_45S_50ntFlanks@10677.10710.34|GCGGTTCCGGCGGCGTCCGGTGAGCTCTCGCTGG</td><td>>hg38_21_+_8433172_8446622_RNA45SN1_45S_50ntFlanks</td><td>10677</td><td>10710</td><td>34</td><td>GCGGTTCCGGCGGCGTCCGGTGAGCTCTCGCTGG</td><td>5277.2</td><td>-4.4</td><td>0.7</td><td>-6.5</td><td>7.2E-11</td><td>1.9E-10</td><td>MB231/MB468</td><td>NUC</td></tr>
<tr><td>>hg38_21_+_8433172_8446622_RNA45SN1_45S_50ntFlanks@10781.10814.34|AGGTGAACAGCCTCTGGCATGTTGGAACAATGTA</td><td>>hg38_21_+_8433172_8446622_RNA45SN1_45S_50ntFlanks</td><td>10781</td><td>10814</td><td>34</td><td>AGGTGAACAGCCTCTGGCATGTTGGAACAATGTA</td><td>330.3</td><td>-8.4</td><td>1.3</td><td>-6.5</td><td>8.7E-11</td><td>1.9E-10</td><td>BT20/MB468</td><td>NUC</td></tr>
<tr><td>>MI0000101|hsa-mir-99a&WithFlank&21|+|16539083|16539175@19.38.20(+1C)&[MIMAT0000097&hsa-miR-99a-5p&offsets|0|-2(+1C);m-64&21|+|16539101|16539121&offsets|0|-1(+1C)]|AACCCGTAGATCCGATCTTGC</td><td>>MI0000101|hsa-mir-99a&WithFlank&21|+|16539083|16539175</td><td>19</td><td>38</td><td>20(+1C)</td><td>AACCCGTAGATCCGATCTTGC</td><td>135.9</td><td>-10.2</td><td>1.5</td><td>-6.8</td><td>7.8E-12</td><td>1.9E-10</td><td>BT20/MB231</td><td>NUC</td></tr>
<tr><td>>hg38_21_+_8433172_8446622_RNA45SN1_45S_50ntFlanks@9198.9218.21|GTTCTCTCGGGGCCACGCGCG</td><td>>hg38_21_+_8433172_8446622_RNA45SN1_45S_50ntFlanks</td><td>9198</td><td>9218</td><td>21</td><td>GTTCTCTCGGGGCCACGCGCG</td><td>268.4</td><td>-11.0</td><td>1.7</td><td>-6.5</td><td>7.2E-11</td><td>1.9E-10</td><td>MB231/MB468</td><td>NUC</td></tr>
<tr><td>>hg38_21_+_8433172_8446622_RNA45SN1_45S_50ntFlanks@10796.10816.21|GGCATGTTGGAACAATGTAGG</td><td>>hg38_21_+_8433172_8446622_RNA45SN1_45S_50ntFlanks</td><td>10796</td><td>10816</td><td>21</td><td>GGCATGTTGGAACAATGTAGG</td><td>76.9</td><td>-10.1</td><td>1.6</td><td>-6.5</td><td>6.8E-11</td><td>1.9E-10</td><td>MB231/MB468</td><td>MP</td></tr>
<tr><td>>hg38_21_+_8433172_8446622_RNA45SN1_45S_50ntFlanks@12909.12930.22|CCGCACGTTCGTGGGGAACCTG</td><td>>hg38_21_+_8433172_8446622_RNA45SN1_45S_50ntFlanks</td><td>12909</td><td>12930</td><td>22</td><td>CCGCACGTTCGTGGGGAACCTG</td><td>104.1</td><td>-10.6</td><td>1.6</td><td>-6.5</td><td>6.8E-11</td><td>1.9E-10</td><td>MB231/MB468</td><td>MP</td></tr>
<tr><td>>hg38_21_+_8433172_8446622_RNA45SN1_45S_50ntFlanks@9459.9500.42|CCGGGGAGGTGGAGCACGAGCGCACGTGTTAGGACCCGAAAG</td><td>>hg38_21_+_8433172_8446622_RNA45SN1_45S_50ntFlanks</td><td>9459</td><td>9500</td><td>42</td><td>CCGGGGAGGTGGAGCACGAGCGCACGTGTTAGGACCCGAAAG</td><td>177.6</td><td>-11.3</td><td>1.7</td><td>-6.5</td><td>8.2E-11</td><td>1.9E-10</td><td>MB231/MB468</td><td>MITO</td></tr>
<tr><td>>hg38_21_+_8433172_8446622_RNA45SN1_45S_50ntFlanks@4328.4352.25|CGGTAATTCCAGCTCCAATAGCGTA</td><td>>hg38_21_+_8433172_8446622_RNA45SN1_45S_50ntFlanks</td><td>4328</td><td>4352</td><td>25</td><td>CGGTAATTCCAGCTCCAATAGCGTA</td><td>112.4</td><td>-10.7</td><td>1.6</td><td>-6.5</td><td>6.8E-11</td><td>1.9E-10</td><td>MB231/MB468</td><td>MP</td></tr>
<tr><td>>hg38_21_+_8433172_8446622_RNA45SN1_45S_50ntFlanks@9739.9764.26|AAACGATCTCAACCTATTCTCAAACT</td><td>>hg38_21_+_8433172_8446622_RNA45SN1_45S_50ntFlanks</td><td>9739</td><td>9764</td><td>26</td><td>AAACGATCTCAACCTATTCTCAAACT</td><td>94.5</td><td>-10.4</td><td>1.6</td><td>-6.5</td><td>6.8E-11</td><td>1.9E-10</td><td>MB231/MB468</td><td>MP</td></tr>
<tr><td>>hg38_21_+_8433172_8446622_RNA45SN1_45S_50ntFlanks@12412.12437.26|ATCCTTCGATGTCGGCTCTTCCTATC</td><td>>hg38_21_+_8433172_8446622_RNA45SN1_45S_50ntFlanks</td><td>12412</td><td>12437</td><td>26</td><td>ATCCTTCGATGTCGGCTCTTCCTATC</td><td>200.0</td><td>-9.7</td><td>1.5</td><td>-6.6</td><td>5.3E-11</td><td>1.9E-10</td><td>BT20/MB468</td><td>MITO</td></tr>
<tr><td>>hg38_21_+_8433172_8446622_RNA45SN1_45S_50ntFlanks@10393.10435.43|CAGCAGTTGAACATGGGTCAGTCGGTCCTGAGAGATGGGCGAG</td><td>>hg38_21_+_8433172_8446622_RNA45SN1_45S_50ntFlanks</td><td>10393</td><td>10435</td><td>43</td><td>CAGCAGTTGAACATGGGTCAGTCGGTCCTGAGAGATGGGCGAG</td><td>206.0</td><td>-11.6</td><td>1.8</td><td>-6.5</td><td>6.8E-11</td><td>1.9E-10</td><td>MB231/MB468</td><td>MP</td></tr>
<tr><td>>hg38_21_+_8433172_8446622_RNA45SN1_45S_50ntFlanks@11573.11591.19|CGACTTAGAACTGGTGCGG</td><td>>hg38_21_+_8433172_8446622_RNA45SN1_45S_50ntFlanks</td><td>11573</td><td>11591</td><td>19</td><td>CGACTTAGAACTGGTGCGG</td><td>1012.3</td><td>-9.2</td><td>1.4</td><td>-6.5</td><td>6.8E-11</td><td>1.9E-10</td><td>MB231/MB468</td><td>MP</td></tr>
<tr><td>>hg38_21_+_8433172_8446622_RNA45SN1_45S_50ntFlanks@5084.5106.23|CGAACGAGACTCTGGCATGCTAA</td><td>>hg38_21_+_8433172_8446622_RNA45SN1_45S_50ntFlanks</td><td>5084</td><td>5106</td><td>23</td><td>CGAACGAGACTCTGGCATGCTAA</td><td>238.3</td><td>-11.8</td><td>1.8</td><td>-6.5</td><td>6.8E-11</td><td>1.9E-10</td><td>MB231/MB468</td><td>MP</td></tr>
<tr><td>>hg38_21_+_8433172_8446622_RNA45SN1_45S_50ntFlanks@11575.11600.26|ACTTAGAACTGGTGCGGACCAGGGGA</td><td>>hg38_21_+_8433172_8446622_RNA45SN1_45S_50ntFlanks</td><td>11575</td><td>11600</td><td>26</td><td>ACTTAGAACTGGTGCGGACCAGGGGA</td><td>1482.0</td><td>-9.7</td><td>1.5</td><td>-6.5</td><td>8.7E-11</td><td>1.9E-10</td><td>BT20/MB468</td><td>NUC</td></tr>
<tr><td>>hg38_21_+_8433172_8446622_RNA45SN1_45S_50ntFlanks@8179.8208.30|CTGATCGAGGCCCAGCCCGTGGACGGTGTG</td><td>>hg38_21_+_8433172_8446622_RNA45SN1_45S_50ntFlanks</td><td>8179</td><td>8208</td><td>30</td><td>CTGATCGAGGCCCAGCCCGTGGACGGTGTG</td><td>651.1</td><td>-6.9</td><td>1.1</td><td>-6.5</td><td>7.2E-11</td><td>1.9E-10</td><td>MB231/MB468</td><td>NUC</td></tr>
<tr><td>>hg38_21_+_8433172_8446622_RNA45SN1_45S_50ntFlanks@11729.11752.24|TAACTATGACTCTCTTAAGGTAGC</td><td>>hg38_21_+_8433172_8446622_RNA45SN1_45S_50ntFlanks</td><td>11729</td><td>11752</td><td>24</td><td>TAACTATGACTCTCTTAAGGTAGC</td><td>1014.4</td><td>-3.1</td><td>0.5</td><td>-6.5</td><td>7.3E-11</td><td>1.9E-10</td><td>MB231/MB468</td><td>CYTO</td></tr>
<tr><td>>hg38_21_+_8433172_8446622_RNA45SN1_45S_50ntFlanks@11452.11474.23|CTCCTCGGCGCGCGGCGGCGGCG</td><td>>hg38_21_+_8433172_8446622_RNA45SN1_45S_50ntFlanks</td><td>11452</td><td>11474</td><td>23</td><td>CTCCTCGGCGCGCGGCGGCGGCG</td><td>82.3</td><td>-10.2</td><td>1.6</td><td>-6.5</td><td>6.9E-11</td><td>1.9E-10</td><td>MB231/MB468</td><td>MP</td></tr>
<tr><td>>hg38_21_+_8433172_8446622_RNA45SN1_45S_50ntFlanks@12307.12326.20|ACGATCCTTCTGACCTTTTG</td><td>>hg38_21_+_8433172_8446622_RNA45SN1_45S_50ntFlanks</td><td>12307</td><td>12326</td><td>20</td><td>ACGATCCTTCTGACCTTTTG</td><td>67.9</td><td>-10.0</td><td>1.5</td><td>-6.5</td><td>6.9E-11</td><td>1.9E-10</td><td>MB231/MB468</td><td>MP</td></tr>
<tr><td>>trna10_AspGTC_12_-_125424193_125424264@38.73.36,trna12_AspGTC_12_-_125411891_125411962@38.73.36,trna144_AspGTC_6_-_27551236_27551307@38.73.36,trna38_AspGTC_17_-_8125556_8125627@38.73.36,trna45_AspGTC_6_+_27447453_27447524@38.73.36,trna48_AspGTC_6_+_27471523_27471594@38.73.36,trna4_AspGTC_12_+_96429799_96429870@38.73.36,trna69_AspGTC_1_-_161440205_161440276@38.73.36,trna72_AspGTC_1_-_161432824_161432895@38.73.36,trna75_AspGTC_1_-_161425414_161425485@38.73.36,trna78_AspGTC_1_-_161418033_161418104@38.73.36,trna81_AspGTC_1_-_161410615_161410686@38.73.36|CGCGGGAGACCGGGGTTCGATTCCCCGACGGGGAGC</td><td>>trna10_AspGTC_12_-_125424193_125424264</td><td>38</td><td>73</td><td>36</td><td>CGCGGGAGACCGGGGTTCGATTCCCCGACGGGGAGC</td><td>4947.0</td><td>5.5</td><td>0.8</td><td>6.5</td><td>7.3E-11</td><td>1.9E-10</td><td>MB231/MB468</td><td>CYTO</td></tr>
<tr><td>>MI0000273|hsa-mir-183&WithFlank&7|-|129774899|129775020@33.53.21&[MIMAT0000261&hsa-miR-183-5p&offsets|0|-1;m-41&7|-|129774966|129774987&offsets|-1|-2]|TATGGCACTGGTAGAATTCAC</td><td>>MI0000273|hsa-mir-183&WithFlank&7|-|129774899|129775020</td><td>33</td><td>53</td><td>21</td><td>TATGGCACTGGTAGAATTCAC</td><td>614.5</td><td>8.7</td><td>1.3</td><td>6.5</td><td>6.5E-11</td><td>1.9E-10</td><td>BT20/MB468</td><td>CYTO</td></tr>
</table>

<table>
<tr><td>>hg38_21_+_8433172_8446622_RNA45SN1_45S_50ntFlanks@6658.6681.24|TAGCGGTGGATCACTCGGCTCGTG</td><td>>hg38_21_+_8433172_8446622_RNA45SN1_45S_50ntFlanks</td><td>6658</td><td>6681</td><td>24</td><td>TAGCGGTGGATCACTCGGCTCGTG</td><td>136.1</td><td>-4.4</td><td>0.7</td><td>-6.5</td><td>7.3E-11</td><td>1.9E-10</td><td>MB231/MB468</td><td>CYTO</td></tr>
<tr><td>>hg38_21_+_8433172_8446622_RNA45SN1_45S_50ntFlanks@10916.10933.18|CCGCGGCTGGACGAGGCG</td><td>>hg38_21_+_8433172_8446622_RNA45SN1_45S_50ntFlanks</td><td>10916</td><td>10933</td><td>18</td><td>CCGCGGCTGGACGAGGCG</td><td>711.9</td><td>-9.2</td><td>1.4</td><td>-6.5</td><td>7.3E-11</td><td>1.9E-10</td><td>MB231/MB468</td><td>NUC</td></tr>
<tr><td>>hg38_21_+_8433172_8446622_RNA45SN1_45S_50ntFlanks@6746.6772.27|CACTTCGAACGCACTTGCGGCCCCGGG</td><td>>hg38_21_+_8433172_8446622_RNA45SN1_45S_50ntFlanks</td><td>6746</td><td>6772</td><td>27</td><td>CACTTCGAACGCACTTGCGGCCCCGGG</td><td>135.6</td><td>-9.2</td><td>1.4</td><td>-6.6</td><td>5.3E-11</td><td>1.9E-10</td><td>BT20/MB468</td><td>MITO</td></tr>
<tr><td>>hg38_21_+_8433172_8446622_RNA45SN1_45S_50ntFlanks@7988.8013.26|CAGACGTGGCGACCCGCTGAATTTAA</td><td>>hg38_21_+_8433172_8446622_RNA45SN1_45S_50ntFlanks</td><td>7988</td><td>8013</td><td>26</td><td>CAGACGTGGCGACCCGCTGAATTTAA</td><td>712.6</td><td>-4.5</td><td>0.7</td><td>-6.5</td><td>8.3E-11</td><td>1.9E-10</td><td>MB231/MB468</td><td>MITO</td></tr>
<tr><td>>hg38_21_+_8433172_8446622_RNA45SN1_45S_50ntFlanks@10761.10785.25|ATATCCGCAGCAGGTCTCCAAGGTG</td><td>>hg38_21_+_8433172_8446622_RNA45SN1_45S_50ntFlanks</td><td>10761</td><td>10785</td><td>25</td><td>ATATCCGCAGCAGGTCTCCAAGGTG</td><td>214.0</td><td>-7.5</td><td>1.2</td><td>-6.5</td><td>7.3E-11</td><td>1.9E-10</td><td>MB231/MB468</td><td>NUC</td></tr>
<tr><td>>hg38_21_+_8433172_8446622_RNA45SN1_45S_50ntFlanks@8155.8175.21|CGCTCGTGGGGGGCCCAAGTC</td><td>>hg38_21_+_8433172_8446622_RNA45SN1_45S_50ntFlanks</td><td>8155</td><td>8175</td><td>21</td><td>CGCTCGTGGGGGGCCCAAGTC</td><td>269.4</td><td>-10.2</td><td>1.5</td><td>-6.6</td><td>5.4E-11</td><td>1.9E-10</td><td>BT20/MB468</td><td>MITO</td></tr>
<tr><td>>hg38_21_+_8433172_8446622_RNA45SN1_45S_50ntFlanks@10357.10378.22|CTTTGAAGGCCGAAGTGGAGAA</td><td>>hg38_21_+_8433172_8446622_RNA45SN1_45S_50ntFlanks</td><td>10357</td><td>10378</td><td>22</td><td>CTTTGAAGGCCGAAGTGGAGAA</td><td>328.6</td><td>-8.3</td><td>1.3</td><td>-6.5</td><td>8.8E-11</td><td>1.9E-10</td><td>BT20/MB468</td><td>NUC</td></tr>
<tr><td>>trna101_AlaAGC_6_-_28831462_28831533@54.75.22,trna102_AlaAGC_6_-_28806221_28806292@54.75.22,trna65_AlaAGC_6_+_28574933_28575004@54.75.22|TCAATCCCCGGCACCTCCACCA</td><td>>trna101_AlaAGC_6_-_28831462_28831533</td><td>54</td><td>75</td><td>22</td><td>TCAATCCCCGGCACCTCCACCA</td><td>407.5</td><td>9.9</td><td>1.5</td><td>6.5</td><td>6.5E-11</td><td>1.9E-10</td><td>BT20/MB468</td><td>CYTO</td></tr>
<tr><td>>hg38_1_-_228634819_228635039_RNA5S12_5S_50ntFlanks@148.169.22|GGAATACCGGGTGCTGTAGGCT</td><td>>hg38_1_-_228634819_228635039_RNA5S12_5S_50ntFlanks</td><td>148</td><td>169</td><td>22</td><td>GGAATACCGGGTGCTGTAGGCT</td><td>3319.4</td><td>3.8</td><td>0.6</td><td>6.5</td><td>6.5E-11</td><td>1.9E-10</td><td>BT20/MB468</td><td>CYTO</td></tr>
<tr><td>>hg38_21_+_8433172_8446622_RNA45SN1_45S_50ntFlanks@11682.11703.22|AATGTCAAAGTGAAGAAATTCA</td><td>>hg38_21_+_8433172_8446622_RNA45SN1_45S_50ntFlanks</td><td>11682</td><td>11703</td><td>22</td><td>AATGTCAAAGTGAAGAAATTCA</td><td>94.3</td><td>-10.4</td><td>1.6</td><td>-6.5</td><td>6.9E-11</td><td>1.9E-10</td><td>MB231/MB468</td><td>MP</td></tr>
<tr><td>>hg38_21_+_8433172_8446622_RNA45SN1_45S_50ntFlanks@12002.12043.42|TGCGGGCCGCCGGTGAAATACCACTACTCTGATCGTTTTTTC</td><td>>hg38_21_+_8433172_8446622_RNA45SN1_45S_50ntFlanks</td><td>12002</td><td>12043</td><td>42</td><td>TGCGGGCCGCCGGTGAAATACCACTACTCTGATCGTTTTTTC</td><td>528.9</td><td>-11.4</td><td>1.7</td><td>-6.5</td><td>6.6E-11</td><td>1.9E-10</td><td>BT20/MB468</td><td>CYTO</td></tr>
<tr><td>>hg38_21_+_8433172_8446622_RNA45SN1_45S_50ntFlanks@8338.8373.36|GTACCGTAAGGGAAAGTTGAAAAGAACTTTGAAGAG</td><td>>hg38_21_+_8433172_8446622_RNA45SN1_45S_50ntFlanks</td><td>8338</td><td>8373</td><td>36</td><td>GTACCGTAAGGGAAAGTTGAAAAGAACTTTGAAGAG</td><td>256.7</td><td>11.8</td><td>1.7</td><td>6.9</td><td>5.3E-12</td><td>1.9E-10</td><td>BT20/MB468</td><td>MP</td></tr>
<tr><td>>hg38_21_+_8433172_8446622_RNA45SN1_45S_50ntFlanks@9941.9961.21|AGGACGGTGGCCATGGAAGTC</td><td>>hg38_21_+_8433172_8446622_RNA45SN1_45S_50ntFlanks</td><td>9941</td><td>9961</td><td>21</td><td>AGGACGGTGGCCATGGAAGTC</td><td>491.5</td><td>12.7</td><td>1.8</td><td>6.9</td><td>5.3E-12</td><td>1.9E-10</td><td>BT20/MB468</td><td>MP</td></tr>
<tr><td>>MI0000101|hsa-mir-99a&WithFlank&21|+|16539083|16539175@18.39.22&[MIMAT0000097&hsa-miR-99a-5p&offsets|-1|-1;m-64&21|+|16539101|16539121&offsets|-1|0]|AAACCCGTAGATCCGATCTTGT</td><td>>MI0000101|hsa-mir-99a&WithFlank&21|+|16539083|16539175</td><td>18</td><td>39</td><td>22</td><td>AAACCCGTAGATCCGATCTTGT</td><td>222.8</td><td>-10.9</td><td>1.6</td><td>-6.9</td><td>5.3E-12</td><td>1.9E-10</td><td>BT20/MB468</td><td>MP</td></tr>
<tr><td>>hg38_21_+_8433172_8446622_RNA45SN1_45S_50ntFlanks@11886.11905.20|TGAGCTTGACTCTAGTCTGG</td><td>>hg38_21_+_8433172_8446622_RNA45SN1_45S_50ntFlanks</td><td>11886</td><td>11905</td><td>20</td><td>TGAGCTTGACTCTAGTCTGG</td><td>493.2</td><td>-8.0</td><td>1.2</td><td>-6.5</td><td>6.6E-11</td><td>1.9E-10</td><td>BT20/MB468</td><td>CYTO</td></tr>
<tr><td>>MI0003776|hsa-mir-320b-1&WithFlank&1|+|116671740|116671823@48.67.20(+1U);MI0003839|hsa-mir-320b-2&WithFlank&1|-|224257034|224257116@47.66.20(+1U);MI0000542|hsa-mir-320a&WithFlank&8|-|22244960|22245043@48.67.20(+1U)&[MIMAT0005792&hsa-miR-320b&offsets|0|-2(+1U);m-183&1|+|116671787|116671806&offsets|0|0(+1U)];[MIMAT0005792_1&hsa-miR-320b&offsets|0|-2(+1U);m-183&1|-|224257051|224257070&offsets|0|0(+1U)];[MIMAT0000510&hsa-miR-320a-3p&offsets|0|-2(+1U);m-60&8|-|22244975|22244996&offsets|0|-2(+1U)]|AAAAGCTGGGTTGAGAGGGCT</td><td>>MI0003776|hsa-mir-320b-1&WithFlank&1|+|116671740|116671823</td><td>48</td><td>67</td><td>20(+1U)</td><td>AAAAGCTGGGTTGAGAGGGCT</td><td>93.3</td><td>2.4</td><td>0.4</td><td>6.5</td><td>8.9E-11</td><td>1.9E-10</td><td>BT20/MB468</td><td>NUC</td></tr>
<tr><td>>hg38_21_+_8433172_8446622_RNA45SN1_45S_50ntFlanks@4003.4020.18|ATAACCTCGGGCCGATCG</td><td>>hg38_21_+_8433172_8446622_RNA45SN1_45S_50ntFlanks</td><td>4003</td><td>4020</td><td>18</td><td>ATAACCTCGGGCCGATCG</td><td>6126.1</td><td>-11.0</td><td>1.7</td><td>-6.5</td><td>8.9E-11</td><td>1.9E-10</td><td>BT20/MB468</td><td>NUC</td></tr>
<tr><td>>hg38_21_+_8433172_8446622_RNA45SN1_45S_50ntFlanks@4163.4181.19|CATCCAAGGAAGGCAGCAG</td><td>>hg38_21_+_8433172_8446622_RNA45SN1_45S_50ntFlanks</td><td>4163</td><td>4181</td><td>19</td><td>CATCCAAGGAAGGCAGCAG</td><td>139.9</td><td>-11.0</td><td>1.7</td><td>-6.5</td><td>7.0E-11</td><td>1.9E-10</td><td>MB231/MB468</td><td>MP</td></tr>
<tr><td>>hg38_21_+_8433172_8446622_RNA45SN1_45S_50ntFlanks@4380.4408.29|CGTAGTTGGATCTTGGGAGCGGGCGGGCG</td><td>>hg38_21_+_8433172_8446622_RNA45SN1_45S_50ntFlanks</td><td>4380</td><td>4408</td><td>29</td><td>CGTAGTTGGATCTTGGGAGCGGGCGGGCG</td><td>633.1</td><td>-7.1</td><td>1.1</td><td>-6.5</td><td>8.9E-11</td><td>1.9E-10</td><td>BT20/MB468</td><td>NUC</td></tr>
<tr><td>>MI0000270|hsa-mir-181b-1&WithFlank&1|-|198858867|198858988@42.63.22(+1C);MI0000683|hsa-mir-181b-2&WithFlank&9|+|124693704|124693804@22.43.22(+1C)&[MIMAT0000257&hsa-miR-181b-5p&offsets|0|-1(+1C);m-73&1|-|198858925|198858947&offsets|0|-1(+1C)];[MIMAT0000257_1&hsa-miR-181b-5p&offsets|0|-1(+1C);m-72&9|+|124693725|124693747&offsets|0|-1(+1C)]|AACATTCATTGCTGTCGGTGGGC</td><td>>MI0000270|hsa-mir-181b-1&WithFlank&1|-|198858867|198858988</td><td>42</td><td>63</td><td>22(+1C)</td><td>AACATTCATTGCTGTCGGTGGGC</td><td>359.8</td><td>5.4</td><td>0.8</td><td>6.5</td><td>7.4E-11</td><td>1.9E-10</td><td>MB231/MB468</td><td>NUC</td></tr>
<tr><td>>hg38_21_+_8433172_8446622_RNA45SN1_45S_50ntFlanks@9202.9221.20|TCTCGGGGCCACGCGCGCGT</td><td>>hg38_21_+_8433172_8446622_RNA45SN1_45S_50ntFlanks</td><td>9202</td><td>9221</td><td>20</td><td>TCTCGGGGCCACGCGCGCGT</td><td>91.4</td><td>-10.4</td><td>1.6</td><td>-6.5</td><td>7.0E-11</td><td>1.9E-10</td><td>MB231/MB468</td><td>MP</td></tr>
<tr><td>>hg38_1_-_228634819_228635039_RNA5S12_5S_50ntFlanks@142.171.30|CGCCTGGGAATACCGGGTGCTGTAGGCTTT</td><td>>hg38_1_-_228634819_228635039_RNA5S12_5S_50ntFlanks</td><td>142</td><td>171</td><td>30</td><td>CGCCTGGGAATACCGGGTGCTGTAGGCTTT</td><td>1649.9</td><td>4.7</td><td>0.7</td><td>6.5</td><td>6.6E-11</td><td>1.9E-10</td><td>BT20/MB468</td><td>CYTO</td></tr>
<tr><td>>MI0000462|hsa-mir-152&WithFlank&17|-|48037155|48037253@60.80.21(+1U)&[MIMAT0000438&hsa-miR-152-3p&offsets|0|0(+1U);m-97&17|-|48037174|48037194&offsets|0|0(+1U)]|TCAGTGCATGACAGAACTTGGT</td><td>>MI0000462|hsa-mir-152&WithFlank&17|-|48037155|48037253</td><td>60</td><td>80</td><td>21(+1U)</td><td>TCAGTGCATGACAGAACTTGGT</td><td>283.0</td><td>11.4</td><td>1.8</td><td>6.5</td><td>8.4E-11</td><td>1.9E-10</td><td>MB231/MB468</td><td>MITO</td></tr>
<tr><td>>hg38_21_+_8433172_8446622_RNA45SN1_45S_50ntFlanks@12893.12926.34|CCTCGCCCGTCACGCACCGCACGTTCGTGGGGAA</td><td>>hg38_21_+_8433172_8446622_RNA45SN1_45S_50ntFlanks</td><td>12893</td><td>12926</td><td>34</td><td>CCTCGCCCGTCACGCACCGCACGTTCGTGGGGAA</td><td>159.6</td><td>-9.4</td><td>1.4</td><td>-6.6</td><td>5.4E-11</td><td>1.9E-10</td><td>BT20/MB468</td><td>MITO</td></tr>
<tr><td>>hg38_21_+_8433172_8446622_RNA45SN1_45S_50ntFlanks@11701.11728.28|TCAATGAAGCGCGGGTAAACGGCGGGAG</td><td>>hg38_21_+_8433172_8446622_RNA45SN1_45S_50ntFlanks</td><td>11701</td><td>11728</td><td>28</td><td>TCAATGAAGCGCGGGTAAACGGCGGGAG</td><td>180.7</td><td>-7.8</td><td>1.2</td><td>-6.5</td><td>7.4E-11</td><td>1.9E-10</td><td>MB231/MB468</td><td>NUC</td></tr>
<tr><td>>MI0000810|hsa-mir-135b&WithFlank&1|-|205448296|205448404@22.44.23&[MIMAT0000758&hsa-miR-135b-5p&offsets|0|0;m-321&1|-|205448361|205448383&offsets|0|0]|TATGGCTTTTCATTCCTATGTGA</td><td>>MI0000810|hsa-mir-135b&WithFlank&1|-|205448296|205448404</td><td>22</td><td>44</td><td>23</td><td>TATGGCTTTTCATTCCTATGTGA</td><td>703.1</td><td>4.3</td><td>0.7</td><td>6.5</td><td>7.4E-11</td><td>1.9E-10</td><td>MB231/MB468</td><td>NUC</td></tr>
<tr><td>>hg38_21_+_8433172_8446622_RNA45SN1_45S_50ntFlanks@3984.4024.41|GCGGCTTTGGTGACTCTAGATAACCTCGGGCCGATCGCACG</td><td>>hg38_21_+_8433172_8446622_RNA45SN1_45S_50ntFlanks</td><td>3984</td><td>4024</td><td>41</td><td>GCGGCTTTGGTGACTCTAGATAACCTCGGGCCGATCGCACG</td><td>262.9</td><td>-8.3</td><td>1.3</td><td>-6.5</td><td>7.4E-11</td><td>2.0E-10</td><td>MB231/MB468</td><td>NUC</td></tr>
<tr><td>>hg38_21_+_8433172_8446622_RNA45SN1_45S_50ntFlanks@4159.4182.24|ACCACATCCAAGGAAGGCAGCAGG</td><td>>hg38_21_+_8433172_8446622_RNA45SN1_45S_50ntFlanks</td><td>4159</td><td>4182</td><td>24</td><td>ACCACATCCAAGGAAGGCAGCAGG</td><td>148.9</td><td>-11.1</td><td>1.7</td><td>-6.5</td><td>7.0E-11</td><td>2.0E-10</td><td>MB231/MB468</td><td>MP</td></tr>
<tr><td>>hg38_21_+_8433172_8446622_RNA45SN1_45S_50ntFlanks@3785.3805.21|TGAAACTGCGAATGGCTCATT</td><td>>hg38_21_+_8433172_8446622_RNA45SN1_45S_50ntFlanks</td><td>3785</td><td>3805</td><td>21</td><td>TGAAACTGCGAATGGCTCATT</td><td>139.2</td><td>-11.0</td><td>1.7</td><td>-6.5</td><td>7.0E-11</td><td>2.0E-10</td><td>MB231/MB468</td><td>MP</td></tr>
<tr><td>>hg38_21_+_8433172_8446622_RNA45SN1_45S_50ntFlanks@11874.11897.24|AGAAGACCCTGTTGAGCTTGACTC</td><td>>hg38_21_+_8433172_8446622_RNA45SN1_45S_50ntFlanks</td><td>11874</td><td>11897</td><td>24</td><td>AGAAGACCCTGTTGAGCTTGACTC</td><td>935.2</td><td>-5.0</td><td>0.8</td><td>-6.5</td><td>8.5E-11</td><td>2.0E-10</td><td>MB231/MB468</td><td>MITO</td></tr>
<tr><td>>trna10_GluCTC_3_-_103879524_103879595@16.37.22,trna116_GluCTC_1_-_145399233_145399304@16.37.22,trna25_GluCTC_2_-_71273488_71273560@16.37.22,trna59_GluCTC_1_+_249168447_249168518@16.37.22,trna71_GluCTC_1_-_161439189_161439260@16.37.22,trna74_GluCTC_1_-_161431809_161431880@16.37.22,trna77_GluCTC_1_-_161424398_161424469@16.37.22,trna77_GluCTC_6_+_28949976_28950047@16.37.22,trna80_GluCTC_1_-_161417018_161417089@16.37.22,trna87_GluCTC_6_-_126101393_126101464@16.37.22|TGGTTAGGATTCGGCGCTCTCA</td><td>>trna10_GluCTC_3_-_103879524_103879595</td><td>16</td><td>37</td><td>22</td><td>TGGTTAGGATTCGGCGCTCTCA</td><td>258.1</td><td>6.9</td><td>1.1</td><td>6.5</td><td>6.7E-11</td><td>2.0E-10</td><td>BT20/MB468</td><td>CYTO</td></tr>
<tr><td>>hg38_21_+_8433172_8446622_RNA45SN1_45S_50ntFlanks@12405.12424.20|CTTTTTGATCCTTCGATGTC</td><td>>hg38_21_+_8433172_8446622_RNA45SN1_45S_50ntFlanks</td><td>12405</td><td>12424</td><td>20</td><td>CTTTTTGATCCTTCGATGTC</td><td>618.6</td><td>-8.1</td><td>1.2</td><td>-6.5</td><td>7.4E-11</td><td>2.0E-10</td><td>MB231/MB468</td><td>NUC</td></tr>
<tr><td>>hg38_21_+_8433172_8446622_RNA45SN1_45S_50ntFlanks@5473.5507.35|CGGCCCTGGCGGAGCGCTGAGAAGACGGTCGAACT</td><td>>hg38_21_+_8433172_8446622_RNA45SN1_45S_50ntFlanks</td><td>5473</td><td>5507</td><td>35</td><td>CGGCCCTGGCGGAGCGCTGAGAAGACGGTCGAACT</td><td>232.5</td><td>5.6</td><td>0.9</td><td>6.5</td><td>7.5E-11</td><td>2.0E-10</td><td>MB231/MB468</td><td>CYTO</td></tr>
<tr><td>>hg38_21_+_8433172_8446622_RNA45SN1_45S_50ntFlanks@6786.6807.22|TACGCCTGTCTGAGCGTCGCTT</td><td>>hg38_21_+_8433172_8446622_RNA45SN1_45S_50ntFlanks</td><td>6786</td><td>6807</td><td>22</td><td>TACGCCTGTCTGAGCGTCGCTT</td><td>780.9</td><td>5.2</td><td>0.8</td><td>6.6</td><td>5.5E-11</td><td>2.0E-10</td><td>BT20/MB468</td><td>MITO</td></tr>
<tr><td>>MI0000065|hsa-let-7d&WithFlank&9|+|94178828|94178926@14.34.21&[MIMAT0000065&hsa-let-7d-5p&offsets|0|-1;m-90&9|+|94178841|94178862&offsets|0|-1]|AGAGGTAGTAGGTTGCATAGT</td><td>>MI0000065|hsa-let-7d&WithFlank&9|+|94178828|94178926</td><td>14</td><td>34</td><td>21</td><td>AGAGGTAGTAGGTTGCATAGT</td><td>977.2</td><td>6.0</td><td>0.9</td><td>6.5</td><td>8.5E-11</td><td>2.0E-10</td><td>MB231/MB468</td><td>MITO</td></tr>
<tr><td>>hg38_21_+_8433172_8446622_RNA45SN1_45S_50ntFlanks@8229.8251.23|CGCGCCGGGCCCGGGTCTTCCCG</td><td>>hg38_21_+_8433172_8446622_RNA45SN1_45S_50ntFlanks</td><td>8229</td><td>8251</td><td>23</td><td>CGCGCCGGGCCCGGGTCTTCCCG</td><td>909.3</td><td>9.6</td><td>1.5</td><td>6.5</td><td>7.5E-11</td><td>2.0E-10</td><td>MB231/MB468</td><td>CYTO</td></tr>
<tr><td>>MI0000253|hsa-mir-148a&WithFlank&7|-|25949913|25949992@50.73.24&[MIMAT0000243&hsa-miR-148a-3p&offsets|0|+2;m-7&7|-|25949922|25949943&offsets|0|+2]|TCAGTGCACTACAGAACTTTGTCT</td><td>>MI0000253|hsa-mir-148a&WithFlank&7|-|25949913|25949992</td><td>50</td><td>73</td><td>24</td><td>TCAGTGCACTACAGAACTTTGTCT</td><td>121.4</td><td>3.4</td><td>0.5</td><td>6.5</td><td>9.0E-11</td><td>2.0E-10</td><td>BT20/MB468</td><td>NUC</td></tr>
<tr><td>>MI0016887|hsa-mir-4521&WithFlank&17|+|8186939|8187010@10.33.24(+1C)&[MIMAT0019058&hsa-miR-4521&offsets|0|+2(+1C);m-765&17|+|8186948|8186969&offsets|0|+2(+1C)]|GCTAAGGAAGTCCTGTGCTCAGTTC</td><td>>MI0016887|hsa-mir-4521&WithFlank&17|+|8186939|8187010</td><td>10</td><td>33</td><td>24(+1C)</td><td>GCTAAGGAAGTCCTGTGCTCAGTTC</td><td>438.3</td><td>9.2</td><td>1.4</td><td>6.5</td><td>6.7E-11</td><td>2.0E-10</td><td>BT20/MB468</td><td>CYTO</td></tr>
<tr><td>>hg38_21_+_8433172_8446622_RNA45SN1_45S_50ntFlanks@5473.5505.33|CGGCCCTGGCGGAGCGCTGAGAAGACGGTCGAA</td><td>>hg38_21_+_8433172_8446622_RNA45SN1_45S_50ntFlanks</td><td>5473</td><td>5505</td><td>33</td><td>CGGCCCTGGCGGAGCGCTGAGAAGACGGTCGAA</td><td>217.5</td><td>5.4</td><td>0.8</td><td>6.5</td><td>7.5E-11</td><td>2.0E-10</td><td>MB231/MB468</td><td>CYTO</td></tr>
<tr><td>>hg38_21_+_8433172_8446622_RNA45SN1_45S_50ntFlanks@7975.8016.42|CGCGACCTCAGATCAGACGTGGCGACCCGCTGAATTTAAGCA</td><td>>hg38_21_+_8433172_8446622_RNA45SN1_45S_50ntFlanks</td><td>7975</td><td>8016</td><td>42</td><td>CGCGACCTCAGATCAGACGTGGCGACCCGCTGAATTTAAGCA</td><td>6370.2</td><td>6.2</td><td>0.9</td><td>6.5</td><td>6.7E-11</td><td>2.0E-10</td><td>BT20/MB468</td><td>CYTO</td></tr>
<tr><td>>hg38_21_+_8433172_8446622_RNA45SN1_45S_50ntFlanks@3846.3874.29|CTTGGATAACTGTGGTAATTCTAGAGCTA</td><td>>hg38_21_+_8433172_8446622_RNA45SN1_45S_50ntFlanks</td><td>3846</td><td>3874</td><td>29</td><td>CTTGGATAACTGTGGTAATTCTAGAGCTA</td><td>353.5</td><td>-8.4</td><td>1.3</td><td>-6.5</td><td>8.5E-11</td><td>2.0E-10</td><td>MB231/MB468</td><td>MITO</td></tr>
<tr><td>>hg38_21_+_8433172_8446622_RNA45SN1_45S_50ntFlanks@10771.10788.18|CAGGTCTCCAAGGTGAAC</td><td>>hg38_21_+_8433172_8446622_RNA45SN1_45S_50ntFlanks</td><td>10771</td><td>10788</td><td>18</td><td>CAGGTCTCCAAGGTGAAC</td><td>83.8</td><td>-10.3</td><td>1.6</td><td>-6.5</td><td>7.1E-11</td><td>2.0E-10</td><td>MB231/MB468</td><td>MP</td></tr>
<tr><td>>MI0000809|hsa-mir-151a&WithFlank&8|-|140732558|140732659@53.75.23&[MIMAT0000757&hsa-miR-151a-3p&offsets|0|+2;m-30&8|-|140732587|140732607&offsets|0|+2]|CTAGACTGAAGCTCCTTGAGGAC</td><td>>MI0000809|hsa-mir-151a&WithFlank&8|-|140732558|140732659</td><td>53</td><td>75</td><td>23</td><td>CTAGACTGAAGCTCCTTGAGGAC</td><td>200.9</td><td>10.9</td><td>1.7</td><td>6.5</td><td>8.6E-11</td><td>2.0E-10</td><td>MB231/MB468</td><td>MITO</td></tr>
<tr><td>>hg38_21_+_8433172_8446622_RNA45SN1_45S_50ntFlanks@6664.6681.18|TGGATCACTCGGCTCGTG</td><td>>hg38_21_+_8433172_8446622_RNA45SN1_45S_50ntFlanks</td><td>6664</td><td>6681</td><td>18</td><td>TGGATCACTCGGCTCGTG</td><td>766.3</td><td>-4.0</td><td>0.6</td><td>-6.5</td><td>6.8E-11</td><td>2.0E-10</td><td>BT20/MB468</td><td>CYTO</td></tr>
</table>

<table>
<tr><th></th><th></th><th></th><th></th><th></th><th></th><th></th><th></th><th></th><th></th><th></th><th></th><th></th><th></th></tr>
<tr><td>>hg38_21_+_8433172_8446622_RNA45SN1_45S_50ntFlanks@10664.10711.48|CTTGGAAAGCGTCGCGGTTCCGGCGGCGTCCGGTGAGCTCTCGCTGGC</td><td>>hg38_21_+_8433172_8446622_RNA45SN1_45S_50ntFlanks</td><td>10664</td><td>10711</td><td>48</td><td>CTTGGAAAGCGTCGCGGTTCCGGCGGCGTCCGGTGAGCTCTCGCTGGC</td><td>776.1</td><td>-12.6</td><td>1.9</td><td>-6.5</td><td>7.5E-11</td><td>2.0E-10</td><td>MB231/MB468</td><td>NUC</td></tr>
<tr><td>>hg38_21_+_8433172_8446622_RNA45SN1_45S_50ntFlanks@12417.12443.27|TCGATGTCGGCTCTTCCTATCATTGTG</td><td>>hg38_21_+_8433172_8446622_RNA45SN1_45S_50ntFlanks</td><td>12417</td><td>12443</td><td>27</td><td>TCGATGTCGGCTCTTCCTATCATTGTG</td><td>7318.2</td><td>-5.5</td><td>0.8</td><td>-6.5</td><td>8.6E-11</td><td>2.0E-10</td><td>MB231/MB468</td><td>MITO</td></tr>
<tr><td>>hg38_21_+_8433172_8446622_RNA45SN1_45S_50ntFlanks@11593.11611.19|CCAGGGGAATCCGACTGTT</td><td>>hg38_21_+_8433172_8446622_RNA45SN1_45S_50ntFlanks</td><td>11593</td><td>11611</td><td>19</td><td>CCAGGGGAATCCGACTGTT</td><td>1934.5</td><td>-3.3</td><td>0.5</td><td>-6.6</td><td>5.6E-11</td><td>2.0E-10</td><td>BT20/MB468</td><td>MITO</td></tr>
<tr><td>>MI0000433|hsa-let-7g&WithFlank&3|-|52268272|52268367@11.30.20(+1C)&[MIMAT0000414&hsa-let-7g-5p&offsets|0|-2(+1C);m-58&3|-|52268336|52268357&offsets|0|-2(+1C)]|TGAGGTAGTAGTTTGTACAGC</td><td>>MI0000433|hsa-let-7g&WithFlank&3|-|52268272|52268367</td><td>11</td><td>30</td><td>20(+1C)</td><td>TGAGGTAGTAGTTTGTACAGC</td><td>215.2</td><td>2.8</td><td>0.4</td><td>6.5</td><td>9.1E-11</td><td>2.0E-10</td><td>BT20/MB468</td><td>NUC</td></tr>
<tr><td>>hg38_21_+_8433172_8446622_RNA45SN1_45S_50ntFlanks@3815.3834.20|ATGGTTCCTTTGGTCGCTCG</td><td>>hg38_21_+_8433172_8446622_RNA45SN1_45S_50ntFlanks</td><td>3815</td><td>3834</td><td>20</td><td>ATGGTTCCTTTGGTCGCTCG</td><td>89.5</td><td>-10.3</td><td>1.6</td><td>-6.5</td><td>7.2E-11</td><td>2.0E-10</td><td>MB231/MB468</td><td>MP</td></tr>
<tr><td>>hg38_21_+_8433172_8446622_RNA45SN1_45S_50ntFlanks@475.497.23|CTTCGTGATCGATGTGGTGACGT</td><td>>hg38_21_+_8433172_8446622_RNA45SN1_45S_50ntFlanks</td><td>475</td><td>497</td><td>23</td><td>CTTCGTGATCGATGTGGTGACGT</td><td>11670.1</td><td>7.4</td><td>1.1</td><td>6.5</td><td>8.7E-11</td><td>2.0E-10</td><td>MB231/MB468</td><td>MITO</td></tr>
<tr><td>>MI0000071|hsa-mir-17&WithFlank&13|+|91350599|91350694@20.41.22(+1A)&[MIMAT0000070&hsa-miR-17-5p&offsets|0|-1(+1A);m-88&13|+|91350618|91350640&offsets|0|-1(+1A)]|CAAAGTGCTTACAGTGCAGGTAA</td><td>>MI0000071|hsa-mir-17&WithFlank&13|+|91350599|91350694</td><td>20</td><td>41</td><td>22(+1A)</td><td>CAAAGTGCTTACAGTGCAGGTAA</td><td>574.8</td><td>4.9</td><td>0.7</td><td>6.5</td><td>7.6E-11</td><td>2.0E-10</td><td>MB231/MB468</td><td>NUC</td></tr>
<tr><td>>MI0000289|hsa-mir-181a-1&WithFlank&1|-|198859038|198859159@30.51.22(+1C);MI0000269|hsa-mir-181a-2&WithFlank&9|+|124692436|124692557@45.66.22(+1C)&[MIMAT0000256&hsa-miR-181a-5p&offsets|0|-1(+1C);m-32&1|-|198859108|198859130&offsets|0|-1(+1C)];[MIMAT0000256_1&hsa-miR-181a-5p&offsets|0|-1(+1C);m-32&9|+|124692480|124692502&offsets|0|-1(+1C)]|AACATTCAACGCTGTCGGTGAGC</td><td>>MI0000289|hsa-mir-181a-1&WithFlank&1|-|198859038|198859159</td><td>30</td><td>51</td><td>22(+1C)</td><td>AACATTCAACGCTGTCGGTGAGC</td><td>577.1</td><td>6.2</td><td>1.0</td><td>6.5</td><td>7.6E-11</td><td>2.0E-10</td><td>MB231/MB468</td><td>NUC</td></tr>
<tr><td>>hg38_21_+_8433172_8446622_RNA45SN1_45S_50ntFlanks@10387.10424.38|TGTGAACAGCAGTTGAACATGGGTCAGTCGGTCCTGAG</td><td>>hg38_21_+_8433172_8446622_RNA45SN1_45S_50ntFlanks</td><td>10387</td><td>10424</td><td>38</td><td>TGTGAACAGCAGTTGAACATGGGTCAGTCGGTCCTGAG</td><td>1633.1</td><td>4.9</td><td>0.8</td><td>6.5</td><td>7.6E-11</td><td>2.0E-10</td><td>MB231/MB468</td><td>NUC</td></tr>
<tr><td>>hg38_21_+_8433172_8446622_RNA45SN1_45S_50ntFlanks@10663.10710.48|CCTTGGAAAGCGTCGCGGTTCCGGCGGCGTCCGGTGAGCTCTCGCTGG</td><td>>hg38_21_+_8433172_8446622_RNA45SN1_45S_50ntFlanks</td><td>10663</td><td>10710</td><td>48</td><td>CCTTGGAAAGCGTCGCGGTTCCGGCGGCGTCCGGTGAGCTCTCGCTGG</td><td>399.8</td><td>-10.7</td><td>1.6</td><td>-6.6</td><td>5.6E-11</td><td>2.0E-10</td><td>BT20/MB468</td><td>MITO</td></tr>
<tr><td>>hg38_21_+_8433172_8446622_RNA45SN1_45S_50ntFlanks@12409.12448.40|TTGATCCTTCGATGTCGGCTCTTCCTATCATTGTGAAGCA</td><td>>hg38_21_+_8433172_8446622_RNA45SN1_45S_50ntFlanks</td><td>12409</td><td>12448</td><td>40</td><td>TTGATCCTTCGATGTCGGCTCTTCCTATCATTGTGAAGCA</td><td>228.1</td><td>5.2</td><td>0.8</td><td>6.5</td><td>7.7E-11</td><td>2.0E-10</td><td>MB231/MB468</td><td>NUC</td></tr>
<tr><td>>hg38_21_+_8433172_8446622_RNA45SN1_45S_50ntFlanks@4734.4766.33|AACGAAAGTCGGAGGTTCGAAGACGATCAGATA</td><td>>hg38_21_+_8433172_8446622_RNA45SN1_45S_50ntFlanks</td><td>4734</td><td>4766</td><td>33</td><td>AACGAAAGTCGGAGGTTCGAAGACGATCAGATA</td><td>221.3</td><td>5.3</td><td>0.8</td><td>6.5</td><td>7.7E-11</td><td>2.0E-10</td><td>MB231/MB468</td><td>CYTO</td></tr>
<tr><td>>MI0001446|hsa-mir-424&WithFlank&X|-|134546608|134546717@54.74.21&[MIMAT0004749&hsa-miR-424-3p&offsets|0|0;m-305&X|-|134546644|134546664&offsets|0|0]|CAAAACGTGAGGCGCTGCTAT</td><td>>MI0001446|hsa-mir-424&WithFlank&X|-|134546608|134546717</td><td>54</td><td>74</td><td>21</td><td>CAAAACGTGAGGCGCTGCTAT</td><td>432.1</td><td>7.3</td><td>1.1</td><td>6.5</td><td>7.7E-11</td><td>2.0E-10</td><td>MB231/MB468</td><td>NUC</td></tr>
<tr><td>>MI0000471|hsa-mir-126&WithFlank&9|+|136670596|136670692@58.78.21(+1A)&[MIMAT0000445&hsa-miR-126-3p&offsets|0|-1(+1A);m-40&9|+|136670653|136670674&offsets|0|-1(+1A)]|TCGTACCGTGAGTAATAATGCA</td><td>>MI0000471|hsa-mir-126&WithFlank&9|+|136670596|136670692</td><td>58</td><td>78</td><td>21(+1A)</td><td>TCGTACCGTGAGTAATAATGCA</td><td>252.6</td><td>8.2</td><td>1.3</td><td>6.5</td><td>9.3E-11</td><td>2.0E-10</td><td>BT20/MB468</td><td>NUC</td></tr>
<tr><td>>hg38_21_+_8433172_8446622_RNA45SN1_45S_50ntFlanks@10731.10757.27|AGGGTGTAAATCTCGCGCCGGGCCGTA</td><td>>hg38_21_+_8433172_8446622_RNA45SN1_45S_50ntFlanks</td><td>10731</td><td>10757</td><td>27</td><td>AGGGTGTAAATCTCGCGCCGGGCCGTA</td><td>622.1</td><td>-9.1</td><td>1.4</td><td>-6.5</td><td>9.3E-11</td><td>2.0E-10</td><td>BT20/MB468</td><td>NUC</td></tr>
<tr><td>>MI0000750|hsa-mir-26a-2&WithFlank&12|-|57824603|57824698@20.41.22(+1C);MI0000083|hsa-mir-26a-1&WithFlank&3|+|37969398|37969486@16.37.22(+1C)&[MIMAT0000082&hsa-miR-26a-5p&offsets|0|0(+1C);m-33&12|-|57824658|57824679&offsets|0|0(+1C)];[MIMAT0000082_1&hsa-miR-26a-5p&offsets|0|0(+1C);m-34&3|+|37969413|37969434&offsets|0|0(+1C)]|TTCAAGTAATCCAGGATAGGCTC</td><td>>MI0000750|hsa-mir-26a-2&WithFlank&12|-|57824603|57824698</td><td>20</td><td>41</td><td>22(+1C)</td><td>TTCAAGTAATCCAGGATAGGCTC</td><td>117.8</td><td>3.0</td><td>0.5</td><td>6.5</td><td>9.3E-11</td><td>2.0E-10</td><td>BT20/MB468</td><td>NUC</td></tr>
<tr><td>>hg38_21_+_8433172_8446622_RNA45SN1_45S_50ntFlanks@12510.12530.21|TCGTGAGACAGGTTAGTTTTA</td><td>>hg38_21_+_8433172_8446622_RNA45SN1_45S_50ntFlanks</td><td>12510</td><td>12530</td><td>21</td><td>TCGTGAGACAGGTTAGTTTTA</td><td>82.5</td><td>-10.2</td><td>1.6</td><td>-6.5</td><td>7.3E-11</td><td>2.0E-10</td><td>MB231/MB468</td><td>MP</td></tr>
<tr><td>>hg38_21_+_8433172_8446622_RNA45SN1_45S_50ntFlanks@4557.4577.21|CAGCTAGGAATAATGGAATAG</td><td>>hg38_21_+_8433172_8446622_RNA45SN1_45S_50ntFlanks</td><td>4557</td><td>4577</td><td>21</td><td>CAGCTAGGAATAATGGAATAG</td><td>75.1</td><td>-10.1</td><td>1.5</td><td>-6.5</td><td>7.3E-11</td><td>2.0E-10</td><td>MB231/MB468</td><td>MP</td></tr>
<tr><td>>hg38_1_-_228634819_228635039_RNA5S12_5S_50ntFlanks@89.123.35|CTGATCTCGGAAGCTAAGCAGGGTCGGGCCTGGTT</td><td>>hg38_1_-_228634819_228635039_RNA5S12_5S_50ntFlanks</td><td>89</td><td>123</td><td>35</td><td>CTGATCTCGGAAGCTAAGCAGGGTCGGGCCTGGTT</td><td>635.1</td><td>7.1</td><td>1.1</td><td>6.5</td><td>7.8E-11</td><td>2.0E-10</td><td>MB231/MB468</td><td>CYTO</td></tr>
<tr><td>>trna13_ValCAC_19_-_4724647_4724719@1.34.34|GTTTCCGTAGTGTAGCGGTTATCACATTCGCCTC</td><td>>trna13_ValCAC_19_-_4724647_4724719</td><td>1</td><td>34</td><td>34</td><td>GTTTCCGTAGTGTAGCGGTTATCACATTCGCCTC</td><td>224.5</td><td>11.0</td><td>1.7</td><td>6.5</td><td>8.8E-11</td><td>2.0E-10</td><td>MB231/MB468</td><td>MITO</td></tr>
<tr><td>>trnaMT_MetCAT_MT_+_4402_4469@54.71.18|ACCCTTCCCGTACTACCA</td><td>>trnaMT_MetCAT_MT_+_4402_4469</td><td>54</td><td>71</td><td>18</td><td>ACCCTTCCCGTACTACCA</td><td>828.0</td><td>8.6</td><td>1.3</td><td>6.5</td><td>6.9E-11</td><td>2.0E-10</td><td>BT20/MB468</td><td>CYTO</td></tr>
<tr><td>>hg38_21_+_8433172_8446622_RNA45SN1_45S_50ntFlanks@9507.9533.27|AACTATGCCTGGGCAGGGCGAAGCCAG</td><td>>hg38_21_+_8433172_8446622_RNA45SN1_45S_50ntFlanks</td><td>9507</td><td>9533</td><td>27</td><td>AACTATGCCTGGGCAGGGCGAAGCCAG</td><td>153.9</td><td>-11.1</td><td>1.7</td><td>-6.5</td><td>7.3E-11</td><td>2.0E-10</td><td>MB231/MB468</td><td>MP</td></tr>
<tr><td>>hg38_21_+_8433172_8446622_RNA45SN1_45S_50ntFlanks@11729.11749.21|TAACTATGACTCTCTTAAGGT</td><td>>hg38_21_+_8433172_8446622_RNA45SN1_45S_50ntFlanks</td><td>11729</td><td>11749</td><td>21</td><td>TAACTATGACTCTCTTAAGGT</td><td>1543.3</td><td>-7.6</td><td>1.2</td><td>-6.5</td><td>7.3E-11</td><td>2.0E-10</td><td>MB231/MB468</td><td>MP</td></tr>
<tr><td>>hg38_1_-_228634819_228635039_RNA5S12_5S_50ntFlanks@72.92.21|AACGCGCCCGATCTCGTCTGA</td><td>>hg38_1_-_228634819_228635039_RNA5S12_5S_50ntFlanks</td><td>72</td><td>92</td><td>21</td><td>AACGCGCCCGATCTCGTCTGA</td><td>586.1</td><td>-7.1</td><td>1.1</td><td>-6.5</td><td>9.4E-11</td><td>2.0E-10</td><td>BT20/MB468</td><td>NUC</td></tr>
<tr><td>>hg38_21_+_8433172_8446622_RNA45SN1_45S_50ntFlanks@10856.10875.20|AGGATTGGCTCTAAGGGCTG</td><td>>hg38_21_+_8433172_8446622_RNA45SN1_45S_50ntFlanks</td><td>10856</td><td>10875</td><td>20</td><td>AGGATTGGCTCTAAGGGCTG</td><td>77.8</td><td>-10.1</td><td>1.6</td><td>-6.5</td><td>7.3E-11</td><td>2.0E-10</td><td>MB231/MB468</td><td>MP</td></tr>
<tr><td>>MI0003763|hsa-mir-767&WithFlank&X|-|152393415|152393535@33.55.23&[MIMAT0003882&hsa-miR-767-5p&offsets|0|0;m-788&X|-|152393481|152393503&offsets|0|0]|TGCACCATGGTTGTCTGAGCATG</td><td>>MI0003763|hsa-mir-767&WithFlank&X|-|152393415|152393535</td><td>33</td><td>55</td><td>23</td><td>TGCACCATGGTTGTCTGAGCATG</td><td>81.2</td><td>-10.2</td><td>1.6</td><td>-6.5</td><td>7.4E-11</td><td>2.0E-10</td><td>MB231/MB468</td><td>MP</td></tr>
<tr><td>>hg38_21_+_8433172_8446622_RNA45SN1_45S_50ntFlanks@12838.12861.24|GTGCGGAGTGCCCTTCGTCCTGGG</td><td>>hg38_21_+_8433172_8446622_RNA45SN1_45S_50ntFlanks</td><td>12838</td><td>12861</td><td>24</td><td>GTGCGGAGTGCCCTTCGTCCTGGG</td><td>185.2</td><td>-11.4</td><td>1.7</td><td>-6.5</td><td>7.4E-11</td><td>2.0E-10</td><td>MB231/MB468</td><td>MP</td></tr>
<tr><td>>hg38_21_+_8433172_8446622_RNA45SN1_45S_50ntFlanks@13002.13023.22|CGATCTATTGAAAGTCAGCCCT</td><td>>hg38_21_+_8433172_8446622_RNA45SN1_45S_50ntFlanks</td><td>13002</td><td>13023</td><td>22</td><td>CGATCTATTGAAAGTCAGCCCT</td><td>231.2</td><td>-9.9</td><td>1.5</td><td>-6.6</td><td>5.7E-11</td><td>2.1E-10</td><td>BT20/MB468</td><td>MITO</td></tr>
<tr><td>>hg38_MT_+_1621_3279_RNR2_16S_50ntFlanks@713.740.28|GCATAAGCCTGCGTCAGATTAAAACACT</td><td>>hg38_MT_+_1621_3279_RNR2_16S_50ntFlanks</td><td>713</td><td>740</td><td>28</td><td>GCATAAGCCTGCGTCAGATTAAAACACT</td><td>118.2</td><td>10.1</td><td>1.6</td><td>6.5</td><td>7.4E-11</td><td>2.1E-10</td><td>MB231/MB468</td><td>MP</td></tr>
<tr><td>>hg38_21_+_8433172_8446622_RNA45SN1_45S_50ntFlanks@11641.11677.37|CGGCGGGTGTTGACGCGATGTGATTTCTGCCCAGTGC</td><td>>hg38_21_+_8433172_8446622_RNA45SN1_45S_50ntFlanks</td><td>11641</td><td>11677</td><td>37</td><td>CGGCGGGTGTTGACGCGATGTGATTTCTGCCCAGTGC</td><td>297.6</td><td>-11.3</td><td>1.6</td><td>-6.9</td><td>5.7E-12</td><td>2.1E-10</td><td>BT20/MB468</td><td>MP</td></tr>
<tr><td>>hg38_21_+_8433172_8446622_RNA45SN1_45S_50ntFlanks@10234.10262.29|CGGACGCTACGCCGCGACGAGTAGGAGGG</td><td>>hg38_21_+_8433172_8446622_RNA45SN1_45S_50ntFlanks</td><td>10234</td><td>10262</td><td>29</td><td>CGGACGCTACGCCGCGACGAGTAGGAGGG</td><td>262.6</td><td>-8.1</td><td>1.2</td><td>-6.5</td><td>8.9E-11</td><td>2.1E-10</td><td>MB231/MB468</td><td>MITO</td></tr>
<tr><td>>am_trnaMT_ValTAC_MT_+_1602_1670@28.66.39|AACTTACACTTAGGAGATTTCAACTTAACTTGACCGCTC</td><td>>am_trnaMT_ValTAC_MT_+_1602_1670</td><td>28</td><td>66</td><td>39</td><td>AACTTACACTTAGGAGATTTCAACTTAACTTGACCGCTC</td><td>89.7</td><td>9.7</td><td>1.5</td><td>6.5</td><td>7.4E-11</td><td>2.1E-10</td><td>MB231/MB468</td><td>MP</td></tr>
<tr><td>>MI0006444|hsa-mir-1307&WithFlank&10|-|103394247|103394407@86.108.23(+1A)&[MIMAT0005951&hsa-miR-1307-3p&offsets|0|+1(+1A);m-66&10|-|103394300|103394322&offsets|0|0(+1A)]|ACTCGGCGTGGCGTCGGTCGTGGA</td><td>>MI0006444|hsa-mir-1307&WithFlank&10|-|103394247|103394407</td><td>86</td><td>108</td><td>23(+1A)</td><td>ACTCGGCGTGGCGTCGGTCGTGGA</td><td>727.4</td><td>8.7</td><td>1.3</td><td>6.5</td><td>7.0E-11</td><td>2.1E-10</td><td>BT20/MB468</td><td>CYTO</td></tr>
<tr><td>>hg38_21_+_8433172_8446622_RNA45SN1_45S_50ntFlanks@10408.10426.19|GGTCAGTCGGTCCTGAGAG</td><td>>hg38_21_+_8433172_8446622_RNA45SN1_45S_50ntFlanks</td><td>10408</td><td>10426</td><td>19</td><td>GGTCAGTCGGTCCTGAGAG</td><td>180.3</td><td>-9.6</td><td>1.5</td><td>-6.5</td><td>5.8E-11</td><td>2.1E-10</td><td>BT20/MB468</td><td>MITO</td></tr>
<tr><td>>hg38_21_+_8433172_8446622_RNA45SN1_45S_50ntFlanks@11491.11516.26|CCGCGGGCCGGTCCCCCCCGCCGGGT</td><td>>hg38_21_+_8433172_8446622_RNA45SN1_45S_50ntFlanks</td><td>11491</td><td>11516</td><td>26</td><td>CCGCGGGCCGGTCCCCCCCGCCGGGT</td><td>143.0</td><td>-11.0</td><td>1.7</td><td>-6.5</td><td>7.4E-11</td><td>2.1E-10</td><td>MB231/MB468</td><td>MP</td></tr>
<tr><td>>am_trna2_LeuTAA_X_-_55207755_55207829@1.18.18,trna4_LeuTAA_20_-_55933428_55933502@1.18.18,trna60_LeuTAA_1_-_238106956_238107030@1.18.18,trna9_LeuTAA_2_+_117781417_117781491@1.18.18,trnaMT_LeuTAA_MT_+_3230_3304@1.18.18|GTTAAGATGGCAGAGCCC</td><td>>am_trna2_LeuTAA_X_-_55207755_55207829</td><td>1</td><td>18</td><td>18</td><td>GTTAAGATGGCAGAGCCC</td><td>501.8</td><td>10.0</td><td>1.5</td><td>6.5</td><td>7.1E-11</td><td>2.1E-10</td><td>BT20/MB468</td><td>CYTO</td></tr>
<tr><td>>hg38_21_+_8433172_8446622_RNA45SN1_45S_50ntFlanks@7989.8011.23|AGACGTGGCGACCCGCTGAATTT</td><td>>hg38_21_+_8433172_8446622_RNA45SN1_45S_50ntFlanks</td><td>7989</td><td>8011</td><td>23</td><td>AGACGTGGCGACCCGCTGAATTT</td><td>1621.3</td><td>5.2</td><td>0.8</td><td>6.5</td><td>7.1E-11</td><td>2.1E-10</td><td>BT20/MB468</td><td>CYTO</td></tr>
<tr><td>>hg38_21_+_8433172_8446622_RNA45SN1_45S_50ntFlanks@10516.10533.18|GAGTGGCGGAGATGGGCG</td><td>>hg38_21_+_8433172_8446622_RNA45SN1_45S_50ntFlanks</td><td>10516</td><td>10533</td><td>18</td><td>GAGTGGCGGAGATGGGCG</td><td>1017.1</td><td>-9.8</td><td>1.5</td><td>-6.5</td><td>9.5E-11</td><td>2.1E-10</td><td>BT20/MB468</td><td>NUC</td></tr>
<tr><td>>hg38_21_+_8433172_8446622_RNA45SN1_45S_50ntFlanks@11783.11813.31|CATGAATGGATGAACGAGATTCCCACTGTCC</td><td>>hg38_21_+_8433172_8446622_RNA45SN1_45S_50ntFlanks</td><td>11783</td><td>11813</td><td>31</td><td>CATGAATGGATGAACGAGATTCCCACTGTCC</td><td>108.7</td><td>-10.6</td><td>1.6</td><td>-6.5</td><td>7.5E-11</td><td>2.1E-10</td><td>MB231/MB468</td><td>MP</td></tr>
<tr><td>>hg38_21_+_8433172_8446622_RNA45SN1_45S_50ntFlanks@5421.5442.22|CGATTGGATGGTTTAGTGAGGC</td><td>>hg38_21_+_8433172_8446622_RNA45SN1_45S_50ntFlanks</td><td>5421</td><td>5442</td><td>22</td><td>CGATTGGATGGTTTAGTGAGGC</td><td>113.8</td><td>-10.7</td><td>1.6</td><td>-6.5</td><td>7.5E-11</td><td>2.1E-10</td><td>MB231/MB468</td><td>MP</td></tr>
<tr><td>>hg38_21_+_8433172_8446622_RNA45SN1_45S_50ntFlanks@4722.4757.36|TCATTAATCAAGAACGAAAGTCGGAGGTTCGAAGAC</td><td>>hg38_21_+_8433172_8446622_RNA45SN1_45S_50ntFlanks</td><td>4722</td><td>4757</td><td>36</td><td>TCATTAATCAAGAACGAAAGTCGGAGGTTCGAAGAC</td><td>152.8</td><td>-11.1</td><td>1.7</td><td>-6.5</td><td>7.5E-11</td><td>2.1E-10</td><td>MB231/MB468</td><td>MP</td></tr>
<tr><td>>hg38_21_+_8433172_8446622_RNA45SN1_45S_50ntFlanks@12223.12249.27|CAGAAACCTCCCGTGGAGCAGAAGGGC</td><td>>hg38_21_+_8433172_8446622_RNA45SN1_45S_50ntFlanks</td><td>12223</td><td>12249</td><td>27</td><td>CAGAAACCTCCCGTGGAGCAGAAGGGC</td><td>77.3</td><td>-10.1</td><td>1.6</td><td>-6.5</td><td>7.5E-11</td><td>2.1E-10</td><td>MB231/MB468</td><td>MP</td></tr>
<tr><td>>hg38_21_+_8433172_8446622_RNA45SN1_45S_50ntFlanks@7975.8001.27|CGCGACCTCAGATCAGACGTGGCGACC</td><td>>hg38_21_+_8433172_8446622_RNA45SN1_45S_50ntFlanks</td><td>7975</td><td>8001</td><td>27</td><td>CGCGACCTCAGATCAGACGTGGCGACC</td><td>345.9</td><td>4.4</td><td>0.7</td><td>6.5</td><td>7.9E-11</td><td>2.1E-10</td><td>MB231/MB468</td><td>NUC</td></tr>
<tr><td>>hg38_21_+_8433172_8446622_RNA45SN1_45S_50ntFlanks@9534.9565.32|AGGAAACTCTGGTGGAGGTCCGTAGCGGTCCT</td><td>>hg38_21_+_8433172_8446622_RNA45SN1_45S_50ntFlanks</td><td>9534</td><td>9565</td><td>32</td><td>AGGAAACTCTGGTGGAGGTCCGTAGCGGTCCT</td><td>1336.7</td><td>-3.1</td><td>0.5</td><td>-6.5</td><td>7.9E-11</td><td>2.1E-10</td><td>MB231/MB468</td><td>CYTO</td></tr>
<tr><td>>hg38_21_+_8433172_8446622_RNA45SN1_45S_50ntFlanks@12221.12248.28|GACAGAAACCTCCCGTGGAGCAGAAGGG</td><td>>hg38_21_+_8433172_8446622_RNA45SN1_45S_50ntFlanks</td><td>12221</td><td>12248</td><td>28</td><td>GACAGAAACCTCCCGTGGAGCAGAAGGG</td><td>83.2</td><td>-10.2</td><td>1.6</td><td>-6.5</td><td>7.5E-11</td><td>2.1E-10</td><td>MB231/MB468</td><td>MP</td></tr>
<tr><td>>hg38_MT_+_1621_3279_RNR2_16S_50ntFlanks@51.74.24|GCTAAACCTAGCCCCAAACCCACT</td><td>>hg38_MT_+_1621_3279_RNR2_16S_50ntFlanks</td><td>51</td><td>74</td><td>24</td><td>GCTAAACCTAGCCCCAAACCCACT</td><td>1022.1</td><td>4.7</td><td>0.7</td><td>6.5</td><td>7.9E-11</td><td>2.1E-10</td><td>MB231/MB468</td><td>NUC</td></tr>
<tr><td>>hg38_1_-_228634819_228635039_RNA5S12_5S_50ntFlanks@154.171.18|CCGGGTGCTGTAGGCTTT</td><td>>hg38_1_-_228634819_228635039_RNA5S12_5S_50ntFlanks</td><td>154</td><td>171</td><td>18</td><td>CCGGGTGCTGTAGGCTTT</td><td>22567.2</td><td>-5.4</td><td>0.8</td><td>-6.5</td><td>7.5E-11</td><td>2.1E-10</td><td>MB231/MB468</td><td>MP</td></tr>
<tr><td>>hg38_21_+_8433172_8446622_RNA45SN1_45S_50ntFlanks@4003.4024.22|ATAACCTCGGGCCGATCGCACG</td><td>>hg38_21_+_8433172_8446622_RNA45SN1_45S_50ntFlanks</td><td>4003</td><td>4024</td><td>22</td><td>ATAACCTCGGGCCGATCGCACG</td><td>1988.1</td><td>-6.9</td><td>1.1</td><td>-6.5</td><td>5.8E-11</td><td>2.1E-10</td><td>BT20/MB468</td><td>MITO</td></tr>
</table>

<table>
<tr><td>>hg38_21_+_8433172_8446622_RNA45SN1_45S_50ntFlanks@11873.11899.27|AAGAAGACCCTGTTGAGCTTGACTCTA</td><td>>hg38_21_+_8433172_8446622_RNA45SN1_45S_50ntFlanks</td><td>11873</td><td>11899</td><td>27</td><td>AAGAAGACCCTGTTGAGCTTGACTCTA</td><td>367.0</td><td>-4.5</td><td>0.7</td><td>-6.5</td><td>8.0E-11</td><td>2.1E-10</td><td>MB231/MB468</td><td>CYTO</td></tr>
<tr><td>>hg38_21_+_8433172_8446622_RNA45SN1_45S_50ntFlanks@11571.11603.33|GCCGACTTAGAACTGGTGCGGACCAGGGGAATC</td><td>>hg38_21_+_8433172_8446622_RNA45SN1_45S_50ntFlanks</td><td>11571</td><td>11603</td><td>33</td><td>GCCGACTTAGAACTGGTGCGGACCAGGGGAATC</td><td>1451.5</td><td>-5.4</td><td>0.8</td><td>-6.5</td><td>9.6E-11</td><td>2.1E-10</td><td>BT20/MB468</td><td>NUC</td></tr>
<tr><td>>hg38_21_+_8433172_8446622_RNA45SN1_45S_50ntFlanks@5186.5226.41|CAATAACAGGTCTGTGATGCCCTTAGATGTCCGGGGCTGCA</td><td>>hg38_21_+_8433172_8446622_RNA45SN1_45S_50ntFlanks</td><td>5186</td><td>5226</td><td>41</td><td>CAATAACAGGTCTGTGATGCCCTTAGATGTCCGGGGCTGCA</td><td>311.7</td><td>-10.4</td><td>1.6</td><td>-6.5</td><td>5.9E-11</td><td>2.1E-10</td><td>BT20/MB468</td><td>MITO</td></tr>
<tr><td>>MI0000266|hsa-mir-10a&WithFlank&17|-|48579832|48579953@28.49.22&[MIMAT0000253&hsa-miR-10a-5p&offsets|0|-1;m-18&17|-|48579905|48579926&offsets|0|0]|TACCCTGTAGATCCGAATTTGT</td><td>>MI0000266|hsa-mir-10a&WithFlank&17|-|48579832|48579953</td><td>28</td><td>49</td><td>22</td><td>TACCCTGTAGATCCGAATTTGT</td><td>5289.6</td><td>8.1</td><td>1.2</td><td>6.5</td><td>7.6E-11</td><td>2.1E-10</td><td>MB231/MB468</td><td>MP</td></tr>
<tr><td>>hg38_21_+_8433172_8446622_RNA45SN1_45S_50ntFlanks@6646.6665.20|TCGTACGACTCTTAGCGGTG</td><td>>hg38_21_+_8433172_8446622_RNA45SN1_45S_50ntFlanks</td><td>6646</td><td>6665</td><td>20</td><td>TCGTACGACTCTTAGCGGTG</td><td>63905.2</td><td>5.4</td><td>0.8</td><td>6.9</td><td>5.8E-12</td><td>2.1E-10</td><td>BT20/MB468</td><td>MP</td></tr>
<tr><td>>trna10_AspGTC_12_-_125424193_125424264@1.30.30,trna12_AspGTC_12_-_125411891_125411962@1.30.30,trna38_AspGTC_17_-_8125556_8125627@1.30.30,trna45_AspGTC_6_+_27447453_27447524@1.30.30,trna48_AspGTC_6_+_27471523_27471594@1.30.30,trna4_AspGTC_12_+_96429799_96429870@1.30.30,trna69_AspGTC_1_-_161440205_161440276@1.30.30,trna72_AspGTC_1_-_161432824_161432895@1.30.30,trna75_AspGTC_1_-_161425414_161425485@1.30.30,trna78_AspGTC_1_-_161418033_161418104@1.30.30,trna81_AspGTC_1_-_161410615_161410686@1.30.30|TCCTCGTTAGTATAGTGGTGAGTATCCCCG</td><td>>trna10_AspGTC_12_-_125424193_125424264</td><td>1</td><td>30</td><td>30</td><td>TCCTCGTTAGTATAGTGGTGAGTATCCCCG</td><td>225.3</td><td>-10.9</td><td>1.6</td><td>-6.9</td><td>5.8E-12</td><td>2.1E-10</td><td>BT20/MB468</td><td>MP</td></tr>
<tr><td>>hg38_21_+_8433172_8446622_RNA45SN1_45S_50ntFlanks@10392.10422.31|ACAGCAGTTGAACATGGGTCAGTCGGTCCTG</td><td>>hg38_21_+_8433172_8446622_RNA45SN1_45S_50ntFlanks</td><td>10392</td><td>10422</td><td>31</td><td>ACAGCAGTTGAACATGGGTCAGTCGGTCCTG</td><td>654.0</td><td>-7.1</td><td>1.1</td><td>-6.5</td><td>7.6E-11</td><td>2.1E-10</td><td>MB231/MB468</td><td>MP</td></tr>
<tr><td>>hg38_21_+_8433172_8446622_RNA45SN1_45S_50ntFlanks@11566.11584.19|TAGCAGCCGACTTAGAACT</td><td>>hg38_21_+_8433172_8446622_RNA45SN1_45S_50ntFlanks</td><td>11566</td><td>11584</td><td>19</td><td>TAGCAGCCGACTTAGAACT</td><td>942.9</td><td>5.7</td><td>0.9</td><td>6.5</td><td>8.0E-11</td><td>2.1E-10</td><td>MB231/MB468</td><td>NUC</td></tr>
<tr><td>>hg38_21_+_8433172_8446622_RNA45SN1_45S_50ntFlanks@4002.4042.41|GATAACCTCGGGCCGATCGCACGCCCCCCGTGGCGGCGACG</td><td>>hg38_21_+_8433172_8446622_RNA45SN1_45S_50ntFlanks</td><td>4002</td><td>4042</td><td>41</td><td>GATAACCTCGGGCCGATCGCACGCCCCCCGTGGCGGCGACG</td><td>1274.2</td><td>-4.9</td><td>0.8</td><td>-6.5</td><td>8.1E-11</td><td>2.1E-10</td><td>MB231/MB468</td><td>CYTO</td></tr>
<tr><td>>hg38_21_+_8433172_8446622_RNA45SN1_45S_50ntFlanks@9565.9596.32|TGACGTGCAAATCGGTCGTCCGACCTGGGTAT</td><td>>hg38_21_+_8433172_8446622_RNA45SN1_45S_50ntFlanks</td><td>9565</td><td>9596</td><td>32</td><td>TGACGTGCAAATCGGTCGTCCGACCTGGGTAT</td><td>7025.0</td><td>-4.7</td><td>0.7</td><td>-6.5</td><td>8.1E-11</td><td>2.1E-10</td><td>MB231/MB468</td><td>CYTO</td></tr>
<tr><td>>hg38_21_+_8433172_8446622_RNA45SN1_45S_50ntFlanks@10342.10368.27|ATATTCAAACGAGAACTTTGAAGGCCG</td><td>>hg38_21_+_8433172_8446622_RNA45SN1_45S_50ntFlanks</td><td>10342</td><td>10368</td><td>27</td><td>ATATTCAAACGAGAACTTTGAAGGCCG</td><td>130.1</td><td>-10.9</td><td>1.7</td><td>-6.5</td><td>7.6E-11</td><td>2.1E-10</td><td>MB231/MB468</td><td>MP</td></tr>
<tr><td>>hg38_21_+_8433172_8446622_RNA45SN1_45S_50ntFlanks@12617.12654.38|TGAGGAGCCAATGGGGCGAAGCTACCATCTGTGGGATT</td><td>>hg38_21_+_8433172_8446622_RNA45SN1_45S_50ntFlanks</td><td>12617</td><td>12654</td><td>38</td><td>TGAGGAGCCAATGGGGCGAAGCTACCATCTGTGGGATT</td><td>956.3</td><td>-9.0</td><td>1.4</td><td>-6.5</td><td>7.6E-11</td><td>2.1E-10</td><td>MB231/MB468</td><td>MP</td></tr>
<tr><td>>hg38_21_+_8433172_8446622_RNA45SN1_45S_50ntFlanks@3835.3859.25|CTCCTCTCCTACTTGGATAACTGTG</td><td>>hg38_21_+_8433172_8446622_RNA45SN1_45S_50ntFlanks</td><td>3835</td><td>3859</td><td>25</td><td>CTCCTCTCCTACTTGGATAACTGTG</td><td>99.9</td><td>-10.5</td><td>1.6</td><td>-6.5</td><td>7.6E-11</td><td>2.1E-10</td><td>MB231/MB468</td><td>MP</td></tr>
<tr><td>>hg38_21_+_8433172_8446622_RNA45SN1_45S_50ntFlanks@8356.8378.23|GAAAAGAACTTTGAAGAGAGAGT</td><td>>hg38_21_+_8433172_8446622_RNA45SN1_45S_50ntFlanks</td><td>8356</td><td>8378</td><td>23</td><td>GAAAAGAACTTTGAAGAGAGAGT</td><td>860.1</td><td>-5.0</td><td>0.8</td><td>-6.5</td><td>9.2E-11</td><td>2.1E-10</td><td>MB231/MB468</td><td>MITO</td></tr>
<tr><td>>hg38_21_+_8433172_8446622_RNA45SN1_45S_50ntFlanks@9858.9901.44|CGGGATGAACCGAACGCCGGGTTAAGGCGCCCGATGCCGACGCT</td><td>>hg38_21_+_8433172_8446622_RNA45SN1_45S_50ntFlanks</td><td>9858</td><td>9901</td><td>44</td><td>CGGGATGAACCGAACGCCGGGTTAAGGCGCCCGATGCCGACGCT</td><td>289.1</td><td>-12.0</td><td>1.8</td><td>-6.5</td><td>9.2E-11</td><td>2.1E-10</td><td>MB231/MB468</td><td>MITO</td></tr>
<tr><td>>Antisense_hg38_21_-_8433172_8446622_RNA45SN1_45S_50ntFlanks@7674.7692.19|CTCTCTCTCTCTCCCCC</td><td>>Antisense_hg38_21_-_8433172_8446622_RNA45SN1_45S_50ntFlanks</td><td>7674</td><td>7692</td><td>19</td><td>CTCTCTCTCTCTCCCCC</td><td>402.7</td><td>-6.8</td><td>1.0</td><td>-6.5</td><td>8.1E-11</td><td>2.1E-10</td><td>MB231/MB468</td><td>CYTO</td></tr>
<tr><td>>hg38_21_+_8433172_8446622_RNA45SN1_45S_50ntFlanks@11027.11045.19|CTCTCTCTCTCTCTCCCCC</td><td>>hg38_21_+_8433172_8446622_RNA45SN1_45S_50ntFlanks</td><td>11027</td><td>11045</td><td>19</td><td>CTCTCTCTCTCTCCCCC</td><td>402.7</td><td>-6.8</td><td>1.0</td><td>-6.5</td><td>8.1E-11</td><td>2.1E-10</td><td>MB231/MB468</td><td>CYTO</td></tr>
<tr><td>>hg38_21_+_8433172_8446622_RNA45SN1_45S_50ntFlanks@9929.9958.30|GATATAGACAGCAGGACGGTGGCCATGGAA</td><td>>hg38_21_+_8433172_8446622_RNA45SN1_45S_50ntFlanks</td><td>9929</td><td>9958</td><td>30</td><td>GATATAGACAGCAGGACGGTGGCCATGGAA</td><td>95.6</td><td>-10.4</td><td>1.6</td><td>-6.5</td><td>7.7E-11</td><td>2.1E-10</td><td>MB231/MB468</td><td>MP</td></tr>
<tr><td>>hg38_21_+_8433172_8446622_RNA45SN1_45S_50ntFlanks@12405.12429.25|CTTTTTGATCCTTCGATGTCGGCTC</td><td>>hg38_21_+_8433172_8446622_RNA45SN1_45S_50ntFlanks</td><td>12405</td><td>12429</td><td>25</td><td>CTTTTTGATCCTTCGATGTCGGCTC</td><td>1766.5</td><td>-8.8</td><td>1.4</td><td>-6.5</td><td>9.2E-11</td><td>2.1E-10</td><td>MB231/MB468</td><td>MITO</td></tr>
<tr><td>>hg38_21_+_8433172_8446622_RNA45SN1_45S_50ntFlanks@9736.9753.18|CCGAAACGATCTCAACCT</td><td>>hg38_21_+_8433172_8446622_RNA45SN1_45S_50ntFlanks</td><td>9736</td><td>9753</td><td>18</td><td>CCGAAACGATCTCAACCT</td><td>445.1</td><td>-8.6</td><td>1.3</td><td>-6.5</td><td>9.2E-11</td><td>2.1E-10</td><td>MB231/MB468</td><td>MITO</td></tr>
<tr><td>>hg38_21_+_8433172_8446622_RNA45SN1_45S_50ntFlanks@11564.11588.25|CCTAGCAGCCGACTTAGAACTGGTG</td><td>>hg38_21_+_8433172_8446622_RNA45SN1_45S_50ntFlanks</td><td>11564</td><td>11588</td><td>25</td><td>CCTAGCAGCCGACTTAGAACTGGTG</td><td>1505.0</td><td>-6.8</td><td>1.0</td><td>-6.5</td><td>9.2E-11</td><td>2.1E-10</td><td>MB231/MB468</td><td>MITO</td></tr>
<tr><td>>MI0000255|hsa-mir-30d&WithFlank&8|-|134804870|134804951@12.35.24(+1C)&[MIMAT0000245&hsa-miR-30d-5p&offsets|0|+2(+1C);m-21&8|-|134804919|134804940&offsets|0|+2(+1C)]|TGTAAACATCCCCGACTGGAAGCTC</td><td>>MI0000255|hsa-mir-30d&WithFlank&8|-|134804870|134804951</td><td>12</td><td>35</td><td>24(+1C)</td><td>TGTAAACATCCCCGACTGGAAGCTC</td><td>343.9</td><td>9.3</td><td>1.4</td><td>6.5</td><td>7.3E-11</td><td>2.1E-10</td><td>BT20/MB468</td><td>CYTO</td></tr>
<tr><td>>hg38_21_+_8433172_8446622_RNA45SN1_45S_50ntFlanks@11999.12033.35|CCCTGCGGGCCGCCGGTGAAATACCACTACTCTGA</td><td>>hg38_21_+_8433172_8446622_RNA45SN1_45S_50ntFlanks</td><td>11999</td><td>12033</td><td>35</td><td>CCCTGCGGGCCGCCGGTGAAATACCACTACTCTGA</td><td>1607.8</td><td>-9.9</td><td>1.5</td><td>-6.5</td><td>9.8E-11</td><td>2.1E-10</td><td>BT20/MB468</td><td>NUC</td></tr>
<tr><td>>hg38_21_+_8433172_8446622_RNA45SN1_45S_50ntFlanks@10389.10408.20|TGAACAGCAGTTGAACATGG</td><td>>hg38_21_+_8433172_8446622_RNA45SN1_45S_50ntFlanks</td><td>10389</td><td>10408</td><td>20</td><td>TGAACAGCAGTTGAACATGG</td><td>200.1</td><td>-5.3</td><td>0.8</td><td>-6.5</td><td>8.1E-11</td><td>2.1E-10</td><td>MB231/MB468</td><td>CYTO</td></tr>
<tr><td>>MI0000456|hsa-mir-140&WithFlank&16|+|69933075|69933186@68.90.23&[MIMAT0004597&hsa-miR-140-3p&offsets|0|+2;m-45&16|+|69933143|69933164&offsets|-1|0]|TACCACAGGGTAGAACCACGGAC</td><td>>MI0000456|hsa-mir-140&WithFlank&16|+|69933075|69933186</td><td>68</td><td>90</td><td>23</td><td>TACCACAGGGTAGAACCACGGAC</td><td>368.6</td><td>8.4</td><td>1.3</td><td>6.5</td><td>8.1E-11</td><td>2.1E-10</td><td>MB231/MB468</td><td>NUC</td></tr>
<tr><td>>hg38_21_+_8433172_8446622_RNA45SN1_45S_50ntFlanks@6647.6672.26|CGTACGACTCTTAGCGGTGGATCACT</td><td>>hg38_21_+_8433172_8446622_RNA45SN1_45S_50ntFlanks</td><td>6647</td><td>6672</td><td>26</td><td>CGTACGACTCTTAGCGGTGGATCACT</td><td>671.4</td><td>4.4</td><td>0.7</td><td>6.5</td><td>8.1E-11</td><td>2.1E-10</td><td>MB231/MB468</td><td>NUC</td></tr>
<tr><td>>hg38_21_+_8433172_8446622_RNA45SN1_45S_50ntFlanks@11452.11469.18|CTCCTCGGCGCGCGGCGG</td><td>>hg38_21_+_8433172_8446622_RNA45SN1_45S_50ntFlanks</td><td>11452</td><td>11469</td><td>18</td><td>CTCCTCGGCGCGCGGCGG</td><td>80.0</td><td>-10.2</td><td>1.6</td><td>-6.5</td><td>7.7E-11</td><td>2.1E-10</td><td>MB231/MB468</td><td>MP</td></tr>
<tr><td>>hg38_21_+_8433172_8446622_RNA45SN1_45S_50ntFlanks@11934.11968.35|ATAAGTGGGAGGCCCCCGGCGCCCCCCCGGTGTCC</td><td>>hg38_21_+_8433172_8446622_RNA45SN1_45S_50ntFlanks</td><td>11934</td><td>11968</td><td>35</td><td>ATAAGTGGGAGGCCCCCGGCGCCCCCCCGGTGTCC</td><td>76.4</td><td>-10.1</td><td>1.6</td><td>-6.5</td><td>7.7E-11</td><td>2.1E-10</td><td>MB231/MB468</td><td>MP</td></tr>
<tr><td>>hg38_21_+_8433172_8446622_RNA45SN1_45S_50ntFlanks@12675.12703.29|CAGAATCCCGCCCAGGCGGAACGATACGG</td><td>>hg38_21_+_8433172_8446622_RNA45SN1_45S_50ntFlanks</td><td>12675</td><td>12703</td><td>29</td><td>CAGAATCCCGCCCAGGCGGAACGATACGG</td><td>126.6</td><td>-10.8</td><td>1.7</td><td>-6.5</td><td>7.7E-11</td><td>2.1E-10</td><td>MB231/MB468</td><td>MP</td></tr>
<tr><td>>hg38_21_+_8433172_8446622_RNA45SN1_45S_50ntFlanks@9505.9522.18|TGAACTATGCCTGGGCAG</td><td>>hg38_21_+_8433172_8446622_RNA45SN1_45S_50ntFlanks</td><td>9505</td><td>9522</td><td>18</td><td>TGAACTATGCCTGGGCAG</td><td>87.4</td><td>-10.3</td><td>1.6</td><td>-6.5</td><td>7.7E-11</td><td>2.1E-10</td><td>MB231/MB468</td><td>MP</td></tr>
<tr><td>>hg38_21_+_8433172_8446622_RNA45SN1_45S_50ntFlanks@11562.11588.27|CGCCTAGCAGCCGACTTAGAACTGGTG</td><td>>hg38_21_+_8433172_8446622_RNA45SN1_45S_50ntFlanks</td><td>11562</td><td>11588</td><td>27</td><td>CGCCTAGCAGCCGACTTAGAACTGGTG</td><td>326.5</td><td>-3.0</td><td>0.5</td><td>-6.5</td><td>7.3E-11</td><td>2.1E-10</td><td>BT20/MB468</td><td>CYTO</td></tr>
<tr><td>>MI0000086|hsa-mir-28&WithFlank&3|+|188688775|188688872@60.82.23&[MIMAT0004502&hsa-miR-28-3p&offsets|0|+1;m-28&3|+|188688834|188688855&offsets|0|+1]|CACTAGATTGTGAGCTCCTGGAG</td><td>>MI0000086|hsa-mir-28&WithFlank&3|+|188688775|188688872</td><td>60</td><td>82</td><td>23</td><td>CACTAGATTGTGAGCTCCTGGAG</td><td>357.6</td><td>9.0</td><td>1.4</td><td>6.5</td><td>8.2E-11</td><td>2.1E-10</td><td>MB231/MB468</td><td>CYTO</td></tr>
<tr><td>>hg38_21_+_8433172_8446622_RNA45SN1_45S_50ntFlanks@12476.12496.21|CCACTAATAGGGAACGTGAGC</td><td>>hg38_21_+_8433172_8446622_RNA45SN1_45S_50ntFlanks</td><td>12476</td><td>12496</td><td>21</td><td>CCACTAATAGGGAACGTGAGC</td><td>2266.9</td><td>-4.0</td><td>0.6</td><td>-6.5</td><td>8.2E-11</td><td>2.1E-10</td><td>MB231/MB468</td><td>NUC</td></tr>
<tr><td>>hg38_MT_+_598_1651_RNR1_12S_50ntFlanks@634.665.32|AACCTCACCACCTCTTGCTCAGCCTATATACC</td><td>>hg38_MT_+_598_1651_RNR1_12S_50ntFlanks</td><td>634</td><td>665</td><td>32</td><td>AACCTCACCACCTCTTGCTCAGCCTATATACC</td><td>983.3</td><td>10.5</td><td>1.6</td><td>6.5</td><td>6.0E-11</td><td>2.1E-10</td><td>BT20/MB468</td><td>MITO</td></tr>
<tr><td>>hg38_21_+_8433172_8446622_RNA45SN1_45S_50ntFlanks@9122.9140.19|CCGCGCGCGGGTCGGGGGG</td><td>>hg38_21_+_8433172_8446622_RNA45SN1_45S_50ntFlanks</td><td>9122</td><td>9140</td><td>19</td><td>CCGCGCGCGGGTCGGGGGG</td><td>186.1</td><td>-9.6</td><td>1.5</td><td>-6.5</td><td>6.0E-11</td><td>2.2E-10</td><td>BT20/MB468</td><td>MITO</td></tr>
<tr><td>>MI0000749|hsa-mir-30e&WithFlank&1|+|40754349|40754452@24.46.23&[MIMAT0000692&hsa-miR-30e-5p&offsets|+1|+2;m-24&1|+|40754371|40754394&offsets|+1|0]|GTAAACATCCTTGACTGGAAGCT</td><td>>MI0000749|hsa-mir-30e&WithFlank&1|+|40754349|40754452</td><td>24</td><td>46</td><td>23</td><td>GTAAACATCCTTGACTGGAAGCT</td><td>174.7</td><td>5.1</td><td>0.8</td><td>6.5</td><td>8.2E-11</td><td>2.2E-10</td><td>MB231/MB468</td><td>NUC</td></tr>
<tr><td>>hg38_21_+_8433172_8446622_RNA45SN1_45S_50ntFlanks@8176.8201.26|CTTCTGATCGAGGCCCAGCCCGTGGA</td><td>>hg38_21_+_8433172_8446622_RNA45SN1_45S_50ntFlanks</td><td>8176</td><td>8201</td><td>26</td><td>CTTCTGATCGAGGCCCAGCCCGTGGA</td><td>485.9</td><td>-3.6</td><td>0.6</td><td>-6.5</td><td>8.2E-11</td><td>2.2E-10</td><td>MB231/MB468</td><td>NUC</td></tr>
<tr><td>>hg38_21_+_8433172_8446622_RNA45SN1_45S_50ntFlanks@9568.9592.25|CGTGCAAATCGGTCGTCCGACCTGG</td><td>>hg38_21_+_8433172_8446622_RNA45SN1_45S_50ntFlanks</td><td>9568</td><td>9592</td><td>25</td><td>CGTGCAAATCGGTCGTCCGACCTGG</td><td>656.8</td><td>-8.2</td><td>1.3</td><td>-6.5</td><td>9.4E-11</td><td>2.2E-10</td><td>MB231/MB468</td><td>MITO</td></tr>
<tr><td>>hg38_21_+_8433172_8446622_RNA45SN1_45S_50ntFlanks@12073.12093.21|CCGAGGGGCTCTCGCTTCTGG</td><td>>hg38_21_+_8433172_8446622_RNA45SN1_45S_50ntFlanks</td><td>12073</td><td>12093</td><td>21</td><td>CCGAGGGGCTCTCGCTTCTGG</td><td>255.7</td><td>-5.5</td><td>0.8</td><td>-6.5</td><td>8.3E-11</td><td>2.2E-10</td><td>MB231/MB468</td><td>CYTO</td></tr>
<tr><td>>hg38_21_+_8433172_8446622_RNA45SN1_45S_50ntFlanks@12632.12657.26|GCGAAGCTACCATCTGTGGGATTATG</td><td>>hg38_21_+_8433172_8446622_RNA45SN1_45S_50ntFlanks</td><td>12632</td><td>12657</td><td>26</td><td>GCGAAGCTACCATCTGTGGGATTATG</td><td>854.0</td><td>-7.0</td><td>1.1</td><td>-6.5</td><td>9.4E-11</td><td>2.2E-10</td><td>MB231/MB468</td><td>MITO</td></tr>
<tr><td>>hg38_21_+_8433172_8446622_RNA45SN1_45S_50ntFlanks@12419.12455.37|GATGTCGGCTCTTCCTATCATTGTGAAGCAGAATTCA</td><td>>hg38_21_+_8433172_8446622_RNA45SN1_45S_50ntFlanks</td><td>12419</td><td>12455</td><td>37</td><td>GATGTCGGCTCTTCCTATCATTGTGAAGCAGAATTCA</td><td>1579.7</td><td>-6.8</td><td>1.0</td><td>-6.5</td><td>9.9E-11</td><td>2.2E-10</td><td>BT20/MB468</td><td>NUC</td></tr>
<tr><td>>hg38_21_+_8433172_8446622_RNA45SN1_45S_50ntFlanks@9546.9571.26|TGGAGGTCCGTAGCGGTCCTGACGTG</td><td>>hg38_21_+_8433172_8446622_RNA45SN1_45S_50ntFlanks</td><td>9546</td><td>9571</td><td>26</td><td>TGGAGGTCCGTAGCGGTCCTGACGTG</td><td>1404.9</td><td>-5.6</td><td>0.9</td><td>-6.5</td><td>9.9E-11</td><td>2.2E-10</td><td>BT20/MB468</td><td>NUC</td></tr>
<tr><td>>hg38_21_+_8433172_8446622_RNA45SN1_45S_50ntFlanks@11873.11894.22|AAGAAGACCCTGTTGAGCTTGA</td><td>>hg38_21_+_8433172_8446622_RNA45SN1_45S_50ntFlanks</td><td>11873</td><td>11894</td><td>22</td><td>AAGAAGACCCTGTTGAGCTTGA</td><td>210.0</td><td>-10.8</td><td>1.6</td><td>-6.9</td><td>6.0E-12</td><td>2.2E-10</td><td>BT20/MB468</td><td>MP</td></tr>
<tr><td>>hg38_MT_+_598_1651_RNR1_12S_50ntFlanks@425.455.31|ACGAAAGTGGCTTTAACATATCTGAACACAC</td><td>>hg38_MT_+_598_1651_RNR1_12S_50ntFlanks</td><td>425</td><td>455</td><td>31</td><td>ACGAAAGTGGCTTTAACATATCTGAACACAC</td><td>936.9</td><td>8.3</td><td>1.3</td><td>6.5</td><td>6.1E-11</td><td>2.2E-10</td><td>BT20/MB468</td><td>MITO</td></tr>
<tr><td>>hg38_21_+_8433172_8446622_RNA45SN1_45S_50ntFlanks@7985.8007.23|GATCAGACGTGGCGACCCGCTGA</td><td>>hg38_21_+_8433172_8446622_RNA45SN1_45S_50ntFlanks</td><td>7985</td><td>8007</td><td>23</td><td>GATCAGACGTGGCGACCCGCTGA</td><td>680.0</td><td>-8.5</td><td>1.3</td><td>-6.5</td><td>9.4E-11</td><td>2.2E-10</td><td>MB231/MB468</td><td>MITO</td></tr>
<tr><td>>MI0000264|hsa-mir-7-2&WithFlank&15|+|88611819|88611940@38.59.22(+1C);MI0000265|hsa-mir-7-3&WithFlank&19|+|4770664|4770785@37.58.22(+1C);MI0000263|hsa-mir-7-1&WithFlank&9|-|83969742|83969863@30.51.22(+1C)&[MIMAT0000252&hsa-miR-7-5p&offsets|0|-2(+1C);m-400&15|+|88611856|88611879&offsets|0|-2(+1C)];[MIMAT0000252_1&hsa-miR-7-5p&offsets|0|-2(+1C);m-402&19|+|4770700|4770723&offsets|0|-2(+1C)];[MIMAT0000252_2&hsa-miR-7-5p&offsets|0|-2(+1C);m-401&9|-|83969811|83969834&offsets|0|-2(+1C)]|TGGAAGACTAGTGATTTTGTTGC</td><td>>MI0000264|hsa-mir-7-2&WithFlank&15|+|88611819|88611940</td><td>38</td><td>59</td><td>22(+1C)</td><td>TGGAAGACTAGTGATTTTGTTGC</td><td>886.3</td><td>6.6</td><td>1.0</td><td>6.5</td><td>9.4E-11</td><td>2.2E-10</td><td>MB231/MB468</td><td>MITO</td></tr>
<tr><td>>hg38_21_+_8433172_8446622_RNA45SN1_45S_50ntFlanks@5444.5467.24|CTCGGATCGGCCCCGCCGGGGTCG</td><td>>hg38_21_+_8433172_8446622_RNA45SN1_45S_50ntFlanks</td><td>5444</td><td>5467</td><td>24</td><td>CTCGGATCGGCCCCGCCGGGGTCG</td><td>999.2</td><td>-4.4</td><td>0.7</td><td>-6.5</td><td>7.4E-11</td><td>2.2E-10</td><td>BT20/MB468</td><td>CYTO</td></tr>
<tr><td>>hg38_21_+_8433172_8446622_RNA45SN1_45S_50ntFlanks@11729.11747.19|TAACTATGACTCTCTTAAG</td><td>>hg38_21_+_8433172_8446622_RNA45SN1_45S_50ntFlanks</td><td>11729</td><td>11747</td><td>19</td><td>TAACTATGACTCTCTTAAG</td><td>460.0</td><td>-8.3</td><td>1.3</td><td>-6.5</td><td>7.8E-11</td><td>2.2E-10</td><td>MB231/MB468</td><td>MP</td></tr>
<tr><td>>trnaMT_ValTAC_MT_+_1602_1670@47.72.26|TCAACTTAACTTGACCGCTCTGACCA</td><td>>trnaMT_ValTAC_MT_+_1602_1670</td><td>47</td><td>72</td><td>26</td><td>TCAACTTAACTTGACCGCTCTGACCA</td><td>1155.9</td><td>5.4</td><td>0.8</td><td>6.5</td><td>8.3E-11</td><td>2.2E-10</td><td>MB231/MB468</td><td>NUC</td></tr>
<tr><td>>hg38_21_+_8433172_8446622_RNA45SN1_45S_50ntFlanks@9133.9151.19|TCGGGGGGCGGGGCGGACT</td><td>>hg38_21_+_8433172_8446622_RNA45SN1_45S_50ntFlanks</td><td>9133</td><td>9151</td><td>19</td><td>TCGGGGGGCGGGGCGGACT</td><td>118.6</td><td>-10.8</td><td>1.7</td><td>-6.5</td><td>7.9E-11</td><td>2.2E-10</td><td>MB231/MB468</td><td>MP</td></tr>
</table>

<table>
<tr><th></th><th></th><th></th><th></th><th></th><th></th><th></th><th></th><th></th><th></th><th></th><th></th><th></th><th></th></tr>
<tr><td>>MI0000767|hsa-mir-365a&WithFlank&16|+|14309279|14309377@62.82.21;MI0000769|hsa-mir-365b&WithFlank&17|+|31575405|31575527@74.94.21&[MIMAT0000710&hsa-miR-365a-3p&offsets|0|-1;m-176&16|+|14309340|14309361&offsets|0|-1];[MIMAT0022834&hsa-miR-365b-3p&offsets|0|-1;m-175&17|+|31575478|31575499&offsets|0|-1]|TAATGCCCCTAAAAATCCTTA</td><td>>MI0000767|hsa-mir-365a&WithFlank&16|+|14309279|14309377</td><td>62</td><td>82</td><td>21</td><td>TAATGCCCCTAAAAATCCTTA</td><td>440.9</td><td>4.8</td><td>0.7</td><td>6.8</td><td>9.0E-12</td><td>2.2E-10</td><td>BT20/MB231</td><td>NUC</td></tr>
<tr><td>>hg38_21_+_8433172_8446622_RNA45SN1_45S_50ntFlanks@10358.10383.26|TTTGAAGGCCGAAGTGGAGAAGGGTT</td><td>>hg38_21_+_8433172_8446622_RNA45SN1_45S_50ntFlanks</td><td>10358</td><td>10383</td><td>26</td><td>TTTGAAGGCCGAAGTGGAGAAGGGTT</td><td>365.4</td><td>-8.5</td><td>1.3</td><td>-6.5</td><td>1.0E-10</td><td>2.2E-10</td><td>BT20/MB468</td><td>NUC</td></tr>
<tr><td>>hg38_21_+_8433172_8446622_RNA45SN1_45S_50ntFlanks@10906.10932.27|TGGGCGCGCGCCGCGGCTGGACGAGGC</td><td>>hg38_21_+_8433172_8446622_RNA45SN1_45S_50ntFlanks</td><td>10906</td><td>10932</td><td>27</td><td>TGGGCGCGCGCCGCGGCTGGACGAGGC</td><td>100.5</td><td>-10.5</td><td>1.6</td><td>-6.5</td><td>7.9E-11</td><td>2.2E-10</td><td>MB231/MB468</td><td>MP</td></tr>
<tr><td>>hg38_21_+_8433172_8446622_RNA45SN1_45S_50ntFlanks@10726.10751.26|GGGAGAGGGTGTAAATCTCGCGCCGG</td><td>>hg38_21_+_8433172_8446622_RNA45SN1_45S_50ntFlanks</td><td>10726</td><td>10751</td><td>26</td><td>GGGAGAGGGTGTAAATCTCGCGCCGG</td><td>113.4</td><td>-10.7</td><td>1.6</td><td>-6.5</td><td>7.9E-11</td><td>2.2E-10</td><td>MB231/MB468</td><td>MP</td></tr>
<tr><td>>hg38_21_+_8433172_8446622_RNA45SN1_45S_50ntFlanks@4555.4577.23|CGCAGCTAGGAATAATGGAATAG</td><td>>hg38_21_+_8433172_8446622_RNA45SN1_45S_50ntFlanks</td><td>4555</td><td>4577</td><td>23</td><td>CGCAGCTAGGAATAATGGAATAG</td><td>94.7</td><td>-10.4</td><td>1.6</td><td>-6.5</td><td>7.9E-11</td><td>2.2E-10</td><td>MB231/MB468</td><td>MP</td></tr>
<tr><td>>hg38_21_+_8433172_8446622_RNA45SN1_45S_50ntFlanks@11999.12043.45|CCCTGCGGGCCGCCGGTGAAATACCACTACTCTGATCGTTTTTTC</td><td>>hg38_21_+_8433172_8446622_RNA45SN1_45S_50ntFlanks</td><td>11999</td><td>12043</td><td>45</td><td>CCCTGCGGGCCGCCGGTGAAATACCACTACTCTGATCGTTTTTTC</td><td>534.3</td><td>-11.4</td><td>1.7</td><td>-6.5</td><td>7.5E-11</td><td>2.2E-10</td><td>BT20/MB468</td><td>CYTO</td></tr>
<tr><td>>am_trna116_GluCTC_1_-_145399233_145399304@1.26.26,trna59_GluCTC_1_+_249168447_249168518@1.26.26,trna71_GluCTC_1_-_161439189_161439260@1.26.26,trna74_GluCTC_1_-_161431809_161431880@1.26.26,trna77_GluCTC_1_-_161424398_161424469@1.26.26,trna77_GluCTC_6_+_28949976_28950047@1.26.26,trna80_GluCTC_1_-_161417018_161417089@1.26.26,trna87_GluCTC_6_-_126101393_126101464@1.26.26|TCCCTGGTGGTCTAGTGGTTAGGATT</td><td>>am_trna116_GluCTC_1_-_145399233_145399304</td><td>1</td><td>26</td><td>26</td><td>TCCCTGGTGGTCTAGTGGTTAGGATT</td><td>7859.2</td><td>3.9</td><td>0.6</td><td>6.5</td><td>7.5E-11</td><td>2.2E-10</td><td>BT20/MB468</td><td>CYTO</td></tr>
<tr><td>>hg38_21_+_8433172_8446622_RNA45SN1_45S_50ntFlanks@12677.12703.27|GAATCCCGCCCAGGCGGAACGATACGG</td><td>>hg38_21_+_8433172_8446622_RNA45SN1_45S_50ntFlanks</td><td>12677</td><td>12703</td><td>27</td><td>GAATCCCGCCCAGGCGGAACGATACGG</td><td>340.7</td><td>-11.5</td><td>1.7</td><td>-6.9</td><td>6.1E-12</td><td>2.2E-10</td><td>BT20/MB468</td><td>MP</td></tr>
<tr><td>>hg38_21_+_8433172_8446622_RNA45SN1_45S_50ntFlanks@11575.11600.26|ACTTAGAACTGGTGCGGACCAGGGGA</td><td>>hg38_21_+_8433172_8446622_RNA45SN1_45S_50ntFlanks</td><td>11575</td><td>11600</td><td>26</td><td>ACTTAGAACTGGTGCGGACCAGGGGA</td><td>724.7</td><td>-6.6</td><td>1.0</td><td>-6.5</td><td>8.4E-11</td><td>2.2E-10</td><td>MB231/MB468</td><td>NUC</td></tr>
<tr><td>>hg38_21_+_8433172_8446622_RNA45SN1_45S_50ntFlanks@5334.5363.30|ATGAACGAGGAATTCCCAGTAAGTGCGGGT</td><td>>hg38_21_+_8433172_8446622_RNA45SN1_45S_50ntFlanks</td><td>5334</td><td>5363</td><td>30</td><td>ATGAACGAGGAATTCCCAGTAAGTGCGGGT</td><td>138.2</td><td>-11.0</td><td>1.7</td><td>-6.5</td><td>7.9E-11</td><td>2.2E-10</td><td>MB231/MB468</td><td>MP</td></tr>
<tr><td>>hg38_21_+_8433172_8446622_RNA45SN1_45S_50ntFlanks@12992.13012.21|TCCCTCGCTGCGATCTATTGA</td><td>>hg38_21_+_8433172_8446622_RNA45SN1_45S_50ntFlanks</td><td>12992</td><td>13012</td><td>21</td><td>TCCCTCGCTGCGATCTATTGA</td><td>120.0</td><td>-9.0</td><td>1.4</td><td>-6.5</td><td>6.1E-11</td><td>2.2E-10</td><td>BT20/MB468</td><td>MITO</td></tr>
<tr><td>>hg38_21_+_8433172_8446622_RNA45SN1_45S_50ntFlanks@6661.6682.22|CGGTGGATCACTCGGCTCGTGC</td><td>>hg38_21_+_8433172_8446622_RNA45SN1_45S_50ntFlanks</td><td>6661</td><td>6682</td><td>22</td><td>CGGTGGATCACTCGGCTCGTGC</td><td>175.3</td><td>4.6</td><td>0.7</td><td>6.5</td><td>8.4E-11</td><td>2.2E-10</td><td>MB231/MB468</td><td>NUC</td></tr>
<tr><td>>hg38_21_+_8433172_8446622_RNA45SN1_45S_50ntFlanks@3802.3834.33|CATTAAATCAGTTATGGTTCCTTTGGTCGCTCG</td><td>>hg38_21_+_8433172_8446622_RNA45SN1_45S_50ntFlanks</td><td>3802</td><td>3834</td><td>33</td><td>CATTAAATCAGTTATGGTTCCTTTGGTCGCTCG</td><td>1462.2</td><td>-9.1</td><td>1.4</td><td>-6.5</td><td>8.4E-11</td><td>2.2E-10</td><td>MB231/MB468</td><td>NUC</td></tr>
<tr><td>>am_trna11_ArgACG_3_-_45730491_45730563@39.56.18,trna138_ArgACG_6_-_27638344_27638416@39.56.18,trna156_ArgACG_6_-_27181623_27181695@39.56.18,trna36_ArgACG_6_+_27182952_27183024@39.56.18|TCAGAAGATTCTAGGTTC</td><td>>am_trna11_ArgACG_3_-_45730491_45730563</td><td>39</td><td>56</td><td>18</td><td>TCAGAAGATTCTAGGTTC</td><td>124.0</td><td>-6.5</td><td>1.0</td><td>-6.5</td><td>7.5E-11</td><td>2.2E-10</td><td>BT20/MB468</td><td>CYTO</td></tr>
<tr><td>>hg38_21_+_8433172_8446622_RNA45SN1_45S_50ntFlanks@10362.10383.22|AAGGCCGAAGTGGAGAAGGGTT</td><td>>hg38_21_+_8433172_8446622_RNA45SN1_45S_50ntFlanks</td><td>10362</td><td>10383</td><td>22</td><td>AAGGCCGAAGTGGAGAAGGGTT</td><td>1998.6</td><td>-4.8</td><td>0.7</td><td>-6.5</td><td>7.5E-11</td><td>2.2E-10</td><td>BT20/MB468</td><td>CYTO</td></tr>
<tr><td>>am_trnaMT_ValTAC_MT_+_1602_1670@49.67.19|AACTTAACTTGACCGCTCT</td><td>>am_trnaMT_ValTAC_MT_+_1602_1670</td><td>49</td><td>67</td><td>19</td><td>AACTTAACTTGACCGCTCT</td><td>2445.2</td><td>6.7</td><td>1.0</td><td>6.5</td><td>6.2E-11</td><td>2.2E-10</td><td>BT20/MB468</td><td>MITO</td></tr>
<tr><td>>hg38_21_+_8433172_8446622_RNA45SN1_45S_50ntFlanks@11604.11623.20|CGACTGTTTAATTAAAACAA</td><td>>hg38_21_+_8433172_8446622_RNA45SN1_45S_50ntFlanks</td><td>11604</td><td>11623</td><td>20</td><td>CGACTGTTTAATTAAAACAA</td><td>5512.8</td><td>4.6</td><td>0.7</td><td>6.5</td><td>6.2E-11</td><td>2.2E-10</td><td>BT20/MB468</td><td>MITO</td></tr>
<tr><td>>hg38_21_+_8433172_8446622_RNA45SN1_45S_50ntFlanks@12412.12437.26|ATCCTTCGATGTCGGCTCTTCCTATC</td><td>>hg38_21_+_8433172_8446622_RNA45SN1_45S_50ntFlanks</td><td>12412</td><td>12437</td><td>26</td><td>ATCCTTCGATGTCGGCTCTTCCTATC</td><td>174.1</td><td>-9.8</td><td>1.5</td><td>-6.5</td><td>7.6E-11</td><td>2.2E-10</td><td>BT20/MB468</td><td>CYTO</td></tr>
<tr><td>>hg38_21_+_8433172_8446622_RNA45SN1_45S_50ntFlanks@6719.6736.18|TGAATTGCAGGACACATT</td><td>>hg38_21_+_8433172_8446622_RNA45SN1_45S_50ntFlanks</td><td>6719</td><td>6736</td><td>18</td><td>TGAATTGCAGGACACATT</td><td>2120.3</td><td>-6.6</td><td>1.0</td><td>-6.5</td><td>8.5E-11</td><td>2.2E-10</td><td>MB231/MB468</td><td>CYTO</td></tr>
<tr><td>>MI0000270|hsa-mir-181b-1&WithFlank&1|-|198858867|198858988@42.65.24;MI0000683|hsa-mir-181b-2&WithFlank&9|+|124693704|124693804@22.45.24&[MIMAT0000257&hsa-miR-181b-5p&offsets|0|+1;m-73&1|-|198858925|198858947&offsets|0|+1];[MIMAT0000257_1&hsa-miR-181b-5p&offsets|0|+1;m-72&9|+|124693725|124693747&offsets|0|+1]|AACATTCATTGCTGTCGGTGGGTT</td><td>>MI0000270|hsa-mir-181b-1&WithFlank&1|-|198858867|198858988</td><td>42</td><td>65</td><td>24</td><td>AACATTCATTGCTGTCGGTGGGTT</td><td>2782.0</td><td>6.2</td><td>1.0</td><td>6.5</td><td>9.6E-11</td><td>2.2E-10</td><td>MB231/MB468</td><td>MITO</td></tr>
<tr><td>>MI0000261|hsa-mir-139&WithFlank&11|-|72615057|72615136@13.34.22(+1C)&[MIMAT0000250&hsa-miR-139-5p&offsets|0|-1(+1C);m-186&11|-|72615102|72615124&offsets|0|-1(+1C)]|TCTACAGTGCACGTGTCTCCAGC</td><td>>MI0000261|hsa-mir-139&WithFlank&11|-|72615057|72615136</td><td>13</td><td>34</td><td>22(+1C)</td><td>TCTACAGTGCACGTGTCTCCAGC</td><td>241.1</td><td>11.1</td><td>1.7</td><td>6.5</td><td>9.6E-11</td><td>2.2E-10</td><td>MB231/MB468</td><td>MITO</td></tr>
<tr><td>>hg38_21_+_8433172_8446622_RNA45SN1_45S_50ntFlanks@10346.10369.24|TCAAACGAGAACTTTGAAGGCCGA</td><td>>hg38_21_+_8433172_8446622_RNA45SN1_45S_50ntFlanks</td><td>10346</td><td>10369</td><td>24</td><td>TCAAACGAGAACTTTGAAGGCCGA</td><td>1238.5</td><td>-4.4</td><td>0.7</td><td>-6.5</td><td>9.6E-11</td><td>2.2E-10</td><td>MB231/MB468</td><td>MITO</td></tr>
<tr><td>>MI0000750|hsa-mir-26a-2&WithFlank&12|-|57824603|57824698@20.39.20;MI0000083|hsa-mir-26a-1&WithFlank&3|+|37969398|37969486@16.35.20&[MIMAT0000082&hsa-miR-26a-5p&offsets|0|-2;m-33&12|-|57824658|57824679&offsets|0|-2];[MIMAT0000082_1&hsa-miR-26a-5p&offsets|0|-2;m-34&3|+|37969413|37969434&offsets|0|-2]|TTCAAGTAATCCAGGATAGG</td><td>>MI0000750|hsa-mir-26a-2&WithFlank&12|-|57824603|57824698</td><td>20</td><td>39</td><td>20</td><td>TTCAAGTAATCCAGGATAGG</td><td>652.5</td><td>6.0</td><td>0.9</td><td>6.5</td><td>8.5E-11</td><td>2.2E-10</td><td>MB231/MB468</td><td>CYTO</td></tr>
<tr><td>>am_trna111_HisGTG_1_-_147774845_147774916@-1G.22.23,trna118_HisGTG_1_-_145396881_145396952@-1G.22.23,trna16_HisGTG_1_+_146544773_146544844@-1G.22.23,trna1_HisGTG_15_+_45493349_45493420@-1G.22.23,trna21_HisGTG_1_+_147753471_147753542@-1G.22.23,trna33_HisGTG_6_+_27125906_27125977@-1G.22.23,trna7_HisGTG_9_-_14433938_14434009@-1G.22.23,trna8_HisGTG_15_-_45492611_45492682@-1G.22.23,trna9_HisGTG_15_-_45490804_45490875@-1G.22.23|GGCCGTGATCGTATAGTGGTTAG</td><td>>am_trna111_HisGTG_1_-_147774845_147774916</td><td>-1G</td><td>22</td><td>23</td><td>GGCCGTGATCGTATAGTGGTTAG</td><td>140.3</td><td>10.4</td><td>1.6</td><td>6.5</td><td>8.0E-11</td><td>2.2E-10</td><td>MB231/MB468</td><td>MP</td></tr>
<tr><td>>hg38_21_+_8433172_8446622_RNA45SN1_45S_50ntFlanks@12625.12661.37|CAATGGGGCGAAGCTACCATCTGTGGGATTATGACTG</td><td>>hg38_21_+_8433172_8446622_RNA45SN1_45S_50ntFlanks</td><td>12625</td><td>12661</td><td>37</td><td>CAATGGGGCGAAGCTACCATCTGTGGGATTATGACTG</td><td>347.3</td><td>-3.7</td><td>0.6</td><td>-6.5</td><td>8.5E-11</td><td>2.2E-10</td><td>MB231/MB468</td><td>NUC</td></tr>
<tr><td>>hg38_21_+_8433172_8446622_RNA45SN1_45S_50ntFlanks@10103.10124.22|CGCGCGTGTGGTGTGCGTCGGA</td><td>>hg38_21_+_8433172_8446622_RNA45SN1_45S_50ntFlanks</td><td>10103</td><td>10124</td><td>22</td><td>CGCGCGTGTGGTGTGCGTCGGA</td><td>76.1</td><td>-10.1</td><td>1.6</td><td>-6.5</td><td>8.0E-11</td><td>2.2E-10</td><td>MB231/MB468</td><td>MP</td></tr>
<tr><td>>hg38_21_+_8433172_8446622_RNA45SN1_45S_50ntFlanks@10351.10381.31|CGAGAACTTTGAAGGCCGAAGTGGAGAAGGG</td><td>>hg38_21_+_8433172_8446622_RNA45SN1_45S_50ntFlanks</td><td>10351</td><td>10381</td><td>31</td><td>CGAGAACTTTGAAGGCCGAAGTGGAGAAGGG</td><td>581.1</td><td>-3.8</td><td>0.6</td><td>-6.5</td><td>7.6E-11</td><td>2.2E-10</td><td>BT20/MB468</td><td>CYTO</td></tr>
<tr><td>>MI0000440|hsa-mir-27b&WithFlank&9|+|95085439|95085547@67.88.22&[MIMAT0000419&hsa-miR-27b-3p&offsets|0|+1;m-39&9|+|95085505|95085525&offsets|0|+1]|TTCACAGTGGCTAAGTTCTGCA</td><td>>MI0000440|hsa-mir-27b&WithFlank&9|+|95085439|95085547</td><td>67</td><td>88</td><td>22</td><td>TTCACAGTGGCTAAGTTCTGCA</td><td>1452.4</td><td>2.4</td><td>0.4</td><td>6.5</td><td>1.0E-10</td><td>2.2E-10</td><td>BT20/MB468</td><td>NUC</td></tr>
<tr><td>>hg38_21_+_8433172_8446622_RNA45SN1_45S_50ntFlanks@4557.4587.31|CAGCTAGGAATAATGGAATAGGACCGCGGTT</td><td>>hg38_21_+_8433172_8446622_RNA45SN1_45S_50ntFlanks</td><td>4557</td><td>4587</td><td>31</td><td>CAGCTAGGAATAATGGAATAGGACCGCGGTT</td><td>344.8</td><td>-3.8</td><td>0.6</td><td>-6.5</td><td>8.5E-11</td><td>2.2E-10</td><td>MB231/MB468</td><td>CYTO</td></tr>
<tr><td>>hg38_21_+_8433172_8446622_RNA45SN1_45S_50ntFlanks@10410.10435.26|TCAGTCGGTCCTGAGAGATGGGCGAG</td><td>>hg38_21_+_8433172_8446622_RNA45SN1_45S_50ntFlanks</td><td>10410</td><td>10435</td><td>26</td><td>TCAGTCGGTCCTGAGAGATGGGCGAG</td><td>233.6</td><td>-7.9</td><td>1.2</td><td>-6.5</td><td>7.6E-11</td><td>2.2E-10</td><td>BT20/MB468</td><td>CYTO</td></tr>
<tr><td>>hg38_21_+_8433172_8446622_RNA45SN1_45S_50ntFlanks@5475.5516.42|GCCCTGGCGGAGCGCTGAGAAGACGGTCGAACTTGACTATCT</td><td>>hg38_21_+_8433172_8446622_RNA45SN1_45S_50ntFlanks</td><td>5475</td><td>5516</td><td>42</td><td>GCCCTGGCGGAGCGCTGAGAAGACGGTCGAACTTGACTATCT</td><td>188.7</td><td>11.3</td><td>1.6</td><td>6.9</td><td>6.2E-12</td><td>2.2E-10</td><td>BT20/MB468</td><td>MP</td></tr>
<tr><td>>hg38_21_+_8433172_8446622_RNA45SN1_45S_50ntFlanks@6786.6806.21|TACGCCTGTCTGAGCGTCGCT</td><td>>hg38_21_+_8433172_8446622_RNA45SN1_45S_50ntFlanks</td><td>6786</td><td>6806</td><td>21</td><td>TACGCCTGTCTGAGCGTCGCT</td><td>5699.2</td><td>5.9</td><td>0.9</td><td>6.5</td><td>6.3E-11</td><td>2.2E-10</td><td>BT20/MB468</td><td>MITO</td></tr>
<tr><td>>hg38_21_+_8433172_8446622_RNA45SN1_45S_50ntFlanks@10396.10419.24|CAGTTGAACATGGGTCAGTCGGTC</td><td>>hg38_21_+_8433172_8446622_RNA45SN1_45S_50ntFlanks</td><td>10396</td><td>10419</td><td>24</td><td>CAGTTGAACATGGGTCAGTCGGTC</td><td>111.2</td><td>-10.7</td><td>1.6</td><td>-6.5</td><td>8.1E-11</td><td>2.2E-10</td><td>MB231/MB468</td><td>MP</td></tr>
<tr><td>>hg38_21_+_8433172_8446622_RNA45SN1_45S_50ntFlanks@10386.10409.24|ATGTGAACAGCAGTTGAACATGGG</td><td>>hg38_21_+_8433172_8446622_RNA45SN1_45S_50ntFlanks</td><td>10386</td><td>10409</td><td>24</td><td>ATGTGAACAGCAGTTGAACATGGG</td><td>441.5</td><td>-12.7</td><td>1.9</td><td>-6.5</td><td>8.1E-11</td><td>2.2E-10</td><td>MB231/MB468</td><td>MP</td></tr>
<tr><td>>hg38_21_+_8433172_8446622_RNA45SN1_45S_50ntFlanks@5476.5516.41|CCCTGGCGGAGCGCTGAGAAGACGGTCGAACTTGACTATCT</td><td>>hg38_21_+_8433172_8446622_RNA45SN1_45S_50ntFlanks</td><td>5476</td><td>5516</td><td>41</td><td>CCCTGGCGGAGCGCTGAGAAGACGGTCGAACTTGACTATCT</td><td>175.2</td><td>11.2</td><td>1.6</td><td>6.9</td><td>6.3E-12</td><td>2.2E-10</td><td>BT20/MB468</td><td>MP</td></tr>
<tr><td>>hg38_21_+_8433172_8446622_RNA45SN1_45S_50ntFlanks@11874.11905.32|AGAAGACCCTGTTGAGCTTGACTCTAGTCTGG</td><td>>hg38_21_+_8433172_8446622_RNA45SN1_45S_50ntFlanks</td><td>11874</td><td>11905</td><td>32</td><td>AGAAGACCCTGTTGAGCTTGACTCTAGTCTGG</td><td>209.1</td><td>-7.0</td><td>1.1</td><td>-6.5</td><td>7.7E-11</td><td>2.2E-10</td><td>BT20/MB468</td><td>CYTO</td></tr>
<tr><td>>hg38_21_+_8433172_8446622_RNA45SN1_45S_50ntFlanks@11210.11232.23|GGCGACTCTGGACGCGAGCCGGG</td><td>>hg38_21_+_8433172_8446622_RNA45SN1_45S_50ntFlanks</td><td>11210</td><td>11232</td><td>23</td><td>GGCGACTCTGGACGCGAGCCGGG</td><td>1533.0</td><td>-9.2</td><td>1.4</td><td>-6.5</td><td>8.5E-11</td><td>2.2E-10</td><td>MB231/MB468</td><td>NUC</td></tr>
<tr><td>>hg38_21_+_8433172_8446622_RNA45SN1_45S_50ntFlanks@11768.11792.25|CTAATTAGTGACGCGCATGAATGGA</td><td>>hg38_21_+_8433172_8446622_RNA45SN1_45S_50ntFlanks</td><td>11768</td><td>11792</td><td>25</td><td>CTAATTAGTGACGCGCATGAATGGA</td><td>417.3</td><td>-4.2</td><td>0.6</td><td>-6.5</td><td>6.3E-11</td><td>2.2E-10</td><td>BT20/MB468</td><td>MITO</td></tr>
<tr><td>>hg38_21_+_8433172_8446622_RNA45SN1_45S_50ntFlanks@12478.12498.21|ACTAATAGGGAACGTGAGCTG</td><td>>hg38_21_+_8433172_8446622_RNA45SN1_45S_50ntFlanks</td><td>12478</td><td>12498</td><td>21</td><td>ACTAATAGGGAACGTGAGCTG</td><td>1742.3</td><td>-7.8</td><td>1.2</td><td>-6.5</td><td>1.0E-10</td><td>2.2E-10</td><td>BT20/MB468</td><td>NUC</td></tr>
<tr><td>>hg38_21_+_8433172_8446622_RNA45SN1_45S_50ntFlanks@4919.4939.21|CCACCAGGAGTGGAGCCTGCG</td><td>>hg38_21_+_8433172_8446622_RNA45SN1_45S_50ntFlanks</td><td>4919</td><td>4939</td><td>21</td><td>CCACCAGGAGTGGAGCCTGCG</td><td>384.6</td><td>-8.1</td><td>1.2</td><td>-6.5</td><td>8.1E-11</td><td>2.2E-10</td><td>MB231/MB468</td><td>MP</td></tr>
<tr><td>>hg38_21_+_8433172_8446622_RNA45SN1_45S_50ntFlanks@9858.9898.41|CGGGATGAACCGAACGCCGGGTTAAGGCGCCCGATGCCGAC</td><td>>hg38_21_+_8433172_8446622_RNA45SN1_45S_50ntFlanks</td><td>9858</td><td>9898</td><td>41</td><td>CGGGATGAACCGAACGCCGGGTTAAGGCGCCCGATGCCGAC</td><td>302.9</td><td>-12.0</td><td>1.9</td><td>-6.5</td><td>9.8E-11</td><td>2.2E-10</td><td>MB231/MB468</td><td>MITO</td></tr>
<tr><td>>hg38_MT_+_1621_3279_RNR2_16S_50ntFlanks@621.652.32|ATATAACTGAACTCCTCACACCCAATTGGACC</td><td>>hg38_MT_+_1621_3279_RNR2_16S_50ntFlanks</td><td>621</td><td>652</td><td>32</td><td>ATATAACTGAACTCCTCACACCCAATTGGACC</td><td>357.0</td><td>9.9</td><td>1.5</td><td>6.5</td><td>6.3E-11</td><td>2.2E-10</td><td>BT20/MB468</td><td>MITO</td></tr>
<tr><td>>MI0000298|hsa-mir-221&WithFlank&X|-|45746151|45746272@31.53.23&[MIMAT0004568&hsa-miR-221-5p&offsets|0|+1;m-405&X|-|45746220|45746242&offsets|0|0]|ACCTGGCATACAATGTAGATTTC</td><td>>MI0000298|hsa-mir-221&WithFlank&X|-|45746151|45746272</td><td>31</td><td>53</td><td>23</td><td>ACCTGGCATACAATGTAGATTTC</td><td>530.5</td><td>5.9</td><td>0.9</td><td>6.5</td><td>9.8E-11</td><td>2.2E-10</td><td>MB231/MB468</td><td>MITO</td></tr>
</table>

| | | | | | | | | | | | | | |
|---|---|---|---|---|---|---|---|---|---|---|---|---|---|
| >hg38_21_+_8433172_8446622_RNA45SN1_45S_50ntFlanks@11874.11893.20\|AGAAGACCCTGTTGAGCTTG | >hg38_21_+_8433172_8446622_RNA45SN1_45S_50ntFlanks | 11874 | 11893 | 20 | AGAAGACCCTGTTGAGCTTG | 1619.0 | -4.0 | 0.6 | -6.5 | 6.3E-11 | 2.2E-10 | BT20/MB468 | MITO |
| >hg38_21_+_8433172_8446622_RNA45SN1_45S_50ntFlanks@4723.4755.33\|CATTAATCAAGAACGAAAGTCGGAGGTTCGAAG | >hg38_21_+_8433172_8446622_RNA45SN1_45S_50ntFlanks | 4723 | 4755 | 33 | CATTAATCAAGAACGAAAGTCGGAGGTTCGAAG | 118.7 | -9.0 | 1.4 | -6.5 | 6.3E-11 | 2.2E-10 | BT20/MB468 | MITO |
| >MI0000102\|hsa-mir-100&WithFlank&11\|-\|122152223\|122152314@19.39.21(+1U)&[MIMAT0000098&hsa-miR-100-5p&offsets\|0\|-1(+1U);m-27&11\|-\|122152276\|122152296&offsets\|0\|0(+1U)]\|AACCCGTAGATCCGAACTTGTT | >MI0000102\|hsa-mir-100&WithFlank&11\|-\|122152223\|122152314 | 19 | 39 | 21(+1U) | AACCCGTAGATCCGAACTTGTT | 6359.6 | 11.2 | 1.7 | 6.5 | 8.1E-11 | 2.2E-10 | MB231/MB468 | MP |
| >hg38_21_+_8433172_8446622_RNA45SN1_45S_50ntFlanks@10345.10382.38\|TTCAAACGAGAACTTTGAAGGCCGAAGTGGAGAAGGGT | >hg38_21_+_8433172_8446622_RNA45SN1_45S_50ntFlanks | 10345 | 10382 | 38 | TTCAAACGAGAACTTTGAAGGCCGAAGTGGAGAAGGGT | 528.0 | 6.7 | 1.0 | 6.5 | 8.6E-11 | 2.3E-10 | MB231/MB468 | CYTO |
| >hg38_21_+_8433172_8446622_RNA45SN1_45S_50ntFlanks@9930.9961.32\|ATATAGACAGCAGGACGGTGGCCATGGAAGTC | >hg38_21_+_8433172_8446622_RNA45SN1_45S_50ntFlanks | 9930 | 9961 | 32 | ATATAGACAGCAGGACGGTGGCCATGGAAGTC | 108.3 | -10.6 | 1.6 | -6.5 | 8.2E-11 | 2.3E-10 | MB231/MB468 | MP |
| >MI0000077\|hsa-mir-21&WithFlank&17\|+\|59841260\|59841343@14.34.21(+1U)&[MIMAT0000076&hsa-miR-21-5p&offsets\|0\|-1(+1U);m-1&17\|+\|59841273\|59841294&offsets\|0\|-1(+1U)]\|TAGCTTATCAGACTGATGTTGT | >MI0000077\|hsa-mir-21&WithFlank&17\|+\|59841260\|59841343 | 14 | 34 | 21(+1U) | TAGCTTATCAGACTGATGTTGT | 731.2 | 6.3 | 1.0 | 6.5 | 6.4E-11 | 2.3E-10 | BT20/MB468 | MITO |
| >hg38_21_+_8433172_8446622_RNA45SN1_45S_50ntFlanks@9738.9758.21\|GAAACGATCTCAACCTATTCT | >hg38_21_+_8433172_8446622_RNA45SN1_45S_50ntFlanks | 9738 | 9758 | 21 | GAAACGATCTCAACCTATTCT | 2019.5 | -4.5 | 0.7 | -6.5 | 8.6E-11 | 2.3E-10 | MB231/MB468 | NUC |
| >hg38_21_+_8433172_8446622_RNA45SN1_45S_50ntFlanks@8169.8212.44\|CCAAGTCCTTCTGATCGAGGCCCAGCCCGTGGACGGTGTGAGGC | >hg38_21_+_8433172_8446622_RNA45SN1_45S_50ntFlanks | 8169 | 8212 | 44 | CCAAGTCCTTCTGATCGAGGCCCAGCCCGTGGACGGTGTGAGGC | 763.4 | -8.6 | 1.3 | -6.5 | 9.9E-11 | 2.3E-10 | MB231/MB468 | MITO |
| >hg38_21_+_8433172_8446622_RNA45SN1_45S_50ntFlanks@5073.5112.40\|AATTCCGATAACGAACGAGACTCTGGCATGCTAACTAGTT | >hg38_21_+_8433172_8446622_RNA45SN1_45S_50ntFlanks | 5073 | 5112 | 40 | AATTCCGATAACGAACGAGACTCTGGCATGCTAACTAGTT | 374.2 | -6.5 | 1.0 | -6.5 | 9.9E-11 | 2.3E-10 | MB231/MB468 | MITO |
| >MI0000746\|hsa-mir-99b&WithFlank&19\|+\|51692606\|51692687@51.70.20(+1U)&[MIMAT0004678&hsa-miR-99b-3p&offsets\|0\|-2(+1U);m-151&19\|+\|51692656\|51692678&offsets\|0\|-3(+1U)]\|CAAGCTCGTGTCTGTGGGTCT | >MI0000746\|hsa-mir-99b&WithFlank&19\|+\|51692606\|51692687 | 51 | 70 | 20(+1U) | CAAGCTCGTGTCTGTGGGTCT | 65.8 | 3.4 | 0.5 | 6.5 | 1.0E-10 | 2.3E-10 | BT20/MB468 | NUC |
| >hg38_21_+_8433172_8446622_RNA45SN1_45S_50ntFlanks@10771.10796.26\|CAGGTCTCCAAGGTGAACAGCCTCTG | >hg38_21_+_8433172_8446622_RNA45SN1_45S_50ntFlanks | 10771 | 10796 | 26 | CAGGTCTCCAAGGTGAACAGCCTCTG | 278.9 | -3.4 | 0.5 | -6.5 | 6.4E-11 | 2.3E-10 | BT20/MB468 | MITO |
| >hg38_21_+_8433172_8446622_RNA45SN1_45S_50ntFlanks@4738.4759.22\|AAAGTCGGAGGTTCGAAGACGA | >hg38_21_+_8433172_8446622_RNA45SN1_45S_50ntFlanks | 4738 | 4759 | 22 | AAAGTCGGAGGTTCGAAGACGA | 644.3 | -5.4 | 0.8 | -6.5 | 9.9E-11 | 2.3E-10 | MB231/MB468 | MITO |
| >hg38_21_+_8433172_8446622_RNA45SN1_45S_50ntFlanks@8159.8178.20\|CGTGGGGGGCCCAAGTCCTT | >hg38_21_+_8433172_8446622_RNA45SN1_45S_50ntFlanks | 8159 | 8178 | 20 | CGTGGGGGGCCCAAGTCCTT | 777.3 | -9.5 | 1.5 | -6.5 | 9.9E-11 | 2.3E-10 | MB231/MB468 | MITO |
| >hg38_21_+_8433172_8446622_RNA45SN1_45S_50ntFlanks@6721.6738.18\|AATTGCAGGACACATTGA | >hg38_21_+_8433172_8446622_RNA45SN1_45S_50ntFlanks | 6721 | 6738 | 18 | AATTGCAGGACACATTGA | 174.4 | -9.8 | 1.5 | -6.5 | 7.8E-11 | 2.3E-10 | BT20/MB468 | CYTO |
| >hg38_21_+_8433172_8446622_RNA45SN1_45S_50ntFlanks@9760.9786.27\|AAACTTTAAATGGGTAAGAAGCCCGGC | >hg38_21_+_8433172_8446622_RNA45SN1_45S_50ntFlanks | 9760 | 9786 | 27 | AAACTTTAAATGGGTAAGAAGCCCGGC | 377.6 | -12.4 | 1.9 | -6.5 | 8.2E-11 | 2.3E-10 | MB231/MB468 | MP |
| >hg38_21_+_8433172_8446622_RNA45SN1_45S_50ntFlanks@5473.5515.43\|CGGCCCTGGCGGAGCGCTGAGAAGACGGTCGAACTTGACTATC | >hg38_21_+_8433172_8446622_RNA45SN1_45S_50ntFlanks | 5473 | 5515 | 43 | CGGCCCTGGCGGAGCGCTGAGAAGACGGTCGAACTTGACTATC | 881.5 | 6.1 | 0.9 | 6.5 | 8.7E-11 | 2.3E-10 | MB231/MB468 | CYTO |
| >hg38_21_+_8433172_8446622_RNA45SN1_45S_50ntFlanks@3845.3866.22\|ACTTGGATAACTGTGGTAATTC | >hg38_21_+_8433172_8446622_RNA45SN1_45S_50ntFlanks | 3845 | 3866 | 22 | ACTTGGATAACTGTGGTAATTC | 306.1 | 4.3 | 0.7 | 6.5 | 8.8E-11 | 2.3E-10 | MB231/MB468 | CYTO |
| >trna152_ValCAC_6_-_27248049_27248121@1.33.33\|GCTTCTGTAGTGTAGTGGTTATCACGTTCGCCT | >trna152_ValCAC_6_-_27248049_27248121 | 1 | 33 | 33 | GCTTCTGTAGTGTAGTGGTTATCACGTTCGCCT | 1273.0 | 9.5 | 1.5 | 6.5 | 8.8E-11 | 2.3E-10 | MB231/MB468 | NUC |
| >hg38_21_+_8433172_8446622_RNA45SN1_45S_50ntFlanks@10234.10273.40\|CGGACGCTACGCCGCGACGAGTAGGAGGGCCGCTGCGGTG | >hg38_21_+_8433172_8446622_RNA45SN1_45S_50ntFlanks | 10234 | 10273 | 40 | CGGACGCTACGCCGCGACGAGTAGGAGGGCCGCTGCGGTG | 455.5 | -6.7 | 1.0 | -6.5 | 1.0E-10 | 2.3E-10 | MB231/MB468 | MITO |
| >hg38_21_+_8433172_8446622_RNA45SN1_45S_50ntFlanks@6650.6689.40\|ACGACTCTTAGCGGTGGATCACTCGGCTCGTGCGTCGATG | >hg38_21_+_8433172_8446622_RNA45SN1_45S_50ntFlanks | 6650 | 6689 | 40 | ACGACTCTTAGCGGTGGATCACTCGGCTCGTGCGTCGATG | 15416.4 | -4.4 | 0.7 | -6.5 | 1.0E-10 | 2.3E-10 | MB231/MB468 | MITO |
| >MI0000061\|hsa-let-7a-2&WithFlank&11\|-\|122146516\|122146599@11.32.22(+1A);MI0000062\|hsa-let-7a-3&WithFlank&22\|+\|46112743\|46112828@10.31.22(+1A);MI0000060\|hsa-let-7a-1&WithFlank&9\|+\|94175951\|94176042@12.33.22(+1A)&[MIMAT0000062&hsa-let-7a-5p&offsets\|0\|0(+1A);m-6&11\|-\|122146568\|122146589&offsets\|0\|0(+1A)];[MIMAT0000062_1&hsa-let-7a-5p&offsets\|0\|0(+1A);m-5&22\|+\|46112752\|46112773&offsets\|0\|0(+1A)];[MIMAT0000062_2&hsa-let-7a-5p&offsets\|0\|0(+1A);m-4&9\|+\|94175962\|94175983&offsets\|0\|0(+1A)]\|TGAGGTAGTAGGTTGTATAGTTA | >MI0000061\|hsa-let-7a-2&WithFlank&11\|-\|122146516\|122146599 | 11 | 32 | 22(+1A) | TGAGGTAGTAGGTTGTATAGTTA | 1089.5 | 5.5 | 0.8 | 6.5 | 8.8E-11 | 2.3E-10 | MB231/MB468 | CYTO |
| >hg38_21_+_8433172_8446622_RNA45SN1_45S_50ntFlanks@12249.12273.25\|CAAAAGCTCGCTTGATCTTGATTTT | >hg38_21_+_8433172_8446622_RNA45SN1_45S_50ntFlanks | 12249 | 12273 | 25 | CAAAAGCTCGCTTGATCTTGATTTT | 489.2 | -11.0 | 1.7 | -6.5 | 6.5E-11 | 2.3E-10 | BT20/MB468 | MITO |
| >hg38_21_+_8433172_8446622_RNA45SN1_45S_50ntFlanks@3793.3814.22\|CGAATGGCTCATTAAATCAGTT | >hg38_21_+_8433172_8446622_RNA45SN1_45S_50ntFlanks | 3793 | 3814 | 22 | CGAATGGCTCATTAAATCAGTT | 185.7 | -10.6 | 1.5 | -6.9 | 6.5E-12 | 2.3E-10 | BT20/MB468 | MP |
| >hg38_21_+_8433172_8446622_RNA45SN1_45S_50ntFlanks@5445.5467.23\|TCGGATCGGCCCCGCCGGGGTCG | >hg38_21_+_8433172_8446622_RNA45SN1_45S_50ntFlanks | 5445 | 5467 | 23 | TCGGATCGGCCCCGCCGGGGTCG | 336.5 | -8.4 | 1.3 | -6.5 | 1.1E-10 | 2.3E-10 | BT20/MB468 | NUC |
| >MI0002468\|hsa-mir-484&WithFlank&16\|+\|15643288\|15643378@14.35.22(+1U)&[MIMAT0002174&hsa-miR-484&offsets\|0\|0(+1U);m-127&16\|+\|15643301\|15643321&offsets\|0\|+1(+1U)]\|TCAGGCTCAGTCCCCTCCCGATT | >MI0002468\|hsa-mir-484&WithFlank&16\|+\|15643288\|15643378 | 14 | 35 | 22(+1U) | TCAGGCTCAGTCCCCTCCCGATT | 433.8 | 5.2 | 0.8 | 6.5 | 8.9E-11 | 2.3E-10 | MB231/MB468 | CYTO |
| >hg38_21_+_8433172_8446622_RNA45SN1_45S_50ntFlanks@10404.10435.32\|CATGGGTCAGTCGGTCCTGAGAGATGGGCGAG | >hg38_21_+_8433172_8446622_RNA45SN1_45S_50ntFlanks | 10404 | 10435 | 32 | CATGGGTCAGTCGGTCCTGAGAGATGGGCGAG | 424.1 | -8.1 | 1.3 | -6.5 | 8.9E-11 | 2.3E-10 | MB231/MB468 | NUC |
| >hg38_21_+_8433172_8446622_RNA45SN1_45S_50ntFlanks@12942.12970.29\|TTCGTAGACGACCTGCTTCTGGGTCGGGG | >hg38_21_+_8433172_8446622_RNA45SN1_45S_50ntFlanks | 12942 | 12970 | 29 | TTCGTAGACGACCTGCTTCTGGGTCGGGG | 137.6 | -11.0 | 1.7 | -6.5 | 8.4E-11 | 2.3E-10 | MB231/MB468 | MP |
| >hg38_21_+_8433172_8446622_RNA45SN1_45S_50ntFlanks@12458.12488.31\|AAGCGTTGGATTGTTCACCCACTAATAGGGA | >hg38_21_+_8433172_8446622_RNA45SN1_45S_50ntFlanks | 12458 | 12488 | 31 | AAGCGTTGGATTGTTCACCCACTAATAGGGA | 351.8 | -8.4 | 1.3 | -6.5 | 1.1E-10 | 2.3E-10 | BT20/MB468 | NUC |
| >hg38_21_+_8433172_8446622_RNA45SN1_45S_50ntFlanks@12643.12661.19\|ATCTGTGGGATTATGACTG | >hg38_21_+_8433172_8446622_RNA45SN1_45S_50ntFlanks | 12643 | 12661 | 19 | ATCTGTGGGATTATGACTG | 123.6 | -9.3 | 1.4 | -6.5 | 8.0E-11 | 2.3E-10 | BT20/MB468 | CYTO |
| >MI0000285\|hsa-mir-205&WithFlank&1\|+\|209432127\|209432248@40.63.24&[MIMAT0000266&hsa-miR-205-5p&offsets\|0\|+2;m-351&1\|+\|209432166\|209432187&offsets\|0\|+2]\|TCCTTCATTCCACCGGAGTCTGTC | >MI0000285\|hsa-mir-205&WithFlank&1\|+\|209432127\|209432248 | 40 | 63 | 24 | TCCTTCATTCCACCGGAGTCTGTC | 590.9 | 7.6 | 1.2 | 6.5 | 8.0E-11 | 2.3E-10 | BT20/MB468 | CYTO |
| >MI0000542\|hsa-mir-320a&WithFlank&8\|-\|22244960\|22245043@48.69.22(+1U)&[MIMAT0000510&hsa-miR-320a-3p&offsets\|0\|0(+1U);m-60&8\|-\|22244975\|22244996&offsets\|0\|0(+1U)]\|AAAAGCTGGGTTGAGAGGGCGAT | >MI0000542\|hsa-mir-320a&WithFlank&8\|-\|22244960\|22245043 | 48 | 69 | 22(+1U) | AAAAGCTGGGTTGAGAGGGCGAT | 3667.9 | 5.7 | 0.9 | 6.5 | 8.9E-11 | 2.3E-10 | MB231/MB468 | NUC |
| >trna119_LysCTT_1_-_145395522_145395594@1.31.31,trna11_LysCTT_5_-_180648979_180649051@1.31.31,trna13_LysCTT_6_+_26556774_26556846@1.31.31,trna32_LysCTT_16_-_3207406_3207478@1.31.31,trna7_LysCTT_16_+_3225692_3225764@1.31.31,trna9_LysCTT_5_+_180634755_180634827@1.31.31\|GCCCGGCTAGCTCAGTCGGTAGAGCATGAGA | >trna119_LysCTT_1_-_145395522_145395594 | 1 | 31 | 31 | GCCCGGCTAGCTCAGTCGGTAGAGCATGAGA | 11059.7 | -4.5 | 0.7 | -6.5 | 8.0E-11 | 2.3E-10 | BT20/MB468 | CYTO |
| >hg38_21_+_8433172_8446622_RNA45SN1_45S_50ntFlanks@9738.9759.22\|GAAACGATCTCAACCTATTCTC | >hg38_21_+_8433172_8446622_RNA45SN1_45S_50ntFlanks | 9738 | 9759 | 22 | GAAACGATCTCAACCTATTCTC | 902.1 | -4.4 | 0.7 | -6.5 | 9.0E-11 | 2.3E-10 | MB231/MB468 | CYTO |
| >trna130_GlnTTG_6_-_27763640_27763711@1.29.29,trna173_GlnTTG_6_-_26311975_26312046@1.29.29,trna174_GlnTTG_6_-_26311424_26311495@1.29.29\|GGCCCCATGGTGTAATGGTTAGCACTCTG | >trna130_GlnTTG_6_-_27763640_27763711 | 1 | 29 | 29 | GGCCCCATGGTGTAATGGTTAGCACTCTG | 122.0 | -10.8 | 1.7 | -6.5 | 8.5E-11 | 2.3E-10 | MB231/MB468 | MP |
| >hg38_21_+_8433172_8446622_RNA45SN1_45S_50ntFlanks@5472.5509.38\|ACGGCCCTGGCGGAGCGCTGAGAAGACGGTCGAACTTG | >hg38_21_+_8433172_8446622_RNA45SN1_45S_50ntFlanks | 5472 | 5509 | 38 | ACGGCCCTGGCGGAGCGCTGAGAAGACGGTCGAACTTG | 4677.9 | 5.5 | 0.9 | 6.5 | 9.0E-11 | 2.3E-10 | MB231/MB468 | CYTO |
| >hg38_21_+_8433172_8446622_RNA45SN1_45S_50ntFlanks@10351.10376.26\|CGAGAACTTTGAAGGCCGAAGTGGAG | >hg38_21_+_8433172_8446622_RNA45SN1_45S_50ntFlanks | 10351 | 10376 | 26 | CGAGAACTTTGAAGGCCGAAGTGGAG | 1205.9 | -4.5 | 0.7 | -6.5 | 8.0E-11 | 2.3E-10 | BT20/MB468 | CYTO |
| >hg38_21_+_8433172_8446622_RNA45SN1_45S_50ntFlanks@11588.11611.24\|GCGGACCAGGGGAATCCGACTGTT | >hg38_21_+_8433172_8446622_RNA45SN1_45S_50ntFlanks | 11588 | 11611 | 24 | GCGGACCAGGGGAATCCGACTGTT | 664.7 | 3.9 | 0.6 | 6.5 | 6.6E-11 | 2.4E-10 | BT20/MB468 | MITO |
| >hg38_21_+_8433172_8446622_RNA45SN1_45S_50ntFlanks@12072.12110.39\|CCCGAGGGGCTCTCGCTTCTGGCGCCAAGCGCCCGGCCG | >hg38_21_+_8433172_8446622_RNA45SN1_45S_50ntFlanks | 12072 | 12110 | 39 | CCCGAGGGGCTCTCGCTTCTGGCGCCAAGCGCCCGGCCG | 400.7 | -8.2 | 1.3 | -6.5 | 1.0E-10 | 2.4E-10 | MB231/MB468 | MITO |
| >hg38_21_+_8433172_8446622_RNA45SN1_45S_50ntFlanks@7984.8006.23\|AGATCAGACGTGGCGACCCGCTG | >hg38_21_+_8433172_8446622_RNA45SN1_45S_50ntFlanks | 7984 | 8006 | 23 | AGATCAGACGTGGCGACCCGCTG | 1295.8 | -6.4 | 1.0 | -6.5 | 9.1E-11 | 2.4E-10 | MB231/MB468 | CYTO |
| >am_trna129_MetCAT_6_-_27870271_27870342@34.51.18,trna142_MetCAT_6_-_27560600_27560671@34.51.18,trna150_MetCAT_6_-_27300764_27300835@34.51.18,trna169_MetCAT_6_-_26330529_26330600@34.51.18,trna171_MetCAT_6_-_26313352_26313423@34.51.18,trna20_MetCAT_17_-_80452597_80452668@34.51.18,trna2_MetCAT_6_+_26286754_26286825@34.51.18,trna32_MetCAT_1_+_153643726_153643797@34.51.18,trna61_MetCAT_6_+_27745664_27745735@34.51.18\|ATAACCCAGAGGTCGATG | >am_trna129_MetCAT_6_-_27870271_27870342 | 34 | 51 | 18 | ATAACCCAGAGGTCGATG | 406.5 | -11.7 | 1.7 | -6.9 | 6.7E-12 | 2.4E-10 | BT20/MB468 | MP |
| >trna111_HisGTG_1_-_147774845_147774916@1.32.32,trna118_HisGTG_1_-_145396881_145396952@1.32.32,trna16_HisGTG_1_+_146544773_146544844@1.32.32,trna1_HisGTG_15_+_45493349_45493420@1.32.32,trna21_HisGTG_1_+_147753471_147753542@1.32.32,trna33_HisGTG_6_+_27125906_27125977@1.32.32,trna7_HisGTG_9_-_14433938_14434009@1.32.32,trna8_HisGTG_15_-_45492611_45492682@1.32.32,trna9_HisGTG_15_-_45490804_45490875@1.32.32\|GCCGTGATCGTATAGTGGTTAGTACTCTGCGT | >trna111_HisGTG_1_-_147774845_147774916 | 1 | 32 | 32 | GCCGTGATCGTATAGTGGTTAGTACTCTGCGT | 731.4 | 10.5 | 1.6 | 6.5 | 6.7E-11 | 2.4E-10 | BT20/MB468 | MITO |

<table>
<tr><td></td><td></td><td></td><td></td><td></td><td></td><td></td><td></td><td></td><td></td><td></td><td></td><td></td><td></td></tr>
<tr><td>>hg38_21_+_8433172_8446622_RNA45SN1_45S_50ntFlanks@4683.4707.25|CAAGACGGACCAGAGCGAAAGCATT</td><td>>hg38_21_+_8433172_8446622_RNA45SN1_45S_50ntFlanks</td><td>4683</td><td>4707</td><td>25</td><td>CAAGACGGACCAGAGCGAAAGCATT</td><td>850.9</td><td>-8.7</td><td>1.3</td><td>-6.5</td><td>8.6E-11</td><td>2.4E-10</td><td>MB231/MB468</td><td>MP</td></tr>
<tr><td>>hg38_21_+_8433172_8446622_RNA45SN1_45S_50ntFlanks@9602.9622.21|CGAAAGACTAATCGAACCATC</td><td>>hg38_21_+_8433172_8446622_RNA45SN1_45S_50ntFlanks</td><td>9602</td><td>9622</td><td>21</td><td>CGAAAGACTAATCGAACCATC</td><td>588.3</td><td>-4.1</td><td>0.6</td><td>-6.5</td><td>9.1E-11</td><td>2.4E-10</td><td>MB231/MB468</td><td>CYTO</td></tr>
<tr><td>>MI0005756|hsa-mir-934&WithFlank&X|+|136550872|136550966@21.40.20(+1A)&[MIMAT0004977&hsa-miR-934&offsets|0|-2(+1A);m-943&X|+|136550892|136550913&offsets|0|-2(+1A)]|TGTCTACTACTGGAGACACTA</td><td>>MI0005756|hsa-mir-934&WithFlank&X|+|136550872|136550966</td><td>21</td><td>40</td><td>20(+1A)</td><td>TGTCTACTACTGGAGACACTA</td><td>147.4</td><td>-11.1</td><td>1.7</td><td>-6.5</td><td>8.6E-11</td><td>2.4E-10</td><td>MB231/MB468</td><td>MP</td></tr>
<tr><td>>hg38_21_+_8433172_8446622_RNA45SN1_45S_50ntFlanks@12628.12651.24|TGGGGCGAAGCTACCATCTGTGGG</td><td>>hg38_21_+_8433172_8446622_RNA45SN1_45S_50ntFlanks</td><td>12628</td><td>12651</td><td>24</td><td>TGGGGCGAAGCTACCATCTGTGGG</td><td>94.0</td><td>-10.4</td><td>1.6</td><td>-6.5</td><td>8.6E-11</td><td>2.4E-10</td><td>MB231/MB468</td><td>MP</td></tr>
<tr><td>>MI0000266|hsa-mir-10a&WithFlank&17|-|48579832|48579953@28.48.21&[MIMAT0000253&hsa-miR-10a-5p&offsets|0|-2;m-18&17|-|48579905|48579926&offsets|0|-1]|TACCCTGTAGATCCGAATTTG</td><td>>MI0000266|hsa-mir-10a&WithFlank&17|-|48579832|48579953</td><td>28</td><td>48</td><td>21</td><td>TACCCTGTAGATCCGAATTTG</td><td>502.6</td><td>12.7</td><td>1.9</td><td>6.9</td><td>6.7E-12</td><td>2.4E-10</td><td>BT20/MB468</td><td>MP</td></tr>
<tr><td>>hg38_21_+_8433172_8446622_RNA45SN1_45S_50ntFlanks@9534.9553.20|AGGAAACTCTGGTGGAGGTC</td><td>>hg38_21_+_8433172_8446622_RNA45SN1_45S_50ntFlanks</td><td>9534</td><td>9553</td><td>20</td><td>AGGAAACTCTGGTGGAGGTC</td><td>36034.9</td><td>-7.2</td><td>1.0</td><td>-6.9</td><td>6.7E-12</td><td>2.4E-10</td><td>BT20/MB468</td><td>MP</td></tr>
<tr><td>>hg38_21_+_8433172_8446622_RNA45SN1_45S_50ntFlanks@7986.8010.25|ATCAGACGTGGCGACCCGCTGAATT</td><td>>hg38_21_+_8433172_8446622_RNA45SN1_45S_50ntFlanks</td><td>7986</td><td>8010</td><td>25</td><td>ATCAGACGTGGCGACCCGCTGAATT</td><td>359.2</td><td>-11.6</td><td>1.7</td><td>-6.9</td><td>6.7E-12</td><td>2.4E-10</td><td>BT20/MB468</td><td>MP</td></tr>
<tr><td>>hg38_21_+_8433172_8446622_RNA45SN1_45S_50ntFlanks@6667.6698.32|ATCACTCGGCTCGTGCGTCGATGAAGAACGCA</td><td>>hg38_21_+_8433172_8446622_RNA45SN1_45S_50ntFlanks</td><td>6667</td><td>6698</td><td>32</td><td>ATCACTCGGCTCGTGCGTCGATGAAGAACGCA</td><td>69.8</td><td>-10.0</td><td>1.5</td><td>-6.5</td><td>8.6E-11</td><td>2.4E-10</td><td>MB231/MB468</td><td>MP</td></tr>
<tr><td>>hg38_21_+_8433172_8446622_RNA45SN1_45S_50ntFlanks@11667.11691.25|CTGCCCAGTGCTCTGAATGTCAAAG</td><td>>hg38_21_+_8433172_8446622_RNA45SN1_45S_50ntFlanks</td><td>11667</td><td>11691</td><td>25</td><td>CTGCCCAGTGCTCTGAATGTCAAAG</td><td>486.0</td><td>6.8</td><td>1.0</td><td>6.5</td><td>9.2E-11</td><td>2.4E-10</td><td>MB231/MB468</td><td>CYTO</td></tr>
<tr><td>>hg38_21_+_8433172_8446622_RNA45SN1_45S_50ntFlanks@12151.12169.19|GTGGGGAGTTTGACTGGGG</td><td>>hg38_21_+_8433172_8446622_RNA45SN1_45S_50ntFlanks</td><td>12151</td><td>12169</td><td>19</td><td>GTGGGGAGTTTGACTGGGG</td><td>88.4</td><td>-10.3</td><td>1.6</td><td>-6.5</td><td>8.6E-11</td><td>2.4E-10</td><td>MB231/MB468</td><td>MP</td></tr>
<tr><td>>am_MI0000809|hsa-mir-151a&WithFlank&8|-|140732558|140732659@53.73.21(+1U)&[MIMAT0000757&hsa-miR-151a-3p&offsets|0|0(+1U);m-30&8|-|140732587|140732607&offsets|0|0(+1U)]|CTAGACTGAAGCTCCTTGAGGT</td><td>>am_MI0000809|hsa-mir-151a&WithFlank&8|-|140732558|140732659</td><td>53</td><td>73</td><td>21(+1U)</td><td>CTAGACTGAAGCTCCTTGAGGT</td><td>7155.6</td><td>6.0</td><td>0.9</td><td>6.9</td><td>6.8E-12</td><td>2.4E-10</td><td>BT20/MB468</td><td>MP</td></tr>
<tr><td>>hg38_21_+_8433172_8446622_RNA45SN1_45S_50ntFlanks@10716.10734.19|GAAAATCCGGGGGAGAGGG</td><td>>hg38_21_+_8433172_8446622_RNA45SN1_45S_50ntFlanks</td><td>10716</td><td>10734</td><td>19</td><td>GAAAATCCGGGGGAGAGGG</td><td>115.8</td><td>-10.7</td><td>1.7</td><td>-6.5</td><td>8.7E-11</td><td>2.4E-10</td><td>MB231/MB468</td><td>MP</td></tr>
<tr><td>>MI0000066|hsa-let-7e&WithFlank&19|+|51692780|51692870@14.31.18&[MIMAT0000066&hsa-let-7e-5p&offsets|0|-4;m-50&19|+|51692793|51692814&offsets|0|-4]|TGAGGTAGGAGGTTGTAT</td><td>>MI0000066|hsa-let-7e&WithFlank&19|+|51692780|51692870</td><td>14</td><td>31</td><td>18</td><td>TGAGGTAGGAGGTTGTAT</td><td>1645.6</td><td>9.3</td><td>1.4</td><td>6.5</td><td>8.2E-11</td><td>2.4E-10</td><td>BT20/MB468</td><td>CYTO</td></tr>
<tr><td>>hg38_21_+_8433172_8446622_RNA45SN1_45S_50ntFlanks@6777.6807.31|TCCCGGGGCTACGCCTGTCTGAGCGTCGCTT</td><td>>hg38_21_+_8433172_8446622_RNA45SN1_45S_50ntFlanks</td><td>6777</td><td>6807</td><td>31</td><td>TCCCGGGGCTACGCCTGTCTGAGCGTCGCTT</td><td>480.6</td><td>4.5</td><td>0.7</td><td>6.5</td><td>6.7E-11</td><td>2.4E-10</td><td>BT20/MB468</td><td>MITO</td></tr>
<tr><td>>am_trna11_ProAGG_16_+_3241989_3242060@1.27.27,trna12_ProAGG_6_+_26555498_26555569@1.27.27,trna12_ProTGG_11_-_75946869_75946940@1.27.27,trna14_ProTGG_5_-_180615854_180615925@1.27.27,trna22_ProAGG_14_-_21081560_21081631@1.27.27,trna23_ProAGG_14_-_21077495_21077566@1.27.27,trna28_ProTGG_16_-_3234133_3234204@1.27.27,trna29_ProAGG_16_-_3232635_3232706@1.27.27,trna2_ProAGG_7_+_128423504_128423575@1.27.27,trna30_ProCGG_6_+_27059521_27059592@1.27.27,trna37_ProCGG_17_-_8126151_8126222@1.27.27,trna3_ProTGG_16_+_3208923_3208994@1.27.27,trna52_ProCGG_1_+_167683962_167684033@1.27.27,trna65_ProAGG_1_-_167684725_167684796@1.27.27,trna6_ProCGG_16_+_3222049_3222120@1.27.27,trna6_ProTGG_14_+_21152175_21152246@1.27.27,trna8_ProTGG_16_+_3238094_3238165@1.27.27,trna9_ProAGG_11_+_75946557_75946628@1.27.27,trna9_ProAGG_16_+_3239634_3239705@1.27.27|GGCTCGTTGGTCTAGGGGTATGATTCT</td><td>>am_trna11_ProAGG_16_+_3241989_3242060</td><td>1</td><td>27</td><td>27</td><td>GGCTCGTTGGTCTAGGGGTATGATTCT</td><td>702.1</td><td>-9.2</td><td>1.4</td><td>-6.5</td><td>9.2E-11</td><td>2.4E-10</td><td>MB231/MB468</td><td>NUC</td></tr>
<tr><td>>trna111_HisGTG_1_-_147774845_147774916@-1T.33.34,trna118_HisGTG_1_-_145396881_145396952@-1T.33.34,trna16_HisGTG_1_+_146544773_146544844@-1T.33.34,trna1_HisGTG_15_+_45493349_45493420@-1T.33.34,trna21_HisGTG_1_+_147753471_147753542@-1T.33.34,trna33_HisGTG_6_+_27125906_27125977@-1T.33.34,trna7_HisGTG_9_-_14433938_14434009@-1T.33.34,trna8_HisGTG_15_-_45492611_45492682@-1T.33.34,trna9_HisGTG_15_-_45490804_45490875@-1T.33.34|TGCCGTGATCGTATAGTGGTTAGTACTCTGCGTT</td><td>>trna111_HisGTG_1_-_147774845_147774916</td><td>-1T</td><td>33</td><td>34</td><td>TGCCGTGATCGTATAGTGGTTAGTACTCTGCGTT</td><td>1042.1</td><td>6.6</td><td>1.0</td><td>6.5</td><td>1.0E-10</td><td>2.4E-10</td><td>MB231/MB468</td><td>MITO</td></tr>
<tr><td>>hg38_21_+_8433172_8446622_RNA45SN1_45S_50ntFlanks@4734.4766.33|AACGAAAGTCGGAGGTTCGAAGACGATCAGATA</td><td>>hg38_21_+_8433172_8446622_RNA45SN1_45S_50ntFlanks</td><td>4734</td><td>4766</td><td>33</td><td>AACGAAAGTCGGAGGTTCGAAGACGATCAGATA</td><td>744.1</td><td>7.4</td><td>1.1</td><td>6.5</td><td>8.2E-11</td><td>2.4E-10</td><td>BT20/MB468</td><td>CYTO</td></tr>
<tr><td>>hg38_21_+_8433172_8446622_RNA45SN1_45S_50ntFlanks@9534.9555.22|AGGAAACTCTGGTGGAGGTCCG</td><td>>hg38_21_+_8433172_8446622_RNA45SN1_45S_50ntFlanks</td><td>9534</td><td>9555</td><td>22</td><td>AGGAAACTCTGGTGGAGGTCCG</td><td>223.4</td><td>-10.1</td><td>1.6</td><td>-6.5</td><td>8.2E-11</td><td>2.4E-10</td><td>BT20/MB468</td><td>CYTO</td></tr>
<tr><td>>hg38_21_+_8433172_8446622_RNA45SN1_45S_50ntFlanks@12033.12055.23|ATCGTTTTTTCACTGACCCGGTG</td><td>>hg38_21_+_8433172_8446622_RNA45SN1_45S_50ntFlanks</td><td>12033</td><td>12055</td><td>23</td><td>ATCGTTTTTTCACTGACCCGGTG</td><td>133.6</td><td>-9.2</td><td>1.4</td><td>-6.5</td><td>6.8E-11</td><td>2.4E-10</td><td>BT20/MB468</td><td>MITO</td></tr>
<tr><td>>hg38_21_+_8433172_8446622_RNA45SN1_45S_50ntFlanks@3782.3800.19|CAGTGAAACTGCGAATGGC</td><td>>hg38_21_+_8433172_8446622_RNA45SN1_45S_50ntFlanks</td><td>3782</td><td>3800</td><td>19</td><td>CAGTGAAACTGCGAATGGC</td><td>765.9</td><td>-5.2</td><td>0.8</td><td>-6.5</td><td>9.2E-11</td><td>2.4E-10</td><td>MB231/MB468</td><td>NUC</td></tr>
<tr><td>>hg38_21_+_8433172_8446622_RNA45SN1_45S_50ntFlanks@12983.13001.19|CAGAGCAGCTCCCTCGCTG</td><td>>hg38_21_+_8433172_8446622_RNA45SN1_45S_50ntFlanks</td><td>12983</td><td>13001</td><td>19</td><td>CAGAGCAGCTCCCTCGCTG</td><td>144.7</td><td>-9.5</td><td>1.5</td><td>-6.5</td><td>8.3E-11</td><td>2.4E-10</td><td>BT20/MB468</td><td>CYTO</td></tr>
<tr><td>>hg38_21_+_8433172_8446622_RNA45SN1_45S_50ntFlanks@11573.11593.21|CGACTTAGAACTGGTGCGGAC</td><td>>hg38_21_+_8433172_8446622_RNA45SN1_45S_50ntFlanks</td><td>11573</td><td>11593</td><td>21</td><td>CGACTTAGAACTGGTGCGGAC</td><td>804.4</td><td>-7.3</td><td>1.1</td><td>-6.5</td><td>1.1E-10</td><td>2.4E-10</td><td>BT20/MB468</td><td>NUC</td></tr>
<tr><td>>hg38_21_+_8433172_8446622_RNA45SN1_45S_50ntFlanks@12629.12654.26|GGGGCGAAGCTACCATCTGTGGGATT</td><td>>hg38_21_+_8433172_8446622_RNA45SN1_45S_50ntFlanks</td><td>12629</td><td>12654</td><td>26</td><td>GGGGCGAAGCTACCATCTGTGGGATT</td><td>94.1</td><td>-10.4</td><td>1.6</td><td>-6.5</td><td>8.7E-11</td><td>2.4E-10</td><td>MB231/MB468</td><td>MP</td></tr>
<tr><td>>hg38_21_+_8433172_8446622_RNA45SN1_45S_50ntFlanks@9806.9841.36|CGTGGAATGCGAGTGCCTAGTGGGCCACTTTTGGTA</td><td>>hg38_21_+_8433172_8446622_RNA45SN1_45S_50ntFlanks</td><td>9806</td><td>9841</td><td>36</td><td>CGTGGAATGCGAGTGCCTAGTGGGCCACTTTTGGTA</td><td>77.1</td><td>-10.1</td><td>1.6</td><td>-6.5</td><td>8.7E-11</td><td>2.4E-10</td><td>MB231/MB468</td><td>MP</td></tr>
<tr><td>>hg38_21_+_8433172_8446622_RNA45SN1_45S_50ntFlanks@9572.9603.32|CAAATCGGTCGTCCGACCTGGGTATAGGGGCG</td><td>>hg38_21_+_8433172_8446622_RNA45SN1_45S_50ntFlanks</td><td>9572</td><td>9603</td><td>32</td><td>CAAATCGGTCGTCCGACCTGGGTATAGGGGCG</td><td>1895.6</td><td>-4.0</td><td>0.6</td><td>-6.5</td><td>9.2E-11</td><td>2.4E-10</td><td>MB231/MB468</td><td>NUC</td></tr>
<tr><td>>hg38_21_+_8433172_8446622_RNA45SN1_45S_50ntFlanks@4922.4940.19|CCAGGAGTGGAGCCTGCGG</td><td>>hg38_21_+_8433172_8446622_RNA45SN1_45S_50ntFlanks</td><td>4922</td><td>4940</td><td>19</td><td>CCAGGAGTGGAGCCTGCGG</td><td>287.4</td><td>-7.9</td><td>1.2</td><td>-6.5</td><td>8.7E-11</td><td>2.4E-10</td><td>MB231/MB468</td><td>MP</td></tr>
<tr><td>>hg38_21_+_8433172_8446622_RNA45SN1_45S_50ntFlanks@11767.11794.28|TCTAATTAGTGACGCGCATGAATGGATG</td><td>>hg38_21_+_8433172_8446622_RNA45SN1_45S_50ntFlanks</td><td>11767</td><td>11794</td><td>28</td><td>TCTAATTAGTGACGCGCATGAATGGATG</td><td>1075.3</td><td>-2.5</td><td>0.4</td><td>-6.5</td><td>8.3E-11</td><td>2.4E-10</td><td>BT20/MB468</td><td>CYTO</td></tr>
<tr><td>>hg38_21_+_8433172_8446622_RNA45SN1_45S_50ntFlanks@10922.10942.21|CTGGACGAGGCGCCGCCGCCC</td><td>>hg38_21_+_8433172_8446622_RNA45SN1_45S_50ntFlanks</td><td>10922</td><td>10942</td><td>21</td><td>CTGGACGAGGCGCCGCCGCCC</td><td>105.5</td><td>-10.6</td><td>1.6</td><td>-6.5</td><td>8.7E-11</td><td>2.4E-10</td><td>MB231/MB468</td><td>MP</td></tr>
<tr><td>>MI0000285|hsa-mir-205&WithFlank&1|+|209432127|209432248@40.59.20(+1C)&[MIMAT0000266&hsa-miR-205-5p&offsets|0|-2(+1C);m-351&1|+|209432166|209432187&offsets|0|-2(+1C)]|TCCTTCATTCCACCGGAGTCC</td><td>>MI0000285|hsa-mir-205&WithFlank&1|+|209432127|209432248</td><td>40</td><td>59</td><td>20(+1C)</td><td>TCCTTCATTCCACCGGAGTCC</td><td>1029.6</td><td>6.6</td><td>1.0</td><td>6.5</td><td>6.8E-11</td><td>2.4E-10</td><td>BT20/MB468</td><td>MITO</td></tr>
<tr><td>>hg38_21_+_8433172_8446622_RNA45SN1_45S_50ntFlanks@11570.11592.23|AGCCGACTTAGAACTGGTGCGGA</td><td>>hg38_21_+_8433172_8446622_RNA45SN1_45S_50ntFlanks</td><td>11570</td><td>11592</td><td>23</td><td>AGCCGACTTAGAACTGGTGCGGA</td><td>238.4</td><td>-5.3</td><td>0.8</td><td>-6.5</td><td>9.3E-11</td><td>2.4E-10</td><td>MB231/MB468</td><td>CYTO</td></tr>
<tr><td>>hg38_21_+_8433172_8446622_RNA45SN1_45S_50ntFlanks@12686.12703.18|CCAGGCGGAACGATACGG</td><td>>hg38_21_+_8433172_8446622_RNA45SN1_45S_50ntFlanks</td><td>12686</td><td>12703</td><td>18</td><td>CCAGGCGGAACGATACGG</td><td>122.8</td><td>-4.9</td><td>0.8</td><td>-6.5</td><td>9.3E-11</td><td>2.4E-10</td><td>MB231/MB468</td><td>CYTO</td></tr>
<tr><td>>hg38_21_+_8433172_8446622_RNA45SN1_45S_50ntFlanks@3835.3871.37|CTCCTCTCCTACTTGGATAACTGTGGTAATTCTAGAG</td><td>>hg38_21_+_8433172_8446622_RNA45SN1_45S_50ntFlanks</td><td>3835</td><td>3871</td><td>37</td><td>CTCCTCTCCTACTTGGATAACTGTGGTAATTCTAGAG</td><td>2163.6</td><td>-10.4</td><td>1.6</td><td>-6.5</td><td>1.1E-10</td><td>2.4E-10</td><td>BT20/MB468</td><td>NUC</td></tr>
<tr><td>>hg38_21_+_8433172_8446622_RNA45SN1_45S_50ntFlanks@7993.8011.19|GTGGCGACCCGCTGAATTT</td><td>>hg38_21_+_8433172_8446622_RNA45SN1_45S_50ntFlanks</td><td>7993</td><td>8011</td><td>19</td><td>GTGGCGACCCGCTGAATTT</td><td>79.8</td><td>2.2</td><td>0.3</td><td>6.5</td><td>1.1E-10</td><td>2.4E-10</td><td>BT20/MB468</td><td>NUC</td></tr>
<tr><td>>hg38_21_+_8433172_8446622_RNA45SN1_45S_50ntFlanks@10885.10902.18|GGCTGGGGCGCGAAGCGG</td><td>>hg38_21_+_8433172_8446622_RNA45SN1_45S_50ntFlanks</td><td>10885</td><td>10902</td><td>18</td><td>GGCTGGGGCGCGAAGCGG</td><td>127.4</td><td>-5.9</td><td>0.9</td><td>-6.5</td><td>9.3E-11</td><td>2.4E-10</td><td>MB231/MB468</td><td>CYTO</td></tr>
<tr><td>>hg38_21_+_8433172_8446622_RNA45SN1_45S_50ntFlanks@10914.10935.22|CGCCGCGGCTGGACGAGGCGCC</td><td>>hg38_21_+_8433172_8446622_RNA45SN1_45S_50ntFlanks</td><td>10914</td><td>10935</td><td>22</td><td>CGCCGCGGCTGGACGAGGCGCC</td><td>250.7</td><td>-11.1</td><td>1.6</td><td>-6.9</td><td>6.9E-12</td><td>2.4E-10</td><td>BT20/MB468</td><td>MP</td></tr>
<tr><td>>MI0001445|hsa-mir-423&WithFlank&17|+|30117073|30117178@58.80.23(+1A)&[MIMAT0001340&hsa-miR-423-3p&offsets|-1|-1(+1A);m-100&17|+|30117131|30117153&offsets|-1|-1(+1A)]|AAGCTCGGTCTGAGGCCCCTCAGA</td><td>>MI0001445|hsa-mir-423&WithFlank&17|+|30117073|30117178</td><td>58</td><td>80</td><td>23(+1A)</td><td>AAGCTCGGTCTGAGGCCCCTCAGA</td><td>382.1</td><td>8.4</td><td>1.3</td><td>6.5</td><td>6.8E-11</td><td>2.4E-10</td><td>BT20/MB468</td><td>MITO</td></tr>
<tr><td>>hg38_21_+_8433172_8446622_RNA45SN1_45S_50ntFlanks@9383.9409.27|CTCGCCGGCCGAGGTGGGATCCCGAGG</td><td>>hg38_21_+_8433172_8446622_RNA45SN1_45S_50ntFlanks</td><td>9383</td><td>9409</td><td>27</td><td>CTCGCCGGCCGAGGTGGGATCCCGAGG</td><td>88.8</td><td>-10.3</td><td>1.6</td><td>-6.5</td><td>8.8E-11</td><td>2.4E-10</td><td>MB231/MB468</td><td>MP</td></tr>
<tr><td>>hg38_21_+_8433172_8446622_RNA45SN1_45S_50ntFlanks@4733.4759.27|GAACGAAAGTCGGAGGTTCGAAGACGA</td><td>>hg38_21_+_8433172_8446622_RNA45SN1_45S_50ntFlanks</td><td>4733</td><td>4759</td><td>27</td><td>GAACGAAAGTCGGAGGTTCGAAGACGA</td><td>916.2</td><td>-4.5</td><td>0.7</td><td>-6.5</td><td>8.8E-11</td><td>2.4E-10</td><td>MB231/MB468</td><td>MP</td></tr>
<tr><td>>hg38_21_+_8433172_8446622_RNA45SN1_45S_50ntFlanks@3873.3899.27|TAATACATGCCGACGGGCGCTGACCCC</td><td>>hg38_21_+_8433172_8446622_RNA45SN1_45S_50ntFlanks</td><td>3873</td><td>3899</td><td>27</td><td>TAATACATGCCGACGGGCGCTGACCCC</td><td>437.0</td><td>-3.6</td><td>0.6</td><td>-6.5</td><td>8.4E-11</td><td>2.4E-10</td><td>BT20/MB468</td><td>CYTO</td></tr>
<tr><td>>hg38_21_+_8433172_8446622_RNA45SN1_45S_50ntFlanks@3782.3800.19|CAGTGAAACTGCGAATGGC</td><td>>hg38_21_+_8433172_8446622_RNA45SN1_45S_50ntFlanks</td><td>3782</td><td>3800</td><td>19</td><td>CAGTGAAACTGCGAATGGC</td><td>1326.0</td><td>-7.5</td><td>1.2</td><td>-6.5</td><td>1.1E-10</td><td>2.4E-10</td><td>MB231/MB468</td><td>MITO</td></tr>
<tr><td>>MI0000441|hsa-mir-30b&WithFlank&8|-|134800514|134800613@23.44.22&[MIMAT0000420&hsa-miR-30b-5p&offsets|0|0;m-76&8|-|134800570|134800591&offsets|0|0]|TGTAAACATCCTACACTCAGCT</td><td>>MI0000441|hsa-mir-30b&WithFlank&8|-|134800514|134800613</td><td>23</td><td>44</td><td>22</td><td>TGTAAACATCCTACACTCAGCT</td><td>9792.0</td><td>8.1</td><td>1.3</td><td>6.5</td><td>1.1E-10</td><td>2.4E-10</td><td>MB231/MB468</td><td>MITO</td></tr>
<tr><td>>MI0001519|hsa-mir-20b&WithFlank&X|-|134169803|134169883@12.34.23&[MIMAT0001413&hsa-miR-20b-5p&offsets|0|0;m-241&X|-|134169850|134169872&offsets|0|0]|CAAAGTGCTCATAGTGCAGGTAG</td><td>>MI0001519|hsa-mir-20b&WithFlank&X|-|134169803|134169883</td><td>12</td><td>34</td><td>23</td><td>CAAAGTGCTCATAGTGCAGGTAG</td><td>81.2</td><td>-10.2</td><td>1.6</td><td>-6.5</td><td>8.8E-11</td><td>2.4E-10</td><td>MB231/MB468</td><td>MP</td></tr>
<tr><td>>hg38_21_+_8433172_8446622_RNA45SN1_45S_50ntFlanks@7985.8016.32|GATCAGACGTGGCGACCCGCTGAATTTAAGCA</td><td>>hg38_21_+_8433172_8446622_RNA45SN1_45S_50ntFlanks</td><td>7985</td><td>8016</td><td>32</td><td>GATCAGACGTGGCGACCCGCTGAATTTAAGCA</td><td>176.1</td><td>-9.6</td><td>1.5</td><td>-6.5</td><td>6.9E-11</td><td>2.4E-10</td><td>BT20/MB468</td><td>MITO</td></tr>
<tr><td>>hg38_21_+_8433172_8446622_RNA45SN1_45S_50ntFlanks@12046.12068.23|TGACCCGGTGAGGCGGGGGGGCG</td><td>>hg38_21_+_8433172_8446622_RNA45SN1_45S_50ntFlanks</td><td>12046</td><td>12068</td><td>23</td><td>TGACCCGGTGAGGCGGGGGGGCG</td><td>94.1</td><td>-10.4</td><td>1.6</td><td>-6.5</td><td>8.9E-11</td><td>2.4E-10</td><td>MB231/MB468</td><td>MP</td></tr>
</table>

| | | | | | | | | | | | | | |
|---|---|---|---|---|---|---|---|---|---|---|---|---|---|
| >hg38_21_+_8433172_8446622_RNA45SN1_45S_50ntFlanks@10411.10429.19\|CAGTCGGTCCTGAGAGATG | >hg38_21_+_8433172_8446622_RNA45SN1_45S_50ntFlanks | 10411 | 10429 | 19 | CAGTCGGTCCTGAGAGATG | 13418.4 | -5.8 | 0.9 | -6.5 | 9.4E-11 | 2.4E-10 | MB231/MB468 | CYTO |
| >hg38_21_+_8433172_8446622_RNA45SN1_45S_50ntFlanks@8799.8823.25\|CCTCGGGAGGGCGCGCGGGTCGGGG | >hg38_21_+_8433172_8446622_RNA45SN1_45S_50ntFlanks | 8799 | 8823 | 25 | CCTCGGGAGGGCGCGCGGGTCGGGG | 166.0 | -6.3 | 1.0 | -6.5 | 9.4E-11 | 2.4E-10 | MB231/MB468 | CYTO |
| >trna9_ArgTCT_1_+_94313129_94313213@1.29.29\|GGCTCCGTGGCGCAATGGATAGCGCATTG | >trna9_ArgTCT_1_+_94313129_94313213 | 1 | 29 | 29 | GGCTCCGTGGCGCAATGGATAGCGCATTG | 364.2 | -7.0 | 1.1 | -6.5 | 9.4E-11 | 2.4E-10 | MB231/MB468 | CYTO |
| >hg38_21_+_8433172_8446622_RNA45SN1_45S_50ntFlanks@6772.6803.32\|GTTCCTCCCGGGGCTACGCCTGTCTGAGCGTC | >hg38_21_+_8433172_8446622_RNA45SN1_45S_50ntFlanks | 6772 | 6803 | 32 | GTTCCTCCCGGGGCTACGCCTGTCTGAGCGTC | 193.2 | -7.4 | 1.1 | -6.5 | 9.4E-11 | 2.4E-10 | MB231/MB468 | NUC |
| >hg38_21_+_8433172_8446622_RNA45SN1_45S_50ntFlanks@10392.10425.34\|ACAGCAGTTGAACATGGGTCAGTCGGTCCTGAGA | >hg38_21_+_8433172_8446622_RNA45SN1_45S_50ntFlanks | 10392 | 10425 | 34 | ACAGCAGTTGAACATGGGTCAGTCGGTCCTGAGA | 443.9 | -8.8 | 1.4 | -6.4 | 1.1E-10 | 2.4E-10 | BT20/MB468 | NUC |
| >hg38_21_+_8433172_8446622_RNA45SN1_45S_50ntFlanks@4047.4064.18\|ATTCGAACGTCTGCCCTA | >hg38_21_+_8433172_8446622_RNA45SN1_45S_50ntFlanks | 4047 | 4064 | 18 | ATTCGAACGTCTGCCCTA | 298.1 | -6.3 | 1.0 | -6.5 | 9.4E-11 | 2.4E-10 | MB231/MB468 | NUC |
| >hg38_21_+_8433172_8446622_RNA45SN1_45S_50ntFlanks@8380.8402.23\|CAAGAGGGCGTGAAACCGTTAAG | >hg38_21_+_8433172_8446622_RNA45SN1_45S_50ntFlanks | 8380 | 8402 | 23 | CAAGAGGGCGTGAAACCGTTAAG | 624.5 | -8.1 | 1.3 | -6.5 | 9.4E-11 | 2.4E-10 | MB231/MB468 | NUC |
| >hg38_21_+_8433172_8446622_RNA45SN1_45S_50ntFlanks@8380.8400.21\|CAAGAGGGCGTGAAACCGTTA | >hg38_21_+_8433172_8446622_RNA45SN1_45S_50ntFlanks | 8380 | 8400 | 21 | CAAGAGGGCGTGAAACCGTTA | 1554.5 | -6.8 | 1.0 | -6.5 | 6.9E-11 | 2.4E-10 | BT20/MB468 | MITO |
| >hg38_21_+_8433172_8446622_RNA45SN1_45S_50ntFlanks@10574.10608.35\|AAGCCGGCGGGAGCCCCGGGGAGAGTTCTCTTTTC | >hg38_21_+_8433172_8446622_RNA45SN1_45S_50ntFlanks | 10574 | 10608 | 35 | AAGCCGGCGGGAGCCCCGGGGAGAGTTCTCTTTTC | 1249.9 | -8.7 | 1.3 | -6.5 | 9.4E-11 | 2.4E-10 | MB231/MB468 | NUC |
| >MI0000077\|hsa-mir-21&WithFlank&17\|+\|59841260\|59841343@14.35.22(+1U)&[MIMAT0000076&hsa-miR-21-5p&offsets\|0\|0(+1U);m-1&17\|+\|59841273\|59841294&offsets\|0\|0(+1U)]\|TAGCTTATCAGACTGATGTTGAT | >MI0000077\|hsa-mir-21&WithFlank&17\|+\|59841260\|59841343 | 14 | 35 | 22(+1U) | TAGCTTATCAGACTGATGTTGAT | 114544.2 | 6.8 | 1.0 | 6.5 | 1.1E-10 | 2.5E-10 | MB231/MB468 | MITO |
| >hg38_21_+_8433172_8446622_RNA45SN1_45S_50ntFlanks@11766.11786.21\|ATCTAATTAGTGACGCGCATG | >hg38_21_+_8433172_8446622_RNA45SN1_45S_50ntFlanks | 11766 | 11786 | 21 | ATCTAATTAGTGACGCGCATG | 151.2 | -9.5 | 1.5 | -6.5 | 8.5E-11 | 2.5E-10 | BT20/MB468 | CYTO |
| >hg38_21_+_8433172_8446622_RNA45SN1_45S_50ntFlanks@9760.9788.29\|AAACTTTAAATGGGTAAGAAGCCCGGCTC | >hg38_21_+_8433172_8446622_RNA45SN1_45S_50ntFlanks | 9760 | 9788 | 29 | AAACTTTAAATGGGTAAGAAGCCCGGCTC | 130.9 | -10.9 | 1.7 | -6.5 | 9.0E-11 | 2.5E-10 | MB231/MB468 | MP |
| >trnaMT_MetCAT_MT_+_4402_4469@33.71.39\|TACCCCGAAAATGTTGGTTATACCCTTCCCGTACTACCA | >trnaMT_MetCAT_MT_+_4402_4469 | 33 | 71 | 39 | TACCCCGAAAATGTTGGTTATACCCTTCCCGTACTACCA | 394.7 | 8.6 | 1.3 | 6.5 | 9.5E-11 | 2.5E-10 | MB231/MB468 | NUC |
| >hg38_21_+_8433172_8446622_RNA45SN1_45S_50ntFlanks@11718.11738.21\|AACGGCGGGAGTAACTATGAC | >hg38_21_+_8433172_8446622_RNA45SN1_45S_50ntFlanks | 11718 | 11738 | 21 | AACGGCGGGAGTAACTATGAC | 502.5 | -8.3 | 1.3 | -6.5 | 9.0E-11 | 2.5E-10 | MB231/MB468 | MP |
| >hg38_21_+_8433172_8446622_RNA45SN1_45S_50ntFlanks@10922.10943.22\|CTGGACGAGGCGCCGCCGCCCC | >hg38_21_+_8433172_8446622_RNA45SN1_45S_50ntFlanks | 10922 | 10943 | 22 | CTGGACGAGGCGCCGCCGCCCC | 128.1 | -10.9 | 1.7 | -6.5 | 9.0E-11 | 2.5E-10 | MB231/MB468 | MP |
| >hg38_21_+_8433172_8446622_RNA45SN1_45S_50ntFlanks@8380.8403.24\|CAAGAGGGCGTGAAACCGTTAAGA | >hg38_21_+_8433172_8446622_RNA45SN1_45S_50ntFlanks | 8380 | 8403 | 24 | CAAGAGGGCGTGAAACCGTTAAGA | 2296.1 | -7.6 | 1.2 | -6.5 | 9.0E-11 | 2.5E-10 | MB231/MB468 | MP |
| >hg38_21_+_8433172_8446622_RNA45SN1_45S_50ntFlanks@4283.4314.32\|CTTTAACGAGGATCCATTGGAGGGCAAGTCTG | >hg38_21_+_8433172_8446622_RNA45SN1_45S_50ntFlanks | 4283 | 4314 | 32 | CTTTAACGAGGATCCATTGGAGGGCAAGTCTG | 171.5 | -9.5 | 1.5 | -6.5 | 7.0E-11 | 2.5E-10 | BT20/MB468 | MITO |
| >MI0000253\|hsa-mir-148a&WithFlank&7\|-\|25949913\|25949992@50.70.21(+1G)&[MIMAT0000243&hsa-miR-148a-3p&offsets\|0\|-1(+1G);m-7&7\|-\|25949922\|25949943&offsets\|0\|-1(+1G)]\|TCAGTGCACTACAGAACTTTGG | >MI0000253\|hsa-mir-148a&WithFlank&7\|-\|25949913\|25949992 | 50 | 70 | 21(+1G) | TCAGTGCACTACAGAACTTTGG | 3448.1 | 5.9 | 0.9 | 6.9 | 6.0E-12 | 2.5E-10 | BT20/MB231 | CYTO |
| >am_trna13_LysCTT_14_-_58706613_58706685@31.51.21,trna2_LysCTT_15_+_79152904_79152976@31.51.21,trna4_LysCTT_18_-_43669271_43669343@31.51.21\|ACTCTTAATCCCAGGGTCGTG | >am_trna13_LysCTT_14_-_58706613_58706685 | 31 | 51 | 21 | ACTCTTAATCCCAGGGTCGTG | 560.8 | -8.0 | 1.2 | -6.5 | 8.6E-11 | 2.5E-10 | BT20/MB468 | CYTO |
| >hg38_21_+_8433172_8446622_RNA45SN1_45S_50ntFlanks@10843.10861.19\|TAACTTCGGGATAAGGATT | >hg38_21_+_8433172_8446622_RNA45SN1_45S_50ntFlanks | 10843 | 10861 | 19 | TAACTTCGGGATAAGGATT | 84.3 | -10.3 | 1.6 | -6.5 | 9.0E-11 | 2.5E-10 | MB231/MB468 | MP |
| >hg38_21_+_8433172_8446622_RNA45SN1_45S_50ntFlanks@9504.9526.23\|GTGAACTATGCCTGGGCAGGGCG | >hg38_21_+_8433172_8446622_RNA45SN1_45S_50ntFlanks | 9504 | 9526 | 23 | GTGAACTATGCCTGGGCAGGGCG | 21813.7 | -9.5 | 1.4 | -6.9 | 7.1E-12 | 2.5E-10 | BT20/MB468 | MP |
| >hg38_21_+_8433172_8446622_RNA45SN1_45S_50ntFlanks@3854.3874.21\|ACTGTGGTAATTCTAGAGCTA | >hg38_21_+_8433172_8446622_RNA45SN1_45S_50ntFlanks | 3854 | 3874 | 21 | ACTGTGGTAATTCTAGAGCTA | 485.9 | -8.5 | 1.3 | -6.5 | 1.1E-10 | 2.5E-10 | MB231/MB468 | MITO |
| >hg38_21_+_8433172_8446622_RNA45SN1_45S_50ntFlanks@10427.10456.30\|ATGGGCGAGCGCCGTTCCGAAGGGACGGGC | >hg38_21_+_8433172_8446622_RNA45SN1_45S_50ntFlanks | 10427 | 10456 | 30 | ATGGGCGAGCGCCGTTCCGAAGGGACGGGC | 1499.2 | -3.3 | 0.5 | -6.5 | 8.6E-11 | 2.5E-10 | BT20/MB468 | CYTO |
| >hg38_21_+_8433172_8446622_RNA45SN1_45S_50ntFlanks@13006.13028.23\|CTATTGAAAGTCAGCCCTCGACA | >hg38_21_+_8433172_8446622_RNA45SN1_45S_50ntFlanks | 13006 | 13028 | 23 | CTATTGAAAGTCAGCCCTCGACA | 1141.3 | -8.7 | 1.3 | -6.5 | 8.6E-11 | 2.5E-10 | BT20/MB468 | CYTO |
| >MI0000734\|hsa-mir-106b&WithFlank&7\|-\|100093987\|100094080@58.78.21(+1A)&[MIMAT0004672&hsa-miR-106b-3p&offsets\|0\|-1(+1A);m-87&7\|-\|100094003\|100094023&offsets\|0\|0(+1A)]\|CCGCACTGTGGGTACTTGCTGA | >MI0000734\|hsa-mir-106b&WithFlank&7\|-\|100093987\|100094080 | 58 | 78 | 21(+1A) | CCGCACTGTGGGTACTTGCTGA | 108.0 | 1.8 | 0.3 | 6.4 | 1.1E-10 | 2.5E-10 | BT20/MB468 | NUC |
| >hg38_21_+_8433172_8446622_RNA45SN1_45S_50ntFlanks@12926.12953.28\|ACCTGGCGCTAAACCATTCGTAGACGAC | >hg38_21_+_8433172_8446622_RNA45SN1_45S_50ntFlanks | 12926 | 12953 | 28 | ACCTGGCGCTAAACCATTCGTAGACGAC | 79.3 | -10.2 | 1.6 | -6.5 | 9.1E-11 | 2.5E-10 | MB231/MB468 | MP |
| >hg38_21_+_8433172_8446622_RNA45SN1_45S_50ntFlanks@12432.12455.24\|CCTATCATTGTGAAGCAGAATTCA | >hg38_21_+_8433172_8446622_RNA45SN1_45S_50ntFlanks | 12432 | 12455 | 24 | CCTATCATTGTGAAGCAGAATTCA | 437.0 | -7.9 | 1.2 | -6.5 | 9.6E-11 | 2.5E-10 | MB231/MB468 | NUC |
| >hg38_21_+_8433172_8446622_RNA45SN1_45S_50ntFlanks@11209.11238.30\|CGGCGACTCTGGACGCGAGCCGGGCCCTTC | >hg38_21_+_8433172_8446622_RNA45SN1_45S_50ntFlanks | 11209 | 11238 | 30 | CGGCGACTCTGGACGCGAGCCGGGCCCTTC | 182.3 | -6.3 | 1.0 | -6.5 | 9.6E-11 | 2.5E-10 | MB231/MB468 | CYTO |
| >hg38_21_+_8433172_8446622_RNA45SN1_45S_50ntFlanks@11759.11777.19\|CCTCGTCATCTAATTAGTG | >hg38_21_+_8433172_8446622_RNA45SN1_45S_50ntFlanks | 11759 | 11777 | 19 | CCTCGTCATCTAATTAGTG | 176.3 | -7.7 | 1.2 | -6.5 | 9.6E-11 | 2.5E-10 | MB231/MB468 | NUC |
| >hg38_21_+_8433172_8446622_RNA45SN1_45S_50ntFlanks@11782.11806.25\|GCATGAATGGATGAACGAGATTCCC | >hg38_21_+_8433172_8446622_RNA45SN1_45S_50ntFlanks | 11782 | 11806 | 25 | GCATGAATGGATGAACGAGATTCCC | 81.4 | -10.2 | 1.6 | -6.5 | 9.1E-11 | 2.5E-10 | MB231/MB468 | MP |
| >hg38_21_+_8433172_8446622_RNA45SN1_45S_50ntFlanks@3984.4003.20\|GCGGCTTTGGTGACTCTAGA | >hg38_21_+_8433172_8446622_RNA45SN1_45S_50ntFlanks | 3984 | 4003 | 20 | GCGGCTTTGGTGACTCTAGA | 106.8 | -10.6 | 1.6 | -6.5 | 9.1E-11 | 2.5E-10 | MB231/MB468 | MP |
| >hg38_21_+_8433172_8446622_RNA45SN1_45S_50ntFlanks@5421.5441.21\|CGATTGGATGGTTTAGTGAGG | >hg38_21_+_8433172_8446622_RNA45SN1_45S_50ntFlanks | 5421 | 5441 | 21 | CGATTGGATGGTTTAGTGAGG | 119.8 | -10.8 | 1.7 | -6.5 | 9.1E-11 | 2.5E-10 | MB231/MB468 | MP |
| >hg38_21_+_8433172_8446622_RNA45SN1_45S_50ntFlanks@12234.12262.29\|CGTGGAGCAGAAGGGCAAAAGCTCGCTTG | >hg38_21_+_8433172_8446622_RNA45SN1_45S_50ntFlanks | 12234 | 12262 | 29 | CGTGGAGCAGAAGGGCAAAAGCTCGCTTG | 75.5 | -10.1 | 1.6 | -6.5 | 9.1E-11 | 2.5E-10 | MB231/MB468 | MP |
| >hg38_21_+_8433172_8446622_RNA45SN1_45S_50ntFlanks@10408.10427.20\|GGTCAGTCGGTCCTGAGAGA | >hg38_21_+_8433172_8446622_RNA45SN1_45S_50ntFlanks | 10408 | 10427 | 20 | GGTCAGTCGGTCCTGAGAGA | 364.0 | -8.7 | 1.3 | -6.4 | 1.2E-10 | 2.5E-10 | BT20/MB468 | NUC |
| >hg38_21_+_8433172_8446622_RNA45SN1_45S_50ntFlanks@10827.10844.18\|CGGCAAGCCGGATCCGTA | >hg38_21_+_8433172_8446622_RNA45SN1_45S_50ntFlanks | 10827 | 10844 | 18 | CGGCAAGCCGGATCCGTA | 89.2 | -10.3 | 1.6 | -6.5 | 9.2E-11 | 2.5E-10 | MB231/MB468 | MP |
| >hg38_21_+_8433172_8446622_RNA45SN1_45S_50ntFlanks@4731.4756.26\|AAGAACGAAAGTCGGAGGTTCGAAGA | >hg38_21_+_8433172_8446622_RNA45SN1_45S_50ntFlanks | 4731 | 4756 | 26 | AAGAACGAAAGTCGGAGGTTCGAAGA | 360.8 | 6.6 | 1.0 | 6.5 | 8.7E-11 | 2.5E-10 | BT20/MB468 | CYTO |
| >hg38_21_+_8433172_8446622_RNA45SN1_45S_50ntFlanks@10783.10822.40\|GTGAACAGCCTCTGGCATGTTGGAACAATGTAGGTAAGGG | >hg38_21_+_8433172_8446622_RNA45SN1_45S_50ntFlanks | 10783 | 10822 | 40 | GTGAACAGCCTCTGGCATGTTGGAACAATGTAGGTAAGGG | 122.8 | -10.8 | 1.7 | -6.5 | 9.2E-11 | 2.5E-10 | MB231/MB468 | MP |
| >trnaMT_GluTTC_MT_-_14674_14742@40.65.26,trnalookalike8_GluTTC_5_-_93905172_93905240@40.65.26\|TTGGTCGTGGTTGTAGTCCGTGCGAG | >trnaMT_GluTTC_MT_-_14674_14742 | 40 | 65 | 26 | TTGGTCGTGGTTGTAGTCCGTGCGAG | 966.3 | 8.2 | 1.3 | 6.5 | 7.2E-11 | 2.5E-10 | BT20/MB468 | MITO |
| >hg38_21_+_8433172_8446622_RNA45SN1_45S_50ntFlanks@4588.4616.29\|CTATTTTGTTGGTTTTCGGAACTGAGGCC | >hg38_21_+_8433172_8446622_RNA45SN1_45S_50ntFlanks | 4588 | 4616 | 29 | CTATTTTGTTGGTTTTCGGAACTGAGGCC | 2015.6 | -7.4 | 1.1 | -6.5 | 9.7E-11 | 2.5E-10 | MB231/MB468 | NUC |
| >hg38_21_+_8433172_8446622_RNA45SN1_45S_50ntFlanks@12408.12448.41\|TTTGATCCTTCGATGTCGGCTCTTCCTATCATTGTGAAGCA | >hg38_21_+_8433172_8446622_RNA45SN1_45S_50ntFlanks | 12408 | 12448 | 41 | TTTGATCCTTCGATGTCGGCTCTTCCTATCATTGTGAAGCA | 346.0 | 9.3 | 1.4 | 6.5 | 9.8E-11 | 2.5E-10 | MB231/MB468 | CYTO |
| >hg38_21_+_8433172_8446622_RNA45SN1_45S_50ntFlanks@6652.6676.25\|GACTCTTAGCGGTGGATCACTCGGC | >hg38_21_+_8433172_8446622_RNA45SN1_45S_50ntFlanks | 6652 | 6676 | 25 | GACTCTTAGCGGTGGATCACTCGGC | 12941.6 | 9.1 | 1.4 | 6.5 | 8.8E-11 | 2.5E-10 | BT20/MB468 | CYTO |
| >hg38_21_+_8433172_8446622_RNA45SN1_45S_50ntFlanks@12081.12107.27\|CTCTCGCTTCTGGCGCCAAGCGCCCGG | >hg38_21_+_8433172_8446622_RNA45SN1_45S_50ntFlanks | 12081 | 12107 | 27 | CTCTCGCTTCTGGCGCCAAGCGCCCGG | 531.6 | -5.8 | 0.9 | -6.5 | 8.8E-11 | 2.5E-10 | BT20/MB468 | CYTO |
| >trna10_AspGTC_12_-_125424193_125424264@37.74.38,trna12_AspGTC_12_-_125411891_125411962@37.74.38,trna144_AspGTC_6_-_27551236_27551307@37.74.38,trna38_AspGTC_17_-_8125556_8125627@37.74.38,trna45_AspGTC_6_+_27447453_27447524@37.74.38,trna48_AspGTC_6_+_27471523_27471594@37.74.38,trna4_AspGTC_12_+_96429799_96429870@37.74.38,trna69_AspGTC_1_-_161440205_161440276@37.74.38,trna72_AspGTC_1_-_161432824_161432895@37.74.38,trna75_AspGTC_1_-_161425414_161425485@37.74.38,trna78_AspGTC_1_-_161418033_161418104@37.74.38,trna81_AspGTC_1_-_161410615_161410686@37.74.38\|ACGCGGGAGACCGGGGTTCGATTCCCCGACGGGGAGCC | >trna10_AspGTC_12_-_125424193_125424264 | 37 | 74 | 38 | ACGCGGGAGACCGGGGTTCGATTCCCCGACGGGGAGCC | 1478.6 | 8.4 | 1.3 | 6.5 | 9.8E-11 | 2.5E-10 | MB231/MB468 | NUC |
| >hg38_21_+_8433172_8446622_RNA45SN1_45S_50ntFlanks@10841.10861.21\|CGTAACTTCGGGATAAGGATT | >hg38_21_+_8433172_8446622_RNA45SN1_45S_50ntFlanks | 10841 | 10861 | 21 | CGTAACTTCGGGATAAGGATT | 857.2 | -3.7 | 0.6 | -6.5 | 9.8E-11 | 2.5E-10 | MB231/MB468 | NUC |
| >hg38_21_+_8433172_8446622_RNA45SN1_45S_50ntFlanks@7979.8003.25\|ACCTCAGATCAGACGTGGCGACCCG | >hg38_21_+_8433172_8446622_RNA45SN1_45S_50ntFlanks | 7979 | 8003 | 25 | ACCTCAGATCAGACGTGGCGACCCG | 1289.1 | -4.9 | 0.8 | -6.5 | 9.8E-11 | 2.5E-10 | MB231/MB468 | NUC |
| >hg38_21_+_8433172_8446622_RNA45SN1_45S_50ntFlanks@10778.10818.41\|CCAAGGTGAACAGCCTCTGGCATGTTGGAACAATGTAGGTA | >hg38_21_+_8433172_8446622_RNA45SN1_45S_50ntFlanks | 10778 | 10818 | 41 | CCAAGGTGAACAGCCTCTGGCATGTTGGAACAATGTAGGTA | 511.2 | -6.8 | 1.1 | -6.5 | 1.1E-10 | 2.5E-10 | MB231/MB468 | MITO |
| >hg38_21_+_8433172_8446622_RNA45SN1_45S_50ntFlanks@11572.11613.42\|CCGACTTAGAACTGGTGCGGACCAGGGGAATCCGACTGTTTA | >hg38_21_+_8433172_8446622_RNA45SN1_45S_50ntFlanks | 11572 | 11613 | 42 | CCGACTTAGAACTGGTGCGGACCAGGGGAATCCGACTGTTTA | 1107.9 | -4.7 | 0.7 | -6.4 | 1.2E-10 | 2.5E-10 | BT20/MB468 | NUC |

<table>
<tr><th></th><th></th><th></th><th></th><th></th><th></th><th></th><th></th><th></th><th></th><th></th><th></th><th></th><th></th></tr>
<tr><td>>hg38_21_+_8433172_8446622_RNA45SN1_45S_50ntFlanks@4241.4264.24|CTCTTTCGAGGCCCTGTAATTGGA</td><td>>hg38_21_+_8433172_8446622_RNA45SN1_45S_50ntFlanks</td><td>4241</td><td>4264</td><td>24</td><td>CTCTTTCGAGGCCCTGTAATTGGA</td><td>118.9</td><td>-5.9</td><td>0.9</td><td>-6.5</td><td>9.8E-11</td><td>2.5E-10</td><td>MB231/MB468</td><td>CYTO</td></tr>
<tr><td>>am_trna116_GluCTC_1_-_145399233_145399304@6.28.23,trna59_GluCTC_1_+_249168447_249168518@6.28.23,trna71_GluCTC_1_-_161439189_161439260@6.28.23,trna74_GluCTC_1_-_161431809_161431880@6.28.23,trna77_GluCTC_1_-_161424398_161424469@6.28.23,trna77_GluCTC_6_+_28949976_28950047@6.28.23,trna80_GluCTC_1_-_161417018_161417089@6.28.23,trna87_GluCTC_6_-_126101393_126101464@6.28.23|GGTGGTCTAGTGGTTAGGATTCG</td><td>>am_trna116_GluCTC_1_-_145399233_145399304</td><td>6</td><td>28</td><td>23</td><td>GGTGGTCTAGTGGTTAGGATTCG</td><td>111.3</td><td>-10.7</td><td>1.6</td><td>-6.5</td><td>9.3E-11</td><td>2.5E-10</td><td>MB231/MB468</td><td>MP</td></tr>
<tr><td>>MI0000812|hsa-mir-331&WithFlank&12|+|95308414|95308519@67.87.21(+1U)&[MIMAT0000760&hsa-miR-331-3p&offsets|0|0(+1U);m-192&12|+|95308480|95308501&offsets|0|-1(+1U)]|GCCCCTGGGCCTATCCTAGAAT</td><td>>MI0000812|hsa-mir-331&WithFlank&12|+|95308414|95308519</td><td>67</td><td>87</td><td>21(+1U)</td><td>GCCCCTGGGCCTATCCTAGAAT</td><td>808.6</td><td>7.4</td><td>1.1</td><td>6.5</td><td>8.8E-11</td><td>2.6E-10</td><td>BT20/MB468</td><td>CYTO</td></tr>
<tr><td>>hg38_21_+_8433172_8446622_RNA45SN1_45S_50ntFlanks@12934.12962.29|CTAAACCATTCGTAGACGACCTGCTTCTG</td><td>>hg38_21_+_8433172_8446622_RNA45SN1_45S_50ntFlanks</td><td>12934</td><td>12962</td><td>29</td><td>CTAAACCATTCGTAGACGACCTGCTTCTG</td><td>902.6</td><td>-6.8</td><td>1.1</td><td>-6.5</td><td>9.9E-11</td><td>2.6E-10</td><td>MB231/MB468</td><td>NUC</td></tr>
<tr><td>>hg38_21_+_8433172_8446622_RNA45SN1_45S_50ntFlanks@7975.8002.28|CGCGACCTCAGATCAGACGTGGCGACCC</td><td>>hg38_21_+_8433172_8446622_RNA45SN1_45S_50ntFlanks</td><td>7975</td><td>8002</td><td>28</td><td>CGCGACCTCAGATCAGACGTGGCGACCC</td><td>771.4</td><td>3.6</td><td>0.6</td><td>6.5</td><td>9.9E-11</td><td>2.6E-10</td><td>MB231/MB468</td><td>NUC</td></tr>
<tr><td>>hg38_21_+_8433172_8446622_RNA45SN1_45S_50ntFlanks@10109.10126.18|TGTGGTGTGCGTCGGAGG</td><td>>hg38_21_+_8433172_8446622_RNA45SN1_45S_50ntFlanks</td><td>10109</td><td>10126</td><td>18</td><td>TGTGGTGTGCGTCGGAGG</td><td>101.1</td><td>-10.5</td><td>1.6</td><td>-6.5</td><td>9.3E-11</td><td>2.6E-10</td><td>MB231/MB468</td><td>MP</td></tr>
<tr><td>>MI0000266|hsa-mir-10a&WithFlank&17|-|48579832|48579953@28.48.21(+1A)&[MIMAT0000253&hsa-miR-10a-5p&offsets|0|-2(+1A);m-18&17|-|48579905|48579926&offsets|0|-1(+1A)]|TACCCTGTAGATCCGAATTTGA</td><td>>MI0000266|hsa-mir-10a&WithFlank&17|-|48579832|48579953</td><td>28</td><td>48</td><td>21(+1A)</td><td>TACCCTGTAGATCCGAATTTGA</td><td>431.7</td><td>12.5</td><td>1.8</td><td>6.9</td><td>7.3E-12</td><td>2.6E-10</td><td>BT20/MB468</td><td>MP</td></tr>
<tr><td>>hg38_21_+_8433172_8446622_RNA45SN1_45S_50ntFlanks@11766.11786.21|ATCTAATTAGTGACGCGCATG</td><td>>hg38_21_+_8433172_8446622_RNA45SN1_45S_50ntFlanks</td><td>11766</td><td>11786</td><td>21</td><td>ATCTAATTAGTGACGCGCATG</td><td>178.7</td><td>-6.7</td><td>1.0</td><td>-6.5</td><td>9.9E-11</td><td>2.6E-10</td><td>MB231/MB468</td><td>CYTO</td></tr>
<tr><td>>hg38_21_+_8433172_8446622_RNA45SN1_45S_50ntFlanks@4069.4097.29|CTTTCGATGGTAGTCGCCGTGCCTACCAT</td><td>>hg38_21_+_8433172_8446622_RNA45SN1_45S_50ntFlanks</td><td>4069</td><td>4097</td><td>29</td><td>CTTTCGATGGTAGTCGCCGTGCCTACCAT</td><td>69.8</td><td>-10.0</td><td>1.5</td><td>-6.5</td><td>9.4E-11</td><td>2.6E-10</td><td>MB231/MB468</td><td>MP</td></tr>
<tr><td>>hg38_21_+_8433172_8446622_RNA45SN1_45S_50ntFlanks@12712.12751.40|CGGAGCCTCGGTTGGCCTCGGATAGCCGGTCCCCCGCCTG</td><td>>hg38_21_+_8433172_8446622_RNA45SN1_45S_50ntFlanks</td><td>12712</td><td>12751</td><td>40</td><td>CGGAGCCTCGGTTGGCCTCGGATAGCCGGTCCCCCGCCTG</td><td>846.9</td><td>-12.8</td><td>1.9</td><td>-6.8</td><td>7.4E-12</td><td>2.6E-10</td><td>BT20/MB468</td><td>MP</td></tr>
<tr><td>>hg38_21_+_8433172_8446622_RNA45SN1_45S_50ntFlanks@9120.9140.21|TGCCGCGCGCGGGTCGGGGGG</td><td>>hg38_21_+_8433172_8446622_RNA45SN1_45S_50ntFlanks</td><td>9120</td><td>9140</td><td>21</td><td>TGCCGCGCGCGGGTCGGGGGG</td><td>118.9</td><td>-10.8</td><td>1.7</td><td>-6.5</td><td>9.4E-11</td><td>2.6E-10</td><td>MB231/MB468</td><td>MP</td></tr>
<tr><td>>MI0000239|hsa-mir-197&WithFlank&1|+|109598887|109598973@54.74.21&[MIMAT0000227&hsa-miR-197-3p&offsets|0|-1;m-99&1|+|109598940|109598961&offsets|0|-1]|TTCACCACCTTCTCCACCCAG</td><td>>MI0000239|hsa-mir-197&WithFlank&1|+|109598887|109598973</td><td>54</td><td>74</td><td>21</td><td>TTCACCACCTTCTCCACCCAG</td><td>127.8</td><td>2.7</td><td>0.4</td><td>6.4</td><td>1.2E-10</td><td>2.6E-10</td><td>BT20/MB468</td><td>NUC</td></tr>
<tr><td>>hg38_21_+_8433172_8446622_RNA45SN1_45S_50ntFlanks@9604.9624.21|AAAGACTAATCGAACCATCTA</td><td>>hg38_21_+_8433172_8446622_RNA45SN1_45S_50ntFlanks</td><td>9604</td><td>9624</td><td>21</td><td>AAAGACTAATCGAACCATCTA</td><td>177.1</td><td>-11.3</td><td>1.7</td><td>-6.5</td><td>9.4E-11</td><td>2.6E-10</td><td>MB231/MB468</td><td>MP</td></tr>
<tr><td>>hg38_21_+_8433172_8446622_RNA45SN1_45S_50ntFlanks@12141.12179.39|ACAGTGCCAGGTGGGGAGTTTGACTGGGGCGGTACACCT</td><td>>hg38_21_+_8433172_8446622_RNA45SN1_45S_50ntFlanks</td><td>12141</td><td>12179</td><td>39</td><td>ACAGTGCCAGGTGGGGAGTTTGACTGGGGCGGTACACCT</td><td>163.7</td><td>-9.4</td><td>1.5</td><td>-6.5</td><td>7.3E-11</td><td>2.6E-10</td><td>BT20/MB468</td><td>MITO</td></tr>
<tr><td>>trna13_ValCAC_19_-_4724647_4724719@1.29.29|GTTTCCGTAGTGTAGCGGTTATCACATTC</td><td>>trna13_ValCAC_19_-_4724647_4724719</td><td>1</td><td>29</td><td>29</td><td>GTTTCCGTAGTGTAGCGGTTATCACATTC</td><td>455.2</td><td>-11.9</td><td>1.7</td><td>-6.8</td><td>7.5E-12</td><td>2.6E-10</td><td>BT20/MB468</td><td>MP</td></tr>
<tr><td>>hg38_21_+_8433172_8446622_RNA45SN1_45S_50ntFlanks@8379.8402.24|TCAAGAGGGCGTGAAACCGTTAAG</td><td>>hg38_21_+_8433172_8446622_RNA45SN1_45S_50ntFlanks</td><td>8379</td><td>8402</td><td>24</td><td>TCAAGAGGGCGTGAAACCGTTAAG</td><td>856.6</td><td>-8.6</td><td>1.3</td><td>-6.5</td><td>9.5E-11</td><td>2.6E-10</td><td>MB231/MB468</td><td>MP</td></tr>
<tr><td>>hg38_21_+_8433172_8446622_RNA45SN1_45S_50ntFlanks@10794.10816.23|CTGGCATGTTGGAACAATGTAGG</td><td>>hg38_21_+_8433172_8446622_RNA45SN1_45S_50ntFlanks</td><td>10794</td><td>10816</td><td>23</td><td>CTGGCATGTTGGAACAATGTAGG</td><td>616.8</td><td>-3.6</td><td>0.6</td><td>-6.5</td><td>9.0E-11</td><td>2.6E-10</td><td>BT20/MB468</td><td>CYTO</td></tr>
<tr><td>>MI0000253|hsa-mir-148a&WithFlank&7|-|25949913|25949992@50.70.21(+1C)&[MIMAT0000243&hsa-miR-148a-3p&offsets|0|-1(+1C);m-7&7|-|25949922|25949943&offsets|0|-1(+1C)]|TCAGTGCACTACAGAACTTTGC</td><td>>MI0000253|hsa-mir-148a&WithFlank&7|-|25949913|25949992</td><td>50</td><td>70</td><td>21(+1C)</td><td>TCAGTGCACTACAGAACTTTGC</td><td>17101.5</td><td>5.3</td><td>0.8</td><td>6.9</td><td>6.4E-12</td><td>2.6E-10</td><td>BT20/MB231</td><td>CYTO</td></tr>
<tr><td>>MI0000299|hsa-mir-222&WithFlank&X|-|45747009|45747130@75.97.23&[MIMAT0000279&hsa-miR-222-3p&offsets|0|+2;m-140&X|-|45747033|45747056&offsets|0|-1]|AGCTACATCTGGCTACTGGGTCT</td><td>>MI0000299|hsa-mir-222&WithFlank&X|-|45747009|45747130</td><td>75</td><td>97</td><td>23</td><td>AGCTACATCTGGCTACTGGGTCT</td><td>6048.3</td><td>5.4</td><td>0.8</td><td>6.5</td><td>1.0E-10</td><td>2.6E-10</td><td>MB231/MB468</td><td>NUC</td></tr>
<tr><td>>MI0000264|hsa-mir-7-2&WithFlank&15|+|88611819|88611940@38.58.21;MI0000265|hsa-mir-7-3&WithFlank&19|+|4770664|4770785@37.57.21;MI0000263|hsa-mir-7-1&WithFlank&9|-|83969742|83969863@30.50.21&[MIMAT0000252&hsa-miR-7-5p&offsets|0|-3;m-400&15|+|88611856|88611879&offsets|0|-3];[MIMAT0000252_1&hsa-miR-7-5p&offsets|0|-3;m-402&19|+|4770700|4770723&offsets|0|-3];[MIMAT0000252_2&hsa-miR-7-5p&offsets|0|-3;m-401&9|-|83969811|83969834&offsets|0|-3]|TGGAAGACTAGTGATTTTGTT</td><td>>MI0000264|hsa-mir-7-2&WithFlank&15|+|88611819|88611940</td><td>38</td><td>58</td><td>21</td><td>TGGAAGACTAGTGATTTTGTT</td><td>1809.0</td><td>4.2</td><td>0.6</td><td>6.5</td><td>9.5E-11</td><td>2.6E-10</td><td>MB231/MB468</td><td>MP</td></tr>
<tr><td>>hg38_21_+_8433172_8446622_RNA45SN1_45S_50ntFlanks@10798.10822.25|CATGTTGGAACAATGTAGGTAAGGG</td><td>>hg38_21_+_8433172_8446622_RNA45SN1_45S_50ntFlanks</td><td>10798</td><td>10822</td><td>25</td><td>CATGTTGGAACAATGTAGGTAAGGG</td><td>205.7</td><td>-4.0</td><td>0.6</td><td>-6.5</td><td>1.0E-10</td><td>2.6E-10</td><td>MB231/MB468</td><td>NUC</td></tr>
<tr><td>>hg38_21_+_8433172_8446622_RNA45SN1_45S_50ntFlanks@5472.5497.26|ACGGCCCTGGCGGAGCGCTGAGAAGA</td><td>>hg38_21_+_8433172_8446622_RNA45SN1_45S_50ntFlanks</td><td>5472</td><td>5497</td><td>26</td><td>ACGGCCCTGGCGGAGCGCTGAGAAGA</td><td>629.6</td><td>5.0</td><td>0.8</td><td>6.5</td><td>1.0E-10</td><td>2.6E-10</td><td>MB231/MB468</td><td>CYTO</td></tr>
<tr><td>>hg38_21_+_8433172_8446622_RNA45SN1_45S_50ntFlanks@7968.8009.42|TCCGAGACGCGACCTCAGATCAGACGTGGCGACCCGCTGAAT</td><td>>hg38_21_+_8433172_8446622_RNA45SN1_45S_50ntFlanks</td><td>7968</td><td>8009</td><td>42</td><td>TCCGAGACGCGACCTCAGATCAGACGTGGCGACCCGCTGAAT</td><td>388.9</td><td>5.5</td><td>0.8</td><td>6.5</td><td>1.0E-10</td><td>2.6E-10</td><td>MB231/MB468</td><td>NUC</td></tr>
<tr><td>>hg38_21_+_8433172_8446622_RNA45SN1_45S_50ntFlanks@3787.3812.26|AAACTGCGAATGGCTCATTAAATCAG</td><td>>hg38_21_+_8433172_8446622_RNA45SN1_45S_50ntFlanks</td><td>3787</td><td>3812</td><td>26</td><td>AAACTGCGAATGGCTCATTAAATCAG</td><td>675.4</td><td>-11.5</td><td>1.8</td><td>-6.5</td><td>7.4E-11</td><td>2.6E-10</td><td>BT20/MB468</td><td>MITO</td></tr>
<tr><td>>hg38_21_+_8433172_8446622_RNA45SN1_45S_50ntFlanks@12939.12966.28|CCATTCGTAGACGACCTGCTTCTGGGTC</td><td>>hg38_21_+_8433172_8446622_RNA45SN1_45S_50ntFlanks</td><td>12939</td><td>12966</td><td>28</td><td>CCATTCGTAGACGACCTGCTTCTGGGTC</td><td>259.2</td><td>-7.7</td><td>1.2</td><td>-6.5</td><td>9.1E-11</td><td>2.6E-10</td><td>BT20/MB468</td><td>CYTO</td></tr>
<tr><td>>hg38_21_+_8433172_8446622_RNA45SN1_45S_50ntFlanks@11190.11223.34|CCCGGGGGGCCGGCGGCGGCGGCGACTCTGGACG</td><td>>hg38_21_+_8433172_8446622_RNA45SN1_45S_50ntFlanks</td><td>11190</td><td>11223</td><td>34</td><td>CCCGGGGGGCCGGCGGCGGCGGCGACTCTGGACG</td><td>187.3</td><td>-11.4</td><td>1.8</td><td>-6.5</td><td>9.5E-11</td><td>2.6E-10</td><td>MB231/MB468</td><td>MP</td></tr>
<tr><td>>hg38_21_+_8433172_8446622_RNA45SN1_45S_50ntFlanks@8191.8212.22|CAGCCCGTGGACGGTGTGAGGC</td><td>>hg38_21_+_8433172_8446622_RNA45SN1_45S_50ntFlanks</td><td>8191</td><td>8212</td><td>22</td><td>CAGCCCGTGGACGGTGTGAGGC</td><td>921.5</td><td>-8.4</td><td>1.3</td><td>-6.4</td><td>1.1E-10</td><td>2.6E-10</td><td>MB231/MB468</td><td>MITO</td></tr>
<tr><td>>trna111_HisGTG_1_-_147774845_147774916@-1T.26.27,trna118_HisGTG_1_-_145396881_145396952@-1T.26.27,trna16_HisGTG_1_+_146544773_146544844@-1T.26.27,trna1_HisGTG_15_+_45493349_45493420@-1T.26.27,trna21_HisGTG_1_+_147753471_147753542@-1T.26.27,trna33_HisGTG_6_+_27125906_27125977@-1T.26.27,trna7_HisGTG_9_-_14433938_14434009@-1T.26.27,trna8_HisGTG_15_-_45492611_45492682@-1T.26.27,trna9_HisGTG_15_-_45490804_45490875@-1T.26.27|TGCCGTGATCGTATAGTGGTTAGTACT</td><td>>trna111_HisGTG_1_-_147774845_147774916</td><td>-1T</td><td>26</td><td>27</td><td>TGCCGTGATCGTATAGTGGTTAGTACT</td><td>801.4</td><td>6.0</td><td>0.9</td><td>6.5</td><td>7.4E-11</td><td>2.6E-10</td><td>BT20/MB468</td><td>MITO</td></tr>
<tr><td>>MI0000102|hsa-mir-100&WithFlank&11|-|122152223|122152314@19.40.22&[MIMAT0000098&hsa-miR-100-5p&offsets|0|0;m-27&11|-|122152276|122152296&offsets|0|+1]|AACCCGTAGATCCGAACTTGTG</td><td>>MI0000102|hsa-mir-100&WithFlank&11|-|122152223|122152314</td><td>19</td><td>40</td><td>22</td><td>AACCCGTAGATCCGAACTTGTG</td><td>2289.8</td><td>6.7</td><td>1.0</td><td>6.4</td><td>1.2E-10</td><td>2.6E-10</td><td>BT20/MB468</td><td>NUC</td></tr>
<tr><td>>hg38_21_+_8433172_8446622_RNA45SN1_45S_50ntFlanks@7988.8018.31|CAGACGTGGCGACCCGCTGAATTTAAGCATA</td><td>>hg38_21_+_8433172_8446622_RNA45SN1_45S_50ntFlanks</td><td>7988</td><td>8018</td><td>31</td><td>CAGACGTGGCGACCCGCTGAATTTAAGCATA</td><td>1088.2</td><td>-4.5</td><td>0.7</td><td>-6.4</td><td>1.1E-10</td><td>2.6E-10</td><td>MB231/MB468</td><td>MITO</td></tr>
<tr><td>>hg38_21_+_8433172_8446622_RNA45SN1_45S_50ntFlanks@5488.5515.28|GCTGAGAAGACGGTCGAACTTGACTATC</td><td>>hg38_21_+_8433172_8446622_RNA45SN1_45S_50ntFlanks</td><td>5488</td><td>5515</td><td>28</td><td>GCTGAGAAGACGGTCGAACTTGACTATC</td><td>312.5</td><td>5.4</td><td>0.8</td><td>6.5</td><td>1.0E-10</td><td>2.6E-10</td><td>MB231/MB468</td><td>CYTO</td></tr>
<tr><td>>hg38_21_+_8433172_8446622_RNA45SN1_45S_50ntFlanks@5419.5445.27|ACCGATTGGATGGTTTAGTGAGGCCCT</td><td>>hg38_21_+_8433172_8446622_RNA45SN1_45S_50ntFlanks</td><td>5419</td><td>5445</td><td>27</td><td>ACCGATTGGATGGTTTAGTGAGGCCCT</td><td>119.5</td><td>-10.8</td><td>1.7</td><td>-6.5</td><td>9.6E-11</td><td>2.6E-10</td><td>MB231/MB468</td><td>MP</td></tr>
<tr><td>>trna10_GlyTCC_17_+_8124866_8124937@1.30.30|GCGTTGGTGGTATAGTGGTAAGCATAGCTG</td><td>>trna10_GlyTCC_17_+_8124866_8124937</td><td>1</td><td>30</td><td>30</td><td>GCGTTGGTGGTATAGTGGTAAGCATAGCTG</td><td>290.7</td><td>-8.4</td><td>1.3</td><td>-6.5</td><td>1.0E-10</td><td>2.6E-10</td><td>MB231/MB468</td><td>NUC</td></tr>
<tr><td>>hg38_21_+_8433172_8446622_RNA45SN1_45S_50ntFlanks@5484.5514.31|GAGCGCTGAGAAGACGGTCGAACTTGACTAT</td><td>>hg38_21_+_8433172_8446622_RNA45SN1_45S_50ntFlanks</td><td>5484</td><td>5514</td><td>31</td><td>GAGCGCTGAGAAGACGGTCGAACTTGACTAT</td><td>3742.3</td><td>-5.5</td><td>0.8</td><td>-6.5</td><td>7.5E-11</td><td>2.6E-10</td><td>BT20/MB468</td><td>MITO</td></tr>
<tr><td>>hg38_21_+_8433172_8446622_RNA45SN1_45S_50ntFlanks@9534.9572.39|AGGAAACTCTGGTGGAGGTCCGTAGCGGTCCTGACGTGC</td><td>>hg38_21_+_8433172_8446622_RNA45SN1_45S_50ntFlanks</td><td>9534</td><td>9572</td><td>39</td><td>AGGAAACTCTGGTGGAGGTCCGTAGCGGTCCTGACGTGC</td><td>1532.0</td><td>-7.9</td><td>1.2</td><td>-6.5</td><td>9.6E-11</td><td>2.6E-10</td><td>MB231/MB468</td><td>MP</td></tr>
<tr><td>>hg38_21_+_8433172_8446622_RNA45SN1_45S_50ntFlanks@3899.3919.21|CCTTCGCGGGGGGGATGCGTG</td><td>>hg38_21_+_8433172_8446622_RNA45SN1_45S_50ntFlanks</td><td>3899</td><td>3919</td><td>21</td><td>CCTTCGCGGGGGGGATGCGTG</td><td>4366.9</td><td>-11.0</td><td>1.7</td><td>-6.4</td><td>1.2E-10</td><td>2.6E-10</td><td>BT20/MB468</td><td>NUC</td></tr>
<tr><td>>hg38_21_+_8433172_8446622_RNA45SN1_45S_50ntFlanks@5073.5108.36|AATTCCGATAACGAACGAGACTCTGGCATGCTAACT</td><td>>hg38_21_+_8433172_8446622_RNA45SN1_45S_50ntFlanks</td><td>5073</td><td>5108</td><td>36</td><td>AATTCCGATAACGAACGAGACTCTGGCATGCTAACT</td><td>704.5</td><td>-5.5</td><td>0.9</td><td>-6.4</td><td>1.2E-10</td><td>2.6E-10</td><td>BT20/MB468</td><td>NUC</td></tr>
<tr><td>>hg38_21_+_8433172_8446622_RNA45SN1_45S_50ntFlanks@9554.9582.29|CGTAGCGGTCCTGACGTGCAAATCGGTCG</td><td>>hg38_21_+_8433172_8446622_RNA45SN1_45S_50ntFlanks</td><td>9554</td><td>9582</td><td>29</td><td>CGTAGCGGTCCTGACGTGCAAATCGGTCG</td><td>1309.1</td><td>-5.2</td><td>0.8</td><td>-6.5</td><td>1.0E-10</td><td>2.6E-10</td><td>MB231/MB468</td><td>NUC</td></tr>
<tr><td>>hg38_21_+_8433172_8446622_RNA45SN1_45S_50ntFlanks@9545.9569.25|GTGGAGGTCCGTAGCGGTCCTGACG</td><td>>hg38_21_+_8433172_8446622_RNA45SN1_45S_50ntFlanks</td><td>9545</td><td>9569</td><td>25</td><td>GTGGAGGTCCGTAGCGGTCCTGACG</td><td>684.8</td><td>-8.4</td><td>1.3</td><td>-6.4</td><td>1.2E-10</td><td>2.6E-10</td><td>MB231/MB468</td><td>MITO</td></tr>
<tr><td>>hg38_21_+_8433172_8446622_RNA45SN1_45S_50ntFlanks@11670.11693.24|CCCAGTGCTCTGAATGTCAAAGTG</td><td>>hg38_21_+_8433172_8446622_RNA45SN1_45S_50ntFlanks</td><td>11670</td><td>11693</td><td>24</td><td>CCCAGTGCTCTGAATGTCAAAGTG</td><td>1086.5</td><td>-3.7</td><td>0.6</td><td>-6.5</td><td>1.0E-10</td><td>2.6E-10</td><td>MB231/MB468</td><td>NUC</td></tr>
<tr><td>>hg38_21_+_8433172_8446622_RNA45SN1_45S_50ntFlanks@11703.11738.36|AATGAAGCGCGGGTAAACGGCGGGAGTAACTATGAC</td><td>>hg38_21_+_8433172_8446622_RNA45SN1_45S_50ntFlanks</td><td>11703</td><td>11738</td><td>36</td><td>AATGAAGCGCGGGTAAACGGCGGGAGTAACTATGAC</td><td>74.5</td><td>-10.1</td><td>1.6</td><td>-6.5</td><td>9.7E-11</td><td>2.6E-10</td><td>MB231/MB468</td><td>MP</td></tr>
<tr><td>>hg38_21_+_8433172_8446622_RNA45SN1_45S_50ntFlanks@8194.8211.18|CCCGTGGACGGTGTGAGG</td><td>>hg38_21_+_8433172_8446622_RNA45SN1_45S_50ntFlanks</td><td>8194</td><td>8211</td><td>18</td><td>CCCGTGGACGGTGTGAGG</td><td>1760.3</td><td>-7.6</td><td>1.2</td><td>-6.5</td><td>1.0E-10</td><td>2.7E-10</td><td>MB231/MB468</td><td>NUC</td></tr>
<tr><td>>hg38_21_+_8433172_8446622_RNA45SN1_45S_50ntFlanks@8182.8208.27|ATCGAGGCCCAGCCCGTGGACGGTGTG</td><td>>hg38_21_+_8433172_8446622_RNA45SN1_45S_50ntFlanks</td><td>8182</td><td>8208</td><td>27</td><td>ATCGAGGCCCAGCCCGTGGACGGTGTG</td><td>653.6</td><td>-8.8</td><td>1.4</td><td>-6.5</td><td>9.7E-11</td><td>2.7E-10</td><td>MB231/MB468</td><td>MP</td></tr>
<tr><td>>hg38_21_+_8433172_8446622_RNA45SN1_45S_50ntFlanks@11564.11605.42|CCTAGCAGCCGACTTAGAACTGGTGCGGACCAGGGGAATCCG</td><td>>hg38_21_+_8433172_8446622_RNA45SN1_45S_50ntFlanks</td><td>11564</td><td>11605</td><td>42</td><td>CCTAGCAGCCGACTTAGAACTGGTGCGGACCAGGGGAATCCG</td><td>595.6</td><td>-8.9</td><td>1.4</td><td>-6.5</td><td>1.0E-10</td><td>2.7E-10</td><td>MB231/MB468</td><td>NUC</td></tr>
<tr><td>>hg38_21_+_8433172_8446622_RNA45SN1_45S_50ntFlanks@4764.4783.20|ATACCGTCGTAGTTCCGACC</td><td>>hg38_21_+_8433172_8446622_RNA45SN1_45S_50ntFlanks</td><td>4764</td><td>4783</td><td>20</td><td>ATACCGTCGTAGTTCCGACC</td><td>83.3</td><td>-10.2</td><td>1.6</td><td>-6.5</td><td>9.8E-11</td><td>2.7E-10</td><td>MB231/MB468</td><td>MP</td></tr>
</table>

<table>
<tr><td>>MI0000471|hsa-mir-126&WithFlank&9|+|136670596|136670692@58.78.21(+1U)&[MIMAT0000445&hsa-miR-126-3p&offsets|0|-1(+1U);m-40&9|+|136670653|136670674&offsets|0|-1(+1U)]|TCGTACCGTGAGTAATAATGCT</td><td>>MI0000471|hsa-mir-126&WithFlank&9|+|136670596|136670692</td><td>58</td><td>78</td><td>21(+1U)</td><td>TCGTACCGTGAGTAATAATGCT</td><td>180.2</td><td>10.7</td><td>1.7</td><td>6.5</td><td>9.8E-11</td><td>2.7E-10</td><td>MB231/MB468</td><td>MP</td></tr>
<tr><td>>hg38_21_+_8433172_8446622_RNA45SN1_45S_50ntFlanks@8177.8195.19|TTCTGATCGAGGCCCAGCC</td><td>>hg38_21_+_8433172_8446622_RNA45SN1_45S_50ntFlanks</td><td>8177</td><td>8195</td><td>19</td><td>TTCTGATCGAGGCCCAGCC</td><td>937.6</td><td>4.8</td><td>0.8</td><td>6.5</td><td>1.0E-10</td><td>2.7E-10</td><td>MB231/MB468</td><td>CYTO</td></tr>
<tr><td>>hg38_21_+_8433172_8446622_RNA45SN1_45S_50ntFlanks@12430.12455.26|TTCCTATCATTGTGAAGCAGAATTCA</td><td>>hg38_21_+_8433172_8446622_RNA45SN1_45S_50ntFlanks</td><td>12430</td><td>12455</td><td>26</td><td>TTCCTATCATTGTGAAGCAGAATTCA</td><td>77.9</td><td>-10.1</td><td>1.6</td><td>-6.5</td><td>9.8E-11</td><td>2.7E-10</td><td>MB231/MB468</td><td>MP</td></tr>
<tr><td>>hg38_21_+_8433172_8446622_RNA45SN1_45S_50ntFlanks@9790.9809.20|CTGGCGTGGAGCCGGGCGTG</td><td>>hg38_21_+_8433172_8446622_RNA45SN1_45S_50ntFlanks</td><td>9790</td><td>9809</td><td>20</td><td>CTGGCGTGGAGCCGGGCGTG</td><td>105.7</td><td>-10.6</td><td>1.6</td><td>-6.5</td><td>9.8E-11</td><td>2.7E-10</td><td>MB231/MB468</td><td>MP</td></tr>
<tr><td>>hg38_21_+_8433172_8446622_RNA45SN1_45S_50ntFlanks@8176.8199.24|CTTCTGATCGAGGCCCAGCCCGTG</td><td>>hg38_21_+_8433172_8446622_RNA45SN1_45S_50ntFlanks</td><td>8176</td><td>8199</td><td>24</td><td>CTTCTGATCGAGGCCCAGCCCGTG</td><td>546.6</td><td>-6.8</td><td>1.0</td><td>-6.5</td><td>9.8E-11</td><td>2.7E-10</td><td>MB231/MB468</td><td>MP</td></tr>
<tr><td>>hg38_21_+_8433172_8446622_RNA45SN1_45S_50ntFlanks@5186.5209.24|CAATAACAGGTCTGTGATGCCCTT</td><td>>hg38_21_+_8433172_8446622_RNA45SN1_45S_50ntFlanks</td><td>5186</td><td>5209</td><td>24</td><td>CAATAACAGGTCTGTGATGCCCTT</td><td>73.1</td><td>-10.1</td><td>1.6</td><td>-6.5</td><td>9.8E-11</td><td>2.7E-10</td><td>MB231/MB468</td><td>MP</td></tr>
<tr><td>>hg38_21_+_8433172_8446622_RNA45SN1_45S_50ntFlanks@11602.11620.19|TCCGACTGTTTAATTAAAA</td><td>>hg38_21_+_8433172_8446622_RNA45SN1_45S_50ntFlanks</td><td>11602</td><td>11620</td><td>19</td><td>TCCGACTGTTTAATTAAAA</td><td>2879.6</td><td>-10.4</td><td>1.6</td><td>-6.4</td><td>1.2E-10</td><td>2.7E-10</td><td>BT20/MB468</td><td>NUC</td></tr>
<tr><td>>hg38_21_+_8433172_8446622_RNA45SN1_45S_50ntFlanks@10349.10376.28|AACGAGAACTTTGAAGGCCGAAGTGGAG</td><td>>hg38_21_+_8433172_8446622_RNA45SN1_45S_50ntFlanks</td><td>10349</td><td>10376</td><td>28</td><td>AACGAGAACTTTGAAGGCCGAAGTGGAG</td><td>2400.8</td><td>-8.1</td><td>1.3</td><td>-6.4</td><td>1.2E-10</td><td>2.7E-10</td><td>BT20/MB468</td><td>NUC</td></tr>
<tr><td>>MI0000103|hsa-mir-101-1&WithFlank&1|-|65058428|65058514@53.73.21(+1U);MI0000739|hsa-mir-101-2&WithFlank&9|+|4850291|4850381@55.75.21(+1U)&[MIMAT0000099&hsa-miR-101-3p&offsets|0|0(+1U);m-17&1|-|65058443|65058462&offsets|0|+1(+1U)];[MIMAT0000099_1&hsa-miR-101-3p&offsets|0|0(+1U);m-16&9|+|4850345|4850364&offsets|0|+1(+1U)]|TACAGTACTGTGATAACTGAAT</td><td>>MI0000103|hsa-mir-101-1&WithFlank&1|-|65058428|65058514</td><td>53</td><td>73</td><td>21(+1U)</td><td>TACAGTACTGTGATAACTGAAT</td><td>790.3</td><td>7.6</td><td>1.2</td><td>6.5</td><td>1.0E-10</td><td>2.7E-10</td><td>MB231/MB468</td><td>NUC</td></tr>
<tr><td>>hg38_21_+_8433172_8446622_RNA45SN1_45S_50ntFlanks@3776.3802.27|CCGGTACAGTGAAACTGCGAATGGCTC</td><td>>hg38_21_+_8433172_8446622_RNA45SN1_45S_50ntFlanks</td><td>3776</td><td>3802</td><td>27</td><td>CCGGTACAGTGAAACTGCGAATGGCTC</td><td>96.2</td><td>-10.5</td><td>1.6</td><td>-6.5</td><td>9.9E-11</td><td>2.7E-10</td><td>MB231/MB468</td><td>MP</td></tr>
<tr><td>>hg38_21_+_8433172_8446622_RNA45SN1_45S_50ntFlanks@8189.8206.18|CCCAGCCCGTGGACGGTG</td><td>>hg38_21_+_8433172_8446622_RNA45SN1_45S_50ntFlanks</td><td>8189</td><td>8206</td><td>18</td><td>CCCAGCCCGTGGACGGTG</td><td>184.8</td><td>-10.6</td><td>1.6</td><td>-6.8</td><td>7.8E-12</td><td>2.7E-10</td><td>BT20/MB468</td><td>MP</td></tr>
<tr><td>>hg38_21_+_8433172_8446622_RNA45SN1_45S_50ntFlanks@9806.9823.18|CGTGGAATGCGAGTGCCT</td><td>>hg38_21_+_8433172_8446622_RNA45SN1_45S_50ntFlanks</td><td>9806</td><td>9823</td><td>18</td><td>CGTGGAATGCGAGTGCCT</td><td>88.2</td><td>-10.3</td><td>1.6</td><td>-6.5</td><td>9.9E-11</td><td>2.7E-10</td><td>MB231/MB468</td><td>MP</td></tr>
<tr><td>>hg38_21_+_8433172_8446622_RNA45SN1_45S_50ntFlanks@11641.11676.36|CGGCGGGTGTTGACGCGATGTGATTTCTGCCCAGTG</td><td>>hg38_21_+_8433172_8446622_RNA45SN1_45S_50ntFlanks</td><td>11641</td><td>11676</td><td>36</td><td>CGGCGGGTGTTGACGCGATGTGATTTCTGCCCAGTG</td><td>210.9</td><td>-9.8</td><td>1.5</td><td>-6.5</td><td>7.7E-11</td><td>2.7E-10</td><td>BT20/MB468</td><td>MITO</td></tr>
<tr><td>>hg38_21_+_8433172_8446622_RNA45SN1_45S_50ntFlanks@13007.13027.21|TATTGAAAGTCAGCCCTCGAC</td><td>>hg38_21_+_8433172_8446622_RNA45SN1_45S_50ntFlanks</td><td>13007</td><td>13027</td><td>21</td><td>TATTGAAAGTCAGCCCTCGAC</td><td>532.5</td><td>-8.8</td><td>1.4</td><td>-6.4</td><td>1.3E-10</td><td>2.7E-10</td><td>BT20/MB468</td><td>NUC</td></tr>
<tr><td>>hg38_21_+_8433172_8446622_RNA45SN1_45S_50ntFlanks@4759.4793.35|ATCAGATACCGTCGTAGTTCCGACCATAAACGATG</td><td>>hg38_21_+_8433172_8446622_RNA45SN1_45S_50ntFlanks</td><td>4759</td><td>4793</td><td>35</td><td>ATCAGATACCGTCGTAGTTCCGACCATAAACGATG</td><td>449.6</td><td>-11.9</td><td>1.7</td><td>-6.8</td><td>7.9E-12</td><td>2.7E-10</td><td>BT20/MB468</td><td>MP</td></tr>
<tr><td>>hg38_21_+_8433172_8446622_RNA45SN1_45S_50ntFlanks@12459.12476.18|AGCGTTGGATTGTTCACC</td><td>>hg38_21_+_8433172_8446622_RNA45SN1_45S_50ntFlanks</td><td>12459</td><td>12476</td><td>18</td><td>AGCGTTGGATTGTTCACC</td><td>3574.7</td><td>-4.3</td><td>0.7</td><td>-6.5</td><td>1.1E-10</td><td>2.7E-10</td><td>MB231/MB468</td><td>CYTO</td></tr>
<tr><td>>MI0000088|hsa-mir-30a&WithFlank&6|-|71403545|71403627@13.35.23(+2U)&[MIMAT0000087&hsa-miR-30a-5p&offsets|+1|+2(+2U);m-15&6|-|71403595|71403616&offsets|+1|+2(+2U)]|GTAAACATCCTCGACTGGAAGCTTT</td><td>>MI0000088|hsa-mir-30a&WithFlank&6|-|71403545|71403627</td><td>13</td><td>35</td><td>23(+2U)</td><td>GTAAACATCCTCGACTGGAAGCTTT</td><td>98.3</td><td>9.8</td><td>1.5</td><td>6.5</td><td>9.9E-11</td><td>2.7E-10</td><td>MB231/MB468</td><td>MP</td></tr>
<tr><td>>hg38_21_+_8433172_8446622_RNA45SN1_45S_50ntFlanks@3813.3830.18|TTATGGTTCCTTTGGTCG</td><td>>hg38_21_+_8433172_8446622_RNA45SN1_45S_50ntFlanks</td><td>3813</td><td>3830</td><td>18</td><td>TTATGGTTCCTTTGGTCG</td><td>72.1</td><td>-10.0</td><td>1.6</td><td>-6.5</td><td>9.9E-11</td><td>2.7E-10</td><td>MB231/MB468</td><td>MP</td></tr>
<tr><td>>hg38_21_+_8433172_8446622_RNA45SN1_45S_50ntFlanks@6745.6771.27|ACACTTCGAACGCACTTGCGGCCCCGG</td><td>>hg38_21_+_8433172_8446622_RNA45SN1_45S_50ntFlanks</td><td>6745</td><td>6771</td><td>27</td><td>ACACTTCGAACGCACTTGCGGCCCCGG</td><td>2164.4</td><td>-6.6</td><td>1.0</td><td>-6.5</td><td>9.4E-11</td><td>2.7E-10</td><td>BT20/MB468</td><td>CYTO</td></tr>
<tr><td>>MI0005756|hsa-mir-934&WithFlank&X|+|136550872|136550966@21.42.22&[MIMAT0004977&hsa-miR-934&offsets|0|0;m-943&X|+|136550892|136550913&offsets|0|0]|TGTCTACTACTGGAGACACTGG</td><td>>MI0005756|hsa-mir-934&WithFlank&X|+|136550872|136550966</td><td>21</td><td>42</td><td>22</td><td>TGTCTACTACTGGAGACACTGG</td><td>1172.1</td><td>13.5</td><td>1.9</td><td>7.1</td><td>1.7E-12</td><td>2.7E-10</td><td>BT20/MB231</td><td>MP</td></tr>
<tr><td>>trna2_GlyGCC_21_-_18827107_18827177@1.28.28,trna35_GlyGCC_1_+_161413094_161413164@1.28.28,trna37_GlyGCC_1_+_161420467_161420537@1.28.28,trna39_GlyGCC_1_+_161427898_161427968@1.28.28,trna41_GlyGCC_1_+_161435258_161435328@1.28.28|GCATGGGTGGTTCAGTGGTAGAATTCTC</td><td>>trna2_GlyGCC_21_-_18827107_18827177</td><td>1</td><td>28</td><td>28</td><td>GCATGGGTGGTTCAGTGGTAGAATTCTC</td><td>5294.5</td><td>7.5</td><td>1.1</td><td>7.1</td><td>1.7E-12</td><td>2.7E-10</td><td>BT20/MB231</td><td>MP</td></tr>
<tr><td>>MI0016887|hsa-mir-4521&WithFlank&17|+|8186939|8187010@13.34.22&[MIMAT0019058&hsa-miR-4521&offsets|+3|+3;m-765&17|+|8186948|8186969&offsets|+3|+3]|AAGGAAGTCCTGTGCTCAGTTT</td><td>>MI0016887|hsa-mir-4521&WithFlank&17|+|8186939|8187010</td><td>13</td><td>34</td><td>22</td><td>AAGGAAGTCCTGTGCTCAGTTT</td><td>115.1</td><td>9.7</td><td>1.5</td><td>6.4</td><td>1.3E-10</td><td>2.7E-10</td><td>BT20/MB468</td><td>NUC</td></tr>
<tr><td>>hg38_21_+_8433172_8446622_RNA45SN1_45S_50ntFlanks@4733.4765.33|GAACGAAAGTCGGAGGTTCGAAGACGATCAGAT</td><td>>hg38_21_+_8433172_8446622_RNA45SN1_45S_50ntFlanks</td><td>4733</td><td>4765</td><td>33</td><td>GAACGAAAGTCGGAGGTTCGAAGACGATCAGAT</td><td>264.7</td><td>4.5</td><td>0.7</td><td>6.5</td><td>1.1E-10</td><td>2.7E-10</td><td>MB231/MB468</td><td>NUC</td></tr>
<tr><td>>hg38_21_+_8433172_8446622_RNA45SN1_45S_50ntFlanks@12605.12634.30|TATGTGCTTGGCTGAGGAGCCAATGGGGCG</td><td>>hg38_21_+_8433172_8446622_RNA45SN1_45S_50ntFlanks</td><td>12605</td><td>12634</td><td>30</td><td>TATGTGCTTGGCTGAGGAGCCAATGGGGCG</td><td>305.5</td><td>-8.0</td><td>1.2</td><td>-6.5</td><td>1.1E-10</td><td>2.7E-10</td><td>MB231/MB468</td><td>NUC</td></tr>
<tr><td>>hg38_21_+_8433172_8446622_RNA45SN1_45S_50ntFlanks@9712.9735.24|AGCGAATGATTAGAGGTCTTGGGG</td><td>>hg38_21_+_8433172_8446622_RNA45SN1_45S_50ntFlanks</td><td>9712</td><td>9735</td><td>24</td><td>AGCGAATGATTAGAGGTCTTGGGG</td><td>84.1</td><td>-10.3</td><td>1.6</td><td>-6.5</td><td>1.0E-10</td><td>2.7E-10</td><td>MB231/MB468</td><td>MP</td></tr>
<tr><td>>hg38_21_+_8433172_8446622_RNA45SN1_45S_50ntFlanks@4372.4398.27|AAAAAGCTCGTAGTTGGATCTTGGGAG</td><td>>hg38_21_+_8433172_8446622_RNA45SN1_45S_50ntFlanks</td><td>4372</td><td>4398</td><td>27</td><td>AAAAAGCTCGTAGTTGGATCTTGGGAG</td><td>268.3</td><td>-8.3</td><td>1.3</td><td>-6.5</td><td>1.1E-10</td><td>2.7E-10</td><td>MB231/MB468</td><td>NUC</td></tr>
<tr><td>>MI0000650|hsa-mir-200c&WithFlank&12|+|6963693|6963772@50.69.20(+1A)&[MIMAT0000617&hsa-miR-200c-3p&offsets|0|-3(+1A);m-54&12|+|6963742|6963764&offsets|0|-3(+1A)]|TAATACTGCCGGGTAATGATA</td><td>>MI0000650|hsa-mir-200c&WithFlank&12|+|6963693|6963772</td><td>50</td><td>69</td><td>20(+1A)</td><td>TAATACTGCCGGGTAATGATA</td><td>383.6</td><td>11.9</td><td>1.7</td><td>7.1</td><td>1.7E-12</td><td>2.7E-10</td><td>BT20/MB231</td><td>MP</td></tr>
<tr><td>>hg38_21_+_8433172_8446622_RNA45SN1_45S_50ntFlanks@4738.4768.31|AAAGTCGGAGGTTCGAAGACGATCAGATACC</td><td>>hg38_21_+_8433172_8446622_RNA45SN1_45S_50ntFlanks</td><td>4738</td><td>4768</td><td>31</td><td>AAAGTCGGAGGTTCGAAGACGATCAGATACC</td><td>1156.8</td><td>8.4</td><td>1.3</td><td>6.5</td><td>1.1E-10</td><td>2.7E-10</td><td>MB231/MB468</td><td>CYTO</td></tr>
<tr><td>>hg38_21_+_8433172_8446622_RNA45SN1_45S_50ntFlanks@10350.10383.34|ACGAGAACTTTGAAGGCCGAAGTGGAGAAGGGTT</td><td>>hg38_21_+_8433172_8446622_RNA45SN1_45S_50ntFlanks</td><td>10350</td><td>10383</td><td>34</td><td>ACGAGAACTTTGAAGGCCGAAGTGGAGAAGGGTT</td><td>4538.9</td><td>-4.0</td><td>0.6</td><td>-6.5</td><td>7.8E-11</td><td>2.7E-10</td><td>BT20/MB468</td><td>MITO</td></tr>
<tr><td>>trna11_GluTTC_15_-_26327381_26327452@1.34.34,trna3_GluTTC_13_-_45492062_45492133@1.34.34|TCCCACATGGTCTAGCGGTTAGGATTCCTGGTTT</td><td>>trna11_GluTTC_15_-_26327381_26327452</td><td>1</td><td>34</td><td>34</td><td>TCCCACATGGTCTAGCGGTTAGGATTCCTGGTTT</td><td>234.5</td><td>11.1</td><td>1.7</td><td>6.4</td><td>1.2E-10</td><td>2.7E-10</td><td>MB231/MB468</td><td>MITO</td></tr>
<tr><td>>hg38_21_+_8433172_8446622_RNA45SN1_45S_50ntFlanks@6732.6774.43|ACATTGATCATCGACACTTCGAACGCACTTGCGGCCCCGGGTT</td><td>>hg38_21_+_8433172_8446622_RNA45SN1_45S_50ntFlanks</td><td>6732</td><td>6774</td><td>43</td><td>ACATTGATCATCGACACTTCGAACGCACTTGCGGCCCCGGGTT</td><td>1906.7</td><td>7.6</td><td>1.2</td><td>6.5</td><td>7.8E-11</td><td>2.7E-10</td><td>BT20/MB468</td><td>MITO</td></tr>
<tr><td>>hg38_21_+_8433172_8446622_RNA45SN1_45S_50ntFlanks@10678.10710.33|CGGTTCCGGCGGCGTCCGGTGAGCTCTCGCTGG</td><td>>hg38_21_+_8433172_8446622_RNA45SN1_45S_50ntFlanks</td><td>10678</td><td>10710</td><td>33</td><td>CGGTTCCGGCGGCGTCCGGTGAGCTCTCGCTGG</td><td>324239.3</td><td>-6.3</td><td>1.0</td><td>-6.5</td><td>7.8E-11</td><td>2.7E-10</td><td>BT20/MB468</td><td>MITO</td></tr>
<tr><td>>hg38_21_+_8433172_8446622_RNA45SN1_45S_50ntFlanks@12640.12670.31|ACCATCTGTGGGATTATGACTGAACGCCTCT</td><td>>hg38_21_+_8433172_8446622_RNA45SN1_45S_50ntFlanks</td><td>12640</td><td>12670</td><td>31</td><td>ACCATCTGTGGGATTATGACTGAACGCCTCT</td><td>495.4</td><td>7.6</td><td>1.2</td><td>6.5</td><td>1.1E-10</td><td>2.7E-10</td><td>MB231/MB468</td><td>CYTO</td></tr>
<tr><td>>hg38_21_+_8433172_8446622_RNA45SN1_45S_50ntFlanks@3738.3756.19|TCTCAAAGATTAAGCCATG</td><td>>hg38_21_+_8433172_8446622_RNA45SN1_45S_50ntFlanks</td><td>3738</td><td>3756</td><td>19</td><td>TCTCAAAGATTAAGCCATG</td><td>196.7</td><td>-9.7</td><td>1.5</td><td>-6.5</td><td>7.8E-11</td><td>2.7E-10</td><td>BT20/MB468</td><td>MITO</td></tr>
<tr><td>>hg38_21_+_8433172_8446622_RNA45SN1_45S_50ntFlanks@9543.9571.29|TGGTGGAGGTCCGTAGCGGTCCTGACGTG</td><td>>hg38_21_+_8433172_8446622_RNA45SN1_45S_50ntFlanks</td><td>9543</td><td>9571</td><td>29</td><td>TGGTGGAGGTCCGTAGCGGTCCTGACGTG</td><td>526.0</td><td>-3.4</td><td>0.5</td><td>-6.5</td><td>1.1E-10</td><td>2.7E-10</td><td>MB231/MB468</td><td>NUC</td></tr>
<tr><td>>hg38_21_+_8433172_8446622_RNA45SN1_45S_50ntFlanks@12388.12434.47|CGTTCATAGCGACGTCGCTTTTTGATCCTTCGATGTCGGCTCTTCCT</td><td>>hg38_21_+_8433172_8446622_RNA45SN1_45S_50ntFlanks</td><td>12388</td><td>12434</td><td>47</td><td>CGTTCATAGCGACGTCGCTTTTTGATCCTTCGATGTCGGCTCTTCCT</td><td>784.7</td><td>6.9</td><td>1.1</td><td>6.5</td><td>1.1E-10</td><td>2.7E-10</td><td>MB231/MB468</td><td>CYTO</td></tr>
<tr><td>>hg38_21_+_8433172_8446622_RNA45SN1_45S_50ntFlanks@9309.9335.27|GGAGTCTAACACGTGCGCGAGTCGGGG</td><td>>hg38_21_+_8433172_8446622_RNA45SN1_45S_50ntFlanks</td><td>9309</td><td>9335</td><td>27</td><td>GGAGTCTAACACGTGCGCGAGTCGGGG</td><td>108.6</td><td>-10.6</td><td>1.6</td><td>-6.5</td><td>1.0E-10</td><td>2.7E-10</td><td>MB231/MB468</td><td>MP</td></tr>
<tr><td>>hg38_21_+_8433172_8446622_RNA45SN1_45S_50ntFlanks@11667.11691.25|CTGCCCAGTGCTCTGAATGTCAAAG</td><td>>hg38_21_+_8433172_8446622_RNA45SN1_45S_50ntFlanks</td><td>11667</td><td>11691</td><td>25</td><td>CTGCCCAGTGCTCTGAATGTCAAAG</td><td>561.2</td><td>7.2</td><td>1.1</td><td>6.5</td><td>9.6E-11</td><td>2.8E-10</td><td>BT20/MB468</td><td>CYTO</td></tr>
<tr><td>>hg38_21_+_8433172_8446622_RNA45SN1_45S_50ntFlanks@10844.10865.22|AACTTCGGGATAAGGATTGGCT</td><td>>hg38_21_+_8433172_8446622_RNA45SN1_45S_50ntFlanks</td><td>10844</td><td>10865</td><td>22</td><td>AACTTCGGGATAAGGATTGGCT</td><td>501.9</td><td>-8.4</td><td>1.3</td><td>-6.5</td><td>1.1E-10</td><td>2.8E-10</td><td>MB231/MB468</td><td>NUC</td></tr>
<tr><td>>trnaMT_IleGAT_MT_+_4263_4331@44.71.28|TAGGAGCTTAAACCCCCTTATTTCTACC</td><td>>trnaMT_IleGAT_MT_+_4263_4331</td><td>44</td><td>71</td><td>28</td><td>TAGGAGCTTAAACCCCCTTATTTCTACC</td><td>98.2</td><td>9.8</td><td>1.5</td><td>6.5</td><td>1.0E-10</td><td>2.8E-10</td><td>MB231/MB468</td><td>MP</td></tr>
<tr><td>>hg38_21_+_8433172_8446622_RNA45SN1_45S_50ntFlanks@11847.11875.29|AACGGGCTTGGCGGAATCAGCGGGGAAAG</td><td>>hg38_21_+_8433172_8446622_RNA45SN1_45S_50ntFlanks</td><td>11847</td><td>11875</td><td>29</td><td>AACGGGCTTGGCGGAATCAGCGGGGAAAG</td><td>71.1</td><td>-10.0</td><td>1.5</td><td>-6.5</td><td>1.0E-10</td><td>2.8E-10</td><td>MB231/MB468</td><td>MP</td></tr>
<tr><td>>hg38_21_+_8433172_8446622_RNA45SN1_45S_50ntFlanks@10409.10429.21|GTCAGTCGGTCCTGAGAGATG</td><td>>hg38_21_+_8433172_8446622_RNA45SN1_45S_50ntFlanks</td><td>10409</td><td>10429</td><td>21</td><td>GTCAGTCGGTCCTGAGAGATG</td><td>528.6</td><td>-4.8</td><td>0.8</td><td>-6.4</td><td>1.2E-10</td><td>2.8E-10</td><td>MB231/MB468</td><td>MITO</td></tr>
<tr><td>>hg38_21_+_8433172_8446622_RNA45SN1_45S_50ntFlanks@7983.8014.32|CAGATCAGACGTGGCGACCCGCTGAATTTAAG</td><td>>hg38_21_+_8433172_8446622_RNA45SN1_45S_50ntFlanks</td><td>7983</td><td>8014</td><td>32</td><td>CAGATCAGACGTGGCGACCCGCTGAATTTAAG</td><td>76.8</td><td>-10.1</td><td>1.6</td><td>-6.5</td><td>1.0E-10</td><td>2.8E-10</td><td>MB231/MB468</td><td>MP</td></tr>
<tr><td>>hg38_21_+_8433172_8446622_RNA45SN1_45S_50ntFlanks@11197.11227.31|GGCCGGCGGCGGCGGCGACTCTGGACGCGAG</td><td>>hg38_21_+_8433172_8446622_RNA45SN1_45S_50ntFlanks</td><td>11197</td><td>11227</td><td>31</td><td>GGCCGGCGGCGGCGGCGACTCTGGACGCGAG</td><td>89.2</td><td>-10.3</td><td>1.6</td><td>-6.5</td><td>1.0E-10</td><td>2.8E-10</td><td>MB231/MB468</td><td>MP</td></tr>
<tr><td>>hg38_21_+_8433172_8446622_RNA45SN1_45S_50ntFlanks@8191.8211.21|CAGCCCGTGGACGGTGTGAGG</td><td>>hg38_21_+_8433172_8446622_RNA45SN1_45S_50ntFlanks</td><td>8191</td><td>8211</td><td>21</td><td>CAGCCCGTGGACGGTGTGAGG</td><td>435.3</td><td>-8.8</td><td>1.4</td><td>-6.4</td><td>1.2E-10</td><td>2.8E-10</td><td>MB231/MB468</td><td>MITO</td></tr>
<tr><td>>hg38_21_+_8433172_8446622_RNA45SN1_45S_50ntFlanks@10403.10433.31|ACATGGGTCAGTCGGTCCTGAGAGATGGGCG</td><td>>hg38_21_+_8433172_8446622_RNA45SN1_45S_50ntFlanks</td><td>10403</td><td>10433</td><td>31</td><td>ACATGGGTCAGTCGGTCCTGAGAGATGGGCG</td><td>97.1</td><td>-10.5</td><td>1.6</td><td>-6.5</td><td>1.0E-10</td><td>2.8E-10</td><td>MB231/MB468</td><td>MP</td></tr>
<tr><td>>hg38_21_+_8433172_8446622_RNA45SN1_45S_50ntFlanks@8176.8197.22|CTTCTGATCGAGGCCCAGCCCG</td><td>>hg38_21_+_8433172_8446622_RNA45SN1_45S_50ntFlanks</td><td>8176</td><td>8197</td><td>22</td><td>CTTCTGATCGAGGCCCAGCCCG</td><td>1040.4</td><td>-8.7</td><td>1.3</td><td>-6.5</td><td>1.0E-10</td><td>2.8E-10</td><td>MB231/MB468</td><td>MP</td></tr>
<tr><td>>hg38_21_+_8433172_8446622_RNA45SN1_45S_50ntFlanks@11565.11585.21|CTAGCAGCCGACTTAGAACTG</td><td>>hg38_21_+_8433172_8446622_RNA45SN1_45S_50ntFlanks</td><td>11565</td><td>11585</td><td>21</td><td>CTAGCAGCCGACTTAGAACTG</td><td>418.2</td><td>-6.7</td><td>1.0</td><td>-6.4</td><td>1.3E-10</td><td>2.8E-10</td><td>BT20/MB468</td><td>NUC</td></tr>
<tr><td>>MI0000746|hsa-mir-99b&WithFlank&19|+|51692606|51692687@51.69.19&[MIMAT0004678&hsa-miR-99b-3p&offsets|0|-3;m-151&19|+|51692656|51692678&offsets|0|-4]|CAAGCTCGTGTCTGTGGGT</td><td>>MI0000746|hsa-mir-99b&WithFlank&19|+|51692606|51692687</td><td>51</td><td>69</td><td>19</td><td>CAAGCTCGTGTCTGTGGGT</td><td>492.2</td><td>8.5</td><td>1.3</td><td>6.5</td><td>9.7E-11</td><td>2.8E-10</td><td>BT20/MB468</td><td>CYTO</td></tr>
<tr><td>>hg38_21_+_8433172_8446622_RNA45SN1_45S_50ntFlanks@5084.5102.19|CGAACGAGACTCTGGCATG</td><td>>hg38_21_+_8433172_8446622_RNA45SN1_45S_50ntFlanks</td><td>5084</td><td>5102</td><td>19</td><td>CGAACGAGACTCTGGCATG</td><td>1376.1</td><td>-8.5</td><td>1.3</td><td>-6.4</td><td>1.3E-10</td><td>2.8E-10</td><td>BT20/MB468</td><td>NUC</td></tr>
</table>

<table>
<tr><td>>am_trna111_HisGTG_1_-_147774845_147774916@3.29.27,trna118_HisGTG_1_-_145396881_145396952@3.29.27,trna16_HisGTG_1_+_146544773_146544844@3.29.27,trna1_HisGTG_15_+_45493349_45493420@3.29.27,trna21_HisGTG_1_+_147753471_147753542@3.29.27,trna33_HisGTG_6_+_27125906_27125977@3.29.27,trna7_HisGTG_9_-_14433938_14434009@3.29.27,trna8_HisGTG_15_-_45492611_45492682@3.29.27,trna9_HisGTG_15_-_45490804_45490875@3.29.27|CGTGATCGTATAGTGGTTAGTACTCTG</td><td>>am_trna111_HisGTG_1_-_147774845_147774916</td><td>3</td><td>29</td><td>27</td><td>CGTGATCGTATAGTGGTTAGTACTCTG</td><td>277.8</td><td>-6.6</td><td>1.0</td><td>-6.5</td><td>1.1E-10</td><td>2.8E-10</td><td>MB231/MB468</td><td>CYTO</td></tr>
<tr><td>>hg38_21_+_8433172_8446622_RNA45SN1_45S_50ntFlanks@7988.8009.22|CAGACGTGGCGACCCGCTGAAT</td><td>>hg38_21_+_8433172_8446622_RNA45SN1_45S_50ntFlanks</td><td>7988</td><td>8009</td><td>22</td><td>CAGACGTGGCGACCCGCTGAAT</td><td>3434.6</td><td>-5.0</td><td>0.8</td><td>-6.4</td><td>1.2E-10</td><td>2.8E-10</td><td>MB231/MB468</td><td>MITO</td></tr>
<tr><td>>hg38_21_+_8433172_8446622_RNA45SN1_45S_50ntFlanks@6743.6771.29|CGACACTTCGAACGCACTTGCGGCCCCGG</td><td>>hg38_21_+_8433172_8446622_RNA45SN1_45S_50ntFlanks</td><td>6743</td><td>6771</td><td>29</td><td>CGACACTTCGAACGCACTTGCGGCCCCGG</td><td>1108.5</td><td>-6.4</td><td>1.0</td><td>-6.5</td><td>1.1E-10</td><td>2.8E-10</td><td>MB231/MB468</td><td>CYTO</td></tr>
<tr><td>>MI0000255|hsa-mir-30d&WithFlank&8|-|134804870|134804951@12.34.23(+1A)&[MIMAT0000245&hsa-miR-30d-5p&offsets|0|+1(+1A);m-21&8|-|134804919|134804940&offsets|0|+1(+1A)]|TGTAAACATCCCCGACTGGAAGCA</td><td>>MI0000255|hsa-mir-30d&WithFlank&8|-|134804870|134804951</td><td>12</td><td>34</td><td>23(+1A)</td><td>TGTAAACATCCCCGACTGGAAGCA</td><td>4494.2</td><td>6.6</td><td>1.0</td><td>6.4</td><td>1.2E-10</td><td>2.8E-10</td><td>MB231/MB468</td><td>MITO</td></tr>
<tr><td>>hg38_21_+_8433172_8446622_RNA45SN1_45S_50ntFlanks@11885.11920.36|TTGAGCTTGACTCTAGTCTGGCACGGTGAAGAGACA</td><td>>hg38_21_+_8433172_8446622_RNA45SN1_45S_50ntFlanks</td><td>11885</td><td>11920</td><td>36</td><td>TTGAGCTTGACTCTAGTCTGGCACGGTGAAGAGACA</td><td>84.8</td><td>-10.3</td><td>1.6</td><td>-6.5</td><td>1.0E-10</td><td>2.8E-10</td><td>MB231/MB468</td><td>MP</td></tr>
<tr><td>>hg38_21_+_8433172_8446622_RNA45SN1_45S_50ntFlanks@6669.6698.30|CACTCGGCTCGTGCGTCGATGAAGAACGCA</td><td>>hg38_21_+_8433172_8446622_RNA45SN1_45S_50ntFlanks</td><td>6669</td><td>6698</td><td>30</td><td>CACTCGGCTCGTGCGTCGATGAAGAACGCA</td><td>72.5</td><td>-10.0</td><td>1.6</td><td>-6.5</td><td>1.0E-10</td><td>2.8E-10</td><td>MB231/MB468</td><td>MP</td></tr>
<tr><td>>hg38_21_+_8433172_8446622_RNA45SN1_45S_50ntFlanks@13002.13027.26|CGATCTATTGAAAGTCAGCCCTCGAC</td><td>>hg38_21_+_8433172_8446622_RNA45SN1_45S_50ntFlanks</td><td>13002</td><td>13027</td><td>26</td><td>CGATCTATTGAAAGTCAGCCCTCGAC</td><td>9280.8</td><td>-9.7</td><td>1.5</td><td>-6.5</td><td>1.1E-10</td><td>2.8E-10</td><td>MB231/MB468</td><td>NUC</td></tr>
<tr><td>>hg38_21_+_8433172_8446622_RNA45SN1_45S_50ntFlanks@3854.3899.46|ACTGTGGTAATTCTAGAGCTAATACATGCCGACGGGCGCTGACCCC</td><td>>hg38_21_+_8433172_8446622_RNA45SN1_45S_50ntFlanks</td><td>3854</td><td>3899</td><td>46</td><td>ACTGTGGTAATTCTAGAGCTAATACATGCCGACGGGCGCTGACCCC</td><td>225.5</td><td>-10.8</td><td>1.7</td><td>-6.5</td><td>1.1E-10</td><td>2.8E-10</td><td>MB231/MB468</td><td>NUC</td></tr>
<tr><td>>hg38_21_+_8433172_8446622_RNA45SN1_45S_50ntFlanks@12412.12446.35|ATCCTTCGATGTCGGCTCTTCCTATCATTGTGAAG</td><td>>hg38_21_+_8433172_8446622_RNA45SN1_45S_50ntFlanks</td><td>12412</td><td>12446</td><td>35</td><td>ATCCTTCGATGTCGGCTCTTCCTATCATTGTGAAG</td><td>110.3</td><td>-8.9</td><td>1.4</td><td>-6.5</td><td>8.0E-11</td><td>2.8E-10</td><td>BT20/MB468</td><td>MITO</td></tr>
<tr><td>>hg38_21_+_8433172_8446622_RNA45SN1_45S_50ntFlanks@12636.12661.26|AGCTACCATCTGTGGGATTATGACTG</td><td>>hg38_21_+_8433172_8446622_RNA45SN1_45S_50ntFlanks</td><td>12636</td><td>12661</td><td>26</td><td>AGCTACCATCTGTGGGATTATGACTG</td><td>1494.0</td><td>-8.0</td><td>1.2</td><td>-6.4</td><td>1.3E-10</td><td>2.8E-10</td><td>BT20/MB468</td><td>NUC</td></tr>
<tr><td>>hg38_21_+_8433172_8446622_RNA45SN1_45S_50ntFlanks@3872.3898.27|CTAATACATGCCGACGGGCGCTGACCC</td><td>>hg38_21_+_8433172_8446622_RNA45SN1_45S_50ntFlanks</td><td>3872</td><td>3898</td><td>27</td><td>CTAATACATGCCGACGGGCGCTGACCC</td><td>1626.2</td><td>-8.3</td><td>1.3</td><td>-6.5</td><td>8.0E-11</td><td>2.8E-10</td><td>BT20/MB468</td><td>MITO</td></tr>
<tr><td>>hg38_21_+_8433172_8446622_RNA45SN1_45S_50ntFlanks@9311.9334.24|AGTCTAACACGTGCGCGAGTCGGG</td><td>>hg38_21_+_8433172_8446622_RNA45SN1_45S_50ntFlanks</td><td>9311</td><td>9334</td><td>24</td><td>AGTCTAACACGTGCGCGAGTCGGG</td><td>415.3</td><td>-7.1</td><td>1.1</td><td>-6.5</td><td>1.0E-10</td><td>2.8E-10</td><td>MB231/MB468</td><td>MP</td></tr>
<tr><td>>MI0000476|hsa-mir-138-1&WithFlank&3|+|44114206|44114316@29.52.24&[MIMAT0000430_1&hsa-miR-138-5p&offsets|0|+1;m-497&3|+|44114234|44114256&offsets|0|+1]|AGCTGGTGTTGTGAATCAGGCCGT</td><td>>MI0000476|hsa-mir-138-1&WithFlank&3|+|44114206|44114316</td><td>29</td><td>52</td><td>24</td><td>AGCTGGTGTTGTGAATCAGGCCGT</td><td>248.0</td><td>11.2</td><td>1.7</td><td>6.4</td><td>1.2E-10</td><td>2.8E-10</td><td>MB231/MB468</td><td>MITO</td></tr>
<tr><td>>hg38_21_+_8433172_8446622_RNA45SN1_45S_50ntFlanks@9199.9222.24|TTCTCTCGGGGCCACGCGCGCGTC</td><td>>hg38_21_+_8433172_8446622_RNA45SN1_45S_50ntFlanks</td><td>9199</td><td>9222</td><td>24</td><td>TTCTCTCGGGGCCACGCGCGCGTC</td><td>102.3</td><td>-10.5</td><td>1.6</td><td>-6.5</td><td>1.0E-10</td><td>2.8E-10</td><td>MB231/MB468</td><td>MP</td></tr>
<tr><td>>MI0000816|hsa-mir-335&WithFlank&7|+|130496105|130496210@58.78.21(+1U)&[MIMAT0004703&hsa-miR-335-3p&offsets|0|-1(+1U);m-159&7|+|130496162|130496183&offsets|0|-1(+1U)]|TTTTTCATTATTGCTCCTGACT</td><td>>MI0000816|hsa-mir-335&WithFlank&7|+|130496105|130496210</td><td>58</td><td>78</td><td>21(+1U)</td><td>TTTTTCATTATTGCTCCTGACT</td><td>1313.7</td><td>6.7</td><td>1.0</td><td>6.4</td><td>1.2E-10</td><td>2.8E-10</td><td>MB231/MB468</td><td>MITO</td></tr>
<tr><td>>hg38_21_+_8433172_8446622_RNA45SN1_45S_50ntFlanks@7974.8004.31|ACGCGACCTCAGATCAGACGTGGCGACCCGC</td><td>>hg38_21_+_8433172_8446622_RNA45SN1_45S_50ntFlanks</td><td>7974</td><td>8004</td><td>31</td><td>ACGCGACCTCAGATCAGACGTGGCGACCCGC</td><td>677.3</td><td>4.8</td><td>0.7</td><td>6.5</td><td>1.1E-10</td><td>2.8E-10</td><td>MB231/MB468</td><td>CYTO</td></tr>
<tr><td>>trna119_LysCTT_1_-_145395522_145395594@1.33.33,trna11_LysCTT_5_-_180648979_180649051@1.33.33,trna13_LysCTT_6_+_26556774_26556846@1.33.33,trna7_LysCTT_16_+_3225692_3225764@1.33.33,trna9_LysCTT_5_+_180634755_180634827@1.33.33|GCCCGGCTAGCTCAGTCGGTAGAGCATGAGACT</td><td>>trna119_LysCTT_1_-_145395522_145395594</td><td>1</td><td>33</td><td>33</td><td>GCCCGGCTAGCTCAGTCGGTAGAGCATGAGACT</td><td>8141.5</td><td>6.1</td><td>0.9</td><td>6.4</td><td>1.2E-10</td><td>2.8E-10</td><td>MB231/MB468</td><td>MITO</td></tr>
<tr><td>>hg38_21_+_8433172_8446622_RNA45SN1_45S_50ntFlanks@12635.12658.24|AAGCTACCATCTGTGGGATTATGA</td><td>>hg38_21_+_8433172_8446622_RNA45SN1_45S_50ntFlanks</td><td>12635</td><td>12658</td><td>24</td><td>AAGCTACCATCTGTGGGATTATGA</td><td>78.8</td><td>-10.2</td><td>1.6</td><td>-6.5</td><td>1.0E-10</td><td>2.8E-10</td><td>MB231/MB468</td><td>MP</td></tr>
<tr><td>>MI0000255|hsa-mir-30d&WithFlank&8|-|134804870|134804951@12.35.24(+4U)&[MIMAT0000245&hsa-miR-30d-5p&offsets|0|+2(+4U);m-21&8|-|134804919|134804940&offsets|0|+2(+4U)]|TGTAAACATCCCCGACTGGAAGCTTTTT</td><td>>MI0000255|hsa-mir-30d&WithFlank&8|-|134804870|134804951</td><td>12</td><td>35</td><td>24(+4U)</td><td>TGTAAACATCCCCGACTGGAAGCTTTTT</td><td>177.3</td><td>10.7</td><td>1.7</td><td>6.4</td><td>1.2E-10</td><td>2.8E-10</td><td>MB231/MB468</td><td>MITO</td></tr>
<tr><td>>hg38_21_+_8433172_8446622_RNA45SN1_45S_50ntFlanks@10365.10388.24|GCCGAAGTGGAGAAGGGTTCCATG</td><td>>hg38_21_+_8433172_8446622_RNA45SN1_45S_50ntFlanks</td><td>10365</td><td>10388</td><td>24</td><td>GCCGAAGTGGAGAAGGGTTCCATG</td><td>83.3</td><td>-10.2</td><td>1.6</td><td>-6.5</td><td>1.0E-10</td><td>2.8E-10</td><td>MB231/MB468</td><td>MP</td></tr>
<tr><td>>hg38_21_+_8433172_8446622_RNA45SN1_45S_50ntFlanks@9539.9563.25|ACTCTGGTGGAGGTCCGTAGCGGTC</td><td>>hg38_21_+_8433172_8446622_RNA45SN1_45S_50ntFlanks</td><td>9539</td><td>9563</td><td>25</td><td>ACTCTGGTGGAGGTCCGTAGCGGTC</td><td>499.9</td><td>-8.3</td><td>1.3</td><td>-6.5</td><td>1.1E-10</td><td>2.8E-10</td><td>MB231/MB468</td><td>NUC</td></tr>
<tr><td>>hg38_21_+_8433172_8446622_RNA45SN1_45S_50ntFlanks@10808.10828.21|CAATGTAGGTAAGGGAAGTCG</td><td>>hg38_21_+_8433172_8446622_RNA45SN1_45S_50ntFlanks</td><td>10808</td><td>10828</td><td>21</td><td>CAATGTAGGTAAGGGAAGTCG</td><td>1147.6</td><td>-8.1</td><td>1.2</td><td>-6.5</td><td>8.1E-11</td><td>2.8E-10</td><td>BT20/MB468</td><td>MITO</td></tr>
<tr><td>>hg38_21_+_8433172_8446622_RNA45SN1_45S_50ntFlanks@4002.4042.41|GATAACCTCGGGCCGATCGCACGCCCCCCGTGGCGGCGACG</td><td>>hg38_21_+_8433172_8446622_RNA45SN1_45S_50ntFlanks</td><td>4002</td><td>4042</td><td>41</td><td>GATAACCTCGGGCCGATCGCACGCCCCCCGTGGCGGCGACG</td><td>780.6</td><td>-8.7</td><td>1.3</td><td>-6.5</td><td>1.1E-10</td><td>2.8E-10</td><td>MB231/MB468</td><td>NUC</td></tr>
<tr><td>>hg38_21_+_8433172_8446622_RNA45SN1_45S_50ntFlanks@8737.8756.20|CCCGTCCGTCCGTCCGTCCG</td><td>>hg38_21_+_8433172_8446622_RNA45SN1_45S_50ntFlanks</td><td>8737</td><td>8756</td><td>20</td><td>CCCGTCCGTCCGTCCGTCCG</td><td>76.5</td><td>-10.1</td><td>1.6</td><td>-6.5</td><td>1.0E-10</td><td>2.8E-10</td><td>MB231/MB468</td><td>MP</td></tr>
<tr><td>>hg38_21_+_8433172_8446622_RNA45SN1_45S_50ntFlanks@3899.3919.21|CCTTCGCGGGGGGGATGCGTG</td><td>>hg38_21_+_8433172_8446622_RNA45SN1_45S_50ntFlanks</td><td>3899</td><td>3919</td><td>21</td><td>CCTTCGCGGGGGGGATGCGTG</td><td>2245.5</td><td>-9.6</td><td>1.5</td><td>-6.5</td><td>1.1E-10</td><td>2.8E-10</td><td>MB231/MB468</td><td>NUC</td></tr>
<tr><td>>hg38_21_+_8433172_8446622_RNA45SN1_45S_50ntFlanks@10869.10889.21|AGGGCTGGGTCGGTCGGGCTG</td><td>>hg38_21_+_8433172_8446622_RNA45SN1_45S_50ntFlanks</td><td>10869</td><td>10889</td><td>21</td><td>AGGGCTGGGTCGGTCGGGCTG</td><td>1604.1</td><td>-9.5</td><td>1.5</td><td>-6.5</td><td>1.1E-10</td><td>2.8E-10</td><td>MB231/MB468</td><td>NUC</td></tr>
<tr><td>>hg38_21_+_8433172_8446622_RNA45SN1_45S_50ntFlanks@8191.8220.30|CAGCCCGTGGACGGTGTGAGGCCGGTAGCG</td><td>>hg38_21_+_8433172_8446622_RNA45SN1_45S_50ntFlanks</td><td>8191</td><td>8220</td><td>30</td><td>CAGCCCGTGGACGGTGTGAGGCCGGTAGCG</td><td>129.2</td><td>-10.9</td><td>1.7</td><td>-6.5</td><td>1.0E-10</td><td>2.8E-10</td><td>MB231/MB468</td><td>MP</td></tr>
<tr><td>>hg38_21_+_8433172_8446622_RNA45SN1_45S_50ntFlanks@9805.9832.28|GCGTGGAATGCGAGTGCCTAGTGGGCCA</td><td>>hg38_21_+_8433172_8446622_RNA45SN1_45S_50ntFlanks</td><td>9805</td><td>9832</td><td>28</td><td>GCGTGGAATGCGAGTGCCTAGTGGGCCA</td><td>135.2</td><td>-10.9</td><td>1.7</td><td>-6.5</td><td>1.0E-10</td><td>2.9E-10</td><td>MB231/MB468</td><td>MP</td></tr>
<tr><td>>MI0000077|hsa-mir-21&WithFlank&17|+|59841260|59841343@14.37.24&[MIMAT0000076&hsa-miR-21-5p&offsets|0|+2;m-1&17|+|59841273|59841294&offsets|0|+2]|TAGCTTATCAGACTGATGTTGACT</td><td>>MI0000077|hsa-mir-21&WithFlank&17|+|59841260|59841343</td><td>14</td><td>37</td><td>24</td><td>TAGCTTATCAGACTGATGTTGACT</td><td>82862.2</td><td>5.8</td><td>0.9</td><td>6.5</td><td>1.1E-10</td><td>2.9E-10</td><td>MB231/MB468</td><td>CYTO</td></tr>
<tr><td>>hg38_21_+_8433172_8446622_RNA45SN1_45S_50ntFlanks@10772.10806.35|AGGTCTCCAAGGTGAACAGCCTCTGGCATGTTGGA</td><td>>hg38_21_+_8433172_8446622_RNA45SN1_45S_50ntFlanks</td><td>10772</td><td>10806</td><td>35</td><td>AGGTCTCCAAGGTGAACAGCCTCTGGCATGTTGGA</td><td>82.4</td><td>-10.2</td><td>1.6</td><td>-6.5</td><td>1.0E-10</td><td>2.9E-10</td><td>MB231/MB468</td><td>MP</td></tr>
<tr><td>>hg38_21_+_8433172_8446622_RNA45SN1_45S_50ntFlanks@11564.11599.36|CCTAGCAGCCGACTTAGAACTGGTGCGGACCAGGGG</td><td>>hg38_21_+_8433172_8446622_RNA45SN1_45S_50ntFlanks</td><td>11564</td><td>11599</td><td>36</td><td>CCTAGCAGCCGACTTAGAACTGGTGCGGACCAGGGG</td><td>102.4</td><td>-10.5</td><td>1.6</td><td>-6.5</td><td>1.1E-10</td><td>2.9E-10</td><td>MB231/MB468</td><td>MP</td></tr>
<tr><td>>hg38_21_+_8433172_8446622_RNA45SN1_45S_50ntFlanks@6651.6693.43|CGACTCTTAGCGGTGGATCACTCGGCTCGTGCGTCGATGAAGA</td><td>>hg38_21_+_8433172_8446622_RNA45SN1_45S_50ntFlanks</td><td>6651</td><td>6693</td><td>43</td><td>CGACTCTTAGCGGTGGATCACTCGGCTCGTGCGTCGATGAAGA</td><td>2626.9</td><td>6.1</td><td>0.9</td><td>6.5</td><td>9.9E-11</td><td>2.9E-10</td><td>BT20/MB468</td><td>CYTO</td></tr>
<tr><td>>hg38_21_+_8433172_8446622_RNA45SN1_45S_50ntFlanks@475.513.39|CTTCGTGATCGATGTGGTGACGTCGTGCTCTCCCGGGCC</td><td>>hg38_21_+_8433172_8446622_RNA45SN1_45S_50ntFlanks</td><td>475</td><td>513</td><td>39</td><td>CTTCGTGATCGATGTGGTGACGTCGTGCTCTCCCGGGCC</td><td>621.5</td><td>8.6</td><td>1.3</td><td>6.5</td><td>8.2E-11</td><td>2.9E-10</td><td>BT20/MB468</td><td>MITO</td></tr>
<tr><td>>hg38_21_+_8433172_8446622_RNA45SN1_45S_50ntFlanks@7975.8020.46|CGCGACCTCAGATCAGACGTGGCGACCCGCTGAATTTAAGCATATT</td><td>>hg38_21_+_8433172_8446622_RNA45SN1_45S_50ntFlanks</td><td>7975</td><td>8020</td><td>46</td><td>CGCGACCTCAGATCAGACGTGGCGACCCGCTGAATTTAAGCATATT</td><td>1893.4</td><td>7.2</td><td>1.1</td><td>6.5</td><td>8.2E-11</td><td>2.9E-10</td><td>BT20/MB468</td><td>MITO</td></tr>
<tr><td>>hg38_21_+_8433172_8446622_RNA45SN1_45S_50ntFlanks@3785.3812.28|TGAAACTGCGAATGGCTCATTAAATCAG</td><td>>hg38_21_+_8433172_8446622_RNA45SN1_45S_50ntFlanks</td><td>3785</td><td>3812</td><td>28</td><td>TGAAACTGCGAATGGCTCATTAAATCAG</td><td>401.0</td><td>-10.7</td><td>1.7</td><td>-6.5</td><td>8.2E-11</td><td>2.9E-10</td><td>BT20/MB468</td><td>MITO</td></tr>
<tr><td>>hg38_21_+_8433172_8446622_RNA45SN1_45S_50ntFlanks@4853.4883.31|AAGTCTTTGGGTTCCGGGGGGAGTATGGTTG</td><td>>hg38_21_+_8433172_8446622_RNA45SN1_45S_50ntFlanks</td><td>4853</td><td>4883</td><td>31</td><td>AAGTCTTTGGGTTCCGGGGGGAGTATGGTTG</td><td>176.1</td><td>-9.5</td><td>1.5</td><td>-6.5</td><td>8.2E-11</td><td>2.9E-10</td><td>BT20/MB468</td><td>MITO</td></tr>
<tr><td>>hg38_21_+_8433172_8446622_RNA45SN1_45S_50ntFlanks@10788.10817.30|CAGCCTCTGGCATGTTGGAACAATGTAGGT</td><td>>hg38_21_+_8433172_8446622_RNA45SN1_45S_50ntFlanks</td><td>10788</td><td>10817</td><td>30</td><td>CAGCCTCTGGCATGTTGGAACAATGTAGGT</td><td>518.0</td><td>-8.2</td><td>1.3</td><td>-6.5</td><td>1.1E-10</td><td>2.9E-10</td><td>MB231/MB468</td><td>MP</td></tr>
<tr><td>>MI0003205|hsa-mir-532&WithFlank&X|+|50003142|50003244@27.47.21&[MIMAT0002888&hsa-miR-532-5p&offsets|+1|0;m-65&X|+|50003167|50003188&offsets|+1|0]|ATGCCTTGAGTGTAGGACCGT</td><td>>MI0003205|hsa-mir-532&WithFlank&X|+|50003142|50003244</td><td>27</td><td>47</td><td>21</td><td>ATGCCTTGAGTGTAGGACCGT</td><td>236.2</td><td>5.9</td><td>0.9</td><td>6.5</td><td>1.0E-10</td><td>2.9E-10</td><td>BT20/MB468</td><td>CYTO</td></tr>
<tr><td>>hg38_21_+_8433172_8446622_RNA45SN1_45S_50ntFlanks@11739.11765.27|TCTCTTAAGGTAGCCAAATGCCTCGTC</td><td>>hg38_21_+_8433172_8446622_RNA45SN1_45S_50ntFlanks</td><td>11739</td><td>11765</td><td>27</td><td>TCTCTTAAGGTAGCCAAATGCCTCGTC</td><td>81.1</td><td>-10.2</td><td>1.6</td><td>-6.5</td><td>1.1E-10</td><td>2.9E-10</td><td>MB231/MB468</td><td>MP</td></tr>
<tr><td>>MI0000085|hsa-mir-27a&WithFlank&19|-|13836434|13836523@16.37.22&[MIMAT0004501&hsa-miR-27a-5p&offsets|0|0;m-270&19|-|13836487|13836508&offsets|0|0]|AGGGCTTAGCTGCTTGTGAGCA</td><td>>MI0000085|hsa-mir-27a&WithFlank&19|-|13836434|13836523</td><td>16</td><td>37</td><td>22</td><td>AGGGCTTAGCTGCTTGTGAGCA</td><td>1714.9</td><td>6.0</td><td>0.9</td><td>6.5</td><td>1.1E-10</td><td>2.9E-10</td><td>MB231/MB468</td><td>CYTO</td></tr>
<tr><td>>hg38_21_+_8433172_8446622_RNA45SN1_45S_50ntFlanks@10866.10903.38|CTAAGGGCTGGGTCGGTCGGGCTGGGGCGCGAAGCGGG</td><td>>hg38_21_+_8433172_8446622_RNA45SN1_45S_50ntFlanks</td><td>10866</td><td>10903</td><td>38</td><td>CTAAGGGCTGGGTCGGTCGGGCTGGGGCGCGAAGCGGG</td><td>86.4</td><td>-10.3</td><td>1.6</td><td>-6.5</td><td>1.1E-10</td><td>2.9E-10</td><td>MB231/MB468</td><td>MP</td></tr>
<tr><td>>am_MI0003581|hsa-mir-574&WithFlank&4|+|38868026|38868133@31.54.24&[MIMAT0004795&hsa-miR-574-5p&offsets|0|+1;m-213&4|+|38868056|38868078&offsets|0|+1]|TGAGTGTGTGTGTGTGAGTGTGTG</td><td>>am_MI0003581|hsa-mir-574&WithFlank&4|+|38868026|38868133</td><td>31</td><td>54</td><td>24</td><td>TGAGTGTGTGTGTGTGAGTGTGTG</td><td>328.3</td><td>4.6</td><td>0.7</td><td>6.4</td><td>1.1E-10</td><td>2.9E-10</td><td>MB231/MB468</td><td>NUC</td></tr>
<tr><td>>trnaMT_GluTTC_MT_-_14674_14742@37.72.36,trnalookalike8_GluTTC_5_-_93905172_93905240@37.72.36|TCATTGGTCGTGGTTGTAGTCCGTGCGAGAATACCA</td><td>>trnaMT_GluTTC_MT_-_14674_14742</td><td>37</td><td>72</td><td>36</td><td>TCATTGGTCGTGGTTGTAGTCCGTGCGAGAATACCA</td><td>10656.8</td><td>5.8</td><td>0.9</td><td>6.4</td><td>1.1E-10</td><td>2.9E-10</td><td>MB231/MB468</td><td>NUC</td></tr>
<tr><td>>hg38_21_+_8433172_8446622_RNA45SN1_45S_50ntFlanks@10807.10831.25|ACAATGTAGGTAAGGGAAGTCGGCA</td><td>>hg38_21_+_8433172_8446622_RNA45SN1_45S_50ntFlanks</td><td>10807</td><td>10831</td><td>25</td><td>ACAATGTAGGTAAGGGAAGTCGGCA</td><td>74.7</td><td>-10.1</td><td>1.6</td><td>-6.5</td><td>1.1E-10</td><td>2.9E-10</td><td>MB231/MB468</td><td>MP</td></tr>
<tr><td>>hg38_21_+_8433172_8446622_RNA45SN1_45S_50ntFlanks@8176.8204.29|CTTCTGATCGAGGCCCAGCCCGTGGACGG</td><td>>hg38_21_+_8433172_8446622_RNA45SN1_45S_50ntFlanks</td><td>8176</td><td>8204</td><td>29</td><td>CTTCTGATCGAGGCCCAGCCCGTGGACGG</td><td>3465.9</td><td>-7.7</td><td>1.2</td><td>-6.5</td><td>1.1E-10</td><td>2.9E-10</td><td>MB231/MB468</td><td>MP</td></tr>
<tr><td>>hg38_21_+_8433172_8446622_RNA45SN1_45S_50ntFlanks@8025.8042.18|AGCGGAGGAGAAGAAACT</td><td>>hg38_21_+_8433172_8446622_RNA45SN1_45S_50ntFlanks</td><td>8025</td><td>8042</td><td>18</td><td>AGCGGAGGAGAAGAAACT</td><td>189.2</td><td>5.1</td><td>0.8</td><td>6.4</td><td>1.1E-10</td><td>2.9E-10</td><td>MB231/MB468</td><td>NUC</td></tr>
</table>

<table>
<tr><th></th><th></th><th></th><th></th><th></th><th></th><th></th><th></th><th></th><th></th><th></th><th></th><th></th><th></th></tr>
<tr><td>>hg38_21_+_8433172_8446622_RNA45SN1_45S_50ntFlanks@12453.12480.28|TCACCAAGCGTTGGATTGTTCACCCACT</td><td>>hg38_21_+_8433172_8446622_RNA45SN1_45S_50ntFlanks</td><td>12453</td><td>12480</td><td>28</td><td>TCACCAAGCGTTGGATTGTTCACCCACT</td><td>223.7</td><td>7.9</td><td>1.2</td><td>6.4</td><td>1.1E-10</td><td>2.9E-10</td><td>MB231/MB468</td><td>CYTO</td></tr>
<tr><td>>hg38_21_+_8433172_8446622_RNA45SN1_45S_50ntFlanks@8357.8375.19|AAAAGAACTTTGAAGAGAG</td><td>>hg38_21_+_8433172_8446622_RNA45SN1_45S_50ntFlanks</td><td>8357</td><td>8375</td><td>19</td><td>AAAAGAACTTTGAAGAGAG</td><td>1818.9</td><td>-4.4</td><td>0.7</td><td>-6.4</td><td>1.1E-10</td><td>2.9E-10</td><td>MB231/MB468</td><td>CYTO</td></tr>
<tr><td>>hg38_21_+_8433172_8446622_RNA45SN1_45S_50ntFlanks@11920.11967.48|ATGAGAGGTGTAGAATAAGTGGGAGGCCCCCGGCGCCCCCCCGGTGTC</td><td>>hg38_21_+_8433172_8446622_RNA45SN1_45S_50ntFlanks</td><td>11920</td><td>11967</td><td>48</td><td>ATGAGAGGTGTAGAATAAGTGGGAGGCCCCCGGCGCCCCCCCGGTGTC</td><td>181.5</td><td>-11.3</td><td>1.8</td><td>-6.4</td><td>1.3E-10</td><td>2.9E-10</td><td>MB231/MB468</td><td>MITO</td></tr>
<tr><td>>hg38_21_+_8433172_8446622_RNA45SN1_45S_50ntFlanks@9198.9221.24|GTTCTCTCGGGGCCACGCGCGCGT</td><td>>hg38_21_+_8433172_8446622_RNA45SN1_45S_50ntFlanks</td><td>9198</td><td>9221</td><td>24</td><td>GTTCTCTCGGGGCCACGCGCGCGT</td><td>87.6</td><td>-10.3</td><td>1.6</td><td>-6.5</td><td>1.1E-10</td><td>2.9E-10</td><td>MB231/MB468</td><td>MP</td></tr>
<tr><td>>hg38_21_+_8433172_8446622_RNA45SN1_45S_50ntFlanks@10425.10453.29|AGATGGGCGAGCGCCGTTCCGAAGGGACG</td><td>>hg38_21_+_8433172_8446622_RNA45SN1_45S_50ntFlanks</td><td>10425</td><td>10453</td><td>29</td><td>AGATGGGCGAGCGCCGTTCCGAAGGGACG</td><td>516.4</td><td>-6.5</td><td>1.0</td><td>-6.4</td><td>1.1E-10</td><td>2.9E-10</td><td>MB231/MB468</td><td>NUC</td></tr>
<tr><td>>MI0000746|hsa-mir-99b&WithFlank&19|+|51692606|51692687@13.33.21(+1C)&[MIMAT0000689&hsa-miR-99b-5p&offsets|0|-1(+1C);m-10&19|+|51692618|51692639&offsets|0|-1(+1C)]|CACCCGTAGAACCGACCTTGCC</td><td>>MI0000746|hsa-mir-99b&WithFlank&19|+|51692606|51692687</td><td>13</td><td>33</td><td>21(+1C)</td><td>CACCCGTAGAACCGACCTTGCC</td><td>1498.1</td><td>8.3</td><td>1.3</td><td>6.5</td><td>1.0E-10</td><td>2.9E-10</td><td>BT20/MB468</td><td>CYTO</td></tr>
<tr><td>>MI0003780|hsa-mir-1296&WithFlank&10|-|63372951|63373054@22.43.22&[MIMAT0005794&hsa-miR-1296-5p&offsets|0|0;m-304&10|-|63373012|63373033&offsets|0|0]|TTAGGGCCCTGGCTCCATCTCC</td><td>>MI0003780|hsa-mir-1296&WithFlank&10|-|63372951|63373054</td><td>22</td><td>43</td><td>22</td><td>TTAGGGCCCTGGCTCCATCTCC</td><td>602.0</td><td>9.8</td><td>1.5</td><td>6.5</td><td>1.0E-10</td><td>2.9E-10</td><td>BT20/MB468</td><td>CYTO</td></tr>
<tr><td>>hg38_21_+_8433172_8446622_RNA45SN1_45S_50ntFlanks@9534.9551.18|AGGAAACTCTGGTGGAGG</td><td>>hg38_21_+_8433172_8446622_RNA45SN1_45S_50ntFlanks</td><td>9534</td><td>9551</td><td>18</td><td>AGGAAACTCTGGTGGAGG</td><td>3583.0</td><td>-5.5</td><td>0.8</td><td>-6.8</td><td>8.5E-12</td><td>2.9E-10</td><td>BT20/MB468</td><td>MP</td></tr>
<tr><td>>hg38_21_+_8433172_8446622_RNA45SN1_45S_50ntFlanks@10784.10816.33|TGAACAGCCTCTGGCATGTTGGAACAATGTAGG</td><td>>hg38_21_+_8433172_8446622_RNA45SN1_45S_50ntFlanks</td><td>10784</td><td>10816</td><td>33</td><td>TGAACAGCCTCTGGCATGTTGGAACAATGTAGG</td><td>72.4</td><td>-10.0</td><td>1.6</td><td>-6.5</td><td>1.1E-10</td><td>2.9E-10</td><td>MB231/MB468</td><td>MP</td></tr>
<tr><td>>hg38_21_+_8433172_8446622_RNA45SN1_45S_50ntFlanks@12626.12649.24|AATGGGGCGAAGCTACCATCTGTG</td><td>>hg38_21_+_8433172_8446622_RNA45SN1_45S_50ntFlanks</td><td>12626</td><td>12649</td><td>24</td><td>AATGGGGCGAAGCTACCATCTGTG</td><td>255.7</td><td>-11.1</td><td>1.6</td><td>-6.8</td><td>8.5E-12</td><td>2.9E-10</td><td>BT20/MB468</td><td>MP</td></tr>
<tr><td>>am_MI0003581|hsa-mir-574&WithFlank&4|+|38868026|38868133@31.54.24&[MIMAT0004795&hsa-miR-574-5p&offsets|0|+1;m-213&4|+|38868056|38868078&offsets|0|+1]|TGAGTGTGTGTGTGTGAGTGTGTG</td><td>>am_MI0003581|hsa-mir-574&WithFlank&4|+|38868026|38868133</td><td>31</td><td>54</td><td>24</td><td>TGAGTGTGTGTGTGTGAGTGTGTG</td><td>433.5</td><td>6.0</td><td>0.9</td><td>6.4</td><td>1.3E-10</td><td>2.9E-10</td><td>MB231/MB468</td><td>MITO</td></tr>
<tr><td>>MI0000767|hsa-mir-365a&WithFlank&16|+|14309279|14309377@62.81.20(+2U);MI0000769|hsa-mir-365b&WithFlank&17|+|31575405|31575527@74.93.20(+2U)&[MIMAT0000710&hsa-miR-365a-3p&offsets|0|-2(+2U);m-176&16|+|14309340|14309361&offsets|0|-2(+2U)];[MIMAT0022834&hsa-miR-365b-3p&offsets|0|-2(+2U);m-175&17|+|31575478|31575499&offsets|0|-2(+2U)]|TAATGCCCCTAAAAATCCTTTT</td><td>>MI0000767|hsa-mir-365a&WithFlank&16|+|14309279|14309377</td><td>62</td><td>81</td><td>20(+2U)</td><td>TAATGCCCCTAAAAATCCTTTT</td><td>321.1</td><td>8.0</td><td>1.2</td><td>6.5</td><td>8.4E-11</td><td>2.9E-10</td><td>BT20/MB468</td><td>MITO</td></tr>
<tr><td>>hg38_21_+_8433172_8446622_RNA45SN1_45S_50ntFlanks@4443.4466.24|CCCTTGCCTCTCGGCGCCCCCTCG</td><td>>hg38_21_+_8433172_8446622_RNA45SN1_45S_50ntFlanks</td><td>4443</td><td>4466</td><td>24</td><td>CCCTTGCCTCTCGGCGCCCCCTCG</td><td>129.8</td><td>-9.1</td><td>1.4</td><td>-6.5</td><td>8.4E-11</td><td>2.9E-10</td><td>BT20/MB468</td><td>MITO</td></tr>
<tr><td>>hg38_21_+_8433172_8446622_RNA45SN1_45S_50ntFlanks@11667.11701.35|CTGCCCAGTGCTCTGAATGTCAAAGTGAAGAAATT</td><td>>hg38_21_+_8433172_8446622_RNA45SN1_45S_50ntFlanks</td><td>11667</td><td>11701</td><td>35</td><td>CTGCCCAGTGCTCTGAATGTCAAAGTGAAGAAATT</td><td>158.4</td><td>6.5</td><td>1.0</td><td>6.4</td><td>1.1E-10</td><td>2.9E-10</td><td>MB231/MB468</td><td>CYTO</td></tr>
<tr><td>>hg38_21_+_8433172_8446622_RNA45SN1_45S_50ntFlanks@10783.10813.31|GTGAACAGCCTCTGGCATGTTGGAACAATGT</td><td>>hg38_21_+_8433172_8446622_RNA45SN1_45S_50ntFlanks</td><td>10783</td><td>10813</td><td>31</td><td>GTGAACAGCCTCTGGCATGTTGGAACAATGT</td><td>120.1</td><td>-10.8</td><td>1.7</td><td>-6.5</td><td>1.1E-10</td><td>2.9E-10</td><td>MB231/MB468</td><td>MP</td></tr>
<tr><td>>hg38_21_+_8433172_8446622_RNA45SN1_45S_50ntFlanks@11212.11252.41|CGACTCTGGACGCGAGCCGGGCCCTTCCCGTGGATCGCCCC</td><td>>hg38_21_+_8433172_8446622_RNA45SN1_45S_50ntFlanks</td><td>11212</td><td>11252</td><td>41</td><td>CGACTCTGGACGCGAGCCGGGCCCTTCCCGTGGATCGCCCC</td><td>968.1</td><td>-5.9</td><td>0.9</td><td>-6.4</td><td>1.3E-10</td><td>2.9E-10</td><td>MB231/MB468</td><td>MITO</td></tr>
<tr><td>>hg38_21_+_8433172_8446622_RNA45SN1_45S_50ntFlanks@10841.10858.18|CGTAACTTCGGGATAAGG</td><td>>hg38_21_+_8433172_8446622_RNA45SN1_45S_50ntFlanks</td><td>10841</td><td>10858</td><td>18</td><td>CGTAACTTCGGGATAAGG</td><td>1909.7</td><td>-7.1</td><td>1.1</td><td>-6.4</td><td>1.1E-10</td><td>2.9E-10</td><td>MB231/MB468</td><td>CYTO</td></tr>
<tr><td>>hg38_21_+_8433172_8446622_RNA45SN1_45S_50ntFlanks@4560.4587.28|CTAGGAATAATGGAATAGGACCGCGGTT</td><td>>hg38_21_+_8433172_8446622_RNA45SN1_45S_50ntFlanks</td><td>4560</td><td>4587</td><td>28</td><td>CTAGGAATAATGGAATAGGACCGCGGTT</td><td>150.7</td><td>-9.3</td><td>1.4</td><td>-6.5</td><td>8.5E-11</td><td>2.9E-10</td><td>BT20/MB468</td><td>MITO</td></tr>
<tr><td>>hg38_21_+_8433172_8446622_RNA45SN1_45S_50ntFlanks@3831.3852.22|CTCGCTCCTCTCCTACTTGGAT</td><td>>hg38_21_+_8433172_8446622_RNA45SN1_45S_50ntFlanks</td><td>3831</td><td>3852</td><td>22</td><td>CTCGCTCCTCTCCTACTTGGAT</td><td>634.9</td><td>-9.2</td><td>1.4</td><td>-6.4</td><td>1.3E-10</td><td>2.9E-10</td><td>MB231/MB468</td><td>MITO</td></tr>
<tr><td>>hg38_21_+_8433172_8446622_RNA45SN1_45S_50ntFlanks@6644.6668.25|CCTCGTACGACTCTTAGCGGTGGAT</td><td>>hg38_21_+_8433172_8446622_RNA45SN1_45S_50ntFlanks</td><td>6644</td><td>6668</td><td>25</td><td>CCTCGTACGACTCTTAGCGGTGGAT</td><td>821.0</td><td>5.8</td><td>0.9</td><td>6.5</td><td>8.5E-11</td><td>2.9E-10</td><td>BT20/MB468</td><td>MITO</td></tr>
<tr><td>>am_MI0000809|hsa-mir-151a&WithFlank&8|-|140732558|140732659@53.71.19&[MIMAT0000757&hsa-miR-151a-3p&offsets|0|-2;m-30&8|-|140732587|140732607&offsets|0|-2]|CTAGACTGAAGCTCCTTGA</td><td>>am_MI0000809|hsa-mir-151a&WithFlank&8|-|140732558|140732659</td><td>53</td><td>71</td><td>19</td><td>CTAGACTGAAGCTCCTTGA</td><td>262.4</td><td>7.5</td><td>1.2</td><td>6.4</td><td>1.2E-10</td><td>3.0E-10</td><td>MB231/MB468</td><td>CYTO</td></tr>
<tr><td>>hg38_21_+_8433172_8446622_RNA45SN1_45S_50ntFlanks@9623.9649.27|TAGTAGCTGGTTCCCTCCGAAGTTTCC</td><td>>hg38_21_+_8433172_8446622_RNA45SN1_45S_50ntFlanks</td><td>9623</td><td>9649</td><td>27</td><td>TAGTAGCTGGTTCCCTCCGAAGTTTCC</td><td>1108.8</td><td>-8.4</td><td>1.3</td><td>-6.4</td><td>1.3E-10</td><td>3.0E-10</td><td>MB231/MB468</td><td>MITO</td></tr>
<tr><td>>hg38_21_+_8433172_8446622_RNA45SN1_45S_50ntFlanks@11876.11899.24|AAGACCCTGTTGAGCTTGACTCTA</td><td>>hg38_21_+_8433172_8446622_RNA45SN1_45S_50ntFlanks</td><td>11876</td><td>11899</td><td>24</td><td>AAGACCCTGTTGAGCTTGACTCTA</td><td>586.6</td><td>-6.5</td><td>1.0</td><td>-6.4</td><td>1.3E-10</td><td>3.0E-10</td><td>MB231/MB468</td><td>MITO</td></tr>
<tr><td>>hg38_21_+_8433172_8446622_RNA45SN1_45S_50ntFlanks@12249.12272.24|CAAAAGCTCGCTTGATCTTGATTT</td><td>>hg38_21_+_8433172_8446622_RNA45SN1_45S_50ntFlanks</td><td>12249</td><td>12272</td><td>24</td><td>CAAAAGCTCGCTTGATCTTGATTT</td><td>3140.0</td><td>-9.7</td><td>1.5</td><td>-6.4</td><td>1.3E-10</td><td>3.0E-10</td><td>MB231/MB468</td><td>MITO</td></tr>
<tr><td>>hg38_21_+_8433172_8446622_RNA45SN1_45S_50ntFlanks@10404.10433.30|CATGGGTCAGTCGGTCCTGAGAGATGGGCG</td><td>>hg38_21_+_8433172_8446622_RNA45SN1_45S_50ntFlanks</td><td>10404</td><td>10433</td><td>30</td><td>CATGGGTCAGTCGGTCCTGAGAGATGGGCG</td><td>158.5</td><td>-4.3</td><td>0.7</td><td>-6.4</td><td>1.2E-10</td><td>3.0E-10</td><td>MB231/MB468</td><td>CYTO</td></tr>
<tr><td>>hg38_21_+_8433172_8446622_RNA45SN1_45S_50ntFlanks@8198.8226.29|TGGACGGTGTGAGGCCGGTAGCGGCCCCC</td><td>>hg38_21_+_8433172_8446622_RNA45SN1_45S_50ntFlanks</td><td>8198</td><td>8226</td><td>29</td><td>TGGACGGTGTGAGGCCGGTAGCGGCCCCC</td><td>163.9</td><td>-4.4</td><td>0.7</td><td>-6.4</td><td>1.2E-10</td><td>3.0E-10</td><td>MB231/MB468</td><td>CYTO</td></tr>
<tr><td>>hg38_21_+_8433172_8446622_RNA45SN1_45S_50ntFlanks@3874.3898.25|AATACATGCCGACGGGCGCTGACCC</td><td>>hg38_21_+_8433172_8446622_RNA45SN1_45S_50ntFlanks</td><td>3874</td><td>3898</td><td>25</td><td>AATACATGCCGACGGGCGCTGACCC</td><td>3845.6</td><td>-5.1</td><td>0.8</td><td>-6.5</td><td>1.0E-10</td><td>3.0E-10</td><td>BT20/MB468</td><td>CYTO</td></tr>
<tr><td>>hg38_21_+_8433172_8446622_RNA45SN1_45S_50ntFlanks@12419.12444.26|GATGTCGGCTCTTCCTATCATTGTGA</td><td>>hg38_21_+_8433172_8446622_RNA45SN1_45S_50ntFlanks</td><td>12419</td><td>12444</td><td>26</td><td>GATGTCGGCTCTTCCTATCATTGTGA</td><td>2550.8</td><td>4.0</td><td>0.6</td><td>6.4</td><td>1.2E-10</td><td>3.0E-10</td><td>MB231/MB468</td><td>CYTO</td></tr>
<tr><td>>hg38_1_-_228634819_228635039_RNA5S12_5S_50ntFlanks@141.171.31|CCGCCTGGGAATACCGGGTGCTGTAGGCTTT</td><td>>hg38_1_-_228634819_228635039_RNA5S12_5S_50ntFlanks</td><td>141</td><td>171</td><td>31</td><td>CCGCCTGGGAATACCGGGTGCTGTAGGCTTT</td><td>1914.9</td><td>4.9</td><td>0.8</td><td>6.4</td><td>1.2E-10</td><td>3.0E-10</td><td>MB231/MB468</td><td>NUC</td></tr>
<tr><td>>hg38_21_+_8433172_8446622_RNA45SN1_45S_50ntFlanks@11906.11924.19|CACGGTGAAGAGACATGAG</td><td>>hg38_21_+_8433172_8446622_RNA45SN1_45S_50ntFlanks</td><td>11906</td><td>11924</td><td>19</td><td>CACGGTGAAGAGACATGAG</td><td>78.6</td><td>-10.2</td><td>1.6</td><td>-6.5</td><td>1.1E-10</td><td>3.0E-10</td><td>MB231/MB468</td><td>MP</td></tr>
<tr><td>>hg38_21_+_8433172_8446622_RNA45SN1_45S_50ntFlanks@3992.4013.22|GGTGACTCTAGATAACCTCGGG</td><td>>hg38_21_+_8433172_8446622_RNA45SN1_45S_50ntFlanks</td><td>3992</td><td>4013</td><td>22</td><td>GGTGACTCTAGATAACCTCGGG</td><td>122.4</td><td>-10.8</td><td>1.7</td><td>-6.5</td><td>1.1E-10</td><td>3.0E-10</td><td>MB231/MB468</td><td>MP</td></tr>
<tr><td>>hg38_MT_+_1621_3279_RNR2_16S_50ntFlanks@1209.1231.23|CAGAACCCAACCTCCGAGCAGTA</td><td>>hg38_MT_+_1621_3279_RNR2_16S_50ntFlanks</td><td>1209</td><td>1231</td><td>23</td><td>CAGAACCCAACCTCCGAGCAGTA</td><td>163.1</td><td>-11.2</td><td>1.7</td><td>-6.5</td><td>1.1E-10</td><td>3.0E-10</td><td>MB231/MB468</td><td>MP</td></tr>
<tr><td>>hg38_21_+_8433172_8446622_RNA45SN1_45S_50ntFlanks@6654.6686.33|CTCTTAGCGGTGGATCACTCGGCTCGTGCGTCG</td><td>>hg38_21_+_8433172_8446622_RNA45SN1_45S_50ntFlanks</td><td>6654</td><td>6686</td><td>33</td><td>CTCTTAGCGGTGGATCACTCGGCTCGTGCGTCG</td><td>109.2</td><td>-10.6</td><td>1.6</td><td>-6.5</td><td>1.1E-10</td><td>3.0E-10</td><td>MB231/MB468</td><td>MP</td></tr>
<tr><td>>hg38_21_+_8433172_8446622_RNA45SN1_45S_50ntFlanks@485.517.33|GATGTGGTGACGTCGTGCTCTCCCGGGCCGGGT</td><td>>hg38_21_+_8433172_8446622_RNA45SN1_45S_50ntFlanks</td><td>485</td><td>517</td><td>33</td><td>GATGTGGTGACGTCGTGCTCTCCCGGGCCGGGT</td><td>366.1</td><td>12.3</td><td>1.8</td><td>6.8</td><td>8.7E-12</td><td>3.0E-10</td><td>BT20/MB468</td><td>MP</td></tr>
<tr><td>>hg38_21_+_8433172_8446622_RNA45SN1_45S_50ntFlanks@13002.13025.24|CGATCTATTGAAAGTCAGCCCTCG</td><td>>hg38_21_+_8433172_8446622_RNA45SN1_45S_50ntFlanks</td><td>13002</td><td>13025</td><td>24</td><td>CGATCTATTGAAAGTCAGCCCTCG</td><td>251.3</td><td>-11.1</td><td>1.6</td><td>-6.8</td><td>8.8E-12</td><td>3.0E-10</td><td>BT20/MB468</td><td>MP</td></tr>
<tr><td>>hg38_21_+_8433172_8446622_RNA45SN1_45S_50ntFlanks@9027.9066.40|CGTCTCCTCGTGGGGGGGCCGGGCCACCCCTCCCACGGCG</td><td>>hg38_21_+_8433172_8446622_RNA45SN1_45S_50ntFlanks</td><td>9027</td><td>9066</td><td>40</td><td>CGTCTCCTCGTGGGGGGGCCGGGCCACCCCTCCCACGGCG</td><td>231.9</td><td>-10.9</td><td>1.6</td><td>-6.8</td><td>8.8E-12</td><td>3.0E-10</td><td>BT20/MB468</td><td>MP</td></tr>
<tr><td>>hg38_21_+_8433172_8446622_RNA45SN1_45S_50ntFlanks@13009.13028.20|TTGAAAGTCAGCCCTCGACA</td><td>>hg38_21_+_8433172_8446622_RNA45SN1_45S_50ntFlanks</td><td>13009</td><td>13028</td><td>20</td><td>TTGAAAGTCAGCCCTCGACA</td><td>346.5</td><td>-6.9</td><td>1.1</td><td>-6.4</td><td>1.2E-10</td><td>3.0E-10</td><td>MB231/MB468</td><td>CYTO</td></tr>
<tr><td>>hg38_21_+_8433172_8446622_RNA45SN1_45S_50ntFlanks@5084.5119.36|CGAACGAGACTCTGGCATGCTAACTAGTTACGCGAC</td><td>>hg38_21_+_8433172_8446622_RNA45SN1_45S_50ntFlanks</td><td>5084</td><td>5119</td><td>36</td><td>CGAACGAGACTCTGGCATGCTAACTAGTTACGCGAC</td><td>274.6</td><td>-12.0</td><td>1.9</td><td>-6.5</td><td>1.1E-10</td><td>3.0E-10</td><td>MB231/MB468</td><td>MP</td></tr>
<tr><td>>hg38_21_+_8433172_8446622_RNA45SN1_45S_50ntFlanks@9554.9572.19|CGTAGCGGTCCTGACGTGC</td><td>>hg38_21_+_8433172_8446622_RNA45SN1_45S_50ntFlanks</td><td>9554</td><td>9572</td><td>19</td><td>CGTAGCGGTCCTGACGTGC</td><td>8590.5</td><td>-6.9</td><td>1.1</td><td>-6.5</td><td>1.1E-10</td><td>3.0E-10</td><td>MB231/MB468</td><td>MP</td></tr>
<tr><td>>MI0000483|hsa-mir-186&WithFlank&1|-|71067625|71067722@21.41.21(+1A)&[MIMAT0000456&hsa-miR-186-5p&offsets|0|-1(+1A);m-84&1|-|71067681|71067702&offsets|0|-1(+1A)]|CAAAGAATTCTCCTTTTGGGCA</td><td>>MI0000483|hsa-mir-186&WithFlank&1|-|71067625|71067722</td><td>21</td><td>41</td><td>21(+1A)</td><td>CAAAGAATTCTCCTTTTGGGCA</td><td>473.6</td><td>8.8</td><td>1.4</td><td>6.5</td><td>1.0E-10</td><td>3.0E-10</td><td>BT20/MB468</td><td>CYTO</td></tr>
<tr><td>>hg38_21_+_8433172_8446622_RNA45SN1_45S_50ntFlanks@12003.12030.28|GCGGGCCGCCGGTGAAATACCACTACTC</td><td>>hg38_21_+_8433172_8446622_RNA45SN1_45S_50ntFlanks</td><td>12003</td><td>12030</td><td>28</td><td>GCGGGCCGCCGGTGAAATACCACTACTC</td><td>1301.0</td><td>-7.9</td><td>1.2</td><td>-6.4</td><td>1.4E-10</td><td>3.0E-10</td><td>BT20/MB468</td><td>NUC</td></tr>
<tr><td>>hg38_21_+_8433172_8446622_RNA45SN1_45S_50ntFlanks@7989.8008.20|AGACGTGGCGACCCGCTGAA</td><td>>hg38_21_+_8433172_8446622_RNA45SN1_45S_50ntFlanks</td><td>7989</td><td>8008</td><td>20</td><td>AGACGTGGCGACCCGCTGAA</td><td>13719.2</td><td>-5.1</td><td>0.8</td><td>-6.4</td><td>1.3E-10</td><td>3.0E-10</td><td>MB231/MB468</td><td>MITO</td></tr>
<tr><td>>hg38_21_+_8433172_8446622_RNA45SN1_45S_50ntFlanks@10350.10378.29|ACGAGAACTTTGAAGGCCGAAGTGGAGAA</td><td>>hg38_21_+_8433172_8446622_RNA45SN1_45S_50ntFlanks</td><td>10350</td><td>10378</td><td>29</td><td>ACGAGAACTTTGAAGGCCGAAGTGGAGAA</td><td>1138.4</td><td>-7.7</td><td>1.2</td><td>-6.5</td><td>1.1E-10</td><td>3.0E-10</td><td>MB231/MB468</td><td>MP</td></tr>
<tr><td>>hg38_21_+_8433172_8446622_RNA45SN1_45S_50ntFlanks@12995.13038.44|CTCGCTGCGATCTATTGAAAGTCAGCCCTCGACACAAGGGTTTG</td><td>>hg38_21_+_8433172_8446622_RNA45SN1_45S_50ntFlanks</td><td>12995</td><td>13038</td><td>44</td><td>CTCGCTGCGATCTATTGAAAGTCAGCCCTCGACACAAGGGTTTG</td><td>1886.0</td><td>8.1</td><td>1.3</td><td>6.5</td><td>1.0E-10</td><td>3.0E-10</td><td>BT20/MB468</td><td>CYTO</td></tr>
<tr><td>>MI0000342|hsa-mir-200b&WithFlank&1|+|1167098|1167204@27.48.22&[MIMAT0004571&hsa-miR-200b-5p&offsets|0|0;m-312&1|+|1167124|1167145&offsets|0|0]|CATCTTACTGGGCAGCATTGGA</td><td>>MI0000342|hsa-mir-200b&WithFlank&1|+|1167098|1167204</td><td>27</td><td>48</td><td>22</td><td>CATCTTACTGGGCAGCATTGGA</td><td>78.4</td><td>-10.2</td><td>1.6</td><td>-6.5</td><td>1.1E-10</td><td>3.0E-10</td><td>MB231/MB468</td><td>MP</td></tr>
<tr><td>>hg38_21_+_8433172_8446622_RNA45SN1_45S_50ntFlanks@10344.10373.30|ATTCAAACGAGAACTTTGAAGGCCGAAGTG</td><td>>hg38_21_+_8433172_8446622_RNA45SN1_45S_50ntFlanks</td><td>10344</td><td>10373</td><td>30</td><td>ATTCAAACGAGAACTTTGAAGGCCGAAGTG</td><td>202.5</td><td>3.8</td><td>0.6</td><td>6.4</td><td>1.2E-10</td><td>3.0E-10</td><td>MB231/MB468</td><td>NUC</td></tr>
<tr><td>>hg38_21_+_8433172_8446622_RNA45SN1_45S_50ntFlanks@11452.11472.21|CTCCTCGGCGCGCGGCGGCGG</td><td>>hg38_21_+_8433172_8446622_RNA45SN1_45S_50ntFlanks</td><td>11452</td><td>11472</td><td>21</td><td>CTCCTCGGCGCGCGGCGGCGG</td><td>554.7</td><td>-7.4</td><td>1.1</td><td>-6.5</td><td>1.0E-10</td><td>3.0E-10</td><td>BT20/MB468</td><td>CYTO</td></tr>
<tr><td>>hg38_21_+_8433172_8446622_RNA45SN1_45S_50ntFlanks@9605.9622.18|AAGACTAATCGAACCATC</td><td>>hg38_21_+_8433172_8446622_RNA45SN1_45S_50ntFlanks</td><td>9605</td><td>9622</td><td>18</td><td>AAGACTAATCGAACCATC</td><td>1490.5</td><td>-8.5</td><td>1.3</td><td>-6.4</td><td>1.4E-10</td><td>3.0E-10</td><td>BT20/MB468</td><td>NUC</td></tr>
<tr><td>>hg38_21_+_8433172_8446622_RNA45SN1_45S_50ntFlanks@12935.12958.24|TAAACCATTCGTAGACGACCTGCT</td><td>>hg38_21_+_8433172_8446622_RNA45SN1_45S_50ntFlanks</td><td>12935</td><td>12958</td><td>24</td><td>TAAACCATTCGTAGACGACCTGCT</td><td>183.8</td><td>-3.1</td><td>0.5</td><td>-6.4</td><td>1.2E-10</td><td>3.0E-10</td><td>MB231/MB468</td><td>NUC</td></tr>
<tr><td>>hg38_21_+_8433172_8446622_RNA45SN1_45S_50ntFlanks@4731.4758.28|AAGAACGAAAGTCGGAGGTTCGAAGACG</td><td>>hg38_21_+_8433172_8446622_RNA45SN1_45S_50ntFlanks</td><td>4731</td><td>4758</td><td>28</td><td>AAGAACGAAAGTCGGAGGTTCGAAGACG</td><td>8387.6</td><td>-4.0</td><td>0.6</td><td>-6.4</td><td>1.2E-10</td><td>3.0E-10</td><td>MB231/MB468</td><td>NUC</td></tr>
<tr><td>>hg38_21_+_8433172_8446622_RNA45SN1_45S_50ntFlanks@4006.4025.20|ACCTCGGGCCGATCGCACGC</td><td>>hg38_21_+_8433172_8446622_RNA45SN1_45S_50ntFlanks</td><td>4006</td><td>4025</td><td>20</td><td>ACCTCGGGCCGATCGCACGC</td><td>800.6</td><td>-8.5</td><td>1.3</td><td>-6.4</td><td>1.3E-10</td><td>3.0E-10</td><td>MB231/MB468</td><td>MITO</td></tr>
</table>

<table>
<tr><th></th><th></th><th></th><th></th><th></th><th></th><th></th><th></th><th></th><th></th><th></th><th></th><th></th><th></th></tr>
<tr><td>>hg38_21_+_8433172_8446622_RNA45SN1_45S_50ntFlanks@7979.7996.18|ACCTCAGATCAGACGTGG</td><td>>hg38_21_+_8433172_8446622_RNA45SN1_45S_50ntFlanks</td><td>7979</td><td>7996</td><td>18</td><td>ACCTCAGATCAGACGTGG</td><td>241.1</td><td>-6.7</td><td>1.0</td><td>-6.4</td><td>1.2E-10</td><td>3.0E-10</td><td>MB231/MB468</td><td>CYTO</td></tr>
<tr><td>>hg38_21_+_8433172_8446622_RNA45SN1_45S_50ntFlanks@3927.3959.33|CAGATCAAAACCAACCCGGTCAGCCCCTCTCCG</td><td>>hg38_21_+_8433172_8446622_RNA45SN1_45S_50ntFlanks</td><td>3927</td><td>3959</td><td>33</td><td>CAGATCAAAACCAACCCGGTCAGCCCCTCTCCG</td><td>442.3</td><td>-8.8</td><td>1.4</td><td>-6.4</td><td>1.4E-10</td><td>3.0E-10</td><td>BT20/MB468</td><td>NUC</td></tr>
<tr><td>>MI0000465|hsa-mir-191&WithFlank&3|-|49020612|49020715@22.45.24&[MIMAT0000440&hsa-miR-191-5p&offsets|0|+1;m-48&3|-|49020672|49020694&offsets|0|+1]|CAACGGAATCCCAAAAGCAGCTGT</td><td>>MI0000465|hsa-mir-191&WithFlank&3|-|49020612|49020715</td><td>22</td><td>45</td><td>24</td><td>CAACGGAATCCCAAAAGCAGCTGT</td><td>3684.1</td><td>6.7</td><td>1.0</td><td>6.4</td><td>1.2E-10</td><td>3.0E-10</td><td>MB231/MB468</td><td>CYTO</td></tr>
<tr><td>>hg38_21_+_8433172_8446622_RNA45SN1_45S_50ntFlanks@8170.8190.21|CAAGTCCTTCTGATCGAGGCC</td><td>>hg38_21_+_8433172_8446622_RNA45SN1_45S_50ntFlanks</td><td>8170</td><td>8190</td><td>21</td><td>CAAGTCCTTCTGATCGAGGCC</td><td>84.5</td><td>-10.3</td><td>1.6</td><td>-6.5</td><td>1.1E-10</td><td>3.0E-10</td><td>MB231/MB468</td><td>MP</td></tr>
<tr><td>>hg38_21_+_8433172_8446622_RNA45SN1_45S_50ntFlanks@3837.3871.35|CCTCTCCTACTTGGATAACTGTGGTAATTCTAGAG</td><td>>hg38_21_+_8433172_8446622_RNA45SN1_45S_50ntFlanks</td><td>3837</td><td>3871</td><td>35</td><td>CCTCTCCTACTTGGATAACTGTGGTAATTCTAGAG</td><td>470.2</td><td>-4.3</td><td>0.7</td><td>-6.4</td><td>1.2E-10</td><td>3.0E-10</td><td>MB231/MB468</td><td>NUC</td></tr>
<tr><td>>hg38_21_+_8433172_8446622_RNA45SN1_45S_50ntFlanks@11930.11958.29|TAGAATAAGTGGGAGGCCCCCGGCGCCCC</td><td>>hg38_21_+_8433172_8446622_RNA45SN1_45S_50ntFlanks</td><td>11930</td><td>11958</td><td>29</td><td>TAGAATAAGTGGGAGGCCCCCGGCGCCCC</td><td>78.9</td><td>-10.2</td><td>1.6</td><td>-6.5</td><td>1.1E-10</td><td>3.0E-10</td><td>MB231/MB468</td><td>MP</td></tr>
<tr><td>>hg38_21_+_8433172_8446622_RNA45SN1_45S_50ntFlanks@3838.3871.34|CTCTCCTACTTGGATAACTGTGGTAATTCTAGAG</td><td>>hg38_21_+_8433172_8446622_RNA45SN1_45S_50ntFlanks</td><td>3838</td><td>3871</td><td>34</td><td>CTCTCCTACTTGGATAACTGTGGTAATTCTAGAG</td><td>540.4</td><td>-3.5</td><td>0.5</td><td>-6.4</td><td>1.2E-10</td><td>3.0E-10</td><td>MB231/MB468</td><td>NUC</td></tr>
<tr><td>>hg38_21_+_8433172_8446622_RNA45SN1_45S_50ntFlanks@10788.10807.20|CAGCCTCTGGCATGTTGGAA</td><td>>hg38_21_+_8433172_8446622_RNA45SN1_45S_50ntFlanks</td><td>10788</td><td>10807</td><td>20</td><td>CAGCCTCTGGCATGTTGGAA</td><td>251.0</td><td>-10.3</td><td>1.6</td><td>-6.5</td><td>1.1E-10</td><td>3.0E-10</td><td>BT20/MB468</td><td>CYTO</td></tr>
<tr><td>>MI0000542|hsa-mir-320a&WithFlank&8|-|22244960|22245043@48.69.22(+1C)&[MIMAT0000510&hsa-miR-320a-3p&offsets|0|0(+1C);m-60&8|-|22244975|22244996&offsets|0|0(+1C)]|AAAAGCTGGGTTGAGAGGGCGAC</td><td>>MI0000542|hsa-mir-320a&WithFlank&8|-|22244960|22245043</td><td>48</td><td>69</td><td>22(+1C)</td><td>AAAAGCTGGGTTGAGAGGGCGAC</td><td>298.5</td><td>4.3</td><td>0.7</td><td>6.4</td><td>1.4E-10</td><td>3.0E-10</td><td>BT20/MB468</td><td>NUC</td></tr>
<tr><td>>hg38_21_+_8433172_8446622_RNA45SN1_45S_50ntFlanks@10866.10892.27|CTAAGGGCTGGGTCGGTCGGGCTGGGG</td><td>>hg38_21_+_8433172_8446622_RNA45SN1_45S_50ntFlanks</td><td>10866</td><td>10892</td><td>27</td><td>CTAAGGGCTGGGTCGGTCGGGCTGGGG</td><td>1169.6</td><td>-9.5</td><td>1.5</td><td>-6.4</td><td>1.4E-10</td><td>3.0E-10</td><td>BT20/MB468</td><td>NUC</td></tr>
<tr><td>>hg38_21_+_8433172_8446622_RNA45SN1_45S_50ntFlanks@4555.4572.18|CGCAGCTAGGAATAATGG</td><td>>hg38_21_+_8433172_8446622_RNA45SN1_45S_50ntFlanks</td><td>4555</td><td>4572</td><td>18</td><td>CGCAGCTAGGAATAATGG</td><td>2205.8</td><td>-10.0</td><td>1.6</td><td>-6.4</td><td>1.4E-10</td><td>3.0E-10</td><td>BT20/MB468</td><td>NUC</td></tr>
<tr><td>>MI0000285|hsa-mir-205&WithFlank&1|+|209432127|209432248@40.60.21&[MIMAT0000266&hsa-miR-205-5p&offsets|0|-1;m-351&1|+|209432166|209432187&offsets|0|-1]|TCCTTCATTCCACCGGAGTCT</td><td>>MI0000285|hsa-mir-205&WithFlank&1|+|209432127|209432248</td><td>40</td><td>60</td><td>21</td><td>TCCTTCATTCCACCGGAGTCT</td><td>6622.2</td><td>6.3</td><td>1.0</td><td>6.5</td><td>8.7E-11</td><td>3.0E-10</td><td>BT20/MB468</td><td>MITO</td></tr>
<tr><td>>hg38_21_+_8433172_8446622_RNA45SN1_45S_50ntFlanks@11876.11895.20|AAGACCCTGTTGAGCTTGAC</td><td>>hg38_21_+_8433172_8446622_RNA45SN1_45S_50ntFlanks</td><td>11876</td><td>11895</td><td>20</td><td>AAGACCCTGTTGAGCTTGAC</td><td>76.6</td><td>-10.1</td><td>1.6</td><td>-6.4</td><td>1.1E-10</td><td>3.0E-10</td><td>MB231/MB468</td><td>MP</td></tr>
<tr><td>>hg38_21_+_8433172_8446622_RNA45SN1_45S_50ntFlanks@10014.10039.26|AAAATGGATGGCGCTGGAGCGTCGGG</td><td>>hg38_21_+_8433172_8446622_RNA45SN1_45S_50ntFlanks</td><td>10014</td><td>10039</td><td>26</td><td>AAAATGGATGGCGCTGGAGCGTCGGG</td><td>192.2</td><td>-11.4</td><td>1.8</td><td>-6.4</td><td>1.1E-10</td><td>3.0E-10</td><td>MB231/MB468</td><td>MP</td></tr>
<tr><td>>hg38_21_+_8433172_8446622_RNA45SN1_45S_50ntFlanks@12073.12095.23|CCGAGGGGCTCTCGCTTCTGGCG</td><td>>hg38_21_+_8433172_8446622_RNA45SN1_45S_50ntFlanks</td><td>12073</td><td>12095</td><td>23</td><td>CCGAGGGGCTCTCGCTTCTGGCG</td><td>1201.6</td><td>-8.1</td><td>1.3</td><td>-6.5</td><td>8.7E-11</td><td>3.0E-10</td><td>BT20/MB468</td><td>MITO</td></tr>
<tr><td>>hg38_21_+_8433172_8446622_RNA45SN1_45S_50ntFlanks@6719.6746.28|TGAATTGCAGGACACATTGATCATCGAC</td><td>>hg38_21_+_8433172_8446622_RNA45SN1_45S_50ntFlanks</td><td>6719</td><td>6746</td><td>28</td><td>TGAATTGCAGGACACATTGATCATCGAC</td><td>784.0</td><td>-8.2</td><td>1.3</td><td>-6.4</td><td>1.2E-10</td><td>3.0E-10</td><td>MB231/MB468</td><td>NUC</td></tr>
<tr><td>>hg38_21_+_8433172_8446622_RNA45SN1_45S_50ntFlanks@4685.4707.23|AGACGGACCAGAGCGAAAGCATT</td><td>>hg38_21_+_8433172_8446622_RNA45SN1_45S_50ntFlanks</td><td>4685</td><td>4707</td><td>23</td><td>AGACGGACCAGAGCGAAAGCATT</td><td>756.1</td><td>-7.2</td><td>1.1</td><td>-6.4</td><td>1.3E-10</td><td>3.0E-10</td><td>MB231/MB468</td><td>MITO</td></tr>
<tr><td>>hg38_MT_+_1621_3279_RNR2_16S_50ntFlanks@51.87.37|GCTAAACCTAGCCCCAAACCCACTCCACCTTACTACC</td><td>>hg38_MT_+_1621_3279_RNR2_16S_50ntFlanks</td><td>51</td><td>87</td><td>37</td><td>GCTAAACCTAGCCCCAAACCCACTCCACCTTACTACC</td><td>6706.4</td><td>6.3</td><td>1.0</td><td>6.5</td><td>8.8E-11</td><td>3.0E-10</td><td>BT20/MB468</td><td>MITO</td></tr>
<tr><td>>MI0005544|hsa-mir-147b&WithFlank&15|+|45433044|45433135@55.73.19&[MIMAT0004928&hsa-miR-147b-3p&offsets|0|-2;m-575&15|+|45433098|45433118&offsets|0|-2]|GTGTGCGGAAATGCTTCTG</td><td>>MI0005544|hsa-mir-147b&WithFlank&15|+|45433044|45433135</td><td>55</td><td>73</td><td>19</td><td>GTGTGCGGAAATGCTTCTG</td><td>102.2</td><td>9.9</td><td>1.5</td><td>6.4</td><td>1.1E-10</td><td>3.1E-10</td><td>MB231/MB468</td><td>MP</td></tr>
<tr><td>>hg38_21_+_8433172_8446622_RNA45SN1_45S_50ntFlanks@6735.6771.37|TTGATCATCGACACTTCGAACGCACTTGCGGCCCCGG</td><td>>hg38_21_+_8433172_8446622_RNA45SN1_45S_50ntFlanks</td><td>6735</td><td>6771</td><td>37</td><td>TTGATCATCGACACTTCGAACGCACTTGCGGCCCCGG</td><td>472.8</td><td>-4.9</td><td>0.8</td><td>-6.4</td><td>1.2E-10</td><td>3.1E-10</td><td>MB231/MB468</td><td>NUC</td></tr>
<tr><td>>hg38_21_+_8433172_8446622_RNA45SN1_45S_50ntFlanks@11315.11334.20|CCACCCCACGTCTCGTCGCG</td><td>>hg38_21_+_8433172_8446622_RNA45SN1_45S_50ntFlanks</td><td>11315</td><td>11334</td><td>20</td><td>CCACCCCACGTCTCGTCGCG</td><td>79.2</td><td>-10.2</td><td>1.6</td><td>-6.4</td><td>1.1E-10</td><td>3.1E-10</td><td>MB231/MB468</td><td>MP</td></tr>
<tr><td>>hg38_21_+_8433172_8446622_RNA45SN1_45S_50ntFlanks@8171.8188.18|AAGTCCTTCTGATCGAGG</td><td>>hg38_21_+_8433172_8446622_RNA45SN1_45S_50ntFlanks</td><td>8171</td><td>8188</td><td>18</td><td>AAGTCCTTCTGATCGAGG</td><td>202.6</td><td>-10.7</td><td>1.6</td><td>-6.8</td><td>9.0E-12</td><td>3.1E-10</td><td>BT20/MB468</td><td>MP</td></tr>
<tr><td>>trna116_GluCTC_1_-_145399233_145399304@1.31.31,trna59_GluCTC_1_+_249168447_249168518@1.31.31,trna71_GluCTC_1_-_161439189_161439260@1.31.31,trna74_GluCTC_1_-_161431809_161431880@1.31.31,trna77_GluCTC_1_-_161424398_161424469@1.31.31,trna77_GluCTC_6_+_28949976_28950047@1.31.31,trna80_GluCTC_1_-_161417018_161417089@1.31.31,trna87_GluCTC_6_-_126101393_126101464@1.31.31|TCCCTGGTGGTCTAGTGGTTAGGATTCGGCG</td><td>>trna116_GluCTC_1_-_145399233_145399304</td><td>1</td><td>31</td><td>31</td><td>TCCCTGGTGGTCTAGTGGTTAGGATTCGGCG</td><td>236876.1</td><td>-4.2</td><td>0.6</td><td>-6.5</td><td>1.1E-10</td><td>3.1E-10</td><td>BT20/MB468</td><td>CYTO</td></tr>
<tr><td>>hg38_21_+_8433172_8446622_RNA45SN1_45S_50ntFlanks@4558.4587.30|AGCTAGGAATAATGGAATAGGACCGCGGTT</td><td>>hg38_21_+_8433172_8446622_RNA45SN1_45S_50ntFlanks</td><td>4558</td><td>4587</td><td>30</td><td>AGCTAGGAATAATGGAATAGGACCGCGGTT</td><td>141.6</td><td>-11.0</td><td>1.7</td><td>-6.4</td><td>1.1E-10</td><td>3.1E-10</td><td>MB231/MB468</td><td>MP</td></tr>
<tr><td>>hg38_21_+_8433172_8446622_RNA45SN1_45S_50ntFlanks@3991.4023.33|TGGTGACTCTAGATAACCTCGGGCCGATCGCAC</td><td>>hg38_21_+_8433172_8446622_RNA45SN1_45S_50ntFlanks</td><td>3991</td><td>4023</td><td>33</td><td>TGGTGACTCTAGATAACCTCGGGCCGATCGCAC</td><td>745.6</td><td>3.9</td><td>0.6</td><td>6.5</td><td>1.1E-10</td><td>3.1E-10</td><td>BT20/MB468</td><td>CYTO</td></tr>
<tr><td>>hg38_21_+_8433172_8446622_RNA45SN1_45S_50ntFlanks@10393.10410.18|CAGCAGTTGAACATGGGT</td><td>>hg38_21_+_8433172_8446622_RNA45SN1_45S_50ntFlanks</td><td>10393</td><td>10410</td><td>18</td><td>CAGCAGTTGAACATGGGT</td><td>2203.2</td><td>-4.7</td><td>0.7</td><td>-6.5</td><td>1.1E-10</td><td>3.1E-10</td><td>BT20/MB468</td><td>CYTO</td></tr>
<tr><td>>hg38_21_+_8433172_8446622_RNA45SN1_45S_50ntFlanks@10788.10816.29|CAGCCTCTGGCATGTTGGAACAATGTAGG</td><td>>hg38_21_+_8433172_8446622_RNA45SN1_45S_50ntFlanks</td><td>10788</td><td>10816</td><td>29</td><td>CAGCCTCTGGCATGTTGGAACAATGTAGG</td><td>1264.5</td><td>-7.7</td><td>1.2</td><td>-6.4</td><td>1.4E-10</td><td>3.1E-10</td><td>BT20/MB468</td><td>NUC</td></tr>
<tr><td>>hg38_21_+_8433172_8446622_RNA45SN1_45S_50ntFlanks@12934.12955.22|CTAAACCATTCGTAGACGACCT</td><td>>hg38_21_+_8433172_8446622_RNA45SN1_45S_50ntFlanks</td><td>12934</td><td>12955</td><td>22</td><td>CTAAACCATTCGTAGACGACCT</td><td>921.6</td><td>-6.7</td><td>1.0</td><td>-6.4</td><td>1.2E-10</td><td>3.1E-10</td><td>MB231/MB468</td><td>NUC</td></tr>
<tr><td>>hg38_21_+_8433172_8446622_RNA45SN1_45S_50ntFlanks@11718.11738.21|AACGGCGGGAGTAACTATGAC</td><td>>hg38_21_+_8433172_8446622_RNA45SN1_45S_50ntFlanks</td><td>11718</td><td>11738</td><td>21</td><td>AACGGCGGGAGTAACTATGAC</td><td>613.5</td><td>-8.2</td><td>1.3</td><td>-6.4</td><td>1.4E-10</td><td>3.1E-10</td><td>MB231/MB468</td><td>MITO</td></tr>
<tr><td>>hg38_21_+_8433172_8446622_RNA45SN1_45S_50ntFlanks@10128.10146.19|CGGCGGCGGCGGCGGCGGC</td><td>>hg38_21_+_8433172_8446622_RNA45SN1_45S_50ntFlanks</td><td>10128</td><td>10146</td><td>19</td><td>CGGCGGCGGCGGCGGCGGC</td><td>99.4</td><td>-10.5</td><td>1.6</td><td>-6.4</td><td>1.1E-10</td><td>3.1E-10</td><td>MB231/MB468</td><td>MP</td></tr>
<tr><td>>hg38_21_+_8433172_8446622_RNA45SN1_45S_50ntFlanks@8845.8863.19|CGGCGGCGGCGGCGGCGGC</td><td>>hg38_21_+_8433172_8446622_RNA45SN1_45S_50ntFlanks</td><td>8845</td><td>8863</td><td>19</td><td>CGGCGGCGGCGGCGGCGGC</td><td>99.4</td><td>-10.5</td><td>1.6</td><td>-6.4</td><td>1.1E-10</td><td>3.1E-10</td><td>MB231/MB468</td><td>MP</td></tr>
<tr><td>>hg38_21_+_8433172_8446622_RNA45SN1_45S_50ntFlanks@10679.10710.32|GGTTCCGGCGGCGTCCGGTGAGCTCTCGCTGG</td><td>>hg38_21_+_8433172_8446622_RNA45SN1_45S_50ntFlanks</td><td>10679</td><td>10710</td><td>32</td><td>GGTTCCGGCGGCGTCCGGTGAGCTCTCGCTGG</td><td>28754.9</td><td>-8.3</td><td>1.3</td><td>-6.4</td><td>1.2E-10</td><td>3.1E-10</td><td>MB231/MB468</td><td>CYTO</td></tr>
<tr><td>>hg38_21_+_8433172_8446622_RNA45SN1_45S_50ntFlanks@11784.11807.24|ATGAATGGATGAACGAGATTCCCA</td><td>>hg38_21_+_8433172_8446622_RNA45SN1_45S_50ntFlanks</td><td>11784</td><td>11807</td><td>24</td><td>ATGAATGGATGAACGAGATTCCCA</td><td>142.5</td><td>-11.0</td><td>1.7</td><td>-6.4</td><td>1.1E-10</td><td>3.1E-10</td><td>MB231/MB468</td><td>MP</td></tr>
<tr><td>>hg38_21_+_8433172_8446622_RNA45SN1_45S_50ntFlanks@3782.3805.24|CAGTGAAACTGCGAATGGCTCATT</td><td>>hg38_21_+_8433172_8446622_RNA45SN1_45S_50ntFlanks</td><td>3782</td><td>3805</td><td>24</td><td>CAGTGAAACTGCGAATGGCTCATT</td><td>1308.0</td><td>-6.3</td><td>1.0</td><td>-6.4</td><td>1.4E-10</td><td>3.1E-10</td><td>MB231/MB468</td><td>MITO</td></tr>
<tr><td>>am_trna128_GlyGCC_6_-_27870686_27870756@3.35.33,trna18_GlyGCC_16_+_70822597_70822667@3.35.33,trna19_GlyGCC_16_+_70823410_70823480@3.35.33,trna19_GlyGCC_2_-_157257659_157257729@3.35.33,trna24_GlyGCC_16_-_70812942_70813012@3.35.33,trna25_GlyGCC_16_-_70812114_70812184@3.35.33,trna5_GlyGCC_17_+_8029064_8029134@3.35.33,trna68_GlyGCC_1_-_161493637_161493707@3.35.33|ATTGGTGGTTCAGTGGTAGAATTCTCGCCTGCC</td><td>>am_trna128_GlyGCC_6_-_27870686_27870756</td><td>3</td><td>35</td><td>33</td><td>ATTGGTGGTTCAGTGGTAGAATTCTCGCCTGCC</td><td>1218.6</td><td>5.3</td><td>0.8</td><td>6.4</td><td>1.4E-10</td><td>3.1E-10</td><td>MB231/MB468</td><td>MITO</td></tr>
<tr><td>>hg38_21_+_8433172_8446622_RNA45SN1_45S_50ntFlanks@12149.12166.18|AGGTGGGGAGTTTGACTG</td><td>>hg38_21_+_8433172_8446622_RNA45SN1_45S_50ntFlanks</td><td>12149</td><td>12166</td><td>18</td><td>AGGTGGGGAGTTTGACTG</td><td>78.5</td><td>-10.2</td><td>1.6</td><td>-6.4</td><td>1.1E-10</td><td>3.1E-10</td><td>MB231/MB468</td><td>MP</td></tr>
<tr><td>>trnaMT_HisGTG_MT_+_12138_12206@46.72.27|GAGGCTTACGACCCCTTATTTACCCCA</td><td>>trnaMT_HisGTG_MT_+_12138_12206</td><td>46</td><td>72</td><td>27</td><td>GAGGCTTACGACCCCTTATTTACCCCA</td><td>101.1</td><td>9.9</td><td>1.5</td><td>6.4</td><td>1.1E-10</td><td>3.1E-10</td><td>MB231/MB468</td><td>MP</td></tr>
<tr><td>>hg38_21_+_8433172_8446622_RNA45SN1_45S_50ntFlanks@12415.12436.22|CTTCGATGTCGGCTCTTCCTAT</td><td>>hg38_21_+_8433172_8446622_RNA45SN1_45S_50ntFlanks</td><td>12415</td><td>12436</td><td>22</td><td>CTTCGATGTCGGCTCTTCCTAT</td><td>4631.9</td><td>-3.8</td><td>0.6</td><td>-6.4</td><td>1.2E-10</td><td>3.1E-10</td><td>MB231/MB468</td><td>CYTO</td></tr>
<tr><td>>MI0005559|hsa-mir-744&WithFlank&17|+|12081893|12082002@17.34.18&[MIMAT0004945&hsa-miR-744-5p&offsets|0|-4;m-172&17|+|12081909|12081929&offsets|0|-3]|TGCGGGGCTAGGGCTAAC</td><td>>MI0005559|hsa-mir-744&WithFlank&17|+|12081893|12082002</td><td>17</td><td>34</td><td>18</td><td>TGCGGGGCTAGGGCTAAC</td><td>405.0</td><td>9.4</td><td>1.5</td><td>6.5</td><td>8.9E-11</td><td>3.1E-10</td><td>BT20/MB468</td><td>MITO</td></tr>
<tr><td>>hg38_21_+_8433172_8446622_RNA45SN1_45S_50ntFlanks@11588.11612.25|GCGGACCAGGGGAATCCGACTGTTT</td><td>>hg38_21_+_8433172_8446622_RNA45SN1_45S_50ntFlanks</td><td>11588</td><td>11612</td><td>25</td><td>GCGGACCAGGGGAATCCGACTGTTT</td><td>279.1</td><td>3.4</td><td>0.5</td><td>6.4</td><td>1.2E-10</td><td>3.1E-10</td><td>MB231/MB468</td><td>CYTO</td></tr>
<tr><td>>hg38_21_+_8433172_8446622_RNA45SN1_45S_50ntFlanks@5484.5526.43|GAGCGCTGAGAAGACGGTCGAACTTGACTATCTAGAGGAAGTA</td><td>>hg38_21_+_8433172_8446622_RNA45SN1_45S_50ntFlanks</td><td>5484</td><td>5526</td><td>43</td><td>GAGCGCTGAGAAGACGGTCGAACTTGACTATCTAGAGGAAGTA</td><td>201.3</td><td>-10.6</td><td>1.6</td><td>-6.4</td><td>1.2E-10</td><td>3.1E-10</td><td>MB231/MB468</td><td>NUC</td></tr>
<tr><td>>hg38_21_+_8433172_8446622_RNA45SN1_45S_50ntFlanks@12003.12031.29|GCGGGCCGCCGGTGAAATACCACTACTCT</td><td>>hg38_21_+_8433172_8446622_RNA45SN1_45S_50ntFlanks</td><td>12003</td><td>12031</td><td>29</td><td>GCGGGCCGCCGGTGAAATACCACTACTCT</td><td>682.7</td><td>-8.3</td><td>1.3</td><td>-6.4</td><td>1.1E-10</td><td>3.1E-10</td><td>MB231/MB468</td><td>MP</td></tr>
<tr><td>>hg38_21_+_8433172_8446622_RNA45SN1_45S_50ntFlanks@9758.9787.30|TCAAACTTTAAATGGGTAAGAAGCCCGGCT</td><td>>hg38_21_+_8433172_8446622_RNA45SN1_45S_50ntFlanks</td><td>9758</td><td>9787</td><td>30</td><td>TCAAACTTTAAATGGGTAAGAAGCCCGGCT</td><td>222.1</td><td>-11.7</td><td>1.8</td><td>-6.4</td><td>1.2E-10</td><td>3.1E-10</td><td>MB231/MB468</td><td>MP</td></tr>
<tr><td>>hg38_21_+_8433172_8446622_RNA45SN1_45S_50ntFlanks@12414.12448.35|CCTTCGATGTCGGCTCTTCCTATCATTGTGAAGCA</td><td>>hg38_21_+_8433172_8446622_RNA45SN1_45S_50ntFlanks</td><td>12414</td><td>12448</td><td>35</td><td>CCTTCGATGTCGGCTCTTCCTATCATTGTGAAGCA</td><td>329.0</td><td>6.6</td><td>1.0</td><td>6.5</td><td>1.1E-10</td><td>3.1E-10</td><td>BT20/MB468</td><td>CYTO</td></tr>
<tr><td>>hg38_21_+_8433172_8446622_RNA45SN1_45S_50ntFlanks@11207.11232.26|GGCGGCGACTCTGGACGCGAGCCGGG</td><td>>hg38_21_+_8433172_8446622_RNA45SN1_45S_50ntFlanks</td><td>11207</td><td>11232</td><td>26</td><td>GGCGGCGACTCTGGACGCGAGCCGGG</td><td>592.7</td><td>-8.1</td><td>1.3</td><td>-6.4</td><td>1.2E-10</td><td>3.1E-10</td><td>MB231/MB468</td><td>NUC</td></tr>
<tr><td>>hg38_21_+_8433172_8446622_RNA45SN1_45S_50ntFlanks@5470.5493.24|CCACGGCCCTGGCGGAGCGCTGAG</td><td>>hg38_21_+_8433172_8446622_RNA45SN1_45S_50ntFlanks</td><td>5470</td><td>5493</td><td>24</td><td>CCACGGCCCTGGCGGAGCGCTGAG</td><td>75.5</td><td>-10.1</td><td>1.6</td><td>-6.4</td><td>1.2E-10</td><td>3.1E-10</td><td>MB231/MB468</td><td>MP</td></tr>
<tr><td>>hg38_21_+_8433172_8446622_RNA45SN1_45S_50ntFlanks@11669.11706.38|GCCCAGTGCTCTGAATGTCAAAGTGAAGAAATTCAATG</td><td>>hg38_21_+_8433172_8446622_RNA45SN1_45S_50ntFlanks</td><td>11669</td><td>11706</td><td>38</td><td>GCCCAGTGCTCTGAATGTCAAAGTGAAGAAATTCAATG</td><td>488.0</td><td>4.9</td><td>0.8</td><td>6.4</td><td>1.2E-10</td><td>3.1E-10</td><td>MB231/MB468</td><td>NUC</td></tr>
<tr><td>>hg38_21_+_8433172_8446622_RNA45SN1_45S_50ntFlanks@10101.10127.27|CGCGCGCGTGTGGTGTGCGTCGGAGGG</td><td>>hg38_21_+_8433172_8446622_RNA45SN1_45S_50ntFlanks</td><td>10101</td><td>10127</td><td>27</td><td>CGCGCGCGTGTGGTGTGCGTCGGAGGG</td><td>1496.7</td><td>-10.2</td><td>1.6</td><td>-6.4</td><td>1.5E-10</td><td>3.1E-10</td><td>BT20/MB468</td><td>NUC</td></tr>
<tr><td>>trna5_AspGTC_12_+_98897281_98897352@37.74.38|ACGCGGGAGACCGGGGTTCAATTCCCCGACGGGGAGCC</td><td>>trna5_AspGTC_12_+_98897281_98897352</td><td>37</td><td>74</td><td>38</td><td>ACGCGGGAGACCGGGGTTCAATTCCCCGACGGGGAGCC</td><td>3664.2</td><td>-7.8</td><td>1.1</td><td>-6.8</td><td>7.8E-12</td><td>3.1E-10</td><td>BT20/MB231</td><td>CYTO</td></tr>
</table>

<table>
<tr><td>>hg38_21_+_8433172_8446622_RNA45SN1_45S_50ntFlanks@12407.12431.25|TTTTGATCCTTCGATGTCGGCTCTT</td><td>>hg38_21_+_8433172_8446622_RNA45SN1_45S_50ntFlanks</td><td>12407</td><td>12431</td><td>25</td><td>TTTTGATCCTTCGATGTCGGCTCTT</td><td>677.5</td><td>-5.2</td><td>0.8</td><td>-6.4</td><td>1.4E-10</td><td>3.1E-10</td><td>MB231/MB468</td><td>MITO</td></tr>
<tr><td>>hg38_21_+_8433172_8446622_RNA45SN1_45S_50ntFlanks@10866.10900.35|CTAAGGGCTGGGTCGGTCGGGCTGGGGCGCGAAGC</td><td>>hg38_21_+_8433172_8446622_RNA45SN1_45S_50ntFlanks</td><td>10866</td><td>10900</td><td>35</td><td>CTAAGGGCTGGGTCGGTCGGGCTGGGGCGCGAAGC</td><td>74.4</td><td>-10.1</td><td>1.6</td><td>-6.4</td><td>1.2E-10</td><td>3.1E-10</td><td>MB231/MB468</td><td>MP</td></tr>
<tr><td>>hg38_21_+_8433172_8446622_RNA45SN1_45S_50ntFlanks@10466.10483.18|CGTTGCCCTCGGCCGATC</td><td>>hg38_21_+_8433172_8446622_RNA45SN1_45S_50ntFlanks</td><td>10466</td><td>10483</td><td>18</td><td>CGTTGCCCTCGGCCGATC</td><td>77.7</td><td>-10.1</td><td>1.6</td><td>-6.4</td><td>1.2E-10</td><td>3.1E-10</td><td>MB231/MB468</td><td>MP</td></tr>
<tr><td>>hg38_21_+_8433172_8446622_RNA45SN1_45S_50ntFlanks@12654.12675.22|TATGACTGAACGCCTCTAAGTC</td><td>>hg38_21_+_8433172_8446622_RNA45SN1_45S_50ntFlanks</td><td>12654</td><td>12675</td><td>22</td><td>TATGACTGAACGCCTCTAAGTC</td><td>6126.2</td><td>-7.7</td><td>1.2</td><td>-6.5</td><td>1.1E-10</td><td>3.1E-10</td><td>BT20/MB468</td><td>CYTO</td></tr>
<tr><td>>hg38_21_+_8433172_8446622_RNA45SN1_45S_50ntFlanks@11641.11678.38|CGGCGGGTGTTGACGCGATGTGATTTCTGCCCAGTGCT</td><td>>hg38_21_+_8433172_8446622_RNA45SN1_45S_50ntFlanks</td><td>11641</td><td>11678</td><td>38</td><td>CGGCGGGTGTTGACGCGATGTGATTTCTGCCCAGTGCT</td><td>270.5</td><td>-10.2</td><td>1.6</td><td>-6.5</td><td>9.1E-11</td><td>3.2E-10</td><td>BT20/MB468</td><td>MITO</td></tr>
<tr><td>>hg38_21_+_8433172_8446622_RNA45SN1_45S_50ntFlanks@5472.5500.29|ACGGCCCTGGCGGAGCGCTGAGAAGACGG</td><td>>hg38_21_+_8433172_8446622_RNA45SN1_45S_50ntFlanks</td><td>5472</td><td>5500</td><td>29</td><td>ACGGCCCTGGCGGAGCGCTGAGAAGACGG</td><td>454.4</td><td>5.7</td><td>0.9</td><td>6.5</td><td>9.1E-11</td><td>3.2E-10</td><td>BT20/MB468</td><td>MITO</td></tr>
<tr><td>>hg38_21_+_8433172_8446622_RNA45SN1_45S_50ntFlanks@8836.8864.29|CGGCGGTGGCGGCGGCGGCGGCGGCGGCG</td><td>>hg38_21_+_8433172_8446622_RNA45SN1_45S_50ntFlanks</td><td>8836</td><td>8864</td><td>29</td><td>CGGCGGTGGCGGCGGCGGCGGCGGCGGCG</td><td>258.5</td><td>-10.1</td><td>1.6</td><td>-6.5</td><td>9.1E-11</td><td>3.2E-10</td><td>BT20/MB468</td><td>MITO</td></tr>
<tr><td>>hg38_21_+_8433172_8446622_RNA45SN1_45S_50ntFlanks@4472.4498.27|CTTAGCTGAGTGTCCCGCGGGGCCCGA</td><td>>hg38_21_+_8433172_8446622_RNA45SN1_45S_50ntFlanks</td><td>4472</td><td>4498</td><td>27</td><td>CTTAGCTGAGTGTCCCGCGGGGCCCGA</td><td>119.2</td><td>-10.8</td><td>1.7</td><td>-6.4</td><td>1.2E-10</td><td>3.2E-10</td><td>MB231/MB468</td><td>MP</td></tr>
<tr><td>>hg38_21_+_8433172_8446622_RNA45SN1_45S_50ntFlanks@6645.6685.41|CTCGTACGACTCTTAGCGGTGGATCACTCGGCTCGTGCGTC</td><td>>hg38_21_+_8433172_8446622_RNA45SN1_45S_50ntFlanks</td><td>6645</td><td>6685</td><td>41</td><td>CTCGTACGACTCTTAGCGGTGGATCACTCGGCTCGTGCGTC</td><td>931.6</td><td>8.2</td><td>1.3</td><td>6.5</td><td>1.1E-10</td><td>3.2E-10</td><td>BT20/MB468</td><td>CYTO</td></tr>
<tr><td>>hg38_21_+_8433172_8446622_RNA45SN1_45S_50ntFlanks@10466.10487.22|CGTTGCCCTCGGCCGATCGAAA</td><td>>hg38_21_+_8433172_8446622_RNA45SN1_45S_50ntFlanks</td><td>10466</td><td>10487</td><td>22</td><td>CGTTGCCCTCGGCCGATCGAAA</td><td>74.1</td><td>-10.1</td><td>1.6</td><td>-6.4</td><td>1.2E-10</td><td>3.2E-10</td><td>MB231/MB468</td><td>MP</td></tr>
<tr><td>>hg38_21_+_8433172_8446622_RNA45SN1_45S_50ntFlanks@10792.10825.34|CTCTGGCATGTTGGAACAATGTAGGTAAGGGAAG</td><td>>hg38_21_+_8433172_8446622_RNA45SN1_45S_50ntFlanks</td><td>10792</td><td>10825</td><td>34</td><td>CTCTGGCATGTTGGAACAATGTAGGTAAGGGAAG</td><td>78.1</td><td>-10.2</td><td>1.6</td><td>-6.4</td><td>1.2E-10</td><td>3.2E-10</td><td>MB231/MB468</td><td>MP</td></tr>
<tr><td>>hg38_21_+_8433172_8446622_RNA45SN1_45S_50ntFlanks@9564.9594.31|CTGACGTGCAAATCGGTCGTCCGACCTGGGT</td><td>>hg38_21_+_8433172_8446622_RNA45SN1_45S_50ntFlanks</td><td>9564</td><td>9594</td><td>31</td><td>CTGACGTGCAAATCGGTCGTCCGACCTGGGT</td><td>1838.2</td><td>-6.9</td><td>1.1</td><td>-6.4</td><td>1.4E-10</td><td>3.2E-10</td><td>MB231/MB468</td><td>MITO</td></tr>
<tr><td>>hg38_21_+_8433172_8446622_RNA45SN1_45S_50ntFlanks@10548.10576.29|GTGCGGTAACGCGACCGATCCCGGAGAAG</td><td>>hg38_21_+_8433172_8446622_RNA45SN1_45S_50ntFlanks</td><td>10548</td><td>10576</td><td>29</td><td>GTGCGGTAACGCGACCGATCCCGGAGAAG</td><td>949.0</td><td>5.8</td><td>0.9</td><td>6.5</td><td>9.2E-11</td><td>3.2E-10</td><td>BT20/MB468</td><td>MITO</td></tr>
<tr><td>>hg38_21_+_8433172_8446622_RNA45SN1_45S_50ntFlanks@10116.10133.18|TGCGTCGGAGGGCGGCGG</td><td>>hg38_21_+_8433172_8446622_RNA45SN1_45S_50ntFlanks</td><td>10116</td><td>10133</td><td>18</td><td>TGCGTCGGAGGGCGGCGG</td><td>105.7</td><td>-9.0</td><td>1.4</td><td>-6.5</td><td>1.1E-10</td><td>3.2E-10</td><td>BT20/MB468</td><td>CYTO</td></tr>
<tr><td>>hg38_21_+_8433172_8446622_RNA45SN1_45S_50ntFlanks@4729.4752.24|TCAAGAACGAAAGTCGGAGGTTCG</td><td>>hg38_21_+_8433172_8446622_RNA45SN1_45S_50ntFlanks</td><td>4729</td><td>4752</td><td>24</td><td>TCAAGAACGAAAGTCGGAGGTTCG</td><td>112.5</td><td>-8.9</td><td>1.4</td><td>-6.5</td><td>9.2E-11</td><td>3.2E-10</td><td>BT20/MB468</td><td>MITO</td></tr>
<tr><td>>MI0000102|hsa-mir-100&WithFlank&11|-|122152223|122152314@19.36.18&[MIMAT0000098&hsa-miR-100-5p&offsets|0|-4;m-27&11|-|122152276|122152296&offsets|0|-3]|AACCCGTAGATCCGAACT</td><td>>MI0000102|hsa-mir-100&WithFlank&11|-|122152223|122152314</td><td>19</td><td>36</td><td>18</td><td>AACCCGTAGATCCGAACT</td><td>427.5</td><td>-6.4</td><td>0.9</td><td>-6.8</td><td>1.1E-11</td><td>3.2E-10</td><td>BT20/MB231</td><td>MITO</td></tr>
<tr><td>>hg38_21_+_8433172_8446622_RNA45SN1_45S_50ntFlanks@12415.12435.21|CTTCGATGTCGGCTCTTCCTA</td><td>>hg38_21_+_8433172_8446622_RNA45SN1_45S_50ntFlanks</td><td>12415</td><td>12435</td><td>21</td><td>CTTCGATGTCGGCTCTTCCTA</td><td>2612.2</td><td>-3.8</td><td>0.6</td><td>-6.4</td><td>1.3E-10</td><td>3.2E-10</td><td>MB231/MB468</td><td>CYTO</td></tr>
<tr><td>>hg38_21_+_8433172_8446622_RNA45SN1_45S_50ntFlanks@11930.11967.38|TAGAATAAGTGGGAGGCCCCCGGCGCCCCCCCGGTGTC</td><td>>hg38_21_+_8433172_8446622_RNA45SN1_45S_50ntFlanks</td><td>11930</td><td>11967</td><td>38</td><td>TAGAATAAGTGGGAGGCCCCCGGCGCCCCCCCGGTGTC</td><td>197.2</td><td>-11.4</td><td>1.8</td><td>-6.4</td><td>1.4E-10</td><td>3.2E-10</td><td>MB231/MB468</td><td>MITO</td></tr>
<tr><td>>hg38_21_+_8433172_8446622_RNA45SN1_45S_50ntFlanks@11580.11602.23|GAACTGGTGCGGACCAGGGGAAT</td><td>>hg38_21_+_8433172_8446622_RNA45SN1_45S_50ntFlanks</td><td>11580</td><td>11602</td><td>23</td><td>GAACTGGTGCGGACCAGGGGAAT</td><td>400.1</td><td>-3.9</td><td>0.6</td><td>-6.4</td><td>1.3E-10</td><td>3.2E-10</td><td>MB231/MB468</td><td>NUC</td></tr>
<tr><td>>MI0000471|hsa-mir-126&WithFlank&9|+|136670596|136670692@58.77.20&[MIMAT0000445&hsa-miR-126-3p&offsets|0|-2;m-40&9|+|136670653|136670674&offsets|0|-2]|TCGTACCGTGAGTAATAATG</td><td>>MI0000471|hsa-mir-126&WithFlank&9|+|136670596|136670692</td><td>58</td><td>77</td><td>20</td><td>TCGTACCGTGAGTAATAATG</td><td>349.6</td><td>8.4</td><td>1.3</td><td>6.4</td><td>1.3E-10</td><td>3.2E-10</td><td>MB231/MB468</td><td>NUC</td></tr>
<tr><td>>MI0000077|hsa-mir-21&WithFlank&17|+|59841260|59841343@14.34.21(+2C)&[MIMAT0000076&hsa-miR-21-5p&offsets|0|-1(+2C);m-1&17|+|59841273|59841294&offsets|0|-1(+2C)]|TAGCTTATCAGACTGATGTTGCC</td><td>>MI0000077|hsa-mir-21&WithFlank&17|+|59841260|59841343</td><td>14</td><td>34</td><td>21(+2C)</td><td>TAGCTTATCAGACTGATGTTGCC</td><td>98.5</td><td>2.0</td><td>0.3</td><td>6.4</td><td>1.5E-10</td><td>3.2E-10</td><td>BT20/MB468</td><td>NUC</td></tr>
<tr><td>>hg38_21_+_8433172_8446622_RNA45SN1_45S_50ntFlanks@3873.3899.27|TAATACATGCCGACGGGCGCTGACCCC</td><td>>hg38_21_+_8433172_8446622_RNA45SN1_45S_50ntFlanks</td><td>3873</td><td>3899</td><td>27</td><td>TAATACATGCCGACGGGCGCTGACCCC</td><td>623.4</td><td>-8.3</td><td>1.3</td><td>-6.4</td><td>1.4E-10</td><td>3.2E-10</td><td>MB231/MB468</td><td>MITO</td></tr>
<tr><td>>trna10_IleTAT_19_-_39902808_39902900@33.52.20,trna29_IleTAT_6_+_26988125_26988218@33.52.20,trna55_IleTAT_6_+_27599200_27599293@33.52.20,trna5_IleTAT_2_+_43037676_43037768@33.52.20,trna63_IleTAT_6_+_28505367_28505460@33.52.20|CTTATAATGCCGAGGTTGTG</td><td>>trna10_IleTAT_19_-_39902808_39902900</td><td>33</td><td>52</td><td>20</td><td>CTTATAATGCCGAGGTTGTG</td><td>100.6</td><td>-10.5</td><td>1.6</td><td>-6.4</td><td>1.2E-10</td><td>3.2E-10</td><td>MB231/MB468</td><td>MP</td></tr>
<tr><td>>MI0000298|hsa-mir-221&WithFlank&X|-|45746151|45746272@71.91.21(+1A)&[MIMAT0000278&hsa-miR-221-3p&offsets|0|-2(+1A);m-91&X|-|45746181|45746202&offsets|0|-1(+1A)]|AGCTACATTGTCTGCTGGGTTA</td><td>>MI0000298|hsa-mir-221&WithFlank&X|-|45746151|45746272</td><td>71</td><td>91</td><td>21(+1A)</td><td>AGCTACATTGTCTGCTGGGTTA</td><td>591.3</td><td>7.6</td><td>1.2</td><td>6.4</td><td>1.1E-10</td><td>3.2E-10</td><td>BT20/MB468</td><td>CYTO</td></tr>
<tr><td>>MI0000264|hsa-mir-7-2&WithFlank&15|+|88611819|88611940@38.60.23;MI0000265|hsa-mir-7-3&WithFlank&19|+|4770664|4770785@37.59.23;MI0000263|hsa-mir-7-1&WithFlank&9|-|83969742|83969863@30.52.23&[MIMAT0000252&hsa-miR-7-5p&offsets|0|-1;m-400&15|+|88611856|88611879&offsets|0|-1];[MIMAT0000252_1&hsa-miR-7-5p&offsets|0|-1;m-402&19|+|4770700|4770723&offsets|0|-1];[MIMAT0000252_2&hsa-miR-7-5p&offsets|0|-1;m-401&9|-|83969811|83969834&offsets|0|-1]|TGGAAGACTAGTGATTTTGTTGT</td><td>>MI0000264|hsa-mir-7-2&WithFlank&15|+|88611819|88611940</td><td>38</td><td>60</td><td>23</td><td>TGGAAGACTAGTGATTTTGTTGT</td><td>6064.7</td><td>6.1</td><td>0.9</td><td>6.4</td><td>1.4E-10</td><td>3.2E-10</td><td>MB231/MB468</td><td>MITO</td></tr>
<tr><td>>hg38_21_+_8433172_8446622_RNA45SN1_45S_50ntFlanks@11821.11841.21|CTATCCAGCGAAACCACAGCC</td><td>>hg38_21_+_8433172_8446622_RNA45SN1_45S_50ntFlanks</td><td>11821</td><td>11841</td><td>21</td><td>CTATCCAGCGAAACCACAGCC</td><td>84.2</td><td>-10.3</td><td>1.6</td><td>-6.4</td><td>1.2E-10</td><td>3.2E-10</td><td>MB231/MB468</td><td>MP</td></tr>
<tr><td>>hg38_21_+_8433172_8446622_RNA45SN1_45S_50ntFlanks@10921.10946.26|GCTGGACGAGGCGCCGCCGCCCCCCC</td><td>>hg38_21_+_8433172_8446622_RNA45SN1_45S_50ntFlanks</td><td>10921</td><td>10946</td><td>26</td><td>GCTGGACGAGGCGCCGCCGCCCCCCC</td><td>286.6</td><td>-8.1</td><td>1.3</td><td>-6.4</td><td>1.4E-10</td><td>3.2E-10</td><td>MB231/MB468</td><td>MITO</td></tr>
<tr><td>>hg38_21_+_8433172_8446622_RNA45SN1_45S_50ntFlanks@4663.4690.28|GTGAAATTCTTGGACCGGCGCAAGACGG</td><td>>hg38_21_+_8433172_8446622_RNA45SN1_45S_50ntFlanks</td><td>4663</td><td>4690</td><td>28</td><td>GTGAAATTCTTGGACCGGCGCAAGACGG</td><td>86.9</td><td>-10.3</td><td>1.6</td><td>-6.4</td><td>1.2E-10</td><td>3.2E-10</td><td>MB231/MB468</td><td>MP</td></tr>
<tr><td>>hg38_21_+_8433172_8446622_RNA45SN1_45S_50ntFlanks@11641.11659.19|CGGCGGGTGTTGACGCGAT</td><td>>hg38_21_+_8433172_8446622_RNA45SN1_45S_50ntFlanks</td><td>11641</td><td>11659</td><td>19</td><td>CGGCGGGTGTTGACGCGAT</td><td>2581.8</td><td>-9.8</td><td>1.4</td><td>-6.8</td><td>9.5E-12</td><td>3.2E-10</td><td>BT20/MB468</td><td>MP</td></tr>
<tr><td>>hg38_21_+_8433172_8446622_RNA45SN1_45S_50ntFlanks@3838.3871.34|CTCTCCTACTTGGATAACTGTGGTAATTCTAGAG</td><td>>hg38_21_+_8433172_8446622_RNA45SN1_45S_50ntFlanks</td><td>3838</td><td>3871</td><td>34</td><td>CTCTCCTACTTGGATAACTGTGGTAATTCTAGAG</td><td>170.7</td><td>-11.3</td><td>1.8</td><td>-6.4</td><td>1.2E-10</td><td>3.2E-10</td><td>MB231/MB468</td><td>MP</td></tr>
<tr><td>>trna116_GluCTC_1_-_145399233_145399304@38.70.33,trna134_GluTTC_1_-_16861774_16861845@38.70.33,trna71_GluCTC_1_-_161439189_161439260@38.70.33,trna74_GluCTC_1_-_161431809_161431880@38.70.33,trna77_GluCTC_1_-_161424398_161424469@38.70.33,trna77_GluCTC_6_+_28949976_28950047@38.70.33,trna80_GluCTC_1_-_161417018_161417089@38.70.33,trna84_GluTTC_1_-_161391883_161391954@38.70.33,trna87_GluCTC_6_-_126101393_126101464@38.70.33|CCGCCGCGGCCCGGGTTCGATTCCCGGTCAGGG</td><td>>trna116_GluCTC_1_-_145399233_145399304</td><td>38</td><td>70</td><td>33</td><td>CCGCCGCGGCCCGGGTTCGATTCCCGGTCAGGG</td><td>88.4</td><td>-10.3</td><td>1.6</td><td>-6.4</td><td>1.2E-10</td><td>3.2E-10</td><td>MB231/MB468</td><td>MP</td></tr>
<tr><td>>hg38_21_+_8433172_8446622_RNA45SN1_45S_50ntFlanks@12456.12486.31|CCAAGCGTTGGATTGTTCACCCACTAATAGG</td><td>>hg38_21_+_8433172_8446622_RNA45SN1_45S_50ntFlanks</td><td>12456</td><td>12486</td><td>31</td><td>CCAAGCGTTGGATTGTTCACCCACTAATAGG</td><td>7013.8</td><td>-10.0</td><td>1.6</td><td>-6.4</td><td>1.2E-10</td><td>3.2E-10</td><td>MB231/MB468</td><td>MP</td></tr>
<tr><td>>MI0000542|hsa-mir-320a&WithFlank&8|-|22244960|22245043@48.70.23(+1U)&[MIMAT0000510&hsa-miR-320a-3p&offsets|0|+1(+1U);m-60&8|-|22244975|22244996&offsets|0|+1(+1U)]|AAAAGCTGGGTTGAGAGGGCGAAT</td><td>>MI0000542|hsa-mir-320a&WithFlank&8|-|22244960|22245043</td><td>48</td><td>70</td><td>23(+1U)</td><td>AAAAGCTGGGTTGAGAGGGCGAAT</td><td>1933.8</td><td>4.8</td><td>0.8</td><td>6.4</td><td>1.3E-10</td><td>3.2E-10</td><td>MB231/MB468</td><td>NUC</td></tr>
<tr><td>>hg38_21_+_8433172_8446622_RNA45SN1_45S_50ntFlanks@11824.11844.21|TCCAGCGAAACCACAGCCAAG</td><td>>hg38_21_+_8433172_8446622_RNA45SN1_45S_50ntFlanks</td><td>11824</td><td>11844</td><td>21</td><td>TCCAGCGAAACCACAGCCAAG</td><td>89.1</td><td>-10.3</td><td>1.6</td><td>-6.4</td><td>1.2E-10</td><td>3.2E-10</td><td>MB231/MB468</td><td>MP</td></tr>
<tr><td>>hg38_21_+_8433172_8446622_RNA45SN1_45S_50ntFlanks@6653.6678.26|ACTCTTAGCGGTGGATCACTCGGCTC</td><td>>hg38_21_+_8433172_8446622_RNA45SN1_45S_50ntFlanks</td><td>6653</td><td>6678</td><td>26</td><td>ACTCTTAGCGGTGGATCACTCGGCTC</td><td>754.6</td><td>5.0</td><td>0.8</td><td>6.5</td><td>9.4E-11</td><td>3.2E-10</td><td>BT20/MB468</td><td>MITO</td></tr>
<tr><td>>hg38_21_+_8433172_8446622_RNA45SN1_45S_50ntFlanks@11561.11589.29|GCGCCTAGCAGCCGACTTAGAACTGGTGC</td><td>>hg38_21_+_8433172_8446622_RNA45SN1_45S_50ntFlanks</td><td>11561</td><td>11589</td><td>29</td><td>GCGCCTAGCAGCCGACTTAGAACTGGTGC</td><td>141.5</td><td>-4.2</td><td>0.7</td><td>-6.4</td><td>1.3E-10</td><td>3.2E-10</td><td>MB231/MB468</td><td>CYTO</td></tr>
<tr><td>>hg38_21_+_8433172_8446622_RNA45SN1_45S_50ntFlanks@10783.10818.36|GTGAACAGCCTCTGGCATGTTGGAACAATGTAGGTA</td><td>>hg38_21_+_8433172_8446622_RNA45SN1_45S_50ntFlanks</td><td>10783</td><td>10818</td><td>36</td><td>GTGAACAGCCTCTGGCATGTTGGAACAATGTAGGTA</td><td>1532.0</td><td>-9.8</td><td>1.5</td><td>-6.4</td><td>1.5E-10</td><td>3.3E-10</td><td>BT20/MB468</td><td>NUC</td></tr>
<tr><td>>hg38_21_+_8433172_8446622_RNA45SN1_45S_50ntFlanks@6788.6807.20|CGCCTGTCTGAGCGTCGCTT</td><td>>hg38_21_+_8433172_8446622_RNA45SN1_45S_50ntFlanks</td><td>6788</td><td>6807</td><td>20</td><td>CGCCTGTCTGAGCGTCGCTT</td><td>1481.2</td><td>3.8</td><td>0.6</td><td>6.4</td><td>1.3E-10</td><td>3.3E-10</td><td>MB231/MB468</td><td>NUC</td></tr>
<tr><td>>hg38_21_+_8433172_8446622_RNA45SN1_45S_50ntFlanks@11270.11294.25|CGGCCGCCCCCGGGGAGCCCGGCGG</td><td>>hg38_21_+_8433172_8446622_RNA45SN1_45S_50ntFlanks</td><td>11270</td><td>11294</td><td>25</td><td>CGGCCGCCCCCGGGGAGCCCGGCGG</td><td>130.7</td><td>-10.9</td><td>1.7</td><td>-6.4</td><td>1.2E-10</td><td>3.3E-10</td><td>MB231/MB468</td><td>MP</td></tr>
<tr><td>>am_trnaMT_SerGCT_MT_+_12207_12265@11.38.28|ACAAGAACTGCTAACTCATGCCCCCATG</td><td>>am_trnaMT_SerGCT_MT_+_12207_12265</td><td>11</td><td>38</td><td>28</td><td>ACAAGAACTGCTAACTCATGCCCCCATG</td><td>1151.4</td><td>-10.3</td><td>1.6</td><td>-6.4</td><td>1.5E-10</td><td>3.3E-10</td><td>BT20/MB468</td><td>NUC</td></tr>
<tr><td>>hg38_21_+_8433172_8446622_RNA45SN1_45S_50ntFlanks@11579.11611.33|AGAACTGGTGCGGACCAGGGGAATCCGACTGTT</td><td>>hg38_21_+_8433172_8446622_RNA45SN1_45S_50ntFlanks</td><td>11579</td><td>11611</td><td>33</td><td>AGAACTGGTGCGGACCAGGGGAATCCGACTGTT</td><td>867.9</td><td>-3.6</td><td>0.6</td><td>-6.4</td><td>1.5E-10</td><td>3.3E-10</td><td>BT20/MB468</td><td>NUC</td></tr>
<tr><td>>hg38_21_+_8433172_8446622_RNA45SN1_45S_50ntFlanks@8144.8163.20|CTCCCCGGCGCCGCTCGTGG</td><td>>hg38_21_+_8433172_8446622_RNA45SN1_45S_50ntFlanks</td><td>8144</td><td>8163</td><td>20</td><td>CTCCCCGGCGCCGCTCGTGG</td><td>77.1</td><td>-10.1</td><td>1.6</td><td>-6.4</td><td>1.2E-10</td><td>3.3E-10</td><td>MB231/MB468</td><td>MP</td></tr>
<tr><td>>hg38_21_+_8433172_8446622_RNA45SN1_45S_50ntFlanks@13009.13028.20|TTGAAAGTCAGCCCTCGACA</td><td>>hg38_21_+_8433172_8446622_RNA45SN1_45S_50ntFlanks</td><td>13009</td><td>13028</td><td>20</td><td>TTGAAAGTCAGCCCTCGACA</td><td>94.5</td><td>-10.4</td><td>1.6</td><td>-6.4</td><td>1.2E-10</td><td>3.3E-10</td><td>MB231/MB468</td><td>MP</td></tr>
<tr><td>>MI0000102|hsa-mir-100&WithFlank&11|-|122152223|122152314@19.40.22(+2U)&[MIMAT0000098&hsa-miR-100-5p&offsets|0|0(+2U);m-27&11|-|122152276|122152296&offsets|0|+1(+2U)]|AACCCGTAGATCCGAACTTGTGTT</td><td>>MI0000102|hsa-mir-100&WithFlank&11|-|122152223|122152314</td><td>19</td><td>40</td><td>22(+2U)</td><td>AACCCGTAGATCCGAACTTGTGTT</td><td>418.3</td><td>-12.1</td><td>1.7</td><td>-7.0</td><td>2.1E-12</td><td>3.3E-10</td><td>BT20/MB231</td><td>MP</td></tr>
<tr><td>>hg38_21_+_8433172_8446622_RNA45SN1_45S_50ntFlanks@9572.9595.24|CAAATCGGTCGTCCGACCTGGGTA</td><td>>hg38_21_+_8433172_8446622_RNA45SN1_45S_50ntFlanks</td><td>9572</td><td>9595</td><td>24</td><td>CAAATCGGTCGTCCGACCTGGGTA</td><td>568.7</td><td>-5.5</td><td>0.9</td><td>-6.4</td><td>1.3E-10</td><td>3.3E-10</td><td>MB231/MB468</td><td>NUC</td></tr>
</table>

| | | | | | | | | | | | | | |
|---|---|---|---|---|---|---|---|---|---|---|---|---|---|
| >MI0000279\|hsa-mir-196a-2&WithFlank&12\|+\|53991732\|53991853@31.51.21;MI0000238\|hsa-mir-196a-1&WithFlank&17\|-\|48632484\|48632565@13.33.21&[MIMAT0000226&hsa-miR-196a-5p&offsets\|0\|-1;m-261&12\|+\|53991762\|53991783&offsets\|0\|-1];[MIMAT0000226_1&hsa-miR-196a-5p&offsets\|0\|-1;m-268&17\|-\|48632532\|48632553&offsets\|0\|-1]\|TAGGTAGTTTCATGTTGTTGG | >MI0000279\|hsa-mir-196a-2&WithFlank&12\|+\|53991732\|53991853 | 31 | 51 | 21 | TAGGTAGTTTCATGTTGTTGG | 441.1 | 12.1 | 1.7 | 7.0 | 2.1E-12 | 3.3E-10 | BT20/MB231 | MP |
| >hg38_21_+_8433172_8446622_RNA45SN1_45S_50ntFlanks@12244.12268.25\|AAGGGCAAAAGCTCGCTTGATCTTG | >hg38_21_+_8433172_8446622_RNA45SN1_45S_50ntFlanks | 12244 | 12268 | 25 | AAGGGCAAAAGCTCGCTTGATCTTG | 427.3 | -4.7 | 0.7 | -6.4 | 1.4E-10 | 3.3E-10 | MB231/MB468 | MITO |
| >MI0000746\|hsa-mir-99b&WithFlank&19\|+\|51692606\|51692687@13.32.20(+1U)&[MIMAT0000689&hsa-miR-99b-5p&offsets\|0\|-2(+1U);m-10&19\|+\|51692618\|51692639&offsets\|0\|-2(+1U)]\|CACCCGTAGAACCGACCTTGT | >MI0000746\|hsa-mir-99b&WithFlank&19\|+\|51692606\|51692687 | 13 | 32 | 20(+1U) | CACCCGTAGAACCGACCTTGT | 1547.5 | 4.2 | 0.7 | 6.4 | 1.3E-10 | 3.3E-10 | MB231/MB468 | NUC |
| >hg38_21_+_8433172_8446622_RNA45SN1_45S_50ntFlanks@11933.11969.37\|AATAAGTGGGAGGCCCCCGGCGCCCCCCCGGTGTCCC | >hg38_21_+_8433172_8446622_RNA45SN1_45S_50ntFlanks | 11933 | 11969 | 37 | AATAAGTGGGAGGCCCCCGGCGCCCCCCCGGTGTCCC | 166.6 | -11.2 | 1.7 | -6.4 | 1.4E-10 | 3.3E-10 | MB231/MB468 | MITO |
| >hg38_21_+_8433172_8446622_RNA45SN1_45S_50ntFlanks@3854.3881.28\|ACTGTGGTAATTCTAGAGCTAATACATG | >hg38_21_+_8433172_8446622_RNA45SN1_45S_50ntFlanks | 3854 | 3881 | 28 | ACTGTGGTAATTCTAGAGCTAATACATG | 1830.1 | -5.7 | 0.9 | -6.4 | 1.3E-10 | 3.3E-10 | MB231/MB468 | CYTO |
| >hg38_21_+_8433172_8446622_RNA45SN1_45S_50ntFlanks@10410.10442.33\|TCAGTCGGTCCTGAGAGATGGGCGAGCGCCGTT | >hg38_21_+_8433172_8446622_RNA45SN1_45S_50ntFlanks | 10410 | 10442 | 33 | TCAGTCGGTCCTGAGAGATGGGCGAGCGCCGTT | 330.9 | -3.6 | 0.6 | -6.4 | 1.3E-10 | 3.3E-10 | MB231/MB468 | NUC |
| >hg38_21_+_8433172_8446622_RNA45SN1_45S_50ntFlanks@11580.11603.24\|GAACTGGTGCGGACCAGGGGAATC | >hg38_21_+_8433172_8446622_RNA45SN1_45S_50ntFlanks | 11580 | 11603 | 24 | GAACTGGTGCGGACCAGGGGAATC | 5652.5 | -8.8 | 1.4 | -6.4 | 1.5E-10 | 3.3E-10 | BT20/MB468 | NUC |
| >hg38_21_+_8433172_8446622_RNA45SN1_45S_50ntFlanks@11567.11600.34\|AGCAGCCGACTTAGAACTGGTGCGGACCAGGGGA | >hg38_21_+_8433172_8446622_RNA45SN1_45S_50ntFlanks | 11567 | 11600 | 34 | AGCAGCCGACTTAGAACTGGTGCGGACCAGGGGA | 1405.8 | -5.8 | 0.9 | -6.4 | 1.5E-10 | 3.3E-10 | BT20/MB468 | NUC |
| >hg38_21_+_8433172_8446622_RNA45SN1_45S_50ntFlanks@4995.5034.40\|ACAGATTGATAGCTCTTTCTCGATTCCGTGGGTGGTGGTG | >hg38_21_+_8433172_8446622_RNA45SN1_45S_50ntFlanks | 4995 | 5034 | 40 | ACAGATTGATAGCTCTTTCTCGATTCCGTGGGTGGTGGTG | 174.8 | -9.5 | 1.5 | -6.5 | 9.5E-11 | 3.3E-10 | BT20/MB468 | MITO |
| >hg38_21_+_8433172_8446622_RNA45SN1_45S_50ntFlanks@12995.13039.45\|CTCGCTGCGATCTATTGAAAGTCAGCCCTCGACACAAGGGTTTGT | >hg38_21_+_8433172_8446622_RNA45SN1_45S_50ntFlanks | 12995 | 13039 | 45 | CTCGCTGCGATCTATTGAAAGTCAGCCCTCGACACAAGGGTTTGT | 181195.3 | 6.8 | 1.1 | 6.5 | 9.5E-11 | 3.3E-10 | BT20/MB468 | MITO |
| >hg38_21_+_8433172_8446622_RNA45SN1_45S_50ntFlanks@11752.11780.29\|CCAAATGCCTCGTCATCTAATTAGTGACG | >hg38_21_+_8433172_8446622_RNA45SN1_45S_50ntFlanks | 11752 | 11780 | 29 | CCAAATGCCTCGTCATCTAATTAGTGACG | 77.2 | -10.1 | 1.6 | -6.4 | 1.2E-10 | 3.3E-10 | MB231/MB468 | MP |
| >hg38_21_+_8433172_8446622_RNA45SN1_45S_50ntFlanks@12644.12661.18\|TCTGTGGGATTATGACTG | >hg38_21_+_8433172_8446622_RNA45SN1_45S_50ntFlanks | 12644 | 12661 | 18 | TCTGTGGGATTATGACTG | 2948.4 | -9.2 | 1.4 | -6.4 | 1.5E-10 | 3.3E-10 | MB231/MB468 | MITO |
| >hg38_21_+_8433172_8446622_RNA45SN1_45S_50ntFlanks@6745.6762.18\|ACACTTCGAACGCACTTG | >hg38_21_+_8433172_8446622_RNA45SN1_45S_50ntFlanks | 6745 | 6762 | 18 | ACACTTCGAACGCACTTG | 259.7 | -5.0 | 0.8 | -6.4 | 1.3E-10 | 3.3E-10 | MB231/MB468 | CYTO |
| >hg38_21_+_8433172_8446622_RNA45SN1_45S_50ntFlanks@6670.6699.30\|ACTCGGCTCGTGCGTCGATGAAGAACGCAG | >hg38_21_+_8433172_8446622_RNA45SN1_45S_50ntFlanks | 6670 | 6699 | 30 | ACTCGGCTCGTGCGTCGATGAAGAACGCAG | 3458.2 | -5.5 | 0.9 | -6.4 | 1.5E-10 | 3.3E-10 | MB231/MB468 | MITO |
| >hg38_21_+_8433172_8446622_RNA45SN1_45S_50ntFlanks@8380.8406.27\|CAAGAGGGCGTGAAACCGTTAAGAGGT | >hg38_21_+_8433172_8446622_RNA45SN1_45S_50ntFlanks | 8380 | 8406 | 27 | CAAGAGGGCGTGAAACCGTTAAGAGGT | 1226.1 | -7.6 | 1.2 | -6.4 | 1.5E-10 | 3.3E-10 | BT20/MB468 | NUC |
| >hg38_21_+_8433172_8446622_RNA45SN1_45S_50ntFlanks@12652.12675.24\|ATTATGACTGAACGCCTCTAAGTC | >hg38_21_+_8433172_8446622_RNA45SN1_45S_50ntFlanks | 12652 | 12675 | 24 | ATTATGACTGAACGCCTCTAAGTC | 294.4 | -8.4 | 1.3 | -6.4 | 1.5E-10 | 3.3E-10 | BT20/MB468 | NUC |
| >MI0000065\|hsa-let-7d&WithFlank&9\|+\|94178828\|94178926@14.34.21(+1C)&[MIMAT0000065&hsa-let-7d-5p&offsets\|0\|-1(+1C);m-90&9\|+\|94178841\|94178862&offsets\|0\|-1(+1C)]\|AGAGGTAGTAGGTTGCATAGTC | >MI0000065\|hsa-let-7d&WithFlank&9\|+\|94178828\|94178926 | 14 | 34 | 21(+1C) | AGAGGTAGTAGGTTGCATAGTC | 547.4 | 2.9 | 0.5 | 6.4 | 1.5E-10 | 3.3E-10 | BT20/MB468 | NUC |
| >hg38_21_+_8433172_8446622_RNA45SN1_45S_50ntFlanks@10186.10208.23\|TCCTCCCCTCCTCCCGCCCACGC | >hg38_21_+_8433172_8446622_RNA45SN1_45S_50ntFlanks | 10186 | 10208 | 23 | TCCTCCCCTCCTCCCGCCCACGC | 1028.5 | -9.6 | 1.5 | -6.4 | 1.5E-10 | 3.3E-10 | BT20/MB468 | NUC |
| >MI0000065\|hsa-let-7d&WithFlank&9\|+\|94178828\|94178926@68.88.21(+1C)&[MIMAT0004484&hsa-let-7d-3p&offsets\|0\|-1(+1C);m-98&9\|+\|94178895\|94178916&offsets\|0\|-1(+1C)]\|CTATACGACCTGCTGCCTTTCC | >MI0000065\|hsa-let-7d&WithFlank&9\|+\|94178828\|94178926 | 68 | 88 | 21(+1C) | CTATACGACCTGCTGCCTTTCC | 358.1 | 4.8 | 0.8 | 6.4 | 1.3E-10 | 3.3E-10 | MB231/MB468 | NUC |
| >hg38_21_+_8433172_8446622_RNA45SN1_45S_50ntFlanks@11843.11868.26\|AGGGAACGGGCTTGGCGGAATCAGCG | >hg38_21_+_8433172_8446622_RNA45SN1_45S_50ntFlanks | 11843 | 11868 | 26 | AGGGAACGGGCTTGGCGGAATCAGCG | 80.7 | -10.2 | 1.6 | -6.4 | 1.2E-10 | 3.3E-10 | MB231/MB468 | MP |
| >hg38_21_+_8433172_8446622_RNA45SN1_45S_50ntFlanks@9352.9378.27\|CCGTGGCGCAATGAAGGTGAAGGCCGG | >hg38_21_+_8433172_8446622_RNA45SN1_45S_50ntFlanks | 9352 | 9378 | 27 | CCGTGGCGCAATGAAGGTGAAGGCCGG | 167.7 | -11.3 | 1.7 | -6.4 | 1.2E-10 | 3.3E-10 | MB231/MB468 | MP |
| >MI0000433\|hsa-let-7g&WithFlank&3\|-\|52268272\|52268367@11.32.22&[MIMAT0000414&hsa-let-7g-5p&offsets\|0\|0;m-58&3\|-\|52268336\|52268357&offsets\|0\|0]\|TGAGGTAGTAGTTTGTACAGTT | >MI0000433\|hsa-let-7g&WithFlank&3\|-\|52268272\|52268367 | 11 | 32 | 22 | TGAGGTAGTAGTTTGTACAGTT | 30015.3 | 4.8 | 0.7 | 6.4 | 1.3E-10 | 3.3E-10 | MB231/MB468 | CYTO |
| >hg38_21_+_8433172_8446622_RNA45SN1_45S_50ntFlanks@11670.11688.19\|CCCAGTGCTCTGAATGTCA | >hg38_21_+_8433172_8446622_RNA45SN1_45S_50ntFlanks | 11670 | 11688 | 19 | CCCAGTGCTCTGAATGTCA | 899.0 | -4.5 | 0.7 | -6.4 | 1.3E-10 | 3.3E-10 | MB231/MB468 | NUC |
| >hg38_21_+_8433172_8446622_RNA45SN1_45S_50ntFlanks@12437.12455.19\|CATTGTGAAGCAGAATTCA | >hg38_21_+_8433172_8446622_RNA45SN1_45S_50ntFlanks | 12437 | 12455 | 19 | CATTGTGAAGCAGAATTCA | 18516.6 | -11.0 | 1.6 | -6.8 | 9.9E-12 | 3.3E-10 | BT20/MB468 | MP |
| >trna111_HisGTG_1_-_147774845_147774916@-1A.26.27,trna118_HisGTG_1_-_145396881_145396952@-1A.26.27,trna16_HisGTG_1_+_146544773_146544844@-1A.26.27,trna1_HisGTG_15_+_45493349_45493420@-1A.26.27,trna21_HisGTG_1_+_147753471_147753542@-1A.26.27,trna33_HisGTG_6_+_27125906_27125977@-1A.26.27,trna7_HisGTG_9_-_14433938_14434009@-1A.26.27,trna8_HisGTG_15_-_45492611_45492682@-1A.26.27,trna9_HisGTG_15_-_45490804_45490875@-1A.26.27\|AGCCGTGATCGTATAGTGGTTAGTACT | >trna111_HisGTG_1_-_147774845_147774916 | -1A | 26 | 27 | AGCCGTGATCGTATAGTGGTTAGTACT | 640.9 | 5.7 | 0.9 | 6.4 | 1.3E-10 | 3.4E-10 | MB231/MB468 | NUC |
| >hg38_21_+_8433172_8446622_RNA45SN1_45S_50ntFlanks@10407.10429.23\|GGGTCAGTCGGTCCTGAGAGATG | >hg38_21_+_8433172_8446622_RNA45SN1_45S_50ntFlanks | 10407 | 10429 | 23 | GGGTCAGTCGGTCCTGAGAGATG | 968.6 | -9.5 | 1.5 | -6.4 | 1.6E-10 | 3.4E-10 | BT20/MB468 | NUC |
| >hg38_21_+_8433172_8446622_RNA45SN1_45S_50ntFlanks@11895.11916.22\|CTCTAGTCTGGCACGGTGAAGA | >hg38_21_+_8433172_8446622_RNA45SN1_45S_50ntFlanks | 11895 | 11916 | 22 | CTCTAGTCTGGCACGGTGAAGA | 289.1 | -8.4 | 1.3 | -6.4 | 1.6E-10 | 3.4E-10 | BT20/MB468 | NUC |
| >hg38_21_+_8433172_8446622_RNA45SN1_45S_50ntFlanks@12347.12365.19\|AGAAAAGTTACCACAGGGA | >hg38_21_+_8433172_8446622_RNA45SN1_45S_50ntFlanks | 12347 | 12365 | 19 | AGAAAAGTTACCACAGGGA | 362.8 | -11.6 | 1.7 | -6.8 | 1.0E-11 | 3.4E-10 | BT20/MB468 | MP |
| >hg38_21_+_8433172_8446622_RNA45SN1_45S_50ntFlanks@8182.8212.31\|ATCGAGGCCCAGCCCGTGGACGGTGTGAGGC | >hg38_21_+_8433172_8446622_RNA45SN1_45S_50ntFlanks | 8182 | 8212 | 31 | ATCGAGGCCCAGCCCGTGGACGGTGTGAGGC | 378.1 | -11.6 | 1.7 | -6.8 | 1.0E-11 | 3.4E-10 | BT20/MB468 | MP |
| >hg38_21_+_8433172_8446622_RNA45SN1_45S_50ntFlanks@4583.4608.26\|CGGTTCTATTTTGTTGGTTTTCGGAA | >hg38_21_+_8433172_8446622_RNA45SN1_45S_50ntFlanks | 4583 | 4608 | 26 | CGGTTCTATTTTGTTGGTTTTCGGAA | 91.1 | -10.4 | 1.6 | -6.4 | 1.3E-10 | 3.4E-10 | MB231/MB468 | MP |
| >hg38_21_+_8433172_8446622_RNA45SN1_45S_50ntFlanks@8104.8140.37\|CCGCGGCGGGGCGCGGGACATGTGGCGTACGGAAGAC | >hg38_21_+_8433172_8446622_RNA45SN1_45S_50ntFlanks | 8104 | 8140 | 37 | CCGCGGCGGGGCGCGGGACATGTGGCGTACGGAAGAC | 11996.9 | -9.5 | 1.5 | -6.4 | 1.3E-10 | 3.4E-10 | MB231/MB468 | MP |
| >hg38_21_+_8433172_8446622_RNA45SN1_45S_50ntFlanks@10350.10388.39\|ACGAGAACTTTGAAGGCCGAAGTGGAGAAGGGTTCCATG | >hg38_21_+_8433172_8446622_RNA45SN1_45S_50ntFlanks | 10350 | 10388 | 39 | ACGAGAACTTTGAAGGCCGAAGTGGAGAAGGGTTCCATG | 3068.3 | -5.1 | 0.8 | -6.4 | 1.3E-10 | 3.4E-10 | MB231/MB468 | CYTO |
| >trna111_HisGTG_1_-_147774845_147774916@-1G.36.37,trna118_HisGTG_1_-_145396881_145396952@-1G.36.37,trna16_HisGTG_1_+_146544773_146544844@-1G.36.37,trna1_HisGTG_15_+_45493349_45493420@-1G.36.37,trna21_HisGTG_1_+_147753471_147753542@-1G.36.37,trna33_HisGTG_6_+_27125906_27125977@-1G.36.37,trna7_HisGTG_9_-_14433938_14434009@-1G.36.37,trna8_HisGTG_15_-_45492611_45492682@-1G.36.37,trna9_HisGTG_15_-_45490804_45490875@-1G.36.37\|GGCCGTGATCGTATAGTGGTTAGTACTCTGCGTTGTG | >trna111_HisGTG_1_-_147774845_147774916 | -1G | 36 | 37 | GGCCGTGATCGTATAGTGGTTAGTACTCTGCGTTGTG | 494.0 | 9.2 | 1.4 | 6.4 | 1.5E-10 | 3.4E-10 | MB231/MB468 | MITO |
| >hg38_21_+_8433172_8446622_RNA45SN1_45S_50ntFlanks@9624.9658.35\|AGTAGCTGGTTCCCTCCGAAGTTTCCCTCAGGATA | >hg38_21_+_8433172_8446622_RNA45SN1_45S_50ntFlanks | 9624 | 9658 | 35 | AGTAGCTGGTTCCCTCCGAAGTTTCCCTCAGGATA | 474.5 | -6.3 | 1.0 | -6.4 | 1.5E-10 | 3.4E-10 | MB231/MB468 | MITO |
| >hg38_21_+_8433172_8446622_RNA45SN1_45S_50ntFlanks@12288.12318.31\|ACCGTGAAAGCGGGGCCTCACGATCCTTCTG | >hg38_21_+_8433172_8446622_RNA45SN1_45S_50ntFlanks | 12288 | 12318 | 31 | ACCGTGAAAGCGGGGCCTCACGATCCTTCTG | 84.6 | -10.3 | 1.6 | -6.4 | 1.3E-10 | 3.4E-10 | MB231/MB468 | MP |
| >hg38_21_+_8433172_8446622_RNA45SN1_45S_50ntFlanks@12992.13015.24\|TCCCTCGCTGCGATCTATTGAAAG | >hg38_21_+_8433172_8446622_RNA45SN1_45S_50ntFlanks | 12992 | 13015 | 24 | TCCCTCGCTGCGATCTATTGAAAG | 844.2 | -6.8 | 1.1 | -6.4 | 1.5E-10 | 3.4E-10 | MB231/MB468 | MITO |
| >am_trna128_GlyGCC_6_-_27870686_27870756@1.35.35,trna18_GlyGCC_16_+_70822597_70822667@1.35.35,trna19_GlyGCC_16_+_70823410_70823480@1.35.35,trna19_GlyGCC_2_-_157257659_157257729@1.35.35,trna24_GlyGCC_16_-_70812942_70813012@1.35.35,trna25_GlyGCC_16_-_70812114_70812184@1.35.35,trna5_GlyGCC_17_+_8029064_8029134@1.35.35,trna68_GlyGCC_1_-_161493637_161493707@1.35.35\|GCATTGGTGGTTCAGTGGTAGAATTCTCGCCTGCC | >am_trna128_GlyGCC_6_-_27870686_27870756 | 1 | 35 | 35 | GCATTGGTGGTTCAGTGGTAGAATTCTCGCCTGCC | 4906.4 | 6.3 | 1.0 | 6.4 | 1.5E-10 | 3.4E-10 | MB231/MB468 | MITO |
| >hg38_21_+_8433172_8446622_RNA45SN1_45S_50ntFlanks@10458.10496.39\|ATGGCCTCCGTTGCCCTCGGCCGATCGAAAGGGAGTCGG | >hg38_21_+_8433172_8446622_RNA45SN1_45S_50ntFlanks | 10458 | 10496 | 39 | ATGGCCTCCGTTGCCCTCGGCCGATCGAAAGGGAGTCGG | 88.0 | -10.3 | 1.6 | -6.4 | 1.3E-10 | 3.4E-10 | MB231/MB468 | MP |
| >hg38_21_+_8433172_8446622_RNA45SN1_45S_50ntFlanks@5227.5244.18\|CGCGCGCTACACTGACTG | >hg38_21_+_8433172_8446622_RNA45SN1_45S_50ntFlanks | 5227 | 5244 | 18 | CGCGCGCTACACTGACTG | 1548.1 | -12.7 | 2.0 | -6.5 | 9.8E-11 | 3.4E-10 | BT20/MB468 | MITO |
| >hg38_21_+_8433172_8446622_RNA45SN1_45S_50ntFlanks@476.503.28\|TTCGTGATCGATGTGGTGACGTCGTGCT | >hg38_21_+_8433172_8446622_RNA45SN1_45S_50ntFlanks | 476 | 503 | 28 | TTCGTGATCGATGTGGTGACGTCGTGCT | 6545.0 | 4.4 | 0.7 | 6.4 | 1.3E-10 | 3.4E-10 | MB231/MB468 | NUC |
| >hg38_21_+_8433172_8446622_RNA45SN1_45S_50ntFlanks@11755.11780.26\|AATGCCTCGTCATCTAATTAGTGACG | >hg38_21_+_8433172_8446622_RNA45SN1_45S_50ntFlanks | 11755 | 11780 | 26 | AATGCCTCGTCATCTAATTAGTGACG | 87.3 | -10.3 | 1.6 | -6.4 | 1.3E-10 | 3.4E-10 | MB231/MB468 | MP |
| >MI0000746\|hsa-mir-99b&WithFlank&19\|+\|51692606\|51692687@13.32.20&[MIMAT0000689&hsa-miR-99b-5p&offsets\|0\|-2;m-10&19\|+\|51692618\|51692639&offsets\|0\|-2]\|CACCCGTAGAACCGACCTTG | >MI0000746\|hsa-mir-99b&WithFlank&19\|+\|51692606\|51692687 | 13 | 32 | 20 | CACCCGTAGAACCGACCTTG | 2622.2 | 6.7 | 1.0 | 6.8 | 1.0E-11 | 3.4E-10 | BT20/MB468 | MP |

<table>
<tr><td>>am_MI0003581|hsa-mir-574&WithFlank&4|+|38868026|38868133@31.53.23&[MIMAT0004795&hsa-miR-574-5p&offsets|0|0;m-213&4|+|38868056|38868078&offsets|0|0]|TGAGTGTGTGTGTGTGAGTGTGT</td><td>>am_MI0003581|hsa-mir-574&WithFlank&4|+|38868026|38868133</td><td>31</td><td>53</td><td>23</td><td>TGAGTGTGTGTGTGTGAGTGTGT</td><td>788.8</td><td>8.4</td><td>1.3</td><td>6.4</td><td>1.3E-10</td><td>3.4E-10</td><td>MB231/MB468</td><td>CYTO</td></tr>
<tr><td>>hg38_21_+_8433172_8446622_RNA45SN1_45S_50ntFlanks@13004.13023.20|ATCTATTGAAAGTCAGCCCT</td><td>>hg38_21_+_8433172_8446622_RNA45SN1_45S_50ntFlanks</td><td>13004</td><td>13023</td><td>20</td><td>ATCTATTGAAAGTCAGCCCT</td><td>87.2</td><td>-10.3</td><td>1.6</td><td>-6.4</td><td>1.3E-10</td><td>3.4E-10</td><td>MB231/MB468</td><td>MP</td></tr>
<tr><td>>hg38_21_+_8433172_8446622_RNA45SN1_45S_50ntFlanks@9504.9521.18|GTGAACTATGCCTGGGCA</td><td>>hg38_21_+_8433172_8446622_RNA45SN1_45S_50ntFlanks</td><td>9504</td><td>9521</td><td>18</td><td>GTGAACTATGCCTGGGCA</td><td>1522.2</td><td>-8.9</td><td>1.4</td><td>-6.4</td><td>1.3E-10</td><td>3.4E-10</td><td>MB231/MB468</td><td>MP</td></tr>
<tr><td>>hg38_21_+_8433172_8446622_RNA45SN1_45S_50ntFlanks@12324.12354.31|TTGGGTTTTAAGCAGGAGGTGTCAGAAAAGT</td><td>>hg38_21_+_8433172_8446622_RNA45SN1_45S_50ntFlanks</td><td>12324</td><td>12354</td><td>31</td><td>TTGGGTTTTAAGCAGGAGGTGTCAGAAAAGT</td><td>87.1</td><td>-10.3</td><td>1.6</td><td>-6.4</td><td>1.3E-10</td><td>3.4E-10</td><td>MB231/MB468</td><td>MP</td></tr>
<tr><td>>hg38_21_+_8433172_8446622_RNA45SN1_45S_50ntFlanks@11857.11886.30|GCGGAATCAGCGGGGAAAGAAGACCCTGTT</td><td>>hg38_21_+_8433172_8446622_RNA45SN1_45S_50ntFlanks</td><td>11857</td><td>11886</td><td>30</td><td>GCGGAATCAGCGGGGAAAGAAGACCCTGTT</td><td>84.7</td><td>-10.3</td><td>1.6</td><td>-6.4</td><td>1.3E-10</td><td>3.4E-10</td><td>MB231/MB468</td><td>MP</td></tr>
<tr><td>>hg38_21_+_8433172_8446622_RNA45SN1_45S_50ntFlanks@2959.2979.21|CGACGTGTGGCGTGGGTCGAC</td><td>>hg38_21_+_8433172_8446622_RNA45SN1_45S_50ntFlanks</td><td>2959</td><td>2979</td><td>21</td><td>CGACGTGTGGCGTGGGTCGAC</td><td>105.6</td><td>-10.6</td><td>1.6</td><td>-6.4</td><td>1.3E-10</td><td>3.4E-10</td><td>MB231/MB468</td><td>MP</td></tr>
<tr><td>>hg38_21_+_8433172_8446622_RNA45SN1_45S_50ntFlanks@9573.9601.29|AAATCGGTCGTCCGACCTGGGTATAGGGG</td><td>>hg38_21_+_8433172_8446622_RNA45SN1_45S_50ntFlanks</td><td>9573</td><td>9601</td><td>29</td><td>AAATCGGTCGTCCGACCTGGGTATAGGGG</td><td>455.9</td><td>-4.2</td><td>0.7</td><td>-6.4</td><td>1.3E-10</td><td>3.4E-10</td><td>MB231/MB468</td><td>MP</td></tr>
<tr><td>>am_MI0006405|hsa-mir-1268a&WithFlank&15|-|22225272|22225335@11.28.18;MI0016748|hsa-mir-1268b&WithFlank&17|+|80098822|80098883@10.27.18&[MIMAT0005922&hsa-miR-1268a&offsets|0|0;m-386&15|-|22225308|22225325&offsets|0|0];[MIMAT0018925&hsa-miR-1268b&offsets|0|-2;m-350&17|+|80098831|80098848&offsets|0|0]|CGGGCGTGGTGGTGGGGG</td><td>>am_MI0006405|hsa-mir-1268a&WithFlank&15|-|22225272|22225335</td><td>11</td><td>28</td><td>18</td><td>CGGGCGTGGTGGTGGGGG</td><td>310.0</td><td>9.4</td><td>1.5</td><td>6.4</td><td>1.2E-10</td><td>3.4E-10</td><td>BT20/MB468</td><td>CYTO</td></tr>
<tr><td>>hg38_21_+_8433172_8446622_RNA45SN1_45S_50ntFlanks@13148.13165.18|CGGGCGTGGTGGTGGGGG</td><td>>hg38_21_+_8433172_8446622_RNA45SN1_45S_50ntFlanks</td><td>13148</td><td>13165</td><td>18</td><td>CGGGCGTGGTGGTGGGGG</td><td>310.0</td><td>9.4</td><td>1.5</td><td>6.4</td><td>1.2E-10</td><td>3.4E-10</td><td>BT20/MB468</td><td>CYTO</td></tr>
<tr><td>>hg38_21_+_8433172_8446622_RNA45SN1_45S_50ntFlanks@9566.9597.32|GACGTGCAAATCGGTCGTCCGACCTGGGTATA</td><td>>hg38_21_+_8433172_8446622_RNA45SN1_45S_50ntFlanks</td><td>9566</td><td>9597</td><td>32</td><td>GACGTGCAAATCGGTCGTCCGACCTGGGTATA</td><td>690.1</td><td>-6.8</td><td>1.1</td><td>-6.4</td><td>1.3E-10</td><td>3.4E-10</td><td>MB231/MB468</td><td>MP</td></tr>
<tr><td>>hg38_21_+_8433172_8446622_RNA45SN1_45S_50ntFlanks@9737.9754.18|CGAAACGATCTCAACCTA</td><td>>hg38_21_+_8433172_8446622_RNA45SN1_45S_50ntFlanks</td><td>9737</td><td>9754</td><td>18</td><td>CGAAACGATCTCAACCTA</td><td>159.9</td><td>-4.7</td><td>0.7</td><td>-6.4</td><td>1.3E-10</td><td>3.4E-10</td><td>MB231/MB468</td><td>CYTO</td></tr>
<tr><td>>trna116_GluCTC_1_-_145399233_145399304@32.73.42,trna71_GluCTC_1_-_161439189_161439260@32.73.42,trna74_GluCTC_1_-_161431809_161431880@32.73.42,trna77_GluCTC_1_-_161424398_161424469@32.73.42,trna77_GluCTC_6_+_28949976_28950047@32.73.42,trna80_GluCTC_1_-_161417018_161417089@32.73.42,trna87_GluCTC_6_-_126101393_126101464@32.73.42|CTCTCACCGCCGCGGCCCGGGTTCGATTCCCGGTCAGGGAAC</td><td>>trna116_GluCTC_1_-_145399233_145399304</td><td>32</td><td>73</td><td>42</td><td>CTCTCACCGCCGCGGCCCGGGTTCGATTCCCGGTCAGGGAAC</td><td>1613.4</td><td>-8.2</td><td>1.3</td><td>-6.4</td><td>1.3E-10</td><td>3.4E-10</td><td>MB231/MB468</td><td>CYTO</td></tr>
<tr><td>>MI0006444|hsa-mir-1307&WithFlank&10|-|103394247|103394407@87.107.21(+1A)&[MIMAT0005951&hsa-miR-1307-3p&offsets|+1|0(+1A);m-66&10|-|103394300|103394322&offsets|+1|-1(+1A)]|CTCGGCGTGGCGTCGGTCGTGA</td><td>>MI0006444|hsa-mir-1307&WithFlank&10|-|103394247|103394407</td><td>87</td><td>107</td><td>21(+1A)</td><td>CTCGGCGTGGCGTCGGTCGTGA</td><td>241.1</td><td>7.2</td><td>1.1</td><td>6.4</td><td>1.3E-10</td><td>3.4E-10</td><td>MB231/MB468</td><td>CYTO</td></tr>
<tr><td>>hg38_21_+_8433172_8446622_RNA45SN1_45S_50ntFlanks@12710.12745.36|CGCGGAGCCTCGGTTGGCCTCGGATAGCCGGTCCCC</td><td>>hg38_21_+_8433172_8446622_RNA45SN1_45S_50ntFlanks</td><td>12710</td><td>12745</td><td>36</td><td>CGCGGAGCCTCGGTTGGCCTCGGATAGCCGGTCCCC</td><td>175.1</td><td>-9.5</td><td>1.5</td><td>-6.5</td><td>9.9E-11</td><td>3.4E-10</td><td>BT20/MB468</td><td>MITO</td></tr>
<tr><td>>hg38_21_+_8433172_8446622_RNA45SN1_45S_50ntFlanks@4069.4089.21|CTTTCGATGGTAGTCGCCGTG</td><td>>hg38_21_+_8433172_8446622_RNA45SN1_45S_50ntFlanks</td><td>4069</td><td>4089</td><td>21</td><td>CTTTCGATGGTAGTCGCCGTG</td><td>233.9</td><td>-7.2</td><td>1.1</td><td>-6.4</td><td>1.2E-10</td><td>3.4E-10</td><td>BT20/MB468</td><td>CYTO</td></tr>
<tr><td>>hg38_21_+_8433172_8446622_RNA45SN1_45S_50ntFlanks@9471.9488.18|AGCACGAGCGCACGTGTT</td><td>>hg38_21_+_8433172_8446622_RNA45SN1_45S_50ntFlanks</td><td>9471</td><td>9488</td><td>18</td><td>AGCACGAGCGCACGTGTT</td><td>176.8</td><td>-11.3</td><td>1.8</td><td>-6.4</td><td>1.3E-10</td><td>3.4E-10</td><td>MB231/MB468</td><td>MP</td></tr>
<tr><td>>hg38_21_+_8433172_8446622_RNA45SN1_45S_50ntFlanks@12625.12652.28|CAATGGGGCGAAGCTACCATCTGTGGGA</td><td>>hg38_21_+_8433172_8446622_RNA45SN1_45S_50ntFlanks</td><td>12625</td><td>12652</td><td>28</td><td>CAATGGGGCGAAGCTACCATCTGTGGGA</td><td>300.1</td><td>-11.3</td><td>1.7</td><td>-6.8</td><td>1.0E-11</td><td>3.4E-10</td><td>BT20/MB468</td><td>MP</td></tr>
<tr><td>>hg38_21_+_8433172_8446622_RNA45SN1_45S_50ntFlanks@12658.12683.26|ACTGAACGCCTCTAAGTCAGAATCCC</td><td>>hg38_21_+_8433172_8446622_RNA45SN1_45S_50ntFlanks</td><td>12658</td><td>12683</td><td>26</td><td>ACTGAACGCCTCTAAGTCAGAATCCC</td><td>87.6</td><td>-10.3</td><td>1.6</td><td>-6.4</td><td>1.3E-10</td><td>3.4E-10</td><td>MB231/MB468</td><td>MP</td></tr>
<tr><td>>hg38_21_+_8433172_8446622_RNA45SN1_45S_50ntFlanks@11576.11593.18|CTTAGAACTGGTGCGGAC</td><td>>hg38_21_+_8433172_8446622_RNA45SN1_45S_50ntFlanks</td><td>11576</td><td>11593</td><td>18</td><td>CTTAGAACTGGTGCGGAC</td><td>795.3</td><td>-8.5</td><td>1.3</td><td>-6.4</td><td>1.3E-10</td><td>3.4E-10</td><td>MB231/MB468</td><td>MP</td></tr>
<tr><td>>hg38_21_+_8433172_8446622_RNA45SN1_45S_50ntFlanks@7973.8005.33|GACGCGACCTCAGATCAGACGTGGCGACCCGCT</td><td>>hg38_21_+_8433172_8446622_RNA45SN1_45S_50ntFlanks</td><td>7973</td><td>8005</td><td>33</td><td>GACGCGACCTCAGATCAGACGTGGCGACCCGCT</td><td>212.8</td><td>3.4</td><td>0.5</td><td>6.4</td><td>1.4E-10</td><td>3.4E-10</td><td>MB231/MB468</td><td>CYTO</td></tr>
<tr><td>>hg38_21_+_8433172_8446622_RNA45SN1_45S_50ntFlanks@6719.6743.25|TGAATTGCAGGACACATTGATCATC</td><td>>hg38_21_+_8433172_8446622_RNA45SN1_45S_50ntFlanks</td><td>6719</td><td>6743</td><td>25</td><td>TGAATTGCAGGACACATTGATCATC</td><td>3646.1</td><td>-6.1</td><td>0.9</td><td>-6.8</td><td>1.0E-11</td><td>3.4E-10</td><td>BT20/MB468</td><td>MP</td></tr>
<tr><td>>hg38_21_+_8433172_8446622_RNA45SN1_45S_50ntFlanks@9166.9187.22|CCGGGCGGGTCGCGCCGTCGGG</td><td>>hg38_21_+_8433172_8446622_RNA45SN1_45S_50ntFlanks</td><td>9166</td><td>9187</td><td>22</td><td>CCGGGCGGGTCGCGCCGTCGGG</td><td>531.2</td><td>-12.1</td><td>1.8</td><td>-6.8</td><td>1.0E-11</td><td>3.4E-10</td><td>BT20/MB468</td><td>MP</td></tr>
<tr><td>>hg38_21_+_8433172_8446622_RNA45SN1_45S_50ntFlanks@12418.12449.32|CGATGTCGGCTCTTCCTATCATTGTGAAGCAG</td><td>>hg38_21_+_8433172_8446622_RNA45SN1_45S_50ntFlanks</td><td>12418</td><td>12449</td><td>32</td><td>CGATGTCGGCTCTTCCTATCATTGTGAAGCAG</td><td>990.5</td><td>-5.7</td><td>0.9</td><td>-6.4</td><td>1.6E-10</td><td>3.5E-10</td><td>BT20/MB468</td><td>NUC</td></tr>
<tr><td>>am_trna10_ValCAC_5_-_180649395_180649467@1.29.29,trna12_ValAAC_5_-_180645270_180645342@1.29.29,trna132_ValAAC_6_-_27721179_27721251@1.29.29,trna136_ValAAC_6_-_27648885_27648957@1.29.29,trna139_ValAAC_6_-_27618707_27618779@1.29.29,trna18_ValCAC_5_-_180529253_180529325@1.29.29,trna2_ValAAC_3_+_169490018_169490090@1.29.29,trna2_ValCAC_5_+_180524070_180524142@1.29.29,trna4_ValAAC_5_+_180591154_180591226@1.29.29,trna5_ValAAC_5_+_180596610_180596682@1.29.29,trna6_ValCAC_5_+_180600650_180600722@1.29.29,trna85_ValCAC_1_-_161369490_161369562@1.29.29,trna90_ValCAC_1_-_149684088_149684161@1.29.29,trna98_ValCAC_1_-_149298555_149298627@1.29.29,trna9_ValCAC_6_+_26538282_26538354@1.29.29|GTTTCCGTAGTGTAGTGGTTATCACGTTC</td><td>>am_trna10_ValCAC_5_-_180649395_180649467</td><td>1</td><td>29</td><td>29</td><td>GTTTCCGTAGTGTAGTGGTTATCACGTTC</td><td>23599.6</td><td>-6.0</td><td>0.9</td><td>-6.4</td><td>1.4E-10</td><td>3.5E-10</td><td>MB231/MB468</td><td>CYTO</td></tr>
<tr><td>>MI0000272|hsa-mir-182&WithFlank&7|-|129770377|129770498@29.50.22&[MIMAT0000259&hsa-miR-182-5p&offsets|0|-2;m-22&7|-|129770449|129770470&offsets|0|0]|TTTGGCAATGGTAGAACTCACA</td><td>>MI0000272|hsa-mir-182&WithFlank&7|-|129770377|129770498</td><td>29</td><td>50</td><td>22</td><td>TTTGGCAATGGTAGAACTCACA</td><td>3247.2</td><td>3.4</td><td>0.5</td><td>6.8</td><td>1.4E-11</td><td>3.5E-10</td><td>BT20/MB231</td><td>NUC</td></tr>
<tr><td>>hg38_21_+_8433172_8446622_RNA45SN1_45S_50ntFlanks@5490.5525.36|TGAGAAGACGGTCGAACTTGACTATCTAGAGGAAGT</td><td>>hg38_21_+_8433172_8446622_RNA45SN1_45S_50ntFlanks</td><td>5490</td><td>5525</td><td>36</td><td>TGAGAAGACGGTCGAACTTGACTATCTAGAGGAAGT</td><td>778.9</td><td>13.4</td><td>2.0</td><td>6.8</td><td>1.0E-11</td><td>3.5E-10</td><td>BT20/MB468</td><td>MP</td></tr>
<tr><td>>MI0000456|hsa-mir-140&WithFlank&16|+|69933075|69933186@68.90.23(+1U)&[MIMAT0004597&hsa-miR-140-3p&offsets|0|+2(+1U);m-45&16|+|69933143|69933164&offsets|-1|0(+1U)]|TACCACAGGGTAGAACCACGGACT</td><td>>MI0000456|hsa-mir-140&WithFlank&16|+|69933075|69933186</td><td>68</td><td>90</td><td>23(+1U)</td><td>TACCACAGGGTAGAACCACGGACT</td><td>205.1</td><td>10.9</td><td>1.7</td><td>6.4</td><td>1.5E-10</td><td>3.5E-10</td><td>MB231/MB468</td><td>MITO</td></tr>
<tr><td>>hg38_21_+_8433172_8446622_RNA45SN1_45S_50ntFlanks@10798.10818.21|CATGTTGGAACAATGTAGGTA</td><td>>hg38_21_+_8433172_8446622_RNA45SN1_45S_50ntFlanks</td><td>10798</td><td>10818</td><td>21</td><td>CATGTTGGAACAATGTAGGTA</td><td>830.5</td><td>-4.4</td><td>0.7</td><td>-6.5</td><td>1.0E-10</td><td>3.5E-10</td><td>BT20/MB468</td><td>MITO</td></tr>
<tr><td>>hg38_21_+_8433172_8446622_RNA45SN1_45S_50ntFlanks@11930.11948.19|TAGAATAAGTGGGAGGCCC</td><td>>hg38_21_+_8433172_8446622_RNA45SN1_45S_50ntFlanks</td><td>11930</td><td>11948</td><td>19</td><td>TAGAATAAGTGGGAGGCCC</td><td>107.6</td><td>-8.8</td><td>1.4</td><td>-6.5</td><td>1.0E-10</td><td>3.5E-10</td><td>BT20/MB468</td><td>MITO</td></tr>
<tr><td>>hg38_21_+_8433172_8446622_RNA45SN1_45S_50ntFlanks@9505.9527.23|TGAACTATGCCTGGGCAGGGCGA</td><td>>hg38_21_+_8433172_8446622_RNA45SN1_45S_50ntFlanks</td><td>9505</td><td>9527</td><td>23</td><td>TGAACTATGCCTGGGCAGGGCGA</td><td>138.9</td><td>-11.0</td><td>1.7</td><td>-6.4</td><td>1.3E-10</td><td>3.5E-10</td><td>MB231/MB468</td><td>MP</td></tr>
<tr><td>>hg38_21_+_8433172_8446622_RNA45SN1_45S_50ntFlanks@11210.11254.45|GGCGACTCTGGACGCGAGCCGGGCCCTTCCCGTGGATCGCCCCAG</td><td>>hg38_21_+_8433172_8446622_RNA45SN1_45S_50ntFlanks</td><td>11210</td><td>11254</td><td>45</td><td>GGCGACTCTGGACGCGAGCCGGGCCCTTCCCGTGGATCGCCCCAG</td><td>782.1</td><td>-6.8</td><td>1.1</td><td>-6.4</td><td>1.4E-10</td><td>3.5E-10</td><td>MB231/MB468</td><td>CYTO</td></tr>
<tr><td>>hg38_21_+_8433172_8446622_RNA45SN1_45S_50ntFlanks@11161.11189.29|CACGGTCCCCCGCGAGGGGGGCCCGGGCA</td><td>>hg38_21_+_8433172_8446622_RNA45SN1_45S_50ntFlanks</td><td>11161</td><td>11189</td><td>29</td><td>CACGGTCCCCCGCGAGGGGGGCCCGGGCA</td><td>271.0</td><td>-10.2</td><td>1.6</td><td>-6.5</td><td>1.0E-10</td><td>3.5E-10</td><td>BT20/MB468</td><td>MITO</td></tr>
<tr><td>>hg38_21_+_8433172_8446622_RNA45SN1_45S_50ntFlanks@10678.10698.21|CGGTTCCGGCGGCGTCCGGTG</td><td>>hg38_21_+_8433172_8446622_RNA45SN1_45S_50ntFlanks</td><td>10678</td><td>10698</td><td>21</td><td>CGGTTCCGGCGGCGTCCGGTG</td><td>116.5</td><td>-10.7</td><td>1.7</td><td>-6.4</td><td>1.3E-10</td><td>3.5E-10</td><td>MB231/MB468</td><td>MP</td></tr>
<tr><td>>am_trna128_GlyGCC_6_-_27870686_27870756@10.29.20,trna133_GlyCCC_1_-_16872434_16872504@10.29.20,trna17_GluTTC_2_-_204229131_204229201@10.29.20,trna18_GlyGCC_16_+_70822597_70822667@10.29.20,trna19_GlyGCC_16_+_70823410_70823480@10.29.20,trna19_GlyGCC_2_-_157257659_157257729@10.29.20,trna24_GlyGCC_16_-_70812942_70813012@10.29.20,trna25_GlyGCC_16_-_70812114_70812184@10.29.20,trna2_GlyGCC_21_-_18827107_18827177@10.29.20,trna35_GlyGCC_1_+_161413094_161413164@10.29.20,trna37_GlyGCC_1_+_161420467_161420537@10.29.20,trna39_GlyGCC_1_+_161427898_161427968@10.29.20,trna41_GlyGCC_1_+_161435258_161435328@10.29.20,trna4_GlyCCC_1_+_17188416_17188486@10.29.20,trna5_GlyGCC_17_+_8029064_8029134@10.29.20,trna68_GlyGCC_1_-_161493637_161493707@10.29.20,trna91_GlyCCC_1_-_149680210_149680280@10.29.20,trna99_ValCAC_1_-_149294666_149294736@10.29.20|GTTCAGTGGTAGAATTCTCG</td><td>>am_trna128_GlyGCC_6_-_27870686_27870756</td><td>10</td><td>29</td><td>20</td><td>GTTCAGTGGTAGAATTCTCG</td><td>163.7</td><td>-11.2</td><td>1.7</td><td>-6.4</td><td>1.3E-10</td><td>3.5E-10</td><td>MB231/MB468</td><td>MP</td></tr>
<tr><td>>hg38_21_+_8433172_8446622_RNA45SN1_45S_50ntFlanks@10360.10390.31|TGAAGGCCGAAGTGGAGAAGGGTTCCATGTG</td><td>>hg38_21_+_8433172_8446622_RNA45SN1_45S_50ntFlanks</td><td>10360</td><td>10390</td><td>31</td><td>TGAAGGCCGAAGTGGAGAAGGGTTCCATGTG</td><td>96.9</td><td>-10.5</td><td>1.6</td><td>-6.4</td><td>1.3E-10</td><td>3.5E-10</td><td>MB231/MB468</td><td>MP</td></tr>
<tr><td>>MI0000465|hsa-mir-191&WithFlank&3|-|49020612|49020715@22.43.22(+1U)&[MIMAT0000440&hsa-miR-191-5p&offsets|0|-1(+1U);m-48&3|-|49020672|49020694&offsets|0|-1(+1U)]|CAACGGAATCCCAAAAGCAGCTT</td><td>>MI0000465|hsa-mir-191&WithFlank&3|-|49020612|49020715</td><td>22</td><td>43</td><td>22(+1U)</td><td>CAACGGAATCCCAAAAGCAGCTT</td><td>343.6</td><td>4.6</td><td>0.7</td><td>6.4</td><td>1.4E-10</td><td>3.5E-10</td><td>MB231/MB468</td><td>NUC</td></tr>
<tr><td>>hg38_21_+_8433172_8446622_RNA45SN1_45S_50ntFlanks@3822.3852.31|CTTTGGTCGCTCGCTCCTCTCCTACTTGGAT</td><td>>hg38_21_+_8433172_8446622_RNA45SN1_45S_50ntFlanks</td><td>3822</td><td>3852</td><td>31</td><td>CTTTGGTCGCTCGCTCCTCTCCTACTTGGAT</td><td>900.2</td><td>-9.6</td><td>1.5</td><td>-6.4</td><td>1.6E-10</td><td>3.5E-10</td><td>BT20/MB468</td><td>NUC</td></tr>
</table>

<table>
<tr><td>>trna16_LeuAAG_16_+_22308461_22308542@14.32.19,trna16_LeuAAG_5_-_180601044_180601125@14.32.19,trna19_LeuAAG_5_-_180524474_180524555@14.32.19,trna1_LeuAAG_14_+_21078291_21078372@14.32.19,trna2_LeuTAG_14_+_21093529_21093610@14.32.19,trna3_LeuAAG_5_+_180528840_180528921@14.32.19,trna42_LeuTAG_17_-_8023632_8023713@14.32.19,trna78_LeuAAG_6_+_28956779_28956860@14.32.19,trna7_LeuAAG_5_+_180614701_180614782@14.32.19,trna98_LeuAAG_6_-_28911399_28911480@14.32.19|AGCGGTCTAAGGCGCTGGA</td><td>>trna16_LeuAAG_16_+_22308461_22308542</td><td>14</td><td>32</td><td>19</td><td>AGCGGTCTAAGGCGCTGGA</td><td>93.2</td><td>-10.4</td><td>1.6</td><td>-6.4</td><td>1.3E-10</td><td>3.5E-10</td><td>MB231/MB468</td><td>MP</td></tr>
<tr><td>>hg38_21_+_8433172_8446622_RNA45SN1_45S_50ntFlanks@11455.11472.18|CTCGGCGCGCGGCGGCGG</td><td>>hg38_21_+_8433172_8446622_RNA45SN1_45S_50ntFlanks</td><td>11455</td><td>11472</td><td>18</td><td>CTCGGCGCGCGGCGGCGG</td><td>92.2</td><td>-10.4</td><td>1.6</td><td>-6.4</td><td>1.3E-10</td><td>3.5E-10</td><td>MB231/MB468</td><td>MP</td></tr>
<tr><td>>hg38_21_+_8433172_8446622_RNA45SN1_45S_50ntFlanks@9821.9850.30|CCTAGTGGGCCACTTTTGGTAAGCAGAACT</td><td>>hg38_21_+_8433172_8446622_RNA45SN1_45S_50ntFlanks</td><td>9821</td><td>9850</td><td>30</td><td>CCTAGTGGGCCACTTTTGGTAAGCAGAACT</td><td>606.8</td><td>-3.7</td><td>0.6</td><td>-6.4</td><td>1.4E-10</td><td>3.5E-10</td><td>MB231/MB468</td><td>NUC</td></tr>
<tr><td>>MI0000078|hsa-mir-22&WithFlank&17|-|1713897|1713993@59.79.21(+1A)&[MIMAT0000077&hsa-miR-22-3p&offsets|0|-1(+1A);m-2&17|-|1713914|1713935&offsets|0|-1(+1A)]|AAGCTGCCAGTTGAAGAACTGA</td><td>>MI0000078|hsa-mir-22&WithFlank&17|-|1713897|1713993</td><td>59</td><td>79</td><td>21(+1A)</td><td>AAGCTGCCAGTTGAAGAACTGA</td><td>175.6</td><td>8.5</td><td>1.3</td><td>6.4</td><td>1.4E-10</td><td>3.5E-10</td><td>MB231/MB468</td><td>NUC</td></tr>
<tr><td>>hg38_21_+_8433172_8446622_RNA45SN1_45S_50ntFlanks@12942.12964.23|TTCGTAGACGACCTGCTTCTGGG</td><td>>hg38_21_+_8433172_8446622_RNA45SN1_45S_50ntFlanks</td><td>12942</td><td>12964</td><td>23</td><td>TTCGTAGACGACCTGCTTCTGGG</td><td>190.6</td><td>-6.0</td><td>0.9</td><td>-6.4</td><td>1.4E-10</td><td>3.5E-10</td><td>MB231/MB468</td><td>CYTO</td></tr>
<tr><td>>MI0000082|hsa-mir-25&WithFlank&7|-|100093554|100093649@58.77.20(+1A)&[MIMAT0000081&hsa-miR-25-3p&offsets|0|-2(+1A);m-23&7|-|100093571|100093592&offsets|0|-2(+1A)]|CATTGCACTTGTCTCGGTCTA</td><td>>MI0000082|hsa-mir-25&WithFlank&7|-|100093554|100093649</td><td>58</td><td>77</td><td>20(+1A)</td><td>CATTGCACTTGTCTCGGTCTA</td><td>189.9</td><td>2.0</td><td>0.3</td><td>6.4</td><td>1.7E-10</td><td>3.5E-10</td><td>BT20/MB468</td><td>NUC</td></tr>
<tr><td>>MI0000298|hsa-mir-221&WithFlank&X|-|45746151|45746272@71.92.22&[MIMAT0000278&hsa-miR-221-3p&offsets|0|-1;m-91&X|-|45746181|45746202&offsets|0|0]|AGCTACATTGTCTGCTGGGTTT</td><td>>MI0000298|hsa-mir-221&WithFlank&X|-|45746151|45746272</td><td>71</td><td>92</td><td>22</td><td>AGCTACATTGTCTGCTGGGTTT</td><td>38613.8</td><td>4.7</td><td>0.7</td><td>6.4</td><td>1.3E-10</td><td>3.5E-10</td><td>MB231/MB468</td><td>MP</td></tr>
<tr><td>>MI0033423|hsa-mir-10399&WithFlank&7|-|139044094|139044163@43.63.21(+1U)&[MIMAT0041630&hsa-miR-10399-3p&offsets|-1|-1(+1U);TJU_CMC_MD2.ID03061.3p-miR&offsets|+3|+2(+1U);m-2657&7|-|139044103|139044124&offsets|+3|+2(+1U)]|TCTCTCGGACAAGCTGTAGGTT</td><td>>MI0033423|hsa-mir-10399&WithFlank&7|-|139044094|139044163</td><td>43</td><td>63</td><td>21(+1U)</td><td>TCTCTCGGACAAGCTGTAGGTT</td><td>155.0</td><td>10.5</td><td>1.6</td><td>6.4</td><td>1.6E-10</td><td>3.5E-10</td><td>MB231/MB468</td><td>MITO</td></tr>
<tr><td>>hg38_21_+_8433172_8446622_RNA45SN1_45S_50ntFlanks@4726.4752.27|TAATCAAGAACGAAAGTCGGAGGTTCG</td><td>>hg38_21_+_8433172_8446622_RNA45SN1_45S_50ntFlanks</td><td>4726</td><td>4752</td><td>27</td><td>TAATCAAGAACGAAAGTCGGAGGTTCG</td><td>199.8</td><td>-9.7</td><td>1.5</td><td>-6.5</td><td>1.0E-10</td><td>3.5E-10</td><td>BT20/MB468</td><td>MITO</td></tr>
<tr><td>>MI0000077|hsa-mir-21&WithFlank&17|+|59841260|59841343@14.36.23(+1A)&[MIMAT0000076&hsa-miR-21-5p&offsets|0|+1(+1A);m-1&17|+|59841273|59841294&offsets|0|+1(+1A)]|TAGCTTATCAGACTGATGTTGACA</td><td>>MI0000077|hsa-mir-21&WithFlank&17|+|59841260|59841343</td><td>14</td><td>36</td><td>23(+1A)</td><td>TAGCTTATCAGACTGATGTTGACA</td><td>81091.3</td><td>5.5</td><td>0.9</td><td>6.4</td><td>1.6E-10</td><td>3.5E-10</td><td>MB231/MB468</td><td>MITO</td></tr>
<tr><td>>hg38_21_+_8433172_8446622_RNA45SN1_45S_50ntFlanks@3987.4025.39|GCTTTGGTGACTCTAGATAACCTCGGGCCGATCGCACGC</td><td>>hg38_21_+_8433172_8446622_RNA45SN1_45S_50ntFlanks</td><td>3987</td><td>4025</td><td>39</td><td>GCTTTGGTGACTCTAGATAACCTCGGGCCGATCGCACGC</td><td>152.7</td><td>-11.1</td><td>1.7</td><td>-6.4</td><td>1.3E-10</td><td>3.5E-10</td><td>MB231/MB468</td><td>MP</td></tr>
<tr><td>>hg38_21_+_8433172_8446622_RNA45SN1_45S_50ntFlanks@12003.12029.27|GCGGGCCGCCGGTGAAATACCACTACT</td><td>>hg38_21_+_8433172_8446622_RNA45SN1_45S_50ntFlanks</td><td>12003</td><td>12029</td><td>27</td><td>GCGGGCCGCCGGTGAAATACCACTACT</td><td>584.4</td><td>-4.9</td><td>0.8</td><td>-6.4</td><td>1.7E-10</td><td>3.6E-10</td><td>BT20/MB468</td><td>NUC</td></tr>
<tr><td>>trnaMT_HisGTG_MT_+_12138_12206@19.55.37|AAACATCAGATTGTGAATCTGACAACAGAGGCTTACG</td><td>>trnaMT_HisGTG_MT_+_12138_12206</td><td>19</td><td>55</td><td>37</td><td>AAACATCAGATTGTGAATCTGACAACAGAGGCTTACG</td><td>582.5</td><td>10.5</td><td>1.6</td><td>6.5</td><td>1.0E-10</td><td>3.6E-10</td><td>BT20/MB468</td><td>MITO</td></tr>
<tr><td>>hg38_21_+_8433172_8446622_RNA45SN1_45S_50ntFlanks@12006.12032.27|GGCCGCCGGTGAAATACCACTACTCTG</td><td>>hg38_21_+_8433172_8446622_RNA45SN1_45S_50ntFlanks</td><td>12006</td><td>12032</td><td>27</td><td>GGCCGCCGGTGAAATACCACTACTCTG</td><td>1391.9</td><td>-7.5</td><td>1.2</td><td>-6.4</td><td>1.6E-10</td><td>3.6E-10</td><td>MB231/MB468</td><td>MITO</td></tr>
<tr><td>>hg38_21_+_8433172_8446622_RNA45SN1_45S_50ntFlanks@11923.11954.32|AGAGGTGTAGAATAAGTGGGAGGCCCCCGGCG</td><td>>hg38_21_+_8433172_8446622_RNA45SN1_45S_50ntFlanks</td><td>11923</td><td>11954</td><td>32</td><td>AGAGGTGTAGAATAAGTGGGAGGCCCCCGGCG</td><td>76.1</td><td>-10.1</td><td>1.6</td><td>-6.4</td><td>1.3E-10</td><td>3.6E-10</td><td>MB231/MB468</td><td>MP</td></tr>
<tr><td>>hg38_21_+_8433172_8446622_RNA45SN1_45S_50ntFlanks@12446.12477.32|GCAGAATTCACCAAGCGTTGGATTGTTCACCC</td><td>>hg38_21_+_8433172_8446622_RNA45SN1_45S_50ntFlanks</td><td>12446</td><td>12477</td><td>32</td><td>GCAGAATTCACCAAGCGTTGGATTGTTCACCC</td><td>281.1</td><td>4.7</td><td>0.7</td><td>6.4</td><td>1.4E-10</td><td>3.6E-10</td><td>MB231/MB468</td><td>CYTO</td></tr>
<tr><td>>MI0000066|hsa-let-7e&WithFlank&19|+|51692780|51692870@14.34.21(+1C)&[MIMAT0000066&hsa-let-7e-5p&offsets|0|-1(+1C);m-50&19|+|51692793|51692814&offsets|0|-1(+1C)]|TGAGGTAGGAGGTTGTATAGTC</td><td>>MI0000066|hsa-let-7e&WithFlank&19|+|51692780|51692870</td><td>14</td><td>34</td><td>21(+1C)</td><td>TGAGGTAGGAGGTTGTATAGTC</td><td>656.1</td><td>3.5</td><td>0.5</td><td>6.4</td><td>1.4E-10</td><td>3.6E-10</td><td>MB231/MB468</td><td>NUC</td></tr>
<tr><td>>hg38_21_+_8433172_8446622_RNA45SN1_45S_50ntFlanks@12938.12972.35|ACCATTCGTAGACGACCTGCTTCTGGGTCGGGGTT</td><td>>hg38_21_+_8433172_8446622_RNA45SN1_45S_50ntFlanks</td><td>12938</td><td>12972</td><td>35</td><td>ACCATTCGTAGACGACCTGCTTCTGGGTCGGGGTT</td><td>82.6</td><td>-10.2</td><td>1.6</td><td>-6.4</td><td>1.3E-10</td><td>3.6E-10</td><td>MB231/MB468</td><td>MP</td></tr>
<tr><td>>hg38_21_+_8433172_8446622_RNA45SN1_45S_50ntFlanks@12157.12183.27|AGTTTGACTGGGGCGGTACACCTGTCA</td><td>>hg38_21_+_8433172_8446622_RNA45SN1_45S_50ntFlanks</td><td>12157</td><td>12183</td><td>27</td><td>AGTTTGACTGGGGCGGTACACCTGTCA</td><td>145.0</td><td>-9.3</td><td>1.4</td><td>-6.5</td><td>1.0E-10</td><td>3.6E-10</td><td>BT20/MB468</td><td>MITO</td></tr>
<tr><td>>hg38_21_+_8433172_8446622_RNA45SN1_45S_50ntFlanks@10351.10371.21|CGAGAACTTTGAAGGCCGAAG</td><td>>hg38_21_+_8433172_8446622_RNA45SN1_45S_50ntFlanks</td><td>10351</td><td>10371</td><td>21</td><td>CGAGAACTTTGAAGGCCGAAG</td><td>2032.8</td><td>-8.0</td><td>1.3</td><td>-6.4</td><td>1.7E-10</td><td>3.6E-10</td><td>BT20/MB468</td><td>NUC</td></tr>
<tr><td>>hg38_21_+_8433172_8446622_RNA45SN1_45S_50ntFlanks@12637.12656.20|GCTACCATCTGTGGGATTAT</td><td>>hg38_21_+_8433172_8446622_RNA45SN1_45S_50ntFlanks</td><td>12637</td><td>12656</td><td>20</td><td>GCTACCATCTGTGGGATTAT</td><td>79.0</td><td>-10.2</td><td>1.6</td><td>-6.4</td><td>1.3E-10</td><td>3.6E-10</td><td>MB231/MB468</td><td>MP</td></tr>
<tr><td>>hg38_21_+_8433172_8446622_RNA45SN1_45S_50ntFlanks@8528.8551.24|CTTCCCCCGCCGCCCCTCCTCCTC</td><td>>hg38_21_+_8433172_8446622_RNA45SN1_45S_50ntFlanks</td><td>8528</td><td>8551</td><td>24</td><td>CTTCCCCCGCCGCCCCTCCTCCTC</td><td>128.5</td><td>-10.9</td><td>1.7</td><td>-6.4</td><td>1.3E-10</td><td>3.6E-10</td><td>MB231/MB468</td><td>MP</td></tr>
<tr><td>>hg38_21_+_8433172_8446622_RNA45SN1_45S_50ntFlanks@10425.10456.32|AGATGGGCGAGCGCCGTTCCGAAGGGACGGGC</td><td>>hg38_21_+_8433172_8446622_RNA45SN1_45S_50ntFlanks</td><td>10425</td><td>10456</td><td>32</td><td>AGATGGGCGAGCGCCGTTCCGAAGGGACGGGC</td><td>812.2</td><td>-3.4</td><td>0.5</td><td>-6.4</td><td>1.3E-10</td><td>3.6E-10</td><td>BT20/MB468</td><td>CYTO</td></tr>
<tr><td>>MI0000077|hsa-mir-21&WithFlank&17|+|59841260|59841343@14.34.21(+3C)&[MIMAT0000076&hsa-miR-21-5p&offsets|0|-1(+3C);m-1&17|+|59841273|59841294&offsets|0|-1(+3C)]|TAGCTTATCAGACTGATGTTGCCC</td><td>>MI0000077|hsa-mir-21&WithFlank&17|+|59841260|59841343</td><td>14</td><td>34</td><td>21(+3C)</td><td>TAGCTTATCAGACTGATGTTGCCC</td><td>235.9</td><td>11.1</td><td>1.7</td><td>6.4</td><td>1.6E-10</td><td>3.6E-10</td><td>MB231/MB468</td><td>MITO</td></tr>
<tr><td>>hg38_21_+_8433172_8446622_RNA45SN1_45S_50ntFlanks@10787.10818.32|ACAGCCTCTGGCATGTTGGAACAATGTAGGTA</td><td>>hg38_21_+_8433172_8446622_RNA45SN1_45S_50ntFlanks</td><td>10787</td><td>10818</td><td>32</td><td>ACAGCCTCTGGCATGTTGGAACAATGTAGGTA</td><td>377.5</td><td>-3.9</td><td>0.6</td><td>-6.4</td><td>1.4E-10</td><td>3.6E-10</td><td>MB231/MB468</td><td>NUC</td></tr>
<tr><td>>hg38_21_+_8433172_8446622_RNA45SN1_45S_50ntFlanks@4121.4146.26|CAGGGTTCGATTCCGGAGAGGGAGCC</td><td>>hg38_21_+_8433172_8446622_RNA45SN1_45S_50ntFlanks</td><td>4121</td><td>4146</td><td>26</td><td>CAGGGTTCGATTCCGGAGAGGGAGCC</td><td>68.6</td><td>-10.0</td><td>1.6</td><td>-6.4</td><td>1.3E-10</td><td>3.6E-10</td><td>MB231/MB468</td><td>MP</td></tr>
<tr><td>>hg38_21_+_8433172_8446622_RNA45SN1_45S_50ntFlanks@10729.10757.29|AGAGGGTGTAAATCTCGCGCCGGGCCGTA</td><td>>hg38_21_+_8433172_8446622_RNA45SN1_45S_50ntFlanks</td><td>10729</td><td>10757</td><td>29</td><td>AGAGGGTGTAAATCTCGCGCCGGGCCGTA</td><td>229.0</td><td>-7.6</td><td>1.2</td><td>-6.4</td><td>1.4E-10</td><td>3.6E-10</td><td>MB231/MB468</td><td>NUC</td></tr>
<tr><td>>hg38_21_+_8433172_8446622_RNA45SN1_45S_50ntFlanks@12488.12525.38|AACGTGAGCTGGGTTTAGACCGTCGTGAGACAGGTTAG</td><td>>hg38_21_+_8433172_8446622_RNA45SN1_45S_50ntFlanks</td><td>12488</td><td>12525</td><td>38</td><td>AACGTGAGCTGGGTTTAGACCGTCGTGAGACAGGTTAG</td><td>544.1</td><td>-9.3</td><td>1.4</td><td>-6.4</td><td>1.4E-10</td><td>3.6E-10</td><td>MB231/MB468</td><td>NUC</td></tr>
<tr><td>>hg38_21_+_8433172_8446622_RNA45SN1_45S_50ntFlanks@4003.4022.20|ATAACCTCGGGCCGATCGCA</td><td>>hg38_21_+_8433172_8446622_RNA45SN1_45S_50ntFlanks</td><td>4003</td><td>4022</td><td>20</td><td>ATAACCTCGGGCCGATCGCA</td><td>665.5</td><td>-3.5</td><td>0.5</td><td>-6.4</td><td>1.4E-10</td><td>3.6E-10</td><td>MB231/MB468</td><td>CYTO</td></tr>
<tr><td>>MI0003137|hsa-mir-193b&WithFlank&16|+|14303961|14304055@57.78.22&[MIMAT0002819&hsa-miR-193b-3p&offsets|0|0;m-166&16|+|14304017|14304038&offsets|0|0]|AACTGGCCCTCAAAGTCCCGCT</td><td>>MI0003137|hsa-mir-193b&WithFlank&16|+|14303961|14304055</td><td>57</td><td>78</td><td>22</td><td>AACTGGCCCTCAAAGTCCCGCT</td><td>199.4</td><td>3.9</td><td>0.6</td><td>6.4</td><td>1.4E-10</td><td>3.6E-10</td><td>MB231/MB468</td><td>NUC</td></tr>
<tr><td>>hg38_21_+_8433172_8446622_RNA45SN1_45S_50ntFlanks@6663.6687.25|GTGGATCACTCGGCTCGTGCGTCGA</td><td>>hg38_21_+_8433172_8446622_RNA45SN1_45S_50ntFlanks</td><td>6663</td><td>6687</td><td>25</td><td>GTGGATCACTCGGCTCGTGCGTCGA</td><td>103.3</td><td>-10.6</td><td>1.6</td><td>-6.4</td><td>1.3E-10</td><td>3.6E-10</td><td>MB231/MB468</td><td>MP</td></tr>
<tr><td>>hg38_21_+_8433172_8446622_RNA45SN1_45S_50ntFlanks@12476.12500.25|CCACTAATAGGGAACGTGAGCTGGG</td><td>>hg38_21_+_8433172_8446622_RNA45SN1_45S_50ntFlanks</td><td>12476</td><td>12500</td><td>25</td><td>CCACTAATAGGGAACGTGAGCTGGG</td><td>757.6</td><td>-5.6</td><td>0.9</td><td>-6.4</td><td>1.4E-10</td><td>3.6E-10</td><td>MB231/MB468</td><td>CYTO</td></tr>
<tr><td>>hg38_21_+_8433172_8446622_RNA45SN1_45S_50ntFlanks@9622.9652.31|CTAGTAGCTGGTTCCCTCCGAAGTTTCCCTC</td><td>>hg38_21_+_8433172_8446622_RNA45SN1_45S_50ntFlanks</td><td>9622</td><td>9652</td><td>31</td><td>CTAGTAGCTGGTTCCCTCCGAAGTTTCCCTC</td><td>329.3</td><td>-3.7</td><td>0.6</td><td>-6.5</td><td>1.0E-10</td><td>3.6E-10</td><td>BT20/MB468</td><td>MITO</td></tr>
<tr><td>>hg38_21_+_8433172_8446622_RNA45SN1_45S_50ntFlanks@10668.10706.39|GAAAGCGTCGCGGTTCCGGCGGCGTCCGGTGAGCTCTCG</td><td>>hg38_21_+_8433172_8446622_RNA45SN1_45S_50ntFlanks</td><td>10668</td><td>10706</td><td>39</td><td>GAAAGCGTCGCGGTTCCGGCGGCGTCCGGTGAGCTCTCG</td><td>259.6</td><td>-11.9</td><td>1.9</td><td>-6.4</td><td>1.3E-10</td><td>3.6E-10</td><td>MB231/MB468</td><td>MP</td></tr>
<tr><td>>hg38_21_+_8433172_8446622_RNA45SN1_45S_50ntFlanks@4614.4633.20|GCCATGATTAAGAGGGACGG</td><td>>hg38_21_+_8433172_8446622_RNA45SN1_45S_50ntFlanks</td><td>4614</td><td>4633</td><td>20</td><td>GCCATGATTAAGAGGGACGG</td><td>128.6</td><td>-10.9</td><td>1.7</td><td>-6.4</td><td>1.4E-10</td><td>3.6E-10</td><td>MB231/MB468</td><td>MP</td></tr>
<tr><td>>hg38_21_+_8433172_8446622_RNA45SN1_45S_50ntFlanks@5495.5526.32|AGACGGTCGAACTTGACTATCTAGAGGAAGTA</td><td>>hg38_21_+_8433172_8446622_RNA45SN1_45S_50ntFlanks</td><td>5495</td><td>5526</td><td>32</td><td>AGACGGTCGAACTTGACTATCTAGAGGAAGTA</td><td>78.3</td><td>-10.2</td><td>1.6</td><td>-6.4</td><td>1.4E-10</td><td>3.6E-10</td><td>MB231/MB468</td><td>MP</td></tr>
<tr><td>>hg38_21_+_8433172_8446622_RNA45SN1_45S_50ntFlanks@11875.11893.19|GAAGACCCTGTTGAGCTTG</td><td>>hg38_21_+_8433172_8446622_RNA45SN1_45S_50ntFlanks</td><td>11875</td><td>11893</td><td>19</td><td>GAAGACCCTGTTGAGCTTG</td><td>922.4</td><td>-3.7</td><td>0.6</td><td>-6.4</td><td>1.3E-10</td><td>3.6E-10</td><td>BT20/MB468</td><td>CYTO</td></tr>
<tr><td>>hg38_21_+_8433172_8446622_RNA45SN1_45S_50ntFlanks@12415.12432.18|CTTCGATGTCGGCTCTTC</td><td>>hg38_21_+_8433172_8446622_RNA45SN1_45S_50ntFlanks</td><td>12415</td><td>12432</td><td>18</td><td>CTTCGATGTCGGCTCTTC</td><td>6156.4</td><td>-3.4</td><td>0.5</td><td>-6.4</td><td>1.3E-10</td><td>3.6E-10</td><td>BT20/MB468</td><td>CYTO</td></tr>
<tr><td>>hg38_21_+_8433172_8446622_RNA45SN1_45S_50ntFlanks@11589.11623.35|CGGACCAGGGGAATCCGACTGTTTAATTAAAACAA</td><td>>hg38_21_+_8433172_8446622_RNA45SN1_45S_50ntFlanks</td><td>11589</td><td>11623</td><td>35</td><td>CGGACCAGGGGAATCCGACTGTTTAATTAAAACAA</td><td>2450.8</td><td>9.4</td><td>1.4</td><td>6.5</td><td>1.1E-10</td><td>3.6E-10</td><td>BT20/MB468</td><td>MITO</td></tr>
<tr><td>>hg38_21_+_8433172_8446622_RNA45SN1_45S_50ntFlanks@12627.12661.35|ATGGGGCGAAGCTACCATCTGTGGGATTATGACTG</td><td>>hg38_21_+_8433172_8446622_RNA45SN1_45S_50ntFlanks</td><td>12627</td><td>12661</td><td>35</td><td>ATGGGGCGAAGCTACCATCTGTGGGATTATGACTG</td><td>3428.6</td><td>-7.2</td><td>1.1</td><td>-6.5</td><td>1.1E-10</td><td>3.6E-10</td><td>BT20/MB468</td><td>MITO</td></tr>
<tr><td>>hg38_21_+_8433172_8446622_RNA45SN1_45S_50ntFlanks@10410.10442.33|TCAGTCGGTCCTGAGAGATGGGCGAGCGCCGTT</td><td>>hg38_21_+_8433172_8446622_RNA45SN1_45S_50ntFlanks</td><td>10410</td><td>10442</td><td>33</td><td>TCAGTCGGTCCTGAGAGATGGGCGAGCGCCGTT</td><td>353.1</td><td>-6.7</td><td>1.0</td><td>-6.4</td><td>1.4E-10</td><td>3.6E-10</td><td>MB231/MB468</td><td>MP</td></tr>
<tr><td>>hg38_21_+_8433172_8446622_RNA45SN1_45S_50ntFlanks@12957.12979.23|CTTCTGGGTCGGGGTTTCGTACG</td><td>>hg38_21_+_8433172_8446622_RNA45SN1_45S_50ntFlanks</td><td>12957</td><td>12979</td><td>23</td><td>CTTCTGGGTCGGGGTTTCGTACG</td><td>603.3</td><td>-5.7</td><td>0.9</td><td>-6.4</td><td>1.3E-10</td><td>3.6E-10</td><td>BT20/MB468</td><td>CYTO</td></tr>
<tr><td>>hg38_21_+_8433172_8446622_RNA45SN1_45S_50ntFlanks@10574.10612.39|AAGCCGGCGGGAGCCCCGGGGAGAGTTCTCTTTTCTTTG</td><td>>hg38_21_+_8433172_8446622_RNA45SN1_45S_50ntFlanks</td><td>10574</td><td>10612</td><td>39</td><td>AAGCCGGCGGGAGCCCCGGGGAGAGTTCTCTTTTCTTTG</td><td>259.1</td><td>-5.8</td><td>0.9</td><td>-6.4</td><td>1.4E-10</td><td>3.6E-10</td><td>MB231/MB468</td><td>CYTO</td></tr>
<tr><td>>hg38_21_+_8433172_8446622_RNA45SN1_45S_50ntFlanks@11208.11227.20|GCGGCGACTCTGGACGCGAG</td><td>>hg38_21_+_8433172_8446622_RNA45SN1_45S_50ntFlanks</td><td>11208</td><td>11227</td><td>20</td><td>GCGGCGACTCTGGACGCGAG</td><td>3003.9</td><td>-8.4</td><td>1.3</td><td>-6.4</td><td>1.4E-10</td><td>3.6E-10</td><td>MB231/MB468</td><td>CYTO</td></tr>
<tr><td>>hg38_21_+_8433172_8446622_RNA45SN1_45S_50ntFlanks@10845.10863.19|ACTTCGGGATAAGGATTGG</td><td>>hg38_21_+_8433172_8446622_RNA45SN1_45S_50ntFlanks</td><td>10845</td><td>10863</td><td>19</td><td>ACTTCGGGATAAGGATTGG</td><td>1141.4</td><td>-6.4</td><td>1.0</td><td>-6.4</td><td>1.3E-10</td><td>3.6E-10</td><td>BT20/MB468</td><td>CYTO</td></tr>
<tr><td>>MI0000434|hsa-let-7i&WithFlank&12|+|62603680|62603775@12.33.22(+2U)&[MIMAT0000415&hsa-let-7i-5p&offsets|0|0(+2U);m-49&12|+|62603691|62603712&offsets|0|0(+2U)]|TGAGGTAGTAGTTTGTGCTGTTTT</td><td>>MI0000434|hsa-let-7i&WithFlank&12|+|62603680|62603775</td><td>12</td><td>33</td><td>22(+2U)</td><td>TGAGGTAGTAGTTTGTGCTGTTTT</td><td>235.2</td><td>6.5</td><td>1.0</td><td>6.4</td><td>1.4E-10</td><td>3.6E-10</td><td>MB231/MB468</td><td>NUC</td></tr>
</table>

| | | | | | | | | | | | | | |
|---|---|---|---|---|---|---|---|---|---|---|---|---|---|
| >hg38_21_+_8433172_8446622_RNA45SN1_45S_50ntFlanks@11768.11790.23\|CTAATTAGTGACGCGCATGAATG | >hg38_21_+_8433172_8446622_RNA45SN1_45S_50ntFlanks | 11768 | 11790 | 23 | CTAATTAGTGACGCGCATGAATG | 4075.5 | -10.5 | 1.6 | -6.4 | 1.7E-10 | 3.6E-10 | BT20/MB468 | NUC |
| >trna111_HisGTG_1_-_147774845_147774916@-1G.36.37,trna118_HisGTG_1_-_145396881_145396952@-1G.36.37,trna16_HisGTG_1_+_146544773_146544844@-1G.36.37,trna1_HisGTG_15_+_45493349_45493420@-1G.36.37,trna21_HisGTG_1_+_147753471_147753542@-1G.36.37,trna33_HisGTG_6_+_27125906_27125977@-1G.36.37,trna7_HisGTG_9_-_14433938_14434009@-1G.36.37,trna8_HisGTG_15_-_45492611_45492682@-1G.36.37,trna9_HisGTG_15_-_45490804_45490875@-1G.36.37\|GGCCGTGATCGTATAGTGGTTAGTACTCTGCGTTGTG | >trna111_HisGTG_1_-_147774845_147774916 | -1G | 36 | 37 | GGCCGTGATCGTATAGTGGTTAGTACTCTGCGTTGTG | 174.8 | 10.7 | 1.7 | 6.4 | 1.4E-10 | 3.6E-10 | MB231/MB468 | MP |
| >hg38_21_+_8433172_8446622_RNA45SN1_45S_50ntFlanks@12479.12498.20\|CTAATAGGGAACGTGAGCTG | >hg38_21_+_8433172_8446622_RNA45SN1_45S_50ntFlanks | 12479 | 12498 | 20 | CTAATAGGGAACGTGAGCTG | 657.0 | -6.9 | 1.1 | -6.4 | 1.4E-10 | 3.6E-10 | MB231/MB468 | MP |
| >trna16_ValTAC_11_-_59318460_59318532@1.34.34\|GGTTCCATAGTGTAGCGGTTATCACGTCTGCTTT | >trna16_ValTAC_11_-_59318460_59318532 | 1 | 34 | 34 | GGTTCCATAGTGTAGCGGTTATCACGTCTGCTTT | 244.5 | 11.2 | 1.7 | 6.4 | 1.4E-10 | 3.6E-10 | MB231/MB468 | MP |
| >hg38_21_+_8433172_8446622_RNA45SN1_45S_50ntFlanks@8196.8214.19\|CGTGGACGGTGTGAGGCCG | >hg38_21_+_8433172_8446622_RNA45SN1_45S_50ntFlanks | 8196 | 8214 | 19 | CGTGGACGGTGTGAGGCCG | 825.4 | -5.1 | 0.8 | -6.4 | 1.6E-10 | 3.6E-10 | MB231/MB468 | MITO |
| >MI0000465\|hsa-mir-191&WithFlank&3\|-\|49020612\|49020715@22.42.21(+1C)&[MIMAT0000440&hsa-miR-191-5p&offsets\|0\|-2(+1C);m-48&3\|-\|49020672\|49020694&offsets\|0\|-2(+1C)]\|CAACGGAATCCCAAAAGCAGCC | >MI0000465\|hsa-mir-191&WithFlank&3\|-\|49020612\|49020715 | 22 | 42 | 21(+1C) | CAACGGAATCCCAAAAGCAGCC | 904.2 | 4.4 | 0.7 | 6.4 | 1.4E-10 | 3.7E-10 | MB231/MB468 | NUC |
| >hg38_21_+_8433172_8446622_RNA45SN1_45S_50ntFlanks@5487.5510.24\|CGCTGAGAAGACGGTCGAACTTGA | >hg38_21_+_8433172_8446622_RNA45SN1_45S_50ntFlanks | 5487 | 5510 | 24 | CGCTGAGAAGACGGTCGAACTTGA | 119.4 | -9.0 | 1.4 | -6.5 | 1.1E-10 | 3.7E-10 | BT20/MB468 | MITO |
| >hg38_21_+_8433172_8446622_RNA45SN1_45S_50ntFlanks@11727.11751.25\|AGTAACTATGACTCTCTTAAGGTAG | >hg38_21_+_8433172_8446622_RNA45SN1_45S_50ntFlanks | 11727 | 11751 | 25 | AGTAACTATGACTCTCTTAAGGTAG | 2600.8 | -10.2 | 1.6 | -6.4 | 1.7E-10 | 3.7E-10 | BT20/MB468 | NUC |
| >MI0000266\|hsa-mir-10a&WithFlank&17\|-\|48579832\|48579953@29.50.22&[MIMAT0000253&hsa-miR-10a-5p&offsets\|+1\|0;m-18&17\|-\|48579905\|48579926&offsets\|+1\|+1]\|ACCCTGTAGATCCGAATTTGTG | >MI0000266\|hsa-mir-10a&WithFlank&17\|-\|48579832\|48579953 | 29 | 50 | 22 | ACCCTGTAGATCCGAATTTGTG | 2199.3 | 10.1 | 1.5 | 6.8 | 1.1E-11 | 3.7E-10 | BT20/MB468 | MP |
| >hg38_21_+_8433172_8446622_RNA45SN1_45S_50ntFlanks@10411.10435.25\|CAGTCGGTCCTGAGAGATGGGCGAG | >hg38_21_+_8433172_8446622_RNA45SN1_45S_50ntFlanks | 10411 | 10435 | 25 | CAGTCGGTCCTGAGAGATGGGCGAG | 2126.0 | -10.2 | 1.5 | -6.8 | 1.1E-11 | 3.7E-10 | BT20/MB468 | MP |
| >hg38_21_+_8433172_8446622_RNA45SN1_45S_50ntFlanks@11569.11586.18\|CAGCCGACTTAGAACTGG | >hg38_21_+_8433172_8446622_RNA45SN1_45S_50ntFlanks | 11569 | 11586 | 18 | CAGCCGACTTAGAACTGG | 1568.4 | -4.6 | 0.7 | -6.8 | 1.1E-11 | 3.7E-10 | BT20/MB468 | MP |
| >hg38_21_+_8433172_8446622_RNA45SN1_45S_50ntFlanks@4000.4024.25\|TAGATAACCTCGGGCCGATCGCACG | >hg38_21_+_8433172_8446622_RNA45SN1_45S_50ntFlanks | 4000 | 4024 | 25 | TAGATAACCTCGGGCCGATCGCACG | 699.9 | -8.2 | 1.3 | -6.4 | 1.6E-10 | 3.7E-10 | MB231/MB468 | MITO |
| >hg38_21_+_8433172_8446622_RNA45SN1_45S_50ntFlanks@10014.10036.23\|AAAATGGATGGCGCTGGAGCGTC | >hg38_21_+_8433172_8446622_RNA45SN1_45S_50ntFlanks | 10014 | 10036 | 23 | AAAATGGATGGCGCTGGAGCGTC | 245.4 | -11.8 | 1.8 | -6.4 | 1.4E-10 | 3.7E-10 | MB231/MB468 | MP |
| >hg38_21_+_8433172_8446622_RNA45SN1_45S_50ntFlanks@5490.5512.23\|TGAGAAGACGGTCGAACTTGACT | >hg38_21_+_8433172_8446622_RNA45SN1_45S_50ntFlanks | 5490 | 5512 | 23 | TGAGAAGACGGTCGAACTTGACT | 1632.7 | 9.7 | 1.5 | 6.4 | 1.3E-10 | 3.7E-10 | BT20/MB468 | CYTO |
| >hg38_21_+_8433172_8446622_RNA45SN1_45S_50ntFlanks@7974.7994.21\|ACGCGACCTCAGATCAGACGT | >hg38_21_+_8433172_8446622_RNA45SN1_45S_50ntFlanks | 7974 | 7994 | 21 | ACGCGACCTCAGATCAGACGT | 10305.6 | 5.2 | 0.8 | 6.5 | 1.1E-10 | 3.7E-10 | BT20/MB468 | MITO |
| >hg38_21_+_8433172_8446622_RNA45SN1_45S_50ntFlanks@12997.13039.43\|CGCTGCGATCTATTGAAAGTCAGCCCTCGACACAAGGGTTTGT | >hg38_21_+_8433172_8446622_RNA45SN1_45S_50ntFlanks | 12997 | 13039 | 43 | CGCTGCGATCTATTGAAAGTCAGCCCTCGACACAAGGGTTTGT | 7025.1 | 5.4 | 0.8 | 6.4 | 1.4E-10 | 3.7E-10 | MB231/MB468 | MP |
| >trnaMT_PheGAA_MT_+_577_647@35.67.33\|GAAAATGTTTAGACGGGCTCACATCACCCCATA | >trnaMT_PheGAA_MT_+_577_647 | 35 | 67 | 33 | GAAAATGTTTAGACGGGCTCACATCACCCCATA | 115.6 | 10.1 | 1.6 | 6.4 | 1.4E-10 | 3.7E-10 | MB231/MB468 | MP |
| >hg38_21_+_8433172_8446622_RNA45SN1_45S_50ntFlanks@4723.4759.37\|CATTAATCAAGAACGAAAGTCGGAGGTTCGAAGACGA | >hg38_21_+_8433172_8446622_RNA45SN1_45S_50ntFlanks | 4723 | 4759 | 37 | CATTAATCAAGAACGAAAGTCGGAGGTTCGAAGACGA | 2017.8 | -6.7 | 1.0 | -6.5 | 1.1E-10 | 3.7E-10 | BT20/MB468 | MITO |
| >hg38_21_+_8433172_8446622_RNA45SN1_45S_50ntFlanks@10357.10388.32\|CTTTGAAGGCCGAAGTGGAGAAGGGTTCCATG | >hg38_21_+_8433172_8446622_RNA45SN1_45S_50ntFlanks | 10357 | 10388 | 32 | CTTTGAAGGCCGAAGTGGAGAAGGGTTCCATG | 395.0 | -8.2 | 1.3 | -6.4 | 1.6E-10 | 3.7E-10 | MB231/MB468 | MITO |
| >hg38_21_+_8433172_8446622_RNA45SN1_45S_50ntFlanks@4499.4519.21\|AGCGTTTACTTTGAAAAAATT | >hg38_21_+_8433172_8446622_RNA45SN1_45S_50ntFlanks | 4499 | 4519 | 21 | AGCGTTTACTTTGAAAAAATT | 624.0 | 5.3 | 0.8 | 6.4 | 1.3E-10 | 3.7E-10 | BT20/MB468 | CYTO |
| >hg38_21_+_8433172_8446622_RNA45SN1_45S_50ntFlanks@8342.8379.38\|CGTAAGGGAAAGTTGAAAAGAACTTTGAAGAGAGAGTT | >hg38_21_+_8433172_8446622_RNA45SN1_45S_50ntFlanks | 8342 | 8379 | 38 | CGTAAGGGAAAGTTGAAAAGAACTTTGAAGAGAGAGTT | 1903.3 | 7.4 | 1.2 | 6.4 | 1.5E-10 | 3.7E-10 | MB231/MB468 | CYTO |
| >hg38_21_+_8433172_8446622_RNA45SN1_45S_50ntFlanks@4657.4680.24\|CTAGAGGTGAAATTCTTGGACCGG | >hg38_21_+_8433172_8446622_RNA45SN1_45S_50ntFlanks | 4657 | 4680 | 24 | CTAGAGGTGAAATTCTTGGACCGG | 246.9 | -5.5 | 0.9 | -6.4 | 1.5E-10 | 3.7E-10 | MB231/MB468 | CYTO |
| >hg38_21_+_8433172_8446622_RNA45SN1_45S_50ntFlanks@10411.10453.43\|CAGTCGGTCCTGAGAGATGGGCGAGCGCCGTTCCGAAGGGACG | >hg38_21_+_8433172_8446622_RNA45SN1_45S_50ntFlanks | 10411 | 10453 | 43 | CAGTCGGTCCTGAGAGATGGGCGAGCGCCGTTCCGAAGGGACG | 838.5 | -7.5 | 1.2 | -6.4 | 1.4E-10 | 3.7E-10 | MB231/MB468 | MP |
| >hg38_21_+_8433172_8446622_RNA45SN1_45S_50ntFlanks@11876.11908.33\|AAGACCCTGTTGAGCTTGACTCTAGTCTGGCAC | >hg38_21_+_8433172_8446622_RNA45SN1_45S_50ntFlanks | 11876 | 11908 | 33 | AAGACCCTGTTGAGCTTGACTCTAGTCTGGCAC | 105.0 | -10.6 | 1.6 | -6.4 | 1.4E-10 | 3.7E-10 | MB231/MB468 | MP |
| >hg38_21_+_8433172_8446622_RNA45SN1_45S_50ntFlanks@11894.11920.27\|ACTCTAGTCTGGCACGGTGAAGAGACA | >hg38_21_+_8433172_8446622_RNA45SN1_45S_50ntFlanks | 11894 | 11920 | 27 | ACTCTAGTCTGGCACGGTGAAGAGACA | 80.1 | -10.2 | 1.6 | -6.4 | 1.4E-10 | 3.7E-10 | MB231/MB468 | MP |
| >hg38_21_+_8433172_8446622_RNA45SN1_45S_50ntFlanks@11212.11232.21\|CGACTCTGGACGCGAGCCGGG | >hg38_21_+_8433172_8446622_RNA45SN1_45S_50ntFlanks | 11212 | 11232 | 21 | CGACTCTGGACGCGAGCCGGG | 3719.7 | -8.4 | 1.3 | -6.4 | 1.5E-10 | 3.7E-10 | MB231/MB468 | CYTO |
| >hg38_21_+_8433172_8446622_RNA45SN1_45S_50ntFlanks@11661.11680.20\|TGATTTCTGCCCAGTGCTCT | >hg38_21_+_8433172_8446622_RNA45SN1_45S_50ntFlanks | 11661 | 11680 | 20 | TGATTTCTGCCCAGTGCTCT | 109.0 | -10.6 | 1.7 | -6.4 | 1.4E-10 | 3.7E-10 | MB231/MB468 | MP |
| >MI0000087\|hsa-mir-29a&WithFlank&7\|-\|130876741\|130876816@47.68.22(+1U)&[MIMAT0000086&hsa-miR-29a-3p&offsets\|-1\|-1(+1U);m-26&7\|-\|130876749\|130876769&offsets\|-1\|0(+1U)]\|CTAGCACCATCTGAAATCGGTTT | >MI0000087\|hsa-mir-29a&WithFlank&7\|-\|130876741\|130876816 | 47 | 68 | 22(+1U) | CTAGCACCATCTGAAATCGGTTT | 1550.8 | 10.1 | 1.6 | 6.4 | 1.6E-10 | 3.7E-10 | MB231/MB468 | MITO |
| >hg38_21_+_8433172_8446622_RNA45SN1_45S_50ntFlanks@6663.6699.37\|GTGGATCACTCGGCTCGTGCGTCGATGAAGAACGCAG | >hg38_21_+_8433172_8446622_RNA45SN1_45S_50ntFlanks | 6663 | 6699 | 37 | GTGGATCACTCGGCTCGTGCGTCGATGAAGAACGCAG | 921.5 | -6.7 | 1.1 | -6.4 | 1.6E-10 | 3.7E-10 | MB231/MB468 | MITO |
| >hg38_21_+_8433172_8446622_RNA45SN1_45S_50ntFlanks@10291.10315.25\|CGCGGGCCCGGGTGGAGCCGCCGCA | >hg38_21_+_8433172_8446622_RNA45SN1_45S_50ntFlanks | 10291 | 10315 | 25 | CGCGGGCCCGGGTGGAGCCGCCGCA | 252.5 | -11.8 | 1.8 | -6.4 | 1.4E-10 | 3.7E-10 | MB231/MB468 | MP |
| >MI0005561\|hsa-mir-877&WithFlank&6\|+\|30584326\|30584423@7.29.23&[MIMAT0004949&hsa-miR-877-5p&offsets\|0\|+3;m-382&6\|+\|30584332\|30584354&offsets\|0\|0]\|GTAGAGGAGATGGCGCAGGGGAC | >MI0005561\|hsa-mir-877&WithFlank&6\|+\|30584326\|30584423 | 7 | 29 | 23 | GTAGAGGAGATGGCGCAGGGGAC | 517.0 | 8.5 | 1.3 | 6.4 | 1.3E-10 | 3.7E-10 | BT20/MB468 | CYTO |
| >hg38_21_+_8433172_8446622_RNA45SN1_45S_50ntFlanks@6743.6770.28\|CGACACTTCGAACGCACTTGCGGCCCCG | >hg38_21_+_8433172_8446622_RNA45SN1_45S_50ntFlanks | 6743 | 6770 | 28 | CGACACTTCGAACGCACTTGCGGCCCCG | 2226.7 | -7.3 | 1.1 | -6.4 | 1.3E-10 | 3.7E-10 | BT20/MB468 | CYTO |
| >hg38_21_+_8433172_8446622_RNA45SN1_45S_50ntFlanks@9027.9059.33\|CGTCTCCTCGTGGGGGGGCCGGGCCACCCCTCC | >hg38_21_+_8433172_8446622_RNA45SN1_45S_50ntFlanks | 9027 | 9059 | 33 | CGTCTCCTCGTGGGGGGGCCGGGCCACCCCTCC | 243.9 | -11.0 | 1.6 | -6.8 | 1.1E-11 | 3.7E-10 | BT20/MB468 | MP |
| >hg38_21_+_8433172_8446622_RNA45SN1_45S_50ntFlanks@8357.8377.21\|AAAAGAACTTTGAAGAGAGAG | >hg38_21_+_8433172_8446622_RNA45SN1_45S_50ntFlanks | 8357 | 8377 | 21 | AAAAGAACTTTGAAGAGAGAG | 2010.8 | -6.4 | 1.0 | -6.4 | 1.5E-10 | 3.7E-10 | MB231/MB468 | CYTO |
| >hg38_21_+_8433172_8446622_RNA45SN1_45S_50ntFlanks@10850.10868.19\|GGGATAAGGATTGGCTCTA | >hg38_21_+_8433172_8446622_RNA45SN1_45S_50ntFlanks | 10850 | 10868 | 19 | GGGATAAGGATTGGCTCTA | 394.4 | -8.6 | 1.3 | -6.4 | 1.8E-10 | 3.8E-10 | BT20/MB468 | NUC |
| >hg38_21_+_8433172_8446622_RNA45SN1_45S_50ntFlanks@6666.6688.23\|GATCACTCGGCTCGTGCGTCGAT | >hg38_21_+_8433172_8446622_RNA45SN1_45S_50ntFlanks | 6666 | 6688 | 23 | GATCACTCGGCTCGTGCGTCGAT | 175.9 | -10.5 | 1.6 | -6.8 | 1.1E-11 | 3.8E-10 | BT20/MB468 | MP |
| >hg38_21_+_8433172_8446622_RNA45SN1_45S_50ntFlanks@9352.9380.29\|CCGTGGCGCAATGAAGGTGAAGGCCGGCG | >hg38_21_+_8433172_8446622_RNA45SN1_45S_50ntFlanks | 9352 | 9380 | 29 | CCGTGGCGCAATGAAGGTGAAGGCCGGCG | 456.5 | -11.9 | 1.8 | -6.8 | 1.1E-11 | 3.8E-10 | BT20/MB468 | MP |
| >hg38_21_+_8433172_8446622_RNA45SN1_45S_50ntFlanks@12456.12498.43\|CCAAGCGTTGGATTGTTCACCCACTAATAGGGAACGTGAGCTG | >hg38_21_+_8433172_8446622_RNA45SN1_45S_50ntFlanks | 12456 | 12498 | 43 | CCAAGCGTTGGATTGTTCACCCACTAATAGGGAACGTGAGCTG | 284.0 | -11.9 | 1.9 | -6.4 | 1.7E-10 | 3.8E-10 | MB231/MB468 | MITO |
| >trna1_GlnCTG_6_+_18836402_18836473@30.70.41,trna3_GlnCTG_17_+_8023070_8023141@30.70.41,trna49_GlnCTG_6_+_27487308_27487379@30.70.41,trna7_GlnCTG_15_-_66161400_66161471@30.70.41,trna99_GlnCTG_6_-_28909378_28909449@30.70.41\|GACTCTGAATCCAGCGATCCGAGTTCAAATCTCGGTGGAAC | >trna1_GlnCTG_6_+_18836402_18836473 | 30 | 70 | 41 | GACTCTGAATCCAGCGATCCGAGTTCAAATCTCGGTGGAAC | 271.0 | -7.3 | 1.1 | -6.4 | 1.5E-10 | 3.8E-10 | MB231/MB468 | CYTO |
| >hg38_21_+_8433172_8446622_RNA45SN1_45S_50ntFlanks@6773.6803.31\|TTCCTCCCGGGGCTACGCCTGTCTGAGCGTC | >hg38_21_+_8433172_8446622_RNA45SN1_45S_50ntFlanks | 6773 | 6803 | 31 | TTCCTCCCGGGGCTACGCCTGTCTGAGCGTC | 851.2 | -5.8 | 0.9 | -6.4 | 1.5E-10 | 3.8E-10 | MB231/MB468 | CYTO |
| >hg38_21_+_8433172_8446622_RNA45SN1_45S_50ntFlanks@10788.10816.29\|CAGCCTCTGGCATGTTGGAACAATGTAGG | >hg38_21_+_8433172_8446622_RNA45SN1_45S_50ntFlanks | 10788 | 10816 | 29 | CAGCCTCTGGCATGTTGGAACAATGTAGG | 541.1 | -8.4 | 1.3 | -6.4 | 1.4E-10 | 3.8E-10 | MB231/MB468 | MP |
| >hg38_21_+_8433172_8446622_RNA45SN1_45S_50ntFlanks@9534.9572.39\|AGGAAACTCTGGTGGAGGTCCGTAGCGGTCCTGACGTGC | >hg38_21_+_8433172_8446622_RNA45SN1_45S_50ntFlanks | 9534 | 9572 | 39 | AGGAAACTCTGGTGGAGGTCCGTAGCGGTCCTGACGTGC | 3062.2 | -3.3 | 0.5 | -6.4 | 1.3E-10 | 3.8E-10 | BT20/MB468 | CYTO |
| >MI0000471\|hsa-mir-126&WithFlank&9\|+\|136670596\|136670692@59.78.20&[MIMAT0000445&hsa-miR-126-3p&offsets\|+1\|-1;m-40&9\|+\|136670653\|136670674&offsets\|+1\|-1]\|CGTACCGTGAGTAATAATGC | >MI0000471\|hsa-mir-126&WithFlank&9\|+\|136670596\|136670692 | 59 | 78 | 20 | CGTACCGTGAGTAATAATGC | 120.8 | 7.4 | 1.2 | 6.4 | 1.8E-10 | 3.8E-10 | BT20/MB468 | NUC |
| >hg38_21_+_8433172_8446622_RNA45SN1_45S_50ntFlanks@11234.11252.19\|CCTTCCCGTGGATCGCCCC | >hg38_21_+_8433172_8446622_RNA45SN1_45S_50ntFlanks | 11234 | 11252 | 19 | CCTTCCCGTGGATCGCCCC | 78.3 | -10.2 | 1.6 | -6.4 | 1.4E-10 | 3.8E-10 | MB231/MB468 | MP |
| >hg38_21_+_8433172_8446622_RNA45SN1_45S_50ntFlanks@4006.4024.19\|ACCTCGGGCCGATCGCACG | >hg38_21_+_8433172_8446622_RNA45SN1_45S_50ntFlanks | 4006 | 4024 | 19 | ACCTCGGGCCGATCGCACG | 774.8 | -8.5 | 1.3 | -6.4 | 1.4E-10 | 3.8E-10 | MB231/MB468 | MP |
| >am_MI0015825\|hsa-mir-378c&WithFlank&10\|-\|130962582\|130962674@17.36.20;MI0000786\|hsa-mir-378a&WithFlank&5\|+\|149732819\|149732896@49.68.20&[MIMAT0016847&hsa-miR-378c&offsets\|0\|-5;m-130&10\|-\|130962639\|130962658&offsets\|0\|0];[MIMAT0000732&hsa-miR-378a-3p&offsets\|0\|-2;m-44&5\|+\|149732867\|149732888&offsets\|0\|-2]\|ACTGGACTTGGAGTCAGAAG | >am_MI0015825\|hsa-mir-378c&WithFlank&10\|-\|130962582\|130962674 | 17 | 36 | 20 | ACTGGACTTGGAGTCAGAAG | 251.2 | 4.1 | 0.6 | 6.4 | 1.5E-10 | 3.8E-10 | MB231/MB468 | NUC |
| >hg38_21_+_8433172_8446622_RNA45SN1_45S_50ntFlanks@9457.9494.38\|CGCCGGGGAGGTGGAGCACGAGCGCACGTGTTAGGACC | >hg38_21_+_8433172_8446622_RNA45SN1_45S_50ntFlanks | 9457 | 9494 | 38 | CGCCGGGGAGGTGGAGCACGAGCGCACGTGTTAGGACC | 140.9 | -11.0 | 1.7 | -6.4 | 1.4E-10 | 3.8E-10 | MB231/MB468 | MP |

<table>
<tr><td>>MI0000253|hsa-mir-148a&WithFlank&7|-|25949913|25949992@50.71.22(+1U)&[MIMAT0000243&hsa-miR-148a-3p&offsets|0|0(+1U);m-7&7|-|25949922|25949943&offsets|0|0(+1U)]|TCAGTGCACTACAGAACTTTGTT</td><td>>MI0000253|hsa-mir-148a&WithFlank&7|-|25949913|25949992</td><td>50</td><td>71</td><td>22(+1U)</td><td>TCAGTGCACTACAGAACTTTGTT</td><td>728.9</td><td>4.8</td><td>0.7</td><td>6.7</td><td>1.6E-11</td><td>3.8E-10</td><td>BT20/MB231</td><td>NUC</td></tr>
<tr><td>>hg38_21_+_8433172_8446622_RNA45SN1_45S_50ntFlanks@12934.12961.28|CTAAACCATTCGTAGACGACCTGCTTCT</td><td>>hg38_21_+_8433172_8446622_RNA45SN1_45S_50ntFlanks</td><td>12934</td><td>12961</td><td>28</td><td>CTAAACCATTCGTAGACGACCTGCTTCT</td><td>404.7</td><td>-6.2</td><td>1.0</td><td>-6.4</td><td>1.7E-10</td><td>3.8E-10</td><td>MB231/MB468</td><td>MITO</td></tr>
<tr><td>>hg38_21_+_8433172_8446622_RNA45SN1_45S_50ntFlanks@6731.6752.22|CACATTGATCATCGACACTTCG</td><td>>hg38_21_+_8433172_8446622_RNA45SN1_45S_50ntFlanks</td><td>6731</td><td>6752</td><td>22</td><td>CACATTGATCATCGACACTTCG</td><td>421.3</td><td>-3.1</td><td>0.5</td><td>-6.4</td><td>1.5E-10</td><td>3.8E-10</td><td>MB231/MB468</td><td>NUC</td></tr>
<tr><td>>hg38_21_+_8433172_8446622_RNA45SN1_45S_50ntFlanks@7975.8013.39|CGCGACCTCAGATCAGACGTGGCGACCCGCTGAATTTAA</td><td>>hg38_21_+_8433172_8446622_RNA45SN1_45S_50ntFlanks</td><td>7975</td><td>8013</td><td>39</td><td>CGCGACCTCAGATCAGACGTGGCGACCCGCTGAATTTAA</td><td>10625.7</td><td>4.5</td><td>0.7</td><td>6.4</td><td>1.5E-10</td><td>3.8E-10</td><td>MB231/MB468</td><td>NUC</td></tr>
<tr><td>>hg38_21_+_8433172_8446622_RNA45SN1_45S_50ntFlanks@4734.4759.26|AACGAAAGTCGGAGGTTCGAAGACGA</td><td>>hg38_21_+_8433172_8446622_RNA45SN1_45S_50ntFlanks</td><td>4734</td><td>4759</td><td>26</td><td>AACGAAAGTCGGAGGTTCGAAGACGA</td><td>2132.2</td><td>-4.4</td><td>0.7</td><td>-6.4</td><td>1.4E-10</td><td>3.8E-10</td><td>MB231/MB468</td><td>MP</td></tr>
<tr><td>>hg38_21_+_8433172_8446622_RNA45SN1_45S_50ntFlanks@11784.11805.22|ATGAATGGATGAACGAGATTCC</td><td>>hg38_21_+_8433172_8446622_RNA45SN1_45S_50ntFlanks</td><td>11784</td><td>11805</td><td>22</td><td>ATGAATGGATGAACGAGATTCC</td><td>554.2</td><td>-3.3</td><td>0.5</td><td>-6.4</td><td>1.5E-10</td><td>3.8E-10</td><td>MB231/MB468</td><td>NUC</td></tr>
<tr><td>>hg38_21_+_8433172_8446622_RNA45SN1_45S_50ntFlanks@10408.10437.30|GGTCAGTCGGTCCTGAGAGATGGGCGAGCG</td><td>>hg38_21_+_8433172_8446622_RNA45SN1_45S_50ntFlanks</td><td>10408</td><td>10437</td><td>30</td><td>GGTCAGTCGGTCCTGAGAGATGGGCGAGCG</td><td>128.7</td><td>-7.0</td><td>1.1</td><td>-6.4</td><td>1.3E-10</td><td>3.8E-10</td><td>BT20/MB468</td><td>CYTO</td></tr>
<tr><td>>hg38_21_+_8433172_8446622_RNA45SN1_45S_50ntFlanks@12892.12934.43|CCCTCGCCCGTCACGCACCGCACGTTCGTGGGGAACCTGGCGC</td><td>>hg38_21_+_8433172_8446622_RNA45SN1_45S_50ntFlanks</td><td>12892</td><td>12934</td><td>43</td><td>CCCTCGCCCGTCACGCACCGCACGTTCGTGGGGAACCTGGCGC</td><td>138.4</td><td>-9.2</td><td>1.4</td><td>-6.5</td><td>1.1E-10</td><td>3.8E-10</td><td>BT20/MB468</td><td>MITO</td></tr>
<tr><td>>hg38_21_+_8433172_8446622_RNA45SN1_45S_50ntFlanks@4882.4903.22|TGCAAAGCTGAAACTTAAAGGA</td><td>>hg38_21_+_8433172_8446622_RNA45SN1_45S_50ntFlanks</td><td>4882</td><td>4903</td><td>22</td><td>TGCAAAGCTGAAACTTAAAGGA</td><td>421.1</td><td>8.6</td><td>1.3</td><td>6.4</td><td>1.3E-10</td><td>3.8E-10</td><td>BT20/MB468</td><td>CYTO</td></tr>
<tr><td>>hg38_21_+_8433172_8446622_RNA45SN1_45S_50ntFlanks@3874.3894.21|AATACATGCCGACGGGCGCTG</td><td>>hg38_21_+_8433172_8446622_RNA45SN1_45S_50ntFlanks</td><td>3874</td><td>3894</td><td>21</td><td>AATACATGCCGACGGGCGCTG</td><td>72.5</td><td>-10.0</td><td>1.6</td><td>-6.4</td><td>1.4E-10</td><td>3.8E-10</td><td>MB231/MB468</td><td>MP</td></tr>
<tr><td>>MI0000440|hsa-mir-27b&WithFlank&9|+|95085439|95085547@70.87.18&[MIMAT0000419&hsa-miR-27b-3p&offsets|+3|0;m-39&9|+|95085505|95085525&offsets|+3|0]|ACAGTGGCTAAGTTCTGC</td><td>>MI0000440|hsa-mir-27b&WithFlank&9|+|95085439|95085547</td><td>70</td><td>87</td><td>18</td><td>ACAGTGGCTAAGTTCTGC</td><td>309.3</td><td>12.0</td><td>1.8</td><td>6.8</td><td>1.2E-11</td><td>3.8E-10</td><td>BT20/MB468</td><td>MP</td></tr>
<tr><td>>hg38_21_+_8433172_8446622_RNA45SN1_45S_50ntFlanks@6661.6701.41|CGGTGGATCACTCGGCTCGTGCGTCGATGAAGAACGCAGCT</td><td>>hg38_21_+_8433172_8446622_RNA45SN1_45S_50ntFlanks</td><td>6661</td><td>6701</td><td>41</td><td>CGGTGGATCACTCGGCTCGTGCGTCGATGAAGAACGCAGCT</td><td>1056.4</td><td>6.5</td><td>1.0</td><td>6.4</td><td>1.4E-10</td><td>3.8E-10</td><td>BT20/MB468</td><td>CYTO</td></tr>
<tr><td>>hg38_21_+_8433172_8446622_RNA45SN1_45S_50ntFlanks@10103.10126.24|CGCGCGTGTGGTGTGCGTCGGAGG</td><td>>hg38_21_+_8433172_8446622_RNA45SN1_45S_50ntFlanks</td><td>10103</td><td>10126</td><td>24</td><td>CGCGCGTGTGGTGTGCGTCGGAGG</td><td>92.6</td><td>-10.4</td><td>1.6</td><td>-6.4</td><td>1.4E-10</td><td>3.8E-10</td><td>MB231/MB468</td><td>MP</td></tr>
<tr><td>>hg38_21_+_8433172_8446622_RNA45SN1_45S_50ntFlanks@11209.11250.42|CGGCGACTCTGGACGCGAGCCGGGCCCTTCCCGTGGATCGCC</td><td>>hg38_21_+_8433172_8446622_RNA45SN1_45S_50ntFlanks</td><td>11209</td><td>11250</td><td>42</td><td>CGGCGACTCTGGACGCGAGCCGGGCCCTTCCCGTGGATCGCC</td><td>373.7</td><td>-6.5</td><td>1.0</td><td>-6.4</td><td>1.5E-10</td><td>3.9E-10</td><td>MB231/MB468</td><td>CYTO</td></tr>
<tr><td>>hg38_21_+_8433172_8446622_RNA45SN1_45S_50ntFlanks@3705.3725.21|TACCTGGTTGATCCTGCCAGT</td><td>>hg38_21_+_8433172_8446622_RNA45SN1_45S_50ntFlanks</td><td>3705</td><td>3725</td><td>21</td><td>TACCTGGTTGATCCTGCCAGT</td><td>1351.6</td><td>4.0</td><td>0.6</td><td>6.4</td><td>1.8E-10</td><td>3.9E-10</td><td>BT20/MB468</td><td>NUC</td></tr>
<tr><td>>hg38_21_+_8433172_8446622_RNA45SN1_45S_50ntFlanks@8110.8143.34|CGGGGCGCGGGACATGTGGCGTACGGAAGACCCG</td><td>>hg38_21_+_8433172_8446622_RNA45SN1_45S_50ntFlanks</td><td>8110</td><td>8143</td><td>34</td><td>CGGGGCGCGGGACATGTGGCGTACGGAAGACCCG</td><td>222.8</td><td>-10.9</td><td>1.6</td><td>-6.8</td><td>1.2E-11</td><td>3.9E-10</td><td>BT20/MB468</td><td>MP</td></tr>
<tr><td>>hg38_21_+_8433172_8446622_RNA45SN1_45S_50ntFlanks@11703.11728.26|AATGAAGCGCGGGTAAACGGCGGGAG</td><td>>hg38_21_+_8433172_8446622_RNA45SN1_45S_50ntFlanks</td><td>11703</td><td>11728</td><td>26</td><td>AATGAAGCGCGGGTAAACGGCGGGAG</td><td>1678.7</td><td>-6.7</td><td>1.0</td><td>-6.4</td><td>1.1E-10</td><td>3.9E-10</td><td>BT20/MB468</td><td>MITO</td></tr>
<tr><td>>hg38_21_+_8433172_8446622_RNA45SN1_45S_50ntFlanks@4370.4394.25|TTAAAAAGCTCGTAGTTGGATCTTG</td><td>>hg38_21_+_8433172_8446622_RNA45SN1_45S_50ntFlanks</td><td>4370</td><td>4394</td><td>25</td><td>TTAAAAAGCTCGTAGTTGGATCTTG</td><td>78.9</td><td>-10.2</td><td>1.6</td><td>-6.4</td><td>1.5E-10</td><td>3.9E-10</td><td>MB231/MB468</td><td>MP</td></tr>
<tr><td>>hg38_21_+_8433172_8446622_RNA45SN1_45S_50ntFlanks@3985.4024.40|CGGCTTTGGTGACTCTAGATAACCTCGGGCCGATCGCACG</td><td>>hg38_21_+_8433172_8446622_RNA45SN1_45S_50ntFlanks</td><td>3985</td><td>4024</td><td>40</td><td>CGGCTTTGGTGACTCTAGATAACCTCGGGCCGATCGCACG</td><td>1298.6</td><td>-8.6</td><td>1.3</td><td>-6.4</td><td>1.1E-10</td><td>3.9E-10</td><td>BT20/MB468</td><td>MITO</td></tr>
<tr><td>>trna14_LysTTT_11_-_59327808_59327880@1.35.35,trna2_LysTTT_17_+_8022473_8022545@1.35.35,trna54_LysTTT_1_+_204475655_204475727@1.35.35,trna5_LysTTT_11_+_59323902_59323974@1.35.35,trna62_LysTTT_1_-_204476158_204476230@1.35.35,trna76_LysTTT_6_+_28918806_28918878@1.35.35|GCCCGGATAGCTCAGTCGGTAGAGCATCAGACTTT</td><td>>trna14_LysTTT_11_-_59327808_59327880</td><td>1</td><td>35</td><td>35</td><td>GCCCGGATAGCTCAGTCGGTAGAGCATCAGACTTT</td><td>500.3</td><td>9.9</td><td>1.5</td><td>6.4</td><td>1.4E-10</td><td>3.9E-10</td><td>BT20/MB468</td><td>CYTO</td></tr>
<tr><td>>hg38_21_+_8433172_8446622_RNA45SN1_45S_50ntFlanks@5074.5092.19|ATTCCGATAACGAACGAGA</td><td>>hg38_21_+_8433172_8446622_RNA45SN1_45S_50ntFlanks</td><td>5074</td><td>5092</td><td>19</td><td>ATTCCGATAACGAACGAGA</td><td>183.4</td><td>-11.4</td><td>1.8</td><td>-6.4</td><td>1.5E-10</td><td>3.9E-10</td><td>MB231/MB468</td><td>MP</td></tr>
<tr><td>>hg38_21_+_8433172_8446622_RNA45SN1_45S_50ntFlanks@9605.9622.18|AAGACTAATCGAACCATC</td><td>>hg38_21_+_8433172_8446622_RNA45SN1_45S_50ntFlanks</td><td>9605</td><td>9622</td><td>18</td><td>AAGACTAATCGAACCATC</td><td>454.8</td><td>-6.3</td><td>1.0</td><td>-6.4</td><td>1.7E-10</td><td>3.9E-10</td><td>MB231/MB468</td><td>MITO</td></tr>
<tr><td>>hg38_21_+_8433172_8446622_RNA45SN1_45S_50ntFlanks@11703.11740.38|AATGAAGCGCGGGTAAACGGCGGGAGTAACTATGACTC</td><td>>hg38_21_+_8433172_8446622_RNA45SN1_45S_50ntFlanks</td><td>11703</td><td>11740</td><td>38</td><td>AATGAAGCGCGGGTAAACGGCGGGAGTAACTATGACTC</td><td>495.8</td><td>-4.4</td><td>0.7</td><td>-6.4</td><td>1.5E-10</td><td>3.9E-10</td><td>MB231/MB468</td><td>NUC</td></tr>
<tr><td>>hg38_21_+_8433172_8446622_RNA45SN1_45S_50ntFlanks@6742.6772.31|TCGACACTTCGAACGCACTTGCGGCCCCGGG</td><td>>hg38_21_+_8433172_8446622_RNA45SN1_45S_50ntFlanks</td><td>6742</td><td>6772</td><td>31</td><td>TCGACACTTCGAACGCACTTGCGGCCCCGGG</td><td>124.8</td><td>-10.8</td><td>1.7</td><td>-6.4</td><td>1.5E-10</td><td>3.9E-10</td><td>MB231/MB468</td><td>MP</td></tr>
<tr><td>>hg38_21_+_8433172_8446622_RNA45SN1_45S_50ntFlanks@12073.12090.18|CCGAGGGGCTCTCGCTTC</td><td>>hg38_21_+_8433172_8446622_RNA45SN1_45S_50ntFlanks</td><td>12073</td><td>12090</td><td>18</td><td>CCGAGGGGCTCTCGCTTC</td><td>661.4</td><td>-5.9</td><td>0.9</td><td>-6.4</td><td>1.5E-10</td><td>3.9E-10</td><td>MB231/MB468</td><td>CYTO</td></tr>
<tr><td>>MI0000061|hsa-let-7a-2&WithFlank&11|-|122146516|122146599@11.31.21(+1A);MI0000062|hsa-let-7a-3&WithFlank&22|+|46112743|46112828@10.30.21(+1A);MI0000060|hsa-let-7a-1&WithFlank&9|+|94175951|94176042@12.32.21(+1A)&[MIMAT0000062&hsa-let-7a-5p&offsets|0|-1(+1A);m-6&11|-|122146568|122146589&offsets|0|-1(+1A)];[MIMAT0000062_1&hsa-let-7a-5p&offsets|0|-1(+1A);m-5&22|+|46112752|46112773&offsets|0|-1(+1A)];[MIMAT0000062_2&hsa-let-7a-5p&offsets|0|-1(+1A);m-4&9|+|94175962|94175983&offsets|0|-1(+1A)]|TGAGGTAGTAGGTTGTATAGTA</td><td>>MI0000061|hsa-let-7a-2&WithFlank&11|-|122146516|122146599</td><td>11</td><td>31</td><td>21(+1A)</td><td>TGAGGTAGTAGGTTGTATAGTA</td><td>1946.8</td><td>6.3</td><td>1.0</td><td>6.4</td><td>1.7E-10</td><td>3.9E-10</td><td>MB231/MB468</td><td>MITO</td></tr>
<tr><td>>hg38_21_+_8433172_8446622_RNA45SN1_45S_50ntFlanks@10808.10829.22|CAATGTAGGTAAGGGAAGTCGG</td><td>>hg38_21_+_8433172_8446622_RNA45SN1_45S_50ntFlanks</td><td>10808</td><td>10829</td><td>22</td><td>CAATGTAGGTAAGGGAAGTCGG</td><td>612.3</td><td>-8.6</td><td>1.3</td><td>-6.4</td><td>1.5E-10</td><td>3.9E-10</td><td>MB231/MB468</td><td>NUC</td></tr>
<tr><td>>hg38_1_-_228634819_228635039_RNA5S12_5S_50ntFlanks@131.171.41|GGATGGGAGACCGCCTGGGAATACCGGGTGCTGTAGGCTTT</td><td>>hg38_1_-_228634819_228635039_RNA5S12_5S_50ntFlanks</td><td>131</td><td>171</td><td>41</td><td>GGATGGGAGACCGCCTGGGAATACCGGGTGCTGTAGGCTTT</td><td>2530.7</td><td>9.3</td><td>1.4</td><td>6.4</td><td>1.4E-10</td><td>3.9E-10</td><td>BT20/MB468</td><td>CYTO</td></tr>
<tr><td>>hg38_21_+_8433172_8446622_RNA45SN1_45S_50ntFlanks@10471.10488.18|CCCTCGGCCGATCGAAAG</td><td>>hg38_21_+_8433172_8446622_RNA45SN1_45S_50ntFlanks</td><td>10471</td><td>10488</td><td>18</td><td>CCCTCGGCCGATCGAAAG</td><td>1788.2</td><td>-4.9</td><td>0.8</td><td>-6.4</td><td>1.6E-10</td><td>3.9E-10</td><td>MB231/MB468</td><td>NUC</td></tr>
<tr><td>>hg38_21_+_8433172_8446622_RNA45SN1_45S_50ntFlanks@9534.9575.42|AGGAAACTCTGGTGGAGGTCCGTAGCGGTCCTGACGTGCAAA</td><td>>hg38_21_+_8433172_8446622_RNA45SN1_45S_50ntFlanks</td><td>9534</td><td>9575</td><td>42</td><td>AGGAAACTCTGGTGGAGGTCCGTAGCGGTCCTGACGTGCAAA</td><td>400.6</td><td>-8.4</td><td>1.3</td><td>-6.4</td><td>1.7E-10</td><td>3.9E-10</td><td>MB231/MB468</td><td>MITO</td></tr>
<tr><td>>hg38_21_+_8433172_8446622_RNA45SN1_45S_50ntFlanks@12893.12925.33|CCTCGCCCGTCACGCACCGCACGTTCGTGGGGA</td><td>>hg38_21_+_8433172_8446622_RNA45SN1_45S_50ntFlanks</td><td>12893</td><td>12925</td><td>33</td><td>CCTCGCCCGTCACGCACCGCACGTTCGTGGGGA</td><td>339.3</td><td>-11.5</td><td>1.7</td><td>-6.8</td><td>1.2E-11</td><td>3.9E-10</td><td>BT20/MB468</td><td>MP</td></tr>
<tr><td>>hg38_21_+_8433172_8446622_RNA45SN1_45S_50ntFlanks@9341.9368.28|CACGAAAGCCGCCGTGGCGCAATGAAGG</td><td>>hg38_21_+_8433172_8446622_RNA45SN1_45S_50ntFlanks</td><td>9341</td><td>9368</td><td>28</td><td>CACGAAAGCCGCCGTGGCGCAATGAAGG</td><td>88.5</td><td>-10.3</td><td>1.6</td><td>-6.4</td><td>1.5E-10</td><td>3.9E-10</td><td>MB231/MB468</td><td>MP</td></tr>
<tr><td>>hg38_21_+_8433172_8446622_RNA45SN1_45S_50ntFlanks@12636.12660.25|AGCTACCATCTGTGGGATTATGACT</td><td>>hg38_21_+_8433172_8446622_RNA45SN1_45S_50ntFlanks</td><td>12636</td><td>12660</td><td>25</td><td>AGCTACCATCTGTGGGATTATGACT</td><td>264.8</td><td>-6.4</td><td>1.0</td><td>-6.4</td><td>1.5E-10</td><td>3.9E-10</td><td>MB231/MB468</td><td>MP</td></tr>
<tr><td>>hg38_1_-_228634819_228635039_RNA5S12_5S_50ntFlanks@138.170.33|AGACCGCCTGGGAATACCGGGTGCTGTAGGCTT</td><td>>hg38_1_-_228634819_228635039_RNA5S12_5S_50ntFlanks</td><td>138</td><td>170</td><td>33</td><td>AGACCGCCTGGGAATACCGGGTGCTGTAGGCTT</td><td>1345.0</td><td>5.3</td><td>0.8</td><td>6.4</td><td>1.6E-10</td><td>3.9E-10</td><td>MB231/MB468</td><td>NUC</td></tr>
<tr><td>>MI0001733|hsa-mir-452&WithFlank&X|-|151959622|151959718@59.80.22&[MIMAT0001636&hsa-miR-452-3p&offsets|-5|-5;m-457&X|-|151959639|151959660&offsets|0|0]|TCAGTCTCATCTGCAAAGAAGT</td><td>>MI0001733|hsa-mir-452&WithFlank&X|-|151959622|151959718</td><td>59</td><td>80</td><td>22</td><td>TCAGTCTCATCTGCAAAGAAGT</td><td>121.2</td><td>10.1</td><td>1.6</td><td>6.4</td><td>1.5E-10</td><td>3.9E-10</td><td>MB231/MB468</td><td>MP</td></tr>
<tr><td>>hg38_21_+_8433172_8446622_RNA45SN1_45S_50ntFlanks@11970.11989.20|CGCGAGGGGCCCGGGGCGGG</td><td>>hg38_21_+_8433172_8446622_RNA45SN1_45S_50ntFlanks</td><td>11970</td><td>11989</td><td>20</td><td>CGCGAGGGGCCCGGGGCGGG</td><td>113.5</td><td>-10.7</td><td>1.7</td><td>-6.4</td><td>1.5E-10</td><td>3.9E-10</td><td>MB231/MB468</td><td>MP</td></tr>
<tr><td>>hg38_21_+_8433172_8446622_RNA45SN1_45S_50ntFlanks@11562.11591.30|CGCCTAGCAGCCGACTTAGAACTGGTGCGG</td><td>>hg38_21_+_8433172_8446622_RNA45SN1_45S_50ntFlanks</td><td>11562</td><td>11591</td><td>30</td><td>CGCCTAGCAGCCGACTTAGAACTGGTGCGG</td><td>244.3</td><td>-8.0</td><td>1.2</td><td>-6.4</td><td>1.7E-10</td><td>3.9E-10</td><td>MB231/MB468</td><td>MITO</td></tr>
<tr><td>>hg38_21_+_8433172_8446622_RNA45SN1_45S_50ntFlanks@11784.11820.37|ATGAATGGATGAACGAGATTCCCACTGTCCCTACCTA</td><td>>hg38_21_+_8433172_8446622_RNA45SN1_45S_50ntFlanks</td><td>11784</td><td>11820</td><td>37</td><td>ATGAATGGATGAACGAGATTCCCACTGTCCCTACCTA</td><td>86.3</td><td>-10.3</td><td>1.6</td><td>-6.4</td><td>1.5E-10</td><td>3.9E-10</td><td>MB231/MB468</td><td>MP</td></tr>
<tr><td>>hg38_21_+_8433172_8446622_RNA45SN1_45S_50ntFlanks@10821.10840.20|GGAAGTCGGCAAGCCGGATC</td><td>>hg38_21_+_8433172_8446622_RNA45SN1_45S_50ntFlanks</td><td>10821</td><td>10840</td><td>20</td><td>GGAAGTCGGCAAGCCGGATC</td><td>3243.8</td><td>-4.8</td><td>0.7</td><td>-6.4</td><td>1.4E-10</td><td>3.9E-10</td><td>BT20/MB468</td><td>CYTO</td></tr>
<tr><td>>MI0000809|hsa-mir-151a&WithFlank&8|-|140732558|140732659@51.74.24&[MIMAT0000757&hsa-miR-151a-3p&offsets|-2|+1;m-30&8|-|140732587|140732607&offsets|-2|+1]|TACTAGACTGAAGCTCCTTGAGGA</td><td>>MI0000809|hsa-mir-151a&WithFlank&8|-|140732558|140732659</td><td>51</td><td>74</td><td>24</td><td>TACTAGACTGAAGCTCCTTGAGGA</td><td>345.1</td><td>7.9</td><td>1.2</td><td>6.4</td><td>1.4E-10</td><td>3.9E-10</td><td>BT20/MB468</td><td>CYTO</td></tr>
<tr><td>>hg38_21_+_8433172_8446622_RNA45SN1_45S_50ntFlanks@9205.9223.19|CGGGGCCACGCGCGCGTCC</td><td>>hg38_21_+_8433172_8446622_RNA45SN1_45S_50ntFlanks</td><td>9205</td><td>9223</td><td>19</td><td>CGGGGCCACGCGCGCGTCC</td><td>109.6</td><td>-10.6</td><td>1.7</td><td>-6.4</td><td>1.5E-10</td><td>3.9E-10</td><td>MB231/MB468</td><td>MP</td></tr>
<tr><td>>MI0000087|hsa-mir-29a&WithFlank&7|-|130876741|130876816@47.68.22(+1U)&[MIMAT0000086&hsa-miR-29a-3p&offsets|-1|-1(+1U);m-26&7|-|130876749|130876769&offsets|-1|0(+1U)]|CTAGCACCATCTGAAATCGGTTT</td><td>>MI0000087|hsa-mir-29a&WithFlank&7|-|130876741|130876816</td><td>47</td><td>68</td><td>22(+1U)</td><td>CTAGCACCATCTGAAATCGGTTT</td><td>979.6</td><td>10.7</td><td>1.7</td><td>6.4</td><td>1.1E-10</td><td>3.9E-10</td><td>BT20/MB468</td><td>MITO</td></tr>
<tr><td>>hg38_21_+_8433172_8446622_RNA45SN1_45S_50ntFlanks@11919.11952.34|CATGAGAGGTGTAGAATAAGTGGGAGGCCCCCGG</td><td>>hg38_21_+_8433172_8446622_RNA45SN1_45S_50ntFlanks</td><td>11919</td><td>11952</td><td>34</td><td>CATGAGAGGTGTAGAATAAGTGGGAGGCCCCCGG</td><td>79.5</td><td>-10.2</td><td>1.6</td><td>-6.4</td><td>1.5E-10</td><td>3.9E-10</td><td>MB231/MB468</td><td>MP</td></tr>
<tr><td>>MI0000477|hsa-mir-146a&WithFlank&5|+|160485346|160485456@27.46.20(+1C)&[MIMAT0000449&hsa-miR-146a-5p&offsets|0|-2(+1C);m-129&5|+|160485372|160485393&offsets|0|-2(+1C)]|TGAGAACTGAATTCCATGGGC</td><td>>MI0000477|hsa-mir-146a&WithFlank&5|+|160485346|160485456</td><td>27</td><td>46</td><td>20(+1C)</td><td>TGAGAACTGAATTCCATGGGC</td><td>270.4</td><td>11.3</td><td>1.8</td><td>6.4</td><td>1.8E-10</td><td>3.9E-10</td><td>MB231/MB468</td><td>MITO</td></tr>
<tr><td>>hg38_21_+_8433172_8446622_RNA45SN1_45S_50ntFlanks@11728.11753.26|GTAACTATGACTCTCTTAAGGTAGCC</td><td>>hg38_21_+_8433172_8446622_RNA45SN1_45S_50ntFlanks</td><td>11728</td><td>11753</td><td>26</td><td>GTAACTATGACTCTCTTAAGGTAGCC</td><td>88.8</td><td>-10.3</td><td>1.6</td><td>-6.4</td><td>1.5E-10</td><td>3.9E-10</td><td>MB231/MB468</td><td>MP</td></tr>
<tr><td>>hg38_21_+_8433172_8446622_RNA45SN1_45S_50ntFlanks@3990.4013.24|TTGGTGACTCTAGATAACCTCGGG</td><td>>hg38_21_+_8433172_8446622_RNA45SN1_45S_50ntFlanks</td><td>3990</td><td>4013</td><td>24</td><td>TTGGTGACTCTAGATAACCTCGGG</td><td>87.6</td><td>-10.3</td><td>1.6</td><td>-6.4</td><td>1.5E-10</td><td>3.9E-10</td><td>MB231/MB468</td><td>MP</td></tr>
</table>

<table>
<tr><td>>trna111_HisGTG_1_-_147774845_147774916@-1T.35.36,trna118_HisGTG_1_-_145396881_145396952@-1T.35.36,trna16_HisGTG_1_+_146544773_146544844@-1T.35.36,trna1_HisGTG_15_+_45493349_45493420@-1T.35.36,trna21_HisGTG_1_+_147753471_147753542@-1T.35.36,trna33_HisGTG_6_+_27125906_27125977@-1T.35.36,trna7_HisGTG_9_-_14433938_14434009@-1T.35.36,trna8_HisGTG_15_-_45492611_45492682@-1T.35.36,trna9_HisGTG_15_-_45490804_45490875@-1T.35.36|TGCCGTGATCGTATAGTGGTTAGTACTCTGCGTTGT</td><td>>trna111_HisGTG_1_-_147774845_147774916</td><td>-1T</td><td>35</td><td>36</td><td>TGCCGTGATCGTATAGTGGTTAGTACTCTGCGTTGT</td><td>203.4</td><td>10.9</td><td>1.7</td><td>6.4</td><td>1.8E-10</td><td>4.0E-10</td><td>MB231/MB468</td><td>MITO</td></tr>
<tr><td>>hg38_21_+_8433172_8446622_RNA45SN1_45S_50ntFlanks@10412.10430.19|AGTCGGTCCTGAGAGATGG</td><td>>hg38_21_+_8433172_8446622_RNA45SN1_45S_50ntFlanks</td><td>10412</td><td>10430</td><td>19</td><td>AGTCGGTCCTGAGAGATGG</td><td>180.5</td><td>-7.3</td><td>1.1</td><td>-6.4</td><td>1.6E-10</td><td>4.0E-10</td><td>MB231/MB468</td><td>NUC</td></tr>
<tr><td>>hg38_21_+_8433172_8446622_RNA45SN1_45S_50ntFlanks@12941.12964.24|ATTCGTAGACGACCTGCTTCTGGG</td><td>>hg38_21_+_8433172_8446622_RNA45SN1_45S_50ntFlanks</td><td>12941</td><td>12964</td><td>24</td><td>ATTCGTAGACGACCTGCTTCTGGG</td><td>372.1</td><td>-8.5</td><td>1.3</td><td>-6.4</td><td>1.9E-10</td><td>4.0E-10</td><td>BT20/MB468</td><td>NUC</td></tr>
<tr><td>>hg38_21_+_8433172_8446622_RNA45SN1_45S_50ntFlanks@12652.12675.24|ATTATGACTGAACGCCTCTAAGTC</td><td>>hg38_21_+_8433172_8446622_RNA45SN1_45S_50ntFlanks</td><td>12652</td><td>12675</td><td>24</td><td>ATTATGACTGAACGCCTCTAAGTC</td><td>225.5</td><td>-7.1</td><td>1.1</td><td>-6.4</td><td>1.6E-10</td><td>4.0E-10</td><td>MB231/MB468</td><td>CYTO</td></tr>
<tr><td>>hg38_21_+_8433172_8446622_RNA45SN1_45S_50ntFlanks@9537.9563.27|AAACTCTGGTGGAGGTCCGTAGCGGTC</td><td>>hg38_21_+_8433172_8446622_RNA45SN1_45S_50ntFlanks</td><td>9537</td><td>9563</td><td>27</td><td>AAACTCTGGTGGAGGTCCGTAGCGGTC</td><td>212.3</td><td>-5.0</td><td>0.8</td><td>-6.4</td><td>1.6E-10</td><td>4.0E-10</td><td>MB231/MB468</td><td>CYTO</td></tr>
<tr><td>>hg38_21_+_8433172_8446622_RNA45SN1_45S_50ntFlanks@10873.10891.19|CTGGGTCGGTCGGGCTGGG</td><td>>hg38_21_+_8433172_8446622_RNA45SN1_45S_50ntFlanks</td><td>10873</td><td>10891</td><td>19</td><td>CTGGGTCGGTCGGGCTGGG</td><td>755.2</td><td>-4.2</td><td>0.7</td><td>-6.4</td><td>1.4E-10</td><td>4.0E-10</td><td>BT20/MB468</td><td>CYTO</td></tr>
<tr><td>>hg38_21_+_8433172_8446622_RNA45SN1_45S_50ntFlanks@10787.10825.39|ACAGCCTCTGGCATGTTGGAACAATGTAGGTAAGGGAAG</td><td>>hg38_21_+_8433172_8446622_RNA45SN1_45S_50ntFlanks</td><td>10787</td><td>10825</td><td>39</td><td>ACAGCCTCTGGCATGTTGGAACAATGTAGGTAAGGGAAG</td><td>122.1</td><td>-10.8</td><td>1.7</td><td>-6.4</td><td>1.5E-10</td><td>4.0E-10</td><td>MB231/MB468</td><td>MP</td></tr>
<tr><td>>hg38_21_+_8433172_8446622_RNA45SN1_45S_50ntFlanks@5489.5532.44|CTGAGAAGACGGTCGAACTTGACTATCTAGAGGAAGTAAAAGTC</td><td>>hg38_21_+_8433172_8446622_RNA45SN1_45S_50ntFlanks</td><td>5489</td><td>5532</td><td>44</td><td>CTGAGAAGACGGTCGAACTTGACTATCTAGAGGAAGTAAAAGTC</td><td>662.0</td><td>6.0</td><td>0.9</td><td>6.4</td><td>1.6E-10</td><td>4.0E-10</td><td>MB231/MB468</td><td>CYTO</td></tr>
<tr><td>>hg38_21_+_8433172_8446622_RNA45SN1_45S_50ntFlanks@10411.10432.22|CAGTCGGTCCTGAGAGATGGGC</td><td>>hg38_21_+_8433172_8446622_RNA45SN1_45S_50ntFlanks</td><td>10411</td><td>10432</td><td>22</td><td>CAGTCGGTCCTGAGAGATGGGC</td><td>1294.3</td><td>-7.6</td><td>1.2</td><td>-6.4</td><td>1.8E-10</td><td>4.0E-10</td><td>MB231/MB468</td><td>MITO</td></tr>
<tr><td>>hg38_21_+_8433172_8446622_RNA45SN1_45S_50ntFlanks@12456.12476.21|CCAAGCGTTGGATTGTTCACC</td><td>>hg38_21_+_8433172_8446622_RNA45SN1_45S_50ntFlanks</td><td>12456</td><td>12476</td><td>21</td><td>CCAAGCGTTGGATTGTTCACC</td><td>36765.3</td><td>-5.5</td><td>0.9</td><td>-6.4</td><td>1.6E-10</td><td>4.0E-10</td><td>MB231/MB468</td><td>CYTO</td></tr>
<tr><td>>hg38_21_+_8433172_8446622_RNA45SN1_45S_50ntFlanks@12438.12455.18|ATTGTGAAGCAGAATTCA</td><td>>hg38_21_+_8433172_8446622_RNA45SN1_45S_50ntFlanks</td><td>12438</td><td>12455</td><td>18</td><td>ATTGTGAAGCAGAATTCA</td><td>2542.9</td><td>-9.1</td><td>1.4</td><td>-6.4</td><td>1.8E-10</td><td>4.0E-10</td><td>MB231/MB468</td><td>MITO</td></tr>
<tr><td>>hg38_21_+_8433172_8446622_RNA45SN1_45S_50ntFlanks@4082.4101.20|TCGCCGTGCCTACCATGGTG</td><td>>hg38_21_+_8433172_8446622_RNA45SN1_45S_50ntFlanks</td><td>4082</td><td>4101</td><td>20</td><td>TCGCCGTGCCTACCATGGTG</td><td>113.1</td><td>-10.7</td><td>1.7</td><td>-6.4</td><td>1.5E-10</td><td>4.0E-10</td><td>MB231/MB468</td><td>MP</td></tr>
<tr><td>>hg38_21_+_8433172_8446622_RNA45SN1_45S_50ntFlanks@12423.12445.23|TCGGCTCTTCCTATCATTGTGAA</td><td>>hg38_21_+_8433172_8446622_RNA45SN1_45S_50ntFlanks</td><td>12423</td><td>12445</td><td>23</td><td>TCGGCTCTTCCTATCATTGTGAA</td><td>493.6</td><td>-8.7</td><td>1.4</td><td>-6.4</td><td>1.9E-10</td><td>4.0E-10</td><td>BT20/MB468</td><td>NUC</td></tr>
<tr><td>>hg38_21_+_8433172_8446622_RNA45SN1_45S_50ntFlanks@4048.4066.19|TTCGAACGTCTGCCCTATC</td><td>>hg38_21_+_8433172_8446622_RNA45SN1_45S_50ntFlanks</td><td>4048</td><td>4066</td><td>19</td><td>TTCGAACGTCTGCCCTATC</td><td>1001.5</td><td>-7.5</td><td>1.2</td><td>-6.4</td><td>1.9E-10</td><td>4.0E-10</td><td>BT20/MB468</td><td>NUC</td></tr>
<tr><td>>hg38_21_+_8433172_8446622_RNA45SN1_45S_50ntFlanks@7972.8011.40|AGACGCGACCTCAGATCAGACGTGGCGACCCGCTGAATTT</td><td>>hg38_21_+_8433172_8446622_RNA45SN1_45S_50ntFlanks</td><td>7972</td><td>8011</td><td>40</td><td>AGACGCGACCTCAGATCAGACGTGGCGACCCGCTGAATTT</td><td>370.2</td><td>5.0</td><td>0.8</td><td>6.4</td><td>1.6E-10</td><td>4.0E-10</td><td>MB231/MB468</td><td>NUC</td></tr>
<tr><td>>MI0003820|hsa-mir-454&WithFlank&17|-|59137752|59137878@70.94.25&[MIMAT0003885&hsa-miR-454-3p&offsets|0|+2;m-272&17|-|59137787|59137809&offsets|0|+2]|TAGTGCAATATTGCTTATAGGGTTT</td><td>>MI0003820|hsa-mir-454&WithFlank&17|-|59137752|59137878</td><td>70</td><td>94</td><td>25</td><td>TAGTGCAATATTGCTTATAGGGTTT</td><td>313.7</td><td>8.2</td><td>1.3</td><td>6.4</td><td>1.4E-10</td><td>4.0E-10</td><td>BT20/MB468</td><td>CYTO</td></tr>
<tr><td>>hg38_21_+_8433172_8446622_RNA45SN1_45S_50ntFlanks@4976.4994.19|CCGGACACGGACAGGATTG</td><td>>hg38_21_+_8433172_8446622_RNA45SN1_45S_50ntFlanks</td><td>4976</td><td>4994</td><td>19</td><td>CCGGACACGGACAGGATTG</td><td>416.4</td><td>-8.3</td><td>1.3</td><td>-6.4</td><td>1.5E-10</td><td>4.0E-10</td><td>MB231/MB468</td><td>MP</td></tr>
<tr><td>>hg38_21_+_8433172_8446622_RNA45SN1_45S_50ntFlanks@8194.8215.22|CCCGTGGACGGTGTGAGGCCGG</td><td>>hg38_21_+_8433172_8446622_RNA45SN1_45S_50ntFlanks</td><td>8194</td><td>8215</td><td>22</td><td>CCCGTGGACGGTGTGAGGCCGG</td><td>3032.8</td><td>-7.8</td><td>1.2</td><td>-6.4</td><td>1.6E-10</td><td>4.0E-10</td><td>MB231/MB468</td><td>NUC</td></tr>
<tr><td>>hg38_21_+_8433172_8446622_RNA45SN1_45S_50ntFlanks@9602.9623.22|CGAAAGACTAATCGAACCATCT</td><td>>hg38_21_+_8433172_8446622_RNA45SN1_45S_50ntFlanks</td><td>9602</td><td>9623</td><td>22</td><td>CGAAAGACTAATCGAACCATCT</td><td>108.7</td><td>-10.6</td><td>1.7</td><td>-6.4</td><td>1.5E-10</td><td>4.0E-10</td><td>MB231/MB468</td><td>MP</td></tr>
<tr><td>>hg38_21_+_8433172_8446622_RNA45SN1_45S_50ntFlanks@5470.5514.45|CCACGGCCCTGGCGGAGCGCTGAGAAGACGGTCGAACTTGACTAT</td><td>>hg38_21_+_8433172_8446622_RNA45SN1_45S_50ntFlanks</td><td>5470</td><td>5514</td><td>45</td><td>CCACGGCCCTGGCGGAGCGCTGAGAAGACGGTCGAACTTGACTAT</td><td>668.2</td><td>-9.1</td><td>1.4</td><td>-6.4</td><td>1.4E-10</td><td>4.0E-10</td><td>BT20/MB468</td><td>CYTO</td></tr>
<tr><td>>hg38_21_+_8433172_8446622_RNA45SN1_45S_50ntFlanks@12419.12437.19|GATGTCGGCTCTTCCTATC</td><td>>hg38_21_+_8433172_8446622_RNA45SN1_45S_50ntFlanks</td><td>12419</td><td>12437</td><td>19</td><td>GATGTCGGCTCTTCCTATC</td><td>1230.7</td><td>2.4</td><td>0.4</td><td>6.4</td><td>1.2E-10</td><td>4.0E-10</td><td>BT20/MB468</td><td>MITO</td></tr>
<tr><td>>hg38_21_+_8433172_8446622_RNA45SN1_45S_50ntFlanks@11729.11768.40|TAACTATGACTCTCTTAAGGTAGCCAAATGCCTCGTCATC</td><td>>hg38_21_+_8433172_8446622_RNA45SN1_45S_50ntFlanks</td><td>11729</td><td>11768</td><td>40</td><td>TAACTATGACTCTCTTAAGGTAGCCAAATGCCTCGTCATC</td><td>82.5</td><td>-10.2</td><td>1.6</td><td>-6.4</td><td>1.5E-10</td><td>4.0E-10</td><td>MB231/MB468</td><td>MP</td></tr>
<tr><td>>hg38_21_+_8433172_8446622_RNA45SN1_45S_50ntFlanks@12405.12430.26|CTTTTTGATCCTTCGATGTCGGCTCT</td><td>>hg38_21_+_8433172_8446622_RNA45SN1_45S_50ntFlanks</td><td>12405</td><td>12430</td><td>26</td><td>CTTTTTGATCCTTCGATGTCGGCTCT</td><td>1071.9</td><td>-6.7</td><td>1.1</td><td>-6.4</td><td>1.6E-10</td><td>4.0E-10</td><td>MB231/MB468</td><td>NUC</td></tr>
<tr><td>>hg38_21_+_8433172_8446622_RNA45SN1_45S_50ntFlanks@7975.8000.26|CGCGACCTCAGATCAGACGTGGCGAC</td><td>>hg38_21_+_8433172_8446622_RNA45SN1_45S_50ntFlanks</td><td>7975</td><td>8000</td><td>26</td><td>CGCGACCTCAGATCAGACGTGGCGAC</td><td>260.1</td><td>5.1</td><td>0.8</td><td>6.4</td><td>1.6E-10</td><td>4.0E-10</td><td>MB231/MB468</td><td>CYTO</td></tr>
<tr><td>>hg38_21_+_8433172_8446622_RNA45SN1_45S_50ntFlanks@11729.11759.31|TAACTATGACTCTCTTAAGGTAGCCAAATGC</td><td>>hg38_21_+_8433172_8446622_RNA45SN1_45S_50ntFlanks</td><td>11729</td><td>11759</td><td>31</td><td>TAACTATGACTCTCTTAAGGTAGCCAAATGC</td><td>145.8</td><td>-11.1</td><td>1.7</td><td>-6.4</td><td>1.5E-10</td><td>4.0E-10</td><td>MB231/MB468</td><td>MP</td></tr>
<tr><td>>hg38_MT_+_1621_3279_RNR2_16S_50ntFlanks@732.759.28|TAAAACACTGAACTGACAATTAACAGCC</td><td>>hg38_MT_+_1621_3279_RNR2_16S_50ntFlanks</td><td>732</td><td>759</td><td>28</td><td>TAAAACACTGAACTGACAATTAACAGCC</td><td>184.0</td><td>10.7</td><td>1.7</td><td>6.4</td><td>1.5E-10</td><td>4.0E-10</td><td>MB231/MB468</td><td>MP</td></tr>
<tr><td>>hg38_21_+_8433172_8446622_RNA45SN1_45S_50ntFlanks@9858.9879.22|CGGGATGAACCGAACGCCGGGT</td><td>>hg38_21_+_8433172_8446622_RNA45SN1_45S_50ntFlanks</td><td>9858</td><td>9879</td><td>22</td><td>CGGGATGAACCGAACGCCGGGT</td><td>184.7</td><td>-11.4</td><td>1.8</td><td>-6.4</td><td>1.5E-10</td><td>4.0E-10</td><td>MB231/MB468</td><td>MP</td></tr>
<tr><td>>hg38_21_+_8433172_8446622_RNA45SN1_45S_50ntFlanks@5236.5263.28|CACTGACTGGCTCAGCGTGTGCCTACCC</td><td>>hg38_21_+_8433172_8446622_RNA45SN1_45S_50ntFlanks</td><td>5236</td><td>5263</td><td>28</td><td>CACTGACTGGCTCAGCGTGTGCCTACCC</td><td>83.7</td><td>-10.3</td><td>1.6</td><td>-6.4</td><td>1.5E-10</td><td>4.0E-10</td><td>MB231/MB468</td><td>MP</td></tr>
<tr><td>>hg38_21_+_8433172_8446622_RNA45SN1_45S_50ntFlanks@3705.3744.40|TACCTGGTTGATCCTGCCAGTAGCATATGCTTGTCTCAAA</td><td>>hg38_21_+_8433172_8446622_RNA45SN1_45S_50ntFlanks</td><td>3705</td><td>3744</td><td>40</td><td>TACCTGGTTGATCCTGCCAGTAGCATATGCTTGTCTCAAA</td><td>2623.2</td><td>8.3</td><td>1.3</td><td>6.4</td><td>1.4E-10</td><td>4.0E-10</td><td>BT20/MB468</td><td>CYTO</td></tr>
<tr><td>>hg38_21_+_8433172_8446622_RNA45SN1_45S_50ntFlanks@6746.6770.25|CACTTCGAACGCACTTGCGGCCCCG</td><td>>hg38_21_+_8433172_8446622_RNA45SN1_45S_50ntFlanks</td><td>6746</td><td>6770</td><td>25</td><td>CACTTCGAACGCACTTGCGGCCCCG</td><td>1434.1</td><td>-8.2</td><td>1.3</td><td>-6.4</td><td>1.4E-10</td><td>4.0E-10</td><td>BT20/MB468</td><td>CYTO</td></tr>
<tr><td>>hg38_21_+_8433172_8446622_RNA45SN1_45S_50ntFlanks@10402.10435.34|AACATGGGTCAGTCGGTCCTGAGAGATGGGCGAG</td><td>>hg38_21_+_8433172_8446622_RNA45SN1_45S_50ntFlanks</td><td>10402</td><td>10435</td><td>34</td><td>AACATGGGTCAGTCGGTCCTGAGAGATGGGCGAG</td><td>890.8</td><td>-6.0</td><td>0.9</td><td>-6.4</td><td>1.6E-10</td><td>4.0E-10</td><td>MB231/MB468</td><td>CYTO</td></tr>
<tr><td>>hg38_21_+_8433172_8446622_RNA45SN1_45S_50ntFlanks@10420.10453.34|CTGAGAGATGGGCGAGCGCCGTTCCGAAGGGACG</td><td>>hg38_21_+_8433172_8446622_RNA45SN1_45S_50ntFlanks</td><td>10420</td><td>10453</td><td>34</td><td>CTGAGAGATGGGCGAGCGCCGTTCCGAAGGGACG</td><td>4220.3</td><td>-10.8</td><td>1.7</td><td>-6.4</td><td>1.9E-10</td><td>4.0E-10</td><td>BT20/MB468</td><td>NUC</td></tr>
<tr><td>>hg38_21_+_8433172_8446622_RNA45SN1_45S_50ntFlanks@6753.6770.18|AACGCACTTGCGGCCCCG</td><td>>hg38_21_+_8433172_8446622_RNA45SN1_45S_50ntFlanks</td><td>6753</td><td>6770</td><td>18</td><td>AACGCACTTGCGGCCCCG</td><td>264.7</td><td>-11.1</td><td>1.6</td><td>-6.8</td><td>1.2E-11</td><td>4.1E-10</td><td>BT20/MB468</td><td>MP</td></tr>
<tr><td>>hg38_21_+_8433172_8446622_RNA45SN1_45S_50ntFlanks@3985.4004.20|CGGCTTTGGTGACTCTAGAT</td><td>>hg38_21_+_8433172_8446622_RNA45SN1_45S_50ntFlanks</td><td>3985</td><td>4004</td><td>20</td><td>CGGCTTTGGTGACTCTAGAT</td><td>275.5</td><td>-10.2</td><td>1.6</td><td>-6.4</td><td>1.2E-10</td><td>4.1E-10</td><td>BT20/MB468</td><td>MITO</td></tr>
<tr><td>>hg38_21_+_8433172_8446622_RNA45SN1_45S_50ntFlanks@12624.12651.28|CCAATGGGGCGAAGCTACCATCTGTGGG</td><td>>hg38_21_+_8433172_8446622_RNA45SN1_45S_50ntFlanks</td><td>12624</td><td>12651</td><td>28</td><td>CCAATGGGGCGAAGCTACCATCTGTGGG</td><td>137.6</td><td>-11.0</td><td>1.7</td><td>-6.4</td><td>1.5E-10</td><td>4.1E-10</td><td>MB231/MB468</td><td>MP</td></tr>
<tr><td>>hg38_21_+_8433172_8446622_RNA45SN1_45S_50ntFlanks@11573.11603.31|CGACTTAGAACTGGTGCGGACCAGGGGAATC</td><td>>hg38_21_+_8433172_8446622_RNA45SN1_45S_50ntFlanks</td><td>11573</td><td>11603</td><td>31</td><td>CGACTTAGAACTGGTGCGGACCAGGGGAATC</td><td>908.4</td><td>-6.6</td><td>1.0</td><td>-6.4</td><td>1.8E-10</td><td>4.1E-10</td><td>MB231/MB468</td><td>MITO</td></tr>
<tr><td>>hg38_21_+_8433172_8446622_RNA45SN1_45S_50ntFlanks@12605.12625.21|TATGTGCTTGGCTGAGGAGCC</td><td>>hg38_21_+_8433172_8446622_RNA45SN1_45S_50ntFlanks</td><td>12605</td><td>12625</td><td>21</td><td>TATGTGCTTGGCTGAGGAGCC</td><td>7529.9</td><td>-4.6</td><td>0.7</td><td>-6.4</td><td>1.6E-10</td><td>4.1E-10</td><td>MB231/MB468</td><td>CYTO</td></tr>
<tr><td>>hg38_21_+_8433172_8446622_RNA45SN1_45S_50ntFlanks@12222.12241.20|ACAGAAACCTCCCGTGGAGC</td><td>>hg38_21_+_8433172_8446622_RNA45SN1_45S_50ntFlanks</td><td>12222</td><td>12241</td><td>20</td><td>ACAGAAACCTCCCGTGGAGC</td><td>127.1</td><td>-10.9</td><td>1.7</td><td>-6.4</td><td>1.5E-10</td><td>4.1E-10</td><td>MB231/MB468</td><td>MP</td></tr>
<tr><td>>hg38_21_+_8433172_8446622_RNA45SN1_45S_50ntFlanks@12076.12096.21|AGGGGCTCTCGCTTCTGGCGC</td><td>>hg38_21_+_8433172_8446622_RNA45SN1_45S_50ntFlanks</td><td>12076</td><td>12096</td><td>21</td><td>AGGGGCTCTCGCTTCTGGCGC</td><td>128.2</td><td>-10.9</td><td>1.7</td><td>-6.4</td><td>1.5E-10</td><td>4.1E-10</td><td>MB231/MB468</td><td>MP</td></tr>
<tr><td>>MI0000542|hsa-mir-320a&WithFlank&8|-|22244960|22245043@48.69.22(+1U)&[MIMAT0000510&hsa-miR-320a-3p&offsets|0|0(+1U);m-60&8|-|22244975|22244996&offsets|0|0(+1U)]|AAAAGCTGGGTTGAGAGGGCGAT</td><td>>MI0000542|hsa-mir-320a&WithFlank&8|-|22244960|22245043</td><td>48</td><td>69</td><td>22(+1U)</td><td>AAAAGCTGGGTTGAGAGGGCGAT</td><td>10570.5</td><td>6.9</td><td>1.0</td><td>6.8</td><td>1.3E-11</td><td>4.1E-10</td><td>BT20/MB468</td><td>MP</td></tr>
<tr><td>>hg38_21_+_8433172_8446622_RNA45SN1_45S_50ntFlanks@11540.11584.45|CGCGGCGCCTCGCCTCGGCCGGCGCCTAGCAGCCGACTTAGAACT</td><td>>hg38_21_+_8433172_8446622_RNA45SN1_45S_50ntFlanks</td><td>11540</td><td>11584</td><td>45</td><td>CGCGGCGCCTCGCCTCGGCCGGCGCCTAGCAGCCGACTTAGAACT</td><td>4348.6</td><td>7.7</td><td>1.2</td><td>6.4</td><td>1.6E-10</td><td>4.1E-10</td><td>MB231/MB468</td><td>CYTO</td></tr>
<tr><td>>MI0000077|hsa-mir-21&WithFlank&17|+|59841260|59841343@14.35.22(+1A)&[MIMAT0000076&hsa-miR-21-5p&offsets|0|0(+1A);m-1&17|+|59841273|59841294&offsets|0|0(+1A)]|TAGCTTATCAGACTGATGTTGAA</td><td>>MI0000077|hsa-mir-21&WithFlank&17|+|59841260|59841343</td><td>14</td><td>35</td><td>22(+1A)</td><td>TAGCTTATCAGACTGATGTTGAA</td><td>19071.1</td><td>6.5</td><td>1.0</td><td>6.4</td><td>1.8E-10</td><td>4.1E-10</td><td>MB231/MB468</td><td>MITO</td></tr>
<tr><td>>hg38_21_+_8433172_8446622_RNA45SN1_45S_50ntFlanks@8112.8142.31|GGGCGCGGGACATGTGGCGTACGGAAGACCC</td><td>>hg38_21_+_8433172_8446622_RNA45SN1_45S_50ntFlanks</td><td>8112</td><td>8142</td><td>31</td><td>GGGCGCGGGACATGTGGCGTACGGAAGACCC</td><td>72.9</td><td>-10.1</td><td>1.6</td><td>-6.4</td><td>1.6E-10</td><td>4.1E-10</td><td>MB231/MB468</td><td>MP</td></tr>
<tr><td>>hg38_21_+_8433172_8446622_RNA45SN1_45S_50ntFlanks@9572.9604.33|CAAATCGGTCGTCCGACCTGGGTATAGGGGCGA</td><td>>hg38_21_+_8433172_8446622_RNA45SN1_45S_50ntFlanks</td><td>9572</td><td>9604</td><td>33</td><td>CAAATCGGTCGTCCGACCTGGGTATAGGGGCGA</td><td>501.4</td><td>-6.5</td><td>1.0</td><td>-6.4</td><td>1.8E-10</td><td>4.1E-10</td><td>MB231/MB468</td><td>MITO</td></tr>
<tr><td>>hg38_21_+_8433172_8446622_RNA45SN1_45S_50ntFlanks@12943.12972.30|TCGTAGACGACCTGCTTCTGGGTCGGGGTT</td><td>>hg38_21_+_8433172_8446622_RNA45SN1_45S_50ntFlanks</td><td>12943</td><td>12972</td><td>30</td><td>TCGTAGACGACCTGCTTCTGGGTCGGGGTT</td><td>177.0</td><td>-5.1</td><td>0.8</td><td>-6.4</td><td>1.6E-10</td><td>4.1E-10</td><td>MB231/MB468</td><td>CYTO</td></tr>
<tr><td>>hg38_21_+_8433172_8446622_RNA45SN1_45S_50ntFlanks@11718.11736.19|AACGGCGGGAGTAACTATG</td><td>>hg38_21_+_8433172_8446622_RNA45SN1_45S_50ntFlanks</td><td>11718</td><td>11736</td><td>19</td><td>AACGGCGGGAGTAACTATG</td><td>1826.4</td><td>-5.2</td><td>0.8</td><td>-6.4</td><td>1.6E-10</td><td>4.1E-10</td><td>MB231/MB468</td><td>NUC</td></tr>
<tr><td>>MI0000487|hsa-mir-193a&WithFlank&17|+|31559990|31560089@27.48.22&[MIMAT0004614&hsa-miR-193a-5p&offsets|0|0;m-123&17|+|31560016|31560037&offsets|0|0]|TGGGTCTTTGCGGGCGAGATGA</td><td>>MI0000487|hsa-mir-193a&WithFlank&17|+|31559990|31560089</td><td>27</td><td>48</td><td>22</td><td>TGGGTCTTTGCGGGCGAGATGA</td><td>607.8</td><td>10.0</td><td>1.6</td><td>6.4</td><td>1.2E-10</td><td>4.1E-10</td><td>BT20/MB468</td><td>MITO</td></tr>
<tr><td>>hg38_21_+_8433172_8446622_RNA45SN1_45S_50ntFlanks@12087.12110.24|CTTCTGGCGCCAAGCGCCCGGCCG</td><td>>hg38_21_+_8433172_8446622_RNA45SN1_45S_50ntFlanks</td><td>12087</td><td>12110</td><td>24</td><td>CTTCTGGCGCCAAGCGCCCGGCCG</td><td>126.2</td><td>-10.8</td><td>1.7</td><td>-6.4</td><td>1.6E-10</td><td>4.1E-10</td><td>MB231/MB468</td><td>MP</td></tr>
<tr><td>>trnaMT_PheGAA_MT_+_577_647@36.56.21|AAAATGTTTAGACGGGCTCAC</td><td>>trnaMT_PheGAA_MT_+_577_647</td><td>36</td><td>56</td><td>21</td><td>AAAATGTTTAGACGGGCTCAC</td><td>63.2</td><td>6.5</td><td>1.0</td><td>6.4</td><td>1.9E-10</td><td>4.1E-10</td><td>BT20/MB468</td><td>NUC</td></tr>
<tr><td>>hg38_21_+_8433172_8446622_RNA45SN1_45S_50ntFlanks@12958.12992.35|TTCTGGGTCGGGGTTTCGTACGTAGCAGAGCAGCT</td><td>>hg38_21_+_8433172_8446622_RNA45SN1_45S_50ntFlanks</td><td>12958</td><td>12992</td><td>35</td><td>TTCTGGGTCGGGGTTTCGTACGTAGCAGAGCAGCT</td><td>330.6</td><td>7.9</td><td>1.2</td><td>6.4</td><td>1.6E-10</td><td>4.1E-10</td><td>MB231/MB468</td><td>CYTO</td></tr>
<tr><td>>hg38_21_+_8433172_8446622_RNA45SN1_45S_50ntFlanks@8359.8378.20|AAGAACTTTGAAGAGAGAGT</td><td>>hg38_21_+_8433172_8446622_RNA45SN1_45S_50ntFlanks</td><td>8359</td><td>8378</td><td>20</td><td>AAGAACTTTGAAGAGAGAGT</td><td>665.3</td><td>-6.0</td><td>0.9</td><td>-6.4</td><td>2.0E-10</td><td>4.1E-10</td><td>BT20/MB468</td><td>NUC</td></tr>
</table>

<table>
<tr><td>>MI0006444|hsa-mir-1307&WithFlank&10|-|103394247|103394407@87.108.22(+1A)&[MIMAT0005951&hsa-miR-1307-3p&offsets|+1|+1(+1A);m-66&10|-|103394300|103394322&offsets|+1|0(+1A)]|CTCGGCGTGGCGTCGGTCGTGGA</td><td>>MI0006444|hsa-mir-1307&WithFlank&10|-|103394247|103394407</td><td>87</td><td>108</td><td>22(+1A)</td><td>CTCGGCGTGGCGTCGGTCGTGGA</td><td>293.3</td><td>7.8</td><td>1.2</td><td>6.4</td><td>1.5E-10</td><td>4.1E-10</td><td>BT20/MB468</td><td>CYTO</td></tr>
<tr><td>>MI0015866|hsa-mir-4326&WithFlank&20|+|63286802|63286872@17.40.24&[MIMAT0016888&hsa-miR-4326&offsets|0|+4;m-339&20|+|63286818|63286841&offsets|0|0]|TGTTCCTCTGTCTCCCAGACTCTG</td><td>>MI0015866|hsa-mir-4326&WithFlank&20|+|63286802|63286872</td><td>17</td><td>40</td><td>24</td><td>TGTTCCTCTGTCTCCCAGACTCTG</td><td>262.8</td><td>6.5</td><td>1.0</td><td>6.4</td><td>1.7E-10</td><td>4.1E-10</td><td>MB231/MB468</td><td>NUC</td></tr>
<tr><td>>MI0000102|hsa-mir-100&WithFlank&11|-|122152223|122152314@19.40.22(+3U)&[MIMAT0000098&hsa-miR-100-5p&offsets|0|0(+3U);m-27&11|-|122152276|122152296&offsets|0|+1(+3U)]|AACCCGTAGATCCGAACTTGTGTTT</td><td>>MI0000102|hsa-mir-100&WithFlank&11|-|122152223|122152314</td><td>19</td><td>40</td><td>22(+3U)</td><td>AACCCGTAGATCCGAACTTGTGTTT</td><td>139.1</td><td>-10.2</td><td>1.5</td><td>-6.7</td><td>1.8E-11</td><td>4.2E-10</td><td>BT20/MB231</td><td>NUC</td></tr>
<tr><td>>hg38_21_+_8433172_8446622_RNA45SN1_45S_50ntFlanks@4760.4780.21|TCAGATACCGTCGTAGTTCCG</td><td>>hg38_21_+_8433172_8446622_RNA45SN1_45S_50ntFlanks</td><td>4760</td><td>4780</td><td>21</td><td>TCAGATACCGTCGTAGTTCCG</td><td>5074.6</td><td>-6.8</td><td>1.1</td><td>-6.4</td><td>1.7E-10</td><td>4.2E-10</td><td>MB231/MB468</td><td>CYTO</td></tr>
<tr><td>>hg38_21_+_8433172_8446622_RNA45SN1_45S_50ntFlanks@10362.10388.27|AAGGCCGAAGTGGAGAAGGGTTCCATG</td><td>>hg38_21_+_8433172_8446622_RNA45SN1_45S_50ntFlanks</td><td>10362</td><td>10388</td><td>27</td><td>AAGGCCGAAGTGGAGAAGGGTTCCATG</td><td>4264.1</td><td>-9.9</td><td>1.6</td><td>-6.4</td><td>1.7E-10</td><td>4.2E-10</td><td>MB231/MB468</td><td>NUC</td></tr>
<tr><td>>hg38_21_+_8433172_8446622_RNA45SN1_45S_50ntFlanks@10379.10424.46|GGGTTCCATGTGAACAGCAGTTGAACATGGGTCAGTCGGTCCTGAG</td><td>>hg38_21_+_8433172_8446622_RNA45SN1_45S_50ntFlanks</td><td>10379</td><td>10424</td><td>46</td><td>GGGTTCCATGTGAACAGCAGTTGAACATGGGTCAGTCGGTCCTGAG</td><td>195.7</td><td>11.4</td><td>1.7</td><td>6.8</td><td>1.3E-11</td><td>4.2E-10</td><td>BT20/MB468</td><td>MP</td></tr>
<tr><td>>hg38_21_+_8433172_8446622_RNA45SN1_45S_50ntFlanks@12934.12959.26|CTAAACCATTCGTAGACGACCTGCTT</td><td>>hg38_21_+_8433172_8446622_RNA45SN1_45S_50ntFlanks</td><td>12934</td><td>12959</td><td>26</td><td>CTAAACCATTCGTAGACGACCTGCTT</td><td>172.8</td><td>-10.5</td><td>1.6</td><td>-6.8</td><td>1.3E-11</td><td>4.2E-10</td><td>BT20/MB468</td><td>MP</td></tr>
<tr><td>>hg38_1_-_228634819_228635039_RNA5S12_5S_50ntFlanks@153.171.19|ACCGGGTGCTGTAGGCTTT</td><td>>hg38_1_-_228634819_228635039_RNA5S12_5S_50ntFlanks</td><td>153</td><td>171</td><td>19</td><td>ACCGGGTGCTGTAGGCTTT</td><td>18630.9</td><td>6.2</td><td>1.0</td><td>6.4</td><td>1.2E-10</td><td>4.2E-10</td><td>BT20/MB468</td><td>MITO</td></tr>
<tr><td>>hg38_21_+_8433172_8446622_RNA45SN1_45S_50ntFlanks@12073.12092.20|CCGAGGGGCTCTCGCTTCTG</td><td>>hg38_21_+_8433172_8446622_RNA45SN1_45S_50ntFlanks</td><td>12073</td><td>12092</td><td>20</td><td>CCGAGGGGCTCTCGCTTCTG</td><td>930.1</td><td>-5.1</td><td>0.8</td><td>-6.4</td><td>1.7E-10</td><td>4.2E-10</td><td>MB231/MB468</td><td>CYTO</td></tr>
<tr><td>>hg38_21_+_8433172_8446622_RNA45SN1_45S_50ntFlanks@11569.11606.38|CAGCCGACTTAGAACTGGTGCGGACCAGGGGAATCCGA</td><td>>hg38_21_+_8433172_8446622_RNA45SN1_45S_50ntFlanks</td><td>11569</td><td>11606</td><td>38</td><td>CAGCCGACTTAGAACTGGTGCGGACCAGGGGAATCCGA</td><td>142.0</td><td>-11.0</td><td>1.7</td><td>-6.4</td><td>1.6E-10</td><td>4.2E-10</td><td>MB231/MB468</td><td>MP</td></tr>
<tr><td>>hg38_21_+_8433172_8446622_RNA45SN1_45S_50ntFlanks@6700.6720.21|CTAGCTGCGAGAATTAATGTG</td><td>>hg38_21_+_8433172_8446622_RNA45SN1_45S_50ntFlanks</td><td>6700</td><td>6720</td><td>21</td><td>CTAGCTGCGAGAATTAATGTG</td><td>14533.4</td><td>-6.0</td><td>0.9</td><td>-6.8</td><td>1.3E-11</td><td>4.2E-10</td><td>BT20/MB468</td><td>MP</td></tr>
<tr><td>>MI0000783|hsa-mir-375&WithFlank&2|-|219001639|219001714@46.67.22&[MIMAT0000728&hsa-miR-375-3p&offsets|0|0;m-92&2|-|219001648|219001669&offsets|0|0]|TTTGTTCGTTCGGCTCGCGTGA</td><td>>MI0000783|hsa-mir-375&WithFlank&2|-|219001639|219001714</td><td>46</td><td>67</td><td>22</td><td>TTTGTTCGTTCGGCTCGCGTGA</td><td>228.9</td><td>7.3</td><td>1.1</td><td>6.8</td><td>1.1E-11</td><td>4.2E-10</td><td>BT20/MB231</td><td>CYTO</td></tr>
<tr><td>>hg38_21_+_8433172_8446622_RNA45SN1_45S_50ntFlanks@5187.5205.19|AATAACAGGTCTGTGATGC</td><td>>hg38_21_+_8433172_8446622_RNA45SN1_45S_50ntFlanks</td><td>5187</td><td>5205</td><td>19</td><td>AATAACAGGTCTGTGATGC</td><td>197.4</td><td>3.1</td><td>0.5</td><td>6.4</td><td>1.7E-10</td><td>4.2E-10</td><td>MB231/MB468</td><td>CYTO</td></tr>
<tr><td>>hg38_21_+_8433172_8446622_RNA45SN1_45S_50ntFlanks@11785.11804.20|TGAATGGATGAACGAGATTC</td><td>>hg38_21_+_8433172_8446622_RNA45SN1_45S_50ntFlanks</td><td>11785</td><td>11804</td><td>20</td><td>TGAATGGATGAACGAGATTC</td><td>6564.8</td><td>-6.0</td><td>0.9</td><td>-6.8</td><td>1.3E-11</td><td>4.2E-10</td><td>BT20/MB468</td><td>MP</td></tr>
<tr><td>>MI0000446|hsa-mir-125b-1&WithFlank&11|-|122099751|122099850@21.42.22(+1A);MI0000470|hsa-mir-125b-2&WithFlank&21|+|16590231|16590331@23.44.22(+1A)&[MIMAT0000423&hsa-miR-125b-5p&offsets|0|0(+1A);m-67&11|-|122099809|122099830&offsets|0|0(+1A)];[MIMAT0000423_1&hsa-miR-125b-5p&offsets|0|0(+1A);m-68&21|+|16590253|16590274&offsets|0|0(+1A)]|TCCCTGAGACCCTAACTTGTGAA</td><td>>MI0000446|hsa-mir-125b-1&WithFlank&11|-|122099751|122099850</td><td>21</td><td>42</td><td>22(+1A)</td><td>TCCCTGAGACCCTAACTTGTGAA</td><td>5812.4</td><td>-6.7</td><td>1.0</td><td>-6.8</td><td>1.5E-11</td><td>4.2E-10</td><td>BT20/MB231</td><td>MITO</td></tr>
<tr><td>>hg38_21_+_8433172_8446622_RNA45SN1_45S_50ntFlanks@4367.4389.23|CAGTTAAAAAGCTCGTAGTTGGA</td><td>>hg38_21_+_8433172_8446622_RNA45SN1_45S_50ntFlanks</td><td>4367</td><td>4389</td><td>23</td><td>CAGTTAAAAAGCTCGTAGTTGGA</td><td>1388.1</td><td>-6.4</td><td>1.0</td><td>-6.4</td><td>1.5E-10</td><td>4.2E-10</td><td>BT20/MB468</td><td>CYTO</td></tr>
<tr><td>>hg38_21_+_8433172_8446622_RNA45SN1_45S_50ntFlanks@9564.9588.25|CTGACGTGCAAATCGGTCGTCCGAC</td><td>>hg38_21_+_8433172_8446622_RNA45SN1_45S_50ntFlanks</td><td>9564</td><td>9588</td><td>25</td><td>CTGACGTGCAAATCGGTCGTCCGAC</td><td>84.3</td><td>-10.3</td><td>1.6</td><td>-6.4</td><td>1.6E-10</td><td>4.2E-10</td><td>MB231/MB468</td><td>MP</td></tr>
<tr><td>>hg38_21_+_8433172_8446622_RNA45SN1_45S_50ntFlanks@12927.12951.25|CCTGGCGCTAAACCATTCGTAGACG</td><td>>hg38_21_+_8433172_8446622_RNA45SN1_45S_50ntFlanks</td><td>12927</td><td>12951</td><td>25</td><td>CCTGGCGCTAAACCATTCGTAGACG</td><td>533.1</td><td>-8.4</td><td>1.3</td><td>-6.4</td><td>1.9E-10</td><td>4.2E-10</td><td>MB231/MB468</td><td>MITO</td></tr>
<tr><td>>MI0000101|hsa-mir-99a&WithFlank&21|+|16539083|16539175@19.40.22(+1U)&[MIMAT0000097&hsa-miR-99a-5p&offsets|0|0(+1U);m-64&21|+|16539101|16539121&offsets|0|+1(+1U)]|AACCCGTAGATCCGATCTTGTGT</td><td>>MI0000101|hsa-mir-99a&WithFlank&21|+|16539083|16539175</td><td>19</td><td>40</td><td>22(+1U)</td><td>AACCCGTAGATCCGATCTTGTGT</td><td>145.6</td><td>-10.3</td><td>1.5</td><td>-6.7</td><td>1.8E-11</td><td>4.2E-10</td><td>BT20/MB231</td><td>NUC</td></tr>
<tr><td>>am_trnaMT_LysTTT_MT_+_8295_8364@28.52.25,trnalookalike7_LysTTT_1_+_568844_568913@28.52.25|TTTTAAGTTAAAGATTAAGAGAACC</td><td>>am_trnaMT_LysTTT_MT_+_8295_8364</td><td>28</td><td>52</td><td>25</td><td>TTTTAAGTTAAAGATTAAGAGAACC</td><td>711.4</td><td>8.7</td><td>1.4</td><td>6.4</td><td>1.5E-10</td><td>4.2E-10</td><td>BT20/MB468</td><td>CYTO</td></tr>
<tr><td>>hg38_21_+_8433172_8446622_RNA45SN1_45S_50ntFlanks@11718.11739.22|AACGGCGGGAGTAACTATGACT</td><td>>hg38_21_+_8433172_8446622_RNA45SN1_45S_50ntFlanks</td><td>11718</td><td>11739</td><td>22</td><td>AACGGCGGGAGTAACTATGACT</td><td>86.6</td><td>-10.3</td><td>1.6</td><td>-6.4</td><td>1.6E-10</td><td>4.2E-10</td><td>MB231/MB468</td><td>MP</td></tr>
<tr><td>>hg38_21_+_8433172_8446622_RNA45SN1_45S_50ntFlanks@11575.11593.19|ACTTAGAACTGGTGCGGAC</td><td>>hg38_21_+_8433172_8446622_RNA45SN1_45S_50ntFlanks</td><td>11575</td><td>11593</td><td>19</td><td>ACTTAGAACTGGTGCGGAC</td><td>1453.0</td><td>-8.9</td><td>1.4</td><td>-6.4</td><td>1.6E-10</td><td>4.2E-10</td><td>MB231/MB468</td><td>MP</td></tr>
<tr><td>>hg38_21_+_8433172_8446622_RNA45SN1_45S_50ntFlanks@10808.10830.23|CAATGTAGGTAAGGGAAGTCGGC</td><td>>hg38_21_+_8433172_8446622_RNA45SN1_45S_50ntFlanks</td><td>10808</td><td>10830</td><td>23</td><td>CAATGTAGGTAAGGGAAGTCGGC</td><td>866.7</td><td>-9.4</td><td>1.5</td><td>-6.4</td><td>2.0E-10</td><td>4.2E-10</td><td>BT20/MB468</td><td>NUC</td></tr>
<tr><td>>MI0003591|hsa-mir-584&WithFlank&5|-|149062307|149062415@22.40.19&[MIMAT0003249&hsa-miR-584-5p&offsets|0|-3;m-167&5|-|149062374|149062394&offsets|0|-2]|TTATGGTTTGCCTGGGACT</td><td>>MI0003591|hsa-mir-584&WithFlank&5|-|149062307|149062415</td><td>22</td><td>40</td><td>19</td><td>TTATGGTTTGCCTGGGACT</td><td>1422.1</td><td>6.7</td><td>1.0</td><td>6.8</td><td>1.3E-11</td><td>4.2E-10</td><td>BT20/MB468</td><td>MP</td></tr>
<tr><td>>MI0001641|hsa-mir-429&WithFlank&1|+|1168999|1169093@57.78.22&[MIMAT0001536&hsa-miR-429&offsets|0|0;m-207&1|+|1169055|1169076&offsets|0|0]|TAATACTGTCTGGTAAAACCGT</td><td>>MI0001641|hsa-mir-429&WithFlank&1|+|1168999|1169093</td><td>57</td><td>78</td><td>22</td><td>TAATACTGTCTGGTAAAACCGT</td><td>1734.7</td><td>-6.0</td><td>0.9</td><td>-6.8</td><td>1.3E-11</td><td>4.2E-10</td><td>BT20/MB468</td><td>MP</td></tr>
<tr><td>>hg38_21_+_8433172_8446622_RNA45SN1_45S_50ntFlanks@6776.6802.27|CTCCCGGGGCTACGCCTGTCTGAGCGT</td><td>>hg38_21_+_8433172_8446622_RNA45SN1_45S_50ntFlanks</td><td>6776</td><td>6802</td><td>27</td><td>CTCCCGGGGCTACGCCTGTCTGAGCGT</td><td>1218.3</td><td>3.6</td><td>0.6</td><td>6.4</td><td>1.2E-10</td><td>4.2E-10</td><td>BT20/MB468</td><td>MITO</td></tr>
<tr><td>>hg38_21_+_8433172_8446622_RNA45SN1_45S_50ntFlanks@7973.7994.22|GACGCGACCTCAGATCAGACGT</td><td>>hg38_21_+_8433172_8446622_RNA45SN1_45S_50ntFlanks</td><td>7973</td><td>7994</td><td>22</td><td>GACGCGACCTCAGATCAGACGT</td><td>1758.6</td><td>5.1</td><td>0.8</td><td>6.4</td><td>1.2E-10</td><td>4.2E-10</td><td>BT20/MB468</td><td>MITO</td></tr>
<tr><td>>hg38_21_+_8433172_8446622_RNA45SN1_45S_50ntFlanks@10458.10480.23|ATGGCCTCCGTTGCCCTCGGCCG</td><td>>hg38_21_+_8433172_8446622_RNA45SN1_45S_50ntFlanks</td><td>10458</td><td>10480</td><td>23</td><td>ATGGCCTCCGTTGCCCTCGGCCG</td><td>142.1</td><td>-11.0</td><td>1.7</td><td>-6.4</td><td>1.6E-10</td><td>4.3E-10</td><td>MB231/MB468</td><td>MP</td></tr>
<tr><td>>hg38_21_+_8433172_8446622_RNA45SN1_45S_50ntFlanks@11573.11605.33|CGACTTAGAACTGGTGCGGACCAGGGGAATCCG</td><td>>hg38_21_+_8433172_8446622_RNA45SN1_45S_50ntFlanks</td><td>11573</td><td>11605</td><td>33</td><td>CGACTTAGAACTGGTGCGGACCAGGGGAATCCG</td><td>600.3</td><td>-6.7</td><td>1.1</td><td>-6.4</td><td>1.9E-10</td><td>4.3E-10</td><td>MB231/MB468</td><td>MITO</td></tr>
<tr><td>>hg38_21_+_8433172_8446622_RNA45SN1_45S_50ntFlanks@8475.8492.18|CCCGGCGGATCTTTCCCG</td><td>>hg38_21_+_8433172_8446622_RNA45SN1_45S_50ntFlanks</td><td>8475</td><td>8492</td><td>18</td><td>CCCGGCGGATCTTTCCCG</td><td>238.1</td><td>-10.0</td><td>1.6</td><td>-6.4</td><td>1.2E-10</td><td>4.3E-10</td><td>BT20/MB468</td><td>MITO</td></tr>
<tr><td>>MI0000264|hsa-mir-7-2&WithFlank&15|+|88611819|88611940@38.58.21;MI0000265|hsa-mir-7-3&WithFlank&19|+|4770664|4770785@37.57.21;MI0000263|hsa-mir-7-1&WithFlank&9|-|83969742|83969863@30.50.21&[MIMAT0000252&hsa-miR-7-5p&offsets|0|-3;m-400&15|+|88611856|88611879&offsets|0|-3];[MIMAT0000252_1&hsa-miR-7-5p&offsets|0|-3;m-402&19|+|4770700|4770723&offsets|0|-3];[MIMAT0000252_2&hsa-miR-7-5p&offsets|0|-3;m-401&9|-|83969811|83969834&offsets|0|-3]|TGGAAGACTAGTGATTTTGTT</td><td>>MI0000264|hsa-mir-7-2&WithFlank&15|+|88611819|88611940</td><td>38</td><td>58</td><td>21</td><td>TGGAAGACTAGTGATTTTGTT</td><td>1290.5</td><td>6.5</td><td>1.0</td><td>6.4</td><td>1.9E-10</td><td>4.3E-10</td><td>MB231/MB468</td><td>MITO</td></tr>
<tr><td>>hg38_21_+_8433172_8446622_RNA45SN1_45S_50ntFlanks@11876.11909.34|AAGACCCTGTTGAGCTTGACTCTAGTCTGGCACG</td><td>>hg38_21_+_8433172_8446622_RNA45SN1_45S_50ntFlanks</td><td>11876</td><td>11909</td><td>34</td><td>AAGACCCTGTTGAGCTTGACTCTAGTCTGGCACG</td><td>525.8</td><td>-8.1</td><td>1.3</td><td>-6.4</td><td>1.9E-10</td><td>4.3E-10</td><td>MB231/MB468</td><td>MITO</td></tr>
<tr><td>>hg38_21_+_8433172_8446622_RNA45SN1_45S_50ntFlanks@4982.5003.22|ACGGACAGGATTGACAGATTGA</td><td>>hg38_21_+_8433172_8446622_RNA45SN1_45S_50ntFlanks</td><td>4982</td><td>5003</td><td>22</td><td>ACGGACAGGATTGACAGATTGA</td><td>457.3</td><td>-12.6</td><td>2.0</td><td>-6.4</td><td>1.9E-10</td><td>4.3E-10</td><td>MB231/MB468</td><td>MITO</td></tr>
<tr><td>>am_MI0000809|hsa-mir-151a&WithFlank&8|-|140732558|140732659@53.72.20(+1U)&[MIMAT0000757&hsa-miR-151a-3p&offsets|0|-1(+1U);m-30&8|-|140732587|140732607&offsets|0|-1(+1U)]|CTAGACTGAAGCTCCTTGAGT</td><td>>am_MI0000809|hsa-mir-151a&WithFlank&8|-|140732558|140732659</td><td>53</td><td>72</td><td>20(+1U)</td><td>CTAGACTGAAGCTCCTTGAGT</td><td>2647.0</td><td>4.7</td><td>0.7</td><td>6.4</td><td>1.6E-10</td><td>4.3E-10</td><td>MB231/MB468</td><td>MP</td></tr>
<tr><td>>hg38_21_+_8433172_8446622_RNA45SN1_45S_50ntFlanks@11641.11665.25|CGGCGGGTGTTGACGCGATGTGATT</td><td>>hg38_21_+_8433172_8446622_RNA45SN1_45S_50ntFlanks</td><td>11641</td><td>11665</td><td>25</td><td>CGGCGGGTGTTGACGCGATGTGATT</td><td>6960.2</td><td>-10.9</td><td>1.7</td><td>-6.4</td><td>2.0E-10</td><td>4.3E-10</td><td>BT20/MB468</td><td>NUC</td></tr>
<tr><td>>hg38_21_+_8433172_8446622_RNA45SN1_45S_50ntFlanks@10841.10875.35|CGTAACTTCGGGATAAGGATTGGCTCTAAGGGCTG</td><td>>hg38_21_+_8433172_8446622_RNA45SN1_45S_50ntFlanks</td><td>10841</td><td>10875</td><td>35</td><td>CGTAACTTCGGGATAAGGATTGGCTCTAAGGGCTG</td><td>887.0</td><td>-4.7</td><td>0.7</td><td>-6.4</td><td>1.7E-10</td><td>4.3E-10</td><td>MB231/MB468</td><td>CYTO</td></tr>
<tr><td>>hg38_21_+_8433172_8446622_RNA45SN1_45S_50ntFlanks@8176.8209.34|CTTCTGATCGAGGCCCAGCCCGTGGACGGTGTGA</td><td>>hg38_21_+_8433172_8446622_RNA45SN1_45S_50ntFlanks</td><td>8176</td><td>8209</td><td>34</td><td>CTTCTGATCGAGGCCCAGCCCGTGGACGGTGTGA</td><td>160.7</td><td>3.5</td><td>0.5</td><td>6.4</td><td>1.7E-10</td><td>4.3E-10</td><td>MB231/MB468</td><td>CYTO</td></tr>
<tr><td>>hg38_1_-_228634819_228635039_RNA5S12_5S_50ntFlanks@51.84.34|GTCTACGGCCATACCACCCTGAACGCGCCCGATC</td><td>>hg38_1_-_228634819_228635039_RNA5S12_5S_50ntFlanks</td><td>51</td><td>84</td><td>34</td><td>GTCTACGGCCATACCACCCTGAACGCGCCCGATC</td><td>5543.4</td><td>5.2</td><td>0.8</td><td>6.4</td><td>1.3E-10</td><td>4.3E-10</td><td>BT20/MB468</td><td>MITO</td></tr>
<tr><td>>hg38_21_+_8433172_8446622_RNA45SN1_45S_50ntFlanks@4069.4093.25|CTTTCGATGGTAGTCGCCGTGCCTA</td><td>>hg38_21_+_8433172_8446622_RNA45SN1_45S_50ntFlanks</td><td>4069</td><td>4093</td><td>25</td><td>CTTTCGATGGTAGTCGCCGTGCCTA</td><td>139.2</td><td>-5.8</td><td>0.9</td><td>-6.4</td><td>1.7E-10</td><td>4.3E-10</td><td>MB231/MB468</td><td>CYTO</td></tr>
<tr><td>>hg38_21_+_8433172_8446622_RNA45SN1_45S_50ntFlanks@7991.8016.26|ACGTGGCGACCCGCTGAATTTAAGCA</td><td>>hg38_21_+_8433172_8446622_RNA45SN1_45S_50ntFlanks</td><td>7991</td><td>8016</td><td>26</td><td>ACGTGGCGACCCGCTGAATTTAAGCA</td><td>2064.2</td><td>-3.5</td><td>0.6</td><td>-6.4</td><td>1.9E-10</td><td>4.3E-10</td><td>MB231/MB468</td><td>MITO</td></tr>
<tr><td>>hg38_21_+_8433172_8446622_RNA45SN1_45S_50ntFlanks@3854.3898.45|ACTGTGGTAATTCTAGAGCTAATACATGCCGACGGGCGCTGACCC</td><td>>hg38_21_+_8433172_8446622_RNA45SN1_45S_50ntFlanks</td><td>3854</td><td>3898</td><td>45</td><td>ACTGTGGTAATTCTAGAGCTAATACATGCCGACGGGCGCTGACCC</td><td>509.8</td><td>-8.7</td><td>1.4</td><td>-6.4</td><td>1.5E-10</td><td>4.3E-10</td><td>BT20/MB468</td><td>CYTO</td></tr>
<tr><td>>hg38_21_+_8433172_8446622_RNA45SN1_45S_50ntFlanks@12959.12990.32|TCTGGGTCGGGGTTTCGTACGTAGCAGAGCAG</td><td>>hg38_21_+_8433172_8446622_RNA45SN1_45S_50ntFlanks</td><td>12959</td><td>12990</td><td>32</td><td>TCTGGGTCGGGGTTTCGTACGTAGCAGAGCAG</td><td>284.6</td><td>-7.6</td><td>1.2</td><td>-6.4</td><td>1.7E-10</td><td>4.3E-10</td><td>MB231/MB468</td><td>NUC</td></tr>
<tr><td>>MI0000114|hsa-mir-107&WithFlank&10|-|89592741|89592833@56.76.21;MI0000108|hsa-mir-103a-2&WithFlank&20|+|3917488|3917577@54.74.21;MI0000109|hsa-mir-103a-1&WithFlank&5|-|168560890|168560979@54.74.21&[MIMAT0000104&hsa-miR-107&offsets|0|-2;m-78&10|-|89592758|89592778&offsets|0|0];[MIMAT0000101&hsa-miR-103a-3p&offsets|0|-2;m-12&20|+|3917541|3917563&offsets|0|-2];[MIMAT0000101_1&hsa-miR-103a-3p&offsets|0|-2;m-13&5|-|168560904|168560926&offsets|0|-2;m-44220&5|-|168560908|168560926&offsets|0|+2]|AGCAGCATTGTACAGGGCTAT</td><td>>MI0000114|hsa-mir-107&WithFlank&10|-|89592741|89592833</td><td>56</td><td>76</td><td>21</td><td>AGCAGCATTGTACAGGGCTAT</td><td>423.6</td><td>6.4</td><td>1.0</td><td>6.4</td><td>1.7E-10</td><td>4.3E-10</td><td>MB231/MB468</td><td>CYTO</td></tr>
<tr><td>>hg38_21_+_8433172_8446622_RNA45SN1_45S_50ntFlanks@5236.5267.32|CACTGACTGGCTCAGCGTGTGCCTACCCTACG</td><td>>hg38_21_+_8433172_8446622_RNA45SN1_45S_50ntFlanks</td><td>5236</td><td>5267</td><td>32</td><td>CACTGACTGGCTCAGCGTGTGCCTACCCTACG</td><td>464.2</td><td>-8.2</td><td>1.3</td><td>-6.4</td><td>1.6E-10</td><td>4.3E-10</td><td>MB231/MB468</td><td>MP</td></tr>
</table>

<table>
<tr><th></th><th></th><th></th><th></th><th></th><th></th><th></th><th></th><th></th><th></th><th></th><th></th><th></th><th></th></tr>
<tr><td>>hg38_21_+_8433172_8446622_RNA45SN1_45S_50ntFlanks@9903.9928.26|ATCAGACCCCAGAAAAGGTGTTGGTT</td><td>>hg38_21_+_8433172_8446622_RNA45SN1_45S_50ntFlanks</td><td>9903</td><td>9928</td><td>26</td><td>ATCAGACCCCAGAAAAGGTGTTGGTT</td><td>177.5</td><td>-9.6</td><td>1.5</td><td>-6.4</td><td>1.3E-10</td><td>4.3E-10</td><td>BT20/MB468</td><td>MITO</td></tr>
<tr><td>>MI0003560|hsa-mir-92b&WithFlank&1|+|155195171|155195278@67.86.20&[MIMAT0003218&hsa-miR-92b-3p&offsets|0|-2;m-115&1|+|155195237|155195258&offsets|0|-2]|TATTGCACTCGTCCCGGCCT</td><td>>MI0003560|hsa-mir-92b&WithFlank&1|+|155195171|155195278</td><td>67</td><td>86</td><td>20</td><td>TATTGCACTCGTCCCGGCCT</td><td>107.7</td><td>3.5</td><td>0.5</td><td>6.4</td><td>2.0E-10</td><td>4.3E-10</td><td>BT20/MB468</td><td>NUC</td></tr>
<tr><td>>hg38_21_+_8433172_8446622_RNA45SN1_45S_50ntFlanks@9570.9603.34|TGCAAATCGGTCGTCCGACCTGGGTATAGGGGCG</td><td>>hg38_21_+_8433172_8446622_RNA45SN1_45S_50ntFlanks</td><td>9570</td><td>9603</td><td>34</td><td>TGCAAATCGGTCGTCCGACCTGGGTATAGGGGCG</td><td>768.9</td><td>-5.0</td><td>0.8</td><td>-6.4</td><td>2.0E-10</td><td>4.3E-10</td><td>BT20/MB468</td><td>NUC</td></tr>
<tr><td>>hg38_21_+_8433172_8446622_RNA45SN1_45S_50ntFlanks@3777.3801.25|CGGTACAGTGAAACTGCGAATGGCT</td><td>>hg38_21_+_8433172_8446622_RNA45SN1_45S_50ntFlanks</td><td>3777</td><td>3801</td><td>25</td><td>CGGTACAGTGAAACTGCGAATGGCT</td><td>1067.0</td><td>-5.5</td><td>0.9</td><td>-6.4</td><td>1.7E-10</td><td>4.3E-10</td><td>MB231/MB468</td><td>NUC</td></tr>
<tr><td>>hg38_21_+_8433172_8446622_RNA45SN1_45S_50ntFlanks@12459.12477.19|AGCGTTGGATTGTTCACCC</td><td>>hg38_21_+_8433172_8446622_RNA45SN1_45S_50ntFlanks</td><td>12459</td><td>12477</td><td>19</td><td>AGCGTTGGATTGTTCACCC</td><td>613.8</td><td>-4.2</td><td>0.7</td><td>-6.4</td><td>1.7E-10</td><td>4.3E-10</td><td>MB231/MB468</td><td>NUC</td></tr>
<tr><td>>hg38_21_+_8433172_8446622_RNA45SN1_45S_50ntFlanks@7984.8015.32|AGATCAGACGTGGCGACCCGCTGAATTTAAGC</td><td>>hg38_21_+_8433172_8446622_RNA45SN1_45S_50ntFlanks</td><td>7984</td><td>8015</td><td>32</td><td>AGATCAGACGTGGCGACCCGCTGAATTTAAGC</td><td>64.0</td><td>-9.9</td><td>1.5</td><td>-6.4</td><td>1.6E-10</td><td>4.3E-10</td><td>MB231/MB468</td><td>MP</td></tr>
<tr><td>>hg38_21_+_8433172_8446622_RNA45SN1_45S_50ntFlanks@12627.12651.25|ATGGGGCGAAGCTACCATCTGTGGG</td><td>>hg38_21_+_8433172_8446622_RNA45SN1_45S_50ntFlanks</td><td>12627</td><td>12651</td><td>25</td><td>ATGGGGCGAAGCTACCATCTGTGGG</td><td>134.7</td><td>-9.2</td><td>1.4</td><td>-6.4</td><td>1.3E-10</td><td>4.3E-10</td><td>BT20/MB468</td><td>MITO</td></tr>
<tr><td>>hg38_21_+_8433172_8446622_RNA45SN1_45S_50ntFlanks@6651.6695.45|CGACTCTTAGCGGTGGATCACTCGGCTCGTGCGTCGATGAAGAAC</td><td>>hg38_21_+_8433172_8446622_RNA45SN1_45S_50ntFlanks</td><td>6651</td><td>6695</td><td>45</td><td>CGACTCTTAGCGGTGGATCACTCGGCTCGTGCGTCGATGAAGAAC</td><td>25088.3</td><td>6.2</td><td>1.0</td><td>6.4</td><td>1.3E-10</td><td>4.3E-10</td><td>BT20/MB468</td><td>MITO</td></tr>
<tr><td>>trna129_MetCAT_6_-_27870271_27870342@31.51.21,trna142_MetCAT_6_-_27560600_27560671@31.51.21,trna150_MetCAT_6_-_27300764_27300835@31.51.21,trna169_MetCAT_6_-_26330529_26330600@31.51.21,trna171_MetCAT_6_-_26313352_26313423@31.51.21,trna20_MetCAT_17_-_80452597_80452668@31.51.21,trna2_MetCAT_6_+_26286754_26286825@31.51.21,trna32_MetCAT_1_+_153643726_153643797@31.51.21,trna61_MetCAT_6_+_27745664_27745735@31.51.21|CCCATAACCCAGAGGTCGATG</td><td>>trna129_MetCAT_6_-_27870271_27870342</td><td>31</td><td>51</td><td>21</td><td>CCCATAACCCAGAGGTCGATG</td><td>258.5</td><td>-10.1</td><td>1.6</td><td>-6.4</td><td>1.3E-10</td><td>4.3E-10</td><td>BT20/MB468</td><td>MITO</td></tr>
<tr><td>>hg38_21_+_8433172_8446622_RNA45SN1_45S_50ntFlanks@4297.4321.25|CATTGGAGGGCAAGTCTGGTGCCAG</td><td>>hg38_21_+_8433172_8446622_RNA45SN1_45S_50ntFlanks</td><td>4297</td><td>4321</td><td>25</td><td>CATTGGAGGGCAAGTCTGGTGCCAG</td><td>213.8</td><td>-9.8</td><td>1.5</td><td>-6.4</td><td>1.3E-10</td><td>4.4E-10</td><td>BT20/MB468</td><td>MITO</td></tr>
<tr><td>>hg38_21_+_8433172_8446622_RNA45SN1_45S_50ntFlanks@9454.9503.50|CCGCGCCGGGGAGGTGGAGCACGAGCGCACGTGTTAGGACCCGAAAGATG</td><td>>hg38_21_+_8433172_8446622_RNA45SN1_45S_50ntFlanks</td><td>9454</td><td>9503</td><td>50</td><td>CCGCGCCGGGGAGGTGGAGCACGAGCGCACGTGTTAGGACCCGAAAGATG</td><td>3961.4</td><td>-7.8</td><td>1.2</td><td>-6.4</td><td>1.3E-10</td><td>4.4E-10</td><td>BT20/MB468</td><td>MITO</td></tr>
<tr><td>>hg38_21_+_8433172_8446622_RNA45SN1_45S_50ntFlanks@6779.6802.24|CCGGGGCTACGCCTGTCTGAGCGT</td><td>>hg38_21_+_8433172_8446622_RNA45SN1_45S_50ntFlanks</td><td>6779</td><td>6802</td><td>24</td><td>CCGGGGCTACGCCTGTCTGAGCGT</td><td>895.7</td><td>4.0</td><td>0.6</td><td>6.4</td><td>1.3E-10</td><td>4.4E-10</td><td>BT20/MB468</td><td>MITO</td></tr>
<tr><td>>hg38_21_+_8433172_8446622_RNA45SN1_45S_50ntFlanks@3860.3898.39|GTAATTCTAGAGCTAATACATGCCGACGGGCGCTGACCC</td><td>>hg38_21_+_8433172_8446622_RNA45SN1_45S_50ntFlanks</td><td>3860</td><td>3898</td><td>39</td><td>GTAATTCTAGAGCTAATACATGCCGACGGGCGCTGACCC</td><td>136.0</td><td>-9.2</td><td>1.4</td><td>-6.4</td><td>1.3E-10</td><td>4.4E-10</td><td>BT20/MB468</td><td>MITO</td></tr>
<tr><td>>hg38_21_+_8433172_8446622_RNA45SN1_45S_50ntFlanks@10317.10341.25|GTGCAGATCTTGGTGGTAGTAGCAA</td><td>>hg38_21_+_8433172_8446622_RNA45SN1_45S_50ntFlanks</td><td>10317</td><td>10341</td><td>25</td><td>GTGCAGATCTTGGTGGTAGTAGCAA</td><td>80.4</td><td>-10.2</td><td>1.6</td><td>-6.4</td><td>1.7E-10</td><td>4.4E-10</td><td>MB231/MB468</td><td>MP</td></tr>
<tr><td>>hg38_21_+_8433172_8446622_RNA45SN1_45S_50ntFlanks@11210.11254.45|GGCGACTCTGGACGCGAGCCGGGCCCTTCCCGTGGATCGCCCCAG</td><td>>hg38_21_+_8433172_8446622_RNA45SN1_45S_50ntFlanks</td><td>11210</td><td>11254</td><td>45</td><td>GGCGACTCTGGACGCGAGCCGGGCCCTTCCCGTGGATCGCCCCAG</td><td>181.5</td><td>-11.4</td><td>1.8</td><td>-6.4</td><td>1.7E-10</td><td>4.4E-10</td><td>MB231/MB468</td><td>MP</td></tr>
<tr><td>>hg38_21_+_8433172_8446622_RNA45SN1_45S_50ntFlanks@9305.9338.34|CCAAGGAGTCTAACACGTGCGCGAGTCGGGGGCT</td><td>>hg38_21_+_8433172_8446622_RNA45SN1_45S_50ntFlanks</td><td>9305</td><td>9338</td><td>34</td><td>CCAAGGAGTCTAACACGTGCGCGAGTCGGGGGCT</td><td>3682.2</td><td>-9.5</td><td>1.5</td><td>-6.4</td><td>2.0E-10</td><td>4.4E-10</td><td>MB231/MB468</td><td>MITO</td></tr>
<tr><td>>hg38_21_+_8433172_8446622_RNA45SN1_45S_50ntFlanks@12043.12063.21|CACTGACCCGGTGAGGCGGGG</td><td>>hg38_21_+_8433172_8446622_RNA45SN1_45S_50ntFlanks</td><td>12043</td><td>12063</td><td>21</td><td>CACTGACCCGGTGAGGCGGGG</td><td>87.9</td><td>-10.3</td><td>1.6</td><td>-6.4</td><td>1.7E-10</td><td>4.4E-10</td><td>MB231/MB468</td><td>MP</td></tr>
<tr><td>>hg38_21_+_8433172_8446622_RNA45SN1_45S_50ntFlanks@12419.12438.20|GATGTCGGCTCTTCCTATCA</td><td>>hg38_21_+_8433172_8446622_RNA45SN1_45S_50ntFlanks</td><td>12419</td><td>12438</td><td>20</td><td>GATGTCGGCTCTTCCTATCA</td><td>992.7</td><td>4.3</td><td>0.7</td><td>6.4</td><td>1.3E-10</td><td>4.4E-10</td><td>BT20/MB468</td><td>MITO</td></tr>
<tr><td>>hg38_21_+_8433172_8446622_RNA45SN1_45S_50ntFlanks@10664.10706.43|CTTGGAAAGCGTCGCGGTTCCGGCGGCGTCCGGTGAGCTCTCG</td><td>>hg38_21_+_8433172_8446622_RNA45SN1_45S_50ntFlanks</td><td>10664</td><td>10706</td><td>43</td><td>CTTGGAAAGCGTCGCGGTTCCGGCGGCGTCCGGTGAGCTCTCG</td><td>3325.1</td><td>-10.7</td><td>1.7</td><td>-6.4</td><td>2.1E-10</td><td>4.4E-10</td><td>BT20/MB468</td><td>NUC</td></tr>
<tr><td>>hg38_21_+_8433172_8446622_RNA45SN1_45S_50ntFlanks@10411.10434.24|CAGTCGGTCCTGAGAGATGGGCGA</td><td>>hg38_21_+_8433172_8446622_RNA45SN1_45S_50ntFlanks</td><td>10411</td><td>10434</td><td>24</td><td>CAGTCGGTCCTGAGAGATGGGCGA</td><td>120.7</td><td>-10.8</td><td>1.7</td><td>-6.4</td><td>1.7E-10</td><td>4.4E-10</td><td>MB231/MB468</td><td>MP</td></tr>
<tr><td>>hg38_21_+_8433172_8446622_RNA45SN1_45S_50ntFlanks@6761.6806.46|TGCGGCCCCGGGTTCCTCCCGGGGCTACGCCTGTCTGAGCGTCGCT</td><td>>hg38_21_+_8433172_8446622_RNA45SN1_45S_50ntFlanks</td><td>6761</td><td>6806</td><td>46</td><td>TGCGGCCCCGGGTTCCTCCCGGGGCTACGCCTGTCTGAGCGTCGCT</td><td>1205.5</td><td>7.4</td><td>1.2</td><td>6.4</td><td>1.3E-10</td><td>4.4E-10</td><td>BT20/MB468</td><td>MITO</td></tr>
<tr><td>>hg38_21_+_8433172_8446622_RNA45SN1_45S_50ntFlanks@10839.10865.27|TCCGTAACTTCGGGATAAGGATTGGCT</td><td>>hg38_21_+_8433172_8446622_RNA45SN1_45S_50ntFlanks</td><td>10839</td><td>10865</td><td>27</td><td>TCCGTAACTTCGGGATAAGGATTGGCT</td><td>459.7</td><td>-6.4</td><td>1.0</td><td>-6.4</td><td>1.8E-10</td><td>4.4E-10</td><td>MB231/MB468</td><td>NUC</td></tr>
<tr><td>>hg38_21_+_8433172_8446622_RNA45SN1_45S_50ntFlanks@8184.8223.40|CGAGGCCCAGCCCGTGGACGGTGTGAGGCCGGTAGCGGCC</td><td>>hg38_21_+_8433172_8446622_RNA45SN1_45S_50ntFlanks</td><td>8184</td><td>8223</td><td>40</td><td>CGAGGCCCAGCCCGTGGACGGTGTGAGGCCGGTAGCGGCC</td><td>423.7</td><td>-6.8</td><td>1.1</td><td>-6.4</td><td>1.8E-10</td><td>4.4E-10</td><td>MB231/MB468</td><td>NUC</td></tr>
<tr><td>>MI0000065|hsa-let-7d&WithFlank&9|+|94178828|94178926@68.89.22&[MIMAT0004484&hsa-let-7d-3p&offsets|0|0;m-98&9|+|94178895|94178916&offsets|0|0]|CTATACGACCTGCTGCCTTTCT</td><td>>MI0000065|hsa-let-7d&WithFlank&9|+|94178828|94178926</td><td>68</td><td>89</td><td>22</td><td>CTATACGACCTGCTGCCTTTCT</td><td>3062.2</td><td>5.9</td><td>0.9</td><td>6.4</td><td>2.0E-10</td><td>4.4E-10</td><td>MB231/MB468</td><td>MITO</td></tr>
<tr><td>>hg38_21_+_8433172_8446622_RNA45SN1_45S_50ntFlanks@6743.6772.30|CGACACTTCGAACGCACTTGCGGCCCCGGG</td><td>>hg38_21_+_8433172_8446622_RNA45SN1_45S_50ntFlanks</td><td>6743</td><td>6772</td><td>30</td><td>CGACACTTCGAACGCACTTGCGGCCCCGGG</td><td>308.8</td><td>-8.5</td><td>1.3</td><td>-6.4</td><td>1.8E-10</td><td>4.4E-10</td><td>MB231/MB468</td><td>NUC</td></tr>
<tr><td>>hg38_21_+_8433172_8446622_RNA45SN1_45S_50ntFlanks@12404.12437.34|GCTTTTTGATCCTTCGATGTCGGCTCTTCCTATC</td><td>>hg38_21_+_8433172_8446622_RNA45SN1_45S_50ntFlanks</td><td>12404</td><td>12437</td><td>34</td><td>GCTTTTTGATCCTTCGATGTCGGCTCTTCCTATC</td><td>871.8</td><td>-6.1</td><td>1.0</td><td>-6.4</td><td>2.1E-10</td><td>4.4E-10</td><td>BT20/MB468</td><td>NUC</td></tr>
<tr><td>>MI0000446|hsa-mir-125b-1&WithFlank&11|-|122099751|122099850@21.41.21(+1G);MI0000470|hsa-mir-125b-2&WithFlank&21|+|16590231|16590331@23.43.21(+1G)&[MIMAT0000423&hsa-miR-125b-5p&offsets|0|-1(+1G);m-67&11|-|122099809|122099830&offsets|0|-1(+1G)];[MIMAT0000423_1&hsa-miR-125b-5p&offsets|0|-1(+1G);m-68&21|+|16590253|16590274&offsets|0|-1(+1G)]|TCCCTGAGACCCTAACTTGTGG</td><td>>MI0000446|hsa-mir-125b-1&WithFlank&11|-|122099751|122099850</td><td>21</td><td>41</td><td>21(+1G)</td><td>TCCCTGAGACCCTAACTTGTGG</td><td>441.8</td><td>-12.2</td><td>1.7</td><td>-7.0</td><td>3.0E-12</td><td>4.4E-10</td><td>BT20/MB231</td><td>MP</td></tr>
<tr><td>>hg38_21_+_8433172_8446622_RNA45SN1_45S_50ntFlanks@9861.9899.39|GATGAACCGAACGCCGGGTTAAGGCGCCCGATGCCGACG</td><td>>hg38_21_+_8433172_8446622_RNA45SN1_45S_50ntFlanks</td><td>9861</td><td>9899</td><td>39</td><td>GATGAACCGAACGCCGGGTTAAGGCGCCCGATGCCGACG</td><td>95.9</td><td>-10.4</td><td>1.6</td><td>-6.4</td><td>1.7E-10</td><td>4.4E-10</td><td>MB231/MB468</td><td>MP</td></tr>
<tr><td>>hg38_21_+_8433172_8446622_RNA45SN1_45S_50ntFlanks@11581.11600.20|AACTGGTGCGGACCAGGGGA</td><td>>hg38_21_+_8433172_8446622_RNA45SN1_45S_50ntFlanks</td><td>11581</td><td>11600</td><td>20</td><td>AACTGGTGCGGACCAGGGGA</td><td>820.1</td><td>-4.6</td><td>0.7</td><td>-6.4</td><td>1.6E-10</td><td>4.4E-10</td><td>BT20/MB468</td><td>CYTO</td></tr>
<tr><td>>hg38_21_+_8433172_8446622_RNA45SN1_45S_50ntFlanks@3854.3881.28|ACTGTGGTAATTCTAGAGCTAATACATG</td><td>>hg38_21_+_8433172_8446622_RNA45SN1_45S_50ntFlanks</td><td>3854</td><td>3881</td><td>28</td><td>ACTGTGGTAATTCTAGAGCTAATACATG</td><td>3289.8</td><td>-10.4</td><td>1.6</td><td>-6.4</td><td>2.1E-10</td><td>4.4E-10</td><td>BT20/MB468</td><td>NUC</td></tr>
<tr><td>>hg38_21_+_8433172_8446622_RNA45SN1_45S_50ntFlanks@7976.8009.34|GCGACCTCAGATCAGACGTGGCGACCCGCTGAAT</td><td>>hg38_21_+_8433172_8446622_RNA45SN1_45S_50ntFlanks</td><td>7976</td><td>8009</td><td>34</td><td>GCGACCTCAGATCAGACGTGGCGACCCGCTGAAT</td><td>591.0</td><td>8.4</td><td>1.3</td><td>6.4</td><td>1.6E-10</td><td>4.4E-10</td><td>BT20/MB468</td><td>CYTO</td></tr>
<tr><td>>trna111_HisGTG_1_-_147774845_147774916@-1T.20.21,trna118_HisGTG_1_-_145396881_145396952@-1T.20.21,trna16_HisGTG_1_+_146544773_146544844@-1T.20.21,trna1_HisGTG_15_+_45493349_45493420@-1T.20.21,trna21_HisGTG_1_+_147753471_147753542@-1T.20.21,trna33_HisGTG_6_+_27125906_27125977@-1T.20.21,trna7_HisGTG_9_-_14433938_14434009@-1T.20.21,trna8_HisGTG_15_-_45492611_45492682@-1T.20.21,trna9_HisGTG_15_-_45490804_45490875@-1T.20.21|TGCCGTGATCGTATAGTGGTT</td><td>>trna111_HisGTG_1_-_147774845_147774916</td><td>-1T</td><td>20</td><td>21</td><td>TGCCGTGATCGTATAGTGGTT</td><td>163.3</td><td>10.6</td><td>1.7</td><td>6.4</td><td>1.7E-10</td><td>4.4E-10</td><td>MB231/MB468</td><td>MP</td></tr>
<tr><td>>hg38_21_+_8433172_8446622_RNA45SN1_45S_50ntFlanks@12456.12499.44|CCAAGCGTTGGATTGTTCACCCACTAATAGGGAACGTGAGCTGG</td><td>>hg38_21_+_8433172_8446622_RNA45SN1_45S_50ntFlanks</td><td>12456</td><td>12499</td><td>44</td><td>CCAAGCGTTGGATTGTTCACCCACTAATAGGGAACGTGAGCTGG</td><td>156.0</td><td>-11.2</td><td>1.7</td><td>-6.4</td><td>1.7E-10</td><td>4.4E-10</td><td>MB231/MB468</td><td>MP</td></tr>
<tr><td>>hg38_21_+_8433172_8446622_RNA45SN1_45S_50ntFlanks@11583.11611.29|CTGGTGCGGACCAGGGGAATCCGACTGTT</td><td>>hg38_21_+_8433172_8446622_RNA45SN1_45S_50ntFlanks</td><td>11583</td><td>11611</td><td>29</td><td>CTGGTGCGGACCAGGGGAATCCGACTGTT</td><td>2026.8</td><td>-3.0</td><td>0.5</td><td>-6.4</td><td>1.8E-10</td><td>4.4E-10</td><td>MB231/MB468</td><td>NUC</td></tr>
<tr><td>>hg38_21_+_8433172_8446622_RNA45SN1_45S_50ntFlanks@11668.11691.24|TGCCCAGTGCTCTGAATGTCAAAG</td><td>>hg38_21_+_8433172_8446622_RNA45SN1_45S_50ntFlanks</td><td>11668</td><td>11691</td><td>24</td><td>TGCCCAGTGCTCTGAATGTCAAAG</td><td>590.7</td><td>6.4</td><td>1.0</td><td>6.4</td><td>1.8E-10</td><td>4.4E-10</td><td>MB231/MB468</td><td>CYTO</td></tr>
<tr><td>>hg38_21_+_8433172_8446622_RNA45SN1_45S_50ntFlanks@9266.9285.20|CGACGTCGGCTACCCACCCG</td><td>>hg38_21_+_8433172_8446622_RNA45SN1_45S_50ntFlanks</td><td>9266</td><td>9285</td><td>20</td><td>CGACGTCGGCTACCCACCCG</td><td>475.4</td><td>-5.0</td><td>0.8</td><td>-6.4</td><td>2.0E-10</td><td>4.4E-10</td><td>MB231/MB468</td><td>MITO</td></tr>
<tr><td>>hg38_21_+_8433172_8446622_RNA45SN1_45S_50ntFlanks@4027.4045.19|CCCCGTGGCGGCGACGACC</td><td>>hg38_21_+_8433172_8446622_RNA45SN1_45S_50ntFlanks</td><td>4027</td><td>4045</td><td>19</td><td>CCCCGTGGCGGCGACGACC</td><td>273.9</td><td>-10.2</td><td>1.6</td><td>-6.4</td><td>1.3E-10</td><td>4.4E-10</td><td>BT20/MB468</td><td>MITO</td></tr>
<tr><td>>trnaMT_PheGAA_MT_+_577_647@36.65.30|AAAATGTTTAGACGGGCTCACATCACCCCA</td><td>>trnaMT_PheGAA_MT_+_577_647</td><td>36</td><td>65</td><td>30</td><td>AAAATGTTTAGACGGGCTCACATCACCCCA</td><td>85.9</td><td>9.6</td><td>1.5</td><td>6.4</td><td>1.7E-10</td><td>4.5E-10</td><td>MB231/MB468</td><td>MP</td></tr>
<tr><td>>hg38_21_+_8433172_8446622_RNA45SN1_45S_50ntFlanks@10134.10157.24|CGGCGGCGGCGGCGGGGGTGTGGG</td><td>>hg38_21_+_8433172_8446622_RNA45SN1_45S_50ntFlanks</td><td>10134</td><td>10157</td><td>24</td><td>CGGCGGCGGCGGCGGGGGTGTGGG</td><td>111.0</td><td>-10.7</td><td>1.7</td><td>-6.4</td><td>1.7E-10</td><td>4.5E-10</td><td>MB231/MB468</td><td>MP</td></tr>
<tr><td>>hg38_21_+_8433172_8446622_RNA45SN1_45S_50ntFlanks@10597.10617.21|AGTTCTCTTTTCTTTGTGAAG</td><td>>hg38_21_+_8433172_8446622_RNA45SN1_45S_50ntFlanks</td><td>10597</td><td>10617</td><td>21</td><td>AGTTCTCTTTTCTTTGTGAAG</td><td>111.8</td><td>-9.8</td><td>1.5</td><td>-6.4</td><td>1.8E-10</td><td>4.5E-10</td><td>MB231/MB468</td><td>NUC</td></tr>
<tr><td>>hg38_21_+_8433172_8446622_RNA45SN1_45S_50ntFlanks@9860.9899.40|GGATGAACCGAACGCCGGGTTAAGGCGCCCGATGCCGACG</td><td>>hg38_21_+_8433172_8446622_RNA45SN1_45S_50ntFlanks</td><td>9860</td><td>9899</td><td>40</td><td>GGATGAACCGAACGCCGGGTTAAGGCGCCCGATGCCGACG</td><td>116.6</td><td>-10.7</td><td>1.7</td><td>-6.4</td><td>1.7E-10</td><td>4.5E-10</td><td>MB231/MB468</td><td>MP</td></tr>
<tr><td>>hg38_21_+_8433172_8446622_RNA45SN1_45S_50ntFlanks@10404.10433.30|CATGGGTCAGTCGGTCCTGAGAGATGGGCG</td><td>>hg38_21_+_8433172_8446622_RNA45SN1_45S_50ntFlanks</td><td>10404</td><td>10433</td><td>30</td><td>CATGGGTCAGTCGGTCCTGAGAGATGGGCG</td><td>556.1</td><td>-9.1</td><td>1.4</td><td>-6.4</td><td>2.1E-10</td><td>4.5E-10</td><td>BT20/MB468</td><td>NUC</td></tr>
<tr><td>>hg38_21_+_8433172_8446622_RNA45SN1_45S_50ntFlanks@10392.10411.20|ACAGCAGTTGAACATGGGTC</td><td>>hg38_21_+_8433172_8446622_RNA45SN1_45S_50ntFlanks</td><td>10392</td><td>10411</td><td>20</td><td>ACAGCAGTTGAACATGGGTC</td><td>502.9</td><td>-4.6</td><td>0.7</td><td>-6.4</td><td>1.8E-10</td><td>4.5E-10</td><td>MB231/MB468</td><td>NUC</td></tr>
<tr><td>>hg38_21_+_8433172_8446622_RNA45SN1_45S_50ntFlanks@8174.8216.43|TCCTTCTGATCGAGGCCCAGCCCGTGGACGGTGTGAGGCCGGT</td><td>>hg38_21_+_8433172_8446622_RNA45SN1_45S_50ntFlanks</td><td>8174</td><td>8216</td><td>43</td><td>TCCTTCTGATCGAGGCCCAGCCCGTGGACGGTGTGAGGCCGGT</td><td>393.1</td><td>5.5</td><td>0.9</td><td>6.4</td><td>1.3E-10</td><td>4.5E-10</td><td>BT20/MB468</td><td>MITO</td></tr>
<tr><td>>hg38_21_+_8433172_8446622_RNA45SN1_45S_50ntFlanks@4733.4751.19|GAACGAAAGTCGGAGGTTC</td><td>>hg38_21_+_8433172_8446622_RNA45SN1_45S_50ntFlanks</td><td>4733</td><td>4751</td><td>19</td><td>GAACGAAAGTCGGAGGTTC</td><td>475.6</td><td>-4.2</td><td>0.7</td><td>-6.4</td><td>2.1E-10</td><td>4.5E-10</td><td>BT20/MB468</td><td>NUC</td></tr>
<tr><td>>hg38_21_+_8433172_8446622_RNA45SN1_45S_50ntFlanks@4161.4186.26|CACATCCAAGGAAGGCAGCAGGCGCG</td><td>>hg38_21_+_8433172_8446622_RNA45SN1_45S_50ntFlanks</td><td>4161</td><td>4186</td><td>26</td><td>CACATCCAAGGAAGGCAGCAGGCGCG</td><td>111.9</td><td>-10.7</td><td>1.7</td><td>-6.4</td><td>1.7E-10</td><td>4.5E-10</td><td>MB231/MB468</td><td>MP</td></tr>
</table>

| | | | | | | | | | | | | | |
|---|---|---|---|---|---|---|---|---|---|---|---|---|---|
| >trna10_AspGTC_12_-_125424193_125424264@1.30.30,trna12_AspGTC_12_-_125411891_125411962@1.30.30,trna38_AspGTC_17_-_8125556_8125627@1.30.30,trna45_AspGTC_6_+_27447453_27447524@1.30.30,trna48_AspGTC_6_+_27471523_27471594@1.30.30,trna4_AspGTC_12_+_96429799_96429870@1.30.30,trna69_AspGTC_1_-_161440205_161440276@1.30.30,trna72_AspGTC_1_-_161432824_161432895@1.30.30,trna75_AspGTC_1_-_161425414_161425485@1.30.30,trna78_AspGTC_1_-_161418033_161418104@1.30.30,trna81_AspGTC_1_-_161410615_161410686@1.30.30\|TCCTCGTTAGTATAGTGGTGAGTATCCCCG | >trna10_AspGTC_12_-_125424193_125424264 | 1 | 30 | 30 | TCCTCGTTAGTATAGTGGTGAGTATCCCCG | 974.1 | -5.1 | 0.8 | -6.4 | 1.8E-10 | 4.5E-10 | MB231/MB468 | CYTO |
| >MI0003129\|hsa-mir-146b&WithFlank&10\|+\|102436506\|102436590@15.38.24&[MIMAT0002809&hsa-miR-146b-5p&offsets\|0\|+1;m-70&10\|+\|102436520\|102436543&offsets\|0\|0]\|TGAGAACTGAATTCCATAGGCTGT | >MI0003129\|hsa-mir-146b&WithFlank&10\|+\|102436506\|102436590 | 15 | 38 | 24 | TGAGAACTGAATTCCATAGGCTGT | 334.8 | 3.7 | 0.6 | 6.4 | 1.8E-10 | 4.5E-10 | MB231/MB468 | NUC |
| >hg38_21_+_8433172_8446622_RNA45SN1_45S_50ntFlanks@12912.12933.22\|CACGTTCGTGGGGAACCTGGCG | >hg38_21_+_8433172_8446622_RNA45SN1_45S_50ntFlanks | 12912 | 12933 | 22 | CACGTTCGTGGGGAACCTGGCG | 1542.9 | -9.3 | 1.5 | -6.4 | 1.7E-10 | 4.5E-10 | MB231/MB468 | MP |
| >hg38_21_+_8433172_8446622_RNA45SN1_45S_50ntFlanks@3882.3899.18\|CCGACGGGCGCTGACCCC | >hg38_21_+_8433172_8446622_RNA45SN1_45S_50ntFlanks | 3882 | 3899 | 18 | CCGACGGGCGCTGACCCC | 2194.2 | -5.7 | 0.9 | -6.4 | 1.8E-10 | 4.5E-10 | MB231/MB468 | CYTO |
| >hg38_21_+_8433172_8446622_RNA45SN1_45S_50ntFlanks@11317.11334.18\|ACCCCACGTCTCGTCGCG | >hg38_21_+_8433172_8446622_RNA45SN1_45S_50ntFlanks | 11317 | 11334 | 18 | ACCCCACGTCTCGTCGCG | 311.5 | -8.5 | 1.3 | -6.4 | 1.8E-10 | 4.5E-10 | MB231/MB468 | NUC |
| >hg38_21_+_8433172_8446622_RNA45SN1_45S_50ntFlanks@10355.10374.20\|AACTTTGAAGGCCGAAGTGG | >hg38_21_+_8433172_8446622_RNA45SN1_45S_50ntFlanks | 10355 | 10374 | 20 | AACTTTGAAGGCCGAAGTGG | 165.3 | -11.2 | 1.8 | -6.4 | 1.7E-10 | 4.5E-10 | MB231/MB468 | MP |
| >hg38_21_+_8433172_8446622_RNA45SN1_45S_50ntFlanks@12626.12647.22\|AATGGGGCGAAGCTACCATCTG | >hg38_21_+_8433172_8446622_RNA45SN1_45S_50ntFlanks | 12626 | 12647 | 22 | AATGGGGCGAAGCTACCATCTG | 90.0 | -10.4 | 1.6 | -6.4 | 1.7E-10 | 4.5E-10 | MB231/MB468 | MP |
| >hg38_21_+_8433172_8446622_RNA45SN1_45S_50ntFlanks@3711.3737.27\|GTTGATCCTGCCAGTAGCATATGCTTG | >hg38_21_+_8433172_8446622_RNA45SN1_45S_50ntFlanks | 3711 | 3737 | 27 | GTTGATCCTGCCAGTAGCATATGCTTG | 1621.5 | -7.6 | 1.2 | -6.4 | 1.8E-10 | 4.5E-10 | MB231/MB468 | NUC |
| >hg38_21_+_8433172_8446622_RNA45SN1_45S_50ntFlanks@8380.8399.20\|CAAGAGGGCGTGAAACCGTT | >hg38_21_+_8433172_8446622_RNA45SN1_45S_50ntFlanks | 8380 | 8399 | 20 | CAAGAGGGCGTGAAACCGTT | 510.1 | -7.4 | 1.2 | -6.4 | 1.6E-10 | 4.5E-10 | BT20/MB468 | CYTO |
| >hg38_21_+_8433172_8446622_RNA45SN1_45S_50ntFlanks@11564.11588.25\|CCTAGCAGCCGACTTAGAACTGGTG | >hg38_21_+_8433172_8446622_RNA45SN1_45S_50ntFlanks | 11564 | 11588 | 25 | CCTAGCAGCCGACTTAGAACTGGTG | 1801.1 | -9.8 | 1.5 | -6.4 | 2.1E-10 | 4.5E-10 | BT20/MB468 | NUC |
| >hg38_21_+_8433172_8446622_RNA45SN1_45S_50ntFlanks@9939.9978.40\|GCAGGACGGTGGCCATGGAAGTCGGAATCCGCTAAGGAGT | >hg38_21_+_8433172_8446622_RNA45SN1_45S_50ntFlanks | 9939 | 9978 | 40 | GCAGGACGGTGGCCATGGAAGTCGGAATCCGCTAAGGAGT | 148.1 | 11.0 | 1.6 | 6.8 | 1.4E-11 | 4.5E-10 | BT20/MB468 | MP |
| >MI0000102\|hsa-mir-100&WithFlank&11\|-\|122152223\|122152314@23.40.18&[MIMAT0000098&hsa-miR-100-5p&offsets\|+4\|0;m-27&11\|-\|122152276\|122152296&offsets\|+4\|+1]\|CGTAGATCCGAACTTGTG | >MI0000102\|hsa-mir-100&WithFlank&11\|-\|122152223\|122152314 | 23 | 40 | 18 | CGTAGATCCGAACTTGTG | 128.2 | 10.2 | 1.6 | 6.4 | 1.7E-10 | 4.5E-10 | MB231/MB468 | MP |
| >hg38_21_+_8433172_8446622_RNA45SN1_45S_50ntFlanks@10880.10902.23\|GGTCGGGCTGGGGCGCGAAGCGG | >hg38_21_+_8433172_8446622_RNA45SN1_45S_50ntFlanks | 10880 | 10902 | 23 | GGTCGGGCTGGGGCGCGAAGCGG | 109.6 | -10.6 | 1.7 | -6.4 | 1.7E-10 | 4.5E-10 | MB231/MB468 | MP |
| >hg38_21_+_8433172_8446622_RNA45SN1_45S_50ntFlanks@6666.6698.33\|GATCACTCGGCTCGTGCGTCGATGAAGAACGCA | >hg38_21_+_8433172_8446622_RNA45SN1_45S_50ntFlanks | 6666 | 6698 | 33 | GATCACTCGGCTCGTGCGTCGATGAAGAACGCA | 72.7 | -10.0 | 1.6 | -6.4 | 1.7E-10 | 4.5E-10 | MB231/MB468 | MP |
| >MI0000542\|hsa-mir-320a&WithFlank&8\|-\|22244960\|22245043@48.68.21(+2U)&[MIMAT0000510&hsa-miR-320a-3p&offsets\|0\|-1(+2U);m-60&8\|-\|22244975\|22244996&offsets\|0\|-1(+2U)]\|AAAAGCTGGGTTGAGAGGGCGTT | >MI0000542\|hsa-mir-320a&WithFlank&8\|-\|22244960\|22245043 | 48 | 68 | 21(+2U) | AAAAGCTGGGTTGAGAGGGCGTT | 776.7 | 8.3 | 1.3 | 6.4 | 1.8E-10 | 4.5E-10 | MB231/MB468 | CYTO |
| >hg38_21_+_8433172_8446622_RNA45SN1_45S_50ntFlanks@7977.8015.39\|CGACCTCAGATCAGACGTGGCGACCCGCTGAATTTAAGC | >hg38_21_+_8433172_8446622_RNA45SN1_45S_50ntFlanks | 7977 | 8015 | 39 | CGACCTCAGATCAGACGTGGCGACCCGCTGAATTTAAGC | 266.3 | 7.3 | 1.1 | 6.4 | 1.8E-10 | 4.5E-10 | MB231/MB468 | CYTO |
| >hg38_21_+_8433172_8446622_RNA45SN1_45S_50ntFlanks@12408.12436.29\|TTTGATCCTTCGATGTCGGCTCTTCCTAT | >hg38_21_+_8433172_8446622_RNA45SN1_45S_50ntFlanks | 12408 | 12436 | 29 | TTTGATCCTTCGATGTCGGCTCTTCCTAT | 635.7 | -4.1 | 0.6 | -6.4 | 1.8E-10 | 4.5E-10 | MB231/MB468 | NUC |
| >trnaMT_GluTTC_MT_-_14674_14742@35.72.38,trnalookalike8_GluTTC_5_-_93905172_93905240@35.72.38\|TATCATTGGTCGTGGTTGTAGTCCGTGCGAGAATACCA | >trnaMT_GluTTC_MT_-_14674_14742 | 35 | 72 | 38 | TATCATTGGTCGTGGTTGTAGTCCGTGCGAGAATACCA | 1051.4 | 7.6 | 1.2 | 6.4 | 1.6E-10 | 4.5E-10 | BT20/MB468 | CYTO |
| >hg38_21_+_8433172_8446622_RNA45SN1_45S_50ntFlanks@4367.4397.31\|CAGTTAAAAAGCTCGTAGTTGGATCTTGGGA | >hg38_21_+_8433172_8446622_RNA45SN1_45S_50ntFlanks | 4367 | 4397 | 31 | CAGTTAAAAAGCTCGTAGTTGGATCTTGGGA | 84.2 | -10.3 | 1.6 | -6.4 | 1.7E-10 | 4.5E-10 | MB231/MB468 | MP |
| >am_trna143_LysTTT_6_-_27559593_27559665@6.24.19,trna14_LysTTT_11_-_59327808_59327880@6.24.19,trna2_LysTTT_17_+_8022473_8022545@6.24.19,trna54_LysTTT_1_+_204475655_204475727@6.24.19,trna5_LysTTT_11_+_59323902_59323974@6.24.19,trna62_LysTTT_1_-_204476158_204476230@6.24.19,trna76_LysTTT_6_+_28918806_28918878@6.24.19\|GATAGCTCAGTCGGTAGAG | >am_trna143_LysTTT_6_-_27559593_27559665 | 6 | 24 | 19 | GATAGCTCAGTCGGTAGAG | 439.0 | -11.1 | 1.7 | -6.4 | 1.6E-10 | 4.5E-10 | BT20/MB468 | CYTO |
| >hg38_21_+_8433172_8446622_RNA45SN1_45S_50ntFlanks@11589.11612.24\|CGGACCAGGGGAATCCGACTGTTT | >hg38_21_+_8433172_8446622_RNA45SN1_45S_50ntFlanks | 11589 | 11612 | 24 | CGGACCAGGGGAATCCGACTGTTT | 485.1 | -4.2 | 0.7 | -6.4 | 2.0E-10 | 4.5E-10 | MB231/MB468 | MITO |
| >hg38_21_+_8433172_8446622_RNA45SN1_45S_50ntFlanks@6650.6685.36\|ACGACTCTTAGCGGTGGATCACTCGGCTCGTGCGTC | >hg38_21_+_8433172_8446622_RNA45SN1_45S_50ntFlanks | 6650 | 6685 | 36 | ACGACTCTTAGCGGTGGATCACTCGGCTCGTGCGTC | 75.8 | 2.6 | 0.4 | 6.3 | 2.2E-10 | 4.6E-10 | BT20/MB468 | NUC |
| >hg38_21_+_8433172_8446622_RNA45SN1_45S_50ntFlanks@12397.12414.18\|CGACGTCGCTTTTTGATC | >hg38_21_+_8433172_8446622_RNA45SN1_45S_50ntFlanks | 12397 | 12414 | 18 | CGACGTCGCTTTTTGATC | 3046.6 | -4.9 | 0.8 | -6.4 | 2.0E-10 | 4.6E-10 | MB231/MB468 | MITO |
| >hg38_21_+_8433172_8446622_RNA45SN1_45S_50ntFlanks@8168.8214.47\|CCCAAGTCCTTCTGATCGAGGCCCAGCCCGTGGACGGTGTGAGGCCG | >hg38_21_+_8433172_8446622_RNA45SN1_45S_50ntFlanks | 8168 | 8214 | 47 | CCCAAGTCCTTCTGATCGAGGCCCAGCCCGTGGACGGTGTGAGGCCG | 581.1 | -7.4 | 1.2 | -6.4 | 2.0E-10 | 4.6E-10 | MB231/MB468 | MITO |
| >hg38_21_+_8433172_8446622_RNA45SN1_45S_50ntFlanks@9129.9149.21\|CGGGTCGGGGGGCGGGGCGGA | >hg38_21_+_8433172_8446622_RNA45SN1_45S_50ntFlanks | 9129 | 9149 | 21 | CGGGTCGGGGGGCGGGGCGGA | 131.3 | -9.1 | 1.4 | -6.4 | 1.4E-10 | 4.6E-10 | BT20/MB468 | MITO |
| >hg38_21_+_8433172_8446622_RNA45SN1_45S_50ntFlanks@10830.10858.29\|CAAGCCGGATCCGTAACTTCGGGATAAGG | >hg38_21_+_8433172_8446622_RNA45SN1_45S_50ntFlanks | 10830 | 10858 | 29 | CAAGCCGGATCCGTAACTTCGGGATAAGG | 466.8 | -8.0 | 1.2 | -6.4 | 1.6E-10 | 4.6E-10 | BT20/MB468 | CYTO |
| >hg38_21_+_8433172_8446622_RNA45SN1_45S_50ntFlanks@12008.12041.34\|CCGCCGGTGAAATACCACTACTCTGATCGTTTTT | >hg38_21_+_8433172_8446622_RNA45SN1_45S_50ntFlanks | 12008 | 12041 | 34 | CCGCCGGTGAAATACCACTACTCTGATCGTTTTT | 321.2 | -7.4 | 1.2 | -6.4 | 1.8E-10 | 4.6E-10 | MB231/MB468 | CYTO |
| >hg38_21_+_8433172_8446622_RNA45SN1_45S_50ntFlanks@11211.11254.44\|GCGACTCTGGACGCGAGCCGGGCCCTTCCCGTGGATCGCCCCAG | >hg38_21_+_8433172_8446622_RNA45SN1_45S_50ntFlanks | 11211 | 11254 | 44 | GCGACTCTGGACGCGAGCCGGGCCCTTCCCGTGGATCGCCCCAG | 96.2 | -10.5 | 1.6 | -6.4 | 1.7E-10 | 4.6E-10 | MB231/MB468 | MP |
| >hg38_21_+_8433172_8446622_RNA45SN1_45S_50ntFlanks@12934.12970.37\|CTAAACCATTCGTAGACGACCTGCTTCTGGGTCGGGG | >hg38_21_+_8433172_8446622_RNA45SN1_45S_50ntFlanks | 12934 | 12970 | 37 | CTAAACCATTCGTAGACGACCTGCTTCTGGGTCGGGG | 112.4 | -8.9 | 1.4 | -6.4 | 1.4E-10 | 4.6E-10 | BT20/MB468 | MITO |
| >hg38_21_+_8433172_8446622_RNA45SN1_45S_50ntFlanks@10423.10457.35\|AGAGATGGGCGAGCGCCGTTCCGAAGGGACGGGCG | >hg38_21_+_8433172_8446622_RNA45SN1_45S_50ntFlanks | 10423 | 10457 | 35 | AGAGATGGGCGAGCGCCGTTCCGAAGGGACGGGCG | 748.9 | -5.5 | 0.9 | -6.4 | 1.8E-10 | 4.6E-10 | MB231/MB468 | CYTO |
| >hg38_21_+_8433172_8446622_RNA45SN1_45S_50ntFlanks@10074.10092.19\|ACGGGAGCGGCGGGGGCGG | >hg38_21_+_8433172_8446622_RNA45SN1_45S_50ntFlanks | 10074 | 10092 | 19 | ACGGGAGCGGCGGGGGCGG | 240.7 | -11.0 | 1.6 | -6.8 | 1.5E-11 | 4.6E-10 | BT20/MB468 | MP |
| >hg38_21_+_8433172_8446622_RNA45SN1_45S_50ntFlanks@6733.6753.21\|CATTGATCATCGACACTTCGA | >hg38_21_+_8433172_8446622_RNA45SN1_45S_50ntFlanks | 6733 | 6753 | 21 | CATTGATCATCGACACTTCGA | 916.3 | -7.1 | 1.1 | -6.4 | 1.8E-10 | 4.6E-10 | MB231/MB468 | MP |
| >hg38_21_+_8433172_8446622_RNA45SN1_45S_50ntFlanks@10351.10391.41\|CGAGAACTTTGAAGGCCGAAGTGGAGAAGGGTTCCATGTGA | >hg38_21_+_8433172_8446622_RNA45SN1_45S_50ntFlanks | 10351 | 10391 | 41 | CGAGAACTTTGAAGGCCGAAGTGGAGAAGGGTTCCATGTGA | 1664.2 | -7.0 | 1.1 | -6.4 | 2.1E-10 | 4.6E-10 | MB231/MB468 | MITO |
| >hg38_21_+_8433172_8446622_RNA45SN1_45S_50ntFlanks@6719.6741.23\|TGAATTGCAGGACACATTGATCA | >hg38_21_+_8433172_8446622_RNA45SN1_45S_50ntFlanks | 6719 | 6741 | 23 | TGAATTGCAGGACACATTGATCA | 338.2 | -7.9 | 1.2 | -6.4 | 2.1E-10 | 4.6E-10 | MB231/MB468 | MITO |
| >hg38_21_+_8433172_8446622_RNA45SN1_45S_50ntFlanks@10869.10890.22\|AGGGCTGGGTCGGTCGGGCTGG | >hg38_21_+_8433172_8446622_RNA45SN1_45S_50ntFlanks | 10869 | 10890 | 22 | AGGGCTGGGTCGGTCGGGCTGG | 523.0 | -8.4 | 1.3 | -6.4 | 2.1E-10 | 4.6E-10 | MB231/MB468 | MITO |
| >hg38_21_+_8433172_8446622_RNA45SN1_45S_50ntFlanks@4738.4762.25\|AAAGTCGGAGGTTCGAAGACGATCA | >hg38_21_+_8433172_8446622_RNA45SN1_45S_50ntFlanks | 4738 | 4762 | 25 | AAAGTCGGAGGTTCGAAGACGATCA | 244.0 | 5.8 | 0.9 | 6.4 | 1.9E-10 | 4.6E-10 | MB231/MB468 | CYTO |
| >hg38_21_+_8433172_8446622_RNA45SN1_45S_50ntFlanks@10843.10862.20\|TAACTTCGGGATAAGGATTG | >hg38_21_+_8433172_8446622_RNA45SN1_45S_50ntFlanks | 10843 | 10862 | 20 | TAACTTCGGGATAAGGATTG | 304.5 | -6.0 | 0.9 | -6.4 | 1.9E-10 | 4.6E-10 | MB231/MB468 | NUC |
| >hg38_21_+_8433172_8446622_RNA45SN1_45S_50ntFlanks@12633.12655.23\|CGAAGCTACCATCTGTGGGATTA | >hg38_21_+_8433172_8446622_RNA45SN1_45S_50ntFlanks | 12633 | 12655 | 23 | CGAAGCTACCATCTGTGGGATTA | 69.9 | -10.0 | 1.6 | -6.4 | 1.8E-10 | 4.6E-10 | MB231/MB468 | MP |
| >hg38_21_+_8433172_8446622_RNA45SN1_45S_50ntFlanks@9573.9592.20\|AAATCGGTCGTCCGACCTGG | >hg38_21_+_8433172_8446622_RNA45SN1_45S_50ntFlanks | 9573 | 9592 | 20 | AAATCGGTCGTCCGACCTGG | 900.1 | -9.1 | 1.4 | -6.4 | 1.8E-10 | 4.6E-10 | MB231/MB468 | MP |
| >trna10_GlyTCC_17_+_8124866_8124937@31.51.21,trna117_GlyTCC_1_-_145397864_145397935@31.51.21,trna2_GlyTCC_19_+_4724082_4724153@31.51.21,trna45_GlyTCC_1_+_161500903_161500974@31.51.21,trna70_GlyTCC_1_-_161439547_161439618@31.51.21,trna73_GlyTCC_1_-_161432166_161432237@31.51.21,trna76_GlyTCC_1_-_161424756_161424827@31.51.21,trna79_GlyTCC_1_-_161417375_161417446@31.51.21,trna82_GlyTCC_1_-_161409961_161410032@31.51.21\|CCTTCCAAGCAGTTGACCCGG | >trna10_GlyTCC_17_+_8124866_8124937 | 31 | 51 | 21 | CCTTCCAAGCAGTTGACCCGG | 132.3 | -10.9 | 1.7 | -6.4 | 1.8E-10 | 4.6E-10 | MB231/MB468 | MP |
| >hg38_21_+_8433172_8446622_RNA45SN1_45S_50ntFlanks@11849.11872.24\|CGGGCTTGGCGGAATCAGCGGGGA | >hg38_21_+_8433172_8446622_RNA45SN1_45S_50ntFlanks | 11849 | 11872 | 24 | CGGGCTTGGCGGAATCAGCGGGGA | 599.6 | -3.7 | 0.6 | -6.4 | 1.4E-10 | 4.6E-10 | BT20/MB468 | MITO |
| >trnaMT_SerGCT_MT_+_12207_12265@1.18.18\|GAGAAAGCTCACAAGAAC | >trnaMT_SerGCT_MT_+_12207_12265 | 1 | 18 | 18 | GAGAAAGCTCACAAGAAC | 5652.4 | 6.7 | 1.1 | 6.4 | 1.9E-10 | 4.6E-10 | MB231/MB468 | NUC |
| >hg38_21_+_8433172_8446622_RNA45SN1_45S_50ntFlanks@3849.3866.18\|GGATAACTGTGGTAATTC | >hg38_21_+_8433172_8446622_RNA45SN1_45S_50ntFlanks | 3849 | 3866 | 18 | GGATAACTGTGGTAATTC | 320.0 | 4.3 | 0.7 | 6.4 | 1.9E-10 | 4.7E-10 | MB231/MB468 | CYTO |
| >hg38_21_+_8433172_8446622_RNA45SN1_45S_50ntFlanks@10412.10430.19\|AGTCGGTCCTGAGAGATGG | >hg38_21_+_8433172_8446622_RNA45SN1_45S_50ntFlanks | 10412 | 10430 | 19 | AGTCGGTCCTGAGAGATGG | 291.4 | -7.1 | 1.1 | -6.4 | 1.7E-10 | 4.7E-10 | BT20/MB468 | CYTO |
| >hg38_21_+_8433172_8446622_RNA45SN1_45S_50ntFlanks@12547.12569.23\|TTGTTGCCATGGTAATCCTGCTC | >hg38_21_+_8433172_8446622_RNA45SN1_45S_50ntFlanks | 12547 | 12569 | 23 | TTGTTGCCATGGTAATCCTGCTC | 75.7 | -10.1 | 1.6 | -6.4 | 1.8E-10 | 4.7E-10 | MB231/MB468 | MP |

| | | | | | | | | | | | | | |
|---|---|---|---|---|---|---|---|---|---|---|---|---|---|
| >hg38_21_+_8433172_8446622_RNA45SN1_45S_50ntFlanks@4724.4751.28\|ATTAATCAAGAACGAAAGTCGGAGGTTC | >hg38_21_+_8433172_8446622_RNA45SN1_45S_50ntFlanks | 4724 | 4751 | 28 | ATTAATCAAGAACGAAAGTCGGAGGTTC | 674.9 | -6.5 | 1.0 | -6.4 | 1.9E-10 | 4.7E-10 | MB231/MB468 | NUC |
| >hg38_21_+_8433172_8446622_RNA45SN1_45S_50ntFlanks@10571.10589.19\|GAGAAGCCGGCGGGAGCCC | >hg38_21_+_8433172_8446622_RNA45SN1_45S_50ntFlanks | 10571 | 10589 | 19 | GAGAAGCCGGCGGGAGCCC | 180.1 | -7.8 | 1.2 | -6.4 | 1.9E-10 | 4.7E-10 | MB231/MB468 | NUC |
| >hg38_21_+_8433172_8446622_RNA45SN1_45S_50ntFlanks@4514.4537.24\|AAAATTAGAGTGTTCAAAGCAGGC | >hg38_21_+_8433172_8446622_RNA45SN1_45S_50ntFlanks | 4514 | 4537 | 24 | AAAATTAGAGTGTTCAAAGCAGGC | 78.5 | -10.2 | 1.6 | -6.4 | 1.8E-10 | 4.7E-10 | MB231/MB468 | MP |
| >trna116_GluCTC_1_-_145399233_145399304@37.74.38,trna134_GluTTC_1_-_16861774_16861845@37.74.38,trna71_GluCTC_1_-_161439189_161439260@37.74.38,trna74_GluCTC_1_-_161431809_161431880@37.74.38,trna77_GluCTC_1_-_161424398_161424469@37.74.38,trna77_GluCTC_6_+_28949976_28950047@37.74.38,trna80_GluCTC_1_-_161417018_161417089@37.74.38,trna84_GluTTC_1_-_161391883_161391954@37.74.38,trna87_GluCTC_6_-_126101393_126101464@37.74.38\|ACCGCCGCGGCCCGGGTTCGATTCCCGGTCAGGGAACC | >trna116_GluCTC_1_-_145399233_145399304 | 37 | 74 | 38 | ACCGCCGCGGCCCGGGTTCGATTCCCGGTCAGGGAACC | 23178.1 | 9.7 | 1.5 | 6.4 | 1.9E-10 | 4.7E-10 | MB231/MB468 | CYTO |
| >MI0000077\|hsa-mir-21&WithFlank&17\|+\|59841260\|59841343@14.35.22(+1G)&[MIMAT0000076&hsa-miR-21-5p&offsets\|0\|0(+1G);m-1&17\|+\|59841273\|59841294&offsets\|0\|0(+1G)]\|TAGCTTATCAGACTGATGTTGAG | >MI0000077\|hsa-mir-21&WithFlank&17\|+\|59841260\|59841343 | 14 | 35 | 22(+1G) | TAGCTTATCAGACTGATGTTGAG | 382.3 | 5.4 | 0.8 | 6.4 | 1.4E-10 | 4.7E-10 | BT20/MB468 | MITO |
| >trnaMT_SerGCT_MT_+_12207_12265@14.45.32\|AGAACTGCTAACTCATGCCCCCATGTCTAACA | >trnaMT_SerGCT_MT_+_12207_12265 | 14 | 45 | 32 | AGAACTGCTAACTCATGCCCCCATGTCTAACA | 852.8 | 5.5 | 0.9 | 6.4 | 1.4E-10 | 4.7E-10 | BT20/MB468 | MITO |
| >trna111_HisGTG_1_-_147774845_147774916@-1T.26.27,trna118_HisGTG_1_-_145396881_145396952@-1T.26.27,trna16_HisGTG_1_+_146544773_146544844@-1T.26.27,trna1_HisGTG_15_+_45493349_45493420@-1T.26.27,trna21_HisGTG_1_+_147753471_147753542@-1T.26.27,trna33_HisGTG_6_+_27125906_27125977@-1T.26.27,trna7_HisGTG_9_-_14433938_14434009@-1T.26.27,trna8_HisGTG_15_-_45492611_45492682@-1T.26.27,trna9_HisGTG_15_-_45490804_45490875@-1T.26.27\|TGCCGTGATCGTATAGTGGTTAGTACT | >trna111_HisGTG_1_-_147774845_147774916 | -1T | 26 | 27 | TGCCGTGATCGTATAGTGGTTAGTACT | 3624.5 | 4.9 | 0.8 | 6.4 | 1.9E-10 | 4.7E-10 | MB231/MB468 | CYTO |
| >MI0000084\|hsa-mir-26b&WithFlank&2\|+\|218402640\|218402728@18.39.22(+1U)&[MIMAT0000083&hsa-miR-26b-5p&offsets\|0\|+1(+1U);m-53&2\|+\|218402657\|218402678&offsets\|0\|0(+1U)]\|TTCAAGTAATTCAGGATAGGTTT | >MI0000084\|hsa-mir-26b&WithFlank&2\|+\|218402640\|218402728 | 18 | 39 | 22(+1U) | TTCAAGTAATTCAGGATAGGTTT | 243.5 | 5.2 | 0.8 | 6.4 | 1.9E-10 | 4.7E-10 | MB231/MB468 | NUC |
| >hg38_21_+_8433172_8446622_RNA45SN1_45S_50ntFlanks@11847.11872.26\|AACGGGCTTGGCGGAATCAGCGGGGA | >hg38_21_+_8433172_8446622_RNA45SN1_45S_50ntFlanks | 11847 | 11872 | 26 | AACGGGCTTGGCGGAATCAGCGGGGA | 1036.9 | -9.4 | 1.5 | -6.3 | 2.2E-10 | 4.7E-10 | BT20/MB468 | NUC |
| >hg38_21_+_8433172_8446622_RNA45SN1_45S_50ntFlanks@10830.10858.29\|CAAGCCGGATCCGTAACTTCGGGATAAGG | >hg38_21_+_8433172_8446622_RNA45SN1_45S_50ntFlanks | 10830 | 10858 | 29 | CAAGCCGGATCCGTAACTTCGGGATAAGG | 183.2 | -7.3 | 1.1 | -6.4 | 1.9E-10 | 4.7E-10 | MB231/MB468 | NUC |
| >hg38_21_+_8433172_8446622_RNA45SN1_45S_50ntFlanks@4050.4069.20\|CGAACGTCTGCCCTATCAAC | >hg38_21_+_8433172_8446622_RNA45SN1_45S_50ntFlanks | 4050 | 4069 | 20 | CGAACGTCTGCCCTATCAAC | 388.1 | 6.2 | 1.0 | 6.4 | 1.9E-10 | 4.7E-10 | MB231/MB468 | CYTO |
| >hg38_21_+_8433172_8446622_RNA45SN1_45S_50ntFlanks@12154.12180.27\|GGGAGTTTGACTGGGGCGGTACACCTG | >hg38_21_+_8433172_8446622_RNA45SN1_45S_50ntFlanks | 12154 | 12180 | 27 | GGGAGTTTGACTGGGGCGGTACACCTG | 113.8 | -8.9 | 1.4 | -6.4 | 1.4E-10 | 4.7E-10 | BT20/MB468 | MITO |
| >hg38_21_+_8433172_8446622_RNA45SN1_45S_50ntFlanks@12324.12345.22\|TTGGGTTTTAAGCAGGAGGTGT | >hg38_21_+_8433172_8446622_RNA45SN1_45S_50ntFlanks | 12324 | 12345 | 22 | TTGGGTTTTAAGCAGGAGGTGT | 841.8 | -6.1 | 1.0 | -6.4 | 1.9E-10 | 4.7E-10 | MB231/MB468 | CYTO |
| >hg38_21_+_8433172_8446622_RNA45SN1_45S_50ntFlanks@9568.9586.19\|CGTGCAAATCGGTCGTCCG | >hg38_21_+_8433172_8446622_RNA45SN1_45S_50ntFlanks | 9568 | 9586 | 19 | CGTGCAAATCGGTCGTCCG | 356.4 | -7.6 | 1.2 | -6.4 | 1.7E-10 | 4.7E-10 | BT20/MB468 | CYTO |
| >hg38_21_+_8433172_8446622_RNA45SN1_45S_50ntFlanks@8388.8405.18\|CGTGAAACCGTTAAGAGG | >hg38_21_+_8433172_8446622_RNA45SN1_45S_50ntFlanks | 8388 | 8405 | 18 | CGTGAAACCGTTAAGAGG | 215.1 | -10.8 | 1.6 | -6.7 | 1.5E-11 | 4.7E-10 | BT20/MB468 | MP |
| >hg38_21_+_8433172_8446622_RNA45SN1_45S_50ntFlanks@10587.10608.22\|CCCCGGGGAGAGTTCTCTTTTC | >hg38_21_+_8433172_8446622_RNA45SN1_45S_50ntFlanks | 10587 | 10608 | 22 | CCCCGGGGAGAGTTCTCTTTTC | 287.7 | -6.9 | 1.1 | -6.4 | 1.9E-10 | 4.7E-10 | MB231/MB468 | CYTO |
| >trna119_LysCTT_1_-_145395522_145395594@4.36.33,trna11_LysCTT_5_-_180648979_180649051@4.36.33,trna13_LysCTT_6_+_26556774_26556846@4.36.33,trna7_LysCTT_16_+_3225692_3225764@4.36.33,trna9_LysCTT_5_+_180634755_180634827@4.36.33\|CGGCTAGCTCAGTCGGTAGAGCATGAGACTCTT | >trna119_LysCTT_1_-_145395522_145395594 | 4 | 36 | 33 | CGGCTAGCTCAGTCGGTAGAGCATGAGACTCTT | 626.1 | 7.5 | 1.2 | 6.4 | 1.9E-10 | 4.7E-10 | MB231/MB468 | NUC |
| >MI0003137\|hsa-mir-193b&WithFlank&16\|+\|14303961\|14304055@57.78.22(+1A)&[MIMAT0002819&hsa-miR-193b-3p&offsets\|0\|0(+1A);m-166&16\|+\|14304017\|14304038&offsets\|0\|0(+1A)]\|AACTGGCCCTCAAAGTCCCGCTA | >MI0003137\|hsa-mir-193b&WithFlank&16\|+\|14303961\|14304055 | 57 | 78 | 22(+1A) | AACTGGCCCTCAAAGTCCCGCTA | 218.5 | 8.4 | 1.2 | 6.7 | 2.0E-11 | 4.7E-10 | BT20/MB231 | NUC |
| >hg38_21_+_8433172_8446622_RNA45SN1_45S_50ntFlanks@3986.4023.38\|GGCTTTGGTGACTCTAGATAACCTCGGGCCGATCGCAC | >hg38_21_+_8433172_8446622_RNA45SN1_45S_50ntFlanks | 3986 | 4023 | 38 | GGCTTTGGTGACTCTAGATAACCTCGGGCCGATCGCAC | 1478.8 | 6.6 | 1.0 | 6.4 | 1.9E-10 | 4.7E-10 | MB231/MB468 | CYTO |
| >hg38_21_+_8433172_8446622_RNA45SN1_45S_50ntFlanks@10771.10807.37\|CAGGTCTCCAAGGTGAACAGCCTCTGGCATGTTGGAA | >hg38_21_+_8433172_8446622_RNA45SN1_45S_50ntFlanks | 10771 | 10807 | 37 | CAGGTCTCCAAGGTGAACAGCCTCTGGCATGTTGGAA | 177.9 | -7.7 | 1.2 | -6.4 | 1.9E-10 | 4.7E-10 | MB231/MB468 | NUC |
| >hg38_21_+_8433172_8446622_RNA45SN1_45S_50ntFlanks@3838.3859.22\|CTCTCCTACTTGGATAACTGTG | >hg38_21_+_8433172_8446622_RNA45SN1_45S_50ntFlanks | 3838 | 3859 | 22 | CTCTCCTACTTGGATAACTGTG | 168.4 | -11.3 | 1.8 | -6.4 | 1.8E-10 | 4.7E-10 | MB231/MB468 | MP |
| >hg38_21_+_8433172_8446622_RNA45SN1_45S_50ntFlanks@6738.6772.35\|ATCATCGACACTTCGAACGCACTTGCGGCCCCGGG | >hg38_21_+_8433172_8446622_RNA45SN1_45S_50ntFlanks | 6738 | 6772 | 35 | ATCATCGACACTTCGAACGCACTTGCGGCCCCGGG | 680.7 | -12.5 | 1.9 | -6.7 | 1.5E-11 | 4.7E-10 | BT20/MB468 | MP |
| >am_trna128_GlyGCC_6_-_27870686_27870756@1.36.36,trna18_GlyGCC_16_+_70822597_70822667@1.36.36,trna19_GlyGCC_16_+_70823410_70823480@1.36.36,trna19_GlyGCC_2_-_157257659_157257729@1.36.36,trna24_GlyGCC_16_-_70812942_70813012@1.36.36,trna25_GlyGCC_16_-_70812114_70812184@1.36.36,trna5_GlyGCC_17_+_8029064_8029134@1.36.36,trna68_GlyGCC_1_-_161493637_161493707@1.36.36\|GCATTGGTGGTTCAGTGGTAGAATTCTCGCCTGCCA | >am_trna128_GlyGCC_6_-_27870686_27870756 | 1 | 36 | 36 | GCATTGGTGGTTCAGTGGTAGAATTCTCGCCTGCCA | 432.5 | 5.6 | 0.9 | 6.4 | 1.9E-10 | 4.7E-10 | MB231/MB468 | NUC |
| >hg38_21_+_8433172_8446622_RNA45SN1_45S_50ntFlanks@6789.6807.19\|GCCTGTCTGAGCGTCGCTT | >hg38_21_+_8433172_8446622_RNA45SN1_45S_50ntFlanks | 6789 | 6807 | 19 | GCCTGTCTGAGCGTCGCTT | 428.7 | 4.8 | 0.8 | 6.4 | 1.7E-10 | 4.7E-10 | BT20/MB468 | CYTO |
| >hg38_21_+_8433172_8446622_RNA45SN1_45S_50ntFlanks@11885.11912.28\|TTGAGCTTGACTCTAGTCTGGCACGGTG | >hg38_21_+_8433172_8446622_RNA45SN1_45S_50ntFlanks | 11885 | 11912 | 28 | TTGAGCTTGACTCTAGTCTGGCACGGTG | 124.1 | -9.3 | 1.5 | -6.4 | 1.7E-10 | 4.8E-10 | BT20/MB468 | CYTO |
| >hg38_21_+_8433172_8446622_RNA45SN1_45S_50ntFlanks@10922.10939.18\|CTGGACGAGGCGCCGCCG | >hg38_21_+_8433172_8446622_RNA45SN1_45S_50ntFlanks | 10922 | 10939 | 18 | CTGGACGAGGCGCCGCCG | 1154.8 | -7.9 | 1.2 | -6.4 | 1.9E-10 | 4.8E-10 | MB231/MB468 | CYTO |
| >hg38_21_+_8433172_8446622_RNA45SN1_45S_50ntFlanks@4889.4916.28\|CTGAAACTTAAAGGAATTGACGGAAGGG | >hg38_21_+_8433172_8446622_RNA45SN1_45S_50ntFlanks | 4889 | 4916 | 28 | CTGAAACTTAAAGGAATTGACGGAAGGG | 71.9 | -10.0 | 1.6 | -6.4 | 1.8E-10 | 4.8E-10 | MB231/MB468 | MP |
| >MI0000077\|hsa-mir-21&WithFlank&17\|+\|59841260\|59841343@14.32.19&[MIMAT0000076&hsa-miR-21-5p&offsets\|0\|-3;m-1&17\|+\|59841273\|59841294&offsets\|0\|-3]\|TAGCTTATCAGACTGATGT | >MI0000077\|hsa-mir-21&WithFlank&17\|+\|59841260\|59841343 | 14 | 32 | 19 | TAGCTTATCAGACTGATGT | 770.9 | 7.4 | 1.2 | 6.4 | 1.4E-10 | 4.8E-10 | BT20/MB468 | MITO |
| >hg38_21_+_8433172_8446622_RNA45SN1_45S_50ntFlanks@11914.11948.35\|AGAGACATGAGAGGTGTAGAATAAGTGGGAGGCCC | >hg38_21_+_8433172_8446622_RNA45SN1_45S_50ntFlanks | 11914 | 11948 | 35 | AGAGACATGAGAGGTGTAGAATAAGTGGGAGGCCC | 132.1 | -10.9 | 1.7 | -6.4 | 1.8E-10 | 4.8E-10 | MB231/MB468 | MP |
| >hg38_21_+_8433172_8446622_RNA45SN1_45S_50ntFlanks@11728.11751.24\|GTAACTATGACTCTCTTAAGGTAG | >hg38_21_+_8433172_8446622_RNA45SN1_45S_50ntFlanks | 11728 | 11751 | 24 | GTAACTATGACTCTCTTAAGGTAG | 110.5 | -10.7 | 1.7 | -6.4 | 1.8E-10 | 4.8E-10 | MB231/MB468 | MP |
| >hg38_MT_+_1621_3279_RNR2_16S_50ntFlanks@718.740.23\|AGCCTGCGTCAGATTAAAACACT | >hg38_MT_+_1621_3279_RNR2_16S_50ntFlanks | 718 | 740 | 23 | AGCCTGCGTCAGATTAAAACACT | 113.3 | 10.0 | 1.6 | 6.3 | 2.2E-10 | 4.8E-10 | MB231/MB468 | MITO |
| >hg38_21_+_8433172_8446622_RNA45SN1_45S_50ntFlanks@12437.12455.19\|CATTGTGAAGCAGAATTCA | >hg38_21_+_8433172_8446622_RNA45SN1_45S_50ntFlanks | 12437 | 12455 | 19 | CATTGTGAAGCAGAATTCA | 16373.6 | -10.4 | 1.6 | -6.4 | 1.7E-10 | 4.8E-10 | BT20/MB468 | CYTO |
| >MI0000433\|hsa-let-7g&WithFlank&3\|-\|52268272\|52268367@11.30.20(+1C)&[MIMAT0000414&hsa-let-7g-5p&offsets\|0\|-2(+1C);m-58&3\|-\|52268336\|52268357&offsets\|0\|-2(+1C)]\|TGAGGTAGTAGTTTGTACAGC | >MI0000433\|hsa-let-7g&WithFlank&3\|-\|52268272\|52268367 | 11 | 30 | 20(+1C) | TGAGGTAGTAGTTTGTACAGC | 249.9 | 5.4 | 0.8 | 6.4 | 1.9E-10 | 4.8E-10 | MB231/MB468 | CYTO |
| >hg38_21_+_8433172_8446622_RNA45SN1_45S_50ntFlanks@11564.11590.27\|CCTAGCAGCCGACTTAGAACTGGTGCG | >hg38_21_+_8433172_8446622_RNA45SN1_45S_50ntFlanks | 11564 | 11590 | 27 | CCTAGCAGCCGACTTAGAACTGGTGCG | 965.5 | -9.0 | 1.4 | -6.4 | 1.8E-10 | 4.8E-10 | MB231/MB468 | MP |
| >hg38_21_+_8433172_8446622_RNA45SN1_45S_50ntFlanks@6690.6718.29\|AAGAACGCAGCTAGCTGCGAGAATTAATG | >hg38_21_+_8433172_8446622_RNA45SN1_45S_50ntFlanks | 6690 | 6718 | 29 | AAGAACGCAGCTAGCTGCGAGAATTAATG | 173.9 | -9.5 | 1.5 | -6.4 | 1.4E-10 | 4.8E-10 | BT20/MB468 | MITO |
| >hg38_21_+_8433172_8446622_RNA45SN1_45S_50ntFlanks@12006.12031.26\|GGCCGCCGGTGAAATACCACTACTCT | >hg38_21_+_8433172_8446622_RNA45SN1_45S_50ntFlanks | 12006 | 12031 | 26 | GGCCGCCGGTGAAATACCACTACTCT | 542.4 | -4.9 | 0.8 | -6.3 | 2.2E-10 | 4.8E-10 | MB231/MB468 | MITO |
| >hg38_21_+_8433172_8446622_RNA45SN1_45S_50ntFlanks@10407.10426.20\|GGGTCAGTCGGTCCTGAGAG | >hg38_21_+_8433172_8446622_RNA45SN1_45S_50ntFlanks | 10407 | 10426 | 20 | GGGTCAGTCGGTCCTGAGAG | 89.9 | -10.4 | 1.6 | -6.4 | 1.8E-10 | 4.8E-10 | MB231/MB468 | MP |
| >hg38_21_+_8433172_8446622_RNA45SN1_45S_50ntFlanks@11027.11050.24\|CTCTCTCTCTCTCTCCCCCGCTCC | >hg38_21_+_8433172_8446622_RNA45SN1_45S_50ntFlanks | 11027 | 11050 | 24 | CTCTCTCTCTCTCTCCCCCGCTCC | 90.2 | -10.4 | 1.6 | -6.4 | 1.8E-10 | 4.8E-10 | MB231/MB468 | MP |
| >am_trnaMT_ArgTCG_MT_+_10405_10469@28.48.21\|ATTTCGACTCATTAAATTATG | >am_trnaMT_ArgTCG_MT_+_10405_10469 | 28 | 48 | 21 | ATTTCGACTCATTAAATTATG | 1590.5 | -9.2 | 1.5 | -6.3 | 2.2E-10 | 4.8E-10 | MB231/MB468 | MITO |
| >hg38_21_+_8433172_8446622_RNA45SN1_45S_50ntFlanks@6772.6806.35\|GTTCCTCCCGGGGCTACGCCTGTCTGAGCGTCGCT | >hg38_21_+_8433172_8446622_RNA45SN1_45S_50ntFlanks | 6772 | 6806 | 35 | GTTCCTCCCGGGGCTACGCCTGTCTGAGCGTCGCT | 7934.5 | -4.1 | 0.6 | -6.3 | 2.2E-10 | 4.8E-10 | MB231/MB468 | MITO |
| >hg38_21_+_8433172_8446622_RNA45SN1_45S_50ntFlanks@11117.11134.18\|GCAGGGGCCGGCGGCGGC | >hg38_21_+_8433172_8446622_RNA45SN1_45S_50ntFlanks | 11117 | 11134 | 18 | GCAGGGGCCGGCGGCGGC | 97.9 | -10.5 | 1.6 | -6.4 | 1.8E-10 | 4.8E-10 | MB231/MB468 | MP |
| >hg38_21_+_8433172_8446622_RNA45SN1_45S_50ntFlanks@4073.4095.23\|CGATGGTAGTCGCCGTGCCTACC | >hg38_21_+_8433172_8446622_RNA45SN1_45S_50ntFlanks | 4073 | 4095 | 23 | CGATGGTAGTCGCCGTGCCTACC | 89.5 | -10.3 | 1.6 | -6.4 | 1.8E-10 | 4.8E-10 | MB231/MB468 | MP |
| >hg38_21_+_8433172_8446622_RNA45SN1_45S_50ntFlanks@11604.11621.18\|CGACTGTTTAATTAAAAC | >hg38_21_+_8433172_8446622_RNA45SN1_45S_50ntFlanks | 11604 | 11621 | 18 | CGACTGTTTAATTAAAAC | 651.3 | -4.1 | 0.6 | -6.3 | 2.2E-10 | 4.8E-10 | MB231/MB468 | MITO |
| >hg38_21_+_8433172_8446622_RNA45SN1_45S_50ntFlanks@12671.12703.33\|AAGTCAGAATCCCGCCCAGGCGGAACGATACGG | >hg38_21_+_8433172_8446622_RNA45SN1_45S_50ntFlanks | 12671 | 12703 | 33 | AAGTCAGAATCCCGCCCAGGCGGAACGATACGG | 129.8 | -10.9 | 1.7 | -6.4 | 1.8E-10 | 4.8E-10 | MB231/MB468 | MP |

| | | | | | | | | | | | | | |
|---|---|---|---|---|---|---|---|---|---|---|---|---|---|
| >hg38_21_+_8433172_8446622_RNA45SN1_45S_50ntFlanks@7979.7999.21\|ACCTCAGATCAGACGTGGCGA | >hg38_21_+_8433172_8446622_RNA45SN1_45S_50ntFlanks | 7979 | 7999 | 21 | ACCTCAGATCAGACGTGGCGA | 235.0 | -6.6 | 1.0 | -6.4 | 2.0E-10 | 4.9E-10 | MB231/MB468 | CYTO |
| >hg38_21_+_8433172_8446622_RNA45SN1_45S_50ntFlanks@8343.8375.33\|GTAAGGGAAAGTTGAAAAGAACTTTGAAGAGAG | >hg38_21_+_8433172_8446622_RNA45SN1_45S_50ntFlanks | 8343 | 8375 | 33 | GTAAGGGAAAGTTGAAAAGAACTTTGAAGAGAG | 1113.0 | 10.9 | 1.6 | 6.7 | 1.5E-11 | 4.9E-10 | BT20/MB468 | MP |
| >hg38_21_+_8433172_8446622_RNA45SN1_45S_50ntFlanks@4884.4907.24\|CAAAGCTGAAACTTAAAGGAATTG | >hg38_21_+_8433172_8446622_RNA45SN1_45S_50ntFlanks | 4884 | 4907 | 24 | CAAAGCTGAAACTTAAAGGAATTG | 168.9 | -10.5 | 1.6 | -6.7 | 1.5E-11 | 4.9E-10 | BT20/MB468 | MP |
| >hg38_21_+_8433172_8446622_RNA45SN1_45S_50ntFlanks@12243.12262.20\|GAAGGGCAAAAGCTCGCTTG | >hg38_21_+_8433172_8446622_RNA45SN1_45S_50ntFlanks | 12243 | 12262 | 20 | GAAGGGCAAAAGCTCGCTTG | 88.2 | -10.3 | 1.6 | -6.4 | 1.9E-10 | 4.9E-10 | MB231/MB468 | MP |
| >hg38_21_+_8433172_8446622_RNA45SN1_45S_50ntFlanks@4727.4765.39\|AATCAAGAACGAAAGTCGGAGGTTCGAAGACGATCAGAT | >hg38_21_+_8433172_8446622_RNA45SN1_45S_50ntFlanks | 4727 | 4765 | 39 | AATCAAGAACGAAAGTCGGAGGTTCGAAGACGATCAGAT | 429.4 | 5.2 | 0.8 | 6.4 | 1.4E-10 | 4.9E-10 | BT20/MB468 | MITO |
| >hg38_21_+_8433172_8446622_RNA45SN1_45S_50ntFlanks@12306.12323.18\|CACGATCCTTCTGACCTT | >hg38_21_+_8433172_8446622_RNA45SN1_45S_50ntFlanks | 12306 | 12323 | 18 | CACGATCCTTCTGACCTT | 3908.3 | -6.5 | 1.0 | -6.4 | 1.4E-10 | 4.9E-10 | BT20/MB468 | MITO |
| >hg38_21_+_8433172_8446622_RNA45SN1_45S_50ntFlanks@12416.12452.37\|TTCGATGTCGGCTCTTCCTATCATTGTGAAGCAGAAT | >hg38_21_+_8433172_8446622_RNA45SN1_45S_50ntFlanks | 12416 | 12452 | 37 | TTCGATGTCGGCTCTTCCTATCATTGTGAAGCAGAAT | 514.6 | 6.9 | 1.1 | 6.4 | 2.0E-10 | 4.9E-10 | MB231/MB468 | CYTO |
| >hg38_21_+_8433172_8446622_RNA45SN1_45S_50ntFlanks@5470.5491.22\|CCACGGCCCTGGCGGAGCGCTG | >hg38_21_+_8433172_8446622_RNA45SN1_45S_50ntFlanks | 5470 | 5491 | 22 | CCACGGCCCTGGCGGAGCGCTG | 85.9 | -10.3 | 1.6 | -6.4 | 1.9E-10 | 4.9E-10 | MB231/MB468 | MP |
| >hg38_21_+_8433172_8446622_RNA45SN1_45S_50ntFlanks@4976.4999.24\|CCGGACACGGACAGGATTGACAGA | >hg38_21_+_8433172_8446622_RNA45SN1_45S_50ntFlanks | 4976 | 4999 | 24 | CCGGACACGGACAGGATTGACAGA | 720.5 | -4.0 | 0.6 | -6.4 | 1.9E-10 | 4.9E-10 | MB231/MB468 | MP |
| >hg38_21_+_8433172_8446622_RNA45SN1_45S_50ntFlanks@11829.11852.24\|CGAAACCACAGCCAAGGGAACGGG | >hg38_21_+_8433172_8446622_RNA45SN1_45S_50ntFlanks | 11829 | 11852 | 24 | CGAAACCACAGCCAAGGGAACGGG | 71.6 | -10.0 | 1.6 | -6.4 | 1.9E-10 | 4.9E-10 | MB231/MB468 | MP |
| >hg38_21_+_8433172_8446622_RNA45SN1_45S_50ntFlanks@10411.10456.46\|CAGTCGGTCCTGAGAGATGGGCGAGCGCCGTTCCGAAGGGACGGGC | >hg38_21_+_8433172_8446622_RNA45SN1_45S_50ntFlanks | 10411 | 10456 | 46 | CAGTCGGTCCTGAGAGATGGGCGAGCGCCGTTCCGAAGGGACGGGC | 1545.7 | -8.8 | 1.4 | -6.3 | 2.2E-10 | 4.9E-10 | MB231/MB468 | MITO |
| >MI0000782\|hsa-mir-374a&WithFlank&X\|-\|74287280\|74287363@18.38.21&[MIMAT0000727&hsa-miR-374a-5p&offsets\|0\|-1;m-184&X\|-\|74287325\|74287346&offsets\|0\|-1]\|TTATAATACAACCTGATAAGT | >MI0000782\|hsa-mir-374a&WithFlank&X\|-\|74287280\|74287363 | 18 | 38 | 21 | TTATAATACAACCTGATAAGT | 247.6 | 5.1 | 0.8 | 6.4 | 2.0E-10 | 4.9E-10 | MB231/MB468 | NUC |
| >hg38_21_+_8433172_8446622_RNA45SN1_45S_50ntFlanks@11781.11805.25\|CGCATGAATGGATGAACGAGATTCC | >hg38_21_+_8433172_8446622_RNA45SN1_45S_50ntFlanks | 11781 | 11805 | 25 | CGCATGAATGGATGAACGAGATTCC | 407.3 | -6.2 | 1.0 | -6.3 | 2.2E-10 | 4.9E-10 | MB231/MB468 | MITO |
| >hg38_21_+_8433172_8446622_RNA45SN1_45S_50ntFlanks@10684.10708.25\|CGGCGGCGTCCGGTGAGCTCTCGCT | >hg38_21_+_8433172_8446622_RNA45SN1_45S_50ntFlanks | 10684 | 10708 | 25 | CGGCGGCGTCCGGTGAGCTCTCGCT | 686.9 | -3.6 | 0.6 | -6.4 | 2.0E-10 | 4.9E-10 | MB231/MB468 | CYTO |
| >hg38_21_+_8433172_8446622_RNA45SN1_45S_50ntFlanks@5485.5512.28\|AGCGCTGAGAAGACGGTCGAACTTGACT | >hg38_21_+_8433172_8446622_RNA45SN1_45S_50ntFlanks | 5485 | 5512 | 28 | AGCGCTGAGAAGACGGTCGAACTTGACT | 964.7 | -7.0 | 1.1 | -6.4 | 1.9E-10 | 4.9E-10 | MB231/MB468 | MP |
| >hg38_21_+_8433172_8446622_RNA45SN1_45S_50ntFlanks@12841.12862.22\|CGGAGTGCCCTTCGTCCTGGGA | >hg38_21_+_8433172_8446622_RNA45SN1_45S_50ntFlanks | 12841 | 12862 | 22 | CGGAGTGCCCTTCGTCCTGGGA | 132.5 | -9.1 | 1.4 | -6.4 | 1.5E-10 | 4.9E-10 | BT20/MB468 | MITO |
| >hg38_21_+_8433172_8446622_RNA45SN1_45S_50ntFlanks@485.531.47\|GATGTGGTGACGTCGTGCTCTCCCGGGCCGGGTCCGAGCCGCGACGG | >hg38_21_+_8433172_8446622_RNA45SN1_45S_50ntFlanks | 485 | 531 | 47 | GATGTGGTGACGTCGTGCTCTCCCGGGCCGGGTCCGAGCCGCGACGG | 256.5 | 10.8 | 1.7 | 6.3 | 2.3E-10 | 4.9E-10 | BT20/MB468 | NUC |
| >hg38_21_+_8433172_8446622_RNA45SN1_45S_50ntFlanks@4297.4323.27\|CATTGGAGGGCAAGTCTGGTGCCAGCA | >hg38_21_+_8433172_8446622_RNA45SN1_45S_50ntFlanks | 4297 | 4323 | 27 | CATTGGAGGGCAAGTCTGGTGCCAGCA | 80.8 | -10.2 | 1.6 | -6.4 | 1.9E-10 | 4.9E-10 | MB231/MB468 | MP |
| >hg38_21_+_8433172_8446622_RNA45SN1_45S_50ntFlanks@11920.11968.49\|ATGAGAGGTGTAGAATAAGTGGGAGGCCCCCGGCGCCCCCCCGGTGTCC | >hg38_21_+_8433172_8446622_RNA45SN1_45S_50ntFlanks | 11920 | 11968 | 49 | ATGAGAGGTGTAGAATAAGTGGGAGGCCCCCGGCGCCCCCCCGGTGTCC | 257.3 | -11.0 | 1.7 | -6.4 | 2.0E-10 | 4.9E-10 | MB231/MB468 | NUC |
| >am_MI0005763\|hsa-mir-941-1&WithFlank&20\|+\|63919443\|63919526@53.74.22(+1U);MI0005764\|hsa-mir-941-2&WithFlank&20\|+\|63919499\|63919582@53.74.22(+1U);MI0005765\|hsa-mir-941-3&WithFlank&20\|+\|63919555\|63919638@53.74.22(+1U);MI0005766\|hsa-mir-941-4&WithFlank&20\|+\|63919750\|63919833@53.74.22(+1U);MI0031520\|hsa-mir-941-5&WithFlank&20\|+\|63919862\|63919945@53.74.22(+1U)&[MIMAT0004984&hsa-miR-941&offsets\|0\|-1(+1U);m-263&20\|+\|63919495\|63919517&offsets\|0\|-1(+1U)];[MIMAT0004984_1&hsa-miR-941&offsets\|0\|-1(+1U);m-263&20\|+\|63919551\|63919573&offsets\|0\|-1(+1U)];[MIMAT0004984_2&hsa-miR-941&offsets\|0\|-1(+1U);m-263&20\|+\|63919607\|63919629&offsets\|0\|-1(+1U)];[MIMAT0004984_3&hsa-miR-941&offsets\|0\|-1(+1U);m-263&20\|+\|63919802\|63919824&offsets\|0\|-1(+1U)];[MIMAT0004984_4&hsa-miR-941&offsets\|0\|-1(+1U);m-263&20\|+\|63919914\|63919936&offsets\|0\|-1(+1U)]\|CACCCGGCTGTGTGCACATGTGT | >am_MI0005763\|hsa-mir-941-1&WithFlank&20\|+\|63919443\|63919526 | 53 | 74 | 22(+1U) | CACCCGGCTGTGTGCACATGTGT | 1483.2 | 5.1 | 0.8 | 6.7 | 1.6E-11 | 4.9E-10 | BT20/MB468 | MP |
| >hg38_21_+_8433172_8446622_RNA45SN1_45S_50ntFlanks@10571.10594.24\|GAGAAGCCGGCGGGAGCCCCGGGG | >hg38_21_+_8433172_8446622_RNA45SN1_45S_50ntFlanks | 10571 | 10594 | 24 | GAGAAGCCGGCGGGAGCCCCGGGG | 87.2 | -10.3 | 1.6 | -6.4 | 1.9E-10 | 4.9E-10 | MB231/MB468 | MP |
| >hg38_21_+_8433172_8446622_RNA45SN1_45S_50ntFlanks@3769.3799.31\|CGCACGGCCGGTACAGTGAAACTGCGAATGG | >hg38_21_+_8433172_8446622_RNA45SN1_45S_50ntFlanks | 3769 | 3799 | 31 | CGCACGGCCGGTACAGTGAAACTGCGAATGG | 83.2 | -10.2 | 1.6 | -6.4 | 1.9E-10 | 4.9E-10 | MB231/MB468 | MP |
| >hg38_21_+_8433172_8446622_RNA45SN1_45S_50ntFlanks@9308.9334.27\|AGGAGTCTAACACGTGCGCGAGTCGGG | >hg38_21_+_8433172_8446622_RNA45SN1_45S_50ntFlanks | 9308 | 9334 | 27 | AGGAGTCTAACACGTGCGCGAGTCGGG | 756.6 | -8.9 | 1.4 | -6.4 | 1.9E-10 | 4.9E-10 | MB231/MB468 | MP |
| >hg38_21_+_8433172_8446622_RNA45SN1_45S_50ntFlanks@9822.9850.29\|CTAGTGGGCCACTTTTGGTAAGCAGAACT | >hg38_21_+_8433172_8446622_RNA45SN1_45S_50ntFlanks | 9822 | 9850 | 29 | CTAGTGGGCCACTTTTGGTAAGCAGAACT | 767.4 | -6.8 | 1.1 | -6.3 | 2.2E-10 | 4.9E-10 | MB231/MB468 | MITO |
| >trna119_LysCTT_1_-_145395522_145395594@4.35.32,trna11_LysCTT_5_-_180648979_180649051@4.35.32,trna13_LysCTT_6_+_26556774_26556846@4.35.32,trna7_LysCTT_16_+_3225692_3225764@4.35.32,trna9_LysCTT_5_+_180634755_180634827@4.35.32\|CGGCTAGCTCAGTCGGTAGAGCATGAGACTCT | >trna119_LysCTT_1_-_145395522_145395594 | 4 | 35 | 32 | CGGCTAGCTCAGTCGGTAGAGCATGAGACTCT | 5712.5 | -5.1 | 0.8 | -6.7 | 1.8E-11 | 5.0E-10 | BT20/MB231 | MITO |
| >hg38_21_+_8433172_8446622_RNA45SN1_45S_50ntFlanks@7980.8009.30\|CCTCAGATCAGACGTGGCGACCCGCTGAAT | >hg38_21_+_8433172_8446622_RNA45SN1_45S_50ntFlanks | 7980 | 8009 | 30 | CCTCAGATCAGACGTGGCGACCCGCTGAAT | 782.8 | -4.1 | 0.6 | -6.3 | 2.2E-10 | 5.0E-10 | MB231/MB468 | MITO |
| >hg38_21_+_8433172_8446622_RNA45SN1_45S_50ntFlanks@6656.6688.33\|CTTAGCGGTGGATCACTCGGCTCGTGCGTCGAT | >hg38_21_+_8433172_8446622_RNA45SN1_45S_50ntFlanks | 6656 | 6688 | 33 | CTTAGCGGTGGATCACTCGGCTCGTGCGTCGAT | 94.4 | -10.4 | 1.6 | -6.4 | 1.9E-10 | 5.0E-10 | MB231/MB468 | MP |
| >hg38_21_+_8433172_8446622_RNA45SN1_45S_50ntFlanks@11784.11804.21\|ATGAATGGATGAACGAGATTC | >hg38_21_+_8433172_8446622_RNA45SN1_45S_50ntFlanks | 11784 | 11804 | 21 | ATGAATGGATGAACGAGATTC | 4900.8 | -5.6 | 0.9 | -6.4 | 1.8E-10 | 5.0E-10 | BT20/MB468 | CYTO |
| >hg38_21_+_8433172_8446622_RNA45SN1_45S_50ntFlanks@12159.12179.21\|TTTGACTGGGGCGGTACACCT | >hg38_21_+_8433172_8446622_RNA45SN1_45S_50ntFlanks | 12159 | 12179 | 21 | TTTGACTGGGGCGGTACACCT | 817.1 | -8.3 | 1.3 | -6.4 | 2.0E-10 | 5.0E-10 | MB231/MB468 | NUC |
| >MI0000298\|hsa-mir-221&WithFlank&X\|-\|45746151\|45746272@71.90.20&[MIMAT0000278&hsa-miR-221-3p&offsets\|0\|-3;m-91&X\|-\|45746181\|45746202&offsets\|0\|-2]\|AGCTACATTGTCTGCTGGGT | >MI0000298\|hsa-mir-221&WithFlank&X\|-\|45746151\|45746272 | 71 | 90 | 20 | AGCTACATTGTCTGCTGGGT | 432.0 | 6.7 | 1.1 | 6.4 | 2.0E-10 | 5.0E-10 | MB231/MB468 | CYTO |
| >hg38_21_+_8433172_8446622_RNA45SN1_45S_50ntFlanks@4686.4707.22\|GACGGACCAGAGCGAAAGCATT | >hg38_21_+_8433172_8446622_RNA45SN1_45S_50ntFlanks | 4686 | 4707 | 22 | GACGGACCAGAGCGAAAGCATT | 81.2 | -10.2 | 1.6 | -6.4 | 1.9E-10 | 5.0E-10 | MB231/MB468 | MP |
| >MI0000086\|hsa-mir-28&WithFlank&3\|+\|188688775\|188688872@60.81.22&[MIMAT0004502&hsa-miR-28-3p&offsets\|0\|0;m-28&3\|+\|188688834\|188688855&offsets\|0\|0]\|CACTAGATTGTGAGCTCCTGGA | >MI0000086\|hsa-mir-28&WithFlank&3\|+\|188688775\|188688872 | 60 | 81 | 22 | CACTAGATTGTGAGCTCCTGGA | 3052.8 | 4.8 | 0.8 | 6.4 | 1.9E-10 | 5.0E-10 | MB231/MB468 | MP |
| >hg38_21_+_8433172_8446622_RNA45SN1_45S_50ntFlanks@3991.4020.30\|TGGTGACTCTAGATAACCTCGGGCCGATCG | >hg38_21_+_8433172_8446622_RNA45SN1_45S_50ntFlanks | 3991 | 4020 | 30 | TGGTGACTCTAGATAACCTCGGGCCGATCG | 72.5 | -10.0 | 1.6 | -6.4 | 1.9E-10 | 5.0E-10 | MB231/MB468 | MP |
| >hg38_21_+_8433172_8446622_RNA45SN1_45S_50ntFlanks@12822.12840.19\|CCGGGACCGGGGTCCGGTG | >hg38_21_+_8433172_8446622_RNA45SN1_45S_50ntFlanks | 12822 | 12840 | 19 | CCGGGACCGGGGTCCGGTG | 322.3 | -12.2 | 1.9 | -6.4 | 1.9E-10 | 5.0E-10 | MB231/MB468 | MP |
| >hg38_21_+_8433172_8446622_RNA45SN1_45S_50ntFlanks@11641.11680.40\|CGGCGGGTGTTGACGCGATGTGATTTCTGCCCAGTGCTCT | >hg38_21_+_8433172_8446622_RNA45SN1_45S_50ntFlanks | 11641 | 11680 | 40 | CGGCGGGTGTTGACGCGATGTGATTTCTGCCCAGTGCTCT | 330.5 | -12.2 | 1.9 | -6.4 | 1.9E-10 | 5.0E-10 | MB231/MB468 | MP |
| >hg38_21_+_8433172_8446622_RNA45SN1_45S_50ntFlanks@6648.6667.20\|GTACGACTCTTAGCGGTGGA | >hg38_21_+_8433172_8446622_RNA45SN1_45S_50ntFlanks | 6648 | 6667 | 20 | GTACGACTCTTAGCGGTGGA | 100.1 | 2.6 | 0.4 | 6.3 | 2.4E-10 | 5.0E-10 | BT20/MB468 | NUC |
| >MI0006444\|hsa-mir-1307&WithFlank&10\|-\|103394247\|103394407@86.109.24&[MIMAT0005951&hsa-miR-1307-3p&offsets\|0\|+2;m-66&10\|-\|103394300\|103394322&offsets\|0\|+1]\|ACTCGGCGTGGCGTCGGTCGTGGT | >MI0006444\|hsa-mir-1307&WithFlank&10\|-\|103394247\|103394407 | 86 | 109 | 24 | ACTCGGCGTGGCGTCGGTCGTGGT | 452.9 | 7.0 | 1.1 | 6.4 | 2.0E-10 | 5.0E-10 | MB231/MB468 | CYTO |
| >trna116_GluCTC_1_-_145399233_145399304@31.70.40,trna71_GluCTC_1_-_161439189_161439260@31.70.40,trna74_GluCTC_1_-_161431809_161431880@31.70.40,trna77_GluCTC_1_-_161424398_161424469@31.70.40,trna77_GluCTC_6_+_28949976_28950047@31.70.40,trna80_GluCTC_1_-_161417018_161417089@31.70.40,trna87_GluCTC_6_-_126101393_126101464@31.70.40\|GCTCTCACCGCCGCGGCCCGGGTTCGATTCCCGGTCAGGG | >trna116_GluCTC_1_-_145399233_145399304 | 31 | 70 | 40 | GCTCTCACCGCCGCGGCCCGGGTTCGATTCCCGGTCAGGG | 218.0 | -6.5 | 1.0 | -6.4 | 2.0E-10 | 5.0E-10 | MB231/MB468 | CYTO |
| >am_trna116_GluCTC_1_-_145399233_145399304@11.29.19,trna25_GluCTC_2_-_71273488_71273560@11.29.19,trna59_GluCTC_1_+_249168447_249168518@11.29.19,trna71_GluCTC_1_-_161439189_161439260@11.29.19,trna74_GluCTC_1_-_161431809_161431880@11.29.19,trna77_GluCTC_1_-_161424398_161424469@11.29.19,trna77_GluCTC_6_+_28949976_28950047@11.29.19,trna80_GluCTC_1_-_161417018_161417089@11.29.19,trna87_GluCTC_6_-_126101393_126101464@11.29.19\|TCTAGTGGTTAGGATTCGG | >am_trna116_GluCTC_1_-_145399233_145399304 | 11 | 29 | 19 | TCTAGTGGTTAGGATTCGG | 98.9 | -10.5 | 1.6 | -6.4 | 1.9E-10 | 5.0E-10 | MB231/MB468 | MP |
| >hg38_21_+_8433172_8446622_RNA45SN1_45S_50ntFlanks@13007.13039.33\|TATTGAAAGTCAGCCCTCGACACAAGGGTTTGT | >hg38_21_+_8433172_8446622_RNA45SN1_45S_50ntFlanks | 13007 | 13039 | 33 | TATTGAAAGTCAGCCCTCGACACAAGGGTTTGT | 2700.4 | 5.7 | 0.9 | 6.4 | 1.8E-10 | 5.0E-10 | BT20/MB468 | CYTO |
| >hg38_21_+_8433172_8446622_RNA45SN1_45S_50ntFlanks@10841.10864.24\|CGTAACTTCGGGATAAGGATTGGC | >hg38_21_+_8433172_8446622_RNA45SN1_45S_50ntFlanks | 10841 | 10864 | 24 | CGTAACTTCGGGATAAGGATTGGC | 1309.7 | -7.5 | 1.2 | -6.3 | 2.2E-10 | 5.0E-10 | MB231/MB468 | MITO |

<table>
<tr><td>>hg38_21_+_8433172_8446622_RNA45SN1_45S_50ntFlanks@12637.12660.24|GCTACCATCTGTGGGATTATGACT</td><td>>hg38_21_+_8433172_8446622_RNA45SN1_45S_50ntFlanks</td><td>12637</td><td>12660</td><td>24</td><td>GCTACCATCTGTGGGATTATGACT</td><td>123.2</td><td>-10.8</td><td>1.7</td><td>-6.4</td><td>1.9E-10</td><td>5.0E-10</td><td>MB231/MB468</td><td>MP</td></tr>
<tr><td>>hg38_21_+_8433172_8446622_RNA45SN1_45S_50ntFlanks@11575.11600.26|ACTTAGAACTGGTGCGGACCAGGGGA</td><td>>hg38_21_+_8433172_8446622_RNA45SN1_45S_50ntFlanks</td><td>11575</td><td>11600</td><td>26</td><td>ACTTAGAACTGGTGCGGACCAGGGGA</td><td>817.1</td><td>-4.9</td><td>0.8</td><td>-6.4</td><td>1.8E-10</td><td>5.0E-10</td><td>BT20/MB468</td><td>CYTO</td></tr>
<tr><td>>hg38_21_+_8433172_8446622_RNA45SN1_45S_50ntFlanks@6651.6695.45|CGACTCTTAGCGGTGGATCACTCGGCTCGTGCGTCGATGAAGAAC</td><td>>hg38_21_+_8433172_8446622_RNA45SN1_45S_50ntFlanks</td><td>6651</td><td>6695</td><td>45</td><td>CGACTCTTAGCGGTGGATCACTCGGCTCGTGCGTCGATGAAGAAC</td><td>3548.0</td><td>7.4</td><td>1.2</td><td>6.4</td><td>1.8E-10</td><td>5.0E-10</td><td>BT20/MB468</td><td>CYTO</td></tr>
<tr><td>>hg38_21_+_8433172_8446622_RNA45SN1_45S_50ntFlanks@475.494.20|CTTCGTGATCGATGTGGTGA</td><td>>hg38_21_+_8433172_8446622_RNA45SN1_45S_50ntFlanks</td><td>475</td><td>494</td><td>20</td><td>CTTCGTGATCGATGTGGTGA</td><td>147.7</td><td>10.4</td><td>1.6</td><td>6.3</td><td>2.3E-10</td><td>5.0E-10</td><td>MB231/MB468</td><td>MITO</td></tr>
<tr><td>>hg38_21_+_8433172_8446622_RNA45SN1_45S_50ntFlanks@3985.4008.24|CGGCTTTGGTGACTCTAGATAACC</td><td>>hg38_21_+_8433172_8446622_RNA45SN1_45S_50ntFlanks</td><td>3985</td><td>4008</td><td>24</td><td>CGGCTTTGGTGACTCTAGATAACC</td><td>135.1</td><td>-10.9</td><td>1.7</td><td>-6.4</td><td>1.9E-10</td><td>5.0E-10</td><td>MB231/MB468</td><td>MP</td></tr>
<tr><td>>hg38_21_+_8433172_8446622_RNA45SN1_45S_50ntFlanks@10782.10800.19|GGTGAACAGCCTCTGGCAT</td><td>>hg38_21_+_8433172_8446622_RNA45SN1_45S_50ntFlanks</td><td>10782</td><td>10800</td><td>19</td><td>GGTGAACAGCCTCTGGCAT</td><td>76.3</td><td>-10.1</td><td>1.6</td><td>-6.4</td><td>1.9E-10</td><td>5.0E-10</td><td>MB231/MB468</td><td>MP</td></tr>
<tr><td>>hg38_21_+_8433172_8446622_RNA45SN1_45S_50ntFlanks@10514.10533.20|CGGAGTGGCGGAGATGGGCG</td><td>>hg38_21_+_8433172_8446622_RNA45SN1_45S_50ntFlanks</td><td>10514</td><td>10533</td><td>20</td><td>CGGAGTGGCGGAGATGGGCG</td><td>207.4</td><td>-5.2</td><td>0.8</td><td>-6.4</td><td>2.0E-10</td><td>5.0E-10</td><td>MB231/MB468</td><td>CYTO</td></tr>
<tr><td>>hg38_21_+_8433172_8446622_RNA45SN1_45S_50ntFlanks@12999.13039.41|CTGCGATCTATTGAAAGTCAGCCCTCGACACAAGGGTTTGT</td><td>>hg38_21_+_8433172_8446622_RNA45SN1_45S_50ntFlanks</td><td>12999</td><td>13039</td><td>41</td><td>CTGCGATCTATTGAAAGTCAGCCCTCGACACAAGGGTTTGT</td><td>11940.0</td><td>5.5</td><td>0.9</td><td>6.4</td><td>2.0E-10</td><td>5.0E-10</td><td>MB231/MB468</td><td>CYTO</td></tr>
<tr><td>>hg38_1_-_228634819_228635039_RNA5S12_5S_50ntFlanks@142.170.29|CGCCTGGGAATACCGGGTGCTGTAGGCTT</td><td>>hg38_1_-_228634819_228635039_RNA5S12_5S_50ntFlanks</td><td>142</td><td>170</td><td>29</td><td>CGCCTGGGAATACCGGGTGCTGTAGGCTT</td><td>788.5</td><td>3.7</td><td>0.6</td><td>6.4</td><td>2.0E-10</td><td>5.0E-10</td><td>MB231/MB468</td><td>NUC</td></tr>
<tr><td>>hg38_21_+_8433172_8446622_RNA45SN1_45S_50ntFlanks@12418.12444.27|CGATGTCGGCTCTTCCTATCATTGTGA</td><td>>hg38_21_+_8433172_8446622_RNA45SN1_45S_50ntFlanks</td><td>12418</td><td>12444</td><td>27</td><td>CGATGTCGGCTCTTCCTATCATTGTGA</td><td>3603.7</td><td>-3.9</td><td>0.6</td><td>-6.4</td><td>2.0E-10</td><td>5.0E-10</td><td>MB231/MB468</td><td>NUC</td></tr>
<tr><td>>hg38_21_+_8433172_8446622_RNA45SN1_45S_50ntFlanks@4731.4748.18|AAGAACGAAAGTCGGAGG</td><td>>hg38_21_+_8433172_8446622_RNA45SN1_45S_50ntFlanks</td><td>4731</td><td>4748</td><td>18</td><td>AAGAACGAAAGTCGGAGG</td><td>992.5</td><td>-4.6</td><td>0.7</td><td>-6.4</td><td>1.9E-10</td><td>5.0E-10</td><td>MB231/MB468</td><td>MP</td></tr>
<tr><td>>hg38_21_+_8433172_8446622_RNA45SN1_45S_50ntFlanks@11670.11701.32|CCCAGTGCTCTGAATGTCAAAGTGAAGAAATT</td><td>>hg38_21_+_8433172_8446622_RNA45SN1_45S_50ntFlanks</td><td>11670</td><td>11701</td><td>32</td><td>CCCAGTGCTCTGAATGTCAAAGTGAAGAAATT</td><td>334.2</td><td>-6.6</td><td>1.0</td><td>-6.4</td><td>1.9E-10</td><td>5.0E-10</td><td>MB231/MB468</td><td>MP</td></tr>
<tr><td>>hg38_21_+_8433172_8446622_RNA45SN1_45S_50ntFlanks@5152.5185.34|AGGGACAAGTGGCGTTCAGCCACCCGAGATTGAG</td><td>>hg38_21_+_8433172_8446622_RNA45SN1_45S_50ntFlanks</td><td>5152</td><td>5185</td><td>34</td><td>AGGGACAAGTGGCGTTCAGCCACCCGAGATTGAG</td><td>856.0</td><td>-11.8</td><td>1.8</td><td>-6.4</td><td>1.5E-10</td><td>5.0E-10</td><td>BT20/MB468</td><td>MITO</td></tr>
<tr><td>>hg38_21_+_8433172_8446622_RNA45SN1_45S_50ntFlanks@11919.11967.49|CATGAGAGGTGTAGAATAAGTGGGAGGCCCCCGGCGCCCCCCCGGTGTC</td><td>>hg38_21_+_8433172_8446622_RNA45SN1_45S_50ntFlanks</td><td>11919</td><td>11967</td><td>49</td><td>CATGAGAGGTGTAGAATAAGTGGGAGGCCCCCGGCGCCCCCCCGGTGTC</td><td>399.2</td><td>-11.6</td><td>1.8</td><td>-6.4</td><td>2.0E-10</td><td>5.0E-10</td><td>MB231/MB468</td><td>NUC</td></tr>
<tr><td>>hg38_21_+_8433172_8446622_RNA45SN1_45S_50ntFlanks@8380.8405.26|CAAGAGGGCGTGAAACCGTTAAGAGG</td><td>>hg38_21_+_8433172_8446622_RNA45SN1_45S_50ntFlanks</td><td>8380</td><td>8405</td><td>26</td><td>CAAGAGGGCGTGAAACCGTTAAGAGG</td><td>4135.9</td><td>-5.3</td><td>0.8</td><td>-6.7</td><td>1.6E-11</td><td>5.0E-10</td><td>BT20/MB468</td><td>MP</td></tr>
<tr><td>>hg38_21_+_8433172_8446622_RNA45SN1_45S_50ntFlanks@12415.12432.18|CTTCGATGTCGGCTCTTC</td><td>>hg38_21_+_8433172_8446622_RNA45SN1_45S_50ntFlanks</td><td>12415</td><td>12432</td><td>18</td><td>CTTCGATGTCGGCTCTTC</td><td>5975.2</td><td>-4.0</td><td>0.6</td><td>-6.4</td><td>2.0E-10</td><td>5.0E-10</td><td>MB231/MB468</td><td>NUC</td></tr>
<tr><td>>hg38_21_+_8433172_8446622_RNA45SN1_45S_50ntFlanks@9563.9596.34|CCTGACGTGCAAATCGGTCGTCCGACCTGGGTAT</td><td>>hg38_21_+_8433172_8446622_RNA45SN1_45S_50ntFlanks</td><td>9563</td><td>9596</td><td>34</td><td>CCTGACGTGCAAATCGGTCGTCCGACCTGGGTAT</td><td>1183.8</td><td>-5.9</td><td>0.9</td><td>-6.4</td><td>2.0E-10</td><td>5.0E-10</td><td>MB231/MB468</td><td>CYTO</td></tr>
<tr><td>>MI0000082|hsa-mir-25&WithFlank&7|-|100093554|100093649@58.78.21&[MIMAT0000081&hsa-miR-25-3p&offsets|0|-1;m-23&7|-|100093571|100093592&offsets|0|-1]|CATTGCACTTGTCTCGGTCTG</td><td>>MI0000082|hsa-mir-25&WithFlank&7|-|100093554|100093649</td><td>58</td><td>78</td><td>21</td><td>CATTGCACTTGTCTCGGTCTG</td><td>3442.2</td><td>4.8</td><td>0.8</td><td>6.3</td><td>2.3E-10</td><td>5.0E-10</td><td>MB231/MB468</td><td>MITO</td></tr>
<tr><td>>hg38_21_+_8433172_8446622_RNA45SN1_45S_50ntFlanks@12045.12063.19|CTGACCCGGTGAGGCGGGG</td><td>>hg38_21_+_8433172_8446622_RNA45SN1_45S_50ntFlanks</td><td>12045</td><td>12063</td><td>19</td><td>CTGACCCGGTGAGGCGGGG</td><td>96.9</td><td>-10.5</td><td>1.6</td><td>-6.4</td><td>1.9E-10</td><td>5.0E-10</td><td>MB231/MB468</td><td>MP</td></tr>
<tr><td>>hg38_21_+_8433172_8446622_RNA45SN1_45S_50ntFlanks@6721.6747.27|AATTGCAGGACACATTGATCATCGACA</td><td>>hg38_21_+_8433172_8446622_RNA45SN1_45S_50ntFlanks</td><td>6721</td><td>6747</td><td>27</td><td>AATTGCAGGACACATTGATCATCGACA</td><td>3256.9</td><td>-7.0</td><td>1.1</td><td>-6.3</td><td>2.3E-10</td><td>5.0E-10</td><td>MB231/MB468</td><td>MITO</td></tr>
<tr><td>>hg38_21_+_8433172_8446622_RNA45SN1_45S_50ntFlanks@10412.10431.20|AGTCGGTCCTGAGAGATGGG</td><td>>hg38_21_+_8433172_8446622_RNA45SN1_45S_50ntFlanks</td><td>10412</td><td>10431</td><td>20</td><td>AGTCGGTCCTGAGAGATGGG</td><td>153.1</td><td>-6.6</td><td>1.0</td><td>-6.4</td><td>1.8E-10</td><td>5.0E-10</td><td>BT20/MB468</td><td>CYTO</td></tr>
<tr><td>>hg38_21_+_8433172_8446622_RNA45SN1_45S_50ntFlanks@11211.11231.21|GCGACTCTGGACGCGAGCCGG</td><td>>hg38_21_+_8433172_8446622_RNA45SN1_45S_50ntFlanks</td><td>11211</td><td>11231</td><td>21</td><td>GCGACTCTGGACGCGAGCCGG</td><td>157.2</td><td>-6.2</td><td>1.0</td><td>-6.4</td><td>2.0E-10</td><td>5.0E-10</td><td>MB231/MB468</td><td>CYTO</td></tr>
<tr><td>>hg38_21_+_8433172_8446622_RNA45SN1_45S_50ntFlanks@11817.11844.28|CCTACTATCCAGCGAAACCACAGCCAAG</td><td>>hg38_21_+_8433172_8446622_RNA45SN1_45S_50ntFlanks</td><td>11817</td><td>11844</td><td>28</td><td>CCTACTATCCAGCGAAACCACAGCCAAG</td><td>72.8</td><td>-10.0</td><td>1.6</td><td>-6.4</td><td>1.9E-10</td><td>5.1E-10</td><td>MB231/MB468</td><td>MP</td></tr>
<tr><td>>hg38_21_+_8433172_8446622_RNA45SN1_45S_50ntFlanks@8844.8867.24|GCGGCGGCGGCGGCGGCGGCGGGA</td><td>>hg38_21_+_8433172_8446622_RNA45SN1_45S_50ntFlanks</td><td>8844</td><td>8867</td><td>24</td><td>GCGGCGGCGGCGGCGGCGGCGGGA</td><td>206.6</td><td>-9.8</td><td>1.5</td><td>-6.4</td><td>1.5E-10</td><td>5.1E-10</td><td>BT20/MB468</td><td>MITO</td></tr>
<tr><td>>hg38_21_+_8433172_8446622_RNA45SN1_45S_50ntFlanks@10842.10865.24|GTAACTTCGGGATAAGGATTGGCT</td><td>>hg38_21_+_8433172_8446622_RNA45SN1_45S_50ntFlanks</td><td>10842</td><td>10865</td><td>24</td><td>GTAACTTCGGGATAAGGATTGGCT</td><td>315.6</td><td>-6.4</td><td>1.0</td><td>-6.4</td><td>2.0E-10</td><td>5.1E-10</td><td>MB231/MB468</td><td>NUC</td></tr>
<tr><td>>MI0000477|hsa-mir-146a&WithFlank&5|+|160485346|160485456@27.44.18&[MIMAT0000449&hsa-miR-146a-5p&offsets|0|-4;m-129&5|+|160485372|160485393&offsets|0|-4]|TGAGAACTGAATTCCATG</td><td>>MI0000477|hsa-mir-146a&WithFlank&5|+|160485346|160485456</td><td>27</td><td>44</td><td>18</td><td>TGAGAACTGAATTCCATG</td><td>815.2</td><td>-13.1</td><td>1.9</td><td>-7.0</td><td>3.4E-12</td><td>5.1E-10</td><td>BT20/MB231</td><td>MP</td></tr>
<tr><td>>hg38_21_+_8433172_8446622_RNA45SN1_45S_50ntFlanks@6651.6669.19|CGACTCTTAGCGGTGGATC</td><td>>hg38_21_+_8433172_8446622_RNA45SN1_45S_50ntFlanks</td><td>6651</td><td>6669</td><td>19</td><td>CGACTCTTAGCGGTGGATC</td><td>5213.0</td><td>2.3</td><td>0.4</td><td>6.3</td><td>2.4E-10</td><td>5.1E-10</td><td>BT20/MB468</td><td>NUC</td></tr>
<tr><td>>hg38_21_+_8433172_8446622_RNA45SN1_45S_50ntFlanks@10116.10134.19|TGCGTCGGAGGGCGGCGGC</td><td>>hg38_21_+_8433172_8446622_RNA45SN1_45S_50ntFlanks</td><td>10116</td><td>10134</td><td>19</td><td>TGCGTCGGAGGGCGGCGGC</td><td>133.2</td><td>-9.1</td><td>1.4</td><td>-6.4</td><td>1.5E-10</td><td>5.1E-10</td><td>BT20/MB468</td><td>MITO</td></tr>
<tr><td>>hg38_21_+_8433172_8446622_RNA45SN1_45S_50ntFlanks@12076.12095.20|AGGGGCTCTCGCTTCTGGCG</td><td>>hg38_21_+_8433172_8446622_RNA45SN1_45S_50ntFlanks</td><td>12076</td><td>12095</td><td>20</td><td>AGGGGCTCTCGCTTCTGGCG</td><td>2886.1</td><td>-7.2</td><td>1.1</td><td>-6.3</td><td>2.3E-10</td><td>5.1E-10</td><td>MB231/MB468</td><td>MITO</td></tr>
<tr><td>>hg38_21_+_8433172_8446622_RNA45SN1_45S_50ntFlanks@4445.4472.28|CTTGCCTCTCGGCGCCCCCTCGATGCTC</td><td>>hg38_21_+_8433172_8446622_RNA45SN1_45S_50ntFlanks</td><td>4445</td><td>4472</td><td>28</td><td>CTTGCCTCTCGGCGCCCCCTCGATGCTC</td><td>622.4</td><td>-9.0</td><td>1.4</td><td>-6.4</td><td>2.1E-10</td><td>5.1E-10</td><td>MB231/MB468</td><td>NUC</td></tr>
<tr><td>>hg38_21_+_8433172_8446622_RNA45SN1_45S_50ntFlanks@4570.4587.18|TGGAATAGGACCGCGGTT</td><td>>hg38_21_+_8433172_8446622_RNA45SN1_45S_50ntFlanks</td><td>4570</td><td>4587</td><td>18</td><td>TGGAATAGGACCGCGGTT</td><td>293.3</td><td>-8.0</td><td>1.3</td><td>-6.3</td><td>2.3E-10</td><td>5.1E-10</td><td>MB231/MB468</td><td>MITO</td></tr>
<tr><td>>hg38_21_+_8433172_8446622_RNA45SN1_45S_50ntFlanks@12621.12654.34|GAGCCAATGGGGCGAAGCTACCATCTGTGGGATT</td><td>>hg38_21_+_8433172_8446622_RNA45SN1_45S_50ntFlanks</td><td>12621</td><td>12654</td><td>34</td><td>GAGCCAATGGGGCGAAGCTACCATCTGTGGGATT</td><td>280.2</td><td>-3.2</td><td>0.5</td><td>-6.4</td><td>1.5E-10</td><td>5.1E-10</td><td>BT20/MB468</td><td>MITO</td></tr>
<tr><td>>MI0000077|hsa-mir-21&WithFlank&17|+|59841260|59841343@14.34.21(+1G)&[MIMAT0000076&hsa-miR-21-5p&offsets|0|-1(+1G);m-1&17|+|59841273|59841294&offsets|0|-1(+1G)]|TAGCTTATCAGACTGATGTTGG</td><td>>MI0000077|hsa-mir-21&WithFlank&17|+|59841260|59841343</td><td>14</td><td>34</td><td>21(+1G)</td><td>TAGCTTATCAGACTGATGTTGG</td><td>1026.4</td><td>5.0</td><td>0.8</td><td>6.3</td><td>2.4E-10</td><td>5.1E-10</td><td>BT20/MB468</td><td>NUC</td></tr>
<tr><td>>am_trna146_GlnCTG_6_-_27515531_27515602@1.29.29,trna1_GlnCTG_6_+_18836402_18836473@1.29.29,trna3_GlnCTG_17_+_8023070_8023141@1.29.29,trna42_GlnCTG_6_+_27263212_27263283@1.29.29,trna49_GlnCTG_6_+_27487308_27487379@1.29.29,trna7_GlnCTG_15_-_66161400_66161471@1.29.29,trna99_GlnCTG_6_-_28909378_28909449@1.29.29|GGTTCCATGGTGTAATGGTTAGCACTCTG</td><td>>am_trna146_GlnCTG_6_-_27515531_27515602</td><td>1</td><td>29</td><td>29</td><td>GGTTCCATGGTGTAATGGTTAGCACTCTG</td><td>3194.7</td><td>-5.6</td><td>0.9</td><td>-6.3</td><td>2.3E-10</td><td>5.1E-10</td><td>MB231/MB468</td><td>MITO</td></tr>
<tr><td>>hg38_21_+_8433172_8446622_RNA45SN1_45S_50ntFlanks@9738.9770.33|GAAACGATCTCAACCTATTCTCAAACTTTAAAT</td><td>>hg38_21_+_8433172_8446622_RNA45SN1_45S_50ntFlanks</td><td>9738</td><td>9770</td><td>33</td><td>GAAACGATCTCAACCTATTCTCAAACTTTAAAT</td><td>129.4</td><td>-10.9</td><td>1.7</td><td>-6.4</td><td>2.0E-10</td><td>5.1E-10</td><td>MB231/MB468</td><td>MP</td></tr>
<tr><td>>hg38_21_+_8433172_8446622_RNA45SN1_45S_50ntFlanks@13004.13027.24|ATCTATTGAAAGTCAGCCCTCGAC</td><td>>hg38_21_+_8433172_8446622_RNA45SN1_45S_50ntFlanks</td><td>13004</td><td>13027</td><td>24</td><td>ATCTATTGAAAGTCAGCCCTCGAC</td><td>164.7</td><td>-10.4</td><td>1.6</td><td>-6.7</td><td>1.6E-11</td><td>5.1E-10</td><td>BT20/MB468</td><td>MP</td></tr>
<tr><td>>hg38_21_+_8433172_8446622_RNA45SN1_45S_50ntFlanks@4283.4321.39|CTTTAACGAGGATCCATTGGAGGGCAAGTCTGGTGCCAG</td><td>>hg38_21_+_8433172_8446622_RNA45SN1_45S_50ntFlanks</td><td>4283</td><td>4321</td><td>39</td><td>CTTTAACGAGGATCCATTGGAGGGCAAGTCTGGTGCCAG</td><td>109.0</td><td>-10.6</td><td>1.7</td><td>-6.4</td><td>2.0E-10</td><td>5.1E-10</td><td>MB231/MB468</td><td>MP</td></tr>
<tr><td>>hg38_21_+_8433172_8446622_RNA45SN1_45S_50ntFlanks@10397.10424.28|AGTTGAACATGGGTCAGTCGGTCCTGAG</td><td>>hg38_21_+_8433172_8446622_RNA45SN1_45S_50ntFlanks</td><td>10397</td><td>10424</td><td>28</td><td>AGTTGAACATGGGTCAGTCGGTCCTGAG</td><td>399.8</td><td>4.9</td><td>0.8</td><td>6.4</td><td>2.1E-10</td><td>5.1E-10</td><td>MB231/MB468</td><td>NUC</td></tr>
<tr><td>>hg38_21_+_8433172_8446622_RNA45SN1_45S_50ntFlanks@4129.4162.34|GATTCCGGAGAGGGAGCCTGAGAAACGGCTACCA</td><td>>hg38_21_+_8433172_8446622_RNA45SN1_45S_50ntFlanks</td><td>4129</td><td>4162</td><td>34</td><td>GATTCCGGAGAGGGAGCCTGAGAAACGGCTACCA</td><td>148.5</td><td>-9.3</td><td>1.5</td><td>-6.4</td><td>1.5E-10</td><td>5.1E-10</td><td>BT20/MB468</td><td>MITO</td></tr>
<tr><td>>hg38_21_+_8433172_8446622_RNA45SN1_45S_50ntFlanks@11190.11220.31|CCCGGGGGGCCGGCGGCGGCGGCGACTCTGG</td><td>>hg38_21_+_8433172_8446622_RNA45SN1_45S_50ntFlanks</td><td>11190</td><td>11220</td><td>31</td><td>CCCGGGGGGCCGGCGGCGGCGGCGACTCTGG</td><td>127.0</td><td>-10.9</td><td>1.7</td><td>-6.4</td><td>2.0E-10</td><td>5.1E-10</td><td>MB231/MB468</td><td>MP</td></tr>
<tr><td>>hg38_21_+_8433172_8446622_RNA45SN1_45S_50ntFlanks@11704.11725.22|ATGAAGCGCGGGTAAACGGCGG</td><td>>hg38_21_+_8433172_8446622_RNA45SN1_45S_50ntFlanks</td><td>11704</td><td>11725</td><td>22</td><td>ATGAAGCGCGGGTAAACGGCGG</td><td>100.7</td><td>-10.5</td><td>1.7</td><td>-6.4</td><td>2.0E-10</td><td>5.1E-10</td><td>MB231/MB468</td><td>MP</td></tr>
<tr><td>>hg38_21_+_8433172_8446622_RNA45SN1_45S_50ntFlanks@5084.5121.38|CGAACGAGACTCTGGCATGCTAACTAGTTACGCGACCC</td><td>>hg38_21_+_8433172_8446622_RNA45SN1_45S_50ntFlanks</td><td>5084</td><td>5121</td><td>38</td><td>CGAACGAGACTCTGGCATGCTAACTAGTTACGCGACCC</td><td>299.1</td><td>-12.1</td><td>1.9</td><td>-6.4</td><td>2.0E-10</td><td>5.1E-10</td><td>MB231/MB468</td><td>MP</td></tr>
<tr><td>>trna139_ValAAC_6_-_27618707_27618779@36.56.21|CACGCGAAAGGTCCCTGGATC</td><td>>trna139_ValAAC_6_-_27618707_27618779</td><td>36</td><td>56</td><td>21</td><td>CACGCGAAAGGTCCCTGGATC</td><td>437.7</td><td>12.5</td><td>1.9</td><td>6.7</td><td>1.6E-11</td><td>5.1E-10</td><td>BT20/MB468</td><td>MP</td></tr>
<tr><td>>hg38_21_+_8433172_8446622_RNA45SN1_45S_50ntFlanks@9201.9223.23|CTCTCGGGGCCACGCGCGCGTCC</td><td>>hg38_21_+_8433172_8446622_RNA45SN1_45S_50ntFlanks</td><td>9201</td><td>9223</td><td>23</td><td>CTCTCGGGGCCACGCGCGCGTCC</td><td>230.6</td><td>-10.9</td><td>1.6</td><td>-6.7</td><td>1.7E-11</td><td>5.1E-10</td><td>BT20/MB468</td><td>MP</td></tr>
<tr><td>>hg38_21_+_8433172_8446622_RNA45SN1_45S_50ntFlanks@12412.12435.24|ATCCTTCGATGTCGGCTCTTCCTA</td><td>>hg38_21_+_8433172_8446622_RNA45SN1_45S_50ntFlanks</td><td>12412</td><td>12435</td><td>24</td><td>ATCCTTCGATGTCGGCTCTTCCTA</td><td>227.9</td><td>-10.1</td><td>1.6</td><td>-6.4</td><td>1.8E-10</td><td>5.1E-10</td><td>BT20/MB468</td><td>CYTO</td></tr>
<tr><td>>hg38_21_+_8433172_8446622_RNA45SN1_45S_50ntFlanks@10355.10383.29|AACTTTGAAGGCCGAAGTGGAGAAGGGTT</td><td>>hg38_21_+_8433172_8446622_RNA45SN1_45S_50ntFlanks</td><td>10355</td><td>10383</td><td>29</td><td>AACTTTGAAGGCCGAAGTGGAGAAGGGTT</td><td>1171.4</td><td>-9.5</td><td>1.5</td><td>-6.3</td><td>2.4E-10</td><td>5.1E-10</td><td>BT20/MB468</td><td>NUC</td></tr>
<tr><td>>hg38_21_+_8433172_8446622_RNA45SN1_45S_50ntFlanks@6742.6772.31|TCGACACTTCGAACGCACTTGCGGCCCCGGG</td><td>>hg38_21_+_8433172_8446622_RNA45SN1_45S_50ntFlanks</td><td>6742</td><td>6772</td><td>31</td><td>TCGACACTTCGAACGCACTTGCGGCCCCGGG</td><td>408.3</td><td>-8.1</td><td>1.3</td><td>-6.3</td><td>2.3E-10</td><td>5.2E-10</td><td>MB231/MB468</td><td>MITO</td></tr>
<tr><td>>hg38_21_+_8433172_8446622_RNA45SN1_45S_50ntFlanks@10841.10863.23|CGTAACTTCGGGATAAGGATTGG</td><td>>hg38_21_+_8433172_8446622_RNA45SN1_45S_50ntFlanks</td><td>10841</td><td>10863</td><td>23</td><td>CGTAACTTCGGGATAAGGATTGG</td><td>824.6</td><td>-8.2</td><td>1.3</td><td>-6.3</td><td>2.3E-10</td><td>5.2E-10</td><td>MB231/MB468</td><td>MITO</td></tr>
<tr><td>>hg38_21_+_8433172_8446622_RNA45SN1_45S_50ntFlanks@11641.11664.24|CGGCGGGTGTTGACGCGATGTGAT</td><td>>hg38_21_+_8433172_8446622_RNA45SN1_45S_50ntFlanks</td><td>11641</td><td>11664</td><td>24</td><td>CGGCGGGTGTTGACGCGATGTGAT</td><td>963.3</td><td>-7.0</td><td>1.1</td><td>-6.4</td><td>2.1E-10</td><td>5.2E-10</td><td>MB231/MB468</td><td>NUC</td></tr>
<tr><td>>MI0000285|hsa-mir-205&WithFlank&1|+|209432127|209432248@40.62.23(+1U)&[MIMAT0000266&hsa-miR-205-5p&offsets|0|+1(+1U);m-351&1|+|209432166|209432187&offsets|0|+1(+1U)]|TCCTTCATTCCACCGGAGTCTGTT</td><td>>MI0000285|hsa-mir-205&WithFlank&1|+|209432127|209432248</td><td>40</td><td>62</td><td>23(+1U)</td><td>TCCTTCATTCCACCGGAGTCTGTT</td><td>1916.8</td><td>6.7</td><td>1.0</td><td>6.4</td><td>1.5E-10</td><td>5.2E-10</td><td>BT20/MB468</td><td>MITO</td></tr>
</table>

<table>
<tr><th></th><th></th><th></th><th></th><th></th><th></th><th></th><th></th><th></th><th></th><th></th><th></th><th></th><th></th></tr>
<tr><td>>MI0001445|hsa-mir-423&WithFlank&17|+|30117073|30117178@59.81.23&[MIMAT0001340&hsa-miR-423-3p&offsets|0|0;m-100&17|+|30117131|30117153&offsets|0|0]|AGCTCGGTCTGAGGCCCCTCAGT</td><td>>MI0001445|hsa-mir-423&WithFlank&17|+|30117073|30117178</td><td>59</td><td>81</td><td>23</td><td>AGCTCGGTCTGAGGCCCCTCAGT</td><td>21647.0</td><td>5.9</td><td>0.9</td><td>6.3</td><td>2.3E-10</td><td>5.2E-10</td><td>MB231/MB468</td><td>MITO</td></tr>
<tr><td>>hg38_21_+_8433172_8446622_RNA45SN1_45S_50ntFlanks@10832.10862.31|AGCCGGATCCGTAACTTCGGGATAAGGATTG</td><td>>hg38_21_+_8433172_8446622_RNA45SN1_45S_50ntFlanks</td><td>10832</td><td>10862</td><td>31</td><td>AGCCGGATCCGTAACTTCGGGATAAGGATTG</td><td>512.4</td><td>-6.6</td><td>1.0</td><td>-6.4</td><td>2.1E-10</td><td>5.2E-10</td><td>MB231/MB468</td><td>CYTO</td></tr>
<tr><td>>hg38_21_+_8433172_8446622_RNA45SN1_45S_50ntFlanks@11886.11905.20|TGAGCTTGACTCTAGTCTGG</td><td>>hg38_21_+_8433172_8446622_RNA45SN1_45S_50ntFlanks</td><td>11886</td><td>11905</td><td>20</td><td>TGAGCTTGACTCTAGTCTGG</td><td>584.4</td><td>-7.6</td><td>1.2</td><td>-6.4</td><td>2.1E-10</td><td>5.2E-10</td><td>MB231/MB468</td><td>CYTO</td></tr>
<tr><td>>hg38_21_+_8433172_8446622_RNA45SN1_45S_50ntFlanks@3913.3937.25|ATGCGTGCATTTATCAGATCAAAAC</td><td>>hg38_21_+_8433172_8446622_RNA45SN1_45S_50ntFlanks</td><td>3913</td><td>3937</td><td>25</td><td>ATGCGTGCATTTATCAGATCAAAAC</td><td>166.0</td><td>-11.2</td><td>1.8</td><td>-6.4</td><td>2.0E-10</td><td>5.2E-10</td><td>MB231/MB468</td><td>MP</td></tr>
<tr><td>>hg38_21_+_8433172_8446622_RNA45SN1_45S_50ntFlanks@10787.10816.30|ACAGCCTCTGGCATGTTGGAACAATGTAGG</td><td>>hg38_21_+_8433172_8446622_RNA45SN1_45S_50ntFlanks</td><td>10787</td><td>10816</td><td>30</td><td>ACAGCCTCTGGCATGTTGGAACAATGTAGG</td><td>1094.2</td><td>-9.4</td><td>1.5</td><td>-6.3</td><td>2.5E-10</td><td>5.2E-10</td><td>BT20/MB468</td><td>NUC</td></tr>
<tr><td>>MI0003591|hsa-mir-584&WithFlank&5|-|149062307|149062415@22.43.22(+1U)&[MIMAT0003249&hsa-miR-584-5p&offsets|0|0(+1U);m-167&5|-|149062374|149062394&offsets|0|+1(+1U)]|TTATGGTTTGCCTGGGACTGAGT</td><td>>MI0003591|hsa-mir-584&WithFlank&5|-|149062307|149062415</td><td>22</td><td>43</td><td>22(+1U)</td><td>TTATGGTTTGCCTGGGACTGAGT</td><td>197.5</td><td>10.8</td><td>1.7</td><td>6.3</td><td>2.3E-10</td><td>5.2E-10</td><td>MB231/MB468</td><td>MITO</td></tr>
<tr><td>>trnaMT_SerGCT_MT_+_12207_12265@10.38.29|CACAAGAACTGCTAACTCATGCCCCCATG</td><td>>trnaMT_SerGCT_MT_+_12207_12265</td><td>10</td><td>38</td><td>29</td><td>CACAAGAACTGCTAACTCATGCCCCCATG</td><td>1096.4</td><td>-7.0</td><td>1.1</td><td>-6.3</td><td>2.3E-10</td><td>5.2E-10</td><td>MB231/MB468</td><td>MITO</td></tr>
<tr><td>>hg38_21_+_8433172_8446622_RNA45SN1_45S_50ntFlanks@8741.8758.18|TCCGTCCGTCCGTCCGTC</td><td>>hg38_21_+_8433172_8446622_RNA45SN1_45S_50ntFlanks</td><td>8741</td><td>8758</td><td>18</td><td>TCCGTCCGTCCGTCCGTC</td><td>136.0</td><td>-11.0</td><td>1.7</td><td>-6.4</td><td>2.0E-10</td><td>5.2E-10</td><td>MB231/MB468</td><td>MP</td></tr>
<tr><td>>MI0000065|hsa-let-7d&WithFlank&9|+|94178828|94178926@14.33.20(+1C)&[MIMAT0000065&hsa-let-7d-5p&offsets|0|-2(+1C);m-90&9|+|94178841|94178862&offsets|0|-2(+1C)]|AGAGGTAGTAGGTTGCATAGC</td><td>>MI0000065|hsa-let-7d&WithFlank&9|+|94178828|94178926</td><td>14</td><td>33</td><td>20(+1C)</td><td>AGAGGTAGTAGGTTGCATAGC</td><td>77.7</td><td>2.8</td><td>0.4</td><td>6.3</td><td>2.5E-10</td><td>5.2E-10</td><td>BT20/MB468</td><td>NUC</td></tr>
<tr><td>>hg38_21_+_8433172_8446622_RNA45SN1_45S_50ntFlanks@5467.5484.18|GGCCCACGGCCCTGGCGG</td><td>>hg38_21_+_8433172_8446622_RNA45SN1_45S_50ntFlanks</td><td>5467</td><td>5484</td><td>18</td><td>GGCCCACGGCCCTGGCGG</td><td>186.3</td><td>-10.6</td><td>1.6</td><td>-6.7</td><td>1.7E-11</td><td>5.2E-10</td><td>BT20/MB468</td><td>MP</td></tr>
<tr><td>>hg38_21_+_8433172_8446622_RNA45SN1_45S_50ntFlanks@5473.5490.18|CGGCCCTGGCGGAGCGCT</td><td>>hg38_21_+_8433172_8446622_RNA45SN1_45S_50ntFlanks</td><td>5473</td><td>5490</td><td>18</td><td>CGGCCCTGGCGGAGCGCT</td><td>1088.0</td><td>8.4</td><td>1.3</td><td>6.4</td><td>1.9E-10</td><td>5.2E-10</td><td>BT20/MB468</td><td>CYTO</td></tr>
<tr><td>>hg38_21_+_8433172_8446622_RNA45SN1_45S_50ntFlanks@9940.9980.41|CAGGACGGTGGCCATGGAAGTCGGAATCCGCTAAGGAGTGT</td><td>>hg38_21_+_8433172_8446622_RNA45SN1_45S_50ntFlanks</td><td>9940</td><td>9980</td><td>41</td><td>CAGGACGGTGGCCATGGAAGTCGGAATCCGCTAAGGAGTGT</td><td>144.9</td><td>10.9</td><td>1.6</td><td>6.7</td><td>1.7E-11</td><td>5.2E-10</td><td>BT20/MB468</td><td>MP</td></tr>
<tr><td>>hg38_21_+_8433172_8446622_RNA45SN1_45S_50ntFlanks@5526.5549.24|AAAAGTCGTAACAAGGTTTCCGTA</td><td>>hg38_21_+_8433172_8446622_RNA45SN1_45S_50ntFlanks</td><td>5526</td><td>5549</td><td>24</td><td>AAAAGTCGTAACAAGGTTTCCGTA</td><td>95.1</td><td>-10.4</td><td>1.6</td><td>-6.4</td><td>2.0E-10</td><td>5.2E-10</td><td>MB231/MB468</td><td>MP</td></tr>
<tr><td>>hg38_21_+_8433172_8446622_RNA45SN1_45S_50ntFlanks@8080.8104.25|AGAGCCCAGCGCCGAATCCCCGCCC</td><td>>hg38_21_+_8433172_8446622_RNA45SN1_45S_50ntFlanks</td><td>8080</td><td>8104</td><td>25</td><td>AGAGCCCAGCGCCGAATCCCCGCCC</td><td>118.9</td><td>-10.8</td><td>1.7</td><td>-6.4</td><td>2.0E-10</td><td>5.2E-10</td><td>MB231/MB468</td><td>MP</td></tr>
<tr><td>>hg38_21_+_8433172_8446622_RNA45SN1_45S_50ntFlanks@12617.12661.45|TGAGGAGCCAATGGGGCGAAGCTACCATCTGTGGGATTATGACTG</td><td>>hg38_21_+_8433172_8446622_RNA45SN1_45S_50ntFlanks</td><td>12617</td><td>12661</td><td>45</td><td>TGAGGAGCCAATGGGGCGAAGCTACCATCTGTGGGATTATGACTG</td><td>605.8</td><td>-7.7</td><td>1.2</td><td>-6.3</td><td>2.4E-10</td><td>5.2E-10</td><td>MB231/MB468</td><td>MITO</td></tr>
<tr><td>>hg38_21_+_8433172_8446622_RNA45SN1_45S_50ntFlanks@4853.4879.27|AAGTCTTTGGGTTCCGGGGGGAGTATG</td><td>>hg38_21_+_8433172_8446622_RNA45SN1_45S_50ntFlanks</td><td>4853</td><td>4879</td><td>27</td><td>AAGTCTTTGGGTTCCGGGGGGAGTATG</td><td>68.5</td><td>-10.0</td><td>1.6</td><td>-6.4</td><td>2.0E-10</td><td>5.2E-10</td><td>MB231/MB468</td><td>MP</td></tr>
<tr><td>>hg38_21_+_8433172_8446622_RNA45SN1_45S_50ntFlanks@11581.11606.26|AACTGGTGCGGACCAGGGGAATCCGA</td><td>>hg38_21_+_8433172_8446622_RNA45SN1_45S_50ntFlanks</td><td>11581</td><td>11606</td><td>26</td><td>AACTGGTGCGGACCAGGGGAATCCGA</td><td>319.4</td><td>-6.4</td><td>1.0</td><td>-6.3</td><td>2.5E-10</td><td>5.2E-10</td><td>BT20/MB468</td><td>NUC</td></tr>
<tr><td>>hg38_21_+_8433172_8446622_RNA45SN1_45S_50ntFlanks@11209.11252.44|CGGCGACTCTGGACGCGAGCCGGGCCCTTCCCGTGGATCGCCCC</td><td>>hg38_21_+_8433172_8446622_RNA45SN1_45S_50ntFlanks</td><td>11209</td><td>11252</td><td>44</td><td>CGGCGACTCTGGACGCGAGCCGGGCCCTTCCCGTGGATCGCCCC</td><td>897.2</td><td>-7.1</td><td>1.1</td><td>-6.3</td><td>2.4E-10</td><td>5.2E-10</td><td>MB231/MB468</td><td>MITO</td></tr>
<tr><td>>MI0000736|hsa-mir-30c-1&WithFlank&1|+|40757278|40757378@23.45.23;MI0000254|hsa-mir-30c-2&WithFlank&6|-|71376954|71377037@13.35.23&[MIMAT0000244&hsa-miR-30c-5p&offsets|0|0;m-57&1|+|40757300|40757323&offsets|0|-1];[MIMAT0000244_1&hsa-miR-30c-5p&offsets|0|0;m-56&6|-|71377002|71377025&offsets|0|-1]|TGTAAACATCCTACACTCTCAGC</td><td>>MI0000736|hsa-mir-30c-1&WithFlank&1|+|40757278|40757378</td><td>23</td><td>45</td><td>23</td><td>TGTAAACATCCTACACTCTCAGC</td><td>40718.6</td><td>6.0</td><td>0.9</td><td>6.3</td><td>2.4E-10</td><td>5.2E-10</td><td>MB231/MB468</td><td>MITO</td></tr>
<tr><td>>hg38_21_+_8433172_8446622_RNA45SN1_45S_50ntFlanks@11821.11842.22|CTATCCAGCGAAACCACAGCCA</td><td>>hg38_21_+_8433172_8446622_RNA45SN1_45S_50ntFlanks</td><td>11821</td><td>11842</td><td>22</td><td>CTATCCAGCGAAACCACAGCCA</td><td>296.0</td><td>-11.3</td><td>1.7</td><td>-6.7</td><td>1.7E-11</td><td>5.2E-10</td><td>BT20/MB468</td><td>MP</td></tr>
<tr><td>>hg38_21_+_8433172_8446622_RNA45SN1_45S_50ntFlanks@8342.8376.35|CGTAAGGGAAAGTTGAAAAGAACTTTGAAGAGAGA</td><td>>hg38_21_+_8433172_8446622_RNA45SN1_45S_50ntFlanks</td><td>8342</td><td>8376</td><td>35</td><td>CGTAAGGGAAAGTTGAAAAGAACTTTGAAGAGAGA</td><td>332.1</td><td>7.9</td><td>1.2</td><td>6.4</td><td>1.9E-10</td><td>5.2E-10</td><td>BT20/MB468</td><td>CYTO</td></tr>
<tr><td>>MI0000266|hsa-mir-10a&WithFlank&17|-|48579832|48579953@29.51.23&[MIMAT0000253&hsa-miR-10a-5p&offsets|+1|+1;m-18&17|-|48579905|48579926&offsets|+1|+2]|ACCCTGTAGATCCGAATTTGTGT</td><td>>MI0000266|hsa-mir-10a&WithFlank&17|-|48579832|48579953</td><td>29</td><td>51</td><td>23</td><td>ACCCTGTAGATCCGAATTTGTGT</td><td>150.0</td><td>11.0</td><td>1.6</td><td>6.7</td><td>1.7E-11</td><td>5.2E-10</td><td>BT20/MB468</td><td>MP</td></tr>
<tr><td>>hg38_21_+_8433172_8446622_RNA45SN1_45S_50ntFlanks@9562.9596.35|TCCTGACGTGCAAATCGGTCGTCCGACCTGGGTAT</td><td>>hg38_21_+_8433172_8446622_RNA45SN1_45S_50ntFlanks</td><td>9562</td><td>9596</td><td>35</td><td>TCCTGACGTGCAAATCGGTCGTCCGACCTGGGTAT</td><td>1622.9</td><td>-8.9</td><td>1.4</td><td>-6.4</td><td>2.1E-10</td><td>5.3E-10</td><td>MB231/MB468</td><td>NUC</td></tr>
<tr><td>>hg38_21_+_8433172_8446622_RNA45SN1_45S_50ntFlanks@12623.12654.32|GCCAATGGGGCGAAGCTACCATCTGTGGGATT</td><td>>hg38_21_+_8433172_8446622_RNA45SN1_45S_50ntFlanks</td><td>12623</td><td>12654</td><td>32</td><td>GCCAATGGGGCGAAGCTACCATCTGTGGGATT</td><td>424.5</td><td>-3.6</td><td>0.6</td><td>-6.4</td><td>1.6E-10</td><td>5.3E-10</td><td>BT20/MB468</td><td>MITO</td></tr>
<tr><td>>MI0003643|hsa-mir-629&WithFlank&15|-|70079366|70079474@28.48.21(+1A)&[MIMAT0004810&hsa-miR-629-5p&offsets|0|0(+1A);m-150&15|-|70079427|70079447&offsets|0|0(+1A)]|TGGGTTTACGTTGGGAGAACTA</td><td>>MI0003643|hsa-mir-629&WithFlank&15|-|70079366|70079474</td><td>28</td><td>48</td><td>21(+1A)</td><td>TGGGTTTACGTTGGGAGAACTA</td><td>405.0</td><td>12.4</td><td>1.8</td><td>6.7</td><td>1.7E-11</td><td>5.3E-10</td><td>BT20/MB468</td><td>MP</td></tr>
<tr><td>>hg38_21_+_8433172_8446622_RNA45SN1_45S_50ntFlanks@4555.4573.19|CGCAGCTAGGAATAATGGA</td><td>>hg38_21_+_8433172_8446622_RNA45SN1_45S_50ntFlanks</td><td>4555</td><td>4573</td><td>19</td><td>CGCAGCTAGGAATAATGGA</td><td>506.9</td><td>-6.5</td><td>1.0</td><td>-6.4</td><td>2.1E-10</td><td>5.3E-10</td><td>MB231/MB468</td><td>NUC</td></tr>
<tr><td>>hg38_21_+_8433172_8446622_RNA45SN1_45S_50ntFlanks@11872.11898.27|AAAGAAGACCCTGTTGAGCTTGACTCT</td><td>>hg38_21_+_8433172_8446622_RNA45SN1_45S_50ntFlanks</td><td>11872</td><td>11898</td><td>27</td><td>AAAGAAGACCCTGTTGAGCTTGACTCT</td><td>67.1</td><td>-9.9</td><td>1.6</td><td>-6.4</td><td>2.0E-10</td><td>5.3E-10</td><td>MB231/MB468</td><td>MP</td></tr>
<tr><td>>hg38_21_+_8433172_8446622_RNA45SN1_45S_50ntFlanks@3777.3794.18|CGGTACAGTGAAACTGCG</td><td>>hg38_21_+_8433172_8446622_RNA45SN1_45S_50ntFlanks</td><td>3777</td><td>3794</td><td>18</td><td>CGGTACAGTGAAACTGCG</td><td>311.1</td><td>-8.0</td><td>1.3</td><td>-6.4</td><td>2.0E-10</td><td>5.3E-10</td><td>MB231/MB468</td><td>MP</td></tr>
<tr><td>>hg38_21_+_8433172_8446622_RNA45SN1_45S_50ntFlanks@11873.11892.20|AAGAAGACCCTGTTGAGCTT</td><td>>hg38_21_+_8433172_8446622_RNA45SN1_45S_50ntFlanks</td><td>11873</td><td>11892</td><td>20</td><td>AAGAAGACCCTGTTGAGCTT</td><td>1495.7</td><td>-7.7</td><td>1.2</td><td>-6.3</td><td>2.5E-10</td><td>5.3E-10</td><td>BT20/MB468</td><td>NUC</td></tr>
<tr><td>>hg38_21_+_8433172_8446622_RNA45SN1_45S_50ntFlanks@11670.11702.33|CCCAGTGCTCTGAATGTCAAAGTGAAGAAATTC</td><td>>hg38_21_+_8433172_8446622_RNA45SN1_45S_50ntFlanks</td><td>11670</td><td>11702</td><td>33</td><td>CCCAGTGCTCTGAATGTCAAAGTGAAGAAATTC</td><td>1203.8</td><td>-7.2</td><td>1.1</td><td>-6.4</td><td>2.0E-10</td><td>5.3E-10</td><td>MB231/MB468</td><td>MP</td></tr>
<tr><td>>hg38_21_+_8433172_8446622_RNA45SN1_45S_50ntFlanks@5471.5516.46|CACGGCCCTGGCGGAGCGCTGAGAAGACGGTCGAACTTGACTATCT</td><td>>hg38_21_+_8433172_8446622_RNA45SN1_45S_50ntFlanks</td><td>5471</td><td>5516</td><td>46</td><td>CACGGCCCTGGCGGAGCGCTGAGAAGACGGTCGAACTTGACTATCT</td><td>9018.1</td><td>-5.6</td><td>0.8</td><td>-6.8</td><td>1.3E-11</td><td>5.3E-10</td><td>BT20/MB231</td><td>CYTO</td></tr>
<tr><td>>hg38_21_+_8433172_8446622_RNA45SN1_45S_50ntFlanks@12399.12435.37|ACGTCGCTTTTTGATCCTTCGATGTCGGCTCTTCCTA</td><td>>hg38_21_+_8433172_8446622_RNA45SN1_45S_50ntFlanks</td><td>12399</td><td>12435</td><td>37</td><td>ACGTCGCTTTTTGATCCTTCGATGTCGGCTCTTCCTA</td><td>284.6</td><td>-6.6</td><td>1.0</td><td>-6.3</td><td>2.4E-10</td><td>5.3E-10</td><td>MB231/MB468</td><td>MITO</td></tr>
<tr><td>>hg38_21_+_8433172_8446622_RNA45SN1_45S_50ntFlanks@4538.4559.22|CCGAGCCGCCTGGATACCGCAG</td><td>>hg38_21_+_8433172_8446622_RNA45SN1_45S_50ntFlanks</td><td>4538</td><td>4559</td><td>22</td><td>CCGAGCCGCCTGGATACCGCAG</td><td>131.4</td><td>-9.1</td><td>1.4</td><td>-6.4</td><td>1.6E-10</td><td>5.3E-10</td><td>BT20/MB468</td><td>MITO</td></tr>
<tr><td>>hg38_21_+_8433172_8446622_RNA45SN1_45S_50ntFlanks@6663.6696.34|GTGGATCACTCGGCTCGTGCGTCGATGAAGAACG</td><td>>hg38_21_+_8433172_8446622_RNA45SN1_45S_50ntFlanks</td><td>6663</td><td>6696</td><td>34</td><td>GTGGATCACTCGGCTCGTGCGTCGATGAAGAACG</td><td>303.5</td><td>-6.1</td><td>1.0</td><td>-6.4</td><td>2.0E-10</td><td>5.3E-10</td><td>MB231/MB468</td><td>MP</td></tr>
<tr><td>>hg38_21_+_8433172_8446622_RNA45SN1_45S_50ntFlanks@11882.11912.31|CTGTTGAGCTTGACTCTAGTCTGGCACGGTG</td><td>>hg38_21_+_8433172_8446622_RNA45SN1_45S_50ntFlanks</td><td>11882</td><td>11912</td><td>31</td><td>CTGTTGAGCTTGACTCTAGTCTGGCACGGTG</td><td>92.2</td><td>-10.4</td><td>1.6</td><td>-6.4</td><td>2.0E-10</td><td>5.3E-10</td><td>MB231/MB468</td><td>MP</td></tr>
<tr><td>>hg38_21_+_8433172_8446622_RNA45SN1_45S_50ntFlanks@3705.3723.19|TACCTGGTTGATCCTGCCA</td><td>>hg38_21_+_8433172_8446622_RNA45SN1_45S_50ntFlanks</td><td>3705</td><td>3723</td><td>19</td><td>TACCTGGTTGATCCTGCCA</td><td>1410.9</td><td>4.4</td><td>0.7</td><td>6.4</td><td>2.1E-10</td><td>5.3E-10</td><td>MB231/MB468</td><td>CYTO</td></tr>
<tr><td>>hg38_21_+_8433172_8446622_RNA45SN1_45S_50ntFlanks@9122.9140.19|CCGCGCGCGGGTCGGGGGG</td><td>>hg38_21_+_8433172_8446622_RNA45SN1_45S_50ntFlanks</td><td>9122</td><td>9140</td><td>19</td><td>CCGCGCGCGGGTCGGGGGG</td><td>234.5</td><td>-11.0</td><td>1.6</td><td>-6.7</td><td>1.7E-11</td><td>5.3E-10</td><td>BT20/MB468</td><td>MP</td></tr>
<tr><td>>MI0000272|hsa-mir-182&WithFlank&7|-|129770377|129770498@29.54.26(+1U)&[MIMAT0000259&hsa-miR-182-5p&offsets|0|+2(+1U);m-22&7|-|129770449|129770470&offsets|0|+4(+1U)]|TTTGGCAATGGTAGAACTCACACTGGT</td><td>>MI0000272|hsa-mir-182&WithFlank&7|-|129770377|129770498</td><td>29</td><td>54</td><td>26(+1U)</td><td>TTTGGCAATGGTAGAACTCACACTGGT</td><td>193.5</td><td>2.1</td><td>0.3</td><td>6.3</td><td>2.5E-10</td><td>5.3E-10</td><td>BT20/MB468</td><td>NUC</td></tr>
<tr><td>>MI0000650|hsa-mir-200c&WithFlank&12|+|6963693|6963772@49.72.24&[MIMAT0000617&hsa-miR-200c-3p&offsets|-1|0;m-54&12|+|6963742|6963764&offsets|-1|0]|CTAATACTGCCGGGTAATGATGGA</td><td>>MI0000650|hsa-mir-200c&WithFlank&12|+|6963693|6963772</td><td>49</td><td>72</td><td>24</td><td>CTAATACTGCCGGGTAATGATGGA</td><td>154.0</td><td>2.3</td><td>0.4</td><td>6.3</td><td>2.5E-10</td><td>5.3E-10</td><td>BT20/MB468</td><td>NUC</td></tr>
<tr><td>>trna146_GlnCTG_6_-_27515531_27515602@17.53.37,trna1_GlnCTG_6_+_18836402_18836473@17.53.37,trna3_GlnCTG_17_+_8023070_8023141@17.53.37,trna49_GlnCTG_6_+_27487308_27487379@17.53.37,trna7_GlnCTG_15_-_66161400_66161471@17.53.37,trna99_GlnCTG_6_-_28909378_28909449@17.53.37|GGTTAGCACTCTGGACTCTGAATCCAGCGATCCGAGT</td><td>>trna146_GlnCTG_6_-_27515531_27515602</td><td>17</td><td>53</td><td>37</td><td>GGTTAGCACTCTGGACTCTGAATCCAGCGATCCGAGT</td><td>282.0</td><td>9.3</td><td>1.5</td><td>6.4</td><td>2.1E-10</td><td>5.3E-10</td><td>MB231/MB468</td><td>CYTO</td></tr>
<tr><td>>MI0003513|hsa-mir-455&WithFlank&9|+|114209428|114209535@22.45.24&[MIMAT0003150&hsa-miR-455-5p&offsets|0|+2;m-232&9|+|114209449|114209470&offsets|0|+2]|TATGTGCCTTTGGACTACATCGTG</td><td>>MI0003513|hsa-mir-455&WithFlank&9|+|114209428|114209535</td><td>22</td><td>45</td><td>24</td><td>TATGTGCCTTTGGACTACATCGTG</td><td>106.1</td><td>10.0</td><td>1.6</td><td>6.4</td><td>2.0E-10</td><td>5.3E-10</td><td>MB231/MB468</td><td>MP</td></tr>
<tr><td>>hg38_21_+_8433172_8446622_RNA45SN1_45S_50ntFlanks@11581.11602.22|AACTGGTGCGGACCAGGGGAAT</td><td>>hg38_21_+_8433172_8446622_RNA45SN1_45S_50ntFlanks</td><td>11581</td><td>11602</td><td>22</td><td>AACTGGTGCGGACCAGGGGAAT</td><td>1442.7</td><td>-9.2</td><td>1.4</td><td>-6.4</td><td>2.0E-10</td><td>5.3E-10</td><td>MB231/MB468</td><td>MP</td></tr>
<tr><td>>MI0000264|hsa-mir-7-2&WithFlank&15|+|88611819|88611940@38.60.23;MI0000265|hsa-mir-7-3&WithFlank&19|+|4770664|4770785@37.59.23;MI0000263|hsa-mir-7-1&WithFlank&9|-|83969742|83969863@30.52.23&[MIMAT0000252&hsa-miR-7-5p&offsets|0|-1;m-400&15|+|88611856|88611879&offsets|0|-1];[MIMAT0000252_1&hsa-miR-7-5p&offsets|0|-1;m-402&19|+|4770700|4770723&offsets|0|-1];[MIMAT0000252_2&hsa-miR-7-5p&offsets|0|-1;m-401&9|-|83969811|83969834&offsets|0|-1]|TGGAAGACTAGTGATTTTGTTGT</td><td>>MI0000264|hsa-mir-7-2&WithFlank&15|+|88611819|88611940</td><td>38</td><td>60</td><td>23</td><td>TGGAAGACTAGTGATTTTGTTGT</td><td>2646.7</td><td>3.4</td><td>0.5</td><td>6.3</td><td>2.5E-10</td><td>5.3E-10</td><td>BT20/MB468</td><td>NUC</td></tr>
<tr><td>>hg38_21_+_8433172_8446622_RNA45SN1_45S_50ntFlanks@8168.8194.27|CCCAAGTCCTTCTGATCGAGGCCCAGC</td><td>>hg38_21_+_8433172_8446622_RNA45SN1_45S_50ntFlanks</td><td>8168</td><td>8194</td><td>27</td><td>CCCAAGTCCTTCTGATCGAGGCCCAGC</td><td>79.5</td><td>-10.2</td><td>1.6</td><td>-6.4</td><td>2.0E-10</td><td>5.3E-10</td><td>MB231/MB468</td><td>MP</td></tr>
<tr><td>>MI0000456|hsa-mir-140&WithFlank&16|+|69933075|69933186@68.90.23(+1U)&[MIMAT0004597&hsa-miR-140-3p&offsets|0|+2(+1U);m-45&16|+|69933143|69933164&offsets|-1|0(+1U)]|TACCACAGGGTAGAACCACGGACT</td><td>>MI0000456|hsa-mir-140&WithFlank&16|+|69933075|69933186</td><td>68</td><td>90</td><td>23(+1U)</td><td>TACCACAGGGTAGAACCACGGACT</td><td>200.9</td><td>7.4</td><td>1.2</td><td>6.3</td><td>2.2E-10</td><td>5.3E-10</td><td>MB231/MB468</td><td>NUC</td></tr>
<tr><td>>hg38_21_+_8433172_8446622_RNA45SN1_45S_50ntFlanks@3853.3898.46|AACTGTGGTAATTCTAGAGCTAATACATGCCGACGGGCGCTGACCC</td><td>>hg38_21_+_8433172_8446622_RNA45SN1_45S_50ntFlanks</td><td>3853</td><td>3898</td><td>46</td><td>AACTGTGGTAATTCTAGAGCTAATACATGCCGACGGGCGCTGACCC</td><td>764.7</td><td>-9.7</td><td>1.5</td><td>-6.3</td><td>2.2E-10</td><td>5.3E-10</td><td>MB231/MB468</td><td>NUC</td></tr>
<tr><td>>hg38_21_+_8433172_8446622_RNA45SN1_45S_50ntFlanks@10014.10041.28|AAAATGGATGGCGCTGGAGCGTCGGGCC</td><td>>hg38_21_+_8433172_8446622_RNA45SN1_45S_50ntFlanks</td><td>10014</td><td>10041</td><td>28</td><td>AAAATGGATGGCGCTGGAGCGTCGGGCC</td><td>107.0</td><td>-10.6</td><td>1.7</td><td>-6.4</td><td>2.0E-10</td><td>5.3E-10</td><td>MB231/MB468</td><td>MP</td></tr>
</table>

| | | | | | | | | | | | | | |
|---|---|---|---|---|---|---|---|---|---|---|---|---|---|
| >hg38_21_+_8433172_8446622_RNA45SN1_45S_50ntFlanks@475.518.44\|CTTCGTGATCGATGTGGTGACGTCGTGCTCTCCCGGGCCGGGTC | >hg38_21_+_8433172_8446622_RNA45SN1_45S_50ntFlanks | 475 | 518 | 44 | CTTCGTGATCGATGTGGTGACGTCGTGCTCTCCCGGGCCGGGTC | 2638.9 | 8.0 | 1.2 | 6.4 | 1.6E-10 | 5.3E-10 | BT20/MB468 | MITO |
| >hg38_21_+_8433172_8446622_RNA45SN1_45S_50ntFlanks@4724.4748.25\|ATTAATCAAGAACGAAAGTCGGAGG | >hg38_21_+_8433172_8446622_RNA45SN1_45S_50ntFlanks | 4724 | 4748 | 25 | ATTAATCAAGAACGAAAGTCGGAGG | 892.3 | -8.6 | 1.3 | -6.4 | 2.1E-10 | 5.3E-10 | MB231/MB468 | MP |
| >hg38_21_+_8433172_8446622_RNA45SN1_45S_50ntFlanks@9166.9188.23\|CCGGGCGGGTCGCGCCGTCGGGC | >hg38_21_+_8433172_8446622_RNA45SN1_45S_50ntFlanks | 9166 | 9188 | 23 | CCGGGCGGGTCGCGCCGTCGGGC | 117.4 | -10.7 | 1.7 | -6.4 | 2.1E-10 | 5.4E-10 | MB231/MB468 | MP |
| >hg38_21_+_8433172_8446622_RNA45SN1_45S_50ntFlanks@11190.11209.20\|CCCGGGGGGCCGGCGGCGGC | >hg38_21_+_8433172_8446622_RNA45SN1_45S_50ntFlanks | 11190 | 11209 | 20 | CCCGGGGGGCCGGCGGCGGC | 145.3 | -11.0 | 1.7 | -6.4 | 2.1E-10 | 5.4E-10 | MB231/MB468 | MP |
| >hg38_21_+_8433172_8446622_RNA45SN1_45S_50ntFlanks@9720.9749.30\|ATTAGAGGTCTTGGGGCCGAAACGATCTCA | >hg38_21_+_8433172_8446622_RNA45SN1_45S_50ntFlanks | 9720 | 9749 | 30 | ATTAGAGGTCTTGGGGCCGAAACGATCTCA | 62.6 | -9.8 | 1.5 | -6.4 | 2.1E-10 | 5.4E-10 | MB231/MB468 | MP |
| >MI0000298\|hsa-mir-221&WithFlank&X\|-\|45746151\|45746272@71.91.21(+1C)&[MIMAT0000278&hsa-miR-221-3p&offsets\|0\|-2(+1C);m-91&X\|-\|45746181\|45746202&offsets\|0\|-1(+1C)]\|AGCTACATTGTCTGCTGGGTTC | >MI0000298\|hsa-mir-221&WithFlank&X\|-\|45746151\|45746272 | 71 | 91 | 21(+1C) | AGCTACATTGTCTGCTGGGTTC | 6632.8 | 5.2 | 0.8 | 6.4 | 2.1E-10 | 5.4E-10 | MB231/MB468 | MP |
| >hg38_21_+_8433172_8446622_RNA45SN1_45S_50ntFlanks@12488.12508.21\|AACGTGAGCTGGGTTTAGACC | >hg38_21_+_8433172_8446622_RNA45SN1_45S_50ntFlanks | 12488 | 12508 | 21 | AACGTGAGCTGGGTTTAGACC | 149.8 | -11.1 | 1.7 | -6.4 | 2.1E-10 | 5.4E-10 | MB231/MB468 | MP |
| >MI0005559\|hsa-mir-744&WithFlank&17\|+\|12081893\|12082002@17.37.21&[MIMAT0004945&hsa-miR-744-5p&offsets\|0\|-1;m-172&17\|+\|12081909\|12081929&offsets\|0\|0]\|TGCGGGGCTAGGGCTAACAGC | >MI0005559\|hsa-mir-744&WithFlank&17\|+\|12081893\|12082002 | 17 | 37 | 21 | TGCGGGGCTAGGGCTAACAGC | 874.6 | 4.5 | 0.7 | 6.3 | 2.2E-10 | 5.4E-10 | MB231/MB468 | NUC |
| >hg38_21_+_8433172_8446622_RNA45SN1_45S_50ntFlanks@9736.9757.22\|CCGAAACGATCTCAACCTATTC | >hg38_21_+_8433172_8446622_RNA45SN1_45S_50ntFlanks | 9736 | 9757 | 22 | CCGAAACGATCTCAACCTATTC | 143.6 | -7.5 | 1.2 | -6.3 | 2.2E-10 | 5.4E-10 | MB231/MB468 | NUC |
| >hg38_21_+_8433172_8446622_RNA45SN1_45S_50ntFlanks@3843.3871.29\|CTACTTGGATAACTGTGGTAATTCTAGAG | >hg38_21_+_8433172_8446622_RNA45SN1_45S_50ntFlanks | 3843 | 3871 | 29 | CTACTTGGATAACTGTGGTAATTCTAGAG | 825.2 | -7.3 | 1.2 | -6.3 | 2.4E-10 | 5.4E-10 | MB231/MB468 | MITO |
| >hg38_21_+_8433172_8446622_RNA45SN1_45S_50ntFlanks@9570.9598.29\|TGCAAATCGGTCGTCCGACCTGGGTATAG | >hg38_21_+_8433172_8446622_RNA45SN1_45S_50ntFlanks | 9570 | 9598 | 29 | TGCAAATCGGTCGTCCGACCTGGGTATAG | 66.9 | -9.9 | 1.6 | -6.4 | 2.1E-10 | 5.4E-10 | MB231/MB468 | MP |
| >hg38_21_+_8433172_8446622_RNA45SN1_45S_50ntFlanks@3802.3834.33\|CATTAAATCAGTTATGGTTCCTTTGGTCGCTCG | >hg38_21_+_8433172_8446622_RNA45SN1_45S_50ntFlanks | 3802 | 3834 | 33 | CATTAAATCAGTTATGGTTCCTTTGGTCGCTCG | 91.3 | -10.4 | 1.6 | -6.4 | 2.1E-10 | 5.4E-10 | MB231/MB468 | MP |
| >hg38_21_+_8433172_8446622_RNA45SN1_45S_50ntFlanks@11569.11611.43\|CAGCCGACTTAGAACTGGTGCGGACCAGGGGAATCCGACTGTT | >hg38_21_+_8433172_8446622_RNA45SN1_45S_50ntFlanks | 11569 | 11611 | 43 | CAGCCGACTTAGAACTGGTGCGGACCAGGGGAATCCGACTGTT | 3705.4 | -6.3 | 1.0 | -6.3 | 2.4E-10 | 5.4E-10 | MB231/MB468 | MITO |
| >hg38_21_+_8433172_8446622_RNA45SN1_45S_50ntFlanks@6739.6771.33\|TCATCGACACTTCGAACGCACTTGCGGCCCCGG | >hg38_21_+_8433172_8446622_RNA45SN1_45S_50ntFlanks | 6739 | 6771 | 33 | TCATCGACACTTCGAACGCACTTGCGGCCCCGG | 717.4 | -9.1 | 1.4 | -6.3 | 2.6E-10 | 5.4E-10 | BT20/MB468 | NUC |
| >hg38_21_+_8433172_8446622_RNA45SN1_45S_50ntFlanks@5083.5119.37\|ACGAACGAGACTCTGGCATGCTAACTAGTTACGCGAC | >hg38_21_+_8433172_8446622_RNA45SN1_45S_50ntFlanks | 5083 | 5119 | 37 | ACGAACGAGACTCTGGCATGCTAACTAGTTACGCGAC | 142.1 | -11.0 | 1.7 | -6.4 | 2.1E-10 | 5.4E-10 | MB231/MB468 | MP |
| >hg38_21_+_8433172_8446622_RNA45SN1_45S_50ntFlanks@10916.10939.24\|CCGCGGCTGGACGAGGCGCCGCCG | >hg38_21_+_8433172_8446622_RNA45SN1_45S_50ntFlanks | 10916 | 10939 | 24 | CCGCGGCTGGACGAGGCGCCGCCG | 114.4 | -10.7 | 1.7 | -6.4 | 2.1E-10 | 5.4E-10 | MB231/MB468 | MP |
| >MI0000078\|hsa-mir-22&WithFlank&17\|-\|1713897\|1713993@59.79.21(+1C)&[MIMAT0000077&hsa-miR-22-3p&offsets\|0\|-1(+1C);m-2&17\|-\|1713914\|1713935&offsets\|0\|-1(+1C)]\|AAGCTGCCAGTTGAAGAACTGC | >MI0000078\|hsa-mir-22&WithFlank&17\|-\|1713897\|1713993 | 59 | 79 | 21(+1C) | AAGCTGCCAGTTGAAGAACTGC | 411.2 | 6.1 | 1.0 | 6.3 | 2.2E-10 | 5.4E-10 | MB231/MB468 | CYTO |
| >hg38_21_+_8433172_8446622_RNA45SN1_45S_50ntFlanks@3776.3814.39\|CCGGTACAGTGAAACTGCGAATGGCTCATTAAATCAGTT | >hg38_21_+_8433172_8446622_RNA45SN1_45S_50ntFlanks | 3776 | 3814 | 39 | CCGGTACAGTGAAACTGCGAATGGCTCATTAAATCAGTT | 435.6 | -6.9 | 1.1 | -6.3 | 2.6E-10 | 5.4E-10 | BT20/MB468 | NUC |
| >MI0000086\|hsa-mir-28&WithFlank&3\|+\|188688775\|188688872@60.80.21&[MIMAT0004502&hsa-miR-28-3p&offsets\|0\|-1;m-28&3\|+\|188688834\|188688855&offsets\|0\|-1]\|CACTAGATTGTGAGCTCCTGG | >MI0000086\|hsa-mir-28&WithFlank&3\|+\|188688775\|188688872 | 60 | 80 | 21 | CACTAGATTGTGAGCTCCTGG | 1527.8 | 6.0 | 0.9 | 6.7 | 1.8E-11 | 5.4E-10 | BT20/MB468 | MP |
| >hg38_21_+_8433172_8446622_RNA45SN1_45S_50ntFlanks@4759.4779.21\|ATCAGATACCGTCGTAGTTCC | >hg38_21_+_8433172_8446622_RNA45SN1_45S_50ntFlanks | 4759 | 4779 | 21 | ATCAGATACCGTCGTAGTTCC | 316.4 | -11.4 | 1.7 | -6.7 | 1.8E-11 | 5.4E-10 | BT20/MB468 | MP |
| >hg38_21_+_8433172_8446622_RNA45SN1_45S_50ntFlanks@12072.12108.37\|CCCGAGGGGCTCTCGCTTCTGGCGCCAAGCGCCCGGC | >hg38_21_+_8433172_8446622_RNA45SN1_45S_50ntFlanks | 12072 | 12108 | 37 | CCCGAGGGGCTCTCGCTTCTGGCGCCAAGCGCCCGGC | 82.0 | -10.2 | 1.6 | -6.4 | 2.1E-10 | 5.4E-10 | MB231/MB468 | MP |
| >MI0000285\|hsa-mir-205&WithFlank&1\|+\|209432127\|209432248@40.60.21(+1A)&[MIMAT0000266&hsa-miR-205-5p&offsets\|0\|-1(+1A);m-351&1\|+\|209432166\|209432187&offsets\|0\|-1(+1A)]\|TCCTTCATTCCACCGGAGTCTA | >MI0000285\|hsa-mir-205&WithFlank&1\|+\|209432127\|209432248 | 40 | 60 | 21(+1A) | TCCTTCATTCCACCGGAGTCTA | 6487.7 | 6.7 | 1.1 | 6.4 | 1.6E-10 | 5.4E-10 | BT20/MB468 | MITO |
| >hg38_21_+_8433172_8446622_RNA45SN1_45S_50ntFlanks@9760.9789.30\|AAACTTTAAATGGGTAAGAAGCCCGGCTCG | >hg38_21_+_8433172_8446622_RNA45SN1_45S_50ntFlanks | 9760 | 9789 | 30 | AAACTTTAAATGGGTAAGAAGCCCGGCTCG | 105.8 | -10.6 | 1.7 | -6.4 | 2.1E-10 | 5.4E-10 | MB231/MB468 | MP |
| >am_trna10_ValCAC_5_-_180649395_180649467@1.33.33,trna12_ValAAC_5_-_180645270_180645342@1.33.33,trna132_ValAAC_6_-_27721179_27721251@1.33.33,trna136_ValAAC_6_-_27648885_27648957@1.33.33,trna139_ValAAC_6_-_27618707_27618779@1.33.33,trna18_ValCAC_5_-_180529253_180529325@1.33.33,trna2_ValAAC_3_+_169490018_169490090@1.33.33,trna2_ValCAC_5_+_180524070_180524142@1.33.33,trna4_ValAAC_5_+_180591154_180591226@1.33.33,trna5_ValAAC_5_+_180596610_180596682@1.33.33,trna6_ValCAC_5_+_180600650_180600722@1.33.33,trna85_ValCAC_1_-_161369490_161369562@1.33.33,trna90_ValCAC_1_-_149684088_149684161@1.33.33,trna98_ValCAC_1_-_149298555_149298627@1.33.33,trna9_ValCAC_6_+_26538282_26538354@1.33.33\|GTTTCCGTAGTGTAGTGGTTATCACGTTCGCCT | >am_trna10_ValCAC_5_-_180649395_180649467 | 1 | 33 | 33 | GTTTCCGTAGTGTAGTGGTTATCACGTTCGCCT | 1998.6 | 7.8 | 1.2 | 6.3 | 2.2E-10 | 5.4E-10 | MB231/MB468 | NUC |
| >hg38_21_+_8433172_8446622_RNA45SN1_45S_50ntFlanks@6722.6744.23\|ATTGCAGGACACATTGATCATCG | >hg38_21_+_8433172_8446622_RNA45SN1_45S_50ntFlanks | 6722 | 6744 | 23 | ATTGCAGGACACATTGATCATCG | 670.8 | -8.4 | 1.3 | -6.4 | 2.1E-10 | 5.4E-10 | MB231/MB468 | MP |
| >hg38_21_+_8433172_8446622_RNA45SN1_45S_50ntFlanks@9794.9820.27\|CGTGGAGCCGGGCGTGGAATGCGAGTG | >hg38_21_+_8433172_8446622_RNA45SN1_45S_50ntFlanks | 9794 | 9820 | 27 | CGTGGAGCCGGGCGTGGAATGCGAGTG | 125.1 | -10.8 | 1.7 | -6.4 | 2.1E-10 | 5.4E-10 | MB231/MB468 | MP |
| >hg38_21_+_8433172_8446622_RNA45SN1_45S_50ntFlanks@4284.4317.34\|TTTAACGAGGATCCATTGGAGGGCAAGTCTGGTG | >hg38_21_+_8433172_8446622_RNA45SN1_45S_50ntFlanks | 4284 | 4317 | 34 | TTTAACGAGGATCCATTGGAGGGCAAGTCTGGTG | 163.8 | -11.2 | 1.8 | -6.4 | 2.1E-10 | 5.4E-10 | MB231/MB468 | MP |
| >hg38_21_+_8433172_8446622_RNA45SN1_45S_50ntFlanks@5492.5525.34\|AGAAGACGGTCGAACTTGACTATCTAGAGGAAGT | >hg38_21_+_8433172_8446622_RNA45SN1_45S_50ntFlanks | 5492 | 5525 | 34 | AGAAGACGGTCGAACTTGACTATCTAGAGGAAGT | 626.7 | -4.6 | 0.7 | -6.3 | 2.5E-10 | 5.5E-10 | MB231/MB468 | MITO |
| >hg38_21_+_8433172_8446622_RNA45SN1_45S_50ntFlanks@9573.9597.25\|AAATCGGTCGTCCGACCTGGGTATA | >hg38_21_+_8433172_8446622_RNA45SN1_45S_50ntFlanks | 9573 | 9597 | 25 | AAATCGGTCGTCCGACCTGGGTATA | 4569.7 | -7.8 | 1.2 | -6.4 | 1.6E-10 | 5.5E-10 | BT20/MB468 | MITO |
| >trna10_LysCTT_16_+_3241501_3241573@1.29.29,trna13_LysCTT_14_-_58706613_58706685@1.29.29,trna2_LysCTT_15_+_79152904_79152976@1.29.29\|GCCCGGCTAGCTCAGTCGGTAGAGCATGG | >trna10_LysCTT_16_+_3241501_3241573 | 1 | 29 | 29 | GCCCGGCTAGCTCAGTCGGTAGAGCATGG | 1149.5 | -5.2 | 0.8 | -6.3 | 2.2E-10 | 5.5E-10 | MB231/MB468 | CYTO |
| >hg38_21_+_8433172_8446622_RNA45SN1_45S_50ntFlanks@10735.10767.33\|TGTAAATCTCGCGCCGGGCCGTACCCATATCCG | >hg38_21_+_8433172_8446622_RNA45SN1_45S_50ntFlanks | 10735 | 10767 | 33 | TGTAAATCTCGCGCCGGGCCGTACCCATATCCG | 496.1 | -7.0 | 1.1 | -6.3 | 2.2E-10 | 5.5E-10 | MB231/MB468 | NUC |
| >hg38_21_+_8433172_8446622_RNA45SN1_45S_50ntFlanks@8189.8212.24\|CCCAGCCCGTGGACGGTGTGAGGC | >hg38_21_+_8433172_8446622_RNA45SN1_45S_50ntFlanks | 8189 | 8212 | 24 | CCCAGCCCGTGGACGGTGTGAGGC | 163.4 | -9.4 | 1.5 | -6.4 | 1.6E-10 | 5.5E-10 | BT20/MB468 | MITO |
| >hg38_21_+_8433172_8446622_RNA45SN1_45S_50ntFlanks@12937.12962.26\|AACCATTCGTAGACGACCTGCTTCTG | >hg38_21_+_8433172_8446622_RNA45SN1_45S_50ntFlanks | 12937 | 12962 | 26 | AACCATTCGTAGACGACCTGCTTCTG | 843.2 | -9.2 | 1.5 | -6.3 | 2.6E-10 | 5.5E-10 | BT20/MB468 | NUC |
| >hg38_21_+_8433172_8446622_RNA45SN1_45S_50ntFlanks@8105.8143.39\|CGCGGCGGGGCGCGGGACATGTGGCGTACGGAAGACCCG | >hg38_21_+_8433172_8446622_RNA45SN1_45S_50ntFlanks | 8105 | 8143 | 39 | CGCGGCGGGGCGCGGGACATGTGGCGTACGGAAGACCCG | 18862.4 | -11.5 | 1.8 | -6.3 | 2.6E-10 | 5.5E-10 | BT20/MB468 | NUC |
| >hg38_21_+_8433172_8446622_RNA45SN1_45S_50ntFlanks@5168.5185.18\|CAGCCACCCGAGATTGAG | >hg38_21_+_8433172_8446622_RNA45SN1_45S_50ntFlanks | 5168 | 5185 | 18 | CAGCCACCCGAGATTGAG | 200.8 | -11.5 | 1.8 | -6.4 | 2.1E-10 | 5.5E-10 | MB231/MB468 | MP |
| >hg38_21_+_8433172_8446622_RNA45SN1_45S_50ntFlanks@10766.10797.32\|CGCAGCAGGTCTCCAAGGTGAACAGCCTCTGG | >hg38_21_+_8433172_8446622_RNA45SN1_45S_50ntFlanks | 10766 | 10797 | 32 | CGCAGCAGGTCTCCAAGGTGAACAGCCTCTGG | 122.6 | -6.5 | 1.0 | -6.4 | 2.0E-10 | 5.5E-10 | BT20/MB468 | CYTO |
| >hg38_21_+_8433172_8446622_RNA45SN1_45S_50ntFlanks@11768.11803.36\|CTAATTAGTGACGCGCATGAATGGATGAACGAGATT | >hg38_21_+_8433172_8446622_RNA45SN1_45S_50ntFlanks | 11768 | 11803 | 36 | CTAATTAGTGACGCGCATGAATGGATGAACGAGATT | 1382.1 | -5.9 | 0.9 | -6.3 | 2.2E-10 | 5.5E-10 | MB231/MB468 | CYTO |
| >hg38_21_+_8433172_8446622_RNA45SN1_45S_50ntFlanks@11567.11584.18\|AGCAGCCGACTTAGAACT | >hg38_21_+_8433172_8446622_RNA45SN1_45S_50ntFlanks | 11567 | 11584 | 18 | AGCAGCCGACTTAGAACT | 3386.0 | 5.0 | 0.8 | 6.3 | 2.2E-10 | 5.5E-10 | MB231/MB468 | NUC |
| >hg38_21_+_8433172_8446622_RNA45SN1_45S_50ntFlanks@10808.10827.20\|CAATGTAGGTAAGGGAAGTC | >hg38_21_+_8433172_8446622_RNA45SN1_45S_50ntFlanks | 10808 | 10827 | 20 | CAATGTAGGTAAGGGAAGTC | 570.0 | -7.4 | 1.2 | -6.3 | 2.2E-10 | 5.5E-10 | MB231/MB468 | NUC |
| >hg38_21_+_8433172_8446622_RNA45SN1_45S_50ntFlanks@12615.12654.40\|GCTGAGGAGCCAATGGGGCGAAGCTACCATCTGTGGGATT | >hg38_21_+_8433172_8446622_RNA45SN1_45S_50ntFlanks | 12615 | 12654 | 40 | GCTGAGGAGCCAATGGGGCGAAGCTACCATCTGTGGGATT | 7468.5 | -5.1 | 0.8 | -6.4 | 1.7E-10 | 5.5E-10 | BT20/MB468 | MITO |
| >hg38_21_+_8433172_8446622_RNA45SN1_45S_50ntFlanks@8171.8212.42\|AAGTCCTTCTGATCGAGGCCCAGCCCGTGGACGGTGTGAGGC | >hg38_21_+_8433172_8446622_RNA45SN1_45S_50ntFlanks | 8171 | 8212 | 42 | AAGTCCTTCTGATCGAGGCCCAGCCCGTGGACGGTGTGAGGC | 3022.4 | -5.6 | 0.9 | -6.3 | 2.3E-10 | 5.6E-10 | MB231/MB468 | NUC |
| >hg38_21_+_8433172_8446622_RNA45SN1_45S_50ntFlanks@8182.8206.25\|ATCGAGGCCCAGCCCGTGGACGGTG | >hg38_21_+_8433172_8446622_RNA45SN1_45S_50ntFlanks | 8182 | 8206 | 25 | ATCGAGGCCCAGCCCGTGGACGGTG | 570.9 | -8.1 | 1.3 | -6.3 | 2.5E-10 | 5.6E-10 | MB231/MB468 | MITO |
| >hg38_21_+_8433172_8446622_RNA45SN1_45S_50ntFlanks@11569.11593.25\|CAGCCGACTTAGAACTGGTGCGGAC | >hg38_21_+_8433172_8446622_RNA45SN1_45S_50ntFlanks | 11569 | 11593 | 25 | CAGCCGACTTAGAACTGGTGCGGAC | 327.1 | -7.9 | 1.2 | -6.3 | 2.5E-10 | 5.6E-10 | MB231/MB468 | MITO |
| >hg38_21_+_8433172_8446622_RNA45SN1_45S_50ntFlanks@10816.10840.25\|GTAAGGGAAGTCGGCAAGCCGGATC | >hg38_21_+_8433172_8446622_RNA45SN1_45S_50ntFlanks | 10816 | 10840 | 25 | GTAAGGGAAGTCGGCAAGCCGGATC | 665.0 | -9.0 | 1.4 | -6.3 | 2.6E-10 | 5.6E-10 | BT20/MB468 | NUC |

| | | | | | | | | | | | | | |
|---|---|---|---|---|---|---|---|---|---|---|---|---|---|
| >trna130_GlnTTG_6_-_27763640_27763711@30.52.23,trna16_GlnTTG_17_+_47269890_47269961@30.52.23,trna173_GlnTTG_6_-_26311975_26312046@30.52.23,trna174_GlnTTG_6_-_26311424_26311495@30.52.23\|GACTTTGAATCCAGCGATCCGAG | >trna130_GlnTTG_6_-_27763640_27763711 | 30 | 52 | 23 | GACTTTGAATCCAGCGATCCGAG | 120.3 | -10.8 | 1.7 | -6.4 | 2.1E-10 | 5.6E-10 | MB231/MB468 | MP |
| >hg38_1_-_228634819_228635039_RNA5S12_5S_50ntFlanks@136.170.35\|GGAGACCGCCTGGGAATACCGGGTGCTGTAGGCTT | >hg38_1_-_228634819_228635039_RNA5S12_5S_50ntFlanks | 136 | 170 | 35 | GGAGACCGCCTGGGAATACCGGGTGCTGTAGGCTT | 1571.0 | 4.3 | 0.7 | 6.3 | 2.3E-10 | 5.6E-10 | MB231/MB468 | NUC |
| >hg38_21_+_8433172_8446622_RNA45SN1_45S_50ntFlanks@12141.12166.26\|ACAGTGCCAGGTGGGGAGTTTGACTG | >hg38_21_+_8433172_8446622_RNA45SN1_45S_50ntFlanks | 12141 | 12166 | 26 | ACAGTGCCAGGTGGGGAGTTTGACTG | 83.8 | -10.3 | 1.6 | -6.4 | 2.1E-10 | 5.6E-10 | MB231/MB468 | MP |
| >hg38_21_+_8433172_8446622_RNA45SN1_45S_50ntFlanks@4760.4779.20\|TCAGATACCGTCGTAGTTCC | >hg38_21_+_8433172_8446622_RNA45SN1_45S_50ntFlanks | 4760 | 4779 | 20 | TCAGATACCGTCGTAGTTCC | 162.3 | -9.4 | 1.5 | -6.4 | 1.7E-10 | 5.6E-10 | BT20/MB468 | MITO |
| >hg38_21_+_8433172_8446622_RNA45SN1_45S_50ntFlanks@3899.3921.23\|CCTTCGCGGGGGGGATGCGTGCA | >hg38_21_+_8433172_8446622_RNA45SN1_45S_50ntFlanks | 3899 | 3921 | 23 | CCTTCGCGGGGGGGATGCGTGCA | 212.0 | -10.8 | 1.6 | -6.7 | 1.8E-11 | 5.6E-10 | BT20/MB468 | MP |
| >MI0000102\|hsa-mir-100&WithFlank&11\|-\|122152223\|122152314@19.38.20(+2A)&[MIMAT0000098&hsa-miR-100-5p&offsets\|0\|-2(+2A);m-27&11\|-\|122152276\|122152296&offsets\|0\|-1(+2A)]\|AACCCGTAGATCCGAACTTGAA | >MI0000102\|hsa-mir-100&WithFlank&11\|-\|122152223\|122152314 | 19 | 38 | 20(+2A) | AACCCGTAGATCCGAACTTGAA | 112.3 | 10.0 | 1.6 | 6.3 | 2.2E-10 | 5.6E-10 | MB231/MB468 | MP |
| >hg38_21_+_8433172_8446622_RNA45SN1_45S_50ntFlanks@12627.12662.36\|ATGGGGCGAAGCTACCATCTGTGGGATTATGACTGA | >hg38_21_+_8433172_8446622_RNA45SN1_45S_50ntFlanks | 12627 | 12662 | 36 | ATGGGGCGAAGCTACCATCTGTGGGATTATGACTGA | 343.5 | -8.2 | 1.3 | -6.3 | 2.5E-10 | 5.6E-10 | MB231/MB468 | MITO |
| >hg38_21_+_8433172_8446622_RNA45SN1_45S_50ntFlanks@10782.10817.36\|GGTGAACAGCCTCTGGCATGTTGGAACAATGTAGGT | >hg38_21_+_8433172_8446622_RNA45SN1_45S_50ntFlanks | 10782 | 10817 | 36 | GGTGAACAGCCTCTGGCATGTTGGAACAATGTAGGT | 67.3 | -9.9 | 1.6 | -6.3 | 2.2E-10 | 5.6E-10 | MB231/MB468 | MP |
| >hg38_21_+_8433172_8446622_RNA45SN1_45S_50ntFlanks@4008.4042.35\|CTCGGGCCGATCGCACGCCCCCCGTGGCGGCGACG | >hg38_21_+_8433172_8446622_RNA45SN1_45S_50ntFlanks | 4008 | 4042 | 35 | CTCGGGCCGATCGCACGCCCCCCGTGGCGGCGACG | 338.6 | -3.6 | 0.6 | -6.4 | 2.0E-10 | 5.6E-10 | BT20/MB468 | CYTO |
| >hg38_21_+_8433172_8446622_RNA45SN1_45S_50ntFlanks@10498.10516.19\|TTCAGATCCCCGAATCCGG | >hg38_21_+_8433172_8446622_RNA45SN1_45S_50ntFlanks | 10498 | 10516 | 19 | TTCAGATCCCCGAATCCGG | 421.6 | -5.9 | 0.9 | -6.4 | 2.0E-10 | 5.6E-10 | BT20/MB468 | CYTO |
| >MI0000746\|hsa-mir-99b&WithFlank&19\|+\|51692606\|51692687@13.32.20(+2U)&[MIMAT0000689&hsa-miR-99b-5p&offsets\|0\|-2(+2U);m-10&19\|+\|51692618\|51692639&offsets\|0\|-2(+2U)]\|CACCCGTAGAACCGACCTTGTT | >MI0000746\|hsa-mir-99b&WithFlank&19\|+\|51692606\|51692687 | 13 | 32 | 20(+2U) | CACCCGTAGAACCGACCTTGTT | 393.1 | 12.0 | 1.7 | 6.9 | 3.9E-12 | 5.6E-10 | BT20/MB231 | MP |
| >hg38_21_+_8433172_8446622_RNA45SN1_45S_50ntFlanks@10781.10806.26\|AGGTGAACAGCCTCTGGCATGTTGGA | >hg38_21_+_8433172_8446622_RNA45SN1_45S_50ntFlanks | 10781 | 10806 | 26 | AGGTGAACAGCCTCTGGCATGTTGGA | 233.2 | -10.9 | 1.6 | -6.7 | 1.9E-11 | 5.6E-10 | BT20/MB468 | MP |
| >hg38_21_+_8433172_8446622_RNA45SN1_45S_50ntFlanks@10818.10840.23\|AAGGGAAGTCGGCAAGCCGGATC | >hg38_21_+_8433172_8446622_RNA45SN1_45S_50ntFlanks | 10818 | 10840 | 23 | AAGGGAAGTCGGCAAGCCGGATC | 242.2 | -3.8 | 0.6 | -6.3 | 2.3E-10 | 5.6E-10 | MB231/MB468 | NUC |
| >hg38_21_+_8433172_8446622_RNA45SN1_45S_50ntFlanks@10678.10696.19\|CGGTTCCGGCGGCGTCCGG | >hg38_21_+_8433172_8446622_RNA45SN1_45S_50ntFlanks | 10678 | 10696 | 19 | CGGTTCCGGCGGCGTCCGG | 220.6 | -10.9 | 1.6 | -6.7 | 1.9E-11 | 5.6E-10 | BT20/MB468 | MP |
| >hg38_21_+_8433172_8446622_RNA45SN1_45S_50ntFlanks@12472.12496.25\|TCACCCACTAATAGGGAACGTGAGC | >hg38_21_+_8433172_8446622_RNA45SN1_45S_50ntFlanks | 12472 | 12496 | 25 | TCACCCACTAATAGGGAACGTGAGC | 355.1 | 8.2 | 1.3 | 6.4 | 1.7E-10 | 5.7E-10 | BT20/MB468 | MITO |
| >hg38_21_+_8433172_8446622_RNA45SN1_45S_50ntFlanks@4974.4994.21\|GCCCGGACACGGACAGGATTG | >hg38_21_+_8433172_8446622_RNA45SN1_45S_50ntFlanks | 4974 | 4994 | 21 | GCCCGGACACGGACAGGATTG | 92.0 | -10.4 | 1.6 | -6.3 | 2.2E-10 | 5.7E-10 | MB231/MB468 | MP |
| >trna11_GluTTC_15_-_26327381_26327452@41.75.35,trna3_GluTTC_13_-_45492062_45492133@41.75.35\|AGGCGGCCCGGGTTCGACTCCCGGTGTGGGAACCA | >trna11_GluTTC_15_-_26327381_26327452 | 41 | 75 | 35 | AGGCGGCCCGGGTTCGACTCCCGGTGTGGGAACCA | 382.1 | 6.9 | 1.1 | 6.4 | 1.7E-10 | 5.7E-10 | BT20/MB468 | MITO |
| >hg38_21_+_8433172_8446622_RNA45SN1_45S_50ntFlanks@10771.10797.27\|CAGGTCTCCAAGGTGAACAGCCTCTGG | >hg38_21_+_8433172_8446622_RNA45SN1_45S_50ntFlanks | 10771 | 10797 | 27 | CAGGTCTCCAAGGTGAACAGCCTCTGG | 120.8 | -9.0 | 1.4 | -6.4 | 1.7E-10 | 5.7E-10 | BT20/MB468 | MITO |
| >hg38_21_+_8433172_8446622_RNA45SN1_45S_50ntFlanks@10806.10840.35\|AACAATGTAGGTAAGGGAAGTCGGCAAGCCGGATC | >hg38_21_+_8433172_8446622_RNA45SN1_45S_50ntFlanks | 10806 | 10840 | 35 | AACAATGTAGGTAAGGGAAGTCGGCAAGCCGGATC | 730.3 | -3.9 | 0.6 | -6.4 | 1.7E-10 | 5.7E-10 | BT20/MB468 | MITO |
| >hg38_21_+_8433172_8446622_RNA45SN1_45S_50ntFlanks@12159.12183.25\|TTTGACTGGGGCGGTACACCTGTCA | >hg38_21_+_8433172_8446622_RNA45SN1_45S_50ntFlanks | 12159 | 12183 | 25 | TTTGACTGGGGCGGTACACCTGTCA | 196.9 | -9.7 | 1.5 | -6.4 | 1.7E-10 | 5.7E-10 | BT20/MB468 | MITO |
| >MI0000483\|hsa-mir-186&WithFlank&1\|-\|71067625\|71067722@21.43.23&[MIMAT0000456&hsa-miR-186-5p&offsets\|0\|+1;m-84&1\|-\|71067681\|71067702&offsets\|0\|+1]\|CAAAGAATTCTCCTTTTGGGCTT | >MI0000483\|hsa-mir-186&WithFlank&1\|-\|71067625\|71067722 | 21 | 43 | 23 | CAAAGAATTCTCCTTTTGGGCTT | 355.6 | 4.0 | 0.6 | 6.3 | 2.7E-10 | 5.7E-10 | BT20/MB468 | NUC |
| >hg38_21_+_8433172_8446622_RNA45SN1_45S_50ntFlanks@10392.10411.20\|ACAGCAGTTGAACATGGGTC | >hg38_21_+_8433172_8446622_RNA45SN1_45S_50ntFlanks | 10392 | 10411 | 20 | ACAGCAGTTGAACATGGGTC | 990.4 | -9.4 | 1.5 | -6.3 | 2.7E-10 | 5.7E-10 | BT20/MB468 | NUC |
| >hg38_21_+_8433172_8446622_RNA45SN1_45S_50ntFlanks@9544.9563.20\|GGTGGAGGTCCGTAGCGGTC | >hg38_21_+_8433172_8446622_RNA45SN1_45S_50ntFlanks | 9544 | 9563 | 20 | GGTGGAGGTCCGTAGCGGTC | 1045.2 | -9.5 | 1.5 | -6.3 | 2.7E-10 | 5.7E-10 | BT20/MB468 | NUC |
| >hg38_21_+_8433172_8446622_RNA45SN1_45S_50ntFlanks@9533.9566.34\|GAGGAAACTCTGGTGGAGGTCCGTAGCGGTCCTG | >hg38_21_+_8433172_8446622_RNA45SN1_45S_50ntFlanks | 9533 | 9566 | 34 | GAGGAAACTCTGGTGGAGGTCCGTAGCGGTCCTG | 608.4 | -8.3 | 1.3 | -6.3 | 2.3E-10 | 5.7E-10 | MB231/MB468 | NUC |
| >hg38_21_+_8433172_8446622_RNA45SN1_45S_50ntFlanks@10735.10752.18\|TGTAAATCTCGCGCCGGG | >hg38_21_+_8433172_8446622_RNA45SN1_45S_50ntFlanks | 10735 | 10752 | 18 | TGTAAATCTCGCGCCGGG | 127.0 | -9.1 | 1.4 | -6.4 | 1.7E-10 | 5.7E-10 | BT20/MB468 | MITO |
| >hg38_21_+_8433172_8446622_RNA45SN1_45S_50ntFlanks@11323.11347.25\|CGTCTCGTCGCGCGCGCGTCCGCTG | >hg38_21_+_8433172_8446622_RNA45SN1_45S_50ntFlanks | 11323 | 11347 | 25 | CGTCTCGTCGCGCGCGCGTCCGCTG | 5494.1 | 7.7 | 1.2 | 6.4 | 1.7E-10 | 5.7E-10 | BT20/MB468 | MITO |
| >hg38_21_+_8433172_8446622_RNA45SN1_45S_50ntFlanks@11575.11603.29\|ACTTAGAACTGGTGCGGACCAGGGGAATC | >hg38_21_+_8433172_8446622_RNA45SN1_45S_50ntFlanks | 11575 | 11603 | 29 | ACTTAGAACTGGTGCGGACCAGGGGAATC | 4401.3 | -6.7 | 1.0 | -6.7 | 1.9E-11 | 5.7E-10 | BT20/MB468 | MP |
| >hg38_21_+_8433172_8446622_RNA45SN1_45S_50ntFlanks@12672.12703.32\|AGTCAGAATCCCGCCCAGGCGGAACGATACGG | >hg38_21_+_8433172_8446622_RNA45SN1_45S_50ntFlanks | 12672 | 12703 | 32 | AGTCAGAATCCCGCCCAGGCGGAACGATACGG | 210.6 | -10.8 | 1.6 | -6.7 | 1.9E-11 | 5.7E-10 | BT20/MB468 | MP |
| >hg38_21_+_8433172_8446622_RNA45SN1_45S_50ntFlanks@10234.10270.37\|CGGACGCTACGCCGCGACGAGTAGGAGGGCCGCTGCG | >hg38_21_+_8433172_8446622_RNA45SN1_45S_50ntFlanks | 10234 | 10270 | 37 | CGGACGCTACGCCGCGACGAGTAGGAGGGCCGCTGCG | 73.9 | -10.1 | 1.6 | -6.3 | 2.2E-10 | 5.7E-10 | MB231/MB468 | MP |
| >hg38_21_+_8433172_8446622_RNA45SN1_45S_50ntFlanks@12307.12325.19\|ACGATCCTTCTGACCTTTT | >hg38_21_+_8433172_8446622_RNA45SN1_45S_50ntFlanks | 12307 | 12325 | 19 | ACGATCCTTCTGACCTTTT | 60.8 | -9.8 | 1.5 | -6.3 | 2.2E-10 | 5.7E-10 | MB231/MB468 | MP |
| >MI0000266\|hsa-mir-10a&WithFlank&17\|-\|48579832\|48579953@28.48.21(+1A)&[MIMAT0000253&hsa-miR-10a-5p&offsets\|0\|-2(+1A);m-18&17\|-\|48579905\|48579926&offsets\|0\|-1(+1A)]\|TACCCTGTAGATCCGAATTTGA | >MI0000266\|hsa-mir-10a&WithFlank&17\|-\|48579832\|48579953 | 28 | 48 | 21(+1A) | TACCCTGTAGATCCGAATTTGA | 178.3 | 10.7 | 1.7 | 6.3 | 2.6E-10 | 5.7E-10 | MB231/MB468 | MITO |
| >hg38_21_+_8433172_8446622_RNA45SN1_45S_50ntFlanks@4660.4690.31\|GAGGTGAAATTCTTGGACCGGCGCAAGACGG | >hg38_21_+_8433172_8446622_RNA45SN1_45S_50ntFlanks | 4660 | 4690 | 31 | GAGGTGAAATTCTTGGACCGGCGCAAGACGG | 93.7 | -10.4 | 1.6 | -6.3 | 2.2E-10 | 5.7E-10 | MB231/MB468 | MP |
| >hg38_21_+_8433172_8446622_RNA45SN1_45S_50ntFlanks@9566.9597.32\|GACGTGCAAATCGGTCGTCCGACCTGGGTATA | >hg38_21_+_8433172_8446622_RNA45SN1_45S_50ntFlanks | 9566 | 9597 | 32 | GACGTGCAAATCGGTCGTCCGACCTGGGTATA | 1405.4 | -3.7 | 0.6 | -6.3 | 2.3E-10 | 5.7E-10 | MB231/MB468 | NUC |
| >hg38_21_+_8433172_8446622_RNA45SN1_45S_50ntFlanks@10391.10435.45\|AACAGCAGTTGAACATGGGTCAGTCGGTCCTGAGAGATGGGCGAG | >hg38_21_+_8433172_8446622_RNA45SN1_45S_50ntFlanks | 10391 | 10435 | 45 | AACAGCAGTTGAACATGGGTCAGTCGGTCCTGAGAGATGGGCGAG | 151.0 | -11.1 | 1.7 | -6.3 | 2.2E-10 | 5.7E-10 | MB231/MB468 | MP |
| >hg38_21_+_8433172_8446622_RNA45SN1_45S_50ntFlanks@10808.10836.29\|CAATGTAGGTAAGGGAAGTCGGCAAGCCG | >hg38_21_+_8433172_8446622_RNA45SN1_45S_50ntFlanks | 10808 | 10836 | 29 | CAATGTAGGTAAGGGAAGTCGGCAAGCCG | 363.0 | -8.0 | 1.3 | -6.3 | 2.6E-10 | 5.7E-10 | MB231/MB468 | MITO |
| >hg38_21_+_8433172_8446622_RNA45SN1_45S_50ntFlanks@10771.10796.26\|CAGGTCTCCAAGGTGAACAGCCTCTG | >hg38_21_+_8433172_8446622_RNA45SN1_45S_50ntFlanks | 10771 | 10796 | 26 | CAGGTCTCCAAGGTGAACAGCCTCTG | 619.7 | -6.4 | 1.0 | -6.3 | 2.6E-10 | 5.7E-10 | MB231/MB468 | MITO |
| >hg38_21_+_8433172_8446622_RNA45SN1_45S_50ntFlanks@9339.9367.29\|CGCACGAAAGCCGCCGTGGCGCAATGAAG | >hg38_21_+_8433172_8446622_RNA45SN1_45S_50ntFlanks | 9339 | 9367 | 29 | CGCACGAAAGCCGCCGTGGCGCAATGAAG | 257.7 | -10.1 | 1.6 | -6.4 | 1.7E-10 | 5.7E-10 | BT20/MB468 | MITO |
| >hg38_21_+_8433172_8446622_RNA45SN1_45S_50ntFlanks@10348.10382.35\|AAACGAGAACTTTGAAGGCCGAAGTGGAGAAGGGT | >hg38_21_+_8433172_8446622_RNA45SN1_45S_50ntFlanks | 10348 | 10382 | 35 | AAACGAGAACTTTGAAGGCCGAAGTGGAGAAGGGT | 902.0 | 3.5 | 0.6 | 6.4 | 2.1E-10 | 5.7E-10 | BT20/MB468 | CYTO |
| >hg38_21_+_8433172_8446622_RNA45SN1_45S_50ntFlanks@4046.4067.22\|CATTCGAACGTCTGCCCTATCA | >hg38_21_+_8433172_8446622_RNA45SN1_45S_50ntFlanks | 4046 | 4067 | 22 | CATTCGAACGTCTGCCCTATCA | 258.6 | -10.1 | 1.6 | -6.4 | 1.7E-10 | 5.7E-10 | BT20/MB468 | MITO |
| >hg38_21_+_8433172_8446622_RNA45SN1_45S_50ntFlanks@4760.4782.23\|TCAGATACCGTCGTAGTTCCGAC | >hg38_21_+_8433172_8446622_RNA45SN1_45S_50ntFlanks | 4760 | 4782 | 23 | TCAGATACCGTCGTAGTTCCGAC | 1126.0 | -8.0 | 1.3 | -6.4 | 2.1E-10 | 5.8E-10 | BT20/MB468 | CYTO |
| >hg38_21_+_8433172_8446622_RNA45SN1_45S_50ntFlanks@8169.8188.20\|CCAAGTCCTTCTGATCGAGG | >hg38_21_+_8433172_8446622_RNA45SN1_45S_50ntFlanks | 8169 | 8188 | 20 | CCAAGTCCTTCTGATCGAGG | 174.8 | -10.5 | 1.6 | -6.7 | 1.9E-11 | 5.8E-10 | BT20/MB468 | MP |
| >hg38_21_+_8433172_8446622_RNA45SN1_45S_50ntFlanks@9804.9832.29\|GGCGTGGAATGCGAGTGCCTAGTGGGCCA | >hg38_21_+_8433172_8446622_RNA45SN1_45S_50ntFlanks | 9804 | 9832 | 29 | GGCGTGGAATGCGAGTGCCTAGTGGGCCA | 96.6 | -10.5 | 1.6 | -6.3 | 2.2E-10 | 5.8E-10 | MB231/MB468 | MP |
| >hg38_21_+_8433172_8446622_RNA45SN1_45S_50ntFlanks@11767.11791.25\|TCTAATTAGTGACGCGCATGAATGG | >hg38_21_+_8433172_8446622_RNA45SN1_45S_50ntFlanks | 11767 | 11791 | 25 | TCTAATTAGTGACGCGCATGAATGG | 408.4 | -8.1 | 1.3 | -6.3 | 2.3E-10 | 5.8E-10 | MB231/MB468 | NUC |
| >hg38_21_+_8433172_8446622_RNA45SN1_45S_50ntFlanks@11622.11640.19\|AAAGCATCGCGAAGGCCCG | >hg38_21_+_8433172_8446622_RNA45SN1_45S_50ntFlanks | 11622 | 11640 | 19 | AAAGCATCGCGAAGGCCCG | 1461.0 | -7.6 | 1.2 | -6.3 | 2.2E-10 | 5.8E-10 | MB231/MB468 | MP |
| >hg38_21_+_8433172_8446622_RNA45SN1_45S_50ntFlanks@6646.6682.37\|TCGTACGACTCTTAGCGGTGGATCACTCGGCTCGTGC | >hg38_21_+_8433172_8446622_RNA45SN1_45S_50ntFlanks | 6646 | 6682 | 37 | TCGTACGACTCTTAGCGGTGGATCACTCGGCTCGTGC | 1461.0 | 5.9 | 0.9 | 6.4 | 1.7E-10 | 5.8E-10 | BT20/MB468 | MITO |
| >hg38_21_+_8433172_8446622_RNA45SN1_45S_50ntFlanks@476.493.18\|TTCGTGATCGATGTGGTG | >hg38_21_+_8433172_8446622_RNA45SN1_45S_50ntFlanks | 476 | 493 | 18 | TTCGTGATCGATGTGGTG | 373.2 | 3.5 | 0.5 | 6.3 | 2.3E-10 | 5.8E-10 | MB231/MB468 | NUC |
| >hg38_21_+_8433172_8446622_RNA45SN1_45S_50ntFlanks@8196.8215.20\|CGTGGACGGTGTGAGGCCGG | >hg38_21_+_8433172_8446622_RNA45SN1_45S_50ntFlanks | 8196 | 8215 | 20 | CGTGGACGGTGTGAGGCCGG | 3311.7 | -5.7 | 0.9 | -6.7 | 1.9E-11 | 5.8E-10 | BT20/MB468 | MP |
| >hg38_21_+_8433172_8446622_RNA45SN1_45S_50ntFlanks@8812.8835.24\|CGCGGGTCGGGGCGGCGGCGGCGG | >hg38_21_+_8433172_8446622_RNA45SN1_45S_50ntFlanks | 8812 | 8835 | 24 | CGCGGGTCGGGGCGGCGGCGGCGG | 122.4 | -10.8 | 1.7 | -6.3 | 2.2E-10 | 5.8E-10 | MB231/MB468 | MP |
| >hg38_21_+_8433172_8446622_RNA45SN1_45S_50ntFlanks@6745.6772.28\|ACACTTCGAACGCACTTGCGGCCCCGGG | >hg38_21_+_8433172_8446622_RNA45SN1_45S_50ntFlanks | 6745 | 6772 | 28 | ACACTTCGAACGCACTTGCGGCCCCGGG | 779.4 | -7.5 | 1.2 | -6.4 | 1.7E-10 | 5.8E-10 | BT20/MB468 | MITO |
| >hg38_21_+_8433172_8446622_RNA45SN1_45S_50ntFlanks@10405.10424.20\|ATGGGTCAGTCGGTCCTGAG | >hg38_21_+_8433172_8446622_RNA45SN1_45S_50ntFlanks | 10405 | 10424 | 20 | ATGGGTCAGTCGGTCCTGAG | 13332.1 | 6.3 | 0.9 | 6.7 | 1.9E-11 | 5.8E-10 | BT20/MB468 | MP |
| >hg38_21_+_8433172_8446622_RNA45SN1_45S_50ntFlanks@12108.12125.18\|CCGCGCGCCGGCCGGGCG | >hg38_21_+_8433172_8446622_RNA45SN1_45S_50ntFlanks | 12108 | 12125 | 18 | CCGCGCGCCGGCCGGGCG | 219.7 | -10.9 | 1.6 | -6.7 | 1.9E-11 | 5.8E-10 | BT20/MB468 | MP |

| | | | | | | | | | | | | | |
|---|---|---|---|---|---|---|---|---|---|---|---|---|---|
| >hg38_21_+_8433172_8446622_RNA45SN1_45S_50ntFlanks@11208.11233.26\|GCGGCGACTCTGGACGCGAGCCGGGC | >hg38_21_+_8433172_8446622_RNA45SN1_45S_50ntFlanks | 11208 | 11233 | 26 | GCGGCGACTCTGGACGCGAGCCGGGC | 6015.0 | -4.5 | 0.7 | -6.4 | 2.1E-10 | 5.8E-10 | BT20/MB468 | CYTO |
| >hg38_21_+_8433172_8446622_RNA45SN1_45S_50ntFlanks@7981.7998.18\|CTCAGATCAGACGTGGCG | >hg38_21_+_8433172_8446622_RNA45SN1_45S_50ntFlanks | 7981 | 7998 | 18 | CTCAGATCAGACGTGGCG | 316.7 | -8.3 | 1.3 | -6.3 | 2.8E-10 | 5.8E-10 | BT20/MB468 | NUC |
| >MI0000734\|hsa-mir-106b&WithFlank&7\|-\|100093987\|100094080@58.78.21(+1A)&[MIMAT0004672&hsa-miR-106b-3p&offsets\|0\|-1(+1A);m-87&7\|-\|100094003\|100094023&offsets\|0\|0(+1A)]\|CCGCACTGTGGGTACTTGCTGA | >MI0000734\|hsa-mir-106b&WithFlank&7\|-\|100093987\|100094080 | 58 | 78 | 21(+1A) | CCGCACTGTGGGTACTTGCTGA | 453.4 | 6.5 | 1.0 | 6.3 | 2.4E-10 | 5.8E-10 | MB231/MB468 | CYTO |
| >hg38_21_+_8433172_8446622_RNA45SN1_45S_50ntFlanks@10818.10840.23\|AAGGGAAGTCGGCAAGCCGGATC | >hg38_21_+_8433172_8446622_RNA45SN1_45S_50ntFlanks | 10818 | 10840 | 23 | AAGGGAAGTCGGCAAGCCGGATC | 477.1 | -6.9 | 1.1 | -6.3 | 2.8E-10 | 5.8E-10 | BT20/MB468 | NUC |
| >hg38_21_+_8433172_8446622_RNA45SN1_45S_50ntFlanks@6699.6719.21\|GCTAGCTGCGAGAATTAATGT | >hg38_21_+_8433172_8446622_RNA45SN1_45S_50ntFlanks | 6699 | 6719 | 21 | GCTAGCTGCGAGAATTAATGT | 3145.5 | -4.8 | 0.8 | -6.3 | 2.4E-10 | 5.8E-10 | MB231/MB468 | NUC |
| >hg38_21_+_8433172_8446622_RNA45SN1_45S_50ntFlanks@11886.11906.21\|TGAGCTTGACTCTAGTCTGGC | >hg38_21_+_8433172_8446622_RNA45SN1_45S_50ntFlanks | 11886 | 11906 | 21 | TGAGCTTGACTCTAGTCTGGC | 89.6 | -10.3 | 1.6 | -6.3 | 2.3E-10 | 5.8E-10 | MB231/MB468 | MP |
| >hg38_21_+_8433172_8446622_RNA45SN1_45S_50ntFlanks@4224.4241.18\|AAAATAACAATACAGGAC | >hg38_21_+_8433172_8446622_RNA45SN1_45S_50ntFlanks | 4224 | 4241 | 18 | AAAATAACAATACAGGAC | 73.9 | -10.1 | 1.6 | -6.3 | 2.3E-10 | 5.8E-10 | MB231/MB468 | MP |
| >MI0000816\|hsa-mir-335&WithFlank&7\|+\|130496105\|130496210@58.79.22&[MIMAT0004703&hsa-miR-335-3p&offsets\|0\|0;m-159&7\|+\|130496162\|130496183&offsets\|0\|0]\|TTTTTCATTATTGCTCCTGACC | >MI0000816\|hsa-mir-335&WithFlank&7\|+\|130496105\|130496210 | 58 | 79 | 22 | TTTTTCATTATTGCTCCTGACC | 2840.5 | 6.1 | 1.0 | 6.3 | 2.7E-10 | 5.8E-10 | MB231/MB468 | MITO |
| >am_trnaMT_ValTAC_MT_+_1602_1670@49.68.20\|AACTTAACTTGACCGCTCTG | >am_trnaMT_ValTAC_MT_+_1602_1670 | 49 | 68 | 20 | AACTTAACTTGACCGCTCTG | 3240.6 | -6.5 | 1.0 | -6.3 | 2.4E-10 | 5.8E-10 | MB231/MB468 | NUC |
| >hg38_21_+_8433172_8446622_RNA45SN1_45S_50ntFlanks@5484.5526.43\|GAGCGCTGAGAAGACGGTCGAACTTGACTATCTAGAGGAAGTA | >hg38_21_+_8433172_8446622_RNA45SN1_45S_50ntFlanks | 5484 | 5526 | 43 | GAGCGCTGAGAAGACGGTCGAACTTGACTATCTAGAGGAAGTA | 189.4 | -11.4 | 1.8 | -6.3 | 2.3E-10 | 5.8E-10 | MB231/MB468 | MP |
| >am_trnaMT_PheGAA_MT_+_577_647@40.58.19\|TGTTTAGACGGGCTCACAT | >am_trnaMT_PheGAA_MT_+_577_647 | 40 | 58 | 19 | TGTTTAGACGGGCTCACAT | 494.4 | 5.1 | 0.8 | 6.3 | 2.7E-10 | 5.8E-10 | MB231/MB468 | MITO |
| >MI0000264\|hsa-mir-7-2&WithFlank&15\|+\|88611819\|88611940@38.59.22;MI0000265\|hsa-mir-7-3&WithFlank&19\|+\|4770664\|4770785@37.58.22;MI0000263\|hsa-mir-7-1&WithFlank&9\|-\|83969742\|83969863@30.51.22&[MIMAT0000252&hsa-miR-7-5p&offsets\|0\|-2;m-400&15\|+\|88611856\|88611879&offsets\|0\|-2];[MIMAT0000252_1&hsa-miR-7-5p&offsets\|0\|-2;m-402&19\|+\|4770700\|4770723&offsets\|0\|-2];[MIMAT0000252_2&hsa-miR-7-5p&offsets\|0\|-2;m-401&9\|-\|83969811\|83969834&offsets\|0\|-2]\|TGGAAGACTAGTGATTTTGTTG | >MI0000264\|hsa-mir-7-2&WithFlank&15\|+\|88611819\|88611940 | 38 | 59 | 22 | TGGAAGACTAGTGATTTTGTTG | 1647.0 | 3.7 | 0.6 | 6.3 | 2.3E-10 | 5.8E-10 | MB231/MB468 | MP |
| >hg38_21_+_8433172_8446622_RNA45SN1_45S_50ntFlanks@6719.6738.20\|TGAATTGCAGGACACATTGA | >hg38_21_+_8433172_8446622_RNA45SN1_45S_50ntFlanks | 6719 | 6738 | 20 | TGAATTGCAGGACACATTGA | 207.3 | -7.5 | 1.2 | -6.3 | 2.4E-10 | 5.9E-10 | MB231/MB468 | NUC |
| >hg38_21_+_8433172_8446622_RNA45SN1_45S_50ntFlanks@8814.8843.30\|CGGGTCGGGGCGGCGGCGGCGGCGGCGGTG | >hg38_21_+_8433172_8446622_RNA45SN1_45S_50ntFlanks | 8814 | 8843 | 30 | CGGGTCGGGGCGGCGGCGGCGGCGGCGGTG | 82.6 | -10.2 | 1.6 | -6.3 | 2.3E-10 | 5.9E-10 | MB231/MB468 | MP |
| >MI0000088\|hsa-mir-30a&WithFlank&6\|-\|71403545\|71403627@54.73.20(+3U)&[MIMAT0000088&hsa-miR-30a-3p&offsets\|+1\|-1(+3U);m-38&6\|-\|71403554\|71403575&offsets\|+1\|-1(+3U)]\|TTTCAGTCGGATGTTTGCAGTTT | >MI0000088\|hsa-mir-30a&WithFlank&6\|-\|71403545\|71403627 | 54 | 73 | 20(+3U) | TTTCAGTCGGATGTTTGCAGTTT | 97.6 | 9.8 | 1.5 | 6.3 | 2.3E-10 | 5.9E-10 | MB231/MB468 | MP |
| >hg38_21_+_8433172_8446622_RNA45SN1_45S_50ntFlanks@10410.10434.25\|TCAGTCGGTCCTGAGAGATGGGCGA | >hg38_21_+_8433172_8446622_RNA45SN1_45S_50ntFlanks | 10410 | 10434 | 25 | TCAGTCGGTCCTGAGAGATGGGCGA | 88.5 | -10.3 | 1.6 | -6.3 | 2.3E-10 | 5.9E-10 | MB231/MB468 | MP |
| >hg38_21_+_8433172_8446622_RNA45SN1_45S_50ntFlanks@12150.12181.32\|GGTGGGGAGTTTGACTGGGGCGGTACACCTGT | >hg38_21_+_8433172_8446622_RNA45SN1_45S_50ntFlanks | 12150 | 12181 | 32 | GGTGGGGAGTTTGACTGGGGCGGTACACCTGT | 73.1 | -10.1 | 1.6 | -6.3 | 2.3E-10 | 5.9E-10 | MB231/MB468 | MP |
| >MI0000440\|hsa-mir-27b&WithFlank&9\|+\|95085439\|95085547@67.86.20(+1A)&[MIMAT0000419&hsa-miR-27b-3p&offsets\|0\|-1(+1A);m-39&9\|+\|95085505\|95085525&offsets\|0\|-1(+1A)]\|TTCACAGTGGCTAAGTTCTGA | >MI0000440\|hsa-mir-27b&WithFlank&9\|+\|95085439\|95085547 | 67 | 86 | 20(+1A) | TTCACAGTGGCTAAGTTCTGA | 2975.3 | 6.5 | 1.0 | 6.3 | 2.7E-10 | 5.9E-10 | MB231/MB468 | MITO |
| >hg38_21_+_8433172_8446622_RNA45SN1_45S_50ntFlanks@7987.8012.26\|TCAGACGTGGCGACCCGCTGAATTTA | >hg38_21_+_8433172_8446622_RNA45SN1_45S_50ntFlanks | 7987 | 8012 | 26 | TCAGACGTGGCGACCCGCTGAATTTA | 269.4 | -8.1 | 1.3 | -6.3 | 2.7E-10 | 5.9E-10 | MB231/MB468 | MITO |
| >hg38_21_+_8433172_8446622_RNA45SN1_45S_50ntFlanks@4785.4803.19\|TAAACGATGCCGACCGGCG | >hg38_21_+_8433172_8446622_RNA45SN1_45S_50ntFlanks | 4785 | 4803 | 19 | TAAACGATGCCGACCGGCG | 96.6 | -10.5 | 1.6 | -6.3 | 2.3E-10 | 5.9E-10 | MB231/MB468 | MP |
| >hg38_21_+_8433172_8446622_RNA45SN1_45S_50ntFlanks@10380.10424.45\|GGTTCCATGTGAACAGCAGTTGAACATGGGTCAGTCGGTCCTGAG | >hg38_21_+_8433172_8446622_RNA45SN1_45S_50ntFlanks | 10380 | 10424 | 45 | GGTTCCATGTGAACAGCAGTTGAACATGGGTCAGTCGGTCCTGAG | 3174.0 | 7.9 | 1.2 | 6.4 | 1.8E-10 | 5.9E-10 | BT20/MB468 | MITO |
| >hg38_21_+_8433172_8446622_RNA45SN1_45S_50ntFlanks@4546.4563.18\|CCTGGATACCGCAGCTAG | >hg38_21_+_8433172_8446622_RNA45SN1_45S_50ntFlanks | 4546 | 4563 | 18 | CCTGGATACCGCAGCTAG | 91.7 | -10.4 | 1.6 | -6.3 | 2.3E-10 | 5.9E-10 | MB231/MB468 | MP |
| >hg38_21_+_8433172_8446622_RNA45SN1_45S_50ntFlanks@12157.12180.24\|AGTTTGACTGGGGCGGTACACCTG | >hg38_21_+_8433172_8446622_RNA45SN1_45S_50ntFlanks | 12157 | 12180 | 24 | AGTTTGACTGGGGCGGTACACCTG | 1136.9 | -6.2 | 1.0 | -6.4 | 1.8E-10 | 5.9E-10 | BT20/MB468 | MITO |
| >hg38_21_+_8433172_8446622_RNA45SN1_45S_50ntFlanks@9178.9195.18\|CGCCGTCGGGCCCGGGGG | >hg38_21_+_8433172_8446622_RNA45SN1_45S_50ntFlanks | 9178 | 9195 | 18 | CGCCGTCGGGCCCGGGGG | 402.2 | -8.5 | 1.3 | -6.3 | 2.7E-10 | 5.9E-10 | MB231/MB468 | MITO |
| >am_MI0000809\|hsa-mir-151a&WithFlank&8\|-\|140732558\|140732659@53.72.20(+2U)&[MIMAT0000757&hsa-miR-151a-3p&offsets\|0\|-1(+2U);m-30&8\|-\|140732587\|140732607&offsets\|0\|-1(+2U)]\|CTAGACTGAAGCTCCTTGAGTT | >am_MI0000809\|hsa-mir-151a&WithFlank&8\|-\|140732558\|140732659 | 53 | 72 | 20(+2U) | CTAGACTGAAGCTCCTTGAGTT | 197.6 | 10.8 | 1.7 | 6.3 | 2.7E-10 | 5.9E-10 | MB231/MB468 | MITO |
| >hg38_21_+_8433172_8446622_RNA45SN1_45S_50ntFlanks@12488.12525.38\|AACGTGAGCTGGGTTTAGACCGTCGTGAGACAGGTTAG | >hg38_21_+_8433172_8446622_RNA45SN1_45S_50ntFlanks | 12488 | 12525 | 38 | AACGTGAGCTGGGTTTAGACCGTCGTGAGACAGGTTAG | 75.5 | -10.1 | 1.6 | -6.3 | 2.3E-10 | 5.9E-10 | MB231/MB468 | MP |
| >hg38_21_+_8433172_8446622_RNA45SN1_45S_50ntFlanks@10361.10383.23\|GAAGGCCGAAGTGGAGAAGGGTT | >hg38_21_+_8433172_8446622_RNA45SN1_45S_50ntFlanks | 10361 | 10383 | 23 | GAAGGCCGAAGTGGAGAAGGGTT | 351.6 | -3.2 | 0.5 | -6.4 | 2.1E-10 | 5.9E-10 | BT20/MB468 | CYTO |
| >hg38_21_+_8433172_8446622_RNA45SN1_45S_50ntFlanks@12657.12674.18\|GACTGAACGCCTCTAAGT | >hg38_21_+_8433172_8446622_RNA45SN1_45S_50ntFlanks | 12657 | 12674 | 18 | GACTGAACGCCTCTAAGT | 2066.5 | -9.9 | 1.6 | -6.3 | 2.8E-10 | 5.9E-10 | BT20/MB468 | NUC |
| >hg38_21_+_8433172_8446622_RNA45SN1_45S_50ntFlanks@6734.6773.40\|ATTGATCATCGACACTTCGAACGCACTTGCGGCCCCGGGT | >hg38_21_+_8433172_8446622_RNA45SN1_45S_50ntFlanks | 6734 | 6773 | 40 | ATTGATCATCGACACTTCGAACGCACTTGCGGCCCCGGGT | 1340.0 | 5.0 | 0.8 | 6.3 | 2.7E-10 | 5.9E-10 | MB231/MB468 | MITO |
| >hg38_21_+_8433172_8446622_RNA45SN1_45S_50ntFlanks@9306.9337.32\|CAAGGAGTCTAACACGTGCGCGAGTCGGGGGC | >hg38_21_+_8433172_8446622_RNA45SN1_45S_50ntFlanks | 9306 | 9337 | 32 | CAAGGAGTCTAACACGTGCGCGAGTCGGGGGC | 327.2 | -8.1 | 1.3 | -6.3 | 2.7E-10 | 5.9E-10 | MB231/MB468 | MITO |
| >hg38_21_+_8433172_8446622_RNA45SN1_45S_50ntFlanks@9605.9622.18\|AAGACTAATCGAACCATC | >hg38_21_+_8433172_8446622_RNA45SN1_45S_50ntFlanks | 9605 | 9622 | 18 | AAGACTAATCGAACCATC | 676.4 | -5.2 | 0.8 | -6.3 | 2.4E-10 | 5.9E-10 | MB231/MB468 | NUC |
| >hg38_21_+_8433172_8446622_RNA45SN1_45S_50ntFlanks@3790.3814.25\|CTGCGAATGGCTCATTAAATCAGTT | >hg38_21_+_8433172_8446622_RNA45SN1_45S_50ntFlanks | 3790 | 3814 | 25 | CTGCGAATGGCTCATTAAATCAGTT | 99.3 | -10.5 | 1.7 | -6.3 | 2.3E-10 | 5.9E-10 | MB231/MB468 | MP |
| >hg38_21_+_8433172_8446622_RNA45SN1_45S_50ntFlanks@7974.8011.38\|ACGCGACCTCAGATCAGACGTGGCGACCCGCTGAATTT | >hg38_21_+_8433172_8446622_RNA45SN1_45S_50ntFlanks | 7974 | 8011 | 38 | ACGCGACCTCAGATCAGACGTGGCGACCCGCTGAATTT | 16840.1 | 4.3 | 0.7 | 6.3 | 2.4E-10 | 5.9E-10 | MB231/MB468 | NUC |
| >hg38_21_+_8433172_8446622_RNA45SN1_45S_50ntFlanks@3782.3812.31\|CAGTGAAACTGCGAATGGCTCATTAAATCAG | >hg38_21_+_8433172_8446622_RNA45SN1_45S_50ntFlanks | 3782 | 3812 | 31 | CAGTGAAACTGCGAATGGCTCATTAAATCAG | 571.4 | -8.9 | 1.4 | -6.3 | 2.4E-10 | 5.9E-10 | MB231/MB468 | NUC |
| >hg38_21_+_8433172_8446622_RNA45SN1_45S_50ntFlanks@11821.11846.26\|CTATCCAGCGAAACCACAGCCAAGGG | >hg38_21_+_8433172_8446622_RNA45SN1_45S_50ntFlanks | 11821 | 11846 | 26 | CTATCCAGCGAAACCACAGCCAAGGG | 99.0 | -10.5 | 1.7 | -6.3 | 2.3E-10 | 5.9E-10 | MB231/MB468 | MP |
| >hg38_21_+_8433172_8446622_RNA45SN1_45S_50ntFlanks@9796.9820.25\|TGGAGCCGGGCGTGGAATGCGAGTG | >hg38_21_+_8433172_8446622_RNA45SN1_45S_50ntFlanks | 9796 | 9820 | 25 | TGGAGCCGGGCGTGGAATGCGAGTG | 118.1 | -10.7 | 1.7 | -6.3 | 2.3E-10 | 5.9E-10 | MB231/MB468 | MP |
| >hg38_1_-_228634819_228635039_RNA5S12_5S_50ntFlanks@127.169.43\|ACTTGGATGGGAGACCGCCTGGGAATACCGGGTGCTGTAGGCT | >hg38_1_-_228634819_228635039_RNA5S12_5S_50ntFlanks | 127 | 169 | 43 | ACTTGGATGGGAGACCGCCTGGGAATACCGGGTGCTGTAGGCT | 425.4 | 9.6 | 1.5 | 6.4 | 1.8E-10 | 5.9E-10 | BT20/MB468 | MITO |
| >hg38_21_+_8433172_8446622_RNA45SN1_45S_50ntFlanks@4287.4307.21\|AACGAGGATCCATTGGAGGGC | >hg38_21_+_8433172_8446622_RNA45SN1_45S_50ntFlanks | 4287 | 4307 | 21 | AACGAGGATCCATTGGAGGGC | 94.2 | -10.4 | 1.6 | -6.3 | 2.3E-10 | 5.9E-10 | MB231/MB468 | MP |
| >hg38_21_+_8433172_8446622_RNA45SN1_45S_50ntFlanks@6741.6770.30\|ATCGACACTTCGAACGCACTTGCGGCCCCG | >hg38_21_+_8433172_8446622_RNA45SN1_45S_50ntFlanks | 6741 | 6770 | 30 | ATCGACACTTCGAACGCACTTGCGGCCCCG | 1122.6 | -7.2 | 1.1 | -6.3 | 2.4E-10 | 5.9E-10 | MB231/MB468 | NUC |
| >MI0003560\|hsa-mir-92b&WithFlank&1\|+\|155195171\|155195278@67.88.22(+1U)&[MIMAT0003218&hsa-miR-92b-3p&offsets\|0\|0(+1U);m-115&1\|+\|155195237\|155195258&offsets\|0\|0(+1U)]\|TATTGCACTCGTCCCGGCCTCCT | >MI0003560\|hsa-mir-92b&WithFlank&1\|+\|155195171\|155195278 | 67 | 88 | 22(+1U) | TATTGCACTCGTCCCGGCCTCCT | 855.8 | 9.4 | 1.5 | 6.4 | 1.8E-10 | 5.9E-10 | BT20/MB468 | MITO |
| >hg38_21_+_8433172_8446622_RNA45SN1_45S_50ntFlanks@9564.9595.32\|CTGACGTGCAAATCGGTCGTCCGACCTGGGTA | >hg38_21_+_8433172_8446622_RNA45SN1_45S_50ntFlanks | 9564 | 9595 | 32 | CTGACGTGCAAATCGGTCGTCCGACCTGGGTA | 6575.8 | -7.6 | 1.2 | -6.3 | 2.7E-10 | 6.0E-10 | MB231/MB468 | MITO |
| >hg38_21_+_8433172_8446622_RNA45SN1_45S_50ntFlanks@10713.10736.24\|CTTGAAAATCCGGGGGAGAGGGTG | >hg38_21_+_8433172_8446622_RNA45SN1_45S_50ntFlanks | 10713 | 10736 | 24 | CTTGAAAATCCGGGGGAGAGGGTG | 122.4 | -9.0 | 1.4 | -6.4 | 1.8E-10 | 6.0E-10 | BT20/MB468 | MITO |
| >hg38_21_+_8433172_8446622_RNA45SN1_45S_50ntFlanks@12412.12432.21\|ATCCTTCGATGTCGGCTCTTC | >hg38_21_+_8433172_8446622_RNA45SN1_45S_50ntFlanks | 12412 | 12432 | 21 | ATCCTTCGATGTCGGCTCTTC | 193.3 | -10.7 | 1.6 | -6.7 | 2.0E-11 | 6.0E-10 | BT20/MB468 | MP |
| >hg38_21_+_8433172_8446622_RNA45SN1_45S_50ntFlanks@12074.12095.22\|CGAGGGGCTCTCGCTTCTGGCG | >hg38_21_+_8433172_8446622_RNA45SN1_45S_50ntFlanks | 12074 | 12095 | 22 | CGAGGGGCTCTCGCTTCTGGCG | 1303.8 | -4.2 | 0.7 | -6.3 | 2.4E-10 | 6.0E-10 | MB231/MB468 | CYTO |
| >trnaMT_PheGAA_MT_+_577_647@35.57.23\|GAAAATGTTTAGACGGGCTCACA | >trnaMT_PheGAA_MT_+_577_647 | 35 | 57 | 23 | GAAAATGTTTAGACGGGCTCACA | 839.6 | 4.6 | 0.7 | 6.3 | 2.7E-10 | 6.0E-10 | MB231/MB468 | MITO |
| >hg38_21_+_8433172_8446622_RNA45SN1_45S_50ntFlanks@12457.12478.22\|CAAGCGTTGGATTGTTCACCCA | >hg38_21_+_8433172_8446622_RNA45SN1_45S_50ntFlanks | 12457 | 12478 | 22 | CAAGCGTTGGATTGTTCACCCA | 4669.0 | -5.1 | 0.8 | -6.3 | 2.4E-10 | 6.0E-10 | MB231/MB468 | CYTO |
| >hg38_21_+_8433172_8446622_RNA45SN1_45S_50ntFlanks@4685.4722.38\|AGACGGACCAGAGCGAAAGCATTTGCCAAGAATGTTTT | >hg38_21_+_8433172_8446622_RNA45SN1_45S_50ntFlanks | 4685 | 4722 | 38 | AGACGGACCAGAGCGAAAGCATTTGCCAAGAATGTTTT | 701.2 | -6.1 | 1.0 | -6.3 | 2.7E-10 | 6.0E-10 | MB231/MB468 | MITO |
| >hg38_21_+_8433172_8446622_RNA45SN1_45S_50ntFlanks@12932.12956.25\|CGCTAAACCATTCGTAGACGACCTG | >hg38_21_+_8433172_8446622_RNA45SN1_45S_50ntFlanks | 12932 | 12956 | 25 | CGCTAAACCATTCGTAGACGACCTG | 2224.3 | -8.0 | 1.3 | -6.3 | 2.9E-10 | 6.0E-10 | BT20/MB468 | NUC |

| | | | | | | | | | | | | | |
|---|---|---|---|---|---|---|---|---|---|---|---|---|---|
| >am_trna128_GlyGCC_6_-_27870686_27870756@1.33.33,trna18_GlyGCC_16_+_70822597_70822667@1.33.33,trna19_GlyGCC_16_+_70823410_70823480@1.33.33,trna19_GlyGCC_2_-_157257659_157257729@1.33.33,trna24_GlyGCC_16_-_70812942_70813012@1.33.33,trna25_GlyGCC_16_-_70812114_70812184@1.33.33,trna5_GlyGCC_17_+_8029064_8029134@1.33.33,trna68_GlyGCC_1_-_161493637_161493707@1.33.33\|GCATTGGTGGTTCAGTGGTAGAATTCTCGCCTG | >am_trna128_GlyGCC_6_-_27870686_27870756 | 1 | 33 | 33 | GCATTGGTGGTTCAGTGGTAGAATTCTCGCCTG | 2120.3 | 5.1 | 0.8 | 6.4 | 1.8E-10 | 6.0E-10 | BT20/MB468 | MITO |
| >hg38_21_+_8433172_8446622_RNA45SN1_45S_50ntFlanks@4734.4752.19\|AACGAAAGTCGGAGGTTCG | >hg38_21_+_8433172_8446622_RNA45SN1_45S_50ntFlanks | 4734 | 4752 | 19 | AACGAAAGTCGGAGGTTCG | 463.7 | -5.0 | 0.8 | -6.3 | 2.7E-10 | 6.0E-10 | MB231/MB468 | MITO |
| >MI0000285\|hsa-mir-205&WithFlank&1\|+\|209432127\|209432248@40.59.20&[MIMAT0000266&hsa-miR-205-5p&offsets\|0\|-2;m-351&1\|+\|209432166\|209432187&offsets\|0\|-2]\|TCCTTCATTCCACCGGAGTC | >MI0000285\|hsa-mir-205&WithFlank&1\|+\|209432127\|209432248 | 40 | 59 | 20 | TCCTTCATTCCACCGGAGTC | 900.2 | 9.0 | 1.4 | 6.3 | 2.2E-10 | 6.0E-10 | BT20/MB468 | CYTO |
| >MI0000433\|hsa-let-7g&WithFlank&3\|-\|52268272\|52268367@11.31.21(+1G)&[MIMAT0000414&hsa-let-7g-5p&offsets\|0\|-1(+1G);m-58&3\|-\|52268336\|52268357&offsets\|0\|-1(+1G)]\|TGAGGTAGTAGTTTGTACAGTG | >MI0000433\|hsa-let-7g&WithFlank&3\|-\|52268272\|52268367 | 11 | 31 | 21(+1G) | TGAGGTAGTAGTTTGTACAGTG | 1575.6 | 5.4 | 0.9 | 6.3 | 2.2E-10 | 6.0E-10 | BT20/MB468 | CYTO |
| >hg38_21_+_8433172_8446622_RNA45SN1_45S_50ntFlanks@9534.9564.31\|AGGAAACTCTGGTGGAGGTCCGTAGCGGTCC | >hg38_21_+_8433172_8446622_RNA45SN1_45S_50ntFlanks | 9534 | 9564 | 31 | AGGAAACTCTGGTGGAGGTCCGTAGCGGTCC | 1463.4 | -4.2 | 0.7 | -6.3 | 2.2E-10 | 6.0E-10 | BT20/MB468 | CYTO |
| >hg38_21_+_8433172_8446622_RNA45SN1_45S_50ntFlanks@12045.12071.27\|CTGACCCGGTGAGGCGGGGGGGCGAGC | >hg38_21_+_8433172_8446622_RNA45SN1_45S_50ntFlanks | 12045 | 12071 | 27 | CTGACCCGGTGAGGCGGGGGGGCGAGC | 433.8 | -8.7 | 1.4 | -6.3 | 2.9E-10 | 6.0E-10 | BT20/MB468 | NUC |
| >hg38_21_+_8433172_8446622_RNA45SN1_45S_50ntFlanks@10926.10946.21\|ACGAGGCGCCGCCGCCCCCCC | >hg38_21_+_8433172_8446622_RNA45SN1_45S_50ntFlanks | 10926 | 10946 | 21 | ACGAGGCGCCGCCGCCCCCCC | 1447.9 | -13.6 | 2.0 | -6.7 | 2.0E-11 | 6.0E-10 | BT20/MB468 | MP |
| >hg38_21_+_8433172_8446622_RNA45SN1_45S_50ntFlanks@5414.5445.32\|CTACTACCGATTGGATGGTTTAGTGAGGCCCT | >hg38_21_+_8433172_8446622_RNA45SN1_45S_50ntFlanks | 5414 | 5445 | 32 | CTACTACCGATTGGATGGTTTAGTGAGGCCCT | 201.1 | -9.7 | 1.5 | -6.4 | 1.8E-10 | 6.0E-10 | BT20/MB468 | MITO |
| >hg38_21_+_8433172_8446622_RNA45SN1_45S_50ntFlanks@11602.11620.19\|TCCGACTGTTTAATTAAAA | >hg38_21_+_8433172_8446622_RNA45SN1_45S_50ntFlanks | 11602 | 11620 | 19 | TCCGACTGTTTAATTAAAA | 1419.1 | -4.1 | 0.7 | -6.3 | 2.5E-10 | 6.0E-10 | MB231/MB468 | NUC |
| >hg38_21_+_8433172_8446622_RNA45SN1_45S_50ntFlanks@10687.10706.20\|CGGCGTCCGGTGAGCTCTCG | >hg38_21_+_8433172_8446622_RNA45SN1_45S_50ntFlanks | 10687 | 10706 | 20 | CGGCGTCCGGTGAGCTCTCG | 5503.1 | -9.5 | 1.5 | -6.3 | 2.5E-10 | 6.0E-10 | MB231/MB468 | NUC |
| >hg38_21_+_8433172_8446622_RNA45SN1_45S_50ntFlanks@12624.12661.38\|CCAATGGGGCGAAGCTACCATCTGTGGGATTATGACTG | >hg38_21_+_8433172_8446622_RNA45SN1_45S_50ntFlanks | 12624 | 12661 | 38 | CCAATGGGGCGAAGCTACCATCTGTGGGATTATGACTG | 384.4 | -2.9 | 0.5 | -6.3 | 2.2E-10 | 6.1E-10 | BT20/MB468 | CYTO |
| >MI0000446\|hsa-mir-125b-1&WithFlank&11\|-\|122099751\|122099850@61.80.20(+2U)&[MIMAT0004592&hsa-miR-125b-1-3p&offsets\|0\|-2(+2U);m-485&11\|-\|122099769\|122099790&offsets\|0\|-2(+2U)]\|ACGGGTTAGGCTCTTGGGAGTT | >MI0000446\|hsa-mir-125b-1&WithFlank&11\|-\|122099751\|122099850 | 61 | 80 | 20(+2U) | ACGGGTTAGGCTCTTGGGAGTT | 100.2 | 9.9 | 1.6 | 6.3 | 2.3E-10 | 6.1E-10 | MB231/MB468 | MP |
| >hg38_21_+_8433172_8446622_RNA45SN1_45S_50ntFlanks@11573.11600.28\|CGACTTAGAACTGGTGCGGACCAGGGGA | >hg38_21_+_8433172_8446622_RNA45SN1_45S_50ntFlanks | 11573 | 11600 | 28 | CGACTTAGAACTGGTGCGGACCAGGGGA | 1515.5 | -7.6 | 1.2 | -6.3 | 2.9E-10 | 6.1E-10 | BT20/MB468 | NUC |
| >hg38_21_+_8433172_8446622_RNA45SN1_45S_50ntFlanks@11643.11663.21\|GCGGGTGTTGACGCGATGTGA | >hg38_21_+_8433172_8446622_RNA45SN1_45S_50ntFlanks | 11643 | 11663 | 21 | GCGGGTGTTGACGCGATGTGA | 201.9 | 5.3 | 0.8 | 6.3 | 2.5E-10 | 6.1E-10 | MB231/MB468 | NUC |
| >trnaMT_SerGCT_MT_+_12207_12265@10.38.29\|CACAAGAACTGCTAACTCATGCCCCCATG | >trnaMT_SerGCT_MT_+_12207_12265 | 10 | 38 | 29 | CACAAGAACTGCTAACTCATGCCCCCATG | 1385.2 | -9.0 | 1.4 | -6.3 | 2.5E-10 | 6.1E-10 | MB231/MB468 | NUC |
| >hg38_21_+_8433172_8446622_RNA45SN1_45S_50ntFlanks@10236.10262.27\|GACGCTACGCCGCGACGAGTAGGAGGG | >hg38_21_+_8433172_8446622_RNA45SN1_45S_50ntFlanks | 10236 | 10262 | 27 | GACGCTACGCCGCGACGAGTAGGAGGG | 109.7 | -10.6 | 1.7 | -6.3 | 2.4E-10 | 6.1E-10 | MB231/MB468 | MP |
| >hg38_21_+_8433172_8446622_RNA45SN1_45S_50ntFlanks@4975.4996.22\|CCCGGACACGGACAGGATTGAC | >hg38_21_+_8433172_8446622_RNA45SN1_45S_50ntFlanks | 4975 | 4996 | 22 | CCCGGACACGGACAGGATTGAC | 82.6 | -10.2 | 1.6 | -6.3 | 2.4E-10 | 6.1E-10 | MB231/MB468 | MP |
| >hg38_21_+_8433172_8446622_RNA45SN1_45S_50ntFlanks@9790.9818.29\|CTGGCGTGGAGCCGGGCGTGGAATGCGAG | >hg38_21_+_8433172_8446622_RNA45SN1_45S_50ntFlanks | 9790 | 9818 | 29 | CTGGCGTGGAGCCGGGCGTGGAATGCGAG | 105.1 | -10.6 | 1.7 | -6.3 | 2.4E-10 | 6.1E-10 | MB231/MB468 | MP |
| >hg38_1_-_228634819_228635039_RNA5S12_5S_50ntFlanks@134.171.38\|TGGGAGACCGCCTGGGAATACCGGGTGCTGTAGGCTTT | >hg38_1_-_228634819_228635039_RNA5S12_5S_50ntFlanks | 134 | 171 | 38 | TGGGAGACCGCCTGGGAATACCGGGTGCTGTAGGCTTT | 15626.7 | 6.5 | 1.0 | 6.4 | 1.8E-10 | 6.1E-10 | BT20/MB468 | MITO |
| >hg38_21_+_8433172_8446622_RNA45SN1_45S_50ntFlanks@7986.8010.25\|ATCAGACGTGGCGACCCGCTGAATT | >hg38_21_+_8433172_8446622_RNA45SN1_45S_50ntFlanks | 7986 | 8010 | 25 | ATCAGACGTGGCGACCCGCTGAATT | 182.2 | -9.6 | 1.5 | -6.4 | 1.8E-10 | 6.1E-10 | BT20/MB468 | MITO |
| >hg38_21_+_8433172_8446622_RNA45SN1_45S_50ntFlanks@3873.3898.26\|TAATACATGCCGACGGGCGCTGACCC | >hg38_21_+_8433172_8446622_RNA45SN1_45S_50ntFlanks | 3873 | 3898 | 26 | TAATACATGCCGACGGGCGCTGACCC | 1917.5 | -8.2 | 1.3 | -6.4 | 1.9E-10 | 6.1E-10 | BT20/MB468 | MITO |
| >MI0001445\|hsa-mir-423&WithFlank&17\|+\|30117073\|30117178@59.79.21&[MIMAT0001340&hsa-miR-423-3p&offsets\|0\|-2;m-100&17\|+\|30117131\|30117153&offsets\|0\|-2]\|AGCTCGGTCTGAGGCCCCTCA | >MI0001445\|hsa-mir-423&WithFlank&17\|+\|30117073\|30117178 | 59 | 79 | 21 | AGCTCGGTCTGAGGCCCCTCA | 1683.9 | 9.1 | 1.4 | 6.3 | 2.2E-10 | 6.1E-10 | BT20/MB468 | CYTO |
| >trnaMT_GluTTC_MT_-_14674_14742@40.57.18,trnalookalike8_GluTTC_5_-_93905172_93905240@40.57.18\|TTGGTCGTGGTTGTAGTC | >trnaMT_GluTTC_MT_-_14674_14742 | 40 | 57 | 18 | TTGGTCGTGGTTGTAGTC | 439.6 | 6.6 | 1.0 | 6.3 | 2.2E-10 | 6.1E-10 | BT20/MB468 | CYTO |
| >hg38_21_+_8433172_8446622_RNA45SN1_45S_50ntFlanks@4772.4793.22\|GTAGTTCCGACCATAAACGATG | >hg38_21_+_8433172_8446622_RNA45SN1_45S_50ntFlanks | 4772 | 4793 | 22 | GTAGTTCCGACCATAAACGATG | 748.7 | -9.3 | 1.5 | -6.3 | 2.9E-10 | 6.1E-10 | BT20/MB468 | NUC |
| >MI0000089\|hsa-mir-31&WithFlank&9\|-\|21512109\|21512191@14.34.21&[MIMAT0000089&hsa-miR-31-5p&offsets\|0\|0;m-360&9\|-\|21512157\|21512178&offsets\|0\|-1]\|AGGCAAGATGCTGGCATAGCT | >MI0000089\|hsa-mir-31&WithFlank&9\|-\|21512109\|21512191 | 14 | 34 | 21 | AGGCAAGATGCTGGCATAGCT | 168.8 | 4.2 | 0.7 | 6.3 | 2.9E-10 | 6.1E-10 | BT20/MB468 | NUC |
| >hg38_21_+_8433172_8446622_RNA45SN1_45S_50ntFlanks@475.499.25\|CTTCGTGATCGATGTGGTGACGTCG | >hg38_21_+_8433172_8446622_RNA45SN1_45S_50ntFlanks | 475 | 499 | 25 | CTTCGTGATCGATGTGGTGACGTCG | 1908.7 | 5.9 | 0.9 | 6.3 | 2.5E-10 | 6.1E-10 | MB231/MB468 | NUC |
| >hg38_21_+_8433172_8446622_RNA45SN1_45S_50ntFlanks@11315.11336.22\|CCACCCCACGTCTCGTCGCGCG | >hg38_21_+_8433172_8446622_RNA45SN1_45S_50ntFlanks | 11315 | 11336 | 22 | CCACCCCACGTCTCGTCGCGCG | 1281.5 | -9.4 | 1.5 | -6.3 | 2.5E-10 | 6.1E-10 | MB231/MB468 | NUC |
| >MI0003177\|hsa-mir-522&WithFlank&19\|+\|53751205\|53751303@60.81.22&[MIMAT0002868&hsa-miR-522-3p&offsets\|0\|0;m-909&19\|+\|53751264\|53751285&offsets\|0\|0]\|AAAATGGTTCCCTTTAGAGTGT | >MI0003177\|hsa-mir-522&WithFlank&19\|+\|53751205\|53751303 | 60 | 81 | 22 | AAAATGGTTCCCTTTAGAGTGT | 130.9 | 10.3 | 1.6 | 6.3 | 2.4E-10 | 6.1E-10 | MB231/MB468 | MP |
| >hg38_21_+_8433172_8446622_RNA45SN1_45S_50ntFlanks@12159.12179.21\|TTTGACTGGGGCGGTACACCT | >hg38_21_+_8433172_8446622_RNA45SN1_45S_50ntFlanks | 12159 | 12179 | 21 | TTTGACTGGGGCGGTACACCT | 1057.4 | -7.6 | 1.2 | -6.3 | 2.5E-10 | 6.1E-10 | MB231/MB468 | CYTO |
| >hg38_21_+_8433172_8446622_RNA45SN1_45S_50ntFlanks@9341.9370.30\|CACGAAAGCCGCCGTGGCGCAATGAAGGTG | >hg38_21_+_8433172_8446622_RNA45SN1_45S_50ntFlanks | 9341 | 9370 | 30 | CACGAAAGCCGCCGTGGCGCAATGAAGGTG | 485.9 | -12.0 | 1.8 | -6.7 | 2.1E-11 | 6.1E-10 | BT20/MB468 | MP |
| >hg38_21_+_8433172_8446622_RNA45SN1_45S_50ntFlanks@11576.11604.29\|CTTAGAACTGGTGCGGACCAGGGGAATCC | >hg38_21_+_8433172_8446622_RNA45SN1_45S_50ntFlanks | 11576 | 11604 | 29 | CTTAGAACTGGTGCGGACCAGGGGAATCC | 392.6 | -8.8 | 1.4 | -6.3 | 2.9E-10 | 6.1E-10 | BT20/MB468 | NUC |
| >hg38_21_+_8433172_8446622_RNA45SN1_45S_50ntFlanks@12641.12659.19\|CCATCTGTGGGATTATGAC | >hg38_21_+_8433172_8446622_RNA45SN1_45S_50ntFlanks | 12641 | 12659 | 19 | CCATCTGTGGGATTATGAC | 3703.8 | -5.9 | 0.9 | -6.4 | 1.9E-10 | 6.1E-10 | BT20/MB468 | MITO |
| >MI0003628\|hsa-mir-615&WithFlank&12\|+\|54033944\|54034051@67.86.20&[MIMAT0003283&hsa-miR-615-3p&offsets\|0\|-2;m-495&12\|+\|54034010\|54034030&offsets\|0\|-1]\|TCCGAGCCTGGGTCTCCCTC | >MI0003628\|hsa-mir-615&WithFlank&12\|+\|54033944\|54034051 | 67 | 86 | 20 | TCCGAGCCTGGGTCTCCCTC | 936.3 | 7.4 | 1.2 | 6.3 | 2.2E-10 | 6.2E-10 | BT20/MB468 | CYTO |
| >hg38_21_+_8433172_8446622_RNA45SN1_45S_50ntFlanks@8384.8402.19\|AGGGCGTGAAACCGTTAAG | >hg38_21_+_8433172_8446622_RNA45SN1_45S_50ntFlanks | 8384 | 8402 | 19 | AGGGCGTGAAACCGTTAAG | 551.6 | -6.8 | 1.1 | -6.3 | 2.4E-10 | 6.2E-10 | MB231/MB468 | MP |
| >hg38_21_+_8433172_8446622_RNA45SN1_45S_50ntFlanks@12940.12973.34\|CATTCGTAGACGACCTGCTTCTGGGTCGGGGTTT | >hg38_21_+_8433172_8446622_RNA45SN1_45S_50ntFlanks | 12940 | 12973 | 34 | CATTCGTAGACGACCTGCTTCTGGGTCGGGGTTT | 159.8 | -6.7 | 1.1 | -6.3 | 2.5E-10 | 6.2E-10 | MB231/MB468 | CYTO |
| >hg38_21_+_8433172_8446622_RNA45SN1_45S_50ntFlanks@5478.5499.22\|CTGGCGGAGCGCTGAGAAGACG | >hg38_21_+_8433172_8446622_RNA45SN1_45S_50ntFlanks | 5478 | 5499 | 22 | CTGGCGGAGCGCTGAGAAGACG | 550.5 | -5.2 | 0.8 | -6.3 | 2.4E-10 | 6.2E-10 | MB231/MB468 | MP |
| >hg38_21_+_8433172_8446622_RNA45SN1_45S_50ntFlanks@12553.12580.28\|CCATGGTAATCCTGCTCAGTACGAGAGG | >hg38_21_+_8433172_8446622_RNA45SN1_45S_50ntFlanks | 12553 | 12580 | 28 | CCATGGTAATCCTGCTCAGTACGAGAGG | 182.3 | -11.3 | 1.8 | -6.3 | 2.8E-10 | 6.2E-10 | MB231/MB468 | MITO |
| >hg38_21_+_8433172_8446622_RNA45SN1_45S_50ntFlanks@4727.4762.36\|AATCAAGAACGAAAGTCGGAGGTTCGAAGACGATCA | >hg38_21_+_8433172_8446622_RNA45SN1_45S_50ntFlanks | 4727 | 4762 | 36 | AATCAAGAACGAAAGTCGGAGGTTCGAAGACGATCA | 849.6 | 10.8 | 1.7 | 6.3 | 2.2E-10 | 6.2E-10 | BT20/MB468 | CYTO |
| >hg38_21_+_8433172_8446622_RNA45SN1_45S_50ntFlanks@3861.3898.38\|TAATTCTAGAGCTAATACATGCCGACGGGCGCTGACCC | >hg38_21_+_8433172_8446622_RNA45SN1_45S_50ntFlanks | 3861 | 3898 | 38 | TAATTCTAGAGCTAATACATGCCGACGGGCGCTGACCC | 391.7 | -5.9 | 0.9 | -6.3 | 2.5E-10 | 6.2E-10 | MB231/MB468 | CYTO |
| >hg38_21_+_8433172_8446622_RNA45SN1_45S_50ntFlanks@12992.13011.20\|TCCCTCGCTGCGATCTATTG | >hg38_21_+_8433172_8446622_RNA45SN1_45S_50ntFlanks | 12992 | 13011 | 20 | TCCCTCGCTGCGATCTATTG | 104.7 | -8.8 | 1.4 | -6.4 | 1.9E-10 | 6.2E-10 | BT20/MB468 | MITO |
| >hg38_21_+_8433172_8446622_RNA45SN1_45S_50ntFlanks@4006.4024.19\|ACCTCGGGCCGATCGCACG | >hg38_21_+_8433172_8446622_RNA45SN1_45S_50ntFlanks | 4006 | 4024 | 19 | ACCTCGGGCCGATCGCACG | 1501.7 | -3.9 | 0.6 | -6.3 | 2.5E-10 | 6.2E-10 | MB231/MB468 | CYTO |
| >hg38_21_+_8433172_8446622_RNA45SN1_45S_50ntFlanks@10391.10430.40\|AACAGCAGTTGAACATGGGTCAGTCGGTCCTGAGAGATGG | >hg38_21_+_8433172_8446622_RNA45SN1_45S_50ntFlanks | 10391 | 10430 | 40 | AACAGCAGTTGAACATGGGTCAGTCGGTCCTGAGAGATGG | 724.5 | -5.8 | 0.9 | -6.3 | 2.5E-10 | 6.2E-10 | MB231/MB468 | CYTO |
| >hg38_21_+_8433172_8446622_RNA45SN1_45S_50ntFlanks@7975.8000.26\|CGCGACCTCAGATCAGACGTGGCGAC | >hg38_21_+_8433172_8446622_RNA45SN1_45S_50ntFlanks | 7975 | 8000 | 26 | CGCGACCTCAGATCAGACGTGGCGAC | 3013.8 | 6.2 | 1.0 | 6.4 | 1.9E-10 | 6.2E-10 | BT20/MB468 | MITO |
| >hg38_21_+_8433172_8446622_RNA45SN1_45S_50ntFlanks@3984.4024.41\|GCGGCTTTGGTGACTCTAGATAACCTCGGGCCGATCGCACG | >hg38_21_+_8433172_8446622_RNA45SN1_45S_50ntFlanks | 3984 | 4024 | 41 | GCGGCTTTGGTGACTCTAGATAACCTCGGGCCGATCGCACG | 521.8 | -9.0 | 1.4 | -6.3 | 3.0E-10 | 6.2E-10 | BT20/MB468 | NUC |
| >hg38_21_+_8433172_8446622_RNA45SN1_45S_50ntFlanks@11784.11803.20\|ATGAATGGATGAACGAGATT | >hg38_21_+_8433172_8446622_RNA45SN1_45S_50ntFlanks | 11784 | 11803 | 20 | ATGAATGGATGAACGAGATT | 548.1 | -6.4 | 1.0 | -6.3 | 2.8E-10 | 6.2E-10 | MB231/MB468 | MITO |
| >hg38_21_+_8433172_8446622_RNA45SN1_45S_50ntFlanks@4472.4492.21\|CTTAGCTGAGTGTCCCGCGGG | >hg38_21_+_8433172_8446622_RNA45SN1_45S_50ntFlanks | 4472 | 4492 | 21 | CTTAGCTGAGTGTCCCGCGGG | 487.9 | -6.9 | 1.1 | -6.3 | 3.0E-10 | 6.2E-10 | BT20/MB468 | NUC |
| >hg38_21_+_8433172_8446622_RNA45SN1_45S_50ntFlanks@11570.11592.23\|AGCCGACTTAGAACTGGTGCGGA | >hg38_21_+_8433172_8446622_RNA45SN1_45S_50ntFlanks | 11570 | 11592 | 23 | AGCCGACTTAGAACTGGTGCGGA | 393.3 | -4.8 | 0.8 | -6.3 | 2.8E-10 | 6.2E-10 | MB231/MB468 | MITO |
| >hg38_21_+_8433172_8446622_RNA45SN1_45S_50ntFlanks@10351.10381.31\|CGAGAACTTTGAAGGCCGAAGTGGAGAAGGG | >hg38_21_+_8433172_8446622_RNA45SN1_45S_50ntFlanks | 10351 | 10381 | 31 | CGAGAACTTTGAAGGCCGAAGTGGAGAAGGG | 1433.9 | -6.8 | 1.1 | -6.3 | 2.5E-10 | 6.2E-10 | MB231/MB468 | NUC |
| >hg38_21_+_8433172_8446622_RNA45SN1_45S_50ntFlanks@6650.6668.19\|ACGACTCTTAGCGGTGGAT | >hg38_21_+_8433172_8446622_RNA45SN1_45S_50ntFlanks | 6650 | 6668 | 19 | ACGACTCTTAGCGGTGGAT | 559.6 | 3.0 | 0.5 | 6.3 | 3.0E-10 | 6.2E-10 | BT20/MB468 | NUC |

| | | | | | | | | | | | | | |
|---|---|---|---|---|---|---|---|---|---|---|---|---|---|
| >MI0000253\|hsa-mir-148a&WithFlank&7\|-\|25949913\|25949992@50.71.22(+1A)&[MIMAT0000243&hsa-miR-148a-3p&offsets\|0\|0(+1A);m-7&7\|-\|25949922\|25949943&offsets\|0\|0(+1A)]\|TCAGTGCACTACAGAACTTTGTA | >MI0000253\|hsa-mir-148a&WithFlank&7\|-\|25949913\|25949992 | 50 | 71 | 22(+1A) | TCAGTGCACTACAGAACTTTGTA | 456.6 | 8.1 | 1.3 | 6.3 | 2.2E-10 | 6.2E-10 | BT20/MB468 | CYTO |
| >hg38_21_+_8433172_8446622_RNA45SN1_45S_50ntFlanks@11821.11842.22\|CTATCCAGCGAAACCACAGCCA | >hg38_21_+_8433172_8446622_RNA45SN1_45S_50ntFlanks | 11821 | 11842 | 22 | CTATCCAGCGAAACCACAGCCA | 174.2 | -9.5 | 1.5 | -6.4 | 1.9E-10 | 6.2E-10 | BT20/MB468 | MITO |
| >hg38_21_+_8433172_8446622_RNA45SN1_45S_50ntFlanks@12259.12289.31\|CTTGATCTTGATTTTCAGTACGAATACAGAC | >hg38_21_+_8433172_8446622_RNA45SN1_45S_50ntFlanks | 12259 | 12289 | 31 | CTTGATCTTGATTTTCAGTACGAATACAGAC | 85.6 | -10.3 | 1.6 | -6.3 | 2.4E-10 | 6.2E-10 | MB231/MB468 | MP |
| >hg38_21_+_8433172_8446622_RNA45SN1_45S_50ntFlanks@10843.10865.23\|TAACTTCGGGATAAGGATTGGCT | >hg38_21_+_8433172_8446622_RNA45SN1_45S_50ntFlanks | 10843 | 10865 | 23 | TAACTTCGGGATAAGGATTGGCT | 2015.8 | -5.3 | 0.8 | -6.3 | 2.2E-10 | 6.2E-10 | BT20/MB468 | CYTO |
| >MI0000088\|hsa-mir-30a&WithFlank&6\|-\|71403545\|71403627@54.71.18&[MIMAT0000088&hsa-miR-30a-3p&offsets\|+1\|-3;m-38&6\|-\|71403554\|71403575&offsets\|+1\|-3]\|TTTCAGTCGGATGTTTGC | >MI0000088\|hsa-mir-30a&WithFlank&6\|-\|71403545\|71403627 | 54 | 71 | 18 | TTTCAGTCGGATGTTTGC | 1206.8 | 5.9 | 0.9 | 6.3 | 2.6E-10 | 6.2E-10 | MB231/MB468 | NUC |
| >hg38_21_+_8433172_8446622_RNA45SN1_45S_50ntFlanks@3705.3740.36\|TACCTGGTTGATCCTGCCAGTAGCATATGCTTGTCT | >hg38_21_+_8433172_8446622_RNA45SN1_45S_50ntFlanks | 3705 | 3740 | 36 | TACCTGGTTGATCCTGCCAGTAGCATATGCTTGTCT | 314.1 | 7.4 | 1.2 | 6.3 | 2.5E-10 | 6.2E-10 | MB231/MB468 | CYTO |
| >hg38_21_+_8433172_8446622_RNA45SN1_45S_50ntFlanks@9029.9058.30\|TCTCCTCGTGGGGGGGCCGGGCCACCCCTC | >hg38_21_+_8433172_8446622_RNA45SN1_45S_50ntFlanks | 9029 | 9058 | 30 | TCTCCTCGTGGGGGGGCCGGGCCACCCCTC | 151.0 | -9.3 | 1.5 | -6.4 | 1.9E-10 | 6.2E-10 | BT20/MB468 | MITO |
| >hg38_21_+_8433172_8446622_RNA45SN1_45S_50ntFlanks@12636.12662.27\|AGCTACCATCTGTGGGATTATGACTGA | >hg38_21_+_8433172_8446622_RNA45SN1_45S_50ntFlanks | 12636 | 12662 | 27 | AGCTACCATCTGTGGGATTATGACTGA | 84.8 | -10.3 | 1.6 | -6.3 | 2.4E-10 | 6.2E-10 | MB231/MB468 | MP |
| >hg38_21_+_8433172_8446622_RNA45SN1_45S_50ntFlanks@4102.4120.19\|ACCACGGGTGACGGGGAAT | >hg38_21_+_8433172_8446622_RNA45SN1_45S_50ntFlanks | 4102 | 4120 | 19 | ACCACGGGTGACGGGGAAT | 577.7 | -9.3 | 1.5 | -6.3 | 3.0E-10 | 6.2E-10 | BT20/MB468 | NUC |
| >hg38_21_+_8433172_8446622_RNA45SN1_45S_50ntFlanks@4445.4470.26\|CTTGCCTCTCGGCGCCCCCTCGATGC | >hg38_21_+_8433172_8446622_RNA45SN1_45S_50ntFlanks | 4445 | 4470 | 26 | CTTGCCTCTCGGCGCCCCCTCGATGC | 711.5 | -9.4 | 1.5 | -6.3 | 3.0E-10 | 6.3E-10 | BT20/MB468 | NUC |
| >hg38_21_+_8433172_8446622_RNA45SN1_45S_50ntFlanks@6666.6699.34\|GATCACTCGGCTCGTGCGTCGATGAAGAACGCAG | >hg38_21_+_8433172_8446622_RNA45SN1_45S_50ntFlanks | 6666 | 6699 | 34 | GATCACTCGGCTCGTGCGTCGATGAAGAACGCAG | 237.9 | -11.0 | 1.6 | -6.7 | 2.1E-11 | 6.3E-10 | BT20/MB468 | MP |
| >hg38_21_+_8433172_8446622_RNA45SN1_45S_50ntFlanks@8354.8390.37\|TTGAAAAGAACTTTGAAGAGAGAGTTCAAGAGGGCGT | >hg38_21_+_8433172_8446622_RNA45SN1_45S_50ntFlanks | 8354 | 8390 | 37 | TTGAAAAGAACTTTGAAGAGAGAGTTCAAGAGGGCGT | 148.0 | 11.0 | 1.6 | 6.7 | 2.1E-11 | 6.3E-10 | BT20/MB468 | MP |
| >hg38_21_+_8433172_8446622_RNA45SN1_45S_50ntFlanks@4560.4577.18\|CTAGGAATAATGGAATAG | >hg38_21_+_8433172_8446622_RNA45SN1_45S_50ntFlanks | 4560 | 4577 | 18 | CTAGGAATAATGGAATAG | 188.0 | -10.6 | 1.6 | -6.7 | 2.1E-11 | 6.3E-10 | BT20/MB468 | MP |
| >hg38_21_+_8433172_8446622_RNA45SN1_45S_50ntFlanks@12152.12169.18\|TGGGGAGTTTGACTGGGG | >hg38_21_+_8433172_8446622_RNA45SN1_45S_50ntFlanks | 12152 | 12169 | 18 | TGGGGAGTTTGACTGGGG | 72.2 | -10.0 | 1.6 | -6.3 | 2.4E-10 | 6.3E-10 | MB231/MB468 | MP |
| >hg38_21_+_8433172_8446622_RNA45SN1_45S_50ntFlanks@4594.4614.21\|TGTTGGTTTTCGGAACTGAGG | >hg38_21_+_8433172_8446622_RNA45SN1_45S_50ntFlanks | 4594 | 4614 | 21 | TGTTGGTTTTCGGAACTGAGG | 78.7 | -10.2 | 1.6 | -6.3 | 2.4E-10 | 6.3E-10 | MB231/MB468 | MP |
| >hg38_21_+_8433172_8446622_RNA45SN1_45S_50ntFlanks@10351.10384.34\|CGAGAACTTTGAAGGCCGAAGTGGAGAAGGGTTC | >hg38_21_+_8433172_8446622_RNA45SN1_45S_50ntFlanks | 10351 | 10384 | 34 | CGAGAACTTTGAAGGCCGAAGTGGAGAAGGGTTC | 1987.6 | -5.0 | 0.8 | -6.4 | 1.9E-10 | 6.3E-10 | BT20/MB468 | MITO |
| >hg38_21_+_8433172_8446622_RNA45SN1_45S_50ntFlanks@10912.10929.18\|CGCGCCGCGGCTGGACGA | >hg38_21_+_8433172_8446622_RNA45SN1_45S_50ntFlanks | 10912 | 10929 | 18 | CGCGCCGCGGCTGGACGA | 71.1 | -10.0 | 1.6 | -6.3 | 2.5E-10 | 6.3E-10 | MB231/MB468 | MP |
| >hg38_21_+_8433172_8446622_RNA45SN1_45S_50ntFlanks@12162.12182.21\|GACTGGGGCGGTACACCTGTC | >hg38_21_+_8433172_8446622_RNA45SN1_45S_50ntFlanks | 12162 | 12182 | 21 | GACTGGGGCGGTACACCTGTC | 79.0 | -10.2 | 1.6 | -6.3 | 2.5E-10 | 6.3E-10 | MB231/MB468 | MP |
| >MI0001448\|hsa-mir-425&WithFlank&3\|-\|49020142\|49020240@20.43.24(+1C)&[MIMAT0003393&hsa-miR-425-5p&offsets\|0\|+1(+1C);m-95&3\|-\|49020200\|49020221&offsets\|0\|+2(+1C)]\|AATGACACGATCACTCCCGTTGAGC | >MI0001448\|hsa-mir-425&WithFlank&3\|-\|49020142\|49020240 | 20 | 43 | 24(+1C) | AATGACACGATCACTCCCGTTGAGC | 772.0 | 6.8 | 1.1 | 6.3 | 2.9E-10 | 6.3E-10 | MB231/MB468 | MITO |
| >MI0000746\|hsa-mir-99b&WithFlank&19\|+\|51692606\|51692687@13.32.20(+1A)&[MIMAT0000689&hsa-miR-99b-5p&offsets\|0\|-2(+1A);m-10&19\|+\|51692618\|51692639&offsets\|0\|-2(+1A)]\|CACCCGTAGAACCGACCTTGA | >MI0000746\|hsa-mir-99b&WithFlank&19\|+\|51692606\|51692687 | 13 | 32 | 20(+1A) | CACCCGTAGAACCGACCTTGA | 403.2 | 4.3 | 0.7 | 6.3 | 2.6E-10 | 6.3E-10 | MB231/MB468 | NUC |
| >hg38_21_+_8433172_8446622_RNA45SN1_45S_50ntFlanks@12073.12098.26\|CCGAGGGGCTCTCGCTTCTGGCGCCA | >hg38_21_+_8433172_8446622_RNA45SN1_45S_50ntFlanks | 12073 | 12098 | 26 | CCGAGGGGCTCTCGCTTCTGGCGCCA | 225.3 | -7.6 | 1.2 | -6.3 | 2.6E-10 | 6.3E-10 | MB231/MB468 | NUC |
| >hg38_21_+_8433172_8446622_RNA45SN1_45S_50ntFlanks@11783.11806.24\|CATGAATGGATGAACGAGATTCCC | >hg38_21_+_8433172_8446622_RNA45SN1_45S_50ntFlanks | 11783 | 11806 | 24 | CATGAATGGATGAACGAGATTCCC | 1601.9 | -4.3 | 0.7 | -6.4 | 1.9E-10 | 6.3E-10 | BT20/MB468 | MITO |
| >hg38_21_+_8433172_8446622_RNA45SN1_45S_50ntFlanks@10830.10851.22\|CAAGCCGGATCCGTAACTTCGG | >hg38_21_+_8433172_8446622_RNA45SN1_45S_50ntFlanks | 10830 | 10851 | 22 | CAAGCCGGATCCGTAACTTCGG | 430.9 | -8.1 | 1.3 | -6.3 | 2.5E-10 | 6.3E-10 | MB231/MB468 | MP |
| >MI0000088\|hsa-mir-30a&WithFlank&6\|-\|71403545\|71403627@12.33.22&[MIMAT0000087&hsa-miR-30a-5p&offsets\|0\|0;m-15&6\|-\|71403595\|71403616&offsets\|0\|0]\|TGTAAACATCCTCGACTGGAAG | >MI0000088\|hsa-mir-30a&WithFlank&6\|-\|71403545\|71403627 | 12 | 33 | 22 | TGTAAACATCCTCGACTGGAAG | 3546.5 | 6.2 | 1.0 | 6.3 | 2.9E-10 | 6.3E-10 | MB231/MB468 | MITO |
| >hg38_21_+_8433172_8446622_RNA45SN1_45S_50ntFlanks@10186.10207.22\|TCCTCCCCTCCTCCCGCCCACG | >hg38_21_+_8433172_8446622_RNA45SN1_45S_50ntFlanks | 10186 | 10207 | 22 | TCCTCCCCTCCTCCCGCCCACG | 439.6 | -7.2 | 1.1 | -6.3 | 2.6E-10 | 6.3E-10 | MB231/MB468 | CYTO |
| >hg38_21_+_8433172_8446622_RNA45SN1_45S_50ntFlanks@6784.6804.21\|GCTACGCCTGTCTGAGCGTCG | >hg38_21_+_8433172_8446622_RNA45SN1_45S_50ntFlanks | 6784 | 6804 | 21 | GCTACGCCTGTCTGAGCGTCG | 95.1 | -10.4 | 1.6 | -6.3 | 2.5E-10 | 6.3E-10 | MB231/MB468 | MP |
| >hg38_21_+_8433172_8446622_RNA45SN1_45S_50ntFlanks@4103.4129.27\|CCACGGGTGACGGGGAATCAGGGTTCG | >hg38_21_+_8433172_8446622_RNA45SN1_45S_50ntFlanks | 4103 | 4129 | 27 | CCACGGGTGACGGGGAATCAGGGTTCG | 69.4 | -10.0 | 1.6 | -6.3 | 2.5E-10 | 6.4E-10 | MB231/MB468 | MP |
| >hg38_21_+_8433172_8446622_RNA45SN1_45S_50ntFlanks@6661.6695.35\|CGGTGGATCACTCGGCTCGTGCGTCGATGAAGAAC | >hg38_21_+_8433172_8446622_RNA45SN1_45S_50ntFlanks | 6661 | 6695 | 35 | CGGTGGATCACTCGGCTCGTGCGTCGATGAAGAAC | 465.8 | 2.9 | 0.5 | 6.3 | 3.0E-10 | 6.4E-10 | BT20/MB468 | NUC |
| >hg38_21_+_8433172_8446622_RNA45SN1_45S_50ntFlanks@9467.9494.28\|GTGGAGCACGAGCGCACGTGTTAGGACC | >hg38_21_+_8433172_8446622_RNA45SN1_45S_50ntFlanks | 9467 | 9494 | 28 | GTGGAGCACGAGCGCACGTGTTAGGACC | 88.0 | -10.3 | 1.6 | -6.3 | 2.5E-10 | 6.4E-10 | MB231/MB468 | MP |
| >hg38_21_+_8433172_8446622_RNA45SN1_45S_50ntFlanks@13005.13025.21\|TCTATTGAAAGTCAGCCCTCG | >hg38_21_+_8433172_8446622_RNA45SN1_45S_50ntFlanks | 13005 | 13025 | 21 | TCTATTGAAAGTCAGCCCTCG | 306.6 | -7.6 | 1.2 | -6.3 | 2.3E-10 | 6.4E-10 | BT20/MB468 | CYTO |
| >hg38_21_+_8433172_8446622_RNA45SN1_45S_50ntFlanks@12247.12268.22\|GGCAAAAGCTCGCTTGATCTTG | >hg38_21_+_8433172_8446622_RNA45SN1_45S_50ntFlanks | 12247 | 12268 | 22 | GGCAAAAGCTCGCTTGATCTTG | 94.0 | -10.4 | 1.6 | -6.3 | 2.5E-10 | 6.4E-10 | MB231/MB468 | MP |
| >MI0000803\|hsa-mir-330&WithFlank&19\|-\|45638988\|45639093@64.86.23(+1U)&[MIMAT0000751&hsa-miR-330-3p&offsets\|+1\|+1(+1U);m-347&19\|-\|45639009\|45639030&offsets\|0\|+1(+1U)]\|CAAAGCACACGGCCTGCAGAGAGT | >MI0000803\|hsa-mir-330&WithFlank&19\|-\|45638988\|45639093 | 64 | 86 | 23(+1U) | CAAAGCACACGGCCTGCAGAGAGT | 452.1 | 9.4 | 1.5 | 6.3 | 2.3E-10 | 6.4E-10 | BT20/MB468 | CYTO |
| >hg38_21_+_8433172_8446622_RNA45SN1_45S_50ntFlanks@9574.9597.24\|AATCGGTCGTCCGACCTGGGTATA | >hg38_21_+_8433172_8446622_RNA45SN1_45S_50ntFlanks | 9574 | 9597 | 24 | AATCGGTCGTCCGACCTGGGTATA | 3955.8 | -7.1 | 1.1 | -6.3 | 2.6E-10 | 6.4E-10 | MB231/MB468 | CYTO |
| >hg38_21_+_8433172_8446622_RNA45SN1_45S_50ntFlanks@4723.4757.35\|CATTAATCAAGAACGAAAGTCGGAGGTTCGAAGAC | >hg38_21_+_8433172_8446622_RNA45SN1_45S_50ntFlanks | 4723 | 4757 | 35 | CATTAATCAAGAACGAAAGTCGGAGGTTCGAAGAC | 170.7 | -9.5 | 1.5 | -6.4 | 1.9E-10 | 6.4E-10 | BT20/MB468 | MITO |
| >hg38_21_+_8433172_8446622_RNA45SN1_45S_50ntFlanks@6733.6772.40\|CATTGATCATCGACACTTCGAACGCACTTGCGGCCCCGGG | >hg38_21_+_8433172_8446622_RNA45SN1_45S_50ntFlanks | 6733 | 6772 | 40 | CATTGATCATCGACACTTCGAACGCACTTGCGGCCCCGGG | 1570.7 | -4.6 | 0.7 | -6.3 | 2.9E-10 | 6.4E-10 | MB231/MB468 | MITO |
| >MI0000285\|hsa-mir-205&WithFlank&1\|+\|209432127\|209432248@40.59.20&[MIMAT0000266&hsa-miR-205-5p&offsets\|0\|-2;m-351&1\|+\|209432166\|209432187&offsets\|0\|-2]\|TCCTTCATTCCACCGGAGTC | >MI0000285\|hsa-mir-205&WithFlank&1\|+\|209432127\|209432248 | 40 | 59 | 20 | TCCTTCATTCCACCGGAGTC | 117.8 | -10.7 | 1.7 | -6.3 | 2.5E-10 | 6.4E-10 | MB231/MB468 | MP |
| >hg38_21_+_8433172_8446622_RNA45SN1_45S_50ntFlanks@9534.9551.18\|AGGAAACTCTGGTGGAGG | >hg38_21_+_8433172_8446622_RNA45SN1_45S_50ntFlanks | 9534 | 9551 | 18 | AGGAAACTCTGGTGGAGG | 1140.8 | -8.3 | 1.3 | -6.3 | 2.9E-10 | 6.4E-10 | MB231/MB468 | MITO |
| >hg38_21_+_8433172_8446622_RNA45SN1_45S_50ntFlanks@4764.4782.19\|ATACCGTCGTAGTTCCGAC | >hg38_21_+_8433172_8446622_RNA45SN1_45S_50ntFlanks | 4764 | 4782 | 19 | ATACCGTCGTAGTTCCGAC | 100.5 | -10.5 | 1.7 | -6.3 | 2.5E-10 | 6.4E-10 | MB231/MB468 | MP |
| >MI0000650\|hsa-mir-200c&WithFlank&12\|+\|6963693\|6963772@50.70.21(+1U)&[MIMAT0000617&hsa-miR-200c-3p&offsets\|0\|-2(+1U);m-54&12\|+\|6963742\|6963764&offsets\|0\|-2(+1U)]\|TAATACTGCCGGGTAATGATGT | >MI0000650\|hsa-mir-200c&WithFlank&12\|+\|6963693\|6963772 | 50 | 70 | 21(+1U) | TAATACTGCCGGGTAATGATGT | 432.4 | 12.1 | 1.7 | 6.9 | 4.5E-12 | 6.4E-10 | BT20/MB231 | MP |
| >MI0000811\|hsa-mir-148b&WithFlank&12\|+\|54337210\|54337320@69.90.22(+1U)&[MIMAT0000759&hsa-miR-148b-3p&offsets\|0\|0(+1U);m-77&12\|+\|54337278\|54337299&offsets\|0\|0(+1U)]\|TCAGTGCATCACAGAACTTTGTT | >MI0000811\|hsa-mir-148b&WithFlank&12\|+\|54337210\|54337320 | 69 | 90 | 22(+1U) | TCAGTGCATCACAGAACTTTGTT | 5369.5 | 6.0 | 0.9 | 6.3 | 2.6E-10 | 6.4E-10 | MB231/MB468 | CYTO |
| >hg38_21_+_8433172_8446622_RNA45SN1_45S_50ntFlanks@9534.9567.34\|AGGAAACTCTGGTGGAGGTCCGTAGCGGTCCTGA | >hg38_21_+_8433172_8446622_RNA45SN1_45S_50ntFlanks | 9534 | 9567 | 34 | AGGAAACTCTGGTGGAGGTCCGTAGCGGTCCTGA | 4776.2 | -10.8 | 1.7 | -6.3 | 3.1E-10 | 6.4E-10 | BT20/MB468 | NUC |
| >hg38_21_+_8433172_8446622_RNA45SN1_45S_50ntFlanks@10430.10456.27\|GGCGAGCGCCGTTCCGAAGGGACGGGC | >hg38_21_+_8433172_8446622_RNA45SN1_45S_50ntFlanks | 10430 | 10456 | 27 | GGCGAGCGCCGTTCCGAAGGGACGGGC | 74.5 | -10.1 | 1.6 | -6.3 | 2.5E-10 | 6.4E-10 | MB231/MB468 | MP |
| >MI0000253\|hsa-mir-148a&WithFlank&7\|-\|25949913\|25949992@50.72.23&[MIMAT0000243&hsa-miR-148a-3p&offsets\|0\|+1;m-7&7\|-\|25949922\|25949943&offsets\|0\|+1]\|TCAGTGCACTACAGAACTTTGTC | >MI0000253\|hsa-mir-148a&WithFlank&7\|-\|25949913\|25949992 | 50 | 72 | 23 | TCAGTGCACTACAGAACTTTGTC | 2160.4 | 4.6 | 0.7 | 6.7 | 2.8E-11 | 6.4E-10 | BT20/MB231 | NUC |
| >hg38_21_+_8433172_8446622_RNA45SN1_45S_50ntFlanks@10873.10890.18\|CTGGGTCGGTCGGGCTGG | >hg38_21_+_8433172_8446622_RNA45SN1_45S_50ntFlanks | 10873 | 10890 | 18 | CTGGGTCGGTCGGGCTGG | 736.0 | -4.9 | 0.8 | -6.3 | 2.6E-10 | 6.4E-10 | MB231/MB468 | CYTO |
| >hg38_21_+_8433172_8446622_RNA45SN1_45S_50ntFlanks@8380.8403.24\|CAAGAGGGCGTGAAACCGTTAAGA | >hg38_21_+_8433172_8446622_RNA45SN1_45S_50ntFlanks | 8380 | 8403 | 24 | CAAGAGGGCGTGAAACCGTTAAGA | 1335.3 | -5.1 | 0.8 | -6.4 | 2.0E-10 | 6.5E-10 | BT20/MB468 | MITO |
| >hg38_21_+_8433172_8446622_RNA45SN1_45S_50ntFlanks@6647.6683.37\|CGTACGACTCTTAGCGGTGGATCACTCGGCTCGTGCG | >hg38_21_+_8433172_8446622_RNA45SN1_45S_50ntFlanks | 6647 | 6683 | 37 | CGTACGACTCTTAGCGGTGGATCACTCGGCTCGTGCG | 320.1 | 8.2 | 1.3 | 6.3 | 2.6E-10 | 6.5E-10 | MB231/MB468 | NUC |
| >hg38_21_+_8433172_8446622_RNA45SN1_45S_50ntFlanks@4163.4186.24\|CATCCAAGGAAGGCAGCAGGCGCG | >hg38_21_+_8433172_8446622_RNA45SN1_45S_50ntFlanks | 4163 | 4186 | 24 | CATCCAAGGAAGGCAGCAGGCGCG | 1525.0 | -9.7 | 1.5 | -6.3 | 3.1E-10 | 6.5E-10 | BT20/MB468 | NUC |
| >am_trnaMT_LysTTT_MT_+_8295_8364@29.52.24,trnalookalike7_LysTTT_1_+_568844_568913@29.52.24\|TTTAAGTTAAAGATTAAGAGAACC | >am_trnaMT_LysTTT_MT_+_8295_8364 | 29 | 52 | 24 | TTTAAGTTAAAGATTAAGAGAACC | 728.3 | 8.7 | 1.4 | 6.3 | 2.3E-10 | 6.5E-10 | BT20/MB468 | CYTO |
| >hg38_21_+_8433172_8446622_RNA45SN1_45S_50ntFlanks@12638.12657.20\|CTACCATCTGTGGGATTATG | >hg38_21_+_8433172_8446622_RNA45SN1_45S_50ntFlanks | 12638 | 12657 | 20 | CTACCATCTGTGGGATTATG | 1536.2 | -9.4 | 1.5 | -6.3 | 3.0E-10 | 6.5E-10 | MB231/MB468 | MITO |
| >hg38_21_+_8433172_8446622_RNA45SN1_45S_50ntFlanks@12949.12988.40\|ACGACCTGCTTCTGGGTCGGGGTTTCGTACGTAGCAGAGC | >hg38_21_+_8433172_8446622_RNA45SN1_45S_50ntFlanks | 12949 | 12988 | 40 | ACGACCTGCTTCTGGGTCGGGGTTTCGTACGTAGCAGAGC | 1083.7 | -9.5 | 1.5 | -6.3 | 3.1E-10 | 6.5E-10 | BT20/MB468 | NUC |

<table>
<tr><th></th><th></th><th></th><th></th><th></th><th></th><th></th><th></th><th></th><th></th><th></th><th></th><th></th><th></th></tr>
<tr><td>>am_trnaMT_HisGTG_MT_+_12138_12206@19.51.33|AAACATCAGATTGTGAATCTGACAACAGAGGCT</td><td>>am_trnaMT_HisGTG_MT_+_12138_12206</td><td>19</td><td>51</td><td>33</td><td>AAACATCAGATTGTGAATCTGACAACAGAGGCT</td><td>1493.8</td><td>8.1</td><td>1.3</td><td>6.4</td><td>2.0E-10</td><td>6.5E-10</td><td>BT20/MB468</td><td>MITO</td></tr>
<tr><td>>hg38_21_+_8433172_8446622_RNA45SN1_45S_50ntFlanks@8184.8223.40|CGAGGCCCAGCCCGTGGACGGTGTGAGGCCGGTAGCGGCC</td><td>>hg38_21_+_8433172_8446622_RNA45SN1_45S_50ntFlanks</td><td>8184</td><td>8223</td><td>40</td><td>CGAGGCCCAGCCCGTGGACGGTGTGAGGCCGGTAGCGGCC</td><td>266.3</td><td>-7.9</td><td>1.2</td><td>-6.3</td><td>3.0E-10</td><td>6.5E-10</td><td>MB231/MB468</td><td>MITO</td></tr>
<tr><td>>hg38_21_+_8433172_8446622_RNA45SN1_45S_50ntFlanks@10681.10712.32|TTCCGGCGGCGTCCGGTGAGCTCTCGCTGGCC</td><td>>hg38_21_+_8433172_8446622_RNA45SN1_45S_50ntFlanks</td><td>10681</td><td>10712</td><td>32</td><td>TTCCGGCGGCGTCCGGTGAGCTCTCGCTGGCC</td><td>2446.8</td><td>-5.0</td><td>0.8</td><td>-6.3</td><td>2.6E-10</td><td>6.5E-10</td><td>MB231/MB468</td><td>CYTO</td></tr>
<tr><td>>hg38_21_+_8433172_8446622_RNA45SN1_45S_50ntFlanks@4663.4689.27|GTGAAATTCTTGGACCGGCGCAAGACG</td><td>>hg38_21_+_8433172_8446622_RNA45SN1_45S_50ntFlanks</td><td>4663</td><td>4689</td><td>27</td><td>GTGAAATTCTTGGACCGGCGCAAGACG</td><td>97.8</td><td>-10.5</td><td>1.7</td><td>-6.3</td><td>2.5E-10</td><td>6.5E-10</td><td>MB231/MB468</td><td>MP</td></tr>
<tr><td>>hg38_21_+_8433172_8446622_RNA45SN1_45S_50ntFlanks@10511.10534.24|ATCCGGAGTGGCGGAGATGGGCGC</td><td>>hg38_21_+_8433172_8446622_RNA45SN1_45S_50ntFlanks</td><td>10511</td><td>10534</td><td>24</td><td>ATCCGGAGTGGCGGAGATGGGCGC</td><td>200.2</td><td>-10.7</td><td>1.6</td><td>-6.7</td><td>2.2E-11</td><td>6.5E-10</td><td>BT20/MB468</td><td>MP</td></tr>
<tr><td>>hg38_21_+_8433172_8446622_RNA45SN1_45S_50ntFlanks@12157.12180.24|AGTTTGACTGGGGCGGTACACCTG</td><td>>hg38_21_+_8433172_8446622_RNA45SN1_45S_50ntFlanks</td><td>12157</td><td>12180</td><td>24</td><td>AGTTTGACTGGGGCGGTACACCTG</td><td>3210.9</td><td>-9.7</td><td>1.4</td><td>-6.7</td><td>2.2E-11</td><td>6.5E-10</td><td>BT20/MB468</td><td>MP</td></tr>
<tr><td>>hg38_21_+_8433172_8446622_RNA45SN1_45S_50ntFlanks@12324.12356.33|TTGGGTTTTAAGCAGGAGGTGTCAGAAAAGTTA</td><td>>hg38_21_+_8433172_8446622_RNA45SN1_45S_50ntFlanks</td><td>12324</td><td>12356</td><td>33</td><td>TTGGGTTTTAAGCAGGAGGTGTCAGAAAAGTTA</td><td>315.5</td><td>-11.4</td><td>1.7</td><td>-6.7</td><td>2.2E-11</td><td>6.5E-10</td><td>BT20/MB468</td><td>MP</td></tr>
<tr><td>>hg38_21_+_8433172_8446622_RNA45SN1_45S_50ntFlanks@7985.8003.19|GATCAGACGTGGCGACCCG</td><td>>hg38_21_+_8433172_8446622_RNA45SN1_45S_50ntFlanks</td><td>7985</td><td>8003</td><td>19</td><td>GATCAGACGTGGCGACCCG</td><td>207.8</td><td>-10.0</td><td>1.6</td><td>-6.3</td><td>2.4E-10</td><td>6.5E-10</td><td>BT20/MB468</td><td>CYTO</td></tr>
<tr><td>>hg38_21_+_8433172_8446622_RNA45SN1_45S_50ntFlanks@12181.12200.20|TCAAACGGTAACGCAGGTGT</td><td>>hg38_21_+_8433172_8446622_RNA45SN1_45S_50ntFlanks</td><td>12181</td><td>12200</td><td>20</td><td>TCAAACGGTAACGCAGGTGT</td><td>132.3</td><td>-10.9</td><td>1.7</td><td>-6.3</td><td>2.5E-10</td><td>6.5E-10</td><td>MB231/MB468</td><td>MP</td></tr>
<tr><td>>hg38_21_+_8433172_8446622_RNA45SN1_45S_50ntFlanks@12934.12971.38|CTAAACCATTCGTAGACGACCTGCTTCTGGGTCGGGGT</td><td>>hg38_21_+_8433172_8446622_RNA45SN1_45S_50ntFlanks</td><td>12934</td><td>12971</td><td>38</td><td>CTAAACCATTCGTAGACGACCTGCTTCTGGGTCGGGGT</td><td>141.1</td><td>-7.1</td><td>1.1</td><td>-6.3</td><td>2.7E-10</td><td>6.5E-10</td><td>MB231/MB468</td><td>CYTO</td></tr>
<tr><td>>hg38_21_+_8433172_8446622_RNA45SN1_45S_50ntFlanks@475.518.44|CTTCGTGATCGATGTGGTGACGTCGTGCTCTCCCGGGCCGGGTC</td><td>>hg38_21_+_8433172_8446622_RNA45SN1_45S_50ntFlanks</td><td>475</td><td>518</td><td>44</td><td>CTTCGTGATCGATGTGGTGACGTCGTGCTCTCCCGGGCCGGGTC</td><td>293.7</td><td>9.6</td><td>1.5</td><td>6.3</td><td>2.4E-10</td><td>6.5E-10</td><td>BT20/MB468</td><td>CYTO</td></tr>
<tr><td>>hg38_21_+_8433172_8446622_RNA45SN1_45S_50ntFlanks@9381.9416.36|CGCTCGCCGGCCGAGGTGGGATCCCGAGGCCTCTCC</td><td>>hg38_21_+_8433172_8446622_RNA45SN1_45S_50ntFlanks</td><td>9381</td><td>9416</td><td>36</td><td>CGCTCGCCGGCCGAGGTGGGATCCCGAGGCCTCTCC</td><td>112.8</td><td>-10.7</td><td>1.7</td><td>-6.3</td><td>2.5E-10</td><td>6.5E-10</td><td>MB231/MB468</td><td>MP</td></tr>
<tr><td>>hg38_21_+_8433172_8446622_RNA45SN1_45S_50ntFlanks@4445.4470.26|CTTGCCTCTCGGCGCCCCCTCGATGC</td><td>>hg38_21_+_8433172_8446622_RNA45SN1_45S_50ntFlanks</td><td>4445</td><td>4470</td><td>26</td><td>CTTGCCTCTCGGCGCCCCCTCGATGC</td><td>133.4</td><td>-9.1</td><td>1.4</td><td>-6.4</td><td>2.0E-10</td><td>6.5E-10</td><td>BT20/MB468</td><td>MITO</td></tr>
<tr><td>>hg38_21_+_8433172_8446622_RNA45SN1_45S_50ntFlanks@4220.4242.23|ACGAAAAATAACAATACAGGACT</td><td>>hg38_21_+_8433172_8446622_RNA45SN1_45S_50ntFlanks</td><td>4220</td><td>4242</td><td>23</td><td>ACGAAAAATAACAATACAGGACT</td><td>2410.0</td><td>-7.5</td><td>1.2</td><td>-6.3</td><td>2.6E-10</td><td>6.6E-10</td><td>MB231/MB468</td><td>MP</td></tr>
<tr><td>>hg38_21_+_8433172_8446622_RNA45SN1_45S_50ntFlanks@6671.6692.22|CTCGGCTCGTGCGTCGATGAAG</td><td>>hg38_21_+_8433172_8446622_RNA45SN1_45S_50ntFlanks</td><td>6671</td><td>6692</td><td>22</td><td>CTCGGCTCGTGCGTCGATGAAG</td><td>206.8</td><td>-10.8</td><td>1.6</td><td>-6.7</td><td>2.2E-11</td><td>6.6E-10</td><td>BT20/MB468</td><td>MP</td></tr>
<tr><td>>hg38_21_+_8433172_8446622_RNA45SN1_45S_50ntFlanks@12970.12990.21|GTTTCGTACGTAGCAGAGCAG</td><td>>hg38_21_+_8433172_8446622_RNA45SN1_45S_50ntFlanks</td><td>12970</td><td>12990</td><td>21</td><td>GTTTCGTACGTAGCAGAGCAG</td><td>266.9</td><td>-7.5</td><td>1.2</td><td>-6.3</td><td>2.7E-10</td><td>6.6E-10</td><td>MB231/MB468</td><td>NUC</td></tr>
<tr><td>>hg38_21_+_8433172_8446622_RNA45SN1_45S_50ntFlanks@12939.12973.35|CCATTCGTAGACGACCTGCTTCTGGGTCGGGGTTT</td><td>>hg38_21_+_8433172_8446622_RNA45SN1_45S_50ntFlanks</td><td>12939</td><td>12973</td><td>35</td><td>CCATTCGTAGACGACCTGCTTCTGGGTCGGGGTTT</td><td>79.8</td><td>-10.2</td><td>1.6</td><td>-6.3</td><td>2.6E-10</td><td>6.6E-10</td><td>MB231/MB468</td><td>MP</td></tr>
<tr><td>>hg38_21_+_8433172_8446622_RNA45SN1_45S_50ntFlanks@11667.11687.21|CTGCCCAGTGCTCTGAATGTC</td><td>>hg38_21_+_8433172_8446622_RNA45SN1_45S_50ntFlanks</td><td>11667</td><td>11687</td><td>21</td><td>CTGCCCAGTGCTCTGAATGTC</td><td>699.5</td><td>5.1</td><td>0.8</td><td>6.3</td><td>2.4E-10</td><td>6.6E-10</td><td>BT20/MB468</td><td>CYTO</td></tr>
<tr><td>>hg38_21_+_8433172_8446622_RNA45SN1_45S_50ntFlanks@3705.3736.32|TACCTGGTTGATCCTGCCAGTAGCATATGCTT</td><td>>hg38_21_+_8433172_8446622_RNA45SN1_45S_50ntFlanks</td><td>3705</td><td>3736</td><td>32</td><td>TACCTGGTTGATCCTGCCAGTAGCATATGCTT</td><td>297.2</td><td>3.4</td><td>0.5</td><td>6.3</td><td>3.2E-10</td><td>6.6E-10</td><td>BT20/MB468</td><td>NUC</td></tr>
<tr><td>>MI0000077|hsa-mir-21&WithFlank&17|+|59841260|59841343@14.35.22(+1U)&[MIMAT0000076&hsa-miR-21-5p&offsets|0|0(+1U);m-1&17|+|59841273|59841294&offsets|0|0(+1U)]|TAGCTTATCAGACTGATGTTGAT</td><td>>MI0000077|hsa-mir-21&WithFlank&17|+|59841260|59841343</td><td>14</td><td>35</td><td>22(+1U)</td><td>TAGCTTATCAGACTGATGTTGAT</td><td>24089.4</td><td>4.4</td><td>0.7</td><td>6.3</td><td>3.2E-10</td><td>6.6E-10</td><td>BT20/MB468</td><td>NUC</td></tr>
<tr><td>>trna2_GlyGCC_21_-_18827107_18827177@1.37.37,trna35_GlyGCC_1_+_161413094_161413164@1.37.37,trna37_GlyGCC_1_+_161420467_161420537@1.37.37,trna39_GlyGCC_1_+_161427898_161427968@1.37.37,trna41_GlyGCC_1_+_161435258_161435328@1.37.37|GCATGGGTGGTTCAGTGGTAGAATTCTCGCCTGCCAC</td><td>>trna2_GlyGCC_21_-_18827107_18827177</td><td>1</td><td>37</td><td>37</td><td>GCATGGGTGGTTCAGTGGTAGAATTCTCGCCTGCCAC</td><td>182.5</td><td>7.9</td><td>1.2</td><td>6.7</td><td>1.7E-11</td><td>6.6E-10</td><td>BT20/MB231</td><td>CYTO</td></tr>
<tr><td>>MI0000809|hsa-mir-151a&WithFlank&8|-|140732558|140732659@51.72.22(+1U)&[MIMAT0000757&hsa-miR-151a-3p&offsets|-2|-1(+1U);m-30&8|-|140732587|140732607&offsets|-2|-1(+1U)]|TACTAGACTGAAGCTCCTTGAGT</td><td>>MI0000809|hsa-mir-151a&WithFlank&8|-|140732558|140732659</td><td>51</td><td>72</td><td>22(+1U)</td><td>TACTAGACTGAAGCTCCTTGAGT</td><td>423.2</td><td>8.2</td><td>1.3</td><td>6.4</td><td>2.0E-10</td><td>6.6E-10</td><td>BT20/MB468</td><td>MITO</td></tr>
<tr><td>>hg38_21_+_8433172_8446622_RNA45SN1_45S_50ntFlanks@10821.10862.42|GGAAGTCGGCAAGCCGGATCCGTAACTTCGGGATAAGGATTG</td><td>>hg38_21_+_8433172_8446622_RNA45SN1_45S_50ntFlanks</td><td>10821</td><td>10862</td><td>42</td><td>GGAAGTCGGCAAGCCGGATCCGTAACTTCGGGATAAGGATTG</td><td>179.4</td><td>-11.3</td><td>1.8</td><td>-6.3</td><td>3.1E-10</td><td>6.7E-10</td><td>MB231/MB468</td><td>MITO</td></tr>
<tr><td>>hg38_21_+_8433172_8446622_RNA45SN1_45S_50ntFlanks@9343.9371.29|CGAAAGCCGCCGTGGCGCAATGAAGGTGA</td><td>>hg38_21_+_8433172_8446622_RNA45SN1_45S_50ntFlanks</td><td>9343</td><td>9371</td><td>29</td><td>CGAAAGCCGCCGTGGCGCAATGAAGGTGA</td><td>68.5</td><td>-10.0</td><td>1.6</td><td>-6.3</td><td>2.6E-10</td><td>6.7E-10</td><td>MB231/MB468</td><td>MP</td></tr>
<tr><td>>hg38_21_+_8433172_8446622_RNA45SN1_45S_50ntFlanks@4739.4759.21|AAGTCGGAGGTTCGAAGACGA</td><td>>hg38_21_+_8433172_8446622_RNA45SN1_45S_50ntFlanks</td><td>4739</td><td>4759</td><td>21</td><td>AAGTCGGAGGTTCGAAGACGA</td><td>358.1</td><td>-6.9</td><td>1.1</td><td>-6.3</td><td>2.6E-10</td><td>6.7E-10</td><td>MB231/MB468</td><td>MP</td></tr>
<tr><td>>hg38_21_+_8433172_8446622_RNA45SN1_45S_50ntFlanks@10871.10892.22|GGCTGGGTCGGTCGGGCTGGGG</td><td>>hg38_21_+_8433172_8446622_RNA45SN1_45S_50ntFlanks</td><td>10871</td><td>10892</td><td>22</td><td>GGCTGGGTCGGTCGGGCTGGGG</td><td>368.8</td><td>-8.2</td><td>1.3</td><td>-6.3</td><td>2.6E-10</td><td>6.7E-10</td><td>MB231/MB468</td><td>MP</td></tr>
<tr><td>>hg38_21_+_8433172_8446622_RNA45SN1_45S_50ntFlanks@10391.10435.45|AACAGCAGTTGAACATGGGTCAGTCGGTCCTGAGAGATGGGCGAG</td><td>>hg38_21_+_8433172_8446622_RNA45SN1_45S_50ntFlanks</td><td>10391</td><td>10435</td><td>45</td><td>AACAGCAGTTGAACATGGGTCAGTCGGTCCTGAGAGATGGGCGAG</td><td>585.5</td><td>-6.5</td><td>1.0</td><td>-6.3</td><td>2.7E-10</td><td>6.7E-10</td><td>MB231/MB468</td><td>CYTO</td></tr>
<tr><td>>hg38_21_+_8433172_8446622_RNA45SN1_45S_50ntFlanks@9570.9594.25|TGCAAATCGGTCGTCCGACCTGGGT</td><td>>hg38_21_+_8433172_8446622_RNA45SN1_45S_50ntFlanks</td><td>9570</td><td>9594</td><td>25</td><td>TGCAAATCGGTCGTCCGACCTGGGT</td><td>809.2</td><td>-8.6</td><td>1.4</td><td>-6.3</td><td>3.1E-10</td><td>6.7E-10</td><td>MB231/MB468</td><td>MITO</td></tr>
<tr><td>>hg38_21_+_8433172_8446622_RNA45SN1_45S_50ntFlanks@12407.12436.30|TTTTGATCCTTCGATGTCGGCTCTTCCTAT</td><td>>hg38_21_+_8433172_8446622_RNA45SN1_45S_50ntFlanks</td><td>12407</td><td>12436</td><td>30</td><td>TTTTGATCCTTCGATGTCGGCTCTTCCTAT</td><td>12443.7</td><td>-11.4</td><td>1.8</td><td>-6.3</td><td>3.2E-10</td><td>6.7E-10</td><td>BT20/MB468</td><td>NUC</td></tr>
<tr><td>>hg38_21_+_8433172_8446622_RNA45SN1_45S_50ntFlanks@5163.5185.23|GCGTTCAGCCACCCGAGATTGAG</td><td>>hg38_21_+_8433172_8446622_RNA45SN1_45S_50ntFlanks</td><td>5163</td><td>5185</td><td>23</td><td>GCGTTCAGCCACCCGAGATTGAG</td><td>190.6</td><td>-11.4</td><td>1.8</td><td>-6.3</td><td>2.6E-10</td><td>6.7E-10</td><td>MB231/MB468</td><td>MP</td></tr>
<tr><td>>MI0000067|hsa-let-7f-1&WithFlank&9|+|94176341|94176439@13.32.20(+1C);MI0000068|hsa-let-7f-2&WithFlank&X|-|53557186|53557280@14.33.20(+1C)&[MIMAT0000067&hsa-let-7f-5p&offsets|0|-2(+1C);m-20&9|+|94176353|94176374&offsets|0|-2(+1C)];[MIMAT0000067_1&hsa-let-7f-5p&offsets|0|-2(+1C);m-19&X|-|53557246|53557267&offsets|0|-2(+1C)]|TGAGGTAGTAGATTGTATAGC</td><td>>MI0000067|hsa-let-7f-1&WithFlank&9|+|94176341|94176439</td><td>13</td><td>32</td><td>20(+1C)</td><td>TGAGGTAGTAGATTGTATAGC</td><td>463.0</td><td>4.6</td><td>0.7</td><td>6.3</td><td>2.7E-10</td><td>6.7E-10</td><td>MB231/MB468</td><td>CYTO</td></tr>
<tr><td>>hg38_21_+_8433172_8446622_RNA45SN1_45S_50ntFlanks@4297.4321.25|CATTGGAGGGCAAGTCTGGTGCCAG</td><td>>hg38_21_+_8433172_8446622_RNA45SN1_45S_50ntFlanks</td><td>4297</td><td>4321</td><td>25</td><td>CATTGGAGGGCAAGTCTGGTGCCAG</td><td>370.0</td><td>-5.1</td><td>0.8</td><td>-6.3</td><td>2.4E-10</td><td>6.7E-10</td><td>BT20/MB468</td><td>CYTO</td></tr>
<tr><td>>hg38_21_+_8433172_8446622_RNA45SN1_45S_50ntFlanks@6721.6770.50|AATTGCAGGACACATTGATCATCGACACTTCGAACGCACTTGCGGCCCCG</td><td>>hg38_21_+_8433172_8446622_RNA45SN1_45S_50ntFlanks</td><td>6721</td><td>6770</td><td>50</td><td>AATTGCAGGACACATTGATCATCGACACTTCGAACGCACTTGCGGCCCCG</td><td>2984.3</td><td>-9.7</td><td>1.5</td><td>-6.3</td><td>3.1E-10</td><td>6.7E-10</td><td>MB231/MB468</td><td>MITO</td></tr>
<tr><td>>MI0000465|hsa-mir-191&WithFlank&3|-|49020612|49020715@23.44.22&[MIMAT0000440&hsa-miR-191-5p&offsets|+1|0;m-48&3|-|49020672|49020694&offsets|+1|0]|AACGGAATCCCAAAAGCAGCTG</td><td>>MI0000465|hsa-mir-191&WithFlank&3|-|49020612|49020715</td><td>23</td><td>44</td><td>22</td><td>AACGGAATCCCAAAAGCAGCTG</td><td>327.2</td><td>4.7</td><td>0.7</td><td>6.3</td><td>2.8E-10</td><td>6.7E-10</td><td>MB231/MB468</td><td>NUC</td></tr>
<tr><td>>hg38_21_+_8433172_8446622_RNA45SN1_45S_50ntFlanks@10466.10484.19|CGTTGCCCTCGGCCGATCG</td><td>>hg38_21_+_8433172_8446622_RNA45SN1_45S_50ntFlanks</td><td>10466</td><td>10484</td><td>19</td><td>CGTTGCCCTCGGCCGATCG</td><td>84.1</td><td>-10.3</td><td>1.6</td><td>-6.3</td><td>2.6E-10</td><td>6.8E-10</td><td>MB231/MB468</td><td>MP</td></tr>
<tr><td>>hg38_21_+_8433172_8446622_RNA45SN1_45S_50ntFlanks@6719.6741.23|TGAATTGCAGGACACATTGATCA</td><td>>hg38_21_+_8433172_8446622_RNA45SN1_45S_50ntFlanks</td><td>6719</td><td>6741</td><td>23</td><td>TGAATTGCAGGACACATTGATCA</td><td>314.2</td><td>-3.8</td><td>0.6</td><td>-6.3</td><td>2.8E-10</td><td>6.8E-10</td><td>MB231/MB468</td><td>CYTO</td></tr>
<tr><td>>hg38_21_+_8433172_8446622_RNA45SN1_45S_50ntFlanks@12008.12036.29|CCGCCGGTGAAATACCACTACTCTGATCG</td><td>>hg38_21_+_8433172_8446622_RNA45SN1_45S_50ntFlanks</td><td>12008</td><td>12036</td><td>29</td><td>CCGCCGGTGAAATACCACTACTCTGATCG</td><td>194.8</td><td>-5.2</td><td>0.8</td><td>-6.3</td><td>2.8E-10</td><td>6.8E-10</td><td>MB231/MB468</td><td>CYTO</td></tr>
<tr><td>>MI0000077|hsa-mir-21&WithFlank&17|+|59841260|59841343@14.36.23(+1G)&[MIMAT0000076&hsa-miR-21-5p&offsets|0|+1(+1G);m-1&17|+|59841273|59841294&offsets|0|+1(+1G)]|TAGCTTATCAGACTGATGTTGACG</td><td>>MI0000077|hsa-mir-21&WithFlank&17|+|59841260|59841343</td><td>14</td><td>36</td><td>23(+1G)</td><td>TAGCTTATCAGACTGATGTTGACG</td><td>3082.2</td><td>3.5</td><td>0.6</td><td>6.3</td><td>3.3E-10</td><td>6.8E-10</td><td>BT20/MB468</td><td>NUC</td></tr>
<tr><td>>MI0000093|hsa-mir-92a-1&WithFlank&13|+|91351308|91351397@54.74.21(+1G);MI0000094|hsa-mir-92a-2&WithFlank&X|-|134169532|134169618@54.74.21(+1G)&[MIMAT0000092&hsa-miR-92a-3p&offsets|0|-1(+1G);m-9&13|+|91351361|91351382&offsets|0|-1(+1G)];[MIMAT0000092_1&hsa-miR-92a-3p&offsets|0|-1(+1G);m-11&X|-|134169544|134169565&offsets|0|-1(+1G)]|TATTGCACTTGTCCCGGCCTGG</td><td>>MI0000093|hsa-mir-92a-1&WithFlank&13|+|91351308|91351397</td><td>54</td><td>74</td><td>21(+1G)</td><td>TATTGCACTTGTCCCGGCCTGG</td><td>1386.4</td><td>4.0</td><td>0.6</td><td>6.3</td><td>2.8E-10</td><td>6.8E-10</td><td>MB231/MB468</td><td>NUC</td></tr>
<tr><td>>hg38_21_+_8433172_8446622_RNA45SN1_45S_50ntFlanks@11808.11827.20|CTGTCCCTACCTACTATCCA</td><td>>hg38_21_+_8433172_8446622_RNA45SN1_45S_50ntFlanks</td><td>11808</td><td>11827</td><td>20</td><td>CTGTCCCTACCTACTATCCA</td><td>149.1</td><td>-11.1</td><td>1.8</td><td>-6.3</td><td>2.6E-10</td><td>6.8E-10</td><td>MB231/MB468</td><td>MP</td></tr>
<tr><td>>hg38_21_+_8433172_8446622_RNA45SN1_45S_50ntFlanks@11188.11206.19|CACCCGGGGGGCCGGCGGC</td><td>>hg38_21_+_8433172_8446622_RNA45SN1_45S_50ntFlanks</td><td>11188</td><td>11206</td><td>19</td><td>CACCCGGGGGGCCGGCGGC</td><td>112.2</td><td>-10.7</td><td>1.7</td><td>-6.3</td><td>2.7E-10</td><td>6.8E-10</td><td>MB231/MB468</td><td>MP</td></tr>
<tr><td>>hg38_21_+_8433172_8446622_RNA45SN1_45S_50ntFlanks@11568.11591.24|GCAGCCGACTTAGAACTGGTGCGG</td><td>>hg38_21_+_8433172_8446622_RNA45SN1_45S_50ntFlanks</td><td>11568</td><td>11591</td><td>24</td><td>GCAGCCGACTTAGAACTGGTGCGG</td><td>157.9</td><td>-4.7</td><td>0.7</td><td>-6.3</td><td>2.8E-10</td><td>6.8E-10</td><td>MB231/MB468</td><td>CYTO</td></tr>
<tr><td>>hg38_1_-_228634819_228635039_RNA5S12_5S_50ntFlanks@51.87.37|GTCTACGGCCATACCACCCTGAACGCGCCCGATCTCG</td><td>>hg38_1_-_228634819_228635039_RNA5S12_5S_50ntFlanks</td><td>51</td><td>87</td><td>37</td><td>GTCTACGGCCATACCACCCTGAACGCGCCCGATCTCG</td><td>944.0</td><td>5.8</td><td>0.9</td><td>6.4</td><td>2.1E-10</td><td>6.8E-10</td><td>BT20/MB468</td><td>MITO</td></tr>
<tr><td>>MI0000301|hsa-mir-224&WithFlank&X|-|151958572|151958664@14.37.24&[MIMAT0000281&hsa-miR-224-5p&offsets|+1|0;m-238&X|-|151958630|151958651&offsets|0|+2]|CAAGTCACTAGTGGTTCCGTTTAG</td><td>>MI0000301|hsa-mir-224&WithFlank&X|-|151958572|151958664</td><td>14</td><td>37</td><td>24</td><td>CAAGTCACTAGTGGTTCCGTTTAG</td><td>1135.3</td><td>9.5</td><td>1.5</td><td>6.3</td><td>3.1E-10</td><td>6.8E-10</td><td>MB231/MB468</td><td>MITO</td></tr>
<tr><td>>MI0000078|hsa-mir-22&WithFlank&17|-|1713897|1713993@59.80.22(+1A)&[MIMAT0000077&hsa-miR-22-3p&offsets|0|0(+1A);m-2&17|-|1713914|1713935&offsets|0|0(+1A)]|AAGCTGCCAGTTGAAGAACTGTA</td><td>>MI0000078|hsa-mir-22&WithFlank&17|-|1713897|1713993</td><td>59</td><td>80</td><td>22(+1A)</td><td>AAGCTGCCAGTTGAAGAACTGTA</td><td>734.8</td><td>8.8</td><td>1.4</td><td>6.3</td><td>2.5E-10</td><td>6.8E-10</td><td>BT20/MB468</td><td>CYTO</td></tr>
<tr><td>>hg38_21_+_8433172_8446622_RNA45SN1_45S_50ntFlanks@10391.10418.28|AACAGCAGTTGAACATGGGTCAGTCGGT</td><td>>hg38_21_+_8433172_8446622_RNA45SN1_45S_50ntFlanks</td><td>10391</td><td>10418</td><td>28</td><td>AACAGCAGTTGAACATGGGTCAGTCGGT</td><td>370.4</td><td>-8.5</td><td>1.4</td><td>-6.3</td><td>3.3E-10</td><td>6.8E-10</td><td>BT20/MB468</td><td>NUC</td></tr>
</table>

| | | | | | | | | | | | | | |
|---|---|---|---|---|---|---|---|---|---|---|---|---|---|
| >hg38_21_+_8433172_8446622_RNA45SN1_45S_50ntFlanks@8423.8467.45\|CGCAGTCCGCCCGGAGGATTCAACCCGGCGGCGGGTCCGGCCGTG | >hg38_21_+_8433172_8446622_RNA45SN1_45S_50ntFlanks | 8423 | 8467 | 45 | CGCAGTCCGCCCGGAGGATTCAACCCGGCGGCGGGTCCGGCCGTG | 206.1 | -11.6 | 1.8 | -6.3 | 2.7E-10 | 6.8E-10 | MB231/MB468 | MP |
| >MI0000093\|hsa-mir-92a-1&WithFlank&13\|+\|91351308\|91351397@54.74.21(+1A);MI0000094\|hsa-mir-92a-2&WithFlank&X\|-\|134169532\|134169618@54.74.21(+1A)&[MIMAT0000092&hsa-miR-92a-3p&offsets\|0\|-1(+1A);m-9&13\|+\|91351361\|91351382&offsets\|0\|-1(+1A)];[MIMAT0000092_1&hsa-miR-92a-3p&offsets\|0\|-1(+1A);m-11&X\|-\|134169544\|134169565&offsets\|0\|-1(+1A)]\|TATTGCACTTGTCCCGGCCTGA | >MI0000093\|hsa-mir-92a-1&WithFlank&13\|+\|91351308\|91351397 | 54 | 74 | 21(+1A) | TATTGCACTTGTCCCGGCCTGA | 2123.7 | 3.9 | 0.6 | 6.3 | 2.8E-10 | 6.8E-10 | MB231/MB468 | NUC |
| >trna3_ArgTCT_11_+_59318767_59318852@1.29.29,trna4_ArgTCT_17_+_8024243_8024330@1.29.29,trna4_ArgTCT_9_-_131102355_131102445@1.29.29,trna52_ArgTCT_6_+_27529963_27530049@1.29.29\|GGCTCTGTGGCGCAATGGATAGCGCATTG | >trna3_ArgTCT_11_+_59318767_59318852 | 1 | 29 | 29 | GGCTCTGTGGCGCAATGGATAGCGCATTG | 123.3 | -10.8 | 1.7 | -6.3 | 2.7E-10 | 6.8E-10 | MB231/MB468 | MP |
| >hg38_21_+_8433172_8446622_RNA45SN1_45S_50ntFlanks@10808.10831.24\|CAATGTAGGTAAGGGAAGTCGGCA | >hg38_21_+_8433172_8446622_RNA45SN1_45S_50ntFlanks | 10808 | 10831 | 24 | CAATGTAGGTAAGGGAAGTCGGCA | 201.9 | -7.9 | 1.3 | -6.3 | 2.8E-10 | 6.8E-10 | MB231/MB468 | NUC |
| >hg38_21_+_8433172_8446622_RNA45SN1_45S_50ntFlanks@10133.10153.21\|GCGGCGGCGGCGGCGGGGGTG | >hg38_21_+_8433172_8446622_RNA45SN1_45S_50ntFlanks | 10133 | 10153 | 21 | GCGGCGGCGGCGGCGGGGGTG | 117.1 | -10.7 | 1.7 | -6.3 | 2.7E-10 | 6.8E-10 | MB231/MB468 | MP |
| >hg38_21_+_8433172_8446622_RNA45SN1_45S_50ntFlanks@3840.3859.20\|CTCCTACTTGGATAACTGTG | >hg38_21_+_8433172_8446622_RNA45SN1_45S_50ntFlanks | 3840 | 3859 | 20 | CTCCTACTTGGATAACTGTG | 302.7 | -12.1 | 1.9 | -6.3 | 2.7E-10 | 6.9E-10 | MB231/MB468 | MP |
| >MI0000279\|hsa-mir-196a-2&WithFlank&12\|+\|53991732\|53991853@31.52.22(+1U);MI0000238\|hsa-mir-196a-1&WithFlank&17\|-\|48632484\|48632565@13.34.22(+1U)&[MIMAT0000226&hsa-miR-196a-5p&offsets\|0\|0(+1U);m-261&12\|+\|53991762\|53991783&offsets\|0\|0(+1U)];[MIMAT0000226_1&hsa-miR-196a-5p&offsets\|0\|0(+1U);m-268&17\|-\|48632532\|48632553&offsets\|0\|0(+1U)]\|TAGGTAGTTTCATGTTGTTGGGT | >MI0000279\|hsa-mir-196a-2&WithFlank&12\|+\|53991732\|53991853 | 31 | 52 | 22(+1U) | TAGGTAGTTTCATGTTGTTGGGT | 389.1 | 11.9 | 1.7 | 6.9 | 4.9E-12 | 6.9E-10 | BT20/MB231 | MP |
| >hg38_21_+_8433172_8446622_RNA45SN1_45S_50ntFlanks@12746.12765.20\|CGCCTGTCCCCGCCGGCGGG | >hg38_21_+_8433172_8446622_RNA45SN1_45S_50ntFlanks | 12746 | 12765 | 20 | CGCCTGTCCCCGCCGGCGGG | 209.4 | -11.6 | 1.8 | -6.3 | 2.7E-10 | 6.9E-10 | MB231/MB468 | MP |
| >MI0000255\|hsa-mir-30d&WithFlank&8\|-\|134804870\|134804951@12.34.23(+1G)&[MIMAT0000245&hsa-miR-30d-5p&offsets\|0\|+1(+1G);m-21&8\|-\|134804919\|134804940&offsets\|0\|+1(+1G)]\|TGTAAACATCCCCGACTGGAAGCG | >MI0000255\|hsa-mir-30d&WithFlank&8\|-\|134804870\|134804951 | 12 | 34 | 23(+1G) | TGTAAACATCCCCGACTGGAAGCG | 661.3 | 3.9 | 0.6 | 6.3 | 3.3E-10 | 6.9E-10 | BT20/MB468 | NUC |
| >hg38_21_+_8433172_8446622_RNA45SN1_45S_50ntFlanks@5525.5548.24\|TAAAAGTCGTAACAAGGTTTCCGT | >hg38_21_+_8433172_8446622_RNA45SN1_45S_50ntFlanks | 5525 | 5548 | 24 | TAAAAGTCGTAACAAGGTTTCCGT | 77.8 | -10.1 | 1.6 | -6.3 | 2.7E-10 | 6.9E-10 | MB231/MB468 | MP |
| >hg38_21_+_8433172_8446622_RNA45SN1_45S_50ntFlanks@9579.9597.19\|GTCGTCCGACCTGGGTATA | >hg38_21_+_8433172_8446622_RNA45SN1_45S_50ntFlanks | 9579 | 9597 | 19 | GTCGTCCGACCTGGGTATA | 74.1 | -10.1 | 1.6 | -6.3 | 2.7E-10 | 6.9E-10 | MB231/MB468 | MP |
| >hg38_21_+_8433172_8446622_RNA45SN1_45S_50ntFlanks@4494.4511.18\|CCCGAAGCGTTTACTTTG | >hg38_21_+_8433172_8446622_RNA45SN1_45S_50ntFlanks | 4494 | 4511 | 18 | CCCGAAGCGTTTACTTTG | 234.9 | -11.7 | 1.9 | -6.3 | 2.7E-10 | 6.9E-10 | MB231/MB468 | MP |
| >hg38_21_+_8433172_8446622_RNA45SN1_45S_50ntFlanks@7982.8005.24\|TCAGATCAGACGTGGCGACCCGCT | >hg38_21_+_8433172_8446622_RNA45SN1_45S_50ntFlanks | 7982 | 8005 | 24 | TCAGATCAGACGTGGCGACCCGCT | 377.4 | -7.9 | 1.2 | -6.3 | 2.7E-10 | 6.9E-10 | MB231/MB468 | MP |
| >hg38_21_+_8433172_8446622_RNA45SN1_45S_50ntFlanks@12000.12032.33\|CCTGCGGGCCGCCGGTGAAATACCACTACTCTG | >hg38_21_+_8433172_8446622_RNA45SN1_45S_50ntFlanks | 12000 | 12032 | 33 | CCTGCGGGCCGCCGGTGAAATACCACTACTCTG | 427.8 | -3.6 | 0.6 | -6.4 | 2.1E-10 | 6.9E-10 | BT20/MB468 | MITO |
| >MI0000750\|hsa-mir-26a-2&WithFlank&12\|-\|57824603\|57824698@23.41.19;MI0000083\|hsa-mir-26a-1&WithFlank&3\|+\|37969398\|37969486@19.37.19&[MIMAT0000082&hsa-miR-26a-5p&offsets\|+3\|0;m-33&12\|-\|57824658\|57824679&offsets\|+3\|0];[MIMAT0000082_1&hsa-miR-26a-5p&offsets\|+3\|0;m-34&3\|+\|37969413\|37969434&offsets\|+3\|0]\|AAGTAATCCAGGATAGGCT | >MI0000750\|hsa-mir-26a-2&WithFlank&12\|-\|57824603\|57824698 | 23 | 41 | 19 | AAGTAATCCAGGATAGGCT | 195.4 | 11.4 | 1.7 | 6.7 | 2.4E-11 | 6.9E-10 | BT20/MB468 | MP |
| >hg38_21_+_8433172_8446622_RNA45SN1_45S_50ntFlanks@8153.8188.36\|GCCGCTCGTGGGGGGCCCAAGTCCTTCTGATCGAGG | >hg38_21_+_8433172_8446622_RNA45SN1_45S_50ntFlanks | 8153 | 8188 | 36 | GCCGCTCGTGGGGGGCCCAAGTCCTTCTGATCGAGG | 76.8 | -10.1 | 1.6 | -6.3 | 2.7E-10 | 6.9E-10 | MB231/MB468 | MP |
| >hg38_21_+_8433172_8446622_RNA45SN1_45S_50ntFlanks@3811.3834.24\|AGTTATGGTTCCTTTGGTCGCTCG | >hg38_21_+_8433172_8446622_RNA45SN1_45S_50ntFlanks | 3811 | 3834 | 24 | AGTTATGGTTCCTTTGGTCGCTCG | 475.0 | -8.0 | 1.3 | -6.3 | 2.8E-10 | 6.9E-10 | MB231/MB468 | NUC |
| >hg38_21_+_8433172_8446622_RNA45SN1_45S_50ntFlanks@11564.11605.42\|CCTAGCAGCCGACTTAGAACTGGTGCGGACCAGGGGAATCCG | >hg38_21_+_8433172_8446622_RNA45SN1_45S_50ntFlanks | 11564 | 11605 | 42 | CCTAGCAGCCGACTTAGAACTGGTGCGGACCAGGGGAATCCG | 251.7 | -11.8 | 1.9 | -6.3 | 2.7E-10 | 6.9E-10 | MB231/MB468 | MP |
| >hg38_21_+_8433172_8446622_RNA45SN1_45S_50ntFlanks@11190.11208.19\|CCCGGGGGGCCGGCGGCGG | >hg38_21_+_8433172_8446622_RNA45SN1_45S_50ntFlanks | 11190 | 11208 | 19 | CCCGGGGGGCCGGCGGCGG | 332.8 | -11.5 | 1.7 | -6.7 | 2.4E-11 | 6.9E-10 | BT20/MB468 | MP |
| >hg38_21_+_8433172_8446622_RNA45SN1_45S_50ntFlanks@4512.4529.18\|AAAAAATTAGAGTGTTCA | >hg38_21_+_8433172_8446622_RNA45SN1_45S_50ntFlanks | 4512 | 4529 | 18 | AAAAAATTAGAGTGTTCA | 186.6 | -10.6 | 1.6 | -6.7 | 2.4E-11 | 6.9E-10 | BT20/MB468 | MP |
| >trna119_LysCTT_1_-_145395522_145395594@5.35.31,trna11_LysCTT_5_-_180648979_180649051@5.35.31,trna13_LysCTT_6_+_26556774_26556846@5.35.31,trna7_LysCTT_16_+_3225692_3225764@5.35.31,trna9_LysCTT_5_+_180634755_180634827@5.35.31\|GGCTAGCTCAGTCGGTAGAGCATGAGACTCT | >trna119_LysCTT_1_-_145395522_145395594 | 5 | 35 | 31 | GGCTAGCTCAGTCGGTAGAGCATGAGACTCT | 2041.2 | -5.1 | 0.8 | -6.7 | 2.5E-11 | 6.9E-10 | BT20/MB231 | MITO |
| >hg38_21_+_8433172_8446622_RNA45SN1_45S_50ntFlanks@10725.10755.31\|GGGGAGAGGGTGTAAATCTCGCGCCGGGCCG | >hg38_21_+_8433172_8446622_RNA45SN1_45S_50ntFlanks | 10725 | 10755 | 31 | GGGGAGAGGGTGTAAATCTCGCGCCGGGCCG | 138.2 | -11.0 | 1.7 | -6.3 | 2.7E-10 | 7.0E-10 | MB231/MB468 | MP |
| >MI0000102\|hsa-mir-100&WithFlank&11\|-\|122152223\|122152314@19.40.22&[MIMAT0000098&hsa-miR-100-5p&offsets\|0\|0;m-27&11\|-\|122152276\|122152296&offsets\|0\|+1]\|AACCCGTAGATCCGAACTTGTG | >MI0000102\|hsa-mir-100&WithFlank&11\|-\|122152223\|122152314 | 19 | 40 | 22 | AACCCGTAGATCCGAACTTGTG | 599454.6 | -6.3 | 0.9 | -6.6 | 3.1E-11 | 7.0E-10 | BT20/MB231 | NUC |
| >trna111_HisGTG_1_-_147774845_147774916@-1G.35.36,trna118_HisGTG_1_-_145396881_145396952@-1G.35.36,trna16_HisGTG_1_+_146544773_146544844@-1G.35.36,trna1_HisGTG_15_+_45493349_45493420@-1G.35.36,trna21_HisGTG_1_+_147753471_147753542@-1G.35.36,trna33_HisGTG_6_+_27125906_27125977@-1G.35.36,trna7_HisGTG_9_-_14433938_14434009@-1G.35.36,trna8_HisGTG_15_-_45492611_45492682@-1G.35.36,trna9_HisGTG_15_-_45490804_45490875@-1G.35.36\|GGCCGTGATCGTATAGTGGTTAGTACTCTGCGTTGT | >trna111_HisGTG_1_-_147774845_147774916 | -1G | 35 | 36 | GGCCGTGATCGTATAGTGGTTAGTACTCTGCGTTGT | 1209.9 | 9.9 | 1.6 | 6.3 | 3.2E-10 | 7.0E-10 | MB231/MB468 | MITO |
| >hg38_21_+_8433172_8446622_RNA45SN1_45S_50ntFlanks@5485.5503.19\|AGCGCTGAGAAGACGGTCG | >hg38_21_+_8433172_8446622_RNA45SN1_45S_50ntFlanks | 5485 | 5503 | 19 | AGCGCTGAGAAGACGGTCG | 245.3 | -11.0 | 1.6 | -6.7 | 2.4E-11 | 7.0E-10 | BT20/MB468 | MP |
| >hg38_21_+_8433172_8446622_RNA45SN1_45S_50ntFlanks@10798.10826.29\|CATGTTGGAACAATGTAGGTAAGGGAAGT | >hg38_21_+_8433172_8446622_RNA45SN1_45S_50ntFlanks | 10798 | 10826 | 29 | CATGTTGGAACAATGTAGGTAAGGGAAGT | 702.0 | -8.3 | 1.3 | -6.3 | 2.7E-10 | 7.0E-10 | MB231/MB468 | MP |
| >hg38_21_+_8433172_8446622_RNA45SN1_45S_50ntFlanks@6652.6693.42\|GACTCTTAGCGGTGGATCACTCGGCTCGTGCGTCGATGAAGA | >hg38_21_+_8433172_8446622_RNA45SN1_45S_50ntFlanks | 6652 | 6693 | 42 | GACTCTTAGCGGTGGATCACTCGGCTCGTGCGTCGATGAAGA | 534.8 | 5.9 | 0.9 | 6.3 | 2.5E-10 | 7.0E-10 | BT20/MB468 | CYTO |
| >Antisense_hg38_21_-_8433172_8446622_RNA45SN1_45S_50ntFlanks@7674.7692.19\|CTCTCTCTCTCTCTCCCCC | >Antisense_hg38_21_-_8433172_8446622_RNA45SN1_45S_50ntFlanks | 7674 | 7692 | 19 | CTCTCTCTCTCTCTCCCCC | 556.5 | -8.8 | 1.4 | -6.3 | 2.9E-10 | 7.0E-10 | MB231/MB468 | NUC |
| >hg38_21_+_8433172_8446622_RNA45SN1_45S_50ntFlanks@11027.11045.19\|CTCTCTCTCTCTCTCCCCC | >hg38_21_+_8433172_8446622_RNA45SN1_45S_50ntFlanks | 11027 | 11045 | 19 | CTCTCTCTCTCTCTCCCCC | 556.5 | -8.8 | 1.4 | -6.3 | 2.9E-10 | 7.0E-10 | MB231/MB468 | NUC |
| >hg38_21_+_8433172_8446622_RNA45SN1_45S_50ntFlanks@3802.3837.36\|CATTAAATCAGTTATGGTTCCTTTGGTCGCTCGCTC | >hg38_21_+_8433172_8446622_RNA45SN1_45S_50ntFlanks | 3802 | 3837 | 36 | CATTAAATCAGTTATGGTTCCTTTGGTCGCTCGCTC | 1203.3 | -9.6 | 1.5 | -6.3 | 3.4E-10 | 7.0E-10 | BT20/MB468 | NUC |
| >hg38_21_+_8433172_8446622_RNA45SN1_45S_50ntFlanks@4449.4471.23\|CCTCTCGGCGCCCCCTCGATGCT | >hg38_21_+_8433172_8446622_RNA45SN1_45S_50ntFlanks | 4449 | 4471 | 23 | CCTCTCGGCGCCCCCTCGATGCT | 2851.9 | -10.2 | 1.6 | -6.3 | 3.4E-10 | 7.0E-10 | BT20/MB468 | NUC |
| >hg38_21_+_8433172_8446622_RNA45SN1_45S_50ntFlanks@6779.6799.21\|CCGGGGCTACGCCTGTCTGAG | >hg38_21_+_8433172_8446622_RNA45SN1_45S_50ntFlanks | 6779 | 6799 | 21 | CCGGGGCTACGCCTGTCTGAG | 111.2 | -10.7 | 1.7 | -6.3 | 2.7E-10 | 7.0E-10 | MB231/MB468 | MP |
| >hg38_21_+_8433172_8446622_RNA45SN1_45S_50ntFlanks@10760.10785.26\|CATATCCGCAGCAGGTCTCCAAGGTG | >hg38_21_+_8433172_8446622_RNA45SN1_45S_50ntFlanks | 10760 | 10785 | 26 | CATATCCGCAGCAGGTCTCCAAGGTG | 134.9 | -9.2 | 1.4 | -6.4 | 2.1E-10 | 7.0E-10 | BT20/MB468 | MITO |
| >hg38_21_+_8433172_8446622_RNA45SN1_45S_50ntFlanks@6651.6678.28\|CGACTCTTAGCGGTGGATCACTCGGCTC | >hg38_21_+_8433172_8446622_RNA45SN1_45S_50ntFlanks | 6651 | 6678 | 28 | CGACTCTTAGCGGTGGATCACTCGGCTC | 4072.7 | 3.2 | 0.5 | 6.3 | 2.9E-10 | 7.0E-10 | MB231/MB468 | CYTO |
| >hg38_21_+_8433172_8446622_RNA45SN1_45S_50ntFlanks@11483.11502.20\|CGGAGGGGCCGCGGGCCGGT | >hg38_21_+_8433172_8446622_RNA45SN1_45S_50ntFlanks | 11483 | 11502 | 20 | CGGAGGGGCCGCGGGCCGGT | 99.6 | -10.5 | 1.7 | -6.3 | 2.7E-10 | 7.0E-10 | MB231/MB468 | MP |
| >hg38_21_+_8433172_8446622_RNA45SN1_45S_50ntFlanks@7979.8007.29\|ACCTCAGATCAGACGTGGCGACCCGCTGA | >hg38_21_+_8433172_8446622_RNA45SN1_45S_50ntFlanks | 7979 | 8007 | 29 | ACCTCAGATCAGACGTGGCGACCCGCTGA | 241.7 | -3.7 | 0.6 | -6.3 | 2.9E-10 | 7.0E-10 | MB231/MB468 | NUC |
| >hg38_21_+_8433172_8446622_RNA45SN1_45S_50ntFlanks@12627.12647.21\|ATGGGGCGAAGCTACCATCTG | >hg38_21_+_8433172_8446622_RNA45SN1_45S_50ntFlanks | 12627 | 12647 | 21 | ATGGGGCGAAGCTACCATCTG | 240.8 | -11.0 | 1.6 | -6.7 | 2.4E-11 | 7.0E-10 | BT20/MB468 | MP |
| >MI0000477\|hsa-mir-146a&WithFlank&5\|+\|160485346\|160485456@26.47.22&[MIMAT0000449&hsa-miR-146a-5p&offsets\|-1\|-1;m-129&5\|+\|160485372\|160485393&offsets\|-1\|-1]\|TTGAGAACTGAATTCCATGGGT | >MI0000477\|hsa-mir-146a&WithFlank&5\|+\|160485346\|160485456 | 26 | 47 | 22 | TTGAGAACTGAATTCCATGGGT | 436.8 | -12.2 | 1.8 | -6.9 | 5.1E-12 | 7.0E-10 | BT20/MB231 | MP |
| >hg38_21_+_8433172_8446622_RNA45SN1_45S_50ntFlanks@12633.12652.20\|CGAAGCTACCATCTGTGGGA | >hg38_21_+_8433172_8446622_RNA45SN1_45S_50ntFlanks | 12633 | 12652 | 20 | CGAAGCTACCATCTGTGGGA | 578.8 | -3.4 | 0.5 | -6.4 | 2.1E-10 | 7.0E-10 | BT20/MB468 | MITO |
| >hg38_21_+_8433172_8446622_RNA45SN1_45S_50ntFlanks@12222.12248.27\|ACAGAAACCTCCCGTGGAGCAGAAGGG | >hg38_21_+_8433172_8446622_RNA45SN1_45S_50ntFlanks | 12222 | 12248 | 27 | ACAGAAACCTCCCGTGGAGCAGAAGGG | 94.9 | -10.4 | 1.7 | -6.3 | 2.7E-10 | 7.0E-10 | MB231/MB468 | MP |
| >hg38_21_+_8433172_8446622_RNA45SN1_45S_50ntFlanks@12633.12653.21\|CGAAGCTACCATCTGTGGGAT | >hg38_21_+_8433172_8446622_RNA45SN1_45S_50ntFlanks | 12633 | 12653 | 21 | CGAAGCTACCATCTGTGGGAT | 2390.6 | -7.1 | 1.1 | -6.3 | 2.9E-10 | 7.0E-10 | MB231/MB468 | CYTO |
| >hg38_21_+_8433172_8446622_RNA45SN1_45S_50ntFlanks@10400.10424.25\|TGAACATGGGTCAGTCGGTCCTGAG | >hg38_21_+_8433172_8446622_RNA45SN1_45S_50ntFlanks | 10400 | 10424 | 25 | TGAACATGGGTCAGTCGGTCCTGAG | 1279.8 | 7.1 | 1.1 | 6.3 | 2.9E-10 | 7.0E-10 | MB231/MB468 | CYTO |
| >hg38_21_+_8433172_8446622_RNA45SN1_45S_50ntFlanks@10357.10390.34\|CTTTGAAGGCCGAAGTGGAGAAGGGTTCCATGTG | >hg38_21_+_8433172_8446622_RNA45SN1_45S_50ntFlanks | 10357 | 10390 | 34 | CTTTGAAGGCCGAAGTGGAGAAGGGTTCCATGTG | 647.1 | -5.0 | 0.8 | -6.3 | 2.9E-10 | 7.0E-10 | MB231/MB468 | NUC |

| | | | | | | | | | | | | | |
|---|---|---|---|---|---|---|---|---|---|---|---|---|---|
| >hg38_21_+_8433172_8446622_RNA45SN1_45S_50ntFlanks@9454.9495.42\|CCGCGCCGGGGAGGTGGAGCACGAGCGCACGTGTTAGGACCC | >hg38_21_+_8433172_8446622_RNA45SN1_45S_50ntFlanks | 9454 | 9495 | 42 | CCGCGCCGGGGAGGTGGAGCACGAGCGCACGTGTTAGGACCC | 130.6 | -10.9 | 1.7 | -6.3 | 2.8E-10 | 7.0E-10 | MB231/MB468 | MP |
| >MI0000087\|hsa-mir-29a&WithFlank&7\|-\|130876741\|130876816@47.67.21(+1C)&[MIMAT0000086&hsa-miR-29a-3p&offsets\|-1\|-2(+1C);m-26&7\|-\|130876749\|130876769&offsets\|-1\|-1(+1C)]\|CTAGCACCATCTGAAATCGGTC | >MI0000087\|hsa-mir-29a&WithFlank&7\|-\|130876741\|130876816 | 47 | 67 | 21(+1C) | CTAGCACCATCTGAAATCGGTC | 482.3 | 7.4 | 1.2 | 6.3 | 2.9E-10 | 7.1E-10 | MB231/MB468 | CYTO |
| >hg38_21_+_8433172_8446622_RNA45SN1_45S_50ntFlanks@9166.9197.32\|CCGGGCGGGTCGCGCCGTCGGGCCCGGGGGAG | >hg38_21_+_8433172_8446622_RNA45SN1_45S_50ntFlanks | 9166 | 9197 | 32 | CCGGGCGGGTCGCGCCGTCGGGCCCGGGGGAG | 1316.0 | -9.6 | 1.4 | -6.7 | 2.4E-11 | 7.1E-10 | BT20/MB468 | MP |
| >am_trna2_ArgCCT_16_+_3202901_3202973@14.31.18,trna3_ArgCCT_7_+_139025446_139025518@14.31.18\|AATGGATAAGGCATTGGC | >am_trna2_ArgCCT_16_+_3202901_3202973 | 14 | 31 | 18 | AATGGATAAGGCATTGGC | 175.0 | -4.6 | 0.7 | -6.3 | 2.9E-10 | 7.1E-10 | MB231/MB468 | CYTO |
| >hg38_21_+_8433172_8446622_RNA45SN1_45S_50ntFlanks@11686.11717.32\|TCAAAGTGAAGAAATTCAATGAAGCGCGGGTA | >hg38_21_+_8433172_8446622_RNA45SN1_45S_50ntFlanks | 11686 | 11717 | 32 | TCAAAGTGAAGAAATTCAATGAAGCGCGGGTA | 248.1 | -7.3 | 1.2 | -6.3 | 2.9E-10 | 7.1E-10 | MB231/MB468 | CYTO |
| >hg38_MT_+_598_1651_RNR1_12S_50ntFlanks@447.480.34\|TGAACACACAATAGCTAAGACCCAAACTGGGATT | >hg38_MT_+_598_1651_RNR1_12S_50ntFlanks | 447 | 480 | 34 | TGAACACACAATAGCTAAGACCCAAACTGGGATT | 241.9 | 11.1 | 1.8 | 6.3 | 3.2E-10 | 7.1E-10 | MB231/MB468 | MITO |
| >MI0000088\|hsa-mir-30a&WithFlank&6\|-\|71403545\|71403627@54.73.20(+1A)&[MIMAT0000088&hsa-miR-30a-3p&offsets\|+1\|-1(+1A);m-38&6\|-\|71403554\|71403575&offsets\|+1\|-1(+1A)]\|TTTCAGTCGGATGTTTGCAGA | >MI0000088\|hsa-mir-30a&WithFlank&6\|-\|71403545\|71403627 | 54 | 73 | 20(+1A) | TTTCAGTCGGATGTTTGCAGA | 194.3 | 10.8 | 1.7 | 6.3 | 3.3E-10 | 7.1E-10 | MB231/MB468 | MITO |
| >hg38_21_+_8433172_8446622_RNA45SN1_45S_50ntFlanks@10349.10382.34\|AACGAGAACTTTGAAGGCCGAAGTGGAGAAGGGT | >hg38_21_+_8433172_8446622_RNA45SN1_45S_50ntFlanks | 10349 | 10382 | 34 | AACGAGAACTTTGAAGGCCGAAGTGGAGAAGGGT | 1338.7 | 2.6 | 0.4 | 6.3 | 2.6E-10 | 7.1E-10 | BT20/MB468 | CYTO |
| >hg38_1_-_228634819_228635039_RNA5S12_5S_50ntFlanks@86.103.18\|CGTCTGATCTCGGAAGCT | >hg38_1_-_228634819_228635039_RNA5S12_5S_50ntFlanks | 86 | 103 | 18 | CGTCTGATCTCGGAAGCT | 90.6 | -10.4 | 1.6 | -6.3 | 2.8E-10 | 7.1E-10 | MB231/MB468 | MP |
| >hg38_21_+_8433172_8446622_RNA45SN1_45S_50ntFlanks@4385.4404.20\|TTGGATCTTGGGAGCGGGCG | >hg38_21_+_8433172_8446622_RNA45SN1_45S_50ntFlanks | 4385 | 4404 | 20 | TTGGATCTTGGGAGCGGGCG | 92.0 | -10.4 | 1.6 | -6.3 | 2.8E-10 | 7.1E-10 | MB231/MB468 | MP |
| >MI0000542\|hsa-mir-320a&WithFlank&8\|-\|22244960\|22245043@48.70.23&[MIMAT0000510&hsa-miR-320a-3p&offsets\|0\|+1;m-60&8\|-\|22244975\|22244996&offsets\|0\|+1]\|AAAAGCTGGGTTGAGAGGGCGAA | >MI0000542\|hsa-mir-320a&WithFlank&8\|-\|22244960\|22245043 | 48 | 70 | 23 | AAAAGCTGGGTTGAGAGGGCGAA | 2857.9 | 4.7 | 0.7 | 6.3 | 2.9E-10 | 7.1E-10 | MB231/MB468 | NUC |
| >hg38_21_+_8433172_8446622_RNA45SN1_45S_50ntFlanks@13006.13025.20\|CTATTGAAAGTCAGCCCTCG | >hg38_21_+_8433172_8446622_RNA45SN1_45S_50ntFlanks | 13006 | 13025 | 20 | CTATTGAAAGTCAGCCCTCG | 1014.2 | -8.4 | 1.3 | -6.3 | 2.6E-10 | 7.1E-10 | BT20/MB468 | CYTO |
| >MI0000440\|hsa-mir-27b&WithFlank&9\|+\|95085439\|95085547@67.88.22&[MIMAT0000419&hsa-miR-27b-3p&offsets\|0\|+1;m-39&9\|+\|95085505\|95085525&offsets\|0\|+1]\|TTCACAGTGGCTAAGTTCTGCA | >MI0000440\|hsa-mir-27b&WithFlank&9\|+\|95085439\|95085547 | 67 | 88 | 22 | TTCACAGTGGCTAAGTTCTGCA | 3797.6 | 5.6 | 0.9 | 6.3 | 3.3E-10 | 7.1E-10 | MB231/MB468 | MITO |
| >am_trna119_LysCTT_1_-_145395522_145395594@7.34.28,trna11_LysCTT_5_-_180648979_180649051@7.34.28,trna128_LysCTT_1_-_55423542_55423614@7.34.28,trna13_LysCTT_6_+_26556774_26556846@7.34.28,trna6_LysCTT_19_-_52425396_52425468@7.34.28,trna7_LysCTT_16_+_3225692_3225764@7.34.28,trna9_LysCTT_5_+_180634755_180634827@7.34.28\|CTAGCTCAGTCGGTAGAGCATGAGACTC | >am_trna119_LysCTT_1_-_145395522_145395594 | 7 | 34 | 28 | CTAGCTCAGTCGGTAGAGCATGAGACTC | 322.5 | -6.0 | 0.9 | -6.7 | 2.6E-11 | 7.1E-10 | BT20/MB231 | MITO |
| >hg38_21_+_8433172_8446622_RNA45SN1_45S_50ntFlanks@8191.8208.18\|CAGCCCGTGGACGGTGTG | >hg38_21_+_8433172_8446622_RNA45SN1_45S_50ntFlanks | 8191 | 8208 | 18 | CAGCCCGTGGACGGTGTG | 120.6 | -10.8 | 1.7 | -6.3 | 2.8E-10 | 7.2E-10 | MB231/MB468 | MP |
| >MI0000067\|hsa-let-7f-1&WithFlank&9\|+\|94176341\|94176439@13.33.21(+1G);MI0000068\|hsa-let-7f-2&WithFlank&X\|-\|53557186\|53557280@14.34.21(+1G)&[MIMAT0000067&hsa-let-7f-5p&offsets\|0\|-1(+1G);m-20&9\|+\|94176353\|94176374&offsets\|0\|-1(+1G)];[MIMAT0000067_1&hsa-let-7f-5p&offsets\|0\|-1(+1G);m-19&X\|-\|53557246\|53557267&offsets\|0\|-1(+1G)]\|TGAGGTAGTAGATTGTATAGTG | >MI0000067\|hsa-let-7f-1&WithFlank&9\|+\|94176341\|94176439 | 13 | 33 | 21(+1G) | TGAGGTAGTAGATTGTATAGTG | 417.7 | 5.4 | 0.8 | 6.3 | 2.2E-10 | 7.2E-10 | BT20/MB468 | MITO |
| >hg38_21_+_8433172_8446622_RNA45SN1_45S_50ntFlanks@12633.12653.21\|CGAAGCTACCATCTGTGGGAT | >hg38_21_+_8433172_8446622_RNA45SN1_45S_50ntFlanks | 12633 | 12653 | 21 | CGAAGCTACCATCTGTGGGAT | 14186.4 | -9.1 | 1.4 | -6.7 | 2.5E-11 | 7.2E-10 | BT20/MB468 | MP |
| >hg38_21_+_8433172_8446622_RNA45SN1_45S_50ntFlanks@4731.4757.27\|AAGAACGAAAGTCGGAGGTTCGAAGAC | >hg38_21_+_8433172_8446622_RNA45SN1_45S_50ntFlanks | 4731 | 4757 | 27 | AAGAACGAAAGTCGGAGGTTCGAAGAC | 887.1 | 4.3 | 0.7 | 6.3 | 2.6E-10 | 7.2E-10 | BT20/MB468 | CYTO |
| >hg38_21_+_8433172_8446622_RNA45SN1_45S_50ntFlanks@10840.10868.29\|CCGTAACTTCGGGATAAGGATTGGCTCTA | >hg38_21_+_8433172_8446622_RNA45SN1_45S_50ntFlanks | 10840 | 10868 | 29 | CCGTAACTTCGGGATAAGGATTGGCTCTA | 617.7 | -8.4 | 1.3 | -6.3 | 2.8E-10 | 7.2E-10 | MB231/MB468 | MP |
| >hg38_21_+_8433172_8446622_RNA45SN1_45S_50ntFlanks@11236.11254.19\|TTCCCGTGGATCGCCCCAG | >hg38_21_+_8433172_8446622_RNA45SN1_45S_50ntFlanks | 11236 | 11254 | 19 | TTCCCGTGGATCGCCCCAG | 80.9 | -10.2 | 1.6 | -6.3 | 2.8E-10 | 7.2E-10 | MB231/MB468 | MP |
| >MI0001446\|hsa-mir-424&WithFlank&X\|-\|134546608\|134546717@54.75.22&[MIMAT0004749&hsa-miR-424-3p&offsets\|0\|+1;m-305&X\|-\|134546644\|134546664&offsets\|0\|+1]\|CAAAACGTGAGGCGCTGCTATA | >MI0001446\|hsa-mir-424&WithFlank&X\|-\|134546608\|134546717 | 54 | 75 | 22 | CAAAACGTGAGGCGCTGCTATA | 207.4 | 7.7 | 1.2 | 6.3 | 3.0E-10 | 7.2E-10 | MB231/MB468 | NUC |
| >hg38_21_+_8433172_8446622_RNA45SN1_45S_50ntFlanks@9029.9066.38\|TCTCCTCGTGGGGGGGCCGGGCCACCCCTCCCACGGCG | >hg38_21_+_8433172_8446622_RNA45SN1_45S_50ntFlanks | 9029 | 9066 | 38 | TCTCCTCGTGGGGGGGCCGGGCCACCCCTCCCACGGCG | 68.7 | -10.0 | 1.6 | -6.3 | 2.8E-10 | 7.2E-10 | MB231/MB468 | MP |
| >MI0000234\|hsa-mir-192&WithFlank&11\|-\|64891131\|64891252@30.50.21&[MIMAT0000222&hsa-miR-192-5p&offsets\|0\|0;m-59&11\|-\|64891203\|64891223&offsets\|0\|0]\|CTGACCTATGAATTGACAGCC | >MI0000234\|hsa-mir-192&WithFlank&11\|-\|64891131\|64891252 | 30 | 50 | 21 | CTGACCTATGAATTGACAGCC | 903.4 | 4.4 | 0.7 | 6.3 | 3.0E-10 | 7.2E-10 | MB231/MB468 | NUC |
| >MI0000440\|hsa-mir-27b&WithFlank&9\|+\|95085439\|95085547@67.85.19(+1A)&[MIMAT0000419&hsa-miR-27b-3p&offsets\|0\|-2(+1A);m-39&9\|+\|95085505\|95085525&offsets\|0\|-2(+1A)]\|TTCACAGTGGCTAAGTTCTA | >MI0000440\|hsa-mir-27b&WithFlank&9\|+\|95085439\|95085547 | 67 | 85 | 19(+1A) | TTCACAGTGGCTAAGTTCTA | 571.9 | 6.3 | 1.0 | 6.3 | 3.3E-10 | 7.2E-10 | MB231/MB468 | MITO |
| >hg38_21_+_8433172_8446622_RNA45SN1_45S_50ntFlanks@4759.4784.26\|ATCAGATACCGTCGTAGTTCCGACCA | >hg38_21_+_8433172_8446622_RNA45SN1_45S_50ntFlanks | 4759 | 4784 | 26 | ATCAGATACCGTCGTAGTTCCGACCA | 175.2 | -10.5 | 1.6 | -6.7 | 2.5E-11 | 7.2E-10 | BT20/MB468 | MP |
| >hg38_21_+_8433172_8446622_RNA45SN1_45S_50ntFlanks@9580.9597.18\|TCGTCCGACCTGGGTATA | >hg38_21_+_8433172_8446622_RNA45SN1_45S_50ntFlanks | 9580 | 9597 | 18 | TCGTCCGACCTGGGTATA | 126.8 | -9.1 | 1.4 | -6.3 | 2.2E-10 | 7.2E-10 | BT20/MB468 | MITO |
| >hg38_21_+_8433172_8446622_RNA45SN1_45S_50ntFlanks@4159.4178.20\|ACCACATCCAAGGAAGGCAG | >hg38_21_+_8433172_8446622_RNA45SN1_45S_50ntFlanks | 4159 | 4178 | 20 | ACCACATCCAAGGAAGGCAG | 96.9 | -10.5 | 1.7 | -6.3 | 2.8E-10 | 7.2E-10 | MB231/MB468 | MP |
| >hg38_21_+_8433172_8446622_RNA45SN1_45S_50ntFlanks@6652.6684.33\|GACTCTTAGCGGTGGATCACTCGGCTCGTGCGT | >hg38_21_+_8433172_8446622_RNA45SN1_45S_50ntFlanks | 6652 | 6684 | 33 | GACTCTTAGCGGTGGATCACTCGGCTCGTGCGT | 2341.4 | 4.9 | 0.8 | 6.3 | 3.0E-10 | 7.2E-10 | MB231/MB468 | NUC |
| >hg38_21_+_8433172_8446622_RNA45SN1_45S_50ntFlanks@9565.9597.33\|TGACGTGCAAATCGGTCGTCCGACCTGGGTATA | >hg38_21_+_8433172_8446622_RNA45SN1_45S_50ntFlanks | 9565 | 9597 | 33 | TGACGTGCAAATCGGTCGTCCGACCTGGGTATA | 1404.1 | -7.1 | 1.1 | -6.3 | 2.8E-10 | 7.2E-10 | MB231/MB468 | MP |
| >hg38_21_+_8433172_8446622_RNA45SN1_45S_50ntFlanks@10351.10373.23\|CGAGAACTTTGAAGGCCGAAGTG | >hg38_21_+_8433172_8446622_RNA45SN1_45S_50ntFlanks | 10351 | 10373 | 23 | CGAGAACTTTGAAGGCCGAAGTG | 3041.9 | -7.9 | 1.3 | -6.3 | 2.8E-10 | 7.2E-10 | MB231/MB468 | MP |
| >hg38_21_+_8433172_8446622_RNA45SN1_45S_50ntFlanks@4732.4761.30\|AGAACGAAAGTCGGAGGTTCGAAGACGATC | >hg38_21_+_8433172_8446622_RNA45SN1_45S_50ntFlanks | 4732 | 4761 | 30 | AGAACGAAAGTCGGAGGTTCGAAGACGATC | 1938.9 | 5.8 | 0.9 | 6.3 | 2.6E-10 | 7.3E-10 | BT20/MB468 | CYTO |
| >hg38_21_+_8433172_8446622_RNA45SN1_45S_50ntFlanks@12936.12959.24\|AAACCATTCGTAGACGACCTGCTT | >hg38_21_+_8433172_8446622_RNA45SN1_45S_50ntFlanks | 12936 | 12959 | 24 | AAACCATTCGTAGACGACCTGCTT | 1076.3 | -4.3 | 0.7 | -6.3 | 3.0E-10 | 7.3E-10 | MB231/MB468 | NUC |
| >hg38_21_+_8433172_8446622_RNA45SN1_45S_50ntFlanks@8176.8214.39\|CTTCTGATCGAGGCCCAGCCCGTGGACGGTGTGAGGCCG | >hg38_21_+_8433172_8446622_RNA45SN1_45S_50ntFlanks | 8176 | 8214 | 39 | CTTCTGATCGAGGCCCAGCCCGTGGACGGTGTGAGGCCG | 2627.3 | -8.2 | 1.3 | -6.3 | 2.8E-10 | 7.3E-10 | MB231/MB468 | MP |
| >hg38_21_+_8433172_8446622_RNA45SN1_45S_50ntFlanks@11571.11601.31\|GCCGACTTAGAACTGGTGCGGACCAGGGGAA | >hg38_21_+_8433172_8446622_RNA45SN1_45S_50ntFlanks | 11571 | 11601 | 31 | GCCGACTTAGAACTGGTGCGGACCAGGGGAA | 1351.4 | -3.7 | 0.6 | -6.3 | 3.0E-10 | 7.3E-10 | MB231/MB468 | NUC |
| >hg38_21_+_8433172_8446622_RNA45SN1_45S_50ntFlanks@8353.8390.38\|GTTGAAAAGAACTTTGAAGAGAGAGTTCAAGAGGGCGT | >hg38_21_+_8433172_8446622_RNA45SN1_45S_50ntFlanks | 8353 | 8390 | 38 | GTTGAAAAGAACTTTGAAGAGAGAGTTCAAGAGGGCGT | 141.9 | 10.9 | 1.6 | 6.7 | 2.5E-11 | 7.3E-10 | BT20/MB468 | MP |
| >hg38_21_+_8433172_8446622_RNA45SN1_45S_50ntFlanks@5473.5501.29\|CGGCCCTGGCGGAGCGCTGAGAAGACGGT | >hg38_21_+_8433172_8446622_RNA45SN1_45S_50ntFlanks | 5473 | 5501 | 29 | CGGCCCTGGCGGAGCGCTGAGAAGACGGT | 1645.6 | 6.5 | 1.0 | 6.3 | 3.0E-10 | 7.3E-10 | MB231/MB468 | CYTO |
| >hg38_21_+_8433172_8446622_RNA45SN1_45S_50ntFlanks@4731.4766.36\|AAGAACGAAAGTCGGAGGTTCGAAGACGATCAGATA | >hg38_21_+_8433172_8446622_RNA45SN1_45S_50ntFlanks | 4731 | 4766 | 36 | AAGAACGAAAGTCGGAGGTTCGAAGACGATCAGATA | 1517.8 | 6.9 | 1.1 | 6.3 | 3.0E-10 | 7.3E-10 | MB231/MB468 | CYTO |
| >hg38_21_+_8433172_8446622_RNA45SN1_45S_50ntFlanks@7988.8015.28\|CAGACGTGGCGACCCGCTGAATTTAAGC | >hg38_21_+_8433172_8446622_RNA45SN1_45S_50ntFlanks | 7988 | 8015 | 28 | CAGACGTGGCGACCCGCTGAATTTAAGC | 299.0 | -7.8 | 1.2 | -6.3 | 2.9E-10 | 7.3E-10 | MB231/MB468 | MP |
| >hg38_21_+_8433172_8446622_RNA45SN1_45S_50ntFlanks@12632.12654.23\|GCGAAGCTACCATCTGTGGGATT | >hg38_21_+_8433172_8446622_RNA45SN1_45S_50ntFlanks | 12632 | 12654 | 23 | GCGAAGCTACCATCTGTGGGATT | 887.4 | -5.7 | 0.9 | -6.3 | 3.0E-10 | 7.3E-10 | MB231/MB468 | CYTO |
| >am_trna2_GlyGCC_21_-_18827107_18827177@1.25.25,trna35_GlyGCC_1_+_161413094_161413164@1.25.25,trna37_GlyGCC_1_+_161420467_161420537@1.25.25,trna39_GlyGCC_1_+_161427898_161427968@1.25.25,trna41_GlyGCC_1_+_161435258_161435328@1.25.25\|GCATGGGTGGTTCAGTGGTAGAATT | >am_trna2_GlyGCC_21_-_18827107_18827177 | 1 | 25 | 25 | GCATGGGTGGTTCAGTGGTAGAATT | 2595.4 | 3.9 | 0.6 | 6.3 | 2.7E-10 | 7.3E-10 | BT20/MB468 | CYTO |
| >MI0000456\|hsa-mir-140&WithFlank&16\|+\|69933075\|69933186@69.87.19(+1A)&[MIMAT0004597&hsa-miR-140-3p&offsets\|+1\|-1(+1A);m-45&16\|+\|69933143\|69933164&offsets\|0\|-3(+1A)]\|ACCACAGGGTAGAACCACGA | >MI0000456\|hsa-mir-140&WithFlank&16\|+\|69933075\|69933186 | 69 | 87 | 19(+1A) | ACCACAGGGTAGAACCACGA | 98.0 | 9.8 | 1.6 | 6.3 | 2.9E-10 | 7.3E-10 | MB231/MB468 | MP |
| >trna17_ValTAC_11_-_59318102_59318174@1.30.30,trna4_ValTAC_X_-_18693029_18693101@1.30.30\|GGTTCCATAGTGTAGTGGTTATCACGTCTG | >trna17_ValTAC_11_-_59318102_59318174 | 1 | 30 | 30 | GGTTCCATAGTGTAGTGGTTATCACGTCTG | 240.6 | -8.1 | 1.3 | -6.3 | 3.0E-10 | 7.3E-10 | MB231/MB468 | NUC |
| >hg38_21_+_8433172_8446622_RNA45SN1_45S_50ntFlanks@6719.6752.34\|TGAATTGCAGGACACATTGATCATCGACACTTCG | >hg38_21_+_8433172_8446622_RNA45SN1_45S_50ntFlanks | 6719 | 6752 | 34 | TGAATTGCAGGACACATTGATCATCGACACTTCG | 546.2 | -8.7 | 1.4 | -6.3 | 2.9E-10 | 7.3E-10 | MB231/MB468 | MP |
| >hg38_21_+_8433172_8446622_RNA45SN1_45S_50ntFlanks@11641.11674.34\|CGGCGGGTGTTGACGCGATGTGATTTCTGCCCAG | >hg38_21_+_8433172_8446622_RNA45SN1_45S_50ntFlanks | 11641 | 11674 | 34 | CGGCGGGTGTTGACGCGATGTGATTTCTGCCCAG | 271.8 | -11.2 | 1.7 | -6.7 | 2.6E-11 | 7.3E-10 | BT20/MB468 | MP |
| >hg38_21_+_8433172_8446622_RNA45SN1_45S_50ntFlanks@9352.9374.23\|CCGTGGCGCAATGAAGGTGAAGG | >hg38_21_+_8433172_8446622_RNA45SN1_45S_50ntFlanks | 9352 | 9374 | 23 | CCGTGGCGCAATGAAGGTGAAGG | 89.0 | -10.3 | 1.6 | -6.3 | 2.9E-10 | 7.3E-10 | MB231/MB468 | MP |
| >trna11_GluTTC_15_-_26327381_26327452@1.33.33,trna3_GluTTC_13_-_45492062_45492133@1.33.33\|TCCCACATGGTCTAGCGGTTAGGATTCCTGGTT | >trna11_GluTTC_15_-_26327381_26327452 | 1 | 33 | 33 | TCCCACATGGTCTAGCGGTTAGGATTCCTGGTT | 228.2 | 3.6 | 0.6 | 6.3 | 3.5E-10 | 7.3E-10 | BT20/MB468 | NUC |
| >hg38_21_+_8433172_8446622_RNA45SN1_45S_50ntFlanks@9738.9757.20\|GAAACGATCTCAACCTATTC | >hg38_21_+_8433172_8446622_RNA45SN1_45S_50ntFlanks | 9738 | 9757 | 20 | GAAACGATCTCAACCTATTC | 231.5 | -9.9 | 1.6 | -6.3 | 2.2E-10 | 7.3E-10 | BT20/MB468 | MITO |

| | | | | | | | | | | | | | |
|---|---|---|---|---|---|---|---|---|---|---|---|---|---|
| >MI0000067\|hsa-let-7f-1&WithFlank&9\|+\|94176341\|94176439@13.33.21;MI0000068\|hsa-let-7f-2&WithFlank&X\|-\|53557186\|53557280@14.34.21&[MIMAT0000067&hsa-let-7f-5p&offsets\|0\|-1;m-20&9\|+\|94176353\|94176374&offsets\|0\|-1];[MIMAT0000067_1&hsa-let-7f-5p&offsets\|0\|-1;m-19&X\|-\|53557246\|53557267&offsets\|0\|-1]\|TGAGGTAGTAGATTGTATAGT | >MI0000067\|hsa-let-7f-1&WithFlank&9\|+\|94176341\|94176439 | 13 | 33 | 21 | TGAGGTAGTAGATTGTATAGT | 4757.7 | 5.3 | 0.8 | 6.3 | 3.4E-10 | 7.3E-10 | MB231/MB468 | MITO |
| >hg38_21_+_8433172_8446622_RNA45SN1_45S_50ntFlanks@9604.9621.18\|AAAGACTAATCGAACCAT | >hg38_21_+_8433172_8446622_RNA45SN1_45S_50ntFlanks | 9604 | 9621 | 18 | AAAGACTAATCGAACCAT | 321.4 | -3.5 | 0.6 | -6.3 | 3.0E-10 | 7.4E-10 | MB231/MB468 | CYTO |
| >hg38_21_+_8433172_8446622_RNA45SN1_45S_50ntFlanks@5489.5526.38\|CTGAGAAGACGGTCGAACTTGACTATCTAGAGGAAGTA | >hg38_21_+_8433172_8446622_RNA45SN1_45S_50ntFlanks | 5489 | 5526 | 38 | CTGAGAAGACGGTCGAACTTGACTATCTAGAGGAAGTA | 245.4 | -7.7 | 1.2 | -6.3 | 3.4E-10 | 7.4E-10 | MB231/MB468 | MITO |
| >hg38_21_+_8433172_8446622_RNA45SN1_45S_50ntFlanks@10107.10132.26\|CGTGTGGTGTGCGTCGGAGGGCGGCG | >hg38_21_+_8433172_8446622_RNA45SN1_45S_50ntFlanks | 10107 | 10132 | 26 | CGTGTGGTGTGCGTCGGAGGGCGGCG | 75.7 | -10.1 | 1.6 | -6.3 | 2.9E-10 | 7.4E-10 | MB231/MB468 | MP |
| >hg38_21_+_8433172_8446622_RNA45SN1_45S_50ntFlanks@11569.11599.31\|CAGCCGACTTAGAACTGGTGCGGACCAGGGG | >hg38_21_+_8433172_8446622_RNA45SN1_45S_50ntFlanks | 11569 | 11599 | 31 | CAGCCGACTTAGAACTGGTGCGGACCAGGGG | 526.7 | -8.2 | 1.3 | -6.3 | 2.9E-10 | 7.4E-10 | MB231/MB468 | MP |
| >hg38_21_+_8433172_8446622_RNA45SN1_45S_50ntFlanks@12633.12661.29\|CGAAGCTACCATCTGTGGGATTATGACTG | >hg38_21_+_8433172_8446622_RNA45SN1_45S_50ntFlanks | 12633 | 12661 | 29 | CGAAGCTACCATCTGTGGGATTATGACTG | 480.7 | -3.3 | 0.5 | -6.3 | 2.7E-10 | 7.4E-10 | BT20/MB468 | CYTO |
| >hg38_21_+_8433172_8446622_RNA45SN1_45S_50ntFlanks@4472.4495.24\|CTTAGCTGAGTGTCCCGCGGGGCC | >hg38_21_+_8433172_8446622_RNA45SN1_45S_50ntFlanks | 4472 | 4495 | 24 | CTTAGCTGAGTGTCCCGCGGGGCC | 86.1 | -10.3 | 1.6 | -6.3 | 2.9E-10 | 7.4E-10 | MB231/MB468 | MP |
| >hg38_21_+_8433172_8446622_RNA45SN1_45S_50ntFlanks@6721.6741.21\|AATTGCAGGACACATTGATCA | >hg38_21_+_8433172_8446622_RNA45SN1_45S_50ntFlanks | 6721 | 6741 | 21 | AATTGCAGGACACATTGATCA | 302.3 | -7.9 | 1.3 | -6.3 | 3.4E-10 | 7.4E-10 | MB231/MB468 | MITO |
| >hg38_21_+_8433172_8446622_RNA45SN1_45S_50ntFlanks@8182.8223.42\|ATCGAGGCCCAGCCCGTGGACGGTGTGAGGCCGGTAGCGGCC | >hg38_21_+_8433172_8446622_RNA45SN1_45S_50ntFlanks | 8182 | 8223 | 42 | ATCGAGGCCCAGCCCGTGGACGGTGTGAGGCCGGTAGCGGCC | 136.6 | -9.2 | 1.4 | -6.3 | 2.3E-10 | 7.4E-10 | BT20/MB468 | MITO |
| >MI0000298\|hsa-mir-221&WithFlank&X\|-\|45746151\|45746272@71.91.21(+1A)&[MIMAT0000278&hsa-miR-221-3p&offsets\|0\|-2(+1A);m-91&X\|-\|45746181\|45746202&offsets\|0\|-1(+1A)]\|AGCTACATTGTCTGCTGGGTTA | >MI0000298\|hsa-mir-221&WithFlank&X\|-\|45746151\|45746272 | 71 | 91 | 21(+1A) | AGCTACATTGTCTGCTGGGTTA | 1655.0 | 5.0 | 0.8 | 6.3 | 2.9E-10 | 7.4E-10 | MB231/MB468 | MP |
| >hg38_21_+_8433172_8446622_RNA45SN1_45S_50ntFlanks@9027.9064.38\|CGTCTCCTCGTGGGGGGGCCGGGCCACCCCTCCCACGG | >hg38_21_+_8433172_8446622_RNA45SN1_45S_50ntFlanks | 9027 | 9064 | 38 | CGTCTCCTCGTGGGGGGGCCGGGCCACCCCTCCCACGG | 467.4 | -8.7 | 1.4 | -6.3 | 3.6E-10 | 7.4E-10 | BT20/MB468 | NUC |
| >hg38_21_+_8433172_8446622_RNA45SN1_45S_50ntFlanks@11561.11580.20\|GCGCCTAGCAGCCGACTTAG | >hg38_21_+_8433172_8446622_RNA45SN1_45S_50ntFlanks | 11561 | 11580 | 20 | GCGCCTAGCAGCCGACTTAG | 147.5 | -9.3 | 1.5 | -6.3 | 2.3E-10 | 7.4E-10 | BT20/MB468 | MITO |
| >hg38_21_+_8433172_8446622_RNA45SN1_45S_50ntFlanks@7978.8009.32\|GACCTCAGATCAGACGTGGCGACCCGCTGAAT | >hg38_21_+_8433172_8446622_RNA45SN1_45S_50ntFlanks | 7978 | 8009 | 32 | GACCTCAGATCAGACGTGGCGACCCGCTGAAT | 475.0 | 7.5 | 1.2 | 6.3 | 3.0E-10 | 7.4E-10 | MB231/MB468 | CYTO |
| >hg38_21_+_8433172_8446622_RNA45SN1_45S_50ntFlanks@9534.9552.19\|AGGAAACTCTGGTGGAGGT | >hg38_21_+_8433172_8446622_RNA45SN1_45S_50ntFlanks | 9534 | 9552 | 19 | AGGAAACTCTGGTGGAGGT | 2924.1 | -4.6 | 0.7 | -6.3 | 2.7E-10 | 7.4E-10 | BT20/MB468 | CYTO |
| >MI0000065\|hsa-let-7d&WithFlank&9\|+\|94178828\|94178926@14.35.22&[MIMAT0000065&hsa-let-7d-5p&offsets\|0\|0;m-90&9\|+\|94178841\|94178862&offsets\|0\|0]\|AGAGGTAGTAGGTTGCATAGTT | >MI0000065\|hsa-let-7d&WithFlank&9\|+\|94178828\|94178926 | 14 | 35 | 22 | AGAGGTAGTAGGTTGCATAGTT | 7386.2 | 4.4 | 0.7 | 6.3 | 3.1E-10 | 7.4E-10 | MB231/MB468 | NUC |
| >hg38_21_+_8433172_8446622_RNA45SN1_45S_50ntFlanks@8154.8178.25\|CCGCTCGTGGGGGGCCCAAGTCCTT | >hg38_21_+_8433172_8446622_RNA45SN1_45S_50ntFlanks | 8154 | 8178 | 25 | CCGCTCGTGGGGGGCCCAAGTCCTT | 297.2 | -11.3 | 1.7 | -6.7 | 2.6E-11 | 7.4E-10 | BT20/MB468 | MP |
| >hg38_21_+_8433172_8446622_RNA45SN1_45S_50ntFlanks@4367.4391.25\|CAGTTAAAAAGCTCGTAGTTGGATC | >hg38_21_+_8433172_8446622_RNA45SN1_45S_50ntFlanks | 4367 | 4391 | 25 | CAGTTAAAAAGCTCGTAGTTGGATC | 2517.5 | -10.5 | 1.7 | -6.3 | 3.6E-10 | 7.4E-10 | BT20/MB468 | NUC |
| >hg38_21_+_8433172_8446622_RNA45SN1_45S_50ntFlanks@12510.12546.37\|TCGTGAGACAGGTTAGTTTTACCCTACTGATGATGTG | >hg38_21_+_8433172_8446622_RNA45SN1_45S_50ntFlanks | 12510 | 12546 | 37 | TCGTGAGACAGGTTAGTTTTACCCTACTGATGATGTG | 83.5 | -10.2 | 1.6 | -6.3 | 2.9E-10 | 7.4E-10 | MB231/MB468 | MP |
| >hg38_21_+_8433172_8446622_RNA45SN1_45S_50ntFlanks@8443.8467.25\|CAACCCGGCGGCGGGTCCGGCCGTG | >hg38_21_+_8433172_8446622_RNA45SN1_45S_50ntFlanks | 8443 | 8467 | 25 | CAACCCGGCGGCGGGTCCGGCCGTG | 69.7 | -10.0 | 1.6 | -6.3 | 2.9E-10 | 7.5E-10 | MB231/MB468 | MP |
| >hg38_21_+_8433172_8446622_RNA45SN1_45S_50ntFlanks@12945.12972.28\|GTAGACGACCTGCTTCTGGGTCGGGGTT | >hg38_21_+_8433172_8446622_RNA45SN1_45S_50ntFlanks | 12945 | 12972 | 28 | GTAGACGACCTGCTTCTGGGTCGGGGTT | 72.2 | -10.0 | 1.6 | -6.3 | 2.9E-10 | 7.5E-10 | MB231/MB468 | MP |
| >hg38_21_+_8433172_8446622_RNA45SN1_45S_50ntFlanks@5070.5098.29\|GTTAATTCCGATAACGAACGAGACTCTGG | >hg38_21_+_8433172_8446622_RNA45SN1_45S_50ntFlanks | 5070 | 5098 | 29 | GTTAATTCCGATAACGAACGAGACTCTGG | 392.4 | -12.5 | 2.0 | -6.3 | 2.9E-10 | 7.5E-10 | MB231/MB468 | MP |
| >hg38_21_+_8433172_8446622_RNA45SN1_45S_50ntFlanks@5445.5463.19\|TCGGATCGGCCCCGCCGGG | >hg38_21_+_8433172_8446622_RNA45SN1_45S_50ntFlanks | 5445 | 5463 | 19 | TCGGATCGGCCCCGCCGGG | 78.9 | -10.2 | 1.6 | -6.3 | 2.9E-10 | 7.5E-10 | MB231/MB468 | MP |
| >hg38_21_+_8433172_8446622_RNA45SN1_45S_50ntFlanks@4005.4024.20\|AACCTCGGGCCGATCGCACG | >hg38_21_+_8433172_8446622_RNA45SN1_45S_50ntFlanks | 4005 | 4024 | 20 | AACCTCGGGCCGATCGCACG | 1006.3 | -3.8 | 0.6 | -6.3 | 3.1E-10 | 7.5E-10 | MB231/MB468 | NUC |
| >hg38_21_+_8433172_8446622_RNA45SN1_45S_50ntFlanks@10781.10818.38\|AGGTGAACAGCCTCTGGCATGTTGGAACAATGTAGGTA | >hg38_21_+_8433172_8446622_RNA45SN1_45S_50ntFlanks | 10781 | 10818 | 38 | AGGTGAACAGCCTCTGGCATGTTGGAACAATGTAGGTA | 2166.2 | -4.8 | 0.8 | -6.3 | 2.7E-10 | 7.5E-10 | BT20/MB468 | CYTO |
| >hg38_21_+_8433172_8446622_RNA45SN1_45S_50ntFlanks@11920.11958.39\|ATGAGAGGTGTAGAATAAGTGGGAGGCCCCCGGCGCCCC | >hg38_21_+_8433172_8446622_RNA45SN1_45S_50ntFlanks | 11920 | 11958 | 39 | ATGAGAGGTGTAGAATAAGTGGGAGGCCCCCGGCGCCCC | 122.7 | -10.8 | 1.7 | -6.3 | 3.0E-10 | 7.5E-10 | MB231/MB468 | MP |
| >hg38_21_+_8433172_8446622_RNA45SN1_45S_50ntFlanks@11161.11181.21\|CACGGTCCCCCGCGAGGGGGG | >hg38_21_+_8433172_8446622_RNA45SN1_45S_50ntFlanks | 11161 | 11181 | 21 | CACGGTCCCCCGCGAGGGGGG | 180.7 | -11.4 | 1.8 | -6.3 | 3.0E-10 | 7.5E-10 | MB231/MB468 | MP |
| >hg38_21_+_8433172_8446622_RNA45SN1_45S_50ntFlanks@10396.10424.29\|CAGTTGAACATGGGTCAGTCGGTCCTGAG | >hg38_21_+_8433172_8446622_RNA45SN1_45S_50ntFlanks | 10396 | 10424 | 29 | CAGTTGAACATGGGTCAGTCGGTCCTGAG | 3754.4 | 4.5 | 0.7 | 6.3 | 3.1E-10 | 7.5E-10 | MB231/MB468 | CYTO |
| >am_trnaMT_PheGAA_MT_+_577_647@43.62.20\|TTAGACGGGCTCACATCACC | >am_trnaMT_PheGAA_MT_+_577_647 | 43 | 62 | 20 | TTAGACGGGCTCACATCACC | 233.8 | 4.8 | 0.8 | 6.3 | 3.5E-10 | 7.6E-10 | MB231/MB468 | MITO |
| >hg38_21_+_8433172_8446622_RNA45SN1_45S_50ntFlanks@8176.8215.40\|CTTCTGATCGAGGCCCAGCCCGTGGACGGTGTGAGGCCGG | >hg38_21_+_8433172_8446622_RNA45SN1_45S_50ntFlanks | 8176 | 8215 | 40 | CTTCTGATCGAGGCCCAGCCCGTGGACGGTGTGAGGCCGG | 624.6 | -8.1 | 1.3 | -6.3 | 3.5E-10 | 7.6E-10 | MB231/MB468 | MITO |
| >hg38_21_+_8433172_8446622_RNA45SN1_45S_50ntFlanks@10186.10207.22\|TCCTCCCCTCCTCCCGCCCACG | >hg38_21_+_8433172_8446622_RNA45SN1_45S_50ntFlanks | 10186 | 10207 | 22 | TCCTCCCCTCCTCCCGCCCACG | 2344.3 | -10.3 | 1.6 | -6.3 | 3.7E-10 | 7.6E-10 | BT20/MB468 | NUC |
| >hg38_21_+_8433172_8446622_RNA45SN1_45S_50ntFlanks@12488.12520.33\|AACGTGAGCTGGGTTTAGACCGTCGTGAGACAG | >hg38_21_+_8433172_8446622_RNA45SN1_45S_50ntFlanks | 12488 | 12520 | 33 | AACGTGAGCTGGGTTTAGACCGTCGTGAGACAG | 484.1 | -8.0 | 1.3 | -6.3 | 3.5E-10 | 7.6E-10 | MB231/MB468 | MITO |
| >hg38_MT_+_1621_3279_RNR2_16S_50ntFlanks@1092.1120.29\|GCAAGACGAGAAGACCCTATGGAGCTTTA | >hg38_MT_+_1621_3279_RNR2_16S_50ntFlanks | 1092 | 1120 | 29 | GCAAGACGAGAAGACCCTATGGAGCTTTA | 532.1 | 5.5 | 0.9 | 6.3 | 2.3E-10 | 7.6E-10 | BT20/MB468 | MITO |
| >hg38_21_+_8433172_8446622_RNA45SN1_45S_50ntFlanks@9170.9197.28\|GCGGGTCGCGCCGTCGGGCCCGGGGGAG | >hg38_21_+_8433172_8446622_RNA45SN1_45S_50ntFlanks | 9170 | 9197 | 28 | GCGGGTCGCGCCGTCGGGCCCGGGGGAG | 269.1 | -11.9 | 1.9 | -6.3 | 3.0E-10 | 7.6E-10 | MB231/MB468 | MP |
| >MI0003674\|hsa-mir-653&WithFlank&7\|-\|93482754\|93482861@22.43.22&[MIMAT0003328&hsa-miR-653-5p&offsets\|+3\|+4;m-368&7\|-\|93482819\|93482840&offsets\|0\|0]\|TTGAAACAATCTCTACTGAACC | >MI0003674\|hsa-mir-653&WithFlank&7\|-\|93482754\|93482861 | 22 | 43 | 22 | TTGAAACAATCTCTACTGAACC | 91.4 | 9.7 | 1.5 | 6.3 | 3.0E-10 | 7.6E-10 | MB231/MB468 | MP |
| >hg38_21_+_8433172_8446622_RNA45SN1_45S_50ntFlanks@10384.10405.22\|CCATGTGAACAGCAGTTGAACA | >hg38_21_+_8433172_8446622_RNA45SN1_45S_50ntFlanks | 10384 | 10405 | 22 | CCATGTGAACAGCAGTTGAACA | 229.7 | -5.5 | 0.9 | -6.3 | 3.1E-10 | 7.6E-10 | MB231/MB468 | CYTO |
| >hg38_21_+_8433172_8446622_RNA45SN1_45S_50ntFlanks@11783.11819.37\|CATGAATGGATGAACGAGATTCCCACTGTCCCTACCT | >hg38_21_+_8433172_8446622_RNA45SN1_45S_50ntFlanks | 11783 | 11819 | 37 | CATGAATGGATGAACGAGATTCCCACTGTCCCTACCT | 451.0 | -8.4 | 1.3 | -6.3 | 3.5E-10 | 7.6E-10 | MB231/MB468 | MITO |
| >hg38_21_+_8433172_8446622_RNA45SN1_45S_50ntFlanks@9379.9409.31\|CGCGCTCGCCGGCCGAGGTGGGATCCCGAGG | >hg38_21_+_8433172_8446622_RNA45SN1_45S_50ntFlanks | 9379 | 9409 | 31 | CGCGCTCGCCGGCCGAGGTGGGATCCCGAGG | 99.2 | -10.5 | 1.7 | -6.3 | 3.0E-10 | 7.6E-10 | MB231/MB468 | MP |
| >MI0000440\|hsa-mir-27b&WithFlank&9\|+\|95085439\|95085547@67.86.20(+1G)&[MIMAT0000419&hsa-miR-27b-3p&offsets\|0\|-1(+1G);m-39&9\|+\|95085505\|95085525&offsets\|0\|-1(+1G)]\|TTCACAGTGGCTAAGTTCTGG | >MI0000440\|hsa-mir-27b&WithFlank&9\|+\|95085439\|95085547 | 67 | 86 | 20(+1G) | TTCACAGTGGCTAAGTTCTGG | 432.7 | 7.5 | 1.2 | 6.3 | 2.8E-10 | 7.6E-10 | BT20/MB468 | CYTO |
| >hg38_21_+_8433172_8446622_RNA45SN1_45S_50ntFlanks@11564.11592.29\|CCTAGCAGCCGACTTAGAACTGGTGCGGA | >hg38_21_+_8433172_8446622_RNA45SN1_45S_50ntFlanks | 11564 | 11592 | 29 | CCTAGCAGCCGACTTAGAACTGGTGCGGA | 379.4 | -6.0 | 0.9 | -6.3 | 2.8E-10 | 7.6E-10 | BT20/MB468 | CYTO |
| >hg38_21_+_8433172_8446622_RNA45SN1_45S_50ntFlanks@11452.11475.24\|CTCCTCGGCGCGCGGCGGCGGCGG | >hg38_21_+_8433172_8446622_RNA45SN1_45S_50ntFlanks | 11452 | 11475 | 24 | CTCCTCGGCGCGCGGCGGCGGCGG | 166.0 | -10.5 | 1.6 | -6.7 | 2.7E-11 | 7.6E-10 | BT20/MB468 | MP |
| >hg38_21_+_8433172_8446622_RNA45SN1_45S_50ntFlanks@10871.10901.31\|GGCTGGGTCGGTCGGGCTGGGGCGCGAAGCG | >hg38_21_+_8433172_8446622_RNA45SN1_45S_50ntFlanks | 10871 | 10901 | 31 | GGCTGGGTCGGTCGGGCTGGGGCGCGAAGCG | 80.4 | -10.2 | 1.6 | -6.3 | 3.0E-10 | 7.6E-10 | MB231/MB468 | MP |
| >hg38_21_+_8433172_8446622_RNA45SN1_45S_50ntFlanks@12944.12964.21\|CGTAGACGACCTGCTTCTGGG | >hg38_21_+_8433172_8446622_RNA45SN1_45S_50ntFlanks | 12944 | 12964 | 21 | CGTAGACGACCTGCTTCTGGG | 543.8 | -4.8 | 0.8 | -6.3 | 3.2E-10 | 7.6E-10 | MB231/MB468 | NUC |
| >am_trna106_HisGTG_1_-_149155828_149155899@7.28.22,trna111_HisGTG_1_-_147774845_147774916@7.28.22,trna118_HisGTG_1_-_145396881_145396952@7.28.22,trna16_HisGTG_1_+_146544773_146544844@7.28.22,trna1_HisGTG_15_+_45493349_45493420@7.28.22,trna21_HisGTG_1_+_147753471_147753542@7.28.22,trna33_HisGTG_6_+_27125906_27125977@7.28.22,trna7_HisGTG_9_-_14433938_14434009@7.28.22,trna8_HisGTG_15_-_45492611_45492682@7.28.22,trna9_HisGTG_15_-_45490804_45490875@7.28.22\|ATCGTATAGTGGTTAGTACTCT | >am_trna106_HisGTG_1_-_149155828_149155899 | 7 | 28 | 22 | ATCGTATAGTGGTTAGTACTCT | 398.1 | -8.0 | 1.3 | -6.3 | 3.0E-10 | 7.6E-10 | MB231/MB468 | MP |
| >hg38_21_+_8433172_8446622_RNA45SN1_45S_50ntFlanks@4666.4689.24\|AAATTCTTGGACCGGCGCAAGACG | >hg38_21_+_8433172_8446622_RNA45SN1_45S_50ntFlanks | 4666 | 4689 | 24 | AAATTCTTGGACCGGCGCAAGACG | 97.9 | -10.5 | 1.7 | -6.3 | 3.0E-10 | 7.6E-10 | MB231/MB468 | MP |
| >hg38_21_+_8433172_8446622_RNA45SN1_45S_50ntFlanks@10512.10532.21\|TCCGGAGTGGCGGAGATGGGC | >hg38_21_+_8433172_8446622_RNA45SN1_45S_50ntFlanks | 10512 | 10532 | 21 | TCCGGAGTGGCGGAGATGGGC | 1228.4 | -7.2 | 1.1 | -6.3 | 3.0E-10 | 7.6E-10 | MB231/MB468 | MP |
| >hg38_21_+_8433172_8446622_RNA45SN1_45S_50ntFlanks@12433.12455.23\|CTATCATTGTGAAGCAGAATTCA | >hg38_21_+_8433172_8446622_RNA45SN1_45S_50ntFlanks | 12433 | 12455 | 23 | CTATCATTGTGAAGCAGAATTCA | 159.7 | -9.4 | 1.5 | -6.3 | 2.3E-10 | 7.7E-10 | BT20/MB468 | MITO |
| >hg38_21_+_8433172_8446622_RNA45SN1_45S_50ntFlanks@6657.6686.30\|TTAGCGGTGGATCACTCGGCTCGTGCGTCG | >hg38_21_+_8433172_8446622_RNA45SN1_45S_50ntFlanks | 6657 | 6686 | 30 | TTAGCGGTGGATCACTCGGCTCGTGCGTCG | 100.2 | -10.5 | 1.7 | -6.3 | 3.0E-10 | 7.7E-10 | MB231/MB468 | MP |
| >hg38_21_+_8433172_8446622_RNA45SN1_45S_50ntFlanks@12141.12162.22\|ACAGTGCCAGGTGGGGAGTTTG | >hg38_21_+_8433172_8446622_RNA45SN1_45S_50ntFlanks | 12141 | 12162 | 22 | ACAGTGCCAGGTGGGGAGTTTG | 146.5 | -9.3 | 1.5 | -6.3 | 2.4E-10 | 7.7E-10 | BT20/MB468 | MITO |

| | | | | | | | | | | | | | |
|---|---|---|---|---|---|---|---|---|---|---|---|---|---|
| >hg38_21_+_8433172_8446622_RNA45SN1_45S_50ntFlanks@8353.8391.39\|GTTGAAAAGAACTTTGAAGAGAGAGTTCAAGAGGGCGTG | >hg38_21_+_8433172_8446622_RNA45SN1_45S_50ntFlanks | 8353 | 8391 | 39 | GTTGAAAAGAACTTTGAAGAGAGAGTTCAAGAGGGCGTG | 139.0 | 10.9 | 1.6 | 6.7 | 2.7E-11 | 7.7E-10 | BT20/MB468 | MP |
| >hg38_21_+_8433172_8446622_RNA45SN1_45S_50ntFlanks@9240.9273.34\|GCGGAGCGAGCGCACGGGGTCGGCGGCGACGTCG | >hg38_21_+_8433172_8446622_RNA45SN1_45S_50ntFlanks | 9240 | 9273 | 34 | GCGGAGCGAGCGCACGGGGTCGGCGGCGACGTCG | 182.4 | -10.6 | 1.6 | -6.7 | 2.7E-11 | 7.7E-10 | BT20/MB468 | MP |
| >hg38_21_+_8433172_8446622_RNA45SN1_45S_50ntFlanks@12033.12070.38\|ATCGTTTTTTCACTGACCCGGTGAGGCGGGGGGGCGAG | >hg38_21_+_8433172_8446622_RNA45SN1_45S_50ntFlanks | 12033 | 12070 | 38 | ATCGTTTTTTCACTGACCCGGTGAGGCGGGGGGGCGAG | 185.7 | -9.6 | 1.5 | -6.3 | 2.4E-10 | 7.7E-10 | BT20/MB468 | MITO |
| >hg38_21_+_8433172_8446622_RNA45SN1_45S_50ntFlanks@12499.12517.19\|GGTTTAGACCGTCGTGAGA | >hg38_21_+_8433172_8446622_RNA45SN1_45S_50ntFlanks | 12499 | 12517 | 19 | GGTTTAGACCGTCGTGAGA | 119.6 | -10.8 | 1.7 | -6.3 | 3.0E-10 | 7.7E-10 | MB231/MB468 | MP |
| >hg38_21_+_8433172_8446622_RNA45SN1_45S_50ntFlanks@11568.11586.19\|GCAGCCGACTTAGAACTGG | >hg38_21_+_8433172_8446622_RNA45SN1_45S_50ntFlanks | 11568 | 11586 | 19 | GCAGCCGACTTAGAACTGG | 873.1 | -4.2 | 0.7 | -6.3 | 3.7E-10 | 7.7E-10 | BT20/MB468 | NUC |
| >hg38_21_+_8433172_8446622_RNA45SN1_45S_50ntFlanks@12163.12182.20\|ACTGGGGCGGTACACCTGTC | >hg38_21_+_8433172_8446622_RNA45SN1_45S_50ntFlanks | 12163 | 12182 | 20 | ACTGGGGCGGTACACCTGTC | 4542.9 | -4.7 | 0.7 | -6.3 | 2.4E-10 | 7.7E-10 | BT20/MB468 | MITO |
| >hg38_21_+_8433172_8446622_RNA45SN1_45S_50ntFlanks@10389.10426.38\|TGAACAGCAGTTGAACATGGGTCAGTCGGTCCTGAGAG | >hg38_21_+_8433172_8446622_RNA45SN1_45S_50ntFlanks | 10389 | 10426 | 38 | TGAACAGCAGTTGAACATGGGTCAGTCGGTCCTGAGAG | 128.2 | -10.9 | 1.7 | -6.3 | 3.0E-10 | 7.7E-10 | MB231/MB468 | MP |
| >hg38_21_+_8433172_8446622_RNA45SN1_45S_50ntFlanks@9567.9595.29\|ACGTGCAAATCGGTCGTCCGACCTGGGTA | >hg38_21_+_8433172_8446622_RNA45SN1_45S_50ntFlanks | 9567 | 9595 | 29 | ACGTGCAAATCGGTCGTCCGACCTGGGTA | 3671.9 | -10.5 | 1.7 | -6.3 | 3.7E-10 | 7.7E-10 | BT20/MB468 | NUC |
| >hg38_21_+_8433172_8446622_RNA45SN1_45S_50ntFlanks@9298.9336.39\|ACACGGACCAAGGAGTCTAACACGTGCGCGAGTCGGGGG | >hg38_21_+_8433172_8446622_RNA45SN1_45S_50ntFlanks | 9298 | 9336 | 39 | ACACGGACCAAGGAGTCTAACACGTGCGCGAGTCGGGGG | 66.3 | -9.9 | 1.6 | -6.3 | 3.0E-10 | 7.7E-10 | MB231/MB468 | MP |
| >hg38_21_+_8433172_8446622_RNA45SN1_45S_50ntFlanks@11579.11606.28\|AGAACTGGTGCGGACCAGGGGAATCCGA | >hg38_21_+_8433172_8446622_RNA45SN1_45S_50ntFlanks | 11579 | 11606 | 28 | AGAACTGGTGCGGACCAGGGGAATCCGA | 332.9 | -8.4 | 1.3 | -6.3 | 3.7E-10 | 7.7E-10 | BT20/MB468 | NUC |
| >MI0000298\|hsa-mir-221&WithFlank&X\|-\|45746151\|45746272@71.92.22(+1U)&[MIMAT0000278&hsa-miR-221-3p&offsets\|0\|-1(+1U);m-91&X\|-\|45746181\|45746202&offsets\|0\|0(+1U)]\|AGCTACATTGTCTGCTGGGTTTT | >MI0000298\|hsa-mir-221&WithFlank&X\|-\|45746151\|45746272 | 71 | 92 | 22(+1U) | AGCTACATTGTCTGCTGGGTTTT | 9179.3 | 4.9 | 0.8 | 6.3 | 3.0E-10 | 7.7E-10 | MB231/MB468 | MP |
| >hg38_21_+_8433172_8446622_RNA45SN1_45S_50ntFlanks@5072.5096.25\|TAATTCCGATAACGAACGAGACTCT | >hg38_21_+_8433172_8446622_RNA45SN1_45S_50ntFlanks | 5072 | 5096 | 25 | TAATTCCGATAACGAACGAGACTCT | 379.5 | 9.0 | 1.4 | 6.3 | 3.2E-10 | 7.7E-10 | MB231/MB468 | CYTO |
| >hg38_21_+_8433172_8446622_RNA45SN1_45S_50ntFlanks@9806.9827.22\|CGTGGAATGCGAGTGCCTAGTG | >hg38_21_+_8433172_8446622_RNA45SN1_45S_50ntFlanks | 9806 | 9827 | 22 | CGTGGAATGCGAGTGCCTAGTG | 110.9 | -10.7 | 1.7 | -6.3 | 3.0E-10 | 7.7E-10 | MB231/MB468 | MP |
| >MI0000462\|hsa-mir-152&WithFlank&17\|-\|48037155\|48037253@60.79.20&[MIMAT0000438&hsa-miR-152-3p&offsets\|0\|-1;m-97&17\|-\|48037174\|48037194&offsets\|0\|-1]\|TCAGTGCATGACAGAACTTG | >MI0000462\|hsa-mir-152&WithFlank&17\|-\|48037155\|48037253 | 60 | 79 | 20 | TCAGTGCATGACAGAACTTG | 221.9 | 11.5 | 1.7 | 6.7 | 2.7E-11 | 7.7E-10 | BT20/MB468 | MP |
| >hg38_21_+_8433172_8446622_RNA45SN1_45S_50ntFlanks@10727.10750.24\|GGAGAGGGTGTAAATCTCGCGCCG | >hg38_21_+_8433172_8446622_RNA45SN1_45S_50ntFlanks | 10727 | 10750 | 24 | GGAGAGGGTGTAAATCTCGCGCCG | 316.2 | -11.4 | 1.7 | -6.7 | 2.7E-11 | 7.7E-10 | BT20/MB468 | MP |
| >hg38_21_+_8433172_8446622_RNA45SN1_45S_50ntFlanks@11621.11660.40\|CAAAGCATCGCGAAGGCCCGCGGCGGGTGTTGACGCGATG | >hg38_21_+_8433172_8446622_RNA45SN1_45S_50ntFlanks | 11621 | 11660 | 40 | CAAAGCATCGCGAAGGCCCGCGGCGGGTGTTGACGCGATG | 247.5 | -11.0 | 1.7 | -6.7 | 2.7E-11 | 7.7E-10 | BT20/MB468 | MP |
| >hg38_21_+_8433172_8446622_RNA45SN1_45S_50ntFlanks@5421.5443.23\|CGATTGGATGGTTTAGTGAGGCC | >hg38_21_+_8433172_8446622_RNA45SN1_45S_50ntFlanks | 5421 | 5443 | 23 | CGATTGGATGGTTTAGTGAGGCC | 1314.6 | -6.0 | 0.9 | -6.7 | 2.7E-11 | 7.7E-10 | BT20/MB468 | MP |
| >hg38_21_+_8433172_8446622_RNA45SN1_45S_50ntFlanks@6719.6740.22\|TGAATTGCAGGACACATTGATC | >hg38_21_+_8433172_8446622_RNA45SN1_45S_50ntFlanks | 6719 | 6740 | 22 | TGAATTGCAGGACACATTGATC | 2217.5 | -5.3 | 0.8 | -6.7 | 2.7E-11 | 7.7E-10 | BT20/MB468 | MP |
| >hg38_21_+_8433172_8446622_RNA45SN1_45S_50ntFlanks@5485.5519.35\|AGCGCTGAGAAGACGGTCGAACTTGACTATCTAGA | >hg38_21_+_8433172_8446622_RNA45SN1_45S_50ntFlanks | 5485 | 5519 | 35 | AGCGCTGAGAAGACGGTCGAACTTGACTATCTAGA | 400.5 | -8.0 | 1.3 | -6.3 | 3.6E-10 | 7.7E-10 | MB231/MB468 | MITO |
| >hg38_21_+_8433172_8446622_RNA45SN1_45S_50ntFlanks@10410.10440.31\|TCAGTCGGTCCTGAGAGATGGGCGAGCGCCG | >hg38_21_+_8433172_8446622_RNA45SN1_45S_50ntFlanks | 10410 | 10440 | 31 | TCAGTCGGTCCTGAGAGATGGGCGAGCGCCG | 263.3 | -7.7 | 1.2 | -6.3 | 3.6E-10 | 7.7E-10 | MB231/MB468 | MITO |
| >hg38_21_+_8433172_8446622_RNA45SN1_45S_50ntFlanks@9534.9564.31\|AGGAAACTCTGGTGGAGGTCCGTAGCGGTCC | >hg38_21_+_8433172_8446622_RNA45SN1_45S_50ntFlanks | 9534 | 9564 | 31 | AGGAAACTCTGGTGGAGGTCCGTAGCGGTCC | 959.4 | -8.4 | 1.3 | -6.3 | 3.6E-10 | 7.7E-10 | MB231/MB468 | MITO |
| >hg38_21_+_8433172_8446622_RNA45SN1_45S_50ntFlanks@8184.8210.27\|CGAGGCCCAGCCCGTGGACGGTGTGAG | >hg38_21_+_8433172_8446622_RNA45SN1_45S_50ntFlanks | 8184 | 8210 | 27 | CGAGGCCCAGCCCGTGGACGGTGTGAG | 438.1 | -3.9 | 0.6 | -6.3 | 3.2E-10 | 7.8E-10 | MB231/MB468 | NUC |
| >hg38_21_+_8433172_8446622_RNA45SN1_45S_50ntFlanks@12894.12931.38\|CTCGCCCGTCACGCACCGCACGTTCGTGGGGAACCTGG | >hg38_21_+_8433172_8446622_RNA45SN1_45S_50ntFlanks | 12894 | 12931 | 38 | CTCGCCCGTCACGCACCGCACGTTCGTGGGGAACCTGG | 159.0 | -11.2 | 1.8 | -6.3 | 3.1E-10 | 7.8E-10 | MB231/MB468 | MP |
| >MI0000079\|hsa-mir-23a&WithFlank&19\|-\|13836581\|13836665@51.72.22&[MIMAT0000078&hsa-miR-23a-3p&offsets\|0\|+1;m-31&19\|-\|13836595\|13836615&offsets\|0\|+1]\|ATCACATTGCCAGGGATTTCCA | >MI0000079\|hsa-mir-23a&WithFlank&19\|-\|13836581\|13836665 | 51 | 72 | 22 | ATCACATTGCCAGGGATTTCCA | 452.4 | -11.9 | 1.8 | -6.7 | 2.8E-11 | 7.8E-10 | BT20/MB468 | MP |
| >hg38_21_+_8433172_8446622_RNA45SN1_45S_50ntFlanks@4160.4178.19\|CCACATCCAAGGAAGGCAG | >hg38_21_+_8433172_8446622_RNA45SN1_45S_50ntFlanks | 4160 | 4178 | 19 | CCACATCCAAGGAAGGCAG | 195.7 | -10.7 | 1.6 | -6.7 | 2.8E-11 | 7.8E-10 | BT20/MB468 | MP |
| >hg38_21_+_8433172_8446622_RNA45SN1_45S_50ntFlanks@12273.12318.46\|TCAGTACGAATACAGACCGTGAAAGCGGGGCCTCACGATCCTTCTG | >hg38_21_+_8433172_8446622_RNA45SN1_45S_50ntFlanks | 12273 | 12318 | 46 | TCAGTACGAATACAGACCGTGAAAGCGGGGCCTCACGATCCTTCTG | 169.4 | -11.2 | 1.8 | -6.3 | 3.6E-10 | 7.8E-10 | MB231/MB468 | MITO |
| >hg38_21_+_8433172_8446622_RNA45SN1_45S_50ntFlanks@11670.11699.30\|CCCAGTGCTCTGAATGTCAAAGTGAAGAAA | >hg38_21_+_8433172_8446622_RNA45SN1_45S_50ntFlanks | 11670 | 11699 | 30 | CCCAGTGCTCTGAATGTCAAAGTGAAGAAA | 1396.6 | -9.5 | 1.5 | -6.3 | 3.8E-10 | 7.8E-10 | BT20/MB468 | NUC |
| >trnaMT_LeuTAA_MT_+_3230_3304@16.45.30\|CCCGGTAATCGCATAAAACTTAAAACTTTA | >trnaMT_LeuTAA_MT_+_3230_3304 | 16 | 45 | 30 | CCCGGTAATCGCATAAAACTTAAAACTTTA | 150.0 | -11.1 | 1.8 | -6.3 | 3.1E-10 | 7.8E-10 | MB231/MB468 | MP |
| >hg38_21_+_8433172_8446622_RNA45SN1_45S_50ntFlanks@8194.8214.21\|CCCGTGGACGGTGTGAGGCCG | >hg38_21_+_8433172_8446622_RNA45SN1_45S_50ntFlanks | 8194 | 8214 | 21 | CCCGTGGACGGTGTGAGGCCG | 429.0 | -2.9 | 0.5 | -6.3 | 2.4E-10 | 7.8E-10 | BT20/MB468 | MITO |
| >hg38_21_+_8433172_8446622_RNA45SN1_45S_50ntFlanks@11452.11474.23\|CTCCTCGGCGCGCGGCGGCGGCG | >hg38_21_+_8433172_8446622_RNA45SN1_45S_50ntFlanks | 11452 | 11474 | 23 | CTCCTCGGCGCGCGGCGGCGGCG | 115.7 | -6.1 | 1.0 | -6.3 | 2.9E-10 | 7.8E-10 | BT20/MB468 | CYTO |
| >hg38_21_+_8433172_8446622_RNA45SN1_45S_50ntFlanks@10407.10433.27\|GGGTCAGTCGGTCCTGAGAGATGGGCG | >hg38_21_+_8433172_8446622_RNA45SN1_45S_50ntFlanks | 10407 | 10433 | 27 | GGGTCAGTCGGTCCTGAGAGATGGGCG | 76.2 | -10.1 | 1.6 | -6.3 | 3.1E-10 | 7.8E-10 | MB231/MB468 | MP |
| >hg38_21_+_8433172_8446622_RNA45SN1_45S_50ntFlanks@11226.11254.29\|AGCCGGGCCCTTCCCGTGGATCGCCCCAG | >hg38_21_+_8433172_8446622_RNA45SN1_45S_50ntFlanks | 11226 | 11254 | 29 | AGCCGGGCCCTTCCCGTGGATCGCCCCAG | 83.4 | -10.2 | 1.6 | -6.3 | 3.1E-10 | 7.8E-10 | MB231/MB468 | MP |
| >hg38_21_+_8433172_8446622_RNA45SN1_45S_50ntFlanks@6776.6806.31\|CTCCCGGGGCTACGCCTGTCTGAGCGTCGCT | >hg38_21_+_8433172_8446622_RNA45SN1_45S_50ntFlanks | 6776 | 6806 | 31 | CTCCCGGGGCTACGCCTGTCTGAGCGTCGCT | 1888.8 | 4.5 | 0.7 | 6.3 | 2.9E-10 | 7.8E-10 | BT20/MB468 | CYTO |
| >hg38_21_+_8433172_8446622_RNA45SN1_45S_50ntFlanks@11580.11609.30\|GAACTGGTGCGGACCAGGGGAATCCGACTG | >hg38_21_+_8433172_8446622_RNA45SN1_45S_50ntFlanks | 11580 | 11609 | 30 | GAACTGGTGCGGACCAGGGGAATCCGACTG | 1471.0 | -4.8 | 0.8 | -6.3 | 3.6E-10 | 7.8E-10 | MB231/MB468 | MITO |
| >MI0000465\|hsa-mir-191&WithFlank&3\|-\|49020612\|49020715@22.42.21(+1A)&[MIMAT0000440&hsa-miR-191-5p&offsets\|0\|-2(+1A);m-48&3\|-\|49020672\|49020694&offsets\|0\|-2(+1A)]\|CAACGGAATCCCAAAAGCAGCA | >MI0000465\|hsa-mir-191&WithFlank&3\|-\|49020612\|49020715 | 22 | 42 | 21(+1A) | CAACGGAATCCCAAAAGCAGCA | 458.5 | 4.4 | 0.7 | 6.3 | 3.3E-10 | 7.9E-10 | MB231/MB468 | NUC |
| >am_trna128_GlyGCC_6_-_27870686_27870756@1.33.33,trna18_GlyGCC_16_+_70822597_70822667@1.33.33,trna19_GlyGCC_16_+_70823410_70823480@1.33.33,trna19_GlyGCC_2_-_157257659_157257729@1.33.33,trna24_GlyGCC_16_-_70812942_70813012@1.33.33,trna25_GlyGCC_16_-_70812114_70812184@1.33.33,trna5_GlyGCC_17_+_8029064_8029134@1.33.33,trna68_GlyGCC_1_-_161493637_161493707@1.33.33\|GCATTGGTGGTTCAGTGGTAGAATTCTCGCCTG | >am_trna128_GlyGCC_6_-_27870686_27870756 | 1 | 33 | 33 | GCATTGGTGGTTCAGTGGTAGAATTCTCGCCTG | 1725.8 | 5.8 | 0.9 | 6.3 | 3.3E-10 | 7.9E-10 | MB231/MB468 | NUC |
| >hg38_21_+_8433172_8446622_RNA45SN1_45S_50ntFlanks@12416.12438.23\|TTCGATGTCGGCTCTTCCTATCA | >hg38_21_+_8433172_8446622_RNA45SN1_45S_50ntFlanks | 12416 | 12438 | 23 | TTCGATGTCGGCTCTTCCTATCA | 1236.5 | 4.7 | 0.7 | 6.3 | 3.3E-10 | 7.9E-10 | MB231/MB468 | NUC |
| >hg38_21_+_8433172_8446622_RNA45SN1_45S_50ntFlanks@12456.12482.27\|CCAAGCGTTGGATTGTTCACCCACTAA | >hg38_21_+_8433172_8446622_RNA45SN1_45S_50ntFlanks | 12456 | 12482 | 27 | CCAAGCGTTGGATTGTTCACCCACTAA | 111.4 | -8.9 | 1.4 | -6.3 | 2.4E-10 | 7.9E-10 | BT20/MB468 | MITO |
| >hg38_21_+_8433172_8446622_RNA45SN1_45S_50ntFlanks@5084.5105.22\|CGAACGAGACTCTGGCATGCTA | >hg38_21_+_8433172_8446622_RNA45SN1_45S_50ntFlanks | 5084 | 5105 | 22 | CGAACGAGACTCTGGCATGCTA | 626.2 | -11.4 | 1.8 | -6.3 | 2.4E-10 | 7.9E-10 | BT20/MB468 | MITO |
| >hg38_21_+_8433172_8446622_RNA45SN1_45S_50ntFlanks@8019.8042.24\|TTAGTCAGCGGAGGAGAAGAAACT | >hg38_21_+_8433172_8446622_RNA45SN1_45S_50ntFlanks | 8019 | 8042 | 24 | TTAGTCAGCGGAGGAGAAGAAACT | 213.7 | 8.1 | 1.3 | 6.3 | 3.2E-10 | 7.9E-10 | MB231/MB468 | CYTO |
| >hg38_21_+_8433172_8446622_RNA45SN1_45S_50ntFlanks@9597.9614.18\|AGGGGCGAAAGACTAATC | >hg38_21_+_8433172_8446622_RNA45SN1_45S_50ntFlanks | 9597 | 9614 | 18 | AGGGGCGAAAGACTAATC | 404.1 | -6.6 | 1.0 | -6.3 | 3.1E-10 | 7.9E-10 | MB231/MB468 | MP |
| >trnaMT_HisGTG_MT_+_12138_12206@46.72.27\|GAGGCTTACGACCCCTTATTTACCCCA | >trnaMT_HisGTG_MT_+_12138_12206 | 46 | 72 | 27 | GAGGCTTACGACCCCTTATTTACCCCA | 428.8 | 12.5 | 1.9 | 6.7 | 2.8E-11 | 7.9E-10 | BT20/MB468 | MP |
| >hg38_MT_+_598_1651_RNR1_12S_50ntFlanks@443.480.38\|TATCTGAACACACAATAGCTAAGACCCAAACTGGGATT | >hg38_MT_+_598_1651_RNR1_12S_50ntFlanks | 443 | 480 | 38 | TATCTGAACACACAATAGCTAAGACCCAAACTGGGATT | 730.2 | 5.3 | 0.8 | 6.3 | 3.6E-10 | 7.9E-10 | MB231/MB468 | MITO |
| >hg38_21_+_8433172_8446622_RNA45SN1_45S_50ntFlanks@6645.6678.34\|CTCGTACGACTCTTAGCGGTGGATCACTCGGCTC | >hg38_21_+_8433172_8446622_RNA45SN1_45S_50ntFlanks | 6645 | 6678 | 34 | CTCGTACGACTCTTAGCGGTGGATCACTCGGCTC | 3276.7 | 4.4 | 0.7 | 6.3 | 2.4E-10 | 7.9E-10 | BT20/MB468 | MITO |
| >hg38_21_+_8433172_8446622_RNA45SN1_45S_50ntFlanks@6650.6668.19\|ACGACTCTTAGCGGTGGAT | >hg38_21_+_8433172_8446622_RNA45SN1_45S_50ntFlanks | 6650 | 6668 | 19 | ACGACTCTTAGCGGTGGAT | 395.7 | 3.7 | 0.6 | 6.3 | 3.3E-10 | 7.9E-10 | MB231/MB468 | NUC |
| >hg38_21_+_8433172_8446622_RNA45SN1_45S_50ntFlanks@11454.11477.24\|CCTCGGCGCGCGGCGGCGGCGGCG | >hg38_21_+_8433172_8446622_RNA45SN1_45S_50ntFlanks | 11454 | 11477 | 24 | CCTCGGCGCGCGGCGGCGGCGGCG | 151.2 | -7.3 | 1.2 | -6.3 | 2.9E-10 | 7.9E-10 | BT20/MB468 | CYTO |
| >hg38_21_+_8433172_8446622_RNA45SN1_45S_50ntFlanks@12457.12474.18\|CAAGCGTTGGATTGTTCA | >hg38_21_+_8433172_8446622_RNA45SN1_45S_50ntFlanks | 12457 | 12474 | 18 | CAAGCGTTGGATTGTTCA | 571.0 | -8.3 | 1.3 | -6.3 | 3.1E-10 | 7.9E-10 | MB231/MB468 | MP |
| >hg38_21_+_8433172_8446622_RNA45SN1_45S_50ntFlanks@12459.12477.19\|AGCGTTGGATTGTTCACCC | >hg38_21_+_8433172_8446622_RNA45SN1_45S_50ntFlanks | 12459 | 12477 | 19 | AGCGTTGGATTGTTCACCC | 476.6 | -6.7 | 1.1 | -6.3 | 3.1E-10 | 7.9E-10 | MB231/MB468 | MP |
| >hg38_21_+_8433172_8446622_RNA45SN1_45S_50ntFlanks@4729.4759.31\|TCAAGAACGAAAGTCGGAGGTTCGAAGACGA | >hg38_21_+_8433172_8446622_RNA45SN1_45S_50ntFlanks | 4729 | 4759 | 31 | TCAAGAACGAAAGTCGGAGGTTCGAAGACGA | 1180.5 | -9.4 | 1.5 | -6.3 | 3.8E-10 | 7.9E-10 | BT20/MB468 | NUC |
| >hg38_21_+_8433172_8446622_RNA45SN1_45S_50ntFlanks@12458.12475.18\|AAGCGTTGGATTGTTCAC | >hg38_21_+_8433172_8446622_RNA45SN1_45S_50ntFlanks | 12458 | 12475 | 18 | AAGCGTTGGATTGTTCAC | 492.9 | -8.9 | 1.4 | -6.3 | 3.8E-10 | 7.9E-10 | BT20/MB468 | NUC |
| >hg38_21_+_8433172_8446622_RNA45SN1_45S_50ntFlanks@4241.4275.35\|CTCTTTCGAGGCCCTGTAATTGGAATGAGTCCACT | >hg38_21_+_8433172_8446622_RNA45SN1_45S_50ntFlanks | 4241 | 4275 | 35 | CTCTTTCGAGGCCCTGTAATTGGAATGAGTCCACT | 332.2 | -8.3 | 1.3 | -6.3 | 3.7E-10 | 8.0E-10 | MB231/MB468 | MITO |

<table>
<tr><td></td><td></td><td></td><td></td><td></td><td></td><td></td><td></td><td></td><td></td><td></td><td></td><td></td><td></td></tr>
<tr><td>>hg38_21_+_8433172_8446622_RNA45SN1_45S_50ntFlanks@6645.6677.33|CTCGTACGACTCTTAGCGGTGGATCACTCGGCT</td><td>>hg38_21_+_8433172_8446622_RNA45SN1_45S_50ntFlanks</td><td>6645</td><td>6677</td><td>33</td><td>CTCGTACGACTCTTAGCGGTGGATCACTCGGCT</td><td>274.3</td><td>3.7</td><td>0.6</td><td>6.3</td><td>3.3E-10</td><td>8.0E-10</td><td>MB231/MB468</td><td>CYTO</td></tr>
<tr><td>>trnaMT_GluTTC_MT_-_14674_14742@37.71.35,trnalookalike8_GluTTC_5_-_93905172_93905240@37.71.35|TCATTGGTCGTGGTTGTAGTCCGTGCGAGAATACC</td><td>>trnaMT_GluTTC_MT_-_14674_14742</td><td>37</td><td>71</td><td>35</td><td>TCATTGGTCGTGGTTGTAGTCCGTGCGAGAATACC</td><td>1054.3</td><td>6.7</td><td>1.1</td><td>6.3</td><td>3.3E-10</td><td>8.0E-10</td><td>MB231/MB468</td><td>NUC</td></tr>
<tr><td>>hg38_21_+_8433172_8446622_RNA45SN1_45S_50ntFlanks@4367.4391.25|CAGTTAAAAAGCTCGTAGTTGGATC</td><td>>hg38_21_+_8433172_8446622_RNA45SN1_45S_50ntFlanks</td><td>4367</td><td>4391</td><td>25</td><td>CAGTTAAAAAGCTCGTAGTTGGATC</td><td>1145.3</td><td>-7.5</td><td>1.2</td><td>-6.3</td><td>3.3E-10</td><td>8.0E-10</td><td>MB231/MB468</td><td>NUC</td></tr>
<tr><td>>hg38_21_+_8433172_8446622_RNA45SN1_45S_50ntFlanks@11843.11862.20|AGGGAACGGGCTTGGCGGAA</td><td>>hg38_21_+_8433172_8446622_RNA45SN1_45S_50ntFlanks</td><td>11843</td><td>11862</td><td>20</td><td>AGGGAACGGGCTTGGCGGAA</td><td>84.3</td><td>-10.3</td><td>1.6</td><td>-6.3</td><td>3.2E-10</td><td>8.0E-10</td><td>MB231/MB468</td><td>MP</td></tr>
<tr><td>>MI0000734|hsa-mir-106b&WithFlank&7|-|100093987|100094080@56.77.22&[MIMAT0004672&hsa-miR-106b-3p&offsets|-2|-2;m-87&7|-|100094003|100094023&offsets|-2|-1]|TACCGCACTGTGGGTACTTGCT</td><td>>MI0000734|hsa-mir-106b&WithFlank&7|-|100093987|100094080</td><td>56</td><td>77</td><td>22</td><td>TACCGCACTGTGGGTACTTGCT</td><td>953.9</td><td>9.3</td><td>1.5</td><td>6.3</td><td>2.5E-10</td><td>8.0E-10</td><td>BT20/MB468</td><td>MITO</td></tr>
<tr><td>>hg38_21_+_8433172_8446622_RNA45SN1_45S_50ntFlanks@10794.10830.37|CTGGCATGTTGGAACAATGTAGGTAAGGGAAGTCGGC</td><td>>hg38_21_+_8433172_8446622_RNA45SN1_45S_50ntFlanks</td><td>10794</td><td>10830</td><td>37</td><td>CTGGCATGTTGGAACAATGTAGGTAAGGGAAGTCGGC</td><td>70.1</td><td>-10.0</td><td>1.6</td><td>-6.3</td><td>3.2E-10</td><td>8.0E-10</td><td>MB231/MB468</td><td>MP</td></tr>
<tr><td>>hg38_21_+_8433172_8446622_RNA45SN1_45S_50ntFlanks@10571.10596.26|GAGAAGCCGGCGGGAGCCCCGGGGAG</td><td>>hg38_21_+_8433172_8446622_RNA45SN1_45S_50ntFlanks</td><td>10571</td><td>10596</td><td>26</td><td>GAGAAGCCGGCGGGAGCCCCGGGGAG</td><td>104.4</td><td>-8.8</td><td>1.4</td><td>-6.3</td><td>2.5E-10</td><td>8.0E-10</td><td>BT20/MB468</td><td>MITO</td></tr>
<tr><td>>hg38_21_+_8433172_8446622_RNA45SN1_45S_50ntFlanks@10362.10390.29|AAGGCCGAAGTGGAGAAGGGTTCCATGTG</td><td>>hg38_21_+_8433172_8446622_RNA45SN1_45S_50ntFlanks</td><td>10362</td><td>10390</td><td>29</td><td>AAGGCCGAAGTGGAGAAGGGTTCCATGTG</td><td>788.5</td><td>-6.2</td><td>1.0</td><td>-6.3</td><td>2.5E-10</td><td>8.0E-10</td><td>BT20/MB468</td><td>MITO</td></tr>
<tr><td>>hg38_21_+_8433172_8446622_RNA45SN1_45S_50ntFlanks@11563.11584.22|GCCTAGCAGCCGACTTAGAACT</td><td>>hg38_21_+_8433172_8446622_RNA45SN1_45S_50ntFlanks</td><td>11563</td><td>11584</td><td>22</td><td>GCCTAGCAGCCGACTTAGAACT</td><td>250.0</td><td>6.6</td><td>1.0</td><td>6.3</td><td>3.3E-10</td><td>8.0E-10</td><td>MB231/MB468</td><td>NUC</td></tr>
<tr><td>>MI0000063|hsa-let-7b&WithFlank&22|+|46113680|46113774@12.33.22(+1C)&[MIMAT0000063&hsa-let-7b-5p&offsets|0|0(+1C);m-14&22|+|46113691|46113712&offsets|0|0(+1C)]|TGAGGTAGTAGGTTGTGTGGTTC</td><td>>MI0000063|hsa-let-7b&WithFlank&22|+|46113680|46113774</td><td>12</td><td>33</td><td>22(+1C)</td><td>TGAGGTAGTAGGTTGTGTGGTTC</td><td>192.1</td><td>4.0</td><td>0.6</td><td>6.3</td><td>3.3E-10</td><td>8.0E-10</td><td>MB231/MB468</td><td>NUC</td></tr>
<tr><td>>hg38_21_+_8433172_8446622_RNA45SN1_45S_50ntFlanks@6647.6681.35|CGTACGACTCTTAGCGGTGGATCACTCGGCTCGTG</td><td>>hg38_21_+_8433172_8446622_RNA45SN1_45S_50ntFlanks</td><td>6647</td><td>6681</td><td>35</td><td>CGTACGACTCTTAGCGGTGGATCACTCGGCTCGTG</td><td>7074.9</td><td>-5.3</td><td>0.8</td><td>-6.3</td><td>3.2E-10</td><td>8.0E-10</td><td>MB231/MB468</td><td>MP</td></tr>
<tr><td>>MI0000746|hsa-mir-99b&WithFlank&19|+|51692606|51692687@13.34.22(+2U)&[MIMAT0000689&hsa-miR-99b-5p&offsets|0|0(+2U);m-10&19|+|51692618|51692639&offsets|0|0(+2U)]|CACCCGTAGAACCGACCTTGCGTT</td><td>>MI0000746|hsa-mir-99b&WithFlank&19|+|51692606|51692687</td><td>13</td><td>34</td><td>22(+2U)</td><td>CACCCGTAGAACCGACCTTGCGTT</td><td>165.5</td><td>10.6</td><td>1.7</td><td>6.3</td><td>3.7E-10</td><td>8.0E-10</td><td>MB231/MB468</td><td>MITO</td></tr>
<tr><td>>hg38_21_+_8433172_8446622_RNA45SN1_45S_50ntFlanks@9911.9928.18|CCAGAAAAGGTGTTGGTT</td><td>>hg38_21_+_8433172_8446622_RNA45SN1_45S_50ntFlanks</td><td>9911</td><td>9928</td><td>18</td><td>CCAGAAAAGGTGTTGGTT</td><td>276.7</td><td>-11.2</td><td>1.7</td><td>-6.7</td><td>2.9E-11</td><td>8.1E-10</td><td>BT20/MB468</td><td>MP</td></tr>
<tr><td>>hg38_21_+_8433172_8446622_RNA45SN1_45S_50ntFlanks@11029.11060.32|CTCTCTCTCTCTCCCCCGCTCCCCGTCCTCCC</td><td>>hg38_21_+_8433172_8446622_RNA45SN1_45S_50ntFlanks</td><td>11029</td><td>11060</td><td>32</td><td>CTCTCTCTCTCTCCCCCGCTCCCCGTCCTCCC</td><td>296.4</td><td>-7.8</td><td>1.3</td><td>-6.3</td><td>3.7E-10</td><td>8.1E-10</td><td>MB231/MB468</td><td>MITO</td></tr>
<tr><td>>hg38_21_+_8433172_8446622_RNA45SN1_45S_50ntFlanks@4163.4186.24|CATCCAAGGAAGGCAGCAGGCGCG</td><td>>hg38_21_+_8433172_8446622_RNA45SN1_45S_50ntFlanks</td><td>4163</td><td>4186</td><td>24</td><td>CATCCAAGGAAGGCAGCAGGCGCG</td><td>524.8</td><td>-8.3</td><td>1.3</td><td>-6.3</td><td>3.2E-10</td><td>8.1E-10</td><td>MB231/MB468</td><td>MP</td></tr>
<tr><td>>hg38_21_+_8433172_8446622_RNA45SN1_45S_50ntFlanks@5486.5516.31|GCGCTGAGAAGACGGTCGAACTTGACTATCT</td><td>>hg38_21_+_8433172_8446622_RNA45SN1_45S_50ntFlanks</td><td>5486</td><td>5516</td><td>31</td><td>GCGCTGAGAAGACGGTCGAACTTGACTATCT</td><td>2764.1</td><td>5.7</td><td>0.9</td><td>6.3</td><td>3.0E-10</td><td>8.1E-10</td><td>BT20/MB468</td><td>CYTO</td></tr>
<tr><td>>MI0000077|hsa-mir-21&WithFlank&17|+|59841260|59841343@14.35.22&[MIMAT0000076&hsa-miR-21-5p&offsets|0|0;m-1&17|+|59841273|59841294&offsets|0|0]|TAGCTTATCAGACTGATGTTGA</td><td>>MI0000077|hsa-mir-21&WithFlank&17|+|59841260|59841343</td><td>14</td><td>35</td><td>22</td><td>TAGCTTATCAGACTGATGTTGA</td><td>96383.4</td><td>6.1</td><td>1.0</td><td>6.3</td><td>3.7E-10</td><td>8.1E-10</td><td>MB231/MB468</td><td>MITO</td></tr>
<tr><td>>hg38_21_+_8433172_8446622_RNA45SN1_45S_50ntFlanks@10410.10443.34|TCAGTCGGTCCTGAGAGATGGGCGAGCGCCGTTC</td><td>>hg38_21_+_8433172_8446622_RNA45SN1_45S_50ntFlanks</td><td>10410</td><td>10443</td><td>34</td><td>TCAGTCGGTCCTGAGAGATGGGCGAGCGCCGTTC</td><td>576.7</td><td>-5.3</td><td>0.8</td><td>-6.3</td><td>3.3E-10</td><td>8.1E-10</td><td>MB231/MB468</td><td>CYTO</td></tr>
<tr><td>>hg38_21_+_8433172_8446622_RNA45SN1_45S_50ntFlanks@10678.10705.28|CGGTTCCGGCGGCGTCCGGTGAGCTCTC</td><td>>hg38_21_+_8433172_8446622_RNA45SN1_45S_50ntFlanks</td><td>10678</td><td>10705</td><td>28</td><td>CGGTTCCGGCGGCGTCCGGTGAGCTCTC</td><td>984.2</td><td>-3.9</td><td>0.6</td><td>-6.3</td><td>3.0E-10</td><td>8.1E-10</td><td>BT20/MB468</td><td>CYTO</td></tr>
<tr><td>>hg38_21_+_8433172_8446622_RNA45SN1_45S_50ntFlanks@11581.11613.33|AACTGGTGCGGACCAGGGGAATCCGACTGTTTA</td><td>>hg38_21_+_8433172_8446622_RNA45SN1_45S_50ntFlanks</td><td>11581</td><td>11613</td><td>33</td><td>AACTGGTGCGGACCAGGGGAATCCGACTGTTTA</td><td>576.1</td><td>-4.6</td><td>0.7</td><td>-6.3</td><td>3.2E-10</td><td>8.1E-10</td><td>MB231/MB468</td><td>MP</td></tr>
<tr><td>>hg38_21_+_8433172_8446622_RNA45SN1_45S_50ntFlanks@11873.11912.40|AAGAAGACCCTGTTGAGCTTGACTCTAGTCTGGCACGGTG</td><td>>hg38_21_+_8433172_8446622_RNA45SN1_45S_50ntFlanks</td><td>11873</td><td>11912</td><td>40</td><td>AAGAAGACCCTGTTGAGCTTGACTCTAGTCTGGCACGGTG</td><td>271.4</td><td>-10.2</td><td>1.6</td><td>-6.3</td><td>2.5E-10</td><td>8.1E-10</td><td>BT20/MB468</td><td>MITO</td></tr>
<tr><td>>hg38_21_+_8433172_8446622_RNA45SN1_45S_50ntFlanks@12159.12182.24|TTTGACTGGGGCGGTACACCTGTC</td><td>>hg38_21_+_8433172_8446622_RNA45SN1_45S_50ntFlanks</td><td>12159</td><td>12182</td><td>24</td><td>TTTGACTGGGGCGGTACACCTGTC</td><td>1288.4</td><td>-6.8</td><td>1.1</td><td>-6.3</td><td>2.5E-10</td><td>8.1E-10</td><td>BT20/MB468</td><td>MITO</td></tr>
<tr><td>>hg38_21_+_8433172_8446622_RNA45SN1_45S_50ntFlanks@8423.8442.20|CGCAGTCCGCCCGGAGGATT</td><td>>hg38_21_+_8433172_8446622_RNA45SN1_45S_50ntFlanks</td><td>8423</td><td>8442</td><td>20</td><td>CGCAGTCCGCCCGGAGGATT</td><td>4564.8</td><td>-5.4</td><td>0.9</td><td>-6.3</td><td>2.5E-10</td><td>8.1E-10</td><td>BT20/MB468</td><td>MITO</td></tr>
<tr><td>>hg38_21_+_8433172_8446622_RNA45SN1_45S_50ntFlanks@3984.4013.30|GCGGCTTTGGTGACTCTAGATAACCTCGGG</td><td>>hg38_21_+_8433172_8446622_RNA45SN1_45S_50ntFlanks</td><td>3984</td><td>4013</td><td>30</td><td>GCGGCTTTGGTGACTCTAGATAACCTCGGG</td><td>107.0</td><td>-10.6</td><td>1.7</td><td>-6.3</td><td>3.2E-10</td><td>8.1E-10</td><td>MB231/MB468</td><td>MP</td></tr>
<tr><td>>trnaMT_GluTTC_MT_-_14674_14742@45.72.28,trnalookalike8_GluTTC_5_-_93905172_93905240@45.72.28|CGTGGTTGTAGTCCGTGCGAGAATACCA</td><td>>trnaMT_GluTTC_MT_-_14674_14742</td><td>45</td><td>72</td><td>28</td><td>CGTGGTTGTAGTCCGTGCGAGAATACCA</td><td>832.8</td><td>5.6</td><td>0.9</td><td>6.3</td><td>3.4E-10</td><td>8.1E-10</td><td>MB231/MB468</td><td>NUC</td></tr>
<tr><td>>hg38_21_+_8433172_8446622_RNA45SN1_45S_50ntFlanks@10574.10598.25|AAGCCGGCGGGAGCCCCGGGGAGAG</td><td>>hg38_21_+_8433172_8446622_RNA45SN1_45S_50ntFlanks</td><td>10574</td><td>10598</td><td>25</td><td>AAGCCGGCGGGAGCCCCGGGGAGAG</td><td>80.2</td><td>-10.2</td><td>1.6</td><td>-6.3</td><td>3.2E-10</td><td>8.1E-10</td><td>MB231/MB468</td><td>MP</td></tr>
<tr><td>>hg38_21_+_8433172_8446622_RNA45SN1_45S_50ntFlanks@11792.11809.18|ATGAACGAGATTCCCACT</td><td>>hg38_21_+_8433172_8446622_RNA45SN1_45S_50ntFlanks</td><td>11792</td><td>11809</td><td>18</td><td>ATGAACGAGATTCCCACT</td><td>137.7</td><td>-11.0</td><td>1.7</td><td>-6.3</td><td>3.2E-10</td><td>8.1E-10</td><td>MB231/MB468</td><td>MP</td></tr>
<tr><td>>hg38_21_+_8433172_8446622_RNA45SN1_45S_50ntFlanks@3839.3861.23|TCTCCTACTTGGATAACTGTGGT</td><td>>hg38_21_+_8433172_8446622_RNA45SN1_45S_50ntFlanks</td><td>3839</td><td>3861</td><td>23</td><td>TCTCCTACTTGGATAACTGTGGT</td><td>285.5</td><td>3.6</td><td>0.6</td><td>6.3</td><td>3.3E-10</td><td>8.1E-10</td><td>MB231/MB468</td><td>CYTO</td></tr>
<tr><td>>hg38_21_+_8433172_8446622_RNA45SN1_45S_50ntFlanks@10801.10818.18|GTTGGAACAATGTAGGTA</td><td>>hg38_21_+_8433172_8446622_RNA45SN1_45S_50ntFlanks</td><td>10801</td><td>10818</td><td>18</td><td>GTTGGAACAATGTAGGTA</td><td>759.9</td><td>-6.5</td><td>1.0</td><td>-6.3</td><td>3.3E-10</td><td>8.1E-10</td><td>MB231/MB468</td><td>CYTO</td></tr>
<tr><td>>trnaMT_GluTTC_MT_-_14674_14742@39.58.20,trnalookalike8_GluTTC_5_-_93905172_93905240@39.58.20|ATTGGTCGTGGTTGTAGTCC</td><td>>trnaMT_GluTTC_MT_-_14674_14742</td><td>39</td><td>58</td><td>20</td><td>ATTGGTCGTGGTTGTAGTCC</td><td>555.1</td><td>8.8</td><td>1.4</td><td>6.3</td><td>3.3E-10</td><td>8.2E-10</td><td>MB231/MB468</td><td>CYTO</td></tr>
<tr><td>>MI0001445|hsa-mir-423&WithFlank&17|+|30117073|30117178@23.43.21&[MIMAT0004748&hsa-miR-423-5p&offsets|0|-2;m-141&17|+|30117095|30117117&offsets|0|-2]|TGAGGGGCAGAGAGCGAGACT</td><td>>MI0001445|hsa-mir-423&WithFlank&17|+|30117073|30117178</td><td>23</td><td>43</td><td>21</td><td>TGAGGGGCAGAGAGCGAGACT</td><td>10062.2</td><td>7.0</td><td>1.1</td><td>6.6</td><td>2.9E-11</td><td>8.2E-10</td><td>BT20/MB468</td><td>MP</td></tr>
<tr><td>>hg38_21_+_8433172_8446622_RNA45SN1_45S_50ntFlanks@8836.8864.29|CGGCGGTGGCGGCGGCGGCGGCGGCGGCG</td><td>>hg38_21_+_8433172_8446622_RNA45SN1_45S_50ntFlanks</td><td>8836</td><td>8864</td><td>29</td><td>CGGCGGTGGCGGCGGCGGCGGCGGCGGCG</td><td>228.1</td><td>-10.9</td><td>1.6</td><td>-6.6</td><td>2.9E-11</td><td>8.2E-10</td><td>BT20/MB468</td><td>MP</td></tr>
<tr><td>>hg38_21_+_8433172_8446622_RNA45SN1_45S_50ntFlanks@10768.10793.26|CAGCAGGTCTCCAAGGTGAACAGCCT</td><td>>hg38_21_+_8433172_8446622_RNA45SN1_45S_50ntFlanks</td><td>10768</td><td>10793</td><td>26</td><td>CAGCAGGTCTCCAAGGTGAACAGCCT</td><td>74.0</td><td>-10.1</td><td>1.6</td><td>-6.3</td><td>3.2E-10</td><td>8.2E-10</td><td>MB231/MB468</td><td>MP</td></tr>
<tr><td>>hg38_21_+_8433172_8446622_RNA45SN1_45S_50ntFlanks@10817.10840.24|TAAGGGAAGTCGGCAAGCCGGATC</td><td>>hg38_21_+_8433172_8446622_RNA45SN1_45S_50ntFlanks</td><td>10817</td><td>10840</td><td>24</td><td>TAAGGGAAGTCGGCAAGCCGGATC</td><td>382.0</td><td>-6.0</td><td>1.0</td><td>-6.3</td><td>3.4E-10</td><td>8.2E-10</td><td>MB231/MB468</td><td>NUC</td></tr>
<tr><td>>MI0000469|hsa-mir-125a&WithFlank&19|+|51693248|51693345@21.41.21&[MIMAT0000443&hsa-miR-125a-5p&offsets|0|-3;m-63&19|+|51693268|51693289&offsets|0|-1]|TCCCTGAGACCCTTTAACCTG</td><td>>MI0000469|hsa-mir-125a&WithFlank&19|+|51693248|51693345</td><td>21</td><td>41</td><td>21</td><td>TCCCTGAGACCCTTTAACCTG</td><td>784.4</td><td>8.9</td><td>1.4</td><td>6.3</td><td>2.5E-10</td><td>8.2E-10</td><td>BT20/MB468</td><td>MITO</td></tr>
<tr><td>>MI0000264|hsa-mir-7-2&WithFlank&15|+|88611819|88611940@38.58.21;MI0000265|hsa-mir-7-3&WithFlank&19|+|4770664|4770785@37.57.21;MI0000263|hsa-mir-7-1&WithFlank&9|-|83969742|83969863@30.50.21&[MIMAT0000252&hsa-miR-7-5p&offsets|0|-3;m-400&15|+|88611856|88611879&offsets|0|-3];[MIMAT0000252_1&hsa-miR-7-5p&offsets|0|-3;m-402&19|+|4770700|4770723&offsets|0|-3];[MIMAT0000252_2&hsa-miR-7-5p&offsets|0|-3;m-401&9|-|83969811|83969834&offsets|0|-3]|TGGAAGACTAGTGATTTTGTT</td><td>>MI0000264|hsa-mir-7-2&WithFlank&15|+|88611819|88611940</td><td>38</td><td>58</td><td>21</td><td>TGGAAGACTAGTGATTTTGTT</td><td>310.8</td><td>4.8</td><td>0.8</td><td>6.3</td><td>3.4E-10</td><td>8.2E-10</td><td>MB231/MB468</td><td>CYTO</td></tr>
<tr><td>>am_trnaMT_PheGAA_MT_+_577_647@41.59.19|GTTTAGACGGGCTCACATC</td><td>>am_trnaMT_PheGAA_MT_+_577_647</td><td>41</td><td>59</td><td>19</td><td>GTTTAGACGGGCTCACATC</td><td>833.3</td><td>10.1</td><td>1.6</td><td>6.3</td><td>3.0E-10</td><td>8.2E-10</td><td>BT20/MB468</td><td>CYTO</td></tr>
<tr><td>>hg38_21_+_8433172_8446622_RNA45SN1_45S_50ntFlanks@6742.6771.30|TCGACACTTCGAACGCACTTGCGGCCCCGG</td><td>>hg38_21_+_8433172_8446622_RNA45SN1_45S_50ntFlanks</td><td>6742</td><td>6771</td><td>30</td><td>TCGACACTTCGAACGCACTTGCGGCCCCGG</td><td>200.9</td><td>-4.8</td><td>0.8</td><td>-6.3</td><td>3.4E-10</td><td>8.2E-10</td><td>MB231/MB468</td><td>CYTO</td></tr>
<tr><td>>hg38_21_+_8433172_8446622_RNA45SN1_45S_50ntFlanks@9564.9598.35|CTGACGTGCAAATCGGTCGTCCGACCTGGGTATAG</td><td>>hg38_21_+_8433172_8446622_RNA45SN1_45S_50ntFlanks</td><td>9564</td><td>9598</td><td>35</td><td>CTGACGTGCAAATCGGTCGTCCGACCTGGGTATAG</td><td>780.5</td><td>-6.6</td><td>1.1</td><td>-6.3</td><td>3.8E-10</td><td>8.2E-10</td><td>MB231/MB468</td><td>MITO</td></tr>
<tr><td>>hg38_21_+_8433172_8446622_RNA45SN1_45S_50ntFlanks@12043.12068.26|CACTGACCCGGTGAGGCGGGGGGGCG</td><td>>hg38_21_+_8433172_8446622_RNA45SN1_45S_50ntFlanks</td><td>12043</td><td>12068</td><td>26</td><td>CACTGACCCGGTGAGGCGGGGGGGCG</td><td>94.5</td><td>-10.4</td><td>1.7</td><td>-6.3</td><td>3.3E-10</td><td>8.2E-10</td><td>MB231/MB468</td><td>MP</td></tr>
<tr><td>>hg38_21_+_8433172_8446622_RNA45SN1_45S_50ntFlanks@10783.10801.19|GTGAACAGCCTCTGGCATG</td><td>>hg38_21_+_8433172_8446622_RNA45SN1_45S_50ntFlanks</td><td>10783</td><td>10801</td><td>19</td><td>GTGAACAGCCTCTGGCATG</td><td>416.6</td><td>-6.2</td><td>1.0</td><td>-6.3</td><td>3.4E-10</td><td>8.3E-10</td><td>MB231/MB468</td><td>CYTO</td></tr>
<tr><td>>hg38_21_+_8433172_8446622_RNA45SN1_45S_50ntFlanks@11235.11269.35|CTTCCCGTGGATCGCCCCAGCTGCGGCGGGCGTCG</td><td>>hg38_21_+_8433172_8446622_RNA45SN1_45S_50ntFlanks</td><td>11235</td><td>11269</td><td>35</td><td>CTTCCCGTGGATCGCCCCAGCTGCGGCGGGCGTCG</td><td>208.8</td><td>-10.8</td><td>1.6</td><td>-6.6</td><td>3.0E-11</td><td>8.3E-10</td><td>BT20/MB468</td><td>MP</td></tr>
<tr><td>>hg38_21_+_8433172_8446622_RNA45SN1_45S_50ntFlanks@12676.12708.33|AGAATCCCGCCCAGGCGGAACGATACGGCAGCG</td><td>>hg38_21_+_8433172_8446622_RNA45SN1_45S_50ntFlanks</td><td>12676</td><td>12708</td><td>33</td><td>AGAATCCCGCCCAGGCGGAACGATACGGCAGCG</td><td>797.4</td><td>-8.7</td><td>1.3</td><td>-6.6</td><td>3.0E-11</td><td>8.3E-10</td><td>BT20/MB468</td><td>MP</td></tr>
<tr><td>>hg38_21_+_8433172_8446622_RNA45SN1_45S_50ntFlanks@12874.12891.18|GCTGGAAAGGCGGCCGCC</td><td>>hg38_21_+_8433172_8446622_RNA45SN1_45S_50ntFlanks</td><td>12874</td><td>12891</td><td>18</td><td>GCTGGAAAGGCGGCCGCC</td><td>184.9</td><td>-10.6</td><td>1.6</td><td>-6.6</td><td>3.0E-11</td><td>8.3E-10</td><td>BT20/MB468</td><td>MP</td></tr>
<tr><td>>hg38_21_+_8433172_8446622_RNA45SN1_45S_50ntFlanks@8198.8215.18|TGGACGGTGTGAGGCCGG</td><td>>hg38_21_+_8433172_8446622_RNA45SN1_45S_50ntFlanks</td><td>8198</td><td>8215</td><td>18</td><td>TGGACGGTGTGAGGCCGG</td><td>2306.1</td><td>-7.8</td><td>1.2</td><td>-6.6</td><td>3.0E-11</td><td>8.3E-10</td><td>BT20/MB468</td><td>MP</td></tr>
<tr><td>>hg38_21_+_8433172_8446622_RNA45SN1_45S_50ntFlanks@9031.9058.28|TCCTCGTGGGGGGGCCGGGCCACCCCTC</td><td>>hg38_21_+_8433172_8446622_RNA45SN1_45S_50ntFlanks</td><td>9031</td><td>9058</td><td>28</td><td>TCCTCGTGGGGGGGCCGGGCCACCCCTC</td><td>190.5</td><td>-10.7</td><td>1.6</td><td>-6.6</td><td>3.0E-11</td><td>8.3E-10</td><td>BT20/MB468</td><td>MP</td></tr>
<tr><td>>hg38_21_+_8433172_8446622_RNA45SN1_45S_50ntFlanks@10767.10785.19|GCAGCAGGTCTCCAAGGTG</td><td>>hg38_21_+_8433172_8446622_RNA45SN1_45S_50ntFlanks</td><td>10767</td><td>10785</td><td>19</td><td>GCAGCAGGTCTCCAAGGTG</td><td>122.2</td><td>-9.0</td><td>1.4</td><td>-6.3</td><td>2.6E-10</td><td>8.3E-10</td><td>BT20/MB468</td><td>MITO</td></tr>
<tr><td>>hg38_21_+_8433172_8446622_RNA45SN1_45S_50ntFlanks@3842.3869.28|CCTACTTGGATAACTGTGGTAATTCTAG</td><td>>hg38_21_+_8433172_8446622_RNA45SN1_45S_50ntFlanks</td><td>3842</td><td>3869</td><td>28</td><td>CCTACTTGGATAACTGTGGTAATTCTAG</td><td>111.5</td><td>-10.7</td><td>1.7</td><td>-6.3</td><td>3.3E-10</td><td>8.3E-10</td><td>MB231/MB468</td><td>MP</td></tr>
<tr><td>>MI0000088|hsa-mir-30a&WithFlank&6|-|71403545|71403627@12.31.20&[MIMAT0000087&hsa-miR-30a-5p&offsets|0|-2;m-15&6|-|71403595|71403616&offsets|0|-2]|TGTAAACATCCTCGACTGGA</td><td>>MI0000088|hsa-mir-30a&WithFlank&6|-|71403545|71403627</td><td>12</td><td>31</td><td>20</td><td>TGTAAACATCCTCGACTGGA</td><td>2638.5</td><td>6.2</td><td>1.0</td><td>6.3</td><td>3.8E-10</td><td>8.3E-10</td><td>MB231/MB468</td><td>MITO</td></tr>
<tr><td>>hg38_21_+_8433172_8446622_RNA45SN1_45S_50ntFlanks@7975.7998.24|CGCGACCTCAGATCAGACGTGGCG</td><td>>hg38_21_+_8433172_8446622_RNA45SN1_45S_50ntFlanks</td><td>7975</td><td>7998</td><td>24</td><td>CGCGACCTCAGATCAGACGTGGCG</td><td>624.2</td><td>4.2</td><td>0.7</td><td>6.3</td><td>3.0E-10</td><td>8.3E-10</td><td>BT20/MB468</td><td>CYTO</td></tr>
</table>

| | | | | | | | | | | | | | |
|---|---|---|---|---|---|---|---|---|---|---|---|---|---|
| >hg38_21_+_8433172_8446622_RNA45SN1_45S_50ntFlanks@6722.6745.24\|ATTGCAGGACACATTGATCATCGA | >hg38_21_+_8433172_8446622_RNA45SN1_45S_50ntFlanks | 6722 | 6745 | 24 | ATTGCAGGACACATTGATCATCGA | 249.3 | -7.6 | 1.2 | -6.3 | 3.0E-10 | 8.3E-10 | BT20/MB468 | CYTO |
| >hg38_21_+_8433172_8446622_RNA45SN1_45S_50ntFlanks@3775.3793.19\|GCCGGTACAGTGAAACTGC | >hg38_21_+_8433172_8446622_RNA45SN1_45S_50ntFlanks | 3775 | 3793 | 19 | GCCGGTACAGTGAAACTGC | 157.6 | -11.2 | 1.8 | -6.3 | 3.3E-10 | 8.3E-10 | MB231/MB468 | MP |
| >hg38_21_+_8433172_8446622_RNA45SN1_45S_50ntFlanks@12045.12068.24\|CTGACCCGGTGAGGCGGGGGGGCG | >hg38_21_+_8433172_8446622_RNA45SN1_45S_50ntFlanks | 12045 | 12068 | 24 | CTGACCCGGTGAGGCGGGGGGGCG | 695.8 | -9.4 | 1.5 | -6.3 | 4.0E-10 | 8.3E-10 | BT20/MB468 | NUC |
| >MI0000066\|hsa-let-7e&WithFlank&19\|+\|51692780\|51692870@14.34.21&[MIMAT0000066&hsa-let-7e-5p&offsets\|0\|-1;m-50&19\|+\|51692793\|51692814&offsets\|0\|-1]\|TGAGGTAGGAGGTTGTATAGT | >MI0000066\|hsa-let-7e&WithFlank&19\|+\|51692780\|51692870 | 14 | 34 | 21 | TGAGGTAGGAGGTTGTATAGT | 1182.0 | 3.8 | 0.6 | 6.3 | 3.4E-10 | 8.3E-10 | MB231/MB468 | NUC |
| >hg38_MT_+_598_1651_RNR1_12S_50ntFlanks@367.404.38\|CCAATAAAGCTAAAACTCACCTGAGTTGTAAAAAACTC | >hg38_MT_+_598_1651_RNR1_12S_50ntFlanks | 367 | 404 | 38 | CCAATAAAGCTAAAACTCACCTGAGTTGTAAAAAACTC | 1212.7 | 5.3 | 0.8 | 6.3 | 2.6E-10 | 8.3E-10 | BT20/MB468 | MITO |
| >hg38_21_+_8433172_8446622_RNA45SN1_45S_50ntFlanks@11629.11669.41\|CGCGAAGGCCCGCGGCGGGTGTTGACGCGATGTGATTTCTG | >hg38_21_+_8433172_8446622_RNA45SN1_45S_50ntFlanks | 11629 | 11669 | 41 | CGCGAAGGCCCGCGGCGGGTGTTGACGCGATGTGATTTCTG | 161.7 | -11.2 | 1.8 | -6.3 | 3.3E-10 | 8.3E-10 | MB231/MB468 | MP |
| >hg38_21_+_8433172_8446622_RNA45SN1_45S_50ntFlanks@4759.4782.24\|ATCAGATACCGTCGTAGTTCCGAC | >hg38_21_+_8433172_8446622_RNA45SN1_45S_50ntFlanks | 4759 | 4782 | 24 | ATCAGATACCGTCGTAGTTCCGAC | 2046.2 | -8.0 | 1.3 | -6.3 | 3.4E-10 | 8.3E-10 | MB231/MB468 | CYTO |
| >trna10_LysCTT_16_+_3241501_3241573@5.34.30,trna13_LysCTT_14_-_58706613_58706685@5.34.30,trna2_LysCTT_15_+_79152904_79152976@5.34.30\|GGCTAGCTCAGTCGGTAGAGCATGGGACTC | >trna10_LysCTT_16_+_3241501_3241573 | 5 | 34 | 30 | GGCTAGCTCAGTCGGTAGAGCATGGGACTC | 153.1 | 10.5 | 1.7 | 6.3 | 3.8E-10 | 8.3E-10 | MB231/MB468 | MITO |
| >hg38_21_+_8433172_8446622_RNA45SN1_45S_50ntFlanks@4742.4759.18\|TCGGAGGTTCGAAGACGA | >hg38_21_+_8433172_8446622_RNA45SN1_45S_50ntFlanks | 4742 | 4759 | 18 | TCGGAGGTTCGAAGACGA | 581.2 | -4.5 | 0.7 | -6.3 | 3.3E-10 | 8.3E-10 | MB231/MB468 | MP |
| >hg38_1_-_228634819_228635039_RNA5S12_5S_50ntFlanks@153.171.19\|ACCGGGTGCTGTAGGCTTT | >hg38_1_-_228634819_228635039_RNA5S12_5S_50ntFlanks | 153 | 171 | 19 | ACCGGGTGCTGTAGGCTTT | 7439.4 | 5.1 | 0.8 | 6.3 | 3.5E-10 | 8.3E-10 | MB231/MB468 | NUC |
| >hg38_21_+_8433172_8446622_RNA45SN1_45S_50ntFlanks@11921.11950.30\|TGAGAGGTGTAGAATAAGTGGGAGGCCCCC | >hg38_21_+_8433172_8446622_RNA45SN1_45S_50ntFlanks | 11921 | 11950 | 30 | TGAGAGGTGTAGAATAAGTGGGAGGCCCCC | 690.1 | -8.8 | 1.4 | -6.3 | 3.3E-10 | 8.4E-10 | MB231/MB468 | MP |
| >hg38_21_+_8433172_8446622_RNA45SN1_45S_50ntFlanks@5312.5330.19\|TCGGGGATTGCAATTATTC | >hg38_21_+_8433172_8446622_RNA45SN1_45S_50ntFlanks | 5312 | 5330 | 19 | TCGGGGATTGCAATTATTC | 121.4 | -10.8 | 1.7 | -6.3 | 3.3E-10 | 8.4E-10 | MB231/MB468 | MP |
| >hg38_21_+_8433172_8446622_RNA45SN1_45S_50ntFlanks@12456.12483.28\|CCAAGCGTTGGATTGTTCACCCACTAAT | >hg38_21_+_8433172_8446622_RNA45SN1_45S_50ntFlanks | 12456 | 12483 | 28 | CCAAGCGTTGGATTGTTCACCCACTAAT | 815.3 | -6.1 | 1.0 | -6.3 | 2.6E-10 | 8.4E-10 | BT20/MB468 | MITO |
| >hg38_21_+_8433172_8446622_RNA45SN1_45S_50ntFlanks@9204.9226.23\|TCGGGGCCACGCGCGCGTCCCCC | >hg38_21_+_8433172_8446622_RNA45SN1_45S_50ntFlanks | 9204 | 9226 | 23 | TCGGGGCCACGCGCGCGTCCCCC | 117.0 | -10.7 | 1.7 | -6.3 | 3.3E-10 | 8.4E-10 | MB231/MB468 | MP |
| >hg38_21_+_8433172_8446622_RNA45SN1_45S_50ntFlanks@10786.10829.44\|AACAGCCTCTGGCATGTTGGAACAATGTAGGTAAGGGAAGTCGG | >hg38_21_+_8433172_8446622_RNA45SN1_45S_50ntFlanks | 10786 | 10829 | 44 | AACAGCCTCTGGCATGTTGGAACAATGTAGGTAAGGGAAGTCGG | 138.0 | -11.0 | 1.7 | -6.3 | 3.3E-10 | 8.4E-10 | MB231/MB468 | MP |
| >hg38_21_+_8433172_8446622_RNA45SN1_45S_50ntFlanks@8806.8823.18\|AGGGCGCGCGGGTCGGGG | >hg38_21_+_8433172_8446622_RNA45SN1_45S_50ntFlanks | 8806 | 8823 | 18 | AGGGCGCGCGGGTCGGGG | 82.6 | -10.2 | 1.6 | -6.3 | 3.3E-10 | 8.4E-10 | MB231/MB468 | MP |
| >MI0001445\|hsa-mir-423&WithFlank&17\|+\|30117073\|30117178@59.80.22(+1C)&[MIMAT0001340&hsa-miR-423-3p&offsets\|0\|-1(+1C);m-100&17\|+\|30117131\|30117153&offsets\|0\|-1(+1C)]\|AGCTCGGTCTGAGGCCCCTCAGC | >MI0001445\|hsa-mir-423&WithFlank&17\|+\|30117073\|30117178 | 59 | 80 | 22(+1C) | AGCTCGGTCTGAGGCCCCTCAGC | 3078.7 | 6.2 | 1.0 | 6.3 | 3.9E-10 | 8.4E-10 | MB231/MB468 | MITO |
| >hg38_21_+_8433172_8446622_RNA45SN1_45S_50ntFlanks@12717.12751.35\|CCTCGGTTGGCCTCGGATAGCCGGTCCCCCGCCTG | >hg38_21_+_8433172_8446622_RNA45SN1_45S_50ntFlanks | 12717 | 12751 | 35 | CCTCGGTTGGCCTCGGATAGCCGGTCCCCCGCCTG | 121.7 | -9.0 | 1.4 | -6.3 | 2.6E-10 | 8.4E-10 | BT20/MB468 | MITO |
| >hg38_21_+_8433172_8446622_RNA45SN1_45S_50ntFlanks@8557.8581.25\|CGGAGGGGGCGGGCTCCGGCGGGTG | >hg38_21_+_8433172_8446622_RNA45SN1_45S_50ntFlanks | 8557 | 8581 | 25 | CGGAGGGGGCGGGCTCCGGCGGGTG | 92.3 | -10.4 | 1.7 | -6.3 | 3.3E-10 | 8.4E-10 | MB231/MB468 | MP |
| >MI0000301\|hsa-mir-224&WithFlank&X\|-\|151958572\|151958664@14.36.23&[MIMAT0000281&hsa-miR-224-5p&offsets\|+1\|-1;m-238&X\|-\|151958630\|151958651&offsets\|0\|+1]\|CAAGTCACTAGTGGTTCCGTTTA | >MI0000301\|hsa-mir-224&WithFlank&X\|-\|151958572\|151958664 | 14 | 36 | 23 | CAAGTCACTAGTGGTTCCGTTTA | 270.7 | 11.8 | 1.8 | 6.6 | 3.1E-11 | 8.4E-10 | BT20/MB468 | MP |
| >hg38_21_+_8433172_8446622_RNA45SN1_45S_50ntFlanks@12044.12070.27\|ACTGACCCGGTGAGGCGGGGGGGCGAG | >hg38_21_+_8433172_8446622_RNA45SN1_45S_50ntFlanks | 12044 | 12070 | 27 | ACTGACCCGGTGAGGCGGGGGGGCGAG | 999.0 | -9.2 | 1.4 | -6.6 | 3.1E-11 | 8.4E-10 | BT20/MB468 | MP |
| >hg38_1_-_228634819_228635039_RNA5S12_5S_50ntFlanks@144.170.27\|CCTGGGAATACCGGGTGCTGTAGGCTT | >hg38_1_-_228634819_228635039_RNA5S12_5S_50ntFlanks | 144 | 170 | 27 | CCTGGGAATACCGGGTGCTGTAGGCTT | 3349.7 | 5.2 | 0.8 | 6.3 | 3.5E-10 | 8.4E-10 | MB231/MB468 | NUC |
| >hg38_21_+_8433172_8446622_RNA45SN1_45S_50ntFlanks@10458.10495.38\|ATGGCCTCCGTTGCCCTCGGCCGATCGAAAGGGAGTCG | >hg38_21_+_8433172_8446622_RNA45SN1_45S_50ntFlanks | 10458 | 10495 | 38 | ATGGCCTCCGTTGCCCTCGGCCGATCGAAAGGGAGTCG | 221.0 | -6.7 | 1.1 | -6.3 | 3.5E-10 | 8.4E-10 | MB231/MB468 | CYTO |
| >hg38_21_+_8433172_8446622_RNA45SN1_45S_50ntFlanks@6734.6773.40\|ATTGATCATCGACACTTCGAACGCACTTGCGGCCCCGGGT | >hg38_21_+_8433172_8446622_RNA45SN1_45S_50ntFlanks | 6734 | 6773 | 40 | ATTGATCATCGACACTTCGAACGCACTTGCGGCCCCGGGT | 983.8 | 9.2 | 1.5 | 6.3 | 3.1E-10 | 8.4E-10 | BT20/MB468 | CYTO |
| >hg38_21_+_8433172_8446622_RNA45SN1_45S_50ntFlanks@10458.10475.18\|ATGGCCTCCGTTGCCCTC | >hg38_21_+_8433172_8446622_RNA45SN1_45S_50ntFlanks | 10458 | 10475 | 18 | ATGGCCTCCGTTGCCCTC | 118.8 | -10.8 | 1.7 | -6.3 | 3.3E-10 | 8.4E-10 | MB231/MB468 | MP |
| >hg38_21_+_8433172_8446622_RNA45SN1_45S_50ntFlanks@12542.12567.26\|ATGTGTTGTTGCCATGGTAATCCTGC | >hg38_21_+_8433172_8446622_RNA45SN1_45S_50ntFlanks | 12542 | 12567 | 26 | ATGTGTTGTTGCCATGGTAATCCTGC | 80.6 | -10.2 | 1.6 | -6.3 | 3.3E-10 | 8.4E-10 | MB231/MB468 | MP |
| >hg38_21_+_8433172_8446622_RNA45SN1_45S_50ntFlanks@11321.11342.22\|CACGTCTCGTCGCGCGCGCGTC | >hg38_21_+_8433172_8446622_RNA45SN1_45S_50ntFlanks | 11321 | 11342 | 22 | CACGTCTCGTCGCGCGCGCGTC | 2272.7 | -8.9 | 1.4 | -6.3 | 3.5E-10 | 8.4E-10 | MB231/MB468 | NUC |
| >trnaMT_PheGAA_MT_+_577_647@56.74.19\|CATCACCCCATAAACACCA | >trnaMT_PheGAA_MT_+_577_647 | 56 | 74 | 19 | CATCACCCCATAAACACCA | 1785.5 | 5.9 | 0.9 | 6.3 | 2.6E-10 | 8.4E-10 | BT20/MB468 | MITO |
| >hg38_21_+_8433172_8446622_RNA45SN1_45S_50ntFlanks@10389.10430.42\|TGAACAGCAGTTGAACATGGGTCAGTCGGTCCTGAGAGATGG | >hg38_21_+_8433172_8446622_RNA45SN1_45S_50ntFlanks | 10389 | 10430 | 42 | TGAACAGCAGTTGAACATGGGTCAGTCGGTCCTGAGAGATGG | 140.7 | -11.0 | 1.8 | -6.3 | 3.4E-10 | 8.5E-10 | MB231/MB468 | MP |
| >hg38_1_-_228634819_228635039_RNA5S12_5S_50ntFlanks@139.170.32\|GACCGCCTGGGAATACCGGGTGCTGTAGGCTT | >hg38_1_-_228634819_228635039_RNA5S12_5S_50ntFlanks | 139 | 170 | 32 | GACCGCCTGGGAATACCGGGTGCTGTAGGCTT | 1242.3 | 5.1 | 0.8 | 6.3 | 3.5E-10 | 8.5E-10 | MB231/MB468 | NUC |
| >hg38_21_+_8433172_8446622_RNA45SN1_45S_50ntFlanks@5073.5097.25\|AATTCCGATAACGAACGAGACTCTG | >hg38_21_+_8433172_8446622_RNA45SN1_45S_50ntFlanks | 5073 | 5097 | 25 | AATTCCGATAACGAACGAGACTCTG | 1311.3 | -7.9 | 1.3 | -6.3 | 3.9E-10 | 8.5E-10 | MB231/MB468 | MITO |
| >hg38_21_+_8433172_8446622_RNA45SN1_45S_50ntFlanks@9572.9590.19\|CAAATCGGTCGTCCGACCT | >hg38_21_+_8433172_8446622_RNA45SN1_45S_50ntFlanks | 9572 | 9590 | 19 | CAAATCGGTCGTCCGACCT | 99.6 | -10.5 | 1.7 | -6.3 | 3.4E-10 | 8.5E-10 | MB231/MB468 | MP |
| >hg38_21_+_8433172_8446622_RNA45SN1_45S_50ntFlanks@12623.12657.35\|GCCAATGGGGCGAAGCTACCATCTGTGGGATTATG | >hg38_21_+_8433172_8446622_RNA45SN1_45S_50ntFlanks | 12623 | 12657 | 35 | GCCAATGGGGCGAAGCTACCATCTGTGGGATTATG | 76.2 | -10.1 | 1.6 | -6.3 | 3.4E-10 | 8.5E-10 | MB231/MB468 | MP |
| >hg38_21_+_8433172_8446622_RNA45SN1_45S_50ntFlanks@476.505.30\|TTCGTGATCGATGTGGTGACGTCGTGCTCT | >hg38_21_+_8433172_8446622_RNA45SN1_45S_50ntFlanks | 476 | 505 | 30 | TTCGTGATCGATGTGGTGACGTCGTGCTCT | 1855.7 | 5.9 | 0.9 | 6.3 | 3.9E-10 | 8.5E-10 | MB231/MB468 | MITO |
| >hg38_21_+_8433172_8446622_RNA45SN1_45S_50ntFlanks@12959.12992.34\|TCTGGGTCGGGGTTTCGTACGTAGCAGAGCAGCT | >hg38_21_+_8433172_8446622_RNA45SN1_45S_50ntFlanks | 12959 | 12992 | 34 | TCTGGGTCGGGGTTTCGTACGTAGCAGAGCAGCT | 422.7 | 6.9 | 1.1 | 6.3 | 3.5E-10 | 8.5E-10 | MB231/MB468 | CYTO |
| >MI0000061\|hsa-let-7a-2&WithFlank&11\|-\|122146516\|122146599@11.28.18;MI0000064\|hsa-let-7c&WithFlank&21\|+\|16539822\|16539917@17.34.18;MI0000062\|hsa-let-7a-3&WithFlank&22\|+\|46112743\|46112828@10.27.18;MI0000060\|hsa-let-7a-1&WithFlank&9\|+\|94175951\|94176042@12.29.18&[MIMAT0000062&hsa-let-7a-5p&offsets\|0\|-4;m-6&11\|-\|122146568\|122146589&offsets\|0\|-4];[MIMAT0000064&hsa-let-7c-5p&offsets\|0\|-4;m-37&21\|+\|16539838\|16539859&offsets\|0\|-4];[MIMAT0000062_1&hsa-let-7a-5p&offsets\|0\|-4;m-5&22\|+\|46112752\|46112773&offsets\|0\|-4];[MIMAT0000062_2&hsa-let-7a-5p&offsets\|0\|-4;m-4&9\|+\|94175962\|94175983&offsets\|0\|-4]\|TGAGGTAGTAGGTTGTAT | >MI0000061\|hsa-let-7a-2&WithFlank&11\|-\|122146516\|122146599 | 11 | 28 | 18 | TGAGGTAGTAGGTTGTAT | 1809.6 | 6.1 | 1.0 | 6.3 | 3.9E-10 | 8.5E-10 | MB231/MB468 | MITO |
| >MI0000809\|hsa-mir-151a&WithFlank&8\|-\|140732558\|140732659@51.72.22(+1U)&[MIMAT0000757&hsa-miR-151a-3p&offsets\|-2\|-1(+1U);m-30&8\|-\|140732587\|140732607&offsets\|-2\|-1(+1U)]\|TACTAGACTGAAGCTCCTTGAGT | >MI0000809\|hsa-mir-151a&WithFlank&8\|-\|140732558\|140732659 | 51 | 72 | 22(+1U) | TACTAGACTGAAGCTCCTTGAGT | 80.2 | 2.2 | 0.4 | 6.2 | 4.1E-10 | 8.6E-10 | BT20/MB468 | NUC |
| >hg38_21_+_8433172_8446622_RNA45SN1_45S_50ntFlanks@12996.13016.21\|TCGCTGCGATCTATTGAAAGT | >hg38_21_+_8433172_8446622_RNA45SN1_45S_50ntFlanks | 12996 | 13016 | 21 | TCGCTGCGATCTATTGAAAGT | 386.9 | 5.1 | 0.8 | 6.3 | 3.6E-10 | 8.6E-10 | MB231/MB468 | NUC |
| >hg38_21_+_8433172_8446622_RNA45SN1_45S_50ntFlanks@5073.5102.30\|AATTCCGATAACGAACGAGACTCTGGCATG | >hg38_21_+_8433172_8446622_RNA45SN1_45S_50ntFlanks | 5073 | 5102 | 30 | AATTCCGATAACGAACGAGACTCTGGCATG | 1304.7 | -8.3 | 1.3 | -6.2 | 4.2E-10 | 8.6E-10 | BT20/MB468 | NUC |
| >hg38_21_+_8433172_8446622_RNA45SN1_45S_50ntFlanks@4588.4615.28\|CTATTTTGTTGGTTTTCGGAACTGAGGC | >hg38_21_+_8433172_8446622_RNA45SN1_45S_50ntFlanks | 4588 | 4615 | 28 | CTATTTTGTTGGTTTTCGGAACTGAGGC | 342.8 | -7.8 | 1.2 | -6.3 | 3.6E-10 | 8.6E-10 | MB231/MB468 | NUC |
| >hg38_21_+_8433172_8446622_RNA45SN1_45S_50ntFlanks@10572.10598.27\|AGAAGCCGGCGGGAGCCCCGGGGAGAG | >hg38_21_+_8433172_8446622_RNA45SN1_45S_50ntFlanks | 10572 | 10598 | 27 | AGAAGCCGGCGGGAGCCCCGGGGAGAG | 79.1 | -10.2 | 1.6 | -6.3 | 3.4E-10 | 8.6E-10 | MB231/MB468 | MP |
| >hg38_21_+_8433172_8446622_RNA45SN1_45S_50ntFlanks@3994.4013.20\|TGACTCTAGATAACCTCGGG | >hg38_21_+_8433172_8446622_RNA45SN1_45S_50ntFlanks | 3994 | 4013 | 20 | TGACTCTAGATAACCTCGGG | 516.4 | -7.3 | 1.2 | -6.2 | 4.2E-10 | 8.6E-10 | BT20/MB468 | NUC |
| >hg38_1_-_228634819_228635039_RNA5S12_5S_50ntFlanks@133.168.36\|ATGGGAGACCGCCTGGGAATACCGGGTGCTGTAGGC | >hg38_1_-_228634819_228635039_RNA5S12_5S_50ntFlanks | 133 | 168 | 36 | ATGGGAGACCGCCTGGGAATACCGGGTGCTGTAGGC | 300.0 | 4.8 | 0.8 | 6.3 | 3.2E-10 | 8.7E-10 | BT20/MB468 | CYTO |
| >MI0006444\|hsa-mir-1307&WithFlank&10\|-\|103394247\|103394407@87.109.23&[MIMAT0005951&hsa-miR-1307-3p&offsets\|+1\|+2;m-66&10\|-\|103394300\|103394322&offsets\|+1\|+1]\|CTCGGCGTGGCGTCGGTCGTGGT | >MI0006444\|hsa-mir-1307&WithFlank&10\|-\|103394247\|103394407 | 87 | 109 | 23 | CTCGGCGTGGCGTCGGTCGTGGT | 253.0 | 4.5 | 0.7 | 6.3 | 3.6E-10 | 8.7E-10 | MB231/MB468 | NUC |
| >am_MI0015825\|hsa-mir-378c&WithFlank&10\|-\|130962582\|130962674@17.37.21&[MIMAT0016847&hsa-miR-378c&offsets\|0\|-4;m-130&10\|-\|130962639\|130962658&offsets\|0\|+1]\|ACTGGACTTGGAGTCAGAAGA | >am_MI0015825\|hsa-mir-378c&WithFlank&10\|-\|130962582\|130962674 | 17 | 37 | 21 | ACTGGACTTGGAGTCAGAAGA | 193.0 | 3.8 | 0.6 | 6.3 | 3.6E-10 | 8.7E-10 | MB231/MB468 | NUC |
| >hg38_21_+_8433172_8446622_RNA45SN1_45S_50ntFlanks@10784.10818.35\|TGAACAGCCTCTGGCATGTTGGAACAATGTAGGTA | >hg38_21_+_8433172_8446622_RNA45SN1_45S_50ntFlanks | 10784 | 10818 | 35 | TGAACAGCCTCTGGCATGTTGGAACAATGTAGGTA | 462.5 | -8.4 | 1.3 | -6.3 | 3.4E-10 | 8.7E-10 | MB231/MB468 | MP |
| >hg38_21_+_8433172_8446622_RNA45SN1_45S_50ntFlanks@11583.11613.31\|CTGGTGCGGACCAGGGGAATCCGACTGTTTA | >hg38_21_+_8433172_8446622_RNA45SN1_45S_50ntFlanks | 11583 | 11613 | 31 | CTGGTGCGGACCAGGGGAATCCGACTGTTTA | 874.4 | -4.8 | 0.8 | -6.3 | 4.0E-10 | 8.7E-10 | MB231/MB468 | MITO |
| >hg38_21_+_8433172_8446622_RNA45SN1_45S_50ntFlanks@6734.6770.37\|ATTGATCATCGACACTTCGAACGCACTTGCGGCCCCG | >hg38_21_+_8433172_8446622_RNA45SN1_45S_50ntFlanks | 6734 | 6770 | 37 | ATTGATCATCGACACTTCGAACGCACTTGCGGCCCCG | 922.6 | -8.8 | 1.4 | -6.3 | 3.5E-10 | 8.7E-10 | MB231/MB468 | MP |

<table>
<tr><td></td><td></td><td></td><td></td><td></td><td></td><td></td><td></td><td></td><td></td><td></td><td></td><td></td><td></td></tr>
<tr><td>>hg38_21_+_8433172_8446622_RNA45SN1_45S_50ntFlanks@5472.5508.37|ACGGCCCTGGCGGAGCGCTGAGAAGACGGTCGAACTT</td><td>>hg38_21_+_8433172_8446622_RNA45SN1_45S_50ntFlanks</td><td>5472</td><td>5508</td><td>37</td><td>ACGGCCCTGGCGGAGCGCTGAGAAGACGGTCGAACTT</td><td>387.8</td><td>7.9</td><td>1.3</td><td>6.3</td><td>3.6E-10</td><td>8.7E-10</td><td>MB231/MB468</td><td>CYTO</td></tr>
<tr><td>>MI0000114|hsa-mir-107&WithFlank&10|-|89592741|89592833@56.76.21;MI0000108|hsa-mir-103a-2&WithFlank&20|+|3917488|3917577@54.74.21;MI0000109|hsa-mir-103a-1&WithFlank&5|-|168560890|168560979@54.74.21&[MIMAT0000104&hsa-miR-107&offsets|0|-2;m-78&10|-|89592758|89592778&offsets|0|0];[MIMAT0000101&hsa-miR-103a-3p&offsets|0|-2;m-12&20|+|3917541|3917563&offsets|0|-2];[MIMAT0000101_1&hsa-miR-103a-3p&offsets|0|-2;m-13&5|-|168560904|168560926&offsets|0|-2;m-44220&5|-|168560908|168560926&offsets|0|+2]|AGCAGCATTGTACAGGGCTAT</td><td>>MI0000114|hsa-mir-107&WithFlank&10|-|89592741|89592833</td><td>56</td><td>76</td><td>21</td><td>AGCAGCATTGTACAGGGCTAT</td><td>1349.2</td><td>6.8</td><td>1.1</td><td>6.3</td><td>4.0E-10</td><td>8.7E-10</td><td>MB231/MB468</td><td>MITO</td></tr>
<tr><td>>am_trna10_ValCAC_5_-_180649395_180649467@1.34.34,trna18_ValCAC_5_-_180529253_180529325@1.34.34,trna2_ValCAC_5_+_180524070_180524142@1.34.34,trna6_ValCAC_5_+_180600650_180600722@1.34.34,trna85_ValCAC_1_-_161369490_161369562@1.34.34,trna90_ValCAC_1_-_149684088_149684161@1.34.34,trna98_ValCAC_1_-_149298555_149298627@1.34.34,trna9_ValCAC_6_+_26538282_26538354@1.34.34|GTTTCCGTAGTGTAGTGGTTATCACGTTCGCCTC</td><td>>am_trna10_ValCAC_5_-_180649395_180649467</td><td>1</td><td>34</td><td>34</td><td>GTTTCCGTAGTGTAGTGGTTATCACGTTCGCCTC</td><td>1166.0</td><td>10.3</td><td>1.6</td><td>6.3</td><td>3.6E-10</td><td>8.7E-10</td><td>MB231/MB468</td><td>NUC</td></tr>
<tr><td>>hg38_21_+_8433172_8446622_RNA45SN1_45S_50ntFlanks@10798.10816.19|CATGTTGGAACAATGTAGG</td><td>>hg38_21_+_8433172_8446622_RNA45SN1_45S_50ntFlanks</td><td>10798</td><td>10816</td><td>19</td><td>CATGTTGGAACAATGTAGG</td><td>447.9</td><td>-4.3</td><td>0.7</td><td>-6.3</td><td>3.2E-10</td><td>8.7E-10</td><td>BT20/MB468</td><td>CYTO</td></tr>
<tr><td>>hg38_21_+_8433172_8446622_RNA45SN1_45S_50ntFlanks@12418.12441.24|CGATGTCGGCTCTTCCTATCATTG</td><td>>hg38_21_+_8433172_8446622_RNA45SN1_45S_50ntFlanks</td><td>12418</td><td>12441</td><td>24</td><td>CGATGTCGGCTCTTCCTATCATTG</td><td>1285.1</td><td>-4.5</td><td>0.7</td><td>-6.3</td><td>2.7E-10</td><td>8.7E-10</td><td>BT20/MB468</td><td>MITO</td></tr>
<tr><td>>MI0000811|hsa-mir-148b&WithFlank&12|+|54337210|54337320@69.90.22&[MIMAT0000759&hsa-miR-148b-3p&offsets|0|0;m-77&12|+|54337278|54337299&offsets|0|0]|TCAGTGCATCACAGAACTTTGT</td><td>>MI0000811|hsa-mir-148b&WithFlank&12|+|54337210|54337320</td><td>69</td><td>90</td><td>22</td><td>TCAGTGCATCACAGAACTTTGT</td><td>1561.9</td><td>2.8</td><td>0.5</td><td>6.2</td><td>4.2E-10</td><td>8.7E-10</td><td>BT20/MB468</td><td>NUC</td></tr>
<tr><td>>hg38_21_+_8433172_8446622_RNA45SN1_45S_50ntFlanks@10781.10809.29|AGGTGAACAGCCTCTGGCATGTTGGAACA</td><td>>hg38_21_+_8433172_8446622_RNA45SN1_45S_50ntFlanks</td><td>10781</td><td>10809</td><td>29</td><td>AGGTGAACAGCCTCTGGCATGTTGGAACA</td><td>331.3</td><td>-6.2</td><td>1.0</td><td>-6.3</td><td>4.1E-10</td><td>8.7E-10</td><td>MB231/MB468</td><td>MITO</td></tr>
<tr><td>>hg38_21_+_8433172_8446622_RNA45SN1_45S_50ntFlanks@11665.11687.23|TTCTGCCCAGTGCTCTGAATGTC</td><td>>hg38_21_+_8433172_8446622_RNA45SN1_45S_50ntFlanks</td><td>11665</td><td>11687</td><td>23</td><td>TTCTGCCCAGTGCTCTGAATGTC</td><td>294.2</td><td>3.7</td><td>0.6</td><td>6.3</td><td>3.7E-10</td><td>8.8E-10</td><td>MB231/MB468</td><td>NUC</td></tr>
<tr><td>>hg38_21_+_8433172_8446622_RNA45SN1_45S_50ntFlanks@10679.10710.32|GGTTCCGGCGGCGTCCGGTGAGCTCTCGCTGG</td><td>>hg38_21_+_8433172_8446622_RNA45SN1_45S_50ntFlanks</td><td>10679</td><td>10710</td><td>32</td><td>GGTTCCGGCGGCGTCCGGTGAGCTCTCGCTGG</td><td>85417.8</td><td>-10.2</td><td>1.6</td><td>-6.2</td><td>4.2E-10</td><td>8.8E-10</td><td>BT20/MB468</td><td>NUC</td></tr>
<tr><td>>hg38_21_+_8433172_8446622_RNA45SN1_45S_50ntFlanks@8189.8211.23|CCCAGCCCGTGGACGGTGTGAGG</td><td>>hg38_21_+_8433172_8446622_RNA45SN1_45S_50ntFlanks</td><td>8189</td><td>8211</td><td>23</td><td>CCCAGCCCGTGGACGGTGTGAGG</td><td>174.0</td><td>-11.3</td><td>1.8</td><td>-6.3</td><td>3.5E-10</td><td>8.8E-10</td><td>MB231/MB468</td><td>MP</td></tr>
<tr><td>>MI0000077|hsa-mir-21&WithFlank&17|+|59841260|59841343@15.36.22(+1A)&[MIMAT0000076&hsa-miR-21-5p&offsets|+1|+1(+1A);m-1&17|+|59841273|59841294&offsets|+1|+1(+1A)]|AGCTTATCAGACTGATGTTGACA</td><td>>MI0000077|hsa-mir-21&WithFlank&17|+|59841260|59841343</td><td>15</td><td>36</td><td>22(+1A)</td><td>AGCTTATCAGACTGATGTTGACA</td><td>530.8</td><td>8.3</td><td>1.3</td><td>6.3</td><td>3.2E-10</td><td>8.8E-10</td><td>BT20/MB468</td><td>CYTO</td></tr>
<tr><td>>hg38_21_+_8433172_8446622_RNA45SN1_45S_50ntFlanks@4297.4332.36|CATTGGAGGGCAAGTCTGGTGCCAGCAGCCGCGGTA</td><td>>hg38_21_+_8433172_8446622_RNA45SN1_45S_50ntFlanks</td><td>4297</td><td>4332</td><td>36</td><td>CATTGGAGGGCAAGTCTGGTGCCAGCAGCCGCGGTA</td><td>106.9</td><td>-10.6</td><td>1.7</td><td>-6.3</td><td>3.5E-10</td><td>8.8E-10</td><td>MB231/MB468</td><td>MP</td></tr>
<tr><td>>am_trna111_HisGTG_1_-_147774845_147774916@2.29.28,trna118_HisGTG_1_-_145396881_145396952@2.29.28,trna16_HisGTG_1_+_146544773_146544844@2.29.28,trna1_HisGTG_15_+_45493349_45493420@2.29.28,trna21_HisGTG_1_+_147753471_147753542@2.29.28,trna33_HisGTG_6_+_27125906_27125977@2.29.28,trna7_HisGTG_9_-_14433938_14434009@2.29.28,trna8_HisGTG_15_-_45492611_45492682@2.29.28,trna9_HisGTG_15_-_45490804_45490875@2.29.28|CCGTGATCGTATAGTGGTTAGTACTCTG</td><td>>am_trna111_HisGTG_1_-_147774845_147774916</td><td>2</td><td>29</td><td>28</td><td>CCGTGATCGTATAGTGGTTAGTACTCTG</td><td>633.6</td><td>-5.5</td><td>0.9</td><td>-6.3</td><td>3.5E-10</td><td>8.8E-10</td><td>MB231/MB468</td><td>MP</td></tr>
<tr><td>>hg38_21_+_8433172_8446622_RNA45SN1_45S_50ntFlanks@9339.9371.33|CGCACGAAAGCCGCCGTGGCGCAATGAAGGTGA</td><td>>hg38_21_+_8433172_8446622_RNA45SN1_45S_50ntFlanks</td><td>9339</td><td>9371</td><td>33</td><td>CGCACGAAAGCCGCCGTGGCGCAATGAAGGTGA</td><td>354.9</td><td>-11.6</td><td>1.7</td><td>-6.6</td><td>3.2E-11</td><td>8.8E-10</td><td>BT20/MB468</td><td>MP</td></tr>
<tr><td>>hg38_21_+_8433172_8446622_RNA45SN1_45S_50ntFlanks@12073.12097.25|CCGAGGGGCTCTCGCTTCTGGCGCC</td><td>>hg38_21_+_8433172_8446622_RNA45SN1_45S_50ntFlanks</td><td>12073</td><td>12097</td><td>25</td><td>CCGAGGGGCTCTCGCTTCTGGCGCC</td><td>778.4</td><td>-8.6</td><td>1.4</td><td>-6.3</td><td>3.5E-10</td><td>8.8E-10</td><td>MB231/MB468</td><td>MP</td></tr>
<tr><td>>hg38_21_+_8433172_8446622_RNA45SN1_45S_50ntFlanks@6700.6719.20|CTAGCTGCGAGAATTAATGT</td><td>>hg38_21_+_8433172_8446622_RNA45SN1_45S_50ntFlanks</td><td>6700</td><td>6719</td><td>20</td><td>CTAGCTGCGAGAATTAATGT</td><td>855.3</td><td>-9.2</td><td>1.5</td><td>-6.2</td><td>4.3E-10</td><td>8.8E-10</td><td>BT20/MB468</td><td>NUC</td></tr>
<tr><td>>MI0000063|hsa-let-7b&WithFlank&22|+|46113680|46113774@12.32.21&[MIMAT0000063&hsa-let-7b-5p&offsets|0|-1;m-14&22|+|46113691|46113712&offsets|0|-1]|TGAGGTAGTAGGTTGTGTGGT</td><td>>MI0000063|hsa-let-7b&WithFlank&22|+|46113680|46113774</td><td>12</td><td>32</td><td>21</td><td>TGAGGTAGTAGGTTGTGTGGT</td><td>1562.6</td><td>5.1</td><td>0.8</td><td>6.3</td><td>3.6E-10</td><td>8.8E-10</td><td>MB231/MB468</td><td>CYTO</td></tr>
<tr><td>>trnaMT_ValTAC_MT_+_1602_1670@53.72.20|TAACTTGACCGCTCTGACCA</td><td>>trnaMT_ValTAC_MT_+_1602_1670</td><td>53</td><td>72</td><td>20</td><td>TAACTTGACCGCTCTGACCA</td><td>270.0</td><td>8.1</td><td>1.3</td><td>6.3</td><td>3.6E-10</td><td>8.8E-10</td><td>MB231/MB468</td><td>CYTO</td></tr>
<tr><td>>hg38_21_+_8433172_8446622_RNA45SN1_45S_50ntFlanks@12931.12953.23|GCGCTAAACCATTCGTAGACGAC</td><td>>hg38_21_+_8433172_8446622_RNA45SN1_45S_50ntFlanks</td><td>12931</td><td>12953</td><td>23</td><td>GCGCTAAACCATTCGTAGACGAC</td><td>65.1</td><td>-9.9</td><td>1.6</td><td>-6.3</td><td>3.5E-10</td><td>8.8E-10</td><td>MB231/MB468</td><td>MP</td></tr>
<tr><td>>MI0000087|hsa-mir-29a&WithFlank&7|-|130876741|130876816@48.69.22(+1C)&[MIMAT0000086&hsa-miR-29a-3p&offsets|0|0(+1C);m-26&7|-|130876749|130876769&offsets|0|+1(+1C)]|TAGCACCATCTGAAATCGGTTAC</td><td>>MI0000087|hsa-mir-29a&WithFlank&7|-|130876741|130876816</td><td>48</td><td>69</td><td>22(+1C)</td><td>TAGCACCATCTGAAATCGGTTAC</td><td>120.1</td><td>10.1</td><td>1.6</td><td>6.3</td><td>3.5E-10</td><td>8.8E-10</td><td>MB231/MB468</td><td>MP</td></tr>
<tr><td>>hg38_21_+_8433172_8446622_RNA45SN1_45S_50ntFlanks@12436.12454.19|TCATTGTGAAGCAGAATTC</td><td>>hg38_21_+_8433172_8446622_RNA45SN1_45S_50ntFlanks</td><td>12436</td><td>12454</td><td>19</td><td>TCATTGTGAAGCAGAATTC</td><td>1125.4</td><td>-9.3</td><td>1.5</td><td>-6.3</td><td>3.2E-10</td><td>8.8E-10</td><td>BT20/MB468</td><td>CYTO</td></tr>
<tr><td>>hg38_21_+_8433172_8446622_RNA45SN1_45S_50ntFlanks@12957.12982.26|CTTCTGGGTCGGGGTTTCGTACGTAG</td><td>>hg38_21_+_8433172_8446622_RNA45SN1_45S_50ntFlanks</td><td>12957</td><td>12982</td><td>26</td><td>CTTCTGGGTCGGGGTTTCGTACGTAG</td><td>1863.3</td><td>-6.3</td><td>1.0</td><td>-6.3</td><td>3.6E-10</td><td>8.8E-10</td><td>MB231/MB468</td><td>CYTO</td></tr>
<tr><td>>hg38_21_+_8433172_8446622_RNA45SN1_45S_50ntFlanks@12625.12667.43|CAATGGGGCGAAGCTACCATCTGTGGGATTATGACTGAACGCC</td><td>>hg38_21_+_8433172_8446622_RNA45SN1_45S_50ntFlanks</td><td>12625</td><td>12667</td><td>43</td><td>CAATGGGGCGAAGCTACCATCTGTGGGATTATGACTGAACGCC</td><td>522.1</td><td>8.8</td><td>1.4</td><td>6.3</td><td>3.7E-10</td><td>8.8E-10</td><td>MB231/MB468</td><td>CYTO</td></tr>
<tr><td>>hg38_21_+_8433172_8446622_RNA45SN1_45S_50ntFlanks@9554.9596.43|CGTAGCGGTCCTGACGTGCAAATCGGTCGTCCGACCTGGGTAT</td><td>>hg38_21_+_8433172_8446622_RNA45SN1_45S_50ntFlanks</td><td>9554</td><td>9596</td><td>43</td><td>CGTAGCGGTCCTGACGTGCAAATCGGTCGTCCGACCTGGGTAT</td><td>151.7</td><td>-11.1</td><td>1.8</td><td>-6.3</td><td>3.5E-10</td><td>8.9E-10</td><td>MB231/MB468</td><td>MP</td></tr>
<tr><td>>hg38_21_+_8433172_8446622_RNA45SN1_45S_50ntFlanks@12990.13013.24|GCTCCCTCGCTGCGATCTATTGAA</td><td>>hg38_21_+_8433172_8446622_RNA45SN1_45S_50ntFlanks</td><td>12990</td><td>13013</td><td>24</td><td>GCTCCCTCGCTGCGATCTATTGAA</td><td>2415.3</td><td>-5.0</td><td>0.8</td><td>-6.3</td><td>4.1E-10</td><td>8.9E-10</td><td>MB231/MB468</td><td>MITO</td></tr>
<tr><td>>hg38_21_+_8433172_8446622_RNA45SN1_45S_50ntFlanks@12646.12664.19|TGTGGGATTATGACTGAAC</td><td>>hg38_21_+_8433172_8446622_RNA45SN1_45S_50ntFlanks</td><td>12646</td><td>12664</td><td>19</td><td>TGTGGGATTATGACTGAAC</td><td>146.9</td><td>10.9</td><td>1.6</td><td>6.6</td><td>3.2E-11</td><td>8.9E-10</td><td>BT20/MB468</td><td>MP</td></tr>
<tr><td>>MI0000465|hsa-mir-191&WithFlank&3|-|49020612|49020715@22.42.21&[MIMAT0000440&hsa-miR-191-5p&offsets|0|-2;m-48&3|-|49020672|49020694&offsets|0|-2]|CAACGGAATCCCAAAAGCAGC</td><td>>MI0000465|hsa-mir-191&WithFlank&3|-|49020612|49020715</td><td>22</td><td>42</td><td>21</td><td>CAACGGAATCCCAAAAGCAGC</td><td>716.8</td><td>4.5</td><td>0.7</td><td>6.3</td><td>3.7E-10</td><td>8.9E-10</td><td>MB231/MB468</td><td>NUC</td></tr>
<tr><td>>hg38_21_+_8433172_8446622_RNA45SN1_45S_50ntFlanks@5468.5514.47|GCCCACGGCCCTGGCGGAGCGCTGAGAAGACGGTCGAACTTGACTAT</td><td>>hg38_21_+_8433172_8446622_RNA45SN1_45S_50ntFlanks</td><td>5468</td><td>5514</td><td>47</td><td>GCCCACGGCCCTGGCGGAGCGCTGAGAAGACGGTCGAACTTGACTAT</td><td>225.3</td><td>-10.8</td><td>1.7</td><td>-6.3</td><td>3.7E-10</td><td>8.9E-10</td><td>MB231/MB468</td><td>NUC</td></tr>
<tr><td>>hg38_21_+_8433172_8446622_RNA45SN1_45S_50ntFlanks@10391.10426.36|AACAGCAGTTGAACATGGGTCAGTCGGTCCTGAGAG</td><td>>hg38_21_+_8433172_8446622_RNA45SN1_45S_50ntFlanks</td><td>10391</td><td>10426</td><td>36</td><td>AACAGCAGTTGAACATGGGTCAGTCGGTCCTGAGAG</td><td>532.8</td><td>-6.6</td><td>1.0</td><td>-6.3</td><td>3.7E-10</td><td>8.9E-10</td><td>MB231/MB468</td><td>NUC</td></tr>
<tr><td>>trnaMT_MetCAT_MT_+_4402_4469@36.71.36|CCCGAAAATGTTGGTTATACCCTTCCCGTACTACCA</td><td>>trnaMT_MetCAT_MT_+_4402_4469</td><td>36</td><td>71</td><td>36</td><td>CCCGAAAATGTTGGTTATACCCTTCCCGTACTACCA</td><td>706.0</td><td>8.6</td><td>1.4</td><td>6.3</td><td>3.7E-10</td><td>8.9E-10</td><td>MB231/MB468</td><td>NUC</td></tr>
<tr><td>>hg38_21_+_8433172_8446622_RNA45SN1_45S_50ntFlanks@10664.10706.43|CTTGGAAAGCGTCGCGGTTCCGGCGGCGTCCGGTGAGCTCTCG</td><td>>hg38_21_+_8433172_8446622_RNA45SN1_45S_50ntFlanks</td><td>10664</td><td>10706</td><td>43</td><td>CTTGGAAAGCGTCGCGGTTCCGGCGGCGTCCGGTGAGCTCTCG</td><td>1648.7</td><td>-9.1</td><td>1.5</td><td>-6.3</td><td>3.7E-10</td><td>8.9E-10</td><td>MB231/MB468</td><td>NUC</td></tr>
<tr><td>>hg38_21_+_8433172_8446622_RNA45SN1_45S_50ntFlanks@6721.6757.37|AATTGCAGGACACATTGATCATCGACACTTCGAACGC</td><td>>hg38_21_+_8433172_8446622_RNA45SN1_45S_50ntFlanks</td><td>6721</td><td>6757</td><td>37</td><td>AATTGCAGGACACATTGATCATCGACACTTCGAACGC</td><td>579.6</td><td>-6.8</td><td>1.1</td><td>-6.2</td><td>4.1E-10</td><td>8.9E-10</td><td>MB231/MB468</td><td>MITO</td></tr>
<tr><td>>MI0000255|hsa-mir-30d&WithFlank&8|-|134804870|134804951@12.31.20&[MIMAT0000245&hsa-miR-30d-5p&offsets|0|-2;m-21&8|-|134804919|134804940&offsets|0|-2]|TGTAAACATCCCCGACTGGA</td><td>>MI0000255|hsa-mir-30d&WithFlank&8|-|134804870|134804951</td><td>12</td><td>31</td><td>20</td><td>TGTAAACATCCCCGACTGGA</td><td>810.5</td><td>6.7</td><td>1.1</td><td>6.3</td><td>3.7E-10</td><td>8.9E-10</td><td>MB231/MB468</td><td>CYTO</td></tr>
<tr><td>>hg38_21_+_8433172_8446622_RNA45SN1_45S_50ntFlanks@4102.4127.26|ACCACGGGTGACGGGGAATCAGGGTT</td><td>>hg38_21_+_8433172_8446622_RNA45SN1_45S_50ntFlanks</td><td>4102</td><td>4127</td><td>26</td><td>ACCACGGGTGACGGGGAATCAGGGTT</td><td>216.4</td><td>-2.8</td><td>0.4</td><td>-6.3</td><td>3.3E-10</td><td>8.9E-10</td><td>BT20/MB468</td><td>CYTO</td></tr>
<tr><td>>hg38_21_+_8433172_8446622_RNA45SN1_45S_50ntFlanks@12395.12418.24|AGCGACGTCGCTTTTTGATCCTTC</td><td>>hg38_21_+_8433172_8446622_RNA45SN1_45S_50ntFlanks</td><td>12395</td><td>12418</td><td>24</td><td>AGCGACGTCGCTTTTTGATCCTTC</td><td>490.0</td><td>6.8</td><td>1.1</td><td>6.3</td><td>3.7E-10</td><td>8.9E-10</td><td>MB231/MB468</td><td>CYTO</td></tr>
<tr><td>>hg38_21_+_8433172_8446622_RNA45SN1_45S_50ntFlanks@8104.8138.35|CCGCGGCGGGGCGCGGGACATGTGGCGTACGGAAG</td><td>>hg38_21_+_8433172_8446622_RNA45SN1_45S_50ntFlanks</td><td>8104</td><td>8138</td><td>35</td><td>CCGCGGCGGGGCGCGGGACATGTGGCGTACGGAAG</td><td>4646.6</td><td>-8.6</td><td>1.4</td><td>-6.3</td><td>3.6E-10</td><td>8.9E-10</td><td>MB231/MB468</td><td>MP</td></tr>
<tr><td>>hg38_21_+_8433172_8446622_RNA45SN1_45S_50ntFlanks@9575.9595.21|ATCGGTCGTCCGACCTGGGTA</td><td>>hg38_21_+_8433172_8446622_RNA45SN1_45S_50ntFlanks</td><td>9575</td><td>9595</td><td>21</td><td>ATCGGTCGTCCGACCTGGGTA</td><td>1366.7</td><td>-8.9</td><td>1.4</td><td>-6.2</td><td>4.2E-10</td><td>8.9E-10</td><td>MB231/MB468</td><td>MITO</td></tr>
<tr><td>>hg38_21_+_8433172_8446622_RNA45SN1_45S_50ntFlanks@10420.10453.34|CTGAGAGATGGGCGAGCGCCGTTCCGAAGGGACG</td><td>>hg38_21_+_8433172_8446622_RNA45SN1_45S_50ntFlanks</td><td>10420</td><td>10453</td><td>34</td><td>CTGAGAGATGGGCGAGCGCCGTTCCGAAGGGACG</td><td>395.6</td><td>-8.1</td><td>1.3</td><td>-6.3</td><td>3.6E-10</td><td>9.0E-10</td><td>MB231/MB468</td><td>MP</td></tr>
<tr><td>>hg38_21_+_8433172_8446622_RNA45SN1_45S_50ntFlanks@10820.10840.21|GGGAAGTCGGCAAGCCGGATC</td><td>>hg38_21_+_8433172_8446622_RNA45SN1_45S_50ntFlanks</td><td>10820</td><td>10840</td><td>21</td><td>GGGAAGTCGGCAAGCCGGATC</td><td>503.5</td><td>-2.9</td><td>0.5</td><td>-6.3</td><td>2.8E-10</td><td>9.0E-10</td><td>BT20/MB468</td><td>MITO</td></tr>
<tr><td>>hg38_21_+_8433172_8446622_RNA45SN1_45S_50ntFlanks@8636.8658.23|CGGCGACCGGCCGCCGCCGGGCG</td><td>>hg38_21_+_8433172_8446622_RNA45SN1_45S_50ntFlanks</td><td>8636</td><td>8658</td><td>23</td><td>CGGCGACCGGCCGCCGCCGGGCG</td><td>417.3</td><td>-10.8</td><td>1.7</td><td>-6.3</td><td>2.8E-10</td><td>9.0E-10</td><td>BT20/MB468</td><td>MITO</td></tr>
<tr><td>>hg38_21_+_8433172_8446622_RNA45SN1_45S_50ntFlanks@10410.10433.24|TCAGTCGGTCCTGAGAGATGGGCG</td><td>>hg38_21_+_8433172_8446622_RNA45SN1_45S_50ntFlanks</td><td>10410</td><td>10433</td><td>24</td><td>TCAGTCGGTCCTGAGAGATGGGCG</td><td>382.5</td><td>-7.7</td><td>1.2</td><td>-6.3</td><td>3.8E-10</td><td>9.0E-10</td><td>MB231/MB468</td><td>NUC</td></tr>
<tr><td>>hg38_21_+_8433172_8446622_RNA45SN1_45S_50ntFlanks@485.532.48|GATGTGGTGACGTCGTGCTCTCCCGGGCCGGGTCCGAGCCGCGACGGG</td><td>>hg38_21_+_8433172_8446622_RNA45SN1_45S_50ntFlanks</td><td>485</td><td>532</td><td>48</td><td>GATGTGGTGACGTCGTGCTCTCCCGGGCCGGGTCCGAGCCGCGACGGG</td><td>3142.7</td><td>11.1</td><td>1.8</td><td>6.3</td><td>3.8E-10</td><td>9.0E-10</td><td>MB231/MB468</td><td>NUC</td></tr>
<tr><td>>hg38_1_-_228634819_228635039_RNA5S12_5S_50ntFlanks@134.168.35|TGGGAGACCGCCTGGGAATACCGGGTGCTGTAGGC</td><td>>hg38_1_-_228634819_228635039_RNA5S12_5S_50ntFlanks</td><td>134</td><td>168</td><td>35</td><td>TGGGAGACCGCCTGGGAATACCGGGTGCTGTAGGC</td><td>695.4</td><td>6.8</td><td>1.1</td><td>6.3</td><td>3.7E-10</td><td>9.0E-10</td><td>MB231/MB468</td><td>CYTO</td></tr>
<tr><td>>hg38_21_+_8433172_8446622_RNA45SN1_45S_50ntFlanks@12251.12280.30|AAAGCTCGCTTGATCTTGATTTTCAGTACG</td><td>>hg38_21_+_8433172_8446622_RNA45SN1_45S_50ntFlanks</td><td>12251</td><td>12280</td><td>30</td><td>AAAGCTCGCTTGATCTTGATTTTCAGTACG</td><td>219.5</td><td>5.6</td><td>0.9</td><td>6.3</td><td>3.7E-10</td><td>9.0E-10</td><td>MB231/MB468</td><td>CYTO</td></tr>
</table>

<table>
<tr><td>>hg38_21_+_8433172_8446622_RNA45SN1_45S_50ntFlanks@11573.11605.33|CGACTTAGAACTGGTGCGGACCAGGGGAATCCG</td><td>>hg38_21_+_8433172_8446622_RNA45SN1_45S_50ntFlanks</td><td>11573</td><td>11605</td><td>33</td><td>CGACTTAGAACTGGTGCGGACCAGGGGAATCCG</td><td>491.6</td><td>-3.0</td><td>0.5</td><td>-6.3</td><td>3.7E-10</td><td>9.0E-10</td><td>MB231/MB468</td><td>CYTO</td></tr>
<tr><td>>hg38_21_+_8433172_8446622_RNA45SN1_45S_50ntFlanks@12203.12222.20|TAAGGCGAGCTCAGGGAGGA</td><td>>hg38_21_+_8433172_8446622_RNA45SN1_45S_50ntFlanks</td><td>12203</td><td>12222</td><td>20</td><td>TAAGGCGAGCTCAGGGAGGA</td><td>95.2</td><td>-10.4</td><td>1.7</td><td>-6.3</td><td>3.6E-10</td><td>9.0E-10</td><td>MB231/MB468</td><td>MP</td></tr>
<tr><td>>hg38_21_+_8433172_8446622_RNA45SN1_45S_50ntFlanks@11579.11605.27|AGAACTGGTGCGGACCAGGGGAATCCG</td><td>>hg38_21_+_8433172_8446622_RNA45SN1_45S_50ntFlanks</td><td>11579</td><td>11605</td><td>27</td><td>AGAACTGGTGCGGACCAGGGGAATCCG</td><td>1687.3</td><td>-7.0</td><td>1.1</td><td>-6.3</td><td>3.8E-10</td><td>9.0E-10</td><td>MB231/MB468</td><td>NUC</td></tr>
<tr><td>>MI0000255|hsa-mir-30d&WithFlank&8|-|134804870|134804951@12.35.24(+1U)&[MIMAT0000245&hsa-miR-30d-5p&offsets|0|+2(+1U);m-21&8|-|134804919|134804940&offsets|0|+2(+1U)]|TGTAAACATCCCCGACTGGAAGCTT</td><td>>MI0000255|hsa-mir-30d&WithFlank&8|-|134804870|134804951</td><td>12</td><td>35</td><td>24(+1U)</td><td>TGTAAACATCCCCGACTGGAAGCTT</td><td>447.7</td><td>2.9</td><td>0.5</td><td>6.2</td><td>4.4E-10</td><td>9.0E-10</td><td>BT20/MB468</td><td>NUC</td></tr>
<tr><td>>MI0000283|hsa-mir-203a&WithFlank&14|+|104117399|104117520@71.92.22&[MIMAT0000264&hsa-miR-203a-3p&offsets|0|0;m-79&14|+|104117469|104117490&offsets|0|0]|GTGAAATGTTTAGGACCACTAG</td><td>>MI0000283|hsa-mir-203a&WithFlank&14|+|104117399|104117520</td><td>71</td><td>92</td><td>22</td><td>GTGAAATGTTTAGGACCACTAG</td><td>1227.6</td><td>8.9</td><td>1.4</td><td>6.6</td><td>4.0E-11</td><td>9.1E-10</td><td>BT20/MB231</td><td>NUC</td></tr>
<tr><td>>hg38_21_+_8433172_8446622_RNA45SN1_45S_50ntFlanks@11925.11948.24|AGGTGTAGAATAAGTGGGAGGCCC</td><td>>hg38_21_+_8433172_8446622_RNA45SN1_45S_50ntFlanks</td><td>11925</td><td>11948</td><td>24</td><td>AGGTGTAGAATAAGTGGGAGGCCC</td><td>328.7</td><td>-7.9</td><td>1.3</td><td>-6.3</td><td>3.6E-10</td><td>9.1E-10</td><td>MB231/MB468</td><td>MP</td></tr>
<tr><td>>hg38_21_+_8433172_8446622_RNA45SN1_45S_50ntFlanks@12184.12201.18|AACGGTAACGCAGGTGTC</td><td>>hg38_21_+_8433172_8446622_RNA45SN1_45S_50ntFlanks</td><td>12184</td><td>12201</td><td>18</td><td>AACGGTAACGCAGGTGTC</td><td>84.8</td><td>-10.3</td><td>1.6</td><td>-6.3</td><td>3.6E-10</td><td>9.1E-10</td><td>MB231/MB468</td><td>MP</td></tr>
<tr><td>>hg38_21_+_8433172_8446622_RNA45SN1_45S_50ntFlanks@4201.4219.19|CCGACCCGGGGAGGTAGTG</td><td>>hg38_21_+_8433172_8446622_RNA45SN1_45S_50ntFlanks</td><td>4201</td><td>4219</td><td>19</td><td>CCGACCCGGGGAGGTAGTG</td><td>160.2</td><td>-9.4</td><td>1.5</td><td>-6.3</td><td>2.8E-10</td><td>9.1E-10</td><td>BT20/MB468</td><td>MITO</td></tr>
<tr><td>>hg38_21_+_8433172_8446622_RNA45SN1_45S_50ntFlanks@11535.11560.26|TTCCGCGCGGCGCCTCGCCTCGGCCG</td><td>>hg38_21_+_8433172_8446622_RNA45SN1_45S_50ntFlanks</td><td>11535</td><td>11560</td><td>26</td><td>TTCCGCGCGGCGCCTCGCCTCGGCCG</td><td>78.4</td><td>-10.2</td><td>1.6</td><td>-6.3</td><td>3.6E-10</td><td>9.1E-10</td><td>MB231/MB468</td><td>MP</td></tr>
<tr><td>>hg38_21_+_8433172_8446622_RNA45SN1_45S_50ntFlanks@9471.9506.36|AGCACGAGCGCACGTGTTAGGACCCGAAAGATGGTG</td><td>>hg38_21_+_8433172_8446622_RNA45SN1_45S_50ntFlanks</td><td>9471</td><td>9506</td><td>36</td><td>AGCACGAGCGCACGTGTTAGGACCCGAAAGATGGTG</td><td>100.4</td><td>-10.5</td><td>1.7</td><td>-6.3</td><td>3.6E-10</td><td>9.1E-10</td><td>MB231/MB468</td><td>MP</td></tr>
<tr><td>>hg38_21_+_8433172_8446622_RNA45SN1_45S_50ntFlanks@6658.6683.26|TAGCGGTGGATCACTCGGCTCGTGCG</td><td>>hg38_21_+_8433172_8446622_RNA45SN1_45S_50ntFlanks</td><td>6658</td><td>6683</td><td>26</td><td>TAGCGGTGGATCACTCGGCTCGTGCG</td><td>331.0</td><td>4.5</td><td>0.7</td><td>6.3</td><td>3.8E-10</td><td>9.1E-10</td><td>MB231/MB468</td><td>NUC</td></tr>
<tr><td>>hg38_21_+_8433172_8446622_RNA45SN1_45S_50ntFlanks@8357.8386.30|AAAAGAACTTTGAAGAGAGAGTTCAAGAGG</td><td>>hg38_21_+_8433172_8446622_RNA45SN1_45S_50ntFlanks</td><td>8357</td><td>8386</td><td>30</td><td>AAAAGAACTTTGAAGAGAGAGTTCAAGAGG</td><td>533.2</td><td>-8.5</td><td>1.4</td><td>-6.3</td><td>3.6E-10</td><td>9.1E-10</td><td>MB231/MB468</td><td>MP</td></tr>
<tr><td>>am_MI0000809|hsa-mir-151a&WithFlank&8|-|140732558|140732659@53.70.18&[MIMAT0000757&hsa-miR-151a-3p&offsets|0|-3;m-30&8|-|140732587|140732607&offsets|0|-3]|CTAGACTGAAGCTCCTTG</td><td>>am_MI0000809|hsa-mir-151a&WithFlank&8|-|140732558|140732659</td><td>53</td><td>70</td><td>18</td><td>CTAGACTGAAGCTCCTTG</td><td>3103.3</td><td>6.6</td><td>1.0</td><td>6.6</td><td>3.3E-11</td><td>9.1E-10</td><td>BT20/MB468</td><td>MP</td></tr>
<tr><td>>hg38_21_+_8433172_8446622_RNA45SN1_45S_50ntFlanks@11919.11954.36|CATGAGAGGTGTAGAATAAGTGGGAGGCCCCCGGCG</td><td>>hg38_21_+_8433172_8446622_RNA45SN1_45S_50ntFlanks</td><td>11919</td><td>11954</td><td>36</td><td>CATGAGAGGTGTAGAATAAGTGGGAGGCCCCCGGCG</td><td>108.5</td><td>-10.6</td><td>1.7</td><td>-6.3</td><td>3.6E-10</td><td>9.1E-10</td><td>MB231/MB468</td><td>MP</td></tr>
<tr><td>>hg38_21_+_8433172_8446622_RNA45SN1_45S_50ntFlanks@8176.8206.31|CTTCTGATCGAGGCCCAGCCCGTGGACGGTG</td><td>>hg38_21_+_8433172_8446622_RNA45SN1_45S_50ntFlanks</td><td>8176</td><td>8206</td><td>31</td><td>CTTCTGATCGAGGCCCAGCCCGTGGACGGTG</td><td>1364.5</td><td>-7.0</td><td>1.1</td><td>-6.3</td><td>3.6E-10</td><td>9.1E-10</td><td>MB231/MB468</td><td>MP</td></tr>
<tr><td>>hg38_21_+_8433172_8446622_RNA45SN1_45S_50ntFlanks@12126.12151.26|CGACCCGCTCCGGGGACAGTGCCAGG</td><td>>hg38_21_+_8433172_8446622_RNA45SN1_45S_50ntFlanks</td><td>12126</td><td>12151</td><td>26</td><td>CGACCCGCTCCGGGGACAGTGCCAGG</td><td>174.6</td><td>-10.5</td><td>1.6</td><td>-6.6</td><td>3.3E-11</td><td>9.1E-10</td><td>BT20/MB468</td><td>MP</td></tr>
<tr><td>>hg38_21_+_8433172_8446622_RNA45SN1_45S_50ntFlanks@12717.12742.26|CCTCGGTTGGCCTCGGATAGCCGGTC</td><td>>hg38_21_+_8433172_8446622_RNA45SN1_45S_50ntFlanks</td><td>12717</td><td>12742</td><td>26</td><td>CCTCGGTTGGCCTCGGATAGCCGGTC</td><td>288.5</td><td>-10.3</td><td>1.6</td><td>-6.3</td><td>2.8E-10</td><td>9.1E-10</td><td>BT20/MB468</td><td>MITO</td></tr>
<tr><td>>hg38_21_+_8433172_8446622_RNA45SN1_45S_50ntFlanks@12971.12989.19|TTTCGTACGTAGCAGAGCA</td><td>>hg38_21_+_8433172_8446622_RNA45SN1_45S_50ntFlanks</td><td>12971</td><td>12989</td><td>19</td><td>TTTCGTACGTAGCAGAGCA</td><td>1112.7</td><td>-9.1</td><td>1.5</td><td>-6.2</td><td>4.4E-10</td><td>9.1E-10</td><td>BT20/MB468</td><td>NUC</td></tr>
<tr><td>>hg38_21_+_8433172_8446622_RNA45SN1_45S_50ntFlanks@11668.11687.20|TGCCCAGTGCTCTGAATGTC</td><td>>hg38_21_+_8433172_8446622_RNA45SN1_45S_50ntFlanks</td><td>11668</td><td>11687</td><td>20</td><td>TGCCCAGTGCTCTGAATGTC</td><td>996.7</td><td>4.6</td><td>0.7</td><td>6.3</td><td>3.8E-10</td><td>9.1E-10</td><td>MB231/MB468</td><td>NUC</td></tr>
<tr><td>>hg38_1_-_228634819_228635039_RNA5S12_5S_50ntFlanks@86.112.27|CGTCTGATCTCGGAAGCTAAGCAGGGT</td><td>>hg38_1_-_228634819_228635039_RNA5S12_5S_50ntFlanks</td><td>86</td><td>112</td><td>27</td><td>CGTCTGATCTCGGAAGCTAAGCAGGGT</td><td>466.0</td><td>-6.9</td><td>1.1</td><td>-6.2</td><td>4.4E-10</td><td>9.1E-10</td><td>BT20/MB468</td><td>NUC</td></tr>
<tr><td>>hg38_21_+_8433172_8446622_RNA45SN1_45S_50ntFlanks@8105.8136.32|CGCGGCGGGGCGCGGGACATGTGGCGTACGGA</td><td>>hg38_21_+_8433172_8446622_RNA45SN1_45S_50ntFlanks</td><td>8105</td><td>8136</td><td>32</td><td>CGCGGCGGGGCGCGGGACATGTGGCGTACGGA</td><td>3432.6</td><td>-8.8</td><td>1.4</td><td>-6.3</td><td>3.6E-10</td><td>9.1E-10</td><td>MB231/MB468</td><td>MP</td></tr>
<tr><td>>hg38_21_+_8433172_8446622_RNA45SN1_45S_50ntFlanks@10352.10369.18|GAGAACTTTGAAGGCCGA</td><td>>hg38_21_+_8433172_8446622_RNA45SN1_45S_50ntFlanks</td><td>10352</td><td>10369</td><td>18</td><td>GAGAACTTTGAAGGCCGA</td><td>424.0</td><td>4.6</td><td>0.7</td><td>6.3</td><td>3.8E-10</td><td>9.2E-10</td><td>MB231/MB468</td><td>CYTO</td></tr>
<tr><td>>MI0003560|hsa-mir-92b&WithFlank&1|+|155195171|155195278@67.87.21(+1U)&[MIMAT0003218&hsa-miR-92b-3p&offsets|0|-1(+1U);m-115&1|+|155195237|155195258&offsets|0|-1(+1U)]|TATTGCACTCGTCCCGGCCTCT</td><td>>MI0003560|hsa-mir-92b&WithFlank&1|+|155195171|155195278</td><td>67</td><td>87</td><td>21(+1U)</td><td>TATTGCACTCGTCCCGGCCTCT</td><td>1768.4</td><td>6.0</td><td>1.0</td><td>6.3</td><td>3.7E-10</td><td>9.2E-10</td><td>MB231/MB468</td><td>MP</td></tr>
<tr><td>>hg38_21_+_8433172_8446622_RNA45SN1_45S_50ntFlanks@5073.5105.33|AATTCCGATAACGAACGAGACTCTGGCATGCTA</td><td>>hg38_21_+_8433172_8446622_RNA45SN1_45S_50ntFlanks</td><td>5073</td><td>5105</td><td>33</td><td>AATTCCGATAACGAACGAGACTCTGGCATGCTA</td><td>215.3</td><td>-11.6</td><td>1.9</td><td>-6.3</td><td>3.7E-10</td><td>9.2E-10</td><td>MB231/MB468</td><td>MP</td></tr>
<tr><td>>hg38_21_+_8433172_8446622_RNA45SN1_45S_50ntFlanks@6782.6801.20|GGGCTACGCCTGTCTGAGCG</td><td>>hg38_21_+_8433172_8446622_RNA45SN1_45S_50ntFlanks</td><td>6782</td><td>6801</td><td>20</td><td>GGGCTACGCCTGTCTGAGCG</td><td>68.1</td><td>-10.0</td><td>1.6</td><td>-6.3</td><td>3.7E-10</td><td>9.2E-10</td><td>MB231/MB468</td><td>MP</td></tr>
<tr><td>>hg38_21_+_8433172_8446622_RNA45SN1_45S_50ntFlanks@6671.6698.28|CTCGGCTCGTGCGTCGATGAAGAACGCA</td><td>>hg38_21_+_8433172_8446622_RNA45SN1_45S_50ntFlanks</td><td>6671</td><td>6698</td><td>28</td><td>CTCGGCTCGTGCGTCGATGAAGAACGCA</td><td>8170.0</td><td>-6.8</td><td>1.1</td><td>-6.3</td><td>2.9E-10</td><td>9.2E-10</td><td>BT20/MB468</td><td>MITO</td></tr>
<tr><td>>hg38_21_+_8433172_8446622_RNA45SN1_45S_50ntFlanks@10393.10420.28|CAGCAGTTGAACATGGGTCAGTCGGTCC</td><td>>hg38_21_+_8433172_8446622_RNA45SN1_45S_50ntFlanks</td><td>10393</td><td>10420</td><td>28</td><td>CAGCAGTTGAACATGGGTCAGTCGGTCC</td><td>489.9</td><td>-8.1</td><td>1.3</td><td>-6.2</td><td>4.3E-10</td><td>9.2E-10</td><td>MB231/MB468</td><td>MITO</td></tr>
<tr><td>>hg38_21_+_8433172_8446622_RNA45SN1_45S_50ntFlanks@9033.9077.45|CTCGTGGGGGGGCCGGGCCACCCCTCCCACGGCGCGACCGCTCTC</td><td>>hg38_21_+_8433172_8446622_RNA45SN1_45S_50ntFlanks</td><td>9033</td><td>9077</td><td>45</td><td>CTCGTGGGGGGGCCGGGCCACCCCTCCCACGGCGCGACCGCTCTC</td><td>450.0</td><td>-8.8</td><td>1.4</td><td>-6.2</td><td>4.3E-10</td><td>9.2E-10</td><td>MB231/MB468</td><td>MITO</td></tr>
<tr><td>>am_trna111_HisGTG_1_-_147774845_147774916@-1G.23.24,trna118_HisGTG_1_-_145396881_145396952@-1G.23.24,trna16_HisGTG_1_+_146544773_146544844@-1G.23.24,trna1_HisGTG_15_+_45493349_45493420@-1G.23.24,trna21_HisGTG_1_+_147753471_147753542@-1G.23.24,trna33_HisGTG_6_+_27125906_27125977@-1G.23.24,trna7_HisGTG_9_-_14433938_14434009@-1G.23.24,trna8_HisGTG_15_-_45492611_45492682@-1G.23.24,trna9_HisGTG_15_-_45490804_45490875@-1G.23.24|GGCCGTGATCGTATAGTGGTTAGT</td><td>>am_trna111_HisGTG_1_-_147774845_147774916</td><td>-1G</td><td>23</td><td>24</td><td>GGCCGTGATCGTATAGTGGTTAGT</td><td>1083.5</td><td>9.6</td><td>1.5</td><td>6.2</td><td>4.3E-10</td><td>9.2E-10</td><td>MB231/MB468</td><td>MITO</td></tr>
<tr><td>>hg38_21_+_8433172_8446622_RNA45SN1_45S_50ntFlanks@11115.11132.18|CGGCAGGGGCCGGCGGCG</td><td>>hg38_21_+_8433172_8446622_RNA45SN1_45S_50ntFlanks</td><td>11115</td><td>11132</td><td>18</td><td>CGGCAGGGGCCGGCGGCG</td><td>252.5</td><td>-11.1</td><td>1.7</td><td>-6.6</td><td>3.4E-11</td><td>9.2E-10</td><td>BT20/MB468</td><td>MP</td></tr>
<tr><td>>hg38_21_+_8433172_8446622_RNA45SN1_45S_50ntFlanks@11974.11998.25|AGGGGCCCGGGGCGGGGTCCGCCGG</td><td>>hg38_21_+_8433172_8446622_RNA45SN1_45S_50ntFlanks</td><td>11974</td><td>11998</td><td>25</td><td>AGGGGCCCGGGGCGGGGTCCGCCGG</td><td>221.8</td><td>-10.9</td><td>1.6</td><td>-6.6</td><td>3.4E-11</td><td>9.2E-10</td><td>BT20/MB468</td><td>MP</td></tr>
<tr><td>>hg38_21_+_8433172_8446622_RNA45SN1_45S_50ntFlanks@8191.8214.24|CAGCCCGTGGACGGTGTGAGGCCG</td><td>>hg38_21_+_8433172_8446622_RNA45SN1_45S_50ntFlanks</td><td>8191</td><td>8214</td><td>24</td><td>CAGCCCGTGGACGGTGTGAGGCCG</td><td>1019.1</td><td>-8.7</td><td>1.4</td><td>-6.3</td><td>3.7E-10</td><td>9.2E-10</td><td>MB231/MB468</td><td>MP</td></tr>
<tr><td>>MI0000542|hsa-mir-320a&WithFlank&8|-|22244960|22245043@48.70.23(+1C)&[MIMAT0000510&hsa-miR-320a-3p&offsets|0|+1(+1C);m-60&8|-|22244975|22244996&offsets|0|+1(+1C)]|AAAAGCTGGGTTGAGAGGGCGAAC</td><td>>MI0000542|hsa-mir-320a&WithFlank&8|-|22244960|22245043</td><td>48</td><td>70</td><td>23(+1C)</td><td>AAAAGCTGGGTTGAGAGGGCGAAC</td><td>362.3</td><td>7.7</td><td>1.2</td><td>6.3</td><td>3.8E-10</td><td>9.2E-10</td><td>MB231/MB468</td><td>CYTO</td></tr>
<tr><td>>MI0000095|hsa-mir-93&WithFlank&7|-|100093762|100093853@17.40.24&[MIMAT0000093&hsa-miR-93-5p&offsets|0|+1;m-29&7|-|100093815|100093837&offsets|0|+1]|CAAAGTGCTGTTCGTGCAGGTAGT</td><td>>MI0000095|hsa-mir-93&WithFlank&7|-|100093762|100093853</td><td>17</td><td>40</td><td>24</td><td>CAAAGTGCTGTTCGTGCAGGTAGT</td><td>1261.2</td><td>5.6</td><td>0.9</td><td>6.3</td><td>2.9E-10</td><td>9.2E-10</td><td>BT20/MB468</td><td>MITO</td></tr>
<tr><td>>am_MI0003639|hsa-mir-625&WithFlank&14|+|65471096|65471192@58.78.21(+1U)&[MIMAT0004808&hsa-miR-625-3p&offsets|0|-1(+1U);m-137&14|+|65471153|65471174&offsets|0|-1(+1U)]|GACTATAGAACTTTCCCCCTCT</td><td>>am_MI0003639|hsa-mir-625&WithFlank&14|+|65471096|65471192</td><td>58</td><td>78</td><td>21(+1U)</td><td>GACTATAGAACTTTCCCCCTCT</td><td>137.3</td><td>10.3</td><td>1.7</td><td>6.2</td><td>4.3E-10</td><td>9.3E-10</td><td>MB231/MB468</td><td>MITO</td></tr>
<tr><td>>MI0000298|hsa-mir-221&WithFlank&X|-|45746151|45746272@71.93.23&[MIMAT0000278&hsa-miR-221-3p&offsets|0|0;m-91&X|-|45746181|45746202&offsets|0|+1]|AGCTACATTGTCTGCTGGGTTTC</td><td>>MI0000298|hsa-mir-221&WithFlank&X|-|45746151|45746272</td><td>71</td><td>93</td><td>23</td><td>AGCTACATTGTCTGCTGGGTTTC</td><td>37081.0</td><td>5.6</td><td>0.9</td><td>6.3</td><td>3.8E-10</td><td>9.3E-10</td><td>MB231/MB468</td><td>CYTO</td></tr>
<tr><td>>hg38_21_+_8433172_8446622_RNA45SN1_45S_50ntFlanks@8355.8373.19|TGAAAAGAACTTTGAAGAG</td><td>>hg38_21_+_8433172_8446622_RNA45SN1_45S_50ntFlanks</td><td>8355</td><td>8373</td><td>19</td><td>TGAAAAGAACTTTGAAGAG</td><td>2075.5</td><td>9.9</td><td>1.5</td><td>6.6</td><td>3.4E-11</td><td>9.3E-10</td><td>BT20/MB468</td><td>MP</td></tr>
<tr><td>>hg38_21_+_8433172_8446622_RNA45SN1_45S_50ntFlanks@12636.12661.26|AGCTACCATCTGTGGGATTATGACTG</td><td>>hg38_21_+_8433172_8446622_RNA45SN1_45S_50ntFlanks</td><td>12636</td><td>12661</td><td>26</td><td>AGCTACCATCTGTGGGATTATGACTG</td><td>741.2</td><td>-4.2</td><td>0.7</td><td>-6.3</td><td>3.9E-10</td><td>9.3E-10</td><td>MB231/MB468</td><td>NUC</td></tr>
<tr><td>>hg38_21_+_8433172_8446622_RNA45SN1_45S_50ntFlanks@10781.10806.26|AGGTGAACAGCCTCTGGCATGTTGGA</td><td>>hg38_21_+_8433172_8446622_RNA45SN1_45S_50ntFlanks</td><td>10781</td><td>10806</td><td>26</td><td>AGGTGAACAGCCTCTGGCATGTTGGA</td><td>390.7</td><td>-7.2</td><td>1.1</td><td>-6.3</td><td>3.8E-10</td><td>9.3E-10</td><td>MB231/MB468</td><td>CYTO</td></tr>
<tr><td>>hg38_21_+_8433172_8446622_RNA45SN1_45S_50ntFlanks@11875.11893.19|GAAGACCCTGTTGAGCTTG</td><td>>hg38_21_+_8433172_8446622_RNA45SN1_45S_50ntFlanks</td><td>11875</td><td>11893</td><td>19</td><td>GAAGACCCTGTTGAGCTTG</td><td>1085.3</td><td>-4.2</td><td>0.7</td><td>-6.3</td><td>3.9E-10</td><td>9.3E-10</td><td>MB231/MB468</td><td>CYTO</td></tr>
<tr><td>>hg38_21_+_8433172_8446622_RNA45SN1_45S_50ntFlanks@12459.12478.20|AGCGTTGGATTGTTCACCCA</td><td>>hg38_21_+_8433172_8446622_RNA45SN1_45S_50ntFlanks</td><td>12459</td><td>12478</td><td>20</td><td>AGCGTTGGATTGTTCACCCA</td><td>464.4</td><td>-8.4</td><td>1.3</td><td>-6.3</td><td>3.7E-10</td><td>9.3E-10</td><td>MB231/MB468</td><td>MP</td></tr>
<tr><td>>hg38_21_+_8433172_8446622_RNA45SN1_45S_50ntFlanks@8176.8203.28|CTTCTGATCGAGGCCCAGCCCGTGGACG</td><td>>hg38_21_+_8433172_8446622_RNA45SN1_45S_50ntFlanks</td><td>8176</td><td>8203</td><td>28</td><td>CTTCTGATCGAGGCCCAGCCCGTGGACG</td><td>3960.1</td><td>-4.6</td><td>0.7</td><td>-6.3</td><td>3.9E-10</td><td>9.3E-10</td><td>MB231/MB468</td><td>NUC</td></tr>
<tr><td>>hg38_21_+_8433172_8446622_RNA45SN1_45S_50ntFlanks@11037.11061.25|CTCTCCCCCGCTCCCCGTCCTCCCC</td><td>>hg38_21_+_8433172_8446622_RNA45SN1_45S_50ntFlanks</td><td>11037</td><td>11061</td><td>25</td><td>CTCTCCCCCGCTCCCCGTCCTCCCC</td><td>362.6</td><td>-8.7</td><td>1.4</td><td>-6.3</td><td>3.9E-10</td><td>9.3E-10</td><td>MB231/MB468</td><td>NUC</td></tr>
<tr><td>>MI0000082|hsa-mir-25&WithFlank&7|-|100093554|100093649@58.79.22&[MIMAT0000081&hsa-miR-25-3p&offsets|0|0;m-23&7|-|100093571|100093592&offsets|0|0]|CATTGCACTTGTCTCGGTCTGA</td><td>>MI0000082|hsa-mir-25&WithFlank&7|-|100093554|100093649</td><td>58</td><td>79</td><td>22</td><td>CATTGCACTTGTCTCGGTCTGA</td><td>21035.3</td><td>5.1</td><td>0.8</td><td>6.2</td><td>4.3E-10</td><td>9.3E-10</td><td>MB231/MB468</td><td>MITO</td></tr>
<tr><td>>MI0000088|hsa-mir-30a&WithFlank&6|-|71403545|71403627@12.33.22(+1U)&[MIMAT0000087&hsa-miR-30a-5p&offsets|0|0(+1U);m-15&6|-|71403595|71403616&offsets|0|0(+1U)]|TGTAAACATCCTCGACTGGAAGT</td><td>>MI0000088|hsa-mir-30a&WithFlank&6|-|71403545|71403627</td><td>12</td><td>33</td><td>22(+1U)</td><td>TGTAAACATCCTCGACTGGAAGT</td><td>8305.6</td><td>6.9</td><td>1.1</td><td>6.2</td><td>4.3E-10</td><td>9.3E-10</td><td>MB231/MB468</td><td>MITO</td></tr>
<tr><td>>hg38_21_+_8433172_8446622_RNA45SN1_45S_50ntFlanks@4898.4918.21|AAAGGAATTGACGGAAGGGCA</td><td>>hg38_21_+_8433172_8446622_RNA45SN1_45S_50ntFlanks</td><td>4898</td><td>4918</td><td>21</td><td>AAAGGAATTGACGGAAGGGCA</td><td>179.9</td><td>-11.4</td><td>1.8</td><td>-6.3</td><td>3.7E-10</td><td>9.3E-10</td><td>MB231/MB468</td><td>MP</td></tr>
<tr><td>>hg38_21_+_8433172_8446622_RNA45SN1_45S_50ntFlanks@4449.4466.18|CCTCTCGGCGCCCCCTCG</td><td>>hg38_21_+_8433172_8446622_RNA45SN1_45S_50ntFlanks</td><td>4449</td><td>4466</td><td>18</td><td>CCTCTCGGCGCCCCCTCG</td><td>76.4</td><td>-10.1</td><td>1.6</td><td>-6.3</td><td>3.7E-10</td><td>9.3E-10</td><td>MB231/MB468</td><td>MP</td></tr>
</table>

<table>
<tr><td>>hg38_21_+_8433172_8446622_RNA45SN1_45S_50ntFlanks@6719.6751.33|TGAATTGCAGGACACATTGATCATCGACACTTC</td><td>>hg38_21_+_8433172_8446622_RNA45SN1_45S_50ntFlanks</td><td>6719</td><td>6751</td><td>33</td><td>TGAATTGCAGGACACATTGATCATCGACACTTC</td><td>643.5</td><td>-6.3</td><td>1.0</td><td>-6.2</td><td>4.4E-10</td><td>9.4E-10</td><td>MB231/MB468</td><td>MITO</td></tr>
<tr><td>>hg38_21_+_8433172_8446622_RNA45SN1_45S_50ntFlanks@12431.12455.25|TCCTATCATTGTGAAGCAGAATTCA</td><td>>hg38_21_+_8433172_8446622_RNA45SN1_45S_50ntFlanks</td><td>12431</td><td>12455</td><td>25</td><td>TCCTATCATTGTGAAGCAGAATTCA</td><td>311.6</td><td>-6.0</td><td>1.0</td><td>-6.3</td><td>3.9E-10</td><td>9.4E-10</td><td>MB231/MB468</td><td>NUC</td></tr>
<tr><td>>MI0000102|hsa-mir-100&WithFlank&11|-|122152223|122152314@19.38.20&[MIMAT0000098&hsa-miR-100-5p&offsets|0|-2;m-27&11|-|122152276|122152296&offsets|0|-1]|AACCCGTAGATCCGAACTTG</td><td>>MI0000102|hsa-mir-100&WithFlank&11|-|122152223|122152314</td><td>19</td><td>38</td><td>20</td><td>AACCCGTAGATCCGAACTTG</td><td>35392.8</td><td>12.4</td><td>2.0</td><td>6.3</td><td>3.7E-10</td><td>9.4E-10</td><td>MB231/MB468</td><td>MP</td></tr>
<tr><td>>hg38_21_+_8433172_8446622_RNA45SN1_45S_50ntFlanks@9562.9597.36|TCCTGACGTGCAAATCGGTCGTCCGACCTGGGTATA</td><td>>hg38_21_+_8433172_8446622_RNA45SN1_45S_50ntFlanks</td><td>9562</td><td>9597</td><td>36</td><td>TCCTGACGTGCAAATCGGTCGTCCGACCTGGGTATA</td><td>835.7</td><td>-8.5</td><td>1.4</td><td>-6.3</td><td>3.9E-10</td><td>9.4E-10</td><td>MB231/MB468</td><td>NUC</td></tr>
<tr><td>>hg38_21_+_8433172_8446622_RNA45SN1_45S_50ntFlanks@12159.12181.23|TTTGACTGGGGCGGTACACCTGT</td><td>>hg38_21_+_8433172_8446622_RNA45SN1_45S_50ntFlanks</td><td>12159</td><td>12181</td><td>23</td><td>TTTGACTGGGGCGGTACACCTGT</td><td>406.4</td><td>-7.8</td><td>1.2</td><td>-6.3</td><td>3.9E-10</td><td>9.4E-10</td><td>MB231/MB468</td><td>NUC</td></tr>
<tr><td>>hg38_21_+_8433172_8446622_RNA45SN1_45S_50ntFlanks@12657.12674.18|GACTGAACGCCTCTAAGT</td><td>>hg38_21_+_8433172_8446622_RNA45SN1_45S_50ntFlanks</td><td>12657</td><td>12674</td><td>18</td><td>GACTGAACGCCTCTAAGT</td><td>305.1</td><td>-7.8</td><td>1.2</td><td>-6.2</td><td>4.4E-10</td><td>9.4E-10</td><td>MB231/MB468</td><td>MITO</td></tr>
<tr><td>>hg38_21_+_8433172_8446622_RNA45SN1_45S_50ntFlanks@10389.10429.41|TGAACAGCAGTTGAACATGGGTCAGTCGGTCCTGAGAGATG</td><td>>hg38_21_+_8433172_8446622_RNA45SN1_45S_50ntFlanks</td><td>10389</td><td>10429</td><td>41</td><td>TGAACAGCAGTTGAACATGGGTCAGTCGGTCCTGAGAGATG</td><td>1121.1</td><td>-5.8</td><td>0.9</td><td>-6.3</td><td>4.0E-10</td><td>9.4E-10</td><td>MB231/MB468</td><td>NUC</td></tr>
<tr><td>>hg38_21_+_8433172_8446622_RNA45SN1_45S_50ntFlanks@10393.10426.34|CAGCAGTTGAACATGGGTCAGTCGGTCCTGAGAG</td><td>>hg38_21_+_8433172_8446622_RNA45SN1_45S_50ntFlanks</td><td>10393</td><td>10426</td><td>34</td><td>CAGCAGTTGAACATGGGTCAGTCGGTCCTGAGAG</td><td>600.5</td><td>-4.9</td><td>0.8</td><td>-6.3</td><td>4.0E-10</td><td>9.4E-10</td><td>MB231/MB468</td><td>NUC</td></tr>
<tr><td>>hg38_21_+_8433172_8446622_RNA45SN1_45S_50ntFlanks@10420.10456.37|CTGAGAGATGGGCGAGCGCCGTTCCGAAGGGACGGGC</td><td>>hg38_21_+_8433172_8446622_RNA45SN1_45S_50ntFlanks</td><td>10420</td><td>10456</td><td>37</td><td>CTGAGAGATGGGCGAGCGCCGTTCCGAAGGGACGGGC</td><td>527.0</td><td>-6.7</td><td>1.1</td><td>-6.3</td><td>3.8E-10</td><td>9.4E-10</td><td>MB231/MB468</td><td>MP</td></tr>
<tr><td>>MI0000255|hsa-mir-30d&WithFlank&8|-|134804870|134804951@12.33.22&[MIMAT0000245&hsa-miR-30d-5p&offsets|0|0;m-21&8|-|134804919|134804940&offsets|0|0]|TGTAAACATCCCCGACTGGAAG</td><td>>MI0000255|hsa-mir-30d&WithFlank&8|-|134804870|134804951</td><td>12</td><td>33</td><td>22</td><td>TGTAAACATCCCCGACTGGAAG</td><td>1481.0</td><td>5.3</td><td>0.8</td><td>6.2</td><td>4.4E-10</td><td>9.4E-10</td><td>MB231/MB468</td><td>MITO</td></tr>
<tr><td>>hg38_21_+_8433172_8446622_RNA45SN1_45S_50ntFlanks@6647.6678.32|CGTACGACTCTTAGCGGTGGATCACTCGGCTC</td><td>>hg38_21_+_8433172_8446622_RNA45SN1_45S_50ntFlanks</td><td>6647</td><td>6678</td><td>32</td><td>CGTACGACTCTTAGCGGTGGATCACTCGGCTC</td><td>2969.5</td><td>4.5</td><td>0.7</td><td>6.3</td><td>2.9E-10</td><td>9.4E-10</td><td>BT20/MB468</td><td>MITO</td></tr>
<tr><td>>hg38_1_-_228634819_228635039_RNA5S12_5S_50ntFlanks@146.170.25|TGGGAATACCGGGTGCTGTAGGCTT</td><td>>hg38_1_-_228634819_228635039_RNA5S12_5S_50ntFlanks</td><td>146</td><td>170</td><td>25</td><td>TGGGAATACCGGGTGCTGTAGGCTT</td><td>1438.2</td><td>4.4</td><td>0.7</td><td>6.3</td><td>4.0E-10</td><td>9.4E-10</td><td>MB231/MB468</td><td>NUC</td></tr>
<tr><td>>hg38_21_+_8433172_8446622_RNA45SN1_45S_50ntFlanks@8381.8402.22|AAGAGGGCGTGAAACCGTTAAG</td><td>>hg38_21_+_8433172_8446622_RNA45SN1_45S_50ntFlanks</td><td>8381</td><td>8402</td><td>22</td><td>AAGAGGGCGTGAAACCGTTAAG</td><td>1074.7</td><td>-4.7</td><td>0.8</td><td>-6.3</td><td>3.9E-10</td><td>9.4E-10</td><td>MB231/MB468</td><td>CYTO</td></tr>
<tr><td>>hg38_21_+_8433172_8446622_RNA45SN1_45S_50ntFlanks@10381.10424.44|GTTCCATGTGAACAGCAGTTGAACATGGGTCAGTCGGTCCTGAG</td><td>>hg38_21_+_8433172_8446622_RNA45SN1_45S_50ntFlanks</td><td>10381</td><td>10424</td><td>44</td><td>GTTCCATGTGAACAGCAGTTGAACATGGGTCAGTCGGTCCTGAG</td><td>629.2</td><td>8.2</td><td>1.3</td><td>6.3</td><td>3.5E-10</td><td>9.4E-10</td><td>BT20/MB468</td><td>CYTO</td></tr>
<tr><td>>hg38_21_+_8433172_8446622_RNA45SN1_45S_50ntFlanks@10131.10157.27|CGGCGGCGGCGGCGGCGGGGGTGTGGG</td><td>>hg38_21_+_8433172_8446622_RNA45SN1_45S_50ntFlanks</td><td>10131</td><td>10157</td><td>27</td><td>CGGCGGCGGCGGCGGCGGGGGTGTGGG</td><td>110.2</td><td>-10.6</td><td>1.7</td><td>-6.3</td><td>3.8E-10</td><td>9.4E-10</td><td>MB231/MB468</td><td>MP</td></tr>
<tr><td>>hg38_21_+_8433172_8446622_RNA45SN1_45S_50ntFlanks@8436.8467.32|GAGGATTCAACCCGGCGGCGGGTCCGGCCGTG</td><td>>hg38_21_+_8433172_8446622_RNA45SN1_45S_50ntFlanks</td><td>8436</td><td>8467</td><td>32</td><td>GAGGATTCAACCCGGCGGCGGGTCCGGCCGTG</td><td>80.6</td><td>-10.2</td><td>1.6</td><td>-6.3</td><td>3.8E-10</td><td>9.5E-10</td><td>MB231/MB468</td><td>MP</td></tr>
<tr><td>>trna11_GluTTC_15_-_26327381_26327452@1.29.29,trna3_GluTTC_13_-_45492062_45492133@1.29.29|TCCCACATGGTCTAGCGGTTAGGATTCCT</td><td>>trna11_GluTTC_15_-_26327381_26327452</td><td>1</td><td>29</td><td>29</td><td>TCCCACATGGTCTAGCGGTTAGGATTCCT</td><td>2813.6</td><td>-4.3</td><td>0.7</td><td>-6.2</td><td>4.4E-10</td><td>9.5E-10</td><td>MB231/MB468</td><td>MITO</td></tr>
<tr><td>>MI0000077|hsa-mir-21&WithFlank&17|+|59841260|59841343@14.34.21(+2C)&[MIMAT0000076&hsa-miR-21-5p&offsets|0|-1(+2C);m-1&17|+|59841273|59841294&offsets|0|-1(+2C)]|TAGCTTATCAGACTGATGTTGCC</td><td>>MI0000077|hsa-mir-21&WithFlank&17|+|59841260|59841343</td><td>14</td><td>34</td><td>21(+2C)</td><td>TAGCTTATCAGACTGATGTTGCC</td><td>1293.3</td><td>6.9</td><td>1.1</td><td>6.3</td><td>3.9E-10</td><td>9.5E-10</td><td>MB231/MB468</td><td>CYTO</td></tr>
<tr><td>>MI0000736|hsa-mir-30c-1&WithFlank&1|+|40757278|40757378@23.44.22(+1A);MI0000254|hsa-mir-30c-2&WithFlank&6|-|71376954|71377037@13.34.22(+1A)&[MIMAT0000244&hsa-miR-30c-5p&offsets|0|-1(+1A);m-57&1|+|40757300|40757323&offsets|0|-2(+1A)];[MIMAT0000244_1&hsa-miR-30c-5p&offsets|0|-1(+1A);m-56&6|-|71377002|71377025&offsets|0|-2(+1A)]|TGTAAACATCCTACACTCTCAGA</td><td>>MI0000736|hsa-mir-30c-1&WithFlank&1|+|40757278|40757378</td><td>23</td><td>44</td><td>22(+1A)</td><td>TGTAAACATCCTACACTCTCAGA</td><td>661.3</td><td>5.7</td><td>0.9</td><td>6.3</td><td>3.9E-10</td><td>9.5E-10</td><td>MB231/MB468</td><td>CYTO</td></tr>
<tr><td>>hg38_21_+_8433172_8446622_RNA45SN1_45S_50ntFlanks@6647.6670.24|CGTACGACTCTTAGCGGTGGATCA</td><td>>hg38_21_+_8433172_8446622_RNA45SN1_45S_50ntFlanks</td><td>6647</td><td>6670</td><td>24</td><td>CGTACGACTCTTAGCGGTGGATCA</td><td>490.6</td><td>-2.7</td><td>0.4</td><td>-6.2</td><td>4.6E-10</td><td>9.5E-10</td><td>BT20/MB468</td><td>NUC</td></tr>
<tr><td>>hg38_21_+_8433172_8446622_RNA45SN1_45S_50ntFlanks@10377.10424.48|AAGGGTTCCATGTGAACAGCAGTTGAACATGGGTCAGTCGGTCCTGAG</td><td>>hg38_21_+_8433172_8446622_RNA45SN1_45S_50ntFlanks</td><td>10377</td><td>10424</td><td>48</td><td>AAGGGTTCCATGTGAACAGCAGTTGAACATGGGTCAGTCGGTCCTGAG</td><td>4588.8</td><td>7.7</td><td>1.2</td><td>6.3</td><td>3.0E-10</td><td>9.5E-10</td><td>BT20/MB468</td><td>MITO</td></tr>
<tr><td>>hg38_21_+_8433172_8446622_RNA45SN1_45S_50ntFlanks@5472.5506.35|ACGGCCCTGGCGGAGCGCTGAGAAGACGGTCGAAC</td><td>>hg38_21_+_8433172_8446622_RNA45SN1_45S_50ntFlanks</td><td>5472</td><td>5506</td><td>35</td><td>ACGGCCCTGGCGGAGCGCTGAGAAGACGGTCGAAC</td><td>640.3</td><td>6.5</td><td>1.0</td><td>6.3</td><td>4.0E-10</td><td>9.5E-10</td><td>MB231/MB468</td><td>NUC</td></tr>
<tr><td>>trna119_LysCTT_1_-_145395522_145395594@4.33.30,trna11_LysCTT_5_-_180648979_180649051@4.33.30,trna13_LysCTT_6_+_26556774_26556846@4.33.30,trna7_LysCTT_16_+_3225692_3225764@4.33.30,trna9_LysCTT_5_+_180634755_180634827@4.33.30|CGGCTAGCTCAGTCGGTAGAGCATGAGACT</td><td>>trna119_LysCTT_1_-_145395522_145395594</td><td>4</td><td>33</td><td>30</td><td>CGGCTAGCTCAGTCGGTAGAGCATGAGACT</td><td>1810.0</td><td>8.3</td><td>1.3</td><td>6.3</td><td>4.0E-10</td><td>9.5E-10</td><td>MB231/MB468</td><td>NUC</td></tr>
<tr><td>>hg38_21_+_8433172_8446622_RNA45SN1_45S_50ntFlanks@12589.12610.22|GTTCAGACATTTGGTGTATGTG</td><td>>hg38_21_+_8433172_8446622_RNA45SN1_45S_50ntFlanks</td><td>12589</td><td>12610</td><td>22</td><td>GTTCAGACATTTGGTGTATGTG</td><td>267.3</td><td>-11.1</td><td>1.7</td><td>-6.6</td><td>3.5E-11</td><td>9.5E-10</td><td>BT20/MB468</td><td>MP</td></tr>
<tr><td>>hg38_21_+_8433172_8446622_RNA45SN1_45S_50ntFlanks@12416.12448.33|TTCGATGTCGGCTCTTCCTATCATTGTGAAGCA</td><td>>hg38_21_+_8433172_8446622_RNA45SN1_45S_50ntFlanks</td><td>12416</td><td>12448</td><td>33</td><td>TTCGATGTCGGCTCTTCCTATCATTGTGAAGCA</td><td>2085.2</td><td>5.9</td><td>0.9</td><td>6.3</td><td>3.5E-10</td><td>9.5E-10</td><td>BT20/MB468</td><td>CYTO</td></tr>
<tr><td>>hg38_21_+_8433172_8446622_RNA45SN1_45S_50ntFlanks@8169.8199.31|CCAAGTCCTTCTGATCGAGGCCCAGCCCGTG</td><td>>hg38_21_+_8433172_8446622_RNA45SN1_45S_50ntFlanks</td><td>8169</td><td>8199</td><td>31</td><td>CCAAGTCCTTCTGATCGAGGCCCAGCCCGTG</td><td>70.0</td><td>-10.0</td><td>1.6</td><td>-6.3</td><td>3.8E-10</td><td>9.6E-10</td><td>MB231/MB468</td><td>MP</td></tr>
<tr><td>>hg38_21_+_8433172_8446622_RNA45SN1_45S_50ntFlanks@10869.10902.34|AGGGCTGGGTCGGTCGGGCTGGGGCGCGAAGCGG</td><td>>hg38_21_+_8433172_8446622_RNA45SN1_45S_50ntFlanks</td><td>10869</td><td>10902</td><td>34</td><td>AGGGCTGGGTCGGTCGGGCTGGGGCGCGAAGCGG</td><td>6942.7</td><td>-10.8</td><td>1.7</td><td>-6.2</td><td>4.6E-10</td><td>9.6E-10</td><td>BT20/MB468</td><td>NUC</td></tr>
<tr><td>>hg38_21_+_8433172_8446622_RNA45SN1_45S_50ntFlanks@4734.4758.25|AACGAAAGTCGGAGGTTCGAAGACG</td><td>>hg38_21_+_8433172_8446622_RNA45SN1_45S_50ntFlanks</td><td>4734</td><td>4758</td><td>25</td><td>AACGAAAGTCGGAGGTTCGAAGACG</td><td>1965.2</td><td>-3.1</td><td>0.5</td><td>-6.3</td><td>4.0E-10</td><td>9.6E-10</td><td>MB231/MB468</td><td>NUC</td></tr>
<tr><td>>hg38_21_+_8433172_8446622_RNA45SN1_45S_50ntFlanks@4161.4184.24|CACATCCAAGGAAGGCAGCAGGCG</td><td>>hg38_21_+_8433172_8446622_RNA45SN1_45S_50ntFlanks</td><td>4161</td><td>4184</td><td>24</td><td>CACATCCAAGGAAGGCAGCAGGCG</td><td>190.6</td><td>-7.4</td><td>1.2</td><td>-6.3</td><td>4.0E-10</td><td>9.6E-10</td><td>MB231/MB468</td><td>NUC</td></tr>
<tr><td>>hg38_21_+_8433172_8446622_RNA45SN1_45S_50ntFlanks@12456.12484.29|CCAAGCGTTGGATTGTTCACCCACTAATA</td><td>>hg38_21_+_8433172_8446622_RNA45SN1_45S_50ntFlanks</td><td>12456</td><td>12484</td><td>29</td><td>CCAAGCGTTGGATTGTTCACCCACTAATA</td><td>6660.2</td><td>-7.9</td><td>1.3</td><td>-6.3</td><td>3.5E-10</td><td>9.6E-10</td><td>BT20/MB468</td><td>CYTO</td></tr>
<tr><td>>hg38_21_+_8433172_8446622_RNA45SN1_45S_50ntFlanks@10389.10409.21|TGAACAGCAGTTGAACATGGG</td><td>>hg38_21_+_8433172_8446622_RNA45SN1_45S_50ntFlanks</td><td>10389</td><td>10409</td><td>21</td><td>TGAACAGCAGTTGAACATGGG</td><td>1300.8</td><td>-8.6</td><td>1.4</td><td>-6.3</td><td>4.0E-10</td><td>9.6E-10</td><td>MB231/MB468</td><td>NUC</td></tr>
<tr><td>>hg38_21_+_8433172_8446622_RNA45SN1_45S_50ntFlanks@10808.10844.37|CAATGTAGGTAAGGGAAGTCGGCAAGCCGGATCCGTA</td><td>>hg38_21_+_8433172_8446622_RNA45SN1_45S_50ntFlanks</td><td>10808</td><td>10844</td><td>37</td><td>CAATGTAGGTAAGGGAAGTCGGCAAGCCGGATCCGTA</td><td>178.3</td><td>-7.7</td><td>1.2</td><td>-6.3</td><td>4.0E-10</td><td>9.6E-10</td><td>MB231/MB468</td><td>NUC</td></tr>
<tr><td>>hg38_21_+_8433172_8446622_RNA45SN1_45S_50ntFlanks@9862.9879.18|ATGAACCGAACGCCGGGT</td><td>>hg38_21_+_8433172_8446622_RNA45SN1_45S_50ntFlanks</td><td>9862</td><td>9879</td><td>18</td><td>ATGAACCGAACGCCGGGT</td><td>283.0</td><td>-12.0</td><td>1.9</td><td>-6.3</td><td>3.8E-10</td><td>9.6E-10</td><td>MB231/MB468</td><td>MP</td></tr>
<tr><td>>MI0006407|hsa-mir-1270&WithFlank&19|-|20399266|20399360@19.41.23&[MIMAT0005924&hsa-miR-1270&offsets|0|0;m-459&19|-|20399320|20399342&offsets|0|0]|CTGGAGATATGGAAGAGCTGTGT</td><td>>MI0006407|hsa-mir-1270&WithFlank&19|-|20399266|20399360</td><td>19</td><td>41</td><td>23</td><td>CTGGAGATATGGAAGAGCTGTGT</td><td>104.0</td><td>9.9</td><td>1.6</td><td>6.3</td><td>3.8E-10</td><td>9.6E-10</td><td>MB231/MB468</td><td>MP</td></tr>
<tr><td>>hg38_21_+_8433172_8446622_RNA45SN1_45S_50ntFlanks@4375.4398.24|AAGCTCGTAGTTGGATCTTGGGAG</td><td>>hg38_21_+_8433172_8446622_RNA45SN1_45S_50ntFlanks</td><td>4375</td><td>4398</td><td>24</td><td>AAGCTCGTAGTTGGATCTTGGGAG</td><td>147.5</td><td>-11.1</td><td>1.8</td><td>-6.3</td><td>3.8E-10</td><td>9.6E-10</td><td>MB231/MB468</td><td>MP</td></tr>
<tr><td>>hg38_21_+_8433172_8446622_RNA45SN1_45S_50ntFlanks@10358.10383.26|TTTGAAGGCCGAAGTGGAGAAGGGTT</td><td>>hg38_21_+_8433172_8446622_RNA45SN1_45S_50ntFlanks</td><td>10358</td><td>10383</td><td>26</td><td>TTTGAAGGCCGAAGTGGAGAAGGGTT</td><td>189.5</td><td>-3.1</td><td>0.5</td><td>-6.3</td><td>4.1E-10</td><td>9.6E-10</td><td>MB231/MB468</td><td>NUC</td></tr>
<tr><td>>hg38_21_+_8433172_8446622_RNA45SN1_45S_50ntFlanks@10808.10827.20|CAATGTAGGTAAGGGAAGTC</td><td>>hg38_21_+_8433172_8446622_RNA45SN1_45S_50ntFlanks</td><td>10808</td><td>10827</td><td>20</td><td>CAATGTAGGTAAGGGAAGTC</td><td>616.6</td><td>-5.9</td><td>0.9</td><td>-6.3</td><td>4.0E-10</td><td>9.6E-10</td><td>MB231/MB468</td><td>CYTO</td></tr>
<tr><td>>hg38_21_+_8433172_8446622_RNA45SN1_45S_50ntFlanks@12418.12455.38|CGATGTCGGCTCTTCCTATCATTGTGAAGCAGAATTCA</td><td>>hg38_21_+_8433172_8446622_RNA45SN1_45S_50ntFlanks</td><td>12418</td><td>12455</td><td>38</td><td>CGATGTCGGCTCTTCCTATCATTGTGAAGCAGAATTCA</td><td>816.9</td><td>-7.6</td><td>1.2</td><td>-6.3</td><td>3.8E-10</td><td>9.6E-10</td><td>MB231/MB468</td><td>MP</td></tr>
<tr><td>>hg38_21_+_8433172_8446622_RNA45SN1_45S_50ntFlanks@8472.8497.26|CGGCCCGGCGGATCTTTCCCGCCCCC</td><td>>hg38_21_+_8433172_8446622_RNA45SN1_45S_50ntFlanks</td><td>8472</td><td>8497</td><td>26</td><td>CGGCCCGGCGGATCTTTCCCGCCCCC</td><td>275.3</td><td>-11.2</td><td>1.7</td><td>-6.6</td><td>3.6E-11</td><td>9.6E-10</td><td>BT20/MB468</td><td>MP</td></tr>
<tr><td>>hg38_21_+_8433172_8446622_RNA45SN1_45S_50ntFlanks@12405.12446.42|CTTTTTGATCCTTCGATGTCGGCTCTTCCTATCATTGTGAAG</td><td>>hg38_21_+_8433172_8446622_RNA45SN1_45S_50ntFlanks</td><td>12405</td><td>12446</td><td>42</td><td>CTTTTTGATCCTTCGATGTCGGCTCTTCCTATCATTGTGAAG</td><td>212.8</td><td>-11.6</td><td>1.9</td><td>-6.3</td><td>3.9E-10</td><td>9.7E-10</td><td>MB231/MB468</td><td>MP</td></tr>
<tr><td>>hg38_21_+_8433172_8446622_RNA45SN1_45S_50ntFlanks@12083.12107.25|CTCGCTTCTGGCGCCAAGCGCCCGG</td><td>>hg38_21_+_8433172_8446622_RNA45SN1_45S_50ntFlanks</td><td>12083</td><td>12107</td><td>25</td><td>CTCGCTTCTGGCGCCAAGCGCCCGG</td><td>256.4</td><td>-6.8</td><td>1.1</td><td>-6.3</td><td>4.0E-10</td><td>9.7E-10</td><td>MB231/MB468</td><td>CYTO</td></tr>
<tr><td>>hg38_21_+_8433172_8446622_RNA45SN1_45S_50ntFlanks@8380.8403.24|CAAGAGGGCGTGAAACCGTTAAGA</td><td>>hg38_21_+_8433172_8446622_RNA45SN1_45S_50ntFlanks</td><td>8380</td><td>8403</td><td>24</td><td>CAAGAGGGCGTGAAACCGTTAAGA</td><td>645.8</td><td>-4.3</td><td>0.7</td><td>-6.3</td><td>4.0E-10</td><td>9.7E-10</td><td>MB231/MB468</td><td>CYTO</td></tr>
<tr><td>>MI0000342|hsa-mir-200b&WithFlank&1|+|1167098|1167204@63.85.23(+1C)&[MIMAT0000318&hsa-miR-200b-3p&offsets|0|+1(+1C);m-93&1|+|1167160|1167182&offsets|0|0(+1C)]|TAATACTGCCTGGTAATGATGACC</td><td>>MI0000342|hsa-mir-200b&WithFlank&1|+|1167098|1167204</td><td>63</td><td>85</td><td>23(+1C)</td><td>TAATACTGCCTGGTAATGATGACC</td><td>157.4</td><td>-10.4</td><td>1.6</td><td>-6.6</td><td>3.6E-11</td><td>9.7E-10</td><td>BT20/MB468</td><td>MP</td></tr>
<tr><td>>MI0000234|hsa-mir-192&WithFlank&11|-|64891131|64891252@30.49.20(+1U)&[MIMAT0000222&hsa-miR-192-5p&offsets|0|-1(+1U);m-59&11|-|64891203|64891223&offsets|0|-1(+1U)]|CTGACCTATGAATTGACAGCT</td><td>>MI0000234|hsa-mir-192&WithFlank&11|-|64891131|64891252</td><td>30</td><td>49</td><td>20(+1U)</td><td>CTGACCTATGAATTGACAGCT</td><td>296.1</td><td>4.6</td><td>0.7</td><td>6.3</td><td>4.1E-10</td><td>9.7E-10</td><td>MB231/MB468</td><td>NUC</td></tr>
<tr><td>>hg38_21_+_8433172_8446622_RNA45SN1_45S_50ntFlanks@4202.4219.18|CGACCCGGGGAGGTAGTG</td><td>>hg38_21_+_8433172_8446622_RNA45SN1_45S_50ntFlanks</td><td>4202</td><td>4219</td><td>18</td><td>CGACCCGGGGAGGTAGTG</td><td>166.6</td><td>-9.5</td><td>1.5</td><td>-6.3</td><td>3.0E-10</td><td>9.7E-10</td><td>BT20/MB468</td><td>MITO</td></tr>
<tr><td>>hg38_21_+_8433172_8446622_RNA45SN1_45S_50ntFlanks@12627.12656.30|ATGGGGCGAAGCTACCATCTGTGGGATTAT</td><td>>hg38_21_+_8433172_8446622_RNA45SN1_45S_50ntFlanks</td><td>12627</td><td>12656</td><td>30</td><td>ATGGGGCGAAGCTACCATCTGTGGGATTAT</td><td>216.9</td><td>-9.9</td><td>1.6</td><td>-6.3</td><td>3.0E-10</td><td>9.7E-10</td><td>BT20/MB468</td><td>MITO</td></tr>
<tr><td>>hg38_21_+_8433172_8446622_RNA45SN1_45S_50ntFlanks@12163.12181.19|ACTGGGGCGGTACACCTGT</td><td>>hg38_21_+_8433172_8446622_RNA45SN1_45S_50ntFlanks</td><td>12163</td><td>12181</td><td>19</td><td>ACTGGGGCGGTACACCTGT</td><td>4406.6</td><td>-9.9</td><td>1.5</td><td>-6.6</td><td>3.6E-11</td><td>9.7E-10</td><td>BT20/MB468</td><td>MP</td></tr>
<tr><td>>hg38_21_+_8433172_8446622_RNA45SN1_45S_50ntFlanks@8190.8212.23|CCAGCCCGTGGACGGTGTGAGGC</td><td>>hg38_21_+_8433172_8446622_RNA45SN1_45S_50ntFlanks</td><td>8190</td><td>8212</td><td>23</td><td>CCAGCCCGTGGACGGTGTGAGGC</td><td>1292.6</td><td>-4.7</td><td>0.7</td><td>-6.6</td><td>3.6E-11</td><td>9.7E-10</td><td>BT20/MB468</td><td>MP</td></tr>
</table>

<table>
<tr><th></th><th></th><th></th><th></th><th></th><th></th><th></th><th></th><th></th><th></th><th></th><th></th><th></th><th></th></tr>
<tr><td>>hg38_21_+_8433172_8446622_RNA45SN1_45S_50ntFlanks@12222.12249.28|ACAGAAACCTCCCGTGGAGCAGAAGGGC</td><td>>hg38_21_+_8433172_8446622_RNA45SN1_45S_50ntFlanks</td><td>12222</td><td>12249</td><td>28</td><td>ACAGAAACCTCCCGTGGAGCAGAAGGGC</td><td>76.1</td><td>-10.1</td><td>1.6</td><td>-6.3</td><td>3.9E-10</td><td>9.7E-10</td><td>MB231/MB468</td><td>MP</td></tr>
<tr><td>>hg38_21_+_8433172_8446622_RNA45SN1_45S_50ntFlanks@4380.4408.29|CGTAGTTGGATCTTGGGAGCGGGCGGGCG</td><td>>hg38_21_+_8433172_8446622_RNA45SN1_45S_50ntFlanks</td><td>4380</td><td>4408</td><td>29</td><td>CGTAGTTGGATCTTGGGAGCGGGCGGGCG</td><td>252.2</td><td>-7.7</td><td>1.2</td><td>-6.2</td><td>4.5E-10</td><td>9.7E-10</td><td>MB231/MB468</td><td>MITO</td></tr>
<tr><td>>hg38_21_+_8433172_8446622_RNA45SN1_45S_50ntFlanks@11733.11751.19|TATGACTCTCTTAAGGTAG</td><td>>hg38_21_+_8433172_8446622_RNA45SN1_45S_50ntFlanks</td><td>11733</td><td>11751</td><td>19</td><td>TATGACTCTCTTAAGGTAG</td><td>244.9</td><td>-4.6</td><td>0.7</td><td>-6.3</td><td>4.0E-10</td><td>9.7E-10</td><td>MB231/MB468</td><td>CYTO</td></tr>
<tr><td>>MI0000239|hsa-mir-197&WithFlank&1|+|109598887|109598973@54.72.19&[MIMAT0000227&hsa-miR-197-3p&offsets|0|-3;m-99&1|+|109598940|109598961&offsets|0|-3]|TTCACCACCTTCTCCACCC</td><td>>MI0000239|hsa-mir-197&WithFlank&1|+|109598887|109598973</td><td>54</td><td>72</td><td>19</td><td>TTCACCACCTTCTCCACCC</td><td>107.9</td><td>3.0</td><td>0.5</td><td>6.2</td><td>4.7E-10</td><td>9.7E-10</td><td>BT20/MB468</td><td>NUC</td></tr>
<tr><td>>trna10_ValCAC_5_-_180649395_180649467@35.54.20,trna12_ValAAC_5_-_180645270_180645342@35.54.20,trna132_ValAAC_6_-_27721179_27721251@35.54.20,trna13_ValCAC_19_-_4724647_4724719@35.54.20,trna152_ValCAC_6_-_27248049_27248121@35.54.20,trna157_ValCAC_6_-_27173867_27173939@35.54.20,trna15_ValAAC_5_-_180615416_180615488@35.54.20,trna18_ValCAC_5_-_180529253_180529325@35.54.20,trna2_ValAAC_3_+_169490018_169490090@35.54.20,trna2_ValCAC_5_+_180524070_180524142@35.54.20,trna37_ValAAC_6_+_27203288_27203360@35.54.20,trna4_ValAAC_5_+_180591154_180591226@35.54.20,trna5_ValAAC_5_+_180596610_180596682@35.54.20,trna6_ValCAC_5_+_180600650_180600722@35.54.20,trna85_ValCAC_1_-_161369490_161369562@35.54.20,trna98_ValCAC_1_-_149298555_149298627@35.54.20,trna9_ValCAC_6_+_26538282_26538354@35.54.20|ACACGCGAAAGGTCCCCGGT</td><td>>trna10_ValCAC_5_-_180649395_180649467</td><td>35</td><td>54</td><td>20</td><td>ACACGCGAAAGGTCCCCGGT</td><td>902.1</td><td>-9.5</td><td>1.5</td><td>-6.3</td><td>3.9E-10</td><td>9.8E-10</td><td>MB231/MB468</td><td>MP</td></tr>
<tr><td>>hg38_21_+_8433172_8446622_RNA45SN1_45S_50ntFlanks@10684.10710.27|CGGCGGCGTCCGGTGAGCTCTCGCTGG</td><td>>hg38_21_+_8433172_8446622_RNA45SN1_45S_50ntFlanks</td><td>10684</td><td>10710</td><td>27</td><td>CGGCGGCGTCCGGTGAGCTCTCGCTGG</td><td>3934.5</td><td>-9.3</td><td>1.5</td><td>-6.3</td><td>3.9E-10</td><td>9.8E-10</td><td>MB231/MB468</td><td>MP</td></tr>
<tr><td>>hg38_21_+_8433172_8446622_RNA45SN1_45S_50ntFlanks@12418.12456.39|CGATGTCGGCTCTTCCTATCATTGTGAAGCAGAATTCAC</td><td>>hg38_21_+_8433172_8446622_RNA45SN1_45S_50ntFlanks</td><td>12418</td><td>12456</td><td>39</td><td>CGATGTCGGCTCTTCCTATCATTGTGAAGCAGAATTCAC</td><td>1021.3</td><td>-4.4</td><td>0.7</td><td>-6.3</td><td>3.6E-10</td><td>9.8E-10</td><td>BT20/MB468</td><td>CYTO</td></tr>
<tr><td>>hg38_21_+_8433172_8446622_RNA45SN1_45S_50ntFlanks@11207.11232.26|GGCGGCGACTCTGGACGCGAGCCGGG</td><td>>hg38_21_+_8433172_8446622_RNA45SN1_45S_50ntFlanks</td><td>11207</td><td>11232</td><td>26</td><td>GGCGGCGACTCTGGACGCGAGCCGGG</td><td>870.7</td><td>-4.2</td><td>0.7</td><td>-6.3</td><td>3.6E-10</td><td>9.8E-10</td><td>BT20/MB468</td><td>CYTO</td></tr>
<tr><td>>hg38_21_+_8433172_8446622_RNA45SN1_45S_50ntFlanks@7974.8013.40|ACGCGACCTCAGATCAGACGTGGCGACCCGCTGAATTTAA</td><td>>hg38_21_+_8433172_8446622_RNA45SN1_45S_50ntFlanks</td><td>7974</td><td>8013</td><td>40</td><td>ACGCGACCTCAGATCAGACGTGGCGACCCGCTGAATTTAA</td><td>2143.2</td><td>4.8</td><td>0.8</td><td>6.2</td><td>4.1E-10</td><td>9.8E-10</td><td>MB231/MB468</td><td>NUC</td></tr>
<tr><td>>MI0000255|hsa-mir-30d&WithFlank&8|-|134804870|134804951@12.35.24(+1A)&[MIMAT0000245&hsa-miR-30d-5p&offsets|0|+2(+1A);m-21&8|-|134804919|134804940&offsets|0|+2(+1A)]|TGTAAACATCCCCGACTGGAAGCTA</td><td>>MI0000255|hsa-mir-30d&WithFlank&8|-|134804870|134804951</td><td>12</td><td>35</td><td>24(+1A)</td><td>TGTAAACATCCCCGACTGGAAGCTA</td><td>213.6</td><td>6.1</td><td>1.0</td><td>6.2</td><td>4.1E-10</td><td>9.8E-10</td><td>MB231/MB468</td><td>NUC</td></tr>
<tr><td>>hg38_21_+_8433172_8446622_RNA45SN1_45S_50ntFlanks@3782.3800.19|CAGTGAAACTGCGAATGGC</td><td>>hg38_21_+_8433172_8446622_RNA45SN1_45S_50ntFlanks</td><td>3782</td><td>3800</td><td>19</td><td>CAGTGAAACTGCGAATGGC</td><td>1420.7</td><td>-6.7</td><td>1.1</td><td>-6.3</td><td>3.9E-10</td><td>9.8E-10</td><td>MB231/MB468</td><td>MP</td></tr>
<tr><td>>hg38_21_+_8433172_8446622_RNA45SN1_45S_50ntFlanks@4722.4753.32|TCATTAATCAAGAACGAAAGTCGGAGGTTCGA</td><td>>hg38_21_+_8433172_8446622_RNA45SN1_45S_50ntFlanks</td><td>4722</td><td>4753</td><td>32</td><td>TCATTAATCAAGAACGAAAGTCGGAGGTTCGA</td><td>127.1</td><td>-9.1</td><td>1.4</td><td>-6.3</td><td>3.1E-10</td><td>9.8E-10</td><td>BT20/MB468</td><td>MITO</td></tr>
<tr><td>>hg38_21_+_8433172_8446622_RNA45SN1_45S_50ntFlanks@3854.3881.28|ACTGTGGTAATTCTAGAGCTAATACATG</td><td>>hg38_21_+_8433172_8446622_RNA45SN1_45S_50ntFlanks</td><td>3854</td><td>3881</td><td>28</td><td>ACTGTGGTAATTCTAGAGCTAATACATG</td><td>1560.6</td><td>-5.8</td><td>0.9</td><td>-6.3</td><td>3.6E-10</td><td>9.8E-10</td><td>BT20/MB468</td><td>CYTO</td></tr>
<tr><td>>hg38_21_+_8433172_8446622_RNA45SN1_45S_50ntFlanks@12648.12675.28|TGGGATTATGACTGAACGCCTCTAAGTC</td><td>>hg38_21_+_8433172_8446622_RNA45SN1_45S_50ntFlanks</td><td>12648</td><td>12675</td><td>28</td><td>TGGGATTATGACTGAACGCCTCTAAGTC</td><td>82.1</td><td>-10.2</td><td>1.6</td><td>-6.3</td><td>3.9E-10</td><td>9.8E-10</td><td>MB231/MB468</td><td>MP</td></tr>
<tr><td>>hg38_21_+_8433172_8446622_RNA45SN1_45S_50ntFlanks@3873.3899.27|TAATACATGCCGACGGGCGCTGACCCC</td><td>>hg38_21_+_8433172_8446622_RNA45SN1_45S_50ntFlanks</td><td>3873</td><td>3899</td><td>27</td><td>TAATACATGCCGACGGGCGCTGACCCC</td><td>507.7</td><td>-4.1</td><td>0.6</td><td>-6.3</td><td>4.1E-10</td><td>9.9E-10</td><td>MB231/MB468</td><td>CYTO</td></tr>
<tr><td>>MI0000066|hsa-let-7e&WithFlank&19|+|51692780|51692870@14.36.23&[MIMAT0000066&hsa-let-7e-5p&offsets|0|+1;m-50&19|+|51692793|51692814&offsets|0|+1]|TGAGGTAGGAGGTTGTATAGTTG</td><td>>MI0000066|hsa-let-7e&WithFlank&19|+|51692780|51692870</td><td>14</td><td>36</td><td>23</td><td>TGAGGTAGGAGGTTGTATAGTTG</td><td>161.9</td><td>1.8</td><td>0.3</td><td>6.2</td><td>4.8E-10</td><td>9.9E-10</td><td>BT20/MB468</td><td>NUC</td></tr>
<tr><td>>hg38_21_+_8433172_8446622_RNA45SN1_45S_50ntFlanks@10481.10499.19|ATCGAAAGGGAGTCGGGTT</td><td>>hg38_21_+_8433172_8446622_RNA45SN1_45S_50ntFlanks</td><td>10481</td><td>10499</td><td>19</td><td>ATCGAAAGGGAGTCGGGTT</td><td>92.2</td><td>-10.4</td><td>1.7</td><td>-6.3</td><td>3.9E-10</td><td>9.9E-10</td><td>MB231/MB468</td><td>MP</td></tr>
<tr><td>>hg38_21_+_8433172_8446622_RNA45SN1_45S_50ntFlanks@11208.11252.45|GCGGCGACTCTGGACGCGAGCCGGGCCCTTCCCGTGGATCGCCCC</td><td>>hg38_21_+_8433172_8446622_RNA45SN1_45S_50ntFlanks</td><td>11208</td><td>11252</td><td>45</td><td>GCGGCGACTCTGGACGCGAGCCGGGCCCTTCCCGTGGATCGCCCC</td><td>894.3</td><td>-5.0</td><td>0.8</td><td>-6.2</td><td>4.1E-10</td><td>9.9E-10</td><td>MB231/MB468</td><td>CYTO</td></tr>
<tr><td>>hg38_21_+_8433172_8446622_RNA45SN1_45S_50ntFlanks@12892.12932.41|CCCTCGCCCGTCACGCACCGCACGTTCGTGGGGAACCTGGC</td><td>>hg38_21_+_8433172_8446622_RNA45SN1_45S_50ntFlanks</td><td>12892</td><td>12932</td><td>41</td><td>CCCTCGCCCGTCACGCACCGCACGTTCGTGGGGAACCTGGC</td><td>3649.7</td><td>-5.1</td><td>0.8</td><td>-6.3</td><td>3.1E-10</td><td>9.9E-10</td><td>BT20/MB468</td><td>MITO</td></tr>
<tr><td>>MI0000071|hsa-mir-17&WithFlank&13|+|91350599|91350694@20.41.22(+1A)&[MIMAT0000070&hsa-miR-17-5p&offsets|0|-1(+1A);m-88&13|+|91350618|91350640&offsets|0|-1(+1A)]|CAAAGTGCTTACAGTGCAGGTAA</td><td>>MI0000071|hsa-mir-17&WithFlank&13|+|91350599|91350694</td><td>20</td><td>41</td><td>22(+1A)</td><td>CAAAGTGCTTACAGTGCAGGTAA</td><td>830.2</td><td>5.6</td><td>0.9</td><td>6.3</td><td>3.6E-10</td><td>9.9E-10</td><td>BT20/MB468</td><td>CYTO</td></tr>
<tr><td>>hg38_21_+_8433172_8446622_RNA45SN1_45S_50ntFlanks@12072.12093.22|CCCGAGGGGCTCTCGCTTCTGG</td><td>>hg38_21_+_8433172_8446622_RNA45SN1_45S_50ntFlanks</td><td>12072</td><td>12093</td><td>22</td><td>CCCGAGGGGCTCTCGCTTCTGG</td><td>288.1</td><td>-5.0</td><td>0.8</td><td>-6.2</td><td>4.1E-10</td><td>9.9E-10</td><td>MB231/MB468</td><td>CYTO</td></tr>
<tr><td>>hg38_21_+_8433172_8446622_RNA45SN1_45S_50ntFlanks@10808.10826.19|CAATGTAGGTAAGGGAAGT</td><td>>hg38_21_+_8433172_8446622_RNA45SN1_45S_50ntFlanks</td><td>10808</td><td>10826</td><td>19</td><td>CAATGTAGGTAAGGGAAGT</td><td>1394.6</td><td>-6.7</td><td>1.1</td><td>-6.3</td><td>3.1E-10</td><td>9.9E-10</td><td>BT20/MB468</td><td>MITO</td></tr>
<tr><td>>hg38_21_+_8433172_8446622_RNA45SN1_45S_50ntFlanks@11209.11234.26|CGGCGACTCTGGACGCGAGCCGGGCC</td><td>>hg38_21_+_8433172_8446622_RNA45SN1_45S_50ntFlanks</td><td>11209</td><td>11234</td><td>26</td><td>CGGCGACTCTGGACGCGAGCCGGGCC</td><td>1047.3</td><td>-6.7</td><td>1.1</td><td>-6.2</td><td>4.2E-10</td><td>9.9E-10</td><td>MB231/MB468</td><td>NUC</td></tr>
<tr><td>>hg38_21_+_8433172_8446622_RNA45SN1_45S_50ntFlanks@12656.12674.19|TGACTGAACGCCTCTAAGT</td><td>>hg38_21_+_8433172_8446622_RNA45SN1_45S_50ntFlanks</td><td>12656</td><td>12674</td><td>19</td><td>TGACTGAACGCCTCTAAGT</td><td>986.0</td><td>-7.3</td><td>1.2</td><td>-6.2</td><td>4.8E-10</td><td>9.9E-10</td><td>BT20/MB468</td><td>NUC</td></tr>
<tr><td>>hg38_21_+_8433172_8446622_RNA45SN1_45S_50ntFlanks@3738.3757.20|TCTCAAAGATTAAGCCATGC</td><td>>hg38_21_+_8433172_8446622_RNA45SN1_45S_50ntFlanks</td><td>3738</td><td>3757</td><td>20</td><td>TCTCAAAGATTAAGCCATGC</td><td>71.3</td><td>-10.0</td><td>1.6</td><td>-6.3</td><td>4.0E-10</td><td>9.9E-10</td><td>MB231/MB468</td><td>MP</td></tr>
<tr><td>>hg38_21_+_8433172_8446622_RNA45SN1_45S_50ntFlanks@9569.9596.28|GTGCAAATCGGTCGTCCGACCTGGGTAT</td><td>>hg38_21_+_8433172_8446622_RNA45SN1_45S_50ntFlanks</td><td>9569</td><td>9596</td><td>28</td><td>GTGCAAATCGGTCGTCCGACCTGGGTAT</td><td>437.1</td><td>-8.6</td><td>1.4</td><td>-6.2</td><td>4.8E-10</td><td>9.9E-10</td><td>BT20/MB468</td><td>NUC</td></tr>
<tr><td>>hg38_21_+_8433172_8446622_RNA45SN1_45S_50ntFlanks@11873.11892.20|AAGAAGACCCTGTTGAGCTT</td><td>>hg38_21_+_8433172_8446622_RNA45SN1_45S_50ntFlanks</td><td>11873</td><td>11892</td><td>20</td><td>AAGAAGACCCTGTTGAGCTT</td><td>690.8</td><td>-8.1</td><td>1.3</td><td>-6.2</td><td>4.6E-10</td><td>1.0E-09</td><td>MB231/MB468</td><td>MITO</td></tr>
<tr><td>>hg38_21_+_8433172_8446622_RNA45SN1_45S_50ntFlanks@10844.10868.25|AACTTCGGGATAAGGATTGGCTCTA</td><td>>hg38_21_+_8433172_8446622_RNA45SN1_45S_50ntFlanks</td><td>10844</td><td>10868</td><td>25</td><td>AACTTCGGGATAAGGATTGGCTCTA</td><td>84.1</td><td>-10.3</td><td>1.6</td><td>-6.3</td><td>4.0E-10</td><td>1.0E-09</td><td>MB231/MB468</td><td>MP</td></tr>
<tr><td>>hg38_21_+_8433172_8446622_RNA45SN1_45S_50ntFlanks@9141.9165.25|CGGGGCGGACTGTCCCCAGTGCGCC</td><td>>hg38_21_+_8433172_8446622_RNA45SN1_45S_50ntFlanks</td><td>9141</td><td>9165</td><td>25</td><td>CGGGGCGGACTGTCCCCAGTGCGCC</td><td>358.0</td><td>-11.6</td><td>1.7</td><td>-6.6</td><td>3.7E-11</td><td>1.0E-09</td><td>BT20/MB468</td><td>MP</td></tr>
<tr><td>>hg38_21_+_8433172_8446622_RNA45SN1_45S_50ntFlanks@9572.9597.26|CAAATCGGTCGTCCGACCTGGGTATA</td><td>>hg38_21_+_8433172_8446622_RNA45SN1_45S_50ntFlanks</td><td>9572</td><td>9597</td><td>26</td><td>CAAATCGGTCGTCCGACCTGGGTATA</td><td>11779.2</td><td>-10.7</td><td>1.6</td><td>-6.6</td><td>3.7E-11</td><td>1.0E-09</td><td>BT20/MB468</td><td>MP</td></tr>
<tr><td>>hg38_21_+_8433172_8446622_RNA45SN1_45S_50ntFlanks@4449.4472.24|CCTCTCGGCGCCCCCTCGATGCTC</td><td>>hg38_21_+_8433172_8446622_RNA45SN1_45S_50ntFlanks</td><td>4449</td><td>4472</td><td>24</td><td>CCTCTCGGCGCCCCCTCGATGCTC</td><td>630.7</td><td>-5.7</td><td>0.9</td><td>-6.2</td><td>4.2E-10</td><td>1.0E-09</td><td>MB231/MB468</td><td>CYTO</td></tr>
<tr><td>>hg38_21_+_8433172_8446622_RNA45SN1_45S_50ntFlanks@10683.10706.24|CCGGCGGCGTCCGGTGAGCTCTCG</td><td>>hg38_21_+_8433172_8446622_RNA45SN1_45S_50ntFlanks</td><td>10683</td><td>10706</td><td>24</td><td>CCGGCGGCGTCCGGTGAGCTCTCG</td><td>7629.8</td><td>-5.5</td><td>0.9</td><td>-6.3</td><td>3.7E-10</td><td>1.0E-09</td><td>BT20/MB468</td><td>CYTO</td></tr>
<tr><td>>hg38_21_+_8433172_8446622_RNA45SN1_45S_50ntFlanks@8357.8383.27|AAAAGAACTTTGAAGAGAGAGTTCAAG</td><td>>hg38_21_+_8433172_8446622_RNA45SN1_45S_50ntFlanks</td><td>8357</td><td>8383</td><td>27</td><td>AAAAGAACTTTGAAGAGAGAGTTCAAG</td><td>79.1</td><td>-10.2</td><td>1.6</td><td>-6.3</td><td>4.0E-10</td><td>1.0E-09</td><td>MB231/MB468</td><td>MP</td></tr>
<tr><td>>hg38_21_+_8433172_8446622_RNA45SN1_45S_50ntFlanks@9534.9552.19|AGGAAACTCTGGTGGAGGT</td><td>>hg38_21_+_8433172_8446622_RNA45SN1_45S_50ntFlanks</td><td>9534</td><td>9552</td><td>19</td><td>AGGAAACTCTGGTGGAGGT</td><td>7945.8</td><td>-10.8</td><td>1.7</td><td>-6.2</td><td>4.9E-10</td><td>1.0E-09</td><td>BT20/MB468</td><td>NUC</td></tr>
<tr><td>>hg38_21_+_8433172_8446622_RNA45SN1_45S_50ntFlanks@12681.12708.28|CCCGCCCAGGCGGAACGATACGGCAGCG</td><td>>hg38_21_+_8433172_8446622_RNA45SN1_45S_50ntFlanks</td><td>12681</td><td>12708</td><td>28</td><td>CCCGCCCAGGCGGAACGATACGGCAGCG</td><td>87.7</td><td>-10.3</td><td>1.6</td><td>-6.3</td><td>4.0E-10</td><td>1.0E-09</td><td>MB231/MB468</td><td>MP</td></tr>
<tr><td>>hg38_21_+_8433172_8446622_RNA45SN1_45S_50ntFlanks@5490.5512.23|TGAGAAGACGGTCGAACTTGACT</td><td>>hg38_21_+_8433172_8446622_RNA45SN1_45S_50ntFlanks</td><td>5490</td><td>5512</td><td>23</td><td>TGAGAAGACGGTCGAACTTGACT</td><td>464.5</td><td>7.6</td><td>1.2</td><td>6.2</td><td>4.2E-10</td><td>1.0E-09</td><td>MB231/MB468</td><td>CYTO</td></tr>
<tr><td>>hg38_21_+_8433172_8446622_RNA45SN1_45S_50ntFlanks@3777.3801.25|CGGTACAGTGAAACTGCGAATGGCT</td><td>>hg38_21_+_8433172_8446622_RNA45SN1_45S_50ntFlanks</td><td>3777</td><td>3801</td><td>25</td><td>CGGTACAGTGAAACTGCGAATGGCT</td><td>671.4</td><td>-4.6</td><td>0.7</td><td>-6.3</td><td>3.1E-10</td><td>1.0E-09</td><td>BT20/MB468</td><td>MITO</td></tr>
<tr><td>>hg38_21_+_8433172_8446622_RNA45SN1_45S_50ntFlanks@4963.4994.32|AACCTCACCCGGCCCGGACACGGACAGGATTG</td><td>>hg38_21_+_8433172_8446622_RNA45SN1_45S_50ntFlanks</td><td>4963</td><td>4994</td><td>32</td><td>AACCTCACCCGGCCCGGACACGGACAGGATTG</td><td>96.2</td><td>-10.5</td><td>1.7</td><td>-6.3</td><td>4.0E-10</td><td>1.0E-09</td><td>MB231/MB468</td><td>MP</td></tr>
<tr><td>>hg38_21_+_8433172_8446622_RNA45SN1_45S_50ntFlanks@12909.12933.25|CCGCACGTTCGTGGGGAACCTGGCG</td><td>>hg38_21_+_8433172_8446622_RNA45SN1_45S_50ntFlanks</td><td>12909</td><td>12933</td><td>25</td><td>CCGCACGTTCGTGGGGAACCTGGCG</td><td>479.2</td><td>-8.2</td><td>1.3</td><td>-6.3</td><td>4.0E-10</td><td>1.0E-09</td><td>MB231/MB468</td><td>MP</td></tr>
<tr><td>>hg38_21_+_8433172_8446622_RNA45SN1_45S_50ntFlanks@13000.13028.29|TGCGATCTATTGAAAGTCAGCCCTCGACA</td><td>>hg38_21_+_8433172_8446622_RNA45SN1_45S_50ntFlanks</td><td>13000</td><td>13028</td><td>29</td><td>TGCGATCTATTGAAAGTCAGCCCTCGACA</td><td>360.6</td><td>-8.5</td><td>1.4</td><td>-6.2</td><td>4.9E-10</td><td>1.0E-09</td><td>BT20/MB468</td><td>NUC</td></tr>
<tr><td>>hg38_21_+_8433172_8446622_RNA45SN1_45S_50ntFlanks@12151.12182.32|GTGGGGAGTTTGACTGGGGCGGTACACCTGTC</td><td>>hg38_21_+_8433172_8446622_RNA45SN1_45S_50ntFlanks</td><td>12151</td><td>12182</td><td>32</td><td>GTGGGGAGTTTGACTGGGGCGGTACACCTGTC</td><td>250.4</td><td>-4.8</td><td>0.8</td><td>-6.2</td><td>4.2E-10</td><td>1.0E-09</td><td>MB231/MB468</td><td>CYTO</td></tr>
<tr><td>>MI0000088|hsa-mir-30a&WithFlank&6|-|71403545|71403627@54.73.20&[MIMAT0000088&hsa-miR-30a-3p&offsets|+1|-1;m-38&6|-|71403554|71403575&offsets|+1|-1]|TTTCAGTCGGATGTTTGCAG</td><td>>MI0000088|hsa-mir-30a&WithFlank&6|-|71403545|71403627</td><td>54</td><td>73</td><td>20</td><td>TTTCAGTCGGATGTTTGCAG</td><td>143.4</td><td>10.4</td><td>1.7</td><td>6.2</td><td>4.7E-10</td><td>1.0E-09</td><td>MB231/MB468</td><td>MITO</td></tr>
<tr><td>>hg38_21_+_8433172_8446622_RNA45SN1_45S_50ntFlanks@7975.7992.18|CGCGACCTCAGATCAGAC</td><td>>hg38_21_+_8433172_8446622_RNA45SN1_45S_50ntFlanks</td><td>7975</td><td>7992</td><td>18</td><td>CGCGACCTCAGATCAGAC</td><td>186601.5</td><td>5.9</td><td>0.9</td><td>6.3</td><td>3.2E-10</td><td>1.0E-09</td><td>BT20/MB468</td><td>MITO</td></tr>
<tr><td>>hg38_21_+_8433172_8446622_RNA45SN1_45S_50ntFlanks@12433.12455.23|CTATCATTGTGAAGCAGAATTCA</td><td>>hg38_21_+_8433172_8446622_RNA45SN1_45S_50ntFlanks</td><td>12433</td><td>12455</td><td>23</td><td>CTATCATTGTGAAGCAGAATTCA</td><td>132.9</td><td>-9.4</td><td>1.5</td><td>-6.3</td><td>3.7E-10</td><td>1.0E-09</td><td>BT20/MB468</td><td>CYTO</td></tr>
<tr><td>>hg38_21_+_8433172_8446622_RNA45SN1_45S_50ntFlanks@9567.9594.28|ACGTGCAAATCGGTCGTCCGACCTGGGT</td><td>>hg38_21_+_8433172_8446622_RNA45SN1_45S_50ntFlanks</td><td>9567</td><td>9594</td><td>28</td><td>ACGTGCAAATCGGTCGTCCGACCTGGGT</td><td>638.5</td><td>-5.6</td><td>0.9</td><td>-6.2</td><td>4.2E-10</td><td>1.0E-09</td><td>MB231/MB468</td><td>CYTO</td></tr>
<tr><td>>MI0000734|hsa-mir-106b&WithFlank&7|-|100093987|100094080@58.78.21&[MIMAT0004672&hsa-miR-106b-3p&offsets|0|-1;m-87&7|-|100094003|100094023&offsets|0|0]|CCGCACTGTGGGTACTTGCTG</td><td>>MI0000734|hsa-mir-106b&WithFlank&7|-|100093987|100094080</td><td>58</td><td>78</td><td>21</td><td>CCGCACTGTGGGTACTTGCTG</td><td>700.2</td><td>5.1</td><td>0.8</td><td>6.2</td><td>4.7E-10</td><td>1.0E-09</td><td>MB231/MB468</td><td>MITO</td></tr>
<tr><td>>hg38_21_+_8433172_8446622_RNA45SN1_45S_50ntFlanks@7975.8011.37|CGCGACCTCAGATCAGACGTGGCGACCCGCTGAATTT</td><td>>hg38_21_+_8433172_8446622_RNA45SN1_45S_50ntFlanks</td><td>7975</td><td>8011</td><td>37</td><td>CGCGACCTCAGATCAGACGTGGCGACCCGCTGAATTT</td><td>125783.4</td><td>4.2</td><td>0.7</td><td>6.2</td><td>4.3E-10</td><td>1.0E-09</td><td>MB231/MB468</td><td>NUC</td></tr>
</table>

| | | | | | | | | | | | | | |
|---|---|---|---|---|---|---|---|---|---|---|---|---|---|
| >MI0005756\|hsa-mir-934&WithFlank&X\|+\|136550872\|136550966@21.41.21(+1A)&[MIMAT0004977&hsa-miR-934&offsets\|0\|-1(+1A);m-943&X\|+\|136550892\|136550913&offsets\|0\|-1(+1A)]\|TGTCTACTACTGGAGACACTGA | >MI0005756\|hsa-mir-934&WithFlank&X\|+\|136550872\|136550966 | 21 | 41 | 21(+1A) | TGTCTACTACTGGAGACACTGA | 196.4 | -11.5 | 1.8 | -6.3 | 4.1E-10 | 1.0E-09 | MB231/MB468 | MP |
| >hg38_21_+_8433172_8446622_RNA45SN1_45S_50ntFlanks@12711.12743.33\|GCGGAGCCTCGGTTGGCCTCGGATAGCCGGTCC | >hg38_21_+_8433172_8446622_RNA45SN1_45S_50ntFlanks | 12711 | 12743 | 33 | GCGGAGCCTCGGTTGGCCTCGGATAGCCGGTCC | 61.6 | -9.8 | 1.6 | -6.3 | 4.1E-10 | 1.0E-09 | MB231/MB468 | MP |
| >hg38_21_+_8433172_8446622_RNA45SN1_45S_50ntFlanks@6751.6770.20\|CGAACGCACTTGCGGCCCCG | >hg38_21_+_8433172_8446622_RNA45SN1_45S_50ntFlanks | 6751 | 6770 | 20 | CGAACGCACTTGCGGCCCCG | 114.7 | -8.9 | 1.4 | -6.3 | 3.2E-10 | 1.0E-09 | BT20/MB468 | MITO |
| >hg38_21_+_8433172_8446622_RNA45SN1_45S_50ntFlanks@12274.12291.18\|CAGTACGAATACAGACCG | >hg38_21_+_8433172_8446622_RNA45SN1_45S_50ntFlanks | 12274 | 12291 | 18 | CAGTACGAATACAGACCG | 64.2 | -9.9 | 1.6 | -6.3 | 4.1E-10 | 1.0E-09 | MB231/MB468 | MP |
| >MI0003643\|hsa-mir-629&WithFlank&15\|-\|70079366\|70079474@28.47.20(+1A)&[MIMAT0004810&hsa-miR-629-5p&offsets\|0\|-1(+1A);m-150&15\|-\|70079427\|70079447&offsets\|0\|-1(+1A)]\|TGGGTTTACGTTGGGAGAACA | >MI0003643\|hsa-mir-629&WithFlank&15\|-\|70079366\|70079474 | 28 | 47 | 20(+1A) | TGGGTTTACGTTGGGAGAACA | 368.6 | 12.3 | 1.9 | 6.6 | 3.8E-11 | 1.0E-09 | BT20/MB468 | MP |
| >hg38_21_+_8433172_8446622_RNA45SN1_45S_50ntFlanks@12405.12438.34\|CTTTTTGATCCTTCGATGTCGGCTCTTCCTATCA | >hg38_21_+_8433172_8446622_RNA45SN1_45S_50ntFlanks | 12405 | 12438 | 34 | CTTTTTGATCCTTCGATGTCGGCTCTTCCTATCA | 3404.5 | -8.5 | 1.4 | -6.2 | 5.0E-10 | 1.0E-09 | BT20/MB468 | NUC |
| >MI0005561\|hsa-mir-877&WithFlank&6\|+\|30584326\|30584423@7.29.23(+1U)&[MIMAT0004949&hsa-miR-877-5p&offsets\|0\|+3(+1U);m-382&6\|+\|30584332\|30584354&offsets\|0\|0(+1U)]\|GTAGAGGAGATGGCGCAGGGGACT | >MI0005561\|hsa-mir-877&WithFlank&6\|+\|30584326\|30584423 | 7 | 29 | 23(+1U) | GTAGAGGAGATGGCGCAGGGGACT | 405.4 | 8.4 | 1.3 | 6.3 | 3.8E-10 | 1.0E-09 | BT20/MB468 | CYTO |
| >hg38_21_+_8433172_8446622_RNA45SN1_45S_50ntFlanks@9539.9571.33\|ACTCTGGTGGAGGTCCGTAGCGGTCCTGACGTG | >hg38_21_+_8433172_8446622_RNA45SN1_45S_50ntFlanks | 9539 | 9571 | 33 | ACTCTGGTGGAGGTCCGTAGCGGTCCTGACGTG | 7377.1 | -8.9 | 1.4 | -6.2 | 5.0E-10 | 1.0E-09 | BT20/MB468 | NUC |
| >hg38_21_+_8433172_8446622_RNA45SN1_45S_50ntFlanks@3985.4025.41\|CGGCTTTGGTGACTCTAGATAACCTCGGGCCGATCGCACGC | >hg38_21_+_8433172_8446622_RNA45SN1_45S_50ntFlanks | 3985 | 4025 | 41 | CGGCTTTGGTGACTCTAGATAACCTCGGGCCGATCGCACGC | 159.0 | -11.2 | 1.8 | -6.3 | 4.1E-10 | 1.0E-09 | MB231/MB468 | MP |
| >hg38_21_+_8433172_8446622_RNA45SN1_45S_50ntFlanks@10778.10818.41\|CCAAGGTGAACAGCCTCTGGCATGTTGGAACAATGTAGGTA | >hg38_21_+_8433172_8446622_RNA45SN1_45S_50ntFlanks | 10778 | 10818 | 41 | CCAAGGTGAACAGCCTCTGGCATGTTGGAACAATGTAGGTA | 138.3 | -11.0 | 1.8 | -6.3 | 4.1E-10 | 1.0E-09 | MB231/MB468 | MP |
| >hg38_21_+_8433172_8446622_RNA45SN1_45S_50ntFlanks@11702.11728.27\|CAATGAAGCGCGGGTAAACGGCGGGAG | >hg38_21_+_8433172_8446622_RNA45SN1_45S_50ntFlanks | 11702 | 11728 | 27 | CAATGAAGCGCGGGTAAACGGCGGGAG | 272.6 | -7.5 | 1.2 | -6.2 | 4.3E-10 | 1.0E-09 | MB231/MB468 | NUC |
| >hg38_21_+_8433172_8446622_RNA45SN1_45S_50ntFlanks@11729.11761.33\|TAACTATGACTCTCTTAAGGTAGCCAAATGCCT | >hg38_21_+_8433172_8446622_RNA45SN1_45S_50ntFlanks | 11729 | 11761 | 33 | TAACTATGACTCTCTTAAGGTAGCCAAATGCCT | 122.7 | -9.0 | 1.4 | -6.3 | 3.2E-10 | 1.0E-09 | BT20/MB468 | MITO |
| >MI0000254\|hsa-mir-30c-2&WithFlank&6\|-\|71376954\|71377037@53.75.23&[MIMAT0004550&hsa-miR-30c-2-3p&offsets\|0\|+1;m-206&6\|-\|71376964\|71376985&offsets\|0\|+1]\|CTGGGAGAAGGCTGTTTACTCTT | >MI0000254\|hsa-mir-30c-2&WithFlank&6\|-\|71376954\|71377037 | 53 | 75 | 23 | CTGGGAGAAGGCTGTTTACTCTT | 288.9 | 6.0 | 1.0 | 6.2 | 4.3E-10 | 1.0E-09 | MB231/MB468 | CYTO |
| >hg38_21_+_8433172_8446622_RNA45SN1_45S_50ntFlanks@12419.12456.38\|GATGTCGGCTCTTCCTATCATTGTGAAGCAGAATTCAC | >hg38_21_+_8433172_8446622_RNA45SN1_45S_50ntFlanks | 12419 | 12456 | 38 | GATGTCGGCTCTTCCTATCATTGTGAAGCAGAATTCAC | 1235.6 | 3.9 | 0.6 | 6.2 | 4.3E-10 | 1.0E-09 | MB231/MB468 | CYTO |
| >hg38_21_+_8433172_8446622_RNA45SN1_45S_50ntFlanks@12457.12477.21\|CAAGCGTTGGATTGTTCACCC | >hg38_21_+_8433172_8446622_RNA45SN1_45S_50ntFlanks | 12457 | 12477 | 21 | CAAGCGTTGGATTGTTCACCC | 1324.8 | -7.3 | 1.2 | -6.3 | 4.1E-10 | 1.0E-09 | MB231/MB468 | MP |
| >hg38_21_+_8433172_8446622_RNA45SN1_45S_50ntFlanks@11540.11560.21\|CGCGGCGCCTCGCCTCGGCCG | >hg38_21_+_8433172_8446622_RNA45SN1_45S_50ntFlanks | 11540 | 11560 | 21 | CGCGGCGCCTCGCCTCGGCCG | 238.7 | -11.0 | 1.7 | -6.6 | 3.9E-11 | 1.0E-09 | BT20/MB468 | MP |
| >hg38_21_+_8433172_8446622_RNA45SN1_45S_50ntFlanks@4005.4024.20\|AACCTCGGGCCGATCGCACG | >hg38_21_+_8433172_8446622_RNA45SN1_45S_50ntFlanks | 4005 | 4024 | 20 | AACCTCGGGCCGATCGCACG | 498.7 | -3.9 | 0.6 | -6.3 | 3.2E-10 | 1.0E-09 | BT20/MB468 | MITO |
| >hg38_21_+_8433172_8446622_RNA45SN1_45S_50ntFlanks@11255.11282.28\|CTGCGGCGGGCGTCGCGGCCGCCCCCGG | >hg38_21_+_8433172_8446622_RNA45SN1_45S_50ntFlanks | 11255 | 11282 | 28 | CTGCGGCGGGCGTCGCGGCCGCCCCCGG | 99.8 | -10.5 | 1.7 | -6.2 | 4.1E-10 | 1.0E-09 | MB231/MB468 | MP |
| >hg38_21_+_8433172_8446622_RNA45SN1_45S_50ntFlanks@4738.4755.18\|AAAGTCGGAGGTTCGAAG | >hg38_21_+_8433172_8446622_RNA45SN1_45S_50ntFlanks | 4738 | 4755 | 18 | AAAGTCGGAGGTTCGAAG | 348.5 | -3.8 | 0.6 | -6.2 | 4.1E-10 | 1.0E-09 | MB231/MB468 | MP |
| >hg38_21_+_8433172_8446622_RNA45SN1_45S_50ntFlanks@11707.11740.34\|AAGCGCGGGTAAACGGCGGGAGTAACTATGACTC | >hg38_21_+_8433172_8446622_RNA45SN1_45S_50ntFlanks | 11707 | 11740 | 34 | AAGCGCGGGTAAACGGCGGGAGTAACTATGACTC | 749.0 | -9.3 | 1.5 | -6.2 | 5.0E-10 | 1.0E-09 | BT20/MB468 | NUC |
| >hg38_1_-_228634819_228635039_RNA5S12_5S_50ntFlanks@104.131.28\|AAGCAGGGTCGGGCCTGGTTAGTACTTG | >hg38_1_-_228634819_228635039_RNA5S12_5S_50ntFlanks | 104 | 131 | 28 | AAGCAGGGTCGGGCCTGGTTAGTACTTG | 531.7 | -6.8 | 1.1 | -6.2 | 5.0E-10 | 1.0E-09 | BT20/MB468 | NUC |
| >hg38_21_+_8433172_8446622_RNA45SN1_45S_50ntFlanks@10843.10866.24\|TAACTTCGGGATAAGGATTGGCTC | >hg38_21_+_8433172_8446622_RNA45SN1_45S_50ntFlanks | 10843 | 10866 | 24 | TAACTTCGGGATAAGGATTGGCTC | 415.3 | -8.7 | 1.4 | -6.2 | 5.0E-10 | 1.0E-09 | BT20/MB468 | NUC |
| >hg38_21_+_8433172_8446622_RNA45SN1_45S_50ntFlanks@8155.8181.27\|CGCTCGTGGGGGGCCCAAGTCCTTCTG | >hg38_21_+_8433172_8446622_RNA45SN1_45S_50ntFlanks | 8155 | 8181 | 27 | CGCTCGTGGGGGGCCCAAGTCCTTCTG | 3045.7 | -4.8 | 0.8 | -6.3 | 3.2E-10 | 1.0E-09 | BT20/MB468 | MITO |
| >MI0000061\|hsa-let-7a-2&WithFlank&11\|-\|122146516\|122146599@11.33.23;MI0000062\|hsa-let-7a-3&WithFlank&22\|+\|46112743\|46112828@10.32.23;MI0000060\|hsa-let-7a-1&WithFlank&9\|+\|94175951\|94176042@12.34.23&[MIMAT0000062&hsa-let-7a-5p&offsets\|0\|+1;m-6&11\|-\|122146568\|122146589&offsets\|0\|+1];[MIMAT0000062_1&hsa-let-7a-5p&offsets\|0\|+1;m-5&22\|+\|46112752\|46112773&offsets\|0\|+1];[MIMAT0000062_2&hsa-let-7a-5p&offsets\|0\|+1;m-4&9\|+\|94175962\|94175983&offsets\|0\|+1]\|TGAGGTAGTAGGTTGTATAGTTT | >MI0000061\|hsa-let-7a-2&WithFlank&11\|-\|122146516\|122146599 | 11 | 33 | 23 | TGAGGTAGTAGGTTGTATAGTTT | 4324.1 | 5.0 | 0.8 | 6.2 | 4.1E-10 | 1.0E-09 | MB231/MB468 | MP |
| >hg38_21_+_8433172_8446622_RNA45SN1_45S_50ntFlanks@5471.5514.44\|CACGGCCCTGGCGGAGCGCTGAGAAGACGGTCGAACTTGACTAT | >hg38_21_+_8433172_8446622_RNA45SN1_45S_50ntFlanks | 5471 | 5514 | 44 | CACGGCCCTGGCGGAGCGCTGAGAAGACGGTCGAACTTGACTAT | 1627.8 | -5.8 | 0.9 | -6.2 | 4.4E-10 | 1.0E-09 | MB231/MB468 | NUC |
| >hg38_21_+_8433172_8446622_RNA45SN1_45S_50ntFlanks@10681.10706.26\|TTCCGGCGGCGTCCGGTGAGCTCTCG | >hg38_21_+_8433172_8446622_RNA45SN1_45S_50ntFlanks | 10681 | 10706 | 26 | TTCCGGCGGCGTCCGGTGAGCTCTCG | 4073.2 | -7.7 | 1.2 | -6.2 | 4.4E-10 | 1.0E-09 | MB231/MB468 | NUC |
| >hg38_21_+_8433172_8446622_RNA45SN1_45S_50ntFlanks@9571.9597.27\|GCAAATCGGTCGTCCGACCTGGGTATA | >hg38_21_+_8433172_8446622_RNA45SN1_45S_50ntFlanks | 9571 | 9597 | 27 | GCAAATCGGTCGTCCGACCTGGGTATA | 800.5 | -6.7 | 1.1 | -6.2 | 4.8E-10 | 1.0E-09 | MB231/MB468 | MITO |
| >hg38_21_+_8433172_8446622_RNA45SN1_45S_50ntFlanks@11633.11660.28\|AAGGCCCGCGGCGGGTGTTGACGCGATG | >hg38_21_+_8433172_8446622_RNA45SN1_45S_50ntFlanks | 11633 | 11660 | 28 | AAGGCCCGCGGCGGGTGTTGACGCGATG | 238.5 | -11.8 | 1.9 | -6.2 | 4.2E-10 | 1.0E-09 | MB231/MB468 | MP |
| >hg38_21_+_8433172_8446622_RNA45SN1_45S_50ntFlanks@12149.12181.33\|AGGTGGGGAGTTTGACTGGGGCGGTACACCTGT | >hg38_21_+_8433172_8446622_RNA45SN1_45S_50ntFlanks | 12149 | 12181 | 33 | AGGTGGGGAGTTTGACTGGGGCGGTACACCTGT | 74.1 | -10.1 | 1.6 | -6.2 | 4.2E-10 | 1.0E-09 | MB231/MB468 | MP |
| >hg38_21_+_8433172_8446622_RNA45SN1_45S_50ntFlanks@10351.10391.41\|CGAGAACTTTGAAGGCCGAAGTGGAGAAGGGTTCCATGTGA | >hg38_21_+_8433172_8446622_RNA45SN1_45S_50ntFlanks | 10351 | 10391 | 41 | CGAGAACTTTGAAGGCCGAAGTGGAGAAGGGTTCCATGTGA | 731.2 | -4.2 | 0.7 | -6.3 | 3.3E-10 | 1.0E-09 | BT20/MB468 | MITO |
| >hg38_21_+_8433172_8446622_RNA45SN1_45S_50ntFlanks@4683.4707.25\|CAAGACGGACCAGAGCGAAAGCATT | >hg38_21_+_8433172_8446622_RNA45SN1_45S_50ntFlanks | 4683 | 4707 | 25 | CAAGACGGACCAGAGCGAAAGCATT | 1082.6 | -7.9 | 1.3 | -6.2 | 5.1E-10 | 1.0E-09 | BT20/MB468 | NUC |
| >hg38_21_+_8433172_8446622_RNA45SN1_45S_50ntFlanks@10357.10383.27\|CTTTGAAGGCCGAAGTGGAGAAGGGTT | >hg38_21_+_8433172_8446622_RNA45SN1_45S_50ntFlanks | 10357 | 10383 | 27 | CTTTGAAGGCCGAAGTGGAGAAGGGTT | 2146.2 | -9.9 | 1.6 | -6.2 | 5.1E-10 | 1.0E-09 | BT20/MB468 | NUC |
| >hg38_21_+_8433172_8446622_RNA45SN1_45S_50ntFlanks@12289.12323.35\|CCGTGAAAGCGGGGCCTCACGATCCTTCTGACCTT | >hg38_21_+_8433172_8446622_RNA45SN1_45S_50ntFlanks | 12289 | 12323 | 35 | CCGTGAAAGCGGGGCCTCACGATCCTTCTGACCTT | 194.3 | -7.9 | 1.3 | -6.2 | 4.4E-10 | 1.0E-09 | MB231/MB468 | NUC |
| >hg38_21_+_8433172_8446622_RNA45SN1_45S_50ntFlanks@12407.12432.26\|TTTTGATCCTTCGATGTCGGCTCTTC | >hg38_21_+_8433172_8446622_RNA45SN1_45S_50ntFlanks | 12407 | 12432 | 26 | TTTTGATCCTTCGATGTCGGCTCTTC | 538.7 | -6.7 | 1.1 | -6.2 | 4.2E-10 | 1.0E-09 | MB231/MB468 | MP |
| >hg38_21_+_8433172_8446622_RNA45SN1_45S_50ntFlanks@4103.4128.26\|CCACGGGTGACGGGGAATCAGGGTTC | >hg38_21_+_8433172_8446622_RNA45SN1_45S_50ntFlanks | 4103 | 4128 | 26 | CCACGGGTGACGGGGAATCAGGGTTC | 68.8 | -10.0 | 1.6 | -6.2 | 4.2E-10 | 1.0E-09 | MB231/MB468 | MP |
| >hg38_21_+_8433172_8446622_RNA45SN1_45S_50ntFlanks@8357.8388.32\|AAAAGAACTTTGAAGAGAGAGTTCAAGAGGGC | >hg38_21_+_8433172_8446622_RNA45SN1_45S_50ntFlanks | 8357 | 8388 | 32 | AAAAGAACTTTGAAGAGAGAGTTCAAGAGGGC | 323.5 | -7.8 | 1.3 | -6.2 | 4.2E-10 | 1.0E-09 | MB231/MB468 | MP |
| >MI0000077\|hsa-mir-21&WithFlank&17\|+\|59841260\|59841343@14.32.19(+1C)&[MIMAT0000076&hsa-miR-21-5p&offsets\|0\|-3(+1C);m-1&17\|+\|59841273\|59841294&offsets\|0\|-3(+1C)]\|TAGCTTATCAGACTGATGTC | >MI0000077\|hsa-mir-21&WithFlank&17\|+\|59841260\|59841343 | 14 | 32 | 19(+1C) | TAGCTTATCAGACTGATGTC | 367.6 | 9.9 | 1.6 | 6.3 | 3.8E-10 | 1.0E-09 | BT20/MB468 | CYTO |
| >am_trna162_MetCAT_6_-_26758550_26758622@31.51.21,trna164_MetCAT_6_-_26735574_26735646@31.51.21,trna20_MetCAT_16_+_71460396_71460468@31.51.21,trna21_MetCAT_6_+_26701712_26701784@31.51.21,trna27_MetCAT_6_+_26766444_26766516@31.51.21,trna75_MetCAT_6_+_28912352_28912424@31.51.21,trna97_MetCAT_6_-_28921042_28921114@31.51.21\|TCTCATAATCTGAAGGTCCTG | >am_trna162_MetCAT_6_-_26758550_26758622 | 31 | 51 | 21 | TCTCATAATCTGAAGGTCCTG | 171.1 | -10.5 | 1.6 | -6.6 | 3.9E-11 | 1.0E-09 | BT20/MB468 | MP |
| >hg38_21_+_8433172_8446622_RNA45SN1_45S_50ntFlanks@5472.5500.29\|ACGGCCCTGGCGGAGCGCTGAGAAGACGG | >hg38_21_+_8433172_8446622_RNA45SN1_45S_50ntFlanks | 5472 | 5500 | 29 | ACGGCCCTGGCGGAGCGCTGAGAAGACGG | 2354.3 | 6.4 | 1.0 | 6.3 | 3.8E-10 | 1.0E-09 | BT20/MB468 | CYTO |
| >hg38_21_+_8433172_8446622_RNA45SN1_45S_50ntFlanks@7973.8013.41\|GACGCGACCTCAGATCAGACGTGGCGACCCGCTGAATTTAA | >hg38_21_+_8433172_8446622_RNA45SN1_45S_50ntFlanks | 7973 | 8013 | 41 | GACGCGACCTCAGATCAGACGTGGCGACCCGCTGAATTTAA | 585.1 | 5.1 | 0.8 | 6.2 | 4.4E-10 | 1.0E-09 | MB231/MB468 | NUC |
| >hg38_21_+_8433172_8446622_RNA45SN1_45S_50ntFlanks@5334.5364.31\|ATGAACGAGGAATTCCCAGTAAGTGCGGGTC | >hg38_21_+_8433172_8446622_RNA45SN1_45S_50ntFlanks | 5334 | 5364 | 31 | ATGAACGAGGAATTCCCAGTAAGTGCGGGTC | 130.3 | -10.9 | 1.7 | -6.2 | 4.2E-10 | 1.0E-09 | MB231/MB468 | MP |
| >hg38_21_+_8433172_8446622_RNA45SN1_45S_50ntFlanks@12634.12661.28\|GAAGCTACCATCTGTGGGATTATGACTG | >hg38_21_+_8433172_8446622_RNA45SN1_45S_50ntFlanks | 12634 | 12661 | 28 | GAAGCTACCATCTGTGGGATTATGACTG | 1340.0 | -9.1 | 1.5 | -6.2 | 4.2E-10 | 1.0E-09 | MB231/MB468 | MP |
| >hg38_21_+_8433172_8446622_RNA45SN1_45S_50ntFlanks@12415.12435.21\|CTTCGATGTCGGCTCTTCCTA | >hg38_21_+_8433172_8446622_RNA45SN1_45S_50ntFlanks | 12415 | 12435 | 21 | CTTCGATGTCGGCTCTTCCTA | 5822.1 | -5.6 | 0.9 | -6.2 | 4.4E-10 | 1.0E-09 | MB231/MB468 | NUC |
| >hg38_21_+_8433172_8446622_RNA45SN1_45S_50ntFlanks@12419.12444.26\|GATGTCGGCTCTTCCTATCATTGTGA | >hg38_21_+_8433172_8446622_RNA45SN1_45S_50ntFlanks | 12419 | 12444 | 26 | GATGTCGGCTCTTCCTATCATTGTGA | 395.6 | 3.8 | 0.6 | 6.3 | 3.3E-10 | 1.0E-09 | BT20/MB468 | MITO |
| >hg38_21_+_8433172_8446622_RNA45SN1_45S_50ntFlanks@4002.4026.25\|GATAACCTCGGGCCGATCGCACGCC | >hg38_21_+_8433172_8446622_RNA45SN1_45S_50ntFlanks | 4002 | 4026 | 25 | GATAACCTCGGGCCGATCGCACGCC | 362.9 | -6.7 | 1.1 | -6.2 | 4.4E-10 | 1.0E-09 | MB231/MB468 | CYTO |
| >hg38_21_+_8433172_8446622_RNA45SN1_45S_50ntFlanks@12413.12434.22\|TCCTTCGATGTCGGCTCTTCCT | >hg38_21_+_8433172_8446622_RNA45SN1_45S_50ntFlanks | 12413 | 12434 | 22 | TCCTTCGATGTCGGCTCTTCCT | 64.6 | -9.9 | 1.6 | -6.2 | 4.2E-10 | 1.0E-09 | MB231/MB468 | MP |

<table>
<tr><td>>MI0000264|hsa-mir-7-2&WithFlank&15|+|88611819|88611940@38.60.23(+1C);MI0000265|hsa-mir-7-3&WithFlank&19|+|4770664|4770785@37.59.23(+1C);MI0000263|hsa-mir-7-1&WithFlank&9|-|83969742|83969863@30.52.23(+1C)&[MIMAT0000252&hsa-miR-7-5p&offsets|0|-1(+1C);m-400&15|+|88611856|88611879&offsets|0|-1(+1C)];[MIMAT0000252_1&hsa-miR-7-5p&offsets|0|-1(+1C);m-402&19|+|4770700|4770723&offsets|0|-1(+1C)];[MIMAT0000252_2&hsa-miR-7-5p&offsets|0|-1(+1C);m-401&9|-|83969811|83969834&offsets|0|-1(+1C)]|TGGAAGACTAGTGATTTTGTTGTC</td><td>>MI0000264|hsa-mir-7-2&WithFlank&15|+|88611819|88611940</td><td>38</td><td>60</td><td>23(+1C)</td><td>TGGAAGACTAGTGATTTTGTTGTC</td><td>3127.2</td><td>3.6</td><td>0.6</td><td>6.2</td><td>5.1E-10</td><td>1.0E-09</td><td>BT20/MB468</td><td>NUC</td></tr>
<tr><td>>trna32_LysCTT_16_-_3207406_3207478@3.33.31|CCGGCTAGCTCAGTCGGTAGAGCATGAGACC</td><td>>trna32_LysCTT_16_-_3207406_3207478</td><td>3</td><td>33</td><td>31</td><td>CCGGCTAGCTCAGTCGGTAGAGCATGAGACC</td><td>177.0</td><td>10.7</td><td>1.7</td><td>6.2</td><td>4.9E-10</td><td>1.0E-09</td><td>MB231/MB468</td><td>MITO</td></tr>
<tr><td>>hg38_21_+_8433172_8446622_RNA45SN1_45S_50ntFlanks@13002.13019.18|CGATCTATTGAAAGTCAG</td><td>>hg38_21_+_8433172_8446622_RNA45SN1_45S_50ntFlanks</td><td>13002</td><td>13019</td><td>18</td><td>CGATCTATTGAAAGTCAG</td><td>102.1</td><td>-10.5</td><td>1.7</td><td>-6.2</td><td>4.2E-10</td><td>1.1E-09</td><td>MB231/MB468</td><td>MP</td></tr>
<tr><td>>trnaMT_LeuTAG_MT_+_12266_12336@43.74.32|AAATTTTGGTGCAACTCCAAATAAAAGTACCA</td><td>>trnaMT_LeuTAG_MT_+_12266_12336</td><td>43</td><td>74</td><td>32</td><td>AAATTTTGGTGCAACTCCAAATAAAAGTACCA</td><td>560.5</td><td>-5.3</td><td>0.9</td><td>-6.2</td><td>4.9E-10</td><td>1.1E-09</td><td>MB231/MB468</td><td>MITO</td></tr>
<tr><td>>hg38_21_+_8433172_8446622_RNA45SN1_45S_50ntFlanks@11610.11640.31|TTTAATTAAAACAAAGCATCGCGAAGGCCCG</td><td>>hg38_21_+_8433172_8446622_RNA45SN1_45S_50ntFlanks</td><td>11610</td><td>11640</td><td>31</td><td>TTTAATTAAAACAAAGCATCGCGAAGGCCCG</td><td>62.0</td><td>-9.8</td><td>1.6</td><td>-6.2</td><td>4.2E-10</td><td>1.1E-09</td><td>MB231/MB468</td><td>MP</td></tr>
<tr><td>>MI0000469|hsa-mir-125a&WithFlank&19|+|51693248|51693345@21.43.23&[MIMAT0000443&hsa-miR-125a-5p&offsets|0|-1;m-63&19|+|51693268|51693289&offsets|0|+1]|TCCCTGAGACCCTTTAACCTGTG</td><td>>MI0000469|hsa-mir-125a&WithFlank&19|+|51693248|51693345</td><td>21</td><td>43</td><td>23</td><td>TCCCTGAGACCCTTTAACCTGTG</td><td>9491.9</td><td>5.3</td><td>0.9</td><td>6.2</td><td>4.4E-10</td><td>1.1E-09</td><td>MB231/MB468</td><td>CYTO</td></tr>
<tr><td>>hg38_21_+_8433172_8446622_RNA45SN1_45S_50ntFlanks@3899.3921.23|CCTTCGCGGGGGGGATGCGTGCA</td><td>>hg38_21_+_8433172_8446622_RNA45SN1_45S_50ntFlanks</td><td>3899</td><td>3921</td><td>23</td><td>CCTTCGCGGGGGGGATGCGTGCA</td><td>149.1</td><td>-9.3</td><td>1.5</td><td>-6.3</td><td>3.3E-10</td><td>1.1E-09</td><td>BT20/MB468</td><td>MITO</td></tr>
<tr><td>>hg38_21_+_8433172_8446622_RNA45SN1_45S_50ntFlanks@11124.11146.23|CCGGCGGCGGCCCGCCGCGGGGC</td><td>>hg38_21_+_8433172_8446622_RNA45SN1_45S_50ntFlanks</td><td>11124</td><td>11146</td><td>23</td><td>CCGGCGGCGGCCCGCCGCGGGGC</td><td>155.3</td><td>-6.9</td><td>1.1</td><td>-6.3</td><td>3.9E-10</td><td>1.1E-09</td><td>BT20/MB468</td><td>CYTO</td></tr>
<tr><td>>trna111_HisGTG_1_-_147774845_147774916@-1T.33.34,trna118_HisGTG_1_-_145396881_145396952@-1T.33.34,trna16_HisGTG_1_+_146544773_146544844@-1T.33.34,trna1_HisGTG_15_+_45493349_45493420@-1T.33.34,trna21_HisGTG_1_+_147753471_147753542@-1T.33.34,trna33_HisGTG_6_+_27125906_27125977@-1T.33.34,trna7_HisGTG_9_-_14433938_14434009@-1T.33.34,trna8_HisGTG_15_-_45492611_45492682@-1T.33.34,trna9_HisGTG_15_-_45490804_45490875@-1T.33.34|TGCCGTGATCGTATAGTGGTTAGTACTCTGCGTT</td><td>>trna111_HisGTG_1_-_147774845_147774916</td><td>-1T</td><td>33</td><td>34</td><td>TGCCGTGATCGTATAGTGGTTAGTACTCTGCGTT</td><td>639.6</td><td>5.5</td><td>0.9</td><td>6.2</td><td>4.2E-10</td><td>1.1E-09</td><td>MB231/MB468</td><td>MP</td></tr>
<tr><td>>hg38_21_+_8433172_8446622_RNA45SN1_45S_50ntFlanks@4666.4695.30|AAATTCTTGGACCGGCGCAAGACGGACCAG</td><td>>hg38_21_+_8433172_8446622_RNA45SN1_45S_50ntFlanks</td><td>4666</td><td>4695</td><td>30</td><td>AAATTCTTGGACCGGCGCAAGACGGACCAG</td><td>69.3</td><td>-10.0</td><td>1.6</td><td>-6.2</td><td>4.2E-10</td><td>1.1E-09</td><td>MB231/MB468</td><td>MP</td></tr>
<tr><td>>hg38_21_+_8433172_8446622_RNA45SN1_45S_50ntFlanks@6652.6672.21|GACTCTTAGCGGTGGATCACT</td><td>>hg38_21_+_8433172_8446622_RNA45SN1_45S_50ntFlanks</td><td>6652</td><td>6672</td><td>21</td><td>GACTCTTAGCGGTGGATCACT</td><td>1419.1</td><td>6.2</td><td>1.0</td><td>6.3</td><td>3.3E-10</td><td>1.1E-09</td><td>BT20/MB468</td><td>MITO</td></tr>
<tr><td>>MI0000439|hsa-mir-23b&WithFlank&9|+|95085202|95085310@67.86.20(+1C)&[MIMAT0000418&hsa-miR-23b-3p&offsets|+3|0(+1C);m-42&9|+|95085265|95085285&offsets|+3|+2(+1C)]|ACATTGCCAGGGATTACCACC</td><td>>MI0000439|hsa-mir-23b&WithFlank&9|+|95085202|95085310</td><td>67</td><td>86</td><td>20(+1C)</td><td>ACATTGCCAGGGATTACCACC</td><td>210.0</td><td>11.5</td><td>1.7</td><td>6.6</td><td>4.0E-11</td><td>1.1E-09</td><td>BT20/MB468</td><td>MP</td></tr>
<tr><td>>MI0003137|hsa-mir-193b&WithFlank&16|+|14303961|14304055@19.40.22&[MIMAT0004767&hsa-miR-193b-5p&offsets|-1|-1;m-343&16|+|14303980|14304001&offsets|-1|-1]|TCGGGGTTTTGAGGGCGAGATG</td><td>>MI0003137|hsa-mir-193b&WithFlank&16|+|14303961|14304055</td><td>19</td><td>40</td><td>22</td><td>TCGGGGTTTTGAGGGCGAGATG</td><td>167.9</td><td>11.1</td><td>1.7</td><td>6.6</td><td>4.0E-11</td><td>1.1E-09</td><td>BT20/MB468</td><td>MP</td></tr>
<tr><td>>hg38_21_+_8433172_8446622_RNA45SN1_45S_50ntFlanks@10535.10564.30|CGCGAGGCGTCCAGTGCGGTAACGCGACCG</td><td>>hg38_21_+_8433172_8446622_RNA45SN1_45S_50ntFlanks</td><td>10535</td><td>10564</td><td>30</td><td>CGCGAGGCGTCCAGTGCGGTAACGCGACCG</td><td>167.5</td><td>-10.5</td><td>1.6</td><td>-6.6</td><td>4.0E-11</td><td>1.1E-09</td><td>BT20/MB468</td><td>MP</td></tr>
<tr><td>>hg38_21_+_8433172_8446622_RNA45SN1_45S_50ntFlanks@9534.9555.22|AGGAAACTCTGGTGGAGGTCCG</td><td>>hg38_21_+_8433172_8446622_RNA45SN1_45S_50ntFlanks</td><td>9534</td><td>9555</td><td>22</td><td>AGGAAACTCTGGTGGAGGTCCG</td><td>614.0</td><td>-12.3</td><td>1.9</td><td>-6.6</td><td>4.0E-11</td><td>1.1E-09</td><td>BT20/MB468</td><td>MP</td></tr>
<tr><td>>trnaMT_LeuTAG_MT_+_12266_12336@42.74.33|AAAATTTTGGTGCAACTCCAAATAAAAGTACCA</td><td>>trnaMT_LeuTAG_MT_+_12266_12336</td><td>42</td><td>74</td><td>33</td><td>AAAATTTTGGTGCAACTCCAAATAAAAGTACCA</td><td>194.6</td><td>-9.7</td><td>1.5</td><td>-6.3</td><td>3.3E-10</td><td>1.1E-09</td><td>BT20/MB468</td><td>MITO</td></tr>
<tr><td>>MI0000440|hsa-mir-27b&WithFlank&9|+|95085439|95085547@67.87.21(+1G)&[MIMAT0000419&hsa-miR-27b-3p&offsets|0|0(+1G);m-39&9|+|95085505|95085525&offsets|0|0(+1G)]|TTCACAGTGGCTAAGTTCTGCG</td><td>>MI0000440|hsa-mir-27b&WithFlank&9|+|95085439|95085547</td><td>67</td><td>87</td><td>21(+1G)</td><td>TTCACAGTGGCTAAGTTCTGCG</td><td>462.3</td><td>6.7</td><td>1.1</td><td>6.3</td><td>3.9E-10</td><td>1.1E-09</td><td>BT20/MB468</td><td>CYTO</td></tr>
<tr><td>>hg38_21_+_8433172_8446622_RNA45SN1_45S_50ntFlanks@6671.6688.18|CTCGGCTCGTGCGTCGAT</td><td>>hg38_21_+_8433172_8446622_RNA45SN1_45S_50ntFlanks</td><td>6671</td><td>6688</td><td>18</td><td>CTCGGCTCGTGCGTCGAT</td><td>219.3</td><td>-10.9</td><td>1.6</td><td>-6.6</td><td>4.0E-11</td><td>1.1E-09</td><td>BT20/MB468</td><td>MP</td></tr>
<tr><td>>hg38_21_+_8433172_8446622_RNA45SN1_45S_50ntFlanks@4683.4722.40|CAAGACGGACCAGAGCGAAAGCATTTGCCAAGAATGTTTT</td><td>>hg38_21_+_8433172_8446622_RNA45SN1_45S_50ntFlanks</td><td>4683</td><td>4722</td><td>40</td><td>CAAGACGGACCAGAGCGAAAGCATTTGCCAAGAATGTTTT</td><td>363.5</td><td>-6.9</td><td>1.1</td><td>-6.2</td><td>5.2E-10</td><td>1.1E-09</td><td>BT20/MB468</td><td>NUC</td></tr>
<tr><td>>hg38_21_+_8433172_8446622_RNA45SN1_45S_50ntFlanks@9568.9592.25|CGTGCAAATCGGTCGTCCGACCTGG</td><td>>hg38_21_+_8433172_8446622_RNA45SN1_45S_50ntFlanks</td><td>9568</td><td>9592</td><td>25</td><td>CGTGCAAATCGGTCGTCCGACCTGG</td><td>948.0</td><td>-7.6</td><td>1.2</td><td>-6.3</td><td>3.9E-10</td><td>1.1E-09</td><td>BT20/MB468</td><td>CYTO</td></tr>
<tr><td>>hg38_21_+_8433172_8446622_RNA45SN1_45S_50ntFlanks@11768.11790.23|CTAATTAGTGACGCGCATGAATG</td><td>>hg38_21_+_8433172_8446622_RNA45SN1_45S_50ntFlanks</td><td>11768</td><td>11790</td><td>23</td><td>CTAATTAGTGACGCGCATGAATG</td><td>1789.3</td><td>-6.2</td><td>1.0</td><td>-6.3</td><td>3.9E-10</td><td>1.1E-09</td><td>BT20/MB468</td><td>CYTO</td></tr>
<tr><td>>hg38_21_+_8433172_8446622_RNA45SN1_45S_50ntFlanks@12539.12587.49|ATGATGTGTTGTTGCCATGGTAATCCTGCTCAGTACGAGAGGAACCGCA</td><td>>hg38_21_+_8433172_8446622_RNA45SN1_45S_50ntFlanks</td><td>12539</td><td>12587</td><td>49</td><td>ATGATGTGTTGTTGCCATGGTAATCCTGCTCAGTACGAGAGGAACCGCA</td><td>549.4</td><td>-11.2</td><td>1.8</td><td>-6.3</td><td>3.3E-10</td><td>1.1E-09</td><td>BT20/MB468</td><td>MITO</td></tr>
<tr><td>>hg38_21_+_8433172_8446622_RNA45SN1_45S_50ntFlanks@10872.10901.30|GCTGGGTCGGTCGGGCTGGGGCGCGAAGCG</td><td>>hg38_21_+_8433172_8446622_RNA45SN1_45S_50ntFlanks</td><td>10872</td><td>10901</td><td>30</td><td>GCTGGGTCGGTCGGGCTGGGGCGCGAAGCG</td><td>89.6</td><td>-10.3</td><td>1.7</td><td>-6.2</td><td>4.3E-10</td><td>1.1E-09</td><td>MB231/MB468</td><td>MP</td></tr>
<tr><td>>hg38_21_+_8433172_8446622_RNA45SN1_45S_50ntFlanks@10806.10830.25|AACAATGTAGGTAAGGGAAGTCGGC</td><td>>hg38_21_+_8433172_8446622_RNA45SN1_45S_50ntFlanks</td><td>10806</td><td>10830</td><td>25</td><td>AACAATGTAGGTAAGGGAAGTCGGC</td><td>156.5</td><td>-2.6</td><td>0.4</td><td>-6.3</td><td>3.9E-10</td><td>1.1E-09</td><td>BT20/MB468</td><td>CYTO</td></tr>
<tr><td>>hg38_21_+_8433172_8446622_RNA45SN1_45S_50ntFlanks@9742.9765.24|CGATCTCAACCTATTCTCAAACTT</td><td>>hg38_21_+_8433172_8446622_RNA45SN1_45S_50ntFlanks</td><td>9742</td><td>9765</td><td>24</td><td>CGATCTCAACCTATTCTCAAACTT</td><td>363.6</td><td>-10.6</td><td>1.7</td><td>-6.3</td><td>3.4E-10</td><td>1.1E-09</td><td>BT20/MB468</td><td>MITO</td></tr>
<tr><td>>hg38_21_+_8433172_8446622_RNA45SN1_45S_50ntFlanks@6648.6683.36|GTACGACTCTTAGCGGTGGATCACTCGGCTCGTGCG</td><td>>hg38_21_+_8433172_8446622_RNA45SN1_45S_50ntFlanks</td><td>6648</td><td>6683</td><td>36</td><td>GTACGACTCTTAGCGGTGGATCACTCGGCTCGTGCG</td><td>376.1</td><td>9.7</td><td>1.6</td><td>6.2</td><td>4.5E-10</td><td>1.1E-09</td><td>MB231/MB468</td><td>NUC</td></tr>
<tr><td>>hg38_21_+_8433172_8446622_RNA45SN1_45S_50ntFlanks@10389.10422.34|TGAACAGCAGTTGAACATGGGTCAGTCGGTCCTG</td><td>>hg38_21_+_8433172_8446622_RNA45SN1_45S_50ntFlanks</td><td>10389</td><td>10422</td><td>34</td><td>TGAACAGCAGTTGAACATGGGTCAGTCGGTCCTG</td><td>502.0</td><td>-8.9</td><td>1.4</td><td>-6.2</td><td>5.2E-10</td><td>1.1E-09</td><td>BT20/MB468</td><td>NUC</td></tr>
<tr><td>>hg38_21_+_8433172_8446622_RNA45SN1_45S_50ntFlanks@10807.10840.34|ACAATGTAGGTAAGGGAAGTCGGCAAGCCGGATC</td><td>>hg38_21_+_8433172_8446622_RNA45SN1_45S_50ntFlanks</td><td>10807</td><td>10840</td><td>34</td><td>ACAATGTAGGTAAGGGAAGTCGGCAAGCCGGATC</td><td>282.9</td><td>-2.8</td><td>0.5</td><td>-6.2</td><td>4.5E-10</td><td>1.1E-09</td><td>MB231/MB468</td><td>CYTO</td></tr>
<tr><td>>hg38_21_+_8433172_8446622_RNA45SN1_45S_50ntFlanks@9339.9379.41|CGCACGAAAGCCGCCGTGGCGCAATGAAGGTGAAGGCCGGC</td><td>>hg38_21_+_8433172_8446622_RNA45SN1_45S_50ntFlanks</td><td>9339</td><td>9379</td><td>41</td><td>CGCACGAAAGCCGCCGTGGCGCAATGAAGGTGAAGGCCGGC</td><td>80.0</td><td>-10.2</td><td>1.6</td><td>-6.2</td><td>4.3E-10</td><td>1.1E-09</td><td>MB231/MB468</td><td>MP</td></tr>
<tr><td>>hg38_21_+_8433172_8446622_RNA45SN1_45S_50ntFlanks@6746.6771.26|CACTTCGAACGCACTTGCGGCCCCGG</td><td>>hg38_21_+_8433172_8446622_RNA45SN1_45S_50ntFlanks</td><td>6746</td><td>6771</td><td>26</td><td>CACTTCGAACGCACTTGCGGCCCCGG</td><td>751.4</td><td>-8.7</td><td>1.4</td><td>-6.2</td><td>5.2E-10</td><td>1.1E-09</td><td>BT20/MB468</td><td>NUC</td></tr>
<tr><td>>hg38_21_+_8433172_8446622_RNA45SN1_45S_50ntFlanks@9788.9820.33|CGCTGGCGTGGAGCCGGGCGTGGAATGCGAGTG</td><td>>hg38_21_+_8433172_8446622_RNA45SN1_45S_50ntFlanks</td><td>9788</td><td>9820</td><td>33</td><td>CGCTGGCGTGGAGCCGGGCGTGGAATGCGAGTG</td><td>749.7</td><td>-8.6</td><td>1.4</td><td>-6.2</td><td>5.0E-10</td><td>1.1E-09</td><td>MB231/MB468</td><td>MITO</td></tr>
<tr><td>>hg38_21_+_8433172_8446622_RNA45SN1_45S_50ntFlanks@12914.12933.20|CGTTCGTGGGGAACCTGGCG</td><td>>hg38_21_+_8433172_8446622_RNA45SN1_45S_50ntFlanks</td><td>12914</td><td>12933</td><td>20</td><td>CGTTCGTGGGGAACCTGGCG</td><td>692.3</td><td>-8.1</td><td>1.3</td><td>-6.2</td><td>5.0E-10</td><td>1.1E-09</td><td>MB231/MB468</td><td>MITO</td></tr>
<tr><td>>hg38_21_+_8433172_8446622_RNA45SN1_45S_50ntFlanks@10680.10710.31|GTTCCGGCGGCGTCCGGTGAGCTCTCGCTGG</td><td>>hg38_21_+_8433172_8446622_RNA45SN1_45S_50ntFlanks</td><td>10680</td><td>10710</td><td>31</td><td>GTTCCGGCGGCGTCCGGTGAGCTCTCGCTGG</td><td>16227.8</td><td>-5.4</td><td>0.9</td><td>-6.3</td><td>3.4E-10</td><td>1.1E-09</td><td>BT20/MB468</td><td>MITO</td></tr>
<tr><td>>hg38_21_+_8433172_8446622_RNA45SN1_45S_50ntFlanks@3839.3879.41|TCTCCTACTTGGATAACTGTGGTAATTCTAGAGCTAATACA</td><td>>hg38_21_+_8433172_8446622_RNA45SN1_45S_50ntFlanks</td><td>3839</td><td>3879</td><td>41</td><td>TCTCCTACTTGGATAACTGTGGTAATTCTAGAGCTAATACA</td><td>308.8</td><td>7.1</td><td>1.1</td><td>6.2</td><td>4.5E-10</td><td>1.1E-09</td><td>MB231/MB468</td><td>CYTO</td></tr>
<tr><td>>hg38_21_+_8433172_8446622_RNA45SN1_45S_50ntFlanks@11572.11601.30|CCGACTTAGAACTGGTGCGGACCAGGGGAA</td><td>>hg38_21_+_8433172_8446622_RNA45SN1_45S_50ntFlanks</td><td>11572</td><td>11601</td><td>30</td><td>CCGACTTAGAACTGGTGCGGACCAGGGGAA</td><td>13225.9</td><td>-5.4</td><td>0.9</td><td>-6.2</td><td>4.5E-10</td><td>1.1E-09</td><td>MB231/MB468</td><td>CYTO</td></tr>
<tr><td>>hg38_21_+_8433172_8446622_RNA45SN1_45S_50ntFlanks@9311.9335.25|AGTCTAACACGTGCGCGAGTCGGGG</td><td>>hg38_21_+_8433172_8446622_RNA45SN1_45S_50ntFlanks</td><td>9311</td><td>9335</td><td>25</td><td>AGTCTAACACGTGCGCGAGTCGGGG</td><td>464.1</td><td>-8.6</td><td>1.4</td><td>-6.2</td><td>4.3E-10</td><td>1.1E-09</td><td>MB231/MB468</td><td>MP</td></tr>
<tr><td>>hg38_21_+_8433172_8446622_RNA45SN1_45S_50ntFlanks@8403.8420.18|AGGTAAACGGGTGGGGTC</td><td>>hg38_21_+_8433172_8446622_RNA45SN1_45S_50ntFlanks</td><td>8403</td><td>8420</td><td>18</td><td>AGGTAAACGGGTGGGGTC</td><td>82.1</td><td>-10.2</td><td>1.6</td><td>-6.2</td><td>4.3E-10</td><td>1.1E-09</td><td>MB231/MB468</td><td>MP</td></tr>
<tr><td>>hg38_21_+_8433172_8446622_RNA45SN1_45S_50ntFlanks@12836.12871.36|CGGTGCGGAGTGCCCTTCGTCCTGGGAAACGGGGCG</td><td>>hg38_21_+_8433172_8446622_RNA45SN1_45S_50ntFlanks</td><td>12836</td><td>12871</td><td>36</td><td>CGGTGCGGAGTGCCCTTCGTCCTGGGAAACGGGGCG</td><td>601.6</td><td>-8.3</td><td>1.3</td><td>-6.2</td><td>4.3E-10</td><td>1.1E-09</td><td>MB231/MB468</td><td>MP</td></tr>
<tr><td>>hg38_21_+_8433172_8446622_RNA45SN1_45S_50ntFlanks@11787.11806.20|AATGGATGAACGAGATTCCC</td><td>>hg38_21_+_8433172_8446622_RNA45SN1_45S_50ntFlanks</td><td>11787</td><td>11806</td><td>20</td><td>AATGGATGAACGAGATTCCC</td><td>335.5</td><td>-11.5</td><td>1.7</td><td>-6.6</td><td>4.1E-11</td><td>1.1E-09</td><td>BT20/MB468</td><td>MP</td></tr>
<tr><td>>hg38_21_+_8433172_8446622_RNA45SN1_45S_50ntFlanks@6731.6757.27|CACATTGATCATCGACACTTCGAACGC</td><td>>hg38_21_+_8433172_8446622_RNA45SN1_45S_50ntFlanks</td><td>6731</td><td>6757</td><td>27</td><td>CACATTGATCATCGACACTTCGAACGC</td><td>86.5</td><td>-10.3</td><td>1.7</td><td>-6.2</td><td>4.3E-10</td><td>1.1E-09</td><td>MB231/MB468</td><td>MP</td></tr>
<tr><td>>hg38_21_+_8433172_8446622_RNA45SN1_45S_50ntFlanks@9788.9816.29|CGCTGGCGTGGAGCCGGGCGTGGAATGCG</td><td>>hg38_21_+_8433172_8446622_RNA45SN1_45S_50ntFlanks</td><td>9788</td><td>9816</td><td>29</td><td>CGCTGGCGTGGAGCCGGGCGTGGAATGCG</td><td>75.6</td><td>-10.1</td><td>1.6</td><td>-6.2</td><td>4.4E-10</td><td>1.1E-09</td><td>MB231/MB468</td><td>MP</td></tr>
<tr><td>>hg38_21_+_8433172_8446622_RNA45SN1_45S_50ntFlanks@7979.8003.25|ACCTCAGATCAGACGTGGCGACCCG</td><td>>hg38_21_+_8433172_8446622_RNA45SN1_45S_50ntFlanks</td><td>7979</td><td>8003</td><td>25</td><td>ACCTCAGATCAGACGTGGCGACCCG</td><td>779.0</td><td>-5.1</td><td>0.8</td><td>-6.2</td><td>5.1E-10</td><td>1.1E-09</td><td>MB231/MB468</td><td>MITO</td></tr>
<tr><td>>hg38_21_+_8433172_8446622_RNA45SN1_45S_50ntFlanks@10690.10712.23|CGTCCGGTGAGCTCTCGCTGGCC</td><td>>hg38_21_+_8433172_8446622_RNA45SN1_45S_50ntFlanks</td><td>10690</td><td>10712</td><td>23</td><td>CGTCCGGTGAGCTCTCGCTGGCC</td><td>127.9</td><td>-10.9</td><td>1.7</td><td>-6.2</td><td>4.4E-10</td><td>1.1E-09</td><td>MB231/MB468</td><td>MP</td></tr>
<tr><td>>MI0000446|hsa-mir-125b-1&WithFlank&11|-|122099751|122099850@21.41.21(+1G);MI0000470|hsa-mir-125b-2&WithFlank&21|+|16590231|16590331@23.43.21(+1G)&[MIMAT0000423&hsa-miR-125b-5p&offsets|0|-1(+1G);m-67&11|-|122099809|122099830&offsets|0|-1(+1G)];[MIMAT0000423_1&hsa-miR-125b-5p&offsets|0|-1(+1G);m-68&21|+|16590253|16590274&offsets|0|-1(+1G)]|TCCCTGAGACCCTAACTTGTGG</td><td>>MI0000446|hsa-mir-125b-1&WithFlank&11|-|122099751|122099850</td><td>21</td><td>41</td><td>21(+1G)</td><td>TCCCTGAGACCCTAACTTGTGG</td><td>292.9</td><td>9.1</td><td>1.5</td><td>6.2</td><td>4.5E-10</td><td>1.1E-09</td><td>MB231/MB468</td><td>CYTO</td></tr>
<tr><td>>hg38_21_+_8433172_8446622_RNA45SN1_45S_50ntFlanks@4367.4391.25|CAGTTAAAAAGCTCGTAGTTGGATC</td><td>>hg38_21_+_8433172_8446622_RNA45SN1_45S_50ntFlanks</td><td>4367</td><td>4391</td><td>25</td><td>CAGTTAAAAAGCTCGTAGTTGGATC</td><td>1440.8</td><td>-3.7</td><td>0.6</td><td>-6.3</td><td>4.0E-10</td><td>1.1E-09</td><td>BT20/MB468</td><td>CYTO</td></tr>
</table>

<table>
<tr><td>>hg38_21_+_8433172_8446622_RNA45SN1_45S_50ntFlanks@9502.9526.25|TGGTGAACTATGCCTGGGCAGGGCG</td><td>>hg38_21_+_8433172_8446622_RNA45SN1_45S_50ntFlanks</td><td>9502</td><td>9526</td><td>25</td><td>TGGTGAACTATGCCTGGGCAGGGCG</td><td>85.1</td><td>-10.3</td><td>1.6</td><td>-6.2</td><td>4.4E-10</td><td>1.1E-09</td><td>MB231/MB468</td><td>MP</td></tr>
<tr><td>>hg38_21_+_8433172_8446622_RNA45SN1_45S_50ntFlanks@11207.11231.25|GGCGGCGACTCTGGACGCGAGCCGG</td><td>>hg38_21_+_8433172_8446622_RNA45SN1_45S_50ntFlanks</td><td>11207</td><td>11231</td><td>25</td><td>GGCGGCGACTCTGGACGCGAGCCGG</td><td>86.0</td><td>-10.3</td><td>1.6</td><td>-6.2</td><td>4.4E-10</td><td>1.1E-09</td><td>MB231/MB468</td><td>MP</td></tr>
<tr><td>>hg38_21_+_8433172_8446622_RNA45SN1_45S_50ntFlanks@10794.10819.26|CTGGCATGTTGGAACAATGTAGGTAA</td><td>>hg38_21_+_8433172_8446622_RNA45SN1_45S_50ntFlanks</td><td>10794</td><td>10819</td><td>26</td><td>CTGGCATGTTGGAACAATGTAGGTAA</td><td>62.9</td><td>-9.8</td><td>1.6</td><td>-6.2</td><td>4.4E-10</td><td>1.1E-09</td><td>MB231/MB468</td><td>MP</td></tr>
<tr><td>>hg38_1_-_228634819_228635039_RNA5S12_5S_50ntFlanks@51.95.45|GTCTACGGCCATACCACCCTGAACGCGCCCGATCTCGTCTGATCT</td><td>>hg38_1_-_228634819_228635039_RNA5S12_5S_50ntFlanks</td><td>51</td><td>95</td><td>45</td><td>GTCTACGGCCATACCACCCTGAACGCGCCCGATCTCGTCTGATCT</td><td>247.7</td><td>5.8</td><td>0.9</td><td>6.2</td><td>4.5E-10</td><td>1.1E-09</td><td>MB231/MB468</td><td>CYTO</td></tr>
<tr><td>>hg38_21_+_8433172_8446622_RNA45SN1_45S_50ntFlanks@6739.6771.33|TCATCGACACTTCGAACGCACTTGCGGCCCCGG</td><td>>hg38_21_+_8433172_8446622_RNA45SN1_45S_50ntFlanks</td><td>6739</td><td>6771</td><td>33</td><td>TCATCGACACTTCGAACGCACTTGCGGCCCCGG</td><td>841.5</td><td>-3.7</td><td>0.6</td><td>-6.3</td><td>3.4E-10</td><td>1.1E-09</td><td>BT20/MB468</td><td>MITO</td></tr>
<tr><td>>hg38_21_+_8433172_8446622_RNA45SN1_45S_50ntFlanks@11567.11605.39|AGCAGCCGACTTAGAACTGGTGCGGACCAGGGGAATCCG</td><td>>hg38_21_+_8433172_8446622_RNA45SN1_45S_50ntFlanks</td><td>11567</td><td>11605</td><td>39</td><td>AGCAGCCGACTTAGAACTGGTGCGGACCAGGGGAATCCG</td><td>162.2</td><td>-11.2</td><td>1.8</td><td>-6.2</td><td>4.4E-10</td><td>1.1E-09</td><td>MB231/MB468</td><td>MP</td></tr>
<tr><td>>hg38_21_+_8433172_8446622_RNA45SN1_45S_50ntFlanks@11820.11842.23|ACTATCCAGCGAAACCACAGCCA</td><td>>hg38_21_+_8433172_8446622_RNA45SN1_45S_50ntFlanks</td><td>11820</td><td>11842</td><td>23</td><td>ACTATCCAGCGAAACCACAGCCA</td><td>80.4</td><td>-10.2</td><td>1.6</td><td>-6.2</td><td>4.4E-10</td><td>1.1E-09</td><td>MB231/MB468</td><td>MP</td></tr>
<tr><td>>trna111_HisGTG_1_-_147774845_147774916@-1T.24.25,trna118_HisGTG_1_-_145396881_145396952@-1T.24.25,trna16_HisGTG_1_+_146544773_146544844@-1T.24.25,trna1_HisGTG_15_+_45493349_45493420@-1T.24.25,trna21_HisGTG_1_+_147753471_147753542@-1T.24.25,trna33_HisGTG_6_+_27125906_27125977@-1T.24.25,trna7_HisGTG_9_-_14433938_14434009@-1T.24.25,trna8_HisGTG_15_-_45492611_45492682@-1T.24.25,trna9_HisGTG_15_-_45490804_45490875@-1T.24.25|TGCCGTGATCGTATAGTGGTTAGTA</td><td>>trna111_HisGTG_1_-_147774845_147774916</td><td>-1T</td><td>24</td><td>25</td><td>TGCCGTGATCGTATAGTGGTTAGTA</td><td>340.0</td><td>4.8</td><td>0.8</td><td>6.2</td><td>4.6E-10</td><td>1.1E-09</td><td>MB231/MB468</td><td>NUC</td></tr>
<tr><td>>hg38_21_+_8433172_8446622_RNA45SN1_45S_50ntFlanks@10823.10862.40|AAGTCGGCAAGCCGGATCCGTAACTTCGGGATAAGGATTG</td><td>>hg38_21_+_8433172_8446622_RNA45SN1_45S_50ntFlanks</td><td>10823</td><td>10862</td><td>40</td><td>AAGTCGGCAAGCCGGATCCGTAACTTCGGGATAAGGATTG</td><td>166.3</td><td>-9.7</td><td>1.5</td><td>-6.3</td><td>4.0E-10</td><td>1.1E-09</td><td>BT20/MB468</td><td>CYTO</td></tr>
<tr><td>>hg38_21_+_8433172_8446622_RNA45SN1_45S_50ntFlanks@12345.12365.21|TCAGAAAAGTTACCACAGGGA</td><td>>hg38_21_+_8433172_8446622_RNA45SN1_45S_50ntFlanks</td><td>12345</td><td>12365</td><td>21</td><td>TCAGAAAAGTTACCACAGGGA</td><td>76.1</td><td>-10.1</td><td>1.6</td><td>-6.2</td><td>4.4E-10</td><td>1.1E-09</td><td>MB231/MB468</td><td>MP</td></tr>
<tr><td>>hg38_21_+_8433172_8446622_RNA45SN1_45S_50ntFlanks@4892.4910.19|AAACTTAAAGGAATTGACG</td><td>>hg38_21_+_8433172_8446622_RNA45SN1_45S_50ntFlanks</td><td>4892</td><td>4910</td><td>19</td><td>AAACTTAAAGGAATTGACG</td><td>82.6</td><td>-10.2</td><td>1.6</td><td>-6.2</td><td>4.4E-10</td><td>1.1E-09</td><td>MB231/MB468</td><td>MP</td></tr>
<tr><td>>hg38_21_+_8433172_8446622_RNA45SN1_45S_50ntFlanks@3876.3898.23|TACATGCCGACGGGCGCTGACCC</td><td>>hg38_21_+_8433172_8446622_RNA45SN1_45S_50ntFlanks</td><td>3876</td><td>3898</td><td>23</td><td>TACATGCCGACGGGCGCTGACCC</td><td>1106.7</td><td>-5.1</td><td>0.8</td><td>-6.2</td><td>4.7E-10</td><td>1.1E-09</td><td>MB231/MB468</td><td>NUC</td></tr>
<tr><td>>hg38_21_+_8433172_8446622_RNA45SN1_45S_50ntFlanks@3985.4023.39|CGGCTTTGGTGACTCTAGATAACCTCGGGCCGATCGCAC</td><td>>hg38_21_+_8433172_8446622_RNA45SN1_45S_50ntFlanks</td><td>3985</td><td>4023</td><td>39</td><td>CGGCTTTGGTGACTCTAGATAACCTCGGGCCGATCGCAC</td><td>114.9</td><td>-10.7</td><td>1.7</td><td>-6.2</td><td>4.4E-10</td><td>1.1E-09</td><td>MB231/MB468</td><td>MP</td></tr>
<tr><td>>hg38_21_+_8433172_8446622_RNA45SN1_45S_50ntFlanks@3985.4026.42|CGGCTTTGGTGACTCTAGATAACCTCGGGCCGATCGCACGCC</td><td>>hg38_21_+_8433172_8446622_RNA45SN1_45S_50ntFlanks</td><td>3985</td><td>4026</td><td>42</td><td>CGGCTTTGGTGACTCTAGATAACCTCGGGCCGATCGCACGCC</td><td>1742.1</td><td>-7.5</td><td>1.2</td><td>-6.2</td><td>4.6E-10</td><td>1.1E-09</td><td>MB231/MB468</td><td>CYTO</td></tr>
<tr><td>>hg38_21_+_8433172_8446622_RNA45SN1_45S_50ntFlanks@12893.12930.38|CCTCGCCCGTCACGCACCGCACGTTCGTGGGGAACCTG</td><td>>hg38_21_+_8433172_8446622_RNA45SN1_45S_50ntFlanks</td><td>12893</td><td>12930</td><td>38</td><td>CCTCGCCCGTCACGCACCGCACGTTCGTGGGGAACCTG</td><td>1598.8</td><td>-9.5</td><td>1.5</td><td>-6.2</td><td>4.5E-10</td><td>1.1E-09</td><td>MB231/MB468</td><td>MP</td></tr>
<tr><td>>hg38_21_+_8433172_8446622_RNA45SN1_45S_50ntFlanks@12640.12668.29|ACCATCTGTGGGATTATGACTGAACGCCT</td><td>>hg38_21_+_8433172_8446622_RNA45SN1_45S_50ntFlanks</td><td>12640</td><td>12668</td><td>29</td><td>ACCATCTGTGGGATTATGACTGAACGCCT</td><td>162.6</td><td>4.8</td><td>0.8</td><td>6.2</td><td>4.7E-10</td><td>1.1E-09</td><td>MB231/MB468</td><td>NUC</td></tr>
<tr><td>>hg38_21_+_8433172_8446622_RNA45SN1_45S_50ntFlanks@12072.12106.35|CCCGAGGGGCTCTCGCTTCTGGCGCCAAGCGCCCG</td><td>>hg38_21_+_8433172_8446622_RNA45SN1_45S_50ntFlanks</td><td>12072</td><td>12106</td><td>35</td><td>CCCGAGGGGCTCTCGCTTCTGGCGCCAAGCGCCCG</td><td>737.3</td><td>-6.3</td><td>1.0</td><td>-6.2</td><td>5.2E-10</td><td>1.1E-09</td><td>MB231/MB468</td><td>MITO</td></tr>
<tr><td>>MI0000650|hsa-mir-200c&WithFlank&12|+|6963693|6963772@50.71.22(+1U)&[MIMAT0000617&hsa-miR-200c-3p&offsets|0|-1(+1U);m-54&12|+|6963742|6963764&offsets|0|-1(+1U)]|TAATACTGCCGGGTAATGATGGT</td><td>>MI0000650|hsa-mir-200c&WithFlank&12|+|6963693|6963772</td><td>50</td><td>71</td><td>22(+1U)</td><td>TAATACTGCCGGGTAATGATGGT</td><td>1699.5</td><td>8.6</td><td>1.3</td><td>6.8</td><td>8.2E-12</td><td>1.1E-09</td><td>BT20/MB231</td><td>MP</td></tr>
<tr><td>>hg38_21_+_8433172_8446622_RNA45SN1_45S_50ntFlanks@12656.12675.20|TGACTGAACGCCTCTAAGTC</td><td>>hg38_21_+_8433172_8446622_RNA45SN1_45S_50ntFlanks</td><td>12656</td><td>12675</td><td>20</td><td>TGACTGAACGCCTCTAAGTC</td><td>310.8</td><td>-8.2</td><td>1.3</td><td>-6.2</td><td>5.2E-10</td><td>1.1E-09</td><td>MB231/MB468</td><td>MITO</td></tr>
<tr><td>>hg38_MT_+_1621_3279_RNR2_16S_50ntFlanks@1349.1388.40|ACAATAGGGTTTACGACCTCGATGTTGGATCAGGACATCC</td><td>>hg38_MT_+_1621_3279_RNR2_16S_50ntFlanks</td><td>1349</td><td>1388</td><td>40</td><td>ACAATAGGGTTTACGACCTCGATGTTGGATCAGGACATCC</td><td>524.7</td><td>5.3</td><td>0.9</td><td>6.2</td><td>5.2E-10</td><td>1.1E-09</td><td>MB231/MB468</td><td>MITO</td></tr>
<tr><td>>hg38_21_+_8433172_8446622_RNA45SN1_45S_50ntFlanks@9180.9197.18|CCGTCGGGCCCGGGGGAG</td><td>>hg38_21_+_8433172_8446622_RNA45SN1_45S_50ntFlanks</td><td>9180</td><td>9197</td><td>18</td><td>CCGTCGGGCCCGGGGGAG</td><td>490.1</td><td>-8.7</td><td>1.4</td><td>-6.2</td><td>4.7E-10</td><td>1.1E-09</td><td>MB231/MB468</td><td>NUC</td></tr>
<tr><td>>MI0000264|hsa-mir-7-2&WithFlank&15|+|88611819|88611940@38.60.23;MI0000265|hsa-mir-7-3&WithFlank&19|+|4770664|4770785@37.59.23;MI0000263|hsa-mir-7-1&WithFlank&9|-|83969742|83969863@30.52.23&[MIMAT0000252&hsa-miR-7-5p&offsets|0|-1;m-400&15|+|88611856|88611879&offsets|0|-1];[MIMAT0000252_1&hsa-miR-7-5p&offsets|0|-1;m-402&19|+|4770700|4770723&offsets|0|-1];[MIMAT0000252_2&hsa-miR-7-5p&offsets|0|-1;m-401&9|-|83969811|83969834&offsets|0|-1]|TGGAAGACTAGTGATTTTGTTGT</td><td>>MI0000264|hsa-mir-7-2&WithFlank&15|+|88611819|88611940</td><td>38</td><td>60</td><td>23</td><td>TGGAAGACTAGTGATTTTGTTGT</td><td>8197.4</td><td>3.5</td><td>0.6</td><td>6.2</td><td>4.5E-10</td><td>1.1E-09</td><td>MB231/MB468</td><td>MP</td></tr>
<tr><td>>hg38_21_+_8433172_8446622_RNA45SN1_45S_50ntFlanks@10727.10763.37|GGAGAGGGTGTAAATCTCGCGCCGGGCCGTACCCATA</td><td>>hg38_21_+_8433172_8446622_RNA45SN1_45S_50ntFlanks</td><td>10727</td><td>10763</td><td>37</td><td>GGAGAGGGTGTAAATCTCGCGCCGGGCCGTACCCATA</td><td>78.5</td><td>-10.2</td><td>1.6</td><td>-6.2</td><td>4.5E-10</td><td>1.1E-09</td><td>MB231/MB468</td><td>MP</td></tr>
<tr><td>>hg38_21_+_8433172_8446622_RNA45SN1_45S_50ntFlanks@9716.9737.22|AATGATTAGAGGTCTTGGGGCC</td><td>>hg38_21_+_8433172_8446622_RNA45SN1_45S_50ntFlanks</td><td>9716</td><td>9737</td><td>22</td><td>AATGATTAGAGGTCTTGGGGCC</td><td>775.8</td><td>-8.3</td><td>1.3</td><td>-6.2</td><td>4.5E-10</td><td>1.1E-09</td><td>MB231/MB468</td><td>MP</td></tr>
<tr><td>>hg38_21_+_8433172_8446622_RNA45SN1_45S_50ntFlanks@11233.11250.18|CCCTTCCCGTGGATCGCC</td><td>>hg38_21_+_8433172_8446622_RNA45SN1_45S_50ntFlanks</td><td>11233</td><td>11250</td><td>18</td><td>CCCTTCCCGTGGATCGCC</td><td>1449.0</td><td>-8.2</td><td>1.3</td><td>-6.2</td><td>4.7E-10</td><td>1.1E-09</td><td>MB231/MB468</td><td>CYTO</td></tr>
<tr><td>>MI0000477|hsa-mir-146a&WithFlank&5|+|160485346|160485456@27.47.21(+1C)&[MIMAT0000449&hsa-miR-146a-5p&offsets|0|-1(+1C);m-129&5|+|160485372|160485393&offsets|0|-1(+1C)]|TGAGAACTGAATTCCATGGGTC</td><td>>MI0000477|hsa-mir-146a&WithFlank&5|+|160485346|160485456</td><td>27</td><td>47</td><td>21(+1C)</td><td>TGAGAACTGAATTCCATGGGTC</td><td>29227.5</td><td>-10.6</td><td>1.6</td><td>-6.8</td><td>8.4E-12</td><td>1.1E-09</td><td>BT20/MB231</td><td>MP</td></tr>
<tr><td>>hg38_21_+_8433172_8446622_RNA45SN1_45S_50ntFlanks@11665.11691.27|TTCTGCCCAGTGCTCTGAATGTCAAAG</td><td>>hg38_21_+_8433172_8446622_RNA45SN1_45S_50ntFlanks</td><td>11665</td><td>11691</td><td>27</td><td>TTCTGCCCAGTGCTCTGAATGTCAAAG</td><td>306.4</td><td>6.5</td><td>1.0</td><td>6.2</td><td>4.1E-10</td><td>1.1E-09</td><td>BT20/MB468</td><td>CYTO</td></tr>
<tr><td>>hg38_21_+_8433172_8446622_RNA45SN1_45S_50ntFlanks@10808.10831.24|CAATGTAGGTAAGGGAAGTCGGCA</td><td>>hg38_21_+_8433172_8446622_RNA45SN1_45S_50ntFlanks</td><td>10808</td><td>10831</td><td>24</td><td>CAATGTAGGTAAGGGAAGTCGGCA</td><td>229.2</td><td>-4.6</td><td>0.7</td><td>-6.2</td><td>4.7E-10</td><td>1.1E-09</td><td>MB231/MB468</td><td>CYTO</td></tr>
<tr><td>>hg38_21_+_8433172_8446622_RNA45SN1_45S_50ntFlanks@6702.6720.19|AGCTGCGAGAATTAATGTG</td><td>>hg38_21_+_8433172_8446622_RNA45SN1_45S_50ntFlanks</td><td>6702</td><td>6720</td><td>19</td><td>AGCTGCGAGAATTAATGTG</td><td>1701.2</td><td>-4.1</td><td>0.6</td><td>-6.2</td><td>4.5E-10</td><td>1.1E-09</td><td>MB231/MB468</td><td>MP</td></tr>
<tr><td>>hg38_21_+_8433172_8446622_RNA45SN1_45S_50ntFlanks@12488.12520.33|AACGTGAGCTGGGTTTAGACCGTCGTGAGACAG</td><td>>hg38_21_+_8433172_8446622_RNA45SN1_45S_50ntFlanks</td><td>12488</td><td>12520</td><td>33</td><td>AACGTGAGCTGGGTTTAGACCGTCGTGAGACAG</td><td>229.3</td><td>-6.1</td><td>1.0</td><td>-6.2</td><td>4.7E-10</td><td>1.1E-09</td><td>MB231/MB468</td><td>CYTO</td></tr>
<tr><td>>hg38_21_+_8433172_8446622_RNA45SN1_45S_50ntFlanks@6649.6678.30|TACGACTCTTAGCGGTGGATCACTCGGCTC</td><td>>hg38_21_+_8433172_8446622_RNA45SN1_45S_50ntFlanks</td><td>6649</td><td>6678</td><td>30</td><td>TACGACTCTTAGCGGTGGATCACTCGGCTC</td><td>2297.5</td><td>4.0</td><td>0.6</td><td>6.3</td><td>3.5E-10</td><td>1.1E-09</td><td>BT20/MB468</td><td>MITO</td></tr>
<tr><td>>MI0000736|hsa-mir-30c-1&WithFlank&1|+|40757278|40757378@23.42.20;MI0000254|hsa-mir-30c-2&WithFlank&6|-|71376954|71377037@13.32.20&[MIMAT0000244&hsa-miR-30c-5p&offsets|0|-3;m-57&1|+|40757300|40757323&offsets|0|-4];[MIMAT0000244_1&hsa-miR-30c-5p&offsets|0|-3;m-56&6|-|71377002|71377025&offsets|0|-4]|TGTAAACATCCTACACTCTC</td><td>>MI0000736|hsa-mir-30c-1&WithFlank&1|+|40757278|40757378</td><td>23</td><td>42</td><td>20</td><td>TGTAAACATCCTACACTCTC</td><td>2013.4</td><td>4.8</td><td>0.8</td><td>6.2</td><td>4.7E-10</td><td>1.1E-09</td><td>MB231/MB468</td><td>NUC</td></tr>
<tr><td>>MI0000101|hsa-mir-99a&WithFlank&21|+|16539083|16539175@19.40.22(+1A)&[MIMAT0000097&hsa-miR-99a-5p&offsets|0|0(+1A);m-64&21|+|16539101|16539121&offsets|0|+1(+1A)]|AACCCGTAGATCCGATCTTGTGA</td><td>>MI0000101|hsa-mir-99a&WithFlank&21|+|16539083|16539175</td><td>19</td><td>40</td><td>22(+1A)</td><td>AACCCGTAGATCCGATCTTGTGA</td><td>2145.8</td><td>-9.5</td><td>1.5</td><td>-6.2</td><td>5.5E-10</td><td>1.1E-09</td><td>BT20/MB468</td><td>NUC</td></tr>
<tr><td>>hg38_21_+_8433172_8446622_RNA45SN1_45S_50ntFlanks@12635.12656.22|AAGCTACCATCTGTGGGATTAT</td><td>>hg38_21_+_8433172_8446622_RNA45SN1_45S_50ntFlanks</td><td>12635</td><td>12656</td><td>22</td><td>AAGCTACCATCTGTGGGATTAT</td><td>160.8</td><td>-10.4</td><td>1.6</td><td>-6.6</td><td>4.3E-11</td><td>1.1E-09</td><td>BT20/MB468</td><td>MP</td></tr>
<tr><td>>hg38_21_+_8433172_8446622_RNA45SN1_45S_50ntFlanks@12083.12106.24|CTCGCTTCTGGCGCCAAGCGCCCG</td><td>>hg38_21_+_8433172_8446622_RNA45SN1_45S_50ntFlanks</td><td>12083</td><td>12106</td><td>24</td><td>CTCGCTTCTGGCGCCAAGCGCCCG</td><td>274.4</td><td>-7.0</td><td>1.1</td><td>-6.2</td><td>4.7E-10</td><td>1.1E-09</td><td>MB231/MB468</td><td>CYTO</td></tr>
<tr><td>>MI0000283|hsa-mir-203a&WithFlank&14|+|104117399|104117520@72.90.19&[MIMAT0000264&hsa-miR-203a-3p&offsets|+1|-2;m-79&14|+|104117469|104117490&offsets|+1|-2]|TGAAATGTTTAGGACCACT</td><td>>MI0000283|hsa-mir-203a&WithFlank&14|+|104117399|104117520</td><td>72</td><td>90</td><td>19</td><td>TGAAATGTTTAGGACCACT</td><td>369.0</td><td>9.3</td><td>1.5</td><td>6.3</td><td>3.5E-10</td><td>1.1E-09</td><td>BT20/MB468</td><td>MITO</td></tr>
<tr><td>>hg38_21_+_8433172_8446622_RNA45SN1_45S_50ntFlanks@11593.11612.20|CCAGGGGAATCCGACTGTTT</td><td>>hg38_21_+_8433172_8446622_RNA45SN1_45S_50ntFlanks</td><td>11593</td><td>11612</td><td>20</td><td>CCAGGGGAATCCGACTGTTT</td><td>1380.4</td><td>-4.8</td><td>0.8</td><td>-6.2</td><td>4.7E-10</td><td>1.1E-09</td><td>MB231/MB468</td><td>CYTO</td></tr>
<tr><td>>hg38_21_+_8433172_8446622_RNA45SN1_45S_50ntFlanks@11603.11620.18|CCGACTGTTTAATTAAAA</td><td>>hg38_21_+_8433172_8446622_RNA45SN1_45S_50ntFlanks</td><td>11603</td><td>11620</td><td>18</td><td>CCGACTGTTTAATTAAAA</td><td>331.3</td><td>-6.2</td><td>1.0</td><td>-6.2</td><td>4.5E-10</td><td>1.1E-09</td><td>MB231/MB468</td><td>MP</td></tr>
<tr><td>>hg38_21_+_8433172_8446622_RNA45SN1_45S_50ntFlanks@11078.11103.26|CCGCGTGGGGGCGGCGGCGGGGGGAG</td><td>>hg38_21_+_8433172_8446622_RNA45SN1_45S_50ntFlanks</td><td>11078</td><td>11103</td><td>26</td><td>CCGCGTGGGGGCGGCGGCGGGGGGAG</td><td>401.7</td><td>-8.6</td><td>1.4</td><td>-6.2</td><td>5.5E-10</td><td>1.1E-09</td><td>BT20/MB468</td><td>NUC</td></tr>
<tr><td>>MI0005756|hsa-mir-934&WithFlank&X|+|136550872|136550966@21.40.20&[MIMAT0004977&hsa-miR-934&offsets|0|-2;m-943&X|+|136550892|136550913&offsets|0|-2]|TGTCTACTACTGGAGACACT</td><td>>MI0005756|hsa-mir-934&WithFlank&X|+|136550872|136550966</td><td>21</td><td>40</td><td>20</td><td>TGTCTACTACTGGAGACACT</td><td>126.4</td><td>-10.8</td><td>1.7</td><td>-6.2</td><td>4.5E-10</td><td>1.1E-09</td><td>MB231/MB468</td><td>MP</td></tr>
<tr><td>>hg38_21_+_8433172_8446622_RNA45SN1_45S_50ntFlanks@11670.11704.35|CCCAGTGCTCTGAATGTCAAAGTGAAGAAATTCAA</td><td>>hg38_21_+_8433172_8446622_RNA45SN1_45S_50ntFlanks</td><td>11670</td><td>11704</td><td>35</td><td>CCCAGTGCTCTGAATGTCAAAGTGAAGAAATTCAA</td><td>101.7</td><td>-10.5</td><td>1.7</td><td>-6.2</td><td>4.6E-10</td><td>1.1E-09</td><td>MB231/MB468</td><td>MP</td></tr>
<tr><td>>hg38_21_+_8433172_8446622_RNA45SN1_45S_50ntFlanks@12942.12990.49|TTCGTAGACGACCTGCTTCTGGGTCGGGGTTTCGTACGTAGCAGAGCAG</td><td>>hg38_21_+_8433172_8446622_RNA45SN1_45S_50ntFlanks</td><td>12942</td><td>12990</td><td>49</td><td>TTCGTAGACGACCTGCTTCTGGGTCGGGGTTTCGTACGTAGCAGAGCAG</td><td>264.5</td><td>-11.0</td><td>1.8</td><td>-6.2</td><td>4.8E-10</td><td>1.1E-09</td><td>MB231/MB468</td><td>NUC</td></tr>
<tr><td>>hg38_21_+_8433172_8446622_RNA45SN1_45S_50ntFlanks@10735.10757.23|TGTAAATCTCGCGCCGGGCCGTA</td><td>>hg38_21_+_8433172_8446622_RNA45SN1_45S_50ntFlanks</td><td>10735</td><td>10757</td><td>23</td><td>TGTAAATCTCGCGCCGGGCCGTA</td><td>1391.8</td><td>-9.7</td><td>1.6</td><td>-6.2</td><td>5.5E-10</td><td>1.1E-09</td><td>BT20/MB468</td><td>NUC</td></tr>
<tr><td>>hg38_21_+_8433172_8446622_RNA45SN1_45S_50ntFlanks@12003.12043.41|GCGGGCCGCCGGTGAAATACCACTACTCTGATCGTTTTTTC</td><td>>hg38_21_+_8433172_8446622_RNA45SN1_45S_50ntFlanks</td><td>12003</td><td>12043</td><td>41</td><td>GCGGGCCGCCGGTGAAATACCACTACTCTGATCGTTTTTTC</td><td>283.5</td><td>-12.0</td><td>1.9</td><td>-6.2</td><td>4.6E-10</td><td>1.1E-09</td><td>MB231/MB468</td><td>MP</td></tr>
<tr><td>>hg38_21_+_8433172_8446622_RNA45SN1_45S_50ntFlanks@4297.4318.22|CATTGGAGGGCAAGTCTGGTGC</td><td>>hg38_21_+_8433172_8446622_RNA45SN1_45S_50ntFlanks</td><td>4297</td><td>4318</td><td>22</td><td>CATTGGAGGGCAAGTCTGGTGC</td><td>392.9</td><td>-8.4</td><td>1.4</td><td>-6.2</td><td>5.5E-10</td><td>1.1E-09</td><td>BT20/MB468</td><td>NUC</td></tr>
</table>

| | | | | | | | | | | | | | |
|---|---|---|---|---|---|---|---|---|---|---|---|---|---|
| >hg38_21_+_8433172_8446622_RNA45SN1_45S_50ntFlanks@11731.11751.21\|ACTATGACTCTCTTAAGGTAG | >hg38_21_+_8433172_8446622_RNA45SN1_45S_50ntFlanks | 11731 | 11751 | 21 | ACTATGACTCTCTTAAGGTAG | 901.0 | -9.2 | 1.5 | -6.2 | 5.5E-10 | 1.1E-09 | BT20/MB468 | NUC |
| >hg38_21_+_8433172_8446622_RNA45SN1_45S_50ntFlanks@11564.11611.48\|CCTAGCAGCCGACTTAGAACTGGTGCGGACCAGGGGAATCCGACTGTT | >hg38_21_+_8433172_8446622_RNA45SN1_45S_50ntFlanks | 11564 | 11611 | 48 | CCTAGCAGCCGACTTAGAACTGGTGCGGACCAGGGGAATCCGACTGTT | 3000.8 | -7.2 | 1.2 | -6.2 | 5.5E-10 | 1.1E-09 | BT20/MB468 | NUC |
| >hg38_21_+_8433172_8446622_RNA45SN1_45S_50ntFlanks@12076.12108.33\|AGGGGCTCTCGCTTCTGGCGCCAAGCGCCCGGC | >hg38_21_+_8433172_8446622_RNA45SN1_45S_50ntFlanks | 12076 | 12108 | 33 | AGGGGCTCTCGCTTCTGGCGCCAAGCGCCCGGC | 63.2 | -9.8 | 1.6 | -6.2 | 4.6E-10 | 1.1E-09 | MB231/MB468 | MP |
| >hg38_21_+_8433172_8446622_RNA45SN1_45S_50ntFlanks@3777.3810.34\|CGGTACAGTGAAACTGCGAATGGCTCATTAAATC | >hg38_21_+_8433172_8446622_RNA45SN1_45S_50ntFlanks | 3777 | 3810 | 34 | CGGTACAGTGAAACTGCGAATGGCTCATTAAATC | 302.9 | -3.9 | 0.6 | -6.2 | 4.8E-10 | 1.1E-09 | MB231/MB468 | NUC |
| >hg38_21_+_8433172_8446622_RNA45SN1_45S_50ntFlanks@10870.10902.33\|GGGCTGGGTCGGTCGGGCTGGGGCGCGAAGCGG | >hg38_21_+_8433172_8446622_RNA45SN1_45S_50ntFlanks | 10870 | 10902 | 33 | GGGCTGGGTCGGTCGGGCTGGGGCGCGAAGCGG | 68.8 | -10.0 | 1.6 | -6.2 | 4.6E-10 | 1.1E-09 | MB231/MB468 | MP |
| >hg38_21_+_8433172_8446622_RNA45SN1_45S_50ntFlanks@10684.10710.27\|CGGCGGCGTCCGGTGAGCTCTCGCTGG | >hg38_21_+_8433172_8446622_RNA45SN1_45S_50ntFlanks | 10684 | 10710 | 27 | CGGCGGCGTCCGGTGAGCTCTCGCTGG | 7780.1 | -4.6 | 0.7 | -6.3 | 3.6E-10 | 1.1E-09 | BT20/MB468 | MITO |
| >hg38_21_+_8433172_8446622_RNA45SN1_45S_50ntFlanks@4657.4689.33\|CTAGAGGTGAAATTCTTGGACCGGCGCAAGACG | >hg38_21_+_8433172_8446622_RNA45SN1_45S_50ntFlanks | 4657 | 4689 | 33 | CTAGAGGTGAAATTCTTGGACCGGCGCAAGACG | 10812.2 | -8.2 | 1.3 | -6.3 | 3.6E-10 | 1.1E-09 | BT20/MB468 | MITO |
| >hg38_21_+_8433172_8446622_RNA45SN1_45S_50ntFlanks@11729.11764.36\|TAACTATGACTCTCTTAAGGTAGCCAAATGCCTCGT | >hg38_21_+_8433172_8446622_RNA45SN1_45S_50ntFlanks | 11729 | 11764 | 36 | TAACTATGACTCTCTTAAGGTAGCCAAATGCCTCGT | 1026.5 | -9.6 | 1.5 | -6.2 | 5.6E-10 | 1.1E-09 | BT20/MB468 | NUC |
| >hg38_21_+_8433172_8446622_RNA45SN1_45S_50ntFlanks@4220.4241.22\|ACGAAAAATAACAATACAGGAC | >hg38_21_+_8433172_8446622_RNA45SN1_45S_50ntFlanks | 4220 | 4241 | 22 | ACGAAAAATAACAATACAGGAC | 148.6 | -10.3 | 1.6 | -6.6 | 4.4E-11 | 1.1E-09 | BT20/MB468 | MP |
| >hg38_21_+_8433172_8446622_RNA45SN1_45S_50ntFlanks@11641.11671.31\|CGGCGGGTGTTGACGCGATGTGATTTCTGCC | >hg38_21_+_8433172_8446622_RNA45SN1_45S_50ntFlanks | 11641 | 11671 | 31 | CGGCGGGTGTTGACGCGATGTGATTTCTGCC | 280.5 | -11.2 | 1.7 | -6.6 | 4.4E-11 | 1.1E-09 | BT20/MB468 | MP |
| >hg38_21_+_8433172_8446622_RNA45SN1_45S_50ntFlanks@12694.12711.18\|AACGATACGGCAGCGCCG | >hg38_21_+_8433172_8446622_RNA45SN1_45S_50ntFlanks | 12694 | 12711 | 18 | AACGATACGGCAGCGCCG | 138.4 | -11.0 | 1.8 | -6.2 | 4.6E-10 | 1.1E-09 | MB231/MB468 | MP |
| >hg38_21_+_8433172_8446622_RNA45SN1_45S_50ntFlanks@8194.8230.37\|CCCGTGGACGGTGTGAGGCCGGTAGCGGCCCCCGGCG | >hg38_21_+_8433172_8446622_RNA45SN1_45S_50ntFlanks | 8194 | 8230 | 37 | CCCGTGGACGGTGTGAGGCCGGTAGCGGCCCCCGGCG | 68.1 | -10.0 | 1.6 | -6.2 | 4.6E-10 | 1.1E-09 | MB231/MB468 | MP |
| >hg38_21_+_8433172_8446622_RNA45SN1_45S_50ntFlanks@11890.11912.23\|CTTGACTCTAGTCTGGCACGGTG | >hg38_21_+_8433172_8446622_RNA45SN1_45S_50ntFlanks | 11890 | 11912 | 23 | CTTGACTCTAGTCTGGCACGGTG | 1051.1 | -5.0 | 0.8 | -6.6 | 4.4E-11 | 1.2E-09 | BT20/MB468 | MP |
| >hg38_21_+_8433172_8446622_RNA45SN1_45S_50ntFlanks@12438.12455.18\|ATTGTGAAGCAGAATTCA | >hg38_21_+_8433172_8446622_RNA45SN1_45S_50ntFlanks | 12438 | 12455 | 18 | ATTGTGAAGCAGAATTCA | 5278.3 | -10.2 | 1.6 | -6.6 | 4.4E-11 | 1.2E-09 | BT20/MB468 | MP |
| >hg38_21_+_8433172_8446622_RNA45SN1_45S_50ntFlanks@10794.10817.24\|CTGGCATGTTGGAACAATGTAGGT | >hg38_21_+_8433172_8446622_RNA45SN1_45S_50ntFlanks | 10794 | 10817 | 24 | CTGGCATGTTGGAACAATGTAGGT | 453.9 | -8.8 | 1.4 | -6.2 | 5.6E-10 | 1.2E-09 | BT20/MB468 | NUC |
| >hg38_21_+_8433172_8446622_RNA45SN1_45S_50ntFlanks@12959.12982.24\|TCTGGGTCGGGGTTTCGTACGTAG | >hg38_21_+_8433172_8446622_RNA45SN1_45S_50ntFlanks | 12959 | 12982 | 24 | TCTGGGTCGGGGTTTCGTACGTAG | 657.8 | -7.7 | 1.2 | -6.2 | 4.9E-10 | 1.2E-09 | MB231/MB468 | NUC |
| >MI0000737\|hsa-mir-200a&WithFlank&1\|+\|1167857\|1167958@23.43.21&[MIMAT0001620&hsa-miR-200a-5p&offsets\|+1\|0;m-154&1\|+\|1167878\|1167899&offsets\|+1\|0]\|ATCTTACCGGACAGTGCTGGA | >MI0000737\|hsa-mir-200a&WithFlank&1\|+\|1167857\|1167958 | 23 | 43 | 21 | ATCTTACCGGACAGTGCTGGA | 163.0 | -10.4 | 1.6 | -6.6 | 4.5E-11 | 1.2E-09 | BT20/MB468 | MP |
| >hg38_21_+_8433172_8446622_RNA45SN1_45S_50ntFlanks@10385.10406.22\|CATGTGAACAGCAGTTGAACAT | >hg38_21_+_8433172_8446622_RNA45SN1_45S_50ntFlanks | 10385 | 10406 | 22 | CATGTGAACAGCAGTTGAACAT | 108.5 | -10.6 | 1.7 | -6.2 | 4.7E-10 | 1.2E-09 | MB231/MB468 | MP |
| >hg38_21_+_8433172_8446622_RNA45SN1_45S_50ntFlanks@12412.12444.33\|ATCCTTCGATGTCGGCTCTTCCTATCATTGTGA | >hg38_21_+_8433172_8446622_RNA45SN1_45S_50ntFlanks | 12412 | 12444 | 33 | ATCCTTCGATGTCGGCTCTTCCTATCATTGTGA | 102.0 | -8.8 | 1.4 | -6.3 | 3.7E-10 | 1.2E-09 | BT20/MB468 | MITO |
| >hg38_21_+_8433172_8446622_RNA45SN1_45S_50ntFlanks@3769.3792.24\|CGCACGGCCGGTACAGTGAAACTG | >hg38_21_+_8433172_8446622_RNA45SN1_45S_50ntFlanks | 3769 | 3792 | 24 | CGCACGGCCGGTACAGTGAAACTG | 1085.0 | -9.0 | 1.4 | -6.2 | 5.4E-10 | 1.2E-09 | MB231/MB468 | MITO |
| >hg38_21_+_8433172_8446622_RNA45SN1_45S_50ntFlanks@6722.6740.19\|ATTGCAGGACACATTGATC | >hg38_21_+_8433172_8446622_RNA45SN1_45S_50ntFlanks | 6722 | 6740 | 19 | ATTGCAGGACACATTGATC | 744.1 | -6.6 | 1.1 | -6.2 | 4.7E-10 | 1.2E-09 | MB231/MB468 | MP |
| >hg38_21_+_8433172_8446622_RNA45SN1_45S_50ntFlanks@5484.5522.39\|GAGCGCTGAGAAGACGGTCGAACTTGACTATCTAGAGGA | >hg38_21_+_8433172_8446622_RNA45SN1_45S_50ntFlanks | 5484 | 5522 | 39 | GAGCGCTGAGAAGACGGTCGAACTTGACTATCTAGAGGA | 113.8 | -10.7 | 1.7 | -6.2 | 4.7E-10 | 1.2E-09 | MB231/MB468 | MP |
| >am_MI0000809\|hsa-mir-151a&WithFlank&8\|-\|140732558\|140732659@17.37.21&[MIMAT0004697&hsa-miR-151a-5p&offsets\|0\|0;m-75&8\|-\|140732623\|140732643&offsets\|0\|0]\|TCGAGGAGCTCACAGTCTAGT | >am_MI0000809\|hsa-mir-151a&WithFlank&8\|-\|140732558\|140732659 | 17 | 37 | 21 | TCGAGGAGCTCACAGTCTAGT | 906.0 | 6.6 | 1.1 | 6.2 | 5.4E-10 | 1.2E-09 | MB231/MB468 | MITO |
| >hg38_21_+_8433172_8446622_RNA45SN1_45S_50ntFlanks@8841.8859.19\|GTGGCGGCGGCGGCGGCGG | >hg38_21_+_8433172_8446622_RNA45SN1_45S_50ntFlanks | 8841 | 8859 | 19 | GTGGCGGCGGCGGCGGCGG | 86.6 | -10.3 | 1.7 | -6.2 | 4.7E-10 | 1.2E-09 | MB231/MB468 | MP |
| >hg38_21_+_8433172_8446622_RNA45SN1_45S_50ntFlanks@4857.4876.20\|CTTTGGGTTCCGGGGGGAGT | >hg38_21_+_8433172_8446622_RNA45SN1_45S_50ntFlanks | 4857 | 4876 | 20 | CTTTGGGTTCCGGGGGGAGT | 370.5 | 9.6 | 1.5 | 6.2 | 4.9E-10 | 1.2E-09 | MB231/MB468 | CYTO |
| >trna129_MetCAT_6_-_27870271_27870342@31.51.21,trna142_MetCAT_6_-_27560600_27560671@31.51.21,trna150_MetCAT_6_-_27300764_27300835@31.51.21,trna169_MetCAT_6_-_26330529_26330600@31.51.21,trna171_MetCAT_6_-_26313352_26313423@31.51.21,trna20_MetCAT_17_-_80452597_80452668@31.51.21,trna2_MetCAT_6_+_26286754_26286825@31.51.21,trna32_MetCAT_1_+_153643726_153643797@31.51.21,trna61_MetCAT_6_+_27745664_27745735@31.51.21\|CCCATAACCCAGAGGTCGATG | >trna129_MetCAT_6_-_27870271_27870342 | 31 | 51 | 21 | CCCATAACCCAGAGGTCGATG | 1323.4 | -9.2 | 1.4 | -6.6 | 4.5E-11 | 1.2E-09 | BT20/MB468 | MP |
| >trna111_HisGTG_1_-_147774845_147774916@-1A.33.34,trna118_HisGTG_1_-_145396881_145396952@-1A.33.34,trna16_HisGTG_1_+_146544773_146544844@-1A.33.34,trna1_HisGTG_15_+_45493349_45493420@-1A.33.34,trna21_HisGTG_1_+_147753471_147753542@-1A.33.34,trna33_HisGTG_6_+_27125906_27125977@-1A.33.34,trna7_HisGTG_9_-_14433938_14434009@-1A.33.34,trna8_HisGTG_15_-_45492611_45492682@-1A.33.34,trna9_HisGTG_15_-_45490804_45490875@-1A.33.34\|AGCCGTGATCGTATAGTGGTTAGTACTCTGCGTT | >trna111_HisGTG_1_-_147774845_147774916 | -1A | 33 | 34 | AGCCGTGATCGTATAGTGGTTAGTACTCTGCGTT | 133.2 | 10.3 | 1.7 | 6.2 | 4.7E-10 | 1.2E-09 | MB231/MB468 | MP |
| >hg38_21_+_8433172_8446622_RNA45SN1_45S_50ntFlanks@11934.11952.19\|ATAAGTGGGAGGCCCCCGG | >hg38_21_+_8433172_8446622_RNA45SN1_45S_50ntFlanks | 11934 | 11952 | 19 | ATAAGTGGGAGGCCCCCGG | 77.9 | -10.1 | 1.6 | -6.2 | 4.7E-10 | 1.2E-09 | MB231/MB468 | MP |
| >hg38_21_+_8433172_8446622_RNA45SN1_45S_50ntFlanks@12324.12348.25\|TTGGGTTTTAAGCAGGAGGTGTCAG | >hg38_21_+_8433172_8446622_RNA45SN1_45S_50ntFlanks | 12324 | 12348 | 25 | TTGGGTTTTAAGCAGGAGGTGTCAG | 786.2 | -7.6 | 1.2 | -6.2 | 4.3E-10 | 1.2E-09 | BT20/MB468 | CYTO |
| >hg38_21_+_8433172_8446622_RNA45SN1_45S_50ntFlanks@6667.6699.33\|ATCACTCGGCTCGTGCGTCGATGAAGAACGCAG | >hg38_21_+_8433172_8446622_RNA45SN1_45S_50ntFlanks | 6667 | 6699 | 33 | ATCACTCGGCTCGTGCGTCGATGAAGAACGCAG | 231.1 | -10.9 | 1.7 | -6.6 | 4.5E-11 | 1.2E-09 | BT20/MB468 | MP |
| >MI0000086\|hsa-mir-28&WithFlank&3\|+\|188688775\|188688872@60.82.23&[MIMAT0004502&hsa-miR-28-3p&offsets\|0\|+1;m-28&3\|+\|188688834\|188688855&offsets\|0\|+1]\|CACTAGATTGTGAGCTCCTGGAG | >MI0000086\|hsa-mir-28&WithFlank&3\|+\|188688775\|188688872 | 60 | 82 | 23 | CACTAGATTGTGAGCTCCTGGAG | 161.8 | 10.6 | 1.7 | 6.2 | 5.4E-10 | 1.2E-09 | MB231/MB468 | MITO |
| >MI0001445\|hsa-mir-423&WithFlank&17\|+\|30117073\|30117178@58.80.23&[MIMAT0001340&hsa-miR-423-3p&offsets\|-1\|-1;m-100&17\|+\|30117131\|30117153&offsets\|-1\|-1]\|AAGCTCGGTCTGAGGCCCCTCAG | >MI0001445\|hsa-mir-423&WithFlank&17\|+\|30117073\|30117178 | 58 | 80 | 23 | AAGCTCGGTCTGAGGCCCCTCAG | 433.0 | 8.2 | 1.3 | 6.2 | 4.3E-10 | 1.2E-09 | BT20/MB468 | CYTO |
| >hg38_21_+_8433172_8446622_RNA45SN1_45S_50ntFlanks@7979.7996.18\|ACCTCAGATCAGACGTGG | >hg38_21_+_8433172_8446622_RNA45SN1_45S_50ntFlanks | 7979 | 7996 | 18 | ACCTCAGATCAGACGTGG | 205.1 | -7.7 | 1.2 | -6.2 | 4.3E-10 | 1.2E-09 | BT20/MB468 | CYTO |
| >hg38_21_+_8433172_8446622_RNA45SN1_45S_50ntFlanks@12907.12934.28\|CACCGCACGTTCGTGGGGAACCTGGCGC | >hg38_21_+_8433172_8446622_RNA45SN1_45S_50ntFlanks | 12907 | 12934 | 28 | CACCGCACGTTCGTGGGGAACCTGGCGC | 74.8 | -10.1 | 1.6 | -6.2 | 4.7E-10 | 1.2E-09 | MB231/MB468 | MP |
| >hg38_21_+_8433172_8446622_RNA45SN1_45S_50ntFlanks@8198.8216.19\|TGGACGGTGTGAGGCCGGT | >hg38_21_+_8433172_8446622_RNA45SN1_45S_50ntFlanks | 8198 | 8216 | 19 | TGGACGGTGTGAGGCCGGT | 174.9 | -10.5 | 1.6 | -6.6 | 4.5E-11 | 1.2E-09 | BT20/MB468 | MP |
| >hg38_21_+_8433172_8446622_RNA45SN1_45S_50ntFlanks@12624.12661.38\|CCAATGGGGCGAAGCTACCATCTGTGGGATTATGACTG | >hg38_21_+_8433172_8446622_RNA45SN1_45S_50ntFlanks | 12624 | 12661 | 38 | CCAATGGGGCGAAGCTACCATCTGTGGGATTATGACTG | 5551.2 | -4.9 | 0.8 | -6.3 | 3.7E-10 | 1.2E-09 | BT20/MB468 | MITO |
| >MI0000254\|hsa-mir-30c-2&WithFlank&6\|-\|71376954\|71377037@53.73.21(+1C)&[MIMAT0004550&hsa-miR-30c-2-3p&offsets\|0\|-1(+1C);m-206&6\|-\|71376964\|71376985&offsets\|0\|-1(+1C)]\|CTGGGAGAAGGCTGTTTACTCC | >MI0000254\|hsa-mir-30c-2&WithFlank&6\|-\|71376954\|71377037 | 53 | 73 | 21(+1C) | CTGGGAGAAGGCTGTTTACTCC | 323.1 | 6.6 | 1.1 | 6.2 | 4.9E-10 | 1.2E-09 | MB231/MB468 | CYTO |
| >hg38_21_+_8433172_8446622_RNA45SN1_45S_50ntFlanks@12654.12671.18\|TATGACTGAACGCCTCTA | >hg38_21_+_8433172_8446622_RNA45SN1_45S_50ntFlanks | 12654 | 12671 | 18 | TATGACTGAACGCCTCTA | 117.8 | -6.5 | 1.0 | -6.2 | 4.3E-10 | 1.2E-09 | BT20/MB468 | CYTO |
| >hg38_21_+_8433172_8446622_RNA45SN1_45S_50ntFlanks@4760.4778.19\|TCAGATACCGTCGTAGTTC | >hg38_21_+_8433172_8446622_RNA45SN1_45S_50ntFlanks | 4760 | 4778 | 19 | TCAGATACCGTCGTAGTTC | 549.1 | -6.3 | 1.0 | -6.2 | 5.0E-10 | 1.2E-09 | MB231/MB468 | NUC |
| >hg38_21_+_8433172_8446622_RNA45SN1_45S_50ntFlanks@475.511.37\|CTTCGTGATCGATGTGGTGACGTCGTGCTCTCCCGGG | >hg38_21_+_8433172_8446622_RNA45SN1_45S_50ntFlanks | 475 | 511 | 37 | CTTCGTGATCGATGTGGTGACGTCGTGCTCTCCCGGG | 1461.9 | 8.9 | 1.4 | 6.2 | 4.3E-10 | 1.2E-09 | BT20/MB468 | CYTO |
| >hg38_21_+_8433172_8446622_RNA45SN1_45S_50ntFlanks@12932.12956.25\|CGCTAAACCATTCGTAGACGACCTG | >hg38_21_+_8433172_8446622_RNA45SN1_45S_50ntFlanks | 12932 | 12956 | 25 | CGCTAAACCATTCGTAGACGACCTG | 1345.4 | -8.3 | 1.3 | -6.2 | 5.5E-10 | 1.2E-09 | MB231/MB468 | MITO |
| >hg38_21_+_8433172_8446622_RNA45SN1_45S_50ntFlanks@10128.10153.26\|CGGCGGCGGCGGCGGCGGCGGGGGTG | >hg38_21_+_8433172_8446622_RNA45SN1_45S_50ntFlanks | 10128 | 10153 | 26 | CGGCGGCGGCGGCGGCGGCGGGGGTG | 210.4 | -10.8 | 1.6 | -6.6 | 4.6E-11 | 1.2E-09 | BT20/MB468 | MP |
| >hg38_21_+_8433172_8446622_RNA45SN1_45S_50ntFlanks@6738.6756.19\|ATCATCGACACTTCGAACG | >hg38_21_+_8433172_8446622_RNA45SN1_45S_50ntFlanks | 6738 | 6756 | 19 | ATCATCGACACTTCGAACG | 736.0 | -7.9 | 1.3 | -6.2 | 4.9E-10 | 1.2E-09 | MB231/MB468 | CYTO |
| >hg38_1_-_228634819_228635039_RNA5S12_5S_50ntFlanks@135.170.36\|GGGAGACCGCCTGGGAATACCGGGTGCTGTAGGCTT | >hg38_1_-_228634819_228635039_RNA5S12_5S_50ntFlanks | 135 | 170 | 36 | GGGAGACCGCCTGGGAATACCGGGTGCTGTAGGCTT | 20660.5 | 4.2 | 0.7 | 6.2 | 5.0E-10 | 1.2E-09 | MB231/MB468 | NUC |
| >hg38_21_+_8433172_8446622_RNA45SN1_45S_50ntFlanks@6719.6749.31\|TGAATTGCAGGACACATTGATCATCGACACT | >hg38_21_+_8433172_8446622_RNA45SN1_45S_50ntFlanks | 6719 | 6749 | 31 | TGAATTGCAGGACACATTGATCATCGACACT | 606.2 | -6.2 | 1.0 | -6.2 | 5.5E-10 | 1.2E-09 | MB231/MB468 | MITO |
| >hg38_21_+_8433172_8446622_RNA45SN1_45S_50ntFlanks@10684.10710.27\|CGGCGGCGTCCGGTGAGCTCTCGCTGG | >hg38_21_+_8433172_8446622_RNA45SN1_45S_50ntFlanks | 10684 | 10710 | 27 | CGGCGGCGTCCGGTGAGCTCTCGCTGG | 25381.1 | -9.6 | 1.5 | -6.2 | 5.8E-10 | 1.2E-09 | BT20/MB468 | NUC |
| >hg38_21_+_8433172_8446622_RNA45SN1_45S_50ntFlanks@8324.8355.32\|CCGATAGTCAACAAGTACCGTAAGGGAAAGTT | >hg38_21_+_8433172_8446622_RNA45SN1_45S_50ntFlanks | 8324 | 8355 | 32 | CCGATAGTCAACAAGTACCGTAAGGGAAAGTT | 66.0 | -9.9 | 1.6 | -6.2 | 4.8E-10 | 1.2E-09 | MB231/MB468 | MP |
| >hg38_21_+_8433172_8446622_RNA45SN1_45S_50ntFlanks@9567.9585.19\|ACGTGCAAATCGGTCGTCC | >hg38_21_+_8433172_8446622_RNA45SN1_45S_50ntFlanks | 9567 | 9585 | 19 | ACGTGCAAATCGGTCGTCC | 74.6 | -10.1 | 1.6 | -6.2 | 4.8E-10 | 1.2E-09 | MB231/MB468 | MP |
| >trna4_ArgTCG_15_+_89878304_89878376@31.53.23\|ACTTCGGATCAGAAGATTGCAGG | >trna4_ArgTCG_15_+_89878304_89878376 | 31 | 53 | 23 | ACTTCGGATCAGAAGATTGCAGG | 172.3 | -11.3 | 1.8 | -6.2 | 4.8E-10 | 1.2E-09 | MB231/MB468 | MP |

<table>
<tr><td>>hg38_21_+_8433172_8446622_RNA45SN1_45S_50ntFlanks@11561.11586.26|GCGCCTAGCAGCCGACTTAGAACTGG</td><td>>hg38_21_+_8433172_8446622_RNA45SN1_45S_50ntFlanks</td><td>11561</td><td>11586</td><td>26</td><td>GCGCCTAGCAGCCGACTTAGAACTGG</td><td>2551.0</td><td>-7.1</td><td>1.1</td><td>-6.2</td><td>5.5E-10</td><td>1.2E-09</td><td>MB231/MB468</td><td>MITO</td></tr>
<tr><td>>hg38_21_+_8433172_8446622_RNA45SN1_45S_50ntFlanks@9738.9759.22|GAAACGATCTCAACCTATTCTC</td><td>>hg38_21_+_8433172_8446622_RNA45SN1_45S_50ntFlanks</td><td>9738</td><td>9759</td><td>22</td><td>GAAACGATCTCAACCTATTCTC</td><td>1617.8</td><td>-7.8</td><td>1.3</td><td>-6.2</td><td>5.8E-10</td><td>1.2E-09</td><td>BT20/MB468</td><td>NUC</td></tr>
<tr><td>>MI0000063|hsa-let-7b&WithFlank&22|+|46113680|46113774@12.33.22(+2A)&[MIMAT0000063&hsa-let-7b-5p&offsets|0|0(+2A);m-14&22|+|46113691|46113712&offsets|0|0(+2A)]|TGAGGTAGTAGGTTGTGTGGTTAA</td><td>>MI0000063|hsa-let-7b&WithFlank&22|+|46113680|46113774</td><td>12</td><td>33</td><td>22(+2A)</td><td>TGAGGTAGTAGGTTGTGTGGTTAA</td><td>506.9</td><td>8.4</td><td>1.4</td><td>6.2</td><td>4.4E-10</td><td>1.2E-09</td><td>BT20/MB468</td><td>CYTO</td></tr>
<tr><td>>hg38_21_+_8433172_8446622_RNA45SN1_45S_50ntFlanks@4297.4319.23|CATTGGAGGGCAAGTCTGGTGCC</td><td>>hg38_21_+_8433172_8446622_RNA45SN1_45S_50ntFlanks</td><td>4297</td><td>4319</td><td>23</td><td>CATTGGAGGGCAAGTCTGGTGCC</td><td>381.9</td><td>-8.3</td><td>1.3</td><td>-6.2</td><td>5.5E-10</td><td>1.2E-09</td><td>MB231/MB468</td><td>MITO</td></tr>
<tr><td>>hg38_21_+_8433172_8446622_RNA45SN1_45S_50ntFlanks@10238.10255.18|CGCTACGCCGCGACGAGT</td><td>>hg38_21_+_8433172_8446622_RNA45SN1_45S_50ntFlanks</td><td>10238</td><td>10255</td><td>18</td><td>CGCTACGCCGCGACGAGT</td><td>74.7</td><td>7.1</td><td>1.1</td><td>6.2</td><td>5.8E-10</td><td>1.2E-09</td><td>BT20/MB468</td><td>NUC</td></tr>
<tr><td>>MI0003560|hsa-mir-92b&WithFlank&1|+|155195171|155195278@67.86.20(+1U)&[MIMAT0003218&hsa-miR-92b-3p&offsets|0|-2(+1U);m-115&1|+|155195237|155195258&offsets|0|-2(+1U)]|TATTGCACTCGTCCCGGCCTT</td><td>>MI0003560|hsa-mir-92b&WithFlank&1|+|155195171|155195278</td><td>67</td><td>86</td><td>20(+1U)</td><td>TATTGCACTCGTCCCGGCCTT</td><td>250.3</td><td>6.4</td><td>1.0</td><td>6.2</td><td>5.0E-10</td><td>1.2E-09</td><td>MB231/MB468</td><td>NUC</td></tr>
<tr><td>>hg38_21_+_8433172_8446622_RNA45SN1_45S_50ntFlanks@3899.3927.29|CCTTCGCGGGGGGGATGCGTGCATTTATC</td><td>>hg38_21_+_8433172_8446622_RNA45SN1_45S_50ntFlanks</td><td>3899</td><td>3927</td><td>29</td><td>CCTTCGCGGGGGGGATGCGTGCATTTATC</td><td>383.1</td><td>-8.3</td><td>1.3</td><td>-6.2</td><td>5.1E-10</td><td>1.2E-09</td><td>MB231/MB468</td><td>NUC</td></tr>
<tr><td>>hg38_21_+_8433172_8446622_RNA45SN1_45S_50ntFlanks@6747.6770.24|ACTTCGAACGCACTTGCGGCCCCG</td><td>>hg38_21_+_8433172_8446622_RNA45SN1_45S_50ntFlanks</td><td>6747</td><td>6770</td><td>24</td><td>ACTTCGAACGCACTTGCGGCCCCG</td><td>307.2</td><td>-7.7</td><td>1.2</td><td>-6.2</td><td>5.0E-10</td><td>1.2E-09</td><td>MB231/MB468</td><td>NUC</td></tr>
<tr><td>>MI0000103|hsa-mir-101-1&WithFlank&1|-|65058428|65058514@52.72.21;MI0000739|hsa-mir-101-2&WithFlank&9|+|4850291|4850381@54.74.21&[MIMAT0000099&hsa-miR-101-3p&offsets|-1|-1;m-17&1|-|65058443|65058462&offsets|-1|0];[MIMAT0000099_1&hsa-miR-101-3p&offsets|-1|-1;m-16&9|+|4850345|4850364&offsets|-1|0]|GTACAGTACTGTGATAACTGA</td><td>>MI0000103|hsa-mir-101-1&WithFlank&1|-|65058428|65058514</td><td>52</td><td>72</td><td>21</td><td>GTACAGTACTGTGATAACTGA</td><td>475.1</td><td>5.2</td><td>0.8</td><td>6.2</td><td>5.1E-10</td><td>1.2E-09</td><td>MB231/MB468</td><td>NUC</td></tr>
<tr><td>>hg38_21_+_8433172_8446622_RNA45SN1_45S_50ntFlanks@12420.12441.22|ATGTCGGCTCTTCCTATCATTG</td><td>>hg38_21_+_8433172_8446622_RNA45SN1_45S_50ntFlanks</td><td>12420</td><td>12441</td><td>22</td><td>ATGTCGGCTCTTCCTATCATTG</td><td>7352.5</td><td>-4.5</td><td>0.7</td><td>-6.2</td><td>5.1E-10</td><td>1.2E-09</td><td>MB231/MB468</td><td>NUC</td></tr>
<tr><td>>hg38_21_+_8433172_8446622_RNA45SN1_45S_50ntFlanks@11583.11600.18|CTGGTGCGGACCAGGGGA</td><td>>hg38_21_+_8433172_8446622_RNA45SN1_45S_50ntFlanks</td><td>11583</td><td>11600</td><td>18</td><td>CTGGTGCGGACCAGGGGA</td><td>540.7</td><td>-8.4</td><td>1.3</td><td>-6.2</td><td>4.8E-10</td><td>1.2E-09</td><td>MB231/MB468</td><td>MP</td></tr>
<tr><td>>hg38_21_+_8433172_8446622_RNA45SN1_45S_50ntFlanks@10471.10497.27|CCCTCGGCCGATCGAAAGGGAGTCGGG</td><td>>hg38_21_+_8433172_8446622_RNA45SN1_45S_50ntFlanks</td><td>10471</td><td>10497</td><td>27</td><td>CCCTCGGCCGATCGAAAGGGAGTCGGG</td><td>102.8</td><td>-10.5</td><td>1.7</td><td>-6.2</td><td>4.8E-10</td><td>1.2E-09</td><td>MB231/MB468</td><td>MP</td></tr>
<tr><td>>hg38_21_+_8433172_8446622_RNA45SN1_45S_50ntFlanks@9123.9152.30|CGCGCGCGGGTCGGGGGGCGGGGCGGACTG</td><td>>hg38_21_+_8433172_8446622_RNA45SN1_45S_50ntFlanks</td><td>9123</td><td>9152</td><td>30</td><td>CGCGCGCGGGTCGGGGGGCGGGGCGGACTG</td><td>111.1</td><td>-10.7</td><td>1.7</td><td>-6.2</td><td>4.8E-10</td><td>1.2E-09</td><td>MB231/MB468</td><td>MP</td></tr>
<tr><td>>hg38_21_+_8433172_8446622_RNA45SN1_45S_50ntFlanks@12526.12559.34|TTTTACCCTACTGATGATGTGTTGTTGCCATGGT</td><td>>hg38_21_+_8433172_8446622_RNA45SN1_45S_50ntFlanks</td><td>12526</td><td>12559</td><td>34</td><td>TTTTACCCTACTGATGATGTGTTGTTGCCATGGT</td><td>69.8</td><td>-10.0</td><td>1.6</td><td>-6.2</td><td>4.8E-10</td><td>1.2E-09</td><td>MB231/MB468</td><td>MP</td></tr>
<tr><td>>hg38_21_+_8433172_8446622_RNA45SN1_45S_50ntFlanks@7980.7998.19|CCTCAGATCAGACGTGGCG</td><td>>hg38_21_+_8433172_8446622_RNA45SN1_45S_50ntFlanks</td><td>7980</td><td>7998</td><td>19</td><td>CCTCAGATCAGACGTGGCG</td><td>181.7</td><td>-5.0</td><td>0.8</td><td>-6.2</td><td>5.0E-10</td><td>1.2E-09</td><td>MB231/MB468</td><td>CYTO</td></tr>
<tr><td>>hg38_21_+_8433172_8446622_RNA45SN1_45S_50ntFlanks@12717.12747.31|CCTCGGTTGGCCTCGGATAGCCGGTCCCCCG</td><td>>hg38_21_+_8433172_8446622_RNA45SN1_45S_50ntFlanks</td><td>12717</td><td>12747</td><td>31</td><td>CCTCGGTTGGCCTCGGATAGCCGGTCCCCCG</td><td>181.3</td><td>-10.5</td><td>1.7</td><td>-6.2</td><td>5.1E-10</td><td>1.2E-09</td><td>MB231/MB468</td><td>NUC</td></tr>
<tr><td>>MI0000650|hsa-mir-200c&WithFlank&12|+|6963693|6963772@50.71.22(+1C)&[MIMAT0000617&hsa-miR-200c-3p&offsets|0|-1(+1C);m-54&12|+|6963742|6963764&offsets|0|-1(+1C)]|TAATACTGCCGGGTAATGATGGC</td><td>>MI0000650|hsa-mir-200c&WithFlank&12|+|6963693|6963772</td><td>50</td><td>71</td><td>22(+1C)</td><td>TAATACTGCCGGGTAATGATGGC</td><td>141.3</td><td>5.2</td><td>0.8</td><td>6.2</td><td>5.9E-10</td><td>1.2E-09</td><td>BT20/MB468</td><td>NUC</td></tr>
<tr><td>>hg38_21_+_8433172_8446622_RNA45SN1_45S_50ntFlanks@12416.12443.28|TTCGATGTCGGCTCTTCCTATCATTGTG</td><td>>hg38_21_+_8433172_8446622_RNA45SN1_45S_50ntFlanks</td><td>12416</td><td>12443</td><td>28</td><td>TTCGATGTCGGCTCTTCCTATCATTGTG</td><td>967.3</td><td>-3.8</td><td>0.6</td><td>-6.2</td><td>5.9E-10</td><td>1.2E-09</td><td>BT20/MB468</td><td>NUC</td></tr>
<tr><td>>hg38_21_+_8433172_8446622_RNA45SN1_45S_50ntFlanks@11315.11343.29|CCACCCCACGTCTCGTCGCGCGCGCGTCC</td><td>>hg38_21_+_8433172_8446622_RNA45SN1_45S_50ntFlanks</td><td>11315</td><td>11343</td><td>29</td><td>CCACCCCACGTCTCGTCGCGCGCGCGTCC</td><td>1006.2</td><td>-8.8</td><td>1.4</td><td>-6.2</td><td>5.1E-10</td><td>1.2E-09</td><td>MB231/MB468</td><td>NUC</td></tr>
<tr><td>>hg38_21_+_8433172_8446622_RNA45SN1_45S_50ntFlanks@10473.10490.18|CTCGGCCGATCGAAAGGG</td><td>>hg38_21_+_8433172_8446622_RNA45SN1_45S_50ntFlanks</td><td>10473</td><td>10490</td><td>18</td><td>CTCGGCCGATCGAAAGGG</td><td>373.4</td><td>-8.5</td><td>1.4</td><td>-6.2</td><td>5.9E-10</td><td>1.2E-09</td><td>BT20/MB468</td><td>NUC</td></tr>
<tr><td>>hg38_21_+_8433172_8446622_RNA45SN1_45S_50ntFlanks@9202.9224.23|TCTCGGGGCCACGCGCGCGTCCC</td><td>>hg38_21_+_8433172_8446622_RNA45SN1_45S_50ntFlanks</td><td>9202</td><td>9224</td><td>23</td><td>TCTCGGGGCCACGCGCGCGTCCC</td><td>227.4</td><td>-10.9</td><td>1.7</td><td>-6.6</td><td>4.7E-11</td><td>1.2E-09</td><td>BT20/MB468</td><td>MP</td></tr>
<tr><td>>hg38_21_+_8433172_8446622_RNA45SN1_45S_50ntFlanks@11874.11898.25|AGAAGACCCTGTTGAGCTTGACTCT</td><td>>hg38_21_+_8433172_8446622_RNA45SN1_45S_50ntFlanks</td><td>11874</td><td>11898</td><td>25</td><td>AGAAGACCCTGTTGAGCTTGACTCT</td><td>2458.7</td><td>-4.8</td><td>0.8</td><td>-6.2</td><td>5.1E-10</td><td>1.2E-09</td><td>MB231/MB468</td><td>NUC</td></tr>
<tr><td>>hg38_21_+_8433172_8446622_RNA45SN1_45S_50ntFlanks@4661.4696.36|AGGTGAAATTCTTGGACCGGCGCAAGACGGACCAGA</td><td>>hg38_21_+_8433172_8446622_RNA45SN1_45S_50ntFlanks</td><td>4661</td><td>4696</td><td>36</td><td>AGGTGAAATTCTTGGACCGGCGCAAGACGGACCAGA</td><td>503.1</td><td>10.4</td><td>1.7</td><td>6.2</td><td>4.5E-10</td><td>1.2E-09</td><td>BT20/MB468</td><td>CYTO</td></tr>
<tr><td>>MI0000542|hsa-mir-320a&WithFlank&8|-|22244960|22245043@48.71.24(+1C)&[MIMAT0000510&hsa-miR-320a-3p&offsets|0|+2(+1C);m-60&8|-|22244975|22244996&offsets|0|+2(+1C)]|AAAAGCTGGGTTGAGAGGGCGAAAC</td><td>>MI0000542|hsa-mir-320a&WithFlank&8|-|22244960|22245043</td><td>48</td><td>71</td><td>24(+1C)</td><td>AAAAGCTGGGTTGAGAGGGCGAAAC</td><td>163.0</td><td>11.1</td><td>1.7</td><td>6.6</td><td>4.7E-11</td><td>1.2E-09</td><td>BT20/MB468</td><td>MP</td></tr>
<tr><td>>hg38_21_+_8433172_8446622_RNA45SN1_45S_50ntFlanks@10326.10352.27|TTGGTGGTAGTAGCAAATATTCAAACG</td><td>>hg38_21_+_8433172_8446622_RNA45SN1_45S_50ntFlanks</td><td>10326</td><td>10352</td><td>27</td><td>TTGGTGGTAGTAGCAAATATTCAAACG</td><td>160.3</td><td>11.1</td><td>1.7</td><td>6.6</td><td>4.7E-11</td><td>1.2E-09</td><td>BT20/MB468</td><td>MP</td></tr>
<tr><td>>hg38_21_+_8433172_8446622_RNA45SN1_45S_50ntFlanks@4738.4761.24|AAAGTCGGAGGTTCGAAGACGATC</td><td>>hg38_21_+_8433172_8446622_RNA45SN1_45S_50ntFlanks</td><td>4738</td><td>4761</td><td>24</td><td>AAAGTCGGAGGTTCGAAGACGATC</td><td>1394.0</td><td>4.1</td><td>0.7</td><td>6.2</td><td>5.1E-10</td><td>1.2E-09</td><td>MB231/MB468</td><td>CYTO</td></tr>
<tr><td>>hg38_21_+_8433172_8446622_RNA45SN1_45S_50ntFlanks@5124.5145.22|GAGCGGTCGGCGTCCCCCAACT</td><td>>hg38_21_+_8433172_8446622_RNA45SN1_45S_50ntFlanks</td><td>5124</td><td>5145</td><td>22</td><td>GAGCGGTCGGCGTCCCCCAACT</td><td>335.7</td><td>8.4</td><td>1.4</td><td>6.2</td><td>5.1E-10</td><td>1.2E-09</td><td>MB231/MB468</td><td>CYTO</td></tr>
<tr><td>>hg38_21_+_8433172_8446622_RNA45SN1_45S_50ntFlanks@10357.10383.27|CTTTGAAGGCCGAAGTGGAGAAGGGTT</td><td>>hg38_21_+_8433172_8446622_RNA45SN1_45S_50ntFlanks</td><td>10357</td><td>10383</td><td>27</td><td>CTTTGAAGGCCGAAGTGGAGAAGGGTT</td><td>288.3</td><td>-3.0</td><td>0.5</td><td>-6.2</td><td>4.5E-10</td><td>1.2E-09</td><td>BT20/MB468</td><td>CYTO</td></tr>
<tr><td>>hg38_21_+_8433172_8446622_RNA45SN1_45S_50ntFlanks@4367.4394.28|CAGTTAAAAAGCTCGTAGTTGGATCTTG</td><td>>hg38_21_+_8433172_8446622_RNA45SN1_45S_50ntFlanks</td><td>4367</td><td>4394</td><td>28</td><td>CAGTTAAAAAGCTCGTAGTTGGATCTTG</td><td>101.8</td><td>-8.8</td><td>1.4</td><td>-6.3</td><td>3.8E-10</td><td>1.2E-09</td><td>BT20/MB468</td><td>MITO</td></tr>
<tr><td>>hg38_21_+_8433172_8446622_RNA45SN1_45S_50ntFlanks@12033.12052.20|ATCGTTTTTTCACTGACCCG</td><td>>hg38_21_+_8433172_8446622_RNA45SN1_45S_50ntFlanks</td><td>12033</td><td>12052</td><td>20</td><td>ATCGTTTTTTCACTGACCCG</td><td>131.5</td><td>-9.1</td><td>1.5</td><td>-6.3</td><td>3.8E-10</td><td>1.2E-09</td><td>BT20/MB468</td><td>MITO</td></tr>
<tr><td>>MI0000446|hsa-mir-125b-1&WithFlank&11|-|122099751|122099850@21.42.22(+1A);MI0000470|hsa-mir-125b-2&WithFlank&21|+|16590231|16590331@23.44.22(+1A)&[MIMAT0000423&hsa-miR-125b-5p&offsets|0|0(+1A);m-67&11|-|122099809|122099830&offsets|0|0(+1A)];[MIMAT0000423_1&hsa-miR-125b-5p&offsets|0|0(+1A);m-68&21|+|16590253|16590274&offsets|0|0(+1A)]|TCCCTGAGACCCTAACTTGTGAA</td><td>>MI0000446|hsa-mir-125b-1&WithFlank&11|-|122099751|122099850</td><td>21</td><td>42</td><td>22(+1A)</td><td>TCCCTGAGACCCTAACTTGTGAA</td><td>554.2</td><td>6.4</td><td>1.0</td><td>6.2</td><td>4.5E-10</td><td>1.2E-09</td><td>BT20/MB468</td><td>CYTO</td></tr>
<tr><td>>hg38_21_+_8433172_8446622_RNA45SN1_45S_50ntFlanks@10408.10432.25|GGTCAGTCGGTCCTGAGAGATGGGC</td><td>>hg38_21_+_8433172_8446622_RNA45SN1_45S_50ntFlanks</td><td>10408</td><td>10432</td><td>25</td><td>GGTCAGTCGGTCCTGAGAGATGGGC</td><td>604.6</td><td>-7.4</td><td>1.2</td><td>-6.2</td><td>4.5E-10</td><td>1.2E-09</td><td>BT20/MB468</td><td>CYTO</td></tr>
<tr><td>>hg38_21_+_8433172_8446622_RNA45SN1_45S_50ntFlanks@4726.4753.28|TAATCAAGAACGAAAGTCGGAGGTTCGA</td><td>>hg38_21_+_8433172_8446622_RNA45SN1_45S_50ntFlanks</td><td>4726</td><td>4753</td><td>28</td><td>TAATCAAGAACGAAAGTCGGAGGTTCGA</td><td>69.6</td><td>-10.0</td><td>1.6</td><td>-6.2</td><td>4.9E-10</td><td>1.2E-09</td><td>MB231/MB468</td><td>MP</td></tr>
<tr><td>>hg38_21_+_8433172_8446622_RNA45SN1_45S_50ntFlanks@9005.9026.22|CTCCCCCGCGGGGGCGCGCCGG</td><td>>hg38_21_+_8433172_8446622_RNA45SN1_45S_50ntFlanks</td><td>9005</td><td>9026</td><td>22</td><td>CTCCCCCGCGGGGGCGCGCCGG</td><td>88.1</td><td>-10.3</td><td>1.7</td><td>-6.2</td><td>4.9E-10</td><td>1.2E-09</td><td>MB231/MB468</td><td>MP</td></tr>
<tr><td>>hg38_21_+_8433172_8446622_RNA45SN1_45S_50ntFlanks@10349.10383.35|AACGAGAACTTTGAAGGCCGAAGTGGAGAAGGGTT</td><td>>hg38_21_+_8433172_8446622_RNA45SN1_45S_50ntFlanks</td><td>10349</td><td>10383</td><td>35</td><td>AACGAGAACTTTGAAGGCCGAAGTGGAGAAGGGTT</td><td>4022.1</td><td>-2.2</td><td>0.3</td><td>-6.2</td><td>4.5E-10</td><td>1.2E-09</td><td>BT20/MB468</td><td>CYTO</td></tr>
<tr><td>>hg38_21_+_8433172_8446622_RNA45SN1_45S_50ntFlanks@9146.9165.20|CGGACTGTCCCCAGTGCGCC</td><td>>hg38_21_+_8433172_8446622_RNA45SN1_45S_50ntFlanks</td><td>9146</td><td>9165</td><td>20</td><td>CGGACTGTCCCCAGTGCGCC</td><td>480.1</td><td>-8.1</td><td>1.3</td><td>-6.2</td><td>5.0E-10</td><td>1.2E-09</td><td>MB231/MB468</td><td>MP</td></tr>
<tr><td>>MI0000270|hsa-mir-181b-1&WithFlank&1|-|198858867|198858988@42.63.22(+1A);MI0000683|hsa-mir-181b-2&WithFlank&9|+|124693704|124693804@22.43.22(+1A)&[MIMAT0000257&hsa-miR-181b-5p&offsets|0|-1(+1A);m-73&1|-|198858925|198858947&offsets|0|-1(+1A)];[MIMAT0000257_1&hsa-miR-181b-5p&offsets|0|-1(+1A);m-72&9|+|124693725|124693747&offsets|0|-1(+1A)]|AACATTCATTGCTGTCGGTGGGA</td><td>>MI0000270|hsa-mir-181b-1&WithFlank&1|-|198858867|198858988</td><td>42</td><td>63</td><td>22(+1A)</td><td>AACATTCATTGCTGTCGGTGGGA</td><td>278.7</td><td>4.5</td><td>0.7</td><td>6.2</td><td>5.2E-10</td><td>1.2E-09</td><td>MB231/MB468</td><td>NUC</td></tr>
<tr><td>>am_trnaMT_SerGCT_MT_+_12207_12265@18.38.21|CTGCTAACTCATGCCCCCATG</td><td>>am_trnaMT_SerGCT_MT_+_12207_12265</td><td>18</td><td>38</td><td>21</td><td>CTGCTAACTCATGCCCCCATG</td><td>106.3</td><td>-10.6</td><td>1.7</td><td>-6.2</td><td>5.0E-10</td><td>1.2E-09</td><td>MB231/MB468</td><td>MP</td></tr>
<tr><td>>hg38_21_+_8433172_8446622_RNA45SN1_45S_50ntFlanks@12417.12441.25|TCGATGTCGGCTCTTCCTATCATTG</td><td>>hg38_21_+_8433172_8446622_RNA45SN1_45S_50ntFlanks</td><td>12417</td><td>12441</td><td>25</td><td>TCGATGTCGGCTCTTCCTATCATTG</td><td>2107.5</td><td>-5.1</td><td>0.8</td><td>-6.2</td><td>5.7E-10</td><td>1.2E-09</td><td>MB231/MB468</td><td>MITO</td></tr>
<tr><td>>hg38_21_+_8433172_8446622_RNA45SN1_45S_50ntFlanks@4297.4314.18|CATTGGAGGGCAAGTCTG</td><td>>hg38_21_+_8433172_8446622_RNA45SN1_45S_50ntFlanks</td><td>4297</td><td>4314</td><td>18</td><td>CATTGGAGGGCAAGTCTG</td><td>1104.6</td><td>-7.5</td><td>1.2</td><td>-6.2</td><td>6.0E-10</td><td>1.2E-09</td><td>BT20/MB468</td><td>NUC</td></tr>
<tr><td>>am_trnaMT_ValTAC_MT_+_1602_1670@51.68.18|CTTAACTTGACCGCTCTG</td><td>>am_trnaMT_ValTAC_MT_+_1602_1670</td><td>51</td><td>68</td><td>18</td><td>CTTAACTTGACCGCTCTG</td><td>9515.5</td><td>-5.6</td><td>0.9</td><td>-6.3</td><td>3.9E-10</td><td>1.2E-09</td><td>BT20/MB468</td><td>MITO</td></tr>
<tr><td>>am_trna128_GlyGCC_6_-_27870686_27870756@2.31.30,trna133_GlyCCC_1_-_16872434_16872504@2.31.30,trna18_GlyGCC_16_+_70822597_70822667@2.31.30,trna19_GlyGCC_16_+_70823410_70823480@2.31.30,trna19_GlyGCC_2_-_157257659_157257729@2.31.30,trna24_GlyGCC_16_-_70812942_70813012@2.31.30,trna25_GlyGCC_16_-_70812114_70812184@2.31.30,trna4_GlyCCC_1_+_17188416_17188486@2.31.30,trna5_GlyGCC_17_+_8029064_8029134@2.31.30,trna68_GlyGCC_1_-_161493637_161493707@2.31.30|CATTGGTGGTTCAGTGGTAGAATTCTCGCC</td><td>>am_trna128_GlyGCC_6_-_27870686_27870756</td><td>2</td><td>31</td><td>30</td><td>CATTGGTGGTTCAGTGGTAGAATTCTCGCC</td><td>146.4</td><td>10.4</td><td>1.7</td><td>6.2</td><td>5.0E-10</td><td>1.2E-09</td><td>MB231/MB468</td><td>MP</td></tr>
<tr><td>>hg38_21_+_8433172_8446622_RNA45SN1_45S_50ntFlanks@10547.10564.18|AGTGCGGTAACGCGACCG</td><td>>hg38_21_+_8433172_8446622_RNA45SN1_45S_50ntFlanks</td><td>10547</td><td>10564</td><td>18</td><td>AGTGCGGTAACGCGACCG</td><td>102.9</td><td>-10.5</td><td>1.7</td><td>-6.2</td><td>5.0E-10</td><td>1.2E-09</td><td>MB231/MB468</td><td>MP</td></tr>
<tr><td>>hg38_21_+_8433172_8446622_RNA45SN1_45S_50ntFlanks@11921.11957.37|TGAGAGGTGTAGAATAAGTGGGAGGCCCCCGGCGCCC</td><td>>hg38_21_+_8433172_8446622_RNA45SN1_45S_50ntFlanks</td><td>11921</td><td>11957</td><td>37</td><td>TGAGAGGTGTAGAATAAGTGGGAGGCCCCCGGCGCCC</td><td>133.4</td><td>-10.9</td><td>1.8</td><td>-6.2</td><td>5.0E-10</td><td>1.2E-09</td><td>MB231/MB468</td><td>MP</td></tr>
<tr><td>>MI0000746|hsa-mir-99b&WithFlank&19|+|51692606|51692687@51.73.23&[MIMAT0004678&hsa-miR-99b-3p&offsets|0|+1;m-151&19|+|51692656|51692678&offsets|0|0]|CAAGCTCGTGTCTGTGGGTCCGT</td><td>>MI0000746|hsa-mir-99b&WithFlank&19|+|51692606|51692687</td><td>51</td><td>73</td><td>23</td><td>CAAGCTCGTGTCTGTGGGTCCGT</td><td>574.2</td><td>5.2</td><td>0.8</td><td>6.2</td><td>5.2E-10</td><td>1.2E-09</td><td>MB231/MB468</td><td>CYTO</td></tr>
<tr><td>>hg38_21_+_8433172_8446622_RNA45SN1_45S_50ntFlanks@12248.12268.21|GCAAAAGCTCGCTTGATCTTG</td><td>>hg38_21_+_8433172_8446622_RNA45SN1_45S_50ntFlanks</td><td>12248</td><td>12268</td><td>21</td><td>GCAAAAGCTCGCTTGATCTTG</td><td>77.8</td><td>-10.1</td><td>1.6</td><td>-6.2</td><td>5.0E-10</td><td>1.2E-09</td><td>MB231/MB468</td><td>MP</td></tr>
</table>

<table>
<tr><td></td><td></td><td></td><td></td><td></td><td></td><td></td><td></td><td></td><td></td><td></td><td></td><td></td><td></td></tr>
<tr><td>>hg38_21_+_8433172_8446622_RNA45SN1_45S_50ntFlanks@12583.12604.22|CCGCAGGTTCAGACATTTGGTG</td><td>>hg38_21_+_8433172_8446622_RNA45SN1_45S_50ntFlanks</td><td>12583</td><td>12604</td><td>22</td><td>CCGCAGGTTCAGACATTTGGTG</td><td>160.3</td><td>-10.4</td><td>1.6</td><td>-6.6</td><td>4.8E-11</td><td>1.2E-09</td><td>BT20/MB468</td><td>MP</td></tr>
<tr><td>>hg38_21_+_8433172_8446622_RNA45SN1_45S_50ntFlanks@11821.11856.36|CTATCCAGCGAAACCACAGCCAAGGGAACGGGCTTG</td><td>>hg38_21_+_8433172_8446622_RNA45SN1_45S_50ntFlanks</td><td>11821</td><td>11856</td><td>36</td><td>CTATCCAGCGAAACCACAGCCAAGGGAACGGGCTTG</td><td>218.6</td><td>-10.9</td><td>1.7</td><td>-6.6</td><td>4.8E-11</td><td>1.2E-09</td><td>BT20/MB468</td><td>MP</td></tr>
<tr><td>>hg38_21_+_8433172_8446622_RNA45SN1_45S_50ntFlanks@12946.12988.43|TAGACGACCTGCTTCTGGGTCGGGGTTTCGTACGTAGCAGAGC</td><td>>hg38_21_+_8433172_8446622_RNA45SN1_45S_50ntFlanks</td><td>12946</td><td>12988</td><td>43</td><td>TAGACGACCTGCTTCTGGGTCGGGGTTTCGTACGTAGCAGAGC</td><td>924.1</td><td>4.6</td><td>0.7</td><td>6.2</td><td>5.2E-10</td><td>1.2E-09</td><td>MB231/MB468</td><td>CYTO</td></tr>
<tr><td>>hg38_21_+_8433172_8446622_RNA45SN1_45S_50ntFlanks@4732.4761.30|AGAACGAAAGTCGGAGGTTCGAAGACGATC</td><td>>hg38_21_+_8433172_8446622_RNA45SN1_45S_50ntFlanks</td><td>4732</td><td>4761</td><td>30</td><td>AGAACGAAAGTCGGAGGTTCGAAGACGATC</td><td>1424.7</td><td>5.0</td><td>0.8</td><td>6.2</td><td>5.2E-10</td><td>1.2E-09</td><td>MB231/MB468</td><td>CYTO</td></tr>
<tr><td>>hg38_21_+_8433172_8446622_RNA45SN1_45S_50ntFlanks@12033.12070.38|ATCGTTTTTTCACTGACCCGGTGAGGCGGGGGGGCGAG</td><td>>hg38_21_+_8433172_8446622_RNA45SN1_45S_50ntFlanks</td><td>12033</td><td>12070</td><td>38</td><td>ATCGTTTTTTCACTGACCCGGTGAGGCGGGGGGGCGAG</td><td>82.2</td><td>-10.2</td><td>1.6</td><td>-6.2</td><td>5.0E-10</td><td>1.2E-09</td><td>MB231/MB468</td><td>MP</td></tr>
<tr><td>>hg38_21_+_8433172_8446622_RNA45SN1_45S_50ntFlanks@11118.11145.28|CAGGGGCCGGCGGCGGCCCGCCGCGGGG</td><td>>hg38_21_+_8433172_8446622_RNA45SN1_45S_50ntFlanks</td><td>11118</td><td>11145</td><td>28</td><td>CAGGGGCCGGCGGCGGCCCGCCGCGGGG</td><td>145.1</td><td>-9.3</td><td>1.5</td><td>-6.3</td><td>3.9E-10</td><td>1.2E-09</td><td>BT20/MB468</td><td>MITO</td></tr>
<tr><td>>hg38_21_+_8433172_8446622_RNA45SN1_45S_50ntFlanks@12008.12036.29|CCGCCGGTGAAATACCACTACTCTGATCG</td><td>>hg38_21_+_8433172_8446622_RNA45SN1_45S_50ntFlanks</td><td>12008</td><td>12036</td><td>29</td><td>CCGCCGGTGAAATACCACTACTCTGATCG</td><td>1033.0</td><td>-8.5</td><td>1.4</td><td>-6.2</td><td>5.8E-10</td><td>1.2E-09</td><td>MB231/MB468</td><td>MITO</td></tr>
<tr><td>>am_trna119_LysCTT_1_-_145395522_145395594@13.33.21,trna11_LysCTT_5_-_180648979_180649051@13.33.21,trna128_LysCTT_1_-_55423542_55423614@13.33.21,trna13_LysCTT_6_+_26556774_26556846@13.33.21,trna6_LysCTT_19_-_52425396_52425468@13.33.21,trna7_LysCTT_16_+_3225692_3225764@13.33.21,trna9_LysCTT_5_+_180634755_180634827@13.33.21|CAGTCGGTAGAGCATGAGACT</td><td>>am_trna119_LysCTT_1_-_145395522_145395594</td><td>13</td><td>33</td><td>21</td><td>CAGTCGGTAGAGCATGAGACT</td><td>311.5</td><td>7.0</td><td>1.1</td><td>6.2</td><td>5.3E-10</td><td>1.2E-09</td><td>MB231/MB468</td><td>NUC</td></tr>
<tr><td>>hg38_21_+_8433172_8446622_RNA45SN1_45S_50ntFlanks@11575.11594.20|ACTTAGAACTGGTGCGGACC</td><td>>hg38_21_+_8433172_8446622_RNA45SN1_45S_50ntFlanks</td><td>11575</td><td>11594</td><td>20</td><td>ACTTAGAACTGGTGCGGACC</td><td>606.6</td><td>-3.9</td><td>0.6</td><td>-6.2</td><td>5.8E-10</td><td>1.2E-09</td><td>MB231/MB468</td><td>MITO</td></tr>
<tr><td>>MI0003760|hsa-mir-671&WithFlank&7|+|151238415|151238544@74.94.21&[MIMAT0004819&hsa-miR-671-3p&offsets|0|0;m-323&7|+|151238488|151238508&offsets|0|0]|TCCGGTTCTCAGGGCTCCACC</td><td>>MI0003760|hsa-mir-671&WithFlank&7|+|151238415|151238544</td><td>74</td><td>94</td><td>21</td><td>TCCGGTTCTCAGGGCTCCACC</td><td>469.8</td><td>6.7</td><td>1.1</td><td>6.2</td><td>4.6E-10</td><td>1.2E-09</td><td>BT20/MB468</td><td>CYTO</td></tr>
<tr><td>>hg38_21_+_8433172_8446622_RNA45SN1_45S_50ntFlanks@12008.12031.24|CCGCCGGTGAAATACCACTACTCT</td><td>>hg38_21_+_8433172_8446622_RNA45SN1_45S_50ntFlanks</td><td>12008</td><td>12031</td><td>24</td><td>CCGCCGGTGAAATACCACTACTCT</td><td>1155.0</td><td>-8.5</td><td>1.4</td><td>-6.2</td><td>5.3E-10</td><td>1.2E-09</td><td>MB231/MB468</td><td>NUC</td></tr>
<tr><td>>hg38_21_+_8433172_8446622_RNA45SN1_45S_50ntFlanks@8356.8378.23|GAAAAGAACTTTGAAGAGAGAGT</td><td>>hg38_21_+_8433172_8446622_RNA45SN1_45S_50ntFlanks</td><td>8356</td><td>8378</td><td>23</td><td>GAAAAGAACTTTGAAGAGAGAGT</td><td>7337.1</td><td>-6.6</td><td>1.0</td><td>-6.6</td><td>4.9E-11</td><td>1.2E-09</td><td>BT20/MB468</td><td>MP</td></tr>
<tr><td>>hg38_21_+_8433172_8446622_RNA45SN1_45S_50ntFlanks@8179.8206.28|CTGATCGAGGCCCAGCCCGTGGACGGTG</td><td>>hg38_21_+_8433172_8446622_RNA45SN1_45S_50ntFlanks</td><td>8179</td><td>8206</td><td>28</td><td>CTGATCGAGGCCCAGCCCGTGGACGGTG</td><td>312.6</td><td>-7.8</td><td>1.3</td><td>-6.2</td><td>5.8E-10</td><td>1.2E-09</td><td>MB231/MB468</td><td>MITO</td></tr>
<tr><td>>MI0003513|hsa-mir-455&WithFlank&9|+|114209428|114209535@24.44.21&[MIMAT0003150&hsa-miR-455-5p&offsets|+2|+1;m-232&9|+|114209449|114209470&offsets|+2|+1]|TGTGCCTTTGGACTACATCGT</td><td>>MI0003513|hsa-mir-455&WithFlank&9|+|114209428|114209535</td><td>24</td><td>44</td><td>21</td><td>TGTGCCTTTGGACTACATCGT</td><td>81.9</td><td>9.6</td><td>1.5</td><td>6.2</td><td>5.1E-10</td><td>1.2E-09</td><td>MB231/MB468</td><td>MP</td></tr>
<tr><td>>trna116_GluCTC_1_-_145399233_145399304@37.75.39,trna134_GluTTC_1_-_16861774_16861845@37.75.39,trna71_GluCTC_1_-_161439189_161439260@37.75.39,trna74_GluCTC_1_-_161431809_161431880@37.75.39,trna77_GluCTC_1_-_161424398_161424469@37.75.39,trna77_GluCTC_6_+_28949976_28950047@37.75.39,trna80_GluCTC_1_-_161417018_161417089@37.75.39,trna84_GluTTC_1_-_161391883_161391954@37.75.39,trna87_GluCTC_6_-_126101393_126101464@37.75.39|ACCGCCGCGGCCCGGGTTCGATTCCCGGTCAGGGAACCA</td><td>>trna116_GluCTC_1_-_145399233_145399304</td><td>37</td><td>75</td><td>39</td><td>ACCGCCGCGGCCCGGGTTCGATTCCCGGTCAGGGAACCA</td><td>30816.6</td><td>8.4</td><td>1.3</td><td>6.2</td><td>5.2E-10</td><td>1.2E-09</td><td>MB231/MB468</td><td>CYTO</td></tr>
<tr><td>>hg38_MT_+_1621_3279_RNR2_16S_50ntFlanks@51.72.22|GCTAAACCTAGCCCCAAACCCA</td><td>>hg38_MT_+_1621_3279_RNR2_16S_50ntFlanks</td><td>51</td><td>72</td><td>22</td><td>GCTAAACCTAGCCCCAAACCCA</td><td>5945.0</td><td>6.1</td><td>1.0</td><td>6.3</td><td>3.9E-10</td><td>1.2E-09</td><td>BT20/MB468</td><td>MITO</td></tr>
<tr><td>>am_trnaMT_ProTGG_MT_-_15956_16023@52.70.19|AAAGACTTTTTCTCTGACC</td><td>>am_trnaMT_ProTGG_MT_-_15956_16023</td><td>52</td><td>70</td><td>19</td><td>AAAGACTTTTTCTCTGACC</td><td>597.7</td><td>8.2</td><td>1.3</td><td>6.2</td><td>5.3E-10</td><td>1.2E-09</td><td>MB231/MB468</td><td>NUC</td></tr>
<tr><td>>hg38_21_+_8433172_8446622_RNA45SN1_45S_50ntFlanks@10392.10432.41|ACAGCAGTTGAACATGGGTCAGTCGGTCCTGAGAGATGGGC</td><td>>hg38_21_+_8433172_8446622_RNA45SN1_45S_50ntFlanks</td><td>10392</td><td>10432</td><td>41</td><td>ACAGCAGTTGAACATGGGTCAGTCGGTCCTGAGAGATGGGC</td><td>144.8</td><td>-11.0</td><td>1.8</td><td>-6.2</td><td>5.1E-10</td><td>1.2E-09</td><td>MB231/MB468</td><td>MP</td></tr>
<tr><td>>hg38_21_+_8433172_8446622_RNA45SN1_45S_50ntFlanks@10461.10480.20|GCCTCCGTTGCCCTCGGCCG</td><td>>hg38_21_+_8433172_8446622_RNA45SN1_45S_50ntFlanks</td><td>10461</td><td>10480</td><td>20</td><td>GCCTCCGTTGCCCTCGGCCG</td><td>356.2</td><td>-8.4</td><td>1.4</td><td>-6.2</td><td>6.1E-10</td><td>1.2E-09</td><td>BT20/MB468</td><td>NUC</td></tr>
<tr><td>>hg38_21_+_8433172_8446622_RNA45SN1_45S_50ntFlanks@10876.10901.26|GGTCGGTCGGGCTGGGGCGCGAAGCG</td><td>>hg38_21_+_8433172_8446622_RNA45SN1_45S_50ntFlanks</td><td>10876</td><td>10901</td><td>26</td><td>GGTCGGTCGGGCTGGGGCGCGAAGCG</td><td>83.9</td><td>-10.3</td><td>1.6</td><td>-6.2</td><td>5.1E-10</td><td>1.2E-09</td><td>MB231/MB468</td><td>MP</td></tr>
<tr><td>>hg38_21_+_8433172_8446622_RNA45SN1_45S_50ntFlanks@10680.10706.27|GTTCCGGCGGCGTCCGGTGAGCTCTCG</td><td>>hg38_21_+_8433172_8446622_RNA45SN1_45S_50ntFlanks</td><td>10680</td><td>10706</td><td>27</td><td>GTTCCGGCGGCGTCCGGTGAGCTCTCG</td><td>11596.1</td><td>-5.3</td><td>0.9</td><td>-6.2</td><td>4.6E-10</td><td>1.2E-09</td><td>BT20/MB468</td><td>CYTO</td></tr>
<tr><td>>hg38_21_+_8433172_8446622_RNA45SN1_45S_50ntFlanks@6719.6737.19|TGAATTGCAGGACACATTG</td><td>>hg38_21_+_8433172_8446622_RNA45SN1_45S_50ntFlanks</td><td>6719</td><td>6737</td><td>19</td><td>TGAATTGCAGGACACATTG</td><td>1972.8</td><td>-6.7</td><td>1.1</td><td>-6.2</td><td>4.6E-10</td><td>1.2E-09</td><td>BT20/MB468</td><td>CYTO</td></tr>
<tr><td>>MI0000749|hsa-mir-30e&WithFlank&1|+|40754349|40754452@65.85.21(+1U)&[MIMAT0000693&hsa-miR-30e-3p&offsets|0|-1(+1U);m-25&1|+|40754413|40754434&offsets|0|-1(+1U)]|CTTTCAGTCGGATGTTTACAGT</td><td>>MI0000749|hsa-mir-30e&WithFlank&1|+|40754349|40754452</td><td>65</td><td>85</td><td>21(+1U)</td><td>CTTTCAGTCGGATGTTTACAGT</td><td>1330.3</td><td>4.2</td><td>0.7</td><td>6.3</td><td>4.0E-10</td><td>1.2E-09</td><td>BT20/MB468</td><td>MITO</td></tr>
<tr><td>>hg38_21_+_8433172_8446622_RNA45SN1_45S_50ntFlanks@10468.10497.30|TTGCCCTCGGCCGATCGAAAGGGAGTCGGG</td><td>>hg38_21_+_8433172_8446622_RNA45SN1_45S_50ntFlanks</td><td>10468</td><td>10497</td><td>30</td><td>TTGCCCTCGGCCGATCGAAAGGGAGTCGGG</td><td>530.3</td><td>-4.6</td><td>0.7</td><td>-6.2</td><td>5.3E-10</td><td>1.3E-09</td><td>MB231/MB468</td><td>CYTO</td></tr>
<tr><td>>hg38_1_-_228634819_228635039_RNA5S12_5S_50ntFlanks@148.170.23|GGAATACCGGGTGCTGTAGGCTT</td><td>>hg38_1_-_228634819_228635039_RNA5S12_5S_50ntFlanks</td><td>148</td><td>170</td><td>23</td><td>GGAATACCGGGTGCTGTAGGCTT</td><td>4212.1</td><td>-4.1</td><td>0.7</td><td>-6.2</td><td>5.9E-10</td><td>1.3E-09</td><td>MB231/MB468</td><td>MITO</td></tr>
<tr><td>>hg38_21_+_8433172_8446622_RNA45SN1_45S_50ntFlanks@12539.12571.33|ATGATGTGTTGTTGCCATGGTAATCCTGCTCAG</td><td>>hg38_21_+_8433172_8446622_RNA45SN1_45S_50ntFlanks</td><td>12539</td><td>12571</td><td>33</td><td>ATGATGTGTTGTTGCCATGGTAATCCTGCTCAG</td><td>85.0</td><td>-10.3</td><td>1.7</td><td>-6.2</td><td>5.1E-10</td><td>1.3E-09</td><td>MB231/MB468</td><td>MP</td></tr>
<tr><td>>hg38_21_+_8433172_8446622_RNA45SN1_45S_50ntFlanks@11669.11687.19|GCCCAGTGCTCTGAATGTC</td><td>>hg38_21_+_8433172_8446622_RNA45SN1_45S_50ntFlanks</td><td>11669</td><td>11687</td><td>19</td><td>GCCCAGTGCTCTGAATGTC</td><td>980.4</td><td>4.2</td><td>0.7</td><td>6.2</td><td>5.4E-10</td><td>1.3E-09</td><td>MB231/MB468</td><td>NUC</td></tr>
<tr><td>>hg38_21_+_8433172_8446622_RNA45SN1_45S_50ntFlanks@4370.4398.29|TTAAAAAGCTCGTAGTTGGATCTTGGGAG</td><td>>hg38_21_+_8433172_8446622_RNA45SN1_45S_50ntFlanks</td><td>4370</td><td>4398</td><td>29</td><td>TTAAAAAGCTCGTAGTTGGATCTTGGGAG</td><td>396.3</td><td>-8.0</td><td>1.3</td><td>-6.2</td><td>5.4E-10</td><td>1.3E-09</td><td>MB231/MB468</td><td>NUC</td></tr>
<tr><td>>hg38_21_+_8433172_8446622_RNA45SN1_45S_50ntFlanks@3815.3852.38|ATGGTTCCTTTGGTCGCTCGCTCCTCTCCTACTTGGAT</td><td>>hg38_21_+_8433172_8446622_RNA45SN1_45S_50ntFlanks</td><td>3815</td><td>3852</td><td>38</td><td>ATGGTTCCTTTGGTCGCTCGCTCCTCTCCTACTTGGAT</td><td>790.3</td><td>-7.5</td><td>1.2</td><td>-6.2</td><td>6.2E-10</td><td>1.3E-09</td><td>BT20/MB468</td><td>NUC</td></tr>
<tr><td>>hg38_1_-_228634819_228635039_RNA5S12_5S_50ntFlanks@144.171.28|CCTGGGAATACCGGGTGCTGTAGGCTTT</td><td>>hg38_1_-_228634819_228635039_RNA5S12_5S_50ntFlanks</td><td>144</td><td>171</td><td>28</td><td>CCTGGGAATACCGGGTGCTGTAGGCTTT</td><td>5023.9</td><td>5.6</td><td>0.9</td><td>6.2</td><td>5.4E-10</td><td>1.3E-09</td><td>MB231/MB468</td><td>NUC</td></tr>
<tr><td>>hg38_21_+_8433172_8446622_RNA45SN1_45S_50ntFlanks@11571.11603.33|GCCGACTTAGAACTGGTGCGGACCAGGGGAATC</td><td>>hg38_21_+_8433172_8446622_RNA45SN1_45S_50ntFlanks</td><td>11571</td><td>11603</td><td>33</td><td>GCCGACTTAGAACTGGTGCGGACCAGGGGAATC</td><td>759.5</td><td>-2.9</td><td>0.5</td><td>-6.2</td><td>5.4E-10</td><td>1.3E-09</td><td>MB231/MB468</td><td>NUC</td></tr>
<tr><td>>hg38_21_+_8433172_8446622_RNA45SN1_45S_50ntFlanks@9739.9758.20|AAACGATCTCAACCTATTCT</td><td>>hg38_21_+_8433172_8446622_RNA45SN1_45S_50ntFlanks</td><td>9739</td><td>9758</td><td>20</td><td>AAACGATCTCAACCTATTCT</td><td>2526.5</td><td>-3.8</td><td>0.6</td><td>-6.2</td><td>5.3E-10</td><td>1.3E-09</td><td>MB231/MB468</td><td>CYTO</td></tr>
<tr><td>>hg38_21_+_8433172_8446622_RNA45SN1_45S_50ntFlanks@3831.3860.30|CTCGCTCCTCTCCTACTTGGATAACTGTGG</td><td>>hg38_21_+_8433172_8446622_RNA45SN1_45S_50ntFlanks</td><td>3831</td><td>3860</td><td>30</td><td>CTCGCTCCTCTCCTACTTGGATAACTGTGG</td><td>118.6</td><td>-10.8</td><td>1.7</td><td>-6.2</td><td>5.1E-10</td><td>1.3E-09</td><td>MB231/MB468</td><td>MP</td></tr>
<tr><td>>hg38_21_+_8433172_8446622_RNA45SN1_45S_50ntFlanks@12003.12033.31|GCGGGCCGCCGGTGAAATACCACTACTCTGA</td><td>>hg38_21_+_8433172_8446622_RNA45SN1_45S_50ntFlanks</td><td>12003</td><td>12033</td><td>31</td><td>GCGGGCCGCCGGTGAAATACCACTACTCTGA</td><td>1235.1</td><td>4.0</td><td>0.6</td><td>6.3</td><td>4.0E-10</td><td>1.3E-09</td><td>BT20/MB468</td><td>MITO</td></tr>
<tr><td>>am_trna116_GluCTC_1_-_145399233_145399304@11.28.18,trna25_GluCTC_2_-_71273488_71273560@11.28.18,trna59_GluCTC_1_+_249168447_249168518@11.28.18,trna71_GluCTC_1_-_161439189_161439260@11.28.18,trna74_GluCTC_1_-_161431809_161431880@11.28.18,trna77_GluCTC_1_-_161424398_161424469@11.28.18,trna77_GluCTC_6_+_28949976_28950047@11.28.18,trna80_GluCTC_1_-_161417018_161417089@11.28.18,trna87_GluCTC_6_-_126101393_126101464@11.28.18|TCTAGTGGTTAGGATTCG</td><td>>am_trna116_GluCTC_1_-_145399233_145399304</td><td>11</td><td>28</td><td>18</td><td>TCTAGTGGTTAGGATTCG</td><td>153.3</td><td>-11.1</td><td>1.8</td><td>-6.2</td><td>5.1E-10</td><td>1.3E-09</td><td>MB231/MB468</td><td>MP</td></tr>
<tr><td>>am_trna2_GlyGCC_21_-_18827107_18827177@1.27.27,trna35_GlyGCC_1_+_161413094_161413164@1.27.27,trna37_GlyGCC_1_+_161420467_161420537@1.27.27,trna39_GlyGCC_1_+_161427898_161427968@1.27.27,trna41_GlyGCC_1_+_161435258_161435328@1.27.27|GCATGGGTGGTTCAGTGGTAGAATTCT</td><td>>am_trna2_GlyGCC_21_-_18827107_18827177</td><td>1</td><td>27</td><td>27</td><td>GCATGGGTGGTTCAGTGGTAGAATTCT</td><td>3490.0</td><td>6.3</td><td>1.0</td><td>6.3</td><td>4.0E-10</td><td>1.3E-09</td><td>BT20/MB468</td><td>MITO</td></tr>
<tr><td>>hg38_21_+_8433172_8446622_RNA45SN1_45S_50ntFlanks@11782.11804.23|GCATGAATGGATGAACGAGATTC</td><td>>hg38_21_+_8433172_8446622_RNA45SN1_45S_50ntFlanks</td><td>11782</td><td>11804</td><td>23</td><td>GCATGAATGGATGAACGAGATTC</td><td>357.6</td><td>-6.5</td><td>1.0</td><td>-6.2</td><td>5.1E-10</td><td>1.3E-09</td><td>MB231/MB468</td><td>MP</td></tr>
<tr><td>>trnaMT_PheGAA_MT_+_577_647@56.74.19|CATCACCCCATAAACACCA</td><td>>trnaMT_PheGAA_MT_+_577_647</td><td>56</td><td>74</td><td>19</td><td>CATCACCCCATAAACACCA</td><td>804.2</td><td>8.5</td><td>1.4</td><td>6.2</td><td>4.7E-10</td><td>1.3E-09</td><td>BT20/MB468</td><td>CYTO</td></tr>
<tr><td>>hg38_21_+_8433172_8446622_RNA45SN1_45S_50ntFlanks@11671.11698.28|CCAGTGCTCTGAATGTCAAAGTGAAGAA</td><td>>hg38_21_+_8433172_8446622_RNA45SN1_45S_50ntFlanks</td><td>11671</td><td>11698</td><td>28</td><td>CCAGTGCTCTGAATGTCAAAGTGAAGAA</td><td>434.9</td><td>-8.6</td><td>1.4</td><td>-6.2</td><td>6.2E-10</td><td>1.3E-09</td><td>BT20/MB468</td><td>NUC</td></tr>
<tr><td>>hg38_21_+_8433172_8446622_RNA45SN1_45S_50ntFlanks@7994.8012.19|TGGCGACCCGCTGAATTTA</td><td>>hg38_21_+_8433172_8446622_RNA45SN1_45S_50ntFlanks</td><td>7994</td><td>8012</td><td>19</td><td>TGGCGACCCGCTGAATTTA</td><td>71.0</td><td>-10.0</td><td>1.6</td><td>-6.2</td><td>5.1E-10</td><td>1.3E-09</td><td>MB231/MB468</td><td>MP</td></tr>
<tr><td>>hg38_21_+_8433172_8446622_RNA45SN1_45S_50ntFlanks@12408.12436.29|TTTGATCCTTCGATGTCGGCTCTTCCTAT</td><td>>hg38_21_+_8433172_8446622_RNA45SN1_45S_50ntFlanks</td><td>12408</td><td>12436</td><td>29</td><td>TTTGATCCTTCGATGTCGGCTCTTCCTAT</td><td>307.8</td><td>-4.4</td><td>0.7</td><td>-6.2</td><td>5.9E-10</td><td>1.3E-09</td><td>MB231/MB468</td><td>MITO</td></tr>
<tr><td>>hg38_21_+_8433172_8446622_RNA45SN1_45S_50ntFlanks@8168.8212.45|CCCAAGTCCTTCTGATCGAGGCCCAGCCCGTGGACGGTGTGAGGC</td><td>>hg38_21_+_8433172_8446622_RNA45SN1_45S_50ntFlanks</td><td>8168</td><td>8212</td><td>45</td><td>CCCAAGTCCTTCTGATCGAGGCCCAGCCCGTGGACGGTGTGAGGC</td><td>1099.0</td><td>-7.0</td><td>1.1</td><td>-6.2</td><td>5.9E-10</td><td>1.3E-09</td><td>MB231/MB468</td><td>MITO</td></tr>
<tr><td>>hg38_1_-_228634819_228635039_RNA5S12_5S_50ntFlanks@131.169.39|GGATGGGAGACCGCCTGGGAATACCGGGTGCTGTAGGCT</td><td>>hg38_1_-_228634819_228635039_RNA5S12_5S_50ntFlanks</td><td>131</td><td>169</td><td>39</td><td>GGATGGGAGACCGCCTGGGAATACCGGGTGCTGTAGGCT</td><td>1370.7</td><td>9.3</td><td>1.5</td><td>6.2</td><td>4.7E-10</td><td>1.3E-09</td><td>BT20/MB468</td><td>CYTO</td></tr>
</table>

<table>
<tr><td>>trna111_HisGTG_1_-_147774845_147774916@1.36.36,trna118_HisGTG_1_-_145396881_145396952@1.36.36,trna16_HisGTG_1_+_146544773_146544844@1.36.36,trna1_HisGTG_15_+_45493349_45493420@1.36.36,trna21_HisGTG_1_+_147753471_147753542@1.36.36,trna33_HisGTG_6_+_27125906_27125977@1.36.36,trna7_HisGTG_9_-_14433938_14434009@1.36.36,trna8_HisGTG_15_-_45492611_45492682@1.36.36,trna9_HisGTG_15_-_45490804_45490875@1.36.36|GCCGTGATCGTATAGTGGTTAGTACTCTGCGTTGTG</td><td>>trna111_HisGTG_1_-_147774845_147774916</td><td>1</td><td>36</td><td>36</td><td>GCCGTGATCGTATAGTGGTTAGTACTCTGCGTTGTG</td><td>1089.7</td><td>9.4</td><td>1.5</td><td>6.2</td><td>4.7E-10</td><td>1.3E-09</td><td>BT20/MB468</td><td>CYTO</td></tr>
<tr><td>>hg38_21_+_8433172_8446622_RNA45SN1_45S_50ntFlanks@13002.13021.20|CGATCTATTGAAAGTCAGCC</td><td>>hg38_21_+_8433172_8446622_RNA45SN1_45S_50ntFlanks</td><td>13002</td><td>13021</td><td>20</td><td>CGATCTATTGAAAGTCAGCC</td><td>70.6</td><td>-10.0</td><td>1.6</td><td>-6.2</td><td>5.2E-10</td><td>1.3E-09</td><td>MB231/MB468</td><td>MP</td></tr>
<tr><td>>hg38_21_+_8433172_8446622_RNA45SN1_45S_50ntFlanks@5482.5499.18|CGGAGCGCTGAGAAGACG</td><td>>hg38_21_+_8433172_8446622_RNA45SN1_45S_50ntFlanks</td><td>5482</td><td>5499</td><td>18</td><td>CGGAGCGCTGAGAAGACG</td><td>413.4</td><td>-8.1</td><td>1.3</td><td>-6.2</td><td>6.0E-10</td><td>1.3E-09</td><td>MB231/MB468</td><td>MITO</td></tr>
<tr><td>>hg38_21_+_8433172_8446622_RNA45SN1_45S_50ntFlanks@12133.12151.19|CTCCGGGGACAGTGCCAGG</td><td>>hg38_21_+_8433172_8446622_RNA45SN1_45S_50ntFlanks</td><td>12133</td><td>12151</td><td>19</td><td>CTCCGGGGACAGTGCCAGG</td><td>180.4</td><td>-10.6</td><td>1.6</td><td>-6.6</td><td>5.0E-11</td><td>1.3E-09</td><td>BT20/MB468</td><td>MP</td></tr>
<tr><td>>hg38_21_+_8433172_8446622_RNA45SN1_45S_50ntFlanks@9569.9597.29|GTGCAAATCGGTCGTCCGACCTGGGTATA</td><td>>hg38_21_+_8433172_8446622_RNA45SN1_45S_50ntFlanks</td><td>9569</td><td>9597</td><td>29</td><td>GTGCAAATCGGTCGTCCGACCTGGGTATA</td><td>1127.8</td><td>-6.7</td><td>1.1</td><td>-6.2</td><td>5.2E-10</td><td>1.3E-09</td><td>MB231/MB468</td><td>MP</td></tr>
<tr><td>>hg38_21_+_8433172_8446622_RNA45SN1_45S_50ntFlanks@4752.4782.31|GAAGACGATCAGATACCGTCGTAGTTCCGAC</td><td>>hg38_21_+_8433172_8446622_RNA45SN1_45S_50ntFlanks</td><td>4752</td><td>4782</td><td>31</td><td>GAAGACGATCAGATACCGTCGTAGTTCCGAC</td><td>475.3</td><td>-11.0</td><td>1.8</td><td>-6.3</td><td>4.0E-10</td><td>1.3E-09</td><td>BT20/MB468</td><td>MITO</td></tr>
<tr><td>>hg38_21_+_8433172_8446622_RNA45SN1_45S_50ntFlanks@10402.10424.23|AACATGGGTCAGTCGGTCCTGAG</td><td>>hg38_21_+_8433172_8446622_RNA45SN1_45S_50ntFlanks</td><td>10402</td><td>10424</td><td>23</td><td>AACATGGGTCAGTCGGTCCTGAG</td><td>6484.7</td><td>-3.7</td><td>0.6</td><td>-6.2</td><td>6.3E-10</td><td>1.3E-09</td><td>BT20/MB468</td><td>NUC</td></tr>
<tr><td>>hg38_21_+_8433172_8446622_RNA45SN1_45S_50ntFlanks@11583.11603.21|CTGGTGCGGACCAGGGGAATC</td><td>>hg38_21_+_8433172_8446622_RNA45SN1_45S_50ntFlanks</td><td>11583</td><td>11603</td><td>21</td><td>CTGGTGCGGACCAGGGGAATC</td><td>2323.1</td><td>-3.9</td><td>0.6</td><td>-6.2</td><td>4.8E-10</td><td>1.3E-09</td><td>BT20/MB468</td><td>CYTO</td></tr>
<tr><td>>hg38_21_+_8433172_8446622_RNA45SN1_45S_50ntFlanks@4043.4063.21|ACCCATTCGAACGTCTGCCCT</td><td>>hg38_21_+_8433172_8446622_RNA45SN1_45S_50ntFlanks</td><td>4043</td><td>4063</td><td>21</td><td>ACCCATTCGAACGTCTGCCCT</td><td>250.5</td><td>-10.1</td><td>1.6</td><td>-6.3</td><td>4.1E-10</td><td>1.3E-09</td><td>BT20/MB468</td><td>MITO</td></tr>
<tr><td>>hg38_21_+_8433172_8446622_RNA45SN1_45S_50ntFlanks@4206.4223.18|CCGGGGAGGTAGTGACGA</td><td>>hg38_21_+_8433172_8446622_RNA45SN1_45S_50ntFlanks</td><td>4206</td><td>4223</td><td>18</td><td>CCGGGGAGGTAGTGACGA</td><td>94.4</td><td>-10.4</td><td>1.7</td><td>-6.2</td><td>5.2E-10</td><td>1.3E-09</td><td>MB231/MB468</td><td>MP</td></tr>
<tr><td>>hg38_21_+_8433172_8446622_RNA45SN1_45S_50ntFlanks@10350.10384.35|ACGAGAACTTTGAAGGCCGAAGTGGAGAAGGGTTC</td><td>>hg38_21_+_8433172_8446622_RNA45SN1_45S_50ntFlanks</td><td>10350</td><td>10384</td><td>35</td><td>ACGAGAACTTTGAAGGCCGAAGTGGAGAAGGGTTC</td><td>4084.8</td><td>-8.1</td><td>1.3</td><td>-6.2</td><td>5.2E-10</td><td>1.3E-09</td><td>MB231/MB468</td><td>MP</td></tr>
<tr><td>>hg38_21_+_8433172_8446622_RNA45SN1_45S_50ntFlanks@7974.8005.32|ACGCGACCTCAGATCAGACGTGGCGACCCGCT</td><td>>hg38_21_+_8433172_8446622_RNA45SN1_45S_50ntFlanks</td><td>7974</td><td>8005</td><td>32</td><td>ACGCGACCTCAGATCAGACGTGGCGACCCGCT</td><td>11489.7</td><td>4.7</td><td>0.8</td><td>6.2</td><td>6.3E-10</td><td>1.3E-09</td><td>BT20/MB468</td><td>NUC</td></tr>
<tr><td>>am_trna128_GlyGCC_6_-_27870686_27870756@1.26.26,trna133_GlyCCC_1_-_16872434_16872504@1.26.26,trna18_GlyGCC_16_+_70822597_70822667@1.26.26,trna19_GlyGCC_16_+_70823410_70823480@1.26.26,trna19_GlyGCC_2_-_157257659_157257729@1.26.26,trna24_GlyGCC_16_-_70812942_70813012@1.26.26,trna25_GlyGCC_16_-_70812114_70812184@1.26.26,trna4_GlyCCC_1_+_17188416_17188486@1.26.26,trna5_GlyGCC_17_+_8029064_8029134@1.26.26,trna68_GlyGCC_1_-_161493637_161493707@1.26.26|GCATTGGTGGTTCAGTGGTAGAATTC</td><td>>am_trna128_GlyGCC_6_-_27870686_27870756</td><td>1</td><td>26</td><td>26</td><td>GCATTGGTGGTTCAGTGGTAGAATTC</td><td>978.7</td><td>-4.9</td><td>0.8</td><td>-6.2</td><td>6.3E-10</td><td>1.3E-09</td><td>BT20/MB468</td><td>NUC</td></tr>
<tr><td>>hg38_21_+_8433172_8446622_RNA45SN1_45S_50ntFlanks@4385.4405.21|TTGGATCTTGGGAGCGGGCGG</td><td>>hg38_21_+_8433172_8446622_RNA45SN1_45S_50ntFlanks</td><td>4385</td><td>4405</td><td>21</td><td>TTGGATCTTGGGAGCGGGCGG</td><td>87.9</td><td>-10.3</td><td>1.7</td><td>-6.2</td><td>5.2E-10</td><td>1.3E-09</td><td>MB231/MB468</td><td>MP</td></tr>
<tr><td>>hg38_21_+_8433172_8446622_RNA45SN1_45S_50ntFlanks@11694.11728.35|AAGAAATTCAATGAAGCGCGGGTAAACGGCGGGAG</td><td>>hg38_21_+_8433172_8446622_RNA45SN1_45S_50ntFlanks</td><td>11694</td><td>11728</td><td>35</td><td>AAGAAATTCAATGAAGCGCGGGTAAACGGCGGGAG</td><td>858.3</td><td>-9.5</td><td>1.5</td><td>-6.2</td><td>6.3E-10</td><td>1.3E-09</td><td>BT20/MB468</td><td>NUC</td></tr>
<tr><td>>hg38_21_+_8433172_8446622_RNA45SN1_45S_50ntFlanks@476.509.34|TTCGTGATCGATGTGGTGACGTCGTGCTCTCCCG</td><td>>hg38_21_+_8433172_8446622_RNA45SN1_45S_50ntFlanks</td><td>476</td><td>509</td><td>34</td><td>TTCGTGATCGATGTGGTGACGTCGTGCTCTCCCG</td><td>8108.9</td><td>4.3</td><td>0.7</td><td>6.2</td><td>5.5E-10</td><td>1.3E-09</td><td>MB231/MB468</td><td>NUC</td></tr>
<tr><td>>hg38_21_+_8433172_8446622_RNA45SN1_45S_50ntFlanks@5524.5549.26|GTAAAAGTCGTAACAAGGTTTCCGTA</td><td>>hg38_21_+_8433172_8446622_RNA45SN1_45S_50ntFlanks</td><td>5524</td><td>5549</td><td>26</td><td>GTAAAAGTCGTAACAAGGTTTCCGTA</td><td>80.0</td><td>-10.2</td><td>1.6</td><td>-6.2</td><td>5.2E-10</td><td>1.3E-09</td><td>MB231/MB468</td><td>MP</td></tr>
<tr><td>>hg38_21_+_8433172_8446622_RNA45SN1_45S_50ntFlanks@12654.12674.21|TATGACTGAACGCCTCTAAGT</td><td>>hg38_21_+_8433172_8446622_RNA45SN1_45S_50ntFlanks</td><td>12654</td><td>12674</td><td>21</td><td>TATGACTGAACGCCTCTAAGT</td><td>446.2</td><td>-6.3</td><td>1.0</td><td>-6.2</td><td>6.1E-10</td><td>1.3E-09</td><td>MB231/MB468</td><td>MITO</td></tr>
<tr><td>>hg38_21_+_8433172_8446622_RNA45SN1_45S_50ntFlanks@6646.6685.40|TCGTACGACTCTTAGCGGTGGATCACTCGGCTCGTGCGTC</td><td>>hg38_21_+_8433172_8446622_RNA45SN1_45S_50ntFlanks</td><td>6646</td><td>6685</td><td>40</td><td>TCGTACGACTCTTAGCGGTGGATCACTCGGCTCGTGCGTC</td><td>5534.8</td><td>4.7</td><td>0.8</td><td>6.2</td><td>6.4E-10</td><td>1.3E-09</td><td>BT20/MB468</td><td>NUC</td></tr>
<tr><td>>trna111_HisGTG_1_-_147774845_147774916@-1A.33.34,trna118_HisGTG_1_-_145396881_145396952@-1A.33.34,trna16_HisGTG_1_+_146544773_146544844@-1A.33.34,trna1_HisGTG_15_+_45493349_45493420@-1A.33.34,trna21_HisGTG_1_+_147753471_147753542@-1A.33.34,trna33_HisGTG_6_+_27125906_27125977@-1A.33.34,trna7_HisGTG_9_-_14433938_14434009@-1A.33.34,trna8_HisGTG_15_-_45492611_45492682@-1A.33.34,trna9_HisGTG_15_-_45490804_45490875@-1A.33.34|AGCCGTGATCGTATAGTGGTTAGTACTCTGCGTT</td><td>>trna111_HisGTG_1_-_147774845_147774916</td><td>-1A</td><td>33</td><td>34</td><td>AGCCGTGATCGTATAGTGGTTAGTACTCTGCGTT</td><td>201.8</td><td>10.9</td><td>1.8</td><td>6.2</td><td>6.1E-10</td><td>1.3E-09</td><td>MB231/MB468</td><td>MITO</td></tr>
<tr><td>>trnaMT_PheGAA_MT_+_577_647@36.54.19|AAAATGTTTAGACGGGCTC</td><td>>trnaMT_PheGAA_MT_+_577_647</td><td>36</td><td>54</td><td>19</td><td>AAAATGTTTAGACGGGCTC</td><td>828.1</td><td>8.7</td><td>1.4</td><td>6.2</td><td>4.1E-10</td><td>1.3E-09</td><td>BT20/MB468</td><td>MITO</td></tr>
<tr><td>>hg38_21_+_8433172_8446622_RNA45SN1_45S_50ntFlanks@9454.9495.42|CCGCGCCGGGGAGGTGGAGCACGAGCGCACGTGTTAGGACCC</td><td>>hg38_21_+_8433172_8446622_RNA45SN1_45S_50ntFlanks</td><td>9454</td><td>9495</td><td>42</td><td>CCGCGCCGGGGAGGTGGAGCACGAGCGCACGTGTTAGGACCC</td><td>137.3</td><td>-9.2</td><td>1.5</td><td>-6.2</td><td>4.1E-10</td><td>1.3E-09</td><td>BT20/MB468</td><td>MITO</td></tr>
<tr><td>>hg38_21_+_8433172_8446622_RNA45SN1_45S_50ntFlanks@9575.9594.20|ATCGGTCGTCCGACCTGGGT</td><td>>hg38_21_+_8433172_8446622_RNA45SN1_45S_50ntFlanks</td><td>9575</td><td>9594</td><td>20</td><td>ATCGGTCGTCCGACCTGGGT</td><td>179.0</td><td>-9.6</td><td>1.5</td><td>-6.2</td><td>4.1E-10</td><td>1.3E-09</td><td>BT20/MB468</td><td>MITO</td></tr>
<tr><td>>hg38_21_+_8433172_8446622_RNA45SN1_45S_50ntFlanks@10737.10755.19|TAAATCTCGCGCCGGGCCG</td><td>>hg38_21_+_8433172_8446622_RNA45SN1_45S_50ntFlanks</td><td>10737</td><td>10755</td><td>19</td><td>TAAATCTCGCGCCGGGCCG</td><td>769.5</td><td>-4.1</td><td>0.7</td><td>-6.2</td><td>4.1E-10</td><td>1.3E-09</td><td>BT20/MB468</td><td>MITO</td></tr>
<tr><td>>hg38_21_+_8433172_8446622_RNA45SN1_45S_50ntFlanks@10781.10818.38|AGGTGAACAGCCTCTGGCATGTTGGAACAATGTAGGTA</td><td>>hg38_21_+_8433172_8446622_RNA45SN1_45S_50ntFlanks</td><td>10781</td><td>10818</td><td>38</td><td>AGGTGAACAGCCTCTGGCATGTTGGAACAATGTAGGTA</td><td>2040.8</td><td>-9.8</td><td>1.6</td><td>-6.2</td><td>6.4E-10</td><td>1.3E-09</td><td>BT20/MB468</td><td>NUC</td></tr>
<tr><td>>hg38_1_-_228634819_228635039_RNA5S12_5S_50ntFlanks@84.101.18|CTCGTCTGATCTCGGAAG</td><td>>hg38_1_-_228634819_228635039_RNA5S12_5S_50ntFlanks</td><td>84</td><td>101</td><td>18</td><td>CTCGTCTGATCTCGGAAG</td><td>2879.8</td><td>-8.3</td><td>1.3</td><td>-6.2</td><td>6.4E-10</td><td>1.3E-09</td><td>BT20/MB468</td><td>NUC</td></tr>
<tr><td>>hg38_21_+_8433172_8446622_RNA45SN1_45S_50ntFlanks@8169.8186.18|CCAAGTCCTTCTGATCGA</td><td>>hg38_21_+_8433172_8446622_RNA45SN1_45S_50ntFlanks</td><td>8169</td><td>8186</td><td>18</td><td>CCAAGTCCTTCTGATCGA</td><td>332.4</td><td>-4.1</td><td>0.7</td><td>-6.2</td><td>5.6E-10</td><td>1.3E-09</td><td>MB231/MB468</td><td>NUC</td></tr>
<tr><td>>hg38_21_+_8433172_8446622_RNA45SN1_45S_50ntFlanks@9790.9830.41|CTGGCGTGGAGCCGGGCGTGGAATGCGAGTGCCTAGTGGGC</td><td>>hg38_21_+_8433172_8446622_RNA45SN1_45S_50ntFlanks</td><td>9790</td><td>9830</td><td>41</td><td>CTGGCGTGGAGCCGGGCGTGGAATGCGAGTGCCTAGTGGGC</td><td>113.3</td><td>-10.7</td><td>1.7</td><td>-6.2</td><td>5.3E-10</td><td>1.3E-09</td><td>MB231/MB468</td><td>MP</td></tr>
<tr><td>>hg38_21_+_8433172_8446622_RNA45SN1_45S_50ntFlanks@4145.4168.24|CCTGAGAAACGGCTACCACATCCA</td><td>>hg38_21_+_8433172_8446622_RNA45SN1_45S_50ntFlanks</td><td>4145</td><td>4168</td><td>24</td><td>CCTGAGAAACGGCTACCACATCCA</td><td>93.7</td><td>-10.4</td><td>1.7</td><td>-6.2</td><td>5.3E-10</td><td>1.3E-09</td><td>MB231/MB468</td><td>MP</td></tr>
<tr><td>>hg38_21_+_8433172_8446622_RNA45SN1_45S_50ntFlanks@5420.5444.25|CCGATTGGATGGTTTAGTGAGGCCC</td><td>>hg38_21_+_8433172_8446622_RNA45SN1_45S_50ntFlanks</td><td>5420</td><td>5444</td><td>25</td><td>CCGATTGGATGGTTTAGTGAGGCCC</td><td>157.0</td><td>-9.4</td><td>1.5</td><td>-6.2</td><td>4.1E-10</td><td>1.3E-09</td><td>BT20/MB468</td><td>MITO</td></tr>
<tr><td>>hg38_21_+_8433172_8446622_RNA45SN1_45S_50ntFlanks@11729.11752.24|TAACTATGACTCTCTTAAGGTAGC</td><td>>hg38_21_+_8433172_8446622_RNA45SN1_45S_50ntFlanks</td><td>11729</td><td>11752</td><td>24</td><td>TAACTATGACTCTCTTAAGGTAGC</td><td>650.5</td><td>-6.5</td><td>1.1</td><td>-6.2</td><td>6.1E-10</td><td>1.3E-09</td><td>MB231/MB468</td><td>MITO</td></tr>
<tr><td>>hg38_21_+_8433172_8446622_RNA45SN1_45S_50ntFlanks@12567.12604.38|CTCAGTACGAGAGGAACCGCAGGTTCAGACATTTGGTG</td><td>>hg38_21_+_8433172_8446622_RNA45SN1_45S_50ntFlanks</td><td>12567</td><td>12604</td><td>38</td><td>CTCAGTACGAGAGGAACCGCAGGTTCAGACATTTGGTG</td><td>87.3</td><td>-10.3</td><td>1.7</td><td>-6.2</td><td>5.3E-10</td><td>1.3E-09</td><td>MB231/MB468</td><td>MP</td></tr>
<tr><td>>hg38_21_+_8433172_8446622_RNA45SN1_45S_50ntFlanks@13007.13025.19|TATTGAAAGTCAGCCCTCG</td><td>>hg38_21_+_8433172_8446622_RNA45SN1_45S_50ntFlanks</td><td>13007</td><td>13025</td><td>19</td><td>TATTGAAAGTCAGCCCTCG</td><td>125.0</td><td>-9.1</td><td>1.4</td><td>-6.2</td><td>4.2E-10</td><td>1.3E-09</td><td>BT20/MB468</td><td>MITO</td></tr>
<tr><td>>hg38_21_+_8433172_8446622_RNA45SN1_45S_50ntFlanks@11781.11798.18|CGCATGAATGGATGAACG</td><td>>hg38_21_+_8433172_8446622_RNA45SN1_45S_50ntFlanks</td><td>11781</td><td>11798</td><td>18</td><td>CGCATGAATGGATGAACG</td><td>780.5</td><td>-5.0</td><td>0.8</td><td>-6.2</td><td>4.9E-10</td><td>1.3E-09</td><td>BT20/MB468</td><td>CYTO</td></tr>
<tr><td>>MI0000283|hsa-mir-203a&WithFlank&14|+|104117399|104117520@72.92.21&[MIMAT0000264&hsa-miR-203a-3p&offsets|+1|0;m-79&14|+|104117469|104117490&offsets|+1|0]|TGAAATGTTTAGGACCACTAG</td><td>>MI0000283|hsa-mir-203a&WithFlank&14|+|104117399|104117520</td><td>72</td><td>92</td><td>21</td><td>TGAAATGTTTAGGACCACTAG</td><td>1420.1</td><td>9.0</td><td>1.4</td><td>6.5</td><td>5.8E-11</td><td>1.3E-09</td><td>BT20/MB231</td><td>NUC</td></tr>
<tr><td>>hg38_21_+_8433172_8446622_RNA45SN1_45S_50ntFlanks@12258.12284.27|GCTTGATCTTGATTTTCAGTACGAATA</td><td>>hg38_21_+_8433172_8446622_RNA45SN1_45S_50ntFlanks</td><td>12258</td><td>12284</td><td>27</td><td>GCTTGATCTTGATTTTCAGTACGAATA</td><td>1467.6</td><td>-9.6</td><td>1.6</td><td>-6.2</td><td>6.4E-10</td><td>1.3E-09</td><td>BT20/MB468</td><td>NUC</td></tr>
<tr><td>>hg38_21_+_8433172_8446622_RNA45SN1_45S_50ntFlanks@475.496.22|CTTCGTGATCGATGTGGTGACG</td><td>>hg38_21_+_8433172_8446622_RNA45SN1_45S_50ntFlanks</td><td>475</td><td>496</td><td>22</td><td>CTTCGTGATCGATGTGGTGACG</td><td>4022.2</td><td>6.4</td><td>1.0</td><td>6.2</td><td>6.1E-10</td><td>1.3E-09</td><td>MB231/MB468</td><td>MITO</td></tr>
<tr><td>>hg38_21_+_8433172_8446622_RNA45SN1_45S_50ntFlanks@3837.3860.24|CCTCTCCTACTTGGATAACTGTGG</td><td>>hg38_21_+_8433172_8446622_RNA45SN1_45S_50ntFlanks</td><td>3837</td><td>3860</td><td>24</td><td>CCTCTCCTACTTGGATAACTGTGG</td><td>822.9</td><td>-7.4</td><td>1.2</td><td>-6.2</td><td>6.1E-10</td><td>1.3E-09</td><td>MB231/MB468</td><td>MITO</td></tr>
<tr><td>>hg38_21_+_8433172_8446622_RNA45SN1_45S_50ntFlanks@4453.4473.21|TCGGCGCCCCCTCGATGCTCT</td><td>>hg38_21_+_8433172_8446622_RNA45SN1_45S_50ntFlanks</td><td>4453</td><td>4473</td><td>21</td><td>TCGGCGCCCCCTCGATGCTCT</td><td>128.9</td><td>10.2</td><td>1.7</td><td>6.2</td><td>6.2E-10</td><td>1.3E-09</td><td>MB231/MB468</td><td>MITO</td></tr>
<tr><td>>hg38_21_+_8433172_8446622_RNA45SN1_45S_50ntFlanks@10905.10925.21|CTGGGCGCGCGCCGCGGCTGG</td><td>>hg38_21_+_8433172_8446622_RNA45SN1_45S_50ntFlanks</td><td>10905</td><td>10925</td><td>21</td><td>CTGGGCGCGCGCCGCGGCTGG</td><td>307.3</td><td>-11.3</td><td>1.7</td><td>-6.6</td><td>5.2E-11</td><td>1.3E-09</td><td>BT20/MB468</td><td>MP</td></tr>
<tr><td>>hg38_21_+_8433172_8446622_RNA45SN1_45S_50ntFlanks@12712.12730.19|CGGAGCCTCGGTTGGCCTC</td><td>>hg38_21_+_8433172_8446622_RNA45SN1_45S_50ntFlanks</td><td>12712</td><td>12730</td><td>19</td><td>CGGAGCCTCGGTTGGCCTC</td><td>145.5</td><td>-11.0</td><td>1.8</td><td>-6.2</td><td>5.3E-10</td><td>1.3E-09</td><td>MB231/MB468</td><td>MP</td></tr>
<tr><td>>hg38_21_+_8433172_8446622_RNA45SN1_45S_50ntFlanks@11233.11255.23|CCCTTCCCGTGGATCGCCCCAGC</td><td>>hg38_21_+_8433172_8446622_RNA45SN1_45S_50ntFlanks</td><td>11233</td><td>11255</td><td>23</td><td>CCCTTCCCGTGGATCGCCCCAGC</td><td>151.4</td><td>-7.5</td><td>1.2</td><td>-6.2</td><td>5.6E-10</td><td>1.3E-09</td><td>MB231/MB468</td><td>NUC</td></tr>
<tr><td>>hg38_21_+_8433172_8446622_RNA45SN1_45S_50ntFlanks@12259.12284.26|CTTGATCTTGATTTTCAGTACGAATA</td><td>>hg38_21_+_8433172_8446622_RNA45SN1_45S_50ntFlanks</td><td>12259</td><td>12284</td><td>26</td><td>CTTGATCTTGATTTTCAGTACGAATA</td><td>833.4</td><td>-4.4</td><td>0.7</td><td>-6.2</td><td>5.6E-10</td><td>1.3E-09</td><td>MB231/MB468</td><td>NUC</td></tr>
<tr><td>>hg38_21_+_8433172_8446622_RNA45SN1_45S_50ntFlanks@12942.12965.24|TTCGTAGACGACCTGCTTCTGGGT</td><td>>hg38_21_+_8433172_8446622_RNA45SN1_45S_50ntFlanks</td><td>12942</td><td>12965</td><td>24</td><td>TTCGTAGACGACCTGCTTCTGGGT</td><td>380.4</td><td>-8.2</td><td>1.3</td><td>-6.2</td><td>5.4E-10</td><td>1.3E-09</td><td>MB231/MB468</td><td>MP</td></tr>
<tr><td>>hg38_21_+_8433172_8446622_RNA45SN1_45S_50ntFlanks@9027.9058.32|CGTCTCCTCGTGGGGGGGCCGGGCCACCCCTC</td><td>>hg38_21_+_8433172_8446622_RNA45SN1_45S_50ntFlanks</td><td>9027</td><td>9058</td><td>32</td><td>CGTCTCCTCGTGGGGGGGCCGGGCCACCCCTC</td><td>225.9</td><td>-10.9</td><td>1.7</td><td>-6.6</td><td>5.2E-11</td><td>1.3E-09</td><td>BT20/MB468</td><td>MP</td></tr>
<tr><td>>hg38_21_+_8433172_8446622_RNA45SN1_45S_50ntFlanks@10682.10706.25|TCCGGCGGCGTCCGGTGAGCTCTCG</td><td>>hg38_21_+_8433172_8446622_RNA45SN1_45S_50ntFlanks</td><td>10682</td><td>10706</td><td>25</td><td>TCCGGCGGCGTCCGGTGAGCTCTCG</td><td>1885.3</td><td>-5.0</td><td>0.8</td><td>-6.2</td><td>4.9E-10</td><td>1.3E-09</td><td>BT20/MB468</td><td>CYTO</td></tr>
<tr><td>>trnaMT_PheGAA_MT_+_577_647@36.65.30|AAAATGTTTAGACGGGCTCACATCACCCCA</td><td>>trnaMT_PheGAA_MT_+_577_647</td><td>36</td><td>65</td><td>30</td><td>AAAATGTTTAGACGGGCTCACATCACCCCA</td><td>572.4</td><td>12.9</td><td>2.0</td><td>6.6</td><td>5.2E-11</td><td>1.3E-09</td><td>BT20/MB468</td><td>MP</td></tr>
<tr><td>>hg38_21_+_8433172_8446622_RNA45SN1_45S_50ntFlanks@4683.4722.40|CAAGACGGACCAGAGCGAAAGCATTTGCCAAGAATGTTTT</td><td>>hg38_21_+_8433172_8446622_RNA45SN1_45S_50ntFlanks</td><td>4683</td><td>4722</td><td>40</td><td>CAAGACGGACCAGAGCGAAAGCATTTGCCAAGAATGTTTT</td><td>198.8</td><td>-11.5</td><td>1.9</td><td>-6.2</td><td>5.4E-10</td><td>1.3E-09</td><td>MB231/MB468</td><td>MP</td></tr>
<tr><td>>hg38_21_+_8433172_8446622_RNA45SN1_45S_50ntFlanks@12456.12499.44|CCAAGCGTTGGATTGTTCACCCACTAATAGGGAACGTGAGCTGG</td><td>>hg38_21_+_8433172_8446622_RNA45SN1_45S_50ntFlanks</td><td>12456</td><td>12499</td><td>44</td><td>CCAAGCGTTGGATTGTTCACCCACTAATAGGGAACGTGAGCTGG</td><td>706.9</td><td>-9.1</td><td>1.5</td><td>-6.2</td><td>6.2E-10</td><td>1.3E-09</td><td>MB231/MB468</td><td>MITO</td></tr>
</table>

<table>
<tr><th></th><th></th><th></th><th></th><th></th><th></th><th></th><th></th><th></th><th></th><th></th><th></th><th></th><th></th><th></th></tr>
<tr><td>>hg38_21_+_8433172_8446622_RNA45SN1_45S_50ntFlanks@11580.11605.26|GAACTGGTGCGGACCAGGGGAATCCG</td><td>>hg38_21_+_8433172_8446622_RNA45SN1_45S_50ntFlanks</td><td>11580</td><td>11605</td><td>26</td><td>GAACTGGTGCGGACCAGGGGAATCCG</td><td>2842.7</td><td>-10.0</td><td>1.6</td><td>-6.2</td><td>6.5E-10</td><td>1.3E-09</td><td>BT20/MB468</td><td>NUC</td></tr>
<tr><td>>hg38_21_+_8433172_8446622_RNA45SN1_45S_50ntFlanks@12456.12491.36|CCAAGCGTTGGATTGTTCACCCACTAATAGGGAACG</td><td>>hg38_21_+_8433172_8446622_RNA45SN1_45S_50ntFlanks</td><td>12456</td><td>12491</td><td>36</td><td>CCAAGCGTTGGATTGTTCACCCACTAATAGGGAACG</td><td>224.3</td><td>-5.7</td><td>0.9</td><td>-6.2</td><td>5.6E-10</td><td>1.3E-09</td><td>MB231/MB468</td><td>CYTO</td></tr>
<tr><td>>hg38_21_+_8433172_8446622_RNA45SN1_45S_50ntFlanks@5237.5268.32|ACTGACTGGCTCAGCGTGTGCCTACCCTACGC</td><td>>hg38_21_+_8433172_8446622_RNA45SN1_45S_50ntFlanks</td><td>5237</td><td>5268</td><td>32</td><td>ACTGACTGGCTCAGCGTGTGCCTACCCTACGC</td><td>277.8</td><td>7.1</td><td>1.2</td><td>6.2</td><td>5.6E-10</td><td>1.3E-09</td><td>MB231/MB468</td><td>CYTO</td></tr>
<tr><td>>hg38_21_+_8433172_8446622_RNA45SN1_45S_50ntFlanks@8476.8497.22|CCGGCGGATCTTTCCCGCCCCC</td><td>>hg38_21_+_8433172_8446622_RNA45SN1_45S_50ntFlanks</td><td>8476</td><td>8497</td><td>22</td><td>CCGGCGGATCTTTCCCGCCCCC</td><td>132.2</td><td>-10.9</td><td>1.8</td><td>-6.2</td><td>5.4E-10</td><td>1.3E-09</td><td>MB231/MB468</td><td>MP</td></tr>
<tr><td>>trna1_GlnCTG_6_+_18836402_18836473@58.75.18,trna3_GlnCTG_17_+_8023070_8023141@58.75.18,trna42_GlnCTG_6_+_27263212_27263283@58.75.18,trna49_GlnCTG_6_+_27487308_27487379@58.75.18,trna7_GlnCTG_15_-_66161400_66161471@58.75.18,trna99_GlnCTG_6_-_28909378_28909449@58.75.18|ATCTCGGTGGAACCTCCA</td><td>>trna1_GlnCTG_6_+_18836402_18836473</td><td>58</td><td>75</td><td>18</td><td>ATCTCGGTGGAACCTCCA</td><td>172.4</td><td>-9.5</td><td>1.5</td><td>-6.2</td><td>4.2E-10</td><td>1.3E-09</td><td>BT20/MB468</td><td>MITO</td></tr>
<tr><td>>MI0000465|hsa-mir-191&WithFlank&3|-|49020612|49020715@22.43.22(+1A)&[MIMAT0000440&hsa-miR-191-5p&offsets|0|-1(+1A);m-48&3|-|49020672|49020694&offsets|0|-1(+1A)]|CAACGGAATCCCAAAAGCAGCTA</td><td>>MI0000465|hsa-mir-191&WithFlank&3|-|49020612|49020715</td><td>22</td><td>43</td><td>22(+1A)</td><td>CAACGGAATCCCAAAAGCAGCTA</td><td>1229.3</td><td>6.2</td><td>1.0</td><td>6.2</td><td>5.6E-10</td><td>1.3E-09</td><td>MB231/MB468</td><td>CYTO</td></tr>
<tr><td>>hg38_21_+_8433172_8446622_RNA45SN1_45S_50ntFlanks@10404.10430.27|CATGGGTCAGTCGGTCCTGAGAGATGG</td><td>>hg38_21_+_8433172_8446622_RNA45SN1_45S_50ntFlanks</td><td>10404</td><td>10430</td><td>27</td><td>CATGGGTCAGTCGGTCCTGAGAGATGG</td><td>496.3</td><td>-8.0</td><td>1.3</td><td>-6.2</td><td>6.2E-10</td><td>1.3E-09</td><td>MB231/MB468</td><td>MITO</td></tr>
<tr><td>>hg38_21_+_8433172_8446622_RNA45SN1_45S_50ntFlanks@4567.4587.21|TAATGGAATAGGACCGCGGTT</td><td>>hg38_21_+_8433172_8446622_RNA45SN1_45S_50ntFlanks</td><td>4567</td><td>4587</td><td>21</td><td>TAATGGAATAGGACCGCGGTT</td><td>431.2</td><td>-8.1</td><td>1.3</td><td>-6.2</td><td>6.2E-10</td><td>1.3E-09</td><td>MB231/MB468</td><td>MITO</td></tr>
<tr><td>>hg38_21_+_8433172_8446622_RNA45SN1_45S_50ntFlanks@4007.4024.18|CCTCGGGCCGATCGCACG</td><td>>hg38_21_+_8433172_8446622_RNA45SN1_45S_50ntFlanks</td><td>4007</td><td>4024</td><td>18</td><td>CCTCGGGCCGATCGCACG</td><td>705.6</td><td>-7.2</td><td>1.2</td><td>-6.2</td><td>6.5E-10</td><td>1.3E-09</td><td>BT20/MB468</td><td>NUC</td></tr>
<tr><td>>hg38_21_+_8433172_8446622_RNA45SN1_45S_50ntFlanks@9736.9767.32|CCGAAACGATCTCAACCTATTCTCAAACTTTA</td><td>>hg38_21_+_8433172_8446622_RNA45SN1_45S_50ntFlanks</td><td>9736</td><td>9767</td><td>32</td><td>CCGAAACGATCTCAACCTATTCTCAAACTTTA</td><td>56.6</td><td>-9.7</td><td>1.6</td><td>-6.2</td><td>5.4E-10</td><td>1.3E-09</td><td>MB231/MB468</td><td>MP</td></tr>
<tr><td>>hg38_21_+_8433172_8446622_RNA45SN1_45S_50ntFlanks@10369.10407.39|AAGTGGAGAAGGGTTCCATGTGAACAGCAGTTGAACATG</td><td>>hg38_21_+_8433172_8446622_RNA45SN1_45S_50ntFlanks</td><td>10369</td><td>10407</td><td>39</td><td>AAGTGGAGAAGGGTTCCATGTGAACAGCAGTTGAACATG</td><td>1122.6</td><td>-5.8</td><td>0.9</td><td>-6.2</td><td>5.6E-10</td><td>1.3E-09</td><td>MB231/MB468</td><td>CYTO</td></tr>
<tr><td>>hg38_21_+_8433172_8446622_RNA45SN1_45S_50ntFlanks@6652.6676.25|GACTCTTAGCGGTGGATCACTCGGC</td><td>>hg38_21_+_8433172_8446622_RNA45SN1_45S_50ntFlanks</td><td>6652</td><td>6676</td><td>25</td><td>GACTCTTAGCGGTGGATCACTCGGC</td><td>1467.4</td><td>5.6</td><td>0.9</td><td>6.2</td><td>5.6E-10</td><td>1.3E-09</td><td>MB231/MB468</td><td>CYTO</td></tr>
<tr><td>>MI0000078|hsa-mir-22&WithFlank&17|-|1713897|1713993@59.81.23&[MIMAT0000077&hsa-miR-22-3p&offsets|0|+1;m-2&17|-|1713914|1713935&offsets|0|+1]|AAGCTGCCAGTTGAAGAACTGTT</td><td>>MI0000078|hsa-mir-22&WithFlank&17|-|1713897|1713993</td><td>59</td><td>81</td><td>23</td><td>AAGCTGCCAGTTGAAGAACTGTT</td><td>73.3</td><td>6.7</td><td>1.1</td><td>6.2</td><td>6.6E-10</td><td>1.3E-09</td><td>BT20/MB468</td><td>NUC</td></tr>
<tr><td>>hg38_21_+_8433172_8446622_RNA45SN1_45S_50ntFlanks@11739.11768.30|TCTCTTAAGGTAGCCAAATGCCTCGTCATC</td><td>>hg38_21_+_8433172_8446622_RNA45SN1_45S_50ntFlanks</td><td>11739</td><td>11768</td><td>30</td><td>TCTCTTAAGGTAGCCAAATGCCTCGTCATC</td><td>124.3</td><td>-7.0</td><td>1.1</td><td>-6.2</td><td>5.0E-10</td><td>1.3E-09</td><td>BT20/MB468</td><td>CYTO</td></tr>
<tr><td>>hg38_21_+_8433172_8446622_RNA45SN1_45S_50ntFlanks@4322.4339.18|CAGCCGCGGTAATTCCAG</td><td>>hg38_21_+_8433172_8446622_RNA45SN1_45S_50ntFlanks</td><td>4322</td><td>4339</td><td>18</td><td>CAGCCGCGGTAATTCCAG</td><td>125.5</td><td>-9.1</td><td>1.5</td><td>-6.2</td><td>4.3E-10</td><td>1.3E-09</td><td>BT20/MB468</td><td>MITO</td></tr>
<tr><td>>hg38_21_+_8433172_8446622_RNA45SN1_45S_50ntFlanks@10798.10823.26|CATGTTGGAACAATGTAGGTAAGGGA</td><td>>hg38_21_+_8433172_8446622_RNA45SN1_45S_50ntFlanks</td><td>10798</td><td>10823</td><td>26</td><td>CATGTTGGAACAATGTAGGTAAGGGA</td><td>565.2</td><td>-8.2</td><td>1.3</td><td>-6.2</td><td>5.4E-10</td><td>1.3E-09</td><td>MB231/MB468</td><td>MP</td></tr>
<tr><td>>trna5_AspGTC_12_+_98897281_98897352@39.74.36|GCGGGAGACCGGGGTTCAATTCCCCGACGGGGAGCC</td><td>>trna5_AspGTC_12_+_98897281_98897352</td><td>39</td><td>74</td><td>36</td><td>GCGGGAGACCGGGGTTCAATTCCCCGACGGGGAGCC</td><td>638.9</td><td>7.3</td><td>1.2</td><td>6.2</td><td>5.6E-10</td><td>1.3E-09</td><td>MB231/MB468</td><td>CYTO</td></tr>
<tr><td>>hg38_21_+_8433172_8446622_RNA45SN1_45S_50ntFlanks@3785.3802.18|TGAAACTGCGAATGGCTC</td><td>>hg38_21_+_8433172_8446622_RNA45SN1_45S_50ntFlanks</td><td>3785</td><td>3802</td><td>18</td><td>TGAAACTGCGAATGGCTC</td><td>353.4</td><td>-8.4</td><td>1.4</td><td>-6.2</td><td>6.6E-10</td><td>1.3E-09</td><td>BT20/MB468</td><td>NUC</td></tr>
<tr><td>>hg38_21_+_8433172_8446622_RNA45SN1_45S_50ntFlanks@4724.4752.29|ATTAATCAAGAACGAAAGTCGGAGGTTCG</td><td>>hg38_21_+_8433172_8446622_RNA45SN1_45S_50ntFlanks</td><td>4724</td><td>4752</td><td>29</td><td>ATTAATCAAGAACGAAAGTCGGAGGTTCG</td><td>963.4</td><td>-5.0</td><td>0.8</td><td>-6.2</td><td>5.6E-10</td><td>1.3E-09</td><td>MB231/MB468</td><td>CYTO</td></tr>
<tr><td>>hg38_21_+_8433172_8446622_RNA45SN1_45S_50ntFlanks@11565.11603.39|CTAGCAGCCGACTTAGAACTGGTGCGGACCAGGGGAATC</td><td>>hg38_21_+_8433172_8446622_RNA45SN1_45S_50ntFlanks</td><td>11565</td><td>11603</td><td>39</td><td>CTAGCAGCCGACTTAGAACTGGTGCGGACCAGGGGAATC</td><td>722.0</td><td>-8.3</td><td>1.3</td><td>-6.2</td><td>6.3E-10</td><td>1.3E-09</td><td>MB231/MB468</td><td>MITO</td></tr>
<tr><td>>hg38_21_+_8433172_8446622_RNA45SN1_45S_50ntFlanks@11769.11791.23|TAATTAGTGACGCGCATGAATGG</td><td>>hg38_21_+_8433172_8446622_RNA45SN1_45S_50ntFlanks</td><td>11769</td><td>11791</td><td>23</td><td>TAATTAGTGACGCGCATGAATGG</td><td>1980.1</td><td>-7.0</td><td>1.1</td><td>-6.2</td><td>5.7E-10</td><td>1.3E-09</td><td>MB231/MB468</td><td>NUC</td></tr>
<tr><td>>hg38_21_+_8433172_8446622_RNA45SN1_45S_50ntFlanks@11255.11281.27|CTGCGGCGGGCGTCGCGGCCGCCCCCG</td><td>>hg38_21_+_8433172_8446622_RNA45SN1_45S_50ntFlanks</td><td>11255</td><td>11281</td><td>27</td><td>CTGCGGCGGGCGTCGCGGCCGCCCCCG</td><td>116.5</td><td>-9.0</td><td>1.4</td><td>-6.2</td><td>4.3E-10</td><td>1.3E-09</td><td>BT20/MB468</td><td>MITO</td></tr>
<tr><td>>MI0000434|hsa-let-7i&WithFlank&12|+|62603680|62603775@12.32.21(+1A)&[MIMAT0000415&hsa-let-7i-5p&offsets|0|-1(+1A);m-49&12|+|62603691|62603712&offsets|0|-1(+1A)]|TGAGGTAGTAGTTTGTGCTGTA</td><td>>MI0000434|hsa-let-7i&WithFlank&12|+|62603680|62603775</td><td>12</td><td>32</td><td>21(+1A)</td><td>TGAGGTAGTAGTTTGTGCTGTA</td><td>937.1</td><td>2.6</td><td>0.4</td><td>6.2</td><td>6.6E-10</td><td>1.3E-09</td><td>BT20/MB468</td><td>NUC</td></tr>
<tr><td>>hg38_21_+_8433172_8446622_RNA45SN1_45S_50ntFlanks@4734.4752.19|AACGAAAGTCGGAGGTTCG</td><td>>hg38_21_+_8433172_8446622_RNA45SN1_45S_50ntFlanks</td><td>4734</td><td>4752</td><td>19</td><td>AACGAAAGTCGGAGGTTCG</td><td>740.6</td><td>-6.6</td><td>1.1</td><td>-6.2</td><td>5.5E-10</td><td>1.3E-09</td><td>MB231/MB468</td><td>MP</td></tr>
<tr><td>>hg38_21_+_8433172_8446622_RNA45SN1_45S_50ntFlanks@5444.5467.24|CTCGGATCGGCCCCGCCGGGGTCG</td><td>>hg38_21_+_8433172_8446622_RNA45SN1_45S_50ntFlanks</td><td>5444</td><td>5467</td><td>24</td><td>CTCGGATCGGCCCCGCCGGGGTCG</td><td>695.1</td><td>-7.7</td><td>1.2</td><td>-6.2</td><td>4.3E-10</td><td>1.3E-09</td><td>BT20/MB468</td><td>MITO</td></tr>
<tr><td>>hg38_21_+_8433172_8446622_RNA45SN1_45S_50ntFlanks@12997.13039.43|CGCTGCGATCTATTGAAAGTCAGCCCTCGACACAAGGGTTTGT</td><td>>hg38_21_+_8433172_8446622_RNA45SN1_45S_50ntFlanks</td><td>12997</td><td>13039</td><td>43</td><td>CGCTGCGATCTATTGAAAGTCAGCCCTCGACACAAGGGTTTGT</td><td>145497.7</td><td>7.8</td><td>1.3</td><td>6.2</td><td>5.0E-10</td><td>1.3E-09</td><td>BT20/MB468</td><td>CYTO</td></tr>
<tr><td>>hg38_21_+_8433172_8446622_RNA45SN1_45S_50ntFlanks@6671.6689.19|CTCGGCTCGTGCGTCGATG</td><td>>hg38_21_+_8433172_8446622_RNA45SN1_45S_50ntFlanks</td><td>6671</td><td>6689</td><td>19</td><td>CTCGGCTCGTGCGTCGATG</td><td>691.5</td><td>-12.5</td><td>1.9</td><td>-6.6</td><td>5.3E-11</td><td>1.3E-09</td><td>BT20/MB468</td><td>MP</td></tr>
<tr><td>>hg38_21_+_8433172_8446622_RNA45SN1_45S_50ntFlanks@11038.11060.23|TCTCCCCCGCTCCCCGTCCTCCC</td><td>>hg38_21_+_8433172_8446622_RNA45SN1_45S_50ntFlanks</td><td>11038</td><td>11060</td><td>23</td><td>TCTCCCCCGCTCCCCGTCCTCCC</td><td>342.7</td><td>-6.0</td><td>1.0</td><td>-6.2</td><td>5.7E-10</td><td>1.3E-09</td><td>MB231/MB468</td><td>CYTO</td></tr>
<tr><td>>hg38_21_+_8433172_8446622_RNA45SN1_45S_50ntFlanks@4003.4042.40|ATAACCTCGGGCCGATCGCACGCCCCCCGTGGCGGCGACG</td><td>>hg38_21_+_8433172_8446622_RNA45SN1_45S_50ntFlanks</td><td>4003</td><td>4042</td><td>40</td><td>ATAACCTCGGGCCGATCGCACGCCCCCCGTGGCGGCGACG</td><td>110.2</td><td>-10.6</td><td>1.7</td><td>-6.2</td><td>5.5E-10</td><td>1.3E-09</td><td>MB231/MB468</td><td>MP</td></tr>
<tr><td>>hg38_21_+_8433172_8446622_RNA45SN1_45S_50ntFlanks@12324.12365.42|TTGGGTTTTAAGCAGGAGGTGTCAGAAAAGTTACCACAGGGA</td><td>>hg38_21_+_8433172_8446622_RNA45SN1_45S_50ntFlanks</td><td>12324</td><td>12365</td><td>42</td><td>TTGGGTTTTAAGCAGGAGGTGTCAGAAAAGTTACCACAGGGA</td><td>169.6</td><td>-11.3</td><td>1.8</td><td>-6.2</td><td>5.5E-10</td><td>1.3E-09</td><td>MB231/MB468</td><td>MP</td></tr>
<tr><td>>hg38_21_+_8433172_8446622_RNA45SN1_45S_50ntFlanks@9929.9946.18|GATATAGACAGCAGGACG</td><td>>hg38_21_+_8433172_8446622_RNA45SN1_45S_50ntFlanks</td><td>9929</td><td>9946</td><td>18</td><td>GATATAGACAGCAGGACG</td><td>449.5</td><td>-6.9</td><td>1.1</td><td>-6.2</td><td>5.5E-10</td><td>1.4E-09</td><td>MB231/MB468</td><td>MP</td></tr>
<tr><td>>hg38_21_+_8433172_8446622_RNA45SN1_45S_50ntFlanks@11124.11146.23|CCGGCGGCGGCCCGCCGCGGGGC</td><td>>hg38_21_+_8433172_8446622_RNA45SN1_45S_50ntFlanks</td><td>11124</td><td>11146</td><td>23</td><td>CCGGCGGCGGCCCGCCGCGGGGC</td><td>176.9</td><td>-11.3</td><td>1.8</td><td>-6.2</td><td>5.5E-10</td><td>1.4E-09</td><td>MB231/MB468</td><td>MP</td></tr>
<tr><td>>hg38_21_+_8433172_8446622_RNA45SN1_45S_50ntFlanks@12074.12108.35|CGAGGGGCTCTCGCTTCTGGCGCCAAGCGCCCGGC</td><td>>hg38_21_+_8433172_8446622_RNA45SN1_45S_50ntFlanks</td><td>12074</td><td>12108</td><td>35</td><td>CGAGGGGCTCTCGCTTCTGGCGCCAAGCGCCCGGC</td><td>81.8</td><td>-10.2</td><td>1.6</td><td>-6.2</td><td>5.5E-10</td><td>1.4E-09</td><td>MB231/MB468</td><td>MP</td></tr>
<tr><td>>trnaMT_LeuTAA_MT_+_3230_3304@11.44.34|CAGAGCCCGGTAATCGCATAAAACTTAAAACTTT</td><td>>trnaMT_LeuTAA_MT_+_3230_3304</td><td>11</td><td>44</td><td>34</td><td>CAGAGCCCGGTAATCGCATAAAACTTAAAACTTT</td><td>148.9</td><td>-10.2</td><td>1.6</td><td>-6.2</td><td>5.8E-10</td><td>1.4E-09</td><td>MB231/MB468</td><td>NUC</td></tr>
<tr><td>>hg38_21_+_8433172_8446622_RNA45SN1_45S_50ntFlanks@5421.5445.25|CGATTGGATGGTTTAGTGAGGCCCT</td><td>>hg38_21_+_8433172_8446622_RNA45SN1_45S_50ntFlanks</td><td>5421</td><td>5445</td><td>25</td><td>CGATTGGATGGTTTAGTGAGGCCCT</td><td>573.7</td><td>-7.3</td><td>1.2</td><td>-6.2</td><td>5.1E-10</td><td>1.4E-09</td><td>BT20/MB468</td><td>CYTO</td></tr>
<tr><td>>MI0000446|hsa-mir-125b-1&WithFlank&11|-|122099751|122099850@61.81.21(+1A)&[MIMAT0004592&hsa-miR-125b-1-3p&offsets|0|-1(+1A);m-485&11|-|122099769|122099790&offsets|0|-1(+1A)]|ACGGGTTAGGCTCTTGGGAGCA</td><td>>MI0000446|hsa-mir-125b-1&WithFlank&11|-|122099751|122099850</td><td>61</td><td>81</td><td>21(+1A)</td><td>ACGGGTTAGGCTCTTGGGAGCA</td><td>127.6</td><td>10.2</td><td>1.6</td><td>6.2</td><td>5.5E-10</td><td>1.4E-09</td><td>MB231/MB468</td><td>MP</td></tr>
<tr><td>>hg38_21_+_8433172_8446622_RNA45SN1_45S_50ntFlanks@3985.4011.27|CGGCTTTGGTGACTCTAGATAACCTCG</td><td>>hg38_21_+_8433172_8446622_RNA45SN1_45S_50ntFlanks</td><td>3985</td><td>4011</td><td>27</td><td>CGGCTTTGGTGACTCTAGATAACCTCG</td><td>89.0</td><td>-10.3</td><td>1.7</td><td>-6.2</td><td>5.6E-10</td><td>1.4E-09</td><td>MB231/MB468</td><td>MP</td></tr>
<tr><td>>hg38_21_+_8433172_8446622_RNA45SN1_45S_50ntFlanks@9240.9283.44|GCGGAGCGAGCGCACGGGGTCGGCGGCGACGTCGGCTACCCACC</td><td>>hg38_21_+_8433172_8446622_RNA45SN1_45S_50ntFlanks</td><td>9240</td><td>9283</td><td>44</td><td>GCGGAGCGAGCGCACGGGGTCGGCGGCGACGTCGGCTACCCACC</td><td>168.7</td><td>-11.2</td><td>1.8</td><td>-6.2</td><td>6.4E-10</td><td>1.4E-09</td><td>MB231/MB468</td><td>MITO</td></tr>
<tr><td>>hg38_21_+_8433172_8446622_RNA45SN1_45S_50ntFlanks@7974.8006.33|ACGCGACCTCAGATCAGACGTGGCGACCCGCTG</td><td>>hg38_21_+_8433172_8446622_RNA45SN1_45S_50ntFlanks</td><td>7974</td><td>8006</td><td>33</td><td>ACGCGACCTCAGATCAGACGTGGCGACCCGCTG</td><td>4076.5</td><td>-4.7</td><td>0.8</td><td>-6.2</td><td>5.6E-10</td><td>1.4E-09</td><td>MB231/MB468</td><td>MP</td></tr>
<tr><td>>hg38_21_+_8433172_8446622_RNA45SN1_45S_50ntFlanks@10819.10839.21|AGGGAAGTCGGCAAGCCGGAT</td><td>>hg38_21_+_8433172_8446622_RNA45SN1_45S_50ntFlanks</td><td>10819</td><td>10839</td><td>21</td><td>AGGGAAGTCGGCAAGCCGGAT</td><td>478.7</td><td>-8.7</td><td>1.4</td><td>-6.2</td><td>6.8E-10</td><td>1.4E-09</td><td>BT20/MB468</td><td>NUC</td></tr>
<tr><td>>hg38_21_+_8433172_8446622_RNA45SN1_45S_50ntFlanks@7974.8008.35|ACGCGACCTCAGATCAGACGTGGCGACCCGCTGAA</td><td>>hg38_21_+_8433172_8446622_RNA45SN1_45S_50ntFlanks</td><td>7974</td><td>8008</td><td>35</td><td>ACGCGACCTCAGATCAGACGTGGCGACCCGCTGAA</td><td>9231.9</td><td>4.5</td><td>0.7</td><td>6.2</td><td>5.8E-10</td><td>1.4E-09</td><td>MB231/MB468</td><td>CYTO</td></tr>
<tr><td>>hg38_21_+_8433172_8446622_RNA45SN1_45S_50ntFlanks@10844.10865.22|AACTTCGGGATAAGGATTGGCT</td><td>>hg38_21_+_8433172_8446622_RNA45SN1_45S_50ntFlanks</td><td>10844</td><td>10865</td><td>22</td><td>AACTTCGGGATAAGGATTGGCT</td><td>601.3</td><td>-4.2</td><td>0.7</td><td>-6.2</td><td>5.1E-10</td><td>1.4E-09</td><td>BT20/MB468</td><td>CYTO</td></tr>
<tr><td>>hg38_21_+_8433172_8446622_RNA45SN1_45S_50ntFlanks@4160.4180.21|CCACATCCAAGGAAGGCAGCA</td><td>>hg38_21_+_8433172_8446622_RNA45SN1_45S_50ntFlanks</td><td>4160</td><td>4180</td><td>21</td><td>CCACATCCAAGGAAGGCAGCA</td><td>107.2</td><td>-10.6</td><td>1.7</td><td>-6.2</td><td>5.6E-10</td><td>1.4E-09</td><td>MB231/MB468</td><td>MP</td></tr>
<tr><td>>hg38_21_+_8433172_8446622_RNA45SN1_45S_50ntFlanks@11727.11748.22|AGTAACTATGACTCTCTTAAGG</td><td>>hg38_21_+_8433172_8446622_RNA45SN1_45S_50ntFlanks</td><td>11727</td><td>11748</td><td>22</td><td>AGTAACTATGACTCTCTTAAGG</td><td>424.1</td><td>3.8</td><td>0.6</td><td>6.2</td><td>5.8E-10</td><td>1.4E-09</td><td>MB231/MB468</td><td>CYTO</td></tr>
<tr><td>>hg38_21_+_8433172_8446622_RNA45SN1_45S_50ntFlanks@11671.11703.33|CCAGTGCTCTGAATGTCAAAGTGAAGAAATTCA</td><td>>hg38_21_+_8433172_8446622_RNA45SN1_45S_50ntFlanks</td><td>11671</td><td>11703</td><td>33</td><td>CCAGTGCTCTGAATGTCAAAGTGAAGAAATTCA</td><td>2809.3</td><td>-10.1</td><td>1.6</td><td>-6.2</td><td>6.8E-10</td><td>1.4E-09</td><td>BT20/MB468</td><td>NUC</td></tr>
<tr><td>>hg38_21_+_8433172_8446622_RNA45SN1_45S_50ntFlanks@7978.8011.34|GACCTCAGATCAGACGTGGCGACCCGCTGAATTT</td><td>>hg38_21_+_8433172_8446622_RNA45SN1_45S_50ntFlanks</td><td>7978</td><td>8011</td><td>34</td><td>GACCTCAGATCAGACGTGGCGACCCGCTGAATTT</td><td>315.5</td><td>7.8</td><td>1.3</td><td>6.2</td><td>5.2E-10</td><td>1.4E-09</td><td>BT20/MB468</td><td>CYTO</td></tr>
<tr><td>>hg38_21_+_8433172_8446622_RNA45SN1_45S_50ntFlanks@13004.13029.26|ATCTATTGAAAGTCAGCCCTCGACAC</td><td>>hg38_21_+_8433172_8446622_RNA45SN1_45S_50ntFlanks</td><td>13004</td><td>13029</td><td>26</td><td>ATCTATTGAAAGTCAGCCCTCGACAC</td><td>293.7</td><td>-6.7</td><td>1.1</td><td>-6.2</td><td>5.9E-10</td><td>1.4E-09</td><td>MB231/MB468</td><td>CYTO</td></tr>
<tr><td>>trna11_GluTTC_15_-_26327381_26327452@1.36.36,trna3_GluTTC_13_-_45492062_45492133@1.36.36|TCCCACATGGTCTAGCGGTTAGGATTCCTGGTTTTC</td><td>>trna11_GluTTC_15_-_26327381_26327452</td><td>1</td><td>36</td><td>36</td><td>TCCCACATGGTCTAGCGGTTAGGATTCCTGGTTTTC</td><td>705.1</td><td>9.0</td><td>1.4</td><td>6.2</td><td>5.2E-10</td><td>1.4E-09</td><td>BT20/MB468</td><td>CYTO</td></tr>
<tr><td>>hg38_21_+_8433172_8446622_RNA45SN1_45S_50ntFlanks@9929.9952.24|GATATAGACAGCAGGACGGTGGCC</td><td>>hg38_21_+_8433172_8446622_RNA45SN1_45S_50ntFlanks</td><td>9929</td><td>9952</td><td>24</td><td>GATATAGACAGCAGGACGGTGGCC</td><td>1372.2</td><td>-7.5</td><td>1.2</td><td>-6.2</td><td>5.7E-10</td><td>1.4E-09</td><td>MB231/MB468</td><td>MP</td></tr>
<tr><td>>hg38_21_+_8433172_8446622_RNA45SN1_45S_50ntFlanks@3988.4020.33|CTTTGGTGACTCTAGATAACCTCGGGCCGATCG</td><td>>hg38_21_+_8433172_8446622_RNA45SN1_45S_50ntFlanks</td><td>3988</td><td>4020</td><td>33</td><td>CTTTGGTGACTCTAGATAACCTCGGGCCGATCG</td><td>64.6</td><td>-9.9</td><td>1.6</td><td>-6.2</td><td>5.7E-10</td><td>1.4E-09</td><td>MB231/MB468</td><td>MP</td></tr>
<tr><td>>hg38_21_+_8433172_8446622_RNA45SN1_45S_50ntFlanks@10393.10419.27|CAGCAGTTGAACATGGGTCAGTCGGTC</td><td>>hg38_21_+_8433172_8446622_RNA45SN1_45S_50ntFlanks</td><td>10393</td><td>10419</td><td>27</td><td>CAGCAGTTGAACATGGGTCAGTCGGTC</td><td>1371.3</td><td>-9.5</td><td>1.5</td><td>-6.2</td><td>6.9E-10</td><td>1.4E-09</td><td>BT20/MB468</td><td>NUC</td></tr>
</table>

| | | | | | | | | | | | | | |
|---|---|---|---|---|---|---|---|---|---|---|---|---|---|
| >hg38_21_+_8433172_8446622_RNA45SN1_45S_50ntFlanks@11581.11623.43\|AACTGGTGCGGACCAGGGGAATCCGACTGTTTAATTAAAACAA | >hg38_21_+_8433172_8446622_RNA45SN1_45S_50ntFlanks | 11581 | 11623 | 43 | AACTGGTGCGGACCAGGGGAATCCGACTGTTTAATTAAAACAA | 670.0 | 6.4 | 1.0 | 6.2 | 4.5E-10 | 1.4E-09 | BT20/MB468 | MITO |
| >hg38_21_+_8433172_8446622_RNA45SN1_45S_50ntFlanks@5467.5498.32\|GGCCCACGGCCCTGGCGGAGCGCTGAGAAGAC | >hg38_21_+_8433172_8446622_RNA45SN1_45S_50ntFlanks | 5467 | 5498 | 32 | GGCCCACGGCCCTGGCGGAGCGCTGAGAAGAC | 184.1 | -11.4 | 1.8 | -6.2 | 5.7E-10 | 1.4E-09 | MB231/MB468 | MP |
| >MI0000465\|hsa-mir-191&WithFlank&3\|-\|49020612\|49020715@22.43.22&[MIMAT0000440&hsa-miR-191-5p&offsets\|0\|-1;m-48&3\|-\|49020672\|49020694&offsets\|0\|-1]\|CAACGGAATCCCAAAAGCAGCT | >MI0000465\|hsa-mir-191&WithFlank&3\|-\|49020612\|49020715 | 22 | 43 | 22 | CAACGGAATCCCAAAAGCAGCT | 7746.7 | 4.4 | 0.7 | 6.2 | 6.0E-10 | 1.4E-09 | MB231/MB468 | NUC |
| >hg38_21_+_8433172_8446622_RNA45SN1_45S_50ntFlanks@11920.11957.38\|ATGAGAGGTGTAGAATAAGTGGGAGGCCCCCGGCGCCC | >hg38_21_+_8433172_8446622_RNA45SN1_45S_50ntFlanks | 11920 | 11957 | 38 | ATGAGAGGTGTAGAATAAGTGGGAGGCCCCCGGCGCCC | 132.5 | -10.9 | 1.8 | -6.2 | 5.7E-10 | 1.4E-09 | MB231/MB468 | MP |
| >hg38_21_+_8433172_8446622_RNA45SN1_45S_50ntFlanks@11092.11116.25\|CGGCGGGGGGAGAAGGGTCGGGGCG | >hg38_21_+_8433172_8446622_RNA45SN1_45S_50ntFlanks | 11092 | 11116 | 25 | CGGCGGGGGGAGAAGGGTCGGGGCG | 96.3 | -10.5 | 1.7 | -6.2 | 5.7E-10 | 1.4E-09 | MB231/MB468 | MP |
| >trnaMT_GluTTC_MT_-_14674_14742@39.66.28,trnalookalike8_GluTTC_5_-_93905172_93905240@39.66.28\|ATTGGTCGTGGTTGTAGTCCGTGCGAGA | >trnaMT_GluTTC_MT_-_14674_14742 | 39 | 66 | 28 | ATTGGTCGTGGTTGTAGTCCGTGCGAGA | 893.7 | 5.9 | 0.9 | 6.6 | 5.6E-11 | 1.4E-09 | BT20/MB468 | MP |
| >hg38_21_+_8433172_8446622_RNA45SN1_45S_50ntFlanks@10829.10851.23\|GCAAGCCGGATCCGTAACTTCGG | >hg38_21_+_8433172_8446622_RNA45SN1_45S_50ntFlanks | 10829 | 10851 | 23 | GCAAGCCGGATCCGTAACTTCGG | 176.8 | -5.9 | 0.9 | -6.2 | 5.7E-10 | 1.4E-09 | MB231/MB468 | MP |
| >hg38_21_+_8433172_8446622_RNA45SN1_45S_50ntFlanks@12158.12180.23\|GTTTGACTGGGGCGGTACACCTG | >hg38_21_+_8433172_8446622_RNA45SN1_45S_50ntFlanks | 12158 | 12180 | 23 | GTTTGACTGGGGCGGTACACCTG | 127.0 | -9.1 | 1.5 | -6.2 | 4.5E-10 | 1.4E-09 | BT20/MB468 | MITO |
| >hg38_21_+_8433172_8446622_RNA45SN1_45S_50ntFlanks@11561.11603.43\|GCGCCTAGCAGCCGACTTAGAACTGGTGCGGACCAGGGGAATC | >hg38_21_+_8433172_8446622_RNA45SN1_45S_50ntFlanks | 11561 | 11603 | 43 | GCGCCTAGCAGCCGACTTAGAACTGGTGCGGACCAGGGGAATC | 3446.1 | -7.8 | 1.3 | -6.2 | 6.0E-10 | 1.4E-09 | MB231/MB468 | NUC |
| >hg38_21_+_8433172_8446622_RNA45SN1_45S_50ntFlanks@10680.10708.29\|GTTCCGGCGGCGTCCGGTGAGCTCTCGCT | >hg38_21_+_8433172_8446622_RNA45SN1_45S_50ntFlanks | 10680 | 10708 | 29 | GTTCCGGCGGCGTCCGGTGAGCTCTCGCT | 16187.5 | -7.6 | 1.2 | -6.2 | 5.9E-10 | 1.4E-09 | MB231/MB468 | CYTO |
| >MI0003834\|hsa-mir-769&WithFlank&19\|+\|46018926\|46019055@36.56.21(+1C)&[MIMAT0003886&hsa-miR-769-5p&offsets\|0\|-1(+1C);m-216&19\|+\|46018961\|46018982&offsets\|0\|-1(+1C)]\|TGAGACCTCTGGGTTCTGAGCC | >MI0003834\|hsa-mir-769&WithFlank&19\|+\|46018926\|46019055 | 36 | 56 | 21(+1C) | TGAGACCTCTGGGTTCTGAGCC | 342.5 | 9.4 | 1.5 | 6.2 | 5.2E-10 | 1.4E-09 | BT20/MB468 | CYTO |
| >hg38_21_+_8433172_8446622_RNA45SN1_45S_50ntFlanks@10869.10894.26\|AGGGCTGGGTCGGTCGGGCTGGGGCG | >hg38_21_+_8433172_8446622_RNA45SN1_45S_50ntFlanks | 10869 | 10894 | 26 | AGGGCTGGGTCGGTCGGGCTGGGGCG | 1410.5 | -9.7 | 1.6 | -6.2 | 6.9E-10 | 1.4E-09 | BT20/MB468 | NUC |
| >trnaMT_MetCAT_MT_+_4402_4469@50.71.22\|TTATACCCTTCCCGTACTACCA | >trnaMT_MetCAT_MT_+_4402_4469 | 50 | 71 | 22 | TTATACCCTTCCCGTACTACCA | 743.2 | 7.2 | 1.2 | 6.2 | 5.9E-10 | 1.4E-09 | MB231/MB468 | CYTO |
| >am_trna2_GlyGCC_21_-_18827107_18827177@1.20.20,trna35_GlyGCC_1_+_161413094_161413164@1.20.20,trna37_GlyGCC_1_+_161420467_161420537@1.20.20,trna39_GlyGCC_1_+_161427898_161427968@1.20.20,trna41_GlyGCC_1_+_161435258_161435328@1.20.20\|GCATGGGTGGTTCAGTGGTA | >am_trna2_GlyGCC_21_-_18827107_18827177 | 1 | 20 | 20 | GCATGGGTGGTTCAGTGGTA | 1484.6 | 5.6 | 0.9 | 6.2 | 5.3E-10 | 1.4E-09 | BT20/MB468 | CYTO |
| >hg38_21_+_8433172_8446622_RNA45SN1_45S_50ntFlanks@11641.11670.30\|CGGCGGGTGTTGACGCGATGTGATTTCTGC | >hg38_21_+_8433172_8446622_RNA45SN1_45S_50ntFlanks | 11641 | 11670 | 30 | CGGCGGGTGTTGACGCGATGTGATTTCTGC | 1133.8 | -6.7 | 1.1 | -6.2 | 6.0E-10 | 1.4E-09 | MB231/MB468 | CYTO |
| >hg38_21_+_8433172_8446622_RNA45SN1_45S_50ntFlanks@6666.6696.31\|GATCACTCGGCTCGTGCGTCGATGAAGAACG | >hg38_21_+_8433172_8446622_RNA45SN1_45S_50ntFlanks | 6666 | 6696 | 31 | GATCACTCGGCTCGTGCGTCGATGAAGAACG | 142.7 | -10.2 | 1.6 | -6.6 | 5.6E-11 | 1.4E-09 | BT20/MB468 | MP |
| >am_trna116_GluCTC_1_-_145399233_145399304@1.28.28,trna59_GluCTC_1_+_249168447_249168518@1.28.28,trna71_GluCTC_1_-_161439189_161439260@1.28.28,trna74_GluCTC_1_-_161431809_161431880@1.28.28,trna77_GluCTC_1_-_161424398_161424469@1.28.28,trna77_GluCTC_6_+_28949976_28950047@1.28.28,trna80_GluCTC_1_-_161417018_161417089@1.28.28,trna87_GluCTC_6_-_126101393_126101464@1.28.28\|TCCCTGGTGGTCTAGTGGTTAGGATTCG | >am_trna116_GluCTC_1_-_145399233_145399304 | 1 | 28 | 28 | TCCCTGGTGGTCTAGTGGTTAGGATTCG | 18530.1 | -5.6 | 0.9 | -6.2 | 6.0E-10 | 1.4E-09 | MB231/MB468 | NUC |
| >hg38_21_+_8433172_8446622_RNA45SN1_45S_50ntFlanks@11913.11950.38\|AAGAGACATGAGAGGTGTAGAATAAGTGGGAGGCCCCC | >hg38_21_+_8433172_8446622_RNA45SN1_45S_50ntFlanks | 11913 | 11950 | 38 | AAGAGACATGAGAGGTGTAGAATAAGTGGGAGGCCCCC | 141.8 | -11.0 | 1.8 | -6.2 | 5.8E-10 | 1.4E-09 | MB231/MB468 | MP |
| >trnaMT_PheGAA_MT_+_577_647@1.20.20\|GTTTATGTAGCTTACCTCCT | >trnaMT_PheGAA_MT_+_577_647 | 1 | 20 | 20 | GTTTATGTAGCTTACCTCCT | 288.6 | 6.3 | 1.0 | 6.2 | 6.1E-10 | 1.4E-09 | MB231/MB468 | NUC |
| >hg38_MT_+_1621_3279_RNR2_16S_50ntFlanks@51.85.35\|GCTAAACCTAGCCCCAAACCCACTCCACCTTACTA | >hg38_MT_+_1621_3279_RNR2_16S_50ntFlanks | 51 | 85 | 35 | GCTAAACCTAGCCCCAAACCCACTCCACCTTACTA | 2962.3 | 6.5 | 1.0 | 6.2 | 4.5E-10 | 1.4E-09 | BT20/MB468 | MITO |
| >hg38_21_+_8433172_8446622_RNA45SN1_45S_50ntFlanks@12219.12236.18\|AGGACAGAAACCTCCCGT | >hg38_21_+_8433172_8446622_RNA45SN1_45S_50ntFlanks | 12219 | 12236 | 18 | AGGACAGAAACCTCCCGT | 2640.6 | 11.0 | 1.8 | 6.2 | 4.5E-10 | 1.4E-09 | BT20/MB468 | MITO |
| >hg38_21_+_8433172_8446622_RNA45SN1_45S_50ntFlanks@11792.11820.29\|ATGAACGAGATTCCCACTGTCCCTACCTA | >hg38_21_+_8433172_8446622_RNA45SN1_45S_50ntFlanks | 11792 | 11820 | 29 | ATGAACGAGATTCCCACTGTCCCTACCTA | 211.8 | -10.8 | 1.6 | -6.6 | 5.6E-11 | 1.4E-09 | BT20/MB468 | MP |
| >hg38_21_+_8433172_8446622_RNA45SN1_45S_50ntFlanks@11604.11626.23\|CGACTGTTTAATTAAAACAAAGC | >hg38_21_+_8433172_8446622_RNA45SN1_45S_50ntFlanks | 11604 | 11626 | 23 | CGACTGTTTAATTAAAACAAAGC | 485.2 | 5.1 | 0.8 | 6.2 | 4.5E-10 | 1.4E-09 | BT20/MB468 | MITO |
| >hg38_21_+_8433172_8446622_RNA45SN1_45S_50ntFlanks@9545.9567.23\|GTGGAGGTCCGTAGCGGTCCTGA | >hg38_21_+_8433172_8446622_RNA45SN1_45S_50ntFlanks | 9545 | 9567 | 23 | GTGGAGGTCCGTAGCGGTCCTGA | 433.2 | -7.5 | 1.2 | -6.2 | 5.3E-10 | 1.4E-09 | BT20/MB468 | CYTO |
| >MI0003599\|hsa-mir-589&WithFlank&7\|-\|5495813\|5495923@30.49.20&[MIMAT0004799&hsa-miR-589-5p&offsets\|0\|-2;m-162&7\|-\|5495874\|5495894&offsets\|0\|-1]\|TGAGAACCACGTCTGCTCTG | >MI0003599\|hsa-mir-589&WithFlank&7\|-\|5495813\|5495923 | 30 | 49 | 20 | TGAGAACCACGTCTGCTCTG | 88.0 | 9.7 | 1.6 | 6.2 | 5.8E-10 | 1.4E-09 | MB231/MB468 | MP |
| >hg38_21_+_8433172_8446622_RNA45SN1_45S_50ntFlanks@6649.6695.47\|TACGACTCTTAGCGGTGGATCACTCGGCTCGTGCGTCGATGAAGAAC | >hg38_21_+_8433172_8446622_RNA45SN1_45S_50ntFlanks | 6649 | 6695 | 47 | TACGACTCTTAGCGGTGGATCACTCGGCTCGTGCGTCGATGAAGAAC | 1153.3 | 6.1 | 1.0 | 6.2 | 4.5E-10 | 1.4E-09 | BT20/MB468 | MITO |
| >hg38_21_+_8433172_8446622_RNA45SN1_45S_50ntFlanks@9167.9187.21\|CGGGCGGGTCGCGCCGTCGGG | >hg38_21_+_8433172_8446622_RNA45SN1_45S_50ntFlanks | 9167 | 9187 | 21 | CGGGCGGGTCGCGCCGTCGGG | 192.5 | -9.7 | 1.6 | -6.2 | 4.5E-10 | 1.4E-09 | BT20/MB468 | MITO |
| >hg38_21_+_8433172_8446622_RNA45SN1_45S_50ntFlanks@6647.6665.19\|CGTACGACTCTTAGCGGTG | >hg38_21_+_8433172_8446622_RNA45SN1_45S_50ntFlanks | 6647 | 6665 | 19 | CGTACGACTCTTAGCGGTG | 15563.7 | 5.1 | 0.8 | 6.6 | 5.7E-11 | 1.4E-09 | BT20/MB468 | MP |
| >hg38_21_+_8433172_8446622_RNA45SN1_45S_50ntFlanks@4823.4844.22\|ATGACCCGCCGGGCAGCTTCCG | >hg38_21_+_8433172_8446622_RNA45SN1_45S_50ntFlanks | 4823 | 4844 | 22 | ATGACCCGCCGGGCAGCTTCCG | 566.6 | 5.1 | 0.8 | 6.2 | 5.3E-10 | 1.4E-09 | BT20/MB468 | CYTO |
| >trna119_LysCTT_1_-_145395522_145395594@1.36.36,trna11_LysCTT_5_-_180648979_180649051@1.36.36,trna13_LysCTT_6_+_26556774_26556846@1.36.36,trna7_LysCTT_16_+_3225692_3225764@1.36.36,trna9_LysCTT_5_+_180634755_180634827@1.36.36\|GCCCGGCTAGCTCAGTCGGTAGAGCATGAGACTCTT | >trna119_LysCTT_1_-_145395522_145395594 | 1 | 36 | 36 | GCCCGGCTAGCTCAGTCGGTAGAGCATGAGACTCTT | 1020.8 | 5.5 | 0.9 | 6.2 | 6.0E-10 | 1.4E-09 | MB231/MB468 | CYTO |
| >hg38_21_+_8433172_8446622_RNA45SN1_45S_50ntFlanks@10830.10848.19\|CAAGCCGGATCCGTAACTT | >hg38_21_+_8433172_8446622_RNA45SN1_45S_50ntFlanks | 10830 | 10848 | 19 | CAAGCCGGATCCGTAACTT | 170.7 | -7.0 | 1.1 | -6.2 | 5.4E-10 | 1.4E-09 | BT20/MB468 | CYTO |
| >hg38_1_-_228634819_228635039_RNA5S12_5S_50ntFlanks@151.171.21\|ATACCGGGTGCTGTAGGCTTT | >hg38_1_-_228634819_228635039_RNA5S12_5S_50ntFlanks | 151 | 171 | 21 | ATACCGGGTGCTGTAGGCTTT | 12580.7 | -4.8 | 0.8 | -6.2 | 5.8E-10 | 1.4E-09 | MB231/MB468 | MP |
| >hg38_21_+_8433172_8446622_RNA45SN1_45S_50ntFlanks@10239.10262.24\|GCTACGCCGCGACGAGTAGGAGGG | >hg38_21_+_8433172_8446622_RNA45SN1_45S_50ntFlanks | 10239 | 10262 | 24 | GCTACGCCGCGACGAGTAGGAGGG | 85.6 | -10.3 | 1.7 | -6.2 | 5.8E-10 | 1.4E-09 | MB231/MB468 | MP |
| >hg38_21_+_8433172_8446622_RNA45SN1_45S_50ntFlanks@4027.4046.20\|CCCCGTGGCGGCGACGACCC | >hg38_21_+_8433172_8446622_RNA45SN1_45S_50ntFlanks | 4027 | 4046 | 20 | CCCCGTGGCGGCGACGACCC | 78.8 | -10.2 | 1.6 | -6.2 | 5.8E-10 | 1.4E-09 | MB231/MB468 | MP |
| >hg38_21_+_8433172_8446622_RNA45SN1_45S_50ntFlanks@10903.10932.30\|GGCTGGGCGCGCGCCGCGGCTGGACGAGGC | >hg38_21_+_8433172_8446622_RNA45SN1_45S_50ntFlanks | 10903 | 10932 | 30 | GGCTGGGCGCGCGCCGCGGCTGGACGAGGC | 84.0 | -10.3 | 1.7 | -6.2 | 5.8E-10 | 1.4E-09 | MB231/MB468 | MP |
| >hg38_21_+_8433172_8446622_RNA45SN1_45S_50ntFlanks@10401.10429.29\|GAACATGGGTCAGTCGGTCCTGAGAGATG | >hg38_21_+_8433172_8446622_RNA45SN1_45S_50ntFlanks | 10401 | 10429 | 29 | GAACATGGGTCAGTCGGTCCTGAGAGATG | 1944.4 | -5.8 | 0.9 | -6.2 | 7.1E-10 | 1.4E-09 | BT20/MB468 | NUC |
| >hg38_21_+_8433172_8446622_RNA45SN1_45S_50ntFlanks@11203.11220.18\|CGGCGGCGGCGACTCTGG | >hg38_21_+_8433172_8446622_RNA45SN1_45S_50ntFlanks | 11203 | 11220 | 18 | CGGCGGCGGCGACTCTGG | 70.6 | -10.0 | 1.6 | -6.2 | 5.9E-10 | 1.4E-09 | MB231/MB468 | MP |
| >hg38_21_+_8433172_8446622_RNA45SN1_45S_50ntFlanks@12656.12674.19\|TGACTGAACGCCTCTAAGT | >hg38_21_+_8433172_8446622_RNA45SN1_45S_50ntFlanks | 12656 | 12674 | 19 | TGACTGAACGCCTCTAAGT | 493.6 | -3.7 | 0.6 | -6.2 | 6.1E-10 | 1.4E-09 | MB231/MB468 | NUC |
| >MI0000299\|hsa-mir-222&WithFlank&X\|-\|45747009\|45747130@75.94.20(+1C)&[MIMAT0000279&hsa-miR-222-3p&offsets\|0\|-1(+1C);m-140&X\|-\|45747033\|45747056&offsets\|0\|-4(+1C)]\|AGCTACATCTGGCTACTGGGC | >MI0000299\|hsa-mir-222&WithFlank&X\|-\|45747009\|45747130 | 75 | 94 | 20(+1C) | AGCTACATCTGGCTACTGGGC | 448.8 | 6.3 | 1.0 | 6.2 | 6.8E-10 | 1.4E-09 | MB231/MB468 | MITO |
| >hg38_21_+_8433172_8446622_RNA45SN1_45S_50ntFlanks@6645.6671.27\|CTCGTACGACTCTTAGCGGTGGATCAC | >hg38_21_+_8433172_8446622_RNA45SN1_45S_50ntFlanks | 6645 | 6671 | 27 | CTCGTACGACTCTTAGCGGTGGATCAC | 459.5 | 4.2 | 0.7 | 6.2 | 4.6E-10 | 1.4E-09 | BT20/MB468 | MITO |
| >hg38_21_+_8433172_8446622_RNA45SN1_45S_50ntFlanks@10399.10429.31\|TTGAACATGGGTCAGTCGGTCCTGAGAGATG | >hg38_21_+_8433172_8446622_RNA45SN1_45S_50ntFlanks | 10399 | 10429 | 31 | TTGAACATGGGTCAGTCGGTCCTGAGAGATG | 122.2 | -10.8 | 1.7 | -6.2 | 5.9E-10 | 1.4E-09 | MB231/MB468 | MP |
| >hg38_21_+_8433172_8446622_RNA45SN1_45S_50ntFlanks@10402.10426.25\|AACATGGGTCAGTCGGTCCTGAGAG | >hg38_21_+_8433172_8446622_RNA45SN1_45S_50ntFlanks | 10402 | 10426 | 25 | AACATGGGTCAGTCGGTCCTGAGAG | 919.3 | -8.4 | 1.4 | -6.2 | 5.9E-10 | 1.4E-09 | MB231/MB468 | MP |
| >hg38_21_+_8433172_8446622_RNA45SN1_45S_50ntFlanks@11729.11764.36\|TAACTATGACTCTCTTAAGGTAGCCAAATGCCTCGT | >hg38_21_+_8433172_8446622_RNA45SN1_45S_50ntFlanks | 11729 | 11764 | 36 | TAACTATGACTCTCTTAAGGTAGCCAAATGCCTCGT | 74.9 | -10.1 | 1.6 | -6.2 | 5.9E-10 | 1.4E-09 | MB231/MB468 | MP |
| >hg38_21_+_8433172_8446622_RNA45SN1_45S_50ntFlanks@9741.9765.25\|ACGATCTCAACCTATTCTCAAACTT | >hg38_21_+_8433172_8446622_RNA45SN1_45S_50ntFlanks | 9741 | 9765 | 25 | ACGATCTCAACCTATTCTCAAACTT | 69.1 | -10.0 | 1.6 | -6.2 | 5.9E-10 | 1.4E-09 | MB231/MB468 | MP |
| >hg38_21_+_8433172_8446622_RNA45SN1_45S_50ntFlanks@12081.12107.27\|CTCTCGCTTCTGGCGCCAAGCGCCCGG | >hg38_21_+_8433172_8446622_RNA45SN1_45S_50ntFlanks | 12081 | 12107 | 27 | CTCTCGCTTCTGGCGCCAAGCGCCCGG | 236.3 | -5.7 | 0.9 | -6.2 | 6.2E-10 | 1.4E-09 | MB231/MB468 | NUC |
| >hg38_21_+_8433172_8446622_RNA45SN1_45S_50ntFlanks@8157.8179.23\|CTCGTGGGGGGCCCAAGTCCTTC | >hg38_21_+_8433172_8446622_RNA45SN1_45S_50ntFlanks | 8157 | 8179 | 23 | CTCGTGGGGGGCCCAAGTCCTTC | 80.8 | -10.2 | 1.6 | -6.2 | 5.9E-10 | 1.4E-09 | MB231/MB468 | MP |
| >hg38_21_+_8433172_8446622_RNA45SN1_45S_50ntFlanks@12942.12965.24\|TTCGTAGACGACCTGCTTCTGGGT | >hg38_21_+_8433172_8446622_RNA45SN1_45S_50ntFlanks | 12942 | 12965 | 24 | TTCGTAGACGACCTGCTTCTGGGT | 349.9 | -6.8 | 1.1 | -6.2 | 5.4E-10 | 1.4E-09 | BT20/MB468 | CYTO |
| >hg38_21_+_8433172_8446622_RNA45SN1_45S_50ntFlanks@12419.12445.27\|GATGTCGGCTCTTCCTATCATTGTGAA | >hg38_21_+_8433172_8446622_RNA45SN1_45S_50ntFlanks | 12419 | 12445 | 27 | GATGTCGGCTCTTCCTATCATTGTGAA | 874.0 | 3.6 | 0.6 | 6.2 | 6.1E-10 | 1.4E-09 | MB231/MB468 | CYTO |
| >hg38_21_+_8433172_8446622_RNA45SN1_45S_50ntFlanks@11581.11612.32\|AACTGGTGCGGACCAGGGGAATCCGACTGTTT | >hg38_21_+_8433172_8446622_RNA45SN1_45S_50ntFlanks | 11581 | 11612 | 32 | AACTGGTGCGGACCAGGGGAATCCGACTGTTT | 2006.7 | -4.5 | 0.7 | -6.2 | 7.1E-10 | 1.4E-09 | BT20/MB468 | NUC |

| | | | | | | | | | | | | | |
|---|---|---|---|---|---|---|---|---|---|---|---|---|---|
| >hg38_21_+_8433172_8446622_RNA45SN1_45S_50ntFlanks@10420.10442.23\|CTGAGAGATGGGCGAGCGCCGTT | >hg38_21_+_8433172_8446622_RNA45SN1_45S_50ntFlanks | 10420 | 10442 | 23 | CTGAGAGATGGGCGAGCGCCGTT | 130.5 | -10.9 | 1.8 | -6.2 | 5.9E-10 | 1.4E-09 | MB231/MB468 | MP |
| >hg38_21_+_8433172_8446622_RNA45SN1_45S_50ntFlanks@7983.8001.19\|CAGATCAGACGTGGCGACC | >hg38_21_+_8433172_8446622_RNA45SN1_45S_50ntFlanks | 7983 | 8001 | 19 | CAGATCAGACGTGGCGACC | 101.8 | -10.5 | 1.7 | -6.2 | 5.9E-10 | 1.4E-09 | MB231/MB468 | MP |
| >MI0000285\|hsa-mir-205&WithFlank&1\|+\|209432127\|209432248@40.59.20(+1A)&[MIMAT0000266&hsa-miR-205-5p&offsets\|0\|-2(+1A);m-351&1\|+\|209432166\|209432187&offsets\|0\|-2(+1A)]\|TCCTTCATTCCACCGGAGTCA | >MI0000285\|hsa-mir-205&WithFlank&1\|+\|209432127\|209432248 | 40 | 59 | 20(+1A) | TCCTTCATTCCACCGGAGTCA | 648.5 | 6.4 | 1.0 | 6.2 | 4.6E-10 | 1.4E-09 | BT20/MB468 | MITO |
| >hg38_21_+_8433172_8446622_RNA45SN1_45S_50ntFlanks@5074.5093.20\|ATTCCGATAACGAACGAGAC | >hg38_21_+_8433172_8446622_RNA45SN1_45S_50ntFlanks | 5074 | 5093 | 20 | ATTCCGATAACGAACGAGAC | 246.0 | -11.8 | 1.9 | -6.2 | 5.9E-10 | 1.4E-09 | MB231/MB468 | MP |
| >hg38_21_+_8433172_8446622_RNA45SN1_45S_50ntFlanks@4770.4793.24\|TCGTAGTTCCGACCATAAACGATG | >hg38_21_+_8433172_8446622_RNA45SN1_45S_50ntFlanks | 4770 | 4793 | 24 | TCGTAGTTCCGACCATAAACGATG | 320.6 | -8.0 | 1.3 | -6.2 | 5.9E-10 | 1.5E-09 | MB231/MB468 | MP |
| >hg38_21_+_8433172_8446622_RNA45SN1_45S_50ntFlanks@12654.12671.18\|TATGACTGAACGCCTCTA | >hg38_21_+_8433172_8446622_RNA45SN1_45S_50ntFlanks | 12654 | 12671 | 18 | TATGACTGAACGCCTCTA | 563.8 | -8.8 | 1.4 | -6.2 | 7.2E-10 | 1.5E-09 | BT20/MB468 | NUC |
| >hg38_21_+_8433172_8446622_RNA45SN1_45S_50ntFlanks@8106.8143.38\|GCGGCGGGGCGCGGGACATGTGGCGTACGGAAGACCCG | >hg38_21_+_8433172_8446622_RNA45SN1_45S_50ntFlanks | 8106 | 8143 | 38 | GCGGCGGGGCGCGGGACATGTGGCGTACGGAAGACCCG | 383.6 | -8.4 | 1.4 | -6.2 | 7.2E-10 | 1.5E-09 | BT20/MB468 | NUC |
| >hg38_21_+_8433172_8446622_RNA45SN1_45S_50ntFlanks@11835.11872.38\|CACAGCCAAGGGAACGGGCTTGGCGGAATCAGCGGGGA | >hg38_21_+_8433172_8446622_RNA45SN1_45S_50ntFlanks | 11835 | 11872 | 38 | CACAGCCAAGGGAACGGGCTTGGCGGAATCAGCGGGGA | 94.1 | -10.4 | 1.7 | -6.2 | 6.0E-10 | 1.5E-09 | MB231/MB468 | MP |
| >hg38_21_+_8433172_8446622_RNA45SN1_45S_50ntFlanks@3846.3873.28\|CTTGGATAACTGTGGTAATTCTAGAGCT | >hg38_21_+_8433172_8446622_RNA45SN1_45S_50ntFlanks | 3846 | 3873 | 28 | CTTGGATAACTGTGGTAATTCTAGAGCT | 232.9 | -6.1 | 1.0 | -6.2 | 6.0E-10 | 1.5E-09 | MB231/MB468 | MP |
| >trna134_GluTTC_1_-_16861774_16861845@1.31.31,trna5_GluTTC_1_+_17199078_17199149@1.31.31,trna84_GluTTC_1_-_161391883_161391954@1.31.31,trna94_GluTTC_1_-_149664355_149664427@1.31.31\|TCCCTGGTGGTCTAGTGGCTAGGATTCGGCG | >trna134_GluTTC_1_-_16861774_16861845 | 1 | 31 | 31 | TCCCTGGTGGTCTAGTGGCTAGGATTCGGCG | 5521.3 | -5.4 | 0.9 | -6.2 | 6.2E-10 | 1.5E-09 | MB231/MB468 | CYTO |
| >hg38_21_+_8433172_8446622_RNA45SN1_45S_50ntFlanks@4047.4073.27\|ATTCGAACGTCTGCCCTATCAACTTTC | >hg38_21_+_8433172_8446622_RNA45SN1_45S_50ntFlanks | 4047 | 4073 | 27 | ATTCGAACGTCTGCCCTATCAACTTTC | 219.2 | -3.5 | 0.6 | -6.2 | 6.3E-10 | 1.5E-09 | MB231/MB468 | NUC |
| >hg38_21_+_8433172_8446622_RNA45SN1_45S_50ntFlanks@4538.4559.22\|CCGAGCCGCCTGGATACCGCAG | >hg38_21_+_8433172_8446622_RNA45SN1_45S_50ntFlanks | 4538 | 4559 | 22 | CCGAGCCGCCTGGATACCGCAG | 82.4 | -10.2 | 1.7 | -6.2 | 6.0E-10 | 1.5E-09 | MB231/MB468 | MP |
| >hg38_21_+_8433172_8446622_RNA45SN1_45S_50ntFlanks@8198.8225.28\|TGGACGGTGTGAGGCCGGTAGCGGCCCC | >hg38_21_+_8433172_8446622_RNA45SN1_45S_50ntFlanks | 8198 | 8225 | 28 | TGGACGGTGTGAGGCCGGTAGCGGCCCC | 803.3 | -9.1 | 1.5 | -6.2 | 7.2E-10 | 1.5E-09 | BT20/MB468 | NUC |
| >hg38_21_+_8433172_8446622_RNA45SN1_45S_50ntFlanks@9462.9488.27\|GGGAGGTGGAGCACGAGCGCACGTGTT | >hg38_21_+_8433172_8446622_RNA45SN1_45S_50ntFlanks | 9462 | 9488 | 27 | GGGAGGTGGAGCACGAGCGCACGTGTT | 99.3 | -10.5 | 1.7 | -6.2 | 6.0E-10 | 1.5E-09 | MB231/MB468 | MP |
| >hg38_21_+_8433172_8446622_RNA45SN1_45S_50ntFlanks@10683.10706.24\|CCGGCGGCGTCCGGTGAGCTCTCG | >hg38_21_+_8433172_8446622_RNA45SN1_45S_50ntFlanks | 10683 | 10706 | 24 | CCGGCGGCGTCCGGTGAGCTCTCG | 16888.9 | -9.7 | 1.6 | -6.2 | 6.9E-10 | 1.5E-09 | MB231/MB468 | MITO |
| >hg38_21_+_8433172_8446622_RNA45SN1_45S_50ntFlanks@9739.9760.22\|AAACGATCTCAACCTATTCTCA | >hg38_21_+_8433172_8446622_RNA45SN1_45S_50ntFlanks | 9739 | 9760 | 22 | AAACGATCTCAACCTATTCTCA | 1170.1 | -7.5 | 1.2 | -6.2 | 6.9E-10 | 1.5E-09 | MB231/MB468 | MITO |
| >hg38_21_+_8433172_8446622_RNA45SN1_45S_50ntFlanks@11576.11612.37\|CTTAGAACTGGTGCGGACCAGGGGAATCCGACTGTTT | >hg38_21_+_8433172_8446622_RNA45SN1_45S_50ntFlanks | 11576 | 11612 | 37 | CTTAGAACTGGTGCGGACCAGGGGAATCCGACTGTTT | 103.4 | -10.6 | 1.7 | -6.2 | 6.0E-10 | 1.5E-09 | MB231/MB468 | MP |
| >hg38_21_+_8433172_8446622_RNA45SN1_45S_50ntFlanks@4886.4917.32\|AAGCTGAAACTTAAAGGAATTGACGGAAGGGC | >hg38_21_+_8433172_8446622_RNA45SN1_45S_50ntFlanks | 4886 | 4917 | 32 | AAGCTGAAACTTAAAGGAATTGACGGAAGGGC | 63.9 | -9.9 | 1.6 | -6.2 | 6.0E-10 | 1.5E-09 | MB231/MB468 | MP |
| >hg38_21_+_8433172_8446622_RNA45SN1_45S_50ntFlanks@5497.5532.36\|ACGGTCGAACTTGACTATCTAGAGGAAGTAAAAGTC | >hg38_21_+_8433172_8446622_RNA45SN1_45S_50ntFlanks | 5497 | 5532 | 36 | ACGGTCGAACTTGACTATCTAGAGGAAGTAAAAGTC | 472.0 | 6.7 | 1.1 | 6.2 | 5.5E-10 | 1.5E-09 | BT20/MB468 | CYTO |
| >hg38_21_+_8433172_8446622_RNA45SN1_45S_50ntFlanks@10425.10457.33\|AGATGGGCGAGCGCCGTTCCGAAGGGACGGGCG | >hg38_21_+_8433172_8446622_RNA45SN1_45S_50ntFlanks | 10425 | 10457 | 33 | AGATGGGCGAGCGCCGTTCCGAAGGGACGGGCG | 253.2 | -8.2 | 1.3 | -6.2 | 6.3E-10 | 1.5E-09 | MB231/MB468 | NUC |
| >hg38_21_+_8433172_8446622_RNA45SN1_45S_50ntFlanks@11572.11601.30\|CCGACTTAGAACTGGTGCGGACCAGGGGAA | >hg38_21_+_8433172_8446622_RNA45SN1_45S_50ntFlanks | 11572 | 11601 | 30 | CCGACTTAGAACTGGTGCGGACCAGGGGAA | 15642.8 | -5.6 | 0.9 | -6.5 | 5.9E-11 | 1.5E-09 | BT20/MB468 | MP |
| >MI0000088\|hsa-mir-30a&WithFlank&6\|-\|71403545\|71403627@12.35.24&[MIMAT0000087&hsa-miR-30a-5p&offsets\|0\|+2;m-15&6\|-\|71403595\|71403616&offsets\|0\|+2]\|TGTAAACATCCTCGACTGGAAGCT | >MI0000088\|hsa-mir-30a&WithFlank&6\|-\|71403545\|71403627 | 12 | 35 | 24 | TGTAAACATCCTCGACTGGAAGCT | 228526.5 | 4.2 | 0.7 | 6.2 | 6.0E-10 | 1.5E-09 | MB231/MB468 | MP |
| >hg38_21_+_8433172_8446622_RNA45SN1_45S_50ntFlanks@11671.11707.37\|CCAGTGCTCTGAATGTCAAAGTGAAGAAATTCAATGA | >hg38_21_+_8433172_8446622_RNA45SN1_45S_50ntFlanks | 11671 | 11707 | 37 | CCAGTGCTCTGAATGTCAAAGTGAAGAAATTCAATGA | 1096.0 | -7.4 | 1.2 | -6.2 | 7.3E-10 | 1.5E-09 | BT20/MB468 | NUC |
| >hg38_21_+_8433172_8446622_RNA45SN1_45S_50ntFlanks@10840.10868.29\|CCGTAACTTCGGGATAAGGATTGGCTCTA | >hg38_21_+_8433172_8446622_RNA45SN1_45S_50ntFlanks | 10840 | 10868 | 29 | CCGTAACTTCGGGATAAGGATTGGCTCTA | 2399.0 | -9.9 | 1.6 | -6.2 | 7.3E-10 | 1.5E-09 | BT20/MB468 | NUC |
| >hg38_21_+_8433172_8446622_RNA45SN1_45S_50ntFlanks@10725.10752.28\|GGGGAGAGGGTGTAAATCTCGCGCCGGG | >hg38_21_+_8433172_8446622_RNA45SN1_45S_50ntFlanks | 10725 | 10752 | 28 | GGGGAGAGGGTGTAAATCTCGCGCCGGG | 134.3 | -10.9 | 1.8 | -6.2 | 6.0E-10 | 1.5E-09 | MB231/MB468 | MP |
| >trna111_HisGTG_1_-_147774845_147774916@-1T.26.27,trna118_HisGTG_1_-_145396881_145396952@-1T.26.27,trna16_HisGTG_1_+_146544773_146544844@-1T.26.27,trna1_HisGTG_15_+_45493349_45493420@-1T.26.27,trna21_HisGTG_1_+_147753471_147753542@-1T.26.27,trna33_HisGTG_6_+_27125906_27125977@-1T.26.27,trna7_HisGTG_9_-_14433938_14434009@-1T.26.27,trna8_HisGTG_15_-_45492611_45492682@-1T.26.27,trna9_HisGTG_15_-_45490804_45490875@-1T.26.27\|TGCCGTGATCGTATAGTGGTTAGTACT | >trna111_HisGTG_1_-_147774845_147774916 | -1T | 26 | 27 | TGCCGTGATCGTATAGTGGTTAGTACT | 4989.4 | 5.4 | 0.9 | 6.2 | 6.3E-10 | 1.5E-09 | MB231/MB468 | NUC |
| >MI0000088\|hsa-mir-30a&WithFlank&6\|-\|71403545\|71403627@53.72.20&[MIMAT0000088&hsa-miR-30a-3p&offsets\|0\|-2;m-38&6\|-\|71403554\|71403575&offsets\|0\|-2]\|CTTTCAGTCGGATGTTTGCA | >MI0000088\|hsa-mir-30a&WithFlank&6\|-\|71403545\|71403627 | 53 | 72 | 20 | CTTTCAGTCGGATGTTTGCA | 145.6 | 10.4 | 1.7 | 6.2 | 6.9E-10 | 1.5E-09 | MB231/MB468 | MITO |
| >hg38_21_+_8433172_8446622_RNA45SN1_45S_50ntFlanks@12300.12319.20\|GGGCCTCACGATCCTTCTGA | >hg38_21_+_8433172_8446622_RNA45SN1_45S_50ntFlanks | 12300 | 12319 | 20 | GGGCCTCACGATCCTTCTGA | 598.3 | -9.2 | 1.5 | -6.2 | 6.9E-10 | 1.5E-09 | MB231/MB468 | MITO |
| >hg38_21_+_8433172_8446622_RNA45SN1_45S_50ntFlanks@8168.8186.19\|CCCAAGTCCTTCTGATCGA | >hg38_21_+_8433172_8446622_RNA45SN1_45S_50ntFlanks | 8168 | 8186 | 19 | CCCAAGTCCTTCTGATCGA | 1409.6 | -8.8 | 1.4 | -6.2 | 7.0E-10 | 1.5E-09 | MB231/MB468 | MITO |
| >hg38_21_+_8433172_8446622_RNA45SN1_45S_50ntFlanks@3903.3938.36\|CGCGGGGGGGATGCGTGCATTTATCAGATCAAAACC | >hg38_21_+_8433172_8446622_RNA45SN1_45S_50ntFlanks | 3903 | 3938 | 36 | CGCGGGGGGGATGCGTGCATTTATCAGATCAAAACC | 175.7 | -11.3 | 1.8 | -6.2 | 6.1E-10 | 1.5E-09 | MB231/MB468 | MP |
| >hg38_21_+_8433172_8446622_RNA45SN1_45S_50ntFlanks@4995.5026.32\|ACAGATTGATAGCTCTTTCTCGATTCCGTGGG | >hg38_21_+_8433172_8446622_RNA45SN1_45S_50ntFlanks | 4995 | 5026 | 32 | ACAGATTGATAGCTCTTTCTCGATTCCGTGGG | 95.7 | -10.4 | 1.7 | -6.2 | 6.1E-10 | 1.5E-09 | MB231/MB468 | MP |
| >hg38_21_+_8433172_8446622_RNA45SN1_45S_50ntFlanks@3911.3937.27\|GGATGCGTGCATTTATCAGATCAAAAC | >hg38_21_+_8433172_8446622_RNA45SN1_45S_50ntFlanks | 3911 | 3937 | 27 | GGATGCGTGCATTTATCAGATCAAAAC | 95.0 | -10.4 | 1.7 | -6.2 | 6.1E-10 | 1.5E-09 | MB231/MB468 | MP |
| >hg38_21_+_8433172_8446622_RNA45SN1_45S_50ntFlanks@8154.8184.31\|CCGCTCGTGGGGGGCCCAAGTCCTTCTGATC | >hg38_21_+_8433172_8446622_RNA45SN1_45S_50ntFlanks | 8154 | 8184 | 31 | CCGCTCGTGGGGGGCCCAAGTCCTTCTGATC | 541.4 | -6.2 | 1.0 | -6.2 | 7.0E-10 | 1.5E-09 | MB231/MB468 | MITO |
| >MI0000811\|hsa-mir-148b&WithFlank&12\|+\|54337210\|54337320@69.90.22&[MIMAT0000759&hsa-miR-148b-3p&offsets\|0\|0;m-77&12\|+\|54337278\|54337299&offsets\|0\|0]\|TCAGTGCATCACAGAACTTTGT | >MI0000811\|hsa-mir-148b&WithFlank&12\|+\|54337210\|54337320 | 69 | 90 | 22 | TCAGTGCATCACAGAACTTTGT | 6814.1 | 6.1 | 1.0 | 6.2 | 6.3E-10 | 1.5E-09 | MB231/MB468 | CYTO |
| >hg38_21_+_8433172_8446622_RNA45SN1_45S_50ntFlanks@11783.11807.25\|CATGAATGGATGAACGAGATTCCCA | >hg38_21_+_8433172_8446622_RNA45SN1_45S_50ntFlanks | 11783 | 11807 | 25 | CATGAATGGATGAACGAGATTCCCA | 392.9 | -11.7 | 1.8 | -6.5 | 6.0E-11 | 1.5E-09 | BT20/MB468 | MP |
| >MI0000749\|hsa-mir-30e&WithFlank&1\|+\|40754349\|40754452@23.45.23(+1C)&[MIMAT0000692&hsa-miR-30e-5p&offsets\|0\|+1(+1C);m-24&1\|+\|40754371\|40754394&offsets\|0\|-1(+1C)]\|TGTAAACATCCTTGACTGGAAGCC | >MI0000749\|hsa-mir-30e&WithFlank&1\|+\|40754349\|40754452 | 23 | 45 | 23(+1C) | TGTAAACATCCTTGACTGGAAGCC | 511.3 | 5.0 | 0.8 | 6.2 | 6.4E-10 | 1.5E-09 | MB231/MB468 | NUC |
| >hg38_21_+_8433172_8446622_RNA45SN1_45S_50ntFlanks@11999.12034.36\|CCCTGCGGGCCGCCGGTGAAATACCACTACTCTGAT | >hg38_21_+_8433172_8446622_RNA45SN1_45S_50ntFlanks | 11999 | 12034 | 36 | CCCTGCGGGCCGCCGGTGAAATACCACTACTCTGAT | 309.5 | -8.0 | 1.3 | -6.2 | 6.4E-10 | 1.5E-09 | MB231/MB468 | NUC |
| >hg38_21_+_8433172_8446622_RNA45SN1_45S_50ntFlanks@7965.7983.19\|CCCTCCGAGACGCGACCTC | >hg38_21_+_8433172_8446622_RNA45SN1_45S_50ntFlanks | 7965 | 7983 | 19 | CCCTCCGAGACGCGACCTC | 196.0 | 4.9 | 0.8 | 6.2 | 6.4E-10 | 1.5E-09 | MB231/MB468 | NUC |
| >hg38_21_+_8433172_8446622_RNA45SN1_45S_50ntFlanks@6733.6773.41\|CATTGATCATCGACACTTCGAACGCACTTGCGGCCCCGGGT | >hg38_21_+_8433172_8446622_RNA45SN1_45S_50ntFlanks | 6733 | 6773 | 41 | CATTGATCATCGACACTTCGAACGCACTTGCGGCCCCGGGT | 378.1 | 7.0 | 1.1 | 6.2 | 5.6E-10 | 1.5E-09 | BT20/MB468 | CYTO |
| >hg38_21_+_8433172_8446622_RNA45SN1_45S_50ntFlanks@12523.12546.24\|TAGTTTTACCCTACTGATGATGTG | >hg38_21_+_8433172_8446622_RNA45SN1_45S_50ntFlanks | 12523 | 12546 | 24 | TAGTTTTACCCTACTGATGATGTG | 69.8 | -10.0 | 1.6 | -6.2 | 6.1E-10 | 1.5E-09 | MB231/MB468 | MP |
| >hg38_21_+_8433172_8446622_RNA45SN1_45S_50ntFlanks@12628.12653.26\|TGGGGCGAAGCTACCATCTGTGGGAT | >hg38_21_+_8433172_8446622_RNA45SN1_45S_50ntFlanks | 12628 | 12653 | 26 | TGGGGCGAAGCTACCATCTGTGGGAT | 954.7 | -6.1 | 1.0 | -6.2 | 6.3E-10 | 1.5E-09 | MB231/MB468 | CYTO |
| >hg38_21_+_8433172_8446622_RNA45SN1_45S_50ntFlanks@12654.12674.21\|TATGACTGAACGCCTCTAAGT | >hg38_21_+_8433172_8446622_RNA45SN1_45S_50ntFlanks | 12654 | 12674 | 21 | TATGACTGAACGCCTCTAAGT | 4943.4 | -10.5 | 1.7 | -6.2 | 7.4E-10 | 1.5E-09 | BT20/MB468 | NUC |
| >hg38_21_+_8433172_8446622_RNA45SN1_45S_50ntFlanks@4995.5012.18\|ACAGATTGATAGCTCTTT | >hg38_21_+_8433172_8446622_RNA45SN1_45S_50ntFlanks | 4995 | 5012 | 18 | ACAGATTGATAGCTCTTT | 888.6 | -7.5 | 1.2 | -6.2 | 6.1E-10 | 1.5E-09 | MB231/MB468 | MP |
| >hg38_21_+_8433172_8446622_RNA45SN1_45S_50ntFlanks@6721.6742.22\|AATTGCAGGACACATTGATCAT | >hg38_21_+_8433172_8446622_RNA45SN1_45S_50ntFlanks | 6721 | 6742 | 22 | AATTGCAGGACACATTGATCAT | 728.0 | -6.7 | 1.1 | -6.2 | 6.5E-10 | 1.5E-09 | MB231/MB468 | NUC |
| >trna13_ValCAC_19_-_4724647_4724719@1.33.33\|GTTTCCGTAGTGTAGCGGTTATCACATTCGCCT | >trna13_ValCAC_19_-_4724647_4724719 | 1 | 33 | 33 | GTTTCCGTAGTGTAGCGGTTATCACATTCGCCT | 647.2 | 10.0 | 1.6 | 6.2 | 6.4E-10 | 1.5E-09 | MB231/MB468 | CYTO |
| >hg38_21_+_8433172_8446622_RNA45SN1_45S_50ntFlanks@12054.12071.18\|TGAGGCGGGGGGGCGAGC | >hg38_21_+_8433172_8446622_RNA45SN1_45S_50ntFlanks | 12054 | 12071 | 18 | TGAGGCGGGGGGGCGAGC | 122.8 | -9.0 | 1.5 | -6.2 | 4.8E-10 | 1.5E-09 | BT20/MB468 | MITO |
| >trna119_LysCTT_1_-_145395522_145395594@2.35.34,trna11_LysCTT_5_-_180648979_180649051@2.35.34,trna13_LysCTT_6_+_26556774_26556846@2.35.34,trna7_LysCTT_16_+_3225692_3225764@2.35.34,trna9_LysCTT_5_+_180634755_180634827@2.35.34\|CCCGGCTAGCTCAGTCGGTAGAGCATGAGACTCT | >trna119_LysCTT_1_-_145395522_145395594 | 2 | 35 | 34 | CCCGGCTAGCTCAGTCGGTAGAGCATGAGACTCT | 177.3 | 10.7 | 1.7 | 6.2 | 7.1E-10 | 1.5E-09 | MB231/MB468 | MITO |
| >MI0000089\|hsa-mir-31&WithFlank&9\|-\|21512109\|21512191@14.35.22&[MIMAT0000089&hsa-miR-31-5p&offsets\|0\|+1;m-360&9\|-\|21512157\|21512178&offsets\|0\|0]\|AGGCAAGATGCTGGCATAGCTG | >MI0000089\|hsa-mir-31&WithFlank&9\|-\|21512109\|21512191 | 14 | 35 | 22 | AGGCAAGATGCTGGCATAGCTG | 126.6 | -10.8 | 1.8 | -6.2 | 6.2E-10 | 1.5E-09 | MB231/MB468 | MP |

<table>
<tr><td>>hg38_21_+_8433172_8446622_RNA45SN1_45S_50ntFlanks@10395.10424.30|GCAGTTGAACATGGGTCAGTCGGTCCTGAG</td><td>>hg38_21_+_8433172_8446622_RNA45SN1_45S_50ntFlanks</td><td>10395</td><td>10424</td><td>30</td><td>GCAGTTGAACATGGGTCAGTCGGTCCTGAG</td><td>1088.3</td><td>4.6</td><td>0.7</td><td>6.2</td><td>5.6E-10</td><td>1.5E-09</td><td>BT20/MB468</td><td>CYTO</td></tr>
<tr><td>>hg38_21_+_8433172_8446622_RNA45SN1_45S_50ntFlanks@12418.12436.19|CGATGTCGGCTCTTCCTAT</td><td>>hg38_21_+_8433172_8446622_RNA45SN1_45S_50ntFlanks</td><td>12418</td><td>12436</td><td>19</td><td>CGATGTCGGCTCTTCCTAT</td><td>16852.9</td><td>-5.0</td><td>0.8</td><td>-6.2</td><td>5.6E-10</td><td>1.5E-09</td><td>BT20/MB468</td><td>CYTO</td></tr>
<tr><td>>MI0000809|hsa-mir-151a&WithFlank&8|-|140732558|140732659@53.74.22&[MIMAT0000757&hsa-miR-151a-3p&offsets|0|+1;m-30&8|-|140732587|140732607&offsets|0|+1]|CTAGACTGAAGCTCCTTGAGGA</td><td>>MI0000809|hsa-mir-151a&WithFlank&8|-|140732558|140732659</td><td>53</td><td>74</td><td>22</td><td>CTAGACTGAAGCTCCTTGAGGA</td><td>12017.4</td><td>5.1</td><td>0.8</td><td>6.5</td><td>6.1E-11</td><td>1.5E-09</td><td>BT20/MB468</td><td>MP</td></tr>
<tr><td>>hg38_21_+_8433172_8446622_RNA45SN1_45S_50ntFlanks@6645.6667.23|CTCGTACGACTCTTAGCGGTGGA</td><td>>hg38_21_+_8433172_8446622_RNA45SN1_45S_50ntFlanks</td><td>6645</td><td>6667</td><td>23</td><td>CTCGTACGACTCTTAGCGGTGGA</td><td>1813.9</td><td>4.7</td><td>0.7</td><td>6.5</td><td>6.1E-11</td><td>1.5E-09</td><td>BT20/MB468</td><td>MP</td></tr>
<tr><td>>hg38_21_+_8433172_8446622_RNA45SN1_45S_50ntFlanks@9938.9980.43|AGCAGGACGGTGGCCATGGAAGTCGGAATCCGCTAAGGAGTGT</td><td>>hg38_21_+_8433172_8446622_RNA45SN1_45S_50ntFlanks</td><td>9938</td><td>9980</td><td>43</td><td>AGCAGGACGGTGGCCATGGAAGTCGGAATCCGCTAAGGAGTGT</td><td>176.7</td><td>11.2</td><td>1.7</td><td>6.5</td><td>6.1E-11</td><td>1.5E-09</td><td>BT20/MB468</td><td>MP</td></tr>
<tr><td>>hg38_21_+_8433172_8446622_RNA45SN1_45S_50ntFlanks@4259.4278.20|ATTGGAATGAGTCCACTTTA</td><td>>hg38_21_+_8433172_8446622_RNA45SN1_45S_50ntFlanks</td><td>4259</td><td>4278</td><td>20</td><td>ATTGGAATGAGTCCACTTTA</td><td>215.0</td><td>-10.8</td><td>1.7</td><td>-6.5</td><td>6.1E-11</td><td>1.5E-09</td><td>BT20/MB468</td><td>MP</td></tr>
<tr><td>>hg38_21_+_8433172_8446622_RNA45SN1_45S_50ntFlanks@6748.6771.24|CTTCGAACGCACTTGCGGCCCCGG</td><td>>hg38_21_+_8433172_8446622_RNA45SN1_45S_50ntFlanks</td><td>6748</td><td>6771</td><td>24</td><td>CTTCGAACGCACTTGCGGCCCCGG</td><td>110.2</td><td>-6.1</td><td>1.0</td><td>-6.2</td><td>5.7E-10</td><td>1.5E-09</td><td>BT20/MB468</td><td>CYTO</td></tr>
<tr><td>>MI0000102|hsa-mir-100&WithFlank&11|-|122152223|122152314@19.38.20(+1A)&[MIMAT0000098&hsa-miR-100-5p&offsets|0|-2(+1A);m-27&11|-|122152276|122152296&offsets|0|-1(+1A)]|AACCCGTAGATCCGAACTTGA</td><td>>MI0000102|hsa-mir-100&WithFlank&11|-|122152223|122152314</td><td>19</td><td>38</td><td>20(+1A)</td><td>AACCCGTAGATCCGAACTTGA</td><td>11880.1</td><td>-7.9</td><td>1.2</td><td>-6.8</td><td>1.2E-11</td><td>1.5E-09</td><td>BT20/MB231</td><td>MP</td></tr>
<tr><td>>hg38_21_+_8433172_8446622_RNA45SN1_45S_50ntFlanks@10826.10858.33|TCGGCAAGCCGGATCCGTAACTTCGGGATAAGG</td><td>>hg38_21_+_8433172_8446622_RNA45SN1_45S_50ntFlanks</td><td>10826</td><td>10858</td><td>33</td><td>TCGGCAAGCCGGATCCGTAACTTCGGGATAAGG</td><td>524.0</td><td>-8.1</td><td>1.3</td><td>-6.2</td><td>6.4E-10</td><td>1.5E-09</td><td>MB231/MB468</td><td>CYTO</td></tr>
<tr><td>>hg38_MT_+_1621_3279_RNR2_16S_50ntFlanks@1182.1216.35|AAAATTTCGGTTGGGGCGACCTCGGAGCAGAACCC</td><td>>hg38_MT_+_1621_3279_RNR2_16S_50ntFlanks</td><td>1182</td><td>1216</td><td>35</td><td>AAAATTTCGGTTGGGGCGACCTCGGAGCAGAACCC</td><td>584.2</td><td>8.6</td><td>1.4</td><td>6.2</td><td>4.9E-10</td><td>1.5E-09</td><td>BT20/MB468</td><td>MITO</td></tr>
<tr><td>>hg38_21_+_8433172_8446622_RNA45SN1_45S_50ntFlanks@11881.11904.24|CCTGTTGAGCTTGACTCTAGTCTG</td><td>>hg38_21_+_8433172_8446622_RNA45SN1_45S_50ntFlanks</td><td>11881</td><td>11904</td><td>24</td><td>CCTGTTGAGCTTGACTCTAGTCTG</td><td>787.9</td><td>-4.2</td><td>0.7</td><td>-6.2</td><td>4.9E-10</td><td>1.5E-09</td><td>BT20/MB468</td><td>MITO</td></tr>
<tr><td>>hg38_21_+_8433172_8446622_RNA45SN1_45S_50ntFlanks@5078.5099.22|CGATAACGAACGAGACTCTGGC</td><td>>hg38_21_+_8433172_8446622_RNA45SN1_45S_50ntFlanks</td><td>5078</td><td>5099</td><td>22</td><td>CGATAACGAACGAGACTCTGGC</td><td>177.4</td><td>-11.3</td><td>1.8</td><td>-6.2</td><td>6.2E-10</td><td>1.5E-09</td><td>MB231/MB468</td><td>MP</td></tr>
<tr><td>>hg38_21_+_8433172_8446622_RNA45SN1_45S_50ntFlanks@10391.10433.43|AACAGCAGTTGAACATGGGTCAGTCGGTCCTGAGAGATGGGCG</td><td>>hg38_21_+_8433172_8446622_RNA45SN1_45S_50ntFlanks</td><td>10391</td><td>10433</td><td>43</td><td>AACAGCAGTTGAACATGGGTCAGTCGGTCCTGAGAGATGGGCG</td><td>715.5</td><td>-7.4</td><td>1.2</td><td>-6.2</td><td>7.2E-10</td><td>1.5E-09</td><td>MB231/MB468</td><td>MITO</td></tr>
<tr><td>>MI0000065|hsa-let-7d&WithFlank&9|+|94178828|94178926@14.31.18&[MIMAT0000065&hsa-let-7d-5p&offsets|0|-4;m-90&9|+|94178841|94178862&offsets|0|-4]|AGAGGTAGTAGGTTGCAT</td><td>>MI0000065|hsa-let-7d&WithFlank&9|+|94178828|94178926</td><td>14</td><td>31</td><td>18</td><td>AGAGGTAGTAGGTTGCAT</td><td>1259.9</td><td>9.5</td><td>1.5</td><td>6.2</td><td>5.7E-10</td><td>1.5E-09</td><td>BT20/MB468</td><td>CYTO</td></tr>
<tr><td>>hg38_21_+_8433172_8446622_RNA45SN1_45S_50ntFlanks@4894.4917.24|ACTTAAAGGAATTGACGGAAGGGC</td><td>>hg38_21_+_8433172_8446622_RNA45SN1_45S_50ntFlanks</td><td>4894</td><td>4917</td><td>24</td><td>ACTTAAAGGAATTGACGGAAGGGC</td><td>125.1</td><td>-10.8</td><td>1.8</td><td>-6.2</td><td>6.3E-10</td><td>1.5E-09</td><td>MB231/MB468</td><td>MP</td></tr>
<tr><td>>MI0000266|hsa-mir-10a&WithFlank&17|-|48579832|48579953@28.48.21&[MIMAT0000253&hsa-miR-10a-5p&offsets|0|-2;m-18&17|-|48579905|48579926&offsets|0|-1]|TACCCTGTAGATCCGAATTTG</td><td>>MI0000266|hsa-mir-10a&WithFlank&17|-|48579832|48579953</td><td>28</td><td>48</td><td>21</td><td>TACCCTGTAGATCCGAATTTG</td><td>145.9</td><td>10.4</td><td>1.7</td><td>6.2</td><td>7.2E-10</td><td>1.5E-09</td><td>MB231/MB468</td><td>MITO</td></tr>
<tr><td>>hg38_21_+_8433172_8446622_RNA45SN1_45S_50ntFlanks@11561.11603.43|GCGCCTAGCAGCCGACTTAGAACTGGTGCGGACCAGGGGAATC</td><td>>hg38_21_+_8433172_8446622_RNA45SN1_45S_50ntFlanks</td><td>11561</td><td>11603</td><td>43</td><td>GCGCCTAGCAGCCGACTTAGAACTGGTGCGGACCAGGGGAATC</td><td>4282.5</td><td>-5.3</td><td>0.9</td><td>-6.2</td><td>4.9E-10</td><td>1.5E-09</td><td>BT20/MB468</td><td>MITO</td></tr>
<tr><td>>hg38_21_+_8433172_8446622_RNA45SN1_45S_50ntFlanks@12417.12448.32|TCGATGTCGGCTCTTCCTATCATTGTGAAGCA</td><td>>hg38_21_+_8433172_8446622_RNA45SN1_45S_50ntFlanks</td><td>12417</td><td>12448</td><td>32</td><td>TCGATGTCGGCTCTTCCTATCATTGTGAAGCA</td><td>2665.2</td><td>4.5</td><td>0.7</td><td>6.2</td><td>6.5E-10</td><td>1.5E-09</td><td>MB231/MB468</td><td>CYTO</td></tr>
<tr><td>>hg38_21_+_8433172_8446622_RNA45SN1_45S_50ntFlanks@10346.10390.45|TCAAACGAGAACTTTGAAGGCCGAAGTGGAGAAGGGTTCCATGTG</td><td>>hg38_21_+_8433172_8446622_RNA45SN1_45S_50ntFlanks</td><td>10346</td><td>10390</td><td>45</td><td>TCAAACGAGAACTTTGAAGGCCGAAGTGGAGAAGGGTTCCATGTG</td><td>1074.7</td><td>-8.1</td><td>1.3</td><td>-6.2</td><td>7.6E-10</td><td>1.5E-09</td><td>BT20/MB468</td><td>NUC</td></tr>
<tr><td>>MI0000342|hsa-mir-200b&WithFlank&1|+|1167098|1167204@63.85.23&[MIMAT0000318&hsa-miR-200b-3p&offsets|0|+1;m-93&1|+|1167160|1167182&offsets|0|0]|TAATACTGCCTGGTAATGATGAC</td><td>>MI0000342|hsa-mir-200b&WithFlank&1|+|1167098|1167204</td><td>63</td><td>85</td><td>23</td><td>TAATACTGCCTGGTAATGATGAC</td><td>2136.6</td><td>-5.0</td><td>0.8</td><td>-6.5</td><td>6.2E-11</td><td>1.5E-09</td><td>BT20/MB468</td><td>MP</td></tr>
<tr><td>>hg38_21_+_8433172_8446622_RNA45SN1_45S_50ntFlanks@6738.6770.33|ATCATCGACACTTCGAACGCACTTGCGGCCCCG</td><td>>hg38_21_+_8433172_8446622_RNA45SN1_45S_50ntFlanks</td><td>6738</td><td>6770</td><td>33</td><td>ATCATCGACACTTCGAACGCACTTGCGGCCCCG</td><td>2066.9</td><td>-7.8</td><td>1.3</td><td>-6.2</td><td>6.5E-10</td><td>1.5E-09</td><td>MB231/MB468</td><td>CYTO</td></tr>
<tr><td>>MI0000803|hsa-mir-330&WithFlank&19|-|45638988|45639093@64.86.23(+1U)&[MIMAT0000751&hsa-miR-330-3p&offsets|+1|+1(+1U);m-347&19|-|45639009|45639030&offsets|0|+1(+1U)]|CAAAGCACACGGCCTGCAGAGAGT</td><td>>MI0000803|hsa-mir-330&WithFlank&19|-|45638988|45639093</td><td>64</td><td>86</td><td>23(+1U)</td><td>CAAAGCACACGGCCTGCAGAGAGT</td><td>272.5</td><td>8.3</td><td>1.4</td><td>6.2</td><td>6.5E-10</td><td>1.5E-09</td><td>MB231/MB468</td><td>CYTO</td></tr>
<tr><td>>hg38_21_+_8433172_8446622_RNA45SN1_45S_50ntFlanks@10344.10373.30|ATTCAAACGAGAACTTTGAAGGCCGAAGTG</td><td>>hg38_21_+_8433172_8446622_RNA45SN1_45S_50ntFlanks</td><td>10344</td><td>10373</td><td>30</td><td>ATTCAAACGAGAACTTTGAAGGCCGAAGTG</td><td>513.3</td><td>6.3</td><td>1.0</td><td>6.2</td><td>6.5E-10</td><td>1.5E-09</td><td>MB231/MB468</td><td>CYTO</td></tr>
<tr><td>>hg38_21_+_8433172_8446622_RNA45SN1_45S_50ntFlanks@6665.6689.25|GGATCACTCGGCTCGTGCGTCGATG</td><td>>hg38_21_+_8433172_8446622_RNA45SN1_45S_50ntFlanks</td><td>6665</td><td>6689</td><td>25</td><td>GGATCACTCGGCTCGTGCGTCGATG</td><td>66.9</td><td>-9.9</td><td>1.6</td><td>-6.2</td><td>6.3E-10</td><td>1.5E-09</td><td>MB231/MB468</td><td>MP</td></tr>
<tr><td>>hg38_21_+_8433172_8446622_RNA45SN1_45S_50ntFlanks@6721.6741.21|AATTGCAGGACACATTGATCA</td><td>>hg38_21_+_8433172_8446622_RNA45SN1_45S_50ntFlanks</td><td>6721</td><td>6741</td><td>21</td><td>AATTGCAGGACACATTGATCA</td><td>658.0</td><td>-7.7</td><td>1.3</td><td>-6.2</td><td>6.6E-10</td><td>1.5E-09</td><td>MB231/MB468</td><td>NUC</td></tr>
<tr><td>>hg38_21_+_8433172_8446622_RNA45SN1_45S_50ntFlanks@11208.11229.22|GCGGCGACTCTGGACGCGAGCC</td><td>>hg38_21_+_8433172_8446622_RNA45SN1_45S_50ntFlanks</td><td>11208</td><td>11229</td><td>22</td><td>GCGGCGACTCTGGACGCGAGCC</td><td>1139.3</td><td>-4.8</td><td>0.8</td><td>-6.2</td><td>6.5E-10</td><td>1.5E-09</td><td>MB231/MB468</td><td>CYTO</td></tr>
<tr><td>>hg38_21_+_8433172_8446622_RNA45SN1_45S_50ntFlanks@9316.9338.23|AACACGTGCGCGAGTCGGGGGCT</td><td>>hg38_21_+_8433172_8446622_RNA45SN1_45S_50ntFlanks</td><td>9316</td><td>9338</td><td>23</td><td>AACACGTGCGCGAGTCGGGGGCT</td><td>484.7</td><td>8.5</td><td>1.4</td><td>6.2</td><td>6.6E-10</td><td>1.5E-09</td><td>MB231/MB468</td><td>CYTO</td></tr>
<tr><td>>hg38_21_+_8433172_8446622_RNA45SN1_45S_50ntFlanks@8194.8214.21|CCCGTGGACGGTGTGAGGCCG</td><td>>hg38_21_+_8433172_8446622_RNA45SN1_45S_50ntFlanks</td><td>8194</td><td>8214</td><td>21</td><td>CCCGTGGACGGTGTGAGGCCG</td><td>1086.9</td><td>-3.7</td><td>0.6</td><td>-6.2</td><td>6.6E-10</td><td>1.5E-09</td><td>MB231/MB468</td><td>CYTO</td></tr>
<tr><td>>hg38_MT_+_1621_3279_RNR2_16S_50ntFlanks@1370.1396.27|ATGTTGGATCAGGACATCCCGATGGTG</td><td>>hg38_MT_+_1621_3279_RNR2_16S_50ntFlanks</td><td>1370</td><td>1396</td><td>27</td><td>ATGTTGGATCAGGACATCCCGATGGTG</td><td>722.5</td><td>5.5</td><td>0.9</td><td>6.2</td><td>5.0E-10</td><td>1.5E-09</td><td>BT20/MB468</td><td>MITO</td></tr>
<tr><td>>hg38_21_+_8433172_8446622_RNA45SN1_45S_50ntFlanks@12510.12527.18|TCGTGAGACAGGTTAGTT</td><td>>hg38_21_+_8433172_8446622_RNA45SN1_45S_50ntFlanks</td><td>12510</td><td>12527</td><td>18</td><td>TCGTGAGACAGGTTAGTT</td><td>1175.3</td><td>-8.6</td><td>1.4</td><td>-6.2</td><td>5.8E-10</td><td>1.5E-09</td><td>BT20/MB468</td><td>CYTO</td></tr>
<tr><td>>hg38_21_+_8433172_8446622_RNA45SN1_45S_50ntFlanks@10914.10931.18|CGCCGCGGCTGGACGAGG</td><td>>hg38_21_+_8433172_8446622_RNA45SN1_45S_50ntFlanks</td><td>10914</td><td>10931</td><td>18</td><td>CGCCGCGGCTGGACGAGG</td><td>1298.9</td><td>-9.4</td><td>1.5</td><td>-6.2</td><td>6.7E-10</td><td>1.5E-09</td><td>MB231/MB468</td><td>NUC</td></tr>
<tr><td>>hg38_21_+_8433172_8446622_RNA45SN1_45S_50ntFlanks@3876.3899.24|TACATGCCGACGGGCGCTGACCCC</td><td>>hg38_21_+_8433172_8446622_RNA45SN1_45S_50ntFlanks</td><td>3876</td><td>3899</td><td>24</td><td>TACATGCCGACGGGCGCTGACCCC</td><td>263.3</td><td>-4.7</td><td>0.8</td><td>-6.2</td><td>7.3E-10</td><td>1.5E-09</td><td>MB231/MB468</td><td>MITO</td></tr>
<tr><td>>hg38_MT_+_1621_3279_RNR2_16S_50ntFlanks@51.91.41|GCTAAACCTAGCCCCAAACCCACTCCACCTTACTACCAGAC</td><td>>hg38_MT_+_1621_3279_RNR2_16S_50ntFlanks</td><td>51</td><td>91</td><td>41</td><td>GCTAAACCTAGCCCCAAACCCACTCCACCTTACTACCAGAC</td><td>1110.4</td><td>5.4</td><td>0.9</td><td>6.2</td><td>5.0E-10</td><td>1.5E-09</td><td>BT20/MB468</td><td>MITO</td></tr>
<tr><td>>hg38_21_+_8433172_8446622_RNA45SN1_45S_50ntFlanks@12258.12288.31|GCTTGATCTTGATTTTCAGTACGAATACAGA</td><td>>hg38_21_+_8433172_8446622_RNA45SN1_45S_50ntFlanks</td><td>12258</td><td>12288</td><td>31</td><td>GCTTGATCTTGATTTTCAGTACGAATACAGA</td><td>113.1</td><td>-10.7</td><td>1.7</td><td>-6.2</td><td>6.4E-10</td><td>1.5E-09</td><td>MB231/MB468</td><td>MP</td></tr>
<tr><td>>hg38_21_+_8433172_8446622_RNA45SN1_45S_50ntFlanks@12712.12757.46|CGGAGCCTCGGTTGGCCTCGGATAGCCGGTCCCCCGCCTGTCCCCG</td><td>>hg38_21_+_8433172_8446622_RNA45SN1_45S_50ntFlanks</td><td>12712</td><td>12757</td><td>46</td><td>CGGAGCCTCGGTTGGCCTCGGATAGCCGGTCCCCCGCCTGTCCCCG</td><td>225.2</td><td>-11.7</td><td>1.9</td><td>-6.2</td><td>6.4E-10</td><td>1.5E-09</td><td>MB231/MB468</td><td>MP</td></tr>
<tr><td>>hg38_21_+_8433172_8446622_RNA45SN1_45S_50ntFlanks@5003.5034.32|ATAGCTCTTTCTCGATTCCGTGGGTGGTGGTG</td><td>>hg38_21_+_8433172_8446622_RNA45SN1_45S_50ntFlanks</td><td>5003</td><td>5034</td><td>32</td><td>ATAGCTCTTTCTCGATTCCGTGGGTGGTGGTG</td><td>118.6</td><td>-10.8</td><td>1.7</td><td>-6.2</td><td>6.4E-10</td><td>1.5E-09</td><td>MB231/MB468</td><td>MP</td></tr>
<tr><td>>hg38_21_+_8433172_8446622_RNA45SN1_45S_50ntFlanks@10872.10889.18|GCTGGGTCGGTCGGGCTG</td><td>>hg38_21_+_8433172_8446622_RNA45SN1_45S_50ntFlanks</td><td>10872</td><td>10889</td><td>18</td><td>GCTGGGTCGGTCGGGCTG</td><td>75.2</td><td>-10.1</td><td>1.6</td><td>-6.2</td><td>6.4E-10</td><td>1.6E-09</td><td>MB231/MB468</td><td>MP</td></tr>
<tr><td>>hg38_21_+_8433172_8446622_RNA45SN1_45S_50ntFlanks@9786.9821.36|CTCGCTGGCGTGGAGCCGGGCGTGGAATGCGAGTGC</td><td>>hg38_21_+_8433172_8446622_RNA45SN1_45S_50ntFlanks</td><td>9786</td><td>9821</td><td>36</td><td>CTCGCTGGCGTGGAGCCGGGCGTGGAATGCGAGTGC</td><td>72.7</td><td>-10.0</td><td>1.6</td><td>-6.2</td><td>6.4E-10</td><td>1.6E-09</td><td>MB231/MB468</td><td>MP</td></tr>
<tr><td>>MI0000446|hsa-mir-125b-1&WithFlank&11|-|122099751|122099850@21.39.19(+1C);MI0000470|hsa-mir-125b-2&WithFlank&21|+|16590231|16590331@23.41.19(+1C)&[MIMAT0000423&hsa-miR-125b-5p&offsets|0|-3(+1C);m-67&11|-|122099809|122099830&offsets|0|-3(+1C)];[MIMAT0000423_1&hsa-miR-125b-5p&offsets|0|-3(+1C);m-68&21|+|16590253|16590274&offsets|0|-3(+1C)]|TCCCTGAGACCCTAACTTGC</td><td>>MI0000446|hsa-mir-125b-1&WithFlank&11|-|122099751|122099850</td><td>21</td><td>39</td><td>19(+1C)</td><td>TCCCTGAGACCCTAACTTGC</td><td>253.4</td><td>6.8</td><td>1.1</td><td>6.2</td><td>6.7E-10</td><td>1.6E-09</td><td>MB231/MB468</td><td>NUC</td></tr>
<tr><td>>hg38_21_+_8433172_8446622_RNA45SN1_45S_50ntFlanks@4047.4065.19|ATTCGAACGTCTGCCCTAT</td><td>>hg38_21_+_8433172_8446622_RNA45SN1_45S_50ntFlanks</td><td>4047</td><td>4065</td><td>19</td><td>ATTCGAACGTCTGCCCTAT</td><td>521.0</td><td>-3.1</td><td>0.5</td><td>-6.2</td><td>6.6E-10</td><td>1.6E-09</td><td>MB231/MB468</td><td>CYTO</td></tr>
<tr><td>>hg38_21_+_8433172_8446622_RNA45SN1_45S_50ntFlanks@3993.4026.34|GTGACTCTAGATAACCTCGGGCCGATCGCACGCC</td><td>>hg38_21_+_8433172_8446622_RNA45SN1_45S_50ntFlanks</td><td>3993</td><td>4026</td><td>34</td><td>GTGACTCTAGATAACCTCGGGCCGATCGCACGCC</td><td>644.6</td><td>-4.2</td><td>0.7</td><td>-6.2</td><td>5.0E-10</td><td>1.6E-09</td><td>BT20/MB468</td><td>MITO</td></tr>
<tr><td>>hg38_21_+_8433172_8446622_RNA45SN1_45S_50ntFlanks@9738.9759.22|GAAACGATCTCAACCTATTCTC</td><td>>hg38_21_+_8433172_8446622_RNA45SN1_45S_50ntFlanks</td><td>9738</td><td>9759</td><td>22</td><td>GAAACGATCTCAACCTATTCTC</td><td>2887.3</td><td>-5.7</td><td>0.9</td><td>-6.2</td><td>5.0E-10</td><td>1.6E-09</td><td>BT20/MB468</td><td>MITO</td></tr>
<tr><td>>hg38_21_+_8433172_8446622_RNA45SN1_45S_50ntFlanks@8170.8203.34|CAAGTCCTTCTGATCGAGGCCCAGCCCGTGGACG</td><td>>hg38_21_+_8433172_8446622_RNA45SN1_45S_50ntFlanks</td><td>8170</td><td>8203</td><td>34</td><td>CAAGTCCTTCTGATCGAGGCCCAGCCCGTGGACG</td><td>396.3</td><td>-11.7</td><td>1.8</td><td>-6.5</td><td>6.3E-11</td><td>1.6E-09</td><td>BT20/MB468</td><td>MP</td></tr>
<tr><td>>hg38_21_+_8433172_8446622_RNA45SN1_45S_50ntFlanks@10458.10481.24|ATGGCCTCCGTTGCCCTCGGCCGA</td><td>>hg38_21_+_8433172_8446622_RNA45SN1_45S_50ntFlanks</td><td>10458</td><td>10481</td><td>24</td><td>ATGGCCTCCGTTGCCCTCGGCCGA</td><td>156.1</td><td>-11.1</td><td>1.8</td><td>-6.2</td><td>6.4E-10</td><td>1.6E-09</td><td>MB231/MB468</td><td>MP</td></tr>
<tr><td>>hg38_21_+_8433172_8446622_RNA45SN1_45S_50ntFlanks@9861.9878.18|GATGAACCGAACGCCGGG</td><td>>hg38_21_+_8433172_8446622_RNA45SN1_45S_50ntFlanks</td><td>9861</td><td>9878</td><td>18</td><td>GATGAACCGAACGCCGGG</td><td>94.3</td><td>-10.4</td><td>1.7</td><td>-6.2</td><td>6.4E-10</td><td>1.6E-09</td><td>MB231/MB468</td><td>MP</td></tr>
<tr><td>>hg38_21_+_8433172_8446622_RNA45SN1_45S_50ntFlanks@12072.12106.35|CCCGAGGGGCTCTCGCTTCTGGCGCCAAGCGCCCG</td><td>>hg38_21_+_8433172_8446622_RNA45SN1_45S_50ntFlanks</td><td>12072</td><td>12106</td><td>35</td><td>CCCGAGGGGCTCTCGCTTCTGGCGCCAAGCGCCCG</td><td>93.7</td><td>-10.4</td><td>1.7</td><td>-6.2</td><td>6.4E-10</td><td>1.6E-09</td><td>MB231/MB468</td><td>MP</td></tr>
<tr><td>>hg38_21_+_8433172_8446622_RNA45SN1_45S_50ntFlanks@7979.7999.21|ACCTCAGATCAGACGTGGCGA</td><td>>hg38_21_+_8433172_8446622_RNA45SN1_45S_50ntFlanks</td><td>7979</td><td>7999</td><td>21</td><td>ACCTCAGATCAGACGTGGCGA</td><td>199.1</td><td>-7.3</td><td>1.2</td><td>-6.2</td><td>5.9E-10</td><td>1.6E-09</td><td>BT20/MB468</td><td>CYTO</td></tr>
<tr><td>>hg38_21_+_8433172_8446622_RNA45SN1_45S_50ntFlanks@11316.11334.19|CACCCCACGTCTCGTCGCG</td><td>>hg38_21_+_8433172_8446622_RNA45SN1_45S_50ntFlanks</td><td>11316</td><td>11334</td><td>19</td><td>CACCCCACGTCTCGTCGCG</td><td>180.4</td><td>-10.6</td><td>1.6</td><td>-6.5</td><td>6.3E-11</td><td>1.6E-09</td><td>BT20/MB468</td><td>MP</td></tr>
<tr><td>>hg38_21_+_8433172_8446622_RNA45SN1_45S_50ntFlanks@12324.12347.24|TTGGGTTTTAAGCAGGAGGTGTCA</td><td>>hg38_21_+_8433172_8446622_RNA45SN1_45S_50ntFlanks</td><td>12324</td><td>12347</td><td>24</td><td>TTGGGTTTTAAGCAGGAGGTGTCA</td><td>146.3</td><td>-6.8</td><td>1.1</td><td>-6.2</td><td>5.9E-10</td><td>1.6E-09</td><td>BT20/MB468</td><td>CYTO</td></tr>
<tr><td>>hg38_21_+_8433172_8446622_RNA45SN1_45S_50ntFlanks@12480.12497.18|TAATAGGGAACGTGAGCT</td><td>>hg38_21_+_8433172_8446622_RNA45SN1_45S_50ntFlanks</td><td>12480</td><td>12497</td><td>18</td><td>TAATAGGGAACGTGAGCT</td><td>2098.3</td><td>4.6</td><td>0.7</td><td>6.2</td><td>5.9E-10</td><td>1.6E-09</td><td>BT20/MB468</td><td>CYTO</td></tr>
<tr><td>>hg38_21_+_8433172_8446622_RNA45SN1_45S_50ntFlanks@4259.4277.19|ATTGGAATGAGTCCACTTT</td><td>>hg38_21_+_8433172_8446622_RNA45SN1_45S_50ntFlanks</td><td>4259</td><td>4277</td><td>19</td><td>ATTGGAATGAGTCCACTTT</td><td>235.8</td><td>-11.0</td><td>1.7</td><td>-6.5</td><td>6.4E-11</td><td>1.6E-09</td><td>BT20/MB468</td><td>MP</td></tr>
<tr><td>>hg38_21_+_8433172_8446622_RNA45SN1_45S_50ntFlanks@9180.9197.18|CCGTCGGGCCCGGGGGAG</td><td>>hg38_21_+_8433172_8446622_RNA45SN1_45S_50ntFlanks</td><td>9180</td><td>9197</td><td>18</td><td>CCGTCGGGCCCGGGGGAG</td><td>957.3</td><td>-9.7</td><td>1.6</td><td>-6.1</td><td>7.8E-10</td><td>1.6E-09</td><td>BT20/MB468</td><td>NUC</td></tr>
<tr><td>>hg38_21_+_8433172_8446622_RNA45SN1_45S_50ntFlanks@11362.11379.18|TCGGGCGGCGGCGGTCGG</td><td>>hg38_21_+_8433172_8446622_RNA45SN1_45S_50ntFlanks</td><td>11362</td><td>11379</td><td>18</td><td>TCGGGCGGCGGCGGTCGG</td><td>72.6</td><td>-10.0</td><td>1.6</td><td>-6.2</td><td>6.5E-10</td><td>1.6E-09</td><td>MB231/MB468</td><td>MP</td></tr>
<tr><td>>hg38_21_+_8433172_8446622_RNA45SN1_45S_50ntFlanks@6789.6807.19|GCCTGTCTGAGCGTCGCTT</td><td>>hg38_21_+_8433172_8446622_RNA45SN1_45S_50ntFlanks</td><td>6789</td><td>6807</td><td>19</td><td>GCCTGTCTGAGCGTCGCTT</td><td>582.8</td><td>3.9</td><td>0.6</td><td>6.2</td><td>6.8E-10</td><td>1.6E-09</td><td>MB231/MB468</td><td>NUC</td></tr>
</table>

| | | | | | | | | | | | | | |
|---|---|---|---|---|---|---|---|---|---|---|---|---|---|
| >hg38_21_+_8433172_8446622_RNA45SN1_45S_50ntFlanks@12125.12146.22\|GCGACCCGCTCCGGGGACAGTG | >hg38_21_+_8433172_8446622_RNA45SN1_45S_50ntFlanks | 12125 | 12146 | 22 | GCGACCCGCTCCGGGGACAGTG | 257.1 | -11.1 | 1.7 | -6.5 | 6.4E-11 | 1.6E-09 | BT20/MB468 | MP |
| >MI0000746\|hsa-mir-99b&WithFlank&19\|+\|51692606\|51692687@13.34.22(+1U)&[MIMAT0000689&hsa-miR-99b-5p&offsets\|0\|0(+1U);m-10&19\|+\|51692618\|51692639&offsets\|0\|0(+1U)]\|CACCCGTAGAACCGACCTTGCGT | >MI0000746\|hsa-mir-99b&WithFlank&19\|+\|51692606\|51692687 | 13 | 34 | 22(+1U) | CACCCGTAGAACCGACCTTGCGT | 1563.2 | 5.5 | 0.9 | 6.2 | 7.4E-10 | 1.6E-09 | MB231/MB468 | MITO |
| >hg38_21_+_8433172_8446622_RNA45SN1_45S_50ntFlanks@12456.12474.19\|CCAAGCGTTGGATTGTTCA | >hg38_21_+_8433172_8446622_RNA45SN1_45S_50ntFlanks | 12456 | 12474 | 19 | CCAAGCGTTGGATTGTTCA | 7524.0 | -10.4 | 1.6 | -6.5 | 6.4E-11 | 1.6E-09 | BT20/MB468 | MP |
| >hg38_21_+_8433172_8446622_RNA45SN1_45S_50ntFlanks@4657.4689.33\|CTAGAGGTGAAATTCTTGGACCGGCGCAAGACG | >hg38_21_+_8433172_8446622_RNA45SN1_45S_50ntFlanks | 4657 | 4689 | 33 | CTAGAGGTGAAATTCTTGGACCGGCGCAAGACG | 4731.7 | -6.2 | 1.0 | -6.2 | 6.7E-10 | 1.6E-09 | MB231/MB468 | CYTO |
| >hg38_21_+_8433172_8446622_RNA45SN1_45S_50ntFlanks@475.498.24\|CTTCGTGATCGATGTGGTGACGTC | >hg38_21_+_8433172_8446622_RNA45SN1_45S_50ntFlanks | 475 | 498 | 24 | CTTCGTGATCGATGTGGTGACGTC | 34387.1 | 5.3 | 0.9 | 6.2 | 6.8E-10 | 1.6E-09 | MB231/MB468 | NUC |
| >MI0000438\|hsa-mir-15b&WithFlank&3\|+\|160404582\|160404691@26.47.22&[MIMAT0000417&hsa-miR-15b-5p&offsets\|0\|0;m-110&3\|+\|160404607\|160404628&offsets\|0\|0]\|TAGCAGCACATCATGGTTTACA | >MI0000438\|hsa-mir-15b&WithFlank&3\|+\|160404582\|160404691 | 26 | 47 | 22 | TAGCAGCACATCATGGTTTACA | 2490.4 | 4.3 | 0.7 | 6.2 | 6.8E-10 | 1.6E-09 | MB231/MB468 | NUC |
| >hg38_21_+_8433172_8446622_RNA45SN1_45S_50ntFlanks@6651.6669.19\|CGACTCTTAGCGGTGGATC | >hg38_21_+_8433172_8446622_RNA45SN1_45S_50ntFlanks | 6651 | 6669 | 19 | CGACTCTTAGCGGTGGATC | 98132.7 | 4.5 | 0.7 | 6.2 | 5.1E-10 | 1.6E-09 | BT20/MB468 | MITO |
| >hg38_21_+_8433172_8446622_RNA45SN1_45S_50ntFlanks@12934.12958.25\|CTAAACCATTCGTAGACGACCTGCT | >hg38_21_+_8433172_8446622_RNA45SN1_45S_50ntFlanks | 12934 | 12958 | 25 | CTAAACCATTCGTAGACGACCTGCT | 517.0 | -6.1 | 1.0 | -6.2 | 7.5E-10 | 1.6E-09 | MB231/MB468 | MITO |
| >MI0000266\|hsa-mir-10a&WithFlank&17\|-\|48579832\|48579953@28.48.21(+1C)&[MIMAT0000253&hsa-miR-10a-5p&offsets\|0\|-2(+1C);m-18&17\|-\|48579905\|48579926&offsets\|0\|-1(+1C)]\|TACCCTGTAGATCCGAATTTGC | >MI0000266\|hsa-mir-10a&WithFlank&17\|-\|48579832\|48579953 | 28 | 48 | 21(+1C) | TACCCTGTAGATCCGAATTTGC | 859.0 | 9.3 | 1.5 | 6.2 | 6.5E-10 | 1.6E-09 | MB231/MB468 | MP |
| >trnaMT_ValTAC_MT_+_1602_1670@41.72.32\|GAGATTTCAACTTAACTTGACCGCTCTGACCA | >trnaMT_ValTAC_MT_+_1602_1670 | 41 | 72 | 32 | GAGATTTCAACTTAACTTGACCGCTCTGACCA | 463.3 | 3.7 | 0.6 | 6.2 | 6.8E-10 | 1.6E-09 | MB231/MB468 | NUC |
| >hg38_21_+_8433172_8446622_RNA45SN1_45S_50ntFlanks@12605.12634.30\|TATGTGCTTGGCTGAGGAGCCAATGGGGCG | >hg38_21_+_8433172_8446622_RNA45SN1_45S_50ntFlanks | 12605 | 12634 | 30 | TATGTGCTTGGCTGAGGAGCCAATGGGGCG | 644.6 | -9.1 | 1.5 | -6.1 | 7.9E-10 | 1.6E-09 | BT20/MB468 | NUC |
| >hg38_21_+_8433172_8446622_RNA45SN1_45S_50ntFlanks@9714.9735.22\|CGAATGATTAGAGGTCTTGGGG | >hg38_21_+_8433172_8446622_RNA45SN1_45S_50ntFlanks | 9714 | 9735 | 22 | CGAATGATTAGAGGTCTTGGGG | 555.7 | -8.7 | 1.4 | -6.1 | 7.9E-10 | 1.6E-09 | BT20/MB468 | NUC |
| >hg38_21_+_8433172_8446622_RNA45SN1_45S_50ntFlanks@11781.11805.25\|CGCATGAATGGATGAACGAGATTCC | >hg38_21_+_8433172_8446622_RNA45SN1_45S_50ntFlanks | 11781 | 11805 | 25 | CGCATGAATGGATGAACGAGATTCC | 415.7 | -3.9 | 0.6 | -6.2 | 6.0E-10 | 1.6E-09 | BT20/MB468 | CYTO |
| >hg38_21_+_8433172_8446622_RNA45SN1_45S_50ntFlanks@12456.12475.20\|CCAAGCGTTGGATTGTTCAC | >hg38_21_+_8433172_8446622_RNA45SN1_45S_50ntFlanks | 12456 | 12475 | 20 | CCAAGCGTTGGATTGTTCAC | 26472.1 | -8.2 | 1.3 | -6.2 | 6.8E-10 | 1.6E-09 | MB231/MB468 | CYTO |
| >hg38_21_+_8433172_8446622_RNA45SN1_45S_50ntFlanks@5497.5532.36\|ACGGTCGAACTTGACTATCTAGAGGAAGTAAAAGTC | >hg38_21_+_8433172_8446622_RNA45SN1_45S_50ntFlanks | 5497 | 5532 | 36 | ACGGTCGAACTTGACTATCTAGAGGAAGTAAAAGTC | 841.6 | 7.3 | 1.2 | 6.2 | 6.8E-10 | 1.6E-09 | MB231/MB468 | CYTO |
| >hg38_21_+_8433172_8446622_RNA45SN1_45S_50ntFlanks@3793.3810.18\|CGAATGGCTCATTAAATC | >hg38_21_+_8433172_8446622_RNA45SN1_45S_50ntFlanks | 3793 | 3810 | 18 | CGAATGGCTCATTAAATC | 250.4 | -10.1 | 1.6 | -6.2 | 5.1E-10 | 1.6E-09 | BT20/MB468 | MITO |
| >am_trna128_GlyGCC_6_-_27870686_27870756@1.34.34,trna18_GlyGCC_16_+_70822597_70822667@1.34.34,trna19_GlyGCC_16_+_70823410_70823480@1.34.34,trna19_GlyGCC_2_-_157257659_157257729@1.34.34,trna24_GlyGCC_16_-_70812942_70813012@1.34.34,trna25_GlyGCC_16_-_70812114_70812184@1.34.34,trna5_GlyGCC_17_+_8029064_8029134@1.34.34,trna68_GlyGCC_1_-_161493637_161493707@1.34.34\|GCATTGGTGGTTCAGTGGTAGAATTCTCGCCTGC | >am_trna128_GlyGCC_6_-_27870686_27870756 | 1 | 34 | 34 | GCATTGGTGGTTCAGTGGTAGAATTCTCGCCTGC | 2884.7 | 6.2 | 1.0 | 6.2 | 7.5E-10 | 1.6E-09 | MB231/MB468 | MITO |
| >hg38_21_+_8433172_8446622_RNA45SN1_45S_50ntFlanks@5484.5519.36\|GAGCGCTGAGAAGACGGTCGAACTTGACTATCTAGA | >hg38_21_+_8433172_8446622_RNA45SN1_45S_50ntFlanks | 5484 | 5519 | 36 | GAGCGCTGAGAAGACGGTCGAACTTGACTATCTAGA | 77.3 | -10.1 | 1.6 | -6.2 | 6.6E-10 | 1.6E-09 | MB231/MB468 | MP |
| >hg38_21_+_8433172_8446622_RNA45SN1_45S_50ntFlanks@11655.11687.33\|GCGATGTGATTTCTGCCCAGTGCTCTGAATGTC | >hg38_21_+_8433172_8446622_RNA45SN1_45S_50ntFlanks | 11655 | 11687 | 33 | GCGATGTGATTTCTGCCCAGTGCTCTGAATGTC | 388.8 | 9.3 | 1.5 | 6.2 | 5.1E-10 | 1.6E-09 | BT20/MB468 | MITO |
| >hg38_21_+_8433172_8446622_RNA45SN1_45S_50ntFlanks@3839.3868.30\|TCTCCTACTTGGATAACTGTGGTAATTCTA | >hg38_21_+_8433172_8446622_RNA45SN1_45S_50ntFlanks | 3839 | 3868 | 30 | TCTCCTACTTGGATAACTGTGGTAATTCTA | 582.5 | 3.8 | 0.6 | 6.2 | 6.8E-10 | 1.6E-09 | MB231/MB468 | CYTO |
| >hg38_21_+_8433172_8446622_RNA45SN1_45S_50ntFlanks@9168.9197.30\|GGGCGGGTCGCGCCGTCGGGCCCGGGGGAG | >hg38_21_+_8433172_8446622_RNA45SN1_45S_50ntFlanks | 9168 | 9197 | 30 | GGGCGGGTCGCGCCGTCGGGCCCGGGGGAG | 203.0 | -11.5 | 1.9 | -6.2 | 6.6E-10 | 1.6E-09 | MB231/MB468 | MP |
| >hg38_21_+_8433172_8446622_RNA45SN1_45S_50ntFlanks@11998.12032.35\|GCCCTGCGGGCCGCCGGTGAAATACCACTACTCTG | >hg38_21_+_8433172_8446622_RNA45SN1_45S_50ntFlanks | 11998 | 12032 | 35 | GCCCTGCGGGCCGCCGGTGAAATACCACTACTCTG | 78.5 | -10.2 | 1.6 | -6.2 | 6.6E-10 | 1.6E-09 | MB231/MB468 | MP |
| >hg38_21_+_8433172_8446622_RNA45SN1_45S_50ntFlanks@4730.4771.42\|CAAGAACGAAAGTCGGAGGTTCGAAGACGATCAGATACCGTC | >hg38_21_+_8433172_8446622_RNA45SN1_45S_50ntFlanks | 4730 | 4771 | 42 | CAAGAACGAAAGTCGGAGGTTCGAAGACGATCAGATACCGTC | 170.0 | 6.5 | 1.1 | 6.2 | 6.8E-10 | 1.6E-09 | MB231/MB468 | CYTO |
| >hg38_21_+_8433172_8446622_RNA45SN1_45S_50ntFlanks@12894.12926.33\|CTCGCCCGTCACGCACCGCACGTTCGTGGGGAA | >hg38_21_+_8433172_8446622_RNA45SN1_45S_50ntFlanks | 12894 | 12926 | 33 | CTCGCCCGTCACGCACCGCACGTTCGTGGGGAA | 85.4 | -10.3 | 1.7 | -6.2 | 6.6E-10 | 1.6E-09 | MB231/MB468 | MP |
| >hg38_21_+_8433172_8446622_RNA45SN1_45S_50ntFlanks@9567.9592.26\|ACGTGCAAATCGGTCGTCCGACCTGG | >hg38_21_+_8433172_8446622_RNA45SN1_45S_50ntFlanks | 9567 | 9592 | 26 | ACGTGCAAATCGGTCGTCCGACCTGG | 1368.5 | -7.9 | 1.3 | -6.2 | 6.0E-10 | 1.6E-09 | BT20/MB468 | CYTO |
| >hg38_21_+_8433172_8446622_RNA45SN1_45S_50ntFlanks@12411.12432.22\|GATCCTTCGATGTCGGCTCTTC | >hg38_21_+_8433172_8446622_RNA45SN1_45S_50ntFlanks | 12411 | 12432 | 22 | GATCCTTCGATGTCGGCTCTTC | 274.5 | -10.4 | 1.7 | -6.2 | 6.1E-10 | 1.6E-09 | BT20/MB468 | CYTO |
| >MI0000285\|hsa-mir-205&WithFlank&1\|+\|209432127\|209432248@40.62.23(+1U)&[MIMAT0000266&hsa-miR-205-5p&offsets\|0\|+1(+1U);m-351&1\|+\|209432166\|209432187&offsets\|0\|+1(+1U)]\|TCCTTCATTCCACCGGAGTCTGTT | >MI0000285\|hsa-mir-205&WithFlank&1\|+\|209432127\|209432248 | 40 | 62 | 23(+1U) | TCCTTCATTCCACCGGAGTCTGTT | 126.7 | -10.8 | 1.8 | -6.2 | 6.6E-10 | 1.6E-09 | MB231/MB468 | MP |
| >hg38_21_+_8433172_8446622_RNA45SN1_45S_50ntFlanks@10802.10830.29\|TTGGAACAATGTAGGTAAGGGAAGTCGGC | >hg38_21_+_8433172_8446622_RNA45SN1_45S_50ntFlanks | 10802 | 10830 | 29 | TTGGAACAATGTAGGTAAGGGAAGTCGGC | 74.5 | -10.1 | 1.6 | -6.2 | 6.7E-10 | 1.6E-09 | MB231/MB468 | MP |
| >am_trna128_GlyGCC_6_-_27870686_27870756@7.29.23,trna133_GlyCCC_1_-_16872434_16872504@7.29.23,trna17_GluTTC_2_-_204229131_204229201@7.29.23,trna18_GlyGCC_16_+_70822597_70822667@7.29.23,trna19_GlyGCC_16_+_70823410_70823480@7.29.23,trna19_GlyGCC_2_-_157257659_157257729@7.29.23,trna24_GlyGCC_16_-_70812942_70813012@7.29.23,trna25_GlyGCC_16_-_70812114_70812184@7.29.23,trna2_GlyGCC_21_-_18827107_18827177@7.29.23,trna35_GlyGCC_1_+_161413094_161413164@7.29.23,trna37_GlyGCC_1_+_161420467_161420537@7.29.23,trna39_GlyGCC_1_+_161427898_161427968@7.29.23,trna41_GlyGCC_1_+_161435258_161435328@7.29.23,trna4_GlyCCC_1_+_17188416_17188486@7.29.23,trna5_GlyGCC_17_+_8029064_8029134@7.29.23,trna68_GlyGCC_1_-_161493637_161493707@7.29.23,trna91_GlyCCC_1_-_149680210_149680280@7.29.23,trna99_ValCAC_1_-_149294666_149294736@7.29.23\|GTGGTTCAGTGGTAGAATTCTCG | >am_trna128_GlyGCC_6_-_27870686_27870756 | 7 | 29 | 23 | GTGGTTCAGTGGTAGAATTCTCG | 90.8 | -10.4 | 1.7 | -6.2 | 6.7E-10 | 1.6E-09 | MB231/MB468 | MP |
| >hg38_21_+_8433172_8446622_RNA45SN1_45S_50ntFlanks@10392.10419.28\|ACAGCAGTTGAACATGGGTCAGTCGGTC | >hg38_21_+_8433172_8446622_RNA45SN1_45S_50ntFlanks | 10392 | 10419 | 28 | ACAGCAGTTGAACATGGGTCAGTCGGTC | 566.8 | -8.3 | 1.3 | -6.2 | 6.7E-10 | 1.6E-09 | MB231/MB468 | MP |
| >hg38_21_+_8433172_8446622_RNA45SN1_45S_50ntFlanks@6651.6696.46\|CGACTCTTAGCGGTGGATCACTCGGCTCGTGCGTCGATGAAGAACG | >hg38_21_+_8433172_8446622_RNA45SN1_45S_50ntFlanks | 6651 | 6696 | 46 | CGACTCTTAGCGGTGGATCACTCGGCTCGTGCGTCGATGAAGAACG | 6368.4 | 4.9 | 0.8 | 6.2 | 6.9E-10 | 1.6E-09 | MB231/MB468 | CYTO |
| >hg38_21_+_8433172_8446622_RNA45SN1_45S_50ntFlanks@9565.9595.31\|TGACGTGCAAATCGGTCGTCCGACCTGGGTA | >hg38_21_+_8433172_8446622_RNA45SN1_45S_50ntFlanks | 9565 | 9595 | 31 | TGACGTGCAAATCGGTCGTCCGACCTGGGTA | 463.3 | -6.4 | 1.0 | -6.2 | 6.7E-10 | 1.6E-09 | MB231/MB468 | MP |
| >hg38_21_+_8433172_8446622_RNA45SN1_45S_50ntFlanks@10107.10125.19\|CGTGTGGTGTGCGTCGGAG | >hg38_21_+_8433172_8446622_RNA45SN1_45S_50ntFlanks | 10107 | 10125 | 19 | CGTGTGGTGTGCGTCGGAG | 551.9 | -7.2 | 1.2 | -6.2 | 6.9E-10 | 1.6E-09 | MB231/MB468 | CYTO |
| >hg38_21_+_8433172_8446622_RNA45SN1_45S_50ntFlanks@6761.6806.46\|TGCGGCCCCGGGTTCCTCCCGGGGCTACGCCTGTCTGAGCGTCGCT | >hg38_21_+_8433172_8446622_RNA45SN1_45S_50ntFlanks | 6761 | 6806 | 46 | TGCGGCCCCGGGTTCCTCCCGGGGCTACGCCTGTCTGAGCGTCGCT | 632.5 | 9.3 | 1.5 | 6.2 | 6.9E-10 | 1.6E-09 | MB231/MB468 | CYTO |
| >hg38_21_+_8433172_8446622_RNA45SN1_45S_50ntFlanks@12405.12430.26\|CTTTTTGATCCTTCGATGTCGGCTCT | >hg38_21_+_8433172_8446622_RNA45SN1_45S_50ntFlanks | 12405 | 12430 | 26 | CTTTTTGATCCTTCGATGTCGGCTCT | 2263.8 | -8.0 | 1.3 | -6.1 | 8.1E-10 | 1.6E-09 | BT20/MB468 | NUC |
| >hg38_21_+_8433172_8446622_RNA45SN1_45S_50ntFlanks@10471.10495.25\|CCCTCGGCCGATCGAAAGGGAGTCG | >hg38_21_+_8433172_8446622_RNA45SN1_45S_50ntFlanks | 10471 | 10495 | 25 | CCCTCGGCCGATCGAAAGGGAGTCG | 321.9 | -7.9 | 1.3 | -6.2 | 7.7E-10 | 1.6E-09 | MB231/MB468 | MITO |
| >hg38_21_+_8433172_8446622_RNA45SN1_45S_50ntFlanks@9545.9573.29\|GTGGAGGTCCGTAGCGGTCCTGACGTGCA | >hg38_21_+_8433172_8446622_RNA45SN1_45S_50ntFlanks | 9545 | 9573 | 29 | GTGGAGGTCCGTAGCGGTCCTGACGTGCA | 1518.8 | -6.6 | 1.1 | -6.2 | 7.7E-10 | 1.6E-09 | MB231/MB468 | MITO |
| >hg38_21_+_8433172_8446622_RNA45SN1_45S_50ntFlanks@10731.10752.22\|AGGGTGTAAATCTCGCGCCGGG | >hg38_21_+_8433172_8446622_RNA45SN1_45S_50ntFlanks | 10731 | 10752 | 22 | AGGGTGTAAATCTCGCGCCGGG | 111.1 | -8.9 | 1.4 | -6.2 | 5.3E-10 | 1.6E-09 | BT20/MB468 | MITO |
| >hg38_21_+_8433172_8446622_RNA45SN1_45S_50ntFlanks@10781.10801.21\|AGGTGAACAGCCTCTGGCATG | >hg38_21_+_8433172_8446622_RNA45SN1_45S_50ntFlanks | 10781 | 10801 | 21 | AGGTGAACAGCCTCTGGCATG | 587.3 | -8.8 | 1.4 | -6.1 | 8.1E-10 | 1.6E-09 | BT20/MB468 | NUC |
| >hg38_21_+_8433172_8446622_RNA45SN1_45S_50ntFlanks@11565.11585.21\|CTAGCAGCCGACTTAGAACTG | >hg38_21_+_8433172_8446622_RNA45SN1_45S_50ntFlanks | 11565 | 11585 | 21 | CTAGCAGCCGACTTAGAACTG | 206.6 | -2.9 | 0.5 | -6.2 | 6.2E-10 | 1.6E-09 | BT20/MB468 | CYTO |
| >hg38_21_+_8433172_8446622_RNA45SN1_45S_50ntFlanks@4552.4572.21\|TACCGCAGCTAGGAATAATGG | >hg38_21_+_8433172_8446622_RNA45SN1_45S_50ntFlanks | 4552 | 4572 | 21 | TACCGCAGCTAGGAATAATGG | 244.7 | -6.5 | 1.1 | -6.1 | 8.1E-10 | 1.6E-09 | BT20/MB468 | NUC |
| >hg38_21_+_8433172_8446622_RNA45SN1_45S_50ntFlanks@3975.4002.28\|CGGGCGCCGGCGGCTTTGGTGACTCTAG | >hg38_21_+_8433172_8446622_RNA45SN1_45S_50ntFlanks | 3975 | 4002 | 28 | CGGGCGCCGGCGGCTTTGGTGACTCTAG | 169.8 | -9.5 | 1.5 | -6.2 | 5.3E-10 | 1.6E-09 | BT20/MB468 | MITO |
| >hg38_21_+_8433172_8446622_RNA45SN1_45S_50ntFlanks@4919.4941.23\|CCACCAGGAGTGGAGCCTGCGGC | >hg38_21_+_8433172_8446622_RNA45SN1_45S_50ntFlanks | 4919 | 4941 | 23 | CCACCAGGAGTGGAGCCTGCGGC | 660.1 | -8.8 | 1.4 | -6.2 | 6.8E-10 | 1.6E-09 | MB231/MB468 | MP |

| | | | | | | | | | | | | | |
|---|---|---|---|---|---|---|---|---|---|---|---|---|---|
| >hg38_21_+_8433172_8446622_RNA45SN1_45S_50ntFlanks@12943.12972.30\|TCGTAGACGACCTGCTTCTGGGTCGGGGTT | >hg38_21_+_8433172_8446622_RNA45SN1_45S_50ntFlanks | 12943 | 12972 | 30 | TCGTAGACGACCTGCTTCTGGGTCGGGGTT | 165.4 | -10.5 | 1.6 | -6.5 | 6.7E-11 | 1.6E-09 | BT20/MB468 | MP |
| >hg38_21_+_8433172_8446622_RNA45SN1_45S_50ntFlanks@4585.4622.38\|GTTCTATTTTGTTGGTTTTCGGAACTGAGGCCATGATT | >hg38_21_+_8433172_8446622_RNA45SN1_45S_50ntFlanks | 4585 | 4622 | 38 | GTTCTATTTTGTTGGTTTTCGGAACTGAGGCCATGATT | 505.6 | 7.6 | 1.2 | 6.2 | 7.0E-10 | 1.6E-09 | MB231/MB468 | CYTO |
| >hg38_21_+_8433172_8446622_RNA45SN1_45S_50ntFlanks@9545.9564.20\|GTGGAGGTCCGTAGCGGTCC | >hg38_21_+_8433172_8446622_RNA45SN1_45S_50ntFlanks | 9545 | 9564 | 20 | GTGGAGGTCCGTAGCGGTCC | 1055.9 | -6.9 | 1.1 | -6.2 | 7.1E-10 | 1.6E-09 | MB231/MB468 | NUC |
| >hg38_21_+_8433172_8446622_RNA45SN1_45S_50ntFlanks@11561.11599.39\|GCGCCTAGCAGCCGACTTAGAACTGGTGCGGACCAGGGG | >hg38_21_+_8433172_8446622_RNA45SN1_45S_50ntFlanks | 11561 | 11599 | 39 | GCGCCTAGCAGCCGACTTAGAACTGGTGCGGACCAGGGG | 336.9 | -8.2 | 1.3 | -6.1 | 8.2E-10 | 1.6E-09 | BT20/MB468 | NUC |
| >hg38_21_+_8433172_8446622_RNA45SN1_45S_50ntFlanks@11561.11587.27\|GCGCCTAGCAGCCGACTTAGAACTGGT | >hg38_21_+_8433172_8446622_RNA45SN1_45S_50ntFlanks | 11561 | 11587 | 27 | GCGCCTAGCAGCCGACTTAGAACTGGT | 382.9 | -8.0 | 1.3 | -6.2 | 6.8E-10 | 1.6E-09 | MB231/MB468 | MP |
| >hg38_21_+_8433172_8446622_RNA45SN1_45S_50ntFlanks@3850.3881.32\|GATAACTGTGGTAATTCTAGAGCTAATACATG | >hg38_21_+_8433172_8446622_RNA45SN1_45S_50ntFlanks | 3850 | 3881 | 32 | GATAACTGTGGTAATTCTAGAGCTAATACATG | 486.9 | -8.0 | 1.3 | -6.1 | 7.8E-10 | 1.7E-09 | MB231/MB468 | MITO |
| >MI0000266\|hsa-mir-10a&WithFlank&17\|-\|48579832\|48579953@28.48.21(+1C)&[MIMAT0000253&hsa-miR-10a-5p&offsets\|0\|-2(+1C);m-18&17\|-\|48579905\|48579926&offsets\|0\|-1(+1C)]\|TACCCTGTAGATCCGAATTTGC | >MI0000266\|hsa-mir-10a&WithFlank&17\|-\|48579832\|48579953 | 28 | 48 | 21(+1C) | TACCCTGTAGATCCGAATTTGC | 2009.6 | 11.4 | 1.7 | 6.5 | 6.8E-11 | 1.7E-09 | BT20/MB468 | MP |
| >MI0001448\|hsa-mir-425&WithFlank&3\|-\|49020142\|49020240@20.43.24&[MIMAT0003393&hsa-miR-425-5p&offsets\|0\|+1;m-95&3\|-\|49020200\|49020221&offsets\|0\|+2]\|AATGACACGATCACTCCCGTTGAG | >MI0001448\|hsa-mir-425&WithFlank&3\|-\|49020142\|49020240 | 20 | 43 | 24 | AATGACACGATCACTCCCGTTGAG | 1251.6 | 5.8 | 0.9 | 6.1 | 7.9E-10 | 1.7E-09 | MB231/MB468 | MITO |
| >hg38_21_+_8433172_8446622_RNA45SN1_45S_50ntFlanks@3785.3810.26\|TGAAACTGCGAATGGCTCATTAAATC | >hg38_21_+_8433172_8446622_RNA45SN1_45S_50ntFlanks | 3785 | 3810 | 26 | TGAAACTGCGAATGGCTCATTAAATC | 435.0 | -8.1 | 1.3 | -6.1 | 7.9E-10 | 1.7E-09 | MB231/MB468 | MITO |
| >hg38_21_+_8433172_8446622_RNA45SN1_45S_50ntFlanks@12076.12110.35\|AGGGGCTCTCGCTTCTGGCGCCAAGCGCCCGGCCG | >hg38_21_+_8433172_8446622_RNA45SN1_45S_50ntFlanks | 12076 | 12110 | 35 | AGGGGCTCTCGCTTCTGGCGCCAAGCGCCCGGCCG | 83.6 | -10.2 | 1.7 | -6.2 | 6.9E-10 | 1.7E-09 | MB231/MB468 | MP |
| >hg38_MT_+_598_1651_RNR1_12S_50ntFlanks@968.1004.37\|ACATGGTAAGTGTACTGGAAAGTGCACTTGGACGAAC | >hg38_MT_+_598_1651_RNR1_12S_50ntFlanks | 968 | 1004 | 37 | ACATGGTAAGTGTACTGGAAAGTGCACTTGGACGAAC | 178.6 | 8.4 | 1.4 | 6.2 | 7.2E-10 | 1.7E-09 | MB231/MB468 | NUC |
| >MI0005559\|hsa-mir-744&WithFlank&17\|+\|12081893\|12082002@17.37.21&[MIMAT0004945&hsa-miR-744-5p&offsets\|0\|-1;m-172&17\|+\|12081909\|12081929&offsets\|0\|0]\|TGCGGGGCTAGGGCTAACAGC | >MI0005559\|hsa-mir-744&WithFlank&17\|+\|12081893\|12082002 | 17 | 37 | 21 | TGCGGGGCTAGGGCTAACAGC | 2896.0 | 6.3 | 1.0 | 6.5 | 6.8E-11 | 1.7E-09 | BT20/MB468 | MP |
| >hg38_21_+_8433172_8446622_RNA45SN1_45S_50ntFlanks@12499.12518.20\|GGTTTAGACCGTCGTGAGAC | >hg38_21_+_8433172_8446622_RNA45SN1_45S_50ntFlanks | 12499 | 12518 | 20 | GGTTTAGACCGTCGTGAGAC | 199.0 | -10.7 | 1.6 | -6.5 | 6.8E-11 | 1.7E-09 | BT20/MB468 | MP |
| >hg38_21_+_8433172_8446622_RNA45SN1_45S_50ntFlanks@8381.8400.20\|AAGAGGGCGTGAAACCGTTA | >hg38_21_+_8433172_8446622_RNA45SN1_45S_50ntFlanks | 8381 | 8400 | 20 | AAGAGGGCGTGAAACCGTTA | 237.5 | -7.3 | 1.2 | -6.2 | 7.2E-10 | 1.7E-09 | MB231/MB468 | NUC |
| >hg38_21_+_8433172_8446622_RNA45SN1_45S_50ntFlanks@3975.4002.28\|CGGGCGCCGGCGGCTTTGGTGACTCTAG | >hg38_21_+_8433172_8446622_RNA45SN1_45S_50ntFlanks | 3975 | 4002 | 28 | CGGGCGCCGGCGGCTTTGGTGACTCTAG | 106.4 | -10.6 | 1.7 | -6.2 | 6.9E-10 | 1.7E-09 | MB231/MB468 | MP |
| >hg38_21_+_8433172_8446622_RNA45SN1_45S_50ntFlanks@12716.12745.30\|GCCTCGGTTGGCCTCGGATAGCCGGTCCCC | >hg38_21_+_8433172_8446622_RNA45SN1_45S_50ntFlanks | 12716 | 12745 | 30 | GCCTCGGTTGGCCTCGGATAGCCGGTCCCC | 107.0 | -10.6 | 1.7 | -6.2 | 6.9E-10 | 1.7E-09 | MB231/MB468 | MP |
| >hg38_21_+_8433172_8446622_RNA45SN1_45S_50ntFlanks@4727.4759.33\|AATCAAGAACGAAAGTCGGAGGTTCGAAGACGA | >hg38_21_+_8433172_8446622_RNA45SN1_45S_50ntFlanks | 4727 | 4759 | 33 | AATCAAGAACGAAAGTCGGAGGTTCGAAGACGA | 505.3 | -4.4 | 0.7 | -6.2 | 6.9E-10 | 1.7E-09 | MB231/MB468 | MP |
| >hg38_21_+_8433172_8446622_RNA45SN1_45S_50ntFlanks@11768.11786.19\|CTAATTAGTGACGCGCATG | >hg38_21_+_8433172_8446622_RNA45SN1_45S_50ntFlanks | 11768 | 11786 | 19 | CTAATTAGTGACGCGCATG | 1108.6 | -8.3 | 1.3 | -6.2 | 6.3E-10 | 1.7E-09 | BT20/MB468 | CYTO |
| >MI0000734\|hsa-mir-106b&WithFlank&7\|-\|100093987\|100094080@58.78.21&[MIMAT0004672&hsa-miR-106b-3p&offsets\|0\|-1;m-87&7\|-\|100094003\|100094023&offsets\|0\|0]\|CCGCACTGTGGGTACTTGCTG | >MI0000734\|hsa-mir-106b&WithFlank&7\|-\|100093987\|100094080 | 58 | 78 | 21 | CCGCACTGTGGGTACTTGCTG | 602.3 | 2.2 | 0.4 | 6.1 | 8.3E-10 | 1.7E-09 | BT20/MB468 | NUC |
| >hg38_21_+_8433172_8446622_RNA45SN1_45S_50ntFlanks@11630.11662.33\|GCGAAGGCCCGCGGCGGGTGTTGACGCGATGTG | >hg38_21_+_8433172_8446622_RNA45SN1_45S_50ntFlanks | 11630 | 11662 | 33 | GCGAAGGCCCGCGGCGGGTGTTGACGCGATGTG | 159.8 | -10.4 | 1.6 | -6.5 | 6.9E-11 | 1.7E-09 | BT20/MB468 | MP |
| >hg38_21_+_8433172_8446622_RNA45SN1_45S_50ntFlanks@9031.9058.28\|TCCTCGTGGGGGGGCCGGGCCACCCCTC | >hg38_21_+_8433172_8446622_RNA45SN1_45S_50ntFlanks | 9031 | 9058 | 28 | TCCTCGTGGGGGGGCCGGGCCACCCCTC | 112.1 | -8.9 | 1.4 | -6.2 | 5.4E-10 | 1.7E-09 | BT20/MB468 | MITO |
| >hg38_21_+_8433172_8446622_RNA45SN1_45S_50ntFlanks@12418.12437.20\|CGATGTCGGCTCTTCCTATC | >hg38_21_+_8433172_8446622_RNA45SN1_45S_50ntFlanks | 12418 | 12437 | 20 | CGATGTCGGCTCTTCCTATC | 4495.7 | -3.3 | 0.5 | -6.2 | 5.4E-10 | 1.7E-09 | BT20/MB468 | MITO |
| >MI0000093\|hsa-mir-92a-1&WithFlank&13\|+\|91351308\|91351397@54.76.23&[MIMAT0000092&hsa-miR-92a-3p&offsets\|0\|+1;m-9&13\|+\|91351361\|91351382&offsets\|0\|+1]\|TATTGCACTTGTCCCGGCCTGTT | >MI0000093\|hsa-mir-92a-1&WithFlank&13\|+\|91351308\|91351397 | 54 | 76 | 23 | TATTGCACTTGTCCCGGCCTGTT | 6196.3 | 5.4 | 0.9 | 6.1 | 8.0E-10 | 1.7E-09 | MB231/MB468 | MITO |
| >am_MI0006370\|hsa-mir-1303&WithFlank&5\|+\|154685770\|154685867@57.79.23&[MIMAT0005891&hsa-miR-1303&offsets\|-1\|0;m-516&5\|+\|154685826\|154685848&offsets\|0\|0]\|TTTTAGAGACGGGGTCTTGCTCT | >am_MI0006370\|hsa-mir-1303&WithFlank&5\|+\|154685770\|154685867 | 57 | 79 | 23 | TTTTAGAGACGGGGTCTTGCTCT | 309.3 | 7.8 | 1.3 | 6.2 | 5.4E-10 | 1.7E-09 | BT20/MB468 | MITO |
| >MI0003190\|hsa-mir-505&WithFlank&X\|-\|139924142\|139924237@57.80.24&[MIMAT0002876&hsa-miR-505-3p&offsets\|0\|+2;m-196&X\|-\|139924160\|139924181&offsets\|0\|+2]\|CGTCAACACTTGCTGGTTTCCTCT | >MI0003190\|hsa-mir-505&WithFlank&X\|-\|139924142\|139924237 | 57 | 80 | 24 | CGTCAACACTTGCTGGTTTCCTCT | 66.5 | 2.5 | 0.4 | 6.1 | 8.4E-10 | 1.7E-09 | BT20/MB468 | NUC |
| >MI0000108\|hsa-mir-103a-2&WithFlank&20\|+\|3917488\|3917577@54.76.23;MI0000109\|hsa-mir-103a-1&WithFlank&5\|-\|168560890\|168560979@54.76.23&[MIMAT0000101&hsa-miR-103a-3p&offsets\|0\|0;m-12&20\|+\|3917541\|3917563&offsets\|0\|0];[MIMAT0000101_1&hsa-miR-103a-3p&offsets\|0\|0;m-13&5\|-\|168560904\|168560926&offsets\|0\|0;m-44220&5\|-\|168560908\|168560926&offsets\|0\|+4]\|AGCAGCATTGTACAGGGCTATGA | >MI0000108\|hsa-mir-103a-2&WithFlank&20\|+\|3917488\|3917577 | 54 | 76 | 23 | AGCAGCATTGTACAGGGCTATGA | 31974.1 | 5.9 | 1.0 | 6.1 | 8.0E-10 | 1.7E-09 | MB231/MB468 | MITO |
| >MI0000284\|hsa-mir-204&WithFlank&9\|-\|70809969\|70810090@39.60.22&[MIMAT0000265&hsa-miR-204-5p&offsets\|0\|0;m-296&9\|-\|70810031\|70810052&offsets\|0\|0]\|TTCCCTTTGTCATCCTATGCCT | >MI0000284\|hsa-mir-204&WithFlank&9\|-\|70809969\|70810090 | 39 | 60 | 22 | TTCCCTTTGTCATCCTATGCCT | 130.4 | 10.2 | 1.7 | 6.1 | 8.0E-10 | 1.7E-09 | MB231/MB468 | MITO |
| >am_trna111_HisGTG_1_-_147774845_147774916@4.29.26,trna118_HisGTG_1_-_145396881_145396952@4.29.26,trna16_HisGTG_1_+_146544773_146544844@4.29.26,trna1_HisGTG_15_+_45493349_45493420@4.29.26,trna21_HisGTG_1_+_147753471_147753542@4.29.26,trna33_HisGTG_6_+_27125906_27125977@4.29.26,trna7_HisGTG_9_-_14433938_14434009@4.29.26,trna8_HisGTG_15_-_45492611_45492682@4.29.26,trna9_HisGTG_15_-_45490804_45490875@4.29.26\|GTGATCGTATAGTGGTTAGTACTCTG | >am_trna111_HisGTG_1_-_147774845_147774916 | 4 | 29 | 26 | GTGATCGTATAGTGGTTAGTACTCTG | 69.2 | -10.0 | 1.6 | -6.2 | 7.0E-10 | 1.7E-09 | MB231/MB468 | MP |
| >MI0000266\|hsa-mir-10a&WithFlank&17\|-\|48579832\|48579953@29.48.20(+1C)&[MIMAT0000253&hsa-miR-10a-5p&offsets\|+1\|-2(+1C);m-18&17\|-\|48579905\|48579926&offsets\|+1\|-1(+1C)]\|ACCCTGTAGATCCGAATTTGC | >MI0000266\|hsa-mir-10a&WithFlank&17\|-\|48579832\|48579953 | 29 | 48 | 20(+1C) | ACCCTGTAGATCCGAATTTGC | 214.3 | 11.5 | 1.8 | 6.5 | 7.0E-11 | 1.7E-09 | BT20/MB468 | MP |
| >hg38_21_+_8433172_8446622_RNA45SN1_45S_50ntFlanks@12283.12306.24\|TACAGACCGTGAAAGCGGGGCCTC | >hg38_21_+_8433172_8446622_RNA45SN1_45S_50ntFlanks | 12283 | 12306 | 24 | TACAGACCGTGAAAGCGGGGCCTC | 95.4 | -10.4 | 1.7 | -6.2 | 7.0E-10 | 1.7E-09 | MB231/MB468 | MP |
| >hg38_21_+_8433172_8446622_RNA45SN1_45S_50ntFlanks@4995.5024.30\|ACAGATTGATAGCTCTTTCTCGATTCCGTG | >hg38_21_+_8433172_8446622_RNA45SN1_45S_50ntFlanks | 4995 | 5024 | 30 | ACAGATTGATAGCTCTTTCTCGATTCCGTG | 3028.2 | -7.0 | 1.1 | -6.2 | 7.3E-10 | 1.7E-09 | MB231/MB468 | NUC |
| >hg38_21_+_8433172_8446622_RNA45SN1_45S_50ntFlanks@4726.4759.34\|TAATCAAGAACGAAAGTCGGAGGTTCGAAGACGA | >hg38_21_+_8433172_8446622_RNA45SN1_45S_50ntFlanks | 4726 | 4759 | 34 | TAATCAAGAACGAAAGTCGGAGGTTCGAAGACGA | 1388.7 | -8.9 | 1.4 | -6.2 | 7.0E-10 | 1.7E-09 | MB231/MB468 | MP |
| >hg38_21_+_8433172_8446622_RNA45SN1_45S_50ntFlanks@10808.10842.35\|CAATGTAGGTAAGGGAAGTCGGCAAGCCGGATCCG | >hg38_21_+_8433172_8446622_RNA45SN1_45S_50ntFlanks | 10808 | 10842 | 35 | CAATGTAGGTAAGGGAAGTCGGCAAGCCGGATCCG | 70.3 | -10.0 | 1.6 | -6.2 | 7.0E-10 | 1.7E-09 | MB231/MB468 | MP |
| >hg38_21_+_8433172_8446622_RNA45SN1_45S_50ntFlanks@4027.4059.33\|CCCCGTGGCGGCGACGACCCATTCGAACGTCTG | >hg38_21_+_8433172_8446622_RNA45SN1_45S_50ntFlanks | 4027 | 4059 | 33 | CCCCGTGGCGGCGACGACCCATTCGAACGTCTG | 82.8 | -10.2 | 1.7 | -6.2 | 7.0E-10 | 1.7E-09 | MB231/MB468 | MP |
| >MI0000433\|hsa-let-7g&WithFlank&3\|-\|52268272\|52268367@11.31.21(+1G)&[MIMAT0000414&hsa-let-7g-5p&offsets\|0\|-1(+1G);m-58&3\|-\|52268336\|52268357&offsets\|0\|-1(+1G)]\|TGAGGTAGTAGTTTGTACAGTG | >MI0000433\|hsa-let-7g&WithFlank&3\|-\|52268272\|52268367 | 11 | 31 | 21(+1G) | TGAGGTAGTAGTTTGTACAGTG | 1110.5 | 4.9 | 0.8 | 6.2 | 7.3E-10 | 1.7E-09 | MB231/MB468 | NUC |
| >hg38_21_+_8433172_8446622_RNA45SN1_45S_50ntFlanks@10760.10785.26\|CATATCCGCAGCAGGTCTCCAAGGTG | >hg38_21_+_8433172_8446622_RNA45SN1_45S_50ntFlanks | 10760 | 10785 | 26 | CATATCCGCAGCAGGTCTCCAAGGTG | 158.1 | -7.1 | 1.2 | -6.2 | 7.3E-10 | 1.7E-09 | MB231/MB468 | NUC |
| >MI0000108\|hsa-mir-103a-2&WithFlank&20\|+\|3917488\|3917577@54.75.22(+1U);MI0000109\|hsa-mir-103a-1&WithFlank&5\|-\|168560890\|168560979@54.75.22(+1U)&[MIMAT0000101&hsa-miR-103a-3p&offsets\|0\|-1(+1U);m-12&20\|+\|3917541\|3917563&offsets\|0\|-1(+1U)];[MIMAT0000101_1&hsa-miR-103a-3p&offsets\|0\|-1(+1U);m-13&5\|-\|168560904\|168560926&offsets\|0\|-1(+1U);m-44220&5\|-\|168560908\|168560926&offsets\|0\|+3(+1U)]\|AGCAGCATTGTACAGGGCTATGT | >MI0000108\|hsa-mir-103a-2&WithFlank&20\|+\|3917488\|3917577 | 54 | 75 | 22(+1U) | AGCAGCATTGTACAGGGCTATGT | 1981.1 | 6.5 | 1.1 | 6.1 | 8.0E-10 | 1.7E-09 | MB231/MB468 | MITO |
| >MI0000102\|hsa-mir-100&WithFlank&11\|-\|122152223\|122152314@19.40.22&[MIMAT0000098&hsa-miR-100-5p&offsets\|0\|0;m-27&11\|-\|122152276\|122152296&offsets\|0\|+1]\|AACCCGTAGATCCGAACTTGTG | >MI0000102\|hsa-mir-100&WithFlank&11\|-\|122152223\|122152314 | 19 | 40 | 22 | AACCCGTAGATCCGAACTTGTG | 883976.2 | -7.7 | 1.1 | -6.8 | 1.3E-11 | 1.7E-09 | BT20/MB231 | MP |
| >hg38_21_+_8433172_8446622_RNA45SN1_45S_50ntFlanks@11792.11810.19\|ATGAACGAGATTCCCACTG | >hg38_21_+_8433172_8446622_RNA45SN1_45S_50ntFlanks | 11792 | 11810 | 19 | ATGAACGAGATTCCCACTG | 215.8 | -7.7 | 1.2 | -6.1 | 8.1E-10 | 1.7E-09 | MB231/MB468 | MITO |
| >hg38_21_+_8433172_8446622_RNA45SN1_45S_50ntFlanks@12892.12925.34\|CCCTCGCCCGTCACGCACCGCACGTTCGTGGGGA | >hg38_21_+_8433172_8446622_RNA45SN1_45S_50ntFlanks | 12892 | 12925 | 34 | CCCTCGCCCGTCACGCACCGCACGTTCGTGGGGA | 170.4 | -10.5 | 1.6 | -6.5 | 7.0E-11 | 1.7E-09 | BT20/MB468 | MP |
| >hg38_21_+_8433172_8446622_RNA45SN1_45S_50ntFlanks@6648.6695.48\|GTACGACTCTTAGCGGTGGATCACTCGGCTCGTGCGTCGATGAAGAAC | >hg38_21_+_8433172_8446622_RNA45SN1_45S_50ntFlanks | 6648 | 6695 | 48 | GTACGACTCTTAGCGGTGGATCACTCGGCTCGTGCGTCGATGAAGAAC | 404.5 | 7.2 | 1.2 | 6.2 | 7.2E-10 | 1.7E-09 | MB231/MB468 | CYTO |
| >hg38_21_+_8433172_8446622_RNA45SN1_45S_50ntFlanks@3782.3814.33\|CAGTGAAACTGCGAATGGCTCATTAAATCAGTT | >hg38_21_+_8433172_8446622_RNA45SN1_45S_50ntFlanks | 3782 | 3814 | 33 | CAGTGAAACTGCGAATGGCTCATTAAATCAGTT | 164.3 | -9.4 | 1.5 | -6.2 | 5.5E-10 | 1.7E-09 | BT20/MB468 | MITO |
| >hg38_21_+_8433172_8446622_RNA45SN1_45S_50ntFlanks@4046.4064.19\|CATTCGAACGTCTGCCCTA | >hg38_21_+_8433172_8446622_RNA45SN1_45S_50ntFlanks | 4046 | 4064 | 19 | CATTCGAACGTCTGCCCTA | 480.7 | -7.0 | 1.1 | -6.1 | 8.1E-10 | 1.7E-09 | MB231/MB468 | MITO |
| >hg38_21_+_8433172_8446622_RNA45SN1_45S_50ntFlanks@12457.12475.19\|CAAGCGTTGGATTGTTCAC | >hg38_21_+_8433172_8446622_RNA45SN1_45S_50ntFlanks | 12457 | 12475 | 19 | CAAGCGTTGGATTGTTCAC | 922.3 | -8.1 | 1.3 | -6.1 | 8.1E-10 | 1.7E-09 | MB231/MB468 | MITO |
| >hg38_21_+_8433172_8446622_RNA45SN1_45S_50ntFlanks@12447.12473.27\|CAGAATTCACCAAGCGTTGGATTGTTC | >hg38_21_+_8433172_8446622_RNA45SN1_45S_50ntFlanks | 12447 | 12473 | 27 | CAGAATTCACCAAGCGTTGGATTGTTC | 633.7 | -4.3 | 0.7 | -6.2 | 7.0E-10 | 1.7E-09 | MB231/MB468 | MP |

| | | | | | | | | | | | | | |
|---|---|---|---|---|---|---|---|---|---|---|---|---|---|
| >hg38_21_+_8433172_8446622_RNA45SN1_45S_50ntFlanks@3899.3936.38\|CCTTCGCGGGGGGGATGCGTGCATTTATCAGATCAAAA | >hg38_21_+_8433172_8446622_RNA45SN1_45S_50ntFlanks | 3899 | 3936 | 38 | CCTTCGCGGGGGGGATGCGTGCATTTATCAGATCAAAA | 166.4 | -11.2 | 1.8 | -6.2 | 7.1E-10 | 1.7E-09 | MB231/MB468 | MP |
| >hg38_21_+_8433172_8446622_RNA45SN1_45S_50ntFlanks@6649.6689.41\|TACGACTCTTAGCGGTGGATCACTCGGCTCGTGCGTCGATG | >hg38_21_+_8433172_8446622_RNA45SN1_45S_50ntFlanks | 6649 | 6689 | 41 | TACGACTCTTAGCGGTGGATCACTCGGCTCGTGCGTCGATG | 11390.9 | -4.5 | 0.7 | -6.1 | 8.1E-10 | 1.7E-09 | MB231/MB468 | MITO |
| >MI0000103\|hsa-mir-101-1&WithFlank&1\|-\|65058428\|65058514@53.74.22;MI0000739\|hsa-mir-101-2&WithFlank&9\|+\|4850291\|4850381@55.76.22&[MIMAT0000099&hsa-miR-101-3p&offsets\|0\|+1;m-17&1\|-\|65058443\|65058462&offsets\|0\|+2];[MIMAT0000099_1&hsa-miR-101-3p&offsets\|0\|+1;m-16&9\|+\|4850345\|4850364&offsets\|0\|+2]\|TACAGTACTGTGATAACTGAAG | >MI0000103\|hsa-mir-101-1&WithFlank&1\|-\|65058428\|65058514 | 53 | 74 | 22 | TACAGTACTGTGATAACTGAAG | 1731.9 | 5.8 | 0.9 | 6.2 | 7.4E-10 | 1.7E-09 | MB231/MB468 | NUC |
| >hg38_21_+_8433172_8446622_RNA45SN1_45S_50ntFlanks@12002.12032.31\|TGCGGGCCGCCGGTGAAATACCACTACTCTG | >hg38_21_+_8433172_8446622_RNA45SN1_45S_50ntFlanks | 12002 | 12032 | 31 | TGCGGGCCGCCGGTGAAATACCACTACTCTG | 5715.6 | -7.7 | 1.3 | -6.2 | 7.4E-10 | 1.7E-09 | MB231/MB468 | NUC |
| >hg38_21_+_8433172_8446622_RNA45SN1_45S_50ntFlanks@10350.10371.22\|ACGAGAACTTTGAAGGCCGAAG | >hg38_21_+_8433172_8446622_RNA45SN1_45S_50ntFlanks | 10350 | 10371 | 22 | ACGAGAACTTTGAAGGCCGAAG | 3383.9 | -4.4 | 0.7 | -6.2 | 7.1E-10 | 1.7E-09 | MB231/MB468 | MP |
| >hg38_21_+_8433172_8446622_RNA45SN1_45S_50ntFlanks@8356.8377.22\|GAAAAGAACTTTGAAGAGAGAG | >hg38_21_+_8433172_8446622_RNA45SN1_45S_50ntFlanks | 8356 | 8377 | 22 | GAAAAGAACTTTGAAGAGAGAG | 1339.6 | -7.9 | 1.3 | -6.1 | 8.5E-10 | 1.7E-09 | BT20/MB468 | NUC |
| >trnaMT_GluTTC_MT_-_14674_14742@34.72.39,trnalookalike8_GluTTC_5_-_93905172_93905240@34.72.39\|ATATCATTGGTCGTGGTTGTAGTCCGTGCGAGAATACCA | >trnaMT_GluTTC_MT_-_14674_14742 | 34 | 72 | 39 | ATATCATTGGTCGTGGTTGTAGTCCGTGCGAGAATACCA | 7666.9 | 5.3 | 0.9 | 6.2 | 5.6E-10 | 1.7E-09 | BT20/MB468 | MITO |
| >hg38_21_+_8433172_8446622_RNA45SN1_45S_50ntFlanks@10403.10431.29\|ACATGGGTCAGTCGGTCCTGAGAGATGGG | >hg38_21_+_8433172_8446622_RNA45SN1_45S_50ntFlanks | 10403 | 10431 | 29 | ACATGGGTCAGTCGGTCCTGAGAGATGGG | 102.2 | -10.5 | 1.7 | -6.2 | 7.1E-10 | 1.7E-09 | MB231/MB468 | MP |
| >hg38_21_+_8433172_8446622_RNA45SN1_45S_50ntFlanks@10384.10405.22\|CCATGTGAACAGCAGTTGAACA | >hg38_21_+_8433172_8446622_RNA45SN1_45S_50ntFlanks | 10384 | 10405 | 22 | CCATGTGAACAGCAGTTGAACA | 929.5 | -7.8 | 1.3 | -6.2 | 7.1E-10 | 1.7E-09 | MB231/MB468 | MP |
| >hg38_21_+_8433172_8446622_RNA45SN1_45S_50ntFlanks@11581.11609.29\|AACTGGTGCGGACCAGGGGAATCCGACTG | >hg38_21_+_8433172_8446622_RNA45SN1_45S_50ntFlanks | 11581 | 11609 | 29 | AACTGGTGCGGACCAGGGGAATCCGACTG | 1565.4 | -2.5 | 0.4 | -6.2 | 6.5E-10 | 1.7E-09 | BT20/MB468 | CYTO |
| >hg38_21_+_8433172_8446622_RNA45SN1_45S_50ntFlanks@3993.4031.39\|GTGACTCTAGATAACCTCGGGCCGATCGCACGCCCCCCG | >hg38_21_+_8433172_8446622_RNA45SN1_45S_50ntFlanks | 3993 | 4031 | 39 | GTGACTCTAGATAACCTCGGGCCGATCGCACGCCCCCCG | 647.8 | 9.0 | 1.5 | 6.2 | 7.4E-10 | 1.7E-09 | MB231/MB468 | CYTO |
| >hg38_21_+_8433172_8446622_RNA45SN1_45S_50ntFlanks@9462.9494.33\|GGGAGGTGGAGCACGAGCGCACGTGTTAGGACC | >hg38_21_+_8433172_8446622_RNA45SN1_45S_50ntFlanks | 9462 | 9494 | 33 | GGGAGGTGGAGCACGAGCGCACGTGTTAGGACC | 108.4 | -10.6 | 1.7 | -6.2 | 7.1E-10 | 1.7E-09 | MB231/MB468 | MP |
| >hg38_21_+_8433172_8446622_RNA45SN1_45S_50ntFlanks@10694.10711.18\|CGGTGAGCTCTCGCTGGC | >hg38_21_+_8433172_8446622_RNA45SN1_45S_50ntFlanks | 10694 | 10711 | 18 | CGGTGAGCTCTCGCTGGC | 497.5 | -4.1 | 0.7 | -6.2 | 5.6E-10 | 1.7E-09 | BT20/MB468 | MITO |
| >MI0000102\|hsa-mir-100&WithFlank&11\|-\|122152223\|122152314@19.42.24&[MIMAT0000098&hsa-miR-100-5p&offsets\|0\|+2;m-27&11\|-\|122152276\|122152296&offsets\|0\|+3]\|AACCCGTAGATCCGAACTTGTGGT | >MI0000102\|hsa-mir-100&WithFlank&11\|-\|122152223\|122152314 | 19 | 42 | 24 | AACCCGTAGATCCGAACTTGTGGT | 99.2 | 9.9 | 1.6 | 6.2 | 7.2E-10 | 1.7E-09 | MB231/MB468 | MP |
| >hg38_21_+_8433172_8446622_RNA45SN1_45S_50ntFlanks@10403.10430.28\|ACATGGGTCAGTCGGTCCTGAGAGATGG | >hg38_21_+_8433172_8446622_RNA45SN1_45S_50ntFlanks | 10403 | 10430 | 28 | ACATGGGTCAGTCGGTCCTGAGAGATGG | 160.6 | -6.4 | 1.0 | -6.2 | 6.5E-10 | 1.7E-09 | BT20/MB468 | CYTO |
| >hg38_21_+_8433172_8446622_RNA45SN1_45S_50ntFlanks@11214.11234.21\|ACTCTGGACGCGAGCCGGGCC | >hg38_21_+_8433172_8446622_RNA45SN1_45S_50ntFlanks | 11214 | 11234 | 21 | ACTCTGGACGCGAGCCGGGCC | 200.9 | -6.5 | 1.1 | -6.2 | 6.5E-10 | 1.7E-09 | BT20/MB468 | CYTO |
| >hg38_21_+_8433172_8446622_RNA45SN1_45S_50ntFlanks@9570.9592.23\|TGCAAATCGGTCGTCCGACCTGG | >hg38_21_+_8433172_8446622_RNA45SN1_45S_50ntFlanks | 9570 | 9592 | 23 | TGCAAATCGGTCGTCCGACCTGG | 179.5 | -6.8 | 1.1 | -6.2 | 6.5E-10 | 1.7E-09 | BT20/MB468 | CYTO |
| >hg38_21_+_8433172_8446622_RNA45SN1_45S_50ntFlanks@11575.11593.19\|ACTTAGAACTGGTGCGGAC | >hg38_21_+_8433172_8446622_RNA45SN1_45S_50ntFlanks | 11575 | 11593 | 19 | ACTTAGAACTGGTGCGGAC | 891.6 | -6.9 | 1.1 | -6.1 | 8.2E-10 | 1.7E-09 | MB231/MB468 | MITO |
| >hg38_21_+_8433172_8446622_RNA45SN1_45S_50ntFlanks@11729.11747.19\|TAACTATGACTCTCTTAAG | >hg38_21_+_8433172_8446622_RNA45SN1_45S_50ntFlanks | 11729 | 11747 | 19 | TAACTATGACTCTCTTAAG | 2881.3 | -7.1 | 1.2 | -6.1 | 8.3E-10 | 1.7E-09 | MB231/MB468 | MITO |
| >MI0000273\|hsa-mir-183&WithFlank&7\|-\|129774899\|129775020@34.53.20(+1C)&[MIMAT0000261&hsa-miR-183-5p&offsets\|+1\|-1(+1C);m-41&7\|-\|129774966\|129774987&offsets\|0\|-2(+1C)]\|ATGGCACTGGTAGAATTCACC | >MI0000273\|hsa-mir-183&WithFlank&7\|-\|129774899\|129775020 | 34 | 53 | 20(+1C) | ATGGCACTGGTAGAATTCACC | 161.9 | 2.6 | 0.4 | 6.1 | 8.6E-10 | 1.7E-09 | BT20/MB468 | NUC |
| >hg38_21_+_8433172_8446622_RNA45SN1_45S_50ntFlanks@10392.10430.39\|ACAGCAGTTGAACATGGGTCAGTCGGTCCTGAGAGATGG | >hg38_21_+_8433172_8446622_RNA45SN1_45S_50ntFlanks | 10392 | 10430 | 39 | ACAGCAGTTGAACATGGGTCAGTCGGTCCTGAGAGATGG | 697.0 | -8.9 | 1.4 | -6.2 | 7.2E-10 | 1.7E-09 | MB231/MB468 | MP |
| >MI0000749\|hsa-mir-30e&WithFlank&1\|+\|40754349\|40754452@65.85.21(+2U)&[MIMAT0000693&hsa-miR-30e-3p&offsets\|0\|-1(+2U);m-25&1\|+\|40754413\|40754434&offsets\|0\|-1(+2U)]\|CTTTCAGTCGGATGTTTACAGTT | >MI0000749\|hsa-mir-30e&WithFlank&1\|+\|40754349\|40754452 | 65 | 85 | 21(+2U) | CTTTCAGTCGGATGTTTACAGTT | 644.8 | 5.1 | 0.8 | 6.2 | 7.4E-10 | 1.7E-09 | MB231/MB468 | CYTO |
| >MI0000061\|hsa-let-7a-2&WithFlank&11\|-\|122146516\|122146599@56.77.22&[MIMAT0010195&hsa-let-7a-2-3p&offsets\|0\|0;m-426&11\|-\|122146523\|122146544&offsets\|0\|0]\|CTGTACAGCCTCCTAGCTTTCC | >MI0000061\|hsa-let-7a-2&WithFlank&11\|-\|122146516\|122146599 | 56 | 77 | 22 | CTGTACAGCCTCCTAGCTTTCC | 99.8 | 9.9 | 1.6 | 6.2 | 7.2E-10 | 1.7E-09 | MB231/MB468 | MP |
| >hg38_21_+_8433172_8446622_RNA45SN1_45S_50ntFlanks@11573.11592.20\|CGACTTAGAACTGGTGCGGA | >hg38_21_+_8433172_8446622_RNA45SN1_45S_50ntFlanks | 11573 | 11592 | 20 | CGACTTAGAACTGGTGCGGA | 825.5 | -7.1 | 1.2 | -6.1 | 8.7E-10 | 1.7E-09 | BT20/MB468 | NUC |
| >hg38_21_+_8433172_8446622_RNA45SN1_45S_50ntFlanks@12405.12442.38\|CTTTTTGATCCTTCGATGTCGGCTCTTCCTATCATTGT | >hg38_21_+_8433172_8446622_RNA45SN1_45S_50ntFlanks | 12405 | 12442 | 38 | CTTTTTGATCCTTCGATGTCGGCTCTTCCTATCATTGT | 1609.0 | -5.4 | 0.9 | -6.2 | 7.5E-10 | 1.7E-09 | MB231/MB468 | NUC |
| >hg38_21_+_8433172_8446622_RNA45SN1_45S_50ntFlanks@11580.11601.22\|GAACTGGTGCGGACCAGGGGAA | >hg38_21_+_8433172_8446622_RNA45SN1_45S_50ntFlanks | 11580 | 11601 | 22 | GAACTGGTGCGGACCAGGGGAA | 1344.3 | -4.8 | 0.8 | -6.2 | 6.6E-10 | 1.7E-09 | BT20/MB468 | CYTO |
| >hg38_21_+_8433172_8446622_RNA45SN1_45S_50ntFlanks@4472.4492.21\|CTTAGCTGAGTGTCCCGCGGG | >hg38_21_+_8433172_8446622_RNA45SN1_45S_50ntFlanks | 4472 | 4492 | 21 | CTTAGCTGAGTGTCCCGCGGG | 234.7 | -10.0 | 1.6 | -6.2 | 5.6E-10 | 1.7E-09 | BT20/MB468 | MITO |
| >trna152_ValCAC_6_-_27248049_27248121@1.29.29\|GCTTCTGTAGTGTAGTGGTTATCACGTTC | >trna152_ValCAC_6_-_27248049_27248121 | 1 | 29 | 29 | GCTTCTGTAGTGTAGTGGTTATCACGTTC | 10151.0 | -5.8 | 0.9 | -6.2 | 7.2E-10 | 1.7E-09 | MB231/MB468 | MP |
| >MI0000809\|hsa-mir-151a&WithFlank&8\|-\|140732558\|140732659@51.72.22(+1U)&[MIMAT0000757&hsa-miR-151a-3p&offsets\|-2\|-1(+1U);m-30&8\|-\|140732587\|140732607&offsets\|-2\|-1(+1U)]\|TACTAGACTGAAGCTCCTTGAGT | >MI0000809\|hsa-mir-151a&WithFlank&8\|-\|140732558\|140732659 | 51 | 72 | 22(+1U) | TACTAGACTGAAGCTCCTTGAGT | 449.4 | 4.7 | 0.8 | 6.2 | 7.2E-10 | 1.7E-09 | MB231/MB468 | MP |
| >hg38_21_+_8433172_8446622_RNA45SN1_45S_50ntFlanks@6771.6806.36\|GGTTCCTCCCGGGGCTACGCCTGTCTGAGCGTCGCT | >hg38_21_+_8433172_8446622_RNA45SN1_45S_50ntFlanks | 6771 | 6806 | 36 | GGTTCCTCCCGGGGCTACGCCTGTCTGAGCGTCGCT | 3224.3 | -4.0 | 0.7 | -6.1 | 8.3E-10 | 1.7E-09 | MB231/MB468 | MITO |
| >trna2_GlyGCC_21_-_18827107_18827177@1.31.31,trna35_GlyGCC_1_+_161413094_161413164@1.31.31,trna37_GlyGCC_1_+_161420467_161420537@1.31.31,trna39_GlyGCC_1_+_161427898_161427968@1.31.31,trna41_GlyGCC_1_+_161435258_161435328@1.31.31\|GCATGGGTGGTTCAGTGGTAGAATTCTCGCC | >trna2_GlyGCC_21_-_18827107_18827177 | 1 | 31 | 31 | GCATGGGTGGTTCAGTGGTAGAATTCTCGCC | 80519.5 | 8.3 | 1.3 | 6.5 | 7.2E-11 | 1.7E-09 | BT20/MB468 | MP |
| >hg38_21_+_8433172_8446622_RNA45SN1_45S_50ntFlanks@10830.10851.22\|CAAGCCGGATCCGTAACTTCGG | >hg38_21_+_8433172_8446622_RNA45SN1_45S_50ntFlanks | 10830 | 10851 | 22 | CAAGCCGGATCCGTAACTTCGG | 264.8 | -2.9 | 0.5 | -6.2 | 6.6E-10 | 1.7E-09 | BT20/MB468 | CYTO |
| >hg38_21_+_8433172_8446622_RNA45SN1_45S_50ntFlanks@12605.12626.22\|TATGTGCTTGGCTGAGGAGCCA | >hg38_21_+_8433172_8446622_RNA45SN1_45S_50ntFlanks | 12605 | 12626 | 22 | TATGTGCTTGGCTGAGGAGCCA | 5398.7 | -6.8 | 1.1 | -6.2 | 7.5E-10 | 1.7E-09 | MB231/MB468 | CYTO |
| >hg38_21_+_8433172_8446622_RNA45SN1_45S_50ntFlanks@10398.10424.27\|GTTGAACATGGGTCAGTCGGTCCTGAG | >hg38_21_+_8433172_8446622_RNA45SN1_45S_50ntFlanks | 10398 | 10424 | 27 | GTTGAACATGGGTCAGTCGGTCCTGAG | 1867.1 | 4.9 | 0.8 | 6.2 | 7.6E-10 | 1.8E-09 | MB231/MB468 | NUC |
| >hg38_21_+_8433172_8446622_RNA45SN1_45S_50ntFlanks@4554.4574.21\|CCGCAGCTAGGAATAATGGAA | >hg38_21_+_8433172_8446622_RNA45SN1_45S_50ntFlanks | 4554 | 4574 | 21 | CCGCAGCTAGGAATAATGGAA | 262.3 | -8.2 | 1.3 | -6.1 | 8.8E-10 | 1.8E-09 | BT20/MB468 | NUC |
| >MI0000750\|hsa-mir-26a-2&WithFlank&12\|-\|57824603\|57824698@20.41.22(+2U);MI0000083\|hsa-mir-26a-1&WithFlank&3\|+\|37969398\|37969486@16.37.22(+2U)&[MIMAT0000082&hsa-miR-26a-5p&offsets\|0\|0(+2U);m-33&12\|-\|57824658\|57824679&offsets\|0\|0(+2U)];[MIMAT0000082_1&hsa-miR-26a-5p&offsets\|0\|0(+2U);m-34&3\|+\|37969413\|37969434&offsets\|0\|0(+2U)]\|TTCAAGTAATCCAGGATAGGCTTT | >MI0000750\|hsa-mir-26a-2&WithFlank&12\|-\|57824603\|57824698 | 20 | 41 | 22(+2U) | TTCAAGTAATCCAGGATAGGCTTT | 2607.2 | 4.9 | 0.8 | 6.2 | 7.3E-10 | 1.8E-09 | MB231/MB468 | MP |
| >hg38_21_+_8433172_8446622_RNA45SN1_45S_50ntFlanks@9554.9573.20\|CGTAGCGGTCCTGACGTGCA | >hg38_21_+_8433172_8446622_RNA45SN1_45S_50ntFlanks | 9554 | 9573 | 20 | CGTAGCGGTCCTGACGTGCA | 12635.7 | -3.6 | 0.6 | -6.2 | 7.6E-10 | 1.8E-09 | MB231/MB468 | NUC |
| >hg38_21_+_8433172_8446622_RNA45SN1_45S_50ntFlanks@10820.10840.21\|GGGAAGTCGGCAAGCCGGATC | >hg38_21_+_8433172_8446622_RNA45SN1_45S_50ntFlanks | 10820 | 10840 | 21 | GGGAAGTCGGCAAGCCGGATC | 2508.6 | -7.4 | 1.2 | -6.2 | 7.3E-10 | 1.8E-09 | MB231/MB468 | MP |
| >hg38_21_+_8433172_8446622_RNA45SN1_45S_50ntFlanks@10725.10755.31\|GGGGAGAGGGTGTAAATCTCGCGCCGGGCCG | >hg38_21_+_8433172_8446622_RNA45SN1_45S_50ntFlanks | 10725 | 10755 | 31 | GGGGAGAGGGTGTAAATCTCGCGCCGGGCCG | 219.7 | -6.4 | 1.0 | -6.2 | 7.5E-10 | 1.8E-09 | MB231/MB468 | CYTO |
| >hg38_21_+_8433172_8446622_RNA45SN1_45S_50ntFlanks@4585.4616.32\|GTTCTATTTTGTTGGTTTTCGGAACTGAGGCC | >hg38_21_+_8433172_8446622_RNA45SN1_45S_50ntFlanks | 4585 | 4616 | 32 | GTTCTATTTTGTTGGTTTTCGGAACTGAGGCC | 245.9 | 5.6 | 0.9 | 6.2 | 7.5E-10 | 1.8E-09 | MB231/MB468 | CYTO |
| >hg38_21_+_8433172_8446622_RNA45SN1_45S_50ntFlanks@11319.11344.26\|CCCACGTCTCGTCGCGCGCGCGTCCG | >hg38_21_+_8433172_8446622_RNA45SN1_45S_50ntFlanks | 11319 | 11344 | 26 | CCCACGTCTCGTCGCGCGCGCGTCCG | 2531.1 | -6.5 | 1.0 | -6.2 | 6.7E-10 | 1.8E-09 | BT20/MB468 | CYTO |
| >MI0000734\|hsa-mir-106b&WithFlank&7\|-\|100093987\|100094080@58.79.22&[MIMAT0004672&hsa-miR-106b-3p&offsets\|0\|0;m-87&7\|-\|100094003\|100094023&offsets\|0\|+1]\|CCGCACTGTGGGTACTTGCTGC | >MI0000734\|hsa-mir-106b&WithFlank&7\|-\|100093987\|100094080 | 58 | 79 | 22 | CCGCACTGTGGGTACTTGCTGC | 1924.4 | 4.4 | 0.7 | 6.1 | 8.4E-10 | 1.8E-09 | MB231/MB468 | MITO |
| >hg38_21_+_8433172_8446622_RNA45SN1_45S_50ntFlanks@4045.4062.18\|CCATTCGAACGTCTGCCC | >hg38_21_+_8433172_8446622_RNA45SN1_45S_50ntFlanks | 4045 | 4062 | 18 | CCATTCGAACGTCTGCCC | 619.9 | -8.2 | 1.3 | -6.1 | 8.4E-10 | 1.8E-09 | MB231/MB468 | MITO |
| >hg38_21_+_8433172_8446622_RNA45SN1_45S_50ntFlanks@5075.5098.24\|TTCCGATAACGAACGAGACTCTGG | >hg38_21_+_8433172_8446622_RNA45SN1_45S_50ntFlanks | 5075 | 5098 | 24 | TTCCGATAACGAACGAGACTCTGG | 303.7 | -12.1 | 2.0 | -6.2 | 7.3E-10 | 1.8E-09 | MB231/MB468 | MP |
| >hg38_21_+_8433172_8446622_RNA45SN1_45S_50ntFlanks@9554.9597.44\|CGTAGCGGTCCTGACGTGCAAATCGGTCGTCCGACCTGGGTATA | >hg38_21_+_8433172_8446622_RNA45SN1_45S_50ntFlanks | 9554 | 9597 | 44 | CGTAGCGGTCCTGACGTGCAAATCGGTCGTCCGACCTGGGTATA | 280.7 | -12.0 | 1.9 | -6.2 | 7.3E-10 | 1.8E-09 | MB231/MB468 | MP |
| >hg38_21_+_8433172_8446622_RNA45SN1_45S_50ntFlanks@6745.6771.27\|ACACTTCGAACGCACTTGCGGCCCCGG | >hg38_21_+_8433172_8446622_RNA45SN1_45S_50ntFlanks | 6745 | 6771 | 27 | ACACTTCGAACGCACTTGCGGCCCCGG | 3200.7 | -10.1 | 1.6 | -6.1 | 8.8E-10 | 1.8E-09 | BT20/MB468 | NUC |
| >hg38_21_+_8433172_8446622_RNA45SN1_45S_50ntFlanks@11188.11207.20\|CACCCGGGGGGCCGGCGGCG | >hg38_21_+_8433172_8446622_RNA45SN1_45S_50ntFlanks | 11188 | 11207 | 20 | CACCCGGGGGGCCGGCGGCG | 225.0 | -7.6 | 1.2 | -6.2 | 7.7E-10 | 1.8E-09 | MB231/MB468 | NUC |

<table>
<tr><td>>hg38_21_+_8433172_8446622_RNA45SN1_45S_50ntFlanks@3802.3837.36|CATTAAATCAGTTATGGTTCCTTTGGTCGCTCGCTC</td><td>>hg38_21_+_8433172_8446622_RNA45SN1_45S_50ntFlanks</td><td>3802</td><td>3837</td><td>36</td><td>CATTAAATCAGTTATGGTTCCTTTGGTCGCTCGCTC</td><td>596.2</td><td>-8.1</td><td>1.3</td><td>-6.2</td><td>7.7E-10</td><td>1.8E-09</td><td>MB231/MB468</td><td>NUC</td></tr>
<tr><td>>hg38_21_+_8433172_8446622_RNA45SN1_45S_50ntFlanks@4129.4162.34|GATTCCGGAGAGGGAGCCTGAGAAACGGCTACCA</td><td>>hg38_21_+_8433172_8446622_RNA45SN1_45S_50ntFlanks</td><td>4129</td><td>4162</td><td>34</td><td>GATTCCGGAGAGGGAGCCTGAGAAACGGCTACCA</td><td>144.2</td><td>-11.0</td><td>1.8</td><td>-6.2</td><td>7.3E-10</td><td>1.8E-09</td><td>MB231/MB468</td><td>MP</td></tr>
<tr><td>>hg38_21_+_8433172_8446622_RNA45SN1_45S_50ntFlanks@4560.4587.28|CTAGGAATAATGGAATAGGACCGCGGTT</td><td>>hg38_21_+_8433172_8446622_RNA45SN1_45S_50ntFlanks</td><td>4560</td><td>4587</td><td>28</td><td>CTAGGAATAATGGAATAGGACCGCGGTT</td><td>669.0</td><td>-5.8</td><td>0.9</td><td>-6.2</td><td>7.6E-10</td><td>1.8E-09</td><td>MB231/MB468</td><td>CYTO</td></tr>
<tr><td>>hg38_21_+_8433172_8446622_RNA45SN1_45S_50ntFlanks@10840.10868.29|CCGTAACTTCGGGATAAGGATTGGCTCTA</td><td>>hg38_21_+_8433172_8446622_RNA45SN1_45S_50ntFlanks</td><td>10840</td><td>10868</td><td>29</td><td>CCGTAACTTCGGGATAAGGATTGGCTCTA</td><td>1168.1</td><td>-6.7</td><td>1.1</td><td>-6.2</td><td>7.7E-10</td><td>1.8E-09</td><td>MB231/MB468</td><td>NUC</td></tr>
<tr><td>>hg38_21_+_8433172_8446622_RNA45SN1_45S_50ntFlanks@4225.4242.18|AAATAACAATACAGGACT</td><td>>hg38_21_+_8433172_8446622_RNA45SN1_45S_50ntFlanks</td><td>4225</td><td>4242</td><td>18</td><td>AAATAACAATACAGGACT</td><td>334.1</td><td>-3.4</td><td>0.5</td><td>-6.2</td><td>5.8E-10</td><td>1.8E-09</td><td>BT20/MB468</td><td>MITO</td></tr>
<tr><td>>MI0000456|hsa-mir-140&WithFlank&16|+|69933075|69933186@29.50.22&[MIMAT0000431&hsa-miR-140-5p&offsets|0|0;m-173&16|+|69933103|69933124&offsets|0|0]|CAGTGGTTTTACCCTATGGTAG</td><td>>MI0000456|hsa-mir-140&WithFlank&16|+|69933075|69933186</td><td>29</td><td>50</td><td>22</td><td>CAGTGGTTTTACCCTATGGTAG</td><td>83.0</td><td>6.6</td><td>1.1</td><td>6.1</td><td>8.9E-10</td><td>1.8E-09</td><td>BT20/MB468</td><td>NUC</td></tr>
<tr><td>>MI0000737|hsa-mir-200a&WithFlank&1|+|1167857|1167958@60.80.21(+1C)&[MIMAT0000682&hsa-miR-200a-3p&offsets|0|-1(+1C);m-142&1|+|1167916|1167938&offsets|0|-2(+1C)]|TAACACTGTCTGGTAACGATGC</td><td>>MI0000737|hsa-mir-200a&WithFlank&1|+|1167857|1167958</td><td>60</td><td>80</td><td>21(+1C)</td><td>TAACACTGTCTGGTAACGATGC</td><td>162.1</td><td>-10.4</td><td>1.6</td><td>-6.5</td><td>7.4E-11</td><td>1.8E-09</td><td>BT20/MB468</td><td>MP</td></tr>
<tr><td>>MI0000279|hsa-mir-196a-2&WithFlank&12|+|53991732|53991853@31.51.21;MI0000238|hsa-mir-196a-1&WithFlank&17|-|48632484|48632565@13.33.21&[MIMAT0000226&hsa-miR-196a-5p&offsets|0|-1;m-261&12|+|53991762|53991783&offsets|0|-1];[MIMAT0000226_1&hsa-miR-196a-5p&offsets|0|-1;m-268&17|-|48632532|48632553&offsets|0|-1]|TAGGTAGTTTCATGTTGTTGG</td><td>>MI0000279|hsa-mir-196a-2&WithFlank&12|+|53991732|53991853</td><td>31</td><td>51</td><td>21</td><td>TAGGTAGTTTCATGTTGTTGG</td><td>339.8</td><td>2.4</td><td>0.4</td><td>6.1</td><td>8.9E-10</td><td>1.8E-09</td><td>BT20/MB468</td><td>NUC</td></tr>
<tr><td>>hg38_21_+_8433172_8446622_RNA45SN1_45S_50ntFlanks@6734.6756.23|ATTGATCATCGACACTTCGAACG</td><td>>hg38_21_+_8433172_8446622_RNA45SN1_45S_50ntFlanks</td><td>6734</td><td>6756</td><td>23</td><td>ATTGATCATCGACACTTCGAACG</td><td>776.1</td><td>3.2</td><td>0.5</td><td>6.2</td><td>5.8E-10</td><td>1.8E-09</td><td>BT20/MB468</td><td>MITO</td></tr>
<tr><td>>MI0000272|hsa-mir-182&WithFlank&7|-|129770377|129770498@29.50.22(+1U)&[MIMAT0000259&hsa-miR-182-5p&offsets|0|-2(+1U);m-22&7|-|129770449|129770470&offsets|0|0(+1U)]|TTTGGCAATGGTAGAACTCACAT</td><td>>MI0000272|hsa-mir-182&WithFlank&7|-|129770377|129770498</td><td>29</td><td>50</td><td>22(+1U)</td><td>TTTGGCAATGGTAGAACTCACAT</td><td>324.0</td><td>9.2</td><td>1.5</td><td>6.2</td><td>5.8E-10</td><td>1.8E-09</td><td>BT20/MB468</td><td>MITO</td></tr>
<tr><td>>hg38_21_+_8433172_8446622_RNA45SN1_45S_50ntFlanks@8379.8400.22|TCAAGAGGGCGTGAAACCGTTA</td><td>>hg38_21_+_8433172_8446622_RNA45SN1_45S_50ntFlanks</td><td>8379</td><td>8400</td><td>22</td><td>TCAAGAGGGCGTGAAACCGTTA</td><td>254.9</td><td>-7.8</td><td>1.3</td><td>-6.1</td><td>8.5E-10</td><td>1.8E-09</td><td>MB231/MB468</td><td>MITO</td></tr>
<tr><td>>hg38_21_+_8433172_8446622_RNA45SN1_45S_50ntFlanks@4592.4616.25|TTTGTTGGTTTTCGGAACTGAGGCC</td><td>>hg38_21_+_8433172_8446622_RNA45SN1_45S_50ntFlanks</td><td>4592</td><td>4616</td><td>25</td><td>TTTGTTGGTTTTCGGAACTGAGGCC</td><td>379.6</td><td>-8.0</td><td>1.3</td><td>-6.2</td><td>7.4E-10</td><td>1.8E-09</td><td>MB231/MB468</td><td>MP</td></tr>
<tr><td>>trna10_LysCTT_16_+_3241501_3241573@1.30.30,trna13_LysCTT_14_-_58706613_58706685@1.30.30,trna2_LysCTT_15_+_79152904_79152976@1.30.30|GCCCGGCTAGCTCAGTCGGTAGAGCATGGG</td><td>>trna10_LysCTT_16_+_3241501_3241573</td><td>1</td><td>30</td><td>30</td><td>GCCCGGCTAGCTCAGTCGGTAGAGCATGGG</td><td>100.8</td><td>-10.5</td><td>1.7</td><td>-6.2</td><td>7.4E-10</td><td>1.8E-09</td><td>MB231/MB468</td><td>MP</td></tr>
<tr><td>>hg38_21_+_8433172_8446622_RNA45SN1_45S_50ntFlanks@6647.6664.18|CGTACGACTCTTAGCGGT</td><td>>hg38_21_+_8433172_8446622_RNA45SN1_45S_50ntFlanks</td><td>6647</td><td>6664</td><td>18</td><td>CGTACGACTCTTAGCGGT</td><td>2070.7</td><td>3.3</td><td>0.5</td><td>6.1</td><td>7.8E-10</td><td>1.8E-09</td><td>MB231/MB468</td><td>NUC</td></tr>
<tr><td>>MI0000082|hsa-mir-25&WithFlank&7|-|100093554|100093649@58.78.21(+1G)&[MIMAT0000081&hsa-miR-25-3p&offsets|0|-1(+1G);m-23&7|-|100093571|100093592&offsets|0|-1(+1G)]|CATTGCACTTGTCTCGGTCTGG</td><td>>MI0000082|hsa-mir-25&WithFlank&7|-|100093554|100093649</td><td>58</td><td>78</td><td>21(+1G)</td><td>CATTGCACTTGTCTCGGTCTGG</td><td>937.5</td><td>4.2</td><td>0.7</td><td>6.1</td><td>7.8E-10</td><td>1.8E-09</td><td>MB231/MB468</td><td>NUC</td></tr>
<tr><td>>hg38_21_+_8433172_8446622_RNA45SN1_45S_50ntFlanks@9567.9588.22|ACGTGCAAATCGGTCGTCCGAC</td><td>>hg38_21_+_8433172_8446622_RNA45SN1_45S_50ntFlanks</td><td>9567</td><td>9588</td><td>22</td><td>ACGTGCAAATCGGTCGTCCGAC</td><td>231.9</td><td>-5.1</td><td>0.8</td><td>-6.2</td><td>7.7E-10</td><td>1.8E-09</td><td>MB231/MB468</td><td>CYTO</td></tr>
<tr><td>>MI0000088|hsa-mir-30a&WithFlank&6|-|71403545|71403627@12.35.24(+2U)&[MIMAT0000087&hsa-miR-30a-5p&offsets|0|+2(+2U);m-15&6|-|71403595|71403616&offsets|0|+2(+2U)]|TGTAAACATCCTCGACTGGAAGCTTT</td><td>>MI0000088|hsa-mir-30a&WithFlank&6|-|71403545|71403627</td><td>12</td><td>35</td><td>24(+2U)</td><td>TGTAAACATCCTCGACTGGAAGCTTT</td><td>2941.2</td><td>-5.5</td><td>0.8</td><td>-6.5</td><td>6.5E-11</td><td>1.8E-09</td><td>BT20/MB231</td><td>MITO</td></tr>
<tr><td>>hg38_21_+_8433172_8446622_RNA45SN1_45S_50ntFlanks@11565.11582.18|CTAGCAGCCGACTTAGAA</td><td>>hg38_21_+_8433172_8446622_RNA45SN1_45S_50ntFlanks</td><td>11565</td><td>11582</td><td>18</td><td>CTAGCAGCCGACTTAGAA</td><td>419.0</td><td>-8.5</td><td>1.4</td><td>-6.1</td><td>8.9E-10</td><td>1.8E-09</td><td>BT20/MB468</td><td>NUC</td></tr>
<tr><td>>hg38_21_+_8433172_8446622_RNA45SN1_45S_50ntFlanks@12084.12108.25|TCGCTTCTGGCGCCAAGCGCCCGGC</td><td>>hg38_21_+_8433172_8446622_RNA45SN1_45S_50ntFlanks</td><td>12084</td><td>12108</td><td>25</td><td>TCGCTTCTGGCGCCAAGCGCCCGGC</td><td>163.7</td><td>-11.2</td><td>1.8</td><td>-6.2</td><td>7.4E-10</td><td>1.8E-09</td><td>MB231/MB468</td><td>MP</td></tr>
<tr><td>>trna10_AspGTC_12_-_125424193_125424264@1.30.30,trna12_AspGTC_12_-_125411891_125411962@1.30.30,trna38_AspGTC_17_-_8125556_8125627@1.30.30,trna45_AspGTC_6_+_27447453_27447524@1.30.30,trna48_AspGTC_6_+_27471523_27471594@1.30.30,trna4_AspGTC_12_+_96429799_96429870@1.30.30,trna69_AspGTC_1_-_161440205_161440276@1.30.30,trna72_AspGTC_1_-_161432824_161432895@1.30.30,trna75_AspGTC_1_-_161425414_161425485@1.30.30,trna78_AspGTC_1_-_161418033_161418104@1.30.30,trna81_AspGTC_1_-_161410615_161410686@1.30.30|TCCTCGTTAGTATAGTGGTGAGTATCCCCG</td><td>>trna10_AspGTC_12_-_125424193_125424264</td><td>1</td><td>30</td><td>30</td><td>TCCTCGTTAGTATAGTGGTGAGTATCCCCG</td><td>611.3</td><td>-7.9</td><td>1.3</td><td>-6.1</td><td>7.8E-10</td><td>1.8E-09</td><td>MB231/MB468</td><td>NUC</td></tr>
<tr><td>>hg38_21_+_8433172_8446622_RNA45SN1_45S_50ntFlanks@6651.6694.44|CGACTCTTAGCGGTGGATCACTCGGCTCGTGCGTCGATGAAGAA</td><td>>hg38_21_+_8433172_8446622_RNA45SN1_45S_50ntFlanks</td><td>6651</td><td>6694</td><td>44</td><td>CGACTCTTAGCGGTGGATCACTCGGCTCGTGCGTCGATGAAGAA</td><td>930.3</td><td>5.2</td><td>0.9</td><td>6.2</td><td>7.7E-10</td><td>1.8E-09</td><td>MB231/MB468</td><td>CYTO</td></tr>
<tr><td>>trna10_LysCTT_16_+_3241501_3241573@4.35.32,trna13_LysCTT_14_-_58706613_58706685@4.35.32,trna2_LysCTT_15_+_79152904_79152976@4.35.32|CGGCTAGCTCAGTCGGTAGAGCATGGGACTCT</td><td>>trna10_LysCTT_16_+_3241501_3241573</td><td>4</td><td>35</td><td>32</td><td>CGGCTAGCTCAGTCGGTAGAGCATGGGACTCT</td><td>1745.5</td><td>7.5</td><td>1.2</td><td>6.1</td><td>8.6E-10</td><td>1.8E-09</td><td>MB231/MB468</td><td>MITO</td></tr>
<tr><td>>hg38_21_+_8433172_8446622_RNA45SN1_45S_50ntFlanks@8171.8214.44|AAGTCCTTCTGATCGAGGCCCAGCCCGTGGACGGTGTGAGGCCG</td><td>>hg38_21_+_8433172_8446622_RNA45SN1_45S_50ntFlanks</td><td>8171</td><td>8214</td><td>44</td><td>AAGTCCTTCTGATCGAGGCCCAGCCCGTGGACGGTGTGAGGCCG</td><td>1274.6</td><td>-5.5</td><td>0.9</td><td>-6.1</td><td>8.6E-10</td><td>1.8E-09</td><td>MB231/MB468</td><td>MITO</td></tr>
<tr><td>>hg38_21_+_8433172_8446622_RNA45SN1_45S_50ntFlanks@11453.11477.25|TCCTCGGCGCGCGGCGGCGGCGGCG</td><td>>hg38_21_+_8433172_8446622_RNA45SN1_45S_50ntFlanks</td><td>11453</td><td>11477</td><td>25</td><td>TCCTCGGCGCGCGGCGGCGGCGGCG</td><td>278.6</td><td>-6.5</td><td>1.1</td><td>-6.2</td><td>7.7E-10</td><td>1.8E-09</td><td>MB231/MB468</td><td>CYTO</td></tr>
<tr><td>>hg38_21_+_8433172_8446622_RNA45SN1_45S_50ntFlanks@12991.13025.35|CTCCCTCGCTGCGATCTATTGAAAGTCAGCCCTCG</td><td>>hg38_21_+_8433172_8446622_RNA45SN1_45S_50ntFlanks</td><td>12991</td><td>13025</td><td>35</td><td>CTCCCTCGCTGCGATCTATTGAAAGTCAGCCCTCG</td><td>121.1</td><td>-9.0</td><td>1.5</td><td>-6.2</td><td>5.9E-10</td><td>1.8E-09</td><td>BT20/MB468</td><td>MITO</td></tr>
<tr><td>>MI0000482|hsa-mir-185&WithFlank&22|+|20033133|20033226@21.42.22&[MIMAT0000455&hsa-miR-185-5p&offsets|0|0;m-145&22|+|20033153|20033174&offsets|0|0]|TGGAGAGAAAGGCAGTTCCTGA</td><td>>MI0000482|hsa-mir-185&WithFlank&22|+|20033133|20033226</td><td>21</td><td>42</td><td>22</td><td>TGGAGAGAAAGGCAGTTCCTGA</td><td>690.2</td><td>3.3</td><td>0.5</td><td>6.1</td><td>7.9E-10</td><td>1.8E-09</td><td>MB231/MB468</td><td>NUC</td></tr>
<tr><td>>hg38_21_+_8433172_8446622_RNA45SN1_45S_50ntFlanks@3846.3868.23|CTTGGATAACTGTGGTAATTCTA</td><td>>hg38_21_+_8433172_8446622_RNA45SN1_45S_50ntFlanks</td><td>3846</td><td>3868</td><td>23</td><td>CTTGGATAACTGTGGTAATTCTA</td><td>988.5</td><td>-6.9</td><td>1.1</td><td>-6.1</td><td>8.6E-10</td><td>1.8E-09</td><td>MB231/MB468</td><td>MITO</td></tr>
<tr><td>>hg38_21_+_8433172_8446622_RNA45SN1_45S_50ntFlanks@9534.9562.29|AGGAAACTCTGGTGGAGGTCCGTAGCGGT</td><td>>hg38_21_+_8433172_8446622_RNA45SN1_45S_50ntFlanks</td><td>9534</td><td>9562</td><td>29</td><td>AGGAAACTCTGGTGGAGGTCCGTAGCGGT</td><td>1502.2</td><td>-8.4</td><td>1.4</td><td>-6.1</td><td>8.6E-10</td><td>1.8E-09</td><td>MB231/MB468</td><td>MITO</td></tr>
<tr><td>>hg38_21_+_8433172_8446622_RNA45SN1_45S_50ntFlanks@9941.9963.23|AGGACGGTGGCCATGGAAGTCGG</td><td>>hg38_21_+_8433172_8446622_RNA45SN1_45S_50ntFlanks</td><td>9941</td><td>9963</td><td>23</td><td>AGGACGGTGGCCATGGAAGTCGG</td><td>982.3</td><td>13.7</td><td>2.1</td><td>6.5</td><td>7.5E-11</td><td>1.8E-09</td><td>BT20/MB468</td><td>MP</td></tr>
<tr><td>>trna13_ValCAC_19_-_4724647_4724719@1.34.34|GTTTCCGTAGTGTAGCGGTTATCACATTCGCCTC</td><td>>trna13_ValCAC_19_-_4724647_4724719</td><td>1</td><td>34</td><td>34</td><td>GTTTCCGTAGTGTAGCGGTTATCACATTCGCCTC</td><td>472.1</td><td>10.4</td><td>1.7</td><td>6.1</td><td>7.9E-10</td><td>1.8E-09</td><td>MB231/MB468</td><td>NUC</td></tr>
<tr><td>>hg38_21_+_8433172_8446622_RNA45SN1_45S_50ntFlanks@8194.8214.21|CCCGTGGACGGTGTGAGGCCG</td><td>>hg38_21_+_8433172_8446622_RNA45SN1_45S_50ntFlanks</td><td>8194</td><td>8214</td><td>21</td><td>CCCGTGGACGGTGTGAGGCCG</td><td>945.3</td><td>-8.1</td><td>1.3</td><td>-6.1</td><td>8.6E-10</td><td>1.8E-09</td><td>MB231/MB468</td><td>MITO</td></tr>
<tr><td>>hg38_21_+_8433172_8446622_RNA45SN1_45S_50ntFlanks@11705.11727.23|TGAAGCGCGGGTAAACGGCGGGA</td><td>>hg38_21_+_8433172_8446622_RNA45SN1_45S_50ntFlanks</td><td>11705</td><td>11727</td><td>23</td><td>TGAAGCGCGGGTAAACGGCGGGA</td><td>83.6</td><td>-10.2</td><td>1.7</td><td>-6.2</td><td>7.5E-10</td><td>1.8E-09</td><td>MB231/MB468</td><td>MP</td></tr>
<tr><td>>hg38_21_+_8433172_8446622_RNA45SN1_45S_50ntFlanks@12940.12972.33|CATTCGTAGACGACCTGCTTCTGGGTCGGGGTT</td><td>>hg38_21_+_8433172_8446622_RNA45SN1_45S_50ntFlanks</td><td>12940</td><td>12972</td><td>33</td><td>CATTCGTAGACGACCTGCTTCTGGGTCGGGGTT</td><td>676.5</td><td>-7.4</td><td>1.2</td><td>-6.1</td><td>7.8E-10</td><td>1.8E-09</td><td>MB231/MB468</td><td>CYTO</td></tr>
<tr><td>>MI0003591|hsa-mir-584&WithFlank&5|-|149062307|149062415@22.41.20&[MIMAT0003249&hsa-miR-584-5p&offsets|0|-2;m-167&5|-|149062374|149062394&offsets|0|-1]|TTATGGTTTGCCTGGGACTG</td><td>>MI0003591|hsa-mir-584&WithFlank&5|-|149062307|149062415</td><td>22</td><td>41</td><td>20</td><td>TTATGGTTTGCCTGGGACTG</td><td>1115.5</td><td>6.0</td><td>0.9</td><td>6.5</td><td>7.5E-11</td><td>1.8E-09</td><td>BT20/MB468</td><td>MP</td></tr>
<tr><td>>MI0003643|hsa-mir-629&WithFlank&15|-|70079366|70079474@28.49.22(+1A)&[MIMAT0004810&hsa-miR-629-5p&offsets|0|+1(+1A);m-150&15|-|70079427|70079447&offsets|0|+1(+1A)]|TGGGTTTACGTTGGGAGAACTTA</td><td>>MI0003643|hsa-mir-629&WithFlank&15|-|70079366|70079474</td><td>28</td><td>49</td><td>22(+1A)</td><td>TGGGTTTACGTTGGGAGAACTTA</td><td>345.6</td><td>12.2</td><td>1.9</td><td>6.5</td><td>7.6E-11</td><td>1.8E-09</td><td>BT20/MB468</td><td>MP</td></tr>
<tr><td>>MI0000101|hsa-mir-99a&WithFlank&21|+|16539083|16539175@19.40.22(+1U)&[MIMAT0000097&hsa-miR-99a-5p&offsets|0|0(+1U);m-64&21|+|16539101|16539121&offsets|0|+1(+1U)]|AACCCGTAGATCCGATCTTGTGT</td><td>>MI0000101|hsa-mir-99a&WithFlank&21|+|16539083|16539175</td><td>19</td><td>40</td><td>22(+1U)</td><td>AACCCGTAGATCCGATCTTGTGT</td><td>199.6</td><td>-10.7</td><td>1.6</td><td>-6.5</td><td>7.6E-11</td><td>1.8E-09</td><td>BT20/MB468</td><td>MP</td></tr>
<tr><td>>hg38_21_+_8433172_8446622_RNA45SN1_45S_50ntFlanks@11888.11905.18|AGCTTGACTCTAGTCTGG</td><td>>hg38_21_+_8433172_8446622_RNA45SN1_45S_50ntFlanks</td><td>11888</td><td>11905</td><td>18</td><td>AGCTTGACTCTAGTCTGG</td><td>153.6</td><td>-10.3</td><td>1.6</td><td>-6.5</td><td>7.6E-11</td><td>1.8E-09</td><td>BT20/MB468</td><td>MP</td></tr>
<tr><td>>hg38_21_+_8433172_8446622_RNA45SN1_45S_50ntFlanks@5522.5550.29|AAGTAAAAGTCGTAACAAGGTTTCCGTAG</td><td>>hg38_21_+_8433172_8446622_RNA45SN1_45S_50ntFlanks</td><td>5522</td><td>5550</td><td>29</td><td>AAGTAAAAGTCGTAACAAGGTTTCCGTAG</td><td>88.0</td><td>-10.3</td><td>1.7</td><td>-6.2</td><td>7.5E-10</td><td>1.8E-09</td><td>MB231/MB468</td><td>MP</td></tr>
<tr><td>>hg38_21_+_8433172_8446622_RNA45SN1_45S_50ntFlanks@11843.11872.30|AGGGAACGGGCTTGGCGGAATCAGCGGGGA</td><td>>hg38_21_+_8433172_8446622_RNA45SN1_45S_50ntFlanks</td><td>11843</td><td>11872</td><td>30</td><td>AGGGAACGGGCTTGGCGGAATCAGCGGGGA</td><td>581.8</td><td>-8.8</td><td>1.4</td><td>-6.1</td><td>9.1E-10</td><td>1.8E-09</td><td>BT20/MB468</td><td>NUC</td></tr>
<tr><td>>MI0000542|hsa-mir-320a&WithFlank&8|-|22244960|22245043@48.69.22(+1C)&[MIMAT0000510&hsa-miR-320a-3p&offsets|0|0(+1C);m-60&8|-|22244975|22244996&offsets|0|0(+1C)]|AAAAGCTGGGTTGAGAGGGCGAC</td><td>>MI0000542|hsa-mir-320a&WithFlank&8|-|22244960|22245043</td><td>48</td><td>69</td><td>22(+1C)</td><td>AAAAGCTGGGTTGAGAGGGCGAC</td><td>322.5</td><td>5.5</td><td>0.9</td><td>6.1</td><td>7.9E-10</td><td>1.8E-09</td><td>MB231/MB468</td><td>NUC</td></tr>
<tr><td>>hg38_21_+_8433172_8446622_RNA45SN1_45S_50ntFlanks@12417.12455.39|TCGATGTCGGCTCTTCCTATCATTGTGAAGCAGAATTCA</td><td>>hg38_21_+_8433172_8446622_RNA45SN1_45S_50ntFlanks</td><td>12417</td><td>12455</td><td>39</td><td>TCGATGTCGGCTCTTCCTATCATTGTGAAGCAGAATTCA</td><td>3809.0</td><td>-10.8</td><td>1.8</td><td>-6.1</td><td>9.1E-10</td><td>1.8E-09</td><td>BT20/MB468</td><td>NUC</td></tr>
<tr><td>>trna134_GluTTC_1_-_16861774_16861845@1.31.31,trna5_GluTTC_1_+_17199078_17199149@1.31.31,trna84_GluTTC_1_-_161391883_161391954@1.31.31,trna94_GluTTC_1_-_149664355_149664427@1.31.31|TCCCTGGTGGTCTAGTGGCTAGGATTCGGCG</td><td>>trna134_GluTTC_1_-_16861774_16861845</td><td>1</td><td>31</td><td>31</td><td>TCCCTGGTGGTCTAGTGGCTAGGATTCGGCG</td><td>544.2</td><td>-5.6</td><td>0.9</td><td>-6.1</td><td>8.7E-10</td><td>1.8E-09</td><td>MB231/MB468</td><td>MITO</td></tr>
<tr><td>>hg38_21_+_8433172_8446622_RNA45SN1_45S_50ntFlanks@9852.9878.27|GCGCTGCGGGATGAACCGAACGCCGGG</td><td>>hg38_21_+_8433172_8446622_RNA45SN1_45S_50ntFlanks</td><td>9852</td><td>9878</td><td>27</td><td>GCGCTGCGGGATGAACCGAACGCCGGG</td><td>129.5</td><td>-10.9</td><td>1.8</td><td>-6.2</td><td>7.5E-10</td><td>1.8E-09</td><td>MB231/MB468</td><td>MP</td></tr>
</table>

<table>
<tr><td>>hg38_21_+_8433172_8446622_RNA45SN1_45S_50ntFlanks@8192.8214.23|AGCCCGTGGACGGTGTGAGGCCG</td><td>>hg38_21_+_8433172_8446622_RNA45SN1_45S_50ntFlanks</td><td>8192</td><td>8214</td><td>23</td><td>AGCCCGTGGACGGTGTGAGGCCG</td><td>628.6</td><td>-9.1</td><td>1.5</td><td>-6.1</td><td>9.1E-10</td><td>1.8E-09</td><td>BT20/MB468</td><td>NUC</td></tr>
<tr><td>>hg38_21_+_8433172_8446622_RNA45SN1_45S_50ntFlanks@10797.10818.22|GCATGTTGGAACAATGTAGGTA</td><td>>hg38_21_+_8433172_8446622_RNA45SN1_45S_50ntFlanks</td><td>10797</td><td>10818</td><td>22</td><td>GCATGTTGGAACAATGTAGGTA</td><td>1369.3</td><td>-7.8</td><td>1.3</td><td>-6.1</td><td>9.1E-10</td><td>1.8E-09</td><td>BT20/MB468</td><td>NUC</td></tr>
<tr><td>>hg38_21_+_8433172_8446622_RNA45SN1_45S_50ntFlanks@10375.10406.32|AGAAGGGTTCCATGTGAACAGCAGTTGAACAT</td><td>>hg38_21_+_8433172_8446622_RNA45SN1_45S_50ntFlanks</td><td>10375</td><td>10406</td><td>32</td><td>AGAAGGGTTCCATGTGAACAGCAGTTGAACAT</td><td>299.9</td><td>-5.4</td><td>0.9</td><td>-6.1</td><td>7.8E-10</td><td>1.8E-09</td><td>MB231/MB468</td><td>CYTO</td></tr>
<tr><td>>hg38_21_+_8433172_8446622_RNA45SN1_45S_50ntFlanks@10786.10820.35|AACAGCCTCTGGCATGTTGGAACAATGTAGGTAAG</td><td>>hg38_21_+_8433172_8446622_RNA45SN1_45S_50ntFlanks</td><td>10786</td><td>10820</td><td>35</td><td>AACAGCCTCTGGCATGTTGGAACAATGTAGGTAAG</td><td>482.9</td><td>-6.4</td><td>1.0</td><td>-6.2</td><td>7.6E-10</td><td>1.8E-09</td><td>MB231/MB468</td><td>MP</td></tr>
<tr><td>>hg38_21_+_8433172_8446622_RNA45SN1_45S_50ntFlanks@12605.12626.22|TATGTGCTTGGCTGAGGAGCCA</td><td>>hg38_21_+_8433172_8446622_RNA45SN1_45S_50ntFlanks</td><td>12605</td><td>12626</td><td>22</td><td>TATGTGCTTGGCTGAGGAGCCA</td><td>860.0</td><td>-4.8</td><td>0.8</td><td>-6.2</td><td>5.9E-10</td><td>1.8E-09</td><td>BT20/MB468</td><td>MITO</td></tr>
<tr><td>>hg38_21_+_8433172_8446622_RNA45SN1_45S_50ntFlanks@10784.10818.35|TGAACAGCCTCTGGCATGTTGGAACAATGTAGGTA</td><td>>hg38_21_+_8433172_8446622_RNA45SN1_45S_50ntFlanks</td><td>10784</td><td>10818</td><td>35</td><td>TGAACAGCCTCTGGCATGTTGGAACAATGTAGGTA</td><td>557.4</td><td>-4.0</td><td>0.7</td><td>-6.1</td><td>7.8E-10</td><td>1.8E-09</td><td>MB231/MB468</td><td>CYTO</td></tr>
<tr><td>>am_trna128_GlyGCC_6_-_27870686_27870756@1.32.32,trna133_GlyCCC_1_-_16872434_16872504@1.32.32,trna18_GlyGCC_16_+_70822597_70822667@1.32.32,trna19_GlyGCC_16_+_70823410_70823480@1.32.32,trna19_GlyGCC_2_-_157257659_157257729@1.32.32,trna24_GlyGCC_16_-_70812942_70813012@1.32.32,trna25_GlyGCC_16_-_70812114_70812184@1.32.32,trna4_GlyCCC_1_+_17188416_17188486@1.32.32,trna5_GlyGCC_17_+_8029064_8029134@1.32.32,trna68_GlyGCC_1_-_161493637_161493707@1.32.32|GCATTGGTGGTTCAGTGGTAGAATTCTCGCCT</td><td>>am_trna128_GlyGCC_6_-_27870686_27870756</td><td>1</td><td>32</td><td>32</td><td>GCATTGGTGGTTCAGTGGTAGAATTCTCGCCT</td><td>13108.6</td><td>6.3</td><td>1.0</td><td>6.2</td><td>7.6E-10</td><td>1.8E-09</td><td>MB231/MB468</td><td>MP</td></tr>
<tr><td>>hg38_21_+_8433172_8446622_RNA45SN1_45S_50ntFlanks@12686.12708.23|CCAGGCGGAACGATACGGCAGCG</td><td>>hg38_21_+_8433172_8446622_RNA45SN1_45S_50ntFlanks</td><td>12686</td><td>12708</td><td>23</td><td>CCAGGCGGAACGATACGGCAGCG</td><td>123.6</td><td>-9.0</td><td>1.5</td><td>-6.2</td><td>5.9E-10</td><td>1.8E-09</td><td>BT20/MB468</td><td>MITO</td></tr>
<tr><td>>hg38_21_+_8433172_8446622_RNA45SN1_45S_50ntFlanks@11641.11678.38|CGGCGGGTGTTGACGCGATGTGATTTCTGCCCAGTGCT</td><td>>hg38_21_+_8433172_8446622_RNA45SN1_45S_50ntFlanks</td><td>11641</td><td>11678</td><td>38</td><td>CGGCGGGTGTTGACGCGATGTGATTTCTGCCCAGTGCT</td><td>208.2</td><td>-10.8</td><td>1.7</td><td>-6.5</td><td>7.7E-11</td><td>1.8E-09</td><td>BT20/MB468</td><td>MP</td></tr>
<tr><td>>hg38_21_+_8433172_8446622_RNA45SN1_45S_50ntFlanks@9034.9059.26|TCGTGGGGGGGCCGGGCCACCCCTCC</td><td>>hg38_21_+_8433172_8446622_RNA45SN1_45S_50ntFlanks</td><td>9034</td><td>9059</td><td>26</td><td>TCGTGGGGGGGCCGGGCCACCCCTCC</td><td>240.6</td><td>-11.0</td><td>1.7</td><td>-6.5</td><td>7.7E-11</td><td>1.8E-09</td><td>BT20/MB468</td><td>MP</td></tr>
<tr><td>>trna2_GlyGCC_21_-_18827107_18827177@1.31.31,trna35_GlyGCC_1_+_161413094_161413164@1.31.31,trna37_GlyGCC_1_+_161420467_161420537@1.31.31,trna39_GlyGCC_1_+_161427898_161427968@1.31.31,trna41_GlyGCC_1_+_161435258_161435328@1.31.31|GCATGGGTGGTTCAGTGGTAGAATTCTCGCC</td><td>>trna2_GlyGCC_21_-_18827107_18827177</td><td>1</td><td>31</td><td>31</td><td>GCATGGGTGGTTCAGTGGTAGAATTCTCGCC</td><td>29096.1</td><td>6.5</td><td>1.0</td><td>6.5</td><td>6.7E-11</td><td>1.8E-09</td><td>BT20/MB231</td><td>MITO</td></tr>
<tr><td>>hg38_21_+_8433172_8446622_RNA45SN1_45S_50ntFlanks@10291.10328.38|CGCGGGCCCGGGTGGAGCCGCCGCAGGTGCAGATCTTG</td><td>>hg38_21_+_8433172_8446622_RNA45SN1_45S_50ntFlanks</td><td>10291</td><td>10328</td><td>38</td><td>CGCGGGCCCGGGTGGAGCCGCCGCAGGTGCAGATCTTG</td><td>65.2</td><td>-9.9</td><td>1.6</td><td>-6.2</td><td>7.6E-10</td><td>1.8E-09</td><td>MB231/MB468</td><td>MP</td></tr>
<tr><td>>hg38_21_+_8433172_8446622_RNA45SN1_45S_50ntFlanks@12631.12654.24|GGCGAAGCTACCATCTGTGGGATT</td><td>>hg38_21_+_8433172_8446622_RNA45SN1_45S_50ntFlanks</td><td>12631</td><td>12654</td><td>24</td><td>GGCGAAGCTACCATCTGTGGGATT</td><td>535.1</td><td>-8.7</td><td>1.4</td><td>-6.1</td><td>9.2E-10</td><td>1.8E-09</td><td>BT20/MB468</td><td>NUC</td></tr>
<tr><td>>hg38_21_+_8433172_8446622_RNA45SN1_45S_50ntFlanks@7991.8008.18|ACGTGGCGACCCGCTGAA</td><td>>hg38_21_+_8433172_8446622_RNA45SN1_45S_50ntFlanks</td><td>7991</td><td>8008</td><td>18</td><td>ACGTGGCGACCCGCTGAA</td><td>32070.4</td><td>-6.8</td><td>1.0</td><td>-6.5</td><td>7.8E-11</td><td>1.8E-09</td><td>BT20/MB468</td><td>MP</td></tr>
<tr><td>>MI0000089|hsa-mir-31&WithFlank&9|-|21512109|21512191@14.35.22(+1A)&[MIMAT0000089&hsa-miR-31-5p&offsets|0|+1(+1A);m-360&9|-|21512157|21512178&offsets|0|0(+1A)]|AGGCAAGATGCTGGCATAGCTGA</td><td>>MI0000089|hsa-mir-31&WithFlank&9|-|21512109|21512191</td><td>14</td><td>35</td><td>22(+1A)</td><td>AGGCAAGATGCTGGCATAGCTGA</td><td>81.4</td><td>-10.2</td><td>1.7</td><td>-6.2</td><td>7.6E-10</td><td>1.8E-09</td><td>MB231/MB468</td><td>MP</td></tr>
<tr><td>>hg38_21_+_8433172_8446622_RNA45SN1_45S_50ntFlanks@4729.4748.20|TCAAGAACGAAAGTCGGAGG</td><td>>hg38_21_+_8433172_8446622_RNA45SN1_45S_50ntFlanks</td><td>4729</td><td>4748</td><td>20</td><td>TCAAGAACGAAAGTCGGAGG</td><td>352.8</td><td>-4.3</td><td>0.7</td><td>-6.2</td><td>7.7E-10</td><td>1.8E-09</td><td>MB231/MB468</td><td>MP</td></tr>
<tr><td>>hg38_21_+_8433172_8446622_RNA45SN1_45S_50ntFlanks@8352.8391.40|AGTTGAAAAGAACTTTGAAGAGAGAGTTCAAGAGGGCGTG</td><td>>hg38_21_+_8433172_8446622_RNA45SN1_45S_50ntFlanks</td><td>8352</td><td>8391</td><td>40</td><td>AGTTGAAAAGAACTTTGAAGAGAGAGTTCAAGAGGGCGTG</td><td>129.6</td><td>10.8</td><td>1.7</td><td>6.5</td><td>7.8E-11</td><td>1.8E-09</td><td>BT20/MB468</td><td>MP</td></tr>
<tr><td>>hg38_21_+_8433172_8446622_RNA45SN1_45S_50ntFlanks@8179.8197.19|CTGATCGAGGCCCAGCCCG</td><td>>hg38_21_+_8433172_8446622_RNA45SN1_45S_50ntFlanks</td><td>8179</td><td>8197</td><td>19</td><td>CTGATCGAGGCCCAGCCCG</td><td>140.7</td><td>-10.2</td><td>1.6</td><td>-6.5</td><td>7.8E-11</td><td>1.8E-09</td><td>BT20/MB468</td><td>MP</td></tr>
<tr><td>>hg38_21_+_8433172_8446622_RNA45SN1_45S_50ntFlanks@3882.3915.34|CCGACGGGCGCTGACCCCCTTCGCGGGGGGGATG</td><td>>hg38_21_+_8433172_8446622_RNA45SN1_45S_50ntFlanks</td><td>3882</td><td>3915</td><td>34</td><td>CCGACGGGCGCTGACCCCCTTCGCGGGGGGGATG</td><td>451.9</td><td>-8.6</td><td>1.4</td><td>-6.1</td><td>9.2E-10</td><td>1.8E-09</td><td>BT20/MB468</td><td>NUC</td></tr>
<tr><td>>MI0000342|hsa-mir-200b&WithFlank&1|+|1167098|1167204@63.85.23(+1U)&[MIMAT0000318&hsa-miR-200b-3p&offsets|0|+1(+1U);m-93&1|+|1167160|1167182&offsets|0|0(+1U)]|TAATACTGCCTGGTAATGATGACT</td><td>>MI0000342|hsa-mir-200b&WithFlank&1|+|1167098|1167204</td><td>63</td><td>85</td><td>23(+1U)</td><td>TAATACTGCCTGGTAATGATGACT</td><td>971.3</td><td>-4.6</td><td>0.8</td><td>-6.2</td><td>7.7E-10</td><td>1.8E-09</td><td>MB231/MB468</td><td>MP</td></tr>
<tr><td>>hg38_21_+_8433172_8446622_RNA45SN1_45S_50ntFlanks@3899.3938.40|CCTTCGCGGGGGGGATGCGTGCATTTATCAGATCAAAACC</td><td>>hg38_21_+_8433172_8446622_RNA45SN1_45S_50ntFlanks</td><td>3899</td><td>3938</td><td>40</td><td>CCTTCGCGGGGGGGATGCGTGCATTTATCAGATCAAAACC</td><td>607.8</td><td>-11.3</td><td>1.8</td><td>-6.2</td><td>6.0E-10</td><td>1.8E-09</td><td>BT20/MB468</td><td>MITO</td></tr>
<tr><td>>hg38_21_+_8433172_8446622_RNA45SN1_45S_50ntFlanks@5473.5499.27|CGGCCCTGGCGGAGCGCTGAGAAGACG</td><td>>hg38_21_+_8433172_8446622_RNA45SN1_45S_50ntFlanks</td><td>5473</td><td>5499</td><td>27</td><td>CGGCCCTGGCGGAGCGCTGAGAAGACG</td><td>3448.1</td><td>-4.8</td><td>0.8</td><td>-6.2</td><td>7.7E-10</td><td>1.8E-09</td><td>MB231/MB468</td><td>MP</td></tr>
<tr><td>>MI0000266|hsa-mir-10a&WithFlank&17|-|48579832|48579953@28.51.24&[MIMAT0000253&hsa-miR-10a-5p&offsets|0|+1;m-18&17|-|48579905|48579926&offsets|0|+2]|TACCCTGTAGATCCGAATTTGTGT</td><td>>MI0000266|hsa-mir-10a&WithFlank&17|-|48579832|48579953</td><td>28</td><td>51</td><td>24</td><td>TACCCTGTAGATCCGAATTTGTGT</td><td>200.6</td><td>11.4</td><td>1.8</td><td>6.5</td><td>7.8E-11</td><td>1.9E-09</td><td>BT20/MB468</td><td>MP</td></tr>
<tr><td>>hg38_21_+_8433172_8446622_RNA45SN1_45S_50ntFlanks@12633.12659.27|CGAAGCTACCATCTGTGGGATTATGAC</td><td>>hg38_21_+_8433172_8446622_RNA45SN1_45S_50ntFlanks</td><td>12633</td><td>12659</td><td>27</td><td>CGAAGCTACCATCTGTGGGATTATGAC</td><td>2017.4</td><td>-9.8</td><td>1.6</td><td>-6.1</td><td>9.3E-10</td><td>1.9E-09</td><td>BT20/MB468</td><td>NUC</td></tr>
<tr><td>>hg38_21_+_8433172_8446622_RNA45SN1_45S_50ntFlanks@4724.4761.38|ATTAATCAAGAACGAAAGTCGGAGGTTCGAAGACGATC</td><td>>hg38_21_+_8433172_8446622_RNA45SN1_45S_50ntFlanks</td><td>4724</td><td>4761</td><td>38</td><td>ATTAATCAAGAACGAAAGTCGGAGGTTCGAAGACGATC</td><td>495.8</td><td>-4.8</td><td>0.8</td><td>-6.1</td><td>8.9E-10</td><td>1.9E-09</td><td>MB231/MB468</td><td>MITO</td></tr>
<tr><td>>hg38_21_+_8433172_8446622_RNA45SN1_45S_50ntFlanks@12072.12090.19|CCCGAGGGGCTCTCGCTTC</td><td>>hg38_21_+_8433172_8446622_RNA45SN1_45S_50ntFlanks</td><td>12072</td><td>12090</td><td>19</td><td>CCCGAGGGGCTCTCGCTTC</td><td>262.8</td><td>-7.8</td><td>1.3</td><td>-6.1</td><td>8.9E-10</td><td>1.9E-09</td><td>MB231/MB468</td><td>MITO</td></tr>
<tr><td>>hg38_21_+_8433172_8446622_RNA45SN1_45S_50ntFlanks@4045.4063.19|CCATTCGAACGTCTGCCCT</td><td>>hg38_21_+_8433172_8446622_RNA45SN1_45S_50ntFlanks</td><td>4045</td><td>4063</td><td>19</td><td>CCATTCGAACGTCTGCCCT</td><td>1090.8</td><td>-6.7</td><td>1.1</td><td>-6.2</td><td>6.1E-10</td><td>1.9E-09</td><td>BT20/MB468</td><td>MITO</td></tr>
<tr><td>>hg38_21_+_8433172_8446622_RNA45SN1_45S_50ntFlanks@10130.10153.24|GCGGCGGCGGCGGCGGCGGGGGTG</td><td>>hg38_21_+_8433172_8446622_RNA45SN1_45S_50ntFlanks</td><td>10130</td><td>10153</td><td>24</td><td>GCGGCGGCGGCGGCGGCGGGGGTG</td><td>89.6</td><td>-10.3</td><td>1.7</td><td>-6.1</td><td>7.8E-10</td><td>1.9E-09</td><td>MB231/MB468</td><td>MP</td></tr>
<tr><td>>hg38_21_+_8433172_8446622_RNA45SN1_45S_50ntFlanks@10408.10432.25|GGTCAGTCGGTCCTGAGAGATGGGC</td><td>>hg38_21_+_8433172_8446622_RNA45SN1_45S_50ntFlanks</td><td>10408</td><td>10432</td><td>25</td><td>GGTCAGTCGGTCCTGAGAGATGGGC</td><td>726.3</td><td>-5.6</td><td>0.9</td><td>-6.1</td><td>8.0E-10</td><td>1.9E-09</td><td>MB231/MB468</td><td>CYTO</td></tr>
<tr><td>>hg38_21_+_8433172_8446622_RNA45SN1_45S_50ntFlanks@10234.10274.41|CGGACGCTACGCCGCGACGAGTAGGAGGGCCGCTGCGGTGA</td><td>>hg38_21_+_8433172_8446622_RNA45SN1_45S_50ntFlanks</td><td>10234</td><td>10274</td><td>41</td><td>CGGACGCTACGCCGCGACGAGTAGGAGGGCCGCTGCGGTGA</td><td>98.1</td><td>-10.5</td><td>1.7</td><td>-6.1</td><td>7.8E-10</td><td>1.9E-09</td><td>MB231/MB468</td><td>MP</td></tr>
<tr><td>>hg38_21_+_8433172_8446622_RNA45SN1_45S_50ntFlanks@11586.11612.27|GTGCGGACCAGGGGAATCCGACTGTTT</td><td>>hg38_21_+_8433172_8446622_RNA45SN1_45S_50ntFlanks</td><td>11586</td><td>11612</td><td>27</td><td>GTGCGGACCAGGGGAATCCGACTGTTT</td><td>1825.8</td><td>-7.1</td><td>1.2</td><td>-6.1</td><td>7.8E-10</td><td>1.9E-09</td><td>MB231/MB468</td><td>MP</td></tr>
<tr><td>>trnaMT_HisGTG_MT_+_12138_12206@19.55.37|AAACATCAGATTGTGAATCTGACAACAGAGGCTTACG</td><td>>trnaMT_HisGTG_MT_+_12138_12206</td><td>19</td><td>55</td><td>37</td><td>AAACATCAGATTGTGAATCTGACAACAGAGGCTTACG</td><td>355.9</td><td>12.2</td><td>1.9</td><td>6.5</td><td>7.9E-11</td><td>1.9E-09</td><td>BT20/MB468</td><td>MP</td></tr>
<tr><td>>hg38_21_+_8433172_8446622_RNA45SN1_45S_50ntFlanks@4047.4066.20|ATTCGAACGTCTGCCCTATC</td><td>>hg38_21_+_8433172_8446622_RNA45SN1_45S_50ntFlanks</td><td>4047</td><td>4066</td><td>20</td><td>ATTCGAACGTCTGCCCTATC</td><td>3820.4</td><td>-5.7</td><td>0.9</td><td>-6.2</td><td>6.1E-10</td><td>1.9E-09</td><td>BT20/MB468</td><td>MITO</td></tr>
<tr><td>>hg38_21_+_8433172_8446622_RNA45SN1_45S_50ntFlanks@10680.10712.33|GTTCCGGCGGCGTCCGGTGAGCTCTCGCTGGCC</td><td>>hg38_21_+_8433172_8446622_RNA45SN1_45S_50ntFlanks</td><td>10680</td><td>10712</td><td>33</td><td>GTTCCGGCGGCGTCCGGTGAGCTCTCGCTGGCC</td><td>14351.8</td><td>-9.2</td><td>1.5</td><td>-6.1</td><td>9.4E-10</td><td>1.9E-09</td><td>BT20/MB468</td><td>NUC</td></tr>
<tr><td>>hg38_21_+_8433172_8446622_RNA45SN1_45S_50ntFlanks@11704.11740.37|ATGAAGCGCGGGTAAACGGCGGGAGTAACTATGACTC</td><td>>hg38_21_+_8433172_8446622_RNA45SN1_45S_50ntFlanks</td><td>11704</td><td>11740</td><td>37</td><td>ATGAAGCGCGGGTAAACGGCGGGAGTAACTATGACTC</td><td>355.5</td><td>-4.4</td><td>0.7</td><td>-6.1</td><td>8.2E-10</td><td>1.9E-09</td><td>MB231/MB468</td><td>NUC</td></tr>
<tr><td>>MI0000760|hsa-mir-361&WithFlank&X|-|85903630|85903713@52.75.24&[MIMAT0004682&hsa-miR-361-3p&offsets|+1|+2;m-124&X|-|85903642|85903663&offsets|+1|+3]|CCCCCAGGTGTGATTCTGATTTGC</td><td>>MI0000760|hsa-mir-361&WithFlank&X|-|85903630|85903713</td><td>52</td><td>75</td><td>24</td><td>CCCCCAGGTGTGATTCTGATTTGC</td><td>491.5</td><td>4.4</td><td>0.7</td><td>6.1</td><td>8.2E-10</td><td>1.9E-09</td><td>MB231/MB468</td><td>NUC</td></tr>
<tr><td>>hg38_21_+_8433172_8446622_RNA45SN1_45S_50ntFlanks@5472.5500.29|ACGGCCCTGGCGGAGCGCTGAGAAGACGG</td><td>>hg38_21_+_8433172_8446622_RNA45SN1_45S_50ntFlanks</td><td>5472</td><td>5500</td><td>29</td><td>ACGGCCCTGGCGGAGCGCTGAGAAGACGG</td><td>414.3</td><td>3.9</td><td>0.6</td><td>6.1</td><td>8.3E-10</td><td>1.9E-09</td><td>MB231/MB468</td><td>NUC</td></tr>
<tr><td>>MI0002470|hsa-mir-486-1&WithFlank&8|-|41660435|41660514@10.31.22;MI0023622|hsa-mir-486-2&WithFlank&8|+|41660438|41660513@7.28.22&[MIMAT0002177&hsa-miR-486-5p&offsets|0|0;m-107&8|-|41660484|41660505&offsets|0|0];[MIMAT0002177_1&hsa-miR-486-5p&offsets|0|0;TJU_CMC_MD2.ID03138.5p-miR&offsets|0|-2;m-106&8|+|41660444|41660465&offsets|0|0]|TCCTGTACTGAGCTGCCCCGAG</td><td>>MI0002470|hsa-mir-486-1&WithFlank&8|-|41660435|41660514</td><td>10</td><td>31</td><td>22</td><td>TCCTGTACTGAGCTGCCCCGAG</td><td>286.0</td><td>-11.6</td><td>1.7</td><td>-6.7</td><td>1.5E-11</td><td>1.9E-09</td><td>BT20/MB231</td><td>MP</td></tr>
<tr><td>>hg38_21_+_8433172_8446622_RNA45SN1_45S_50ntFlanks@12327.12346.20|GGTTTTAAGCAGGAGGTGTC</td><td>>hg38_21_+_8433172_8446622_RNA45SN1_45S_50ntFlanks</td><td>12327</td><td>12346</td><td>20</td><td>GGTTTTAAGCAGGAGGTGTC</td><td>162.7</td><td>-4.6</td><td>0.8</td><td>-6.1</td><td>8.1E-10</td><td>1.9E-09</td><td>MB231/MB468</td><td>CYTO</td></tr>
<tr><td>>hg38_21_+_8433172_8446622_RNA45SN1_45S_50ntFlanks@12405.12438.34|CTTTTTGATCCTTCGATGTCGGCTCTTCCTATCA</td><td>>hg38_21_+_8433172_8446622_RNA45SN1_45S_50ntFlanks</td><td>12405</td><td>12438</td><td>34</td><td>CTTTTTGATCCTTCGATGTCGGCTCTTCCTATCA</td><td>101.0</td><td>-10.5</td><td>1.7</td><td>-6.1</td><td>7.9E-10</td><td>1.9E-09</td><td>MB231/MB468</td><td>MP</td></tr>
<tr><td>>hg38_21_+_8433172_8446622_RNA45SN1_45S_50ntFlanks@10384.10430.47|CCATGTGAACAGCAGTTGAACATGGGTCAGTCGGTCCTGAGAGATGG</td><td>>hg38_21_+_8433172_8446622_RNA45SN1_45S_50ntFlanks</td><td>10384</td><td>10430</td><td>47</td><td>CCATGTGAACAGCAGTTGAACATGGGTCAGTCGGTCCTGAGAGATGG</td><td>410.7</td><td>-11.6</td><td>1.9</td><td>-6.1</td><td>8.3E-10</td><td>1.9E-09</td><td>MB231/MB468</td><td>NUC</td></tr>
<tr><td>>hg38_21_+_8433172_8446622_RNA45SN1_45S_50ntFlanks@12249.12269.21|CAAAAGCTCGCTTGATCTTGA</td><td>>hg38_21_+_8433172_8446622_RNA45SN1_45S_50ntFlanks</td><td>12249</td><td>12269</td><td>21</td><td>CAAAAGCTCGCTTGATCTTGA</td><td>401.3</td><td>-10.7</td><td>1.7</td><td>-6.2</td><td>6.2E-10</td><td>1.9E-09</td><td>BT20/MB468</td><td>MITO</td></tr>
<tr><td>>hg38_21_+_8433172_8446622_RNA45SN1_45S_50ntFlanks@8179.8210.32|CTGATCGAGGCCCAGCCCGTGGACGGTGTGAG</td><td>>hg38_21_+_8433172_8446622_RNA45SN1_45S_50ntFlanks</td><td>8179</td><td>8210</td><td>32</td><td>CTGATCGAGGCCCAGCCCGTGGACGGTGTGAG</td><td>343.1</td><td>-4.7</td><td>0.8</td><td>-6.1</td><td>8.3E-10</td><td>1.9E-09</td><td>MB231/MB468</td><td>NUC</td></tr>
<tr><td>>hg38_1_-_228634819_228635039_RNA5S12_5S_50ntFlanks@126.170.45|TACTTGGATGGGAGACCGCCTGGGAATACCGGGTGCTGTAGGCTT</td><td>>hg38_1_-_228634819_228635039_RNA5S12_5S_50ntFlanks</td><td>126</td><td>170</td><td>45</td><td>TACTTGGATGGGAGACCGCCTGGGAATACCGGGTGCTGTAGGCTT</td><td>1250.7</td><td>7.7</td><td>1.2</td><td>6.2</td><td>6.2E-10</td><td>1.9E-09</td><td>BT20/MB468</td><td>MITO</td></tr>
<tr><td>>hg38_21_+_8433172_8446622_RNA45SN1_45S_50ntFlanks@10914.10939.26|CGCCGCGGCTGGACGAGGCGCCGCCG</td><td>>hg38_21_+_8433172_8446622_RNA45SN1_45S_50ntFlanks</td><td>10914</td><td>10939</td><td>26</td><td>CGCCGCGGCTGGACGAGGCGCCGCCG</td><td>105.9</td><td>-8.8</td><td>1.4</td><td>-6.2</td><td>6.2E-10</td><td>1.9E-09</td><td>BT20/MB468</td><td>MITO</td></tr>
<tr><td>>hg38_21_+_8433172_8446622_RNA45SN1_45S_50ntFlanks@8153.8178.26|GCCGCTCGTGGGGGGCCCAAGTCCTT</td><td>>hg38_21_+_8433172_8446622_RNA45SN1_45S_50ntFlanks</td><td>8153</td><td>8178</td><td>26</td><td>GCCGCTCGTGGGGGGCCCAAGTCCTT</td><td>244.2</td><td>-10.0</td><td>1.6</td><td>-6.2</td><td>6.2E-10</td><td>1.9E-09</td><td>BT20/MB468</td><td>MITO</td></tr>
<tr><td>>hg38_21_+_8433172_8446622_RNA45SN1_45S_50ntFlanks@4007.4024.18|CCTCGGGCCGATCGCACG</td><td>>hg38_21_+_8433172_8446622_RNA45SN1_45S_50ntFlanks</td><td>4007</td><td>4024</td><td>18</td><td>CCTCGGGCCGATCGCACG</td><td>957.7</td><td>-6.8</td><td>1.1</td><td>-6.1</td><td>7.9E-10</td><td>1.9E-09</td><td>MB231/MB468</td><td>MP</td></tr>
</table>

<table>
<tr><td>>hg38_21_+_8433172_8446622_RNA45SN1_45S_50ntFlanks@6653.6700.48|ACTCTTAGCGGTGGATCACTCGGCTCGTGCGTCGATGAAGAACGCAGC</td><td>>hg38_21_+_8433172_8446622_RNA45SN1_45S_50ntFlanks</td><td>6653</td><td>6700</td><td>48</td><td>ACTCTTAGCGGTGGATCACTCGGCTCGTGCGTCGATGAAGAACGCAGC</td><td>299.2</td><td>12.0</td><td>1.8</td><td>6.5</td><td>8.1E-11</td><td>1.9E-09</td><td>BT20/MB468</td><td>MP</td></tr>
<tr><td>>hg38_21_+_8433172_8446622_RNA45SN1_45S_50ntFlanks@12615.12657.43|GCTGAGGAGCCAATGGGGCGAAGCTACCATCTGTGGGATTATG</td><td>>hg38_21_+_8433172_8446622_RNA45SN1_45S_50ntFlanks</td><td>12615</td><td>12657</td><td>43</td><td>GCTGAGGAGCCAATGGGGCGAAGCTACCATCTGTGGGATTATG</td><td>132.4</td><td>-10.9</td><td>1.8</td><td>-6.1</td><td>8.0E-10</td><td>1.9E-09</td><td>MB231/MB468</td><td>MP</td></tr>
<tr><td>>MI0000102|hsa-mir-100&WithFlank&11|-|122152223|122152314@19.38.20(+1C)&[MIMAT0000098&hsa-miR-100-5p&offsets|0|-2(+1C);m-27&11|-|122152276|122152296&offsets|0|-1(+1C)]|AACCCGTAGATCCGAACTTGC</td><td>>MI0000102|hsa-mir-100&WithFlank&11|-|122152223|122152314</td><td>19</td><td>38</td><td>20(+1C)</td><td>AACCCGTAGATCCGAACTTGC</td><td>47855.1</td><td>-7.7</td><td>1.1</td><td>-6.7</td><td>1.6E-11</td><td>1.9E-09</td><td>BT20/MB231</td><td>MP</td></tr>
<tr><td>>MI0000476|hsa-mir-138-1&WithFlank&3|+|44114206|44114316@29.53.25&[MIMAT0000430_1&hsa-miR-138-5p&offsets|0|+2;m-497&3|+|44114234|44114256&offsets|0|+2]|AGCTGGTGTTGTGAATCAGGCCGTT</td><td>>MI0000476|hsa-mir-138-1&WithFlank&3|+|44114206|44114316</td><td>29</td><td>53</td><td>25</td><td>AGCTGGTGTTGTGAATCAGGCCGTT</td><td>318.9</td><td>-11.7</td><td>1.7</td><td>-6.7</td><td>1.6E-11</td><td>1.9E-09</td><td>BT20/MB231</td><td>MP</td></tr>
<tr><td>>hg38_21_+_8433172_8446622_RNA45SN1_45S_50ntFlanks@10771.10806.36|CAGGTCTCCAAGGTGAACAGCCTCTGGCATGTTGGA</td><td>>hg38_21_+_8433172_8446622_RNA45SN1_45S_50ntFlanks</td><td>10771</td><td>10806</td><td>36</td><td>CAGGTCTCCAAGGTGAACAGCCTCTGGCATGTTGGA</td><td>978.7</td><td>-7.0</td><td>1.1</td><td>-6.1</td><td>8.4E-10</td><td>1.9E-09</td><td>MB231/MB468</td><td>NUC</td></tr>
<tr><td>>hg38_21_+_8433172_8446622_RNA45SN1_45S_50ntFlanks@12419.12438.20|GATGTCGGCTCTTCCTATCA</td><td>>hg38_21_+_8433172_8446622_RNA45SN1_45S_50ntFlanks</td><td>12419</td><td>12438</td><td>20</td><td>GATGTCGGCTCTTCCTATCA</td><td>1880.6</td><td>4.2</td><td>0.7</td><td>6.1</td><td>8.4E-10</td><td>1.9E-09</td><td>MB231/MB468</td><td>NUC</td></tr>
<tr><td>>hg38_21_+_8433172_8446622_RNA45SN1_45S_50ntFlanks@6656.6680.25|CTTAGCGGTGGATCACTCGGCTCGT</td><td>>hg38_21_+_8433172_8446622_RNA45SN1_45S_50ntFlanks</td><td>6656</td><td>6680</td><td>25</td><td>CTTAGCGGTGGATCACTCGGCTCGT</td><td>131.1</td><td>-5.5</td><td>0.9</td><td>-6.1</td><td>8.2E-10</td><td>1.9E-09</td><td>MB231/MB468</td><td>CYTO</td></tr>
<tr><td>>hg38_21_+_8433172_8446622_RNA45SN1_45S_50ntFlanks@11702.11728.27|CAATGAAGCGCGGGTAAACGGCGGGAG</td><td>>hg38_21_+_8433172_8446622_RNA45SN1_45S_50ntFlanks</td><td>11702</td><td>11728</td><td>27</td><td>CAATGAAGCGCGGGTAAACGGCGGGAG</td><td>106.8</td><td>-8.8</td><td>1.4</td><td>-6.2</td><td>6.3E-10</td><td>1.9E-09</td><td>BT20/MB468</td><td>MITO</td></tr>
<tr><td>>hg38_21_+_8433172_8446622_RNA45SN1_45S_50ntFlanks@4724.4767.44|ATTAATCAAGAACGAAAGTCGGAGGTTCGAAGACGATCAGATAC</td><td>>hg38_21_+_8433172_8446622_RNA45SN1_45S_50ntFlanks</td><td>4724</td><td>4767</td><td>44</td><td>ATTAATCAAGAACGAAAGTCGGAGGTTCGAAGACGATCAGATAC</td><td>531.2</td><td>8.7</td><td>1.4</td><td>6.1</td><td>8.2E-10</td><td>1.9E-09</td><td>MB231/MB468</td><td>CYTO</td></tr>
<tr><td>>MI0000469|hsa-mir-125a&WithFlank&19|+|51693248|51693345@21.44.24&[MIMAT0000443&hsa-miR-125a-5p&offsets|0|0;m-63&19|+|51693268|51693289&offsets|0|+2]|TCCCTGAGACCCTTTAACCTGTGA</td><td>>MI0000469|hsa-mir-125a&WithFlank&19|+|51693248|51693345</td><td>21</td><td>44</td><td>24</td><td>TCCCTGAGACCCTTTAACCTGTGA</td><td>2363.0</td><td>5.1</td><td>0.8</td><td>6.1</td><td>8.2E-10</td><td>1.9E-09</td><td>MB231/MB468</td><td>CYTO</td></tr>
<tr><td>>MI0000111|hsa-mir-105-1&WithFlank&X|-|152392213|152392305@19.40.22;MI0000112|hsa-mir-105-2&WithFlank&X|-|152394406|152394498@19.40.22&[MIMAT0000102&hsa-miR-105-5p&offsets|0|-1;m-735&X|-|152392266|152392287&offsets|0|0];[MIMAT0000102_1&hsa-miR-105-5p&offsets|0|-1;m-735&X|-|152394459|152394480&offsets|0|0]|TCAAATGCTCAGACTCCTGTGG</td><td>>MI0000111|hsa-mir-105-1&WithFlank&X|-|152392213|152392305</td><td>19</td><td>40</td><td>22</td><td>TCAAATGCTCAGACTCCTGTGG</td><td>325.3</td><td>11.7</td><td>1.7</td><td>6.7</td><td>1.6E-11</td><td>1.9E-09</td><td>BT20/MB231</td><td>MP</td></tr>
<tr><td>>MI0000782|hsa-mir-374a&WithFlank&X|-|74287280|74287363@18.39.22&[MIMAT0000727&hsa-miR-374a-5p&offsets|0|0;m-184&X|-|74287325|74287346&offsets|0|0]|TTATAATACAACCTGATAAGTG</td><td>>MI0000782|hsa-mir-374a&WithFlank&X|-|74287280|74287363</td><td>18</td><td>39</td><td>22</td><td>TTATAATACAACCTGATAAGTG</td><td>351.5</td><td>5.0</td><td>0.8</td><td>6.1</td><td>8.4E-10</td><td>1.9E-09</td><td>MB231/MB468</td><td>NUC</td></tr>
<tr><td>>hg38_21_+_8433172_8446622_RNA45SN1_45S_50ntFlanks@8475.8492.18|CCCGGCGGATCTTTCCCG</td><td>>hg38_21_+_8433172_8446622_RNA45SN1_45S_50ntFlanks</td><td>8475</td><td>8492</td><td>18</td><td>CCCGGCGGATCTTTCCCG</td><td>161.7</td><td>-10.4</td><td>1.6</td><td>-6.5</td><td>8.2E-11</td><td>1.9E-09</td><td>BT20/MB468</td><td>MP</td></tr>
<tr><td>>MI0000283|hsa-mir-203a&WithFlank&14|+|104117399|104117520@71.90.20&[MIMAT0000264&hsa-miR-203a-3p&offsets|0|-2;m-79&14|+|104117469|104117490&offsets|0|-2]|GTGAAATGTTTAGGACCACT</td><td>>MI0000283|hsa-mir-203a&WithFlank&14|+|104117399|104117520</td><td>71</td><td>90</td><td>20</td><td>GTGAAATGTTTAGGACCACT</td><td>322.1</td><td>11.7</td><td>1.7</td><td>6.7</td><td>1.6E-11</td><td>1.9E-09</td><td>BT20/MB231</td><td>MP</td></tr>
<tr><td>>MI0000294|hsa-mir-218-1&WithFlank&4|+|20528269|20528390@31.52.22(+1U);MI0000295|hsa-mir-218-2&WithFlank&5|-|168768140|168768261@31.52.22(+1U)&[MIMAT0000275&hsa-miR-218-5p&offsets|0|+1(+1U);m-185&4|+|20528299|20528319&offsets|0|+1(+1U)];[MIMAT0000275_1&hsa-miR-218-5p&offsets|0|+1(+1U);m-187&5|-|168768211|168768231&offsets|0|+1(+1U)]|TTGTGCTTGATCTAACCATGTGT</td><td>>MI0000294|hsa-mir-218-1&WithFlank&4|+|20528269|20528390</td><td>31</td><td>52</td><td>22(+1U)</td><td>TTGTGCTTGATCTAACCATGTGT</td><td>359.5</td><td>-11.9</td><td>1.8</td><td>-6.7</td><td>1.6E-11</td><td>1.9E-09</td><td>BT20/MB231</td><td>MP</td></tr>
<tr><td>>hg38_21_+_8433172_8446622_RNA45SN1_45S_50ntFlanks@11919.11967.49|CATGAGAGGTGTAGAATAAGTGGGAGGCCCCCGGCGCCCCCCCGGTGTC</td><td>>hg38_21_+_8433172_8446622_RNA45SN1_45S_50ntFlanks</td><td>11919</td><td>11967</td><td>49</td><td>CATGAGAGGTGTAGAATAAGTGGGAGGCCCCCGGCGCCCCCCCGGTGTC</td><td>121.4</td><td>-9.2</td><td>1.5</td><td>-6.2</td><td>7.3E-10</td><td>1.9E-09</td><td>BT20/MB468</td><td>CYTO</td></tr>
<tr><td>>hg38_21_+_8433172_8446622_RNA45SN1_45S_50ntFlanks@8109.8139.31|GCGGGGCGCGGGACATGTGGCGTACGGAAGA</td><td>>hg38_21_+_8433172_8446622_RNA45SN1_45S_50ntFlanks</td><td>8109</td><td>8139</td><td>31</td><td>GCGGGGCGCGGGACATGTGGCGTACGGAAGA</td><td>84.6</td><td>-10.3</td><td>1.7</td><td>-6.1</td><td>8.0E-10</td><td>1.9E-09</td><td>MB231/MB468</td><td>MP</td></tr>
<tr><td>>trna116_GluCTC_1_-_145399233_145399304@1.37.37,trna59_GluCTC_1_+_249168447_249168518@1.37.37,trna71_GluCTC_1_-_161439189_161439260@1.37.37,trna74_GluCTC_1_-_161431809_161431880@1.37.37,trna77_GluCTC_1_-_161424398_161424469@1.37.37,trna77_GluCTC_6_+_28949976_28950047@1.37.37,trna80_GluCTC_1_-_161417018_161417089@1.37.37,trna87_GluCTC_6_-_126101393_126101464@1.37.37|TCCCTGGTGGTCTAGTGGTTAGGATTCGGCGCTCTCA</td><td>>trna116_GluCTC_1_-_145399233_145399304</td><td>1</td><td>37</td><td>37</td><td>TCCCTGGTGGTCTAGTGGTTAGGATTCGGCGCTCTCA</td><td>479.4</td><td>8.3</td><td>1.4</td><td>6.2</td><td>7.3E-10</td><td>1.9E-09</td><td>BT20/MB468</td><td>CYTO</td></tr>
<tr><td>>hg38_21_+_8433172_8446622_RNA45SN1_45S_50ntFlanks@11770.11794.25|AATTAGTGACGCGCATGAATGGATG</td><td>>hg38_21_+_8433172_8446622_RNA45SN1_45S_50ntFlanks</td><td>11770</td><td>11794</td><td>25</td><td>AATTAGTGACGCGCATGAATGGATG</td><td>1520.4</td><td>3.7</td><td>0.6</td><td>6.1</td><td>8.3E-10</td><td>1.9E-09</td><td>MB231/MB468</td><td>CYTO</td></tr>
<tr><td>>trnaMT_PheGAA_MT_+_577_647@35.57.23|GAAAATGTTTAGACGGGCTCACA</td><td>>trnaMT_PheGAA_MT_+_577_647</td><td>35</td><td>57</td><td>23</td><td>GAAAATGTTTAGACGGGCTCACA</td><td>1972.5</td><td>5.9</td><td>0.9</td><td>6.5</td><td>8.2E-11</td><td>1.9E-09</td><td>BT20/MB468</td><td>MP</td></tr>
<tr><td>>MI0000255|hsa-mir-30d&WithFlank&8|-|134804870|134804951@12.33.22(+1U)&[MIMAT0000245&hsa-miR-30d-5p&offsets|0|0(+1U);m-21&8|-|134804919|134804940&offsets|0|0(+1U)]|TGTAAACATCCCCGACTGGAAGT</td><td>>MI0000255|hsa-mir-30d&WithFlank&8|-|134804870|134804951</td><td>12</td><td>33</td><td>22(+1U)</td><td>TGTAAACATCCCCGACTGGAAGT</td><td>3303.3</td><td>5.4</td><td>0.9</td><td>6.1</td><td>9.3E-10</td><td>1.9E-09</td><td>MB231/MB468</td><td>MITO</td></tr>
<tr><td>>hg38_21_+_8433172_8446622_RNA45SN1_45S_50ntFlanks@12420.12456.37|ATGTCGGCTCTTCCTATCATTGTGAAGCAGAATTCAC</td><td>>hg38_21_+_8433172_8446622_RNA45SN1_45S_50ntFlanks</td><td>12420</td><td>12456</td><td>37</td><td>ATGTCGGCTCTTCCTATCATTGTGAAGCAGAATTCAC</td><td>810.3</td><td>-2.8</td><td>0.5</td><td>-6.2</td><td>7.4E-10</td><td>1.9E-09</td><td>BT20/MB468</td><td>CYTO</td></tr>
<tr><td>>trna32_LysCTT_16_-_3207406_3207478@1.33.33|GCCCGGCTAGCTCAGTCGGTAGAGCATGAGACC</td><td>>trna32_LysCTT_16_-_3207406_3207478</td><td>1</td><td>33</td><td>33</td><td>GCCCGGCTAGCTCAGTCGGTAGAGCATGAGACC</td><td>1308.0</td><td>6.4</td><td>1.0</td><td>6.1</td><td>9.3E-10</td><td>1.9E-09</td><td>MB231/MB468</td><td>MITO</td></tr>
<tr><td>>hg38_21_+_8433172_8446622_RNA45SN1_45S_50ntFlanks@10350.10380.31|ACGAGAACTTTGAAGGCCGAAGTGGAGAAGG</td><td>>hg38_21_+_8433172_8446622_RNA45SN1_45S_50ntFlanks</td><td>10350</td><td>10380</td><td>31</td><td>ACGAGAACTTTGAAGGCCGAAGTGGAGAAGG</td><td>373.1</td><td>-2.8</td><td>0.5</td><td>-6.1</td><td>8.5E-10</td><td>1.9E-09</td><td>MB231/MB468</td><td>NUC</td></tr>
<tr><td>>trna10_AspGTC_12_-_125424193_125424264@32.73.42,trna12_AspGTC_12_-_125411891_125411962@32.73.42,trna144_AspGTC_6_-_27551236_27551307@32.73.42,trna38_AspGTC_17_-_8125556_8125627@32.73.42,trna45_AspGTC_6_+_27447453_27447524@32.73.42,trna48_AspGTC_6_+_27471523_27471594@32.73.42,trna4_AspGTC_12_+_96429799_96429870@32.73.42,trna69_AspGTC_1_-_161440205_161440276@32.73.42,trna72_AspGTC_1_-_161432824_161432895@32.73.42,trna75_AspGTC_1_-_161425414_161425485@32.73.42,trna78_AspGTC_1_-_161418033_161418104@32.73.42,trna81_AspGTC_1_-_161410615_161410686@32.73.42|CTGTCACGCGGGAGACCGGGGTTCGATTCCCCGACGGGGAGC</td><td>>trna10_AspGTC_12_-_125424193_125424264</td><td>32</td><td>73</td><td>42</td><td>CTGTCACGCGGGAGACCGGGGTTCGATTCCCCGACGGGGAGC</td><td>76.4</td><td>-10.1</td><td>1.6</td><td>-6.1</td><td>8.1E-10</td><td>1.9E-09</td><td>MB231/MB468</td><td>MP</td></tr>
<tr><td>>hg38_21_+_8433172_8446622_RNA45SN1_45S_50ntFlanks@11907.11929.23|ACGGTGAAGAGACATGAGAGGTG</td><td>>hg38_21_+_8433172_8446622_RNA45SN1_45S_50ntFlanks</td><td>11907</td><td>11929</td><td>23</td><td>ACGGTGAAGAGACATGAGAGGTG</td><td>164.5</td><td>-7.2</td><td>1.2</td><td>-6.1</td><td>8.5E-10</td><td>1.9E-09</td><td>MB231/MB468</td><td>NUC</td></tr>
<tr><td>>hg38_21_+_8433172_8446622_RNA45SN1_45S_50ntFlanks@7974.8005.32|ACGCGACCTCAGATCAGACGTGGCGACCCGCT</td><td>>hg38_21_+_8433172_8446622_RNA45SN1_45S_50ntFlanks</td><td>7974</td><td>8005</td><td>32</td><td>ACGCGACCTCAGATCAGACGTGGCGACCCGCT</td><td>1239.5</td><td>3.3</td><td>0.5</td><td>6.1</td><td>8.4E-10</td><td>1.9E-09</td><td>MB231/MB468</td><td>CYTO</td></tr>
<tr><td>>hg38_21_+_8433172_8446622_RNA45SN1_45S_50ntFlanks@11564.11601.38|CCTAGCAGCCGACTTAGAACTGGTGCGGACCAGGGGAA</td><td>>hg38_21_+_8433172_8446622_RNA45SN1_45S_50ntFlanks</td><td>11564</td><td>11601</td><td>38</td><td>CCTAGCAGCCGACTTAGAACTGGTGCGGACCAGGGGAA</td><td>1633.2</td><td>-5.0</td><td>0.8</td><td>-6.1</td><td>8.4E-10</td><td>1.9E-09</td><td>MB231/MB468</td><td>CYTO</td></tr>
<tr><td>>hg38_21_+_8433172_8446622_RNA45SN1_45S_50ntFlanks@9534.9559.26|AGGAAACTCTGGTGGAGGTCCGTAGC</td><td>>hg38_21_+_8433172_8446622_RNA45SN1_45S_50ntFlanks</td><td>9534</td><td>9559</td><td>26</td><td>AGGAAACTCTGGTGGAGGTCCGTAGC</td><td>312.1</td><td>-8.0</td><td>1.3</td><td>-6.1</td><td>9.3E-10</td><td>1.9E-09</td><td>MB231/MB468</td><td>MITO</td></tr>
<tr><td>>am_trnaMT_IleGAT_MT_+_4263_4331@1.33.33|AGAAATATGTCTGATAAAAGAGTTACTTTGATA</td><td>>am_trnaMT_IleGAT_MT_+_4263_4331</td><td>1</td><td>33</td><td>33</td><td>AGAAATATGTCTGATAAAAGAGTTACTTTGATA</td><td>154.9</td><td>10.5</td><td>1.7</td><td>6.1</td><td>9.3E-10</td><td>1.9E-09</td><td>MB231/MB468</td><td>MITO</td></tr>
<tr><td>>hg38_21_+_8433172_8446622_RNA45SN1_45S_50ntFlanks@5492.5509.18|AGAAGACGGTCGAACTTG</td><td>>hg38_21_+_8433172_8446622_RNA45SN1_45S_50ntFlanks</td><td>5492</td><td>5509</td><td>18</td><td>AGAAGACGGTCGAACTTG</td><td>497.9</td><td>-7.8</td><td>1.3</td><td>-6.1</td><td>9.3E-10</td><td>1.9E-09</td><td>MB231/MB468</td><td>MITO</td></tr>
<tr><td>>MI0000077|hsa-mir-21&WithFlank&17|+|59841260|59841343@14.34.21(+1G)&[MIMAT0000076&hsa-miR-21-5p&offsets|0|-1(+1G);m-1&17|+|59841273|59841294&offsets|0|-1(+1G)]|TAGCTTATCAGACTGATGTTGG</td><td>>MI0000077|hsa-mir-21&WithFlank&17|+|59841260|59841343</td><td>14</td><td>34</td><td>21(+1G)</td><td>TAGCTTATCAGACTGATGTTGG</td><td>662.2</td><td>5.4</td><td>0.9</td><td>6.2</td><td>6.4E-10</td><td>1.9E-09</td><td>BT20/MB468</td><td>MITO</td></tr>
<tr><td>>MI0000805|hsa-mir-342&WithFlank&14|+|100109649|100109759@67.90.24&[MIMAT0000753&hsa-miR-342-3p&offsets|0|+1;m-128&14|+|100109715|100109737&offsets|0|+1]|TCTCACACAGAAATCGCACCCGTC</td><td>>MI0000805|hsa-mir-342&WithFlank&14|+|100109649|100109759</td><td>67</td><td>90</td><td>24</td><td>TCTCACACAGAAATCGCACCCGTC</td><td>540.7</td><td>5.3</td><td>0.9</td><td>6.1</td><td>8.4E-10</td><td>2.0E-09</td><td>MB231/MB468</td><td>CYTO</td></tr>
<tr><td>>MI0000088|hsa-mir-30a&WithFlank&6|-|71403545|71403627@54.74.21(+1A)&[MIMAT0000088&hsa-miR-30a-3p&offsets|+1|0(+1A);m-38&6|-|71403554|71403575&offsets|+1|0(+1A)]|TTTCAGTCGGATGTTTGCAGCA</td><td>>MI0000088|hsa-mir-30a&WithFlank&6|-|71403545|71403627</td><td>54</td><td>74</td><td>21(+1A)</td><td>TTTCAGTCGGATGTTTGCAGCA</td><td>165.2</td><td>10.6</td><td>1.7</td><td>6.1</td><td>9.3E-10</td><td>2.0E-09</td><td>MB231/MB468</td><td>MITO</td></tr>
<tr><td>>hg38_21_+_8433172_8446622_RNA45SN1_45S_50ntFlanks@3885.3915.31|ACGGGCGCTGACCCCCTTCGCGGGGGGGATG</td><td>>hg38_21_+_8433172_8446622_RNA45SN1_45S_50ntFlanks</td><td>3885</td><td>3915</td><td>31</td><td>ACGGGCGCTGACCCCCTTCGCGGGGGGGATG</td><td>290.8</td><td>-7.6</td><td>1.2</td><td>-6.1</td><td>9.3E-10</td><td>2.0E-09</td><td>MB231/MB468</td><td>MITO</td></tr>
<tr><td>>hg38_21_+_8433172_8446622_RNA45SN1_45S_50ntFlanks@11641.11668.28|CGGCGGGTGTTGACGCGATGTGATTTCT</td><td>>hg38_21_+_8433172_8446622_RNA45SN1_45S_50ntFlanks</td><td>11641</td><td>11668</td><td>28</td><td>CGGCGGGTGTTGACGCGATGTGATTTCT</td><td>542.6</td><td>-6.5</td><td>1.1</td><td>-6.1</td><td>8.6E-10</td><td>2.0E-09</td><td>MB231/MB468</td><td>NUC</td></tr>
<tr><td>>hg38_21_+_8433172_8446622_RNA45SN1_45S_50ntFlanks@3835.3862.28|CTCCTCTCCTACTTGGATAACTGTGGTA</td><td>>hg38_21_+_8433172_8446622_RNA45SN1_45S_50ntFlanks</td><td>3835</td><td>3862</td><td>28</td><td>CTCCTCTCCTACTTGGATAACTGTGGTA</td><td>513.9</td><td>-8.6</td><td>1.4</td><td>-6.1</td><td>9.8E-10</td><td>2.0E-09</td><td>BT20/MB468</td><td>NUC</td></tr>
<tr><td>>hg38_21_+_8433172_8446622_RNA45SN1_45S_50ntFlanks@5421.5466.46|CGATTGGATGGTTTAGTGAGGCCCTCGGATCGGCCCCGCCGGGGTC</td><td>>hg38_21_+_8433172_8446622_RNA45SN1_45S_50ntFlanks</td><td>5421</td><td>5466</td><td>46</td><td>CGATTGGATGGTTTAGTGAGGCCCTCGGATCGGCCCCGCCGGGGTC</td><td>103.1</td><td>-10.6</td><td>1.7</td><td>-6.1</td><td>8.2E-10</td><td>2.0E-09</td><td>MB231/MB468</td><td>MP</td></tr>
<tr><td>>hg38_21_+_8433172_8446622_RNA45SN1_45S_50ntFlanks@6698.6718.21|AGCTAGCTGCGAGAATTAATG</td><td>>hg38_21_+_8433172_8446622_RNA45SN1_45S_50ntFlanks</td><td>6698</td><td>6718</td><td>21</td><td>AGCTAGCTGCGAGAATTAATG</td><td>26135.1</td><td>-11.5</td><td>1.9</td><td>-6.1</td><td>9.8E-10</td><td>2.0E-09</td><td>BT20/MB468</td><td>NUC</td></tr>
<tr><td>>hg38_21_+_8433172_8446622_RNA45SN1_45S_50ntFlanks@4722.4757.36|TCATTAATCAAGAACGAAAGTCGGAGGTTCGAAGAC</td><td>>hg38_21_+_8433172_8446622_RNA45SN1_45S_50ntFlanks</td><td>4722</td><td>4757</td><td>36</td><td>TCATTAATCAAGAACGAAAGTCGGAGGTTCGAAGAC</td><td>230.5</td><td>-5.8</td><td>0.9</td><td>-6.1</td><td>8.6E-10</td><td>2.0E-09</td><td>MB231/MB468</td><td>NUC</td></tr>
<tr><td>>hg38_21_+_8433172_8446622_RNA45SN1_45S_50ntFlanks@10250.10273.24|ACGAGTAGGAGGGCCGCTGCGGTG</td><td>>hg38_21_+_8433172_8446622_RNA45SN1_45S_50ntFlanks</td><td>10250</td><td>10273</td><td>24</td><td>ACGAGTAGGAGGGCCGCTGCGGTG</td><td>122.9</td><td>-10.8</td><td>1.8</td><td>-6.1</td><td>8.2E-10</td><td>2.0E-09</td><td>MB231/MB468</td><td>MP</td></tr>
</table>

<table>
<tr><td>>hg38_21_+_8433172_8446622_RNA45SN1_45S_50ntFlanks@10683.10708.26|CCGGCGGCGTCCGGTGAGCTCTCGCT</td><td>>hg38_21_+_8433172_8446622_RNA45SN1_45S_50ntFlanks</td><td>10683</td><td>10708</td><td>26</td><td>CCGGCGGCGTCCGGTGAGCTCTCGCT</td><td>537.6</td><td>-4.6</td><td>0.8</td><td>-6.1</td><td>9.4E-10</td><td>2.0E-09</td><td>MB231/MB468</td><td>MITO</td></tr>
<tr><td>>hg38_21_+_8433172_8446622_RNA45SN1_45S_50ntFlanks@12940.12969.30|CATTCGTAGACGACCTGCTTCTGGGTCGGG</td><td>>hg38_21_+_8433172_8446622_RNA45SN1_45S_50ntFlanks</td><td>12940</td><td>12969</td><td>30</td><td>CATTCGTAGACGACCTGCTTCTGGGTCGGG</td><td>132.5</td><td>-7.0</td><td>1.1</td><td>-6.1</td><td>8.5E-10</td><td>2.0E-09</td><td>MB231/MB468</td><td>CYTO</td></tr>
<tr><td>>hg38_21_+_8433172_8446622_RNA45SN1_45S_50ntFlanks@12941.12966.26|ATTCGTAGACGACCTGCTTCTGGGTC</td><td>>hg38_21_+_8433172_8446622_RNA45SN1_45S_50ntFlanks</td><td>12941</td><td>12966</td><td>26</td><td>ATTCGTAGACGACCTGCTTCTGGGTC</td><td>331.4</td><td>-8.4</td><td>1.4</td><td>-6.1</td><td>9.4E-10</td><td>2.0E-09</td><td>MB231/MB468</td><td>MITO</td></tr>
<tr><td>>hg38_21_+_8433172_8446622_RNA45SN1_45S_50ntFlanks@11578.11603.26|TAGAACTGGTGCGGACCAGGGGAATC</td><td>>hg38_21_+_8433172_8446622_RNA45SN1_45S_50ntFlanks</td><td>11578</td><td>11603</td><td>26</td><td>TAGAACTGGTGCGGACCAGGGGAATC</td><td>688.3</td><td>-8.1</td><td>1.3</td><td>-6.1</td><td>9.5E-10</td><td>2.0E-09</td><td>MB231/MB468</td><td>MITO</td></tr>
<tr><td>>MI0000102|hsa-mir-100&WithFlank&11|-|122152223|122152314@19.39.21&[MIMAT0000098&hsa-miR-100-5p&offsets|0|-1;m-27&11|-|122152276|122152296&offsets|0|0]|AACCCGTAGATCCGAACTTGT</td><td>>MI0000102|hsa-mir-100&WithFlank&11|-|122152223|122152314</td><td>19</td><td>39</td><td>21</td><td>AACCCGTAGATCCGAACTTGT</td><td>533.8</td><td>6.0</td><td>1.0</td><td>6.1</td><td>9.9E-10</td><td>2.0E-09</td><td>BT20/MB468</td><td>NUC</td></tr>
<tr><td>>hg38_21_+_8433172_8446622_RNA45SN1_45S_50ntFlanks@5077.5112.36|CCGATAACGAACGAGACTCTGGCATGCTAACTAGTT</td><td>>hg38_21_+_8433172_8446622_RNA45SN1_45S_50ntFlanks</td><td>5077</td><td>5112</td><td>36</td><td>CCGATAACGAACGAGACTCTGGCATGCTAACTAGTT</td><td>141.6</td><td>-11.0</td><td>1.8</td><td>-6.1</td><td>8.2E-10</td><td>2.0E-09</td><td>MB231/MB468</td><td>MP</td></tr>
<tr><td>>hg38_21_+_8433172_8446622_RNA45SN1_45S_50ntFlanks@12159.12176.18|TTTGACTGGGGCGGTACA</td><td>>hg38_21_+_8433172_8446622_RNA45SN1_45S_50ntFlanks</td><td>12159</td><td>12176</td><td>18</td><td>TTTGACTGGGGCGGTACA</td><td>167.2</td><td>-10.5</td><td>1.6</td><td>-6.5</td><td>8.4E-11</td><td>2.0E-09</td><td>BT20/MB468</td><td>MP</td></tr>
<tr><td>>hg38_21_+_8433172_8446622_RNA45SN1_45S_50ntFlanks@4660.4691.32|GAGGTGAAATTCTTGGACCGGCGCAAGACGGA</td><td>>hg38_21_+_8433172_8446622_RNA45SN1_45S_50ntFlanks</td><td>4660</td><td>4691</td><td>32</td><td>GAGGTGAAATTCTTGGACCGGCGCAAGACGGA</td><td>90.6</td><td>-10.4</td><td>1.7</td><td>-6.1</td><td>8.2E-10</td><td>2.0E-09</td><td>MB231/MB468</td><td>MP</td></tr>
<tr><td>>hg38_21_+_8433172_8446622_RNA45SN1_45S_50ntFlanks@4243.4264.22|CTTTCGAGGCCCTGTAATTGGA</td><td>>hg38_21_+_8433172_8446622_RNA45SN1_45S_50ntFlanks</td><td>4243</td><td>4264</td><td>22</td><td>CTTTCGAGGCCCTGTAATTGGA</td><td>364.3</td><td>-8.0</td><td>1.3</td><td>-6.1</td><td>9.5E-10</td><td>2.0E-09</td><td>MB231/MB468</td><td>MITO</td></tr>
<tr><td>>hg38_21_+_8433172_8446622_RNA45SN1_45S_50ntFlanks@6731.6756.26|CACATTGATCATCGACACTTCGAACG</td><td>>hg38_21_+_8433172_8446622_RNA45SN1_45S_50ntFlanks</td><td>6731</td><td>6756</td><td>26</td><td>CACATTGATCATCGACACTTCGAACG</td><td>1108.1</td><td>-4.5</td><td>0.7</td><td>-6.1</td><td>9.5E-10</td><td>2.0E-09</td><td>MB231/MB468</td><td>MITO</td></tr>
<tr><td>>hg38_21_+_8433172_8446622_RNA45SN1_45S_50ntFlanks@12072.12090.19|CCCGAGGGGCTCTCGCTTC</td><td>>hg38_21_+_8433172_8446622_RNA45SN1_45S_50ntFlanks</td><td>12072</td><td>12090</td><td>19</td><td>CCCGAGGGGCTCTCGCTTC</td><td>113.6</td><td>-8.9</td><td>1.4</td><td>-6.2</td><td>6.5E-10</td><td>2.0E-09</td><td>BT20/MB468</td><td>MITO</td></tr>
<tr><td>>hg38_21_+_8433172_8446622_RNA45SN1_45S_50ntFlanks@10444.10462.19|CGAAGGGACGGGCGATGGC</td><td>>hg38_21_+_8433172_8446622_RNA45SN1_45S_50ntFlanks</td><td>10444</td><td>10462</td><td>19</td><td>CGAAGGGACGGGCGATGGC</td><td>1061.0</td><td>-9.3</td><td>1.5</td><td>-6.1</td><td>1.0E-09</td><td>2.0E-09</td><td>BT20/MB468</td><td>NUC</td></tr>
<tr><td>>hg38_21_+_8433172_8446622_RNA45SN1_45S_50ntFlanks@10786.10817.32|AACAGCCTCTGGCATGTTGGAACAATGTAGGT</td><td>>hg38_21_+_8433172_8446622_RNA45SN1_45S_50ntFlanks</td><td>10786</td><td>10817</td><td>32</td><td>AACAGCCTCTGGCATGTTGGAACAATGTAGGT</td><td>979.7</td><td>-7.3</td><td>1.2</td><td>-6.1</td><td>1.0E-09</td><td>2.0E-09</td><td>BT20/MB468</td><td>NUC</td></tr>
<tr><td>>hg38_21_+_8433172_8446622_RNA45SN1_45S_50ntFlanks@4390.4407.18|TCTTGGGAGCGGGCGGGC</td><td>>hg38_21_+_8433172_8446622_RNA45SN1_45S_50ntFlanks</td><td>4390</td><td>4407</td><td>18</td><td>TCTTGGGAGCGGGCGGGC</td><td>81.5</td><td>-10.2</td><td>1.7</td><td>-6.1</td><td>8.3E-10</td><td>2.0E-09</td><td>MB231/MB468</td><td>MP</td></tr>
<tr><td>>hg38_21_+_8433172_8446622_RNA45SN1_45S_50ntFlanks@10771.10801.31|CAGGTCTCCAAGGTGAACAGCCTCTGGCATG</td><td>>hg38_21_+_8433172_8446622_RNA45SN1_45S_50ntFlanks</td><td>10771</td><td>10801</td><td>31</td><td>CAGGTCTCCAAGGTGAACAGCCTCTGGCATG</td><td>87.0</td><td>-10.3</td><td>1.7</td><td>-6.1</td><td>8.3E-10</td><td>2.0E-09</td><td>MB231/MB468</td><td>MP</td></tr>
<tr><td>>hg38_21_+_8433172_8446622_RNA45SN1_45S_50ntFlanks@12635.12657.23|AAGCTACCATCTGTGGGATTATG</td><td>>hg38_21_+_8433172_8446622_RNA45SN1_45S_50ntFlanks</td><td>12635</td><td>12657</td><td>23</td><td>AAGCTACCATCTGTGGGATTATG</td><td>2018.8</td><td>-7.3</td><td>1.1</td><td>-6.5</td><td>8.5E-11</td><td>2.0E-09</td><td>BT20/MB468</td><td>MP</td></tr>
<tr><td>>hg38_21_+_8433172_8446622_RNA45SN1_45S_50ntFlanks@4046.4065.20|CATTCGAACGTCTGCCCTAT</td><td>>hg38_21_+_8433172_8446622_RNA45SN1_45S_50ntFlanks</td><td>4046</td><td>4065</td><td>20</td><td>CATTCGAACGTCTGCCCTAT</td><td>58.7</td><td>-9.7</td><td>1.6</td><td>-6.1</td><td>8.3E-10</td><td>2.0E-09</td><td>MB231/MB468</td><td>MP</td></tr>
<tr><td>>MI0000270|hsa-mir-181b-1&WithFlank&1|-|198858867|198858988@42.64.23;MI0000683|hsa-mir-181b-2&WithFlank&9|+|124693704|124693804@22.44.23&[MIMAT0000257&hsa-miR-181b-5p&offsets|0|0;m-73&1|-|198858925|198858947&offsets|0|0];[MIMAT0000257_1&hsa-miR-181b-5p&offsets|0|0;m-72&9|+|124693725|124693747&offsets|0|0]|AACATTCATTGCTGTCGGTGGGT</td><td>>MI0000270|hsa-mir-181b-1&WithFlank&1|-|198858867|198858988</td><td>42</td><td>64</td><td>23</td><td>AACATTCATTGCTGTCGGTGGGT</td><td>2361.6</td><td>5.7</td><td>0.9</td><td>6.1</td><td>9.6E-10</td><td>2.0E-09</td><td>MB231/MB468</td><td>MITO</td></tr>
<tr><td>>hg38_21_+_8433172_8446622_RNA45SN1_45S_50ntFlanks@10362.10384.23|AAGGCCGAAGTGGAGAAGGGTTC</td><td>>hg38_21_+_8433172_8446622_RNA45SN1_45S_50ntFlanks</td><td>10362</td><td>10384</td><td>23</td><td>AAGGCCGAAGTGGAGAAGGGTTC</td><td>1071.7</td><td>-8.0</td><td>1.3</td><td>-6.1</td><td>8.7E-10</td><td>2.0E-09</td><td>MB231/MB468</td><td>NUC</td></tr>
<tr><td>>am_trnaMT_TyrGTA_MT_-_5826_5891@13.32.20|GAGTGAAGCATTGGACTGTA</td><td>>am_trnaMT_TyrGTA_MT_-_5826_5891</td><td>13</td><td>32</td><td>20</td><td>GAGTGAAGCATTGGACTGTA</td><td>519.6</td><td>7.3</td><td>1.2</td><td>6.2</td><td>6.5E-10</td><td>2.0E-09</td><td>BT20/MB468</td><td>MITO</td></tr>
<tr><td>>MI0000063|hsa-let-7b&WithFlank&22|+|46113680|46113774@12.31.20(+1C)&[MIMAT0000063&hsa-let-7b-5p&offsets|0|-2(+1C);m-14&22|+|46113691|46113712&offsets|0|-2(+1C)]|TGAGGTAGTAGGTTGTGTGGC</td><td>>MI0000063|hsa-let-7b&WithFlank&22|+|46113680|46113774</td><td>12</td><td>31</td><td>20(+1C)</td><td>TGAGGTAGTAGGTTGTGTGGC</td><td>278.3</td><td>2.5</td><td>0.4</td><td>6.1</td><td>1.0E-09</td><td>2.0E-09</td><td>BT20/MB468</td><td>NUC</td></tr>
<tr><td>>hg38_21_+_8433172_8446622_RNA45SN1_45S_50ntFlanks@10711.10736.26|CCCTTGAAAATCCGGGGGAGAGGGTG</td><td>>hg38_21_+_8433172_8446622_RNA45SN1_45S_50ntFlanks</td><td>10711</td><td>10736</td><td>26</td><td>CCCTTGAAAATCCGGGGGAGAGGGTG</td><td>85.1</td><td>-10.3</td><td>1.7</td><td>-6.1</td><td>8.4E-10</td><td>2.0E-09</td><td>MB231/MB468</td><td>MP</td></tr>
<tr><td>>hg38_21_+_8433172_8446622_RNA45SN1_45S_50ntFlanks@11540.11580.41|CGCGGCGCCTCGCCTCGGCCGGCGCCTAGCAGCCGACTTAG</td><td>>hg38_21_+_8433172_8446622_RNA45SN1_45S_50ntFlanks</td><td>11540</td><td>11580</td><td>41</td><td>CGCGGCGCCTCGCCTCGGCCGGCGCCTAGCAGCCGACTTAG</td><td>2629.4</td><td>-6.8</td><td>1.1</td><td>-6.1</td><td>1.0E-09</td><td>2.0E-09</td><td>BT20/MB468</td><td>NUC</td></tr>
<tr><td>>hg38_21_+_8433172_8446622_RNA45SN1_45S_50ntFlanks@12477.12499.23|CACTAATAGGGAACGTGAGCTGG</td><td>>hg38_21_+_8433172_8446622_RNA45SN1_45S_50ntFlanks</td><td>12477</td><td>12499</td><td>23</td><td>CACTAATAGGGAACGTGAGCTGG</td><td>321.0</td><td>-3.8</td><td>0.6</td><td>-6.1</td><td>8.8E-10</td><td>2.0E-09</td><td>MB231/MB468</td><td>NUC</td></tr>
<tr><td>>hg38_21_+_8433172_8446622_RNA45SN1_45S_50ntFlanks@10683.10711.29|CCGGCGGCGTCCGGTGAGCTCTCGCTGGC</td><td>>hg38_21_+_8433172_8446622_RNA45SN1_45S_50ntFlanks</td><td>10683</td><td>10711</td><td>29</td><td>CCGGCGGCGTCCGGTGAGCTCTCGCTGGC</td><td>1064.5</td><td>-6.8</td><td>1.1</td><td>-6.1</td><td>8.8E-10</td><td>2.0E-09</td><td>MB231/MB468</td><td>NUC</td></tr>
<tr><td>>hg38_21_+_8433172_8446622_RNA45SN1_45S_50ntFlanks@10826.10865.40|TCGGCAAGCCGGATCCGTAACTTCGGGATAAGGATTGGCT</td><td>>hg38_21_+_8433172_8446622_RNA45SN1_45S_50ntFlanks</td><td>10826</td><td>10865</td><td>40</td><td>TCGGCAAGCCGGATCCGTAACTTCGGGATAAGGATTGGCT</td><td>97.1</td><td>-10.5</td><td>1.7</td><td>-6.1</td><td>8.4E-10</td><td>2.0E-09</td><td>MB231/MB468</td><td>MP</td></tr>
<tr><td>>MI0000736|hsa-mir-30c-1&WithFlank&1|+|40757278|40757378@23.42.20;MI0000254|hsa-mir-30c-2&WithFlank&6|-|71376954|71377037@13.32.20&[MIMAT0000244&hsa-miR-30c-5p&offsets|0|-3;m-57&1|+|40757300|40757323&offsets|0|-4];[MIMAT0000244_1&hsa-miR-30c-5p&offsets|0|-3;m-56&6|-|71377002|71377025&offsets|0|-4]|TGTAAACATCCTACACTCTC</td><td>>MI0000736|hsa-mir-30c-1&WithFlank&1|+|40757278|40757378</td><td>23</td><td>42</td><td>20</td><td>TGTAAACATCCTACACTCTC</td><td>696.0</td><td>5.7</td><td>0.9</td><td>6.1</td><td>8.7E-10</td><td>2.0E-09</td><td>MB231/MB468</td><td>CYTO</td></tr>
<tr><td>>hg38_21_+_8433172_8446622_RNA45SN1_45S_50ntFlanks@6671.6698.28|CTCGGCTCGTGCGTCGATGAAGAACGCA</td><td>>hg38_21_+_8433172_8446622_RNA45SN1_45S_50ntFlanks</td><td>6671</td><td>6698</td><td>28</td><td>CTCGGCTCGTGCGTCGATGAAGAACGCA</td><td>1494.9</td><td>-9.8</td><td>1.6</td><td>-6.1</td><td>1.0E-09</td><td>2.0E-09</td><td>BT20/MB468</td><td>NUC</td></tr>
<tr><td>>MI0003820|hsa-mir-454&WithFlank&17|-|59137752|59137878@70.95.26&[MIMAT0003885&hsa-miR-454-3p&offsets|0|+3;m-272&17|-|59137787|59137809&offsets|0|+3]|TAGTGCAATATTGCTTATAGGGTTTT</td><td>>MI0003820|hsa-mir-454&WithFlank&17|-|59137752|59137878</td><td>70</td><td>95</td><td>26</td><td>TAGTGCAATATTGCTTATAGGGTTTT</td><td>91.0</td><td>9.7</td><td>1.6</td><td>6.1</td><td>8.4E-10</td><td>2.0E-09</td><td>MB231/MB468</td><td>MP</td></tr>
<tr><td>>hg38_21_+_8433172_8446622_RNA45SN1_45S_50ntFlanks@11602.11621.20|TCCGACTGTTTAATTAAAAC</td><td>>hg38_21_+_8433172_8446622_RNA45SN1_45S_50ntFlanks</td><td>11602</td><td>11621</td><td>20</td><td>TCCGACTGTTTAATTAAAAC</td><td>284.0</td><td>-6.1</td><td>1.0</td><td>-6.1</td><td>8.5E-10</td><td>2.0E-09</td><td>MB231/MB468</td><td>MP</td></tr>
<tr><td>>hg38_21_+_8433172_8446622_RNA45SN1_45S_50ntFlanks@12283.12309.27|TACAGACCGTGAAAGCGGGGCCTCACG</td><td>>hg38_21_+_8433172_8446622_RNA45SN1_45S_50ntFlanks</td><td>12283</td><td>12309</td><td>27</td><td>TACAGACCGTGAAAGCGGGGCCTCACG</td><td>67.0</td><td>-9.9</td><td>1.6</td><td>-6.1</td><td>8.5E-10</td><td>2.0E-09</td><td>MB231/MB468</td><td>MP</td></tr>
<tr><td>>hg38_21_+_8433172_8446622_RNA45SN1_45S_50ntFlanks@9166.9194.29|CCGGGCGGGTCGCGCCGTCGGGCCCGGGG</td><td>>hg38_21_+_8433172_8446622_RNA45SN1_45S_50ntFlanks</td><td>9166</td><td>9194</td><td>29</td><td>CCGGGCGGGTCGCGCCGTCGGGCCCGGGG</td><td>84.3</td><td>-10.3</td><td>1.7</td><td>-6.1</td><td>8.5E-10</td><td>2.0E-09</td><td>MB231/MB468</td><td>MP</td></tr>
<tr><td>>hg38_21_+_8433172_8446622_RNA45SN1_45S_50ntFlanks@4069.4101.33|CTTTCGATGGTAGTCGCCGTGCCTACCATGGTG</td><td>>hg38_21_+_8433172_8446622_RNA45SN1_45S_50ntFlanks</td><td>4069</td><td>4101</td><td>33</td><td>CTTTCGATGGTAGTCGCCGTGCCTACCATGGTG</td><td>454.3</td><td>-3.7</td><td>0.6</td><td>-6.1</td><td>8.5E-10</td><td>2.0E-09</td><td>MB231/MB468</td><td>MP</td></tr>
<tr><td>>hg38_21_+_8433172_8446622_RNA45SN1_45S_50ntFlanks@483.519.37|TCGATGTGGTGACGTCGTGCTCTCCCGGGCCGGGTCC</td><td>>hg38_21_+_8433172_8446622_RNA45SN1_45S_50ntFlanks</td><td>483</td><td>519</td><td>37</td><td>TCGATGTGGTGACGTCGTGCTCTCCCGGGCCGGGTCC</td><td>279.1</td><td>5.2</td><td>0.9</td><td>6.1</td><td>8.9E-10</td><td>2.0E-09</td><td>MB231/MB468</td><td>NUC</td></tr>
<tr><td>>hg38_21_+_8433172_8446622_RNA45SN1_45S_50ntFlanks@8194.8215.22|CCCGTGGACGGTGTGAGGCCGG</td><td>>hg38_21_+_8433172_8446622_RNA45SN1_45S_50ntFlanks</td><td>8194</td><td>8215</td><td>22</td><td>CCCGTGGACGGTGTGAGGCCGG</td><td>729.8</td><td>-3.8</td><td>0.6</td><td>-6.2</td><td>6.6E-10</td><td>2.0E-09</td><td>BT20/MB468</td><td>MITO</td></tr>
<tr><td>>hg38_21_+_8433172_8446622_RNA45SN1_45S_50ntFlanks@6651.6674.24|CGACTCTTAGCGGTGGATCACTCG</td><td>>hg38_21_+_8433172_8446622_RNA45SN1_45S_50ntFlanks</td><td>6651</td><td>6674</td><td>24</td><td>CGACTCTTAGCGGTGGATCACTCG</td><td>20011.4</td><td>4.8</td><td>0.8</td><td>6.2</td><td>6.7E-10</td><td>2.0E-09</td><td>BT20/MB468</td><td>MITO</td></tr>
<tr><td>>MI0000816|hsa-mir-335&WithFlank&7|+|130496105|130496210@57.79.23&[MIMAT0004703&hsa-miR-335-3p&offsets|-1|0;m-159&7|+|130496162|130496183&offsets|-1|0]|GTTTTTCATTATTGCTCCTGACC</td><td>>MI0000816|hsa-mir-335&WithFlank&7|+|130496105|130496210</td><td>57</td><td>79</td><td>23</td><td>GTTTTTCATTATTGCTCCTGACC</td><td>328.0</td><td>6.2</td><td>1.0</td><td>6.1</td><td>8.8E-10</td><td>2.0E-09</td><td>MB231/MB468</td><td>CYTO</td></tr>
<tr><td>>hg38_21_+_8433172_8446622_RNA45SN1_45S_50ntFlanks@11561.11581.21|GCGCCTAGCAGCCGACTTAGA</td><td>>hg38_21_+_8433172_8446622_RNA45SN1_45S_50ntFlanks</td><td>11561</td><td>11581</td><td>21</td><td>GCGCCTAGCAGCCGACTTAGA</td><td>133.3</td><td>-9.1</td><td>1.5</td><td>-6.2</td><td>6.7E-10</td><td>2.0E-09</td><td>BT20/MB468</td><td>MITO</td></tr>
<tr><td>>hg38_21_+_8433172_8446622_RNA45SN1_45S_50ntFlanks@10421.10453.33|TGAGAGATGGGCGAGCGCCGTTCCGAAGGGACG</td><td>>hg38_21_+_8433172_8446622_RNA45SN1_45S_50ntFlanks</td><td>10421</td><td>10453</td><td>33</td><td>TGAGAGATGGGCGAGCGCCGTTCCGAAGGGACG</td><td>822.9</td><td>-8.1</td><td>1.3</td><td>-6.1</td><td>9.0E-10</td><td>2.0E-09</td><td>MB231/MB468</td><td>NUC</td></tr>
<tr><td>>hg38_21_+_8433172_8446622_RNA45SN1_45S_50ntFlanks@12638.12659.22|CTACCATCTGTGGGATTATGAC</td><td>>hg38_21_+_8433172_8446622_RNA45SN1_45S_50ntFlanks</td><td>12638</td><td>12659</td><td>22</td><td>CTACCATCTGTGGGATTATGAC</td><td>442.4</td><td>-9.0</td><td>1.5</td><td>-6.1</td><td>1.0E-09</td><td>2.0E-09</td><td>BT20/MB468</td><td>NUC</td></tr>
<tr><td>>MI0000440|hsa-mir-27b&WithFlank&9|+|95085439|95085547@67.87.21&[MIMAT0000419&hsa-miR-27b-3p&offsets|0|0;m-39&9|+|95085505|95085525&offsets|0|0]|TTCACAGTGGCTAAGTTCTGC</td><td>>MI0000440|hsa-mir-27b&WithFlank&9|+|95085439|95085547</td><td>67</td><td>87</td><td>21</td><td>TTCACAGTGGCTAAGTTCTGC</td><td>18282.6</td><td>5.4</td><td>0.9</td><td>6.1</td><td>8.8E-10</td><td>2.0E-09</td><td>MB231/MB468</td><td>CYTO</td></tr>
<tr><td>>trna2_GlyGCC_21_-_18827107_18827177@1.30.30,trna35_GlyGCC_1_+_161413094_161413164@1.30.30,trna37_GlyGCC_1_+_161420467_161420537@1.30.30,trna39_GlyGCC_1_+_161427898_161427968@1.30.30,trna41_GlyGCC_1_+_161435258_161435328@1.30.30|GCATGGGTGGTTCAGTGGTAGAATTCTCGC</td><td>>trna2_GlyGCC_21_-_18827107_18827177</td><td>1</td><td>30</td><td>30</td><td>GCATGGGTGGTTCAGTGGTAGAATTCTCGC</td><td>4024.8</td><td>-6.0</td><td>1.0</td><td>-6.1</td><td>9.0E-10</td><td>2.0E-09</td><td>MB231/MB468</td><td>NUC</td></tr>
<tr><td>>hg38_21_+_8433172_8446622_RNA45SN1_45S_50ntFlanks@5236.5267.32|CACTGACTGGCTCAGCGTGTGCCTACCCTACG</td><td>>hg38_21_+_8433172_8446622_RNA45SN1_45S_50ntFlanks</td><td>5236</td><td>5267</td><td>32</td><td>CACTGACTGGCTCAGCGTGTGCCTACCCTACG</td><td>249.4</td><td>-3.1</td><td>0.5</td><td>-6.1</td><td>9.0E-10</td><td>2.0E-09</td><td>MB231/MB468</td><td>NUC</td></tr>
<tr><td>>hg38_21_+_8433172_8446622_RNA45SN1_45S_50ntFlanks@11999.12042.44|CCCTGCGGGCCGCCGGTGAAATACCACTACTCTGATCGTTTTTT</td><td>>hg38_21_+_8433172_8446622_RNA45SN1_45S_50ntFlanks</td><td>11999</td><td>12042</td><td>44</td><td>CCCTGCGGGCCGCCGGTGAAATACCACTACTCTGATCGTTTTTT</td><td>334.1</td><td>-11.3</td><td>1.9</td><td>-6.1</td><td>9.0E-10</td><td>2.0E-09</td><td>MB231/MB468</td><td>NUC</td></tr>
<tr><td>>hg38_21_+_8433172_8446622_RNA45SN1_45S_50ntFlanks@4762.4780.19|AGATACCGTCGTAGTTCCG</td><td>>hg38_21_+_8433172_8446622_RNA45SN1_45S_50ntFlanks</td><td>4762</td><td>4780</td><td>19</td><td>AGATACCGTCGTAGTTCCG</td><td>729.5</td><td>-4.0</td><td>0.6</td><td>-6.1</td><td>8.8E-10</td><td>2.0E-09</td><td>MB231/MB468</td><td>CYTO</td></tr>
<tr><td>>hg38_21_+_8433172_8446622_RNA45SN1_45S_50ntFlanks@3840.3877.38|CTCCTACTTGGATAACTGTGGTAATTCTAGAGCTAATA</td><td>>hg38_21_+_8433172_8446622_RNA45SN1_45S_50ntFlanks</td><td>3840</td><td>3877</td><td>38</td><td>CTCCTACTTGGATAACTGTGGTAATTCTAGAGCTAATA</td><td>568.0</td><td>7.3</td><td>1.2</td><td>6.1</td><td>7.8E-10</td><td>2.0E-09</td><td>BT20/MB468</td><td>CYTO</td></tr>
<tr><td>>trnaMT_GluTTC_MT_-_14674_14742@39.70.32,trnalookalike8_GluTTC_5_-_93905172_93905240@39.70.32|ATTGGTCGTGGTTGTAGTCCGTGCGAGAATAC</td><td>>trnaMT_GluTTC_MT_-_14674_14742</td><td>39</td><td>70</td><td>32</td><td>ATTGGTCGTGGTTGTAGTCCGTGCGAGAATAC</td><td>730.0</td><td>10.2</td><td>1.6</td><td>6.2</td><td>6.7E-10</td><td>2.1E-09</td><td>BT20/MB468</td><td>MITO</td></tr>
<tr><td>>am_trnaMT_ArgTCG_MT_+_10405_10469@28.48.21|ATTTCGACTCATTAAATTATG</td><td>>am_trnaMT_ArgTCG_MT_+_10405_10469</td><td>28</td><td>48</td><td>21</td><td>ATTTCGACTCATTAAATTATG</td><td>5302.2</td><td>-11.7</td><td>1.9</td><td>-6.1</td><td>1.0E-09</td><td>2.1E-09</td><td>BT20/MB468</td><td>NUC</td></tr>
<tr><td>>hg38_21_+_8433172_8446622_RNA45SN1_45S_50ntFlanks@5489.5516.28|CTGAGAAGACGGTCGAACTTGACTATCT</td><td>>hg38_21_+_8433172_8446622_RNA45SN1_45S_50ntFlanks</td><td>5489</td><td>5516</td><td>28</td><td>CTGAGAAGACGGTCGAACTTGACTATCT</td><td>27625.8</td><td>6.1</td><td>1.0</td><td>6.1</td><td>7.8E-10</td><td>2.1E-09</td><td>BT20/MB468</td><td>CYTO</td></tr>
</table>

<table>
<tr><td>>am_MI0000239|hsa-mir-197&WithFlank&1|+|109598887|109598973@54.71.18(+1U)&[MIMAT0000227&hsa-miR-197-3p&offsets|0|-4(+1U);m-99&1|+|109598940|109598961&offsets|0|-4(+1U)]|TTCACCACCTTCTCCACCT</td><td>>am_MI0000239|hsa-mir-197&WithFlank&1|+|109598887|109598973</td><td>54</td><td>71</td><td>18(+1U)</td><td>TTCACCACCTTCTCCACCT</td><td>71.7</td><td>4.0</td><td>0.6</td><td>6.1</td><td>1.0E-09</td><td>2.1E-09</td><td>BT20/MB468</td><td>NUC</td></tr>
<tr><td>>hg38_21_+_8433172_8446622_RNA45SN1_45S_50ntFlanks@11630.11660.31|GCGAAGGCCCGCGGCGGGTGTTGACGCGATG</td><td>>hg38_21_+_8433172_8446622_RNA45SN1_45S_50ntFlanks</td><td>11630</td><td>11660</td><td>31</td><td>GCGAAGGCCCGCGGCGGGTGTTGACGCGATG</td><td>256.5</td><td>-11.1</td><td>1.7</td><td>-6.5</td><td>8.9E-11</td><td>2.1E-09</td><td>BT20/MB468</td><td>MP</td></tr>
<tr><td>>hg38_21_+_8433172_8446622_RNA45SN1_45S_50ntFlanks@3593.3610.18|CCCGTCCGTCCGTCCGTC</td><td>>hg38_21_+_8433172_8446622_RNA45SN1_45S_50ntFlanks</td><td>3593</td><td>3610</td><td>18</td><td>CCCGTCCGTCCGTCCGTC</td><td>268.0</td><td>-11.1</td><td>1.7</td><td>-6.5</td><td>8.9E-11</td><td>2.1E-09</td><td>BT20/MB468</td><td>MP</td></tr>
<tr><td>>hg38_21_+_8433172_8446622_RNA45SN1_45S_50ntFlanks@8737.8754.18|CCCGTCCGTCCGTCCGTC</td><td>>hg38_21_+_8433172_8446622_RNA45SN1_45S_50ntFlanks</td><td>8737</td><td>8754</td><td>18</td><td>CCCGTCCGTCCGTCCGTC</td><td>268.0</td><td>-11.1</td><td>1.7</td><td>-6.5</td><td>8.9E-11</td><td>2.1E-09</td><td>BT20/MB468</td><td>MP</td></tr>
<tr><td>>hg38_21_+_8433172_8446622_RNA45SN1_45S_50ntFlanks@4047.4068.22|ATTCGAACGTCTGCCCTATCAA</td><td>>hg38_21_+_8433172_8446622_RNA45SN1_45S_50ntFlanks</td><td>4047</td><td>4068</td><td>22</td><td>ATTCGAACGTCTGCCCTATCAA</td><td>244.9</td><td>-10.0</td><td>1.6</td><td>-6.2</td><td>6.8E-10</td><td>2.1E-09</td><td>BT20/MB468</td><td>MITO</td></tr>
<tr><td>>hg38_21_+_8433172_8446622_RNA45SN1_45S_50ntFlanks@6653.6676.24|ACTCTTAGCGGTGGATCACTCGGC</td><td>>hg38_21_+_8433172_8446622_RNA45SN1_45S_50ntFlanks</td><td>6653</td><td>6676</td><td>24</td><td>ACTCTTAGCGGTGGATCACTCGGC</td><td>399.2</td><td>5.3</td><td>0.9</td><td>6.1</td><td>9.1E-10</td><td>2.1E-09</td><td>MB231/MB468</td><td>NUC</td></tr>
<tr><td>>hg38_21_+_8433172_8446622_RNA45SN1_45S_50ntFlanks@6648.6669.22|GTACGACTCTTAGCGGTGGATC</td><td>>hg38_21_+_8433172_8446622_RNA45SN1_45S_50ntFlanks</td><td>6648</td><td>6669</td><td>22</td><td>GTACGACTCTTAGCGGTGGATC</td><td>1579.4</td><td>4.2</td><td>0.7</td><td>6.1</td><td>7.9E-10</td><td>2.1E-09</td><td>BT20/MB468</td><td>CYTO</td></tr>
<tr><td>>hg38_21_+_8433172_8446622_RNA45SN1_45S_50ntFlanks@11586.11605.20|GTGCGGACCAGGGGAATCCG</td><td>>hg38_21_+_8433172_8446622_RNA45SN1_45S_50ntFlanks</td><td>11586</td><td>11605</td><td>20</td><td>GTGCGGACCAGGGGAATCCG</td><td>322.1</td><td>-6.7</td><td>1.1</td><td>-6.1</td><td>8.9E-10</td><td>2.1E-09</td><td>MB231/MB468</td><td>CYTO</td></tr>
<tr><td>>hg38_21_+_8433172_8446622_RNA45SN1_45S_50ntFlanks@475.516.42|CTTCGTGATCGATGTGGTGACGTCGTGCTCTCCCGGGCCGGG</td><td>>hg38_21_+_8433172_8446622_RNA45SN1_45S_50ntFlanks</td><td>475</td><td>516</td><td>42</td><td>CTTCGTGATCGATGTGGTGACGTCGTGCTCTCCCGGGCCGGG</td><td>3783.0</td><td>8.0</td><td>1.3</td><td>6.2</td><td>6.8E-10</td><td>2.1E-09</td><td>BT20/MB468</td><td>MITO</td></tr>
<tr><td>>hg38_21_+_8433172_8446622_RNA45SN1_45S_50ntFlanks@11161.11178.18|CACGGTCCCCCGCGAGGG</td><td>>hg38_21_+_8433172_8446622_RNA45SN1_45S_50ntFlanks</td><td>11161</td><td>11178</td><td>18</td><td>CACGGTCCCCCGCGAGGG</td><td>3469.9</td><td>-10.1</td><td>1.7</td><td>-6.1</td><td>1.0E-09</td><td>2.1E-09</td><td>BT20/MB468</td><td>NUC</td></tr>
<tr><td>>hg38_21_+_8433172_8446622_RNA45SN1_45S_50ntFlanks@11704.11740.37|ATGAAGCGCGGGTAAACGGCGGGAGTAACTATGACTC</td><td>>hg38_21_+_8433172_8446622_RNA45SN1_45S_50ntFlanks</td><td>11704</td><td>11740</td><td>37</td><td>ATGAAGCGCGGGTAAACGGCGGGAGTAACTATGACTC</td><td>689.6</td><td>-9.0</td><td>1.5</td><td>-6.1</td><td>1.0E-09</td><td>2.1E-09</td><td>BT20/MB468</td><td>NUC</td></tr>
<tr><td>>hg38_21_+_8433172_8446622_RNA45SN1_45S_50ntFlanks@5492.5524.33|AGAAGACGGTCGAACTTGACTATCTAGAGGAAG</td><td>>hg38_21_+_8433172_8446622_RNA45SN1_45S_50ntFlanks</td><td>5492</td><td>5524</td><td>33</td><td>AGAAGACGGTCGAACTTGACTATCTAGAGGAAG</td><td>291.1</td><td>-6.4</td><td>1.0</td><td>-6.1</td><td>8.7E-10</td><td>2.1E-09</td><td>MB231/MB468</td><td>MP</td></tr>
<tr><td>>hg38_21_+_8433172_8446622_RNA45SN1_45S_50ntFlanks@4760.4782.23|TCAGATACCGTCGTAGTTCCGAC</td><td>>hg38_21_+_8433172_8446622_RNA45SN1_45S_50ntFlanks</td><td>4760</td><td>4782</td><td>23</td><td>TCAGATACCGTCGTAGTTCCGAC</td><td>1357.4</td><td>-6.3</td><td>1.0</td><td>-6.1</td><td>9.0E-10</td><td>2.1E-09</td><td>MB231/MB468</td><td>CYTO</td></tr>
<tr><td>>hg38_21_+_8433172_8446622_RNA45SN1_45S_50ntFlanks@12997.13036.40|CGCTGCGATCTATTGAAAGTCAGCCCTCGACACAAGGGTT</td><td>>hg38_21_+_8433172_8446622_RNA45SN1_45S_50ntFlanks</td><td>12997</td><td>13036</td><td>40</td><td>CGCTGCGATCTATTGAAAGTCAGCCCTCGACACAAGGGTT</td><td>2531.3</td><td>6.9</td><td>1.1</td><td>6.2</td><td>6.9E-10</td><td>2.1E-09</td><td>BT20/MB468</td><td>MITO</td></tr>
<tr><td>>hg38_21_+_8433172_8446622_RNA45SN1_45S_50ntFlanks@12407.12437.31|TTTTGATCCTTCGATGTCGGCTCTTCCTATC</td><td>>hg38_21_+_8433172_8446622_RNA45SN1_45S_50ntFlanks</td><td>12407</td><td>12437</td><td>31</td><td>TTTTGATCCTTCGATGTCGGCTCTTCCTATC</td><td>1442.7</td><td>-4.3</td><td>0.7</td><td>-6.1</td><td>1.0E-09</td><td>2.1E-09</td><td>MB231/MB468</td><td>MITO</td></tr>
<tr><td>>hg38_21_+_8433172_8446622_RNA45SN1_45S_50ntFlanks@8109.8136.28|GCGGGGCGCGGGACATGTGGCGTACGGA</td><td>>hg38_21_+_8433172_8446622_RNA45SN1_45S_50ntFlanks</td><td>8109</td><td>8136</td><td>28</td><td>GCGGGGCGCGGGACATGTGGCGTACGGA</td><td>140.3</td><td>-11.0</td><td>1.8</td><td>-6.1</td><td>8.8E-10</td><td>2.1E-09</td><td>MB231/MB468</td><td>MP</td></tr>
<tr><td>>MI0000102|hsa-mir-100&WithFlank&11|-|122152223|122152314@19.37.19(+1A)&[MIMAT0000098&hsa-miR-100-5p&offsets|0|-3(+1A);m-27&11|-|122152276|122152296&offsets|0|-2(+1A)]|AACCCGTAGATCCGAACTTA</td><td>>MI0000102|hsa-mir-100&WithFlank&11|-|122152223|122152314</td><td>19</td><td>37</td><td>19(+1A)</td><td>AACCCGTAGATCCGAACTTA</td><td>2565.2</td><td>-7.0</td><td>1.1</td><td>-6.5</td><td>7.8E-11</td><td>2.1E-09</td><td>BT20/MB231</td><td>MITO</td></tr>
<tr><td>>hg38_21_+_8433172_8446622_RNA45SN1_45S_50ntFlanks@11580.11607.28|GAACTGGTGCGGACCAGGGGAATCCGAC</td><td>>hg38_21_+_8433172_8446622_RNA45SN1_45S_50ntFlanks</td><td>11580</td><td>11607</td><td>28</td><td>GAACTGGTGCGGACCAGGGGAATCCGAC</td><td>623.2</td><td>-4.0</td><td>0.7</td><td>-6.1</td><td>1.0E-09</td><td>2.1E-09</td><td>MB231/MB468</td><td>MITO</td></tr>
<tr><td>>hg38_21_+_8433172_8446622_RNA45SN1_45S_50ntFlanks@10350.10374.25|ACGAGAACTTTGAAGGCCGAAGTGG</td><td>>hg38_21_+_8433172_8446622_RNA45SN1_45S_50ntFlanks</td><td>10350</td><td>10374</td><td>25</td><td>ACGAGAACTTTGAAGGCCGAAGTGG</td><td>5020.0</td><td>-6.5</td><td>1.0</td><td>-6.5</td><td>9.1E-11</td><td>2.1E-09</td><td>BT20/MB468</td><td>MP</td></tr>
<tr><td>>hg38_21_+_8433172_8446622_RNA45SN1_45S_50ntFlanks@6645.6667.23|CTCGTACGACTCTTAGCGGTGGA</td><td>>hg38_21_+_8433172_8446622_RNA45SN1_45S_50ntFlanks</td><td>6645</td><td>6667</td><td>23</td><td>CTCGTACGACTCTTAGCGGTGGA</td><td>1099.2</td><td>3.6</td><td>0.6</td><td>6.1</td><td>9.3E-10</td><td>2.1E-09</td><td>MB231/MB468</td><td>NUC</td></tr>
<tr><td>>hg38_21_+_8433172_8446622_RNA45SN1_45S_50ntFlanks@9739.9764.26|AAACGATCTCAACCTATTCTCAAACT</td><td>>hg38_21_+_8433172_8446622_RNA45SN1_45S_50ntFlanks</td><td>9739</td><td>9764</td><td>26</td><td>AAACGATCTCAACCTATTCTCAAACT</td><td>543.4</td><td>-4.6</td><td>0.8</td><td>-6.1</td><td>1.0E-09</td><td>2.1E-09</td><td>MB231/MB468</td><td>MITO</td></tr>
<tr><td>>hg38_21_+_8433172_8446622_RNA45SN1_45S_50ntFlanks@12626.12649.24|AATGGGGCGAAGCTACCATCTGTG</td><td>>hg38_21_+_8433172_8446622_RNA45SN1_45S_50ntFlanks</td><td>12626</td><td>12649</td><td>24</td><td>AATGGGGCGAAGCTACCATCTGTG</td><td>134.5</td><td>-9.2</td><td>1.5</td><td>-6.2</td><td>6.9E-10</td><td>2.1E-09</td><td>BT20/MB468</td><td>MITO</td></tr>
<tr><td>>am_MI0000809|hsa-mir-151a&WithFlank&8|-|140732558|140732659@53.72.20(+1A)&[MIMAT0000757&hsa-miR-151a-3p&offsets|0|-1(+1A);m-30&8|-|140732587|140732607&offsets|0|-1(+1A)]|CTAGACTGAAGCTCCTTGAGA</td><td>>am_MI0000809|hsa-mir-151a&WithFlank&8|-|140732558|140732659</td><td>53</td><td>72</td><td>20(+1A)</td><td>CTAGACTGAAGCTCCTTGAGA</td><td>12691.4</td><td>6.0</td><td>0.9</td><td>6.5</td><td>9.1E-11</td><td>2.1E-09</td><td>BT20/MB468</td><td>MP</td></tr>
<tr><td>>hg38_21_+_8433172_8446622_RNA45SN1_45S_50ntFlanks@4007.4025.19|CCTCGGGCCGATCGCACGC</td><td>>hg38_21_+_8433172_8446622_RNA45SN1_45S_50ntFlanks</td><td>4007</td><td>4025</td><td>19</td><td>CCTCGGGCCGATCGCACGC</td><td>516.8</td><td>-8.3</td><td>1.4</td><td>-6.1</td><td>1.0E-09</td><td>2.1E-09</td><td>MB231/MB468</td><td>MITO</td></tr>
<tr><td>>hg38_21_+_8433172_8446622_RNA45SN1_45S_50ntFlanks@6698.6718.21|AGCTAGCTGCGAGAATTAATG</td><td>>hg38_21_+_8433172_8446622_RNA45SN1_45S_50ntFlanks</td><td>6698</td><td>6718</td><td>21</td><td>AGCTAGCTGCGAGAATTAATG</td><td>6748.8</td><td>-4.2</td><td>0.7</td><td>-6.1</td><td>9.2E-10</td><td>2.1E-09</td><td>MB231/MB468</td><td>CYTO</td></tr>
<tr><td>>MI0000734|hsa-mir-106b&WithFlank&7|-|100093987|100094080@58.78.21(+1U)&[MIMAT0004672&hsa-miR-106b-3p&offsets|0|-1(+1U);m-87&7|-|100094003|100094023&offsets|0|0(+1U)]|CCGCACTGTGGGTACTTGCTGT</td><td>>MI0000734|hsa-mir-106b&WithFlank&7|-|100093987|100094080</td><td>58</td><td>78</td><td>21(+1U)</td><td>CCGCACTGTGGGTACTTGCTGT</td><td>662.1</td><td>5.1</td><td>0.8</td><td>6.1</td><td>1.0E-09</td><td>2.1E-09</td><td>MB231/MB468</td><td>MITO</td></tr>
<tr><td>>hg38_21_+_8433172_8446622_RNA45SN1_45S_50ntFlanks@11667.11687.21|CTGCCCAGTGCTCTGAATGTC</td><td>>hg38_21_+_8433172_8446622_RNA45SN1_45S_50ntFlanks</td><td>11667</td><td>11687</td><td>21</td><td>CTGCCCAGTGCTCTGAATGTC</td><td>429.6</td><td>3.6</td><td>0.6</td><td>6.1</td><td>9.4E-10</td><td>2.1E-09</td><td>MB231/MB468</td><td>NUC</td></tr>
<tr><td>>hg38_21_+_8433172_8446622_RNA45SN1_45S_50ntFlanks@11233.11250.18|CCCTTCCCGTGGATCGCC</td><td>>hg38_21_+_8433172_8446622_RNA45SN1_45S_50ntFlanks</td><td>11233</td><td>11250</td><td>18</td><td>CCCTTCCCGTGGATCGCC</td><td>165.5</td><td>-10.5</td><td>1.6</td><td>-6.5</td><td>9.2E-11</td><td>2.1E-09</td><td>BT20/MB468</td><td>MP</td></tr>
<tr><td>>hg38_21_+_8433172_8446622_RNA45SN1_45S_50ntFlanks@9564.9598.35|CTGACGTGCAAATCGGTCGTCCGACCTGGGTATAG</td><td>>hg38_21_+_8433172_8446622_RNA45SN1_45S_50ntFlanks</td><td>9564</td><td>9598</td><td>35</td><td>CTGACGTGCAAATCGGTCGTCCGACCTGGGTATAG</td><td>91.4</td><td>-10.4</td><td>1.7</td><td>-6.1</td><td>8.9E-10</td><td>2.1E-09</td><td>MB231/MB468</td><td>MP</td></tr>
<tr><td>>hg38_21_+_8433172_8446622_RNA45SN1_45S_50ntFlanks@12415.12455.41|CTTCGATGTCGGCTCTTCCTATCATTGTGAAGCAGAATTCA</td><td>>hg38_21_+_8433172_8446622_RNA45SN1_45S_50ntFlanks</td><td>12415</td><td>12455</td><td>41</td><td>CTTCGATGTCGGCTCTTCCTATCATTGTGAAGCAGAATTCA</td><td>170.5</td><td>-11.3</td><td>1.8</td><td>-6.1</td><td>8.9E-10</td><td>2.1E-09</td><td>MB231/MB468</td><td>MP</td></tr>
<tr><td>>hg38_21_+_8433172_8446622_RNA45SN1_45S_50ntFlanks@5045.5072.28|CTTAGTTGGTGGAGCGATTTGTCTGGTT</td><td>>hg38_21_+_8433172_8446622_RNA45SN1_45S_50ntFlanks</td><td>5045</td><td>5072</td><td>28</td><td>CTTAGTTGGTGGAGCGATTTGTCTGGTT</td><td>114.3</td><td>-10.7</td><td>1.7</td><td>-6.1</td><td>8.9E-10</td><td>2.1E-09</td><td>MB231/MB468</td><td>MP</td></tr>
<tr><td>>hg38_21_+_8433172_8446622_RNA45SN1_45S_50ntFlanks@10821.10838.18|GGAAGTCGGCAAGCCGGA</td><td>>hg38_21_+_8433172_8446622_RNA45SN1_45S_50ntFlanks</td><td>10821</td><td>10838</td><td>18</td><td>GGAAGTCGGCAAGCCGGA</td><td>774.6</td><td>-4.3</td><td>0.7</td><td>-6.1</td><td>9.3E-10</td><td>2.1E-09</td><td>MB231/MB468</td><td>CYTO</td></tr>
<tr><td>>am_trnaMT_LysTTT_MT_+_8295_8364@35.52.18,trnalookalike7_LysTTT_1_+_568844_568913@35.52.18|TTAAAGATTAAGAGAACC</td><td>>am_trnaMT_LysTTT_MT_+_8295_8364</td><td>35</td><td>52</td><td>18</td><td>TTAAAGATTAAGAGAACC</td><td>365.1</td><td>9.2</td><td>1.5</td><td>6.1</td><td>8.1E-10</td><td>2.1E-09</td><td>BT20/MB468</td><td>CYTO</td></tr>
<tr><td>>hg38_21_+_8433172_8446622_RNA45SN1_45S_50ntFlanks@5081.5102.22|TAACGAACGAGACTCTGGCATG</td><td>>hg38_21_+_8433172_8446622_RNA45SN1_45S_50ntFlanks</td><td>5081</td><td>5102</td><td>22</td><td>TAACGAACGAGACTCTGGCATG</td><td>117.1</td><td>-10.7</td><td>1.8</td><td>-6.1</td><td>9.0E-10</td><td>2.1E-09</td><td>MB231/MB468</td><td>MP</td></tr>
<tr><td>>hg38_21_+_8433172_8446622_RNA45SN1_45S_50ntFlanks@3999.4024.26|CTAGATAACCTCGGGCCGATCGCACG</td><td>>hg38_21_+_8433172_8446622_RNA45SN1_45S_50ntFlanks</td><td>3999</td><td>4024</td><td>26</td><td>CTAGATAACCTCGGGCCGATCGCACG</td><td>299.0</td><td>-7.8</td><td>1.3</td><td>-6.1</td><td>1.0E-09</td><td>2.1E-09</td><td>MB231/MB468</td><td>MITO</td></tr>
<tr><td>>hg38_21_+_8433172_8446622_RNA45SN1_45S_50ntFlanks@5078.5117.40|CGATAACGAACGAGACTCTGGCATGCTAACTAGTTACGCG</td><td>>hg38_21_+_8433172_8446622_RNA45SN1_45S_50ntFlanks</td><td>5078</td><td>5117</td><td>40</td><td>CGATAACGAACGAGACTCTGGCATGCTAACTAGTTACGCG</td><td>254.3</td><td>3.4</td><td>0.6</td><td>6.1</td><td>9.3E-10</td><td>2.2E-09</td><td>MB231/MB468</td><td>CYTO</td></tr>
<tr><td>>hg38_21_+_8433172_8446622_RNA45SN1_45S_50ntFlanks@6728.6760.33|GGACACATTGATCATCGACACTTCGAACGCACT</td><td>>hg38_21_+_8433172_8446622_RNA45SN1_45S_50ntFlanks</td><td>6728</td><td>6760</td><td>33</td><td>GGACACATTGATCATCGACACTTCGAACGCACT</td><td>137.6</td><td>7.3</td><td>1.2</td><td>6.1</td><td>9.3E-10</td><td>2.2E-09</td><td>MB231/MB468</td><td>CYTO</td></tr>
<tr><td>>hg38_21_+_8433172_8446622_RNA45SN1_45S_50ntFlanks@10689.10709.21|GCGTCCGGTGAGCTCTCGCTG</td><td>>hg38_21_+_8433172_8446622_RNA45SN1_45S_50ntFlanks</td><td>10689</td><td>10709</td><td>21</td><td>GCGTCCGGTGAGCTCTCGCTG</td><td>894.7</td><td>-8.2</td><td>1.3</td><td>-6.1</td><td>9.5E-10</td><td>2.2E-09</td><td>MB231/MB468</td><td>NUC</td></tr>
<tr><td>>hg38_21_+_8433172_8446622_RNA45SN1_45S_50ntFlanks@8357.8376.20|AAAAGAACTTTGAAGAGAGA</td><td>>hg38_21_+_8433172_8446622_RNA45SN1_45S_50ntFlanks</td><td>8357</td><td>8376</td><td>20</td><td>AAAAGAACTTTGAAGAGAGA</td><td>2664.1</td><td>-7.7</td><td>1.3</td><td>-6.1</td><td>9.0E-10</td><td>2.2E-09</td><td>MB231/MB468</td><td>MP</td></tr>
<tr><td>>hg38_21_+_8433172_8446622_RNA45SN1_45S_50ntFlanks@12945.12964.20|GTAGACGACCTGCTTCTGGG</td><td>>hg38_21_+_8433172_8446622_RNA45SN1_45S_50ntFlanks</td><td>12945</td><td>12964</td><td>20</td><td>GTAGACGACCTGCTTCTGGG</td><td>115.0</td><td>-10.7</td><td>1.7</td><td>-6.1</td><td>9.0E-10</td><td>2.2E-09</td><td>MB231/MB468</td><td>MP</td></tr>
<tr><td>>hg38_21_+_8433172_8446622_RNA45SN1_45S_50ntFlanks@11027.11053.27|CTCTCTCTCTCTCTCCCCCGCTCCCCG</td><td>>hg38_21_+_8433172_8446622_RNA45SN1_45S_50ntFlanks</td><td>11027</td><td>11053</td><td>27</td><td>CTCTCTCTCTCTCTCCCCCGCTCCCCG</td><td>2294.2</td><td>-9.6</td><td>1.6</td><td>-6.1</td><td>9.5E-10</td><td>2.2E-09</td><td>MB231/MB468</td><td>NUC</td></tr>
<tr><td>>hg38_21_+_8433172_8446622_RNA45SN1_45S_50ntFlanks@7977.8006.30|CGACCTCAGATCAGACGTGGCGACCCGCTG</td><td>>hg38_21_+_8433172_8446622_RNA45SN1_45S_50ntFlanks</td><td>7977</td><td>8006</td><td>30</td><td>CGACCTCAGATCAGACGTGGCGACCCGCTG</td><td>1677.1</td><td>-8.7</td><td>1.4</td><td>-6.1</td><td>9.1E-10</td><td>2.2E-09</td><td>MB231/MB468</td><td>MP</td></tr>
<tr><td>>MI0000446|hsa-mir-125b-1&WithFlank&11|-|122099751|122099850@61.80.20(+1U)&[MIMAT0004592&hsa-miR-125b-1-3p&offsets|0|-2(+1U);m-485&11|-|122099769|122099790&offsets|0|-2(+1U)]|ACGGGTTAGGCTCTTGGGAGT</td><td>>MI0000446|hsa-mir-125b-1&WithFlank&11|-|122099751|122099850</td><td>61</td><td>80</td><td>20(+1U)</td><td>ACGGGTTAGGCTCTTGGGAGT</td><td>135.8</td><td>10.3</td><td>1.7</td><td>6.1</td><td>1.0E-09</td><td>2.2E-09</td><td>MB231/MB468</td><td>MITO</td></tr>
<tr><td>>hg38_21_+_8433172_8446622_RNA45SN1_45S_50ntFlanks@3846.3868.23|CTTGGATAACTGTGGTAATTCTA</td><td>>hg38_21_+_8433172_8446622_RNA45SN1_45S_50ntFlanks</td><td>3846</td><td>3868</td><td>23</td><td>CTTGGATAACTGTGGTAATTCTA</td><td>944.1</td><td>-7.4</td><td>1.2</td><td>-6.1</td><td>9.1E-10</td><td>2.2E-09</td><td>MB231/MB468</td><td>MP</td></tr>
<tr><td>>hg38_21_+_8433172_8446622_RNA45SN1_45S_50ntFlanks@10393.10433.41|CAGCAGTTGAACATGGGTCAGTCGGTCCTGAGAGATGGGCG</td><td>>hg38_21_+_8433172_8446622_RNA45SN1_45S_50ntFlanks</td><td>10393</td><td>10433</td><td>41</td><td>CAGCAGTTGAACATGGGTCAGTCGGTCCTGAGAGATGGGCG</td><td>631.4</td><td>-5.1</td><td>0.8</td><td>-6.1</td><td>1.0E-09</td><td>2.2E-09</td><td>MB231/MB468</td><td>MITO</td></tr>
<tr><td>>hg38_21_+_8433172_8446622_RNA45SN1_45S_50ntFlanks@11217.11250.34|CTGGACGCGAGCCGGGCCCTTCCCGTGGATCGCC</td><td>>hg38_21_+_8433172_8446622_RNA45SN1_45S_50ntFlanks</td><td>11217</td><td>11250</td><td>34</td><td>CTGGACGCGAGCCGGGCCCTTCCCGTGGATCGCC</td><td>418.3</td><td>-6.6</td><td>1.1</td><td>-6.1</td><td>9.4E-10</td><td>2.2E-09</td><td>MB231/MB468</td><td>CYTO</td></tr>
<tr><td>>hg38_21_+_8433172_8446622_RNA45SN1_45S_50ntFlanks@8355.8390.36|TGAAAAGAACTTTGAAGAGAGAGTTCAAGAGGGCGT</td><td>>hg38_21_+_8433172_8446622_RNA45SN1_45S_50ntFlanks</td><td>8355</td><td>8390</td><td>36</td><td>TGAAAAGAACTTTGAAGAGAGAGTTCAAGAGGGCGT</td><td>119.5</td><td>10.6</td><td>1.6</td><td>6.5</td><td>9.4E-11</td><td>2.2E-09</td><td>BT20/MB468</td><td>MP</td></tr>
<tr><td>>MI0000477|hsa-mir-146a&WithFlank&5|+|160485346|160485456@27.47.21(+1G)&[MIMAT0000449&hsa-miR-146a-5p&offsets|0|-1(+1G);m-129&5|+|160485372|160485393&offsets|0|-1(+1G)]|TGAGAACTGAATTCCATGGGTG</td><td>>MI0000477|hsa-mir-146a&WithFlank&5|+|160485346|160485456</td><td>27</td><td>47</td><td>21(+1G)</td><td>TGAGAACTGAATTCCATGGGTG</td><td>291.3</td><td>-11.3</td><td>1.7</td><td>-6.5</td><td>9.9E-11</td><td>2.2E-09</td><td>BT20/MB231</td><td>NUC</td></tr>
<tr><td>>hg38_21_+_8433172_8446622_RNA45SN1_45S_50ntFlanks@7985.8018.34|GATCAGACGTGGCGACCCGCTGAATTTAAGCATA</td><td>>hg38_21_+_8433172_8446622_RNA45SN1_45S_50ntFlanks</td><td>7985</td><td>8018</td><td>34</td><td>GATCAGACGTGGCGACCCGCTGAATTTAAGCATA</td><td>148.2</td><td>-9.3</td><td>1.5</td><td>-6.2</td><td>7.2E-10</td><td>2.2E-09</td><td>BT20/MB468</td><td>MITO</td></tr>
<tr><td>>hg38_21_+_8433172_8446622_RNA45SN1_45S_50ntFlanks@12490.12518.29|CGTGAGCTGGGTTTAGACCGTCGTGAGAC</td><td>>hg38_21_+_8433172_8446622_RNA45SN1_45S_50ntFlanks</td><td>12490</td><td>12518</td><td>29</td><td>CGTGAGCTGGGTTTAGACCGTCGTGAGAC</td><td>87.0</td><td>-10.3</td><td>1.7</td><td>-6.1</td><td>9.1E-10</td><td>2.2E-09</td><td>MB231/MB468</td><td>MP</td></tr>
</table>

<table>
<tr><td>>hg38_21_+_8433172_8446622_RNA45SN1_45S_50ntFlanks@12476.12500.25|CCACTAATAGGGAACGTGAGCTGGG</td><td>>hg38_21_+_8433172_8446622_RNA45SN1_45S_50ntFlanks</td><td>12476</td><td>12500</td><td>25</td><td>CCACTAATAGGGAACGTGAGCTGGG</td><td>310.6</td><td>-7.7</td><td>1.3</td><td>-6.1</td><td>1.1E-09</td><td>2.2E-09</td><td>MB231/MB468</td><td>MITO</td></tr>
<tr><td>>hg38_21_+_8433172_8446622_RNA45SN1_45S_50ntFlanks@12457.12475.19|CAAGCGTTGGATTGTTCAC</td><td>>hg38_21_+_8433172_8446622_RNA45SN1_45S_50ntFlanks</td><td>12457</td><td>12475</td><td>19</td><td>CAAGCGTTGGATTGTTCAC</td><td>1709.1</td><td>-6.5</td><td>1.1</td><td>-6.1</td><td>9.5E-10</td><td>2.2E-09</td><td>MB231/MB468</td><td>CYTO</td></tr>
<tr><td>>hg38_21_+_8433172_8446622_RNA45SN1_45S_50ntFlanks@6719.6741.23|TGAATTGCAGGACACATTGATCA</td><td>>hg38_21_+_8433172_8446622_RNA45SN1_45S_50ntFlanks</td><td>6719</td><td>6741</td><td>23</td><td>TGAATTGCAGGACACATTGATCA</td><td>503.6</td><td>-6.2</td><td>1.0</td><td>-6.1</td><td>9.7E-10</td><td>2.2E-09</td><td>MB231/MB468</td><td>NUC</td></tr>
<tr><td>>hg38_21_+_8433172_8446622_RNA45SN1_45S_50ntFlanks@10349.10382.34|AACGAGAACTTTGAAGGCCGAAGTGGAGAAGGGT</td><td>>hg38_21_+_8433172_8446622_RNA45SN1_45S_50ntFlanks</td><td>10349</td><td>10382</td><td>34</td><td>AACGAGAACTTTGAAGGCCGAAGTGGAGAAGGGT</td><td>1189.3</td><td>-4.0</td><td>0.7</td><td>-6.1</td><td>1.1E-09</td><td>2.2E-09</td><td>BT20/MB468</td><td>NUC</td></tr>
<tr><td>>hg38_21_+_8433172_8446622_RNA45SN1_45S_50ntFlanks@10350.10388.39|ACGAGAACTTTGAAGGCCGAAGTGGAGAAGGGTTCCATG</td><td>>hg38_21_+_8433172_8446622_RNA45SN1_45S_50ntFlanks</td><td>10350</td><td>10388</td><td>39</td><td>ACGAGAACTTTGAAGGCCGAAGTGGAGAAGGGTTCCATG</td><td>2723.6</td><td>-9.3</td><td>1.5</td><td>-6.1</td><td>9.2E-10</td><td>2.2E-09</td><td>MB231/MB468</td><td>MP</td></tr>
<tr><td>>hg38_21_+_8433172_8446622_RNA45SN1_45S_50ntFlanks@11793.11810.18|TGAACGAGATTCCCACTG</td><td>>hg38_21_+_8433172_8446622_RNA45SN1_45S_50ntFlanks</td><td>11793</td><td>11810</td><td>18</td><td>TGAACGAGATTCCCACTG</td><td>96.9</td><td>-10.5</td><td>1.7</td><td>-6.1</td><td>9.2E-10</td><td>2.2E-09</td><td>MB231/MB468</td><td>MP</td></tr>
<tr><td>>hg38_21_+_8433172_8446622_RNA45SN1_45S_50ntFlanks@10357.10383.27|CTTTGAAGGCCGAAGTGGAGAAGGGTT</td><td>>hg38_21_+_8433172_8446622_RNA45SN1_45S_50ntFlanks</td><td>10357</td><td>10383</td><td>27</td><td>CTTTGAAGGCCGAAGTGGAGAAGGGTT</td><td>1032.0</td><td>-6.5</td><td>1.1</td><td>-6.1</td><td>9.7E-10</td><td>2.2E-09</td><td>MB231/MB468</td><td>NUC</td></tr>
<tr><td>>hg38_21_+_8433172_8446622_RNA45SN1_45S_50ntFlanks@3961.3983.23|CCCCGGCCGGGGGGCGGGCGCCG</td><td>>hg38_21_+_8433172_8446622_RNA45SN1_45S_50ntFlanks</td><td>3961</td><td>3983</td><td>23</td><td>CCCCGGCCGGGGGGCGGGCGCCG</td><td>81.5</td><td>-10.2</td><td>1.7</td><td>-6.1</td><td>9.2E-10</td><td>2.2E-09</td><td>MB231/MB468</td><td>MP</td></tr>
<tr><td>>hg38_21_+_8433172_8446622_RNA45SN1_45S_50ntFlanks@11572.11605.34|CCGACTTAGAACTGGTGCGGACCAGGGGAATCCG</td><td>>hg38_21_+_8433172_8446622_RNA45SN1_45S_50ntFlanks</td><td>11572</td><td>11605</td><td>34</td><td>CCGACTTAGAACTGGTGCGGACCAGGGGAATCCG</td><td>2780.3</td><td>-4.6</td><td>0.7</td><td>-6.1</td><td>9.5E-10</td><td>2.2E-09</td><td>MB231/MB468</td><td>CYTO</td></tr>
<tr><td>>hg38_21_+_8433172_8446622_RNA45SN1_45S_50ntFlanks@11076.11103.28|CGCCGCGTGGGGGCGGCGGCGGGGGGAG</td><td>>hg38_21_+_8433172_8446622_RNA45SN1_45S_50ntFlanks</td><td>11076</td><td>11103</td><td>28</td><td>CGCCGCGTGGGGGCGGCGGCGGGGGGAG</td><td>237.6</td><td>-10.0</td><td>1.6</td><td>-6.2</td><td>7.3E-10</td><td>2.2E-09</td><td>BT20/MB468</td><td>MITO</td></tr>
<tr><td>>trna10_AspGTC_12_-_125424193_125424264@32.73.42,trna12_AspGTC_12_-_125411891_125411962@32.73.42,trna144_AspGTC_6_-_27551236_27551307@32.73.42,trna38_AspGTC_17_-_8125556_8125627@32.73.42,trna45_AspGTC_6_+_27447453_27447524@32.73.42,trna48_AspGTC_6_+_27471523_27471594@32.73.42,trna4_AspGTC_12_+_96429799_96429870@32.73.42,trna69_AspGTC_1_-_161440205_161440276@32.73.42,trna72_AspGTC_1_-_161432824_161432895@32.73.42,trna75_AspGTC_1_-_161425414_161425485@32.73.42,trna78_AspGTC_1_-_161418033_161418104@32.73.42,trna81_AspGTC_1_-_161410615_161410686@32.73.42|CTGTCACGCGGGAGACCGGGGTTCGATTCCCCGACGGGGAGC</td><td>>trna10_AspGTC_12_-_125424193_125424264</td><td>32</td><td>73</td><td>42</td><td>CTGTCACGCGGGAGACCGGGGTTCGATTCCCCGACGGGGAGC</td><td>391.4</td><td>-6.8</td><td>1.1</td><td>-6.1</td><td>9.6E-10</td><td>2.2E-09</td><td>MB231/MB468</td><td>CYTO</td></tr>
<tr><td>>hg38_21_+_8433172_8446622_RNA45SN1_45S_50ntFlanks@6708.6731.24|GAGAATTAATGTGAATTGCAGGAC</td><td>>hg38_21_+_8433172_8446622_RNA45SN1_45S_50ntFlanks</td><td>6708</td><td>6731</td><td>24</td><td>GAGAATTAATGTGAATTGCAGGAC</td><td>141.1</td><td>5.0</td><td>0.8</td><td>6.1</td><td>9.7E-10</td><td>2.2E-09</td><td>MB231/MB468</td><td>NUC</td></tr>
<tr><td>>hg38_21_+_8433172_8446622_RNA45SN1_45S_50ntFlanks@8168.8185.18|CCCAAGTCCTTCTGATCG</td><td>>hg38_21_+_8433172_8446622_RNA45SN1_45S_50ntFlanks</td><td>8168</td><td>8185</td><td>18</td><td>CCCAAGTCCTTCTGATCG</td><td>1249.5</td><td>-9.5</td><td>1.6</td><td>-6.1</td><td>1.1E-09</td><td>2.2E-09</td><td>BT20/MB468</td><td>NUC</td></tr>
<tr><td>>hg38_21_+_8433172_8446622_RNA45SN1_45S_50ntFlanks@11785.11804.20|TGAATGGATGAACGAGATTC</td><td>>hg38_21_+_8433172_8446622_RNA45SN1_45S_50ntFlanks</td><td>11785</td><td>11804</td><td>20</td><td>TGAATGGATGAACGAGATTC</td><td>7448.9</td><td>-6.0</td><td>1.0</td><td>-6.1</td><td>8.4E-10</td><td>2.2E-09</td><td>BT20/MB468</td><td>CYTO</td></tr>
<tr><td>>hg38_21_+_8433172_8446622_RNA45SN1_45S_50ntFlanks@8177.8196.20|TTCTGATCGAGGCCCAGCCC</td><td>>hg38_21_+_8433172_8446622_RNA45SN1_45S_50ntFlanks</td><td>8177</td><td>8196</td><td>20</td><td>TTCTGATCGAGGCCCAGCCC</td><td>162.8</td><td>3.8</td><td>0.6</td><td>6.1</td><td>9.7E-10</td><td>2.2E-09</td><td>MB231/MB468</td><td>CYTO</td></tr>
<tr><td>>MI0003205|hsa-mir-532&WithFlank&X|+|50003142|50003244@26.47.22&[MIMAT0002888&hsa-miR-532-5p&offsets|0|0;m-65&X|+|50003167|50003188&offsets|0|0]|CATGCCTTGAGTGTAGGACCGT</td><td>>MI0003205|hsa-mir-532&WithFlank&X|+|50003142|50003244</td><td>26</td><td>47</td><td>22</td><td>CATGCCTTGAGTGTAGGACCGT</td><td>2804.0</td><td>3.7</td><td>0.6</td><td>6.1</td><td>9.9E-10</td><td>2.2E-09</td><td>MB231/MB468</td><td>NUC</td></tr>
<tr><td>>MI0000255|hsa-mir-30d&WithFlank&8|-|134804870|134804951@12.35.24(+1U)&[MIMAT0000245&hsa-miR-30d-5p&offsets|0|+2(+1U);m-21&8|-|134804919|134804940&offsets|0|+2(+1U)]|TGTAAACATCCCCGACTGGAAGCTT</td><td>>MI0000255|hsa-mir-30d&WithFlank&8|-|134804870|134804951</td><td>12</td><td>35</td><td>24(+1U)</td><td>TGTAAACATCCCCGACTGGAAGCTT</td><td>1277.7</td><td>4.5</td><td>0.7</td><td>6.1</td><td>9.4E-10</td><td>2.2E-09</td><td>MB231/MB468</td><td>MP</td></tr>
<tr><td>>hg38_21_+_8433172_8446622_RNA45SN1_45S_50ntFlanks@4687.4723.37|ACGGACCAGAGCGAAAGCATTTGCCAAGAATGTTTTC</td><td>>hg38_21_+_8433172_8446622_RNA45SN1_45S_50ntFlanks</td><td>4687</td><td>4723</td><td>37</td><td>ACGGACCAGAGCGAAAGCATTTGCCAAGAATGTTTTC</td><td>126.6</td><td>-10.8</td><td>1.8</td><td>-6.1</td><td>9.4E-10</td><td>2.2E-09</td><td>MB231/MB468</td><td>MP</td></tr>
<tr><td>>hg38_21_+_8433172_8446622_RNA45SN1_45S_50ntFlanks@6734.6756.23|ATTGATCATCGACACTTCGAACG</td><td>>hg38_21_+_8433172_8446622_RNA45SN1_45S_50ntFlanks</td><td>6734</td><td>6756</td><td>23</td><td>ATTGATCATCGACACTTCGAACG</td><td>471.3</td><td>-4.2</td><td>0.7</td><td>-6.1</td><td>9.4E-10</td><td>2.2E-09</td><td>MB231/MB468</td><td>MP</td></tr>
<tr><td>>MI0000465|hsa-mir-191&WithFlank&3|-|49020612|49020715@22.43.22&[MIMAT0000440&hsa-miR-191-5p&offsets|0|-1;m-48&3|-|49020672|49020694&offsets|0|-1]|CAACGGAATCCCAAAAGCAGCT</td><td>>MI0000465|hsa-mir-191&WithFlank&3|-|49020612|49020715</td><td>22</td><td>43</td><td>22</td><td>CAACGGAATCCCAAAAGCAGCT</td><td>4450.7</td><td>5.4</td><td>0.9</td><td>6.1</td><td>9.7E-10</td><td>2.2E-09</td><td>MB231/MB468</td><td>CYTO</td></tr>
<tr><td>>hg38_21_+_8433172_8446622_RNA45SN1_45S_50ntFlanks@10391.10429.39|AACAGCAGTTGAACATGGGTCAGTCGGTCCTGAGAGATG</td><td>>hg38_21_+_8433172_8446622_RNA45SN1_45S_50ntFlanks</td><td>10391</td><td>10429</td><td>39</td><td>AACAGCAGTTGAACATGGGTCAGTCGGTCCTGAGAGATG</td><td>1107.4</td><td>-6.1</td><td>1.0</td><td>-6.1</td><td>9.4E-10</td><td>2.2E-09</td><td>MB231/MB468</td><td>MP</td></tr>
<tr><td>>hg38_21_+_8433172_8446622_RNA45SN1_45S_50ntFlanks@12152.12181.30|TGGGGAGTTTGACTGGGGCGGTACACCTGT</td><td>>hg38_21_+_8433172_8446622_RNA45SN1_45S_50ntFlanks</td><td>12152</td><td>12181</td><td>30</td><td>TGGGGAGTTTGACTGGGGCGGTACACCTGT</td><td>153.3</td><td>-10.3</td><td>1.6</td><td>-6.5</td><td>9.7E-11</td><td>2.2E-09</td><td>BT20/MB468</td><td>MP</td></tr>
<tr><td>>MI0003820|hsa-mir-454&WithFlank&17|-|59137752|59137878@70.93.24&[MIMAT0003885&hsa-miR-454-3p&offsets|0|+1;m-272&17|-|59137787|59137809&offsets|0|+1]|TAGTGCAATATTGCTTATAGGGTT</td><td>>MI0003820|hsa-mir-454&WithFlank&17|-|59137752|59137878</td><td>70</td><td>93</td><td>24</td><td>TAGTGCAATATTGCTTATAGGGTT</td><td>791.6</td><td>6.2</td><td>1.0</td><td>6.1</td><td>1.1E-09</td><td>2.2E-09</td><td>MB231/MB468</td><td>MITO</td></tr>
<tr><td>>hg38_21_+_8433172_8446622_RNA45SN1_45S_50ntFlanks@11235.11252.18|CTTCCCGTGGATCGCCCC</td><td>>hg38_21_+_8433172_8446622_RNA45SN1_45S_50ntFlanks</td><td>11235</td><td>11252</td><td>18</td><td>CTTCCCGTGGATCGCCCC</td><td>4352.5</td><td>-8.2</td><td>1.3</td><td>-6.1</td><td>1.1E-09</td><td>2.2E-09</td><td>BT20/MB468</td><td>NUC</td></tr>
<tr><td>>hg38_21_+_8433172_8446622_RNA45SN1_45S_50ntFlanks@12678.12705.28|AATCCCGCCCAGGCGGAACGATACGGCA</td><td>>hg38_21_+_8433172_8446622_RNA45SN1_45S_50ntFlanks</td><td>12678</td><td>12705</td><td>28</td><td>AATCCCGCCCAGGCGGAACGATACGGCA</td><td>70.0</td><td>-10.0</td><td>1.6</td><td>-6.1</td><td>9.4E-10</td><td>2.3E-09</td><td>MB231/MB468</td><td>MP</td></tr>
<tr><td>>hg38_21_+_8433172_8446622_RNA45SN1_45S_50ntFlanks@5485.5506.22|AGCGCTGAGAAGACGGTCGAAC</td><td>>hg38_21_+_8433172_8446622_RNA45SN1_45S_50ntFlanks</td><td>5485</td><td>5506</td><td>22</td><td>AGCGCTGAGAAGACGGTCGAAC</td><td>1977.8</td><td>-8.8</td><td>1.4</td><td>-6.1</td><td>9.4E-10</td><td>2.3E-09</td><td>MB231/MB468</td><td>MP</td></tr>
<tr><td>>hg38_21_+_8433172_8446622_RNA45SN1_45S_50ntFlanks@5482.5499.18|CGGAGCGCTGAGAAGACG</td><td>>hg38_21_+_8433172_8446622_RNA45SN1_45S_50ntFlanks</td><td>5482</td><td>5499</td><td>18</td><td>CGGAGCGCTGAGAAGACG</td><td>451.7</td><td>-7.0</td><td>1.2</td><td>-6.1</td><td>1.1E-09</td><td>2.3E-09</td><td>BT20/MB468</td><td>NUC</td></tr>
<tr><td>>hg38_21_+_8433172_8446622_RNA45SN1_45S_50ntFlanks@12938.12959.22|ACCATTCGTAGACGACCTGCTT</td><td>>hg38_21_+_8433172_8446622_RNA45SN1_45S_50ntFlanks</td><td>12938</td><td>12959</td><td>22</td><td>ACCATTCGTAGACGACCTGCTT</td><td>524.2</td><td>-6.8</td><td>1.1</td><td>-6.1</td><td>1.1E-09</td><td>2.3E-09</td><td>BT20/MB468</td><td>NUC</td></tr>
<tr><td>>trnaMT_MetCAT_MT_+_4402_4469@52.70.19|ATACCCTTCCCGTACTACC</td><td>>trnaMT_MetCAT_MT_+_4402_4469</td><td>52</td><td>70</td><td>19</td><td>ATACCCTTCCCGTACTACC</td><td>2479.3</td><td>15.3</td><td>2.5</td><td>6.1</td><td>9.8E-10</td><td>2.3E-09</td><td>MB231/MB468</td><td>CYTO</td></tr>
<tr><td>>MI0000457|hsa-mir-141&WithFlank&12|+|6964091|6964197@65.85.21&[MIMAT0000432&hsa-miR-141-3p&offsets|0|-1;m-134&12|+|6964155|6964175&offsets|0|0]|TAACACTGTCTGGTAAAGATG</td><td>>MI0000457|hsa-mir-141&WithFlank&12|+|6964091|6964197</td><td>65</td><td>85</td><td>21</td><td>TAACACTGTCTGGTAAAGATG</td><td>730.8</td><td>7.9</td><td>1.3</td><td>6.1</td><td>8.6E-10</td><td>2.3E-09</td><td>BT20/MB468</td><td>CYTO</td></tr>
<tr><td>>hg38_21_+_8433172_8446622_RNA45SN1_45S_50ntFlanks@8848.8867.20|CGGCGGCGGCGGCGGCGGGA</td><td>>hg38_21_+_8433172_8446622_RNA45SN1_45S_50ntFlanks</td><td>8848</td><td>8867</td><td>20</td><td>CGGCGGCGGCGGCGGCGGGA</td><td>236.2</td><td>-11.0</td><td>1.7</td><td>-6.5</td><td>9.8E-11</td><td>2.3E-09</td><td>BT20/MB468</td><td>MP</td></tr>
<tr><td>>hg38_21_+_8433172_8446622_RNA45SN1_45S_50ntFlanks@12646.12666.21|TGTGGGATTATGACTGAACGC</td><td>>hg38_21_+_8433172_8446622_RNA45SN1_45S_50ntFlanks</td><td>12646</td><td>12666</td><td>21</td><td>TGTGGGATTATGACTGAACGC</td><td>224.2</td><td>11.6</td><td>1.8</td><td>6.5</td><td>9.8E-11</td><td>2.3E-09</td><td>BT20/MB468</td><td>MP</td></tr>
<tr><td>>hg38_21_+_8433172_8446622_RNA45SN1_45S_50ntFlanks@6653.6683.31|ACTCTTAGCGGTGGATCACTCGGCTCGTGCG</td><td>>hg38_21_+_8433172_8446622_RNA45SN1_45S_50ntFlanks</td><td>6653</td><td>6683</td><td>31</td><td>ACTCTTAGCGGTGGATCACTCGGCTCGTGCG</td><td>4525.1</td><td>5.2</td><td>0.8</td><td>6.1</td><td>1.0E-09</td><td>2.3E-09</td><td>MB231/MB468</td><td>NUC</td></tr>
<tr><td>>hg38_21_+_8433172_8446622_RNA45SN1_45S_50ntFlanks@8113.8143.31|GGCGCGGGACATGTGGCGTACGGAAGACCCG</td><td>>hg38_21_+_8433172_8446622_RNA45SN1_45S_50ntFlanks</td><td>8113</td><td>8143</td><td>31</td><td>GGCGCGGGACATGTGGCGTACGGAAGACCCG</td><td>3146.4</td><td>-9.5</td><td>1.5</td><td>-6.5</td><td>9.9E-11</td><td>2.3E-09</td><td>BT20/MB468</td><td>MP</td></tr>
<tr><td>>hg38_21_+_8433172_8446622_RNA45SN1_45S_50ntFlanks@8323.8354.32|ACCGATAGTCAACAAGTACCGTAAGGGAAAGT</td><td>>hg38_21_+_8433172_8446622_RNA45SN1_45S_50ntFlanks</td><td>8323</td><td>8354</td><td>32</td><td>ACCGATAGTCAACAAGTACCGTAAGGGAAAGT</td><td>76.8</td><td>-10.1</td><td>1.7</td><td>-6.1</td><td>9.5E-10</td><td>2.3E-09</td><td>MB231/MB468</td><td>MP</td></tr>
<tr><td>>MI0000234|hsa-mir-192&WithFlank&11|-|64891131|64891252@30.51.22(+1U)&[MIMAT0000222&hsa-miR-192-5p&offsets|0|+1(+1U);m-59&11|-|64891203|64891223&offsets|0|+1(+1U)]|CTGACCTATGAATTGACAGCCAT</td><td>>MI0000234|hsa-mir-192&WithFlank&11|-|64891131|64891252</td><td>30</td><td>51</td><td>22(+1U)</td><td>CTGACCTATGAATTGACAGCCAT</td><td>452.1</td><td>5.1</td><td>0.8</td><td>6.1</td><td>8.6E-10</td><td>2.3E-09</td><td>BT20/MB468</td><td>CYTO</td></tr>
<tr><td>>hg38_21_+_8433172_8446622_RNA45SN1_45S_50ntFlanks@9545.9572.28|GTGGAGGTCCGTAGCGGTCCTGACGTGC</td><td>>hg38_21_+_8433172_8446622_RNA45SN1_45S_50ntFlanks</td><td>9545</td><td>9572</td><td>28</td><td>GTGGAGGTCCGTAGCGGTCCTGACGTGC</td><td>4651.2</td><td>-8.2</td><td>1.4</td><td>-6.1</td><td>1.1E-09</td><td>2.3E-09</td><td>BT20/MB468</td><td>NUC</td></tr>
<tr><td>>hg38_21_+_8433172_8446622_RNA45SN1_45S_50ntFlanks@12405.12444.40|CTTTTTGATCCTTCGATGTCGGCTCTTCCTATCATTGTGA</td><td>>hg38_21_+_8433172_8446622_RNA45SN1_45S_50ntFlanks</td><td>12405</td><td>12444</td><td>40</td><td>CTTTTTGATCCTTCGATGTCGGCTCTTCCTATCATTGTGA</td><td>133.8</td><td>-10.9</td><td>1.8</td><td>-6.1</td><td>9.5E-10</td><td>2.3E-09</td><td>MB231/MB468</td><td>MP</td></tr>
<tr><td>>am_trna162_MetCAT_6_-_26758550_26758622@31.51.21,trna164_MetCAT_6_-_26735574_26735646@31.51.21,trna20_MetCAT_16_+_71460396_71460468@31.51.21,trna21_MetCAT_6_+_26701712_26701784@31.51.21,trna27_MetCAT_6_+_26766444_26766516@31.51.21,trna75_MetCAT_6_+_28912352_28912424@31.51.21,trna97_MetCAT_6_-_28921042_28921114@31.51.21|TCTCATAATCTGAAGGTCCTG</td><td>>am_trna162_MetCAT_6_-_26758550_26758622</td><td>31</td><td>51</td><td>21</td><td>TCTCATAATCTGAAGGTCCTG</td><td>114.7</td><td>-8.9</td><td>1.5</td><td>-6.2</td><td>7.5E-10</td><td>2.3E-09</td><td>BT20/MB468</td><td>MITO</td></tr>
<tr><td>>hg38_21_+_8433172_8446622_RNA45SN1_45S_50ntFlanks@6671.6689.19|CTCGGCTCGTGCGTCGATG</td><td>>hg38_21_+_8433172_8446622_RNA45SN1_45S_50ntFlanks</td><td>6671</td><td>6689</td><td>19</td><td>CTCGGCTCGTGCGTCGATG</td><td>151.3</td><td>-9.3</td><td>1.5</td><td>-6.2</td><td>7.5E-10</td><td>2.3E-09</td><td>BT20/MB468</td><td>MITO</td></tr>
<tr><td>>hg38_21_+_8433172_8446622_RNA45SN1_45S_50ntFlanks@5059.5077.19|CGATTTGTCTGGTTAATTC</td><td>>hg38_21_+_8433172_8446622_RNA45SN1_45S_50ntFlanks</td><td>5059</td><td>5077</td><td>19</td><td>CGATTTGTCTGGTTAATTC</td><td>81.1</td><td>-10.2</td><td>1.7</td><td>-6.1</td><td>9.5E-10</td><td>2.3E-09</td><td>MB231/MB468</td><td>MP</td></tr>
<tr><td>>MI0000088|hsa-mir-30a&WithFlank&6|-|71403545|71403627@12.35.24(+2U)&[MIMAT0000087&hsa-miR-30a-5p&offsets|0|+2(+2U);m-15&6|-|71403595|71403616&offsets|0|+2(+2U)]|TGTAAACATCCTCGACTGGAAGCTTT</td><td>>MI0000088|hsa-mir-30a&WithFlank&6|-|71403545|71403627</td><td>12</td><td>35</td><td>24(+2U)</td><td>TGTAAACATCCTCGACTGGAAGCTTT</td><td>2876.2</td><td>8.3</td><td>1.4</td><td>6.1</td><td>1.1E-09</td><td>2.3E-09</td><td>MB231/MB468</td><td>MITO</td></tr>
<tr><td>>hg38_MT_+_1621_3279_RNR2_16S_50ntFlanks@1439.1477.39|ACGTGATCTGAGTTCAGACCGGAGTAATCCAGGTCGGTT</td><td>>hg38_MT_+_1621_3279_RNR2_16S_50ntFlanks</td><td>1439</td><td>1477</td><td>39</td><td>ACGTGATCTGAGTTCAGACCGGAGTAATCCAGGTCGGTT</td><td>590.6</td><td>5.1</td><td>0.8</td><td>6.2</td><td>7.5E-10</td><td>2.3E-09</td><td>BT20/MB468</td><td>MITO</td></tr>
<tr><td>>hg38_21_+_8433172_8446622_RNA45SN1_45S_50ntFlanks@4759.4781.23|ATCAGATACCGTCGTAGTTCCGA</td><td>>hg38_21_+_8433172_8446622_RNA45SN1_45S_50ntFlanks</td><td>4759</td><td>4781</td><td>23</td><td>ATCAGATACCGTCGTAGTTCCGA</td><td>150.6</td><td>-9.3</td><td>1.5</td><td>-6.2</td><td>7.5E-10</td><td>2.3E-09</td><td>BT20/MB468</td><td>MITO</td></tr>
<tr><td>>hg38_21_+_8433172_8446622_RNA45SN1_45S_50ntFlanks@12157.12174.18|AGTTTGACTGGGGCGGTA</td><td>>hg38_21_+_8433172_8446622_RNA45SN1_45S_50ntFlanks</td><td>12157</td><td>12174</td><td>18</td><td>AGTTTGACTGGGGCGGTA</td><td>70.0</td><td>-10.0</td><td>1.6</td><td>-6.1</td><td>9.6E-10</td><td>2.3E-09</td><td>MB231/MB468</td><td>MP</td></tr>
<tr><td>>trnaMT_ValTAC_MT_+_1602_1670@54.72.19|AACTTGACCGCTCTGACCA</td><td>>trnaMT_ValTAC_MT_+_1602_1670</td><td>54</td><td>72</td><td>19</td><td>AACTTGACCGCTCTGACCA</td><td>3029.9</td><td>11.2</td><td>1.8</td><td>6.1</td><td>8.7E-10</td><td>2.3E-09</td><td>BT20/MB468</td><td>CYTO</td></tr>
<tr><td>>hg38_21_+_8433172_8446622_RNA45SN1_45S_50ntFlanks@4822.4844.23|CATGACCCGCCGGGCAGCTTCCG</td><td>>hg38_21_+_8433172_8446622_RNA45SN1_45S_50ntFlanks</td><td>4822</td><td>4844</td><td>23</td><td>CATGACCCGCCGGGCAGCTTCCG</td><td>391.5</td><td>5.9</td><td>1.0</td><td>6.1</td><td>9.9E-10</td><td>2.3E-09</td><td>MB231/MB468</td><td>CYTO</td></tr>
<tr><td>>hg38_21_+_8433172_8446622_RNA45SN1_45S_50ntFlanks@11454.11471.18|CCTCGGCGCGCGGCGGCG</td><td>>hg38_21_+_8433172_8446622_RNA45SN1_45S_50ntFlanks</td><td>11454</td><td>11471</td><td>18</td><td>CCTCGGCGCGCGGCGGCG</td><td>195.9</td><td>-6.1</td><td>1.0</td><td>-6.1</td><td>9.9E-10</td><td>2.3E-09</td><td>MB231/MB468</td><td>CYTO</td></tr>
<tr><td>>hg38_21_+_8433172_8446622_RNA45SN1_45S_50ntFlanks@4975.4994.20|CCCGGACACGGACAGGATTG</td><td>>hg38_21_+_8433172_8446622_RNA45SN1_45S_50ntFlanks</td><td>4975</td><td>4994</td><td>20</td><td>CCCGGACACGGACAGGATTG</td><td>523.3</td><td>-11.1</td><td>1.8</td><td>-6.2</td><td>7.6E-10</td><td>2.3E-09</td><td>BT20/MB468</td><td>MITO</td></tr>
<tr><td>>hg38_21_+_8433172_8446622_RNA45SN1_45S_50ntFlanks@11199.11233.35|CCGGCGGCGGCGGCGACTCTGGACGCGAGCCGGGC</td><td>>hg38_21_+_8433172_8446622_RNA45SN1_45S_50ntFlanks</td><td>11199</td><td>11233</td><td>35</td><td>CCGGCGGCGGCGGCGACTCTGGACGCGAGCCGGGC</td><td>188.7</td><td>-7.8</td><td>1.3</td><td>-6.1</td><td>1.0E-09</td><td>2.3E-09</td><td>MB231/MB468</td><td>NUC</td></tr>
</table>

| | | | | | | | | | | | | | |
|---|---|---|---|---|---|---|---|---|---|---|---|---|---|
| >hg38_21_+_8433172_8446622_RNA45SN1_45S_50ntFlanks@12997.13035.39\|CGCTGCGATCTATTGAAAGTCAGCCCTCGACACAAGGGT | >hg38_21_+_8433172_8446622_RNA45SN1_45S_50ntFlanks | 12997 | 13035 | 39 | CGCTGCGATCTATTGAAAGTCAGCCCTCGACACAAGGGT | 882.1 | 8.5 | 1.4 | 6.1 | 1.0E-09 | 2.3E-09 | MB231/MB468 | CYTO |
| >hg38_21_+_8433172_8446622_RNA45SN1_45S_50ntFlanks@8171.8197.27\|AAGTCCTTCTGATCGAGGCCCAGCCCG | >hg38_21_+_8433172_8446622_RNA45SN1_45S_50ntFlanks | 8171 | 8197 | 27 | AAGTCCTTCTGATCGAGGCCCAGCCCG | 2099.7 | -10.2 | 1.7 | -6.1 | 1.2E-09 | 2.3E-09 | BT20/MB468 | NUC |
| >hg38_21_+_8433172_8446622_RNA45SN1_45S_50ntFlanks@4003.4042.40\|ATAACCTCGGGCCGATCGCACGCCCCCCGTGGCGGCGACG | >hg38_21_+_8433172_8446622_RNA45SN1_45S_50ntFlanks | 4003 | 4042 | 40 | ATAACCTCGGGCCGATCGCACGCCCCCCGTGGCGGCGACG | 4556.6 | -10.1 | 1.7 | -6.1 | 1.2E-09 | 2.3E-09 | BT20/MB468 | NUC |
| >hg38_21_+_8433172_8446622_RNA45SN1_45S_50ntFlanks@8672.8699.28\|GCGGTGCGCCGCGACCGGCTCCGGGACG | >hg38_21_+_8433172_8446622_RNA45SN1_45S_50ntFlanks | 8672 | 8699 | 28 | GCGGTGCGCCGCGACCGGCTCCGGGACG | 83.0 | -10.2 | 1.7 | -6.1 | 9.6E-10 | 2.3E-09 | MB231/MB468 | MP |
| >hg38_21_+_8433172_8446622_RNA45SN1_45S_50ntFlanks@10683.10705.23\|CCGGCGGCGTCCGGTGAGCTCTC | >hg38_21_+_8433172_8446622_RNA45SN1_45S_50ntFlanks | 10683 | 10705 | 23 | CCGGCGGCGTCCGGTGAGCTCTC | 676.1 | -3.0 | 0.5 | -6.1 | 8.8E-10 | 2.3E-09 | BT20/MB468 | CYTO |
| >hg38_21_+_8433172_8446622_RNA45SN1_45S_50ntFlanks@10402.10431.30\|AACATGGGTCAGTCGGTCCTGAGAGATGGG | >hg38_21_+_8433172_8446622_RNA45SN1_45S_50ntFlanks | 10402 | 10431 | 30 | AACATGGGTCAGTCGGTCCTGAGAGATGGG | 5105.6 | -10.4 | 1.7 | -6.1 | 1.2E-09 | 2.3E-09 | BT20/MB468 | NUC |
| >hg38_21_+_8433172_8446622_RNA45SN1_45S_50ntFlanks@6730.6752.23\|ACACATTGATCATCGACACTTCG | >hg38_21_+_8433172_8446622_RNA45SN1_45S_50ntFlanks | 6730 | 6752 | 23 | ACACATTGATCATCGACACTTCG | 357.0 | -8.3 | 1.4 | -6.1 | 1.2E-09 | 2.3E-09 | BT20/MB468 | NUC |
| >hg38_21_+_8433172_8446622_RNA45SN1_45S_50ntFlanks@12938.12960.23\|ACCATTCGTAGACGACCTGCTTC | >hg38_21_+_8433172_8446622_RNA45SN1_45S_50ntFlanks | 12938 | 12960 | 23 | ACCATTCGTAGACGACCTGCTTC | 370.1 | 3.9 | 0.6 | 6.1 | 8.8E-10 | 2.3E-09 | BT20/MB468 | CYTO |
| >hg38_21_+_8433172_8446622_RNA45SN1_45S_50ntFlanks@4296.4315.20\|CCATTGGAGGGCAAGTCTGG | >hg38_21_+_8433172_8446622_RNA45SN1_45S_50ntFlanks | 4296 | 4315 | 20 | CCATTGGAGGGCAAGTCTGG | 61.6 | -9.8 | 1.6 | -6.1 | 9.7E-10 | 2.3E-09 | MB231/MB468 | MP |
| >hg38_21_+_8433172_8446622_RNA45SN1_45S_50ntFlanks@12635.12668.34\|AAGCTACCATCTGTGGGATTATGACTGAACGCCT | >hg38_21_+_8433172_8446622_RNA45SN1_45S_50ntFlanks | 12635 | 12668 | 34 | AAGCTACCATCTGTGGGATTATGACTGAACGCCT | 224.1 | 6.9 | 1.1 | 6.1 | 1.0E-09 | 2.3E-09 | MB231/MB468 | CYTO |
| >hg38_21_+_8433172_8446622_RNA45SN1_45S_50ntFlanks@11587.11623.37\|TGCGGACCAGGGGAATCCGACTGTTTAATTAAAACAA | >hg38_21_+_8433172_8446622_RNA45SN1_45S_50ntFlanks | 11587 | 11623 | 37 | TGCGGACCAGGGGAATCCGACTGTTTAATTAAAACAA | 199.9 | 11.4 | 1.8 | 6.5 | 1.0E-10 | 2.3E-09 | BT20/MB468 | MP |
| >hg38_21_+_8433172_8446622_RNA45SN1_45S_50ntFlanks@12933.12965.33\|GCTAAACCATTCGTAGACGACCTGCTTCTGGGT | >hg38_21_+_8433172_8446622_RNA45SN1_45S_50ntFlanks | 12933 | 12965 | 33 | GCTAAACCATTCGTAGACGACCTGCTTCTGGGT | 137.2 | -11.0 | 1.8 | -6.1 | 9.7E-10 | 2.3E-09 | MB231/MB468 | MP |
| >trnaMT_GluTTC_MT_-_14674_14742@32.72.41,trnalookalike8_GluTTC_5_-_93905172_93905240@32.72.41\|TCATATCATTGGTCGTGGTTGTAGTCCGTGCGAGAATACCA | >trnaMT_GluTTC_MT_-_14674_14742 | 32 | 72 | 41 | TCATATCATTGGTCGTGGTTGTAGTCCGTGCGAGAATACCA | 1125.6 | 7.6 | 1.3 | 6.1 | 1.0E-09 | 2.3E-09 | MB231/MB468 | CYTO |
| >hg38_21_+_8433172_8446622_RNA45SN1_45S_50ntFlanks@8177.8210.34\|TTCTGATCGAGGCCCAGCCCGTGGACGGTGTGAG | >hg38_21_+_8433172_8446622_RNA45SN1_45S_50ntFlanks | 8177 | 8210 | 34 | TTCTGATCGAGGCCCAGCCCGTGGACGGTGTGAG | 166.8 | 3.5 | 0.6 | 6.1 | 1.0E-09 | 2.3E-09 | MB231/MB468 | CYTO |
| >MI0000266\|hsa-mir-10a&WithFlank&17\|-\|48579832\|48579953@29.48.20(+1C)&[MIMAT0000253&hsa-miR-10a-5p&offsets\|+1\|-2(+1C);m-18&17\|-\|48579905\|48579926&offsets\|+1\|-1(+1C)]\|ACCCTGTAGATCCGAATTTGC | >MI0000266\|hsa-mir-10a&WithFlank&17\|-\|48579832\|48579953 | 29 | 48 | 20(+1C) | ACCCTGTAGATCCGAATTTGC | 141.3 | 10.4 | 1.7 | 6.1 | 1.1E-09 | 2.3E-09 | MB231/MB468 | MITO |
| >hg38_21_+_8433172_8446622_RNA45SN1_45S_50ntFlanks@13008.13025.18\|ATTGAAAGTCAGCCCTCG | >hg38_21_+_8433172_8446622_RNA45SN1_45S_50ntFlanks | 13008 | 13025 | 18 | ATTGAAAGTCAGCCCTCG | 2364.2 | -10.0 | 1.6 | -6.1 | 1.2E-09 | 2.3E-09 | BT20/MB468 | NUC |
| >hg38_21_+_8433172_8446622_RNA45SN1_45S_50ntFlanks@6777.6802.26\|TCCCGGGGCTACGCCTGTCTGAGCGT | >hg38_21_+_8433172_8446622_RNA45SN1_45S_50ntFlanks | 6777 | 6802 | 26 | TCCCGGGGCTACGCCTGTCTGAGCGT | 468.3 | 5.0 | 0.8 | 6.2 | 7.7E-10 | 2.3E-09 | BT20/MB468 | MITO |
| >hg38_21_+_8433172_8446622_RNA45SN1_45S_50ntFlanks@10726.10750.25\|GGGAGAGGGTGTAAATCTCGCGCCG | >hg38_21_+_8433172_8446622_RNA45SN1_45S_50ntFlanks | 10726 | 10750 | 25 | GGGAGAGGGTGTAAATCTCGCGCCG | 137.0 | -11.0 | 1.8 | -6.1 | 9.8E-10 | 2.3E-09 | MB231/MB468 | MP |
| >trnaMT_GluTTC_MT_-_14674_14742@39.71.33,trnalookalike8_GluTTC_5_-_93905172_93905240@39.71.33\|ATTGGTCGTGGTTGTAGTCCGTGCGAGAATACC | >trnaMT_GluTTC_MT_-_14674_14742 | 39 | 71 | 33 | ATTGGTCGTGGTTGTAGTCCGTGCGAGAATACC | 1193.7 | 4.5 | 0.7 | 6.1 | 1.1E-09 | 2.3E-09 | MB231/MB468 | MITO |
| >hg38_21_+_8433172_8446622_RNA45SN1_45S_50ntFlanks@12539.12587.49\|ATGATGTGTTGTTGCCATGGTAATCCTGCTCAGTACGAGAGGAACCGCA | >hg38_21_+_8433172_8446622_RNA45SN1_45S_50ntFlanks | 12539 | 12587 | 49 | ATGATGTGTTGTTGCCATGGTAATCCTGCTCAGTACGAGAGGAACCGCA | 177.1 | -9.8 | 1.6 | -6.1 | 8.9E-10 | 2.3E-09 | BT20/MB468 | CYTO |
| >MI0000102\|hsa-mir-100&WithFlank&11\|-\|122152223\|122152314@19.40.22(+1U)&[MIMAT0000098&hsa-miR-100-5p&offsets\|0\|0(+1U);m-27&11\|-\|122152276\|122152296&offsets\|0\|+1(+1U)]\|AACCCGTAGATCCGAACTTGTGT | >MI0000102\|hsa-mir-100&WithFlank&11\|-\|122152223\|122152314 | 19 | 40 | 22(+1U) | AACCCGTAGATCCGAACTTGTGT | 45997.4 | -6.6 | 1.0 | -6.5 | 1.1E-10 | 2.3E-09 | BT20/MB231 | NUC |
| >hg38_21_+_8433172_8446622_RNA45SN1_45S_50ntFlanks@5420.5441.22\|CCGATTGGATGGTTTAGTGAGG | >hg38_21_+_8433172_8446622_RNA45SN1_45S_50ntFlanks | 5420 | 5441 | 22 | CCGATTGGATGGTTTAGTGAGG | 71.3 | -10.0 | 1.6 | -6.1 | 9.8E-10 | 2.3E-09 | MB231/MB468 | MP |
| >hg38_21_+_8433172_8446622_RNA45SN1_45S_50ntFlanks@12124.12146.23\|CGCGACCCGCTCCGGGGACAGTG | >hg38_21_+_8433172_8446622_RNA45SN1_45S_50ntFlanks | 12124 | 12146 | 23 | CGCGACCCGCTCCGGGGACAGTG | 226.5 | -10.9 | 1.7 | -6.5 | 1.0E-10 | 2.3E-09 | BT20/MB468 | MP |
| >am_trnaMT_HisGTG_MT_+_12138_12206@26.45.20\|AGATTGTGAATCTGACAACA | >am_trnaMT_HisGTG_MT_+_12138_12206 | 26 | 45 | 20 | AGATTGTGAATCTGACAACA | 907.1 | 5.9 | 1.0 | 6.2 | 7.7E-10 | 2.3E-09 | BT20/MB468 | MITO |
| >MI0000746\|hsa-mir-99b&WithFlank&19\|+\|51692606\|51692687@13.31.19&[MIMAT0000689&hsa-miR-99b-5p&offsets\|0\|-3;m-10&19\|+\|51692618\|51692639&offsets\|0\|-3]\|CACCCGTAGAACCGACCTT | >MI0000746\|hsa-mir-99b&WithFlank&19\|+\|51692606\|51692687 | 13 | 31 | 19 | CACCCGTAGAACCGACCTT | 509.8 | 9.5 | 1.5 | 6.1 | 9.0E-10 | 2.3E-09 | BT20/MB468 | CYTO |
| >hg38_21_+_8433172_8446622_RNA45SN1_45S_50ntFlanks@11665.11703.39\|TTCTGCCCAGTGCTCTGAATGTCAAAGTGAAGAAATTCA | >hg38_21_+_8433172_8446622_RNA45SN1_45S_50ntFlanks | 11665 | 11703 | 39 | TTCTGCCCAGTGCTCTGAATGTCAAAGTGAAGAAATTCA | 320.2 | 6.2 | 1.0 | 6.1 | 1.0E-09 | 2.4E-09 | MB231/MB468 | CYTO |
| >hg38_21_+_8433172_8446622_RNA45SN1_45S_50ntFlanks@6743.6770.28\|CGACACTTCGAACGCACTTGCGGCCCCG | >hg38_21_+_8433172_8446622_RNA45SN1_45S_50ntFlanks | 6743 | 6770 | 28 | CGACACTTCGAACGCACTTGCGGCCCCG | 1165.9 | -7.2 | 1.2 | -6.1 | 1.0E-09 | 2.4E-09 | MB231/MB468 | NUC |
| >hg38_21_+_8433172_8446622_RNA45SN1_45S_50ntFlanks@12223.12240.18\|CAGAAACCTCCCGTGGAG | >hg38_21_+_8433172_8446622_RNA45SN1_45S_50ntFlanks | 12223 | 12240 | 18 | CAGAAACCTCCCGTGGAG | 71.6 | -10.0 | 1.6 | -6.1 | 9.9E-10 | 2.4E-09 | MB231/MB468 | MP |
| >hg38_21_+_8433172_8446622_RNA45SN1_45S_50ntFlanks@11572.11591.20\|CCGACTTAGAACTGGTGCGG | >hg38_21_+_8433172_8446622_RNA45SN1_45S_50ntFlanks | 11572 | 11591 | 20 | CCGACTTAGAACTGGTGCGG | 1902.3 | -3.7 | 0.6 | -6.1 | 7.8E-10 | 2.4E-09 | BT20/MB468 | MITO |
| >hg38_1_-_228634819_228635039_RNA5S12_5S_50ntFlanks@130.170.41\|TGGATGGGAGACCGCCTGGGAATACCGGGTGCTGTAGGCTT | >hg38_1_-_228634819_228635039_RNA5S12_5S_50ntFlanks | 130 | 170 | 41 | TGGATGGGAGACCGCCTGGGAATACCGGGTGCTGTAGGCTT | 4830.4 | 6.7 | 1.0 | 6.5 | 1.0E-10 | 2.4E-09 | BT20/MB468 | MP |
| >hg38_21_+_8433172_8446622_RNA45SN1_45S_50ntFlanks@6734.6774.41\|ATTGATCATCGACACTTCGAACGCACTTGCGGCCCCGGGTT | >hg38_21_+_8433172_8446622_RNA45SN1_45S_50ntFlanks | 6734 | 6774 | 41 | ATTGATCATCGACACTTCGAACGCACTTGCGGCCCCGGGTT | 147.6 | 11.0 | 1.7 | 6.5 | 1.0E-10 | 2.4E-09 | BT20/MB468 | MP |
| >hg38_21_+_8433172_8446622_RNA45SN1_45S_50ntFlanks@11999.12043.45\|CCCTGCGGGCCGCCGGTGAAATACCACTACTCTGATCGTTTTTTC | >hg38_21_+_8433172_8446622_RNA45SN1_45S_50ntFlanks | 11999 | 12043 | 45 | CCCTGCGGGCCGCCGGTGAAATACCACTACTCTGATCGTTTTTTC | 317.6 | -12.2 | 2.0 | -6.1 | 9.9E-10 | 2.4E-09 | MB231/MB468 | MP |
| >am_trna128_GlyGCC_6_-_27870686_27870756@19.37.19,trna19_GlyGCC_16_+_70823410_70823480@19.37.19,trna19_GlyGCC_2_-_157257659_157257729@19.37.19,trna24_GlyGCC_16_-_70812942_70813012@19.37.19,trna25_GlyGCC_16_-_70812114_70812184@19.37.19,trna2_GlyGCC_21_-_18827107_18827177@19.37.19,trna35_GlyGCC_1_+_161413094_161413164@19.37.19,trna37_GlyGCC_1_+_161420467_161420537@19.37.19,trna39_GlyGCC_1_+_161427898_161427968@19.37.19,trna41_GlyGCC_1_+_161435258_161435328@19.37.19,trna5_GlyGCC_17_+_8029064_8029134@19.37.19,trna68_GlyGCC_1_-_161493637_161493707@19.37.19\|TAGAATTCTCGCCTGCCAC | >am_trna128_GlyGCC_6_-_27870686_27870756 | 19 | 37 | 19 | TAGAATTCTCGCCTGCCAC | 1234.8 | 9.1 | 1.5 | 6.1 | 9.0E-10 | 2.4E-09 | BT20/MB468 | CYTO |
| >hg38_21_+_8433172_8446622_RNA45SN1_45S_50ntFlanks@4223.4242.20\|AAAAATAACAATACAGGACT | >hg38_21_+_8433172_8446622_RNA45SN1_45S_50ntFlanks | 4223 | 4242 | 20 | AAAAATAACAATACAGGACT | 429.8 | -7.8 | 1.3 | -6.1 | 1.1E-09 | 2.4E-09 | MB231/MB468 | MITO |
| >hg38_21_+_8433172_8446622_RNA45SN1_45S_50ntFlanks@6740.6758.19\|CATCGACACTTCGAACGCA | >hg38_21_+_8433172_8446622_RNA45SN1_45S_50ntFlanks | 6740 | 6758 | 19 | CATCGACACTTCGAACGCA | 82.2 | -10.2 | 1.7 | -6.1 | 9.9E-10 | 2.4E-09 | MB231/MB468 | MP |
| >hg38_21_+_8433172_8446622_RNA45SN1_45S_50ntFlanks@10663.10711.49\|CCTTGGAAAGCGTCGCGGTTCCGGCGGCGTCCGGTGAGCTCTCGCTGGC | >hg38_21_+_8433172_8446622_RNA45SN1_45S_50ntFlanks | 10663 | 10711 | 49 | CCTTGGAAAGCGTCGCGGTTCCGGCGGCGTCCGGTGAGCTCTCGCTGGC | 245.0 | -7.7 | 1.3 | -6.1 | 1.0E-09 | 2.4E-09 | MB231/MB468 | CYTO |
| >trna116_GluCTC_1_-_145399233_145399304@32.50.19,trna59_GluCTC_1_+_249168447_249168518@32.50.19,trna71_GluCTC_1_-_161439189_161439260@32.50.19,trna74_GluCTC_1_-_161431809_161431880@32.50.19,trna77_GluCTC_1_-_161424398_161424469@32.50.19,trna77_GluCTC_6_+_28949976_28950047@32.50.19,trna80_GluCTC_1_-_161417018_161417089@32.50.19,trna87_GluCTC_6_-_126101393_126101464@32.50.19\|CTCTCACCGCCGCGGCCCG | >trna116_GluCTC_1_-_145399233_145399304 | 32 | 50 | 19 | CTCTCACCGCCGCGGCCCG | 323.1 | -11.3 | 1.9 | -6.1 | 1.0E-09 | 2.4E-09 | MB231/MB468 | NUC |
| >hg38_21_+_8433172_8446622_RNA45SN1_45S_50ntFlanks@11316.11344.29\|CACCCCACGTCTCGTCGCGCGCGCGTCCG | >hg38_21_+_8433172_8446622_RNA45SN1_45S_50ntFlanks | 11316 | 11344 | 29 | CACCCCACGTCTCGTCGCGCGCGCGTCCG | 79226.4 | -12.1 | 2.0 | -6.1 | 1.2E-09 | 2.4E-09 | BT20/MB468 | NUC |
| >hg38_21_+_8433172_8446622_RNA45SN1_45S_50ntFlanks@12444.12474.31\|AAGCAGAATTCACCAAGCGTTGGATTGTTCA | >hg38_21_+_8433172_8446622_RNA45SN1_45S_50ntFlanks | 12444 | 12474 | 31 | AAGCAGAATTCACCAAGCGTTGGATTGTTCA | 89.5 | -10.3 | 1.7 | -6.1 | 1.0E-09 | 2.4E-09 | MB231/MB468 | MP |
| >hg38_21_+_8433172_8446622_RNA45SN1_45S_50ntFlanks@10348.10367.20\|AAACGAGAACTTTGAAGGCC | >hg38_21_+_8433172_8446622_RNA45SN1_45S_50ntFlanks | 10348 | 10367 | 20 | AAACGAGAACTTTGAAGGCC | 496.0 | -4.0 | 0.7 | -6.1 | 1.2E-09 | 2.4E-09 | BT20/MB468 | NUC |
| >hg38_21_+_8433172_8446622_RNA45SN1_45S_50ntFlanks@7984.8005.22\|AGATCAGACGTGGCGACCCGCT | >hg38_21_+_8433172_8446622_RNA45SN1_45S_50ntFlanks | 7984 | 8005 | 22 | AGATCAGACGTGGCGACCCGCT | 1026.3 | -8.4 | 1.4 | -6.1 | 1.0E-09 | 2.4E-09 | MB231/MB468 | MP |
| >hg38_21_+_8433172_8446622_RNA45SN1_45S_50ntFlanks@10841.10875.35\|CGTAACTTCGGGATAAGGATTGGCTCTAAGGGCTG | >hg38_21_+_8433172_8446622_RNA45SN1_45S_50ntFlanks | 10841 | 10875 | 35 | CGTAACTTCGGGATAAGGATTGGCTCTAAGGGCTG | 2004.1 | -9.8 | 1.6 | -6.1 | 1.2E-09 | 2.4E-09 | BT20/MB468 | NUC |
| >hg38_21_+_8433172_8446622_RNA45SN1_45S_50ntFlanks@9454.9500.47\|CCGCGCCGGGGAGGTGGAGCACGAGCGCACGTGTTAGGACCCGAAAG | >hg38_21_+_8433172_8446622_RNA45SN1_45S_50ntFlanks | 9454 | 9500 | 47 | CCGCGCCGGGGAGGTGGAGCACGAGCGCACGTGTTAGGACCCGAAAG | 170.2 | -11.3 | 1.8 | -6.1 | 1.0E-09 | 2.4E-09 | MB231/MB468 | MP |
| >hg38_21_+_8433172_8446622_RNA45SN1_45S_50ntFlanks@5515.5544.30\|CTAGAGGAAGTAAAAGTCGTAACAAGGTTT | >hg38_21_+_8433172_8446622_RNA45SN1_45S_50ntFlanks | 5515 | 5544 | 30 | CTAGAGGAAGTAAAAGTCGTAACAAGGTTT | 68.2 | -10.0 | 1.6 | -6.1 | 1.0E-09 | 2.4E-09 | MB231/MB468 | MP |
| >MI0000102\|hsa-mir-100&WithFlank&11\|-\|122152223\|122152314@19.38.20&[MIMAT0000098&hsa-miR-100-5p&offsets\|0\|-2;m-27&11\|-\|122152276\|122152296&offsets\|0\|-1]\|AACCCGTAGATCCGAACTTG | >MI0000102\|hsa-mir-100&WithFlank&11\|-\|122152223\|122152314 | 19 | 38 | 20 | AACCCGTAGATCCGAACTTG | 113.1 | 9.7 | 1.6 | 6.1 | 1.2E-09 | 2.4E-09 | BT20/MB468 | NUC |
| >hg38_21_+_8433172_8446622_RNA45SN1_45S_50ntFlanks@13006.13024.19\|CTATTGAAAGTCAGCCCTC | >hg38_21_+_8433172_8446622_RNA45SN1_45S_50ntFlanks | 13006 | 13024 | 19 | CTATTGAAAGTCAGCCCTC | 148.6 | -6.5 | 1.1 | -6.1 | 9.2E-10 | 2.4E-09 | BT20/MB468 | CYTO |
| >hg38_21_+_8433172_8446622_RNA45SN1_45S_50ntFlanks@3781.3801.21\|ACAGTGAAACTGCGAATGGCT | >hg38_21_+_8433172_8446622_RNA45SN1_45S_50ntFlanks | 3781 | 3801 | 21 | ACAGTGAAACTGCGAATGGCT | 607.3 | -5.3 | 0.9 | -6.1 | 1.2E-09 | 2.4E-09 | MB231/MB468 | MITO |

| | | | | | | | | | | | | | |
|---|---|---|---|---|---|---|---|---|---|---|---|---|---|
| >hg38_21_+_8433172_8446622_RNA45SN1_45S_50ntFlanks@6730.6772.43\|ACACATTGATCATCGACACTTCGAACGCACTTGCGGCCCCGGG | >hg38_21_+_8433172_8446622_RNA45SN1_45S_50ntFlanks | 6730 | 6772 | 43 | ACACATTGATCATCGACACTTCGAACGCACTTGCGGCCCCGGG | 435.7 | -6.4 | 1.0 | -6.1 | 1.2E-09 | 2.4E-09 | MB231/MB468 | MITO |
| >hg38_21_+_8433172_8446622_RNA45SN1_45S_50ntFlanks@10391.10433.43\|AACAGCAGTTGAACATGGGTCAGTCGGTCCTGAGAGATGGGCG | >hg38_21_+_8433172_8446622_RNA45SN1_45S_50ntFlanks | 10391 | 10433 | 43 | AACAGCAGTTGAACATGGGTCAGTCGGTCCTGAGAGATGGGCG | 162.5 | -11.2 | 1.8 | -6.1 | 1.0E-09 | 2.4E-09 | MB231/MB468 | MP |
| >hg38_21_+_8433172_8446622_RNA45SN1_45S_50ntFlanks@11787.11804.18\|AATGGATGAACGAGATTC | >hg38_21_+_8433172_8446622_RNA45SN1_45S_50ntFlanks | 11787 | 11804 | 18 | AATGGATGAACGAGATTC | 3468.1 | -5.3 | 0.9 | -6.1 | 9.2E-10 | 2.4E-09 | BT20/MB468 | CYTO |
| >hg38_21_+_8433172_8446622_RNA45SN1_45S_50ntFlanks@3841.3859.19\|TCCTACTTGGATAACTGTG | >hg38_21_+_8433172_8446622_RNA45SN1_45S_50ntFlanks | 3841 | 3859 | 19 | TCCTACTTGGATAACTGTG | 123.0 | -10.8 | 1.8 | -6.1 | 1.0E-09 | 2.4E-09 | MB231/MB468 | MP |
| >hg38_21_+_8433172_8446622_RNA45SN1_45S_50ntFlanks@10351.10372.22\|CGAGAACTTTGAAGGCCGAAGT | >hg38_21_+_8433172_8446622_RNA45SN1_45S_50ntFlanks | 10351 | 10372 | 22 | CGAGAACTTTGAAGGCCGAAGT | 1370.5 | -4.9 | 0.8 | -6.1 | 1.0E-09 | 2.4E-09 | MB231/MB468 | MP |
| >hg38_21_+_8433172_8446622_RNA45SN1_45S_50ntFlanks@12004.12035.32\|CGGGCCGCCGGTGAAATACCACTACTCTGATC | >hg38_21_+_8433172_8446622_RNA45SN1_45S_50ntFlanks | 12004 | 12035 | 32 | CGGGCCGCCGGTGAAATACCACTACTCTGATC | 434.6 | 5.3 | 0.8 | 6.5 | 6.3E-11 | 2.4E-09 | BT20/MB231 | CYTO |
| >MI0000449\|hsa-mir-132&WithFlank&17\|-\|2049902\|2050014@65.86.22(+1U)&[MIMAT0000426&hsa-miR-132-3p&offsets\|0\|0(+1U);m-138&17\|-\|2049929\|2049950&offsets\|0\|0(+1U)]\|TAACAGTCTACAGCCATGGTCGT | >MI0000449\|hsa-mir-132&WithFlank&17\|-\|2049902\|2050014 | 65 | 86 | 22(+1U) | TAACAGTCTACAGCCATGGTCGT | 124.9 | 10.2 | 1.7 | 6.1 | 1.2E-09 | 2.4E-09 | MB231/MB468 | MITO |
| >hg38_21_+_8433172_8446622_RNA45SN1_45S_50ntFlanks@12324.12358.35\|TTGGGTTTTAAGCAGGAGGTGTCAGAAAAGTTACC | >hg38_21_+_8433172_8446622_RNA45SN1_45S_50ntFlanks | 12324 | 12358 | 35 | TTGGGTTTTAAGCAGGAGGTGTCAGAAAAGTTACC | 86.2 | -10.3 | 1.7 | -6.1 | 1.0E-09 | 2.4E-09 | MB231/MB468 | MP |
| >hg38_21_+_8433172_8446622_RNA45SN1_45S_50ntFlanks@12417.12436.20\|TCGATGTCGGCTCTTCCTAT | >hg38_21_+_8433172_8446622_RNA45SN1_45S_50ntFlanks | 12417 | 12436 | 20 | TCGATGTCGGCTCTTCCTAT | 1062.4 | -8.4 | 1.4 | -6.1 | 1.0E-09 | 2.4E-09 | MB231/MB468 | MP |
| >hg38_21_+_8433172_8446622_RNA45SN1_45S_50ntFlanks@11882.11905.24\|CTGTTGAGCTTGACTCTAGTCTGG | >hg38_21_+_8433172_8446622_RNA45SN1_45S_50ntFlanks | 11882 | 11905 | 24 | CTGTTGAGCTTGACTCTAGTCTGG | 114.4 | -6.8 | 1.1 | -6.1 | 9.3E-10 | 2.4E-09 | BT20/MB468 | CYTO |
| >MI0033423\|hsa-mir-10399&WithFlank&7\|-\|139044094\|139044163@44.63.20(+1U)&[MIMAT0041630&hsa-miR-10399-3p&offsets\|0\|-1(+1U);TJU_CMC_MD2.ID03061.3p-miR&offsets\|+4\|+2(+1U);m-2657&7\|-\|139044103\|139044124&offsets\|+4\|+2(+1U)]\|CTCTCGGACAAGCTGTAGGTT | >MI0033423\|hsa-mir-10399&WithFlank&7\|-\|139044094\|139044163 | 44 | 63 | 20(+1U) | CTCTCGGACAAGCTGTAGGTT | 86.6 | 9.7 | 1.6 | 6.1 | 1.0E-09 | 2.4E-09 | MB231/MB468 | MP |
| >hg38_21_+_8433172_8446622_RNA45SN1_45S_50ntFlanks@12004.12032.29\|CGGGCCGCCGGTGAAATACCACTACTCTG | >hg38_21_+_8433172_8446622_RNA45SN1_45S_50ntFlanks | 12004 | 12032 | 29 | CGGGCCGCCGGTGAAATACCACTACTCTG | 1554.6 | -7.2 | 1.2 | -6.1 | 1.0E-09 | 2.4E-09 | MB231/MB468 | MP |
| >hg38_21_+_8433172_8446622_RNA45SN1_45S_50ntFlanks@9738.9763.26\|GAAACGATCTCAACCTATTCTCAAAC | >hg38_21_+_8433172_8446622_RNA45SN1_45S_50ntFlanks | 9738 | 9763 | 26 | GAAACGATCTCAACCTATTCTCAAAC | 463.7 | -4.5 | 0.7 | -6.1 | 1.2E-09 | 2.4E-09 | MB231/MB468 | MITO |
| >hg38_21_+_8433172_8446622_RNA45SN1_45S_50ntFlanks@10865.10891.27\|TCTAAGGGCTGGGTCGGTCGGGCTGGG | >hg38_21_+_8433172_8446622_RNA45SN1_45S_50ntFlanks | 10865 | 10891 | 27 | TCTAAGGGCTGGGTCGGTCGGGCTGGG | 62.6 | -9.8 | 1.6 | -6.1 | 1.0E-09 | 2.4E-09 | MB231/MB468 | MP |
| >hg38_21_+_8433172_8446622_RNA45SN1_45S_50ntFlanks@11321.11344.24\|CACGTCTCGTCGCGCGCGCGTCCG | >hg38_21_+_8433172_8446622_RNA45SN1_45S_50ntFlanks | 11321 | 11344 | 24 | CACGTCTCGTCGCGCGCGCGTCCG | 11411.3 | -9.7 | 1.6 | -6.1 | 1.1E-09 | 2.4E-09 | MB231/MB468 | NUC |
| >hg38_21_+_8433172_8446622_RNA45SN1_45S_50ntFlanks@12712.12757.46\|CGGAGCCTCGGTTGGCCTCGGATAGCCGGTCCCCCGCCTGTCCCCG | >hg38_21_+_8433172_8446622_RNA45SN1_45S_50ntFlanks | 12712 | 12757 | 46 | CGGAGCCTCGGTTGGCCTCGGATAGCCGGTCCCCCGCCTGTCCCCG | 189.7 | -9.7 | 1.6 | -6.1 | 8.1E-10 | 2.4E-09 | BT20/MB468 | MITO |
| >hg38_21_+_8433172_8446622_RNA45SN1_45S_50ntFlanks@6721.6760.40\|AATTGCAGGACACATTGATCATCGACACTTCGAACGCACT | >hg38_21_+_8433172_8446622_RNA45SN1_45S_50ntFlanks | 6721 | 6760 | 40 | AATTGCAGGACACATTGATCATCGACACTTCGAACGCACT | 2676.2 | -8.5 | 1.4 | -6.1 | 1.2E-09 | 2.4E-09 | BT20/MB468 | NUC |
| >hg38_21_+_8433172_8446622_RNA45SN1_45S_50ntFlanks@9540.9569.30\|CTCTGGTGGAGGTCCGTAGCGGTCCTGACG | >hg38_21_+_8433172_8446622_RNA45SN1_45S_50ntFlanks | 9540 | 9569 | 30 | CTCTGGTGGAGGTCCGTAGCGGTCCTGACG | 627.2 | -7.2 | 1.2 | -6.1 | 1.2E-09 | 2.4E-09 | BT20/MB468 | NUC |
| >hg38_21_+_8433172_8446622_RNA45SN1_45S_50ntFlanks@6646.6685.40\|TCGTACGACTCTTAGCGGTGGATCACTCGGCTCGTGCGTC | >hg38_21_+_8433172_8446622_RNA45SN1_45S_50ntFlanks | 6646 | 6685 | 40 | TCGTACGACTCTTAGCGGTGGATCACTCGGCTCGTGCGTC | 29974.4 | 5.8 | 1.0 | 6.1 | 8.1E-10 | 2.4E-09 | BT20/MB468 | MITO |
| >MI0000469\|hsa-mir-125a&WithFlank&19\|+\|51693248\|51693345@21.42.22(+1A)&[MIMAT0000443&hsa-miR-125a-5p&offsets\|0\|-2(+1A);m-63&19\|+\|51693268\|51693289&offsets\|0\|0(+1A)]\|TCCCTGAGACCCTTTAACCTGTA | >MI0000469\|hsa-mir-125a&WithFlank&19\|+\|51693248\|51693345 | 21 | 42 | 22(+1A) | TCCCTGAGACCCTTTAACCTGTA | 1088.9 | 5.6 | 0.9 | 6.1 | 1.1E-09 | 2.4E-09 | MB231/MB468 | CYTO |
| >hg38_21_+_8433172_8446622_RNA45SN1_45S_50ntFlanks@6719.6739.21\|TGAATTGCAGGACACATTGAT | >hg38_21_+_8433172_8446622_RNA45SN1_45S_50ntFlanks | 6719 | 6739 | 21 | TGAATTGCAGGACACATTGAT | 768.3 | -5.7 | 0.9 | -6.1 | 1.1E-09 | 2.5E-09 | MB231/MB468 | CYTO |
| >MI0000650\|hsa-mir-200c&WithFlank&12\|+\|6963693\|6963772@50.72.23&[MIMAT0000617&hsa-miR-200c-3p&offsets\|0\|0;m-54&12\|+\|6963742\|6963764&offsets\|0\|0]\|TAATACTGCCGGGTAATGATGGA | >MI0000650\|hsa-mir-200c&WithFlank&12\|+\|6963693\|6963772 | 50 | 72 | 23 | TAATACTGCCGGGTAATGATGGA | 88450.6 | 5.7 | 0.9 | 6.5 | 1.1E-10 | 2.5E-09 | BT20/MB468 | MP |
| >hg38_21_+_8433172_8446622_RNA45SN1_45S_50ntFlanks@4283.4307.25\|CTTTAACGAGGATCCATTGGAGGGC | >hg38_21_+_8433172_8446622_RNA45SN1_45S_50ntFlanks | 4283 | 4307 | 25 | CTTTAACGAGGATCCATTGGAGGGC | 171.8 | -4.7 | 0.8 | -6.1 | 1.1E-09 | 2.5E-09 | MB231/MB468 | CYTO |
| >MI0000098\|hsa-mir-96&WithFlank&7\|-\|129774686\|129774775@15.37.23&[MIMAT0000095&hsa-miR-96-5p&offsets\|0\|0;m-248&7\|-\|129774739\|129774761&offsets\|0\|0]\|TTTGGCACTAGCACATTTTTGCT | >MI0000098\|hsa-mir-96&WithFlank&7\|-\|129774686\|129774775 | 15 | 37 | 23 | TTTGGCACTAGCACATTTTTGCT | 1016.9 | 4.2 | 0.7 | 6.1 | 1.1E-09 | 2.5E-09 | MB231/MB468 | NUC |
| >MI0000108\|hsa-mir-103a-2&WithFlank&20\|+\|3917488\|3917577@54.75.22;MI0000109\|hsa-mir-103a-1&WithFlank&5\|-\|168560890\|168560979@54.75.22&[MIMAT0000101&hsa-miR-103a-3p&offsets\|0\|-1;m-12&20\|+\|3917541\|3917563&offsets\|0\|-1];[MIMAT0000101_1&hsa-miR-103a-3p&offsets\|0\|-1;m-13&5\|-\|168560904\|168560926&offsets\|0\|-1;m-44220&5\|-\|168560908\|168560926&offsets\|0\|+3]\|AGCAGCATTGTACAGGGCTATG | >MI0000108\|hsa-mir-103a-2&WithFlank&20\|+\|3917488\|3917577 | 54 | 75 | 22 | AGCAGCATTGTACAGGGCTATG | 6456.8 | 5.4 | 0.9 | 6.1 | 1.2E-09 | 2.5E-09 | MB231/MB468 | MITO |
| >hg38_21_+_8433172_8446622_RNA45SN1_45S_50ntFlanks@5444.5465.22\|CTCGGATCGGCCCCGCCGGGGT | >hg38_21_+_8433172_8446622_RNA45SN1_45S_50ntFlanks | 5444 | 5465 | 22 | CTCGGATCGGCCCCGCCGGGGT | 491.6 | -7.9 | 1.3 | -6.1 | 1.2E-09 | 2.5E-09 | MB231/MB468 | MITO |
| >hg38_21_+_8433172_8446622_RNA45SN1_45S_50ntFlanks@6750.6770.21\|TCGAACGCACTTGCGGCCCCG | >hg38_21_+_8433172_8446622_RNA45SN1_45S_50ntFlanks | 6750 | 6770 | 21 | TCGAACGCACTTGCGGCCCCG | 181.9 | -10.6 | 1.6 | -6.5 | 1.1E-10 | 2.5E-09 | BT20/MB468 | MP |
| >hg38_21_+_8433172_8446622_RNA45SN1_45S_50ntFlanks@10385.10404.20\|CATGTGAACAGCAGTTGAAC | >hg38_21_+_8433172_8446622_RNA45SN1_45S_50ntFlanks | 10385 | 10404 | 20 | CATGTGAACAGCAGTTGAAC | 108.4 | -10.6 | 1.7 | -6.1 | 1.0E-09 | 2.5E-09 | MB231/MB468 | MP |
| >hg38_21_+_8433172_8446622_RNA45SN1_45S_50ntFlanks@10785.10818.34\|GAACAGCCTCTGGCATGTTGGAACAATGTAGGTA | >hg38_21_+_8433172_8446622_RNA45SN1_45S_50ntFlanks | 10785 | 10818 | 34 | GAACAGCCTCTGGCATGTTGGAACAATGTAGGTA | 69.4 | -10.0 | 1.6 | -6.1 | 1.0E-09 | 2.5E-09 | MB231/MB468 | MP |
| >hg38_21_+_8433172_8446622_RNA45SN1_45S_50ntFlanks@4732.4761.30\|AGAACGAAAGTCGGAGGTTCGAAGACGATC | >hg38_21_+_8433172_8446622_RNA45SN1_45S_50ntFlanks | 4732 | 4761 | 30 | AGAACGAAAGTCGGAGGTTCGAAGACGATC | 377.8 | 3.1 | 0.5 | 6.1 | 8.1E-10 | 2.5E-09 | BT20/MB468 | MITO |
| >am_MI0005763\|hsa-mir-941-1&WithFlank&20\|+\|63919443\|63919526@53.74.22(+1A);MI0005764\|hsa-mir-941-2&WithFlank&20\|+\|63919499\|63919582@53.74.22(+1A);MI0005765\|hsa-mir-941-3&WithFlank&20\|+\|63919555\|63919638@53.74.22(+1A);MI0005766\|hsa-mir-941-4&WithFlank&20\|+\|63919750\|63919833@53.74.22(+1A);MI0031520\|hsa-mir-941-5&WithFlank&20\|+\|63919862\|63919945@53.74.22(+1A)&[MIMAT0004984&hsa-miR-941&offsets\|0\|-1(+1A);m-263&20\|+\|63919495\|63919517&offsets\|0\|-1(+1A)];[MIMAT0004984_1&hsa-miR-941&offsets\|0\|-1(+1A);m-263&20\|+\|63919551\|63919573&offsets\|0\|-1(+1A)];[MIMAT0004984_2&hsa-miR-941&offsets\|0\|-1(+1A);m-263&20\|+\|63919607\|63919629&offsets\|0\|-1(+1A)];[MIMAT0004984_3&hsa-miR-941&offsets\|0\|-1(+1A);m-263&20\|+\|63919802\|63919824&offsets\|0\|-1(+1A)];[MIMAT0004984_4&hsa-miR-941&offsets\|0\|-1(+1A);m-263&20\|+\|63919914\|63919936&offsets\|0\|-1(+1A)]\|CACCCGGCTGTGTGCACATGTGA | >am_MI0005763\|hsa-mir-941-1&WithFlank&20\|+\|63919443\|63919526 | 53 | 74 | 22(+1A) | CACCCGGCTGTGTGCACATGTGA | 151.3 | 2.3 | 0.4 | 6.1 | 1.2E-09 | 2.5E-09 | BT20/MB468 | NUC |
| >hg38_21_+_8433172_8446622_RNA45SN1_45S_50ntFlanks@8380.8401.22\|CAAGAGGGCGTGAAACCGTTAA | >hg38_21_+_8433172_8446622_RNA45SN1_45S_50ntFlanks | 8380 | 8401 | 22 | CAAGAGGGCGTGAAACCGTTAA | 453.2 | -7.1 | 1.2 | -6.1 | 9.4E-10 | 2.5E-09 | BT20/MB468 | CYTO |
| >hg38_21_+_8433172_8446622_RNA45SN1_45S_50ntFlanks@11769.11803.35\|TAATTAGTGACGCGCATGAATGGATGAACGAGATT | >hg38_21_+_8433172_8446622_RNA45SN1_45S_50ntFlanks | 11769 | 11803 | 35 | TAATTAGTGACGCGCATGAATGGATGAACGAGATT | 1861.9 | -7.7 | 1.3 | -6.1 | 1.2E-09 | 2.5E-09 | BT20/MB468 | NUC |
| >hg38_21_+_8433172_8446622_RNA45SN1_45S_50ntFlanks@10317.10336.20\|GTGCAGATCTTGGTGGTAGT | >hg38_21_+_8433172_8446622_RNA45SN1_45S_50ntFlanks | 10317 | 10336 | 20 | GTGCAGATCTTGGTGGTAGT | 157.2 | 11.0 | 1.7 | 6.5 | 1.1E-10 | 2.5E-09 | BT20/MB468 | MP |
| >hg38_21_+_8433172_8446622_RNA45SN1_45S_50ntFlanks@12405.12439.35\|CTTTTTGATCCTTCGATGTCGGCTCTTCCTATCAT | >hg38_21_+_8433172_8446622_RNA45SN1_45S_50ntFlanks | 12405 | 12439 | 35 | CTTTTTGATCCTTCGATGTCGGCTCTTCCTATCAT | 596.2 | -4.1 | 0.7 | -6.1 | 1.2E-09 | 2.5E-09 | MB231/MB468 | MITO |
| >hg38_21_+_8433172_8446622_RNA45SN1_45S_50ntFlanks@11588.11623.36\|GCGGACCAGGGGAATCCGACTGTTTAATTAAAACAA | >hg38_21_+_8433172_8446622_RNA45SN1_45S_50ntFlanks | 11588 | 11623 | 36 | GCGGACCAGGGGAATCCGACTGTTTAATTAAAACAA | 235.6 | 11.6 | 1.8 | 6.5 | 1.1E-10 | 2.5E-09 | BT20/MB468 | MP |
| >hg38_21_+_8433172_8446622_RNA45SN1_45S_50ntFlanks@12071.12093.23\|CCCCGAGGGGCTCTCGCTTCTGG | >hg38_21_+_8433172_8446622_RNA45SN1_45S_50ntFlanks | 12071 | 12093 | 23 | CCCCGAGGGGCTCTCGCTTCTGG | 273.2 | -5.5 | 0.9 | -6.1 | 1.1E-09 | 2.5E-09 | MB231/MB468 | CYTO |
| >hg38_21_+_8433172_8446622_RNA45SN1_45S_50ntFlanks@10427.10456.30\|ATGGGCGAGCGCCGTTCCGAAGGGACGGGC | >hg38_21_+_8433172_8446622_RNA45SN1_45S_50ntFlanks | 10427 | 10456 | 30 | ATGGGCGAGCGCCGTTCCGAAGGGACGGGC | 886.1 | -7.0 | 1.1 | -6.1 | 1.0E-09 | 2.5E-09 | MB231/MB468 | MP |
| >hg38_21_+_8433172_8446622_RNA45SN1_45S_50ntFlanks@11188.11225.38\|CACCCGGGGGGCCGGCGGCGGCGGCGACTCTGGACGCG | >hg38_21_+_8433172_8446622_RNA45SN1_45S_50ntFlanks | 11188 | 11225 | 38 | CACCCGGGGGGCCGGCGGCGGCGGCGACTCTGGACGCG | 787.0 | -9.2 | 1.5 | -6.1 | 1.3E-09 | 2.5E-09 | BT20/MB468 | NUC |
| >hg38_21_+_8433172_8446622_RNA45SN1_45S_50ntFlanks@8118.8136.19\|GGGACATGTGGCGTACGGA | >hg38_21_+_8433172_8446622_RNA45SN1_45S_50ntFlanks | 8118 | 8136 | 19 | GGGACATGTGGCGTACGGA | 3424.0 | 15.1 | 2.3 | 6.7 | 2.1E-11 | 2.5E-09 | BT20/MB231 | MP |
| >MI0000650\|hsa-mir-200c&WithFlank&12\|+\|6963693\|6963772@50.72.23&[MIMAT0000617&hsa-miR-200c-3p&offsets\|0\|0;m-54&12\|+\|6963742\|6963764&offsets\|0\|0]\|TAATACTGCCGGGTAATGATGGA | >MI0000650\|hsa-mir-200c&WithFlank&12\|+\|6963693\|6963772 | 50 | 72 | 23 | TAATACTGCCGGGTAATGATGGA | 3522.5 | -4.7 | 0.8 | -6.1 | 1.0E-09 | 2.5E-09 | MB231/MB468 | MP |
| >hg38_21_+_8433172_8446622_RNA45SN1_45S_50ntFlanks@9624.9658.35\|AGTAGCTGGTTCCCTCCGAAGTTTCCCTCAGGATA | >hg38_21_+_8433172_8446622_RNA45SN1_45S_50ntFlanks | 9624 | 9658 | 35 | AGTAGCTGGTTCCCTCCGAAGTTTCCCTCAGGATA | 68.0 | -10.0 | 1.6 | -6.1 | 1.0E-09 | 2.5E-09 | MB231/MB468 | MP |
| >hg38_21_+_8433172_8446622_RNA45SN1_45S_50ntFlanks@6744.6771.28\|GACACTTCGAACGCACTTGCGGCCCCGG | >hg38_21_+_8433172_8446622_RNA45SN1_45S_50ntFlanks | 6744 | 6771 | 28 | GACACTTCGAACGCACTTGCGGCCCCGG | 741.5 | -4.1 | 0.7 | -6.1 | 9.5E-10 | 2.5E-09 | BT20/MB468 | CYTO |
| >hg38_21_+_8433172_8446622_RNA45SN1_45S_50ntFlanks@12905.12930.26\|CGCACCGCACGTTCGTGGGGAACCTG | >hg38_21_+_8433172_8446622_RNA45SN1_45S_50ntFlanks | 12905 | 12930 | 26 | CGCACCGCACGTTCGTGGGGAACCTG | 67.0 | -9.9 | 1.6 | -6.1 | 1.0E-09 | 2.5E-09 | MB231/MB468 | MP |

| | | | | | | | | | | | | | |
|---|---|---|---|---|---|---|---|---|---|---|---|---|---|
| >hg38_21_+_8433172_8446622_RNA45SN1_45S_50ntFlanks@8380.8407.28\|CAAGAGGGCGTGAAACCGTTAAGAGGTA | >hg38_21_+_8433172_8446622_RNA45SN1_45S_50ntFlanks | 8380 | 8407 | 28 | CAAGAGGGCGTGAAACCGTTAAGAGGTA | 580.4 | -5.3 | 0.9 | -6.1 | 1.0E-09 | 2.5E-09 | MB231/MB468 | MP |
| >hg38_21_+_8433172_8446622_RNA45SN1_45S_50ntFlanks@485.516.32\|GATGTGGTGACGTCGTGCTCTCCCGGGCCGGG | >hg38_21_+_8433172_8446622_RNA45SN1_45S_50ntFlanks | 485 | 516 | 32 | GATGTGGTGACGTCGTGCTCTCCCGGGCCGGG | 6983.8 | 13.4 | 2.2 | 6.1 | 1.1E-09 | 2.5E-09 | MB231/MB468 | NUC |
| >hg38_21_+_8433172_8446622_RNA45SN1_45S_50ntFlanks@3987.4023.37\|GCTTTGGTGACTCTAGATAACCTCGGGCCGATCGCAC | >hg38_21_+_8433172_8446622_RNA45SN1_45S_50ntFlanks | 3987 | 4023 | 37 | GCTTTGGTGACTCTAGATAACCTCGGGCCGATCGCAC | 677.8 | 6.1 | 1.0 | 6.1 | 1.1E-09 | 2.5E-09 | MB231/MB468 | CYTO |
| >hg38_21_+_8433172_8446622_RNA45SN1_45S_50ntFlanks@12707.12742.36\|CGCCGCGGAGCCTCGGTTGGCCTCGGATAGCCGGTC | >hg38_21_+_8433172_8446622_RNA45SN1_45S_50ntFlanks | 12707 | 12742 | 36 | CGCCGCGGAGCCTCGGTTGGCCTCGGATAGCCGGTC | 64.7 | -9.9 | 1.6 | -6.1 | 1.1E-09 | 2.5E-09 | MB231/MB468 | MP |
| >hg38_21_+_8433172_8446622_RNA45SN1_45S_50ntFlanks@12994.13017.24\|CCTCGCTGCGATCTATTGAAAGTC | >hg38_21_+_8433172_8446622_RNA45SN1_45S_50ntFlanks | 12994 | 13017 | 24 | CCTCGCTGCGATCTATTGAAAGTC | 183.3 | 3.3 | 0.5 | 6.1 | 1.1E-09 | 2.5E-09 | MB231/MB468 | CYTO |
| >hg38_21_+_8433172_8446622_RNA45SN1_45S_50ntFlanks@6722.6743.22\|ATTGCAGGACACATTGATCATC | >hg38_21_+_8433172_8446622_RNA45SN1_45S_50ntFlanks | 6722 | 6743 | 22 | ATTGCAGGACACATTGATCATC | 401.2 | -4.1 | 0.7 | -6.1 | 1.2E-09 | 2.5E-09 | MB231/MB468 | MITO |
| >hg38_21_+_8433172_8446622_RNA45SN1_45S_50ntFlanks@10873.10892.20\|CTGGGTCGGTCGGGCTGGGG | >hg38_21_+_8433172_8446622_RNA45SN1_45S_50ntFlanks | 10873 | 10892 | 20 | CTGGGTCGGTCGGGCTGGGG | 896.5 | -8.2 | 1.4 | -6.1 | 1.1E-09 | 2.5E-09 | MB231/MB468 | NUC |
| >hg38_21_+_8433172_8446622_RNA45SN1_45S_50ntFlanks@10786.10821.36\|AACAGCCTCTGGCATGTTGGAACAATGTAGGTAAGG | >hg38_21_+_8433172_8446622_RNA45SN1_45S_50ntFlanks | 10786 | 10821 | 36 | AACAGCCTCTGGCATGTTGGAACAATGTAGGTAAGG | 1375.8 | -5.0 | 0.8 | -6.1 | 1.2E-09 | 2.5E-09 | MB231/MB468 | MITO |
| >hg38_21_+_8433172_8446622_RNA45SN1_45S_50ntFlanks@11729.11754.26\|TAACTATGACTCTCTTAAGGTAGCCA | >hg38_21_+_8433172_8446622_RNA45SN1_45S_50ntFlanks | 11729 | 11754 | 26 | TAACTATGACTCTCTTAAGGTAGCCA | 199.4 | -9.9 | 1.6 | -6.1 | 9.6E-10 | 2.5E-09 | BT20/MB468 | CYTO |
| >MI0000440\|hsa-mir-27b&WithFlank&9\|+\|95085439\|95085547@67.86.20(+1U)&[MIMAT0000419&hsa-miR-27b-3p&offsets\|0\|-1(+1U);m-39&9\|+\|95085505\|95085525&offsets\|0\|-1(+1U)]\|TTCACAGTGGCTAAGTTCTGT | >MI0000440\|hsa-mir-27b&WithFlank&9\|+\|95085439\|95085547 | 67 | 86 | 20(+1U) | TTCACAGTGGCTAAGTTCTGT | 3182.8 | 5.6 | 0.9 | 6.1 | 1.1E-09 | 2.5E-09 | MB231/MB468 | CYTO |
| >hg38_21_+_8433172_8446622_RNA45SN1_45S_50ntFlanks@10781.10807.27\|AGGTGAACAGCCTCTGGCATGTTGGAA | >hg38_21_+_8433172_8446622_RNA45SN1_45S_50ntFlanks | 10781 | 10807 | 27 | AGGTGAACAGCCTCTGGCATGTTGGAA | 111.4 | -8.9 | 1.4 | -6.1 | 8.3E-10 | 2.5E-09 | BT20/MB468 | MITO |
| >hg38_21_+_8433172_8446622_RNA45SN1_45S_50ntFlanks@10118.10138.21\|CGTCGGAGGGCGGCGGCGGCG | >hg38_21_+_8433172_8446622_RNA45SN1_45S_50ntFlanks | 10118 | 10138 | 21 | CGTCGGAGGGCGGCGGCGGCG | 123.5 | -10.8 | 1.8 | -6.1 | 1.1E-09 | 2.5E-09 | MB231/MB468 | MP |
| >hg38_21_+_8433172_8446622_RNA45SN1_45S_50ntFlanks@11729.11765.37\|TAACTATGACTCTCTTAAGGTAGCCAAATGCCTCGTC | >hg38_21_+_8433172_8446622_RNA45SN1_45S_50ntFlanks | 11729 | 11765 | 37 | TAACTATGACTCTCTTAAGGTAGCCAAATGCCTCGTC | 8715.4 | -6.5 | 1.1 | -6.1 | 9.7E-10 | 2.5E-09 | BT20/MB468 | CYTO |
| >hg38_21_+_8433172_8446622_RNA45SN1_45S_50ntFlanks@12625.12661.37\|CAATGGGGCGAAGCTACCATCTGTGGGATTATGACTG | >hg38_21_+_8433172_8446622_RNA45SN1_45S_50ntFlanks | 12625 | 12661 | 37 | CAATGGGGCGAAGCTACCATCTGTGGGATTATGACTG | 2473.5 | -7.2 | 1.2 | -6.1 | 1.2E-09 | 2.5E-09 | MB231/MB468 | MITO |
| >hg38_21_+_8433172_8446622_RNA45SN1_45S_50ntFlanks@11661.11682.22\|TGATTTCTGCCCAGTGCTCTGA | >hg38_21_+_8433172_8446622_RNA45SN1_45S_50ntFlanks | 11661 | 11682 | 22 | TGATTTCTGCCCAGTGCTCTGA | 302.0 | -7.6 | 1.3 | -6.1 | 1.2E-09 | 2.5E-09 | MB231/MB468 | MITO |
| >hg38_21_+_8433172_8446622_RNA45SN1_45S_50ntFlanks@8717.8736.20\|GGGAAGGTGGCTCGGGGGGC | >hg38_21_+_8433172_8446622_RNA45SN1_45S_50ntFlanks | 8717 | 8736 | 20 | GGGAAGGTGGCTCGGGGGGC | 90.1 | -10.4 | 1.7 | -6.1 | 1.1E-09 | 2.5E-09 | MB231/MB468 | MP |
| >hg38_21_+_8433172_8446622_RNA45SN1_45S_50ntFlanks@10929.10946.18\|AGGCGCCGCCGCCCCCCC | >hg38_21_+_8433172_8446622_RNA45SN1_45S_50ntFlanks | 10929 | 10946 | 18 | AGGCGCCGCCGCCCCCCC | 198.5 | -11.5 | 1.9 | -6.1 | 1.1E-09 | 2.5E-09 | MB231/MB468 | MP |
| >am_trna16_TyrGTA_14_-_21131351_21131444@1.25.25,trna17_TyrGTA_14_-_21128117_21128210@1.25.25,trna17_TyrGTA_6_+_26595102_26595190@1.25.25,trna18_TyrGTA_14_-_21125623_21125716@1.25.25,trna19_TyrGTA_14_-_21121258_21121351@1.25.25,trna4_TyrGTA_8_+_67025602_67025694@1.25.25,trna5_TyrGTA_14_+_21151432_21151520@1.25.25,trna5_TyrGTA_8_+_67026223_67026311@1.25.25\|CCTTCGATAGCTCAGCTGGTAGAGC | >am_trna16_TyrGTA_14_-_21131351_21131444 | 1 | 25 | 25 | CCTTCGATAGCTCAGCTGGTAGAGC | 1420.2 | 10.0 | 1.6 | 6.1 | 9.7E-10 | 2.5E-09 | BT20/MB468 | CYTO |
| >MI0000279\|hsa-mir-196a-2&WithFlank&12\|+\|53991732\|53991853@31.51.21(+1A);MI0000238\|hsa-mir-196a-1&WithFlank&17\|-\|48632484\|48632565@13.33.21(+1A)&[MIMAT0000226&hsa-miR-196a-5p&offsets\|0\|-1(+1A);m-261&12\|+\|53991762\|53991783&offsets\|0\|-1(+1A)];[MIMAT0000226_1&hsa-miR-196a-5p&offsets\|0\|-1(+1A);m-268&17\|-\|48632532\|48632553&offsets\|0\|-1(+1A)]\|TAGGTAGTTTCATGTTGTTGGA | >MI0000279\|hsa-mir-196a-2&WithFlank&12\|+\|53991732\|53991853 | 31 | 51 | 21(+1A) | TAGGTAGTTTCATGTTGTTGGA | 402.6 | 12.0 | 1.8 | 6.7 | 2.2E-11 | 2.5E-09 | BT20/MB231 | MP |
| >MI0000102\|hsa-mir-100&WithFlank&11\|-\|122152223\|122152314@22.40.19&[MIMAT0000098&hsa-miR-100-5p&offsets\|+3\|0;m-27&11\|-\|122152276\|122152296&offsets\|+3\|+1]\|CCGTAGATCCGAACTTGTG | >MI0000102\|hsa-mir-100&WithFlank&11\|-\|122152223\|122152314 | 22 | 40 | 19 | CCGTAGATCCGAACTTGTG | 675.6 | -12.8 | 1.9 | -6.7 | 2.2E-11 | 2.5E-09 | BT20/MB231 | MP |
| >MI0000294\|hsa-mir-218-1&WithFlank&4\|+\|20528269\|20528390@31.52.22;MI0000295\|hsa-mir-218-2&WithFlank&5\|-\|168768140\|168768261@31.52.22&[MIMAT0000275&hsa-miR-218-5p&offsets\|0\|+1;m-185&4\|+\|20528299\|20528319&offsets\|0\|+1];[MIMAT0000275_1&hsa-miR-218-5p&offsets\|0\|+1;m-187&5\|-\|168768211\|168768231&offsets\|0\|+1]\|TTGTGCTTGATCTAACCATGTG | >MI0000294\|hsa-mir-218-1&WithFlank&4\|+\|20528269\|20528390 | 31 | 52 | 22 | TTGTGCTTGATCTAACCATGTG | 348.8 | -11.8 | 1.8 | -6.7 | 2.3E-11 | 2.5E-09 | BT20/MB231 | MP |
| >hg38_21_+_8433172_8446622_RNA45SN1_45S_50ntFlanks@12413.12443.31\|TCCTTCGATGTCGGCTCTTCCTATCATTGTG | >hg38_21_+_8433172_8446622_RNA45SN1_45S_50ntFlanks | 12413 | 12443 | 31 | TCCTTCGATGTCGGCTCTTCCTATCATTGTG | 1681.2 | -7.8 | 1.3 | -6.1 | 1.3E-09 | 2.5E-09 | BT20/MB468 | NUC |
| >hg38_21_+_8433172_8446622_RNA45SN1_45S_50ntFlanks@12456.12496.41\|CCAAGCGTTGGATTGTTCACCCACTAATAGGGAACGTGAGC | >hg38_21_+_8433172_8446622_RNA45SN1_45S_50ntFlanks | 12456 | 12496 | 41 | CCAAGCGTTGGATTGTTCACCCACTAATAGGGAACGTGAGC | 1974.8 | -8.2 | 1.3 | -6.1 | 1.3E-09 | 2.5E-09 | BT20/MB468 | NUC |
| >hg38_21_+_8433172_8446622_RNA45SN1_45S_50ntFlanks@12635.12654.20\|AAGCTACCATCTGTGGGATT | >hg38_21_+_8433172_8446622_RNA45SN1_45S_50ntFlanks | 12635 | 12654 | 20 | AAGCTACCATCTGTGGGATT | 746.7 | -6.3 | 1.0 | -6.1 | 1.1E-09 | 2.5E-09 | MB231/MB468 | NUC |
| >hg38_21_+_8433172_8446622_RNA45SN1_45S_50ntFlanks@3975.4005.31\|CGGGCGCCGGCGGCTTTGGTGACTCTAGATA | >hg38_21_+_8433172_8446622_RNA45SN1_45S_50ntFlanks | 3975 | 4005 | 31 | CGGGCGCCGGCGGCTTTGGTGACTCTAGATA | 122.1 | -10.8 | 1.8 | -6.1 | 1.1E-09 | 2.5E-09 | MB231/MB468 | MP |
| >hg38_21_+_8433172_8446622_RNA45SN1_45S_50ntFlanks@10386.10424.39\|ATGTGAACAGCAGTTGAACATGGGTCAGTCGGTCCTGAG | >hg38_21_+_8433172_8446622_RNA45SN1_45S_50ntFlanks | 10386 | 10424 | 39 | ATGTGAACAGCAGTTGAACATGGGTCAGTCGGTCCTGAG | 1941.9 | 7.5 | 1.2 | 6.1 | 9.7E-10 | 2.5E-09 | BT20/MB468 | CYTO |
| >trnaMT_HisGTG_MT_+_12138_12206@32.71.40\|TGAATCTGACAACAGAGGCTTACGACCCCTTATTTACCCC | >trnaMT_HisGTG_MT_+_12138_12206 | 32 | 71 | 40 | TGAATCTGACAACAGAGGCTTACGACCCCTTATTTACCCC | 1345.5 | 7.4 | 1.2 | 6.1 | 1.1E-09 | 2.5E-09 | MB231/MB468 | CYTO |
| >hg38_21_+_8433172_8446622_RNA45SN1_45S_50ntFlanks@9574.9591.18\|AATCGGTCGTCCGACCTG | >hg38_21_+_8433172_8446622_RNA45SN1_45S_50ntFlanks | 9574 | 9591 | 18 | AATCGGTCGTCCGACCTG | 106.8 | -10.6 | 1.7 | -6.1 | 1.1E-09 | 2.5E-09 | MB231/MB468 | MP |
| >hg38_21_+_8433172_8446622_RNA45SN1_45S_50ntFlanks@12717.12745.29\|CCTCGGTTGGCCTCGGATAGCCGGTCCCC | >hg38_21_+_8433172_8446622_RNA45SN1_45S_50ntFlanks | 12717 | 12745 | 29 | CCTCGGTTGGCCTCGGATAGCCGGTCCCC | 126.0 | -9.1 | 1.5 | -6.1 | 8.4E-10 | 2.5E-09 | BT20/MB468 | MITO |
| >hg38_21_+_8433172_8446622_RNA45SN1_45S_50ntFlanks@12044.12067.24\|ACTGACCCGGTGAGGCGGGGGGGC | >hg38_21_+_8433172_8446622_RNA45SN1_45S_50ntFlanks | 12044 | 12067 | 24 | ACTGACCCGGTGAGGCGGGGGGGC | 92.3 | -10.4 | 1.7 | -6.1 | 1.1E-09 | 2.5E-09 | MB231/MB468 | MP |
| >hg38_21_+_8433172_8446622_RNA45SN1_45S_50ntFlanks@3838.3860.23\|CTCTCCTACTTGGATAACTGTGG | >hg38_21_+_8433172_8446622_RNA45SN1_45S_50ntFlanks | 3838 | 3860 | 23 | CTCTCCTACTTGGATAACTGTGG | 1137.9 | -7.0 | 1.2 | -6.1 | 1.2E-09 | 2.5E-09 | MB231/MB468 | MITO |
| >hg38_21_+_8433172_8446622_RNA45SN1_45S_50ntFlanks@6741.6771.31\|ATCGACACTTCGAACGCACTTGCGGCCCCGG | >hg38_21_+_8433172_8446622_RNA45SN1_45S_50ntFlanks | 6741 | 6771 | 31 | ATCGACACTTCGAACGCACTTGCGGCCCCGG | 998.3 | -6.6 | 1.1 | -6.1 | 1.1E-09 | 2.5E-09 | MB231/MB468 | NUC |
| >MI0003560\|hsa-mir-92b&WithFlank&1\|+\|155195171\|155195278@67.87.21(+1U)&[MIMAT0003218&hsa-miR-92b-3p&offsets\|0\|-1(+1U);m-115&1\|+\|155195237\|155195258&offsets\|0\|-1(+1U)]\|TATTGCACTCGTCCCGGCCTCT | >MI0003560\|hsa-mir-92b&WithFlank&1\|+\|155195171\|155195278 | 67 | 87 | 21(+1U) | TATTGCACTCGTCCCGGCCTCT | 3293.0 | 7.4 | 1.2 | 6.4 | 1.1E-10 | 2.6E-09 | BT20/MB468 | MP |
| >hg38_21_+_8433172_8446622_RNA45SN1_45S_50ntFlanks@8168.8187.20\|CCCAAGTCCTTCTGATCGAG | >hg38_21_+_8433172_8446622_RNA45SN1_45S_50ntFlanks | 8168 | 8187 | 20 | CCCAAGTCCTTCTGATCGAG | 511.1 | -8.0 | 1.3 | -6.1 | 1.2E-09 | 2.6E-09 | MB231/MB468 | MITO |
| >hg38_21_+_8433172_8446622_RNA45SN1_45S_50ntFlanks@11876.11893.18\|AAGACCCTGTTGAGCTTG | >hg38_21_+_8433172_8446622_RNA45SN1_45S_50ntFlanks | 11876 | 11893 | 18 | AAGACCCTGTTGAGCTTG | 1508.5 | -6.9 | 1.1 | -6.1 | 1.1E-09 | 2.6E-09 | MB231/MB468 | NUC |
| >hg38_21_+_8433172_8446622_RNA45SN1_45S_50ntFlanks@8380.8407.28\|CAAGAGGGCGTGAAACCGTTAAGAGGTA | >hg38_21_+_8433172_8446622_RNA45SN1_45S_50ntFlanks | 8380 | 8407 | 28 | CAAGAGGGCGTGAAACCGTTAAGAGGTA | 761.4 | -6.5 | 1.1 | -6.1 | 1.2E-09 | 2.6E-09 | MB231/MB468 | MITO |
| >MI0000078\|hsa-mir-22&WithFlank&17\|-\|1713897\|1713993@21.41.21(+1U)&[MIMAT0004495&hsa-miR-22-5p&offsets\|0\|-1(+1U);m-217&17\|-\|1713953\|1713973&offsets\|0\|0(+1U)]\|AGTTCTTCAGTGGCAAGCTTTT | >MI0000078\|hsa-mir-22&WithFlank&17\|-\|1713897\|1713993 | 21 | 41 | 21(+1U) | AGTTCTTCAGTGGCAAGCTTTT | 87.3 | 9.7 | 1.6 | 6.1 | 1.1E-09 | 2.6E-09 | MB231/MB468 | MP |
| >hg38_21_+_8433172_8446622_RNA45SN1_45S_50ntFlanks@3843.3872.30\|CTACTTGGATAACTGTGGTAATTCTAGAGC | >hg38_21_+_8433172_8446622_RNA45SN1_45S_50ntFlanks | 3843 | 3872 | 30 | CTACTTGGATAACTGTGGTAATTCTAGAGC | 291.4 | -6.3 | 1.0 | -6.1 | 1.1E-09 | 2.6E-09 | MB231/MB468 | MP |
| >hg38_21_+_8433172_8446622_RNA45SN1_45S_50ntFlanks@3782.3799.18\|CAGTGAAACTGCGAATGG | >hg38_21_+_8433172_8446622_RNA45SN1_45S_50ntFlanks | 3782 | 3799 | 18 | CAGTGAAACTGCGAATGG | 324.1 | -7.7 | 1.3 | -6.1 | 1.1E-09 | 2.6E-09 | MB231/MB468 | MP |
| >hg38_21_+_8433172_8446622_RNA45SN1_45S_50ntFlanks@4557.4574.18\|CAGCTAGGAATAATGGAA | >hg38_21_+_8433172_8446622_RNA45SN1_45S_50ntFlanks | 4557 | 4574 | 18 | CAGCTAGGAATAATGGAA | 138.1 | -10.2 | 1.6 | -6.4 | 1.1E-10 | 2.6E-09 | BT20/MB468 | MP |
| >hg38_21_+_8433172_8446622_RNA45SN1_45S_50ntFlanks@7984.8008.25\|AGATCAGACGTGGCGACCCGCTGAA | >hg38_21_+_8433172_8446622_RNA45SN1_45S_50ntFlanks | 7984 | 8008 | 25 | AGATCAGACGTGGCGACCCGCTGAA | 4141.4 | -5.8 | 0.9 | -6.4 | 1.1E-10 | 2.6E-09 | BT20/MB468 | MP |
| >hg38_21_+_8433172_8446622_RNA45SN1_45S_50ntFlanks@11792.11810.19\|ATGAACGAGATTCCCACTG | >hg38_21_+_8433172_8446622_RNA45SN1_45S_50ntFlanks | 11792 | 11810 | 19 | ATGAACGAGATTCCCACTG | 92.3 | -8.6 | 1.4 | -6.1 | 8.5E-10 | 2.6E-09 | BT20/MB468 | MITO |
| >hg38_21_+_8433172_8446622_RNA45SN1_45S_50ntFlanks@11534.11551.18\|GTTCCGCGCGGCGCCTCG | >hg38_21_+_8433172_8446622_RNA45SN1_45S_50ntFlanks | 11534 | 11551 | 18 | GTTCCGCGCGGCGCCTCG | 1572.6 | -6.1 | 1.0 | -6.1 | 1.1E-09 | 2.6E-09 | MB231/MB468 | CYTO |
| >hg38_21_+_8433172_8446622_RNA45SN1_45S_50ntFlanks@9794.9831.38\|CGTGGAGCCGGGCGTGGAATGCGAGTGCCTAGTGGGCC | >hg38_21_+_8433172_8446622_RNA45SN1_45S_50ntFlanks | 9794 | 9831 | 38 | CGTGGAGCCGGGCGTGGAATGCGAGTGCCTAGTGGGCC | 62.0 | -9.8 | 1.6 | -6.1 | 1.1E-09 | 2.6E-09 | MB231/MB468 | MP |
| >hg38_21_+_8433172_8446622_RNA45SN1_45S_50ntFlanks@6730.6753.24\|ACACATTGATCATCGACACTTCGA | >hg38_21_+_8433172_8446622_RNA45SN1_45S_50ntFlanks | 6730 | 6753 | 24 | ACACATTGATCATCGACACTTCGA | 60.2 | -9.8 | 1.6 | -6.1 | 1.1E-09 | 2.6E-09 | MB231/MB468 | MP |
| >hg38_21_+_8433172_8446622_RNA45SN1_45S_50ntFlanks@12396.12414.19\|GCGACGTCGCTTTTTGATC | >hg38_21_+_8433172_8446622_RNA45SN1_45S_50ntFlanks | 12396 | 12414 | 19 | GCGACGTCGCTTTTTGATC | 3414.3 | -5.1 | 0.8 | -6.1 | 1.3E-09 | 2.6E-09 | BT20/MB468 | NUC |
| >hg38_21_+_8433172_8446622_RNA45SN1_45S_50ntFlanks@11315.11336.22\|CCACCCCACGTCTCGTCGCGCG | >hg38_21_+_8433172_8446622_RNA45SN1_45S_50ntFlanks | 11315 | 11336 | 22 | CCACCCCACGTCTCGTCGCGCG | 610.7 | -7.5 | 1.2 | -6.1 | 1.1E-09 | 2.6E-09 | MB231/MB468 | CYTO |
| >MI0000811\|hsa-mir-148b&WithFlank&12\|+\|54337210\|54337320@69.91.23&[MIMAT0000759&hsa-miR-148b-3p&offsets\|0\|+1;m-77&12\|+\|54337278\|54337299&offsets\|0\|+1]\|TCAGTGCATCACAGAACTTTGTC | >MI0000811\|hsa-mir-148b&WithFlank&12\|+\|54337210\|54337320 | 69 | 91 | 23 | TCAGTGCATCACAGAACTTTGTC | 138.0 | 2.9 | 0.5 | 6.1 | 1.3E-09 | 2.6E-09 | BT20/MB468 | NUC |

<table>
<tr><td></td><td></td><td></td><td></td><td></td><td></td><td></td><td></td><td></td><td></td><td></td><td></td><td></td><td></td></tr>
<tr><td>>hg38_1_-_228634819_228635039_RNA5S12_5S_50ntFlanks@131.171.41|GGATGGGAGACCGCCTGGGAATACCGGGTGCTGTAGGCTTT</td><td>>hg38_1_-_228634819_228635039_RNA5S12_5S_50ntFlanks</td><td>131</td><td>171</td><td>41</td><td>GGATGGGAGACCGCCTGGGAATACCGGGTGCTGTAGGCTTT</td><td>711.2</td><td>5.8</td><td>1.0</td><td>6.1</td><td>1.1E-09</td><td>2.6E-09</td><td>MB231/MB468</td><td>NUC</td></tr>
<tr><td>>hg38_21_+_8433172_8446622_RNA45SN1_45S_50ntFlanks@12526.12549.24|TTTTACCCTACTGATGATGTGTTG</td><td>>hg38_21_+_8433172_8446622_RNA45SN1_45S_50ntFlanks</td><td>12526</td><td>12549</td><td>24</td><td>TTTTACCCTACTGATGATGTGTTG</td><td>209.3</td><td>-10.8</td><td>1.7</td><td>-6.4</td><td>1.2E-10</td><td>2.6E-09</td><td>BT20/MB468</td><td>MP</td></tr>
<tr><td>>hg38_21_+_8433172_8446622_RNA45SN1_45S_50ntFlanks@9501.9523.23|ATGGTGAACTATGCCTGGGCAGG</td><td>>hg38_21_+_8433172_8446622_RNA45SN1_45S_50ntFlanks</td><td>9501</td><td>9523</td><td>23</td><td>ATGGTGAACTATGCCTGGGCAGG</td><td>85.7</td><td>-10.3</td><td>1.7</td><td>-6.1</td><td>1.1E-09</td><td>2.6E-09</td><td>MB231/MB468</td><td>MP</td></tr>
<tr><td>>hg38_21_+_8433172_8446622_RNA45SN1_45S_50ntFlanks@10534.10567.34|CCGCGAGGCGTCCAGTGCGGTAACGCGACCGATC</td><td>>hg38_21_+_8433172_8446622_RNA45SN1_45S_50ntFlanks</td><td>10534</td><td>10567</td><td>34</td><td>CCGCGAGGCGTCCAGTGCGGTAACGCGACCGATC</td><td>691.7</td><td>-8.5</td><td>1.3</td><td>-6.4</td><td>1.2E-10</td><td>2.6E-09</td><td>BT20/MB468</td><td>MP</td></tr>
<tr><td>>hg38_21_+_8433172_8446622_RNA45SN1_45S_50ntFlanks@12413.12432.20|TCCTTCGATGTCGGCTCTTC</td><td>>hg38_21_+_8433172_8446622_RNA45SN1_45S_50ntFlanks</td><td>12413</td><td>12432</td><td>20</td><td>TCCTTCGATGTCGGCTCTTC</td><td>1091.8</td><td>-4.1</td><td>0.7</td><td>-6.1</td><td>1.2E-09</td><td>2.6E-09</td><td>MB231/MB468</td><td>NUC</td></tr>
<tr><td>>hg38_21_+_8433172_8446622_RNA45SN1_45S_50ntFlanks@9533.9552.20|GAGGAAACTCTGGTGGAGGT</td><td>>hg38_21_+_8433172_8446622_RNA45SN1_45S_50ntFlanks</td><td>9533</td><td>9552</td><td>20</td><td>GAGGAAACTCTGGTGGAGGT</td><td>395.9</td><td>-3.8</td><td>0.6</td><td>-6.1</td><td>1.2E-09</td><td>2.6E-09</td><td>MB231/MB468</td><td>NUC</td></tr>
<tr><td>>MI0000098|hsa-mir-96&WithFlank&7|-|129774686|129774775@15.36.22(+1A)&[MIMAT0000095&hsa-miR-96-5p&offsets|0|-1(+1A);m-248&7|-|129774739|129774761&offsets|0|-1(+1A)]|TTTGGCACTAGCACATTTTTGCA</td><td>>MI0000098|hsa-mir-96&WithFlank&7|-|129774686|129774775</td><td>15</td><td>36</td><td>22(+1A)</td><td>TTTGGCACTAGCACATTTTTGCA</td><td>390.9</td><td>6.3</td><td>1.0</td><td>6.5</td><td>6.9E-11</td><td>2.6E-09</td><td>BT20/MB231</td><td>CYTO</td></tr>
<tr><td>>hg38_21_+_8433172_8446622_RNA45SN1_45S_50ntFlanks@10349.10372.24|AACGAGAACTTTGAAGGCCGAAGT</td><td>>hg38_21_+_8433172_8446622_RNA45SN1_45S_50ntFlanks</td><td>10349</td><td>10372</td><td>24</td><td>AACGAGAACTTTGAAGGCCGAAGT</td><td>604.1</td><td>-2.5</td><td>0.4</td><td>-6.1</td><td>1.3E-09</td><td>2.6E-09</td><td>BT20/MB468</td><td>NUC</td></tr>
<tr><td>>hg38_21_+_8433172_8446622_RNA45SN1_45S_50ntFlanks@10393.10431.39|CAGCAGTTGAACATGGGTCAGTCGGTCCTGAGAGATGGG</td><td>>hg38_21_+_8433172_8446622_RNA45SN1_45S_50ntFlanks</td><td>10393</td><td>10431</td><td>39</td><td>CAGCAGTTGAACATGGGTCAGTCGGTCCTGAGAGATGGG</td><td>697.4</td><td>-9.2</td><td>1.5</td><td>-6.1</td><td>1.3E-09</td><td>2.6E-09</td><td>BT20/MB468</td><td>NUC</td></tr>
<tr><td>>hg38_21_+_8433172_8446622_RNA45SN1_45S_50ntFlanks@10914.10945.32|CGCCGCGGCTGGACGAGGCGCCGCCGCCCCCC</td><td>>hg38_21_+_8433172_8446622_RNA45SN1_45S_50ntFlanks</td><td>10914</td><td>10945</td><td>32</td><td>CGCCGCGGCTGGACGAGGCGCCGCCGCCCCCC</td><td>152.4</td><td>-10.3</td><td>1.6</td><td>-6.4</td><td>1.2E-10</td><td>2.6E-09</td><td>BT20/MB468</td><td>MP</td></tr>
<tr><td>>hg38_21_+_8433172_8446622_RNA45SN1_45S_50ntFlanks@5077.5115.39|CCGATAACGAACGAGACTCTGGCATGCTAACTAGTTACG</td><td>>hg38_21_+_8433172_8446622_RNA45SN1_45S_50ntFlanks</td><td>5077</td><td>5115</td><td>39</td><td>CCGATAACGAACGAGACTCTGGCATGCTAACTAGTTACG</td><td>95.0</td><td>-10.4</td><td>1.7</td><td>-6.1</td><td>1.1E-09</td><td>2.6E-09</td><td>MB231/MB468</td><td>MP</td></tr>
<tr><td>>hg38_21_+_8433172_8446622_RNA45SN1_45S_50ntFlanks@6733.6772.40|CATTGATCATCGACACTTCGAACGCACTTGCGGCCCCGGG</td><td>>hg38_21_+_8433172_8446622_RNA45SN1_45S_50ntFlanks</td><td>6733</td><td>6772</td><td>40</td><td>CATTGATCATCGACACTTCGAACGCACTTGCGGCCCCGGG</td><td>278.5</td><td>-12.0</td><td>2.0</td><td>-6.1</td><td>1.1E-09</td><td>2.6E-09</td><td>MB231/MB468</td><td>MP</td></tr>
<tr><td>>hg38_21_+_8433172_8446622_RNA45SN1_45S_50ntFlanks@9569.9596.28|GTGCAAATCGGTCGTCCGACCTGGGTAT</td><td>>hg38_21_+_8433172_8446622_RNA45SN1_45S_50ntFlanks</td><td>9569</td><td>9596</td><td>28</td><td>GTGCAAATCGGTCGTCCGACCTGGGTAT</td><td>1093.9</td><td>-6.4</td><td>1.0</td><td>-6.1</td><td>1.3E-09</td><td>2.6E-09</td><td>MB231/MB468</td><td>MITO</td></tr>
<tr><td>>hg38_21_+_8433172_8446622_RNA45SN1_45S_50ntFlanks@4725.4758.34|TTAATCAAGAACGAAAGTCGGAGGTTCGAAGACG</td><td>>hg38_21_+_8433172_8446622_RNA45SN1_45S_50ntFlanks</td><td>4725</td><td>4758</td><td>34</td><td>TTAATCAAGAACGAAAGTCGGAGGTTCGAAGACG</td><td>936.3</td><td>-3.6</td><td>0.6</td><td>-6.1</td><td>8.7E-10</td><td>2.6E-09</td><td>BT20/MB468</td><td>MITO</td></tr>
<tr><td>>hg38_21_+_8433172_8446622_RNA45SN1_45S_50ntFlanks@12044.12071.28|ACTGACCCGGTGAGGCGGGGGGGCGAGC</td><td>>hg38_21_+_8433172_8446622_RNA45SN1_45S_50ntFlanks</td><td>12044</td><td>12071</td><td>28</td><td>ACTGACCCGGTGAGGCGGGGGGGCGAGC</td><td>626.3</td><td>-7.9</td><td>1.3</td><td>-6.1</td><td>1.2E-09</td><td>2.6E-09</td><td>MB231/MB468</td><td>NUC</td></tr>
<tr><td>>hg38_21_+_8433172_8446622_RNA45SN1_45S_50ntFlanks@4046.4063.18|CATTCGAACGTCTGCCCT</td><td>>hg38_21_+_8433172_8446622_RNA45SN1_45S_50ntFlanks</td><td>4046</td><td>4063</td><td>18</td><td>CATTCGAACGTCTGCCCT</td><td>3462.4</td><td>-4.3</td><td>0.7</td><td>-6.1</td><td>1.0E-09</td><td>2.6E-09</td><td>BT20/MB468</td><td>CYTO</td></tr>
<tr><td>>MI0000749|hsa-mir-30e&WithFlank&1|+|40754349|40754452@65.85.21(+1U)&[MIMAT0000693&hsa-miR-30e-3p&offsets|0|-1(+1U);m-25&1|+|40754413|40754434&offsets|0|-1(+1U)]|CTTTCAGTCGGATGTTTACAGT</td><td>>MI0000749|hsa-mir-30e&WithFlank&1|+|40754349|40754452</td><td>65</td><td>85</td><td>21(+1U)</td><td>CTTTCAGTCGGATGTTTACAGT</td><td>3695.6</td><td>3.6</td><td>0.6</td><td>6.1</td><td>1.2E-09</td><td>2.6E-09</td><td>MB231/MB468</td><td>NUC</td></tr>
<tr><td>>hg38_21_+_8433172_8446622_RNA45SN1_45S_50ntFlanks@10346.10383.38|TCAAACGAGAACTTTGAAGGCCGAAGTGGAGAAGGGTT</td><td>>hg38_21_+_8433172_8446622_RNA45SN1_45S_50ntFlanks</td><td>10346</td><td>10383</td><td>38</td><td>TCAAACGAGAACTTTGAAGGCCGAAGTGGAGAAGGGTT</td><td>1048.0</td><td>-4.6</td><td>0.8</td><td>-6.1</td><td>1.3E-09</td><td>2.6E-09</td><td>MB231/MB468</td><td>MITO</td></tr>
<tr><td>>trna14_LysTTT_11_-_59327808_59327880@1.36.36,trna2_LysTTT_17_+_8022473_8022545@1.36.36,trna54_LysTTT_1_+_204475655_204475727@1.36.36,trna5_LysTTT_11_+_59323902_59323974@1.36.36,trna62_LysTTT_1_-_204476158_204476230@1.36.36,trna76_LysTTT_6_+_28918806_28918878@1.36.36|GCCCGGATAGCTCAGTCGGTAGAGCATCAGACTTTT</td><td>>trna14_LysTTT_11_-_59327808_59327880</td><td>1</td><td>36</td><td>36</td><td>GCCCGGATAGCTCAGTCGGTAGAGCATCAGACTTTT</td><td>770.3</td><td>8.8</td><td>1.4</td><td>6.1</td><td>1.0E-09</td><td>2.6E-09</td><td>BT20/MB468</td><td>CYTO</td></tr>
<tr><td>>hg38_21_+_8433172_8446622_RNA45SN1_45S_50ntFlanks@3899.3938.40|CCTTCGCGGGGGGGATGCGTGCATTTATCAGATCAAAACC</td><td>>hg38_21_+_8433172_8446622_RNA45SN1_45S_50ntFlanks</td><td>3899</td><td>3938</td><td>40</td><td>CCTTCGCGGGGGGGATGCGTGCATTTATCAGATCAAAACC</td><td>439.0</td><td>-11.9</td><td>1.8</td><td>-6.4</td><td>1.2E-10</td><td>2.6E-09</td><td>BT20/MB468</td><td>MP</td></tr>
<tr><td>>hg38_21_+_8433172_8446622_RNA45SN1_45S_50ntFlanks@9240.9272.33|GCGGAGCGAGCGCACGGGGTCGGCGGCGACGTC</td><td>>hg38_21_+_8433172_8446622_RNA45SN1_45S_50ntFlanks</td><td>9240</td><td>9272</td><td>33</td><td>GCGGAGCGAGCGCACGGGGTCGGCGGCGACGTC</td><td>141.3</td><td>-10.2</td><td>1.6</td><td>-6.4</td><td>1.2E-10</td><td>2.6E-09</td><td>BT20/MB468</td><td>MP</td></tr>
<tr><td>>hg38_21_+_8433172_8446622_RNA45SN1_45S_50ntFlanks@10347.10374.28|CAAACGAGAACTTTGAAGGCCGAAGTGG</td><td>>hg38_21_+_8433172_8446622_RNA45SN1_45S_50ntFlanks</td><td>10347</td><td>10374</td><td>28</td><td>CAAACGAGAACTTTGAAGGCCGAAGTGG</td><td>629.2</td><td>-8.9</td><td>1.5</td><td>-6.1</td><td>1.3E-09</td><td>2.6E-09</td><td>BT20/MB468</td><td>NUC</td></tr>
<tr><td>>hg38_21_+_8433172_8446622_RNA45SN1_45S_50ntFlanks@4721.4761.41|TTCATTAATCAAGAACGAAAGTCGGAGGTTCGAAGACGATC</td><td>>hg38_21_+_8433172_8446622_RNA45SN1_45S_50ntFlanks</td><td>4721</td><td>4761</td><td>41</td><td>TTCATTAATCAAGAACGAAAGTCGGAGGTTCGAAGACGATC</td><td>245.6</td><td>6.8</td><td>1.1</td><td>6.1</td><td>1.2E-09</td><td>2.6E-09</td><td>MB231/MB468</td><td>CYTO</td></tr>
<tr><td>>MI0003591|hsa-mir-584&WithFlank&5|-|149062307|149062415@22.44.23(+1U)&[MIMAT0003249&hsa-miR-584-5p&offsets|0|+1(+1U);m-167&5|-|149062374|149062394&offsets|0|+2(+1U)]|TTATGGTTTGCCTGGGACTGAGGT</td><td>>MI0003591|hsa-mir-584&WithFlank&5|-|149062307|149062415</td><td>22</td><td>44</td><td>23(+1U)</td><td>TTATGGTTTGCCTGGGACTGAGGT</td><td>110.8</td><td>10.0</td><td>1.6</td><td>6.1</td><td>1.1E-09</td><td>2.6E-09</td><td>MB231/MB468</td><td>MP</td></tr>
<tr><td>>MI0000093|hsa-mir-92a-1&WithFlank&13|+|91351308|91351397@54.75.22;MI0000094|hsa-mir-92a-2&WithFlank&X|-|134169532|134169618@54.75.22&[MIMAT0000092&hsa-miR-92a-3p&offsets|0|0;m-9&13|+|91351361|91351382&offsets|0|0];[MIMAT0000092_1&hsa-miR-92a-3p&offsets|0|0;m-11&X|-|134169544|134169565&offsets|0|0]|TATTGCACTTGTCCCGGCCTGT</td><td>>MI0000093|hsa-mir-92a-1&WithFlank&13|+|91351308|91351397</td><td>54</td><td>75</td><td>22</td><td>TATTGCACTTGTCCCGGCCTGT</td><td>76808.1</td><td>4.0</td><td>0.7</td><td>6.1</td><td>1.2E-09</td><td>2.6E-09</td><td>MB231/MB468</td><td>NUC</td></tr>
<tr><td>>hg38_21_+_8433172_8446622_RNA45SN1_45S_50ntFlanks@10408.10457.50|GGTCAGTCGGTCCTGAGAGATGGGCGAGCGCCGTTCCGAAGGGACGGGCG</td><td>>hg38_21_+_8433172_8446622_RNA45SN1_45S_50ntFlanks</td><td>10408</td><td>10457</td><td>50</td><td>GGTCAGTCGGTCCTGAGAGATGGGCGAGCGCCGTTCCGAAGGGACGGGCG</td><td>208.9</td><td>-10.7</td><td>1.8</td><td>-6.1</td><td>1.2E-09</td><td>2.6E-09</td><td>MB231/MB468</td><td>NUC</td></tr>
<tr><td>>hg38_21_+_8433172_8446622_RNA45SN1_45S_50ntFlanks@9720.9746.27|ATTAGAGGTCTTGGGGCCGAAACGATC</td><td>>hg38_21_+_8433172_8446622_RNA45SN1_45S_50ntFlanks</td><td>9720</td><td>9746</td><td>27</td><td>ATTAGAGGTCTTGGGGCCGAAACGATC</td><td>134.5</td><td>-10.9</td><td>1.8</td><td>-6.1</td><td>1.1E-09</td><td>2.6E-09</td><td>MB231/MB468</td><td>MP</td></tr>
<tr><td>>MI0000737|hsa-mir-200a&WithFlank&1|+|1167857|1167958@60.81.22(+1A)&[MIMAT0000682&hsa-miR-200a-3p&offsets|0|0(+1A);m-142&1|+|1167916|1167938&offsets|0|-1(+1A)]|TAACACTGTCTGGTAACGATGTA</td><td>>MI0000737|hsa-mir-200a&WithFlank&1|+|1167857|1167958</td><td>60</td><td>81</td><td>22(+1A)</td><td>TAACACTGTCTGGTAACGATGTA</td><td>237.1</td><td>-11.0</td><td>1.7</td><td>-6.4</td><td>1.2E-10</td><td>2.6E-09</td><td>BT20/MB468</td><td>MP</td></tr>
<tr><td>>MI0000734|hsa-mir-106b&WithFlank&7|-|100093987|100094080@58.77.20&[MIMAT0004672&hsa-miR-106b-3p&offsets|0|-2;m-87&7|-|100094003|100094023&offsets|0|-1]|CCGCACTGTGGGTACTTGCT</td><td>>MI0000734|hsa-mir-106b&WithFlank&7|-|100093987|100094080</td><td>58</td><td>77</td><td>20</td><td>CCGCACTGTGGGTACTTGCT</td><td>298.4</td><td>2.0</td><td>0.3</td><td>6.1</td><td>1.3E-09</td><td>2.6E-09</td><td>BT20/MB468</td><td>NUC</td></tr>
<tr><td>>hg38_21_+_8433172_8446622_RNA45SN1_45S_50ntFlanks@6721.6749.29|AATTGCAGGACACATTGATCATCGACACT</td><td>>hg38_21_+_8433172_8446622_RNA45SN1_45S_50ntFlanks</td><td>6721</td><td>6749</td><td>29</td><td>AATTGCAGGACACATTGATCATCGACACT</td><td>435.2</td><td>-4.1</td><td>0.7</td><td>-6.1</td><td>1.3E-09</td><td>2.6E-09</td><td>MB231/MB468</td><td>MITO</td></tr>
<tr><td>>hg38_21_+_8433172_8446622_RNA45SN1_45S_50ntFlanks@6734.6770.37|ATTGATCATCGACACTTCGAACGCACTTGCGGCCCCG</td><td>>hg38_21_+_8433172_8446622_RNA45SN1_45S_50ntFlanks</td><td>6734</td><td>6770</td><td>37</td><td>ATTGATCATCGACACTTCGAACGCACTTGCGGCCCCG</td><td>1187.1</td><td>-3.6</td><td>0.6</td><td>-6.1</td><td>1.2E-09</td><td>2.6E-09</td><td>MB231/MB468</td><td>NUC</td></tr>
<tr><td>>hg38_21_+_8433172_8446622_RNA45SN1_45S_50ntFlanks@9556.9582.27|TAGCGGTCCTGACGTGCAAATCGGTCG</td><td>>hg38_21_+_8433172_8446622_RNA45SN1_45S_50ntFlanks</td><td>9556</td><td>9582</td><td>27</td><td>TAGCGGTCCTGACGTGCAAATCGGTCG</td><td>343.0</td><td>-7.7</td><td>1.3</td><td>-6.1</td><td>1.3E-09</td><td>2.6E-09</td><td>MB231/MB468</td><td>MITO</td></tr>
<tr><td>>hg38_21_+_8433172_8446622_RNA45SN1_45S_50ntFlanks@9166.9197.32|CCGGGCGGGTCGCGCCGTCGGGCCCGGGGGAG</td><td>>hg38_21_+_8433172_8446622_RNA45SN1_45S_50ntFlanks</td><td>9166</td><td>9197</td><td>32</td><td>CCGGGCGGGTCGCGCCGTCGGGCCCGGGGGAG</td><td>569.0</td><td>-8.5</td><td>1.4</td><td>-6.1</td><td>1.2E-09</td><td>2.6E-09</td><td>MB231/MB468</td><td>NUC</td></tr>
<tr><td>>hg38_21_+_8433172_8446622_RNA45SN1_45S_50ntFlanks@4153.4175.23|ACGGCTACCACATCCAAGGAAGG</td><td>>hg38_21_+_8433172_8446622_RNA45SN1_45S_50ntFlanks</td><td>4153</td><td>4175</td><td>23</td><td>ACGGCTACCACATCCAAGGAAGG</td><td>78.9</td><td>-10.2</td><td>1.7</td><td>-6.1</td><td>1.1E-09</td><td>2.7E-09</td><td>MB231/MB468</td><td>MP</td></tr>
<tr><td>>hg38_21_+_8433172_8446622_RNA45SN1_45S_50ntFlanks@4729.4759.31|TCAAGAACGAAAGTCGGAGGTTCGAAGACGA</td><td>>hg38_21_+_8433172_8446622_RNA45SN1_45S_50ntFlanks</td><td>4729</td><td>4759</td><td>31</td><td>TCAAGAACGAAAGTCGGAGGTTCGAAGACGA</td><td>1126.4</td><td>-6.7</td><td>1.1</td><td>-6.1</td><td>1.1E-09</td><td>2.7E-09</td><td>MB231/MB468</td><td>MP</td></tr>
<tr><td>>hg38_21_+_8433172_8446622_RNA45SN1_45S_50ntFlanks@9457.9493.37|CGCCGGGGAGGTGGAGCACGAGCGCACGTGTTAGGAC</td><td>>hg38_21_+_8433172_8446622_RNA45SN1_45S_50ntFlanks</td><td>9457</td><td>9493</td><td>37</td><td>CGCCGGGGAGGTGGAGCACGAGCGCACGTGTTAGGAC</td><td>97.2</td><td>-10.5</td><td>1.7</td><td>-6.1</td><td>1.1E-09</td><td>2.7E-09</td><td>MB231/MB468</td><td>MP</td></tr>
<tr><td>>hg38_21_+_8433172_8446622_RNA45SN1_45S_50ntFlanks@9545.9569.25|GTGGAGGTCCGTAGCGGTCCTGACG</td><td>>hg38_21_+_8433172_8446622_RNA45SN1_45S_50ntFlanks</td><td>9545</td><td>9569</td><td>25</td><td>GTGGAGGTCCGTAGCGGTCCTGACG</td><td>443.9</td><td>-8.6</td><td>1.4</td><td>-6.1</td><td>1.4E-09</td><td>2.7E-09</td><td>BT20/MB468</td><td>NUC</td></tr>
<tr><td>>hg38_21_+_8433172_8446622_RNA45SN1_45S_50ntFlanks@4128.4159.32|CGATTCCGGAGAGGGAGCCTGAGAAACGGCTA</td><td>>hg38_21_+_8433172_8446622_RNA45SN1_45S_50ntFlanks</td><td>4128</td><td>4159</td><td>32</td><td>CGATTCCGGAGAGGGAGCCTGAGAAACGGCTA</td><td>90.1</td><td>-10.4</td><td>1.7</td><td>-6.1</td><td>1.1E-09</td><td>2.7E-09</td><td>MB231/MB468</td><td>MP</td></tr>
<tr><td>>hg38_21_+_8433172_8446622_RNA45SN1_45S_50ntFlanks@9467.9486.20|GTGGAGCACGAGCGCACGTG</td><td>>hg38_21_+_8433172_8446622_RNA45SN1_45S_50ntFlanks</td><td>9467</td><td>9486</td><td>20</td><td>GTGGAGCACGAGCGCACGTG</td><td>213.4</td><td>-11.6</td><td>1.9</td><td>-6.1</td><td>1.1E-09</td><td>2.7E-09</td><td>MB231/MB468</td><td>MP</td></tr>
<tr><td>>trnaMT_MetCAT_MT_+_4402_4469@53.71.19|TACCCTTCCCGTACTACCA</td><td>>trnaMT_MetCAT_MT_+_4402_4469</td><td>53</td><td>71</td><td>19</td><td>TACCCTTCCCGTACTACCA</td><td>765.6</td><td>9.2</td><td>1.5</td><td>6.1</td><td>1.2E-09</td><td>2.7E-09</td><td>MB231/MB468</td><td>CYTO</td></tr>
<tr><td>>hg38_21_+_8433172_8446622_RNA45SN1_45S_50ntFlanks@10391.10409.19|AACAGCAGTTGAACATGGG</td><td>>hg38_21_+_8433172_8446622_RNA45SN1_45S_50ntFlanks</td><td>10391</td><td>10409</td><td>19</td><td>AACAGCAGTTGAACATGGG</td><td>742.8</td><td>-8.8</td><td>1.4</td><td>-6.4</td><td>1.2E-10</td><td>2.7E-09</td><td>BT20/MB468</td><td>MP</td></tr>
<tr><td>>hg38_21_+_8433172_8446622_RNA45SN1_45S_50ntFlanks@10431.10456.26|GCGAGCGCCGTTCCGAAGGGACGGGC</td><td>>hg38_21_+_8433172_8446622_RNA45SN1_45S_50ntFlanks</td><td>10431</td><td>10456</td><td>26</td><td>GCGAGCGCCGTTCCGAAGGGACGGGC</td><td>201.9</td><td>-10.7</td><td>1.7</td><td>-6.4</td><td>1.2E-10</td><td>2.7E-09</td><td>BT20/MB468</td><td>MP</td></tr>
<tr><td>>hg38_21_+_8433172_8446622_RNA45SN1_45S_50ntFlanks@11847.11872.26|AACGGGCTTGGCGGAATCAGCGGGGA</td><td>>hg38_21_+_8433172_8446622_RNA45SN1_45S_50ntFlanks</td><td>11847</td><td>11872</td><td>26</td><td>AACGGGCTTGGCGGAATCAGCGGGGA</td><td>1660.4</td><td>-4.8</td><td>0.7</td><td>-6.4</td><td>1.2E-10</td><td>2.7E-09</td><td>BT20/MB468</td><td>MP</td></tr>
<tr><td>>hg38_21_+_8433172_8446622_RNA45SN1_45S_50ntFlanks@9567.9598.32|ACGTGCAAATCGGTCGTCCGACCTGGGTATAG</td><td>>hg38_21_+_8433172_8446622_RNA45SN1_45S_50ntFlanks</td><td>9567</td><td>9598</td><td>32</td><td>ACGTGCAAATCGGTCGTCCGACCTGGGTATAG</td><td>57.4</td><td>-9.7</td><td>1.6</td><td>-6.1</td><td>1.1E-09</td><td>2.7E-09</td><td>MB231/MB468</td><td>MP</td></tr>
<tr><td>>hg38_21_+_8433172_8446622_RNA45SN1_45S_50ntFlanks@9568.9594.27|CGTGCAAATCGGTCGTCCGACCTGGGT</td><td>>hg38_21_+_8433172_8446622_RNA45SN1_45S_50ntFlanks</td><td>9568</td><td>9594</td><td>27</td><td>CGTGCAAATCGGTCGTCCGACCTGGGT</td><td>701.7</td><td>-8.9</td><td>1.5</td><td>-6.1</td><td>1.4E-09</td><td>2.7E-09</td><td>BT20/MB468</td><td>NUC</td></tr>
<tr><td>>hg38_21_+_8433172_8446622_RNA45SN1_45S_50ntFlanks@8169.8212.44|CCAAGTCCTTCTGATCGAGGCCCAGCCCGTGGACGGTGTGAGGC</td><td>>hg38_21_+_8433172_8446622_RNA45SN1_45S_50ntFlanks</td><td>8169</td><td>8212</td><td>44</td><td>CCAAGTCCTTCTGATCGAGGCCCAGCCCGTGGACGGTGTGAGGC</td><td>828.2</td><td>-5.5</td><td>0.9</td><td>-6.1</td><td>1.2E-09</td><td>2.7E-09</td><td>MB231/MB468</td><td>NUC</td></tr>
<tr><td>>hg38_21_+_8433172_8446622_RNA45SN1_45S_50ntFlanks@11921.11958.38|TGAGAGGTGTAGAATAAGTGGGAGGCCCCCGGCGCCCC</td><td>>hg38_21_+_8433172_8446622_RNA45SN1_45S_50ntFlanks</td><td>11921</td><td>11958</td><td>38</td><td>TGAGAGGTGTAGAATAAGTGGGAGGCCCCCGGCGCCCC</td><td>114.8</td><td>-10.7</td><td>1.8</td><td>-6.1</td><td>1.1E-09</td><td>2.7E-09</td><td>MB231/MB468</td><td>MP</td></tr>
</table>

<table>
<tr><th></th><th></th><th></th><th></th><th></th><th></th><th></th><th></th><th></th><th></th><th></th><th></th><th></th><th></th></tr>
<tr><td>>hg38_1_-_228634819_228635039_RNA5S12_5S_50ntFlanks@127.171.45|ACTTGGATGGGAGACCGCCTGGGAATACCGGGTGCTGTAGGCTTT</td><td>>hg38_1_-_228634819_228635039_RNA5S12_5S_50ntFlanks</td><td>127</td><td>171</td><td>45</td><td>ACTTGGATGGGAGACCGCCTGGGAATACCGGGTGCTGTAGGCTTT</td><td>186.4</td><td>11.3</td><td>1.8</td><td>6.4</td><td>1.2E-10</td><td>2.7E-09</td><td>BT20/MB468</td><td>MP</td></tr>
<tr><td>>hg38_21_+_8433172_8446622_RNA45SN1_45S_50ntFlanks@10905.10926.22|CTGGGCGCGCGCCGCGGCTGGA</td><td>>hg38_21_+_8433172_8446622_RNA45SN1_45S_50ntFlanks</td><td>10905</td><td>10926</td><td>22</td><td>CTGGGCGCGCGCCGCGGCTGGA</td><td>267.8</td><td>-11.1</td><td>1.7</td><td>-6.4</td><td>1.2E-10</td><td>2.7E-09</td><td>BT20/MB468</td><td>MP</td></tr>
<tr><td>>MI0000062|hsa-let-7a-3&WithFlank&22|+|46112743|46112828@58.78.21(+1U);MI0000060|hsa-let-7a-1&WithFlank&9|+|94175951|94176042@63.83.21(+1U)&[MIMAT0004481&hsa-let-7a-3p&offsets|0|0(+1U);m-180&22|+|46112800|46112821&offsets|0|-1(+1U)];[MIMAT0004481_1&hsa-let-7a-3p&offsets|0|0(+1U);m-181&9|+|94176013|94176034&offsets|0|-1(+1U)]|CTATACAATCTACTGTCTTTCT</td><td>>MI0000062|hsa-let-7a-3&WithFlank&22|+|46112743|46112828</td><td>58</td><td>78</td><td>21(+1U)</td><td>CTATACAATCTACTGTCTTTCT</td><td>1386.6</td><td>6.3</td><td>1.0</td><td>6.1</td><td>1.0E-09</td><td>2.7E-09</td><td>BT20/MB468</td><td>CYTO</td></tr>
<tr><td>>MI0001733|hsa-mir-452&WithFlank&X|-|151959622|151959718@59.80.22&[MIMAT0001636&hsa-miR-452-3p&offsets|-5|-5;m-457&X|-|151959639|151959660&offsets|0|0]|TCAGTCTCATCTGCAAAGAAGT</td><td>>MI0001733|hsa-mir-452&WithFlank&X|-|151959622|151959718</td><td>59</td><td>80</td><td>22</td><td>TCAGTCTCATCTGCAAAGAAGT</td><td>366.3</td><td>12.3</td><td>1.9</td><td>6.4</td><td>1.2E-10</td><td>2.7E-09</td><td>BT20/MB468</td><td>MP</td></tr>
<tr><td>>am_trna162_MetCAT_6_-_26758550_26758622@31.53.23,trna164_MetCAT_6_-_26735574_26735646@31.53.23,trna20_MetCAT_16_+_71460396_71460468@31.53.23,trna21_MetCAT_6_+_26701712_26701784@31.53.23,trna27_MetCAT_6_+_26766444_26766516@31.53.23,trna75_MetCAT_6_+_28912352_28912424@31.53.23,trna97_MetCAT_6_-_28921042_28921114@31.53.23|TCTCATAATCTGAAGGTCCTGAG</td><td>>am_trna162_MetCAT_6_-_26758550_26758622</td><td>31</td><td>53</td><td>23</td><td>TCTCATAATCTGAAGGTCCTGAG</td><td>171.3</td><td>-10.5</td><td>1.6</td><td>-6.4</td><td>1.2E-10</td><td>2.7E-09</td><td>BT20/MB468</td><td>MP</td></tr>
<tr><td>>MI0000088|hsa-mir-30a&WithFlank&6|-|71403545|71403627@54.75.22&[MIMAT0000088&hsa-miR-30a-3p&offsets|+1|+1;m-38&6|-|71403554|71403575&offsets|+1|+1]|TTTCAGTCGGATGTTTGCAGCT</td><td>>MI0000088|hsa-mir-30a&WithFlank&6|-|71403545|71403627</td><td>54</td><td>75</td><td>22</td><td>TTTCAGTCGGATGTTTGCAGCT</td><td>709.7</td><td>9.3</td><td>1.5</td><td>6.1</td><td>1.3E-09</td><td>2.7E-09</td><td>MB231/MB468</td><td>MITO</td></tr>
<tr><td>>hg38_21_+_8433172_8446622_RNA45SN1_45S_50ntFlanks@3991.4026.36|TGGTGACTCTAGATAACCTCGGGCCGATCGCACGCC</td><td>>hg38_21_+_8433172_8446622_RNA45SN1_45S_50ntFlanks</td><td>3991</td><td>4026</td><td>36</td><td>TGGTGACTCTAGATAACCTCGGGCCGATCGCACGCC</td><td>604.2</td><td>-4.5</td><td>0.7</td><td>-6.1</td><td>1.2E-09</td><td>2.7E-09</td><td>MB231/MB468</td><td>NUC</td></tr>
<tr><td>>hg38_21_+_8433172_8446622_RNA45SN1_45S_50ntFlanks@10713.10732.20|CTTGAAAATCCGGGGGAGAG</td><td>>hg38_21_+_8433172_8446622_RNA45SN1_45S_50ntFlanks</td><td>10713</td><td>10732</td><td>20</td><td>CTTGAAAATCCGGGGGAGAG</td><td>87.5</td><td>-10.3</td><td>1.7</td><td>-6.1</td><td>1.1E-09</td><td>2.7E-09</td><td>MB231/MB468</td><td>MP</td></tr>
<tr><td>>MI0003513|hsa-mir-455&WithFlank&9|+|114209428|114209535@23.44.22&[MIMAT0003150&hsa-miR-455-5p&offsets|+1|+1;m-232&9|+|114209449|114209470&offsets|+1|+1]|ATGTGCCTTTGGACTACATCGT</td><td>>MI0003513|hsa-mir-455&WithFlank&9|+|114209428|114209535</td><td>23</td><td>44</td><td>22</td><td>ATGTGCCTTTGGACTACATCGT</td><td>81.6</td><td>9.6</td><td>1.6</td><td>6.1</td><td>1.1E-09</td><td>2.7E-09</td><td>MB231/MB468</td><td>MP</td></tr>
<tr><td>>hg38_21_+_8433172_8446622_RNA45SN1_45S_50ntFlanks@11208.11252.45|GCGGCGACTCTGGACGCGAGCCGGGCCCTTCCCGTGGATCGCCCC</td><td>>hg38_21_+_8433172_8446622_RNA45SN1_45S_50ntFlanks</td><td>11208</td><td>11252</td><td>45</td><td>GCGGCGACTCTGGACGCGAGCCGGGCCCTTCCCGTGGATCGCCCC</td><td>1002.2</td><td>-8.6</td><td>1.4</td><td>-6.1</td><td>1.3E-09</td><td>2.7E-09</td><td>MB231/MB468</td><td>MITO</td></tr>
<tr><td>>MI0000342|hsa-mir-200b&WithFlank&1|+|1167098|1167204@64.85.22(+1U)&[MIMAT0000318&hsa-miR-200b-3p&offsets|+1|+1(+1U);m-93&1|+|1167160|1167182&offsets|+1|0(+1U)]|AATACTGCCTGGTAATGATGACT</td><td>>MI0000342|hsa-mir-200b&WithFlank&1|+|1167098|1167204</td><td>64</td><td>85</td><td>22(+1U)</td><td>AATACTGCCTGGTAATGATGACT</td><td>883.0</td><td>-8.9</td><td>1.5</td><td>-6.1</td><td>1.4E-09</td><td>2.7E-09</td><td>BT20/MB468</td><td>NUC</td></tr>
<tr><td>>hg38_21_+_8433172_8446622_RNA45SN1_45S_50ntFlanks@10546.10564.19|CAGTGCGGTAACGCGACCG</td><td>>hg38_21_+_8433172_8446622_RNA45SN1_45S_50ntFlanks</td><td>10546</td><td>10564</td><td>19</td><td>CAGTGCGGTAACGCGACCG</td><td>114.3</td><td>-10.7</td><td>1.8</td><td>-6.1</td><td>1.2E-09</td><td>2.7E-09</td><td>MB231/MB468</td><td>MP</td></tr>
<tr><td>>MI0000477|hsa-mir-146a&WithFlank&5|+|160485346|160485456@27.46.20(+1C)&[MIMAT0000449&hsa-miR-146a-5p&offsets|0|-2(+1C);m-129&5|+|160485372|160485393&offsets|0|-2(+1C)]|TGAGAACTGAATTCCATGGGC</td><td>>MI0000477|hsa-mir-146a&WithFlank&5|+|160485346|160485456</td><td>27</td><td>46</td><td>20(+1C)</td><td>TGAGAACTGAATTCCATGGGC</td><td>223.3</td><td>-10.9</td><td>1.7</td><td>-6.4</td><td>1.3E-10</td><td>2.7E-09</td><td>BT20/MB231</td><td>NUC</td></tr>
<tr><td>>hg38_21_+_8433172_8446622_RNA45SN1_45S_50ntFlanks@9548.9573.26|GAGGTCCGTAGCGGTCCTGACGTGCA</td><td>>hg38_21_+_8433172_8446622_RNA45SN1_45S_50ntFlanks</td><td>9548</td><td>9573</td><td>26</td><td>GAGGTCCGTAGCGGTCCTGACGTGCA</td><td>685.8</td><td>-4.7</td><td>0.8</td><td>-6.1</td><td>1.4E-09</td><td>2.7E-09</td><td>BT20/MB468</td><td>NUC</td></tr>
<tr><td>>hg38_21_+_8433172_8446622_RNA45SN1_45S_50ntFlanks@11918.11948.31|ACATGAGAGGTGTAGAATAAGTGGGAGGCCC</td><td>>hg38_21_+_8433172_8446622_RNA45SN1_45S_50ntFlanks</td><td>11918</td><td>11948</td><td>31</td><td>ACATGAGAGGTGTAGAATAAGTGGGAGGCCC</td><td>953.0</td><td>-8.0</td><td>1.3</td><td>-6.1</td><td>1.2E-09</td><td>2.7E-09</td><td>MB231/MB468</td><td>CYTO</td></tr>
<tr><td>>hg38_21_+_8433172_8446622_RNA45SN1_45S_50ntFlanks@6772.6803.32|GTTCCTCCCGGGGCTACGCCTGTCTGAGCGTC</td><td>>hg38_21_+_8433172_8446622_RNA45SN1_45S_50ntFlanks</td><td>6772</td><td>6803</td><td>32</td><td>GTTCCTCCCGGGGCTACGCCTGTCTGAGCGTC</td><td>229.0</td><td>-4.5</td><td>0.7</td><td>-6.1</td><td>1.2E-09</td><td>2.7E-09</td><td>MB231/MB468</td><td>CYTO</td></tr>
<tr><td>>hg38_21_+_8433172_8446622_RNA45SN1_45S_50ntFlanks@9790.9831.42|CTGGCGTGGAGCCGGGCGTGGAATGCGAGTGCCTAGTGGGCC</td><td>>hg38_21_+_8433172_8446622_RNA45SN1_45S_50ntFlanks</td><td>9790</td><td>9831</td><td>42</td><td>CTGGCGTGGAGCCGGGCGTGGAATGCGAGTGCCTAGTGGGCC</td><td>131.9</td><td>-10.9</td><td>1.8</td><td>-6.1</td><td>1.2E-09</td><td>2.7E-09</td><td>MB231/MB468</td><td>MP</td></tr>
<tr><td>>am_MI0000809|hsa-mir-151a&WithFlank&8|-|140732558|140732659@17.37.21&[MIMAT0004697&hsa-miR-151a-5p&offsets|0|0;m-75&8|-|140732623|140732643&offsets|0|0]|TCGAGGAGCTCACAGTCTAGT</td><td>>am_MI0000809|hsa-mir-151a&WithFlank&8|-|140732558|140732659</td><td>17</td><td>37</td><td>21</td><td>TCGAGGAGCTCACAGTCTAGT</td><td>879.3</td><td>7.8</td><td>1.3</td><td>6.1</td><td>1.2E-09</td><td>2.7E-09</td><td>MB231/MB468</td><td>CYTO</td></tr>
<tr><td>>MI0000085|hsa-mir-27a&WithFlank&19|-|13836434|13836523@57.76.20(+2U)&[MIMAT0000084&hsa-miR-27a-3p&offsets|0|-1(+2U);m-43&19|-|13836447|13836467&offsets|0|-1(+2U)]|TTCACAGTGGCTAAGTTCCGTT</td><td>>MI0000085|hsa-mir-27a&WithFlank&19|-|13836434|13836523</td><td>57</td><td>76</td><td>20(+2U)</td><td>TTCACAGTGGCTAAGTTCCGTT</td><td>890.4</td><td>4.3</td><td>0.7</td><td>6.1</td><td>1.1E-09</td><td>2.7E-09</td><td>BT20/MB468</td><td>CYTO</td></tr>
<tr><td>>hg38_21_+_8433172_8446622_RNA45SN1_45S_50ntFlanks@6660.6706.47|GCGGTGGATCACTCGGCTCGTGCGTCGATGAAGAACGCAGCTAGCTG</td><td>>hg38_21_+_8433172_8446622_RNA45SN1_45S_50ntFlanks</td><td>6660</td><td>6706</td><td>47</td><td>GCGGTGGATCACTCGGCTCGTGCGTCGATGAAGAACGCAGCTAGCTG</td><td>1004.4</td><td>4.9</td><td>0.8</td><td>6.1</td><td>9.1E-10</td><td>2.8E-09</td><td>BT20/MB468</td><td>MITO</td></tr>
<tr><td>>MI0000734|hsa-mir-106b&WithFlank&7|-|100093987|100094080@58.76.19&[MIMAT0004672&hsa-miR-106b-3p&offsets|0|-3;m-87&7|-|100094003|100094023&offsets|0|-2]|CCGCACTGTGGGTACTTGC</td><td>>MI0000734|hsa-mir-106b&WithFlank&7|-|100093987|100094080</td><td>58</td><td>76</td><td>19</td><td>CCGCACTGTGGGTACTTGC</td><td>247.0</td><td>4.8</td><td>0.8</td><td>6.1</td><td>1.2E-09</td><td>2.8E-09</td><td>MB231/MB468</td><td>NUC</td></tr>
<tr><td>>hg38_21_+_8433172_8446622_RNA45SN1_45S_50ntFlanks@5487.5523.37|CGCTGAGAAGACGGTCGAACTTGACTATCTAGAGGAA</td><td>>hg38_21_+_8433172_8446622_RNA45SN1_45S_50ntFlanks</td><td>5487</td><td>5523</td><td>37</td><td>CGCTGAGAAGACGGTCGAACTTGACTATCTAGAGGAA</td><td>83.2</td><td>-10.2</td><td>1.7</td><td>-6.1</td><td>1.2E-09</td><td>2.8E-09</td><td>MB231/MB468</td><td>MP</td></tr>
<tr><td>>hg38_21_+_8433172_8446622_RNA45SN1_45S_50ntFlanks@10873.10890.18|CTGGGTCGGTCGGGCTGG</td><td>>hg38_21_+_8433172_8446622_RNA45SN1_45S_50ntFlanks</td><td>10873</td><td>10890</td><td>18</td><td>CTGGGTCGGTCGGGCTGG</td><td>437.1</td><td>-8.1</td><td>1.3</td><td>-6.1</td><td>1.2E-09</td><td>2.8E-09</td><td>MB231/MB468</td><td>MP</td></tr>
<tr><td>>hg38_21_+_8433172_8446622_RNA45SN1_45S_50ntFlanks@3780.3801.22|TACAGTGAAACTGCGAATGGCT</td><td>>hg38_21_+_8433172_8446622_RNA45SN1_45S_50ntFlanks</td><td>3780</td><td>3801</td><td>22</td><td>TACAGTGAAACTGCGAATGGCT</td><td>796.8</td><td>-8.4</td><td>1.4</td><td>-6.1</td><td>1.3E-09</td><td>2.8E-09</td><td>MB231/MB468</td><td>MITO</td></tr>
<tr><td>>trnaMT_PheGAA_MT_+_577_647@36.59.24|AAAATGTTTAGACGGGCTCACATC</td><td>>trnaMT_PheGAA_MT_+_577_647</td><td>36</td><td>59</td><td>24</td><td>AAAATGTTTAGACGGGCTCACATC</td><td>1003.1</td><td>7.8</td><td>1.3</td><td>6.1</td><td>9.2E-10</td><td>2.8E-09</td><td>BT20/MB468</td><td>MITO</td></tr>
<tr><td>>hg38_21_+_8433172_8446622_RNA45SN1_45S_50ntFlanks@8472.8496.25|CGGCCCGGCGGATCTTTCCCGCCCC</td><td>>hg38_21_+_8433172_8446622_RNA45SN1_45S_50ntFlanks</td><td>8472</td><td>8496</td><td>25</td><td>CGGCCCGGCGGATCTTTCCCGCCCC</td><td>116.8</td><td>-9.0</td><td>1.5</td><td>-6.1</td><td>9.2E-10</td><td>2.8E-09</td><td>BT20/MB468</td><td>MITO</td></tr>
<tr><td>>MI0000100|hsa-mir-98&WithFlank&X|-|53556217|53556347@28.49.22&[MIMAT0000096&hsa-miR-98-5p&offsets|0|0;m-143&X|-|53556299|53556320&offsets|0|0]|TGAGGTAGTAAGTTGTATTGTT</td><td>>MI0000100|hsa-mir-98&WithFlank&X|-|53556217|53556347</td><td>28</td><td>49</td><td>22</td><td>TGAGGTAGTAAGTTGTATTGTT</td><td>4145.3</td><td>3.1</td><td>0.5</td><td>6.1</td><td>1.2E-09</td><td>2.8E-09</td><td>MB231/MB468</td><td>NUC</td></tr>
<tr><td>>MI0000736|hsa-mir-30c-1&WithFlank&1|+|40757278|40757378@23.46.24(+1A);MI0000254|hsa-mir-30c-2&WithFlank&6|-|71376954|71377037@13.36.24(+1A)&[MIMAT0000244&hsa-miR-30c-5p&offsets|0|+1(+1A);m-57&1|+|40757300|40757323&offsets|0|0(+1A)];[MIMAT0000244_1&hsa-miR-30c-5p&offsets|0|+1(+1A);m-56&6|-|71377002|71377025&offsets|0|0(+1A)]|TGTAAACATCCTACACTCTCAGCTA</td><td>>MI0000736|hsa-mir-30c-1&WithFlank&1|+|40757278|40757378</td><td>23</td><td>46</td><td>24(+1A)</td><td>TGTAAACATCCTACACTCTCAGCTA</td><td>311.7</td><td>3.9</td><td>0.6</td><td>6.1</td><td>1.2E-09</td><td>2.8E-09</td><td>MB231/MB468</td><td>NUC</td></tr>
<tr><td>>hg38_21_+_8433172_8446622_RNA45SN1_45S_50ntFlanks@4370.4396.27|TTAAAAAGCTCGTAGTTGGATCTTGGG</td><td>>hg38_21_+_8433172_8446622_RNA45SN1_45S_50ntFlanks</td><td>4370</td><td>4396</td><td>27</td><td>TTAAAAAGCTCGTAGTTGGATCTTGGG</td><td>141.8</td><td>-7.0</td><td>1.1</td><td>-6.1</td><td>1.2E-09</td><td>2.8E-09</td><td>MB231/MB468</td><td>NUC</td></tr>
<tr><td>>hg38_21_+_8433172_8446622_RNA45SN1_45S_50ntFlanks@10457.10497.41|GATGGCCTCCGTTGCCCTCGGCCGATCGAAAGGGAGTCGGG</td><td>>hg38_21_+_8433172_8446622_RNA45SN1_45S_50ntFlanks</td><td>10457</td><td>10497</td><td>41</td><td>GATGGCCTCCGTTGCCCTCGGCCGATCGAAAGGGAGTCGGG</td><td>143.2</td><td>-9.3</td><td>1.5</td><td>-6.1</td><td>9.2E-10</td><td>2.8E-09</td><td>BT20/MB468</td><td>MITO</td></tr>
<tr><td>>hg38_21_+_8433172_8446622_RNA45SN1_45S_50ntFlanks@10729.10749.21|AGAGGGTGTAAATCTCGCGCC</td><td>>hg38_21_+_8433172_8446622_RNA45SN1_45S_50ntFlanks</td><td>10729</td><td>10749</td><td>21</td><td>AGAGGGTGTAAATCTCGCGCC</td><td>69.8</td><td>-10.0</td><td>1.6</td><td>-6.1</td><td>1.2E-09</td><td>2.8E-09</td><td>MB231/MB468</td><td>MP</td></tr>
<tr><td>>MI0000077|hsa-mir-21&WithFlank&17|+|59841260|59841343@14.34.21(+1G)&[MIMAT0000076&hsa-miR-21-5p&offsets|0|-1(+1G);m-1&17|+|59841273|59841294&offsets|0|-1(+1G)]|TAGCTTATCAGACTGATGTTGG</td><td>>MI0000077|hsa-mir-21&WithFlank&17|+|59841260|59841343</td><td>14</td><td>34</td><td>21(+1G)</td><td>TAGCTTATCAGACTGATGTTGG</td><td>6131.3</td><td>6.8</td><td>1.1</td><td>6.1</td><td>1.1E-09</td><td>2.8E-09</td><td>BT20/MB468</td><td>CYTO</td></tr>
<tr><td>>hg38_21_+_8433172_8446622_RNA45SN1_45S_50ntFlanks@12285.12306.22|CAGACCGTGAAAGCGGGGCCTC</td><td>>hg38_21_+_8433172_8446622_RNA45SN1_45S_50ntFlanks</td><td>12285</td><td>12306</td><td>22</td><td>CAGACCGTGAAAGCGGGGCCTC</td><td>100.2</td><td>-10.5</td><td>1.7</td><td>-6.1</td><td>1.2E-09</td><td>2.8E-09</td><td>MB231/MB468</td><td>MP</td></tr>
<tr><td>>hg38_21_+_8433172_8446622_RNA45SN1_45S_50ntFlanks@6721.6757.37|AATTGCAGGACACATTGATCATCGACACTTCGAACGC</td><td>>hg38_21_+_8433172_8446622_RNA45SN1_45S_50ntFlanks</td><td>6721</td><td>6757</td><td>37</td><td>AATTGCAGGACACATTGATCATCGACACTTCGAACGC</td><td>911.2</td><td>-9.4</td><td>1.6</td><td>-6.1</td><td>1.4E-09</td><td>2.8E-09</td><td>BT20/MB468</td><td>NUC</td></tr>
<tr><td>>trnaMT_LeuTAA_MT_+_3230_3304@16.44.29|CCCGGTAATCGCATAAAACTTAAAACTTT</td><td>>trnaMT_LeuTAA_MT_+_3230_3304</td><td>16</td><td>44</td><td>29</td><td>CCCGGTAATCGCATAAAACTTAAAACTTT</td><td>1513.0</td><td>-9.3</td><td>1.5</td><td>-6.1</td><td>1.2E-09</td><td>2.8E-09</td><td>MB231/MB468</td><td>MP</td></tr>
<tr><td>>hg38_21_+_8433172_8446622_RNA45SN1_45S_50ntFlanks@12822.12856.35|CCGGGACCGGGGTCCGGTGCGGAGTGCCCTTCGTC</td><td>>hg38_21_+_8433172_8446622_RNA45SN1_45S_50ntFlanks</td><td>12822</td><td>12856</td><td>35</td><td>CCGGGACCGGGGTCCGGTGCGGAGTGCCCTTCGTC</td><td>146.2</td><td>-10.3</td><td>1.6</td><td>-6.4</td><td>1.3E-10</td><td>2.8E-09</td><td>BT20/MB468</td><td>MP</td></tr>
<tr><td>>hg38_21_+_8433172_8446622_RNA45SN1_45S_50ntFlanks@12072.12097.26|CCCGAGGGGCTCTCGCTTCTGGCGCC</td><td>>hg38_21_+_8433172_8446622_RNA45SN1_45S_50ntFlanks</td><td>12072</td><td>12097</td><td>26</td><td>CCCGAGGGGCTCTCGCTTCTGGCGCC</td><td>928.5</td><td>-6.4</td><td>1.1</td><td>-6.1</td><td>1.4E-09</td><td>2.8E-09</td><td>MB231/MB468</td><td>MITO</td></tr>
<tr><td>>hg38_21_+_8433172_8446622_RNA45SN1_45S_50ntFlanks@12934.12955.22|CTAAACCATTCGTAGACGACCT</td><td>>hg38_21_+_8433172_8446622_RNA45SN1_45S_50ntFlanks</td><td>12934</td><td>12955</td><td>22</td><td>CTAAACCATTCGTAGACGACCT</td><td>549.9</td><td>-7.1</td><td>1.2</td><td>-6.1</td><td>1.1E-09</td><td>2.8E-09</td><td>BT20/MB468</td><td>CYTO</td></tr>
<tr><td>>hg38_21_+_8433172_8446622_RNA45SN1_45S_50ntFlanks@13022.13039.18|CTCGACACAAGGGTTTGT</td><td>>hg38_21_+_8433172_8446622_RNA45SN1_45S_50ntFlanks</td><td>13022</td><td>13039</td><td>18</td><td>CTCGACACAAGGGTTTGT</td><td>1001.4</td><td>3.8</td><td>0.6</td><td>6.1</td><td>1.1E-09</td><td>2.8E-09</td><td>BT20/MB468</td><td>CYTO</td></tr>
<tr><td>>hg38_21_+_8433172_8446622_RNA45SN1_45S_50ntFlanks@9533.9562.30|GAGGAAACTCTGGTGGAGGTCCGTAGCGGT</td><td>>hg38_21_+_8433172_8446622_RNA45SN1_45S_50ntFlanks</td><td>9533</td><td>9562</td><td>30</td><td>GAGGAAACTCTGGTGGAGGTCCGTAGCGGT</td><td>698.8</td><td>-4.9</td><td>0.8</td><td>-6.1</td><td>1.3E-09</td><td>2.8E-09</td><td>MB231/MB468</td><td>NUC</td></tr>
<tr><td>>hg38_21_+_8433172_8446622_RNA45SN1_45S_50ntFlanks@8381.8400.20|AAGAGGGCGTGAAACCGTTA</td><td>>hg38_21_+_8433172_8446622_RNA45SN1_45S_50ntFlanks</td><td>8381</td><td>8400</td><td>20</td><td>AAGAGGGCGTGAAACCGTTA</td><td>444.3</td><td>-4.3</td><td>0.7</td><td>-6.1</td><td>9.3E-10</td><td>2.8E-09</td><td>BT20/MB468</td><td>MITO</td></tr>
<tr><td>>MI0000087|hsa-mir-29a&WithFlank&7|-|130876741|130876816@47.68.22(+1C)&[MIMAT0000086&hsa-miR-29a-3p&offsets|-1|-1(+1C);m-26&7|-|130876749|130876769&offsets|-1|0(+1C)]|CTAGCACCATCTGAAATCGGTTC</td><td>>MI0000087|hsa-mir-29a&WithFlank&7|-|130876741|130876816</td><td>47</td><td>68</td><td>22(+1C)</td><td>CTAGCACCATCTGAAATCGGTTC</td><td>362.9</td><td>12.3</td><td>1.9</td><td>6.4</td><td>1.3E-10</td><td>2.8E-09</td><td>BT20/MB468</td><td>MP</td></tr>
<tr><td>>hg38_21_+_8433172_8446622_RNA45SN1_45S_50ntFlanks@3992.4024.33|GGTGACTCTAGATAACCTCGGGCCGATCGCACG</td><td>>hg38_21_+_8433172_8446622_RNA45SN1_45S_50ntFlanks</td><td>3992</td><td>4024</td><td>33</td><td>GGTGACTCTAGATAACCTCGGGCCGATCGCACG</td><td>534.0</td><td>-4.6</td><td>0.8</td><td>-6.1</td><td>1.4E-09</td><td>2.8E-09</td><td>BT20/MB468</td><td>NUC</td></tr>
<tr><td>>hg38_21_+_8433172_8446622_RNA45SN1_45S_50ntFlanks@12072.12108.37|CCCGAGGGGCTCTCGCTTCTGGCGCCAAGCGCCCGGC</td><td>>hg38_21_+_8433172_8446622_RNA45SN1_45S_50ntFlanks</td><td>12072</td><td>12108</td><td>37</td><td>CCCGAGGGGCTCTCGCTTCTGGCGCCAAGCGCCCGGC</td><td>263.3</td><td>-7.6</td><td>1.3</td><td>-6.1</td><td>1.4E-09</td><td>2.8E-09</td><td>MB231/MB468</td><td>MITO</td></tr>
</table>

<table>
<tr><th></th><th></th><th></th><th></th><th></th><th></th><th></th><th></th><th></th><th></th><th></th><th></th><th></th><th></th></tr>
<tr><td>>hg38_21_+_8433172_8446622_RNA45SN1_45S_50ntFlanks@5070.5097.28|GTTAATTCCGATAACGAACGAGACTCTG</td><td>>hg38_21_+_8433172_8446622_RNA45SN1_45S_50ntFlanks</td><td>5070</td><td>5097</td><td>28</td><td>GTTAATTCCGATAACGAACGAGACTCTG</td><td>202.3</td><td>-11.5</td><td>1.9</td><td>-6.1</td><td>1.2E-09</td><td>2.8E-09</td><td>MB231/MB468</td><td>MP</td></tr>
<tr><td>>hg38_21_+_8433172_8446622_RNA45SN1_45S_50ntFlanks@7975.8006.32|CGCGACCTCAGATCAGACGTGGCGACCCGCTG</td><td>>hg38_21_+_8433172_8446622_RNA45SN1_45S_50ntFlanks</td><td>7975</td><td>8006</td><td>32</td><td>CGCGACCTCAGATCAGACGTGGCGACCCGCTG</td><td>70771.4</td><td>4.3</td><td>0.7</td><td>6.4</td><td>1.3E-10</td><td>2.8E-09</td><td>BT20/MB231</td><td>NUC</td></tr>
<tr><td>>hg38_21_+_8433172_8446622_RNA45SN1_45S_50ntFlanks@5472.5492.21|ACGGCCCTGGCGGAGCGCTGA</td><td>>hg38_21_+_8433172_8446622_RNA45SN1_45S_50ntFlanks</td><td>5472</td><td>5492</td><td>21</td><td>ACGGCCCTGGCGGAGCGCTGA</td><td>306.3</td><td>6.2</td><td>1.0</td><td>6.1</td><td>1.2E-09</td><td>2.8E-09</td><td>MB231/MB468</td><td>CYTO</td></tr>
<tr><td>>hg38_21_+_8433172_8446622_RNA45SN1_45S_50ntFlanks@11805.11827.23|CCACTGTCCCTACCTACTATCCA</td><td>>hg38_21_+_8433172_8446622_RNA45SN1_45S_50ntFlanks</td><td>11805</td><td>11827</td><td>23</td><td>CCACTGTCCCTACCTACTATCCA</td><td>164.2</td><td>-7.6</td><td>1.3</td><td>-6.1</td><td>1.3E-09</td><td>2.8E-09</td><td>MB231/MB468</td><td>NUC</td></tr>
<tr><td>>hg38_21_+_8433172_8446622_RNA45SN1_45S_50ntFlanks@6737.6756.20|GATCATCGACACTTCGAACG</td><td>>hg38_21_+_8433172_8446622_RNA45SN1_45S_50ntFlanks</td><td>6737</td><td>6756</td><td>20</td><td>GATCATCGACACTTCGAACG</td><td>141.7</td><td>-10.2</td><td>1.6</td><td>-6.4</td><td>1.3E-10</td><td>2.8E-09</td><td>BT20/MB468</td><td>MP</td></tr>
<tr><td>>hg38_21_+_8433172_8446622_RNA45SN1_45S_50ntFlanks@6745.6762.18|ACACTTCGAACGCACTTG</td><td>>hg38_21_+_8433172_8446622_RNA45SN1_45S_50ntFlanks</td><td>6745</td><td>6762</td><td>18</td><td>ACACTTCGAACGCACTTG</td><td>140.0</td><td>-10.2</td><td>1.6</td><td>-6.4</td><td>1.3E-10</td><td>2.8E-09</td><td>BT20/MB468</td><td>MP</td></tr>
<tr><td>>am_trna119_LysCTT_1_-_145395522_145395594@13.33.21,trna11_LysCTT_5_-_180648979_180649051@13.33.21,trna128_LysCTT_1_-_55423542_55423614@13.33.21,trna13_LysCTT_6_+_26556774_26556846@13.33.21,trna6_LysCTT_19_-_52425396_52425468@13.33.21,trna7_LysCTT_16_+_3225692_3225764@13.33.21,trna9_LysCTT_5_+_180634755_180634827@13.33.21|CAGTCGGTAGAGCATGAGACT</td><td>>am_trna119_LysCTT_1_-_145395522_145395594</td><td>13</td><td>33</td><td>21</td><td>CAGTCGGTAGAGCATGAGACT</td><td>917.6</td><td>7.5</td><td>1.2</td><td>6.1</td><td>1.4E-09</td><td>2.9E-09</td><td>MB231/MB468</td><td>MITO</td></tr>
<tr><td>>hg38_21_+_8433172_8446622_RNA45SN1_45S_50ntFlanks@3853.3898.46|AACTGTGGTAATTCTAGAGCTAATACATGCCGACGGGCGCTGACCC</td><td>>hg38_21_+_8433172_8446622_RNA45SN1_45S_50ntFlanks</td><td>3853</td><td>3898</td><td>46</td><td>AACTGTGGTAATTCTAGAGCTAATACATGCCGACGGGCGCTGACCC</td><td>192.2</td><td>-9.9</td><td>1.6</td><td>-6.1</td><td>1.1E-09</td><td>2.9E-09</td><td>BT20/MB468</td><td>CYTO</td></tr>
<tr><td>>hg38_21_+_8433172_8446622_RNA45SN1_45S_50ntFlanks@475.507.33|CTTCGTGATCGATGTGGTGACGTCGTGCTCTCC</td><td>>hg38_21_+_8433172_8446622_RNA45SN1_45S_50ntFlanks</td><td>475</td><td>507</td><td>33</td><td>CTTCGTGATCGATGTGGTGACGTCGTGCTCTCC</td><td>1668.7</td><td>9.0</td><td>1.5</td><td>6.1</td><td>1.1E-09</td><td>2.9E-09</td><td>BT20/MB468</td><td>CYTO</td></tr>
<tr><td>>MI0000255|hsa-mir-30d&WithFlank&8|-|134804870|134804951@12.31.20&[MIMAT0000245&hsa-miR-30d-5p&offsets|0|-2;m-21&8|-|134804919|134804940&offsets|0|-2]|TGTAAACATCCCCGACTGGA</td><td>>MI0000255|hsa-mir-30d&WithFlank&8|-|134804870|134804951</td><td>12</td><td>31</td><td>20</td><td>TGTAAACATCCCCGACTGGA</td><td>3111.6</td><td>6.6</td><td>1.0</td><td>6.4</td><td>1.3E-10</td><td>2.9E-09</td><td>BT20/MB468</td><td>MP</td></tr>
<tr><td>>MI0000239|hsa-mir-197&WithFlank&1|+|109598887|109598973@54.74.21(+1U)&[MIMAT0000227&hsa-miR-197-3p&offsets|0|-1(+1U);m-99&1|+|109598940|109598961&offsets|0|-1(+1U)]|TTCACCACCTTCTCCACCCAGT</td><td>>MI0000239|hsa-mir-197&WithFlank&1|+|109598887|109598973</td><td>54</td><td>74</td><td>21(+1U)</td><td>TTCACCACCTTCTCCACCCAGT</td><td>566.9</td><td>6.7</td><td>1.1</td><td>6.1</td><td>1.3E-09</td><td>2.9E-09</td><td>MB231/MB468</td><td>CYTO</td></tr>
<tr><td>>hg38_21_+_8433172_8446622_RNA45SN1_45S_50ntFlanks@10393.10429.37|CAGCAGTTGAACATGGGTCAGTCGGTCCTGAGAGATG</td><td>>hg38_21_+_8433172_8446622_RNA45SN1_45S_50ntFlanks</td><td>10393</td><td>10429</td><td>37</td><td>CAGCAGTTGAACATGGGTCAGTCGGTCCTGAGAGATG</td><td>1884.2</td><td>-5.1</td><td>0.8</td><td>-6.1</td><td>1.4E-09</td><td>2.9E-09</td><td>MB231/MB468</td><td>MITO</td></tr>
<tr><td>>hg38_21_+_8433172_8446622_RNA45SN1_45S_50ntFlanks@4727.4759.33|AATCAAGAACGAAAGTCGGAGGTTCGAAGACGA</td><td>>hg38_21_+_8433172_8446622_RNA45SN1_45S_50ntFlanks</td><td>4727</td><td>4759</td><td>33</td><td>AATCAAGAACGAAAGTCGGAGGTTCGAAGACGA</td><td>496.9</td><td>-4.7</td><td>0.8</td><td>-6.1</td><td>1.4E-09</td><td>2.9E-09</td><td>MB231/MB468</td><td>MITO</td></tr>
<tr><td>>hg38_21_+_8433172_8446622_RNA45SN1_45S_50ntFlanks@4732.4759.28|AGAACGAAAGTCGGAGGTTCGAAGACGA</td><td>>hg38_21_+_8433172_8446622_RNA45SN1_45S_50ntFlanks</td><td>4732</td><td>4759</td><td>28</td><td>AGAACGAAAGTCGGAGGTTCGAAGACGA</td><td>609.5</td><td>-4.9</td><td>0.8</td><td>-6.1</td><td>1.4E-09</td><td>2.9E-09</td><td>MB231/MB468</td><td>MITO</td></tr>
<tr><td>>hg38_21_+_8433172_8446622_RNA45SN1_45S_50ntFlanks@9573.9597.25|AAATCGGTCGTCCGACCTGGGTATA</td><td>>hg38_21_+_8433172_8446622_RNA45SN1_45S_50ntFlanks</td><td>9573</td><td>9597</td><td>25</td><td>AAATCGGTCGTCCGACCTGGGTATA</td><td>1080.5</td><td>-9.3</td><td>1.5</td><td>-6.0</td><td>1.5E-09</td><td>2.9E-09</td><td>BT20/MB468</td><td>NUC</td></tr>
<tr><td>>hg38_21_+_8433172_8446622_RNA45SN1_45S_50ntFlanks@5484.5515.32|GAGCGCTGAGAAGACGGTCGAACTTGACTATC</td><td>>hg38_21_+_8433172_8446622_RNA45SN1_45S_50ntFlanks</td><td>5484</td><td>5515</td><td>32</td><td>GAGCGCTGAGAAGACGGTCGAACTTGACTATC</td><td>377.9</td><td>-8.0</td><td>1.3</td><td>-6.1</td><td>1.2E-09</td><td>2.9E-09</td><td>MB231/MB468</td><td>MP</td></tr>
<tr><td>>MI0000093|hsa-mir-92a-1&WithFlank&13|+|91351308|91351397@54.74.21(+1C);MI0000094|hsa-mir-92a-2&WithFlank&X|-|134169532|134169618@54.74.21(+1C)&[MIMAT0000092&hsa-miR-92a-3p&offsets|0|-1(+1C);m-9&13|+|91351361|91351382&offsets|0|-1(+1C)];[MIMAT0000092_1&hsa-miR-92a-3p&offsets|0|-1(+1C);m-11&X|-|134169544|134169565&offsets|0|-1(+1C)]|TATTGCACTTGTCCCGGCCTGC</td><td>>MI0000093|hsa-mir-92a-1&WithFlank&13|+|91351308|91351397</td><td>54</td><td>74</td><td>21(+1C)</td><td>TATTGCACTTGTCCCGGCCTGC</td><td>8637.9</td><td>3.9</td><td>0.6</td><td>6.1</td><td>1.3E-09</td><td>2.9E-09</td><td>MB231/MB468</td><td>NUC</td></tr>
<tr><td>>hg38_21_+_8433172_8446622_RNA45SN1_45S_50ntFlanks@10101.10124.24|CGCGCGCGTGTGGTGTGCGTCGGA</td><td>>hg38_21_+_8433172_8446622_RNA45SN1_45S_50ntFlanks</td><td>10101</td><td>10124</td><td>24</td><td>CGCGCGCGTGTGGTGTGCGTCGGA</td><td>236.7</td><td>-11.0</td><td>1.7</td><td>-6.4</td><td>1.3E-10</td><td>2.9E-09</td><td>BT20/MB468</td><td>MP</td></tr>
<tr><td>>MI0000462|hsa-mir-152&WithFlank&17|-|48037155|48037253@60.81.22(+1U)&[MIMAT0000438&hsa-miR-152-3p&offsets|0|+1(+1U);m-97&17|-|48037174|48037194&offsets|0|+1(+1U)]|TCAGTGCATGACAGAACTTGGGT</td><td>>MI0000462|hsa-mir-152&WithFlank&17|-|48037155|48037253</td><td>60</td><td>81</td><td>22(+1U)</td><td>TCAGTGCATGACAGAACTTGGGT</td><td>473.6</td><td>9.1</td><td>1.5</td><td>6.1</td><td>1.2E-09</td><td>2.9E-09</td><td>MB231/MB468</td><td>MP</td></tr>
<tr><td>>hg38_21_+_8433172_8446622_RNA45SN1_45S_50ntFlanks@4732.4765.34|AGAACGAAAGTCGGAGGTTCGAAGACGATCAGAT</td><td>>hg38_21_+_8433172_8446622_RNA45SN1_45S_50ntFlanks</td><td>4732</td><td>4765</td><td>34</td><td>AGAACGAAAGTCGGAGGTTCGAAGACGATCAGAT</td><td>1963.2</td><td>7.3</td><td>1.2</td><td>6.1</td><td>1.1E-09</td><td>2.9E-09</td><td>BT20/MB468</td><td>CYTO</td></tr>
<tr><td>>hg38_21_+_8433172_8446622_RNA45SN1_45S_50ntFlanks@11569.11602.34|CAGCCGACTTAGAACTGGTGCGGACCAGGGGAAT</td><td>>hg38_21_+_8433172_8446622_RNA45SN1_45S_50ntFlanks</td><td>11569</td><td>11602</td><td>34</td><td>CAGCCGACTTAGAACTGGTGCGGACCAGGGGAAT</td><td>1666.9</td><td>-3.6</td><td>0.6</td><td>-6.1</td><td>1.1E-09</td><td>2.9E-09</td><td>BT20/MB468</td><td>CYTO</td></tr>
<tr><td>>hg38_21_+_8433172_8446622_RNA45SN1_45S_50ntFlanks@6747.6770.24|ACTTCGAACGCACTTGCGGCCCCG</td><td>>hg38_21_+_8433172_8446622_RNA45SN1_45S_50ntFlanks</td><td>6747</td><td>6770</td><td>24</td><td>ACTTCGAACGCACTTGCGGCCCCG</td><td>674.1</td><td>-7.4</td><td>1.2</td><td>-6.1</td><td>1.1E-09</td><td>2.9E-09</td><td>BT20/MB468</td><td>CYTO</td></tr>
<tr><td>>hg38_21_+_8433172_8446622_RNA45SN1_45S_50ntFlanks@11918.11959.42|ACATGAGAGGTGTAGAATAAGTGGGAGGCCCCCGGCGCCCCC</td><td>>hg38_21_+_8433172_8446622_RNA45SN1_45S_50ntFlanks</td><td>11918</td><td>11959</td><td>42</td><td>ACATGAGAGGTGTAGAATAAGTGGGAGGCCCCCGGCGCCCCC</td><td>145.2</td><td>-9.5</td><td>1.6</td><td>-6.1</td><td>1.1E-09</td><td>2.9E-09</td><td>BT20/MB468</td><td>CYTO</td></tr>
<tr><td>>hg38_21_+_8433172_8446622_RNA45SN1_45S_50ntFlanks@3846.3867.22|CTTGGATAACTGTGGTAATTCT</td><td>>hg38_21_+_8433172_8446622_RNA45SN1_45S_50ntFlanks</td><td>3846</td><td>3867</td><td>22</td><td>CTTGGATAACTGTGGTAATTCT</td><td>73.2</td><td>-10.1</td><td>1.7</td><td>-6.1</td><td>1.2E-09</td><td>2.9E-09</td><td>MB231/MB468</td><td>MP</td></tr>
<tr><td>>hg38_21_+_8433172_8446622_RNA45SN1_45S_50ntFlanks@11584.11612.29|TGGTGCGGACCAGGGGAATCCGACTGTTT</td><td>>hg38_21_+_8433172_8446622_RNA45SN1_45S_50ntFlanks</td><td>11584</td><td>11612</td><td>29</td><td>TGGTGCGGACCAGGGGAATCCGACTGTTT</td><td>711.0</td><td>5.3</td><td>0.9</td><td>6.1</td><td>9.6E-10</td><td>2.9E-09</td><td>BT20/MB468</td><td>MITO</td></tr>
<tr><td>>hg38_21_+_8433172_8446622_RNA45SN1_45S_50ntFlanks@5472.5513.42|ACGGCCCTGGCGGAGCGCTGAGAAGACGGTCGAACTTGACTA</td><td>>hg38_21_+_8433172_8446622_RNA45SN1_45S_50ntFlanks</td><td>5472</td><td>5513</td><td>42</td><td>ACGGCCCTGGCGGAGCGCTGAGAAGACGGTCGAACTTGACTA</td><td>1895.7</td><td>5.4</td><td>0.9</td><td>6.1</td><td>1.3E-09</td><td>2.9E-09</td><td>MB231/MB468</td><td>CYTO</td></tr>
<tr><td>>hg38_21_+_8433172_8446622_RNA45SN1_45S_50ntFlanks@6753.6770.18|AACGCACTTGCGGCCCCG</td><td>>hg38_21_+_8433172_8446622_RNA45SN1_45S_50ntFlanks</td><td>6753</td><td>6770</td><td>18</td><td>AACGCACTTGCGGCCCCG</td><td>144.2</td><td>-6.8</td><td>1.1</td><td>-6.1</td><td>1.1E-09</td><td>2.9E-09</td><td>BT20/MB468</td><td>CYTO</td></tr>
<tr><td>>hg38_21_+_8433172_8446622_RNA45SN1_45S_50ntFlanks@12003.12034.32|GCGGGCCGCCGGTGAAATACCACTACTCTGAT</td><td>>hg38_21_+_8433172_8446622_RNA45SN1_45S_50ntFlanks</td><td>12003</td><td>12034</td><td>32</td><td>GCGGGCCGCCGGTGAAATACCACTACTCTGAT</td><td>945.7</td><td>5.4</td><td>0.9</td><td>6.1</td><td>1.3E-09</td><td>2.9E-09</td><td>MB231/MB468</td><td>NUC</td></tr>
<tr><td>>hg38_21_+_8433172_8446622_RNA45SN1_45S_50ntFlanks@12045.12068.24|CTGACCCGGTGAGGCGGGGGGGCG</td><td>>hg38_21_+_8433172_8446622_RNA45SN1_45S_50ntFlanks</td><td>12045</td><td>12068</td><td>24</td><td>CTGACCCGGTGAGGCGGGGGGGCG</td><td>356.6</td><td>-8.2</td><td>1.4</td><td>-6.1</td><td>1.3E-09</td><td>2.9E-09</td><td>MB231/MB468</td><td>NUC</td></tr>
<tr><td>>hg38_21_+_8433172_8446622_RNA45SN1_45S_50ntFlanks@10834.10867.34|CCGGATCCGTAACTTCGGGATAAGGATTGGCTCT</td><td>>hg38_21_+_8433172_8446622_RNA45SN1_45S_50ntFlanks</td><td>10834</td><td>10867</td><td>34</td><td>CCGGATCCGTAACTTCGGGATAAGGATTGGCTCT</td><td>603.1</td><td>3.9</td><td>0.6</td><td>6.1</td><td>1.1E-09</td><td>2.9E-09</td><td>BT20/MB468</td><td>CYTO</td></tr>
<tr><td>>hg38_21_+_8433172_8446622_RNA45SN1_45S_50ntFlanks@6648.6669.22|GTACGACTCTTAGCGGTGGATC</td><td>>hg38_21_+_8433172_8446622_RNA45SN1_45S_50ntFlanks</td><td>6648</td><td>6669</td><td>22</td><td>GTACGACTCTTAGCGGTGGATC</td><td>9517.4</td><td>4.1</td><td>0.7</td><td>6.1</td><td>9.7E-10</td><td>2.9E-09</td><td>BT20/MB468</td><td>MITO</td></tr>
<tr><td>>hg38_21_+_8433172_8446622_RNA45SN1_45S_50ntFlanks@10388.10410.23|GTGAACAGCAGTTGAACATGGGT</td><td>>hg38_21_+_8433172_8446622_RNA45SN1_45S_50ntFlanks</td><td>10388</td><td>10410</td><td>23</td><td>GTGAACAGCAGTTGAACATGGGT</td><td>1821.8</td><td>-9.6</td><td>1.6</td><td>-6.0</td><td>1.5E-09</td><td>2.9E-09</td><td>BT20/MB468</td><td>NUC</td></tr>
<tr><td>>hg38_21_+_8433172_8446622_RNA45SN1_45S_50ntFlanks@9720.9737.18|ATTAGAGGTCTTGGGGCC</td><td>>hg38_21_+_8433172_8446622_RNA45SN1_45S_50ntFlanks</td><td>9720</td><td>9737</td><td>18</td><td>ATTAGAGGTCTTGGGGCC</td><td>140.8</td><td>-7.2</td><td>1.2</td><td>-6.1</td><td>1.1E-09</td><td>2.9E-09</td><td>BT20/MB468</td><td>CYTO</td></tr>
<tr><td>>MI0000085|hsa-mir-27a&WithFlank&19|-|13836434|13836523@57.76.20(+1U)&[MIMAT0000084&hsa-miR-27a-3p&offsets|0|-1(+1U);m-43&19|-|13836447|13836467&offsets|0|-1(+1U)]|TTCACAGTGGCTAAGTTCCGT</td><td>>MI0000085|hsa-mir-27a&WithFlank&19|-|13836434|13836523</td><td>57</td><td>76</td><td>20(+1U)</td><td>TTCACAGTGGCTAAGTTCCGT</td><td>2361.5</td><td>4.8</td><td>0.8</td><td>6.1</td><td>1.1E-09</td><td>2.9E-09</td><td>BT20/MB468</td><td>CYTO</td></tr>
<tr><td>>hg38_21_+_8433172_8446622_RNA45SN1_45S_50ntFlanks@4575.4610.36|TAGGACCGCGGTTCTATTTTGTTGGTTTTCGGAACT</td><td>>hg38_21_+_8433172_8446622_RNA45SN1_45S_50ntFlanks</td><td>4575</td><td>4610</td><td>36</td><td>TAGGACCGCGGTTCTATTTTGTTGGTTTTCGGAACT</td><td>354.5</td><td>6.0</td><td>1.0</td><td>6.1</td><td>1.3E-09</td><td>2.9E-09</td><td>MB231/MB468</td><td>CYTO</td></tr>
<tr><td>>hg38_21_+_8433172_8446622_RNA45SN1_45S_50ntFlanks@3705.3743.39|TACCTGGTTGATCCTGCCAGTAGCATATGCTTGTCTCAA</td><td>>hg38_21_+_8433172_8446622_RNA45SN1_45S_50ntFlanks</td><td>3705</td><td>3743</td><td>39</td><td>TACCTGGTTGATCCTGCCAGTAGCATATGCTTGTCTCAA</td><td>1364.1</td><td>4.2</td><td>0.7</td><td>6.1</td><td>9.7E-10</td><td>2.9E-09</td><td>BT20/MB468</td><td>MITO</td></tr>
<tr><td>>hg38_21_+_8433172_8446622_RNA45SN1_45S_50ntFlanks@6661.6706.46|CGGTGGATCACTCGGCTCGTGCGTCGATGAAGAACGCAGCTAGCTG</td><td>>hg38_21_+_8433172_8446622_RNA45SN1_45S_50ntFlanks</td><td>6661</td><td>6706</td><td>46</td><td>CGGTGGATCACTCGGCTCGTGCGTCGATGAAGAACGCAGCTAGCTG</td><td>13147.7</td><td>4.8</td><td>0.8</td><td>6.1</td><td>9.7E-10</td><td>2.9E-09</td><td>BT20/MB468</td><td>MITO</td></tr>
<tr><td>>MI0000283|hsa-mir-203a&WithFlank&14|+|104117399|104117520@72.90.19&[MIMAT0000264&hsa-miR-203a-3p&offsets|+1|-2;m-79&14|+|104117469|104117490&offsets|+1|-2]|TGAAATGTTTAGGACCACT</td><td>>MI0000283|hsa-mir-203a&WithFlank&14|+|104117399|104117520</td><td>72</td><td>90</td><td>19</td><td>TGAAATGTTTAGGACCACT</td><td>292.8</td><td>11.5</td><td>1.7</td><td>6.7</td><td>2.7E-11</td><td>2.9E-09</td><td>BT20/MB231</td><td>MP</td></tr>
<tr><td>>hg38_21_+_8433172_8446622_RNA45SN1_45S_50ntFlanks@7980.8004.25|CCTCAGATCAGACGTGGCGACCCGC</td><td>>hg38_21_+_8433172_8446622_RNA45SN1_45S_50ntFlanks</td><td>7980</td><td>8004</td><td>25</td><td>CCTCAGATCAGACGTGGCGACCCGC</td><td>595.6</td><td>-8.0</td><td>1.3</td><td>-6.1</td><td>1.4E-09</td><td>2.9E-09</td><td>MB231/MB468</td><td>MITO</td></tr>
<tr><td>>hg38_21_+_8433172_8446622_RNA45SN1_45S_50ntFlanks@7975.8004.30|CGCGACCTCAGATCAGACGTGGCGACCCGC</td><td>>hg38_21_+_8433172_8446622_RNA45SN1_45S_50ntFlanks</td><td>7975</td><td>8004</td><td>30</td><td>CGCGACCTCAGATCAGACGTGGCGACCCGC</td><td>5407.7</td><td>4.8</td><td>0.8</td><td>6.1</td><td>1.3E-09</td><td>2.9E-09</td><td>MB231/MB468</td><td>CYTO</td></tr>
<tr><td>>hg38_21_+_8433172_8446622_RNA45SN1_45S_50ntFlanks@6651.6668.18|CGACTCTTAGCGGTGGAT</td><td>>hg38_21_+_8433172_8446622_RNA45SN1_45S_50ntFlanks</td><td>6651</td><td>6668</td><td>18</td><td>CGACTCTTAGCGGTGGAT</td><td>4836.0</td><td>2.9</td><td>0.5</td><td>6.0</td><td>1.5E-09</td><td>2.9E-09</td><td>BT20/MB468</td><td>NUC</td></tr>
<tr><td>>hg38_21_+_8433172_8446622_RNA45SN1_45S_50ntFlanks@6649.6677.29|TACGACTCTTAGCGGTGGATCACTCGGCT</td><td>>hg38_21_+_8433172_8446622_RNA45SN1_45S_50ntFlanks</td><td>6649</td><td>6677</td><td>29</td><td>TACGACTCTTAGCGGTGGATCACTCGGCT</td><td>2630.9</td><td>4.0</td><td>0.6</td><td>6.1</td><td>9.8E-10</td><td>2.9E-09</td><td>BT20/MB468</td><td>MITO</td></tr>
<tr><td>>am_trnaMT_ValTAC_MT_+_1602_1670@33.52.20|ACACTTAGGAGATTTCAACT</td><td>>am_trnaMT_ValTAC_MT_+_1602_1670</td><td>33</td><td>52</td><td>20</td><td>ACACTTAGGAGATTTCAACT</td><td>509.5</td><td>9.7</td><td>1.6</td><td>6.1</td><td>1.1E-09</td><td>2.9E-09</td><td>BT20/MB468</td><td>CYTO</td></tr>
<tr><td>>hg38_21_+_8433172_8446622_RNA45SN1_45S_50ntFlanks@12456.12496.41|CCAAGCGTTGGATTGTTCACCCACTAATAGGGAACGTGAGC</td><td>>hg38_21_+_8433172_8446622_RNA45SN1_45S_50ntFlanks</td><td>12456</td><td>12496</td><td>41</td><td>CCAAGCGTTGGATTGTTCACCCACTAATAGGGAACGTGAGC</td><td>842.7</td><td>-5.3</td><td>0.9</td><td>-6.1</td><td>1.4E-09</td><td>2.9E-09</td><td>MB231/MB468</td><td>MITO</td></tr>
<tr><td>>MI0000465|hsa-mir-191&WithFlank&3|-|49020612|49020715@22.43.22(+1U)&[MIMAT0000440&hsa-miR-191-5p&offsets|0|-1(+1U);m-48&3|-|49020672|49020694&offsets|0|-1(+1U)]|CAACGGAATCCCAAAAGCAGCTT</td><td>>MI0000465|hsa-mir-191&WithFlank&3|-|49020612|49020715</td><td>22</td><td>43</td><td>22(+1U)</td><td>CAACGGAATCCCAAAAGCAGCTT</td><td>295.2</td><td>6.6</td><td>1.1</td><td>6.1</td><td>1.3E-09</td><td>2.9E-09</td><td>MB231/MB468</td><td>CYTO</td></tr>
<tr><td>>hg38_21_+_8433172_8446622_RNA45SN1_45S_50ntFlanks@6738.6771.34|ATCATCGACACTTCGAACGCACTTGCGGCCCCGG</td><td>>hg38_21_+_8433172_8446622_RNA45SN1_45S_50ntFlanks</td><td>6738</td><td>6771</td><td>34</td><td>ATCATCGACACTTCGAACGCACTTGCGGCCCCGG</td><td>3141.0</td><td>-8.8</td><td>1.4</td><td>-6.1</td><td>1.3E-09</td><td>2.9E-09</td><td>MB231/MB468</td><td>NUC</td></tr>
<tr><td>>hg38_21_+_8433172_8446622_RNA45SN1_45S_50ntFlanks@4727.4752.26|AATCAAGAACGAAAGTCGGAGGTTCG</td><td>>hg38_21_+_8433172_8446622_RNA45SN1_45S_50ntFlanks</td><td>4727</td><td>4752</td><td>26</td><td>AATCAAGAACGAAAGTCGGAGGTTCG</td><td>442.2</td><td>-6.3</td><td>1.0</td><td>-6.1</td><td>1.2E-09</td><td>2.9E-09</td><td>MB231/MB468</td><td>MP</td></tr>
<tr><td>>hg38_21_+_8433172_8446622_RNA45SN1_45S_50ntFlanks@13008.13025.18|ATTGAAAGTCAGCCCTCG</td><td>>hg38_21_+_8433172_8446622_RNA45SN1_45S_50ntFlanks</td><td>13008</td><td>13025</td><td>18</td><td>ATTGAAAGTCAGCCCTCG</td><td>520.0</td><td>-7.8</td><td>1.3</td><td>-6.1</td><td>1.4E-09</td><td>2.9E-09</td><td>MB231/MB468</td><td>MITO</td></tr>
</table>

<table>
<tr><th></th><th></th><th></th><th></th><th></th><th></th><th></th><th></th><th></th><th></th><th></th><th></th><th></th><th></th></tr>
<tr><td>>MI0000095|hsa-mir-93&WithFlank&7|-|100093762|100093853@17.39.23(+1A)&[MIMAT0000093&hsa-miR-93-5p&offsets|0|0(+1A);m-29&7|-|100093815|100093837&offsets|0|0(+1A)]|CAAAGTGCTGTTCGTGCAGGTAGA</td><td>>MI0000095|hsa-mir-93&WithFlank&7|-|100093762|100093853</td><td>17</td><td>39</td><td>23(+1A)</td><td>CAAAGTGCTGTTCGTGCAGGTAGA</td><td>1601.1</td><td>4.2</td><td>0.7</td><td>6.1</td><td>1.3E-09</td><td>2.9E-09</td><td>MB231/MB468</td><td>NUC</td></tr>
<tr><td>>hg38_21_+_8433172_8446622_RNA45SN1_45S_50ntFlanks@12727.12747.21|CCTCGGATAGCCGGTCCCCCG</td><td>>hg38_21_+_8433172_8446622_RNA45SN1_45S_50ntFlanks</td><td>12727</td><td>12747</td><td>21</td><td>CCTCGGATAGCCGGTCCCCCG</td><td>94.5</td><td>-10.4</td><td>1.7</td><td>-6.1</td><td>1.3E-09</td><td>3.0E-09</td><td>MB231/MB468</td><td>MP</td></tr>
<tr><td>>hg38_21_+_8433172_8446622_RNA45SN1_45S_50ntFlanks@12997.13039.43|CGCTGCGATCTATTGAAAGTCAGCCCTCGACACAAGGGTTTGT</td><td>>hg38_21_+_8433172_8446622_RNA45SN1_45S_50ntFlanks</td><td>12997</td><td>13039</td><td>43</td><td>CGCTGCGATCTATTGAAAGTCAGCCCTCGACACAAGGGTTTGT</td><td>107419.6</td><td>6.2</td><td>1.0</td><td>6.1</td><td>9.8E-10</td><td>3.0E-09</td><td>BT20/MB468</td><td>MITO</td></tr>
<tr><td>>hg38_21_+_8433172_8446622_RNA45SN1_45S_50ntFlanks@6671.6692.22|CTCGGCTCGTGCGTCGATGAAG</td><td>>hg38_21_+_8433172_8446622_RNA45SN1_45S_50ntFlanks</td><td>6671</td><td>6692</td><td>22</td><td>CTCGGCTCGTGCGTCGATGAAG</td><td>193.7</td><td>-9.7</td><td>1.6</td><td>-6.1</td><td>9.9E-10</td><td>3.0E-09</td><td>BT20/MB468</td><td>MITO</td></tr>
<tr><td>>hg38_21_+_8433172_8446622_RNA45SN1_45S_50ntFlanks@3988.4010.23|CTTTGGTGACTCTAGATAACCTC</td><td>>hg38_21_+_8433172_8446622_RNA45SN1_45S_50ntFlanks</td><td>3988</td><td>4010</td><td>23</td><td>CTTTGGTGACTCTAGATAACCTC</td><td>416.4</td><td>4.1</td><td>0.7</td><td>6.1</td><td>1.1E-09</td><td>3.0E-09</td><td>BT20/MB468</td><td>CYTO</td></tr>
<tr><td>>MI0000736|hsa-mir-30c-1&WithFlank&1|+|40757278|40757378@23.45.23;MI0000254|hsa-mir-30c-2&WithFlank&6|-|71376954|71377037@13.35.23&[MIMAT0000244&hsa-miR-30c-5p&offsets|0|0;m-57&1|+|40757300|40757323&offsets|0|-1];[MIMAT0000244_1&hsa-miR-30c-5p&offsets|0|0;m-56&6|-|71377002|71377025&offsets|0|-1]|TGTAAACATCCTACACTCTCAGC</td><td>>MI0000736|hsa-mir-30c-1&WithFlank&1|+|40757278|40757378</td><td>23</td><td>45</td><td>23</td><td>TGTAAACATCCTACACTCTCAGC</td><td>8536.6</td><td>4.9</td><td>0.8</td><td>6.1</td><td>9.9E-10</td><td>3.0E-09</td><td>BT20/MB468</td><td>MITO</td></tr>
<tr><td>>hg38_21_+_8433172_8446622_RNA45SN1_45S_50ntFlanks@8423.8459.37|CGCAGTCCGCCCGGAGGATTCAACCCGGCGGCGGGTC</td><td>>hg38_21_+_8433172_8446622_RNA45SN1_45S_50ntFlanks</td><td>8423</td><td>8459</td><td>37</td><td>CGCAGTCCGCCCGGAGGATTCAACCCGGCGGCGGGTC</td><td>94.8</td><td>-10.4</td><td>1.7</td><td>-6.1</td><td>1.3E-09</td><td>3.0E-09</td><td>MB231/MB468</td><td>MP</td></tr>
<tr><td>>hg38_21_+_8433172_8446622_RNA45SN1_45S_50ntFlanks@4499.4520.22|AGCGTTTACTTTGAAAAAATTA</td><td>>hg38_21_+_8433172_8446622_RNA45SN1_45S_50ntFlanks</td><td>4499</td><td>4520</td><td>22</td><td>AGCGTTTACTTTGAAAAAATTA</td><td>1266.1</td><td>5.7</td><td>0.9</td><td>6.1</td><td>1.1E-09</td><td>3.0E-09</td><td>BT20/MB468</td><td>CYTO</td></tr>
<tr><td>>hg38_21_+_8433172_8446622_RNA45SN1_45S_50ntFlanks@12957.12985.29|CTTCTGGGTCGGGGTTTCGTACGTAGCAG</td><td>>hg38_21_+_8433172_8446622_RNA45SN1_45S_50ntFlanks</td><td>12957</td><td>12985</td><td>29</td><td>CTTCTGGGTCGGGGTTTCGTACGTAGCAG</td><td>384.1</td><td>-7.7</td><td>1.3</td><td>-6.1</td><td>1.3E-09</td><td>3.0E-09</td><td>MB231/MB468</td><td>NUC</td></tr>
<tr><td>>hg38_21_+_8433172_8446622_RNA45SN1_45S_50ntFlanks@4008.4042.35|CTCGGGCCGATCGCACGCCCCCCGTGGCGGCGACG</td><td>>hg38_21_+_8433172_8446622_RNA45SN1_45S_50ntFlanks</td><td>4008</td><td>4042</td><td>35</td><td>CTCGGGCCGATCGCACGCCCCCCGTGGCGGCGACG</td><td>343.0</td><td>-6.3</td><td>1.0</td><td>-6.1</td><td>1.4E-09</td><td>3.0E-09</td><td>MB231/MB468</td><td>MITO</td></tr>
<tr><td>>hg38_21_+_8433172_8446622_RNA45SN1_45S_50ntFlanks@10794.10826.33|CTGGCATGTTGGAACAATGTAGGTAAGGGAAGT</td><td>>hg38_21_+_8433172_8446622_RNA45SN1_45S_50ntFlanks</td><td>10794</td><td>10826</td><td>33</td><td>CTGGCATGTTGGAACAATGTAGGTAAGGGAAGT</td><td>213.8</td><td>-4.9</td><td>0.8</td><td>-6.1</td><td>1.3E-09</td><td>3.0E-09</td><td>MB231/MB468</td><td>CYTO</td></tr>
<tr><td>>hg38_21_+_8433172_8446622_RNA45SN1_45S_50ntFlanks@9572.9594.23|CAAATCGGTCGTCCGACCTGGGT</td><td>>hg38_21_+_8433172_8446622_RNA45SN1_45S_50ntFlanks</td><td>9572</td><td>9594</td><td>23</td><td>CAAATCGGTCGTCCGACCTGGGT</td><td>134.3</td><td>-7.3</td><td>1.2</td><td>-6.1</td><td>1.3E-09</td><td>3.0E-09</td><td>MB231/MB468</td><td>NUC</td></tr>
<tr><td>>hg38_21_+_8433172_8446622_RNA45SN1_45S_50ntFlanks@6654.6681.28|CTCTTAGCGGTGGATCACTCGGCTCGTG</td><td>>hg38_21_+_8433172_8446622_RNA45SN1_45S_50ntFlanks</td><td>6654</td><td>6681</td><td>28</td><td>CTCTTAGCGGTGGATCACTCGGCTCGTG</td><td>1056.1</td><td>-5.9</td><td>1.0</td><td>-6.1</td><td>1.3E-09</td><td>3.0E-09</td><td>MB231/MB468</td><td>CYTO</td></tr>
<tr><td>>hg38_21_+_8433172_8446622_RNA45SN1_45S_50ntFlanks@12949.12982.34|ACGACCTGCTTCTGGGTCGGGGTTTCGTACGTAG</td><td>>hg38_21_+_8433172_8446622_RNA45SN1_45S_50ntFlanks</td><td>12949</td><td>12982</td><td>34</td><td>ACGACCTGCTTCTGGGTCGGGGTTTCGTACGTAG</td><td>83.6</td><td>-10.2</td><td>1.7</td><td>-6.1</td><td>1.3E-09</td><td>3.0E-09</td><td>MB231/MB468</td><td>MP</td></tr>
<tr><td>>hg38_21_+_8433172_8446622_RNA45SN1_45S_50ntFlanks@10777.10818.42|TCCAAGGTGAACAGCCTCTGGCATGTTGGAACAATGTAGGTA</td><td>>hg38_21_+_8433172_8446622_RNA45SN1_45S_50ntFlanks</td><td>10777</td><td>10818</td><td>42</td><td>TCCAAGGTGAACAGCCTCTGGCATGTTGGAACAATGTAGGTA</td><td>129.3</td><td>-10.9</td><td>1.8</td><td>-6.1</td><td>1.3E-09</td><td>3.0E-09</td><td>MB231/MB468</td><td>MP</td></tr>
<tr><td>>hg38_21_+_8433172_8446622_RNA45SN1_45S_50ntFlanks@11670.11692.23|CCCAGTGCTCTGAATGTCAAAGT</td><td>>hg38_21_+_8433172_8446622_RNA45SN1_45S_50ntFlanks</td><td>11670</td><td>11692</td><td>23</td><td>CCCAGTGCTCTGAATGTCAAAGT</td><td>468.1</td><td>3.4</td><td>0.6</td><td>6.1</td><td>9.9E-10</td><td>3.0E-09</td><td>BT20/MB468</td><td>MITO</td></tr>
<tr><td>>hg38_21_+_8433172_8446622_RNA45SN1_45S_50ntFlanks@9739.9767.29|AAACGATCTCAACCTATTCTCAAACTTTA</td><td>>hg38_21_+_8433172_8446622_RNA45SN1_45S_50ntFlanks</td><td>9739</td><td>9767</td><td>29</td><td>AAACGATCTCAACCTATTCTCAAACTTTA</td><td>557.9</td><td>-8.0</td><td>1.3</td><td>-6.1</td><td>1.3E-09</td><td>3.0E-09</td><td>MB231/MB468</td><td>MP</td></tr>
<tr><td>>MI0000289|hsa-mir-181a-1&WithFlank&1|-|198859038|198859159@30.54.25(+1U);MI0000269|hsa-mir-181a-2&WithFlank&9|+|124692436|124692557@45.69.25(+1U)&[MIMAT0000256&hsa-miR-181a-5p&offsets|0|+2(+1U);m-32&1|-|198859108|198859130&offsets|0|+2(+1U)];[MIMAT0000256_1&hsa-miR-181a-5p&offsets|0|+2(+1U);m-32&9|+|124692480|124692502&offsets|0|+2(+1U)]|AACATTCAACGCTGTCGGTGAGTTTT</td><td>>MI0000289|hsa-mir-181a-1&WithFlank&1|-|198859038|198859159</td><td>30</td><td>54</td><td>25(+1U)</td><td>AACATTCAACGCTGTCGGTGAGTTTT</td><td>1137.0</td><td>10.2</td><td>1.7</td><td>6.0</td><td>1.5E-09</td><td>3.0E-09</td><td>MB231/MB468</td><td>MITO</td></tr>
<tr><td>>MI0000065|hsa-let-7d&WithFlank&9|+|94178828|94178926@14.35.22&[MIMAT0000065&hsa-let-7d-5p&offsets|0|0;m-90&9|+|94178841|94178862&offsets|0|0]|AGAGGTAGTAGGTTGCATAGTT</td><td>>MI0000065|hsa-let-7d&WithFlank&9|+|94178828|94178926</td><td>14</td><td>35</td><td>22</td><td>AGAGGTAGTAGGTTGCATAGTT</td><td>5150.5</td><td>5.5</td><td>0.9</td><td>6.0</td><td>1.5E-09</td><td>3.0E-09</td><td>MB231/MB468</td><td>MITO</td></tr>
<tr><td>>hg38_21_+_8433172_8446622_RNA45SN1_45S_50ntFlanks@12258.12284.27|GCTTGATCTTGATTTTCAGTACGAATA</td><td>>hg38_21_+_8433172_8446622_RNA45SN1_45S_50ntFlanks</td><td>12258</td><td>12284</td><td>27</td><td>GCTTGATCTTGATTTTCAGTACGAATA</td><td>1166.6</td><td>-7.4</td><td>1.2</td><td>-6.0</td><td>1.5E-09</td><td>3.0E-09</td><td>MB231/MB468</td><td>MITO</td></tr>
<tr><td>>trna112_GlnCTG_1_-_147737382_147737453@30.50.21,trna131_GlnCTG_6_-_27759135_27759206@30.50.21,trna146_GlnCTG_6_-_27515531_27515602@30.50.21,trna15_GlnCTG_1_+_145963304_145963375@30.50.21,trna19_GlnCTG_1_+_147505038_147505109@30.50.21,trna1_GlnCTG_6_+_18836402_18836473@30.50.21,trna28_GlnCTG_1_+_149186125_149186196@30.50.21,trna3_GlnCTG_17_+_8023070_8023141@30.50.21,trna49_GlnCTG_6_+_27487308_27487379@30.50.21,trna7_GlnCTG_15_-_66161400_66161471@30.50.21,trna99_GlnCTG_6_-_28909378_28909449@30.50.21|GACTCTGAATCCAGCGATCCG</td><td>>trna112_GlnCTG_1_-_147737382_147737453</td><td>30</td><td>50</td><td>21</td><td>GACTCTGAATCCAGCGATCCG</td><td>276.7</td><td>-6.5</td><td>1.1</td><td>-6.1</td><td>1.3E-09</td><td>3.0E-09</td><td>MB231/MB468</td><td>CYTO</td></tr>
<tr><td>>am_trna119_LysCTT_1_-_145395522_145395594@7.35.29,trna11_LysCTT_5_-_180648979_180649051@7.35.29,trna128_LysCTT_1_-_55423542_55423614@7.35.29,trna13_LysCTT_6_+_26556774_26556846@7.35.29,trna6_LysCTT_19_-_52425396_52425468@7.35.29,trna7_LysCTT_16_+_3225692_3225764@7.35.29,trna9_LysCTT_5_+_180634755_180634827@7.35.29|CTAGCTCAGTCGGTAGAGCATGAGACTCT</td><td>>am_trna119_LysCTT_1_-_145395522_145395594</td><td>7</td><td>35</td><td>29</td><td>CTAGCTCAGTCGGTAGAGCATGAGACTCT</td><td>968.6</td><td>-5.1</td><td>0.8</td><td>-6.5</td><td>1.1E-10</td><td>3.0E-09</td><td>BT20/MB231</td><td>MITO</td></tr>
<tr><td>>MI0000465|hsa-mir-191&WithFlank&3|-|49020612|49020715@22.44.23(+1C)&[MIMAT0000440&hsa-miR-191-5p&offsets|0|0(+1C);m-48&3|-|49020672|49020694&offsets|0|0(+1C)]|CAACGGAATCCCAAAAGCAGCTGC</td><td>>MI0000465|hsa-mir-191&WithFlank&3|-|49020612|49020715</td><td>22</td><td>44</td><td>23(+1C)</td><td>CAACGGAATCCCAAAAGCAGCTGC</td><td>410.9</td><td>6.5</td><td>1.1</td><td>6.1</td><td>1.3E-09</td><td>3.0E-09</td><td>MB231/MB468</td><td>CYTO</td></tr>
<tr><td>>hg38_21_+_8433172_8446622_RNA45SN1_45S_50ntFlanks@11208.11239.32|GCGGCGACTCTGGACGCGAGCCGGGCCCTTCC</td><td>>hg38_21_+_8433172_8446622_RNA45SN1_45S_50ntFlanks</td><td>11208</td><td>11239</td><td>32</td><td>GCGGCGACTCTGGACGCGAGCCGGGCCCTTCC</td><td>70.0</td><td>-10.0</td><td>1.6</td><td>-6.1</td><td>1.3E-09</td><td>3.0E-09</td><td>MB231/MB468</td><td>MP</td></tr>
<tr><td>>hg38_21_+_8433172_8446622_RNA45SN1_45S_50ntFlanks@10351.10389.39|CGAGAACTTTGAAGGCCGAAGTGGAGAAGGGTTCCATGT</td><td>>hg38_21_+_8433172_8446622_RNA45SN1_45S_50ntFlanks</td><td>10351</td><td>10389</td><td>39</td><td>CGAGAACTTTGAAGGCCGAAGTGGAGAAGGGTTCCATGT</td><td>196.3</td><td>-9.7</td><td>1.6</td><td>-6.1</td><td>1.0E-09</td><td>3.0E-09</td><td>BT20/MB468</td><td>MITO</td></tr>
<tr><td>>hg38_21_+_8433172_8446622_RNA45SN1_45S_50ntFlanks@12437.12455.19|CATTGTGAAGCAGAATTCA</td><td>>hg38_21_+_8433172_8446622_RNA45SN1_45S_50ntFlanks</td><td>12437</td><td>12455</td><td>19</td><td>CATTGTGAAGCAGAATTCA</td><td>6302.5</td><td>-7.5</td><td>1.2</td><td>-6.1</td><td>1.0E-09</td><td>3.0E-09</td><td>BT20/MB468</td><td>MITO</td></tr>
<tr><td>>am_trnaMT_PheGAA_MT_+_577_647@44.62.19|TAGACGGGCTCACATCACC</td><td>>am_trnaMT_PheGAA_MT_+_577_647</td><td>44</td><td>62</td><td>19</td><td>TAGACGGGCTCACATCACC</td><td>2217.5</td><td>6.2</td><td>1.0</td><td>6.1</td><td>1.0E-09</td><td>3.0E-09</td><td>BT20/MB468</td><td>MITO</td></tr>
<tr><td>>hg38_21_+_8433172_8446622_RNA45SN1_45S_50ntFlanks@11104.11131.28|AAGGGTCGGGGCGGCAGGGGCCGGCGGC</td><td>>hg38_21_+_8433172_8446622_RNA45SN1_45S_50ntFlanks</td><td>11104</td><td>11131</td><td>28</td><td>AAGGGTCGGGGCGGCAGGGGCCGGCGGC</td><td>496.0</td><td>-8.6</td><td>1.4</td><td>-6.0</td><td>1.5E-09</td><td>3.0E-09</td><td>BT20/MB468</td><td>NUC</td></tr>
<tr><td>>hg38_21_+_8433172_8446622_RNA45SN1_45S_50ntFlanks@10116.10137.22|TGCGTCGGAGGGCGGCGGCGGC</td><td>>hg38_21_+_8433172_8446622_RNA45SN1_45S_50ntFlanks</td><td>10116</td><td>10137</td><td>22</td><td>TGCGTCGGAGGGCGGCGGCGGC</td><td>141.6</td><td>-11.0</td><td>1.8</td><td>-6.1</td><td>1.3E-09</td><td>3.0E-09</td><td>MB231/MB468</td><td>MP</td></tr>
<tr><td>>hg38_21_+_8433172_8446622_RNA45SN1_45S_50ntFlanks@10797.10823.27|GCATGTTGGAACAATGTAGGTAAGGGA</td><td>>hg38_21_+_8433172_8446622_RNA45SN1_45S_50ntFlanks</td><td>10797</td><td>10823</td><td>27</td><td>GCATGTTGGAACAATGTAGGTAAGGGA</td><td>778.5</td><td>-8.3</td><td>1.4</td><td>-6.1</td><td>1.3E-09</td><td>3.0E-09</td><td>MB231/MB468</td><td>MP</td></tr>
<tr><td>>MI0000465|hsa-mir-191&WithFlank&3|-|49020612|49020715@22.44.23&[MIMAT0000440&hsa-miR-191-5p&offsets|0|0;m-48&3|-|49020672|49020694&offsets|0|0]|CAACGGAATCCCAAAAGCAGCTG</td><td>>MI0000465|hsa-mir-191&WithFlank&3|-|49020612|49020715</td><td>22</td><td>44</td><td>23</td><td>CAACGGAATCCCAAAAGCAGCTG</td><td>10182.0</td><td>5.7</td><td>0.9</td><td>6.1</td><td>1.3E-09</td><td>3.0E-09</td><td>MB231/MB468</td><td>CYTO</td></tr>
<tr><td>>hg38_21_+_8433172_8446622_RNA45SN1_45S_50ntFlanks@5527.5549.23|AAAGTCGTAACAAGGTTTCCGTA</td><td>>hg38_21_+_8433172_8446622_RNA45SN1_45S_50ntFlanks</td><td>5527</td><td>5549</td><td>23</td><td>AAAGTCGTAACAAGGTTTCCGTA</td><td>242.4</td><td>-5.2</td><td>0.9</td><td>-6.1</td><td>1.3E-09</td><td>3.0E-09</td><td>MB231/MB468</td><td>CYTO</td></tr>
<tr><td>>hg38_21_+_8433172_8446622_RNA45SN1_45S_50ntFlanks@12159.12178.20|TTTGACTGGGGCGGTACACC</td><td>>hg38_21_+_8433172_8446622_RNA45SN1_45S_50ntFlanks</td><td>12159</td><td>12178</td><td>20</td><td>TTTGACTGGGGCGGTACACC</td><td>1065.1</td><td>-9.4</td><td>1.6</td><td>-6.0</td><td>1.5E-09</td><td>3.0E-09</td><td>BT20/MB468</td><td>NUC</td></tr>
<tr><td>>hg38_21_+_8433172_8446622_RNA45SN1_45S_50ntFlanks@6728.6758.31|GGACACATTGATCATCGACACTTCGAACGCA</td><td>>hg38_21_+_8433172_8446622_RNA45SN1_45S_50ntFlanks</td><td>6728</td><td>6758</td><td>31</td><td>GGACACATTGATCATCGACACTTCGAACGCA</td><td>152.0</td><td>3.5</td><td>0.6</td><td>6.1</td><td>1.3E-09</td><td>3.0E-09</td><td>MB231/MB468</td><td>CYTO</td></tr>
<tr><td>>MI0003205|hsa-mir-532&WithFlank&X|+|50003142|50003244@26.46.21(+1C)&[MIMAT0002888&hsa-miR-532-5p&offsets|0|-1(+1C);m-65&X|+|50003167|50003188&offsets|0|-1(+1C)]|CATGCCTTGAGTGTAGGACCGC</td><td>>MI0003205|hsa-mir-532&WithFlank&X|+|50003142|50003244</td><td>26</td><td>46</td><td>21(+1C)</td><td>CATGCCTTGAGTGTAGGACCGC</td><td>292.3</td><td>3.6</td><td>0.6</td><td>6.1</td><td>1.3E-09</td><td>3.0E-09</td><td>MB231/MB468</td><td>NUC</td></tr>
<tr><td>>hg38_21_+_8433172_8446622_RNA45SN1_45S_50ntFlanks@7976.8010.35|GCGACCTCAGATCAGACGTGGCGACCCGCTGAATT</td><td>>hg38_21_+_8433172_8446622_RNA45SN1_45S_50ntFlanks</td><td>7976</td><td>8010</td><td>35</td><td>GCGACCTCAGATCAGACGTGGCGACCCGCTGAATT</td><td>644.0</td><td>5.1</td><td>0.8</td><td>6.1</td><td>1.4E-09</td><td>3.0E-09</td><td>MB231/MB468</td><td>NUC</td></tr>
<tr><td>>hg38_21_+_8433172_8446622_RNA45SN1_45S_50ntFlanks@7978.8010.33|GACCTCAGATCAGACGTGGCGACCCGCTGAATT</td><td>>hg38_21_+_8433172_8446622_RNA45SN1_45S_50ntFlanks</td><td>7978</td><td>8010</td><td>33</td><td>GACCTCAGATCAGACGTGGCGACCCGCTGAATT</td><td>224.9</td><td>4.1</td><td>0.7</td><td>6.1</td><td>1.4E-09</td><td>3.0E-09</td><td>MB231/MB468</td><td>NUC</td></tr>
<tr><td>>hg38_21_+_8433172_8446622_RNA45SN1_45S_50ntFlanks@8116.8141.26|GCGGGACATGTGGCGTACGGAAGACC</td><td>>hg38_21_+_8433172_8446622_RNA45SN1_45S_50ntFlanks</td><td>8116</td><td>8141</td><td>26</td><td>GCGGGACATGTGGCGTACGGAAGACC</td><td>1103.5</td><td>13.4</td><td>2.0</td><td>6.7</td><td>2.8E-11</td><td>3.0E-09</td><td>BT20/MB231</td><td>MP</td></tr>
<tr><td>>MI0000272|hsa-mir-182&WithFlank&7|-|129770377|129770498@29.53.25(+1C)&[MIMAT0000259&hsa-miR-182-5p&offsets|0|+1(+1C);m-22&7|-|129770449|129770470&offsets|0|+3(+1C)]|TTTGGCAATGGTAGAACTCACACTGC</td><td>>MI0000272|hsa-mir-182&WithFlank&7|-|129770377|129770498</td><td>29</td><td>53</td><td>25(+1C)</td><td>TTTGGCAATGGTAGAACTCACACTGC</td><td>76.9</td><td>3.2</td><td>0.5</td><td>6.0</td><td>1.5E-09</td><td>3.0E-09</td><td>BT20/MB468</td><td>NUC</td></tr>
<tr><td>>hg38_21_+_8433172_8446622_RNA45SN1_45S_50ntFlanks@10140.10158.19|CGGCGGCGGGGGTGTGGGG</td><td>>hg38_21_+_8433172_8446622_RNA45SN1_45S_50ntFlanks</td><td>10140</td><td>10158</td><td>19</td><td>CGGCGGCGGGGGTGTGGGG</td><td>73.3</td><td>-10.1</td><td>1.7</td><td>-6.1</td><td>1.3E-09</td><td>3.0E-09</td><td>MB231/MB468</td><td>MP</td></tr>
<tr><td>>MI0003591|hsa-mir-584&WithFlank&5|-|149062307|149062415@22.41.20(+1U)&[MIMAT0003249&hsa-miR-584-5p&offsets|0|-2(+1U);m-167&5|-|149062374|149062394&offsets|0|-1(+1U)]|TTATGGTTTGCCTGGGACTGT</td><td>>MI0003591|hsa-mir-584&WithFlank&5|-|149062307|149062415</td><td>22</td><td>41</td><td>20(+1U)</td><td>TTATGGTTTGCCTGGGACTGT</td><td>1587.4</td><td>6.8</td><td>1.1</td><td>6.4</td><td>1.4E-10</td><td>3.0E-09</td><td>BT20/MB468</td><td>MP</td></tr>
<tr><td>>hg38_21_+_8433172_8446622_RNA45SN1_45S_50ntFlanks@4153.4184.32|ACGGCTACCACATCCAAGGAAGGCAGCAGGCG</td><td>>hg38_21_+_8433172_8446622_RNA45SN1_45S_50ntFlanks</td><td>4153</td><td>4184</td><td>32</td><td>ACGGCTACCACATCCAAGGAAGGCAGCAGGCG</td><td>78.1</td><td>-10.2</td><td>1.7</td><td>-6.1</td><td>1.3E-09</td><td>3.0E-09</td><td>MB231/MB468</td><td>MP</td></tr>
</table>

<table>
<tr><td>>hg38_21_+_8433172_8446622_RNA45SN1_45S_50ntFlanks@6670.6699.30|ACTCGGCTCGTGCGTCGATGAAGAACGCAG</td><td>>hg38_21_+_8433172_8446622_RNA45SN1_45S_50ntFlanks</td><td>6670</td><td>6699</td><td>30</td><td>ACTCGGCTCGTGCGTCGATGAAGAACGCAG</td><td>1053.1</td><td>-7.0</td><td>1.2</td><td>-6.1</td><td>1.3E-09</td><td>3.0E-09</td><td>MB231/MB468</td><td>MP</td></tr>
<tr><td>>hg38_1_-_228634819_228635039_RNA5S12_5S_50ntFlanks@150.171.22|AATACCGGGTGCTGTAGGCTTT</td><td>>hg38_1_-_228634819_228635039_RNA5S12_5S_50ntFlanks</td><td>150</td><td>171</td><td>22</td><td>AATACCGGGTGCTGTAGGCTTT</td><td>4200.2</td><td>4.2</td><td>0.7</td><td>6.1</td><td>1.2E-09</td><td>3.0E-09</td><td>BT20/MB468</td><td>CYTO</td></tr>
<tr><td>>hg38_21_+_8433172_8446622_RNA45SN1_45S_50ntFlanks@12396.12417.22|GCGACGTCGCTTTTTGATCCTT</td><td>>hg38_21_+_8433172_8446622_RNA45SN1_45S_50ntFlanks</td><td>12396</td><td>12417</td><td>22</td><td>GCGACGTCGCTTTTTGATCCTT</td><td>131.9</td><td>-10.9</td><td>1.8</td><td>-6.1</td><td>1.3E-09</td><td>3.0E-09</td><td>MB231/MB468</td><td>MP</td></tr>
<tr><td>>hg38_21_+_8433172_8446622_RNA45SN1_45S_50ntFlanks@10402.10431.30|AACATGGGTCAGTCGGTCCTGAGAGATGGG</td><td>>hg38_21_+_8433172_8446622_RNA45SN1_45S_50ntFlanks</td><td>10402</td><td>10431</td><td>30</td><td>AACATGGGTCAGTCGGTCCTGAGAGATGGG</td><td>934.4</td><td>-8.2</td><td>1.4</td><td>-6.0</td><td>1.5E-09</td><td>3.0E-09</td><td>MB231/MB468</td><td>MITO</td></tr>
<tr><td>>hg38_21_+_8433172_8446622_RNA45SN1_45S_50ntFlanks@12669.12703.35|CTAAGTCAGAATCCCGCCCAGGCGGAACGATACGG</td><td>>hg38_21_+_8433172_8446622_RNA45SN1_45S_50ntFlanks</td><td>12669</td><td>12703</td><td>35</td><td>CTAAGTCAGAATCCCGCCCAGGCGGAACGATACGG</td><td>85.0</td><td>-10.3</td><td>1.7</td><td>-6.1</td><td>1.3E-09</td><td>3.0E-09</td><td>MB231/MB468</td><td>MP</td></tr>
<tr><td>>hg38_21_+_8433172_8446622_RNA45SN1_45S_50ntFlanks@6647.6683.37|CGTACGACTCTTAGCGGTGGATCACTCGGCTCGTGCG</td><td>>hg38_21_+_8433172_8446622_RNA45SN1_45S_50ntFlanks</td><td>6647</td><td>6683</td><td>37</td><td>CGTACGACTCTTAGCGGTGGATCACTCGGCTCGTGCG</td><td>59.0</td><td>4.6</td><td>0.8</td><td>6.0</td><td>1.6E-09</td><td>3.0E-09</td><td>BT20/MB468</td><td>NUC</td></tr>
<tr><td>>hg38_21_+_8433172_8446622_RNA45SN1_45S_50ntFlanks@8170.8194.25|CAAGTCCTTCTGATCGAGGCCCAGC</td><td>>hg38_21_+_8433172_8446622_RNA45SN1_45S_50ntFlanks</td><td>8170</td><td>8194</td><td>25</td><td>CAAGTCCTTCTGATCGAGGCCCAGC</td><td>161.0</td><td>-10.4</td><td>1.6</td><td>-6.4</td><td>1.4E-10</td><td>3.0E-09</td><td>BT20/MB468</td><td>MP</td></tr>
<tr><td>>hg38_21_+_8433172_8446622_RNA45SN1_45S_50ntFlanks@5484.5518.35|GAGCGCTGAGAAGACGGTCGAACTTGACTATCTAG</td><td>>hg38_21_+_8433172_8446622_RNA45SN1_45S_50ntFlanks</td><td>5484</td><td>5518</td><td>35</td><td>GAGCGCTGAGAAGACGGTCGAACTTGACTATCTAG</td><td>280.4</td><td>-7.9</td><td>1.3</td><td>-6.1</td><td>1.4E-09</td><td>3.1E-09</td><td>MB231/MB468</td><td>NUC</td></tr>
<tr><td>>hg38_21_+_8433172_8446622_RNA45SN1_45S_50ntFlanks@12025.12055.31|CTACTCTGATCGTTTTTTCACTGACCCGGTG</td><td>>hg38_21_+_8433172_8446622_RNA45SN1_45S_50ntFlanks</td><td>12025</td><td>12055</td><td>31</td><td>CTACTCTGATCGTTTTTTCACTGACCCGGTG</td><td>127.8</td><td>10.7</td><td>1.7</td><td>6.4</td><td>1.4E-10</td><td>3.1E-09</td><td>BT20/MB468</td><td>MP</td></tr>
<tr><td>>hg38_21_+_8433172_8446622_RNA45SN1_45S_50ntFlanks@10781.10807.27|AGGTGAACAGCCTCTGGCATGTTGGAA</td><td>>hg38_21_+_8433172_8446622_RNA45SN1_45S_50ntFlanks</td><td>10781</td><td>10807</td><td>27</td><td>AGGTGAACAGCCTCTGGCATGTTGGAA</td><td>207.0</td><td>-10.8</td><td>1.7</td><td>-6.4</td><td>1.4E-10</td><td>3.1E-09</td><td>BT20/MB468</td><td>MP</td></tr>
<tr><td>>hg38_21_+_8433172_8446622_RNA45SN1_45S_50ntFlanks@10826.10848.23|TCGGCAAGCCGGATCCGTAACTT</td><td>>hg38_21_+_8433172_8446622_RNA45SN1_45S_50ntFlanks</td><td>10826</td><td>10848</td><td>23</td><td>TCGGCAAGCCGGATCCGTAACTT</td><td>232.0</td><td>-10.2</td><td>1.7</td><td>-6.1</td><td>1.2E-09</td><td>3.1E-09</td><td>BT20/MB468</td><td>CYTO</td></tr>
<tr><td>>MI0000255|hsa-mir-30d&WithFlank&8|-|134804870|134804951@12.35.24(+3U)&[MIMAT0000245&hsa-miR-30d-5p&offsets|0|+2(+3U);m-21&8|-|134804919|134804940&offsets|0|+2(+3U)]|TGTAAACATCCCCGACTGGAAGCTTTT</td><td>>MI0000255|hsa-mir-30d&WithFlank&8|-|134804870|134804951</td><td>12</td><td>35</td><td>24(+3U)</td><td>TGTAAACATCCCCGACTGGAAGCTTTT</td><td>264.9</td><td>8.3</td><td>1.4</td><td>6.1</td><td>1.3E-09</td><td>3.1E-09</td><td>MB231/MB468</td><td>CYTO</td></tr>
<tr><td>>hg38_21_+_8433172_8446622_RNA45SN1_45S_50ntFlanks@10794.10825.32|CTGGCATGTTGGAACAATGTAGGTAAGGGAAG</td><td>>hg38_21_+_8433172_8446622_RNA45SN1_45S_50ntFlanks</td><td>10794</td><td>10825</td><td>32</td><td>CTGGCATGTTGGAACAATGTAGGTAAGGGAAG</td><td>386.0</td><td>-3.5</td><td>0.6</td><td>-6.1</td><td>1.0E-09</td><td>3.1E-09</td><td>BT20/MB468</td><td>MITO</td></tr>
<tr><td>>hg38_21_+_8433172_8446622_RNA45SN1_45S_50ntFlanks@11452.11477.26|CTCCTCGGCGCGCGGCGGCGGCGG</td><td>>hg38_21_+_8433172_8446622_RNA45SN1_45S_50ntFlanks</td><td>11452</td><td>11477</td><td>26</td><td>CTCCTCGGCGCGCGGCGGCGGCGG</td><td>140.5</td><td>-10.2</td><td>1.6</td><td>-6.4</td><td>1.4E-10</td><td>3.1E-09</td><td>BT20/MB468</td><td>MP</td></tr>
<tr><td>>hg38_21_+_8433172_8446622_RNA45SN1_45S_50ntFlanks@6773.6807.35|TTCCTCCCGGGGCTACGCCTGTCTGAGCGTCGCTT</td><td>>hg38_21_+_8433172_8446622_RNA45SN1_45S_50ntFlanks</td><td>6773</td><td>6807</td><td>35</td><td>TTCCTCCCGGGGCTACGCCTGTCTGAGCGTCGCTT</td><td>777.2</td><td>-3.9</td><td>0.6</td><td>-6.0</td><td>1.6E-09</td><td>3.1E-09</td><td>BT20/MB468</td><td>NUC</td></tr>
<tr><td>>hg38_21_+_8433172_8446622_RNA45SN1_45S_50ntFlanks@9568.9594.27|CGTGCAAATCGGTCGTCCGACCTGGGT</td><td>>hg38_21_+_8433172_8446622_RNA45SN1_45S_50ntFlanks</td><td>9568</td><td>9594</td><td>27</td><td>CGTGCAAATCGGTCGTCCGACCTGGGT</td><td>541.3</td><td>-5.6</td><td>0.9</td><td>-6.1</td><td>1.4E-09</td><td>3.1E-09</td><td>MB231/MB468</td><td>CYTO</td></tr>
<tr><td>>trna111_HisGTG_1_-_147774845_147774916@-1A.23.24,trna118_HisGTG_1_-_145396881_145396952@-1A.23.24,trna16_HisGTG_1_+_146544773_146544844@-1A.23.24,trna1_HisGTG_15_+_45493349_45493420@-1A.23.24,trna21_HisGTG_1_+_147753471_147753542@-1A.23.24,trna33_HisGTG_6_+_27125906_27125977@-1A.23.24,trna7_HisGTG_9_-_14433938_14434009@-1A.23.24,trna8_HisGTG_15_-_45492611_45492682@-1A.23.24,trna9_HisGTG_15_-_45490804_45490875@-1A.23.24|AGCCGTGATCGTATAGTGGTTAGT</td><td>>trna111_HisGTG_1_-_147774845_147774916</td><td>-1A</td><td>23</td><td>24</td><td>AGCCGTGATCGTATAGTGGTTAGT</td><td>117.7</td><td>10.1</td><td>1.7</td><td>6.1</td><td>1.3E-09</td><td>3.1E-09</td><td>MB231/MB468</td><td>MP</td></tr>
<tr><td>>hg38_21_+_8433172_8446622_RNA45SN1_45S_50ntFlanks@9597.9615.19|AGGGGCGAAAGACTAATCG</td><td>>hg38_21_+_8433172_8446622_RNA45SN1_45S_50ntFlanks</td><td>9597</td><td>9615</td><td>19</td><td>AGGGGCGAAAGACTAATCG</td><td>273.8</td><td>-7.8</td><td>1.3</td><td>-6.1</td><td>1.3E-09</td><td>3.1E-09</td><td>MB231/MB468</td><td>MP</td></tr>
<tr><td>>hg38_21_+_8433172_8446622_RNA45SN1_45S_50ntFlanks@10872.10892.21|GCTGGGTCGGTCGGGCTGGGG</td><td>>hg38_21_+_8433172_8446622_RNA45SN1_45S_50ntFlanks</td><td>10872</td><td>10892</td><td>21</td><td>GCTGGGTCGGTCGGGCTGGGG</td><td>339.9</td><td>-8.0</td><td>1.3</td><td>-6.1</td><td>1.3E-09</td><td>3.1E-09</td><td>MB231/MB468</td><td>MP</td></tr>
<tr><td>>hg38_21_+_8433172_8446622_RNA45SN1_45S_50ntFlanks@10389.10411.23|TGAACAGCAGTTGAACATGGGTC</td><td>>hg38_21_+_8433172_8446622_RNA45SN1_45S_50ntFlanks</td><td>10389</td><td>10411</td><td>23</td><td>TGAACAGCAGTTGAACATGGGTC</td><td>474.9</td><td>-8.5</td><td>1.4</td><td>-6.0</td><td>1.6E-09</td><td>3.1E-09</td><td>BT20/MB468</td><td>NUC</td></tr>
<tr><td>>hg38_21_+_8433172_8446622_RNA45SN1_45S_50ntFlanks@10361.10390.30|GAAGGCCGAAGTGGAGAAGGGTTCCATGTG</td><td>>hg38_21_+_8433172_8446622_RNA45SN1_45S_50ntFlanks</td><td>10361</td><td>10390</td><td>30</td><td>GAAGGCCGAAGTGGAGAAGGGTTCCATGTG</td><td>626.7</td><td>-8.4</td><td>1.4</td><td>-6.1</td><td>1.3E-09</td><td>3.1E-09</td><td>MB231/MB468</td><td>MP</td></tr>
<tr><td>>MI0000767|hsa-mir-365a&WithFlank&16|+|14309279|14309377@62.81.20;MI0000769|hsa-mir-365b&WithFlank&17|+|31575405|31575527@74.93.20&[MIMAT0000710&hsa-miR-365a-3p&offsets|0|-2;m-176&16|+|14309340|14309361&offsets|0|-2];[MIMAT0022834&hsa-miR-365b-3p&offsets|0|-2;m-175&17|+|31575478|31575499&offsets|0|-2]|TAATGCCCCTAAAAATCCTT</td><td>>MI0000767|hsa-mir-365a&WithFlank&16|+|14309279|14309377</td><td>62</td><td>81</td><td>20</td><td>TAATGCCCCTAAAAATCCTT</td><td>99.1</td><td>4.7</td><td>0.8</td><td>6.0</td><td>1.6E-09</td><td>3.1E-09</td><td>BT20/MB468</td><td>NUC</td></tr>
<tr><td>>MI0000809|hsa-mir-151a&WithFlank&8|-|140732558|140732659@51.72.22&[MIMAT0000757&hsa-miR-151a-3p&offsets|-2|-1;m-30&8|-|140732587|140732607&offsets|-2|-1]|TACTAGACTGAAGCTCCTTGAG</td><td>>MI0000809|hsa-mir-151a&WithFlank&8|-|140732558|140732659</td><td>51</td><td>72</td><td>22</td><td>TACTAGACTGAAGCTCCTTGAG</td><td>278.5</td><td>5.6</td><td>0.9</td><td>6.1</td><td>1.4E-09</td><td>3.1E-09</td><td>MB231/MB468</td><td>CYTO</td></tr>
<tr><td>>hg38_21_+_8433172_8446622_RNA45SN1_45S_50ntFlanks@10664.10710.47|CTTGGAAAGCGTCGCGGTTCCGGCGGCGTCCGGTGAGCTCTCGCTGG</td><td>>hg38_21_+_8433172_8446622_RNA45SN1_45S_50ntFlanks</td><td>10664</td><td>10710</td><td>47</td><td>CTTGGAAAGCGTCGCGGTTCCGGCGGCGTCCGGTGAGCTCTCGCTGG</td><td>420.0</td><td>-7.0</td><td>1.2</td><td>-6.1</td><td>1.4E-09</td><td>3.1E-09</td><td>MB231/MB468</td><td>CYTO</td></tr>
<tr><td>>MI0000087|hsa-mir-29a&WithFlank&7|-|130876741|130876816@48.66.19&[MIMAT0000086&hsa-miR-29a-3p&offsets|0|-3;m-26&7|-|130876749|130876769&offsets|0|-2]|TAGCACCATCTGAAATCGG</td><td>>MI0000087|hsa-mir-29a&WithFlank&7|-|130876741|130876816</td><td>48</td><td>66</td><td>19</td><td>TAGCACCATCTGAAATCGG</td><td>140.7</td><td>10.4</td><td>1.7</td><td>6.0</td><td>1.5E-09</td><td>3.1E-09</td><td>MB231/MB468</td><td>MITO</td></tr>
<tr><td>>hg38_21_+_8433172_8446622_RNA45SN1_45S_50ntFlanks@9575.9592.18|ATCGGTCGTCCGACCTGG</td><td>>hg38_21_+_8433172_8446622_RNA45SN1_45S_50ntFlanks</td><td>9575</td><td>9592</td><td>18</td><td>ATCGGTCGTCCGACCTGG</td><td>344.7</td><td>-11.5</td><td>1.8</td><td>-6.4</td><td>1.4E-10</td><td>3.1E-09</td><td>BT20/MB468</td><td>MP</td></tr>
<tr><td>>hg38_21_+_8433172_8446622_RNA45SN1_45S_50ntFlanks@12445.12475.31|AGCAGAATTCACCAAGCGTTGGATTGTTCAC</td><td>>hg38_21_+_8433172_8446622_RNA45SN1_45S_50ntFlanks</td><td>12445</td><td>12475</td><td>31</td><td>AGCAGAATTCACCAAGCGTTGGATTGTTCAC</td><td>100.9</td><td>-10.5</td><td>1.7</td><td>-6.1</td><td>1.3E-09</td><td>3.1E-09</td><td>MB231/MB468</td><td>MP</td></tr>
<tr><td>>hg38_21_+_8433172_8446622_RNA45SN1_45S_50ntFlanks@9939.9980.42|GCAGGACGGTGGCCATGGAAGTCGGAATCCGCTAAGGAGTGT</td><td>>hg38_21_+_8433172_8446622_RNA45SN1_45S_50ntFlanks</td><td>9939</td><td>9980</td><td>42</td><td>GCAGGACGGTGGCCATGGAAGTCGGAATCCGCTAAGGAGTGT</td><td>274.3</td><td>11.8</td><td>1.8</td><td>6.4</td><td>1.4E-10</td><td>3.1E-09</td><td>BT20/MB468</td><td>MP</td></tr>
<tr><td>>hg38_21_+_8433172_8446622_RNA45SN1_45S_50ntFlanks@4099.4119.21|GTGACCACGGGTGACGGGGAA</td><td>>hg38_21_+_8433172_8446622_RNA45SN1_45S_50ntFlanks</td><td>4099</td><td>4119</td><td>21</td><td>GTGACCACGGGTGACGGGGAA</td><td>64.3</td><td>-9.9</td><td>1.6</td><td>-6.1</td><td>1.3E-09</td><td>3.1E-09</td><td>MB231/MB468</td><td>MP</td></tr>
<tr><td>>hg38_21_+_8433172_8446622_RNA45SN1_45S_50ntFlanks@10384.10419.36|CCATGTGAACAGCAGTTGAACATGGGTCAGTCGGTC</td><td>>hg38_21_+_8433172_8446622_RNA45SN1_45S_50ntFlanks</td><td>10384</td><td>10419</td><td>36</td><td>CCATGTGAACAGCAGTTGAACATGGGTCAGTCGGTC</td><td>99.0</td><td>-8.9</td><td>1.5</td><td>-6.1</td><td>1.2E-09</td><td>3.1E-09</td><td>BT20/MB468</td><td>CYTO</td></tr>
<tr><td>>trna119_LysCTT_1_-_145395522_145395594@4.34.31,trna11_LysCTT_5_-_180648979_180649051@4.34.31,trna13_LysCTT_6_+_26556774_26556846@4.34.31,trna7_LysCTT_16_+_3225692_3225764@4.34.31,trna9_LysCTT_5_+_180634755_180634827@4.34.31|CGGCTAGCTCAGTCGGTAGAGCATGAGACTC</td><td>>trna119_LysCTT_1_-_145395522_145395594</td><td>4</td><td>34</td><td>31</td><td>CGGCTAGCTCAGTCGGTAGAGCATGAGACTC</td><td>681.6</td><td>6.9</td><td>1.1</td><td>6.0</td><td>1.5E-09</td><td>3.1E-09</td><td>MB231/MB468</td><td>MITO</td></tr>
<tr><td>>hg38_21_+_8433172_8446622_RNA45SN1_45S_50ntFlanks@11828.11856.29|GCGAAACCACAGCCAAGGGAACGGGCTTG</td><td>>hg38_21_+_8433172_8446622_RNA45SN1_45S_50ntFlanks</td><td>11828</td><td>11856</td><td>29</td><td>GCGAAACCACAGCCAAGGGAACGGGCTTG</td><td>89.7</td><td>-10.3</td><td>1.7</td><td>-6.1</td><td>1.3E-09</td><td>3.1E-09</td><td>MB231/MB468</td><td>MP</td></tr>
<tr><td>>hg38_21_+_8433172_8446622_RNA45SN1_45S_50ntFlanks@10735.10752.18|TGTAAATCTCGCGCCGGG</td><td>>hg38_21_+_8433172_8446622_RNA45SN1_45S_50ntFlanks</td><td>10735</td><td>10752</td><td>18</td><td>TGTAAATCTCGCGCCGGG</td><td>137.1</td><td>-10.2</td><td>1.6</td><td>-6.4</td><td>1.4E-10</td><td>3.1E-09</td><td>BT20/MB468</td><td>MP</td></tr>
<tr><td>>hg38_21_+_8433172_8446622_RNA45SN1_45S_50ntFlanks@8348.8373.26|GGAAAGTTGAAAAGAACTTTGAAGAG</td><td>>hg38_21_+_8433172_8446622_RNA45SN1_45S_50ntFlanks</td><td>8348</td><td>8373</td><td>26</td><td>GGAAAGTTGAAAAGAACTTTGAAGAG</td><td>1262.4</td><td>10.4</td><td>1.6</td><td>6.4</td><td>1.4E-10</td><td>3.1E-09</td><td>BT20/MB468</td><td>MP</td></tr>
<tr><td>>hg38_21_+_8433172_8446622_RNA45SN1_45S_50ntFlanks@9381.9418.38|CGCTCGCCGGCCGAGGTGGGATCCCGAGGCCTCTCCAG</td><td>>hg38_21_+_8433172_8446622_RNA45SN1_45S_50ntFlanks</td><td>9381</td><td>9418</td><td>38</td><td>CGCTCGCCGGCCGAGGTGGGATCCCGAGGCCTCTCCAG</td><td>74.9</td><td>-10.1</td><td>1.7</td><td>-6.1</td><td>1.3E-09</td><td>3.1E-09</td><td>MB231/MB468</td><td>MP</td></tr>
<tr><td>>hg38_21_+_8433172_8446622_RNA45SN1_45S_50ntFlanks@9031.9059.29|TCCTCGTGGGGGGGCCGGGCCACCCCTCC</td><td>>hg38_21_+_8433172_8446622_RNA45SN1_45S_50ntFlanks</td><td>9031</td><td>9059</td><td>29</td><td>TCCTCGTGGGGGGGCCGGGCCACCCCTCC</td><td>165.2</td><td>-10.4</td><td>1.6</td><td>-6.4</td><td>1.4E-10</td><td>3.1E-09</td><td>BT20/MB468</td><td>MP</td></tr>
<tr><td>>hg38_21_+_8433172_8446622_RNA45SN1_45S_50ntFlanks@5084.5110.27|CGAACGAGACTCTGGCATGCTAACTAG</td><td>>hg38_21_+_8433172_8446622_RNA45SN1_45S_50ntFlanks</td><td>5084</td><td>5110</td><td>27</td><td>CGAACGAGACTCTGGCATGCTAACTAG</td><td>1282.9</td><td>-5.2</td><td>0.9</td><td>-6.1</td><td>1.4E-09</td><td>3.1E-09</td><td>MB231/MB468</td><td>NUC</td></tr>
<tr><td>>hg38_21_+_8433172_8446622_RNA45SN1_45S_50ntFlanks@8171.8213.43|AAGTCCTTCTGATCGAGGCCCAGCCCGTGGACGGTGTGAGGCC</td><td>>hg38_21_+_8433172_8446622_RNA45SN1_45S_50ntFlanks</td><td>8171</td><td>8213</td><td>43</td><td>AAGTCCTTCTGATCGAGGCCCAGCCCGTGGACGGTGTGAGGCC</td><td>172.6</td><td>-11.3</td><td>1.9</td><td>-6.1</td><td>1.3E-09</td><td>3.1E-09</td><td>MB231/MB468</td><td>MP</td></tr>
<tr><td>>hg38_21_+_8433172_8446622_RNA45SN1_45S_50ntFlanks@11874.11907.34|AGAAGACCCTGTTGAGCTTGACTCTAGTCTGGCA</td><td>>hg38_21_+_8433172_8446622_RNA45SN1_45S_50ntFlanks</td><td>11874</td><td>11907</td><td>34</td><td>AGAAGACCCTGTTGAGCTTGACTCTAGTCTGGCA</td><td>157.0</td><td>-7.1</td><td>1.2</td><td>-6.1</td><td>1.4E-09</td><td>3.1E-09</td><td>MB231/MB468</td><td>NUC</td></tr>
<tr><td>>hg38_21_+_8433172_8446622_RNA45SN1_45S_50ntFlanks@12251.12289.39|AAAGCTCGCTTGATCTTGATTTTCAGTACGAATACAGAC</td><td>>hg38_21_+_8433172_8446622_RNA45SN1_45S_50ntFlanks</td><td>12251</td><td>12289</td><td>39</td><td>AAAGCTCGCTTGATCTTGATTTTCAGTACGAATACAGAC</td><td>469.7</td><td>6.3</td><td>1.0</td><td>6.1</td><td>1.4E-09</td><td>3.1E-09</td><td>MB231/MB468</td><td>CYTO</td></tr>
<tr><td>>hg38_21_+_8433172_8446622_RNA45SN1_45S_50ntFlanks@12990.13012.23|GCTCCCTCGCTGCGATCTATTGA</td><td>>hg38_21_+_8433172_8446622_RNA45SN1_45S_50ntFlanks</td><td>12990</td><td>13012</td><td>23</td><td>GCTCCCTCGCTGCGATCTATTGA</td><td>2945.5</td><td>-10.3</td><td>1.7</td><td>-6.0</td><td>1.6E-09</td><td>3.1E-09</td><td>BT20/MB468</td><td>NUC</td></tr>
<tr><td>>hg38_21_+_8433172_8446622_RNA45SN1_45S_50ntFlanks@12073.12093.21|CCGAGGGGCTCTCGCTTCTGG</td><td>>hg38_21_+_8433172_8446622_RNA45SN1_45S_50ntFlanks</td><td>12073</td><td>12093</td><td>21</td><td>CCGAGGGGCTCTCGCTTCTGG</td><td>224.8</td><td>-10.9</td><td>1.7</td><td>-6.4</td><td>1.5E-10</td><td>3.1E-09</td><td>BT20/MB468</td><td>MP</td></tr>
<tr><td>>hg38_21_+_8433172_8446622_RNA45SN1_45S_50ntFlanks@11767.11804.38|TCTAATTAGTGACGCGCATGAATGGATGAACGAGATTC</td><td>>hg38_21_+_8433172_8446622_RNA45SN1_45S_50ntFlanks</td><td>11767</td><td>11804</td><td>38</td><td>TCTAATTAGTGACGCGCATGAATGGATGAACGAGATTC</td><td>82.6</td><td>-10.2</td><td>1.7</td><td>-6.1</td><td>1.3E-09</td><td>3.1E-09</td><td>MB231/MB468</td><td>MP</td></tr>
<tr><td>>hg38_21_+_8433172_8446622_RNA45SN1_45S_50ntFlanks@10731.10749.19|AGGGTGTAAATCTCGCGCC</td><td>>hg38_21_+_8433172_8446622_RNA45SN1_45S_50ntFlanks</td><td>10731</td><td>10749</td><td>19</td><td>AGGGTGTAAATCTCGCGCC</td><td>85.1</td><td>-10.3</td><td>1.7</td><td>-6.1</td><td>1.3E-09</td><td>3.1E-09</td><td>MB231/MB468</td><td>MP</td></tr>
<tr><td>>hg38_21_+_8433172_8446622_RNA45SN1_45S_50ntFlanks@4130.4162.33|ATTCCGGAGAGGGAGCCTGAGAAACGGCTACCA</td><td>>hg38_21_+_8433172_8446622_RNA45SN1_45S_50ntFlanks</td><td>4130</td><td>4162</td><td>33</td><td>ATTCCGGAGAGGGAGCCTGAGAAACGGCTACCA</td><td>809.4</td><td>-4.0</td><td>0.7</td><td>-6.1</td><td>1.4E-09</td><td>3.1E-09</td><td>MB231/MB468</td><td>CYTO</td></tr>
<tr><td>>hg38_21_+_8433172_8446622_RNA45SN1_45S_50ntFlanks@12932.12960.29|CGCTAAACCATTCGTAGACGACCTGCTTC</td><td>>hg38_21_+_8433172_8446622_RNA45SN1_45S_50ntFlanks</td><td>12932</td><td>12960</td><td>29</td><td>CGCTAAACCATTCGTAGACGACCTGCTTC</td><td>428.6</td><td>3.7</td><td>0.6</td><td>6.1</td><td>1.2E-09</td><td>3.1E-09</td><td>BT20/MB468</td><td>CYTO</td></tr>
<tr><td>>hg38_21_+_8433172_8446622_RNA45SN1_45S_50ntFlanks@12836.12859.24|CGGTGCGGAGTGCCCTTCGTCCTG</td><td>>hg38_21_+_8433172_8446622_RNA45SN1_45S_50ntFlanks</td><td>12836</td><td>12859</td><td>24</td><td>CGGTGCGGAGTGCCCTTCGTCCTG</td><td>109.0</td><td>-10.6</td><td>1.8</td><td>-6.1</td><td>1.3E-09</td><td>3.1E-09</td><td>MB231/MB468</td><td>MP</td></tr>
</table>

| | | | | | | | | | | | | | |
|---|---|---|---|---|---|---|---|---|---|---|---|---|---|
| >hg38_21_+_8433172_8446622_RNA45SN1_45S_50ntFlanks@10786.10816.31\|AACAGCCTCTGGCATGTTGGAACAATGTAGG | >hg38_21_+_8433172_8446622_RNA45SN1_45S_50ntFlanks | 10786 | 10816 | 31 | AACAGCCTCTGGCATGTTGGAACAATGTAGG | 1018.9 | -7.2 | 1.2 | -6.1 | 1.3E-09 | 3.1E-09 | MB231/MB468 | MP |
| >hg38_21_+_8433172_8446622_RNA45SN1_45S_50ntFlanks@11552.11591.40\|CCTCGGCCGGCGCCTAGCAGCCGACTTAGAACTGGTGCGG | >hg38_21_+_8433172_8446622_RNA45SN1_45S_50ntFlanks | 11552 | 11591 | 40 | CCTCGGCCGGCGCCTAGCAGCCGACTTAGAACTGGTGCGG | 424.8 | -5.6 | 0.9 | -6.1 | 1.4E-09 | 3.1E-09 | MB231/MB468 | CYTO |
| >MI0000077\|hsa-mir-21&WithFlank&17\|+\|59841260\|59841343@14.36.23&[MIMAT0000076&hsa-miR-21-5p&offsets\|0\|+1;m-1&17\|+\|59841273\|59841294&offsets\|0\|+1]\|TAGCTTATCAGACTGATGTTGAC | >MI0000077\|hsa-mir-21&WithFlank&17\|+\|59841260\|59841343 | 14 | 36 | 23 | TAGCTTATCAGACTGATGTTGAC | 448061.7 | 5.8 | 1.0 | 6.1 | 1.4E-09 | 3.1E-09 | MB231/MB468 | CYTO |
| >hg38_21_+_8433172_8446622_RNA45SN1_45S_50ntFlanks@10513.10533.21\|CCGGAGTGGCGGAGATGGGCG | >hg38_21_+_8433172_8446622_RNA45SN1_45S_50ntFlanks | 10513 | 10533 | 21 | CCGGAGTGGCGGAGATGGGCG | 1005.6 | -6.1 | 1.0 | -6.1 | 1.4E-09 | 3.1E-09 | MB231/MB468 | CYTO |
| >hg38_21_+_8433172_8446622_RNA45SN1_45S_50ntFlanks@11874.11897.24\|AGAAGACCCTGTTGAGCTTGACTC | >hg38_21_+_8433172_8446622_RNA45SN1_45S_50ntFlanks | 11874 | 11897 | 24 | AGAAGACCCTGTTGAGCTTGACTC | 937.7 | -4.2 | 0.7 | -6.1 | 1.4E-09 | 3.2E-09 | MB231/MB468 | CYTO |
| >hg38_21_+_8433172_8446622_RNA45SN1_45S_50ntFlanks@9504.9527.24\|GTGAACTATGCCTGGGCAGGGCGA | >hg38_21_+_8433172_8446622_RNA45SN1_45S_50ntFlanks | 9504 | 9527 | 24 | GTGAACTATGCCTGGGCAGGGCGA | 373.3 | -7.1 | 1.2 | -6.1 | 1.2E-09 | 3.2E-09 | BT20/MB468 | CYTO |
| >hg38_21_+_8433172_8446622_RNA45SN1_45S_50ntFlanks@12419.12437.19\|GATGTCGGCTCTTCCTATC | >hg38_21_+_8433172_8446622_RNA45SN1_45S_50ntFlanks | 12419 | 12437 | 19 | GATGTCGGCTCTTCCTATC | 9060.4 | 3.4 | 0.6 | 6.1 | 1.4E-09 | 3.2E-09 | MB231/MB468 | CYTO |
| >hg38_21_+_8433172_8446622_RNA45SN1_45S_50ntFlanks@4760.4777.18\|TCAGATACCGTCGTAGTT | >hg38_21_+_8433172_8446622_RNA45SN1_45S_50ntFlanks | 4760 | 4777 | 18 | TCAGATACCGTCGTAGTT | 163.8 | -5.1 | 0.8 | -6.1 | 1.4E-09 | 3.2E-09 | MB231/MB468 | CYTO |
| >hg38_21_+_8433172_8446622_RNA45SN1_45S_50ntFlanks@10393.10430.38\|CAGCAGTTGAACATGGGTCAGTCGGTCCTGAGAGATGG | >hg38_21_+_8433172_8446622_RNA45SN1_45S_50ntFlanks | 10393 | 10430 | 38 | CAGCAGTTGAACATGGGTCAGTCGGTCCTGAGAGATGG | 1022.9 | -6.8 | 1.1 | -6.1 | 1.4E-09 | 3.2E-09 | MB231/MB468 | NUC |
| >hg38_21_+_8433172_8446622_RNA45SN1_45S_50ntFlanks@12503.12521.19\|TAGACCGTCGTGAGACAGG | >hg38_21_+_8433172_8446622_RNA45SN1_45S_50ntFlanks | 12503 | 12521 | 19 | TAGACCGTCGTGAGACAGG | 116.7 | -10.7 | 1.8 | -6.1 | 1.4E-09 | 3.2E-09 | MB231/MB468 | MP |
| >hg38_21_+_8433172_8446622_RNA45SN1_45S_50ntFlanks@12417.12435.19\|TCGATGTCGGCTCTTCCTA | >hg38_21_+_8433172_8446622_RNA45SN1_45S_50ntFlanks | 12417 | 12435 | 19 | TCGATGTCGGCTCTTCCTA | 5363.0 | -4.4 | 0.7 | -6.1 | 1.2E-09 | 3.2E-09 | BT20/MB468 | CYTO |
| >hg38_21_+_8433172_8446622_RNA45SN1_45S_50ntFlanks@11572.11603.32\|CCGACTTAGAACTGGTGCGGACCAGGGGAATC | >hg38_21_+_8433172_8446622_RNA45SN1_45S_50ntFlanks | 11572 | 11603 | 32 | CCGACTTAGAACTGGTGCGGACCAGGGGAATC | 9170.2 | -5.2 | 0.8 | -6.4 | 1.5E-10 | 3.2E-09 | BT20/MB468 | MP |
| >hg38_21_+_8433172_8446622_RNA45SN1_45S_50ntFlanks@12434.12455.22\|TATCATTGTGAAGCAGAATTCA | >hg38_21_+_8433172_8446622_RNA45SN1_45S_50ntFlanks | 12434 | 12455 | 22 | TATCATTGTGAAGCAGAATTCA | 232.6 | -7.6 | 1.2 | -6.1 | 1.4E-09 | 3.2E-09 | MB231/MB468 | MP |
| >MI0000086\|hsa-mir-28&WithFlank&3\|+\|188688775\|188688872@62.82.21(+1A)&[MIMAT0004502&hsa-miR-28-3p&offsets\|+2\|+1(+1A);m-28&3\|+\|188688834\|188688855&offsets\|+2\|+1(+1A)]\|CTAGATTGTGAGCTCCTGGAGA | >MI0000086\|hsa-mir-28&WithFlank&3\|+\|188688775\|188688872 | 62 | 82 | 21(+1A) | CTAGATTGTGAGCTCCTGGAGA | 129.9 | 10.8 | 1.7 | 6.4 | 1.5E-10 | 3.2E-09 | BT20/MB468 | MP |
| >hg38_21_+_8433172_8446622_RNA45SN1_45S_50ntFlanks@12826.12847.22\|GACCGGGGTCCGGTGCGGAGTG | >hg38_21_+_8433172_8446622_RNA45SN1_45S_50ntFlanks | 12826 | 12847 | 22 | GACCGGGGTCCGGTGCGGAGTG | 68.4 | -10.0 | 1.6 | -6.1 | 1.4E-09 | 3.2E-09 | MB231/MB468 | MP |
| >hg38_21_+_8433172_8446622_RNA45SN1_45S_50ntFlanks@8381.8403.23\|AAGAGGGCGTGAAACCGTTAAGA | >hg38_21_+_8433172_8446622_RNA45SN1_45S_50ntFlanks | 8381 | 8403 | 23 | AAGAGGGCGTGAAACCGTTAAGA | 860.7 | -8.4 | 1.4 | -6.0 | 1.6E-09 | 3.2E-09 | MB231/MB468 | MITO |
| >MI0000063\|hsa-let-7b&WithFlank&22\|+\|46113680\|46113774@12.32.21(+1A)&[MIMAT0000063&hsa-let-7b-5p&offsets\|0\|-1(+1A);m-14&22\|+\|46113691\|46113712&offsets\|0\|-1(+1A)]\|TGAGGTAGTAGGTTGTGTGGTA | >MI0000063\|hsa-let-7b&WithFlank&22\|+\|46113680\|46113774 | 12 | 32 | 21(+1A) | TGAGGTAGTAGGTTGTGTGGTA | 2218.7 | 3.3 | 0.5 | 6.4 | 1.5E-10 | 3.2E-09 | BT20/MB231 | NUC |
| >hg38_21_+_8433172_8446622_RNA45SN1_45S_50ntFlanks@11571.11613.43\|GCCGACTTAGAACTGGTGCGGACCAGGGGAATCCGACTGTTTA | >hg38_21_+_8433172_8446622_RNA45SN1_45S_50ntFlanks | 11571 | 11613 | 43 | GCCGACTTAGAACTGGTGCGGACCAGGGGAATCCGACTGTTTA | 817.1 | 5.4 | 0.9 | 6.1 | 1.1E-09 | 3.2E-09 | BT20/MB468 | MITO |
| >hg38_21_+_8433172_8446622_RNA45SN1_45S_50ntFlanks@12202.12222.21\|CTAAGGCGAGCTCAGGGAGGA | >hg38_21_+_8433172_8446622_RNA45SN1_45S_50ntFlanks | 12202 | 12222 | 21 | CTAAGGCGAGCTCAGGGAGGA | 313.0 | -10.4 | 1.7 | -6.1 | 1.1E-09 | 3.2E-09 | BT20/MB468 | MITO |
| >trna14_LysTTT_11_-_59327808_59327880@1.31.31,trna2_LysTTT_17_+_8022473_8022545@1.31.31,trna54_LysTTT_1_+_204475655_204475727@1.31.31,trna5_LysTTT_11_+_59323902_59323974@1.31.31,trna62_LysTTT_1_-_204476158_204476230@1.31.31,trna76_LysTTT_6_+_28918806_28918878@1.31.31\|GCCCGGATAGCTCAGTCGGTAGAGCATCAGA | >trna14_LysTTT_11_-_59327808_59327880 | 1 | 31 | 31 | GCCCGGATAGCTCAGTCGGTAGAGCATCAGA | 278.7 | -11.2 | 1.7 | -6.4 | 1.5E-10 | 3.2E-09 | BT20/MB468 | MP |
| >hg38_21_+_8433172_8446622_RNA45SN1_45S_50ntFlanks@5071.5093.23\|TTAATTCCGATAACGAACGAGAC | >hg38_21_+_8433172_8446622_RNA45SN1_45S_50ntFlanks | 5071 | 5093 | 23 | TTAATTCCGATAACGAACGAGAC | 193.3 | -11.5 | 1.9 | -6.1 | 1.4E-09 | 3.2E-09 | MB231/MB468 | MP |
| >hg38_21_+_8433172_8446622_RNA45SN1_45S_50ntFlanks@11829.11856.28\|CGAAACCACAGCCAAGGGAACGGGCTTG | >hg38_21_+_8433172_8446622_RNA45SN1_45S_50ntFlanks | 11829 | 11856 | 28 | CGAAACCACAGCCAAGGGAACGGGCTTG | 208.3 | -10.8 | 1.7 | -6.4 | 1.5E-10 | 3.2E-09 | BT20/MB468 | MP |
| >MI0000088\|hsa-mir-30a&WithFlank&6\|-\|71403545\|71403627@12.34.23&[MIMAT0000087&hsa-miR-30a-5p&offsets\|0\|+1;m-15&6\|-\|71403595\|71403616&offsets\|0\|+1]\|TGTAAACATCCTCGACTGGAAGC | >MI0000088\|hsa-mir-30a&WithFlank&6\|-\|71403545\|71403627 | 12 | 34 | 23 | TGTAAACATCCTCGACTGGAAGC | 6887.9 | 5.6 | 0.9 | 6.1 | 1.4E-09 | 3.2E-09 | MB231/MB468 | CYTO |
| >MI0000750\|hsa-mir-26a-2&WithFlank&12\|-\|57824603\|57824698@20.40.21;MI0000083\|hsa-mir-26a-1&WithFlank&3\|+\|37969398\|37969486@16.36.21&[MIMAT0000082&hsa-miR-26a-5p&offsets\|0\|-1;m-33&12\|-\|57824658\|57824679&offsets\|0\|-1];[MIMAT0000082_1&hsa-miR-26a-5p&offsets\|0\|-1;m-34&3\|+\|37969413\|37969434&offsets\|0\|-1]\|TTCAAGTAATCCAGGATAGGC | >MI0000750\|hsa-mir-26a-2&WithFlank&12\|-\|57824603\|57824698 | 20 | 40 | 21 | TTCAAGTAATCCAGGATAGGC | 1271.5 | 6.0 | 1.0 | 6.1 | 1.4E-09 | 3.2E-09 | MB231/MB468 | CYTO |
| >hg38_21_+_8433172_8446622_RNA45SN1_45S_50ntFlanks@12997.13038.42\|CGCTGCGATCTATTGAAAGTCAGCCCTCGACACAAGGGTTTG | >hg38_21_+_8433172_8446622_RNA45SN1_45S_50ntFlanks | 12997 | 13038 | 42 | CGCTGCGATCTATTGAAAGTCAGCCCTCGACACAAGGGTTTG | 74.0 | 9.4 | 1.6 | 6.1 | 1.4E-09 | 3.2E-09 | MB231/MB468 | MP |
| >hg38_21_+_8433172_8446622_RNA45SN1_45S_50ntFlanks@9339.9370.32\|CGCACGAAAGCCGCCGTGGCGCAATGAAGGTG | >hg38_21_+_8433172_8446622_RNA45SN1_45S_50ntFlanks | 9339 | 9370 | 32 | CGCACGAAAGCCGCCGTGGCGCAATGAAGGTG | 6857.5 | -10.4 | 1.6 | -6.4 | 1.5E-10 | 3.2E-09 | BT20/MB468 | MP |
| >hg38_21_+_8433172_8446622_RNA45SN1_45S_50ntFlanks@9504.9522.19\|GTGAACTATGCCTGGGCAG | >hg38_21_+_8433172_8446622_RNA45SN1_45S_50ntFlanks | 9504 | 9522 | 19 | GTGAACTATGCCTGGGCAG | 1048.0 | -6.9 | 1.1 | -6.1 | 1.4E-09 | 3.2E-09 | MB231/MB468 | NUC |
| >MI0000542\|hsa-mir-320a&WithFlank&8\|-\|22244960\|22245043@48.69.22(+1U)&[MIMAT0000510&hsa-miR-320a-3p&offsets\|0\|0(+1U);m-60&8\|-\|22244975\|22244996&offsets\|0\|0(+1U)]\|AAAAGCTGGGTTGAGAGGGCGAT | >MI0000542\|hsa-mir-320a&WithFlank&8\|-\|22244960\|22245043 | 48 | 69 | 22(+1U) | AAAAGCTGGGTTGAGAGGGCGAT | 769.3 | 6.0 | 1.0 | 6.0 | 1.6E-09 | 3.2E-09 | MB231/MB468 | MITO |
| >hg38_21_+_8433172_8446622_RNA45SN1_45S_50ntFlanks@9201.9224.24\|CTCTCGGGGCCACGCGCGCGTCCC | >hg38_21_+_8433172_8446622_RNA45SN1_45S_50ntFlanks | 9201 | 9224 | 24 | CTCTCGGGGCCACGCGCGCGTCCC | 171.9 | -10.5 | 1.6 | -6.4 | 1.5E-10 | 3.2E-09 | BT20/MB468 | MP |
| >MI0000469\|hsa-mir-125a&WithFlank&19\|+\|51693248\|51693345@21.42.22(+1U)&[MIMAT0000443&hsa-miR-125a-5p&offsets\|0\|-2(+1U);m-63&19\|+\|51693268\|51693289&offsets\|0\|0(+1U)]\|TCCCTGAGACCCTTTAACCTGTT | >MI0000469\|hsa-mir-125a&WithFlank&19\|+\|51693248\|51693345 | 21 | 42 | 22(+1U) | TCCCTGAGACCCTTTAACCTGTT | 509.6 | 8.6 | 1.4 | 6.1 | 1.1E-09 | 3.2E-09 | BT20/MB468 | MITO |
| >hg38_21_+_8433172_8446622_RNA45SN1_45S_50ntFlanks@11873.11907.35\|AAGAAGACCCTGTTGAGCTTGACTCTAGTCTGGCA | >hg38_21_+_8433172_8446622_RNA45SN1_45S_50ntFlanks | 11873 | 11907 | 35 | AAGAAGACCCTGTTGAGCTTGACTCTAGTCTGGCA | 410.7 | -4.0 | 0.7 | -6.1 | 1.1E-09 | 3.2E-09 | BT20/MB468 | MITO |
| >MI0000478\|hsa-mir-149&WithFlank&2\|+\|240455995\|240456095@21.43.23&[MIMAT0000450&hsa-miR-149-5p&offsets\|0\|0;m-195&2\|+\|240456015\|240456037&offsets\|0\|0]\|TCTGGCTCCGTGTCTTCACTCCC | >MI0000478\|hsa-mir-149&WithFlank&2\|+\|240455995\|240456095 | 21 | 43 | 23 | TCTGGCTCCGTGTCTTCACTCCC | 127.7 | 10.2 | 1.7 | 6.0 | 1.6E-09 | 3.2E-09 | MB231/MB468 | MITO |
| >hg38_1_-_228634819_228635039_RNA5S12_5S_50ntFlanks@84.101.18\|CTCGTCTGATCTCGGAAG | >hg38_1_-_228634819_228635039_RNA5S12_5S_50ntFlanks | 84 | 101 | 18 | CTCGTCTGATCTCGGAAG | 1969.4 | -4.4 | 0.7 | -6.1 | 1.2E-09 | 3.2E-09 | BT20/MB468 | CYTO |
| >hg38_21_+_8433172_8446622_RNA45SN1_45S_50ntFlanks@9574.9595.22\|AATCGGTCGTCCGACCTGGGTA | >hg38_21_+_8433172_8446622_RNA45SN1_45S_50ntFlanks | 9574 | 9595 | 22 | AATCGGTCGTCCGACCTGGGTA | 8008.1 | -6.8 | 1.1 | -6.4 | 1.5E-10 | 3.2E-09 | BT20/MB468 | MP |
| >hg38_21_+_8433172_8446622_RNA45SN1_45S_50ntFlanks@12244.12280.37\|AAGGGCAAAAGCTCGCTTGATCTTGATTTTCAGTACG | >hg38_21_+_8433172_8446622_RNA45SN1_45S_50ntFlanks | 12244 | 12280 | 37 | AAGGGCAAAAGCTCGCTTGATCTTGATTTTCAGTACG | 138.3 | 5.6 | 0.9 | 6.1 | 1.4E-09 | 3.2E-09 | MB231/MB468 | CYTO |
| >MI0000102\|hsa-mir-100&WithFlank&11\|-\|122152223\|122152314@19.40.22(+1U)&[MIMAT0000098&hsa-miR-100-5p&offsets\|0\|0(+1U);m-27&11\|-\|122152276\|122152296&offsets\|0\|+1(+1U)]\|AACCCGTAGATCCGAACTTGTGT | >MI0000102\|hsa-mir-100&WithFlank&11\|-\|122152223\|122152314 | 19 | 40 | 22(+1U) | AACCCGTAGATCCGAACTTGTGT | 137.3 | 9.9 | 1.6 | 6.0 | 1.6E-09 | 3.2E-09 | BT20/MB468 | NUC |
| >MI0000063\|hsa-let-7b&WithFlank&22\|+\|46113680\|46113774@12.32.21(+1A)&[MIMAT0000063&hsa-let-7b-5p&offsets\|0\|-1(+1A);m-14&22\|+\|46113691\|46113712&offsets\|0\|-1(+1A)]\|TGAGGTAGTAGGTTGTGTGGTA | >MI0000063\|hsa-let-7b&WithFlank&22\|+\|46113680\|46113774 | 12 | 32 | 21(+1A) | TGAGGTAGTAGGTTGTGTGGTA | 519.8 | 5.6 | 0.9 | 6.1 | 1.4E-09 | 3.2E-09 | MB231/MB468 | CYTO |
| >trnaMT_GluTTC_MT_-_14674_14742@40.57.18,trnalookalike8_GluTTC_5_-_93905172_93905240@40.57.18\|TTGGTCGTGGTTGTAGTC | >trnaMT_GluTTC_MT_-_14674_14742 | 40 | 57 | 18 | TTGGTCGTGGTTGTAGTC | 354.8 | 6.0 | 1.0 | 6.1 | 1.4E-09 | 3.2E-09 | MB231/MB468 | CYTO |
| >MI0000088\|hsa-mir-30a&WithFlank&6\|-\|71403545\|71403627@13.35.23&[MIMAT0000087&hsa-miR-30a-5p&offsets\|+1\|+2;m-15&6\|-\|71403595\|71403616&offsets\|+1\|+2]\|GTAAACATCCTCGACTGGAAGCT | >MI0000088\|hsa-mir-30a&WithFlank&6\|-\|71403545\|71403627 | 13 | 35 | 23 | GTAAACATCCTCGACTGGAAGCT | 466.1 | 6.5 | 1.1 | 6.1 | 1.4E-09 | 3.2E-09 | MB231/MB468 | CYTO |
| >hg38_21_+_8433172_8446622_RNA45SN1_45S_50ntFlanks@11876.11904.29\|AAGACCCTGTTGAGCTTGACTCTAGTCTG | >hg38_21_+_8433172_8446622_RNA45SN1_45S_50ntFlanks | 11876 | 11904 | 29 | AAGACCCTGTTGAGCTTGACTCTAGTCTG | 185.0 | -9.6 | 1.6 | -6.1 | 1.1E-09 | 3.2E-09 | BT20/MB468 | MITO |
| >hg38_21_+_8433172_8446622_RNA45SN1_45S_50ntFlanks@11686.11728.43\|TCAAAGTGAAGAAATTCAATGAAGCGCGGGTAAACGGCGGGAG | >hg38_21_+_8433172_8446622_RNA45SN1_45S_50ntFlanks | 11686 | 11728 | 43 | TCAAAGTGAAGAAATTCAATGAAGCGCGGGTAAACGGCGGGAG | 161.9 | -11.1 | 1.8 | -6.0 | 1.6E-09 | 3.2E-09 | MB231/MB468 | MITO |
| >MI0000438\|hsa-mir-15b&WithFlank&3\|+\|160404582\|160404691@26.46.21&[MIMAT0000417&hsa-miR-15b-5p&offsets\|0\|-1;m-110&3\|+\|160404607\|160404628&offsets\|0\|-1]\|TAGCAGCACATCATGGTTTAC | >MI0000438\|hsa-mir-15b&WithFlank&3\|+\|160404582\|160404691 | 26 | 46 | 21 | TAGCAGCACATCATGGTTTAC | 519.7 | 4.5 | 0.7 | 6.0 | 1.4E-09 | 3.2E-09 | MB231/MB468 | NUC |
| >hg38_21_+_8433172_8446622_RNA45SN1_45S_50ntFlanks@5472.5497.26\|ACGGCCCTGGCGGAGCGCTGAGAAGA | >hg38_21_+_8433172_8446622_RNA45SN1_45S_50ntFlanks | 5472 | 5497 | 26 | ACGGCCCTGGCGGAGCGCTGAGAAGA | 2146.6 | 7.1 | 1.2 | 6.1 | 1.3E-09 | 3.2E-09 | BT20/MB468 | CYTO |
| >hg38_21_+_8433172_8446622_RNA45SN1_45S_50ntFlanks@7981.8008.28\|CTCAGATCAGACGTGGCGACCCGCTGAA | >hg38_21_+_8433172_8446622_RNA45SN1_45S_50ntFlanks | 7981 | 8008 | 28 | CTCAGATCAGACGTGGCGACCCGCTGAA | 543.5 | -4.1 | 0.7 | -6.0 | 1.5E-09 | 3.2E-09 | MB231/MB468 | NUC |
| >trna5_AspGTC_12_+_98897281_98897352@37.74.38\|ACGCGGGAGACCGGGGTTCAATTCCCCGACGGGGAGCC | >trna5_AspGTC_12_+_98897281_98897352 | 37 | 74 | 38 | ACGCGGGAGACCGGGGTTCAATTCCCCGACGGGGAGCC | 6175.1 | 16.6 | 2.7 | 6.1 | 1.4E-09 | 3.2E-09 | MB231/MB468 | CYTO |
| >hg38_21_+_8433172_8446622_RNA45SN1_45S_50ntFlanks@11999.12032.34\|CCCTGCGGGCCGCCGGTGAAATACCACTACTCTG | >hg38_21_+_8433172_8446622_RNA45SN1_45S_50ntFlanks | 11999 | 12032 | 34 | CCCTGCGGGCCGCCGGTGAAATACCACTACTCTG | 2498.0 | -9.4 | 1.5 | -6.4 | 1.5E-10 | 3.2E-09 | BT20/MB468 | MP |
| >hg38_21_+_8433172_8446622_RNA45SN1_45S_50ntFlanks@12159.12180.22\|TTTGACTGGGGCGGTACACCTG | >hg38_21_+_8433172_8446622_RNA45SN1_45S_50ntFlanks | 12159 | 12180 | 22 | TTTGACTGGGGCGGTACACCTG | 6078.7 | -8.3 | 1.3 | -6.4 | 1.5E-10 | 3.2E-09 | BT20/MB468 | MP |

| | | | | | | | | | | | | | |
|---|---|---|---|---|---|---|---|---|---|---|---|---|---|
| >trna130_GlnTTG_6_-_27763640_27763711@1.29.29,trna173_GlnTTG_6_-_26311975_26312046@1.29.29,trna174_GlnTTG_6_-_26311424_26311495@1.29.29\|GGCCCCATGGTGTAATGGTTAGCACTCTG | >trna130_GlnTTG_6_-_27763640_27763711 | 1 | 29 | 29 | GGCCCCATGGTGTAATGGTTAGCACTCTG | 128.6 | -9.1 | 1.5 | -6.1 | 1.1E-09 | 3.2E-09 | BT20/MB468 | MITO |
| >hg38_21_+_8433172_8446622_RNA45SN1_45S_50ntFlanks@10839.10868.30\|TCCGTAACTTCGGGATAAGGATTGGCTCTA | >hg38_21_+_8433172_8446622_RNA45SN1_45S_50ntFlanks | 10839 | 10868 | 30 | TCCGTAACTTCGGGATAAGGATTGGCTCTA | 373.8 | -6.0 | 1.0 | -6.0 | 1.5E-09 | 3.2E-09 | MB231/MB468 | NUC |
| >am_MI0003772\|hsa-mir-151b&WithFlank&14\|-\|100109413\|100109520@66.85.20(+1C);MI0000809\|hsa-mir-151a&WithFlank&8\|-\|140732558\|140732659@17.36.20(+1C)&[MIMAT0010214&hsa-miR-151b&offsets\|0\|+2(+1C);m-179&14\|-\|100109436\|100109455&offsets\|0\|0(+1C)];[MIMAT0004697&hsa-miR-151a-5p&offsets\|0\|-1(+1C);m-75&8\|-\|140732623\|140732643&offsets\|0\|-1(+1C)]\|TCGAGGAGCTCACAGTCTAGC | >am_MI0003772\|hsa-mir-151b&WithFlank&14\|-\|100109413\|100109520 | 66 | 85 | 20(+1C) | TCGAGGAGCTCACAGTCTAGC | 117.7 | 6.9 | 1.1 | 6.0 | 1.7E-09 | 3.3E-09 | BT20/MB468 | NUC |
| >hg38_21_+_8433172_8446622_RNA45SN1_45S_50ntFlanks@5472.5498.27\|ACGGCCCTGGCGGAGCGCTGAGAAGAC | >hg38_21_+_8433172_8446622_RNA45SN1_45S_50ntFlanks | 5472 | 5498 | 27 | ACGGCCCTGGCGGAGCGCTGAGAAGAC | 3906.9 | 6.7 | 1.1 | 6.1 | 1.3E-09 | 3.3E-09 | BT20/MB468 | CYTO |
| >hg38_21_+_8433172_8446622_RNA45SN1_45S_50ntFlanks@12996.13014.19\|TCGCTGCGATCTATTGAAA | >hg38_21_+_8433172_8446622_RNA45SN1_45S_50ntFlanks | 12996 | 13014 | 19 | TCGCTGCGATCTATTGAAA | 1642.9 | 4.1 | 0.7 | 6.1 | 1.3E-09 | 3.3E-09 | BT20/MB468 | CYTO |
| >hg38_21_+_8433172_8446622_RNA45SN1_45S_50ntFlanks@10819.10837.19\|AGGGAAGTCGGCAAGCCGG | >hg38_21_+_8433172_8446622_RNA45SN1_45S_50ntFlanks | 10819 | 10837 | 19 | AGGGAAGTCGGCAAGCCGG | 623.8 | -9.2 | 1.5 | -6.0 | 1.7E-09 | 3.3E-09 | BT20/MB468 | NUC |
| >hg38_21_+_8433172_8446622_RNA45SN1_45S_50ntFlanks@6719.6743.25\|TGAATTGCAGGACACATTGATCATC | >hg38_21_+_8433172_8446622_RNA45SN1_45S_50ntFlanks | 6719 | 6743 | 25 | TGAATTGCAGGACACATTGATCATC | 5425.0 | -4.8 | 0.8 | -6.1 | 1.4E-09 | 3.3E-09 | MB231/MB468 | CYTO |
| >MI0003763\|hsa-mir-767&WithFlank&X\|-\|152393415\|152393535@33.52.20&[MIMAT0003882&hsa-miR-767-5p&offsets\|0\|-3;m-788&X\|-\|152393481\|152393503&offsets\|0\|-3]\|TGCACCATGGTTGTCTGAGC | >MI0003763\|hsa-mir-767&WithFlank&X\|-\|152393415\|152393535 | 33 | 52 | 20 | TGCACCATGGTTGTCTGAGC | 263.1 | 7.5 | 1.2 | 6.1 | 1.3E-09 | 3.3E-09 | BT20/MB468 | CYTO |
| >MI0016005\|hsa-mir-3615&WithFlank&17\|+\|74748607\|74748705@57.78.22(+2U)&[MIMAT0017994&hsa-miR-3615&offsets\|0\|+1(+2U);m-271&17\|+\|74748663\|74748684&offsets\|0\|0(+2U)]\|TCTCTCGGCTCCTCGCGGCTCGTT | >MI0016005\|hsa-mir-3615&WithFlank&17\|+\|74748607\|74748705 | 57 | 78 | 22(+2U) | TCTCTCGGCTCCTCGCGGCTCGTT | 248.3 | 4.6 | 0.8 | 6.0 | 1.5E-09 | 3.3E-09 | MB231/MB468 | NUC |
| >hg38_21_+_8433172_8446622_RNA45SN1_45S_50ntFlanks@10738.10755.18\|AAATCTCGCGCCGGGCCG | >hg38_21_+_8433172_8446622_RNA45SN1_45S_50ntFlanks | 10738 | 10755 | 18 | AAATCTCGCGCCGGGCCG | 665.5 | -4.8 | 0.8 | -6.0 | 1.6E-09 | 3.3E-09 | MB231/MB468 | MITO |
| >hg38_21_+_8433172_8446622_RNA45SN1_45S_50ntFlanks@9602.9622.21\|CGAAAGACTAATCGAACCATC | >hg38_21_+_8433172_8446622_RNA45SN1_45S_50ntFlanks | 9602 | 9622 | 21 | CGAAAGACTAATCGAACCATC | 394.7 | -8.6 | 1.4 | -6.0 | 1.7E-09 | 3.3E-09 | BT20/MB468 | NUC |
| >hg38_21_+_8433172_8446622_RNA45SN1_45S_50ntFlanks@12459.12477.19\|AGCGTTGGATTGTTCACCC | >hg38_21_+_8433172_8446622_RNA45SN1_45S_50ntFlanks | 12459 | 12477 | 19 | AGCGTTGGATTGTTCACCC | 1205.6 | -9.3 | 1.5 | -6.0 | 1.7E-09 | 3.3E-09 | BT20/MB468 | NUC |
| >hg38_21_+_8433172_8446622_RNA45SN1_45S_50ntFlanks@11769.11803.35\|TAATTAGTGACGCGCATGAATGGATGAACGAGATT | >hg38_21_+_8433172_8446622_RNA45SN1_45S_50ntFlanks | 11769 | 11803 | 35 | TAATTAGTGACGCGCATGAATGGATGAACGAGATT | 941.9 | -4.0 | 0.7 | -6.0 | 1.5E-09 | 3.3E-09 | MB231/MB468 | NUC |
| >hg38_21_+_8433172_8446622_RNA45SN1_45S_50ntFlanks@3846.3869.24\|CTTGGATAACTGTGGTAATTCTAG | >hg38_21_+_8433172_8446622_RNA45SN1_45S_50ntFlanks | 3846 | 3869 | 24 | CTTGGATAACTGTGGTAATTCTAG | 1013.9 | -8.7 | 1.4 | -6.0 | 1.6E-09 | 3.3E-09 | MB231/MB468 | MITO |
| >hg38_21_+_8433172_8446622_RNA45SN1_45S_50ntFlanks@11318.11338.21\|CCCCACGTCTCGTCGCGCGCG | >hg38_21_+_8433172_8446622_RNA45SN1_45S_50ntFlanks | 11318 | 11338 | 21 | CCCCACGTCTCGTCGCGCGCG | 739.6 | -5.9 | 1.0 | -6.1 | 1.3E-09 | 3.3E-09 | BT20/MB468 | CYTO |
| >MI0000089\|hsa-mir-31&WithFlank&9\|-\|21512109\|21512191@14.36.23&[MIMAT0000089&hsa-miR-31-5p&offsets\|0\|+2;m-360&9\|-\|21512157\|21512178&offsets\|0\|+1]\|AGGCAAGATGCTGGCATAGCTGT | >MI0000089\|hsa-mir-31&WithFlank&9\|-\|21512109\|21512191 | 14 | 36 | 23 | AGGCAAGATGCTGGCATAGCTGT | 494.3 | 4.3 | 0.7 | 6.0 | 1.7E-09 | 3.3E-09 | BT20/MB468 | NUC |
| >trnaMT_MetCAT_MT_+_4402_4469@52.70.19\|ATACCCTTCCCGTACTACC | >trnaMT_MetCAT_MT_+_4402_4469 | 52 | 70 | 19 | ATACCCTTCCCGTACTACC | 655.2 | 8.3 | 1.4 | 6.0 | 1.5E-09 | 3.3E-09 | MB231/MB468 | NUC |
| >hg38_21_+_8433172_8446622_RNA45SN1_45S_50ntFlanks@8737.8758.22\|CCCGTCCGTCCGTCCGTCCGTC | >hg38_21_+_8433172_8446622_RNA45SN1_45S_50ntFlanks | 8737 | 8758 | 22 | CCCGTCCGTCCGTCCGTCCGTC | 264.7 | -11.1 | 1.7 | -6.4 | 1.6E-10 | 3.3E-09 | BT20/MB468 | MP |
| >hg38_21_+_8433172_8446622_RNA45SN1_45S_50ntFlanks@485.524.40\|GATGTGGTGACGTCGTGCTCTCCCGGGCCGGGTCCGAGCC | >hg38_21_+_8433172_8446622_RNA45SN1_45S_50ntFlanks | 485 | 524 | 40 | GATGTGGTGACGTCGTGCTCTCCCGGGCCGGGTCCGAGCC | 324.6 | 12.1 | 1.9 | 6.4 | 1.6E-10 | 3.3E-09 | BT20/MB468 | MP |
| >hg38_21_+_8433172_8446622_RNA45SN1_45S_50ntFlanks@12567.12587.21\|CTCAGTACGAGAGGAACCGCA | >hg38_21_+_8433172_8446622_RNA45SN1_45S_50ntFlanks | 12567 | 12587 | 21 | CTCAGTACGAGAGGAACCGCA | 336.0 | -6.9 | 1.1 | -6.1 | 1.3E-09 | 3.3E-09 | BT20/MB468 | CYTO |
| >hg38_21_+_8433172_8446622_RNA45SN1_45S_50ntFlanks@10383.10424.42\|TCCATGTGAACAGCAGTTGAACATGGGTCAGTCGGTCCTGAG | >hg38_21_+_8433172_8446622_RNA45SN1_45S_50ntFlanks | 10383 | 10424 | 42 | TCCATGTGAACAGCAGTTGAACATGGGTCAGTCGGTCCTGAG | 1283.1 | 7.6 | 1.3 | 6.1 | 1.3E-09 | 3.3E-09 | BT20/MB468 | CYTO |
| >MI0000093\|hsa-mir-92a-1&WithFlank&13\|+\|91351308\|91351397@54.74.21;MI0000094\|hsa-mir-92a-2&WithFlank&X\|-\|134169532\|134169618@54.74.21&[MIMAT0000092&hsa-miR-92a-3p&offsets\|0\|-1;m-9&13\|+\|91351361\|91351382&offsets\|0\|-1];[MIMAT0000092_1&hsa-miR-92a-3p&offsets\|0\|-1;m-11&X\|-\|134169544\|134169565&offsets\|0\|-1]\|TATTGCACTTGTCCCGGCCTG | >MI0000093\|hsa-mir-92a-1&WithFlank&13\|+\|91351308\|91351397 | 54 | 74 | 21 | TATTGCACTTGTCCCGGCCTG | 1169.9 | 4.2 | 0.7 | 6.0 | 1.5E-09 | 3.3E-09 | MB231/MB468 | NUC |
| >trna103_AsnGTT_1_-_149230570_149230643@29.55.27,trna107_AsnGTT_1_-_148760356_148760429@29.55.27,trna108_AsnGTT_1_-_148598314_148598387@29.55.27,trna1_AsnGTT_19_+_1383562_1383635@29.55.27,trna26_AsnGTT_1_+_148248115_148248188@29.55.27,trna31_AsnGTT_17_-_36908034_36908107@29.55.27,trna47_AsnGTT_1_+_161510031_161510104@29.55.27,trna4_AsnGTT_10_-_22518438_22518511@29.55.27,trna7_AsnGTT_13_-_31248101_31248174@29.55.27,trna83_AsnGTT_1_-_161397867_161397940@29.55.27\|TCGGCTGTTAACCGAAAGGTTGGTGGT | >trna103_AsnGTT_1_-_149230570_149230643 | 29 | 55 | 27 | TCGGCTGTTAACCGAAAGGTTGGTGGT | 638.1 | 10.4 | 1.7 | 6.1 | 1.3E-09 | 3.3E-09 | BT20/MB468 | CYTO |
| >hg38_21_+_8433172_8446622_RNA45SN1_45S_50ntFlanks@9180.9210.31\|CCGTCGGGCCCGGGGGAGGTTCTCTCGGGGC | >hg38_21_+_8433172_8446622_RNA45SN1_45S_50ntFlanks | 9180 | 9210 | 31 | CCGTCGGGCCCGGGGGAGGTTCTCTCGGGGC | 559.9 | -8.2 | 1.4 | -6.0 | 1.6E-09 | 3.3E-09 | MB231/MB468 | MITO |
| >hg38_21_+_8433172_8446622_RNA45SN1_45S_50ntFlanks@12405.12433.29\|CTTTTTGATCCTTCGATGTCGGCTCTTCC | >hg38_21_+_8433172_8446622_RNA45SN1_45S_50ntFlanks | 12405 | 12433 | 29 | CTTTTTGATCCTTCGATGTCGGCTCTTCC | 3966.9 | -4.8 | 0.8 | -6.1 | 1.3E-09 | 3.3E-09 | BT20/MB468 | CYTO |
| >hg38_21_+_8433172_8446622_RNA45SN1_45S_50ntFlanks@11359.11383.25\|CGGTCGGGCGGCGGCGGTCGGCGGG | >hg38_21_+_8433172_8446622_RNA45SN1_45S_50ntFlanks | 11359 | 11383 | 25 | CGGTCGGGCGGCGGCGGTCGGCGGG | 91.3 | -10.4 | 1.7 | -6.1 | 1.4E-09 | 3.3E-09 | MB231/MB468 | MP |
| >hg38_21_+_8433172_8446622_RNA45SN1_45S_50ntFlanks@10402.10420.19\|AACATGGGTCAGTCGGTCC | >hg38_21_+_8433172_8446622_RNA45SN1_45S_50ntFlanks | 10402 | 10420 | 19 | AACATGGGTCAGTCGGTCC | 2435.5 | -8.5 | 1.4 | -6.0 | 1.6E-09 | 3.3E-09 | MB231/MB468 | MITO |
| >hg38_21_+_8433172_8446622_RNA45SN1_45S_50ntFlanks@10349.10384.36\|AACGAGAACTTTGAAGGCCGAAGTGGAGAAGGGTTC | >hg38_21_+_8433172_8446622_RNA45SN1_45S_50ntFlanks | 10349 | 10384 | 36 | AACGAGAACTTTGAAGGCCGAAGTGGAGAAGGGTTC | 1965.9 | -4.3 | 0.7 | -6.0 | 1.5E-09 | 3.3E-09 | MB231/MB468 | NUC |
| >MI0000434\|hsa-let-7i&WithFlank&12\|+\|62603680\|62603775@12.33.22(+1C)&[MIMAT0000415&hsa-let-7i-5p&offsets\|0\|0(+1C);m-49&12\|+\|62603691\|62603712&offsets\|0\|0(+1C)]\|TGAGGTAGTAGTTTGTGCTGTTC | >MI0000434\|hsa-let-7i&WithFlank&12\|+\|62603680\|62603775 | 12 | 33 | 22(+1C) | TGAGGTAGTAGTTTGTGCTGTTC | 381.5 | 7.9 | 1.3 | 6.1 | 1.1E-09 | 3.3E-09 | BT20/MB468 | MITO |
| >hg38_21_+_8433172_8446622_RNA45SN1_45S_50ntFlanks@10411.10440.30\|CAGTCGGTCCTGAGAGATGGGCGAGCGCCG | >hg38_21_+_8433172_8446622_RNA45SN1_45S_50ntFlanks | 10411 | 10440 | 30 | CAGTCGGTCCTGAGAGATGGGCGAGCGCCG | 236.4 | -6.6 | 1.1 | -6.1 | 1.3E-09 | 3.3E-09 | BT20/MB468 | CYTO |
| >hg38_21_+_8433172_8446622_RNA45SN1_45S_50ntFlanks@4260.4278.19\|TTGGAATGAGTCCACTTTA | >hg38_21_+_8433172_8446622_RNA45SN1_45S_50ntFlanks | 4260 | 4278 | 19 | TTGGAATGAGTCCACTTTA | 149.4 | -10.3 | 1.6 | -6.4 | 1.6E-10 | 3.4E-09 | BT20/MB468 | MP |
| >trna116_GluCTC_1_-_145399233_145399304@1.32.32,trna59_GluCTC_1_+_249168447_249168518@1.32.32,trna71_GluCTC_1_-_161439189_161439260@1.32.32,trna74_GluCTC_1_-_161431809_161431880@1.32.32,trna77_GluCTC_1_-_161424398_161424469@1.32.32,trna77_GluCTC_6_+_28949976_28950047@1.32.32,trna80_GluCTC_1_-_161417018_161417089@1.32.32,trna87_GluCTC_6_-_126101393_126101464@1.32.32\|TCCCTGGTGGTCTAGTGGTTAGGATTCGGCGC | >trna116_GluCTC_1_-_145399233_145399304 | 1 | 32 | 32 | TCCCTGGTGGTCTAGTGGTTAGGATTCGGCGC | 1353.0 | 5.4 | 0.9 | 6.0 | 1.6E-09 | 3.4E-09 | MB231/MB468 | MITO |
| >hg38_21_+_8433172_8446622_RNA45SN1_45S_50ntFlanks@10396.10435.40\|CAGTTGAACATGGGTCAGTCGGTCCTGAGAGATGGGCGAG | >hg38_21_+_8433172_8446622_RNA45SN1_45S_50ntFlanks | 10396 | 10435 | 40 | CAGTTGAACATGGGTCAGTCGGTCCTGAGAGATGGGCGAG | 122.0 | -10.8 | 1.8 | -6.1 | 1.4E-09 | 3.4E-09 | MB231/MB468 | MP |
| >hg38_21_+_8433172_8446622_RNA45SN1_45S_50ntFlanks@4027.4044.18\|CCCCGTGGCGGCGACGAC | >hg38_21_+_8433172_8446622_RNA45SN1_45S_50ntFlanks | 4027 | 4044 | 18 | CCCCGTGGCGGCGACGAC | 155.6 | -9.4 | 1.5 | -6.1 | 1.1E-09 | 3.4E-09 | BT20/MB468 | MITO |
| >hg38_21_+_8433172_8446622_RNA45SN1_45S_50ntFlanks@10871.10902.32\|GGCTGGGTCGGTCGGGCTGGGGCGCGAAGCGG | >hg38_21_+_8433172_8446622_RNA45SN1_45S_50ntFlanks | 10871 | 10902 | 32 | GGCTGGGTCGGTCGGGCTGGGGCGCGAAGCGG | 182.1 | -7.3 | 1.2 | -6.0 | 1.5E-09 | 3.4E-09 | MB231/MB468 | NUC |
| >hg38_21_+_8433172_8446622_RNA45SN1_45S_50ntFlanks@12942.12973.32\|TTCGTAGACGACCTGCTTCTGGGTCGGGGTTT | >hg38_21_+_8433172_8446622_RNA45SN1_45S_50ntFlanks | 12942 | 12973 | 32 | TTCGTAGACGACCTGCTTCTGGGTCGGGGTTT | 365.7 | -7.0 | 1.2 | -6.0 | 1.5E-09 | 3.4E-09 | MB231/MB468 | CYTO |
| >MI0000066\|hsa-let-7e&WithFlank&19\|+\|51692780\|51692870@14.32.19&[MIMAT0000066&hsa-let-7e-5p&offsets\|0\|-3;m-50&19\|+\|51692793\|51692814&offsets\|0\|-3]\|TGAGGTAGGAGGTTGTATA | >MI0000066\|hsa-let-7e&WithFlank&19\|+\|51692780\|51692870 | 14 | 32 | 19 | TGAGGTAGGAGGTTGTATA | 1146.0 | 9.1 | 1.5 | 6.1 | 1.3E-09 | 3.4E-09 | BT20/MB468 | CYTO |
| >hg38_21_+_8433172_8446622_RNA45SN1_45S_50ntFlanks@10095.10117.23\|CGCGCGCGCGCGCGTGTGGTGTG | >hg38_21_+_8433172_8446622_RNA45SN1_45S_50ntFlanks | 10095 | 10117 | 23 | CGCGCGCGCGCGCGTGTGGTGTG | 133.4 | -10.1 | 1.6 | -6.4 | 1.6E-10 | 3.4E-09 | BT20/MB468 | MP |
| >hg38_21_+_8433172_8446622_RNA45SN1_45S_50ntFlanks@5519.5545.27\|AGGAAGTAAAAGTCGTAACAAGGTTTC | >hg38_21_+_8433172_8446622_RNA45SN1_45S_50ntFlanks | 5519 | 5545 | 27 | AGGAAGTAAAAGTCGTAACAAGGTTTC | 107.5 | -10.6 | 1.8 | -6.1 | 1.4E-09 | 3.4E-09 | MB231/MB468 | MP |
| >hg38_21_+_8433172_8446622_RNA45SN1_45S_50ntFlanks@10905.10925.21\|CTGGGCGCGCGCCGCGGCTGG | >hg38_21_+_8433172_8446622_RNA45SN1_45S_50ntFlanks | 10905 | 10925 | 21 | CTGGGCGCGCGCCGCGGCTGG | 114.2 | -8.9 | 1.5 | -6.1 | 1.1E-09 | 3.4E-09 | BT20/MB468 | MITO |
| >hg38_21_+_8433172_8446622_RNA45SN1_45S_50ntFlanks@10349.10388.40\|AACGAGAACTTTGAAGGCCGAAGTGGAGAAGGGTTCCATG | >hg38_21_+_8433172_8446622_RNA45SN1_45S_50ntFlanks | 10349 | 10388 | 40 | AACGAGAACTTTGAAGGCCGAAGTGGAGAAGGGTTCCATG | 1778.5 | -4.9 | 0.8 | -6.0 | 1.7E-09 | 3.4E-09 | MB231/MB468 | MITO |
| >MI0001641\|hsa-mir-429&WithFlank&1\|+\|1168999\|1169093@57.78.22&[MIMAT0001536&hsa-miR-429&offsets\|0\|0;m-207&1\|+\|1169055\|1169076&offsets\|0\|0]\|TAATACTGTCTGGTAAAACCGT | >MI0001641\|hsa-mir-429&WithFlank&1\|+\|1168999\|1169093 | 57 | 78 | 22 | TAATACTGTCTGGTAAAACCGT | 2539.6 | -5.5 | 0.9 | -6.1 | 1.4E-09 | 3.4E-09 | MB231/MB468 | MP |
| >hg38_21_+_8433172_8446622_RNA45SN1_45S_50ntFlanks@4005.4024.20\|AACCTCGGGCCGATCGCACG | >hg38_21_+_8433172_8446622_RNA45SN1_45S_50ntFlanks | 4005 | 4024 | 20 | AACCTCGGGCCGATCGCACG | 1367.9 | -3.0 | 0.5 | -6.0 | 1.5E-09 | 3.4E-09 | MB231/MB468 | CYTO |
| >hg38_21_+_8433172_8446622_RNA45SN1_45S_50ntFlanks@11581.11609.29\|AACTGGTGCGGACCAGGGGAATCCGACTG | >hg38_21_+_8433172_8446622_RNA45SN1_45S_50ntFlanks | 11581 | 11609 | 29 | AACTGGTGCGGACCAGGGGAATCCGACTG | 1127.1 | -3.2 | 0.5 | -6.0 | 1.5E-09 | 3.4E-09 | MB231/MB468 | NUC |

| | | | | | | | | | | | | | |
|---|---|---|---|---|---|---|---|---|---|---|---|---|---|
| >MI0000298\|hsa-mir-221&WithFlank&X\|-\|45746151\|45746272@71.92.22(+1A)&[MIMAT0000278&hsa-miR-221-3p&offsets\|0\|-1(+1A);m-91&X\|-\|45746181\|45746202&offsets\|0\|0(+1A)]\|AGCTACATTGTCTGCTGGGTTTA | >MI0000298\|hsa-mir-221&WithFlank&X\|-\|45746151\|45746272 | 71 | 92 | 22(+1A) | AGCTACATTGTCTGCTGGGTTTA | 1015.3 | 5.6 | 0.9 | 6.0 | 1.7E-09 | 3.4E-09 | MB231/MB468 | MITO |
| >hg38_21_+_8433172_8446622_RNA45SN1_45S_50ntFlanks@3787.3810.24\|AAACTGCGAATGGCTCATTAAATC | >hg38_21_+_8433172_8446622_RNA45SN1_45S_50ntFlanks | 3787 | 3810 | 24 | AAACTGCGAATGGCTCATTAAATC | 1212.5 | -3.9 | 0.7 | -6.0 | 1.5E-09 | 3.4E-09 | MB231/MB468 | NUC |
| >hg38_21_+_8433172_8446622_RNA45SN1_45S_50ntFlanks@10350.10391.42\|ACGAGAACTTTGAAGGCCGAAGTGGAGAAGGGTTCCATGTGA | >hg38_21_+_8433172_8446622_RNA45SN1_45S_50ntFlanks | 10350 | 10391 | 42 | ACGAGAACTTTGAAGGCCGAAGTGGAGAAGGGTTCCATGTGA | 1087.4 | -9.7 | 1.6 | -6.0 | 1.7E-09 | 3.4E-09 | BT20/MB468 | NUC |
| >hg38_21_+_8433172_8446622_RNA45SN1_45S_50ntFlanks@12044.12066.23\|ACTGACCCGGTGAGGCGGGGGGG | >hg38_21_+_8433172_8446622_RNA45SN1_45S_50ntFlanks | 12044 | 12066 | 23 | ACTGACCCGGTGAGGCGGGGGGG | 176.5 | -9.6 | 1.6 | -6.1 | 1.1E-09 | 3.4E-09 | BT20/MB468 | MITO |
| >hg38_21_+_8433172_8446622_RNA45SN1_45S_50ntFlanks@12838.12856.19\|GTGCGGAGTGCCCTTCGTC | >hg38_21_+_8433172_8446622_RNA45SN1_45S_50ntFlanks | 12838 | 12856 | 19 | GTGCGGAGTGCCCTTCGTC | 123.6 | -10.8 | 1.8 | -6.0 | 1.5E-09 | 3.4E-09 | MB231/MB468 | MP |
| >hg38_21_+_8433172_8446622_RNA45SN1_45S_50ntFlanks@12597.12614.18\|ATTTGGTGTATGTGCTTG | >hg38_21_+_8433172_8446622_RNA45SN1_45S_50ntFlanks | 12597 | 12614 | 18 | ATTTGGTGTATGTGCTTG | 471.3 | -8.3 | 1.4 | -6.0 | 1.7E-09 | 3.4E-09 | MB231/MB468 | MITO |
| >hg38_21_+_8433172_8446622_RNA45SN1_45S_50ntFlanks@11050.11075.26\|CCCGTCCTCCCCCCTCCCCGGGGGAG | >hg38_21_+_8433172_8446622_RNA45SN1_45S_50ntFlanks | 11050 | 11075 | 26 | CCCGTCCTCCCCCCTCCCCGGGGGAG | 108.2 | -6.8 | 1.1 | -6.1 | 1.3E-09 | 3.4E-09 | BT20/MB468 | CYTO |
| >hg38_21_+_8433172_8446622_RNA45SN1_45S_50ntFlanks@11671.11704.34\|CCAGTGCTCTGAATGTCAAAGTGAAGAAATTCAA | >hg38_21_+_8433172_8446622_RNA45SN1_45S_50ntFlanks | 11671 | 11704 | 34 | CCAGTGCTCTGAATGTCAAAGTGAAGAAATTCAA | 858.6 | -3.9 | 0.6 | -6.0 | 1.5E-09 | 3.4E-09 | MB231/MB468 | NUC |
| >MI0003188\|hsa-mir-503&WithFlank&X\|-\|134546322\|134546404@12.30.19&[MIMAT0002874&hsa-miR-503-5p&offsets\|0\|-4;m-297&X\|-\|134546371\|134546393&offsets\|0\|-4]\|TAGCAGCGGGAACAGTTCT | >MI0003188\|hsa-mir-503&WithFlank&X\|-\|134546322\|134546404 | 12 | 30 | 19 | TAGCAGCGGGAACAGTTCT | 264.9 | 11.8 | 1.8 | 6.4 | 1.6E-10 | 3.4E-09 | BT20/MB468 | MP |
| >hg38_21_+_8433172_8446622_RNA45SN1_45S_50ntFlanks@11539.11584.46\|GCGCGGCGCCTCGCCTCGGCCGGCGCCTAGCAGCCGACTTAGAACT | >hg38_21_+_8433172_8446622_RNA45SN1_45S_50ntFlanks | 11539 | 11584 | 46 | GCGCGGCGCCTCGCCTCGGCCGGCGCCTAGCAGCCGACTTAGAACT | 808.2 | 9.0 | 1.5 | 6.0 | 1.5E-09 | 3.4E-09 | MB231/MB468 | CYTO |
| >hg38_21_+_8433172_8446622_RNA45SN1_45S_50ntFlanks@6647.6664.18\|CGTACGACTCTTAGCGGT | >hg38_21_+_8433172_8446622_RNA45SN1_45S_50ntFlanks | 6647 | 6664 | 18 | CGTACGACTCTTAGCGGT | 1401.2 | 3.9 | 0.6 | 6.0 | 1.5E-09 | 3.4E-09 | MB231/MB468 | CYTO |
| >hg38_21_+_8433172_8446622_RNA45SN1_45S_50ntFlanks@11730.11751.22\|AACTATGACTCTCTTAAGGTAG | >hg38_21_+_8433172_8446622_RNA45SN1_45S_50ntFlanks | 11730 | 11751 | 22 | AACTATGACTCTCTTAAGGTAG | 70.6 | -10.0 | 1.7 | -6.0 | 1.5E-09 | 3.4E-09 | MB231/MB468 | MP |
| >hg38_21_+_8433172_8446622_RNA45SN1_45S_50ntFlanks@12627.12659.33\|ATGGGGCGAAGCTACCATCTGTGGGATTATGAC | >hg38_21_+_8433172_8446622_RNA45SN1_45S_50ntFlanks | 12627 | 12659 | 33 | ATGGGGCGAAGCTACCATCTGTGGGATTATGAC | 5962.5 | -8.0 | 1.3 | -6.1 | 1.2E-09 | 3.4E-09 | BT20/MB468 | MITO |
| >hg38_21_+_8433172_8446622_RNA45SN1_45S_50ntFlanks@7989.8016.28\|AGACGTGGCGACCCGCTGAATTTAAGCA | >hg38_21_+_8433172_8446622_RNA45SN1_45S_50ntFlanks | 7989 | 8016 | 28 | AGACGTGGCGACCCGCTGAATTTAAGCA | 987.3 | 6.6 | 1.1 | 6.0 | 1.5E-09 | 3.5E-09 | MB231/MB468 | CYTO |
| >hg38_21_+_8433172_8446622_RNA45SN1_45S_50ntFlanks@9839.9857.19\|GTAAGCAGAACTGGCGCTG | >hg38_21_+_8433172_8446622_RNA45SN1_45S_50ntFlanks | 9839 | 9857 | 19 | GTAAGCAGAACTGGCGCTG | 76.1 | -10.1 | 1.7 | -6.0 | 1.5E-09 | 3.5E-09 | MB231/MB468 | MP |
| >hg38_21_+_8433172_8446622_RNA45SN1_45S_50ntFlanks@6698.6720.23\|AGCTAGCTGCGAGAATTAATGTG | >hg38_21_+_8433172_8446622_RNA45SN1_45S_50ntFlanks | 6698 | 6720 | 23 | AGCTAGCTGCGAGAATTAATGTG | 34552.4 | -5.2 | 0.9 | -6.0 | 1.6E-09 | 3.5E-09 | MB231/MB468 | NUC |
| >hg38_21_+_8433172_8446622_RNA45SN1_45S_50ntFlanks@5494.5514.21\|AAGACGGTCGAACTTGACTAT | >hg38_21_+_8433172_8446622_RNA45SN1_45S_50ntFlanks | 5494 | 5514 | 21 | AAGACGGTCGAACTTGACTAT | 156.5 | -9.4 | 1.5 | -6.1 | 1.2E-09 | 3.5E-09 | BT20/MB468 | MITO |
| >MI0000301\|hsa-mir-224&WithFlank&X\|-\|151958572\|151958664@14.35.22&[MIMAT0000281&hsa-miR-224-5p&offsets\|+1\|-2;m-238&X\|-\|151958630\|151958651&offsets\|0\|0]\|CAAGTCACTAGTGGTTCCGTTT | >MI0000301\|hsa-mir-224&WithFlank&X\|-\|151958572\|151958664 | 14 | 35 | 22 | CAAGTCACTAGTGGTTCCGTTT | 1251.8 | 8.6 | 1.4 | 6.0 | 1.7E-09 | 3.5E-09 | MB231/MB468 | MITO |
| >MI0000088\|hsa-mir-30a&WithFlank&6\|-\|71403545\|71403627@12.34.23(+1C)&[MIMAT0000087&hsa-miR-30a-5p&offsets\|0\|+1(+1C);m-15&6\|-\|71403595\|71403616&offsets\|0\|+1(+1C)]\|TGTAAACATCCTCGACTGGAAGCC | >MI0000088\|hsa-mir-30a&WithFlank&6\|-\|71403545\|71403627 | 12 | 34 | 23(+1C) | TGTAAACATCCTCGACTGGAAGCC | 39080.9 | 4.4 | 0.7 | 6.0 | 1.5E-09 | 3.5E-09 | MB231/MB468 | MP |
| >hg38_21_+_8433172_8446622_RNA45SN1_45S_50ntFlanks@11616.11640.25\|TAAAACAAAGCATCGCGAAGGCCCG | >hg38_21_+_8433172_8446622_RNA45SN1_45S_50ntFlanks | 11616 | 11640 | 25 | TAAAACAAAGCATCGCGAAGGCCCG | 131.2 | -10.1 | 1.6 | -6.4 | 1.7E-10 | 3.5E-09 | BT20/MB468 | MP |
| >hg38_21_+_8433172_8446622_RNA45SN1_45S_50ntFlanks@10391.10425.35\|AACAGCAGTTGAACATGGGTCAGTCGGTCCTGAGA | >hg38_21_+_8433172_8446622_RNA45SN1_45S_50ntFlanks | 10391 | 10425 | 35 | AACAGCAGTTGAACATGGGTCAGTCGGTCCTGAGA | 77.8 | -10.1 | 1.7 | -6.0 | 1.5E-09 | 3.5E-09 | MB231/MB468 | MP |
| >hg38_21_+_8433172_8446622_RNA45SN1_45S_50ntFlanks@12932.12965.34\|CGCTAAACCATTCGTAGACGACCTGCTTCTGGGT | >hg38_21_+_8433172_8446622_RNA45SN1_45S_50ntFlanks | 12932 | 12965 | 34 | CGCTAAACCATTCGTAGACGACCTGCTTCTGGGT | 88.5 | -10.3 | 1.7 | -6.0 | 1.5E-09 | 3.5E-09 | MB231/MB468 | MP |
| >MI0000441\|hsa-mir-30b&WithFlank&8\|-\|134800514\|134800613@23.43.21(+1C)&[MIMAT0000420&hsa-miR-30b-5p&offsets\|0\|-1(+1C);m-76&8\|-\|134800570\|134800591&offsets\|0\|-1(+1C)]\|TGTAAACATCCTACACTCAGCC | >MI0000441\|hsa-mir-30b&WithFlank&8\|-\|134800514\|134800613 | 23 | 43 | 21(+1C) | TGTAAACATCCTACACTCAGCC | 307.9 | 4.6 | 0.8 | 6.0 | 1.8E-09 | 3.5E-09 | BT20/MB468 | NUC |
| >hg38_21_+_8433172_8446622_RNA45SN1_45S_50ntFlanks@10404.10424.21\|CATGGGTCAGTCGGTCCTGAG | >hg38_21_+_8433172_8446622_RNA45SN1_45S_50ntFlanks | 10404 | 10424 | 21 | CATGGGTCAGTCGGTCCTGAG | 6995.8 | 4.7 | 0.7 | 6.4 | 1.7E-10 | 3.5E-09 | BT20/MB468 | MP |
| >hg38_21_+_8433172_8446622_RNA45SN1_45S_50ntFlanks@11320.11344.25\|CCACGTCTCGTCGCGCGCGCGTCCG | >hg38_21_+_8433172_8446622_RNA45SN1_45S_50ntFlanks | 11320 | 11344 | 25 | CCACGTCTCGTCGCGCGCGCGTCCG | 7179.2 | -9.4 | 1.6 | -6.0 | 1.6E-09 | 3.5E-09 | MB231/MB468 | NUC |
| >hg38_21_+_8433172_8446622_RNA45SN1_45S_50ntFlanks@12589.12616.28\|GTTCAGACATTTGGTGTATGTGCTTGGC | >hg38_21_+_8433172_8446622_RNA45SN1_45S_50ntFlanks | 12589 | 12616 | 28 | GTTCAGACATTTGGTGTATGTGCTTGGC | 79.3 | -10.2 | 1.7 | -6.0 | 1.5E-09 | 3.5E-09 | MB231/MB468 | MP |
| >hg38_21_+_8433172_8446622_RNA45SN1_45S_50ntFlanks@10680.10705.26\|GTTCCGGCGGCGTCCGGTGAGCTCTC | >hg38_21_+_8433172_8446622_RNA45SN1_45S_50ntFlanks | 10680 | 10705 | 26 | GTTCCGGCGGCGTCCGGTGAGCTCTC | 690.7 | -8.1 | 1.3 | -6.0 | 1.5E-09 | 3.5E-09 | MB231/MB468 | MP |
| >hg38_21_+_8433172_8446622_RNA45SN1_45S_50ntFlanks@8115.8143.29\|CGCGGGACATGTGGCGTACGGAAGACCCG | >hg38_21_+_8433172_8446622_RNA45SN1_45S_50ntFlanks | 8115 | 8143 | 29 | CGCGGGACATGTGGCGTACGGAAGACCCG | 14064.7 | -8.6 | 1.3 | -6.4 | 1.7E-10 | 3.5E-09 | BT20/MB468 | MP |
| >hg38_21_+_8433172_8446622_RNA45SN1_45S_50ntFlanks@12259.12280.22\|CTTGATCTTGATTTTCAGTACG | >hg38_21_+_8433172_8446622_RNA45SN1_45S_50ntFlanks | 12259 | 12280 | 22 | CTTGATCTTGATTTTCAGTACG | 243.4 | -6.1 | 1.0 | -6.0 | 1.5E-09 | 3.5E-09 | MB231/MB468 | MP |
| >hg38_21_+_8433172_8446622_RNA45SN1_45S_50ntFlanks@9033.9057.25\|CTCGTGGGGGGGCCGGGCCACCCCT | >hg38_21_+_8433172_8446622_RNA45SN1_45S_50ntFlanks | 9033 | 9057 | 25 | CTCGTGGGGGGGCCGGGCCACCCCT | 163.5 | -9.4 | 1.6 | -6.1 | 1.2E-09 | 3.5E-09 | BT20/MB468 | MITO |
| >hg38_21_+_8433172_8446622_RNA45SN1_45S_50ntFlanks@12926.12953.28\|ACCTGGCGCTAAACCATTCGTAGACGAC | >hg38_21_+_8433172_8446622_RNA45SN1_45S_50ntFlanks | 12926 | 12953 | 28 | ACCTGGCGCTAAACCATTCGTAGACGAC | 361.7 | -6.0 | 1.0 | -6.0 | 1.7E-09 | 3.5E-09 | MB231/MB468 | MITO |
| >hg38_21_+_8433172_8446622_RNA45SN1_45S_50ntFlanks@12406.12443.38\|TTTTTGATCCTTCGATGTCGGCTCTTCCTATCATTGTG | >hg38_21_+_8433172_8446622_RNA45SN1_45S_50ntFlanks | 12406 | 12443 | 38 | TTTTTGATCCTTCGATGTCGGCTCTTCCTATCATTGTG | 1880.3 | -4.5 | 0.7 | -6.0 | 1.6E-09 | 3.5E-09 | MB231/MB468 | NUC |
| >MI0000477\|hsa-mir-146a&WithFlank&5\|+\|160485346\|160485456@27.49.23(+1A)&[MIMAT0000449&hsa-miR-146a-5p&offsets\|0\|+1(+1A);m-129&5\|+\|160485372\|160485393&offsets\|0\|+1(+1A)]\|TGAGAACTGAATTCCATGGGTTGA | >MI0000477\|hsa-mir-146a&WithFlank&5\|+\|160485346\|160485456 | 27 | 49 | 23(+1A) | TGAGAACTGAATTCCATGGGTTGA | 300.6 | -11.6 | 1.8 | -6.6 | 3.3E-11 | 3.5E-09 | BT20/MB231 | MP |
| >hg38_21_+_8433172_8446622_RNA45SN1_45S_50ntFlanks@9572.9603.32\|CAAATCGGTCGTCCGACCTGGGTATAGGGGCG | >hg38_21_+_8433172_8446622_RNA45SN1_45S_50ntFlanks | 9572 | 9603 | 32 | CAAATCGGTCGTCCGACCTGGGTATAGGGGCG | 3806.8 | -8.1 | 1.3 | -6.0 | 1.8E-09 | 3.5E-09 | BT20/MB468 | NUC |
| >hg38_21_+_8433172_8446622_RNA45SN1_45S_50ntFlanks@12712.12741.30\|CGGAGCCTCGGTTGGCCTCGGATAGCCGGT | >hg38_21_+_8433172_8446622_RNA45SN1_45S_50ntFlanks | 12712 | 12741 | 30 | CGGAGCCTCGGTTGGCCTCGGATAGCCGGT | 159.0 | -10.4 | 1.6 | -6.4 | 1.7E-10 | 3.5E-09 | BT20/MB468 | MP |
| >hg38_21_+_8433172_8446622_RNA45SN1_45S_50ntFlanks@10432.10453.22\|CGAGCGCCGTTCCGAAGGGACG | >hg38_21_+_8433172_8446622_RNA45SN1_45S_50ntFlanks | 10432 | 10453 | 22 | CGAGCGCCGTTCCGAAGGGACG | 130.7 | -9.1 | 1.5 | -6.1 | 1.2E-09 | 3.5E-09 | BT20/MB468 | MITO |
| >hg38_21_+_8433172_8446622_RNA45SN1_45S_50ntFlanks@5157.5185.29\|CAAGTGGCGTTCAGCCACCCGAGATTGAG | >hg38_21_+_8433172_8446622_RNA45SN1_45S_50ntFlanks | 5157 | 5185 | 29 | CAAGTGGCGTTCAGCCACCCGAGATTGAG | 107.0 | -10.6 | 1.8 | -6.0 | 1.5E-09 | 3.5E-09 | MB231/MB468 | MP |
| >hg38_21_+_8433172_8446622_RNA45SN1_45S_50ntFlanks@10404.10429.26\|CATGGGTCAGTCGGTCCTGAGAGATG | >hg38_21_+_8433172_8446622_RNA45SN1_45S_50ntFlanks | 10404 | 10429 | 26 | CATGGGTCAGTCGGTCCTGAGAGATG | 1230.3 | -2.9 | 0.5 | -6.1 | 1.4E-09 | 3.5E-09 | BT20/MB468 | CYTO |
| >trnaMT_PheGAA_MT_+_577_647@22.57.36\|AAAGCAATACACTGAAAATGTTTAGACGGGCTCACA | >trnaMT_PheGAA_MT_+_577_647 | 22 | 57 | 36 | AAAGCAATACACTGAAAATGTTTAGACGGGCTCACA | 396.0 | 9.9 | 1.6 | 6.1 | 1.2E-09 | 3.5E-09 | BT20/MB468 | MITO |
| >MI0000434\|hsa-let-7i&WithFlank&12\|+\|62603680\|62603775@12.33.22(+1A)&[MIMAT0000415&hsa-let-7i-5p&offsets\|0\|0(+1A);m-49&12\|+\|62603691\|62603712&offsets\|0\|0(+1A)]\|TGAGGTAGTAGTTTGTGCTGTTA | >MI0000434\|hsa-let-7i&WithFlank&12\|+\|62603680\|62603775 | 12 | 33 | 22(+1A) | TGAGGTAGTAGTTTGTGCTGTTA | 1312.1 | 1.8 | 0.3 | 6.0 | 1.8E-09 | 3.5E-09 | BT20/MB468 | NUC |
| >hg38_21_+_8433172_8446622_RNA45SN1_45S_50ntFlanks@12892.12931.40\|CCCTCGCCCGTCACGCACCGCACGTTCGTGGGGAACCTGG | >hg38_21_+_8433172_8446622_RNA45SN1_45S_50ntFlanks | 12892 | 12931 | 40 | CCCTCGCCCGTCACGCACCGCACGTTCGTGGGGAACCTGG | 121.3 | -10.8 | 1.8 | -6.0 | 1.5E-09 | 3.5E-09 | MB231/MB468 | MP |
| >hg38_21_+_8433172_8446622_RNA45SN1_45S_50ntFlanks@9573.9598.26\|AAATCGGTCGTCCGACCTGGGTATAG | >hg38_21_+_8433172_8446622_RNA45SN1_45S_50ntFlanks | 9573 | 9598 | 26 | AAATCGGTCGTCCGACCTGGGTATAG | 96.4 | -8.7 | 1.4 | -6.1 | 1.2E-09 | 3.5E-09 | BT20/MB468 | MITO |
| >hg38_21_+_8433172_8446622_RNA45SN1_45S_50ntFlanks@13008.13027.20\|ATTGAAAGTCAGCCCTCGAC | >hg38_21_+_8433172_8446622_RNA45SN1_45S_50ntFlanks | 13008 | 13027 | 20 | ATTGAAAGTCAGCCCTCGAC | 306.1 | -7.4 | 1.2 | -6.0 | 1.6E-09 | 3.5E-09 | MB231/MB468 | NUC |
| >hg38_21_+_8433172_8446622_RNA45SN1_45S_50ntFlanks@10738.10757.20\|AAATCTCGCGCCGGGCCGTA | >hg38_21_+_8433172_8446622_RNA45SN1_45S_50ntFlanks | 10738 | 10757 | 20 | AAATCTCGCGCCGGGCCGTA | 1248.8 | -7.4 | 1.2 | -6.0 | 1.8E-09 | 3.5E-09 | BT20/MB468 | NUC |
| >hg38_21_+_8433172_8446622_RNA45SN1_45S_50ntFlanks@4297.4317.21\|CATTGGAGGGCAAGTCTGGTG | >hg38_21_+_8433172_8446622_RNA45SN1_45S_50ntFlanks | 4297 | 4317 | 21 | CATTGGAGGGCAAGTCTGGTG | 939.3 | -8.4 | 1.4 | -6.0 | 1.7E-09 | 3.5E-09 | MB231/MB468 | MITO |
| >hg38_21_+_8433172_8446622_RNA45SN1_45S_50ntFlanks@10914.10936.23\|CGCCGCGGCTGGACGAGGCGCCG | >hg38_21_+_8433172_8446622_RNA45SN1_45S_50ntFlanks | 10914 | 10936 | 23 | CGCCGCGGCTGGACGAGGCGCCG | 435.1 | -8.5 | 1.4 | -6.0 | 1.8E-09 | 3.5E-09 | BT20/MB468 | NUC |
| >hg38_21_+_8433172_8446622_RNA45SN1_45S_50ntFlanks@4762.4780.19\|AGATACCGTCGTAGTTCCG | >hg38_21_+_8433172_8446622_RNA45SN1_45S_50ntFlanks | 4762 | 4780 | 19 | AGATACCGTCGTAGTTCCG | 867.1 | -7.4 | 1.2 | -6.0 | 1.8E-09 | 3.5E-09 | BT20/MB468 | NUC |
| >MI0005756\|hsa-mir-934&WithFlank&X\|+\|136550872\|136550966@21.42.22(+1A)&[MIMAT0004977&hsa-miR-934&offsets\|0\|0(+1A);m-943&X\|+\|136550892\|136550913&offsets\|0\|0(+1A)]\|TGTCTACTACTGGAGACACTGGA | >MI0005756\|hsa-mir-934&WithFlank&X\|+\|136550872\|136550966 | 21 | 42 | 22(+1A) | TGTCTACTACTGGAGACACTGGA | 398.8 | 12.0 | 1.8 | 6.6 | 3.4E-11 | 3.6E-09 | BT20/MB231 | MP |
| >MI0000750\|hsa-mir-26a-2&WithFlank&12\|-\|57824603\|57824698@20.41.22;MI0000083\|hsa-mir-26a-1&WithFlank&3\|+\|37969398\|37969486@16.37.22&[MIMAT0000082&hsa-miR-26a-5p&offsets\|0\|0;m-33&12\|-\|57824658\|57824679&offsets\|0\|0];[MIMAT0000082_1&hsa-miR-26a-5p&offsets\|0\|0;m-34&3\|+\|37969413\|37969434&offsets\|0\|0]\|TTCAAGTAATCCAGGATAGGCT | >MI0000750\|hsa-mir-26a-2&WithFlank&12\|-\|57824603\|57824698 | 20 | 41 | 22 | TTCAAGTAATCCAGGATAGGCT | 63674.0 | 5.4 | 0.9 | 6.0 | 1.6E-09 | 3.6E-09 | MB231/MB468 | CYTO |

| | | | | | | | | | | | | | |
|---|---|---|---|---|---|---|---|---|---|---|---|---|---|
| >hg38_21_+_8433172_8446622_RNA45SN1_45S_50ntFlanks@11729.11749.21\|TAACTATGACTCTCTTAAGGT | >hg38_21_+_8433172_8446622_RNA45SN1_45S_50ntFlanks | 11729 | 11749 | 21 | TAACTATGACTCTCTTAAGGT | 1135.5 | -3.0 | 0.5 | -6.1 | 1.2E-09 | 3.6E-09 | BT20/MB468 | MITO |
| >hg38_21_+_8433172_8446622_RNA45SN1_45S_50ntFlanks@4064.4101.38\|ATCAACTTTCGATGGTAGTCGCCGTGCCTACCATGGTG | >hg38_21_+_8433172_8446622_RNA45SN1_45S_50ntFlanks | 4064 | 4101 | 38 | ATCAACTTTCGATGGTAGTCGCCGTGCCTACCATGGTG | 110.9 | -8.9 | 1.5 | -6.1 | 1.2E-09 | 3.6E-09 | BT20/MB468 | MITO |
| >hg38_21_+_8433172_8446622_RNA45SN1_45S_50ntFlanks@10798.10823.26\|CATGTTGGAACAATGTAGGTAAGGGA | >hg38_21_+_8433172_8446622_RNA45SN1_45S_50ntFlanks | 10798 | 10823 | 26 | CATGTTGGAACAATGTAGGTAAGGGA | 151.1 | -4.3 | 0.7 | -6.0 | 1.6E-09 | 3.6E-09 | MB231/MB468 | CYTO |
| >hg38_21_+_8433172_8446622_RNA45SN1_45S_50ntFlanks@10669.10706.38\|AAAGCGTCGCGGTTCCGGCGGCGTCCGGTGAGCTCTCG | >hg38_21_+_8433172_8446622_RNA45SN1_45S_50ntFlanks | 10669 | 10706 | 38 | AAAGCGTCGCGGTTCCGGCGGCGTCCGGTGAGCTCTCG | 224.9 | -11.7 | 1.9 | -6.0 | 1.5E-09 | 3.6E-09 | MB231/MB468 | MP |
| >hg38_21_+_8433172_8446622_RNA45SN1_45S_50ntFlanks@8352.8390.39\|AGTTGAAAAGAACTTTGAAGAGAGAGTTCAAGAGGGCGT | >hg38_21_+_8433172_8446622_RNA45SN1_45S_50ntFlanks | 8352 | 8390 | 39 | AGTTGAAAAGAACTTTGAAGAGAGAGTTCAAGAGGGCGT | 113.9 | 10.6 | 1.7 | 6.4 | 1.7E-10 | 3.6E-09 | BT20/MB468 | MP |
| >hg38_21_+_8433172_8446622_RNA45SN1_45S_50ntFlanks@7985.8016.32\|GATCAGACGTGGCGACCCGCTGAATTTAAGCA | >hg38_21_+_8433172_8446622_RNA45SN1_45S_50ntFlanks | 7985 | 8016 | 32 | GATCAGACGTGGCGACCCGCTGAATTTAAGCA | 160.3 | -10.4 | 1.6 | -6.4 | 1.7E-10 | 3.6E-09 | BT20/MB468 | MP |
| >hg38_21_+_8433172_8446622_RNA45SN1_45S_50ntFlanks@11783.11817.35\|CATGAATGGATGAACGAGATTCCCACTGTCCCTAC | >hg38_21_+_8433172_8446622_RNA45SN1_45S_50ntFlanks | 11783 | 11817 | 35 | CATGAATGGATGAACGAGATTCCCACTGTCCCTAC | 80.0 | -10.2 | 1.7 | -6.0 | 1.5E-09 | 3.6E-09 | MB231/MB468 | MP |
| >hg38_21_+_8433172_8446622_RNA45SN1_45S_50ntFlanks@7975.7998.24\|CGCGACCTCAGATCAGACGTGGCG | >hg38_21_+_8433172_8446622_RNA45SN1_45S_50ntFlanks | 7975 | 7998 | 24 | CGCGACCTCAGATCAGACGTGGCG | 462.9 | 4.4 | 0.7 | 6.0 | 1.8E-09 | 3.6E-09 | BT20/MB468 | NUC |
| >hg38_21_+_8433172_8446622_RNA45SN1_45S_50ntFlanks@5056.5082.27\|GAGCGATTTGTCTGGTTAATTCCGATA | >hg38_21_+_8433172_8446622_RNA45SN1_45S_50ntFlanks | 5056 | 5082 | 27 | GAGCGATTTGTCTGGTTAATTCCGATA | 570.6 | -5.5 | 0.9 | -6.0 | 1.8E-09 | 3.6E-09 | BT20/MB468 | NUC |
| >hg38_21_+_8433172_8446622_RNA45SN1_45S_50ntFlanks@11670.11700.31\|CCCAGTGCTCTGAATGTCAAAGTGAAGAAAT | >hg38_21_+_8433172_8446622_RNA45SN1_45S_50ntFlanks | 11670 | 11700 | 31 | CCCAGTGCTCTGAATGTCAAAGTGAAGAAAT | 986.1 | -6.4 | 1.1 | -6.0 | 1.6E-09 | 3.6E-09 | MB231/MB468 | NUC |
| >hg38_21_+_8433172_8446622_RNA45SN1_45S_50ntFlanks@12157.12179.23\|AGTTTGACTGGGGCGGTACACCT | >hg38_21_+_8433172_8446622_RNA45SN1_45S_50ntFlanks | 12157 | 12179 | 23 | AGTTTGACTGGGGCGGTACACCT | 257.2 | -6.5 | 1.1 | -6.1 | 1.4E-09 | 3.6E-09 | BT20/MB468 | CYTO |
| >MI0006444\|hsa-mir-1307&WithFlank&10\|-\|103394247\|103394407@86.109.24&[MIMAT0005951&hsa-miR-1307-3p&offsets\|0\|+2;m-66&10\|-\|103394300\|103394322&offsets\|0\|+1]\|ACTCGGCGTGGCGTCGGTCGTGGT | >MI0006444\|hsa-mir-1307&WithFlank&10\|-\|103394247\|103394407 | 86 | 109 | 24 | ACTCGGCGTGGCGTCGGTCGTGGT | 333.7 | 5.3 | 0.9 | 6.0 | 1.6E-09 | 3.6E-09 | MB231/MB468 | NUC |
| >hg38_21_+_8433172_8446622_RNA45SN1_45S_50ntFlanks@10826.10865.40\|TCGGCAAGCCGGATCCGTAACTTCGGGATAAGGATTGGCT | >hg38_21_+_8433172_8446622_RNA45SN1_45S_50ntFlanks | 10826 | 10865 | 40 | TCGGCAAGCCGGATCCGTAACTTCGGGATAAGGATTGGCT | 98.2 | -8.7 | 1.4 | -6.1 | 1.2E-09 | 3.6E-09 | BT20/MB468 | MITO |
| >hg38_21_+_8433172_8446622_RNA45SN1_45S_50ntFlanks@11579.11609.31\|AGAACTGGTGCGGACCAGGGGAATCCGACTG | >hg38_21_+_8433172_8446622_RNA45SN1_45S_50ntFlanks | 11579 | 11609 | 31 | AGAACTGGTGCGGACCAGGGGAATCCGACTG | 1101.8 | -5.1 | 0.8 | -6.0 | 1.8E-09 | 3.6E-09 | MB231/MB468 | MITO |
| >hg38_21_+_8433172_8446622_RNA45SN1_45S_50ntFlanks@6741.6772.32\|ATCGACACTTCGAACGCACTTGCGGCCCCGGG | >hg38_21_+_8433172_8446622_RNA45SN1_45S_50ntFlanks | 6741 | 6772 | 32 | ATCGACACTTCGAACGCACTTGCGGCCCCGGG | 821.8 | -9.1 | 1.5 | -6.0 | 1.9E-09 | 3.6E-09 | BT20/MB468 | NUC |
| >hg38_21_+_8433172_8446622_RNA45SN1_45S_50ntFlanks@11586.11612.27\|GTGCGGACCAGGGGAATCCGACTGTTT | >hg38_21_+_8433172_8446622_RNA45SN1_45S_50ntFlanks | 11586 | 11612 | 27 | GTGCGGACCAGGGGAATCCGACTGTTT | 1123.8 | -3.7 | 0.6 | -6.0 | 1.6E-09 | 3.6E-09 | MB231/MB468 | NUC |
| >hg38_21_+_8433172_8446622_RNA45SN1_45S_50ntFlanks@11562.11591.30\|CGCCTAGCAGCCGACTTAGAACTGGTGCGG | >hg38_21_+_8433172_8446622_RNA45SN1_45S_50ntFlanks | 11562 | 11591 | 30 | CGCCTAGCAGCCGACTTAGAACTGGTGCGG | 132.9 | -5.7 | 0.9 | -6.0 | 1.6E-09 | 3.6E-09 | MB231/MB468 | CYTO |
| >MI0000082\|hsa-mir-25&WithFlank&7\|-\|100093554\|100093649@58.77.20(+1A)&[MIMAT0000081&hsa-miR-25-3p&offsets\|0\|-2(+1A);m-23&7\|-\|100093571\|100093592&offsets\|0\|-2(+1A)]\|CATTGCACTTGTCTCGGTCTA | >MI0000082\|hsa-mir-25&WithFlank&7\|-\|100093554\|100093649 | 58 | 77 | 20(+1A) | CATTGCACTTGTCTCGGTCTA | 492.2 | 5.3 | 0.9 | 6.0 | 1.8E-09 | 3.6E-09 | MB231/MB468 | MITO |
| >MI0033672\|hsa-mir-10525&WithFlank&7\|+\|74595962\|74596032@45.66.22&[MIMAT0041995&hsa-miR-10525-3p&offsets\|0\|+1]\|TGACTATGATGTGCACCTGATT | >MI0033672\|hsa-mir-10525&WithFlank&7\|+\|74595962\|74596032 | 45 | 66 | 22 | TGACTATGATGTGCACCTGATT | 429.7 | 4.8 | 0.8 | 6.0 | 1.6E-09 | 3.6E-09 | MB231/MB468 | NUC |
| >hg38_21_+_8433172_8446622_RNA45SN1_45S_50ntFlanks@11886.11905.20\|TGAGCTTGACTCTAGTCTGG | >hg38_21_+_8433172_8446622_RNA45SN1_45S_50ntFlanks | 11886 | 11905 | 20 | TGAGCTTGACTCTAGTCTGG | 167.2 | -9.5 | 1.6 | -6.1 | 1.2E-09 | 3.6E-09 | BT20/MB468 | MITO |
| >MI0000477\|hsa-mir-146a&WithFlank&5\|+\|160485346\|160485456@27.48.22(+1U)&[MIMAT0000449&hsa-miR-146a-5p&offsets\|0\|0(+1U);m-129&5\|+\|160485372\|160485393&offsets\|0\|0(+1U)]\|TGAGAACTGAATTCCATGGGTTT | >MI0000477\|hsa-mir-146a&WithFlank&5\|+\|160485346\|160485456 | 27 | 48 | 22(+1U) | TGAGAACTGAATTCCATGGGTTT | 12268.2 | -9.9 | 1.5 | -6.6 | 3.5E-11 | 3.6E-09 | BT20/MB231 | MP |
| >hg38_MT_+_1621_3279_RNR2_16S_50ntFlanks@716.740.25\|TAAGCCTGCGTCAGATTAAAACACT | >hg38_MT_+_1621_3279_RNR2_16S_50ntFlanks | 716 | 740 | 25 | TAAGCCTGCGTCAGATTAAAACACT | 94.5 | 9.8 | 1.6 | 6.0 | 1.8E-09 | 3.6E-09 | MB231/MB468 | MITO |
| >hg38_21_+_8433172_8446622_RNA45SN1_45S_50ntFlanks@11564.11603.40\|CCTAGCAGCCGACTTAGAACTGGTGCGGACCAGGGGAATC | >hg38_21_+_8433172_8446622_RNA45SN1_45S_50ntFlanks | 11564 | 11603 | 40 | CCTAGCAGCCGACTTAGAACTGGTGCGGACCAGGGGAATC | 7129.7 | -5.0 | 0.8 | -6.1 | 1.2E-09 | 3.7E-09 | BT20/MB468 | MITO |
| >hg38_21_+_8433172_8446622_RNA45SN1_45S_50ntFlanks@8105.8132.28\|CGCGGCGGGGCGCGGGACATGTGGCGTA | >hg38_21_+_8433172_8446622_RNA45SN1_45S_50ntFlanks | 8105 | 8132 | 28 | CGCGGCGGGGCGCGGGACATGTGGCGTA | 101.0 | -10.5 | 1.7 | -6.0 | 1.6E-09 | 3.7E-09 | MB231/MB468 | MP |
| >hg38_21_+_8433172_8446622_RNA45SN1_45S_50ntFlanks@11212.11229.18\|CGACTCTGGACGCGAGCC | >hg38_21_+_8433172_8446622_RNA45SN1_45S_50ntFlanks | 11212 | 11229 | 18 | CGACTCTGGACGCGAGCC | 276.7 | -7.6 | 1.3 | -6.0 | 1.6E-09 | 3.7E-09 | MB231/MB468 | MP |
| >hg38_21_+_8433172_8446622_RNA45SN1_45S_50ntFlanks@6733.6771.39\|CATTGATCATCGACACTTCGAACGCACTTGCGGCCCCGG | >hg38_21_+_8433172_8446622_RNA45SN1_45S_50ntFlanks | 6733 | 6771 | 39 | CATTGATCATCGACACTTCGAACGCACTTGCGGCCCCGG | 6049.9 | -8.7 | 1.4 | -6.0 | 1.9E-09 | 3.7E-09 | BT20/MB468 | NUC |
| >hg38_21_+_8433172_8446622_RNA45SN1_45S_50ntFlanks@476.504.29\|TTCGTGATCGATGTGGTGACGTCGTGCTC | >hg38_21_+_8433172_8446622_RNA45SN1_45S_50ntFlanks | 476 | 504 | 29 | TTCGTGATCGATGTGGTGACGTCGTGCTC | 367.1 | 7.5 | 1.2 | 6.0 | 1.6E-09 | 3.7E-09 | MB231/MB468 | CYTO |
| >hg38_21_+_8433172_8446622_RNA45SN1_45S_50ntFlanks@3782.3810.29\|CAGTGAAACTGCGAATGGCTCATTAAATC | >hg38_21_+_8433172_8446622_RNA45SN1_45S_50ntFlanks | 3782 | 3810 | 29 | CAGTGAAACTGCGAATGGCTCATTAAATC | 551.2 | -6.6 | 1.1 | -6.0 | 1.8E-09 | 3.7E-09 | MB231/MB468 | MITO |
| >hg38_21_+_8433172_8446622_RNA45SN1_45S_50ntFlanks@11670.11703.34\|CCCAGTGCTCTGAATGTCAAAGTGAAGAAATTCA | >hg38_21_+_8433172_8446622_RNA45SN1_45S_50ntFlanks | 11670 | 11703 | 34 | CCCAGTGCTCTGAATGTCAAAGTGAAGAAATTCA | 909.7 | -6.3 | 1.0 | -6.0 | 1.8E-09 | 3.7E-09 | MB231/MB468 | MITO |
| >hg38_21_+_8433172_8446622_RNA45SN1_45S_50ntFlanks@9304.9338.35\|ACCAAGGAGTCTAACACGTGCGCGAGTCGGGGGCT | >hg38_21_+_8433172_8446622_RNA45SN1_45S_50ntFlanks | 9304 | 9338 | 35 | ACCAAGGAGTCTAACACGTGCGCGAGTCGGGGGCT | 143.1 | -7.0 | 1.2 | -6.0 | 1.7E-09 | 3.7E-09 | MB231/MB468 | NUC |
| >MI0000301\|hsa-mir-224&WithFlank&X\|-\|151958572\|151958664@14.36.23&[MIMAT0000281&hsa-miR-224-5p&offsets\|+1\|-1;m-238&X\|-\|151958630\|151958651&offsets\|0\|+1]\|CAAGTCACTAGTGGTTCCGTTTA | >MI0000301\|hsa-mir-224&WithFlank&X\|-\|151958572\|151958664 | 14 | 36 | 23 | CAAGTCACTAGTGGTTCCGTTTA | 362.1 | 7.9 | 1.3 | 6.0 | 1.7E-09 | 3.7E-09 | MB231/MB468 | NUC |
| >hg38_21_+_8433172_8446622_RNA45SN1_45S_50ntFlanks@10680.10712.33\|GTTCCGGCGGCGTCCGGTGAGCTCTCGCTGGCC | >hg38_21_+_8433172_8446622_RNA45SN1_45S_50ntFlanks | 10680 | 10712 | 33 | GTTCCGGCGGCGTCCGGTGAGCTCTCGCTGGCC | 4738.1 | -5.0 | 0.8 | -6.1 | 1.4E-09 | 3.7E-09 | BT20/MB468 | CYTO |
| >hg38_21_+_8433172_8446622_RNA45SN1_45S_50ntFlanks@12285.12317.33\|CAGACCGTGAAAGCGGGGCCTCACGATCCTTCT | >hg38_21_+_8433172_8446622_RNA45SN1_45S_50ntFlanks | 12285 | 12317 | 33 | CAGACCGTGAAAGCGGGGCCTCACGATCCTTCT | 192.6 | -9.7 | 1.6 | -6.1 | 1.2E-09 | 3.7E-09 | BT20/MB468 | MITO |
| >hg38_21_+_8433172_8446622_RNA45SN1_45S_50ntFlanks@10685.10706.22\|GGCGGCGTCCGGTGAGCTCTCG | >hg38_21_+_8433172_8446622_RNA45SN1_45S_50ntFlanks | 10685 | 10706 | 22 | GGCGGCGTCCGGTGAGCTCTCG | 215.2 | -4.8 | 0.8 | -6.0 | 1.6E-09 | 3.7E-09 | MB231/MB468 | CYTO |
| >hg38_21_+_8433172_8446622_RNA45SN1_45S_50ntFlanks@4728.4753.26\|ATCAAGAACGAAAGTCGGAGGTTCGA | >hg38_21_+_8433172_8446622_RNA45SN1_45S_50ntFlanks | 4728 | 4753 | 26 | ATCAAGAACGAAAGTCGGAGGTTCGA | 140.9 | -10.2 | 1.6 | -6.4 | 1.8E-10 | 3.7E-09 | BT20/MB468 | MP |
| >trna10_AspGTC_12_-_125424193_125424264@40.74.35,trna12_AspGTC_12_-_125411891_125411962@40.74.35,trna144_AspGTC_6_-_27551236_27551307@40.74.35,trna38_AspGTC_17_-_8125556_8125627@40.74.35,trna45_AspGTC_6_+_27447453_27447524@40.74.35,trna48_AspGTC_6_+_27471523_27471594@40.74.35,trna4_AspGTC_12_+_96429799_96429870@40.74.35,trna69_AspGTC_1_-_161440205_161440276@40.74.35,trna72_AspGTC_1_-_161432824_161432895@40.74.35,trna75_AspGTC_1_-_161425414_161425485@40.74.35,trna78_AspGTC_1_-_161418033_161418104@40.74.35,trna81_AspGTC_1_-_161410615_161410686@40.74.35\|CGGGAGACCGGGGTTCGATTCCCCGACGGGGAGCC | >trna10_AspGTC_12_-_125424193_125424264 | 40 | 74 | 35 | CGGGAGACCGGGGTTCGATTCCCCGACGGGGAGCC | 638.1 | 7.5 | 1.3 | 6.0 | 1.6E-09 | 3.7E-09 | MB231/MB468 | CYTO |
| >hg38_21_+_8433172_8446622_RNA45SN1_45S_50ntFlanks@6646.6667.22\|TCGTACGACTCTTAGCGGTGGA | >hg38_21_+_8433172_8446622_RNA45SN1_45S_50ntFlanks | 6646 | 6667 | 22 | TCGTACGACTCTTAGCGGTGGA | 2786.8 | 3.4 | 0.6 | 6.0 | 1.7E-09 | 3.7E-09 | MB231/MB468 | NUC |
| >hg38_21_+_8433172_8446622_RNA45SN1_45S_50ntFlanks@4472.4496.25\|CTTAGCTGAGTGTCCCGCGGGGCCC | >hg38_21_+_8433172_8446622_RNA45SN1_45S_50ntFlanks | 4472 | 4496 | 25 | CTTAGCTGAGTGTCCCGCGGGGCCC | 306.3 | -6.8 | 1.1 | -6.1 | 1.4E-09 | 3.7E-09 | BT20/MB468 | CYTO |
| >hg38_21_+_8433172_8446622_RNA45SN1_45S_50ntFlanks@11646.11676.31\|GGTGTTGACGCGATGTGATTTCTGCCCAGTG | >hg38_21_+_8433172_8446622_RNA45SN1_45S_50ntFlanks | 11646 | 11676 | 31 | GGTGTTGACGCGATGTGATTTCTGCCCAGTG | 101.4 | -10.5 | 1.7 | -6.0 | 1.6E-09 | 3.7E-09 | MB231/MB468 | MP |
| >hg38_21_+_8433172_8446622_RNA45SN1_45S_50ntFlanks@10351.10387.37\|CGAGAACTTTGAAGGCCGAAGTGGAGAAGGGTTCCAT | >hg38_21_+_8433172_8446622_RNA45SN1_45S_50ntFlanks | 10351 | 10387 | 37 | CGAGAACTTTGAAGGCCGAAGTGGAGAAGGGTTCCAT | 329.9 | -7.9 | 1.3 | -6.0 | 1.8E-09 | 3.7E-09 | MB231/MB468 | MITO |
| >hg38_21_+_8433172_8446622_RNA45SN1_45S_50ntFlanks@476.510.35\|TTCGTGATCGATGTGGTGACGTCGTGCTCTCCCGG | >hg38_21_+_8433172_8446622_RNA45SN1_45S_50ntFlanks | 476 | 510 | 35 | TTCGTGATCGATGTGGTGACGTCGTGCTCTCCCGG | 996.5 | 7.2 | 1.2 | 6.1 | 1.2E-09 | 3.7E-09 | BT20/MB468 | MITO |
| >hg38_21_+_8433172_8446622_RNA45SN1_45S_50ntFlanks@6773.6799.27\|TTCCTCCCGGGGCTACGCCTGTCTGAG | >hg38_21_+_8433172_8446622_RNA45SN1_45S_50ntFlanks | 6773 | 6799 | 27 | TTCCTCCCGGGGCTACGCCTGTCTGAG | 571.8 | -8.6 | 1.4 | -6.0 | 1.9E-09 | 3.7E-09 | BT20/MB468 | NUC |
| >MI0000767\|hsa-mir-365a&WithFlank&16\|+\|14309279\|14309377@62.84.23;MI0000769\|hsa-mir-365b&WithFlank&17\|+\|31575405\|31575527@74.96.23&[MIMAT0000710&hsa-miR-365a-3p&offsets\|0\|+1;m-176&16\|+\|14309340\|14309361&offsets\|0\|+1];[MIMAT0022834&hsa-miR-365b-3p&offsets\|0\|+1;m-175&17\|+\|31575478\|31575499&offsets\|0\|+1]\|TAATGCCCCTAAAAATCCTTATT | >MI0000767\|hsa-mir-365a&WithFlank&16\|+\|14309279\|14309377 | 62 | 84 | 23 | TAATGCCCCTAAAAATCCTTATT | 579.8 | 3.5 | 0.6 | 6.4 | 1.7E-10 | 3.7E-09 | BT20/MB231 | NUC |
| >hg38_21_+_8433172_8446622_RNA45SN1_45S_50ntFlanks@3861.3898.38\|TAATTCTAGAGCTAATACATGCCGACGGGCGCTGACCC | >hg38_21_+_8433172_8446622_RNA45SN1_45S_50ntFlanks | 3861 | 3898 | 38 | TAATTCTAGAGCTAATACATGCCGACGGGCGCTGACCC | 667.3 | -9.3 | 1.6 | -6.0 | 1.9E-09 | 3.7E-09 | BT20/MB468 | NUC |

| | | | | | | | | | | | | | |
|---|---|---|---|---|---|---|---|---|---|---|---|---|---|
| >hg38_21_+_8433172_8446622_RNA45SN1_45S_50ntFlanks@7973.8008.36\|GACGCGACCTCAGATCAGACGTGGCGACCCGCTGAA | >hg38_21_+_8433172_8446622_RNA45SN1_45S_50ntFlanks | 7973 | 8008 | 36 | GACGCGACCTCAGATCAGACGTGGCGACCCGCTGAA | 1055.7 | 4.3 | 0.7 | 6.0 | 1.7E-09 | 3.7E-09 | MB231/MB468 | CYTO |
| >hg38_21_+_8433172_8446622_RNA45SN1_45S_50ntFlanks@11200.11232.33\|CGGCGGCGGCGGCGACTCTGGACGCGAGCCGGG | >hg38_21_+_8433172_8446622_RNA45SN1_45S_50ntFlanks | 11200 | 11232 | 33 | CGGCGGCGGCGGCGACTCTGGACGCGAGCCGGG | 1528.2 | -7.1 | 1.2 | -6.0 | 1.8E-09 | 3.7E-09 | MB231/MB468 | MITO |
| >hg38_21_+_8433172_8446622_RNA45SN1_45S_50ntFlanks@3854.3898.45\|ACTGTGGTAATTCTAGAGCTAATACATGCCGACGGGCGCTGACCC | >hg38_21_+_8433172_8446622_RNA45SN1_45S_50ntFlanks | 3854 | 3898 | 45 | ACTGTGGTAATTCTAGAGCTAATACATGCCGACGGGCGCTGACCC | 352.8 | -12.3 | 2.0 | -6.0 | 1.6E-09 | 3.7E-09 | MB231/MB468 | MP |
| >MI0000088\|hsa-mir-30a&WithFlank&6\|-\|71403545\|71403627@12.31.20&[MIMAT0000087&hsa-miR-30a-5p&offsets\|0\|-2;m-15&6\|-\|71403595\|71403616&offsets\|0\|-2]\|TGTAAACATCCTCGACTGGA | >MI0000088\|hsa-mir-30a&WithFlank&6\|-\|71403545\|71403627 | 12 | 31 | 20 | TGTAAACATCCTCGACTGGA | 742.4 | 6.1 | 1.0 | 6.0 | 1.7E-09 | 3.7E-09 | MB231/MB468 | CYTO |
| >hg38_21_+_8433172_8446622_RNA45SN1_45S_50ntFlanks@12435.12454.20\|ATCATTGTGAAGCAGAATTC | >hg38_21_+_8433172_8446622_RNA45SN1_45S_50ntFlanks | 12435 | 12454 | 20 | ATCATTGTGAAGCAGAATTC | 364.0 | -10.8 | 1.8 | -6.0 | 1.5E-09 | 3.7E-09 | BT20/MB468 | CYTO |
| >MI0000066\|hsa-let-7e&WithFlank&19\|+\|51692780\|51692870@14.34.21(+1G)&[MIMAT0000066&hsa-let-7e-5p&offsets\|0\|-1(+1G);m-50&19\|+\|51692793\|51692814&offsets\|0\|-1(+1G)]\|TGAGGTAGGAGGTTGTATAGTG | >MI0000066\|hsa-let-7e&WithFlank&19\|+\|51692780\|51692870 | 14 | 34 | 21(+1G) | TGAGGTAGGAGGTTGTATAGTG | 809.0 | 7.8 | 1.3 | 6.0 | 1.5E-09 | 3.7E-09 | BT20/MB468 | CYTO |
| >hg38_21_+_8433172_8446622_RNA45SN1_45S_50ntFlanks@9202.9222.21\|TCTCGGGGCCACGCGCGCGTC | >hg38_21_+_8433172_8446622_RNA45SN1_45S_50ntFlanks | 9202 | 9222 | 21 | TCTCGGGGCCACGCGCGCGTC | 914.2 | -8.8 | 1.4 | -6.4 | 1.8E-10 | 3.7E-09 | BT20/MB468 | MP |
| >hg38_21_+_8433172_8446622_RNA45SN1_45S_50ntFlanks@12490.12521.32\|CGTGAGCTGGGTTTAGACCGTCGTGAGACAGG | >hg38_21_+_8433172_8446622_RNA45SN1_45S_50ntFlanks | 12490 | 12521 | 32 | CGTGAGCTGGGTTTAGACCGTCGTGAGACAGG | 142.7 | -9.2 | 1.5 | -6.1 | 1.3E-09 | 3.7E-09 | BT20/MB468 | MITO |
| >hg38_21_+_8433172_8446622_RNA45SN1_45S_50ntFlanks@9556.9581.26\|TAGCGGTCCTGACGTGCAAATCGGTC | >hg38_21_+_8433172_8446622_RNA45SN1_45S_50ntFlanks | 9556 | 9581 | 26 | TAGCGGTCCTGACGTGCAAATCGGTC | 548.6 | -6.8 | 1.1 | -6.0 | 1.9E-09 | 3.7E-09 | BT20/MB468 | NUC |
| >hg38_21_+_8433172_8446622_RNA45SN1_45S_50ntFlanks@9352.9379.28\|CCGTGGCGCAATGAAGGTGAAGGCCGGC | >hg38_21_+_8433172_8446622_RNA45SN1_45S_50ntFlanks | 9352 | 9379 | 28 | CCGTGGCGCAATGAAGGTGAAGGCCGGC | 205.7 | -10.8 | 1.7 | -6.4 | 1.8E-10 | 3.7E-09 | BT20/MB468 | MP |
| >am_MI0000809\|hsa-mir-151a&WithFlank&8\|-\|140732558\|140732659@53.73.21&[MIMAT0000757&hsa-miR-151a-3p&offsets\|0\|0;m-30&8\|-\|140732587\|140732607&offsets\|0\|0]\|CTAGACTGAAGCTCCTTGAGG | >am_MI0000809\|hsa-mir-151a&WithFlank&8\|-\|140732558\|140732659 | 53 | 73 | 21 | CTAGACTGAAGCTCCTTGAGG | 20277.7 | 3.0 | 0.5 | 6.0 | 1.9E-09 | 3.8E-09 | BT20/MB468 | NUC |
| >hg38_21_+_8433172_8446622_RNA45SN1_45S_50ntFlanks@8327.8354.28\|ATAGTCAACAAGTACCGTAAGGGAAAGT | >hg38_21_+_8433172_8446622_RNA45SN1_45S_50ntFlanks | 8327 | 8354 | 28 | ATAGTCAACAAGTACCGTAAGGGAAAGT | 74.0 | -10.1 | 1.7 | -6.0 | 1.6E-09 | 3.8E-09 | MB231/MB468 | MP |
| >hg38_1_-_228634819_228635039_RNA5S12_5S_50ntFlanks@134.172.39\|TGGGAGACCGCCTGGGAATACCGGGTGCTGTAGGCTTTT | >hg38_1_-_228634819_228635039_RNA5S12_5S_50ntFlanks | 134 | 172 | 39 | TGGGAGACCGCCTGGGAATACCGGGTGCTGTAGGCTTTT | 821.8 | 7.1 | 1.2 | 6.0 | 1.7E-09 | 3.8E-09 | MB231/MB468 | NUC |
| >MI0000283\|hsa-mir-203a&WithFlank&14\|+\|104117399\|104117520@72.92.21(+1U)&[MIMAT0000264&hsa-miR-203a-3p&offsets\|+1\|0(+1U);m-79&14\|+\|104117469\|104117490&offsets\|+1\|0(+1U)]\|TGAAATGTTTAGGACCACTAGT | >MI0000283\|hsa-mir-203a&WithFlank&14\|+\|104117399\|104117520 | 72 | 92 | 21(+1U) | TGAAATGTTTAGGACCACTAGT | 678.1 | 9.7 | 1.6 | 6.1 | 1.3E-09 | 3.8E-09 | BT20/MB468 | MITO |
| >MI0000456\|hsa-mir-140&WithFlank&16\|+\|69933075\|69933186@29.50.22&[MIMAT0000431&hsa-miR-140-5p&offsets\|0\|0;m-173&16\|+\|69933103\|69933124&offsets\|0\|0]\|CAGTGGTTTTACCCTATGGTAG | >MI0000456\|hsa-mir-140&WithFlank&16\|+\|69933075\|69933186 | 29 | 50 | 22 | CAGTGGTTTTACCCTATGGTAG | 1116.5 | 5.3 | 0.9 | 6.0 | 1.6E-09 | 3.8E-09 | MB231/MB468 | MP |
| >hg38_21_+_8433172_8446622_RNA45SN1_45S_50ntFlanks@10841.10875.35\|CGTAACTTCGGGATAAGGATTGGCTCTAAGGGCTG | >hg38_21_+_8433172_8446622_RNA45SN1_45S_50ntFlanks | 10841 | 10875 | 35 | CGTAACTTCGGGATAAGGATTGGCTCTAAGGGCTG | 758.6 | -4.6 | 0.8 | -6.0 | 1.5E-09 | 3.8E-09 | BT20/MB468 | CYTO |
| >hg38_21_+_8433172_8446622_RNA45SN1_45S_50ntFlanks@8325.8356.32\|CGATAGTCAACAAGTACCGTAAGGGAAAGTTG | >hg38_21_+_8433172_8446622_RNA45SN1_45S_50ntFlanks | 8325 | 8356 | 32 | CGATAGTCAACAAGTACCGTAAGGGAAAGTTG | 207.6 | -6.8 | 1.1 | -6.0 | 1.5E-09 | 3.8E-09 | BT20/MB468 | CYTO |
| >hg38_21_+_8433172_8446622_RNA45SN1_45S_50ntFlanks@8114.8133.20\|GCGCGGGACATGTGGCGTAC | >hg38_21_+_8433172_8446622_RNA45SN1_45S_50ntFlanks | 8114 | 8133 | 20 | GCGCGGGACATGTGGCGTAC | 88.2 | -10.3 | 1.7 | -6.0 | 1.6E-09 | 3.8E-09 | MB231/MB468 | MP |
| >am_trna116_GluCTC_1_-_145399233_145399304@1.26.26,trna59_GluCTC_1_+_249168447_249168518@1.26.26,trna71_GluCTC_1_-_161439189_161439260@1.26.26,trna74_GluCTC_1_-_161431809_161431880@1.26.26,trna77_GluCTC_1_-_161424398_161424469@1.26.26,trna77_GluCTC_6_+_28949976_28950047@1.26.26,trna80_GluCTC_1_-_161417018_161417089@1.26.26,trna87_GluCTC_6_-_126101393_126101464@1.26.26\|TCCCTGGTGGTCTAGTGGTTAGGATT | >am_trna116_GluCTC_1_-_145399233_145399304 | 1 | 26 | 26 | TCCCTGGTGGTCTAGTGGTTAGGATT | 1440.7 | -4.4 | 0.7 | -6.0 | 1.9E-09 | 3.8E-09 | BT20/MB468 | NUC |
| >hg38_21_+_8433172_8446622_RNA45SN1_45S_50ntFlanks@4122.4157.36\|AGGGTTCGATTCCGGAGAGGGAGCCTGAGAAACGGC | >hg38_21_+_8433172_8446622_RNA45SN1_45S_50ntFlanks | 4122 | 4157 | 36 | AGGGTTCGATTCCGGAGAGGGAGCCTGAGAAACGGC | 83.5 | -10.2 | 1.7 | -6.0 | 1.6E-09 | 3.8E-09 | MB231/MB468 | MP |
| >hg38_21_+_8433172_8446622_RNA45SN1_45S_50ntFlanks@6645.6677.33\|CTCGTACGACTCTTAGCGGTGGATCACTCGGCT | >hg38_21_+_8433172_8446622_RNA45SN1_45S_50ntFlanks | 6645 | 6677 | 33 | CTCGTACGACTCTTAGCGGTGGATCACTCGGCT | 3639.0 | 4.2 | 0.7 | 6.1 | 1.3E-09 | 3.8E-09 | BT20/MB468 | MITO |
| >hg38_21_+_8433172_8446622_RNA45SN1_45S_50ntFlanks@11661.11678.18\|TGATTTCTGCCCAGTGCT | >hg38_21_+_8433172_8446622_RNA45SN1_45S_50ntFlanks | 11661 | 11678 | 18 | TGATTTCTGCCCAGTGCT | 364.5 | -7.9 | 1.3 | -6.0 | 1.5E-09 | 3.8E-09 | BT20/MB468 | CYTO |
| >hg38_21_+_8433172_8446622_RNA45SN1_45S_50ntFlanks@10346.10374.29\|TCAAACGAGAACTTTGAAGGCCGAAGTGG | >hg38_21_+_8433172_8446622_RNA45SN1_45S_50ntFlanks | 10346 | 10374 | 29 | TCAAACGAGAACTTTGAAGGCCGAAGTGG | 1309.9 | -3.2 | 0.5 | -6.1 | 1.3E-09 | 3.8E-09 | BT20/MB468 | MITO |
| >hg38_21_+_8433172_8446622_RNA45SN1_45S_50ntFlanks@11663.11685.23\|ATTTCTGCCCAGTGCTCTGAATG | >hg38_21_+_8433172_8446622_RNA45SN1_45S_50ntFlanks | 11663 | 11685 | 23 | ATTTCTGCCCAGTGCTCTGAATG | 1547.0 | -7.9 | 1.3 | -6.0 | 1.6E-09 | 3.8E-09 | MB231/MB468 | MP |
| >hg38_21_+_8433172_8446622_RNA45SN1_45S_50ntFlanks@10727.10751.25\|GGAGAGGGTGTAAATCTCGCGCCGG | >hg38_21_+_8433172_8446622_RNA45SN1_45S_50ntFlanks | 10727 | 10751 | 25 | GGAGAGGGTGTAAATCTCGCGCCGG | 242.4 | -11.0 | 1.7 | -6.4 | 1.8E-10 | 3.8E-09 | BT20/MB468 | MP |
| >hg38_21_+_8433172_8446622_RNA45SN1_45S_50ntFlanks@11729.11754.26\|TAACTATGACTCTCTTAAGGTAGCCA | >hg38_21_+_8433172_8446622_RNA45SN1_45S_50ntFlanks | 11729 | 11754 | 26 | TAACTATGACTCTCTTAAGGTAGCCA | 276.0 | -11.2 | 1.8 | -6.4 | 1.8E-10 | 3.8E-09 | BT20/MB468 | MP |
| >hg38_21_+_8433172_8446622_RNA45SN1_45S_50ntFlanks@4736.4758.23\|CGAAAGTCGGAGGTTCGAAGACG | >hg38_21_+_8433172_8446622_RNA45SN1_45S_50ntFlanks | 4736 | 4758 | 23 | CGAAAGTCGGAGGTTCGAAGACG | 721.3 | -3.9 | 0.7 | -6.0 | 1.7E-09 | 3.8E-09 | MB231/MB468 | NUC |
| >hg38_21_+_8433172_8446622_RNA45SN1_45S_50ntFlanks@4008.4025.18\|CTCGGGCCGATCGCACGC | >hg38_21_+_8433172_8446622_RNA45SN1_45S_50ntFlanks | 4008 | 4025 | 18 | CTCGGGCCGATCGCACGC | 400.6 | -5.2 | 0.9 | -6.0 | 1.9E-09 | 3.8E-09 | MB231/MB468 | MITO |
| >hg38_21_+_8433172_8446622_RNA45SN1_45S_50ntFlanks@10248.10285.38\|CGACGAGTAGGAGGGCCGCTGCGGTGAGCCTTGAAGCC | >hg38_21_+_8433172_8446622_RNA45SN1_45S_50ntFlanks | 10248 | 10285 | 38 | CGACGAGTAGGAGGGCCGCTGCGGTGAGCCTTGAAGCC | 128.1 | 10.7 | 1.7 | 6.4 | 1.8E-10 | 3.8E-09 | BT20/MB468 | MP |
| >hg38_MT_+_1621_3279_RNR2_16S_50ntFlanks@488.518.31\|GAACAGCTCTTTGGACACTAGGAAAAAACCT | >hg38_MT_+_1621_3279_RNR2_16S_50ntFlanks | 488 | 518 | 31 | GAACAGCTCTTTGGACACTAGGAAAAAACCT | 174.4 | 4.7 | 0.8 | 6.0 | 1.9E-09 | 3.8E-09 | MB231/MB468 | MITO |
| >hg38_21_+_8433172_8446622_RNA45SN1_45S_50ntFlanks@10468.10497.30\|TTGCCCTCGGCCGATCGAAAGGGAGTCGGG | >hg38_21_+_8433172_8446622_RNA45SN1_45S_50ntFlanks | 10468 | 10497 | 30 | TTGCCCTCGGCCGATCGAAAGGGAGTCGGG | 341.9 | -7.7 | 1.3 | -6.0 | 1.9E-09 | 3.8E-09 | MB231/MB468 | MITO |
| >hg38_21_+_8433172_8446622_RNA45SN1_45S_50ntFlanks@10409.10429.21\|GTCAGTCGGTCCTGAGAGATG | >hg38_21_+_8433172_8446622_RNA45SN1_45S_50ntFlanks | 10409 | 10429 | 21 | GTCAGTCGGTCCTGAGAGATG | 707.7 | -7.0 | 1.2 | -6.0 | 1.6E-09 | 3.8E-09 | MB231/MB468 | MP |
| >trna116_GluCTC_1_-_145399233_145399304@1.40.40,trna59_GluCTC_1_+_249168447_249168518@1.40.40,trna71_GluCTC_1_-_161439189_161439260@1.40.40,trna74_GluCTC_1_-_161431809_161431880@1.40.40,trna77_GluCTC_1_-_161424398_161424469@1.40.40,trna77_GluCTC_6_+_28949976_28950047@1.40.40,trna80_GluCTC_1_-_161417018_161417089@1.40.40,trna87_GluCTC_6_-_126101393_126101464@1.40.40\|TCCCTGGTGGTCTAGTGGTTAGGATTCGGCGCTCTCACCG | >trna116_GluCTC_1_-_145399233_145399304 | 1 | 40 | 40 | TCCCTGGTGGTCTAGTGGTTAGGATTCGGCGCTCTCACCG | 706.9 | 9.2 | 1.5 | 6.1 | 1.3E-09 | 3.8E-09 | BT20/MB468 | MITO |
| >hg38_21_+_8433172_8446622_RNA45SN1_45S_50ntFlanks@5059.5080.22\|CGATTTGTCTGGTTAATTCCGA | >hg38_21_+_8433172_8446622_RNA45SN1_45S_50ntFlanks | 5059 | 5080 | 22 | CGATTTGTCTGGTTAATTCCGA | 159.4 | -11.2 | 1.9 | -6.0 | 1.6E-09 | 3.8E-09 | MB231/MB468 | MP |
| >hg38_21_+_8433172_8446622_RNA45SN1_45S_50ntFlanks@6722.6744.23\|ATTGCAGGACACATTGATCATCG | >hg38_21_+_8433172_8446622_RNA45SN1_45S_50ntFlanks | 6722 | 6744 | 23 | ATTGCAGGACACATTGATCATCG | 650.0 | -7.9 | 1.3 | -6.0 | 1.9E-09 | 3.8E-09 | MB231/MB468 | MITO |
| >hg38_21_+_8433172_8446622_RNA45SN1_45S_50ntFlanks@11569.11611.43\|CAGCCGACTTAGAACTGGTGCGGACCAGGGGAATCCGACTGTT | >hg38_21_+_8433172_8446622_RNA45SN1_45S_50ntFlanks | 11569 | 11611 | 43 | CAGCCGACTTAGAACTGGTGCGGACCAGGGGAATCCGACTGTT | 1229.2 | -6.4 | 1.1 | -6.0 | 2.0E-09 | 3.8E-09 | BT20/MB468 | NUC |
| >hg38_21_+_8433172_8446622_RNA45SN1_45S_50ntFlanks@10786.10813.28\|AACAGCCTCTGGCATGTTGGAACAATGT | >hg38_21_+_8433172_8446622_RNA45SN1_45S_50ntFlanks | 10786 | 10813 | 28 | AACAGCCTCTGGCATGTTGGAACAATGT | 204.9 | -10.8 | 1.7 | -6.4 | 1.9E-10 | 3.8E-09 | BT20/MB468 | MP |
| >hg38_21_+_8433172_8446622_RNA45SN1_45S_50ntFlanks@9032.9056.25\|CCTCGTGGGGGGGCCGGGCCACCCC | >hg38_21_+_8433172_8446622_RNA45SN1_45S_50ntFlanks | 9032 | 9056 | 25 | CCTCGTGGGGGGGCCGGGCCACCCC | 156.6 | -10.4 | 1.6 | -6.4 | 1.9E-10 | 3.8E-09 | BT20/MB468 | MP |
| >hg38_21_+_8433172_8446622_RNA45SN1_45S_50ntFlanks@6738.6758.21\|ATCATCGACACTTCGAACGCA | >hg38_21_+_8433172_8446622_RNA45SN1_45S_50ntFlanks | 6738 | 6758 | 21 | ATCATCGACACTTCGAACGCA | 265.4 | -6.6 | 1.1 | -6.0 | 1.7E-09 | 3.8E-09 | MB231/MB468 | CYTO |
| >hg38_21_+_8433172_8446622_RNA45SN1_45S_50ntFlanks@7990.8007.18\|GACGTGGCGACCCGCTGA | >hg38_21_+_8433172_8446622_RNA45SN1_45S_50ntFlanks | 7990 | 8007 | 18 | GACGTGGCGACCCGCTGA | 796.6 | -8.3 | 1.4 | -6.0 | 1.6E-09 | 3.8E-09 | MB231/MB468 | MP |
| >hg38_21_+_8433172_8446622_RNA45SN1_45S_50ntFlanks@5524.5550.27\|GTAAAAGTCGTAACAAGGTTTCCGTAG | >hg38_21_+_8433172_8446622_RNA45SN1_45S_50ntFlanks | 5524 | 5550 | 27 | GTAAAAGTCGTAACAAGGTTTCCGTAG | 291.8 | -5.0 | 0.8 | -6.0 | 1.7E-09 | 3.8E-09 | MB231/MB468 | CYTO |
| >hg38_21_+_8433172_8446622_RNA45SN1_45S_50ntFlanks@12258.12282.25\|GCTTGATCTTGATTTTCAGTACGAA | >hg38_21_+_8433172_8446622_RNA45SN1_45S_50ntFlanks | 12258 | 12282 | 25 | GCTTGATCTTGATTTTCAGTACGAA | 120.8 | -10.8 | 1.8 | -6.0 | 1.7E-09 | 3.8E-09 | MB231/MB468 | MP |
| >trnaMT_PheGAA_MT_+_577_647@35.67.33\|GAAAATGTTTAGACGGGCTCACATCACCCCATA | >trnaMT_PheGAA_MT_+_577_647 | 35 | 67 | 33 | GAAAATGTTTAGACGGGCTCACATCACCCCATA | 466.3 | 12.6 | 2.0 | 6.4 | 1.9E-10 | 3.9E-09 | BT20/MB468 | MP |
| >hg38_21_+_8433172_8446622_RNA45SN1_45S_50ntFlanks@3990.4023.34\|TTGGTGACTCTAGATAACCTCGGGCCGATCGCAC | >hg38_21_+_8433172_8446622_RNA45SN1_45S_50ntFlanks | 3990 | 4023 | 34 | TTGGTGACTCTAGATAACCTCGGGCCGATCGCAC | 522.3 | 3.9 | 0.6 | 6.1 | 1.3E-09 | 3.9E-09 | BT20/MB468 | MITO |
| >hg38_21_+_8433172_8446622_RNA45SN1_45S_50ntFlanks@12396.12417.22\|GCGACGTCGCTTTTTGATCCTT | >hg38_21_+_8433172_8446622_RNA45SN1_45S_50ntFlanks | 12396 | 12417 | 22 | GCGACGTCGCTTTTTGATCCTT | 2229.9 | -7.5 | 1.2 | -6.0 | 1.9E-09 | 3.9E-09 | MB231/MB468 | MITO |
| >trna15_ValAAC_5_-_180615416_180615488@1.33.33\|GTTTCCGTAGTGTAGTGGTCATCACGTTCGCCT | >trna15_ValAAC_5_-_180615416_180615488 | 1 | 33 | 33 | GTTTCCGTAGTGTAGTGGTCATCACGTTCGCCT | 169.2 | 10.6 | 1.8 | 6.0 | 1.7E-09 | 3.9E-09 | MB231/MB468 | MP |
| >hg38_21_+_8433172_8446622_RNA45SN1_45S_50ntFlanks@4082.4099.18\|TCGCCGTGCCTACCATGG | >hg38_21_+_8433172_8446622_RNA45SN1_45S_50ntFlanks | 4082 | 4099 | 18 | TCGCCGTGCCTACCATGG | 70.1 | -10.0 | 1.7 | -6.0 | 1.7E-09 | 3.9E-09 | MB231/MB468 | MP |

<table>
<tr><td>>hg38_21_+_8433172_8446622_RNA45SN1_45S_50ntFlanks@9465.9500.36|AGGTGGAGCACGAGCGCACGTGTTAGGACCCGAAAG</td><td>>hg38_21_+_8433172_8446622_RNA45SN1_45S_50ntFlanks</td><td>9465</td><td>9500</td><td>36</td><td>AGGTGGAGCACGAGCGCACGTGTTAGGACCCGAAAG</td><td>53.5</td><td>-9.6</td><td>1.6</td><td>-6.0</td><td>1.7E-09</td><td>3.9E-09</td><td>MB231/MB468</td><td>MP</td></tr>
<tr><td>>hg38_21_+_8433172_8446622_RNA45SN1_45S_50ntFlanks@9738.9766.29|GAAACGATCTCAACCTATTCTCAAACTTT</td><td>>hg38_21_+_8433172_8446622_RNA45SN1_45S_50ntFlanks</td><td>9738</td><td>9766</td><td>29</td><td>GAAACGATCTCAACCTATTCTCAAACTTT</td><td>1257.4</td><td>-4.6</td><td>0.8</td><td>-6.1</td><td>1.3E-09</td><td>3.9E-09</td><td>BT20/MB468</td><td>MITO</td></tr>
<tr><td>>MI0000439|hsa-mir-23b&WithFlank&9|+|95085202|95085310@65.86.22(+1U)&[MIMAT0000418&hsa-miR-23b-3p&offsets|+1|0(+1U);m-42&9|+|95085265|95085285&offsets|+1|+2(+1U)]|TCACATTGCCAGGGATTACCACT</td><td>>MI0000439|hsa-mir-23b&WithFlank&9|+|95085202|95085310</td><td>65</td><td>86</td><td>22(+1U)</td><td>TCACATTGCCAGGGATTACCACT</td><td>444.3</td><td>6.8</td><td>1.1</td><td>6.0</td><td>1.7E-09</td><td>3.9E-09</td><td>MB231/MB468</td><td>CYTO</td></tr>
<tr><td>>hg38_21_+_8433172_8446622_RNA45SN1_45S_50ntFlanks@10759.10785.27|CCATATCCGCAGCAGGTCTCCAAGGTG</td><td>>hg38_21_+_8433172_8446622_RNA45SN1_45S_50ntFlanks</td><td>10759</td><td>10785</td><td>27</td><td>CCATATCCGCAGCAGGTCTCCAAGGTG</td><td>68.0</td><td>-9.9</td><td>1.7</td><td>-6.0</td><td>1.7E-09</td><td>3.9E-09</td><td>MB231/MB468</td><td>MP</td></tr>
<tr><td>>trna111_HisGTG_1_-_147774845_147774916@1.33.33,trna118_HisGTG_1_-_145396881_145396952@1.33.33,trna16_HisGTG_1_+_146544773_146544844@1.33.33,trna1_HisGTG_15_+_45493349_45493420@1.33.33,trna21_HisGTG_1_+_147753471_147753542@1.33.33,trna33_HisGTG_6_+_27125906_27125977@1.33.33,trna7_HisGTG_9_-_14433938_14434009@1.33.33,trna8_HisGTG_15_-_45492611_45492682@1.33.33,trna9_HisGTG_15_-_45490804_45490875@1.33.33|GCCGTGATCGTATAGTGGTTAGTACTCTGCGTT</td><td>>trna111_HisGTG_1_-_147774845_147774916</td><td>1</td><td>33</td><td>33</td><td>GCCGTGATCGTATAGTGGTTAGTACTCTGCGTT</td><td>1510.8</td><td>5.6</td><td>0.9</td><td>6.0</td><td>1.8E-09</td><td>3.9E-09</td><td>MB231/MB468</td><td>NUC</td></tr>
<tr><td>>hg38_21_+_8433172_8446622_RNA45SN1_45S_50ntFlanks@11562.11602.41|CGCCTAGCAGCCGACTTAGAACTGGTGCGGACCAGGGGAAT</td><td>>hg38_21_+_8433172_8446622_RNA45SN1_45S_50ntFlanks</td><td>11562</td><td>11602</td><td>41</td><td>CGCCTAGCAGCCGACTTAGAACTGGTGCGGACCAGGGGAAT</td><td>517.9</td><td>-6.6</td><td>1.1</td><td>-6.0</td><td>1.9E-09</td><td>3.9E-09</td><td>MB231/MB468</td><td>MITO</td></tr>
<tr><td>>hg38_21_+_8433172_8446622_RNA45SN1_45S_50ntFlanks@12933.12956.24|GCTAAACCATTCGTAGACGACCTG</td><td>>hg38_21_+_8433172_8446622_RNA45SN1_45S_50ntFlanks</td><td>12933</td><td>12956</td><td>24</td><td>GCTAAACCATTCGTAGACGACCTG</td><td>866.7</td><td>-8.0</td><td>1.3</td><td>-6.0</td><td>1.9E-09</td><td>3.9E-09</td><td>MB231/MB468</td><td>MITO</td></tr>
<tr><td>>hg38_21_+_8433172_8446622_RNA45SN1_45S_50ntFlanks@3854.3889.36|ACTGTGGTAATTCTAGAGCTAATACATGCCGACGGG</td><td>>hg38_21_+_8433172_8446622_RNA45SN1_45S_50ntFlanks</td><td>3854</td><td>3889</td><td>36</td><td>ACTGTGGTAATTCTAGAGCTAATACATGCCGACGGG</td><td>593.0</td><td>-6.3</td><td>1.0</td><td>-6.0</td><td>1.8E-09</td><td>3.9E-09</td><td>MB231/MB468</td><td>NUC</td></tr>
<tr><td>>hg38_21_+_8433172_8446622_RNA45SN1_45S_50ntFlanks@12615.12661.47|GCTGAGGAGCCAATGGGGCGAAGCTACCATCTGTGGGATTATGACTG</td><td>>hg38_21_+_8433172_8446622_RNA45SN1_45S_50ntFlanks</td><td>12615</td><td>12661</td><td>47</td><td>GCTGAGGAGCCAATGGGGCGAAGCTACCATCTGTGGGATTATGACTG</td><td>318.4</td><td>-12.2</td><td>2.0</td><td>-6.0</td><td>1.7E-09</td><td>3.9E-09</td><td>MB231/MB468</td><td>MP</td></tr>
<tr><td>>hg38_21_+_8433172_8446622_RNA45SN1_45S_50ntFlanks@9534.9561.28|AGGAAACTCTGGTGGAGGTCCGTAGCGG</td><td>>hg38_21_+_8433172_8446622_RNA45SN1_45S_50ntFlanks</td><td>9534</td><td>9561</td><td>28</td><td>AGGAAACTCTGGTGGAGGTCCGTAGCGG</td><td>2660.7</td><td>-5.2</td><td>0.9</td><td>-6.0</td><td>1.5E-09</td><td>3.9E-09</td><td>BT20/MB468</td><td>CYTO</td></tr>
<tr><td>>MI0000088|hsa-mir-30a&WithFlank&6|-|71403545|71403627@12.32.21(+1A)&[MIMAT0000087&hsa-miR-30a-5p&offsets|0|-1(+1A);m-15&6|-|71403595|71403616&offsets|0|-1(+1A)]|TGTAAACATCCTCGACTGGAAA</td><td>>MI0000088|hsa-mir-30a&WithFlank&6|-|71403545|71403627</td><td>12</td><td>32</td><td>21(+1A)</td><td>TGTAAACATCCTCGACTGGAAA</td><td>336.8</td><td>8.1</td><td>1.3</td><td>6.0</td><td>1.5E-09</td><td>3.9E-09</td><td>BT20/MB468</td><td>CYTO</td></tr>
<tr><td>>hg38_21_+_8433172_8446622_RNA45SN1_45S_50ntFlanks@11729.11766.38|TAACTATGACTCTCTTAAGGTAGCCAAATGCCTCGTCA</td><td>>hg38_21_+_8433172_8446622_RNA45SN1_45S_50ntFlanks</td><td>11729</td><td>11766</td><td>38</td><td>TAACTATGACTCTCTTAAGGTAGCCAAATGCCTCGTCA</td><td>679.8</td><td>-6.2</td><td>1.0</td><td>-6.0</td><td>1.7E-09</td><td>3.9E-09</td><td>MB231/MB468</td><td>CYTO</td></tr>
<tr><td>>hg38_21_+_8433172_8446622_RNA45SN1_45S_50ntFlanks@12990.13015.26|GCTCCCTCGCTGCGATCTATTGAAAG</td><td>>hg38_21_+_8433172_8446622_RNA45SN1_45S_50ntFlanks</td><td>12990</td><td>13015</td><td>26</td><td>GCTCCCTCGCTGCGATCTATTGAAAG</td><td>1988.3</td><td>-9.9</td><td>1.6</td><td>-6.0</td><td>2.0E-09</td><td>3.9E-09</td><td>BT20/MB468</td><td>NUC</td></tr>
<tr><td>>hg38_21_+_8433172_8446622_RNA45SN1_45S_50ntFlanks@10512.10531.20|TCCGGAGTGGCGGAGATGGG</td><td>>hg38_21_+_8433172_8446622_RNA45SN1_45S_50ntFlanks</td><td>10512</td><td>10531</td><td>20</td><td>TCCGGAGTGGCGGAGATGGG</td><td>139.7</td><td>-11.0</td><td>1.8</td><td>-6.0</td><td>1.7E-09</td><td>3.9E-09</td><td>MB231/MB468</td><td>MP</td></tr>
<tr><td>>hg38_21_+_8433172_8446622_RNA45SN1_45S_50ntFlanks@4129.4162.34|GATTCCGGAGAGGGAGCCTGAGAAACGGCTACCA</td><td>>hg38_21_+_8433172_8446622_RNA45SN1_45S_50ntFlanks</td><td>4129</td><td>4162</td><td>34</td><td>GATTCCGGAGAGGGAGCCTGAGAAACGGCTACCA</td><td>533.0</td><td>-3.6</td><td>0.6</td><td>-6.0</td><td>1.7E-09</td><td>3.9E-09</td><td>MB231/MB468</td><td>CYTO</td></tr>
<tr><td>>hg38_21_+_8433172_8446622_RNA45SN1_45S_50ntFlanks@11604.11621.18|CGACTGTTTAATTAAAAC</td><td>>hg38_21_+_8433172_8446622_RNA45SN1_45S_50ntFlanks</td><td>11604</td><td>11621</td><td>18</td><td>CGACTGTTTAATTAAAAC</td><td>761.6</td><td>-6.4</td><td>1.1</td><td>-6.0</td><td>1.7E-09</td><td>3.9E-09</td><td>MB231/MB468</td><td>MP</td></tr>
<tr><td>>trnaMT_PheGAA_MT_+_577_647@35.60.26|GAAAATGTTTAGACGGGCTCACATCA</td><td>>trnaMT_PheGAA_MT_+_577_647</td><td>35</td><td>60</td><td>26</td><td>GAAAATGTTTAGACGGGCTCACATCA</td><td>1681.6</td><td>9.3</td><td>1.5</td><td>6.1</td><td>1.3E-09</td><td>3.9E-09</td><td>BT20/MB468</td><td>MITO</td></tr>
<tr><td>>hg38_21_+_8433172_8446622_RNA45SN1_45S_50ntFlanks@11345.11367.23|CTGGGGGCGGGGAGCGGTCGGGC</td><td>>hg38_21_+_8433172_8446622_RNA45SN1_45S_50ntFlanks</td><td>11345</td><td>11367</td><td>23</td><td>CTGGGGGCGGGGAGCGGTCGGGC</td><td>105.9</td><td>-10.6</td><td>1.8</td><td>-6.0</td><td>1.7E-09</td><td>3.9E-09</td><td>MB231/MB468</td><td>MP</td></tr>
<tr><td>>trnaMT_GluTTC_MT_-_14674_14742@40.63.24,trnalookalike8_GluTTC_5_-_93905172_93905240@40.63.24|TTGGTCGTGGTTGTAGTCCGTGCG</td><td>>trnaMT_GluTTC_MT_-_14674_14742</td><td>40</td><td>63</td><td>24</td><td>TTGGTCGTGGTTGTAGTCCGTGCG</td><td>225.3</td><td>4.3</td><td>0.7</td><td>6.0</td><td>1.8E-09</td><td>3.9E-09</td><td>MB231/MB468</td><td>NUC</td></tr>
<tr><td>>hg38_21_+_8433172_8446622_RNA45SN1_45S_50ntFlanks@4444.4471.28|CCTTGCCTCTCGGCGCCCCCTCGATGCT</td><td>>hg38_21_+_8433172_8446622_RNA45SN1_45S_50ntFlanks</td><td>4444</td><td>4471</td><td>28</td><td>CCTTGCCTCTCGGCGCCCCCTCGATGCT</td><td>350.8</td><td>-6.9</td><td>1.1</td><td>-6.1</td><td>1.3E-09</td><td>3.9E-09</td><td>BT20/MB468</td><td>MITO</td></tr>
<tr><td>>MI0003834|hsa-mir-769&WithFlank&19|+|46018926|46019055@36.57.22&[MIMAT0003886&hsa-miR-769-5p&offsets|0|0;m-216&19|+|46018961|46018982&offsets|0|0]|TGAGACCTCTGGGTTCTGAGCT</td><td>>MI0003834|hsa-mir-769&WithFlank&19|+|46018926|46019055</td><td>36</td><td>57</td><td>22</td><td>TGAGACCTCTGGGTTCTGAGCT</td><td>882.8</td><td>6.0</td><td>1.0</td><td>6.0</td><td>1.8E-09</td><td>3.9E-09</td><td>MB231/MB468</td><td>CYTO</td></tr>
<tr><td>>hg38_21_+_8433172_8446622_RNA45SN1_45S_50ntFlanks@12893.12927.35|CCTCGCCCGTCACGCACCGCACGTTCGTGGGGAAC</td><td>>hg38_21_+_8433172_8446622_RNA45SN1_45S_50ntFlanks</td><td>12893</td><td>12927</td><td>35</td><td>CCTCGCCCGTCACGCACCGCACGTTCGTGGGGAAC</td><td>126.2</td><td>-9.1</td><td>1.5</td><td>-6.1</td><td>1.3E-09</td><td>3.9E-09</td><td>BT20/MB468</td><td>MITO</td></tr>
<tr><td>>hg38_21_+_8433172_8446622_RNA45SN1_45S_50ntFlanks@12007.12043.37|GCCGCCGGTGAAATACCACTACTCTGATCGTTTTTTC</td><td>>hg38_21_+_8433172_8446622_RNA45SN1_45S_50ntFlanks</td><td>12007</td><td>12043</td><td>37</td><td>GCCGCCGGTGAAATACCACTACTCTGATCGTTTTTTC</td><td>6826.7</td><td>-11.2</td><td>1.9</td><td>-6.0</td><td>2.0E-09</td><td>4.0E-09</td><td>BT20/MB468</td><td>NUC</td></tr>
<tr><td>>MI0000455|hsa-mir-138-2&WithFlank&16|+|56858512|56858607@16.36.21;MI0000476|hsa-mir-138-1&WithFlank&3|+|44114206|44114316@29.49.21&[MIMAT0000430&hsa-miR-138-5p&offsets|0|-2;m-509&16|+|56858527|56858549&offsets|0|-2];[MIMAT0000430_1&hsa-miR-138-5p&offsets|0|-2;m-497&3|+|44114234|44114256&offsets|0|-2]|AGCTGGTGTTGTGAATCAGGC</td><td>>MI0000455|hsa-mir-138-2&WithFlank&16|+|56858512|56858607</td><td>16</td><td>36</td><td>21</td><td>AGCTGGTGTTGTGAATCAGGC</td><td>80.2</td><td>9.5</td><td>1.6</td><td>6.0</td><td>1.7E-09</td><td>4.0E-09</td><td>MB231/MB468</td><td>MP</td></tr>
<tr><td>>hg38_21_+_8433172_8446622_RNA45SN1_45S_50ntFlanks@4729.4765.37|TCAAGAACGAAAGTCGGAGGTTCGAAGACGATCAGAT</td><td>>hg38_21_+_8433172_8446622_RNA45SN1_45S_50ntFlanks</td><td>4729</td><td>4765</td><td>37</td><td>TCAAGAACGAAAGTCGGAGGTTCGAAGACGATCAGAT</td><td>389.2</td><td>6.5</td><td>1.1</td><td>6.0</td><td>1.5E-09</td><td>4.0E-09</td><td>BT20/MB468</td><td>CYTO</td></tr>
<tr><td>>hg38_21_+_8433172_8446622_RNA45SN1_45S_50ntFlanks@10395.10424.30|GCAGTTGAACATGGGTCAGTCGGTCCTGAG</td><td>>hg38_21_+_8433172_8446622_RNA45SN1_45S_50ntFlanks</td><td>10395</td><td>10424</td><td>30</td><td>GCAGTTGAACATGGGTCAGTCGGTCCTGAG</td><td>1664.4</td><td>4.9</td><td>0.8</td><td>6.0</td><td>1.8E-09</td><td>4.0E-09</td><td>MB231/MB468</td><td>CYTO</td></tr>
<tr><td>>am_MI0000809|hsa-mir-151a&WithFlank&8|-|140732558|140732659@53.73.21&[MIMAT0000757&hsa-miR-151a-3p&offsets|0|0;m-30&8|-|140732587|140732607&offsets|0|0]|CTAGACTGAAGCTCCTTGAGG</td><td>>am_MI0000809|hsa-mir-151a&WithFlank&8|-|140732558|140732659</td><td>53</td><td>73</td><td>21</td><td>CTAGACTGAAGCTCCTTGAGG</td><td>104342.6</td><td>5.9</td><td>0.9</td><td>6.4</td><td>1.9E-10</td><td>4.0E-09</td><td>BT20/MB468</td><td>MP</td></tr>
<tr><td>>trna10_AspGTC_12_-_125424193_125424264@32.72.41,trna12_AspGTC_12_-_125411891_125411962@32.72.41,trna144_AspGTC_6_-_27551236_27551307@32.72.41,trna38_AspGTC_17_-_8125556_8125627@32.72.41,trna45_AspGTC_6_+_27447453_27447524@32.72.41,trna48_AspGTC_6_+_27471523_27471594@32.72.41,trna4_AspGTC_12_+_96429799_96429870@32.72.41,trna69_AspGTC_1_-_161440205_161440276@32.72.41,trna72_AspGTC_1_-_161432824_161432895@32.72.41,trna75_AspGTC_1_-_161425414_161425485@32.72.41,trna78_AspGTC_1_-_161418033_161418104@32.72.41,trna81_AspGTC_1_-_161410615_161410686@32.72.41|CTGTCACGCGGGAGACCGGGGTTCGATTCCCCGACGGGGAG</td><td>>trna10_AspGTC_12_-_125424193_125424264</td><td>32</td><td>72</td><td>41</td><td>CTGTCACGCGGGAGACCGGGGTTCGATTCCCCGACGGGGAG</td><td>268.8</td><td>-11.2</td><td>1.8</td><td>-6.4</td><td>1.9E-10</td><td>4.0E-09</td><td>BT20/MB468</td><td>MP</td></tr>
<tr><td>>hg38_21_+_8433172_8446622_RNA45SN1_45S_50ntFlanks@6649.6667.19|TACGACTCTTAGCGGTGGA</td><td>>hg38_21_+_8433172_8446622_RNA45SN1_45S_50ntFlanks</td><td>6649</td><td>6667</td><td>19</td><td>TACGACTCTTAGCGGTGGA</td><td>998.1</td><td>5.7</td><td>0.9</td><td>6.1</td><td>1.3E-09</td><td>4.0E-09</td><td>BT20/MB468</td><td>MITO</td></tr>
<tr><td>>hg38_21_+_8433172_8446622_RNA45SN1_45S_50ntFlanks@5421.5444.24|CGATTGGATGGTTTAGTGAGGCCC</td><td>>hg38_21_+_8433172_8446622_RNA45SN1_45S_50ntFlanks</td><td>5421</td><td>5444</td><td>24</td><td>CGATTGGATGGTTTAGTGAGGCCC</td><td>124.5</td><td>-6.0</td><td>1.0</td><td>-6.0</td><td>1.6E-09</td><td>4.0E-09</td><td>BT20/MB468</td><td>CYTO</td></tr>
<tr><td>>hg38_21_+_8433172_8446622_RNA45SN1_45S_50ntFlanks@5523.5550.28|AGTAAAAGTCGTAACAAGGTTTCCGTAG</td><td>>hg38_21_+_8433172_8446622_RNA45SN1_45S_50ntFlanks</td><td>5523</td><td>5550</td><td>28</td><td>AGTAAAAGTCGTAACAAGGTTTCCGTAG</td><td>1059.6</td><td>-8.1</td><td>1.3</td><td>-6.0</td><td>1.8E-09</td><td>4.0E-09</td><td>MB231/MB468</td><td>NUC</td></tr>
<tr><td>>hg38_21_+_8433172_8446622_RNA45SN1_45S_50ntFlanks@12678.12704.27|AATCCCGCCCAGGCGGAACGATACGGC</td><td>>hg38_21_+_8433172_8446622_RNA45SN1_45S_50ntFlanks</td><td>12678</td><td>12704</td><td>27</td><td>AATCCCGCCCAGGCGGAACGATACGGC</td><td>228.4</td><td>-10.9</td><td>1.7</td><td>-6.4</td><td>1.9E-10</td><td>4.0E-09</td><td>BT20/MB468</td><td>MP</td></tr>
<tr><td>>hg38_21_+_8433172_8446622_RNA45SN1_45S_50ntFlanks@6743.6771.29|CGACACTTCGAACGCACTTGCGGCCCCGG</td><td>>hg38_21_+_8433172_8446622_RNA45SN1_45S_50ntFlanks</td><td>6743</td><td>6771</td><td>29</td><td>CGACACTTCGAACGCACTTGCGGCCCCGG</td><td>945.6</td><td>-6.6</td><td>1.1</td><td>-6.0</td><td>1.8E-09</td><td>4.0E-09</td><td>MB231/MB468</td><td>NUC</td></tr>
<tr><td>>hg38_21_+_8433172_8446622_RNA45SN1_45S_50ntFlanks@11876.11893.18|AAGACCCTGTTGAGCTTG</td><td>>hg38_21_+_8433172_8446622_RNA45SN1_45S_50ntFlanks</td><td>11876</td><td>11893</td><td>18</td><td>AAGACCCTGTTGAGCTTG</td><td>3052.3</td><td>-4.6</td><td>0.7</td><td>-6.4</td><td>2.0E-10</td><td>4.0E-09</td><td>BT20/MB468</td><td>MP</td></tr>
<tr><td>>hg38_21_+_8433172_8446622_RNA45SN1_45S_50ntFlanks@11915.11940.26|GAGACATGAGAGGTGTAGAATAAGTG</td><td>>hg38_21_+_8433172_8446622_RNA45SN1_45S_50ntFlanks</td><td>11915</td><td>11940</td><td>26</td><td>GAGACATGAGAGGTGTAGAATAAGTG</td><td>143.0</td><td>10.9</td><td>1.7</td><td>6.4</td><td>2.0E-10</td><td>4.0E-09</td><td>BT20/MB468</td><td>MP</td></tr>
<tr><td>>MI0000650|hsa-mir-200c&WithFlank&12|+|6963693|6963772@50.72.23(+1C)&[MIMAT0000617&hsa-miR-200c-3p&offsets|0|0(+1C);m-54&12|+|6963742|6963764&offsets|0|0(+1C)]|TAATACTGCCGGGTAATGATGGAC</td><td>>MI0000650|hsa-mir-200c&WithFlank&12|+|6963693|6963772</td><td>50</td><td>72</td><td>23(+1C)</td><td>TAATACTGCCGGGTAATGATGGAC</td><td>285.0</td><td>11.5</td><td>1.7</td><td>6.6</td><td>3.9E-11</td><td>4.0E-09</td><td>BT20/MB231</td><td>MP</td></tr>
<tr><td>>trna10_LysCTT_16_+_3241501_3241573@1.36.36,trna13_LysCTT_14_-_58706613_58706685@1.36.36,trna2_LysCTT_15_+_79152904_79152976@1.36.36|GCCCGGCTAGCTCAGTCGGTAGAGCATGGGACTCTT</td><td>>trna10_LysCTT_16_+_3241501_3241573</td><td>1</td><td>36</td><td>36</td><td>GCCCGGCTAGCTCAGTCGGTAGAGCATGGGACTCTT</td><td>924.1</td><td>5.1</td><td>0.8</td><td>6.0</td><td>1.8E-09</td><td>4.0E-09</td><td>MB231/MB468</td><td>CYTO</td></tr>
<tr><td>>hg38_21_+_8433172_8446622_RNA45SN1_45S_50ntFlanks@8178.8202.25|TCTGATCGAGGCCCAGCCCGTGGAC</td><td>>hg38_21_+_8433172_8446622_RNA45SN1_45S_50ntFlanks</td><td>8178</td><td>8202</td><td>25</td><td>TCTGATCGAGGCCCAGCCCGTGGAC</td><td>169.8</td><td>3.4</td><td>0.6</td><td>6.0</td><td>1.8E-09</td><td>4.0E-09</td><td>MB231/MB468</td><td>CYTO</td></tr>
<tr><td>>hg38_21_+_8433172_8446622_RNA45SN1_45S_50ntFlanks@10845.10866.22|ACTTCGGGATAAGGATTGGCTC</td><td>>hg38_21_+_8433172_8446622_RNA45SN1_45S_50ntFlanks</td><td>10845</td><td>10866</td><td>22</td><td>ACTTCGGGATAAGGATTGGCTC</td><td>928.6</td><td>-9.2</td><td>1.5</td><td>-6.0</td><td>2.1E-09</td><td>4.0E-09</td><td>BT20/MB468</td><td>NUC</td></tr>
<tr><td>>hg38_21_+_8433172_8446622_RNA45SN1_45S_50ntFlanks@6682.6699.18|CGTCGATGAAGAACGCAG</td><td>>hg38_21_+_8433172_8446622_RNA45SN1_45S_50ntFlanks</td><td>6682</td><td>6699</td><td>18</td><td>CGTCGATGAAGAACGCAG</td><td>87.3</td><td>-10.3</td><td>1.7</td><td>-6.0</td><td>1.7E-09</td><td>4.0E-09</td><td>MB231/MB468</td><td>MP</td></tr>
<tr><td>>hg38_21_+_8433172_8446622_RNA45SN1_45S_50ntFlanks@11576.11609.34|CTTAGAACTGGTGCGGACCAGGGGAATCCGACTG</td><td>>hg38_21_+_8433172_8446622_RNA45SN1_45S_50ntFlanks</td><td>11576</td><td>11609</td><td>34</td><td>CTTAGAACTGGTGCGGACCAGGGGAATCCGACTG</td><td>580.1</td><td>-4.0</td><td>0.7</td><td>-6.0</td><td>1.6E-09</td><td>4.0E-09</td><td>BT20/MB468</td><td>CYTO</td></tr>
<tr><td>>hg38_MT_+_1621_3279_RNR2_16S_50ntFlanks@1182.1215.34|AAAATTTCGGTTGGGGCGACCTCGGAGCAGAACC</td><td>>hg38_MT_+_1621_3279_RNR2_16S_50ntFlanks</td><td>1182</td><td>1215</td><td>34</td><td>AAAATTTCGGTTGGGGCGACCTCGGAGCAGAACC</td><td>699.0</td><td>8.7</td><td>1.4</td><td>6.1</td><td>1.4E-09</td><td>4.0E-09</td><td>BT20/MB468</td><td>MITO</td></tr>
<tr><td>>hg38_21_+_8433172_8446622_RNA45SN1_45S_50ntFlanks@10512.10533.22|TCCGGAGTGGCGGAGATGGGCG</td><td>>hg38_21_+_8433172_8446622_RNA45SN1_45S_50ntFlanks</td><td>10512</td><td>10533</td><td>22</td><td>TCCGGAGTGGCGGAGATGGGCG</td><td>1692.0</td><td>-8.4</td><td>1.4</td><td>-6.1</td><td>1.4E-09</td><td>4.0E-09</td><td>BT20/MB468</td><td>MITO</td></tr>
<tr><td>>hg38_21_+_8433172_8446622_RNA45SN1_45S_50ntFlanks@11581.11601.21|AACTGGTGCGGACCAGGGGAA</td><td>>hg38_21_+_8433172_8446622_RNA45SN1_45S_50ntFlanks</td><td>11581</td><td>11601</td><td>21</td><td>AACTGGTGCGGACCAGGGGAA</td><td>679.0</td><td>-6.2</td><td>1.0</td><td>-6.0</td><td>1.8E-09</td><td>4.0E-09</td><td>MB231/MB468</td><td>NUC</td></tr>
</table>

| | | | | | | | | | | | | | |
|---|---|---|---|---|---|---|---|---|---|---|---|---|---|
| >hg38_21_+_8433172_8446622_RNA45SN1_45S_50ntFlanks@4591.4616.26\|TTTTGTTGGTTTTCGGAACTGAGGCC | >hg38_21_+_8433172_8446622_RNA45SN1_45S_50ntFlanks | 4591 | 4616 | 26 | TTTTGTTGGTTTTCGGAACTGAGGCC | 493.7 | -8.6 | 1.4 | -6.0 | 2.1E-09 | 4.0E-09 | BT20/MB468 | NUC |
| >hg38_21_+_8433172_8446622_RNA45SN1_45S_50ntFlanks@12616.12657.42\|CTGAGGAGCCAATGGGGCGAAGCTACCATCTGTGGGATTATG | >hg38_21_+_8433172_8446622_RNA45SN1_45S_50ntFlanks | 12616 | 12657 | 42 | CTGAGGAGCCAATGGGGCGAAGCTACCATCTGTGGGATTATG | 724.0 | -8.3 | 1.4 | -6.0 | 2.0E-09 | 4.0E-09 | MB231/MB468 | MITO |
| >hg38_21_+_8433172_8446622_RNA45SN1_45S_50ntFlanks@10402.10433.32\|AACATGGGTCAGTCGGTCCTGAGAGATGGGCG | >hg38_21_+_8433172_8446622_RNA45SN1_45S_50ntFlanks | 10402 | 10433 | 32 | AACATGGGTCAGTCGGTCCTGAGAGATGGGCG | 1123.3 | -8.4 | 1.4 | -6.0 | 1.8E-09 | 4.0E-09 | MB231/MB468 | NUC |
| >hg38_21_+_8433172_8446622_RNA45SN1_45S_50ntFlanks@7975.7997.23\|CGCGACCTCAGATCAGACGTGGC | >hg38_21_+_8433172_8446622_RNA45SN1_45S_50ntFlanks | 7975 | 7997 | 23 | CGCGACCTCAGATCAGACGTGGC | 12068.5 | 5.0 | 0.8 | 6.0 | 1.8E-09 | 4.0E-09 | MB231/MB468 | CYTO |
| >hg38_21_+_8433172_8446622_RNA45SN1_45S_50ntFlanks@12456.12481.26\|CCAAGCGTTGGATTGTTCACCCACTA | >hg38_21_+_8433172_8446622_RNA45SN1_45S_50ntFlanks | 12456 | 12481 | 26 | CCAAGCGTTGGATTGTTCACCCACTA | 1239.9 | -6.3 | 1.1 | -6.0 | 1.8E-09 | 4.0E-09 | MB231/MB468 | CYTO |
| >hg38_21_+_8433172_8446622_RNA45SN1_45S_50ntFlanks@11873.11899.27\|AAGAAGACCCTGTTGAGCTTGACTCTA | >hg38_21_+_8433172_8446622_RNA45SN1_45S_50ntFlanks | 11873 | 11899 | 27 | AAGAAGACCCTGTTGAGCTTGACTCTA | 1007.6 | -9.1 | 1.5 | -6.0 | 2.1E-09 | 4.0E-09 | BT20/MB468 | NUC |
| >MI0000299\|hsa-mir-222&WithFlank&X\|-\|45747009\|45747130@75.99.25&[MIMAT0000279&hsa-miR-222-3p&offsets\|0\|+4;m-140&X\|-\|45747033\|45747056&offsets\|0\|+1]\|AGCTACATCTGGCTACTGGGTCTCT | >MI0000299\|hsa-mir-222&WithFlank&X\|-\|45747009\|45747130 | 75 | 99 | 25 | AGCTACATCTGGCTACTGGGTCTCT | 10522.1 | 4.4 | 0.7 | 6.0 | 1.7E-09 | 4.0E-09 | MB231/MB468 | MP |
| >MI0000465\|hsa-mir-191&WithFlank&3\|-\|49020612\|49020715@22.42.21(+1C)&[MIMAT0000440&hsa-miR-191-5p&offsets\|0\|-2(+1C);m-48&3\|-\|49020672\|49020694&offsets\|0\|-2(+1C)]\|CAACGGAATCCCAAAAGCAGCC | >MI0000465\|hsa-mir-191&WithFlank&3\|-\|49020612\|49020715 | 22 | 42 | 21(+1C) | CAACGGAATCCCAAAAGCAGCC | 897.8 | 5.6 | 0.9 | 6.0 | 2.0E-09 | 4.0E-09 | MB231/MB468 | MITO |
| >hg38_21_+_8433172_8446622_RNA45SN1_45S_50ntFlanks@9554.9584.31\|CGTAGCGGTCCTGACGTGCAAATCGGTCGTC | >hg38_21_+_8433172_8446622_RNA45SN1_45S_50ntFlanks | 9554 | 9584 | 31 | CGTAGCGGTCCTGACGTGCAAATCGGTCGTC | 3047.6 | -3.7 | 0.6 | -6.0 | 1.8E-09 | 4.0E-09 | MB231/MB468 | CYTO |
| >hg38_21_+_8433172_8446622_RNA45SN1_45S_50ntFlanks@12973.13000.28\|TCGTACGTAGCAGAGCAGCTCCCTCGCT | >hg38_21_+_8433172_8446622_RNA45SN1_45S_50ntFlanks | 12973 | 13000 | 28 | TCGTACGTAGCAGAGCAGCTCCCTCGCT | 228.5 | -7.5 | 1.3 | -6.0 | 2.0E-09 | 4.0E-09 | MB231/MB468 | MITO |
| >MI0003760\|hsa-mir-671&WithFlank&7\|+\|151238415\|151238544@74.94.21&[MIMAT0004819&hsa-miR-671-3p&offsets\|0\|0;m-323&7\|+\|151238488\|151238508&offsets\|0\|0]\|TCCGGTTCTCAGGGCTCCACC | >MI0003760\|hsa-mir-671&WithFlank&7\|+\|151238415\|151238544 | 74 | 94 | 21 | TCCGGTTCTCAGGGCTCCACC | 352.9 | 6.1 | 1.0 | 6.0 | 1.8E-09 | 4.0E-09 | MB231/MB468 | CYTO |
| >trna10_LysCTT_16_+_3241501_3241573@1.35.35,trna13_LysCTT_14_-_58706613_58706685@1.35.35,trna2_LysCTT_15_+_79152904_79152976@1.35.35\|GCCCGGCTAGCTCAGTCGGTAGAGCATGGGACTCT | >trna10_LysCTT_16_+_3241501_3241573 | 1 | 35 | 35 | GCCCGGCTAGCTCAGTCGGTAGAGCATGGGACTCT | 5595.1 | 7.2 | 1.2 | 6.1 | 1.4E-09 | 4.0E-09 | BT20/MB468 | MITO |
| >hg38_21_+_8433172_8446622_RNA45SN1_45S_50ntFlanks@12635.12659.25\|AAGCTACCATCTGTGGGATTATGAC | >hg38_21_+_8433172_8446622_RNA45SN1_45S_50ntFlanks | 12635 | 12659 | 25 | AAGCTACCATCTGTGGGATTATGAC | 5625.2 | -5.4 | 0.9 | -6.1 | 1.4E-09 | 4.0E-09 | BT20/MB468 | MITO |
| >trnaMT_SerGCT_MT_+_12207_12265@1.25.25\|GAGAAAGCTCACAAGAACTGCTAAC | >trnaMT_SerGCT_MT_+_12207_12265 | 1 | 25 | 25 | GAGAAAGCTCACAAGAACTGCTAAC | 2851.2 | 5.6 | 0.9 | 6.0 | 1.8E-09 | 4.0E-09 | MB231/MB468 | NUC |
| >trnaMT_GluTTC_MT_-_14674_14742@35.71.37,trnalookalike8_GluTTC_5_-_93905172_93905240@35.71.37\|TATCATTGGTCGTGGTTGTAGTCCGTGCGAGAATACC | >trnaMT_GluTTC_MT_-_14674_14742 | 35 | 71 | 37 | TATCATTGGTCGTGGTTGTAGTCCGTGCGAGAATACC | 624.8 | 6.2 | 1.0 | 6.0 | 1.8E-09 | 4.0E-09 | MB231/MB468 | NUC |
| >hg38_21_+_8433172_8446622_RNA45SN1_45S_50ntFlanks@12893.12935.43\|CCTCGCCCGTCACGCACCGCACGTTCGTGGGGAACCTGGCGCT | >hg38_21_+_8433172_8446622_RNA45SN1_45S_50ntFlanks | 12893 | 12935 | 43 | CCTCGCCCGTCACGCACCGCACGTTCGTGGGGAACCTGGCGCT | 148.2 | -11.1 | 1.8 | -6.0 | 1.7E-09 | 4.0E-09 | MB231/MB468 | MP |
| >MI0000465\|hsa-mir-191&WithFlank&3\|-\|49020612\|49020715@22.43.22(+1A)&[MIMAT0000440&hsa-miR-191-5p&offsets\|0\|-1(+1A);m-48&3\|-\|49020672\|49020694&offsets\|0\|-1(+1A)]\|CAACGGAATCCCAAAAGCAGCTA | >MI0000465\|hsa-mir-191&WithFlank&3\|-\|49020612\|49020715 | 22 | 43 | 22(+1A) | CAACGGAATCCCAAAAGCAGCTA | 831.1 | 4.3 | 0.7 | 6.0 | 1.8E-09 | 4.0E-09 | MB231/MB468 | NUC |
| >trna13_ValCAC_19_-_4724647_4724719@1.34.34\|GTTTCCGTAGTGTAGCGGTTATCACATTCGCCTC | >trna13_ValCAC_19_-_4724647_4724719 | 1 | 34 | 34 | GTTTCCGTAGTGTAGCGGTTATCACATTCGCCTC | 128.3 | 10.2 | 1.7 | 6.0 | 1.7E-09 | 4.0E-09 | MB231/MB468 | MP |
| >hg38_21_+_8433172_8446622_RNA45SN1_45S_50ntFlanks@12710.12730.21\|CGCGGAGCCTCGGTTGGCCTC | >hg38_21_+_8433172_8446622_RNA45SN1_45S_50ntFlanks | 12710 | 12730 | 21 | CGCGGAGCCTCGGTTGGCCTC | 107.6 | -10.6 | 1.8 | -6.0 | 1.7E-09 | 4.0E-09 | MB231/MB468 | MP |
| >hg38_21_+_8433172_8446622_RNA45SN1_45S_50ntFlanks@12456.12482.27\|CCAAGCGTTGGATTGTTCACCCACTAA | >hg38_21_+_8433172_8446622_RNA45SN1_45S_50ntFlanks | 12456 | 12482 | 27 | CCAAGCGTTGGATTGTTCACCCACTAA | 206.1 | -10.8 | 1.7 | -6.4 | 2.0E-10 | 4.1E-09 | BT20/MB468 | MP |
| >hg38_21_+_8433172_8446622_RNA45SN1_45S_50ntFlanks@3899.3918.20\|CCTTCGCGGGGGGGATGCGT | >hg38_21_+_8433172_8446622_RNA45SN1_45S_50ntFlanks | 3899 | 3918 | 20 | CCTTCGCGGGGGGGATGCGT | 174.3 | -10.5 | 1.7 | -6.4 | 2.0E-10 | 4.1E-09 | BT20/MB468 | MP |
| >trnaMT_PheGAA_MT_+_577_647@35.63.29\|GAAAATGTTTAGACGGGCTCACATCACCC | >trnaMT_PheGAA_MT_+_577_647 | 35 | 63 | 29 | GAAAATGTTTAGACGGGCTCACATCACCC | 110.7 | 10.0 | 1.7 | 6.0 | 1.7E-09 | 4.1E-09 | MB231/MB468 | MP |
| >hg38_21_+_8433172_8446622_RNA45SN1_45S_50ntFlanks@5073.5093.21\|AATTCCGATAACGAACGAGAC | >hg38_21_+_8433172_8446622_RNA45SN1_45S_50ntFlanks | 5073 | 5093 | 21 | AATTCCGATAACGAACGAGAC | 503.7 | -12.1 | 1.9 | -6.4 | 2.0E-10 | 4.1E-09 | BT20/MB468 | MP |
| >MI0000471\|hsa-mir-126&WithFlank&9\|+\|136670596\|136670692@58.78.21&[MIMAT0000445&hsa-miR-126-3p&offsets\|0\|-1;m-40&9\|+\|136670653\|136670674&offsets\|0\|-1]\|TCGTACCGTGAGTAATAATGC | >MI0000471\|hsa-mir-126&WithFlank&9\|+\|136670596\|136670692 | 58 | 78 | 21 | TCGTACCGTGAGTAATAATGC | 507.6 | 6.1 | 1.0 | 6.0 | 2.1E-09 | 4.1E-09 | BT20/MB468 | NUC |
| >hg38_21_+_8433172_8446622_RNA45SN1_45S_50ntFlanks@9534.9579.46\|AGGAAACTCTGGTGGAGGTCCGTAGCGGTCCTGACGTGCAAATCGG | >hg38_21_+_8433172_8446622_RNA45SN1_45S_50ntFlanks | 9534 | 9579 | 46 | AGGAAACTCTGGTGGAGGTCCGTAGCGGTCCTGACGTGCAAATCGG | 203.0 | -10.6 | 1.8 | -6.0 | 1.9E-09 | 4.1E-09 | MB231/MB468 | NUC |
| >hg38_21_+_8433172_8446622_RNA45SN1_45S_50ntFlanks@5487.5524.38\|CGCTGAGAAGACGGTCGAACTTGACTATCTAGAGGAAG | >hg38_21_+_8433172_8446622_RNA45SN1_45S_50ntFlanks | 5487 | 5524 | 38 | CGCTGAGAAGACGGTCGAACTTGACTATCTAGAGGAAG | 86.8 | -10.3 | 1.7 | -6.0 | 1.8E-09 | 4.1E-09 | MB231/MB468 | MP |
| >MI0000100\|hsa-mir-98&WithFlank&X\|-\|53556217\|53556347@28.49.22&[MIMAT0000096&hsa-miR-98-5p&offsets\|0\|0;m-143&X\|-\|53556299\|53556320&offsets\|0\|0]\|TGAGGTAGTAAGTTGTATTGTT | >MI0000100\|hsa-mir-98&WithFlank&X\|-\|53556217\|53556347 | 28 | 49 | 22 | TGAGGTAGTAAGTTGTATTGTT | 7797.0 | 4.8 | 0.8 | 6.0 | 1.6E-09 | 4.1E-09 | BT20/MB468 | CYTO |
| >hg38_21_+_8433172_8446622_RNA45SN1_45S_50ntFlanks@11583.11601.19\|CTGGTGCGGACCAGGGGAA | >hg38_21_+_8433172_8446622_RNA45SN1_45S_50ntFlanks | 11583 | 11601 | 19 | CTGGTGCGGACCAGGGGAA | 1532.6 | -4.5 | 0.7 | -6.0 | 1.8E-09 | 4.1E-09 | MB231/MB468 | CYTO |
| >hg38_21_+_8433172_8446622_RNA45SN1_45S_50ntFlanks@4385.4423.39\|TTGGATCTTGGGAGCGGGCGGGCGGTCCGCCGCGAGGCG | >hg38_21_+_8433172_8446622_RNA45SN1_45S_50ntFlanks | 4385 | 4423 | 39 | TTGGATCTTGGGAGCGGGCGGGCGGTCCGCCGCGAGGCG | 84.5 | -10.3 | 1.7 | -6.0 | 1.8E-09 | 4.1E-09 | MB231/MB468 | MP |
| >hg38_21_+_8433172_8446622_RNA45SN1_45S_50ntFlanks@10797.10818.22\|GCATGTTGGAACAATGTAGGTA | >hg38_21_+_8433172_8446622_RNA45SN1_45S_50ntFlanks | 10797 | 10818 | 22 | GCATGTTGGAACAATGTAGGTA | 2165.1 | -7.0 | 1.2 | -6.0 | 2.0E-09 | 4.1E-09 | MB231/MB468 | MITO |
| >hg38_21_+_8433172_8446622_RNA45SN1_45S_50ntFlanks@9624.9648.25\|AGTAGCTGGTTCCCTCCGAAGTTTC | >hg38_21_+_8433172_8446622_RNA45SN1_45S_50ntFlanks | 9624 | 9648 | 25 | AGTAGCTGGTTCCCTCCGAAGTTTC | 132.8 | -10.9 | 1.8 | -6.0 | 1.8E-09 | 4.1E-09 | MB231/MB468 | MP |
| >hg38_21_+_8433172_8446622_RNA45SN1_45S_50ntFlanks@11729.11746.18\|TAACTATGACTCTCTTAA | >hg38_21_+_8433172_8446622_RNA45SN1_45S_50ntFlanks | 11729 | 11746 | 18 | TAACTATGACTCTCTTAA | 62.4 | -9.8 | 1.6 | -6.0 | 1.8E-09 | 4.1E-09 | MB231/MB468 | MP |
| >hg38_21_+_8433172_8446622_RNA45SN1_45S_50ntFlanks@4069.4089.21\|CTTTCGATGGTAGTCGCCGTG | >hg38_21_+_8433172_8446622_RNA45SN1_45S_50ntFlanks | 4069 | 4089 | 21 | CTTTCGATGGTAGTCGCCGTG | 2597.0 | -8.8 | 1.5 | -6.0 | 2.0E-09 | 4.1E-09 | MB231/MB468 | MITO |
| >hg38_21_+_8433172_8446622_RNA45SN1_45S_50ntFlanks@8170.8187.18\|CAAGTCCTTCTGATCGAG | >hg38_21_+_8433172_8446622_RNA45SN1_45S_50ntFlanks | 8170 | 8187 | 18 | CAAGTCCTTCTGATCGAG | 1311.6 | -9.1 | 1.5 | -6.0 | 2.1E-09 | 4.1E-09 | BT20/MB468 | NUC |
| >hg38_21_+_8433172_8446622_RNA45SN1_45S_50ntFlanks@8196.8223.28\|CGTGGACGGTGTGAGGCCGGTAGCGGCC | >hg38_21_+_8433172_8446622_RNA45SN1_45S_50ntFlanks | 8196 | 8223 | 28 | CGTGGACGGTGTGAGGCCGGTAGCGGCC | 636.8 | -3.9 | 0.7 | -6.0 | 1.9E-09 | 4.1E-09 | MB231/MB468 | NUC |
| >hg38_21_+_8433172_8446622_RNA45SN1_45S_50ntFlanks@12071.12096.26\|CCCCGAGGGGCTCTCGCTTCTGGCGC | >hg38_21_+_8433172_8446622_RNA45SN1_45S_50ntFlanks | 12071 | 12096 | 26 | CCCCGAGGGGCTCTCGCTTCTGGCGC | 530.6 | -8.8 | 1.5 | -6.0 | 2.1E-09 | 4.1E-09 | BT20/MB468 | NUC |
| >hg38_21_+_8433172_8446622_RNA45SN1_45S_50ntFlanks@11874.11902.29\|AGAAGACCCTGTTGAGCTTGACTCTAGTC | >hg38_21_+_8433172_8446622_RNA45SN1_45S_50ntFlanks | 11874 | 11902 | 29 | AGAAGACCCTGTTGAGCTTGACTCTAGTC | 288.6 | -5.9 | 1.0 | -6.0 | 2.0E-09 | 4.1E-09 | MB231/MB468 | MITO |
| >hg38_21_+_8433172_8446622_RNA45SN1_45S_50ntFlanks@12488.12518.31\|AACGTGAGCTGGGTTTAGACCGTCGTGAGAC | >hg38_21_+_8433172_8446622_RNA45SN1_45S_50ntFlanks | 12488 | 12518 | 31 | AACGTGAGCTGGGTTTAGACCGTCGTGAGAC | 365.4 | -7.7 | 1.3 | -6.0 | 2.0E-09 | 4.1E-09 | MB231/MB468 | MITO |
| >hg38_21_+_8433172_8446622_RNA45SN1_45S_50ntFlanks@12358.12376.19\|CACAGGGATAACTGGCTTG | >hg38_21_+_8433172_8446622_RNA45SN1_45S_50ntFlanks | 12358 | 12376 | 19 | CACAGGGATAACTGGCTTG | 167.6 | -10.5 | 1.6 | -6.4 | 2.0E-10 | 4.1E-09 | BT20/MB468 | MP |
| >MI0000482\|hsa-mir-185&WithFlank&22\|+\|20033133\|20033226@21.42.22(+1A)&[MIMAT0000455&hsa-miR-185-5p&offsets\|0\|0(+1A);m-145&22\|+\|20033153\|20033174&offsets\|0\|0(+1A)]\|TGGAGAGAAAGGCAGTTCCTGAA | >MI0000482\|hsa-mir-185&WithFlank&22\|+\|20033133\|20033226 | 21 | 42 | 22(+1A) | TGGAGAGAAAGGCAGTTCCTGAA | 97.6 | 2.1 | 0.4 | 6.0 | 2.1E-09 | 4.1E-09 | BT20/MB468 | NUC |
| >MI0000478\|hsa-mir-149&WithFlank&2\|+\|240455995\|240456095@21.42.22&[MIMAT0000450&hsa-miR-149-5p&offsets\|0\|-1;m-195&2\|+\|240456015\|240456037&offsets\|0\|-1]\|TCTGGCTCCGTGTCTTCACTCC | >MI0000478\|hsa-mir-149&WithFlank&2\|+\|240455995\|240456095 | 21 | 42 | 22 | TCTGGCTCCGTGTCTTCACTCC | 122.3 | 10.2 | 1.7 | 6.0 | 2.0E-09 | 4.1E-09 | MB231/MB468 | MITO |
| >hg38_21_+_8433172_8446622_RNA45SN1_45S_50ntFlanks@12936.12961.26\|AAACCATTCGTAGACGACCTGCTTCT | >hg38_21_+_8433172_8446622_RNA45SN1_45S_50ntFlanks | 12936 | 12961 | 26 | AAACCATTCGTAGACGACCTGCTTCT | 2280.5 | -9.7 | 1.6 | -6.0 | 2.1E-09 | 4.1E-09 | BT20/MB468 | NUC |
| >MI0003591\|hsa-mir-584&WithFlank&5\|-\|149062307\|149062415@22.41.20&[MIMAT0003249&hsa-miR-584-5p&offsets\|0\|-2;m-167&5\|-\|149062374\|149062394&offsets\|0\|-1]\|TTATGGTTTGCCTGGGACTG | >MI0003591\|hsa-mir-584&WithFlank&5\|-\|149062307\|149062415 | 22 | 41 | 20 | TTATGGTTTGCCTGGGACTG | 485.3 | 4.2 | 0.7 | 6.0 | 1.8E-09 | 4.1E-09 | MB231/MB468 | MP |
| >hg38_21_+_8433172_8446622_RNA45SN1_45S_50ntFlanks@12993.13013.21\|CCCTCGCTGCGATCTATTGAA | >hg38_21_+_8433172_8446622_RNA45SN1_45S_50ntFlanks | 12993 | 13013 | 21 | CCCTCGCTGCGATCTATTGAA | 3597.5 | -10.1 | 1.7 | -6.0 | 2.1E-09 | 4.1E-09 | BT20/MB468 | NUC |
| >hg38_21_+_8433172_8446622_RNA45SN1_45S_50ntFlanks@11455.11478.24\|CTCGGCGCGCGGCGGCGGCGGCGG | >hg38_21_+_8433172_8446622_RNA45SN1_45S_50ntFlanks | 11455 | 11478 | 24 | CTCGGCGCGCGGCGGCGGCGGCGG | 193.2 | -10.7 | 1.7 | -6.4 | 2.0E-10 | 4.1E-09 | BT20/MB468 | MP |
| >hg38_21_+_8433172_8446622_RNA45SN1_45S_50ntFlanks@7975.8015.41\|CGCGACCTCAGATCAGACGTGGCGACCCGCTGAATTTAAGC | >hg38_21_+_8433172_8446622_RNA45SN1_45S_50ntFlanks | 7975 | 8015 | 41 | CGCGACCTCAGATCAGACGTGGCGACCCGCTGAATTTAAGC | 7761.7 | 5.3 | 0.9 | 6.0 | 1.9E-09 | 4.1E-09 | MB231/MB468 | NUC |
| >hg38_21_+_8433172_8446622_RNA45SN1_45S_50ntFlanks@10411.10433.23\|CAGTCGGTCCTGAGAGATGGGCG | >hg38_21_+_8433172_8446622_RNA45SN1_45S_50ntFlanks | 10411 | 10433 | 23 | CAGTCGGTCCTGAGAGATGGGCG | 952.8 | -5.3 | 0.9 | -6.1 | 1.4E-09 | 4.1E-09 | BT20/MB468 | MITO |
| >hg38_21_+_8433172_8446622_RNA45SN1_45S_50ntFlanks@9740.9758.19\|AACGATCTCAACCTATTCT | >hg38_21_+_8433172_8446622_RNA45SN1_45S_50ntFlanks | 9740 | 9758 | 19 | AACGATCTCAACCTATTCT | 1064.4 | -5.7 | 0.9 | -6.0 | 2.0E-09 | 4.1E-09 | MB231/MB468 | MITO |
| >hg38_21_+_8433172_8446622_RNA45SN1_45S_50ntFlanks@11600.11623.24\|AATCCGACTGTTTAATTAAAACAA | >hg38_21_+_8433172_8446622_RNA45SN1_45S_50ntFlanks | 11600 | 11623 | 24 | AATCCGACTGTTTAATTAAAACAA | 201.6 | 11.4 | 1.8 | 6.4 | 2.1E-10 | 4.1E-09 | BT20/MB468 | MP |
| >MI0000088\|hsa-mir-30a&WithFlank&6\|-\|71403545\|71403627@12.33.22(+1A)&[MIMAT0000087&hsa-miR-30a-5p&offsets\|0\|0(+1A);m-15&6\|-\|71403595\|71403616&offsets\|0\|0(+1A)]\|TGTAAACATCCTCGACTGGAAGA | >MI0000088\|hsa-mir-30a&WithFlank&6\|-\|71403545\|71403627 | 12 | 33 | 22(+1A) | TGTAAACATCCTCGACTGGAAGA | 1514.5 | 7.8 | 1.3 | 6.0 | 1.6E-09 | 4.2E-09 | BT20/MB468 | CYTO |
| >hg38_21_+_8433172_8446622_RNA45SN1_45S_50ntFlanks@9575.9602.28\|ATCGGTCGTCCGACCTGGGTATAGGGGC | >hg38_21_+_8433172_8446622_RNA45SN1_45S_50ntFlanks | 9575 | 9602 | 28 | ATCGGTCGTCCGACCTGGGTATAGGGGC | 193.7 | -10.7 | 1.7 | -6.4 | 2.1E-10 | 4.2E-09 | BT20/MB468 | MP |

<table>
<tr><td>>hg38_21_+_8433172_8446622_RNA45SN1_45S_50ntFlanks@6740.6771.32|CATCGACACTTCGAACGCACTTGCGGCCCCGG</td><td>>hg38_21_+_8433172_8446622_RNA45SN1_45S_50ntFlanks</td><td>6740</td><td>6771</td><td>32</td><td>CATCGACACTTCGAACGCACTTGCGGCCCCGG</td><td>569.7</td><td>-5.5</td><td>0.9</td><td>-6.0</td><td>1.9E-09</td><td>4.2E-09</td><td>MB231/MB468</td><td>CYTO</td></tr>
<tr><td>>hg38_21_+_8433172_8446622_RNA45SN1_45S_50ntFlanks@11577.11602.26|TTAGAACTGGTGCGGACCAGGGGAAT</td><td>>hg38_21_+_8433172_8446622_RNA45SN1_45S_50ntFlanks</td><td>11577</td><td>11602</td><td>26</td><td>TTAGAACTGGTGCGGACCAGGGGAAT</td><td>388.4</td><td>-8.4</td><td>1.4</td><td>-6.0</td><td>2.2E-09</td><td>4.2E-09</td><td>BT20/MB468</td><td>NUC</td></tr>
<tr><td>>hg38_21_+_8433172_8446622_RNA45SN1_45S_50ntFlanks@9134.9152.19|CGGGGGGCGGGGCGGACTG</td><td>>hg38_21_+_8433172_8446622_RNA45SN1_45S_50ntFlanks</td><td>9134</td><td>9152</td><td>19</td><td>CGGGGGGCGGGGCGGACTG</td><td>215.6</td><td>-10.8</td><td>1.7</td><td>-6.4</td><td>2.1E-10</td><td>4.2E-09</td><td>BT20/MB468</td><td>MP</td></tr>
<tr><td>>hg38_21_+_8433172_8446622_RNA45SN1_45S_50ntFlanks@7970.8005.36|CGAGACGCGACCTCAGATCAGACGTGGCGACCCGCT</td><td>>hg38_21_+_8433172_8446622_RNA45SN1_45S_50ntFlanks</td><td>7970</td><td>8005</td><td>36</td><td>CGAGACGCGACCTCAGATCAGACGTGGCGACCCGCT</td><td>220.7</td><td>4.8</td><td>0.8</td><td>6.0</td><td>2.2E-09</td><td>4.2E-09</td><td>BT20/MB468</td><td>NUC</td></tr>
<tr><td>>hg38_21_+_8433172_8446622_RNA45SN1_45S_50ntFlanks@10571.10600.30|GAGAAGCCGGCGGGAGCCCCGGGGAGAGTT</td><td>>hg38_21_+_8433172_8446622_RNA45SN1_45S_50ntFlanks</td><td>10571</td><td>10600</td><td>30</td><td>GAGAAGCCGGCGGGAGCCCCGGGGAGAGTT</td><td>69.4</td><td>-10.0</td><td>1.7</td><td>-6.0</td><td>1.8E-09</td><td>4.2E-09</td><td>MB231/MB468</td><td>MP</td></tr>
<tr><td>>hg38_21_+_8433172_8446622_RNA45SN1_45S_50ntFlanks@6652.6695.44|GACTCTTAGCGGTGGATCACTCGGCTCGTGCGTCGATGAAGAAC</td><td>>hg38_21_+_8433172_8446622_RNA45SN1_45S_50ntFlanks</td><td>6652</td><td>6695</td><td>44</td><td>GACTCTTAGCGGTGGATCACTCGGCTCGTGCGTCGATGAAGAAC</td><td>1910.7</td><td>5.2</td><td>0.9</td><td>6.0</td><td>1.9E-09</td><td>4.2E-09</td><td>MB231/MB468</td><td>NUC</td></tr>
<tr><td>>hg38_21_+_8433172_8446622_RNA45SN1_45S_50ntFlanks@12365.12382.18|ATAACTGGCTTGTGGCGG</td><td>>hg38_21_+_8433172_8446622_RNA45SN1_45S_50ntFlanks</td><td>12365</td><td>12382</td><td>18</td><td>ATAACTGGCTTGTGGCGG</td><td>1061.7</td><td>-7.7</td><td>1.3</td><td>-6.0</td><td>1.6E-09</td><td>4.2E-09</td><td>BT20/MB468</td><td>CYTO</td></tr>
<tr><td>>trna14_LysTTT_11_-_59327808_59327880@1.33.33,trna2_LysTTT_17_+_8022473_8022545@1.33.33,trna54_LysTTT_1_+_204475655_204475727@1.33.33,trna5_LysTTT_11_+_59323902_59323974@1.33.33,trna62_LysTTT_1_-_204476158_204476230@1.33.33,trna76_LysTTT_6_+_28918806_28918878@1.33.33|GCCCGGATAGCTCAGTCGGTAGAGCATCAGACT</td><td>>trna14_LysTTT_11_-_59327808_59327880</td><td>1</td><td>33</td><td>33</td><td>GCCCGGATAGCTCAGTCGGTAGAGCATCAGACT</td><td>2841.7</td><td>6.6</td><td>1.1</td><td>6.1</td><td>1.4E-09</td><td>4.2E-09</td><td>BT20/MB468</td><td>MITO</td></tr>
<tr><td>>hg38_21_+_8433172_8446622_RNA45SN1_45S_50ntFlanks@12306.12326.21|CACGATCCTTCTGACCTTTTG</td><td>>hg38_21_+_8433172_8446622_RNA45SN1_45S_50ntFlanks</td><td>12306</td><td>12326</td><td>21</td><td>CACGATCCTTCTGACCTTTTG</td><td>70.1</td><td>-10.0</td><td>1.7</td><td>-6.0</td><td>1.8E-09</td><td>4.2E-09</td><td>MB231/MB468</td><td>MP</td></tr>
<tr><td>>MI0005756|hsa-mir-934&WithFlank&X|+|136550872|136550966@21.41.21(+1U)&[MIMAT0004977&hsa-miR-934&offsets|0|-1(+1U);m-943&X|+|136550892|136550913&offsets|0|-1(+1U)]|TGTCTACTACTGGAGACACTGT</td><td>>MI0005756|hsa-mir-934&WithFlank&X|+|136550872|136550966</td><td>21</td><td>41</td><td>21(+1U)</td><td>TGTCTACTACTGGAGACACTGT</td><td>95.4</td><td>-10.4</td><td>1.7</td><td>-6.0</td><td>1.8E-09</td><td>4.2E-09</td><td>MB231/MB468</td><td>MP</td></tr>
<tr><td>>am_trna162_MetCAT_6_-_26758550_26758622@31.51.21,trna164_MetCAT_6_-_26735574_26735646@31.51.21,trna20_MetCAT_16_+_71460396_71460468@31.51.21,trna21_MetCAT_6_+_26701712_26701784@31.51.21,trna27_MetCAT_6_+_26766444_26766516@31.51.21,trna75_MetCAT_6_+_28912352_28912424@31.51.21,trna97_MetCAT_6_-_28921042_28921114@31.51.21|TCTCATAATCTGAAGGTCCTG</td><td>>am_trna162_MetCAT_6_-_26758550_26758622</td><td>31</td><td>51</td><td>21</td><td>TCTCATAATCTGAAGGTCCTG</td><td>298.7</td><td>-8.3</td><td>1.4</td><td>-6.0</td><td>1.6E-09</td><td>4.2E-09</td><td>BT20/MB468</td><td>CYTO</td></tr>
<tr><td>>hg38_21_+_8433172_8446622_RNA45SN1_45S_50ntFlanks@10024.10072.49|GCGCTGGAGCGTCGGGCCCATACCCGGCCGTCGCCGGCAGTCGAGAGTG</td><td>>hg38_21_+_8433172_8446622_RNA45SN1_45S_50ntFlanks</td><td>10024</td><td>10072</td><td>49</td><td>GCGCTGGAGCGTCGGGCCCATACCCGGCCGTCGCCGGCAGTCGAGAGTG</td><td>137.4</td><td>-11.0</td><td>1.8</td><td>-6.0</td><td>1.8E-09</td><td>4.2E-09</td><td>MB231/MB468</td><td>MP</td></tr>
<tr><td>>hg38_21_+_8433172_8446622_RNA45SN1_45S_50ntFlanks@6645.6665.21|CTCGTACGACTCTTAGCGGTG</td><td>>hg38_21_+_8433172_8446622_RNA45SN1_45S_50ntFlanks</td><td>6645</td><td>6665</td><td>21</td><td>CTCGTACGACTCTTAGCGGTG</td><td>22101.2</td><td>5.5</td><td>0.9</td><td>6.4</td><td>2.1E-10</td><td>4.2E-09</td><td>BT20/MB468</td><td>MP</td></tr>
<tr><td>>hg38_21_+_8433172_8446622_RNA45SN1_45S_50ntFlanks@6652.6669.18|GACTCTTAGCGGTGGATC</td><td>>hg38_21_+_8433172_8446622_RNA45SN1_45S_50ntFlanks</td><td>6652</td><td>6669</td><td>18</td><td>GACTCTTAGCGGTGGATC</td><td>1825.6</td><td>4.4</td><td>0.7</td><td>6.1</td><td>1.4E-09</td><td>4.2E-09</td><td>BT20/MB468</td><td>MITO</td></tr>
<tr><td>>hg38_21_+_8433172_8446622_RNA45SN1_45S_50ntFlanks@8116.8136.21|GCGGGACATGTGGCGTACGGA</td><td>>hg38_21_+_8433172_8446622_RNA45SN1_45S_50ntFlanks</td><td>8116</td><td>8136</td><td>21</td><td>GCGGGACATGTGGCGTACGGA</td><td>2629.1</td><td>14.7</td><td>2.2</td><td>6.6</td><td>4.1E-11</td><td>4.2E-09</td><td>BT20/MB231</td><td>MP</td></tr>
<tr><td>>hg38_21_+_8433172_8446622_RNA45SN1_45S_50ntFlanks@12414.12437.24|CCTTCGATGTCGGCTCTTCCTATC</td><td>>hg38_21_+_8433172_8446622_RNA45SN1_45S_50ntFlanks</td><td>12414</td><td>12437</td><td>24</td><td>CCTTCGATGTCGGCTCTTCCTATC</td><td>279.4</td><td>4.0</td><td>0.7</td><td>6.0</td><td>1.9E-09</td><td>4.2E-09</td><td>MB231/MB468</td><td>CYTO</td></tr>
<tr><td>>hg38_21_+_8433172_8446622_RNA45SN1_45S_50ntFlanks@12405.12442.38|CTTTTTGATCCTTCGATGTCGGCTCTTCCTATCATTGT</td><td>>hg38_21_+_8433172_8446622_RNA45SN1_45S_50ntFlanks</td><td>12405</td><td>12442</td><td>38</td><td>CTTTTTGATCCTTCGATGTCGGCTCTTCCTATCATTGT</td><td>3432.4</td><td>-10.5</td><td>1.8</td><td>-6.0</td><td>2.2E-09</td><td>4.2E-09</td><td>BT20/MB468</td><td>NUC</td></tr>
<tr><td>>MI0000255|hsa-mir-30d&WithFlank&8|-|134804870|134804951@12.34.23&[MIMAT0000245&hsa-miR-30d-5p&offsets|0|+1;m-21&8|-|134804919|134804940&offsets|0|+1]|TGTAAACATCCCCGACTGGAAGC</td><td>>MI0000255|hsa-mir-30d&WithFlank&8|-|134804870|134804951</td><td>12</td><td>34</td><td>23</td><td>TGTAAACATCCCCGACTGGAAGC</td><td>11416.5</td><td>4.9</td><td>0.8</td><td>6.0</td><td>2.1E-09</td><td>4.2E-09</td><td>MB231/MB468</td><td>MITO</td></tr>
<tr><td>>hg38_21_+_8433172_8446622_RNA45SN1_45S_50ntFlanks@11314.11344.31|CCCACCCCACGTCTCGTCGCGCGCGCGTCCG</td><td>>hg38_21_+_8433172_8446622_RNA45SN1_45S_50ntFlanks</td><td>11314</td><td>11344</td><td>31</td><td>CCCACCCCACGTCTCGTCGCGCGCGCGTCCG</td><td>4959.0</td><td>-10.3</td><td>1.7</td><td>-6.0</td><td>2.2E-09</td><td>4.2E-09</td><td>BT20/MB468</td><td>NUC</td></tr>
<tr><td>>MI0000301|hsa-mir-224&WithFlank&X|-|151958572|151958664@14.34.21(+1C)&[MIMAT0000281&hsa-miR-224-5p&offsets|+1|-3(+1C);m-238&X|-|151958630|151958651&offsets|0|-1(+1C)]|CAAGTCACTAGTGGTTCCGTTC</td><td>>MI0000301|hsa-mir-224&WithFlank&X|-|151958572|151958664</td><td>14</td><td>34</td><td>21(+1C)</td><td>CAAGTCACTAGTGGTTCCGTTC</td><td>151.4</td><td>10.5</td><td>1.7</td><td>6.0</td><td>2.1E-09</td><td>4.2E-09</td><td>MB231/MB468</td><td>MITO</td></tr>
<tr><td>>hg38_21_+_8433172_8446622_RNA45SN1_45S_50ntFlanks@5107.5136.30|CTAGTTACGCGACCCCCGAGCGGTCGGCGT</td><td>>hg38_21_+_8433172_8446622_RNA45SN1_45S_50ntFlanks</td><td>5107</td><td>5136</td><td>30</td><td>CTAGTTACGCGACCCCCGAGCGGTCGGCGT</td><td>120.8</td><td>-10.8</td><td>1.8</td><td>-6.0</td><td>1.8E-09</td><td>4.2E-09</td><td>MB231/MB468</td><td>MP</td></tr>
<tr><td>>hg38_21_+_8433172_8446622_RNA45SN1_45S_50ntFlanks@7990.8011.22|GACGTGGCGACCCGCTGAATTT</td><td>>hg38_21_+_8433172_8446622_RNA45SN1_45S_50ntFlanks</td><td>7990</td><td>8011</td><td>22</td><td>GACGTGGCGACCCGCTGAATTT</td><td>1028.6</td><td>6.4</td><td>1.1</td><td>6.0</td><td>1.9E-09</td><td>4.2E-09</td><td>MB231/MB468</td><td>CYTO</td></tr>
<tr><td>>hg38_21_+_8433172_8446622_RNA45SN1_45S_50ntFlanks@8102.8140.39|CCCCGCGGCGGGGCGCGGGACATGTGGCGTACGGAAGAC</td><td>>hg38_21_+_8433172_8446622_RNA45SN1_45S_50ntFlanks</td><td>8102</td><td>8140</td><td>39</td><td>CCCCGCGGCGGGGCGCGGGACATGTGGCGTACGGAAGAC</td><td>92.0</td><td>-10.4</td><td>1.7</td><td>-6.0</td><td>1.8E-09</td><td>4.2E-09</td><td>MB231/MB468</td><td>MP</td></tr>
<tr><td>>hg38_21_+_8433172_8446622_RNA45SN1_45S_50ntFlanks@10378.10424.47|AGGGTTCCATGTGAACAGCAGTTGAACATGGGTCAGTCGGTCCTGAG</td><td>>hg38_21_+_8433172_8446622_RNA45SN1_45S_50ntFlanks</td><td>10378</td><td>10424</td><td>47</td><td>AGGGTTCCATGTGAACAGCAGTTGAACATGGGTCAGTCGGTCCTGAG</td><td>463.5</td><td>4.8</td><td>0.8</td><td>6.0</td><td>1.9E-09</td><td>4.2E-09</td><td>MB231/MB468</td><td>CYTO</td></tr>
<tr><td>>hg38_21_+_8433172_8446622_RNA45SN1_45S_50ntFlanks@3840.3867.28|CTCCTACTTGGATAACTGTGGTAATTCT</td><td>>hg38_21_+_8433172_8446622_RNA45SN1_45S_50ntFlanks</td><td>3840</td><td>3867</td><td>28</td><td>CTCCTACTTGGATAACTGTGGTAATTCT</td><td>273.5</td><td>3.3</td><td>0.5</td><td>6.0</td><td>1.9E-09</td><td>4.2E-09</td><td>MB231/MB468</td><td>CYTO</td></tr>
<tr><td>>hg38_21_+_8433172_8446622_RNA45SN1_45S_50ntFlanks@9572.9603.32|CAAATCGGTCGTCCGACCTGGGTATAGGGGCG</td><td>>hg38_21_+_8433172_8446622_RNA45SN1_45S_50ntFlanks</td><td>9572</td><td>9603</td><td>32</td><td>CAAATCGGTCGTCCGACCTGGGTATAGGGGCG</td><td>4501.9</td><td>-3.9</td><td>0.6</td><td>-6.0</td><td>1.4E-09</td><td>4.2E-09</td><td>BT20/MB468</td><td>MITO</td></tr>
<tr><td>>MI0000471|hsa-mir-126&WithFlank&9|+|136670596|136670692@58.77.20&[MIMAT0000445&hsa-miR-126-3p&offsets|0|-2;m-40&9|+|136670653|136670674&offsets|0|-2]|TCGTACCGTGAGTAATAATG</td><td>>MI0000471|hsa-mir-126&WithFlank&9|+|136670596|136670692</td><td>58</td><td>77</td><td>20</td><td>TCGTACCGTGAGTAATAATG</td><td>114.0</td><td>5.8</td><td>1.0</td><td>6.0</td><td>2.2E-09</td><td>4.3E-09</td><td>BT20/MB468</td><td>NUC</td></tr>
<tr><td>>hg38_21_+_8433172_8446622_RNA45SN1_45S_50ntFlanks@12939.12961.23|CCATTCGTAGACGACCTGCTTCT</td><td>>hg38_21_+_8433172_8446622_RNA45SN1_45S_50ntFlanks</td><td>12939</td><td>12961</td><td>23</td><td>CCATTCGTAGACGACCTGCTTCT</td><td>224.1</td><td>-3.0</td><td>0.5</td><td>-6.0</td><td>1.9E-09</td><td>4.3E-09</td><td>MB231/MB468</td><td>NUC</td></tr>
<tr><td>>hg38_21_+_8433172_8446622_RNA45SN1_45S_50ntFlanks@4002.4026.25|GATAACCTCGGGCCGATCGCACGCC</td><td>>hg38_21_+_8433172_8446622_RNA45SN1_45S_50ntFlanks</td><td>4002</td><td>4026</td><td>25</td><td>GATAACCTCGGGCCGATCGCACGCC</td><td>304.2</td><td>-6.6</td><td>1.1</td><td>-6.0</td><td>1.7E-09</td><td>4.3E-09</td><td>BT20/MB468</td><td>CYTO</td></tr>
<tr><td>>hg38_21_+_8433172_8446622_RNA45SN1_45S_50ntFlanks@11600.11627.28|AATCCGACTGTTTAATTAAAACAAAGCA</td><td>>hg38_21_+_8433172_8446622_RNA45SN1_45S_50ntFlanks</td><td>11600</td><td>11627</td><td>28</td><td>AATCCGACTGTTTAATTAAAACAAAGCA</td><td>138.1</td><td>10.9</td><td>1.7</td><td>6.4</td><td>2.1E-10</td><td>4.3E-09</td><td>BT20/MB468</td><td>MP</td></tr>
<tr><td>>hg38_21_+_8433172_8446622_RNA45SN1_45S_50ntFlanks@4068.4089.22|ACTTTCGATGGTAGTCGCCGTG</td><td>>hg38_21_+_8433172_8446622_RNA45SN1_45S_50ntFlanks</td><td>4068</td><td>4089</td><td>22</td><td>ACTTTCGATGGTAGTCGCCGTG</td><td>288.4</td><td>-6.0</td><td>1.0</td><td>-6.0</td><td>1.8E-09</td><td>4.3E-09</td><td>MB231/MB468</td><td>MP</td></tr>
<tr><td>>hg38_21_+_8433172_8446622_RNA45SN1_45S_50ntFlanks@6661.6700.40|CGGTGGATCACTCGGCTCGTGCGTCGATGAAGAACGCAGC</td><td>>hg38_21_+_8433172_8446622_RNA45SN1_45S_50ntFlanks</td><td>6661</td><td>6700</td><td>40</td><td>CGGTGGATCACTCGGCTCGTGCGTCGATGAAGAACGCAGC</td><td>385.3</td><td>4.0</td><td>0.7</td><td>6.0</td><td>2.2E-09</td><td>4.3E-09</td><td>BT20/MB468</td><td>NUC</td></tr>
<tr><td>>MI0000077|hsa-mir-21&WithFlank&17|+|59841260|59841343@15.36.22&[MIMAT0000076&hsa-miR-21-5p&offsets|+1|+1;m-1&17|+|59841273|59841294&offsets|+1|+1]|AGCTTATCAGACTGATGTTGAC</td><td>>MI0000077|hsa-mir-21&WithFlank&17|+|59841260|59841343</td><td>15</td><td>36</td><td>22</td><td>AGCTTATCAGACTGATGTTGAC</td><td>1451.0</td><td>5.8</td><td>1.0</td><td>6.0</td><td>2.1E-09</td><td>4.3E-09</td><td>MB231/MB468</td><td>MITO</td></tr>
<tr><td>>hg38_21_+_8433172_8446622_RNA45SN1_45S_50ntFlanks@3788.3810.23|AACTGCGAATGGCTCATTAAATC</td><td>>hg38_21_+_8433172_8446622_RNA45SN1_45S_50ntFlanks</td><td>3788</td><td>3810</td><td>23</td><td>AACTGCGAATGGCTCATTAAATC</td><td>394.9</td><td>-3.5</td><td>0.6</td><td>-6.0</td><td>1.9E-09</td><td>4.3E-09</td><td>MB231/MB468</td><td>NUC</td></tr>
<tr><td>>hg38_21_+_8433172_8446622_RNA45SN1_45S_50ntFlanks@7991.8014.24|ACGTGGCGACCCGCTGAATTTAAG</td><td>>hg38_21_+_8433172_8446622_RNA45SN1_45S_50ntFlanks</td><td>7991</td><td>8014</td><td>24</td><td>ACGTGGCGACCCGCTGAATTTAAG</td><td>2466.5</td><td>-7.5</td><td>1.2</td><td>-6.0</td><td>1.8E-09</td><td>4.3E-09</td><td>MB231/MB468</td><td>MP</td></tr>
<tr><td>>hg38_21_+_8433172_8446622_RNA45SN1_45S_50ntFlanks@11729.11767.39|TAACTATGACTCTCTTAAGGTAGCCAAATGCCTCGTCAT</td><td>>hg38_21_+_8433172_8446622_RNA45SN1_45S_50ntFlanks</td><td>11729</td><td>11767</td><td>39</td><td>TAACTATGACTCTCTTAAGGTAGCCAAATGCCTCGTCAT</td><td>1098.0</td><td>-9.3</td><td>1.5</td><td>-6.4</td><td>2.1E-10</td><td>4.3E-09</td><td>BT20/MB468</td><td>MP</td></tr>
<tr><td>>MI0000093|hsa-mir-92a-1&WithFlank&13|+|91351308|91351397@54.74.21(+1C);MI0000094|hsa-mir-92a-2&WithFlank&X|-|134169532|134169618@54.74.21(+1C)&[MIMAT0000092&hsa-miR-92a-3p&offsets|0|-1(+1C);m-9&13|+|91351361|91351382&offsets|0|-1(+1C)];[MIMAT0000092_1&hsa-miR-92a-3p&offsets|0|-1(+1C);m-11&X|-|134169544|134169565&offsets|0|-1(+1C)]|TATTGCACTTGTCCCGGCCTGC</td><td>>MI0000093|hsa-mir-92a-1&WithFlank&13|+|91351308|91351397</td><td>54</td><td>74</td><td>21(+1C)</td><td>TATTGCACTTGTCCCGGCCTGC</td><td>10852.7</td><td>5.6</td><td>0.9</td><td>6.0</td><td>2.1E-09</td><td>4.3E-09</td><td>MB231/MB468</td><td>MITO</td></tr>
<tr><td>>hg38_21_+_8433172_8446622_RNA45SN1_45S_50ntFlanks@11783.11805.23|CATGAATGGATGAACGAGATTCC</td><td>>hg38_21_+_8433172_8446622_RNA45SN1_45S_50ntFlanks</td><td>11783</td><td>11805</td><td>23</td><td>CATGAATGGATGAACGAGATTCC</td><td>2366.9</td><td>-6.9</td><td>1.2</td><td>-6.0</td><td>1.8E-09</td><td>4.3E-09</td><td>MB231/MB468</td><td>MP</td></tr>
<tr><td>>MI0000743|hsa-mir-34c&WithFlank&11|+|111513433|111513521@19.40.22(+1A)&[MIMAT0000686&hsa-miR-34c-5p&offsets|0|-1(+1A);m-336&11|+|111513451|111513473&offsets|0|-1(+1A)]|AGGCAGTGTAGTTAGCTGATTGA</td><td>>MI0000743|hsa-mir-34c&WithFlank&11|+|111513433|111513521</td><td>19</td><td>40</td><td>22(+1A)</td><td>AGGCAGTGTAGTTAGCTGATTGA</td><td>72.5</td><td>9.4</td><td>1.6</td><td>6.0</td><td>1.9E-09</td><td>4.3E-09</td><td>MB231/MB468</td><td>MP</td></tr>
<tr><td>>hg38_21_+_8433172_8446622_RNA45SN1_45S_50ntFlanks@12139.12162.24|GGACAGTGCCAGGTGGGGAGTTTG</td><td>>hg38_21_+_8433172_8446622_RNA45SN1_45S_50ntFlanks</td><td>12139</td><td>12162</td><td>24</td><td>GGACAGTGCCAGGTGGGGAGTTTG</td><td>74.7</td><td>-10.1</td><td>1.7</td><td>-6.0</td><td>1.9E-09</td><td>4.3E-09</td><td>MB231/MB468</td><td>MP</td></tr>
<tr><td>>MI0000285|hsa-mir-205&WithFlank&1|+|209432127|209432248@40.59.20(+1C)&[MIMAT0000266&hsa-miR-205-5p&offsets|0|-2(+1C);m-351&1|+|209432166|209432187&offsets|0|-2(+1C)]|TCCTTCATTCCACCGGAGTCC</td><td>>MI0000285|hsa-mir-205&WithFlank&1|+|209432127|209432248</td><td>40</td><td>59</td><td>20(+1C)</td><td>TCCTTCATTCCACCGGAGTCC</td><td>485.9</td><td>12.3</td><td>1.9</td><td>6.6</td><td>4.3E-11</td><td>4.3E-09</td><td>BT20/MB231</td><td>MP</td></tr>
<tr><td>>hg38_21_+_8433172_8446622_RNA45SN1_45S_50ntFlanks@12073.12097.25|CCGAGGGGCTCTCGCTTCTGGCGCC</td><td>>hg38_21_+_8433172_8446622_RNA45SN1_45S_50ntFlanks</td><td>12073</td><td>12097</td><td>25</td><td>CCGAGGGGCTCTCGCTTCTGGCGCC</td><td>1631.2</td><td>-4.0</td><td>0.7</td><td>-6.0</td><td>1.5E-09</td><td>4.3E-09</td><td>BT20/MB468</td><td>MITO</td></tr>
<tr><td>>hg38_21_+_8433172_8446622_RNA45SN1_45S_50ntFlanks@9454.9488.35|CCGCGCCGGGGAGGTGGAGCACGAGCGCACGTGTT</td><td>>hg38_21_+_8433172_8446622_RNA45SN1_45S_50ntFlanks</td><td>9454</td><td>9488</td><td>35</td><td>CCGCGCCGGGGAGGTGGAGCACGAGCGCACGTGTT</td><td>148.6</td><td>-7.0</td><td>1.2</td><td>-6.0</td><td>2.0E-09</td><td>4.3E-09</td><td>MB231/MB468</td><td>NUC</td></tr>
<tr><td>>hg38_21_+_8433172_8446622_RNA45SN1_45S_50ntFlanks@12635.12656.22|AAGCTACCATCTGTGGGATTAT</td><td>>hg38_21_+_8433172_8446622_RNA45SN1_45S_50ntFlanks</td><td>12635</td><td>12656</td><td>22</td><td>AAGCTACCATCTGTGGGATTAT</td><td>241.2</td><td>-7.6</td><td>1.3</td><td>-6.0</td><td>1.9E-09</td><td>4.3E-09</td><td>MB231/MB468</td><td>MP</td></tr>
<tr><td>>hg38_21_+_8433172_8446622_RNA45SN1_45S_50ntFlanks@11580.11599.20|GAACTGGTGCGGACCAGGGG</td><td>>hg38_21_+_8433172_8446622_RNA45SN1_45S_50ntFlanks</td><td>11580</td><td>11599</td><td>20</td><td>GAACTGGTGCGGACCAGGGG</td><td>90.4</td><td>-10.4</td><td>1.7</td><td>-6.0</td><td>1.9E-09</td><td>4.3E-09</td><td>MB231/MB468</td><td>MP</td></tr>
</table>

<table>
<tr><th></th><th></th><th></th><th></th><th></th><th></th><th></th><th></th><th></th><th></th><th></th><th></th><th></th><th></th><th></th></tr>
<tr><td>>hg38_21_+_8433172_8446622_RNA45SN1_45S_50ntFlanks@8318.8356.39|ACGAGACCGATAGTCAACAAGTACCGTAAGGGAAAGTTG</td><td>>hg38_21_+_8433172_8446622_RNA45SN1_45S_50ntFlanks</td><td>8318</td><td>8356</td><td>39</td><td>ACGAGACCGATAGTCAACAAGTACCGTAAGGGAAAGTTG</td><td>113.9</td><td>-10.7</td><td>1.8</td><td>-6.0</td><td>1.9E-09</td><td>4.3E-09</td><td>MB231/MB468</td><td>MP</td></tr>
<tr><td>>hg38_21_+_8433172_8446622_RNA45SN1_45S_50ntFlanks@11769.11792.24|TAATTAGTGACGCGCATGAATGGA</td><td>>hg38_21_+_8433172_8446622_RNA45SN1_45S_50ntFlanks</td><td>11769</td><td>11792</td><td>24</td><td>TAATTAGTGACGCGCATGAATGGA</td><td>825.7</td><td>-4.0</td><td>0.7</td><td>-6.0</td><td>1.9E-09</td><td>4.3E-09</td><td>MB231/MB468</td><td>CYTO</td></tr>
<tr><td>>hg38_21_+_8433172_8446622_RNA45SN1_45S_50ntFlanks@10694.10711.18|CGGTGAGCTCTCGCTGGC</td><td>>hg38_21_+_8433172_8446622_RNA45SN1_45S_50ntFlanks</td><td>10694</td><td>10711</td><td>18</td><td>CGGTGAGCTCTCGCTGGC</td><td>1309.9</td><td>-7.2</td><td>1.2</td><td>-6.0</td><td>1.9E-09</td><td>4.3E-09</td><td>MB231/MB468</td><td>MP</td></tr>
<tr><td>>hg38_21_+_8433172_8446622_RNA45SN1_45S_50ntFlanks@3899.3937.39|CCTTCGCGGGGGGGATGCGTGCATTTATCAGATCAAAAC</td><td>>hg38_21_+_8433172_8446622_RNA45SN1_45S_50ntFlanks</td><td>3899</td><td>3937</td><td>39</td><td>CCTTCGCGGGGGGGATGCGTGCATTTATCAGATCAAAAC</td><td>485.3</td><td>-8.6</td><td>1.4</td><td>-6.0</td><td>2.0E-09</td><td>4.3E-09</td><td>MB231/MB468</td><td>NUC</td></tr>
<tr><td>>hg38_21_+_8433172_8446622_RNA45SN1_45S_50ntFlanks@11573.11602.30|CGACTTAGAACTGGTGCGGACCAGGGGAAT</td><td>>hg38_21_+_8433172_8446622_RNA45SN1_45S_50ntFlanks</td><td>11573</td><td>11602</td><td>30</td><td>CGACTTAGAACTGGTGCGGACCAGGGGAAT</td><td>2364.8</td><td>-7.6</td><td>1.3</td><td>-6.0</td><td>2.3E-09</td><td>4.3E-09</td><td>BT20/MB468</td><td>NUC</td></tr>
<tr><td>>hg38_21_+_8433172_8446622_RNA45SN1_45S_50ntFlanks@4069.4104.36|CTTTCGATGGTAGTCGCCGTGCCTACCATGGTGACC</td><td>>hg38_21_+_8433172_8446622_RNA45SN1_45S_50ntFlanks</td><td>4069</td><td>4104</td><td>36</td><td>CTTTCGATGGTAGTCGCCGTGCCTACCATGGTGACC</td><td>243.4</td><td>4.3</td><td>0.7</td><td>6.0</td><td>1.9E-09</td><td>4.3E-09</td><td>MB231/MB468</td><td>CYTO</td></tr>
<tr><td>>hg38_21_+_8433172_8446622_RNA45SN1_45S_50ntFlanks@7984.8008.25|AGATCAGACGTGGCGACCCGCTGAA</td><td>>hg38_21_+_8433172_8446622_RNA45SN1_45S_50ntFlanks</td><td>7984</td><td>8008</td><td>25</td><td>AGATCAGACGTGGCGACCCGCTGAA</td><td>2816.5</td><td>-3.9</td><td>0.6</td><td>-6.0</td><td>1.5E-09</td><td>4.3E-09</td><td>BT20/MB468</td><td>MITO</td></tr>
<tr><td>>hg38_21_+_8433172_8446622_RNA45SN1_45S_50ntFlanks@7975.8008.34|CGCGACCTCAGATCAGACGTGGCGACCCGCTGAA</td><td>>hg38_21_+_8433172_8446622_RNA45SN1_45S_50ntFlanks</td><td>7975</td><td>8008</td><td>34</td><td>CGCGACCTCAGATCAGACGTGGCGACCCGCTGAA</td><td>65193.1</td><td>5.0</td><td>0.8</td><td>6.0</td><td>1.9E-09</td><td>4.3E-09</td><td>MB231/MB468</td><td>CYTO</td></tr>
<tr><td>>hg38_21_+_8433172_8446622_RNA45SN1_45S_50ntFlanks@9352.9379.28|CCGTGGCGCAATGAAGGTGAAGGCCGGC</td><td>>hg38_21_+_8433172_8446622_RNA45SN1_45S_50ntFlanks</td><td>9352</td><td>9379</td><td>28</td><td>CCGTGGCGCAATGAAGGTGAAGGCCGGC</td><td>126.9</td><td>-9.1</td><td>1.5</td><td>-6.0</td><td>1.5E-09</td><td>4.3E-09</td><td>BT20/MB468</td><td>MITO</td></tr>
<tr><td>>hg38_21_+_8433172_8446622_RNA45SN1_45S_50ntFlanks@3979.4002.24|CGCCGGCGGCTTTGGTGACTCTAG</td><td>>hg38_21_+_8433172_8446622_RNA45SN1_45S_50ntFlanks</td><td>3979</td><td>4002</td><td>24</td><td>CGCCGGCGGCTTTGGTGACTCTAG</td><td>215.8</td><td>-9.8</td><td>1.6</td><td>-6.0</td><td>1.5E-09</td><td>4.3E-09</td><td>BT20/MB468</td><td>MITO</td></tr>
<tr><td>>hg38_21_+_8433172_8446622_RNA45SN1_45S_50ntFlanks@3899.3937.39|CCTTCGCGGGGGGGATGCGTGCATTTATCAGATCAAAAC</td><td>>hg38_21_+_8433172_8446622_RNA45SN1_45S_50ntFlanks</td><td>3899</td><td>3937</td><td>39</td><td>CCTTCGCGGGGGGGATGCGTGCATTTATCAGATCAAAAC</td><td>712.4</td><td>-12.6</td><td>2.0</td><td>-6.3</td><td>2.2E-10</td><td>4.3E-09</td><td>BT20/MB468</td><td>MP</td></tr>
<tr><td>>hg38_21_+_8433172_8446622_RNA45SN1_45S_50ntFlanks@10771.10808.38|CAGGTCTCCAAGGTGAACAGCCTCTGGCATGTTGGAAC</td><td>>hg38_21_+_8433172_8446622_RNA45SN1_45S_50ntFlanks</td><td>10771</td><td>10808</td><td>38</td><td>CAGGTCTCCAAGGTGAACAGCCTCTGGCATGTTGGAAC</td><td>97.0</td><td>-10.5</td><td>1.7</td><td>-6.0</td><td>1.9E-09</td><td>4.4E-09</td><td>MB231/MB468</td><td>MP</td></tr>
<tr><td>>trnaMT_GluTTC_MT_-_14674_14742@39.69.31,trnalookalike8_GluTTC_5_-_93905172_93905240@39.69.31|ATTGGTCGTGGTTGTAGTCCGTGCGAGAATA</td><td>>trnaMT_GluTTC_MT_-_14674_14742</td><td>39</td><td>69</td><td>31</td><td>ATTGGTCGTGGTTGTAGTCCGTGCGAGAATA</td><td>848.6</td><td>9.2</td><td>1.5</td><td>6.0</td><td>1.5E-09</td><td>4.4E-09</td><td>BT20/MB468</td><td>MITO</td></tr>
<tr><td>>hg38_21_+_8433172_8446622_RNA45SN1_45S_50ntFlanks@8179.8211.33|CTGATCGAGGCCCAGCCCGTGGACGGTGTGAGG</td><td>>hg38_21_+_8433172_8446622_RNA45SN1_45S_50ntFlanks</td><td>8179</td><td>8211</td><td>33</td><td>CTGATCGAGGCCCAGCCCGTGGACGGTGTGAGG</td><td>117.6</td><td>-9.0</td><td>1.5</td><td>-6.0</td><td>1.5E-09</td><td>4.4E-09</td><td>BT20/MB468</td><td>MITO</td></tr>
<tr><td>>hg38_21_+_8433172_8446622_RNA45SN1_45S_50ntFlanks@4046.4070.25|CATTCGAACGTCTGCCCTATCAACT</td><td>>hg38_21_+_8433172_8446622_RNA45SN1_45S_50ntFlanks</td><td>4046</td><td>4070</td><td>25</td><td>CATTCGAACGTCTGCCCTATCAACT</td><td>193.8</td><td>6.5</td><td>1.1</td><td>6.0</td><td>2.0E-09</td><td>4.4E-09</td><td>MB231/MB468</td><td>CYTO</td></tr>
<tr><td>>hg38_21_+_8433172_8446622_RNA45SN1_45S_50ntFlanks@9459.9500.42|CCGGGGAGGTGGAGCACGAGCGCACGTGTTAGGACCCGAAAG</td><td>>hg38_21_+_8433172_8446622_RNA45SN1_45S_50ntFlanks</td><td>9459</td><td>9500</td><td>42</td><td>CCGGGGAGGTGGAGCACGAGCGCACGTGTTAGGACCCGAAAG</td><td>86.9</td><td>-10.3</td><td>1.7</td><td>-6.0</td><td>1.9E-09</td><td>4.4E-09</td><td>MB231/MB468</td><td>MP</td></tr>
<tr><td>>hg38_21_+_8433172_8446622_RNA45SN1_45S_50ntFlanks@476.499.24|TTCGTGATCGATGTGGTGACGTCG</td><td>>hg38_21_+_8433172_8446622_RNA45SN1_45S_50ntFlanks</td><td>476</td><td>499</td><td>24</td><td>TTCGTGATCGATGTGGTGACGTCG</td><td>2770.5</td><td>5.9</td><td>1.0</td><td>6.0</td><td>2.2E-09</td><td>4.4E-09</td><td>MB231/MB468</td><td>MITO</td></tr>
<tr><td>>hg38_21_+_8433172_8446622_RNA45SN1_45S_50ntFlanks@12633.12660.28|CGAAGCTACCATCTGTGGGATTATGACT</td><td>>hg38_21_+_8433172_8446622_RNA45SN1_45S_50ntFlanks</td><td>12633</td><td>12660</td><td>28</td><td>CGAAGCTACCATCTGTGGGATTATGACT</td><td>388.1</td><td>-6.1</td><td>1.0</td><td>-6.0</td><td>2.2E-09</td><td>4.4E-09</td><td>MB231/MB468</td><td>MITO</td></tr>
<tr><td>>hg38_21_+_8433172_8446622_RNA45SN1_45S_50ntFlanks@12072.12095.24|CCCGAGGGGCTCTCGCTTCTGGCG</td><td>>hg38_21_+_8433172_8446622_RNA45SN1_45S_50ntFlanks</td><td>12072</td><td>12095</td><td>24</td><td>CCCGAGGGGCTCTCGCTTCTGGCG</td><td>2565.2</td><td>-6.5</td><td>1.1</td><td>-6.0</td><td>1.7E-09</td><td>4.4E-09</td><td>BT20/MB468</td><td>CYTO</td></tr>
<tr><td>>hg38_21_+_8433172_8446622_RNA45SN1_45S_50ntFlanks@4753.4782.30|AAGACGATCAGATACCGTCGTAGTTCCGAC</td><td>>hg38_21_+_8433172_8446622_RNA45SN1_45S_50ntFlanks</td><td>4753</td><td>4782</td><td>30</td><td>AAGACGATCAGATACCGTCGTAGTTCCGAC</td><td>443.2</td><td>-3.7</td><td>0.6</td><td>-6.0</td><td>1.7E-09</td><td>4.4E-09</td><td>BT20/MB468</td><td>CYTO</td></tr>
<tr><td>>MI0016887|hsa-mir-4521&WithFlank&17|+|8186939|8187010@14.34.21&[MIMAT0019058&hsa-miR-4521&offsets|+4|+3;m-765&17|+|8186948|8186969&offsets|+4|+3]|AGGAAGTCCTGTGCTCAGTTT</td><td>>MI0016887|hsa-mir-4521&WithFlank&17|+|8186939|8187010</td><td>14</td><td>34</td><td>21</td><td>AGGAAGTCCTGTGCTCAGTTT</td><td>806.9</td><td>10.0</td><td>1.7</td><td>6.0</td><td>1.7E-09</td><td>4.4E-09</td><td>BT20/MB468</td><td>CYTO</td></tr>
<tr><td>>am_MI0005763|hsa-mir-941-1&WithFlank&20|+|63919443|63919526@53.75.23;MI0005764|hsa-mir-941-2&WithFlank&20|+|63919499|63919582@53.75.23;MI0005765|hsa-mir-941-3&WithFlank&20|+|63919555|63919638@53.75.23;MI0005766|hsa-mir-941-4&WithFlank&20|+|63919750|63919833@53.75.23;MI0031520|hsa-mir-941-5&WithFlank&20|+|63919862|63919945@53.75.23&[MIMAT0004984&hsa-miR-941&offsets|0|0;m-263&20|+|63919495|63919517&offsets|0|0];[MIMAT0004984_1&hsa-miR-941&offsets|0|0;m-263&20|+|63919551|63919573&offsets|0|0];[MIMAT0004984_2&hsa-miR-941&offsets|0|0;m-263&20|+|63919607|63919629&offsets|0|0];[MIMAT0004984_3&hsa-miR-941&offsets|0|0;m-263&20|+|63919802|63919824&offsets|0|0];[MIMAT0004984_4&hsa-miR-941&offsets|0|0;m-263&20|+|63919914|63919936&offsets|0|0]|CACCCGGCTGTGTGCACATGTGC</td><td>>am_MI0005763|hsa-mir-941-1&WithFlank&20|+|63919443|63919526</td><td>53</td><td>75</td><td>23</td><td>CACCCGGCTGTGTGCACATGTGC</td><td>5331.7</td><td>4.9</td><td>0.8</td><td>6.3</td><td>2.2E-10</td><td>4.4E-09</td><td>BT20/MB468</td><td>MP</td></tr>
<tr><td>>hg38_21_+_8433172_8446622_RNA45SN1_45S_50ntFlanks@10481.10498.18|ATCGAAAGGGAGTCGGGT</td><td>>hg38_21_+_8433172_8446622_RNA45SN1_45S_50ntFlanks</td><td>10481</td><td>10498</td><td>18</td><td>ATCGAAAGGGAGTCGGGT</td><td>167.2</td><td>-10.5</td><td>1.6</td><td>-6.3</td><td>2.2E-10</td><td>4.4E-09</td><td>BT20/MB468</td><td>MP</td></tr>
<tr><td>>hg38_21_+_8433172_8446622_RNA45SN1_45S_50ntFlanks@11214.11234.21|ACTCTGGACGCGAGCCGGGCC</td><td>>hg38_21_+_8433172_8446622_RNA45SN1_45S_50ntFlanks</td><td>11214</td><td>11234</td><td>21</td><td>ACTCTGGACGCGAGCCGGGCC</td><td>97.3</td><td>-10.5</td><td>1.7</td><td>-6.0</td><td>1.9E-09</td><td>4.4E-09</td><td>MB231/MB468</td><td>MP</td></tr>
<tr><td>>hg38_21_+_8433172_8446622_RNA45SN1_45S_50ntFlanks@11582.11602.21|ACTGGTGCGGACCAGGGGAAT</td><td>>hg38_21_+_8433172_8446622_RNA45SN1_45S_50ntFlanks</td><td>11582</td><td>11602</td><td>21</td><td>ACTGGTGCGGACCAGGGGAAT</td><td>385.3</td><td>-2.6</td><td>0.4</td><td>-6.0</td><td>2.3E-09</td><td>4.4E-09</td><td>BT20/MB468</td><td>NUC</td></tr>
<tr><td>>MI0000446|hsa-mir-125b-1&WithFlank&11|-|122099751|122099850@61.81.21(+1C)&[MIMAT0004592&hsa-miR-125b-1-3p&offsets|0|-1(+1C);m-485&11|-|122099769|122099790&offsets|0|-1(+1C)]|ACGGGTTAGGCTCTTGGGAGCC</td><td>>MI0000446|hsa-mir-125b-1&WithFlank&11|-|122099751|122099850</td><td>61</td><td>81</td><td>21(+1C)</td><td>ACGGGTTAGGCTCTTGGGAGCC</td><td>339.2</td><td>-11.8</td><td>1.8</td><td>-6.6</td><td>4.4E-11</td><td>4.4E-09</td><td>BT20/MB231</td><td>MP</td></tr>
<tr><td>>hg38_21_+_8433172_8446622_RNA45SN1_45S_50ntFlanks@3975.4001.27|CGGGCGCCGGCGGCTTTGGTGACTCTA</td><td>>hg38_21_+_8433172_8446622_RNA45SN1_45S_50ntFlanks</td><td>3975</td><td>4001</td><td>27</td><td>CGGGCGCCGGCGGCTTTGGTGACTCTA</td><td>125.3</td><td>-10.8</td><td>1.8</td><td>-6.0</td><td>1.9E-09</td><td>4.4E-09</td><td>MB231/MB468</td><td>MP</td></tr>
<tr><td>>hg38_21_+_8433172_8446622_RNA45SN1_45S_50ntFlanks@10500.10533.34|CAGATCCCCGAATCCGGAGTGGCGGAGATGGGCG</td><td>>hg38_21_+_8433172_8446622_RNA45SN1_45S_50ntFlanks</td><td>10500</td><td>10533</td><td>34</td><td>CAGATCCCCGAATCCGGAGTGGCGGAGATGGGCG</td><td>166.8</td><td>-10.5</td><td>1.6</td><td>-6.3</td><td>2.2E-10</td><td>4.4E-09</td><td>BT20/MB468</td><td>MP</td></tr>
<tr><td>>hg38_21_+_8433172_8446622_RNA45SN1_45S_50ntFlanks@3833.3860.28|CGCTCCTCTCCTACTTGGATAACTGTGG</td><td>>hg38_21_+_8433172_8446622_RNA45SN1_45S_50ntFlanks</td><td>3833</td><td>3860</td><td>28</td><td>CGCTCCTCTCCTACTTGGATAACTGTGG</td><td>928.2</td><td>-7.0</td><td>1.2</td><td>-6.0</td><td>2.3E-09</td><td>4.4E-09</td><td>BT20/MB468</td><td>NUC</td></tr>
<tr><td>>hg38_21_+_8433172_8446622_RNA45SN1_45S_50ntFlanks@12627.12661.35|ATGGGGCGAAGCTACCATCTGTGGGATTATGACTG</td><td>>hg38_21_+_8433172_8446622_RNA45SN1_45S_50ntFlanks</td><td>12627</td><td>12661</td><td>35</td><td>ATGGGGCGAAGCTACCATCTGTGGGATTATGACTG</td><td>2254.1</td><td>-9.2</td><td>1.5</td><td>-6.3</td><td>2.2E-10</td><td>4.4E-09</td><td>BT20/MB468</td><td>MP</td></tr>
<tr><td>>hg38_21_+_8433172_8446622_RNA45SN1_45S_50ntFlanks@4565.4587.23|AATAATGGAATAGGACCGCGGTT</td><td>>hg38_21_+_8433172_8446622_RNA45SN1_45S_50ntFlanks</td><td>4565</td><td>4587</td><td>23</td><td>AATAATGGAATAGGACCGCGGTT</td><td>1142.6</td><td>-4.3</td><td>0.7</td><td>-6.0</td><td>2.0E-09</td><td>4.4E-09</td><td>MB231/MB468</td><td>NUC</td></tr>
<tr><td>>hg38_21_+_8433172_8446622_RNA45SN1_45S_50ntFlanks@11666.11691.26|TCTGCCCAGTGCTCTGAATGTCAAAG</td><td>>hg38_21_+_8433172_8446622_RNA45SN1_45S_50ntFlanks</td><td>11666</td><td>11691</td><td>26</td><td>TCTGCCCAGTGCTCTGAATGTCAAAG</td><td>331.6</td><td>3.8</td><td>0.6</td><td>6.0</td><td>2.0E-09</td><td>4.4E-09</td><td>MB231/MB468</td><td>CYTO</td></tr>
<tr><td>>hg38_21_+_8433172_8446622_RNA45SN1_45S_50ntFlanks@4068.4095.28|ACTTTCGATGGTAGTCGCCGTGCCTACC</td><td>>hg38_21_+_8433172_8446622_RNA45SN1_45S_50ntFlanks</td><td>4068</td><td>4095</td><td>28</td><td>ACTTTCGATGGTAGTCGCCGTGCCTACC</td><td>237.5</td><td>-6.3</td><td>1.1</td><td>-6.0</td><td>1.9E-09</td><td>4.4E-09</td><td>MB231/MB468</td><td>MP</td></tr>
<tr><td>>hg38_21_+_8433172_8446622_RNA45SN1_45S_50ntFlanks@4729.4751.23|TCAAGAACGAAAGTCGGAGGTTC</td><td>>hg38_21_+_8433172_8446622_RNA45SN1_45S_50ntFlanks</td><td>4729</td><td>4751</td><td>23</td><td>TCAAGAACGAAAGTCGGAGGTTC</td><td>741.8</td><td>-8.9</td><td>1.5</td><td>-6.0</td><td>2.3E-09</td><td>4.4E-09</td><td>BT20/MB468</td><td>NUC</td></tr>
<tr><td>>hg38_21_+_8433172_8446622_RNA45SN1_45S_50ntFlanks@11668.11703.36|TGCCCAGTGCTCTGAATGTCAAAGTGAAGAAATTCA</td><td>>hg38_21_+_8433172_8446622_RNA45SN1_45S_50ntFlanks</td><td>11668</td><td>11703</td><td>36</td><td>TGCCCAGTGCTCTGAATGTCAAAGTGAAGAAATTCA</td><td>653.8</td><td>3.6</td><td>0.6</td><td>6.0</td><td>1.7E-09</td><td>4.4E-09</td><td>BT20/MB468</td><td>CYTO</td></tr>
<tr><td>>MI0000749|hsa-mir-30e&WithFlank&1|+|40754349|40754452@65.85.21(+2U)&[MIMAT0000693&hsa-miR-30e-3p&offsets|0|-1(+2U);m-25&1|+|40754413|40754434&offsets|0|-1(+2U)]|CTTTCAGTCGGATGTTTACAGTT</td><td>>MI0000749|hsa-mir-30e&WithFlank&1|+|40754349|40754452</td><td>65</td><td>85</td><td>21(+2U)</td><td>CTTTCAGTCGGATGTTTACAGTT</td><td>1012.7</td><td>3.6</td><td>0.6</td><td>6.0</td><td>2.0E-09</td><td>4.4E-09</td><td>MB231/MB468</td><td>NUC</td></tr>
<tr><td>>hg38_21_+_8433172_8446622_RNA45SN1_45S_50ntFlanks@12456.12480.25|CCAAGCGTTGGATTGTTCACCCACT</td><td>>hg38_21_+_8433172_8446622_RNA45SN1_45S_50ntFlanks</td><td>12456</td><td>12480</td><td>25</td><td>CCAAGCGTTGGATTGTTCACCCACT</td><td>1197.4</td><td>-5.5</td><td>0.9</td><td>-6.0</td><td>2.0E-09</td><td>4.5E-09</td><td>MB231/MB468</td><td>NUC</td></tr>
<tr><td>>hg38_21_+_8433172_8446622_RNA45SN1_45S_50ntFlanks@4735.4761.27|ACGAAAGTCGGAGGTTCGAAGACGATC</td><td>>hg38_21_+_8433172_8446622_RNA45SN1_45S_50ntFlanks</td><td>4735</td><td>4761</td><td>27</td><td>ACGAAAGTCGGAGGTTCGAAGACGATC</td><td>941.5</td><td>6.2</td><td>1.0</td><td>6.0</td><td>2.0E-09</td><td>4.5E-09</td><td>MB231/MB468</td><td>CYTO</td></tr>
<tr><td>>hg38_21_+_8433172_8446622_RNA45SN1_45S_50ntFlanks@3875.3897.23|ATACATGCCGACGGGCGCTGACC</td><td>>hg38_21_+_8433172_8446622_RNA45SN1_45S_50ntFlanks</td><td>3875</td><td>3897</td><td>23</td><td>ATACATGCCGACGGGCGCTGACC</td><td>249.3</td><td>-4.5</td><td>0.8</td><td>-6.0</td><td>2.2E-09</td><td>4.5E-09</td><td>MB231/MB468</td><td>MITO</td></tr>
<tr><td>>MI0000102|hsa-mir-100&WithFlank&11|-|122152223|122152314@19.38.20&[MIMAT0000098&hsa-miR-100-5p&offsets|0|-2;m-27&11|-|122152276|122152296&offsets|0|-1]|AACCCGTAGATCCGAACTTG</td><td>>MI0000102|hsa-mir-100&WithFlank&11|-|122152223|122152314</td><td>19</td><td>38</td><td>20</td><td>AACCCGTAGATCCGAACTTG</td><td>546.8</td><td>10.1</td><td>1.7</td><td>6.0</td><td>1.5E-09</td><td>4.5E-09</td><td>BT20/MB468</td><td>MITO</td></tr>
<tr><td>>hg38_21_+_8433172_8446622_RNA45SN1_45S_50ntFlanks@11452.11478.27|CTCCTCGGCGCGCGGCGGCGGCGG</td><td>>hg38_21_+_8433172_8446622_RNA45SN1_45S_50ntFlanks</td><td>11452</td><td>11478</td><td>27</td><td>CTCCTCGGCGCGCGGCGGCGGCGGCGG</td><td>849.7</td><td>-7.5</td><td>1.2</td><td>-6.0</td><td>1.8E-09</td><td>4.5E-09</td><td>BT20/MB468</td><td>CYTO</td></tr>
<tr><td>>hg38_21_+_8433172_8446622_RNA45SN1_45S_50ntFlanks@8342.8373.32|CGTAAGGGAAAGTTGAAAAGAACTTTGAAGAG</td><td>>hg38_21_+_8433172_8446622_RNA45SN1_45S_50ntFlanks</td><td>8342</td><td>8373</td><td>32</td><td>CGTAAGGGAAAGTTGAAAAGAACTTTGAAGAG</td><td>645.6</td><td>10.2</td><td>1.6</td><td>6.3</td><td>2.3E-10</td><td>4.5E-09</td><td>BT20/MB468</td><td>MP</td></tr>
<tr><td>>hg38_21_+_8433172_8446622_RNA45SN1_45S_50ntFlanks@10537.10564.28|CGAGGCGTCCAGTGCGGTAACGCGACCG</td><td>>hg38_21_+_8433172_8446622_RNA45SN1_45S_50ntFlanks</td><td>10537</td><td>10564</td><td>28</td><td>CGAGGCGTCCAGTGCGGTAACGCGACCG</td><td>164.1</td><td>-10.4</td><td>1.6</td><td>-6.3</td><td>2.3E-10</td><td>4.5E-09</td><td>BT20/MB468</td><td>MP</td></tr>
<tr><td>>MI0000108|hsa-mir-103a-2&WithFlank&20|+|3917488|3917577@54.76.23;MI0000109|hsa-mir-103a-1&WithFlank&5|-|168560890|168560979@54.76.23&[MIMAT0000101&hsa-miR-103a-3p&offsets|0|0;m-12&20|+|3917541|3917563&offsets|0|0];[MIMAT0000101_1&hsa-miR-103a-3p&offsets|0|0;m-13&5|-|168560904|168560926&offsets|0|0;m-44220&5|-|168560908|168560926&offsets|0|+4]|AGCAGCATTGTACAGGGCTATGA</td><td>>MI0000108|hsa-mir-103a-2&WithFlank&20|+|3917488|3917577</td><td>54</td><td>76</td><td>23</td><td>AGCAGCATTGTACAGGGCTATGA</td><td>12429.1</td><td>5.6</td><td>0.9</td><td>6.0</td><td>2.0E-09</td><td>4.5E-09</td><td>MB231/MB468</td><td>CYTO</td></tr>
<tr><td>>hg38_21_+_8433172_8446622_RNA45SN1_45S_50ntFlanks@7990.8008.19|GACGTGGCGACCCGCTGAA</td><td>>hg38_21_+_8433172_8446622_RNA45SN1_45S_50ntFlanks</td><td>7990</td><td>8008</td><td>19</td><td>GACGTGGCGACCCGCTGAA</td><td>3513.1</td><td>-4.8</td><td>0.8</td><td>-6.0</td><td>2.2E-09</td><td>4.5E-09</td><td>MB231/MB468</td><td>MITO</td></tr>
</table>

| | | | | | | | | | | | | | |
|---|---|---|---|---|---|---|---|---|---|---|---|---|---|
| >hg38_21_+_8433172_8446622_RNA45SN1_45S_50ntFlanks@9930.9968.39\|ATATAGACAGCAGGACGGTGGCCATGGAAGTCGGAATCC | >hg38_21_+_8433172_8446622_RNA45SN1_45S_50ntFlanks | 9930 | 9968 | 39 | ATATAGACAGCAGGACGGTGGCCATGGAAGTCGGAATCC | 114.5 | -10.7 | 1.8 | -6.0 | 1.9E-09 | 4.5E-09 | MB231/MB468 | MP |
| >hg38_21_+_8433172_8446622_RNA45SN1_45S_50ntFlanks@6779.6797.19\|CCGGGGCTACGCCTGTCTG | >hg38_21_+_8433172_8446622_RNA45SN1_45S_50ntFlanks | 6779 | 6797 | 19 | CCGGGGCTACGCCTGTCTG | 78.0 | -10.1 | 1.7 | -6.0 | 1.9E-09 | 4.5E-09 | MB231/MB468 | MP |
| >MI0000115\|hsa-mir-16-2&WithFlank&3\|+\|160404739\|160404831@58.79.22&[MIMAT0004518&hsa-miR-16-2-3p&offsets\|-1\|-1;m-252&3\|+\|160404796\|160404816&offsets\|0\|+1]\|ACCAATATTACTGTGCTGCTTT | >MI0000115\|hsa-mir-16-2&WithFlank&3\|+\|160404739\|160404831 | 58 | 79 | 22 | ACCAATATTACTGTGCTGCTTT | 881.3 | 4.7 | 0.8 | 6.0 | 2.1E-09 | 4.5E-09 | MB231/MB468 | NUC |
| >hg38_21_+_8433172_8446622_RNA45SN1_45S_50ntFlanks@12163.12180.18\|ACTGGGGCGGTACACCTG | >hg38_21_+_8433172_8446622_RNA45SN1_45S_50ntFlanks | 12163 | 12180 | 18 | ACTGGGGCGGTACACCTG | 2232.7 | -7.0 | 1.2 | -6.0 | 1.5E-09 | 4.5E-09 | BT20/MB468 | MITO |
| >hg38_21_+_8433172_8446622_RNA45SN1_45S_50ntFlanks@12626.12652.27\|AATGGGGCGAAGCTACCATCTGTGGGA | >hg38_21_+_8433172_8446622_RNA45SN1_45S_50ntFlanks | 12626 | 12652 | 27 | AATGGGGCGAAGCTACCATCTGTGGGA | 188.7 | -5.4 | 0.9 | -6.0 | 2.1E-09 | 4.5E-09 | MB231/MB468 | NUC |
| >MI0003137\|hsa-mir-193b&WithFlank&16\|+\|14303961\|14304055@20.40.21(+1U)&[MIMAT0004767&hsa-miR-193b-5p&offsets\|0\|-1(+1U);m-343&16\|+\|14303980\|14304001&offsets\|0\|-1(+1U)]\|CGGGGTTTTGAGGGCGAGATGT | >MI0003137\|hsa-mir-193b&WithFlank&16\|+\|14303961\|14304055 | 20 | 40 | 21(+1U) | CGGGGTTTTGAGGGCGAGATGT | 200.1 | 11.4 | 1.8 | 6.3 | 2.3E-10 | 4.5E-09 | BT20/MB468 | MP |
| >hg38_21_+_8433172_8446622_RNA45SN1_45S_50ntFlanks@8325.8356.32\|CGATAGTCAACAAGTACCGTAAGGGAAAGTTG | >hg38_21_+_8433172_8446622_RNA45SN1_45S_50ntFlanks | 8325 | 8356 | 32 | CGATAGTCAACAAGTACCGTAAGGGAAAGTTG | 141.6 | -10.2 | 1.6 | -6.3 | 2.3E-10 | 4.5E-09 | BT20/MB468 | MP |
| >hg38_21_+_8433172_8446622_RNA45SN1_45S_50ntFlanks@11886.11903.18\|TGAGCTTGACTCTAGTCT | >hg38_21_+_8433172_8446622_RNA45SN1_45S_50ntFlanks | 11886 | 11903 | 18 | TGAGCTTGACTCTAGTCT | 296.3 | -7.9 | 1.3 | -6.0 | 1.8E-09 | 4.5E-09 | BT20/MB468 | CYTO |
| >hg38_21_+_8433172_8446622_RNA45SN1_45S_50ntFlanks@7990.8016.27\|GACGTGGCGACCCGCTGAATTTAAGCA | >hg38_21_+_8433172_8446622_RNA45SN1_45S_50ntFlanks | 7990 | 8016 | 27 | GACGTGGCGACCCGCTGAATTTAAGCA | 774.4 | 7.4 | 1.2 | 6.0 | 2.0E-09 | 4.5E-09 | MB231/MB468 | CYTO |
| >hg38_21_+_8433172_8446622_RNA45SN1_45S_50ntFlanks@5473.5498.26\|CGGCCCTGGCGGAGCGCTGAGAAGAC | >hg38_21_+_8433172_8446622_RNA45SN1_45S_50ntFlanks | 5473 | 5498 | 26 | CGGCCCTGGCGGAGCGCTGAGAAGAC | 342.5 | 4.0 | 0.7 | 6.0 | 2.0E-09 | 4.5E-09 | MB231/MB468 | CYTO |
| >MI0000108\|hsa-mir-103a-2&WithFlank&20\|+\|3917488\|3917577@54.75.22(+1U);MI0000109\|hsa-mir-103a-1&WithFlank&5\|-\|168560890\|168560979@54.75.22(+1U)&[MIMAT0000101&hsa-miR-103a-3p&offsets\|0\|-1(+1U);m-12&20\|+\|3917541\|3917563&offsets\|0\|-1(+1U)];[MIMAT0000101_1&hsa-miR-103a-3p&offsets\|0\|-1(+1U);m-13&5\|-\|168560904\|168560926&offsets\|0\|-1(+1U);m-44220&5\|-\|168560908\|168560926&offsets\|0\|+3(+1U)]\|AGCAGCATTGTACAGGGCTATGT | >MI0000108\|hsa-mir-103a-2&WithFlank&20\|+\|3917488\|3917577 | 54 | 75 | 22(+1U) | AGCAGCATTGTACAGGGCTATGT | 348.0 | 3.2 | 0.5 | 6.0 | 2.4E-09 | 4.5E-09 | BT20/MB468 | NUC |
| >am_trnaMT_ValTAC_MT_+_1602_1670@48.67.20\|CAACTTAACTTGACCGCTCT | >am_trnaMT_ValTAC_MT_+_1602_1670 | 48 | 67 | 20 | CAACTTAACTTGACCGCTCT | 21672.7 | 9.5 | 1.6 | 6.0 | 1.8E-09 | 4.5E-09 | BT20/MB468 | CYTO |
| >hg38_21_+_8433172_8446622_RNA45SN1_45S_50ntFlanks@12275.12293.19\|AGTACGAATACAGACCGTG | >hg38_21_+_8433172_8446622_RNA45SN1_45S_50ntFlanks | 12275 | 12293 | 19 | AGTACGAATACAGACCGTG | 119.5 | -10.8 | 1.8 | -6.0 | 2.0E-09 | 4.5E-09 | MB231/MB468 | MP |
| >MI0000440\|hsa-mir-27b&WithFlank&9\|+\|95085439\|95085547@67.85.19(+1A)&[MIMAT0000419&hsa-miR-27b-3p&offsets\|0\|-2(+1A);m-39&9\|+\|95085505\|95085525&offsets\|0\|-2(+1A)]\|TTCACAGTGGCTAAGTTCTA | >MI0000440\|hsa-mir-27b&WithFlank&9\|+\|95085439\|95085547 | 67 | 85 | 19(+1A) | TTCACAGTGGCTAAGTTCTA | 511.1 | 6.2 | 1.0 | 6.0 | 2.0E-09 | 4.5E-09 | MB231/MB468 | CYTO |
| >hg38_21_+_8433172_8446622_RNA45SN1_45S_50ntFlanks@12024.12055.32\|ACTACTCTGATCGTTTTTTCACTGACCCGGTG | >hg38_21_+_8433172_8446622_RNA45SN1_45S_50ntFlanks | 12024 | 12055 | 32 | ACTACTCTGATCGTTTTTTCACTGACCCGGTG | 127.7 | 10.7 | 1.7 | 6.3 | 2.3E-10 | 4.5E-09 | BT20/MB468 | MP |
| >hg38_21_+_8433172_8446622_RNA45SN1_45S_50ntFlanks@12476.12498.23\|CCACTAATAGGGAACGTGAGCTG | >hg38_21_+_8433172_8446622_RNA45SN1_45S_50ntFlanks | 12476 | 12498 | 23 | CCACTAATAGGGAACGTGAGCTG | 2283.7 | -6.1 | 1.0 | -6.0 | 2.0E-09 | 4.5E-09 | MB231/MB468 | CYTO |
| >hg38_21_+_8433172_8446622_RNA45SN1_45S_50ntFlanks@12415.12436.22\|CTTCGATGTCGGCTCTTCCTAT | >hg38_21_+_8433172_8446622_RNA45SN1_45S_50ntFlanks | 12415 | 12436 | 22 | CTTCGATGTCGGCTCTTCCTAT | 23347.4 | -7.3 | 1.2 | -6.0 | 2.4E-09 | 4.5E-09 | BT20/MB468 | NUC |
| >trna111_HisGTG_1_-_147774845_147774916@-1T.22.23,trna118_HisGTG_1_-_145396881_145396952@-1T.22.23,trna16_HisGTG_1_+_146544773_146544844@-1T.22.23,trna1_HisGTG_15_+_45493349_45493420@-1T.22.23,trna21_HisGTG_1_+_147753471_147753542@-1T.22.23,trna33_HisGTG_6_+_27125906_27125977@-1T.22.23,trna7_HisGTG_9_-_14433938_14434009@-1T.22.23,trna8_HisGTG_15_-_45492611_45492682@-1T.22.23,trna9_HisGTG_15_-_45490804_45490875@-1T.22.23\|TGCCGTGATCGTATAGTGGTTAG | >trna111_HisGTG_1_-_147774845_147774916 | -1T | 22 | 23 | TGCCGTGATCGTATAGTGGTTAG | 375.5 | 8.9 | 1.5 | 6.0 | 2.1E-09 | 4.6E-09 | MB231/MB468 | NUC |
| >hg38_21_+_8433172_8446622_RNA45SN1_45S_50ntFlanks@4982.5002.21\|ACGGACAGGATTGACAGATTG | >hg38_21_+_8433172_8446622_RNA45SN1_45S_50ntFlanks | 4982 | 5002 | 21 | ACGGACAGGATTGACAGATTG | 839.7 | -9.7 | 1.6 | -6.0 | 2.3E-09 | 4.6E-09 | MB231/MB468 | MITO |
| >am_MI0003639\|hsa-mir-625&WithFlank&14\|+\|65471096\|65471192@58.78.21&[MIMAT0004808&hsa-miR-625-3p&offsets\|0\|-1;m-137&14\|+\|65471153\|65471174&offsets\|0\|-1]\|GACTATAGAACTTTCCCCCTC | >am_MI0003639\|hsa-mir-625&WithFlank&14\|+\|65471096\|65471192 | 58 | 78 | 21 | GACTATAGAACTTTCCCCCTC | 82.3 | 9.6 | 1.6 | 6.0 | 2.0E-09 | 4.6E-09 | MB231/MB468 | MP |
| >hg38_21_+_8433172_8446622_RNA45SN1_45S_50ntFlanks@11587.11616.30\|TGCGGACCAGGGGAATCCGACTGTTTAATT | >hg38_21_+_8433172_8446622_RNA45SN1_45S_50ntFlanks | 11587 | 11616 | 30 | TGCGGACCAGGGGAATCCGACTGTTTAATT | 179.7 | 4.1 | 0.7 | 6.0 | 2.1E-09 | 4.6E-09 | MB231/MB468 | CYTO |
| >hg38_21_+_8433172_8446622_RNA45SN1_45S_50ntFlanks@7988.8016.29\|CAGACGTGGCGACCCGCTGAATTTAAGCA | >hg38_21_+_8433172_8446622_RNA45SN1_45S_50ntFlanks | 7988 | 8016 | 29 | CAGACGTGGCGACCCGCTGAATTTAAGCA | 771.0 | -4.4 | 0.7 | -6.3 | 2.3E-10 | 4.6E-09 | BT20/MB468 | MP |
| >hg38_21_+_8433172_8446622_RNA45SN1_45S_50ntFlanks@12412.12443.32\|ATCCTTCGATGTCGGCTCTTCCTATCATTGTG | >hg38_21_+_8433172_8446622_RNA45SN1_45S_50ntFlanks | 12412 | 12443 | 32 | ATCCTTCGATGTCGGCTCTTCCTATCATTGTG | 69.1 | -10.0 | 1.7 | -6.0 | 2.0E-09 | 4.6E-09 | MB231/MB468 | MP |
| >hg38_21_+_8433172_8446622_RNA45SN1_45S_50ntFlanks@8170.8201.32\|CAAGTCCTTCTGATCGAGGCCCAGCCCGTGGA | >hg38_21_+_8433172_8446622_RNA45SN1_45S_50ntFlanks | 8170 | 8201 | 32 | CAAGTCCTTCTGATCGAGGCCCAGCCCGTGGA | 614.7 | -8.0 | 1.3 | -6.0 | 2.3E-09 | 4.6E-09 | MB231/MB468 | MITO |
| >trna9_ArgTCT_1_+_94313129_94313213@1.29.29\|GGCTCCGTGGCGCAATGGATAGCGCATTG | >trna9_ArgTCT_1_+_94313129_94313213 | 1 | 29 | 29 | GGCTCCGTGGCGCAATGGATAGCGCATTG | 303.1 | -6.9 | 1.2 | -6.0 | 1.8E-09 | 4.6E-09 | BT20/MB468 | CYTO |
| >MI0000266\|hsa-mir-10a&WithFlank&17\|-\|48579832\|48579953@69.89.21&[MIMAT0004555&hsa-miR-10a-3p&offsets\|0\|-1;m-468&17\|-\|48579865\|48579885&offsets\|0\|0]\|CAAATTCGTATCTAGGGGAAT | >MI0000266\|hsa-mir-10a&WithFlank&17\|-\|48579832\|48579953 | 69 | 89 | 21 | CAAATTCGTATCTAGGGGAAT | 410.6 | 12.4 | 2.0 | 6.3 | 2.3E-10 | 4.6E-09 | BT20/MB468 | MP |
| >hg38_21_+_8433172_8446622_RNA45SN1_45S_50ntFlanks@6721.6753.33\|AATTGCAGGACACATTGATCATCGACACTTCGA | >hg38_21_+_8433172_8446622_RNA45SN1_45S_50ntFlanks | 6721 | 6753 | 33 | AATTGCAGGACACATTGATCATCGACACTTCGA | 496.7 | -5.9 | 1.0 | -6.0 | 2.3E-09 | 4.6E-09 | MB231/MB468 | MITO |
| >hg38_21_+_8433172_8446622_RNA45SN1_45S_50ntFlanks@10230.10252.23\|CCCGCGGACGCTACGCCGCGACG | >hg38_21_+_8433172_8446622_RNA45SN1_45S_50ntFlanks | 10230 | 10252 | 23 | CCCGCGGACGCTACGCCGCGACG | 211.4 | 11.5 | 1.8 | 6.3 | 2.3E-10 | 4.6E-09 | BT20/MB468 | MP |
| >hg38_21_+_8433172_8446622_RNA45SN1_45S_50ntFlanks@12513.12538.26\|TGAGACAGGTTAGTTTTACCCTACTG | >hg38_21_+_8433172_8446622_RNA45SN1_45S_50ntFlanks | 12513 | 12538 | 26 | TGAGACAGGTTAGTTTTACCCTACTG | 208.7 | -10.8 | 1.7 | -6.3 | 2.3E-10 | 4.6E-09 | BT20/MB468 | MP |
| >hg38_21_+_8433172_8446622_RNA45SN1_45S_50ntFlanks@11540.11560.21\|CGCGGCGCCTCGCCTCGGCCG | >hg38_21_+_8433172_8446622_RNA45SN1_45S_50ntFlanks | 11540 | 11560 | 21 | CGCGGCGCCTCGCCTCGGCCG | 351.5 | -8.1 | 1.3 | -6.0 | 2.0E-09 | 4.6E-09 | MB231/MB468 | MP |
| >hg38_1_-_228634819_228635039_RNA5S12_5S_50ntFlanks@153.172.20\|ACCGGGTGCTGTAGGCTTTT | >hg38_1_-_228634819_228635039_RNA5S12_5S_50ntFlanks | 153 | 172 | 20 | ACCGGGTGCTGTAGGCTTTT | 452.8 | 8.6 | 1.4 | 6.0 | 2.1E-09 | 4.6E-09 | MB231/MB468 | CYTO |
| >hg38_21_+_8433172_8446622_RNA45SN1_45S_50ntFlanks@11661.11680.20\|TGATTTCTGCCCAGTGCTCT | >hg38_21_+_8433172_8446622_RNA45SN1_45S_50ntFlanks | 11661 | 11680 | 20 | TGATTTCTGCCCAGTGCTCT | 314.8 | -4.5 | 0.7 | -6.0 | 2.3E-09 | 4.6E-09 | MB231/MB468 | MITO |
| >hg38_21_+_8433172_8446622_RNA45SN1_45S_50ntFlanks@10689.10710.22\|GCGTCCGGTGAGCTCTCGCTGG | >hg38_21_+_8433172_8446622_RNA45SN1_45S_50ntFlanks | 10689 | 10710 | 22 | GCGTCCGGTGAGCTCTCGCTGG | 188.4 | -10.6 | 1.7 | -6.3 | 2.3E-10 | 4.6E-09 | BT20/MB468 | MP |
| >hg38_21_+_8433172_8446622_RNA45SN1_45S_50ntFlanks@6661.6687.27\|CGGTGGATCACTCGGCTCGTGCGTCGA | >hg38_21_+_8433172_8446622_RNA45SN1_45S_50ntFlanks | 6661 | 6687 | 27 | CGGTGGATCACTCGGCTCGTGCGTCGA | 3076.9 | 4.7 | 0.8 | 6.0 | 1.6E-09 | 4.6E-09 | BT20/MB468 | MITO |
| >MI0000736\|hsa-mir-30c-1&WithFlank&1\|+\|40757278\|40757378@23.42.20;MI0000254\|hsa-mir-30c-2&WithFlank&6\|-\|71376954\|71377037@13.32.20&[MIMAT0000244&hsa-miR-30c-5p&offsets\|0\|-3;m-57&1\|+\|40757300\|40757323&offsets\|0\|-4];[MIMAT0000244_1&hsa-miR-30c-5p&offsets\|0\|-3;m-56&6\|-\|71377002\|71377025&offsets\|0\|-4]\|TGTAAACATCCTACACTCTC | >MI0000736\|hsa-mir-30c-1&WithFlank&1\|+\|40757278\|40757378 | 23 | 42 | 20 | TGTAAACATCCTACACTCTC | 2227.6 | 5.5 | 0.9 | 6.0 | 2.3E-09 | 4.6E-09 | MB231/MB468 | MITO |
| >hg38_21_+_8433172_8446622_RNA45SN1_45S_50ntFlanks@10798.10825.28\|CATGTTGGAACAATGTAGGTAAGGGAAG | >hg38_21_+_8433172_8446622_RNA45SN1_45S_50ntFlanks | 10798 | 10825 | 28 | CATGTTGGAACAATGTAGGTAAGGGAAG | 505.4 | -7.7 | 1.3 | -6.0 | 2.3E-09 | 4.6E-09 | MB231/MB468 | MITO |
| >hg38_21_+_8433172_8446622_RNA45SN1_45S_50ntFlanks@11323.11340.18\|CGTCTCGTCGCGCGCGCG | >hg38_21_+_8433172_8446622_RNA45SN1_45S_50ntFlanks | 11323 | 11340 | 18 | CGTCTCGTCGCGCGCGCG | 980.7 | -8.1 | 1.3 | -6.0 | 2.1E-09 | 4.6E-09 | MB231/MB468 | NUC |
| >hg38_21_+_8433172_8446622_RNA45SN1_45S_50ntFlanks@11600.11626.27\|AATCCGACTGTTTAATTAAAACAAAGC | >hg38_21_+_8433172_8446622_RNA45SN1_45S_50ntFlanks | 11600 | 11626 | 27 | AATCCGACTGTTTAATTAAAACAAAGC | 127.8 | 10.7 | 1.7 | 6.3 | 2.4E-10 | 4.6E-09 | BT20/MB468 | MP |
| >hg38_21_+_8433172_8446622_RNA45SN1_45S_50ntFlanks@12002.12029.28\|TGCGGGCCGCCGGTGAAATACCACTACT | >hg38_21_+_8433172_8446622_RNA45SN1_45S_50ntFlanks | 12002 | 12029 | 28 | TGCGGGCCGCCGGTGAAATACCACTACT | 144.6 | -10.3 | 1.6 | -6.3 | 2.4E-10 | 4.6E-09 | BT20/MB468 | MP |
| >hg38_21_+_8433172_8446622_RNA45SN1_45S_50ntFlanks@10350.10381.32\|ACGAGAACTTTGAAGGCCGAAGTGGAGAAGGG | >hg38_21_+_8433172_8446622_RNA45SN1_45S_50ntFlanks | 10350 | 10381 | 32 | ACGAGAACTTTGAAGGCCGAAGTGGAGAAGGG | 896.1 | -6.9 | 1.1 | -6.0 | 2.0E-09 | 4.6E-09 | MB231/MB468 | MP |
| >MI0000809\|hsa-mir-151a&WithFlank&8\|-\|140732558\|140732659@51.72.22&[MIMAT0000757&hsa-miR-151a-3p&offsets\|-2\|-1;m-30&8\|-\|140732587\|140732607&offsets\|-2\|-1]\|TACTAGACTGAAGCTCCTTGAG | >MI0000809\|hsa-mir-151a&WithFlank&8\|-\|140732558\|140732659 | 51 | 72 | 22 | TACTAGACTGAAGCTCCTTGAG | 477.2 | 5.6 | 0.9 | 6.0 | 2.3E-09 | 4.6E-09 | MB231/MB468 | MITO |
| >hg38_21_+_8433172_8446622_RNA45SN1_45S_50ntFlanks@12444.12484.41\|AAGCAGAATTCACCAAGCGTTGGATTGTTCACCCACTAATA | >hg38_21_+_8433172_8446622_RNA45SN1_45S_50ntFlanks | 12444 | 12484 | 41 | AAGCAGAATTCACCAAGCGTTGGATTGTTCACCCACTAATA | 103.5 | -10.6 | 1.8 | -6.0 | 2.0E-09 | 4.6E-09 | MB231/MB468 | MP |
| >hg38_21_+_8433172_8446622_RNA45SN1_45S_50ntFlanks@5379.5410.32\|ATTAAGTCCCTGCCCTTTGTACACACCGCCCG | >hg38_21_+_8433172_8446622_RNA45SN1_45S_50ntFlanks | 5379 | 5410 | 32 | ATTAAGTCCCTGCCCTTTGTACACACCGCCCG | 257.3 | -10.1 | 1.7 | -6.0 | 1.6E-09 | 4.7E-09 | BT20/MB468 | MITO |
| >trnaMT_PheGAA_MT_+_577_647@35.54.20\|GAAAATGTTTAGACGGGCTC | >trnaMT_PheGAA_MT_+_577_647 | 35 | 54 | 20 | GAAAATGTTTAGACGGGCTC | 678.5 | 5.7 | 0.9 | 6.0 | 2.3E-09 | 4.7E-09 | MB231/MB468 | MITO |
| >hg38_21_+_8433172_8446622_RNA45SN1_45S_50ntFlanks@9570.9596.27\|TGCAAATCGGTCGTCCGACCTGGGTAT | >hg38_21_+_8433172_8446622_RNA45SN1_45S_50ntFlanks | 9570 | 9596 | 27 | TGCAAATCGGTCGTCCGACCTGGGTAT | 3377.1 | -7.2 | 1.2 | -6.0 | 1.6E-09 | 4.7E-09 | BT20/MB468 | MITO |
| >trna13_ValCAC_19_-_4724647_4724719@1.34.34\|GTTTCCGTAGTGTAGCGGTTATCACATTCGCCTC | >trna13_ValCAC_19_-_4724647_4724719 | 1 | 34 | 34 | GTTTCCGTAGTGTAGCGGTTATCACATTCGCCTC | 260.5 | -11.1 | 1.8 | -6.3 | 2.2E-10 | 4.7E-09 | BT20/MB231 | NUC |
| >hg38_21_+_8433172_8446622_RNA45SN1_45S_50ntFlanks@11922.11950.29\|GAGAGGTGTAGAATAAGTGGGAGGCCCCC | >hg38_21_+_8433172_8446622_RNA45SN1_45S_50ntFlanks | 11922 | 11950 | 29 | GAGAGGTGTAGAATAAGTGGGAGGCCCCC | 2084.1 | -5.7 | 0.9 | -6.4 | 1.2E-10 | 4.7E-09 | BT20/MB231 | CYTO |
| >hg38_21_+_8433172_8446622_RNA45SN1_45S_50ntFlanks@3777.3801.25\|CGGTACAGTGAAACTGCGAATGGCT | >hg38_21_+_8433172_8446622_RNA45SN1_45S_50ntFlanks | 3777 | 3801 | 25 | CGGTACAGTGAAACTGCGAATGGCT | 1550.7 | -8.4 | 1.4 | -6.0 | 2.3E-09 | 4.7E-09 | MB231/MB468 | MITO |

<table>
<tr><th></th><th></th><th></th><th></th><th></th><th></th><th></th><th></th><th></th><th></th><th></th><th></th><th></th><th></th><th></th></tr>
<tr><td>>hg38_21_+_8433172_8446622_RNA45SN1_45S_50ntFlanks@12947.12988.42|AGACGACCTGCTTCTGGGTCGGGGTTTCGTACGTAGCAGAGC</td><td>>hg38_21_+_8433172_8446622_RNA45SN1_45S_50ntFlanks</td><td>12947</td><td>12988</td><td>42</td><td>AGACGACCTGCTTCTGGGTCGGGGTTTCGTACGTAGCAGAGC</td><td>341.0</td><td>4.1</td><td>0.7</td><td>6.0</td><td>2.1E-09</td><td>4.7E-09</td><td>MB231/MB468</td><td>CYTO</td></tr>
<tr><td>>MI0000750|hsa-mir-26a-2&WithFlank&12|-|57824603|57824698@20.39.20(+2U);MI0000083|hsa-mir-26a-1&WithFlank&3|+|37969398|37969486@16.35.20(+2U)&[MIMAT0000082&hsa-miR-26a-5p&offsets|0|-2(+2U);m-33&12|-|57824658|57824679&offsets|0|-2(+2U)];[MIMAT0000082_1&hsa-miR-26a-5p&offsets|0|-2(+2U);m-34&3|+|37969413|37969434&offsets|0|-2(+2U)]|TTCAAGTAATCCAGGATAGGTT</td><td>>MI0000750|hsa-mir-26a-2&WithFlank&12|-|57824603|57824698</td><td>20</td><td>39</td><td>20(+2U)</td><td>TTCAAGTAATCCAGGATAGGTT</td><td>505.3</td><td>6.5</td><td>1.1</td><td>6.0</td><td>1.8E-09</td><td>4.7E-09</td><td>BT20/MB468</td><td>CYTO</td></tr>
<tr><td>>trna15_ValAAC_5_-_180615416_180615488@1.32.32|GTTTCCGTAGTGTAGTGGTCATCACGTTCGCC</td><td>>trna15_ValAAC_5_-_180615416_180615488</td><td>1</td><td>32</td><td>32</td><td>GTTTCCGTAGTGTAGTGGTCATCACGTTCGCC</td><td>674.7</td><td>7.2</td><td>1.2</td><td>6.0</td><td>1.8E-09</td><td>4.7E-09</td><td>BT20/MB468</td><td>CYTO</td></tr>
<tr><td>>hg38_21_+_8433172_8446622_RNA45SN1_45S_50ntFlanks@5495.5516.22|AGACGGTCGAACTTGACTATCT</td><td>>hg38_21_+_8433172_8446622_RNA45SN1_45S_50ntFlanks</td><td>5495</td><td>5516</td><td>22</td><td>AGACGGTCGAACTTGACTATCT</td><td>1443.2</td><td>9.0</td><td>1.5</td><td>6.0</td><td>1.6E-09</td><td>4.7E-09</td><td>BT20/MB468</td><td>MITO</td></tr>
<tr><td>>hg38_21_+_8433172_8446622_RNA45SN1_45S_50ntFlanks@12182.12211.30|CAAACGGTAACGCAGGTGTCCTAAGGCGAG</td><td>>hg38_21_+_8433172_8446622_RNA45SN1_45S_50ntFlanks</td><td>12182</td><td>12211</td><td>30</td><td>CAAACGGTAACGCAGGTGTCCTAAGGCGAG</td><td>92.2</td><td>-8.6</td><td>1.4</td><td>-6.0</td><td>1.6E-09</td><td>4.7E-09</td><td>BT20/MB468</td><td>MITO</td></tr>
<tr><td>>MI0000542|hsa-mir-320a&WithFlank&8|-|22244960|22245043@48.69.22&[MIMAT0000510&hsa-miR-320a-3p&offsets|0|0;m-60&8|-|22244975|22244996&offsets|0|0]|AAAAGCTGGGTTGAGAGGGCGA</td><td>>MI0000542|hsa-mir-320a&WithFlank&8|-|22244960|22245043</td><td>48</td><td>69</td><td>22</td><td>AAAAGCTGGGTTGAGAGGGCGA</td><td>3711.0</td><td>4.9</td><td>0.8</td><td>6.0</td><td>2.1E-09</td><td>4.7E-09</td><td>MB231/MB468</td><td>NUC</td></tr>
<tr><td>>hg38_21_+_8433172_8446622_RNA45SN1_45S_50ntFlanks@12242.12262.21|AGAAGGGCAAAAGCTCGCTTG</td><td>>hg38_21_+_8433172_8446622_RNA45SN1_45S_50ntFlanks</td><td>12242</td><td>12262</td><td>21</td><td>AGAAGGGCAAAAGCTCGCTTG</td><td>78.2</td><td>-10.2</td><td>1.7</td><td>-6.0</td><td>2.0E-09</td><td>4.7E-09</td><td>MB231/MB468</td><td>MP</td></tr>
<tr><td>>hg38_1_-_228634819_228635039_RNA5S12_5S_50ntFlanks@131.171.41|GGATGGGAGACCGCCTGGGAATACCGGGTGCTGTAGGCTTT</td><td>>hg38_1_-_228634819_228635039_RNA5S12_5S_50ntFlanks</td><td>131</td><td>171</td><td>41</td><td>GGATGGGAGACCGCCTGGGAATACCGGGTGCTGTAGGCTTT</td><td>1046.1</td><td>7.3</td><td>1.2</td><td>6.0</td><td>1.6E-09</td><td>4.7E-09</td><td>BT20/MB468</td><td>MITO</td></tr>
<tr><td>>hg38_21_+_8433172_8446622_RNA45SN1_45S_50ntFlanks@10411.10453.43|CAGTCGGTCCTGAGAGATGGGCGAGCGCCGTTCCGAAGGGACG</td><td>>hg38_21_+_8433172_8446622_RNA45SN1_45S_50ntFlanks</td><td>10411</td><td>10453</td><td>43</td><td>CAGTCGGTCCTGAGAGATGGGCGAGCGCCGTTCCGAAGGGACG</td><td>1305.7</td><td>-5.5</td><td>0.9</td><td>-6.0</td><td>2.1E-09</td><td>4.7E-09</td><td>MB231/MB468</td><td>NUC</td></tr>
<tr><td>>MI0000301|hsa-mir-224&WithFlank&X|-|151958572|151958664@14.34.21&[MIMAT0000281&hsa-miR-224-5p&offsets|+1|-3;m-238&X|-|151958630|151958651&offsets|0|-1]|CAAGTCACTAGTGGTTCCGTT</td><td>>MI0000301|hsa-mir-224&WithFlank&X|-|151958572|151958664</td><td>14</td><td>34</td><td>21</td><td>CAAGTCACTAGTGGTTCCGTT</td><td>98.8</td><td>9.8</td><td>1.6</td><td>6.0</td><td>2.0E-09</td><td>4.7E-09</td><td>MB231/MB468</td><td>MP</td></tr>
<tr><td>>hg38_21_+_8433172_8446622_RNA45SN1_45S_50ntFlanks@12162.12179.18|GACTGGGGCGGTACACCT</td><td>>hg38_21_+_8433172_8446622_RNA45SN1_45S_50ntFlanks</td><td>12162</td><td>12179</td><td>18</td><td>GACTGGGGCGGTACACCT</td><td>271.4</td><td>-10.2</td><td>1.7</td><td>-6.0</td><td>1.6E-09</td><td>4.7E-09</td><td>BT20/MB468</td><td>MITO</td></tr>
<tr><td>>hg38_21_+_8433172_8446622_RNA45SN1_45S_50ntFlanks@5473.5509.37|CGGCCCTGGCGGAGCGCTGAGAAGACGGTCGAACTTG</td><td>>hg38_21_+_8433172_8446622_RNA45SN1_45S_50ntFlanks</td><td>5473</td><td>5509</td><td>37</td><td>CGGCCCTGGCGGAGCGCTGAGAAGACGGTCGAACTTG</td><td>1988.2</td><td>5.7</td><td>0.9</td><td>6.0</td><td>1.9E-09</td><td>4.7E-09</td><td>BT20/MB468</td><td>CYTO</td></tr>
<tr><td>>hg38_21_+_8433172_8446622_RNA45SN1_45S_50ntFlanks@11669.11692.24|GCCCAGTGCTCTGAATGTCAAAGT</td><td>>hg38_21_+_8433172_8446622_RNA45SN1_45S_50ntFlanks</td><td>11669</td><td>11692</td><td>24</td><td>GCCCAGTGCTCTGAATGTCAAAGT</td><td>373.2</td><td>7.0</td><td>1.2</td><td>6.0</td><td>2.1E-09</td><td>4.7E-09</td><td>MB231/MB468</td><td>CYTO</td></tr>
<tr><td>>hg38_21_+_8433172_8446622_RNA45SN1_45S_50ntFlanks@11874.11898.25|AGAAGACCCTGTTGAGCTTGACTCT</td><td>>hg38_21_+_8433172_8446622_RNA45SN1_45S_50ntFlanks</td><td>11874</td><td>11898</td><td>25</td><td>AGAAGACCCTGTTGAGCTTGACTCT</td><td>1651.9</td><td>-8.6</td><td>1.4</td><td>-6.0</td><td>2.0E-09</td><td>4.7E-09</td><td>MB231/MB468</td><td>MP</td></tr>
<tr><td>>MI0000736|hsa-mir-30c-1&WithFlank&1|+|40757278|40757378@23.46.24(+1U);MI0000254|hsa-mir-30c-2&WithFlank&6|-|71376954|71377037@13.36.24(+1U)&[MIMAT0000244&hsa-miR-30c-5p&offsets|0|+1(+1U);m-57&1|+|40757300|40757323&offsets|0|0(+1U)];[MIMAT0000244_1&hsa-miR-30c-5p&offsets|0|+1(+1U);m-56&6|-|71377002|71377025&offsets|0|0(+1U)]|TGTAAACATCCTACACTCTCAGCTT</td><td>>MI0000736|hsa-mir-30c-1&WithFlank&1|+|40757278|40757378</td><td>23</td><td>46</td><td>24(+1U)</td><td>TGTAAACATCCTACACTCTCAGCTT</td><td>1914.7</td><td>6.1</td><td>1.0</td><td>6.0</td><td>2.3E-09</td><td>4.7E-09</td><td>MB231/MB468</td><td>MITO</td></tr>
<tr><td>>hg38_21_+_8433172_8446622_RNA45SN1_45S_50ntFlanks@12444.12484.41|AAGCAGAATTCACCAAGCGTTGGATTGTTCACCCACTAATA</td><td>>hg38_21_+_8433172_8446622_RNA45SN1_45S_50ntFlanks</td><td>12444</td><td>12484</td><td>41</td><td>AAGCAGAATTCACCAAGCGTTGGATTGTTCACCCACTAATA</td><td>249.5</td><td>-10.1</td><td>1.7</td><td>-6.0</td><td>1.6E-09</td><td>4.7E-09</td><td>BT20/MB468</td><td>MITO</td></tr>
<tr><td>>hg38_21_+_8433172_8446622_RNA45SN1_45S_50ntFlanks@11783.11800.18|CATGAATGGATGAACGAG</td><td>>hg38_21_+_8433172_8446622_RNA45SN1_45S_50ntFlanks</td><td>11783</td><td>11800</td><td>18</td><td>CATGAATGGATGAACGAG</td><td>392.0</td><td>-5.4</td><td>0.9</td><td>-6.0</td><td>2.1E-09</td><td>4.7E-09</td><td>MB231/MB468</td><td>CYTO</td></tr>
<tr><td>>hg38_21_+_8433172_8446622_RNA45SN1_45S_50ntFlanks@10781.10817.37|AGGTGAACAGCCTCTGGCATGTTGGAACAATGTAGGT</td><td>>hg38_21_+_8433172_8446622_RNA45SN1_45S_50ntFlanks</td><td>10781</td><td>10817</td><td>37</td><td>AGGTGAACAGCCTCTGGCATGTTGGAACAATGTAGGT</td><td>572.9</td><td>-8.6</td><td>1.4</td><td>-6.0</td><td>2.5E-09</td><td>4.7E-09</td><td>BT20/MB468</td><td>NUC</td></tr>
<tr><td>>hg38_21_+_8433172_8446622_RNA45SN1_45S_50ntFlanks@9567.9595.29|ACGTGCAAATCGGTCGTCCGACCTGGGTA</td><td>>hg38_21_+_8433172_8446622_RNA45SN1_45S_50ntFlanks</td><td>9567</td><td>9595</td><td>29</td><td>ACGTGCAAATCGGTCGTCCGACCTGGGTA</td><td>2339.0</td><td>-4.4</td><td>0.7</td><td>-6.0</td><td>1.6E-09</td><td>4.7E-09</td><td>BT20/MB468</td><td>MITO</td></tr>
<tr><td>>hg38_21_+_8433172_8446622_RNA45SN1_45S_50ntFlanks@7979.8011.33|ACCTCAGATCAGACGTGGCGACCCGCTGAATTT</td><td>>hg38_21_+_8433172_8446622_RNA45SN1_45S_50ntFlanks</td><td>7979</td><td>8011</td><td>33</td><td>ACCTCAGATCAGACGTGGCGACCCGCTGAATTT</td><td>345.5</td><td>3.3</td><td>0.6</td><td>6.0</td><td>2.1E-09</td><td>4.7E-09</td><td>MB231/MB468</td><td>CYTO</td></tr>
<tr><td>>hg38_21_+_8433172_8446622_RNA45SN1_45S_50ntFlanks@12128.12146.19|ACCCGCTCCGGGGACAGTG</td><td>>hg38_21_+_8433172_8446622_RNA45SN1_45S_50ntFlanks</td><td>12128</td><td>12146</td><td>19</td><td>ACCCGCTCCGGGGACAGTG</td><td>137.7</td><td>-9.2</td><td>1.5</td><td>-6.0</td><td>1.6E-09</td><td>4.7E-09</td><td>BT20/MB468</td><td>MITO</td></tr>
<tr><td>>MI0000746|hsa-mir-99b&WithFlank&19|+|51692606|51692687@14.32.19(+1U)&[MIMAT0000689&hsa-miR-99b-5p&offsets|+1|-2(+1U);m-10&19|+|51692618|51692639&offsets|+1|-2(+1U)]|ACCCGTAGAACCGACCTTGT</td><td>>MI0000746|hsa-mir-99b&WithFlank&19|+|51692606|51692687</td><td>14</td><td>32</td><td>19(+1U)</td><td>ACCCGTAGAACCGACCTTGT</td><td>238.7</td><td>5.9</td><td>0.9</td><td>6.4</td><td>1.3E-10</td><td>4.7E-09</td><td>BT20/MB231</td><td>CYTO</td></tr>
<tr><td>>hg38_1_-_228634819_228635039_RNA5S12_5S_50ntFlanks@51.76.26|GTCTACGGCCATACCACCCTGAACGC</td><td>>hg38_1_-_228634819_228635039_RNA5S12_5S_50ntFlanks</td><td>51</td><td>76</td><td>26</td><td>GTCTACGGCCATACCACCCTGAACGC</td><td>767.7</td><td>5.4</td><td>0.9</td><td>6.0</td><td>2.1E-09</td><td>4.8E-09</td><td>MB231/MB468</td><td>CYTO</td></tr>
<tr><td>>hg38_21_+_8433172_8446622_RNA45SN1_45S_50ntFlanks@11918.11960.43|ACATGAGAGGTGTAGAATAAGTGGGAGGCCCCCGGCGCCCCCC</td><td>>hg38_21_+_8433172_8446622_RNA45SN1_45S_50ntFlanks</td><td>11918</td><td>11960</td><td>43</td><td>ACATGAGAGGTGTAGAATAAGTGGGAGGCCCCCGGCGCCCCCC</td><td>116.4</td><td>-5.3</td><td>0.9</td><td>-6.0</td><td>2.1E-09</td><td>4.8E-09</td><td>MB231/MB468</td><td>CYTO</td></tr>
<tr><td>>hg38_21_+_8433172_8446622_RNA45SN1_45S_50ntFlanks@4008.4026.19|CTCGGGCCGATCGCACGCC</td><td>>hg38_21_+_8433172_8446622_RNA45SN1_45S_50ntFlanks</td><td>4008</td><td>4026</td><td>19</td><td>CTCGGGCCGATCGCACGCC</td><td>139.3</td><td>-4.2</td><td>0.7</td><td>-6.0</td><td>2.1E-09</td><td>4.8E-09</td><td>MB231/MB468</td><td>CYTO</td></tr>
<tr><td>>hg38_21_+_8433172_8446622_RNA45SN1_45S_50ntFlanks@11785.11805.21|TGAATGGATGAACGAGATTCC</td><td>>hg38_21_+_8433172_8446622_RNA45SN1_45S_50ntFlanks</td><td>11785</td><td>11805</td><td>21</td><td>TGAATGGATGAACGAGATTCC</td><td>955.7</td><td>-6.6</td><td>1.1</td><td>-6.0</td><td>2.1E-09</td><td>4.8E-09</td><td>MB231/MB468</td><td>MP</td></tr>
<tr><td>>hg38_21_+_8433172_8446622_RNA45SN1_45S_50ntFlanks@9572.9601.30|CAAATCGGTCGTCCGACCTGGGTATAGGGG</td><td>>hg38_21_+_8433172_8446622_RNA45SN1_45S_50ntFlanks</td><td>9572</td><td>9601</td><td>30</td><td>CAAATCGGTCGTCCGACCTGGGTATAGGGG</td><td>638.5</td><td>-2.8</td><td>0.5</td><td>-6.0</td><td>1.6E-09</td><td>4.8E-09</td><td>BT20/MB468</td><td>MITO</td></tr>
<tr><td>>trnaMT_GluTTC_MT_-_14674_14742@39.72.34,trnalookalike8_GluTTC_5_-_93905172_93905240@39.72.34|ATTGGTCGTGGTTGTAGTCCGTGCGAGAATACCA</td><td>>trnaMT_GluTTC_MT_-_14674_14742</td><td>39</td><td>72</td><td>34</td><td>ATTGGTCGTGGTTGTAGTCCGTGCGAGAATACCA</td><td>18647.5</td><td>8.7</td><td>1.5</td><td>6.0</td><td>2.2E-09</td><td>4.8E-09</td><td>MB231/MB468</td><td>CYTO</td></tr>
<tr><td>>hg38_21_+_8433172_8446622_RNA45SN1_45S_50ntFlanks@8349.8373.25|GAAAGTTGAAAAGAACTTTGAAGAG</td><td>>hg38_21_+_8433172_8446622_RNA45SN1_45S_50ntFlanks</td><td>8349</td><td>8373</td><td>25</td><td>GAAAGTTGAAAAGAACTTTGAAGAG</td><td>1082.1</td><td>10.3</td><td>1.6</td><td>6.3</td><td>2.4E-10</td><td>4.8E-09</td><td>BT20/MB468</td><td>MP</td></tr>
<tr><td>>hg38_21_+_8433172_8446622_RNA45SN1_45S_50ntFlanks@9932.9950.19|ATAGACAGCAGGACGGTGG</td><td>>hg38_21_+_8433172_8446622_RNA45SN1_45S_50ntFlanks</td><td>9932</td><td>9950</td><td>19</td><td>ATAGACAGCAGGACGGTGG</td><td>93.5</td><td>-10.4</td><td>1.7</td><td>-6.0</td><td>2.1E-09</td><td>4.8E-09</td><td>MB231/MB468</td><td>MP</td></tr>
<tr><td>>MI0000737|hsa-mir-200a&WithFlank&1|+|1167857|1167958@60.81.22(+1A)&[MIMAT0000682&hsa-miR-200a-3p&offsets|0|0(+1A);m-142&1|+|1167916|1167938&offsets|0|-1(+1A)]|TAACACTGTCTGGTAACGATGTA</td><td>>MI0000737|hsa-mir-200a&WithFlank&1|+|1167857|1167958</td><td>60</td><td>81</td><td>22(+1A)</td><td>TAACACTGTCTGGTAACGATGTA</td><td>335.3</td><td>-8.1</td><td>1.4</td><td>-6.0</td><td>2.1E-09</td><td>4.8E-09</td><td>MB231/MB468</td><td>MP</td></tr>
<tr><td>>MI0001445|hsa-mir-423&WithFlank&17|+|30117073|30117178@58.80.23(+1C)&[MIMAT0001340&hsa-miR-423-3p&offsets|-1|-1(+1C);m-100&17|+|30117131|30117153&offsets|-1|-1(+1C)]|AAGCTCGGTCTGAGGCCCCTCAGC</td><td>>MI0001445|hsa-mir-423&WithFlank&17|+|30117073|30117178</td><td>58</td><td>80</td><td>23(+1C)</td><td>AAGCTCGGTCTGAGGCCCCTCAGC</td><td>473.9</td><td>6.4</td><td>1.1</td><td>6.0</td><td>2.4E-09</td><td>4.8E-09</td><td>MB231/MB468</td><td>MITO</td></tr>
<tr><td>>hg38_21_+_8433172_8446622_RNA45SN1_45S_50ntFlanks@10357.10384.28|CTTTGAAGGCCGAAGTGGAGAAGGGTTC</td><td>>hg38_21_+_8433172_8446622_RNA45SN1_45S_50ntFlanks</td><td>10357</td><td>10384</td><td>28</td><td>CTTTGAAGGCCGAAGTGGAGAAGGGTTC</td><td>217.1</td><td>-5.6</td><td>0.9</td><td>-6.0</td><td>2.2E-09</td><td>4.8E-09</td><td>MB231/MB468</td><td>NUC</td></tr>
<tr><td>>hg38_21_+_8433172_8446622_RNA45SN1_45S_50ntFlanks@12635.12660.26|AAGCTACCATCTGTGGGATTATGACT</td><td>>hg38_21_+_8433172_8446622_RNA45SN1_45S_50ntFlanks</td><td>12635</td><td>12660</td><td>26</td><td>AAGCTACCATCTGTGGGATTATGACT</td><td>839.2</td><td>-6.9</td><td>1.2</td><td>-6.0</td><td>2.1E-09</td><td>4.8E-09</td><td>MB231/MB468</td><td>MP</td></tr>
<tr><td>>hg38_21_+_8433172_8446622_RNA45SN1_45S_50ntFlanks@12872.12891.20|CGGCTGGAAAGGCGGCCGCC</td><td>>hg38_21_+_8433172_8446622_RNA45SN1_45S_50ntFlanks</td><td>12872</td><td>12891</td><td>20</td><td>CGGCTGGAAAGGCGGCCGCC</td><td>103.9</td><td>-10.6</td><td>1.8</td><td>-6.0</td><td>2.1E-09</td><td>4.8E-09</td><td>MB231/MB468</td><td>MP</td></tr>
<tr><td>>hg38_21_+_8433172_8446622_RNA45SN1_45S_50ntFlanks@5467.5494.28|GGCCCACGGCCCTGGCGGAGCGCTGAGA</td><td>>hg38_21_+_8433172_8446622_RNA45SN1_45S_50ntFlanks</td><td>5467</td><td>5494</td><td>28</td><td>GGCCCACGGCCCTGGCGGAGCGCTGAGA</td><td>123.6</td><td>-10.8</td><td>1.8</td><td>-6.0</td><td>2.1E-09</td><td>4.8E-09</td><td>MB231/MB468</td><td>MP</td></tr>
<tr><td>>hg38_21_+_8433172_8446622_RNA45SN1_45S_50ntFlanks@12420.12437.18|ATGTCGGCTCTTCCTATC</td><td>>hg38_21_+_8433172_8446622_RNA45SN1_45S_50ntFlanks</td><td>12420</td><td>12437</td><td>18</td><td>ATGTCGGCTCTTCCTATC</td><td>2201.4</td><td>-2.3</td><td>0.4</td><td>-6.0</td><td>1.7E-09</td><td>4.8E-09</td><td>BT20/MB468</td><td>MITO</td></tr>
<tr><td>>MI0000263|hsa-mir-7-1&WithFlank&9|-|83969742|83969863@30.54.25&[MIMAT0000252_2&hsa-miR-7-5p&offsets|0|+1;m-401&9|-|83969811|83969834&offsets|0|+1]|TGGAAGACTAGTGATTTTGTTGTTT</td><td>>MI0000263|hsa-mir-7-1&WithFlank&9|-|83969742|83969863</td><td>30</td><td>54</td><td>25</td><td>TGGAAGACTAGTGATTTTGTTGTTT</td><td>1725.5</td><td>6.1</td><td>1.0</td><td>6.0</td><td>2.4E-09</td><td>4.8E-09</td><td>MB231/MB468</td><td>MITO</td></tr>
<tr><td>>hg38_21_+_8433172_8446622_RNA45SN1_45S_50ntFlanks@11199.11225.27|CCGGCGGCGGCGGCGACTCTGGACGCG</td><td>>hg38_21_+_8433172_8446622_RNA45SN1_45S_50ntFlanks</td><td>11199</td><td>11225</td><td>27</td><td>CCGGCGGCGGCGGCGACTCTGGACGCG</td><td>2848.5</td><td>-8.9</td><td>1.5</td><td>-6.0</td><td>2.2E-09</td><td>4.8E-09</td><td>MB231/MB468</td><td>NUC</td></tr>
<tr><td>>hg38_21_+_8433172_8446622_RNA45SN1_45S_50ntFlanks@9562.9592.31|TCCTGACGTGCAAATCGGTCGTCCGACCTGG</td><td>>hg38_21_+_8433172_8446622_RNA45SN1_45S_50ntFlanks</td><td>9562</td><td>9592</td><td>31</td><td>TCCTGACGTGCAAATCGGTCGTCCGACCTGG</td><td>299.0</td><td>-8.0</td><td>1.3</td><td>-6.0</td><td>2.5E-09</td><td>4.9E-09</td><td>BT20/MB468</td><td>NUC</td></tr>
<tr><td>>hg38_21_+_8433172_8446622_RNA45SN1_45S_50ntFlanks@5073.5108.36|AATTCCGATAACGAACGAGACTCTGGCATGCTAACT</td><td>>hg38_21_+_8433172_8446622_RNA45SN1_45S_50ntFlanks</td><td>5073</td><td>5108</td><td>36</td><td>AATTCCGATAACGAACGAGACTCTGGCATGCTAACT</td><td>836.7</td><td>-7.4</td><td>1.2</td><td>-6.0</td><td>2.4E-09</td><td>4.9E-09</td><td>MB231/MB468</td><td>MITO</td></tr>
<tr><td>>hg38_21_+_8433172_8446622_RNA45SN1_45S_50ntFlanks@4367.4384.18|CAGTTAAAAAGCTCGTAG</td><td>>hg38_21_+_8433172_8446622_RNA45SN1_45S_50ntFlanks</td><td>4367</td><td>4384</td><td>18</td><td>CAGTTAAAAAGCTCGTAG</td><td>130.5</td><td>-10.1</td><td>1.6</td><td>-6.3</td><td>2.5E-10</td><td>4.9E-09</td><td>BT20/MB468</td><td>MP</td></tr>
<tr><td>>hg38_21_+_8433172_8446622_RNA45SN1_45S_50ntFlanks@12640.12668.29|ACCATCTGTGGGATTATGACTGAACGCCT</td><td>>hg38_21_+_8433172_8446622_RNA45SN1_45S_50ntFlanks</td><td>12640</td><td>12668</td><td>29</td><td>ACCATCTGTGGGATTATGACTGAACGCCT</td><td>310.7</td><td>7.3</td><td>1.2</td><td>6.0</td><td>2.2E-09</td><td>4.9E-09</td><td>MB231/MB468</td><td>CYTO</td></tr>
<tr><td>>hg38_21_+_8433172_8446622_RNA45SN1_45S_50ntFlanks@9032.9057.26|CCTCGTGGGGGGGCCGGGCCACCCCT</td><td>>hg38_21_+_8433172_8446622_RNA45SN1_45S_50ntFlanks</td><td>9032</td><td>9057</td><td>26</td><td>CCTCGTGGGGGGGCCGGGCCACCCCT</td><td>164.0</td><td>-10.4</td><td>1.6</td><td>-6.3</td><td>2.5E-10</td><td>4.9E-09</td><td>BT20/MB468</td><td>MP</td></tr>
<tr><td>>hg38_21_+_8433172_8446622_RNA45SN1_45S_50ntFlanks@9738.9768.31|GAAACGATCTCAACCTATTCTCAAACTTTAA</td><td>>hg38_21_+_8433172_8446622_RNA45SN1_45S_50ntFlanks</td><td>9738</td><td>9768</td><td>31</td><td>GAAACGATCTCAACCTATTCTCAAACTTTAA</td><td>335.2</td><td>-7.6</td><td>1.3</td><td>-6.0</td><td>2.4E-09</td><td>4.9E-09</td><td>MB231/MB468</td><td>MITO</td></tr>
<tr><td>>hg38_21_+_8433172_8446622_RNA45SN1_45S_50ntFlanks@3841.3861.21|TCCTACTTGGATAACTGTGGT</td><td>>hg38_21_+_8433172_8446622_RNA45SN1_45S_50ntFlanks</td><td>3841</td><td>3861</td><td>21</td><td>TCCTACTTGGATAACTGTGGT</td><td>429.1</td><td>4.0</td><td>0.7</td><td>6.0</td><td>1.9E-09</td><td>4.9E-09</td><td>BT20/MB468</td><td>CYTO</td></tr>
<tr><td>>hg38_21_+_8433172_8446622_RNA45SN1_45S_50ntFlanks@6657.6689.33|TTAGCGGTGGATCACTCGGCTCGTGCGTCGATG</td><td>>hg38_21_+_8433172_8446622_RNA45SN1_45S_50ntFlanks</td><td>6657</td><td>6689</td><td>33</td><td>TTAGCGGTGGATCACTCGGCTCGTGCGTCGATG</td><td>368.5</td><td>-8.0</td><td>1.3</td><td>-6.0</td><td>2.1E-09</td><td>4.9E-09</td><td>MB231/MB468</td><td>MP</td></tr>
<tr><td>>hg38_21_+_8433172_8446622_RNA45SN1_45S_50ntFlanks@11210.11254.45|GGCGACTCTGGACGCGAGCCGGGCCCTTCCCGTGGATCGCCCCAG</td><td>>hg38_21_+_8433172_8446622_RNA45SN1_45S_50ntFlanks</td><td>11210</td><td>11254</td><td>45</td><td>GGCGACTCTGGACGCGAGCCGGGCCCTTCCCGTGGATCGCCCCAG</td><td>2258.4</td><td>-9.3</td><td>1.5</td><td>-6.0</td><td>2.2E-09</td><td>4.9E-09</td><td>MB231/MB468</td><td>NUC</td></tr>
<tr><td>>hg38_21_+_8433172_8446622_RNA45SN1_45S_50ntFlanks@12941.12972.32|ATTCGTAGACGACCTGCTTCTGGGTCGGGGTT</td><td>>hg38_21_+_8433172_8446622_RNA45SN1_45S_50ntFlanks</td><td>12941</td><td>12972</td><td>32</td><td>ATTCGTAGACGACCTGCTTCTGGGTCGGGGTT</td><td>251.2</td><td>-7.6</td><td>1.3</td><td>-6.0</td><td>1.9E-09</td><td>4.9E-09</td><td>BT20/MB468</td><td>CYTO</td></tr>
</table>

<table>
<tr><td>>hg38_21_+_8433172_8446622_RNA45SN1_45S_50ntFlanks@11313.11344.32|CCCCACCCCACGTCTCGTCGCGCGCGCGTCCG</td><td>>hg38_21_+_8433172_8446622_RNA45SN1_45S_50ntFlanks</td><td>11313</td><td>11344</td><td>32</td><td>CCCCACCCCACGTCTCGTCGCGCGCGCGTCCG</td><td>588.4</td><td>-7.8</td><td>1.3</td><td>-6.0</td><td>2.2E-09</td><td>4.9E-09</td><td>MB231/MB468</td><td>NUC</td></tr>
<tr><td>>hg38_21_+_8433172_8446622_RNA45SN1_45S_50ntFlanks@12957.12981.25|CTTCTGGGTCGGGGTTTCGTACGTA</td><td>>hg38_21_+_8433172_8446622_RNA45SN1_45S_50ntFlanks</td><td>12957</td><td>12981</td><td>25</td><td>CTTCTGGGTCGGGGTTTCGTACGTA</td><td>944.7</td><td>-9.1</td><td>1.5</td><td>-6.0</td><td>2.5E-09</td><td>4.9E-09</td><td>BT20/MB468</td><td>NUC</td></tr>
<tr><td>>am_trnaMT_GluTTC_MT_-_14674_14742@27.52.26,trnalookalike8_GluTTC_5_-_93905172_93905240@27.52.26|GTTTTTCATATCATTGGTCGTGGTTG</td><td>>am_trnaMT_GluTTC_MT_-_14674_14742</td><td>27</td><td>52</td><td>26</td><td>GTTTTTCATATCATTGGTCGTGGTTG</td><td>368.2</td><td>-7.8</td><td>1.3</td><td>-6.0</td><td>2.1E-09</td><td>4.9E-09</td><td>MB231/MB468</td><td>MP</td></tr>
<tr><td>>hg38_21_+_8433172_8446622_RNA45SN1_45S_50ntFlanks@12182.12209.28|CAAACGGTAACGCAGGTGTCCTAAGGCG</td><td>>hg38_21_+_8433172_8446622_RNA45SN1_45S_50ntFlanks</td><td>12182</td><td>12209</td><td>28</td><td>CAAACGGTAACGCAGGTGTCCTAAGGCG</td><td>89.7</td><td>-10.3</td><td>1.7</td><td>-6.0</td><td>2.1E-09</td><td>4.9E-09</td><td>MB231/MB468</td><td>MP</td></tr>
<tr><td>>MI0000469|hsa-mir-125a&WithFlank&19|+|51693248|51693345@59.78.20(+1U)&[MIMAT0004602&hsa-miR-125a-3p&offsets|0|-2(+1U);m-333&19|+|51693306|51693326&offsets|0|-1(+1U)]|ACAGGTGAGGTTCTTGGGAGT</td><td>>MI0000469|hsa-mir-125a&WithFlank&19|+|51693248|51693345</td><td>59</td><td>78</td><td>20(+1U)</td><td>ACAGGTGAGGTTCTTGGGAGT</td><td>212.0</td><td>11.5</td><td>1.8</td><td>6.3</td><td>2.5E-10</td><td>4.9E-09</td><td>BT20/MB468</td><td>MP</td></tr>
<tr><td>>hg38_21_+_8433172_8446622_RNA45SN1_45S_50ntFlanks@12935.12953.19|TAAACCATTCGTAGACGAC</td><td>>hg38_21_+_8433172_8446622_RNA45SN1_45S_50ntFlanks</td><td>12935</td><td>12953</td><td>19</td><td>TAAACCATTCGTAGACGAC</td><td>493.3</td><td>-5.5</td><td>0.9</td><td>-6.0</td><td>2.2E-09</td><td>4.9E-09</td><td>MB231/MB468</td><td>CYTO</td></tr>
<tr><td>>hg38_21_+_8433172_8446622_RNA45SN1_45S_50ntFlanks@11787.11804.18|AATGGATGAACGAGATTC</td><td>>hg38_21_+_8433172_8446622_RNA45SN1_45S_50ntFlanks</td><td>11787</td><td>11804</td><td>18</td><td>AATGGATGAACGAGATTC</td><td>4092.0</td><td>-4.9</td><td>0.8</td><td>-6.0</td><td>2.2E-09</td><td>4.9E-09</td><td>MB231/MB468</td><td>CYTO</td></tr>
<tr><td>>hg38_21_+_8433172_8446622_RNA45SN1_45S_50ntFlanks@7974.8002.29|ACGCGACCTCAGATCAGACGTGGCGACCC</td><td>>hg38_21_+_8433172_8446622_RNA45SN1_45S_50ntFlanks</td><td>7974</td><td>8002</td><td>29</td><td>ACGCGACCTCAGATCAGACGTGGCGACCC</td><td>506.6</td><td>4.0</td><td>0.7</td><td>6.0</td><td>2.5E-09</td><td>4.9E-09</td><td>BT20/MB468</td><td>NUC</td></tr>
<tr><td>>hg38_21_+_8433172_8446622_RNA45SN1_45S_50ntFlanks@8153.8181.29|GCCGCTCGTGGGGGGCCCAAGTCCTTCTG</td><td>>hg38_21_+_8433172_8446622_RNA45SN1_45S_50ntFlanks</td><td>8153</td><td>8181</td><td>29</td><td>GCCGCTCGTGGGGGGCCCAAGTCCTTCTG</td><td>758.4</td><td>-9.0</td><td>1.5</td><td>-6.0</td><td>2.6E-09</td><td>4.9E-09</td><td>BT20/MB468</td><td>NUC</td></tr>
<tr><td>>hg38_MT_+_1621_3279_RNR2_16S_50ntFlanks@51.88.38|GCTAAACCTAGCCCCAAACCCACTCCACCTTACTACCA</td><td>>hg38_MT_+_1621_3279_RNR2_16S_50ntFlanks</td><td>51</td><td>88</td><td>38</td><td>GCTAAACCTAGCCCCAAACCCACTCCACCTTACTACCA</td><td>1377.9</td><td>6.2</td><td>1.0</td><td>6.0</td><td>2.2E-09</td><td>4.9E-09</td><td>MB231/MB468</td><td>NUC</td></tr>
<tr><td>>hg38_21_+_8433172_8446622_RNA45SN1_45S_50ntFlanks@9546.9572.27|TGGAGGTCCGTAGCGGTCCTGACGTGC</td><td>>hg38_21_+_8433172_8446622_RNA45SN1_45S_50ntFlanks</td><td>9546</td><td>9572</td><td>27</td><td>TGGAGGTCCGTAGCGGTCCTGACGTGC</td><td>373.9</td><td>-3.8</td><td>0.6</td><td>-6.0</td><td>2.6E-09</td><td>4.9E-09</td><td>BT20/MB468</td><td>NUC</td></tr>
<tr><td>>hg38_21_+_8433172_8446622_RNA45SN1_45S_50ntFlanks@11317.11342.26|ACCCCACGTCTCGTCGCGCGCGCGTC</td><td>>hg38_21_+_8433172_8446622_RNA45SN1_45S_50ntFlanks</td><td>11317</td><td>11342</td><td>26</td><td>ACCCCACGTCTCGTCGCGCGCGCGTC</td><td>3855.8</td><td>-9.0</td><td>1.5</td><td>-6.0</td><td>2.2E-09</td><td>4.9E-09</td><td>MB231/MB468</td><td>NUC</td></tr>
<tr><td>>hg38_MT_+_598_1651_RNR1_12S_50ntFlanks@52.83.32|ATAGGTTTGGTCCTAGCCTTTCTATTAGCTCT</td><td>>hg38_MT_+_598_1651_RNR1_12S_50ntFlanks</td><td>52</td><td>83</td><td>32</td><td>ATAGGTTTGGTCCTAGCCTTTCTATTAGCTCT</td><td>292.3</td><td>5.3</td><td>0.9</td><td>6.0</td><td>2.3E-09</td><td>4.9E-09</td><td>MB231/MB468</td><td>NUC</td></tr>
<tr><td>>hg38_21_+_8433172_8446622_RNA45SN1_45S_50ntFlanks@11586.11609.24|GTGCGGACCAGGGGAATCCGACTG</td><td>>hg38_21_+_8433172_8446622_RNA45SN1_45S_50ntFlanks</td><td>11586</td><td>11609</td><td>24</td><td>GTGCGGACCAGGGGAATCCGACTG</td><td>922.8</td><td>-6.8</td><td>1.1</td><td>-6.0</td><td>2.1E-09</td><td>4.9E-09</td><td>MB231/MB468</td><td>MP</td></tr>
<tr><td>>hg38_21_+_8433172_8446622_RNA45SN1_45S_50ntFlanks@10291.10318.28|CGCGGGCCCGGGTGGAGCCGCCGCAGGT</td><td>>hg38_21_+_8433172_8446622_RNA45SN1_45S_50ntFlanks</td><td>10291</td><td>10318</td><td>28</td><td>CGCGGGCCCGGGTGGAGCCGCCGCAGGT</td><td>92.4</td><td>-10.4</td><td>1.7</td><td>-6.0</td><td>2.1E-09</td><td>4.9E-09</td><td>MB231/MB468</td><td>MP</td></tr>
<tr><td>>hg38_21_+_8433172_8446622_RNA45SN1_45S_50ntFlanks@3780.3800.21|TACAGTGAAACTGCGAATGGC</td><td>>hg38_21_+_8433172_8446622_RNA45SN1_45S_50ntFlanks</td><td>3780</td><td>3800</td><td>21</td><td>TACAGTGAAACTGCGAATGGC</td><td>103.8</td><td>-10.6</td><td>1.8</td><td>-6.0</td><td>2.1E-09</td><td>4.9E-09</td><td>MB231/MB468</td><td>MP</td></tr>
<tr><td>>hg38_1_-_228634819_228635039_RNA5S12_5S_50ntFlanks@134.170.37|TGGGAGACCGCCTGGGAATACCGGGTGCTGTAGGCTT</td><td>>hg38_1_-_228634819_228635039_RNA5S12_5S_50ntFlanks</td><td>134</td><td>170</td><td>37</td><td>TGGGAGACCGCCTGGGAATACCGGGTGCTGTAGGCTT</td><td>88259.0</td><td>5.5</td><td>0.9</td><td>6.0</td><td>2.3E-09</td><td>4.9E-09</td><td>MB231/MB468</td><td>NUC</td></tr>
<tr><td>>hg38_21_+_8433172_8446622_RNA45SN1_45S_50ntFlanks@4732.4758.27|AGAACGAAAGTCGGAGGTTCGAAGACG</td><td>>hg38_21_+_8433172_8446622_RNA45SN1_45S_50ntFlanks</td><td>4732</td><td>4758</td><td>27</td><td>AGAACGAAAGTCGGAGGTTCGAAGACG</td><td>2320.3</td><td>-5.3</td><td>0.9</td><td>-6.0</td><td>2.4E-09</td><td>4.9E-09</td><td>MB231/MB468</td><td>MITO</td></tr>
<tr><td>>hg38_21_+_8433172_8446622_RNA45SN1_45S_50ntFlanks@6661.6684.24|CGGTGGATCACTCGGCTCGTGCGT</td><td>>hg38_21_+_8433172_8446622_RNA45SN1_45S_50ntFlanks</td><td>6661</td><td>6684</td><td>24</td><td>CGGTGGATCACTCGGCTCGTGCGT</td><td>405.4</td><td>3.7</td><td>0.6</td><td>6.0</td><td>2.3E-09</td><td>4.9E-09</td><td>MB231/MB468</td><td>NUC</td></tr>
<tr><td>>MI0000736|hsa-mir-30c-1&WithFlank&1|+|40757278|40757378@23.44.22(+1U);MI0000254|hsa-mir-30c-2&WithFlank&6|-|71376954|71377037@13.34.22(+1U)&[MIMAT0000244&hsa-miR-30c-5p&offsets|0|-1(+1U);m-57&1|+|40757300|40757323&offsets|0|-2(+1U)];[MIMAT0000244_1&hsa-miR-30c-5p&offsets|0|-1(+1U);m-56&6|-|71377002|71377025&offsets|0|-2(+1U)]|TGTAAACATCCTACACTCTCAGT</td><td>>MI0000736|hsa-mir-30c-1&WithFlank&1|+|40757278|40757378</td><td>23</td><td>44</td><td>22(+1U)</td><td>TGTAAACATCCTACACTCTCAGT</td><td>12874.3</td><td>6.4</td><td>1.1</td><td>6.0</td><td>2.5E-09</td><td>4.9E-09</td><td>MB231/MB468</td><td>MITO</td></tr>
<tr><td>>hg38_21_+_8433172_8446622_RNA45SN1_45S_50ntFlanks@11255.11272.18|CTGCGGCGGGCGTCGCGG</td><td>>hg38_21_+_8433172_8446622_RNA45SN1_45S_50ntFlanks</td><td>11255</td><td>11272</td><td>18</td><td>CTGCGGCGGGCGTCGCGG</td><td>90.2</td><td>-10.4</td><td>1.7</td><td>-6.0</td><td>2.1E-09</td><td>4.9E-09</td><td>MB231/MB468</td><td>MP</td></tr>
<tr><td>>hg38_21_+_8433172_8446622_RNA45SN1_45S_50ntFlanks@10845.10863.19|ACTTCGGGATAAGGATTGG</td><td>>hg38_21_+_8433172_8446622_RNA45SN1_45S_50ntFlanks</td><td>10845</td><td>10863</td><td>19</td><td>ACTTCGGGATAAGGATTGG</td><td>946.9</td><td>-9.2</td><td>1.5</td><td>-6.0</td><td>2.6E-09</td><td>4.9E-09</td><td>BT20/MB468</td><td>NUC</td></tr>
<tr><td>>hg38_21_+_8433172_8446622_RNA45SN1_45S_50ntFlanks@11782.11804.23|GCATGAATGGATGAACGAGATTC</td><td>>hg38_21_+_8433172_8446622_RNA45SN1_45S_50ntFlanks</td><td>11782</td><td>11804</td><td>23</td><td>GCATGAATGGATGAACGAGATTC</td><td>920.1</td><td>-7.1</td><td>1.2</td><td>-6.0</td><td>2.6E-09</td><td>4.9E-09</td><td>BT20/MB468</td><td>NUC</td></tr>
<tr><td>>hg38_21_+_8433172_8446622_RNA45SN1_45S_50ntFlanks@12003.12043.41|GCGGGCCGCCGGTGAAATACCACTACTCTGATCGTTTTTTC</td><td>>hg38_21_+_8433172_8446622_RNA45SN1_45S_50ntFlanks</td><td>12003</td><td>12043</td><td>41</td><td>GCGGGCCGCCGGTGAAATACCACTACTCTGATCGTTTTTTC</td><td>207.8</td><td>-10.0</td><td>1.7</td><td>-6.0</td><td>2.0E-09</td><td>4.9E-09</td><td>BT20/MB468</td><td>CYTO</td></tr>
<tr><td>>hg38_21_+_8433172_8446622_RNA45SN1_45S_50ntFlanks@6659.6689.31|AGCGGTGGATCACTCGGCTCGTGCGTCGATG</td><td>>hg38_21_+_8433172_8446622_RNA45SN1_45S_50ntFlanks</td><td>6659</td><td>6689</td><td>31</td><td>AGCGGTGGATCACTCGGCTCGTGCGTCGATG</td><td>1053.9</td><td>-6.4</td><td>1.1</td><td>-6.0</td><td>2.5E-09</td><td>5.0E-09</td><td>MB231/MB468</td><td>MITO</td></tr>
<tr><td>>MI0000093|hsa-mir-92a-1&WithFlank&13|+|91351308|91351397@54.76.23(+1U)&[MIMAT0000092&hsa-miR-92a-3p&offsets|0|+1(+1U);m-9&13|+|91351361|91351382&offsets|0|+1(+1U)]|TATTGCACTTGTCCCGGCCTGTTT</td><td>>MI0000093|hsa-mir-92a-1&WithFlank&13|+|91351308|91351397</td><td>54</td><td>76</td><td>23(+1U)</td><td>TATTGCACTTGTCCCGGCCTGTTT</td><td>430.7</td><td>4.0</td><td>0.7</td><td>6.0</td><td>2.3E-09</td><td>5.0E-09</td><td>MB231/MB468</td><td>NUC</td></tr>
<tr><td>>hg38_21_+_8433172_8446622_RNA45SN1_45S_50ntFlanks@10409.10433.25|GTCAGTCGGTCCTGAGAGATGGGCG</td><td>>hg38_21_+_8433172_8446622_RNA45SN1_45S_50ntFlanks</td><td>10409</td><td>10433</td><td>25</td><td>GTCAGTCGGTCCTGAGAGATGGGCG</td><td>318.5</td><td>-4.8</td><td>0.8</td><td>-6.0</td><td>2.2E-09</td><td>5.0E-09</td><td>MB231/MB468</td><td>CYTO</td></tr>
<tr><td>>am_trna128_GlyGCC_6_-_27870686_27870756@1.29.29,trna133_GlyCCC_1_-_16872434_16872504@1.29.29,trna18_GlyGCC_16_+_70822597_70822667@1.29.29,trna19_GlyGCC_16_+_70823410_70823480@1.29.29,trna19_GlyGCC_2_-_157257659_157257729@1.29.29,trna24_GlyGCC_16_-_70812942_70813012@1.29.29,trna25_GlyGCC_16_-_70812114_70812184@1.29.29,trna4_GlyCCC_1_+_17188416_17188486@1.29.29,trna5_GlyGCC_17_+_8029064_8029134@1.29.29,trna68_GlyGCC_1_-_161493637_161493707@1.29.29|GCATTGGTGGTTCAGTGGTAGAATTCTCG</td><td>>am_trna128_GlyGCC_6_-_27870686_27870756</td><td>1</td><td>29</td><td>29</td><td>GCATTGGTGGTTCAGTGGTAGAATTCTCG</td><td>12535.4</td><td>-5.1</td><td>0.8</td><td>-6.3</td><td>2.6E-10</td><td>5.0E-09</td><td>BT20/MB468</td><td>MP</td></tr>
<tr><td>>hg38_21_+_8433172_8446622_RNA45SN1_45S_50ntFlanks@11565.11603.39|CTAGCAGCCGACTTAGAACTGGTGCGGACCAGGGGAATC</td><td>>hg38_21_+_8433172_8446622_RNA45SN1_45S_50ntFlanks</td><td>11565</td><td>11603</td><td>39</td><td>CTAGCAGCCGACTTAGAACTGGTGCGGACCAGGGGAATC</td><td>625.5</td><td>-9.1</td><td>1.5</td><td>-6.0</td><td>2.6E-09</td><td>5.0E-09</td><td>BT20/MB468</td><td>NUC</td></tr>
<tr><td>>MI0000093|hsa-mir-92a-1&WithFlank&13|+|91351308|91351397@54.75.22;MI0000094|hsa-mir-92a-2&WithFlank&X|-|134169532|134169618@54.75.22&[MIMAT0000092&hsa-miR-92a-3p&offsets|0|0;m-9&13|+|91351361|91351382&offsets|0|0];[MIMAT0000092_1&hsa-miR-92a-3p&offsets|0|0;m-11&X|-|134169544|134169565&offsets|0|0]|TATTGCACTTGTCCCGGCCTGT</td><td>>MI0000093|hsa-mir-92a-1&WithFlank&13|+|91351308|91351397</td><td>54</td><td>75</td><td>22</td><td>TATTGCACTTGTCCCGGCCTGT</td><td>70446.8</td><td>5.3</td><td>0.9</td><td>6.0</td><td>2.5E-09</td><td>5.0E-09</td><td>MB231/MB468</td><td>MITO</td></tr>
<tr><td>>MI0000299|hsa-mir-222&WithFlank&X|-|45747009|45747130@75.98.24(+1A)&[MIMAT0000279&hsa-miR-222-3p&offsets|0|+3(+1A);m-140&X|-|45747033|45747056&offsets|0|0(+1A)]|AGCTACATCTGGCTACTGGGTCTCA</td><td>>MI0000299|hsa-mir-222&WithFlank&X|-|45747009|45747130</td><td>75</td><td>98</td><td>24(+1A)</td><td>AGCTACATCTGGCTACTGGGTCTCA</td><td>1343.3</td><td>4.6</td><td>0.8</td><td>6.0</td><td>2.2E-09</td><td>5.0E-09</td><td>MB231/MB468</td><td>MP</td></tr>
<tr><td>>hg38_21_+_8433172_8446622_RNA45SN1_45S_50ntFlanks@11729.11767.39|TAACTATGACTCTCTTAAGGTAGCCAAATGCCTCGTCAT</td><td>>hg38_21_+_8433172_8446622_RNA45SN1_45S_50ntFlanks</td><td>11729</td><td>11767</td><td>39</td><td>TAACTATGACTCTCTTAAGGTAGCCAAATGCCTCGTCAT</td><td>3812.4</td><td>-8.9</td><td>1.5</td><td>-6.0</td><td>2.0E-09</td><td>5.0E-09</td><td>BT20/MB468</td><td>CYTO</td></tr>
<tr><td>>hg38_21_+_8433172_8446622_RNA45SN1_45S_50ntFlanks@6700.6717.18|CTAGCTGCGAGAATTAAT</td><td>>hg38_21_+_8433172_8446622_RNA45SN1_45S_50ntFlanks</td><td>6700</td><td>6717</td><td>18</td><td>CTAGCTGCGAGAATTAAT</td><td>199.4</td><td>-10.7</td><td>1.7</td><td>-6.3</td><td>2.6E-10</td><td>5.0E-09</td><td>BT20/MB468</td><td>MP</td></tr>
<tr><td>>hg38_1_-_228634819_228635039_RNA5S12_5S_50ntFlanks@150.171.22|AATACCGGGTGCTGTAGGCTTT</td><td>>hg38_1_-_228634819_228635039_RNA5S12_5S_50ntFlanks</td><td>150</td><td>171</td><td>22</td><td>AATACCGGGTGCTGTAGGCTTT</td><td>7001.0</td><td>4.7</td><td>0.8</td><td>6.0</td><td>2.3E-09</td><td>5.0E-09</td><td>MB231/MB468</td><td>CYTO</td></tr>
<tr><td>>hg38_21_+_8433172_8446622_RNA45SN1_45S_50ntFlanks@10359.10382.24|TTGAAGGCCGAAGTGGAGAAGGGT</td><td>>hg38_21_+_8433172_8446622_RNA45SN1_45S_50ntFlanks</td><td>10359</td><td>10382</td><td>24</td><td>TTGAAGGCCGAAGTGGAGAAGGGT</td><td>225.2</td><td>5.9</td><td>1.0</td><td>6.0</td><td>2.3E-09</td><td>5.0E-09</td><td>MB231/MB468</td><td>CYTO</td></tr>
<tr><td>>hg38_21_+_8433172_8446622_RNA45SN1_45S_50ntFlanks@7981.8004.24|CTCAGATCAGACGTGGCGACCCGC</td><td>>hg38_21_+_8433172_8446622_RNA45SN1_45S_50ntFlanks</td><td>7981</td><td>8004</td><td>24</td><td>CTCAGATCAGACGTGGCGACCCGC</td><td>640.4</td><td>-7.8</td><td>1.3</td><td>-6.0</td><td>2.5E-09</td><td>5.0E-09</td><td>MB231/MB468</td><td>MITO</td></tr>
<tr><td>>hg38_21_+_8433172_8446622_RNA45SN1_45S_50ntFlanks@11670.11700.31|CCCAGTGCTCTGAATGTCAAAGTGAAGAAAT</td><td>>hg38_21_+_8433172_8446622_RNA45SN1_45S_50ntFlanks</td><td>11670</td><td>11700</td><td>31</td><td>CCCAGTGCTCTGAATGTCAAAGTGAAGAAAT</td><td>2044.3</td><td>-9.6</td><td>1.6</td><td>-6.0</td><td>2.6E-09</td><td>5.1E-09</td><td>BT20/MB468</td><td>NUC</td></tr>
<tr><td>>hg38_21_+_8433172_8446622_RNA45SN1_45S_50ntFlanks@10475.10497.23|CGGCCGATCGAAAGGGAGTCGGG</td><td>>hg38_21_+_8433172_8446622_RNA45SN1_45S_50ntFlanks</td><td>10475</td><td>10497</td><td>23</td><td>CGGCCGATCGAAAGGGAGTCGGG</td><td>241.2</td><td>-11.0</td><td>1.7</td><td>-6.3</td><td>2.6E-10</td><td>5.1E-09</td><td>BT20/MB468</td><td>MP</td></tr>
<tr><td>>hg38_MT_+_598_1651_RNR1_12S_50ntFlanks@52.87.36|ATAGGTTTGGTCCTAGCCTTTCTATTAGCTCTTAGT</td><td>>hg38_MT_+_598_1651_RNR1_12S_50ntFlanks</td><td>52</td><td>87</td><td>36</td><td>ATAGGTTTGGTCCTAGCCTTTCTATTAGCTCTTAGT</td><td>212.0</td><td>3.6</td><td>0.6</td><td>6.0</td><td>2.3E-09</td><td>5.1E-09</td><td>MB231/MB468</td><td>NUC</td></tr>
<tr><td>>hg38_21_+_8433172_8446622_RNA45SN1_45S_50ntFlanks@9526.9572.47|GAAGCCAGAGGAAACTCTGGTGGAGGTCCGTAGCGGTCCTGACGTGC</td><td>>hg38_21_+_8433172_8446622_RNA45SN1_45S_50ntFlanks</td><td>9526</td><td>9572</td><td>47</td><td>GAAGCCAGAGGAAACTCTGGTGGAGGTCCGTAGCGGTCCTGACGTGC</td><td>408.7</td><td>6.3</td><td>1.1</td><td>6.0</td><td>2.3E-09</td><td>5.1E-09</td><td>MB231/MB468</td><td>CYTO</td></tr>
<tr><td>>am_trna162_MetCAT_6_-_26758550_26758622@31.54.24,trna164_MetCAT_6_-_26735574_26735646@31.54.24,trna20_MetCAT_16_+_71460396_71460468@31.54.24,trna21_MetCAT_6_+_26701712_26701784@31.54.24,trna27_MetCAT_6_+_26766444_26766516@31.54.24,trna75_MetCAT_6_+_28912352_28912424@31.54.24,trna97_MetCAT_6_-_28921042_28921114@31.54.24|TCTCATAATCTGAAGGTCCTGAGT</td><td>>am_trna162_MetCAT_6_-_26758550_26758622</td><td>31</td><td>54</td><td>24</td><td>TCTCATAATCTGAAGGTCCTGAGT</td><td>378.9</td><td>-4.8</td><td>0.8</td><td>-6.0</td><td>2.2E-09</td><td>5.1E-09</td><td>MB231/MB468</td><td>MP</td></tr>
<tr><td>>hg38_21_+_8433172_8446622_RNA45SN1_45S_50ntFlanks@10393.10424.32|CAGCAGTTGAACATGGGTCAGTCGGTCCTGAG</td><td>>hg38_21_+_8433172_8446622_RNA45SN1_45S_50ntFlanks</td><td>10393</td><td>10424</td><td>32</td><td>CAGCAGTTGAACATGGGTCAGTCGGTCCTGAG</td><td>15809.7</td><td>4.2</td><td>0.7</td><td>6.0</td><td>1.8E-09</td><td>5.1E-09</td><td>BT20/MB468</td><td>MITO</td></tr>
<tr><td>>hg38_21_+_8433172_8446622_RNA45SN1_45S_50ntFlanks@10377.10424.48|AAGGGTTCCATGTGAACAGCAGTTGAACATGGGTCAGTCGGTCCTGAG</td><td>>hg38_21_+_8433172_8446622_RNA45SN1_45S_50ntFlanks</td><td>10377</td><td>10424</td><td>48</td><td>AAGGGTTCCATGTGAACAGCAGTTGAACATGGGTCAGTCGGTCCTGAG</td><td>425.9</td><td>5.1</td><td>0.9</td><td>6.0</td><td>2.3E-09</td><td>5.1E-09</td><td>MB231/MB468</td><td>CYTO</td></tr>
<tr><td>>hg38_21_+_8433172_8446622_RNA45SN1_45S_50ntFlanks@6656.6679.24|CTTAGCGGTGGATCACTCGGCTCG</td><td>>hg38_21_+_8433172_8446622_RNA45SN1_45S_50ntFlanks</td><td>6656</td><td>6679</td><td>24</td><td>CTTAGCGGTGGATCACTCGGCTCG</td><td>85.8</td><td>-10.3</td><td>1.7</td><td>-6.0</td><td>2.2E-09</td><td>5.1E-09</td><td>MB231/MB468</td><td>MP</td></tr>
</table>

<table>
<tr><td>>hg38_21_+_8433172_8446622_RNA45SN1_45S_50ntFlanks@12405.12441.37|CTTTTTGATCCTTCGATGTCGGCTCTTCCTATCATTG</td><td>>hg38_21_+_8433172_8446622_RNA45SN1_45S_50ntFlanks</td><td>12405</td><td>12441</td><td>37</td><td>CTTTTTGATCCTTCGATGTCGGCTCTTCCTATCATTG</td><td>2184.2</td><td>-6.5</td><td>1.1</td><td>-6.0</td><td>2.0E-09</td><td>5.1E-09</td><td>BT20/MB468</td><td>CYTO</td></tr>
<tr><td>>hg38_21_+_8433172_8446622_RNA45SN1_45S_50ntFlanks@3813.3852.40|TTATGGTTCCTTTGGTCGCTCGCTCCTCTCCTACTTGGAT</td><td>>hg38_21_+_8433172_8446622_RNA45SN1_45S_50ntFlanks</td><td>3813</td><td>3852</td><td>40</td><td>TTATGGTTCCTTTGGTCGCTCGCTCCTCTCCTACTTGGAT</td><td>776.8</td><td>-9.4</td><td>1.6</td><td>-6.0</td><td>2.7E-09</td><td>5.1E-09</td><td>BT20/MB468</td><td>NUC</td></tr>
<tr><td>>hg38_21_+_8433172_8446622_RNA45SN1_45S_50ntFlanks@11583.11602.20|CTGGTGCGGACCAGGGGAAT</td><td>>hg38_21_+_8433172_8446622_RNA45SN1_45S_50ntFlanks</td><td>11583</td><td>11602</td><td>20</td><td>CTGGTGCGGACCAGGGGAAT</td><td>1307.4</td><td>-8.5</td><td>1.4</td><td>-6.0</td><td>2.2E-09</td><td>5.1E-09</td><td>MB231/MB468</td><td>MP</td></tr>
<tr><td>>hg38_21_+_8433172_8446622_RNA45SN1_45S_50ntFlanks@11592.11609.18|ACCAGGGGAATCCGACTG</td><td>>hg38_21_+_8433172_8446622_RNA45SN1_45S_50ntFlanks</td><td>11592</td><td>11609</td><td>18</td><td>ACCAGGGGAATCCGACTG</td><td>175.3</td><td>-10.5</td><td>1.7</td><td>-6.3</td><td>2.6E-10</td><td>5.1E-09</td><td>BT20/MB468</td><td>MP</td></tr>
<tr><td>>hg38_21_+_8433172_8446622_RNA45SN1_45S_50ntFlanks@10412.10433.22|AGTCGGTCCTGAGAGATGGGCG</td><td>>hg38_21_+_8433172_8446622_RNA45SN1_45S_50ntFlanks</td><td>10412</td><td>10433</td><td>22</td><td>AGTCGGTCCTGAGAGATGGGCG</td><td>87.6</td><td>-10.3</td><td>1.7</td><td>-6.0</td><td>2.2E-09</td><td>5.1E-09</td><td>MB231/MB468</td><td>MP</td></tr>
<tr><td>>hg38_21_+_8433172_8446622_RNA45SN1_45S_50ntFlanks@5188.5207.20|ATAACAGGTCTGTGATGCCC</td><td>>hg38_21_+_8433172_8446622_RNA45SN1_45S_50ntFlanks</td><td>5188</td><td>5207</td><td>20</td><td>ATAACAGGTCTGTGATGCCC</td><td>96.9</td><td>-10.5</td><td>1.7</td><td>-6.0</td><td>2.2E-09</td><td>5.1E-09</td><td>MB231/MB468</td><td>MP</td></tr>
<tr><td>>hg38_21_+_8433172_8446622_RNA45SN1_45S_50ntFlanks@475.511.37|CTTCGTGATCGATGTGGTGACGTCGTGCTCTCCCGGG</td><td>>hg38_21_+_8433172_8446622_RNA45SN1_45S_50ntFlanks</td><td>475</td><td>511</td><td>37</td><td>CTTCGTGATCGATGTGGTGACGTCGTGCTCTCCCGGG</td><td>826.3</td><td>7.8</td><td>1.3</td><td>6.0</td><td>2.3E-09</td><td>5.1E-09</td><td>MB231/MB468</td><td>CYTO</td></tr>
<tr><td>>hg38_21_+_8433172_8446622_RNA45SN1_45S_50ntFlanks@5489.5516.28|CTGAGAAGACGGTCGAACTTGACTATCT</td><td>>hg38_21_+_8433172_8446622_RNA45SN1_45S_50ntFlanks</td><td>5489</td><td>5516</td><td>28</td><td>CTGAGAAGACGGTCGAACTTGACTATCT</td><td>14921.3</td><td>4.9</td><td>0.8</td><td>6.0</td><td>2.3E-09</td><td>5.1E-09</td><td>MB231/MB468</td><td>CYTO</td></tr>
<tr><td>>hg38_21_+_8433172_8446622_RNA45SN1_45S_50ntFlanks@8112.8137.26|GGGCGCGGGACATGTGGCGTACGGAA</td><td>>hg38_21_+_8433172_8446622_RNA45SN1_45S_50ntFlanks</td><td>8112</td><td>8137</td><td>26</td><td>GGGCGCGGGACATGTGGCGTACGGAA</td><td>86.8</td><td>-10.3</td><td>1.7</td><td>-6.0</td><td>2.2E-09</td><td>5.1E-09</td><td>MB231/MB468</td><td>MP</td></tr>
<tr><td>>MI0006444|hsa-mir-1307&WithFlank&10|-|103394247|103394407@86.107.22(+1U)&[MIMAT0005951&hsa-miR-1307-3p&offsets|0|0(+1U);m-66&10|-|103394300|103394322&offsets|0|-1(+1U)]|ACTCGGCGTGGCGTCGGTCGTGT</td><td>>MI0006444|hsa-mir-1307&WithFlank&10|-|103394247|103394407</td><td>86</td><td>107</td><td>22(+1U)</td><td>ACTCGGCGTGGCGTCGGTCGTGT</td><td>991.6</td><td>6.0</td><td>1.0</td><td>6.0</td><td>2.6E-09</td><td>5.1E-09</td><td>MB231/MB468</td><td>MITO</td></tr>
<tr><td>>trna10_AspGTC_12_-_125424193_125424264@31.75.45,trna12_AspGTC_12_-_125411891_125411962@31.75.45,trna38_AspGTC_17_-_8125556_8125627@31.75.45,trna45_AspGTC_6_+_27447453_27447524@31.75.45,trna48_AspGTC_6_+_27471523_27471594@31.75.45,trna4_AspGTC_12_+_96429799_96429870@31.75.45,trna69_AspGTC_1_-_161440205_161440276@31.75.45,trna72_AspGTC_1_-_161432824_161432895@31.75.45,trna75_AspGTC_1_-_161425414_161425485@31.75.45,trna78_AspGTC_1_-_161418033_161418104@31.75.45,trna81_AspGTC_1_-_161410615_161410686@31.75.45|CCTGTCACGCGGGAGACCGGGGTTCGATTCCCCGACGGGGAGCCA</td><td>>trna10_AspGTC_12_-_125424193_125424264</td><td>31</td><td>75</td><td>45</td><td>CCTGTCACGCGGGAGACCGGGGTTCGATTCCCCGACGGGGAGCCA</td><td>179.1</td><td>-11.3</td><td>1.9</td><td>-6.0</td><td>2.6E-09</td><td>5.1E-09</td><td>MB231/MB468</td><td>MITO</td></tr>
<tr><td>>hg38_21_+_8433172_8446622_RNA45SN1_45S_50ntFlanks@5473.5500.28|CGGCCCTGGCGGAGCGCTGAGAAGACGG</td><td>>hg38_21_+_8433172_8446622_RNA45SN1_45S_50ntFlanks</td><td>5473</td><td>5500</td><td>28</td><td>CGGCCCTGGCGGAGCGCTGAGAAGACGG</td><td>466.4</td><td>7.6</td><td>1.3</td><td>6.0</td><td>2.3E-09</td><td>5.1E-09</td><td>MB231/MB468</td><td>CYTO</td></tr>
<tr><td>>hg38_21_+_8433172_8446622_RNA45SN1_45S_50ntFlanks@5485.5518.34|AGCGCTGAGAAGACGGTCGAACTTGACTATCTAG</td><td>>hg38_21_+_8433172_8446622_RNA45SN1_45S_50ntFlanks</td><td>5485</td><td>5518</td><td>34</td><td>AGCGCTGAGAAGACGGTCGAACTTGACTATCTAG</td><td>621.5</td><td>-8.1</td><td>1.4</td><td>-6.0</td><td>2.4E-09</td><td>5.1E-09</td><td>MB231/MB468</td><td>NUC</td></tr>
<tr><td>>hg38_21_+_8433172_8446622_RNA45SN1_45S_50ntFlanks@10362.10390.29|AAGGCCGAAGTGGAGAAGGGTTCCATGTG</td><td>>hg38_21_+_8433172_8446622_RNA45SN1_45S_50ntFlanks</td><td>10362</td><td>10390</td><td>29</td><td>AAGGCCGAAGTGGAGAAGGGTTCCATGTG</td><td>2407.9</td><td>-7.1</td><td>1.2</td><td>-6.0</td><td>2.0E-09</td><td>5.1E-09</td><td>BT20/MB468</td><td>CYTO</td></tr>
<tr><td>>hg38_21_+_8433172_8446622_RNA45SN1_45S_50ntFlanks@12942.12966.25|TTCGTAGACGACCTGCTTCTGGGTC</td><td>>hg38_21_+_8433172_8446622_RNA45SN1_45S_50ntFlanks</td><td>12942</td><td>12966</td><td>25</td><td>TTCGTAGACGACCTGCTTCTGGGTC</td><td>999.0</td><td>-8.0</td><td>1.3</td><td>-6.0</td><td>2.0E-09</td><td>5.2E-09</td><td>BT20/MB468</td><td>CYTO</td></tr>
<tr><td>>hg38_1_-_228634819_228635039_RNA5S12_5S_50ntFlanks@143.171.29|GCCTGGGAATACCGGGTGCTGTAGGCTTT</td><td>>hg38_1_-_228634819_228635039_RNA5S12_5S_50ntFlanks</td><td>143</td><td>171</td><td>29</td><td>GCCTGGGAATACCGGGTGCTGTAGGCTTT</td><td>371.5</td><td>7.4</td><td>1.2</td><td>6.0</td><td>2.3E-09</td><td>5.2E-09</td><td>MB231/MB468</td><td>CYTO</td></tr>
<tr><td>>hg38_21_+_8433172_8446622_RNA45SN1_45S_50ntFlanks@4003.4021.19|ATAACCTCGGGCCGATCGC</td><td>>hg38_21_+_8433172_8446622_RNA45SN1_45S_50ntFlanks</td><td>4003</td><td>4021</td><td>19</td><td>ATAACCTCGGGCCGATCGC</td><td>447.4</td><td>-8.6</td><td>1.4</td><td>-5.9</td><td>2.7E-09</td><td>5.2E-09</td><td>BT20/MB468</td><td>NUC</td></tr>
<tr><td>>hg38_21_+_8433172_8446622_RNA45SN1_45S_50ntFlanks@11561.11589.29|GCGCCTAGCAGCCGACTTAGAACTGGTGC</td><td>>hg38_21_+_8433172_8446622_RNA45SN1_45S_50ntFlanks</td><td>11561</td><td>11589</td><td>29</td><td>GCGCCTAGCAGCCGACTTAGAACTGGTGC</td><td>451.7</td><td>-6.2</td><td>1.0</td><td>-6.0</td><td>2.6E-09</td><td>5.2E-09</td><td>MB231/MB468</td><td>MITO</td></tr>
<tr><td>>hg38_21_+_8433172_8446622_RNA45SN1_45S_50ntFlanks@10396.10433.38|CAGTTGAACATGGGTCAGTCGGTCCTGAGAGATGGGCG</td><td>>hg38_21_+_8433172_8446622_RNA45SN1_45S_50ntFlanks</td><td>10396</td><td>10433</td><td>38</td><td>CAGTTGAACATGGGTCAGTCGGTCCTGAGAGATGGGCG</td><td>97.1</td><td>-10.5</td><td>1.8</td><td>-6.0</td><td>2.2E-09</td><td>5.2E-09</td><td>MB231/MB468</td><td>MP</td></tr>
<tr><td>>hg38_21_+_8433172_8446622_RNA45SN1_45S_50ntFlanks@3777.3810.34|CGGTACAGTGAAACTGCGAATGGCTCATTAAATC</td><td>>hg38_21_+_8433172_8446622_RNA45SN1_45S_50ntFlanks</td><td>3777</td><td>3810</td><td>34</td><td>CGGTACAGTGAAACTGCGAATGGCTCATTAAATC</td><td>88.7</td><td>-10.3</td><td>1.7</td><td>-6.0</td><td>2.2E-09</td><td>5.2E-09</td><td>MB231/MB468</td><td>MP</td></tr>
<tr><td>>hg38_21_+_8433172_8446622_RNA45SN1_45S_50ntFlanks@10410.10442.33|TCAGTCGGTCCTGAGAGATGGGCGAGCGCCGTT</td><td>>hg38_21_+_8433172_8446622_RNA45SN1_45S_50ntFlanks</td><td>10410</td><td>10442</td><td>33</td><td>TCAGTCGGTCCTGAGAGATGGGCGAGCGCCGTT</td><td>431.8</td><td>-3.4</td><td>0.6</td><td>-6.0</td><td>2.3E-09</td><td>5.2E-09</td><td>MB231/MB468</td><td>CYTO</td></tr>
<tr><td>>hg38_21_+_8433172_8446622_RNA45SN1_45S_50ntFlanks@3835.3861.27|CTCCTCTCCTACTTGGATAACTGTGGT</td><td>>hg38_21_+_8433172_8446622_RNA45SN1_45S_50ntFlanks</td><td>3835</td><td>3861</td><td>27</td><td>CTCCTCTCCTACTTGGATAACTGTGGT</td><td>1553.1</td><td>-7.8</td><td>1.3</td><td>-6.0</td><td>2.6E-09</td><td>5.2E-09</td><td>MB231/MB468</td><td>MITO</td></tr>
<tr><td>>hg38_21_+_8433172_8446622_RNA45SN1_45S_50ntFlanks@8158.8178.21|TCGTGGGGGGCCCAAGTCCTT</td><td>>hg38_21_+_8433172_8446622_RNA45SN1_45S_50ntFlanks</td><td>8158</td><td>8178</td><td>21</td><td>TCGTGGGGGGCCCAAGTCCTT</td><td>262.3</td><td>-10.1</td><td>1.7</td><td>-6.0</td><td>1.8E-09</td><td>5.2E-09</td><td>BT20/MB468</td><td>MITO</td></tr>
<tr><td>>hg38_21_+_8433172_8446622_RNA45SN1_45S_50ntFlanks@9125.9152.28|CGCGCGGGTCGGGGGGCGGGGCGGACTG</td><td>>hg38_21_+_8433172_8446622_RNA45SN1_45S_50ntFlanks</td><td>9125</td><td>9152</td><td>28</td><td>CGCGCGGGTCGGGGGGCGGGGCGGACTG</td><td>213.6</td><td>-10.8</td><td>1.7</td><td>-6.3</td><td>2.7E-10</td><td>5.2E-09</td><td>BT20/MB468</td><td>MP</td></tr>
<tr><td>>hg38_21_+_8433172_8446622_RNA45SN1_45S_50ntFlanks@10537.10567.31|CGAGGCGTCCAGTGCGGTAACGCGACCGATC</td><td>>hg38_21_+_8433172_8446622_RNA45SN1_45S_50ntFlanks</td><td>10537</td><td>10567</td><td>31</td><td>CGAGGCGTCCAGTGCGGTAACGCGACCGATC</td><td>136.0</td><td>-10.2</td><td>1.6</td><td>-6.3</td><td>2.7E-10</td><td>5.2E-09</td><td>BT20/MB468</td><td>MP</td></tr>
<tr><td>>hg38_21_+_8433172_8446622_RNA45SN1_45S_50ntFlanks@4507.4528.22|CTTTGAAAAAATTAGAGTGTTC</td><td>>hg38_21_+_8433172_8446622_RNA45SN1_45S_50ntFlanks</td><td>4507</td><td>4528</td><td>22</td><td>CTTTGAAAAAATTAGAGTGTTC</td><td>86.5</td><td>-10.3</td><td>1.7</td><td>-6.0</td><td>2.3E-09</td><td>5.2E-09</td><td>MB231/MB468</td><td>MP</td></tr>
<tr><td>>hg38_21_+_8433172_8446622_RNA45SN1_45S_50ntFlanks@10786.10815.30|AACAGCCTCTGGCATGTTGGAACAATGTAG</td><td>>hg38_21_+_8433172_8446622_RNA45SN1_45S_50ntFlanks</td><td>10786</td><td>10815</td><td>30</td><td>AACAGCCTCTGGCATGTTGGAACAATGTAG</td><td>314.7</td><td>-7.5</td><td>1.3</td><td>-6.0</td><td>2.6E-09</td><td>5.2E-09</td><td>MB231/MB468</td><td>MITO</td></tr>
<tr><td>>hg38_21_+_8433172_8446622_RNA45SN1_45S_50ntFlanks@12141.12180.40|ACAGTGCCAGGTGGGGAGTTTGACTGGGGCGGTACACCTG</td><td>>hg38_21_+_8433172_8446622_RNA45SN1_45S_50ntFlanks</td><td>12141</td><td>12180</td><td>40</td><td>ACAGTGCCAGGTGGGGAGTTTGACTGGGGCGGTACACCTG</td><td>109.7</td><td>-8.9</td><td>1.5</td><td>-6.0</td><td>1.8E-09</td><td>5.2E-09</td><td>BT20/MB468</td><td>MITO</td></tr>
<tr><td>>MI0005566|hsa-mir-374b&WithFlank&X|-|74218541|74218624@17.38.22&[MIMAT0004955&hsa-miR-374b-5p&offsets|0|0;m-153&X|-|74218587|74218608&offsets|0|0]|ATATAATACAACCTGCTAAGTG</td><td>>MI0005566|hsa-mir-374b&WithFlank&X|-|74218541|74218624</td><td>17</td><td>38</td><td>22</td><td>ATATAATACAACCTGCTAAGTG</td><td>830.6</td><td>5.2</td><td>0.9</td><td>6.0</td><td>1.8E-09</td><td>5.2E-09</td><td>BT20/MB468</td><td>MITO</td></tr>
<tr><td>>trna119_LysCTT_1_-_145395522_145395594@1.34.34,trna11_LysCTT_5_-_180648979_180649051@1.34.34,trna13_LysCTT_6_+_26556774_26556846@1.34.34,trna7_LysCTT_16_+_3225692_3225764@1.34.34,trna9_LysCTT_5_+_180634755_180634827@1.34.34|GCCCGGCTAGCTCAGTCGGTAGAGCATGAGACTC</td><td>>trna119_LysCTT_1_-_145395522_145395594</td><td>1</td><td>34</td><td>34</td><td>GCCCGGCTAGCTCAGTCGGTAGAGCATGAGACTC</td><td>4954.7</td><td>6.0</td><td>1.0</td><td>6.0</td><td>2.6E-09</td><td>5.2E-09</td><td>MB231/MB468</td><td>MITO</td></tr>
<tr><td>>trna124_ArgTCG_6_-_28510891_28510963@31.53.23,trna19_ArgTCG_17_+_73031208_73031280@31.53.23,trna3_ArgTCG_6_+_26299905_26299977@31.53.23,trna4_ArgTCG_6_+_26323046_26323118@31.53.23|ACTTCGGATCAGAAGATTGAGGG</td><td>>trna124_ArgTCG_6_-_28510891_28510963</td><td>31</td><td>53</td><td>23</td><td>ACTTCGGATCAGAAGATTGAGGG</td><td>114.2</td><td>-10.7</td><td>1.8</td><td>-6.0</td><td>2.3E-09</td><td>5.2E-09</td><td>MB231/MB468</td><td>MP</td></tr>
<tr><td>>hg38_21_+_8433172_8446622_RNA45SN1_45S_50ntFlanks@3930.3952.23|ATCAAAACCAACCCGGTCAGCCC</td><td>>hg38_21_+_8433172_8446622_RNA45SN1_45S_50ntFlanks</td><td>3930</td><td>3952</td><td>23</td><td>ATCAAAACCAACCCGGTCAGCCC</td><td>64.1</td><td>-9.9</td><td>1.7</td><td>-6.0</td><td>2.3E-09</td><td>5.2E-09</td><td>MB231/MB468</td><td>MP</td></tr>
<tr><td>>hg38_21_+_8433172_8446622_RNA45SN1_45S_50ntFlanks@10690.10711.22|CGTCCGGTGAGCTCTCGCTGGC</td><td>>hg38_21_+_8433172_8446622_RNA45SN1_45S_50ntFlanks</td><td>10690</td><td>10711</td><td>22</td><td>CGTCCGGTGAGCTCTCGCTGGC</td><td>349.8</td><td>-11.5</td><td>1.8</td><td>-6.3</td><td>2.7E-10</td><td>5.2E-09</td><td>BT20/MB468</td><td>MP</td></tr>
<tr><td>>hg38_21_+_8433172_8446622_RNA45SN1_45S_50ntFlanks@12605.12625.21|TATGTGCTTGGCTGAGGAGCC</td><td>>hg38_21_+_8433172_8446622_RNA45SN1_45S_50ntFlanks</td><td>12605</td><td>12625</td><td>21</td><td>TATGTGCTTGGCTGAGGAGCC</td><td>2524.5</td><td>-5.4</td><td>0.9</td><td>-6.3</td><td>2.7E-10</td><td>5.2E-09</td><td>BT20/MB468</td><td>MP</td></tr>
<tr><td>>hg38_21_+_8433172_8446622_RNA45SN1_45S_50ntFlanks@8686.8706.21|CCGGCTCCGGGACGGCTGGGA</td><td>>hg38_21_+_8433172_8446622_RNA45SN1_45S_50ntFlanks</td><td>8686</td><td>8706</td><td>21</td><td>CCGGCTCCGGGACGGCTGGGA</td><td>469.1</td><td>9.3</td><td>1.6</td><td>6.0</td><td>2.1E-09</td><td>5.2E-09</td><td>BT20/MB468</td><td>CYTO</td></tr>
<tr><td>>hg38_21_+_8433172_8446622_RNA45SN1_45S_50ntFlanks@10788.10816.29|CAGCCTCTGGCATGTTGGAACAATGTAGG</td><td>>hg38_21_+_8433172_8446622_RNA45SN1_45S_50ntFlanks</td><td>10788</td><td>10816</td><td>29</td><td>CAGCCTCTGGCATGTTGGAACAATGTAGG</td><td>1014.1</td><td>-3.4</td><td>0.6</td><td>-6.0</td><td>2.4E-09</td><td>5.2E-09</td><td>MB231/MB468</td><td>CYTO</td></tr>
<tr><td>>hg38_21_+_8433172_8446622_RNA45SN1_45S_50ntFlanks@3802.3836.35|CATTAAATCAGTTATGGTTCCTTTGGTCGCTCGCT</td><td>>hg38_21_+_8433172_8446622_RNA45SN1_45S_50ntFlanks</td><td>3802</td><td>3836</td><td>35</td><td>CATTAAATCAGTTATGGTTCCTTTGGTCGCTCGCT</td><td>938.7</td><td>-9.2</td><td>1.5</td><td>-5.9</td><td>2.7E-09</td><td>5.2E-09</td><td>BT20/MB468</td><td>NUC</td></tr>
<tr><td>>hg38_21_+_8433172_8446622_RNA45SN1_45S_50ntFlanks@3899.3932.34|CCTTCGCGGGGGGGATGCGTGCATTTATCAGATC</td><td>>hg38_21_+_8433172_8446622_RNA45SN1_45S_50ntFlanks</td><td>3899</td><td>3932</td><td>34</td><td>CCTTCGCGGGGGGGATGCGTGCATTTATCAGATC</td><td>186.1</td><td>-10.6</td><td>1.7</td><td>-6.3</td><td>2.7E-10</td><td>5.2E-09</td><td>BT20/MB468</td><td>MP</td></tr>
<tr><td>>hg38_21_+_8433172_8446622_RNA45SN1_45S_50ntFlanks@12139.12158.20|GGACAGTGCCAGGTGGGGAG</td><td>>hg38_21_+_8433172_8446622_RNA45SN1_45S_50ntFlanks</td><td>12139</td><td>12158</td><td>20</td><td>GGACAGTGCCAGGTGGGGAG</td><td>65.7</td><td>-9.9</td><td>1.7</td><td>-6.0</td><td>2.3E-09</td><td>5.2E-09</td><td>MB231/MB468</td><td>MP</td></tr>
<tr><td>>hg38_21_+_8433172_8446622_RNA45SN1_45S_50ntFlanks@7971.8009.39|GAGACGCGACCTCAGATCAGACGTGGCGACCCGCTGAAT</td><td>>hg38_21_+_8433172_8446622_RNA45SN1_45S_50ntFlanks</td><td>7971</td><td>8009</td><td>39</td><td>GAGACGCGACCTCAGATCAGACGTGGCGACCCGCTGAAT</td><td>443.3</td><td>6.5</td><td>1.1</td><td>6.0</td><td>1.8E-09</td><td>5.2E-09</td><td>BT20/MB468</td><td>MITO</td></tr>
<tr><td>>hg38_21_+_8433172_8446622_RNA45SN1_45S_50ntFlanks@11575.11604.30|ACTTAGAACTGGTGCGGACCAGGGGAATCC</td><td>>hg38_21_+_8433172_8446622_RNA45SN1_45S_50ntFlanks</td><td>11575</td><td>11604</td><td>30</td><td>ACTTAGAACTGGTGCGGACCAGGGGAATCC</td><td>266.3</td><td>-4.2</td><td>0.7</td><td>-6.0</td><td>2.4E-09</td><td>5.2E-09</td><td>MB231/MB468</td><td>NUC</td></tr>
<tr><td>>am_trnaMT_GluTTC_MT_-_14674_14742@27.48.22,trnalookalike8_GluTTC_5_-_93905172_93905240@27.48.22|GTTTTTCATATCATTGGTCGTG</td><td>>am_trnaMT_GluTTC_MT_-_14674_14742</td><td>27</td><td>48</td><td>22</td><td>GTTTTTCATATCATTGGTCGTG</td><td>163.2</td><td>-6.1</td><td>1.0</td><td>-6.0</td><td>2.4E-09</td><td>5.2E-09</td><td>MB231/MB468</td><td>CYTO</td></tr>
<tr><td>>MI0000542|hsa-mir-320a&WithFlank&8|-|22244960|22245043@48.70.23(+1G)&[MIMAT0000510&hsa-miR-320a-3p&offsets|0|+1(+1G);m-60&8|-|22244975|22244996&offsets|0|+1(+1G)]|AAAAGCTGGGTTGAGAGGGCGAAG</td><td>>MI0000542|hsa-mir-320a&WithFlank&8|-|22244960|22245043</td><td>48</td><td>70</td><td>23(+1G)</td><td>AAAAGCTGGGTTGAGAGGGCGAAG</td><td>964.8</td><td>7.6</td><td>1.2</td><td>6.3</td><td>2.7E-10</td><td>5.2E-09</td><td>BT20/MB468</td><td>MP</td></tr>
<tr><td>>hg38_21_+_8433172_8446622_RNA45SN1_45S_50ntFlanks@12626.12661.36|AATGGGGCGAAGCTACCATCTGTGGGATTATGACTG</td><td>>hg38_21_+_8433172_8446622_RNA45SN1_45S_50ntFlanks</td><td>12626</td><td>12661</td><td>36</td><td>AATGGGGCGAAGCTACCATCTGTGGGATTATGACTG</td><td>3887.0</td><td>-4.5</td><td>0.8</td><td>-6.0</td><td>1.8E-09</td><td>5.2E-09</td><td>BT20/MB468</td><td>MITO</td></tr>
<tr><td>>hg38_21_+_8433172_8446622_RNA45SN1_45S_50ntFlanks@11918.11956.39|ACATGAGAGGTGTAGAATAAGTGGGAGGCCCCCGGCGCC</td><td>>hg38_21_+_8433172_8446622_RNA45SN1_45S_50ntFlanks</td><td>11918</td><td>11956</td><td>39</td><td>ACATGAGAGGTGTAGAATAAGTGGGAGGCCCCCGGCGCC</td><td>76.5</td><td>-10.1</td><td>1.7</td><td>-6.0</td><td>2.3E-09</td><td>5.2E-09</td><td>MB231/MB468</td><td>MP</td></tr>
</table>

| | | | | | | | | | | | | | |
|---|---|---|---|---|---|---|---|---|---|---|---|---|---|
| >hg38_21_+_8433172_8446622_RNA45SN1_45S_50ntFlanks@6730.6771.42\|ACACATTGATCATCGACACTTCGAACGCACTTGCGGCCCCGG | >hg38_21_+_8433172_8446622_RNA45SN1_45S_50ntFlanks | 6730 | 6771 | 42 | ACACATTGATCATCGACACTTCGAACGCACTTGCGGCCCCGG | 157.8 | -11.2 | 1.9 | -6.0 | 2.3E-09 | 5.2E-09 | MB231/MB468 | MP |
| >MI0005544\|hsa-mir-147b&WithFlank&15\|+\|45433044\|45433135@55.74.20&[MIMAT0004928&hsa-miR-147b-3p&offsets\|0\|-1;m-575&15\|+\|45433098\|45433118&offsets\|0\|-1]\|GTGTGCGGAAATGCTTCTGC | >MI0005544\|hsa-mir-147b&WithFlank&15\|+\|45433044\|45433135 | 55 | 74 | 20 | GTGTGCGGAAATGCTTCTGC | 78.7 | 9.5 | 1.6 | 6.0 | 2.3E-09 | 5.2E-09 | MB231/MB468 | MP |
| >MI0000809\|hsa-mir-151a&WithFlank&8\|-\|140732558\|140732659@53.74.22(+1A)&[MIMAT0000757&hsa-miR-151a-3p&offsets\|0\|+1(+1A);m-30&8\|-\|140732587\|140732607&offsets\|0\|+1(+1A)]\|CTAGACTGAAGCTCCTTGAGGAA | >MI0000809\|hsa-mir-151a&WithFlank&8\|-\|140732558\|140732659 | 53 | 74 | 22(+1A) | CTAGACTGAAGCTCCTTGAGGAA | 1631.0 | 4.9 | 0.8 | 6.0 | 2.6E-09 | 5.3E-09 | MB231/MB468 | MITO |
| >MI0000077\|hsa-mir-21&WithFlank&17\|+\|59841260\|59841343@14.31.18&[MIMAT0000076&hsa-miR-21-5p&offsets\|0\|-4;m-1&17\|+\|59841273\|59841294&offsets\|0\|-4]\|TAGCTTATCAGACTGATG | >MI0000077\|hsa-mir-21&WithFlank&17\|+\|59841260\|59841343 | 14 | 31 | 18 | TAGCTTATCAGACTGATG | 385.7 | 4.6 | 0.8 | 6.0 | 2.6E-09 | 5.3E-09 | MB231/MB468 | MITO |
| >hg38_21_+_8433172_8446622_RNA45SN1_45S_50ntFlanks@4560.4578.19\|CTAGGAATAATGGAATAGG | >hg38_21_+_8433172_8446622_RNA45SN1_45S_50ntFlanks | 4560 | 4578 | 19 | CTAGGAATAATGGAATAGG | 181.6 | -4.7 | 0.8 | -6.0 | 2.1E-09 | 5.3E-09 | BT20/MB468 | CYTO |
| >hg38_21_+_8433172_8446622_RNA45SN1_45S_50ntFlanks@12626.12652.27\|AATGGGGCGAAGCTACCATCTGTGGGA | >hg38_21_+_8433172_8446622_RNA45SN1_45S_50ntFlanks | 12626 | 12652 | 27 | AATGGGGCGAAGCTACCATCTGTGGGA | 388.8 | -8.2 | 1.4 | -5.9 | 2.8E-09 | 5.3E-09 | BT20/MB468 | NUC |
| >hg38_21_+_8433172_8446622_RNA45SN1_45S_50ntFlanks@6689.6720.32\|GAAGAACGCAGCTAGCTGCGAGAATTAATGTG | >hg38_21_+_8433172_8446622_RNA45SN1_45S_50ntFlanks | 6689 | 6720 | 32 | GAAGAACGCAGCTAGCTGCGAGAATTAATGTG | 97.5 | -8.7 | 1.4 | -6.0 | 1.8E-09 | 5.3E-09 | BT20/MB468 | MITO |
| >hg38_21_+_8433172_8446622_RNA45SN1_45S_50ntFlanks@11719.11740.22\|ACGGCGGGAGTAACTATGACTC | >hg38_21_+_8433172_8446622_RNA45SN1_45S_50ntFlanks | 11719 | 11740 | 22 | ACGGCGGGAGTAACTATGACTC | 832.0 | -9.2 | 1.5 | -5.9 | 2.8E-09 | 5.3E-09 | BT20/MB468 | NUC |
| >hg38_21_+_8433172_8446622_RNA45SN1_45S_50ntFlanks@3777.3814.38\|CGGTACAGTGAAACTGCGAATGGCTCATTAAATCAGTT | >hg38_21_+_8433172_8446622_RNA45SN1_45S_50ntFlanks | 3777 | 3814 | 38 | CGGTACAGTGAAACTGCGAATGGCTCATTAAATCAGTT | 300.9 | -3.6 | 0.6 | -6.0 | 2.4E-09 | 5.3E-09 | MB231/MB468 | NUC |
| >MI0000750\|hsa-mir-26a-2&WithFlank&12\|-\|57824603\|57824698@20.40.21;MI0000083\|hsa-mir-26a-1&WithFlank&3\|+\|37969398\|37969486@16.36.21&[MIMAT0000082&hsa-miR-26a-5p&offsets\|0\|-1;m-33&12\|-\|57824658\|57824679&offsets\|0\|-1];[MIMAT0000082_1&hsa-miR-26a-5p&offsets\|0\|-1;m-34&3\|+\|37969413\|37969434&offsets\|0\|-1]\|TTCAAGTAATCCAGGATAGGC | >MI0000750\|hsa-mir-26a-2&WithFlank&12\|-\|57824603\|57824698 | 20 | 40 | 21 | TTCAAGTAATCCAGGATAGGC | 1361.1 | 5.8 | 1.0 | 6.0 | 2.6E-09 | 5.3E-09 | MB231/MB468 | MITO |
| >MI0001445\|hsa-mir-423&WithFlank&17\|+\|30117073\|30117178@59.81.23(+1U)&[MIMAT0001340&hsa-miR-423-3p&offsets\|0\|0(+1U);m-100&17\|+\|30117131\|30117153&offsets\|0\|0(+1U)]\|AGCTCGGTCTGAGGCCCCTCAGTT | >MI0001445\|hsa-mir-423&WithFlank&17\|+\|30117073\|30117178 | 59 | 81 | 23(+1U) | AGCTCGGTCTGAGGCCCCTCAGTT | 1852.5 | 5.5 | 0.9 | 6.0 | 2.6E-09 | 5.3E-09 | MB231/MB468 | MITO |
| >MI0000087\|hsa-mir-29a&WithFlank&7\|-\|130876741\|130876816@48.68.21(+2U)&[MIMAT0000086&hsa-miR-29a-3p&offsets\|0\|-1(+2U);m-26&7\|-\|130876749\|130876769&offsets\|0\|0(+2U)]\|TAGCACCATCTGAAATCGGTTTT | >MI0000087\|hsa-mir-29a&WithFlank&7\|-\|130876741\|130876816 | 48 | 68 | 21(+2U) | TAGCACCATCTGAAATCGGTTTT | 101.4 | 9.9 | 1.7 | 6.0 | 2.3E-09 | 5.3E-09 | MB231/MB468 | MP |
| >hg38_21_+_8433172_8446622_RNA45SN1_45S_50ntFlanks@12936.12970.35\|AAACCATTCGTAGACGACCTGCTTCTGGGTCGGGG | >hg38_21_+_8433172_8446622_RNA45SN1_45S_50ntFlanks | 12936 | 12970 | 35 | AAACCATTCGTAGACGACCTGCTTCTGGGTCGGGG | 59.2 | -9.8 | 1.6 | -6.0 | 2.3E-09 | 5.3E-09 | MB231/MB468 | MP |
| >MI0000086\|hsa-mir-28&WithFlank&3\|+\|188688775\|188688872@20.40.21&[MIMAT0000085&hsa-miR-28-5p&offsets\|0\|-1;m-133&3\|+\|188688794\|188688815&offsets\|0\|-1]\|AAGGAGCTCACAGTCTATTGA | >MI0000086\|hsa-mir-28&WithFlank&3\|+\|188688775\|188688872 | 20 | 40 | 21 | AAGGAGCTCACAGTCTATTGA | 133.8 | 10.3 | 1.7 | 6.0 | 2.6E-09 | 5.3E-09 | MB231/MB468 | MITO |
| >hg38_21_+_8433172_8446622_RNA45SN1_45S_50ntFlanks@12406.12435.30\|TTTTTGATCCTTCGATGTCGGCTCTTCCTA | >hg38_21_+_8433172_8446622_RNA45SN1_45S_50ntFlanks | 12406 | 12435 | 30 | TTTTTGATCCTTCGATGTCGGCTCTTCCTA | 180.8 | -4.7 | 0.8 | -6.0 | 2.4E-09 | 5.3E-09 | MB231/MB468 | CYTO |
| >hg38_21_+_8433172_8446622_RNA45SN1_45S_50ntFlanks@9716.9735.20\|AATGATTAGAGGTCTTGGGG | >hg38_21_+_8433172_8446622_RNA45SN1_45S_50ntFlanks | 9716 | 9735 | 20 | AATGATTAGAGGTCTTGGGG | 262.0 | -7.7 | 1.3 | -6.0 | 2.3E-09 | 5.3E-09 | MB231/MB468 | MP |
| >hg38_21_+_8433172_8446622_RNA45SN1_45S_50ntFlanks@10726.10747.22\|GGGAGAGGGTGTAAATCTCGCG | >hg38_21_+_8433172_8446622_RNA45SN1_45S_50ntFlanks | 10726 | 10747 | 22 | GGGAGAGGGTGTAAATCTCGCG | 73.7 | -10.1 | 1.7 | -6.0 | 2.3E-09 | 5.3E-09 | MB231/MB468 | MP |
| >hg38_21_+_8433172_8446622_RNA45SN1_45S_50ntFlanks@8105.8143.39\|CGCGGCGGGGCGCGGGACATGTGGCGTACGGAAGACCCG | >hg38_21_+_8433172_8446622_RNA45SN1_45S_50ntFlanks | 8105 | 8143 | 39 | CGCGGCGGGGCGCGGGACATGTGGCGTACGGAAGACCCG | 46530.8 | -7.4 | 1.2 | -6.3 | 2.8E-10 | 5.3E-09 | BT20/MB468 | MP |
| >hg38_21_+_8433172_8446622_RNA45SN1_45S_50ntFlanks@4728.4747.20\|ATCAAGAACGAAAGTCGGAG | >hg38_21_+_8433172_8446622_RNA45SN1_45S_50ntFlanks | 4728 | 4747 | 20 | ATCAAGAACGAAAGTCGGAG | 181.7 | -10.6 | 1.7 | -6.3 | 2.8E-10 | 5.3E-09 | BT20/MB468 | MP |
| >MI0000811\|hsa-mir-148b&WithFlank&12\|+\|54337210\|54337320@69.90.22(+1U)&[MIMAT0000759&hsa-miR-148b-3p&offsets\|0\|0(+1U);m-77&12\|+\|54337278\|54337299&offsets\|0\|0(+1U)]\|TCAGTGCATCACAGAACTTTGTT | >MI0000811\|hsa-mir-148b&WithFlank&12\|+\|54337210\|54337320 | 69 | 90 | 22(+1U) | TCAGTGCATCACAGAACTTTGTT | 5150.0 | 6.2 | 1.0 | 6.0 | 2.7E-09 | 5.3E-09 | MB231/MB468 | MITO |
| >hg38_21_+_8433172_8446622_RNA45SN1_45S_50ntFlanks@13007.13039.33\|TATTGAAAGTCAGCCCTCGACACAAGGGTTTGT | >hg38_21_+_8433172_8446622_RNA45SN1_45S_50ntFlanks | 13007 | 13039 | 33 | TATTGAAAGTCAGCCCTCGACACAAGGGTTTGT | 1242.8 | 4.3 | 0.7 | 6.0 | 2.4E-09 | 5.3E-09 | MB231/MB468 | CYTO |
| >hg38_21_+_8433172_8446622_RNA45SN1_45S_50ntFlanks@11564.11582.19\|CCTAGCAGCCGACTTAGAA | >hg38_21_+_8433172_8446622_RNA45SN1_45S_50ntFlanks | 11564 | 11582 | 19 | CCTAGCAGCCGACTTAGAA | 1125.3 | -5.8 | 1.0 | -6.0 | 2.4E-09 | 5.3E-09 | MB231/MB468 | CYTO |
| >hg38_21_+_8433172_8446622_RNA45SN1_45S_50ntFlanks@10357.10388.32\|CTTTGAAGGCCGAAGTGGAGAAGGGTTCCATG | >hg38_21_+_8433172_8446622_RNA45SN1_45S_50ntFlanks | 10357 | 10388 | 32 | CTTTGAAGGCCGAAGTGGAGAAGGGTTCCATG | 864.1 | -8.4 | 1.4 | -6.0 | 2.4E-09 | 5.3E-09 | MB231/MB468 | NUC |
| >hg38_21_+_8433172_8446622_RNA45SN1_45S_50ntFlanks@10663.10711.49\|CCTTGGAAAGCGTCGCGGTTCCGGCGGCGTCCGGTGAGCTCTCGCTGGC | >hg38_21_+_8433172_8446622_RNA45SN1_45S_50ntFlanks | 10663 | 10711 | 49 | CCTTGGAAAGCGTCGCGGTTCCGGCGGCGTCCGGTGAGCTCTCGCTGGC | 208.9 | -10.0 | 1.7 | -6.0 | 2.1E-09 | 5.3E-09 | BT20/MB468 | CYTO |
| >MI0003778\|hsa-mir-320c-1&WithFlank&18\|+\|21683512\|21683595@49.67.19;MI0008191\|hsa-mir-320c-2&WithFlank&18\|+\|24321669\|24321752@49.67.19&[MIMAT0005793&hsa-miR-320c&offsets\|+1\|0;m-222&18\|+\|21683559\|21683578&offsets\|+1\|0];[MIMAT0005793_1&hsa-miR-320c&offsets\|+1\|0;m-219&18\|+\|24321716\|24321735&offsets\|+1\|0]\|AAAGCTGGGTTGAGAGGGT | >MI0003778\|hsa-mir-320c-1&WithFlank&18\|+\|21683512\|21683595 | 49 | 67 | 19 | AAAGCTGGGTTGAGAGGGT | 190.6 | 11.3 | 1.8 | 6.3 | 2.8E-10 | 5.3E-09 | BT20/MB468 | MP |
| >hg38_21_+_8433172_8446622_RNA45SN1_45S_50ntFlanks@12259.12284.26\|CTTGATCTTGATTTTCAGTACGAATA | >hg38_21_+_8433172_8446622_RNA45SN1_45S_50ntFlanks | 12259 | 12284 | 26 | CTTGATCTTGATTTTCAGTACGAATA | 1754.8 | -7.8 | 1.3 | -5.9 | 2.8E-09 | 5.3E-09 | BT20/MB468 | NUC |
| >MI0000067\|hsa-let-7f-1&WithFlank&9\|+\|94176341\|94176439@13.33.21(+1C);MI0000068\|hsa-let-7f-2&WithFlank&X\|-\|53557186\|53557280@14.34.21(+1C)&[MIMAT0000067&hsa-let-7f-5p&offsets\|0\|-1(+1C);m-20&9\|+\|94176353\|94176374&offsets\|0\|-1(+1C)];[MIMAT0000067_1&hsa-let-7f-5p&offsets\|0\|-1(+1C);m-19&X\|-\|53557246\|53557267&offsets\|0\|-1(+1C)]\|TGAGGTAGTAGATTGTATAGTC | >MI0000067\|hsa-let-7f-1&WithFlank&9\|+\|94176341\|94176439 | 13 | 33 | 21(+1C) | TGAGGTAGTAGATTGTATAGTC | 5811.8 | 4.3 | 0.7 | 6.0 | 2.4E-09 | 5.3E-09 | MB231/MB468 | CYTO |
| >hg38_21_+_8433172_8446622_RNA45SN1_45S_50ntFlanks@11208.11225.18\|GCGGCGACTCTGGACGCG | >hg38_21_+_8433172_8446622_RNA45SN1_45S_50ntFlanks | 11208 | 11225 | 18 | GCGGCGACTCTGGACGCG | 12168.9 | -6.5 | 1.0 | -6.3 | 2.8E-10 | 5.3E-09 | BT20/MB468 | MP |
| >hg38_1_-_228634819_228635039_RNA5S12_5S_50ntFlanks@150.170.21\|AATACCGGGTGCTGTAGGCTT | >hg38_1_-_228634819_228635039_RNA5S12_5S_50ntFlanks | 150 | 170 | 21 | AATACCGGGTGCTGTAGGCTT | 15648.6 | -4.6 | 0.8 | -6.0 | 2.3E-09 | 5.3E-09 | MB231/MB468 | MP |
| >hg38_21_+_8433172_8446622_RNA45SN1_45S_50ntFlanks@11905.11931.27\|GCACGGTGAAGAGACATGAGAGGTGTA | >hg38_21_+_8433172_8446622_RNA45SN1_45S_50ntFlanks | 11905 | 11931 | 27 | GCACGGTGAAGAGACATGAGAGGTGTA | 112.0 | -10.7 | 1.8 | -6.0 | 2.3E-09 | 5.3E-09 | MB231/MB468 | MP |
| >hg38_21_+_8433172_8446622_RNA45SN1_45S_50ntFlanks@11579.11611.33\|AGAACTGGTGCGGACCAGGGGAATCCGACTGTT | >hg38_21_+_8433172_8446622_RNA45SN1_45S_50ntFlanks | 11579 | 11611 | 33 | AGAACTGGTGCGGACCAGGGGAATCCGACTGTT | 2891.7 | -4.5 | 0.8 | -6.0 | 2.7E-09 | 5.3E-09 | MB231/MB468 | MITO |
| >hg38_21_+_8433172_8446622_RNA45SN1_45S_50ntFlanks@11593.11613.21\|CCAGGGGAATCCGACTGTTTA | >hg38_21_+_8433172_8446622_RNA45SN1_45S_50ntFlanks | 11593 | 11613 | 21 | CCAGGGGAATCCGACTGTTTA | 1065.4 | -7.3 | 1.2 | -5.9 | 2.8E-09 | 5.3E-09 | BT20/MB468 | NUC |
| >hg38_21_+_8433172_8446622_RNA45SN1_45S_50ntFlanks@11568.11588.21\|GCAGCCGACTTAGAACTGGTG | >hg38_21_+_8433172_8446622_RNA45SN1_45S_50ntFlanks | 11568 | 11588 | 21 | GCAGCCGACTTAGAACTGGTG | 1452.8 | -5.0 | 0.8 | -6.0 | 2.3E-09 | 5.4E-09 | MB231/MB468 | MP |
| >hg38_21_+_8433172_8446622_RNA45SN1_45S_50ntFlanks@9534.9566.33\|AGGAAACTCTGGTGGAGGTCCGTAGCGGTCCTG | >hg38_21_+_8433172_8446622_RNA45SN1_45S_50ntFlanks | 9534 | 9566 | 33 | AGGAAACTCTGGTGGAGGTCCGTAGCGGTCCTG | 757.4 | -3.0 | 0.5 | -6.0 | 1.9E-09 | 5.4E-09 | BT20/MB468 | MITO |
| >hg38_21_+_8433172_8446622_RNA45SN1_45S_50ntFlanks@9029.9058.30\|TCTCCTCGTGGGGGGGCCGGGCCACCCCTC | >hg38_21_+_8433172_8446622_RNA45SN1_45S_50ntFlanks | 9029 | 9058 | 30 | TCTCCTCGTGGGGGGGCCGGGCCACCCCTC | 59.3 | -9.8 | 1.6 | -6.0 | 2.3E-09 | 5.4E-09 | MB231/MB468 | MP |
| >hg38_21_+_8433172_8446622_RNA45SN1_45S_50ntFlanks@10346.10388.43\|TCAAACGAGAACTTTGAAGGCCGAAGTGGAGAAGGGTTCCATG | >hg38_21_+_8433172_8446622_RNA45SN1_45S_50ntFlanks | 10346 | 10388 | 43 | TCAAACGAGAACTTTGAAGGCCGAAGTGGAGAAGGGTTCCATG | 1157.9 | -7.7 | 1.3 | -5.9 | 2.8E-09 | 5.4E-09 | BT20/MB468 | NUC |
| >hg38_21_+_8433172_8446622_RNA45SN1_45S_50ntFlanks@10361.10391.31\|GAAGGCCGAAGTGGAGAAGGGTTCCATGTGA | >hg38_21_+_8433172_8446622_RNA45SN1_45S_50ntFlanks | 10361 | 10391 | 31 | GAAGGCCGAAGTGGAGAAGGGTTCCATGTGA | 63.2 | -9.8 | 1.6 | -6.0 | 2.3E-09 | 5.4E-09 | MB231/MB468 | MP |
| >hg38_21_+_8433172_8446622_RNA45SN1_45S_50ntFlanks@4734.4771.38\|AACGAAAGTCGGAGGTTCGAAGACGATCAGATACCGTC | >hg38_21_+_8433172_8446622_RNA45SN1_45S_50ntFlanks | 4734 | 4771 | 38 | AACGAAAGTCGGAGGTTCGAAGACGATCAGATACCGTC | 228.8 | 6.8 | 1.1 | 6.0 | 2.4E-09 | 5.4E-09 | MB231/MB468 | CYTO |
| >MI0000760\|hsa-mir-361&WithFlank&X\|-\|85903630\|85903713@12.33.22&[MIMAT0000703&hsa-miR-361-5p&offsets\|0\|0;m-89&X\|-\|85903681\|85903702&offsets\|0\|0]\|TTATCAGAATCTCCAGGGGTAC | >MI0000760\|hsa-mir-361&WithFlank&X\|-\|85903630\|85903713 | 12 | 33 | 22 | TTATCAGAATCTCCAGGGGTAC | 315.2 | 4.5 | 0.8 | 6.0 | 2.5E-09 | 5.4E-09 | MB231/MB468 | NUC |
| >hg38_21_+_8433172_8446622_RNA45SN1_45S_50ntFlanks@10798.10817.20\|CATGTTGGAACAATGTAGGT | >hg38_21_+_8433172_8446622_RNA45SN1_45S_50ntFlanks | 10798 | 10817 | 20 | CATGTTGGAACAATGTAGGT | 457.5 | -6.2 | 1.0 | -5.9 | 2.7E-09 | 5.4E-09 | MB231/MB468 | MITO |
| >hg38_21_+_8433172_8446622_RNA45SN1_45S_50ntFlanks@11582.11611.30\|ACTGGTGCGGACCAGGGGAATCCGACTGTT | >hg38_21_+_8433172_8446622_RNA45SN1_45S_50ntFlanks | 11582 | 11611 | 30 | ACTGGTGCGGACCAGGGGAATCCGACTGTT | 2160.6 | -4.0 | 0.7 | -5.9 | 2.7E-09 | 5.4E-09 | MB231/MB468 | MITO |
| >MI0000098\|hsa-mir-96&WithFlank&7\|-\|129774686\|129774775@15.36.22(+1C)&[MIMAT0000095&hsa-miR-96-5p&offsets\|0\|-1(+1C);m-248&7\|-\|129774739\|129774761&offsets\|0\|-1(+1C)]\|TTTGGCACTAGCACATTTTTGCC | >MI0000098\|hsa-mir-96&WithFlank&7\|-\|129774686\|129774775 | 15 | 36 | 22(+1C) | TTTGGCACTAGCACATTTTTGCC | 411.6 | 5.0 | 0.8 | 6.4 | 1.5E-10 | 5.4E-09 | BT20/MB231 | CYTO |
| >MI0003185\|hsa-mir-501&WithFlank&X\|+\|50009716\|50009811@57.78.22&[MIMAT0004774&hsa-miR-501-3p&offsets\|0\|0;m-168&X\|+\|50009772\|50009793&offsets\|0\|0]\|AATGCACCCGGGCAAGGATTCT | >MI0003185\|hsa-mir-501&WithFlank&X\|+\|50009716\|50009811 | 57 | 78 | 22 | AATGCACCCGGGCAAGGATTCT | 248.7 | 4.0 | 0.7 | 6.0 | 2.5E-09 | 5.4E-09 | MB231/MB468 | NUC |
| >hg38_21_+_8433172_8446622_RNA45SN1_45S_50ntFlanks@9775.9819.45\|AAGAAGCCCGGCTCGCTGGCGTGGAGCCGGGCGTGGAATGCGAGT | >hg38_21_+_8433172_8446622_RNA45SN1_45S_50ntFlanks | 9775 | 9819 | 45 | AAGAAGCCCGGCTCGCTGGCGTGGAGCCGGGCGTGGAATGCGAGT | 262.1 | 11.8 | 1.9 | 6.3 | 2.8E-10 | 5.4E-09 | BT20/MB468 | MP |
| >MI0000101\|hsa-mir-99a&WithFlank&21\|+\|16539083\|16539175@19.39.21(+1A)&[MIMAT0000097&hsa-miR-99a-5p&offsets\|0\|-1(+1A);m-64&21\|+\|16539101\|16539121&offsets\|0\|0(+1A)]\|AACCCGTAGATCCGATCTTGTA | >MI0000101\|hsa-mir-99a&WithFlank&21\|+\|16539083\|16539175 | 19 | 39 | 21(+1A) | AACCCGTAGATCCGATCTTGTA | 155.4 | -10.4 | 1.6 | -6.3 | 2.8E-10 | 5.4E-09 | BT20/MB468 | MP |
| >hg38_21_+_8433172_8446622_RNA45SN1_45S_50ntFlanks@9573.9595.23\|AAATCGGTCGTCCGACCTGGGTA | >hg38_21_+_8433172_8446622_RNA45SN1_45S_50ntFlanks | 9573 | 9595 | 23 | AAATCGGTCGTCCGACCTGGGTA | 3537.1 | -6.5 | 1.0 | -6.3 | 2.8E-10 | 5.4E-09 | BT20/MB468 | MP |
| >hg38_21_+_8433172_8446622_RNA45SN1_45S_50ntFlanks@12488.12522.35\|AACGTGAGCTGGGTTTAGACCGTCGTGAGACAGGT | >hg38_21_+_8433172_8446622_RNA45SN1_45S_50ntFlanks | 12488 | 12522 | 35 | AACGTGAGCTGGGTTTAGACCGTCGTGAGACAGGT | 289.8 | -7.8 | 1.3 | -5.9 | 2.7E-09 | 5.4E-09 | MB231/MB468 | MITO |

<table>
<tr><th></th><th></th><th></th><th></th><th></th><th></th><th></th><th></th><th></th><th></th><th></th><th></th><th></th><th></th></tr>
<tr><td>>hg38_21_+_8433172_8446622_RNA45SN1_45S_50ntFlanks@12415.12443.29|CTTCGATGTCGGCTCTTCCTATCATTGTG</td><td>>hg38_21_+_8433172_8446622_RNA45SN1_45S_50ntFlanks</td><td>12415</td><td>12443</td><td>29</td><td>CTTCGATGTCGGCTCTTCCTATCATTGTG</td><td>12187.0</td><td>-5.4</td><td>0.9</td><td>-5.9</td><td>2.7E-09</td><td>5.4E-09</td><td>MB231/MB468</td><td>MITO</td></tr>
<tr><td>>trnaMT_ProTGG_MT_-_15956_16023@50.71.22|TTAAAGACTTTTTCTCTGACCA</td><td>>trnaMT_ProTGG_MT_-_15956_16023</td><td>50</td><td>71</td><td>22</td><td>TTAAAGACTTTTTCTCTGACCA</td><td>686.5</td><td>8.3</td><td>1.4</td><td>6.0</td><td>2.5E-09</td><td>5.4E-09</td><td>MB231/MB468</td><td>CYTO</td></tr>
<tr><td>>hg38_1_-_228634819_228635039_RNA5S12_5S_50ntFlanks@51.85.35|GTCTACGGCCATACCACCCTGAACGCGCCCGATCT</td><td>>hg38_1_-_228634819_228635039_RNA5S12_5S_50ntFlanks</td><td>51</td><td>85</td><td>35</td><td>GTCTACGGCCATACCACCCTGAACGCGCCCGATCT</td><td>1779.1</td><td>5.0</td><td>0.8</td><td>6.0</td><td>2.5E-09</td><td>5.4E-09</td><td>MB231/MB468</td><td>CYTO</td></tr>
<tr><td>>hg38_21_+_8433172_8446622_RNA45SN1_45S_50ntFlanks@10787.10818.32|ACAGCCTCTGGCATGTTGGAACAATGTAGGTA</td><td>>hg38_21_+_8433172_8446622_RNA45SN1_45S_50ntFlanks</td><td>10787</td><td>10818</td><td>32</td><td>ACAGCCTCTGGCATGTTGGAACAATGTAGGTA</td><td>1138.1</td><td>-4.5</td><td>0.8</td><td>-6.0</td><td>2.5E-09</td><td>5.4E-09</td><td>MB231/MB468</td><td>CYTO</td></tr>
<tr><td>>hg38_21_+_8433172_8446622_RNA45SN1_45S_50ntFlanks@11918.11957.40|ACATGAGAGGTGTAGAATAAGTGGGAGGCCCCCGGCGCCC</td><td>>hg38_21_+_8433172_8446622_RNA45SN1_45S_50ntFlanks</td><td>11918</td><td>11957</td><td>40</td><td>ACATGAGAGGTGTAGAATAAGTGGGAGGCCCCCGGCGCCC</td><td>116.3</td><td>-10.7</td><td>1.8</td><td>-6.0</td><td>2.4E-09</td><td>5.4E-09</td><td>MB231/MB468</td><td>MP</td></tr>
<tr><td>>hg38_21_+_8433172_8446622_RNA45SN1_45S_50ntFlanks@8739.8758.20|CGTCCGTCCGTCCGTCCGTC</td><td>>hg38_21_+_8433172_8446622_RNA45SN1_45S_50ntFlanks</td><td>8739</td><td>8758</td><td>20</td><td>CGTCCGTCCGTCCGTCCGTC</td><td>112.6</td><td>-10.7</td><td>1.8</td><td>-6.0</td><td>2.4E-09</td><td>5.4E-09</td><td>MB231/MB468</td><td>MP</td></tr>
<tr><td>>hg38_21_+_8433172_8446622_RNA45SN1_45S_50ntFlanks@5238.5268.31|CTGACTGGCTCAGCGTGTGCCTACCCTACGC</td><td>>hg38_21_+_8433172_8446622_RNA45SN1_45S_50ntFlanks</td><td>5238</td><td>5268</td><td>31</td><td>CTGACTGGCTCAGCGTGTGCCTACCCTACGC</td><td>217.8</td><td>7.6</td><td>1.3</td><td>6.0</td><td>2.5E-09</td><td>5.4E-09</td><td>MB231/MB468</td><td>CYTO</td></tr>
<tr><td>>hg38_21_+_8433172_8446622_RNA45SN1_45S_50ntFlanks@9173.9197.25|GGTCGCGCCGTCGGGCCCGGGGGAG</td><td>>hg38_21_+_8433172_8446622_RNA45SN1_45S_50ntFlanks</td><td>9173</td><td>9197</td><td>25</td><td>GGTCGCGCCGTCGGGCCCGGGGGAG</td><td>112.4</td><td>-10.7</td><td>1.8</td><td>-6.0</td><td>2.4E-09</td><td>5.4E-09</td><td>MB231/MB468</td><td>MP</td></tr>
<tr><td>>MI0025242|hsa-mir-7706&WithFlank&15|+|85380590|85380668@50.71.22(+1U)&[MIMAT0030021&hsa-miR-7706&offsets|0|-2(+1U);m-390&15|+|85380639|85380662&offsets|0|-2(+1U)]|TGAAGCGCCTGTGCTCTGCCGAT</td><td>>MI0025242|hsa-mir-7706&WithFlank&15|+|85380590|85380668</td><td>50</td><td>71</td><td>22(+1U)</td><td>TGAAGCGCCTGTGCTCTGCCGAT</td><td>275.5</td><td>7.7</td><td>1.3</td><td>6.0</td><td>2.2E-09</td><td>5.4E-09</td><td>BT20/MB468</td><td>CYTO</td></tr>
<tr><td>>hg38_21_+_8433172_8446622_RNA45SN1_45S_50ntFlanks@12126.12149.24|CGACCCGCTCCGGGGACAGTGCCA</td><td>>hg38_21_+_8433172_8446622_RNA45SN1_45S_50ntFlanks</td><td>12126</td><td>12149</td><td>24</td><td>CGACCCGCTCCGGGGACAGTGCCA</td><td>107.7</td><td>-8.8</td><td>1.5</td><td>-6.0</td><td>1.9E-09</td><td>5.4E-09</td><td>BT20/MB468</td><td>MITO</td></tr>
<tr><td>>hg38_21_+_8433172_8446622_RNA45SN1_45S_50ntFlanks@4283.4317.35|CTTTAACGAGGATCCATTGGAGGGCAAGTCTGGTG</td><td>>hg38_21_+_8433172_8446622_RNA45SN1_45S_50ntFlanks</td><td>4283</td><td>4317</td><td>35</td><td>CTTTAACGAGGATCCATTGGAGGGCAAGTCTGGTG</td><td>825.4</td><td>-5.9</td><td>1.0</td><td>-6.0</td><td>1.9E-09</td><td>5.5E-09</td><td>BT20/MB468</td><td>MITO</td></tr>
<tr><td>>hg38_21_+_8433172_8446622_RNA45SN1_45S_50ntFlanks@12973.12990.18|TCGTACGTAGCAGAGCAG</td><td>>hg38_21_+_8433172_8446622_RNA45SN1_45S_50ntFlanks</td><td>12973</td><td>12990</td><td>18</td><td>TCGTACGTAGCAGAGCAG</td><td>6362.2</td><td>-6.8</td><td>1.1</td><td>-6.0</td><td>2.5E-09</td><td>5.5E-09</td><td>MB231/MB468</td><td>CYTO</td></tr>
<tr><td>>trna143_LysTTT_6_-_27559593_27559665@13.30.18,trna149_LysTTT_6_-_27302769_27302841@13.30.18,trna14_LysTTT_11_-_59327808_59327880@13.30.18,trna2_LysTTT_17_+_8022473_8022545@13.30.18,trna54_LysTTT_1_+_204475655_204475727@13.30.18,trna5_LysTTT_11_+_59323902_59323974@13.30.18,trna62_LysTTT_1_-_204476158_204476230@13.30.18,trna76_LysTTT_6_+_28918806_28918878@13.30.18|CAGTCGGTAGAGCATCAG</td><td>>trna143_LysTTT_6_-_27559593_27559665</td><td>13</td><td>30</td><td>18</td><td>CAGTCGGTAGAGCATCAG</td><td>356.5</td><td>-6.4</td><td>1.1</td><td>-6.0</td><td>2.4E-09</td><td>5.5E-09</td><td>MB231/MB468</td><td>MP</td></tr>
<tr><td>>hg38_21_+_8433172_8446622_RNA45SN1_45S_50ntFlanks@6732.6774.43|ACATTGATCATCGACACTTCGAACGCACTTGCGGCCCCGGGTT</td><td>>hg38_21_+_8433172_8446622_RNA45SN1_45S_50ntFlanks</td><td>6732</td><td>6774</td><td>43</td><td>ACATTGATCATCGACACTTCGAACGCACTTGCGGCCCCGGGTT</td><td>438.3</td><td>5.4</td><td>0.9</td><td>6.0</td><td>2.5E-09</td><td>5.5E-09</td><td>MB231/MB468</td><td>NUC</td></tr>
<tr><td>>hg38_21_+_8433172_8446622_RNA45SN1_45S_50ntFlanks@11188.11207.20|CACCCGGGGGGCCGGCGGCG</td><td>>hg38_21_+_8433172_8446622_RNA45SN1_45S_50ntFlanks</td><td>11188</td><td>11207</td><td>20</td><td>CACCCGGGGGGCCGGCGGCG</td><td>228.6</td><td>-10.9</td><td>1.7</td><td>-6.3</td><td>2.9E-10</td><td>5.5E-09</td><td>BT20/MB468</td><td>MP</td></tr>
<tr><td>>hg38_21_+_8433172_8446622_RNA45SN1_45S_50ntFlanks@9129.9152.24|CGGGTCGGGGGGCGGGGCGGACTG</td><td>>hg38_21_+_8433172_8446622_RNA45SN1_45S_50ntFlanks</td><td>9129</td><td>9152</td><td>24</td><td>CGGGTCGGGGGGCGGGGCGGACTG</td><td>423.4</td><td>-8.5</td><td>1.4</td><td>-5.9</td><td>2.9E-09</td><td>5.5E-09</td><td>BT20/MB468</td><td>NUC</td></tr>
<tr><td>>hg38_21_+_8433172_8446622_RNA45SN1_45S_50ntFlanks@12936.12953.18|AAACCATTCGTAGACGAC</td><td>>hg38_21_+_8433172_8446622_RNA45SN1_45S_50ntFlanks</td><td>12936</td><td>12953</td><td>18</td><td>AAACCATTCGTAGACGAC</td><td>494.5</td><td>-7.7</td><td>1.3</td><td>-5.9</td><td>2.7E-09</td><td>5.5E-09</td><td>MB231/MB468</td><td>MITO</td></tr>
<tr><td>>MI0003513|hsa-mir-455&WithFlank&9|+|114209428|114209535@60.80.21(+1C)&[MIMAT0004784&hsa-miR-455-3p&offsets|0|0(+1C);m-135&9|+|114209487|114209508&offsets|0|-1(+1C)]|GCAGTCCATGGGCATATACACC</td><td>>MI0003513|hsa-mir-455&WithFlank&9|+|114209428|114209535</td><td>60</td><td>80</td><td>21(+1C)</td><td>GCAGTCCATGGGCATATACACC</td><td>86.3</td><td>9.7</td><td>1.6</td><td>6.0</td><td>2.4E-09</td><td>5.5E-09</td><td>MB231/MB468</td><td>MP</td></tr>
<tr><td>>hg38_21_+_8433172_8446622_RNA45SN1_45S_50ntFlanks@10539.10571.33|AGGCGTCCAGTGCGGTAACGCGACCGATCCCGG</td><td>>hg38_21_+_8433172_8446622_RNA45SN1_45S_50ntFlanks</td><td>10539</td><td>10571</td><td>33</td><td>AGGCGTCCAGTGCGGTAACGCGACCGATCCCGG</td><td>168.5</td><td>-10.5</td><td>1.7</td><td>-6.3</td><td>2.9E-10</td><td>5.5E-09</td><td>BT20/MB468</td><td>MP</td></tr>
<tr><td>>hg38_21_+_8433172_8446622_RNA45SN1_45S_50ntFlanks@4783.4800.18|CATAAACGATGCCGACCG</td><td>>hg38_21_+_8433172_8446622_RNA45SN1_45S_50ntFlanks</td><td>4783</td><td>4800</td><td>18</td><td>CATAAACGATGCCGACCG</td><td>103.7</td><td>-6.3</td><td>1.1</td><td>-6.0</td><td>2.2E-09</td><td>5.5E-09</td><td>BT20/MB468</td><td>CYTO</td></tr>
<tr><td>>hg38_21_+_8433172_8446622_RNA45SN1_45S_50ntFlanks@8552.8579.28|CTCCCCGGAGGGGGCGGGCTCCGGCGGG</td><td>>hg38_21_+_8433172_8446622_RNA45SN1_45S_50ntFlanks</td><td>8552</td><td>8579</td><td>28</td><td>CTCCCCGGAGGGGGCGGGCTCCGGCGGG</td><td>162.9</td><td>-11.2</td><td>1.9</td><td>-6.0</td><td>2.4E-09</td><td>5.5E-09</td><td>MB231/MB468</td><td>MP</td></tr>
<tr><td>>am_trna119_LysCTT_1_-_145395522_145395594@7.35.29,trna11_LysCTT_5_-_180648979_180649051@7.35.29,trna128_LysCTT_1_-_55423542_55423614@7.35.29,trna13_LysCTT_6_+_26556774_26556846@7.35.29,trna6_LysCTT_19_-_52425396_52425468@7.35.29,trna7_LysCTT_16_+_3225692_3225764@7.35.29,trna9_LysCTT_5_+_180634755_180634827@7.35.29|CTAGCTCAGTCGGTAGAGCATGAGACTCT</td><td>>am_trna119_LysCTT_1_-_145395522_145395594</td><td>7</td><td>35</td><td>29</td><td>CTAGCTCAGTCGGTAGAGCATGAGACTCT</td><td>84.4</td><td>9.6</td><td>1.6</td><td>6.0</td><td>2.4E-09</td><td>5.5E-09</td><td>MB231/MB468</td><td>MP</td></tr>
<tr><td>>hg38_21_+_8433172_8446622_RNA45SN1_45S_50ntFlanks@9534.9560.27|AGGAAACTCTGGTGGAGGTCCGTAGCG</td><td>>hg38_21_+_8433172_8446622_RNA45SN1_45S_50ntFlanks</td><td>9534</td><td>9560</td><td>27</td><td>AGGAAACTCTGGTGGAGGTCCGTAGCG</td><td>1056.3</td><td>-4.9</td><td>0.8</td><td>-6.3</td><td>2.9E-10</td><td>5.5E-09</td><td>BT20/MB468</td><td>MP</td></tr>
<tr><td>>hg38_21_+_8433172_8446622_RNA45SN1_45S_50ntFlanks@5073.5110.38|AATTCCGATAACGAACGAGACTCTGGCATGCTAACTAG</td><td>>hg38_21_+_8433172_8446622_RNA45SN1_45S_50ntFlanks</td><td>5073</td><td>5110</td><td>38</td><td>AATTCCGATAACGAACGAGACTCTGGCATGCTAACTAG</td><td>298.2</td><td>-12.1</td><td>2.0</td><td>-6.0</td><td>2.4E-09</td><td>5.5E-09</td><td>MB231/MB468</td><td>MP</td></tr>
<tr><td>>hg38_21_+_8433172_8446622_RNA45SN1_45S_50ntFlanks@11671.11703.33|CCAGTGCTCTGAATGTCAAAGTGAAGAAATTCA</td><td>>hg38_21_+_8433172_8446622_RNA45SN1_45S_50ntFlanks</td><td>11671</td><td>11703</td><td>33</td><td>CCAGTGCTCTGAATGTCAAAGTGAAGAAATTCA</td><td>419.9</td><td>-4.3</td><td>0.7</td><td>-5.9</td><td>2.8E-09</td><td>5.5E-09</td><td>MB231/MB468</td><td>MITO</td></tr>
<tr><td>>hg38_21_+_8433172_8446622_RNA45SN1_45S_50ntFlanks@11592.11611.20|ACCAGGGGAATCCGACTGTT</td><td>>hg38_21_+_8433172_8446622_RNA45SN1_45S_50ntFlanks</td><td>11592</td><td>11611</td><td>20</td><td>ACCAGGGGAATCCGACTGTT</td><td>576.5</td><td>-6.7</td><td>1.1</td><td>-5.9</td><td>2.9E-09</td><td>5.5E-09</td><td>BT20/MB468</td><td>NUC</td></tr>
<tr><td>>trna11_GluTTC_15_-_26327381_26327452@1.30.30,trna3_GluTTC_13_-_45492062_45492133@1.30.30|TCCCACATGGTCTAGCGGTTAGGATTCCTG</td><td>>trna11_GluTTC_15_-_26327381_26327452</td><td>1</td><td>30</td><td>30</td><td>TCCCACATGGTCTAGCGGTTAGGATTCCTG</td><td>1944.6</td><td>-9.6</td><td>1.6</td><td>-5.9</td><td>2.9E-09</td><td>5.5E-09</td><td>BT20/MB468</td><td>NUC</td></tr>
<tr><td>>hg38_21_+_8433172_8446622_RNA45SN1_45S_50ntFlanks@11576.11600.25|CTTAGAACTGGTGCGGACCAGGGGA</td><td>>hg38_21_+_8433172_8446622_RNA45SN1_45S_50ntFlanks</td><td>11576</td><td>11600</td><td>25</td><td>CTTAGAACTGGTGCGGACCAGGGGA</td><td>485.9</td><td>-7.6</td><td>1.3</td><td>-6.0</td><td>2.5E-09</td><td>5.5E-09</td><td>MB231/MB468</td><td>NUC</td></tr>
<tr><td>>hg38_21_+_8433172_8446622_RNA45SN1_45S_50ntFlanks@6699.6721.23|GCTAGCTGCGAGAATTAATGTGA</td><td>>hg38_21_+_8433172_8446622_RNA45SN1_45S_50ntFlanks</td><td>6699</td><td>6721</td><td>23</td><td>GCTAGCTGCGAGAATTAATGTGA</td><td>1195.3</td><td>-7.3</td><td>1.2</td><td>-5.9</td><td>2.9E-09</td><td>5.5E-09</td><td>BT20/MB468</td><td>NUC</td></tr>
<tr><td>>trna128_GlyGCC_6_-_27870686_27870756@20.38.19,trna19_GlyGCC_16_+_70823410_70823480@20.38.19,trna19_GlyGCC_2_-_157257659_157257729@20.38.19,trna24_GlyGCC_16_-_70812942_70813012@20.38.19,trna25_GlyGCC_16_-_70812114_70812184@20.38.19,trna2_GlyGCC_21_-_18827107_18827177@20.38.19,trna35_GlyGCC_1_+_161413094_161413164@20.38.19,trna37_GlyGCC_1_+_161420467_161420537@20.38.19,trna39_GlyGCC_1_+_161427898_161427968@20.38.19,trna41_GlyGCC_1_+_161435258_161435328@20.38.19,trna5_GlyGCC_17_+_8029064_8029134@20.38.19,trna68_GlyGCC_1_-_161493637_161493707@20.38.19|AGAATTCTCGCCTGCCACG</td><td>>trna128_GlyGCC_6_-_27870686_27870756</td><td>20</td><td>38</td><td>19</td><td>AGAATTCTCGCCTGCCACG</td><td>460.0</td><td>8.1</td><td>1.4</td><td>6.0</td><td>2.2E-09</td><td>5.5E-09</td><td>BT20/MB468</td><td>CYTO</td></tr>
<tr><td>>MI0000061|hsa-let-7a-2&WithFlank&11|-|122146516|122146599@11.32.22(+1A);MI0000062|hsa-let-7a-3&WithFlank&22|+|46112743|46112828@10.31.22(+1A);MI0000060|hsa-let-7a-1&WithFlank&9|+|94175951|94176042@12.33.22(+1A)&[MIMAT0000062&hsa-let-7a-5p&offsets|0|0(+1A);m-6&11|-|122146568|122146589&offsets|0|0(+1A)];[MIMAT0000062_1&hsa-let-7a-5p&offsets|0|0(+1A);m-5&22|+|46112752|46112773&offsets|0|0(+1A)];[MIMAT0000062_2&hsa-let-7a-5p&offsets|0|0(+1A);m-4&9|+|94175962|94175983&offsets|0|0(+1A)]|TGAGGTAGTAGGTTGTATAGTTA</td><td>>MI0000061|hsa-let-7a-2&WithFlank&11|-|122146516|122146599</td><td>11</td><td>32</td><td>22(+1A)</td><td>TGAGGTAGTAGGTTGTATAGTTA</td><td>885.8</td><td>5.4</td><td>0.9</td><td>5.9</td><td>2.8E-09</td><td>5.5E-09</td><td>MB231/MB468</td><td>MITO</td></tr>
<tr><td>>hg38_21_+_8433172_8446622_RNA45SN1_45S_50ntFlanks@4220.4243.24|ACGAAAAATAACAATACAGGACTC</td><td>>hg38_21_+_8433172_8446622_RNA45SN1_45S_50ntFlanks</td><td>4220</td><td>4243</td><td>24</td><td>ACGAAAAATAACAATACAGGACTC</td><td>115.4</td><td>-8.9</td><td>1.5</td><td>-6.0</td><td>1.9E-09</td><td>5.5E-09</td><td>BT20/MB468</td><td>MITO</td></tr>
<tr><td>>hg38_21_+_8433172_8446622_RNA45SN1_45S_50ntFlanks@4129.4162.34|GATTCCGGAGAGGGAGCCTGAGAAACGGCTACCA</td><td>>hg38_21_+_8433172_8446622_RNA45SN1_45S_50ntFlanks</td><td>4129</td><td>4162</td><td>34</td><td>GATTCCGGAGAGGGAGCCTGAGAAACGGCTACCA</td><td>362.1</td><td>-7.8</td><td>1.3</td><td>-5.9</td><td>2.8E-09</td><td>5.6E-09</td><td>MB231/MB468</td><td>MITO</td></tr>
<tr><td>>MI0000088|hsa-mir-30a&WithFlank&6|-|71403545|71403627@12.33.22(+1A)&[MIMAT0000087&hsa-miR-30a-5p&offsets|0|0(+1A);m-15&6|-|71403595|71403616&offsets|0|0(+1A)]|TGTAAACATCCTCGACTGGAAGA</td><td>>MI0000088|hsa-mir-30a&WithFlank&6|-|71403545|71403627</td><td>12</td><td>33</td><td>22(+1A)</td><td>TGTAAACATCCTCGACTGGAAGA</td><td>1698.6</td><td>6.4</td><td>1.1</td><td>5.9</td><td>2.8E-09</td><td>5.6E-09</td><td>MB231/MB468</td><td>MITO</td></tr>
<tr><td>>hg38_21_+_8433172_8446622_RNA45SN1_45S_50ntFlanks@13006.13039.34|CTATTGAAAGTCAGCCCTCGACACAAGGGTTTGT</td><td>>hg38_21_+_8433172_8446622_RNA45SN1_45S_50ntFlanks</td><td>13006</td><td>13039</td><td>34</td><td>CTATTGAAAGTCAGCCCTCGACACAAGGGTTTGT</td><td>4080.1</td><td>5.5</td><td>0.9</td><td>6.0</td><td>1.9E-09</td><td>5.6E-09</td><td>BT20/MB468</td><td>MITO</td></tr>
<tr><td>>hg38_21_+_8433172_8446622_RNA45SN1_45S_50ntFlanks@11682.11700.19|AATGTCAAAGTGAAGAAAT</td><td>>hg38_21_+_8433172_8446622_RNA45SN1_45S_50ntFlanks</td><td>11682</td><td>11700</td><td>19</td><td>AATGTCAAAGTGAAGAAAT</td><td>91.1</td><td>-10.4</td><td>1.7</td><td>-6.0</td><td>2.4E-09</td><td>5.6E-09</td><td>MB231/MB468</td><td>MP</td></tr>
<tr><td>>MI0000462|hsa-mir-152&WithFlank&17|-|48037155|48037253@60.80.21(+1C)&[MIMAT0000438&hsa-miR-152-3p&offsets|0|0(+1C);m-97&17|-|48037174|48037194&offsets|0|0(+1C)]|TCAGTGCATGACAGAACTTGGC</td><td>>MI0000462|hsa-mir-152&WithFlank&17|-|48037155|48037253</td><td>60</td><td>80</td><td>21(+1C)</td><td>TCAGTGCATGACAGAACTTGGC</td><td>93.5</td><td>9.8</td><td>1.6</td><td>6.0</td><td>2.4E-09</td><td>5.6E-09</td><td>MB231/MB468</td><td>MP</td></tr>
<tr><td>>MI0000802|hsa-mir-340&WithFlank&5|-|180015297|180015403@22.43.22&[MIMAT0004692&hsa-miR-340-5p&offsets|0|0;m-220&5|-|180015361|180015382&offsets|0|0]|TTATAAAGCAATGAGACTGATT</td><td>>MI0000802|hsa-mir-340&WithFlank&5|-|180015297|180015403</td><td>22</td><td>43</td><td>22</td><td>TTATAAAGCAATGAGACTGATT</td><td>1486.3</td><td>4.5</td><td>0.8</td><td>6.0</td><td>2.6E-09</td><td>5.6E-09</td><td>MB231/MB468</td><td>NUC</td></tr>
<tr><td>>hg38_21_+_8433172_8446622_RNA45SN1_45S_50ntFlanks@8198.8215.18|TGGACGGTGTGAGGCCGG</td><td>>hg38_21_+_8433172_8446622_RNA45SN1_45S_50ntFlanks</td><td>8198</td><td>8215</td><td>18</td><td>TGGACGGTGTGAGGCCGG</td><td>4475.4</td><td>-7.7</td><td>1.3</td><td>-6.0</td><td>2.5E-09</td><td>5.6E-09</td><td>MB231/MB468</td><td>CYTO</td></tr>
<tr><td>>hg38_21_+_8433172_8446622_RNA45SN1_45S_50ntFlanks@4555.4576.22|CGCAGCTAGGAATAATGGAATA</td><td>>hg38_21_+_8433172_8446622_RNA45SN1_45S_50ntFlanks</td><td>4555</td><td>4576</td><td>22</td><td>CGCAGCTAGGAATAATGGAATA</td><td>225.1</td><td>-4.0</td><td>0.7</td><td>-6.0</td><td>2.6E-09</td><td>5.6E-09</td><td>MB231/MB468</td><td>NUC</td></tr>
<tr><td>>hg38_21_+_8433172_8446622_RNA45SN1_45S_50ntFlanks@8168.8197.30|CCCAAGTCCTTCTGATCGAGGCCCAGCCCG</td><td>>hg38_21_+_8433172_8446622_RNA45SN1_45S_50ntFlanks</td><td>8168</td><td>8197</td><td>30</td><td>CCCAAGTCCTTCTGATCGAGGCCCAGCCCG</td><td>1527.3</td><td>-9.2</td><td>1.5</td><td>-6.0</td><td>2.6E-09</td><td>5.6E-09</td><td>MB231/MB468</td><td>NUC</td></tr>
</table>

<table>
<tr><th></th><th></th><th></th><th></th><th></th><th></th><th></th><th></th><th></th><th></th><th></th><th></th><th></th><th></th></tr>
<tr><td>>hg38_21_+_8433172_8446622_RNA45SN1_45S_50ntFlanks@4657.4689.33|CTAGAGGTGAAATTCTTGGACCGGCGCAAGACG</td><td>>hg38_21_+_8433172_8446622_RNA45SN1_45S_50ntFlanks</td><td>4657</td><td>4689</td><td>33</td><td>CTAGAGGTGAAATTCTTGGACCGGCGCAAGACG</td><td>9086.9</td><td>-10.6</td><td>1.8</td><td>-5.9</td><td>2.9E-09</td><td>5.6E-09</td><td>BT20/MB468</td><td>NUC</td></tr>
<tr><td>>hg38_21_+_8433172_8446622_RNA45SN1_45S_50ntFlanks@8104.8136.33|CCGCGGCGGGGCGCGGGACATGTGGCGTACGGA</td><td>>hg38_21_+_8433172_8446622_RNA45SN1_45S_50ntFlanks</td><td>8104</td><td>8136</td><td>33</td><td>CCGCGGCGGGGCGCGGGACATGTGGCGTACGGA</td><td>592.8</td><td>12.5</td><td>1.9</td><td>6.5</td><td>5.8E-11</td><td>5.6E-09</td><td>BT20/MB231</td><td>MP</td></tr>
<tr><td>>MI0000477|hsa-mir-146a&WithFlank&5|+|160485346|160485456@27.49.23(+1C)&[MIMAT0000449&hsa-miR-146a-5p&offsets|0|+1(+1C);m-129&5|+|160485372|160485393&offsets|0|+1(+1C)]|TGAGAACTGAATTCCATGGGTTGC</td><td>>MI0000477|hsa-mir-146a&WithFlank&5|+|160485346|160485456</td><td>27</td><td>49</td><td>23(+1C)</td><td>TGAGAACTGAATTCCATGGGTTGC</td><td>380.9</td><td>-12.0</td><td>1.8</td><td>-6.5</td><td>5.8E-11</td><td>5.6E-09</td><td>BT20/MB231</td><td>MP</td></tr>
<tr><td>>hg38_21_+_8433172_8446622_RNA45SN1_45S_50ntFlanks@11323.11341.19|CGTCTCGTCGCGCGCGCGT</td><td>>hg38_21_+_8433172_8446622_RNA45SN1_45S_50ntFlanks</td><td>11323</td><td>11341</td><td>19</td><td>CGTCTCGTCGCGCGCGCGT</td><td>228.6</td><td>-7.2</td><td>1.2</td><td>-6.0</td><td>2.6E-09</td><td>5.6E-09</td><td>MB231/MB468</td><td>NUC</td></tr>
<tr><td>>hg38_21_+_8433172_8446622_RNA45SN1_45S_50ntFlanks@12634.12669.36|GAAGCTACCATCTGTGGGATTATGACTGAACGCCTC</td><td>>hg38_21_+_8433172_8446622_RNA45SN1_45S_50ntFlanks</td><td>12634</td><td>12669</td><td>36</td><td>GAAGCTACCATCTGTGGGATTATGACTGAACGCCTC</td><td>293.4</td><td>8.3</td><td>1.4</td><td>6.0</td><td>2.6E-09</td><td>5.6E-09</td><td>MB231/MB468</td><td>CYTO</td></tr>
<tr><td>>trnaMT_PheGAA_MT_+_577_647@35.57.23|GAAAATGTTTAGACGGGCTCACA</td><td>>trnaMT_PheGAA_MT_+_577_647</td><td>35</td><td>57</td><td>23</td><td>GAAAATGTTTAGACGGGCTCACA</td><td>549.8</td><td>3.5</td><td>0.6</td><td>6.0</td><td>2.5E-09</td><td>5.6E-09</td><td>MB231/MB468</td><td>MP</td></tr>
<tr><td>>MI0000542|hsa-mir-320a&WithFlank&8|-|22244960|22245043@50.71.22(+1U)&[MIMAT0000510&hsa-miR-320a-3p&offsets|+2|+2(+1U);m-60&8|-|22244975|22244996&offsets|+2|+2(+1U)]|AAGCTGGGTTGAGAGGGCGAAAT</td><td>>MI0000542|hsa-mir-320a&WithFlank&8|-|22244960|22245043</td><td>50</td><td>71</td><td>22(+1U)</td><td>AAGCTGGGTTGAGAGGGCGAAAT</td><td>180.6</td><td>11.2</td><td>1.8</td><td>6.3</td><td>3.0E-10</td><td>5.6E-09</td><td>BT20/MB468</td><td>MP</td></tr>
<tr><td>>hg38_21_+_8433172_8446622_RNA45SN1_45S_50ntFlanks@5084.5120.37|CGAACGAGACTCTGGCATGCTAACTAGTTACGCGACC</td><td>>hg38_21_+_8433172_8446622_RNA45SN1_45S_50ntFlanks</td><td>5084</td><td>5120</td><td>37</td><td>CGAACGAGACTCTGGCATGCTAACTAGTTACGCGACC</td><td>2079.4</td><td>-10.3</td><td>1.7</td><td>-5.9</td><td>3.0E-09</td><td>5.6E-09</td><td>BT20/MB468</td><td>NUC</td></tr>
<tr><td>>trnaMT_GluTTC_MT_-_14674_14742@29.72.44,trnalookalike8_GluTTC_5_-_93905172_93905240@29.72.44|TTTTCATATCATTGGTCGTGGTTGTAGTCCGTGCGAGAATACCA</td><td>>trnaMT_GluTTC_MT_-_14674_14742</td><td>29</td><td>72</td><td>44</td><td>TTTTCATATCATTGGTCGTGGTTGTAGTCCGTGCGAGAATACCA</td><td>378.2</td><td>8.5</td><td>1.4</td><td>6.0</td><td>2.6E-09</td><td>5.7E-09</td><td>MB231/MB468</td><td>CYTO</td></tr>
<tr><td>>hg38_21_+_8433172_8446622_RNA45SN1_45S_50ntFlanks@11578.11611.34|TAGAACTGGTGCGGACCAGGGGAATCCGACTGTT</td><td>>hg38_21_+_8433172_8446622_RNA45SN1_45S_50ntFlanks</td><td>11578</td><td>11611</td><td>34</td><td>TAGAACTGGTGCGGACCAGGGGAATCCGACTGTT</td><td>636.1</td><td>-3.8</td><td>0.6</td><td>-5.9</td><td>2.8E-09</td><td>5.7E-09</td><td>MB231/MB468</td><td>MITO</td></tr>
<tr><td>>hg38_21_+_8433172_8446622_RNA45SN1_45S_50ntFlanks@12043.12071.29|CACTGACCCGGTGAGGCGGGGGGGCGAGC</td><td>>hg38_21_+_8433172_8446622_RNA45SN1_45S_50ntFlanks</td><td>12043</td><td>12071</td><td>29</td><td>CACTGACCCGGTGAGGCGGGGGGGCGAGC</td><td>70.6</td><td>-10.0</td><td>1.7</td><td>-6.0</td><td>2.5E-09</td><td>5.7E-09</td><td>MB231/MB468</td><td>MP</td></tr>
<tr><td>>hg38_21_+_8433172_8446622_RNA45SN1_45S_50ntFlanks@8844.8867.24|GCGGCGGCGGCGGCGGCGGCGGGA</td><td>>hg38_21_+_8433172_8446622_RNA45SN1_45S_50ntFlanks</td><td>8844</td><td>8867</td><td>24</td><td>GCGGCGGCGGCGGCGGCGGCGGGA</td><td>183.8</td><td>-10.6</td><td>1.7</td><td>-6.3</td><td>3.0E-10</td><td>5.7E-09</td><td>BT20/MB468</td><td>MP</td></tr>
<tr><td>>hg38_21_+_8433172_8446622_RNA45SN1_45S_50ntFlanks@10823.10865.43|AAGTCGGCAAGCCGGATCCGTAACTTCGGGATAAGGATTGGCT</td><td>>hg38_21_+_8433172_8446622_RNA45SN1_45S_50ntFlanks</td><td>10823</td><td>10865</td><td>43</td><td>AAGTCGGCAAGCCGGATCCGTAACTTCGGGATAAGGATTGGCT</td><td>991.5</td><td>-9.0</td><td>1.5</td><td>-6.0</td><td>2.6E-09</td><td>5.7E-09</td><td>MB231/MB468</td><td>NUC</td></tr>
<tr><td>>MI0000746|hsa-mir-99b&WithFlank&19|+|51692606|51692687@13.32.20(+1A)&[MIMAT0000689&hsa-miR-99b-5p&offsets|0|-2(+1A);m-10&19|+|51692618|51692639&offsets|0|-2(+1A)]|CACCCGTAGAACCGACCTTGA</td><td>>MI0000746|hsa-mir-99b&WithFlank&19|+|51692606|51692687</td><td>13</td><td>32</td><td>20(+1A)</td><td>CACCCGTAGAACCGACCTTGA</td><td>617.2</td><td>6.7</td><td>1.1</td><td>6.0</td><td>2.6E-09</td><td>5.7E-09</td><td>MB231/MB468</td><td>CYTO</td></tr>
<tr><td>>hg38_21_+_8433172_8446622_RNA45SN1_45S_50ntFlanks@4732.4764.33|AGAACGAAAGTCGGAGGTTCGAAGACGATCAGA</td><td>>hg38_21_+_8433172_8446622_RNA45SN1_45S_50ntFlanks</td><td>4732</td><td>4764</td><td>33</td><td>AGAACGAAAGTCGGAGGTTCGAAGACGATCAGA</td><td>266.6</td><td>7.0</td><td>1.2</td><td>6.0</td><td>2.6E-09</td><td>5.7E-09</td><td>MB231/MB468</td><td>CYTO</td></tr>
<tr><td>>hg38_21_+_8433172_8446622_RNA45SN1_45S_50ntFlanks@10392.10435.44|ACAGCAGTTGAACATGGGTCAGTCGGTCCTGAGAGATGGGCGAG</td><td>>hg38_21_+_8433172_8446622_RNA45SN1_45S_50ntFlanks</td><td>10392</td><td>10435</td><td>44</td><td>ACAGCAGTTGAACATGGGTCAGTCGGTCCTGAGAGATGGGCGAG</td><td>882.0</td><td>-6.5</td><td>1.1</td><td>-6.0</td><td>2.6E-09</td><td>5.7E-09</td><td>MB231/MB468</td><td>CYTO</td></tr>
<tr><td>>hg38_21_+_8433172_8446622_RNA45SN1_45S_50ntFlanks@12893.12925.33|CCTCGCCCGTCACGCACCGCACGTTCGTGGGGA</td><td>>hg38_21_+_8433172_8446622_RNA45SN1_45S_50ntFlanks</td><td>12893</td><td>12925</td><td>33</td><td>CCTCGCCCGTCACGCACCGCACGTTCGTGGGGA</td><td>124.4</td><td>-9.0</td><td>1.5</td><td>-6.0</td><td>2.0E-09</td><td>5.7E-09</td><td>BT20/MB468</td><td>MITO</td></tr>
<tr><td>>am_trna2_GlyTCC_19_+_4724082_4724153@1.30.30|GCGTTGGTGGTATAGTGGTTAGCATAGCTG</td><td>>am_trna2_GlyTCC_19_+_4724082_4724153</td><td>1</td><td>30</td><td>30</td><td>GCGTTGGTGGTATAGTGGTTAGCATAGCTG</td><td>1273.9</td><td>-5.8</td><td>1.0</td><td>-6.0</td><td>2.6E-09</td><td>5.7E-09</td><td>MB231/MB468</td><td>CYTO</td></tr>
<tr><td>>hg38_21_+_8433172_8446622_RNA45SN1_45S_50ntFlanks@10349.10391.43|AACGAGAACTTTGAAGGCCGAAGTGGAGAAGGGTTCCATGTGA</td><td>>hg38_21_+_8433172_8446622_RNA45SN1_45S_50ntFlanks</td><td>10349</td><td>10391</td><td>43</td><td>AACGAGAACTTTGAAGGCCGAAGTGGAGAAGGGTTCCATGTGA</td><td>380.8</td><td>-5.0</td><td>0.9</td><td>-5.9</td><td>3.0E-09</td><td>5.7E-09</td><td>BT20/MB468</td><td>NUC</td></tr>
<tr><td>>hg38_21_+_8433172_8446622_RNA45SN1_45S_50ntFlanks@485.516.32|GATGTGGTGACGTCGTGCTCTCCCGGGCCGGG</td><td>>hg38_21_+_8433172_8446622_RNA45SN1_45S_50ntFlanks</td><td>485</td><td>516</td><td>32</td><td>GATGTGGTGACGTCGTGCTCTCCCGGGCCGGG</td><td>200.7</td><td>11.4</td><td>1.8</td><td>6.3</td><td>3.0E-10</td><td>5.7E-09</td><td>BT20/MB468</td><td>MP</td></tr>
<tr><td>>hg38_21_+_8433172_8446622_RNA45SN1_45S_50ntFlanks@11115.11133.19|CGGCAGGGGCCGGCGGCGG</td><td>>hg38_21_+_8433172_8446622_RNA45SN1_45S_50ntFlanks</td><td>11115</td><td>11133</td><td>19</td><td>CGGCAGGGGCCGGCGGCGG</td><td>212.3</td><td>-10.8</td><td>1.7</td><td>-6.3</td><td>3.0E-10</td><td>5.7E-09</td><td>BT20/MB468</td><td>MP</td></tr>
<tr><td>>hg38_21_+_8433172_8446622_RNA45SN1_45S_50ntFlanks@5487.5514.28|CGCTGAGAAGACGGTCGAACTTGACTAT</td><td>>hg38_21_+_8433172_8446622_RNA45SN1_45S_50ntFlanks</td><td>5487</td><td>5514</td><td>28</td><td>CGCTGAGAAGACGGTCGAACTTGACTAT</td><td>9669.0</td><td>-6.6</td><td>1.1</td><td>-6.3</td><td>3.0E-10</td><td>5.7E-09</td><td>BT20/MB468</td><td>MP</td></tr>
<tr><td>>hg38_21_+_8433172_8446622_RNA45SN1_45S_50ntFlanks@5446.5466.21|CGGATCGGCCCCGCCGGGGTC</td><td>>hg38_21_+_8433172_8446622_RNA45SN1_45S_50ntFlanks</td><td>5446</td><td>5466</td><td>21</td><td>CGGATCGGCCCCGCCGGGGTC</td><td>12321.9</td><td>-9.0</td><td>1.5</td><td>-5.9</td><td>3.0E-09</td><td>5.7E-09</td><td>BT20/MB468</td><td>NUC</td></tr>
<tr><td>>hg38_21_+_8433172_8446622_RNA45SN1_45S_50ntFlanks@11552.11591.40|CCTCGGCCGGCGCCTAGCAGCCGACTTAGAACTGGTGCGG</td><td>>hg38_21_+_8433172_8446622_RNA45SN1_45S_50ntFlanks</td><td>11552</td><td>11591</td><td>40</td><td>CCTCGGCCGGCGCCTAGCAGCCGACTTAGAACTGGTGCGG</td><td>1165.4</td><td>-6.4</td><td>1.1</td><td>-5.9</td><td>2.9E-09</td><td>5.7E-09</td><td>MB231/MB468</td><td>MITO</td></tr>
<tr><td>>am_trna106_PheGAA_6_-_28775610_28775682@31.52.22,trna109_PheGAA_6_-_28758499_28758571@31.52.22,trna11_PheGAA_12_-_125412389_125412461@31.52.22,trna13_PheGAA_11_-_59333853_59333925@31.52.22,trna14_PheGAA_19_-_1383361_1383433@31.52.22,trna15_PheGAA_11_-_59324970_59325042@31.52.22,trna1_PheGAA_13_-_95201904_95201976@31.52.22,trna72_PheGAA_6_+_28732164_28732236@31.52.22,trna96_PheGAA_6_-_28949449_28949521@31.52.22|ACTGAAGATCTAAAGGTCCCTG</td><td>>am_trna106_PheGAA_6_-_28775610_28775682</td><td>31</td><td>52</td><td>22</td><td>ACTGAAGATCTAAAGGTCCCTG</td><td>122.8</td><td>-10.0</td><td>1.6</td><td>-6.3</td><td>3.1E-10</td><td>5.7E-09</td><td>BT20/MB468</td><td>MP</td></tr>
<tr><td>>MI0000440|hsa-mir-27b&WithFlank&9|+|95085439|95085547@67.87.21(+1U)&[MIMAT0000419&hsa-miR-27b-3p&offsets|0|0(+1U);m-39&9|+|95085505|95085525&offsets|0|0(+1U)]|TTCACAGTGGCTAAGTTCTGCT</td><td>>MI0000440|hsa-mir-27b&WithFlank&9|+|95085439|95085547</td><td>67</td><td>87</td><td>21(+1U)</td><td>TTCACAGTGGCTAAGTTCTGCT</td><td>2104.7</td><td>6.5</td><td>1.1</td><td>5.9</td><td>2.9E-09</td><td>5.7E-09</td><td>MB231/MB468</td><td>MITO</td></tr>
<tr><td>>hg38_21_+_8433172_8446622_RNA45SN1_45S_50ntFlanks@4996.5024.29|CAGATTGATAGCTCTTTCTCGATTCCGTG</td><td>>hg38_21_+_8433172_8446622_RNA45SN1_45S_50ntFlanks</td><td>4996</td><td>5024</td><td>29</td><td>CAGATTGATAGCTCTTTCTCGATTCCGTG</td><td>133.2</td><td>-10.9</td><td>1.8</td><td>-6.0</td><td>2.5E-09</td><td>5.7E-09</td><td>MB231/MB468</td><td>MP</td></tr>
<tr><td>>hg38_21_+_8433172_8446622_RNA45SN1_45S_50ntFlanks@12437.12475.39|CATTGTGAAGCAGAATTCACCAAGCGTTGGATTGTTCAC</td><td>>hg38_21_+_8433172_8446622_RNA45SN1_45S_50ntFlanks</td><td>12437</td><td>12475</td><td>39</td><td>CATTGTGAAGCAGAATTCACCAAGCGTTGGATTGTTCAC</td><td>113.3</td><td>-10.7</td><td>1.8</td><td>-6.0</td><td>2.5E-09</td><td>5.7E-09</td><td>MB231/MB468</td><td>MP</td></tr>
<tr><td>>hg38_21_+_8433172_8446622_RNA45SN1_45S_50ntFlanks@12626.12659.34|AATGGGGCGAAGCTACCATCTGTGGGATTATGAC</td><td>>hg38_21_+_8433172_8446622_RNA45SN1_45S_50ntFlanks</td><td>12626</td><td>12659</td><td>34</td><td>AATGGGGCGAAGCTACCATCTGTGGGATTATGAC</td><td>1088.9</td><td>-6.7</td><td>1.1</td><td>-6.0</td><td>2.6E-09</td><td>5.7E-09</td><td>MB231/MB468</td><td>NUC</td></tr>
<tr><td>>hg38_21_+_8433172_8446622_RNA45SN1_45S_50ntFlanks@11190.11216.27|CCCGGGGGGGCCGGCGGCGGCGGCGACT</td><td>>hg38_21_+_8433172_8446622_RNA45SN1_45S_50ntFlanks</td><td>11190</td><td>11216</td><td>27</td><td>CCCGGGGGGGCCGGCGGCGGCGGCGACT</td><td>106.2</td><td>-10.6</td><td>1.8</td><td>-6.0</td><td>2.5E-09</td><td>5.7E-09</td><td>MB231/MB468</td><td>MP</td></tr>
<tr><td>>MI0005561|hsa-mir-877&WithFlank&6|+|30584326|30584423@7.30.24&[MIMAT0004949&hsa-miR-877-5p&offsets|0|+4;m-382&6|+|30584332|30584354&offsets|0|+1]|GTAGAGGAGATGGCGCAGGGGACA</td><td>>MI0005561|hsa-mir-877&WithFlank&6|+|30584326|30584423</td><td>7</td><td>30</td><td>24</td><td>GTAGAGGAGATGGCGCAGGGGACA</td><td>680.5</td><td>8.7</td><td>1.5</td><td>6.0</td><td>2.3E-09</td><td>5.7E-09</td><td>BT20/MB468</td><td>CYTO</td></tr>
<tr><td>>hg38_21_+_8433172_8446622_RNA45SN1_45S_50ntFlanks@10903.10935.33|GGCTGGGCGCGCGCCGCGGCTGGACGAGGCGCC</td><td>>hg38_21_+_8433172_8446622_RNA45SN1_45S_50ntFlanks</td><td>10903</td><td>10935</td><td>33</td><td>GGCTGGGCGCGCGCCGCGGCTGGACGAGGCGCC</td><td>96.1</td><td>-10.4</td><td>1.8</td><td>-6.0</td><td>2.5E-09</td><td>5.7E-09</td><td>MB231/MB468</td><td>MP</td></tr>
<tr><td>>hg38_21_+_8433172_8446622_RNA45SN1_45S_50ntFlanks@10458.10486.29|ATGGCCTCCGTTGCCCTCGGCCGATCGAA</td><td>>hg38_21_+_8433172_8446622_RNA45SN1_45S_50ntFlanks</td><td>10458</td><td>10486</td><td>29</td><td>ATGGCCTCCGTTGCCCTCGGCCGATCGAA</td><td>715.5</td><td>-9.1</td><td>1.5</td><td>-5.9</td><td>3.0E-09</td><td>5.8E-09</td><td>BT20/MB468</td><td>NUC</td></tr>
<tr><td>>hg38_MT_+_1621_3279_RNR2_16S_50ntFlanks@1349.1389.41|ACAATAGGGTTTACGACCTCGATGTTGGATCAGGACATCCC</td><td>>hg38_MT_+_1621_3279_RNR2_16S_50ntFlanks</td><td>1349</td><td>1389</td><td>41</td><td>ACAATAGGGTTTACGACCTCGATGTTGGATCAGGACATCCC</td><td>86.6</td><td>9.7</td><td>1.6</td><td>6.0</td><td>2.5E-09</td><td>5.8E-09</td><td>MB231/MB468</td><td>MP</td></tr>
<tr><td>>hg38_21_+_8433172_8446622_RNA45SN1_45S_50ntFlanks@9739.9767.29|AAACGATCTCAACCTATTCTCAAACTTTA</td><td>>hg38_21_+_8433172_8446622_RNA45SN1_45S_50ntFlanks</td><td>9739</td><td>9767</td><td>29</td><td>AAACGATCTCAACCTATTCTCAAACTTTA</td><td>1778.1</td><td>-7.9</td><td>1.3</td><td>-5.9</td><td>3.0E-09</td><td>5.8E-09</td><td>BT20/MB468</td><td>NUC</td></tr>
<tr><td>>hg38_21_+_8433172_8446622_RNA45SN1_45S_50ntFlanks@12974.12992.19|CGTACGTAGCAGAGCAGCT</td><td>>hg38_21_+_8433172_8446622_RNA45SN1_45S_50ntFlanks</td><td>12974</td><td>12992</td><td>19</td><td>CGTACGTAGCAGAGCAGCT</td><td>1267.5</td><td>-9.2</td><td>1.6</td><td>-5.9</td><td>3.0E-09</td><td>5.8E-09</td><td>BT20/MB468</td><td>NUC</td></tr>
<tr><td>>hg38_21_+_8433172_8446622_RNA45SN1_45S_50ntFlanks@8111.8141.31|GGGGCGCGGGACATGTGGCGTACGGAAGACC</td><td>>hg38_21_+_8433172_8446622_RNA45SN1_45S_50ntFlanks</td><td>8111</td><td>8141</td><td>31</td><td>GGGGCGCGGGACATGTGGCGTACGGAAGACC</td><td>106.8</td><td>-10.6</td><td>1.8</td><td>-6.0</td><td>2.5E-09</td><td>5.8E-09</td><td>MB231/MB468</td><td>MP</td></tr>
<tr><td>>hg38_21_+_8433172_8446622_RNA45SN1_45S_50ntFlanks@10513.10533.21|CCGGAGTGGCGGAGATGGGCG</td><td>>hg38_21_+_8433172_8446622_RNA45SN1_45S_50ntFlanks</td><td>10513</td><td>10533</td><td>21</td><td>CCGGAGTGGCGGAGATGGGCG</td><td>6359.3</td><td>-10.6</td><td>1.8</td><td>-5.9</td><td>3.0E-09</td><td>5.8E-09</td><td>BT20/MB468</td><td>NUC</td></tr>
<tr><td>>hg38_21_+_8433172_8446622_RNA45SN1_45S_50ntFlanks@4555.4574.20|CGCAGCTAGGAATAATGGAA</td><td>>hg38_21_+_8433172_8446622_RNA45SN1_45S_50ntFlanks</td><td>4555</td><td>4574</td><td>20</td><td>CGCAGCTAGGAATAATGGAA</td><td>539.5</td><td>-8.7</td><td>1.5</td><td>-5.9</td><td>3.0E-09</td><td>5.8E-09</td><td>BT20/MB468</td><td>NUC</td></tr>
<tr><td>>hg38_21_+_8433172_8446622_RNA45SN1_45S_50ntFlanks@3829.3860.32|CGCTCGCTCCTCTCCTACTTGGATAACTGTGG</td><td>>hg38_21_+_8433172_8446622_RNA45SN1_45S_50ntFlanks</td><td>3829</td><td>3860</td><td>32</td><td>CGCTCGCTCCTCTCCTACTTGGATAACTGTGG</td><td>532.8</td><td>-8.5</td><td>1.4</td><td>-5.9</td><td>3.0E-09</td><td>5.8E-09</td><td>BT20/MB468</td><td>NUC</td></tr>
<tr><td>>hg38_21_+_8433172_8446622_RNA45SN1_45S_50ntFlanks@11874.11907.34|AGAAGACCCTGTTGAGCTTGACTCTAGTCTGGCA</td><td>>hg38_21_+_8433172_8446622_RNA45SN1_45S_50ntFlanks</td><td>11874</td><td>11907</td><td>34</td><td>AGAAGACCCTGTTGAGCTTGACTCTAGTCTGGCA</td><td>280.0</td><td>-6.7</td><td>1.1</td><td>-6.0</td><td>2.3E-09</td><td>5.8E-09</td><td>BT20/MB468</td><td>CYTO</td></tr>
<tr><td>>MI0003820|hsa-mir-454&WithFlank&17|-|59137752|59137878@70.94.25&[MIMAT0003885&hsa-miR-454-3p&offsets|0|+2;m-272&17|-|59137787|59137809&offsets|0|+2]|TAGTGCAATATTGCTTATAGGGTTT</td><td>>MI0003820|hsa-mir-454&WithFlank&17|-|59137752|59137878</td><td>70</td><td>94</td><td>25</td><td>TAGTGCAATATTGCTTATAGGGTTT</td><td>1315.7</td><td>7.6</td><td>1.3</td><td>5.9</td><td>2.9E-09</td><td>5.8E-09</td><td>MB231/MB468</td><td>MITO</td></tr>
<tr><td>>hg38_21_+_8433172_8446622_RNA45SN1_45S_50ntFlanks@12477.12500.24|CACTAATAGGGAACGTGAGCTGGG</td><td>>hg38_21_+_8433172_8446622_RNA45SN1_45S_50ntFlanks</td><td>12477</td><td>12500</td><td>24</td><td>CACTAATAGGGAACGTGAGCTGGG</td><td>190.8</td><td>-3.5</td><td>0.6</td><td>-6.0</td><td>2.7E-09</td><td>5.8E-09</td><td>MB231/MB468</td><td>NUC</td></tr>
<tr><td>>hg38_21_+_8433172_8446622_RNA45SN1_45S_50ntFlanks@483.525.43|TCGATGTGGTGACGTCGTGCTCTCCCGGGCCGGGTCCGAGCCG</td><td>>hg38_21_+_8433172_8446622_RNA45SN1_45S_50ntFlanks</td><td>483</td><td>525</td><td>43</td><td>TCGATGTGGTGACGTCGTGCTCTCCCGGGCCGGGTCCGAGCCG</td><td>533.9</td><td>5.5</td><td>0.9</td><td>6.0</td><td>2.7E-09</td><td>5.8E-09</td><td>MB231/MB468</td><td>NUC</td></tr>
<tr><td>>MI0000088|hsa-mir-30a&WithFlank&6|-|71403545|71403627@12.34.23(+1A)&[MIMAT0000087&hsa-miR-30a-5p&offsets|0|+1(+1A);m-15&6|-|71403595|71403616&offsets|0|+1(+1A)]|TGTAAACATCCTCGACTGGAAGCA</td><td>>MI0000088|hsa-mir-30a&WithFlank&6|-|71403545|71403627</td><td>12</td><td>34</td><td>23(+1A)</td><td>TGTAAACATCCTCGACTGGAAGCA</td><td>2329.5</td><td>6.0</td><td>1.0</td><td>6.0</td><td>2.7E-09</td><td>5.8E-09</td><td>MB231/MB468</td><td>CYTO</td></tr>
<tr><td>>MI0000088|hsa-mir-30a&WithFlank&6|-|71403545|71403627@12.34.23(+1G)&[MIMAT0000087&hsa-miR-30a-5p&offsets|0|+1(+1G);m-15&6|-|71403595|71403616&offsets|0|+1(+1G)]|TGTAAACATCCTCGACTGGAAGCG</td><td>>MI0000088|hsa-mir-30a&WithFlank&6|-|71403545|71403627</td><td>12</td><td>34</td><td>23(+1G)</td><td>TGTAAACATCCTCGACTGGAAGCG</td><td>3046.0</td><td>6.5</td><td>1.1</td><td>5.9</td><td>2.9E-09</td><td>5.8E-09</td><td>MB231/MB468</td><td>MITO</td></tr>
</table>

| | | | | | | | | | | | | | |
|---|---|---|---|---|---|---|---|---|---|---|---|---|---|
| >MI0000084\|hsa-mir-26b&WithFlank&2\|+\|218402640\|218402728@18.39.22&[MIMAT0000083&hsa-miR-26b-5p&offsets\|0\|+1;m-53&2\|+\|218402657\|218402678&offsets\|0\|0]\|TTCAAGTAATTCAGGATAGGTT | >MI0000084\|hsa-mir-26b&WithFlank&2\|+\|218402640\|218402728 | 18 | 39 | 22 | TTCAAGTAATTCAGGATAGGTT | 2692.0 | 2.6 | 0.4 | 5.9 | 3.1E-09 | 5.8E-09 | BT20/MB468 | NUC |
| >hg38_21_+_8433172_8446622_RNA45SN1_45S_50ntFlanks@11795.11820.26\|AACGAGATTCCCACTGTCCCTACCTA | >hg38_21_+_8433172_8446622_RNA45SN1_45S_50ntFlanks | 11795 | 11820 | 26 | AACGAGATTCCCACTGTCCCTACCTA | 98.1 | -10.5 | 1.8 | -6.0 | 2.6E-09 | 5.8E-09 | MB231/MB468 | MP |
| >hg38_21_+_8433172_8446622_RNA45SN1_45S_50ntFlanks@3910.3937.28\|GGGATGCGTGCATTTATCAGATCAAAAC | >hg38_21_+_8433172_8446622_RNA45SN1_45S_50ntFlanks | 3910 | 3937 | 28 | GGGATGCGTGCATTTATCAGATCAAAAC | 137.3 | -11.0 | 1.8 | -6.0 | 2.6E-09 | 5.8E-09 | MB231/MB468 | MP |
| >hg38_21_+_8433172_8446622_RNA45SN1_45S_50ntFlanks@12071.12096.26\|CCCCGAGGGGCTCTCGCTTCTGGCGC | >hg38_21_+_8433172_8446622_RNA45SN1_45S_50ntFlanks | 12071 | 12096 | 26 | CCCCGAGGGGCTCTCGCTTCTGGCGC | 172.9 | -11.3 | 1.9 | -6.0 | 2.6E-09 | 5.8E-09 | MB231/MB468 | MP |
| >MI0000066\|hsa-let-7e&WithFlank&19\|+\|51692780\|51692870@14.35.22&[MIMAT0000066&hsa-let-7e-5p&offsets\|0\|0;m-50&19\|+\|51692793\|51692814&offsets\|0\|0]\|TGAGGTAGGAGGTTGTATAGTT | >MI0000066\|hsa-let-7e&WithFlank&19\|+\|51692780\|51692870 | 14 | 35 | 22 | TGAGGTAGGAGGTTGTATAGTT | 6577.4 | 3.4 | 0.6 | 5.9 | 2.7E-09 | 5.8E-09 | MB231/MB468 | NUC |
| >hg38_21_+_8433172_8446622_RNA45SN1_45S_50ntFlanks@12617.12654.38\|TGAGGAGCCAATGGGGCGAAGCTACCATCTGTGGGATT | >hg38_21_+_8433172_8446622_RNA45SN1_45S_50ntFlanks | 12617 | 12654 | 38 | TGAGGAGCCAATGGGGCGAAGCTACCATCTGTGGGATT | 3452.2 | -10.2 | 1.7 | -5.9 | 3.1E-09 | 5.8E-09 | BT20/MB468 | NUC |
| >MI0000239\|hsa-mir-197&WithFlank&1\|+\|109598887\|109598973@54.76.23&[MIMAT0000227&hsa-miR-197-3p&offsets\|0\|+1;m-99&1\|+\|109598940\|109598961&offsets\|0\|+1]\|TTCACCACCTTCTCCACCCAGCA | >MI0000239\|hsa-mir-197&WithFlank&1\|+\|109598887\|109598973 | 54 | 76 | 23 | TTCACCACCTTCTCCACCCAGCA | 334.9 | 9.0 | 1.5 | 6.0 | 2.0E-09 | 5.8E-09 | BT20/MB468 | MITO |
| >hg38_21_+_8433172_8446622_RNA45SN1_45S_50ntFlanks@10411.10430.20\|CAGTCGGTCCTGAGAGATGG | >hg38_21_+_8433172_8446622_RNA45SN1_45S_50ntFlanks | 10411 | 10430 | 20 | CAGTCGGTCCTGAGAGATGG | 4564.2 | -5.7 | 0.9 | -6.3 | 3.1E-10 | 5.9E-09 | BT20/MB468 | MP |
| >hg38_21_+_8433172_8446622_RNA45SN1_45S_50ntFlanks@12045.12072.28\|CTGACCCGGTGAGGCGGGGGGGCGAGCC | >hg38_21_+_8433172_8446622_RNA45SN1_45S_50ntFlanks | 12045 | 12072 | 28 | CTGACCCGGTGAGGCGGGGGGGCGAGCC | 151.4 | -10.3 | 1.6 | -6.3 | 3.1E-10 | 5.9E-09 | BT20/MB468 | MP |
| >MI0000082\|hsa-mir-25&WithFlank&7\|-\|100093554\|100093649@58.79.22(+1A)&[MIMAT0000081&hsa-miR-25-3p&offsets\|0\|0(+1A);m-23&7\|-\|100093571\|100093592&offsets\|0\|0(+1A)]\|CATTGCACTTGTCTCGGTCTGAA | >MI0000082\|hsa-mir-25&WithFlank&7\|-\|100093554\|100093649 | 58 | 79 | 22(+1A) | CATTGCACTTGTCTCGGTCTGAA | 1346.1 | 3.9 | 0.7 | 5.9 | 2.7E-09 | 5.9E-09 | MB231/MB468 | NUC |
| >hg38_21_+_8433172_8446622_RNA45SN1_45S_50ntFlanks@10864.10889.26\|CTCTAAGGGCTGGGTCGGTCGGGCTG | >hg38_21_+_8433172_8446622_RNA45SN1_45S_50ntFlanks | 10864 | 10889 | 26 | CTCTAAGGGCTGGGTCGGTCGGGCTG | 73.8 | -10.1 | 1.7 | -6.0 | 2.6E-09 | 5.9E-09 | MB231/MB468 | MP |
| >hg38_21_+_8433172_8446622_RNA45SN1_45S_50ntFlanks@12633.12656.24\|CGAAGCTACCATCTGTGGGATTAT | >hg38_21_+_8433172_8446622_RNA45SN1_45S_50ntFlanks | 12633 | 12656 | 24 | CGAAGCTACCATCTGTGGGATTAT | 334.9 | -7.7 | 1.3 | -6.0 | 2.6E-09 | 5.9E-09 | MB231/MB468 | MP |
| >hg38_21_+_8433172_8446622_RNA45SN1_45S_50ntFlanks@4367.4402.36\|CAGTTAAAAAGCTCGTAGTTGGATCTTGGGAGCGGG | >hg38_21_+_8433172_8446622_RNA45SN1_45S_50ntFlanks | 4367 | 4402 | 36 | CAGTTAAAAAGCTCGTAGTTGGATCTTGGGAGCGGG | 124.6 | -10.8 | 1.8 | -6.0 | 2.6E-09 | 5.9E-09 | MB231/MB468 | MP |
| >hg38_21_+_8433172_8446622_RNA45SN1_45S_50ntFlanks@5084.5102.19\|CGAACGAGACTCTGGCATG | >hg38_21_+_8433172_8446622_RNA45SN1_45S_50ntFlanks | 5084 | 5102 | 19 | CGAACGAGACTCTGGCATG | 2003.3 | -8.2 | 1.4 | -5.9 | 2.9E-09 | 5.9E-09 | MB231/MB468 | MITO |
| >hg38_21_+_8433172_8446622_RNA45SN1_45S_50ntFlanks@11567.11591.25\|AGCAGCCGACTTAGAACTGGTGCGG | >hg38_21_+_8433172_8446622_RNA45SN1_45S_50ntFlanks | 11567 | 11591 | 25 | AGCAGCCGACTTAGAACTGGTGCGG | 331.3 | -11.5 | 1.8 | -6.3 | 3.1E-10 | 5.9E-09 | BT20/MB468 | MP |
| >hg38_21_+_8433172_8446622_RNA45SN1_45S_50ntFlanks@7975.8001.27\|CGCGACCTCAGATCAGACGTGGCGACC | >hg38_21_+_8433172_8446622_RNA45SN1_45S_50ntFlanks | 7975 | 8001 | 27 | CGCGACCTCAGATCAGACGTGGCGACC | 3441.3 | 5.9 | 1.0 | 6.0 | 2.0E-09 | 5.9E-09 | BT20/MB468 | MITO |
| >hg38_21_+_8433172_8446622_RNA45SN1_45S_50ntFlanks@5500.5526.27\|GTCGAACTTGACTATCTAGAGGAAGTA | >hg38_21_+_8433172_8446622_RNA45SN1_45S_50ntFlanks | 5500 | 5526 | 27 | GTCGAACTTGACTATCTAGAGGAAGTA | 119.2 | -10.0 | 1.6 | -6.3 | 3.1E-10 | 5.9E-09 | BT20/MB468 | MP |
| >hg38_21_+_8433172_8446622_RNA45SN1_45S_50ntFlanks@12035.12053.19\|CGTTTTTTCACTGACCCGG | >hg38_21_+_8433172_8446622_RNA45SN1_45S_50ntFlanks | 12035 | 12053 | 19 | CGTTTTTTCACTGACCCGG | 74.2 | -10.1 | 1.7 | -6.0 | 2.6E-09 | 5.9E-09 | MB231/MB468 | MP |
| >hg38_21_+_8433172_8446622_RNA45SN1_45S_50ntFlanks@11922.11953.32\|GAGAGGTGTAGAATAAGTGGGAGGCCCCCGGC | >hg38_21_+_8433172_8446622_RNA45SN1_45S_50ntFlanks | 11922 | 11953 | 32 | GAGAGGTGTAGAATAAGTGGGAGGCCCCCGGC | 520.8 | 8.1 | 1.4 | 5.9 | 2.7E-09 | 5.9E-09 | MB231/MB468 | CYTO |
| >hg38_1_-_228634819_228635039_RNA5S12_5S_50ntFlanks@86.112.27\|CGTCTGATCTCGGAAGCTAAGCAGGGT | >hg38_1_-_228634819_228635039_RNA5S12_5S_50ntFlanks | 86 | 112 | 27 | CGTCTGATCTCGGAAGCTAAGCAGGGT | 215.5 | -7.2 | 1.2 | -5.9 | 2.7E-09 | 5.9E-09 | MB231/MB468 | NUC |
| >hg38_21_+_8433172_8446622_RNA45SN1_45S_50ntFlanks@4008.4042.35\|CTCGGGCCGATCGCACGCCCCCCGTGGCGGCGACG | >hg38_21_+_8433172_8446622_RNA45SN1_45S_50ntFlanks | 4008 | 4042 | 35 | CTCGGGCCGATCGCACGCCCCCCGTGGCGGCGACG | 381.0 | -6.5 | 1.1 | -5.9 | 3.1E-09 | 5.9E-09 | BT20/MB468 | NUC |
| >hg38_21_+_8433172_8446622_RNA45SN1_45S_50ntFlanks@11785.11807.23\|TGAATGGATGAACGAGATTCCCA | >hg38_21_+_8433172_8446622_RNA45SN1_45S_50ntFlanks | 11785 | 11807 | 23 | TGAATGGATGAACGAGATTCCCA | 95.0 | -10.4 | 1.8 | -6.0 | 2.6E-09 | 5.9E-09 | MB231/MB468 | MP |
| >hg38_21_+_8433172_8446622_RNA45SN1_45S_50ntFlanks@12934.12960.27\|CTAAACCATTCGTAGACGACCTGCTTC | >hg38_21_+_8433172_8446622_RNA45SN1_45S_50ntFlanks | 12934 | 12960 | 27 | CTAAACCATTCGTAGACGACCTGCTTC | 2141.7 | -9.6 | 1.6 | -5.9 | 3.1E-09 | 5.9E-09 | BT20/MB468 | NUC |
| >hg38_21_+_8433172_8446622_RNA45SN1_45S_50ntFlanks@10781.10805.25\|AGGTGAACAGCCTCTGGCATGTTGG | >hg38_21_+_8433172_8446622_RNA45SN1_45S_50ntFlanks | 10781 | 10805 | 25 | AGGTGAACAGCCTCTGGCATGTTGG | 447.2 | -7.7 | 1.3 | -6.0 | 2.3E-09 | 5.9E-09 | BT20/MB468 | CYTO |
| >hg38_21_+_8433172_8446622_RNA45SN1_45S_50ntFlanks@9718.9735.18\|TGATTAGAGGTCTTGGGG | >hg38_21_+_8433172_8446622_RNA45SN1_45S_50ntFlanks | 9718 | 9735 | 18 | TGATTAGAGGTCTTGGGG | 66.4 | -9.9 | 1.7 | -6.0 | 2.6E-09 | 5.9E-09 | MB231/MB468 | MP |
| >hg38_21_+_8433172_8446622_RNA45SN1_45S_50ntFlanks@12488.12522.35\|AACGTGAGCTGGGTTTAGACCGTCGTGAGACAGGT | >hg38_21_+_8433172_8446622_RNA45SN1_45S_50ntFlanks | 12488 | 12522 | 35 | AACGTGAGCTGGGTTTAGACCGTCGTGAGACAGGT | 210.6 | -5.9 | 1.0 | -5.9 | 2.7E-09 | 5.9E-09 | MB231/MB468 | CYTO |
| >hg38_21_+_8433172_8446622_RNA45SN1_45S_50ntFlanks@8191.8214.24\|CAGCCCGTGGACGGTGTGAGGCCG | >hg38_21_+_8433172_8446622_RNA45SN1_45S_50ntFlanks | 8191 | 8214 | 24 | CAGCCCGTGGACGGTGTGAGGCCG | 514.0 | -3.1 | 0.5 | -6.0 | 2.1E-09 | 5.9E-09 | BT20/MB468 | MITO |
| >MI0000077\|hsa-mir-21&WithFlank&17\|+\|59841260\|59841343@14.35.22(+1G)&[MIMAT0000076&hsa-miR-21-5p&offsets\|0\|0(+1G);m-1&17\|+\|59841273\|59841294&offsets\|0\|0(+1G)]\|TAGCTTATCAGACTGATGTTGAG | >MI0000077\|hsa-mir-21&WithFlank&17\|+\|59841260\|59841343 | 14 | 35 | 22(+1G) | TAGCTTATCAGACTGATGTTGAG | 5178.0 | 7.5 | 1.3 | 6.0 | 2.4E-09 | 5.9E-09 | BT20/MB468 | CYTO |
| >trnaMT_ValTAC_MT_+_1602_1670@48.72.25\|CAACTTAACTTGACCGCTCTGACCA | >trnaMT_ValTAC_MT_+_1602_1670 | 48 | 72 | 25 | CAACTTAACTTGACCGCTCTGACCA | 1813.8 | 4.7 | 0.8 | 5.9 | 2.7E-09 | 5.9E-09 | MB231/MB468 | NUC |
| >hg38_21_+_8433172_8446622_RNA45SN1_45S_50ntFlanks@4760.4783.24\|TCAGATACCGTCGTAGTTCCGACC | >hg38_21_+_8433172_8446622_RNA45SN1_45S_50ntFlanks | 4760 | 4783 | 24 | TCAGATACCGTCGTAGTTCCGACC | 777.8 | -8.9 | 1.5 | -5.9 | 3.1E-09 | 5.9E-09 | BT20/MB468 | NUC |
| >hg38_21_+_8433172_8446622_RNA45SN1_45S_50ntFlanks@9546.9576.31\|TGGAGGTCCGTAGCGGTCCTGACGTGCAAAT | >hg38_21_+_8433172_8446622_RNA45SN1_45S_50ntFlanks | 9546 | 9576 | 31 | TGGAGGTCCGTAGCGGTCCTGACGTGCAAAT | 147.5 | 6.1 | 1.0 | 5.9 | 2.7E-09 | 5.9E-09 | MB231/MB468 | CYTO |
| >am_trnaMT_GluTTC_MT_-_14674_14742@27.55.29,trnalookalike8_GluTTC_5_-_93905172_93905240@27.55.29\|GTTTTTCATATCATTGGTCGTGGTTGTAG | >am_trnaMT_GluTTC_MT_-_14674_14742 | 27 | 55 | 29 | GTTTTTCATATCATTGGTCGTGGTTGTAG | 91.8 | -9.2 | 1.6 | -5.9 | 2.7E-09 | 5.9E-09 | MB231/MB468 | CYTO |
| >hg38_21_+_8433172_8446622_RNA45SN1_45S_50ntFlanks@10678.10708.31\|CGGTTCCGGCGGCGTCCGGTGAGCTCTCGCT | >hg38_21_+_8433172_8446622_RNA45SN1_45S_50ntFlanks | 10678 | 10708 | 31 | CGGTTCCGGCGGCGTCCGGTGAGCTCTCGCT | 76450.1 | -8.1 | 1.4 | -5.9 | 3.1E-09 | 5.9E-09 | BT20/MB468 | NUC |
| >hg38_21_+_8433172_8446622_RNA45SN1_45S_50ntFlanks@11875.11912.38\|GAAGACCCTGTTGAGCTTGACTCTAGTCTGGCACGGTG | >hg38_21_+_8433172_8446622_RNA45SN1_45S_50ntFlanks | 11875 | 11912 | 38 | GAAGACCCTGTTGAGCTTGACTCTAGTCTGGCACGGTG | 69.0 | -10.0 | 1.7 | -6.0 | 2.6E-09 | 5.9E-09 | MB231/MB468 | MP |
| >hg38_21_+_8433172_8446622_RNA45SN1_45S_50ntFlanks@11881.11903.23\|CCTGTTGAGCTTGACTCTAGTCT | >hg38_21_+_8433172_8446622_RNA45SN1_45S_50ntFlanks | 11881 | 11903 | 23 | CCTGTTGAGCTTGACTCTAGTCT | 397.4 | -7.8 | 1.3 | -5.9 | 3.0E-09 | 5.9E-09 | MB231/MB468 | MITO |
| >hg38_21_+_8433172_8446622_RNA45SN1_45S_50ntFlanks@12632.12659.28\|GCGAAGCTACCATCTGTGGGATTATGAC | >hg38_21_+_8433172_8446622_RNA45SN1_45S_50ntFlanks | 12632 | 12659 | 28 | GCGAAGCTACCATCTGTGGGATTATGAC | 837.7 | -7.3 | 1.2 | -5.9 | 3.1E-09 | 6.0E-09 | BT20/MB468 | NUC |
| >hg38_21_+_8433172_8446622_RNA45SN1_45S_50ntFlanks@3829.3852.24\|CGCTCGCTCCTCTCCTACTTGGAT | >hg38_21_+_8433172_8446622_RNA45SN1_45S_50ntFlanks | 3829 | 3852 | 24 | CGCTCGCTCCTCTCCTACTTGGAT | 420.4 | -8.7 | 1.5 | -5.9 | 3.1E-09 | 6.0E-09 | BT20/MB468 | NUC |
| >hg38_21_+_8433172_8446622_RNA45SN1_45S_50ntFlanks@7971.8009.39\|GAGACGCGACCTCAGATCAGACGTGGCGACCCGCTGAAT | >hg38_21_+_8433172_8446622_RNA45SN1_45S_50ntFlanks | 7971 | 8009 | 39 | GAGACGCGACCTCAGATCAGACGTGGCGACCCGCTGAAT | 225.2 | 4.1 | 0.7 | 5.9 | 3.2E-09 | 6.0E-09 | BT20/MB468 | NUC |
| >hg38_21_+_8433172_8446622_RNA45SN1_45S_50ntFlanks@11666.11702.37\|TCTGCCCAGTGCTCTGAATGTCAAAGTGAAGAAATTC | >hg38_21_+_8433172_8446622_RNA45SN1_45S_50ntFlanks | 11666 | 11702 | 37 | TCTGCCCAGTGCTCTGAATGTCAAAGTGAAGAAATTC | 2109.7 | 3.7 | 0.6 | 5.9 | 2.7E-09 | 6.0E-09 | MB231/MB468 | CYTO |
| >hg38_21_+_8433172_8446622_RNA45SN1_45S_50ntFlanks@11208.11230.23\|GCGGCGACTCTGGACGCGAGCCG | >hg38_21_+_8433172_8446622_RNA45SN1_45S_50ntFlanks | 11208 | 11230 | 23 | GCGGCGACTCTGGACGCGAGCCG | 1529.7 | -7.0 | 1.2 | -5.9 | 2.8E-09 | 6.0E-09 | MB231/MB468 | NUC |
| >trnaMT_IleGAT_MT_+_4263_4331@43.71.29\|ATAGGAGCTTAAACCCCCTTATTTCTACC | >trnaMT_IleGAT_MT_+_4263_4331 | 43 | 71 | 29 | ATAGGAGCTTAAACCCCCTTATTTCTACC | 69.5 | 9.3 | 1.6 | 6.0 | 2.6E-09 | 6.0E-09 | MB231/MB468 | MP |
| >MI0001445\|hsa-mir-423&WithFlank&17\|+\|30117073\|30117178@59.81.23&[MIMAT0001340&hsa-miR-423-3p&offsets\|0\|0;m-100&17\|+\|30117131\|30117153&offsets\|0\|0]\|AGCTCGGTCTGAGGCCCCTCAGT | >MI0001445\|hsa-mir-423&WithFlank&17\|+\|30117073\|30117178 | 59 | 81 | 23 | AGCTCGGTCTGAGGCCCCTCAGT | 11678.0 | 5.4 | 0.9 | 6.3 | 3.2E-10 | 6.0E-09 | BT20/MB468 | MP |
| >hg38_21_+_8433172_8446622_RNA45SN1_45S_50ntFlanks@8357.8402.46\|AAAAGAACTTTGAAGAGAGAGTTCAAGAGGGCGTGAAACCGTTAAG | >hg38_21_+_8433172_8446622_RNA45SN1_45S_50ntFlanks | 8357 | 8402 | 46 | AAAAGAACTTTGAAGAGAGAGTTCAAGAGGGCGTGAAACCGTTAAG | 91.5 | -10.4 | 1.7 | -6.0 | 2.6E-09 | 6.0E-09 | MB231/MB468 | MP |
| >hg38_21_+_8433172_8446622_RNA45SN1_45S_50ntFlanks@12625.12652.28\|CAATGGGGCGAAGCTACCATCTGTGGGA | >hg38_21_+_8433172_8446622_RNA45SN1_45S_50ntFlanks | 12625 | 12652 | 28 | CAATGGGGCGAAGCTACCATCTGTGGGA | 281.8 | -7.5 | 1.3 | -5.9 | 3.0E-09 | 6.0E-09 | MB231/MB468 | MITO |
| >hg38_21_+_8433172_8446622_RNA45SN1_45S_50ntFlanks@10795.10818.24\|TGGCATGTTGGAACAATGTAGGTA | >hg38_21_+_8433172_8446622_RNA45SN1_45S_50ntFlanks | 10795 | 10818 | 24 | TGGCATGTTGGAACAATGTAGGTA | 428.2 | -6.1 | 1.0 | -5.9 | 2.8E-09 | 6.0E-09 | MB231/MB468 | NUC |
| >hg38_21_+_8433172_8446622_RNA45SN1_45S_50ntFlanks@5077.5117.41\|CCGATAACGAACGAGACTCTGGCATGCTAACTAGTTACGCG | >hg38_21_+_8433172_8446622_RNA45SN1_45S_50ntFlanks | 5077 | 5117 | 41 | CCGATAACGAACGAGACTCTGGCATGCTAACTAGTTACGCG | 125.6 | -10.8 | 1.8 | -6.0 | 2.6E-09 | 6.0E-09 | MB231/MB468 | MP |
| >hg38_21_+_8433172_8446622_RNA45SN1_45S_50ntFlanks@10356.10388.33\|ACTTTGAAGGCCGAAGTGGAGAAGGGTTCCATG | >hg38_21_+_8433172_8446622_RNA45SN1_45S_50ntFlanks | 10356 | 10388 | 33 | ACTTTGAAGGCCGAAGTGGAGAAGGGTTCCATG | 99.2 | -10.5 | 1.8 | -6.0 | 2.6E-09 | 6.0E-09 | MB231/MB468 | MP |
| >hg38_21_+_8433172_8446622_RNA45SN1_45S_50ntFlanks@10926.10945.20\|ACGAGGCGCCGCCGCCCCCC | >hg38_21_+_8433172_8446622_RNA45SN1_45S_50ntFlanks | 10926 | 10945 | 20 | ACGAGGCGCCGCCGCCCCCC | 659.2 | -12.4 | 2.0 | -6.3 | 3.2E-10 | 6.0E-09 | BT20/MB468 | MP |
| >hg38_21_+_8433172_8446622_RNA45SN1_45S_50ntFlanks@12942.12962.21\|TTCGTAGACGACCTGCTTCTG | >hg38_21_+_8433172_8446622_RNA45SN1_45S_50ntFlanks | 12942 | 12962 | 21 | TTCGTAGACGACCTGCTTCTG | 256.0 | -7.7 | 1.3 | -6.0 | 2.6E-09 | 6.0E-09 | MB231/MB468 | MP |
| >hg38_21_+_8433172_8446622_RNA45SN1_45S_50ntFlanks@10353.10376.24\|AGAACTTTGAAGGCCGAAGTGGAG | >hg38_21_+_8433172_8446622_RNA45SN1_45S_50ntFlanks | 10353 | 10376 | 24 | AGAACTTTGAAGGCCGAAGTGGAG | 91.2 | -10.4 | 1.7 | -6.0 | 2.6E-09 | 6.0E-09 | MB231/MB468 | MP |
| >hg38_21_+_8433172_8446622_RNA45SN1_45S_50ntFlanks@9788.9820.33\|CGCTGGCGTGGAGCCGGGCGTGGAATGCGAGTG | >hg38_21_+_8433172_8446622_RNA45SN1_45S_50ntFlanks | 9788 | 9820 | 33 | CGCTGGCGTGGAGCCGGGCGTGGAATGCGAGTG | 594.6 | -8.1 | 1.4 | -6.0 | 2.7E-09 | 6.1E-09 | MB231/MB468 | MP |
| >hg38_21_+_8433172_8446622_RNA45SN1_45S_50ntFlanks@9303.9338.36\|GACCAAGGAGTCTAACACGTGCGCGAGTCGGGGGCT | >hg38_21_+_8433172_8446622_RNA45SN1_45S_50ntFlanks | 9303 | 9338 | 36 | GACCAAGGAGTCTAACACGTGCGCGAGTCGGGGGCT | 288.8 | -7.6 | 1.3 | -6.0 | 2.7E-09 | 6.1E-09 | MB231/MB468 | MP |

| | | | | | | | | | | | | | |
|---|---|---|---|---|---|---|---|---|---|---|---|---|---|
| >hg38_21_+_8433172_8446622_RNA45SN1_45S_50ntFlanks@10738.10756.19\|AAATCTCGCGCCGGGCCGT | >hg38_21_+_8433172_8446622_RNA45SN1_45S_50ntFlanks | 10738 | 10756 | 19 | AAATCTCGCGCCGGGCCGT | 1287.9 | 5.6 | 0.9 | 5.9 | 2.8E-09 | 6.1E-09 | MB231/MB468 | CYTO |
| >hg38_21_+_8433172_8446622_RNA45SN1_45S_50ntFlanks@10402.10432.31\|AACATGGGTCAGTCGGTCCTGAGAGATGGGC | >hg38_21_+_8433172_8446622_RNA45SN1_45S_50ntFlanks | 10402 | 10432 | 31 | AACATGGGTCAGTCGGTCCTGAGAGATGGGC | 306.8 | -7.8 | 1.3 | -6.0 | 2.7E-09 | 6.1E-09 | MB231/MB468 | MP |
| >hg38_21_+_8433172_8446622_RNA45SN1_45S_50ntFlanks@11773.11794.22\|TAGTGACGCGCATGAATGGATG | >hg38_21_+_8433172_8446622_RNA45SN1_45S_50ntFlanks | 11773 | 11794 | 22 | TAGTGACGCGCATGAATGGATG | 386.8 | 3.7 | 0.6 | 5.9 | 2.8E-09 | 6.1E-09 | MB231/MB468 | CYTO |
| >hg38_21_+_8433172_8446622_RNA45SN1_45S_50ntFlanks@11589.11611.23\|CGGACCAGGGGAATCCGACTGTT | >hg38_21_+_8433172_8446622_RNA45SN1_45S_50ntFlanks | 11589 | 11611 | 23 | CGGACCAGGGGAATCCGACTGTT | 1204.1 | -3.4 | 0.6 | -5.9 | 3.2E-09 | 6.1E-09 | BT20/MB468 | NUC |
| >hg38_21_+_8433172_8446622_RNA45SN1_45S_50ntFlanks@12418.12437.20\|CGATGTCGGCTCTTCCTATC | >hg38_21_+_8433172_8446622_RNA45SN1_45S_50ntFlanks | 12418 | 12437 | 20 | CGATGTCGGCTCTTCCTATC | 6516.2 | -3.0 | 0.5 | -6.0 | 2.4E-09 | 6.1E-09 | BT20/MB468 | CYTO |
| >hg38_21_+_8433172_8446622_RNA45SN1_45S_50ntFlanks@6721.6756.36\|AATTGCAGGACACATTGATCATCGACACTTCGAACG | >hg38_21_+_8433172_8446622_RNA45SN1_45S_50ntFlanks | 6721 | 6756 | 36 | AATTGCAGGACACATTGATCATCGACACTTCGAACG | 436.3 | -3.3 | 0.6 | -6.0 | 2.4E-09 | 6.1E-09 | BT20/MB468 | CYTO |
| >hg38_1_-_228634819_228635039_RNA5S12_5S_50ntFlanks@130.169.40\|TGGATGGGAGACCGCCTGGGAATACCGGGTGCTGTAGGCT | >hg38_1_-_228634819_228635039_RNA5S12_5S_50ntFlanks | 130 | 169 | 40 | TGGATGGGAGACCGCCTGGGAATACCGGGTGCTGTAGGCT | 660.4 | 8.3 | 1.4 | 6.0 | 2.1E-09 | 6.1E-09 | BT20/MB468 | MITO |
| >hg38_21_+_8433172_8446622_RNA45SN1_45S_50ntFlanks@12394.12414.21\|TAGCGACGTCGCTTTTTGATC | >hg38_21_+_8433172_8446622_RNA45SN1_45S_50ntFlanks | 12394 | 12414 | 21 | TAGCGACGTCGCTTTTTGATC | 2477.0 | -5.4 | 0.9 | -5.9 | 3.2E-09 | 6.1E-09 | BT20/MB468 | NUC |
| >hg38_21_+_8433172_8446622_RNA45SN1_45S_50ntFlanks@12405.12424.20\|CTTTTTGATCCTTCGATGTC | >hg38_21_+_8433172_8446622_RNA45SN1_45S_50ntFlanks | 12405 | 12424 | 20 | CTTTTTGATCCTTCGATGTC | 2383.4 | -8.6 | 1.4 | -6.0 | 2.4E-09 | 6.1E-09 | BT20/MB468 | CYTO |
| >hg38_21_+_8433172_8446622_RNA45SN1_45S_50ntFlanks@7988.8007.20\|CAGACGTGGCGACCCGCTGA | >hg38_21_+_8433172_8446622_RNA45SN1_45S_50ntFlanks | 7988 | 8007 | 20 | CAGACGTGGCGACCCGCTGA | 1635.8 | -3.8 | 0.6 | -6.0 | 2.1E-09 | 6.1E-09 | BT20/MB468 | MITO |
| >hg38_21_+_8433172_8446622_RNA45SN1_45S_50ntFlanks@4005.4025.21\|AACCTCGGGCCGATCGCACGC | >hg38_21_+_8433172_8446622_RNA45SN1_45S_50ntFlanks | 4005 | 4025 | 21 | AACCTCGGGCCGATCGCACGC | 276.7 | -2.7 | 0.5 | -6.0 | 2.4E-09 | 6.1E-09 | BT20/MB468 | CYTO |
| >hg38_21_+_8433172_8446622_RNA45SN1_45S_50ntFlanks@12709.12745.37\|CCGCGGAGCCTCGGTTGGCCTCGGATAGCCGGTCCCC | >hg38_21_+_8433172_8446622_RNA45SN1_45S_50ntFlanks | 12709 | 12745 | 37 | CCGCGGAGCCTCGGTTGGCCTCGGATAGCCGGTCCCC | 179.3 | -10.6 | 1.7 | -6.3 | 3.3E-10 | 6.1E-09 | BT20/MB468 | MP |
| >hg38_21_+_8433172_8446622_RNA45SN1_45S_50ntFlanks@11766.11791.26\|ATCTAATTAGTGACGCGCATGAATGG | >hg38_21_+_8433172_8446622_RNA45SN1_45S_50ntFlanks | 11766 | 11791 | 26 | ATCTAATTAGTGACGCGCATGAATGG | 120.2 | -10.0 | 1.6 | -6.3 | 3.3E-10 | 6.1E-09 | BT20/MB468 | MP |
| >hg38_21_+_8433172_8446622_RNA45SN1_45S_50ntFlanks@8737.8758.22\|CCCGTCCGTCCGTCCGTCCGTC | >hg38_21_+_8433172_8446622_RNA45SN1_45S_50ntFlanks | 8737 | 8758 | 22 | CCCGTCCGTCCGTCCGTCCGTC | 391.1 | -8.8 | 1.5 | -5.9 | 2.8E-09 | 6.1E-09 | MB231/MB468 | NUC |
| >hg38_21_+_8433172_8446622_RNA45SN1_45S_50ntFlanks@10806.10830.25\|AACAATGTAGGTAAGGGAAGTCGGC | >hg38_21_+_8433172_8446622_RNA45SN1_45S_50ntFlanks | 10806 | 10830 | 25 | AACAATGTAGGTAAGGGAAGTCGGC | 399.3 | -8.3 | 1.4 | -5.9 | 3.2E-09 | 6.1E-09 | BT20/MB468 | NUC |
| >MI0003513\|hsa-mir-455&WithFlank&9\|+\|114209428\|114209535@60.81.22&[MIMAT0004784&hsa-miR-455-3p&offsets\|0\|+1;m-135&9\|+\|114209487\|114209508&offsets\|0\|0]\|GCAGTCCATGGGCATATACACT | >MI0003513\|hsa-mir-455&WithFlank&9\|+\|114209428\|114209535 | 60 | 81 | 22 | GCAGTCCATGGGCATATACACT | 113.7 | 10.0 | 1.7 | 5.9 | 2.7E-09 | 6.1E-09 | MB231/MB468 | MP |
| >trna27_GlyCCC_2_-_70476123_70476193@57.74.18,trna34_GlyCCC_16_-_686736_686806@57.74.18\|TTCCCGGGCGGCGCACCA | >trna27_GlyCCC_2_-_70476123_70476193 | 57 | 74 | 18 | TTCCCGGGCGGCGCACCA | 466.3 | 6.8 | 1.1 | 6.0 | 2.4E-09 | 6.1E-09 | BT20/MB468 | CYTO |
| >hg38_21_+_8433172_8446622_RNA45SN1_45S_50ntFlanks@10782.10818.37\|GGTGAACAGCCTCTGGCATGTTGGAACAATGTAGGTA | >hg38_21_+_8433172_8446622_RNA45SN1_45S_50ntFlanks | 10782 | 10818 | 37 | GGTGAACAGCCTCTGGCATGTTGGAACAATGTAGGTA | 203.2 | -10.7 | 1.7 | -6.3 | 3.3E-10 | 6.1E-09 | BT20/MB468 | MP |
| >hg38_21_+_8433172_8446622_RNA45SN1_45S_50ntFlanks@13006.13027.22\|CTATTGAAAGTCAGCCCTCGAC | >hg38_21_+_8433172_8446622_RNA45SN1_45S_50ntFlanks | 13006 | 13027 | 22 | CTATTGAAAGTCAGCCCTCGAC | 119.7 | -10.0 | 1.6 | -6.3 | 3.3E-10 | 6.1E-09 | BT20/MB468 | MP |
| >hg38_21_+_8433172_8446622_RNA45SN1_45S_50ntFlanks@12405.12439.35\|CTTTTTGATCCTTCGATGTCGGCTCTTCCTATCAT | >hg38_21_+_8433172_8446622_RNA45SN1_45S_50ntFlanks | 12405 | 12439 | 35 | CTTTTTGATCCTTCGATGTCGGCTCTTCCTATCAT | 2442.7 | -9.9 | 1.7 | -5.9 | 3.2E-09 | 6.1E-09 | BT20/MB468 | NUC |
| >hg38_21_+_8433172_8446622_RNA45SN1_45S_50ntFlanks@11768.11794.27\|CTAATTAGTGACGCGCATGAATGGATG | >hg38_21_+_8433172_8446622_RNA45SN1_45S_50ntFlanks | 11768 | 11794 | 27 | CTAATTAGTGACGCGCATGAATGGATG | 3005.8 | -3.5 | 0.6 | -5.9 | 2.8E-09 | 6.1E-09 | MB231/MB468 | CYTO |
| >hg38_21_+_8433172_8446622_RNA45SN1_45S_50ntFlanks@6653.6684.32\|ACTCTTAGCGGTGGATCACTCGGCTCGTGCGT | >hg38_21_+_8433172_8446622_RNA45SN1_45S_50ntFlanks | 6653 | 6684 | 32 | ACTCTTAGCGGTGGATCACTCGGCTCGTGCGT | 402.4 | 4.3 | 0.7 | 5.9 | 2.8E-09 | 6.1E-09 | MB231/MB468 | NUC |
| >hg38_21_+_8433172_8446622_RNA45SN1_45S_50ntFlanks@12405.12438.34\|CTTTTTGATCCTTCGATGTCGGCTCTTCCTATCA | >hg38_21_+_8433172_8446622_RNA45SN1_45S_50ntFlanks | 12405 | 12438 | 34 | CTTTTTGATCCTTCGATGTCGGCTCTTCCTATCA | 1693.1 | -4.6 | 0.8 | -5.9 | 2.8E-09 | 6.1E-09 | MB231/MB468 | NUC |
| >hg38_21_+_8433172_8446622_RNA45SN1_45S_50ntFlanks@6733.6769.37\|CATTGATCATCGACACTTCGAACGCACTTGCGGCCCC | >hg38_21_+_8433172_8446622_RNA45SN1_45S_50ntFlanks | 6733 | 6769 | 37 | CATTGATCATCGACACTTCGAACGCACTTGCGGCCCC | 94.3 | -10.4 | 1.8 | -5.9 | 2.7E-09 | 6.1E-09 | MB231/MB468 | MP |
| >hg38_21_+_8433172_8446622_RNA45SN1_45S_50ntFlanks@9712.9729.18\|AGCGAATGATTAGAGGTC | >hg38_21_+_8433172_8446622_RNA45SN1_45S_50ntFlanks | 9712 | 9729 | 18 | AGCGAATGATTAGAGGTC | 276.8 | -10.2 | 1.7 | -6.0 | 2.1E-09 | 6.2E-09 | BT20/MB468 | MITO |
| >hg38_21_+_8433172_8446622_RNA45SN1_45S_50ntFlanks@10910.10928.19\|CGCGCGCCGCGGCTGGACG | >hg38_21_+_8433172_8446622_RNA45SN1_45S_50ntFlanks | 10910 | 10928 | 19 | CGCGCGCCGCGGCTGGACG | 341.2 | -6.5 | 1.1 | -5.9 | 3.1E-09 | 6.2E-09 | MB231/MB468 | MITO |
| >hg38_21_+_8433172_8446622_RNA45SN1_45S_50ntFlanks@12416.12444.29\|TTCGATGTCGGCTCTTCCTATCATTGTGA | >hg38_21_+_8433172_8446622_RNA45SN1_45S_50ntFlanks | 12416 | 12444 | 29 | TTCGATGTCGGCTCTTCCTATCATTGTGA | 3562.9 | 3.9 | 0.7 | 5.9 | 2.8E-09 | 6.2E-09 | MB231/MB468 | CYTO |
| >hg38_21_+_8433172_8446622_RNA45SN1_45S_50ntFlanks@11712.11742.31\|CGGGTAAACGGCGGGAGTAACTATGACTCTC | >hg38_21_+_8433172_8446622_RNA45SN1_45S_50ntFlanks | 11712 | 11742 | 31 | CGGGTAAACGGCGGGAGTAACTATGACTCTC | 312.5 | -8.1 | 1.4 | -5.9 | 3.3E-09 | 6.2E-09 | BT20/MB468 | NUC |
| >hg38_21_+_8433172_8446622_RNA45SN1_45S_50ntFlanks@5486.5513.28\|GCGCTGAGAAGACGGTCGAACTTGACTA | >hg38_21_+_8433172_8446622_RNA45SN1_45S_50ntFlanks | 5486 | 5513 | 28 | GCGCTGAGAAGACGGTCGAACTTGACTA | 393.2 | -7.9 | 1.3 | -5.9 | 2.7E-09 | 6.2E-09 | MB231/MB468 | MP |
| >hg38_21_+_8433172_8446622_RNA45SN1_45S_50ntFlanks@11569.11594.26\|CAGCCGACTTAGAACTGGTGCGGACC | >hg38_21_+_8433172_8446622_RNA45SN1_45S_50ntFlanks | 11569 | 11594 | 26 | CAGCCGACTTAGAACTGGTGCGGACC | 976.7 | -6.8 | 1.1 | -5.9 | 2.7E-09 | 6.2E-09 | MB231/MB468 | MP |
| >hg38_21_+_8433172_8446622_RNA45SN1_45S_50ntFlanks@4728.4759.32\|ATCAAGAACGAAAGTCGGAGGTTCGAAGACGA | >hg38_21_+_8433172_8446622_RNA45SN1_45S_50ntFlanks | 4728 | 4759 | 32 | ATCAAGAACGAAAGTCGGAGGTTCGAAGACGA | 1189.8 | -8.5 | 1.4 | -5.9 | 2.7E-09 | 6.2E-09 | MB231/MB468 | MP |
| >hg38_21_+_8433172_8446622_RNA45SN1_45S_50ntFlanks@9565.9595.31\|TGACGTGCAAATCGGTCGTCCGACCTGGGTA | >hg38_21_+_8433172_8446622_RNA45SN1_45S_50ntFlanks | 9565 | 9595 | 31 | TGACGTGCAAATCGGTCGTCCGACCTGGGTA | 1719.9 | -5.3 | 0.9 | -5.9 | 3.1E-09 | 6.2E-09 | MB231/MB468 | MITO |
| >hg38_21_+_8433172_8446622_RNA45SN1_45S_50ntFlanks@6659.6683.25\|AGCGGTGGATCACTCGGCTCGTGCG | >hg38_21_+_8433172_8446622_RNA45SN1_45S_50ntFlanks | 6659 | 6683 | 25 | AGCGGTGGATCACTCGGCTCGTGCG | 356.8 | 4.5 | 0.8 | 5.9 | 2.9E-09 | 6.2E-09 | MB231/MB468 | NUC |
| >MI0000440\|hsa-mir-27b&WithFlank&9\|+\|95085439\|95085547@67.87.21(+2U)&[MIMAT0000419&hsa-miR-27b-3p&offsets\|0\|0(+2U);m-39&9\|+\|95085505\|95085525&offsets\|0\|0(+2U)]\|TTCACAGTGGCTAAGTTCTGCTT | >MI0000440\|hsa-mir-27b&WithFlank&9\|+\|95085439\|95085547 | 67 | 87 | 21(+2U) | TTCACAGTGGCTAAGTTCTGCTT | 560.7 | 7.1 | 1.2 | 6.0 | 2.2E-09 | 6.2E-09 | BT20/MB468 | MITO |
| >hg38_21_+_8433172_8446622_RNA45SN1_45S_50ntFlanks@10786.10827.42\|AACAGCCTCTGGCATGTTGGAACAATGTAGGTAAGGGAAGTC | >hg38_21_+_8433172_8446622_RNA45SN1_45S_50ntFlanks | 10786 | 10827 | 42 | AACAGCCTCTGGCATGTTGGAACAATGTAGGTAAGGGAAGTC | 878.7 | -6.7 | 1.1 | -5.9 | 3.1E-09 | 6.2E-09 | MB231/MB468 | MITO |
| >hg38_21_+_8433172_8446622_RNA45SN1_45S_50ntFlanks@11759.11780.22\|CCTCGTCATCTAATTAGTGACG | >hg38_21_+_8433172_8446622_RNA45SN1_45S_50ntFlanks | 11759 | 11780 | 22 | CCTCGTCATCTAATTAGTGACG | 234.5 | -7.5 | 1.3 | -6.0 | 2.5E-09 | 6.2E-09 | BT20/MB468 | CYTO |
| >trna10_LysCTT_16_+_3241501_3241573@1.34.34,trna13_LysCTT_14_-_58706613_58706685@1.34.34,trna2_LysCTT_15_+_79152904_79152976@1.34.34\|GCCCGGCTAGCTCAGTCGGTAGAGCATGGGACTC | >trna10_LysCTT_16_+_3241501_3241573 | 1 | 34 | 34 | GCCCGGCTAGCTCAGTCGGTAGAGCATGGGACTC | 3209.0 | 6.8 | 1.2 | 5.9 | 3.1E-09 | 6.2E-09 | MB231/MB468 | MITO |
| >hg38_21_+_8433172_8446622_RNA45SN1_45S_50ntFlanks@11578.11603.26\|TAGAACTGGTGCGGACCAGGGGAATC | >hg38_21_+_8433172_8446622_RNA45SN1_45S_50ntFlanks | 11578 | 11603 | 26 | TAGAACTGGTGCGGACCAGGGGAATC | 1321.0 | -3.6 | 0.6 | -6.0 | 2.5E-09 | 6.2E-09 | BT20/MB468 | CYTO |
| >hg38_21_+_8433172_8446622_RNA45SN1_45S_50ntFlanks@12404.12435.32\|GCTTTTTGATCCTTCGATGTCGGCTCTTCCTA | >hg38_21_+_8433172_8446622_RNA45SN1_45S_50ntFlanks | 12404 | 12435 | 32 | GCTTTTTGATCCTTCGATGTCGGCTCTTCCTA | 572.1 | -6.8 | 1.1 | -5.9 | 2.9E-09 | 6.2E-09 | MB231/MB468 | NUC |
| >hg38_MT_+_1621_3279_RNR2_16S_50ntFlanks@460.492.33\|TTGTAAATTTAACTGTTAGTCCAAAGAGGAACA | >hg38_MT_+_1621_3279_RNR2_16S_50ntFlanks | 460 | 492 | 33 | TTGTAAATTTAACTGTTAGTCCAAAGAGGAACA | 224.5 | 7.5 | 1.3 | 5.9 | 2.9E-09 | 6.2E-09 | MB231/MB468 | NUC |
| >hg38_21_+_8433172_8446622_RNA45SN1_45S_50ntFlanks@12486.12509.24\|GGAACGTGAGCTGGGTTTAGACCG | >hg38_21_+_8433172_8446622_RNA45SN1_45S_50ntFlanks | 12486 | 12509 | 24 | GGAACGTGAGCTGGGTTTAGACCG | 95.0 | -10.4 | 1.8 | -5.9 | 2.7E-09 | 6.2E-09 | MB231/MB468 | MP |
| >hg38_21_+_8433172_8446622_RNA45SN1_45S_50ntFlanks@4047.4081.35\|ATTCGAACGTCTGCCCTATCAACTTTCGATGGTAG | >hg38_21_+_8433172_8446622_RNA45SN1_45S_50ntFlanks | 4047 | 4081 | 35 | ATTCGAACGTCTGCCCTATCAACTTTCGATGGTAG | 313.2 | -7.7 | 1.3 | -5.9 | 2.9E-09 | 6.2E-09 | MB231/MB468 | NUC |
| >hg38_21_+_8433172_8446622_RNA45SN1_45S_50ntFlanks@9806.9835.30\|CGTGGAATGCGAGTGCCTAGTGGGCCACTT | >hg38_21_+_8433172_8446622_RNA45SN1_45S_50ntFlanks | 9806 | 9835 | 30 | CGTGGAATGCGAGTGCCTAGTGGGCCACTT | 101.6 | -10.5 | 1.8 | -5.9 | 2.7E-09 | 6.2E-09 | MB231/MB468 | MP |
| >hg38_21_+_8433172_8446622_RNA45SN1_45S_50ntFlanks@10788.10816.29\|CAGCCTCTGGCATGTTGGAACAATGTAGG | >hg38_21_+_8433172_8446622_RNA45SN1_45S_50ntFlanks | 10788 | 10816 | 29 | CAGCCTCTGGCATGTTGGAACAATGTAGG | 858.3 | -3.4 | 0.6 | -6.0 | 2.5E-09 | 6.2E-09 | BT20/MB468 | CYTO |
| >hg38_21_+_8433172_8446622_RNA45SN1_45S_50ntFlanks@6721.6746.26\|AATTGCAGGACACATTGATCATCGAC | >hg38_21_+_8433172_8446622_RNA45SN1_45S_50ntFlanks | 6721 | 6746 | 26 | AATTGCAGGACACATTGATCATCGAC | 2042.7 | -5.7 | 1.0 | -5.9 | 2.9E-09 | 6.2E-09 | MB231/MB468 | CYTO |
| >MI0000239\|hsa-mir-197&WithFlank&1\|+\|109598887\|109598973@54.74.21(+1A)&[MIMAT0000227&hsa-miR-197-3p&offsets\|0\|-1(+1A);m-99&1\|+\|109598940\|109598961&offsets\|0\|-1(+1A)]\|TTCACCACCTTCTCCACCCAGA | >MI0000239\|hsa-mir-197&WithFlank&1\|+\|109598887\|109598973 | 54 | 74 | 21(+1A) | TTCACCACCTTCTCCACCCAGA | 653.5 | 3.9 | 0.6 | 6.3 | 3.0E-10 | 6.2E-09 | BT20/MB231 | NUC |
| >hg38_21_+_8433172_8446622_RNA45SN1_45S_50ntFlanks@10794.10821.28\|CTGGCATGTTGGAACAATGTAGGTAAGG | >hg38_21_+_8433172_8446622_RNA45SN1_45S_50ntFlanks | 10794 | 10821 | 28 | CTGGCATGTTGGAACAATGTAGGTAAGG | 255.5 | -7.6 | 1.3 | -5.9 | 2.7E-09 | 6.3E-09 | MB231/MB468 | MP |
| >hg38_21_+_8433172_8446622_RNA45SN1_45S_50ntFlanks@8115.8136.22\|CGCGGGACATGTGGCGTACGGA | >hg38_21_+_8433172_8446622_RNA45SN1_45S_50ntFlanks | 8115 | 8136 | 22 | CGCGGGACATGTGGCGTACGGA | 2624.8 | 14.7 | 2.2 | 6.5 | 6.5E-11 | 6.3E-09 | BT20/MB231 | MP |
| >hg38_21_+_8433172_8446622_RNA45SN1_45S_50ntFlanks@7976.8011.36\|GCGACCTCAGATCAGACGTGGCGACCCGCTGAATTT | >hg38_21_+_8433172_8446622_RNA45SN1_45S_50ntFlanks | 7976 | 8011 | 36 | GCGACCTCAGATCAGACGTGGCGACCCGCTGAATTT | 1329.6 | 5.3 | 0.9 | 5.9 | 2.9E-09 | 6.3E-09 | MB231/MB468 | NUC |
| >hg38_21_+_8433172_8446622_RNA45SN1_45S_50ntFlanks@4046.4066.21\|CATTCGAACGTCTGCCCTATC | >hg38_21_+_8433172_8446622_RNA45SN1_45S_50ntFlanks | 4046 | 4066 | 21 | CATTCGAACGTCTGCCCTATC | 2292.1 | -4.2 | 0.7 | -5.9 | 2.9E-09 | 6.3E-09 | MB231/MB468 | NUC |

| | | | | | | | | | | | | | |
|---|---|---|---|---|---|---|---|---|---|---|---|---|---|
| >hg38_21_+_8433172_8446622_RNA45SN1_45S_50ntFlanks@12324.12344.21\|TTGGGTTTTAAGCAGGAGGTG | >hg38_21_+_8433172_8446622_RNA45SN1_45S_50ntFlanks | 12324 | 12344 | 21 | TTGGGTTTTAAGCAGGAGGTG | 2835.6 | -8.3 | 1.4 | -6.0 | 2.5E-09 | 6.3E-09 | BT20/MB468 | CYTO |
| >hg38_1_-_228634819_228635039_RNA5S12_5S_50ntFlanks@86.112.27\|CGTCTGATCTCGGAAGCTAAGCAGGGT | >hg38_1_-_228634819_228635039_RNA5S12_5S_50ntFlanks | 86 | 112 | 27 | CGTCTGATCTCGGAAGCTAAGCAGGGT | 83.5 | -10.2 | 1.7 | -5.9 | 2.8E-09 | 6.3E-09 | MB231/MB468 | MP |
| >MI0000433\|hsa-let-7g&WithFlank&3\|-\|52268272\|52268367@11.31.21(+1C)&[MIMAT0000414&hsa-let-7g-5p&offsets\|0\|-1(+1C);m-58&3\|-\|52268336\|52268357&offsets\|0\|-1(+1C)]\|TGAGGTAGTAGTTTGTACAGTC | >MI0000433\|hsa-let-7g&WithFlank&3\|-\|52268272\|52268367 | 11 | 31 | 21(+1C) | TGAGGTAGTAGTTTGTACAGTC | 3437.4 | 4.7 | 0.8 | 5.9 | 2.9E-09 | 6.3E-09 | MB231/MB468 | CYTO |
| >hg38_21_+_8433172_8446622_RNA45SN1_45S_50ntFlanks@11209.11243.35\|CGGCGACTCTGGACGCGAGCCGGGCCCTTCCCGTG | >hg38_21_+_8433172_8446622_RNA45SN1_45S_50ntFlanks | 11209 | 11243 | 35 | CGGCGACTCTGGACGCGAGCCGGGCCCTTCCCGTG | 60.0 | -9.8 | 1.6 | -5.9 | 2.8E-09 | 6.3E-09 | MB231/MB468 | MP |
| >hg38_21_+_8433172_8446622_RNA45SN1_45S_50ntFlanks@11805.11827.23\|CCACTGTCCCTACCTACTATCCA | >hg38_21_+_8433172_8446622_RNA45SN1_45S_50ntFlanks | 11805 | 11827 | 23 | CCACTGTCCCTACCTACTATCCA | 98.7 | -8.7 | 1.5 | -6.0 | 2.2E-09 | 6.3E-09 | BT20/MB468 | MITO |
| >hg38_21_+_8433172_8446622_RNA45SN1_45S_50ntFlanks@6651.6676.26\|CGACTCTTAGCGGTGGATCACTCGGC | >hg38_21_+_8433172_8446622_RNA45SN1_45S_50ntFlanks | 6651 | 6676 | 26 | CGACTCTTAGCGGTGGATCACTCGGC | 3961.0 | 5.0 | 0.8 | 5.9 | 2.9E-09 | 6.3E-09 | MB231/MB468 | CYTO |
| >MI0000264\|hsa-mir-7-2&WithFlank&15\|+\|88611819\|88611940@38.59.22(+1C);MI0000265\|hsa-mir-7-3&WithFlank&19\|+\|4770664\|4770785@37.58.22(+1C);MI0000263\|hsa-mir-7-1&WithFlank&9\|-\|83969742\|83969863@30.51.22(+1C)&[MIMAT0000252&hsa-miR-7-5p&offsets\|0\|-2(+1C);m-400&15\|+\|88611856\|88611879&offsets\|0\|-2(+1C)];[MIMAT0000252_1&hsa-miR-7-5p&offsets\|0\|-2(+1C);m-402&19\|+\|4770700\|4770723&offsets\|0\|-2(+1C)];[MIMAT0000252_2&hsa-miR-7-5p&offsets\|0\|-2(+1C);m-401&9\|-\|83969811\|83969834&offsets\|0\|-2(+1C)]\|TGGAAGACTAGTGATTTTGTTGC | >MI0000264\|hsa-mir-7-2&WithFlank&15\|+\|88611819\|88611940 | 38 | 59 | 22(+1C) | TGGAAGACTAGTGATTTTGTTGC | 1252.9 | 3.7 | 0.6 | 5.9 | 2.8E-09 | 6.3E-09 | MB231/MB468 | MP |
| >hg38_21_+_8433172_8446622_RNA45SN1_45S_50ntFlanks@4661.4689.29\|AGGTGAAATTCTTGGACCGGCGCAAGACG | >hg38_21_+_8433172_8446622_RNA45SN1_45S_50ntFlanks | 4661 | 4689 | 29 | AGGTGAAATTCTTGGACCGGCGCAAGACG | 609.0 | -8.5 | 1.4 | -5.9 | 2.8E-09 | 6.3E-09 | MB231/MB468 | MP |
| >hg38_21_+_8433172_8446622_RNA45SN1_45S_50ntFlanks@11480.11500.21\|CGGCGGAGGGGCCGCGGGCCG | >hg38_21_+_8433172_8446622_RNA45SN1_45S_50ntFlanks | 11480 | 11500 | 21 | CGGCGGAGGGGCCGCGGGCCG | 91.3 | -10.4 | 1.7 | -5.9 | 2.8E-09 | 6.3E-09 | MB231/MB468 | MP |
| >trna10_AlaCGC_6_+_26553731_26553802@1.19.19,trna119_AlaCGC_6_-_28641613_28641684@1.19.19,trna120_AlaAGC_6_-_28626014_28626085@1.19.19,trna13_AlaCGC_2_+_157257281_157257352@1.19.19,trna13_AlaTGC_12_-_125406301_125406372@1.19.19,trna66_AlaTGC_6_+_28611222_28611293@1.19.19,trna8_AlaTGC_12_+_125424512_125424583@1.19.19,trna8_AlaTGC_5_+_180633868_180633939@1.19.19\|GGGGATGTAGCTCAGTGGT | >trna10_AlaCGC_6_+_26553731_26553802 | 1 | 19 | 19 | GGGGATGTAGCTCAGTGGT | 1032.8 | 5.3 | 0.9 | 5.9 | 2.9E-09 | 6.3E-09 | MB231/MB468 | CYTO |
| >MI0000255\|hsa-mir-30d&WithFlank&8\|-\|134804870\|134804951@12.35.24(+2U)&[MIMAT0000245&hsa-miR-30d-5p&offsets\|0\|+2(+2U);m-21&8\|-\|134804919\|134804940&offsets\|0\|+2(+2U)]\|TGTAAACATCCCCGACTGGAAGCTTT | >MI0000255\|hsa-mir-30d&WithFlank&8\|-\|134804870\|134804951 | 12 | 35 | 24(+2U) | TGTAAACATCCCCGACTGGAAGCTTT | 744.6 | 7.5 | 1.3 | 5.9 | 3.2E-09 | 6.3E-09 | MB231/MB468 | MITO |
| >hg38_21_+_8433172_8446622_RNA45SN1_45S_50ntFlanks@9736.9765.30\|CCGAAACGATCTCAACCTATTCTCAAACTT | >hg38_21_+_8433172_8446622_RNA45SN1_45S_50ntFlanks | 9736 | 9765 | 30 | CCGAAACGATCTCAACCTATTCTCAAACTT | 1388.6 | -8.4 | 1.4 | -5.9 | 2.9E-09 | 6.3E-09 | MB231/MB468 | NUC |
| >hg38_21_+_8433172_8446622_RNA45SN1_45S_50ntFlanks@4443.4472.30\|CCCTTGCCTCTCGGCGCCCCCTCGATGCTC | >hg38_21_+_8433172_8446622_RNA45SN1_45S_50ntFlanks | 4443 | 4472 | 30 | CCCTTGCCTCTCGGCGCCCCCTCGATGCTC | 655.7 | -7.9 | 1.3 | -6.0 | 2.5E-09 | 6.4E-09 | BT20/MB468 | CYTO |
| >MI0006444\|hsa-mir-1307&WithFlank&10\|-\|103394247\|103394407@86.108.23(+1A)&[MIMAT0005951&hsa-miR-1307-3p&offsets\|0\|+1(+1A);m-66&10\|-\|103394300\|103394322&offsets\|0\|0(+1A)]\|ACTCGGCGTGGCGTCGGTCGTGGA | >MI0006444\|hsa-mir-1307&WithFlank&10\|-\|103394247\|103394407 | 86 | 108 | 23(+1A) | ACTCGGCGTGGCGTCGGTCGTGGA | 341.4 | 7.3 | 1.2 | 5.9 | 2.9E-09 | 6.4E-09 | MB231/MB468 | CYTO |
| >hg38_21_+_8433172_8446622_RNA45SN1_45S_50ntFlanks@3775.3794.20\|GCCGGTACAGTGAAACTGCG | >hg38_21_+_8433172_8446622_RNA45SN1_45S_50ntFlanks | 3775 | 3794 | 20 | GCCGGTACAGTGAAACTGCG | 455.2 | -8.4 | 1.4 | -5.9 | 3.2E-09 | 6.4E-09 | MB231/MB468 | MITO |
| >hg38_21_+_8433172_8446622_RNA45SN1_45S_50ntFlanks@5494.5516.23\|AAGACGGTCGAACTTGACTATCT | >hg38_21_+_8433172_8446622_RNA45SN1_45S_50ntFlanks | 5494 | 5516 | 23 | AAGACGGTCGAACTTGACTATCT | 532.4 | 3.6 | 0.6 | 5.9 | 3.0E-09 | 6.4E-09 | MB231/MB468 | NUC |
| >hg38_21_+_8433172_8446622_RNA45SN1_45S_50ntFlanks@13004.13028.25\|ATCTATTGAAAGTCAGCCCTCGACA | >hg38_21_+_8433172_8446622_RNA45SN1_45S_50ntFlanks | 13004 | 13028 | 25 | ATCTATTGAAAGTCAGCCCTCGACA | 129.4 | -10.1 | 1.6 | -6.3 | 3.5E-10 | 6.4E-09 | BT20/MB468 | MP |
| >hg38_1_-_228634819_228635039_RNA5S12_5S_50ntFlanks@150.170.21\|AATACCGGGTGCTGTAGGCTT | >hg38_1_-_228634819_228635039_RNA5S12_5S_50ntFlanks | 150 | 170 | 21 | AATACCGGGTGCTGTAGGCTT | 5303.1 | 4.6 | 0.8 | 5.9 | 2.9E-09 | 6.4E-09 | MB231/MB468 | CYTO |
| >hg38_21_+_8433172_8446622_RNA45SN1_45S_50ntFlanks@6731.6753.23\|CACATTGATCATCGACACTTCGA | >hg38_21_+_8433172_8446622_RNA45SN1_45S_50ntFlanks | 6731 | 6753 | 23 | CACATTGATCATCGACACTTCGA | 403.2 | -6.5 | 1.1 | -5.9 | 3.4E-09 | 6.4E-09 | BT20/MB468 | NUC |
| >hg38_21_+_8433172_8446622_RNA45SN1_45S_50ntFlanks@3837.3861.25\|CCTCTCCTACTTGGATAACTGTGGT | >hg38_21_+_8433172_8446622_RNA45SN1_45S_50ntFlanks | 3837 | 3861 | 25 | CCTCTCCTACTTGGATAACTGTGGT | 405.1 | -3.5 | 0.6 | -5.9 | 3.0E-09 | 6.4E-09 | MB231/MB468 | NUC |
| >MI0000440\|hsa-mir-27b&WithFlank&9\|+\|95085439\|95085547@67.86.20(+1A)&[MIMAT0000419&hsa-miR-27b-3p&offsets\|0\|-1(+1A);m-39&9\|+\|95085505\|95085525&offsets\|0\|-1(+1A)]\|TTCACAGTGGCTAAGTTCTGA | >MI0000440\|hsa-mir-27b&WithFlank&9\|+\|95085439\|95085547 | 67 | 86 | 20(+1A) | TTCACAGTGGCTAAGTTCTGA | 1241.0 | 5.8 | 1.0 | 5.9 | 2.9E-09 | 6.4E-09 | MB231/MB468 | CYTO |
| >hg38_21_+_8433172_8446622_RNA45SN1_45S_50ntFlanks@9739.9767.29\|AAACGATCTCAACCTATTCTCAAACTTTA | >hg38_21_+_8433172_8446622_RNA45SN1_45S_50ntFlanks | 9739 | 9767 | 29 | AAACGATCTCAACCTATTCTCAAACTTTA | 2493.1 | -3.6 | 0.6 | -6.0 | 2.2E-09 | 6.4E-09 | BT20/MB468 | MITO |
| >hg38_21_+_8433172_8446622_RNA45SN1_45S_50ntFlanks@8238.8260.23\|CCCGGGTCTTCCCGGAGTCGGGT | >hg38_21_+_8433172_8446622_RNA45SN1_45S_50ntFlanks | 8238 | 8260 | 23 | CCCGGGTCTTCCCGGAGTCGGGT | 61.6 | -9.8 | 1.7 | -5.9 | 2.8E-09 | 6.4E-09 | MB231/MB468 | MP |
| >hg38_1_-_228634819_228635039_RNA5S12_5S_50ntFlanks@134.172.39\|TGGGAGACCGCCTGGGAATACCGGGTGCTGTAGGCTTTT | >hg38_1_-_228634819_228635039_RNA5S12_5S_50ntFlanks | 134 | 172 | 39 | TGGGAGACCGCCTGGGAATACCGGGTGCTGTAGGCTTTT | 336.5 | 11.6 | 2.0 | 5.9 | 2.8E-09 | 6.4E-09 | MB231/MB468 | MP |
| >MI0000266\|hsa-mir-10a&WithFlank&17\|-\|48579832\|48579953@29.49.21&[MIMAT0000253&hsa-miR-10a-5p&offsets\|+1\|-1;m-18&17\|-\|48579905\|48579926&offsets\|+1\|0]\|ACCCTGTAGATCCGAATTTGT | >MI0000266\|hsa-mir-10a&WithFlank&17\|-\|48579832\|48579953 | 29 | 49 | 21 | ACCCTGTAGATCCGAATTTGT | 1550.3 | 9.6 | 1.5 | 6.3 | 3.5E-10 | 6.4E-09 | BT20/MB468 | MP |
| >hg38_21_+_8433172_8446622_RNA45SN1_45S_50ntFlanks@3985.4007.23\|CGGCTTTGGTGACTCTAGATAAC | >hg38_21_+_8433172_8446622_RNA45SN1_45S_50ntFlanks | 3985 | 4007 | 23 | CGGCTTTGGTGACTCTAGATAAC | 80.1 | -10.2 | 1.7 | -5.9 | 2.8E-09 | 6.4E-09 | MB231/MB468 | MP |
| >hg38_21_+_8433172_8446622_RNA45SN1_45S_50ntFlanks@6682.6718.37\|CGTCGATGAAGAACGCAGCTAGCTGCGAGAATTAATG | >hg38_21_+_8433172_8446622_RNA45SN1_45S_50ntFlanks | 6682 | 6718 | 37 | CGTCGATGAAGAACGCAGCTAGCTGCGAGAATTAATG | 153.1 | -11.1 | 1.9 | -5.9 | 2.8E-09 | 6.5E-09 | MB231/MB468 | MP |
| >MI0000683\|hsa-mir-181b-2&WithFlank&9\|+\|124693704\|124693804@22.46.25(+1U)&[MIMAT0000257_1&hsa-miR-181b-5p&offsets\|0\|+2(+1U);m-72&9\|+\|124693725\|124693747&offsets\|0\|+2(+1U)]\|AACATTCATTGCTGTCGGTGGGTTTT | >MI0000683\|hsa-mir-181b-2&WithFlank&9\|+\|124693704\|124693804 | 22 | 46 | 25(+1U) | AACATTCATTGCTGTCGGTGGGTTTT | 727.8 | 7.7 | 1.3 | 5.9 | 3.2E-09 | 6.5E-09 | MB231/MB468 | MITO |
| >hg38_1_-_228634819_228635039_RNA5S12_5S_50ntFlanks@140.170.31\|ACCGCCTGGGAATACCGGGTGCTGTAGGCTT | >hg38_1_-_228634819_228635039_RNA5S12_5S_50ntFlanks | 140 | 170 | 31 | ACCGCCTGGGAATACCGGGTGCTGTAGGCTT | 13727.1 | 4.5 | 0.8 | 5.9 | 3.0E-09 | 6.5E-09 | MB231/MB468 | NUC |
| >hg38_21_+_8433172_8446622_RNA45SN1_45S_50ntFlanks@6699.6720.22\|GCTAGCTGCGAGAATTAATGTG | >hg38_21_+_8433172_8446622_RNA45SN1_45S_50ntFlanks | 6699 | 6720 | 22 | GCTAGCTGCGAGAATTAATGTG | 60468.5 | -5.5 | 0.9 | -5.9 | 3.0E-09 | 6.5E-09 | MB231/MB468 | NUC |
| >hg38_21_+_8433172_8446622_RNA45SN1_45S_50ntFlanks@11270.11293.24\|CGGCCGCCCCCGGGGAGCCCGGCG | >hg38_21_+_8433172_8446622_RNA45SN1_45S_50ntFlanks | 11270 | 11293 | 24 | CGGCCGCCCCCGGGGAGCCCGGCG | 1474.2 | -9.6 | 1.5 | -6.3 | 3.5E-10 | 6.5E-09 | BT20/MB468 | MP |
| >hg38_21_+_8433172_8446622_RNA45SN1_45S_50ntFlanks@5268.5295.28\|CCGGCAGGCGCGGGTAACCCGTTGAACC | >hg38_21_+_8433172_8446622_RNA45SN1_45S_50ntFlanks | 5268 | 5295 | 28 | CCGGCAGGCGCGGGTAACCCGTTGAACC | 87.3 | -10.3 | 1.7 | -5.9 | 2.8E-09 | 6.5E-09 | MB231/MB468 | MP |
| >hg38_21_+_8433172_8446622_RNA45SN1_45S_50ntFlanks@11209.11233.25\|CGGCGACTCTGGACGCGAGCCGGGC | >hg38_21_+_8433172_8446622_RNA45SN1_45S_50ntFlanks | 11209 | 11233 | 25 | CGGCGACTCTGGACGCGAGCCGGGC | 1356.3 | -6.7 | 1.1 | -5.9 | 3.0E-09 | 6.5E-09 | MB231/MB468 | NUC |
| >trna10_AspGTC_12_-_125424193_125424264@39.74.36,trna12_AspGTC_12_-_125411891_125411962@39.74.36,trna144_AspGTC_6_-_27551236_27551307@39.74.36,trna38_AspGTC_17_-_8125556_8125627@39.74.36,trna45_AspGTC_6_+_27447453_27447524@39.74.36,trna48_AspGTC_6_+_27471523_27471594@39.74.36,trna4_AspGTC_12_+_96429799_96429870@39.74.36,trna69_AspGTC_1_-_161440205_161440276@39.74.36,trna72_AspGTC_1_-_161432824_161432895@39.74.36,trna75_AspGTC_1_-_161425414_161425485@39.74.36,trna78_AspGTC_1_-_161418033_161418104@39.74.36,trna81_AspGTC_1_-_161410615_161410686@39.74.36\|GCGGGAGACCGGGGTTCGATTCCCCGACGGGGAGCC | >trna10_AspGTC_12_-_125424193_125424264 | 39 | 74 | 36 | GCGGGAGACCGGGGTTCGATTCCCCGACGGGGAGCC | 3910.7 | 6.2 | 1.0 | 5.9 | 3.0E-09 | 6.5E-09 | MB231/MB468 | CYTO |
| >hg38_21_+_8433172_8446622_RNA45SN1_45S_50ntFlanks@4453.4472.20\|TCGGCGCCCCCTCGATGCTC | >hg38_21_+_8433172_8446622_RNA45SN1_45S_50ntFlanks | 4453 | 4472 | 20 | TCGGCGCCCCCTCGATGCTC | 128.1 | -10.1 | 1.6 | -6.3 | 3.5E-10 | 6.5E-09 | BT20/MB468 | MP |
| >hg38_21_+_8433172_8446622_RNA45SN1_45S_50ntFlanks@6651.6680.30\|CGACTCTTAGCGGTGGATCACTCGGCTCGT | >hg38_21_+_8433172_8446622_RNA45SN1_45S_50ntFlanks | 6651 | 6680 | 30 | CGACTCTTAGCGGTGGATCACTCGGCTCGT | 4317.8 | -3.9 | 0.7 | -5.9 | 2.9E-09 | 6.5E-09 | MB231/MB468 | MP |
| >hg38_21_+_8433172_8446622_RNA45SN1_45S_50ntFlanks@5084.5121.38\|CGAACGAGACTCTGGCATGCTAACTAGTTACGCGACCC | >hg38_21_+_8433172_8446622_RNA45SN1_45S_50ntFlanks | 5084 | 5121 | 38 | CGAACGAGACTCTGGCATGCTAACTAGTTACGCGACCC | 1235.1 | -10.0 | 1.7 | -5.9 | 3.5E-09 | 6.5E-09 | BT20/MB468 | NUC |
| >MI0033423\|hsa-mir-10399&WithFlank&7\|-\|139044094\|139044163@7.27.21&[MIMAT0041629&hsa-miR-10399-5p&offsets\|0\|0;TJU_CMC_MD2.ID03061.5p-miR&offsets\|0\|0;m-2644&7\|-\|139044137\|139044157&offsets\|0\|0]\|AATTACAGATTGTCTCAGAGA | >MI0033423\|hsa-mir-10399&WithFlank&7\|-\|139044094\|139044163 | 7 | 27 | 21 | AATTACAGATTGTCTCAGAGA | 84.0 | 9.6 | 1.6 | 5.9 | 2.9E-09 | 6.6E-09 | MB231/MB468 | MP |
| >hg38_21_+_8433172_8446622_RNA45SN1_45S_50ntFlanks@11572.11603.32\|CCGACTTAGAACTGGTGCGGACCAGGGGAATC | >hg38_21_+_8433172_8446622_RNA45SN1_45S_50ntFlanks | 11572 | 11603 | 32 | CCGACTTAGAACTGGTGCGGACCAGGGGAATC | 6032.5 | -4.7 | 0.8 | -5.9 | 3.0E-09 | 6.6E-09 | MB231/MB468 | CYTO |

<table>
<tr><th></th><th></th><th></th><th></th><th></th><th></th><th></th><th></th><th></th><th></th><th></th><th></th><th></th><th></th></tr>
<tr><td>>hg38_21_+_8433172_8446622_RNA45SN1_45S_50ntFlanks@10770.10797.28|GCAGGTCTCCAAGGTGAACAGCCTCTGG</td><td>>hg38_21_+_8433172_8446622_RNA45SN1_45S_50ntFlanks</td><td>10770</td><td>10797</td><td>28</td><td>GCAGGTCTCCAAGGTGAACAGCCTCTGG</td><td>115.1</td><td>-7.1</td><td>1.2</td><td>-5.9</td><td>3.1E-09</td><td>6.6E-09</td><td>MB231/MB468</td><td>NUC</td></tr>
<tr><td>>hg38_21_+_8433172_8446622_RNA45SN1_45S_50ntFlanks@12133.12151.19|CTCCGGGGACAGTGCCAGG</td><td>>hg38_21_+_8433172_8446622_RNA45SN1_45S_50ntFlanks</td><td>12133</td><td>12151</td><td>19</td><td>CTCCGGGGACAGTGCCAGG</td><td>260.0</td><td>-7.6</td><td>1.3</td><td>-5.9</td><td>2.9E-09</td><td>6.6E-09</td><td>MB231/MB468</td><td>MP</td></tr>
<tr><td>>hg38_21_+_8433172_8446622_RNA45SN1_45S_50ntFlanks@5046.5078.33|TTAGTTGGTGGAGCGATTTGTCTGGTTAATTCC</td><td>>hg38_21_+_8433172_8446622_RNA45SN1_45S_50ntFlanks</td><td>5046</td><td>5078</td><td>33</td><td>TTAGTTGGTGGAGCGATTTGTCTGGTTAATTCC</td><td>201.7</td><td>7.7</td><td>1.3</td><td>5.9</td><td>3.0E-09</td><td>6.6E-09</td><td>MB231/MB468</td><td>CYTO</td></tr>
<tr><td>>hg38_21_+_8433172_8446622_RNA45SN1_45S_50ntFlanks@12456.12490.35|CCAAGCGTTGGATTGTTCACCCACTAATAGGGAAC</td><td>>hg38_21_+_8433172_8446622_RNA45SN1_45S_50ntFlanks</td><td>12456</td><td>12490</td><td>35</td><td>CCAAGCGTTGGATTGTTCACCCACTAATAGGGAAC</td><td>1118.7</td><td>-3.9</td><td>0.7</td><td>-5.9</td><td>3.0E-09</td><td>6.6E-09</td><td>MB231/MB468</td><td>CYTO</td></tr>
<tr><td>>hg38_21_+_8433172_8446622_RNA45SN1_45S_50ntFlanks@6732.6771.40|ACATTGATCATCGACACTTCGAACGCACTTGCGGCCCCGG</td><td>>hg38_21_+_8433172_8446622_RNA45SN1_45S_50ntFlanks</td><td>6732</td><td>6771</td><td>40</td><td>ACATTGATCATCGACACTTCGAACGCACTTGCGGCCCCGG</td><td>706.7</td><td>-7.7</td><td>1.3</td><td>-5.9</td><td>2.9E-09</td><td>6.6E-09</td><td>MB231/MB468</td><td>MP</td></tr>
<tr><td>>hg38_21_+_8433172_8446622_RNA45SN1_45S_50ntFlanks@12658.12676.19|ACTGAACGCCTCTAAGTCA</td><td>>hg38_21_+_8433172_8446622_RNA45SN1_45S_50ntFlanks</td><td>12658</td><td>12676</td><td>19</td><td>ACTGAACGCCTCTAAGTCA</td><td>77.0</td><td>-10.1</td><td>1.7</td><td>-5.9</td><td>2.9E-09</td><td>6.6E-09</td><td>MB231/MB468</td><td>MP</td></tr>
<tr><td>>hg38_21_+_8433172_8446622_RNA45SN1_45S_50ntFlanks@3823.3861.39|TTTGGTCGCTCGCTCCTCTCCTACTTGGATAACTGTGGT</td><td>>hg38_21_+_8433172_8446622_RNA45SN1_45S_50ntFlanks</td><td>3823</td><td>3861</td><td>39</td><td>TTTGGTCGCTCGCTCCTCTCCTACTTGGATAACTGTGGT</td><td>306.5</td><td>4.9</td><td>0.8</td><td>5.9</td><td>3.1E-09</td><td>6.6E-09</td><td>MB231/MB468</td><td>CYTO</td></tr>
<tr><td>>hg38_21_+_8433172_8446622_RNA45SN1_45S_50ntFlanks@9534.9568.35|AGGAAACTCTGGTGGAGGTCCGTAGCGGTCCTGAC</td><td>>hg38_21_+_8433172_8446622_RNA45SN1_45S_50ntFlanks</td><td>9534</td><td>9568</td><td>35</td><td>AGGAAACTCTGGTGGAGGTCCGTAGCGGTCCTGAC</td><td>1739.1</td><td>-9.9</td><td>1.7</td><td>-5.9</td><td>3.5E-09</td><td>6.6E-09</td><td>BT20/MB468</td><td>NUC</td></tr>
<tr><td>>hg38_21_+_8433172_8446622_RNA45SN1_45S_50ntFlanks@9738.9762.25|GAAACGATCTCAACCTATTCTCAAA</td><td>>hg38_21_+_8433172_8446622_RNA45SN1_45S_50ntFlanks</td><td>9738</td><td>9762</td><td>25</td><td>GAAACGATCTCAACCTATTCTCAAA</td><td>1077.5</td><td>-8.7</td><td>1.5</td><td>-5.9</td><td>3.3E-09</td><td>6.6E-09</td><td>MB231/MB468</td><td>MITO</td></tr>
<tr><td>>hg38_21_+_8433172_8446622_RNA45SN1_45S_50ntFlanks@4297.4317.21|CATTGGAGGGCAAGTCTGGTG</td><td>>hg38_21_+_8433172_8446622_RNA45SN1_45S_50ntFlanks</td><td>4297</td><td>4317</td><td>21</td><td>CATTGGAGGGCAAGTCTGGTG</td><td>706.7</td><td>-4.7</td><td>0.8</td><td>-5.9</td><td>3.1E-09</td><td>6.6E-09</td><td>MB231/MB468</td><td>CYTO</td></tr>
<tr><td>>hg38_21_+_8433172_8446622_RNA45SN1_45S_50ntFlanks@10731.10757.27|AGGGTGTAAATCTCGCGCCGGGCCGTA</td><td>>hg38_21_+_8433172_8446622_RNA45SN1_45S_50ntFlanks</td><td>10731</td><td>10757</td><td>27</td><td>AGGGTGTAAATCTCGCGCCGGGCCGTA</td><td>306.2</td><td>-7.4</td><td>1.2</td><td>-5.9</td><td>3.1E-09</td><td>6.6E-09</td><td>MB231/MB468</td><td>NUC</td></tr>
<tr><td>>hg38_21_+_8433172_8446622_RNA45SN1_45S_50ntFlanks@12640.12669.30|ACCATCTGTGGGATTATGACTGAACGCCTC</td><td>>hg38_21_+_8433172_8446622_RNA45SN1_45S_50ntFlanks</td><td>12640</td><td>12669</td><td>30</td><td>ACCATCTGTGGGATTATGACTGAACGCCTC</td><td>1114.7</td><td>8.4</td><td>1.4</td><td>5.9</td><td>3.1E-09</td><td>6.6E-09</td><td>MB231/MB468</td><td>CYTO</td></tr>
<tr><td>>hg38_21_+_8433172_8446622_RNA45SN1_45S_50ntFlanks@12404.12436.33|GCTTTTTGATCCTTCGATGTCGGCTCTTCCTAT</td><td>>hg38_21_+_8433172_8446622_RNA45SN1_45S_50ntFlanks</td><td>12404</td><td>12436</td><td>33</td><td>GCTTTTTGATCCTTCGATGTCGGCTCTTCCTAT</td><td>1078.5</td><td>-5.3</td><td>0.9</td><td>-5.9</td><td>3.1E-09</td><td>6.7E-09</td><td>MB231/MB468</td><td>NUC</td></tr>
<tr><td>>hg38_21_+_8433172_8446622_RNA45SN1_45S_50ntFlanks@9567.9584.18|ACGTGCAAATCGGTCGTC</td><td>>hg38_21_+_8433172_8446622_RNA45SN1_45S_50ntFlanks</td><td>9567</td><td>9584</td><td>18</td><td>ACGTGCAAATCGGTCGTC</td><td>684.8</td><td>-6.1</td><td>1.0</td><td>-5.9</td><td>3.1E-09</td><td>6.7E-09</td><td>MB231/MB468</td><td>CYTO</td></tr>
<tr><td>>hg38_21_+_8433172_8446622_RNA45SN1_45S_50ntFlanks@11704.11724.21|ATGAAGCGCGGGTAAACGGCG</td><td>>hg38_21_+_8433172_8446622_RNA45SN1_45S_50ntFlanks</td><td>11704</td><td>11724</td><td>21</td><td>ATGAAGCGCGGGTAAACGGCG</td><td>95.2</td><td>-10.4</td><td>1.8</td><td>-5.9</td><td>2.9E-09</td><td>6.7E-09</td><td>MB231/MB468</td><td>MP</td></tr>
<tr><td>>hg38_21_+_8433172_8446622_RNA45SN1_45S_50ntFlanks@10687.10708.22|CGGCGTCCGGTGAGCTCTCGCT</td><td>>hg38_21_+_8433172_8446622_RNA45SN1_45S_50ntFlanks</td><td>10687</td><td>10708</td><td>22</td><td>CGGCGTCCGGTGAGCTCTCGCT</td><td>5251.6</td><td>-6.2</td><td>1.1</td><td>-5.9</td><td>3.5E-09</td><td>6.7E-09</td><td>BT20/MB468</td><td>NUC</td></tr>
<tr><td>>hg38_21_+_8433172_8446622_RNA45SN1_45S_50ntFlanks@11535.11560.26|TTCCGCGCGGCGCCTCGCCTCGGCCG</td><td>>hg38_21_+_8433172_8446622_RNA45SN1_45S_50ntFlanks</td><td>11535</td><td>11560</td><td>26</td><td>TTCCGCGCGGCGCCTCGCCTCGGCCG</td><td>100.2</td><td>-8.7</td><td>1.5</td><td>-6.0</td><td>2.3E-09</td><td>6.7E-09</td><td>BT20/MB468</td><td>MITO</td></tr>
<tr><td>>trnaMT_GluTTC_MT_-_14674_14742@40.66.27,trnalookalike8_GluTTC_5_-_93905172_93905240@40.66.27|TTGGTCGTGGTTGTAGTCCGTGCGAGA</td><td>>trnaMT_GluTTC_MT_-_14674_14742</td><td>40</td><td>66</td><td>27</td><td>TTGGTCGTGGTTGTAGTCCGTGCGAGA</td><td>435.3</td><td>4.4</td><td>0.8</td><td>5.9</td><td>3.1E-09</td><td>6.7E-09</td><td>MB231/MB468</td><td>NUC</td></tr>
<tr><td>>MI0000088|hsa-mir-30a&WithFlank&6|-|71403545|71403627@53.74.22(+1A)&[MIMAT0000088&hsa-miR-30a-3p&offsets|0|0(+1A);m-38&6|-|71403554|71403575&offsets|0|0(+1A)]|CTTTCAGTCGGATGTTTGCAGCA</td><td>>MI0000088|hsa-mir-30a&WithFlank&6|-|71403545|71403627</td><td>53</td><td>74</td><td>22(+1A)</td><td>CTTTCAGTCGGATGTTTGCAGCA</td><td>610.9</td><td>6.7</td><td>1.1</td><td>5.9</td><td>3.1E-09</td><td>6.7E-09</td><td>MB231/MB468</td><td>CYTO</td></tr>
<tr><td>>hg38_21_+_8433172_8446622_RNA45SN1_45S_50ntFlanks@8175.8202.28|CCTTCTGATCGAGGCCCAGCCCGTGGAC</td><td>>hg38_21_+_8433172_8446622_RNA45SN1_45S_50ntFlanks</td><td>8175</td><td>8202</td><td>28</td><td>CCTTCTGATCGAGGCCCAGCCCGTGGAC</td><td>553.1</td><td>8.0</td><td>1.3</td><td>6.0</td><td>2.7E-09</td><td>6.7E-09</td><td>BT20/MB468</td><td>CYTO</td></tr>
<tr><td>>hg38_21_+_8433172_8446622_RNA45SN1_45S_50ntFlanks@12406.12432.27|TTTTTGATCCTTCGATGTCGGCTCTTC</td><td>>hg38_21_+_8433172_8446622_RNA45SN1_45S_50ntFlanks</td><td>12406</td><td>12432</td><td>27</td><td>TTTTTGATCCTTCGATGTCGGCTCTTC</td><td>550.3</td><td>-8.6</td><td>1.5</td><td>-5.9</td><td>3.5E-09</td><td>6.7E-09</td><td>BT20/MB468</td><td>NUC</td></tr>
<tr><td>>hg38_21_+_8433172_8446622_RNA45SN1_45S_50ntFlanks@8154.8181.28|CCGCTCGTGGGGGGCCCAAGTCCTTCTG</td><td>>hg38_21_+_8433172_8446622_RNA45SN1_45S_50ntFlanks</td><td>8154</td><td>8181</td><td>28</td><td>CCGCTCGTGGGGGGCCCAAGTCCTTCTG</td><td>3712.6</td><td>-10.1</td><td>1.7</td><td>-5.9</td><td>3.6E-09</td><td>6.7E-09</td><td>BT20/MB468</td><td>NUC</td></tr>
<tr><td>>hg38_21_+_8433172_8446622_RNA45SN1_45S_50ntFlanks@10369.10406.38|AAGTGGAGAAGGGTTCCATGTGAACAGCAGTTGAACAT</td><td>>hg38_21_+_8433172_8446622_RNA45SN1_45S_50ntFlanks</td><td>10369</td><td>10406</td><td>38</td><td>AAGTGGAGAAGGGTTCCATGTGAACAGCAGTTGAACAT</td><td>199.6</td><td>-4.6</td><td>0.8</td><td>-5.9</td><td>3.1E-09</td><td>6.7E-09</td><td>MB231/MB468</td><td>CYTO</td></tr>
<tr><td>>am_trna143_LysTTT_6_-_27559593_27559665@7.26.20,trna14_LysTTT_11_-_59327808_59327880@7.26.20,trna2_LysTTT_17_+_8022473_8022545@7.26.20,trna54_LysTTT_1_+_204475655_204475727@7.26.20,trna5_LysTTT_11_+_59323902_59323974@7.26.20,trna62_LysTTT_1_-_204476158_204476230@7.26.20,trna76_LysTTT_6_+_28918806_28918878@7.26.20|ATAGCTCAGTCGGTAGAGCA</td><td>>am_trna143_LysTTT_6_-_27559593_27559665</td><td>7</td><td>26</td><td>20</td><td>ATAGCTCAGTCGGTAGAGCA</td><td>54.0</td><td>-9.6</td><td>1.6</td><td>-5.9</td><td>3.0E-09</td><td>6.7E-09</td><td>MB231/MB468</td><td>MP</td></tr>
<tr><td>>hg38_21_+_8433172_8446622_RNA45SN1_45S_50ntFlanks@6650.6676.27|ACGACTCTTAGCGGTGGATCACTCGGC</td><td>>hg38_21_+_8433172_8446622_RNA45SN1_45S_50ntFlanks</td><td>6650</td><td>6676</td><td>27</td><td>ACGACTCTTAGCGGTGGATCACTCGGC</td><td>3977.8</td><td>9.2</td><td>1.5</td><td>5.9</td><td>2.7E-09</td><td>6.7E-09</td><td>BT20/MB468</td><td>CYTO</td></tr>
<tr><td>>MI0000803|hsa-mir-330&WithFlank&19|-|45638988|45639093@63.86.24(+1U)&[MIMAT0000751&hsa-miR-330-3p&offsets|0|+1(+1U);m-347&19|-|45639009|45639030&offsets|-1|+1(+1U)]|GCAAAGCACACGGCCTGCAGAGAGT</td><td>>MI0000803|hsa-mir-330&WithFlank&19|-|45638988|45639093</td><td>63</td><td>86</td><td>24(+1U)</td><td>GCAAAGCACACGGCCTGCAGAGAGT</td><td>550.7</td><td>4.4</td><td>0.7</td><td>5.9</td><td>3.1E-09</td><td>6.7E-09</td><td>MB231/MB468</td><td>NUC</td></tr>
<tr><td>>hg38_21_+_8433172_8446622_RNA45SN1_45S_50ntFlanks@12008.12042.35|CCGCCGGTGAAATACCACTACTCTGATCGTTTTTT</td><td>>hg38_21_+_8433172_8446622_RNA45SN1_45S_50ntFlanks</td><td>12008</td><td>12042</td><td>35</td><td>CCGCCGGTGAAATACCACTACTCTGATCGTTTTTT</td><td>674.7</td><td>-7.8</td><td>1.3</td><td>-5.9</td><td>3.1E-09</td><td>6.7E-09</td><td>MB231/MB468</td><td>CYTO</td></tr>
<tr><td>>hg38_21_+_8433172_8446622_RNA45SN1_45S_50ntFlanks@9860.9880.21|GGATGAACCGAACGCCGGGTT</td><td>>hg38_21_+_8433172_8446622_RNA45SN1_45S_50ntFlanks</td><td>9860</td><td>9880</td><td>21</td><td>GGATGAACCGAACGCCGGGTT</td><td>162.4</td><td>-11.2</td><td>1.9</td><td>-5.9</td><td>3.0E-09</td><td>6.8E-09</td><td>MB231/MB468</td><td>MP</td></tr>
<tr><td>>trnaMT_GluTTC_MT_-_14674_14742@34.72.39,trnalookalike8_GluTTC_5_-_93905172_93905240@34.72.39|ATATCATTGGTCGTGGTTGTAGTCCGTGCGAGAATACCA</td><td>>trnaMT_GluTTC_MT_-_14674_14742</td><td>34</td><td>72</td><td>39</td><td>ATATCATTGGTCGTGGTTGTAGTCCGTGCGAGAATACCA</td><td>629.8</td><td>7.1</td><td>1.2</td><td>5.9</td><td>2.7E-09</td><td>6.8E-09</td><td>BT20/MB468</td><td>CYTO</td></tr>
<tr><td>>hg38_21_+_8433172_8446622_RNA45SN1_45S_50ntFlanks@3802.3834.33|CATTAAATCAGTTATGGTTCCTTTGGTCGCTCG</td><td>>hg38_21_+_8433172_8446622_RNA45SN1_45S_50ntFlanks</td><td>3802</td><td>3834</td><td>33</td><td>CATTAAATCAGTTATGGTTCCTTTGGTCGCTCG</td><td>2876.7</td><td>-10.1</td><td>1.7</td><td>-5.9</td><td>3.6E-09</td><td>6.8E-09</td><td>BT20/MB468</td><td>NUC</td></tr>
<tr><td>>hg38_21_+_8433172_8446622_RNA45SN1_45S_50ntFlanks@6721.6752.32|AATTGCAGGACACATTGATCATCGACACTTCG</td><td>>hg38_21_+_8433172_8446622_RNA45SN1_45S_50ntFlanks</td><td>6721</td><td>6752</td><td>32</td><td>AATTGCAGGACACATTGATCATCGACACTTCG</td><td>672.1</td><td>-6.9</td><td>1.2</td><td>-5.9</td><td>3.0E-09</td><td>6.8E-09</td><td>MB231/MB468</td><td>MP</td></tr>
<tr><td>>hg38_21_+_8433172_8446622_RNA45SN1_45S_50ntFlanks@6721.6758.38|AATTGCAGGACACATTGATCATCGACACTTCGAACGCA</td><td>>hg38_21_+_8433172_8446622_RNA45SN1_45S_50ntFlanks</td><td>6721</td><td>6758</td><td>38</td><td>AATTGCAGGACACATTGATCATCGACACTTCGAACGCA</td><td>8594.1</td><td>-10.9</td><td>1.8</td><td>-5.9</td><td>3.6E-09</td><td>6.8E-09</td><td>BT20/MB468</td><td>NUC</td></tr>
<tr><td>>hg38_21_+_8433172_8446622_RNA45SN1_45S_50ntFlanks@3853.3881.29|AACTGTGGTAATTCTAGAGCTAATACATG</td><td>>hg38_21_+_8433172_8446622_RNA45SN1_45S_50ntFlanks</td><td>3853</td><td>3881</td><td>29</td><td>AACTGTGGTAATTCTAGAGCTAATACATG</td><td>757.2</td><td>-8.1</td><td>1.4</td><td>-5.9</td><td>3.0E-09</td><td>6.8E-09</td><td>MB231/MB468</td><td>MP</td></tr>
<tr><td>>trnaMT_PheGAA_MT_+_577_647@39.60.22|ATGTTTAGACGGGCTCACATCA</td><td>>trnaMT_PheGAA_MT_+_577_647</td><td>39</td><td>60</td><td>22</td><td>ATGTTTAGACGGGCTCACATCA</td><td>505.7</td><td>7.3</td><td>1.2</td><td>6.0</td><td>2.4E-09</td><td>6.8E-09</td><td>BT20/MB468</td><td>MITO</td></tr>
<tr><td>>hg38_21_+_8433172_8446622_RNA45SN1_45S_50ntFlanks@11534.11560.27|GTTCCGCGCGGCGCCTCGCCTCGGCCG</td><td>>hg38_21_+_8433172_8446622_RNA45SN1_45S_50ntFlanks</td><td>11534</td><td>11560</td><td>27</td><td>GTTCCGCGCGGCGCCTCGCCTCGGCCG</td><td>510.9</td><td>-7.0</td><td>1.2</td><td>-6.0</td><td>2.4E-09</td><td>6.8E-09</td><td>BT20/MB468</td><td>MITO</td></tr>
<tr><td>>MI0000342|hsa-mir-200b&WithFlank&1|+|1167098|1167204@63.84.22(+1A)&[MIMAT0000318&hsa-miR-200b-3p&offsets|0|0(+1A);m-93&1|+|1167160|1167182&offsets|0|-1(+1A)]|TAATACTGCCTGGTAATGATGAA</td><td>>MI0000342|hsa-mir-200b&WithFlank&1|+|1167098|1167204</td><td>63</td><td>84</td><td>22(+1A)</td><td>TAATACTGCCTGGTAATGATGAA</td><td>167.8</td><td>-10.5</td><td>1.7</td><td>-6.3</td><td>3.7E-10</td><td>6.8E-09</td><td>BT20/MB468</td><td>MP</td></tr>
<tr><td>>hg38_21_+_8433172_8446622_RNA45SN1_45S_50ntFlanks@11317.11342.26|ACCCCACGTCTCGTCGCGCGCGCGTC</td><td>>hg38_21_+_8433172_8446622_RNA45SN1_45S_50ntFlanks</td><td>11317</td><td>11342</td><td>26</td><td>ACCCCACGTCTCGTCGCGCGCGCGTC</td><td>179.8</td><td>-10.6</td><td>1.7</td><td>-6.3</td><td>3.7E-10</td><td>6.8E-09</td><td>BT20/MB468</td><td>MP</td></tr>
<tr><td>>MI0001445|hsa-mir-423&WithFlank&17|+|30117073|30117178@23.44.22(+1C)&[MIMAT0004748&hsa-miR-423-5p&offsets|0|-1(+1C);m-141&17|+|30117095|30117117&offsets|0|-1(+1C)]|TGAGGGGCAGAGAGCGAGACTTC</td><td>>MI0001445|hsa-mir-423&WithFlank&17|+|30117073|30117178</td><td>23</td><td>44</td><td>22(+1C)</td><td>TGAGGGGCAGAGAGCGAGACTTC</td><td>242.7</td><td>4.8</td><td>0.8</td><td>5.9</td><td>3.2E-09</td><td>6.8E-09</td><td>MB231/MB468</td><td>NUC</td></tr>
<tr><td>>hg38_21_+_8433172_8446622_RNA45SN1_45S_50ntFlanks@6690.6720.31|AAGAACGCAGCTAGCTGCGAGAATTAATGTG</td><td>>hg38_21_+_8433172_8446622_RNA45SN1_45S_50ntFlanks</td><td>6690</td><td>6720</td><td>31</td><td>AAGAACGCAGCTAGCTGCGAGAATTAATGTG</td><td>914.0</td><td>-6.3</td><td>1.1</td><td>-5.9</td><td>3.2E-09</td><td>6.8E-09</td><td>MB231/MB468</td><td>NUC</td></tr>
<tr><td>>hg38_21_+_8433172_8446622_RNA45SN1_45S_50ntFlanks@11573.11601.29|CGACTTAGAACTGGTGCGGACCAGGGGAA</td><td>>hg38_21_+_8433172_8446622_RNA45SN1_45S_50ntFlanks</td><td>11573</td><td>11601</td><td>29</td><td>CGACTTAGAACTGGTGCGGACCAGGGGAA</td><td>693.7</td><td>-6.2</td><td>1.0</td><td>-5.9</td><td>3.4E-09</td><td>6.8E-09</td><td>MB231/MB468</td><td>MITO</td></tr>
<tr><td>>hg38_21_+_8433172_8446622_RNA45SN1_45S_50ntFlanks@11572.11600.29|CCGACTTAGAACTGGTGCGGACCAGGGGA</td><td>>hg38_21_+_8433172_8446622_RNA45SN1_45S_50ntFlanks</td><td>11572</td><td>11600</td><td>29</td><td>CCGACTTAGAACTGGTGCGGACCAGGGGA</td><td>4187.5</td><td>-3.9</td><td>0.7</td><td>-5.9</td><td>2.7E-09</td><td>6.8E-09</td><td>BT20/MB468</td><td>CYTO</td></tr>
<tr><td>>hg38_21_+_8433172_8446622_RNA45SN1_45S_50ntFlanks@10391.10435.45|AACAGCAGTTGAACATGGGTCAGTCGGTCCTGAGAGATGGGCGAG</td><td>>hg38_21_+_8433172_8446622_RNA45SN1_45S_50ntFlanks</td><td>10391</td><td>10435</td><td>45</td><td>AACAGCAGTTGAACATGGGTCAGTCGGTCCTGAGAGATGGGCGAG</td><td>496.1</td><td>-8.1</td><td>1.4</td><td>-5.9</td><td>2.7E-09</td><td>6.8E-09</td><td>BT20/MB468</td><td>CYTO</td></tr>
<tr><td>>trnaMT_ValTAC_MT_+_1602_1670@50.70.21|ACTTAACTTGACCGCTCTGAC</td><td>>trnaMT_ValTAC_MT_+_1602_1670</td><td>50</td><td>70</td><td>21</td><td>ACTTAACTTGACCGCTCTGAC</td><td>540.6</td><td>9.4</td><td>1.6</td><td>5.9</td><td>2.7E-09</td><td>6.8E-09</td><td>BT20/MB468</td><td>CYTO</td></tr>
<tr><td>>hg38_21_+_8433172_8446622_RNA45SN1_45S_50ntFlanks@9926.9949.24|GTTGATATAGACAGCAGGACGGTG</td><td>>hg38_21_+_8433172_8446622_RNA45SN1_45S_50ntFlanks</td><td>9926</td><td>9949</td><td>24</td><td>GTTGATATAGACAGCAGGACGGTG</td><td>120.3</td><td>10.7</td><td>1.7</td><td>6.3</td><td>3.7E-10</td><td>6.8E-09</td><td>BT20/MB468</td><td>MP</td></tr>
<tr><td>>hg38_21_+_8433172_8446622_RNA45SN1_45S_50ntFlanks@12712.12739.28|CGGAGCCTCGGTTGGCCTCGGATAGCCG</td><td>>hg38_21_+_8433172_8446622_RNA45SN1_45S_50ntFlanks</td><td>12712</td><td>12739</td><td>28</td><td>CGGAGCCTCGGTTGGCCTCGGATAGCCG</td><td>161.5</td><td>-10.4</td><td>1.7</td><td>-6.3</td><td>3.7E-10</td><td>6.8E-09</td><td>BT20/MB468</td><td>MP</td></tr>
<tr><td>>hg38_21_+_8433172_8446622_RNA45SN1_45S_50ntFlanks@11315.11342.28|CCACCCCACGTCTCGTCGCGCGCGCGTC</td><td>>hg38_21_+_8433172_8446622_RNA45SN1_45S_50ntFlanks</td><td>11315</td><td>11342</td><td>28</td><td>CCACCCCACGTCTCGTCGCGCGCGCGTC</td><td>212.3</td><td>-10.8</td><td>1.7</td><td>-6.3</td><td>3.7E-10</td><td>6.8E-09</td><td>BT20/MB468</td><td>MP</td></tr>
<tr><td>>MI0000076|hsa-mir-20a&WithFlank&13|+|91351059|91351141@14.37.24&[MIMAT0000075&hsa-miR-20a-5p&offsets|0|+1;m-96&13|+|91351072|91351094&offsets|0|+1]|TAAAGTGCTTATAGTGCAGGTAGT</td><td>>MI0000076|hsa-mir-20a&WithFlank&13|+|91351059|91351141</td><td>14</td><td>37</td><td>24</td><td>TAAAGTGCTTATAGTGCAGGTAGT</td><td>782.0</td><td>-6.0</td><td>0.9</td><td>-6.5</td><td>7.3E-11</td><td>6.8E-09</td><td>BT20/MB231</td><td>MP</td></tr>
<tr><td>>MI0000102|hsa-mir-100&WithFlank&11|-|122152223|122152314@20.39.20(+1A)&[MIMAT0000098&hsa-miR-100-5p&offsets|+1|-1(+1A);m-27&11|-|122152276|122152296&offsets|+1|0(+1A)]|ACCCGTAGATCCGAACTTGTA</td><td>>MI0000102|hsa-mir-100&WithFlank&11|-|122152223|122152314</td><td>20</td><td>39</td><td>20(+1A)</td><td>ACCCGTAGATCCGAACTTGTA</td><td>288.1</td><td>-11.6</td><td>1.8</td><td>-6.5</td><td>7.3E-11</td><td>6.8E-09</td><td>BT20/MB231</td><td>MP</td></tr>
<tr><td>>trnaMT_MetCAT_MT_+_4402_4469@54.71.18|ACCCTTCCCGTACTACCA</td><td>>trnaMT_MetCAT_MT_+_4402_4469</td><td>54</td><td>71</td><td>18</td><td>ACCCTTCCCGTACTACCA</td><td>586.0</td><td>7.8</td><td>1.3</td><td>5.9</td><td>3.2E-09</td><td>6.8E-09</td><td>MB231/MB468</td><td>CYTO</td></tr>
</table>

<table>
<tr><td>>hg38_21_+_8433172_8446622_RNA45SN1_45S_50ntFlanks@6790.6807.18|CCTGTCTGAGCGTCGCTT</td><td>>hg38_21_+_8433172_8446622_RNA45SN1_45S_50ntFlanks</td><td>6790</td><td>6807</td><td>18</td><td>CCTGTCTGAGCGTCGCTT</td><td>1489.7</td><td>3.6</td><td>0.6</td><td>5.9</td><td>3.2E-09</td><td>6.9E-09</td><td>MB231/MB468</td><td>NUC</td></tr>
<tr><td>>hg38_21_+_8433172_8446622_RNA45SN1_45S_50ntFlanks@11027.11075.49|CTCTCTCTCTCTCTCCCCCGCTCCCCGTCCTCCCCCCTCCCCGGGGGAG</td><td>>hg38_21_+_8433172_8446622_RNA45SN1_45S_50ntFlanks</td><td>11027</td><td>11075</td><td>49</td><td>CTCTCTCTCTCTCTCCCCCGCTCCCCGTCCTCCCCCCTCCCCGGGGGAG</td><td>321.5</td><td>-10.4</td><td>1.7</td><td>-6.0</td><td>2.4E-09</td><td>6.9E-09</td><td>BT20/MB468</td><td>MITO</td></tr>
<tr><td>>hg38_21_+_8433172_8446622_RNA45SN1_45S_50ntFlanks@11666.11703.38|TCTGCCCAGTGCTCTGAATGTCAAAGTGAAGAAATTCA</td><td>>hg38_21_+_8433172_8446622_RNA45SN1_45S_50ntFlanks</td><td>11666</td><td>11703</td><td>38</td><td>TCTGCCCAGTGCTCTGAATGTCAAAGTGAAGAAATTCA</td><td>626.9</td><td>4.0</td><td>0.7</td><td>5.9</td><td>3.2E-09</td><td>6.9E-09</td><td>MB231/MB468</td><td>CYTO</td></tr>
<tr><td>>hg38_21_+_8433172_8446622_RNA45SN1_45S_50ntFlanks@476.499.24|TTCGTGATCGATGTGGTGACGTCG</td><td>>hg38_21_+_8433172_8446622_RNA45SN1_45S_50ntFlanks</td><td>476</td><td>499</td><td>24</td><td>TTCGTGATCGATGTGGTGACGTCG</td><td>8607.0</td><td>5.1</td><td>0.9</td><td>5.9</td><td>3.2E-09</td><td>6.9E-09</td><td>MB231/MB468</td><td>NUC</td></tr>
<tr><td>>hg38_21_+_8433172_8446622_RNA45SN1_45S_50ntFlanks@10853.10872.20|ATAAGGATTGGCTCTAAGGG</td><td>>hg38_21_+_8433172_8446622_RNA45SN1_45S_50ntFlanks</td><td>10853</td><td>10872</td><td>20</td><td>ATAAGGATTGGCTCTAAGGG</td><td>62.9</td><td>-9.8</td><td>1.7</td><td>-5.9</td><td>3.1E-09</td><td>6.9E-09</td><td>MB231/MB468</td><td>MP</td></tr>
<tr><td>>hg38_21_+_8433172_8446622_RNA45SN1_45S_50ntFlanks@10342.10361.20|ATATTCAAACGAGAACTTTG</td><td>>hg38_21_+_8433172_8446622_RNA45SN1_45S_50ntFlanks</td><td>10342</td><td>10361</td><td>20</td><td>ATATTCAAACGAGAACTTTG</td><td>1209.4</td><td>-4.1</td><td>0.7</td><td>-6.3</td><td>3.8E-10</td><td>6.9E-09</td><td>BT20/MB468</td><td>MP</td></tr>
<tr><td>>hg38_21_+_8433172_8446622_RNA45SN1_45S_50ntFlanks@10406.10424.19|TGGGTCAGTCGGTCCTGAG</td><td>>hg38_21_+_8433172_8446622_RNA45SN1_45S_50ntFlanks</td><td>10406</td><td>10424</td><td>19</td><td>TGGGTCAGTCGGTCCTGAG</td><td>1881.7</td><td>5.7</td><td>0.9</td><td>6.3</td><td>3.8E-10</td><td>6.9E-09</td><td>BT20/MB468</td><td>MP</td></tr>
<tr><td>>hg38_21_+_8433172_8446622_RNA45SN1_45S_50ntFlanks@10130.10149.20|GCGGCGGCGGCGGCGGCGGG</td><td>>hg38_21_+_8433172_8446622_RNA45SN1_45S_50ntFlanks</td><td>10130</td><td>10149</td><td>20</td><td>GCGGCGGCGGCGGCGGCGGG</td><td>162.7</td><td>-10.4</td><td>1.7</td><td>-6.3</td><td>3.8E-10</td><td>6.9E-09</td><td>BT20/MB468</td><td>MP</td></tr>
<tr><td>>hg38_21_+_8433172_8446622_RNA45SN1_45S_50ntFlanks@12418.12446.29|CGATGTCGGCTCTTCCTATCATTGTGAAG</td><td>>hg38_21_+_8433172_8446622_RNA45SN1_45S_50ntFlanks</td><td>12418</td><td>12446</td><td>29</td><td>CGATGTCGGCTCTTCCTATCATTGTGAAG</td><td>793.2</td><td>-4.2</td><td>0.7</td><td>-6.3</td><td>3.8E-10</td><td>6.9E-09</td><td>BT20/MB468</td><td>MP</td></tr>
<tr><td>>hg38_21_+_8433172_8446622_RNA45SN1_45S_50ntFlanks@8847.8866.20|GCGGCGGCGGCGGCGGCGGG</td><td>>hg38_21_+_8433172_8446622_RNA45SN1_45S_50ntFlanks</td><td>8847</td><td>8866</td><td>20</td><td>GCGGCGGCGGCGGCGGCGGG</td><td>162.7</td><td>-10.4</td><td>1.7</td><td>-6.3</td><td>3.8E-10</td><td>6.9E-09</td><td>BT20/MB468</td><td>MP</td></tr>
<tr><td>>hg38_21_+_8433172_8446622_RNA45SN1_45S_50ntFlanks@10402.10432.31|AACATGGGTCAGTCGGTCCTGAGAGATGGGC</td><td>>hg38_21_+_8433172_8446622_RNA45SN1_45S_50ntFlanks</td><td>10402</td><td>10432</td><td>31</td><td>AACATGGGTCAGTCGGTCCTGAGAGATGGGC</td><td>573.1</td><td>-6.3</td><td>1.1</td><td>-5.9</td><td>3.2E-09</td><td>7.0E-09</td><td>MB231/MB468</td><td>NUC</td></tr>
<tr><td>>MI0000440|hsa-mir-27b&WithFlank&9|+|95085439|95085547@67.87.21(+1U)&[MIMAT0000419&hsa-miR-27b-3p&offsets|0|0(+1U);m-39&9|+|95085505|95085525&offsets|0|0(+1U)]|TTCACAGTGGCTAAGTTCTGCT</td><td>>MI0000440|hsa-mir-27b&WithFlank&9|+|95085439|95085547</td><td>67</td><td>87</td><td>21(+1U)</td><td>TTCACAGTGGCTAAGTTCTGCT</td><td>290.2</td><td>4.0</td><td>0.7</td><td>5.9</td><td>3.7E-09</td><td>7.0E-09</td><td>BT20/MB468</td><td>NUC</td></tr>
<tr><td>>hg38_21_+_8433172_8446622_RNA45SN1_45S_50ntFlanks@9534.9565.32|AGGAAACTCTGGTGGAGGTCCGTAGCGGTCCT</td><td>>hg38_21_+_8433172_8446622_RNA45SN1_45S_50ntFlanks</td><td>9534</td><td>9565</td><td>32</td><td>AGGAAACTCTGGTGGAGGTCCGTAGCGGTCCT</td><td>5904.3</td><td>-10.4</td><td>1.8</td><td>-5.9</td><td>3.7E-09</td><td>7.0E-09</td><td>BT20/MB468</td><td>NUC</td></tr>
<tr><td>>MI0000285|hsa-mir-205&WithFlank&1|+|209432127|209432248@40.62.23&[MIMAT0000266&hsa-miR-205-5p&offsets|0|+1;m-351&1|+|209432166|209432187&offsets|0|+1]|TCCTTCATTCCACCGGAGTCTGT</td><td>>MI0000285|hsa-mir-205&WithFlank&1|+|209432127|209432248</td><td>40</td><td>62</td><td>23</td><td>TCCTTCATTCCACCGGAGTCTGT</td><td>3390.7</td><td>10.2</td><td>1.6</td><td>6.5</td><td>7.5E-11</td><td>7.0E-09</td><td>BT20/MB231</td><td>MP</td></tr>
<tr><td>>MI0000285|hsa-mir-205&WithFlank&1|+|209432127|209432248@40.60.21(+1U)&[MIMAT0000266&hsa-miR-205-5p&offsets|0|-1(+1U);m-351&1|+|209432166|209432187&offsets|0|-1(+1U)]|TCCTTCATTCCACCGGAGTCTT</td><td>>MI0000285|hsa-mir-205&WithFlank&1|+|209432127|209432248</td><td>40</td><td>60</td><td>21(+1U)</td><td>TCCTTCATTCCACCGGAGTCTT</td><td>1144.2</td><td>6.6</td><td>1.1</td><td>6.0</td><td>2.4E-09</td><td>7.0E-09</td><td>BT20/MB468</td><td>MITO</td></tr>
<tr><td>>hg38_21_+_8433172_8446622_RNA45SN1_45S_50ntFlanks@9027.9044.18|CGTCTCCTCGTGGGGGGG</td><td>>hg38_21_+_8433172_8446622_RNA45SN1_45S_50ntFlanks</td><td>9027</td><td>9044</td><td>18</td><td>CGTCTCCTCGTGGGGGGG</td><td>113.3</td><td>-8.9</td><td>1.5</td><td>-6.0</td><td>2.4E-09</td><td>7.0E-09</td><td>BT20/MB468</td><td>MITO</td></tr>
<tr><td>>MI0001733|hsa-mir-452&WithFlank&X|-|151959622|151959718@20.41.22&[MIMAT0001635&hsa-miR-452-5p&offsets|0|0;m-188&X|-|151959677|151959699&offsets|0|-1]|AACTGTTTGCAGAGGAAACTGA</td><td>>MI0001733|hsa-mir-452&WithFlank&X|-|151959622|151959718</td><td>20</td><td>41</td><td>22</td><td>AACTGTTTGCAGAGGAAACTGA</td><td>409.7</td><td>12.4</td><td>2.0</td><td>6.3</td><td>3.9E-10</td><td>7.0E-09</td><td>BT20/MB468</td><td>MP</td></tr>
<tr><td>>hg38_21_+_8433172_8446622_RNA45SN1_45S_50ntFlanks@11211.11254.44|GCGACTCTGGACGCGAGCCGGGCCCTTCCCGTGGATCGCCCCAG</td><td>>hg38_21_+_8433172_8446622_RNA45SN1_45S_50ntFlanks</td><td>11211</td><td>11254</td><td>44</td><td>GCGACTCTGGACGCGAGCCGGGCCCTTCCCGTGGATCGCCCCAG</td><td>749.1</td><td>-8.5</td><td>1.4</td><td>-5.9</td><td>3.5E-09</td><td>7.0E-09</td><td>MB231/MB468</td><td>MITO</td></tr>
<tr><td>>am_trna17_LeuCAG_16_+_57333863_57333945@1.25.25,trna26_LeuCAG_16_-_57334392_57334474@1.25.25,trna34_LeuCAG_1_+_161411323_161411405@1.25.25,trna36_LeuCAG_1_+_161418741_161418823@1.25.25,trna38_LeuCAG_1_+_161426122_161426204@1.25.25,trna40_LeuCAG_1_+_161433532_161433614@1.25.25,trna42_LeuCAG_1_+_161440913_161440995@1.25.25,trna67_LeuCAG_1_-_161500132_161500214@1.25.25,trna7_LeuCAG_6_+_26521436_26521518@1.25.25|GTCAGGATGGCCGAGCGGTCTAAGG</td><td>>am_trna17_LeuCAG_16_+_57333863_57333945</td><td>1</td><td>25</td><td>25</td><td>GTCAGGATGGCCGAGCGGTCTAAGG</td><td>2424.1</td><td>8.5</td><td>1.4</td><td>5.9</td><td>3.2E-09</td><td>7.0E-09</td><td>MB231/MB468</td><td>CYTO</td></tr>
<tr><td>>hg38_21_+_8433172_8446622_RNA45SN1_45S_50ntFlanks@11537.11560.24|CCGCGCGGCGCCTCGCCTCGGCCG</td><td>>hg38_21_+_8433172_8446622_RNA45SN1_45S_50ntFlanks</td><td>11537</td><td>11560</td><td>24</td><td>CCGCGCGGCGCCTCGCCTCGGCCG</td><td>734.3</td><td>-9.3</td><td>1.6</td><td>-5.9</td><td>3.7E-09</td><td>7.0E-09</td><td>BT20/MB468</td><td>NUC</td></tr>
<tr><td>>hg38_21_+_8433172_8446622_RNA45SN1_45S_50ntFlanks@4722.4759.38|TCATTAATCAAGAACGAAAGTCGGAGGTTCGAAGACGA</td><td>>hg38_21_+_8433172_8446622_RNA45SN1_45S_50ntFlanks</td><td>4722</td><td>4759</td><td>38</td><td>TCATTAATCAAGAACGAAAGTCGGAGGTTCGAAGACGA</td><td>5023.8</td><td>-5.5</td><td>0.9</td><td>-5.9</td><td>3.3E-09</td><td>7.0E-09</td><td>MB231/MB468</td><td>NUC</td></tr>
<tr><td>>MI0000446|hsa-mir-125b-1&WithFlank&11|-|122099751|122099850@21.42.22;MI0000470|hsa-mir-125b-2&WithFlank&21|+|16590231|16590331@23.44.22&[MIMAT0000423&hsa-miR-125b-5p&offsets|0|0;m-67&11|-|122099809|122099830&offsets|0|0];[MIMAT0000423_1&hsa-miR-125b-5p&offsets|0|0;m-68&21|+|16590253|16590274&offsets|0|0]|TCCCTGAGACCCTAACTTGTGA</td><td>>MI0000446|hsa-mir-125b-1&WithFlank&11|-|122099751|122099850</td><td>21</td><td>42</td><td>22</td><td>TCCCTGAGACCCTAACTTGTGA</td><td>15102.1</td><td>5.5</td><td>0.9</td><td>5.9</td><td>3.1E-09</td><td>7.0E-09</td><td>MB231/MB468</td><td>MP</td></tr>
<tr><td>>hg38_21_+_8433172_8446622_RNA45SN1_45S_50ntFlanks@8198.8223.26|TGGACGGTGTGAGGCCGGTAGCGGCC</td><td>>hg38_21_+_8433172_8446622_RNA45SN1_45S_50ntFlanks</td><td>8198</td><td>8223</td><td>26</td><td>TGGACGGTGTGAGGCCGGTAGCGGCC</td><td>796.7</td><td>-6.2</td><td>1.1</td><td>-5.9</td><td>3.3E-09</td><td>7.0E-09</td><td>MB231/MB468</td><td>NUC</td></tr>
<tr><td>>hg38_21_+_8433172_8446622_RNA45SN1_45S_50ntFlanks@10108.10127.20|GTGTGGTGTGCGTCGGAGGG</td><td>>hg38_21_+_8433172_8446622_RNA45SN1_45S_50ntFlanks</td><td>10108</td><td>10127</td><td>20</td><td>GTGTGGTGTGCGTCGGAGGG</td><td>67.9</td><td>-9.9</td><td>1.7</td><td>-5.9</td><td>3.1E-09</td><td>7.0E-09</td><td>MB231/MB468</td><td>MP</td></tr>
<tr><td>>MI0000542|hsa-mir-320a&WithFlank&8|-|22244960|22245043@49.70.22&[MIMAT0000510&hsa-miR-320a-3p&offsets|+1|+1;m-60&8|-|22244975|22244996&offsets|+1|+1]|AAAGCTGGGTTGAGAGGGCGAA</td><td>>MI0000542|hsa-mir-320a&WithFlank&8|-|22244960|22245043</td><td>49</td><td>70</td><td>22</td><td>AAAGCTGGGTTGAGAGGGCGAA</td><td>137.0</td><td>2.3</td><td>0.4</td><td>5.9</td><td>3.7E-09</td><td>7.0E-09</td><td>BT20/MB468</td><td>NUC</td></tr>
<tr><td>>hg38_21_+_8433172_8446622_RNA45SN1_45S_50ntFlanks@12959.12982.24|TCTGGGTCGGGGTTTCGTACGTAG</td><td>>hg38_21_+_8433172_8446622_RNA45SN1_45S_50ntFlanks</td><td>12959</td><td>12982</td><td>24</td><td>TCTGGGTCGGGGTTTCGTACGTAG</td><td>1336.1</td><td>-9.3</td><td>1.6</td><td>-5.9</td><td>3.7E-09</td><td>7.0E-09</td><td>BT20/MB468</td><td>NUC</td></tr>
<tr><td>>hg38_21_+_8433172_8446622_RNA45SN1_45S_50ntFlanks@10919.10936.18|CGGCTGGACGAGGCGCCG</td><td>>hg38_21_+_8433172_8446622_RNA45SN1_45S_50ntFlanks</td><td>10919</td><td>10936</td><td>18</td><td>CGGCTGGACGAGGCGCCG</td><td>60.3</td><td>-9.8</td><td>1.6</td><td>-5.9</td><td>3.1E-09</td><td>7.0E-09</td><td>MB231/MB468</td><td>MP</td></tr>
<tr><td>>hg38_21_+_8433172_8446622_RNA45SN1_45S_50ntFlanks@11781.11810.30|CGCATGAATGGATGAACGAGATTCCCACTG</td><td>>hg38_21_+_8433172_8446622_RNA45SN1_45S_50ntFlanks</td><td>11781</td><td>11810</td><td>30</td><td>CGCATGAATGGATGAACGAGATTCCCACTG</td><td>118.3</td><td>-10.8</td><td>1.8</td><td>-5.9</td><td>3.1E-09</td><td>7.0E-09</td><td>MB231/MB468</td><td>MP</td></tr>
<tr><td>>trnaMT_GlyTCC_MT_+_9991_10058@28.68.41|ACTTCCAATTAACTAGTTTTGACAACATTCAAAAAAGAGTA</td><td>>trnaMT_GlyTCC_MT_+_9991_10058</td><td>28</td><td>68</td><td>41</td><td>ACTTCCAATTAACTAGTTTTGACAACATTCAAAAAAGAGTA</td><td>394.1</td><td>-11.6</td><td>2.0</td><td>-5.9</td><td>3.3E-09</td><td>7.0E-09</td><td>MB231/MB468</td><td>NUC</td></tr>
<tr><td>>hg38_21_+_8433172_8446622_RNA45SN1_45S_50ntFlanks@10905.10925.21|CTGGGCGCGCGCCGCGGCTGG</td><td>>hg38_21_+_8433172_8446622_RNA45SN1_45S_50ntFlanks</td><td>10905</td><td>10925</td><td>21</td><td>CTGGGCGCGCGCCGCGGCTGG</td><td>296.5</td><td>-7.3</td><td>1.2</td><td>-5.9</td><td>2.8E-09</td><td>7.0E-09</td><td>BT20/MB468</td><td>CYTO</td></tr>
<tr><td>>MI0003628|hsa-mir-615&WithFlank&12|+|54033944|54034051@67.85.19(+1U)&[MIMAT0003283&hsa-miR-615-3p&offsets|0|-3(+1U);m-495&12|+|54034010|54034030&offsets|0|-2(+1U)]|TCCGAGCCTGGGTCTCCCTT</td><td>>MI0003628|hsa-mir-615&WithFlank&12|+|54033944|54034051</td><td>67</td><td>85</td><td>19(+1U)</td><td>TCCGAGCCTGGGTCTCCCTT</td><td>151.2</td><td>5.9</td><td>0.9</td><td>6.4</td><td>1.9E-10</td><td>7.0E-09</td><td>BT20/MB231</td><td>CYTO</td></tr>
<tr><td>>MI0000102|hsa-mir-100&WithFlank&11|-|122152223|122152314@19.37.19(+2U)&[MIMAT0000098&hsa-miR-100-5p&offsets|0|-3(+2U);m-27&11|-|122152276|122152296&offsets|0|-2(+2U)]|AACCCGTAGATCCGAACTTTT</td><td>>MI0000102|hsa-mir-100&WithFlank&11|-|122152223|122152314</td><td>19</td><td>37</td><td>19(+2U)</td><td>AACCCGTAGATCCGAACTTTT</td><td>94.0</td><td>9.8</td><td>1.6</td><td>5.9</td><td>3.1E-09</td><td>7.0E-09</td><td>MB231/MB468</td><td>MP</td></tr>
<tr><td>>hg38_21_+_8433172_8446622_RNA45SN1_45S_50ntFlanks@9862.9900.39|ATGAACCGAACGCCGGGTTAAGGCGCCCGATGCCGACGC</td><td>>hg38_21_+_8433172_8446622_RNA45SN1_45S_50ntFlanks</td><td>9862</td><td>9900</td><td>39</td><td>ATGAACCGAACGCCGGGTTAAGGCGCCCGATGCCGACGC</td><td>97.4</td><td>-10.5</td><td>1.8</td><td>-5.9</td><td>3.1E-09</td><td>7.1E-09</td><td>MB231/MB468</td><td>MP</td></tr>
<tr><td>>hg38_21_+_8433172_8446622_RNA45SN1_45S_50ntFlanks@11924.11950.27|GAGGTGTAGAATAAGTGGGAGGCCCCC</td><td>>hg38_21_+_8433172_8446622_RNA45SN1_45S_50ntFlanks</td><td>11924</td><td>11950</td><td>27</td><td>GAGGTGTAGAATAAGTGGGAGGCCCCC</td><td>4464.5</td><td>15.5</td><td>2.4</td><td>6.5</td><td>7.7E-11</td><td>7.1E-09</td><td>BT20/MB231</td><td>MP</td></tr>
<tr><td>>hg38_21_+_8433172_8446622_RNA45SN1_45S_50ntFlanks@6661.6686.26|CGGTGGATCACTCGGCTCGTGCGTCG</td><td>>hg38_21_+_8433172_8446622_RNA45SN1_45S_50ntFlanks</td><td>6661</td><td>6686</td><td>26</td><td>CGGTGGATCACTCGGCTCGTGCGTCG</td><td>1335.9</td><td>-1.6</td><td>0.3</td><td>-5.9</td><td>3.8E-09</td><td>7.1E-09</td><td>BT20/MB468</td><td>NUC</td></tr>
<tr><td>>hg38_21_+_8433172_8446622_RNA45SN1_45S_50ntFlanks@11579.11600.22|AGAACTGGTGCGGACCAGGGGA</td><td>>hg38_21_+_8433172_8446622_RNA45SN1_45S_50ntFlanks</td><td>11579</td><td>11600</td><td>22</td><td>AGAACTGGTGCGGACCAGGGGA</td><td>295.4</td><td>-7.2</td><td>1.2</td><td>-5.9</td><td>3.3E-09</td><td>7.1E-09</td><td>MB231/MB468</td><td>CYTO</td></tr>
<tr><td>>hg38_21_+_8433172_8446622_RNA45SN1_45S_50ntFlanks@8157.8185.29|CTCGTGGGGGGCCCAAGTCCTTCTGATCG</td><td>>hg38_21_+_8433172_8446622_RNA45SN1_45S_50ntFlanks</td><td>8157</td><td>8185</td><td>29</td><td>CTCGTGGGGGGCCCAAGTCCTTCTGATCG</td><td>344.3</td><td>-5.9</td><td>1.0</td><td>-5.9</td><td>3.1E-09</td><td>7.1E-09</td><td>MB231/MB468</td><td>MP</td></tr>
<tr><td>>hg38_21_+_8433172_8446622_RNA45SN1_45S_50ntFlanks@9339.9365.27|CGCACGAAAGCCGCCGTGGCGCAATGA</td><td>>hg38_21_+_8433172_8446622_RNA45SN1_45S_50ntFlanks</td><td>9339</td><td>9365</td><td>27</td><td>CGCACGAAAGCCGCCGTGGCGCAATGA</td><td>157.1</td><td>-10.4</td><td>1.7</td><td>-6.3</td><td>3.9E-10</td><td>7.1E-09</td><td>BT20/MB468</td><td>MP</td></tr>
<tr><td>>hg38_21_+_8433172_8446622_RNA45SN1_45S_50ntFlanks@10410.10442.33|TCAGTCGGTCCTGAGAGATGGGCGAGCGCCGTT</td><td>>hg38_21_+_8433172_8446622_RNA45SN1_45S_50ntFlanks</td><td>10410</td><td>10442</td><td>33</td><td>TCAGTCGGTCCTGAGAGATGGGCGAGCGCCGTT</td><td>445.0</td><td>-5.1</td><td>0.9</td><td>-5.9</td><td>3.6E-09</td><td>7.1E-09</td><td>MB231/MB468</td><td>MITO</td></tr>
<tr><td>>hg38_21_+_8433172_8446622_RNA45SN1_45S_50ntFlanks@5492.5510.19|AGAAGACGGTCGAACTTGA</td><td>>hg38_21_+_8433172_8446622_RNA45SN1_45S_50ntFlanks</td><td>5492</td><td>5510</td><td>19</td><td>AGAAGACGGTCGAACTTGA</td><td>305.4</td><td>-7.6</td><td>1.3</td><td>-5.9</td><td>3.6E-09</td><td>7.1E-09</td><td>MB231/MB468</td><td>MITO</td></tr>
<tr><td>>hg38_21_+_8433172_8446622_RNA45SN1_45S_50ntFlanks@4045.4062.18|CCATTCGAACGTCTGCCC</td><td>>hg38_21_+_8433172_8446622_RNA45SN1_45S_50ntFlanks</td><td>4045</td><td>4062</td><td>18</td><td>CCATTCGAACGTCTGCCC</td><td>1278.3</td><td>-9.0</td><td>1.5</td><td>-5.9</td><td>3.8E-09</td><td>7.1E-09</td><td>BT20/MB468</td><td>NUC</td></tr>
<tr><td>>am_MI0000786|hsa-mir-378a&WithFlank&5|+|149732819|149732896@49.69.21(+1U)&[MIMAT0000732&hsa-miR-378a-3p&offsets|0|-1(+1U);m-44&5|+|149732867|149732888&offsets|0|-1(+1U)]|ACTGGACTTGGAGTCAGAAGGT</td><td>>am_MI0000786|hsa-mir-378a&WithFlank&5|+|149732819|149732896</td><td>49</td><td>69</td><td>21(+1U)</td><td>ACTGGACTTGGAGTCAGAAGGT</td><td>2245.4</td><td>3.2</td><td>0.5</td><td>5.9</td><td>3.3E-09</td><td>7.1E-09</td><td>MB231/MB468</td><td>NUC</td></tr>
<tr><td>>hg38_21_+_8433172_8446622_RNA45SN1_45S_50ntFlanks@3899.3917.19|CCTTCGCGGGGGGGATGCG</td><td>>hg38_21_+_8433172_8446622_RNA45SN1_45S_50ntFlanks</td><td>3899</td><td>3917</td><td>19</td><td>CCTTCGCGGGGGGGATGCG</td><td>150.4</td><td>-10.3</td><td>1.6</td><td>-6.3</td><td>3.9E-10</td><td>7.1E-09</td><td>BT20/MB468</td><td>MP</td></tr>
<tr><td>>hg38_21_+_8433172_8446622_RNA45SN1_45S_50ntFlanks@10402.10419.18|AACATGGGTCAGTCGGTC</td><td>>hg38_21_+_8433172_8446622_RNA45SN1_45S_50ntFlanks</td><td>10402</td><td>10419</td><td>18</td><td>AACATGGGTCAGTCGGTC</td><td>1439.4</td><td>-8.0</td><td>1.3</td><td>-5.9</td><td>3.3E-09</td><td>7.1E-09</td><td>MB231/MB468</td><td>NUC</td></tr>
<tr><td>>hg38_21_+_8433172_8446622_RNA45SN1_45S_50ntFlanks@10353.10388.36|AGAACTTTGAAGGCCGAAGTGGAGAAGGGTTCCATG</td><td>>hg38_21_+_8433172_8446622_RNA45SN1_45S_50ntFlanks</td><td>10353</td><td>10388</td><td>36</td><td>AGAACTTTGAAGGCCGAAGTGGAGAAGGGTTCCATG</td><td>846.9</td><td>-7.6</td><td>1.3</td><td>-5.9</td><td>3.8E-09</td><td>7.1E-09</td><td>BT20/MB468</td><td>NUC</td></tr>
<tr><td>>hg38_21_+_8433172_8446622_RNA45SN1_45S_50ntFlanks@5431.5466.36|GTTTAGTGAGGCCCTCGGATCGGCCCCGCCGGGGTC</td><td>>hg38_21_+_8433172_8446622_RNA45SN1_45S_50ntFlanks</td><td>5431</td><td>5466</td><td>36</td><td>GTTTAGTGAGGCCCTCGGATCGGCCCCGCCGGGGTC</td><td>208.7</td><td>-7.1</td><td>1.2</td><td>-5.9</td><td>3.3E-09</td><td>7.1E-09</td><td>MB231/MB468</td><td>NUC</td></tr>
<tr><td>>hg38_21_+_8433172_8446622_RNA45SN1_45S_50ntFlanks@6722.6758.37|ATTGCAGGACACATTGATCATCGACACTTCGAACGCA</td><td>>hg38_21_+_8433172_8446622_RNA45SN1_45S_50ntFlanks</td><td>6722</td><td>6758</td><td>37</td><td>ATTGCAGGACACATTGATCATCGACACTTCGAACGCA</td><td>287.0</td><td>-4.0</td><td>0.7</td><td>-5.9</td><td>3.4E-09</td><td>7.2E-09</td><td>MB231/MB468</td><td>NUC</td></tr>
</table>

<table>
<tr><th></th><th></th><th></th><th></th><th></th><th></th><th></th><th></th><th></th><th></th><th></th><th></th><th></th><th></th></tr>
<tr><td>>hg38_MT_+_598_1651_RNR1_12S_50ntFlanks@570.611.42|AAGGACCTGGCGGTGCTTCATATCCCTCTAGAGGAGCCTGTT</td><td>>hg38_MT_+_598_1651_RNR1_12S_50ntFlanks</td><td>570</td><td>611</td><td>42</td><td>AAGGACCTGGCGGTGCTTCATATCCCTCTAGAGGAGCCTGTT</td><td>67.6</td><td>9.3</td><td>1.6</td><td>5.9</td><td>3.2E-09</td><td>7.2E-09</td><td>MB231/MB468</td><td>MP</td></tr>
<tr><td>>hg38_21_+_8433172_8446622_RNA45SN1_45S_50ntFlanks@9534.9572.39|AGGAAACTCTGGTGGAGGTCCGTAGCGGTCCTGACGTGC</td><td>>hg38_21_+_8433172_8446622_RNA45SN1_45S_50ntFlanks</td><td>9534</td><td>9572</td><td>39</td><td>AGGAAACTCTGGTGGAGGTCCGTAGCGGTCCTGACGTGC</td><td>2363.3</td><td>-7.1</td><td>1.2</td><td>-5.9</td><td>3.6E-09</td><td>7.2E-09</td><td>MB231/MB468</td><td>MITO</td></tr>
<tr><td>>hg38_21_+_8433172_8446622_RNA45SN1_45S_50ntFlanks@9574.9598.25|AATCGGTCGTCCGACCTGGGTATAG</td><td>>hg38_21_+_8433172_8446622_RNA45SN1_45S_50ntFlanks</td><td>9574</td><td>9598</td><td>25</td><td>AATCGGTCGTCCGACCTGGGTATAG</td><td>434.2</td><td>-6.0</td><td>1.0</td><td>-5.9</td><td>3.6E-09</td><td>7.2E-09</td><td>MB231/MB468</td><td>MITO</td></tr>
<tr><td>>trnaMT_SerGCT_MT_+_12207_12265@1.25.25|GAGAAAGCTCACAAGAACTGCTAAC</td><td>>trnaMT_SerGCT_MT_+_12207_12265</td><td>1</td><td>25</td><td>25</td><td>GAGAAAGCTCACAAGAACTGCTAAC</td><td>3392.2</td><td>7.7</td><td>1.3</td><td>5.9</td><td>2.9E-09</td><td>7.2E-09</td><td>BT20/MB468</td><td>CYTO</td></tr>
<tr><td>>am_trnaMT_PheGAA_MT_+_577_647@42.59.18|TTTAGACGGGCTCACATC</td><td>>am_trnaMT_PheGAA_MT_+_577_647</td><td>42</td><td>59</td><td>18</td><td>TTTAGACGGGCTCACATC</td><td>275.8</td><td>9.0</td><td>1.5</td><td>5.9</td><td>2.9E-09</td><td>7.2E-09</td><td>BT20/MB468</td><td>CYTO</td></tr>
<tr><td>>hg38_21_+_8433172_8446622_RNA45SN1_45S_50ntFlanks@10781.10823.43|AGGTGAACAGCCTCTGGCATGTTGGAACAATGTAGGTAAGGGA</td><td>>hg38_21_+_8433172_8446622_RNA45SN1_45S_50ntFlanks</td><td>10781</td><td>10823</td><td>43</td><td>AGGTGAACAGCCTCTGGCATGTTGGAACAATGTAGGTAAGGGA</td><td>98.7</td><td>-10.5</td><td>1.8</td><td>-5.9</td><td>3.2E-09</td><td>7.2E-09</td><td>MB231/MB468</td><td>MP</td></tr>
<tr><td>>MI0016005|hsa-mir-3615&WithFlank&17|+|74748607|74748705@57.78.22(+2U)&[MIMAT0017994&hsa-miR-3615&offsets|0|+1(+2U);m-271&17|+|74748663|74748684&offsets|0|0(+2U)]|TCTCTCGGCTCCTCGCGGCTCGTT</td><td>>MI0016005|hsa-mir-3615&WithFlank&17|+|74748607|74748705</td><td>57</td><td>78</td><td>22(+2U)</td><td>TCTCTCGGCTCCTCGCGGCTCGTT</td><td>414.4</td><td>7.4</td><td>1.2</td><td>5.9</td><td>3.3E-09</td><td>7.2E-09</td><td>MB231/MB468</td><td>CYTO</td></tr>
<tr><td>>hg38_21_+_8433172_8446622_RNA45SN1_45S_50ntFlanks@475.505.31|CTTCGTGATCGATGTGGTGACGTCGTGCTCT</td><td>>hg38_21_+_8433172_8446622_RNA45SN1_45S_50ntFlanks</td><td>475</td><td>505</td><td>31</td><td>CTTCGTGATCGATGTGGTGACGTCGTGCTCT</td><td>5064.9</td><td>5.8</td><td>1.0</td><td>5.9</td><td>3.6E-09</td><td>7.2E-09</td><td>MB231/MB468</td><td>MITO</td></tr>
<tr><td>>hg38_21_+_8433172_8446622_RNA45SN1_45S_50ntFlanks@12822.12871.50|CCGGGACCGGGGTCCGGTGCGGAGTGCCCTTCGTCCTGGGAAACGGGGCG</td><td>>hg38_21_+_8433172_8446622_RNA45SN1_45S_50ntFlanks</td><td>12822</td><td>12871</td><td>50</td><td>CCGGGACCGGGGTCCGGTGCGGAGTGCCCTTCGTCCTGGGAAACGGGGCG</td><td>173.1</td><td>-9.5</td><td>1.6</td><td>-6.0</td><td>2.5E-09</td><td>7.2E-09</td><td>BT20/MB468</td><td>MITO</td></tr>
<tr><td>>hg38_21_+_8433172_8446622_RNA45SN1_45S_50ntFlanks@5227.5245.19|CGCGCGCTACACTGACTGG</td><td>>hg38_21_+_8433172_8446622_RNA45SN1_45S_50ntFlanks</td><td>5227</td><td>5245</td><td>19</td><td>CGCGCGCTACACTGACTGG</td><td>104.3</td><td>-10.6</td><td>1.8</td><td>-5.9</td><td>3.2E-09</td><td>7.2E-09</td><td>MB231/MB468</td><td>MP</td></tr>
<tr><td>>hg38_21_+_8433172_8446622_RNA45SN1_45S_50ntFlanks@10684.10706.23|CGGCGGCGTCCGGTGAGCTCTCG</td><td>>hg38_21_+_8433172_8446622_RNA45SN1_45S_50ntFlanks</td><td>10684</td><td>10706</td><td>23</td><td>CGGCGGCGTCCGGTGAGCTCTCG</td><td>5053.9</td><td>-4.7</td><td>0.8</td><td>-6.0</td><td>2.5E-09</td><td>7.2E-09</td><td>BT20/MB468</td><td>MITO</td></tr>
<tr><td>>hg38_21_+_8433172_8446622_RNA45SN1_45S_50ntFlanks@10458.10488.31|ATGGCCTCCGTTGCCCTCGGCCGATCGAAAG</td><td>>hg38_21_+_8433172_8446622_RNA45SN1_45S_50ntFlanks</td><td>10458</td><td>10488</td><td>31</td><td>ATGGCCTCCGTTGCCCTCGGCCGATCGAAAG</td><td>1611.8</td><td>-5.3</td><td>0.9</td><td>-5.9</td><td>3.4E-09</td><td>7.2E-09</td><td>MB231/MB468</td><td>NUC</td></tr>
<tr><td>>MI0003513|hsa-mir-455&WithFlank&9|+|114209428|114209535@22.42.21(+1A)&[MIMAT0003150&hsa-miR-455-5p&offsets|0|-1(+1A);m-232&9|+|114209449|114209470&offsets|0|-1(+1A)]|TATGTGCCTTTGGACTACATCA</td><td>>MI0003513|hsa-mir-455&WithFlank&9|+|114209428|114209535</td><td>22</td><td>42</td><td>21(+1A)</td><td>TATGTGCCTTTGGACTACATCA</td><td>84.8</td><td>9.6</td><td>1.6</td><td>5.9</td><td>3.2E-09</td><td>7.2E-09</td><td>MB231/MB468</td><td>MP</td></tr>
<tr><td>>hg38_21_+_8433172_8446622_RNA45SN1_45S_50ntFlanks@11575.11611.37|ACTTAGAACTGGTGCGGACCAGGGGAATCCGACTGTT</td><td>>hg38_21_+_8433172_8446622_RNA45SN1_45S_50ntFlanks</td><td>11575</td><td>11611</td><td>37</td><td>ACTTAGAACTGGTGCGGACCAGGGGAATCCGACTGTT</td><td>1011.1</td><td>-7.0</td><td>1.2</td><td>-5.9</td><td>3.2E-09</td><td>7.2E-09</td><td>MB231/MB468</td><td>MP</td></tr>
<tr><td>>hg38_21_+_8433172_8446622_RNA45SN1_45S_50ntFlanks@12158.12178.21|GTTTGACTGGGGCGGTACACC</td><td>>hg38_21_+_8433172_8446622_RNA45SN1_45S_50ntFlanks</td><td>12158</td><td>12178</td><td>21</td><td>GTTTGACTGGGGCGGTACACC</td><td>71.1</td><td>-10.0</td><td>1.7</td><td>-5.9</td><td>3.2E-09</td><td>7.2E-09</td><td>MB231/MB468</td><td>MP</td></tr>
<tr><td>>MI0000063|hsa-let-7b&WithFlank&22|+|46113680|46113774@12.34.23&[MIMAT0000063&hsa-let-7b-5p&offsets|0|+1;m-14&22|+|46113691|46113712&offsets|0|+1]|TGAGGTAGTAGGTTGTGTGGTTT</td><td>>MI0000063|hsa-let-7b&WithFlank&22|+|46113680|46113774</td><td>12</td><td>34</td><td>23</td><td>TGAGGTAGTAGGTTGTGTGGTTT</td><td>1544.9</td><td>5.2</td><td>0.9</td><td>5.9</td><td>3.7E-09</td><td>7.3E-09</td><td>MB231/MB468</td><td>MITO</td></tr>
<tr><td>>MI0000077|hsa-mir-21&WithFlank&17|+|59841260|59841343@14.34.21&[MIMAT0000076&hsa-miR-21-5p&offsets|0|-1;m-1&17|+|59841273|59841294&offsets|0|-1]|TAGCTTATCAGACTGATGTTG</td><td>>MI0000077|hsa-mir-21&WithFlank&17|+|59841260|59841343</td><td>14</td><td>34</td><td>21</td><td>TAGCTTATCAGACTGATGTTG</td><td>8785.1</td><td>5.4</td><td>0.9</td><td>5.9</td><td>3.7E-09</td><td>7.3E-09</td><td>MB231/MB468</td><td>MITO</td></tr>
<tr><td>>hg38_21_+_8433172_8446622_RNA45SN1_45S_50ntFlanks@8189.8215.27|CCCAGCCCGTGGACGGTGTGAGGCCGG</td><td>>hg38_21_+_8433172_8446622_RNA45SN1_45S_50ntFlanks</td><td>8189</td><td>8215</td><td>27</td><td>CCCAGCCCGTGGACGGTGTGAGGCCGG</td><td>134.6</td><td>-10.2</td><td>1.6</td><td>-6.3</td><td>4.0E-10</td><td>7.3E-09</td><td>BT20/MB468</td><td>MP</td></tr>
<tr><td>>MI0000253|hsa-mir-148a&WithFlank&7|-|25949913|25949992@50.70.21&[MIMAT0000243&hsa-miR-148a-3p&offsets|0|-1;m-7&7|-|25949922|25949943&offsets|0|-1]|TCAGTGCACTACAGAACTTTG</td><td>>MI0000253|hsa-mir-148a&WithFlank&7|-|25949913|25949992</td><td>50</td><td>70</td><td>21</td><td>TCAGTGCACTACAGAACTTTG</td><td>419.8</td><td>8.0</td><td>1.3</td><td>5.9</td><td>2.9E-09</td><td>7.3E-09</td><td>BT20/MB468</td><td>CYTO</td></tr>
<tr><td>>hg38_21_+_8433172_8446622_RNA45SN1_45S_50ntFlanks@12654.12674.21|TATGACTGAACGCCTCTAAGT</td><td>>hg38_21_+_8433172_8446622_RNA45SN1_45S_50ntFlanks</td><td>12654</td><td>12674</td><td>21</td><td>TATGACTGAACGCCTCTAAGT</td><td>870.9</td><td>-5.7</td><td>1.0</td><td>-5.9</td><td>3.4E-09</td><td>7.3E-09</td><td>MB231/MB468</td><td>CYTO</td></tr>
<tr><td>>MI0000465|hsa-mir-191&WithFlank&3|-|49020612|49020715@22.43.22&[MIMAT0000440&hsa-miR-191-5p&offsets|0|-1;m-48&3|-|49020672|49020694&offsets|0|-1]|CAACGGAATCCCAAAAGCAGCT</td><td>>MI0000465|hsa-mir-191&WithFlank&3|-|49020612|49020715</td><td>22</td><td>43</td><td>22</td><td>CAACGGAATCCCAAAAGCAGCT</td><td>5691.4</td><td>5.1</td><td>0.9</td><td>5.9</td><td>3.7E-09</td><td>7.3E-09</td><td>MB231/MB468</td><td>MITO</td></tr>
<tr><td>>hg38_21_+_8433172_8446622_RNA45SN1_45S_50ntFlanks@11564.11590.27|CCTAGCAGCCGACTTAGAACTGGTGCG</td><td>>hg38_21_+_8433172_8446622_RNA45SN1_45S_50ntFlanks</td><td>11564</td><td>11590</td><td>27</td><td>CCTAGCAGCCGACTTAGAACTGGTGCG</td><td>283.8</td><td>-7.9</td><td>1.3</td><td>-5.9</td><td>3.9E-09</td><td>7.3E-09</td><td>BT20/MB468</td><td>NUC</td></tr>
<tr><td>>hg38_21_+_8433172_8446622_RNA45SN1_45S_50ntFlanks@11161.11179.19|CACGGTCCCCCGCGAGGGG</td><td>>hg38_21_+_8433172_8446622_RNA45SN1_45S_50ntFlanks</td><td>11161</td><td>11179</td><td>19</td><td>CACGGTCCCCCGCGAGGGG</td><td>996.9</td><td>-9.1</td><td>1.5</td><td>-6.3</td><td>4.1E-10</td><td>7.3E-09</td><td>BT20/MB468</td><td>MP</td></tr>
<tr><td>>hg38_21_+_8433172_8446622_RNA45SN1_45S_50ntFlanks@4730.4752.23|CAAGAACGAAAGTCGGAGGTTCG</td><td>>hg38_21_+_8433172_8446622_RNA45SN1_45S_50ntFlanks</td><td>4730</td><td>4752</td><td>23</td><td>CAAGAACGAAAGTCGGAGGTTCG</td><td>365.9</td><td>-7.6</td><td>1.3</td><td>-5.9</td><td>3.7E-09</td><td>7.3E-09</td><td>MB231/MB468</td><td>MITO</td></tr>
<tr><td>>hg38_21_+_8433172_8446622_RNA45SN1_45S_50ntFlanks@10458.10487.30|ATGGCCTCCGTTGCCCTCGGCCGATCGAAA</td><td>>hg38_21_+_8433172_8446622_RNA45SN1_45S_50ntFlanks</td><td>10458</td><td>10487</td><td>30</td><td>ATGGCCTCCGTTGCCCTCGGCCGATCGAAA</td><td>796.7</td><td>-6.2</td><td>1.1</td><td>-5.9</td><td>3.7E-09</td><td>7.3E-09</td><td>MB231/MB468</td><td>MITO</td></tr>
<tr><td>>hg38_21_+_8433172_8446622_RNA45SN1_45S_50ntFlanks@13008.13027.20|ATTGAAAGTCAGCCCTCGAC</td><td>>hg38_21_+_8433172_8446622_RNA45SN1_45S_50ntFlanks</td><td>13008</td><td>13027</td><td>20</td><td>ATTGAAAGTCAGCCCTCGAC</td><td>210.1</td><td>-7.6</td><td>1.3</td><td>-5.9</td><td>3.7E-09</td><td>7.3E-09</td><td>MB231/MB468</td><td>MITO</td></tr>
<tr><td>>am_trna60_LeuTAA_1_-_238106956_238107030@27.44.18,trnaMT_LeuTAA_MT_+_3230_3304@27.44.18|CATAAAACTTAAAACTTT</td><td>>am_trna60_LeuTAA_1_-_238106956_238107030</td><td>27</td><td>44</td><td>18</td><td>CATAAAACTTAAAACTTT</td><td>130.8</td><td>-10.1</td><td>1.6</td><td>-6.3</td><td>4.1E-10</td><td>7.3E-09</td><td>BT20/MB468</td><td>MP</td></tr>
<tr><td>>hg38_21_+_8433172_8446622_RNA45SN1_45S_50ntFlanks@11999.12026.28|CCCTGCGGGCCGCCGGTGAAATACCACT</td><td>>hg38_21_+_8433172_8446622_RNA45SN1_45S_50ntFlanks</td><td>11999</td><td>12026</td><td>28</td><td>CCCTGCGGGCCGCCGGTGAAATACCACT</td><td>166.8</td><td>-11.2</td><td>1.9</td><td>-5.9</td><td>3.2E-09</td><td>7.3E-09</td><td>MB231/MB468</td><td>MP</td></tr>
<tr><td>>hg38_21_+_8433172_8446622_RNA45SN1_45S_50ntFlanks@10099.10125.27|CGCGCGCGCGTGTGGTGTGCGTCGGAG</td><td>>hg38_21_+_8433172_8446622_RNA45SN1_45S_50ntFlanks</td><td>10099</td><td>10125</td><td>27</td><td>CGCGCGCGCGTGTGGTGTGCGTCGGAG</td><td>110.3</td><td>-10.6</td><td>1.8</td><td>-5.9</td><td>3.2E-09</td><td>7.4E-09</td><td>MB231/MB468</td><td>MP</td></tr>
<tr><td>>hg38_21_+_8433172_8446622_RNA45SN1_45S_50ntFlanks@9925.9951.27|GGTTGATATAGACAGCAGGACGGTGGC</td><td>>hg38_21_+_8433172_8446622_RNA45SN1_45S_50ntFlanks</td><td>9925</td><td>9951</td><td>27</td><td>GGTTGATATAGACAGCAGGACGGTGGC</td><td>124.3</td><td>10.7</td><td>1.7</td><td>6.3</td><td>4.1E-10</td><td>7.4E-09</td><td>BT20/MB468</td><td>MP</td></tr>
<tr><td>>hg38_21_+_8433172_8446622_RNA45SN1_45S_50ntFlanks@11562.11603.42|CGCCTAGCAGCCGACTTAGAACTGGTGCGGACCAGGGGAATC</td><td>>hg38_21_+_8433172_8446622_RNA45SN1_45S_50ntFlanks</td><td>11562</td><td>11603</td><td>42</td><td>CGCCTAGCAGCCGACTTAGAACTGGTGCGGACCAGGGGAATC</td><td>158.7</td><td>-11.2</td><td>1.9</td><td>-5.9</td><td>3.3E-09</td><td>7.4E-09</td><td>MB231/MB468</td><td>MP</td></tr>
<tr><td>>trna14_LysTTT_11_-_59327808_59327880@1.30.30,trna2_LysTTT_17_+_8022473_8022545@1.30.30,trna54_LysTTT_1_+_204475655_204475727@1.30.30,trna5_LysTTT_11_+_59323902_59323974@1.30.30,trna62_LysTTT_1_-_204476158_204476230@1.30.30,trna76_LysTTT_6_+_28918806_28918878@1.30.30|GCCCGGATAGCTCAGTCGGTAGAGCATCAG</td><td>>trna14_LysTTT_11_-_59327808_59327880</td><td>1</td><td>30</td><td>30</td><td>GCCCGGATAGCTCAGTCGGTAGAGCATCAG</td><td>104.0</td><td>-8.8</td><td>1.5</td><td>-6.0</td><td>2.6E-09</td><td>7.4E-09</td><td>BT20/MB468</td><td>MITO</td></tr>
<tr><td>>hg38_21_+_8433172_8446622_RNA45SN1_45S_50ntFlanks@12413.12436.24|TCCTTCGATGTCGGCTCTTCCTAT</td><td>>hg38_21_+_8433172_8446622_RNA45SN1_45S_50ntFlanks</td><td>12413</td><td>12436</td><td>24</td><td>TCCTTCGATGTCGGCTCTTCCTAT</td><td>450.6</td><td>-7.8</td><td>1.3</td><td>-5.9</td><td>3.7E-09</td><td>7.4E-09</td><td>MB231/MB468</td><td>MITO</td></tr>
<tr><td>>trnaMT_MetCAT_MT_+_4402_4469@34.71.38|ACCCCGAAAATGTTGGTTATACCCTTCCCGTACTACCA</td><td>>trnaMT_MetCAT_MT_+_4402_4469</td><td>34</td><td>71</td><td>38</td><td>ACCCCGAAAATGTTGGTTATACCCTTCCCGTACTACCA</td><td>544.3</td><td>9.8</td><td>1.7</td><td>5.9</td><td>3.5E-09</td><td>7.4E-09</td><td>MB231/MB468</td><td>NUC</td></tr>
<tr><td>>hg38_21_+_8433172_8446622_RNA45SN1_45S_50ntFlanks@11209.11232.24|CGGCGACTCTGGACGCGAGCCGGG</td><td>>hg38_21_+_8433172_8446622_RNA45SN1_45S_50ntFlanks</td><td>11209</td><td>11232</td><td>24</td><td>CGGCGACTCTGGACGCGAGCCGGG</td><td>13749.3</td><td>-6.1</td><td>1.0</td><td>-6.2</td><td>4.1E-10</td><td>7.4E-09</td><td>BT20/MB468</td><td>MP</td></tr>
<tr><td>>MI0000760|hsa-mir-361&WithFlank&X|-|85903630|85903713@51.74.24&[MIMAT0004682&hsa-miR-361-3p&offsets|0|+1;m-124&X|-|85903642|85903663&offsets|0|+2]|TCCCCCAGGTGTGATTCTGATTTG</td><td>>MI0000760|hsa-mir-361&WithFlank&X|-|85903630|85903713</td><td>51</td><td>74</td><td>24</td><td>TCCCCCAGGTGTGATTCTGATTTG</td><td>281.1</td><td>5.6</td><td>0.9</td><td>5.9</td><td>3.8E-09</td><td>7.4E-09</td><td>MB231/MB468</td><td>MITO</td></tr>
<tr><td>>trna119_LysCTT_1_-_145395522_145395594@1.29.29,trna11_LysCTT_5_-_180648979_180649051@1.29.29,trna13_LysCTT_6_+_26556774_26556846@1.29.29,trna32_LysCTT_16_-_3207406_3207478@1.29.29,trna7_LysCTT_16_+_3225692_3225764@1.29.29,trna9_LysCTT_5_+_180634755_180634827@1.29.29|GCCCGGCTAGCTCAGTCGGTAGAGCATGA</td><td>>trna119_LysCTT_1_-_145395522_145395594</td><td>1</td><td>29</td><td>29</td><td>GCCCGGCTAGCTCAGTCGGTAGAGCATGA</td><td>1029.9</td><td>-3.9</td><td>0.7</td><td>-5.9</td><td>3.5E-09</td><td>7.4E-09</td><td>MB231/MB468</td><td>NUC</td></tr>
<tr><td>>hg38_21_+_8433172_8446622_RNA45SN1_45S_50ntFlanks@12932.12972.41|CGCTAAACCATTCGTAGACGACCTGCTTCTGGGTCGGGGTT</td><td>>hg38_21_+_8433172_8446622_RNA45SN1_45S_50ntFlanks</td><td>12932</td><td>12972</td><td>41</td><td>CGCTAAACCATTCGTAGACGACCTGCTTCTGGGTCGGGGTT</td><td>545.8</td><td>-5.5</td><td>0.9</td><td>-5.9</td><td>3.5E-09</td><td>7.4E-09</td><td>MB231/MB468</td><td>NUC</td></tr>
<tr><td>>hg38_21_+_8433172_8446622_RNA45SN1_45S_50ntFlanks@5475.5516.42|GCCCTGGCGGAGCGCTGAGAAGACGGTCGAACTTGACTATCT</td><td>>hg38_21_+_8433172_8446622_RNA45SN1_45S_50ntFlanks</td><td>5475</td><td>5516</td><td>42</td><td>GCCCTGGCGGAGCGCTGAGAAGACGGTCGAACTTGACTATCT</td><td>4041.6</td><td>7.3</td><td>1.2</td><td>6.3</td><td>2.8E-10</td><td>7.5E-09</td><td>BT20/MB231</td><td>MITO</td></tr>
<tr><td>>hg38_21_+_8433172_8446622_RNA45SN1_45S_50ntFlanks@8346.8375.30|AGGGAAAGTTGAAAAGAACTTTGAAGAGAG</td><td>>hg38_21_+_8433172_8446622_RNA45SN1_45S_50ntFlanks</td><td>8346</td><td>8375</td><td>30</td><td>AGGGAAAGTTGAAAAGAACTTTGAAGAGAG</td><td>1557.7</td><td>9.4</td><td>1.5</td><td>6.2</td><td>4.2E-10</td><td>7.5E-09</td><td>BT20/MB468</td><td>MP</td></tr>
<tr><td>>hg38_21_+_8433172_8446622_RNA45SN1_45S_50ntFlanks@10787.10826.40|ACAGCCTCTGGCATGTTGGAACAATGTAGGTAAGGGAAGT</td><td>>hg38_21_+_8433172_8446622_RNA45SN1_45S_50ntFlanks</td><td>10787</td><td>10826</td><td>40</td><td>ACAGCCTCTGGCATGTTGGAACAATGTAGGTAAGGGAAGT</td><td>739.9</td><td>-8.0</td><td>1.4</td><td>-5.9</td><td>3.8E-09</td><td>7.5E-09</td><td>MB231/MB468</td><td>MITO</td></tr>
<tr><td>>hg38_21_+_8433172_8446622_RNA45SN1_45S_50ntFlanks@11576.11606.31|CTTAGAACTGGTGCGGACCAGGGGAATCCGA</td><td>>hg38_21_+_8433172_8446622_RNA45SN1_45S_50ntFlanks</td><td>11576</td><td>11606</td><td>31</td><td>CTTAGAACTGGTGCGGACCAGGGGAATCCGA</td><td>207.2</td><td>-4.9</td><td>0.8</td><td>-5.9</td><td>3.5E-09</td><td>7.5E-09</td><td>MB231/MB468</td><td>CYTO</td></tr>
<tr><td>>hg38_21_+_8433172_8446622_RNA45SN1_45S_50ntFlanks@10392.10432.41|ACAGCAGTTGAACATGGGTCAGTCGGTCCTGAGAGATGGGC</td><td>>hg38_21_+_8433172_8446622_RNA45SN1_45S_50ntFlanks</td><td>10392</td><td>10432</td><td>41</td><td>ACAGCAGTTGAACATGGGTCAGTCGGTCCTGAGAGATGGGC</td><td>452.4</td><td>-5.2</td><td>0.9</td><td>-5.9</td><td>3.5E-09</td><td>7.5E-09</td><td>MB231/MB468</td><td>NUC</td></tr>
<tr><td>>trnaMT_PheGAA_MT_+_577_647@36.62.27|AAAATGTTTAGACGGGCTCACATCACC</td><td>>trnaMT_PheGAA_MT_+_577_647</td><td>36</td><td>62</td><td>27</td><td>AAAATGTTTAGACGGGCTCACATCACC</td><td>562.1</td><td>6.7</td><td>1.1</td><td>5.9</td><td>3.8E-09</td><td>7.5E-09</td><td>MB231/MB468</td><td>MITO</td></tr>
<tr><td>>hg38_21_+_8433172_8446622_RNA45SN1_45S_50ntFlanks@10350.10367.18|ACGAGAACTTTGAAGGCC</td><td>>hg38_21_+_8433172_8446622_RNA45SN1_45S_50ntFlanks</td><td>10350</td><td>10367</td><td>18</td><td>ACGAGAACTTTGAAGGCC</td><td>384.7</td><td>-4.0</td><td>0.7</td><td>-5.9</td><td>3.8E-09</td><td>7.5E-09</td><td>MB231/MB468</td><td>MITO</td></tr>
<tr><td>>hg38_21_+_8433172_8446622_RNA45SN1_45S_50ntFlanks@3837.3871.35|CCTCTCCTACTTGGATAACTGTGGTAATTCTAGAG</td><td>>hg38_21_+_8433172_8446622_RNA45SN1_45S_50ntFlanks</td><td>3837</td><td>3871</td><td>35</td><td>CCTCTCCTACTTGGATAACTGTGGTAATTCTAGAG</td><td>520.4</td><td>-7.7</td><td>1.3</td><td>-5.9</td><td>3.8E-09</td><td>7.5E-09</td><td>MB231/MB468</td><td>MITO</td></tr>
<tr><td>>hg38_21_+_8433172_8446622_RNA45SN1_45S_50ntFlanks@4445.4471.27|CTTGCCTCTCGGCGCCCCCTCGATGCT</td><td>>hg38_21_+_8433172_8446622_RNA45SN1_45S_50ntFlanks</td><td>4445</td><td>4471</td><td>27</td><td>CTTGCCTCTCGGCGCCCCCTCGATGCT</td><td>733.3</td><td>-5.7</td><td>1.0</td><td>-5.9</td><td>3.5E-09</td><td>7.5E-09</td><td>MB231/MB468</td><td>CYTO</td></tr>
<tr><td>>hg38_21_+_8433172_8446622_RNA45SN1_45S_50ntFlanks@10683.10710.28|CCGGCGGCGTCCGGTGAGCTCTCGCTGG</td><td>>hg38_21_+_8433172_8446622_RNA45SN1_45S_50ntFlanks</td><td>10683</td><td>10710</td><td>28</td><td>CCGGCGGCGTCCGGTGAGCTCTCGCTGG</td><td>140.5</td><td>-7.4</td><td>1.3</td><td>-5.9</td><td>3.5E-09</td><td>7.5E-09</td><td>MB231/MB468</td><td>NUC</td></tr>
</table>

| | | | | | | | | | | | | | |
|---|---|---|---|---|---|---|---|---|---|---|---|---|---|
| >hg38_21_+_8433172_8446622_RNA45SN1_45S_50ntFlanks@8343.8378.36\|GTAAGGGAAAGTTGAAAAGAACTTTGAAGAGAGAGT | >hg38_21_+_8433172_8446622_RNA45SN1_45S_50ntFlanks | 8343 | 8378 | 36 | GTAAGGGAAAGTTGAAAAGAACTTTGAAGAGAGAGT | 124.6 | 10.7 | 1.7 | 6.2 | 4.2E-10 | 7.5E-09 | BT20/MB468 | MP |
| >hg38_21_+_8433172_8446622_RNA45SN1_45S_50ntFlanks@5071.5102.32\|TTAATTCCGATAACGAACGAGACTCTGGCATG | >hg38_21_+_8433172_8446622_RNA45SN1_45S_50ntFlanks | 5071 | 5102 | 32 | TTAATTCCGATAACGAACGAGACTCTGGCATG | 442.5 | -8.6 | 1.5 | -5.9 | 3.8E-09 | 7.5E-09 | MB231/MB468 | MITO |
| >MI0000477\|hsa-mir-146a&WithFlank&5\|+\|160485346\|160485456@27.47.21(+1G)&[MIMAT0000449&hsa-miR-146a-5p&offsets\|0\|-1(+1G);m-129&5\|+\|160485372\|160485393&offsets\|0\|-1(+1G)]\|TGAGAACTGAATTCCATGGGTG | >MI0000477\|hsa-mir-146a&WithFlank&5\|+\|160485346\|160485456 | 27 | 47 | 21(+1G) | TGAGAACTGAATTCCATGGGTG | 193.3 | 10.8 | 1.8 | 5.9 | 3.8E-09 | 7.5E-09 | MB231/MB468 | MITO |
| >hg38_21_+_8433172_8446622_RNA45SN1_45S_50ntFlanks@10760.10785.26\|CATATCCGCAGCAGGTCTCCAAGGTG | >hg38_21_+_8433172_8446622_RNA45SN1_45S_50ntFlanks | 10760 | 10785 | 26 | CATATCCGCAGCAGGTCTCCAAGGTG | 310.8 | -7.6 | 1.3 | -5.9 | 3.8E-09 | 7.5E-09 | MB231/MB468 | MITO |
| >hg38_21_+_8433172_8446622_RNA45SN1_45S_50ntFlanks@3996.4024.29\|ACTCTAGATAACCTCGGGCCGATCGCACG | >hg38_21_+_8433172_8446622_RNA45SN1_45S_50ntFlanks | 3996 | 4024 | 29 | ACTCTAGATAACCTCGGGCCGATCGCACG | 725.0 | -6.3 | 1.1 | -5.9 | 3.8E-09 | 7.5E-09 | MB231/MB468 | MITO |
| >trnaMT_ProTGG_MT_-_15956_16023@41.67.27\|ATGGTGGAGTTAAAGACTTTTTCTCTG | >trnaMT_ProTGG_MT_-_15956_16023 | 41 | 67 | 27 | ATGGTGGAGTTAAAGACTTTTTCTCTG | 283.0 | -6.0 | 1.0 | -5.9 | 3.3E-09 | 7.5E-09 | MB231/MB468 | MP |
| >hg38_21_+_8433172_8446622_RNA45SN1_45S_50ntFlanks@6652.6674.23\|GACTCTTAGCGGTGGATCACTCG | >hg38_21_+_8433172_8446622_RNA45SN1_45S_50ntFlanks | 6652 | 6674 | 23 | GACTCTTAGCGGTGGATCACTCG | 1481.3 | 4.9 | 0.8 | 6.0 | 2.6E-09 | 7.5E-09 | BT20/MB468 | MITO |
| >hg38_21_+_8433172_8446622_RNA45SN1_45S_50ntFlanks@10786.10820.35\|AACAGCCTCTGGCATGTTGGAACAATGTAGGTAAG | >hg38_21_+_8433172_8446622_RNA45SN1_45S_50ntFlanks | 10786 | 10820 | 35 | AACAGCCTCTGGCATGTTGGAACAATGTAGGTAAG | 468.9 | -3.3 | 0.5 | -6.0 | 2.6E-09 | 7.5E-09 | BT20/MB468 | MITO |
| >MI0000286\|hsa-mir-210&WithFlank&11\|-\|568083\|568204@72.93.22(+2A)&[MIMAT0000267&hsa-miR-210-3p&offsets\|0\|0(+2A);m-102&11\|-\|568112\|568133&offsets\|0\|0(+2A)]\|CTGTGCGTGTGACAGCGGCTGAAA | >MI0000286\|hsa-mir-210&WithFlank&11\|-\|568083\|568204 | 72 | 93 | 22(+2A) | CTGTGCGTGTGACAGCGGCTGAAA | 775.9 | 5.6 | 1.0 | 5.9 | 3.8E-09 | 7.5E-09 | MB231/MB468 | MITO |
| >hg38_21_+_8433172_8446622_RNA45SN1_45S_50ntFlanks@4560.4589.30\|CTAGGAATAATGGAATAGGACCGCGGTTCT | >hg38_21_+_8433172_8446622_RNA45SN1_45S_50ntFlanks | 4560 | 4589 | 30 | CTAGGAATAATGGAATAGGACCGCGGTTCT | 417.7 | -8.2 | 1.4 | -5.9 | 3.8E-09 | 7.5E-09 | MB231/MB468 | MITO |
| >hg38_21_+_8433172_8446622_RNA45SN1_45S_50ntFlanks@4472.4493.22\|CTTAGCTGAGTGTCCCGCGGGG | >hg38_21_+_8433172_8446622_RNA45SN1_45S_50ntFlanks | 4472 | 4493 | 22 | CTTAGCTGAGTGTCCCGCGGGG | 387.6 | -8.1 | 1.4 | -5.9 | 3.3E-09 | 7.5E-09 | MB231/MB468 | MP |
| >trna111_HisGTG_1_-_147774845_147774916@-1G.33.34,trna118_HisGTG_1_-_145396881_145396952@-1G.33.34,trna16_HisGTG_1_+_146544773_146544844@-1G.33.34,trna1_HisGTG_15_+_45493349_45493420@-1G.33.34,trna21_HisGTG_1_+_147753471_147753542@-1G.33.34,trna33_HisGTG_6_+_27125906_27125977@-1G.33.34,trna7_HisGTG_9_-_14433938_14434009@-1G.33.34,trna8_HisGTG_15_-_45492611_45492682@-1G.33.34,trna9_HisGTG_15_-_45490804_45490875@-1G.33.34\|GGCCGTGATCGTATAGTGGTTAGTACTCTGCGTT | >trna111_HisGTG_1_-_147774845_147774916 | -1G | 33 | 34 | GGCCGTGATCGTATAGTGGTTAGTACTCTGCGTT | 10059.9 | 8.7 | 1.5 | 6.0 | 2.6E-09 | 7.5E-09 | BT20/MB468 | MITO |
| >MI0001445\|hsa-mir-423&WithFlank&17\|+\|30117073\|30117178@59.80.22(+1C)&[MIMAT0001340&hsa-miR-423-3p&offsets\|0\|-1(+1C);m-100&17\|+\|30117131\|30117153&offsets\|0\|-1(+1C)]\|AGCTCGGTCTGAGGCCCCTCAGC | >MI0001445\|hsa-mir-423&WithFlank&17\|+\|30117073\|30117178 | 59 | 80 | 22(+1C) | AGCTCGGTCTGAGGCCCCTCAGC | 1521.0 | 5.6 | 0.9 | 6.2 | 4.2E-10 | 7.6E-09 | BT20/MB468 | MP |
| >MI0000294\|hsa-mir-218-1&WithFlank&4\|+\|20528269\|20528390@31.52.22;MI0000295\|hsa-mir-218-2&WithFlank&5\|-\|168768140\|168768261@31.52.22&[MIMAT0000275&hsa-miR-218-5p&offsets\|0\|+1;m-185&4\|+\|20528299\|20528319&offsets\|0\|+1];[MIMAT0000275_1&hsa-miR-218-5p&offsets\|0\|+1;m-187&5\|-\|168768211\|168768231&offsets\|0\|+1]\|TTGTGCTTGATCTAACCATGTG | >MI0000294\|hsa-mir-218-1&WithFlank&4\|+\|20528269\|20528390 | 31 | 52 | 22 | TTGTGCTTGATCTAACCATGTG | 133.0 | 10.3 | 1.7 | 5.9 | 3.8E-09 | 7.6E-09 | MB231/MB468 | MITO |
| >hg38_21_+_8433172_8446622_RNA45SN1_45S_50ntFlanks@12787.12805.19\|CCGCGCGCGCGGGAGGGCG | >hg38_21_+_8433172_8446622_RNA45SN1_45S_50ntFlanks | 12787 | 12805 | 19 | CCGCGCGCGCGGGAGGGCG | 142.3 | -10.2 | 1.6 | -6.2 | 4.2E-10 | 7.6E-09 | BT20/MB468 | MP |
| >hg38_21_+_8433172_8446622_RNA45SN1_45S_50ntFlanks@10386.10410.25\|ATGTGAACAGCAGTTGAACATGGGT | >hg38_21_+_8433172_8446622_RNA45SN1_45S_50ntFlanks | 10386 | 10410 | 25 | ATGTGAACAGCAGTTGAACATGGGT | 1176.2 | -6.4 | 1.1 | -5.9 | 3.4E-09 | 7.6E-09 | MB231/MB468 | MP |
| >hg38_21_+_8433172_8446622_RNA45SN1_45S_50ntFlanks@10738.10755.18\|AAATCTCGCGCCGGGCCG | >hg38_21_+_8433172_8446622_RNA45SN1_45S_50ntFlanks | 10738 | 10755 | 18 | AAATCTCGCGCCGGGCCG | 4957.3 | -8.1 | 1.4 | -5.9 | 4.0E-09 | 7.6E-09 | BT20/MB468 | NUC |
| >hg38_21_+_8433172_8446622_RNA45SN1_45S_50ntFlanks@12451.12476.26\|ATTCACCAAGCGTTGGATTGTTCACC | >hg38_21_+_8433172_8446622_RNA45SN1_45S_50ntFlanks | 12451 | 12476 | 26 | ATTCACCAAGCGTTGGATTGTTCACC | 231.7 | 4.9 | 0.8 | 5.9 | 3.6E-09 | 7.6E-09 | MB231/MB468 | NUC |
| >hg38_21_+_8433172_8446622_RNA45SN1_45S_50ntFlanks@10393.10420.28\|CAGCAGTTGAACATGGGTCAGTCGGTCC | >hg38_21_+_8433172_8446622_RNA45SN1_45S_50ntFlanks | 10393 | 10420 | 28 | CAGCAGTTGAACATGGGTCAGTCGGTCC | 212.7 | -4.6 | 0.8 | -5.9 | 3.5E-09 | 7.6E-09 | MB231/MB468 | CYTO |
| >hg38_21_+_8433172_8446622_RNA45SN1_45S_50ntFlanks@10781.10803.23\|AGGTGAACAGCCTCTGGCATGTT | >hg38_21_+_8433172_8446622_RNA45SN1_45S_50ntFlanks | 10781 | 10803 | 23 | AGGTGAACAGCCTCTGGCATGTT | 497.0 | -5.8 | 1.0 | -5.9 | 3.5E-09 | 7.6E-09 | MB231/MB468 | CYTO |
| >hg38_21_+_8433172_8446622_RNA45SN1_45S_50ntFlanks@10366.10388.23\|CCGAAGTGGAGAAGGGTTCCATG | >hg38_21_+_8433172_8446622_RNA45SN1_45S_50ntFlanks | 10366 | 10388 | 23 | CCGAAGTGGAGAAGGGTTCCATG | 114.7 | -9.9 | 1.6 | -6.2 | 4.3E-10 | 7.6E-09 | BT20/MB468 | MP |
| >hg38_21_+_8433172_8446622_RNA45SN1_45S_50ntFlanks@5496.5514.19\|GACGGTCGAACTTGACTAT | >hg38_21_+_8433172_8446622_RNA45SN1_45S_50ntFlanks | 5496 | 5514 | 19 | GACGGTCGAACTTGACTAT | 227.7 | -10.9 | 1.7 | -6.2 | 4.3E-10 | 7.6E-09 | BT20/MB468 | MP |
| >hg38_21_+_8433172_8446622_RNA45SN1_45S_50ntFlanks@7980.8009.30\|CCTCAGATCAGACGTGGCGACCCGCTGAAT | >hg38_21_+_8433172_8446622_RNA45SN1_45S_50ntFlanks | 7980 | 8009 | 30 | CCTCAGATCAGACGTGGCGACCCGCTGAAT | 559.7 | -4.5 | 0.8 | -5.9 | 3.4E-09 | 7.6E-09 | MB231/MB468 | MP |
| >hg38_21_+_8433172_8446622_RNA45SN1_45S_50ntFlanks@4995.5021.27\|ACAGATTGATAGCTCTTTCTCGATTCC | >hg38_21_+_8433172_8446622_RNA45SN1_45S_50ntFlanks | 4995 | 5021 | 27 | ACAGATTGATAGCTCTTTCTCGATTCC | 98.5 | -10.5 | 1.8 | -5.9 | 3.4E-09 | 7.6E-09 | MB231/MB468 | MP |
| >hg38_21_+_8433172_8446622_RNA45SN1_45S_50ntFlanks@5459.5499.41\|CCGGGGTCGGCCCACGGCCCTGGCGGAGCGCTGAGAAGACG | >hg38_21_+_8433172_8446622_RNA45SN1_45S_50ntFlanks | 5459 | 5499 | 41 | CCGGGGTCGGCCCACGGCCCTGGCGGAGCGCTGAGAAGACG | 88.0 | -10.3 | 1.7 | -5.9 | 3.4E-09 | 7.6E-09 | MB231/MB468 | MP |
| >hg38_21_+_8433172_8446622_RNA45SN1_45S_50ntFlanks@12588.12614.27\|GGTTCAGACATTTGGTGTATGTGCTTG | >hg38_21_+_8433172_8446622_RNA45SN1_45S_50ntFlanks | 12588 | 12614 | 27 | GGTTCAGACATTTGGTGTATGTGCTTG | 97.9 | -8.7 | 1.5 | -6.0 | 2.7E-09 | 7.6E-09 | BT20/MB468 | MITO |
| >hg38_21_+_8433172_8446622_RNA45SN1_45S_50ntFlanks@10345.10373.29\|TTCAAACGAGAACTTTGAAGGCCGAAGTG | >hg38_21_+_8433172_8446622_RNA45SN1_45S_50ntFlanks | 10345 | 10373 | 29 | TTCAAACGAGAACTTTGAAGGCCGAAGTG | 751.9 | 4.8 | 0.8 | 5.9 | 3.5E-09 | 7.6E-09 | MB231/MB468 | CYTO |
| >hg38_21_+_8433172_8446622_RNA45SN1_45S_50ntFlanks@5073.5098.26\|AATTCCGATAACGAACGAGACTCTGG | >hg38_21_+_8433172_8446622_RNA45SN1_45S_50ntFlanks | 5073 | 5098 | 26 | AATTCCGATAACGAACGAGACTCTGG | 2753.5 | -6.0 | 1.0 | -5.9 | 3.6E-09 | 7.6E-09 | MB231/MB468 | NUC |
| >hg38_21_+_8433172_8446622_RNA45SN1_45S_50ntFlanks@9718.9737.20\|TGATTAGAGGTCTTGGGGCC | >hg38_21_+_8433172_8446622_RNA45SN1_45S_50ntFlanks | 9718 | 9737 | 20 | TGATTAGAGGTCTTGGGGCC | 126.7 | -10.1 | 1.6 | -6.2 | 4.3E-10 | 7.6E-09 | BT20/MB468 | MP |
| >hg38_21_+_8433172_8446622_RNA45SN1_45S_50ntFlanks@9565.9597.33\|TGACGTGCAAATCGGTCGTCCGACCTGGGTATA | >hg38_21_+_8433172_8446622_RNA45SN1_45S_50ntFlanks | 9565 | 9597 | 33 | TGACGTGCAAATCGGTCGTCCGACCTGGGTATA | 8158.5 | -10.6 | 1.8 | -5.9 | 4.1E-09 | 7.7E-09 | BT20/MB468 | NUC |
| >hg38_21_+_8433172_8446622_RNA45SN1_45S_50ntFlanks@8357.8380.24\|AAAAGAACTTTGAAGAGAGAGTTC | >hg38_21_+_8433172_8446622_RNA45SN1_45S_50ntFlanks | 8357 | 8380 | 24 | AAAAGAACTTTGAAGAGAGAGTTC | 162.3 | -10.4 | 1.7 | -6.2 | 4.3E-10 | 7.7E-09 | BT20/MB468 | MP |
| >hg38_21_+_8433172_8446622_RNA45SN1_45S_50ntFlanks@5527.5549.23\|AAAGTCGTAACAAGGTTTCCGTA | >hg38_21_+_8433172_8446622_RNA45SN1_45S_50ntFlanks | 5527 | 5549 | 23 | AAAGTCGTAACAAGGTTTCCGTA | 111.1 | -8.9 | 1.5 | -5.9 | 2.7E-09 | 7.7E-09 | BT20/MB468 | MITO |
| >hg38_21_+_8433172_8446622_RNA45SN1_45S_50ntFlanks@11933.11968.36\|AATAAGTGGGAGGCCCCCGGCGCCCCCCCGGTGTCC | >hg38_21_+_8433172_8446622_RNA45SN1_45S_50ntFlanks | 11933 | 11968 | 36 | AATAAGTGGGAGGCCCCCGGCGCCCCCCCGGTGTCC | 138.9 | -10.2 | 1.6 | -6.2 | 4.3E-10 | 7.7E-09 | BT20/MB468 | MP |
| >MI0000272\|hsa-mir-182&WithFlank&7\|-\|129770377\|129770498@29.48.20(+1U)&[MIMAT0000259&hsa-miR-182-5p&offsets\|0\|-4(+1U);m-22&7\|-\|129770449\|129770470&offsets\|0\|-2(+1U)]\|TTTGGCAATGGTAGAACTCAT | >MI0000272\|hsa-mir-182&WithFlank&7\|-\|129770377\|129770498 | 29 | 48 | 20(+1U) | TTTGGCAATGGTAGAACTCAT | 190.8 | 7.4 | 1.2 | 6.4 | 2.1E-10 | 7.7E-09 | BT20/MB231 | CYTO |
| >hg38_21_+_8433172_8446622_RNA45SN1_45S_50ntFlanks@10351.10390.40\|CGAGAACTTTGAAGGCCGAAGTGGAGAAGGGTTCCATGTG | >hg38_21_+_8433172_8446622_RNA45SN1_45S_50ntFlanks | 10351 | 10390 | 40 | CGAGAACTTTGAAGGCCGAAGTGGAGAAGGGTTCCATGTG | 10157.4 | -5.0 | 0.8 | -5.9 | 2.7E-09 | 7.7E-09 | BT20/MB468 | MITO |
| >hg38_21_+_8433172_8446622_RNA45SN1_45S_50ntFlanks@4047.4065.19\|ATTCGAACGTCTGCCCTAT | >hg38_21_+_8433172_8446622_RNA45SN1_45S_50ntFlanks | 4047 | 4065 | 19 | ATTCGAACGTCTGCCCTAT | 463.6 | -6.1 | 1.0 | -5.9 | 3.6E-09 | 7.7E-09 | MB231/MB468 | NUC |
| >hg38_21_+_8433172_8446622_RNA45SN1_45S_50ntFlanks@3705.3725.21\|TACCTGGTTGATCCTGCCAGT | >hg38_21_+_8433172_8446622_RNA45SN1_45S_50ntFlanks | 3705 | 3725 | 21 | TACCTGGTTGATCCTGCCAGT | 935.8 | 3.9 | 0.7 | 5.9 | 3.6E-09 | 7.7E-09 | MB231/MB468 | CYTO |
| >hg38_21_+_8433172_8446622_RNA45SN1_45S_50ntFlanks@12619.12654.36\|AGGAGCCAATGGGGCGAAGCTACCATCTGTGGGATT | >hg38_21_+_8433172_8446622_RNA45SN1_45S_50ntFlanks | 12619 | 12654 | 36 | AGGAGCCAATGGGGCGAAGCTACCATCTGTGGGATT | 1380.5 | -4.4 | 0.7 | -5.9 | 3.6E-09 | 7.7E-09 | MB231/MB468 | CYTO |
| >MI0000114\|hsa-mir-107&WithFlank&10\|-\|89592741\|89592833@56.76.21(+1A);MI0000108\|hsa-mir-103a-2&WithFlank&20\|+\|3917488\|3917577@54.74.21(+1A);MI0000109\|hsa-mir-103a-1&WithFlank&5\|-\|168560890\|168560979@54.74.21(+1A)&[MIMAT0000104&hsa-miR-107&offsets\|0\|-2(+1A);m-78&10\|-\|89592758\|89592778&offsets\|0\|0(+1A)];[MIMAT0000101&hsa-miR-103a-3p&offsets\|0\|-2(+1A);m-12&20\|+\|3917541\|3917563&offsets\|0\|-2(+1A)];[MIMAT0000101_1&hsa-miR-103a-3p&offsets\|0\|-2(+1A);m-13&5\|-\|168560904\|168560926&offsets\|0\|-2(+1A);m-44220&5\|-\|168560908\|168560926&offsets\|0\|+2(+1A)]\|AGCAGCATTGTACAGGGCTATA | >MI0000114\|hsa-mir-107&WithFlank&10\|-\|89592741\|89592833 | 56 | 76 | 21(+1A) | AGCAGCATTGTACAGGGCTATA | 189.5 | 3.7 | 0.6 | 5.9 | 4.1E-09 | 7.7E-09 | BT20/MB468 | NUC |
| >MI0000542\|hsa-mir-320a&WithFlank&8\|-\|22244960\|22245043@48.68.21&[MIMAT0000510&hsa-miR-320a-3p&offsets\|0\|-1;m-60&8\|-\|22244975\|22244996&offsets\|0\|-1]\|AAAAGCTGGGTTGAGAGGGCG | >MI0000542\|hsa-mir-320a&WithFlank&8\|-\|22244960\|22245043 | 48 | 68 | 21 | AAAAGCTGGGTTGAGAGGGCG | 1169.4 | 2.5 | 0.4 | 5.9 | 4.1E-09 | 7.7E-09 | BT20/MB468 | NUC |
| >hg38_21_+_8433172_8446622_RNA45SN1_45S_50ntFlanks@4723.4752.30\|CATTAATCAAGAACGAAAGTCGGAGGTTCG | >hg38_21_+_8433172_8446622_RNA45SN1_45S_50ntFlanks | 4723 | 4752 | 30 | CATTAATCAAGAACGAAAGTCGGAGGTTCG | 813.8 | -4.8 | 0.8 | -5.9 | 3.1E-09 | 7.7E-09 | BT20/MB468 | CYTO |
| >hg38_21_+_8433172_8446622_RNA45SN1_45S_50ntFlanks@12407.12431.25\|TTTTGATCCTTCGATGTCGGCTCTT | >hg38_21_+_8433172_8446622_RNA45SN1_45S_50ntFlanks | 12407 | 12431 | 25 | TTTTGATCCTTCGATGTCGGCTCTT | 2337.2 | -3.9 | 0.7 | -5.9 | 3.1E-09 | 7.8E-09 | BT20/MB468 | CYTO |
| >hg38_21_+_8433172_8446622_RNA45SN1_45S_50ntFlanks@5084.5109.26\|CGAACGAGACTCTGGCATGCTAACTA | >hg38_21_+_8433172_8446622_RNA45SN1_45S_50ntFlanks | 5084 | 5109 | 26 | CGAACGAGACTCTGGCATGCTAACTA | 117.5 | -10.7 | 1.8 | -5.9 | 3.4E-09 | 7.8E-09 | MB231/MB468 | MP |
| >trna119_LysCTT_1_-_145395522_145395594@4.33.30,trna11_LysCTT_5_-_180648979_180649051@4.33.30,trna13_LysCTT_6_+_26556774_26556846@4.33.30,trna7_LysCTT_16_+_3225692_3225764@4.33.30,trna9_LysCTT_5_+_180634755_180634827@4.33.30\|CGGCTAGCTCAGTCGGTAGAGCATGAGACT | >trna119_LysCTT_1_-_145395522_145395594 | 4 | 33 | 30 | CGGCTAGCTCAGTCGGTAGAGCATGAGACT | 541.4 | 9.3 | 1.6 | 5.9 | 3.6E-09 | 7.8E-09 | MB231/MB468 | CYTO |

<table>
<tr><td>>hg38_21_+_8433172_8446622_RNA45SN1_45S_50ntFlanks@11572.11597.26|CCGACTTAGAACTGGTGCGGACCAGG</td><td>>hg38_21_+_8433172_8446622_RNA45SN1_45S_50ntFlanks</td><td>11572</td><td>11597</td><td>26</td><td>CCGACTTAGAACTGGTGCGGACCAGG</td><td>356.5</td><td>-4.1</td><td>0.7</td><td>-5.9</td><td>4.0E-09</td><td>7.8E-09</td><td>MB231/MB468</td><td>MITO</td></tr>
<tr><td>>hg38_21_+_8433172_8446622_RNA45SN1_45S_50ntFlanks@4000.4022.23|TAGATAACCTCGGGCCGATCGCA</td><td>>hg38_21_+_8433172_8446622_RNA45SN1_45S_50ntFlanks</td><td>4000</td><td>4022</td><td>23</td><td>TAGATAACCTCGGGCCGATCGCA</td><td>276.4</td><td>-6.2</td><td>1.0</td><td>-5.9</td><td>3.7E-09</td><td>7.8E-09</td><td>MB231/MB468</td><td>NUC</td></tr>
<tr><td>>hg38_21_+_8433172_8446622_RNA45SN1_45S_50ntFlanks@9554.9576.23|CGTAGCGGTCCTGACGTGCAAAT</td><td>>hg38_21_+_8433172_8446622_RNA45SN1_45S_50ntFlanks</td><td>9554</td><td>9576</td><td>23</td><td>CGTAGCGGTCCTGACGTGCAAAT</td><td>2553.7</td><td>3.4</td><td>0.6</td><td>5.9</td><td>3.6E-09</td><td>7.8E-09</td><td>MB231/MB468</td><td>CYTO</td></tr>
<tr><td>>hg38_21_+_8433172_8446622_RNA45SN1_45S_50ntFlanks@7980.7999.20|CCTCAGATCAGACGTGGCGA</td><td>>hg38_21_+_8433172_8446622_RNA45SN1_45S_50ntFlanks</td><td>7980</td><td>7999</td><td>20</td><td>CCTCAGATCAGACGTGGCGA</td><td>189.5</td><td>-6.2</td><td>1.0</td><td>-5.9</td><td>3.6E-09</td><td>7.8E-09</td><td>MB231/MB468</td><td>CYTO</td></tr>
<tr><td>>hg38_21_+_8433172_8446622_RNA45SN1_45S_50ntFlanks@11876.11898.23|AAGACCCTGTTGAGCTTGACTCT</td><td>>hg38_21_+_8433172_8446622_RNA45SN1_45S_50ntFlanks</td><td>11876</td><td>11898</td><td>23</td><td>AAGACCCTGTTGAGCTTGACTCT</td><td>1285.2</td><td>-5.0</td><td>0.8</td><td>-5.9</td><td>4.0E-09</td><td>7.9E-09</td><td>MB231/MB468</td><td>MITO</td></tr>
<tr><td>>trna10_GlyTCC_17_+_8124866_8124937@1.29.29|GCGTTGGTGGTATAGTGGTAAGCATAGCT</td><td>>trna10_GlyTCC_17_+_8124866_8124937</td><td>1</td><td>29</td><td>29</td><td>GCGTTGGTGGTATAGTGGTAAGCATAGCT</td><td>341.9</td><td>-3.9</td><td>0.7</td><td>-5.9</td><td>3.2E-09</td><td>7.9E-09</td><td>BT20/MB468</td><td>CYTO</td></tr>
<tr><td>>hg38_21_+_8433172_8446622_RNA45SN1_45S_50ntFlanks@3705.3747.43|TACCTGGTTGATCCTGCCAGTAGCATATGCTTGTCTCAAAGAT</td><td>>hg38_21_+_8433172_8446622_RNA45SN1_45S_50ntFlanks</td><td>3705</td><td>3747</td><td>43</td><td>TACCTGGTTGATCCTGCCAGTAGCATATGCTTGTCTCAAAGAT</td><td>964.0</td><td>8.1</td><td>1.4</td><td>5.9</td><td>3.7E-09</td><td>7.9E-09</td><td>MB231/MB468</td><td>CYTO</td></tr>
<tr><td>>hg38_21_+_8433172_8446622_RNA45SN1_45S_50ntFlanks@9454.9473.20|CCGCGCCGGGGAGGTGGAGC</td><td>>hg38_21_+_8433172_8446622_RNA45SN1_45S_50ntFlanks</td><td>9454</td><td>9473</td><td>20</td><td>CCGCGCCGGGGAGGTGGAGC</td><td>78.6</td><td>-10.2</td><td>1.7</td><td>-5.9</td><td>3.5E-09</td><td>7.9E-09</td><td>MB231/MB468</td><td>MP</td></tr>
<tr><td>>MI0000103|hsa-mir-101-1&WithFlank&1|-|65058428|65058514@52.73.22;MI0000739|hsa-mir-101-2&WithFlank&9|+|4850291|4850381@54.75.22&[MIMAT0000099&hsa-miR-101-3p&offsets|-1|0;m-17&1|-|65058443|65058462&offsets|-1|+1];[MIMAT0000099_1&hsa-miR-101-3p&offsets|-1|0;m-16&9|+|4850345|4850364&offsets|-1|+1]|GTACAGTACTGTGATAACTGAA</td><td>>MI0000103|hsa-mir-101-1&WithFlank&1|-|65058428|65058514</td><td>52</td><td>73</td><td>22</td><td>GTACAGTACTGTGATAACTGAA</td><td>744.2</td><td>5.0</td><td>0.8</td><td>5.9</td><td>3.2E-09</td><td>7.9E-09</td><td>BT20/MB468</td><td>CYTO</td></tr>
<tr><td>>hg38_21_+_8433172_8446622_RNA45SN1_45S_50ntFlanks@9941.9980.40|AGGACGGTGGCCATGGAAGTCGGAATCCGCTAAGGAGTGT</td><td>>hg38_21_+_8433172_8446622_RNA45SN1_45S_50ntFlanks</td><td>9941</td><td>9980</td><td>40</td><td>AGGACGGTGGCCATGGAAGTCGGAATCCGCTAAGGAGTGT</td><td>414.1</td><td>12.4</td><td>2.0</td><td>6.2</td><td>4.5E-10</td><td>7.9E-09</td><td>BT20/MB468</td><td>MP</td></tr>
<tr><td>>hg38_21_+_8433172_8446622_RNA45SN1_45S_50ntFlanks@10805.10825.21|GAACAATGTAGGTAAGGGAAG</td><td>>hg38_21_+_8433172_8446622_RNA45SN1_45S_50ntFlanks</td><td>10805</td><td>10825</td><td>21</td><td>GAACAATGTAGGTAAGGGAAG</td><td>113.9</td><td>-9.9</td><td>1.6</td><td>-6.2</td><td>4.5E-10</td><td>7.9E-09</td><td>BT20/MB468</td><td>MP</td></tr>
<tr><td>>hg38_21_+_8433172_8446622_RNA45SN1_45S_50ntFlanks@4122.4158.37|AGGGTTCGATTCCGGAGAGGGAGCCTGAGAAACGGCT</td><td>>hg38_21_+_8433172_8446622_RNA45SN1_45S_50ntFlanks</td><td>4122</td><td>4158</td><td>37</td><td>AGGGTTCGATTCCGGAGAGGGAGCCTGAGAAACGGCT</td><td>86.0</td><td>-10.3</td><td>1.7</td><td>-5.9</td><td>3.5E-09</td><td>7.9E-09</td><td>MB231/MB468</td><td>MP</td></tr>
<tr><td>>hg38_21_+_8433172_8446622_RNA45SN1_45S_50ntFlanks@9806.9824.19|CGTGGAATGCGAGTGCCTA</td><td>>hg38_21_+_8433172_8446622_RNA45SN1_45S_50ntFlanks</td><td>9806</td><td>9824</td><td>19</td><td>CGTGGAATGCGAGTGCCTA</td><td>91.2</td><td>-10.4</td><td>1.8</td><td>-5.9</td><td>3.5E-09</td><td>7.9E-09</td><td>MB231/MB468</td><td>MP</td></tr>
<tr><td>>hg38_21_+_8433172_8446622_RNA45SN1_45S_50ntFlanks@11879.11904.26|ACCCTGTTGAGCTTGACTCTAGTCTG</td><td>>hg38_21_+_8433172_8446622_RNA45SN1_45S_50ntFlanks</td><td>11879</td><td>11904</td><td>26</td><td>ACCCTGTTGAGCTTGACTCTAGTCTG</td><td>407.0</td><td>-8.3</td><td>1.4</td><td>-5.9</td><td>4.2E-09</td><td>7.9E-09</td><td>BT20/MB468</td><td>NUC</td></tr>
<tr><td>>hg38_21_+_8433172_8446622_RNA45SN1_45S_50ntFlanks@5083.5105.23|ACGAACGAGACTCTGGCATGCTA</td><td>>hg38_21_+_8433172_8446622_RNA45SN1_45S_50ntFlanks</td><td>5083</td><td>5105</td><td>23</td><td>ACGAACGAGACTCTGGCATGCTA</td><td>431.3</td><td>-8.5</td><td>1.4</td><td>-5.9</td><td>3.5E-09</td><td>7.9E-09</td><td>MB231/MB468</td><td>MP</td></tr>
<tr><td>>MI0000478|hsa-mir-149&WithFlank&2|+|240455995|240456095@21.42.22(+1A)&[MIMAT0000450&hsa-miR-149-5p&offsets|0|-1(+1A);m-195&2|+|240456015|240456037&offsets|0|-1(+1A)]|TCTGGCTCCGTGTCTTCACTCCA</td><td>>MI0000478|hsa-mir-149&WithFlank&2|+|240455995|240456095</td><td>21</td><td>42</td><td>22(+1A)</td><td>TCTGGCTCCGTGTCTTCACTCCA</td><td>116.5</td><td>10.6</td><td>1.7</td><td>6.2</td><td>4.5E-10</td><td>7.9E-09</td><td>BT20/MB468</td><td>MP</td></tr>
<tr><td>>trna111_HisGTG_1_-_147774845_147774916@-1G.36.37,trna118_HisGTG_1_-_145396881_145396952@-1G.36.37,trna16_HisGTG_1_+_146544773_146544844@-1G.36.37,trna1_HisGTG_15_+_45493349_45493420@-1G.36.37,trna21_HisGTG_1_+_147753471_147753542@-1G.36.37,trna33_HisGTG_6_+_27125906_27125977@-1G.36.37,trna7_HisGTG_9_-_14433938_14434009@-1G.36.37,trna8_HisGTG_15_-_45492611_45492682@-1G.36.37,trna9_HisGTG_15_-_45490804_45490875@-1G.36.37|GGCCGTGATCGTATAGTGGTTAGTACTCTGCGTTGTG</td><td>>trna111_HisGTG_1_-_147774845_147774916</td><td>-1G</td><td>36</td><td>37</td><td>GGCCGTGATCGTATAGTGGTTAGTACTCTGCGTTGTG</td><td>450.7</td><td>8.3</td><td>1.4</td><td>5.9</td><td>3.7E-09</td><td>8.0E-09</td><td>MB231/MB468</td><td>CYTO</td></tr>
<tr><td>>hg38_21_+_8433172_8446622_RNA45SN1_45S_50ntFlanks@7985.8005.21|GATCAGACGTGGCGACCCGCT</td><td>>hg38_21_+_8433172_8446622_RNA45SN1_45S_50ntFlanks</td><td>7985</td><td>8005</td><td>21</td><td>GATCAGACGTGGCGACCCGCT</td><td>787.1</td><td>-8.0</td><td>1.4</td><td>-5.9</td><td>4.0E-09</td><td>8.0E-09</td><td>MB231/MB468</td><td>MITO</td></tr>
<tr><td>>hg38_21_+_8433172_8446622_RNA45SN1_45S_50ntFlanks@12623.12661.39|GCCAATGGGGCGAAGCTACCATCTGTGGGATTATGACTG</td><td>>hg38_21_+_8433172_8446622_RNA45SN1_45S_50ntFlanks</td><td>12623</td><td>12661</td><td>39</td><td>GCCAATGGGGCGAAGCTACCATCTGTGGGATTATGACTG</td><td>122.1</td><td>-10.8</td><td>1.8</td><td>-5.9</td><td>3.5E-09</td><td>8.0E-09</td><td>MB231/MB468</td><td>MP</td></tr>
<tr><td>>hg38_21_+_8433172_8446622_RNA45SN1_45S_50ntFlanks@475.504.30|CTTCGTGATCGATGTGGTGACGTCGTGCTC</td><td>>hg38_21_+_8433172_8446622_RNA45SN1_45S_50ntFlanks</td><td>475</td><td>504</td><td>30</td><td>CTTCGTGATCGATGTGGTGACGTCGTGCTC</td><td>1557.4</td><td>6.0</td><td>1.0</td><td>6.2</td><td>4.5E-10</td><td>8.0E-09</td><td>BT20/MB468</td><td>MP</td></tr>
<tr><td>>hg38_21_+_8433172_8446622_RNA45SN1_45S_50ntFlanks@8176.8220.45|CTTCTGATCGAGGCCCAGCCCGTGGACGGTGTGAGGCCGGTAGCG</td><td>>hg38_21_+_8433172_8446622_RNA45SN1_45S_50ntFlanks</td><td>8176</td><td>8220</td><td>45</td><td>CTTCTGATCGAGGCCCAGCCCGTGGACGGTGTGAGGCCGGTAGCG</td><td>974.9</td><td>-9.5</td><td>1.6</td><td>-5.9</td><td>4.2E-09</td><td>8.0E-09</td><td>BT20/MB468</td><td>NUC</td></tr>
<tr><td>>hg38_21_+_8433172_8446622_RNA45SN1_45S_50ntFlanks@11641.11674.34|CGGCGGGTGTTGACGCGATGTGATTTCTGCCCAG</td><td>>hg38_21_+_8433172_8446622_RNA45SN1_45S_50ntFlanks</td><td>11641</td><td>11674</td><td>34</td><td>CGGCGGGTGTTGACGCGATGTGATTTCTGCCCAG</td><td>117.3</td><td>-9.0</td><td>1.5</td><td>-5.9</td><td>2.8E-09</td><td>8.0E-09</td><td>BT20/MB468</td><td>MITO</td></tr>
<tr><td>>hg38_21_+_8433172_8446622_RNA45SN1_45S_50ntFlanks@9534.9573.40|AGGAAACTCTGGTGGAGGTCCGTAGCGGTCCTGACGTGCA</td><td>>hg38_21_+_8433172_8446622_RNA45SN1_45S_50ntFlanks</td><td>9534</td><td>9573</td><td>40</td><td>AGGAAACTCTGGTGGAGGTCCGTAGCGGTCCTGACGTGCA</td><td>1346.7</td><td>-6.8</td><td>1.2</td><td>-5.9</td><td>4.1E-09</td><td>8.0E-09</td><td>MB231/MB468</td><td>MITO</td></tr>
<tr><td>>hg38_21_+_8433172_8446622_RNA45SN1_45S_50ntFlanks@10682.10710.29|TCCGGCGGCGTCCGGTGAGCTCTCGCTGG</td><td>>hg38_21_+_8433172_8446622_RNA45SN1_45S_50ntFlanks</td><td>10682</td><td>10710</td><td>29</td><td>TCCGGCGGCGTCCGGTGAGCTCTCGCTGG</td><td>315.0</td><td>-7.6</td><td>1.3</td><td>-5.9</td><td>4.1E-09</td><td>8.0E-09</td><td>MB231/MB468</td><td>MITO</td></tr>
<tr><td>>MI0000079|hsa-mir-23a&WithFlank&19|-|13836581|13836665@52.74.23&[MIMAT0000078&hsa-miR-23a-3p&offsets|+1|+3;m-31&19|-|13836595|13836615&offsets|+1|+3]|TCACATTGCCAGGGATTTCCAAC</td><td>>MI0000079|hsa-mir-23a&WithFlank&19|-|13836581|13836665</td><td>52</td><td>74</td><td>23</td><td>TCACATTGCCAGGGATTTCCAAC</td><td>407.9</td><td>6.7</td><td>1.1</td><td>5.9</td><td>3.7E-09</td><td>8.0E-09</td><td>MB231/MB468</td><td>CYTO</td></tr>
<tr><td>>hg38_21_+_8433172_8446622_RNA45SN1_45S_50ntFlanks@7990.8011.22|GACGTGGCGACCCGCTGAATTT</td><td>>hg38_21_+_8433172_8446622_RNA45SN1_45S_50ntFlanks</td><td>7990</td><td>8011</td><td>22</td><td>GACGTGGCGACCCGCTGAATTT</td><td>840.5</td><td>4.2</td><td>0.7</td><td>5.9</td><td>3.8E-09</td><td>8.0E-09</td><td>MB231/MB468</td><td>NUC</td></tr>
<tr><td>>hg38_21_+_8433172_8446622_RNA45SN1_45S_50ntFlanks@12456.12474.19|CCAAGCGTTGGATTGTTCA</td><td>>hg38_21_+_8433172_8446622_RNA45SN1_45S_50ntFlanks</td><td>12456</td><td>12474</td><td>19</td><td>CCAAGCGTTGGATTGTTCA</td><td>1548.3</td><td>-7.4</td><td>1.3</td><td>-5.9</td><td>3.8E-09</td><td>8.0E-09</td><td>MB231/MB468</td><td>NUC</td></tr>
<tr><td>>hg38_21_+_8433172_8446622_RNA45SN1_45S_50ntFlanks@10316.10341.26|GGTGCAGATCTTGGTGGTAGTAGCAA</td><td>>hg38_21_+_8433172_8446622_RNA45SN1_45S_50ntFlanks</td><td>10316</td><td>10341</td><td>26</td><td>GGTGCAGATCTTGGTGGTAGTAGCAA</td><td>78.6</td><td>-10.2</td><td>1.7</td><td>-5.9</td><td>3.5E-09</td><td>8.0E-09</td><td>MB231/MB468</td><td>MP</td></tr>
<tr><td>>hg38_21_+_8433172_8446622_RNA45SN1_45S_50ntFlanks@10389.10410.22|TGAACAGCAGTTGAACATGGGT</td><td>>hg38_21_+_8433172_8446622_RNA45SN1_45S_50ntFlanks</td><td>10389</td><td>10410</td><td>22</td><td>TGAACAGCAGTTGAACATGGGT</td><td>618.6</td><td>-7.9</td><td>1.3</td><td>-5.9</td><td>4.1E-09</td><td>8.0E-09</td><td>MB231/MB468</td><td>MITO</td></tr>
<tr><td>>hg38_21_+_8433172_8446622_RNA45SN1_45S_50ntFlanks@6646.6683.38|TCGTACGACTCTTAGCGGTGGATCACTCGGCTCGTGCG</td><td>>hg38_21_+_8433172_8446622_RNA45SN1_45S_50ntFlanks</td><td>6646</td><td>6683</td><td>38</td><td>TCGTACGACTCTTAGCGGTGGATCACTCGGCTCGTGCG</td><td>1392.7</td><td>7.4</td><td>1.3</td><td>5.9</td><td>3.2E-09</td><td>8.0E-09</td><td>BT20/MB468</td><td>CYTO</td></tr>
<tr><td>>hg38_21_+_8433172_8446622_RNA45SN1_45S_50ntFlanks@476.508.33|TTCGTGATCGATGTGGTGACGTCGTGCTCTCCC</td><td>>hg38_21_+_8433172_8446622_RNA45SN1_45S_50ntFlanks</td><td>476</td><td>508</td><td>33</td><td>TTCGTGATCGATGTGGTGACGTCGTGCTCTCCC</td><td>364.0</td><td>6.9</td><td>1.2</td><td>5.9</td><td>2.8E-09</td><td>8.1E-09</td><td>BT20/MB468</td><td>MITO</td></tr>
<tr><td>>hg38_21_+_8433172_8446622_RNA45SN1_45S_50ntFlanks@11573.11605.33|CGACTTAGAACTGGTGCGGACCAGGGGAATCCG</td><td>>hg38_21_+_8433172_8446622_RNA45SN1_45S_50ntFlanks</td><td>11573</td><td>11605</td><td>33</td><td>CGACTTAGAACTGGTGCGGACCAGGGGAATCCG</td><td>1383.6</td><td>-9.3</td><td>1.6</td><td>-5.9</td><td>4.3E-09</td><td>8.1E-09</td><td>BT20/MB468</td><td>NUC</td></tr>
<tr><td>>hg38_21_+_8433172_8446622_RNA45SN1_45S_50ntFlanks@12308.12325.18|CGATCCTTCTGACCTTTT</td><td>>hg38_21_+_8433172_8446622_RNA45SN1_45S_50ntFlanks</td><td>12308</td><td>12325</td><td>18</td><td>CGATCCTTCTGACCTTTT</td><td>143.6</td><td>-9.3</td><td>1.6</td><td>-5.9</td><td>2.8E-09</td><td>8.1E-09</td><td>BT20/MB468</td><td>MITO</td></tr>
<tr><td>>hg38_21_+_8433172_8446622_RNA45SN1_45S_50ntFlanks@8839.8864.26|CGGTGGCGGCGGCGGCGGCGGCGGCG</td><td>>hg38_21_+_8433172_8446622_RNA45SN1_45S_50ntFlanks</td><td>8839</td><td>8864</td><td>26</td><td>CGGTGGCGGCGGCGGCGGCGGCGGCG</td><td>179.0</td><td>-10.6</td><td>1.7</td><td>-6.2</td><td>4.6E-10</td><td>8.1E-09</td><td>BT20/MB468</td><td>MP</td></tr>
<tr><td>>hg38_21_+_8433172_8446622_RNA45SN1_45S_50ntFlanks@10411.10433.23|CAGTCGGTCCTGAGAGATGGGCG</td><td>>hg38_21_+_8433172_8446622_RNA45SN1_45S_50ntFlanks</td><td>10411</td><td>10433</td><td>23</td><td>CAGTCGGTCCTGAGAGATGGGCG</td><td>1127.0</td><td>-7.8</td><td>1.3</td><td>-5.9</td><td>3.2E-09</td><td>8.1E-09</td><td>BT20/MB468</td><td>CYTO</td></tr>
<tr><td>>hg38_21_+_8433172_8446622_RNA45SN1_45S_50ntFlanks@12939.12972.34|CCATTCGTAGACGACCTGCTTCTGGGTCGGGGTT</td><td>>hg38_21_+_8433172_8446622_RNA45SN1_45S_50ntFlanks</td><td>12939</td><td>12972</td><td>34</td><td>CCATTCGTAGACGACCTGCTTCTGGGTCGGGGTT</td><td>186.2</td><td>-7.2</td><td>1.2</td><td>-5.9</td><td>3.3E-09</td><td>8.1E-09</td><td>BT20/MB468</td><td>CYTO</td></tr>
<tr><td>>hg38_21_+_8433172_8446622_RNA45SN1_45S_50ntFlanks@4738.4761.24|AAAGTCGGAGGTTCGAAGACGATC</td><td>>hg38_21_+_8433172_8446622_RNA45SN1_45S_50ntFlanks</td><td>4738</td><td>4761</td><td>24</td><td>AAAGTCGGAGGTTCGAAGACGATC</td><td>1229.7</td><td>4.2</td><td>0.7</td><td>5.9</td><td>3.3E-09</td><td>8.1E-09</td><td>BT20/MB468</td><td>CYTO</td></tr>
<tr><td>>hg38_21_+_8433172_8446622_RNA45SN1_45S_50ntFlanks@9736.9757.22|CCGAAACGATCTCAACCTATTC</td><td>>hg38_21_+_8433172_8446622_RNA45SN1_45S_50ntFlanks</td><td>9736</td><td>9757</td><td>22</td><td>CCGAAACGATCTCAACCTATTC</td><td>166.5</td><td>-6.4</td><td>1.1</td><td>-5.9</td><td>3.3E-09</td><td>8.1E-09</td><td>BT20/MB468</td><td>CYTO</td></tr>
<tr><td>>hg38_MT_+_598_1651_RNR1_12S_50ntFlanks@916.946.31|AAGGACATTTAACTAAAACCCCTACGCATTT</td><td>>hg38_MT_+_598_1651_RNR1_12S_50ntFlanks</td><td>916</td><td>946</td><td>31</td><td>AAGGACATTTAACTAAAACCCCTACGCATTT</td><td>1160.7</td><td>8.9</td><td>1.5</td><td>5.9</td><td>2.8E-09</td><td>8.1E-09</td><td>BT20/MB468</td><td>MITO</td></tr>
<tr><td>>MI0000446|hsa-mir-125b-1&WithFlank&11|-|122099751|122099850@61.81.21&[MIMAT0004592&hsa-miR-125b-1-3p&offsets|0|-1;m-485&11|-|122099769|122099790&offsets|0|-1]|ACGGGTTAGGCTCTTGGGAGC</td><td>>MI0000446|hsa-mir-125b-1&WithFlank&11|-|122099751|122099850</td><td>61</td><td>81</td><td>21</td><td>ACGGGTTAGGCTCTTGGGAGC</td><td>114.1</td><td>10.1</td><td>1.7</td><td>5.9</td><td>4.1E-09</td><td>8.1E-09</td><td>MB231/MB468</td><td>MITO</td></tr>
<tr><td>>hg38_21_+_8433172_8446622_RNA45SN1_45S_50ntFlanks@9575.9597.23|ATCGGTCGTCCGACCTGGGTATA</td><td>>hg38_21_+_8433172_8446622_RNA45SN1_45S_50ntFlanks</td><td>9575</td><td>9597</td><td>23</td><td>ATCGGTCGTCCGACCTGGGTATA</td><td>7046.3</td><td>-9.2</td><td>1.6</td><td>-5.9</td><td>4.1E-09</td><td>8.1E-09</td><td>MB231/MB468</td><td>MITO</td></tr>
<tr><td>>trnaMT_GluTTC_MT_-_14674_14742@45.72.28,trnalookalike8_GluTTC_5_-_93905172_93905240@45.72.28|CGTGGTTGTAGTCCGTGCGAGAATACCA</td><td>>trnaMT_GluTTC_MT_-_14674_14742</td><td>45</td><td>72</td><td>28</td><td>CGTGGTTGTAGTCCGTGCGAGAATACCA</td><td>1215.1</td><td>8.4</td><td>1.4</td><td>5.9</td><td>3.8E-09</td><td>8.1E-09</td><td>MB231/MB468</td><td>CYTO</td></tr>
<tr><td>>MI0000750|hsa-mir-26a-2&WithFlank&12|-|57824603|57824698@20.39.20;MI0000083|hsa-mir-26a-1&WithFlank&3|+|37969398|37969486@16.35.20&[MIMAT0000082&hsa-miR-26a-5p&offsets|0|-2;m-33&12|-|57824658|57824679&offsets|0|-2];[MIMAT0000082_1&hsa-miR-26a-5p&offsets|0|-2;m-34&3|+|37969413|37969434&offsets|0|-2]|TTCAAGTAATCCAGGATAGG</td><td>>MI0000750|hsa-mir-26a-2&WithFlank&12|-|57824603|57824698</td><td>20</td><td>39</td><td>20</td><td>TTCAAGTAATCCAGGATAGG</td><td>458.1</td><td>5.8</td><td>1.0</td><td>5.9</td><td>4.1E-09</td><td>8.1E-09</td><td>MB231/MB468</td><td>MITO</td></tr>
<tr><td>>hg38_21_+_8433172_8446622_RNA45SN1_45S_50ntFlanks@12252.12284.33|AAGCTCGCTTGATCTTGATTTTCAGTACGAATA</td><td>>hg38_21_+_8433172_8446622_RNA45SN1_45S_50ntFlanks</td><td>12252</td><td>12284</td><td>33</td><td>AAGCTCGCTTGATCTTGATTTTCAGTACGAATA</td><td>692.3</td><td>-8.4</td><td>1.4</td><td>-5.9</td><td>4.1E-09</td><td>8.1E-09</td><td>MB231/MB468</td><td>MITO</td></tr>
<tr><td>>hg38_21_+_8433172_8446622_RNA45SN1_45S_50ntFlanks@4975.5000.26|CCCGGACACGGACAGGATTGACAGAT</td><td>>hg38_21_+_8433172_8446622_RNA45SN1_45S_50ntFlanks</td><td>4975</td><td>5000</td><td>26</td><td>CCCGGACACGGACAGGATTGACAGAT</td><td>146.0</td><td>-10.3</td><td>1.6</td><td>-6.2</td><td>4.6E-10</td><td>8.1E-09</td><td>BT20/MB468</td><td>MP</td></tr>
<tr><td>>trnaMT_HisGTG_MT_+_12138_12206@19.55.37|AAACATCAGATTGTGAATCTGACAACAGAGGCTTACG</td><td>>trnaMT_HisGTG_MT_+_12138_12206</td><td>19</td><td>55</td><td>37</td><td>AAACATCAGATTGTGAATCTGACAACAGAGGCTTACG</td><td>70.4</td><td>9.4</td><td>1.6</td><td>5.9</td><td>3.6E-09</td><td>8.1E-09</td><td>MB231/MB468</td><td>MP</td></tr>
<tr><td>>hg38_21_+_8433172_8446622_RNA45SN1_45S_50ntFlanks@12405.12440.36|CTTTTTGATCCTTCGATGTCGGCTCTTCCTATCATT</td><td>>hg38_21_+_8433172_8446622_RNA45SN1_45S_50ntFlanks</td><td>12405</td><td>12440</td><td>36</td><td>CTTTTTGATCCTTCGATGTCGGCTCTTCCTATCATT</td><td>78.4</td><td>-10.2</td><td>1.7</td><td>-5.9</td><td>3.6E-09</td><td>8.1E-09</td><td>MB231/MB468</td><td>MP</td></tr>
</table>

| | | | | | | | | | | | | | |
|---|---|---|---|---|---|---|---|---|---|---|---|---|---|
| >hg38_21_+_8433172_8446622_RNA45SN1_45S_50ntFlanks@4726.4759.34\|TAATCAAGAACGAAAGTCGGAGGTTCGAAGACGA | >hg38_21_+_8433172_8446622_RNA45SN1_45S_50ntFlanks | 4726 | 4759 | 34 | TAATCAAGAACGAAAGTCGGAGGTTCGAAGACGA | 1905.3 | -6.8 | 1.2 | -5.9 | 3.8E-09 | 8.1E-09 | MB231/MB468 | NUC |
| >hg38_21_+_8433172_8446622_RNA45SN1_45S_50ntFlanks@9033.9060.28\|CTCGTGGGGGGGCCGGGCCACCCCTCCC | >hg38_21_+_8433172_8446622_RNA45SN1_45S_50ntFlanks | 9033 | 9060 | 28 | CTCGTGGGGGGGCCGGGCCACCCCTCCC | 164.7 | -10.4 | 1.7 | -6.2 | 4.6E-10 | 8.2E-09 | BT20/MB468 | MP |
| >trna111_HisGTG_1_-_147774845_147774916@-1G.34.35,trna118_HisGTG_1_-_145396881_145396952@-1G.34.35,trna16_HisGTG_1_+_146544773_146544844@-1G.34.35,trna1_HisGTG_15_+_45493349_45493420@-1G.34.35,trna21_HisGTG_1_+_147753471_147753542@-1G.34.35,trna33_HisGTG_6_+_27125906_27125977@-1G.34.35,trna7_HisGTG_9_-_14433938_14434009@-1G.34.35,trna8_HisGTG_15_-_45492611_45492682@-1G.34.35,trna9_HisGTG_15_-_45490804_45490875@-1G.34.35\|GGCCGTGATCGTATAGTGGTTAGTACTCTGCGTTG | >trna111_HisGTG_1_-_147774845_147774916 | -1G | 34 | 35 | GGCCGTGATCGTATAGTGGTTAGTACTCTGCGTTG | 2946.7 | 8.7 | 1.5 | 5.9 | 3.3E-09 | 8.2E-09 | BT20/MB468 | CYTO |
| >hg38_21_+_8433172_8446622_RNA45SN1_45S_50ntFlanks@7991.8012.22\|ACGTGGCGACCCGCTGAATTTA | >hg38_21_+_8433172_8446622_RNA45SN1_45S_50ntFlanks | 7991 | 8012 | 22 | ACGTGGCGACCCGCTGAATTTA | 1314.4 | -3.0 | 0.5 | -5.9 | 3.8E-09 | 8.2E-09 | MB231/MB468 | NUC |
| >hg38_21_+_8433172_8446622_RNA45SN1_45S_50ntFlanks@8189.8212.24\|CCCAGCCCGTGGACGGTGTGAGGC | >hg38_21_+_8433172_8446622_RNA45SN1_45S_50ntFlanks | 8189 | 8212 | 24 | CCCAGCCCGTGGACGGTGTGAGGC | 712.7 | -8.3 | 1.3 | -6.2 | 4.7E-10 | 8.2E-09 | BT20/MB468 | MP |
| >hg38_21_+_8433172_8446622_RNA45SN1_45S_50ntFlanks@11787.11805.19\|AATGGATGAACGAGATTCC | >hg38_21_+_8433172_8446622_RNA45SN1_45S_50ntFlanks | 11787 | 11805 | 19 | AATGGATGAACGAGATTCC | 125.8 | -10.1 | 1.6 | -6.2 | 4.7E-10 | 8.2E-09 | BT20/MB468 | MP |
| >hg38_21_+_8433172_8446622_RNA45SN1_45S_50ntFlanks@4283.4316.34\|CTTTAACGAGGATCCATTGGAGGGCAAGTCTGGT | >hg38_21_+_8433172_8446622_RNA45SN1_45S_50ntFlanks | 4283 | 4316 | 34 | CTTTAACGAGGATCCATTGGAGGGCAAGTCTGGT | 256.8 | -7.1 | 1.2 | -5.9 | 3.9E-09 | 8.2E-09 | MB231/MB468 | NUC |
| >hg38_21_+_8433172_8446622_RNA45SN1_45S_50ntFlanks@8351.8391.41\|AAGTTGAAAAGAACTTTGAAGAGAGAGTTCAAGAGGGCGTG | >hg38_21_+_8433172_8446622_RNA45SN1_45S_50ntFlanks | 8351 | 8391 | 41 | AAGTTGAAAAGAACTTTGAAGAGAGAGTTCAAGAGGGCGTG | 104.7 | 10.5 | 1.7 | 6.2 | 4.7E-10 | 8.2E-09 | BT20/MB468 | MP |
| >hg38_21_+_8433172_8446622_RNA45SN1_45S_50ntFlanks@12912.12935.24\|CACGTTCGTGGGGAACCTGGCGCT | >hg38_21_+_8433172_8446622_RNA45SN1_45S_50ntFlanks | 12912 | 12935 | 24 | CACGTTCGTGGGGAACCTGGCGCT | 532.8 | -8.4 | 1.4 | -5.9 | 3.6E-09 | 8.2E-09 | MB231/MB468 | MP |
| >hg38_21_+_8433172_8446622_RNA45SN1_45S_50ntFlanks@10458.10484.27\|ATGGCCTCCGTTGCCCTCGGCCGATCG | >hg38_21_+_8433172_8446622_RNA45SN1_45S_50ntFlanks | 10458 | 10484 | 27 | ATGGCCTCCGTTGCCCTCGGCCGATCG | 201.2 | -10.7 | 1.7 | -6.2 | 4.7E-10 | 8.2E-09 | BT20/MB468 | MP |
| >trna10_GlyTCC_17_+_8124866_8124937@31.70.40,trna117_GlyTCC_1_-_145397864_145397935@31.70.40,trna2_GlyTCC_19_+_4724082_4724153@31.70.40,trna45_GlyTCC_1_+_161500903_161500974@31.70.40,trna70_GlyTCC_1_-_161439547_161439618@31.70.40,trna73_GlyTCC_1_-_161432166_161432237@31.70.40,trna76_GlyTCC_1_-_161424756_161424827@31.70.40,trna79_GlyTCC_1_-_161417375_161417446@31.70.40\|CCTTCCAAGCAGTTGACCCGGGTTCGATTCCCGGCCAACG | >trna10_GlyTCC_17_+_8124866_8124937 | 31 | 70 | 40 | CCTTCCAAGCAGTTGACCCGGGTTCGATTCCCGGCCAACG | 164.0 | -9.7 | 1.6 | -5.9 | 3.3E-09 | 8.2E-09 | BT20/MB468 | CYTO |
| >hg38_21_+_8433172_8446622_RNA45SN1_45S_50ntFlanks@3909.3932.24\|GGGGATGCGTGCATTTATCAGATC | >hg38_21_+_8433172_8446622_RNA45SN1_45S_50ntFlanks | 3909 | 3932 | 24 | GGGGATGCGTGCATTTATCAGATC | 178.0 | -11.3 | 1.9 | -5.9 | 3.6E-09 | 8.2E-09 | MB231/MB468 | MP |
| >hg38_21_+_8433172_8446622_RNA45SN1_45S_50ntFlanks@9532.9572.41\|AGAGGAAACTCTGGTGGAGGTCCGTAGCGGTCCTGACGTGC | >hg38_21_+_8433172_8446622_RNA45SN1_45S_50ntFlanks | 9532 | 9572 | 41 | AGAGGAAACTCTGGTGGAGGTCCGTAGCGGTCCTGACGTGC | 132.9 | 10.8 | 1.7 | 6.2 | 4.7E-10 | 8.2E-09 | BT20/MB468 | MP |
| >hg38_21_+_8433172_8446622_RNA45SN1_45S_50ntFlanks@10869.10890.22\|AGGGCTGGGTCGGTCGGGCTGG | >hg38_21_+_8433172_8446622_RNA45SN1_45S_50ntFlanks | 10869 | 10890 | 22 | AGGGCTGGGTCGGTCGGGCTGG | 1132.8 | -5.3 | 0.9 | -5.9 | 3.8E-09 | 8.2E-09 | MB231/MB468 | CYTO |
| >hg38_21_+_8433172_8446622_RNA45SN1_45S_50ntFlanks@10477.10497.21\|GCCGATCGAAAGGGAGTCGGG | >hg38_21_+_8433172_8446622_RNA45SN1_45S_50ntFlanks | 10477 | 10497 | 21 | GCCGATCGAAAGGGAGTCGGG | 89.2 | -10.3 | 1.8 | -5.9 | 3.7E-09 | 8.2E-09 | MB231/MB468 | MP |
| >MI0000090\|hsa-mir-32&WithFlank&9\|-\|109046223\|109046304@53.73.21&[MIMAT0004505&hsa-miR-32-3p&offsets\|0\|-1;m-518&9\|-\|109046232\|109046252&offsets\|0\|0]\|CAATTTAGTGTGTGTGATATT | >MI0000090\|hsa-mir-32&WithFlank&9\|-\|109046223\|109046304 | 53 | 73 | 21 | CAATTTAGTGTGTGTGATATT | 128.4 | 10.2 | 1.7 | 5.9 | 4.2E-09 | 8.2E-09 | MB231/MB468 | MITO |
| >hg38_21_+_8433172_8446622_RNA45SN1_45S_50ntFlanks@11482.11502.21\|GCGGAGGGGCCGCGGGCCGGT | >hg38_21_+_8433172_8446622_RNA45SN1_45S_50ntFlanks | 11482 | 11502 | 21 | GCGGAGGGGCCGCGGGCCGGT | 86.7 | -10.3 | 1.7 | -5.9 | 3.7E-09 | 8.3E-09 | MB231/MB468 | MP |
| >hg38_21_+_8433172_8446622_RNA45SN1_45S_50ntFlanks@10402.10433.32\|AACATGGGTCAGTCGGTCCTGAGAGATGGGCG | >hg38_21_+_8433172_8446622_RNA45SN1_45S_50ntFlanks | 10402 | 10433 | 32 | AACATGGGTCAGTCGGTCCTGAGAGATGGGCG | 541.0 | -7.7 | 1.3 | -5.9 | 4.2E-09 | 8.3E-09 | MB231/MB468 | MITO |
| >hg38_21_+_8433172_8446622_RNA45SN1_45S_50ntFlanks@12007.12032.26\|GCCGCCGGTGAAATACCACTACTCTG | >hg38_21_+_8433172_8446622_RNA45SN1_45S_50ntFlanks | 12007 | 12032 | 26 | GCCGCCGGTGAAATACCACTACTCTG | 2805.9 | -4.5 | 0.8 | -5.9 | 3.3E-09 | 8.3E-09 | BT20/MB468 | CYTO |
| >hg38_21_+_8433172_8446622_RNA45SN1_45S_50ntFlanks@12251.12271.21\|AAAGCTCGCTTGATCTTGATT | >hg38_21_+_8433172_8446622_RNA45SN1_45S_50ntFlanks | 12251 | 12271 | 21 | AAAGCTCGCTTGATCTTGATT | 183.3 | 4.3 | 0.7 | 5.9 | 3.9E-09 | 8.3E-09 | MB231/MB468 | NUC |
| >hg38_21_+_8433172_8446622_RNA45SN1_45S_50ntFlanks@12526.12557.32\|TTTTACCCTACTGATGATGTGTTGTTGCCATG | >hg38_21_+_8433172_8446622_RNA45SN1_45S_50ntFlanks | 12526 | 12557 | 32 | TTTTACCCTACTGATGATGTGTTGTTGCCATG | 171.0 | -10.5 | 1.7 | -6.2 | 4.7E-10 | 8.3E-09 | BT20/MB468 | MP |
| >hg38_21_+_8433172_8446622_RNA45SN1_45S_50ntFlanks@6646.6695.50\|TCGTACGACTCTTAGCGGTGGATCACTCGGCTCGTGCGTCGATGAAGAAC | >hg38_21_+_8433172_8446622_RNA45SN1_45S_50ntFlanks | 6646 | 6695 | 50 | TCGTACGACTCTTAGCGGTGGATCACTCGGCTCGTGCGTCGATGAAGAAC | 15667.9 | 7.2 | 1.2 | 5.9 | 2.9E-09 | 8.3E-09 | BT20/MB468 | MITO |
| >hg38_21_+_8433172_8446622_RNA45SN1_45S_50ntFlanks@5472.5509.38\|ACGGCCCTGGCGGAGCGCTGAGAAGACGGTCGAACTTG | >hg38_21_+_8433172_8446622_RNA45SN1_45S_50ntFlanks | 5472 | 5509 | 38 | ACGGCCCTGGCGGAGCGCTGAGAAGACGGTCGAACTTG | 1319.2 | 5.6 | 0.9 | 5.9 | 2.9E-09 | 8.3E-09 | BT20/MB468 | MITO |
| >hg38_21_+_8433172_8446622_RNA45SN1_45S_50ntFlanks@8423.8470.48\|CGCAGTCCGCCCGGAGGATTCAACCCGGCGGCGGGTCCGGCCGTGTCG | >hg38_21_+_8433172_8446622_RNA45SN1_45S_50ntFlanks | 8423 | 8470 | 48 | CGCAGTCCGCCCGGAGGATTCAACCCGGCGGCGGGTCCGGCCGTGTCG | 187.8 | -9.6 | 1.6 | -5.9 | 2.9E-09 | 8.3E-09 | BT20/MB468 | MITO |
| >hg38_21_+_8433172_8446622_RNA45SN1_45S_50ntFlanks@11163.11189.27\|CGGTCCCCCGCGAGGGGGGCCCGGGCA | >hg38_21_+_8433172_8446622_RNA45SN1_45S_50ntFlanks | 11163 | 11189 | 27 | CGGTCCCCCGCGAGGGGGGCCCGGGCA | 146.0 | -9.3 | 1.6 | -5.9 | 2.9E-09 | 8.3E-09 | BT20/MB468 | MITO |
| >hg38_21_+_8433172_8446622_RNA45SN1_45S_50ntFlanks@7985.8011.27\|GATCAGACGTGGCGACCCGCTGAATTT | >hg38_21_+_8433172_8446622_RNA45SN1_45S_50ntFlanks | 7985 | 8011 | 27 | GATCAGACGTGGCGACCCGCTGAATTT | 114.9 | -8.9 | 1.5 | -5.9 | 2.9E-09 | 8.3E-09 | BT20/MB468 | MITO |
| >hg38_1_-_228634819_228635039_RNA5S12_5S_50ntFlanks@51.69.19\|GTCTACGGCCATACCACCC | >hg38_1_-_228634819_228635039_RNA5S12_5S_50ntFlanks | 51 | 69 | 19 | GTCTACGGCCATACCACCC | 667.9 | 4.7 | 0.8 | 5.9 | 3.3E-09 | 8.3E-09 | BT20/MB468 | CYTO |
| >hg38_21_+_8433172_8446622_RNA45SN1_45S_50ntFlanks@11581.11601.21\|AACTGGTGCGGACCAGGGGAA | >hg38_21_+_8433172_8446622_RNA45SN1_45S_50ntFlanks | 11581 | 11601 | 21 | AACTGGTGCGGACCAGGGGAA | 1422.0 | -9.3 | 1.6 | -5.9 | 4.4E-09 | 8.3E-09 | BT20/MB468 | NUC |
| >hg38_21_+_8433172_8446622_RNA45SN1_45S_50ntFlanks@8198.8225.28\|TGGACGGTGTGAGGCCGGTAGCGGCCCC | >hg38_21_+_8433172_8446622_RNA45SN1_45S_50ntFlanks | 8198 | 8225 | 28 | TGGACGGTGTGAGGCCGGTAGCGGCCCC | 383.9 | -5.9 | 1.0 | -5.9 | 3.9E-09 | 8.3E-09 | MB231/MB468 | NUC |
| >hg38_21_+_8433172_8446622_RNA45SN1_45S_50ntFlanks@10405.10424.20\|ATGGGTCAGTCGGTCCTGAG | >hg38_21_+_8433172_8446622_RNA45SN1_45S_50ntFlanks | 10405 | 10424 | 20 | ATGGGTCAGTCGGTCCTGAG | 1667.9 | 4.4 | 0.7 | 5.9 | 3.9E-09 | 8.3E-09 | MB231/MB468 | NUC |
| >hg38_21_+_8433172_8446622_RNA45SN1_45S_50ntFlanks@12625.12661.37\|CAATGGGGCGAAGCTACCATCTGTGGGATTATGACTG | >hg38_21_+_8433172_8446622_RNA45SN1_45S_50ntFlanks | 12625 | 12661 | 37 | CAATGGGGCGAAGCTACCATCTGTGGGATTATGACTG | 805.1 | -8.6 | 1.5 | -5.9 | 3.7E-09 | 8.3E-09 | MB231/MB468 | MP |
| >MI0000065\|hsa-let-7d&WithFlank&9\|+\|94178828\|94178926@68.88.21(+1C)&[MIMAT0004484&hsa-let-7d-3p&offsets\|0\|-1(+1C);m-98&9\|+\|94178895\|94178916&offsets\|0\|-1(+1C)]\|CTATACGACCTGCTGCCTTTCC | >MI0000065\|hsa-let-7d&WithFlank&9\|+\|94178828\|94178926 | 68 | 88 | 21(+1C) | CTATACGACCTGCTGCCTTTCC | 274.8 | 6.1 | 1.0 | 5.9 | 3.9E-09 | 8.3E-09 | MB231/MB468 | CYTO |
| >hg38_21_+_8433172_8446622_RNA45SN1_45S_50ntFlanks@10808.10844.37\|CAATGTAGGTAAGGGAAGTCGGCAAGCCGGATCCGTA | >hg38_21_+_8433172_8446622_RNA45SN1_45S_50ntFlanks | 10808 | 10844 | 37 | CAATGTAGGTAAGGGAAGTCGGCAAGCCGGATCCGTA | 175.3 | -4.1 | 0.7 | -5.9 | 3.9E-09 | 8.3E-09 | MB231/MB468 | CYTO |
| >hg38_21_+_8433172_8446622_RNA45SN1_45S_50ntFlanks@10761.10785.25\|ATATCCGCAGCAGGTCTCCAAGGTG | >hg38_21_+_8433172_8446622_RNA45SN1_45S_50ntFlanks | 10761 | 10785 | 25 | ATATCCGCAGCAGGTCTCCAAGGTG | 130.6 | -10.1 | 1.6 | -6.2 | 4.8E-10 | 8.4E-09 | BT20/MB468 | MP |
| >am_MI0000748\|hsa-mir-130b&WithFlank&22\|+\|21653298\|21653391@19.39.21&[MIMAT0004680&hsa-miR-130b-5p&offsets\|0\|0;m-234&22\|+\|21653316\|21653337&offsets\|0\|-1]\|ACTCTTTCCCTGTTGCACTAC | >am_MI0000748\|hsa-mir-130b&WithFlank&22\|+\|21653298\|21653391 | 19 | 39 | 21 | ACTCTTTCCCTGTTGCACTAC | 985.8 | 7.9 | 1.3 | 5.9 | 3.9E-09 | 8.4E-09 | MB231/MB468 | CYTO |
| >trna10_AspGTC_12_-_125424193_125424264@36.74.39,trna12_AspGTC_12_-_125411891_125411962@36.74.39,trna144_AspGTC_6_-_27551236_27551307@36.74.39,trna38_AspGTC_17_-_8125556_8125627@36.74.39,trna45_AspGTC_6_+_27447453_27447524@36.74.39,trna48_AspGTC_6_+_27471523_27471594@36.74.39,trna4_AspGTC_12_+_96429799_96429870@36.74.39,trna69_AspGTC_1_-_161440205_161440276@36.74.39,trna72_AspGTC_1_-_161432824_161432895@36.74.39,trna75_AspGTC_1_-_161425414_161425485@36.74.39,trna78_AspGTC_1_-_161418033_161418104@36.74.39,trna81_AspGTC_1_-_161410615_161410686@36.74.39\|CACGCGGGAGACCGGGGTTCGATTCCCCGACGGGGAGCC | >trna10_AspGTC_12_-_125424193_125424264 | 36 | 74 | 39 | CACGCGGGAGACCGGGGTTCGATTCCCCGACGGGGAGCC | 124.8 | 10.2 | 1.7 | 5.9 | 4.3E-09 | 8.4E-09 | MB231/MB468 | MITO |
| >hg38_21_+_8433172_8446622_RNA45SN1_45S_50ntFlanks@12360.12384.25\|CAGGGATAACTGGCTTGTGGCGGCC | >hg38_21_+_8433172_8446622_RNA45SN1_45S_50ntFlanks | 12360 | 12384 | 25 | CAGGGATAACTGGCTTGTGGCGGCC | 386.0 | -8.2 | 1.4 | -5.9 | 4.5E-09 | 8.4E-09 | BT20/MB468 | NUC |
| >hg38_21_+_8433172_8446622_RNA45SN1_45S_50ntFlanks@9304.9338.35\|ACCAAGGAGTCTAACACGTGCGCGAGTCGGGGGCT | >hg38_21_+_8433172_8446622_RNA45SN1_45S_50ntFlanks | 9304 | 9338 | 35 | ACCAAGGAGTCTAACACGTGCGCGAGTCGGGGGCT | 303.9 | -8.0 | 1.4 | -5.9 | 4.5E-09 | 8.4E-09 | BT20/MB468 | NUC |
| >hg38_21_+_8433172_8446622_RNA45SN1_45S_50ntFlanks@3777.3798.22\|CGGTACAGTGAAACTGCGAATG | >hg38_21_+_8433172_8446622_RNA45SN1_45S_50ntFlanks | 3777 | 3798 | 22 | CGGTACAGTGAAACTGCGAATG | 256.1 | -7.7 | 1.3 | -5.9 | 3.7E-09 | 8.4E-09 | MB231/MB468 | MP |
| >MI0001445\|hsa-mir-423&WithFlank&17\|+\|30117073\|30117178@58.78.21&[MIMAT0001340&hsa-miR-423-3p&offsets\|-1\|-3;m-100&17\|+\|30117131\|30117153&offsets\|-1\|-3]\|AAGCTCGGTCTGAGGCCCCTC | >MI0001445\|hsa-mir-423&WithFlank&17\|+\|30117073\|30117178 | 58 | 78 | 21 | AAGCTCGGTCTGAGGCCCCTC | 718.7 | 4.7 | 0.8 | 5.9 | 4.0E-09 | 8.4E-09 | MB231/MB468 | NUC |
| >hg38_21_+_8433172_8446622_RNA45SN1_45S_50ntFlanks@9597.9622.26\|AGGGGCGAAAGACTAATCGAACCATC | >hg38_21_+_8433172_8446622_RNA45SN1_45S_50ntFlanks | 9597 | 9622 | 26 | AGGGGCGAAAGACTAATCGAACCATC | 368.5 | -11.6 | 1.9 | -6.2 | 4.8E-10 | 8.4E-09 | BT20/MB468 | MP |

<table>
<tr><td>>hg38_21_+_8433172_8446622_RNA45SN1_45S_50ntFlanks@10357.10377.21|CTTTGAAGGCCGAAGTGGAGA</td><td>>hg38_21_+_8433172_8446622_RNA45SN1_45S_50ntFlanks</td><td>10357</td><td>10377</td><td>21</td><td>CTTTGAAGGCCGAAGTGGAGA</td><td>623.6</td><td>-8.6</td><td>1.5</td><td>-5.9</td><td>4.5E-09</td><td>8.4E-09</td><td>BT20/MB468</td><td>NUC</td></tr>
<tr><td>>hg38_21_+_8433172_8446622_RNA45SN1_45S_50ntFlanks@10681.10711.31|TTCCGGCGGCGTCCGGTGAGCTCTCGCTGGC</td><td>>hg38_21_+_8433172_8446622_RNA45SN1_45S_50ntFlanks</td><td>10681</td><td>10711</td><td>31</td><td>TTCCGGCGGCGTCCGGTGAGCTCTCGCTGGC</td><td>2313.5</td><td>-9.9</td><td>1.7</td><td>-5.9</td><td>4.5E-09</td><td>8.4E-09</td><td>BT20/MB468</td><td>NUC</td></tr>
<tr><td>>hg38_1_-_228634819_228635039_RNA5S12_5S_50ntFlanks@85.123.39|TCGTCTGATCTCGGAAGCTAAGCAGGGTCGGGCCTGGTT</td><td>>hg38_1_-_228634819_228635039_RNA5S12_5S_50ntFlanks</td><td>85</td><td>123</td><td>39</td><td>TCGTCTGATCTCGGAAGCTAAGCAGGGTCGGGCCTGGTT</td><td>129.6</td><td>10.8</td><td>1.7</td><td>6.2</td><td>4.9E-10</td><td>8.4E-09</td><td>BT20/MB468</td><td>MP</td></tr>
<tr><td>>hg38_21_+_8433172_8446622_RNA45SN1_45S_50ntFlanks@3985.4003.19|CGGCTTTGGTGACTCTAGA</td><td>>hg38_21_+_8433172_8446622_RNA45SN1_45S_50ntFlanks</td><td>3985</td><td>4003</td><td>19</td><td>CGGCTTTGGTGACTCTAGA</td><td>400.2</td><td>-8.3</td><td>1.4</td><td>-5.9</td><td>4.5E-09</td><td>8.4E-09</td><td>BT20/MB468</td><td>NUC</td></tr>
<tr><td>>hg38_21_+_8433172_8446622_RNA45SN1_45S_50ntFlanks@12417.12447.31|TCGATGTCGGCTCTTCCTATCATTGTGAAGC</td><td>>hg38_21_+_8433172_8446622_RNA45SN1_45S_50ntFlanks</td><td>12417</td><td>12447</td><td>31</td><td>TCGATGTCGGCTCTTCCTATCATTGTGAAGC</td><td>971.0</td><td>4.0</td><td>0.7</td><td>5.9</td><td>3.9E-09</td><td>8.4E-09</td><td>MB231/MB468</td><td>CYTO</td></tr>
<tr><td>>hg38_21_+_8433172_8446622_RNA45SN1_45S_50ntFlanks@12456.12473.18|CCAAGCGTTGGATTGTTC</td><td>>hg38_21_+_8433172_8446622_RNA45SN1_45S_50ntFlanks</td><td>12456</td><td>12473</td><td>18</td><td>CCAAGCGTTGGATTGTTC</td><td>1748.0</td><td>-5.6</td><td>1.0</td><td>-5.9</td><td>4.0E-09</td><td>8.5E-09</td><td>MB231/MB468</td><td>NUC</td></tr>
<tr><td>>hg38_1_-_228634819_228635039_RNA5S12_5S_50ntFlanks@125.171.47|GTACTTGGATGGGAGACCGCCTGGGAATACCGGGTGCTGTAGGCTTT</td><td>>hg38_1_-_228634819_228635039_RNA5S12_5S_50ntFlanks</td><td>125</td><td>171</td><td>47</td><td>GTACTTGGATGGGAGACCGCCTGGGAATACCGGGTGCTGTAGGCTTT</td><td>736.5</td><td>9.3</td><td>1.6</td><td>5.9</td><td>3.9E-09</td><td>8.5E-09</td><td>MB231/MB468</td><td>CYTO</td></tr>
<tr><td>>hg38_21_+_8433172_8446622_RNA45SN1_45S_50ntFlanks@9576.9595.20|TCGGTCGTCCGACCTGGGTA</td><td>>hg38_21_+_8433172_8446622_RNA45SN1_45S_50ntFlanks</td><td>9576</td><td>9595</td><td>20</td><td>TCGGTCGTCCGACCTGGGTA</td><td>228.6</td><td>-10.9</td><td>1.8</td><td>-6.2</td><td>4.9E-10</td><td>8.5E-09</td><td>BT20/MB468</td><td>MP</td></tr>
<tr><td>>hg38_21_+_8433172_8446622_RNA45SN1_45S_50ntFlanks@6651.6700.50|CGACTCTTAGCGGTGGATCACTCGGCTCGTGCGTCGATGAAGAACGCAGC</td><td>>hg38_21_+_8433172_8446622_RNA45SN1_45S_50ntFlanks</td><td>6651</td><td>6700</td><td>50</td><td>CGACTCTTAGCGGTGGATCACTCGGCTCGTGCGTCGATGAAGAACGCAGC</td><td>2734.3</td><td>6.4</td><td>1.1</td><td>5.9</td><td>3.9E-09</td><td>8.5E-09</td><td>MB231/MB468</td><td>CYTO</td></tr>
<tr><td>>MI0000456|hsa-mir-140&WithFlank&16|+|69933075|69933186@69.87.19&[MIMAT0004597&hsa-miR-140-3p&offsets|+1|-1;m-45&16|+|69933143|69933164&offsets|0|-3]|ACCACAGGGTAGAACCACG</td><td>>MI0000456|hsa-mir-140&WithFlank&16|+|69933075|69933186</td><td>69</td><td>87</td><td>19</td><td>ACCACAGGGTAGAACCACG</td><td>71.0</td><td>9.4</td><td>1.6</td><td>5.9</td><td>3.8E-09</td><td>8.5E-09</td><td>MB231/MB468</td><td>MP</td></tr>
<tr><td>>trna17_ValTAC_11_-_59318102_59318174@1.34.34,trna4_ValTAC_X_-_18693029_18693101@1.34.34|GGTTCCATAGTGTAGTGGTTATCACGTCTGCTTT</td><td>>trna17_ValTAC_11_-_59318102_59318174</td><td>1</td><td>34</td><td>34</td><td>GGTTCCATAGTGTAGTGGTTATCACGTCTGCTTT</td><td>160.4</td><td>10.5</td><td>1.8</td><td>5.9</td><td>3.8E-09</td><td>8.5E-09</td><td>MB231/MB468</td><td>MP</td></tr>
<tr><td>>hg38_21_+_8433172_8446622_RNA45SN1_45S_50ntFlanks@9862.9905.44|ATGAACCGAACGCCGGGTTAAGGCGCCCGATGCCGACGCTCATC</td><td>>hg38_21_+_8433172_8446622_RNA45SN1_45S_50ntFlanks</td><td>9862</td><td>9905</td><td>44</td><td>ATGAACCGAACGCCGGGTTAAGGCGCCCGATGCCGACGCTCATC</td><td>60.1</td><td>-9.8</td><td>1.7</td><td>-5.9</td><td>3.8E-09</td><td>8.5E-09</td><td>MB231/MB468</td><td>MP</td></tr>
<tr><td>>hg38_21_+_8433172_8446622_RNA45SN1_45S_50ntFlanks@4439.4471.33|CCGCCCCTTGCCTCTCGGCGCCCCCTCGATGCT</td><td>>hg38_21_+_8433172_8446622_RNA45SN1_45S_50ntFlanks</td><td>4439</td><td>4471</td><td>33</td><td>CCGCCCCTTGCCTCTCGGCGCCCCCTCGATGCT</td><td>65.1</td><td>-9.9</td><td>1.7</td><td>-5.9</td><td>3.8E-09</td><td>8.5E-09</td><td>MB231/MB468</td><td>MP</td></tr>
<tr><td>>hg38_21_+_8433172_8446622_RNA45SN1_45S_50ntFlanks@11999.12038.40|CCCTGCGGGCCGCCGGTGAAATACCACTACTCTGATCGTT</td><td>>hg38_21_+_8433172_8446622_RNA45SN1_45S_50ntFlanks</td><td>11999</td><td>12038</td><td>40</td><td>CCCTGCGGGCCGCCGGTGAAATACCACTACTCTGATCGTT</td><td>258.8</td><td>-7.8</td><td>1.3</td><td>-5.9</td><td>4.0E-09</td><td>8.5E-09</td><td>MB231/MB468</td><td>NUC</td></tr>
<tr><td>>hg38_21_+_8433172_8446622_RNA45SN1_45S_50ntFlanks@9554.9584.31|CGTAGCGGTCCTGACGTGCAAATCGGTCGTC</td><td>>hg38_21_+_8433172_8446622_RNA45SN1_45S_50ntFlanks</td><td>9554</td><td>9584</td><td>31</td><td>CGTAGCGGTCCTGACGTGCAAATCGGTCGTC</td><td>2533.8</td><td>-4.3</td><td>0.7</td><td>-5.9</td><td>3.4E-09</td><td>8.5E-09</td><td>BT20/MB468</td><td>CYTO</td></tr>
<tr><td>>hg38_21_+_8433172_8446622_RNA45SN1_45S_50ntFlanks@8196.8226.31|CGTGGACGGTGTGAGGCCGGTAGCGGCCCCC</td><td>>hg38_21_+_8433172_8446622_RNA45SN1_45S_50ntFlanks</td><td>8196</td><td>8226</td><td>31</td><td>CGTGGACGGTGTGAGGCCGGTAGCGGCCCCC</td><td>612.8</td><td>-3.4</td><td>0.6</td><td>-5.9</td><td>3.4E-09</td><td>8.5E-09</td><td>BT20/MB468</td><td>CYTO</td></tr>
<tr><td>>hg38_21_+_8433172_8446622_RNA45SN1_45S_50ntFlanks@12942.12965.24|TTCGTAGACGACCTGCTTCTGGGT</td><td>>hg38_21_+_8433172_8446622_RNA45SN1_45S_50ntFlanks</td><td>12942</td><td>12965</td><td>24</td><td>TTCGTAGACGACCTGCTTCTGGGT</td><td>423.7</td><td>-4.8</td><td>0.8</td><td>-5.9</td><td>4.0E-09</td><td>8.5E-09</td><td>MB231/MB468</td><td>CYTO</td></tr>
<tr><td>>hg38_21_+_8433172_8446622_RNA45SN1_45S_50ntFlanks@8353.8375.23|GTTGAAAAGAACTTTGAAGAGAG</td><td>>hg38_21_+_8433172_8446622_RNA45SN1_45S_50ntFlanks</td><td>8353</td><td>8375</td><td>23</td><td>GTTGAAAAGAACTTTGAAGAGAG</td><td>4883.6</td><td>10.1</td><td>1.6</td><td>6.2</td><td>4.9E-10</td><td>8.6E-09</td><td>BT20/MB468</td><td>MP</td></tr>
<tr><td>>hg38_21_+_8433172_8446622_RNA45SN1_45S_50ntFlanks@10806.10840.35|AACAATGTAGGTAAGGGAAGTCGGCAAGCCGGATC</td><td>>hg38_21_+_8433172_8446622_RNA45SN1_45S_50ntFlanks</td><td>10806</td><td>10840</td><td>35</td><td>AACAATGTAGGTAAGGGAAGTCGGCAAGCCGGATC</td><td>645.3</td><td>-8.0</td><td>1.4</td><td>-5.9</td><td>3.8E-09</td><td>8.6E-09</td><td>MB231/MB468</td><td>MP</td></tr>
<tr><td>>hg38_21_+_8433172_8446622_RNA45SN1_45S_50ntFlanks@3984.4009.26|GCGGCTTTGGTGACTCTAGATAACCT</td><td>>hg38_21_+_8433172_8446622_RNA45SN1_45S_50ntFlanks</td><td>3984</td><td>4009</td><td>26</td><td>GCGGCTTTGGTGACTCTAGATAACCT</td><td>75.6</td><td>-10.1</td><td>1.7</td><td>-5.9</td><td>3.8E-09</td><td>8.6E-09</td><td>MB231/MB468</td><td>MP</td></tr>
<tr><td>>MI0000812|hsa-mir-331&WithFlank&12|+|95308414|95308519@67.87.21(+1U)&[MIMAT0000760&hsa-miR-331-3p&offsets|0|0(+1U);m-192&12|+|95308480|95308501&offsets|0|-1(+1U)]|GCCCCTGGGCCTATCCTAGAAT</td><td>>MI0000812|hsa-mir-331&WithFlank&12|+|95308414|95308519</td><td>67</td><td>87</td><td>21(+1U)</td><td>GCCCCTGGGCCTATCCTAGAAT</td><td>777.7</td><td>5.8</td><td>1.0</td><td>5.9</td><td>4.4E-09</td><td>8.6E-09</td><td>MB231/MB468</td><td>MITO</td></tr>
<tr><td>>hg38_21_+_8433172_8446622_RNA45SN1_45S_50ntFlanks@11874.11900.27|AGAAGACCCTGTTGAGCTTGACTCTAG</td><td>>hg38_21_+_8433172_8446622_RNA45SN1_45S_50ntFlanks</td><td>11874</td><td>11900</td><td>27</td><td>AGAAGACCCTGTTGAGCTTGACTCTAG</td><td>482.7</td><td>-3.6</td><td>0.6</td><td>-5.9</td><td>3.0E-09</td><td>8.6E-09</td><td>BT20/MB468</td><td>MITO</td></tr>
<tr><td>>hg38_21_+_8433172_8446622_RNA45SN1_45S_50ntFlanks@9569.9597.29|GTGCAAATCGGTCGTCCGACCTGGGTATA</td><td>>hg38_21_+_8433172_8446622_RNA45SN1_45S_50ntFlanks</td><td>9569</td><td>9597</td><td>29</td><td>GTGCAAATCGGTCGTCCGACCTGGGTATA</td><td>1141.3</td><td>-3.0</td><td>0.5</td><td>-5.9</td><td>3.0E-09</td><td>8.6E-09</td><td>BT20/MB468</td><td>MITO</td></tr>
<tr><td>>hg38_21_+_8433172_8446622_RNA45SN1_45S_50ntFlanks@11783.11807.25|CATGAATGGATGAACGAGATTCCCA</td><td>>hg38_21_+_8433172_8446622_RNA45SN1_45S_50ntFlanks</td><td>11783</td><td>11807</td><td>25</td><td>CATGAATGGATGAACGAGATTCCCA</td><td>552.7</td><td>-8.6</td><td>1.5</td><td>-5.9</td><td>3.8E-09</td><td>8.6E-09</td><td>MB231/MB468</td><td>MP</td></tr>
<tr><td>>MI0000077|hsa-mir-21&WithFlank&17|+|59841260|59841343@14.35.22(+1G)&[MIMAT0000076&hsa-miR-21-5p&offsets|0|0(+1G);m-1&17|+|59841273|59841294&offsets|0|0(+1G)]|TAGCTTATCAGACTGATGTTGAG</td><td>>MI0000077|hsa-mir-21&WithFlank&17|+|59841260|59841343</td><td>14</td><td>35</td><td>22(+1G)</td><td>TAGCTTATCAGACTGATGTTGAG</td><td>2315.0</td><td>6.8</td><td>1.2</td><td>5.9</td><td>4.4E-09</td><td>8.6E-09</td><td>MB231/MB468</td><td>MITO</td></tr>
<tr><td>>hg38_21_+_8433172_8446622_RNA45SN1_45S_50ntFlanks@8381.8403.23|AAGAGGGCGTGAAACCGTTAAGA</td><td>>hg38_21_+_8433172_8446622_RNA45SN1_45S_50ntFlanks</td><td>8381</td><td>8403</td><td>23</td><td>AAGAGGGCGTGAAACCGTTAAGA</td><td>315.3</td><td>-6.3</td><td>1.1</td><td>-5.9</td><td>4.6E-09</td><td>8.6E-09</td><td>BT20/MB468</td><td>NUC</td></tr>
<tr><td>>hg38_21_+_8433172_8446622_RNA45SN1_45S_50ntFlanks@4044.4063.20|CCCATTCGAACGTCTGCCCT</td><td>>hg38_21_+_8433172_8446622_RNA45SN1_45S_50ntFlanks</td><td>4044</td><td>4063</td><td>20</td><td>CCCATTCGAACGTCTGCCCT</td><td>144.6</td><td>-9.3</td><td>1.6</td><td>-5.9</td><td>3.0E-09</td><td>8.6E-09</td><td>BT20/MB468</td><td>MITO</td></tr>
<tr><td>>MI0000114|hsa-mir-107&WithFlank&10|-|89592741|89592833@56.76.21(+1A);MI0000108|hsa-mir-103a-2&WithFlank&20|+|3917488|3917577@54.74.21(+1A);MI0000109|hsa-mir-103a-1&WithFlank&5|-|168560890|168560979@54.74.21(+1A)&[MIMAT0000104&hsa-miR-107&offsets|0|-2(+1A);m-78&10|-|89592758|89592778&offsets|0|0(+1A)];[MIMAT0000101&hsa-miR-103a-3p&offsets|0|-2(+1A);m-12&20|+|3917541|3917563&offsets|0|-2(+1A)];[MIMAT0000101_1&hsa-miR-103a-3p&offsets|0|-2(+1A);m-13&5|-|168560904|168560926&offsets|0|-2(+1A);m-44220&5|-|168560908|168560926&offsets|0|+2(+1A)]|AGCAGCATTGTACAGGGCTATA</td><td>>MI0000114|hsa-mir-107&WithFlank&10|-|89592741|89592833</td><td>56</td><td>76</td><td>21(+1A)</td><td>AGCAGCATTGTACAGGGCTATA</td><td>800.2</td><td>6.2</td><td>1.1</td><td>5.9</td><td>4.4E-09</td><td>8.6E-09</td><td>MB231/MB468</td><td>MITO</td></tr>
<tr><td>>hg38_21_+_8433172_8446622_RNA45SN1_45S_50ntFlanks@10402.10426.25|AACATGGGTCAGTCGGTCCTGAGAG</td><td>>hg38_21_+_8433172_8446622_RNA45SN1_45S_50ntFlanks</td><td>10402</td><td>10426</td><td>25</td><td>AACATGGGTCAGTCGGTCCTGAGAG</td><td>613.6</td><td>-6.0</td><td>1.0</td><td>-5.9</td><td>4.4E-09</td><td>8.6E-09</td><td>MB231/MB468</td><td>MITO</td></tr>
<tr><td>>hg38_21_+_8433172_8446622_RNA45SN1_45S_50ntFlanks@12449.12486.38|GAATTCACCAAGCGTTGGATTGTTCACCCACTAATAGG</td><td>>hg38_21_+_8433172_8446622_RNA45SN1_45S_50ntFlanks</td><td>12449</td><td>12486</td><td>38</td><td>GAATTCACCAAGCGTTGGATTGTTCACCCACTAATAGG</td><td>344.3</td><td>7.3</td><td>1.2</td><td>5.9</td><td>4.0E-09</td><td>8.6E-09</td><td>MB231/MB468</td><td>CYTO</td></tr>
<tr><td>>hg38_21_+_8433172_8446622_RNA45SN1_45S_50ntFlanks@475.497.23|CTTCGTGATCGATGTGGTGACGT</td><td>>hg38_21_+_8433172_8446622_RNA45SN1_45S_50ntFlanks</td><td>475</td><td>497</td><td>23</td><td>CTTCGTGATCGATGTGGTGACGT</td><td>805.9</td><td>8.2</td><td>1.4</td><td>5.9</td><td>4.0E-09</td><td>8.6E-09</td><td>MB231/MB468</td><td>CYTO</td></tr>
<tr><td>>MI0000462|hsa-mir-152&WithFlank&17|-|48037155|48037253@60.81.22(+1A)&[MIMAT0000438&hsa-miR-152-3p&offsets|0|+1(+1A);m-97&17|-|48037174|48037194&offsets|0|+1(+1A)]|TCAGTGCATGACAGAACTTGGGA</td><td>>MI0000462|hsa-mir-152&WithFlank&17|-|48037155|48037253</td><td>60</td><td>81</td><td>22(+1A)</td><td>TCAGTGCATGACAGAACTTGGGA</td><td>175.7</td><td>11.2</td><td>1.8</td><td>6.2</td><td>5.0E-10</td><td>8.6E-09</td><td>BT20/MB468</td><td>MP</td></tr>
<tr><td>>hg38_21_+_8433172_8446622_RNA45SN1_45S_50ntFlanks@6740.6770.31|CATCGACACTTCGAACGCACTTGCGGCCCCG</td><td>>hg38_21_+_8433172_8446622_RNA45SN1_45S_50ntFlanks</td><td>6740</td><td>6770</td><td>31</td><td>CATCGACACTTCGAACGCACTTGCGGCCCCG</td><td>1247.6</td><td>-8.3</td><td>1.4</td><td>-5.9</td><td>4.1E-09</td><td>8.6E-09</td><td>MB231/MB468</td><td>NUC</td></tr>
<tr><td>>MI0000266|hsa-mir-10a&WithFlank&17|-|48579832|48579953@28.50.23&[MIMAT0000253&hsa-miR-10a-5p&offsets|0|0;m-18&17|-|48579905|48579926&offsets|0|+1]|TACCCTGTAGATCCGAATTTGTG</td><td>>MI0000266|hsa-mir-10a&WithFlank&17|-|48579832|48579953</td><td>28</td><td>50</td><td>23</td><td>TACCCTGTAGATCCGAATTTGTG</td><td>5869.0</td><td>10.4</td><td>1.7</td><td>6.2</td><td>5.0E-10</td><td>8.6E-09</td><td>BT20/MB468</td><td>MP</td></tr>
<tr><td>>MI0000650|hsa-mir-200c&WithFlank&12|+|6963693|6963772@50.71.22&[MIMAT0000617&hsa-miR-200c-3p&offsets|0|-1;m-54&12|+|6963742|6963764&offsets|0|-1]|TAATACTGCCGGGTAATGATGG</td><td>>MI0000650|hsa-mir-200c&WithFlank&12|+|6963693|6963772</td><td>50</td><td>71</td><td>22</td><td>TAATACTGCCGGGTAATGATGG</td><td>893.5</td><td>-4.7</td><td>0.8</td><td>-5.9</td><td>3.8E-09</td><td>8.6E-09</td><td>MB231/MB468</td><td>MP</td></tr>
<tr><td>>hg38_21_+_8433172_8446622_RNA45SN1_45S_50ntFlanks@12417.12437.21|TCGATGTCGGCTCTTCCTATC</td><td>>hg38_21_+_8433172_8446622_RNA45SN1_45S_50ntFlanks</td><td>12417</td><td>12437</td><td>21</td><td>TCGATGTCGGCTCTTCCTATC</td><td>3559.2</td><td>-3.9</td><td>0.7</td><td>-5.9</td><td>4.4E-09</td><td>8.6E-09</td><td>MB231/MB468</td><td>MITO</td></tr>
<tr><td>>hg38_21_+_8433172_8446622_RNA45SN1_45S_50ntFlanks@3987.4026.40|GCTTTGGTGACTCTAGATAACCTCGGGCCGATCGCACGCC</td><td>>hg38_21_+_8433172_8446622_RNA45SN1_45S_50ntFlanks</td><td>3987</td><td>4026</td><td>40</td><td>GCTTTGGTGACTCTAGATAACCTCGGGCCGATCGCACGCC</td><td>952.1</td><td>-5.5</td><td>0.9</td><td>-5.9</td><td>4.4E-09</td><td>8.6E-09</td><td>MB231/MB468</td><td>MITO</td></tr>
<tr><td>>hg38_21_+_8433172_8446622_RNA45SN1_45S_50ntFlanks@12157.12182.26|AGTTTGACTGGGGCGGTACACCTGTC</td><td>>hg38_21_+_8433172_8446622_RNA45SN1_45S_50ntFlanks</td><td>12157</td><td>12182</td><td>26</td><td>AGTTTGACTGGGGCGGTACACCTGTC</td><td>3426.2</td><td>-8.0</td><td>1.3</td><td>-6.2</td><td>5.0E-10</td><td>8.6E-09</td><td>BT20/MB468</td><td>MP</td></tr>
<tr><td>>hg38_21_+_8433172_8446622_RNA45SN1_45S_50ntFlanks@6661.6700.40|CGGTGGATCACTCGGCTCGTGCGTCGATGAAGAACGCAGC</td><td>>hg38_21_+_8433172_8446622_RNA45SN1_45S_50ntFlanks</td><td>6661</td><td>6700</td><td>40</td><td>CGGTGGATCACTCGGCTCGTGCGTCGATGAAGAACGCAGC</td><td>382.8</td><td>5.1</td><td>0.9</td><td>5.9</td><td>4.1E-09</td><td>8.7E-09</td><td>MB231/MB468</td><td>NUC</td></tr>
<tr><td>>hg38_21_+_8433172_8446622_RNA45SN1_45S_50ntFlanks@3842.3868.27|CCTACTTGGATAACTGTGGTAATTCTA</td><td>>hg38_21_+_8433172_8446622_RNA45SN1_45S_50ntFlanks</td><td>3842</td><td>3868</td><td>27</td><td>CCTACTTGGATAACTGTGGTAATTCTA</td><td>646.9</td><td>-3.9</td><td>0.7</td><td>-5.9</td><td>4.6E-09</td><td>8.7E-09</td><td>BT20/MB468</td><td>NUC</td></tr>
<tr><td>>hg38_21_+_8433172_8446622_RNA45SN1_45S_50ntFlanks@8118.8135.18|GGGACATGTGGCGTACGG</td><td>>hg38_21_+_8433172_8446622_RNA45SN1_45S_50ntFlanks</td><td>8118</td><td>8135</td><td>18</td><td>GGGACATGTGGCGTACGG</td><td>279.6</td><td>11.9</td><td>1.9</td><td>6.2</td><td>5.0E-10</td><td>8.7E-09</td><td>BT20/MB468</td><td>MP</td></tr>
<tr><td>>am_MI0000809|hsa-mir-151a&WithFlank&8|-|140732558|140732659@53.70.18(+3G)&[MIMAT0000757&hsa-miR-151a-3p&offsets|0|-3(+3G);m-30&8|-|140732587|140732607&offsets|0|-3(+3G)]|CTAGACTGAAGCTCCTTGGGG</td><td>>am_MI0000809|hsa-mir-151a&WithFlank&8|-|140732558|140732659</td><td>53</td><td>70</td><td>18(+3G)</td><td>CTAGACTGAAGCTCCTTGGGG</td><td>112.4</td><td>10.0</td><td>1.7</td><td>5.9</td><td>4.4E-09</td><td>8.7E-09</td><td>MB231/MB468</td><td>MITO</td></tr>
<tr><td>>hg38_21_+_8433172_8446622_RNA45SN1_45S_50ntFlanks@11703.11726.24|AATGAAGCGCGGGTAAACGGCGGG</td><td>>hg38_21_+_8433172_8446622_RNA45SN1_45S_50ntFlanks</td><td>11703</td><td>11726</td><td>24</td><td>AATGAAGCGCGGGTAAACGGCGGG</td><td>333.2</td><td>-7.5</td><td>1.3</td><td>-5.9</td><td>4.4E-09</td><td>8.7E-09</td><td>MB231/MB468</td><td>MITO</td></tr>
<tr><td>>hg38_21_+_8433172_8446622_RNA45SN1_45S_50ntFlanks@10866.10901.36|CTAAGGGCTGGGTCGGTCGGGCTGGGGCGCGAAGCG</td><td>>hg38_21_+_8433172_8446622_RNA45SN1_45S_50ntFlanks</td><td>10866</td><td>10901</td><td>36</td><td>CTAAGGGCTGGGTCGGTCGGGCTGGGGCGCGAAGCG</td><td>4094.3</td><td>-10.3</td><td>1.8</td><td>-5.9</td><td>4.6E-09</td><td>8.7E-09</td><td>BT20/MB468</td><td>NUC</td></tr>
<tr><td>>hg38_21_+_8433172_8446622_RNA45SN1_45S_50ntFlanks@11038.11060.23|TCTCCCCCGCTCCCCGTCCTCCC</td><td>>hg38_21_+_8433172_8446622_RNA45SN1_45S_50ntFlanks</td><td>11038</td><td>11060</td><td>23</td><td>TCTCCCCCGCTCCCCGTCCTCCC</td><td>1308.6</td><td>-9.0</td><td>1.5</td><td>-5.9</td><td>4.1E-09</td><td>8.7E-09</td><td>MB231/MB468</td><td>NUC</td></tr>
<tr><td>>MI0000477|hsa-mir-146a&WithFlank&5|+|160485346|160485456@27.48.22&[MIMAT0000449&hsa-miR-146a-5p&offsets|0|0;m-129&5|+|160485372|160485393&offsets|0|0]|TGAGAACTGAATTCCATGGGTT</td><td>>MI0000477|hsa-mir-146a&WithFlank&5|+|160485346|160485456</td><td>27</td><td>48</td><td>22</td><td>TGAGAACTGAATTCCATGGGTT</td><td>167841.0</td><td>-10.6</td><td>1.6</td><td>-6.5</td><td>9.7E-11</td><td>8.7E-09</td><td>BT20/MB231</td><td>MP</td></tr>
<tr><td>>MI0005756|hsa-mir-934&WithFlank&X|+|136550872|136550966@21.40.20(+1A)&[MIMAT0004977&hsa-miR-934&offsets|0|-2(+1A);m-943&X|+|136550892|136550913&offsets|0|-2(+1A)]|TGTCTACTACTGGAGACACTA</td><td>>MI0005756|hsa-mir-934&WithFlank&X|+|136550872|136550966</td><td>21</td><td>40</td><td>20(+1A)</td><td>TGTCTACTACTGGAGACACTA</td><td>605.1</td><td>12.6</td><td>1.9</td><td>6.5</td><td>9.7E-11</td><td>8.7E-09</td><td>BT20/MB231</td><td>MP</td></tr>
</table>

| | | | | | | | | | | | | | |
|---|---|---|---|---|---|---|---|---|---|---|---|---|---|
| >hg38_21_+_8433172_8446622_RNA45SN1_45S_50ntFlanks@3860.3898.39\|GTAATTCTAGAGCTAATACATGCCGACGGGCGCTGACCC | >hg38_21_+_8433172_8446622_RNA45SN1_45S_50ntFlanks | 3860 | 3898 | 39 | GTAATTCTAGAGCTAATACATGCCGACGGGCGCTGACCC | 233.5 | -5.6 | 1.0 | -5.9 | 4.1E-09 | 8.7E-09 | MB231/MB468 | CYTO |
| >hg38_21_+_8433172_8446622_RNA45SN1_45S_50ntFlanks@6664.6681.18\|TGGATCACTCGGCTCGTG | >hg38_21_+_8433172_8446622_RNA45SN1_45S_50ntFlanks | 6664 | 6681 | 18 | TGGATCACTCGGCTCGTG | 888.3 | -5.7 | 1.0 | -5.9 | 4.1E-09 | 8.7E-09 | MB231/MB468 | CYTO |
| >hg38_21_+_8433172_8446622_RNA45SN1_45S_50ntFlanks@10797.10822.26\|GCATGTTGGAACAATGTAGGTAAGGG | >hg38_21_+_8433172_8446622_RNA45SN1_45S_50ntFlanks | 10797 | 10822 | 26 | GCATGTTGGAACAATGTAGGTAAGGG | 381.8 | -5.9 | 1.0 | -5.9 | 4.4E-09 | 8.7E-09 | MB231/MB468 | MITO |
| >MI0000234\|hsa-mir-192&WithFlank&11\|-\|64891131\|64891252@31.51.21(+1U)&[MIMAT0000222&hsa-miR-192-5p&offsets\|+1\|+1(+1U);m-59&11\|-\|64891203\|64891223&offsets\|+1\|+1(+1U)]\|TGACCTATGAATTGACAGCCAT | >MI0000234\|hsa-mir-192&WithFlank&11\|-\|64891131\|64891252 | 31 | 51 | 21(+1U) | TGACCTATGAATTGACAGCCAT | 214.8 | 4.6 | 0.8 | 5.9 | 4.1E-09 | 8.7E-09 | MB231/MB468 | NUC |
| >hg38_21_+_8433172_8446622_RNA45SN1_45S_50ntFlanks@12628.12657.30\|TGGGGCGAAGCTACCATCTGTGGGATTATG | >hg38_21_+_8433172_8446622_RNA45SN1_45S_50ntFlanks | 12628 | 12657 | 30 | TGGGGCGAAGCTACCATCTGTGGGATTATG | 259.8 | -7.4 | 1.3 | -5.9 | 4.1E-09 | 8.7E-09 | MB231/MB468 | NUC |
| >hg38_21_+_8433172_8446622_RNA45SN1_45S_50ntFlanks@10777.10818.42\|TCCAAGGTGAACAGCCTCTGGCATGTTGGAACAATGTAGGTA | >hg38_21_+_8433172_8446622_RNA45SN1_45S_50ntFlanks | 10777 | 10818 | 42 | TCCAAGGTGAACAGCCTCTGGCATGTTGGAACAATGTAGGTA | 442.0 | -6.4 | 1.1 | -5.9 | 4.5E-09 | 8.7E-09 | MB231/MB468 | MITO |
| >MI0000471\|hsa-mir-126&WithFlank&9\|+\|136670596\|136670692@59.78.20(+1A)&[MIMAT0000445&hsa-miR-126-3p&offsets\|+1\|-1(+1A);m-40&9\|+\|136670653\|136670674&offsets\|+1\|-1(+1A)]\|CGTACCGTGAGTAATAATGCA | >MI0000471\|hsa-mir-126&WithFlank&9\|+\|136670596\|136670692 | 59 | 78 | 20(+1A) | CGTACCGTGAGTAATAATGCA | 108.5 | 10.0 | 1.7 | 5.9 | 3.9E-09 | 8.7E-09 | MB231/MB468 | MP |
| >hg38_21_+_8433172_8446622_RNA45SN1_45S_50ntFlanks@3855.3881.27\|CTGTGGTAATTCTAGAGCTAATACATG | >hg38_21_+_8433172_8446622_RNA45SN1_45S_50ntFlanks | 3855 | 3881 | 27 | CTGTGGTAATTCTAGAGCTAATACATG | 172.5 | -9.5 | 1.6 | -5.9 | 3.1E-09 | 8.7E-09 | BT20/MB468 | MITO |
| >hg38_21_+_8433172_8446622_RNA45SN1_45S_50ntFlanks@9308.9333.26\|AGGAGTCTAACACGTGCGCGAGTCGG | >hg38_21_+_8433172_8446622_RNA45SN1_45S_50ntFlanks | 9308 | 9333 | 26 | AGGAGTCTAACACGTGCGCGAGTCGG | 96.4 | -10.5 | 1.8 | -5.9 | 3.9E-09 | 8.7E-09 | MB231/MB468 | MP |
| >hg38_21_+_8433172_8446622_RNA45SN1_45S_50ntFlanks@4443.4460.18\|CCCTTGCCTCTCGGCGCC | >hg38_21_+_8433172_8446622_RNA45SN1_45S_50ntFlanks | 4443 | 4460 | 18 | CCCTTGCCTCTCGGCGCC | 180.2 | -10.6 | 1.7 | -6.2 | 5.1E-10 | 8.7E-09 | BT20/MB468 | MP |
| >hg38_21_+_8433172_8446622_RNA45SN1_45S_50ntFlanks@7987.8012.26\|TCAGACGTGGCGACCCGCTGAATTTA | >hg38_21_+_8433172_8446622_RNA45SN1_45S_50ntFlanks | 7987 | 8012 | 26 | TCAGACGTGGCGACCCGCTGAATTTA | 118.0 | -9.0 | 1.5 | -5.9 | 3.1E-09 | 8.8E-09 | BT20/MB468 | MITO |
| >hg38_21_+_8433172_8446622_RNA45SN1_45S_50ntFlanks@11647.11687.41\|GTGTTGACGCGATGTGATTTCTGCCCAGTGCTCTGAATGTC | >hg38_21_+_8433172_8446622_RNA45SN1_45S_50ntFlanks | 11647 | 11687 | 41 | GTGTTGACGCGATGTGATTTCTGCCCAGTGCTCTGAATGTC | 695.9 | 13.2 | 2.1 | 6.2 | 5.1E-10 | 8.8E-09 | BT20/MB468 | MP |
| >hg38_21_+_8433172_8446622_RNA45SN1_45S_50ntFlanks@10534.10568.35\|CCGCGAGGCGTCCAGTGCGGTAACGCGACCGATCC | >hg38_21_+_8433172_8446622_RNA45SN1_45S_50ntFlanks | 10534 | 10568 | 35 | CCGCGAGGCGTCCAGTGCGGTAACGCGACCGATCC | 128.8 | -10.1 | 1.6 | -6.2 | 5.1E-10 | 8.8E-09 | BT20/MB468 | MP |
| >trnaMT_GluTTC_MT_-_14674_14742@34.71.38,trnalookalike8_GluTTC_5_-_93905172_93905240@34.71.38\|ATATCATTGGTCGTGGTTGTAGTCCGTGCGAGAATACC | >trnaMT_GluTTC_MT_-_14674_14742 | 34 | 71 | 38 | ATATCATTGGTCGTGGTTGTAGTCCGTGCGAGAATACC | 735.1 | 5.4 | 0.9 | 5.9 | 3.1E-09 | 8.8E-09 | BT20/MB468 | MITO |
| >hg38_21_+_8433172_8446622_RNA45SN1_45S_50ntFlanks@12633.12654.22\|CGAAGCTACCATCTGTGGGATT | >hg38_21_+_8433172_8446622_RNA45SN1_45S_50ntFlanks | 12633 | 12654 | 22 | CGAAGCTACCATCTGTGGGATT | 3421.6 | -6.7 | 1.1 | -5.9 | 4.1E-09 | 8.8E-09 | MB231/MB468 | CYTO |
| >hg38_21_+_8433172_8446622_RNA45SN1_45S_50ntFlanks@3858.3881.24\|TGGTAATTCTAGAGCTAATACATG | >hg38_21_+_8433172_8446622_RNA45SN1_45S_50ntFlanks | 3858 | 3881 | 24 | TGGTAATTCTAGAGCTAATACATG | 69.7 | -10.0 | 1.7 | -5.9 | 3.9E-09 | 8.8E-09 | MB231/MB468 | MP |
| >MI0000283\|hsa-mir-203a&WithFlank&14\|+\|104117399\|104117520@71.90.20&[MIMAT0000264&hsa-miR-203a-3p&offsets\|0\|-2;m-79&14\|+\|104117469\|104117490&offsets\|0\|-2]\|GTGAAATGTTTAGGACCACT | >MI0000283\|hsa-mir-203a&WithFlank&14\|+\|104117399\|104117520 | 71 | 90 | 20 | GTGAAATGTTTAGGACCACT | 467.5 | 8.2 | 1.3 | 6.2 | 4.3E-10 | 8.8E-09 | BT20/MB231 | NUC |
| >hg38_21_+_8433172_8446622_RNA45SN1_45S_50ntFlanks@12948.12982.35\|GACGACCTGCTTCTGGGTCGGGGTTTCGTACGTAG | >hg38_21_+_8433172_8446622_RNA45SN1_45S_50ntFlanks | 12948 | 12982 | 35 | GACGACCTGCTTCTGGGTCGGGGTTTCGTACGTAG | 504.5 | -8.4 | 1.4 | -5.9 | 4.7E-09 | 8.8E-09 | BT20/MB468 | NUC |
| >hg38_21_+_8433172_8446622_RNA45SN1_45S_50ntFlanks@8079.8104.26\|AAGAGCCCAGCGCCGAATCCCCGCCC | >hg38_21_+_8433172_8446622_RNA45SN1_45S_50ntFlanks | 8079 | 8104 | 26 | AAGAGCCCAGCGCCGAATCCCCGCCC | 2935.8 | -6.5 | 1.0 | -6.2 | 5.1E-10 | 8.8E-09 | BT20/MB468 | MP |
| >MI0000095\|hsa-mir-93&WithFlank&7\|-\|100093762\|100093853@17.40.24&[MIMAT0000093&hsa-miR-93-5p&offsets\|0\|+1;m-29&7\|-\|100093815\|100093837&offsets\|0\|+1]\|CAAAGTGCTGTTCGTGCAGGTAGT | >MI0000095\|hsa-mir-93&WithFlank&7\|-\|100093762\|100093853 | 17 | 40 | 24 | CAAAGTGCTGTTCGTGCAGGTAGT | 3089.0 | 5.6 | 1.0 | 5.9 | 4.5E-09 | 8.8E-09 | MB231/MB468 | MITO |
| >hg38_21_+_8433172_8446622_RNA45SN1_45S_50ntFlanks@10393.10410.18\|CAGCAGTTGAACATGGGT | >hg38_21_+_8433172_8446622_RNA45SN1_45S_50ntFlanks | 10393 | 10410 | 18 | CAGCAGTTGAACATGGGT | 905.2 | -8.2 | 1.4 | -5.9 | 4.5E-09 | 8.9E-09 | MB231/MB468 | MITO |
| >hg38_21_+_8433172_8446622_RNA45SN1_45S_50ntFlanks@4591.4616.26\|TTTTGTTGGTTTTCGGAACTGAGGCC | >hg38_21_+_8433172_8446622_RNA45SN1_45S_50ntFlanks | 4591 | 4616 | 26 | TTTTGTTGGTTTTCGGAACTGAGGCC | 449.0 | -7.9 | 1.3 | -5.9 | 3.9E-09 | 8.9E-09 | MB231/MB468 | MP |
| >hg38_21_+_8433172_8446622_RNA45SN1_45S_50ntFlanks@4733.4765.33\|GAACGAAAGTCGGAGGTTCGAAGACGATCAGAT | >hg38_21_+_8433172_8446622_RNA45SN1_45S_50ntFlanks | 4733 | 4765 | 33 | GAACGAAAGTCGGAGGTTCGAAGACGATCAGAT | 485.1 | 4.9 | 0.8 | 6.2 | 5.2E-10 | 8.9E-09 | BT20/MB468 | MP |
| >hg38_MT_+_598_1651_RNR1_12S_50ntFlanks@374.404.31\|AGCTAAAACTCACCTGAGTTGTAAAAAACTC | >hg38_MT_+_598_1651_RNR1_12S_50ntFlanks | 374 | 404 | 31 | AGCTAAAACTCACCTGAGTTGTAAAAAACTC | 519.5 | 5.0 | 0.8 | 5.9 | 3.1E-09 | 8.9E-09 | BT20/MB468 | MITO |
| >hg38_21_+_8433172_8446622_RNA45SN1_45S_50ntFlanks@6667.6699.33\|ATCACTCGGCTCGTGCGTCGATGAAGAACGCAG | >hg38_21_+_8433172_8446622_RNA45SN1_45S_50ntFlanks | 6667 | 6699 | 33 | ATCACTCGGCTCGTGCGTCGATGAAGAACGCAG | 1028.6 | -8.1 | 1.4 | -5.9 | 4.5E-09 | 8.9E-09 | MB231/MB468 | MITO |
| >hg38_21_+_8433172_8446622_RNA45SN1_45S_50ntFlanks@11572.11599.28\|CCGACTTAGAACTGGTGCGGACCAGGGG | >hg38_21_+_8433172_8446622_RNA45SN1_45S_50ntFlanks | 11572 | 11599 | 28 | CCGACTTAGAACTGGTGCGGACCAGGGG | 872.6 | -6.9 | 1.2 | -5.9 | 4.0E-09 | 8.9E-09 | MB231/MB468 | MP |
| >hg38_21_+_8433172_8446622_RNA45SN1_45S_50ntFlanks@12306.12328.23\|CACGATCCTTCTGACCTTTTGGG | >hg38_21_+_8433172_8446622_RNA45SN1_45S_50ntFlanks | 12306 | 12328 | 23 | CACGATCCTTCTGACCTTTTGGG | 101.9 | -10.5 | 1.8 | -5.9 | 4.0E-09 | 8.9E-09 | MB231/MB468 | MP |
| >hg38_21_+_8433172_8446622_RNA45SN1_45S_50ntFlanks@11572.11590.19\|CCGACTTAGAACTGGTGCG | >hg38_21_+_8433172_8446622_RNA45SN1_45S_50ntFlanks | 11572 | 11590 | 19 | CCGACTTAGAACTGGTGCG | 5105.7 | -7.3 | 1.2 | -5.9 | 4.0E-09 | 8.9E-09 | MB231/MB468 | MP |
| >trnaMT_PheGAA_MT_+_577_647@36.60.25\|AAAATGTTTAGACGGGCTCACATCA | >trnaMT_PheGAA_MT_+_577_647 | 36 | 60 | 25 | AAAATGTTTAGACGGGCTCACATCA | 741.7 | 9.9 | 1.7 | 5.9 | 3.2E-09 | 8.9E-09 | BT20/MB468 | MITO |
| >hg38_MT_+_1621_3279_RNR2_16S_50ntFlanks@51.82.32\|GCTAAACCTAGCCCCAAACCCACTCCACCTTA | >hg38_MT_+_1621_3279_RNR2_16S_50ntFlanks | 51 | 82 | 32 | GCTAAACCTAGCCCCAAACCCACTCCACCTTA | 3842.2 | 6.0 | 1.0 | 5.9 | 3.2E-09 | 8.9E-09 | BT20/MB468 | MITO |
| >hg38_21_+_8433172_8446622_RNA45SN1_45S_50ntFlanks@11701.11728.28\|TCAATGAAGCGCGGGTAAACGGCGGGAG | >hg38_21_+_8433172_8446622_RNA45SN1_45S_50ntFlanks | 11701 | 11728 | 28 | TCAATGAAGCGCGGGTAAACGGCGGGAG | 97.4 | -8.7 | 1.5 | -5.9 | 3.2E-09 | 8.9E-09 | BT20/MB468 | MITO |
| >trnaMT_ProTGG_MT_-_15956_16023@50.71.22\|TTAAAGACTTTTTCTCTGACCA | >trnaMT_ProTGG_MT_-_15956_16023 | 50 | 71 | 22 | TTAAAGACTTTTTCTCTGACCA | 906.3 | 8.3 | 1.4 | 5.9 | 4.2E-09 | 8.9E-09 | MB231/MB468 | NUC |
| >hg38_21_+_8433172_8446622_RNA45SN1_45S_50ntFlanks@12083.12108.26\|CTCGCTTCTGGCGCCAAGCGCCCGGC | >hg38_21_+_8433172_8446622_RNA45SN1_45S_50ntFlanks | 12083 | 12108 | 26 | CTCGCTTCTGGCGCCAAGCGCCCGGC | 426.4 | -7.7 | 1.3 | -5.9 | 4.0E-09 | 8.9E-09 | MB231/MB468 | MP |
| >trna10_AspGTC_12_-_125424193_125424264@36.74.39,trna12_AspGTC_12_-_125411891_125411962@36.74.39,trna144_AspGTC_6_-_27551236_27551307@36.74.39,trna38_AspGTC_17_-_8125556_8125627@36.74.39,trna45_AspGTC_6_+_27447453_27447524@36.74.39,trna48_AspGTC_6_+_27471523_27471594@36.74.39,trna4_AspGTC_12_+_96429799_96429870@36.74.39,trna69_AspGTC_1_-_161440205_161440276@36.74.39,trna72_AspGTC_1_-_161432824_161432895@36.74.39,trna75_AspGTC_1_-_161425414_161425485@36.74.39,trna78_AspGTC_1_-_161418033_161418104@36.74.39,trna81_AspGTC_1_-_161410615_161410686@36.74.39\|CACGCGGGAGACCGGGGTTCGATTCCCCGACGGGGAGCC | >trna10_AspGTC_12_-_125424193_125424264 | 36 | 74 | 39 | CACGCGGGAGACCGGGGTTCGATTCCCCGACGGGGAGCC | 112.8 | 10.0 | 1.7 | 5.9 | 4.0E-09 | 8.9E-09 | MB231/MB468 | MP |
| >hg38_21_+_8433172_8446622_RNA45SN1_45S_50ntFlanks@12404.12434.31\|GCTTTTTGATCCTTCGATGTCGGCTCTTCCT | >hg38_21_+_8433172_8446622_RNA45SN1_45S_50ntFlanks | 12404 | 12434 | 31 | GCTTTTTGATCCTTCGATGTCGGCTCTTCCT | 2264.2 | -5.6 | 0.9 | -5.9 | 4.8E-09 | 8.9E-09 | BT20/MB468 | NUC |
| >hg38_21_+_8433172_8446622_RNA45SN1_45S_50ntFlanks@3789.3810.22\|ACTGCGAATGGCTCATTAAATC | >hg38_21_+_8433172_8446622_RNA45SN1_45S_50ntFlanks | 3789 | 3810 | 22 | ACTGCGAATGGCTCATTAAATC | 865.7 | -6.6 | 1.1 | -5.9 | 4.6E-09 | 8.9E-09 | MB231/MB468 | MITO |
| >am_trna10_LysCTT_16_+_3241501_3241573@31.51.21,trna119_LysCTT_1_-_145395522_145395594@31.51.21,trna11_LysCTT_5_-_180648979_180649051@31.51.21,trna13_LysCTT_16_+_3246153_3246225@31.51.21,trna13_LysCTT_6_+_26556774_26556846@31.51.21,trna30_LysCTT_16_-_3230555_3230627@31.51.21,trna5_LysCTT_16_+_3214939_3215011@31.51.21,trna7_LysCTT_16_+_3225692_3225764@31.51.21,trna8_LysCTT_1_+_39970195_39970267@31.51.21,trna9_LysCTT_5_+_180634755_180634827@31.51.21\|ACTCTTAATCTCAGGGTCGTG | >am_trna10_LysCTT_16_+_3241501_3241573 | 31 | 51 | 21 | ACTCTTAATCTCAGGGTCGTG | 90.3 | -8.6 | 1.4 | -5.9 | 3.2E-09 | 8.9E-09 | BT20/MB468 | MITO |
| >hg38_21_+_8433172_8446622_RNA45SN1_45S_50ntFlanks@11115.11133.19\|CGGCAGGGGCCGGCGGCGG | >hg38_21_+_8433172_8446622_RNA45SN1_45S_50ntFlanks | 11115 | 11133 | 19 | CGGCAGGGGCCGGCGGCGG | 115.7 | -8.9 | 1.5 | -5.9 | 3.2E-09 | 8.9E-09 | BT20/MB468 | MITO |
| >hg38_21_+_8433172_8446622_RNA45SN1_45S_50ntFlanks@5186.5206.21\|CAATAACAGGTCTGTGATGCC | >hg38_21_+_8433172_8446622_RNA45SN1_45S_50ntFlanks | 5186 | 5206 | 21 | CAATAACAGGTCTGTGATGCC | 418.8 | -10.8 | 1.8 | -5.9 | 3.2E-09 | 8.9E-09 | BT20/MB468 | MITO |
| >trna116_GluCTC_1_-_145399233_145399304@1.37.37,trna59_GluCTC_1_+_249168447_249168518@1.37.37,trna71_GluCTC_1_-_161439189_161439260@1.37.37,trna74_GluCTC_1_-_161431809_161431880@1.37.37,trna77_GluCTC_1_-_161424398_161424469@1.37.37,trna77_GluCTC_6_+_28949976_28950047@1.37.37,trna80_GluCTC_1_-_161417018_161417089@1.37.37,trna87_GluCTC_6_-_126101393_126101464@1.37.37\|TCCCTGGTGGTCTAGTGGTTAGGATTCGGCGCTCTCA | >trna116_GluCTC_1_-_145399233_145399304 | 1 | 37 | 37 | TCCCTGGTGGTCTAGTGGTTAGGATTCGGCGCTCTCA | 466.7 | 8.6 | 1.4 | 5.9 | 3.2E-09 | 8.9E-09 | BT20/MB468 | MITO |
| >hg38_21_+_8433172_8446622_RNA45SN1_45S_50ntFlanks@11235.11269.35\|CTTCCCGTGGATCGCCCCAGCTGCGGCGGGCGTCG | >hg38_21_+_8433172_8446622_RNA45SN1_45S_50ntFlanks | 11235 | 11269 | 35 | CTTCCCGTGGATCGCCCCAGCTGCGGCGGGCGTCG | 3038.7 | -10.3 | 1.8 | -5.9 | 4.8E-09 | 9.0E-09 | BT20/MB468 | NUC |
| >hg38_21_+_8433172_8446622_RNA45SN1_45S_50ntFlanks@11792.11819.28\|ATGAACGAGATTCCCACTGTCCCTACCT | >hg38_21_+_8433172_8446622_RNA45SN1_45S_50ntFlanks | 11792 | 11819 | 28 | ATGAACGAGATTCCCACTGTCCCTACCT | 152.9 | -10.3 | 1.7 | -6.2 | 5.2E-10 | 9.0E-09 | BT20/MB468 | MP |
| >hg38_21_+_8433172_8446622_RNA45SN1_45S_50ntFlanks@10687.10706.20\|CGGCGTCCGGTGAGCTCTCG | >hg38_21_+_8433172_8446622_RNA45SN1_45S_50ntFlanks | 10687 | 10706 | 20 | CGGCGTCCGGTGAGCTCTCG | 4736.7 | -7.0 | 1.2 | -5.9 | 4.2E-09 | 9.0E-09 | MB231/MB468 | CYTO |

| | | | | | | | | | | | | | |
|---|---|---|---|---|---|---|---|---|---|---|---|---|---|
| >hg38_21_+_8433172_8446622_RNA45SN1_45S_50ntFlanks@12417.12436.20\|TCGATGTCGGCTCTTCCTAT | >hg38_21_+_8433172_8446622_RNA45SN1_45S_50ntFlanks | 12417 | 12436 | 20 | TCGATGTCGGCTCTTCCTAT | 24118.1 | -7.7 | 1.3 | -5.9 | 4.8E-09 | 9.0E-09 | BT20/MB468 | NUC |
| >hg38_21_+_8433172_8446622_RNA45SN1_45S_50ntFlanks@7988.8010.23\|CAGACGTGGCGACCCGCTGAATT | >hg38_21_+_8433172_8446622_RNA45SN1_45S_50ntFlanks | 7988 | 8010 | 23 | CAGACGTGGCGACCCGCTGAATT | 317.5 | 3.7 | 0.6 | 5.9 | 3.6E-09 | 9.0E-09 | BT20/MB468 | CYTO |
| >hg38_21_+_8433172_8446622_RNA45SN1_45S_50ntFlanks@10783.10800.18\|GTGAACAGCCTCTGGCAT | >hg38_21_+_8433172_8446622_RNA45SN1_45S_50ntFlanks | 10783 | 10800 | 18 | GTGAACAGCCTCTGGCAT | 65.7 | -9.9 | 1.7 | -5.9 | 4.0E-09 | 9.0E-09 | MB231/MB468 | MP |
| >hg38_21_+_8433172_8446622_RNA45SN1_45S_50ntFlanks@9915.9948.34\|AAAAGGTGTTGGTTGATATAGACAGCAGGACGGT | >hg38_21_+_8433172_8446622_RNA45SN1_45S_50ntFlanks | 9915 | 9948 | 34 | AAAAGGTGTTGGTTGATATAGACAGCAGGACGGT | 104.1 | 10.4 | 1.7 | 6.2 | 5.3E-10 | 9.0E-09 | BT20/MB468 | MP |
| >hg38_21_+_8433172_8446622_RNA45SN1_45S_50ntFlanks@5333.5362.30\|CATGAACGAGGAATTCCCAGTAAGTGCGGG | >hg38_21_+_8433172_8446622_RNA45SN1_45S_50ntFlanks | 5333 | 5362 | 30 | CATGAACGAGGAATTCCCAGTAAGTGCGGG | 149.5 | -11.1 | 1.9 | -5.9 | 4.0E-09 | 9.0E-09 | MB231/MB468 | MP |
| >trna134_GluTTC_1_-_16861774_16861845@1.31.31,trna5_GluTTC_1_+_17199078_17199149@1.31.31,trna84_GluTTC_1_-_161391883_161391954@1.31.31,trna94_GluTTC_1_-_149664355_149664427@1.31.31\|TCCCTGGTGGTCTAGTGGCTAGGATTCGGCG | >trna134_GluTTC_1_-_16861774_16861845 | 1 | 31 | 31 | TCCCTGGTGGTCTAGTGGCTAGGATTCGGCG | 6208.0 | -9.4 | 1.6 | -5.9 | 4.3E-09 | 9.0E-09 | MB231/MB468 | NUC |
| >trna152_ValCAC_6_-_27248049_27248121@1.36.36\|GCTTCTGTAGTGTAGTGGTTATCACGTTCGCCTCAC | >trna152_ValCAC_6_-_27248049_27248121 | 1 | 36 | 36 | GCTTCTGTAGTGTAGTGGTTATCACGTTCGCCTCAC | 126.7 | 10.2 | 1.7 | 5.9 | 4.0E-09 | 9.0E-09 | MB231/MB468 | MP |
| >hg38_21_+_8433172_8446622_RNA45SN1_45S_50ntFlanks@12937.12956.20\|AACCATTCGTAGACGACCTG | >hg38_21_+_8433172_8446622_RNA45SN1_45S_50ntFlanks | 12937 | 12956 | 20 | AACCATTCGTAGACGACCTG | 1710.9 | -3.8 | 0.6 | -5.9 | 3.2E-09 | 9.0E-09 | BT20/MB468 | MITO |
| >hg38_21_+_8433172_8446622_RNA45SN1_45S_50ntFlanks@3852.3874.23\|TAACTGTGGTAATTCTAGAGCTA | >hg38_21_+_8433172_8446622_RNA45SN1_45S_50ntFlanks | 3852 | 3874 | 23 | TAACTGTGGTAATTCTAGAGCTA | 284.6 | -5.9 | 1.0 | -5.9 | 4.0E-09 | 9.0E-09 | MB231/MB468 | MP |
| >hg38_21_+_8433172_8446622_RNA45SN1_45S_50ntFlanks@4241.4258.18\|CTCTTTCGAGGCCCTGTA | >hg38_21_+_8433172_8446622_RNA45SN1_45S_50ntFlanks | 4241 | 4258 | 18 | CTCTTTCGAGGCCCTGTA | 368.1 | -7.4 | 1.3 | -5.9 | 4.3E-09 | 9.0E-09 | MB231/MB468 | NUC |
| >hg38_21_+_8433172_8446622_RNA45SN1_45S_50ntFlanks@10427.10456.30\|ATGGGCGAGCGCCGTTCCGAAGGGACGGGC | >hg38_21_+_8433172_8446622_RNA45SN1_45S_50ntFlanks | 10427 | 10456 | 30 | ATGGGCGAGCGCCGTTCCGAAGGGACGGGC | 1937.8 | -9.6 | 1.6 | -5.9 | 4.8E-09 | 9.1E-09 | BT20/MB468 | NUC |
| >hg38_21_+_8433172_8446622_RNA45SN1_45S_50ntFlanks@9739.9759.21\|AAACGATCTCAACCTATTCTC | >hg38_21_+_8433172_8446622_RNA45SN1_45S_50ntFlanks | 9739 | 9759 | 21 | AAACGATCTCAACCTATTCTC | 1921.1 | -7.4 | 1.3 | -5.9 | 4.8E-09 | 9.1E-09 | BT20/MB468 | NUC |
| >hg38_21_+_8433172_8446622_RNA45SN1_45S_50ntFlanks@10403.10429.27\|ACATGGGTCAGTCGGTCCTGAGAGATG | >hg38_21_+_8433172_8446622_RNA45SN1_45S_50ntFlanks | 10403 | 10429 | 27 | ACATGGGTCAGTCGGTCCTGAGAGATG | 665.7 | -6.6 | 1.1 | -5.9 | 4.0E-09 | 9.1E-09 | MB231/MB468 | MP |
| >hg38_21_+_8433172_8446622_RNA45SN1_45S_50ntFlanks@9484.9510.27\|GTGTTAGGACCCGAAAGATGGTGAACT | >hg38_21_+_8433172_8446622_RNA45SN1_45S_50ntFlanks | 9484 | 9510 | 27 | GTGTTAGGACCCGAAAGATGGTGAACT | 138.9 | 10.9 | 1.7 | 6.2 | 5.3E-10 | 9.1E-09 | BT20/MB468 | MP |
| >hg38_21_+_8433172_8446622_RNA45SN1_45S_50ntFlanks@9104.9121.18\|CCGGCGACGGGGGGGGTG | >hg38_21_+_8433172_8446622_RNA45SN1_45S_50ntFlanks | 9104 | 9121 | 18 | CCGGCGACGGGGGGGGTG | 138.3 | -10.2 | 1.6 | -6.2 | 5.3E-10 | 9.1E-09 | BT20/MB468 | MP |
| >hg38_21_+_8433172_8446622_RNA45SN1_45S_50ntFlanks@12654.12673.20\|TATGACTGAACGCCTCTAAG | >hg38_21_+_8433172_8446622_RNA45SN1_45S_50ntFlanks | 12654 | 12673 | 20 | TATGACTGAACGCCTCTAAG | 290.2 | -7.3 | 1.2 | -5.9 | 4.3E-09 | 9.1E-09 | MB231/MB468 | NUC |
| >hg38_21_+_8433172_8446622_RNA45SN1_45S_50ntFlanks@10727.10755.29\|GGAGAGGGTGTAAATCTCGCGCCGGGCCG | >hg38_21_+_8433172_8446622_RNA45SN1_45S_50ntFlanks | 10727 | 10755 | 29 | GGAGAGGGTGTAAATCTCGCGCCGGGCCG | 1194.5 | -8.7 | 1.4 | -6.2 | 5.3E-10 | 9.1E-09 | BT20/MB468 | MP |
| >hg38_21_+_8433172_8446622_RNA45SN1_45S_50ntFlanks@9788.9821.34\|CGCTGGCGTGGAGCCGGGCGTGGAATGCGAGTGC | >hg38_21_+_8433172_8446622_RNA45SN1_45S_50ntFlanks | 9788 | 9821 | 34 | CGCTGGCGTGGAGCCGGGCGTGGAATGCGAGTGC | 366.0 | -7.8 | 1.3 | -5.9 | 4.6E-09 | 9.1E-09 | MB231/MB468 | MITO |
| >hg38_21_+_8433172_8446622_RNA45SN1_45S_50ntFlanks@10668.10706.39\|GAAAGCGTCGCGGTTCCGGCGGCGTCCGGTGAGCTCTCG | >hg38_21_+_8433172_8446622_RNA45SN1_45S_50ntFlanks | 10668 | 10706 | 39 | GAAAGCGTCGCGGTTCCGGCGGCGTCCGGTGAGCTCTCG | 2605.9 | -10.1 | 1.7 | -5.9 | 4.9E-09 | 9.1E-09 | BT20/MB468 | NUC |
| >hg38_21_+_8433172_8446622_RNA45SN1_45S_50ntFlanks@10682.10706.25\|TCCGGCGGCGTCCGGTGAGCTCTCG | >hg38_21_+_8433172_8446622_RNA45SN1_45S_50ntFlanks | 10682 | 10706 | 25 | TCCGGCGGCGTCCGGTGAGCTCTCG | 1012.0 | -3.5 | 0.6 | -5.9 | 3.2E-09 | 9.1E-09 | BT20/MB468 | MITO |
| >hg38_21_+_8433172_8446622_RNA45SN1_45S_50ntFlanks@9533.9561.29\|GAGGAAACTCTGGTGGAGGTCCGTAGCGG | >hg38_21_+_8433172_8446622_RNA45SN1_45S_50ntFlanks | 9533 | 9561 | 29 | GAGGAAACTCTGGTGGAGGTCCGTAGCGG | 436.5 | -7.9 | 1.3 | -5.9 | 4.1E-09 | 9.1E-09 | MB231/MB468 | MP |
| >MI0000749\|hsa-mir-30e&WithFlank&1\|+\|40754349\|40754452@23.45.23&[MIMAT0000692&hsa-miR-30e-5p&offsets\|0\|+1;m-24&1\|+\|40754371\|40754394&offsets\|0\|-1]\|TGTAAACATCCTTGACTGGAAGC | >MI0000749\|hsa-mir-30e&WithFlank&1\|+\|40754349\|40754452 | 23 | 45 | 23 | TGTAAACATCCTTGACTGGAAGC | 171.6 | 4.9 | 0.8 | 5.9 | 4.3E-09 | 9.1E-09 | MB231/MB468 | NUC |
| >hg38_21_+_8433172_8446622_RNA45SN1_45S_50ntFlanks@12935.12956.22\|TAAACCATTCGTAGACGACCTG | >hg38_21_+_8433172_8446622_RNA45SN1_45S_50ntFlanks | 12935 | 12956 | 22 | TAAACCATTCGTAGACGACCTG | 658.8 | -3.3 | 0.6 | -5.9 | 3.2E-09 | 9.1E-09 | BT20/MB468 | MITO |
| >hg38_21_+_8433172_8446622_RNA45SN1_45S_50ntFlanks@10822.10865.44\|GAAGTCGGCAAGCCGGATCCGTAACTTCGGGATAAGGATTGGCT | >hg38_21_+_8433172_8446622_RNA45SN1_45S_50ntFlanks | 10822 | 10865 | 44 | GAAGTCGGCAAGCCGGATCCGTAACTTCGGGATAAGGATTGGCT | 388.6 | -6.4 | 1.1 | -5.9 | 4.7E-09 | 9.1E-09 | MB231/MB468 | MITO |
| >hg38_21_+_8433172_8446622_RNA45SN1_45S_50ntFlanks@11703.11726.24\|AATGAAGCGCGGGTAAACGGCGGG | >hg38_21_+_8433172_8446622_RNA45SN1_45S_50ntFlanks | 11703 | 11726 | 24 | AATGAAGCGCGGGTAAACGGCGGG | 407.8 | -6.1 | 1.0 | -5.9 | 4.3E-09 | 9.1E-09 | MB231/MB468 | NUC |
| >MI0000082\|hsa-mir-25&WithFlank&7\|-\|100093554\|100093649@58.77.20&[MIMAT0000081&hsa-miR-25-3p&offsets\|0\|-2;m-23&7\|-\|100093571\|100093592&offsets\|0\|-2]\|CATTGCACTTGTCTCGGTCT | >MI0000082\|hsa-mir-25&WithFlank&7\|-\|100093554\|100093649 | 58 | 77 | 20 | CATTGCACTTGTCTCGGTCT | 298.0 | 7.6 | 1.3 | 5.9 | 4.3E-09 | 9.1E-09 | MB231/MB468 | CYTO |
| >hg38_21_+_8433172_8446622_RNA45SN1_45S_50ntFlanks@12992.13017.26\|TCCCTCGCTGCGATCTATTGAAAGTC | >hg38_21_+_8433172_8446622_RNA45SN1_45S_50ntFlanks | 12992 | 13017 | 26 | TCCCTCGCTGCGATCTATTGAAAGTC | 458.1 | 4.0 | 0.7 | 5.9 | 4.3E-09 | 9.1E-09 | MB231/MB468 | CYTO |
| >hg38_21_+_8433172_8446622_RNA45SN1_45S_50ntFlanks@8158.8188.31\|TCGTGGGGGGCCCAAGTCCTTCTGATCGAGG | >hg38_21_+_8433172_8446622_RNA45SN1_45S_50ntFlanks | 8158 | 8188 | 31 | TCGTGGGGGGCCCAAGTCCTTCTGATCGAGG | 391.4 | -4.0 | 0.7 | -5.9 | 3.2E-09 | 9.1E-09 | BT20/MB468 | MITO |
| >hg38_21_+_8433172_8446622_RNA45SN1_45S_50ntFlanks@11210.11233.24\|GGCGACTCTGGACGCGAGCCGGGC | >hg38_21_+_8433172_8446622_RNA45SN1_45S_50ntFlanks | 11210 | 11233 | 24 | GGCGACTCTGGACGCGAGCCGGGC | 289.7 | -7.5 | 1.3 | -5.9 | 4.7E-09 | 9.2E-09 | MB231/MB468 | MITO |
| >hg38_21_+_8433172_8446622_RNA45SN1_45S_50ntFlanks@4003.4042.40\|ATAACCTCGGGCCGATCGCACGCCCCCCGTGGCGGCGACG | >hg38_21_+_8433172_8446622_RNA45SN1_45S_50ntFlanks | 4003 | 4042 | 40 | ATAACCTCGGGCCGATCGCACGCCCCCCGTGGCGGCGACG | 2252.6 | -7.1 | 1.2 | -5.9 | 3.7E-09 | 9.2E-09 | BT20/MB468 | CYTO |
| >hg38_21_+_8433172_8446622_RNA45SN1_45S_50ntFlanks@10806.10840.35\|AACAATGTAGGTAAGGGAAGTCGGCAAGCCGGATC | >hg38_21_+_8433172_8446622_RNA45SN1_45S_50ntFlanks | 10806 | 10840 | 35 | AACAATGTAGGTAAGGGAAGTCGGCAAGCCGGATC | 555.3 | -6.2 | 1.1 | -5.9 | 4.3E-09 | 9.2E-09 | MB231/MB468 | NUC |
| >hg38_21_+_8433172_8446622_RNA45SN1_45S_50ntFlanks@10735.10752.18\|TGTAAATCTCGCGCCGGG | >hg38_21_+_8433172_8446622_RNA45SN1_45S_50ntFlanks | 10735 | 10752 | 18 | TGTAAATCTCGCGCCGGG | 260.8 | -6.5 | 1.1 | -5.9 | 3.7E-09 | 9.2E-09 | BT20/MB468 | CYTO |
| >trna116_GluCTC_1_-_145399233_145399304@30.70.41,trna71_GluCTC_1_-_161439189_161439260@30.70.41,trna74_GluCTC_1_-_161431809_161431880@30.70.41,trna77_GluCTC_1_-_161424398_161424469@30.70.41,trna77_GluCTC_6_+_28949976_28950047@30.70.41,trna80_GluCTC_1_-_161417018_161417089@30.70.41,trna87_GluCTC_6_-_126101393_126101464@30.70.41\|CGCTCTCACCGCCGCGGCCCGGGTTCGATTCCCGGTCAGGG | >trna116_GluCTC_1_-_145399233_145399304 | 30 | 70 | 41 | CGCTCTCACCGCCGCGGCCCGGGTTCGATTCCCGGTCAGGG | 406.5 | -6.9 | 1.2 | -5.9 | 4.3E-09 | 9.2E-09 | MB231/MB468 | CYTO |
| >hg38_21_+_8433172_8446622_RNA45SN1_45S_50ntFlanks@11163.11188.26\|CGGTCCCCCGCGAGGGGGGCCCGGGC | >hg38_21_+_8433172_8446622_RNA45SN1_45S_50ntFlanks | 11163 | 11188 | 26 | CGGTCCCCCGCGAGGGGGGCCCGGGC | 130.2 | -9.1 | 1.5 | -5.9 | 3.3E-09 | 9.2E-09 | BT20/MB468 | MITO |
| >hg38_21_+_8433172_8446622_RNA45SN1_45S_50ntFlanks@3843.3869.27\|CTACTTGGATAACTGTGGTAATTCTAG | >hg38_21_+_8433172_8446622_RNA45SN1_45S_50ntFlanks | 3843 | 3869 | 27 | CTACTTGGATAACTGTGGTAATTCTAG | 185.7 | -9.6 | 1.6 | -5.9 | 3.3E-09 | 9.2E-09 | BT20/MB468 | MITO |
| >hg38_21_+_8433172_8446622_RNA45SN1_45S_50ntFlanks@9862.9901.40\|ATGAACCGAACGCCGGGTTAAGGCGCCCGATGCCGACGCT | >hg38_21_+_8433172_8446622_RNA45SN1_45S_50ntFlanks | 9862 | 9901 | 40 | ATGAACCGAACGCCGGGTTAAGGCGCCCGATGCCGACGCT | 158.7 | -11.1 | 1.9 | -5.9 | 4.7E-09 | 9.2E-09 | MB231/MB468 | MITO |
| >hg38_21_+_8433172_8446622_RNA45SN1_45S_50ntFlanks@9913.9940.28\|AGAAAAGGTGTTGGTTGATATAGACAGC | >hg38_21_+_8433172_8446622_RNA45SN1_45S_50ntFlanks | 9913 | 9940 | 28 | AGAAAAGGTGTTGGTTGATATAGACAGC | 142.1 | 10.9 | 1.8 | 6.2 | 5.4E-10 | 9.2E-09 | BT20/MB468 | MP |
| >hg38_21_+_8433172_8446622_RNA45SN1_45S_50ntFlanks@8356.8380.25\|GAAAAGAACTTTGAAGAGAGAGTTC | >hg38_21_+_8433172_8446622_RNA45SN1_45S_50ntFlanks | 8356 | 8380 | 25 | GAAAAGAACTTTGAAGAGAGAGTTC | 179.5 | -10.6 | 1.7 | -6.2 | 5.4E-10 | 9.2E-09 | BT20/MB468 | MP |
| >MI0000455\|hsa-mir-138-2&WithFlank&16\|+\|56858512\|56858607@16.38.23(+1C);MI0000476\|hsa-mir-138-1&WithFlank&3\|+\|44114206\|44114316@29.51.23(+1C)&[MIMAT0000430&hsa-miR-138-5p&offsets\|0\|0(+1C);m-509&16\|+\|56858527\|56858549&offsets\|0\|0(+1C)];[MIMAT0000430_1&hsa-miR-138-5p&offsets\|0\|0(+1C);m-497&3\|+\|44114234\|44114256&offsets\|0\|0(+1C)]\|AGCTGGTGTTGTGAATCAGGCCGC | >MI0000455\|hsa-mir-138-2&WithFlank&16\|+\|56858512\|56858607 | 16 | 38 | 23(+1C) | AGCTGGTGTTGTGAATCAGGCCGC | 229.0 | -11.2 | 1.7 | -6.5 | 1.0E-10 | 9.2E-09 | BT20/MB231 | MP |
| >MI0000465\|hsa-mir-191&WithFlank&3\|-\|49020612\|49020715@22.42.21(+1A)&[MIMAT0000440&hsa-miR-191-5p&offsets\|0\|-2(+1A);m-48&3\|-\|49020672\|49020694&offsets\|0\|-2(+1A)]\|CAACGGAATCCCAAAAGCAGCA | >MI0000465\|hsa-mir-191&WithFlank&3\|-\|49020612\|49020715 | 22 | 42 | 21(+1A) | CAACGGAATCCCAAAAGCAGCA | 366.1 | 5.7 | 1.0 | 5.9 | 4.3E-09 | 9.2E-09 | MB231/MB468 | CYTO |
| >hg38_21_+_8433172_8446622_RNA45SN1_45S_50ntFlanks@4971.4999.29\|CCGGCCCGGACACGGACAGGATTGACAGA | >hg38_21_+_8433172_8446622_RNA45SN1_45S_50ntFlanks | 4971 | 4999 | 29 | CCGGCCCGGACACGGACAGGATTGACAGA | 1073.6 | -6.9 | 1.2 | -5.9 | 4.7E-09 | 9.2E-09 | MB231/MB468 | MITO |
| >MI0000816\|hsa-mir-335&WithFlank&7\|+\|130496105\|130496210@57.79.23&[MIMAT0004703&hsa-miR-335-3p&offsets\|-1\|0;m-159&7\|+\|130496162\|130496183&offsets\|-1\|0]\|GTTTTTCATTATTGCTCCTGACC | >MI0000816\|hsa-mir-335&WithFlank&7\|+\|130496105\|130496210 | 57 | 79 | 23 | GTTTTTCATTATTGCTCCTGACC | 202.5 | 4.0 | 0.7 | 5.9 | 4.4E-09 | 9.2E-09 | MB231/MB468 | NUC |
| >MI0000061\|hsa-let-7a-2&WithFlank&11\|-\|122146516\|122146599@11.31.21;MI0000062\|hsa-let-7a-3&WithFlank&22\|+\|46112743\|46112828@10.30.21;MI0000060\|hsa-let-7a-1&WithFlank&9\|+\|94175951\|94176042@12.32.21&[MIMAT0000062&hsa-let-7a-5p&offsets\|0\|-1;m-6&11\|-\|122146568\|122146589&offsets\|0\|-1];[MIMAT0000062_1&hsa-let-7a-5p&offsets\|0\|-1;m-5&22\|+\|46112752\|46112773&offsets\|0\|-1];[MIMAT0000062_2&hsa-let-7a-5p&offsets\|0\|-1;m-4&9\|+\|94175962\|94175983&offsets\|0\|-1]\|TGAGGTAGTAGGTTGTATAGT | >MI0000061\|hsa-let-7a-2&WithFlank&11\|-\|122146516\|122146599 | 11 | 31 | 21 | TGAGGTAGTAGGTTGTATAGT | 5676.7 | 4.5 | 0.8 | 5.9 | 4.1E-09 | 9.2E-09 | MB231/MB468 | MP |
| >hg38_21_+_8433172_8446622_RNA45SN1_45S_50ntFlanks@12442.12473.32\|TGAAGCAGAATTCACCAAGCGTTGGATTGTTC | >hg38_21_+_8433172_8446622_RNA45SN1_45S_50ntFlanks | 12442 | 12473 | 32 | TGAAGCAGAATTCACCAAGCGTTGGATTGTTC | 169.8 | -10.5 | 1.7 | -6.2 | 5.5E-10 | 9.3E-09 | BT20/MB468 | MP |
| >MI0000065\|hsa-let-7d&WithFlank&9\|+\|94178828\|94178926@68.88.21(+1C)&[MIMAT0004484&hsa-let-7d-3p&offsets\|0\|-1(+1C);m-98&9\|+\|94178895\|94178916&offsets\|0\|-1(+1C)]\|CTATACGACCTGCTGCCTTTCC | >MI0000065\|hsa-let-7d&WithFlank&9\|+\|94178828\|94178926 | 68 | 88 | 21(+1C) | CTATACGACCTGCTGCCTTTCC | 517.4 | 6.2 | 1.1 | 5.9 | 4.7E-09 | 9.3E-09 | MB231/MB468 | MITO |
| >hg38_21_+_8433172_8446622_RNA45SN1_45S_50ntFlanks@10806.10844.39\|AACAATGTAGGTAAGGGAAGTCGGCAAGCCGGATCCGTA | >hg38_21_+_8433172_8446622_RNA45SN1_45S_50ntFlanks | 10806 | 10844 | 39 | AACAATGTAGGTAAGGGAAGTCGGCAAGCCGGATCCGTA | 301.6 | -7.8 | 1.3 | -5.9 | 4.7E-09 | 9.3E-09 | MB231/MB468 | MITO |

| | | | | | | | | | | | | | |
|---|---|---|---|---|---|---|---|---|---|---|---|---|---|
| >hg38_21_+_8433172_8446622_RNA45SN1_45S_50ntFlanks@10351.10375.25\|CGAGAACTTTGAAGGCCGAAGTGGA | >hg38_21_+_8433172_8446622_RNA45SN1_45S_50ntFlanks | 10351 | 10375 | 25 | CGAGAACTTTGAAGGCCGAAGTGGA | 229.6 | -5.6 | 1.0 | -5.9 | 4.4E-09 | 9.3E-09 | MB231/MB468 | NUC |
| >hg38_21_+_8433172_8446622_RNA45SN1_45S_50ntFlanks@5083.5102.20\|ACGAACGAGACTCTGGCATG | >hg38_21_+_8433172_8446622_RNA45SN1_45S_50ntFlanks | 5083 | 5102 | 20 | ACGAACGAGACTCTGGCATG | 1064.9 | -7.4 | 1.3 | -5.9 | 4.1E-09 | 9.3E-09 | MB231/MB468 | MP |
| >hg38_21_+_8433172_8446622_RNA45SN1_45S_50ntFlanks@11670.11702.33\|CCCAGTGCTCTGAATGTCAAAGTGAAGAAATTC | >hg38_21_+_8433172_8446622_RNA45SN1_45S_50ntFlanks | 11670 | 11702 | 33 | CCCAGTGCTCTGAATGTCAAAGTGAAGAAATTC | 1622.6 | -3.8 | 0.6 | -5.9 | 3.8E-09 | 9.3E-09 | BT20/MB468 | CYTO |
| >hg38_21_+_8433172_8446622_RNA45SN1_45S_50ntFlanks@5175.5197.23\|CCGAGATTGAGCAATAACAGGTC | >hg38_21_+_8433172_8446622_RNA45SN1_45S_50ntFlanks | 5175 | 5197 | 23 | CCGAGATTGAGCAATAACAGGTC | 98.5 | -10.5 | 1.8 | -5.9 | 4.1E-09 | 9.3E-09 | MB231/MB468 | MP |
| >hg38_21_+_8433172_8446622_RNA45SN1_45S_50ntFlanks@8360.8379.20\|AGAACTTTGAAGAGAGAGTT | >hg38_21_+_8433172_8446622_RNA45SN1_45S_50ntFlanks | 8360 | 8379 | 20 | AGAACTTTGAAGAGAGAGTT | 93.2 | -10.4 | 1.8 | -5.9 | 4.1E-09 | 9.3E-09 | MB231/MB468 | MP |
| >hg38_21_+_8433172_8446622_RNA45SN1_45S_50ntFlanks@3777.3814.38\|CGGTACAGTGAAACTGCGAATGGCTCATTAAATCAGTT | >hg38_21_+_8433172_8446622_RNA45SN1_45S_50ntFlanks | 3777 | 3814 | 38 | CGGTACAGTGAAACTGCGAATGGCTCATTAAATCAGTT | 582.8 | -6.8 | 1.2 | -5.8 | 5.0E-09 | 9.3E-09 | BT20/MB468 | NUC |
| >hg38_21_+_8433172_8446622_RNA45SN1_45S_50ntFlanks@8196.8215.20\|CGTGGACGGTGTGAGGCCGG | >hg38_21_+_8433172_8446622_RNA45SN1_45S_50ntFlanks | 8196 | 8215 | 20 | CGTGGACGGTGTGAGGCCGG | 1300.2 | -6.6 | 1.1 | -5.9 | 4.4E-09 | 9.3E-09 | MB231/MB468 | NUC |
| >MI0000270\|hsa-mir-181b-1&WithFlank&1\|-\|198858867\|198858988@42.63.22;MI0000683\|hsa-mir-181b-2&WithFlank&9\|+\|124693704\|124693804@22.43.22&[MIMAT0000257&hsa-miR-181b-5p&offsets\|0\|-1;m-73&1\|-\|198858925\|198858947&offsets\|0\|-1];[MIMAT0000257_1&hsa-miR-181b-5p&offsets\|0\|-1;m-72&9\|+\|124693725\|124693747&offsets\|0\|-1]\|AACATTCATTGCTGTCGGTGGG | >MI0000270\|hsa-mir-181b-1&WithFlank&1\|-\|198858867\|198858988 | 42 | 63 | 22 | AACATTCATTGCTGTCGGTGGG | 448.3 | 4.9 | 0.8 | 5.9 | 4.4E-09 | 9.3E-09 | MB231/MB468 | NUC |
| >MI0000736\|hsa-mir-30c-1&WithFlank&1\|+\|40757278\|40757378@23.45.23(+1G);MI0000254\|hsa-mir-30c-2&WithFlank&6\|-\|71376954\|71377037@13.35.23(+1G)&[MIMAT0000244&hsa-miR-30c-5p&offsets\|0\|0(+1G);m-57&1\|+\|40757300\|40757323&offsets\|0\|-1(+1G)];[MIMAT0000244_1&hsa-miR-30c-5p&offsets\|0\|0(+1G);m-56&6\|-\|71377002\|71377025&offsets\|0\|-1(+1G)]\|TGTAAACATCCTACACTCTCAGCG | >MI0000736\|hsa-mir-30c-1&WithFlank&1\|+\|40757278\|40757378 | 23 | 45 | 23(+1G) | TGTAAACATCCTACACTCTCAGCG | 860.6 | 4.0 | 0.7 | 5.9 | 4.4E-09 | 9.3E-09 | MB231/MB468 | NUC |
| >hg38_21_+_8433172_8446622_RNA45SN1_45S_50ntFlanks@483.507.25\|TCGATGTGGTGACGTCGTGCTCTCC | >hg38_21_+_8433172_8446622_RNA45SN1_45S_50ntFlanks | 483 | 507 | 25 | TCGATGTGGTGACGTCGTGCTCTCC | 690.1 | 4.7 | 0.8 | 5.9 | 4.4E-09 | 9.3E-09 | MB231/MB468 | NUC |
| >hg38_21_+_8433172_8446622_RNA45SN1_45S_50ntFlanks@10729.10750.22\|AGAGGGTGTAAATCTCGCGCCG | >hg38_21_+_8433172_8446622_RNA45SN1_45S_50ntFlanks | 10729 | 10750 | 22 | AGAGGGTGTAAATCTCGCGCCG | 170.3 | -10.5 | 1.7 | -6.2 | 5.5E-10 | 9.3E-09 | BT20/MB468 | MP |
| >hg38_21_+_8433172_8446622_RNA45SN1_45S_50ntFlanks@11575.11601.27\|ACTTAGAACTGGTGCGGACCAGGGGAA | >hg38_21_+_8433172_8446622_RNA45SN1_45S_50ntFlanks | 11575 | 11601 | 27 | ACTTAGAACTGGTGCGGACCAGGGGAA | 1389.3 | -5.0 | 0.8 | -5.9 | 3.8E-09 | 9.3E-09 | BT20/MB468 | CYTO |
| >hg38_21_+_8433172_8446622_RNA45SN1_45S_50ntFlanks@12942.12962.21\|TTCGTAGACGACCTGCTTCTG | >hg38_21_+_8433172_8446622_RNA45SN1_45S_50ntFlanks | 12942 | 12962 | 21 | TTCGTAGACGACCTGCTTCTG | 297.9 | -3.0 | 0.5 | -5.9 | 4.4E-09 | 9.4E-09 | MB231/MB468 | NUC |
| >hg38_21_+_8433172_8446622_RNA45SN1_45S_50ntFlanks@10131.10158.28\|CGGCGGCGGCGGCGGCGGGGGTGTGGGG | >hg38_21_+_8433172_8446622_RNA45SN1_45S_50ntFlanks | 10131 | 10158 | 28 | CGGCGGCGGCGGCGGCGGGGGTGTGGGG | 66.6 | -9.9 | 1.7 | -5.9 | 4.2E-09 | 9.4E-09 | MB231/MB468 | MP |
| >hg38_21_+_8433172_8446622_RNA45SN1_45S_50ntFlanks@4569.4587.19\|ATGGAATAGGACCGCGGTT | >hg38_21_+_8433172_8446622_RNA45SN1_45S_50ntFlanks | 4569 | 4587 | 19 | ATGGAATAGGACCGCGGTT | 299.8 | -6.4 | 1.1 | -5.9 | 4.2E-09 | 9.4E-09 | MB231/MB468 | MP |
| >hg38_21_+_8433172_8446622_RNA45SN1_45S_50ntFlanks@11576.11611.36\|CTTAGAACTGGTGCGGACCAGGGGAATCCGACTGTT | >hg38_21_+_8433172_8446622_RNA45SN1_45S_50ntFlanks | 11576 | 11611 | 36 | CTTAGAACTGGTGCGGACCAGGGGAATCCGACTGTT | 695.5 | -7.0 | 1.2 | -5.9 | 4.2E-09 | 9.4E-09 | MB231/MB468 | MP |
| >trna10_AspGTC_12_-_125424193_125424264@38.72.35,trna12_AspGTC_12_-_125411891_125411962@38.72.35,trna144_AspGTC_6_-_27551236_27551307@38.72.35,trna38_AspGTC_17_-_8125556_8125627@38.72.35,trna45_AspGTC_6_+_27447453_27447524@38.72.35,trna48_AspGTC_6_+_27471523_27471594@38.72.35,trna4_AspGTC_12_+_96429799_96429870@38.72.35,trna69_AspGTC_1_-_161440205_161440276@38.72.35,trna72_AspGTC_1_-_161432824_161432895@38.72.35,trna75_AspGTC_1_-_161425414_161425485@38.72.35,trna78_AspGTC_1_-_161418033_161418104@38.72.35,trna81_AspGTC_1_-_161410615_161410686@38.72.35\|CGCGGGAGACCGGGGTTCGATTCCCCGACGGGGAG | >trna10_AspGTC_12_-_125424193_125424264 | 38 | 72 | 35 | CGCGGGAGACCGGGGTTCGATTCCCCGACGGGGAG | 777.9 | -5.1 | 0.9 | -5.9 | 4.2E-09 | 9.4E-09 | MB231/MB468 | MP |
| >hg38_21_+_8433172_8446622_RNA45SN1_45S_50ntFlanks@9653.9670.18\|AGGATAGCTGGCGCTCTC | >hg38_21_+_8433172_8446622_RNA45SN1_45S_50ntFlanks | 9653 | 9670 | 18 | AGGATAGCTGGCGCTCTC | 63.8 | -9.9 | 1.7 | -5.9 | 4.2E-09 | 9.4E-09 | MB231/MB468 | MP |
| >hg38_21_+_8433172_8446622_RNA45SN1_45S_50ntFlanks@6655.6696.42\|TCTTAGCGGTGGATCACTCGGCTCGTGCGTCGATGAAGAACG | >hg38_21_+_8433172_8446622_RNA45SN1_45S_50ntFlanks | 6655 | 6696 | 42 | TCTTAGCGGTGGATCACTCGGCTCGTGCGTCGATGAAGAACG | 314.4 | 4.2 | 0.7 | 5.9 | 4.5E-09 | 9.4E-09 | MB231/MB468 | NUC |
| >hg38_21_+_8433172_8446622_RNA45SN1_45S_50ntFlanks@12457.12484.28\|CAAGCGTTGGATTGTTCACCCACTAATA | >hg38_21_+_8433172_8446622_RNA45SN1_45S_50ntFlanks | 12457 | 12484 | 28 | CAAGCGTTGGATTGTTCACCCACTAATA | 1432.3 | -8.6 | 1.5 | -5.9 | 4.8E-09 | 9.4E-09 | MB231/MB468 | MITO |
| >hg38_21_+_8433172_8446622_RNA45SN1_45S_50ntFlanks@8175.8202.28\|CCTTCTGATCGAGGCCCAGCCCGTGGAC | >hg38_21_+_8433172_8446622_RNA45SN1_45S_50ntFlanks | 8175 | 8202 | 28 | CCTTCTGATCGAGGCCCAGCCCGTGGAC | 494.5 | 7.5 | 1.3 | 5.9 | 4.4E-09 | 9.4E-09 | MB231/MB468 | CYTO |
| >MI0000465\|hsa-mir-191&WithFlank&3\|-\|49020612\|49020715@22.42.21&[MIMAT0000440&hsa-miR-191-5p&offsets\|0\|-2;m-48&3\|-\|49020672\|49020694&offsets\|0\|-2]\|CAACGGAATCCCAAAAGCAGC | >MI0000465\|hsa-mir-191&WithFlank&3\|-\|49020612\|49020715 | 22 | 42 | 21 | CAACGGAATCCCAAAAGCAGC | 330.5 | 5.4 | 0.9 | 5.9 | 4.8E-09 | 9.4E-09 | MB231/MB468 | MITO |
| >hg38_21_+_8433172_8446622_RNA45SN1_45S_50ntFlanks@9379.9409.31\|CGCGCTCGCCGGCCGAGGTGGGATCCCGAGG | >hg38_21_+_8433172_8446622_RNA45SN1_45S_50ntFlanks | 9379 | 9409 | 31 | CGCGCTCGCCGGCCGAGGTGGGATCCCGAGG | 649.2 | -9.1 | 1.6 | -5.9 | 4.8E-09 | 9.4E-09 | MB231/MB468 | MITO |
| >am_trna128_GlyGCC_6_-_27870686_27870756@1.29.29,trna133_GlyCCC_1_-_16872434_16872504@1.29.29,trna18_GlyGCC_16_+_70822597_70822667@1.29.29,trna19_GlyGCC_16_+_70823410_70823480@1.29.29,trna19_GlyGCC_2_-_157257659_157257729@1.29.29,trna24_GlyGCC_16_-_70812942_70813012@1.29.29,trna25_GlyGCC_16_-_70812114_70812184@1.29.29,trna4_GlyCCC_1_+_17188416_17188486@1.29.29,trna5_GlyGCC_17_+_8029064_8029134@1.29.29,trna68_GlyGCC_1_-_161493637_161493707@1.29.29\|GCATTGGTGGTTCAGTGGTAGAATTCTCG | >am_trna128_GlyGCC_6_-_27870686_27870756 | 1 | 29 | 29 | GCATTGGTGGTTCAGTGGTAGAATTCTCG | 18371.1 | -5.3 | 0.9 | -5.9 | 4.2E-09 | 9.4E-09 | MB231/MB468 | MP |
| >hg38_21_+_8433172_8446622_RNA45SN1_45S_50ntFlanks@10534.10569.36\|CCGCGAGGCGTCCAGTGCGGTAACGCGACCGATCCC | >hg38_21_+_8433172_8446622_RNA45SN1_45S_50ntFlanks | 10534 | 10569 | 36 | CCGCGAGGCGTCCAGTGCGGTAACGCGACCGATCCC | 156.5 | -10.4 | 1.7 | -6.2 | 5.6E-10 | 9.4E-09 | BT20/MB468 | MP |
| >hg38_21_+_8433172_8446622_RNA45SN1_45S_50ntFlanks@9166.9195.30\|CCGGGCGGGTCGCGCCGTCGGGCCCGGGGG | >hg38_21_+_8433172_8446622_RNA45SN1_45S_50ntFlanks | 9166 | 9195 | 30 | CCGGGCGGGTCGCGCCGTCGGGCCCGGGGG | 185.1 | -10.6 | 1.7 | -6.2 | 5.6E-10 | 9.4E-09 | BT20/MB468 | MP |
| >hg38_21_+_8433172_8446622_RNA45SN1_45S_50ntFlanks@10781.10804.24\|AGGTGAACAGCCTCTGGCATGTTG | >hg38_21_+_8433172_8446622_RNA45SN1_45S_50ntFlanks | 10781 | 10804 | 24 | AGGTGAACAGCCTCTGGCATGTTG | 399.6 | -8.3 | 1.4 | -5.9 | 3.8E-09 | 9.4E-09 | BT20/MB468 | CYTO |
| >MI0000809\|hsa-mir-151a&WithFlank&8\|-\|140732558\|140732659@53.74.22(+1A)&[MIMAT0000757&hsa-miR-151a-3p&offsets\|0\|+1(+1A);m-30&8\|-\|140732587\|140732607&offsets\|0\|+1(+1A)]\|CTAGACTGAAGCTCCTTGAGGAA | >MI0000809\|hsa-mir-151a&WithFlank&8\|-\|140732558\|140732659 | 53 | 74 | 22(+1A) | CTAGACTGAAGCTCCTTGAGGAA | 1380.8 | 2.6 | 0.5 | 5.8 | 5.1E-09 | 9.4E-09 | BT20/MB468 | NUC |
| >hg38_21_+_8433172_8446622_RNA45SN1_45S_50ntFlanks@12415.12433.19\|CTTCGATGTCGGCTCTTCC | >hg38_21_+_8433172_8446622_RNA45SN1_45S_50ntFlanks | 12415 | 12433 | 19 | CTTCGATGTCGGCTCTTCC | 2812.9 | -4.0 | 0.7 | -5.8 | 5.1E-09 | 9.4E-09 | BT20/MB468 | NUC |
| >hg38_21_+_8433172_8446622_RNA45SN1_45S_50ntFlanks@10694.10711.18\|CGGTGAGCTCTCGCTGGC | >hg38_21_+_8433172_8446622_RNA45SN1_45S_50ntFlanks | 10694 | 10711 | 18 | CGGTGAGCTCTCGCTGGC | 636.3 | -5.6 | 0.9 | -5.9 | 3.8E-09 | 9.4E-09 | BT20/MB468 | CYTO |
| >hg38_21_+_8433172_8446622_RNA45SN1_45S_50ntFlanks@9534.9568.35\|AGGAAACTCTGGTGGAGGTCCGTAGCGGTCCTGAC | >hg38_21_+_8433172_8446622_RNA45SN1_45S_50ntFlanks | 9534 | 9568 | 35 | AGGAAACTCTGGTGGAGGTCCGTAGCGGTCCTGAC | 912.6 | -5.1 | 0.9 | -5.9 | 4.5E-09 | 9.5E-09 | MB231/MB468 | NUC |
| >hg38_21_+_8433172_8446622_RNA45SN1_45S_50ntFlanks@3985.4003.19\|CGGCTTTGGTGACTCTAGA | >hg38_21_+_8433172_8446622_RNA45SN1_45S_50ntFlanks | 3985 | 4003 | 19 | CGGCTTTGGTGACTCTAGA | 193.2 | -7.0 | 1.2 | -5.9 | 4.5E-09 | 9.5E-09 | MB231/MB468 | NUC |
| >hg38_21_+_8433172_8446622_RNA45SN1_45S_50ntFlanks@12973.12992.20\|TCGTACGTAGCAGAGCAGCT | >hg38_21_+_8433172_8446622_RNA45SN1_45S_50ntFlanks | 12973 | 12992 | 20 | TCGTACGTAGCAGAGCAGCT | 733.0 | -6.0 | 1.0 | -5.9 | 4.9E-09 | 9.5E-09 | MB231/MB468 | MITO |
| >hg38_21_+_8433172_8446622_RNA45SN1_45S_50ntFlanks@6745.6766.22\|ACACTTCGAACGCACTTGCGGC | >hg38_21_+_8433172_8446622_RNA45SN1_45S_50ntFlanks | 6745 | 6766 | 22 | ACACTTCGAACGCACTTGCGGC | 261.6 | -7.9 | 1.3 | -5.9 | 4.9E-09 | 9.5E-09 | MB231/MB468 | MITO |
| >hg38_21_+_8433172_8446622_RNA45SN1_45S_50ntFlanks@11569.11604.36\|CAGCCGACTTAGAACTGGTGCGGACCAGGGGAATCC | >hg38_21_+_8433172_8446622_RNA45SN1_45S_50ntFlanks | 11569 | 11604 | 36 | CAGCCGACTTAGAACTGGTGCGGACCAGGGGAATCC | 76.7 | -10.1 | 1.7 | -5.9 | 4.2E-09 | 9.5E-09 | MB231/MB468 | MP |
| >hg38_21_+_8433172_8446622_RNA45SN1_45S_50ntFlanks@12677.12704.28\|GAATCCCGCCCAGGCGGAACGATACGGC | >hg38_21_+_8433172_8446622_RNA45SN1_45S_50ntFlanks | 12677 | 12704 | 28 | GAATCCCGCCCAGGCGGAACGATACGGC | 74.1 | -10.1 | 1.7 | -5.9 | 4.2E-09 | 9.5E-09 | MB231/MB468 | MP |
| >hg38_21_+_8433172_8446622_RNA45SN1_45S_50ntFlanks@7985.8008.24\|GATCAGACGTGGCGACCCGCTGAA | >hg38_21_+_8433172_8446622_RNA45SN1_45S_50ntFlanks | 7985 | 8008 | 24 | GATCAGACGTGGCGACCCGCTGAA | 2177.2 | -8.5 | 1.5 | -5.9 | 4.9E-09 | 9.5E-09 | MB231/MB468 | MITO |
| >hg38_21_+_8433172_8446622_RNA45SN1_45S_50ntFlanks@8170.8216.47\|CAAGTCCTTCTGATCGAGGCCCAGCCCGTGGACGGTGTGAGGCCGGT | >hg38_21_+_8433172_8446622_RNA45SN1_45S_50ntFlanks | 8170 | 8216 | 47 | CAAGTCCTTCTGATCGAGGCCCAGCCCGTGGACGGTGTGAGGCCGGT | 987.5 | -9.1 | 1.6 | -5.8 | 5.1E-09 | 9.5E-09 | BT20/MB468 | NUC |
| >hg38_21_+_8433172_8446622_RNA45SN1_45S_50ntFlanks@12940.12966.27\|CATTCGTAGACGACCTGCTTCTGGGTC | >hg38_21_+_8433172_8446622_RNA45SN1_45S_50ntFlanks | 12940 | 12966 | 27 | CATTCGTAGACGACCTGCTTCTGGGTC | 416.8 | -6.5 | 1.1 | -5.9 | 4.9E-09 | 9.5E-09 | MB231/MB468 | MITO |
| >MI0000462\|hsa-mir-152&WithFlank&17\|-\|48037155\|48037253@60.80.21(+1A)&[MIMAT0000438&hsa-miR-152-3p&offsets\|0\|0(+1A);m-97&17\|-\|48037174\|48037194&offsets\|0\|0(+1A)]\|TCAGTGCATGACAGAACTTGGA | >MI0000462\|hsa-mir-152&WithFlank&17\|-\|48037155\|48037253 | 60 | 80 | 21(+1A) | TCAGTGCATGACAGAACTTGGA | 167.9 | 11.1 | 1.8 | 6.2 | 5.6E-10 | 9.5E-09 | BT20/MB468 | MP |
| >hg38_21_+_8433172_8446622_RNA45SN1_45S_50ntFlanks@5074.5091.18\|ATTCCGATAACGAACGAG | >hg38_21_+_8433172_8446622_RNA45SN1_45S_50ntFlanks | 5074 | 5091 | 18 | ATTCCGATAACGAACGAG | 131.6 | -10.9 | 1.9 | -5.9 | 4.3E-09 | 9.5E-09 | MB231/MB468 | MP |
| >hg38_21_+_8433172_8446622_RNA45SN1_45S_50ntFlanks@6659.6681.23\|AGCGGTGGATCACTCGGCTCGTG | >hg38_21_+_8433172_8446622_RNA45SN1_45S_50ntFlanks | 6659 | 6681 | 23 | AGCGGTGGATCACTCGGCTCGTG | 513.3 | -3.6 | 0.6 | -5.9 | 4.5E-09 | 9.5E-09 | MB231/MB468 | CYTO |
| >hg38_21_+_8433172_8446622_RNA45SN1_45S_50ntFlanks@3705.3739.35\|TACCTGGTTGATCCTGCCAGTAGCATATGCTTGTC | >hg38_21_+_8433172_8446622_RNA45SN1_45S_50ntFlanks | 3705 | 3739 | 35 | TACCTGGTTGATCCTGCCAGTAGCATATGCTTGTC | 1198.4 | 3.7 | 0.6 | 5.9 | 4.5E-09 | 9.6E-09 | MB231/MB468 | CYTO |
| >hg38_21_+_8433172_8446622_RNA45SN1_45S_50ntFlanks@10350.10377.28\|ACGAGAACTTTGAAGGCCGAAGTGGAGA | >hg38_21_+_8433172_8446622_RNA45SN1_45S_50ntFlanks | 10350 | 10377 | 28 | ACGAGAACTTTGAAGGCCGAAGTGGAGA | 1605.8 | -6.4 | 1.1 | -5.9 | 4.9E-09 | 9.6E-09 | MB231/MB468 | MITO |

| | | | | | | | | | | | | | |
|---|---|---|---|---|---|---|---|---|---|---|---|---|---|
| >hg38_21_+_8433172_8446622_RNA45SN1_45S_50ntFlanks@11894.11912.19\|ACTCTAGTCTGGCACGGTG | >hg38_21_+_8433172_8446622_RNA45SN1_45S_50ntFlanks | 11894 | 11912 | 19 | ACTCTAGTCTGGCACGGTG | 259.0 | -3.7 | 0.6 | -5.9 | 3.4E-09 | 9.6E-09 | BT20/MB468 | MITO |
| >trnaMT_PheGAA_MT_+_577_647@35.55.21\|GAAAATGTTTAGACGGGCTCA | >trnaMT_PheGAA_MT_+_577_647 | 35 | 55 | 21 | GAAAATGTTTAGACGGGCTCA | 361.5 | 4.0 | 0.7 | 5.9 | 4.3E-09 | 9.6E-09 | MB231/MB468 | MP |
| >hg38_21_+_8433172_8446622_RNA45SN1_45S_50ntFlanks@12285.12319.35\|CAGACCGTGAAAGCGGGGCCTCACGATCCTTCTGA | >hg38_21_+_8433172_8446622_RNA45SN1_45S_50ntFlanks | 12285 | 12319 | 35 | CAGACCGTGAAAGCGGGGCCTCACGATCCTTCTGA | 81.9 | -10.2 | 1.7 | -5.9 | 4.3E-09 | 9.6E-09 | MB231/MB468 | MP |
| >trna111_HisGTG_1_-_147774845_147774916@-1T.23.24,trna118_HisGTG_1_-_145396881_145396952@-1T.23.24,trna16_HisGTG_1_+_146544773_146544844@-1T.23.24,trna1_HisGTG_15_+_45493349_45493420@-1T.23.24,trna21_HisGTG_1_+_147753471_147753542@-1T.23.24,trna33_HisGTG_6_+_27125906_27125977@-1T.23.24,trna7_HisGTG_9_-_14433938_14434009@-1T.23.24,trna8_HisGTG_15_-_45492611_45492682@-1T.23.24,trna9_HisGTG_15_-_45490804_45490875@-1T.23.24\|TGCCGTGATCGTATAGTGGTTAGT | >trna111_HisGTG_1_-_147774845_147774916 | -1T | 23 | 24 | TGCCGTGATCGTATAGTGGTTAGT | 4296.9 | -4.5 | 0.7 | -6.2 | 4.7E-10 | 9.6E-09 | BT20/MB231 | NUC |
| >am_trna119_LysCTT_1_-_145395522_145395594@13.30.18,trna11_LysCTT_5_-_180648979_180649051@13.30.18,trna128_LysCTT_1_-_55423542_55423614@13.30.18,trna13_LysCTT_6_+_26556774_26556846@13.30.18,trna32_LysCTT_16_-_3207406_3207478@13.30.18,trna6_LysCTT_19_-_52425396_52425468@13.30.18,trna7_LysCTT_16_+_3225692_3225764@13.30.18,trna9_LysCTT_5_+_180634755_180634827@13.30.18\|CAGTCGGTAGAGCATGAG | >am_trna119_LysCTT_1_-_145395522_145395594 | 13 | 30 | 18 | CAGTCGGTAGAGCATGAG | 493.6 | -6.9 | 1.2 | -5.8 | 5.1E-09 | 9.6E-09 | BT20/MB468 | NUC |
| >hg38_21_+_8433172_8446622_RNA45SN1_45S_50ntFlanks@8170.8188.19\|CAAGTCCTTCTGATCGAGG | >hg38_21_+_8433172_8446622_RNA45SN1_45S_50ntFlanks | 8170 | 8188 | 19 | CAAGTCCTTCTGATCGAGG | 169.1 | -9.5 | 1.6 | -5.9 | 3.4E-09 | 9.6E-09 | BT20/MB468 | MITO |
| >hg38_21_+_8433172_8446622_RNA45SN1_45S_50ntFlanks@11569.11601.33\|CAGCCGACTTAGAACTGGTGCGGACCAGGGGAA | >hg38_21_+_8433172_8446622_RNA45SN1_45S_50ntFlanks | 11569 | 11601 | 33 | CAGCCGACTTAGAACTGGTGCGGACCAGGGGAA | 3594.2 | -3.8 | 0.6 | -5.9 | 3.4E-09 | 9.6E-09 | BT20/MB468 | MITO |
| >hg38_21_+_8433172_8446622_RNA45SN1_45S_50ntFlanks@11766.11790.25\|ATCTAATTAGTGACGCGCATGAATG | >hg38_21_+_8433172_8446622_RNA45SN1_45S_50ntFlanks | 11766 | 11790 | 25 | ATCTAATTAGTGACGCGCATGAATG | 183.3 | -10.6 | 1.7 | -6.2 | 5.7E-10 | 9.6E-09 | BT20/MB468 | MP |
| >hg38_21_+_8433172_8446622_RNA45SN1_45S_50ntFlanks@8191.8211.21\|CAGCCCGTGGACGGTGTGAGG | >hg38_21_+_8433172_8446622_RNA45SN1_45S_50ntFlanks | 8191 | 8211 | 21 | CAGCCCGTGGACGGTGTGAGG | 185.6 | -9.6 | 1.6 | -5.9 | 3.4E-09 | 9.6E-09 | BT20/MB468 | MITO |
| >hg38_21_+_8433172_8446622_RNA45SN1_45S_50ntFlanks@7984.8018.35\|AGATCAGACGTGGCGACCCGCTGAATTTAAGCATA | >hg38_21_+_8433172_8446622_RNA45SN1_45S_50ntFlanks | 7984 | 8018 | 35 | AGATCAGACGTGGCGACCCGCTGAATTTAAGCATA | 83.7 | -10.3 | 1.7 | -5.9 | 4.3E-09 | 9.6E-09 | MB231/MB468 | MP |
| >hg38_21_+_8433172_8446622_RNA45SN1_45S_50ntFlanks@12366.12383.18\|TAACTGGCTTGTGGCGGC | >hg38_21_+_8433172_8446622_RNA45SN1_45S_50ntFlanks | 12366 | 12383 | 18 | TAACTGGCTTGTGGCGGC | 2034.5 | -8.3 | 1.4 | -5.9 | 3.9E-09 | 9.7E-09 | BT20/MB468 | CYTO |
| >hg38_21_+_8433172_8446622_RNA45SN1_45S_50ntFlanks@12133.12150.18\|CTCCGGGGACAGTGCCAG | >hg38_21_+_8433172_8446622_RNA45SN1_45S_50ntFlanks | 12133 | 12150 | 18 | CTCCGGGGACAGTGCCAG | 2098.0 | -8.3 | 1.4 | -5.9 | 3.4E-09 | 9.7E-09 | BT20/MB468 | MITO |
| >hg38_21_+_8433172_8446622_RNA45SN1_45S_50ntFlanks@4068.4089.22\|ACTTTCGATGGTAGTCGCCGTG | >hg38_21_+_8433172_8446622_RNA45SN1_45S_50ntFlanks | 4068 | 4089 | 22 | ACTTTCGATGGTAGTCGCCGTG | 462.5 | -10.9 | 1.9 | -5.9 | 3.4E-09 | 9.7E-09 | BT20/MB468 | MITO |
| >hg38_21_+_8433172_8446622_RNA45SN1_45S_50ntFlanks@11583.11613.31\|CTGGTGCGGACCAGGGGAATCCGACTGTTTA | >hg38_21_+_8433172_8446622_RNA45SN1_45S_50ntFlanks | 11583 | 11613 | 31 | CTGGTGCGGACCAGGGGAATCCGACTGTTTA | 387.6 | -4.8 | 0.8 | -5.9 | 4.3E-09 | 9.7E-09 | MB231/MB468 | MP |
| >hg38_21_+_8433172_8446622_RNA45SN1_45S_50ntFlanks@8737.8758.22\|CCCGTCCGTCCGTCCGTCCGTC | >hg38_21_+_8433172_8446622_RNA45SN1_45S_50ntFlanks | 8737 | 8758 | 22 | CCCGTCCGTCCGTCCGTCCGTC | 747.6 | -9.5 | 1.6 | -5.8 | 5.2E-09 | 9.7E-09 | BT20/MB468 | NUC |
| >hg38_21_+_8433172_8446622_RNA45SN1_45S_50ntFlanks@12347.12376.30\|AGAAAAGTTACCACAGGGATAACTGGCTTG | >hg38_21_+_8433172_8446622_RNA45SN1_45S_50ntFlanks | 12347 | 12376 | 30 | AGAAAAGTTACCACAGGGATAACTGGCTTG | 236.9 | -11.0 | 1.8 | -6.2 | 5.7E-10 | 9.7E-09 | BT20/MB468 | MP |
| >hg38_21_+_8433172_8446622_RNA45SN1_45S_50ntFlanks@11876.11897.22\|AAGACCCTGTTGAGCTTGACTC | >hg38_21_+_8433172_8446622_RNA45SN1_45S_50ntFlanks | 11876 | 11897 | 22 | AAGACCCTGTTGAGCTTGACTC | 396.4 | -7.7 | 1.3 | -5.9 | 4.3E-09 | 9.7E-09 | MB231/MB468 | MP |
| >am_trnaMT_ValTAC_MT_+_1602_1670@48.66.19\|CAACTTAACTTGACCGCTC | >am_trnaMT_ValTAC_MT_+_1602_1670 | 48 | 66 | 19 | CAACTTAACTTGACCGCTC | 1346.2 | 8.9 | 1.5 | 5.9 | 3.9E-09 | 9.7E-09 | BT20/MB468 | CYTO |
| >trnaMT_GluTTC_MT_-_14674_14742@41.71.31,trnalookalike8_GluTTC_5_-_93905172_93905240@41.71.31\|TGGTCGTGGTTGTAGTCCGTGCGAGAATACC | >trnaMT_GluTTC_MT_-_14674_14742 | 41 | 71 | 31 | TGGTCGTGGTTGTAGTCCGTGCGAGAATACC | 569.3 | 5.0 | 0.9 | 5.9 | 4.6E-09 | 9.7E-09 | MB231/MB468 | NUC |
| >am_trna2_GlyTCC_19_+_4724082_4724153@1.29.29\|GCGTTGGTGGTATAGTGGTTAGCATAGCT | >am_trna2_GlyTCC_19_+_4724082_4724153 | 1 | 29 | 29 | GCGTTGGTGGTATAGTGGTTAGCATAGCT | 508.1 | -5.0 | 0.9 | -5.9 | 4.6E-09 | 9.7E-09 | MB231/MB468 | CYTO |
| >hg38_21_+_8433172_8446622_RNA45SN1_45S_50ntFlanks@6650.6677.28\|ACGACTCTTAGCGGTGGATCACTCGGCT | >hg38_21_+_8433172_8446622_RNA45SN1_45S_50ntFlanks | 6650 | 6677 | 28 | ACGACTCTTAGCGGTGGATCACTCGGCT | 468.1 | 3.4 | 0.6 | 5.9 | 3.9E-09 | 9.7E-09 | BT20/MB468 | CYTO |
| >MI0000102\|hsa-mir-100&WithFlank&11\|-\|122152223\|122152314@19.37.19(+1U)&[MIMAT0000098&hsa-miR-100-5p&offsets\|0\|-3(+1U);m-27&11\|-\|122152276\|122152296&offsets\|0\|-2(+1U)]\|AACCCGTAGATCCGAACTTT | >MI0000102\|hsa-mir-100&WithFlank&11\|-\|122152223\|122152314 | 19 | 37 | 19(+1U) | AACCCGTAGATCCGAACTTT | 136.8 | 10.3 | 1.8 | 5.8 | 5.0E-09 | 9.7E-09 | MB231/MB468 | MITO |
| >am_trnaMT_SerGCT_MT_+_12207_12265@12.38.27\|CAAGAACTGCTAACTCATGCCCCCATG | >am_trnaMT_SerGCT_MT_+_12207_12265 | 12 | 38 | 27 | CAAGAACTGCTAACTCATGCCCCCATG | 1053.0 | -4.3 | 0.7 | -5.8 | 5.0E-09 | 9.7E-09 | MB231/MB468 | MITO |
| >MI0000434\|hsa-let-7i&WithFlank&12\|+\|62603680\|62603775@12.33.22(+1U)&[MIMAT0000415&hsa-let-7i-5p&offsets\|0\|0(+1U);m-49&12\|+\|62603691\|62603712&offsets\|0\|0(+1U)]\|TGAGGTAGTAGTTTGTGCTGTTT | >MI0000434\|hsa-let-7i&WithFlank&12\|+\|62603680\|62603775 | 12 | 33 | 22(+1U) | TGAGGTAGTAGTTTGTGCTGTTT | 10332.7 | 5.1 | 0.9 | 5.9 | 4.4E-09 | 9.8E-09 | MB231/MB468 | MP |
| >hg38_21_+_8433172_8446622_RNA45SN1_45S_50ntFlanks@11575.11594.20\|ACTTAGAACTGGTGCGGACC | >hg38_21_+_8433172_8446622_RNA45SN1_45S_50ntFlanks | 11575 | 11594 | 20 | ACTTAGAACTGGTGCGGACC | 1549.0 | -3.2 | 0.5 | -5.9 | 4.0E-09 | 9.8E-09 | BT20/MB468 | CYTO |
| >hg38_21_+_8433172_8446622_RNA45SN1_45S_50ntFlanks@6780.6802.23\|CGGGGCTACGCCTGTCTGAGCGT | >hg38_21_+_8433172_8446622_RNA45SN1_45S_50ntFlanks | 6780 | 6802 | 23 | CGGGGCTACGCCTGTCTGAGCGT | 805.0 | 4.9 | 0.8 | 5.9 | 4.6E-09 | 9.8E-09 | MB231/MB468 | CYTO |
| >hg38_21_+_8433172_8446622_RNA45SN1_45S_50ntFlanks@10411.10429.19\|CAGTCGGTCCTGAGAGATG | >hg38_21_+_8433172_8446622_RNA45SN1_45S_50ntFlanks | 10411 | 10429 | 19 | CAGTCGGTCCTGAGAGATG | 4766.9 | -3.8 | 0.6 | -5.9 | 3.5E-09 | 9.8E-09 | BT20/MB468 | MITO |
| >hg38_21_+_8433172_8446622_RNA45SN1_45S_50ntFlanks@12938.12956.19\|ACCATTCGTAGACGACCTG | >hg38_21_+_8433172_8446622_RNA45SN1_45S_50ntFlanks | 12938 | 12956 | 19 | ACCATTCGTAGACGACCTG | 6225.1 | -10.3 | 1.8 | -5.8 | 5.3E-09 | 9.8E-09 | BT20/MB468 | NUC |
| >hg38_21_+_8433172_8446622_RNA45SN1_45S_50ntFlanks@6652.6681.30\|GACTCTTAGCGGTGGATCACTCGGCTCGTG | >hg38_21_+_8433172_8446622_RNA45SN1_45S_50ntFlanks | 6652 | 6681 | 30 | GACTCTTAGCGGTGGATCACTCGGCTCGTG | 9310.8 | -5.4 | 0.9 | -5.9 | 4.4E-09 | 9.8E-09 | MB231/MB468 | MP |
| >hg38_21_+_8433172_8446622_RNA45SN1_45S_50ntFlanks@12419.12436.18\|GATGTCGGCTCTTCCTAT | >hg38_21_+_8433172_8446622_RNA45SN1_45S_50ntFlanks | 12419 | 12436 | 18 | GATGTCGGCTCTTCCTAT | 5133.6 | -3.2 | 0.6 | -5.9 | 4.7E-09 | 9.8E-09 | MB231/MB468 | NUC |
| >hg38_21_+_8433172_8446622_RNA45SN1_45S_50ntFlanks@4064.4081.18\|ATCAACTTTCGATGGTAG | >hg38_21_+_8433172_8446622_RNA45SN1_45S_50ntFlanks | 4064 | 4081 | 18 | ATCAACTTTCGATGGTAG | 349.2 | -10.5 | 1.8 | -5.9 | 3.5E-09 | 9.8E-09 | BT20/MB468 | MITO |
| >hg38_21_+_8433172_8446622_RNA45SN1_45S_50ntFlanks@11569.11607.39\|CAGCCGACTTAGAACTGGTGCGGACCAGGGGAATCCGAC | >hg38_21_+_8433172_8446622_RNA45SN1_45S_50ntFlanks | 11569 | 11607 | 39 | CAGCCGACTTAGAACTGGTGCGGACCAGGGGAATCCGAC | 93.1 | -10.4 | 1.8 | -5.9 | 4.4E-09 | 9.8E-09 | MB231/MB468 | MP |
| >hg38_21_+_8433172_8446622_RNA45SN1_45S_50ntFlanks@4241.4264.24\|CTCTTTCGAGGCCCTGTAATTGGA | >hg38_21_+_8433172_8446622_RNA45SN1_45S_50ntFlanks | 4241 | 4264 | 24 | CTCTTTCGAGGCCCTGTAATTGGA | 81.7 | -10.2 | 1.7 | -5.9 | 4.4E-09 | 9.8E-09 | MB231/MB468 | MP |
| >hg38_21_+_8433172_8446622_RNA45SN1_45S_50ntFlanks@11534.11553.20\|GTTCCGCGCGGCGCCTCGCC | >hg38_21_+_8433172_8446622_RNA45SN1_45S_50ntFlanks | 11534 | 11553 | 20 | GTTCCGCGCGGCGCCTCGCC | 368.1 | -6.9 | 1.2 | -5.9 | 3.5E-09 | 9.8E-09 | BT20/MB468 | MITO |
| >hg38_21_+_8433172_8446622_RNA45SN1_45S_50ntFlanks@10571.10596.26\|GAGAAGCCGGCGGGAGCCCCGGGGAG | >hg38_21_+_8433172_8446622_RNA45SN1_45S_50ntFlanks | 10571 | 10596 | 26 | GAGAAGCCGGCGGGAGCCCCGGGGAG | 400.1 | -7.5 | 1.3 | -5.9 | 4.7E-09 | 9.8E-09 | MB231/MB468 | NUC |
| >hg38_21_+_8433172_8446622_RNA45SN1_45S_50ntFlanks@6732.6760.29\|ACATTGATCATCGACACTTCGAACGCACT | >hg38_21_+_8433172_8446622_RNA45SN1_45S_50ntFlanks | 6732 | 6760 | 29 | ACATTGATCATCGACACTTCGAACGCACT | 484.4 | 3.7 | 0.6 | 5.9 | 3.5E-09 | 9.8E-09 | BT20/MB468 | MITO |
| >hg38_21_+_8433172_8446622_RNA45SN1_45S_50ntFlanks@7975.8002.28\|CGCGACCTCAGATCAGACGTGGCGACCC | >hg38_21_+_8433172_8446622_RNA45SN1_45S_50ntFlanks | 7975 | 8002 | 28 | CGCGACCTCAGATCAGACGTGGCGACCC | 887.1 | 3.7 | 0.6 | 5.9 | 4.6E-09 | 9.9E-09 | MB231/MB468 | CYTO |
| >hg38_21_+_8433172_8446622_RNA45SN1_45S_50ntFlanks@12072.12095.24\|CCCGAGGGGCTCTCGCTTCTGGCG | >hg38_21_+_8433172_8446622_RNA45SN1_45S_50ntFlanks | 12072 | 12095 | 24 | CCCGAGGGGCTCTCGCTTCTGGCG | 707.3 | -8.4 | 1.4 | -6.2 | 5.9E-10 | 9.9E-09 | BT20/MB468 | MP |
| >hg38_21_+_8433172_8446622_RNA45SN1_45S_50ntFlanks@12943.12966.24\|TCGTAGACGACCTGCTTCTGGGTC | >hg38_21_+_8433172_8446622_RNA45SN1_45S_50ntFlanks | 12943 | 12966 | 24 | TCGTAGACGACCTGCTTCTGGGTC | 75.3 | -10.1 | 1.7 | -5.9 | 4.4E-09 | 9.9E-09 | MB231/MB468 | MP |
| >hg38_21_+_8433172_8446622_RNA45SN1_45S_50ntFlanks@8170.8204.35\|CAAGTCCTTCTGATCGAGGCCCAGCCCGTGGACGG | >hg38_21_+_8433172_8446622_RNA45SN1_45S_50ntFlanks | 8170 | 8204 | 35 | CAAGTCCTTCTGATCGAGGCCCAGCCCGTGGACGG | 258.6 | -11.1 | 1.8 | -6.2 | 5.9E-10 | 9.9E-09 | BT20/MB468 | MP |
| >hg38_21_+_8433172_8446622_RNA45SN1_45S_50ntFlanks@7975.8008.34\|CGCGACCTCAGATCAGACGTGGCGACCCGCTGAA | >hg38_21_+_8433172_8446622_RNA45SN1_45S_50ntFlanks | 7975 | 8008 | 34 | CGCGACCTCAGATCAGACGTGGCGACCCGCTGAA | 528071.5 | 8.3 | 1.4 | 5.9 | 4.0E-09 | 9.9E-09 | BT20/MB468 | CYTO |
| >MI0000102\|hsa-mir-100&WithFlank&11\|-\|122152223\|122152314@19.39.21&[MIMAT0000098&hsa-miR-100-5p&offsets\|0\|-1;m-27&11\|-\|122152276\|122152296&offsets\|0\|0]\|AACCCGTAGATCCGAACTTGT | >MI0000102\|hsa-mir-100&WithFlank&11\|-\|122152223\|122152314 | 19 | 39 | 21 | AACCCGTAGATCCGAACTTGT | 145142.0 | -6.3 | 1.0 | -6.2 | 4.8E-10 | 9.9E-09 | BT20/MB231 | NUC |
| >hg38_21_+_8433172_8446622_RNA45SN1_45S_50ntFlanks@6721.6749.29\|AATTGCAGGACACATTGATCATCGACACT | >hg38_21_+_8433172_8446622_RNA45SN1_45S_50ntFlanks | 6721 | 6749 | 29 | AATTGCAGGACACATTGATCATCGACACT | 372.8 | -3.2 | 0.5 | -5.9 | 4.7E-09 | 9.9E-09 | MB231/MB468 | NUC |
| >MI0000750\|hsa-mir-26a-2&WithFlank&12\|-\|57824603\|57824698@20.40.21(+1A);MI0000083\|hsa-mir-26a-1&WithFlank&3\|+\|37969398\|37969486@16.36.21(+1A)&[MIMAT0000082&hsa-miR-26a-5p&offsets\|0\|-1(+1A);m-33&12\|-\|57824658\|57824679&offsets\|0\|-1(+1A)];[MIMAT0000082_1&hsa-miR-26a-5p&offsets\|0\|-1(+1A);m-34&3\|+\|37969413\|37969434&offsets\|0\|-1(+1A)]\|TTCAAGTAATCCAGGATAGGCA | >MI0000750\|hsa-mir-26a-2&WithFlank&12\|-\|57824603\|57824698 | 20 | 40 | 21(+1A) | TTCAAGTAATCCAGGATAGGCA | 10364.2 | 5.5 | 0.9 | 5.8 | 5.1E-09 | 9.9E-09 | MB231/MB468 | MITO |
| >hg38_21_+_8433172_8446622_RNA45SN1_45S_50ntFlanks@10795.10818.24\|TGGCATGTTGGAACAATGTAGGTA | >hg38_21_+_8433172_8446622_RNA45SN1_45S_50ntFlanks | 10795 | 10818 | 24 | TGGCATGTTGGAACAATGTAGGTA | 1040.9 | -3.6 | 0.6 | -5.9 | 4.0E-09 | 9.9E-09 | BT20/MB468 | CYTO |
| >hg38_21_+_8433172_8446622_RNA45SN1_45S_50ntFlanks@10474.10497.24\|TCGGCCGATCGAAAGGGAGTCGGG | >hg38_21_+_8433172_8446622_RNA45SN1_45S_50ntFlanks | 10474 | 10497 | 24 | TCGGCCGATCGAAAGGGAGTCGGG | 167.7 | -10.5 | 1.7 | -6.2 | 5.9E-10 | 9.9E-09 | BT20/MB468 | MP |
| >hg38_21_+_8433172_8446622_RNA45SN1_45S_50ntFlanks@9570.9595.26\|TGCAAATCGGTCGTCCGACCTGGGTA | >hg38_21_+_8433172_8446622_RNA45SN1_45S_50ntFlanks | 9570 | 9595 | 26 | TGCAAATCGGTCGTCCGACCTGGGTA | 1871.6 | -9.6 | 1.6 | -5.8 | 5.3E-09 | 9.9E-09 | BT20/MB468 | NUC |
| >hg38_21_+_8433172_8446622_RNA45SN1_45S_50ntFlanks@12423.12446.24\|TCGGCTCTTCCTATCATTGTGAAG | >hg38_21_+_8433172_8446622_RNA45SN1_45S_50ntFlanks | 12423 | 12446 | 24 | TCGGCTCTTCCTATCATTGTGAAG | 111.1 | -6.4 | 1.1 | -5.9 | 4.0E-09 | 9.9E-09 | BT20/MB468 | CYTO |
| >hg38_21_+_8433172_8446622_RNA45SN1_45S_50ntFlanks@5003.5034.32\|ATAGCTCTTTCTCGATTCCGTGGGTGGTGGTG | >hg38_21_+_8433172_8446622_RNA45SN1_45S_50ntFlanks | 5003 | 5034 | 32 | ATAGCTCTTTCTCGATTCCGTGGGTGGTGGTG | 347.2 | -10.5 | 1.8 | -5.9 | 3.6E-09 | 1.0E-08 | BT20/MB468 | MITO |

<table>
<tr><th></th><th></th><th></th><th></th><th></th><th></th><th></th><th></th><th></th><th></th><th></th><th></th><th></th><th></th></tr>
<tr><td>>MI0000760|hsa-mir-361&WithFlank&X|-|85903630|85903713@51.75.25&[MIMAT0004682&hsa-miR-361-3p&offsets|0|+2;m-124&X|-|85903642|85903663&offsets|0|+3]|TCCCCCAGGTGTGATTCTGATTTGC</td><td>>MI0000760|hsa-mir-361&WithFlank&X|-|85903630|85903713</td><td>51</td><td>75</td><td>25</td><td>TCCCCCAGGTGTGATTCTGATTTGC</td><td>337.4</td><td>7.2</td><td>1.2</td><td>5.9</td><td>4.7E-09</td><td>1.0E-08</td><td>MB231/MB468</td><td>CYTO</td></tr>
<tr><td>>hg38_21_+_8433172_8446622_RNA45SN1_45S_50ntFlanks@10826.10844.19|TCGGCAAGCCGGATCCGTA</td><td>>hg38_21_+_8433172_8446622_RNA45SN1_45S_50ntFlanks</td><td>10826</td><td>10844</td><td>19</td><td>TCGGCAAGCCGGATCCGTA</td><td>605.3</td><td>-8.6</td><td>1.5</td><td>-5.9</td><td>4.0E-09</td><td>1.0E-08</td><td>BT20/MB468</td><td>CYTO</td></tr>
<tr><td>>hg38_21_+_8433172_8446622_RNA45SN1_45S_50ntFlanks@12412.12436.25|ATCCTTCGATGTCGGCTCTTCCTAT</td><td>>hg38_21_+_8433172_8446622_RNA45SN1_45S_50ntFlanks</td><td>12412</td><td>12436</td><td>25</td><td>ATCCTTCGATGTCGGCTCTTCCTAT</td><td>1356.7</td><td>-8.5</td><td>1.4</td><td>-5.8</td><td>5.2E-09</td><td>1.0E-08</td><td>MB231/MB468</td><td>MITO</td></tr>
<tr><td>>MI0000063|hsa-let-7b&WithFlank&22|+|46113680|46113774@12.33.22(+1A)&[MIMAT0000063&hsa-let-7b-5p&offsets|0|0(+1A);m-14&22|+|46113691|46113712&offsets|0|0(+1A)]|TGAGGTAGTAGGTTGTGTGGTTA</td><td>>MI0000063|hsa-let-7b&WithFlank&22|+|46113680|46113774</td><td>12</td><td>33</td><td>22(+1A)</td><td>TGAGGTAGTAGGTTGTGTGGTTA</td><td>510.5</td><td>4.9</td><td>0.8</td><td>5.9</td><td>4.7E-09</td><td>1.0E-08</td><td>MB231/MB468</td><td>CYTO</td></tr>
<tr><td>>hg38_21_+_8433172_8446622_RNA45SN1_45S_50ntFlanks@11569.11609.41|CAGCCGACTTAGAACTGGTGCGGACCAGGGGAATCCGACTG</td><td>>hg38_21_+_8433172_8446622_RNA45SN1_45S_50ntFlanks</td><td>11569</td><td>11609</td><td>41</td><td>CAGCCGACTTAGAACTGGTGCGGACCAGGGGAATCCGACTG</td><td>505.6</td><td>-5.1</td><td>0.9</td><td>-5.8</td><td>5.4E-09</td><td>1.0E-08</td><td>BT20/MB468</td><td>NUC</td></tr>
<tr><td>>hg38_21_+_8433172_8446622_RNA45SN1_45S_50ntFlanks@3838.3874.37|CTCTCCTACTTGGATAACTGTGGTAATTCTAGAGCTA</td><td>>hg38_21_+_8433172_8446622_RNA45SN1_45S_50ntFlanks</td><td>3838</td><td>3874</td><td>37</td><td>CTCTCCTACTTGGATAACTGTGGTAATTCTAGAGCTA</td><td>190.3</td><td>-3.4</td><td>0.6</td><td>-5.9</td><td>4.8E-09</td><td>1.0E-08</td><td>MB231/MB468</td><td>NUC</td></tr>
<tr><td>>MI0000736|hsa-mir-30c-1&WithFlank&1|+|40757278|40757378@23.44.22(+1A);MI0000254|hsa-mir-30c-2&WithFlank&6|-|71376954|71377037@13.34.22(+1A)&[MIMAT0000244&hsa-miR-30c-5p&offsets|0|-1(+1A);m-57&1|+|40757300|40757323&offsets|0|-2(+1A)];[MIMAT0000244_1&hsa-miR-30c-5p&offsets|0|-1(+1A);m-56&6|-|71377002|71377025&offsets|0|-2(+1A)]|TGTAAACATCCTACACTCTCAGA</td><td>>MI0000736|hsa-mir-30c-1&WithFlank&1|+|40757278|40757378</td><td>23</td><td>44</td><td>22(+1A)</td><td>TGTAAACATCCTACACTCTCAGA</td><td>2591.8</td><td>6.2</td><td>1.1</td><td>5.8</td><td>5.2E-09</td><td>1.0E-08</td><td>MB231/MB468</td><td>MITO</td></tr>
<tr><td>>trna10_AspGTC_12_-_125424193_125424264@31.71.41,trna12_AspGTC_12_-_125411891_125411962@31.71.41,trna38_AspGTC_17_-_8125556_8125627@31.71.41,trna45_AspGTC_6_+_27447453_27447524@31.71.41,trna48_AspGTC_6_+_27471523_27471594@31.71.41,trna4_AspGTC_12_+_96429799_96429870@31.71.41,trna69_AspGTC_1_-_161440205_161440276@31.71.41,trna72_AspGTC_1_-_161432824_161432895@31.71.41,trna75_AspGTC_1_-_161425414_161425485@31.71.41,trna78_AspGTC_1_-_161418033_161418104@31.71.41,trna81_AspGTC_1_-_161410615_161410686@31.71.41|CCTGTCACGCGGGAGACCGGGGTTCGATTCCCCGACGGGGA</td><td>>trna10_AspGTC_12_-_125424193_125424264</td><td>31</td><td>71</td><td>41</td><td>CCTGTCACGCGGGAGACCGGGGTTCGATTCCCCGACGGGGA</td><td>235.6</td><td>-11.7</td><td>2.0</td><td>-5.8</td><td>5.2E-09</td><td>1.0E-08</td><td>MB231/MB468</td><td>MITO</td></tr>
<tr><td>>MI0000746|hsa-mir-99b&WithFlank&19|+|51692606|51692687@13.33.21&[MIMAT0000689&hsa-miR-99b-5p&offsets|0|-1;m-10&19|+|51692618|51692639&offsets|0|-1]|CACCCGTAGAACCGACCTTGC</td><td>>MI0000746|hsa-mir-99b&WithFlank&19|+|51692606|51692687</td><td>13</td><td>33</td><td>21</td><td>CACCCGTAGAACCGACCTTGC</td><td>4718.5</td><td>4.4</td><td>0.7</td><td>5.8</td><td>5.2E-09</td><td>1.0E-08</td><td>MB231/MB468</td><td>MITO</td></tr>
<tr><td>>hg38_21_+_8433172_8446622_RNA45SN1_45S_50ntFlanks@11575.11612.38|ACTTAGAACTGGTGCGGACCAGGGGAATCCGACTGTTT</td><td>>hg38_21_+_8433172_8446622_RNA45SN1_45S_50ntFlanks</td><td>11575</td><td>11612</td><td>38</td><td>ACTTAGAACTGGTGCGGACCAGGGGAATCCGACTGTTT</td><td>1235.9</td><td>-5.4</td><td>0.9</td><td>-5.8</td><td>5.2E-09</td><td>1.0E-08</td><td>MB231/MB468</td><td>MITO</td></tr>
<tr><td>>hg38_21_+_8433172_8446622_RNA45SN1_45S_50ntFlanks@10786.10825.40|AACAGCCTCTGGCATGTTGGAACAATGTAGGTAAGGGAAG</td><td>>hg38_21_+_8433172_8446622_RNA45SN1_45S_50ntFlanks</td><td>10786</td><td>10825</td><td>40</td><td>AACAGCCTCTGGCATGTTGGAACAATGTAGGTAAGGGAAG</td><td>1167.2</td><td>-4.5</td><td>0.8</td><td>-5.9</td><td>3.6E-09</td><td>1.0E-08</td><td>BT20/MB468</td><td>MITO</td></tr>
<tr><td>>trnaMT_GluTTC_MT_-_14674_14742@25.72.48,trnalookalike8_GluTTC_5_-_93905172_93905240@25.72.48|TGGTTTTTCATATCATTGGTCGTGGTTGTAGTCCGTGCGAGAATACCA</td><td>>trnaMT_GluTTC_MT_-_14674_14742</td><td>25</td><td>72</td><td>48</td><td>TGGTTTTTCATATCATTGGTCGTGGTTGTAGTCCGTGCGAGAATACCA</td><td>501.7</td><td>8.8</td><td>1.5</td><td>5.9</td><td>4.7E-09</td><td>1.0E-08</td><td>MB231/MB468</td><td>CYTO</td></tr>
<tr><td>>hg38_MT_+_1621_3279_RNR2_16S_50ntFlanks@51.90.40|GCTAAACCTAGCCCCAAACCCACTCCACCTTACTACCAGA</td><td>>hg38_MT_+_1621_3279_RNR2_16S_50ntFlanks</td><td>51</td><td>90</td><td>40</td><td>GCTAAACCTAGCCCCAAACCCACTCCACCTTACTACCAGA</td><td>766.4</td><td>6.3</td><td>1.1</td><td>5.9</td><td>4.8E-09</td><td>1.0E-08</td><td>MB231/MB468</td><td>NUC</td></tr>
<tr><td>>hg38_21_+_8433172_8446622_RNA45SN1_45S_50ntFlanks@12995.13016.22|CTCGCTGCGATCTATTGAAAGT</td><td>>hg38_21_+_8433172_8446622_RNA45SN1_45S_50ntFlanks</td><td>12995</td><td>13016</td><td>22</td><td>CTCGCTGCGATCTATTGAAAGT</td><td>863.1</td><td>4.7</td><td>0.8</td><td>5.9</td><td>4.8E-09</td><td>1.0E-08</td><td>MB231/MB468</td><td>NUC</td></tr>
<tr><td>>hg38_21_+_8433172_8446622_RNA45SN1_45S_50ntFlanks@9551.9570.20|GTCCGTAGCGGTCCTGACGT</td><td>>hg38_21_+_8433172_8446622_RNA45SN1_45S_50ntFlanks</td><td>9551</td><td>9570</td><td>20</td><td>GTCCGTAGCGGTCCTGACGT</td><td>349.5</td><td>3.5</td><td>0.6</td><td>5.9</td><td>4.8E-09</td><td>1.0E-08</td><td>MB231/MB468</td><td>CYTO</td></tr>
<tr><td>>hg38_21_+_8433172_8446622_RNA45SN1_45S_50ntFlanks@4163.4182.20|CATCCAAGGAAGGCAGCAGG</td><td>>hg38_21_+_8433172_8446622_RNA45SN1_45S_50ntFlanks</td><td>4163</td><td>4182</td><td>20</td><td>CATCCAAGGAAGGCAGCAGG</td><td>602.4</td><td>-5.1</td><td>0.9</td><td>-5.9</td><td>4.8E-09</td><td>1.0E-08</td><td>MB231/MB468</td><td>CYTO</td></tr>
<tr><td>>hg38_21_+_8433172_8446622_RNA45SN1_45S_50ntFlanks@483.506.24|TCGATGTGGTGACGTCGTGCTCTC</td><td>>hg38_21_+_8433172_8446622_RNA45SN1_45S_50ntFlanks</td><td>483</td><td>506</td><td>24</td><td>TCGATGTGGTGACGTCGTGCTCTC</td><td>955.8</td><td>5.2</td><td>0.9</td><td>5.9</td><td>4.8E-09</td><td>1.0E-08</td><td>MB231/MB468</td><td>NUC</td></tr>
<tr><td>>hg38_MT_+_1621_3279_RNR2_16S_50ntFlanks@51.95.45|GCTAAACCTAGCCCCAAACCCACTCCACCTTACTACCAGACAACC</td><td>>hg38_MT_+_1621_3279_RNR2_16S_50ntFlanks</td><td>51</td><td>95</td><td>45</td><td>GCTAAACCTAGCCCCAAACCCACTCCACCTTACTACCAGACAACC</td><td>7364.9</td><td>5.9</td><td>1.0</td><td>5.9</td><td>3.6E-09</td><td>1.0E-08</td><td>BT20/MB468</td><td>MITO</td></tr>
<tr><td>>hg38_21_+_8433172_8446622_RNA45SN1_45S_50ntFlanks@3984.4024.41|GCGGCTTTGGTGACTCTAGATAACCTCGGGCCGATCGCACG</td><td>>hg38_21_+_8433172_8446622_RNA45SN1_45S_50ntFlanks</td><td>3984</td><td>4024</td><td>41</td><td>GCGGCTTTGGTGACTCTAGATAACCTCGGGCCGATCGCACG</td><td>3128.6</td><td>-10.0</td><td>1.6</td><td>-6.2</td><td>6.1E-10</td><td>1.0E-08</td><td>BT20/MB468</td><td>MP</td></tr>
<tr><td>>hg38_21_+_8433172_8446622_RNA45SN1_45S_50ntFlanks@12479.12498.20|CTAATAGGGAACGTGAGCTG</td><td>>hg38_21_+_8433172_8446622_RNA45SN1_45S_50ntFlanks</td><td>12479</td><td>12498</td><td>20</td><td>CTAATAGGGAACGTGAGCTG</td><td>307.3</td><td>-5.9</td><td>1.0</td><td>-5.8</td><td>5.2E-09</td><td>1.0E-08</td><td>MB231/MB468</td><td>MITO</td></tr>
<tr><td>>MI0000067|hsa-let-7f-1&WithFlank&9|+|94176341|94176439@13.34.22;MI0000068|hsa-let-7f-2&WithFlank&X|-|53557186|53557280@14.35.22&[MIMAT0000067&hsa-let-7f-5p&offsets|0|0;m-20&9|+|94176353|94176374&offsets|0|0];[MIMAT0000067_1&hsa-let-7f-5p&offsets|0|0;m-19&X|-|53557246|53557267&offsets|0|0]|TGAGGTAGTAGATTGTATAGTT</td><td>>MI0000067|hsa-let-7f-1&WithFlank&9|+|94176341|94176439</td><td>13</td><td>34</td><td>22</td><td>TGAGGTAGTAGATTGTATAGTT</td><td>56430.1</td><td>5.1</td><td>0.9</td><td>5.8</td><td>5.2E-09</td><td>1.0E-08</td><td>MB231/MB468</td><td>MITO</td></tr>
<tr><td>>hg38_21_+_8433172_8446622_RNA45SN1_45S_50ntFlanks@475.503.29|CTTCGTGATCGATGTGGTGACGTCGTGCT</td><td>>hg38_21_+_8433172_8446622_RNA45SN1_45S_50ntFlanks</td><td>475</td><td>503</td><td>29</td><td>CTTCGTGATCGATGTGGTGACGTCGTGCT</td><td>297.1</td><td>7.1</td><td>1.2</td><td>5.9</td><td>4.8E-09</td><td>1.0E-08</td><td>MB231/MB468</td><td>CYTO</td></tr>
<tr><td>>hg38_21_+_8433172_8446622_RNA45SN1_45S_50ntFlanks@11718.11743.26|AACGGCGGGAGTAACTATGACTCTCT</td><td>>hg38_21_+_8433172_8446622_RNA45SN1_45S_50ntFlanks</td><td>11718</td><td>11743</td><td>26</td><td>AACGGCGGGAGTAACTATGACTCTCT</td><td>383.6</td><td>-6.1</td><td>1.0</td><td>-5.9</td><td>4.6E-09</td><td>1.0E-08</td><td>MB231/MB468</td><td>MP</td></tr>
<tr><td>>hg38_21_+_8433172_8446622_RNA45SN1_45S_50ntFlanks@10408.10427.20|GGTCAGTCGGTCCTGAGAGA</td><td>>hg38_21_+_8433172_8446622_RNA45SN1_45S_50ntFlanks</td><td>10408</td><td>10427</td><td>20</td><td>GGTCAGTCGGTCCTGAGAGA</td><td>177.8</td><td>-7.2</td><td>1.2</td><td>-5.9</td><td>4.9E-09</td><td>1.0E-08</td><td>MB231/MB468</td><td>NUC</td></tr>
<tr><td>>hg38_21_+_8433172_8446622_RNA45SN1_45S_50ntFlanks@11730.11751.22|AACTATGACTCTCTTAAGGTAG</td><td>>hg38_21_+_8433172_8446622_RNA45SN1_45S_50ntFlanks</td><td>11730</td><td>11751</td><td>22</td><td>AACTATGACTCTCTTAAGGTAG</td><td>611.3</td><td>-8.6</td><td>1.5</td><td>-5.8</td><td>5.5E-09</td><td>1.0E-08</td><td>BT20/MB468</td><td>NUC</td></tr>
<tr><td>>MI0000087|hsa-mir-29a&WithFlank&7|-|130876741|130876816@47.68.22(+1U)&[MIMAT0000086&hsa-miR-29a-3p&offsets|-1|-1(+1U);m-26&7|-|130876749|130876769&offsets|-1|0(+1U)]|CTAGCACCATCTGAAATCGGTTT</td><td>>MI0000087|hsa-mir-29a&WithFlank&7|-|130876741|130876816</td><td>47</td><td>68</td><td>22(+1U)</td><td>CTAGCACCATCTGAAATCGGTTT</td><td>212.6</td><td>7.1</td><td>1.2</td><td>5.8</td><td>5.5E-09</td><td>1.0E-08</td><td>BT20/MB468</td><td>NUC</td></tr>
<tr><td>>hg38_21_+_8433172_8446622_RNA45SN1_45S_50ntFlanks@3776.3810.35|CCGGTACAGTGAAACTGCGAATGGCTCATTAAATC</td><td>>hg38_21_+_8433172_8446622_RNA45SN1_45S_50ntFlanks</td><td>3776</td><td>3810</td><td>35</td><td>CCGGTACAGTGAAACTGCGAATGGCTCATTAAATC</td><td>191.6</td><td>-3.4</td><td>0.6</td><td>-5.9</td><td>4.9E-09</td><td>1.0E-08</td><td>MB231/MB468</td><td>NUC</td></tr>
<tr><td>>hg38_21_+_8433172_8446622_RNA45SN1_45S_50ntFlanks@12406.12431.26|TTTTTGATCCTTCGATGTCGGCTCTT</td><td>>hg38_21_+_8433172_8446622_RNA45SN1_45S_50ntFlanks</td><td>12406</td><td>12431</td><td>26</td><td>TTTTTGATCCTTCGATGTCGGCTCTT</td><td>815.8</td><td>-4.5</td><td>0.8</td><td>-5.9</td><td>4.8E-09</td><td>1.0E-08</td><td>MB231/MB468</td><td>CYTO</td></tr>
<tr><td>>MI0000093|hsa-mir-92a-1&WithFlank&13|+|91351308|91351397@54.74.21;MI0000094|hsa-mir-92a-2&WithFlank&X|-|134169532|134169618@54.74.21&[MIMAT0000092&hsa-miR-92a-3p&offsets|0|-1;m-9&13|+|91351361|91351382&offsets|0|-1];[MIMAT0000092_1&hsa-miR-92a-3p&offsets|0|-1;m-11&X|-|134169544|134169565&offsets|0|-1]|TATTGCACTTGTCCCGGCCTG</td><td>>MI0000093|hsa-mir-92a-1&WithFlank&13|+|91351308|91351397</td><td>54</td><td>74</td><td>21</td><td>TATTGCACTTGTCCCGGCCTG</td><td>708.4</td><td>4.8</td><td>0.8</td><td>5.8</td><td>5.3E-09</td><td>1.0E-08</td><td>MB231/MB468</td><td>MITO</td></tr>
<tr><td>>hg38_21_+_8433172_8446622_RNA45SN1_45S_50ntFlanks@11712.11740.29|CGGGTAAACGGCGGGAGTAACTATGACTC</td><td>>hg38_21_+_8433172_8446622_RNA45SN1_45S_50ntFlanks</td><td>11712</td><td>11740</td><td>29</td><td>CGGGTAAACGGCGGGAGTAACTATGACTC</td><td>336.7</td><td>-4.8</td><td>0.8</td><td>-5.9</td><td>4.9E-09</td><td>1.0E-08</td><td>MB231/MB468</td><td>NUC</td></tr>
<tr><td>>am_trna111_HisGTG_1_-_147774845_147774916@-1G.23.24,trna118_HisGTG_1_-_145396881_145396952@-1G.23.24,trna16_HisGTG_1_+_146544773_146544844@-1G.23.24,trna1_HisGTG_15_+_45493349_45493420@-1G.23.24,trna21_HisGTG_1_+_147753471_147753542@-1G.23.24,trna33_HisGTG_6_+_27125906_27125977@-1G.23.24,trna7_HisGTG_9_-_14433938_14434009@-1G.23.24,trna8_HisGTG_15_-_45492611_45492682@-1G.23.24,trna9_HisGTG_15_-_45490804_45490875@-1G.23.24|GGCCGTGATCGTATAGTGGTTAGT</td><td>>am_trna111_HisGTG_1_-_147774845_147774916</td><td>-1G</td><td>23</td><td>24</td><td>GGCCGTGATCGTATAGTGGTTAGT</td><td>856.8</td><td>7.8</td><td>1.3</td><td>5.9</td><td>4.8E-09</td><td>1.0E-08</td><td>MB231/MB468</td><td>CYTO</td></tr>
<tr><td>>hg38_21_+_8433172_8446622_RNA45SN1_45S_50ntFlanks@10346.10368.23|TCAAACGAGAACTTTGAAGGCCG</td><td>>hg38_21_+_8433172_8446622_RNA45SN1_45S_50ntFlanks</td><td>10346</td><td>10368</td><td>23</td><td>TCAAACGAGAACTTTGAAGGCCG</td><td>3109.9</td><td>-3.1</td><td>0.5</td><td>-5.9</td><td>4.8E-09</td><td>1.0E-08</td><td>MB231/MB468</td><td>CYTO</td></tr>
<tr><td>>hg38_21_+_8433172_8446622_RNA45SN1_45S_50ntFlanks@5061.5093.33|ATTTGTCTGGTTAATTCCGATAACGAACGAGAC</td><td>>hg38_21_+_8433172_8446622_RNA45SN1_45S_50ntFlanks</td><td>5061</td><td>5093</td><td>33</td><td>ATTTGTCTGGTTAATTCCGATAACGAACGAGAC</td><td>136.3</td><td>-11.0</td><td>1.9</td><td>-5.9</td><td>4.6E-09</td><td>1.0E-08</td><td>MB231/MB468</td><td>MP</td></tr>
<tr><td>>hg38_21_+_8433172_8446622_RNA45SN1_45S_50ntFlanks@10361.10383.23|GAAGGCCGAAGTGGAGAAGGGTT</td><td>>hg38_21_+_8433172_8446622_RNA45SN1_45S_50ntFlanks</td><td>10361</td><td>10383</td><td>23</td><td>GAAGGCCGAAGTGGAGAAGGGTT</td><td>421.5</td><td>-5.9</td><td>1.0</td><td>-5.8</td><td>5.3E-09</td><td>1.0E-08</td><td>MB231/MB468</td><td>MITO</td></tr>
<tr><td>>hg38_21_+_8433172_8446622_RNA45SN1_45S_50ntFlanks@11322.11344.23|ACGTCTCGTCGCGCGCGCGTCCG</td><td>>hg38_21_+_8433172_8446622_RNA45SN1_45S_50ntFlanks</td><td>11322</td><td>11344</td><td>23</td><td>ACGTCTCGTCGCGCGCGCGTCCG</td><td>9500.7</td><td>-5.6</td><td>1.0</td><td>-5.9</td><td>4.8E-09</td><td>1.0E-08</td><td>MB231/MB468</td><td>CYTO</td></tr>
<tr><td>>MI0000272|hsa-mir-182&WithFlank&7|-|129770377|129770498@29.51.23(+1A)&[MIMAT0000259&hsa-miR-182-5p&offsets|0|-1(+1A);m-22&7|-|129770449|129770470&offsets|0|+1(+1A)]|TTTGGCAATGGTAGAACTCACACA</td><td>>MI0000272|hsa-mir-182&WithFlank&7|-|129770377|129770498</td><td>29</td><td>51</td><td>23(+1A)</td><td>TTTGGCAATGGTAGAACTCACACA</td><td>9922.6</td><td>5.6</td><td>0.9</td><td>6.3</td><td>2.9E-10</td><td>1.0E-08</td><td>BT20/MB231</td><td>CYTO</td></tr>
<tr><td>>MI0000272|hsa-mir-182&WithFlank&7|-|129770377|129770498@29.51.23(+1C)&[MIMAT0000259&hsa-miR-182-5p&offsets|0|-1(+1C);m-22&7|-|129770449|129770470&offsets|0|+1(+1C)]|TTTGGCAATGGTAGAACTCACACC</td><td>>MI0000272|hsa-mir-182&WithFlank&7|-|129770377|129770498</td><td>29</td><td>51</td><td>23(+1C)</td><td>TTTGGCAATGGTAGAACTCACACC</td><td>13348.9</td><td>5.0</td><td>0.8</td><td>6.3</td><td>2.9E-10</td><td>1.0E-08</td><td>BT20/MB231</td><td>CYTO</td></tr>
<tr><td>>hg38_21_+_8433172_8446622_RNA45SN1_45S_50ntFlanks@3813.3850.38|TTATGGTTCCTTTGGTCGCTCGCTCCTCTCCTACTTGG</td><td>>hg38_21_+_8433172_8446622_RNA45SN1_45S_50ntFlanks</td><td>3813</td><td>3850</td><td>38</td><td>TTATGGTTCCTTTGGTCGCTCGCTCCTCTCCTACTTGG</td><td>336.8</td><td>-4.7</td><td>0.8</td><td>-5.8</td><td>4.9E-09</td><td>1.0E-08</td><td>MB231/MB468</td><td>NUC</td></tr>
<tr><td>>hg38_21_+_8433172_8446622_RNA45SN1_45S_50ntFlanks@11217.11234.18|CTGGACGCGAGCCGGGCC</td><td>>hg38_21_+_8433172_8446622_RNA45SN1_45S_50ntFlanks</td><td>11217</td><td>11234</td><td>18</td><td>CTGGACGCGAGCCGGGCC</td><td>2099.7</td><td>-5.9</td><td>1.0</td><td>-5.9</td><td>4.6E-09</td><td>1.0E-08</td><td>MB231/MB468</td><td>MP</td></tr>
<tr><td>>hg38_21_+_8433172_8446622_RNA45SN1_45S_50ntFlanks@6653.6677.25|ACTCTTAGCGGTGGATCACTCGGCT</td><td>>hg38_21_+_8433172_8446622_RNA45SN1_45S_50ntFlanks</td><td>6653</td><td>6677</td><td>25</td><td>ACTCTTAGCGGTGGATCACTCGGCT</td><td>662.8</td><td>3.8</td><td>0.7</td><td>5.9</td><td>3.7E-09</td><td>1.0E-08</td><td>BT20/MB468</td><td>MITO</td></tr>
<tr><td>>hg38_21_+_8433172_8446622_RNA45SN1_45S_50ntFlanks@10589.10608.20|CCGGGGAGAGTTCTCTTTTC</td><td>>hg38_21_+_8433172_8446622_RNA45SN1_45S_50ntFlanks</td><td>10589</td><td>10608</td><td>20</td><td>CCGGGGAGAGTTCTCTTTTC</td><td>59.7</td><td>-9.8</td><td>1.7</td><td>-5.9</td><td>4.6E-09</td><td>1.0E-08</td><td>MB231/MB468</td><td>MP</td></tr>
</table>

<table>
<tr><td>>hg38_21_+_8433172_8446622_RNA45SN1_45S_50ntFlanks@11124.11145.22|CCGGCGGCGGCCCGCCGCGGGG</td><td>>hg38_21_+_8433172_8446622_RNA45SN1_45S_50ntFlanks</td><td>11124</td><td>11145</td><td>22</td><td>CCGGCGGCGGCCCGCCGCGGGG</td><td>191.6</td><td>-11.4</td><td>2.0</td><td>-5.9</td><td>4.6E-09</td><td>1.0E-08</td><td>MB231/MB468</td><td>MP</td></tr>
<tr><td>>hg38_21_+_8433172_8446622_RNA45SN1_45S_50ntFlanks@12937.12955.19|AACCATTCGTAGACGACCT</td><td>>hg38_21_+_8433172_8446622_RNA45SN1_45S_50ntFlanks</td><td>12937</td><td>12955</td><td>19</td><td>AACCATTCGTAGACGACCT</td><td>365.4</td><td>-2.8</td><td>0.5</td><td>-5.9</td><td>4.9E-09</td><td>1.0E-08</td><td>MB231/MB468</td><td>CYTO</td></tr>
<tr><td>>hg38_21_+_8433172_8446622_RNA45SN1_45S_50ntFlanks@8176.8197.22|CTTCTGATCGAGGCCCAGCCCG</td><td>>hg38_21_+_8433172_8446622_RNA45SN1_45S_50ntFlanks</td><td>8176</td><td>8197</td><td>22</td><td>CTTCTGATCGAGGCCCAGCCCG</td><td>956.2</td><td>-3.2</td><td>0.5</td><td>-5.9</td><td>4.9E-09</td><td>1.0E-08</td><td>MB231/MB468</td><td>CYTO</td></tr>
<tr><td>>hg38_21_+_8433172_8446622_RNA45SN1_45S_50ntFlanks@12539.12557.19|ATGATGTGTTGTTGCCATG</td><td>>hg38_21_+_8433172_8446622_RNA45SN1_45S_50ntFlanks</td><td>12539</td><td>12557</td><td>19</td><td>ATGATGTGTTGTTGCCATG</td><td>134.2</td><td>-6.0</td><td>1.0</td><td>-5.9</td><td>4.9E-09</td><td>1.0E-08</td><td>MB231/MB468</td><td>CYTO</td></tr>
<tr><td>>hg38_21_+_8433172_8446622_RNA45SN1_45S_50ntFlanks@9504.9533.30|GTGAACTATGCCTGGGCAGGGCGAAGCCAG</td><td>>hg38_21_+_8433172_8446622_RNA45SN1_45S_50ntFlanks</td><td>9504</td><td>9533</td><td>30</td><td>GTGAACTATGCCTGGGCAGGGCGAAGCCAG</td><td>1396.9</td><td>-9.1</td><td>1.5</td><td>-6.2</td><td>6.2E-10</td><td>1.0E-08</td><td>BT20/MB468</td><td>MP</td></tr>
<tr><td>>hg38_21_+_8433172_8446622_RNA45SN1_45S_50ntFlanks@11913.11949.37|AAGAGACATGAGAGGTGTAGAATAAGTGGGAGGCCCC</td><td>>hg38_21_+_8433172_8446622_RNA45SN1_45S_50ntFlanks</td><td>11913</td><td>11949</td><td>37</td><td>AAGAGACATGAGAGGTGTAGAATAAGTGGGAGGCCCC</td><td>106.1</td><td>-10.6</td><td>1.8</td><td>-5.9</td><td>4.7E-09</td><td>1.0E-08</td><td>MB231/MB468</td><td>MP</td></tr>
<tr><td>>hg38_21_+_8433172_8446622_RNA45SN1_45S_50ntFlanks@4853.4883.31|AAGTCTTTGGGTTCCGGGGGGAGTATGGTTG</td><td>>hg38_21_+_8433172_8446622_RNA45SN1_45S_50ntFlanks</td><td>4853</td><td>4883</td><td>31</td><td>AAGTCTTTGGGTTCCGGGGGGAGTATGGTTG</td><td>226.7</td><td>-4.5</td><td>0.8</td><td>-5.8</td><td>4.9E-09</td><td>1.0E-08</td><td>MB231/MB468</td><td>CYTO</td></tr>
<tr><td>>hg38_21_+_8433172_8446622_RNA45SN1_45S_50ntFlanks@8179.8203.25|CTGATCGAGGCCCAGCCCGTGGACG</td><td>>hg38_21_+_8433172_8446622_RNA45SN1_45S_50ntFlanks</td><td>8179</td><td>8203</td><td>25</td><td>CTGATCGAGGCCCAGCCCGTGGACG</td><td>366.1</td><td>-5.8</td><td>1.0</td><td>-5.8</td><td>5.4E-09</td><td>1.0E-08</td><td>MB231/MB468</td><td>MITO</td></tr>
<tr><td>>hg38_21_+_8433172_8446622_RNA45SN1_45S_50ntFlanks@12670.12703.34|TAAGTCAGAATCCCGCCCAGGCGGAACGATACGG</td><td>>hg38_21_+_8433172_8446622_RNA45SN1_45S_50ntFlanks</td><td>12670</td><td>12703</td><td>34</td><td>TAAGTCAGAATCCCGCCCAGGCGGAACGATACGG</td><td>158.2</td><td>-10.4</td><td>1.7</td><td>-6.2</td><td>6.3E-10</td><td>1.0E-08</td><td>BT20/MB468</td><td>MP</td></tr>
<tr><td>>MI0000471|hsa-mir-126&WithFlank&9|+|136670596|136670692@21.41.21(+1U)&[MIMAT0000444&hsa-miR-126-5p&offsets|0|0(+1U);m-114&9|+|136670616|136670636&offsets|0|0(+1U)]|CATTATTACTTTTGGTACGCGT</td><td>>MI0000471|hsa-mir-126&WithFlank&9|+|136670596|136670692</td><td>21</td><td>41</td><td>21(+1U)</td><td>CATTATTACTTTTGGTACGCGT</td><td>124.6</td><td>10.2</td><td>1.7</td><td>5.9</td><td>4.7E-09</td><td>1.0E-08</td><td>MB231/MB468</td><td>MP</td></tr>
<tr><td>>hg38_21_+_8433172_8446622_RNA45SN1_45S_50ntFlanks@10821.10840.20|GGAAGTCGGCAAGCCGGATC</td><td>>hg38_21_+_8433172_8446622_RNA45SN1_45S_50ntFlanks</td><td>10821</td><td>10840</td><td>20</td><td>GGAAGTCGGCAAGCCGGATC</td><td>5713.9</td><td>-7.7</td><td>1.3</td><td>-5.9</td><td>4.7E-09</td><td>1.0E-08</td><td>MB231/MB468</td><td>MP</td></tr>
<tr><td>>hg38_21_+_8433172_8446622_RNA45SN1_45S_50ntFlanks@6732.6771.40|ACATTGATCATCGACACTTCGAACGCACTTGCGGCCCCGG</td><td>>hg38_21_+_8433172_8446622_RNA45SN1_45S_50ntFlanks</td><td>6732</td><td>6771</td><td>40</td><td>ACATTGATCATCGACACTTCGAACGCACTTGCGGCCCCGG</td><td>3306.2</td><td>-8.3</td><td>1.4</td><td>-5.8</td><td>5.6E-09</td><td>1.1E-08</td><td>BT20/MB468</td><td>NUC</td></tr>
<tr><td>>trna10_LysCTT_16_+_3241501_3241573@1.28.28,trna119_LysCTT_1_-_145395522_145395594@1.28.28,trna11_LysCTT_5_-_180648979_180649051@1.28.28,trna13_LysCTT_14_-_58706613_58706685@1.28.28,trna13_LysCTT_6_+_26556774_26556846@1.28.28,trna2_LysCTT_15_+_79152904_79152976@1.28.28,trna32_LysCTT_16_-_3207406_3207478@1.28.28,trna7_LysCTT_16_+_3225692_3225764@1.28.28,trna9_LysCTT_5_+_180634755_180634827@1.28.28|GCCCGGCTAGCTCAGTCGGTAGAGCATG</td><td>>trna10_LysCTT_16_+_3241501_3241573</td><td>1</td><td>28</td><td>28</td><td>GCCCGGCTAGCTCAGTCGGTAGAGCATG</td><td>3650.9</td><td>-4.2</td><td>0.7</td><td>-5.8</td><td>5.4E-09</td><td>1.1E-08</td><td>MB231/MB468</td><td>MITO</td></tr>
<tr><td>>MI0000298|hsa-mir-221&WithFlank&X|-|45746151|45746272@71.93.23(+1G)&[MIMAT0000278&hsa-miR-221-3p&offsets|0|0(+1G);m-91&X|-|45746181|45746202&offsets|0|+1(+1G)]|AGCTACATTGTCTGCTGGGTTTCG</td><td>>MI0000298|hsa-mir-221&WithFlank&X|-|45746151|45746272</td><td>71</td><td>93</td><td>23(+1G)</td><td>AGCTACATTGTCTGCTGGGTTTCG</td><td>594.9</td><td>5.0</td><td>0.8</td><td>5.8</td><td>5.0E-09</td><td>1.1E-08</td><td>MB231/MB468</td><td>CYTO</td></tr>
<tr><td>>MI0000811|hsa-mir-148b&WithFlank&12|+|54337210|54337320@69.90.22&[MIMAT0000759&hsa-miR-148b-3p&offsets|0|0;m-77&12|+|54337278|54337299&offsets|0|0]|TCAGTGCATCACAGAACTTTGT</td><td>>MI0000811|hsa-mir-148b&WithFlank&12|+|54337210|54337320</td><td>69</td><td>90</td><td>22</td><td>TCAGTGCATCACAGAACTTTGT</td><td>6161.0</td><td>6.2</td><td>1.1</td><td>5.8</td><td>5.4E-09</td><td>1.1E-08</td><td>MB231/MB468</td><td>MITO</td></tr>
<tr><td>>hg38_21_+_8433172_8446622_RNA45SN1_45S_50ntFlanks@4738.4767.30|AAAGTCGGAGGTTCGAAGACGATCAGATAC</td><td>>hg38_21_+_8433172_8446622_RNA45SN1_45S_50ntFlanks</td><td>4738</td><td>4767</td><td>30</td><td>AAAGTCGGAGGTTCGAAGACGATCAGATAC</td><td>1730.5</td><td>8.8</td><td>1.5</td><td>5.9</td><td>4.3E-09</td><td>1.1E-08</td><td>BT20/MB468</td><td>CYTO</td></tr>
<tr><td>>hg38_21_+_8433172_8446622_RNA45SN1_45S_50ntFlanks@10388.10410.23|GTGAACAGCAGTTGAACATGGGT</td><td>>hg38_21_+_8433172_8446622_RNA45SN1_45S_50ntFlanks</td><td>10388</td><td>10410</td><td>23</td><td>GTGAACAGCAGTTGAACATGGGT</td><td>480.8</td><td>-5.1</td><td>0.9</td><td>-5.8</td><td>5.0E-09</td><td>1.1E-08</td><td>MB231/MB468</td><td>CYTO</td></tr>
<tr><td>>MI0000264|hsa-mir-7-2&WithFlank&15|+|88611819|88611940@38.60.23(+1A);MI0000265|hsa-mir-7-3&WithFlank&19|+|4770664|4770785@37.59.23(+1A);MI0000263|hsa-mir-7-1&WithFlank&9|-|83969742|83969863@30.52.23(+1A)&[MIMAT0000252&hsa-miR-7-5p&offsets|0|-1(+1A);m-400&15|+|88611856|88611879&offsets|0|-1(+1A)];[MIMAT0000252_1&hsa-miR-7-5p&offsets|0|-1(+1A);m-402&19|+|4770700|4770723&offsets|0|-1(+1A)];[MIMAT0000252_2&hsa-miR-7-5p&offsets|0|-1(+1A);m-401&9|-|83969811|83969834&offsets|0|-1(+1A)]|TGGAAGACTAGTGATTTTGTTGTA</td><td>>MI0000264|hsa-mir-7-2&WithFlank&15|+|88611819|88611940</td><td>38</td><td>60</td><td>23(+1A)</td><td>TGGAAGACTAGTGATTTTGTTGTA</td><td>1182.0</td><td>5.6</td><td>1.0</td><td>5.8</td><td>5.5E-09</td><td>1.1E-08</td><td>MB231/MB468</td><td>MITO</td></tr>
<tr><td>>hg38_MT_+_1621_3279_RNR2_16S_50ntFlanks@729.759.31|GATTAAAACACTGAACTGACAATTAACAGCC</td><td>>hg38_MT_+_1621_3279_RNR2_16S_50ntFlanks</td><td>729</td><td>759</td><td>31</td><td>GATTAAAACACTGAACTGACAATTAACAGCC</td><td>88.9</td><td>9.7</td><td>1.7</td><td>5.8</td><td>5.5E-09</td><td>1.1E-08</td><td>MB231/MB468</td><td>MITO</td></tr>
<tr><td>>hg38_21_+_8433172_8446622_RNA45SN1_45S_50ntFlanks@12249.12268.20|CAAAAGCTCGCTTGATCTTG</td><td>>hg38_21_+_8433172_8446622_RNA45SN1_45S_50ntFlanks</td><td>12249</td><td>12268</td><td>20</td><td>CAAAAGCTCGCTTGATCTTG</td><td>136.4</td><td>-10.2</td><td>1.6</td><td>-6.2</td><td>6.3E-10</td><td>1.1E-08</td><td>BT20/MB468</td><td>MP</td></tr>
<tr><td>>hg38_21_+_8433172_8446622_RNA45SN1_45S_50ntFlanks@6644.6665.22|CCTCGTACGACTCTTAGCGGTG</td><td>>hg38_21_+_8433172_8446622_RNA45SN1_45S_50ntFlanks</td><td>6644</td><td>6665</td><td>22</td><td>CCTCGTACGACTCTTAGCGGTG</td><td>663.9</td><td>3.6</td><td>0.6</td><td>5.8</td><td>5.1E-09</td><td>1.1E-08</td><td>MB231/MB468</td><td>NUC</td></tr>
<tr><td>>hg38_21_+_8433172_8446622_RNA45SN1_45S_50ntFlanks@6690.6719.30|AAGAACGCAGCTAGCTGCGAGAATTAATGT</td><td>>hg38_21_+_8433172_8446622_RNA45SN1_45S_50ntFlanks</td><td>6690</td><td>6719</td><td>30</td><td>AAGAACGCAGCTAGCTGCGAGAATTAATGT</td><td>105.0</td><td>-8.8</td><td>1.5</td><td>-5.9</td><td>3.8E-09</td><td>1.1E-08</td><td>BT20/MB468</td><td>MITO</td></tr>
<tr><td>>MI0000101|hsa-mir-99a&WithFlank&21|+|16539083|16539175@19.40.22(+1A)&[MIMAT0000097&hsa-miR-99a-5p&offsets|0|0(+1A);m-64&21|+|16539101|16539121&offsets|0|+1(+1A)]|AACCCGTAGATCCGATCTTGTGA</td><td>>MI0000101|hsa-mir-99a&WithFlank&21|+|16539083|16539175</td><td>19</td><td>40</td><td>22(+1A)</td><td>AACCCGTAGATCCGATCTTGTGA</td><td>422.0</td><td>-5.9</td><td>0.9</td><td>-6.3</td><td>4.0E-10</td><td>1.1E-08</td><td>BT20/MB231</td><td>MITO</td></tr>
<tr><td>>hg38_21_+_8433172_8446622_RNA45SN1_45S_50ntFlanks@12360.12384.25|CAGGGATAACTGGCTTGTGGCGGCC</td><td>>hg38_21_+_8433172_8446622_RNA45SN1_45S_50ntFlanks</td><td>12360</td><td>12384</td><td>25</td><td>CAGGGATAACTGGCTTGTGGCGGCC</td><td>266.7</td><td>-7.8</td><td>1.3</td><td>-5.9</td><td>4.7E-09</td><td>1.1E-08</td><td>MB231/MB468</td><td>MP</td></tr>
<tr><td>>hg38_21_+_8433172_8446622_RNA45SN1_45S_50ntFlanks@7991.8015.25|ACGTGGCGACCCGCTGAATTTAAGC</td><td>>hg38_21_+_8433172_8446622_RNA45SN1_45S_50ntFlanks</td><td>7991</td><td>8015</td><td>25</td><td>ACGTGGCGACCCGCTGAATTTAAGC</td><td>222.3</td><td>-4.3</td><td>0.7</td><td>-5.9</td><td>4.7E-09</td><td>1.1E-08</td><td>MB231/MB468</td><td>MP</td></tr>
<tr><td>>hg38_21_+_8433172_8446622_RNA45SN1_45S_50ntFlanks@3992.4026.35|GGTGACTCTAGATAACCTCGGGCCGATCGCACGCC</td><td>>hg38_21_+_8433172_8446622_RNA45SN1_45S_50ntFlanks</td><td>3992</td><td>4026</td><td>35</td><td>GGTGACTCTAGATAACCTCGGGCCGATCGCACGCC</td><td>495.2</td><td>-6.8</td><td>1.2</td><td>-5.8</td><td>5.7E-09</td><td>1.1E-08</td><td>BT20/MB468</td><td>NUC</td></tr>
<tr><td>>hg38_21_+_8433172_8446622_RNA45SN1_45S_50ntFlanks@4099.4127.29|GTGACCACGGGTGACGGGGAATCAGGGTT</td><td>>hg38_21_+_8433172_8446622_RNA45SN1_45S_50ntFlanks</td><td>4099</td><td>4127</td><td>29</td><td>GTGACCACGGGTGACGGGGAATCAGGGTT</td><td>474.4</td><td>-8.4</td><td>1.4</td><td>-5.8</td><td>5.7E-09</td><td>1.1E-08</td><td>BT20/MB468</td><td>NUC</td></tr>
<tr><td>>hg38_21_+_8433172_8446622_RNA45SN1_45S_50ntFlanks@11581.11605.25|AACTGGTGCGGACCAGGGGAATCCG</td><td>>hg38_21_+_8433172_8446622_RNA45SN1_45S_50ntFlanks</td><td>11581</td><td>11605</td><td>25</td><td>AACTGGTGCGGACCAGGGGAATCCG</td><td>3430.9</td><td>-9.9</td><td>1.7</td><td>-5.8</td><td>5.7E-09</td><td>1.1E-08</td><td>BT20/MB468</td><td>NUC</td></tr>
<tr><td>>hg38_21_+_8433172_8446622_RNA45SN1_45S_50ntFlanks@12635.12656.22|AAGCTACCATCTGTGGGATTAT</td><td>>hg38_21_+_8433172_8446622_RNA45SN1_45S_50ntFlanks</td><td>12635</td><td>12656</td><td>22</td><td>AAGCTACCATCTGTGGGATTAT</td><td>320.5</td><td>-8.3</td><td>1.4</td><td>-5.8</td><td>5.5E-09</td><td>1.1E-08</td><td>MB231/MB468</td><td>MITO</td></tr>
<tr><td>>trna111_HisGTG_1_-_147774845_147774916@-1T.23.24,trna118_HisGTG_1_-_145396881_145396952@-1T.23.24,trna16_HisGTG_1_+_146544773_146544844@-1T.23.24,trna1_HisGTG_15_+_45493349_45493420@-1T.23.24,trna21_HisGTG_1_+_147753471_147753542@-1T.23.24,trna33_HisGTG_6_+_27125906_27125977@-1T.23.24,trna7_HisGTG_9_-_14433938_14434009@-1T.23.24,trna8_HisGTG_15_-_45492611_45492682@-1T.23.24,trna9_HisGTG_15_-_45490804_45490875@-1T.23.24|TGCCGTGATCGTATAGTGGTTAGT</td><td>>trna111_HisGTG_1_-_147774845_147774916</td><td>-1T</td><td>23</td><td>24</td><td>TGCCGTGATCGTATAGTGGTTAGT</td><td>5351.6</td><td>-4.8</td><td>0.8</td><td>-6.3</td><td>4.1E-10</td><td>1.1E-08</td><td>BT20/MB231</td><td>MITO</td></tr>
<tr><td>>hg38_21_+_8433172_8446622_RNA45SN1_45S_50ntFlanks@11917.11940.24|GACATGAGAGGTGTAGAATAAGTG</td><td>>hg38_21_+_8433172_8446622_RNA45SN1_45S_50ntFlanks</td><td>11917</td><td>11940</td><td>24</td><td>GACATGAGAGGTGTAGAATAAGTG</td><td>155.5</td><td>11.0</td><td>1.8</td><td>6.2</td><td>6.4E-10</td><td>1.1E-08</td><td>BT20/MB468</td><td>MP</td></tr>
<tr><td>>hg38_21_+_8433172_8446622_RNA45SN1_45S_50ntFlanks@10353.10374.22|AGAACTTTGAAGGCCGAAGTGG</td><td>>hg38_21_+_8433172_8446622_RNA45SN1_45S_50ntFlanks</td><td>10353</td><td>10374</td><td>22</td><td>AGAACTTTGAAGGCCGAAGTGG</td><td>198.3</td><td>-10.7</td><td>1.7</td><td>-6.2</td><td>6.4E-10</td><td>1.1E-08</td><td>BT20/MB468</td><td>MP</td></tr>
<tr><td>>hg38_21_+_8433172_8446622_RNA45SN1_45S_50ntFlanks@10794.10826.33|CTGGCATGTTGGAACAATGTAGGTAAGGGAAGT</td><td>>hg38_21_+_8433172_8446622_RNA45SN1_45S_50ntFlanks</td><td>10794</td><td>10826</td><td>33</td><td>CTGGCATGTTGGAACAATGTAGGTAAGGGAAGT</td><td>386.9</td><td>-2.8</td><td>0.5</td><td>-5.9</td><td>3.8E-09</td><td>1.1E-08</td><td>BT20/MB468</td><td>MITO</td></tr>
<tr><td>>hg38_1_-_228634819_228635039_RNA5S12_5S_50ntFlanks@139.171.33|GACCGCCTGGGAATACCGGGTGCTGTAGGCTTT</td><td>>hg38_1_-_228634819_228635039_RNA5S12_5S_50ntFlanks</td><td>139</td><td>171</td><td>33</td><td>GACCGCCTGGGAATACCGGGTGCTGTAGGCTTT</td><td>325.0</td><td>3.8</td><td>0.7</td><td>5.9</td><td>3.8E-09</td><td>1.1E-08</td><td>BT20/MB468</td><td>MITO</td></tr>
<tr><td>>hg38_21_+_8433172_8446622_RNA45SN1_45S_50ntFlanks@12456.12491.36|CCAAGCGTTGGATTGTTCACCCACTAATAGGGAACG</td><td>>hg38_21_+_8433172_8446622_RNA45SN1_45S_50ntFlanks</td><td>12456</td><td>12491</td><td>36</td><td>CCAAGCGTTGGATTGTTCACCCACTAATAGGGAACG</td><td>240.6</td><td>-11.0</td><td>1.8</td><td>-6.2</td><td>6.4E-10</td><td>1.1E-08</td><td>BT20/MB468</td><td>MP</td></tr>
<tr><td>>hg38_21_+_8433172_8446622_RNA45SN1_45S_50ntFlanks@10404.10422.19|CATGGGTCAGTCGGTCCTG</td><td>>hg38_21_+_8433172_8446622_RNA45SN1_45S_50ntFlanks</td><td>10404</td><td>10422</td><td>19</td><td>CATGGGTCAGTCGGTCCTG</td><td>321.9</td><td>-5.1</td><td>0.9</td><td>-5.8</td><td>5.1E-09</td><td>1.1E-08</td><td>MB231/MB468</td><td>CYTO</td></tr>
<tr><td>>hg38_21_+_8433172_8446622_RNA45SN1_45S_50ntFlanks@10786.10822.37|AACAGCCTCTGGCATGTTGGAACAATGTAGGTAAGGG</td><td>>hg38_21_+_8433172_8446622_RNA45SN1_45S_50ntFlanks</td><td>10786</td><td>10822</td><td>37</td><td>AACAGCCTCTGGCATGTTGGAACAATGTAGGTAAGGG</td><td>556.2</td><td>-5.4</td><td>0.9</td><td>-5.8</td><td>5.1E-09</td><td>1.1E-08</td><td>MB231/MB468</td><td>CYTO</td></tr>
<tr><td>>hg38_21_+_8433172_8446622_RNA45SN1_45S_50ntFlanks@8212.8230.19|CCGGTAGCGGCCCCCGGCG</td><td>>hg38_21_+_8433172_8446622_RNA45SN1_45S_50ntFlanks</td><td>8212</td><td>8230</td><td>19</td><td>CCGGTAGCGGCCCCCGGCG</td><td>117.9</td><td>-10.0</td><td>1.6</td><td>-6.2</td><td>6.4E-10</td><td>1.1E-08</td><td>BT20/MB468</td><td>MP</td></tr>
<tr><td>>hg38_1_-_228634819_228635039_RNA5S12_5S_50ntFlanks@51.82.32|GTCTACGGCCATACCACCCTGAACGCGCCCGA</td><td>>hg38_1_-_228634819_228635039_RNA5S12_5S_50ntFlanks</td><td>51</td><td>82</td><td>32</td><td>GTCTACGGCCATACCACCCTGAACGCGCCCGA</td><td>3346.1</td><td>5.8</td><td>1.0</td><td>5.9</td><td>4.3E-09</td><td>1.1E-08</td><td>BT20/MB468</td><td>CYTO</td></tr>
<tr><td>>hg38_21_+_8433172_8446622_RNA45SN1_45S_50ntFlanks@5489.5514.26|CTGAGAAGACGGTCGAACTTGACTAT</td><td>>hg38_21_+_8433172_8446622_RNA45SN1_45S_50ntFlanks</td><td>5489</td><td>5514</td><td>26</td><td>CTGAGAAGACGGTCGAACTTGACTAT</td><td>714.0</td><td>-5.6</td><td>0.9</td><td>-5.9</td><td>3.8E-09</td><td>1.1E-08</td><td>BT20/MB468</td><td>MITO</td></tr>
<tr><td>>hg38_21_+_8433172_8446622_RNA45SN1_45S_50ntFlanks@4102.4128.27|ACCACGGGTGACGGGGAATCAGGGTTC</td><td>>hg38_21_+_8433172_8446622_RNA45SN1_45S_50ntFlanks</td><td>4102</td><td>4128</td><td>27</td><td>ACCACGGGTGACGGGGAATCAGGGTTC</td><td>176.7</td><td>-5.9</td><td>1.0</td><td>-5.9</td><td>4.8E-09</td><td>1.1E-08</td><td>MB231/MB468</td><td>MP</td></tr>
<tr><td>>hg38_21_+_8433172_8446622_RNA45SN1_45S_50ntFlanks@4593.4616.24|TTGTTGGTTTTCGGAACTGAGGCC</td><td>>hg38_21_+_8433172_8446622_RNA45SN1_45S_50ntFlanks</td><td>4593</td><td>4616</td><td>24</td><td>TTGTTGGTTTTCGGAACTGAGGCC</td><td>285.3</td><td>-6.1</td><td>1.0</td><td>-5.9</td><td>4.8E-09</td><td>1.1E-08</td><td>MB231/MB468</td><td>MP</td></tr>
<tr><td>>hg38_21_+_8433172_8446622_RNA45SN1_45S_50ntFlanks@12488.12520.33|AACGTGAGCTGGGTTTAGACCGTCGTGAGACAG</td><td>>hg38_21_+_8433172_8446622_RNA45SN1_45S_50ntFlanks</td><td>12488</td><td>12520</td><td>33</td><td>AACGTGAGCTGGGTTTAGACCGTCGTGAGACAG</td><td>243.0</td><td>-5.9</td><td>1.0</td><td>-5.8</td><td>5.1E-09</td><td>1.1E-08</td><td>MB231/MB468</td><td>NUC</td></tr>
</table>

<table>
<tr><th></th><th></th><th></th><th></th><th></th><th></th><th></th><th></th><th></th><th></th><th></th><th></th><th></th><th></th></tr>
<tr><td>>MI0000456|hsa-mir-140&WithFlank&16|+|69933075|69933186@68.89.22&[MIMAT0004597&hsa-miR-140-3p&offsets|0|+1;m-45&16|+|69933143|69933164&offsets|-1|-1]|TACCACAGGGTAGAACCACGGA</td><td>>MI0000456|hsa-mir-140&WithFlank&16|+|69933075|69933186</td><td>68</td><td>89</td><td>22</td><td>TACCACAGGGTAGAACCACGGA</td><td>130.1</td><td>5.8</td><td>1.0</td><td>5.8</td><td>5.8E-09</td><td>1.1E-08</td><td>BT20/MB468</td><td>NUC</td></tr>
<tr><td>>hg38_21_+_8433172_8446622_RNA45SN1_45S_50ntFlanks@9534.9567.34|AGGAAACTCTGGTGGAGGTCCGTAGCGGTCCTGA</td><td>>hg38_21_+_8433172_8446622_RNA45SN1_45S_50ntFlanks</td><td>9534</td><td>9567</td><td>34</td><td>AGGAAACTCTGGTGGAGGTCCGTAGCGGTCCTGA</td><td>2432.8</td><td>-8.9</td><td>1.5</td><td>-5.8</td><td>5.1E-09</td><td>1.1E-08</td><td>MB231/MB468</td><td>NUC</td></tr>
<tr><td>>hg38_21_+_8433172_8446622_RNA45SN1_45S_50ntFlanks@12479.12499.21|CTAATAGGGAACGTGAGCTGG</td><td>>hg38_21_+_8433172_8446622_RNA45SN1_45S_50ntFlanks</td><td>12479</td><td>12499</td><td>21</td><td>CTAATAGGGAACGTGAGCTGG</td><td>540.5</td><td>-6.2</td><td>1.1</td><td>-5.8</td><td>5.1E-09</td><td>1.1E-08</td><td>MB231/MB468</td><td>NUC</td></tr>
<tr><td>>hg38_MT_+_1621_3279_RNR2_16S_50ntFlanks@382.416.35|GAGCCTGGTGATAGCTGGTTGTCCAAGATAGAATC</td><td>>hg38_MT_+_1621_3279_RNR2_16S_50ntFlanks</td><td>382</td><td>416</td><td>35</td><td>GAGCCTGGTGATAGCTGGTTGTCCAAGATAGAATC</td><td>243.2</td><td>11.7</td><td>1.9</td><td>6.2</td><td>6.5E-10</td><td>1.1E-08</td><td>BT20/MB468</td><td>MP</td></tr>
<tr><td>>hg38_21_+_8433172_8446622_RNA45SN1_45S_50ntFlanks@12715.12751.37|AGCCTCGGTTGGCCTCGGATAGCCGGTCCCCCGCCTG</td><td>>hg38_21_+_8433172_8446622_RNA45SN1_45S_50ntFlanks</td><td>12715</td><td>12751</td><td>37</td><td>AGCCTCGGTTGGCCTCGGATAGCCGGTCCCCCGCCTG</td><td>197.9</td><td>-11.5</td><td>2.0</td><td>-5.9</td><td>4.8E-09</td><td>1.1E-08</td><td>MB231/MB468</td><td>MP</td></tr>
<tr><td>>hg38_21_+_8433172_8446622_RNA45SN1_45S_50ntFlanks@4442.4471.30|CCCCTTGCCTCTCGGCGCCCCCTCGATGCT</td><td>>hg38_21_+_8433172_8446622_RNA45SN1_45S_50ntFlanks</td><td>4442</td><td>4471</td><td>30</td><td>CCCCTTGCCTCTCGGCGCCCCCTCGATGCT</td><td>635.1</td><td>-9.1</td><td>1.6</td><td>-5.8</td><td>5.8E-09</td><td>1.1E-08</td><td>BT20/MB468</td><td>NUC</td></tr>
<tr><td>>hg38_21_+_8433172_8446622_RNA45SN1_45S_50ntFlanks@10342.10361.20|ATATTCAAACGAGAACTTTG</td><td>>hg38_21_+_8433172_8446622_RNA45SN1_45S_50ntFlanks</td><td>10342</td><td>10361</td><td>20</td><td>ATATTCAAACGAGAACTTTG</td><td>789.1</td><td>-4.3</td><td>0.7</td><td>-5.8</td><td>5.1E-09</td><td>1.1E-08</td><td>MB231/MB468</td><td>CYTO</td></tr>
<tr><td>>hg38_21_+_8433172_8446622_RNA45SN1_45S_50ntFlanks@6651.6690.40|CGACTCTTAGCGGTGGATCACTCGGCTCGTGCGTCGATGA</td><td>>hg38_21_+_8433172_8446622_RNA45SN1_45S_50ntFlanks</td><td>6651</td><td>6690</td><td>40</td><td>CGACTCTTAGCGGTGGATCACTCGGCTCGTGCGTCGATGA</td><td>1578.3</td><td>5.0</td><td>0.9</td><td>5.9</td><td>4.4E-09</td><td>1.1E-08</td><td>BT20/MB468</td><td>CYTO</td></tr>
<tr><td>>trnaMT_PheGAA_MT_+_577_647@35.56.22|GAAAATGTTTAGACGGGCTCAC</td><td>>trnaMT_PheGAA_MT_+_577_647</td><td>35</td><td>56</td><td>22</td><td>GAAAATGTTTAGACGGGCTCAC</td><td>125.8</td><td>7.2</td><td>1.2</td><td>5.8</td><td>5.8E-09</td><td>1.1E-08</td><td>BT20/MB468</td><td>NUC</td></tr>
<tr><td>>hg38_21_+_8433172_8446622_RNA45SN1_45S_50ntFlanks@11209.11253.45|CGGCGACTCTGGACGCGAGCCGGGCCCTTCCCGTGGATCGCCCCA</td><td>>hg38_21_+_8433172_8446622_RNA45SN1_45S_50ntFlanks</td><td>11209</td><td>11253</td><td>45</td><td>CGGCGACTCTGGACGCGAGCCGGGCCCTTCCCGTGGATCGCCCCA</td><td>587.3</td><td>-8.9</td><td>1.5</td><td>-5.8</td><td>5.2E-09</td><td>1.1E-08</td><td>MB231/MB468</td><td>NUC</td></tr>
<tr><td>>hg38_21_+_8433172_8446622_RNA45SN1_45S_50ntFlanks@9202.9220.19|TCTCGGGGCCACGCGCGCG</td><td>>hg38_21_+_8433172_8446622_RNA45SN1_45S_50ntFlanks</td><td>9202</td><td>9220</td><td>19</td><td>TCTCGGGGCCACGCGCGCG</td><td>809.9</td><td>-8.6</td><td>1.4</td><td>-6.2</td><td>6.5E-10</td><td>1.1E-08</td><td>BT20/MB468</td><td>MP</td></tr>
<tr><td>>hg38_21_+_8433172_8446622_RNA45SN1_45S_50ntFlanks@476.506.31|TTCGTGATCGATGTGGTGACGTCGTGCTCTC</td><td>>hg38_21_+_8433172_8446622_RNA45SN1_45S_50ntFlanks</td><td>476</td><td>506</td><td>31</td><td>TTCGTGATCGATGTGGTGACGTCGTGCTCTC</td><td>15024.4</td><td>4.3</td><td>0.7</td><td>5.8</td><td>5.2E-09</td><td>1.1E-08</td><td>MB231/MB468</td><td>NUC</td></tr>
<tr><td>>hg38_21_+_8433172_8446622_RNA45SN1_45S_50ntFlanks@10427.10457.31|ATGGGCGAGCGCCGTTCCGAAGGGACGGGCG</td><td>>hg38_21_+_8433172_8446622_RNA45SN1_45S_50ntFlanks</td><td>10427</td><td>10457</td><td>31</td><td>ATGGGCGAGCGCCGTTCCGAAGGGACGGGCG</td><td>114.4</td><td>-9.9</td><td>1.6</td><td>-6.2</td><td>6.5E-10</td><td>1.1E-08</td><td>BT20/MB468</td><td>MP</td></tr>
<tr><td>>am_trna143_LysTTT_6_-_27559593_27559665@7.24.18,trna14_LysTTT_11_-_59327808_59327880@7.24.18,trna2_LysTTT_17_+_8022473_8022545@7.24.18,trna54_LysTTT_1_+_204475655_204475727@7.24.18,trna5_LysTTT_11_+_59323902_59323974@7.24.18,trna62_LysTTT_1_-_204476158_204476230@7.24.18,trna76_LysTTT_6_+_28918806_28918878@7.24.18|ATAGCTCAGTCGGTAGAG</td><td>>am_trna143_LysTTT_6_-_27559593_27559665</td><td>7</td><td>24</td><td>18</td><td>ATAGCTCAGTCGGTAGAG</td><td>117.9</td><td>-10.0</td><td>1.6</td><td>-6.2</td><td>6.5E-10</td><td>1.1E-08</td><td>BT20/MB468</td><td>MP</td></tr>
<tr><td>>hg38_21_+_8433172_8446622_RNA45SN1_45S_50ntFlanks@10427.10456.30|ATGGGCGAGCGCCGTTCCGAAGGGACGGGC</td><td>>hg38_21_+_8433172_8446622_RNA45SN1_45S_50ntFlanks</td><td>10427</td><td>10456</td><td>30</td><td>ATGGGCGAGCGCCGTTCCGAAGGGACGGGC</td><td>591.2</td><td>-6.4</td><td>1.1</td><td>-5.8</td><td>5.6E-09</td><td>1.1E-08</td><td>MB231/MB468</td><td>MITO</td></tr>
<tr><td>>MI0000434|hsa-let-7i&WithFlank&12|+|62603680|62603775@12.32.21(+1G)&[MIMAT0000415&hsa-let-7i-5p&offsets|0|-1(+1G);m-49&12|+|62603691|62603712&offsets|0|-1(+1G)]|TGAGGTAGTAGTTTGTGCTGTG</td><td>>MI0000434|hsa-let-7i&WithFlank&12|+|62603680|62603775</td><td>12</td><td>32</td><td>21(+1G)</td><td>TGAGGTAGTAGTTTGTGCTGTG</td><td>482.4</td><td>2.3</td><td>0.4</td><td>5.8</td><td>5.8E-09</td><td>1.1E-08</td><td>BT20/MB468</td><td>NUC</td></tr>
<tr><td>>hg38_21_+_8433172_8446622_RNA45SN1_45S_50ntFlanks@10480.10497.18|GATCGAAAGGGAGTCGGG</td><td>>hg38_21_+_8433172_8446622_RNA45SN1_45S_50ntFlanks</td><td>10480</td><td>10497</td><td>18</td><td>GATCGAAAGGGAGTCGGG</td><td>182.3</td><td>-10.6</td><td>1.7</td><td>-6.2</td><td>6.6E-10</td><td>1.1E-08</td><td>BT20/MB468</td><td>MP</td></tr>
<tr><td>>MI0000289|hsa-mir-181a-1&WithFlank&1|-|198859038|198859159@30.52.23;MI0000269|hsa-mir-181a-2&WithFlank&9|+|124692436|124692557@45.67.23&[MIMAT0000256&hsa-miR-181a-5p&offsets|0|0;m-32&1|-|198859108|198859130&offsets|0|0];[MIMAT0000256_1&hsa-miR-181a-5p&offsets|0|0;m-32&9|+|124692480|124692502&offsets|0|0]|AACATTCAACGCTGTCGGTGAGT</td><td>>MI0000289|hsa-mir-181a-1&WithFlank&1|-|198859038|198859159</td><td>30</td><td>52</td><td>23</td><td>AACATTCAACGCTGTCGGTGAGT</td><td>6688.5</td><td>5.7</td><td>1.0</td><td>5.8</td><td>5.6E-09</td><td>1.1E-08</td><td>MB231/MB468</td><td>MITO</td></tr>
<tr><td>>hg38_21_+_8433172_8446622_RNA45SN1_45S_50ntFlanks@11642.11660.19|GGCGGGTGTTGACGCGATG</td><td>>hg38_21_+_8433172_8446622_RNA45SN1_45S_50ntFlanks</td><td>11642</td><td>11660</td><td>19</td><td>GGCGGGTGTTGACGCGATG</td><td>382.1</td><td>-7.8</td><td>1.3</td><td>-5.8</td><td>5.6E-09</td><td>1.1E-08</td><td>MB231/MB468</td><td>MITO</td></tr>
<tr><td>>MI0000085|hsa-mir-27a&WithFlank&19|-|13836434|13836523@57.75.19&[MIMAT0000084&hsa-miR-27a-3p&offsets|0|-2;m-43&19|-|13836447|13836467&offsets|0|-2]|TTCACAGTGGCTAAGTTCC</td><td>>MI0000085|hsa-mir-27a&WithFlank&19|-|13836434|13836523</td><td>57</td><td>75</td><td>19</td><td>TTCACAGTGGCTAAGTTCC</td><td>344.7</td><td>5.3</td><td>0.9</td><td>5.8</td><td>5.1E-09</td><td>1.1E-08</td><td>MB231/MB468</td><td>CYTO</td></tr>
<tr><td>>hg38_21_+_8433172_8446622_RNA45SN1_45S_50ntFlanks@8381.8403.23|AAGAGGGCGTGAAACCGTTAAGA</td><td>>hg38_21_+_8433172_8446622_RNA45SN1_45S_50ntFlanks</td><td>8381</td><td>8403</td><td>23</td><td>AAGAGGGCGTGAAACCGTTAAGA</td><td>162.6</td><td>-3.0</td><td>0.5</td><td>-5.8</td><td>5.2E-09</td><td>1.1E-08</td><td>MB231/MB468</td><td>NUC</td></tr>
<tr><td>>hg38_21_+_8433172_8446622_RNA45SN1_45S_50ntFlanks@3925.3959.35|ATCAGATCAAAACCAACCCGGTCAGCCCCTCTCCG</td><td>>hg38_21_+_8433172_8446622_RNA45SN1_45S_50ntFlanks</td><td>3925</td><td>3959</td><td>35</td><td>ATCAGATCAAAACCAACCCGGTCAGCCCCTCTCCG</td><td>128.5</td><td>-5.5</td><td>0.9</td><td>-5.8</td><td>5.1E-09</td><td>1.1E-08</td><td>MB231/MB468</td><td>CYTO</td></tr>
<tr><td>>hg38_21_+_8433172_8446622_RNA45SN1_45S_50ntFlanks@4005.4026.22|AACCTCGGGCCGATCGCACGCC</td><td>>hg38_21_+_8433172_8446622_RNA45SN1_45S_50ntFlanks</td><td>4005</td><td>4026</td><td>22</td><td>AACCTCGGGCCGATCGCACGCC</td><td>319.9</td><td>-4.0</td><td>0.7</td><td>-5.8</td><td>5.1E-09</td><td>1.1E-08</td><td>MB231/MB468</td><td>CYTO</td></tr>
<tr><td>>hg38_21_+_8433172_8446622_RNA45SN1_45S_50ntFlanks@3979.4002.24|CGCCGGCGGCTTTGGTGACTCTAG</td><td>>hg38_21_+_8433172_8446622_RNA45SN1_45S_50ntFlanks</td><td>3979</td><td>4002</td><td>24</td><td>CGCCGGCGGCTTTGGTGACTCTAG</td><td>96.1</td><td>-10.5</td><td>1.8</td><td>-5.9</td><td>4.9E-09</td><td>1.1E-08</td><td>MB231/MB468</td><td>MP</td></tr>
<tr><td>>hg38_21_+_8433172_8446622_RNA45SN1_45S_50ntFlanks@6739.6772.34|TCATCGACACTTCGAACGCACTTGCGGCCCCGGG</td><td>>hg38_21_+_8433172_8446622_RNA45SN1_45S_50ntFlanks</td><td>6739</td><td>6772</td><td>34</td><td>TCATCGACACTTCGAACGCACTTGCGGCCCCGGG</td><td>128.7</td><td>-10.9</td><td>1.9</td><td>-5.9</td><td>4.9E-09</td><td>1.1E-08</td><td>MB231/MB468</td><td>MP</td></tr>
<tr><td>>MI0003560|hsa-mir-92b&WithFlank&1|+|155195171|155195278@67.88.22&[MIMAT0003218&hsa-miR-92b-3p&offsets|0|0;m-115&1|+|155195237|155195258&offsets|0|0]|TATTGCACTCGTCCCGGCCTCC</td><td>>MI0003560|hsa-mir-92b&WithFlank&1|+|155195171|155195278</td><td>67</td><td>88</td><td>22</td><td>TATTGCACTCGTCCCGGCCTCC</td><td>6945.5</td><td>7.3</td><td>1.2</td><td>6.2</td><td>6.6E-10</td><td>1.1E-08</td><td>BT20/MB468</td><td>MP</td></tr>
<tr><td>>hg38_21_+_8433172_8446622_RNA45SN1_45S_50ntFlanks@6647.6664.18|CGTACGACTCTTAGCGGT</td><td>>hg38_21_+_8433172_8446622_RNA45SN1_45S_50ntFlanks</td><td>6647</td><td>6664</td><td>18</td><td>CGTACGACTCTTAGCGGT</td><td>5761.3</td><td>4.4</td><td>0.7</td><td>6.2</td><td>6.6E-10</td><td>1.1E-08</td><td>BT20/MB468</td><td>MP</td></tr>
<tr><td>>hg38_21_+_8433172_8446622_RNA45SN1_45S_50ntFlanks@8176.8193.18|CTTCTGATCGAGGCCCAG</td><td>>hg38_21_+_8433172_8446622_RNA45SN1_45S_50ntFlanks</td><td>8176</td><td>8193</td><td>18</td><td>CTTCTGATCGAGGCCCAG</td><td>686.8</td><td>-6.6</td><td>1.1</td><td>-5.9</td><td>4.9E-09</td><td>1.1E-08</td><td>MB231/MB468</td><td>MP</td></tr>
<tr><td>>hg38_21_+_8433172_8446622_RNA45SN1_45S_50ntFlanks@6733.6771.39|CATTGATCATCGACACTTCGAACGCACTTGCGGCCCCGG</td><td>>hg38_21_+_8433172_8446622_RNA45SN1_45S_50ntFlanks</td><td>6733</td><td>6771</td><td>39</td><td>CATTGATCATCGACACTTCGAACGCACTTGCGGCCCCGG</td><td>476.7</td><td>-3.5</td><td>0.6</td><td>-5.9</td><td>4.4E-09</td><td>1.1E-08</td><td>BT20/MB468</td><td>CYTO</td></tr>
<tr><td>>hg38_21_+_8433172_8446622_RNA45SN1_45S_50ntFlanks@7976.8008.33|GCGACCTCAGATCAGACGTGGCGACCCGCTGAA</td><td>>hg38_21_+_8433172_8446622_RNA45SN1_45S_50ntFlanks</td><td>7976</td><td>8008</td><td>33</td><td>GCGACCTCAGATCAGACGTGGCGACCCGCTGAA</td><td>1459.0</td><td>3.9</td><td>0.7</td><td>5.9</td><td>3.9E-09</td><td>1.1E-08</td><td>BT20/MB468</td><td>MITO</td></tr>
<tr><td>>hg38_21_+_8433172_8446622_RNA45SN1_45S_50ntFlanks@8191.8213.23|CAGCCCGTGGACGGTGTGAGGCC</td><td>>hg38_21_+_8433172_8446622_RNA45SN1_45S_50ntFlanks</td><td>8191</td><td>8213</td><td>23</td><td>CAGCCCGTGGACGGTGTGAGGCC</td><td>1168.9</td><td>-4.2</td><td>0.7</td><td>-5.8</td><td>5.2E-09</td><td>1.1E-08</td><td>MB231/MB468</td><td>NUC</td></tr>
<tr><td>>hg38_21_+_8433172_8446622_RNA45SN1_45S_50ntFlanks@8118.8139.22|GGGACATGTGGCGTACGGAAGA</td><td>>hg38_21_+_8433172_8446622_RNA45SN1_45S_50ntFlanks</td><td>8118</td><td>8139</td><td>22</td><td>GGGACATGTGGCGTACGGAAGA</td><td>1913.3</td><td>14.2</td><td>2.2</td><td>6.4</td><td>1.2E-10</td><td>1.1E-08</td><td>BT20/MB231</td><td>MP</td></tr>
<tr><td>>hg38_21_+_8433172_8446622_RNA45SN1_45S_50ntFlanks@12616.12654.39|CTGAGGAGCCAATGGGGCGAAGCTACCATCTGTGGGATT</td><td>>hg38_21_+_8433172_8446622_RNA45SN1_45S_50ntFlanks</td><td>12616</td><td>12654</td><td>39</td><td>CTGAGGAGCCAATGGGGCGAAGCTACCATCTGTGGGATT</td><td>3528.7</td><td>-10.2</td><td>1.8</td><td>-5.8</td><td>5.9E-09</td><td>1.1E-08</td><td>BT20/MB468</td><td>NUC</td></tr>
<tr><td>>hg38_21_+_8433172_8446622_RNA45SN1_45S_50ntFlanks@9538.9563.26|AACTCTGGTGGAGGTCCGTAGCGGTC</td><td>>hg38_21_+_8433172_8446622_RNA45SN1_45S_50ntFlanks</td><td>9538</td><td>9563</td><td>26</td><td>AACTCTGGTGGAGGTCCGTAGCGGTC</td><td>274.2</td><td>-7.5</td><td>1.3</td><td>-5.8</td><td>5.2E-09</td><td>1.1E-08</td><td>MB231/MB468</td><td>NUC</td></tr>
<tr><td>>hg38_21_+_8433172_8446622_RNA45SN1_45S_50ntFlanks@12940.12966.27|CATTCGTAGACGACCTGCTTCTGGGTC</td><td>>hg38_21_+_8433172_8446622_RNA45SN1_45S_50ntFlanks</td><td>12940</td><td>12966</td><td>27</td><td>CATTCGTAGACGACCTGCTTCTGGGTC</td><td>1329.3</td><td>-8.0</td><td>1.4</td><td>-5.9</td><td>4.4E-09</td><td>1.1E-08</td><td>BT20/MB468</td><td>CYTO</td></tr>
<tr><td>>hg38_21_+_8433172_8446622_RNA45SN1_45S_50ntFlanks@4498.4524.27|AAGCGTTTACTTTGAAAAAATTAGAGT</td><td>>hg38_21_+_8433172_8446622_RNA45SN1_45S_50ntFlanks</td><td>4498</td><td>4524</td><td>27</td><td>AAGCGTTTACTTTGAAAAAATTAGAGT</td><td>595.1</td><td>5.5</td><td>0.9</td><td>5.9</td><td>4.4E-09</td><td>1.1E-08</td><td>BT20/MB468</td><td>CYTO</td></tr>
<tr><td>>hg38_21_+_8433172_8446622_RNA45SN1_45S_50ntFlanks@10808.10840.33|CAATGTAGGTAAGGGAAGTCGGCAAGCCGGATC</td><td>>hg38_21_+_8433172_8446622_RNA45SN1_45S_50ntFlanks</td><td>10808</td><td>10840</td><td>33</td><td>CAATGTAGGTAAGGGAAGTCGGCAAGCCGGATC</td><td>1306.7</td><td>-5.4</td><td>0.9</td><td>-5.8</td><td>5.2E-09</td><td>1.1E-08</td><td>MB231/MB468</td><td>CYTO</td></tr>
<tr><td>>hg38_21_+_8433172_8446622_RNA45SN1_45S_50ntFlanks@10821.10865.45|GGAAGTCGGCAAGCCGGATCCGTAACTTCGGGATAAGGATTGGCT</td><td>>hg38_21_+_8433172_8446622_RNA45SN1_45S_50ntFlanks</td><td>10821</td><td>10865</td><td>45</td><td>GGAAGTCGGCAAGCCGGATCCGTAACTTCGGGATAAGGATTGGCT</td><td>121.7</td><td>-10.8</td><td>1.8</td><td>-5.9</td><td>4.9E-09</td><td>1.1E-08</td><td>MB231/MB468</td><td>MP</td></tr>
<tr><td>>hg38_21_+_8433172_8446622_RNA45SN1_45S_50ntFlanks@3768.3791.24|ACGCACGGCCGGTACAGTGAAACT</td><td>>hg38_21_+_8433172_8446622_RNA45SN1_45S_50ntFlanks</td><td>3768</td><td>3791</td><td>24</td><td>ACGCACGGCCGGTACAGTGAAACT</td><td>150.9</td><td>5.4</td><td>0.9</td><td>5.8</td><td>5.2E-09</td><td>1.1E-08</td><td>MB231/MB468</td><td>CYTO</td></tr>
<tr><td>>hg38_21_+_8433172_8446622_RNA45SN1_45S_50ntFlanks@12892.12933.42|CCCTCGCCCGTCACGCACCGCACGTTCGTGGGGAACCTGGCG</td><td>>hg38_21_+_8433172_8446622_RNA45SN1_45S_50ntFlanks</td><td>12892</td><td>12933</td><td>42</td><td>CCCTCGCCCGTCACGCACCGCACGTTCGTGGGGAACCTGGCG</td><td>399.4</td><td>-11.7</td><td>1.9</td><td>-6.2</td><td>6.7E-10</td><td>1.1E-08</td><td>BT20/MB468</td><td>MP</td></tr>
<tr><td>>hg38_21_+_8433172_8446622_RNA45SN1_45S_50ntFlanks@12183.12211.29|AAACGGTAACGCAGGTGTCCTAAGGCGAG</td><td>>hg38_21_+_8433172_8446622_RNA45SN1_45S_50ntFlanks</td><td>12183</td><td>12211</td><td>29</td><td>AAACGGTAACGCAGGTGTCCTAAGGCGAG</td><td>96.8</td><td>-8.7</td><td>1.5</td><td>-5.9</td><td>3.9E-09</td><td>1.1E-08</td><td>BT20/MB468</td><td>MITO</td></tr>
<tr><td>>hg38_21_+_8433172_8446622_RNA45SN1_45S_50ntFlanks@12008.12044.37|CCGCCGGTGAAATACCACTACTCTGATCGTTTTTTCA</td><td>>hg38_21_+_8433172_8446622_RNA45SN1_45S_50ntFlanks</td><td>12008</td><td>12044</td><td>37</td><td>CCGCCGGTGAAATACCACTACTCTGATCGTTTTTTCA</td><td>143.7</td><td>-9.3</td><td>1.6</td><td>-5.9</td><td>4.0E-09</td><td>1.1E-08</td><td>BT20/MB468</td><td>MITO</td></tr>
<tr><td>>hg38_21_+_8433172_8446622_RNA45SN1_45S_50ntFlanks@8423.8469.47|CGCAGTCCGCCCGGAGGATTCAACCCGGCGGCGGGTCCGGCCGTGTC</td><td>>hg38_21_+_8433172_8446622_RNA45SN1_45S_50ntFlanks</td><td>8423</td><td>8469</td><td>47</td><td>CGCAGTCCGCCCGGAGGATTCAACCCGGCGGCGGGTCCGGCCGTGTC</td><td>310.1</td><td>-11.2</td><td>1.9</td><td>-5.8</td><td>5.3E-09</td><td>1.1E-08</td><td>MB231/MB468</td><td>NUC</td></tr>
<tr><td>>hg38_21_+_8433172_8446622_RNA45SN1_45S_50ntFlanks@7986.8013.28|ATCAGACGTGGCGACCCGCTGAATTTAA</td><td>>hg38_21_+_8433172_8446622_RNA45SN1_45S_50ntFlanks</td><td>7986</td><td>8013</td><td>28</td><td>ATCAGACGTGGCGACCCGCTGAATTTAA</td><td>127.7</td><td>-10.1</td><td>1.6</td><td>-6.2</td><td>6.7E-10</td><td>1.1E-08</td><td>BT20/MB468</td><td>MP</td></tr>
<tr><td>>hg38_21_+_8433172_8446622_RNA45SN1_45S_50ntFlanks@485.518.34|GATGTGGTGACGTCGTGCTCTCCCGGGCCGGGTC</td><td>>hg38_21_+_8433172_8446622_RNA45SN1_45S_50ntFlanks</td><td>485</td><td>518</td><td>34</td><td>GATGTGGTGACGTCGTGCTCTCCCGGGCCGGGTC</td><td>207.9</td><td>11.4</td><td>1.9</td><td>6.2</td><td>6.7E-10</td><td>1.1E-08</td><td>BT20/MB468</td><td>MP</td></tr>
<tr><td>>hg38_1_-_228634819_228635039_RNA5S12_5S_50ntFlanks@125.170.46|GTACTTGGATGGGAGACCGCCTGGGAATACCGGGTGCTGTAGGCTT</td><td>>hg38_1_-_228634819_228635039_RNA5S12_5S_50ntFlanks</td><td>125</td><td>170</td><td>46</td><td>GTACTTGGATGGGAGACCGCCTGGGAATACCGGGTGCTGTAGGCTT</td><td>995.8</td><td>8.7</td><td>1.5</td><td>5.9</td><td>4.0E-09</td><td>1.1E-08</td><td>BT20/MB468</td><td>MITO</td></tr>
<tr><td>>MI0000102|hsa-mir-100&WithFlank&11|-|122152223|122152314@20.41.22&[MIMAT0000098&hsa-miR-100-5p&offsets|+1|+1;m-27&11|-|122152276|122152296&offsets|+1|+2]|ACCCGTAGATCCGAACTTGTGG</td><td>>MI0000102|hsa-mir-100&WithFlank&11|-|122152223|122152314</td><td>20</td><td>41</td><td>22</td><td>ACCCGTAGATCCGAACTTGTGG</td><td>73.9</td><td>9.4</td><td>1.6</td><td>5.8</td><td>5.0E-09</td><td>1.1E-08</td><td>MB231/MB468</td><td>MP</td></tr>
<tr><td>>MI0000439|hsa-mir-23b&WithFlank&9|+|95085202|95085310@65.86.22(+1U)&[MIMAT0000418&hsa-miR-23b-3p&offsets|+1|0(+1U);m-42&9|+|95085265|95085285&offsets|+1|+2(+1U)]|TCACATTGCCAGGGATTACCACT</td><td>>MI0000439|hsa-mir-23b&WithFlank&9|+|95085202|95085310</td><td>65</td><td>86</td><td>22(+1U)</td><td>TCACATTGCCAGGGATTACCACT</td><td>81.6</td><td>2.9</td><td>0.5</td><td>5.8</td><td>6.0E-09</td><td>1.1E-08</td><td>BT20/MB468</td><td>NUC</td></tr>
</table>

| | | | | | | | | | | | | | |
|---|---|---|---|---|---|---|---|---|---|---|---|---|---|
| >hg38_21_+_8433172_8446622_RNA45SN1_45S_50ntFlanks@11572.11590.19\|CCGACTTAGAACTGGTGCG | >hg38_21_+_8433172_8446622_RNA45SN1_45S_50ntFlanks | 11572 | 11590 | 19 | CCGACTTAGAACTGGTGCG | 2592.1 | -3.3 | 0.6 | -5.8 | 5.2E-09 | 1.1E-08 | MB231/MB468 | CYTO |
| >hg38_21_+_8433172_8446622_RNA45SN1_45S_50ntFlanks@11920.11956.37\|ATGAGAGGTGTAGAATAAGTGGGAGGCCCCCGGCGCC | >hg38_21_+_8433172_8446622_RNA45SN1_45S_50ntFlanks | 11920 | 11956 | 37 | ATGAGAGGTGTAGAATAAGTGGGAGGCCCCCGGCGCC | 370.9 | -8.3 | 1.4 | -5.8 | 6.0E-09 | 1.1E-08 | BT20/MB468 | NUC |
| >MI0000812\|hsa-mir-331&WithFlank&12\|+\|95308414\|95308519@67.87.21(+1U)&[MIMAT0000760&hsa-miR-331-3p&offsets\|0\|0(+1U);m-192&12\|+\|95308480\|95308501&offsets\|0\|-1(+1U)]\|GCCCCTGGGCCTATCCTAGAAT | >MI0000812\|hsa-mir-331&WithFlank&12\|+\|95308414\|95308519 | 67 | 87 | 21(+1U) | GCCCCTGGGCCTATCCTAGAAT | 263.4 | 3.8 | 0.6 | 5.8 | 6.0E-09 | 1.1E-08 | BT20/MB468 | NUC |
| >hg38_21_+_8433172_8446622_RNA45SN1_45S_50ntFlanks@7985.8011.27\|GATCAGACGTGGCGACCCGCTGAATTT | >hg38_21_+_8433172_8446622_RNA45SN1_45S_50ntFlanks | 7985 | 8011 | 27 | GATCAGACGTGGCGACCCGCTGAATTT | 147.1 | -10.3 | 1.7 | -6.2 | 6.8E-10 | 1.1E-08 | BT20/MB468 | MP |
| >hg38_21_+_8433172_8446622_RNA45SN1_45S_50ntFlanks@10409.10430.22\|GTCAGTCGGTCCTGAGAGATGG | >hg38_21_+_8433172_8446622_RNA45SN1_45S_50ntFlanks | 10409 | 10430 | 22 | GTCAGTCGGTCCTGAGAGATGG | 2484.6 | -9.6 | 1.7 | -5.8 | 6.0E-09 | 1.1E-08 | BT20/MB468 | NUC |
| >hg38_21_+_8433172_8446622_RNA45SN1_45S_50ntFlanks@10684.10708.25\|CGGCGGCGTCCGGTGAGCTCTCGCT | >hg38_21_+_8433172_8446622_RNA45SN1_45S_50ntFlanks | 10684 | 10708 | 25 | CGGCGGCGTCCGGTGAGCTCTCGCT | 1982.5 | -5.5 | 0.9 | -5.8 | 6.0E-09 | 1.1E-08 | BT20/MB468 | NUC |
| >hg38_21_+_8433172_8446622_RNA45SN1_45S_50ntFlanks@12410.12448.39\|TGATCCTTCGATGTCGGCTCTTCCTATCATTGTGAAGCA | >hg38_21_+_8433172_8446622_RNA45SN1_45S_50ntFlanks | 12410 | 12448 | 39 | TGATCCTTCGATGTCGGCTCTTCCTATCATTGTGAAGCA | 158.4 | 6.0 | 1.0 | 5.8 | 5.3E-09 | 1.1E-08 | MB231/MB468 | NUC |
| >MI0000542\|hsa-mir-320a&WithFlank&8\|-\|22244960\|22245043@48.71.24(+1U)&[MIMAT0000510&hsa-miR-320a-3p&offsets\|0\|+2(+1U);m-60&8\|-\|22244975\|22244996&offsets\|0\|+2(+1U)]\|AAAAGCTGGGTTGAGAGGGCGAAAT | >MI0000542\|hsa-mir-320a&WithFlank&8\|-\|22244960\|22245043 | 48 | 71 | 24(+1U) | AAAAGCTGGGTTGAGAGGGCGAAAT | 144.4 | 3.2 | 0.5 | 5.8 | 6.0E-09 | 1.1E-08 | BT20/MB468 | NUC |
| >MI0000477\|hsa-mir-146a&WithFlank&5\|+\|160485346\|160485456@27.48.22(+2U)&[MIMAT0000449&hsa-miR-146a-5p&offsets\|0\|0(+2U);m-129&5\|+\|160485372\|160485393&offsets\|0\|0(+2U)]\|TGAGAACTGAATTCCATGGGTTTT | >MI0000477\|hsa-mir-146a&WithFlank&5\|+\|160485346\|160485456 | 27 | 48 | 22(+2U) | TGAGAACTGAATTCCATGGGTTTT | 159.1 | 10.5 | 1.8 | 5.8 | 5.8E-09 | 1.1E-08 | MB231/MB468 | MITO |
| >MI0000270\|hsa-mir-181b-1&WithFlank&1\|-\|198858867\|198858988@42.63.22;MI0000683\|hsa-mir-181b-2&WithFlank&9\|+\|124693704\|124693804@22.43.22&[MIMAT0000257&hsa-miR-181b-5p&offsets\|0\|-1;m-73&1\|-\|198858925\|198858947&offsets\|0\|-1];[MIMAT0000257_1&hsa-miR-181b-5p&offsets\|0\|-1;m-72&9\|+\|124693725\|124693747&offsets\|0\|-1]\|AACATTCATTGCTGTCGGTGGG | >MI0000270\|hsa-mir-181b-1&WithFlank&1\|-\|198858867\|198858988 | 42 | 63 | 22 | AACATTCATTGCTGTCGGTGGG | 400.0 | 5.1 | 0.9 | 5.8 | 5.3E-09 | 1.1E-08 | MB231/MB468 | CYTO |
| >hg38_21_+_8433172_8446622_RNA45SN1_45S_50ntFlanks@6734.6775.42\|ATTGATCATCGACACTTCGAACGCACTTGCGGCCCCGGGTTC | >hg38_21_+_8433172_8446622_RNA45SN1_45S_50ntFlanks | 6734 | 6775 | 42 | ATTGATCATCGACACTTCGAACGCACTTGCGGCCCCGGGTTC | 320.6 | 7.6 | 1.3 | 5.9 | 4.0E-09 | 1.1E-08 | BT20/MB468 | MITO |
| >hg38_21_+_8433172_8446622_RNA45SN1_45S_50ntFlanks@12892.12933.42\|CCCTCGCCCGTCACGCACCGCACGTTCGTGGGGAACCTGGCG | >hg38_21_+_8433172_8446622_RNA45SN1_45S_50ntFlanks | 12892 | 12933 | 42 | CCCTCGCCCGTCACGCACCGCACGTTCGTGGGGAACCTGGCG | 1434.4 | -7.7 | 1.3 | -5.9 | 4.0E-09 | 1.1E-08 | BT20/MB468 | MITO |
| >MI0000342\|hsa-mir-200b&WithFlank&1\|+\|1167098\|1167204@63.82.20(+1A)&[MIMAT0000318&hsa-miR-200b-3p&offsets\|0\|-2(+1A);m-93&1\|+\|1167160\|1167182&offsets\|0\|-3(+1A)]\|TAATACTGCCTGGTAATGATA | >MI0000342\|hsa-mir-200b&WithFlank&1\|+\|1167098\|1167204 | 63 | 82 | 20(+1A) | TAATACTGCCTGGTAATGATA | 117.3 | -10.0 | 1.6 | -6.2 | 6.8E-10 | 1.1E-08 | BT20/MB468 | MP |
| >hg38_21_+_8433172_8446622_RNA45SN1_45S_50ntFlanks@11577.11609.33\|TTAGAACTGGTGCGGACCAGGGGAATCCGACTG | >hg38_21_+_8433172_8446622_RNA45SN1_45S_50ntFlanks | 11577 | 11609 | 33 | TTAGAACTGGTGCGGACCAGGGGAATCCGACTG | 285.9 | -7.8 | 1.3 | -5.8 | 5.0E-09 | 1.1E-08 | MB231/MB468 | MP |
| >hg38_21_+_8433172_8446622_RNA45SN1_45S_50ntFlanks@3835.3861.27\|CTCCTCTCCTACTTGGATAACTGTGGT | >hg38_21_+_8433172_8446622_RNA45SN1_45S_50ntFlanks | 3835 | 3861 | 27 | CTCCTCTCCTACTTGGATAACTGTGGT | 64.5 | -9.9 | 1.7 | -5.8 | 5.0E-09 | 1.1E-08 | MB231/MB468 | MP |
| >MI0000465\|hsa-mir-191&WithFlank&3\|-\|49020612\|49020715@22.42.21&[MIMAT0000440&hsa-miR-191-5p&offsets\|0\|-2;m-48&3\|-\|49020672\|49020694&offsets\|0\|-2]\|CAACGGAATCCCAAAAGCAGC | >MI0000465\|hsa-mir-191&WithFlank&3\|-\|49020612\|49020715 | 22 | 42 | 21 | CAACGGAATCCCAAAAGCAGC | 419.5 | 5.6 | 1.0 | 5.8 | 5.3E-09 | 1.1E-08 | MB231/MB468 | CYTO |
| >hg38_MT_+_1621_3279_RNR2_16S_50ntFlanks@482.504.23\|AAAGAGGAACAGCTCTTTGGACA | >hg38_MT_+_1621_3279_RNR2_16S_50ntFlanks | 482 | 504 | 23 | AAAGAGGAACAGCTCTTTGGACA | 331.9 | 5.0 | 0.8 | 5.9 | 4.1E-09 | 1.1E-08 | BT20/MB468 | MITO |
| >hg38_21_+_8433172_8446622_RNA45SN1_45S_50ntFlanks@8184.8223.40\|CGAGGCCCAGCCCGTGGACGGTGTGAGGCCGGTAGCGGCC | >hg38_21_+_8433172_8446622_RNA45SN1_45S_50ntFlanks | 8184 | 8223 | 40 | CGAGGCCCAGCCCGTGGACGGTGTGAGGCCGGTAGCGGCC | 64.0 | -9.9 | 1.7 | -5.8 | 5.1E-09 | 1.1E-08 | MB231/MB468 | MP |
| >hg38_21_+_8433172_8446622_RNA45SN1_45S_50ntFlanks@5073.5100.28\|AATTCCGATAACGAACGAGACTCTGGCA | >hg38_21_+_8433172_8446622_RNA45SN1_45S_50ntFlanks | 5073 | 5100 | 28 | AATTCCGATAACGAACGAGACTCTGGCA | 120.6 | -10.8 | 1.8 | -5.8 | 5.1E-09 | 1.1E-08 | MB231/MB468 | MP |
| >hg38_21_+_8433172_8446622_RNA45SN1_45S_50ntFlanks@9861.9899.39\|GATGAACCGAACGCCGGGTTAAGGCGCCCGATGCCGACG | >hg38_21_+_8433172_8446622_RNA45SN1_45S_50ntFlanks | 9861 | 9899 | 39 | GATGAACCGAACGCCGGGTTAAGGCGCCCGATGCCGACG | 168.1 | -11.2 | 1.9 | -5.8 | 5.9E-09 | 1.1E-08 | MB231/MB468 | MITO |
| >hg38_21_+_8433172_8446622_RNA45SN1_45S_50ntFlanks@4047.4069.23\|ATTCGAACGTCTGCCCTATCAAC | >hg38_21_+_8433172_8446622_RNA45SN1_45S_50ntFlanks | 4047 | 4069 | 23 | ATTCGAACGTCTGCCCTATCAAC | 490.7 | -6.7 | 1.1 | -5.8 | 6.1E-09 | 1.1E-08 | BT20/MB468 | NUC |
| >hg38_21_+_8433172_8446622_RNA45SN1_45S_50ntFlanks@11729.11750.22\|TAACTATGACTCTCTTAAGGTA | >hg38_21_+_8433172_8446622_RNA45SN1_45S_50ntFlanks | 11729 | 11750 | 22 | TAACTATGACTCTCTTAAGGTA | 574.2 | -6.2 | 1.1 | -5.8 | 5.9E-09 | 1.1E-08 | MB231/MB468 | MITO |
| >MI0016005\|hsa-mir-3615&WithFlank&17\|+\|74748607\|74748705@57.79.23(+1U)&[MIMAT0017994&hsa-miR-3615&offsets\|0\|+2(+1U);m-271&17\|+\|74748663\|74748684&offsets\|0\|+1(+1U)]\|TCTCTCGGCTCCTCGCGGCTCGCT | >MI0016005\|hsa-mir-3615&WithFlank&17\|+\|74748607\|74748705 | 57 | 79 | 23(+1U) | TCTCTCGGCTCCTCGCGGCTCGCT | 255.2 | 6.9 | 1.2 | 5.8 | 5.4E-09 | 1.1E-08 | MB231/MB468 | CYTO |
| >hg38_21_+_8433172_8446622_RNA45SN1_45S_50ntFlanks@10686.10711.26\|GCGGCGTCCGGTGAGCTCTCGCTGGC | >hg38_21_+_8433172_8446622_RNA45SN1_45S_50ntFlanks | 10686 | 10711 | 26 | GCGGCGTCCGGTGAGCTCTCGCTGGC | 676.2 | -4.1 | 0.7 | -5.8 | 5.4E-09 | 1.1E-08 | MB231/MB468 | NUC |
| >hg38_21_+_8433172_8446622_RNA45SN1_45S_50ntFlanks@8472.8492.21\|CGGCCCGGCGGATCTTTCCCG | >hg38_21_+_8433172_8446622_RNA45SN1_45S_50ntFlanks | 8472 | 8492 | 21 | CGGCCCGGCGGATCTTTCCCG | 397.9 | -6.6 | 1.1 | -5.8 | 5.4E-09 | 1.1E-08 | MB231/MB468 | CYTO |
| >hg38_21_+_8433172_8446622_RNA45SN1_45S_50ntFlanks@12949.12990.42\|ACGACCTGCTTCTGGGTCGGGGTTTCGTACGTAGCAGAGCAG | >hg38_21_+_8433172_8446622_RNA45SN1_45S_50ntFlanks | 12949 | 12990 | 42 | ACGACCTGCTTCTGGGTCGGGGTTTCGTACGTAGCAGAGCAG | 865.6 | -8.6 | 1.5 | -5.8 | 5.5E-09 | 1.1E-08 | MB231/MB468 | NUC |
| >hg38_21_+_8433172_8446622_RNA45SN1_45S_50ntFlanks@12627.12661.35\|ATGGGGCGAAGCTACCATCTGTGGGATTATGACTG | >hg38_21_+_8433172_8446622_RNA45SN1_45S_50ntFlanks | 12627 | 12661 | 35 | ATGGGGCGAAGCTACCATCTGTGGGATTATGACTG | 837.6 | -6.2 | 1.1 | -5.8 | 5.5E-09 | 1.1E-08 | MB231/MB468 | NUC |
| >hg38_21_+_8433172_8446622_RNA45SN1_45S_50ntFlanks@7968.7997.30\|TCCGAGACGCGACCTCAGATCAGACGTGGC | >hg38_21_+_8433172_8446622_RNA45SN1_45S_50ntFlanks | 7968 | 7997 | 30 | TCCGAGACGCGACCTCAGATCAGACGTGGC | 167.6 | 4.0 | 0.7 | 5.8 | 5.5E-09 | 1.1E-08 | MB231/MB468 | NUC |
| >hg38_21_+_8433172_8446622_RNA45SN1_45S_50ntFlanks@6653.6696.44\|ACTCTTAGCGGTGGATCACTCGGCTCGTGCGTCGATGAAGAACG | >hg38_21_+_8433172_8446622_RNA45SN1_45S_50ntFlanks | 6653 | 6696 | 44 | ACTCTTAGCGGTGGATCACTCGGCTCGTGCGTCGATGAAGAACG | 238.4 | 5.5 | 0.9 | 5.8 | 5.4E-09 | 1.1E-08 | MB231/MB468 | CYTO |
| >hg38_21_+_8433172_8446622_RNA45SN1_45S_50ntFlanks@10787.10817.31\|ACAGCCTCTGGCATGTTGGAACAATGTAGGT | >hg38_21_+_8433172_8446622_RNA45SN1_45S_50ntFlanks | 10787 | 10817 | 31 | ACAGCCTCTGGCATGTTGGAACAATGTAGGT | 318.6 | -5.8 | 1.0 | -5.8 | 5.9E-09 | 1.1E-08 | MB231/MB468 | MITO |
| >hg38_21_+_8433172_8446622_RNA45SN1_45S_50ntFlanks@13006.13028.23\|CTATTGAAAGTCAGCCCTCGACA | >hg38_21_+_8433172_8446622_RNA45SN1_45S_50ntFlanks | 13006 | 13028 | 23 | CTATTGAAAGTCAGCCCTCGACA | 117.4 | -10.0 | 1.6 | -6.2 | 7.0E-10 | 1.1E-08 | BT20/MB468 | MP |
| >MI0000650\|hsa-mir-200c&WithFlank&12\|+\|6963693\|6963772@50.70.21(+1A)&[MIMAT0000617&hsa-miR-200c-3p&offsets\|0\|-2(+1A);m-54&12\|+\|6963742\|6963764&offsets\|0\|-2(+1A)]\|TAATACTGCCGGGTAATGATGA | >MI0000650\|hsa-mir-200c&WithFlank&12\|+\|6963693\|6963772 | 50 | 70 | 21(+1A) | TAATACTGCCGGGTAATGATGA | 2221.8 | 9.9 | 1.5 | 6.4 | 1.3E-10 | 1.1E-08 | BT20/MB231 | MP |
| >hg38_21_+_8433172_8446622_RNA45SN1_45S_50ntFlanks@6660.6679.20\|GCGGTGGATCACTCGGCTCG | >hg38_21_+_8433172_8446622_RNA45SN1_45S_50ntFlanks | 6660 | 6679 | 20 | GCGGTGGATCACTCGGCTCG | 122.7 | -10.8 | 1.8 | -5.8 | 5.2E-09 | 1.1E-08 | MB231/MB468 | MP |
| >hg38_21_+_8433172_8446622_RNA45SN1_45S_50ntFlanks@11622.11660.39\|AAAGCATCGCGAAGGCCCGCGGCGGGTGTTGACGCGATG | >hg38_21_+_8433172_8446622_RNA45SN1_45S_50ntFlanks | 11622 | 11660 | 39 | AAAGCATCGCGAAGGCCCGCGGCGGGTGTTGACGCGATG | 59.0 | -9.7 | 1.7 | -5.8 | 5.2E-09 | 1.1E-08 | MB231/MB468 | MP |
| >am_trna10_ValCAC_5_-_180649395_180649467@1.32.32,trna12_ValAAC_5_-_180645270_180645342@1.32.32,trna132_ValAAC_6_-_27721179_27721251@1.32.32,trna136_ValAAC_6_-_27648885_27648957@1.32.32,trna139_ValAAC_6_-_27618707_27618779@1.32.32,trna18_ValCAC_5_-_180529253_180529325@1.32.32,trna2_ValAAC_3_+_169490018_169490090@1.32.32,trna2_ValCAC_5_+_180524070_180524142@1.32.32,trna4_ValAAC_5_+_180591154_180591226@1.32.32,trna5_ValAAC_5_+_180596610_180596682@1.32.32,trna6_ValCAC_5_+_180600650_180600722@1.32.32,trna85_ValCAC_1_-_161369490_161369562@1.32.32,trna90_ValCAC_1_-_149684088_149684161@1.32.32,trna98_ValCAC_1_-_149298555_149298627@1.32.32,trna9_ValCAC_6_+_26538282_26538354@1.32.32\|GTTTCCGTAGTGTAGTGGTTATCACGTTCGCC | >am_trna10_ValCAC_5_-_180649395_180649467 | 1 | 32 | 32 | GTTTCCGTAGTGTAGTGGTTATCACGTTCGCC | 1396.4 | 5.7 | 1.0 | 5.8 | 5.2E-09 | 1.1E-08 | MB231/MB468 | MP |
| >hg38_21_+_8433172_8446622_RNA45SN1_45S_50ntFlanks@11787.11806.20\|AATGGATGAACGAGATTCCC | >hg38_21_+_8433172_8446622_RNA45SN1_45S_50ntFlanks | 11787 | 11806 | 20 | AATGGATGAACGAGATTCCC | 482.8 | -8.3 | 1.4 | -5.8 | 5.2E-09 | 1.1E-08 | MB231/MB468 | MP |
| >hg38_21_+_8433172_8446622_RNA45SN1_45S_50ntFlanks@12459.12478.20\|AGCGTTGGATTGTTCACCCA | >hg38_21_+_8433172_8446622_RNA45SN1_45S_50ntFlanks | 12459 | 12478 | 20 | AGCGTTGGATTGTTCACCCA | 500.5 | -6.2 | 1.1 | -5.8 | 5.5E-09 | 1.2E-08 | MB231/MB468 | NUC |
| >hg38_21_+_8433172_8446622_RNA45SN1_45S_50ntFlanks@9716.9735.20\|AATGATTAGAGGTCTTGGGG | >hg38_21_+_8433172_8446622_RNA45SN1_45S_50ntFlanks | 9716 | 9735 | 20 | AATGATTAGAGGTCTTGGGG | 107.3 | -4.6 | 0.8 | -5.8 | 5.5E-09 | 1.2E-08 | MB231/MB468 | CYTO |
| >hg38_21_+_8433172_8446622_RNA45SN1_45S_50ntFlanks@9564.9594.31\|CTGACGTGCAAATCGGTCGTCCGACCTGGGT | >hg38_21_+_8433172_8446622_RNA45SN1_45S_50ntFlanks | 9564 | 9594 | 31 | CTGACGTGCAAATCGGTCGTCCGACCTGGGT | 498.3 | -2.9 | 0.5 | -5.9 | 4.7E-09 | 1.2E-08 | BT20/MB468 | CYTO |
| >am_trnaMT_ValTAC_MT_+_1602_1670@47.67.21\|TCAACTTAACTTGACCGCTCT | >am_trnaMT_ValTAC_MT_+_1602_1670 | 47 | 67 | 21 | TCAACTTAACTTGACCGCTCT | 477.7 | 9.6 | 1.6 | 5.9 | 4.7E-09 | 1.2E-08 | BT20/MB468 | CYTO |
| >hg38_21_+_8433172_8446622_RNA45SN1_45S_50ntFlanks@10389.10435.47\|TGAACAGCAGTTGAACATGGGTCAGTCGGTCCTGAGAGATGGGCGAG | >hg38_21_+_8433172_8446622_RNA45SN1_45S_50ntFlanks | 10389 | 10435 | 47 | TGAACAGCAGTTGAACATGGGTCAGTCGGTCCTGAGAGATGGGCGAG | 206.4 | -10.7 | 1.8 | -5.8 | 5.5E-09 | 1.2E-08 | MB231/MB468 | NUC |
| >hg38_1_-_228634819_228635039_RNA5S12_5S_50ntFlanks@151.169.19\|ATACCGGGTGCTGTAGGCT | >hg38_1_-_228634819_228635039_RNA5S12_5S_50ntFlanks | 151 | 169 | 19 | ATACCGGGTGCTGTAGGCT | 2690.9 | -4.2 | 0.7 | -5.8 | 5.2E-09 | 1.2E-08 | MB231/MB468 | MP |
| >hg38_21_+_8433172_8446622_RNA45SN1_45S_50ntFlanks@6746.6771.26\|CACTTCGAACGCACTTGCGGCCCCGG | >hg38_21_+_8433172_8446622_RNA45SN1_45S_50ntFlanks | 6746 | 6771 | 26 | CACTTCGAACGCACTTGCGGCCCCGG | 1023.5 | -5.8 | 1.0 | -5.9 | 4.2E-09 | 1.2E-08 | BT20/MB468 | MITO |
| >hg38_21_+_8433172_8446622_RNA45SN1_45S_50ntFlanks@11562.11602.41\|CGCCTAGCAGCCGACTTAGAACTGGTGCGGACCAGGGGAAT | >hg38_21_+_8433172_8446622_RNA45SN1_45S_50ntFlanks | 11562 | 11602 | 41 | CGCCTAGCAGCCGACTTAGAACTGGTGCGGACCAGGGGAAT | 136.7 | -11.0 | 1.9 | -5.8 | 5.2E-09 | 1.2E-08 | MB231/MB468 | MP |
| >hg38_21_+_8433172_8446622_RNA45SN1_45S_50ntFlanks@12183.12200.18\|AAACGGTAACGCAGGTGT | >hg38_21_+_8433172_8446622_RNA45SN1_45S_50ntFlanks | 12183 | 12200 | 18 | AAACGGTAACGCAGGTGT | 82.9 | -10.2 | 1.8 | -5.8 | 5.2E-09 | 1.2E-08 | MB231/MB468 | MP |

<table>
<tr><td></td><td></td><td></td><td></td><td></td><td></td><td></td><td></td><td></td><td></td><td></td><td></td><td></td><td></td></tr>
<tr><td>>trnaMT_GluTTC_MT_-_14674_14742@34.71.38,trnalookalike8_GluTTC_5_-_93905172_93905240@34.71.38|ATATCATTGGTCGTGGTTGTAGTCCGTGCGAGAATACC</td><td>>trnaMT_GluTTC_MT_-_14674_14742</td><td>34</td><td>71</td><td>38</td><td>ATATCATTGGTCGTGGTTGTAGTCCGTGCGAGAATACC</td><td>437.4</td><td>6.7</td><td>1.1</td><td>5.8</td><td>5.6E-09</td><td>1.2E-08</td><td>MB231/MB468</td><td>NUC</td></tr>
<tr><td>>hg38_21_+_8433172_8446622_RNA45SN1_45S_50ntFlanks@8117.8136.20|CGGGACATGTGGCGTACGGA</td><td>>hg38_21_+_8433172_8446622_RNA45SN1_45S_50ntFlanks</td><td>8117</td><td>8136</td><td>20</td><td>CGGGACATGTGGCGTACGGA</td><td>903.0</td><td>13.2</td><td>2.0</td><td>6.4</td><td>1.4E-10</td><td>1.2E-08</td><td>BT20/MB231</td><td>MP</td></tr>
<tr><td>>hg38_21_+_8433172_8446622_RNA45SN1_45S_50ntFlanks@3984.4010.27|GCGGCTTTGGTGACTCTAGATAACCTC</td><td>>hg38_21_+_8433172_8446622_RNA45SN1_45S_50ntFlanks</td><td>3984</td><td>4010</td><td>27</td><td>GCGGCTTTGGTGACTCTAGATAACCTC</td><td>634.0</td><td>-8.6</td><td>1.5</td><td>-5.8</td><td>6.3E-09</td><td>1.2E-08</td><td>BT20/MB468</td><td>NUC</td></tr>
<tr><td>>hg38_21_+_8433172_8446622_RNA45SN1_45S_50ntFlanks@12008.12031.24|CCGCCGGTGAAATACCACTACTCT</td><td>>hg38_21_+_8433172_8446622_RNA45SN1_45S_50ntFlanks</td><td>12008</td><td>12031</td><td>24</td><td>CCGCCGGTGAAATACCACTACTCT</td><td>2414.9</td><td>-7.9</td><td>1.4</td><td>-5.8</td><td>6.3E-09</td><td>1.2E-08</td><td>BT20/MB468</td><td>NUC</td></tr>
<tr><td>>hg38_21_+_8433172_8446622_RNA45SN1_45S_50ntFlanks@10827.10865.39|CGGCAAGCCGGATCCGTAACTTCGGGATAAGGATTGGCT</td><td>>hg38_21_+_8433172_8446622_RNA45SN1_45S_50ntFlanks</td><td>10827</td><td>10865</td><td>39</td><td>CGGCAAGCCGGATCCGTAACTTCGGGATAAGGATTGGCT</td><td>70.8</td><td>-10.0</td><td>1.7</td><td>-5.8</td><td>5.2E-09</td><td>1.2E-08</td><td>MB231/MB468</td><td>MP</td></tr>
<tr><td>>MI0000077|hsa-mir-21&WithFlank&17|+|59841260|59841343@14.33.20&[MIMAT0000076&hsa-miR-21-5p&offsets|0|-2;m-1&17|+|59841273|59841294&offsets|0|-2]|TAGCTTATCAGACTGATGTT</td><td>>MI0000077|hsa-mir-21&WithFlank&17|+|59841260|59841343</td><td>14</td><td>33</td><td>20</td><td>TAGCTTATCAGACTGATGTT</td><td>490.8</td><td>4.9</td><td>0.8</td><td>5.8</td><td>6.3E-09</td><td>1.2E-08</td><td>BT20/MB468</td><td>NUC</td></tr>
<tr><td>>hg38_21_+_8433172_8446622_RNA45SN1_45S_50ntFlanks@10866.10884.19|CTAAGGGCTGGGTCGGTCG</td><td>>hg38_21_+_8433172_8446622_RNA45SN1_45S_50ntFlanks</td><td>10866</td><td>10884</td><td>19</td><td>CTAAGGGCTGGGTCGGTCG</td><td>881.5</td><td>-7.1</td><td>1.2</td><td>-5.8</td><td>5.5E-09</td><td>1.2E-08</td><td>MB231/MB468</td><td>CYTO</td></tr>
<tr><td>>hg38_21_+_8433172_8446622_RNA45SN1_45S_50ntFlanks@11781.11803.23|CGCATGAATGGATGAACGAGATT</td><td>>hg38_21_+_8433172_8446622_RNA45SN1_45S_50ntFlanks</td><td>11781</td><td>11803</td><td>23</td><td>CGCATGAATGGATGAACGAGATT</td><td>797.6</td><td>-4.1</td><td>0.7</td><td>-5.9</td><td>4.7E-09</td><td>1.2E-08</td><td>BT20/MB468</td><td>CYTO</td></tr>
<tr><td>>hg38_21_+_8433172_8446622_RNA45SN1_45S_50ntFlanks@11783.11803.21|CATGAATGGATGAACGAGATT</td><td>>hg38_21_+_8433172_8446622_RNA45SN1_45S_50ntFlanks</td><td>11783</td><td>11803</td><td>21</td><td>CATGAATGGATGAACGAGATT</td><td>4045.5</td><td>-4.5</td><td>0.8</td><td>-5.8</td><td>5.5E-09</td><td>1.2E-08</td><td>MB231/MB468</td><td>CYTO</td></tr>
<tr><td>>MI0000065|hsa-let-7d&WithFlank&9|+|94178828|94178926@14.34.21(+1C)&[MIMAT0000065&hsa-let-7d-5p&offsets|0|-1(+1C);m-90&9|+|94178841|94178862&offsets|0|-1(+1C)]|AGAGGTAGTAGGTTGCATAGTC</td><td>>MI0000065|hsa-let-7d&WithFlank&9|+|94178828|94178926</td><td>14</td><td>34</td><td>21(+1C)</td><td>AGAGGTAGTAGGTTGCATAGTC</td><td>726.1</td><td>5.9</td><td>1.0</td><td>5.8</td><td>6.1E-09</td><td>1.2E-08</td><td>MB231/MB468</td><td>MITO</td></tr>
<tr><td>>hg38_21_+_8433172_8446622_RNA45SN1_45S_50ntFlanks@12321.12346.26|CTTTTGGGTTTTAAGCAGGAGGTGTC</td><td>>hg38_21_+_8433172_8446622_RNA45SN1_45S_50ntFlanks</td><td>12321</td><td>12346</td><td>26</td><td>CTTTTGGGTTTTAAGCAGGAGGTGTC</td><td>122.4</td><td>-10.0</td><td>1.6</td><td>-6.2</td><td>7.2E-10</td><td>1.2E-08</td><td>BT20/MB468</td><td>MP</td></tr>
<tr><td>>MI0000264|hsa-mir-7-2&WithFlank&15|+|88611819|88611940@38.60.23(+1C);MI0000265|hsa-mir-7-3&WithFlank&19|+|4770664|4770785@37.59.23(+1C);MI0000263|hsa-mir-7-1&WithFlank&9|-|83969742|83969863@30.52.23(+1C)&[MIMAT0000252&hsa-miR-7-5p&offsets|0|-1(+1C);m-400&15|+|88611856|88611879&offsets|0|-1(+1C)];[MIMAT0000252_1&hsa-miR-7-5p&offsets|0|-1(+1C);m-402&19|+|4770700|4770723&offsets|0|-1(+1C)];[MIMAT0000252_2&hsa-miR-7-5p&offsets|0|-1(+1C);m-401&9|-|83969811|83969834&offsets|0|-1(+1C)]|TGGAAGACTAGTGATTTTGTTGTC</td><td>>MI0000264|hsa-mir-7-2&WithFlank&15|+|88611819|88611940</td><td>38</td><td>60</td><td>23(+1C)</td><td>TGGAAGACTAGTGATTTTGTTGTC</td><td>5663.4</td><td>5.5</td><td>1.0</td><td>5.8</td><td>6.1E-09</td><td>1.2E-08</td><td>MB231/MB468</td><td>MITO</td></tr>
<tr><td>>hg38_21_+_8433172_8446622_RNA45SN1_45S_50ntFlanks@5074.5110.37|ATTCCGATAACGAACGAGACTCTGGCATGCTAACTAG</td><td>>hg38_21_+_8433172_8446622_RNA45SN1_45S_50ntFlanks</td><td>5074</td><td>5110</td><td>37</td><td>ATTCCGATAACGAACGAGACTCTGGCATGCTAACTAG</td><td>603.8</td><td>-8.8</td><td>1.5</td><td>-5.8</td><td>6.3E-09</td><td>1.2E-08</td><td>BT20/MB468</td><td>NUC</td></tr>
<tr><td>>hg38_21_+_8433172_8446622_RNA45SN1_45S_50ntFlanks@3846.3881.36|CTTGGATAACTGTGGTAATTCTAGAGCTAATACATG</td><td>>hg38_21_+_8433172_8446622_RNA45SN1_45S_50ntFlanks</td><td>3846</td><td>3881</td><td>36</td><td>CTTGGATAACTGTGGTAATTCTAGAGCTAATACATG</td><td>814.7</td><td>-7.1</td><td>1.2</td><td>-5.8</td><td>6.1E-09</td><td>1.2E-08</td><td>MB231/MB468</td><td>MITO</td></tr>
<tr><td>>am_trna116_GluCTC_1_-_145399233_145399304@1.29.29,trna59_GluCTC_1_+_249168447_249168518@1.29.29,trna71_GluCTC_1_-_161439189_161439260@1.29.29,trna74_GluCTC_1_-_161431809_161431880@1.29.29,trna77_GluCTC_1_-_161424398_161424469@1.29.29,trna77_GluCTC_6_+_28949976_28950047@1.29.29,trna80_GluCTC_1_-_161417018_161417089@1.29.29,trna87_GluCTC_6_-_126101393_126101464@1.29.29|TCCCTGGTGGTCTAGTGGTTAGGATTCGG</td><td>>am_trna116_GluCTC_1_-_145399233_145399304</td><td>1</td><td>29</td><td>29</td><td>TCCCTGGTGGTCTAGTGGTTAGGATTCGG</td><td>33138.1</td><td>-5.5</td><td>0.9</td><td>-5.8</td><td>5.6E-09</td><td>1.2E-08</td><td>MB231/MB468</td><td>NUC</td></tr>
<tr><td>>hg38_21_+_8433172_8446622_RNA45SN1_45S_50ntFlanks@11573.11601.29|CGACTTAGAACTGGTGCGGACCAGGGGAA</td><td>>hg38_21_+_8433172_8446622_RNA45SN1_45S_50ntFlanks</td><td>11573</td><td>11601</td><td>29</td><td>CGACTTAGAACTGGTGCGGACCAGGGGAA</td><td>893.5</td><td>-3.9</td><td>0.7</td><td>-5.9</td><td>4.8E-09</td><td>1.2E-08</td><td>BT20/MB468</td><td>CYTO</td></tr>
<tr><td>>trnaMT_PheGAA_MT_+_577_647@39.56.18|ATGTTTAGACGGGCTCAC</td><td>>trnaMT_PheGAA_MT_+_577_647</td><td>39</td><td>56</td><td>18</td><td>ATGTTTAGACGGGCTCAC</td><td>3890.8</td><td>5.2</td><td>0.8</td><td>6.2</td><td>7.2E-10</td><td>1.2E-08</td><td>BT20/MB468</td><td>MP</td></tr>
<tr><td>>hg38_21_+_8433172_8446622_RNA45SN1_45S_50ntFlanks@11930.11952.23|TAGAATAAGTGGGAGGCCCCCGG</td><td>>hg38_21_+_8433172_8446622_RNA45SN1_45S_50ntFlanks</td><td>11930</td><td>11952</td><td>23</td><td>TAGAATAAGTGGGAGGCCCCCGG</td><td>165.4</td><td>-10.5</td><td>1.7</td><td>-6.2</td><td>7.2E-10</td><td>1.2E-08</td><td>BT20/MB468</td><td>MP</td></tr>
<tr><td>>hg38_21_+_8433172_8446622_RNA45SN1_45S_50ntFlanks@10351.10388.38|CGAGAACTTTGAAGGCCGAAGTGGAGAAGGGTTCCATG</td><td>>hg38_21_+_8433172_8446622_RNA45SN1_45S_50ntFlanks</td><td>10351</td><td>10388</td><td>38</td><td>CGAGAACTTTGAAGGCCGAAGTGGAGAAGGGTTCCATG</td><td>6789.7</td><td>-5.4</td><td>0.9</td><td>-5.9</td><td>4.8E-09</td><td>1.2E-08</td><td>BT20/MB468</td><td>CYTO</td></tr>
<tr><td>>hg38_21_+_8433172_8446622_RNA45SN1_45S_50ntFlanks@4442.4471.30|CCCCTTGCCTCTCGGCGCCCCCTCGATGCT</td><td>>hg38_21_+_8433172_8446622_RNA45SN1_45S_50ntFlanks</td><td>4442</td><td>4471</td><td>30</td><td>CCCCTTGCCTCTCGGCGCCCCCTCGATGCT</td><td>325.4</td><td>-8.1</td><td>1.4</td><td>-5.8</td><td>5.7E-09</td><td>1.2E-08</td><td>MB231/MB468</td><td>NUC</td></tr>
<tr><td>>MI0000067|hsa-let-7f-1&WithFlank&9|+|94176341|94176439@13.33.21(+1C);MI0000068|hsa-let-7f-2&WithFlank&X|-|53557186|53557280@14.34.21(+1C)&[MIMAT0000067&hsa-let-7f-5p&offsets|0|-1(+1C);m-20&9|+|94176353|94176374&offsets|0|-1(+1C)];[MIMAT0000067_1&hsa-let-7f-5p&offsets|0|-1(+1C);m-19&X|-|53557246|53557267&offsets|0|-1(+1C)]|TGAGGTAGTAGATTGTATAGTC</td><td>>MI0000067|hsa-let-7f-1&WithFlank&9|+|94176341|94176439</td><td>13</td><td>33</td><td>21(+1C)</td><td>TGAGGTAGTAGATTGTATAGTC</td><td>7800.7</td><td>5.3</td><td>0.9</td><td>5.8</td><td>6.1E-09</td><td>1.2E-08</td><td>MB231/MB468</td><td>MITO</td></tr>
<tr><td>>MI0000085|hsa-mir-27a&WithFlank&19|-|13836434|13836523@57.74.18;MI0000440|hsa-mir-27b&WithFlank&9|+|95085439|95085547@67.84.18&[MIMAT0000084&hsa-miR-27a-3p&offsets|0|-3;m-43&19|-|13836447|13836467&offsets|0|-3];[MIMAT0000419&hsa-miR-27b-3p&offsets|0|-3;m-39&9|+|95085505|95085525&offsets|0|-3]|TTCACAGTGGCTAAGTTC</td><td>>MI0000085|hsa-mir-27a&WithFlank&19|-|13836434|13836523</td><td>57</td><td>74</td><td>18</td><td>TTCACAGTGGCTAAGTTC</td><td>407.0</td><td>5.7</td><td>1.0</td><td>5.8</td><td>5.6E-09</td><td>1.2E-08</td><td>MB231/MB468</td><td>CYTO</td></tr>
<tr><td>>hg38_21_+_8433172_8446622_RNA45SN1_45S_50ntFlanks@7982.8006.25|TCAGATCAGACGTGGCGACCCGCTG</td><td>>hg38_21_+_8433172_8446622_RNA45SN1_45S_50ntFlanks</td><td>7982</td><td>8006</td><td>25</td><td>TCAGATCAGACGTGGCGACCCGCTG</td><td>3556.0</td><td>-5.5</td><td>0.9</td><td>-6.2</td><td>7.3E-10</td><td>1.2E-08</td><td>BT20/MB468</td><td>MP</td></tr>
<tr><td>>hg38_21_+_8433172_8446622_RNA45SN1_45S_50ntFlanks@11783.11803.21|CATGAATGGATGAACGAGATT</td><td>>hg38_21_+_8433172_8446622_RNA45SN1_45S_50ntFlanks</td><td>11783</td><td>11803</td><td>21</td><td>CATGAATGGATGAACGAGATT</td><td>1067.3</td><td>-3.5</td><td>0.6</td><td>-5.9</td><td>4.3E-09</td><td>1.2E-08</td><td>BT20/MB468</td><td>MITO</td></tr>
<tr><td>>hg38_21_+_8433172_8446622_RNA45SN1_45S_50ntFlanks@10402.10419.18|AACATGGGTCAGTCGGTC</td><td>>hg38_21_+_8433172_8446622_RNA45SN1_45S_50ntFlanks</td><td>10402</td><td>10419</td><td>18</td><td>AACATGGGTCAGTCGGTC</td><td>1476.5</td><td>-8.8</td><td>1.5</td><td>-5.8</td><td>5.3E-09</td><td>1.2E-08</td><td>MB231/MB468</td><td>MP</td></tr>
<tr><td>>hg38_21_+_8433172_8446622_RNA45SN1_45S_50ntFlanks@10572.10607.36|AGAAGCCGGCGGGAGCCCCGGGGAGAGTTCTCTTTT</td><td>>hg38_21_+_8433172_8446622_RNA45SN1_45S_50ntFlanks</td><td>10572</td><td>10607</td><td>36</td><td>AGAAGCCGGCGGGAGCCCCGGGGAGAGTTCTCTTTT</td><td>470.9</td><td>-6.8</td><td>1.2</td><td>-5.8</td><td>5.6E-09</td><td>1.2E-08</td><td>MB231/MB468</td><td>CYTO</td></tr>
<tr><td>>hg38_21_+_8433172_8446622_RNA45SN1_45S_50ntFlanks@12991.13039.49|CTCCCTCGCTGCGATCTATTGAAAGTCAGCCCTCGACACAAGGGTTTGT</td><td>>hg38_21_+_8433172_8446622_RNA45SN1_45S_50ntFlanks</td><td>12991</td><td>13039</td><td>49</td><td>CTCCCTCGCTGCGATCTATTGAAAGTCAGCCCTCGACACAAGGGTTTGT</td><td>339.2</td><td>-12.3</td><td>2.1</td><td>-5.8</td><td>5.3E-09</td><td>1.2E-08</td><td>MB231/MB468</td><td>MP</td></tr>
<tr><td>>hg38_21_+_8433172_8446622_RNA45SN1_45S_50ntFlanks@11769.11791.23|TAATTAGTGACGCGCATGAATGG</td><td>>hg38_21_+_8433172_8446622_RNA45SN1_45S_50ntFlanks</td><td>11769</td><td>11791</td><td>23</td><td>TAATTAGTGACGCGCATGAATGG</td><td>934.4</td><td>-4.2</td><td>0.7</td><td>-5.9</td><td>4.8E-09</td><td>1.2E-08</td><td>BT20/MB468</td><td>CYTO</td></tr>
<tr><td>>hg38_21_+_8433172_8446622_RNA45SN1_45S_50ntFlanks@6661.6680.20|CGGTGGATCACTCGGCTCGT</td><td>>hg38_21_+_8433172_8446622_RNA45SN1_45S_50ntFlanks</td><td>6661</td><td>6680</td><td>20</td><td>CGGTGGATCACTCGGCTCGT</td><td>1360.9</td><td>3.6</td><td>0.6</td><td>5.9</td><td>4.3E-09</td><td>1.2E-08</td><td>BT20/MB468</td><td>MITO</td></tr>
<tr><td>>MI0000063|hsa-let-7b&WithFlank&22|+|46113680|46113774@12.31.20&[MIMAT0000063&hsa-let-7b-5p&offsets|0|-2;m-14&22|+|46113691|46113712&offsets|0|-2]|TGAGGTAGTAGGTTGTGTGG</td><td>>MI0000063|hsa-let-7b&WithFlank&22|+|46113680|46113774</td><td>12</td><td>31</td><td>20</td><td>TGAGGTAGTAGGTTGTGTGG</td><td>2477.8</td><td>6.0</td><td>1.0</td><td>6.2</td><td>7.3E-10</td><td>1.2E-08</td><td>BT20/MB468</td><td>MP</td></tr>
<tr><td>>hg38_21_+_8433172_8446622_RNA45SN1_45S_50ntFlanks@11199.11234.36|CCGGCGGCGGCGGCGACTCTGGACGCGAGCCGGGCC</td><td>>hg38_21_+_8433172_8446622_RNA45SN1_45S_50ntFlanks</td><td>11199</td><td>11234</td><td>36</td><td>CCGGCGGCGGCGGCGACTCTGGACGCGAGCCGGGCC</td><td>61.7</td><td>-9.8</td><td>1.7</td><td>-5.8</td><td>5.4E-09</td><td>1.2E-08</td><td>MB231/MB468</td><td>MP</td></tr>
<tr><td>>hg38_21_+_8433172_8446622_RNA45SN1_45S_50ntFlanks@5080.5102.23|ATAACGAACGAGACTCTGGCATG</td><td>>hg38_21_+_8433172_8446622_RNA45SN1_45S_50ntFlanks</td><td>5080</td><td>5102</td><td>23</td><td>ATAACGAACGAGACTCTGGCATG</td><td>622.5</td><td>-8.7</td><td>1.5</td><td>-5.8</td><td>5.4E-09</td><td>1.2E-08</td><td>MB231/MB468</td><td>MP</td></tr>
<tr><td>>hg38_21_+_8433172_8446622_RNA45SN1_45S_50ntFlanks@12437.12454.18|CATTGTGAAGCAGAATTC</td><td>>hg38_21_+_8433172_8446622_RNA45SN1_45S_50ntFlanks</td><td>12437</td><td>12454</td><td>18</td><td>CATTGTGAAGCAGAATTC</td><td>2164.8</td><td>-7.0</td><td>1.2</td><td>-5.9</td><td>4.9E-09</td><td>1.2E-08</td><td>BT20/MB468</td><td>CYTO</td></tr>
<tr><td>>trna10_LysCTT_16_+_3241501_3241573@3.33.31,trna13_LysCTT_14_-_58706613_58706685@3.33.31,trna2_LysCTT_15_+_79152904_79152976@3.33.31|CCGGCTAGCTCAGTCGGTAGAGCATGGGACT</td><td>>trna10_LysCTT_16_+_3241501_3241573</td><td>3</td><td>33</td><td>31</td><td>CCGGCTAGCTCAGTCGGTAGAGCATGGGACT</td><td>831.7</td><td>7.3</td><td>1.3</td><td>5.8</td><td>5.7E-09</td><td>1.2E-08</td><td>MB231/MB468</td><td>NUC</td></tr>
<tr><td>>hg38_21_+_8433172_8446622_RNA45SN1_45S_50ntFlanks@5084.5104.21|CGAACGAGACTCTGGCATGCT</td><td>>hg38_21_+_8433172_8446622_RNA45SN1_45S_50ntFlanks</td><td>5084</td><td>5104</td><td>21</td><td>CGAACGAGACTCTGGCATGCT</td><td>628.3</td><td>-5.8</td><td>1.0</td><td>-5.8</td><td>5.4E-09</td><td>1.2E-08</td><td>MB231/MB468</td><td>MP</td></tr>
<tr><td>>hg38_21_+_8433172_8446622_RNA45SN1_45S_50ntFlanks@12997.13036.40|CGCTGCGATCTATTGAAAGTCAGCCCTCGACACAAGGGTT</td><td>>hg38_21_+_8433172_8446622_RNA45SN1_45S_50ntFlanks</td><td>12997</td><td>13036</td><td>40</td><td>CGCTGCGATCTATTGAAAGTCAGCCCTCGACACAAGGGTT</td><td>1466.2</td><td>5.5</td><td>0.9</td><td>5.8</td><td>5.7E-09</td><td>1.2E-08</td><td>MB231/MB468</td><td>NUC</td></tr>
<tr><td>>hg38_21_+_8433172_8446622_RNA45SN1_45S_50ntFlanks@10786.10818.33|AACAGCCTCTGGCATGTTGGAACAATGTAGGTA</td><td>>hg38_21_+_8433172_8446622_RNA45SN1_45S_50ntFlanks</td><td>10786</td><td>10818</td><td>33</td><td>AACAGCCTCTGGCATGTTGGAACAATGTAGGTA</td><td>2877.5</td><td>-9.7</td><td>1.7</td><td>-5.8</td><td>6.5E-09</td><td>1.2E-08</td><td>BT20/MB468</td><td>NUC</td></tr>
<tr><td>>hg38_21_+_8433172_8446622_RNA45SN1_45S_50ntFlanks@7977.8006.30|CGACCTCAGATCAGACGTGGCGACCCGCTG</td><td>>hg38_21_+_8433172_8446622_RNA45SN1_45S_50ntFlanks</td><td>7977</td><td>8006</td><td>30</td><td>CGACCTCAGATCAGACGTGGCGACCCGCTG</td><td>2146.9</td><td>-3.2</td><td>0.5</td><td>-5.9</td><td>4.3E-09</td><td>1.2E-08</td><td>BT20/MB468</td><td>MITO</td></tr>
<tr><td>>MI0000542|hsa-mir-320a&WithFlank&8|-|22244960|22245043@47.68.22&[MIMAT0000510&hsa-miR-320a-3p&offsets|-1|-1;m-60&8|-|22244975|22244996&offsets|-1|-1]|GAAAAGCTGGGTTGAGAGGGCG</td><td>>MI0000542|hsa-mir-320a&WithFlank&8|-|22244960|22245043</td><td>47</td><td>68</td><td>22</td><td>GAAAAGCTGGGTTGAGAGGGCG</td><td>267.0</td><td>4.8</td><td>0.8</td><td>6.3</td><td>3.4E-10</td><td>1.2E-08</td><td>BT20/MB231</td><td>CYTO</td></tr>
<tr><td>>hg38_21_+_8433172_8446622_RNA45SN1_45S_50ntFlanks@6740.6771.32|CATCGACACTTCGAACGCACTTGCGGCCCCGG</td><td>>hg38_21_+_8433172_8446622_RNA45SN1_45S_50ntFlanks</td><td>6740</td><td>6771</td><td>32</td><td>CATCGACACTTCGAACGCACTTGCGGCCCCGG</td><td>2877.7</td><td>-9.8</td><td>1.7</td><td>-5.8</td><td>6.5E-09</td><td>1.2E-08</td><td>BT20/MB468</td><td>NUC</td></tr>
<tr><td>>hg38_21_+_8433172_8446622_RNA45SN1_45S_50ntFlanks@5073.5104.32|AATTCCGATAACGAACGAGACTCTGGCATGCT</td><td>>hg38_21_+_8433172_8446622_RNA45SN1_45S_50ntFlanks</td><td>5073</td><td>5104</td><td>32</td><td>AATTCCGATAACGAACGAGACTCTGGCATGCT</td><td>159.7</td><td>-11.2</td><td>1.9</td><td>-5.8</td><td>5.4E-09</td><td>1.2E-08</td><td>MB231/MB468</td><td>MP</td></tr>
<tr><td>>MI0000071|hsa-mir-17&WithFlank&13|+|91350599|91350694@20.43.24&[MIMAT0000070&hsa-miR-17-5p&offsets|0|+1;m-88&13|+|91350618|91350640&offsets|0|+1]|CAAAGTGCTTACAGTGCAGGTAGT</td><td>>MI0000071|hsa-mir-17&WithFlank&13|+|91350599|91350694</td><td>20</td><td>43</td><td>24</td><td>CAAAGTGCTTACAGTGCAGGTAGT</td><td>3860.3</td><td>6.3</td><td>1.1</td><td>5.8</td><td>6.2E-09</td><td>1.2E-08</td><td>MB231/MB468</td><td>MITO</td></tr>
<tr><td>>MI0000108|hsa-mir-103a-2&WithFlank&20|+|3917488|3917577@54.75.22(+1U);MI0000109|hsa-mir-103a-1&WithFlank&5|-|168560890|168560979@54.75.22(+1U)&[MIMAT0000101&hsa-miR-103a-3p&offsets|0|-1(+1U);m-12&20|+|3917541|3917563&offsets|0|-1(+1U)];[MIMAT0000101_1&hsa-miR-103a-3p&offsets|0|-1(+1U);m-13&5|-|168560904|168560926&offsets|0|-1(+1U);m-44220&5|-|168560908|168560926&offsets|0|+3(+1U)]|AGCAGCATTGTACAGGGCTATGT</td><td>>MI0000108|hsa-mir-103a-2&WithFlank&20|+|3917488|3917577</td><td>54</td><td>75</td><td>22(+1U)</td><td>AGCAGCATTGTACAGGGCTATGT</td><td>681.9</td><td>6.0</td><td>1.0</td><td>5.8</td><td>5.7E-09</td><td>1.2E-08</td><td>MB231/MB468</td><td>CYTO</td></tr>
</table>

| | | | | | | | | | | | | | |
|---|---|---|---|---|---|---|---|---|---|---|---|---|---|
| >hg38_21_+_8433172_8446622_RNA45SN1_45S_50ntFlanks@6660.6680.21\|GCGGTGGATCACTCGGCTCGT | >hg38_21_+_8433172_8446622_RNA45SN1_45S_50ntFlanks | 6660 | 6680 | 21 | GCGGTGGATCACTCGGCTCGT | 272.2 | -6.5 | 1.1 | -5.8 | 5.4E-09 | 1.2E-08 | MB231/MB468 | MP |
| >hg38_21_+_8433172_8446622_RNA45SN1_45S_50ntFlanks@11876.11910.35\|AAGACCCTGTTGAGCTTGACTCTAGTCTGGCACGG | >hg38_21_+_8433172_8446622_RNA45SN1_45S_50ntFlanks | 11876 | 11910 | 35 | AAGACCCTGTTGAGCTTGACTCTAGTCTGGCACGG | 287.2 | -3.0 | 0.5 | -5.9 | 4.3E-09 | 1.2E-08 | BT20/MB468 | MITO |
| >MI0000465\|hsa-mir-191&WithFlank&3\|-\|49020612\|49020715@22.42.21(+1C)&[MIMAT0000440&hsa-miR-191-5p&offsets\|0\|-2(+1C);m-48&3\|-\|49020672\|49020694&offsets\|0\|-2(+1C)]\|CAACGGAATCCCAAAAGCAGCC | >MI0000465\|hsa-mir-191&WithFlank&3\|-\|49020612\|49020715 | 22 | 42 | 21(+1C) | CAACGGAATCCCAAAAGCAGCC | 542.7 | 5.3 | 0.9 | 5.8 | 5.7E-09 | 1.2E-08 | MB231/MB468 | CYTO |
| >trnaMT_ValTAC_MT_+_1602_1670@49.71.23\|AACTTAACTTGACCGCTCTGACC | >trnaMT_ValTAC_MT_+_1602_1670 | 49 | 71 | 23 | AACTTAACTTGACCGCTCTGACC | 234.2 | 6.5 | 1.1 | 5.8 | 5.7E-09 | 1.2E-08 | MB231/MB468 | CYTO |
| >hg38_21_+_8433172_8446622_RNA45SN1_45S_50ntFlanks@10391.10431.41\|AACAGCAGTTGAACATGGGTCAGTCGGTCCTGAGAGATGGG | >hg38_21_+_8433172_8446622_RNA45SN1_45S_50ntFlanks | 10391 | 10431 | 41 | AACAGCAGTTGAACATGGGTCAGTCGGTCCTGAGAGATGGG | 107.6 | -10.6 | 1.8 | -5.8 | 5.4E-09 | 1.2E-08 | MB231/MB468 | MP |
| >MI0000809\|hsa-mir-151a&WithFlank&8\|-\|140732558\|140732659@53.74.22&[MIMAT0000757&hsa-miR-151a-3p&offsets\|0\|+1;m-30&8\|-\|140732587\|140732607&offsets\|0\|+1]\|CTAGACTGAAGCTCCTTGAGGA | >MI0000809\|hsa-mir-151a&WithFlank&8\|-\|140732558\|140732659 | 53 | 74 | 22 | CTAGACTGAAGCTCCTTGAGGA | 6897.1 | 3.6 | 0.6 | 5.8 | 5.4E-09 | 1.2E-08 | MB231/MB468 | MP |
| >hg38_21_+_8433172_8446622_RNA45SN1_45S_50ntFlanks@12632.12659.28\|GCGAAGCTACCATCTGTGGGATTATGAC | >hg38_21_+_8433172_8446622_RNA45SN1_45S_50ntFlanks | 12632 | 12659 | 28 | GCGAAGCTACCATCTGTGGGATTATGAC | 3234.5 | -7.2 | 1.2 | -5.8 | 6.3E-09 | 1.2E-08 | MB231/MB468 | MITO |
| >hg38_21_+_8433172_8446622_RNA45SN1_45S_50ntFlanks@11641.11685.45\|CGGCGGGTGTTGACGCGATGTGATTTCTGCCCAGTGCTCTGAATG | >hg38_21_+_8433172_8446622_RNA45SN1_45S_50ntFlanks | 11641 | 11685 | 45 | CGGCGGGTGTTGACGCGATGTGATTTCTGCCCAGTGCTCTGAATG | 1726.0 | -5.9 | 1.0 | -5.8 | 5.9E-09 | 1.2E-08 | MB231/MB468 | NUC |
| >hg38_21_+_8433172_8446622_RNA45SN1_45S_50ntFlanks@475.517.43\|CTTCGTGATCGATGTGGTGACGTCGTGCTCTCCCGGGCCGGGT | >hg38_21_+_8433172_8446622_RNA45SN1_45S_50ntFlanks | 475 | 517 | 43 | CTTCGTGATCGATGTGGTGACGTCGTGCTCTCCCGGGCCGGGT | 189.4 | 11.3 | 1.8 | 6.2 | 7.5E-10 | 1.2E-08 | BT20/MB468 | MP |
| >hg38_21_+_8433172_8446622_RNA45SN1_45S_50ntFlanks@9565.9595.31\|TGACGTGCAAATCGGTCGTCCGACCTGGGTA | >hg38_21_+_8433172_8446622_RNA45SN1_45S_50ntFlanks | 9565 | 9595 | 31 | TGACGTGCAAATCGGTCGTCCGACCTGGGTA | 1390.5 | -3.9 | 0.7 | -5.8 | 5.9E-09 | 1.2E-08 | MB231/MB468 | NUC |
| >hg38_21_+_8433172_8446622_RNA45SN1_45S_50ntFlanks@10402.10431.30\|AACATGGGTCAGTCGGTCCTGAGAGATGGG | >hg38_21_+_8433172_8446622_RNA45SN1_45S_50ntFlanks | 10402 | 10431 | 30 | AACATGGGTCAGTCGGTCCTGAGAGATGGG | 1135.7 | -5.7 | 1.0 | -5.8 | 5.0E-09 | 1.2E-08 | BT20/MB468 | CYTO |
| >hg38_21_+_8433172_8446622_RNA45SN1_45S_50ntFlanks@4045.4066.22\|CCATTCGAACGTCTGCCCTATC | >hg38_21_+_8433172_8446622_RNA45SN1_45S_50ntFlanks | 4045 | 4066 | 22 | CCATTCGAACGTCTGCCCTATC | 2426.5 | -7.3 | 1.3 | -5.8 | 6.3E-09 | 1.2E-08 | MB231/MB468 | MITO |
| >hg38_21_+_8433172_8446622_RNA45SN1_45S_50ntFlanks@11561.11600.40\|GCGCCTAGCAGCCGACTTAGAACTGGTGCGGACCAGGGGA | >hg38_21_+_8433172_8446622_RNA45SN1_45S_50ntFlanks | 11561 | 11600 | 40 | GCGCCTAGCAGCCGACTTAGAACTGGTGCGGACCAGGGGA | 3461.4 | -9.9 | 1.7 | -5.8 | 6.6E-09 | 1.2E-08 | BT20/MB468 | NUC |
| >hg38_21_+_8433172_8446622_RNA45SN1_45S_50ntFlanks@11564.11600.37\|CCTAGCAGCCGACTTAGAACTGGTGCGGACCAGGGGA | >hg38_21_+_8433172_8446622_RNA45SN1_45S_50ntFlanks | 11564 | 11600 | 37 | CCTAGCAGCCGACTTAGAACTGGTGCGGACCAGGGGA | 1822.9 | -7.5 | 1.3 | -5.8 | 5.5E-09 | 1.2E-08 | MB231/MB468 | MP |
| >hg38_21_+_8433172_8446622_RNA45SN1_45S_50ntFlanks@4160.4184.25\|CCACATCCAAGGAAGGCAGCAGGCG | >hg38_21_+_8433172_8446622_RNA45SN1_45S_50ntFlanks | 4160 | 4184 | 25 | CCACATCCAAGGAAGGCAGCAGGCG | 141.4 | -4.3 | 0.7 | -5.8 | 5.8E-09 | 1.2E-08 | MB231/MB468 | CYTO |
| >trna119_LysCTT_1_-_145395522_145395594@2.33.32,trna11_LysCTT_5_-_180648979_180649051@2.33.32,trna13_LysCTT_6_+_26556774_26556846@2.33.32,trna7_LysCTT_16_+_3225692_3225764@2.33.32,trna9_LysCTT_5_+_180634755_180634827@2.33.32\|CCCGGCTAGCTCAGTCGGTAGAGCATGAGACT | >trna119_LysCTT_1_-_145395522_145395594 | 2 | 33 | 32 | CCCGGCTAGCTCAGTCGGTAGAGCATGAGACT | 384.9 | -5.9 | 0.9 | -6.2 | 4.7E-10 | 1.2E-08 | BT20/MB231 | MITO |
| >MI0000077\|hsa-mir-21&WithFlank&17\|+\|59841260\|59841343@52.72.21&[MIMAT0004494&hsa-miR-21-3p&offsets\|0\|0;m-52&17\|+\|59841311\|59841332&offsets\|0\|-1]\|CAACACCAGTCGATGGGCTGT | >MI0000077\|hsa-mir-21&WithFlank&17\|+\|59841260\|59841343 | 52 | 72 | 21 | CAACACCAGTCGATGGGCTGT | 124.8 | 6.4 | 1.1 | 5.8 | 6.7E-09 | 1.2E-08 | BT20/MB468 | NUC |
| >hg38_21_+_8433172_8446622_RNA45SN1_45S_50ntFlanks@5084.5102.19\|CGAACGAGACTCTGGCATG | >hg38_21_+_8433172_8446622_RNA45SN1_45S_50ntFlanks | 5084 | 5102 | 19 | CGAACGAGACTCTGGCATG | 716.7 | -5.5 | 0.9 | -5.8 | 5.9E-09 | 1.2E-08 | MB231/MB468 | NUC |
| >hg38_21_+_8433172_8446622_RNA45SN1_45S_50ntFlanks@11452.11469.18\|CTCCTCGGCGCGCGGCGG | >hg38_21_+_8433172_8446622_RNA45SN1_45S_50ntFlanks | 11452 | 11469 | 18 | CTCCTCGGCGCGCGGCGG | 298.8 | -8.4 | 1.4 | -5.8 | 5.9E-09 | 1.2E-08 | MB231/MB468 | NUC |
| >hg38_21_+_8433172_8446622_RNA45SN1_45S_50ntFlanks@11234.11252.19\|CCTTCCCGTGGATCGCCCC | >hg38_21_+_8433172_8446622_RNA45SN1_45S_50ntFlanks | 11234 | 11252 | 19 | CCTTCCCGTGGATCGCCCC | 372.4 | -8.3 | 1.4 | -5.8 | 6.7E-09 | 1.2E-08 | BT20/MB468 | NUC |
| >hg38_21_+_8433172_8446622_RNA45SN1_45S_50ntFlanks@11603.11620.18\|CCGACTGTTTAATTAAAA | >hg38_21_+_8433172_8446622_RNA45SN1_45S_50ntFlanks | 11603 | 11620 | 18 | CCGACTGTTTAATTAAAA | 1521.1 | -9.5 | 1.6 | -5.8 | 6.7E-09 | 1.2E-08 | BT20/MB468 | NUC |
| >hg38_21_+_8433172_8446622_RNA45SN1_45S_50ntFlanks@4130.4150.21\|ATTCCGGAGAGGGAGCCTGAG | >hg38_21_+_8433172_8446622_RNA45SN1_45S_50ntFlanks | 4130 | 4150 | 21 | ATTCCGGAGAGGGAGCCTGAG | 207.0 | -3.6 | 0.6 | -5.8 | 6.0E-09 | 1.2E-08 | MB231/MB468 | NUC |
| >hg38_21_+_8433172_8446622_RNA45SN1_45S_50ntFlanks@12459.12478.20\|AGCGTTGGATTGTTCACCCA | >hg38_21_+_8433172_8446622_RNA45SN1_45S_50ntFlanks | 12459 | 12478 | 20 | AGCGTTGGATTGTTCACCCA | 584.7 | -7.8 | 1.4 | -5.8 | 6.4E-09 | 1.2E-08 | MB231/MB468 | MITO |
| >trna116_GluCTC_1_-_145399233_145399304@37.75.39,trna134_GluTTC_1_-_16861774_16861845@37.75.39,trna71_GluCTC_1_-_161439189_161439260@37.75.39,trna74_GluCTC_1_-_161431809_161431880@37.75.39,trna77_GluCTC_1_-_161424398_161424469@37.75.39,trna77_GluCTC_6_+_28949976_28950047@37.75.39,trna80_GluCTC_1_-_161417018_161417089@37.75.39,trna84_GluTTC_1_-_161391883_161391954@37.75.39,trna87_GluCTC_6_-_126101393_126101464@37.75.39\|ACCGCCGCGGCCCGGGTTCGATTCCCGGTCAGGGAACCA | >trna116_GluCTC_1_-_145399233_145399304 | 37 | 75 | 39 | ACCGCCGCGGCCCGGGTTCGATTCCCGGTCAGGGAACCA | 8048.6 | 8.1 | 1.4 | 5.8 | 6.0E-09 | 1.2E-08 | MB231/MB468 | NUC |
| >trna11_GluTTC_15_-_26327381_26327452@1.29.29,trna3_GluTTC_13_-_45492062_45492133@1.29.29\|TCCCACATGGTCTAGCGGTTAGGATTCCT | >trna11_GluTTC_15_-_26327381_26327452 | 1 | 29 | 29 | TCCCACATGGTCTAGCGGTTAGGATTCCT | 1371.9 | -5.6 | 1.0 | -5.8 | 5.6E-09 | 1.2E-08 | MB231/MB468 | MP |
| >trnaMT_PheGAA_MT_+_577_647@56.74.19\|CATCACCCCATAAACACCA | >trnaMT_PheGAA_MT_+_577_647 | 56 | 74 | 19 | CATCACCCCATAAACACCA | 697.0 | 8.0 | 1.4 | 5.8 | 5.9E-09 | 1.2E-08 | MB231/MB468 | CYTO |
| >hg38_21_+_8433172_8446622_RNA45SN1_45S_50ntFlanks@12157.12180.24\|AGTTTGACTGGGGCGGTACACCTG | >hg38_21_+_8433172_8446622_RNA45SN1_45S_50ntFlanks | 12157 | 12180 | 24 | AGTTTGACTGGGGCGGTACACCTG | 465.8 | -6.7 | 1.1 | -5.8 | 5.9E-09 | 1.2E-08 | MB231/MB468 | CYTO |
| >hg38_21_+_8433172_8446622_RNA45SN1_45S_50ntFlanks@11876.11920.45\|AAGACCCTGTTGAGCTTGACTCTAGTCTGGCACGGTGAAGAGACA | >hg38_21_+_8433172_8446622_RNA45SN1_45S_50ntFlanks | 11876 | 11920 | 45 | AAGACCCTGTTGAGCTTGACTCTAGTCTGGCACGGTGAAGAGACA | 370.0 | -6.4 | 1.1 | -5.8 | 6.5E-09 | 1.2E-08 | MB231/MB468 | MITO |
| >hg38_21_+_8433172_8446622_RNA45SN1_45S_50ntFlanks@12622.12661.40\|AGCCAATGGGGCGAAGCTACCATCTGTGGGATTATGACTG | >hg38_21_+_8433172_8446622_RNA45SN1_45S_50ntFlanks | 12622 | 12661 | 40 | AGCCAATGGGGCGAAGCTACCATCTGTGGGATTATGACTG | 480.0 | -6.7 | 1.2 | -5.8 | 6.5E-09 | 1.2E-08 | MB231/MB468 | MITO |
| >hg38_21_+_8433172_8446622_RNA45SN1_45S_50ntFlanks@11784.11806.23\|ATGAATGGATGAACGAGATTCCC | >hg38_21_+_8433172_8446622_RNA45SN1_45S_50ntFlanks | 11784 | 11806 | 23 | ATGAATGGATGAACGAGATTCCC | 817.3 | -6.3 | 1.1 | -5.8 | 6.5E-09 | 1.2E-08 | MB231/MB468 | MITO |
| >hg38_21_+_8433172_8446622_RNA45SN1_45S_50ntFlanks@10681.10708.28\|TTCCGGCGGCGTCCGGTGAGCTCTCGCT | >hg38_21_+_8433172_8446622_RNA45SN1_45S_50ntFlanks | 10681 | 10708 | 28 | TTCCGGCGGCGTCCGGTGAGCTCTCGCT | 372.8 | -6.9 | 1.2 | -5.8 | 5.9E-09 | 1.2E-08 | MB231/MB468 | CYTO |
| >hg38_21_+_8433172_8446622_RNA45SN1_45S_50ntFlanks@3984.4004.21\|GCGGCTTTGGTGACTCTAGAT | >hg38_21_+_8433172_8446622_RNA45SN1_45S_50ntFlanks | 3984 | 4004 | 21 | GCGGCTTTGGTGACTCTAGAT | 180.1 | -9.6 | 1.6 | -5.9 | 4.5E-09 | 1.2E-08 | BT20/MB468 | MITO |
| >hg38_21_+_8433172_8446622_RNA45SN1_45S_50ntFlanks@12488.12519.32\|AACGTGAGCTGGGTTTAGACCGTCGTGAGACA | >hg38_21_+_8433172_8446622_RNA45SN1_45S_50ntFlanks | 12488 | 12519 | 32 | AACGTGAGCTGGGTTTAGACCGTCGTGAGACA | 135.6 | -4.4 | 0.8 | -5.8 | 5.9E-09 | 1.2E-08 | MB231/MB468 | CYTO |
| >hg38_MT_+_598_1651_RNR1_12S_50ntFlanks@52.88.37\|ATAGGTTTGGTCCTAGCCTTTCTATTAGCTCTTAGTA | >hg38_MT_+_598_1651_RNR1_12S_50ntFlanks | 52 | 88 | 37 | ATAGGTTTGGTCCTAGCCTTTCTATTAGCTCTTAGTA | 336.4 | 5.9 | 1.0 | 5.8 | 5.9E-09 | 1.2E-08 | MB231/MB468 | CYTO |
| >hg38_21_+_8433172_8446622_RNA45SN1_45S_50ntFlanks@11886.11912.27\|TGAGCTTGACTCTAGTCTGGCACGGTG | >hg38_21_+_8433172_8446622_RNA45SN1_45S_50ntFlanks | 11886 | 11912 | 27 | TGAGCTTGACTCTAGTCTGGCACGGTG | 201.7 | -10.7 | 1.7 | -6.2 | 7.7E-10 | 1.2E-08 | BT20/MB468 | MP |
| >hg38_21_+_8433172_8446622_RNA45SN1_45S_50ntFlanks@11667.11690.24\|CTGCCCAGTGCTCTGAATGTCAAA | >hg38_21_+_8433172_8446622_RNA45SN1_45S_50ntFlanks | 11667 | 11690 | 24 | CTGCCCAGTGCTCTGAATGTCAAA | 213.9 | 8.7 | 1.5 | 5.8 | 5.1E-09 | 1.3E-08 | BT20/MB468 | CYTO |
| >hg38_21_+_8433172_8446622_RNA45SN1_45S_50ntFlanks@6722.6743.22\|ATTGCAGGACACATTGATCATC | >hg38_21_+_8433172_8446622_RNA45SN1_45S_50ntFlanks | 6722 | 6743 | 22 | ATTGCAGGACACATTGATCATC | 1366.3 | -4.3 | 0.7 | -5.8 | 6.0E-09 | 1.3E-08 | MB231/MB468 | NUC |
| >trna139_ValAAC_6_-_27618707_27618779@36.56.21\|CACGCGAAAGGTCCCTGGATC | >trna139_ValAAC_6_-_27618707_27618779 | 36 | 56 | 21 | CACGCGAAAGGTCCCTGGATC | 280.5 | 11.5 | 1.8 | 6.4 | 1.5E-10 | 1.3E-08 | BT20/MB231 | MP |
| >MI0006444\|hsa-mir-1307&WithFlank&10\|-\|103394247\|103394407@87.108.22&[MIMAT0005951&hsa-miR-1307-3p&offsets\|+1\|+1;m-66&10\|-\|103394300\|103394322&offsets\|+1\|0]\|CTCGGCGTGGCGTCGGTCGTGG | >MI0006444\|hsa-mir-1307&WithFlank&10\|-\|103394247\|103394407 | 87 | 108 | 22 | CTCGGCGTGGCGTCGGTCGTGG | 1656.2 | 4.3 | 0.7 | 5.8 | 6.1E-09 | 1.3E-08 | MB231/MB468 | NUC |
| >hg38_21_+_8433172_8446622_RNA45SN1_45S_50ntFlanks@9554.9573.20\|CGTAGCGGTCCTGACGTGCA | >hg38_21_+_8433172_8446622_RNA45SN1_45S_50ntFlanks | 9554 | 9573 | 20 | CGTAGCGGTCCTGACGTGCA | 6427.5 | -5.7 | 1.0 | -5.8 | 5.7E-09 | 1.3E-08 | MB231/MB468 | MP |
| >hg38_21_+_8433172_8446622_RNA45SN1_45S_50ntFlanks@9180.9198.19\|CCGTCGGGCCCGGGGGAGG | >hg38_21_+_8433172_8446622_RNA45SN1_45S_50ntFlanks | 9180 | 9198 | 19 | CCGTCGGGCCCGGGGGAGG | 83.4 | -10.2 | 1.8 | -5.8 | 5.7E-09 | 1.3E-08 | MB231/MB468 | MP |
| >hg38_21_+_8433172_8446622_RNA45SN1_45S_50ntFlanks@10846.10864.19\|CTTCGGGATAAGGATTGGC | >hg38_21_+_8433172_8446622_RNA45SN1_45S_50ntFlanks | 10846 | 10864 | 19 | CTTCGGGATAAGGATTGGC | 483.5 | -7.5 | 1.3 | -5.8 | 6.1E-09 | 1.3E-08 | MB231/MB468 | NUC |
| >hg38_21_+_8433172_8446622_RNA45SN1_45S_50ntFlanks@10402.10428.27\|AACATGGGTCAGTCGGTCCTGAGAGAT | >hg38_21_+_8433172_8446622_RNA45SN1_45S_50ntFlanks | 10402 | 10428 | 27 | AACATGGGTCAGTCGGTCCTGAGAGAT | 343.1 | -3.4 | 0.6 | -5.8 | 6.1E-09 | 1.3E-08 | MB231/MB468 | NUC |
| >hg38_21_+_8433172_8446622_RNA45SN1_45S_50ntFlanks@3838.3873.36\|CTCTCCTACTTGGATAACTGTGGTAATTCTAGAGCT | >hg38_21_+_8433172_8446622_RNA45SN1_45S_50ntFlanks | 3838 | 3873 | 36 | CTCTCCTACTTGGATAACTGTGGTAATTCTAGAGCT | 134.8 | -10.9 | 1.9 | -5.8 | 5.7E-09 | 1.3E-08 | MB231/MB468 | MP |
| >hg38_21_+_8433172_8446622_RNA45SN1_45S_50ntFlanks@3711.3729.19\|GTTGATCCTGCCAGTAGCA | >hg38_21_+_8433172_8446622_RNA45SN1_45S_50ntFlanks | 3711 | 3729 | 19 | GTTGATCCTGCCAGTAGCA | 667.6 | -3.6 | 0.6 | -5.8 | 6.9E-09 | 1.3E-08 | BT20/MB468 | NUC |
| >hg38_21_+_8433172_8446622_RNA45SN1_45S_50ntFlanks@10316.10340.25\|GGTGCAGATCTTGGTGGTAGTAGCA | >hg38_21_+_8433172_8446622_RNA45SN1_45S_50ntFlanks | 10316 | 10340 | 25 | GGTGCAGATCTTGGTGGTAGTAGCA | 2064.6 | 14.3 | 2.2 | 6.4 | 1.5E-10 | 1.3E-08 | BT20/MB231 | MP |
| >MI0000081\|hsa-mir-24-2&WithFlank&19\|-\|13836281\|13836365@56.77.22(+1U);MI0000080\|hsa-mir-24-1&WithFlank&9\|+\|95086015\|95086094@50.71.22(+1U)&[MIMAT0000080&hsa-miR-24-3p&offsets\|0\|0(+1U);m-36&19\|-\|13836288\|13836310&offsets\|0\|-1(+1U)];[MIMAT0000080_1&hsa-miR-24-3p&offsets\|0\|0(+1U);m-35&9\|+\|95086064\|95086086&offsets\|0\|-1(+1U)]\|TGGCTCAGTTCAGCAGGAACAGT | >MI0000081\|hsa-mir-24-2&WithFlank&19\|-\|13836281\|13836365 | 56 | 77 | 22(+1U) | TGGCTCAGTTCAGCAGGAACAGT | 10634.1 | 5.8 | 1.0 | 5.8 | 6.6E-09 | 1.3E-08 | MB231/MB468 | MITO |
| >trnaMT_GluTTC_MT_-_14674_14742@20.58.39,trnalookalike8_GluTTC_5_-_93905172_93905240@20.58.39\|AACGATGGTTTTTCATATCATTGGTCGTGGTTGTAGTCC | >trnaMT_GluTTC_MT_-_14674_14742 | 20 | 58 | 39 | AACGATGGTTTTTCATATCATTGGTCGTGGTTGTAGTCC | 273.0 | 11.8 | 1.9 | 6.1 | 7.9E-10 | 1.3E-08 | BT20/MB468 | MP |
| >hg38_21_+_8433172_8446622_RNA45SN1_45S_50ntFlanks@3854.3872.19\|ACTGTGGTAATTCTAGAGC | >hg38_21_+_8433172_8446622_RNA45SN1_45S_50ntFlanks | 3854 | 3872 | 19 | ACTGTGGTAATTCTAGAGC | 618.3 | -8.5 | 1.5 | -5.8 | 6.6E-09 | 1.3E-08 | MB231/MB468 | MITO |

<table>
<tr><th></th><th></th><th></th><th></th><th></th><th></th><th></th><th></th><th></th><th></th><th></th><th></th><th></th><th></th></tr>
<tr><td>>MI0000477|hsa-mir-146a&WithFlank&5|+|160485346|160485456@28.48.21&[MIMAT0000449&hsa-miR-146a-5p&offsets|+1|0;m-129&5|+|160485372|160485393&offsets|+1|0]|GAGAACTGAATTCCATGGGTT</td><td>>MI0000477|hsa-mir-146a&WithFlank&5|+|160485346|160485456</td><td>28</td><td>48</td><td>21</td><td>GAGAACTGAATTCCATGGGTT</td><td>114.2</td><td>10.1</td><td>1.7</td><td>5.8</td><td>5.7E-09</td><td>1.3E-08</td><td>MB231/MB468</td><td>MP</td></tr>
<tr><td>>hg38_21_+_8433172_8446622_RNA45SN1_45S_50ntFlanks@4005.4025.21|AACCTCGGGCCGATCGCACGC</td><td>>hg38_21_+_8433172_8446622_RNA45SN1_45S_50ntFlanks</td><td>4005</td><td>4025</td><td>21</td><td>AACCTCGGGCCGATCGCACGC</td><td>668.8</td><td>-8.0</td><td>1.4</td><td>-5.8</td><td>6.6E-09</td><td>1.3E-08</td><td>MB231/MB468</td><td>MITO</td></tr>
<tr><td>>hg38_21_+_8433172_8446622_RNA45SN1_45S_50ntFlanks@8356.8391.36|GAAAAGAACTTTGAAGAGAGAGTTCAAGAGGGCGTG</td><td>>hg38_21_+_8433172_8446622_RNA45SN1_45S_50ntFlanks</td><td>8356</td><td>8391</td><td>36</td><td>GAAAAGAACTTTGAAGAGAGAGTTCAAGAGGGCGTG</td><td>294.0</td><td>-3.1</td><td>0.5</td><td>-5.8</td><td>6.2E-09</td><td>1.3E-08</td><td>MB231/MB468</td><td>NUC</td></tr>
<tr><td>>MI0000542|hsa-mir-320a&WithFlank&8|-|22244960|22245043@48.68.21(+1C)&[MIMAT0000510&hsa-miR-320a-3p&offsets|0|-1(+1C);m-60&8|-|22244975|22244996&offsets|0|-1(+1C)]|AAAAGCTGGGTTGAGAGGGCGC</td><td>>MI0000542|hsa-mir-320a&WithFlank&8|-|22244960|22245043</td><td>48</td><td>68</td><td>21(+1C)</td><td>AAAAGCTGGGTTGAGAGGGCGC</td><td>5620.6</td><td>8.8</td><td>1.4</td><td>6.1</td><td>7.9E-10</td><td>1.3E-08</td><td>BT20/MB468</td><td>MP</td></tr>
<tr><td>>hg38_21_+_8433172_8446622_RNA45SN1_45S_50ntFlanks@12536.12569.34|CTGATGATGTGTTGTTGCCATGGTAATCCTGCTC</td><td>>hg38_21_+_8433172_8446622_RNA45SN1_45S_50ntFlanks</td><td>12536</td><td>12569</td><td>34</td><td>CTGATGATGTGTTGTTGCCATGGTAATCCTGCTC</td><td>75.3</td><td>-10.1</td><td>1.7</td><td>-5.8</td><td>5.8E-09</td><td>1.3E-08</td><td>MB231/MB468</td><td>MP</td></tr>
<tr><td>>hg38_21_+_8433172_8446622_RNA45SN1_45S_50ntFlanks@12071.12106.36|CCCCGAGGGGCTCTCGCTTCTGGCGCCAAGCGCCCG</td><td>>hg38_21_+_8433172_8446622_RNA45SN1_45S_50ntFlanks</td><td>12071</td><td>12106</td><td>36</td><td>CCCCGAGGGGCTCTCGCTTCTGGCGCCAAGCGCCCG</td><td>371.9</td><td>-5.5</td><td>0.9</td><td>-5.8</td><td>6.1E-09</td><td>1.3E-08</td><td>MB231/MB468</td><td>CYTO</td></tr>
<tr><td>>trnaMT_HisGTG_MT_+_12138_12206@18.55.38|AAAACATCAGATTGTGAATCTGACAACAGAGGCTTACG</td><td>>trnaMT_HisGTG_MT_+_12138_12206</td><td>18</td><td>55</td><td>38</td><td>AAAACATCAGATTGTGAATCTGACAACAGAGGCTTACG</td><td>327.5</td><td>12.1</td><td>2.0</td><td>6.1</td><td>7.9E-10</td><td>1.3E-08</td><td>BT20/MB468</td><td>MP</td></tr>
<tr><td>>MI0000102|hsa-mir-100&WithFlank&11|-|122152223|122152314@19.40.22(+2A)&[MIMAT0000098&hsa-miR-100-5p&offsets|0|0(+2A);m-27&11|-|122152276|122152296&offsets|0|+1(+2A)]|AACCCGTAGATCCGAACTTGTGAA</td><td>>MI0000102|hsa-mir-100&WithFlank&11|-|122152223|122152314</td><td>19</td><td>40</td><td>22(+2A)</td><td>AACCCGTAGATCCGAACTTGTGAA</td><td>1777.2</td><td>-6.3</td><td>1.0</td><td>-6.2</td><td>5.0E-10</td><td>1.3E-08</td><td>BT20/MB231</td><td>MITO</td></tr>
<tr><td>>am_trna128_GlyGCC_6_-_27870686_27870756@3.35.33,trna18_GlyGCC_16_+_70822597_70822667@3.35.33,trna19_GlyGCC_16_+_70823410_70823480@3.35.33,trna19_GlyGCC_2_-_157257659_157257729@3.35.33,trna24_GlyGCC_16_-_70812942_70813012@3.35.33,trna25_GlyGCC_16_-_70812114_70812184@3.35.33,trna5_GlyGCC_17_+_8029064_8029134@3.35.33,trna68_GlyGCC_1_-_161493637_161493707@3.35.33|ATTGGTGGTTCAGTGGTAGAATTCTCGCCTGCC</td><td>>am_trna128_GlyGCC_6_-_27870686_27870756</td><td>3</td><td>35</td><td>33</td><td>ATTGGTGGTTCAGTGGTAGAATTCTCGCCTGCC</td><td>1575.9</td><td>-4.6</td><td>0.7</td><td>-6.2</td><td>5.0E-10</td><td>1.3E-08</td><td>BT20/MB231</td><td>MITO</td></tr>
<tr><td>>hg38_21_+_8433172_8446622_RNA45SN1_45S_50ntFlanks@12991.13027.37|CTCCCTCGCTGCGATCTATTGAAAGTCAGCCCTCGAC</td><td>>hg38_21_+_8433172_8446622_RNA45SN1_45S_50ntFlanks</td><td>12991</td><td>13027</td><td>37</td><td>CTCCCTCGCTGCGATCTATTGAAAGTCAGCCCTCGAC</td><td>80.5</td><td>-10.2</td><td>1.8</td><td>-5.8</td><td>5.8E-09</td><td>1.3E-08</td><td>MB231/MB468</td><td>MP</td></tr>
<tr><td>>hg38_21_+_8433172_8446622_RNA45SN1_45S_50ntFlanks@8839.8859.21|CGGTGGCGGCGGCGGCGGCGG</td><td>>hg38_21_+_8433172_8446622_RNA45SN1_45S_50ntFlanks</td><td>8839</td><td>8859</td><td>21</td><td>CGGTGGCGGCGGCGGCGGCGG</td><td>165.3</td><td>-10.4</td><td>1.7</td><td>-6.1</td><td>8.0E-10</td><td>1.3E-08</td><td>BT20/MB468</td><td>MP</td></tr>
<tr><td>>hg38_21_+_8433172_8446622_RNA45SN1_45S_50ntFlanks@6719.6746.28|TGAATTGCAGGACACATTGATCATCGAC</td><td>>hg38_21_+_8433172_8446622_RNA45SN1_45S_50ntFlanks</td><td>6719</td><td>6746</td><td>28</td><td>TGAATTGCAGGACACATTGATCATCGAC</td><td>993.1</td><td>-5.4</td><td>0.9</td><td>-5.8</td><td>6.1E-09</td><td>1.3E-08</td><td>MB231/MB468</td><td>CYTO</td></tr>
<tr><td>>hg38_21_+_8433172_8446622_RNA45SN1_45S_50ntFlanks@10346.10383.38|TCAAACGAGAACTTTGAAGGCCGAAGTGGAGAAGGGTT</td><td>>hg38_21_+_8433172_8446622_RNA45SN1_45S_50ntFlanks</td><td>10346</td><td>10383</td><td>38</td><td>TCAAACGAGAACTTTGAAGGCCGAAGTGGAGAAGGGTT</td><td>372.6</td><td>-4.3</td><td>0.7</td><td>-5.8</td><td>5.8E-09</td><td>1.3E-08</td><td>MB231/MB468</td><td>MP</td></tr>
<tr><td>>hg38_21_+_8433172_8446622_RNA45SN1_45S_50ntFlanks@10396.10431.36|CAGTTGAACATGGGTCAGTCGGTCCTGAGAGATGGG</td><td>>hg38_21_+_8433172_8446622_RNA45SN1_45S_50ntFlanks</td><td>10396</td><td>10431</td><td>36</td><td>CAGTTGAACATGGGTCAGTCGGTCCTGAGAGATGGG</td><td>65.9</td><td>-9.9</td><td>1.7</td><td>-5.8</td><td>5.8E-09</td><td>1.3E-08</td><td>MB231/MB468</td><td>MP</td></tr>
<tr><td>>hg38_21_+_8433172_8446622_RNA45SN1_45S_50ntFlanks@10841.10859.19|CGTAACTTCGGGATAAGGA</td><td>>hg38_21_+_8433172_8446622_RNA45SN1_45S_50ntFlanks</td><td>10841</td><td>10859</td><td>19</td><td>CGTAACTTCGGGATAAGGA</td><td>1202.6</td><td>-6.7</td><td>1.1</td><td>-5.8</td><td>6.1E-09</td><td>1.3E-08</td><td>MB231/MB468</td><td>CYTO</td></tr>
<tr><td>>hg38_21_+_8433172_8446622_RNA45SN1_45S_50ntFlanks@10127.10149.23|GCGGCGGCGGCGGCGGCGGCGGG</td><td>>hg38_21_+_8433172_8446622_RNA45SN1_45S_50ntFlanks</td><td>10127</td><td>10149</td><td>23</td><td>GCGGCGGCGGCGGCGGCGGCGGG</td><td>147.9</td><td>-9.3</td><td>1.6</td><td>-5.9</td><td>4.7E-09</td><td>1.3E-08</td><td>BT20/MB468</td><td>MITO</td></tr>
<tr><td>>hg38_21_+_8433172_8446622_RNA45SN1_45S_50ntFlanks@8844.8866.23|GCGGCGGCGGCGGCGGCGGCGGG</td><td>>hg38_21_+_8433172_8446622_RNA45SN1_45S_50ntFlanks</td><td>8844</td><td>8866</td><td>23</td><td>GCGGCGGCGGCGGCGGCGGCGGG</td><td>147.9</td><td>-9.3</td><td>1.6</td><td>-5.9</td><td>4.7E-09</td><td>1.3E-08</td><td>BT20/MB468</td><td>MITO</td></tr>
<tr><td>>hg38_21_+_8433172_8446622_RNA45SN1_45S_50ntFlanks@4045.4067.23|CCATTCGAACGTCTGCCCTATCA</td><td>>hg38_21_+_8433172_8446622_RNA45SN1_45S_50ntFlanks</td><td>4045</td><td>4067</td><td>23</td><td>CCATTCGAACGTCTGCCCTATCA</td><td>149.9</td><td>-9.3</td><td>1.6</td><td>-5.9</td><td>4.7E-09</td><td>1.3E-08</td><td>BT20/MB468</td><td>MITO</td></tr>
<tr><td>>hg38_21_+_8433172_8446622_RNA45SN1_45S_50ntFlanks@9241.9274.34|CGGAGCGAGCGCACGGGGTCGGCGGCGACGTCGG</td><td>>hg38_21_+_8433172_8446622_RNA45SN1_45S_50ntFlanks</td><td>9241</td><td>9274</td><td>34</td><td>CGGAGCGAGCGCACGGGGTCGGCGGCGACGTCGG</td><td>147.1</td><td>-10.3</td><td>1.7</td><td>-6.1</td><td>8.0E-10</td><td>1.3E-08</td><td>BT20/MB468</td><td>MP</td></tr>
<tr><td>>hg38_21_+_8433172_8446622_RNA45SN1_45S_50ntFlanks@6651.6678.28|CGACTCTTAGCGGTGGATCACTCGGCTC</td><td>>hg38_21_+_8433172_8446622_RNA45SN1_45S_50ntFlanks</td><td>6651</td><td>6678</td><td>28</td><td>CGACTCTTAGCGGTGGATCACTCGGCTC</td><td>77334.4</td><td>4.4</td><td>0.8</td><td>5.9</td><td>4.7E-09</td><td>1.3E-08</td><td>BT20/MB468</td><td>MITO</td></tr>
<tr><td>>MI0000441|hsa-mir-30b&WithFlank&8|-|134800514|134800613@23.43.21&[MIMAT0000420&hsa-miR-30b-5p&offsets|0|-1;m-76&8|-|134800570|134800591&offsets|0|-1]|TGTAAACATCCTACACTCAGC</td><td>>MI0000441|hsa-mir-30b&WithFlank&8|-|134800514|134800613</td><td>23</td><td>43</td><td>21</td><td>TGTAAACATCCTACACTCAGC</td><td>205.8</td><td>8.5</td><td>1.5</td><td>5.8</td><td>6.2E-09</td><td>1.3E-08</td><td>MB231/MB468</td><td>NUC</td></tr>
<tr><td>>hg38_MT_+_1621_3279_RNR2_16S_50ntFlanks@51.92.42|GCTAAACCTAGCCCCAAACCCACTCCACCTTACTACCAGACA</td><td>>hg38_MT_+_1621_3279_RNR2_16S_50ntFlanks</td><td>51</td><td>92</td><td>42</td><td>GCTAAACCTAGCCCCAAACCCACTCCACCTTACTACCAGACA</td><td>1820.0</td><td>5.3</td><td>0.9</td><td>5.9</td><td>4.7E-09</td><td>1.3E-08</td><td>BT20/MB468</td><td>MITO</td></tr>
<tr><td>>hg38_21_+_8433172_8446622_RNA45SN1_45S_50ntFlanks@6734.6774.41|ATTGATCATCGACACTTCGAACGCACTTGCGGCCCCGGGTT</td><td>>hg38_21_+_8433172_8446622_RNA45SN1_45S_50ntFlanks</td><td>6734</td><td>6774</td><td>41</td><td>ATTGATCATCGACACTTCGAACGCACTTGCGGCCCCGGGTT</td><td>1052.1</td><td>5.5</td><td>1.0</td><td>5.8</td><td>6.7E-09</td><td>1.3E-08</td><td>MB231/MB468</td><td>MITO</td></tr>
<tr><td>>hg38_21_+_8433172_8446622_RNA45SN1_45S_50ntFlanks@4243.4275.33|CTTTCGAGGCCCTGTAATTGGAATGAGTCCACT</td><td>>hg38_21_+_8433172_8446622_RNA45SN1_45S_50ntFlanks</td><td>4243</td><td>4275</td><td>33</td><td>CTTTCGAGGCCCTGTAATTGGAATGAGTCCACT</td><td>449.5</td><td>-5.9</td><td>1.0</td><td>-5.8</td><td>6.7E-09</td><td>1.3E-08</td><td>MB231/MB468</td><td>MITO</td></tr>
<tr><td>>hg38_21_+_8433172_8446622_RNA45SN1_45S_50ntFlanks@3853.3874.22|AACTGTGGTAATTCTAGAGCTA</td><td>>hg38_21_+_8433172_8446622_RNA45SN1_45S_50ntFlanks</td><td>3853</td><td>3874</td><td>22</td><td>AACTGTGGTAATTCTAGAGCTA</td><td>450.8</td><td>-6.2</td><td>1.1</td><td>-5.8</td><td>5.8E-09</td><td>1.3E-08</td><td>MB231/MB468</td><td>MP</td></tr>
<tr><td>>hg38_21_+_8433172_8446622_RNA45SN1_45S_50ntFlanks@4760.4786.27|TCAGATACCGTCGTAGTTCCGACCATA</td><td>>hg38_21_+_8433172_8446622_RNA45SN1_45S_50ntFlanks</td><td>4760</td><td>4786</td><td>27</td><td>TCAGATACCGTCGTAGTTCCGACCATA</td><td>290.1</td><td>-3.5</td><td>0.6</td><td>-5.8</td><td>5.3E-09</td><td>1.3E-08</td><td>BT20/MB468</td><td>CYTO</td></tr>
<tr><td>>hg38_21_+_8433172_8446622_RNA45SN1_45S_50ntFlanks@8358.8379.22|AAAGAACTTTGAAGAGAGAGTT</td><td>>hg38_21_+_8433172_8446622_RNA45SN1_45S_50ntFlanks</td><td>8358</td><td>8379</td><td>22</td><td>AAAGAACTTTGAAGAGAGAGTT</td><td>500.7</td><td>-5.8</td><td>1.0</td><td>-5.8</td><td>6.2E-09</td><td>1.3E-08</td><td>MB231/MB468</td><td>CYTO</td></tr>
<tr><td>>MI0000289|hsa-mir-181a-1&WithFlank&1|-|198859038|198859159@30.53.24(+1A);MI0000269|hsa-mir-181a-2&WithFlank&9|+|124692436|124692557@45.68.24(+1A)&[MIMAT0000256&hsa-miR-181a-5p&offsets|0|+1(+1A);m-32&1|-|198859108|198859130&offsets|0|+1(+1A)];[MIMAT0000256_1&hsa-miR-181a-5p&offsets|0|+1(+1A);m-32&9|+|124692480|124692502&offsets|0|+1(+1A)]|AACATTCAACGCTGTCGGTGAGTTA</td><td>>MI0000289|hsa-mir-181a-1&WithFlank&1|-|198859038|198859159</td><td>30</td><td>53</td><td>24(+1A)</td><td>AACATTCAACGCTGTCGGTGAGTTA</td><td>260.3</td><td>-11.4</td><td>1.8</td><td>-6.4</td><td>1.6E-10</td><td>1.3E-08</td><td>BT20/MB231</td><td>MP</td></tr>
<tr><td>>hg38_21_+_8433172_8446622_RNA45SN1_45S_50ntFlanks@8556.8581.26|CCGGAGGGGGCGGGCTCCGGCGGGTG</td><td>>hg38_21_+_8433172_8446622_RNA45SN1_45S_50ntFlanks</td><td>8556</td><td>8581</td><td>26</td><td>CCGGAGGGGGCGGGCTCCGGCGGGTG</td><td>70.0</td><td>-10.0</td><td>1.7</td><td>-5.8</td><td>5.9E-09</td><td>1.3E-08</td><td>MB231/MB468</td><td>MP</td></tr>
<tr><td>>hg38_21_+_8433172_8446622_RNA45SN1_45S_50ntFlanks@11729.11746.18|TAACTATGACTCTCTTAA</td><td>>hg38_21_+_8433172_8446622_RNA45SN1_45S_50ntFlanks</td><td>11729</td><td>11746</td><td>18</td><td>TAACTATGACTCTCTTAA</td><td>405.1</td><td>-8.3</td><td>1.4</td><td>-5.8</td><td>7.1E-09</td><td>1.3E-08</td><td>BT20/MB468</td><td>NUC</td></tr>
<tr><td>>am_trna10_ValCAC_5_-_180649395_180649467@1.33.33,trna12_ValAAC_5_-_180645270_180645342@1.33.33,trna132_ValAAC_6_-_27721179_27721251@1.33.33,trna136_ValAAC_6_-_27648885_27648957@1.33.33,trna139_ValAAC_6_-_27618707_27618779@1.33.33,trna18_ValCAC_5_-_180529253_180529325@1.33.33,trna2_ValAAC_3_+_169490018_169490090@1.33.33,trna2_ValCAC_5_+_180524070_180524142@1.33.33,trna4_ValAAC_5_+_180591154_180591226@1.33.33,trna5_ValAAC_5_+_180596610_180596682@1.33.33,trna6_ValCAC_5_+_180600650_180600722@1.33.33,trna85_ValCAC_1_-_161369490_161369562@1.33.33,trna90_ValCAC_1_-_149684088_149684161@1.33.33,trna98_ValCAC_1_-_149298555_149298627@1.33.33,trna9_ValCAC_6_+_26538282_26538354@1.33.33|GTTTCCGTAGTGTAGTGGTTATCACGTTCGCCT</td><td>>am_trna10_ValCAC_5_-_180649395_180649467</td><td>1</td><td>33</td><td>33</td><td>GTTTCCGTAGTGTAGTGGTTATCACGTTCGCCT</td><td>2001.4</td><td>8.0</td><td>1.4</td><td>5.8</td><td>5.3E-09</td><td>1.3E-08</td><td>BT20/MB468</td><td>CYTO</td></tr>
<tr><td>>MI0000441|hsa-mir-30b&WithFlank&8|-|134800514|134800613@23.44.22&[MIMAT0000420&hsa-miR-30b-5p&offsets|0|0;m-76&8|-|134800570|134800591&offsets|0|0]|TGTAAACATCCTACACTCAGCT</td><td>>MI0000441|hsa-mir-30b&WithFlank&8|-|134800514|134800613</td><td>23</td><td>44</td><td>22</td><td>TGTAAACATCCTACACTCAGCT</td><td>1249.6</td><td>6.0</td><td>1.0</td><td>5.8</td><td>6.2E-09</td><td>1.3E-08</td><td>MB231/MB468</td><td>CYTO</td></tr>
<tr><td>>hg38_21_+_8433172_8446622_RNA45SN1_45S_50ntFlanks@10771.10796.26|CAGGTCTCCAAGGTGAACAGCCTCTG</td><td>>hg38_21_+_8433172_8446622_RNA45SN1_45S_50ntFlanks</td><td>10771</td><td>10796</td><td>26</td><td>CAGGTCTCCAAGGTGAACAGCCTCTG</td><td>523.9</td><td>-8.1</td><td>1.3</td><td>-6.1</td><td>8.1E-10</td><td>1.3E-08</td><td>BT20/MB468</td><td>MP</td></tr>
<tr><td>>MI0000446|hsa-mir-125b-1&WithFlank&11|-|122099751|122099850@21.41.21(+1U);MI0000470|hsa-mir-125b-2&WithFlank&21|+|16590231|16590331@23.43.21(+1U)&[MIMAT0000423&hsa-miR-125b-5p&offsets|0|-1(+1U);m-67&11|-|122099809|122099830&offsets|0|-1(+1U)];[MIMAT0000423_1&hsa-miR-125b-5p&offsets|0|-1(+1U);m-68&21|+|16590253|16590274&offsets|0|-1(+1U)]|TCCCTGAGACCCTAACTTGTGT</td><td>>MI0000446|hsa-mir-125b-1&WithFlank&11|-|122099751|122099850</td><td>21</td><td>41</td><td>21(+1U)</td><td>TCCCTGAGACCCTAACTTGTGT</td><td>1435.9</td><td>9.6</td><td>1.7</td><td>5.8</td><td>6.8E-09</td><td>1.3E-08</td><td>MB231/MB468</td><td>MITO</td></tr>
<tr><td>>MI0000434|hsa-let-7i&WithFlank&12|+|62603680|62603775@12.33.22(+1A)&[MIMAT0000415&hsa-let-7i-5p&offsets|0|0(+1A);m-49&12|+|62603691|62603712&offsets|0|0(+1A)]|TGAGGTAGTAGTTTGTGCTGTTA</td><td>>MI0000434|hsa-let-7i&WithFlank&12|+|62603680|62603775</td><td>12</td><td>33</td><td>22(+1A)</td><td>TGAGGTAGTAGTTTGTGCTGTTA</td><td>1956.3</td><td>4.7</td><td>0.8</td><td>5.8</td><td>6.2E-09</td><td>1.3E-08</td><td>MB231/MB468</td><td>CYTO</td></tr>
<tr><td>>hg38_21_+_8433172_8446622_RNA45SN1_45S_50ntFlanks@4983.5001.19|CGGACAGGATTGACAGATT</td><td>>hg38_21_+_8433172_8446622_RNA45SN1_45S_50ntFlanks</td><td>4983</td><td>5001</td><td>19</td><td>CGGACAGGATTGACAGATT</td><td>339.1</td><td>6.4</td><td>1.1</td><td>5.8</td><td>5.4E-09</td><td>1.3E-08</td><td>BT20/MB468</td><td>CYTO</td></tr>
<tr><td>>hg38_MT_+_1621_3279_RNR2_16S_50ntFlanks@1416.1453.38|GTTTGTTCAACGATTAAAGTCCTACGTGATCTGAGTTC</td><td>>hg38_MT_+_1621_3279_RNR2_16S_50ntFlanks</td><td>1416</td><td>1453</td><td>38</td><td>GTTTGTTCAACGATTAAAGTCCTACGTGATCTGAGTTC</td><td>66.3</td><td>9.3</td><td>1.6</td><td>5.8</td><td>5.9E-09</td><td>1.3E-08</td><td>MB231/MB468</td><td>MP</td></tr>
<tr><td>>hg38_21_+_8433172_8446622_RNA45SN1_45S_50ntFlanks@9004.9026.23|TCTCCCCCGCGGGGGCGCGCCGG</td><td>>hg38_21_+_8433172_8446622_RNA45SN1_45S_50ntFlanks</td><td>9004</td><td>9026</td><td>23</td><td>TCTCCCCCGCGGGGGCGCGCCGG</td><td>3373.7</td><td>-8.7</td><td>1.5</td><td>-5.9</td><td>4.7E-09</td><td>1.3E-08</td><td>BT20/MB468</td><td>MITO</td></tr>
<tr><td>>hg38_21_+_8433172_8446622_RNA45SN1_45S_50ntFlanks@9852.9879.28|GCGCTGCGGGATGAACCGAACGCCGGGT</td><td>>hg38_21_+_8433172_8446622_RNA45SN1_45S_50ntFlanks</td><td>9852</td><td>9879</td><td>28</td><td>GCGCTGCGGGATGAACCGAACGCCGGGT</td><td>98.2</td><td>-10.5</td><td>1.8</td><td>-5.8</td><td>5.9E-09</td><td>1.3E-08</td><td>MB231/MB468</td><td>MP</td></tr>
<tr><td>>hg38_21_+_8433172_8446622_RNA45SN1_45S_50ntFlanks@10393.10430.38|CAGCAGTTGAACATGGGTCAGTCGGTCCTGAGAGATGG</td><td>>hg38_21_+_8433172_8446622_RNA45SN1_45S_50ntFlanks</td><td>10393</td><td>10430</td><td>38</td><td>CAGCAGTTGAACATGGGTCAGTCGGTCCTGAGAGATGG</td><td>2073.3</td><td>-9.7</td><td>1.7</td><td>-5.8</td><td>7.1E-09</td><td>1.3E-08</td><td>BT20/MB468</td><td>NUC</td></tr>
</table>

<table>
<tr><td>>hg38_21_+_8433172_8446622_RNA45SN1_45S_50ntFlanks@9570.9594.25|TGCAAATCGGTCGTCCGACCTGGGT</td><td>>hg38_21_+_8433172_8446622_RNA45SN1_45S_50ntFlanks</td><td>9570</td><td>9594</td><td>25</td><td>TGCAAATCGGTCGTCCGACCTGGGT</td><td>415.5</td><td>-5.5</td><td>1.0</td><td>-5.8</td><td>6.3E-09</td><td>1.3E-08</td><td>MB231/MB468</td><td>CYTO</td></tr>
<tr><td>>hg38_21_+_8433172_8446622_RNA45SN1_45S_50ntFlanks@11999.12034.36|CCCTGCGGGCCGCCGGTGAAATACCACTACTCTGAT</td><td>>hg38_21_+_8433172_8446622_RNA45SN1_45S_50ntFlanks</td><td>11999</td><td>12034</td><td>36</td><td>CCCTGCGGGCCGCCGGTGAAATACCACTACTCTGAT</td><td>646.3</td><td>-8.9</td><td>1.5</td><td>-5.8</td><td>7.2E-09</td><td>1.3E-08</td><td>BT20/MB468</td><td>NUC</td></tr>
<tr><td>>hg38_21_+_8433172_8446622_RNA45SN1_45S_50ntFlanks@10771.10807.37|CAGGTCTCCAAGGTGAACAGCCTCTGGCATGTTGGAA</td><td>>hg38_21_+_8433172_8446622_RNA45SN1_45S_50ntFlanks</td><td>10771</td><td>10807</td><td>37</td><td>CAGGTCTCCAAGGTGAACAGCCTCTGGCATGTTGGAA</td><td>98.5</td><td>-8.7</td><td>1.5</td><td>-5.9</td><td>4.8E-09</td><td>1.3E-08</td><td>BT20/MB468</td><td>MITO</td></tr>
<tr><td>>hg38_21_+_8433172_8446622_RNA45SN1_45S_50ntFlanks@4760.4779.20|TCAGATACCGTCGTAGTTCC</td><td>>hg38_21_+_8433172_8446622_RNA45SN1_45S_50ntFlanks</td><td>4760</td><td>4779</td><td>20</td><td>TCAGATACCGTCGTAGTTCC</td><td>502.6</td><td>-3.1</td><td>0.5</td><td>-5.8</td><td>6.3E-09</td><td>1.3E-08</td><td>MB231/MB468</td><td>CYTO</td></tr>
<tr><td>>MI0000264|hsa-mir-7-2&WithFlank&15|+|88611819|88611940@38.61.24(+1A);MI0000265|hsa-mir-7-3&WithFlank&19|+|4770664|4770785@37.60.24(+1A);MI0000263|hsa-mir-7-1&WithFlank&9|-|83969742|83969863@30.53.24(+1A)&[MIMAT0000252&hsa-miR-7-5p&offsets|0|0(+1A);m-400&15|+|88611856|88611879&offsets|0|0(+1A)];[MIMAT0000252_1&hsa-miR-7-5p&offsets|0|0(+1A);m-402&19|+|4770700|4770723&offsets|0|0(+1A)];[MIMAT0000252_2&hsa-miR-7-5p&offsets|0|0(+1A);m-401&9|-|83969811|83969834&offsets|0|0(+1A)]|TGGAAGACTAGTGATTTTGTTGTTA</td><td>>MI0000264|hsa-mir-7-2&WithFlank&15|+|88611819|88611940</td><td>38</td><td>61</td><td>24(+1A)</td><td>TGGAAGACTAGTGATTTTGTTGTTA</td><td>552.0</td><td>5.8</td><td>1.0</td><td>5.8</td><td>6.9E-09</td><td>1.3E-08</td><td>MB231/MB468</td><td>MITO</td></tr>
<tr><td>>hg38_21_+_8433172_8446622_RNA45SN1_45S_50ntFlanks@3813.3837.25|TTATGGTTCCTTTGGTCGCTCGCTC</td><td>>hg38_21_+_8433172_8446622_RNA45SN1_45S_50ntFlanks</td><td>3813</td><td>3837</td><td>25</td><td>TTATGGTTCCTTTGGTCGCTCGCTC</td><td>319.7</td><td>-7.5</td><td>1.3</td><td>-5.8</td><td>6.9E-09</td><td>1.3E-08</td><td>MB231/MB468</td><td>MITO</td></tr>
<tr><td>>hg38_21_+_8433172_8446622_RNA45SN1_45S_50ntFlanks@4047.4072.26|ATTCGAACGTCTGCCCTATCAACTTT</td><td>>hg38_21_+_8433172_8446622_RNA45SN1_45S_50ntFlanks</td><td>4047</td><td>4072</td><td>26</td><td>ATTCGAACGTCTGCCCTATCAACTTT</td><td>448.5</td><td>-8.4</td><td>1.4</td><td>-5.8</td><td>6.9E-09</td><td>1.3E-08</td><td>MB231/MB468</td><td>MITO</td></tr>
<tr><td>>hg38_21_+_8433172_8446622_RNA45SN1_45S_50ntFlanks@12044.12063.20|ACTGACCCGGTGAGGCGGGG</td><td>>hg38_21_+_8433172_8446622_RNA45SN1_45S_50ntFlanks</td><td>12044</td><td>12063</td><td>20</td><td>ACTGACCCGGTGAGGCGGGG</td><td>254.0</td><td>-11.1</td><td>1.8</td><td>-6.1</td><td>8.2E-10</td><td>1.3E-08</td><td>BT20/MB468</td><td>MP</td></tr>
<tr><td>>hg38_21_+_8433172_8446622_RNA45SN1_45S_50ntFlanks@6667.6696.30|ATCACTCGGCTCGTGCGTCGATGAAGAACG</td><td>>hg38_21_+_8433172_8446622_RNA45SN1_45S_50ntFlanks</td><td>6667</td><td>6696</td><td>30</td><td>ATCACTCGGCTCGTGCGTCGATGAAGAACG</td><td>114.4</td><td>-9.9</td><td>1.6</td><td>-6.1</td><td>8.2E-10</td><td>1.3E-08</td><td>BT20/MB468</td><td>MP</td></tr>
<tr><td>>trnaMT_PheGAA_MT_+_577_647@39.62.24|ATGTTTAGACGGGCTCACATCACC</td><td>>trnaMT_PheGAA_MT_+_577_647</td><td>39</td><td>62</td><td>24</td><td>ATGTTTAGACGGGCTCACATCACC</td><td>736.5</td><td>4.9</td><td>0.8</td><td>5.8</td><td>6.9E-09</td><td>1.3E-08</td><td>MB231/MB468</td><td>MITO</td></tr>
<tr><td>>hg38_21_+_8433172_8446622_RNA45SN1_45S_50ntFlanks@12289.12318.30|CCGTGAAAGCGGGGCCTCACGATCCTTCTG</td><td>>hg38_21_+_8433172_8446622_RNA45SN1_45S_50ntFlanks</td><td>12289</td><td>12318</td><td>30</td><td>CCGTGAAAGCGGGGCCTCACGATCCTTCTG</td><td>66.6</td><td>-9.9</td><td>1.7</td><td>-5.8</td><td>6.0E-09</td><td>1.3E-08</td><td>MB231/MB468</td><td>MP</td></tr>
<tr><td>>hg38_21_+_8433172_8446622_RNA45SN1_45S_50ntFlanks@12627.12658.32|ATGGGGCGAAGCTACCATCTGTGGGATTATGA</td><td>>hg38_21_+_8433172_8446622_RNA45SN1_45S_50ntFlanks</td><td>12627</td><td>12658</td><td>32</td><td>ATGGGGCGAAGCTACCATCTGTGGGATTATGA</td><td>57.9</td><td>-9.7</td><td>1.7</td><td>-5.8</td><td>6.0E-09</td><td>1.3E-08</td><td>MB231/MB468</td><td>MP</td></tr>
<tr><td>>hg38_21_+_8433172_8446622_RNA45SN1_45S_50ntFlanks@6787.6807.21|ACGCCTGTCTGAGCGTCGCTT</td><td>>hg38_21_+_8433172_8446622_RNA45SN1_45S_50ntFlanks</td><td>6787</td><td>6807</td><td>21</td><td>ACGCCTGTCTGAGCGTCGCTT</td><td>706.6</td><td>4.7</td><td>0.8</td><td>5.9</td><td>4.8E-09</td><td>1.3E-08</td><td>BT20/MB468</td><td>MITO</td></tr>
<tr><td>>am_trna162_MetCAT_6_-_26758550_26758622@31.53.23,trna164_MetCAT_6_-_26735574_26735646@31.53.23,trna20_MetCAT_16_+_71460396_71460468@31.53.23,trna21_MetCAT_6_+_26701712_26701784@31.53.23,trna27_MetCAT_6_+_26766444_26766516@31.53.23,trna75_MetCAT_6_+_28912352_28912424@31.53.23,trna97_MetCAT_6_-_28921042_28921114@31.53.23|TCTCATAATCTGAAGGTCCTGAG</td><td>>am_trna162_MetCAT_6_-_26758550_26758622</td><td>31</td><td>53</td><td>23</td><td>TCTCATAATCTGAAGGTCCTGAG</td><td>93.1</td><td>-8.6</td><td>1.5</td><td>-5.9</td><td>4.8E-09</td><td>1.3E-08</td><td>BT20/MB468</td><td>MITO</td></tr>
<tr><td>>hg38_21_+_8433172_8446622_RNA45SN1_45S_50ntFlanks@4560.4579.20|CTAGGAATAATGGAATAGGA</td><td>>hg38_21_+_8433172_8446622_RNA45SN1_45S_50ntFlanks</td><td>4560</td><td>4579</td><td>20</td><td>CTAGGAATAATGGAATAGGA</td><td>449.4</td><td>-3.9</td><td>0.7</td><td>-5.8</td><td>6.3E-09</td><td>1.3E-08</td><td>MB231/MB468</td><td>CYTO</td></tr>
<tr><td>>hg38_21_+_8433172_8446622_RNA45SN1_45S_50ntFlanks@12528.12546.19|TTACCCTACTGATGATGTG</td><td>>hg38_21_+_8433172_8446622_RNA45SN1_45S_50ntFlanks</td><td>12528</td><td>12546</td><td>19</td><td>TTACCCTACTGATGATGTG</td><td>76.3</td><td>-10.1</td><td>1.7</td><td>-5.8</td><td>6.0E-09</td><td>1.3E-08</td><td>MB231/MB468</td><td>MP</td></tr>
<tr><td>>hg38_21_+_8433172_8446622_RNA45SN1_45S_50ntFlanks@10783.10817.35|GTGAACAGCCTCTGGCATGTTGGAACAATGTAGGT</td><td>>hg38_21_+_8433172_8446622_RNA45SN1_45S_50ntFlanks</td><td>10783</td><td>10817</td><td>35</td><td>GTGAACAGCCTCTGGCATGTTGGAACAATGTAGGT</td><td>278.7</td><td>-5.8</td><td>1.0</td><td>-5.8</td><td>6.9E-09</td><td>1.3E-08</td><td>MB231/MB468</td><td>MITO</td></tr>
<tr><td>>hg38_21_+_8433172_8446622_RNA45SN1_45S_50ntFlanks@12728.12747.20|CTCGGATAGCCGGTCCCCCG</td><td>>hg38_21_+_8433172_8446622_RNA45SN1_45S_50ntFlanks</td><td>12728</td><td>12747</td><td>20</td><td>CTCGGATAGCCGGTCCCCCG</td><td>205.9</td><td>-7.5</td><td>1.3</td><td>-5.8</td><td>6.9E-09</td><td>1.3E-08</td><td>MB231/MB468</td><td>MITO</td></tr>
<tr><td>>hg38_21_+_8433172_8446622_RNA45SN1_45S_50ntFlanks@10680.10708.29|GTTCCGGCGGCGTCCGGTGAGCTCTCGCT</td><td>>hg38_21_+_8433172_8446622_RNA45SN1_45S_50ntFlanks</td><td>10680</td><td>10708</td><td>29</td><td>GTTCCGGCGGCGTCCGGTGAGCTCTCGCT</td><td>16444.7</td><td>-4.4</td><td>0.7</td><td>-5.9</td><td>4.8E-09</td><td>1.3E-08</td><td>BT20/MB468</td><td>MITO</td></tr>
<tr><td>>hg38_21_+_8433172_8446622_RNA45SN1_45S_50ntFlanks@11892.11922.31|TGACTCTAGTCTGGCACGGTGAAGAGACATG</td><td>>hg38_21_+_8433172_8446622_RNA45SN1_45S_50ntFlanks</td><td>11892</td><td>11922</td><td>31</td><td>TGACTCTAGTCTGGCACGGTGAAGAGACATG</td><td>433.5</td><td>9.2</td><td>1.6</td><td>5.9</td><td>4.8E-09</td><td>1.3E-08</td><td>BT20/MB468</td><td>MITO</td></tr>
<tr><td>>trna10_AspGTC_12_-_125424193_125424264@38.74.37,trna12_AspGTC_12_-_125411891_125411962@38.74.37,trna144_AspGTC_6_-_27551236_27551307@38.74.37,trna38_AspGTC_17_-_8125556_8125627@38.74.37,trna45_AspGTC_6_+_27447453_27447524@38.74.37,trna48_AspGTC_6_+_27471523_27471594@38.74.37,trna4_AspGTC_12_+_96429799_96429870@38.74.37,trna69_AspGTC_1_-_161440205_161440276@38.74.37,trna72_AspGTC_1_-_161432824_161432895@38.74.37,trna75_AspGTC_1_-_161425414_161425485@38.74.37,trna78_AspGTC_1_-_161418033_161418104@38.74.37,trna81_AspGTC_1_-_161410615_161410686@38.74.37|CGCGGGAGACCGGGGTTCGATTCCCCGACGGGGAGCC</td><td>>trna10_AspGTC_12_-_125424193_125424264</td><td>38</td><td>74</td><td>37</td><td>CGCGGGAGACCGGGGTTCGATTCCCCGACGGGGAGCC</td><td>327.4</td><td>4.6</td><td>0.8</td><td>5.8</td><td>5.4E-09</td><td>1.3E-08</td><td>BT20/MB468</td><td>CYTO</td></tr>
<tr><td>>hg38_21_+_8433172_8446622_RNA45SN1_45S_50ntFlanks@10771.10807.37|CAGGTCTCCAAGGTGAACAGCCTCTGGCATGTTGGAA</td><td>>hg38_21_+_8433172_8446622_RNA45SN1_45S_50ntFlanks</td><td>10771</td><td>10807</td><td>37</td><td>CAGGTCTCCAAGGTGAACAGCCTCTGGCATGTTGGAA</td><td>110.9</td><td>-10.7</td><td>1.8</td><td>-5.8</td><td>6.0E-09</td><td>1.3E-08</td><td>MB231/MB468</td><td>MP</td></tr>
<tr><td>>hg38_21_+_8433172_8446622_RNA45SN1_45S_50ntFlanks@6656.6681.26|CTTAGCGGTGGATCACTCGGCTCGTG</td><td>>hg38_21_+_8433172_8446622_RNA45SN1_45S_50ntFlanks</td><td>6656</td><td>6681</td><td>26</td><td>CTTAGCGGTGGATCACTCGGCTCGTG</td><td>1760.7</td><td>-3.5</td><td>0.6</td><td>-5.8</td><td>6.4E-09</td><td>1.3E-08</td><td>MB231/MB468</td><td>NUC</td></tr>
<tr><td>>hg38_21_+_8433172_8446622_RNA45SN1_45S_50ntFlanks@11885.11902.18|TTGAGCTTGACTCTAGTC</td><td>>hg38_21_+_8433172_8446622_RNA45SN1_45S_50ntFlanks</td><td>11885</td><td>11902</td><td>18</td><td>TTGAGCTTGACTCTAGTC</td><td>68.2</td><td>-10.0</td><td>1.7</td><td>-5.8</td><td>6.0E-09</td><td>1.3E-08</td><td>MB231/MB468</td><td>MP</td></tr>
<tr><td>>am_trnaMT_ValTAC_MT_+_1602_1670@49.68.20|AACTTAACTTGACCGCTCTG</td><td>>am_trnaMT_ValTAC_MT_+_1602_1670</td><td>49</td><td>68</td><td>20</td><td>AACTTAACTTGACCGCTCTG</td><td>510.1</td><td>-6.0</td><td>1.0</td><td>-5.8</td><td>6.0E-09</td><td>1.3E-08</td><td>MB231/MB468</td><td>MP</td></tr>
<tr><td>>trnaMT_SerGCT_MT_+_12207_12265@1.19.19|GAGAAAGCTCACAAGAACT</td><td>>trnaMT_SerGCT_MT_+_12207_12265</td><td>1</td><td>19</td><td>19</td><td>GAGAAAGCTCACAAGAACT</td><td>1298.5</td><td>5.0</td><td>0.9</td><td>5.8</td><td>6.4E-09</td><td>1.3E-08</td><td>MB231/MB468</td><td>NUC</td></tr>
<tr><td>>hg38_21_+_8433172_8446622_RNA45SN1_45S_50ntFlanks@8319.8356.38|CGAGACCGATAGTCAACAAGTACCGTAAGGGAAAGTTG</td><td>>hg38_21_+_8433172_8446622_RNA45SN1_45S_50ntFlanks</td><td>8319</td><td>8356</td><td>38</td><td>CGAGACCGATAGTCAACAAGTACCGTAAGGGAAAGTTG</td><td>523.4</td><td>-4.4</td><td>0.8</td><td>-5.8</td><td>6.4E-09</td><td>1.3E-08</td><td>MB231/MB468</td><td>CYTO</td></tr>
<tr><td>>hg38_21_+_8433172_8446622_RNA45SN1_45S_50ntFlanks@12416.12445.30|TTCGATGTCGGCTCTTCCTATCATTGTGAA</td><td>>hg38_21_+_8433172_8446622_RNA45SN1_45S_50ntFlanks</td><td>12416</td><td>12445</td><td>30</td><td>TTCGATGTCGGCTCTTCCTATCATTGTGAA</td><td>1241.4</td><td>3.7</td><td>0.6</td><td>5.8</td><td>6.4E-09</td><td>1.3E-08</td><td>MB231/MB468</td><td>CYTO</td></tr>
<tr><td>>hg38_21_+_8433172_8446622_RNA45SN1_45S_50ntFlanks@12476.12493.18|CCACTAATAGGGAACGTG</td><td>>hg38_21_+_8433172_8446622_RNA45SN1_45S_50ntFlanks</td><td>12476</td><td>12493</td><td>18</td><td>CCACTAATAGGGAACGTG</td><td>452.9</td><td>-4.9</td><td>0.8</td><td>-5.8</td><td>6.4E-09</td><td>1.3E-08</td><td>MB231/MB468</td><td>CYTO</td></tr>
<tr><td>>MI0000456|hsa-mir-140&WithFlank&16|+|69933075|69933186@69.88.20&[MIMAT0004597&hsa-miR-140-3p&offsets|+1|0;m-45&16|+|69933143|69933164&offsets|0|-2]|ACCACAGGGTAGAACCACGG</td><td>>MI0000456|hsa-mir-140&WithFlank&16|+|69933075|69933186</td><td>69</td><td>88</td><td>20</td><td>ACCACAGGGTAGAACCACGG</td><td>592.3</td><td>6.3</td><td>1.1</td><td>5.8</td><td>6.0E-09</td><td>1.3E-08</td><td>MB231/MB468</td><td>MP</td></tr>
<tr><td>>hg38_21_+_8433172_8446622_RNA45SN1_45S_50ntFlanks@12970.12990.21|GTTTCGTACGTAGCAGAGCAG</td><td>>hg38_21_+_8433172_8446622_RNA45SN1_45S_50ntFlanks</td><td>12970</td><td>12990</td><td>21</td><td>GTTTCGTACGTAGCAGAGCAG</td><td>65.2</td><td>-9.9</td><td>1.7</td><td>-5.8</td><td>6.0E-09</td><td>1.3E-08</td><td>MB231/MB468</td><td>MP</td></tr>
<tr><td>>hg38_21_+_8433172_8446622_RNA45SN1_45S_50ntFlanks@10024.10042.19|GCGCTGGAGCGTCGGGCCC</td><td>>hg38_21_+_8433172_8446622_RNA45SN1_45S_50ntFlanks</td><td>10024</td><td>10042</td><td>19</td><td>GCGCTGGAGCGTCGGGCCC</td><td>352.5</td><td>8.3</td><td>1.4</td><td>5.8</td><td>6.4E-09</td><td>1.3E-08</td><td>MB231/MB468</td><td>CYTO</td></tr>
<tr><td>>hg38_21_+_8433172_8446622_RNA45SN1_45S_50ntFlanks@12407.12432.26|TTTTGATCCTTCGATGTCGGCTCTTC</td><td>>hg38_21_+_8433172_8446622_RNA45SN1_45S_50ntFlanks</td><td>12407</td><td>12432</td><td>26</td><td>TTTTGATCCTTCGATGTCGGCTCTTC</td><td>461.1</td><td>-4.7</td><td>0.8</td><td>-5.8</td><td>7.0E-09</td><td>1.3E-08</td><td>MB231/MB468</td><td>MITO</td></tr>
<tr><td>>MI0000746|hsa-mir-99b&WithFlank&19|+|51692606|51692687@13.34.22&[MIMAT0000689&hsa-miR-99b-5p&offsets|0|0;m-10&19|+|51692618|51692639&offsets|0|0]|CACCCGTAGAACCGACCTTGCG</td><td>>MI0000746|hsa-mir-99b&WithFlank&19|+|51692606|51692687</td><td>13</td><td>34</td><td>22</td><td>CACCCGTAGAACCGACCTTGCG</td><td>13416.7</td><td>4.5</td><td>0.8</td><td>5.8</td><td>7.0E-09</td><td>1.3E-08</td><td>MB231/MB468</td><td>MITO</td></tr>
<tr><td>>MI0000298|hsa-mir-221&WithFlank&X|-|45746151|45746272@71.92.22(+2U)&[MIMAT0000278&hsa-miR-221-3p&offsets|0|-1(+2U);m-91&X|-|45746181|45746202&offsets|0|0(+2U)]|AGCTACATTGTCTGCTGGGTTTTT</td><td>>MI0000298|hsa-mir-221&WithFlank&X|-|45746151|45746272</td><td>71</td><td>92</td><td>22(+2U)</td><td>AGCTACATTGTCTGCTGGGTTTTT</td><td>619.4</td><td>6.8</td><td>1.2</td><td>5.8</td><td>7.0E-09</td><td>1.3E-08</td><td>MB231/MB468</td><td>MITO</td></tr>
<tr><td>>MI0000289|hsa-mir-181a-1&WithFlank&1|-|198859038|198859159@30.49.20;MI0000269|hsa-mir-181a-2&WithFlank&9|+|124692436|124692557@45.64.20&[MIMAT0000256&hsa-miR-181a-5p&offsets|0|-3;m-32&1|-|198859108|198859130&offsets|0|-3];[MIMAT0000256_1&hsa-miR-181a-5p&offsets|0|-3;m-32&9|+|124692480|124692502&offsets|0|-3]|AACATTCAACGCTGTCGGTG</td><td>>MI0000289|hsa-mir-181a-1&WithFlank&1|-|198859038|198859159</td><td>30</td><td>49</td><td>20</td><td>AACATTCAACGCTGTCGGTG</td><td>646.8</td><td>5.4</td><td>0.9</td><td>5.8</td><td>6.5E-09</td><td>1.3E-08</td><td>MB231/MB468</td><td>NUC</td></tr>
<tr><td>>hg38_21_+_8433172_8446622_RNA45SN1_45S_50ntFlanks@8198.8222.25|TGGACGGTGTGAGGCCGGTAGCGGC</td><td>>hg38_21_+_8433172_8446622_RNA45SN1_45S_50ntFlanks</td><td>8198</td><td>8222</td><td>25</td><td>TGGACGGTGTGAGGCCGGTAGCGGC</td><td>274.6</td><td>-7.8</td><td>1.3</td><td>-5.8</td><td>6.1E-09</td><td>1.3E-08</td><td>MB231/MB468</td><td>MP</td></tr>
<tr><td>>hg38_21_+_8433172_8446622_RNA45SN1_45S_50ntFlanks@11161.11189.29|CACGGTCCCCCGCGAGGGGGGCCCGGGCA</td><td>>hg38_21_+_8433172_8446622_RNA45SN1_45S_50ntFlanks</td><td>11161</td><td>11189</td><td>29</td><td>CACGGTCCCCCGCGAGGGGGGCCCGGGCA</td><td>299.7</td><td>-11.3</td><td>1.8</td><td>-6.1</td><td>8.4E-10</td><td>1.3E-08</td><td>BT20/MB468</td><td>MP</td></tr>
<tr><td>>trnaMT_PheGAA_MT_+_577_647@42.74.33|TTTAGACGGGCTCACATCACCCCATAAACACCA</td><td>>trnaMT_PheGAA_MT_+_577_647</td><td>42</td><td>74</td><td>33</td><td>TTTAGACGGGCTCACATCACCCCATAAACACCA</td><td>970.7</td><td>5.9</td><td>1.0</td><td>5.9</td><td>4.9E-09</td><td>1.3E-08</td><td>BT20/MB468</td><td>MITO</td></tr>
<tr><td>>hg38_21_+_8433172_8446622_RNA45SN1_45S_50ntFlanks@11833.11872.40|ACCACAGCCAAGGGAACGGGCTTGGCGGAATCAGCGGGGA</td><td>>hg38_21_+_8433172_8446622_RNA45SN1_45S_50ntFlanks</td><td>11833</td><td>11872</td><td>40</td><td>ACCACAGCCAAGGGAACGGGCTTGGCGGAATCAGCGGGGA</td><td>64.4</td><td>-9.9</td><td>1.7</td><td>-5.8</td><td>6.1E-09</td><td>1.3E-08</td><td>MB231/MB468</td><td>MP</td></tr>
<tr><td>>hg38_21_+_8433172_8446622_RNA45SN1_45S_50ntFlanks@10845.10864.20|ACTTCGGGATAAGGATTGGC</td><td>>hg38_21_+_8433172_8446622_RNA45SN1_45S_50ntFlanks</td><td>10845</td><td>10864</td><td>20</td><td>ACTTCGGGATAAGGATTGGC</td><td>710.8</td><td>-7.8</td><td>1.3</td><td>-5.8</td><td>6.5E-09</td><td>1.3E-08</td><td>MB231/MB468</td><td>NUC</td></tr>
<tr><td>>hg38_21_+_8433172_8446622_RNA45SN1_45S_50ntFlanks@4785.4812.28|TAAACGATGCCGACCGGCGATGCGGCGG</td><td>>hg38_21_+_8433172_8446622_RNA45SN1_45S_50ntFlanks</td><td>4785</td><td>4812</td><td>28</td><td>TAAACGATGCCGACCGGCGATGCGGCGG</td><td>65.2</td><td>-9.9</td><td>1.7</td><td>-5.8</td><td>6.1E-09</td><td>1.3E-08</td><td>MB231/MB468</td><td>MP</td></tr>
<tr><td>>hg38_21_+_8433172_8446622_RNA45SN1_45S_50ntFlanks@4724.4753.30|ATTAATCAAGAACGAAAGTCGGAGGTTCGA</td><td>>hg38_21_+_8433172_8446622_RNA45SN1_45S_50ntFlanks</td><td>4724</td><td>4753</td><td>30</td><td>ATTAATCAAGAACGAAAGTCGGAGGTTCGA</td><td>151.2</td><td>-9.3</td><td>1.6</td><td>-5.9</td><td>4.9E-09</td><td>1.4E-08</td><td>BT20/MB468</td><td>MITO</td></tr>
<tr><td>>hg38_21_+_8433172_8446622_RNA45SN1_45S_50ntFlanks@10786.10822.37|AACAGCCTCTGGCATGTTGGAACAATGTAGGTAAGGG</td><td>>hg38_21_+_8433172_8446622_RNA45SN1_45S_50ntFlanks</td><td>10786</td><td>10822</td><td>37</td><td>AACAGCCTCTGGCATGTTGGAACAATGTAGGTAAGGG</td><td>1171.1</td><td>-3.6</td><td>0.6</td><td>-5.9</td><td>4.9E-09</td><td>1.4E-08</td><td>BT20/MB468</td><td>MITO</td></tr>
<tr><td>>hg38_21_+_8433172_8446622_RNA45SN1_45S_50ntFlanks@11161.11179.19|CACGGTCCCCCGCGAGGGG</td><td>>hg38_21_+_8433172_8446622_RNA45SN1_45S_50ntFlanks</td><td>11161</td><td>11179</td><td>19</td><td>CACGGTCCCCCGCGAGGGG</td><td>441.9</td><td>-7.0</td><td>1.2</td><td>-5.9</td><td>4.9E-09</td><td>1.4E-08</td><td>BT20/MB468</td><td>MITO</td></tr>
<tr><td>>hg38_21_+_8433172_8446622_RNA45SN1_45S_50ntFlanks@12244.12265.22|AAGGGCAAAAGCTCGCTTGATC</td><td>>hg38_21_+_8433172_8446622_RNA45SN1_45S_50ntFlanks</td><td>12244</td><td>12265</td><td>22</td><td>AAGGGCAAAAGCTCGCTTGATC</td><td>179.2</td><td>-6.0</td><td>1.0</td><td>-5.8</td><td>6.1E-09</td><td>1.4E-08</td><td>MB231/MB468</td><td>MP</td></tr>
</table>

<table>
<tr><td></td><td></td><td></td><td></td><td></td><td></td><td></td><td></td><td></td><td></td><td></td><td></td><td></td><td></td></tr>
<tr><td>>hg38_21_+_8433172_8446622_RNA45SN1_45S_50ntFlanks@4445.4470.26|CTTGCCTCTCGGCGCCCCCTCGATGC</td><td>>hg38_21_+_8433172_8446622_RNA45SN1_45S_50ntFlanks</td><td>4445</td><td>4470</td><td>26</td><td>CTTGCCTCTCGGCGCCCCCTCGATGC</td><td>356.5</td><td>-7.8</td><td>1.4</td><td>-5.8</td><td>6.6E-09</td><td>1.4E-08</td><td>MB231/MB468</td><td>NUC</td></tr>
<tr><td>>hg38_21_+_8433172_8446622_RNA45SN1_45S_50ntFlanks@10411.10456.46|CAGTCGGTCCTGAGAGATGGGCGAGCGCCGTTCCGAAGGGACGGGC</td><td>>hg38_21_+_8433172_8446622_RNA45SN1_45S_50ntFlanks</td><td>10411</td><td>10456</td><td>46</td><td>CAGTCGGTCCTGAGAGATGGGCGAGCGCCGTTCCGAAGGGACGGGC</td><td>510.4</td><td>-6.2</td><td>1.1</td><td>-5.8</td><td>6.5E-09</td><td>1.4E-08</td><td>MB231/MB468</td><td>CYTO</td></tr>
<tr><td>>hg38_21_+_8433172_8446622_RNA45SN1_45S_50ntFlanks@11718.11751.34|AACGGCGGGAGTAACTATGACTCTCTTAAGGTAG</td><td>>hg38_21_+_8433172_8446622_RNA45SN1_45S_50ntFlanks</td><td>11718</td><td>11751</td><td>34</td><td>AACGGCGGGAGTAACTATGACTCTCTTAAGGTAG</td><td>713.2</td><td>-5.1</td><td>0.9</td><td>-5.8</td><td>5.5E-09</td><td>1.4E-08</td><td>BT20/MB468</td><td>CYTO</td></tr>
<tr><td>>MI0001445|hsa-mir-423&WithFlank&17|+|30117073|30117178@58.77.20(+1U)&[MIMAT0001340&hsa-miR-423-3p&offsets|-1|-4(+1U);m-100&17|+|30117131|30117153&offsets|-1|-4(+1U)]|AAGCTCGGTCTGAGGCCCCTT</td><td>>MI0001445|hsa-mir-423&WithFlank&17|+|30117073|30117178</td><td>58</td><td>77</td><td>20(+1U)</td><td>AAGCTCGGTCTGAGGCCCCTT</td><td>142.1</td><td>10.9</td><td>1.8</td><td>6.1</td><td>8.5E-10</td><td>1.4E-08</td><td>BT20/MB468</td><td>MP</td></tr>
<tr><td>>hg38_21_+_8433172_8446622_RNA45SN1_45S_50ntFlanks@3705.3739.35|TACCTGGTTGATCCTGCCAGTAGCATATGCTTGTC</td><td>>hg38_21_+_8433172_8446622_RNA45SN1_45S_50ntFlanks</td><td>3705</td><td>3739</td><td>35</td><td>TACCTGGTTGATCCTGCCAGTAGCATATGCTTGTC</td><td>1640.3</td><td>3.3</td><td>0.5</td><td>6.2</td><td>6.7E-10</td><td>1.4E-08</td><td>BT20/MB231</td><td>NUC</td></tr>
<tr><td>>am_trna10_ValCAC_5_-_180649395_180649467@1.29.29,trna12_ValAAC_5_-_180645270_180645342@1.29.29,trna132_ValAAC_6_-_27721179_27721251@1.29.29,trna136_ValAAC_6_-_27648885_27648957@1.29.29,trna139_ValAAC_6_-_27618707_27618779@1.29.29,trna18_ValCAC_5_-_180529253_180529325@1.29.29,trna2_ValAAC_3_+_169490018_169490090@1.29.29,trna2_ValCAC_5_+_180524070_180524142@1.29.29,trna4_ValAAC_5_+_180591154_180591226@1.29.29,trna5_ValAAC_5_+_180596610_180596682@1.29.29,trna6_ValCAC_5_+_180600650_180600722@1.29.29,trna85_ValCAC_1_-_161369490_161369562@1.29.29,trna90_ValCAC_1_-_149684088_149684161@1.29.29,trna98_ValCAC_1_-_149298555_149298627@1.29.29,trna9_ValCAC_6_+_26538282_26538354@1.29.29|GTTTCCGTAGTGTAGTGGTTATCACGTTC</td><td>>am_trna10_ValCAC_5_-_180649395_180649467</td><td>1</td><td>29</td><td>29</td><td>GTTTCCGTAGTGTAGTGGTTATCACGTTC</td><td>19891.5</td><td>-5.3</td><td>0.9</td><td>-5.8</td><td>5.6E-09</td><td>1.4E-08</td><td>BT20/MB468</td><td>CYTO</td></tr>
<tr><td>>hg38_21_+_8433172_8446622_RNA45SN1_45S_50ntFlanks@4972.4999.28|CGGCCCGGACACGGACAGGATTGACAGA</td><td>>hg38_21_+_8433172_8446622_RNA45SN1_45S_50ntFlanks</td><td>4972</td><td>4999</td><td>28</td><td>CGGCCCGGACACGGACAGGATTGACAGA</td><td>468.9</td><td>-5.5</td><td>1.0</td><td>-5.8</td><td>7.1E-09</td><td>1.4E-08</td><td>MB231/MB468</td><td>MITO</td></tr>
<tr><td>>MI0000299|hsa-mir-222&WithFlank&X|-|45747009|45747130@75.98.24(+1G)&[MIMAT0000279&hsa-miR-222-3p&offsets|0|+3(+1G);m-140&X|-|45747033|45747056&offsets|0|0(+1G)]|AGCTACATCTGGCTACTGGGTCTCG</td><td>>MI0000299|hsa-mir-222&WithFlank&X|-|45747009|45747130</td><td>75</td><td>98</td><td>24(+1G)</td><td>AGCTACATCTGGCTACTGGGTCTCG</td><td>721.3</td><td>5.8</td><td>1.0</td><td>5.8</td><td>7.1E-09</td><td>1.4E-08</td><td>MB231/MB468</td><td>MITO</td></tr>
<tr><td>>hg38_21_+_8433172_8446622_RNA45SN1_45S_50ntFlanks@10677.10711.35|GCGGTTCCGGCGGCGTCCGGTGAGCTCTCGCTGGC</td><td>>hg38_21_+_8433172_8446622_RNA45SN1_45S_50ntFlanks</td><td>10677</td><td>10711</td><td>35</td><td>GCGGTTCCGGCGGCGTCCGGTGAGCTCTCGCTGGC</td><td>1384.4</td><td>-4.0</td><td>0.7</td><td>-5.8</td><td>7.1E-09</td><td>1.4E-08</td><td>MB231/MB468</td><td>MITO</td></tr>
<tr><td>>hg38_21_+_8433172_8446622_RNA45SN1_45S_50ntFlanks@10345.10373.29|TTCAAACGAGAACTTTGAAGGCCGAAGTG</td><td>>hg38_21_+_8433172_8446622_RNA45SN1_45S_50ntFlanks</td><td>10345</td><td>10373</td><td>29</td><td>TTCAAACGAGAACTTTGAAGGCCGAAGTG</td><td>368.8</td><td>4.0</td><td>0.7</td><td>5.8</td><td>5.6E-09</td><td>1.4E-08</td><td>BT20/MB468</td><td>CYTO</td></tr>
<tr><td>>hg38_21_+_8433172_8446622_RNA45SN1_45S_50ntFlanks@12417.12455.39|TCGATGTCGGCTCTTCCTATCATTGTGAAGCAGAATTCA</td><td>>hg38_21_+_8433172_8446622_RNA45SN1_45S_50ntFlanks</td><td>12417</td><td>12455</td><td>39</td><td>TCGATGTCGGCTCTTCCTATCATTGTGAAGCAGAATTCA</td><td>1369.9</td><td>-4.9</td><td>0.8</td><td>-5.8</td><td>5.6E-09</td><td>1.4E-08</td><td>BT20/MB468</td><td>CYTO</td></tr>
<tr><td>>hg38_21_+_8433172_8446622_RNA45SN1_45S_50ntFlanks@3775.3794.20|GCCGGTACAGTGAAACTGCG</td><td>>hg38_21_+_8433172_8446622_RNA45SN1_45S_50ntFlanks</td><td>3775</td><td>3794</td><td>20</td><td>GCCGGTACAGTGAAACTGCG</td><td>189.1</td><td>-9.7</td><td>1.7</td><td>-5.8</td><td>5.0E-09</td><td>1.4E-08</td><td>BT20/MB468</td><td>MITO</td></tr>
<tr><td>>hg38_21_+_8433172_8446622_RNA45SN1_45S_50ntFlanks@12420.12448.29|ATGTCGGCTCTTCCTATCATTGTGAAGCA</td><td>>hg38_21_+_8433172_8446622_RNA45SN1_45S_50ntFlanks</td><td>12420</td><td>12448</td><td>29</td><td>ATGTCGGCTCTTCCTATCATTGTGAAGCA</td><td>1115.8</td><td>-4.7</td><td>0.8</td><td>-5.8</td><td>7.4E-09</td><td>1.4E-08</td><td>BT20/MB468</td><td>NUC</td></tr>
<tr><td>>trna10_AspGTC_12_-_125424193_125424264@38.70.33,trna12_AspGTC_12_-_125411891_125411962@38.70.33,trna144_AspGTC_6_-_27551236_27551307@38.70.33,trna38_AspGTC_17_-_8125556_8125627@38.70.33,trna45_AspGTC_6_+_27447453_27447524@38.70.33,trna48_AspGTC_6_+_27471523_27471594@38.70.33,trna4_AspGTC_12_+_96429799_96429870@38.70.33,trna69_AspGTC_1_-_161440205_161440276@38.70.33,trna72_AspGTC_1_-_161432824_161432895@38.70.33,trna75_AspGTC_1_-_161425414_161425485@38.70.33,trna78_AspGTC_1_-_161418033_161418104@38.70.33,trna81_AspGTC_1_-_161410615_161410686@38.70.33|CGCGGGAGACCGGGGTTCGATTCCCCGACGGGG</td><td>>trna10_AspGTC_12_-_125424193_125424264</td><td>38</td><td>70</td><td>33</td><td>CGCGGGAGACCGGGGTTCGATTCCCCGACGGGG</td><td>2187.1</td><td>-5.0</td><td>0.9</td><td>-5.8</td><td>6.2E-09</td><td>1.4E-08</td><td>MB231/MB468</td><td>MP</td></tr>
<tr><td>>am_MI0000809|hsa-mir-151a&WithFlank&8|-|140732558|140732659@53.73.21(+3U)&[MIMAT0000757&hsa-miR-151a-3p&offsets|0|0(+3U);m-30&8|-|140732587|140732607&offsets|0|0(+3U)]|CTAGACTGAAGCTCCTTGAGGTTT</td><td>>am_MI0000809|hsa-mir-151a&WithFlank&8|-|140732558|140732659</td><td>53</td><td>73</td><td>21(+3U)</td><td>CTAGACTGAAGCTCCTTGAGGTTT</td><td>564.8</td><td>8.1</td><td>1.4</td><td>5.8</td><td>7.1E-09</td><td>1.4E-08</td><td>MB231/MB468</td><td>MITO</td></tr>
<tr><td>>hg38_21_+_8433172_8446622_RNA45SN1_45S_50ntFlanks@11326.11346.21|CTCGTCGCGCGCGCGTCCGCT</td><td>>hg38_21_+_8433172_8446622_RNA45SN1_45S_50ntFlanks</td><td>11326</td><td>11346</td><td>21</td><td>CTCGTCGCGCGCGCGTCCGCT</td><td>148.7</td><td>10.4</td><td>1.8</td><td>5.8</td><td>7.1E-09</td><td>1.4E-08</td><td>MB231/MB468</td><td>MITO</td></tr>
<tr><td>>MI0000440|hsa-mir-27b&WithFlank&9|+|95085439|95085547@67.87.21(+2U)&[MIMAT0000419&hsa-miR-27b-3p&offsets|0|0(+2U);m-39&9|+|95085505|95085525&offsets|0|0(+2U)]|TTCACAGTGGCTAAGTTCTGCTT</td><td>>MI0000440|hsa-mir-27b&WithFlank&9|+|95085439|95085547</td><td>67</td><td>87</td><td>21(+2U)</td><td>TTCACAGTGGCTAAGTTCTGCTT</td><td>119.7</td><td>3.9</td><td>0.7</td><td>5.8</td><td>7.4E-09</td><td>1.4E-08</td><td>BT20/MB468</td><td>NUC</td></tr>
<tr><td>>hg38_21_+_8433172_8446622_RNA45SN1_45S_50ntFlanks@10128.10147.20|CGGCGGCGGCGGCGGCGGCG</td><td>>hg38_21_+_8433172_8446622_RNA45SN1_45S_50ntFlanks</td><td>10128</td><td>10147</td><td>20</td><td>CGGCGGCGGCGGCGGCGGCG</td><td>1261.9</td><td>-7.4</td><td>1.2</td><td>-6.1</td><td>8.6E-10</td><td>1.4E-08</td><td>BT20/MB468</td><td>MP</td></tr>
<tr><td>>hg38_21_+_8433172_8446622_RNA45SN1_45S_50ntFlanks@8845.8864.20|CGGCGGCGGCGGCGGCGGCG</td><td>>hg38_21_+_8433172_8446622_RNA45SN1_45S_50ntFlanks</td><td>8845</td><td>8864</td><td>20</td><td>CGGCGGCGGCGGCGGCGGCG</td><td>1261.9</td><td>-7.4</td><td>1.2</td><td>-6.1</td><td>8.6E-10</td><td>1.4E-08</td><td>BT20/MB468</td><td>MP</td></tr>
<tr><td>>hg38_21_+_8433172_8446622_RNA45SN1_45S_50ntFlanks@10922.10945.24|CTGGACGAGGCGCCGCCGCCCCCC</td><td>>hg38_21_+_8433172_8446622_RNA45SN1_45S_50ntFlanks</td><td>10922</td><td>10945</td><td>24</td><td>CTGGACGAGGCGCCGCCGCCCCCC</td><td>1178.2</td><td>-6.8</td><td>1.2</td><td>-5.8</td><td>6.6E-09</td><td>1.4E-08</td><td>MB231/MB468</td><td>CYTO</td></tr>
<tr><td>>hg38_21_+_8433172_8446622_RNA45SN1_45S_50ntFlanks@9738.9764.27|GAAACGATCTCAACCTATTCTCAAACT</td><td>>hg38_21_+_8433172_8446622_RNA45SN1_45S_50ntFlanks</td><td>9738</td><td>9764</td><td>27</td><td>GAAACGATCTCAACCTATTCTCAAACT</td><td>842.4</td><td>-6.9</td><td>1.2</td><td>-5.8</td><td>7.5E-09</td><td>1.4E-08</td><td>BT20/MB468</td><td>NUC</td></tr>
<tr><td>>trna116_GluCTC_1_-_145399233_145399304@36.74.39,trna134_GluTTC_1_-_16861774_16861845@36.74.39,trna71_GluCTC_1_-_161439189_161439260@36.74.39,trna74_GluCTC_1_-_161431809_161431880@36.74.39,trna77_GluCTC_1_-_161424398_161424469@36.74.39,trna77_GluCTC_6_+_28949976_28950047@36.74.39,trna80_GluCTC_1_-_161417018_161417089@36.74.39,trna84_GluTTC_1_-_161391883_161391954@36.74.39,trna87_GluCTC_6_-_126101393_126101464@36.74.39|CACCGCCGCGGCCCGGGTTCGATTCCCGGTCAGGGAACC</td><td>>trna116_GluCTC_1_-_145399233_145399304</td><td>36</td><td>74</td><td>39</td><td>CACCGCCGCGGCCCGGGTTCGATTCCCGGTCAGGGAACC</td><td>4012.0</td><td>7.7</td><td>1.3</td><td>5.8</td><td>6.6E-09</td><td>1.4E-08</td><td>MB231/MB468</td><td>CYTO</td></tr>
<tr><td>>hg38_21_+_8433172_8446622_RNA45SN1_45S_50ntFlanks@475.511.37|CTTCGTGATCGATGTGGTGACGTCGTGCTCTCCCGGG</td><td>>hg38_21_+_8433172_8446622_RNA45SN1_45S_50ntFlanks</td><td>475</td><td>511</td><td>37</td><td>CTTCGTGATCGATGTGGTGACGTCGTGCTCTCCCGGG</td><td>2113.5</td><td>7.9</td><td>1.4</td><td>5.8</td><td>5.0E-09</td><td>1.4E-08</td><td>BT20/MB468</td><td>MITO</td></tr>
<tr><td>>hg38_21_+_8433172_8446622_RNA45SN1_45S_50ntFlanks@6738.6769.32|ATCATCGACACTTCGAACGCACTTGCGGCCCC</td><td>>hg38_21_+_8433172_8446622_RNA45SN1_45S_50ntFlanks</td><td>6738</td><td>6769</td><td>32</td><td>ATCATCGACACTTCGAACGCACTTGCGGCCCC</td><td>149.7</td><td>-10.3</td><td>1.7</td><td>-6.1</td><td>8.6E-10</td><td>1.4E-08</td><td>BT20/MB468</td><td>MP</td></tr>
<tr><td>>hg38_21_+_8433172_8446622_RNA45SN1_45S_50ntFlanks@3984.4026.43|GCGGCTTTGGTGACTCTAGATAACCTCGGGCCGATCGCACGCC</td><td>>hg38_21_+_8433172_8446622_RNA45SN1_45S_50ntFlanks</td><td>3984</td><td>4026</td><td>43</td><td>GCGGCTTTGGTGACTCTAGATAACCTCGGGCCGATCGCACGCC</td><td>741.4</td><td>-8.1</td><td>1.4</td><td>-5.8</td><td>6.6E-09</td><td>1.4E-08</td><td>MB231/MB468</td><td>CYTO</td></tr>
<tr><td>>hg38_21_+_8433172_8446622_RNA45SN1_45S_50ntFlanks@9570.9595.26|TGCAAATCGGTCGTCCGACCTGGGTA</td><td>>hg38_21_+_8433172_8446622_RNA45SN1_45S_50ntFlanks</td><td>9570</td><td>9595</td><td>26</td><td>TGCAAATCGGTCGTCCGACCTGGGTA</td><td>1918.4</td><td>-4.3</td><td>0.7</td><td>-5.8</td><td>5.0E-09</td><td>1.4E-08</td><td>BT20/MB468</td><td>MITO</td></tr>
<tr><td>>hg38_21_+_8433172_8446622_RNA45SN1_45S_50ntFlanks@8079.8104.26|AAGAGCCCAGCGCCGAATCCCCGCCC</td><td>>hg38_21_+_8433172_8446622_RNA45SN1_45S_50ntFlanks</td><td>8079</td><td>8104</td><td>26</td><td>AAGAGCCCAGCGCCGAATCCCCGCCC</td><td>317.2</td><td>-8.1</td><td>1.4</td><td>-5.8</td><td>7.5E-09</td><td>1.4E-08</td><td>BT20/MB468</td><td>NUC</td></tr>
<tr><td>>hg38_21_+_8433172_8446622_RNA45SN1_45S_50ntFlanks@4724.4747.24|ATTAATCAAGAACGAAAGTCGGAG</td><td>>hg38_21_+_8433172_8446622_RNA45SN1_45S_50ntFlanks</td><td>4724</td><td>4747</td><td>24</td><td>ATTAATCAAGAACGAAAGTCGGAG</td><td>128.3</td><td>-10.1</td><td>1.6</td><td>-6.1</td><td>8.7E-10</td><td>1.4E-08</td><td>BT20/MB468</td><td>MP</td></tr>
<tr><td>>hg38_21_+_8433172_8446622_RNA45SN1_45S_50ntFlanks@11564.11581.18|CCTAGCAGCCGACTTAGA</td><td>>hg38_21_+_8433172_8446622_RNA45SN1_45S_50ntFlanks</td><td>11564</td><td>11581</td><td>18</td><td>CCTAGCAGCCGACTTAGA</td><td>1322.1</td><td>-6.6</td><td>1.1</td><td>-5.8</td><td>6.7E-09</td><td>1.4E-08</td><td>MB231/MB468</td><td>NUC</td></tr>
<tr><td>>hg38_MT_+_598_1651_RNR1_12S_50ntFlanks@51.88.38|AATAGGTTTGGTCCTAGCCTTTCTATTAGCTCTTAGTA</td><td>>hg38_MT_+_598_1651_RNR1_12S_50ntFlanks</td><td>51</td><td>88</td><td>38</td><td>AATAGGTTTGGTCCTAGCCTTTCTATTAGCTCTTAGTA</td><td>1563.4</td><td>5.2</td><td>0.9</td><td>5.8</td><td>5.1E-09</td><td>1.4E-08</td><td>BT20/MB468</td><td>MITO</td></tr>
<tr><td>>hg38_21_+_8433172_8446622_RNA45SN1_45S_50ntFlanks@9858.9890.33|CGGGATGAACCGAACGCCGGGTTAAGGCGCCCG</td><td>>hg38_21_+_8433172_8446622_RNA45SN1_45S_50ntFlanks</td><td>9858</td><td>9890</td><td>33</td><td>CGGGATGAACCGAACGCCGGGTTAAGGCGCCCG</td><td>77.8</td><td>-10.1</td><td>1.7</td><td>-5.8</td><td>6.3E-09</td><td>1.4E-08</td><td>MB231/MB468</td><td>MP</td></tr>
<tr><td>>hg38_21_+_8433172_8446622_RNA45SN1_45S_50ntFlanks@12631.12661.31|GGCGAAGCTACCATCTGTGGGATTATGACTG</td><td>>hg38_21_+_8433172_8446622_RNA45SN1_45S_50ntFlanks</td><td>12631</td><td>12661</td><td>31</td><td>GGCGAAGCTACCATCTGTGGGATTATGACTG</td><td>761.2</td><td>-9.2</td><td>1.6</td><td>-5.8</td><td>6.3E-09</td><td>1.4E-08</td><td>MB231/MB468</td><td>MP</td></tr>
<tr><td>>hg38_21_+_8433172_8446622_RNA45SN1_45S_50ntFlanks@12499.12518.20|GGTTTAGACCGTCGTGAGAC</td><td>>hg38_21_+_8433172_8446622_RNA45SN1_45S_50ntFlanks</td><td>12499</td><td>12518</td><td>20</td><td>GGTTTAGACCGTCGTGAGAC</td><td>133.3</td><td>-9.1</td><td>1.6</td><td>-5.8</td><td>5.1E-09</td><td>1.4E-08</td><td>BT20/MB468</td><td>MITO</td></tr>
<tr><td>>hg38_21_+_8433172_8446622_RNA45SN1_45S_50ntFlanks@8342.8375.34|CGTAAGGGAAAGTTGAAAAGAACTTTGAAGAGAG</td><td>>hg38_21_+_8433172_8446622_RNA45SN1_45S_50ntFlanks</td><td>8342</td><td>8375</td><td>34</td><td>CGTAAGGGAAAGTTGAAAAGAACTTTGAAGAGAG</td><td>2111.9</td><td>10.7</td><td>1.7</td><td>6.1</td><td>8.7E-10</td><td>1.4E-08</td><td>BT20/MB468</td><td>MP</td></tr>
<tr><td>>hg38_21_+_8433172_8446622_RNA45SN1_45S_50ntFlanks@4443.4462.20|CCCTTGCCTCTCGGCGCCCC</td><td>>hg38_21_+_8433172_8446622_RNA45SN1_45S_50ntFlanks</td><td>4443</td><td>4462</td><td>20</td><td>CCCTTGCCTCTCGGCGCCCC</td><td>147.6</td><td>-10.3</td><td>1.7</td><td>-6.1</td><td>8.8E-10</td><td>1.4E-08</td><td>BT20/MB468</td><td>MP</td></tr>
<tr><td>>hg38_21_+_8433172_8446622_RNA45SN1_45S_50ntFlanks@9034.9058.25|TCGTGGGGGGGCCGGGCCACCCCTC</td><td>>hg38_21_+_8433172_8446622_RNA45SN1_45S_50ntFlanks</td><td>9034</td><td>9058</td><td>25</td><td>TCGTGGGGGGGCCGGGCCACCCCTC</td><td>197.8</td><td>-10.7</td><td>1.7</td><td>-6.1</td><td>8.7E-10</td><td>1.4E-08</td><td>BT20/MB468</td><td>MP</td></tr>
<tr><td>>hg38_21_+_8433172_8446622_RNA45SN1_45S_50ntFlanks@3837.3869.33|CCTCTCCTACTTGGATAACTGTGGTAATTCTAG</td><td>>hg38_21_+_8433172_8446622_RNA45SN1_45S_50ntFlanks</td><td>3837</td><td>3869</td><td>33</td><td>CCTCTCCTACTTGGATAACTGTGGTAATTCTAG</td><td>155.5</td><td>-6.7</td><td>1.2</td><td>-5.8</td><td>6.7E-09</td><td>1.4E-08</td><td>MB231/MB468</td><td>NUC</td></tr>
<tr><td>>hg38_21_+_8433172_8446622_RNA45SN1_45S_50ntFlanks@11322.11344.23|ACGTCTCGTCGCGCGCGCGTCCG</td><td>>hg38_21_+_8433172_8446622_RNA45SN1_45S_50ntFlanks</td><td>11322</td><td>11344</td><td>23</td><td>ACGTCTCGTCGCGCGCGCGTCCG</td><td>13918.3</td><td>-5.7</td><td>1.0</td><td>-5.8</td><td>6.7E-09</td><td>1.4E-08</td><td>MB231/MB468</td><td>NUC</td></tr>
<tr><td>>hg38_21_+_8433172_8446622_RNA45SN1_45S_50ntFlanks@5444.5467.24|CTCGGATCGGCCCCGCCGGGGTCG</td><td>>hg38_21_+_8433172_8446622_RNA45SN1_45S_50ntFlanks</td><td>5444</td><td>5467</td><td>24</td><td>CTCGGATCGGCCCCGCCGGGGTCG</td><td>1634.9</td><td>-7.1</td><td>1.2</td><td>-5.8</td><td>7.3E-09</td><td>1.4E-08</td><td>MB231/MB468</td><td>MITO</td></tr>
</table>

<table>
<tr><td>>hg38_21_+_8433172_8446622_RNA45SN1_45S_50ntFlanks@10878.10902.25|TCGGTCGGGCTGGGGCGCGAAGCGG</td><td>>hg38_21_+_8433172_8446622_RNA45SN1_45S_50ntFlanks</td><td>10878</td><td>10902</td><td>25</td><td>TCGGTCGGGCTGGGGCGCGAAGCGG</td><td>599.6</td><td>-6.3</td><td>1.1</td><td>-5.8</td><td>6.8E-09</td><td>1.4E-08</td><td>MB231/MB468</td><td>NUC</td></tr>
<tr><td>>hg38_MT_+_1621_3279_RNR2_16S_50ntFlanks@713.749.37|GCATAAGCCTGCGTCAGATTAAAACACTGAACTGACA</td><td>>hg38_MT_+_1621_3279_RNR2_16S_50ntFlanks</td><td>713</td><td>749</td><td>37</td><td>GCATAAGCCTGCGTCAGATTAAAACACTGAACTGACA</td><td>65.9</td><td>9.3</td><td>1.6</td><td>5.8</td><td>6.3E-09</td><td>1.4E-08</td><td>MB231/MB468</td><td>MP</td></tr>
<tr><td>>hg38_21_+_8433172_8446622_RNA45SN1_45S_50ntFlanks@9533.9551.19|GAGGAAACTCTGGTGGAGG</td><td>>hg38_21_+_8433172_8446622_RNA45SN1_45S_50ntFlanks</td><td>9533</td><td>9551</td><td>19</td><td>GAGGAAACTCTGGTGGAGG</td><td>433.8</td><td>-5.3</td><td>0.9</td><td>-5.8</td><td>6.3E-09</td><td>1.4E-08</td><td>MB231/MB468</td><td>MP</td></tr>
<tr><td>>hg38_21_+_8433172_8446622_RNA45SN1_45S_50ntFlanks@4729.4766.38|TCAAGAACGAAAGTCGGAGGTTCGAAGACGATCAGATA</td><td>>hg38_21_+_8433172_8446622_RNA45SN1_45S_50ntFlanks</td><td>4729</td><td>4766</td><td>38</td><td>TCAAGAACGAAAGTCGGAGGTTCGAAGACGATCAGATA</td><td>390.1</td><td>7.1</td><td>1.2</td><td>5.8</td><td>5.7E-09</td><td>1.4E-08</td><td>BT20/MB468</td><td>CYTO</td></tr>
<tr><td>>hg38_21_+_8433172_8446622_RNA45SN1_45S_50ntFlanks@9532.9553.22|AGAGGAAACTCTGGTGGAGGTC</td><td>>hg38_21_+_8433172_8446622_RNA45SN1_45S_50ntFlanks</td><td>9532</td><td>9553</td><td>22</td><td>AGAGGAAACTCTGGTGGAGGTC</td><td>370.6</td><td>4.6</td><td>0.8</td><td>5.8</td><td>6.7E-09</td><td>1.4E-08</td><td>MB231/MB468</td><td>CYTO</td></tr>
<tr><td>>MI0000734|hsa-mir-106b&WithFlank&7|-|100093987|100094080@18.38.21&[MIMAT0000680&hsa-miR-106b-5p&offsets|0|0;m-103&7|-|100094043|100094063&offsets|0|0]|TAAAGTGCTGACAGTGCAGAT</td><td>>MI0000734|hsa-mir-106b&WithFlank&7|-|100093987|100094080</td><td>18</td><td>38</td><td>21</td><td>TAAAGTGCTGACAGTGCAGAT</td><td>1379.9</td><td>6.7</td><td>1.2</td><td>5.8</td><td>7.3E-09</td><td>1.4E-08</td><td>MB231/MB468</td><td>MITO</td></tr>
<tr><td>>MI0000434|hsa-let-7i&WithFlank&12|+|62603680|62603775@12.34.23&[MIMAT0000415&hsa-let-7i-5p&offsets|0|+1;m-49&12|+|62603691|62603712&offsets|0|+1]|TGAGGTAGTAGTTTGTGCTGTTG</td><td>>MI0000434|hsa-let-7i&WithFlank&12|+|62603680|62603775</td><td>12</td><td>34</td><td>23</td><td>TGAGGTAGTAGTTTGTGCTGTTG</td><td>79.7</td><td>2.1</td><td>0.4</td><td>5.8</td><td>7.6E-09</td><td>1.4E-08</td><td>BT20/MB468</td><td>NUC</td></tr>
<tr><td>>hg38_21_+_8433172_8446622_RNA45SN1_45S_50ntFlanks@9533.9566.34|GAGGAAACTCTGGTGGAGGTCCGTAGCGGTCCTG</td><td>>hg38_21_+_8433172_8446622_RNA45SN1_45S_50ntFlanks</td><td>9533</td><td>9566</td><td>34</td><td>GAGGAAACTCTGGTGGAGGTCCGTAGCGGTCCTG</td><td>1213.8</td><td>-9.2</td><td>1.6</td><td>-5.8</td><td>7.6E-09</td><td>1.4E-08</td><td>BT20/MB468</td><td>NUC</td></tr>
<tr><td>>hg38_21_+_8433172_8446622_RNA45SN1_45S_50ntFlanks@4367.4388.22|CAGTTAAAAAGCTCGTAGTTGG</td><td>>hg38_21_+_8433172_8446622_RNA45SN1_45S_50ntFlanks</td><td>4367</td><td>4388</td><td>22</td><td>CAGTTAAAAAGCTCGTAGTTGG</td><td>278.9</td><td>-5.8</td><td>1.0</td><td>-5.8</td><td>6.7E-09</td><td>1.4E-08</td><td>MB231/MB468</td><td>CYTO</td></tr>
<tr><td>>hg38_21_+_8433172_8446622_RNA45SN1_45S_50ntFlanks@6652.6694.43|GACTCTTAGCGGTGGATCACTCGGCTCGTGCGTCGATGAAGAA</td><td>>hg38_21_+_8433172_8446622_RNA45SN1_45S_50ntFlanks</td><td>6652</td><td>6694</td><td>43</td><td>GACTCTTAGCGGTGGATCACTCGGCTCGTGCGTCGATGAAGAA</td><td>304.8</td><td>5.7</td><td>1.0</td><td>5.8</td><td>6.7E-09</td><td>1.4E-08</td><td>MB231/MB468</td><td>CYTO</td></tr>
<tr><td>>hg38_21_+_8433172_8446622_RNA45SN1_45S_50ntFlanks@3987.4023.37|GCTTTGGTGACTCTAGATAACCTCGGGCCGATCGCAC</td><td>>hg38_21_+_8433172_8446622_RNA45SN1_45S_50ntFlanks</td><td>3987</td><td>4023</td><td>37</td><td>GCTTTGGTGACTCTAGATAACCTCGGGCCGATCGCAC</td><td>422.9</td><td>5.2</td><td>0.9</td><td>5.8</td><td>5.1E-09</td><td>1.4E-08</td><td>BT20/MB468</td><td>MITO</td></tr>
<tr><td>>MI0000299|hsa-mir-222&WithFlank&X|-|45747009|45747130@75.94.20(+1C)&[MIMAT0000279&hsa-miR-222-3p&offsets|0|-1(+1C);m-140&X|-|45747033|45747056&offsets|0|-4(+1C)]|AGCTACATCTGGCTACTGGGC</td><td>>MI0000299|hsa-mir-222&WithFlank&X|-|45747009|45747130</td><td>75</td><td>94</td><td>20(+1C)</td><td>AGCTACATCTGGCTACTGGGC</td><td>1071.4</td><td>4.6</td><td>0.8</td><td>5.8</td><td>6.4E-09</td><td>1.4E-08</td><td>MB231/MB468</td><td>MP</td></tr>
<tr><td>>MI0000078|hsa-mir-22&WithFlank&17|-|1713897|1713993@59.79.21(+1A)&[MIMAT0000077&hsa-miR-22-3p&offsets|0|-1(+1A);m-2&17|-|1713914|1713935&offsets|0|-1(+1A)]|AAGCTGCCAGTTGAAGAACTGA</td><td>>MI0000078|hsa-mir-22&WithFlank&17|-|1713897|1713993</td><td>59</td><td>79</td><td>21(+1A)</td><td>AAGCTGCCAGTTGAAGAACTGA</td><td>1313.1</td><td>5.8</td><td>1.0</td><td>5.8</td><td>6.4E-09</td><td>1.4E-08</td><td>MB231/MB468</td><td>MP</td></tr>
<tr><td>>hg38_21_+_8433172_8446622_RNA45SN1_45S_50ntFlanks@4443.4470.28|CCCTTGCCTCTCGGCGCCCCCTCGATGC</td><td>>hg38_21_+_8433172_8446622_RNA45SN1_45S_50ntFlanks</td><td>4443</td><td>4470</td><td>28</td><td>CCCTTGCCTCTCGGCGCCCCCTCGATGC</td><td>145.3</td><td>-6.8</td><td>1.2</td><td>-5.8</td><td>5.8E-09</td><td>1.4E-08</td><td>BT20/MB468</td><td>CYTO</td></tr>
<tr><td>>MI0000285|hsa-mir-205&WithFlank&1|+|209432127|209432248@40.61.22(+1G)&[MIMAT0000266&hsa-miR-205-5p&offsets|0|0(+1G);m-351&1|+|209432166|209432187&offsets|0|0(+1G)]|TCCTTCATTCCACCGGAGTCTGG</td><td>>MI0000285|hsa-mir-205&WithFlank&1|+|209432127|209432248</td><td>40</td><td>61</td><td>22(+1G)</td><td>TCCTTCATTCCACCGGAGTCTGG</td><td>1719.7</td><td>5.6</td><td>1.0</td><td>5.8</td><td>5.1E-09</td><td>1.4E-08</td><td>BT20/MB468</td><td>MITO</td></tr>
<tr><td>>hg38_21_+_8433172_8446622_RNA45SN1_45S_50ntFlanks@8155.8182.28|CGCTCGTGGGGGGCCCAAGTCCTTCTGA</td><td>>hg38_21_+_8433172_8446622_RNA45SN1_45S_50ntFlanks</td><td>8155</td><td>8182</td><td>28</td><td>CGCTCGTGGGGGGCCCAAGTCCTTCTGA</td><td>122.3</td><td>-9.0</td><td>1.5</td><td>-5.8</td><td>5.1E-09</td><td>1.4E-08</td><td>BT20/MB468</td><td>MITO</td></tr>
<tr><td>>hg38_21_+_8433172_8446622_RNA45SN1_45S_50ntFlanks@12934.12972.39|CTAAACCATTCGTAGACGACCTGCTTCTGGGTCGGGGTT</td><td>>hg38_21_+_8433172_8446622_RNA45SN1_45S_50ntFlanks</td><td>12934</td><td>12972</td><td>39</td><td>CTAAACCATTCGTAGACGACCTGCTTCTGGGTCGGGGTT</td><td>1175.9</td><td>-4.3</td><td>0.7</td><td>-5.8</td><td>5.2E-09</td><td>1.4E-08</td><td>BT20/MB468</td><td>MITO</td></tr>
<tr><td>>hg38_21_+_8433172_8446622_RNA45SN1_45S_50ntFlanks@4588.4608.21|CTATTTTGTTGGTTTTCGGAA</td><td>>hg38_21_+_8433172_8446622_RNA45SN1_45S_50ntFlanks</td><td>4588</td><td>4608</td><td>21</td><td>CTATTTTGTTGGTTTTCGGAA</td><td>86.3</td><td>-8.5</td><td>1.5</td><td>-5.8</td><td>5.2E-09</td><td>1.4E-08</td><td>BT20/MB468</td><td>MITO</td></tr>
<tr><td>>hg38_21_+_8433172_8446622_RNA45SN1_45S_50ntFlanks@10392.10431.40|ACAGCAGTTGAACATGGGTCAGTCGGTCCTGAGAGATGGG</td><td>>hg38_21_+_8433172_8446622_RNA45SN1_45S_50ntFlanks</td><td>10392</td><td>10431</td><td>40</td><td>ACAGCAGTTGAACATGGGTCAGTCGGTCCTGAGAGATGGG</td><td>397.8</td><td>-6.5</td><td>1.1</td><td>-5.8</td><td>7.4E-09</td><td>1.4E-08</td><td>MB231/MB468</td><td>MITO</td></tr>
<tr><td>>hg38_21_+_8433172_8446622_RNA45SN1_45S_50ntFlanks@4257.4282.26|TAATTGGAATGAGTCCACTTTAAATC</td><td>>hg38_21_+_8433172_8446622_RNA45SN1_45S_50ntFlanks</td><td>4257</td><td>4282</td><td>26</td><td>TAATTGGAATGAGTCCACTTTAAATC</td><td>63.7</td><td>-9.9</td><td>1.7</td><td>-5.8</td><td>6.4E-09</td><td>1.4E-08</td><td>MB231/MB468</td><td>MP</td></tr>
<tr><td>>hg38_21_+_8433172_8446622_RNA45SN1_45S_50ntFlanks@11315.11338.24|CCACCCCACGTCTCGTCGCGCGCG</td><td>>hg38_21_+_8433172_8446622_RNA45SN1_45S_50ntFlanks</td><td>11315</td><td>11338</td><td>24</td><td>CCACCCCACGTCTCGTCGCGCGCG</td><td>123.2</td><td>-10.0</td><td>1.6</td><td>-6.1</td><td>8.9E-10</td><td>1.4E-08</td><td>BT20/MB468</td><td>MP</td></tr>
<tr><td>>MI0000086|hsa-mir-28&WithFlank&3|+|188688775|188688872@61.82.22(+1U)&[MIMAT0004502&hsa-miR-28-3p&offsets|+1|+1(+1U);m-28&3|+|188688834|188688855&offsets|+1|+1(+1U)]|ACTAGATTGTGAGCTCCTGGAGT</td><td>>MI0000086|hsa-mir-28&WithFlank&3|+|188688775|188688872</td><td>61</td><td>82</td><td>22(+1U)</td><td>ACTAGATTGTGAGCTCCTGGAGT</td><td>192.3</td><td>6.1</td><td>1.0</td><td>5.8</td><td>6.9E-09</td><td>1.4E-08</td><td>MB231/MB468</td><td>NUC</td></tr>
<tr><td>>hg38_21_+_8433172_8446622_RNA45SN1_45S_50ntFlanks@11320.11338.19|CCACGTCTCGTCGCGCGCG</td><td>>hg38_21_+_8433172_8446622_RNA45SN1_45S_50ntFlanks</td><td>11320</td><td>11338</td><td>19</td><td>CCACGTCTCGTCGCGCGCG</td><td>132.2</td><td>-10.1</td><td>1.7</td><td>-6.1</td><td>8.9E-10</td><td>1.4E-08</td><td>BT20/MB468</td><td>MP</td></tr>
<tr><td>>hg38_21_+_8433172_8446622_RNA45SN1_45S_50ntFlanks@5497.5526.30|ACGGTCGAACTTGACTATCTAGAGGAAGTA</td><td>>hg38_21_+_8433172_8446622_RNA45SN1_45S_50ntFlanks</td><td>5497</td><td>5526</td><td>30</td><td>ACGGTCGAACTTGACTATCTAGAGGAAGTA</td><td>497.8</td><td>-6.4</td><td>1.1</td><td>-5.8</td><td>6.4E-09</td><td>1.4E-08</td><td>MB231/MB468</td><td>MP</td></tr>
<tr><td>>hg38_21_+_8433172_8446622_RNA45SN1_45S_50ntFlanks@3987.4005.19|GCTTTGGTGACTCTAGATA</td><td>>hg38_21_+_8433172_8446622_RNA45SN1_45S_50ntFlanks</td><td>3987</td><td>4005</td><td>19</td><td>GCTTTGGTGACTCTAGATA</td><td>268.2</td><td>-6.4</td><td>1.1</td><td>-5.8</td><td>6.4E-09</td><td>1.4E-08</td><td>MB231/MB468</td><td>MP</td></tr>
<tr><td>>hg38_21_+_8433172_8446622_RNA45SN1_45S_50ntFlanks@4729.4759.31|TCAAGAACGAAAGTCGGAGGTTCGAAGACGA</td><td>>hg38_21_+_8433172_8446622_RNA45SN1_45S_50ntFlanks</td><td>4729</td><td>4759</td><td>31</td><td>TCAAGAACGAAAGTCGGAGGTTCGAAGACGA</td><td>334.4</td><td>-3.0</td><td>0.5</td><td>-5.8</td><td>5.2E-09</td><td>1.4E-08</td><td>BT20/MB468</td><td>MITO</td></tr>
<tr><td>>hg38_21_+_8433172_8446622_RNA45SN1_45S_50ntFlanks@10735.10755.21|TGTAAATCTCGCGCCGGGCCG</td><td>>hg38_21_+_8433172_8446622_RNA45SN1_45S_50ntFlanks</td><td>10735</td><td>10755</td><td>21</td><td>TGTAAATCTCGCGCCGGGCCG</td><td>3532.0</td><td>-8.9</td><td>1.5</td><td>-5.8</td><td>6.9E-09</td><td>1.4E-08</td><td>MB231/MB468</td><td>NUC</td></tr>
<tr><td>>hg38_21_+_8433172_8446622_RNA45SN1_45S_50ntFlanks@9941.9963.23|AGGACGGTGGCCATGGAAGTCGG</td><td>>hg38_21_+_8433172_8446622_RNA45SN1_45S_50ntFlanks</td><td>9941</td><td>9963</td><td>23</td><td>AGGACGGTGGCCATGGAAGTCGG</td><td>2932.8</td><td>14.9</td><td>2.3</td><td>6.4</td><td>1.7E-10</td><td>1.4E-08</td><td>BT20/MB231</td><td>MP</td></tr>
<tr><td>>hg38_21_+_8433172_8446622_RNA45SN1_45S_50ntFlanks@12408.12437.30|TTTGATCCTTCGATGTCGGCTCTTCCTATC</td><td>>hg38_21_+_8433172_8446622_RNA45SN1_45S_50ntFlanks</td><td>12408</td><td>12437</td><td>30</td><td>TTTGATCCTTCGATGTCGGCTCTTCCTATC</td><td>464.1</td><td>-4.4</td><td>0.8</td><td>-5.8</td><td>7.7E-09</td><td>1.4E-08</td><td>BT20/MB468</td><td>NUC</td></tr>
<tr><td>>hg38_21_+_8433172_8446622_RNA45SN1_45S_50ntFlanks@10069.10094.26|AGTGGACGGGAGCGGCGGGGGCGGCG</td><td>>hg38_21_+_8433172_8446622_RNA45SN1_45S_50ntFlanks</td><td>10069</td><td>10094</td><td>26</td><td>AGTGGACGGGAGCGGCGGGGGCGGCG</td><td>66.5</td><td>-9.9</td><td>1.7</td><td>-5.8</td><td>6.4E-09</td><td>1.4E-08</td><td>MB231/MB468</td><td>MP</td></tr>
<tr><td>>hg38_21_+_8433172_8446622_RNA45SN1_45S_50ntFlanks@10845.10868.24|ACTTCGGGATAAGGATTGGCTCTA</td><td>>hg38_21_+_8433172_8446622_RNA45SN1_45S_50ntFlanks</td><td>10845</td><td>10868</td><td>24</td><td>ACTTCGGGATAAGGATTGGCTCTA</td><td>1199.9</td><td>-6.3</td><td>1.1</td><td>-5.8</td><td>6.8E-09</td><td>1.4E-08</td><td>MB231/MB468</td><td>CYTO</td></tr>
<tr><td>>hg38_21_+_8433172_8446622_RNA45SN1_45S_50ntFlanks@10834.10867.34|CCGGATCCGTAACTTCGGGATAAGGATTGGCTCT</td><td>>hg38_21_+_8433172_8446622_RNA45SN1_45S_50ntFlanks</td><td>10834</td><td>10867</td><td>34</td><td>CCGGATCCGTAACTTCGGGATAAGGATTGGCTCT</td><td>1285.0</td><td>4.8</td><td>0.8</td><td>5.8</td><td>6.8E-09</td><td>1.4E-08</td><td>MB231/MB468</td><td>CYTO</td></tr>
<tr><td>>hg38_21_+_8433172_8446622_RNA45SN1_45S_50ntFlanks@12348.12376.29|GAAAAGTTACCACAGGGATAACTGGCTTG</td><td>>hg38_21_+_8433172_8446622_RNA45SN1_45S_50ntFlanks</td><td>12348</td><td>12376</td><td>29</td><td>GAAAAGTTACCACAGGGATAACTGGCTTG</td><td>66.0</td><td>-9.9</td><td>1.7</td><td>-5.8</td><td>6.5E-09</td><td>1.4E-08</td><td>MB231/MB468</td><td>MP</td></tr>
<tr><td>>hg38_21_+_8433172_8446622_RNA45SN1_45S_50ntFlanks@11672.11699.28|CAGTGCTCTGAATGTCAAAGTGAAGAAA</td><td>>hg38_21_+_8433172_8446622_RNA45SN1_45S_50ntFlanks</td><td>11672</td><td>11699</td><td>28</td><td>CAGTGCTCTGAATGTCAAAGTGAAGAAA</td><td>375.0</td><td>-6.3</td><td>1.1</td><td>-5.8</td><td>7.8E-09</td><td>1.4E-08</td><td>BT20/MB468</td><td>NUC</td></tr>
<tr><td>>hg38_21_+_8433172_8446622_RNA45SN1_45S_50ntFlanks@6651.6672.22|CGACTCTTAGCGGTGGATCACT</td><td>>hg38_21_+_8433172_8446622_RNA45SN1_45S_50ntFlanks</td><td>6651</td><td>6672</td><td>22</td><td>CGACTCTTAGCGGTGGATCACT</td><td>3765.0</td><td>4.2</td><td>0.7</td><td>5.8</td><td>6.9E-09</td><td>1.4E-08</td><td>MB231/MB468</td><td>NUC</td></tr>
<tr><td>>hg38_21_+_8433172_8446622_RNA45SN1_45S_50ntFlanks@10684.10712.29|CGGCGGCGTCCGGTGAGCTCTCGCTGGCC</td><td>>hg38_21_+_8433172_8446622_RNA45SN1_45S_50ntFlanks</td><td>10684</td><td>10712</td><td>29</td><td>CGGCGGCGTCCGGTGAGCTCTCGCTGGCC</td><td>623.4</td><td>-4.1</td><td>0.7</td><td>-5.8</td><td>6.9E-09</td><td>1.4E-08</td><td>MB231/MB468</td><td>NUC</td></tr>
<tr><td>>hg38_21_+_8433172_8446622_RNA45SN1_45S_50ntFlanks@9362.9380.19|ATGAAGGTGAAGGCCGGCG</td><td>>hg38_21_+_8433172_8446622_RNA45SN1_45S_50ntFlanks</td><td>9362</td><td>9380</td><td>19</td><td>ATGAAGGTGAAGGCCGGCG</td><td>86.2</td><td>-10.3</td><td>1.8</td><td>-5.8</td><td>6.5E-09</td><td>1.4E-08</td><td>MB231/MB468</td><td>MP</td></tr>
<tr><td>>hg38_21_+_8433172_8446622_RNA45SN1_45S_50ntFlanks@11203.11232.30|CGGCGGCGGCGACTCTGGACGCGAGCCGGG</td><td>>hg38_21_+_8433172_8446622_RNA45SN1_45S_50ntFlanks</td><td>11203</td><td>11232</td><td>30</td><td>CGGCGGCGGCGACTCTGGACGCGAGCCGGG</td><td>463.7</td><td>-8.0</td><td>1.4</td><td>-5.8</td><td>6.5E-09</td><td>1.4E-08</td><td>MB231/MB468</td><td>MP</td></tr>
<tr><td>>hg38_21_+_8433172_8446622_RNA45SN1_45S_50ntFlanks@11586.11613.28|GTGCGGACCAGGGGAATCCGACTGTTTA</td><td>>hg38_21_+_8433172_8446622_RNA45SN1_45S_50ntFlanks</td><td>11586</td><td>11613</td><td>28</td><td>GTGCGGACCAGGGGAATCCGACTGTTTA</td><td>417.7</td><td>-4.0</td><td>0.7</td><td>-5.8</td><td>7.5E-09</td><td>1.4E-08</td><td>MB231/MB468</td><td>MITO</td></tr>
<tr><td>>hg38_21_+_8433172_8446622_RNA45SN1_45S_50ntFlanks@6719.6756.38|TGAATTGCAGGACACATTGATCATCGACACTTCGAACG</td><td>>hg38_21_+_8433172_8446622_RNA45SN1_45S_50ntFlanks</td><td>6719</td><td>6756</td><td>38</td><td>TGAATTGCAGGACACATTGATCATCGACACTTCGAACG</td><td>489.1</td><td>-6.3</td><td>1.1</td><td>-5.8</td><td>7.5E-09</td><td>1.4E-08</td><td>MB231/MB468</td><td>MITO</td></tr>
<tr><td>>hg38_21_+_8433172_8446622_RNA45SN1_45S_50ntFlanks@11561.11599.39|GCGCCTAGCAGCCGACTTAGAACTGGTGCGGACCAGGGG</td><td>>hg38_21_+_8433172_8446622_RNA45SN1_45S_50ntFlanks</td><td>11561</td><td>11599</td><td>39</td><td>GCGCCTAGCAGCCGACTTAGAACTGGTGCGGACCAGGGG</td><td>83.9</td><td>-10.3</td><td>1.8</td><td>-5.8</td><td>6.5E-09</td><td>1.4E-08</td><td>MB231/MB468</td><td>MP</td></tr>
<tr><td>>hg38_21_+_8433172_8446622_RNA45SN1_45S_50ntFlanks@12108.12125.18|CCGCGCGCCGGCCGGGCG</td><td>>hg38_21_+_8433172_8446622_RNA45SN1_45S_50ntFlanks</td><td>12108</td><td>12125</td><td>18</td><td>CCGCGCGCCGGCCGGGCG</td><td>117.8</td><td>-9.0</td><td>1.5</td><td>-5.8</td><td>5.2E-09</td><td>1.4E-08</td><td>BT20/MB468</td><td>MITO</td></tr>
<tr><td>>MI0000440|hsa-mir-27b&WithFlank&9|+|95085439|95085547@67.88.22&[MIMAT0000419&hsa-miR-27b-3p&offsets|0|+1;m-39&9|+|95085505|95085525&offsets|0|+1]|TTCACAGTGGCTAAGTTCTGCA</td><td>>MI0000440|hsa-mir-27b&WithFlank&9|+|95085439|95085547</td><td>67</td><td>88</td><td>22</td><td>TTCACAGTGGCTAAGTTCTGCA</td><td>2144.3</td><td>4.8</td><td>0.8</td><td>5.8</td><td>6.9E-09</td><td>1.4E-08</td><td>MB231/MB468</td><td>CYTO</td></tr>
<tr><td>>hg38_21_+_8433172_8446622_RNA45SN1_45S_50ntFlanks@6778.6807.30|CCCGGGGCTACGCCTGTCTGAGCGTCGCTT</td><td>>hg38_21_+_8433172_8446622_RNA45SN1_45S_50ntFlanks</td><td>6778</td><td>6807</td><td>30</td><td>CCCGGGGCTACGCCTGTCTGAGCGTCGCTT</td><td>267.3</td><td>3.7</td><td>0.6</td><td>5.8</td><td>7.0E-09</td><td>1.4E-08</td><td>MB231/MB468</td><td>NUC</td></tr>
<tr><td>>hg38_21_+_8433172_8446622_RNA45SN1_45S_50ntFlanks@10410.10433.24|TCAGTCGGTCCTGAGAGATGGGCG</td><td>>hg38_21_+_8433172_8446622_RNA45SN1_45S_50ntFlanks</td><td>10410</td><td>10433</td><td>24</td><td>TCAGTCGGTCCTGAGAGATGGGCG</td><td>518.6</td><td>-4.0</td><td>0.7</td><td>-5.8</td><td>5.2E-09</td><td>1.4E-08</td><td>BT20/MB468</td><td>MITO</td></tr>
<tr><td>>hg38_21_+_8433172_8446622_RNA45SN1_45S_50ntFlanks@9556.9573.18|TAGCGGTCCTGACGTGCA</td><td>>hg38_21_+_8433172_8446622_RNA45SN1_45S_50ntFlanks</td><td>9556</td><td>9573</td><td>18</td><td>TAGCGGTCCTGACGTGCA</td><td>131.0</td><td>-9.1</td><td>1.6</td><td>-5.8</td><td>5.2E-09</td><td>1.4E-08</td><td>BT20/MB468</td><td>MITO</td></tr>
<tr><td>>hg38_21_+_8433172_8446622_RNA45SN1_45S_50ntFlanks@11568.11603.36|GCAGCCGACTTAGAACTGGTGCGGACCAGGGGAATC</td><td>>hg38_21_+_8433172_8446622_RNA45SN1_45S_50ntFlanks</td><td>11568</td><td>11603</td><td>36</td><td>GCAGCCGACTTAGAACTGGTGCGGACCAGGGGAATC</td><td>1189.5</td><td>-6.9</td><td>1.2</td><td>-5.8</td><td>6.5E-09</td><td>1.4E-08</td><td>MB231/MB468</td><td>MP</td></tr>
<tr><td>>hg38_21_+_8433172_8446622_RNA45SN1_45S_50ntFlanks@12416.12460.45|TTCGATGTCGGCTCTTCCTATCATTGTGAAGCAGAATTCACCAAG</td><td>>hg38_21_+_8433172_8446622_RNA45SN1_45S_50ntFlanks</td><td>12416</td><td>12460</td><td>45</td><td>TTCGATGTCGGCTCTTCCTATCATTGTGAAGCAGAATTCACCAAG</td><td>162.9</td><td>6.5</td><td>1.1</td><td>5.8</td><td>6.9E-09</td><td>1.4E-08</td><td>MB231/MB468</td><td>CYTO</td></tr>
<tr><td>>hg38_21_+_8433172_8446622_RNA45SN1_45S_50ntFlanks@4728.4759.32|ATCAAGAACGAAAGTCGGAGGTTCGAAGACGA</td><td>>hg38_21_+_8433172_8446622_RNA45SN1_45S_50ntFlanks</td><td>4728</td><td>4759</td><td>32</td><td>ATCAAGAACGAAAGTCGGAGGTTCGAAGACGA</td><td>450.9</td><td>-7.6</td><td>1.3</td><td>-5.8</td><td>7.5E-09</td><td>1.4E-08</td><td>MB231/MB468</td><td>MITO</td></tr>
<tr><td>>hg38_21_+_8433172_8446622_RNA45SN1_45S_50ntFlanks@5489.5516.28|CTGAGAAGACGGTCGAACTTGACTATCT</td><td>>hg38_21_+_8433172_8446622_RNA45SN1_45S_50ntFlanks</td><td>5489</td><td>5516</td><td>28</td><td>CTGAGAAGACGGTCGAACTTGACTATCT</td><td>5274.1</td><td>4.1</td><td>0.7</td><td>5.8</td><td>5.3E-09</td><td>1.4E-08</td><td>BT20/MB468</td><td>MITO</td></tr>
<tr><td>>hg38_21_+_8433172_8446622_RNA45SN1_45S_50ntFlanks@11170.11188.19|CCGCGAGGGGGGCCCGGGC</td><td>>hg38_21_+_8433172_8446622_RNA45SN1_45S_50ntFlanks</td><td>11170</td><td>11188</td><td>19</td><td>CCGCGAGGGGGGCCCGGGC</td><td>1129.4</td><td>6.7</td><td>1.2</td><td>5.8</td><td>7.0E-09</td><td>1.4E-08</td><td>MB231/MB468</td><td>NUC</td></tr>
</table>

<table>
<tr><td>>hg38_21_+_8433172_8446622_RNA45SN1_45S_50ntFlanks@7979.8005.27|ACCTCAGATCAGACGTGGCGACCCGCT</td><td>>hg38_21_+_8433172_8446622_RNA45SN1_45S_50ntFlanks</td><td>7979</td><td>8005</td><td>27</td><td>ACCTCAGATCAGACGTGGCGACCCGCT</td><td>323.7</td><td>-5.9</td><td>1.0</td><td>-5.8</td><td>6.5E-09</td><td>1.4E-08</td><td>MB231/MB468</td><td>MP</td></tr>
<tr><td>>hg38_21_+_8433172_8446622_RNA45SN1_45S_50ntFlanks@7988.8017.30|CAGACGTGGCGACCCGCTGAATTTAAGCAT</td><td>>hg38_21_+_8433172_8446622_RNA45SN1_45S_50ntFlanks</td><td>7988</td><td>8017</td><td>30</td><td>CAGACGTGGCGACCCGCTGAATTTAAGCAT</td><td>695.4</td><td>-4.9</td><td>0.8</td><td>-6.1</td><td>9.1E-10</td><td>1.4E-08</td><td>BT20/MB468</td><td>MP</td></tr>
<tr><td>>hg38_21_+_8433172_8446622_RNA45SN1_45S_50ntFlanks@10786.10826.41|AACAGCCTCTGGCATGTTGGAACAATGTAGGTAAGGGAAGT</td><td>>hg38_21_+_8433172_8446622_RNA45SN1_45S_50ntFlanks</td><td>10786</td><td>10826</td><td>41</td><td>AACAGCCTCTGGCATGTTGGAACAATGTAGGTAAGGGAAGT</td><td>1928.9</td><td>-4.7</td><td>0.8</td><td>-5.8</td><td>5.3E-09</td><td>1.4E-08</td><td>BT20/MB468</td><td>MITO</td></tr>
<tr><td>>hg38_21_+_8433172_8446622_RNA45SN1_45S_50ntFlanks@10357.10377.21|CTTTGAAGGCCGAAGTGGAGA</td><td>>hg38_21_+_8433172_8446622_RNA45SN1_45S_50ntFlanks</td><td>10357</td><td>10377</td><td>21</td><td>CTTTGAAGGCCGAAGTGGAGA</td><td>278.9</td><td>-7.6</td><td>1.3</td><td>-5.8</td><td>6.5E-09</td><td>1.4E-08</td><td>MB231/MB468</td><td>MP</td></tr>
<tr><td>>hg38_21_+_8433172_8446622_RNA45SN1_45S_50ntFlanks@12632.12660.29|GCGAAGCTACCATCTGTGGGATTATGACT</td><td>>hg38_21_+_8433172_8446622_RNA45SN1_45S_50ntFlanks</td><td>12632</td><td>12660</td><td>29</td><td>GCGAAGCTACCATCTGTGGGATTATGACT</td><td>94.9</td><td>-10.4</td><td>1.8</td><td>-5.8</td><td>6.6E-09</td><td>1.4E-08</td><td>MB231/MB468</td><td>MP</td></tr>
<tr><td>>trna16_ValTAC_11_-_59318460_59318532@31.53.23,trna17_ValTAC_11_-_59318102_59318174@31.53.23,trna4_ValTAC_X_-_18693029_18693101@31.53.23,trna6_ValTAC_10_-_5895674_5895746@31.53.23|CTTTACACGCAGAAGGTCCTGGG</td><td>>trna16_ValTAC_11_-_59318460_59318532</td><td>31</td><td>53</td><td>23</td><td>CTTTACACGCAGAAGGTCCTGGG</td><td>66.1</td><td>-9.9</td><td>1.7</td><td>-5.8</td><td>6.6E-09</td><td>1.4E-08</td><td>MB231/MB468</td><td>MP</td></tr>
<tr><td>>MI0000266|hsa-mir-10a&WithFlank&17|-|48579832|48579953@28.48.21(+1G)&[MIMAT0000253&hsa-miR-10a-5p&offsets|0|-2(+1G);m-18&17|-|48579905|48579926&offsets|0|-1(+1G)]|TACCCTGTAGATCCGAATTTGG</td><td>>MI0000266|hsa-mir-10a&WithFlank&17|-|48579832|48579953</td><td>28</td><td>48</td><td>21(+1G)</td><td>TACCCTGTAGATCCGAATTTGG</td><td>73.5</td><td>9.4</td><td>1.6</td><td>5.8</td><td>6.6E-09</td><td>1.4E-08</td><td>MB231/MB468</td><td>MP</td></tr>
<tr><td>>hg38_21_+_8433172_8446622_RNA45SN1_45S_50ntFlanks@12948.12982.35|GACGACCTGCTTCTGGGTCGGGGTTTCGTACGTAG</td><td>>hg38_21_+_8433172_8446622_RNA45SN1_45S_50ntFlanks</td><td>12948</td><td>12982</td><td>35</td><td>GACGACCTGCTTCTGGGTCGGGGTTTCGTACGTAG</td><td>267.3</td><td>-4.7</td><td>0.8</td><td>-5.8</td><td>5.9E-09</td><td>1.4E-08</td><td>BT20/MB468</td><td>CYTO</td></tr>
<tr><td>>hg38_21_+_8433172_8446622_RNA45SN1_45S_50ntFlanks@10683.10712.30|CCGGCGGCGTCCGGTGAGCTCTCGCTGGCC</td><td>>hg38_21_+_8433172_8446622_RNA45SN1_45S_50ntFlanks</td><td>10683</td><td>10712</td><td>30</td><td>CCGGCGGCGTCCGGTGAGCTCTCGCTGGCC</td><td>2375.2</td><td>-4.2</td><td>0.7</td><td>-5.8</td><td>7.0E-09</td><td>1.4E-08</td><td>MB231/MB468</td><td>NUC</td></tr>
<tr><td>>hg38_21_+_8433172_8446622_RNA45SN1_45S_50ntFlanks@10869.10904.36|AGGGCTGGGTCGGTCGGGCTGGGGCGCGAAGCGGGG</td><td>>hg38_21_+_8433172_8446622_RNA45SN1_45S_50ntFlanks</td><td>10869</td><td>10904</td><td>36</td><td>AGGGCTGGGTCGGTCGGGCTGGGGCGCGAAGCGGGG</td><td>1728.7</td><td>-9.4</td><td>1.6</td><td>-5.8</td><td>7.9E-09</td><td>1.5E-08</td><td>BT20/MB468</td><td>NUC</td></tr>
<tr><td>>hg38_21_+_8433172_8446622_RNA45SN1_45S_50ntFlanks@12720.12742.23|CGGTTGGCCTCGGATAGCCGGTC</td><td>>hg38_21_+_8433172_8446622_RNA45SN1_45S_50ntFlanks</td><td>12720</td><td>12742</td><td>23</td><td>CGGTTGGCCTCGGATAGCCGGTC</td><td>91.4</td><td>-10.4</td><td>1.8</td><td>-5.8</td><td>6.6E-09</td><td>1.5E-08</td><td>MB231/MB468</td><td>MP</td></tr>
<tr><td>>trnaMT_PheGAA_MT_+_577_647@37.54.18|AAATGTTTAGACGGGCTC</td><td>>trnaMT_PheGAA_MT_+_577_647</td><td>37</td><td>54</td><td>18</td><td>AAATGTTTAGACGGGCTC</td><td>326.8</td><td>9.2</td><td>1.6</td><td>5.8</td><td>5.3E-09</td><td>1.5E-08</td><td>BT20/MB468</td><td>MITO</td></tr>
<tr><td>>hg38_21_+_8433172_8446622_RNA45SN1_45S_50ntFlanks@10546.10566.21|CAGTGCGGTAACGCGACCGAT</td><td>>hg38_21_+_8433172_8446622_RNA45SN1_45S_50ntFlanks</td><td>10546</td><td>10566</td><td>21</td><td>CAGTGCGGTAACGCGACCGAT</td><td>61.4</td><td>-9.8</td><td>1.7</td><td>-5.8</td><td>6.6E-09</td><td>1.5E-08</td><td>MB231/MB468</td><td>MP</td></tr>
<tr><td>>hg38_21_+_8433172_8446622_RNA45SN1_45S_50ntFlanks@9570.9596.27|TGCAAATCGGTCGTCCGACCTGGGTAT</td><td>>hg38_21_+_8433172_8446622_RNA45SN1_45S_50ntFlanks</td><td>9570</td><td>9596</td><td>27</td><td>TGCAAATCGGTCGTCCGACCTGGGTAT</td><td>2012.6</td><td>-8.1</td><td>1.4</td><td>-5.8</td><td>6.0E-09</td><td>1.5E-08</td><td>BT20/MB468</td><td>CYTO</td></tr>
<tr><td>>hg38_21_+_8433172_8446622_RNA45SN1_45S_50ntFlanks@4770.4793.24|TCGTAGTTCCGACCATAAACGATG</td><td>>hg38_21_+_8433172_8446622_RNA45SN1_45S_50ntFlanks</td><td>4770</td><td>4793</td><td>24</td><td>TCGTAGTTCCGACCATAAACGATG</td><td>329.5</td><td>-5.8</td><td>1.0</td><td>-5.8</td><td>7.6E-09</td><td>1.5E-08</td><td>MB231/MB468</td><td>MITO</td></tr>
<tr><td>>MI0000650|hsa-mir-200c&WithFlank&12|+|6963693|6963772@50.72.23(+1A)&[MIMAT0000617&hsa-miR-200c-3p&offsets|0|0(+1A);m-54&12|+|6963742|6963764&offsets|0|0(+1A)]|TAATACTGCCGGGTAATGATGGAA</td><td>>MI0000650|hsa-mir-200c&WithFlank&12|+|6963693|6963772</td><td>50</td><td>72</td><td>23(+1A)</td><td>TAATACTGCCGGGTAATGATGGAA</td><td>246.0</td><td>-5.6</td><td>1.0</td><td>-5.8</td><td>7.1E-09</td><td>1.5E-08</td><td>MB231/MB468</td><td>NUC</td></tr>
<tr><td>>MI0000273|hsa-mir-183&WithFlank&7|-|129774899|129775020@34.53.20(+1C)&[MIMAT0000261&hsa-miR-183-5p&offsets|+1|-1(+1C);m-41&7|-|129774966|129774987&offsets|0|-2(+1C)]|ATGGCACTGGTAGAATTCACC</td><td>>MI0000273|hsa-mir-183&WithFlank&7|-|129774899|129775020</td><td>34</td><td>53</td><td>20(+1C)</td><td>ATGGCACTGGTAGAATTCACC</td><td>491.4</td><td>3.5</td><td>0.6</td><td>6.2</td><td>7.2E-10</td><td>1.5E-08</td><td>BT20/MB231</td><td>NUC</td></tr>
<tr><td>>hg38_21_+_8433172_8446622_RNA45SN1_45S_50ntFlanks@3705.3749.45|TACCTGGTTGATCCTGCCAGTAGCATATGCTTGTCTCAAAGATTA</td><td>>hg38_21_+_8433172_8446622_RNA45SN1_45S_50ntFlanks</td><td>3705</td><td>3749</td><td>45</td><td>TACCTGGTTGATCCTGCCAGTAGCATATGCTTGTCTCAAAGATTA</td><td>279.1</td><td>6.7</td><td>1.2</td><td>5.8</td><td>7.0E-09</td><td>1.5E-08</td><td>MB231/MB468</td><td>CYTO</td></tr>
<tr><td>>hg38_21_+_8433172_8446622_RNA45SN1_45S_50ntFlanks@11027.11075.49|CTCTCTCTCTCTCTCCCCCGCTCCCCGTCCTCCCCCCTCCCCGGGGGAG</td><td>>hg38_21_+_8433172_8446622_RNA45SN1_45S_50ntFlanks</td><td>11027</td><td>11075</td><td>49</td><td>CTCTCTCTCTCTCTCCCCCGCTCCCCGTCCTCCCCCCTCCCCGGGGGAG</td><td>305.0</td><td>-6.6</td><td>1.1</td><td>-5.8</td><td>7.0E-09</td><td>1.5E-08</td><td>MB231/MB468</td><td>CYTO</td></tr>
<tr><td>>hg38_21_+_8433172_8446622_RNA45SN1_45S_50ntFlanks@7991.8019.29|ACGTGGCGACCCGCTGAATTTAAGCATAT</td><td>>hg38_21_+_8433172_8446622_RNA45SN1_45S_50ntFlanks</td><td>7991</td><td>8019</td><td>29</td><td>ACGTGGCGACCCGCTGAATTTAAGCATAT</td><td>307.5</td><td>8.3</td><td>1.4</td><td>5.8</td><td>7.0E-09</td><td>1.5E-08</td><td>MB231/MB468</td><td>CYTO</td></tr>
<tr><td>>hg38_21_+_8433172_8446622_RNA45SN1_45S_50ntFlanks@12624.12661.38|CCAATGGGGCGAAGCTACCATCTGTGGGATTATGACTG</td><td>>hg38_21_+_8433172_8446622_RNA45SN1_45S_50ntFlanks</td><td>12624</td><td>12661</td><td>38</td><td>CCAATGGGGCGAAGCTACCATCTGTGGGATTATGACTG</td><td>4043.0</td><td>-9.9</td><td>1.7</td><td>-5.8</td><td>7.9E-09</td><td>1.5E-08</td><td>BT20/MB468</td><td>NUC</td></tr>
<tr><td>>hg38_21_+_8433172_8446622_RNA45SN1_45S_50ntFlanks@5066.5098.33|TCTGGTTAATTCCGATAACGAACGAGACTCTGG</td><td>>hg38_21_+_8433172_8446622_RNA45SN1_45S_50ntFlanks</td><td>5066</td><td>5098</td><td>33</td><td>TCTGGTTAATTCCGATAACGAACGAGACTCTGG</td><td>182.3</td><td>-11.4</td><td>2.0</td><td>-5.8</td><td>6.6E-09</td><td>1.5E-08</td><td>MB231/MB468</td><td>MP</td></tr>
<tr><td>>am_trna143_LysTTT_6_-_27559593_27559665@6.24.19,trna14_LysTTT_11_-_59327808_59327880@6.24.19,trna2_LysTTT_17_+_8022473_8022545@6.24.19,trna54_LysTTT_1_+_204475655_204475727@6.24.19,trna5_LysTTT_11_+_59323902_59323974@6.24.19,trna62_LysTTT_1_-_204476158_204476230@6.24.19,trna76_LysTTT_6_+_28918806_28918878@6.24.19|GATAGCTCAGTCGGTAGAG</td><td>>am_trna143_LysTTT_6_-_27559593_27559665</td><td>6</td><td>24</td><td>19</td><td>GATAGCTCAGTCGGTAGAG</td><td>145.6</td><td>-10.3</td><td>1.7</td><td>-6.1</td><td>9.2E-10</td><td>1.5E-08</td><td>BT20/MB468</td><td>MP</td></tr>
<tr><td>>hg38_21_+_8433172_8446622_RNA45SN1_45S_50ntFlanks@11208.11235.28|GCGGCGACTCTGGACGCGAGCCGGGCCC</td><td>>hg38_21_+_8433172_8446622_RNA45SN1_45S_50ntFlanks</td><td>11208</td><td>11235</td><td>28</td><td>GCGGCGACTCTGGACGCGAGCCGGGCCC</td><td>269.6</td><td>-6.3</td><td>1.1</td><td>-5.8</td><td>7.0E-09</td><td>1.5E-08</td><td>MB231/MB468</td><td>CYTO</td></tr>
<tr><td>>hg38_21_+_8433172_8446622_RNA45SN1_45S_50ntFlanks@10681.10712.32|TTCCGGCGGCGTCCGGTGAGCTCTCGCTGGCC</td><td>>hg38_21_+_8433172_8446622_RNA45SN1_45S_50ntFlanks</td><td>10681</td><td>10712</td><td>32</td><td>TTCCGGCGGCGTCCGGTGAGCTCTCGCTGGCC</td><td>1301.1</td><td>-3.7</td><td>0.6</td><td>-5.8</td><td>5.4E-09</td><td>1.5E-08</td><td>BT20/MB468</td><td>MITO</td></tr>
<tr><td>>hg38_21_+_8433172_8446622_RNA45SN1_45S_50ntFlanks@4731.4765.35|AAGAACGAAAGTCGGAGGTTCGAAGACGATCAGAT</td><td>>hg38_21_+_8433172_8446622_RNA45SN1_45S_50ntFlanks</td><td>4731</td><td>4765</td><td>35</td><td>AAGAACGAAAGTCGGAGGTTCGAAGACGATCAGAT</td><td>533.7</td><td>3.8</td><td>0.7</td><td>5.8</td><td>7.1E-09</td><td>1.5E-08</td><td>MB231/MB468</td><td>NUC</td></tr>
<tr><td>>hg38_21_+_8433172_8446622_RNA45SN1_45S_50ntFlanks@6661.6701.41|CGGTGGATCACTCGGCTCGTGCGTCGATGAAGAACGCAGCT</td><td>>hg38_21_+_8433172_8446622_RNA45SN1_45S_50ntFlanks</td><td>6661</td><td>6701</td><td>41</td><td>CGGTGGATCACTCGGCTCGTGCGTCGATGAAGAACGCAGCT</td><td>579.4</td><td>4.3</td><td>0.7</td><td>5.8</td><td>7.1E-09</td><td>1.5E-08</td><td>MB231/MB468</td><td>NUC</td></tr>
<tr><td>>hg38_21_+_8433172_8446622_RNA45SN1_45S_50ntFlanks@11199.11229.31|CCGGCGGCGGCGGCGACTCTGGACGCGAGCC</td><td>>hg38_21_+_8433172_8446622_RNA45SN1_45S_50ntFlanks</td><td>11199</td><td>11229</td><td>31</td><td>CCGGCGGCGGCGGCGACTCTGGACGCGAGCC</td><td>244.0</td><td>-2.6</td><td>0.5</td><td>-5.8</td><td>6.0E-09</td><td>1.5E-08</td><td>BT20/MB468</td><td>CYTO</td></tr>
<tr><td>>hg38_21_+_8433172_8446622_RNA45SN1_45S_50ntFlanks@12073.12097.25|CCGAGGGGCTCTCGCTTCTGGCGCC</td><td>>hg38_21_+_8433172_8446622_RNA45SN1_45S_50ntFlanks</td><td>12073</td><td>12097</td><td>25</td><td>CCGAGGGGCTCTCGCTTCTGGCGCC</td><td>4297.6</td><td>-8.1</td><td>1.4</td><td>-5.8</td><td>8.0E-09</td><td>1.5E-08</td><td>BT20/MB468</td><td>NUC</td></tr>
<tr><td>>am_trnaMT_PheGAA_MT_+_577_647@42.62.21|TTTAGACGGGCTCACATCACC</td><td>>am_trnaMT_PheGAA_MT_+_577_647</td><td>42</td><td>62</td><td>21</td><td>TTTAGACGGGCTCACATCACC</td><td>738.0</td><td>8.5</td><td>1.5</td><td>5.8</td><td>6.0E-09</td><td>1.5E-08</td><td>BT20/MB468</td><td>CYTO</td></tr>
<tr><td>>hg38_21_+_8433172_8446622_RNA45SN1_45S_50ntFlanks@4565.4587.23|AATAATGGAATAGGACCGCGGTT</td><td>>hg38_21_+_8433172_8446622_RNA45SN1_45S_50ntFlanks</td><td>4565</td><td>4587</td><td>23</td><td>AATAATGGAATAGGACCGCGGTT</td><td>524.4</td><td>-4.4</td><td>0.8</td><td>-5.8</td><td>7.7E-09</td><td>1.5E-08</td><td>MB231/MB468</td><td>MITO</td></tr>
<tr><td>>MI0000088|hsa-mir-30a&WithFlank&6|-|71403545|71403627@12.35.24(+1A)&[MIMAT0000087&hsa-miR-30a-5p&offsets|0|+2(+1A);m-15&6|-|71403595|71403616&offsets|0|+2(+1A)]|TGTAAACATCCTCGACTGGAAGCTA</td><td>>MI0000088|hsa-mir-30a&WithFlank&6|-|71403545|71403627</td><td>12</td><td>35</td><td>24(+1A)</td><td>TGTAAACATCCTCGACTGGAAGCTA</td><td>438.6</td><td>7.8</td><td>1.4</td><td>5.8</td><td>7.1E-09</td><td>1.5E-08</td><td>MB231/MB468</td><td>NUC</td></tr>
<tr><td>>hg38_21_+_8433172_8446622_RNA45SN1_45S_50ntFlanks@12300.12317.18|GGGCCTCACGATCCTTCT</td><td>>hg38_21_+_8433172_8446622_RNA45SN1_45S_50ntFlanks</td><td>12300</td><td>12317</td><td>18</td><td>GGGCCTCACGATCCTTCT</td><td>226.5</td><td>-9.9</td><td>1.7</td><td>-5.8</td><td>5.4E-09</td><td>1.5E-08</td><td>BT20/MB468</td><td>MITO</td></tr>
<tr><td>>MI0000108|hsa-mir-103a-2&WithFlank&20|+|3917488|3917577@54.75.22;MI0000109|hsa-mir-103a-1&WithFlank&5|-|168560890|168560979@54.75.22&[MIMAT0000101&hsa-miR-103a-3p&offsets|0|-1;m-12&20|+|3917541|3917563&offsets|0|-1];[MIMAT0000101_1&hsa-miR-103a-3p&offsets|0|-1;m-13&5|-|168560904|168560926&offsets|0|-1;m-44220&5|-|168560908|168560926&offsets|0|+3]|AGCAGCATTGTACAGGGCTATG</td><td>>MI0000108|hsa-mir-103a-2&WithFlank&20|+|3917488|3917577</td><td>54</td><td>75</td><td>22</td><td>AGCAGCATTGTACAGGGCTATG</td><td>2768.3</td><td>5.4</td><td>0.9</td><td>5.8</td><td>7.1E-09</td><td>1.5E-08</td><td>MB231/MB468</td><td>CYTO</td></tr>
<tr><td>>hg38_21_+_8433172_8446622_RNA45SN1_45S_50ntFlanks@4973.4999.27|GGCCCGGACACGGACAGGATTGACAGA</td><td>>hg38_21_+_8433172_8446622_RNA45SN1_45S_50ntFlanks</td><td>4973</td><td>4999</td><td>27</td><td>GGCCCGGACACGGACAGGATTGACAGA</td><td>79.9</td><td>-10.2</td><td>1.8</td><td>-5.8</td><td>6.7E-09</td><td>1.5E-08</td><td>MB231/MB468</td><td>MP</td></tr>
<tr><td>>hg38_21_+_8433172_8446622_RNA45SN1_45S_50ntFlanks@10466.10488.23|CGTTGCCCTCGGCCGATCGAAAG</td><td>>hg38_21_+_8433172_8446622_RNA45SN1_45S_50ntFlanks</td><td>10466</td><td>10488</td><td>23</td><td>CGTTGCCCTCGGCCGATCGAAAG</td><td>731.1</td><td>-6.8</td><td>1.2</td><td>-5.8</td><td>7.7E-09</td><td>1.5E-08</td><td>MB231/MB468</td><td>MITO</td></tr>
<tr><td>>MI0000298|hsa-mir-221&WithFlank&X|-|45746151|45746272@71.91.21&[MIMAT0000278&hsa-miR-221-3p&offsets|0|-2;m-91&X|-|45746181|45746202&offsets|0|-1]|AGCTACATTGTCTGCTGGGTT</td><td>>MI0000298|hsa-mir-221&WithFlank&X|-|45746151|45746272</td><td>71</td><td>91</td><td>21</td><td>AGCTACATTGTCTGCTGGGTT</td><td>5178.4</td><td>4.7</td><td>0.8</td><td>5.8</td><td>6.7E-09</td><td>1.5E-08</td><td>MB231/MB468</td><td>MP</td></tr>
<tr><td>>MI0000542|hsa-mir-320a&WithFlank&8|-|22244960|22245043@48.70.23(+1U)&[MIMAT0000510&hsa-miR-320a-3p&offsets|0|+1(+1U);m-60&8|-|22244975|22244996&offsets|0|+1(+1U)]|AAAAGCTGGGTTGAGAGGGCGAAT</td><td>>MI0000542|hsa-mir-320a&WithFlank&8|-|22244960|22245043</td><td>48</td><td>70</td><td>23(+1U)</td><td>AAAAGCTGGGTTGAGAGGGCGAAT</td><td>647.0</td><td>5.6</td><td>1.0</td><td>5.8</td><td>7.7E-09</td><td>1.5E-08</td><td>MB231/MB468</td><td>MITO</td></tr>
<tr><td>>hg38_21_+_8433172_8446622_RNA45SN1_45S_50ntFlanks@11783.11808.26|CATGAATGGATGAACGAGATTCCCAC</td><td>>hg38_21_+_8433172_8446622_RNA45SN1_45S_50ntFlanks</td><td>11783</td><td>11808</td><td>26</td><td>CATGAATGGATGAACGAGATTCCCAC</td><td>79.9</td><td>-10.2</td><td>1.8</td><td>-5.8</td><td>6.7E-09</td><td>1.5E-08</td><td>MB231/MB468</td><td>MP</td></tr>
<tr><td>>hg38_21_+_8433172_8446622_RNA45SN1_45S_50ntFlanks@9552.9571.20|TCCGTAGCGGTCCTGACGTG</td><td>>hg38_21_+_8433172_8446622_RNA45SN1_45S_50ntFlanks</td><td>9552</td><td>9571</td><td>20</td><td>TCCGTAGCGGTCCTGACGTG</td><td>524.2</td><td>-4.7</td><td>0.8</td><td>-5.8</td><td>7.8E-09</td><td>1.5E-08</td><td>MB231/MB468</td><td>MITO</td></tr>
<tr><td>>MI0000301|hsa-mir-224&WithFlank&X|-|151958572|151958664@14.37.24&[MIMAT0000281&hsa-miR-224-5p&offsets|+1|0;m-238&X|-|151958630|151958651&offsets|0|+2]|CAAGTCACTAGTGGTTCCGTTTAG</td><td>>MI0000301|hsa-mir-224&WithFlank&X|-|151958572|151958664</td><td>14</td><td>37</td><td>24</td><td>CAAGTCACTAGTGGTTCCGTTTAG</td><td>640.3</td><td>9.9</td><td>1.6</td><td>6.1</td><td>9.4E-10</td><td>1.5E-08</td><td>BT20/MB468</td><td>MP</td></tr>
<tr><td>>trnaMT_ValTAC_MT_+_1602_1670@46.71.26|TTCAACTTAACTTGACCGCTCTGACC</td><td>>trnaMT_ValTAC_MT_+_1602_1670</td><td>46</td><td>71</td><td>26</td><td>TTCAACTTAACTTGACCGCTCTGACC</td><td>267.9</td><td>4.8</td><td>0.8</td><td>5.8</td><td>7.2E-09</td><td>1.5E-08</td><td>MB231/MB468</td><td>NUC</td></tr>
<tr><td>>hg38_21_+_8433172_8446622_RNA45SN1_45S_50ntFlanks@6721.6746.26|AATTGCAGGACACATTGATCATCGAC</td><td>>hg38_21_+_8433172_8446622_RNA45SN1_45S_50ntFlanks</td><td>6721</td><td>6746</td><td>26</td><td>AATTGCAGGACACATTGATCATCGAC</td><td>1118.9</td><td>-3.9</td><td>0.7</td><td>-5.8</td><td>5.4E-09</td><td>1.5E-08</td><td>BT20/MB468</td><td>MITO</td></tr>
<tr><td>>hg38_21_+_8433172_8446622_RNA45SN1_45S_50ntFlanks@12247.12273.27|GGCAAAAGCTCGCTTGATCTTGATTTT</td><td>>hg38_21_+_8433172_8446622_RNA45SN1_45S_50ntFlanks</td><td>12247</td><td>12273</td><td>27</td><td>GGCAAAAGCTCGCTTGATCTTGATTTT</td><td>307.1</td><td>-8.2</td><td>1.4</td><td>-5.8</td><td>7.8E-09</td><td>1.5E-08</td><td>MB231/MB468</td><td>MITO</td></tr>
<tr><td>>hg38_21_+_8433172_8446622_RNA45SN1_45S_50ntFlanks@10914.10936.23|CGCCGCGGCTGGACGAGGCGCCG</td><td>>hg38_21_+_8433172_8446622_RNA45SN1_45S_50ntFlanks</td><td>10914</td><td>10936</td><td>23</td><td>CGCCGCGGCTGGACGAGGCGCCG</td><td>857.9</td><td>-8.4</td><td>1.4</td><td>-6.1</td><td>9.4E-10</td><td>1.5E-08</td><td>BT20/MB468</td><td>MP</td></tr>
<tr><td>>hg38_21_+_8433172_8446622_RNA45SN1_45S_50ntFlanks@11678.11702.25|TCTGAATGTCAAAGTGAAGAAATTC</td><td>>hg38_21_+_8433172_8446622_RNA45SN1_45S_50ntFlanks</td><td>11678</td><td>11702</td><td>25</td><td>TCTGAATGTCAAAGTGAAGAAATTC</td><td>115.1</td><td>-9.9</td><td>1.6</td><td>-6.1</td><td>9.4E-10</td><td>1.5E-08</td><td>BT20/MB468</td><td>MP</td></tr>
<tr><td>>hg38_21_+_8433172_8446622_RNA45SN1_45S_50ntFlanks@3705.3734.30|TACCTGGTTGATCCTGCCAGTAGCATATGC</td><td>>hg38_21_+_8433172_8446622_RNA45SN1_45S_50ntFlanks</td><td>3705</td><td>3734</td><td>30</td><td>TACCTGGTTGATCCTGCCAGTAGCATATGC</td><td>295.4</td><td>7.8</td><td>1.3</td><td>5.8</td><td>6.1E-09</td><td>1.5E-08</td><td>BT20/MB468</td><td>CYTO</td></tr>
</table>

<table>
<tr><td>>hg38_21_+_8433172_8446622_RNA45SN1_45S_50ntFlanks@12415.12437.23|CTTCGATGTCGGCTCTTCCTATC</td><td>>hg38_21_+_8433172_8446622_RNA45SN1_45S_50ntFlanks</td><td>12415</td><td>12437</td><td>23</td><td>CTTCGATGTCGGCTCTTCCTATC</td><td>5340.3</td><td>-4.5</td><td>0.8</td><td>-5.8</td><td>7.8E-09</td><td>1.5E-08</td><td>MB231/MB468</td><td>MITO</td></tr>
<tr><td>>MI0000816|hsa-mir-335&WithFlank&7|+|130496105|130496210@58.78.21(+1A)&[MIMAT0004703&hsa-miR-335-3p&offsets|0|-1(+1A);m-159&7|+|130496162|130496183&offsets|0|-1(+1A)]|TTTTTCATTATTGCTCCTGACA</td><td>>MI0000816|hsa-mir-335&WithFlank&7|+|130496105|130496210</td><td>58</td><td>78</td><td>21(+1A)</td><td>TTTTTCATTATTGCTCCTGACA</td><td>560.2</td><td>6.3</td><td>1.1</td><td>5.8</td><td>7.8E-09</td><td>1.5E-08</td><td>MB231/MB468</td><td>MITO</td></tr>
<tr><td>>am_MI0000809|hsa-mir-151a&WithFlank&8|-|140732558|140732659@52.73.22&[MIMAT0000757&hsa-miR-151a-3p&offsets|-1|0;m-30&8|-|140732587|140732607&offsets|-1|0]|ACTAGACTGAAGCTCCTTGAGG</td><td>>am_MI0000809|hsa-mir-151a&WithFlank&8|-|140732558|140732659</td><td>52</td><td>73</td><td>22</td><td>ACTAGACTGAAGCTCCTTGAGG</td><td>598.1</td><td>5.0</td><td>0.9</td><td>5.8</td><td>7.8E-09</td><td>1.5E-08</td><td>MB231/MB468</td><td>MITO</td></tr>
<tr><td>>hg38_21_+_8433172_8446622_RNA45SN1_45S_50ntFlanks@12074.12095.22|CGAGGGGCTCTCGCTTCTGGCG</td><td>>hg38_21_+_8433172_8446622_RNA45SN1_45S_50ntFlanks</td><td>12074</td><td>12095</td><td>22</td><td>CGAGGGGCTCTCGCTTCTGGCG</td><td>679.1</td><td>-3.4</td><td>0.6</td><td>-5.8</td><td>5.5E-09</td><td>1.5E-08</td><td>BT20/MB468</td><td>MITO</td></tr>
<tr><td>>MI0000478|hsa-mir-149&WithFlank&2|+|240455995|240456095@21.41.21&[MIMAT0000450&hsa-miR-149-5p&offsets|0|-2;m-195&2|+|240456015|240456037&offsets|0|-2]|TCTGGCTCCGTGTCTTCACTC</td><td>>MI0000478|hsa-mir-149&WithFlank&2|+|240455995|240456095</td><td>21</td><td>41</td><td>21</td><td>TCTGGCTCCGTGTCTTCACTC</td><td>98.6</td><td>4.3</td><td>0.7</td><td>5.8</td><td>8.2E-09</td><td>1.5E-08</td><td>BT20/MB468</td><td>NUC</td></tr>
<tr><td>>MI0000542|hsa-mir-320a&WithFlank&8|-|22244960|22245043@49.70.22(+1U)&[MIMAT0000510&hsa-miR-320a-3p&offsets|+1|+1(+1U);m-60&8|-|22244975|22244996&offsets|+1|+1(+1U)]|AAAGCTGGGTTGAGAGGGCGAAT</td><td>>MI0000542|hsa-mir-320a&WithFlank&8|-|22244960|22245043</td><td>49</td><td>70</td><td>22(+1U)</td><td>AAAGCTGGGTTGAGAGGGCGAAT</td><td>109.0</td><td>2.8</td><td>0.5</td><td>5.8</td><td>8.2E-09</td><td>1.5E-08</td><td>BT20/MB468</td><td>NUC</td></tr>
<tr><td>>hg38_21_+_8433172_8446622_RNA45SN1_45S_50ntFlanks@12071.12106.36|CCCCGAGGGGCTCTCGCTTCTGGCGCCAAGCGCCCG</td><td>>hg38_21_+_8433172_8446622_RNA45SN1_45S_50ntFlanks</td><td>12071</td><td>12106</td><td>36</td><td>CCCCGAGGGGCTCTCGCTTCTGGCGCCAAGCGCCCG</td><td>493.7</td><td>-4.7</td><td>0.8</td><td>-5.8</td><td>7.3E-09</td><td>1.5E-08</td><td>MB231/MB468</td><td>NUC</td></tr>
<tr><td>>hg38_21_+_8433172_8446622_RNA45SN1_45S_50ntFlanks@12081.12110.30|CTCTCGCTTCTGGCGCCAAGCGCCCGGCCG</td><td>>hg38_21_+_8433172_8446622_RNA45SN1_45S_50ntFlanks</td><td>12081</td><td>12110</td><td>30</td><td>CTCTCGCTTCTGGCGCCAAGCGCCCGGCCG</td><td>821.2</td><td>-8.9</td><td>1.5</td><td>-5.8</td><td>8.2E-09</td><td>1.5E-08</td><td>BT20/MB468</td><td>NUC</td></tr>
<tr><td>>hg38_MT_+_1621_3279_RNR2_16S_50ntFlanks@493.518.26|GCTCTTTGGACACTAGGAAAAAACCT</td><td>>hg38_MT_+_1621_3279_RNR2_16S_50ntFlanks</td><td>493</td><td>518</td><td>26</td><td>GCTCTTTGGACACTAGGAAAAAACCT</td><td>809.5</td><td>4.6</td><td>0.8</td><td>5.8</td><td>7.9E-09</td><td>1.5E-08</td><td>MB231/MB468</td><td>MITO</td></tr>
<tr><td>>trna116_GluCTC_1_-_145399233_145399304@3.36.34,trna59_GluCTC_1_+_249168447_249168518@3.36.34,trna71_GluCTC_1_-_161439189_161439260@3.36.34,trna74_GluCTC_1_-_161431809_161431880@3.36.34,trna77_GluCTC_1_-_161424398_161424469@3.36.34,trna77_GluCTC_6_+_28949976_28950047@3.36.34,trna80_GluCTC_1_-_161417018_161417089@3.36.34,trna87_GluCTC_6_-_126101393_126101464@3.36.34|CCTGGTGGTCTAGTGGTTAGGATTCGGCGCTCTC</td><td>>trna116_GluCTC_1_-_145399233_145399304</td><td>3</td><td>36</td><td>34</td><td>CCTGGTGGTCTAGTGGTTAGGATTCGGCGCTCTC</td><td>663.6</td><td>4.7</td><td>0.8</td><td>5.8</td><td>7.9E-09</td><td>1.5E-08</td><td>MB231/MB468</td><td>MITO</td></tr>
<tr><td>>hg38_21_+_8433172_8446622_RNA45SN1_45S_50ntFlanks@10817.10840.24|TAAGGGAAGTCGGCAAGCCGGATC</td><td>>hg38_21_+_8433172_8446622_RNA45SN1_45S_50ntFlanks</td><td>10817</td><td>10840</td><td>24</td><td>TAAGGGAAGTCGGCAAGCCGGATC</td><td>279.2</td><td>-2.4</td><td>0.4</td><td>-5.8</td><td>6.2E-09</td><td>1.5E-08</td><td>BT20/MB468</td><td>CYTO</td></tr>
<tr><td>>trna111_HisGTG_1_-_147774845_147774916@-1G.34.35,trna118_HisGTG_1_-_145396881_145396952@-1G.34.35,trna16_HisGTG_1_+_146544773_146544844@-1G.34.35,trna1_HisGTG_15_+_45493349_45493420@-1G.34.35,trna21_HisGTG_1_+_147753471_147753542@-1G.34.35,trna33_HisGTG_6_+_27125906_27125977@-1G.34.35,trna7_HisGTG_9_-_14433938_14434009@-1G.34.35,trna8_HisGTG_15_-_45492611_45492682@-1G.34.35,trna9_HisGTG_15_-_45490804_45490875@-1G.34.35|GGCCGTGATCGTATAGTGGTTAGTACTCTGCGTTG</td><td>>trna111_HisGTG_1_-_147774845_147774916</td><td>-1G</td><td>34</td><td>35</td><td>GGCCGTGATCGTATAGTGGTTAGTACTCTGCGTTG</td><td>7142.5</td><td>6.0</td><td>1.0</td><td>5.8</td><td>7.9E-09</td><td>1.5E-08</td><td>MB231/MB468</td><td>MITO</td></tr>
<tr><td>>hg38_21_+_8433172_8446622_RNA45SN1_45S_50ntFlanks@11666.11691.26|TCTGCCCAGTGCTCTGAATGTCAAAG</td><td>>hg38_21_+_8433172_8446622_RNA45SN1_45S_50ntFlanks</td><td>11666</td><td>11691</td><td>26</td><td>TCTGCCCAGTGCTCTGAATGTCAAAG</td><td>464.0</td><td>4.5</td><td>0.8</td><td>5.8</td><td>6.2E-09</td><td>1.5E-08</td><td>BT20/MB468</td><td>CYTO</td></tr>
<tr><td>>hg38_21_+_8433172_8446622_RNA45SN1_45S_50ntFlanks@5086.5104.19|AACGAGACTCTGGCATGCT</td><td>>hg38_21_+_8433172_8446622_RNA45SN1_45S_50ntFlanks</td><td>5086</td><td>5104</td><td>19</td><td>AACGAGACTCTGGCATGCT</td><td>228.7</td><td>5.2</td><td>0.9</td><td>5.8</td><td>7.2E-09</td><td>1.5E-08</td><td>MB231/MB468</td><td>CYTO</td></tr>
<tr><td>>MI0000809|hsa-mir-151a&WithFlank&8|-|140732558|140732659@51.72.22(+1A)&[MIMAT0000757&hsa-miR-151a-3p&offsets|-2|-1(+1A);m-30&8|-|140732587|140732607&offsets|-2|-1(+1A)]|TACTAGACTGAAGCTCCTTGAGA</td><td>>MI0000809|hsa-mir-151a&WithFlank&8|-|140732558|140732659</td><td>51</td><td>72</td><td>22(+1A)</td><td>TACTAGACTGAAGCTCCTTGAGA</td><td>356.7</td><td>4.0</td><td>0.7</td><td>5.8</td><td>6.8E-09</td><td>1.5E-08</td><td>MB231/MB468</td><td>MP</td></tr>
<tr><td>>hg38_21_+_8433172_8446622_RNA45SN1_45S_50ntFlanks@8157.8175.19|CTCGTGGGGGGCCCAAGTC</td><td>>hg38_21_+_8433172_8446622_RNA45SN1_45S_50ntFlanks</td><td>8157</td><td>8175</td><td>19</td><td>CTCGTGGGGGGCCCAAGTC</td><td>144.6</td><td>-10.3</td><td>1.7</td><td>-6.1</td><td>9.6E-10</td><td>1.5E-08</td><td>BT20/MB468</td><td>MP</td></tr>
<tr><td>>hg38_21_+_8433172_8446622_RNA45SN1_45S_50ntFlanks@9740.9757.18|AACGATCTCAACCTATTC</td><td>>hg38_21_+_8433172_8446622_RNA45SN1_45S_50ntFlanks</td><td>9740</td><td>9757</td><td>18</td><td>AACGATCTCAACCTATTC</td><td>589.2</td><td>-8.2</td><td>1.4</td><td>-5.8</td><td>8.0E-09</td><td>1.5E-08</td><td>MB231/MB468</td><td>MITO</td></tr>
<tr><td>>hg38_21_+_8433172_8446622_RNA45SN1_45S_50ntFlanks@4163.4182.20|CATCCAAGGAAGGCAGCAGG</td><td>>hg38_21_+_8433172_8446622_RNA45SN1_45S_50ntFlanks</td><td>4163</td><td>4182</td><td>20</td><td>CATCCAAGGAAGGCAGCAGG</td><td>519.1</td><td>-3.8</td><td>0.6</td><td>-5.8</td><td>6.2E-09</td><td>1.5E-08</td><td>BT20/MB468</td><td>CYTO</td></tr>
<tr><td>>hg38_21_+_8433172_8446622_RNA45SN1_45S_50ntFlanks@11876.11917.42|AAGACCCTGTTGAGCTTGACTCTAGTCTGGCACGGTGAAGAG</td><td>>hg38_21_+_8433172_8446622_RNA45SN1_45S_50ntFlanks</td><td>11876</td><td>11917</td><td>42</td><td>AAGACCCTGTTGAGCTTGACTCTAGTCTGGCACGGTGAAGAG</td><td>221.0</td><td>-7.8</td><td>1.4</td><td>-5.8</td><td>8.0E-09</td><td>1.5E-08</td><td>MB231/MB468</td><td>MITO</td></tr>
<tr><td>>hg38_21_+_8433172_8446622_RNA45SN1_45S_50ntFlanks@4759.4776.18|ATCAGATACCGTCGTAGT</td><td>>hg38_21_+_8433172_8446622_RNA45SN1_45S_50ntFlanks</td><td>4759</td><td>4776</td><td>18</td><td>ATCAGATACCGTCGTAGT</td><td>112.3</td><td>-9.9</td><td>1.6</td><td>-6.1</td><td>9.7E-10</td><td>1.5E-08</td><td>BT20/MB468</td><td>MP</td></tr>
<tr><td>>MI0000746|hsa-mir-99b&WithFlank&19|+|51692606|51692687@51.73.23&[MIMAT0004678&hsa-miR-99b-3p&offsets|0|+1;m-151&19|+|51692656|51692678&offsets|0|0]|CAAGCTCGTGTCTGTGGGTCCGT</td><td>>MI0000746|hsa-mir-99b&WithFlank&19|+|51692606|51692687</td><td>51</td><td>73</td><td>23</td><td>CAAGCTCGTGTCTGTGGGTCCGT</td><td>476.0</td><td>4.9</td><td>0.9</td><td>5.8</td><td>7.4E-09</td><td>1.5E-08</td><td>MB231/MB468</td><td>NUC</td></tr>
<tr><td>>hg38_21_+_8433172_8446622_RNA45SN1_45S_50ntFlanks@12407.12441.35|TTTTGATCCTTCGATGTCGGCTCTTCCTATCATTG</td><td>>hg38_21_+_8433172_8446622_RNA45SN1_45S_50ntFlanks</td><td>12407</td><td>12441</td><td>35</td><td>TTTTGATCCTTCGATGTCGGCTCTTCCTATCATTG</td><td>2231.3</td><td>-3.5</td><td>0.6</td><td>-5.8</td><td>7.5E-09</td><td>1.5E-08</td><td>MB231/MB468</td><td>NUC</td></tr>
<tr><td>>hg38_21_+_8433172_8446622_RNA45SN1_45S_50ntFlanks@8170.8211.42|CAAGTCCTTCTGATCGAGGCCCAGCCCGTGGACGGTGTGAGG</td><td>>hg38_21_+_8433172_8446622_RNA45SN1_45S_50ntFlanks</td><td>8170</td><td>8211</td><td>42</td><td>CAAGTCCTTCTGATCGAGGCCCAGCCCGTGGACGGTGTGAGG</td><td>222.5</td><td>-10.9</td><td>1.8</td><td>-6.1</td><td>9.7E-10</td><td>1.5E-08</td><td>BT20/MB468</td><td>MP</td></tr>
<tr><td>>hg38_21_+_8433172_8446622_RNA45SN1_45S_50ntFlanks@8176.8203.28|CTTCTGATCGAGGCCCAGCCCGTGGACG</td><td>>hg38_21_+_8433172_8446622_RNA45SN1_45S_50ntFlanks</td><td>8176</td><td>8203</td><td>28</td><td>CTTCTGATCGAGGCCCAGCCCGTGGACG</td><td>2658.8</td><td>-2.9</td><td>0.5</td><td>-5.8</td><td>6.3E-09</td><td>1.5E-08</td><td>BT20/MB468</td><td>CYTO</td></tr>
<tr><td>>hg38_21_+_8433172_8446622_RNA45SN1_45S_50ntFlanks@9862.9900.39|ATGAACCGAACGCCGGGTTAAGGCGCCCGATGCCGACGC</td><td>>hg38_21_+_8433172_8446622_RNA45SN1_45S_50ntFlanks</td><td>9862</td><td>9900</td><td>39</td><td>ATGAACCGAACGCCGGGTTAAGGCGCCCGATGCCGACGC</td><td>162.4</td><td>-11.1</td><td>1.9</td><td>-5.8</td><td>8.1E-09</td><td>1.5E-08</td><td>MB231/MB468</td><td>MITO</td></tr>
<tr><td>>hg38_21_+_8433172_8446622_RNA45SN1_45S_50ntFlanks@4722.4758.37|TCATTAATCAAGAACGAAAGTCGGAGGTTCGAAGACG</td><td>>hg38_21_+_8433172_8446622_RNA45SN1_45S_50ntFlanks</td><td>4722</td><td>4758</td><td>37</td><td>TCATTAATCAAGAACGAAAGTCGGAGGTTCGAAGACG</td><td>5409.2</td><td>-4.9</td><td>0.8</td><td>-5.8</td><td>6.3E-09</td><td>1.5E-08</td><td>BT20/MB468</td><td>CYTO</td></tr>
<tr><td>>hg38_21_+_8433172_8446622_RNA45SN1_45S_50ntFlanks@10411.10445.35|CAGTCGGTCCTGAGAGATGGGCGAGCGCCGTTCCG</td><td>>hg38_21_+_8433172_8446622_RNA45SN1_45S_50ntFlanks</td><td>10411</td><td>10445</td><td>35</td><td>CAGTCGGTCCTGAGAGATGGGCGAGCGCCGTTCCG</td><td>389.0</td><td>-3.3</td><td>0.6</td><td>-5.8</td><td>5.6E-09</td><td>1.5E-08</td><td>BT20/MB468</td><td>MITO</td></tr>
<tr><td>>hg38_21_+_8433172_8446622_RNA45SN1_45S_50ntFlanks@9930.9948.19|ATATAGACAGCAGGACGGT</td><td>>hg38_21_+_8433172_8446622_RNA45SN1_45S_50ntFlanks</td><td>9930</td><td>9948</td><td>19</td><td>ATATAGACAGCAGGACGGT</td><td>447.0</td><td>-6.4</td><td>1.1</td><td>-5.8</td><td>8.1E-09</td><td>1.5E-08</td><td>MB231/MB468</td><td>MITO</td></tr>
<tr><td>>hg38_21_+_8433172_8446622_RNA45SN1_45S_50ntFlanks@12712.12747.36|CGGAGCCTCGGTTGGCCTCGGATAGCCGGTCCCCCG</td><td>>hg38_21_+_8433172_8446622_RNA45SN1_45S_50ntFlanks</td><td>12712</td><td>12747</td><td>36</td><td>CGGAGCCTCGGTTGGCCTCGGATAGCCGGTCCCCCG</td><td>486.6</td><td>-7.3</td><td>1.3</td><td>-5.8</td><td>5.6E-09</td><td>1.5E-08</td><td>BT20/MB468</td><td>MITO</td></tr>
<tr><td>>hg38_21_+_8433172_8446622_RNA45SN1_45S_50ntFlanks@4588.4614.27|CTATTTTGTTGGTTTTCGGAACTGAGG</td><td>>hg38_21_+_8433172_8446622_RNA45SN1_45S_50ntFlanks</td><td>4588</td><td>4614</td><td>27</td><td>CTATTTTGTTGGTTTTCGGAACTGAGG</td><td>530.7</td><td>-5.4</td><td>0.9</td><td>-5.8</td><td>6.3E-09</td><td>1.5E-08</td><td>BT20/MB468</td><td>CYTO</td></tr>
<tr><td>>hg38_21_+_8433172_8446622_RNA45SN1_45S_50ntFlanks@8354.8375.22|TTGAAAAGAACTTTGAAGAGAG</td><td>>hg38_21_+_8433172_8446622_RNA45SN1_45S_50ntFlanks</td><td>8354</td><td>8375</td><td>22</td><td>TTGAAAAGAACTTTGAAGAGAG</td><td>1483.4</td><td>7.5</td><td>1.2</td><td>6.1</td><td>9.8E-10</td><td>1.5E-08</td><td>BT20/MB468</td><td>MP</td></tr>
<tr><td>>am_trnaMT_PheGAA_MT_+_577_647@42.62.21|TTTAGACGGGCTCACATCACC</td><td>>am_trnaMT_PheGAA_MT_+_577_647</td><td>42</td><td>62</td><td>21</td><td>TTTAGACGGGCTCACATCACC</td><td>240.9</td><td>4.7</td><td>0.8</td><td>5.8</td><td>8.1E-09</td><td>1.5E-08</td><td>MB231/MB468</td><td>MITO</td></tr>
<tr><td>>hg38_21_+_8433172_8446622_RNA45SN1_45S_50ntFlanks@10786.10822.37|AACAGCCTCTGGCATGTTGGAACAATGTAGGTAAGGG</td><td>>hg38_21_+_8433172_8446622_RNA45SN1_45S_50ntFlanks</td><td>10786</td><td>10822</td><td>37</td><td>AACAGCCTCTGGCATGTTGGAACAATGTAGGTAAGGG</td><td>895.5</td><td>-8.9</td><td>1.5</td><td>-5.8</td><td>8.4E-09</td><td>1.5E-08</td><td>BT20/MB468</td><td>NUC</td></tr>
<tr><td>>hg38_21_+_8433172_8446622_RNA45SN1_45S_50ntFlanks@13002.13023.22|CGATCTATTGAAAGTCAGCCCT</td><td>>hg38_21_+_8433172_8446622_RNA45SN1_45S_50ntFlanks</td><td>13002</td><td>13023</td><td>22</td><td>CGATCTATTGAAAGTCAGCCCT</td><td>866.7</td><td>-7.2</td><td>1.2</td><td>-5.8</td><td>7.4E-09</td><td>1.5E-08</td><td>MB231/MB468</td><td>CYTO</td></tr>
<tr><td>>hg38_21_+_8433172_8446622_RNA45SN1_45S_50ntFlanks@11783.11800.18|CATGAATGGATGAACGAG</td><td>>hg38_21_+_8433172_8446622_RNA45SN1_45S_50ntFlanks</td><td>11783</td><td>11800</td><td>18</td><td>CATGAATGGATGAACGAG</td><td>399.1</td><td>-7.5</td><td>1.3</td><td>-5.8</td><td>8.1E-09</td><td>1.5E-08</td><td>MB231/MB468</td><td>MITO</td></tr>
<tr><td>>hg38_21_+_8433172_8446622_RNA45SN1_45S_50ntFlanks@5492.5524.33|AGAAGACGGTCGAACTTGACTATCTAGAGGAAG</td><td>>hg38_21_+_8433172_8446622_RNA45SN1_45S_50ntFlanks</td><td>5492</td><td>5524</td><td>33</td><td>AGAAGACGGTCGAACTTGACTATCTAGAGGAAG</td><td>389.5</td><td>-7.5</td><td>1.3</td><td>-5.8</td><td>8.1E-09</td><td>1.5E-08</td><td>MB231/MB468</td><td>MITO</td></tr>
<tr><td>>hg38_21_+_8433172_8446622_RNA45SN1_45S_50ntFlanks@4003.4025.23|ATAACCTCGGGCCGATCGCACGC</td><td>>hg38_21_+_8433172_8446622_RNA45SN1_45S_50ntFlanks</td><td>4003</td><td>4025</td><td>23</td><td>ATAACCTCGGGCCGATCGCACGC</td><td>3317.1</td><td>-9.7</td><td>1.7</td><td>-5.8</td><td>8.4E-09</td><td>1.5E-08</td><td>BT20/MB468</td><td>NUC</td></tr>
<tr><td>>hg38_21_+_8433172_8446622_RNA45SN1_45S_50ntFlanks@5467.5484.18|GGCCCACGGCCCTGGCGG</td><td>>hg38_21_+_8433172_8446622_RNA45SN1_45S_50ntFlanks</td><td>5467</td><td>5484</td><td>18</td><td>GGCCCACGGCCCTGGCGG</td><td>631.6</td><td>-8.8</td><td>1.5</td><td>-5.8</td><td>8.4E-09</td><td>1.5E-08</td><td>BT20/MB468</td><td>NUC</td></tr>
<tr><td>>hg38_21_+_8433172_8446622_RNA45SN1_45S_50ntFlanks@10731.10755.25|AGGGTGTAAATCTCGCGCCGGGCCG</td><td>>hg38_21_+_8433172_8446622_RNA45SN1_45S_50ntFlanks</td><td>10731</td><td>10755</td><td>25</td><td>AGGGTGTAAATCTCGCGCCGGGCCG</td><td>395.7</td><td>-6.5</td><td>1.1</td><td>-5.8</td><td>7.4E-09</td><td>1.5E-08</td><td>MB231/MB468</td><td>CYTO</td></tr>
<tr><td>>hg38_21_+_8433172_8446622_RNA45SN1_45S_50ntFlanks@3811.3834.24|AGTTATGGTTCCTTTGGTCGCTCG</td><td>>hg38_21_+_8433172_8446622_RNA45SN1_45S_50ntFlanks</td><td>3811</td><td>3834</td><td>24</td><td>AGTTATGGTTCCTTTGGTCGCTCG</td><td>282.2</td><td>-7.8</td><td>1.3</td><td>-5.8</td><td>8.1E-09</td><td>1.5E-08</td><td>MB231/MB468</td><td>MITO</td></tr>
<tr><td>>hg38_21_+_8433172_8446622_RNA45SN1_45S_50ntFlanks@8380.8418.39|CAAGAGGGCGTGAAACCGTTAAGAGGTAAACGGGTGGGG</td><td>>hg38_21_+_8433172_8446622_RNA45SN1_45S_50ntFlanks</td><td>8380</td><td>8418</td><td>39</td><td>CAAGAGGGCGTGAAACCGTTAAGAGGTAAACGGGTGGGG</td><td>110.2</td><td>-10.6</td><td>1.8</td><td>-5.8</td><td>7.0E-09</td><td>1.5E-08</td><td>MB231/MB468</td><td>MP</td></tr>
<tr><td>>MI0000078|hsa-mir-22&WithFlank&17|-|1713897|1713993@59.78.20&[MIMAT0000077&hsa-miR-22-3p&offsets|0|-2;m-2&17|-|1713914|1713935&offsets|0|-2]|AAGCTGCCAGTTGAAGAACT</td><td>>MI0000078|hsa-mir-22&WithFlank&17|-|1713897|1713993</td><td>59</td><td>78</td><td>20</td><td>AAGCTGCCAGTTGAAGAACT</td><td>566.6</td><td>7.9</td><td>1.4</td><td>5.8</td><td>8.2E-09</td><td>1.5E-08</td><td>MB231/MB468</td><td>MITO</td></tr>
<tr><td>>trna10_AspGTC_12_-_125424193_125424264@40.70.31,trna12_AspGTC_12_-_125411891_125411962@40.70.31,trna144_AspGTC_6_-_27551236_27551307@40.70.31,trna38_AspGTC_17_-_8125556_8125627@40.70.31,trna45_AspGTC_6_+_27447453_27447524@40.70.31,trna48_AspGTC_6_+_27471523_27471594@40.70.31,trna4_AspGTC_12_+_96429799_96429870@40.70.31,trna69_AspGTC_1_-_161440205_161440276@40.70.31,trna72_AspGTC_1_-_161432824_161432895@40.70.31,trna75_AspGTC_1_-_161425414_161425485@40.70.31,trna78_AspGTC_1_-_161418033_161418104@40.70.31,trna81_AspGTC_1_-_161410615_161410686@40.70.31|CGGGAGACCGGGGTTCGATTCCCCGACGGGG</td><td>>trna10_AspGTC_12_-_125424193_125424264</td><td>40</td><td>70</td><td>31</td><td>CGGGAGACCGGGGTTCGATTCCCCGACGGGG</td><td>755.2</td><td>-5.1</td><td>0.9</td><td>-5.8</td><td>7.0E-09</td><td>1.5E-08</td><td>MB231/MB468</td><td>MP</td></tr>
<tr><td>>hg38_21_+_8433172_8446622_RNA45SN1_45S_50ntFlanks@9534.9551.18|AGGAAACTCTGGTGGAGG</td><td>>hg38_21_+_8433172_8446622_RNA45SN1_45S_50ntFlanks</td><td>9534</td><td>9551</td><td>18</td><td>AGGAAACTCTGGTGGAGG</td><td>5485.9</td><td>-9.3</td><td>1.6</td><td>-5.8</td><td>7.0E-09</td><td>1.6E-08</td><td>MB231/MB468</td><td>MP</td></tr>
</table>

<table>
<tr><th></th><th></th><th></th><th></th><th></th><th></th><th></th><th></th><th></th><th></th><th></th><th></th><th></th><th></th></tr>
<tr><td>>hg38_21_+_8433172_8446622_RNA45SN1_45S_50ntFlanks@11718.11744.27|AACGGCGGGAGTAACTATGACTCTCTT</td><td>>hg38_21_+_8433172_8446622_RNA45SN1_45S_50ntFlanks</td><td>11718</td><td>11744</td><td>27</td><td>AACGGCGGGAGTAACTATGACTCTCTT</td><td>724.8</td><td>-6.7</td><td>1.2</td><td>-5.8</td><td>7.5E-09</td><td>1.6E-08</td><td>MB231/MB468</td><td>CYTO</td></tr>
<tr><td>>hg38_21_+_8433172_8446622_RNA45SN1_45S_50ntFlanks@6660.6685.26|GCGGTGGATCACTCGGCTCGTGCGTC</td><td>>hg38_21_+_8433172_8446622_RNA45SN1_45S_50ntFlanks</td><td>6660</td><td>6685</td><td>26</td><td>GCGGTGGATCACTCGGCTCGTGCGTC</td><td>432.1</td><td>4.1</td><td>0.7</td><td>5.8</td><td>5.7E-09</td><td>1.6E-08</td><td>BT20/MB468</td><td>MITO</td></tr>
<tr><td>>hg38_21_+_8433172_8446622_RNA45SN1_45S_50ntFlanks@11885.11904.20|TTGAGCTTGACTCTAGTCTG</td><td>>hg38_21_+_8433172_8446622_RNA45SN1_45S_50ntFlanks</td><td>11885</td><td>11904</td><td>20</td><td>TTGAGCTTGACTCTAGTCTG</td><td>408.9</td><td>-5.8</td><td>1.0</td><td>-5.8</td><td>7.6E-09</td><td>1.6E-08</td><td>MB231/MB468</td><td>NUC</td></tr>
<tr><td>>hg38_21_+_8433172_8446622_RNA45SN1_45S_50ntFlanks@5492.5515.24|AGAAGACGGTCGAACTTGACTATC</td><td>>hg38_21_+_8433172_8446622_RNA45SN1_45S_50ntFlanks</td><td>5492</td><td>5515</td><td>24</td><td>AGAAGACGGTCGAACTTGACTATC</td><td>389.2</td><td>-6.1</td><td>1.1</td><td>-5.8</td><td>7.1E-09</td><td>1.6E-08</td><td>MB231/MB468</td><td>MP</td></tr>
<tr><td>>hg38_21_+_8433172_8446622_RNA45SN1_45S_50ntFlanks@8144.8175.32|CTCCCCGGCGCCGCTCGTGGGGGGCCCAAGTC</td><td>>hg38_21_+_8433172_8446622_RNA45SN1_45S_50ntFlanks</td><td>8144</td><td>8175</td><td>32</td><td>CTCCCCGGCGCCGCTCGTGGGGGGCCCAAGTC</td><td>95.0</td><td>-10.4</td><td>1.8</td><td>-5.8</td><td>7.1E-09</td><td>1.6E-08</td><td>MB231/MB468</td><td>MP</td></tr>
<tr><td>>hg38_21_+_8433172_8446622_RNA45SN1_45S_50ntFlanks@11455.11475.21|CTCGGCGCGCGGCGGCGGCGG</td><td>>hg38_21_+_8433172_8446622_RNA45SN1_45S_50ntFlanks</td><td>11455</td><td>11475</td><td>21</td><td>CTCGGCGCGCGGCGGCGGCGG</td><td>131.8</td><td>-10.1</td><td>1.7</td><td>-6.1</td><td>9.9E-10</td><td>1.6E-08</td><td>BT20/MB468</td><td>MP</td></tr>
<tr><td>>hg38_21_+_8433172_8446622_RNA45SN1_45S_50ntFlanks@10663.10706.44|CCTTGGAAAGCGTCGCGGTTCCGGCGGCGTCCGGTGAGCTCTCG</td><td>>hg38_21_+_8433172_8446622_RNA45SN1_45S_50ntFlanks</td><td>10663</td><td>10706</td><td>44</td><td>CCTTGGAAAGCGTCGCGGTTCCGGCGGCGTCCGGTGAGCTCTCG</td><td>780.0</td><td>-8.4</td><td>1.5</td><td>-5.8</td><td>7.6E-09</td><td>1.6E-08</td><td>MB231/MB468</td><td>NUC</td></tr>
<tr><td>>MI0000471|hsa-mir-126&WithFlank&9|+|136670596|136670692@59.77.19(+1U)&[MIMAT0000445&hsa-miR-126-3p&offsets|+1|-2(+1U);m-40&9|+|136670653|136670674&offsets|+1|-2(+1U)]|CGTACCGTGAGTAATAATGT</td><td>>MI0000471|hsa-mir-126&WithFlank&9|+|136670596|136670692</td><td>59</td><td>77</td><td>19(+1U)</td><td>CGTACCGTGAGTAATAATGT</td><td>191.1</td><td>11.3</td><td>1.9</td><td>6.1</td><td>1.0E-09</td><td>1.6E-08</td><td>BT20/MB468</td><td>MP</td></tr>
<tr><td>>am_MI0000239|hsa-mir-197&WithFlank&1|+|109598887|109598973@54.71.18(+1U)&[MIMAT0000227&hsa-miR-197-3p&offsets|0|-4(+1U);m-99&1|+|109598940|109598961&offsets|0|-4(+1U)]|TTCACCACCTTCTCCACCT</td><td>>am_MI0000239|hsa-mir-197&WithFlank&1|+|109598887|109598973</td><td>54</td><td>71</td><td>18(+1U)</td><td>TTCACCACCTTCTCCACCT</td><td>241.0</td><td>3.8</td><td>0.6</td><td>6.1</td><td>7.9E-10</td><td>1.6E-08</td><td>BT20/MB231</td><td>NUC</td></tr>
<tr><td>>hg38_21_+_8433172_8446622_RNA45SN1_45S_50ntFlanks@6646.6685.40|TCGTACGACTCTTAGCGGTGGATCACTCGGCTCGTGCGTC</td><td>>hg38_21_+_8433172_8446622_RNA45SN1_45S_50ntFlanks</td><td>6646</td><td>6685</td><td>40</td><td>TCGTACGACTCTTAGCGGTGGATCACTCGGCTCGTGCGTC</td><td>18906.5</td><td>5.2</td><td>0.8</td><td>6.1</td><td>7.9E-10</td><td>1.6E-08</td><td>BT20/MB231</td><td>NUC</td></tr>
<tr><td>>hg38_21_+_8433172_8446622_RNA45SN1_45S_50ntFlanks@11208.11231.24|GCGGCGACTCTGGACGCGAGCCGG</td><td>>hg38_21_+_8433172_8446622_RNA45SN1_45S_50ntFlanks</td><td>11208</td><td>11231</td><td>24</td><td>GCGGCGACTCTGGACGCGAGCCGG</td><td>2065.1</td><td>-3.8</td><td>0.7</td><td>-5.8</td><td>6.4E-09</td><td>1.6E-08</td><td>BT20/MB468</td><td>CYTO</td></tr>
<tr><td>>hg38_21_+_8433172_8446622_RNA45SN1_45S_50ntFlanks@5003.5022.20|ATAGCTCTTTCTCGATTCCG</td><td>>hg38_21_+_8433172_8446622_RNA45SN1_45S_50ntFlanks</td><td>5003</td><td>5022</td><td>20</td><td>ATAGCTCTTTCTCGATTCCG</td><td>205.8</td><td>-9.8</td><td>1.7</td><td>-5.8</td><td>5.8E-09</td><td>1.6E-08</td><td>BT20/MB468</td><td>MITO</td></tr>
<tr><td>>hg38_21_+_8433172_8446622_RNA45SN1_45S_50ntFlanks@11570.11611.42|AGCCGACTTAGAACTGGTGCGGACCAGGGGAATCCGACTGTT</td><td>>hg38_21_+_8433172_8446622_RNA45SN1_45S_50ntFlanks</td><td>11570</td><td>11611</td><td>42</td><td>AGCCGACTTAGAACTGGTGCGGACCAGGGGAATCCGACTGTT</td><td>1194.1</td><td>-4.9</td><td>0.9</td><td>-5.8</td><td>8.3E-09</td><td>1.6E-08</td><td>MB231/MB468</td><td>MITO</td></tr>
<tr><td>>hg38_21_+_8433172_8446622_RNA45SN1_45S_50ntFlanks@7973.8016.44|GACGCGACCTCAGATCAGACGTGGCGACCCGCTGAATTTAAGCA</td><td>>hg38_21_+_8433172_8446622_RNA45SN1_45S_50ntFlanks</td><td>7973</td><td>8016</td><td>44</td><td>GACGCGACCTCAGATCAGACGTGGCGACCCGCTGAATTTAAGCA</td><td>598.3</td><td>4.9</td><td>0.8</td><td>5.8</td><td>5.8E-09</td><td>1.6E-08</td><td>BT20/MB468</td><td>MITO</td></tr>
<tr><td>>hg38_21_+_8433172_8446622_RNA45SN1_45S_50ntFlanks@12607.12624.18|TGTGCTTGGCTGAGGAGC</td><td>>hg38_21_+_8433172_8446622_RNA45SN1_45S_50ntFlanks</td><td>12607</td><td>12624</td><td>18</td><td>TGTGCTTGGCTGAGGAGC</td><td>622.3</td><td>-4.0</td><td>0.7</td><td>-5.8</td><td>8.6E-09</td><td>1.6E-08</td><td>BT20/MB468</td><td>NUC</td></tr>
<tr><td>>hg38_21_+_8433172_8446622_RNA45SN1_45S_50ntFlanks@6719.6742.24|TGAATTGCAGGACACATTGATCAT</td><td>>hg38_21_+_8433172_8446622_RNA45SN1_45S_50ntFlanks</td><td>6719</td><td>6742</td><td>24</td><td>TGAATTGCAGGACACATTGATCAT</td><td>342.3</td><td>-7.1</td><td>1.2</td><td>-5.8</td><td>6.5E-09</td><td>1.6E-08</td><td>BT20/MB468</td><td>CYTO</td></tr>
<tr><td>>hg38_21_+_8433172_8446622_RNA45SN1_45S_50ntFlanks@12424.12456.33|CGGCTCTTCCTATCATTGTGAAGCAGAATTCAC</td><td>>hg38_21_+_8433172_8446622_RNA45SN1_45S_50ntFlanks</td><td>12424</td><td>12456</td><td>33</td><td>CGGCTCTTCCTATCATTGTGAAGCAGAATTCAC</td><td>393.0</td><td>-8.4</td><td>1.5</td><td>-5.8</td><td>8.6E-09</td><td>1.6E-08</td><td>BT20/MB468</td><td>NUC</td></tr>
<tr><td>>hg38_21_+_8433172_8446622_RNA45SN1_45S_50ntFlanks@9568.9596.29|CGTGCAAATCGGTCGTCCGACCTGGGTAT</td><td>>hg38_21_+_8433172_8446622_RNA45SN1_45S_50ntFlanks</td><td>9568</td><td>9596</td><td>29</td><td>CGTGCAAATCGGTCGTCCGACCTGGGTAT</td><td>3772.3</td><td>-6.8</td><td>1.2</td><td>-5.8</td><td>6.5E-09</td><td>1.6E-08</td><td>BT20/MB468</td><td>CYTO</td></tr>
<tr><td>>hg38_21_+_8433172_8446622_RNA45SN1_45S_50ntFlanks@7975.8002.28|CGCGACCTCAGATCAGACGTGGCGACCC</td><td>>hg38_21_+_8433172_8446622_RNA45SN1_45S_50ntFlanks</td><td>7975</td><td>8002</td><td>28</td><td>CGCGACCTCAGATCAGACGTGGCGACCC</td><td>8151.0</td><td>3.9</td><td>0.6</td><td>6.1</td><td>8.0E-10</td><td>1.6E-08</td><td>BT20/MB231</td><td>NUC</td></tr>
<tr><td>>hg38_21_+_8433172_8446622_RNA45SN1_45S_50ntFlanks@3854.3874.21|ACTGTGGTAATTCTAGAGCTA</td><td>>hg38_21_+_8433172_8446622_RNA45SN1_45S_50ntFlanks</td><td>3854</td><td>3874</td><td>21</td><td>ACTGTGGTAATTCTAGAGCTA</td><td>323.9</td><td>-3.3</td><td>0.6</td><td>-5.8</td><td>7.6E-09</td><td>1.6E-08</td><td>MB231/MB468</td><td>CYTO</td></tr>
<tr><td>>hg38_21_+_8433172_8446622_RNA45SN1_45S_50ntFlanks@10843.10865.23|TAACTTCGGGATAAGGATTGGCT</td><td>>hg38_21_+_8433172_8446622_RNA45SN1_45S_50ntFlanks</td><td>10843</td><td>10865</td><td>23</td><td>TAACTTCGGGATAAGGATTGGCT</td><td>415.6</td><td>-7.8</td><td>1.3</td><td>-5.8</td><td>7.2E-09</td><td>1.6E-08</td><td>MB231/MB468</td><td>MP</td></tr>
<tr><td>>hg38_21_+_8433172_8446622_RNA45SN1_45S_50ntFlanks@11582.11612.31|ACTGGTGCGGACCAGGGGAATCCGACTGTTT</td><td>>hg38_21_+_8433172_8446622_RNA45SN1_45S_50ntFlanks</td><td>11582</td><td>11612</td><td>31</td><td>ACTGGTGCGGACCAGGGGAATCCGACTGTTT</td><td>516.9</td><td>-4.0</td><td>0.7</td><td>-5.8</td><td>7.2E-09</td><td>1.6E-08</td><td>MB231/MB468</td><td>MP</td></tr>
<tr><td>>hg38_21_+_8433172_8446622_RNA45SN1_45S_50ntFlanks@11564.11611.48|CCTAGCAGCCGACTTAGAACTGGTGCGGACCAGGGGAATCCGACTGTT</td><td>>hg38_21_+_8433172_8446622_RNA45SN1_45S_50ntFlanks</td><td>11564</td><td>11611</td><td>48</td><td>CCTAGCAGCCGACTTAGAACTGGTGCGGACCAGGGGAATCCGACTGTT</td><td>7561.4</td><td>-7.0</td><td>1.2</td><td>-5.8</td><td>8.3E-09</td><td>1.6E-08</td><td>MB231/MB468</td><td>MITO</td></tr>
<tr><td>>hg38_21_+_8433172_8446622_RNA45SN1_45S_50ntFlanks@12996.13039.44|TCGCTGCGATCTATTGAAAGTCAGCCCTCGACACAAGGGTTTGT</td><td>>hg38_21_+_8433172_8446622_RNA45SN1_45S_50ntFlanks</td><td>12996</td><td>13039</td><td>44</td><td>TCGCTGCGATCTATTGAAAGTCAGCCCTCGACACAAGGGTTTGT</td><td>126917.7</td><td>6.3</td><td>1.1</td><td>5.8</td><td>5.8E-09</td><td>1.6E-08</td><td>BT20/MB468</td><td>MITO</td></tr>
<tr><td>>hg38_21_+_8433172_8446622_RNA45SN1_45S_50ntFlanks@8171.8212.42|AAGTCCTTCTGATCGAGGCCCAGCCCGTGGACGGTGTGAGGC</td><td>>hg38_21_+_8433172_8446622_RNA45SN1_45S_50ntFlanks</td><td>8171</td><td>8212</td><td>42</td><td>AAGTCCTTCTGATCGAGGCCCAGCCCGTGGACGGTGTGAGGC</td><td>5845.7</td><td>-10.5</td><td>1.8</td><td>-5.8</td><td>8.6E-09</td><td>1.6E-08</td><td>BT20/MB468</td><td>NUC</td></tr>
<tr><td>>hg38_21_+_8433172_8446622_RNA45SN1_45S_50ntFlanks@9597.9618.22|AGGGGCGAAAGACTAATCGAAC</td><td>>hg38_21_+_8433172_8446622_RNA45SN1_45S_50ntFlanks</td><td>9597</td><td>9618</td><td>22</td><td>AGGGGCGAAAGACTAATCGAAC</td><td>224.5</td><td>-7.5</td><td>1.3</td><td>-5.8</td><td>7.2E-09</td><td>1.6E-08</td><td>MB231/MB468</td><td>MP</td></tr>
<tr><td>>hg38_21_+_8433172_8446622_RNA45SN1_45S_50ntFlanks@11569.11591.23|CAGCCGACTTAGAACTGGTGCGG</td><td>>hg38_21_+_8433172_8446622_RNA45SN1_45S_50ntFlanks</td><td>11569</td><td>11591</td><td>23</td><td>CAGCCGACTTAGAACTGGTGCGG</td><td>254.6</td><td>-7.4</td><td>1.3</td><td>-5.8</td><td>8.3E-09</td><td>1.6E-08</td><td>MB231/MB468</td><td>MITO</td></tr>
<tr><td>>hg38_21_+_8433172_8446622_RNA45SN1_45S_50ntFlanks@10873.10904.32|CTGGGTCGGTCGGGCTGGGGCGCGAAGCGGGG</td><td>>hg38_21_+_8433172_8446622_RNA45SN1_45S_50ntFlanks</td><td>10873</td><td>10904</td><td>32</td><td>CTGGGTCGGTCGGGCTGGGGCGCGAAGCGGGG</td><td>988.8</td><td>-9.0</td><td>1.6</td><td>-5.8</td><td>8.6E-09</td><td>1.6E-08</td><td>BT20/MB468</td><td>NUC</td></tr>
<tr><td>>hg38_21_+_8433172_8446622_RNA45SN1_45S_50ntFlanks@11719.11740.22|ACGGCGGGAGTAACTATGACTC</td><td>>hg38_21_+_8433172_8446622_RNA45SN1_45S_50ntFlanks</td><td>11719</td><td>11740</td><td>22</td><td>ACGGCGGGAGTAACTATGACTC</td><td>424.7</td><td>-6.3</td><td>1.1</td><td>-5.8</td><td>7.7E-09</td><td>1.6E-08</td><td>MB231/MB468</td><td>NUC</td></tr>
<tr><td>>hg38_21_+_8433172_8446622_RNA45SN1_45S_50ntFlanks@6777.6802.26|TCCCGGGGCTACGCCTGTCTGAGCGT</td><td>>hg38_21_+_8433172_8446622_RNA45SN1_45S_50ntFlanks</td><td>6777</td><td>6802</td><td>26</td><td>TCCCGGGGCTACGCCTGTCTGAGCGT</td><td>285.7</td><td>4.4</td><td>0.8</td><td>5.8</td><td>7.6E-09</td><td>1.6E-08</td><td>MB231/MB468</td><td>CYTO</td></tr>
<tr><td>>hg38_21_+_8433172_8446622_RNA45SN1_45S_50ntFlanks@12637.12654.18|GCTACCATCTGTGGGATT</td><td>>hg38_21_+_8433172_8446622_RNA45SN1_45S_50ntFlanks</td><td>12637</td><td>12654</td><td>18</td><td>GCTACCATCTGTGGGATT</td><td>251.6</td><td>-5.1</td><td>0.9</td><td>-5.8</td><td>7.6E-09</td><td>1.6E-08</td><td>MB231/MB468</td><td>CYTO</td></tr>
<tr><td>>hg38_21_+_8433172_8446622_RNA45SN1_45S_50ntFlanks@3789.3812.24|ACTGCGAATGGCTCATTAAATCAG</td><td>>hg38_21_+_8433172_8446622_RNA45SN1_45S_50ntFlanks</td><td>3789</td><td>3812</td><td>24</td><td>ACTGCGAATGGCTCATTAAATCAG</td><td>326.2</td><td>-10.4</td><td>1.8</td><td>-5.8</td><td>5.8E-09</td><td>1.6E-08</td><td>BT20/MB468</td><td>MITO</td></tr>
<tr><td>>MI0006383|hsa-mir-1248&WithFlank&3|+|186786666|186786783@10.32.23&[MIMAT0005900&hsa-miR-1248&offsets|0|-4;m-827&3|+|186786675|186786701&offsets|0|-4]|ACCTTCTTGTATAAGCACTGTGC</td><td>>MI0006383|hsa-mir-1248&WithFlank&3|+|186786666|186786783</td><td>10</td><td>32</td><td>23</td><td>ACCTTCTTGTATAAGCACTGTGC</td><td>532.6</td><td>8.2</td><td>1.4</td><td>5.8</td><td>6.6E-09</td><td>1.6E-08</td><td>BT20/MB468</td><td>CYTO</td></tr>
<tr><td>>hg38_21_+_8433172_8446622_RNA45SN1_45S_50ntFlanks@3705.3734.30|TACCTGGTTGATCCTGCCAGTAGCATATGC</td><td>>hg38_21_+_8433172_8446622_RNA45SN1_45S_50ntFlanks</td><td>3705</td><td>3734</td><td>30</td><td>TACCTGGTTGATCCTGCCAGTAGCATATGC</td><td>227.3</td><td>7.2</td><td>1.3</td><td>5.8</td><td>7.7E-09</td><td>1.6E-08</td><td>MB231/MB468</td><td>CYTO</td></tr>
<tr><td>>hg38_21_+_8433172_8446622_RNA45SN1_45S_50ntFlanks@3993.4013.21|GTGACTCTAGATAACCTCGGG</td><td>>hg38_21_+_8433172_8446622_RNA45SN1_45S_50ntFlanks</td><td>3993</td><td>4013</td><td>21</td><td>GTGACTCTAGATAACCTCGGG</td><td>2419.8</td><td>-7.6</td><td>1.3</td><td>-5.8</td><td>8.7E-09</td><td>1.6E-08</td><td>BT20/MB468</td><td>NUC</td></tr>
<tr><td>>hg38_21_+_8433172_8446622_RNA45SN1_45S_50ntFlanks@12394.12419.26|TAGCGACGTCGCTTTTTGATCCTTCG</td><td>>hg38_21_+_8433172_8446622_RNA45SN1_45S_50ntFlanks</td><td>12394</td><td>12419</td><td>26</td><td>TAGCGACGTCGCTTTTTGATCCTTCG</td><td>1782.8</td><td>-7.6</td><td>1.3</td><td>-5.8</td><td>8.7E-09</td><td>1.6E-08</td><td>BT20/MB468</td><td>NUC</td></tr>
<tr><td>>hg38_21_+_8433172_8446622_RNA45SN1_45S_50ntFlanks@6771.6806.36|GGTTCCTCCCGGGGCTACGCCTGTCTGAGCGTCGCT</td><td>>hg38_21_+_8433172_8446622_RNA45SN1_45S_50ntFlanks</td><td>6771</td><td>6806</td><td>36</td><td>GGTTCCTCCCGGGGCTACGCCTGTCTGAGCGTCGCT</td><td>1675.9</td><td>-4.0</td><td>0.7</td><td>-5.8</td><td>7.3E-09</td><td>1.6E-08</td><td>MB231/MB468</td><td>MP</td></tr>
<tr><td>>hg38_21_+_8433172_8446622_RNA45SN1_45S_50ntFlanks@12643.12661.19|ATCTGTGGGATTATGACTG</td><td>>hg38_21_+_8433172_8446622_RNA45SN1_45S_50ntFlanks</td><td>12643</td><td>12661</td><td>19</td><td>ATCTGTGGGATTATGACTG</td><td>1681.0</td><td>-8.6</td><td>1.5</td><td>-5.8</td><td>8.4E-09</td><td>1.6E-08</td><td>MB231/MB468</td><td>MITO</td></tr>
<tr><td>>hg38_21_+_8433172_8446622_RNA45SN1_45S_50ntFlanks@3835.3869.35|CTCCTCTCCTACTTGGATAACTGTGGTAATTCTAG</td><td>>hg38_21_+_8433172_8446622_RNA45SN1_45S_50ntFlanks</td><td>3835</td><td>3869</td><td>35</td><td>CTCCTCTCCTACTTGGATAACTGTGGTAATTCTAG</td><td>294.8</td><td>-7.7</td><td>1.3</td><td>-5.8</td><td>8.4E-09</td><td>1.6E-08</td><td>MB231/MB468</td><td>MITO</td></tr>
<tr><td>>hg38_21_+_8433172_8446622_RNA45SN1_45S_50ntFlanks@10843.10868.26|TAACTTCGGGATAAGGATTGGCTCTA</td><td>>hg38_21_+_8433172_8446622_RNA45SN1_45S_50ntFlanks</td><td>10843</td><td>10868</td><td>26</td><td>TAACTTCGGGATAAGGATTGGCTCTA</td><td>1105.9</td><td>-6.3</td><td>1.1</td><td>-5.8</td><td>7.7E-09</td><td>1.6E-08</td><td>MB231/MB468</td><td>CYTO</td></tr>
<tr><td>>hg38_21_+_8433172_8446622_RNA45SN1_45S_50ntFlanks@12472.12496.25|TCACCCACTAATAGGGAACGTGAGC</td><td>>hg38_21_+_8433172_8446622_RNA45SN1_45S_50ntFlanks</td><td>12472</td><td>12496</td><td>25</td><td>TCACCCACTAATAGGGAACGTGAGC</td><td>97.2</td><td>10.4</td><td>1.7</td><td>6.1</td><td>1.0E-09</td><td>1.6E-08</td><td>BT20/MB468</td><td>MP</td></tr>
<tr><td>>hg38_21_+_8433172_8446622_RNA45SN1_45S_50ntFlanks@8356.8379.24|GAAAAGAACTTTGAAGAGAGAGTT</td><td>>hg38_21_+_8433172_8446622_RNA45SN1_45S_50ntFlanks</td><td>8356</td><td>8379</td><td>24</td><td>GAAAAGAACTTTGAAGAGAGAGTT</td><td>14474.1</td><td>-8.5</td><td>1.4</td><td>-6.1</td><td>1.0E-09</td><td>1.6E-08</td><td>BT20/MB468</td><td>MP</td></tr>
<tr><td>>MI0000809|hsa-mir-151a&WithFlank&8|-|140732558|140732659@53.74.22(+1A)&[MIMAT0000757&hsa-miR-151a-3p&offsets|0|+1(+1A);m-30&8|-|140732587|140732607&offsets|0|+1(+1A)]|CTAGACTGAAGCTCCTTGAGGAA</td><td>>MI0000809|hsa-mir-151a&WithFlank&8|-|140732558|140732659</td><td>53</td><td>74</td><td>22(+1A)</td><td>CTAGACTGAAGCTCCTTGAGGAA</td><td>1876.3</td><td>3.5</td><td>0.6</td><td>5.8</td><td>7.3E-09</td><td>1.6E-08</td><td>MB231/MB468</td><td>MP</td></tr>
<tr><td>>hg38_21_+_8433172_8446622_RNA45SN1_45S_50ntFlanks@3937.3959.23|CCAACCCGGTCAGCCCCTCTCCG</td><td>>hg38_21_+_8433172_8446622_RNA45SN1_45S_50ntFlanks</td><td>3937</td><td>3959</td><td>23</td><td>CCAACCCGGTCAGCCCCTCTCCG</td><td>181.6</td><td>-9.6</td><td>1.6</td><td>-5.8</td><td>5.9E-09</td><td>1.6E-08</td><td>BT20/MB468</td><td>MITO</td></tr>
<tr><td>>hg38_21_+_8433172_8446622_RNA45SN1_45S_50ntFlanks@3899.3918.20|CCTTCGCGGGGGGGATGCGT</td><td>>hg38_21_+_8433172_8446622_RNA45SN1_45S_50ntFlanks</td><td>3899</td><td>3918</td><td>20</td><td>CCTTCGCGGGGGGGATGCGT</td><td>113.8</td><td>-8.9</td><td>1.5</td><td>-5.8</td><td>5.9E-09</td><td>1.6E-08</td><td>BT20/MB468</td><td>MITO</td></tr>
<tr><td>>hg38_21_+_8433172_8446622_RNA45SN1_45S_50ntFlanks@5081.5098.18|TAACGAACGAGACTCTGG</td><td>>hg38_21_+_8433172_8446622_RNA45SN1_45S_50ntFlanks</td><td>5081</td><td>5098</td><td>18</td><td>TAACGAACGAGACTCTGG</td><td>111.2</td><td>-10.7</td><td>1.8</td><td>-5.8</td><td>7.3E-09</td><td>1.6E-08</td><td>MB231/MB468</td><td>MP</td></tr>
<tr><td>>MI0000081|hsa-mir-24-2&WithFlank&19|-|13836281|13836365@56.77.22(+1C);MI0000080|hsa-mir-24-1&WithFlank&9|+|95086015|95086094@50.71.22(+1C)&[MIMAT0000080&hsa-miR-24-3p&offsets|0|0(+1C);m-36&19|-|13836288|13836310&offsets|0|-1(+1C)];[MIMAT0000080_1&hsa-miR-24-3p&offsets|0|0(+1C);m-35&9|+|95086064|95086086&offsets|0|-1(+1C)]|TGGCTCAGTTCAGCAGGAACAGC</td><td>>MI0000081|hsa-mir-24-2&WithFlank&19|-|13836281|13836365</td><td>56</td><td>77</td><td>22(+1C)</td><td>TGGCTCAGTTCAGCAGGAACAGC</td><td>1515.1</td><td>6.1</td><td>1.1</td><td>5.8</td><td>8.5E-09</td><td>1.6E-08</td><td>MB231/MB468</td><td>MITO</td></tr>
<tr><td>>MI0000542|hsa-mir-320a&WithFlank&8|-|22244960|22245043@49.70.22(+1G)&[MIMAT0000510&hsa-miR-320a-3p&offsets|+1|+1(+1G);m-60&8|-|22244975|22244996&offsets|+1|+1(+1G)]|AAAGCTGGGTTGAGAGGGCGAAG</td><td>>MI0000542|hsa-mir-320a&WithFlank&8|-|22244960|22245043</td><td>49</td><td>70</td><td>22(+1G)</td><td>AAAGCTGGGTTGAGAGGGCGAAG</td><td>132.7</td><td>10.8</td><td>1.8</td><td>6.1</td><td>1.0E-09</td><td>1.6E-08</td><td>BT20/MB468</td><td>MP</td></tr>
</table>

| | | | | | | | | | | | | | |
|---|---|---|---|---|---|---|---|---|---|---|---|---|---|
| >hg38_21_+_8433172_8446622_RNA45SN1_45S_50ntFlanks@6647.6676.30\|CGTACGACTCTTAGCGGTGGATCACTCGGC | >hg38_21_+_8433172_8446622_RNA45SN1_45S_50ntFlanks | 6647 | 6676 | 30 | CGTACGACTCTTAGCGGTGGATCACTCGGC | 930.7 | 7.3 | 1.3 | 5.8 | 6.7E-09 | 1.6E-08 | BT20/MB468 | CYTO |
| >MI0000477\|hsa-mir-146a&WithFlank&5\|+\|160485346\|160485456@27.48.22(+1C)&[MIMAT0000449&hsa-miR-146a-5p&offsets\|0\|0(+1C);m-129&5\|+\|160485372\|160485393&offsets\|0\|0(+1C)]\|TGAGAACTGAATTCCATGGGTTC | >MI0000477\|hsa-mir-146a&WithFlank&5\|+\|160485346\|160485456 | 27 | 48 | 22(+1C) | TGAGAACTGAATTCCATGGGTTC | 181.2 | 10.7 | 1.9 | 5.8 | 8.5E-09 | 1.6E-08 | MB231/MB468 | MITO |
| >hg38_21_+_8433172_8446622_RNA45SN1_45S_50ntFlanks@12126.12145.20\|CGACCCGCTCCGGGGACAGT | >hg38_21_+_8433172_8446622_RNA45SN1_45S_50ntFlanks | 12126 | 12145 | 20 | CGACCCGCTCCGGGGACAGT | 141.1 | -10.2 | 1.7 | -6.1 | 1.0E-09 | 1.6E-08 | BT20/MB468 | MP |
| >hg38_21_+_8433172_8446622_RNA45SN1_45S_50ntFlanks@12828.12847.20\|CCGGGGTCCGGTGCGGAGTG | >hg38_21_+_8433172_8446622_RNA45SN1_45S_50ntFlanks | 12828 | 12847 | 20 | CCGGGGTCCGGTGCGGAGTG | 88.9 | -10.3 | 1.8 | -5.8 | 7.4E-09 | 1.6E-08 | MB231/MB468 | MP |
| >hg38_21_+_8433172_8446622_RNA45SN1_45S_50ntFlanks@480.505.26\|TGATCGATGTGGTGACGTCGTGCTCT | >hg38_21_+_8433172_8446622_RNA45SN1_45S_50ntFlanks | 480 | 505 | 26 | TGATCGATGTGGTGACGTCGTGCTCT | 238.3 | 4.8 | 0.8 | 5.8 | 7.9E-09 | 1.6E-08 | MB231/MB468 | NUC |
| >MI0006443\|hsa-mir-1306&WithFlank&22\|+\|20086052\|20086148@21.42.22&[MIMAT0022726&hsa-miR-1306-5p&offsets\|0\|0;m-301&22\|+\|20086072\|20086092&offsets\|0\|+1]\|CCACCTCCCCTGCAAACGTCCA | >MI0006443\|hsa-mir-1306&WithFlank&22\|+\|20086052\|20086148 | 21 | 42 | 22 | CCACCTCCCCTGCAAACGTCCA | 361.8 | 9.0 | 1.5 | 5.8 | 6.7E-09 | 1.6E-08 | BT20/MB468 | CYTO |
| >hg38_21_+_8433172_8446622_RNA45SN1_45S_50ntFlanks@11568.11602.35\|GCAGCCGACTTAGAACTGGTGCGGACCAGGGGAAT | >hg38_21_+_8433172_8446622_RNA45SN1_45S_50ntFlanks | 11568 | 11602 | 35 | GCAGCCGACTTAGAACTGGTGCGGACCAGGGGAAT | 2122.9 | -5.5 | 1.0 | -5.8 | 8.9E-09 | 1.6E-08 | BT20/MB468 | NUC |
| >hg38_21_+_8433172_8446622_RNA45SN1_45S_50ntFlanks@10678.10695.18\|CGGTTCCGGCGGCGTCCG | >hg38_21_+_8433172_8446622_RNA45SN1_45S_50ntFlanks | 10678 | 10695 | 18 | CGGTTCCGGCGGCGTCCG | 84.7 | -8.5 | 1.5 | -5.8 | 6.0E-09 | 1.6E-08 | BT20/MB468 | MITO |
| >hg38_21_+_8433172_8446622_RNA45SN1_45S_50ntFlanks@10816.10840.25\|GTAAGGGAAGTCGGCAAGCCGGATC | >hg38_21_+_8433172_8446622_RNA45SN1_45S_50ntFlanks | 10816 | 10840 | 25 | GTAAGGGAAGTCGGCAAGCCGGATC | 323.1 | -7.6 | 1.3 | -5.8 | 7.4E-09 | 1.6E-08 | MB231/MB468 | MP |
| >hg38_21_+_8433172_8446622_RNA45SN1_45S_50ntFlanks@6646.6683.38\|TCGTACGACTCTTAGCGGTGGATCACTCGGCTCGTGCG | >hg38_21_+_8433172_8446622_RNA45SN1_45S_50ntFlanks | 6646 | 6683 | 38 | TCGTACGACTCTTAGCGGTGGATCACTCGGCTCGTGCG | 387.4 | 3.8 | 0.7 | 5.7 | 8.9E-09 | 1.6E-08 | BT20/MB468 | NUC |
| >hg38_21_+_8433172_8446622_RNA45SN1_45S_50ntFlanks@12433.12454.22\|CTATCATTGTGAAGCAGAATTC | >hg38_21_+_8433172_8446622_RNA45SN1_45S_50ntFlanks | 12433 | 12454 | 22 | CTATCATTGTGAAGCAGAATTC | 684.9 | -6.5 | 1.1 | -5.8 | 7.5E-09 | 1.6E-08 | MB231/MB468 | MP |
| >trnaMT_ValTAC_MT_+_1602_1670@47.71.25\|TCAACTTAACTTGACCGCTCTGACC | >trnaMT_ValTAC_MT_+_1602_1670 | 47 | 71 | 25 | TCAACTTAACTTGACCGCTCTGACC | 313.3 | 5.7 | 1.0 | 5.8 | 8.0E-09 | 1.6E-08 | MB231/MB468 | NUC |
| >MI0003643\|hsa-mir-629&WithFlank&15\|-\|70079366\|70079474@28.47.20&[MIMAT0004810&hsa-miR-629-5p&offsets\|0\|-1;m-150&15\|-\|70079427\|70079447&offsets\|0\|-1]\|TGGGTTTACGTTGGGAGAAC | >MI0003643\|hsa-mir-629&WithFlank&15\|-\|70079366\|70079474 | 28 | 47 | 20 | TGGGTTTACGTTGGGAGAAC | 182.0 | 11.3 | 1.8 | 6.1 | 1.1E-09 | 1.6E-08 | BT20/MB468 | MP |
| >trnaMT_PheGAA_MT_+_577_647@37.54.18\|AAATGTTTAGACGGGCTC | >trnaMT_PheGAA_MT_+_577_647 | 37 | 54 | 18 | AAATGTTTAGACGGGCTC | 166.3 | 11.1 | 1.8 | 6.1 | 1.1E-09 | 1.6E-08 | BT20/MB468 | MP |
| >hg38_21_+_8433172_8446622_RNA45SN1_45S_50ntFlanks@12002.12032.31\|TGCGGGCCGCCGGTGAAATACCACTACTCTG | >hg38_21_+_8433172_8446622_RNA45SN1_45S_50ntFlanks | 12002 | 12032 | 31 | TGCGGGCCGCCGGTGAAATACCACTACTCTG | 3206.9 | -9.2 | 1.5 | -6.1 | 1.1E-09 | 1.6E-08 | BT20/MB468 | MP |
| >hg38_21_+_8433172_8446622_RNA45SN1_45S_50ntFlanks@9533.9570.38\|GAGGAAACTCTGGTGGAGGTCCGTAGCGGTCCTGACGT | >hg38_21_+_8433172_8446622_RNA45SN1_45S_50ntFlanks | 9533 | 9570 | 38 | GAGGAAACTCTGGTGGAGGTCCGTAGCGGTCCTGACGT | 1697.8 | -5.8 | 1.0 | -5.7 | 9.0E-09 | 1.6E-08 | BT20/MB468 | NUC |
| >hg38_21_+_8433172_8446622_RNA45SN1_45S_50ntFlanks@4353.4391.39\|TATTAAAGTTGCTGCAGTTAAAAAGCTCGTAGTTGGATC | >hg38_21_+_8433172_8446622_RNA45SN1_45S_50ntFlanks | 4353 | 4391 | 39 | TATTAAAGTTGCTGCAGTTAAAAAGCTCGTAGTTGGATC | 229.0 | -7.6 | 1.3 | -5.8 | 8.7E-09 | 1.6E-08 | MB231/MB468 | MITO |
| >hg38_21_+_8433172_8446622_RNA45SN1_45S_50ntFlanks@10116.10133.18\|TGCGTCGGAGGGCGGCGG | >hg38_21_+_8433172_8446622_RNA45SN1_45S_50ntFlanks | 10116 | 10133 | 18 | TGCGTCGGAGGGCGGCGG | 126.3 | -10.1 | 1.6 | -6.1 | 1.1E-09 | 1.6E-08 | BT20/MB468 | MP |
| >hg38_21_+_8433172_8446622_RNA45SN1_45S_50ntFlanks@3835.3868.34\|CTCCTCTCCTACTTGGATAACTGTGGTAATTCTA | >hg38_21_+_8433172_8446622_RNA45SN1_45S_50ntFlanks | 3835 | 3868 | 34 | CTCCTCTCCTACTTGGATAACTGTGGTAATTCTA | 563.4 | -3.7 | 0.6 | -5.8 | 8.0E-09 | 1.6E-08 | MB231/MB468 | NUC |
| >hg38_21_+_8433172_8446622_RNA45SN1_45S_50ntFlanks@6732.6770.39\|ACATTGATCATCGACACTTCGAACGCACTTGCGGCCCCG | >hg38_21_+_8433172_8446622_RNA45SN1_45S_50ntFlanks | 6732 | 6770 | 39 | ACATTGATCATCGACACTTCGAACGCACTTGCGGCCCCG | 1572.5 | -4.4 | 0.8 | -5.8 | 8.0E-09 | 1.6E-08 | MB231/MB468 | NUC |
| >hg38_21_+_8433172_8446622_RNA45SN1_45S_50ntFlanks@12638.12659.22\|CTACCATCTGTGGGATTATGAC | >hg38_21_+_8433172_8446622_RNA45SN1_45S_50ntFlanks | 12638 | 12659 | 22 | CTACCATCTGTGGGATTATGAC | 3066.7 | -5.5 | 0.9 | -5.8 | 6.1E-09 | 1.6E-08 | BT20/MB468 | MITO |
| >MI0003834\|hsa-mir-769&WithFlank&19\|+\|46018926\|46019055@36.56.21&[MIMAT0003886&hsa-miR-769-5p&offsets\|0\|-1;m-216&19\|+\|46018961\|46018982&offsets\|0\|-1]\|TGAGACCTCTGGGTTCTGAGC | >MI0003834\|hsa-mir-769&WithFlank&19\|+\|46018926\|46019055 | 36 | 56 | 21 | TGAGACCTCTGGGTTCTGAGC | 416.8 | 7.8 | 1.3 | 5.8 | 6.8E-09 | 1.6E-08 | BT20/MB468 | CYTO |
| >hg38_21_+_8433172_8446622_RNA45SN1_45S_50ntFlanks@11769.11794.26\|TAATTAGTGACGCGCATGAATGGATG | >hg38_21_+_8433172_8446622_RNA45SN1_45S_50ntFlanks | 11769 | 11794 | 26 | TAATTAGTGACGCGCATGAATGGATG | 376.2 | -4.4 | 0.8 | -5.8 | 8.7E-09 | 1.6E-08 | MB231/MB468 | MITO |
| >hg38_21_+_8433172_8446622_RNA45SN1_45S_50ntFlanks@8738.8758.21\|CCGTCCGTCCGTCCGTCCGTC | >hg38_21_+_8433172_8446622_RNA45SN1_45S_50ntFlanks | 8738 | 8758 | 21 | CCGTCCGTCCGTCCGTCCGTC | 297.7 | -11.3 | 1.9 | -6.1 | 1.1E-09 | 1.7E-08 | BT20/MB468 | MP |
| >hg38_21_+_8433172_8446622_RNA45SN1_45S_50ntFlanks@11778.11804.27\|ACGCGCATGAATGGATGAACGAGATTC | >hg38_21_+_8433172_8446622_RNA45SN1_45S_50ntFlanks | 11778 | 11804 | 27 | ACGCGCATGAATGGATGAACGAGATTC | 239.0 | -4.7 | 0.8 | -5.8 | 8.0E-09 | 1.7E-08 | MB231/MB468 | CYTO |
| >MI0000477\|hsa-mir-146a&WithFlank&5\|+\|160485346\|160485456@27.46.20(+1G)&[MIMAT0000449&hsa-miR-146a-5p&offsets\|0\|-2(+1G);m-129&5\|+\|160485372\|160485393&offsets\|0\|-2(+1G)]\|TGAGAACTGAATTCCATGGGG | >MI0000477\|hsa-mir-146a&WithFlank&5\|+\|160485346\|160485456 | 27 | 46 | 20(+1G) | TGAGAACTGAATTCCATGGGG | 149.3 | 10.4 | 1.8 | 5.8 | 7.5E-09 | 1.7E-08 | MB231/MB468 | MP |
| >hg38_21_+_8433172_8446622_RNA45SN1_45S_50ntFlanks@10802.10827.26\|TTGGAACAATGTAGGTAAGGGAAGTC | >hg38_21_+_8433172_8446622_RNA45SN1_45S_50ntFlanks | 10802 | 10827 | 26 | TTGGAACAATGTAGGTAAGGGAAGTC | 239.1 | -7.6 | 1.3 | -5.8 | 7.5E-09 | 1.7E-08 | MB231/MB468 | MP |
| >hg38_21_+_8433172_8446622_RNA45SN1_45S_50ntFlanks@11920.11950.31\|ATGAGAGGTGTAGAATAAGTGGGAGGCCCCC | >hg38_21_+_8433172_8446622_RNA45SN1_45S_50ntFlanks | 11920 | 11950 | 31 | ATGAGAGGTGTAGAATAAGTGGGAGGCCCCC | 400.1 | -8.4 | 1.5 | -5.7 | 9.1E-09 | 1.7E-08 | BT20/MB468 | NUC |
| >hg38_21_+_8433172_8446622_RNA45SN1_45S_50ntFlanks@8194.8211.18\|CCCGTGGACGGTGTGAGG | >hg38_21_+_8433172_8446622_RNA45SN1_45S_50ntFlanks | 8194 | 8211 | 18 | CCCGTGGACGGTGTGAGG | 3458.7 | -10.0 | 1.7 | -5.7 | 9.1E-09 | 1.7E-08 | BT20/MB468 | NUC |
| >hg38_21_+_8433172_8446622_RNA45SN1_45S_50ntFlanks@10411.10445.35\|CAGTCGGTCCTGAGAGATGGGCGAGCGCCGTTCCG | >hg38_21_+_8433172_8446622_RNA45SN1_45S_50ntFlanks | 10411 | 10445 | 35 | CAGTCGGTCCTGAGAGATGGGCGAGCGCCGTTCCG | 612.6 | -8.6 | 1.5 | -5.7 | 9.1E-09 | 1.7E-08 | BT20/MB468 | NUC |
| >trna16_GlnTTG_17_+_47269890_47269961@1.29.29,trna64_GlnTTG_6_+_28557156_28557227@1.29.29,trna84_GlnTTG_6_+_145503859_145503930@1.29.29\|GGTCCCATGGTGTAATGGTTAGCACTCTG | >trna16_GlnTTG_17_+_47269890_47269961 | 1 | 29 | 29 | GGTCCCATGGTGTAATGGTTAGCACTCTG | 1277.8 | -7.5 | 1.3 | -5.8 | 7.6E-09 | 1.7E-08 | MB231/MB468 | MP |
| >hg38_21_+_8433172_8446622_RNA45SN1_45S_50ntFlanks@11876.11912.37\|AAGACCCTGTTGAGCTTGACTCTAGTCTGGCACGGTG | >hg38_21_+_8433172_8446622_RNA45SN1_45S_50ntFlanks | 11876 | 11912 | 37 | AAGACCCTGTTGAGCTTGACTCTAGTCTGGCACGGTG | 167.6 | -10.5 | 1.7 | -6.1 | 1.1E-09 | 1.7E-08 | BT20/MB468 | MP |
| >trnaMT_SerGCT_MT_+_12207_12265@1.22.22\|GAGAAAGCTCACAAGAACTGCT | >trnaMT_SerGCT_MT_+_12207_12265 | 1 | 22 | 22 | GAGAAAGCTCACAAGAACTGCT | 732.1 | 4.3 | 0.8 | 5.8 | 8.1E-09 | 1.7E-08 | MB231/MB468 | NUC |
| >MI0000465\|hsa-mir-191&WithFlank&3\|-\|49020612\|49020715@22.42.21(+1G)&[MIMAT0000440&hsa-miR-191-5p&offsets\|0\|-2(+1G);m-48&3\|-\|49020672\|49020694&offsets\|0\|-2(+1G)]\|CAACGGAATCCCAAAAGCAGCG | >MI0000465\|hsa-mir-191&WithFlank&3\|-\|49020612\|49020715 | 22 | 42 | 21(+1G) | CAACGGAATCCCAAAAGCAGCG | 192.1 | 4.3 | 0.7 | 5.8 | 8.1E-09 | 1.7E-08 | MB231/MB468 | NUC |
| >trnaMT_ValTAC_MT_+_1602_1670@53.72.20\|TAACTTGACCGCTCTGACCA | >trnaMT_ValTAC_MT_+_1602_1670 | 53 | 72 | 20 | TAACTTGACCGCTCTGACCA | 2060.9 | 5.1 | 0.9 | 5.8 | 8.1E-09 | 1.7E-08 | MB231/MB468 | NUC |
| >hg38_21_+_8433172_8446622_RNA45SN1_45S_50ntFlanks@10349.10368.20\|AACGAGAACTTTGAAGGCCG | >hg38_21_+_8433172_8446622_RNA45SN1_45S_50ntFlanks | 10349 | 10368 | 20 | AACGAGAACTTTGAAGGCCG | 2371.3 | -3.6 | 0.6 | -5.8 | 8.1E-09 | 1.7E-08 | MB231/MB468 | CYTO |
| >hg38_21_+_8433172_8446622_RNA45SN1_45S_50ntFlanks@11918.11960.43\|ACATGAGAGGTGTAGAATAAGTGGGAGGCCCCCGGCGCCCCCC | >hg38_21_+_8433172_8446622_RNA45SN1_45S_50ntFlanks | 11918 | 11960 | 43 | ACATGAGAGGTGTAGAATAAGTGGGAGGCCCCCGGCGCCCCCC | 97.3 | -8.9 | 1.5 | -5.8 | 6.9E-09 | 1.7E-08 | BT20/MB468 | CYTO |
| >hg38_21_+_8433172_8446622_RNA45SN1_45S_50ntFlanks@11569.11599.31\|CAGCCGACTTAGAACTGGTGCGGACCAGGGG | >hg38_21_+_8433172_8446622_RNA45SN1_45S_50ntFlanks | 11569 | 11599 | 31 | CAGCCGACTTAGAACTGGTGCGGACCAGGGG | 1300.2 | -7.2 | 1.3 | -5.7 | 9.2E-09 | 1.7E-08 | BT20/MB468 | NUC |
| >hg38_21_+_8433172_8446622_RNA45SN1_45S_50ntFlanks@9307.9330.24\|AAGGAGTCTAACACGTGCGCGAGT | >hg38_21_+_8433172_8446622_RNA45SN1_45S_50ntFlanks | 9307 | 9330 | 24 | AAGGAGTCTAACACGTGCGCGAGT | 575.0 | 7.5 | 1.3 | 5.8 | 8.1E-09 | 1.7E-08 | MB231/MB468 | CYTO |
| >hg38_21_+_8433172_8446622_RNA45SN1_45S_50ntFlanks@9564.9595.32\|CTGACGTGCAAATCGGTCGTCCGACCTGGGTA | >hg38_21_+_8433172_8446622_RNA45SN1_45S_50ntFlanks | 9564 | 9595 | 32 | CTGACGTGCAAATCGGTCGTCCGACCTGGGTA | 6688.4 | -10.4 | 1.8 | -5.7 | 9.2E-09 | 1.7E-08 | BT20/MB468 | NUC |
| >hg38_21_+_8433172_8446622_RNA45SN1_45S_50ntFlanks@11641.11668.28\|CGGCGGGTGTTGACGCGATGTGATTTCT | >hg38_21_+_8433172_8446622_RNA45SN1_45S_50ntFlanks | 11641 | 11668 | 28 | CGGCGGGTGTTGACGCGATGTGATTTCT | 750.5 | -4.3 | 0.7 | -5.8 | 6.2E-09 | 1.7E-08 | BT20/MB468 | MITO |
| >hg38_1_-_228634819_228635039_RNA5S12_5S_50ntFlanks@133.169.37\|ATGGGAGACCGCCTGGGAATACCGGGTGCTGTAGGCT | >hg38_1_-_228634819_228635039_RNA5S12_5S_50ntFlanks | 133 | 169 | 37 | ATGGGAGACCGCCTGGGAATACCGGGTGCTGTAGGCT | 9126.2 | 5.2 | 0.9 | 5.8 | 6.9E-09 | 1.7E-08 | BT20/MB468 | CYTO |
| >hg38_21_+_8433172_8446622_RNA45SN1_45S_50ntFlanks@5492.5510.19\|AGAAGACGGTCGAACTTGA | >hg38_21_+_8433172_8446622_RNA45SN1_45S_50ntFlanks | 5492 | 5510 | 19 | AGAAGACGGTCGAACTTGA | 846.1 | -8.1 | 1.4 | -5.8 | 7.6E-09 | 1.7E-08 | MB231/MB468 | MP |
| >hg38_21_+_8433172_8446622_RNA45SN1_45S_50ntFlanks@6773.6803.31\|TTCCTCCCGGGGCTACGCCTGTCTGAGCGTC | >hg38_21_+_8433172_8446622_RNA45SN1_45S_50ntFlanks | 6773 | 6803 | 31 | TTCCTCCCGGGGCTACGCCTGTCTGAGCGTC | 2994.8 | -4.4 | 0.8 | -5.8 | 6.2E-09 | 1.7E-08 | BT20/MB468 | MITO |
| >hg38_21_+_8433172_8446622_RNA45SN1_45S_50ntFlanks@11209.11227.19\|CGGCGACTCTGGACGCGAG | >hg38_21_+_8433172_8446622_RNA45SN1_45S_50ntFlanks | 11209 | 11227 | 19 | CGGCGACTCTGGACGCGAG | 444.1 | -5.9 | 1.0 | -5.8 | 8.2E-09 | 1.7E-08 | MB231/MB468 | NUC |
| >MI0003832\|hsa-mir-1283-1&WithFlank&19\|+\|53688475\|53688573@21.43.23;MI0006430\|hsa-mir-1283-2&WithFlank&19\|+\|53758226\|53758324@21.43.23&[MIMAT0005799&hsa-miR-1283&offsets\|+1\|+2;m-905&19\|+\|53688495\|53688516&offsets\|0\|+1];[MIMAT0005799_1&hsa-miR-1283&offsets\|+1\|+2;m-905&19\|+\|53758246\|53758267&offsets\|0\|+1]\|CTACAAAGGAAAGCGCTTTCTGT | >MI0003832\|hsa-mir-1283-1&WithFlank&19\|+\|53688475\|53688573 | 21 | 43 | 23 | CTACAAAGGAAAGCGCTTTCTGT | 81.7 | 9.6 | 1.7 | 5.8 | 7.6E-09 | 1.7E-08 | MB231/MB468 | MP |
| >hg38_21_+_8433172_8446622_RNA45SN1_45S_50ntFlanks@4049.4066.18\|TCGAACGTCTGCCCTATC | >hg38_21_+_8433172_8446622_RNA45SN1_45S_50ntFlanks | 4049 | 4066 | 18 | TCGAACGTCTGCCCTATC | 458.0 | -6.6 | 1.1 | -5.7 | 9.2E-09 | 1.7E-08 | BT20/MB468 | NUC |
| >hg38_21_+_8433172_8446622_RNA45SN1_45S_50ntFlanks@6784.6801.18\|GCTACGCCTGTCTGAGCG | >hg38_21_+_8433172_8446622_RNA45SN1_45S_50ntFlanks | 6784 | 6801 | 18 | GCTACGCCTGTCTGAGCG | 383.5 | -7.5 | 1.3 | -5.8 | 7.7E-09 | 1.7E-08 | MB231/MB468 | MP |
| >MI0000289\|hsa-mir-181a-1&WithFlank&1\|-\|198859038\|198859159@30.54.25(+1U);MI0000269\|hsa-mir-181a-2&WithFlank&9\|+\|124692436\|124692557@45.69.25(+1U)&[MIMAT0000256&hsa-miR-181a-5p&offsets\|0\|+2(+1U);m-32&1\|-\|198859108\|198859130&offsets\|0\|+2(+1U)];[MIMAT0000256_1&hsa-miR-181a-5p&offsets\|0\|+2(+1U);m-32&9\|+\|124692480\|124692502&offsets\|0\|+2(+1U)]\|AACATTCAACGCTGTCGGTGAGTTTT | >MI0000289\|hsa-mir-181a-1&WithFlank&1\|-\|198859038\|198859159 | 30 | 54 | 25(+1U) | AACATTCAACGCTGTCGGTGAGTTTT | 1100.2 | -6.8 | 1.1 | -6.2 | 6.7E-10 | 1.7E-08 | BT20/MB231 | MITO |
| >hg38_21_+_8433172_8446622_RNA45SN1_45S_50ntFlanks@5083.5112.30\|ACGAACGAGACTCTGGCATGCTAACTAGTT | >hg38_21_+_8433172_8446622_RNA45SN1_45S_50ntFlanks | 5083 | 5112 | 30 | ACGAACGAGACTCTGGCATGCTAACTAGTT | 598.3 | -7.4 | 1.3 | -5.8 | 7.7E-09 | 1.7E-08 | MB231/MB468 | MP |

<table>
<tr><th></th><th></th><th></th><th></th><th></th><th></th><th></th><th></th><th></th><th></th><th></th><th></th><th></th><th></th></tr>
<tr><td>>hg38_21_+_8433172_8446622_RNA45SN1_45S_50ntFlanks@4297.4319.23|CATTGGAGGGCAAGTCTGGTGCC</td><td>>hg38_21_+_8433172_8446622_RNA45SN1_45S_50ntFlanks</td><td>4297</td><td>4319</td><td>23</td><td>CATTGGAGGGCAAGTCTGGTGCC</td><td>246.8</td><td>-5.6</td><td>1.0</td><td>-5.8</td><td>8.2E-09</td><td>1.7E-08</td><td>MB231/MB468</td><td>NUC</td></tr>
<tr><td>>MI0000273|hsa-mir-183&WithFlank&7|-|129774899|129775020@33.53.21(+1G)&[MIMAT0000261&hsa-miR-183-5p&offsets|0|-1(+1G);m-41&7|-|129774966|129774987&offsets|-1|-2(+1G)]|TATGGCACTGGTAGAATTCACG</td><td>>MI0000273|hsa-mir-183&WithFlank&7|-|129774899|129775020</td><td>33</td><td>53</td><td>21(+1G)</td><td>TATGGCACTGGTAGAATTCACG</td><td>730.0</td><td>6.6</td><td>1.1</td><td>5.8</td><td>7.0E-09</td><td>1.7E-08</td><td>BT20/MB468</td><td>CYTO</td></tr>
<tr><td>>hg38_21_+_8433172_8446622_RNA45SN1_45S_50ntFlanks@9545.9575.31|GTGGAGGTCCGTAGCGGTCCTGACGTGCAAA</td><td>>hg38_21_+_8433172_8446622_RNA45SN1_45S_50ntFlanks</td><td>9545</td><td>9575</td><td>31</td><td>GTGGAGGTCCGTAGCGGTCCTGACGTGCAAA</td><td>883.4</td><td>-8.9</td><td>1.5</td><td>-5.7</td><td>9.3E-09</td><td>1.7E-08</td><td>BT20/MB468</td><td>NUC</td></tr>
<tr><td>>MI0000093|hsa-mir-92a-1&WithFlank&13|+|91351308|91351397@54.74.21(+1A);MI0000094|hsa-mir-92a-2&WithFlank&X|-|134169532|134169618@54.74.21(+1A)&[MIMAT0000092&hsa-miR-92a-3p&offsets|0|-1(+1A);m-9&13|+|91351361|91351382&offsets|0|-1(+1A)];[MIMAT0000092_1&hsa-miR-92a-3p&offsets|0|-1(+1A);m-11&X|-|134169544|134169565&offsets|0|-1(+1A)]|TATTGCACTTGTCCCGGCCTGA</td><td>>MI0000093|hsa-mir-92a-1&WithFlank&13|+|91351308|91351397</td><td>54</td><td>74</td><td>21(+1A)</td><td>TATTGCACTTGTCCCGGCCTGA</td><td>2530.1</td><td>5.4</td><td>0.9</td><td>5.8</td><td>8.9E-09</td><td>1.7E-08</td><td>MB231/MB468</td><td>MITO</td></tr>
<tr><td>>hg38_21_+_8433172_8446622_RNA45SN1_45S_50ntFlanks@9568.9601.34|CGTGCAAATCGGTCGTCCGACCTGGGTATAGGGG</td><td>>hg38_21_+_8433172_8446622_RNA45SN1_45S_50ntFlanks</td><td>9568</td><td>9601</td><td>34</td><td>CGTGCAAATCGGTCGTCCGACCTGGGTATAGGGG</td><td>438.7</td><td>-7.6</td><td>1.3</td><td>-5.8</td><td>8.9E-09</td><td>1.7E-08</td><td>MB231/MB468</td><td>MITO</td></tr>
<tr><td>>hg38_21_+_8433172_8446622_RNA45SN1_45S_50ntFlanks@8176.8206.31|CTTCTGATCGAGGCCCAGCCCGTGGACGGTG</td><td>>hg38_21_+_8433172_8446622_RNA45SN1_45S_50ntFlanks</td><td>8176</td><td>8206</td><td>31</td><td>CTTCTGATCGAGGCCCAGCCCGTGGACGGTG</td><td>2811.6</td><td>-7.7</td><td>1.3</td><td>-5.7</td><td>9.3E-09</td><td>1.7E-08</td><td>BT20/MB468</td><td>NUC</td></tr>
<tr><td>>hg38_21_+_8433172_8446622_RNA45SN1_45S_50ntFlanks@5444.5463.20|CTCGGATCGGCCCCGCCGGG</td><td>>hg38_21_+_8433172_8446622_RNA45SN1_45S_50ntFlanks</td><td>5444</td><td>5463</td><td>20</td><td>CTCGGATCGGCCCCGCCGGG</td><td>85.8</td><td>-10.3</td><td>1.8</td><td>-5.8</td><td>7.7E-09</td><td>1.7E-08</td><td>MB231/MB468</td><td>MP</td></tr>
<tr><td>>hg38_21_+_8433172_8446622_RNA45SN1_45S_50ntFlanks@12709.12727.19|CCGCGGAGCCTCGGTTGGC</td><td>>hg38_21_+_8433172_8446622_RNA45SN1_45S_50ntFlanks</td><td>12709</td><td>12727</td><td>19</td><td>CCGCGGAGCCTCGGTTGGC</td><td>67.0</td><td>-9.9</td><td>1.7</td><td>-5.8</td><td>7.7E-09</td><td>1.7E-08</td><td>MB231/MB468</td><td>MP</td></tr>
<tr><td>>MI0000108|hsa-mir-103a-2&WithFlank&20|+|3917488|3917577@56.76.21;MI0000109|hsa-mir-103a-1&WithFlank&5|-|168560890|168560979@56.76.21&[MIMAT0000101&hsa-miR-103a-3p&offsets|+2|0;m-12&20|+|3917541|3917563&offsets|+2|0];[MIMAT0000101_1&hsa-miR-103a-3p&offsets|+2|0;m-13&5|-|168560904|168560926&offsets|+2|0;m-44220&5|-|168560908|168560926&offsets|+2|+4]|CAGCATTGTACAGGGCTATGA</td><td>>MI0000108|hsa-mir-103a-2&WithFlank&20|+|3917488|3917577</td><td>56</td><td>76</td><td>21</td><td>CAGCATTGTACAGGGCTATGA</td><td>241.3</td><td>4.2</td><td>0.7</td><td>5.8</td><td>8.3E-09</td><td>1.7E-08</td><td>MB231/MB468</td><td>NUC</td></tr>
<tr><td>>hg38_21_+_8433172_8446622_RNA45SN1_45S_50ntFlanks@11630.11662.33|GCGAAGGCCCGCGGCGGGTGTTGACGCGATGTG</td><td>>hg38_21_+_8433172_8446622_RNA45SN1_45S_50ntFlanks</td><td>11630</td><td>11662</td><td>33</td><td>GCGAAGGCCCGCGGCGGGTGTTGACGCGATGTG</td><td>223.0</td><td>-7.3</td><td>1.3</td><td>-5.8</td><td>7.7E-09</td><td>1.7E-08</td><td>MB231/MB468</td><td>MP</td></tr>
<tr><td>>hg38_21_+_8433172_8446622_RNA45SN1_45S_50ntFlanks@6662.6699.38|GGTGGATCACTCGGCTCGTGCGTCGATGAAGAACGCAG</td><td>>hg38_21_+_8433172_8446622_RNA45SN1_45S_50ntFlanks</td><td>6662</td><td>6699</td><td>38</td><td>GGTGGATCACTCGGCTCGTGCGTCGATGAAGAACGCAG</td><td>73.3</td><td>-10.1</td><td>1.7</td><td>-5.8</td><td>7.7E-09</td><td>1.7E-08</td><td>MB231/MB468</td><td>MP</td></tr>
<tr><td>>hg38_21_+_8433172_8446622_RNA45SN1_45S_50ntFlanks@12084.12107.24|TCGCTTCTGGCGCCAAGCGCCCGG</td><td>>hg38_21_+_8433172_8446622_RNA45SN1_45S_50ntFlanks</td><td>12084</td><td>12107</td><td>24</td><td>TCGCTTCTGGCGCCAAGCGCCCGG</td><td>220.0</td><td>-10.9</td><td>1.8</td><td>-6.1</td><td>1.1E-09</td><td>1.7E-08</td><td>BT20/MB468</td><td>MP</td></tr>
<tr><td>>hg38_21_+_8433172_8446622_RNA45SN1_45S_50ntFlanks@9534.9560.27|AGGAAACTCTGGTGGAGGTCCGTAGCG</td><td>>hg38_21_+_8433172_8446622_RNA45SN1_45S_50ntFlanks</td><td>9534</td><td>9560</td><td>27</td><td>AGGAAACTCTGGTGGAGGTCCGTAGCG</td><td>702.6</td><td>-4.4</td><td>0.8</td><td>-5.8</td><td>8.3E-09</td><td>1.7E-08</td><td>MB231/MB468</td><td>NUC</td></tr>
<tr><td>>am_trna106_HisGTG_1_-_149155828_149155899@9.29.21,trna111_HisGTG_1_-_147774845_147774916@9.29.21,trna118_HisGTG_1_-_145396881_145396952@9.29.21,trna16_HisGTG_1_+_146544773_146544844@9.29.21,trna1_HisGTG_15_+_45493349_45493420@9.29.21,trna21_HisGTG_1_+_147753471_147753542@9.29.21,trna33_HisGTG_6_+_27125906_27125977@9.29.21,trna7_HisGTG_9_-_14433938_14434009@9.29.21,trna8_HisGTG_15_-_45492611_45492682@9.29.21,trna9_HisGTG_15_-_45490804_45490875@9.29.21|CGTATAGTGGTTAGTACTCTG</td><td>>am_trna106_HisGTG_1_-_149155828_149155899</td><td>9</td><td>29</td><td>21</td><td>CGTATAGTGGTTAGTACTCTG</td><td>104.4</td><td>-9.8</td><td>1.6</td><td>-6.1</td><td>1.1E-09</td><td>1.7E-08</td><td>BT20/MB468</td><td>MP</td></tr>
<tr><td>>am_MI0000809|hsa-mir-151a&WithFlank&8|-|140732558|140732659@53.73.21(+1G)&[MIMAT0000757&hsa-miR-151a-3p&offsets|0|0(+1G);m-30&8|-|140732587|140732607&offsets|0|0(+1G)]|CTAGACTGAAGCTCCTTGAGGG</td><td>>am_MI0000809|hsa-mir-151a&WithFlank&8|-|140732558|140732659</td><td>53</td><td>73</td><td>21(+1G)</td><td>CTAGACTGAAGCTCCTTGAGGG</td><td>149.6</td><td>3.2</td><td>0.6</td><td>5.7</td><td>9.4E-09</td><td>1.7E-08</td><td>BT20/MB468</td><td>NUC</td></tr>
<tr><td>>hg38_21_+_8433172_8446622_RNA45SN1_45S_50ntFlanks@12366.12392.27|TAACTGGCTTGTGGCGGCCAAGCGTTC</td><td>>hg38_21_+_8433172_8446622_RNA45SN1_45S_50ntFlanks</td><td>12366</td><td>12392</td><td>27</td><td>TAACTGGCTTGTGGCGGCCAAGCGTTC</td><td>58.6</td><td>-9.7</td><td>1.7</td><td>-5.8</td><td>7.8E-09</td><td>1.7E-08</td><td>MB231/MB468</td><td>MP</td></tr>
<tr><td>>hg38_21_+_8433172_8446622_RNA45SN1_45S_50ntFlanks@10388.10407.20|GTGAACAGCAGTTGAACATG</td><td>>hg38_21_+_8433172_8446622_RNA45SN1_45S_50ntFlanks</td><td>10388</td><td>10407</td><td>20</td><td>GTGAACAGCAGTTGAACATG</td><td>206.2</td><td>-5.9</td><td>1.0</td><td>-5.8</td><td>7.8E-09</td><td>1.7E-08</td><td>MB231/MB468</td><td>MP</td></tr>
<tr><td>>hg38_21_+_8433172_8446622_RNA45SN1_45S_50ntFlanks@8182.8223.42|ATCGAGGCCCAGCCCGTGGACGGTGTGAGGCCGGTAGCGGCC</td><td>>hg38_21_+_8433172_8446622_RNA45SN1_45S_50ntFlanks</td><td>8182</td><td>8223</td><td>42</td><td>ATCGAGGCCCAGCCCGTGGACGGTGTGAGGCCGGTAGCGGCC</td><td>97.7</td><td>-10.5</td><td>1.8</td><td>-5.8</td><td>7.8E-09</td><td>1.7E-08</td><td>MB231/MB468</td><td>MP</td></tr>
<tr><td>>hg38_21_+_8433172_8446622_RNA45SN1_45S_50ntFlanks@12709.12745.37|CCGCGGAGCCTCGGTTGGCCTCGGATAGCCGGTCCCC</td><td>>hg38_21_+_8433172_8446622_RNA45SN1_45S_50ntFlanks</td><td>12709</td><td>12745</td><td>37</td><td>CCGCGGAGCCTCGGTTGGCCTCGGATAGCCGGTCCCC</td><td>686.3</td><td>-7.2</td><td>1.2</td><td>-5.8</td><td>6.3E-09</td><td>1.7E-08</td><td>BT20/MB468</td><td>MITO</td></tr>
<tr><td>>hg38_21_+_8433172_8446622_RNA45SN1_45S_50ntFlanks@12994.13013.20|CCTCGCTGCGATCTATTGAA</td><td>>hg38_21_+_8433172_8446622_RNA45SN1_45S_50ntFlanks</td><td>12994</td><td>13013</td><td>20</td><td>CCTCGCTGCGATCTATTGAA</td><td>1391.4</td><td>-6.8</td><td>1.2</td><td>-5.7</td><td>9.0E-09</td><td>1.7E-08</td><td>MB231/MB468</td><td>MITO</td></tr>
<tr><td>>hg38_21_+_8433172_8446622_RNA45SN1_45S_50ntFlanks@12567.12587.21|CTCAGTACGAGAGGAACCGCA</td><td>>hg38_21_+_8433172_8446622_RNA45SN1_45S_50ntFlanks</td><td>12567</td><td>12587</td><td>21</td><td>CTCAGTACGAGAGGAACCGCA</td><td>400.0</td><td>-6.6</td><td>1.1</td><td>-5.8</td><td>8.2E-09</td><td>1.7E-08</td><td>MB231/MB468</td><td>CYTO</td></tr>
<tr><td>>hg38_21_+_8433172_8446622_RNA45SN1_45S_50ntFlanks@8213.8230.18|CGGTAGCGGCCCCCGGCG</td><td>>hg38_21_+_8433172_8446622_RNA45SN1_45S_50ntFlanks</td><td>8213</td><td>8230</td><td>18</td><td>CGGTAGCGGCCCCCGGCG</td><td>108.2</td><td>-9.8</td><td>1.6</td><td>-6.1</td><td>1.1E-09</td><td>1.7E-08</td><td>BT20/MB468</td><td>MP</td></tr>
<tr><td>>hg38_21_+_8433172_8446622_RNA45SN1_45S_50ntFlanks@10665.10706.42|TTGGAAAGCGTCGCGGTTCCGGCGGCGTCCGGTGAGCTCTCG</td><td>>hg38_21_+_8433172_8446622_RNA45SN1_45S_50ntFlanks</td><td>10665</td><td>10706</td><td>42</td><td>TTGGAAAGCGTCGCGGTTCCGGCGGCGTCCGGTGAGCTCTCG</td><td>398.3</td><td>-8.8</td><td>1.5</td><td>-5.7</td><td>9.4E-09</td><td>1.7E-08</td><td>BT20/MB468</td><td>NUC</td></tr>
<tr><td>>hg38_21_+_8433172_8446622_RNA45SN1_45S_50ntFlanks@9915.9949.35|AAAAGGTGTTGGTTGATATAGACAGCAGGACGGTG</td><td>>hg38_21_+_8433172_8446622_RNA45SN1_45S_50ntFlanks</td><td>9915</td><td>9949</td><td>35</td><td>AAAAGGTGTTGGTTGATATAGACAGCAGGACGGTG</td><td>93.8</td><td>10.3</td><td>1.7</td><td>6.1</td><td>1.1E-09</td><td>1.7E-08</td><td>BT20/MB468</td><td>MP</td></tr>
<tr><td>>hg38_21_+_8433172_8446622_RNA45SN1_45S_50ntFlanks@10808.10831.24|CAATGTAGGTAAGGGAAGTCGGCA</td><td>>hg38_21_+_8433172_8446622_RNA45SN1_45S_50ntFlanks</td><td>10808</td><td>10831</td><td>24</td><td>CAATGTAGGTAAGGGAAGTCGGCA</td><td>266.1</td><td>-7.5</td><td>1.3</td><td>-5.8</td><td>7.8E-09</td><td>1.7E-08</td><td>MB231/MB468</td><td>MP</td></tr>
<tr><td>>hg38_21_+_8433172_8446622_RNA45SN1_45S_50ntFlanks@475.496.22|CTTCGTGATCGATGTGGTGACG</td><td>>hg38_21_+_8433172_8446622_RNA45SN1_45S_50ntFlanks</td><td>475</td><td>496</td><td>22</td><td>CTTCGTGATCGATGTGGTGACG</td><td>518.5</td><td>7.7</td><td>1.3</td><td>5.8</td><td>8.3E-09</td><td>1.7E-08</td><td>MB231/MB468</td><td>CYTO</td></tr>
<tr><td>>hg38_21_+_8433172_8446622_RNA45SN1_45S_50ntFlanks@11212.11254.43|CGACTCTGGACGCGAGCCGGGCCCTTCCCGTGGATCGCCCCAG</td><td>>hg38_21_+_8433172_8446622_RNA45SN1_45S_50ntFlanks</td><td>11212</td><td>11254</td><td>43</td><td>CGACTCTGGACGCGAGCCGGGCCCTTCCCGTGGATCGCCCCAG</td><td>425.1</td><td>-6.7</td><td>1.2</td><td>-5.8</td><td>8.3E-09</td><td>1.7E-08</td><td>MB231/MB468</td><td>CYTO</td></tr>
<tr><td>>hg38_21_+_8433172_8446622_RNA45SN1_45S_50ntFlanks@12001.12030.30|CTGCGGGCCGCCGGTGAAATACCACTACTC</td><td>>hg38_21_+_8433172_8446622_RNA45SN1_45S_50ntFlanks</td><td>12001</td><td>12030</td><td>30</td><td>CTGCGGGCCGCCGGTGAAATACCACTACTC</td><td>312.6</td><td>-7.4</td><td>1.3</td><td>-5.7</td><td>9.1E-09</td><td>1.7E-08</td><td>MB231/MB468</td><td>MITO</td></tr>
<tr><td>>hg38_21_+_8433172_8446622_RNA45SN1_45S_50ntFlanks@12431.12455.25|TCCTATCATTGTGAAGCAGAATTCA</td><td>>hg38_21_+_8433172_8446622_RNA45SN1_45S_50ntFlanks</td><td>12431</td><td>12455</td><td>25</td><td>TCCTATCATTGTGAAGCAGAATTCA</td><td>503.6</td><td>-7.7</td><td>1.3</td><td>-5.7</td><td>9.1E-09</td><td>1.7E-08</td><td>MB231/MB468</td><td>MITO</td></tr>
<tr><td>>hg38_21_+_8433172_8446622_RNA45SN1_45S_50ntFlanks@12415.12452.38|CTTCGATGTCGGCTCTTCCTATCATTGTGAAGCAGAAT</td><td>>hg38_21_+_8433172_8446622_RNA45SN1_45S_50ntFlanks</td><td>12415</td><td>12452</td><td>38</td><td>CTTCGATGTCGGCTCTTCCTATCATTGTGAAGCAGAAT</td><td>207.0</td><td>5.2</td><td>0.9</td><td>5.8</td><td>8.3E-09</td><td>1.7E-08</td><td>MB231/MB468</td><td>CYTO</td></tr>
<tr><td>>hg38_21_+_8433172_8446622_RNA45SN1_45S_50ntFlanks@7986.8006.21|ATCAGACGTGGCGACCCGCTG</td><td>>hg38_21_+_8433172_8446622_RNA45SN1_45S_50ntFlanks</td><td>7986</td><td>8006</td><td>21</td><td>ATCAGACGTGGCGACCCGCTG</td><td>1532.9</td><td>-7.3</td><td>1.3</td><td>-5.8</td><td>8.3E-09</td><td>1.7E-08</td><td>MB231/MB468</td><td>CYTO</td></tr>
<tr><td>>hg38_21_+_8433172_8446622_RNA45SN1_45S_50ntFlanks@9457.9503.47|CGCCGGGGAGGTGGAGCACGAGCGCACGTGTTAGGACCCGAAAGATG</td><td>>hg38_21_+_8433172_8446622_RNA45SN1_45S_50ntFlanks</td><td>9457</td><td>9503</td><td>47</td><td>CGCCGGGGAGGTGGAGCACGAGCGCACGTGTTAGGACCCGAAAGATG</td><td>958.8</td><td>-7.7</td><td>1.3</td><td>-5.8</td><td>6.3E-09</td><td>1.7E-08</td><td>BT20/MB468</td><td>MITO</td></tr>
<tr><td>>hg38_21_+_8433172_8446622_RNA45SN1_45S_50ntFlanks@10841.10861.21|CGTAACTTCGGGATAAGGATT</td><td>>hg38_21_+_8433172_8446622_RNA45SN1_45S_50ntFlanks</td><td>10841</td><td>10861</td><td>21</td><td>CGTAACTTCGGGATAAGGATT</td><td>2898.4</td><td>-3.9</td><td>0.7</td><td>-5.8</td><td>7.1E-09</td><td>1.7E-08</td><td>BT20/MB468</td><td>CYTO</td></tr>
<tr><td>>hg38_21_+_8433172_8446622_RNA45SN1_45S_50ntFlanks@8472.8492.21|CGGCCCGGCGGATCTTTCCCG</td><td>>hg38_21_+_8433172_8446622_RNA45SN1_45S_50ntFlanks</td><td>8472</td><td>8492</td><td>21</td><td>CGGCCCGGCGGATCTTTCCCG</td><td>118.0</td><td>-10.0</td><td>1.6</td><td>-6.1</td><td>1.1E-09</td><td>1.7E-08</td><td>BT20/MB468</td><td>MP</td></tr>
<tr><td>>hg38_21_+_8433172_8446622_RNA45SN1_45S_50ntFlanks@11570.11605.36|AGCCGACTTAGAACTGGTGCGGACCAGGGGAATCCG</td><td>>hg38_21_+_8433172_8446622_RNA45SN1_45S_50ntFlanks</td><td>11570</td><td>11605</td><td>36</td><td>AGCCGACTTAGAACTGGTGCGGACCAGGGGAATCCG</td><td>674.5</td><td>-8.6</td><td>1.5</td><td>-5.7</td><td>9.5E-09</td><td>1.7E-08</td><td>BT20/MB468</td><td>NUC</td></tr>
<tr><td>>MI0003513|hsa-mir-455&WithFlank&9|+|114209428|114209535@24.45.22&[MIMAT0003150&hsa-miR-455-5p&offsets|+2|+2;m-232&9|+|114209449|114209470&offsets|+2|+2]|TGTGCCTTTGGACTACATCGTG</td><td>>MI0003513|hsa-mir-455&WithFlank&9|+|114209428|114209535</td><td>24</td><td>45</td><td>22</td><td>TGTGCCTTTGGACTACATCGTG</td><td>69.5</td><td>9.3</td><td>1.6</td><td>5.8</td><td>7.9E-09</td><td>1.7E-08</td><td>MB231/MB468</td><td>MP</td></tr>
<tr><td>>hg38_21_+_8433172_8446622_RNA45SN1_45S_50ntFlanks@10350.10384.35|ACGAGAACTTTGAAGGCCGAAGTGGAGAAGGGTTC</td><td>>hg38_21_+_8433172_8446622_RNA45SN1_45S_50ntFlanks</td><td>10350</td><td>10384</td><td>35</td><td>ACGAGAACTTTGAAGGCCGAAGTGGAGAAGGGTTC</td><td>1942.4</td><td>-6.6</td><td>1.1</td><td>-5.7</td><td>9.1E-09</td><td>1.7E-08</td><td>MB231/MB468</td><td>MITO</td></tr>
<tr><td>>MI0000449|hsa-mir-132&WithFlank&17|-|2049902|2050014@65.85.21&[MIMAT0000426&hsa-miR-132-3p&offsets|0|-1;m-138&17|-|2049929|2049950&offsets|0|-1]|TAACAGTCTACAGCCATGGTC</td><td>>MI0000449|hsa-mir-132&WithFlank&17|-|2049902|2050014</td><td>65</td><td>85</td><td>21</td><td>TAACAGTCTACAGCCATGGTC</td><td>85.3</td><td>9.6</td><td>1.7</td><td>5.8</td><td>7.9E-09</td><td>1.7E-08</td><td>MB231/MB468</td><td>MP</td></tr>
<tr><td>>hg38_21_+_8433172_8446622_RNA45SN1_45S_50ntFlanks@4734.4751.18|AACGAAAGTCGGAGGTTC</td><td>>hg38_21_+_8433172_8446622_RNA45SN1_45S_50ntFlanks</td><td>4734</td><td>4751</td><td>18</td><td>AACGAAAGTCGGAGGTTC</td><td>429.0</td><td>-3.8</td><td>0.7</td><td>-5.7</td><td>9.1E-09</td><td>1.7E-08</td><td>MB231/MB468</td><td>MITO</td></tr>
<tr><td>>hg38_21_+_8433172_8446622_RNA45SN1_45S_50ntFlanks@4738.4765.28|AAAGTCGGAGGTTCGAAGACGATCAGAT</td><td>>hg38_21_+_8433172_8446622_RNA45SN1_45S_50ntFlanks</td><td>4738</td><td>4765</td><td>28</td><td>AAAGTCGGAGGTTCGAAGACGATCAGAT</td><td>712.3</td><td>6.5</td><td>1.1</td><td>5.8</td><td>7.2E-09</td><td>1.7E-08</td><td>BT20/MB468</td><td>CYTO</td></tr>
<tr><td>>hg38_21_+_8433172_8446622_RNA45SN1_45S_50ntFlanks@10692.10711.20|TCCGGTGAGCTCTCGCTGGC</td><td>>hg38_21_+_8433172_8446622_RNA45SN1_45S_50ntFlanks</td><td>10692</td><td>10711</td><td>20</td><td>TCCGGTGAGCTCTCGCTGGC</td><td>63.5</td><td>-9.9</td><td>1.7</td><td>-5.8</td><td>7.9E-09</td><td>1.7E-08</td><td>MB231/MB468</td><td>MP</td></tr>
<tr><td>>hg38_MT_+_598_1651_RNR1_12S_50ntFlanks@52.88.37|ATAGGTTTGGTCCTAGCCTTTCTATTAGCTCTTAGTA</td><td>>hg38_MT_+_598_1651_RNR1_12S_50ntFlanks</td><td>52</td><td>88</td><td>37</td><td>ATAGGTTTGGTCCTAGCCTTTCTATTAGCTCTTAGTA</td><td>27635.4</td><td>5.7</td><td>1.0</td><td>5.8</td><td>6.4E-09</td><td>1.7E-08</td><td>BT20/MB468</td><td>MITO</td></tr>
<tr><td>>am_trnaMT_GluTTC_MT_-_14674_14742@27.48.22,trnalookalike8_GluTTC_5_-_93905172_93905240@27.48.22|GTTTTTCATATCATTGGTCGTG</td><td>>am_trnaMT_GluTTC_MT_-_14674_14742</td><td>27</td><td>48</td><td>22</td><td>GTTTTTCATATCATTGGTCGTG</td><td>158.0</td><td>-7.1</td><td>1.2</td><td>-5.8</td><td>8.5E-09</td><td>1.7E-08</td><td>MB231/MB468</td><td>NUC</td></tr>
<tr><td>>hg38_21_+_8433172_8446622_RNA45SN1_45S_50ntFlanks@3985.4010.26|CGGCTTTGGTGACTCTAGATAACCTC</td><td>>hg38_21_+_8433172_8446622_RNA45SN1_45S_50ntFlanks</td><td>3985</td><td>4010</td><td>26</td><td>CGGCTTTGGTGACTCTAGATAACCTC</td><td>1071.5</td><td>-7.1</td><td>1.2</td><td>-5.7</td><td>9.5E-09</td><td>1.7E-08</td><td>BT20/MB468</td><td>NUC</td></tr>
<tr><td>>hg38_21_+_8433172_8446622_RNA45SN1_45S_50ntFlanks@7974.8017.44|ACGCGACCTCAGATCAGACGTGGCGACCCGCTGAATTTAAGCAT</td><td>>hg38_21_+_8433172_8446622_RNA45SN1_45S_50ntFlanks</td><td>7974</td><td>8017</td><td>44</td><td>ACGCGACCTCAGATCAGACGTGGCGACCCGCTGAATTTAAGCAT</td><td>1689.4</td><td>5.1</td><td>0.9</td><td>5.8</td><td>6.4E-09</td><td>1.7E-08</td><td>BT20/MB468</td><td>MITO</td></tr>
<tr><td>>MI0006407|hsa-mir-1270&WithFlank&19|-|20399266|20399360@19.41.23&[MIMAT0005924&hsa-miR-1270&offsets|0|0;m-459&19|-|20399320|20399342&offsets|0|0]|CTGGAGATATGGAAGAGCTGTGT</td><td>>MI0006407|hsa-mir-1270&WithFlank&19|-|20399266|20399360</td><td>19</td><td>41</td><td>23</td><td>CTGGAGATATGGAAGAGCTGTGT</td><td>323.3</td><td>12.1</td><td>2.0</td><td>6.1</td><td>1.1E-09</td><td>1.7E-08</td><td>BT20/MB468</td><td>MP</td></tr>
</table>

| | | | | | | | | | | | | | |
|---|---|---|---|---|---|---|---|---|---|---|---|---|---|
| >hg38_21_+_8433172_8446622_RNA45SN1_45S_50ntFlanks@10345.10380.36\|TTCAAACGAGAACTTTGAAGGCCGAAGTGGAGAAGG | >hg38_21_+_8433172_8446622_RNA45SN1_45S_50ntFlanks | 10345 | 10380 | 36 | TTCAAACGAGAACTTTGAAGGCCGAAGTGGAGAAGG | 99.9 | 10.4 | 1.7 | 6.1 | 1.1E-09 | 1.7E-08 | BT20/MB468 | MP |
| >hg38_21_+_8433172_8446622_RNA45SN1_45S_50ntFlanks@8170.8215.46\|CAAGTCCTTCTGATCGAGGCCCAGCCCGTGGACGGTGTGAGGCCGG | >hg38_21_+_8433172_8446622_RNA45SN1_45S_50ntFlanks | 8170 | 8215 | 46 | CAAGTCCTTCTGATCGAGGCCCAGCCCGTGGACGGTGTGAGGCCGG | 2245.1 | -7.3 | 1.3 | -5.8 | 8.5E-09 | 1.7E-08 | MB231/MB468 | NUC |
| >hg38_21_+_8433172_8446622_RNA45SN1_45S_50ntFlanks@12804.12821.18\|CGCGTGCCCCGCCGCGCG | >hg38_21_+_8433172_8446622_RNA45SN1_45S_50ntFlanks | 12804 | 12821 | 18 | CGCGTGCCCCGCCGCGCG | 217.6 | -10.8 | 1.8 | -6.1 | 1.1E-09 | 1.7E-08 | BT20/MB468 | MP |
| >trnaMT_ProTGG_MT_-_15956_16023@51.71.21\|TAAAGACTTTTTCTCTGACCA | >trnaMT_ProTGG_MT_-_15956_16023 | 51 | 71 | 21 | TAAAGACTTTTTCTCTGACCA | 682.4 | 5.8 | 1.0 | 5.8 | 8.5E-09 | 1.7E-08 | MB231/MB468 | NUC |
| >hg38_21_+_8433172_8446622_RNA45SN1_45S_50ntFlanks@9738.9767.30\|GAAACGATCTCAACCTATTCTCAAACTTTA | >hg38_21_+_8433172_8446622_RNA45SN1_45S_50ntFlanks | 9738 | 9767 | 30 | GAAACGATCTCAACCTATTCTCAAACTTTA | 932.4 | -8.1 | 1.4 | -5.8 | 8.0E-09 | 1.7E-08 | MB231/MB468 | MP |
| >hg38_21_+_8433172_8446622_RNA45SN1_45S_50ntFlanks@8380.8397.18\|CAAGAGGGCGTGAAACCG | >hg38_21_+_8433172_8446622_RNA45SN1_45S_50ntFlanks | 8380 | 8397 | 18 | CAAGAGGGCGTGAAACCG | 1583.5 | -4.8 | 0.8 | -6.1 | 1.1E-09 | 1.8E-08 | BT20/MB468 | MP |
| >hg38_21_+_8433172_8446622_RNA45SN1_45S_50ntFlanks@12944.12965.22\|CGTAGACGACCTGCTTCTGGGT | >hg38_21_+_8433172_8446622_RNA45SN1_45S_50ntFlanks | 12944 | 12965 | 22 | CGTAGACGACCTGCTTCTGGGT | 138.7 | -4.0 | 0.7 | -5.8 | 8.5E-09 | 1.8E-08 | MB231/MB468 | CYTO |
| >hg38_21_+_8433172_8446622_RNA45SN1_45S_50ntFlanks@6671.6697.27\|CTCGGCTCGTGCGTCGATGAAGAACGC | >hg38_21_+_8433172_8446622_RNA45SN1_45S_50ntFlanks | 6671 | 6697 | 27 | CTCGGCTCGTGCGTCGATGAAGAACGC | 487.9 | -3.5 | 0.6 | -5.8 | 8.6E-09 | 1.8E-08 | MB231/MB468 | NUC |
| >hg38_21_+_8433172_8446622_RNA45SN1_45S_50ntFlanks@12223.12241.19\|CAGAAACCTCCCGTGGAGC | >hg38_21_+_8433172_8446622_RNA45SN1_45S_50ntFlanks | 12223 | 12241 | 19 | CAGAAACCTCCCGTGGAGC | 131.3 | -10.1 | 1.7 | -6.1 | 1.1E-09 | 1.8E-08 | BT20/MB468 | MP |
| >hg38_21_+_8433172_8446622_RNA45SN1_45S_50ntFlanks@8352.8373.22\|AGTTGAAAAGAACTTTGAAGAG | >hg38_21_+_8433172_8446622_RNA45SN1_45S_50ntFlanks | 8352 | 8373 | 22 | AGTTGAAAAGAACTTTGAAGAG | 1653.8 | 9.3 | 1.5 | 6.1 | 1.2E-09 | 1.8E-08 | BT20/MB468 | MP |
| >hg38_21_+_8433172_8446622_RNA45SN1_45S_50ntFlanks@11643.11660.18\|GCGGGTGTTGACGCGATG | >hg38_21_+_8433172_8446622_RNA45SN1_45S_50ntFlanks | 11643 | 11660 | 18 | GCGGGTGTTGACGCGATG | 1369.1 | -6.1 | 1.1 | -5.7 | 9.3E-09 | 1.8E-08 | MB231/MB468 | MITO |
| >hg38_21_+_8433172_8446622_RNA45SN1_45S_50ntFlanks@10729.10757.29\|AGAGGGTGTAAATCTCGCGCCGGGCCGTA | >hg38_21_+_8433172_8446622_RNA45SN1_45S_50ntFlanks | 10729 | 10757 | 29 | AGAGGGTGTAAATCTCGCGCCGGGCCGTA | 463.6 | -8.3 | 1.5 | -5.7 | 9.7E-09 | 1.8E-08 | BT20/MB468 | NUC |
| >hg38_21_+_8433172_8446622_RNA45SN1_45S_50ntFlanks@9714.9732.19\|CGAATGATTAGAGGTCTTG | >hg38_21_+_8433172_8446622_RNA45SN1_45S_50ntFlanks | 9714 | 9732 | 19 | CGAATGATTAGAGGTCTTG | 354.2 | -8.0 | 1.4 | -5.7 | 9.3E-09 | 1.8E-08 | MB231/MB468 | MITO |
| >hg38_21_+_8433172_8446622_RNA45SN1_45S_50ntFlanks@11768.11790.23\|CTAATTAGTGACGCGCATGAATG | >hg38_21_+_8433172_8446622_RNA45SN1_45S_50ntFlanks | 11768 | 11790 | 23 | CTAATTAGTGACGCGCATGAATG | 1979.4 | -6.6 | 1.2 | -5.8 | 8.6E-09 | 1.8E-08 | MB231/MB468 | NUC |
| >hg38_21_+_8433172_8446622_RNA45SN1_45S_50ntFlanks@9156.9187.32\|CCAGTGCGCCCCGGGCGGGTCGCGCCGTCGGG | >hg38_21_+_8433172_8446622_RNA45SN1_45S_50ntFlanks | 9156 | 9187 | 32 | CCAGTGCGCCCCGGGCGGGTCGCGCCGTCGGG | 105.7 | -9.8 | 1.6 | -6.1 | 1.2E-09 | 1.8E-08 | BT20/MB468 | MP |
| >hg38_21_+_8433172_8446622_RNA45SN1_45S_50ntFlanks@7985.8017.33\|GATCAGACGTGGCGACCCGCTGAATTTAAGCAT | >hg38_21_+_8433172_8446622_RNA45SN1_45S_50ntFlanks | 7985 | 8017 | 33 | GATCAGACGTGGCGACCCGCTGAATTTAAGCAT | 167.8 | -10.5 | 1.7 | -6.1 | 1.2E-09 | 1.8E-08 | BT20/MB468 | MP |
| >MI0000737\|hsa-mir-200a&WithFlank&1\|+\|1167857\|1167958@60.83.24&[MIMAT0000682&hsa-miR-200a-3p&offsets\|0\|+2;m-142&1\|+\|1167916\|1167938&offsets\|0\|+1]\|TAACACTGTCTGGTAACGATGTTC | >MI0000737\|hsa-mir-200a&WithFlank&1\|+\|1167857\|1167958 | 60 | 83 | 24 | TAACACTGTCTGGTAACGATGTTC | 174.1 | -10.5 | 1.7 | -6.1 | 1.2E-09 | 1.8E-08 | BT20/MB468 | MP |
| >hg38_21_+_8433172_8446622_RNA45SN1_45S_50ntFlanks@8319.8356.38\|CGAGACCGATAGTCAACAAGTACCGTAAGGGAAAGTTG | >hg38_21_+_8433172_8446622_RNA45SN1_45S_50ntFlanks | 8319 | 8356 | 38 | CGAGACCGATAGTCAACAAGTACCGTAAGGGAAAGTTG | 658.4 | -8.7 | 1.5 | -5.7 | 9.7E-09 | 1.8E-08 | BT20/MB468 | NUC |
| >hg38_21_+_8433172_8446622_RNA45SN1_45S_50ntFlanks@6725.6743.19\|GCAGGACACATTGATCATC | >hg38_21_+_8433172_8446622_RNA45SN1_45S_50ntFlanks | 6725 | 6743 | 19 | GCAGGACACATTGATCATC | 1268.3 | -5.0 | 0.9 | -5.7 | 9.7E-09 | 1.8E-08 | BT20/MB468 | NUC |
| >hg38_21_+_8433172_8446622_RNA45SN1_45S_50ntFlanks@8170.8187.18\|CAAGTCCTTCTGATCGAG | >hg38_21_+_8433172_8446622_RNA45SN1_45S_50ntFlanks | 8170 | 8187 | 18 | CAAGTCCTTCTGATCGAG | 767.5 | -7.9 | 1.4 | -5.7 | 9.4E-09 | 1.8E-08 | MB231/MB468 | MITO |
| >hg38_21_+_8433172_8446622_RNA45SN1_45S_50ntFlanks@12617.12653.37\|TGAGGAGCCAATGGGGCGAAGCTACCATCTGTGGGAT | >hg38_21_+_8433172_8446622_RNA45SN1_45S_50ntFlanks | 12617 | 12653 | 37 | TGAGGAGCCAATGGGGCGAAGCTACCATCTGTGGGAT | 381.0 | -4.8 | 0.8 | -5.8 | 8.6E-09 | 1.8E-08 | MB231/MB468 | CYTO |
| >hg38_21_+_8433172_8446622_RNA45SN1_45S_50ntFlanks@9027.9044.18\|CGTCTCCTCGTGGGGGGG | >hg38_21_+_8433172_8446622_RNA45SN1_45S_50ntFlanks | 9027 | 9044 | 18 | CGTCTCCTCGTGGGGGGG | 216.1 | -10.8 | 1.8 | -6.1 | 1.2E-09 | 1.8E-08 | BT20/MB468 | MP |
| >hg38_21_+_8433172_8446622_RNA45SN1_45S_50ntFlanks@9545.9562.18\|GTGGAGGTCCGTAGCGGT | >hg38_21_+_8433172_8446622_RNA45SN1_45S_50ntFlanks | 9545 | 9562 | 18 | GTGGAGGTCCGTAGCGGT | 326.9 | -7.7 | 1.3 | -5.8 | 8.1E-09 | 1.8E-08 | MB231/MB468 | MP |
| >hg38_21_+_8433172_8446622_RNA45SN1_45S_50ntFlanks@8475.8496.22\|CCCGGCGGATCTTTCCCGCCCC | >hg38_21_+_8433172_8446622_RNA45SN1_45S_50ntFlanks | 8475 | 8496 | 22 | CCCGGCGGATCTTTCCCGCCCC | 103.0 | -10.5 | 1.8 | -5.8 | 8.1E-09 | 1.8E-08 | MB231/MB468 | MP |
| >hg38_21_+_8433172_8446622_RNA45SN1_45S_50ntFlanks@12638.12657.20\|CTACCATCTGTGGGATTATG | >hg38_21_+_8433172_8446622_RNA45SN1_45S_50ntFlanks | 12638 | 12657 | 20 | CTACCATCTGTGGGATTATG | 1073.9 | -8.5 | 1.4 | -6.1 | 1.2E-09 | 1.8E-08 | BT20/MB468 | MP |
| >hg38_21_+_8433172_8446622_RNA45SN1_45S_50ntFlanks@9143.9161.19\|GGGCGGACTGTCCCCAGTG | >hg38_21_+_8433172_8446622_RNA45SN1_45S_50ntFlanks | 9143 | 9161 | 19 | GGGCGGACTGTCCCCAGTG | 115.0 | -9.9 | 1.6 | -6.1 | 1.2E-09 | 1.8E-08 | BT20/MB468 | MP |
| >MI0000750\|hsa-mir-26a-2&WithFlank&12\|-\|57824603\|57824698@20.41.22(+1U);MI0000083\|hsa-mir-26a-1&WithFlank&3\|+\|37969398\|37969486@16.37.22(+1U)&[MIMAT0000082&hsa-miR-26a-5p&offsets\|0\|0(+1U);m-33&12\|-\|57824658\|57824679&offsets\|0\|0(+1U)];[MIMAT0000082_1&hsa-miR-26a-5p&offsets\|0\|0(+1U);m-34&3\|+\|37969413\|37969434&offsets\|0\|0(+1U)]\|TTCAAGTAATCCAGGATAGGCTT | >MI0000750\|hsa-mir-26a-2&WithFlank&12\|-\|57824603\|57824698 | 20 | 41 | 22(+1U) | TTCAAGTAATCCAGGATAGGCTT | 1809.0 | 5.3 | 0.9 | 5.8 | 8.6E-09 | 1.8E-08 | MB231/MB468 | CYTO |
| >hg38_21_+_8433172_8446622_RNA45SN1_45S_50ntFlanks@6651.6686.36\|CGACTCTTAGCGGTGGATCACTCGGCTCGTGCGTCG | >hg38_21_+_8433172_8446622_RNA45SN1_45S_50ntFlanks | 6651 | 6686 | 36 | CGACTCTTAGCGGTGGATCACTCGGCTCGTGCGTCG | 1849.2 | -4.1 | 0.7 | -5.8 | 8.1E-09 | 1.8E-08 | MB231/MB468 | MP |
| >hg38_21_+_8433172_8446622_RNA45SN1_45S_50ntFlanks@3835.3859.25\|CTCCTCTCCTACTTGGATAACTGTG | >hg38_21_+_8433172_8446622_RNA45SN1_45S_50ntFlanks | 3835 | 3859 | 25 | CTCCTCTCCTACTTGGATAACTGTG | 239.7 | -10.0 | 1.7 | -5.8 | 6.6E-09 | 1.8E-08 | BT20/MB468 | MITO |
| >hg38_21_+_8433172_8446622_RNA45SN1_45S_50ntFlanks@12329.12346.18\|TTTTAAGCAGGAGGTGTC | >hg38_21_+_8433172_8446622_RNA45SN1_45S_50ntFlanks | 12329 | 12346 | 18 | TTTTAAGCAGGAGGTGTC | 134.4 | -10.2 | 1.7 | -6.1 | 1.2E-09 | 1.8E-08 | BT20/MB468 | MP |
| >hg38_21_+_8433172_8446622_RNA45SN1_45S_50ntFlanks@12641.12675.35\|CCATCTGTGGGATTATGACTGAACGCCTCTAAGTC | >hg38_21_+_8433172_8446622_RNA45SN1_45S_50ntFlanks | 12641 | 12675 | 35 | CCATCTGTGGGATTATGACTGAACGCCTCTAAGTC | 59.2 | -9.8 | 1.7 | -5.8 | 8.1E-09 | 1.8E-08 | MB231/MB468 | MP |
| >hg38_21_+_8433172_8446622_RNA45SN1_45S_50ntFlanks@4367.4388.22\|CAGTTAAAAAGCTCGTAGTTGG | >hg38_21_+_8433172_8446622_RNA45SN1_45S_50ntFlanks | 4367 | 4388 | 22 | CAGTTAAAAAGCTCGTAGTTGG | 110.6 | -9.9 | 1.6 | -6.1 | 1.2E-09 | 1.8E-08 | BT20/MB468 | MP |
| >hg38_21_+_8433172_8446622_RNA45SN1_45S_50ntFlanks@9534.9558.25\|AGGAAACTCTGGTGGAGGTCCGTAG | >hg38_21_+_8433172_8446622_RNA45SN1_45S_50ntFlanks | 9534 | 9558 | 25 | AGGAAACTCTGGTGGAGGTCCGTAG | 750.5 | -8.7 | 1.4 | -6.1 | 1.2E-09 | 1.8E-08 | BT20/MB468 | MP |
| >hg38_21_+_8433172_8446622_RNA45SN1_45S_50ntFlanks@12635.12661.27\|AAGCTACCATCTGTGGGATTATGACTG | >hg38_21_+_8433172_8446622_RNA45SN1_45S_50ntFlanks | 12635 | 12661 | 27 | AAGCTACCATCTGTGGGATTATGACTG | 4385.6 | -8.1 | 1.4 | -5.7 | 9.8E-09 | 1.8E-08 | BT20/MB468 | NUC |
| >MI0000086\|hsa-mir-28&WithFlank&3\|+\|188688775\|188688872@20.41.22&[MIMAT0000085&hsa-miR-28-5p&offsets\|0\|0;m-133&3\|+\|188688794\|188688815&offsets\|0\|0]\|AAGGAGCTCACAGTCTATTGAG | >MI0000086\|hsa-mir-28&WithFlank&3\|+\|188688775\|188688872 | 20 | 41 | 22 | AAGGAGCTCACAGTCTATTGAG | 270.0 | 6.2 | 1.1 | 5.8 | 8.7E-09 | 1.8E-08 | MB231/MB468 | NUC |
| >hg38_1_-_228634819_228635039_RNA5S12_5S_50ntFlanks@131.170.40\|GGATGGGAGACCGCCTGGGAATACCGGGTGCTGTAGGCTT | >hg38_1_-_228634819_228635039_RNA5S12_5S_50ntFlanks | 131 | 170 | 40 | GGATGGGAGACCGCCTGGGAATACCGGGTGCTGTAGGCTT | 826.8 | 5.2 | 0.9 | 5.8 | 8.8E-09 | 1.8E-08 | MB231/MB468 | NUC |
| >MI0000077\|hsa-mir-21&WithFlank&17\|+\|59841260\|59841343@14.34.21(+1C)&[MIMAT0000076&hsa-miR-21-5p&offsets\|0\|-1(+1C);m-1&17\|+\|59841273\|59841294&offsets\|0\|-1(+1C)]\|TAGCTTATCAGACTGATGTTGC | >MI0000077\|hsa-mir-21&WithFlank&17\|+\|59841260\|59841343 | 14 | 34 | 21(+1C) | TAGCTTATCAGACTGATGTTGC | 79.5 | 5.6 | 1.0 | 5.7 | 9.8E-09 | 1.8E-08 | BT20/MB468 | NUC |
| >hg38_21_+_8433172_8446622_RNA45SN1_45S_50ntFlanks@9791.9819.29\|TGGCGTGGAGCCGGGCGTGGAATGCGAGT | >hg38_21_+_8433172_8446622_RNA45SN1_45S_50ntFlanks | 9791 | 9819 | 29 | TGGCGTGGAGCCGGGCGTGGAATGCGAGT | 202.4 | 11.4 | 1.9 | 6.1 | 1.2E-09 | 1.8E-08 | BT20/MB468 | MP |
| >hg38_21_+_8433172_8446622_RNA45SN1_45S_50ntFlanks@9556.9582.27\|TAGCGGTCCTGACGTGCAAATCGGTCG | >hg38_21_+_8433172_8446622_RNA45SN1_45S_50ntFlanks | 9556 | 9582 | 27 | TAGCGGTCCTGACGTGCAAATCGGTCG | 910.9 | -6.7 | 1.2 | -5.8 | 8.8E-09 | 1.8E-08 | MB231/MB468 | NUC |
| >hg38_21_+_8433172_8446622_RNA45SN1_45S_50ntFlanks@4736.4758.23\|CGAAAGTCGGAGGTTCGAAGACG | >hg38_21_+_8433172_8446622_RNA45SN1_45S_50ntFlanks | 4736 | 4758 | 23 | CGAAAGTCGGAGGTTCGAAGACG | 584.3 | -6.7 | 1.2 | -5.7 | 9.5E-09 | 1.8E-08 | MB231/MB468 | MITO |
| >hg38_21_+_8433172_8446622_RNA45SN1_45S_50ntFlanks@11768.11805.38\|CTAATTAGTGACGCGCATGAATGGATGAACGAGATTCC | >hg38_21_+_8433172_8446622_RNA45SN1_45S_50ntFlanks | 11768 | 11805 | 38 | CTAATTAGTGACGCGCATGAATGGATGAACGAGATTCC | 69.2 | -10.0 | 1.7 | -5.8 | 8.2E-09 | 1.8E-08 | MB231/MB468 | MP |
| >hg38_21_+_8433172_8446622_RNA45SN1_45S_50ntFlanks@12244.12273.30\|AAGGGCAAAAGCTCGCTTGATCTTGATTTT | >hg38_21_+_8433172_8446622_RNA45SN1_45S_50ntFlanks | 12244 | 12273 | 30 | AAGGGCAAAAGCTCGCTTGATCTTGATTTT | 829.8 | -6.7 | 1.2 | -5.7 | 9.5E-09 | 1.8E-08 | MB231/MB468 | MITO |
| >hg38_21_+_8433172_8446622_RNA45SN1_45S_50ntFlanks@10841.10875.35\|CGTAACTTCGGGATAAGGATTGGCTCTAAGGGCTG | >hg38_21_+_8433172_8446622_RNA45SN1_45S_50ntFlanks | 10841 | 10875 | 35 | CGTAACTTCGGGATAAGGATTGGCTCTAAGGGCTG | 1002.9 | -6.6 | 1.2 | -5.8 | 8.8E-09 | 1.8E-08 | MB231/MB468 | NUC |
| >am_trna2_GlyGCC_21_-_18827107_18827177@1.27.27,trna35_GlyGCC_1_+_161413094_161413164@1.27.27,trna37_GlyGCC_1_+_161420467_161420537@1.27.27,trna39_GlyGCC_1_+_161427898_161427968@1.27.27,trna41_GlyGCC_1_+_161435258_161435328@1.27.27\|GCATGGGTGGTTCAGTGGTAGAATTCT | >am_trna2_GlyGCC_21_-_18827107_18827177 | 1 | 27 | 27 | GCATGGGTGGTTCAGTGGTAGAATTCT | 1120.1 | -8.1 | 1.4 | -5.8 | 8.8E-09 | 1.8E-08 | MB231/MB468 | NUC |
| >hg38_21_+_8433172_8446622_RNA45SN1_45S_50ntFlanks@11576.11607.32\|CTTAGAACTGGTGCGGACCAGGGGAATCCGAC | >hg38_21_+_8433172_8446622_RNA45SN1_45S_50ntFlanks | 11576 | 11607 | 32 | CTTAGAACTGGTGCGGACCAGGGGAATCCGAC | 287.7 | -7.9 | 1.4 | -5.8 | 8.2E-09 | 1.8E-08 | MB231/MB468 | MP |
| >hg38_21_+_8433172_8446622_RNA45SN1_45S_50ntFlanks@475.518.44\|CTTCGTGATCGATGTGGTGACGTCGTGCTCTCCCGGGCCGGGTC | >hg38_21_+_8433172_8446622_RNA45SN1_45S_50ntFlanks | 475 | 518 | 44 | CTTCGTGATCGATGTGGTGACGTCGTGCTCTCCCGGGCCGGGTC | 162.3 | 11.1 | 1.8 | 6.1 | 1.2E-09 | 1.8E-08 | BT20/MB468 | MP |
| >hg38_21_+_8433172_8446622_RNA45SN1_45S_50ntFlanks@12459.12478.20\|AGCGTTGGATTGTTCACCCA | >hg38_21_+_8433172_8446622_RNA45SN1_45S_50ntFlanks | 12459 | 12478 | 20 | AGCGTTGGATTGTTCACCCA | 839.2 | -5.8 | 1.0 | -5.8 | 8.7E-09 | 1.8E-08 | MB231/MB468 | CYTO |
| >hg38_21_+_8433172_8446622_RNA45SN1_45S_50ntFlanks@10905.10932.28\|CTGGGCGCGCGCCGCGGCTGGACGAGGC | >hg38_21_+_8433172_8446622_RNA45SN1_45S_50ntFlanks | 10905 | 10932 | 28 | CTGGGCGCGCGCCGCGGCTGGACGAGGC | 536.5 | -8.4 | 1.5 | -5.8 | 8.9E-09 | 1.8E-08 | MB231/MB468 | NUC |
| >hg38_21_+_8433172_8446622_RNA45SN1_45S_50ntFlanks@7984.8003.20\|AGATCAGACGTGGCGACCCG | >hg38_21_+_8433172_8446622_RNA45SN1_45S_50ntFlanks | 7984 | 8003 | 20 | AGATCAGACGTGGCGACCCG | 566.4 | -7.6 | 1.3 | -5.7 | 9.6E-09 | 1.8E-08 | MB231/MB468 | MITO |
| >hg38_21_+_8433172_8446622_RNA45SN1_45S_50ntFlanks@6646.6680.35\|TCGTACGACTCTTAGCGGTGGATCACTCGGCTCGT | >hg38_21_+_8433172_8446622_RNA45SN1_45S_50ntFlanks | 6646 | 6680 | 35 | TCGTACGACTCTTAGCGGTGGATCACTCGGCTCGT | 6558.3 | 4.2 | 0.7 | 5.8 | 7.5E-09 | 1.8E-08 | BT20/MB468 | CYTO |
| >trnaMT_PheGAA_MT_+_577_647@1.19.19\|GTTTATGTAGCTTACCTCC | >trnaMT_PheGAA_MT_+_577_647 | 1 | 19 | 19 | GTTTATGTAGCTTACCTCC | 368.8 | 9.2 | 1.6 | 5.8 | 7.5E-09 | 1.8E-08 | BT20/MB468 | CYTO |
| >hg38_21_+_8433172_8446622_RNA45SN1_45S_50ntFlanks@6700.6717.18\|CTAGCTGCGAGAATTAAT | >hg38_21_+_8433172_8446622_RNA45SN1_45S_50ntFlanks | 6700 | 6717 | 18 | CTAGCTGCGAGAATTAAT | 397.7 | -3.9 | 0.7 | -5.8 | 8.9E-09 | 1.8E-08 | MB231/MB468 | NUC |
| >hg38_21_+_8433172_8446622_RNA45SN1_45S_50ntFlanks@12954.12973.20\|CTGCTTCTGGGTCGGGGTTT | >hg38_21_+_8433172_8446622_RNA45SN1_45S_50ntFlanks | 12954 | 12973 | 20 | CTGCTTCTGGGTCGGGGTTT | 235.7 | -7.6 | 1.3 | -5.8 | 8.9E-09 | 1.8E-08 | MB231/MB468 | NUC |
| >MI0000077\|hsa-mir-21&WithFlank&17\|+\|59841260\|59841343@13.37.25&[MIMAT0000076&hsa-miR-21-5p&offsets\|-1\|+2;m-1&17\|+\|59841273\|59841294&offsets\|-1\|+2]\|GTAGCTTATCAGACTGATGTTGACT | >MI0000077\|hsa-mir-21&WithFlank&17\|+\|59841260\|59841343 | 13 | 37 | 25 | GTAGCTTATCAGACTGATGTTGACT | 345.5 | 7.4 | 1.3 | 5.8 | 8.9E-09 | 1.8E-08 | MB231/MB468 | NUC |

| | | | | | | | | | | | | | |
|---|---|---|---|---|---|---|---|---|---|---|---|---|---|
| >hg38_21_+_8433172_8446622_RNA45SN1_45S_50ntFlanks@12263.12284.22\|ATCTTGATTTTCAGTACGAATA | >hg38_21_+_8433172_8446622_RNA45SN1_45S_50ntFlanks | 12263 | 12284 | 22 | ATCTTGATTTTCAGTACGAATA | 82.2 | -10.2 | 1.8 | -5.8 | 8.3E-09 | 1.8E-08 | MB231/MB468 | MP |
| >hg38_1_-_228634819_228635039_RNA5S12_5S_50ntFlanks@127.169.43\|ACTTGGATGGGAGACCGCCTGGGAATACCGGGTGCTGTAGGCT | >hg38_1_-_228634819_228635039_RNA5S12_5S_50ntFlanks | 127 | 169 | 43 | ACTTGGATGGGAGACCGCCTGGGAATACCGGGTGCTGTAGGCT | 1144.3 | 10.6 | 1.8 | 5.8 | 7.5E-09 | 1.8E-08 | BT20/MB468 | CYTO |
| >hg38_21_+_8433172_8446622_RNA45SN1_45S_50ntFlanks@12656.12675.20\|TGACTGAACGCCTCTAAGTC | >hg38_21_+_8433172_8446622_RNA45SN1_45S_50ntFlanks | 12656 | 12675 | 20 | TGACTGAACGCCTCTAAGTC | 488.9 | -5.6 | 1.0 | -5.8 | 8.8E-09 | 1.8E-08 | MB231/MB468 | CYTO |
| >hg38_21_+_8433172_8446622_RNA45SN1_45S_50ntFlanks@12418.12446.29\|CGATGTCGGCTCTTCCTATCATTGTGAAG | >hg38_21_+_8433172_8446622_RNA45SN1_45S_50ntFlanks | 12418 | 12446 | 29 | CGATGTCGGCTCTTCCTATCATTGTGAAG | 1131.6 | -6.5 | 1.1 | -5.8 | 8.3E-09 | 1.8E-08 | MB231/MB468 | MP |
| >hg38_21_+_8433172_8446622_RNA45SN1_45S_50ntFlanks@4264.4282.19\|AATGAGTCCACTTTAAATC | >hg38_21_+_8433172_8446622_RNA45SN1_45S_50ntFlanks | 4264 | 4282 | 19 | AATGAGTCCACTTTAAATC | 154.9 | -9.4 | 1.6 | -5.8 | 6.7E-09 | 1.8E-08 | BT20/MB468 | MITO |
| >hg38_21_+_8433172_8446622_RNA45SN1_45S_50ntFlanks@10391.10433.43\|AACAGCAGTTGAACATGGGTCAGTCGGTCCTGAGAGATGGGCG | >hg38_21_+_8433172_8446622_RNA45SN1_45S_50ntFlanks | 10391 | 10433 | 43 | AACAGCAGTTGAACATGGGTCAGTCGGTCCTGAGAGATGGGCG | 614.2 | -8.6 | 1.5 | -5.7 | 8.9E-09 | 1.8E-08 | MB231/MB468 | NUC |
| >hg38_21_+_8433172_8446622_RNA45SN1_45S_50ntFlanks@11783.11802.20\|CATGAATGGATGAACGAGAT | >hg38_21_+_8433172_8446622_RNA45SN1_45S_50ntFlanks | 11783 | 11802 | 20 | CATGAATGGATGAACGAGAT | 1117.8 | -6.9 | 1.2 | -5.8 | 8.3E-09 | 1.8E-08 | MB231/MB468 | MP |
| >hg38_21_+_8433172_8446622_RNA45SN1_45S_50ntFlanks@12636.12659.24\|AGCTACCATCTGTGGGATTATGAC | >hg38_21_+_8433172_8446622_RNA45SN1_45S_50ntFlanks | 12636 | 12659 | 24 | AGCTACCATCTGTGGGATTATGAC | 500.0 | -4.4 | 0.8 | -5.7 | 9.0E-09 | 1.8E-08 | MB231/MB468 | NUC |
| >am_trnaMT_GluTTC_MT_-_14674_14742@27.52.26,trnalookalike8_GluTTC_5_-_93905172_93905240@27.52.26\|GTTTTTCATATCATTGGTCGTGGTTG | >am_trnaMT_GluTTC_MT_-_14674_14742 | 27 | 52 | 26 | GTTTTTCATATCATTGGTCGTGGTTG | 302.6 | -7.0 | 1.2 | -5.8 | 8.8E-09 | 1.8E-08 | MB231/MB468 | CYTO |
| >hg38_21_+_8433172_8446622_RNA45SN1_45S_50ntFlanks@11029.11060.32\|CTCTCTCTCTCTCCCCCGCTCCCCGTCCTCCC | >hg38_21_+_8433172_8446622_RNA45SN1_45S_50ntFlanks | 11029 | 11060 | 32 | CTCTCTCTCTCTCCCCCGCTCCCCGTCCTCCC | 563.2 | -8.8 | 1.5 | -5.7 | 9.0E-09 | 1.8E-08 | MB231/MB468 | NUC |
| >MI0006444\|hsa-mir-1307&WithFlank&10\|-\|103394247\|103394407@86.108.23&[MIMAT0005951&hsa-miR-1307-3p&offsets\|0\|+1;m-66&10\|-\|103394300\|103394322&offsets\|0\|0]\|ACTCGGCGTGGCGTCGGTCGTGG | >MI0006444\|hsa-mir-1307&WithFlank&10\|-\|103394247\|103394407 | 86 | 108 | 23 | ACTCGGCGTGGCGTCGGTCGTGG | 1772.5 | 4.6 | 0.8 | 5.7 | 9.7E-09 | 1.8E-08 | MB231/MB468 | MITO |
| >hg38_21_+_8433172_8446622_RNA45SN1_45S_50ntFlanks@11517.11539.23\|CCGCCCCCGGGGCCGCGGTTCCG | >hg38_21_+_8433172_8446622_RNA45SN1_45S_50ntFlanks | 11517 | 11539 | 23 | CCGCCCCCGGGGCCGCGGTTCCG | 609.7 | -7.9 | 1.4 | -5.7 | 9.0E-09 | 1.8E-08 | MB231/MB468 | NUC |
| >hg38_21_+_8433172_8446622_RNA45SN1_45S_50ntFlanks@10427.10457.31\|ATGGGCGAGCGCCGTTCCGAAGGGACGGGCG | >hg38_21_+_8433172_8446622_RNA45SN1_45S_50ntFlanks | 10427 | 10457 | 31 | ATGGGCGAGCGCCGTTCCGAAGGGACGGGCG | 392.9 | -7.0 | 1.2 | -5.8 | 7.6E-09 | 1.8E-08 | BT20/MB468 | CYTO |
| >am_MI0005763\|hsa-mir-941-1&WithFlank&20\|+\|63919443\|63919526@53.74.22(+1U);MI0005764\|hsa-mir-941-2&WithFlank&20\|+\|63919499\|63919582@53.74.22(+1U);MI0005765\|hsa-mir-941-3&WithFlank&20\|+\|63919555\|63919638@53.74.22(+1U);MI0005766\|hsa-mir-941-4&WithFlank&20\|+\|63919750\|63919833@53.74.22(+1U);MI0031520\|hsa-mir-941-5&WithFlank&20\|+\|63919862\|63919945@53.74.22(+1U)&[MIMAT0004984&hsa-miR-941&offsets\|0\|-1(+1U);m-263&20\|+\|63919495\|63919517&offsets\|0\|-1(+1U)];[MIMAT0004984_1&hsa-miR-941&offsets\|0\|-1(+1U);m-263&20\|+\|63919551\|63919573&offsets\|0\|-1(+1U)];[MIMAT0004984_2&hsa-miR-941&offsets\|0\|-1(+1U);m-263&20\|+\|63919607\|63919629&offsets\|0\|-1(+1U)];[MIMAT0004984_3&hsa-miR-941&offsets\|0\|-1(+1U);m-263&20\|+\|63919802\|63919824&offsets\|0\|-1(+1U)];[MIMAT0004984_4&hsa-miR-941&offsets\|0\|-1(+1U);m-263&20\|+\|63919914\|63919936&offsets\|0\|-1(+1U)]\|CACCCGGCTGTGTGCACATGTGT | >am_MI0005763\|hsa-mir-941-1&WithFlank&20\|+\|63919443\|63919526 | 53 | 74 | 22(+1U) | CACCCGGCTGTGTGCACATGTGT | 1128.3 | 4.1 | 0.7 | 5.8 | 8.4E-09 | 1.8E-08 | MB231/MB468 | MP |
| >hg38_21_+_8433172_8446622_RNA45SN1_45S_50ntFlanks@4726.4765.40\|TAATCAAGAACGAAAGTCGGAGGTTCGAAGACGATCAGAT | >hg38_21_+_8433172_8446622_RNA45SN1_45S_50ntFlanks | 4726 | 4765 | 40 | TAATCAAGAACGAAAGTCGGAGGTTCGAAGACGATCAGAT | 505.4 | 5.8 | 1.0 | 5.8 | 8.9E-09 | 1.8E-08 | MB231/MB468 | CYTO |
| >hg38_21_+_8433172_8446622_RNA45SN1_45S_50ntFlanks@11200.11221.22\|CGGCGGCGGCGGCGACTCTGGA | >hg38_21_+_8433172_8446622_RNA45SN1_45S_50ntFlanks | 11200 | 11221 | 22 | CGGCGGCGGCGGCGACTCTGGA | 65.6 | -9.9 | 1.7 | -5.8 | 8.4E-09 | 1.8E-08 | MB231/MB468 | MP |
| >hg38_21_+_8433172_8446622_RNA45SN1_45S_50ntFlanks@12255.12284.30\|CTCGCTTGATCTTGATTTTCAGTACGAATA | >hg38_21_+_8433172_8446622_RNA45SN1_45S_50ntFlanks | 12255 | 12284 | 30 | CTCGCTTGATCTTGATTTTCAGTACGAATA | 110.8 | -10.7 | 1.8 | -5.8 | 8.4E-09 | 1.8E-08 | MB231/MB468 | MP |
| >hg38_21_+_8433172_8446622_RNA45SN1_45S_50ntFlanks@12456.12490.35\|CCAAGCGTTGGATTGTTCACCCACTAATAGGGAAC | >hg38_21_+_8433172_8446622_RNA45SN1_45S_50ntFlanks | 12456 | 12490 | 35 | CCAAGCGTTGGATTGTTCACCCACTAATAGGGAAC | 2202.9 | -4.8 | 0.8 | -5.7 | 9.0E-09 | 1.8E-08 | MB231/MB468 | NUC |
| >hg38_21_+_8433172_8446622_RNA45SN1_45S_50ntFlanks@4661.4689.29\|AGGTGAAATTCTTGGACCGGCGCAAGACG | >hg38_21_+_8433172_8446622_RNA45SN1_45S_50ntFlanks | 4661 | 4689 | 29 | AGGTGAAATTCTTGGACCGGCGCAAGACG | 631.6 | -7.2 | 1.3 | -5.7 | 1.0E-08 | 1.8E-08 | BT20/MB468 | NUC |
| >hg38_21_+_8433172_8446622_RNA45SN1_45S_50ntFlanks@10240.10262.23\|CTACGCCGCGACGAGTAGGAGGG | >hg38_21_+_8433172_8446622_RNA45SN1_45S_50ntFlanks | 10240 | 10262 | 23 | CTACGCCGCGACGAGTAGGAGGG | 275.1 | -7.4 | 1.3 | -5.8 | 8.4E-09 | 1.8E-08 | MB231/MB468 | MP |
| >MI0000750\|hsa-mir-26a-2&WithFlank&12\|-\|57824603\|57824698@20.40.21(+1C);MI0000083\|hsa-mir-26a-1&WithFlank&3\|+\|37969398\|37969486@16.36.21(+1C)&[MIMAT0000082&hsa-miR-26a-5p&offsets\|0\|-1(+1C);m-33&12\|-\|57824658\|57824679&offsets\|0\|-1(+1C)];[MIMAT0000082_1&hsa-miR-26a-5p&offsets\|0\|-1(+1C);m-34&3\|+\|37969413\|37969434&offsets\|0\|-1(+1C)]\|TTCAAGTAATCCAGGATAGGCC | >MI0000750\|hsa-mir-26a-2&WithFlank&12\|-\|57824603\|57824698 | 20 | 40 | 21(+1C) | TTCAAGTAATCCAGGATAGGCC | 16950.6 | 5.3 | 0.9 | 5.7 | 9.8E-09 | 1.8E-08 | MB231/MB468 | MITO |
| >hg38_21_+_8433172_8446622_RNA45SN1_45S_50ntFlanks@6721.6756.36\|AATTGCAGGACACATTGATCATCGACACTTCGAACG | >hg38_21_+_8433172_8446622_RNA45SN1_45S_50ntFlanks | 6721 | 6756 | 36 | AATTGCAGGACACATTGATCATCGACACTTCGAACG | 1353.9 | -6.3 | 1.1 | -5.7 | 9.8E-09 | 1.8E-08 | MB231/MB468 | MITO |
| >trna116_GluCTC_1_-_145399233_145399304@1.36.36,trna59_GluCTC_1_+_249168447_249168518@1.36.36,trna71_GluCTC_1_-_161439189_161439260@1.36.36,trna74_GluCTC_1_-_161431809_161431880@1.36.36,trna77_GluCTC_1_-_161424398_161424469@1.36.36,trna77_GluCTC_6_+_28949976_28950047@1.36.36,trna80_GluCTC_1_-_161417018_161417089@1.36.36,trna87_GluCTC_6_-_126101393_126101464@1.36.36\|TCCCTGGTGGTCTAGTGGTTAGGATTCGGCGCTCTC | >trna116_GluCTC_1_-_145399233_145399304 | 1 | 36 | 36 | TCCCTGGTGGTCTAGTGGTTAGGATTCGGCGCTCTC | 2288.1 | 5.1 | 0.9 | 5.7 | 9.8E-09 | 1.8E-08 | MB231/MB468 | MITO |
| >MI0000071\|hsa-mir-17&WithFlank&13\|+\|91350599\|91350694@20.42.23(+1C)&[MIMAT0000070&hsa-miR-17-5p&offsets\|0\|0(+1C);m-88&13\|+\|91350618\|91350640&offsets\|0\|0(+1C)]\|CAAAGTGCTTACAGTGCAGGTAGC | >MI0000071\|hsa-mir-17&WithFlank&13\|+\|91350599\|91350694 | 20 | 42 | 23(+1C) | CAAAGTGCTTACAGTGCAGGTAGC | 264.1 | -11.4 | 1.8 | -6.3 | 2.3E-10 | 1.8E-08 | BT20/MB231 | MP |
| >hg38_21_+_8433172_8446622_RNA45SN1_45S_50ntFlanks@7975.7997.23\|CGCGACCTCAGATCAGACGTGGC | >hg38_21_+_8433172_8446622_RNA45SN1_45S_50ntFlanks | 7975 | 7997 | 23 | CGCGACCTCAGATCAGACGTGGC | 25188.1 | 3.9 | 0.7 | 5.7 | 1.0E-08 | 1.8E-08 | BT20/MB468 | NUC |
| >hg38_21_+_8433172_8446622_RNA45SN1_45S_50ntFlanks@5186.5204.19\|CAATAACAGGTCTGTGATG | >hg38_21_+_8433172_8446622_RNA45SN1_45S_50ntFlanks | 5186 | 5204 | 19 | CAATAACAGGTCTGTGATG | 585.6 | -5.0 | 0.9 | -5.8 | 8.4E-09 | 1.8E-08 | MB231/MB468 | MP |
| >MI0000063\|hsa-let-7b&WithFlank&22\|+\|46113680\|46113774@12.32.21(+1A)&[MIMAT0000063&hsa-let-7b-5p&offsets\|0\|-1(+1A);m-14&22\|+\|46113691\|46113712&offsets\|0\|-1(+1A)]\|TGAGGTAGTAGGTTGTGTGGTA | >MI0000063\|hsa-let-7b&WithFlank&22\|+\|46113680\|46113774 | 12 | 32 | 21(+1A) | TGAGGTAGTAGGTTGTGTGGTA | 689.5 | 3.2 | 0.6 | 5.7 | 1.0E-08 | 1.8E-08 | BT20/MB468 | NUC |
| >hg38_21_+_8433172_8446622_RNA45SN1_45S_50ntFlanks@9653.9677.25\|AGGATAGCTGGCGCTCTCGCAGACC | >hg38_21_+_8433172_8446622_RNA45SN1_45S_50ntFlanks | 9653 | 9677 | 25 | AGGATAGCTGGCGCTCTCGCAGACC | 1366.2 | -5.2 | 0.9 | -5.7 | 9.8E-09 | 1.8E-08 | MB231/MB468 | MITO |
| >hg38_21_+_8433172_8446622_RNA45SN1_45S_50ntFlanks@8168.8203.36\|CCCAAGTCCTTCTGATCGAGGCCCAGCCCGTGGACG | >hg38_21_+_8433172_8446622_RNA45SN1_45S_50ntFlanks | 8168 | 8203 | 36 | CCCAAGTCCTTCTGATCGAGGCCCAGCCCGTGGACG | 166.7 | -9.5 | 1.6 | -5.8 | 6.9E-09 | 1.9E-08 | BT20/MB468 | MITO |
| >hg38_21_+_8433172_8446622_RNA45SN1_45S_50ntFlanks@11670.11698.29\|CCCAGTGCTCTGAATGTCAAAGTGAAGAA | >hg38_21_+_8433172_8446622_RNA45SN1_45S_50ntFlanks | 11670 | 11698 | 29 | CCCAGTGCTCTGAATGTCAAAGTGAAGAA | 1310.4 | -9.1 | 1.6 | -5.7 | 1.0E-08 | 1.9E-08 | BT20/MB468 | NUC |
| >MI0000809\|hsa-mir-151a&WithFlank&8\|-\|140732558\|140732659@53.74.22(+2U)&[MIMAT0000757&hsa-miR-151a-3p&offsets\|0\|+1(+2U);m-30&8\|-\|140732587\|140732607&offsets\|0\|+1(+2U)]\|CTAGACTGAAGCTCCTTGAGGATT | >MI0000809\|hsa-mir-151a&WithFlank&8\|-\|140732558\|140732659 | 53 | 74 | 22(+2U) | CTAGACTGAAGCTCCTTGAGGATT | 526.7 | 6.9 | 1.2 | 5.7 | 9.9E-09 | 1.9E-08 | MB231/MB468 | MITO |
| >hg38_21_+_8433172_8446622_RNA45SN1_45S_50ntFlanks@12624.12660.37\|CCAATGGGGCGAAGCTACCATCTGTGGGATTATGACT | >hg38_21_+_8433172_8446622_RNA45SN1_45S_50ntFlanks | 12624 | 12660 | 37 | CCAATGGGGCGAAGCTACCATCTGTGGGATTATGACT | 388.6 | -6.5 | 1.1 | -5.7 | 9.9E-09 | 1.9E-08 | MB231/MB468 | MITO |
| >MI0000098\|hsa-mir-96&WithFlank&7\|-\|129774686\|129774775@15.36.22(+1C)&[MIMAT0000095&hsa-miR-96-5p&offsets\|0\|-1(+1C);m-248&7\|-\|129774739\|129774761&offsets\|0\|-1(+1C)]\|TTTGGCACTAGCACATTTTTGCC | >MI0000098\|hsa-mir-96&WithFlank&7\|-\|129774686\|129774775 | 15 | 36 | 22(+1C) | TTTGGCACTAGCACATTTTTGCC | 313.8 | 4.4 | 0.8 | 5.7 | 1.0E-08 | 1.9E-08 | BT20/MB468 | NUC |
| >hg38_21_+_8433172_8446622_RNA45SN1_45S_50ntFlanks@5078.5105.28\|CGATAACGAACGAGACTCTGGCATGCTA | >hg38_21_+_8433172_8446622_RNA45SN1_45S_50ntFlanks | 5078 | 5105 | 28 | CGATAACGAACGAGACTCTGGCATGCTA | 226.1 | -9.9 | 1.7 | -5.8 | 6.9E-09 | 1.9E-08 | BT20/MB468 | MITO |
| >hg38_21_+_8433172_8446622_RNA45SN1_45S_50ntFlanks@11712.11740.29\|CGGGTAAACGGCGGGAGTAACTATGACTC | >hg38_21_+_8433172_8446622_RNA45SN1_45S_50ntFlanks | 11712 | 11740 | 29 | CGGGTAAACGGCGGGAGTAACTATGACTC | 648.4 | -8.8 | 1.5 | -5.7 | 1.0E-08 | 1.9E-08 | BT20/MB468 | NUC |
| >hg38_21_+_8433172_8446622_RNA45SN1_45S_50ntFlanks@6745.6769.25\|ACACTTCGAACGCACTTGCGGCCCC | >hg38_21_+_8433172_8446622_RNA45SN1_45S_50ntFlanks | 6745 | 6769 | 25 | ACACTTCGAACGCACTTGCGGCCCC | 386.4 | -4.8 | 0.8 | -5.7 | 9.0E-09 | 1.9E-08 | MB231/MB468 | CYTO |
| >hg38_1_-_228634819_228635039_RNA5S12_5S_50ntFlanks@84.112.29\|CTCGTCTGATCTCGGAAGCTAAGCAGGGT | >hg38_1_-_228634819_228635039_RNA5S12_5S_50ntFlanks | 84 | 112 | 29 | CTCGTCTGATCTCGGAAGCTAAGCAGGGT | 86.1 | -8.5 | 1.5 | -5.8 | 6.9E-09 | 1.9E-08 | BT20/MB468 | MITO |
| >hg38_21_+_8433172_8446622_RNA45SN1_45S_50ntFlanks@12365.12383.19\|ATAACTGGCTTGTGGCGGC | >hg38_21_+_8433172_8446622_RNA45SN1_45S_50ntFlanks | 12365 | 12383 | 19 | ATAACTGGCTTGTGGCGGC | 296.0 | -6.6 | 1.1 | -5.8 | 7.7E-09 | 1.9E-08 | BT20/MB468 | CYTO |
| >MI0000093\|hsa-mir-92a-1&WithFlank&13\|+\|91351308\|91351397@54.75.22(+1C);MI0000094\|hsa-mir-92a-2&WithFlank&X\|-\|134169532\|134169618@54.75.22(+1C)&[MIMAT0000092&hsa-miR-92a-3p&offsets\|0\|0(+1C);m-9&13\|+\|91351361\|91351382&offsets\|0\|0(+1C)];[MIMAT0000092_1&hsa-miR-92a-3p&offsets\|0\|0(+1C);m-11&X\|-\|134169544\|134169565&offsets\|0\|0(+1C)]\|TATTGCACTTGTCCCGGCCTGTC | >MI0000093\|hsa-mir-92a-1&WithFlank&13\|+\|91351308\|91351397 | 54 | 75 | 22(+1C) | TATTGCACTTGTCCCGGCCTGTC | 956.3 | 5.7 | 1.0 | 5.7 | 9.9E-09 | 1.9E-08 | MB231/MB468 | MITO |

| | | | | | | | | | | | | | |
|---|---|---|---|---|---|---|---|---|---|---|---|---|---|
| >hg38_21_+_8433172_8446622_RNA45SN1_45S_50ntFlanks@10403.10422.20\|ACATGGGTCAGTCGGTCCTG | >hg38_21_+_8433172_8446622_RNA45SN1_45S_50ntFlanks | 10403 | 10422 | 20 | ACATGGGTCAGTCGGTCCTG | 190.9 | -6.1 | 1.1 | -5.8 | 8.6E-09 | 1.9E-08 | MB231/MB468 | MP |
| >hg38_MT_+_1621_3279_RNR2_16S_50ntFlanks@483.518.36\|AAGAGGAACAGCTCTTTGGACACTAGGAAAAAACCT | >hg38_MT_+_1621_3279_RNR2_16S_50ntFlanks | 483 | 518 | 36 | AAGAGGAACAGCTCTTTGGACACTAGGAAAAAACCT | 112.4 | 10.6 | 1.7 | 6.1 | 1.2E-09 | 1.9E-08 | BT20/MB468 | MP |
| >trna2_GlyGCC_21_-_18827107_18827177@1.31.31,trna35_GlyGCC_1_+_161413094_161413164@1.31.31,trna37_GlyGCC_1_+_161420467_161420537@1.31.31,trna39_GlyGCC_1_+_161427898_161427968@1.31.31,trna41_GlyGCC_1_+_161435258_161435328@1.31.31\|GCATGGGTGGTTCAGTGGTAGAATTCTCGCC | >trna2_GlyGCC_21_-_18827107_18827177 | 1 | 31 | 31 | GCATGGGTGGTTCAGTGGTAGAATTCTCGCC | 42180.9 | 5.3 | 0.9 | 6.2 | 5.4E-10 | 1.9E-08 | BT20/MB231 | CYTO |
| >hg38_21_+_8433172_8446622_RNA45SN1_45S_50ntFlanks@5059.5079.21\|CGATTTGTCTGGTTAATTCCG | >hg38_21_+_8433172_8446622_RNA45SN1_45S_50ntFlanks | 5059 | 5079 | 21 | CGATTTGTCTGGTTAATTCCG | 101.2 | -10.5 | 1.8 | -5.8 | 8.6E-09 | 1.9E-08 | MB231/MB468 | MP |
| >hg38_21_+_8433172_8446622_RNA45SN1_45S_50ntFlanks@11667.11702.36\|CTGCCCAGTGCTCTGAATGTCAAAGTGAAGAAATTC | >hg38_21_+_8433172_8446622_RNA45SN1_45S_50ntFlanks | 11667 | 11702 | 36 | CTGCCCAGTGCTCTGAATGTCAAAGTGAAGAAATTC | 388.8 | -3.9 | 0.7 | -5.7 | 1.0E-08 | 1.9E-08 | BT20/MB468 | NUC |
| >hg38_21_+_8433172_8446622_RNA45SN1_45S_50ntFlanks@8354.8379.26\|TTGAAAAGAACTTTGAAGAGAGAGTT | >hg38_21_+_8433172_8446622_RNA45SN1_45S_50ntFlanks | 8354 | 8379 | 26 | TTGAAAAGAACTTTGAAGAGAGAGTT | 81.6 | -10.2 | 1.8 | -5.8 | 8.6E-09 | 1.9E-08 | MB231/MB468 | MP |
| >hg38_21_+_8433172_8446622_RNA45SN1_45S_50ntFlanks@4069.4101.33\|CTTTCGATGGTAGTCGCCGTGCCTACCATGGTG | >hg38_21_+_8433172_8446622_RNA45SN1_45S_50ntFlanks | 4069 | 4101 | 33 | CTTTCGATGGTAGTCGCCGTGCCTACCATGGTG | 209.6 | -7.1 | 1.2 | -5.7 | 9.3E-09 | 1.9E-08 | MB231/MB468 | NUC |
| >trna10_GlyTCC_17_+_8124866_8124937@1.30.30\|GCGTTGGTGGTATAGTGGTAAGCATAGCTG | >trna10_GlyTCC_17_+_8124866_8124937 | 1 | 30 | 30 | GCGTTGGTGGTATAGTGGTAAGCATAGCTG | 136.5 | -9.2 | 1.6 | -5.8 | 7.0E-09 | 1.9E-08 | BT20/MB468 | MITO |
| >hg38_21_+_8433172_8446622_RNA45SN1_45S_50ntFlanks@4546.4573.28\|CCTGGATACCGCAGCTAGGAATAATGGA | >hg38_21_+_8433172_8446622_RNA45SN1_45S_50ntFlanks | 4546 | 4573 | 28 | CCTGGATACCGCAGCTAGGAATAATGGA | 347.2 | -11.5 | 1.9 | -6.1 | 1.2E-09 | 1.9E-08 | BT20/MB468 | MP |
| >hg38_21_+_8433172_8446622_RNA45SN1_45S_50ntFlanks@4727.4767.41\|AATCAAGAACGAAAGTCGGAGGTTCGAAGACGATCAGATAC | >hg38_21_+_8433172_8446622_RNA45SN1_45S_50ntFlanks | 4727 | 4767 | 41 | AATCAAGAACGAAAGTCGGAGGTTCGAAGACGATCAGATAC | 134.1 | 10.8 | 1.8 | 6.1 | 1.3E-09 | 1.9E-08 | BT20/MB468 | MP |
| >hg38_21_+_8433172_8446622_RNA45SN1_45S_50ntFlanks@11323.11344.22\|CGTCTCGTCGCGCGCGCGTCCG | >hg38_21_+_8433172_8446622_RNA45SN1_45S_50ntFlanks | 11323 | 11344 | 22 | CGTCTCGTCGCGCGCGCGTCCG | 7946.9 | -8.0 | 1.3 | -6.1 | 1.3E-09 | 1.9E-08 | BT20/MB468 | MP |
| >hg38_21_+_8433172_8446622_RNA45SN1_45S_50ntFlanks@3815.3850.36\|ATGGTTCCTTTGGTCGCTCGCTCCTCTCCTACTTGG | >hg38_21_+_8433172_8446622_RNA45SN1_45S_50ntFlanks | 3815 | 3850 | 36 | ATGGTTCCTTTGGTCGCTCGCTCCTCTCCTACTTGG | 638.5 | -8.7 | 1.5 | -5.7 | 1.0E-08 | 1.9E-08 | BT20/MB468 | NUC |
| >hg38_21_+_8433172_8446622_RNA45SN1_45S_50ntFlanks@12526.12552.27\|TTTTACCCTACTGATGATGTGTTGTTG | >hg38_21_+_8433172_8446622_RNA45SN1_45S_50ntFlanks | 12526 | 12552 | 27 | TTTTACCCTACTGATGATGTGTTGTTG | 128.0 | -10.1 | 1.7 | -6.1 | 1.3E-09 | 1.9E-08 | BT20/MB468 | MP |
| >hg38_21_+_8433172_8446622_RNA45SN1_45S_50ntFlanks@10726.10748.23\|GGGAGAGGGTGTAAATCTCGCGC | >hg38_21_+_8433172_8446622_RNA45SN1_45S_50ntFlanks | 10726 | 10748 | 23 | GGGAGAGGGTGTAAATCTCGCGC | 70.4 | -10.0 | 1.7 | -5.8 | 8.7E-09 | 1.9E-08 | MB231/MB468 | MP |
| >MI0000477\|hsa-mir-146a&WithFlank&5\|+\|160485346\|160485456@27.47.21(+1A)&[MIMAT0000449&hsa-miR-146a-5p&offsets\|0\|-1(+1A);m-129&5\|+\|160485372\|160485393&offsets\|0\|-1(+1A)]\|TGAGAACTGAATTCCATGGGTA | >MI0000477\|hsa-mir-146a&WithFlank&5\|+\|160485346\|160485456 | 27 | 47 | 21(+1A) | TGAGAACTGAATTCCATGGGTA | 8321.8 | -11.0 | 1.7 | -6.3 | 2.3E-10 | 1.9E-08 | BT20/MB231 | MP |
| >hg38_21_+_8433172_8446622_RNA45SN1_45S_50ntFlanks@8190.8210.21\|CCAGCCCGTGGACGGTGTGAG | >hg38_21_+_8433172_8446622_RNA45SN1_45S_50ntFlanks | 8190 | 8210 | 21 | CCAGCCCGTGGACGGTGTGAG | 59.2 | -9.8 | 1.7 | -5.8 | 8.7E-09 | 1.9E-08 | MB231/MB468 | MP |
| >trna111_HisGTG_1_-_147774845_147774916@-1G.32.33,trna118_HisGTG_1_-_145396881_145396952@-1G.32.33,trna16_HisGTG_1_+_146544773_146544844@-1G.32.33,trna1_HisGTG_15_+_45493349_45493420@-1G.32.33,trna21_HisGTG_1_+_147753471_147753542@-1G.32.33,trna33_HisGTG_6_+_27125906_27125977@-1G.32.33,trna7_HisGTG_9_-_14433938_14434009@-1G.32.33,trna8_HisGTG_15_-_45492611_45492682@-1G.32.33,trna9_HisGTG_15_-_45490804_45490875@-1G.32.33\|GGCCGTGATCGTATAGTGGTTAGTACTCTGCGT | >trna111_HisGTG_1_-_147774845_147774916 | -1G | 32 | 33 | GGCCGTGATCGTATAGTGGTTAGTACTCTGCGT | 3115.6 | 10.2 | 1.8 | 5.8 | 8.7E-09 | 1.9E-08 | MB231/MB468 | MP |
| >hg38_21_+_8433172_8446622_RNA45SN1_45S_50ntFlanks@12992.13011.20\|TCCCTCGCTGCGATCTATTG | >hg38_21_+_8433172_8446622_RNA45SN1_45S_50ntFlanks | 12992 | 13011 | 20 | TCCCTCGCTGCGATCTATTG | 247.8 | -7.3 | 1.3 | -5.7 | 1.0E-08 | 1.9E-08 | MB231/MB468 | MITO |
| >hg38_21_+_8433172_8446622_RNA45SN1_45S_50ntFlanks@5083.5102.20\|ACGAACGAGACTCTGGCATG | >hg38_21_+_8433172_8446622_RNA45SN1_45S_50ntFlanks | 5083 | 5102 | 20 | ACGAACGAGACTCTGGCATG | 723.7 | -6.1 | 1.1 | -5.7 | 1.0E-08 | 1.9E-08 | MB231/MB468 | MITO |
| >hg38_21_+_8433172_8446622_RNA45SN1_45S_50ntFlanks@3843.3868.26\|CTACTTGGATAACTGTGGTAATTCTA | >hg38_21_+_8433172_8446622_RNA45SN1_45S_50ntFlanks | 3843 | 3868 | 26 | CTACTTGGATAACTGTGGTAATTCTA | 792.4 | -7.7 | 1.3 | -5.8 | 8.7E-09 | 1.9E-08 | MB231/MB468 | MP |
| >hg38_21_+_8433172_8446622_RNA45SN1_45S_50ntFlanks@9482.9510.29\|ACGTGTTAGGACCCGAAAGATGGTGAACT | >hg38_21_+_8433172_8446622_RNA45SN1_45S_50ntFlanks | 9482 | 9510 | 29 | ACGTGTTAGGACCCGAAAGATGGTGAACT | 170.4 | 11.2 | 1.8 | 6.1 | 1.3E-09 | 1.9E-08 | BT20/MB468 | MP |
| >hg38_21_+_8433172_8446622_RNA45SN1_45S_50ntFlanks@9307.9336.30\|AAGGAGTCTAACACGTGCGCGAGTCGGGGG | >hg38_21_+_8433172_8446622_RNA45SN1_45S_50ntFlanks | 9307 | 9336 | 30 | AAGGAGTCTAACACGTGCGCGAGTCGGGGG | 1591.4 | -7.0 | 1.2 | -5.8 | 8.7E-09 | 1.9E-08 | MB231/MB468 | MP |
| >MI0000298\|hsa-mir-221&WithFlank&X\|-\|45746151\|45746272@71.92.22(+2U)&[MIMAT0000278&hsa-miR-221-3p&offsets\|0\|-1(+2U);m-91&X\|-\|45746181\|45746202&offsets\|0\|0(+2U)]\|AGCTACATTGTCTGCTGGGTTTTT | >MI0000298\|hsa-mir-221&WithFlank&X\|-\|45746151\|45746272 | 71 | 92 | 22(+2U) | AGCTACATTGTCTGCTGGGTTTTT | 286.3 | 6.6 | 1.2 | 5.7 | 9.2E-09 | 1.9E-08 | MB231/MB468 | CYTO |
| >hg38_21_+_8433172_8446622_RNA45SN1_45S_50ntFlanks@7975.8017.43\|CGCGACCTCAGATCAGACGTGGCGACCCGCTGAATTTAAGCAT | >hg38_21_+_8433172_8446622_RNA45SN1_45S_50ntFlanks | 7975 | 8017 | 43 | CGCGACCTCAGATCAGACGTGGCGACCCGCTGAATTTAAGCAT | 2486.2 | 5.5 | 1.0 | 5.7 | 9.2E-09 | 1.9E-08 | MB231/MB468 | CYTO |
| >hg38_21_+_8433172_8446622_RNA45SN1_45S_50ntFlanks@12324.12357.34\|TTGGGTTTTAAGCAGGAGGTGTCAGAAAAGTTAC | >hg38_21_+_8433172_8446622_RNA45SN1_45S_50ntFlanks | 12324 | 12357 | 34 | TTGGGTTTTAAGCAGGAGGTGTCAGAAAAGTTAC | 67.7 | -9.9 | 1.7 | -5.8 | 8.7E-09 | 1.9E-08 | MB231/MB468 | MP |
| >am_trna2_GlyTCC_19_+_4724082_4724153@1.30.30\|GCGTTGGTGGTATAGTGGTTAGCATAGCTG | >am_trna2_GlyTCC_19_+_4724082_4724153 | 1 | 30 | 30 | GCGTTGGTGGTATAGTGGTTAGCATAGCTG | 116.9 | -9.0 | 1.5 | -5.8 | 7.0E-09 | 1.9E-08 | BT20/MB468 | MITO |
| >hg38_21_+_8433172_8446622_RNA45SN1_45S_50ntFlanks@3782.3805.24\|CAGTGAAACTGCGAATGGCTCATT | >hg38_21_+_8433172_8446622_RNA45SN1_45S_50ntFlanks | 3782 | 3805 | 24 | CAGTGAAACTGCGAATGGCTCATT | 761.9 | -5.6 | 1.0 | -5.8 | 8.7E-09 | 1.9E-08 | MB231/MB468 | MP |
| >hg38_21_+_8433172_8446622_RNA45SN1_45S_50ntFlanks@8154.8175.22\|CCGCTCGTGGGGGGCCCAAGTC | >hg38_21_+_8433172_8446622_RNA45SN1_45S_50ntFlanks | 8154 | 8175 | 22 | CCGCTCGTGGGGGGCCCAAGTC | 178.0 | -10.6 | 1.7 | -6.1 | 1.3E-09 | 1.9E-08 | BT20/MB468 | MP |
| >MI0006444\|hsa-mir-1307&WithFlank&10\|-\|103394247\|103394407@87.107.21(+1A)&[MIMAT0005951&hsa-miR-1307-3p&offsets\|+1\|0(+1A);m-66&10\|-\|103394300\|103394322&offsets\|+1\|-1(+1A)]\|CTCGGCGTGGCGTCGGTCGTGA | >MI0006444\|hsa-mir-1307&WithFlank&10\|-\|103394247\|103394407 | 87 | 107 | 21(+1A) | CTCGGCGTGGCGTCGGTCGTGA | 213.9 | 4.2 | 0.7 | 5.7 | 9.4E-09 | 1.9E-08 | MB231/MB468 | NUC |
| >hg38_21_+_8433172_8446622_RNA45SN1_45S_50ntFlanks@8118.8140.23\|GGGACATGTGGCGTACGGAAGAC | >hg38_21_+_8433172_8446622_RNA45SN1_45S_50ntFlanks | 8118 | 8140 | 23 | GGGACATGTGGCGTACGGAAGAC | 1364.8 | 13.7 | 2.2 | 6.3 | 2.4E-10 | 1.9E-08 | BT20/MB231 | MP |
| >hg38_21_+_8433172_8446622_RNA45SN1_45S_50ntFlanks@11569.11612.44\|CAGCCGACTTAGAACTGGTGCGGACCAGGGGAATCCGACTGTTT | >hg38_21_+_8433172_8446622_RNA45SN1_45S_50ntFlanks | 11569 | 11612 | 44 | CAGCCGACTTAGAACTGGTGCGGACCAGGGGAATCCGACTGTTT | 4233.1 | -6.9 | 1.2 | -5.7 | 1.1E-08 | 1.9E-08 | BT20/MB468 | NUC |
| >MI0000439\|hsa-mir-23b&WithFlank&9\|+\|95085202\|95085310@65.86.22(+1U)&[MIMAT0000418&hsa-miR-23b-3p&offsets\|+1\|0(+1U);m-42&9\|+\|95085265\|95085285&offsets\|+1\|+2(+1U)]\|TCACATTGCCAGGGATTACCACT | >MI0000439\|hsa-mir-23b&WithFlank&9\|+\|95085202\|95085310 | 65 | 86 | 22(+1U) | TCACATTGCCAGGGATTACCACT | 103.1 | 9.9 | 1.7 | 5.8 | 8.7E-09 | 1.9E-08 | MB231/MB468 | MP |
| >MI0000750\|hsa-mir-26a-2&WithFlank&12\|-\|57824603\|57824698@20.40.21(+1A);MI0000083\|hsa-mir-26a-1&WithFlank&3\|+\|37969398\|37969486@16.36.21(+1A)&[MIMAT0000082&hsa-miR-26a-5p&offsets\|0\|-1(+1A);m-33&12\|-\|57824658\|57824679&offsets\|0\|-1(+1A)];[MIMAT0000082_1&hsa-miR-26a-5p&offsets\|0\|-1(+1A);m-34&3\|+\|37969413\|37969434&offsets\|0\|-1(+1A)]\|TTCAAGTAATCCAGGATAGGCA | >MI0000750\|hsa-mir-26a-2&WithFlank&12\|-\|57824603\|57824698 | 20 | 40 | 21(+1A) | TTCAAGTAATCCAGGATAGGCA | 5718.7 | 5.6 | 1.0 | 5.7 | 9.3E-09 | 1.9E-08 | MB231/MB468 | CYTO |
| >hg38_MT_+_1621_3279_RNR2_16S_50ntFlanks@51.94.44\|GCTAAACCTAGCCCCAAACCCACTCCACCTTACTACCAGACAAC | >hg38_MT_+_1621_3279_RNR2_16S_50ntFlanks | 51 | 94 | 44 | GCTAAACCTAGCCCCAAACCCACTCCACCTTACTACCAGACAAC | 3716.4 | 5.4 | 0.9 | 5.8 | 7.1E-09 | 1.9E-08 | BT20/MB468 | MITO |
| >hg38_21_+_8433172_8446622_RNA45SN1_45S_50ntFlanks@12478.12498.21\|ACTAATAGGGAACGTGAGCTG | >hg38_21_+_8433172_8446622_RNA45SN1_45S_50ntFlanks | 12478 | 12498 | 21 | ACTAATAGGGAACGTGAGCTG | 389.5 | -6.4 | 1.1 | -5.8 | 8.8E-09 | 1.9E-08 | MB231/MB468 | MP |
| >hg38_21_+_8433172_8446622_RNA45SN1_45S_50ntFlanks@6645.6685.41\|CTCGTACGACTCTTAGCGGTGGATCACTCGGCTCGTGCGTC | >hg38_21_+_8433172_8446622_RNA45SN1_45S_50ntFlanks | 6645 | 6685 | 41 | CTCGTACGACTCTTAGCGGTGGATCACTCGGCTCGTGCGTC | 817.9 | 4.7 | 0.8 | 5.8 | 7.1E-09 | 1.9E-08 | BT20/MB468 | MITO |
| >hg38_21_+_8433172_8446622_RNA45SN1_45S_50ntFlanks@12995.13036.42\|CTCGCTGCGATCTATTGAAAGTCAGCCCTCGACACAAGGGTT | >hg38_21_+_8433172_8446622_RNA45SN1_45S_50ntFlanks | 12995 | 13036 | 42 | CTCGCTGCGATCTATTGAAAGTCAGCCCTCGACACAAGGGTT | 2589.0 | 7.3 | 1.3 | 5.8 | 7.1E-09 | 1.9E-08 | BT20/MB468 | MITO |
| >hg38_21_+_8433172_8446622_RNA45SN1_45S_50ntFlanks@12635.12654.20\|AAGCTACCATCTGTGGGATT | >hg38_21_+_8433172_8446622_RNA45SN1_45S_50ntFlanks | 12635 | 12654 | 20 | AAGCTACCATCTGTGGGATT | 1230.0 | -7.7 | 1.3 | -5.8 | 7.9E-09 | 1.9E-08 | BT20/MB468 | CYTO |
| >hg38_21_+_8433172_8446622_RNA45SN1_45S_50ntFlanks@10821.10840.20\|GGAAGTCGGCAAGCCGGATC | >hg38_21_+_8433172_8446622_RNA45SN1_45S_50ntFlanks | 10821 | 10840 | 20 | GGAAGTCGGCAAGCCGGATC | 817.6 | -3.4 | 0.6 | -5.8 | 7.1E-09 | 1.9E-08 | BT20/MB468 | MITO |
| >hg38_21_+_8433172_8446622_RNA45SN1_45S_50ntFlanks@6645.6666.22\|CTCGTACGACTCTTAGCGGTGG | >hg38_21_+_8433172_8446622_RNA45SN1_45S_50ntFlanks | 6645 | 6666 | 22 | CTCGTACGACTCTTAGCGGTGG | 14800.5 | 4.4 | 0.8 | 5.8 | 7.1E-09 | 1.9E-08 | BT20/MB468 | MITO |
| >hg38_21_+_8433172_8446622_RNA45SN1_45S_50ntFlanks@12420.12455.36\|ATGTCGGCTCTTCCTATCATTGTGAAGCAGAATTCA | >hg38_21_+_8433172_8446622_RNA45SN1_45S_50ntFlanks | 12420 | 12455 | 36 | ATGTCGGCTCTTCCTATCATTGTGAAGCAGAATTCA | 4500.9 | -5.0 | 0.9 | -5.7 | 9.5E-09 | 1.9E-08 | MB231/MB468 | NUC |
| >hg38_21_+_8433172_8446622_RNA45SN1_45S_50ntFlanks@7974.8009.36\|ACGCGACCTCAGATCAGACGTGGCGACCCGCTGAAT | >hg38_21_+_8433172_8446622_RNA45SN1_45S_50ntFlanks | 7974 | 8009 | 36 | ACGCGACCTCAGATCAGACGTGGCGACCCGCTGAAT | 8430.3 | 3.3 | 0.6 | 5.7 | 9.5E-09 | 1.9E-08 | MB231/MB468 | NUC |
| >hg38_21_+_8433172_8446622_RNA45SN1_45S_50ntFlanks@11739.11758.20\|TCTCTTAAGGTAGCCAAATG | >hg38_21_+_8433172_8446622_RNA45SN1_45S_50ntFlanks | 11739 | 11758 | 20 | TCTCTTAAGGTAGCCAAATG | 460.1 | -5.8 | 1.0 | -5.7 | 9.4E-09 | 1.9E-08 | MB231/MB468 | CYTO |
| >MI0000078\|hsa-mir-22&WithFlank&17\|-\|1713897\|1713993@59.79.21(+1C)&[MIMAT0000077&hsa-miR-22-3p&offsets\|0\|-1(+1C);m-2&17\|-\|1713914\|1713935&offsets\|0\|-1(+1C)]\|AAGCTGCCAGTTGAAGAACTGC | >MI0000078\|hsa-mir-22&WithFlank&17\|-\|1713897\|1713993 | 59 | 79 | 21(+1C) | AAGCTGCCAGTTGAAGAACTGC | 5846.6 | 5.5 | 1.0 | 5.8 | 8.8E-09 | 1.9E-08 | MB231/MB468 | MP |

<table>
<tr><th></th><th></th><th></th><th></th><th></th><th></th><th></th><th></th><th></th><th></th><th></th><th></th><th></th><th></th></tr>
<tr><td>>hg38_21_+_8433172_8446622_RNA45SN1_45S_50ntFlanks@4824.4844.21|TGACCCGCCGGGCAGCTTCCG</td><td>>hg38_21_+_8433172_8446622_RNA45SN1_45S_50ntFlanks</td><td>4824</td><td>4844</td><td>21</td><td>TGACCCGCCGGGCAGCTTCCG</td><td>802.3</td><td>6.2</td><td>1.1</td><td>5.7</td><td>9.4E-09</td><td>1.9E-08</td><td>MB231/MB468</td><td>CYTO</td></tr>
<tr><td>>hg38_21_+_8433172_8446622_RNA45SN1_45S_50ntFlanks@12420.12455.36|ATGTCGGCTCTTCCTATCATTGTGAAGCAGAATTCA</td><td>>hg38_21_+_8433172_8446622_RNA45SN1_45S_50ntFlanks</td><td>12420</td><td>12455</td><td>36</td><td>ATGTCGGCTCTTCCTATCATTGTGAAGCAGAATTCA</td><td>3827.3</td><td>-5.4</td><td>0.9</td><td>-5.8</td><td>7.2E-09</td><td>1.9E-08</td><td>BT20/MB468</td><td>MITO</td></tr>
<tr><td>>MI0000465|hsa-mir-191&WithFlank&3|-|49020612|49020715@22.44.23&[MIMAT0000440&hsa-miR-191-5p&offsets|0|0;m-48&3|-|49020672|49020694&offsets|0|0]|CAACGGAATCCCAAAAGCAGCTG</td><td>>MI0000465|hsa-mir-191&WithFlank&3|-|49020612|49020715</td><td>22</td><td>44</td><td>23</td><td>CAACGGAATCCCAAAAGCAGCTG</td><td>7735.8</td><td>4.7</td><td>0.8</td><td>5.7</td><td>1.0E-08</td><td>1.9E-08</td><td>MB231/MB468</td><td>MITO</td></tr>
<tr><td>>trna111_HisGTG_1_-_147774845_147774916@-1T.23.24,trna118_HisGTG_1_-_145396881_145396952@-1T.23.24,trna16_HisGTG_1_+_146544773_146544844@-1T.23.24,trna1_HisGTG_15_+_45493349_45493420@-1T.23.24,trna21_HisGTG_1_+_147753471_147753542@-1T.23.24,trna33_HisGTG_6_+_27125906_27125977@-1T.23.24,trna7_HisGTG_9_-_14433938_14434009@-1T.23.24,trna8_HisGTG_15_-_45492611_45492682@-1T.23.24,trna9_HisGTG_15_-_45490804_45490875@-1T.23.24|TGCCGTGATCGTATAGTGGTTAGT</td><td>>trna111_HisGTG_1_-_147774845_147774916</td><td>-1T</td><td>23</td><td>24</td><td>TGCCGTGATCGTATAGTGGTTAGT</td><td>366.4</td><td>6.8</td><td>1.2</td><td>5.8</td><td>7.2E-09</td><td>1.9E-08</td><td>BT20/MB468</td><td>MITO</td></tr>
<tr><td>>trnaMT_LeuTAA_MT_+_3230_3304@1.37.37|GTTAAGATGGCAGAGCCCGGTAATCGCATAAAACTTA</td><td>>trnaMT_LeuTAA_MT_+_3230_3304</td><td>1</td><td>37</td><td>37</td><td>GTTAAGATGGCAGAGCCCGGTAATCGCATAAAACTTA</td><td>80.1</td><td>9.5</td><td>1.7</td><td>5.8</td><td>8.9E-09</td><td>1.9E-08</td><td>MB231/MB468</td><td>MP</td></tr>
<tr><td>>am_trnaMT_LysTTT_MT_+_8295_8364@33.52.20,trnalookalike7_LysTTT_1_+_568844_568913@33.52.20|AGTTAAAGATTAAGAGAACC</td><td>>am_trnaMT_LysTTT_MT_+_8295_8364</td><td>33</td><td>52</td><td>20</td><td>AGTTAAAGATTAAGAGAACC</td><td>565.2</td><td>8.2</td><td>1.4</td><td>5.8</td><td>8.1E-09</td><td>1.9E-08</td><td>BT20/MB468</td><td>CYTO</td></tr>
<tr><td>>hg38_21_+_8433172_8446622_RNA45SN1_45S_50ntFlanks@12627.12649.23|ATGGGGCGAAGCTACCATCTGTG</td><td>>hg38_21_+_8433172_8446622_RNA45SN1_45S_50ntFlanks</td><td>12627</td><td>12649</td><td>23</td><td>ATGGGGCGAAGCTACCATCTGTG</td><td>104.0</td><td>-9.8</td><td>1.6</td><td>-6.1</td><td>1.3E-09</td><td>1.9E-08</td><td>BT20/MB468</td><td>MP</td></tr>
<tr><td>>hg38_21_+_8433172_8446622_RNA45SN1_45S_50ntFlanks@12447.12475.29|CAGAATTCACCAAGCGTTGGATTGTTCAC</td><td>>hg38_21_+_8433172_8446622_RNA45SN1_45S_50ntFlanks</td><td>12447</td><td>12475</td><td>29</td><td>CAGAATTCACCAAGCGTTGGATTGTTCAC</td><td>194.2</td><td>-10.7</td><td>1.8</td><td>-6.1</td><td>1.3E-09</td><td>1.9E-08</td><td>BT20/MB468</td><td>MP</td></tr>
<tr><td>>hg38_21_+_8433172_8446622_RNA45SN1_45S_50ntFlanks@4047.4071.25|ATTCGAACGTCTGCCCTATCAACTT</td><td>>hg38_21_+_8433172_8446622_RNA45SN1_45S_50ntFlanks</td><td>4047</td><td>4071</td><td>25</td><td>ATTCGAACGTCTGCCCTATCAACTT</td><td>209.4</td><td>-9.8</td><td>1.7</td><td>-5.8</td><td>7.2E-09</td><td>1.9E-08</td><td>BT20/MB468</td><td>MITO</td></tr>
<tr><td>>hg38_21_+_8433172_8446622_RNA45SN1_45S_50ntFlanks@483.516.34|TCGATGTGGTGACGTCGTGCTCTCCCGGGCCGGG</td><td>>hg38_21_+_8433172_8446622_RNA45SN1_45S_50ntFlanks</td><td>483</td><td>516</td><td>34</td><td>TCGATGTGGTGACGTCGTGCTCTCCCGGGCCGGG</td><td>777.4</td><td>4.4</td><td>0.8</td><td>5.7</td><td>9.6E-09</td><td>1.9E-08</td><td>MB231/MB468</td><td>NUC</td></tr>
<tr><td>>hg38_21_+_8433172_8446622_RNA45SN1_45S_50ntFlanks@7986.8013.28|ATCAGACGTGGCGACCCGCTGAATTTAA</td><td>>hg38_21_+_8433172_8446622_RNA45SN1_45S_50ntFlanks</td><td>7986</td><td>8013</td><td>28</td><td>ATCAGACGTGGCGACCCGCTGAATTTAA</td><td>105.8</td><td>-8.8</td><td>1.5</td><td>-5.8</td><td>7.2E-09</td><td>1.9E-08</td><td>BT20/MB468</td><td>MITO</td></tr>
<tr><td>>hg38_21_+_8433172_8446622_RNA45SN1_45S_50ntFlanks@474.505.32|TCTTCGTGATCGATGTGGTGACGTCGTGCTCT</td><td>>hg38_21_+_8433172_8446622_RNA45SN1_45S_50ntFlanks</td><td>474</td><td>505</td><td>32</td><td>TCTTCGTGATCGATGTGGTGACGTCGTGCTCT</td><td>2601.6</td><td>4.5</td><td>0.8</td><td>5.7</td><td>9.6E-09</td><td>2.0E-08</td><td>MB231/MB468</td><td>NUC</td></tr>
<tr><td>>hg38_21_+_8433172_8446622_RNA45SN1_45S_50ntFlanks@3771.3794.24|CACGGCCGGTACAGTGAAACTGCG</td><td>>hg38_21_+_8433172_8446622_RNA45SN1_45S_50ntFlanks</td><td>3771</td><td>3794</td><td>24</td><td>CACGGCCGGTACAGTGAAACTGCG</td><td>695.4</td><td>-6.5</td><td>1.1</td><td>-5.7</td><td>1.0E-08</td><td>2.0E-08</td><td>MB231/MB468</td><td>MITO</td></tr>
<tr><td>>hg38_21_+_8433172_8446622_RNA45SN1_45S_50ntFlanks@12418.12441.24|CGATGTCGGCTCTTCCTATCATTG</td><td>>hg38_21_+_8433172_8446622_RNA45SN1_45S_50ntFlanks</td><td>12418</td><td>12441</td><td>24</td><td>CGATGTCGGCTCTTCCTATCATTG</td><td>433.1</td><td>-7.8</td><td>1.4</td><td>-5.7</td><td>9.0E-09</td><td>2.0E-08</td><td>MB231/MB468</td><td>MP</td></tr>
<tr><td>>hg38_21_+_8433172_8446622_RNA45SN1_45S_50ntFlanks@4279.4314.36|AATCCTTTAACGAGGATCCATTGGAGGGCAAGTCTG</td><td>>hg38_21_+_8433172_8446622_RNA45SN1_45S_50ntFlanks</td><td>4279</td><td>4314</td><td>36</td><td>AATCCTTTAACGAGGATCCATTGGAGGGCAAGTCTG</td><td>88.7</td><td>-10.3</td><td>1.8</td><td>-5.7</td><td>9.0E-09</td><td>2.0E-08</td><td>MB231/MB468</td><td>MP</td></tr>
<tr><td>>trnaMT_PheGAA_MT_+_577_647@32.55.24|ACTGAAAATGTTTAGACGGGCTCA</td><td>>trnaMT_PheGAA_MT_+_577_647</td><td>32</td><td>55</td><td>24</td><td>ACTGAAAATGTTTAGACGGGCTCA</td><td>128.8</td><td>10.8</td><td>1.8</td><td>6.1</td><td>1.3E-09</td><td>2.0E-08</td><td>BT20/MB468</td><td>MP</td></tr>
<tr><td>>trna119_LysCTT_1_-_145395522_145395594@4.33.30,trna11_LysCTT_5_-_180648979_180649051@4.33.30,trna13_LysCTT_6_+_26556774_26556846@4.33.30,trna7_LysCTT_16_+_3225692_3225764@4.33.30,trna9_LysCTT_5_+_180634755_180634827@4.33.30|CGGCTAGCTCAGTCGGTAGAGCATGAGACT</td><td>>trna119_LysCTT_1_-_145395522_145395594</td><td>4</td><td>33</td><td>30</td><td>CGGCTAGCTCAGTCGGTAGAGCATGAGACT</td><td>516.2</td><td>4.7</td><td>0.8</td><td>5.7</td><td>9.0E-09</td><td>2.0E-08</td><td>MB231/MB468</td><td>MP</td></tr>
<tr><td>>hg38_21_+_8433172_8446622_RNA45SN1_45S_50ntFlanks@8198.8215.18|TGGACGGTGTGAGGCCGG</td><td>>hg38_21_+_8433172_8446622_RNA45SN1_45S_50ntFlanks</td><td>8198</td><td>8215</td><td>18</td><td>TGGACGGTGTGAGGCCGG</td><td>3858.9</td><td>-6.4</td><td>1.1</td><td>-5.8</td><td>8.2E-09</td><td>2.0E-08</td><td>BT20/MB468</td><td>CYTO</td></tr>
<tr><td>>hg38_21_+_8433172_8446622_RNA45SN1_45S_50ntFlanks@10731.10750.20|AGGGTGTAAATCTCGCGCCG</td><td>>hg38_21_+_8433172_8446622_RNA45SN1_45S_50ntFlanks</td><td>10731</td><td>10750</td><td>20</td><td>AGGGTGTAAATCTCGCGCCG</td><td>162.8</td><td>-10.4</td><td>1.7</td><td>-6.1</td><td>1.3E-09</td><td>2.0E-08</td><td>BT20/MB468</td><td>MP</td></tr>
<tr><td>>hg38_21_+_8433172_8446622_RNA45SN1_45S_50ntFlanks@6738.6758.21|ATCATCGACACTTCGAACGCA</td><td>>hg38_21_+_8433172_8446622_RNA45SN1_45S_50ntFlanks</td><td>6738</td><td>6758</td><td>21</td><td>ATCATCGACACTTCGAACGCA</td><td>119.3</td><td>-10.0</td><td>1.6</td><td>-6.1</td><td>1.3E-09</td><td>2.0E-08</td><td>BT20/MB468</td><td>MP</td></tr>
<tr><td>>hg38_21_+_8433172_8446622_RNA45SN1_45S_50ntFlanks@13002.13025.24|CGATCTATTGAAAGTCAGCCCTCG</td><td>>hg38_21_+_8433172_8446622_RNA45SN1_45S_50ntFlanks</td><td>13002</td><td>13025</td><td>24</td><td>CGATCTATTGAAAGTCAGCCCTCG</td><td>86.8</td><td>-8.5</td><td>1.5</td><td>-5.8</td><td>7.4E-09</td><td>2.0E-08</td><td>BT20/MB468</td><td>MITO</td></tr>
<tr><td>>hg38_21_+_8433172_8446622_RNA45SN1_45S_50ntFlanks@10687.10706.20|CGGCGTCCGGTGAGCTCTCG</td><td>>hg38_21_+_8433172_8446622_RNA45SN1_45S_50ntFlanks</td><td>10687</td><td>10706</td><td>20</td><td>CGGCGTCCGGTGAGCTCTCG</td><td>11214.9</td><td>-10.6</td><td>1.8</td><td>-5.7</td><td>1.1E-08</td><td>2.0E-08</td><td>BT20/MB468</td><td>NUC</td></tr>
<tr><td>>hg38_21_+_8433172_8446622_RNA45SN1_45S_50ntFlanks@11573.11600.28|CGACTTAGAACTGGTGCGGACCAGGGGA</td><td>>hg38_21_+_8433172_8446622_RNA45SN1_45S_50ntFlanks</td><td>11573</td><td>11600</td><td>28</td><td>CGACTTAGAACTGGTGCGGACCAGGGGA</td><td>1111.7</td><td>-6.5</td><td>1.1</td><td>-5.7</td><td>9.1E-09</td><td>2.0E-08</td><td>MB231/MB468</td><td>MP</td></tr>
<tr><td>>hg38_21_+_8433172_8446622_RNA45SN1_45S_50ntFlanks@10471.10488.18|CCCTCGGCCGATCGAAAG</td><td>>hg38_21_+_8433172_8446622_RNA45SN1_45S_50ntFlanks</td><td>10471</td><td>10488</td><td>18</td><td>CCCTCGGCCGATCGAAAG</td><td>3439.1</td><td>-10.0</td><td>1.7</td><td>-5.7</td><td>1.1E-08</td><td>2.0E-08</td><td>BT20/MB468</td><td>NUC</td></tr>
<tr><td>>hg38_21_+_8433172_8446622_RNA45SN1_45S_50ntFlanks@10378.10424.47|AGGGTTCCATGTGAACAGCAGTTGAACATGGGTCAGTCGGTCCTGAG</td><td>>hg38_21_+_8433172_8446622_RNA45SN1_45S_50ntFlanks</td><td>10378</td><td>10424</td><td>47</td><td>AGGGTTCCATGTGAACAGCAGTTGAACATGGGTCAGTCGGTCCTGAG</td><td>3251.3</td><td>6.8</td><td>1.2</td><td>5.8</td><td>7.4E-09</td><td>2.0E-08</td><td>BT20/MB468</td><td>MITO</td></tr>
<tr><td>>hg38_21_+_8433172_8446622_RNA45SN1_45S_50ntFlanks@10402.10433.32|AACATGGGTCAGTCGGTCCTGAGAGATGGGCG</td><td>>hg38_21_+_8433172_8446622_RNA45SN1_45S_50ntFlanks</td><td>10402</td><td>10433</td><td>32</td><td>AACATGGGTCAGTCGGTCCTGAGAGATGGGCG</td><td>497.9</td><td>-5.0</td><td>0.9</td><td>-5.8</td><td>8.2E-09</td><td>2.0E-08</td><td>BT20/MB468</td><td>CYTO</td></tr>
<tr><td>>hg38_21_+_8433172_8446622_RNA45SN1_45S_50ntFlanks@3899.3920.22|CCTTCGCGGGGGGGATGCGTGC</td><td>>hg38_21_+_8433172_8446622_RNA45SN1_45S_50ntFlanks</td><td>3899</td><td>3920</td><td>22</td><td>CCTTCGCGGGGGGGATGCGTGC</td><td>808.8</td><td>-5.6</td><td>1.0</td><td>-5.7</td><td>9.6E-09</td><td>2.0E-08</td><td>MB231/MB468</td><td>CYTO</td></tr>
<tr><td>>hg38_21_+_8433172_8446622_RNA45SN1_45S_50ntFlanks@6648.6667.20|GTACGACTCTTAGCGGTGGA</td><td>>hg38_21_+_8433172_8446622_RNA45SN1_45S_50ntFlanks</td><td>6648</td><td>6667</td><td>20</td><td>GTACGACTCTTAGCGGTGGA</td><td>427.8</td><td>6.3</td><td>1.1</td><td>5.8</td><td>8.3E-09</td><td>2.0E-08</td><td>BT20/MB468</td><td>CYTO</td></tr>
<tr><td>>hg38_21_+_8433172_8446622_RNA45SN1_45S_50ntFlanks@11207.11225.19|GGCGGCGACTCTGGACGCG</td><td>>hg38_21_+_8433172_8446622_RNA45SN1_45S_50ntFlanks</td><td>11207</td><td>11225</td><td>19</td><td>GGCGGCGACTCTGGACGCG</td><td>589.5</td><td>-5.4</td><td>0.9</td><td>-5.8</td><td>8.3E-09</td><td>2.0E-08</td><td>BT20/MB468</td><td>CYTO</td></tr>
<tr><td>>hg38_21_+_8433172_8446622_RNA45SN1_45S_50ntFlanks@10663.10709.47|CCTTGGAAAGCGTCGCGGTTCCGGCGGCGTCCGGTGAGCTCTCGCTG</td><td>>hg38_21_+_8433172_8446622_RNA45SN1_45S_50ntFlanks</td><td>10663</td><td>10709</td><td>47</td><td>CCTTGGAAAGCGTCGCGGTTCCGGCGGCGTCCGGTGAGCTCTCGCTG</td><td>110.2</td><td>-7.0</td><td>1.2</td><td>-5.7</td><td>9.7E-09</td><td>2.0E-08</td><td>MB231/MB468</td><td>CYTO</td></tr>
<tr><td>>hg38_21_+_8433172_8446622_RNA45SN1_45S_50ntFlanks@12324.12346.23|TTGGGTTTTAAGCAGGAGGTGTC</td><td>>hg38_21_+_8433172_8446622_RNA45SN1_45S_50ntFlanks</td><td>12324</td><td>12346</td><td>23</td><td>TTGGGTTTTAAGCAGGAGGTGTC</td><td>867.3</td><td>-7.5</td><td>1.3</td><td>-5.8</td><td>7.4E-09</td><td>2.0E-08</td><td>BT20/MB468</td><td>MITO</td></tr>
<tr><td>>hg38_21_+_8433172_8446622_RNA45SN1_45S_50ntFlanks@10784.10823.40|TGAACAGCCTCTGGCATGTTGGAACAATGTAGGTAAGGGA</td><td>>hg38_21_+_8433172_8446622_RNA45SN1_45S_50ntFlanks</td><td>10784</td><td>10823</td><td>40</td><td>TGAACAGCCTCTGGCATGTTGGAACAATGTAGGTAAGGGA</td><td>208.0</td><td>-7.8</td><td>1.4</td><td>-5.7</td><td>1.1E-08</td><td>2.0E-08</td><td>MB231/MB468</td><td>MITO</td></tr>
<tr><td>>hg38_21_+_8433172_8446622_RNA45SN1_45S_50ntFlanks@3991.4024.34|TGGTGACTCTAGATAACCTCGGGCCGATCGCACG</td><td>>hg38_21_+_8433172_8446622_RNA45SN1_45S_50ntFlanks</td><td>3991</td><td>4024</td><td>34</td><td>TGGTGACTCTAGATAACCTCGGGCCGATCGCACG</td><td>361.3</td><td>-2.4</td><td>0.4</td><td>-5.8</td><td>7.5E-09</td><td>2.0E-08</td><td>BT20/MB468</td><td>MITO</td></tr>
<tr><td>>hg38_21_+_8433172_8446622_RNA45SN1_45S_50ntFlanks@12510.12538.29|TCGTGAGACAGGTTAGTTTTACCCTACTG</td><td>>hg38_21_+_8433172_8446622_RNA45SN1_45S_50ntFlanks</td><td>12510</td><td>12538</td><td>29</td><td>TCGTGAGACAGGTTAGTTTTACCCTACTG</td><td>167.5</td><td>-10.5</td><td>1.7</td><td>-6.1</td><td>1.3E-09</td><td>2.0E-08</td><td>BT20/MB468</td><td>MP</td></tr>
<tr><td>>hg38_21_+_8433172_8446622_RNA45SN1_45S_50ntFlanks@5467.5514.48|GGCCCACGGCCCTGGCGGAGCGCTGAGAAGACGGTCGAACTTGACTAT</td><td>>hg38_21_+_8433172_8446622_RNA45SN1_45S_50ntFlanks</td><td>5467</td><td>5514</td><td>48</td><td>GGCCCACGGCCCTGGCGGAGCGCTGAGAAGACGGTCGAACTTGACTAT</td><td>741.4</td><td>-8.3</td><td>1.4</td><td>-5.8</td><td>8.3E-09</td><td>2.0E-08</td><td>BT20/MB468</td><td>CYTO</td></tr>
<tr><td>>hg38_21_+_8433172_8446622_RNA45SN1_45S_50ntFlanks@8144.8165.22|CTCCCCGGCGCCGCTCGTGGGG</td><td>>hg38_21_+_8433172_8446622_RNA45SN1_45S_50ntFlanks</td><td>8144</td><td>8165</td><td>22</td><td>CTCCCCGGCGCCGCTCGTGGGG</td><td>123.8</td><td>-10.8</td><td>1.9</td><td>-5.7</td><td>9.2E-09</td><td>2.0E-08</td><td>MB231/MB468</td><td>MP</td></tr>
<tr><td>>hg38_21_+_8433172_8446622_RNA45SN1_45S_50ntFlanks@9504.9525.22|GTGAACTATGCCTGGGCAGGGC</td><td>>hg38_21_+_8433172_8446622_RNA45SN1_45S_50ntFlanks</td><td>9504</td><td>9525</td><td>22</td><td>GTGAACTATGCCTGGGCAGGGC</td><td>604.8</td><td>-3.8</td><td>0.7</td><td>-5.8</td><td>7.5E-09</td><td>2.0E-08</td><td>BT20/MB468</td><td>MITO</td></tr>
<tr><td>>MI0000342|hsa-mir-200b&WithFlank&1|+|1167098|1167204@64.85.22(+1U)&[MIMAT0000318&hsa-miR-200b-3p&offsets|+1|+1(+1U);m-93&1|+|1167160|1167182&offsets|+1|0(+1U)]|AATACTGCCTGGTAATGATGACT</td><td>>MI0000342|hsa-mir-200b&WithFlank&1|+|1167098|1167204</td><td>64</td><td>85</td><td>22(+1U)</td><td>AATACTGCCTGGTAATGATGACT</td><td>433.0</td><td>-7.7</td><td>1.3</td><td>-5.7</td><td>9.2E-09</td><td>2.0E-08</td><td>MB231/MB468</td><td>MP</td></tr>
<tr><td>>MI0000077|hsa-mir-21&WithFlank&17|+|59841260|59841343@14.33.20(+1U)&[MIMAT0000076&hsa-miR-21-5p&offsets|0|-2(+1U);m-1&17|+|59841273|59841294&offsets|0|-2(+1U)]|TAGCTTATCAGACTGATGTTT</td><td>>MI0000077|hsa-mir-21&WithFlank&17|+|59841260|59841343</td><td>14</td><td>33</td><td>20(+1U)</td><td>TAGCTTATCAGACTGATGTTT</td><td>68.3</td><td>6.6</td><td>1.2</td><td>5.7</td><td>1.1E-08</td><td>2.0E-08</td><td>BT20/MB468</td><td>NUC</td></tr>
<tr><td>>hg38_21_+_8433172_8446622_RNA45SN1_45S_50ntFlanks@11319.11336.18|CCCACGTCTCGTCGCGCG</td><td>>hg38_21_+_8433172_8446622_RNA45SN1_45S_50ntFlanks</td><td>11319</td><td>11336</td><td>18</td><td>CCCACGTCTCGTCGCGCG</td><td>235.6</td><td>-7.2</td><td>1.3</td><td>-5.8</td><td>8.4E-09</td><td>2.0E-08</td><td>BT20/MB468</td><td>CYTO</td></tr>
<tr><td>>hg38_21_+_8433172_8446622_RNA45SN1_45S_50ntFlanks@11573.11613.41|CGACTTAGAACTGGTGCGGACCAGGGGAATCCGACTGTTTA</td><td>>hg38_21_+_8433172_8446622_RNA45SN1_45S_50ntFlanks</td><td>11573</td><td>11613</td><td>41</td><td>CGACTTAGAACTGGTGCGGACCAGGGGAATCCGACTGTTTA</td><td>756.0</td><td>4.9</td><td>0.9</td><td>5.7</td><td>9.8E-09</td><td>2.0E-08</td><td>MB231/MB468</td><td>CYTO</td></tr>
<tr><td>>hg38_21_+_8433172_8446622_RNA45SN1_45S_50ntFlanks@10795.10818.24|TGGCATGTTGGAACAATGTAGGTA</td><td>>hg38_21_+_8433172_8446622_RNA45SN1_45S_50ntFlanks</td><td>10795</td><td>10818</td><td>24</td><td>TGGCATGTTGGAACAATGTAGGTA</td><td>883.1</td><td>-8.9</td><td>1.6</td><td>-5.7</td><td>1.1E-08</td><td>2.0E-08</td><td>BT20/MB468</td><td>NUC</td></tr>
<tr><td>>hg38_21_+_8433172_8446622_RNA45SN1_45S_50ntFlanks@9766.9795.30|TAAATGGGTAAGAAGCCCGGCTCGCTGGCG</td><td>>hg38_21_+_8433172_8446622_RNA45SN1_45S_50ntFlanks</td><td>9766</td><td>9795</td><td>30</td><td>TAAATGGGTAAGAAGCCCGGCTCGCTGGCG</td><td>195.8</td><td>-9.7</td><td>1.7</td><td>-5.8</td><td>7.5E-09</td><td>2.0E-08</td><td>BT20/MB468</td><td>MITO</td></tr>
<tr><td>>hg38_21_+_8433172_8446622_RNA45SN1_45S_50ntFlanks@10352.10371.20|GAGAACTTTGAAGGCCGAAG</td><td>>hg38_21_+_8433172_8446622_RNA45SN1_45S_50ntFlanks</td><td>10352</td><td>10371</td><td>20</td><td>GAGAACTTTGAAGGCCGAAG</td><td>297.8</td><td>4.3</td><td>0.7</td><td>5.7</td><td>1.0E-08</td><td>2.0E-08</td><td>MB231/MB468</td><td>NUC</td></tr>
<tr><td>>MI0000750|hsa-mir-26a-2&WithFlank&12|-|57824603|57824698@20.41.22;MI0000083|hsa-mir-26a-1&WithFlank&3|+|37969398|37969486@16.37.22&[MIMAT0000082&hsa-miR-26a-5p&offsets|0|0;m-33&12|-|57824658|57824679&offsets|0|0];[MIMAT0000082_1&hsa-miR-26a-5p&offsets|0|0;m-34&3|+|37969413|37969434&offsets|0|0]|TTCAAGTAATCCAGGATAGGCT</td><td>>MI0000750|hsa-mir-26a-2&WithFlank&12|-|57824603|57824698</td><td>20</td><td>41</td><td>22</td><td>TTCAAGTAATCCAGGATAGGCT</td><td>107314.5</td><td>5.1</td><td>0.9</td><td>5.7</td><td>1.1E-08</td><td>2.0E-08</td><td>MB231/MB468</td><td>MITO</td></tr>
<tr><td>>MI0000085|hsa-mir-27a&WithFlank&19|-|13836434|13836523@57.74.18;MI0000440|hsa-mir-27b&WithFlank&9|+|95085439|95085547@67.84.18&[MIMAT0000084&hsa-miR-27a-3p&offsets|0|-3;m-43&19|-|13836447|13836467&offsets|0|-3];[MIMAT0000419&hsa-miR-27b-3p&offsets|0|-3;m-39&9|+|95085505|95085525&offsets|0|-3]|TTCACAGTGGCTAAGTTC</td><td>>MI0000085|hsa-mir-27a&WithFlank&19|-|13836434|13836523</td><td>57</td><td>74</td><td>18</td><td>TTCACAGTGGCTAAGTTC</td><td>733.3</td><td>5.5</td><td>1.0</td><td>5.7</td><td>1.1E-08</td><td>2.0E-08</td><td>MB231/MB468</td><td>MITO</td></tr>
<tr><td>>hg38_21_+_8433172_8446622_RNA45SN1_45S_50ntFlanks@12957.12981.25|CTTCTGGGTCGGGGTTTCGTACGTA</td><td>>hg38_21_+_8433172_8446622_RNA45SN1_45S_50ntFlanks</td><td>12957</td><td>12981</td><td>25</td><td>CTTCTGGGTCGGGGTTTCGTACGTA</td><td>453.3</td><td>-7.5</td><td>1.3</td><td>-5.7</td><td>1.0E-08</td><td>2.0E-08</td><td>MB231/MB468</td><td>NUC</td></tr>
</table>

| | | | | | | | | | | | | | |
|---|---|---|---|---|---|---|---|---|---|---|---|---|---|
| >trna10_LysCTT_16_+_3241501_3241573@3.33.31,trna13_LysCTT_14_-_58706613_58706685@3.33.31,trna2_LysCTT_15_+_79152904_79152976@3.33.31\|CCGGCTAGCTCAGTCGGTAGAGCATGGGACT | >trna10_LysCTT_16_+_3241501_3241573 | 3 | 33 | 31 | CCGGCTAGCTCAGTCGGTAGAGCATGGGACT | 807.9 | 8.1 | 1.4 | 5.7 | 9.9E-09 | 2.0E-08 | MB231/MB468 | CYTO |
| >hg38_21_+_8433172_8446622_RNA45SN1_45S_50ntFlanks@10845.10862.18\|ACTTCGGGATAAGGATTG | >hg38_21_+_8433172_8446622_RNA45SN1_45S_50ntFlanks | 10845 | 10862 | 18 | ACTTCGGGATAAGGATTG | 541.8 | -7.7 | 1.3 | -5.7 | 1.0E-08 | 2.0E-08 | MB231/MB468 | NUC |
| >MI0003666\|hsa-mir-651&WithFlank&X\|+\|8126959\|8127067@22.43.22&[MIMAT0003321&hsa-miR-651-5p&offsets\|0\|0;m-399&X\|+\|8126980\|8127001&offsets\|0\|0]\|TTTAGGATAAGCTTGACTTTTG | >MI0003666\|hsa-mir-651&WithFlank&X\|+\|8126959\|8127067 | 22 | 43 | 22 | TTTAGGATAAGCTTGACTTTTG | 188.4 | 5.8 | 1.0 | 5.7 | 1.0E-08 | 2.0E-08 | MB231/MB468 | NUC |
| >hg38_21_+_8433172_8446622_RNA45SN1_45S_50ntFlanks@8169.8197.29\|CCAAGTCCTTCTGATCGAGGCCCAGCCCG | >hg38_21_+_8433172_8446622_RNA45SN1_45S_50ntFlanks | 8169 | 8197 | 29 | CCAAGTCCTTCTGATCGAGGCCCAGCCCG | 625.4 | -8.3 | 1.5 | -5.7 | 1.0E-08 | 2.0E-08 | MB231/MB468 | NUC |
| >trna10_AspGTC_12_-_125424193_125424264@37.74.38,trna12_AspGTC_12_-_125411891_125411962@37.74.38,trna144_AspGTC_6_-_27551236_27551307@37.74.38,trna38_AspGTC_17_-_8125556_8125627@37.74.38,trna45_AspGTC_6_+_27447453_27447524@37.74.38,trna48_AspGTC_6_+_27471523_27471594@37.74.38,trna4_AspGTC_12_+_96429799_96429870@37.74.38,trna69_AspGTC_1_-_161440205_161440276@37.74.38,trna72_AspGTC_1_-_161432824_161432895@37.74.38,trna75_AspGTC_1_-_161425414_161425485@37.74.38,trna78_AspGTC_1_-_161418033_161418104@37.74.38,trna81_AspGTC_1_-_161410615_161410686@37.74.38\|ACGCGGGAGACCGGGGTTCGATTCCCCGACGGGGAGCC | >trna10_AspGTC_12_-_125424193_125424264 | 37 | 74 | 38 | ACGCGGGAGACCGGGGTTCGATTCCCCGACGGGGAGCC | 159.4 | 10.5 | 1.8 | 5.7 | 1.1E-08 | 2.0E-08 | MB231/MB468 | MITO |
| >hg38_21_+_8433172_8446622_RNA45SN1_45S_50ntFlanks@11783.11806.24\|CATGAATGGATGAACGAGATTCCC | >hg38_21_+_8433172_8446622_RNA45SN1_45S_50ntFlanks | 11783 | 11806 | 24 | CATGAATGGATGAACGAGATTCCC | 2115.8 | -6.5 | 1.1 | -5.7 | 9.9E-09 | 2.0E-08 | MB231/MB468 | CYTO |
| >hg38_21_+_8433172_8446622_RNA45SN1_45S_50ntFlanks@10346.10376.31\|TCAAACGAGAACTTTGAAGGCCGAAGTGGAG | >hg38_21_+_8433172_8446622_RNA45SN1_45S_50ntFlanks | 10346 | 10376 | 31 | TCAAACGAGAACTTTGAAGGCCGAAGTGGAG | 305.9 | -6.2 | 1.1 | -5.7 | 9.3E-09 | 2.0E-08 | MB231/MB468 | MP |
| >hg38_21_+_8433172_8446622_RNA45SN1_45S_50ntFlanks@12154.12182.29\|GGGAGTTTGACTGGGGCGGTACACCTGTC | >hg38_21_+_8433172_8446622_RNA45SN1_45S_50ntFlanks | 12154 | 12182 | 29 | GGGAGTTTGACTGGGGCGGTACACCTGTC | 268.3 | -7.3 | 1.3 | -5.7 | 1.1E-08 | 2.0E-08 | MB231/MB468 | MITO |
| >MI0000483\|hsa-mir-186&WithFlank&1\|-\|71067625\|71067722@21.44.24(+1U)&[MIMAT0000456&hsa-miR-186-5p&offsets\|0\|+2(+1U);m-84&1\|-\|71067681\|71067702&offsets\|0\|+2(+1U)]\|CAAAGAATTCTCCTTTTGGGCTTTT | >MI0000483\|hsa-mir-186&WithFlank&1\|-\|71067625\|71067722 | 21 | 44 | 24(+1U) | CAAAGAATTCTCCTTTTGGGCTTTT | 244.5 | -11.3 | 1.8 | -6.3 | 2.6E-10 | 2.0E-08 | BT20/MB231 | MP |
| >MI0000456\|hsa-mir-140&WithFlank&16\|+\|69933075\|69933186@68.89.22(+1U)&[MIMAT0004597&hsa-miR-140-3p&offsets\|0\|+1(+1U);m-45&16\|+\|69933143\|69933164&offsets\|-1\|-1(+1U)]\|TACCACAGGGTAGAACCACGGAT | >MI0000456\|hsa-mir-140&WithFlank&16\|+\|69933075\|69933186 | 68 | 89 | 22(+1U) | TACCACAGGGTAGAACCACGGAT | 84.7 | 9.6 | 1.7 | 5.7 | 9.4E-09 | 2.0E-08 | MB231/MB468 | MP |
| >hg38_21_+_8433172_8446622_RNA45SN1_45S_50ntFlanks@3990.4013.24\|TTGGTGACTCTAGATAACCTCGGG | >hg38_21_+_8433172_8446622_RNA45SN1_45S_50ntFlanks | 3990 | 4013 | 24 | TTGGTGACTCTAGATAACCTCGGG | 544.0 | -8.4 | 1.5 | -5.7 | 1.1E-08 | 2.0E-08 | BT20/MB468 | NUC |
| >hg38_21_+_8433172_8446622_RNA45SN1_45S_50ntFlanks@10395.10429.35\|GCAGTTGAACATGGGTCAGTCGGTCCTGAGAGATG | >hg38_21_+_8433172_8446622_RNA45SN1_45S_50ntFlanks | 10395 | 10429 | 35 | GCAGTTGAACATGGGTCAGTCGGTCCTGAGAGATG | 449.8 | -4.5 | 0.8 | -5.7 | 1.1E-08 | 2.0E-08 | BT20/MB468 | NUC |
| >MI0003820\|hsa-mir-454&WithFlank&17\|-\|59137752\|59137878@70.94.25&[MIMAT0003885&hsa-miR-454-3p&offsets\|0\|+2;m-272&17\|-\|59137787\|59137809&offsets\|0\|+2]\|TAGTGCAATATTGCTTATAGGGTTT | >MI0003820\|hsa-mir-454&WithFlank&17\|-\|59137752\|59137878 | 70 | 94 | 25 | TAGTGCAATATTGCTTATAGGGTTT | 1181.7 | -5.8 | 0.9 | -6.3 | 2.6E-10 | 2.0E-08 | BT20/MB231 | MP |
| >hg38_21_+_8433172_8446622_RNA45SN1_45S_50ntFlanks@5471.5510.40\|CACGGCCCTGGCGGAGCGCTGAGAAGACGGTCGAACTTGA | >hg38_21_+_8433172_8446622_RNA45SN1_45S_50ntFlanks | 5471 | 5510 | 40 | CACGGCCCTGGCGGAGCGCTGAGAAGACGGTCGAACTTGA | 72.9 | -10.1 | 1.8 | -5.7 | 9.4E-09 | 2.0E-08 | MB231/MB468 | MP |
| >hg38_21_+_8433172_8446622_RNA45SN1_45S_50ntFlanks@3843.3874.32\|CTACTTGGATAACTGTGGTAATTCTAGAGCTA | >hg38_21_+_8433172_8446622_RNA45SN1_45S_50ntFlanks | 3843 | 3874 | 32 | CTACTTGGATAACTGTGGTAATTCTAGAGCTA | 194.7 | -10.7 | 1.8 | -6.1 | 1.4E-09 | 2.0E-08 | BT20/MB468 | MP |
| >hg38_21_+_8433172_8446622_RNA45SN1_45S_50ntFlanks@4730.4767.38\|CAAGAACGAAAGTCGGAGGTTCGAAGACGATCAGATAC | >hg38_21_+_8433172_8446622_RNA45SN1_45S_50ntFlanks | 4730 | 4767 | 38 | CAAGAACGAAAGTCGGAGGTTCGAAGACGATCAGATAC | 110.1 | 10.5 | 1.7 | 6.1 | 1.4E-09 | 2.0E-08 | BT20/MB468 | MP |
| >hg38_21_+_8433172_8446622_RNA45SN1_45S_50ntFlanks@10544.10564.21\|TCCAGTGCGGTAACGCGACCG | >hg38_21_+_8433172_8446622_RNA45SN1_45S_50ntFlanks | 10544 | 10564 | 21 | TCCAGTGCGGTAACGCGACCG | 116.4 | -9.9 | 1.6 | -6.1 | 1.4E-09 | 2.0E-08 | BT20/MB468 | MP |
| >hg38_21_+_8433172_8446622_RNA45SN1_45S_50ntFlanks@6699.6718.20\|GCTAGCTGCGAGAATTAATG | >hg38_21_+_8433172_8446622_RNA45SN1_45S_50ntFlanks | 6699 | 6718 | 20 | GCTAGCTGCGAGAATTAATG | 26453.0 | -7.3 | 1.2 | -6.1 | 1.4E-09 | 2.0E-08 | BT20/MB468 | MP |
| >trnaMT_SerGCT_MT_+_12207_12265@1.26.26\|GAGAAAGCTCACAAGAACTGCTAACT | >trnaMT_SerGCT_MT_+_12207_12265 | 1 | 26 | 26 | GAGAAAGCTCACAAGAACTGCTAACT | 1794.0 | 4.2 | 0.7 | 5.7 | 1.0E-08 | 2.1E-08 | MB231/MB468 | NUC |
| >hg38_21_+_8433172_8446622_RNA45SN1_45S_50ntFlanks@11729.11766.38\|TAACTATGACTCTCTTAAGGTAGCCAAATGCCTCGTCA | >hg38_21_+_8433172_8446622_RNA45SN1_45S_50ntFlanks | 11729 | 11766 | 38 | TAACTATGACTCTCTTAAGGTAGCCAAATGCCTCGTCA | 143.6 | -9.3 | 1.6 | -5.8 | 7.7E-09 | 2.1E-08 | BT20/MB468 | MITO |
| >hg38_21_+_8433172_8446622_RNA45SN1_45S_50ntFlanks@9839.9861.23\|GTAAGCAGAACTGGCGCTGCGGG | >hg38_21_+_8433172_8446622_RNA45SN1_45S_50ntFlanks | 9839 | 9861 | 23 | GTAAGCAGAACTGGCGCTGCGGG | 98.0 | -10.5 | 1.8 | -5.7 | 9.5E-09 | 2.1E-08 | MB231/MB468 | MP |
| >hg38_21_+_8433172_8446622_RNA45SN1_45S_50ntFlanks@10684.10708.25\|CGGCGGCGTCCGGTGAGCTCTCGCT | >hg38_21_+_8433172_8446622_RNA45SN1_45S_50ntFlanks | 10684 | 10708 | 25 | CGGCGGCGTCCGGTGAGCTCTCGCT | 541.8 | -4.4 | 0.8 | -5.7 | 9.5E-09 | 2.1E-08 | MB231/MB468 | MP |
| >hg38_21_+_8433172_8446622_RNA45SN1_45S_50ntFlanks@4003.4022.20\|ATAACCTCGGGCCGATCGCA | >hg38_21_+_8433172_8446622_RNA45SN1_45S_50ntFlanks | 4003 | 4022 | 20 | ATAACCTCGGGCCGATCGCA | 4059.9 | -10.0 | 1.8 | -5.7 | 1.1E-08 | 2.1E-08 | BT20/MB468 | NUC |
| >hg38_21_+_8433172_8446622_RNA45SN1_45S_50ntFlanks@10832.10862.31\|AGCCGGATCCGTAACTTCGGGATAAGGATTG | >hg38_21_+_8433172_8446622_RNA45SN1_45S_50ntFlanks | 10832 | 10862 | 31 | AGCCGGATCCGTAACTTCGGGATAAGGATTG | 153.7 | -10.3 | 1.7 | -6.1 | 1.4E-09 | 2.1E-08 | BT20/MB468 | MP |
| >hg38_21_+_8433172_8446622_RNA45SN1_45S_50ntFlanks@11845.11872.28\|GGAACGGGCTTGGCGGAATCAGCGGGGA | >hg38_21_+_8433172_8446622_RNA45SN1_45S_50ntFlanks | 11845 | 11872 | 28 | GGAACGGGCTTGGCGGAATCAGCGGGGA | 2323.7 | -5.7 | 0.9 | -6.1 | 1.4E-09 | 2.1E-08 | BT20/MB468 | MP |
| >hg38_21_+_8433172_8446622_RNA45SN1_45S_50ntFlanks@12630.12653.24\|GGGCGAAGCTACCATCTGTGGGAT | >hg38_21_+_8433172_8446622_RNA45SN1_45S_50ntFlanks | 12630 | 12653 | 24 | GGGCGAAGCTACCATCTGTGGGAT | 121.1 | -10.0 | 1.7 | -6.1 | 1.4E-09 | 2.1E-08 | BT20/MB468 | MP |
| >hg38_21_+_8433172_8446622_RNA45SN1_45S_50ntFlanks@9198.9226.29\|GTTCTCTCGGGGCCACGCGCGCGTCCCCC | >hg38_21_+_8433172_8446622_RNA45SN1_45S_50ntFlanks | 9198 | 9226 | 29 | GTTCTCTCGGGGCCACGCGCGCGTCCCCC | 996.9 | -6.9 | 1.2 | -5.7 | 1.1E-08 | 2.1E-08 | MB231/MB468 | MITO |
| >hg38_21_+_8433172_8446622_RNA45SN1_45S_50ntFlanks@5118.5136.19\|ACCCCCGAGCGGTCGGCGT | >hg38_21_+_8433172_8446622_RNA45SN1_45S_50ntFlanks | 5118 | 5136 | 19 | ACCCCCGAGCGGTCGGCGT | 189.2 | -11.4 | 2.0 | -5.7 | 9.5E-09 | 2.1E-08 | MB231/MB468 | MP |
| >hg38_21_+_8433172_8446622_RNA45SN1_45S_50ntFlanks@10357.10383.27\|CTTTGAAGGCCGAAGTGGAGAAGGGTT | >hg38_21_+_8433172_8446622_RNA45SN1_45S_50ntFlanks | 10357 | 10383 | 27 | CTTTGAAGGCCGAAGTGGAGAAGGGTT | 354.1 | -5.9 | 1.0 | -5.7 | 1.1E-08 | 2.1E-08 | MB231/MB468 | MITO |
| >MI0000447\|hsa-mir-128-1&WithFlank&2\|+\|135665391\|135665484@56.74.19;MI0000727\|hsa-mir-128-2&WithFlank&3\|+\|35744470\|35744565@58.76.19&[MIMAT0000424&hsa-miR-128-3p&offsets\|0\|-2;m-108&2\|+\|135665446\|135665466&offsets\|0\|-2];[MIMAT0000424_1&hsa-miR-128-3p&offsets\|0\|-2;m-117&3\|+\|35744527\|35744547&offsets\|0\|-2]\|TCACAGTGAACCGGTCTCT | >MI0000447\|hsa-mir-128-1&WithFlank&2\|+\|135665391\|135665484 | 56 | 74 | 19 | TCACAGTGAACCGGTCTCT | 211.7 | 4.4 | 0.8 | 5.7 | 1.0E-08 | 2.1E-08 | MB231/MB468 | NUC |
| >trna111_HisGTG_1_-_147774845_147774916@1.30.30,trna118_HisGTG_1_-_145396881_145396952@1.30.30,trna16_HisGTG_1_+_146544773_146544844@1.30.30,trna1_HisGTG_15_+_45493349_45493420@1.30.30,trna21_HisGTG_1_+_147753471_147753542@1.30.30,trna33_HisGTG_6_+_27125906_27125977@1.30.30,trna7_HisGTG_9_-_14433938_14434009@1.30.30,trna8_HisGTG_15_-_45492611_45492682@1.30.30,trna9_HisGTG_15_-_45490804_45490875@1.30.30\|GCCGTGATCGTATAGTGGTTAGTACTCTGC | >trna111_HisGTG_1_-_147774845_147774916 | 1 | 30 | 30 | GCCGTGATCGTATAGTGGTTAGTACTCTGC | 1834.0 | -5.0 | 0.9 | -5.7 | 1.1E-08 | 2.1E-08 | BT20/MB468 | NUC |
| >hg38_21_+_8433172_8446622_RNA45SN1_45S_50ntFlanks@5423.5455.33\|ATTGGATGGTTTAGTGAGGCCCTCGGATCGGCC | >hg38_21_+_8433172_8446622_RNA45SN1_45S_50ntFlanks | 5423 | 5455 | 33 | ATTGGATGGTTTAGTGAGGCCCTCGGATCGGCC | 69.6 | -10.0 | 1.7 | -5.7 | 9.5E-09 | 2.1E-08 | MB231/MB468 | MP |
| >hg38_21_+_8433172_8446622_RNA45SN1_45S_50ntFlanks@10873.10901.29\|CTGGGTCGGTCGGGCTGGGGCGCGAAGCG | >hg38_21_+_8433172_8446622_RNA45SN1_45S_50ntFlanks | 10873 | 10901 | 29 | CTGGGTCGGTCGGGCTGGGGCGCGAAGCG | 833.3 | -8.8 | 1.5 | -5.7 | 1.1E-08 | 2.1E-08 | BT20/MB468 | NUC |
| >hg38_21_+_8433172_8446622_RNA45SN1_45S_50ntFlanks@12626.12659.34\|AATGGGGCGAAGCTACCATCTGTGGGATTATGAC | >hg38_21_+_8433172_8446622_RNA45SN1_45S_50ntFlanks | 12626 | 12659 | 34 | AATGGGGCGAAGCTACCATCTGTGGGATTATGAC | 2177.4 | -9.5 | 1.7 | -5.7 | 1.1E-08 | 2.1E-08 | BT20/MB468 | NUC |
| >hg38_21_+_8433172_8446622_RNA45SN1_45S_50ntFlanks@10826.10844.19\|TCGGCAAGCCGGATCCGTA | >hg38_21_+_8433172_8446622_RNA45SN1_45S_50ntFlanks | 10826 | 10844 | 19 | TCGGCAAGCCGGATCCGTA | 106.2 | -9.8 | 1.6 | -6.1 | 1.4E-09 | 2.1E-08 | BT20/MB468 | MP |
| >hg38_21_+_8433172_8446622_RNA45SN1_45S_50ntFlanks@12157.12182.26\|AGTTTGACTGGGGCGGTACACCTGTC | >hg38_21_+_8433172_8446622_RNA45SN1_45S_50ntFlanks | 12157 | 12182 | 26 | AGTTTGACTGGGGCGGTACACCTGTC | 810.9 | -5.9 | 1.0 | -5.8 | 7.7E-09 | 2.1E-08 | BT20/MB468 | MITO |
| >hg38_21_+_8433172_8446622_RNA45SN1_45S_50ntFlanks@12412.12435.24\|ATCCTTCGATGTCGGCTCTTCCTA | >hg38_21_+_8433172_8446622_RNA45SN1_45S_50ntFlanks | 12412 | 12435 | 24 | ATCCTTCGATGTCGGCTCTTCCTA | 513.1 | -7.8 | 1.4 | -5.7 | 1.1E-08 | 2.1E-08 | MB231/MB468 | MITO |
| >hg38_21_+_8433172_8446622_RNA45SN1_45S_50ntFlanks@4025.4045.21\|CCCCCCGTGGCGGCGACGACC | >hg38_21_+_8433172_8446622_RNA45SN1_45S_50ntFlanks | 4025 | 4045 | 21 | CCCCCCGTGGCGGCGACGACC | 136.5 | -9.2 | 1.6 | -5.8 | 7.8E-09 | 2.1E-08 | BT20/MB468 | MITO |
| >hg38_21_+_8433172_8446622_RNA45SN1_45S_50ntFlanks@12690.12711.22\|GCGGAACGATACGGCAGCGCCG | >hg38_21_+_8433172_8446622_RNA45SN1_45S_50ntFlanks | 12690 | 12711 | 22 | GCGGAACGATACGGCAGCGCCG | 297.4 | -7.5 | 1.3 | -5.7 | 9.6E-09 | 2.1E-08 | MB231/MB468 | MP |
| >hg38_21_+_8433172_8446622_RNA45SN1_45S_50ntFlanks@6645.6664.20\|CTCGTACGACTCTTAGCGGT | >hg38_21_+_8433172_8446622_RNA45SN1_45S_50ntFlanks | 6645 | 6664 | 20 | CTCGTACGACTCTTAGCGGT | 5723.2 | 4.7 | 0.8 | 6.1 | 1.4E-09 | 2.1E-08 | BT20/MB468 | MP |
| >MI0000746\|hsa-mir-99b&WithFlank&19\|+\|51692606\|51692687@13.34.22&[MIMAT0000689&hsa-miR-99b-5p&offsets\|0\|0;m-10&19\|+\|51692618\|51692639&offsets\|0\|0]\|CACCCGTAGAACCGACCTTGCG | >MI0000746\|hsa-mir-99b&WithFlank&19\|+\|51692606\|51692687 | 13 | 34 | 22 | CACCCGTAGAACCGACCTTGCG | 24698.0 | 4.0 | 0.7 | 5.7 | 1.0E-08 | 2.1E-08 | MB231/MB468 | NUC |
| >hg38_1_-_228634819_228635039_RNA5S12_5S_50ntFlanks@51.74.24\|GTCTACGGCCATACCACCCTGAAC | >hg38_1_-_228634819_228635039_RNA5S12_5S_50ntFlanks | 51 | 74 | 24 | GTCTACGGCCATACCACCCTGAAC | 1592.4 | 9.0 | 1.6 | 5.8 | 8.7E-09 | 2.1E-08 | BT20/MB468 | CYTO |
| >hg38_21_+_8433172_8446622_RNA45SN1_45S_50ntFlanks@10500.10533.34\|CAGATCCCCGAATCCGGAGTGGCGGAGATGGGCG | >hg38_21_+_8433172_8446622_RNA45SN1_45S_50ntFlanks | 10500 | 10533 | 34 | CAGATCCCCGAATCCGGAGTGGCGGAGATGGGCG | 84.7 | -8.5 | 1.5 | -5.8 | 7.8E-09 | 2.1E-08 | BT20/MB468 | MITO |

| | | | | | | | | | | | | |
|---|---|---|---|---|---|---|---|---|---|---|---|---|
| >am_MI0005763\|hsa-mir-941-1&WithFlank&20\|+\|63919443\|63919526@53.75.23(+1U);MI0005764\|hsa-mir-941-2&WithFlank&20\|+\|63919499\|63919582@53.75.23(+1U);MI0005765\|hsa-mir-941-3&WithFlank&20\|+\|63919555\|63919638@53.75.23(+1U);MI0005766\|hsa-mir-941-4&WithFlank&20\|+\|63919750\|63919833@53.75.23(+1U);MI0031520\|hsa-mir-941-5&WithFlank&20\|+\|63919862\|63919945@53.75.23(+1U)&[MIMAT0004984&hsa-miR-941&offsets\|0\|0(+1U);m-263&20\|+\|63919495\|63919517&offsets\|0\|0(+1U)];[MIMAT0004984_1&hsa-miR-941&offsets\|0\|0(+1U);m-263&20\|+\|63919551\|63919573&offsets\|0\|0(+1U)];[MIMAT0004984_2&hsa-miR-941&offsets\|0\|0(+1U);m-263&20\|+\|63919607\|63919629&offsets\|0\|0(+1U)];[MIMAT0004984_3&hsa-miR-941&offsets\|0\|0(+1U);m-263&20\|+\|63919802\|63919824&offsets\|0\|0(+1U)];[MIMAT0004984_4&hsa-miR-941&offsets\|0\|0(+1U);m-263&20\|+\|63919914\|63919936&offsets\|0\|0(+1U)]\|CACCCGGCTGTGTGCACATGTGCT | >am_MI0005763\|hsa-mir-941-1&WithFlank&20\|+\|63919443\|63919526 | 53 | 75 | 23(+1U) | CACCCGGCTGTGTGCACATGTGCT | 706.2 | 8.0 | 1.4 | 5.8 | 8.7E-09 | 2.1E-08 | BT20/MB468 | CYTO |
| >hg38_21_+_8433172_8446622_RNA45SN1_45S_50ntFlanks@4724.4752.29\|ATTAATCAAGAACGAAAGTCGGAGGTTCG | >hg38_21_+_8433172_8446622_RNA45SN1_45S_50ntFlanks | 4724 | 4752 | 29 | ATTAATCAAGAACGAAAGTCGGAGGTTCG | 807.8 | -4.5 | 0.8 | -5.8 | 8.7E-09 | 2.1E-08 | BT20/MB468 | CYTO |
| >hg38_21_+_8433172_8446622_RNA45SN1_45S_50ntFlanks@10823.10862.40\|AAGTCGGCAAGCCGGATCCGTAACTTCGGGATAAGGATTG | >hg38_21_+_8433172_8446622_RNA45SN1_45S_50ntFlanks | 10823 | 10862 | 40 | AAGTCGGCAAGCCGGATCCGTAACTTCGGGATAAGGATTG | 197.0 | -6.3 | 1.1 | -5.7 | 1.0E-08 | 2.1E-08 | MB231/MB468 | CYTO |
| >hg38_21_+_8433172_8446622_RNA45SN1_45S_50ntFlanks@12044.12066.23\|ACTGACCCGGTGAGGCGGGGGGG | >hg38_21_+_8433172_8446622_RNA45SN1_45S_50ntFlanks | 12044 | 12066 | 23 | ACTGACCCGGTGAGGCGGGGGGG | 216.6 | -10.8 | 1.8 | -6.1 | 1.4E-09 | 2.1E-08 | BT20/MB468 | MP |
| >hg38_21_+_8433172_8446622_RNA45SN1_45S_50ntFlanks@10402.10420.19\|AACATGGGTCAGTCGGTCC | >hg38_21_+_8433172_8446622_RNA45SN1_45S_50ntFlanks | 10402 | 10420 | 19 | AACATGGGTCAGTCGGTCC | 869.3 | -6.3 | 1.1 | -5.7 | 1.0E-08 | 2.1E-08 | MB231/MB468 | NUC |
| >MI0000093\|hsa-mir-92a-1&WithFlank&13\|+\|91351308\|91351397@54.73.20;MI0000094\|hsa-mir-92a-2&WithFlank&X\|-\|134169532\|134169618@54.73.20&[MIMAT0000092&hsa-miR-92a-3p&offsets\|0\|-2;m-9&13\|+\|91351361\|91351382&offsets\|0\|-2];[MIMAT0000092_1&hsa-miR-92a-3p&offsets\|0\|-2;m-11&X\|-\|134169544\|134169565&offsets\|0\|-2]\|TATTGCACTTGTCCCGGCCT | >MI0000093\|hsa-mir-92a-1&WithFlank&13\|+\|91351308\|91351397 | 54 | 73 | 20 | TATTGCACTTGTCCCGGCCT | 372.6 | 7.4 | 1.3 | 5.7 | 1.0E-08 | 2.1E-08 | MB231/MB468 | CYTO |
| >hg38_21_+_8433172_8446622_RNA45SN1_45S_50ntFlanks@6697.6718.22\|CAGCTAGCTGCGAGAATTAATG | >hg38_21_+_8433172_8446622_RNA45SN1_45S_50ntFlanks | 6697 | 6718 | 22 | CAGCTAGCTGCGAGAATTAATG | 1885.5 | -6.9 | 1.2 | -5.7 | 9.6E-09 | 2.1E-08 | MB231/MB468 | MP |
| >hg38_21_+_8433172_8446622_RNA45SN1_45S_50ntFlanks@11163.11188.26\|CGGTCCCCCGCGAGGGGGGCCCGGGC | >hg38_21_+_8433172_8446622_RNA45SN1_45S_50ntFlanks | 11163 | 11188 | 26 | CGGTCCCCCGCGAGGGGGGCCCGGGC | 333.7 | -11.5 | 1.9 | -6.1 | 1.4E-09 | 2.1E-08 | BT20/MB468 | MP |
| >trna119_LysCTT_1_-_145395522_145395594@1.35.35,trna11_LysCTT_5_-_180648979_180649051@1.35.35,trna13_LysCTT_6_+_26556774_26556846@1.35.35,trna7_LysCTT_16_+_3225692_3225764@1.35.35,trna9_LysCTT_5_+_180634755_180634827@1.35.35\|GCCCGGCTAGCTCAGTCGGTAGAGCATGAGACTCT | >trna119_LysCTT_1_-_145395522_145395594 | 1 | 35 | 35 | GCCCGGCTAGCTCAGTCGGTAGAGCATGAGACTCT | 3786.7 | 5.5 | 1.0 | 5.7 | 1.1E-08 | 2.1E-08 | MB231/MB468 | MITO |
| >hg38_21_+_8433172_8446622_RNA45SN1_45S_50ntFlanks@12482.12500.19\|ATAGGGAACGTGAGCTGGG | >hg38_21_+_8433172_8446622_RNA45SN1_45S_50ntFlanks | 12482 | 12500 | 19 | ATAGGGAACGTGAGCTGGG | 277.3 | -4.6 | 0.8 | -5.7 | 9.7E-09 | 2.1E-08 | MB231/MB468 | MP |
| >hg38_21_+_8433172_8446622_RNA45SN1_45S_50ntFlanks@4759.4793.35\|ATCAGATACCGTCGTAGTTCCGACCATAAACGATG | >hg38_21_+_8433172_8446622_RNA45SN1_45S_50ntFlanks | 4759 | 4793 | 35 | ATCAGATACCGTCGTAGTTCCGACCATAAACGATG | 916.8 | -8.0 | 1.4 | -5.7 | 1.1E-08 | 2.1E-08 | MB231/MB468 | MITO |
| >hg38_21_+_8433172_8446622_RNA45SN1_45S_50ntFlanks@9651.9668.18\|TCAGGATAGCTGGCGCTC | >hg38_21_+_8433172_8446622_RNA45SN1_45S_50ntFlanks | 9651 | 9668 | 18 | TCAGGATAGCTGGCGCTC | 346.2 | -8.4 | 1.5 | -5.7 | 1.2E-08 | 2.1E-08 | BT20/MB468 | NUC |
| >hg38_21_+_8433172_8446622_RNA45SN1_45S_50ntFlanks@9766.9788.23\|TAAATGGGTAAGAAGCCCGGCTC | >hg38_21_+_8433172_8446622_RNA45SN1_45S_50ntFlanks | 9766 | 9788 | 23 | TAAATGGGTAAGAAGCCCGGCTC | 78.8 | -10.2 | 1.8 | -5.7 | 9.7E-09 | 2.1E-08 | MB231/MB468 | MP |
| >hg38_21_+_8433172_8446622_RNA45SN1_45S_50ntFlanks@4731.4763.33\|AAGAACGAAAGTCGGAGGTTCGAAGACGATCAG | >hg38_21_+_8433172_8446622_RNA45SN1_45S_50ntFlanks | 4731 | 4763 | 33 | AAGAACGAAAGTCGGAGGTTCGAAGACGATCAG | 357.5 | 4.3 | 0.7 | 5.7 | 1.0E-08 | 2.1E-08 | MB231/MB468 | CYTO |
| >hg38_21_+_8433172_8446622_RNA45SN1_45S_50ntFlanks@9569.9595.27\|GTGCAAATCGGTCGTCCGACCTGGGTA | >hg38_21_+_8433172_8446622_RNA45SN1_45S_50ntFlanks | 9569 | 9595 | 27 | GTGCAAATCGGTCGTCCGACCTGGGTA | 357.7 | -2.3 | 0.4 | -5.8 | 8.8E-09 | 2.1E-08 | BT20/MB468 | CYTO |
| >MI0000061\|hsa-let-7a-2&WithFlank&11\|-\|122146516\|122146599@11.29.19;MI0000062\|hsa-let-7a-3&WithFlank&22\|+\|46112743\|46112828@10.28.19;MI0000060\|hsa-let-7a-1&WithFlank&9\|+\|94175951\|94176042@12.30.19&[MIMAT0000062&hsa-let-7a-5p&offsets\|0\|-3;m-6&11\|-\|122146568\|122146589&offsets\|0\|-3];[MIMAT0000062_1&hsa-let-7a-5p&offsets\|0\|-3;m-5&22\|+\|46112752\|46112773&offsets\|0\|-3];[MIMAT0000062_2&hsa-let-7a-5p&offsets\|0\|-3;m-4&9\|+\|94175962\|94175983&offsets\|0\|-3]\|TGAGGTAGTAGGTTGTATA | >MI0000061\|hsa-let-7a-2&WithFlank&11\|-\|122146516\|122146599 | 11 | 29 | 19 | TGAGGTAGTAGGTTGTATA | 307.7 | 8.8 | 1.5 | 5.8 | 7.9E-09 | 2.1E-08 | BT20/MB468 | MITO |
| >MI0000272\|hsa-mir-182&WithFlank&7\|-\|129770377\|129770498@29.49.21&[MIMAT0000259&hsa-miR-182-5p&offsets\|0\|-3;m-22&7\|-\|129770449\|129770470&offsets\|0\|-1]\|TTTGGCAATGGTAGAACTCAC | >MI0000272\|hsa-mir-182&WithFlank&7\|-\|129770377\|129770498 | 29 | 49 | 21 | TTTGGCAATGGTAGAACTCAC | 508.7 | 3.6 | 0.6 | 6.1 | 1.1E-09 | 2.1E-08 | BT20/MB231 | NUC |
| >hg38_21_+_8433172_8446622_RNA45SN1_45S_50ntFlanks@8356.8391.36\|GAAAAGAACTTTGAAGAGAGAGTTCAAGAGGGCGTG | >hg38_21_+_8433172_8446622_RNA45SN1_45S_50ntFlanks | 8356 | 8391 | 36 | GAAAAGAACTTTGAAGAGAGAGTTCAAGAGGGCGTG | 586.6 | -4.7 | 0.8 | -5.7 | 1.2E-08 | 2.1E-08 | BT20/MB468 | NUC |
| >hg38_21_+_8433172_8446622_RNA45SN1_45S_50ntFlanks@7989.8018.30\|AGACGTGGCGACCCGCTGAATTTAAGCATA | >hg38_21_+_8433172_8446622_RNA45SN1_45S_50ntFlanks | 7989 | 8018 | 30 | AGACGTGGCGACCCGCTGAATTTAAGCATA | 276.9 | -4.1 | 0.7 | -5.7 | 9.7E-09 | 2.1E-08 | MB231/MB468 | MP |
| >hg38_21_+_8433172_8446622_RNA45SN1_45S_50ntFlanks@12394.12417.24\|TAGCGACGTCGCTTTTTGATCCTT | >hg38_21_+_8433172_8446622_RNA45SN1_45S_50ntFlanks | 12394 | 12417 | 24 | TAGCGACGTCGCTTTTTGATCCTT | 510.7 | -8.5 | 1.5 | -5.7 | 1.2E-08 | 2.1E-08 | BT20/MB468 | NUC |
| >hg38_21_+_8433172_8446622_RNA45SN1_45S_50ntFlanks@12437.12474.38\|CATTGTGAAGCAGAATTCACCAAGCGTTGGATTGTTCA | >hg38_21_+_8433172_8446622_RNA45SN1_45S_50ntFlanks | 12437 | 12474 | 38 | CATTGTGAAGCAGAATTCACCAAGCGTTGGATTGTTCA | 793.6 | -9.4 | 1.6 | -5.7 | 1.2E-08 | 2.1E-08 | BT20/MB468 | NUC |
| >MI0001445\|hsa-mir-423&WithFlank&17\|+\|30117073\|30117178@23.46.24&[MIMAT0004748&hsa-miR-423-5p&offsets\|0\|+1;m-141&17\|+\|30117095\|30117117&offsets\|0\|+1]\|TGAGGGGCAGAGAGCGAGACTTTT | >MI0001445\|hsa-mir-423&WithFlank&17\|+\|30117073\|30117178 | 23 | 46 | 24 | TGAGGGGCAGAGAGCGAGACTTTT | 6398.8 | 6.5 | 1.1 | 6.1 | 1.4E-09 | 2.1E-08 | BT20/MB468 | MP |
| >hg38_21_+_8433172_8446622_RNA45SN1_45S_50ntFlanks@12022.12055.34\|CCACTACTCTGATCGTTTTTTCACTGACCCGGTG | >hg38_21_+_8433172_8446622_RNA45SN1_45S_50ntFlanks | 12022 | 12055 | 34 | CCACTACTCTGATCGTTTTTTCACTGACCCGGTG | 104.7 | 10.5 | 1.7 | 6.1 | 1.4E-09 | 2.1E-08 | BT20/MB468 | MP |
| >hg38_21_+_8433172_8446622_RNA45SN1_45S_50ntFlanks@6672.6700.29\|TCGGCTCGTGCGTCGATGAAGAACGCAGC | >hg38_21_+_8433172_8446622_RNA45SN1_45S_50ntFlanks | 6672 | 6700 | 29 | TCGGCTCGTGCGTCGATGAAGAACGCAGC | 105.0 | 10.5 | 1.7 | 6.1 | 1.4E-09 | 2.1E-08 | BT20/MB468 | MP |
| >hg38_21_+_8433172_8446622_RNA45SN1_45S_50ntFlanks@12191.12211.21\|ACGCAGGTGTCCTAAGGCGAG | >hg38_21_+_8433172_8446622_RNA45SN1_45S_50ntFlanks | 12191 | 12211 | 21 | ACGCAGGTGTCCTAAGGCGAG | 133.2 | -10.1 | 1.7 | -6.1 | 1.4E-09 | 2.1E-08 | BT20/MB468 | MP |
| >hg38_21_+_8433172_8446622_RNA45SN1_45S_50ntFlanks@12251.12279.29\|AAAGCTCGCTTGATCTTGATTTTCAGTAC | >hg38_21_+_8433172_8446622_RNA45SN1_45S_50ntFlanks | 12251 | 12279 | 29 | AAAGCTCGCTTGATCTTGATTTTCAGTAC | 616.7 | 5.2 | 0.9 | 5.7 | 1.0E-08 | 2.1E-08 | MB231/MB468 | CYTO |
| >hg38_21_+_8433172_8446622_RNA45SN1_45S_50ntFlanks@11731.11751.21\|ACTATGACTCTCTTAAGGTAG | >hg38_21_+_8433172_8446622_RNA45SN1_45S_50ntFlanks | 11731 | 11751 | 21 | ACTATGACTCTCTTAAGGTAG | 104.6 | -9.8 | 1.6 | -6.1 | 1.4E-09 | 2.1E-08 | BT20/MB468 | MP |
| >hg38_21_+_8433172_8446622_RNA45SN1_45S_50ntFlanks@10349.10376.28\|AACGAGAACTTTGAAGGCCGAAGTGGAG | >hg38_21_+_8433172_8446622_RNA45SN1_45S_50ntFlanks | 10349 | 10376 | 28 | AACGAGAACTTTGAAGGCCGAAGTGGAG | 1791.5 | -7.5 | 1.3 | -5.7 | 9.8E-09 | 2.1E-08 | MB231/MB468 | MP |
| >hg38_21_+_8433172_8446622_RNA45SN1_45S_50ntFlanks@9533.9564.32\|GAGGAAACTCTGGTGGAGGTCCGTAGCGGTCC | >hg38_21_+_8433172_8446622_RNA45SN1_45S_50ntFlanks | 9533 | 9564 | 32 | GAGGAAACTCTGGTGGAGGTCCGTAGCGGTCC | 875.6 | -4.8 | 0.8 | -5.7 | 1.1E-08 | 2.1E-08 | MB231/MB468 | NUC |
| >trnaMT_ValTAC_MT_+_1602_1670@54.71.18\|AACTTGACCGCTCTGACC | >trnaMT_ValTAC_MT_+_1602_1670 | 54 | 71 | 18 | AACTTGACCGCTCTGACC | 1027.0 | 6.0 | 1.0 | 5.7 | 1.1E-08 | 2.1E-08 | MB231/MB468 | NUC |
| >MI0000746\|hsa-mir-99b&WithFlank&19\|+\|51692606\|51692687@13.33.21(+1A)&[MIMAT0000689&hsa-miR-99b-5p&offsets\|0\|-1(+1A);m-10&19\|+\|51692618\|51692639&offsets\|0\|-1(+1A)]\|CACCCGTAGAACCGACCTTGCA | >MI0000746\|hsa-mir-99b&WithFlank&19\|+\|51692606\|51692687 | 13 | 33 | 21(+1A) | CACCCGTAGAACCGACCTTGCA | 3173.1 | 5.1 | 0.9 | 5.7 | 1.1E-08 | 2.1E-08 | MB231/MB468 | MITO |
| >hg38_21_+_8433172_8446622_RNA45SN1_45S_50ntFlanks@12503.12520.18\|TAGACCGTCGTGAGACAG | >hg38_21_+_8433172_8446622_RNA45SN1_45S_50ntFlanks | 12503 | 12520 | 18 | TAGACCGTCGTGAGACAG | 117.1 | -10.7 | 1.9 | -5.7 | 9.8E-09 | 2.1E-08 | MB231/MB468 | MP |
| >hg38_21_+_8433172_8446622_RNA45SN1_45S_50ntFlanks@5473.5514.42\|CGGCCCTGGCGGAGCGCTGAGAAGACGGTCGAACTTGACTAT | >hg38_21_+_8433172_8446622_RNA45SN1_45S_50ntFlanks | 5473 | 5514 | 42 | CGGCCCTGGCGGAGCGCTGAGAAGACGGTCGAACTTGACTAT | 1179.2 | -4.6 | 0.8 | -5.7 | 1.1E-08 | 2.1E-08 | MB231/MB468 | MITO |
| >MI0000102\|hsa-mir-100&WithFlank&11\|-\|122152223\|122152314@19.40.22(+1A)&[MIMAT0000098&hsa-miR-100-5p&offsets\|0\|0(+1A);m-27&11\|-\|122152276\|122152296&offsets\|0\|+1(+1A)]\|AACCCGTAGATCCGAACTTGTGA | >MI0000102\|hsa-mir-100&WithFlank&11\|-\|122152223\|122152314 | 19 | 40 | 22(+1A) | AACCCGTAGATCCGAACTTGTGA | 37701.7 | -5.9 | 1.0 | -6.1 | 1.1E-09 | 2.1E-08 | BT20/MB231 | NUC |
| >hg38_21_+_8433172_8446622_RNA45SN1_45S_50ntFlanks@11569.11592.24\|CAGCCGACTTAGAACTGGTGCGGA | >hg38_21_+_8433172_8446622_RNA45SN1_45S_50ntFlanks | 11569 | 11592 | 24 | CAGCCGACTTAGAACTGGTGCGGA | 593.1 | -7.8 | 1.4 | -5.7 | 1.1E-08 | 2.1E-08 | MB231/MB468 | MITO |
| >trnaMT_ValTAC_MT_+_1602_1670@43.72.30\|GATTTCAACTTAACTTGACCGCTCTGACCA | >trnaMT_ValTAC_MT_+_1602_1670 | 43 | 72 | 30 | GATTTCAACTTAACTTGACCGCTCTGACCA | 6544.9 | 6.4 | 1.1 | 5.8 | 8.0E-09 | 2.1E-08 | BT20/MB468 | MITO |
| >hg38_21_+_8433172_8446622_RNA45SN1_45S_50ntFlanks@12578.12614.37\|AGGAACCGCAGGTTCAGACATTTGGTGTATGTGCTTG | >hg38_21_+_8433172_8446622_RNA45SN1_45S_50ntFlanks | 12578 | 12614 | 37 | AGGAACCGCAGGTTCAGACATTTGGTGTATGTGCTTG | 65.0 | -9.9 | 1.7 | -5.7 | 9.9E-09 | 2.1E-08 | MB231/MB468 | MP |
| >hg38_21_+_8433172_8446622_RNA45SN1_45S_50ntFlanks@5236.5257.22\|CACTGACTGGCTCAGCGTGTGC | >hg38_21_+_8433172_8446622_RNA45SN1_45S_50ntFlanks | 5236 | 5257 | 22 | CACTGACTGGCTCAGCGTGTGC | 310.3 | -6.2 | 1.1 | -5.7 | 9.9E-09 | 2.2E-08 | MB231/MB468 | MP |
| >hg38_21_+_8433172_8446622_RNA45SN1_45S_50ntFlanks@6653.6701.49\|ACTCTTAGCGGTGGATCACTCGGCTCGTGCGTCGATGAAGAACGCAGCT | >hg38_21_+_8433172_8446622_RNA45SN1_45S_50ntFlanks | 6653 | 6701 | 49 | ACTCTTAGCGGTGGATCACTCGGCTCGTGCGTCGATGAAGAACGCAGCT | 500.2 | 8.2 | 1.4 | 5.8 | 8.1E-09 | 2.2E-08 | BT20/MB468 | MITO |
| >trnaMT_GlnTTG_MT_-_4329_4400@1.30.30,trnalookalike2_GlnTTG_1_-_564879_564950@1.30.30\|TAGGATGGGGTGTGATAGGTGGCACGGAGA | >trnaMT_GlnTTG_MT_-_4329_4400 | 1 | 30 | 30 | TAGGATGGGGTGTGATAGGTGGCACGGAGA | 1868.2 | 6.1 | 1.1 | 5.8 | 8.1E-09 | 2.2E-08 | BT20/MB468 | MITO |
| >hg38_21_+_8433172_8446622_RNA45SN1_45S_50ntFlanks@12147.12179.33\|CCAGGTGGGGAGTTTGACTGGGGCGGTACACCT | >hg38_21_+_8433172_8446622_RNA45SN1_45S_50ntFlanks | 12147 | 12179 | 33 | CCAGGTGGGGAGTTTGACTGGGGCGGTACACCT | 105.1 | -9.8 | 1.6 | -6.0 | 1.5E-09 | 2.2E-08 | BT20/MB468 | MP |
| >hg38_21_+_8433172_8446622_RNA45SN1_45S_50ntFlanks@11677.11702.26\|CTCTGAATGTCAAAGTGAAGAAATTC | >hg38_21_+_8433172_8446622_RNA45SN1_45S_50ntFlanks | 11677 | 11702 | 26 | CTCTGAATGTCAAAGTGAAGAAATTC | 773.3 | -6.2 | 1.1 | -5.7 | 9.9E-09 | 2.2E-08 | MB231/MB468 | MP |

<table>
<tr><th></th><th></th><th></th><th></th><th></th><th></th><th></th><th></th><th></th><th></th><th></th><th></th><th></th><th></th></tr>
<tr><td>>trnaMT_ArgTCG_MT_+_10405_10469@28.68.41|ATTTCGACTCATTAAATTATGATAATCATATTTACCAACCA</td><td>>trnaMT_ArgTCG_MT_+_10405_10469</td><td>28</td><td>68</td><td>41</td><td>ATTTCGACTCATTAAATTATGATAATCATATTTACCAACCA</td><td>138.8</td><td>-9.2</td><td>1.6</td><td>-5.8</td><td>8.1E-09</td><td>2.2E-08</td><td>BT20/MB468</td><td>MITO</td></tr>
<tr><td>>hg38_21_+_8433172_8446622_RNA45SN1_45S_50ntFlanks@7980.7999.20|CCTCAGATCAGACGTGGCGA</td><td>>hg38_21_+_8433172_8446622_RNA45SN1_45S_50ntFlanks</td><td>7980</td><td>7999</td><td>20</td><td>CCTCAGATCAGACGTGGCGA</td><td>159.1</td><td>-6.6</td><td>1.2</td><td>-5.7</td><td>9.0E-09</td><td>2.2E-08</td><td>BT20/MB468</td><td>CYTO</td></tr>
<tr><td>>hg38_21_+_8433172_8446622_RNA45SN1_45S_50ntFlanks@9822.9850.29|CTAGTGGGCCACTTTTGGTAAGCAGAACT</td><td>>hg38_21_+_8433172_8446622_RNA45SN1_45S_50ntFlanks</td><td>9822</td><td>9850</td><td>29</td><td>CTAGTGGGCCACTTTTGGTAAGCAGAACT</td><td>1167.5</td><td>-7.5</td><td>1.3</td><td>-5.7</td><td>1.2E-08</td><td>2.2E-08</td><td>BT20/MB468</td><td>NUC</td></tr>
<tr><td>>MI0000650|hsa-mir-200c&WithFlank&12|+|6963693|6963772@50.72.23(+1A)&[MIMAT0000617&hsa-miR-200c-3p&offsets|0|0(+1A);m-54&12|+|6963742|6963764&offsets|0|0(+1A)]|TAATACTGCCGGGTAATGATGGAA</td><td>>MI0000650|hsa-mir-200c&WithFlank&12|+|6963693|6963772</td><td>50</td><td>72</td><td>23(+1A)</td><td>TAATACTGCCGGGTAATGATGGAA</td><td>3310.0</td><td>4.9</td><td>0.8</td><td>6.0</td><td>1.5E-09</td><td>2.2E-08</td><td>BT20/MB468</td><td>MP</td></tr>
<tr><td>>MI0003602|hsa-mir-590&WithFlank&7|+|74191192|74191300@62.82.21&[MIMAT0004801&hsa-miR-590-3p&offsets|0|0;m-332&7|+|74191253|74191273&offsets|0|0]|TAATTTTATGTATAAGCTAGT</td><td>>MI0003602|hsa-mir-590&WithFlank&7|+|74191192|74191300</td><td>62</td><td>82</td><td>21</td><td>TAATTTTATGTATAAGCTAGT</td><td>227.2</td><td>-11.2</td><td>1.8</td><td>-6.3</td><td>2.8E-10</td><td>2.2E-08</td><td>BT20/MB231</td><td>MP</td></tr>
<tr><td>>hg38_21_+_8433172_8446622_RNA45SN1_45S_50ntFlanks@12163.12181.19|ACTGGGGCGGTACACCTGT</td><td>>hg38_21_+_8433172_8446622_RNA45SN1_45S_50ntFlanks</td><td>12163</td><td>12181</td><td>19</td><td>ACTGGGGCGGTACACCTGT</td><td>2071.6</td><td>-7.2</td><td>1.2</td><td>-5.8</td><td>8.1E-09</td><td>2.2E-08</td><td>BT20/MB468</td><td>MITO</td></tr>
<tr><td>>hg38_21_+_8433172_8446622_RNA45SN1_45S_50ntFlanks@3775.3799.25|GCCGGTACAGTGAAACTGCGAATGG</td><td>>hg38_21_+_8433172_8446622_RNA45SN1_45S_50ntFlanks</td><td>3775</td><td>3799</td><td>25</td><td>GCCGGTACAGTGAAACTGCGAATGG</td><td>144.0</td><td>-9.3</td><td>1.6</td><td>-5.8</td><td>8.1E-09</td><td>2.2E-08</td><td>BT20/MB468</td><td>MITO</td></tr>
<tr><td>>hg38_21_+_8433172_8446622_RNA45SN1_45S_50ntFlanks@8194.8223.30|CCCGTGGACGGTGTGAGGCCGGTAGCGGCC</td><td>>hg38_21_+_8433172_8446622_RNA45SN1_45S_50ntFlanks</td><td>8194</td><td>8223</td><td>30</td><td>CCCGTGGACGGTGTGAGGCCGGTAGCGGCC</td><td>103.4</td><td>-9.8</td><td>1.6</td><td>-6.0</td><td>1.5E-09</td><td>2.2E-08</td><td>BT20/MB468</td><td>MP</td></tr>
<tr><td>>hg38_21_+_8433172_8446622_RNA45SN1_45S_50ntFlanks@6721.6742.22|AATTGCAGGACACATTGATCAT</td><td>>hg38_21_+_8433172_8446622_RNA45SN1_45S_50ntFlanks</td><td>6721</td><td>6742</td><td>22</td><td>AATTGCAGGACACATTGATCAT</td><td>245.8</td><td>-7.5</td><td>1.3</td><td>-5.7</td><td>1.0E-08</td><td>2.2E-08</td><td>MB231/MB468</td><td>MP</td></tr>
<tr><td>>hg38_21_+_8433172_8446622_RNA45SN1_45S_50ntFlanks@3985.4025.41|CGGCTTTGGTGACTCTAGATAACCTCGGGCCGATCGCACGC</td><td>>hg38_21_+_8433172_8446622_RNA45SN1_45S_50ntFlanks</td><td>3985</td><td>4025</td><td>41</td><td>CGGCTTTGGTGACTCTAGATAACCTCGGGCCGATCGCACGC</td><td>159.8</td><td>-9.4</td><td>1.6</td><td>-5.8</td><td>8.1E-09</td><td>2.2E-08</td><td>BT20/MB468</td><td>MITO</td></tr>
<tr><td>>hg38_21_+_8433172_8446622_RNA45SN1_45S_50ntFlanks@12934.12953.20|CTAAACCATTCGTAGACGAC</td><td>>hg38_21_+_8433172_8446622_RNA45SN1_45S_50ntFlanks</td><td>12934</td><td>12953</td><td>20</td><td>CTAAACCATTCGTAGACGAC</td><td>841.5</td><td>-6.3</td><td>1.1</td><td>-5.7</td><td>1.1E-08</td><td>2.2E-08</td><td>MB231/MB468</td><td>NUC</td></tr>
<tr><td>>hg38_21_+_8433172_8446622_RNA45SN1_45S_50ntFlanks@10458.10487.30|ATGGCCTCCGTTGCCCTCGGCCGATCGAAA</td><td>>hg38_21_+_8433172_8446622_RNA45SN1_45S_50ntFlanks</td><td>10458</td><td>10487</td><td>30</td><td>ATGGCCTCCGTTGCCCTCGGCCGATCGAAA</td><td>357.8</td><td>-3.1</td><td>0.5</td><td>-5.8</td><td>8.2E-09</td><td>2.2E-08</td><td>BT20/MB468</td><td>MITO</td></tr>
<tr><td>>hg38_21_+_8433172_8446622_RNA45SN1_45S_50ntFlanks@10390.10424.35|GAACAGCAGTTGAACATGGGTCAGTCGGTCCTGAG</td><td>>hg38_21_+_8433172_8446622_RNA45SN1_45S_50ntFlanks</td><td>10390</td><td>10424</td><td>35</td><td>GAACAGCAGTTGAACATGGGTCAGTCGGTCCTGAG</td><td>670.3</td><td>4.7</td><td>0.8</td><td>5.7</td><td>1.1E-08</td><td>2.2E-08</td><td>MB231/MB468</td><td>NUC</td></tr>
<tr><td>>hg38_21_+_8433172_8446622_RNA45SN1_45S_50ntFlanks@8170.8206.37|CAAGTCCTTCTGATCGAGGCCCAGCCCGTGGACGGTG</td><td>>hg38_21_+_8433172_8446622_RNA45SN1_45S_50ntFlanks</td><td>8170</td><td>8206</td><td>37</td><td>CAAGTCCTTCTGATCGAGGCCCAGCCCGTGGACGGTG</td><td>241.1</td><td>-11.0</td><td>1.8</td><td>-6.0</td><td>1.5E-09</td><td>2.2E-08</td><td>BT20/MB468</td><td>MP</td></tr>
<tr><td>>MI0000067|hsa-let-7f-1&WithFlank&9|+|94176341|94176439@13.32.20(+1C);MI0000068|hsa-let-7f-2&WithFlank&X|-|53557186|53557280@14.33.20(+1C)&[MIMAT0000067&hsa-let-7f-5p&offsets|0|-2(+1C);m-20&9|+|94176353|94176374&offsets|0|-2(+1C)];[MIMAT0000067_1&hsa-let-7f-5p&offsets|0|-2(+1C);m-19&X|-|53557246|53557267&offsets|0|-2(+1C)]|TGAGGTAGTAGATTGTATAGC</td><td>>MI0000067|hsa-let-7f-1&WithFlank&9|+|94176341|94176439</td><td>13</td><td>32</td><td>20(+1C)</td><td>TGAGGTAGTAGATTGTATAGC</td><td>612.4</td><td>5.2</td><td>0.9</td><td>5.7</td><td>1.2E-08</td><td>2.2E-08</td><td>MB231/MB468</td><td>MITO</td></tr>
<tr><td>>am_trna10_LysCTT_16_+_3241501_3241573@11.28.18,trna119_LysCTT_1_-_145395522_145395594@11.28.18,trna11_LysCTT_5_-_180648979_180649051@11.28.18,trna128_LysCTT_1_-_55423542_55423614@11.28.18,trna13_LysCTT_14_-_58706613_58706685@11.28.18,trna13_LysCTT_6_+_26556774_26556846@11.28.18,trna2_LysCTT_15_+_79152904_79152976@11.28.18,trna32_LysCTT_16_-_3207406_3207478@11.28.18,trna4_LysCTT_18_-_43669271_43669343@11.28.18,trna6_LysCTT_19_-_52425396_52425468@11.28.18,trna7_LysCTT_16_+_3225692_3225764@11.28.18,trna9_LysCTT_5_+_180634755_180634827@11.28.18|CTCAGTCGGTAGAGCATG</td><td>>am_trna10_LysCTT_16_+_3241501_3241573</td><td>11</td><td>28</td><td>18</td><td>CTCAGTCGGTAGAGCATG</td><td>803.9</td><td>-3.9</td><td>0.7</td><td>-5.7</td><td>1.0E-08</td><td>2.2E-08</td><td>MB231/MB468</td><td>MP</td></tr>
<tr><td>>hg38_21_+_8433172_8446622_RNA45SN1_45S_50ntFlanks@3802.3829.28|CATTAAATCAGTTATGGTTCCTTTGGTC</td><td>>hg38_21_+_8433172_8446622_RNA45SN1_45S_50ntFlanks</td><td>3802</td><td>3829</td><td>28</td><td>CATTAAATCAGTTATGGTTCCTTTGGTC</td><td>231.4</td><td>-6.6</td><td>1.1</td><td>-5.7</td><td>1.1E-08</td><td>2.2E-08</td><td>MB231/MB468</td><td>NUC</td></tr>
<tr><td>>hg38_1_-_228634819_228635039_RNA5S12_5S_50ntFlanks@124.147.24|AGTACTTGGATGGGAGACCGCCTG</td><td>>hg38_1_-_228634819_228635039_RNA5S12_5S_50ntFlanks</td><td>124</td><td>147</td><td>24</td><td>AGTACTTGGATGGGAGACCGCCTG</td><td>923.3</td><td>-7.3</td><td>1.3</td><td>-5.7</td><td>1.2E-08</td><td>2.2E-08</td><td>BT20/MB468</td><td>NUC</td></tr>
<tr><td>>hg38_21_+_8433172_8446622_RNA45SN1_45S_50ntFlanks@12233.12265.33|CCGTGGAGCAGAAGGGCAAAAGCTCGCTTGATC</td><td>>hg38_21_+_8433172_8446622_RNA45SN1_45S_50ntFlanks</td><td>12233</td><td>12265</td><td>33</td><td>CCGTGGAGCAGAAGGGCAAAAGCTCGCTTGATC</td><td>637.8</td><td>-6.3</td><td>1.1</td><td>-5.7</td><td>1.2E-08</td><td>2.2E-08</td><td>MB231/MB468</td><td>MITO</td></tr>
<tr><td>>hg38_21_+_8433172_8446622_RNA45SN1_45S_50ntFlanks@6645.6668.24|CTCGTACGACTCTTAGCGGTGGAT</td><td>>hg38_21_+_8433172_8446622_RNA45SN1_45S_50ntFlanks</td><td>6645</td><td>6668</td><td>24</td><td>CTCGTACGACTCTTAGCGGTGGAT</td><td>770.5</td><td>3.4</td><td>0.6</td><td>5.7</td><td>1.1E-08</td><td>2.2E-08</td><td>MB231/MB468</td><td>CYTO</td></tr>
<tr><td>>hg38_MT_+_1621_3279_RNR2_16S_50ntFlanks@717.740.24|AAGCCTGCGTCAGATTAAAACACT</td><td>>hg38_MT_+_1621_3279_RNR2_16S_50ntFlanks</td><td>717</td><td>740</td><td>24</td><td>AAGCCTGCGTCAGATTAAAACACT</td><td>80.4</td><td>9.6</td><td>1.7</td><td>5.7</td><td>1.2E-08</td><td>2.2E-08</td><td>MB231/MB468</td><td>MITO</td></tr>
<tr><td>>hg38_21_+_8433172_8446622_RNA45SN1_45S_50ntFlanks@7980.8008.29|CCTCAGATCAGACGTGGCGACCCGCTGAA</td><td>>hg38_21_+_8433172_8446622_RNA45SN1_45S_50ntFlanks</td><td>7980</td><td>8008</td><td>29</td><td>CCTCAGATCAGACGTGGCGACCCGCTGAA</td><td>1888.6</td><td>-3.1</td><td>0.5</td><td>-5.8</td><td>8.2E-09</td><td>2.2E-08</td><td>BT20/MB468</td><td>MITO</td></tr>
<tr><td>>hg38_21_+_8433172_8446622_RNA45SN1_45S_50ntFlanks@6734.6772.39|ATTGATCATCGACACTTCGAACGCACTTGCGGCCCCGGG</td><td>>hg38_21_+_8433172_8446622_RNA45SN1_45S_50ntFlanks</td><td>6734</td><td>6772</td><td>39</td><td>ATTGATCATCGACACTTCGAACGCACTTGCGGCCCCGGG</td><td>124.9</td><td>-10.8</td><td>1.9</td><td>-5.7</td><td>1.0E-08</td><td>2.2E-08</td><td>MB231/MB468</td><td>MP</td></tr>
<tr><td>>MI0000088|hsa-mir-30a&WithFlank&6|-|71403545|71403627@12.33.22(+2A)&[MIMAT0000087&hsa-miR-30a-5p&offsets|0|0(+2A);m-15&6|-|71403595|71403616&offsets|0|0(+2A)]|TGTAAACATCCTCGACTGGAAGAA</td><td>>MI0000088|hsa-mir-30a&WithFlank&6|-|71403545|71403627</td><td>12</td><td>33</td><td>22(+2A)</td><td>TGTAAACATCCTCGACTGGAAGAA</td><td>73.0</td><td>9.4</td><td>1.6</td><td>5.7</td><td>1.0E-08</td><td>2.2E-08</td><td>MB231/MB468</td><td>MP</td></tr>
<tr><td>>hg38_21_+_8433172_8446622_RNA45SN1_45S_50ntFlanks@9310.9338.29|GAGTCTAACACGTGCGCGAGTCGGGGGCT</td><td>>hg38_21_+_8433172_8446622_RNA45SN1_45S_50ntFlanks</td><td>9310</td><td>9338</td><td>29</td><td>GAGTCTAACACGTGCGCGAGTCGGGGGCT</td><td>362.7</td><td>-7.9</td><td>1.4</td><td>-5.7</td><td>1.0E-08</td><td>2.2E-08</td><td>MB231/MB468</td><td>MP</td></tr>
<tr><td>>hg38_21_+_8433172_8446622_RNA45SN1_45S_50ntFlanks@12418.12455.38|CGATGTCGGCTCTTCCTATCATTGTGAAGCAGAATTCA</td><td>>hg38_21_+_8433172_8446622_RNA45SN1_45S_50ntFlanks</td><td>12418</td><td>12455</td><td>38</td><td>CGATGTCGGCTCTTCCTATCATTGTGAAGCAGAATTCA</td><td>8514.1</td><td>-10.9</td><td>1.9</td><td>-5.7</td><td>1.2E-08</td><td>2.2E-08</td><td>BT20/MB468</td><td>NUC</td></tr>
<tr><td>>hg38_21_+_8433172_8446622_RNA45SN1_45S_50ntFlanks@10866.10889.24|CTAAGGGCTGGGTCGGTCGGGCTG</td><td>>hg38_21_+_8433172_8446622_RNA45SN1_45S_50ntFlanks</td><td>10866</td><td>10889</td><td>24</td><td>CTAAGGGCTGGGTCGGTCGGGCTG</td><td>2089.7</td><td>-9.5</td><td>1.7</td><td>-5.7</td><td>1.2E-08</td><td>2.2E-08</td><td>BT20/MB468</td><td>NUC</td></tr>
<tr><td>>hg38_21_+_8433172_8446622_RNA45SN1_45S_50ntFlanks@11575.11603.29|ACTTAGAACTGGTGCGGACCAGGGGAATC</td><td>>hg38_21_+_8433172_8446622_RNA45SN1_45S_50ntFlanks</td><td>11575</td><td>11603</td><td>29</td><td>ACTTAGAACTGGTGCGGACCAGGGGAATC</td><td>2048.5</td><td>-4.3</td><td>0.7</td><td>-5.8</td><td>8.2E-09</td><td>2.2E-08</td><td>BT20/MB468</td><td>MITO</td></tr>
<tr><td>>hg38_21_+_8433172_8446622_RNA45SN1_45S_50ntFlanks@12974.13001.28|CGTACGTAGCAGAGCAGCTCCCTCGCTG</td><td>>hg38_21_+_8433172_8446622_RNA45SN1_45S_50ntFlanks</td><td>12974</td><td>13001</td><td>28</td><td>CGTACGTAGCAGAGCAGCTCCCTCGCTG</td><td>112.3</td><td>-5.9</td><td>1.0</td><td>-5.7</td><td>9.1E-09</td><td>2.2E-08</td><td>BT20/MB468</td><td>CYTO</td></tr>
<tr><td>>hg38_21_+_8433172_8446622_RNA45SN1_45S_50ntFlanks@9306.9338.33|CAAGGAGTCTAACACGTGCGCGAGTCGGGGGCT</td><td>>hg38_21_+_8433172_8446622_RNA45SN1_45S_50ntFlanks</td><td>9306</td><td>9338</td><td>33</td><td>CAAGGAGTCTAACACGTGCGCGAGTCGGGGGCT</td><td>356.2</td><td>-8.1</td><td>1.4</td><td>-5.7</td><td>1.2E-08</td><td>2.2E-08</td><td>BT20/MB468</td><td>NUC</td></tr>
<tr><td>>hg38_21_+_8433172_8446622_RNA45SN1_45S_50ntFlanks@9788.9830.43|CGCTGGCGTGGAGCCGGGCGTGGAATGCGAGTGCCTAGTGGGC</td><td>>hg38_21_+_8433172_8446622_RNA45SN1_45S_50ntFlanks</td><td>9788</td><td>9830</td><td>43</td><td>CGCTGGCGTGGAGCCGGGCGTGGAATGCGAGTGCCTAGTGGGC</td><td>84.2</td><td>-10.3</td><td>1.8</td><td>-5.7</td><td>1.0E-08</td><td>2.2E-08</td><td>MB231/MB468</td><td>MP</td></tr>
<tr><td>>hg38_21_+_8433172_8446622_RNA45SN1_45S_50ntFlanks@10574.10596.23|AAGCCGGCGGGAGCCCCGGGGAG</td><td>>hg38_21_+_8433172_8446622_RNA45SN1_45S_50ntFlanks</td><td>10574</td><td>10596</td><td>23</td><td>AAGCCGGCGGGAGCCCCGGGGAG</td><td>116.3</td><td>-9.9</td><td>1.6</td><td>-6.0</td><td>1.5E-09</td><td>2.2E-08</td><td>BT20/MB468</td><td>MP</td></tr>
<tr><td>>hg38_21_+_8433172_8446622_RNA45SN1_45S_50ntFlanks@12940.12965.26|CATTCGTAGACGACCTGCTTCTGGGT</td><td>>hg38_21_+_8433172_8446622_RNA45SN1_45S_50ntFlanks</td><td>12940</td><td>12965</td><td>26</td><td>CATTCGTAGACGACCTGCTTCTGGGT</td><td>251.5</td><td>-2.7</td><td>0.5</td><td>-5.7</td><td>1.1E-08</td><td>2.2E-08</td><td>MB231/MB468</td><td>CYTO</td></tr>
<tr><td>>hg38_21_+_8433172_8446622_RNA45SN1_45S_50ntFlanks@4202.4222.21|CGACCCGGGGAGGTAGTGACG</td><td>>hg38_21_+_8433172_8446622_RNA45SN1_45S_50ntFlanks</td><td>4202</td><td>4222</td><td>21</td><td>CGACCCGGGGAGGTAGTGACG</td><td>353.4</td><td>-7.6</td><td>1.3</td><td>-5.7</td><td>1.2E-08</td><td>2.2E-08</td><td>MB231/MB468</td><td>MITO</td></tr>
<tr><td>>hg38_21_+_8433172_8446622_RNA45SN1_45S_50ntFlanks@4727.4752.26|AATCAAGAACGAAAGTCGGAGGTTCG</td><td>>hg38_21_+_8433172_8446622_RNA45SN1_45S_50ntFlanks</td><td>4727</td><td>4752</td><td>26</td><td>AATCAAGAACGAAAGTCGGAGGTTCG</td><td>297.2</td><td>-6.1</td><td>1.1</td><td>-5.7</td><td>1.2E-08</td><td>2.2E-08</td><td>BT20/MB468</td><td>NUC</td></tr>
<tr><td>>MI0000087|hsa-mir-29a&WithFlank&7|-|130876741|130876816@48.67.20&[MIMAT0000086&hsa-miR-29a-3p&offsets|0|-2;m-26&7|-|130876749|130876769&offsets|0|-1]|TAGCACCATCTGAAATCGGT</td><td>>MI0000087|hsa-mir-29a&WithFlank&7|-|130876741|130876816</td><td>48</td><td>67</td><td>20</td><td>TAGCACCATCTGAAATCGGT</td><td>538.8</td><td>9.3</td><td>1.6</td><td>5.7</td><td>1.2E-08</td><td>2.2E-08</td><td>MB231/MB468</td><td>MITO</td></tr>
<tr><td>>hg38_21_+_8433172_8446622_RNA45SN1_45S_50ntFlanks@8195.8212.18|CCGTGGACGGTGTGAGGC</td><td>>hg38_21_+_8433172_8446622_RNA45SN1_45S_50ntFlanks</td><td>8195</td><td>8212</td><td>18</td><td>CCGTGGACGGTGTGAGGC</td><td>328.3</td><td>3.3</td><td>0.6</td><td>5.7</td><td>1.1E-08</td><td>2.2E-08</td><td>MB231/MB468</td><td>CYTO</td></tr>
<tr><td>>hg38_21_+_8433172_8446622_RNA45SN1_45S_50ntFlanks@12676.12706.31|AGAATCCCGCCCAGGCGGAACGATACGGCAG</td><td>>hg38_21_+_8433172_8446622_RNA45SN1_45S_50ntFlanks</td><td>12676</td><td>12706</td><td>31</td><td>AGAATCCCGCCCAGGCGGAACGATACGGCAG</td><td>99.1</td><td>-9.7</td><td>1.6</td><td>-6.0</td><td>1.5E-09</td><td>2.2E-08</td><td>BT20/MB468</td><td>MP</td></tr>
<tr><td>>MI0000067|hsa-let-7f-1&WithFlank&9|+|94176341|94176439@13.33.21(+1G);MI0000068|hsa-let-7f-2&WithFlank&X|-|53557186|53557280@14.34.21(+1G)&[MIMAT0000067&hsa-let-7f-5p&offsets|0|-1(+1G);m-20&9|+|94176353|94176374&offsets|0|-1(+1G)];[MIMAT0000067_1&hsa-let-7f-5p&offsets|0|-1(+1G);m-19&X|-|53557246|53557267&offsets|0|-1(+1G)]|TGAGGTAGTAGATTGTATAGTG</td><td>>MI0000067|hsa-let-7f-1&WithFlank&9|+|94176341|94176439</td><td>13</td><td>33</td><td>21(+1G)</td><td>TGAGGTAGTAGATTGTATAGTG</td><td>2823.6</td><td>5.4</td><td>0.9</td><td>5.7</td><td>9.2E-09</td><td>2.2E-08</td><td>BT20/MB468</td><td>CYTO</td></tr>
<tr><td>>hg38_21_+_8433172_8446622_RNA45SN1_45S_50ntFlanks@9574.9597.24|AATCGGTCGTCCGACCTGGGTATA</td><td>>hg38_21_+_8433172_8446622_RNA45SN1_45S_50ntFlanks</td><td>9574</td><td>9597</td><td>24</td><td>AATCGGTCGTCCGACCTGGGTATA</td><td>7953.7</td><td>-7.5</td><td>1.3</td><td>-5.8</td><td>8.3E-09</td><td>2.2E-08</td><td>BT20/MB468</td><td>MITO</td></tr>
<tr><td>>hg38_MT_+_1621_3279_RNR2_16S_50ntFlanks@497.518.22|TTTGGACACTAGGAAAAAACCT</td><td>>hg38_MT_+_1621_3279_RNR2_16S_50ntFlanks</td><td>497</td><td>518</td><td>22</td><td>TTTGGACACTAGGAAAAAACCT</td><td>98.7</td><td>10.4</td><td>1.7</td><td>6.0</td><td>1.5E-09</td><td>2.2E-08</td><td>BT20/MB468</td><td>MP</td></tr>
<tr><td>>hg38_21_+_8433172_8446622_RNA45SN1_45S_50ntFlanks@9759.9786.28|CAAACTTTAAATGGGTAAGAAGCCCGGC</td><td>>hg38_21_+_8433172_8446622_RNA45SN1_45S_50ntFlanks</td><td>9759</td><td>9786</td><td>28</td><td>CAAACTTTAAATGGGTAAGAAGCCCGGC</td><td>386.9</td><td>-11.7</td><td>1.9</td><td>-6.0</td><td>1.5E-09</td><td>2.2E-08</td><td>BT20/MB468</td><td>MP</td></tr>
<tr><td>>hg38_21_+_8433172_8446622_RNA45SN1_45S_50ntFlanks@11234.11252.19|CCTTCCCGTGGATCGCCCC</td><td>>hg38_21_+_8433172_8446622_RNA45SN1_45S_50ntFlanks</td><td>11234</td><td>11252</td><td>19</td><td>CCTTCCCGTGGATCGCCCC</td><td>187.8</td><td>-7.3</td><td>1.3</td><td>-5.7</td><td>1.1E-08</td><td>2.2E-08</td><td>MB231/MB468</td><td>NUC</td></tr>
<tr><td>>trna143_LysTTT_6_-_27559593_27559665@13.30.18,trna149_LysTTT_6_-_27302769_27302841@13.30.18,trna14_LysTTT_11_-_59327808_59327880@13.30.18,trna2_LysTTT_17_+_8022473_8022545@13.30.18,trna54_LysTTT_1_+_204475655_204475727@13.30.18,trna5_LysTTT_11_+_59323902_59323974@13.30.18,trna62_LysTTT_1_-_204476158_204476230@13.30.18,trna76_LysTTT_6_+_28918806_28918878@13.30.18|CAGTCGGTAGAGCATCAG</td><td>>trna143_LysTTT_6_-_27559593_27559665</td><td>13</td><td>30</td><td>18</td><td>CAGTCGGTAGAGCATCAG</td><td>823.4</td><td>-4.5</td><td>0.8</td><td>-5.7</td><td>9.3E-09</td><td>2.2E-08</td><td>BT20/MB468</td><td>CYTO</td></tr>
</table>

<table>
<tr><th></th><th></th><th></th><th></th><th></th><th></th><th></th><th></th><th></th><th></th><th></th><th></th><th></th><th></th></tr>
<tr><td>>hg38_21_+_8433172_8446622_RNA45SN1_45S_50ntFlanks@475.505.31|CTTCGTGATCGATGTGGTGACGTCGTGCTCT</td><td>>hg38_21_+_8433172_8446622_RNA45SN1_45S_50ntFlanks</td><td>475</td><td>505</td><td>31</td><td>CTTCGTGATCGATGTGGTGACGTCGTGCTCT</td><td>72499.3</td><td>4.3</td><td>0.8</td><td>5.7</td><td>1.1E-08</td><td>2.2E-08</td><td>MB231/MB468</td><td>NUC</td></tr>
<tr><td>>hg38_21_+_8433172_8446622_RNA45SN1_45S_50ntFlanks@9563.9595.33|CCTGACGTGCAAATCGGTCGTCCGACCTGGGTA</td><td>>hg38_21_+_8433172_8446622_RNA45SN1_45S_50ntFlanks</td><td>9563</td><td>9595</td><td>33</td><td>CCTGACGTGCAAATCGGTCGTCCGACCTGGGTA</td><td>654.2</td><td>-6.1</td><td>1.1</td><td>-5.7</td><td>1.2E-08</td><td>2.2E-08</td><td>MB231/MB468</td><td>MITO</td></tr>
<tr><td>>hg38_21_+_8433172_8446622_RNA45SN1_45S_50ntFlanks@3900.3936.37|CTTCGCGGGGGGGATGCGTGCATTTATCAGATCAAAA</td><td>>hg38_21_+_8433172_8446622_RNA45SN1_45S_50ntFlanks</td><td>3900</td><td>3936</td><td>37</td><td>CTTCGCGGGGGGGATGCGTGCATTTATCAGATCAAAA</td><td>104.4</td><td>-10.6</td><td>1.8</td><td>-5.7</td><td>1.0E-08</td><td>2.2E-08</td><td>MB231/MB468</td><td>MP</td></tr>
<tr><td>>hg38_21_+_8433172_8446622_RNA45SN1_45S_50ntFlanks@12511.12546.36|CGTGAGACAGGTTAGTTTTACCCTACTGATGATGTG</td><td>>hg38_21_+_8433172_8446622_RNA45SN1_45S_50ntFlanks</td><td>12511</td><td>12546</td><td>36</td><td>CGTGAGACAGGTTAGTTTTACCCTACTGATGATGTG</td><td>51.9</td><td>-9.6</td><td>1.7</td><td>-5.7</td><td>1.0E-08</td><td>2.2E-08</td><td>MB231/MB468</td><td>MP</td></tr>
<tr><td>>hg38_21_+_8433172_8446622_RNA45SN1_45S_50ntFlanks@11667.11691.25|CTGCCCAGTGCTCTGAATGTCAAAG</td><td>>hg38_21_+_8433172_8446622_RNA45SN1_45S_50ntFlanks</td><td>11667</td><td>11691</td><td>25</td><td>CTGCCCAGTGCTCTGAATGTCAAAG</td><td>841.8</td><td>5.1</td><td>0.9</td><td>5.8</td><td>8.4E-09</td><td>2.2E-08</td><td>BT20/MB468</td><td>MITO</td></tr>
<tr><td>>MI0000266|hsa-mir-10a&WithFlank&17|-|48579832|48579953@29.49.21&[MIMAT0000253&hsa-miR-10a-5p&offsets|+1|-1;m-18&17|-|48579905|48579926&offsets|+1|0]|ACCCTGTAGATCCGAATTTGT</td><td>>MI0000266|hsa-mir-10a&WithFlank&17|-|48579832|48579953</td><td>29</td><td>49</td><td>21</td><td>ACCCTGTAGATCCGAATTTGT</td><td>790.8</td><td>7.9</td><td>1.4</td><td>5.7</td><td>1.0E-08</td><td>2.2E-08</td><td>MB231/MB468</td><td>MP</td></tr>
<tr><td>>hg38_21_+_8433172_8446622_RNA45SN1_45S_50ntFlanks@10393.10432.40|CAGCAGTTGAACATGGGTCAGTCGGTCCTGAGAGATGGGC</td><td>>hg38_21_+_8433172_8446622_RNA45SN1_45S_50ntFlanks</td><td>10393</td><td>10432</td><td>40</td><td>CAGCAGTTGAACATGGGTCAGTCGGTCCTGAGAGATGGGC</td><td>83.2</td><td>-10.2</td><td>1.8</td><td>-5.7</td><td>1.0E-08</td><td>2.2E-08</td><td>MB231/MB468</td><td>MP</td></tr>
<tr><td>>hg38_21_+_8433172_8446622_RNA45SN1_45S_50ntFlanks@11923.11948.26|AGAGGTGTAGAATAAGTGGGAGGCCC</td><td>>hg38_21_+_8433172_8446622_RNA45SN1_45S_50ntFlanks</td><td>11923</td><td>11948</td><td>26</td><td>AGAGGTGTAGAATAAGTGGGAGGCCC</td><td>438.7</td><td>12.1</td><td>1.9</td><td>6.3</td><td>2.9E-10</td><td>2.2E-08</td><td>BT20/MB231</td><td>MP</td></tr>
<tr><td>>hg38_21_+_8433172_8446622_RNA45SN1_45S_50ntFlanks@11759.11777.19|CCTCGTCATCTAATTAGTG</td><td>>hg38_21_+_8433172_8446622_RNA45SN1_45S_50ntFlanks</td><td>11759</td><td>11777</td><td>19</td><td>CCTCGTCATCTAATTAGTG</td><td>105.5</td><td>-9.8</td><td>1.6</td><td>-6.0</td><td>1.5E-09</td><td>2.2E-08</td><td>BT20/MB468</td><td>MP</td></tr>
<tr><td>>hg38_21_+_8433172_8446622_RNA45SN1_45S_50ntFlanks@3813.3853.41|TTATGGTTCCTTTGGTCGCTCGCTCCTCTCCTACTTGGATA</td><td>>hg38_21_+_8433172_8446622_RNA45SN1_45S_50ntFlanks</td><td>3813</td><td>3853</td><td>41</td><td>TTATGGTTCCTTTGGTCGCTCGCTCCTCTCCTACTTGGATA</td><td>224.4</td><td>-8.0</td><td>1.4</td><td>-5.7</td><td>1.1E-08</td><td>2.2E-08</td><td>MB231/MB468</td><td>NUC</td></tr>
<tr><td>>hg38_21_+_8433172_8446622_RNA45SN1_45S_50ntFlanks@10787.10818.32|ACAGCCTCTGGCATGTTGGAACAATGTAGGTA</td><td>>hg38_21_+_8433172_8446622_RNA45SN1_45S_50ntFlanks</td><td>10787</td><td>10818</td><td>32</td><td>ACAGCCTCTGGCATGTTGGAACAATGTAGGTA</td><td>1542.2</td><td>-4.4</td><td>0.7</td><td>-6.0</td><td>1.5E-09</td><td>2.2E-08</td><td>BT20/MB468</td><td>MP</td></tr>
<tr><td>>hg38_21_+_8433172_8446622_RNA45SN1_45S_50ntFlanks@11318.11344.27|CCCCACGTCTCGTCGCGCGCGCGTCCG</td><td>>hg38_21_+_8433172_8446622_RNA45SN1_45S_50ntFlanks</td><td>11318</td><td>11344</td><td>27</td><td>CCCCACGTCTCGTCGCGCGCGCGTCCG</td><td>6211.7</td><td>-8.2</td><td>1.4</td><td>-5.7</td><td>1.1E-08</td><td>2.2E-08</td><td>MB231/MB468</td><td>CYTO</td></tr>
<tr><td>>hg38_21_+_8433172_8446622_RNA45SN1_45S_50ntFlanks@4568.4589.22|AATGGAATAGGACCGCGGTTCT</td><td>>hg38_21_+_8433172_8446622_RNA45SN1_45S_50ntFlanks</td><td>4568</td><td>4589</td><td>22</td><td>AATGGAATAGGACCGCGGTTCT</td><td>577.1</td><td>4.7</td><td>0.8</td><td>5.7</td><td>1.1E-08</td><td>2.2E-08</td><td>MB231/MB468</td><td>NUC</td></tr>
<tr><td>>MI0000103|hsa-mir-101-1&WithFlank&1|-|65058428|65058514@53.74.22(+1U);MI0000739|hsa-mir-101-2&WithFlank&9|+|4850291|4850381@55.76.22(+1U)&[MIMAT0000099&hsa-miR-101-3p&offsets|0|+1(+1U);m-17&1|-|65058443|65058462&offsets|0|+2(+1U)];[MIMAT0000099_1&hsa-miR-101-3p&offsets|0|+1(+1U);m-16&9|+|4850345|4850364&offsets|0|+2(+1U)]|TACAGTACTGTGATAACTGAAGT</td><td>>MI0000103|hsa-mir-101-1&WithFlank&1|-|65058428|65058514</td><td>53</td><td>74</td><td>22(+1U)</td><td>TACAGTACTGTGATAACTGAAGT</td><td>542.7</td><td>6.8</td><td>1.2</td><td>5.7</td><td>1.1E-08</td><td>2.2E-08</td><td>MB231/MB468</td><td>NUC</td></tr>
<tr><td>>hg38_21_+_8433172_8446622_RNA45SN1_45S_50ntFlanks@12957.12991.35|CTTCTGGGTCGGGGTTTCGTACGTAGCAGAGCAGC</td><td>>hg38_21_+_8433172_8446622_RNA45SN1_45S_50ntFlanks</td><td>12957</td><td>12991</td><td>35</td><td>CTTCTGGGTCGGGGTTTCGTACGTAGCAGAGCAGC</td><td>58.7</td><td>-9.7</td><td>1.7</td><td>-5.7</td><td>1.0E-08</td><td>2.2E-08</td><td>MB231/MB468</td><td>MP</td></tr>
<tr><td>>hg38_21_+_8433172_8446622_RNA45SN1_45S_50ntFlanks@11821.11850.30|CTATCCAGCGAAACCACAGCCAAGGGAACG</td><td>>hg38_21_+_8433172_8446622_RNA45SN1_45S_50ntFlanks</td><td>11821</td><td>11850</td><td>30</td><td>CTATCCAGCGAAACCACAGCCAAGGGAACG</td><td>66.5</td><td>-9.9</td><td>1.7</td><td>-5.7</td><td>1.0E-08</td><td>2.2E-08</td><td>MB231/MB468</td><td>MP</td></tr>
<tr><td>>am_trnaMT_ArgTCG_MT_+_10405_10469@31.48.18|TCGACTCATTAAATTATG</td><td>>am_trnaMT_ArgTCG_MT_+_10405_10469</td><td>31</td><td>48</td><td>18</td><td>TCGACTCATTAAATTATG</td><td>133.6</td><td>-10.1</td><td>1.7</td><td>-6.0</td><td>1.6E-09</td><td>2.2E-08</td><td>BT20/MB468</td><td>MP</td></tr>
<tr><td>>hg38_21_+_8433172_8446622_RNA45SN1_45S_50ntFlanks@10392.10433.42|ACAGCAGTTGAACATGGGTCAGTCGGTCCTGAGAGATGGGCG</td><td>>hg38_21_+_8433172_8446622_RNA45SN1_45S_50ntFlanks</td><td>10392</td><td>10433</td><td>42</td><td>ACAGCAGTTGAACATGGGTCAGTCGGTCCTGAGAGATGGGCG</td><td>909.0</td><td>-6.7</td><td>1.2</td><td>-5.7</td><td>1.2E-08</td><td>2.3E-08</td><td>MB231/MB468</td><td>MITO</td></tr>
<tr><td>>hg38_21_+_8433172_8446622_RNA45SN1_45S_50ntFlanks@10186.10208.23|TCCTCCCCTCCTCCCGCCCACGC</td><td>>hg38_21_+_8433172_8446622_RNA45SN1_45S_50ntFlanks</td><td>10186</td><td>10208</td><td>23</td><td>TCCTCCCCTCCTCCCGCCCACGC</td><td>238.5</td><td>-6.1</td><td>1.1</td><td>-5.7</td><td>1.1E-08</td><td>2.3E-08</td><td>MB231/MB468</td><td>CYTO</td></tr>
<tr><td>>MI0000077|hsa-mir-21&WithFlank&17|+|59841260|59841343@14.34.21(+1U)&[MIMAT0000076&hsa-miR-21-5p&offsets|0|-1(+1U);m-1&17|+|59841273|59841294&offsets|0|-1(+1U)]|TAGCTTATCAGACTGATGTTGT</td><td>>MI0000077|hsa-mir-21&WithFlank&17|+|59841260|59841343</td><td>14</td><td>34</td><td>21(+1U)</td><td>TAGCTTATCAGACTGATGTTGT</td><td>2961.9</td><td>7.1</td><td>1.2</td><td>5.7</td><td>1.2E-08</td><td>2.3E-08</td><td>MB231/MB468</td><td>MITO</td></tr>
<tr><td>>hg38_21_+_8433172_8446622_RNA45SN1_45S_50ntFlanks@10788.10818.31|CAGCCTCTGGCATGTTGGAACAATGTAGGTA</td><td>>hg38_21_+_8433172_8446622_RNA45SN1_45S_50ntFlanks</td><td>10788</td><td>10818</td><td>31</td><td>CAGCCTCTGGCATGTTGGAACAATGTAGGTA</td><td>2501.3</td><td>-4.6</td><td>0.8</td><td>-6.0</td><td>1.6E-09</td><td>2.3E-08</td><td>BT20/MB468</td><td>MP</td></tr>
<tr><td>>hg38_21_+_8433172_8446622_RNA45SN1_45S_50ntFlanks@11104.11132.29|AAGGGTCGGGGCGGCAGGGGCCGGCGGCG</td><td>>hg38_21_+_8433172_8446622_RNA45SN1_45S_50ntFlanks</td><td>11104</td><td>11132</td><td>29</td><td>AAGGGTCGGGGCGGCAGGGGCCGGCGGCG</td><td>163.2</td><td>-10.4</td><td>1.7</td><td>-6.0</td><td>1.6E-09</td><td>2.3E-08</td><td>BT20/MB468</td><td>MP</td></tr>
<tr><td>>trna10_AspGTC_12_-_125424193_125424264@36.73.38,trna12_AspGTC_12_-_125411891_125411962@36.73.38,trna144_AspGTC_6_-_27551236_27551307@36.73.38,trna38_AspGTC_17_-_8125556_8125627@36.73.38,trna45_AspGTC_6_+_27447453_27447524@36.73.38,trna48_AspGTC_6_+_27471523_27471594@36.73.38,trna4_AspGTC_12_+_96429799_96429870@36.73.38,trna69_AspGTC_1_-_161440205_161440276@36.73.38,trna72_AspGTC_1_-_161432824_161432895@36.73.38,trna75_AspGTC_1_-_161425414_161425485@36.73.38,trna78_AspGTC_1_-_161418033_161418104@36.73.38,trna81_AspGTC_1_-_161410615_161410686@36.73.38|CACGCGGGAGACCGGGGTTCGATTCCCCGACGGGGAGC</td><td>>trna10_AspGTC_12_-_125424193_125424264</td><td>36</td><td>73</td><td>38</td><td>CACGCGGGAGACCGGGGTTCGATTCCCCGACGGGGAGC</td><td>698.3</td><td>4.8</td><td>0.8</td><td>5.7</td><td>9.4E-09</td><td>2.3E-08</td><td>BT20/MB468</td><td>CYTO</td></tr>
<tr><td>>MI0000076|hsa-mir-20a&WithFlank&13|+|91351059|91351141@14.37.24&[MIMAT0000075&hsa-miR-20a-5p&offsets|0|+1;m-96&13|+|91351072|91351094&offsets|0|+1]|TAAAGTGCTTATAGTGCAGGTAGT</td><td>>MI0000076|hsa-mir-20a&WithFlank&13|+|91351059|91351141</td><td>14</td><td>37</td><td>24</td><td>TAAAGTGCTTATAGTGCAGGTAGT</td><td>2934.5</td><td>6.3</td><td>1.1</td><td>5.7</td><td>1.2E-08</td><td>2.3E-08</td><td>MB231/MB468</td><td>MITO</td></tr>
<tr><td>>hg38_21_+_8433172_8446622_RNA45SN1_45S_50ntFlanks@12420.12458.39|ATGTCGGCTCTTCCTATCATTGTGAAGCAGAATTCACCA</td><td>>hg38_21_+_8433172_8446622_RNA45SN1_45S_50ntFlanks</td><td>12420</td><td>12458</td><td>39</td><td>ATGTCGGCTCTTCCTATCATTGTGAAGCAGAATTCACCA</td><td>1260.5</td><td>3.8</td><td>0.7</td><td>5.7</td><td>1.1E-08</td><td>2.3E-08</td><td>MB231/MB468</td><td>CYTO</td></tr>
<tr><td>>hg38_21_+_8433172_8446622_RNA45SN1_45S_50ntFlanks@10458.10497.40|ATGGCCTCCGTTGCCCTCGGCCGATCGAAAGGGAGTCGGG</td><td>>hg38_21_+_8433172_8446622_RNA45SN1_45S_50ntFlanks</td><td>10458</td><td>10497</td><td>40</td><td>ATGGCCTCCGTTGCCCTCGGCCGATCGAAAGGGAGTCGGG</td><td>442.8</td><td>-7.2</td><td>1.3</td><td>-5.7</td><td>9.4E-09</td><td>2.3E-08</td><td>BT20/MB468</td><td>CYTO</td></tr>
<tr><td>>hg38_21_+_8433172_8446622_RNA45SN1_45S_50ntFlanks@4007.4024.18|CCTCGGGCCGATCGCACG</td><td>>hg38_21_+_8433172_8446622_RNA45SN1_45S_50ntFlanks</td><td>4007</td><td>4024</td><td>18</td><td>CCTCGGGCCGATCGCACG</td><td>769.3</td><td>-3.4</td><td>0.6</td><td>-5.7</td><td>9.5E-09</td><td>2.3E-08</td><td>BT20/MB468</td><td>CYTO</td></tr>
<tr><td>>hg38_21_+_8433172_8446622_RNA45SN1_45S_50ntFlanks@6698.6717.20|AGCTAGCTGCGAGAATTAAT</td><td>>hg38_21_+_8433172_8446622_RNA45SN1_45S_50ntFlanks</td><td>6698</td><td>6717</td><td>20</td><td>AGCTAGCTGCGAGAATTAAT</td><td>439.0</td><td>-4.4</td><td>0.8</td><td>-5.7</td><td>1.1E-08</td><td>2.3E-08</td><td>MB231/MB468</td><td>NUC</td></tr>
<tr><td>>hg38_21_+_8433172_8446622_RNA45SN1_45S_50ntFlanks@7983.8003.21|CAGATCAGACGTGGCGACCCG</td><td>>hg38_21_+_8433172_8446622_RNA45SN1_45S_50ntFlanks</td><td>7983</td><td>8003</td><td>21</td><td>CAGATCAGACGTGGCGACCCG</td><td>767.0</td><td>-7.9</td><td>1.4</td><td>-5.7</td><td>1.1E-08</td><td>2.3E-08</td><td>MB231/MB468</td><td>NUC</td></tr>
<tr><td>>MI0000760|hsa-mir-361&WithFlank&X|-|85903630|85903713@12.33.22&[MIMAT0000703&hsa-miR-361-5p&offsets|0|0;m-89&X|-|85903681|85903702&offsets|0|0]|TTATCAGAATCTCCAGGGGTAC</td><td>>MI0000760|hsa-mir-361&WithFlank&X|-|85903630|85903713</td><td>12</td><td>33</td><td>22</td><td>TTATCAGAATCTCCAGGGGTAC</td><td>377.3</td><td>6.1</td><td>1.1</td><td>5.7</td><td>1.1E-08</td><td>2.3E-08</td><td>MB231/MB468</td><td>CYTO</td></tr>
<tr><td>>hg38_21_+_8433172_8446622_RNA45SN1_45S_50ntFlanks@12407.12443.37|TTTTGATCCTTCGATGTCGGCTCTTCCTATCATTGTG</td><td>>hg38_21_+_8433172_8446622_RNA45SN1_45S_50ntFlanks</td><td>12407</td><td>12443</td><td>37</td><td>TTTTGATCCTTCGATGTCGGCTCTTCCTATCATTGTG</td><td>5485.6</td><td>-4.0</td><td>0.7</td><td>-5.7</td><td>1.1E-08</td><td>2.3E-08</td><td>MB231/MB468</td><td>NUC</td></tr>
<tr><td>>hg38_21_+_8433172_8446622_RNA45SN1_45S_50ntFlanks@10771.10818.48|CAGGTCTCCAAGGTGAACAGCCTCTGGCATGTTGGAACAATGTAGGTA</td><td>>hg38_21_+_8433172_8446622_RNA45SN1_45S_50ntFlanks</td><td>10771</td><td>10818</td><td>48</td><td>CAGGTCTCCAAGGTGAACAGCCTCTGGCATGTTGGAACAATGTAGGTA</td><td>1012.1</td><td>-8.4</td><td>1.5</td><td>-5.7</td><td>1.2E-08</td><td>2.3E-08</td><td>MB231/MB468</td><td>MITO</td></tr>
<tr><td>>MI0000433|hsa-let-7g&WithFlank&3|-|52268272|52268367@11.31.21(+1C)&[MIMAT0000414&hsa-let-7g-5p&offsets|0|-1(+1C);m-58&3|-|52268336|52268357&offsets|0|-1(+1C)]|TGAGGTAGTAGTTTGTACAGTC</td><td>>MI0000433|hsa-let-7g&WithFlank&3|-|52268272|52268367</td><td>11</td><td>31</td><td>21(+1C)</td><td>TGAGGTAGTAGTTTGTACAGTC</td><td>4548.1</td><td>5.0</td><td>0.9</td><td>5.7</td><td>1.2E-08</td><td>2.3E-08</td><td>MB231/MB468</td><td>MITO</td></tr>
<tr><td>>hg38_21_+_8433172_8446622_RNA45SN1_45S_50ntFlanks@4003.4042.40|ATAACCTCGGGCCGATCGCACGCCCCCCGTGGCGGCGACG</td><td>>hg38_21_+_8433172_8446622_RNA45SN1_45S_50ntFlanks</td><td>4003</td><td>4042</td><td>40</td><td>ATAACCTCGGGCCGATCGCACGCCCCCCGTGGCGGCGACG</td><td>730.8</td><td>-5.6</td><td>1.0</td><td>-5.8</td><td>8.6E-09</td><td>2.3E-08</td><td>BT20/MB468</td><td>MITO</td></tr>
<tr><td>>hg38_21_+_8433172_8446622_RNA45SN1_45S_50ntFlanks@11577.11601.25|TTAGAACTGGTGCGGACCAGGGGAA</td><td>>hg38_21_+_8433172_8446622_RNA45SN1_45S_50ntFlanks</td><td>11577</td><td>11601</td><td>25</td><td>TTAGAACTGGTGCGGACCAGGGGAA</td><td>143.7</td><td>-10.2</td><td>1.7</td><td>-6.0</td><td>1.6E-09</td><td>2.3E-08</td><td>BT20/MB468</td><td>MP</td></tr>
<tr><td>>hg38_21_+_8433172_8446622_RNA45SN1_45S_50ntFlanks@5488.5514.27|GCTGAGAAGACGGTCGAACTTGACTAT</td><td>>hg38_21_+_8433172_8446622_RNA45SN1_45S_50ntFlanks</td><td>5488</td><td>5514</td><td>27</td><td>GCTGAGAAGACGGTCGAACTTGACTAT</td><td>213.9</td><td>-5.9</td><td>1.0</td><td>-5.7</td><td>1.1E-08</td><td>2.3E-08</td><td>MB231/MB468</td><td>MP</td></tr>
<tr><td>>hg38_21_+_8433172_8446622_RNA45SN1_45S_50ntFlanks@10511.10532.22|ATCCGGAGTGGCGGAGATGGGC</td><td>>hg38_21_+_8433172_8446622_RNA45SN1_45S_50ntFlanks</td><td>10511</td><td>10532</td><td>22</td><td>ATCCGGAGTGGCGGAGATGGGC</td><td>149.3</td><td>-10.3</td><td>1.7</td><td>-6.0</td><td>1.6E-09</td><td>2.3E-08</td><td>BT20/MB468</td><td>MP</td></tr>
<tr><td>>hg38_21_+_8433172_8446622_RNA45SN1_45S_50ntFlanks@10351.10379.29|CGAGAACTTTGAAGGCCGAAGTGGAGAAG</td><td>>hg38_21_+_8433172_8446622_RNA45SN1_45S_50ntFlanks</td><td>10351</td><td>10379</td><td>29</td><td>CGAGAACTTTGAAGGCCGAAGTGGAGAAG</td><td>1244.4</td><td>-6.2</td><td>1.1</td><td>-5.7</td><td>1.2E-08</td><td>2.3E-08</td><td>MB231/MB468</td><td>MITO</td></tr>
<tr><td>>MI0000455|hsa-mir-138-2&WithFlank&16|+|56858512|56858607@16.37.22(+1A);MI0000476|hsa-mir-138-1&WithFlank&3|+|44114206|44114316@29.50.22(+1A)&[MIMAT0000430&hsa-miR-138-5p&offsets|0|-1(+1A);m-509&16|+|56858527|56858549&offsets|0|-1(+1A)];[MIMAT0000430_1&hsa-miR-138-5p&offsets|0|-1(+1A);m-497&3|+|44114234|44114256&offsets|0|-1(+1A)]|AGCTGGTGTTGTGAATCAGGCCA</td><td>>MI0000455|hsa-mir-138-2&WithFlank&16|+|56858512|56858607</td><td>16</td><td>37</td><td>22(+1A)</td><td>AGCTGGTGTTGTGAATCAGGCCA</td><td>65.5</td><td>9.3</td><td>1.6</td><td>5.7</td><td>1.1E-08</td><td>2.3E-08</td><td>MB231/MB468</td><td>MP</td></tr>
<tr><td>>hg38_21_+_8433172_8446622_RNA45SN1_45S_50ntFlanks@11212.11250.39|CGACTCTGGACGCGAGCCGGGCCCTTCCCGTGGATCGCC</td><td>>hg38_21_+_8433172_8446622_RNA45SN1_45S_50ntFlanks</td><td>11212</td><td>11250</td><td>39</td><td>CGACTCTGGACGCGAGCCGGGCCCTTCCCGTGGATCGCC</td><td>435.1</td><td>-5.0</td><td>0.9</td><td>-5.7</td><td>1.2E-08</td><td>2.3E-08</td><td>MB231/MB468</td><td>MITO</td></tr>
<tr><td>>hg38_21_+_8433172_8446622_RNA45SN1_45S_50ntFlanks@10679.10705.27|GGTTCCGGCGGCGTCCGGTGAGCTCTC</td><td>>hg38_21_+_8433172_8446622_RNA45SN1_45S_50ntFlanks</td><td>10679</td><td>10705</td><td>27</td><td>GGTTCCGGCGGCGTCCGGTGAGCTCTC</td><td>471.5</td><td>-4.4</td><td>0.8</td><td>-5.7</td><td>9.6E-09</td><td>2.3E-08</td><td>BT20/MB468</td><td>CYTO</td></tr>
<tr><td>>hg38_21_+_8433172_8446622_RNA45SN1_45S_50ntFlanks@10794.10826.33|CTGGCATGTTGGAACAATGTAGGTAAGGGAAGT</td><td>>hg38_21_+_8433172_8446622_RNA45SN1_45S_50ntFlanks</td><td>10794</td><td>10826</td><td>33</td><td>CTGGCATGTTGGAACAATGTAGGTAAGGGAAGT</td><td>312.1</td><td>-5.8</td><td>1.0</td><td>-5.7</td><td>1.1E-08</td><td>2.3E-08</td><td>MB231/MB468</td><td>NUC</td></tr>
<tr><td>>hg38_21_+_8433172_8446622_RNA45SN1_45S_50ntFlanks@5059.5098.40|CGATTTGTCTGGTTAATTCCGATAACGAACGAGACTCTGG</td><td>>hg38_21_+_8433172_8446622_RNA45SN1_45S_50ntFlanks</td><td>5059</td><td>5098</td><td>40</td><td>CGATTTGTCTGGTTAATTCCGATAACGAACGAGACTCTGG</td><td>528.7</td><td>-6.7</td><td>1.2</td><td>-5.7</td><td>1.2E-08</td><td>2.3E-08</td><td>MB231/MB468</td><td>MITO</td></tr>
</table>

| | | | | | | | | | | | | | |
|---|---|---|---|---|---|---|---|---|---|---|---|---|---|
| >MI0000478\|hsa-mir-149&WithFlank&2\|+\|240455995\|240456095@21.41.21(+1A)&[MIMAT0000450&hsa-miR-149-5p&offsets\|0\|-2(+1A);m-195&2\|+\|240456015\|240456037&offsets\|0\|-2(+1A)]\|TCTGGCTCCGTGTCTTCACTCA | >MI0000478\|hsa-mir-149&WithFlank&2\|+\|240455995\|240456095 | 21 | 41 | 21(+1A) | TCTGGCTCCGTGTCTTCACTCA | 108.2 | 10.5 | 1.7 | 6.0 | 1.6E-09 | 2.3E-08 | BT20/MB468 | MP |
| >hg38_21_+_8433172_8446622_RNA45SN1_45S_50ntFlanks@4297.4319.23\|CATTGGAGGGCAAGTCTGGTGCC | >hg38_21_+_8433172_8446622_RNA45SN1_45S_50ntFlanks | 4297 | 4319 | 23 | CATTGGAGGGCAAGTCTGGTGCC | 243.7 | -3.3 | 0.6 | -5.7 | 9.6E-09 | 2.3E-08 | BT20/MB468 | CYTO |
| >hg38_21_+_8433172_8446622_RNA45SN1_45S_50ntFlanks@10869.10891.23\|AGGGCTGGGTCGGTCGGGCTGGG | >hg38_21_+_8433172_8446622_RNA45SN1_45S_50ntFlanks | 10869 | 10891 | 23 | AGGGCTGGGTCGGTCGGGCTGGG | 1394.3 | -7.3 | 1.3 | -5.7 | 1.3E-08 | 2.3E-08 | BT20/MB468 | NUC |
| >hg38_21_+_8433172_8446622_RNA45SN1_45S_50ntFlanks@6747.6770.24\|ACTTCGAACGCACTTGCGGCCCCG | >hg38_21_+_8433172_8446622_RNA45SN1_45S_50ntFlanks | 6747 | 6770 | 24 | ACTTCGAACGCACTTGCGGCCCCG | 598.5 | -5.7 | 1.0 | -5.8 | 8.7E-09 | 2.3E-08 | BT20/MB468 | MITO |
| >hg38_21_+_8433172_8446622_RNA45SN1_45S_50ntFlanks@4380.4423.44\|CGTAGTTGGATCTTGGGAGCGGGCGGGCGGTCCGCCGCGAGGCG | >hg38_21_+_8433172_8446622_RNA45SN1_45S_50ntFlanks | 4380 | 4423 | 44 | CGTAGTTGGATCTTGGGAGCGGGCGGGCGGTCCGCCGCGAGGCG | 307.2 | -6.3 | 1.1 | -5.7 | 1.2E-08 | 2.3E-08 | MB231/MB468 | NUC |
| >hg38_21_+_8433172_8446622_RNA45SN1_45S_50ntFlanks@4729.4758.30\|TCAAGAACGAAAGTCGGAGGTTCGAAGACG | >hg38_21_+_8433172_8446622_RNA45SN1_45S_50ntFlanks | 4729 | 4758 | 30 | TCAAGAACGAAAGTCGGAGGTTCGAAGACG | 1387.2 | -4.0 | 0.7 | -5.8 | 8.7E-09 | 2.3E-08 | BT20/MB468 | MITO |
| >hg38_21_+_8433172_8446622_RNA45SN1_45S_50ntFlanks@11768.11794.27\|CTAATTAGTGACGCGCATGAATGGATG | >hg38_21_+_8433172_8446622_RNA45SN1_45S_50ntFlanks | 11768 | 11794 | 27 | CTAATTAGTGACGCGCATGAATGGATG | 2039.6 | -4.5 | 0.7 | -6.0 | 1.6E-09 | 2.3E-08 | BT20/MB468 | MP |
| >hg38_21_+_8433172_8446622_RNA45SN1_45S_50ntFlanks@475.516.42\|CTTCGTGATCGATGTGGTGACGTCGTGCTCTCCCGGGCCGGG | >hg38_21_+_8433172_8446622_RNA45SN1_45S_50ntFlanks | 475 | 516 | 42 | CTTCGTGATCGATGTGGTGACGTCGTGCTCTCCCGGGCCGGG | 166.8 | 11.1 | 1.8 | 6.0 | 1.6E-09 | 2.3E-08 | BT20/MB468 | MP |
| >hg38_21_+_8433172_8446622_RNA45SN1_45S_50ntFlanks@6645.6664.20\|CTCGTACGACTCTTAGCGGT | >hg38_21_+_8433172_8446622_RNA45SN1_45S_50ntFlanks | 6645 | 6664 | 20 | CTCGTACGACTCTTAGCGGT | 2522.5 | 4.5 | 0.8 | 5.7 | 1.1E-08 | 2.3E-08 | MB231/MB468 | CYTO |
| >hg38_21_+_8433172_8446622_RNA45SN1_45S_50ntFlanks@7983.8009.27\|CAGATCAGACGTGGCGACCCGCTGAAT | >hg38_21_+_8433172_8446622_RNA45SN1_45S_50ntFlanks | 7983 | 8009 | 27 | CAGATCAGACGTGGCGACCCGCTGAAT | 1488.2 | -4.3 | 0.8 | -5.7 | 1.2E-08 | 2.3E-08 | MB231/MB468 | MITO |
| >hg38_21_+_8433172_8446622_RNA45SN1_45S_50ntFlanks@12393.12419.27\|ATAGCGACGTCGCTTTTTGATCCTTCG | >hg38_21_+_8433172_8446622_RNA45SN1_45S_50ntFlanks | 12393 | 12419 | 27 | ATAGCGACGTCGCTTTTTGATCCTTCG | 56.6 | -9.7 | 1.7 | -5.7 | 1.1E-08 | 2.3E-08 | MB231/MB468 | MP |
| >hg38_21_+_8433172_8446622_RNA45SN1_45S_50ntFlanks@8379.8406.28\|TCAAGAGGGCGTGAAACCGTTAAGAGGT | >hg38_21_+_8433172_8446622_RNA45SN1_45S_50ntFlanks | 8379 | 8406 | 28 | TCAAGAGGGCGTGAAACCGTTAAGAGGT | 190.9 | -4.0 | 0.7 | -5.7 | 1.1E-08 | 2.3E-08 | MB231/MB468 | MP |
| >hg38_21_+_8433172_8446622_RNA45SN1_45S_50ntFlanks@10846.10863.18\|CTTCGGGATAAGGATTGG | >hg38_21_+_8433172_8446622_RNA45SN1_45S_50ntFlanks | 10846 | 10863 | 18 | CTTCGGGATAAGGATTGG | 319.6 | -6.1 | 1.1 | -5.7 | 1.2E-08 | 2.3E-08 | MB231/MB468 | NUC |
| >hg38_21_+_8433172_8446622_RNA45SN1_45S_50ntFlanks@10337.10373.37\|AGCAAATATTCAAACGAGAACTTTGAAGGCCGAAGTG | >hg38_21_+_8433172_8446622_RNA45SN1_45S_50ntFlanks | 10337 | 10373 | 37 | AGCAAATATTCAAACGAGAACTTTGAAGGCCGAAGTG | 97.0 | 10.3 | 1.7 | 6.0 | 1.6E-09 | 2.3E-08 | BT20/MB468 | MP |
| >MI0000087\|hsa-mir-29a&WithFlank&7\|-\|130876741\|130876816@47.69.23&[MIMAT0000086&hsa-miR-29a-3p&offsets\|-1\|0;m-26&7\|-\|130876749\|130876769&offsets\|-1\|+1]\|CTAGCACCATCTGAAATCGGTTA | >MI0000087\|hsa-mir-29a&WithFlank&7\|-\|130876741\|130876816 | 47 | 69 | 23 | CTAGCACCATCTGAAATCGGTTA | 264.0 | 6.1 | 1.1 | 5.7 | 1.3E-08 | 2.3E-08 | BT20/MB468 | NUC |
| >hg38_21_+_8433172_8446622_RNA45SN1_45S_50ntFlanks@4002.4022.21\|GATAACCTCGGGCCGATCGCA | >hg38_21_+_8433172_8446622_RNA45SN1_45S_50ntFlanks | 4002 | 4022 | 21 | GATAACCTCGGGCCGATCGCA | 1267.1 | -6.8 | 1.2 | -5.7 | 1.2E-08 | 2.3E-08 | MB231/MB468 | NUC |
| >hg38_21_+_8433172_8446622_RNA45SN1_45S_50ntFlanks@4733.4761.29\|GAACGAAAGTCGGAGGTTCGAAGACGATC | >hg38_21_+_8433172_8446622_RNA45SN1_45S_50ntFlanks | 4733 | 4761 | 29 | GAACGAAAGTCGGAGGTTCGAAGACGATC | 1265.9 | 6.0 | 1.0 | 5.7 | 1.1E-08 | 2.3E-08 | MB231/MB468 | CYTO |
| >hg38_21_+_8433172_8446622_RNA45SN1_45S_50ntFlanks@10574.10607.34\|AAGCCGGCGGGAGCCCCGGGGAGAGTTCTCTTTT | >hg38_21_+_8433172_8446622_RNA45SN1_45S_50ntFlanks | 10574 | 10607 | 34 | AAGCCGGCGGGAGCCCCGGGGAGAGTTCTCTTTT | 347.2 | -7.3 | 1.3 | -5.7 | 1.2E-08 | 2.3E-08 | MB231/MB468 | NUC |
| >hg38_21_+_8433172_8446622_RNA45SN1_45S_50ntFlanks@8350.8391.42\|AAAGTTGAAAAGAACTTTGAAGAGAGAGTTCAAGAGGGCGTG | >hg38_21_+_8433172_8446622_RNA45SN1_45S_50ntFlanks | 8350 | 8391 | 42 | AAAGTTGAAAAGAACTTTGAAGAGAGAGTTCAAGAGGGCGTG | 98.4 | 10.4 | 1.7 | 6.0 | 1.6E-09 | 2.3E-08 | BT20/MB468 | MP |
| >hg38_21_+_8433172_8446622_RNA45SN1_45S_50ntFlanks@5492.5526.35\|AGAAGACGGTCGAACTTGACTATCTAGAGGAAGTA | >hg38_21_+_8433172_8446622_RNA45SN1_45S_50ntFlanks | 5492 | 5526 | 35 | AGAAGACGGTCGAACTTGACTATCTAGAGGAAGTA | 376.1 | -5.6 | 1.0 | -5.7 | 1.2E-08 | 2.3E-08 | MB231/MB468 | NUC |
| >hg38_21_+_8433172_8446622_RNA45SN1_45S_50ntFlanks@13004.13039.36\|ATCTATTGAAAGTCAGCCCTCGACACAAGGGTTTGT | >hg38_21_+_8433172_8446622_RNA45SN1_45S_50ntFlanks | 13004 | 13039 | 36 | ATCTATTGAAAGTCAGCCCTCGACACAAGGGTTTGT | 203.0 | -9.8 | 1.7 | -5.8 | 8.8E-09 | 2.3E-08 | BT20/MB468 | MITO |
| >hg38_21_+_8433172_8446622_RNA45SN1_45S_50ntFlanks@12570.12588.19\|AGTACGAGAGGAACCGCAG | >hg38_21_+_8433172_8446622_RNA45SN1_45S_50ntFlanks | 12570 | 12588 | 19 | AGTACGAGAGGAACCGCAG | 127.4 | -10.1 | 1.7 | -6.0 | 1.6E-09 | 2.3E-08 | BT20/MB468 | MP |
| >hg38_21_+_8433172_8446622_RNA45SN1_45S_50ntFlanks@12183.12201.19\|AAACGGTAACGCAGGTGTC | >hg38_21_+_8433172_8446622_RNA45SN1_45S_50ntFlanks | 12183 | 12201 | 19 | AAACGGTAACGCAGGTGTC | 71.8 | -10.0 | 1.8 | -5.7 | 1.1E-08 | 2.3E-08 | MB231/MB468 | MP |
| >MI0000746\|hsa-mir-99b&WithFlank&19\|+\|51692606\|51692687@51.73.23&[MIMAT0004678&hsa-miR-99b-3p&offsets\|0\|+1;m-151&19\|+\|51692656\|51692678&offsets\|0\|0]\|CAAGCTCGTGTCTGTGGGTCCGT | >MI0000746\|hsa-mir-99b&WithFlank&19\|+\|51692606\|51692687 | 51 | 73 | 23 | CAAGCTCGTGTCTGTGGGTCCGT | 244.3 | 2.8 | 0.5 | 5.7 | 1.3E-08 | 2.3E-08 | BT20/MB468 | NUC |
| >hg38_21_+_8433172_8446622_RNA45SN1_45S_50ntFlanks@4050.4081.32\|CGAACGTCTGCCCTATCAACTTTCGATGGTAG | >hg38_21_+_8433172_8446622_RNA45SN1_45S_50ntFlanks | 4050 | 4081 | 32 | CGAACGTCTGCCCTATCAACTTTCGATGGTAG | 369.1 | -7.8 | 1.4 | -5.7 | 1.3E-08 | 2.3E-08 | BT20/MB468 | NUC |
| >hg38_21_+_8433172_8446622_RNA45SN1_45S_50ntFlanks@12995.13039.45\|CTCGCTGCGATCTATTGAAAGTCAGCCCTCGACACAAGGGTTTGT | >hg38_21_+_8433172_8446622_RNA45SN1_45S_50ntFlanks | 12995 | 13039 | 45 | CTCGCTGCGATCTATTGAAAGTCAGCCCTCGACACAAGGGTTTGT | 288266.2 | 7.8 | 1.4 | 5.7 | 9.8E-09 | 2.4E-08 | BT20/MB468 | CYTO |
| >MI0000088\|hsa-mir-30a&WithFlank&6\|-\|71403545\|71403627@54.73.20(+1U)&[MIMAT0000088&hsa-miR-30a-3p&offsets\|+1\|-1(+1U);m-38&6\|-\|71403554\|71403575&offsets\|+1\|-1(+1U)]\|TTTCAGTCGGATGTTTGCAGT | >MI0000088\|hsa-mir-30a&WithFlank&6\|-\|71403545\|71403627 | 54 | 73 | 20(+1U) | TTTCAGTCGGATGTTTGCAGT | 2407.4 | 4.5 | 0.8 | 5.7 | 1.1E-08 | 2.4E-08 | MB231/MB468 | MP |
| >hg38_21_+_8433172_8446622_RNA45SN1_45S_50ntFlanks@10921.10946.26\|GCTGGACGAGGCGCCGCCGCCCCCCC | >hg38_21_+_8433172_8446622_RNA45SN1_45S_50ntFlanks | 10921 | 10946 | 26 | GCTGGACGAGGCGCCGCCGCCCCCCC | 370.4 | -8.3 | 1.4 | -5.7 | 1.1E-08 | 2.4E-08 | MB231/MB468 | MP |
| >hg38_21_+_8433172_8446622_RNA45SN1_45S_50ntFlanks@10846.10864.19\|CTTCGGGATAAGGATTGGC | >hg38_21_+_8433172_8446622_RNA45SN1_45S_50ntFlanks | 10846 | 10864 | 19 | CTTCGGGATAAGGATTGGC | 909.3 | -5.4 | 1.0 | -5.7 | 1.2E-08 | 2.4E-08 | MB231/MB468 | CYTO |
| >hg38_21_+_8433172_8446622_RNA45SN1_45S_50ntFlanks@10687.10709.23\|CGGCGTCCGGTGAGCTCTCGCTG | >hg38_21_+_8433172_8446622_RNA45SN1_45S_50ntFlanks | 10687 | 10709 | 23 | CGGCGTCCGGTGAGCTCTCGCTG | 2381.2 | -6.9 | 1.2 | -5.7 | 1.2E-08 | 2.4E-08 | MB231/MB468 | NUC |
| >MI0000063\|hsa-let-7b&WithFlank&22\|+\|46113680\|46113774@12.30.19&[MIMAT0000063&hsa-let-7b-5p&offsets\|0\|-3;m-14&22\|+\|46113691\|46113712&offsets\|0\|-3]\|TGAGGTAGTAGGTTGTGTG | >MI0000063\|hsa-let-7b&WithFlank&22\|+\|46113680\|46113774 | 12 | 30 | 19 | TGAGGTAGTAGGTTGTGTG | 289.9 | 7.6 | 1.3 | 5.7 | 9.9E-09 | 2.4E-08 | BT20/MB468 | CYTO |
| >hg38_21_+_8433172_8446622_RNA45SN1_45S_50ntFlanks@5077.5098.22\|CCGATAACGAACGAGACTCTGG | >hg38_21_+_8433172_8446622_RNA45SN1_45S_50ntFlanks | 5077 | 5098 | 22 | CCGATAACGAACGAGACTCTGG | 517.4 | -8.4 | 1.5 | -5.7 | 1.3E-08 | 2.4E-08 | MB231/MB468 | MITO |
| >hg38_21_+_8433172_8446622_RNA45SN1_45S_50ntFlanks@7984.8005.22\|AGATCAGACGTGGCGACCCGCT | >hg38_21_+_8433172_8446622_RNA45SN1_45S_50ntFlanks | 7984 | 8005 | 22 | AGATCAGACGTGGCGACCCGCT | 221.2 | -6.9 | 1.2 | -5.7 | 1.2E-08 | 2.4E-08 | MB231/MB468 | NUC |
| >hg38_21_+_8433172_8446622_RNA45SN1_45S_50ntFlanks@12083.12107.25\|CTCGCTTCTGGCGCCAAGCGCCCGG | >hg38_21_+_8433172_8446622_RNA45SN1_45S_50ntFlanks | 12083 | 12107 | 25 | CTCGCTTCTGGCGCCAAGCGCCCGG | 862.6 | -7.8 | 1.4 | -5.7 | 1.3E-08 | 2.4E-08 | MB231/MB468 | MITO |
| >hg38_21_+_8433172_8446622_RNA45SN1_45S_50ntFlanks@7979.8008.30\|ACCTCAGATCAGACGTGGCGACCCGCTGAA | >hg38_21_+_8433172_8446622_RNA45SN1_45S_50ntFlanks | 7979 | 8008 | 30 | ACCTCAGATCAGACGTGGCGACCCGCTGAA | 1493.2 | -3.6 | 0.6 | -5.7 | 1.2E-08 | 2.4E-08 | MB231/MB468 | NUC |
| >hg38_21_+_8433172_8446622_RNA45SN1_45S_50ntFlanks@10783.10818.36\|GTGAACAGCCTCTGGCATGTTGGAACAATGTAGGTA | >hg38_21_+_8433172_8446622_RNA45SN1_45S_50ntFlanks | 10783 | 10818 | 36 | GTGAACAGCCTCTGGCATGTTGGAACAATGTAGGTA | 962.8 | -5.6 | 0.9 | -6.0 | 1.7E-09 | 2.4E-08 | BT20/MB468 | MP |
| >hg38_21_+_8433172_8446622_RNA45SN1_45S_50ntFlanks@11210.11227.18\|GGCGACTCTGGACGCGAG | >hg38_21_+_8433172_8446622_RNA45SN1_45S_50ntFlanks | 11210 | 11227 | 18 | GGCGACTCTGGACGCGAG | 315.2 | -8.0 | 1.4 | -5.7 | 9.9E-09 | 2.4E-08 | BT20/MB468 | CYTO |
| >hg38_21_+_8433172_8446622_RNA45SN1_45S_50ntFlanks@11572.11601.30\|CCGACTTAGAACTGGTGCGGACCAGGGGAA | >hg38_21_+_8433172_8446622_RNA45SN1_45S_50ntFlanks | 11572 | 11601 | 30 | CCGACTTAGAACTGGTGCGGACCAGGGGAA | 4406.8 | -3.3 | 0.6 | -5.7 | 9.0E-09 | 2.4E-08 | BT20/MB468 | MITO |
| >am_trna128_GlyGCC_6_-_27870686_27870756@1.35.35,trna18_GlyGCC_16_+_70822597_70822667@1.35.35,trna19_GlyGCC_16_+_70823410_70823480@1.35.35,trna19_GlyGCC_2_-_157257659_157257729@1.35.35,trna24_GlyGCC_16_-_70812942_70813012@1.35.35,trna25_GlyGCC_16_-_70812114_70812184@1.35.35,trna5_GlyGCC_17_+_8029064_8029134@1.35.35,trna68_GlyGCC_1_-_161493637_161493707@1.35.35\|GCATTGGTGGTTCAGTGGTAGAATTCTCGCCTGCC | >am_trna128_GlyGCC_6_-_27870686_27870756 | 1 | 35 | 35 | GCATTGGTGGTTCAGTGGTAGAATTCTCGCCTGCC | 3507.6 | 6.0 | 1.0 | 5.7 | 1.1E-08 | 2.4E-08 | MB231/MB468 | MP |
| >hg38_21_+_8433172_8446622_RNA45SN1_45S_50ntFlanks@9477.9497.21\|AGCGCACGTGTTAGGACCCGA | >hg38_21_+_8433172_8446622_RNA45SN1_45S_50ntFlanks | 9477 | 9497 | 21 | AGCGCACGTGTTAGGACCCGA | 818.1 | 7.4 | 1.3 | 5.7 | 1.2E-08 | 2.4E-08 | MB231/MB468 | NUC |
| >hg38_21_+_8433172_8446622_RNA45SN1_45S_50ntFlanks@3833.3860.28\|CGCTCCTCTCCTACTTGGATAACTGTGG | >hg38_21_+_8433172_8446622_RNA45SN1_45S_50ntFlanks | 3833 | 3860 | 28 | CGCTCCTCTCCTACTTGGATAACTGTGG | 511.3 | -2.3 | 0.4 | -5.7 | 1.2E-08 | 2.4E-08 | MB231/MB468 | NUC |
| >hg38_21_+_8433172_8446622_RNA45SN1_45S_50ntFlanks@6698.6719.22\|AGCTAGCTGCGAGAATTAATGT | >hg38_21_+_8433172_8446622_RNA45SN1_45S_50ntFlanks | 6698 | 6719 | 22 | AGCTAGCTGCGAGAATTAATGT | 1824.1 | -4.3 | 0.8 | -5.7 | 1.2E-08 | 2.4E-08 | MB231/MB468 | NUC |
| >hg38_21_+_8433172_8446622_RNA45SN1_45S_50ntFlanks@6745.6762.18\|ACACTTCGAACGCACTTG | >hg38_21_+_8433172_8446622_RNA45SN1_45S_50ntFlanks | 6745 | 6762 | 18 | ACACTTCGAACGCACTTG | 97.4 | -8.7 | 1.5 | -5.7 | 9.0E-09 | 2.4E-08 | BT20/MB468 | MITO |
| >MI0000063\|hsa-let-7b&WithFlank&22\|+\|46113680\|46113774@12.32.21(+1C)&[MIMAT0000063&hsa-let-7b-5p&offsets\|0\|-1(+1C);m-14&22\|+\|46113691\|46113712&offsets\|0\|-1(+1C)]\|TGAGGTAGTAGGTTGTGTGGTC | >MI0000063\|hsa-let-7b&WithFlank&22\|+\|46113680\|46113774 | 12 | 32 | 21(+1C) | TGAGGTAGTAGGTTGTGTGGTC | 1220.3 | 3.4 | 0.6 | 5.7 | 1.2E-08 | 2.4E-08 | MB231/MB468 | NUC |
| >MI0000088\|hsa-mir-30a&WithFlank&6\|-\|71403545\|71403627@12.32.21&[MIMAT0000087&hsa-miR-30a-5p&offsets\|0\|-1;m-15&6\|-\|71403595\|71403616&offsets\|0\|-1]\|TGTAAACATCCTCGACTGGAA | >MI0000088\|hsa-mir-30a&WithFlank&6\|-\|71403545\|71403627 | 12 | 32 | 21 | TGTAAACATCCTCGACTGGAA | 887.2 | 5.5 | 1.0 | 5.7 | 1.3E-08 | 2.4E-08 | MB231/MB468 | MITO |
| >hg38_21_+_8433172_8446622_RNA45SN1_45S_50ntFlanks@11317.11342.26\|ACCCCACGTCTCGTCGCGCGCGCGTC | >hg38_21_+_8433172_8446622_RNA45SN1_45S_50ntFlanks | 11317 | 11342 | 26 | ACCCCACGTCTCGTCGCGCGCGCGTC | 1768.3 | -7.4 | 1.3 | -5.7 | 1.2E-08 | 2.4E-08 | MB231/MB468 | CYTO |
| >hg38_21_+_8433172_8446622_RNA45SN1_45S_50ntFlanks@10458.10488.31\|ATGGCCTCCGTTGCCCTCGGCCGATCGAAAG | >hg38_21_+_8433172_8446622_RNA45SN1_45S_50ntFlanks | 10458 | 10488 | 31 | ATGGCCTCCGTTGCCCTCGGCCGATCGAAAG | 3080.6 | -10.1 | 1.8 | -5.7 | 1.3E-08 | 2.4E-08 | BT20/MB468 | NUC |
| >trnaMT_ValTAC_MT_+_1602_1670@49.71.23\|AACTTAACTTGACCGCTCTGACC | >trnaMT_ValTAC_MT_+_1602_1670 | 49 | 71 | 23 | AACTTAACTTGACCGCTCTGACC | 890.8 | 8.7 | 1.5 | 5.7 | 1.0E-08 | 2.4E-08 | BT20/MB468 | CYTO |

<table>
<tr><th></th><th></th><th></th><th></th><th></th><th></th><th></th><th></th><th></th><th></th><th></th><th></th><th></th><th></th></tr>
<tr><td>>hg38_21_+_8433172_8446622_RNA45SN1_45S_50ntFlanks@4443.4471.29|CCCTTGCCTCTCGGCGCCCCCTCGATGCT</td><td>>hg38_21_+_8433172_8446622_RNA45SN1_45S_50ntFlanks</td><td>4443</td><td>4471</td><td>29</td><td>CCCTTGCCTCTCGGCGCCCCCTCGATGCT</td><td>1300.7</td><td>-7.9</td><td>1.4</td><td>-5.7</td><td>1.0E-08</td><td>2.4E-08</td><td>BT20/MB468</td><td>CYTO</td></tr>
<tr><td>>hg38_21_+_8433172_8446622_RNA45SN1_45S_50ntFlanks@11661.11685.25|TGATTTCTGCCCAGTGCTCTGAATG</td><td>>hg38_21_+_8433172_8446622_RNA45SN1_45S_50ntFlanks</td><td>11661</td><td>11685</td><td>25</td><td>TGATTTCTGCCCAGTGCTCTGAATG</td><td>6320.1</td><td>-7.1</td><td>1.2</td><td>-5.7</td><td>1.0E-08</td><td>2.4E-08</td><td>BT20/MB468</td><td>CYTO</td></tr>
<tr><td>>hg38_21_+_8433172_8446622_RNA45SN1_45S_50ntFlanks@10844.10865.22|AACTTCGGGATAAGGATTGGCT</td><td>>hg38_21_+_8433172_8446622_RNA45SN1_45S_50ntFlanks</td><td>10844</td><td>10865</td><td>22</td><td>AACTTCGGGATAAGGATTGGCT</td><td>693.9</td><td>-5.4</td><td>0.9</td><td>-5.7</td><td>1.2E-08</td><td>2.4E-08</td><td>MB231/MB468</td><td>CYTO</td></tr>
<tr><td>>hg38_21_+_8433172_8446622_RNA45SN1_45S_50ntFlanks@11672.11710.39|CAGTGCTCTGAATGTCAAAGTGAAGAAATTCAATGAAGC</td><td>>hg38_21_+_8433172_8446622_RNA45SN1_45S_50ntFlanks</td><td>11672</td><td>11710</td><td>39</td><td>CAGTGCTCTGAATGTCAAAGTGAAGAAATTCAATGAAGC</td><td>203.8</td><td>5.9</td><td>1.0</td><td>5.7</td><td>1.2E-08</td><td>2.4E-08</td><td>MB231/MB468</td><td>CYTO</td></tr>
<tr><td>>hg38_21_+_8433172_8446622_RNA45SN1_45S_50ntFlanks@11783.11807.25|CATGAATGGATGAACGAGATTCCCA</td><td>>hg38_21_+_8433172_8446622_RNA45SN1_45S_50ntFlanks</td><td>11783</td><td>11807</td><td>25</td><td>CATGAATGGATGAACGAGATTCCCA</td><td>108.7</td><td>-8.9</td><td>1.5</td><td>-5.7</td><td>9.1E-09</td><td>2.4E-08</td><td>BT20/MB468</td><td>MITO</td></tr>
<tr><td>>MI0000102|hsa-mir-100&WithFlank&11|-|122152223|122152314@19.38.20(+1C)&[MIMAT0000098&hsa-miR-100-5p&offsets|0|-2(+1C);m-27&11|-|122152276|122152296&offsets|0|-1(+1C)]|AACCCGTAGATCCGAACTTGC</td><td>>MI0000102|hsa-mir-100&WithFlank&11|-|122152223|122152314</td><td>19</td><td>38</td><td>20(+1C)</td><td>AACCCGTAGATCCGAACTTGC</td><td>83.4</td><td>9.2</td><td>1.6</td><td>5.7</td><td>1.3E-08</td><td>2.4E-08</td><td>BT20/MB468</td><td>NUC</td></tr>
<tr><td>>MI0000750|hsa-mir-26a-2&WithFlank&12|-|57824603|57824698@20.41.22(+1U);MI0000083|hsa-mir-26a-1&WithFlank&3|+|37969398|37969486@16.37.22(+1U)&[MIMAT0000082&hsa-miR-26a-5p&offsets|0|0(+1U);m-33&12|-|57824658|57824679&offsets|0|0(+1U)];[MIMAT0000082_1&hsa-miR-26a-5p&offsets|0|0(+1U);m-34&3|+|37969413|37969434&offsets|0|0(+1U)]|TTCAAGTAATCCAGGATAGGCTT</td><td>>MI0000750|hsa-mir-26a-2&WithFlank&12|-|57824603|57824698</td><td>20</td><td>41</td><td>22(+1U)</td><td>TTCAAGTAATCCAGGATAGGCTT</td><td>799.5</td><td>2.5</td><td>0.4</td><td>5.7</td><td>1.3E-08</td><td>2.4E-08</td><td>BT20/MB468</td><td>NUC</td></tr>
<tr><td>>hg38_21_+_8433172_8446622_RNA45SN1_45S_50ntFlanks@5071.5098.28|TTAATTCCGATAACGAACGAGACTCTGG</td><td>>hg38_21_+_8433172_8446622_RNA45SN1_45S_50ntFlanks</td><td>5071</td><td>5098</td><td>28</td><td>TTAATTCCGATAACGAACGAGACTCTGG</td><td>2879.5</td><td>-10.3</td><td>1.8</td><td>-5.7</td><td>1.3E-08</td><td>2.4E-08</td><td>BT20/MB468</td><td>NUC</td></tr>
<tr><td>>hg38_21_+_8433172_8446622_RNA45SN1_45S_50ntFlanks@9714.9732.19|CGAATGATTAGAGGTCTTG</td><td>>hg38_21_+_8433172_8446622_RNA45SN1_45S_50ntFlanks</td><td>9714</td><td>9732</td><td>19</td><td>CGAATGATTAGAGGTCTTG</td><td>148.6</td><td>-9.3</td><td>1.6</td><td>-5.7</td><td>9.1E-09</td><td>2.4E-08</td><td>BT20/MB468</td><td>MITO</td></tr>
<tr><td>>hg38_21_+_8433172_8446622_RNA45SN1_45S_50ntFlanks@12202.12220.19|CTAAGGCGAGCTCAGGGAG</td><td>>hg38_21_+_8433172_8446622_RNA45SN1_45S_50ntFlanks</td><td>12202</td><td>12220</td><td>19</td><td>CTAAGGCGAGCTCAGGGAG</td><td>1063.4</td><td>-9.0</td><td>1.6</td><td>-5.7</td><td>1.3E-08</td><td>2.4E-08</td><td>BT20/MB468</td><td>NUC</td></tr>
<tr><td>>hg38_21_+_8433172_8446622_RNA45SN1_45S_50ntFlanks@10457.10481.25|GATGGCCTCCGTTGCCCTCGGCCGA</td><td>>hg38_21_+_8433172_8446622_RNA45SN1_45S_50ntFlanks</td><td>10457</td><td>10481</td><td>25</td><td>GATGGCCTCCGTTGCCCTCGGCCGA</td><td>70.4</td><td>-10.0</td><td>1.8</td><td>-5.7</td><td>1.1E-08</td><td>2.4E-08</td><td>MB231/MB468</td><td>MP</td></tr>
<tr><td>>hg38_21_+_8433172_8446622_RNA45SN1_45S_50ntFlanks@11873.11903.31|AAGAAGACCCTGTTGAGCTTGACTCTAGTCT</td><td>>hg38_21_+_8433172_8446622_RNA45SN1_45S_50ntFlanks</td><td>11873</td><td>11903</td><td>31</td><td>AAGAAGACCCTGTTGAGCTTGACTCTAGTCT</td><td>253.5</td><td>-4.2</td><td>0.7</td><td>-5.7</td><td>1.2E-08</td><td>2.4E-08</td><td>MB231/MB468</td><td>CYTO</td></tr>
<tr><td>>hg38_21_+_8433172_8446622_RNA45SN1_45S_50ntFlanks@11703.11726.24|AATGAAGCGCGGGTAAACGGCGGG</td><td>>hg38_21_+_8433172_8446622_RNA45SN1_45S_50ntFlanks</td><td>11703</td><td>11726</td><td>24</td><td>AATGAAGCGCGGGTAAACGGCGGG</td><td>525.8</td><td>-8.0</td><td>1.4</td><td>-5.7</td><td>1.1E-08</td><td>2.4E-08</td><td>MB231/MB468</td><td>MP</td></tr>
<tr><td>>trnaMT_MetCAT_MT_+_4402_4469@52.71.20|ATACCCTTCCCGTACTACCA</td><td>>trnaMT_MetCAT_MT_+_4402_4469</td><td>52</td><td>71</td><td>20</td><td>ATACCCTTCCCGTACTACCA</td><td>1666.2</td><td>6.9</td><td>1.2</td><td>5.7</td><td>1.2E-08</td><td>2.4E-08</td><td>MB231/MB468</td><td>NUC</td></tr>
<tr><td>>MI0001445|hsa-mir-423&WithFlank&17|+|30117073|30117178@23.44.22&[MIMAT0004748&hsa-miR-423-5p&offsets|0|-1;m-141&17|+|30117095|30117117&offsets|0|-1]|TGAGGGGCAGAGAGCGAGACTT</td><td>>MI0001445|hsa-mir-423&WithFlank&17|+|30117073|30117178</td><td>23</td><td>44</td><td>22</td><td>TGAGGGGCAGAGAGCGAGACTT</td><td>116.2</td><td>2.4</td><td>0.4</td><td>5.7</td><td>1.4E-08</td><td>2.4E-08</td><td>BT20/MB468</td><td>NUC</td></tr>
<tr><td>>trna119_LysCTT_1_-_145395522_145395594@1.32.32,trna11_LysCTT_5_-_180648979_180649051@1.32.32,trna13_LysCTT_6_+_26556774_26556846@1.32.32,trna32_LysCTT_16_-_3207406_3207478@1.32.32,trna7_LysCTT_16_+_3225692_3225764@1.32.32,trna9_LysCTT_5_+_180634755_180634827@1.32.32|GCCCGGCTAGCTCAGTCGGTAGAGCATGAGAC</td><td>>trna119_LysCTT_1_-_145395522_145395594</td><td>1</td><td>32</td><td>32</td><td>GCCCGGCTAGCTCAGTCGGTAGAGCATGAGAC</td><td>556.5</td><td>-4.0</td><td>0.7</td><td>-5.7</td><td>1.4E-08</td><td>2.4E-08</td><td>BT20/MB468</td><td>NUC</td></tr>
<tr><td>>hg38_21_+_8433172_8446622_RNA45SN1_45S_50ntFlanks@4130.4162.33|ATTCCGGAGAGGGAGCCTGAGAAACGGCTACCA</td><td>>hg38_21_+_8433172_8446622_RNA45SN1_45S_50ntFlanks</td><td>4130</td><td>4162</td><td>33</td><td>ATTCCGGAGAGGGAGCCTGAGAAACGGCTACCA</td><td>767.3</td><td>-6.4</td><td>1.1</td><td>-5.7</td><td>1.2E-08</td><td>2.4E-08</td><td>MB231/MB468</td><td>NUC</td></tr>
<tr><td>>hg38_21_+_8433172_8446622_RNA45SN1_45S_50ntFlanks@6651.6693.43|CGACTCTTAGCGGTGGATCACTCGGCTCGTGCGTCGATGAAGA</td><td>>hg38_21_+_8433172_8446622_RNA45SN1_45S_50ntFlanks</td><td>6651</td><td>6693</td><td>43</td><td>CGACTCTTAGCGGTGGATCACTCGGCTCGTGCGTCGATGAAGA</td><td>18986.7</td><td>4.7</td><td>0.8</td><td>5.7</td><td>9.2E-09</td><td>2.4E-08</td><td>BT20/MB468</td><td>MITO</td></tr>
<tr><td>>hg38_MT_+_598_1651_RNR1_12S_50ntFlanks@52.83.32|ATAGGTTTGGTCCTAGCCTTTCTATTAGCTCT</td><td>>hg38_MT_+_598_1651_RNR1_12S_50ntFlanks</td><td>52</td><td>83</td><td>32</td><td>ATAGGTTTGGTCCTAGCCTTTCTATTAGCTCT</td><td>380.6</td><td>6.9</td><td>1.2</td><td>5.7</td><td>1.2E-08</td><td>2.4E-08</td><td>MB231/MB468</td><td>CYTO</td></tr>
<tr><td>>hg38_21_+_8433172_8446622_RNA45SN1_45S_50ntFlanks@10383.10424.42|TCCATGTGAACAGCAGTTGAACATGGGTCAGTCGGTCCTGAG</td><td>>hg38_21_+_8433172_8446622_RNA45SN1_45S_50ntFlanks</td><td>10383</td><td>10424</td><td>42</td><td>TCCATGTGAACAGCAGTTGAACATGGGTCAGTCGGTCCTGAG</td><td>1857.7</td><td>5.2</td><td>0.9</td><td>5.7</td><td>1.2E-08</td><td>2.4E-08</td><td>MB231/MB468</td><td>NUC</td></tr>
<tr><td>>hg38_21_+_8433172_8446622_RNA45SN1_45S_50ntFlanks@4046.4064.19|CATTCGAACGTCTGCCCTA</td><td>>hg38_21_+_8433172_8446622_RNA45SN1_45S_50ntFlanks</td><td>4046</td><td>4064</td><td>19</td><td>CATTCGAACGTCTGCCCTA</td><td>153.2</td><td>-6.7</td><td>1.2</td><td>-5.7</td><td>1.2E-08</td><td>2.4E-08</td><td>MB231/MB468</td><td>NUC</td></tr>
<tr><td>>trna12_TrpCCA_17_+_19411494_19411565@30.52.23,trna168_TrpCCA_6_-_26331672_26331743@30.52.23,trna170_TrpCCA_6_-_26319330_26319401@30.52.23,trna1_TrpCCA_7_+_99067307_99067378@30.52.23,trna39_TrpCCA_17_-_8124187_8124258@30.52.23,trna6_TrpCCA_17_+_8089676_8089747@30.52.23|ACTCCAGATCAGAAGGTTGCGTG</td><td>>trna12_TrpCCA_17_+_19411494_19411565</td><td>30</td><td>52</td><td>23</td><td>ACTCCAGATCAGAAGGTTGCGTG</td><td>112.0</td><td>-9.9</td><td>1.6</td><td>-6.0</td><td>1.7E-09</td><td>2.4E-08</td><td>BT20/MB468</td><td>MP</td></tr>
<tr><td>>hg38_21_+_8433172_8446622_RNA45SN1_45S_50ntFlanks@11209.11232.24|CGGCGACTCTGGACGCGAGCCGGG</td><td>>hg38_21_+_8433172_8446622_RNA45SN1_45S_50ntFlanks</td><td>11209</td><td>11232</td><td>24</td><td>CGGCGACTCTGGACGCGAGCCGGG</td><td>21125.8</td><td>-5.4</td><td>0.9</td><td>-5.7</td><td>1.0E-08</td><td>2.4E-08</td><td>BT20/MB468</td><td>CYTO</td></tr>
<tr><td>>hg38_21_+_8433172_8446622_RNA45SN1_45S_50ntFlanks@12328.12346.19|GTTTTAAGCAGGAGGTGTC</td><td>>hg38_21_+_8433172_8446622_RNA45SN1_45S_50ntFlanks</td><td>12328</td><td>12346</td><td>19</td><td>GTTTTAAGCAGGAGGTGTC</td><td>144.6</td><td>-6.0</td><td>1.1</td><td>-5.7</td><td>1.0E-08</td><td>2.4E-08</td><td>BT20/MB468</td><td>CYTO</td></tr>
<tr><td>>hg38_21_+_8433172_8446622_RNA45SN1_45S_50ntFlanks@5420.5466.47|CCGATTGGATGGTTTAGTGAGGCCCTCGGATCGGCCCCGCCGGGGTC</td><td>>hg38_21_+_8433172_8446622_RNA45SN1_45S_50ntFlanks</td><td>5420</td><td>5466</td><td>47</td><td>CCGATTGGATGGTTTAGTGAGGCCCTCGGATCGGCCCCGCCGGGGTC</td><td>94.9</td><td>-10.4</td><td>1.8</td><td>-5.7</td><td>1.1E-08</td><td>2.4E-08</td><td>MB231/MB468</td><td>MP</td></tr>
<tr><td>>trnaMT_GluTTC_MT_-_14674_14742@39.59.21,trnalookalike8_GluTTC_5_-_93905172_93905240@39.59.21|ATTGGTCGTGGTTGTAGTCCG</td><td>>trnaMT_GluTTC_MT_-_14674_14742</td><td>39</td><td>59</td><td>21</td><td>ATTGGTCGTGGTTGTAGTCCG</td><td>431.3</td><td>9.2</td><td>1.6</td><td>5.7</td><td>9.2E-09</td><td>2.4E-08</td><td>BT20/MB468</td><td>MITO</td></tr>
<tr><td>>hg38_21_+_8433172_8446622_RNA45SN1_45S_50ntFlanks@10843.10872.30|TAACTTCGGGATAAGGATTGGCTCTAAGGG</td><td>>hg38_21_+_8433172_8446622_RNA45SN1_45S_50ntFlanks</td><td>10843</td><td>10872</td><td>30</td><td>TAACTTCGGGATAAGGATTGGCTCTAAGGG</td><td>297.8</td><td>-7.8</td><td>1.4</td><td>-5.7</td><td>1.4E-08</td><td>2.4E-08</td><td>BT20/MB468</td><td>NUC</td></tr>
<tr><td>>hg38_21_+_8433172_8446622_RNA45SN1_45S_50ntFlanks@12071.12097.27|CCCCGAGGGGCTCTCGCTTCTGGCGCC</td><td>>hg38_21_+_8433172_8446622_RNA45SN1_45S_50ntFlanks</td><td>12071</td><td>12097</td><td>27</td><td>CCCCGAGGGGCTCTCGCTTCTGGCGCC</td><td>385.4</td><td>-4.2</td><td>0.7</td><td>-5.7</td><td>1.2E-08</td><td>2.4E-08</td><td>MB231/MB468</td><td>NUC</td></tr>
<tr><td>>hg38_21_+_8433172_8446622_RNA45SN1_45S_50ntFlanks@11199.11224.26|CCGGCGGCGGCGGCGACTCTGGACGC</td><td>>hg38_21_+_8433172_8446622_RNA45SN1_45S_50ntFlanks</td><td>11199</td><td>11224</td><td>26</td><td>CCGGCGGCGGCGGCGACTCTGGACGC</td><td>322.0</td><td>-7.7</td><td>1.4</td><td>-5.7</td><td>1.2E-08</td><td>2.4E-08</td><td>MB231/MB468</td><td>NUC</td></tr>
<tr><td>>hg38_21_+_8433172_8446622_RNA45SN1_45S_50ntFlanks@10349.10388.40|AACGAGAACTTTGAAGGCCGAAGTGGAGAAGGGTTCCATG</td><td>>hg38_21_+_8433172_8446622_RNA45SN1_45S_50ntFlanks</td><td>10349</td><td>10388</td><td>40</td><td>AACGAGAACTTTGAAGGCCGAAGTGGAGAAGGGTTCCATG</td><td>2731.4</td><td>-7.9</td><td>1.4</td><td>-5.7</td><td>1.4E-08</td><td>2.4E-08</td><td>BT20/MB468</td><td>NUC</td></tr>
<tr><td>>hg38_21_+_8433172_8446622_RNA45SN1_45S_50ntFlanks@11579.11612.34|AGAACTGGTGCGGACCAGGGGAATCCGACTGTTT</td><td>>hg38_21_+_8433172_8446622_RNA45SN1_45S_50ntFlanks</td><td>11579</td><td>11612</td><td>34</td><td>AGAACTGGTGCGGACCAGGGGAATCCGACTGTTT</td><td>431.0</td><td>-4.2</td><td>0.7</td><td>-5.7</td><td>1.3E-08</td><td>2.5E-08</td><td>MB231/MB468</td><td>MITO</td></tr>
<tr><td>>hg38_21_+_8433172_8446622_RNA45SN1_45S_50ntFlanks@10808.10837.30|CAATGTAGGTAAGGGAAGTCGGCAAGCCGG</td><td>>hg38_21_+_8433172_8446622_RNA45SN1_45S_50ntFlanks</td><td>10808</td><td>10837</td><td>30</td><td>CAATGTAGGTAAGGGAAGTCGGCAAGCCGG</td><td>372.2</td><td>-7.7</td><td>1.3</td><td>-5.7</td><td>1.1E-08</td><td>2.5E-08</td><td>MB231/MB468</td><td>MP</td></tr>
<tr><td>>hg38_21_+_8433172_8446622_RNA45SN1_45S_50ntFlanks@9533.9561.29|GAGGAAACTCTGGTGGAGGTCCGTAGCGG</td><td>>hg38_21_+_8433172_8446622_RNA45SN1_45S_50ntFlanks</td><td>9533</td><td>9561</td><td>29</td><td>GAGGAAACTCTGGTGGAGGTCCGTAGCGG</td><td>1060.9</td><td>-7.1</td><td>1.2</td><td>-5.7</td><td>1.2E-08</td><td>2.5E-08</td><td>MB231/MB468</td><td>NUC</td></tr>
<tr><td>>hg38_21_+_8433172_8446622_RNA45SN1_45S_50ntFlanks@12419.12449.31|GATGTCGGCTCTTCCTATCATTGTGAAGCAG</td><td>>hg38_21_+_8433172_8446622_RNA45SN1_45S_50ntFlanks</td><td>12419</td><td>12449</td><td>31</td><td>GATGTCGGCTCTTCCTATCATTGTGAAGCAG</td><td>711.3</td><td>3.1</td><td>0.5</td><td>5.7</td><td>1.0E-08</td><td>2.5E-08</td><td>BT20/MB468</td><td>CYTO</td></tr>
<tr><td>>hg38_21_+_8433172_8446622_RNA45SN1_45S_50ntFlanks@11317.11344.28|ACCCCACGTCTCGTCGCGCGCGCGTCCG</td><td>>hg38_21_+_8433172_8446622_RNA45SN1_45S_50ntFlanks</td><td>11317</td><td>11344</td><td>28</td><td>ACCCCACGTCTCGTCGCGCGCGCGTCCG</td><td>49378.5</td><td>-11.3</td><td>2.0</td><td>-5.7</td><td>1.4E-08</td><td>2.5E-08</td><td>BT20/MB468</td><td>NUC</td></tr>
<tr><td>>hg38_21_+_8433172_8446622_RNA45SN1_45S_50ntFlanks@12954.12979.26|CTGCTTCTGGGTCGGGGTTTCGTACG</td><td>>hg38_21_+_8433172_8446622_RNA45SN1_45S_50ntFlanks</td><td>12954</td><td>12979</td><td>26</td><td>CTGCTTCTGGGTCGGGGTTTCGTACG</td><td>453.6</td><td>-8.6</td><td>1.5</td><td>-5.7</td><td>1.4E-08</td><td>2.5E-08</td><td>BT20/MB468</td><td>NUC</td></tr>
<tr><td>>hg38_21_+_8433172_8446622_RNA45SN1_45S_50ntFlanks@11161.11189.29|CACGGTCCCCCGCGAGGGGGGCCCGGGCA</td><td>>hg38_21_+_8433172_8446622_RNA45SN1_45S_50ntFlanks</td><td>11161</td><td>11189</td><td>29</td><td>CACGGTCCCCCGCGAGGGGGGCCCGGGCA</td><td>252.2</td><td>-7.3</td><td>1.3</td><td>-5.7</td><td>1.0E-08</td><td>2.5E-08</td><td>BT20/MB468</td><td>CYTO</td></tr>
<tr><td>>hg38_21_+_8433172_8446622_RNA45SN1_45S_50ntFlanks@11667.11685.19|CTGCCCAGTGCTCTGAATG</td><td>>hg38_21_+_8433172_8446622_RNA45SN1_45S_50ntFlanks</td><td>11667</td><td>11685</td><td>19</td><td>CTGCCCAGTGCTCTGAATG</td><td>936.4</td><td>-4.5</td><td>0.8</td><td>-5.7</td><td>1.3E-08</td><td>2.5E-08</td><td>MB231/MB468</td><td>MITO</td></tr>
<tr><td>>hg38_21_+_8433172_8446622_RNA45SN1_45S_50ntFlanks@7974.8015.42|ACGCGACCTCAGATCAGACGTGGCGACCCGCTGAATTTAAGC</td><td>>hg38_21_+_8433172_8446622_RNA45SN1_45S_50ntFlanks</td><td>7974</td><td>8015</td><td>42</td><td>ACGCGACCTCAGATCAGACGTGGCGACCCGCTGAATTTAAGC</td><td>1066.9</td><td>5.1</td><td>0.9</td><td>5.7</td><td>1.2E-08</td><td>2.5E-08</td><td>MB231/MB468</td><td>NUC</td></tr>
<tr><td>>MI0000273|hsa-mir-183&WithFlank&7|-|129774899|129775020@33.54.22(+1A)&[MIMAT0000261&hsa-miR-183-5p&offsets|0|0(+1A);m-41&7|-|129774966|129774987&offsets|-1|-1(+1A)]|TATGGCACTGGTAGAATTCACTA</td><td>>MI0000273|hsa-mir-183&WithFlank&7|-|129774899|129775020</td><td>33</td><td>54</td><td>22(+1A)</td><td>TATGGCACTGGTAGAATTCACTA</td><td>511.7</td><td>3.3</td><td>0.5</td><td>6.1</td><td>1.3E-09</td><td>2.5E-08</td><td>BT20/MB231</td><td>NUC</td></tr>
<tr><td>>hg38_21_+_8433172_8446622_RNA45SN1_45S_50ntFlanks@7994.8011.18|TGGCGACCCGCTGAATTT</td><td>>hg38_21_+_8433172_8446622_RNA45SN1_45S_50ntFlanks</td><td>7994</td><td>8011</td><td>18</td><td>TGGCGACCCGCTGAATTT</td><td>306.8</td><td>4.8</td><td>0.8</td><td>6.1</td><td>1.3E-09</td><td>2.5E-08</td><td>BT20/MB231</td><td>NUC</td></tr>
<tr><td>>hg38_21_+_8433172_8446622_RNA45SN1_45S_50ntFlanks@6645.6664.20|CTCGTACGACTCTTAGCGGT</td><td>>hg38_21_+_8433172_8446622_RNA45SN1_45S_50ntFlanks</td><td>6645</td><td>6664</td><td>20</td><td>CTCGTACGACTCTTAGCGGT</td><td>2131.3</td><td>3.4</td><td>0.6</td><td>5.7</td><td>1.2E-08</td><td>2.5E-08</td><td>MB231/MB468</td><td>NUC</td></tr>
<tr><td>>hg38_21_+_8433172_8446622_RNA45SN1_45S_50ntFlanks@9549.9570.22|AGGTCCGTAGCGGTCCTGACGT</td><td>>hg38_21_+_8433172_8446622_RNA45SN1_45S_50ntFlanks</td><td>9549</td><td>9570</td><td>22</td><td>AGGTCCGTAGCGGTCCTGACGT</td><td>283.3</td><td>4.6</td><td>0.8</td><td>6.0</td><td>1.7E-09</td><td>2.5E-08</td><td>BT20/MB468</td><td>MP</td></tr>
<tr><td>>hg38_21_+_8433172_8446622_RNA45SN1_45S_50ntFlanks@10405.10424.20|ATGGGTCAGTCGGTCCTGAG</td><td>>hg38_21_+_8433172_8446622_RNA45SN1_45S_50ntFlanks</td><td>10405</td><td>10424</td><td>20</td><td>ATGGGTCAGTCGGTCCTGAG</td><td>18372.3</td><td>6.3</td><td>1.1</td><td>5.7</td><td>1.1E-08</td><td>2.5E-08</td><td>MB231/MB468</td><td>MP</td></tr>
<tr><td>>hg38_21_+_8433172_8446622_RNA45SN1_45S_50ntFlanks@4728.4752.25|ATCAAGAACGAAAGTCGGAGGTTCG</td><td>>hg38_21_+_8433172_8446622_RNA45SN1_45S_50ntFlanks</td><td>4728</td><td>4752</td><td>25</td><td>ATCAAGAACGAAAGTCGGAGGTTCG</td><td>908.8</td><td>-8.1</td><td>1.4</td><td>-5.7</td><td>1.1E-08</td><td>2.5E-08</td><td>MB231/MB468</td><td>MP</td></tr>
<tr><td>>hg38_21_+_8433172_8446622_RNA45SN1_45S_50ntFlanks@8345.8375.31|AAGGGAAAGTTGAAAAGAACTTTGAAGAGAG</td><td>>hg38_21_+_8433172_8446622_RNA45SN1_45S_50ntFlanks</td><td>8345</td><td>8375</td><td>31</td><td>AAGGGAAAGTTGAAAAGAACTTTGAAGAGAG</td><td>1869.9</td><td>10.5</td><td>1.7</td><td>6.0</td><td>1.7E-09</td><td>2.5E-08</td><td>BT20/MB468</td><td>MP</td></tr>
<tr><td>>MI0003643|hsa-mir-629&WithFlank&15|-|70079366|70079474@28.48.21(+1A)&[MIMAT0004810&hsa-miR-629-5p&offsets|0|0(+1A);m-150&15|-|70079427|70079447&offsets|0|0(+1A)]|TGGGTTTACGTTGGGAGAACTA</td><td>>MI0003643|hsa-mir-629&WithFlank&15|-|70079366|70079474</td><td>28</td><td>48</td><td>21(+1A)</td><td>TGGGTTTACGTTGGGAGAACTA</td><td>81.4</td><td>9.6</td><td>1.7</td><td>5.7</td><td>1.1E-08</td><td>2.5E-08</td><td>MB231/MB468</td><td>MP</td></tr>
<tr><td>>hg38_21_+_8433172_8446622_RNA45SN1_45S_50ntFlanks@8184.8221.38|CGAGGCCCAGCCCGTGGACGGTGTGAGGCCGGTAGCGG</td><td>>hg38_21_+_8433172_8446622_RNA45SN1_45S_50ntFlanks</td><td>8184</td><td>8221</td><td>38</td><td>CGAGGCCCAGCCCGTGGACGGTGTGAGGCCGGTAGCGG</td><td>60.5</td><td>-9.8</td><td>1.7</td><td>-5.7</td><td>1.1E-08</td><td>2.5E-08</td><td>MB231/MB468</td><td>MP</td></tr>
</table>

<table>
<tr><td>>hg38_21_+_8433172_8446622_RNA45SN1_45S_50ntFlanks@4724.4764.41|ATTAATCAAGAACGAAAGTCGGAGGTTCGAAGACGATCAGA</td><td>>hg38_21_+_8433172_8446622_RNA45SN1_45S_50ntFlanks</td><td>4724</td><td>4764</td><td>41</td><td>ATTAATCAAGAACGAAAGTCGGAGGTTCGAAGACGATCAGA</td><td>883.7</td><td>-8.3</td><td>1.5</td><td>-5.7</td><td>1.3E-08</td><td>2.5E-08</td><td>MB231/MB468</td><td>MITO</td></tr>
<tr><td>>hg38_21_+_8433172_8446622_RNA45SN1_45S_50ntFlanks@5207.5235.29|CTTAGATGTCCGGGGCTGCACGCGCGCTA</td><td>>hg38_21_+_8433172_8446622_RNA45SN1_45S_50ntFlanks</td><td>5207</td><td>5235</td><td>29</td><td>CTTAGATGTCCGGGGCTGCACGCGCGCTA</td><td>568.3</td><td>-8.9</td><td>1.6</td><td>-5.7</td><td>1.3E-08</td><td>2.5E-08</td><td>MB231/MB468</td><td>MITO</td></tr>
<tr><td>>MI0000449|hsa-mir-132&WithFlank&17|-|2049902|2050014@65.85.21&[MIMAT0000426&hsa-miR-132-3p&offsets|0|-1;m-138&17|-|2049929|2049950&offsets|0|-1]|TAACAGTCTACAGCCATGGTC</td><td>>MI0000449|hsa-mir-132&WithFlank&17|-|2049902|2050014</td><td>65</td><td>85</td><td>21</td><td>TAACAGTCTACAGCCATGGTC</td><td>309.4</td><td>12.0</td><td>2.0</td><td>6.0</td><td>1.7E-09</td><td>2.5E-08</td><td>BT20/MB468</td><td>MP</td></tr>
<tr><td>>hg38_21_+_8433172_8446622_RNA45SN1_45S_50ntFlanks@10843.10862.20|TAACTTCGGGATAAGGATTG</td><td>>hg38_21_+_8433172_8446622_RNA45SN1_45S_50ntFlanks</td><td>10843</td><td>10862</td><td>20</td><td>TAACTTCGGGATAAGGATTG</td><td>825.8</td><td>-7.4</td><td>1.3</td><td>-5.7</td><td>1.0E-08</td><td>2.5E-08</td><td>BT20/MB468</td><td>CYTO</td></tr>
<tr><td>>MI0000446|hsa-mir-125b-1&WithFlank&11|-|122099751|122099850@22.42.21;MI0000470|hsa-mir-125b-2&WithFlank&21|+|16590231|16590331@24.44.21&[MIMAT0000423&hsa-miR-125b-5p&offsets|+1|0;m-67&11|-|122099809|122099830&offsets|+1|0];[MIMAT0000423_1&hsa-miR-125b-5p&offsets|+1|0;m-68&21|+|16590253|16590274&offsets|+1|0]|CCCTGAGACCCTAACTTGTGA</td><td>>MI0000446|hsa-mir-125b-1&WithFlank&11|-|122099751|122099850</td><td>22</td><td>42</td><td>21</td><td>CCCTGAGACCCTAACTTGTGA</td><td>68.1</td><td>9.3</td><td>1.6</td><td>5.7</td><td>1.2E-08</td><td>2.5E-08</td><td>MB231/MB468</td><td>MP</td></tr>
<tr><td>>hg38_21_+_8433172_8446622_RNA45SN1_45S_50ntFlanks@12053.12070.18|GTGAGGCGGGGGGGCGAG</td><td>>hg38_21_+_8433172_8446622_RNA45SN1_45S_50ntFlanks</td><td>12053</td><td>12070</td><td>18</td><td>GTGAGGCGGGGGGGCGAG</td><td>138.0</td><td>-10.2</td><td>1.7</td><td>-6.0</td><td>1.8E-09</td><td>2.5E-08</td><td>BT20/MB468</td><td>MP</td></tr>
<tr><td>>hg38_21_+_8433172_8446622_RNA45SN1_45S_50ntFlanks@11564.11602.39|CCTAGCAGCCGACTTAGAACTGGTGCGGACCAGGGGAAT</td><td>>hg38_21_+_8433172_8446622_RNA45SN1_45S_50ntFlanks</td><td>11564</td><td>11602</td><td>39</td><td>CCTAGCAGCCGACTTAGAACTGGTGCGGACCAGGGGAAT</td><td>2183.3</td><td>-7.8</td><td>1.4</td><td>-5.7</td><td>1.2E-08</td><td>2.5E-08</td><td>MB231/MB468</td><td>MP</td></tr>
<tr><td>>trna10_AspGTC_12_-_125424193_125424264@37.72.36,trna12_AspGTC_12_-_125411891_125411962@37.72.36,trna144_AspGTC_6_-_27551236_27551307@37.72.36,trna38_AspGTC_17_-_8125556_8125627@37.72.36,trna45_AspGTC_6_+_27447453_27447524@37.72.36,trna48_AspGTC_6_+_27471523_27471594@37.72.36,trna4_AspGTC_12_+_96429799_96429870@37.72.36,trna69_AspGTC_1_-_161440205_161440276@37.72.36,trna72_AspGTC_1_-_161432824_161432895@37.72.36,trna75_AspGTC_1_-_161425414_161425485@37.72.36,trna78_AspGTC_1_-_161418033_161418104@37.72.36,trna81_AspGTC_1_-_161410615_161410686@37.72.36|ACGCGGGAGACCGGGGTTCGATTCCCCGACGGGGAG</td><td>>trna10_AspGTC_12_-_125424193_125424264</td><td>37</td><td>72</td><td>36</td><td>ACGCGGGAGACCGGGGTTCGATTCCCCGACGGGGAG</td><td>581.8</td><td>-6.6</td><td>1.2</td><td>-5.7</td><td>1.4E-08</td><td>2.5E-08</td><td>BT20/MB468</td><td>NUC</td></tr>
<tr><td>>hg38_21_+_8433172_8446622_RNA45SN1_45S_50ntFlanks@4997.5024.28|AGATTGATAGCTCTTTCTCGATTCCGTG</td><td>>hg38_21_+_8433172_8446622_RNA45SN1_45S_50ntFlanks</td><td>4997</td><td>5024</td><td>28</td><td>AGATTGATAGCTCTTTCTCGATTCCGTG</td><td>103.1</td><td>-10.6</td><td>1.8</td><td>-5.7</td><td>1.2E-08</td><td>2.5E-08</td><td>MB231/MB468</td><td>MP</td></tr>
<tr><td>>hg38_21_+_8433172_8446622_RNA45SN1_45S_50ntFlanks@9352.9370.19|CCGTGGCGCAATGAAGGTG</td><td>>hg38_21_+_8433172_8446622_RNA45SN1_45S_50ntFlanks</td><td>9352</td><td>9370</td><td>19</td><td>CCGTGGCGCAATGAAGGTG</td><td>2201.4</td><td>-7.9</td><td>1.4</td><td>-5.7</td><td>9.5E-09</td><td>2.5E-08</td><td>BT20/MB468</td><td>MITO</td></tr>
<tr><td>>MI0000441|hsa-mir-30b&WithFlank&8|-|134800514|134800613@23.43.21(+1A)&[MIMAT0000420&hsa-miR-30b-5p&offsets|0|-1(+1A);m-76&8|-|134800570|134800591&offsets|0|-1(+1A)]|TGTAAACATCCTACACTCAGCA</td><td>>MI0000441|hsa-mir-30b&WithFlank&8|-|134800514|134800613</td><td>23</td><td>43</td><td>21(+1A)</td><td>TGTAAACATCCTACACTCAGCA</td><td>192.8</td><td>5.6</td><td>1.0</td><td>5.7</td><td>1.4E-08</td><td>2.5E-08</td><td>BT20/MB468</td><td>NUC</td></tr>
<tr><td>>hg38_21_+_8433172_8446622_RNA45SN1_45S_50ntFlanks@5485.5506.22|AGCGCTGAGAAGACGGTCGAAC</td><td>>hg38_21_+_8433172_8446622_RNA45SN1_45S_50ntFlanks</td><td>5485</td><td>5506</td><td>22</td><td>AGCGCTGAGAAGACGGTCGAAC</td><td>281.7</td><td>-7.5</td><td>1.3</td><td>-5.7</td><td>1.3E-08</td><td>2.5E-08</td><td>MB231/MB468</td><td>MITO</td></tr>
<tr><td>>hg38_21_+_8433172_8446622_RNA45SN1_45S_50ntFlanks@12447.12474.28|CAGAATTCACCAAGCGTTGGATTGTTCA</td><td>>hg38_21_+_8433172_8446622_RNA45SN1_45S_50ntFlanks</td><td>12447</td><td>12474</td><td>28</td><td>CAGAATTCACCAAGCGTTGGATTGTTCA</td><td>109.9</td><td>-10.6</td><td>1.9</td><td>-5.7</td><td>1.2E-08</td><td>2.5E-08</td><td>MB231/MB468</td><td>MP</td></tr>
<tr><td>>hg38_21_+_8433172_8446622_RNA45SN1_45S_50ntFlanks@6732.6770.39|ACATTGATCATCGACACTTCGAACGCACTTGCGGCCCCG</td><td>>hg38_21_+_8433172_8446622_RNA45SN1_45S_50ntFlanks</td><td>6732</td><td>6770</td><td>39</td><td>ACATTGATCATCGACACTTCGAACGCACTTGCGGCCCCG</td><td>1619.7</td><td>-3.0</td><td>0.5</td><td>-5.7</td><td>9.5E-09</td><td>2.5E-08</td><td>BT20/MB468</td><td>MITO</td></tr>
<tr><td>>hg38_21_+_8433172_8446622_RNA45SN1_45S_50ntFlanks@7986.8011.26|ATCAGACGTGGCGACCCGCTGAATTT</td><td>>hg38_21_+_8433172_8446622_RNA45SN1_45S_50ntFlanks</td><td>7986</td><td>8011</td><td>26</td><td>ATCAGACGTGGCGACCCGCTGAATTT</td><td>1110.0</td><td>-8.0</td><td>1.4</td><td>-5.7</td><td>1.4E-08</td><td>2.5E-08</td><td>MB231/MB468</td><td>MITO</td></tr>
<tr><td>>hg38_21_+_8433172_8446622_RNA45SN1_45S_50ntFlanks@6731.6772.42|CACATTGATCATCGACACTTCGAACGCACTTGCGGCCCCGGG</td><td>>hg38_21_+_8433172_8446622_RNA45SN1_45S_50ntFlanks</td><td>6731</td><td>6772</td><td>42</td><td>CACATTGATCATCGACACTTCGAACGCACTTGCGGCCCCGGG</td><td>1297.0</td><td>-6.3</td><td>1.1</td><td>-5.7</td><td>1.4E-08</td><td>2.5E-08</td><td>MB231/MB468</td><td>MITO</td></tr>
<tr><td>>hg38_21_+_8433172_8446622_RNA45SN1_45S_50ntFlanks@11834.11852.19|CCACAGCCAAGGGAACGGG</td><td>>hg38_21_+_8433172_8446622_RNA45SN1_45S_50ntFlanks</td><td>11834</td><td>11852</td><td>19</td><td>CCACAGCCAAGGGAACGGG</td><td>102.1</td><td>-9.8</td><td>1.6</td><td>-6.0</td><td>1.8E-09</td><td>2.5E-08</td><td>BT20/MB468</td><td>MP</td></tr>
<tr><td>>hg38_21_+_8433172_8446622_RNA45SN1_45S_50ntFlanks@12677.12708.32|GAATCCCGCCCAGGCGGAACGATACGGCAGCG</td><td>>hg38_21_+_8433172_8446622_RNA45SN1_45S_50ntFlanks</td><td>12677</td><td>12708</td><td>32</td><td>GAATCCCGCCCAGGCGGAACGATACGGCAGCG</td><td>99.1</td><td>-9.7</td><td>1.6</td><td>-6.0</td><td>1.8E-09</td><td>2.5E-08</td><td>BT20/MB468</td><td>MP</td></tr>
<tr><td>>hg38_21_+_8433172_8446622_RNA45SN1_45S_50ntFlanks@10784.10818.35|TGAACAGCCTCTGGCATGTTGGAACAATGTAGGTA</td><td>>hg38_21_+_8433172_8446622_RNA45SN1_45S_50ntFlanks</td><td>10784</td><td>10818</td><td>35</td><td>TGAACAGCCTCTGGCATGTTGGAACAATGTAGGTA</td><td>177.5</td><td>-3.0</td><td>0.5</td><td>-5.7</td><td>1.3E-08</td><td>2.5E-08</td><td>MB231/MB468</td><td>NUC</td></tr>
<tr><td>>trna124_ArgTCG_6_-_28510891_28510963@31.52.22,trna19_ArgTCG_17_+_73031208_73031280@31.52.22,trna3_ArgTCG_6_+_26299905_26299977@31.52.22,trna4_ArgTCG_6_+_26323046_26323118@31.52.22|ACTTCGGATCAGAAGATTGAGG</td><td>>trna124_ArgTCG_6_-_28510891_28510963</td><td>31</td><td>52</td><td>22</td><td>ACTTCGGATCAGAAGATTGAGG</td><td>91.1</td><td>-10.4</td><td>1.8</td><td>-5.7</td><td>1.2E-08</td><td>2.5E-08</td><td>MB231/MB468</td><td>MP</td></tr>
<tr><td>>trna6_ValTAC_10_-_5895674_5895746@1.30.30|GGTTCCATAGTGTAGTGGTTATCACATCTG</td><td>>trna6_ValTAC_10_-_5895674_5895746</td><td>1</td><td>30</td><td>30</td><td>GGTTCCATAGTGTAGTGGTTATCACATCTG</td><td>160.3</td><td>-6.2</td><td>1.1</td><td>-5.7</td><td>1.1E-08</td><td>2.5E-08</td><td>BT20/MB468</td><td>CYTO</td></tr>
<tr><td>>hg38_21_+_8433172_8446622_RNA45SN1_45S_50ntFlanks@11678.11702.25|TCTGAATGTCAAAGTGAAGAAATTC</td><td>>hg38_21_+_8433172_8446622_RNA45SN1_45S_50ntFlanks</td><td>11678</td><td>11702</td><td>25</td><td>TCTGAATGTCAAAGTGAAGAAATTC</td><td>169.6</td><td>-5.7</td><td>1.0</td><td>-5.7</td><td>1.2E-08</td><td>2.5E-08</td><td>MB231/MB468</td><td>MP</td></tr>
<tr><td>>trna116_GluCTC_1_-_145399233_145399304@34.75.42,trna71_GluCTC_1_-_161439189_161439260@34.75.42,trna74_GluCTC_1_-_161431809_161431880@34.75.42,trna77_GluCTC_1_-_161424398_161424469@34.75.42,trna77_GluCTC_6_+_28949976_28950047@34.75.42,trna80_GluCTC_1_-_161417018_161417089@34.75.42,trna87_GluCTC_6_-_126101393_126101464@34.75.42|CTCACCGCCGCGGCCCGGGTTCGATTCCCGGTCAGGGAACCA</td><td>>trna116_GluCTC_1_-_145399233_145399304</td><td>34</td><td>75</td><td>42</td><td>CTCACCGCCGCGGCCCGGGTTCGATTCCCGGTCAGGGAACCA</td><td>1523.1</td><td>7.7</td><td>1.4</td><td>5.7</td><td>1.2E-08</td><td>2.5E-08</td><td>MB231/MB468</td><td>CYTO</td></tr>
<tr><td>>hg38_21_+_8433172_8446622_RNA45SN1_45S_50ntFlanks@9306.9338.33|CAAGGAGTCTAACACGTGCGCGAGTCGGGGGCT</td><td>>hg38_21_+_8433172_8446622_RNA45SN1_45S_50ntFlanks</td><td>9306</td><td>9338</td><td>33</td><td>CAAGGAGTCTAACACGTGCGCGAGTCGGGGGCT</td><td>166.9</td><td>-6.8</td><td>1.2</td><td>-5.7</td><td>1.3E-08</td><td>2.5E-08</td><td>MB231/MB468</td><td>NUC</td></tr>
<tr><td>>hg38_21_+_8433172_8446622_RNA45SN1_45S_50ntFlanks@10808.10827.20|CAATGTAGGTAAGGGAAGTC</td><td>>hg38_21_+_8433172_8446622_RNA45SN1_45S_50ntFlanks</td><td>10808</td><td>10827</td><td>20</td><td>CAATGTAGGTAAGGGAAGTC</td><td>1106.7</td><td>-9.3</td><td>1.6</td><td>-5.7</td><td>1.4E-08</td><td>2.5E-08</td><td>BT20/MB468</td><td>NUC</td></tr>
<tr><td>>hg38_21_+_8433172_8446622_RNA45SN1_45S_50ntFlanks@10362.10383.22|AAGGCCGAAGTGGAGAAGGGTT</td><td>>hg38_21_+_8433172_8446622_RNA45SN1_45S_50ntFlanks</td><td>10362</td><td>10383</td><td>22</td><td>AAGGCCGAAGTGGAGAAGGGTT</td><td>2592.2</td><td>-5.6</td><td>0.9</td><td>-6.0</td><td>1.8E-09</td><td>2.5E-08</td><td>BT20/MB468</td><td>MP</td></tr>
<tr><td>>hg38_MT_+_598_1651_RNR1_12S_50ntFlanks@606.633.28|CCTGTTCTGTAATCGATAAACCCCGATC</td><td>>hg38_MT_+_598_1651_RNR1_12S_50ntFlanks</td><td>606</td><td>633</td><td>28</td><td>CCTGTTCTGTAATCGATAAACCCCGATC</td><td>307.5</td><td>6.6</td><td>1.2</td><td>5.7</td><td>1.2E-08</td><td>2.5E-08</td><td>MB231/MB468</td><td>CYTO</td></tr>
<tr><td>>hg38_21_+_8433172_8446622_RNA45SN1_45S_50ntFlanks@7975.8003.29|CGCGACCTCAGATCAGACGTGGCGACCCG</td><td>>hg38_21_+_8433172_8446622_RNA45SN1_45S_50ntFlanks</td><td>7975</td><td>8003</td><td>29</td><td>CGCGACCTCAGATCAGACGTGGCGACCCG</td><td>2214.1</td><td>-3.3</td><td>0.6</td><td>-5.7</td><td>1.2E-08</td><td>2.5E-08</td><td>MB231/MB468</td><td>MP</td></tr>
<tr><td>>hg38_21_+_8433172_8446622_RNA45SN1_45S_50ntFlanks@9545.9574.30|GTGGAGGTCCGTAGCGGTCCTGACGTGCAA</td><td>>hg38_21_+_8433172_8446622_RNA45SN1_45S_50ntFlanks</td><td>9545</td><td>9574</td><td>30</td><td>GTGGAGGTCCGTAGCGGTCCTGACGTGCAA</td><td>992.5</td><td>-3.3</td><td>0.6</td><td>-5.7</td><td>1.2E-08</td><td>2.5E-08</td><td>MB231/MB468</td><td>CYTO</td></tr>
<tr><td>>am_trnaMT_SerGCT_MT_+_12207_12265@13.38.26|AAGAACTGCTAACTCATGCCCCCATG</td><td>>am_trnaMT_SerGCT_MT_+_12207_12265</td><td>13</td><td>38</td><td>26</td><td>AAGAACTGCTAACTCATGCCCCCATG</td><td>1016.6</td><td>-4.9</td><td>0.9</td><td>-5.7</td><td>1.3E-08</td><td>2.5E-08</td><td>MB231/MB468</td><td>NUC</td></tr>
<tr><td>>MI0000811|hsa-mir-148b&WithFlank&12|+|54337210|54337320@69.91.23&[MIMAT0000759&hsa-miR-148b-3p&offsets|0|+1;m-77&12|+|54337278|54337299&offsets|0|+1]|TCAGTGCATCACAGAACTTTGTC</td><td>>MI0000811|hsa-mir-148b&WithFlank&12|+|54337210|54337320</td><td>69</td><td>91</td><td>23</td><td>TCAGTGCATCACAGAACTTTGTC</td><td>581.4</td><td>5.7</td><td>1.0</td><td>5.7</td><td>1.3E-08</td><td>2.6E-08</td><td>MB231/MB468</td><td>CYTO</td></tr>
<tr><td>>hg38_21_+_8433172_8446622_RNA45SN1_45S_50ntFlanks@12406.12429.24|TTTTTGATCCTTCGATGTCGGCTC</td><td>>hg38_21_+_8433172_8446622_RNA45SN1_45S_50ntFlanks</td><td>12406</td><td>12429</td><td>24</td><td>TTTTTGATCCTTCGATGTCGGCTC</td><td>299.4</td><td>-7.5</td><td>1.3</td><td>-5.7</td><td>1.4E-08</td><td>2.6E-08</td><td>BT20/MB468</td><td>NUC</td></tr>
<tr><td>>am_trna10_ValCAC_5_-_180649395_180649467@1.23.23,trna12_ValAAC_5_-_180645270_180645342@1.23.23,trna132_ValAAC_6_-_27721179_27721251@1.23.23,trna136_ValAAC_6_-_27648885_27648957@1.23.23,trna139_ValAAC_6_-_27618707_27618779@1.23.23,trna18_ValCAC_5_-_180529253_180529325@1.23.23,trna2_ValAAC_3_+_169490018_169490090@1.23.23,trna2_ValCAC_5_+_180524070_180524142@1.23.23,trna37_ValAAC_6_+_27203288_27203360@1.23.23,trna4_ValAAC_5_+_180591154_180591226@1.23.23,trna5_ValAAC_5_+_180596610_180596682@1.23.23,trna6_ValCAC_5_+_180600650_180600722@1.23.23,trna85_ValCAC_1_-_161369490_161369562@1.23.23,trna90_ValCAC_1_-_149684088_149684161@1.23.23,trna98_ValCAC_1_-_149298555_149298627@1.23.23,trna9_ValCAC_6_+_26538282_26538354@1.23.23|GTTTCCGTAGTGTAGTGGTTATC</td><td>>am_trna10_ValCAC_5_-_180649395_180649467</td><td>1</td><td>23</td><td>23</td><td>GTTTCCGTAGTGTAGTGGTTATC</td><td>1552.5</td><td>-2.9</td><td>0.5</td><td>-5.7</td><td>1.4E-08</td><td>2.6E-08</td><td>BT20/MB468</td><td>NUC</td></tr>
<tr><td>>hg38_21_+_8433172_8446622_RNA45SN1_45S_50ntFlanks@10690.10709.20|CGTCCGGTGAGCTCTCGCTG</td><td>>hg38_21_+_8433172_8446622_RNA45SN1_45S_50ntFlanks</td><td>10690</td><td>10709</td><td>20</td><td>CGTCCGGTGAGCTCTCGCTG</td><td>1061.0</td><td>-4.0</td><td>0.7</td><td>-5.7</td><td>9.7E-09</td><td>2.6E-08</td><td>BT20/MB468</td><td>MITO</td></tr>
<tr><td>>hg38_21_+_8433172_8446622_RNA45SN1_45S_50ntFlanks@9156.9188.33|CCAGTGCGCCCCGGGCGGGTCGCGCCGTCGGGC</td><td>>hg38_21_+_8433172_8446622_RNA45SN1_45S_50ntFlanks</td><td>9156</td><td>9188</td><td>33</td><td>CCAGTGCGCCCCGGGCGGGTCGCGCCGTCGGGC</td><td>1633.3</td><td>9.9</td><td>1.7</td><td>5.7</td><td>1.3E-08</td><td>2.6E-08</td><td>MB231/MB468</td><td>NUC</td></tr>
<tr><td>>MI0000093|hsa-mir-92a-1&WithFlank&13|+|91351308|91351397@54.73.20;MI0000094|hsa-mir-92a-2&WithFlank&X|-|134169532|134169618@54.73.20&[MIMAT0000092&hsa-miR-92a-3p&offsets|0|-2;m-9&13|+|91351361|91351382&offsets|0|-2];[MIMAT0000092_1&hsa-miR-92a-3p&offsets|0|-2;m-11&X|-|134169544|134169565&offsets|0|-2]|TATTGCACTTGTCCCGGCCT</td><td>>MI0000093|hsa-mir-92a-1&WithFlank&13|+|91351308|91351397</td><td>54</td><td>73</td><td>20</td><td>TATTGCACTTGTCCCGGCCT</td><td>289.5</td><td>5.7</td><td>1.0</td><td>5.7</td><td>1.4E-08</td><td>2.6E-08</td><td>MB231/MB468</td><td>MITO</td></tr>
<tr><td>>hg38_21_+_8433172_8446622_RNA45SN1_45S_50ntFlanks@12273.12290.18|TCAGTACGAATACAGACC</td><td>>hg38_21_+_8433172_8446622_RNA45SN1_45S_50ntFlanks</td><td>12273</td><td>12290</td><td>18</td><td>TCAGTACGAATACAGACC</td><td>129.8</td><td>-10.1</td><td>1.7</td><td>-6.0</td><td>1.8E-09</td><td>2.6E-08</td><td>BT20/MB468</td><td>MP</td></tr>
<tr><td>>hg38_MT_+_1621_3279_RNR2_16S_50ntFlanks@1216.1248.33|CAACCTCCGAGCAGTACATGCTAAGACTTCACC</td><td>>hg38_MT_+_1621_3279_RNR2_16S_50ntFlanks</td><td>1216</td><td>1248</td><td>33</td><td>CAACCTCCGAGCAGTACATGCTAAGACTTCACC</td><td>1338.6</td><td>6.9</td><td>1.2</td><td>5.7</td><td>9.8E-09</td><td>2.6E-08</td><td>BT20/MB468</td><td>MITO</td></tr>
<tr><td>>hg38_21_+_8433172_8446622_RNA45SN1_45S_50ntFlanks@12150.12183.34|GGTGGGGAGTTTGACTGGGGCGGTACACCTGTCA</td><td>>hg38_21_+_8433172_8446622_RNA45SN1_45S_50ntFlanks</td><td>12150</td><td>12183</td><td>34</td><td>GGTGGGGAGTTTGACTGGGGCGGTACACCTGTCA</td><td>62.1</td><td>-9.8</td><td>1.7</td><td>-5.7</td><td>1.2E-08</td><td>2.6E-08</td><td>MB231/MB468</td><td>MP</td></tr>
</table>

<table>
<tr><td>>hg38_21_+_8433172_8446622_RNA45SN1_45S_50ntFlanks@10727.10753.27|GGAGAGGGTGTAAATCTCGCGCCGGGC</td><td>>hg38_21_+_8433172_8446622_RNA45SN1_45S_50ntFlanks</td><td>10727</td><td>10753</td><td>27</td><td>GGAGAGGGTGTAAATCTCGCGCCGGGC</td><td>250.3</td><td>-7.6</td><td>1.3</td><td>-5.7</td><td>1.2E-08</td><td>2.6E-08</td><td>MB231/MB468</td><td>MP</td></tr>
<tr><td>>hg38_21_+_8433172_8446622_RNA45SN1_45S_50ntFlanks@12449.12476.28|GAATTCACCAAGCGTTGGATTGTTCACC</td><td>>hg38_21_+_8433172_8446622_RNA45SN1_45S_50ntFlanks</td><td>12449</td><td>12476</td><td>28</td><td>GAATTCACCAAGCGTTGGATTGTTCACC</td><td>400.3</td><td>6.0</td><td>1.1</td><td>5.7</td><td>1.3E-08</td><td>2.6E-08</td><td>MB231/MB468</td><td>CYTO</td></tr>
<tr><td>>hg38_21_+_8433172_8446622_RNA45SN1_45S_50ntFlanks@13007.13025.19|TATTGAAAGTCAGCCCTCG</td><td>>hg38_21_+_8433172_8446622_RNA45SN1_45S_50ntFlanks</td><td>13007</td><td>13025</td><td>19</td><td>TATTGAAAGTCAGCCCTCG</td><td>293.1</td><td>-7.4</td><td>1.3</td><td>-5.7</td><td>1.4E-08</td><td>2.6E-08</td><td>MB231/MB468</td><td>MITO</td></tr>
<tr><td>>hg38_21_+_8433172_8446622_RNA45SN1_45S_50ntFlanks@11920.11956.37|ATGAGAGGTGTAGAATAAGTGGGAGGCCCCCGGCGCC</td><td>>hg38_21_+_8433172_8446622_RNA45SN1_45S_50ntFlanks</td><td>11920</td><td>11956</td><td>37</td><td>ATGAGAGGTGTAGAATAAGTGGGAGGCCCCCGGCGCC</td><td>78.6</td><td>-10.2</td><td>1.8</td><td>-5.7</td><td>1.2E-08</td><td>2.6E-08</td><td>MB231/MB468</td><td>MP</td></tr>
<tr><td>>hg38_21_+_8433172_8446622_RNA45SN1_45S_50ntFlanks@6661.6710.50|CGGTGGATCACTCGGCTCGTGCGTCGATGAAGAACGCAGCTAGCTGCGAG</td><td>>hg38_21_+_8433172_8446622_RNA45SN1_45S_50ntFlanks</td><td>6661</td><td>6710</td><td>50</td><td>CGGTGGATCACTCGGCTCGTGCGTCGATGAAGAACGCAGCTAGCTGCGAG</td><td>3718.2</td><td>6.0</td><td>1.0</td><td>5.7</td><td>9.8E-09</td><td>2.6E-08</td><td>BT20/MB468</td><td>MITO</td></tr>
<tr><td>>hg38_21_+_8433172_8446622_RNA45SN1_45S_50ntFlanks@11581.11602.22|AACTGGTGCGGACCAGGGGAAT</td><td>>hg38_21_+_8433172_8446622_RNA45SN1_45S_50ntFlanks</td><td>11581</td><td>11602</td><td>22</td><td>AACTGGTGCGGACCAGGGGAAT</td><td>899.7</td><td>-5.1</td><td>0.9</td><td>-5.7</td><td>1.4E-08</td><td>2.6E-08</td><td>MB231/MB468</td><td>MITO</td></tr>
<tr><td>>hg38_21_+_8433172_8446622_RNA45SN1_45S_50ntFlanks@9466.9503.38|GGTGGAGCACGAGCGCACGTGTTAGGACCCGAAAGATG</td><td>>hg38_21_+_8433172_8446622_RNA45SN1_45S_50ntFlanks</td><td>9466</td><td>9503</td><td>38</td><td>GGTGGAGCACGAGCGCACGTGTTAGGACCCGAAAGATG</td><td>77.6</td><td>-10.1</td><td>1.8</td><td>-5.7</td><td>1.2E-08</td><td>2.6E-08</td><td>MB231/MB468</td><td>MP</td></tr>
<tr><td>>hg38_21_+_8433172_8446622_RNA45SN1_45S_50ntFlanks@12415.12432.18|CTTCGATGTCGGCTCTTC</td><td>>hg38_21_+_8433172_8446622_RNA45SN1_45S_50ntFlanks</td><td>12415</td><td>12432</td><td>18</td><td>CTTCGATGTCGGCTCTTC</td><td>1905.3</td><td>-8.5</td><td>1.5</td><td>-5.7</td><td>1.2E-08</td><td>2.6E-08</td><td>MB231/MB468</td><td>MP</td></tr>
<tr><td>>hg38_21_+_8433172_8446622_RNA45SN1_45S_50ntFlanks@3840.3879.40|CTCCTACTTGGATAACTGTGGTAATTCTAGAGCTAATACA</td><td>>hg38_21_+_8433172_8446622_RNA45SN1_45S_50ntFlanks</td><td>3840</td><td>3879</td><td>40</td><td>CTCCTACTTGGATAACTGTGGTAATTCTAGAGCTAATACA</td><td>1533.1</td><td>7.6</td><td>1.3</td><td>5.7</td><td>1.1E-08</td><td>2.6E-08</td><td>BT20/MB468</td><td>CYTO</td></tr>
<tr><td>>hg38_21_+_8433172_8446622_RNA45SN1_45S_50ntFlanks@9575.9595.21|ATCGGTCGTCCGACCTGGGTA</td><td>>hg38_21_+_8433172_8446622_RNA45SN1_45S_50ntFlanks</td><td>9575</td><td>9595</td><td>21</td><td>ATCGGTCGTCCGACCTGGGTA</td><td>1078.3</td><td>-7.1</td><td>1.3</td><td>-5.7</td><td>1.3E-08</td><td>2.6E-08</td><td>MB231/MB468</td><td>CYTO</td></tr>
<tr><td>>hg38_21_+_8433172_8446622_RNA45SN1_45S_50ntFlanks@11670.11701.32|CCCAGTGCTCTGAATGTCAAAGTGAAGAAATT</td><td>>hg38_21_+_8433172_8446622_RNA45SN1_45S_50ntFlanks</td><td>11670</td><td>11701</td><td>32</td><td>CCCAGTGCTCTGAATGTCAAAGTGAAGAAATT</td><td>485.7</td><td>-5.8</td><td>1.0</td><td>-5.7</td><td>1.4E-08</td><td>2.6E-08</td><td>MB231/MB468</td><td>MITO</td></tr>
<tr><td>>hg38_21_+_8433172_8446622_RNA45SN1_45S_50ntFlanks@4738.4757.20|AAAGTCGGAGGTTCGAAGAC</td><td>>hg38_21_+_8433172_8446622_RNA45SN1_45S_50ntFlanks</td><td>4738</td><td>4757</td><td>20</td><td>AAAGTCGGAGGTTCGAAGAC</td><td>538.5</td><td>4.0</td><td>0.7</td><td>5.7</td><td>1.3E-08</td><td>2.6E-08</td><td>MB231/MB468</td><td>CYTO</td></tr>
<tr><td>>hg38_21_+_8433172_8446622_RNA45SN1_45S_50ntFlanks@10346.10373.28|TCAAACGAGAACTTTGAAGGCCGAAGTG</td><td>>hg38_21_+_8433172_8446622_RNA45SN1_45S_50ntFlanks</td><td>10346</td><td>10373</td><td>28</td><td>TCAAACGAGAACTTTGAAGGCCGAAGTG</td><td>695.7</td><td>-4.1</td><td>0.7</td><td>-5.7</td><td>1.4E-08</td><td>2.6E-08</td><td>MB231/MB468</td><td>MITO</td></tr>
<tr><td>>hg38_21_+_8433172_8446622_RNA45SN1_45S_50ntFlanks@10402.10428.27|AACATGGGTCAGTCGGTCCTGAGAGAT</td><td>>hg38_21_+_8433172_8446622_RNA45SN1_45S_50ntFlanks</td><td>10402</td><td>10428</td><td>27</td><td>AACATGGGTCAGTCGGTCCTGAGAGAT</td><td>275.0</td><td>-5.9</td><td>1.0</td><td>-5.7</td><td>1.2E-08</td><td>2.6E-08</td><td>MB231/MB468</td><td>MP</td></tr>
<tr><td>>hg38_21_+_8433172_8446622_RNA45SN1_45S_50ntFlanks@12073.12091.19|CCGAGGGGCTCTCGCTTCT</td><td>>hg38_21_+_8433172_8446622_RNA45SN1_45S_50ntFlanks</td><td>12073</td><td>12091</td><td>19</td><td>CCGAGGGGCTCTCGCTTCT</td><td>236.9</td><td>-6.3</td><td>1.1</td><td>-5.7</td><td>1.3E-08</td><td>2.6E-08</td><td>MB231/MB468</td><td>CYTO</td></tr>
<tr><td>>hg38_21_+_8433172_8446622_RNA45SN1_45S_50ntFlanks@11593.11612.20|CCAGGGGAATCCGACTGTTT</td><td>>hg38_21_+_8433172_8446622_RNA45SN1_45S_50ntFlanks</td><td>11593</td><td>11612</td><td>20</td><td>CCAGGGGAATCCGACTGTTT</td><td>991.1</td><td>-6.3</td><td>1.1</td><td>-5.7</td><td>1.3E-08</td><td>2.6E-08</td><td>MB231/MB468</td><td>NUC</td></tr>
<tr><td>>hg38_21_+_8433172_8446622_RNA45SN1_45S_50ntFlanks@8323.8356.34|ACCGATAGTCAACAAGTACCGTAAGGGAAAGTTG</td><td>>hg38_21_+_8433172_8446622_RNA45SN1_45S_50ntFlanks</td><td>8323</td><td>8356</td><td>34</td><td>ACCGATAGTCAACAAGTACCGTAAGGGAAAGTTG</td><td>116.5</td><td>-9.9</td><td>1.7</td><td>-6.0</td><td>1.8E-09</td><td>2.6E-08</td><td>BT20/MB468</td><td>MP</td></tr>
<tr><td>>hg38_21_+_8433172_8446622_RNA45SN1_45S_50ntFlanks@3845.3881.37|ACTTGGATAACTGTGGTAATTCTAGAGCTAATACATG</td><td>>hg38_21_+_8433172_8446622_RNA45SN1_45S_50ntFlanks</td><td>3845</td><td>3881</td><td>37</td><td>ACTTGGATAACTGTGGTAATTCTAGAGCTAATACATG</td><td>260.8</td><td>-7.5</td><td>1.3</td><td>-5.7</td><td>1.4E-08</td><td>2.6E-08</td><td>MB231/MB468</td><td>MITO</td></tr>
<tr><td>>hg38_21_+_8433172_8446622_RNA45SN1_45S_50ntFlanks@5472.5501.30|ACGGCCCTGGCGGAGCGCTGAGAAGACGGT</td><td>>hg38_21_+_8433172_8446622_RNA45SN1_45S_50ntFlanks</td><td>5472</td><td>5501</td><td>30</td><td>ACGGCCCTGGCGGAGCGCTGAGAAGACGGT</td><td>754.4</td><td>3.9</td><td>0.7</td><td>5.7</td><td>1.3E-08</td><td>2.6E-08</td><td>MB231/MB468</td><td>NUC</td></tr>
<tr><td>>hg38_21_+_8433172_8446622_RNA45SN1_45S_50ntFlanks@12242.12272.31|AGAAGGGCAAAAGCTCGCTTGATCTTGATTT</td><td>>hg38_21_+_8433172_8446622_RNA45SN1_45S_50ntFlanks</td><td>12242</td><td>12272</td><td>31</td><td>AGAAGGGCAAAAGCTCGCTTGATCTTGATTT</td><td>998.7</td><td>-5.2</td><td>0.9</td><td>-5.7</td><td>1.4E-08</td><td>2.6E-08</td><td>MB231/MB468</td><td>MITO</td></tr>
<tr><td>>hg38_21_+_8433172_8446622_RNA45SN1_45S_50ntFlanks@11208.11239.32|GCGGCGACTCTGGACGCGAGCCGGGCCCTTCC</td><td>>hg38_21_+_8433172_8446622_RNA45SN1_45S_50ntFlanks</td><td>11208</td><td>11239</td><td>32</td><td>GCGGCGACTCTGGACGCGAGCCGGGCCCTTCC</td><td>268.3</td><td>-7.3</td><td>1.3</td><td>-5.7</td><td>1.4E-08</td><td>2.6E-08</td><td>MB231/MB468</td><td>MITO</td></tr>
<tr><td>>hg38_21_+_8433172_8446622_RNA45SN1_45S_50ntFlanks@11827.11844.18|AGCGAAACCACAGCCAAG</td><td>>hg38_21_+_8433172_8446622_RNA45SN1_45S_50ntFlanks</td><td>11827</td><td>11844</td><td>18</td><td>AGCGAAACCACAGCCAAG</td><td>72.9</td><td>-10.1</td><td>1.8</td><td>-5.7</td><td>1.2E-08</td><td>2.6E-08</td><td>MB231/MB468</td><td>MP</td></tr>
<tr><td>>hg38_21_+_8433172_8446622_RNA45SN1_45S_50ntFlanks@11821.11847.27|CTATCCAGCGAAACCACAGCCAAGGGA</td><td>>hg38_21_+_8433172_8446622_RNA45SN1_45S_50ntFlanks</td><td>11821</td><td>11847</td><td>27</td><td>CTATCCAGCGAAACCACAGCCAAGGGA</td><td>61.4</td><td>-9.8</td><td>1.7</td><td>-5.7</td><td>1.2E-08</td><td>2.6E-08</td><td>MB231/MB468</td><td>MP</td></tr>
<tr><td>>MI0003591|hsa-mir-584&WithFlank&5|-|149062307|149062415@22.42.21&[MIMAT0003249&hsa-miR-584-5p&offsets|0|-1;m-167&5|-|149062374|149062394&offsets|0|0]|TTATGGTTTGCCTGGGACTGA</td><td>>MI0003591|hsa-mir-584&WithFlank&5|-|149062307|149062415</td><td>22</td><td>42</td><td>21</td><td>TTATGGTTTGCCTGGGACTGA</td><td>2281.1</td><td>3.9</td><td>0.7</td><td>5.7</td><td>1.2E-08</td><td>2.6E-08</td><td>MB231/MB468</td><td>MP</td></tr>
<tr><td>>hg38_21_+_8433172_8446622_RNA45SN1_45S_50ntFlanks@12958.12993.36|TTCTGGGTCGGGGTTTCGTACGTAGCAGAGCAGCTC</td><td>>hg38_21_+_8433172_8446622_RNA45SN1_45S_50ntFlanks</td><td>12958</td><td>12993</td><td>36</td><td>TTCTGGGTCGGGGTTTCGTACGTAGCAGAGCAGCTC</td><td>413.9</td><td>7.0</td><td>1.2</td><td>5.7</td><td>1.1E-08</td><td>2.6E-08</td><td>BT20/MB468</td><td>CYTO</td></tr>
<tr><td>>hg38_21_+_8433172_8446622_RNA45SN1_45S_50ntFlanks@9534.9565.32|AGGAAACTCTGGTGGAGGTCCGTAGCGGTCCT</td><td>>hg38_21_+_8433172_8446622_RNA45SN1_45S_50ntFlanks</td><td>9534</td><td>9565</td><td>32</td><td>AGGAAACTCTGGTGGAGGTCCGTAGCGGTCCT</td><td>693.7</td><td>-8.1</td><td>1.4</td><td>-5.7</td><td>1.2E-08</td><td>2.6E-08</td><td>MB231/MB468</td><td>MP</td></tr>
<tr><td>>MI0000471|hsa-mir-126&WithFlank&9|+|136670596|136670692@58.78.21(+1A)&[MIMAT0000445&hsa-miR-126-3p&offsets|0|-1(+1A);m-40&9|+|136670653|136670674&offsets|0|-1(+1A)]|TCGTACCGTGAGTAATAATGCA</td><td>>MI0000471|hsa-mir-126&WithFlank&9|+|136670596|136670692</td><td>58</td><td>78</td><td>21(+1A)</td><td>TCGTACCGTGAGTAATAATGCA</td><td>633.0</td><td>9.0</td><td>1.6</td><td>5.7</td><td>1.4E-08</td><td>2.6E-08</td><td>MB231/MB468</td><td>MITO</td></tr>
<tr><td>>hg38_21_+_8433172_8446622_RNA45SN1_45S_50ntFlanks@10421.10443.23|TGAGAGATGGGCGAGCGCCGTTC</td><td>>hg38_21_+_8433172_8446622_RNA45SN1_45S_50ntFlanks</td><td>10421</td><td>10443</td><td>23</td><td>TGAGAGATGGGCGAGCGCCGTTC</td><td>149.0</td><td>-6.6</td><td>1.2</td><td>-5.7</td><td>1.3E-08</td><td>2.6E-08</td><td>MB231/MB468</td><td>NUC</td></tr>
<tr><td>>hg38_21_+_8433172_8446622_RNA45SN1_45S_50ntFlanks@12418.12443.26|CGATGTCGGCTCTTCCTATCATTGTG</td><td>>hg38_21_+_8433172_8446622_RNA45SN1_45S_50ntFlanks</td><td>12418</td><td>12443</td><td>26</td><td>CGATGTCGGCTCTTCCTATCATTGTG</td><td>2007.5</td><td>-8.6</td><td>1.5</td><td>-5.7</td><td>1.2E-08</td><td>2.6E-08</td><td>MB231/MB468</td><td>MP</td></tr>
<tr><td>>hg38_1_-_228634819_228635039_RNA5S12_5S_50ntFlanks@137.171.35|GAGACCGCCTGGGAATACCGGGTGCTGTAGGCTTT</td><td>>hg38_1_-_228634819_228635039_RNA5S12_5S_50ntFlanks</td><td>137</td><td>171</td><td>35</td><td>GAGACCGCCTGGGAATACCGGGTGCTGTAGGCTTT</td><td>804.9</td><td>4.9</td><td>0.8</td><td>5.7</td><td>1.0E-08</td><td>2.6E-08</td><td>BT20/MB468</td><td>MITO</td></tr>
<tr><td>>hg38_21_+_8433172_8446622_RNA45SN1_45S_50ntFlanks@475.504.30|CTTCGTGATCGATGTGGTGACGTCGTGCTC</td><td>>hg38_21_+_8433172_8446622_RNA45SN1_45S_50ntFlanks</td><td>475</td><td>504</td><td>30</td><td>CTTCGTGATCGATGTGGTGACGTCGTGCTC</td><td>2422.6</td><td>5.6</td><td>1.0</td><td>5.7</td><td>1.4E-08</td><td>2.6E-08</td><td>MB231/MB468</td><td>MITO</td></tr>
<tr><td>>hg38_21_+_8433172_8446622_RNA45SN1_45S_50ntFlanks@12417.12455.39|TCGATGTCGGCTCTTCCTATCATTGTGAAGCAGAATTCA</td><td>>hg38_21_+_8433172_8446622_RNA45SN1_45S_50ntFlanks</td><td>12417</td><td>12455</td><td>39</td><td>TCGATGTCGGCTCTTCCTATCATTGTGAAGCAGAATTCA</td><td>3999.2</td><td>-5.2</td><td>0.9</td><td>-5.7</td><td>1.4E-08</td><td>2.6E-08</td><td>MB231/MB468</td><td>MITO</td></tr>
<tr><td>>hg38_21_+_8433172_8446622_RNA45SN1_45S_50ntFlanks@8194.8224.31|CCCGTGGACGGTGTGAGGCCGGTAGCGGCCC</td><td>>hg38_21_+_8433172_8446622_RNA45SN1_45S_50ntFlanks</td><td>8194</td><td>8224</td><td>31</td><td>CCCGTGGACGGTGTGAGGCCGGTAGCGGCCC</td><td>107.3</td><td>-9.8</td><td>1.6</td><td>-6.0</td><td>1.9E-09</td><td>2.6E-08</td><td>BT20/MB468</td><td>MP</td></tr>
<tr><td>>hg38_21_+_8433172_8446622_RNA45SN1_45S_50ntFlanks@10770.10796.27|GCAGGTCTCCAAGGTGAACAGCCTCTG</td><td>>hg38_21_+_8433172_8446622_RNA45SN1_45S_50ntFlanks</td><td>10770</td><td>10796</td><td>27</td><td>GCAGGTCTCCAAGGTGAACAGCCTCTG</td><td>1160.5</td><td>-7.0</td><td>1.2</td><td>-5.7</td><td>1.5E-08</td><td>2.6E-08</td><td>BT20/MB468</td><td>NUC</td></tr>
<tr><td>>hg38_21_+_8433172_8446622_RNA45SN1_45S_50ntFlanks@12054.12071.18|TGAGGCGGGGGGGCGAGC</td><td>>hg38_21_+_8433172_8446622_RNA45SN1_45S_50ntFlanks</td><td>12054</td><td>12071</td><td>18</td><td>TGAGGCGGGGGGGCGAGC</td><td>161.5</td><td>-6.2</td><td>1.1</td><td>-5.7</td><td>1.3E-08</td><td>2.6E-08</td><td>MB231/MB468</td><td>CYTO</td></tr>
<tr><td>>hg38_21_+_8433172_8446622_RNA45SN1_45S_50ntFlanks@12202.12240.39|CTAAGGCGAGCTCAGGGAGGACAGAAACCTCCCGTGGAG</td><td>>hg38_21_+_8433172_8446622_RNA45SN1_45S_50ntFlanks</td><td>12202</td><td>12240</td><td>39</td><td>CTAAGGCGAGCTCAGGGAGGACAGAAACCTCCCGTGGAG</td><td>153.0</td><td>-11.1</td><td>2.0</td><td>-5.7</td><td>1.2E-08</td><td>2.6E-08</td><td>MB231/MB468</td><td>MP</td></tr>
<tr><td>>hg38_21_+_8433172_8446622_RNA45SN1_45S_50ntFlanks@12616.12653.38|CTGAGGAGCCAATGGGGCGAAGCTACCATCTGTGGGAT</td><td>>hg38_21_+_8433172_8446622_RNA45SN1_45S_50ntFlanks</td><td>12616</td><td>12653</td><td>38</td><td>CTGAGGAGCCAATGGGGCGAAGCTACCATCTGTGGGAT</td><td>755.8</td><td>-4.5</td><td>0.8</td><td>-5.7</td><td>1.0E-08</td><td>2.6E-08</td><td>BT20/MB468</td><td>MITO</td></tr>
<tr><td>>MI0000749|hsa-mir-30e&WithFlank&1|+|40754349|40754452@23.45.23(+1A)&[MIMAT0000692&hsa-miR-30e-5p&offsets|0|+1(+1A);m-24&1|+|40754371|40754394&offsets|0|-1(+1A)]|TGTAAACATCCTTGACTGGAAGCA</td><td>>MI0000749|hsa-mir-30e&WithFlank&1|+|40754349|40754452</td><td>23</td><td>45</td><td>23(+1A)</td><td>TGTAAACATCCTTGACTGGAAGCA</td><td>273.2</td><td>4.9</td><td>0.9</td><td>5.7</td><td>1.3E-08</td><td>2.6E-08</td><td>MB231/MB468</td><td>NUC</td></tr>
<tr><td>>hg38_21_+_8433172_8446622_RNA45SN1_45S_50ntFlanks@11322.11344.23|ACGTCTCGTCGCGCGCGCGTCCG</td><td>>hg38_21_+_8433172_8446622_RNA45SN1_45S_50ntFlanks</td><td>11322</td><td>11344</td><td>23</td><td>ACGTCTCGTCGCGCGCGCGTCCG</td><td>8114.9</td><td>-4.9</td><td>0.9</td><td>-5.7</td><td>1.1E-08</td><td>2.6E-08</td><td>BT20/MB468</td><td>CYTO</td></tr>
<tr><td>>hg38_21_+_8433172_8446622_RNA45SN1_45S_50ntFlanks@10788.10807.20|CAGCCTCTGGCATGTTGGAA</td><td>>hg38_21_+_8433172_8446622_RNA45SN1_45S_50ntFlanks</td><td>10788</td><td>10807</td><td>20</td><td>CAGCCTCTGGCATGTTGGAA</td><td>295.1</td><td>-6.8</td><td>1.2</td><td>-5.7</td><td>1.3E-08</td><td>2.7E-08</td><td>MB231/MB468</td><td>CYTO</td></tr>
<tr><td>>MI0000266|hsa-mir-10a&WithFlank&17|-|48579832|48579953@28.50.23&[MIMAT0000253&hsa-miR-10a-5p&offsets|0|0;m-18&17|-|48579905|48579926&offsets|0|+1]|TACCCTGTAGATCCGAATTTGTG</td><td>>MI0000266|hsa-mir-10a&WithFlank&17|-|48579832|48579953</td><td>28</td><td>50</td><td>23</td><td>TACCCTGTAGATCCGAATTTGTG</td><td>2573.2</td><td>8.5</td><td>1.5</td><td>5.7</td><td>1.2E-08</td><td>2.7E-08</td><td>MB231/MB468</td><td>MP</td></tr>
<tr><td>>hg38_21_+_8433172_8446622_RNA45SN1_45S_50ntFlanks@10713.10735.23|CTTGAAAATCCGGGGGAGAGGGT</td><td>>hg38_21_+_8433172_8446622_RNA45SN1_45S_50ntFlanks</td><td>10713</td><td>10735</td><td>23</td><td>CTTGAAAATCCGGGGGAGAGGGT</td><td>246.8</td><td>-7.3</td><td>1.3</td><td>-5.7</td><td>1.2E-08</td><td>2.7E-08</td><td>MB231/MB468</td><td>MP</td></tr>
<tr><td>>hg38_21_+_8433172_8446622_RNA45SN1_45S_50ntFlanks@8104.8143.40|CCGCGGCGGGGCGCGGGACATGTGGCGTACGGAAGACCCG</td><td>>hg38_21_+_8433172_8446622_RNA45SN1_45S_50ntFlanks</td><td>8104</td><td>8143</td><td>40</td><td>CCGCGGCGGGGCGCGGGACATGTGGCGTACGGAAGACCCG</td><td>22355.8</td><td>-7.7</td><td>1.3</td><td>-6.0</td><td>1.9E-09</td><td>2.7E-08</td><td>BT20/MB468</td><td>MP</td></tr>
<tr><td>>trna27_GlyCCC_2_-_70476123_70476193@1.35.35,trna34_GlyCCC_16_-_686736_686806@1.35.35|GCGCCGCTGGTGTAGTGGTATCATGCAAGATTCCC</td><td>>trna27_GlyCCC_2_-_70476123_70476193</td><td>1</td><td>35</td><td>35</td><td>GCGCCGCTGGTGTAGTGGTATCATGCAAGATTCCC</td><td>77.5</td><td>9.5</td><td>1.7</td><td>5.7</td><td>1.2E-08</td><td>2.7E-08</td><td>MB231/MB468</td><td>MP</td></tr>
<tr><td>>hg38_21_+_8433172_8446622_RNA45SN1_45S_50ntFlanks@10392.10428.37|ACAGCAGTTGAACATGGGTCAGTCGGTCCTGAGAGAT</td><td>>hg38_21_+_8433172_8446622_RNA45SN1_45S_50ntFlanks</td><td>10392</td><td>10428</td><td>37</td><td>ACAGCAGTTGAACATGGGTCAGTCGGTCCTGAGAGAT</td><td>391.0</td><td>-3.9</td><td>0.7</td><td>-5.7</td><td>1.3E-08</td><td>2.7E-08</td><td>MB231/MB468</td><td>NUC</td></tr>
<tr><td>>hg38_21_+_8433172_8446622_RNA45SN1_45S_50ntFlanks@4005.4022.18|AACCTCGGGCCGATCGCA</td><td>>hg38_21_+_8433172_8446622_RNA45SN1_45S_50ntFlanks</td><td>4005</td><td>4022</td><td>18</td><td>AACCTCGGGCCGATCGCA</td><td>861.3</td><td>-4.1</td><td>0.7</td><td>-5.7</td><td>1.3E-08</td><td>2.7E-08</td><td>MB231/MB468</td><td>NUC</td></tr>
<tr><td>>MI0000286|hsa-mir-210&WithFlank&11|-|568083|568204@72.93.22&[MIMAT0000267&hsa-miR-210-3p&offsets|0|0;m-102&11|-|568112|568133&offsets|0|0]|CTGTGCGTGTGACAGCGGCTGA</td><td>>MI0000286|hsa-mir-210&WithFlank&11|-|568083|568204</td><td>72</td><td>93</td><td>22</td><td>CTGTGCGTGTGACAGCGGCTGA</td><td>563.5</td><td>6.6</td><td>1.2</td><td>5.7</td><td>1.4E-08</td><td>2.7E-08</td><td>MB231/MB468</td><td>MITO</td></tr>
<tr><td>>hg38_21_+_8433172_8446622_RNA45SN1_45S_50ntFlanks@10788.10822.35|CAGCCTCTGGCATGTTGGAACAATGTAGGTAAGGG</td><td>>hg38_21_+_8433172_8446622_RNA45SN1_45S_50ntFlanks</td><td>10788</td><td>10822</td><td>35</td><td>CAGCCTCTGGCATGTTGGAACAATGTAGGTAAGGG</td><td>448.9</td><td>-7.6</td><td>1.3</td><td>-5.7</td><td>1.4E-08</td><td>2.7E-08</td><td>MB231/MB468</td><td>MITO</td></tr>
<tr><td>>MI0000076|hsa-mir-20a&WithFlank&13|+|91351059|91351141@14.34.21&[MIMAT0000075&hsa-miR-20a-5p&offsets|0|-2;m-96&13|+|91351072|91351094&offsets|0|-2]|TAAAGTGCTTATAGTGCAGGT</td><td>>MI0000076|hsa-mir-20a&WithFlank&13|+|91351059|91351141</td><td>14</td><td>34</td><td>21</td><td>TAAAGTGCTTATAGTGCAGGT</td><td>2057.9</td><td>6.1</td><td>1.1</td><td>5.7</td><td>1.4E-08</td><td>2.7E-08</td><td>MB231/MB468</td><td>MITO</td></tr>
<tr><td>>hg38_21_+_8433172_8446622_RNA45SN1_45S_50ntFlanks@7973.8010.38|GACGCGACCTCAGATCAGACGTGGCGACCCGCTGAATT</td><td>>hg38_21_+_8433172_8446622_RNA45SN1_45S_50ntFlanks</td><td>7973</td><td>8010</td><td>38</td><td>GACGCGACCTCAGATCAGACGTGGCGACCCGCTGAATT</td><td>2586.5</td><td>3.6</td><td>0.6</td><td>5.7</td><td>1.4E-08</td><td>2.7E-08</td><td>MB231/MB468</td><td>MITO</td></tr>
<tr><td>>hg38_21_+_8433172_8446622_RNA45SN1_45S_50ntFlanks@3835.3862.28|CTCCTCTCCTACTTGGATAACTGTGGTA</td><td>>hg38_21_+_8433172_8446622_RNA45SN1_45S_50ntFlanks</td><td>3835</td><td>3862</td><td>28</td><td>CTCCTCTCCTACTTGGATAACTGTGGTA</td><td>250.9</td><td>-5.5</td><td>1.0</td><td>-5.7</td><td>1.3E-08</td><td>2.7E-08</td><td>MB231/MB468</td><td>NUC</td></tr>
</table>

<table>
<tr><td>>hg38_21_+_8433172_8446622_RNA45SN1_45S_50ntFlanks@10686.10707.22|GCGGCGTCCGGTGAGCTCTCGC</td><td>>hg38_21_+_8433172_8446622_RNA45SN1_45S_50ntFlanks</td><td>10686</td><td>10707</td><td>22</td><td>GCGGCGTCCGGTGAGCTCTCGC</td><td>1258.9</td><td>5.3</td><td>0.9</td><td>5.7</td><td>1.0E-08</td><td>2.7E-08</td><td>BT20/MB468</td><td>MITO</td></tr>
<tr><td>>hg38_21_+_8433172_8446622_RNA45SN1_45S_50ntFlanks@10735.10755.21|TGTAAATCTCGCGCCGGGCCG</td><td>>hg38_21_+_8433172_8446622_RNA45SN1_45S_50ntFlanks</td><td>10735</td><td>10755</td><td>21</td><td>TGTAAATCTCGCGCCGGGCCG</td><td>583.6</td><td>-3.8</td><td>0.7</td><td>-5.7</td><td>1.0E-08</td><td>2.7E-08</td><td>BT20/MB468</td><td>MITO</td></tr>
<tr><td>>am_trna111_HisGTG_1_-_147774845_147774916@-1G.26.27,trna118_HisGTG_1_-_145396881_145396952@-1G.26.27,trna16_HisGTG_1_+_146544773_146544844@-1G.26.27,trna1_HisGTG_15_+_45493349_45493420@-1G.26.27,trna21_HisGTG_1_+_147753471_147753542@-1G.26.27,trna33_HisGTG_6_+_27125906_27125977@-1G.26.27,trna7_HisGTG_9_-_14433938_14434009@-1G.26.27,trna8_HisGTG_15_-_45492611_45492682@-1G.26.27,trna9_HisGTG_15_-_45490804_45490875@-1G.26.27|GGCCGTGATCGTATAGTGGTTAGTACT</td><td>>am_trna111_HisGTG_1_-_147774845_147774916</td><td>-1G</td><td>26</td><td>27</td><td>GGCCGTGATCGTATAGTGGTTAGTACT</td><td>13465.2</td><td>6.7</td><td>1.2</td><td>5.7</td><td>1.3E-08</td><td>2.7E-08</td><td>MB231/MB468</td><td>CYTO</td></tr>
<tr><td>>MI0000298|hsa-mir-221&WithFlank&X|-|45746151|45746272@71.91.21(+1C)&[MIMAT0000278&hsa-miR-221-3p&offsets|0|-2(+1C);m-91&X|-|45746181|45746202&offsets|0|-1(+1C)]|AGCTACATTGTCTGCTGGGTTC</td><td>>MI0000298|hsa-mir-221&WithFlank&X|-|45746151|45746272</td><td>71</td><td>91</td><td>21(+1C)</td><td>AGCTACATTGTCTGCTGGGTTC</td><td>193.7</td><td>2.7</td><td>0.5</td><td>5.7</td><td>1.5E-08</td><td>2.7E-08</td><td>BT20/MB468</td><td>NUC</td></tr>
<tr><td>>hg38_21_+_8433172_8446622_RNA45SN1_45S_50ntFlanks@8158.8188.31|TCGTGGGGGGCCCAAGTCCTTCTGATCGAGG</td><td>>hg38_21_+_8433172_8446622_RNA45SN1_45S_50ntFlanks</td><td>8158</td><td>8188</td><td>31</td><td>TCGTGGGGGGCCCAAGTCCTTCTGATCGAGG</td><td>925.2</td><td>-6.3</td><td>1.1</td><td>-5.7</td><td>1.4E-08</td><td>2.7E-08</td><td>MB231/MB468</td><td>MITO</td></tr>
<tr><td>>hg38_21_+_8433172_8446622_RNA45SN1_45S_50ntFlanks@3838.3872.35|CTCTCCTACTTGGATAACTGTGGTAATTCTAGAGC</td><td>>hg38_21_+_8433172_8446622_RNA45SN1_45S_50ntFlanks</td><td>3838</td><td>3872</td><td>35</td><td>CTCTCCTACTTGGATAACTGTGGTAATTCTAGAGC</td><td>98.1</td><td>-10.5</td><td>1.8</td><td>-5.7</td><td>1.2E-08</td><td>2.7E-08</td><td>MB231/MB468</td><td>MP</td></tr>
<tr><td>>am_MI0000809|hsa-mir-151a&WithFlank&8|-|140732558|140732659@53.72.20&[MIMAT0000757&hsa-miR-151a-3p&offsets|0|-1;m-30&8|-|140732587|140732607&offsets|0|-1]|CTAGACTGAAGCTCCTTGAG</td><td>>am_MI0000809|hsa-mir-151a&WithFlank&8|-|140732558|140732659</td><td>53</td><td>72</td><td>20</td><td>CTAGACTGAAGCTCCTTGAG</td><td>6144.7</td><td>5.9</td><td>1.0</td><td>6.0</td><td>1.9E-09</td><td>2.7E-08</td><td>BT20/MB468</td><td>MP</td></tr>
<tr><td>>hg38_21_+_8433172_8446622_RNA45SN1_45S_50ntFlanks@12488.12521.34|AACGTGAGCTGGGTTTAGACCGTCGTGAGACAGG</td><td>>hg38_21_+_8433172_8446622_RNA45SN1_45S_50ntFlanks</td><td>12488</td><td>12521</td><td>34</td><td>AACGTGAGCTGGGTTTAGACCGTCGTGAGACAGG</td><td>439.3</td><td>-7.2</td><td>1.3</td><td>-5.7</td><td>1.0E-08</td><td>2.7E-08</td><td>BT20/MB468</td><td>MITO</td></tr>
<tr><td>>hg38_21_+_8433172_8446622_RNA45SN1_45S_50ntFlanks@10597.10617.21|AGTTCTCTTTTCTTTGTGAAG</td><td>>hg38_21_+_8433172_8446622_RNA45SN1_45S_50ntFlanks</td><td>10597</td><td>10617</td><td>21</td><td>AGTTCTCTTTTCTTTGTGAAG</td><td>106.1</td><td>-9.8</td><td>1.6</td><td>-6.0</td><td>1.9E-09</td><td>2.7E-08</td><td>BT20/MB468</td><td>MP</td></tr>
<tr><td>>hg38_21_+_8433172_8446622_RNA45SN1_45S_50ntFlanks@10348.10372.25|AAACGAGAACTTTGAAGGCCGAAGT</td><td>>hg38_21_+_8433172_8446622_RNA45SN1_45S_50ntFlanks</td><td>10348</td><td>10372</td><td>25</td><td>AAACGAGAACTTTGAAGGCCGAAGT</td><td>671.7</td><td>2.9</td><td>0.5</td><td>5.7</td><td>1.0E-08</td><td>2.7E-08</td><td>BT20/MB468</td><td>MITO</td></tr>
<tr><td>>hg38_21_+_8433172_8446622_RNA45SN1_45S_50ntFlanks@3873.3899.27|TAATACATGCCGACGGGCGCTGACCCC</td><td>>hg38_21_+_8433172_8446622_RNA45SN1_45S_50ntFlanks</td><td>3873</td><td>3899</td><td>27</td><td>TAATACATGCCGACGGGCGCTGACCCC</td><td>759.2</td><td>-8.5</td><td>1.5</td><td>-5.7</td><td>1.5E-08</td><td>2.7E-08</td><td>BT20/MB468</td><td>NUC</td></tr>
<tr><td>>hg38_21_+_8433172_8446622_RNA45SN1_45S_50ntFlanks@12023.12043.21|CACTACTCTGATCGTTTTTTC</td><td>>hg38_21_+_8433172_8446622_RNA45SN1_45S_50ntFlanks</td><td>12023</td><td>12043</td><td>21</td><td>CACTACTCTGATCGTTTTTTC</td><td>412.3</td><td>-7.7</td><td>1.4</td><td>-5.7</td><td>1.4E-08</td><td>2.7E-08</td><td>MB231/MB468</td><td>MITO</td></tr>
<tr><td>>MI0000434|hsa-let-7i&WithFlank&12|+|62603680|62603775@12.32.21&[MIMAT0000415&hsa-let-7i-5p&offsets|0|-1;m-49&12|+|62603691|62603712&offsets|0|-1]|TGAGGTAGTAGTTTGTGCTGT</td><td>>MI0000434|hsa-let-7i&WithFlank&12|+|62603680|62603775</td><td>12</td><td>32</td><td>21</td><td>TGAGGTAGTAGTTTGTGCTGT</td><td>1711.8</td><td>5.4</td><td>0.9</td><td>5.7</td><td>1.4E-08</td><td>2.7E-08</td><td>MB231/MB468</td><td>MITO</td></tr>
<tr><td>>hg38_21_+_8433172_8446622_RNA45SN1_45S_50ntFlanks@11581.11601.21|AACTGGTGCGGACCAGGGGAA</td><td>>hg38_21_+_8433172_8446622_RNA45SN1_45S_50ntFlanks</td><td>11581</td><td>11601</td><td>21</td><td>AACTGGTGCGGACCAGGGGAA</td><td>1252.5</td><td>-9.2</td><td>1.5</td><td>-6.0</td><td>1.9E-09</td><td>2.7E-08</td><td>BT20/MB468</td><td>MP</td></tr>
<tr><td>>MI0000270|hsa-mir-181b-1&WithFlank&1|-|198858867|198858988@42.64.23;MI0000683|hsa-mir-181b-2&WithFlank&9|+|124693704|124693804@22.44.23&[MIMAT0000257&hsa-miR-181b-5p&offsets|0|0;m-73&1|-|198858925|198858947&offsets|0|0];[MIMAT0000257_1&hsa-miR-181b-5p&offsets|0|0;m-72&9|+|124693725|124693747&offsets|0|0]|AACATTCATTGCTGTCGGTGGGT</td><td>>MI0000270|hsa-mir-181b-1&WithFlank&1|-|198858867|198858988</td><td>42</td><td>64</td><td>23</td><td>AACATTCATTGCTGTCGGTGGGT</td><td>1952.1</td><td>4.8</td><td>0.9</td><td>5.7</td><td>1.3E-08</td><td>2.7E-08</td><td>MB231/MB468</td><td>CYTO</td></tr>
<tr><td>>hg38_21_+_8433172_8446622_RNA45SN1_45S_50ntFlanks@10392.10418.27|ACAGCAGTTGAACATGGGTCAGTCGGT</td><td>>hg38_21_+_8433172_8446622_RNA45SN1_45S_50ntFlanks</td><td>10392</td><td>10418</td><td>27</td><td>ACAGCAGTTGAACATGGGTCAGTCGGT</td><td>758.1</td><td>-8.7</td><td>1.5</td><td>-5.7</td><td>1.5E-08</td><td>2.7E-08</td><td>BT20/MB468</td><td>NUC</td></tr>
<tr><td>>MI0000447|hsa-mir-128-1&WithFlank&2|+|135665391|135665484@56.75.20;MI0000727|hsa-mir-128-2&WithFlank&3|+|35744470|35744565@58.77.20&[MIMAT0000424&hsa-miR-128-3p&offsets|0|-1;m-108&2|+|135665446|135665466&offsets|0|-1];[MIMAT0000424_1&hsa-miR-128-3p&offsets|0|-1;m-117&3|+|35744527|35744547&offsets|0|-1]|TCACAGTGAACCGGTCTCTT</td><td>>MI0000447|hsa-mir-128-1&WithFlank&2|+|135665391|135665484</td><td>56</td><td>75</td><td>20</td><td>TCACAGTGAACCGGTCTCTT</td><td>442.4</td><td>4.1</td><td>0.7</td><td>5.7</td><td>1.3E-08</td><td>2.7E-08</td><td>MB231/MB468</td><td>CYTO</td></tr>
<tr><td>>MI0000101|hsa-mir-99a&WithFlank&21|+|16539083|16539175@19.39.21&[MIMAT0000097&hsa-miR-99a-5p&offsets|0|-1;m-64&21|+|16539101|16539121&offsets|0|0]|AACCCGTAGATCCGATCTTGT</td><td>>MI0000101|hsa-mir-99a&WithFlank&21|+|16539083|16539175</td><td>19</td><td>39</td><td>21</td><td>AACCCGTAGATCCGATCTTGT</td><td>4241.5</td><td>-9.6</td><td>1.7</td><td>-5.7</td><td>1.5E-08</td><td>2.7E-08</td><td>BT20/MB468</td><td>NUC</td></tr>
<tr><td>>hg38_21_+_8433172_8446622_RNA45SN1_45S_50ntFlanks@12404.12436.33|GCTTTTTGATCCTTCGATGTCGGCTCTTCCTAT</td><td>>hg38_21_+_8433172_8446622_RNA45SN1_45S_50ntFlanks</td><td>12404</td><td>12436</td><td>33</td><td>GCTTTTTGATCCTTCGATGTCGGCTCTTCCTAT</td><td>2219.3</td><td>-9.9</td><td>1.8</td><td>-5.7</td><td>1.5E-08</td><td>2.7E-08</td><td>BT20/MB468</td><td>NUC</td></tr>
<tr><td>>hg38_21_+_8433172_8446622_RNA45SN1_45S_50ntFlanks@12475.12497.23|CCCACTAATAGGGAACGTGAGCT</td><td>>hg38_21_+_8433172_8446622_RNA45SN1_45S_50ntFlanks</td><td>12475</td><td>12497</td><td>23</td><td>CCCACTAATAGGGAACGTGAGCT</td><td>406.0</td><td>-4.5</td><td>0.8</td><td>-5.7</td><td>1.5E-08</td><td>2.7E-08</td><td>BT20/MB468</td><td>NUC</td></tr>
<tr><td>>MI0000542|hsa-mir-320a&WithFlank&8|-|22244960|22245043@49.68.20(+1U)&[MIMAT0000510&hsa-miR-320a-3p&offsets|+1|-1(+1U);m-60&8|-|22244975|22244996&offsets|+1|-1(+1U)]|AAAGCTGGGTTGAGAGGGCGT</td><td>>MI0000542|hsa-mir-320a&WithFlank&8|-|22244960|22245043</td><td>49</td><td>68</td><td>20(+1U)</td><td>AAAGCTGGGTTGAGAGGGCGT</td><td>2054.2</td><td>8.4</td><td>1.4</td><td>6.0</td><td>1.9E-09</td><td>2.7E-08</td><td>BT20/MB468</td><td>MP</td></tr>
<tr><td>>hg38_21_+_8433172_8446622_RNA45SN1_45S_50ntFlanks@11783.11816.34|CATGAATGGATGAACGAGATTCCCACTGTCCCTA</td><td>>hg38_21_+_8433172_8446622_RNA45SN1_45S_50ntFlanks</td><td>11783</td><td>11816</td><td>34</td><td>CATGAATGGATGAACGAGATTCCCACTGTCCCTA</td><td>230.9</td><td>-7.8</td><td>1.4</td><td>-5.7</td><td>1.5E-08</td><td>2.7E-08</td><td>MB231/MB468</td><td>MITO</td></tr>
<tr><td>>hg38_21_+_8433172_8446622_RNA45SN1_45S_50ntFlanks@11316.11343.28|CACCCCACGTCTCGTCGCGCGCGCGTCC</td><td>>hg38_21_+_8433172_8446622_RNA45SN1_45S_50ntFlanks</td><td>11316</td><td>11343</td><td>28</td><td>CACCCCACGTCTCGTCGCGCGCGCGTCC</td><td>3717.1</td><td>-8.9</td><td>1.6</td><td>-5.7</td><td>1.4E-08</td><td>2.7E-08</td><td>MB231/MB468</td><td>NUC</td></tr>
<tr><td>>hg38_21_+_8433172_8446622_RNA45SN1_45S_50ntFlanks@10386.10424.39|ATGTGAACAGCAGTTGAACATGGGTCAGTCGGTCCTGAG</td><td>>hg38_21_+_8433172_8446622_RNA45SN1_45S_50ntFlanks</td><td>10386</td><td>10424</td><td>39</td><td>ATGTGAACAGCAGTTGAACATGGGTCAGTCGGTCCTGAG</td><td>1856.4</td><td>4.8</td><td>0.8</td><td>5.7</td><td>1.4E-08</td><td>2.7E-08</td><td>MB231/MB468</td><td>NUC</td></tr>
<tr><td>>hg38_21_+_8433172_8446622_RNA45SN1_45S_50ntFlanks@11807.11828.22|ACTGTCCCTACCTACTATCCAG</td><td>>hg38_21_+_8433172_8446622_RNA45SN1_45S_50ntFlanks</td><td>11807</td><td>11828</td><td>22</td><td>ACTGTCCCTACCTACTATCCAG</td><td>70.5</td><td>-10.0</td><td>1.8</td><td>-5.7</td><td>1.3E-08</td><td>2.7E-08</td><td>MB231/MB468</td><td>MP</td></tr>
<tr><td>>hg38_21_+_8433172_8446622_RNA45SN1_45S_50ntFlanks@10823.10858.36|AAGTCGGCAAGCCGGATCCGTAACTTCGGGATAAGG</td><td>>hg38_21_+_8433172_8446622_RNA45SN1_45S_50ntFlanks</td><td>10823</td><td>10858</td><td>36</td><td>AAGTCGGCAAGCCGGATCCGTAACTTCGGGATAAGG</td><td>66.2</td><td>-9.9</td><td>1.7</td><td>-5.7</td><td>1.3E-08</td><td>2.7E-08</td><td>MB231/MB468</td><td>MP</td></tr>
<tr><td>>hg38_21_+_8433172_8446622_RNA45SN1_45S_50ntFlanks@3845.3881.37|ACTTGGATAACTGTGGTAATTCTAGAGCTAATACATG</td><td>>hg38_21_+_8433172_8446622_RNA45SN1_45S_50ntFlanks</td><td>3845</td><td>3881</td><td>37</td><td>ACTTGGATAACTGTGGTAATTCTAGAGCTAATACATG</td><td>84.2</td><td>-10.3</td><td>1.8</td><td>-5.7</td><td>1.3E-08</td><td>2.7E-08</td><td>MB231/MB468</td><td>MP</td></tr>
<tr><td>>hg38_21_+_8433172_8446622_RNA45SN1_45S_50ntFlanks@7987.8008.22|TCAGACGTGGCGACCCGCTGAA</td><td>>hg38_21_+_8433172_8446622_RNA45SN1_45S_50ntFlanks</td><td>7987</td><td>8008</td><td>22</td><td>TCAGACGTGGCGACCCGCTGAA</td><td>562.5</td><td>-5.4</td><td>1.0</td><td>-5.7</td><td>1.5E-08</td><td>2.7E-08</td><td>MB231/MB468</td><td>MITO</td></tr>
<tr><td>>hg38_21_+_8433172_8446622_RNA45SN1_45S_50ntFlanks@10713.10736.24|CTTGAAAATCCGGGGGAGAGGGTG</td><td>>hg38_21_+_8433172_8446622_RNA45SN1_45S_50ntFlanks</td><td>10713</td><td>10736</td><td>24</td><td>CTTGAAAATCCGGGGGAGAGGGTG</td><td>312.0</td><td>-3.3</td><td>0.6</td><td>-5.7</td><td>1.1E-08</td><td>2.7E-08</td><td>BT20/MB468</td><td>CYTO</td></tr>
<tr><td>>hg38_1_-_228634819_228635039_RNA5S12_5S_50ntFlanks@51.92.42|GTCTACGGCCATACCACCCTGAACGCGCCCGATCTCGTCTGA</td><td>>hg38_1_-_228634819_228635039_RNA5S12_5S_50ntFlanks</td><td>51</td><td>92</td><td>42</td><td>GTCTACGGCCATACCACCCTGAACGCGCCCGATCTCGTCTGA</td><td>2016.9</td><td>8.3</td><td>1.5</td><td>5.7</td><td>1.1E-08</td><td>2.7E-08</td><td>BT20/MB468</td><td>CYTO</td></tr>
<tr><td>>hg38_21_+_8433172_8446622_RNA45SN1_45S_50ntFlanks@10385.10409.25|CATGTGAACAGCAGTTGAACATGGG</td><td>>hg38_21_+_8433172_8446622_RNA45SN1_45S_50ntFlanks</td><td>10385</td><td>10409</td><td>25</td><td>CATGTGAACAGCAGTTGAACATGGG</td><td>184.7</td><td>-10.6</td><td>1.8</td><td>-6.0</td><td>1.9E-09</td><td>2.7E-08</td><td>BT20/MB468</td><td>MP</td></tr>
<tr><td>>hg38_21_+_8433172_8446622_RNA45SN1_45S_50ntFlanks@10345.10382.38|TTCAAACGAGAACTTTGAAGGCCGAAGTGGAGAAGGGT</td><td>>hg38_21_+_8433172_8446622_RNA45SN1_45S_50ntFlanks</td><td>10345</td><td>10382</td><td>38</td><td>TTCAAACGAGAACTTTGAAGGCCGAAGTGGAGAAGGGT</td><td>393.7</td><td>6.5</td><td>1.1</td><td>5.7</td><td>1.1E-08</td><td>2.7E-08</td><td>BT20/MB468</td><td>CYTO</td></tr>
<tr><td>>hg38_21_+_8433172_8446622_RNA45SN1_45S_50ntFlanks@6719.6749.31|TGAATTGCAGGACACATTGATCATCGACACT</td><td>>hg38_21_+_8433172_8446622_RNA45SN1_45S_50ntFlanks</td><td>6719</td><td>6749</td><td>31</td><td>TGAATTGCAGGACACATTGATCATCGACACT</td><td>317.9</td><td>-7.7</td><td>1.4</td><td>-5.7</td><td>1.3E-08</td><td>2.7E-08</td><td>MB231/MB468</td><td>MP</td></tr>
<tr><td>>hg38_21_+_8433172_8446622_RNA45SN1_45S_50ntFlanks@3854.3881.28|ACTGTGGTAATTCTAGAGCTAATACATG</td><td>>hg38_21_+_8433172_8446622_RNA45SN1_45S_50ntFlanks</td><td>3854</td><td>3881</td><td>28</td><td>ACTGTGGTAATTCTAGAGCTAATACATG</td><td>1590.5</td><td>-6.6</td><td>1.2</td><td>-5.7</td><td>1.4E-08</td><td>2.7E-08</td><td>MB231/MB468</td><td>NUC</td></tr>
<tr><td>>hg38_21_+_8433172_8446622_RNA45SN1_45S_50ntFlanks@10840.10865.26|CCGTAACTTCGGGATAAGGATTGGCT</td><td>>hg38_21_+_8433172_8446622_RNA45SN1_45S_50ntFlanks</td><td>10840</td><td>10865</td><td>26</td><td>CCGTAACTTCGGGATAAGGATTGGCT</td><td>910.6</td><td>-4.0</td><td>0.7</td><td>-5.7</td><td>1.0E-08</td><td>2.7E-08</td><td>BT20/MB468</td><td>MITO</td></tr>
<tr><td>>hg38_21_+_8433172_8446622_RNA45SN1_45S_50ntFlanks@7974.8008.35|ACGCGACCTCAGATCAGACGTGGCGACCCGCTGAA</td><td>>hg38_21_+_8433172_8446622_RNA45SN1_45S_50ntFlanks</td><td>7974</td><td>8008</td><td>35</td><td>ACGCGACCTCAGATCAGACGTGGCGACCCGCTGAA</td><td>89725.1</td><td>8.2</td><td>1.4</td><td>5.7</td><td>1.2E-08</td><td>2.7E-08</td><td>BT20/MB468</td><td>CYTO</td></tr>
<tr><td>>trna10_LysCTT_16_+_3241501_3241573@4.34.31,trna13_LysCTT_14_-_58706613_58706685@4.34.31,trna2_LysCTT_15_+_79152904_79152976@4.34.31|CGGCTAGCTCAGTCGGTAGAGCATGGGACTC</td><td>>trna10_LysCTT_16_+_3241501_3241573</td><td>4</td><td>34</td><td>31</td><td>CGGCTAGCTCAGTCGGTAGAGCATGGGACTC</td><td>80.6</td><td>9.6</td><td>1.7</td><td>5.7</td><td>1.3E-08</td><td>2.7E-08</td><td>MB231/MB468</td><td>MP</td></tr>
<tr><td>>hg38_21_+_8433172_8446622_RNA45SN1_45S_50ntFlanks@12940.12969.30|CATTCGTAGACGACCTGCTTCTGGGTCGGG</td><td>>hg38_21_+_8433172_8446622_RNA45SN1_45S_50ntFlanks</td><td>12940</td><td>12969</td><td>30</td><td>CATTCGTAGACGACCTGCTTCTGGGTCGGG</td><td>112.4</td><td>-9.1</td><td>1.6</td><td>-5.7</td><td>1.2E-08</td><td>2.7E-08</td><td>BT20/MB468</td><td>CYTO</td></tr>
<tr><td>>hg38_21_+_8433172_8446622_RNA45SN1_45S_50ntFlanks@12949.12972.24|ACGACCTGCTTCTGGGTCGGGGTT</td><td>>hg38_21_+_8433172_8446622_RNA45SN1_45S_50ntFlanks</td><td>12949</td><td>12972</td><td>24</td><td>ACGACCTGCTTCTGGGTCGGGGTT</td><td>80.0</td><td>-10.2</td><td>1.8</td><td>-5.7</td><td>1.3E-08</td><td>2.7E-08</td><td>MB231/MB468</td><td>MP</td></tr>
<tr><td>>hg38_21_+_8433172_8446622_RNA45SN1_45S_50ntFlanks@10678.10707.30|CGGTTCCGGCGGCGTCCGGTGAGCTCTCGC</td><td>>hg38_21_+_8433172_8446622_RNA45SN1_45S_50ntFlanks</td><td>10678</td><td>10707</td><td>30</td><td>CGGTTCCGGCGGCGTCCGGTGAGCTCTCGC</td><td>370.9</td><td>-4.1</td><td>0.7</td><td>-5.7</td><td>1.4E-08</td><td>2.7E-08</td><td>MB231/MB468</td><td>CYTO</td></tr>
<tr><td>>MI0000298|hsa-mir-221&WithFlank&X|-|45746151|45746272@71.94.24&[MIMAT0000278&hsa-miR-221-3p&offsets|0|+1;m-91&X|-|45746181|45746202&offsets|0|+2]|AGCTACATTGTCTGCTGGGTTTCA</td><td>>MI0000298|hsa-mir-221&WithFlank&X|-|45746151|45746272</td><td>71</td><td>94</td><td>24</td><td>AGCTACATTGTCTGCTGGGTTTCA</td><td>2568.2</td><td>5.0</td><td>0.9</td><td>5.7</td><td>1.5E-08</td><td>2.7E-08</td><td>MB231/MB468</td><td>MITO</td></tr>
<tr><td>>hg38_21_+_8433172_8446622_RNA45SN1_45S_50ntFlanks@5491.5532.42|GAGAAGACGGTCGAACTTGACTATCTAGAGGAAGTAAAAGTC</td><td>>hg38_21_+_8433172_8446622_RNA45SN1_45S_50ntFlanks</td><td>5491</td><td>5532</td><td>42</td><td>GAGAAGACGGTCGAACTTGACTATCTAGAGGAAGTAAAAGTC</td><td>285.1</td><td>11.9</td><td>2.0</td><td>6.0</td><td>2.0E-09</td><td>2.8E-08</td><td>BT20/MB468</td><td>MP</td></tr>
<tr><td>>hg38_21_+_8433172_8446622_RNA45SN1_45S_50ntFlanks@10128.10149.22|CGGCGGCGGCGGCGGCGGCGGG</td><td>>hg38_21_+_8433172_8446622_RNA45SN1_45S_50ntFlanks</td><td>10128</td><td>10149</td><td>22</td><td>CGGCGGCGGCGGCGGCGGCGGG</td><td>1134.9</td><td>-8.7</td><td>1.5</td><td>-6.0</td><td>2.0E-09</td><td>2.8E-08</td><td>BT20/MB468</td><td>MP</td></tr>
<tr><td>>hg38_21_+_8433172_8446622_RNA45SN1_45S_50ntFlanks@8845.8866.22|CGGCGGCGGCGGCGGCGGCGGG</td><td>>hg38_21_+_8433172_8446622_RNA45SN1_45S_50ntFlanks</td><td>8845</td><td>8866</td><td>22</td><td>CGGCGGCGGCGGCGGCGGCGGG</td><td>1134.9</td><td>-8.7</td><td>1.5</td><td>-6.0</td><td>2.0E-09</td><td>2.8E-08</td><td>BT20/MB468</td><td>MP</td></tr>
</table>

| | | | | | | | | | | | | | |
|---|---|---|---|---|---|---|---|---|---|---|---|---|---|
| >MI0000746\|hsa-mir-99b&WithFlank&19\|+\|51692606\|51692687@13.34.22(+1A)&[MIMAT0000689&hsa-miR-99b-5p&offsets\|0\|0(+1A);m-10&19\|+\|51692618\|51692639&offsets\|0\|0(+1A)]\|CACCCGTAGAACCGACCTTGCGA | >MI0000746\|hsa-mir-99b&WithFlank&19\|+\|51692606\|51692687 | 13 | 34 | 22(+1A) | CACCCGTAGAACCGACCTTGCGA | 739.4 | 4.0 | 0.7 | 5.7 | 1.4E-08 | 2.8E-08 | MB231/MB468 | NUC |
| >hg38_21_+_8433172_8446622_RNA45SN1_45S_50ntFlanks@10846.10872.27\|CTTCGGGATAAGGATTGGCTCTAAGGG | >hg38_21_+_8433172_8446622_RNA45SN1_45S_50ntFlanks | 10846 | 10872 | 27 | CTTCGGGATAAGGATTGGCTCTAAGGG | 214.2 | -5.7 | 1.0 | -5.7 | 1.4E-08 | 2.8E-08 | MB231/MB468 | NUC |
| >hg38_21_+_8433172_8446622_RNA45SN1_45S_50ntFlanks@4725.4758.34\|TTAATCAAGAACGAAAGTCGGAGGTTCGAAGACG | >hg38_21_+_8433172_8446622_RNA45SN1_45S_50ntFlanks | 4725 | 4758 | 34 | TTAATCAAGAACGAAAGTCGGAGGTTCGAAGACG | 1824.1 | -7.2 | 1.3 | -5.7 | 1.3E-08 | 2.8E-08 | MB231/MB468 | MP |
| >hg38_21_+_8433172_8446622_RNA45SN1_45S_50ntFlanks@13004.13025.22\|ATCTATTGAAAGTCAGCCCTCG | >hg38_21_+_8433172_8446622_RNA45SN1_45S_50ntFlanks | 13004 | 13025 | 22 | ATCTATTGAAAGTCAGCCCTCG | 107.6 | -9.8 | 1.6 | -6.0 | 2.0E-09 | 2.8E-08 | BT20/MB468 | MP |
| >hg38_21_+_8433172_8446622_RNA45SN1_45S_50ntFlanks@4884.4911.28\|CAAAGCTGAAACTTAAAGGAATTGACGG | >hg38_21_+_8433172_8446622_RNA45SN1_45S_50ntFlanks | 4884 | 4911 | 28 | CAAAGCTGAAACTTAAAGGAATTGACGG | 394.0 | -8.4 | 1.5 | -5.7 | 1.5E-08 | 2.8E-08 | MB231/MB468 | MITO |
| >hg38_21_+_8433172_8446622_RNA45SN1_45S_50ntFlanks@6721.6752.32\|AATTGCAGGACACATTGATCATCGACACTTCG | >hg38_21_+_8433172_8446622_RNA45SN1_45S_50ntFlanks | 6721 | 6752 | 32 | AATTGCAGGACACATTGATCATCGACACTTCG | 1315.7 | -9.1 | 1.6 | -5.7 | 1.6E-08 | 2.8E-08 | BT20/MB468 | NUC |
| >hg38_21_+_8433172_8446622_RNA45SN1_45S_50ntFlanks@12949.12988.40\|ACGACCTGCTTCTGGGTCGGGGTTTCGTACGTAGCAGAGC | >hg38_21_+_8433172_8446622_RNA45SN1_45S_50ntFlanks | 12949 | 12988 | 40 | ACGACCTGCTTCTGGGTCGGGGTTTCGTACGTAGCAGAGC | 554.3 | -3.9 | 0.7 | -5.7 | 1.4E-08 | 2.8E-08 | MB231/MB468 | NUC |
| >hg38_21_+_8433172_8446622_RNA45SN1_45S_50ntFlanks@8179.8211.33\|CTGATCGAGGCCCAGCCCGTGGACGGTGTGAGG | >hg38_21_+_8433172_8446622_RNA45SN1_45S_50ntFlanks | 8179 | 8211 | 33 | CTGATCGAGGCCCAGCCCGTGGACGGTGTGAGG | 249.2 | -7.6 | 1.3 | -5.7 | 1.2E-08 | 2.8E-08 | BT20/MB468 | CYTO |
| >hg38_21_+_8433172_8446622_RNA45SN1_45S_50ntFlanks@12582.12604.23\|ACCGCAGGTTCAGACATTTGGTG | >hg38_21_+_8433172_8446622_RNA45SN1_45S_50ntFlanks | 12582 | 12604 | 23 | ACCGCAGGTTCAGACATTTGGTG | 103.1 | -9.8 | 1.6 | -6.0 | 2.0E-09 | 2.8E-08 | BT20/MB468 | MP |
| >MI0000542\|hsa-mir-320a&WithFlank&8\|-\|22244960\|22245043@48.71.24&[MIMAT0000510&hsa-miR-320a-3p&offsets\|0\|+2;m-60&8\|-\|22244975\|22244996&offsets\|0\|+2]\|AAAAGCTGGGTTGAGAGGGCGAAA | >MI0000542\|hsa-mir-320a&WithFlank&8\|-\|22244960\|22245043 | 48 | 71 | 24 | AAAAGCTGGGTTGAGAGGGCGAAA | 567.7 | 4.6 | 0.8 | 5.7 | 1.4E-08 | 2.8E-08 | MB231/MB468 | NUC |
| >hg38_21_+_8433172_8446622_RNA45SN1_45S_50ntFlanks@11212.11232.21\|CGACTCTGGACGCGAGCCGGG | >hg38_21_+_8433172_8446622_RNA45SN1_45S_50ntFlanks | 11212 | 11232 | 21 | CGACTCTGGACGCGAGCCGGG | 3158.3 | -6.8 | 1.2 | -5.7 | 1.2E-08 | 2.8E-08 | BT20/MB468 | CYTO |
| >hg38_21_+_8433172_8446622_RNA45SN1_45S_50ntFlanks@11316.11338.23\|CACCCCACGTCTCGTCGCGCGCG | >hg38_21_+_8433172_8446622_RNA45SN1_45S_50ntFlanks | 11316 | 11338 | 23 | CACCCCACGTCTCGTCGCGCGCG | 2501.0 | -6.1 | 1.1 | -5.7 | 1.2E-08 | 2.8E-08 | BT20/MB468 | CYTO |
| >hg38_21_+_8433172_8446622_RNA45SN1_45S_50ntFlanks@3769.3799.31\|CGCACGGCCGGTACAGTGAAACTGCGAATGG | >hg38_21_+_8433172_8446622_RNA45SN1_45S_50ntFlanks | 3769 | 3799 | 31 | CGCACGGCCGGTACAGTGAAACTGCGAATGG | 193.0 | -5.3 | 0.9 | -5.7 | 1.4E-08 | 2.8E-08 | MB231/MB468 | NUC |
| >hg38_21_+_8433172_8446622_RNA45SN1_45S_50ntFlanks@10466.10488.23\|CGTTGCCCTCGGCCGATCGAAAG | >hg38_21_+_8433172_8446622_RNA45SN1_45S_50ntFlanks | 10466 | 10488 | 23 | CGTTGCCCTCGGCCGATCGAAAG | 741.5 | -9.0 | 1.6 | -5.7 | 1.6E-08 | 2.8E-08 | BT20/MB468 | NUC |
| >hg38_21_+_8433172_8446622_RNA45SN1_45S_50ntFlanks@3705.3744.40\|TACCTGGTTGATCCTGCCAGTAGCATATGCTTGTCTCAAA | >hg38_21_+_8433172_8446622_RNA45SN1_45S_50ntFlanks | 3705 | 3744 | 40 | TACCTGGTTGATCCTGCCAGTAGCATATGCTTGTCTCAAA | 603.2 | 5.9 | 1.0 | 5.7 | 1.4E-08 | 2.8E-08 | MB231/MB468 | CYTO |
| >hg38_21_+_8433172_8446622_RNA45SN1_45S_50ntFlanks@10663.10712.50\|CCTTGGAAAGCGTCGCGGTTCCGGCGGCGTCCGGTGAGCTCTCGCTGGCC | >hg38_21_+_8433172_8446622_RNA45SN1_45S_50ntFlanks | 10663 | 10712 | 50 | CCTTGGAAAGCGTCGCGGTTCCGGCGGCGTCCGGTGAGCTCTCGCTGGCC | 126.0 | -9.1 | 1.6 | -5.7 | 1.1E-08 | 2.8E-08 | BT20/MB468 | MITO |
| >hg38_21_+_8433172_8446622_RNA45SN1_45S_50ntFlanks@4201.4223.23\|CCGACCCGGGGAGGTAGTGACGA | >hg38_21_+_8433172_8446622_RNA45SN1_45S_50ntFlanks | 4201 | 4223 | 23 | CCGACCCGGGGAGGTAGTGACGA | 513.9 | -8.2 | 1.4 | -5.7 | 1.3E-08 | 2.8E-08 | MB231/MB468 | MP |
| >hg38_21_+_8433172_8446622_RNA45SN1_45S_50ntFlanks@3823.3853.31\|TTTGGTCGCTCGCTCCTCTCCTACTTGGATA | >hg38_21_+_8433172_8446622_RNA45SN1_45S_50ntFlanks | 3823 | 3853 | 31 | TTTGGTCGCTCGCTCCTCTCCTACTTGGATA | 505.6 | -8.6 | 1.5 | -5.7 | 1.6E-08 | 2.8E-08 | BT20/MB468 | NUC |
| >hg38_21_+_8433172_8446622_RNA45SN1_45S_50ntFlanks@7977.8012.36\|CGACCTCAGATCAGACGTGGCGACCCGCTGAATTTA | >hg38_21_+_8433172_8446622_RNA45SN1_45S_50ntFlanks | 7977 | 8012 | 36 | CGACCTCAGATCAGACGTGGCGACCCGCTGAATTTA | 880.4 | 5.3 | 0.9 | 5.7 | 1.4E-08 | 2.8E-08 | MB231/MB468 | CYTO |
| >hg38_21_+_8433172_8446622_RNA45SN1_45S_50ntFlanks@4102.4121.20\|ACCACGGGTGACGGGGAATC | >hg38_21_+_8433172_8446622_RNA45SN1_45S_50ntFlanks | 4102 | 4121 | 20 | ACCACGGGTGACGGGGAATC | 1539.9 | -7.7 | 1.4 | -5.7 | 1.5E-08 | 2.8E-08 | MB231/MB468 | MITO |
| >hg38_21_+_8433172_8446622_RNA45SN1_45S_50ntFlanks@10463.10483.21\|CTCCGTTGCCCTCGGCCGATC | >hg38_21_+_8433172_8446622_RNA45SN1_45S_50ntFlanks | 10463 | 10483 | 21 | CTCCGTTGCCCTCGGCCGATC | 262.2 | -5.0 | 0.9 | -5.7 | 1.4E-08 | 2.8E-08 | MB231/MB468 | CYTO |
| >hg38_21_+_8433172_8446622_RNA45SN1_45S_50ntFlanks@11703.11727.25\|AATGAAGCGCGGGTAAACGGCGGGA | >hg38_21_+_8433172_8446622_RNA45SN1_45S_50ntFlanks | 11703 | 11727 | 25 | AATGAAGCGCGGGTAAACGGCGGGA | 236.9 | -5.5 | 1.0 | -5.7 | 1.4E-08 | 2.8E-08 | MB231/MB468 | NUC |
| >hg38_21_+_8433172_8446622_RNA45SN1_45S_50ntFlanks@9572.9597.26\|CAAATCGGTCGTCCGACCTGGGTATA | >hg38_21_+_8433172_8446622_RNA45SN1_45S_50ntFlanks | 9572 | 9597 | 26 | CAAATCGGTCGTCCGACCTGGGTATA | 1161.3 | -6.2 | 1.1 | -5.7 | 1.4E-08 | 2.8E-08 | MB231/MB468 | NUC |
| >hg38_21_+_8433172_8446622_RNA45SN1_45S_50ntFlanks@10389.10419.31\|TGAACAGCAGTTGAACATGGGTCAGTCGGTC | >hg38_21_+_8433172_8446622_RNA45SN1_45S_50ntFlanks | 10389 | 10419 | 31 | TGAACAGCAGTTGAACATGGGTCAGTCGGTC | 104.9 | -5.8 | 1.0 | -5.7 | 1.2E-08 | 2.8E-08 | BT20/MB468 | CYTO |
| >hg38_21_+_8433172_8446622_RNA45SN1_45S_50ntFlanks@6732.6760.29\|ACATTGATCATCGACACTTCGAACGCACT | >hg38_21_+_8433172_8446622_RNA45SN1_45S_50ntFlanks | 6732 | 6760 | 29 | ACATTGATCATCGACACTTCGAACGCACT | 271.1 | 2.5 | 0.4 | 5.7 | 1.4E-08 | 2.8E-08 | MB231/MB468 | CYTO |
| >hg38_21_+_8433172_8446622_RNA45SN1_45S_50ntFlanks@12991.13039.49\|CTCCCTCGCTGCGATCTATTGAAAGTCAGCCCTCGACACAAGGGTTTGT | >hg38_21_+_8433172_8446622_RNA45SN1_45S_50ntFlanks | 12991 | 13039 | 49 | CTCCCTCGCTGCGATCTATTGAAAGTCAGCCCTCGACACAAGGGTTTGT | 23178.4 | -12.0 | 2.1 | -5.7 | 1.6E-08 | 2.8E-08 | BT20/MB468 | NUC |
| >hg38_21_+_8433172_8446622_RNA45SN1_45S_50ntFlanks@9578.9603.26\|GGTCGTCCGACCTGGGTATAGGGGCG | >hg38_21_+_8433172_8446622_RNA45SN1_45S_50ntFlanks | 9578 | 9603 | 26 | GGTCGTCCGACCTGGGTATAGGGGCG | 466.2 | 9.6 | 1.6 | 6.0 | 2.0E-09 | 2.8E-08 | BT20/MB468 | MP |
| >hg38_21_+_8433172_8446622_RNA45SN1_45S_50ntFlanks@12414.12446.33\|CCTTCGATGTCGGCTCTTCCTATCATTGTGAAG | >hg38_21_+_8433172_8446622_RNA45SN1_45S_50ntFlanks | 12414 | 12446 | 33 | CCTTCGATGTCGGCTCTTCCTATCATTGTGAAG | 385.8 | 3.1 | 0.5 | 5.7 | 1.4E-08 | 2.8E-08 | MB231/MB468 | CYTO |
| >hg38_21_+_8433172_8446622_RNA45SN1_45S_50ntFlanks@12624.12656.33\|CCAATGGGGCGAAGCTACCATCTGTGGGATTAT | >hg38_21_+_8433172_8446622_RNA45SN1_45S_50ntFlanks | 12624 | 12656 | 33 | CCAATGGGGCGAAGCTACCATCTGTGGGATTAT | 102.5 | -9.8 | 1.6 | -6.0 | 2.0E-09 | 2.8E-08 | BT20/MB468 | MP |
| >hg38_21_+_8433172_8446622_RNA45SN1_45S_50ntFlanks@10387.10410.24\|TGTGAACAGCAGTTGAACATGGGT | >hg38_21_+_8433172_8446622_RNA45SN1_45S_50ntFlanks | 10387 | 10410 | 24 | TGTGAACAGCAGTTGAACATGGGT | 1112.7 | -8.9 | 1.6 | -5.7 | 1.6E-08 | 2.8E-08 | BT20/MB468 | NUC |
| >hg38_21_+_8433172_8446622_RNA45SN1_45S_50ntFlanks@11570.11591.22\|AGCCGACTTAGAACTGGTGCGG | >hg38_21_+_8433172_8446622_RNA45SN1_45S_50ntFlanks | 11570 | 11591 | 22 | AGCCGACTTAGAACTGGTGCGG | 399.5 | -5.3 | 0.9 | -5.7 | 1.4E-08 | 2.8E-08 | MB231/MB468 | CYTO |
| >hg38_21_+_8433172_8446622_RNA45SN1_45S_50ntFlanks@9736.9754.19\|CCGAAACGATCTCAACCTA | >hg38_21_+_8433172_8446622_RNA45SN1_45S_50ntFlanks | 9736 | 9754 | 19 | CCGAAACGATCTCAACCTA | 920.6 | -6.4 | 1.1 | -5.7 | 1.4E-08 | 2.8E-08 | MB231/MB468 | CYTO |
| >hg38_MT_+_1621_3279_RNR2_16S_50ntFlanks@51.70.20\|GCTAAACCTAGCCCCAAACC | >hg38_MT_+_1621_3279_RNR2_16S_50ntFlanks | 51 | 70 | 20 | GCTAAACCTAGCCCCAAACC | 2075.9 | 6.2 | 1.1 | 5.7 | 1.5E-08 | 2.8E-08 | MB231/MB468 | MITO |
| >hg38_21_+_8433172_8446622_RNA45SN1_45S_50ntFlanks@9569.9595.27\|GTGCAAATCGGTCGTCCGACCTGGGTA | >hg38_21_+_8433172_8446622_RNA45SN1_45S_50ntFlanks | 9569 | 9595 | 27 | GTGCAAATCGGTCGTCCGACCTGGGTA | 384.6 | -6.1 | 1.1 | -5.7 | 1.3E-08 | 2.8E-08 | MB231/MB468 | MP |
| >hg38_21_+_8433172_8446622_RNA45SN1_45S_50ntFlanks@4567.4587.21\|TAATGGAATAGGACCGCGGTT | >hg38_21_+_8433172_8446622_RNA45SN1_45S_50ntFlanks | 4567 | 4587 | 21 | TAATGGAATAGGACCGCGGTT | 279.5 | -6.4 | 1.1 | -5.7 | 1.3E-08 | 2.8E-08 | MB231/MB468 | MP |
| >hg38_21_+_8433172_8446622_RNA45SN1_45S_50ntFlanks@11537.11560.24\|CCGCGCGGCGCCTCGCCTCGGCCG | >hg38_21_+_8433172_8446622_RNA45SN1_45S_50ntFlanks | 11537 | 11560 | 24 | CCGCGCGGCGCCTCGCCTCGGCCG | 214.2 | -6.3 | 1.1 | -5.7 | 1.2E-08 | 2.8E-08 | BT20/MB468 | CYTO |
| >hg38_21_+_8433172_8446622_RNA45SN1_45S_50ntFlanks@5225.5244.20\|CACGCGCGCTACACTGACTG | >hg38_21_+_8433172_8446622_RNA45SN1_45S_50ntFlanks | 5225 | 5244 | 20 | CACGCGCGCTACACTGACTG | 538.6 | -11.2 | 2.0 | -5.7 | 1.1E-08 | 2.8E-08 | BT20/MB468 | MITO |
| >MI0003776\|hsa-mir-320b-1&WithFlank&1\|+\|116671740\|116671823@48.69.22;MI0003839\|hsa-mir-320b-2&WithFlank&1\|-\|224257034\|224257116@47.68.22&[MIMAT0005792&hsa-miR-320b&offsets\|0\|0;m-183&1\|+\|116671787\|116671806&offsets\|0\|+2];[MIMAT0005792_1&hsa-miR-320b&offsets\|0\|0;m-183&1\|-\|224257051\|224257070&offsets\|0\|+2]\|AAAAGCTGGGTTGAGAGGGCAA | >MI0003776\|hsa-mir-320b-1&WithFlank&1\|+\|116671740\|116671823 | 48 | 69 | 22 | AAAAGCTGGGTTGAGAGGGCAA | 153.4 | 11.0 | 1.8 | 6.0 | 2.0E-09 | 2.8E-08 | BT20/MB468 | MP |
| >hg38_21_+_8433172_8446622_RNA45SN1_45S_50ntFlanks@10681.10710.30\|TTCCGGCGGCGTCCGGTGAGCTCTCGCTGG | >hg38_21_+_8433172_8446622_RNA45SN1_45S_50ntFlanks | 10681 | 10710 | 30 | TTCCGGCGGCGTCCGGTGAGCTCTCGCTGG | 1174.2 | -8.5 | 1.4 | -6.0 | 2.0E-09 | 2.8E-08 | BT20/MB468 | MP |
| >hg38_21_+_8433172_8446622_RNA45SN1_45S_50ntFlanks@10399.10424.26\|TTGAACATGGGTCAGTCGGTCCTGAG | >hg38_21_+_8433172_8446622_RNA45SN1_45S_50ntFlanks | 10399 | 10424 | 26 | TTGAACATGGGTCAGTCGGTCCTGAG | 342.0 | 4.4 | 0.8 | 5.7 | 1.4E-08 | 2.9E-08 | MB231/MB468 | CYTO |
| >hg38_21_+_8433172_8446622_RNA45SN1_45S_50ntFlanks@10546.10570.25\|CAGTGCGGTAACGCGACCGATCCCG | >hg38_21_+_8433172_8446622_RNA45SN1_45S_50ntFlanks | 10546 | 10570 | 25 | CAGTGCGGTAACGCGACCGATCCCG | 331.5 | -7.4 | 1.3 | -5.7 | 1.3E-08 | 2.9E-08 | MB231/MB468 | MP |
| >hg38_21_+_8433172_8446622_RNA45SN1_45S_50ntFlanks@11569.11613.45\|CAGCCGACTTAGAACTGGTGCGGACCAGGGGAATCCGACTGTTTA | >hg38_21_+_8433172_8446622_RNA45SN1_45S_50ntFlanks | 11569 | 11613 | 45 | CAGCCGACTTAGAACTGGTGCGGACCAGGGGAATCCGACTGTTTA | 637.0 | -5.2 | 0.9 | -5.6 | 1.6E-08 | 2.9E-08 | BT20/MB468 | NUC |
| >hg38_21_+_8433172_8446622_RNA45SN1_45S_50ntFlanks@8384.8405.22\|AGGGCGTGAAACCGTTAAGAGG | >hg38_21_+_8433172_8446622_RNA45SN1_45S_50ntFlanks | 8384 | 8405 | 22 | AGGGCGTGAAACCGTTAAGAGG | 507.5 | -4.7 | 0.8 | -5.7 | 1.3E-08 | 2.9E-08 | MB231/MB468 | MP |
| >hg38_21_+_8433172_8446622_RNA45SN1_45S_50ntFlanks@10479.10511.33\|CGATCGAAAGGGAGTCGGGTTCAGATCCCCGAA | >hg38_21_+_8433172_8446622_RNA45SN1_45S_50ntFlanks | 10479 | 10511 | 33 | CGATCGAAAGGGAGTCGGGTTCAGATCCCCGAA | 175.2 | -5.7 | 1.0 | -5.7 | 1.4E-08 | 2.9E-08 | MB231/MB468 | CYTO |
| >hg38_21_+_8433172_8446622_RNA45SN1_45S_50ntFlanks@10458.10484.27\|ATGGCCTCCGTTGCCCTCGGCCGATCG | >hg38_21_+_8433172_8446622_RNA45SN1_45S_50ntFlanks | 10458 | 10484 | 27 | ATGGCCTCCGTTGCCCTCGGCCGATCG | 338.1 | -7.2 | 1.3 | -5.7 | 1.4E-08 | 2.9E-08 | MB231/MB468 | NUC |
| >hg38_21_+_8433172_8446622_RNA45SN1_45S_50ntFlanks@10819.10865.47\|AGGGAAGTCGGCAAGCCGGATCCGTAACTTCGGGATAAGGATTGGCT | >hg38_21_+_8433172_8446622_RNA45SN1_45S_50ntFlanks | 10819 | 10865 | 47 | AGGGAAGTCGGCAAGCCGGATCCGTAACTTCGGGATAAGGATTGGCT | 261.8 | -8.1 | 1.4 | -5.7 | 1.2E-08 | 2.9E-08 | BT20/MB468 | CYTO |
| >hg38_21_+_8433172_8446622_RNA45SN1_45S_50ntFlanks@10806.10823.18\|AACAATGTAGGTAAGGGA | >hg38_21_+_8433172_8446622_RNA45SN1_45S_50ntFlanks | 10806 | 10823 | 18 | AACAATGTAGGTAAGGGA | 488.7 | -5.5 | 1.0 | -5.7 | 1.1E-08 | 2.9E-08 | BT20/MB468 | MITO |
| >hg38_21_+_8433172_8446622_RNA45SN1_45S_50ntFlanks@5035.5052.18\|CATGGCCGTTCTTAGTTG | >hg38_21_+_8433172_8446622_RNA45SN1_45S_50ntFlanks | 5035 | 5052 | 18 | CATGGCCGTTCTTAGTTG | 653.3 | -11.4 | 2.0 | -5.7 | 1.1E-08 | 2.9E-08 | BT20/MB468 | MITO |
| >hg38_21_+_8433172_8446622_RNA45SN1_45S_50ntFlanks@6651.6689.39\|CGACTCTTAGCGGTGGATCACTCGGCTCGTGCGTCGATG | >hg38_21_+_8433172_8446622_RNA45SN1_45S_50ntFlanks | 6651 | 6689 | 39 | CGACTCTTAGCGGTGGATCACTCGGCTCGTGCGTCGATG | 36831.9 | -4.8 | 0.8 | -5.7 | 1.3E-08 | 2.9E-08 | MB231/MB468 | MP |
| >hg38_21_+_8433172_8446622_RNA45SN1_45S_50ntFlanks@3705.3728.24\|TACCTGGTTGATCCTGCCAGTAGC | >hg38_21_+_8433172_8446622_RNA45SN1_45S_50ntFlanks | 3705 | 3728 | 24 | TACCTGGTTGATCCTGCCAGTAGC | 225.6 | 2.7 | 0.5 | 5.6 | 1.6E-08 | 2.9E-08 | BT20/MB468 | NUC |
| >hg38_21_+_8433172_8446622_RNA45SN1_45S_50ntFlanks@3843.3868.26\|CTACTTGGATAACTGTGGTAATTCTA | >hg38_21_+_8433172_8446622_RNA45SN1_45S_50ntFlanks | 3843 | 3868 | 26 | CTACTTGGATAACTGTGGTAATTCTA | 1346.8 | -6.9 | 1.2 | -5.7 | 1.6E-08 | 2.9E-08 | MB231/MB468 | MITO |
| >trna5_SupTTA_4_-_7325838_7325909@33.55.23\|TTAGAGCCAGACTGCCTGGGTTT | >trna5_SupTTA_4_-_7325838_7325909 | 33 | 55 | 23 | TTAGAGCCAGACTGCCTGGGTTT | 225.1 | 11.6 | 1.9 | 6.0 | 2.1E-09 | 2.9E-08 | BT20/MB468 | MP |

<table>
<tr><td>>hg38_21_+_8433172_8446622_RNA45SN1_45S_50ntFlanks@10798.10817.20|CATGTTGGAACAATGTAGGT</td><td>>hg38_21_+_8433172_8446622_RNA45SN1_45S_50ntFlanks</td><td>10798</td><td>10817</td><td>20</td><td>CATGTTGGAACAATGTAGGT</td><td>723.7</td><td>-3.9</td><td>0.7</td><td>-5.7</td><td>1.4E-08</td><td>2.9E-08</td><td>MB231/MB468</td><td>CYTO</td></tr>
<tr><td>>MI0000809|hsa-mir-151a&WithFlank&8|-|140732558|140732659@51.72.22(+1C)&[MIMAT0000757&hsa-miR-151a-3p&offsets|-2|-1(+1C);m-30&8|-|140732587|140732607&offsets|-2|-1(+1C)]|TACTAGACTGAAGCTCCTTGAGC</td><td>>MI0000809|hsa-mir-151a&WithFlank&8|-|140732558|140732659</td><td>51</td><td>72</td><td>22(+1C)</td><td>TACTAGACTGAAGCTCCTTGAGC</td><td>76.7</td><td>9.5</td><td>1.7</td><td>5.7</td><td>1.4E-08</td><td>2.9E-08</td><td>MB231/MB468</td><td>MP</td></tr>
<tr><td>>hg38_21_+_8433172_8446622_RNA45SN1_45S_50ntFlanks@8437.8470.34|AGGATTCAACCCGGCGGCGGGTCCGGCCGTGTCG</td><td>>hg38_21_+_8433172_8446622_RNA45SN1_45S_50ntFlanks</td><td>8437</td><td>8470</td><td>34</td><td>AGGATTCAACCCGGCGGCGGGTCCGGCCGTGTCG</td><td>115.6</td><td>-9.9</td><td>1.7</td><td>-6.0</td><td>2.1E-09</td><td>2.9E-08</td><td>BT20/MB468</td><td>MP</td></tr>
<tr><td>>hg38_21_+_8433172_8446622_RNA45SN1_45S_50ntFlanks@3855.3881.27|CTGTGGTAATTCTAGAGCTAATACATG</td><td>>hg38_21_+_8433172_8446622_RNA45SN1_45S_50ntFlanks</td><td>3855</td><td>3881</td><td>27</td><td>CTGTGGTAATTCTAGAGCTAATACATG</td><td>416.7</td><td>-8.0</td><td>1.4</td><td>-5.7</td><td>1.6E-08</td><td>2.9E-08</td><td>MB231/MB468</td><td>MITO</td></tr>
<tr><td>>hg38_21_+_8433172_8446622_RNA45SN1_45S_50ntFlanks@10366.10390.25|CCGAAGTGGAGAAGGGTTCCATGTG</td><td>>hg38_21_+_8433172_8446622_RNA45SN1_45S_50ntFlanks</td><td>10366</td><td>10390</td><td>25</td><td>CCGAAGTGGAGAAGGGTTCCATGTG</td><td>102.1</td><td>-9.8</td><td>1.6</td><td>-6.0</td><td>2.1E-09</td><td>2.9E-08</td><td>BT20/MB468</td><td>MP</td></tr>
<tr><td>>hg38_21_+_8433172_8446622_RNA45SN1_45S_50ntFlanks@11575.11609.35|ACTTAGAACTGGTGCGGACCAGGGGAATCCGACTG</td><td>>hg38_21_+_8433172_8446622_RNA45SN1_45S_50ntFlanks</td><td>11575</td><td>11609</td><td>35</td><td>ACTTAGAACTGGTGCGGACCAGGGGAATCCGACTG</td><td>1001.2</td><td>-6.3</td><td>1.1</td><td>-5.7</td><td>1.6E-08</td><td>2.9E-08</td><td>MB231/MB468</td><td>MITO</td></tr>
<tr><td>>hg38_21_+_8433172_8446622_RNA45SN1_45S_50ntFlanks@12488.12517.30|AACGTGAGCTGGGTTTAGACCGTCGTGAGA</td><td>>hg38_21_+_8433172_8446622_RNA45SN1_45S_50ntFlanks</td><td>12488</td><td>12517</td><td>30</td><td>AACGTGAGCTGGGTTTAGACCGTCGTGAGA</td><td>66.4</td><td>-9.9</td><td>1.7</td><td>-5.7</td><td>1.4E-08</td><td>2.9E-08</td><td>MB231/MB468</td><td>MP</td></tr>
<tr><td>>hg38_21_+_8433172_8446622_RNA45SN1_45S_50ntFlanks@4047.4072.26|ATTCGAACGTCTGCCCTATCAACTTT</td><td>>hg38_21_+_8433172_8446622_RNA45SN1_45S_50ntFlanks</td><td>4047</td><td>4072</td><td>26</td><td>ATTCGAACGTCTGCCCTATCAACTTT</td><td>185.7</td><td>-9.6</td><td>1.7</td><td>-5.7</td><td>1.1E-08</td><td>2.9E-08</td><td>BT20/MB468</td><td>MITO</td></tr>
<tr><td>>trnaMT_PheGAA_MT_+_577_647@28.55.28|ATACACTGAAAATGTTTAGACGGGCTCA</td><td>>trnaMT_PheGAA_MT_+_577_647</td><td>28</td><td>55</td><td>28</td><td>ATACACTGAAAATGTTTAGACGGGCTCA</td><td>204.4</td><td>11.4</td><td>1.9</td><td>6.0</td><td>2.1E-09</td><td>2.9E-08</td><td>BT20/MB468</td><td>MP</td></tr>
<tr><td>>hg38_21_+_8433172_8446622_RNA45SN1_45S_50ntFlanks@4982.5002.21|ACGGACAGGATTGACAGATTG</td><td>>hg38_21_+_8433172_8446622_RNA45SN1_45S_50ntFlanks</td><td>4982</td><td>5002</td><td>21</td><td>ACGGACAGGATTGACAGATTG</td><td>362.2</td><td>-7.9</td><td>1.4</td><td>-5.7</td><td>1.4E-08</td><td>2.9E-08</td><td>MB231/MB468</td><td>MP</td></tr>
<tr><td>>hg38_21_+_8433172_8446622_RNA45SN1_45S_50ntFlanks@11686.11717.32|TCAAAGTGAAGAAATTCAATGAAGCGCGGGTA</td><td>>hg38_21_+_8433172_8446622_RNA45SN1_45S_50ntFlanks</td><td>11686</td><td>11717</td><td>32</td><td>TCAAAGTGAAGAAATTCAATGAAGCGCGGGTA</td><td>87.5</td><td>-8.5</td><td>1.5</td><td>-5.7</td><td>1.1E-08</td><td>2.9E-08</td><td>BT20/MB468</td><td>MITO</td></tr>
<tr><td>>hg38_21_+_8433172_8446622_RNA45SN1_45S_50ntFlanks@12155.12178.24|GGAGTTTGACTGGGGCGGTACACC</td><td>>hg38_21_+_8433172_8446622_RNA45SN1_45S_50ntFlanks</td><td>12155</td><td>12178</td><td>24</td><td>GGAGTTTGACTGGGGCGGTACACC</td><td>65.5</td><td>-9.9</td><td>1.7</td><td>-5.7</td><td>1.4E-08</td><td>2.9E-08</td><td>MB231/MB468</td><td>MP</td></tr>
<tr><td>>MI0005559|hsa-mir-744&WithFlank&17|+|12081893|12082002@17.38.22&[MIMAT0004945&hsa-miR-744-5p&offsets|0|0;m-172&17|+|12081909|12081929&offsets|0|+1]|TGCGGGGCTAGGGCTAACAGCA</td><td>>MI0005559|hsa-mir-744&WithFlank&17|+|12081893|12082002</td><td>17</td><td>38</td><td>22</td><td>TGCGGGGCTAGGGCTAACAGCA</td><td>255.7</td><td>4.3</td><td>0.8</td><td>5.7</td><td>1.5E-08</td><td>2.9E-08</td><td>MB231/MB468</td><td>NUC</td></tr>
<tr><td>>MI0000066|hsa-let-7e&WithFlank&19|+|51692780|51692870@14.36.23&[MIMAT0000066&hsa-let-7e-5p&offsets|0|+1;m-50&19|+|51692793|51692814&offsets|0|+1]|TGAGGTAGGAGGTTGTATAGTTG</td><td>>MI0000066|hsa-let-7e&WithFlank&19|+|51692780|51692870</td><td>14</td><td>36</td><td>23</td><td>TGAGGTAGGAGGTTGTATAGTTG</td><td>392.8</td><td>6.4</td><td>1.1</td><td>5.7</td><td>1.5E-08</td><td>2.9E-08</td><td>MB231/MB468</td><td>CYTO</td></tr>
<tr><td>>hg38_21_+_8433172_8446622_RNA45SN1_45S_50ntFlanks@11999.12030.32|CCCTGCGGGCCGCCGGTGAAATACCACTACTC</td><td>>hg38_21_+_8433172_8446622_RNA45SN1_45S_50ntFlanks</td><td>11999</td><td>12030</td><td>32</td><td>CCCTGCGGGCCGCCGGTGAAATACCACTACTC</td><td>412.6</td><td>-6.5</td><td>1.1</td><td>-5.7</td><td>1.5E-08</td><td>3.0E-08</td><td>MB231/MB468</td><td>CYTO</td></tr>
<tr><td>>hg38_21_+_8433172_8446622_RNA45SN1_45S_50ntFlanks@9623.9648.26|TAGTAGCTGGTTCCCTCCGAAGTTTC</td><td>>hg38_21_+_8433172_8446622_RNA45SN1_45S_50ntFlanks</td><td>9623</td><td>9648</td><td>26</td><td>TAGTAGCTGGTTCCCTCCGAAGTTTC</td><td>989.9</td><td>-3.6</td><td>0.6</td><td>-5.7</td><td>1.1E-08</td><td>3.0E-08</td><td>BT20/MB468</td><td>MITO</td></tr>
<tr><td>>MI0016746|hsa-mir-548o-2&WithFlank&20|+|38516557|38516638@51.72.22;MI0006402|hsa-mir-548o&WithFlank&7|-|102405736|102405861@93.114.22&[MIMAT0005919&hsa-miR-548o-3p&offsets|0|0;m-431&20|+|38516607|38516628&offsets|0|0];[MIMAT0005919_1&hsa-miR-548o-3p&offsets|0|0;m-430&7|-|102405748|102405769&offsets|0|0]|CCAAAACTGCAGTTACTTTTGC</td><td>>MI0016746|hsa-mir-548o-2&WithFlank&20|+|38516557|38516638</td><td>51</td><td>72</td><td>22</td><td>CCAAAACTGCAGTTACTTTTGC</td><td>229.9</td><td>5.4</td><td>1.0</td><td>5.7</td><td>1.6E-08</td><td>3.0E-08</td><td>MB231/MB468</td><td>MITO</td></tr>
<tr><td>>hg38_21_+_8433172_8446622_RNA45SN1_45S_50ntFlanks@6651.6684.34|CGACTCTTAGCGGTGGATCACTCGGCTCGTGCGT</td><td>>hg38_21_+_8433172_8446622_RNA45SN1_45S_50ntFlanks</td><td>6651</td><td>6684</td><td>34</td><td>CGACTCTTAGCGGTGGATCACTCGGCTCGTGCGT</td><td>10900.1</td><td>3.5</td><td>0.6</td><td>5.7</td><td>1.1E-08</td><td>3.0E-08</td><td>BT20/MB468</td><td>MITO</td></tr>
<tr><td>>hg38_21_+_8433172_8446622_RNA45SN1_45S_50ntFlanks@12704.12742.39|CAGCGCCGCGGAGCCTCGGTTGGCCTCGGATAGCCGGTC</td><td>>hg38_21_+_8433172_8446622_RNA45SN1_45S_50ntFlanks</td><td>12704</td><td>12742</td><td>39</td><td>CAGCGCCGCGGAGCCTCGGTTGGCCTCGGATAGCCGGTC</td><td>56.9</td><td>-9.7</td><td>1.7</td><td>-5.7</td><td>1.4E-08</td><td>3.0E-08</td><td>MB231/MB468</td><td>MP</td></tr>
<tr><td>>hg38_21_+_8433172_8446622_RNA45SN1_45S_50ntFlanks@11808.11830.23|CTGTCCCTACCTACTATCCAGCG</td><td>>hg38_21_+_8433172_8446622_RNA45SN1_45S_50ntFlanks</td><td>11808</td><td>11830</td><td>23</td><td>CTGTCCCTACCTACTATCCAGCG</td><td>316.8</td><td>-7.8</td><td>1.4</td><td>-5.7</td><td>1.4E-08</td><td>3.0E-08</td><td>MB231/MB468</td><td>MP</td></tr>
<tr><td>>hg38_21_+_8433172_8446622_RNA45SN1_45S_50ntFlanks@3920.3952.33|CATTTATCAGATCAAAACCAACCCGGTCAGCCC</td><td>>hg38_21_+_8433172_8446622_RNA45SN1_45S_50ntFlanks</td><td>3920</td><td>3952</td><td>33</td><td>CATTTATCAGATCAAAACCAACCCGGTCAGCCC</td><td>109.4</td><td>-8.9</td><td>1.6</td><td>-5.7</td><td>1.1E-08</td><td>3.0E-08</td><td>BT20/MB468</td><td>MITO</td></tr>
<tr><td>>hg38_21_+_8433172_8446622_RNA45SN1_45S_50ntFlanks@3840.3871.32|CTCCTACTTGGATAACTGTGGTAATTCTAGAG</td><td>>hg38_21_+_8433172_8446622_RNA45SN1_45S_50ntFlanks</td><td>3840</td><td>3871</td><td>32</td><td>CTCCTACTTGGATAACTGTGGTAATTCTAGAG</td><td>653.6</td><td>4.0</td><td>0.7</td><td>5.7</td><td>1.3E-08</td><td>3.0E-08</td><td>BT20/MB468</td><td>CYTO</td></tr>
<tr><td>>MI0000298|hsa-mir-221&WithFlank&X|-|45746151|45746272@71.92.22(+1A)&[MIMAT0000278&hsa-miR-221-3p&offsets|0|-1(+1A);m-91&X|-|45746181|45746202&offsets|0|0(+1A)]|AGCTACATTGTCTGCTGGGTTTA</td><td>>MI0000298|hsa-mir-221&WithFlank&X|-|45746151|45746272</td><td>71</td><td>92</td><td>22(+1A)</td><td>AGCTACATTGTCTGCTGGGTTTA</td><td>1118.0</td><td>5.9</td><td>1.0</td><td>5.7</td><td>1.5E-08</td><td>3.0E-08</td><td>MB231/MB468</td><td>CYTO</td></tr>
<tr><td>>MI0000077|hsa-mir-21&WithFlank&17|+|59841260|59841343@14.39.26(+1U)&[MIMAT0000076&hsa-miR-21-5p&offsets|0|+4(+1U);m-1&17|+|59841273|59841294&offsets|0|+4(+1U)]|TAGCTTATCAGACTGATGTTGACTGTT</td><td>>MI0000077|hsa-mir-21&WithFlank&17|+|59841260|59841343</td><td>14</td><td>39</td><td>26(+1U)</td><td>TAGCTTATCAGACTGATGTTGACTGTT</td><td>270.3</td><td>4.2</td><td>0.7</td><td>5.7</td><td>1.5E-08</td><td>3.0E-08</td><td>MB231/MB468</td><td>NUC</td></tr>
<tr><td>>MI0000542|hsa-mir-320a&WithFlank&8|-|22244960|22245043@48.70.23&[MIMAT0000510&hsa-miR-320a-3p&offsets|0|+1;m-60&8|-|22244975|22244996&offsets|0|+1]|AAAAGCTGGGTTGAGAGGGCGAA</td><td>>MI0000542|hsa-mir-320a&WithFlank&8|-|22244960|22245043</td><td>48</td><td>70</td><td>23</td><td>AAAAGCTGGGTTGAGAGGGCGAA</td><td>595.6</td><td>5.6</td><td>1.0</td><td>5.6</td><td>1.6E-08</td><td>3.0E-08</td><td>MB231/MB468</td><td>MITO</td></tr>
<tr><td>>hg38_21_+_8433172_8446622_RNA45SN1_45S_50ntFlanks@12274.12305.32|CAGTACGAATACAGACCGTGAAAGCGGGGCCT</td><td>>hg38_21_+_8433172_8446622_RNA45SN1_45S_50ntFlanks</td><td>12274</td><td>12305</td><td>32</td><td>CAGTACGAATACAGACCGTGAAAGCGGGGCCT</td><td>193.3</td><td>-9.7</td><td>1.7</td><td>-5.7</td><td>1.1E-08</td><td>3.0E-08</td><td>BT20/MB468</td><td>MITO</td></tr>
<tr><td>>hg38_21_+_8433172_8446622_RNA45SN1_45S_50ntFlanks@7984.8014.31|AGATCAGACGTGGCGACCCGCTGAATTTAAG</td><td>>hg38_21_+_8433172_8446622_RNA45SN1_45S_50ntFlanks</td><td>7984</td><td>8014</td><td>31</td><td>AGATCAGACGTGGCGACCCGCTGAATTTAAG</td><td>208.9</td><td>-7.6</td><td>1.4</td><td>-5.6</td><td>1.6E-08</td><td>3.0E-08</td><td>MB231/MB468</td><td>MITO</td></tr>
<tr><td>>MI0000750|hsa-mir-26a-2&WithFlank&12|-|57824603|57824698@20.40.21(+1C);MI0000083|hsa-mir-26a-1&WithFlank&3|+|37969398|37969486@16.36.21(+1C)&[MIMAT0000082&hsa-miR-26a-5p&offsets|0|-1(+1C);m-33&12|-|57824658|57824679&offsets|0|-1(+1C)];[MIMAT0000082_1&hsa-miR-26a-5p&offsets|0|-1(+1C);m-34&3|+|37969413|37969434&offsets|0|-1(+1C)]|TTCAAGTAATCCAGGATAGGCC</td><td>>MI0000750|hsa-mir-26a-2&WithFlank&12|-|57824603|57824698</td><td>20</td><td>40</td><td>21(+1C)</td><td>TTCAAGTAATCCAGGATAGGCC</td><td>7560.7</td><td>5.2</td><td>0.9</td><td>5.7</td><td>1.5E-08</td><td>3.0E-08</td><td>MB231/MB468</td><td>CYTO</td></tr>
<tr><td>>am_trna2_GlyTCC_19_+_4724082_4724153@1.29.29|GCGTTGGTGGTATAGTGGTTAGCATAGCT</td><td>>am_trna2_GlyTCC_19_+_4724082_4724153</td><td>1</td><td>29</td><td>29</td><td>GCGTTGGTGGTATAGTGGTTAGCATAGCT</td><td>188.4</td><td>-7.3</td><td>1.3</td><td>-5.7</td><td>1.4E-08</td><td>3.0E-08</td><td>MB231/MB468</td><td>MP</td></tr>
<tr><td>>hg38_21_+_8433172_8446622_RNA45SN1_45S_50ntFlanks@11567.11600.34|AGCAGCCGACTTAGAACTGGTGCGGACCAGGGGA</td><td>>hg38_21_+_8433172_8446622_RNA45SN1_45S_50ntFlanks</td><td>11567</td><td>11600</td><td>34</td><td>AGCAGCCGACTTAGAACTGGTGCGGACCAGGGGA</td><td>363.5</td><td>-3.8</td><td>0.7</td><td>-5.7</td><td>1.4E-08</td><td>3.0E-08</td><td>MB231/MB468</td><td>MP</td></tr>
<tr><td>>hg38_21_+_8433172_8446622_RNA45SN1_45S_50ntFlanks@11571.11603.33|GCCGACTTAGAACTGGTGCGGACCAGGGGAATC</td><td>>hg38_21_+_8433172_8446622_RNA45SN1_45S_50ntFlanks</td><td>11571</td><td>11603</td><td>33</td><td>GCCGACTTAGAACTGGTGCGGACCAGGGGAATC</td><td>1460.0</td><td>-4.8</td><td>0.8</td><td>-5.6</td><td>1.6E-08</td><td>3.0E-08</td><td>MB231/MB468</td><td>MITO</td></tr>
<tr><td>>hg38_21_+_8433172_8446622_RNA45SN1_45S_50ntFlanks@6668.6686.19|TCACTCGGCTCGTGCGTCG</td><td>>hg38_21_+_8433172_8446622_RNA45SN1_45S_50ntFlanks</td><td>6668</td><td>6686</td><td>19</td><td>TCACTCGGCTCGTGCGTCG</td><td>225.6</td><td>-7.4</td><td>1.3</td><td>-5.6</td><td>1.6E-08</td><td>3.0E-08</td><td>MB231/MB468</td><td>MITO</td></tr>
<tr><td>>hg38_21_+_8433172_8446622_RNA45SN1_45S_50ntFlanks@3835.3868.34|CTCCTCTCCTACTTGGATAACTGTGGTAATTCTA</td><td>>hg38_21_+_8433172_8446622_RNA45SN1_45S_50ntFlanks</td><td>3835</td><td>3868</td><td>34</td><td>CTCCTCTCCTACTTGGATAACTGTGGTAATTCTA</td><td>83.1</td><td>-10.2</td><td>1.8</td><td>-5.7</td><td>1.4E-08</td><td>3.0E-08</td><td>MB231/MB468</td><td>MP</td></tr>
<tr><td>>hg38_21_+_8433172_8446622_RNA45SN1_45S_50ntFlanks@11683.11701.19|ATGTCAAAGTGAAGAAATT</td><td>>hg38_21_+_8433172_8446622_RNA45SN1_45S_50ntFlanks</td><td>11683</td><td>11701</td><td>19</td><td>ATGTCAAAGTGAAGAAATT</td><td>63.1</td><td>-9.8</td><td>1.7</td><td>-5.7</td><td>1.4E-08</td><td>3.0E-08</td><td>MB231/MB468</td><td>MP</td></tr>
<tr><td>>hg38_21_+_8433172_8446622_RNA45SN1_45S_50ntFlanks@11583.11612.30|CTGGTGCGGACCAGGGGAATCCGACTGTTT</td><td>>hg38_21_+_8433172_8446622_RNA45SN1_45S_50ntFlanks</td><td>11583</td><td>11612</td><td>30</td><td>CTGGTGCGGACCAGGGGAATCCGACTGTTT</td><td>1013.0</td><td>-2.8</td><td>0.5</td><td>-5.7</td><td>1.5E-08</td><td>3.0E-08</td><td>MB231/MB468</td><td>NUC</td></tr>
<tr><td>>hg38_21_+_8433172_8446622_RNA45SN1_45S_50ntFlanks@12045.12071.27|CTGACCCGGTGAGGCGGGGGGCGAGC</td><td>>hg38_21_+_8433172_8446622_RNA45SN1_45S_50ntFlanks</td><td>12045</td><td>12071</td><td>27</td><td>CTGACCCGGTGAGGCGGGGGGCGAGC</td><td>215.9</td><td>-7.1</td><td>1.3</td><td>-5.7</td><td>1.5E-08</td><td>3.0E-08</td><td>MB231/MB468</td><td>NUC</td></tr>
<tr><td>>hg38_21_+_8433172_8446622_RNA45SN1_45S_50ntFlanks@6776.6799.24|CTCCCGGGGCTACGCCTGTCTGAG</td><td>>hg38_21_+_8433172_8446622_RNA45SN1_45S_50ntFlanks</td><td>6776</td><td>6799</td><td>24</td><td>CTCCCGGGGCTACGCCTGTCTGAG</td><td>117.7</td><td>-10.0</td><td>1.7</td><td>-6.0</td><td>2.2E-09</td><td>3.0E-08</td><td>BT20/MB468</td><td>MP</td></tr>
<tr><td>>hg38_21_+_8433172_8446622_RNA45SN1_45S_50ntFlanks@9566.9592.27|GACGTGCAAATCGGTCGTCCGACCTGG</td><td>>hg38_21_+_8433172_8446622_RNA45SN1_45S_50ntFlanks</td><td>9566</td><td>9592</td><td>27</td><td>GACGTGCAAATCGGTCGTCCGACCTGG</td><td>399.4</td><td>-3.3</td><td>0.6</td><td>-5.7</td><td>1.3E-08</td><td>3.0E-08</td><td>BT20/MB468</td><td>CYTO</td></tr>
<tr><td>>hg38_21_+_8433172_8446622_RNA45SN1_45S_50ntFlanks@10458.10489.32|ATGGCCTCCGTTGCCCTCGGCCGATCGAAAGG</td><td>>hg38_21_+_8433172_8446622_RNA45SN1_45S_50ntFlanks</td><td>10458</td><td>10489</td><td>32</td><td>ATGGCCTCCGTTGCCCTCGGCCGATCGAAAGG</td><td>411.3</td><td>-8.3</td><td>1.5</td><td>-5.6</td><td>1.7E-08</td><td>3.0E-08</td><td>BT20/MB468</td><td>NUC</td></tr>
<tr><td>>hg38_21_+_8433172_8446622_RNA45SN1_45S_50ntFlanks@9198.9223.26|GTTCTCTCGGGGCCACGCGCGCGTCC</td><td>>hg38_21_+_8433172_8446622_RNA45SN1_45S_50ntFlanks</td><td>9198</td><td>9223</td><td>26</td><td>GTTCTCTCGGGGCCACGCGCGCGTCC</td><td>540.1</td><td>-8.3</td><td>1.4</td><td>-6.0</td><td>2.2E-09</td><td>3.0E-08</td><td>BT20/MB468</td><td>MP</td></tr>
<tr><td>>hg38_21_+_8433172_8446622_RNA45SN1_45S_50ntFlanks@4367.4406.40|CAGTTAAAAAGCTCGTAGTTGGATCTTGGGAGCGGGCGGG</td><td>>hg38_21_+_8433172_8446622_RNA45SN1_45S_50ntFlanks</td><td>4367</td><td>4406</td><td>40</td><td>CAGTTAAAAAGCTCGTAGTTGGATCTTGGGAGCGGGCGGG</td><td>116.4</td><td>-10.7</td><td>1.9</td><td>-5.7</td><td>1.4E-08</td><td>3.0E-08</td><td>MB231/MB468</td><td>MP</td></tr>
<tr><td>>hg38_21_+_8433172_8446622_RNA45SN1_45S_50ntFlanks@12948.12982.35|GACGACCTGCTTCTGGGTCGGGGTTTCGTACGTAG</td><td>>hg38_21_+_8433172_8446622_RNA45SN1_45S_50ntFlanks</td><td>12948</td><td>12982</td><td>35</td><td>GACGACCTGCTTCTGGGTCGGGGTTTCGTACGTAG</td><td>335.0</td><td>-3.1</td><td>0.6</td><td>-5.7</td><td>1.5E-08</td><td>3.0E-08</td><td>MB231/MB468</td><td>CYTO</td></tr>
<tr><td>>MI0003180|hsa-mir-516a-1&WithFlank&19|+|53756735|53756836@22.44.23;MI0003181|hsa-mir-516a-2&WithFlank&19|+|53761127|53761228@22.44.23&[MIMAT0004770&hsa-miR-516a-5p&offsets|0|0;m-852&19|+|53756756|53756777&offsets|0|+1];[MIMAT0004770_1&hsa-miR-516a-5p&offsets|0|0;m-852&19|+|53761148|53761169&offsets|0|+1]|TTCTCGAGGAAAGAAGCACTTTC</td><td>>MI0003180|hsa-mir-516a-1&WithFlank&19|+|53756735|53756836</td><td>22</td><td>44</td><td>23</td><td>TTCTCGAGGAAAGAAGCACTTTC</td><td>63.1</td><td>9.2</td><td>1.6</td><td>5.7</td><td>1.4E-08</td><td>3.0E-08</td><td>MB231/MB468</td><td>MP</td></tr>
<tr><td>>hg38_21_+_8433172_8446622_RNA45SN1_45S_50ntFlanks@10247.10279.33|GCGACGAGTAGGAGGGCCGCTGCGGTGAGCCTT</td><td>>hg38_21_+_8433172_8446622_RNA45SN1_45S_50ntFlanks</td><td>10247</td><td>10279</td><td>33</td><td>GCGACGAGTAGGAGGGCCGCTGCGGTGAGCCTT</td><td>481.8</td><td>12.2</td><td>2.0</td><td>6.3</td><td>4.0E-10</td><td>3.0E-08</td><td>BT20/MB231</td><td>MP</td></tr>
<tr><td>>trnaMT_LeuTAG_MT_+_12266_12336@40.74.35|CAAAAATTTTGGTGCAACTCCAAATAAAAGTACCA</td><td>>trnaMT_LeuTAG_MT_+_12266_12336</td><td>40</td><td>74</td><td>35</td><td>CAAAAATTTTGGTGCAACTCCAAATAAAAGTACCA</td><td>250.1</td><td>-11.4</td><td>1.8</td><td>-6.3</td><td>4.0E-10</td><td>3.0E-08</td><td>BT20/MB231</td><td>MP</td></tr>
<tr><td>>hg38_21_+_8433172_8446622_RNA45SN1_45S_50ntFlanks@12991.13028.38|CTCCCTCGCTGCGATCTATTGAAAGTCAGCCCTCGACA</td><td>>hg38_21_+_8433172_8446622_RNA45SN1_45S_50ntFlanks</td><td>12991</td><td>13028</td><td>38</td><td>CTCCCTCGCTGCGATCTATTGAAAGTCAGCCCTCGACA</td><td>65.2</td><td>-9.9</td><td>1.7</td><td>-5.7</td><td>1.4E-08</td><td>3.0E-08</td><td>MB231/MB468</td><td>MP</td></tr>
<tr><td>>MI0000746|hsa-mir-99b&WithFlank&19|+|51692606|51692687@51.72.22&[MIMAT0004678&hsa-miR-99b-3p&offsets|0|0;m-151&19|+|51692656|51692678&offsets|0|-1]|CAAGCTCGTGTCTGTGGGTCCG</td><td>>MI0000746|hsa-mir-99b&WithFlank&19|+|51692606|51692687</td><td>51</td><td>72</td><td>22</td><td>CAAGCTCGTGTCTGTGGGTCCG</td><td>628.7</td><td>4.0</td><td>0.7</td><td>5.7</td><td>1.5E-08</td><td>3.0E-08</td><td>MB231/MB468</td><td>CYTO</td></tr>
</table>

| | | | | | | | | | | | | | |
|---|---|---|---|---|---|---|---|---|---|---|---|---|---|
| >hg38_21_+_8433172_8446622_RNA45SN1_45S_50ntFlanks@11820.11856.37\|ACTATCCAGCGAAACCACAGCCAAGGGAACGGGCTTG | >hg38_21_+_8433172_8446622_RNA45SN1_45S_50ntFlanks | 11820 | 11856 | 37 | ACTATCCAGCGAAACCACAGCCAAGGGAACGGGCTTG | 50.5 | -9.5 | 1.7 | -5.7 | 1.4E-08 | 3.0E-08 | MB231/MB468 | MP |
| >hg38_21_+_8433172_8446622_RNA45SN1_45S_50ntFlanks@10687.10708.22\|CGGCGTCCGGTGAGCTCTCGCT | >hg38_21_+_8433172_8446622_RNA45SN1_45S_50ntFlanks | 10687 | 10708 | 22 | CGGCGTCCGGTGAGCTCTCGCT | 933.2 | -8.1 | 1.4 | -5.7 | 1.4E-08 | 3.0E-08 | MB231/MB468 | MP |
| >hg38_21_+_8433172_8446622_RNA45SN1_45S_50ntFlanks@12458.12478.21\|AAGCGTTGGATTGTTCACCCA | >hg38_21_+_8433172_8446622_RNA45SN1_45S_50ntFlanks | 12458 | 12478 | 21 | AAGCGTTGGATTGTTCACCCA | 1940.0 | -5.1 | 0.9 | -5.6 | 1.7E-08 | 3.0E-08 | BT20/MB468 | NUC |
| >hg38_21_+_8433172_8446622_RNA45SN1_45S_50ntFlanks@3853.3881.29\|AACTGTGGTAATTCTAGAGCTAATACATG | >hg38_21_+_8433172_8446622_RNA45SN1_45S_50ntFlanks | 3853 | 3881 | 29 | AACTGTGGTAATTCTAGAGCTAATACATG | 1000.9 | -8.9 | 1.6 | -5.6 | 1.7E-08 | 3.0E-08 | BT20/MB468 | NUC |
| >hg38_21_+_8433172_8446622_RNA45SN1_45S_50ntFlanks@9339.9366.28\|CGCACGAAAGCCGCCGTGGCGCAATGAA | >hg38_21_+_8433172_8446622_RNA45SN1_45S_50ntFlanks | 9339 | 9366 | 28 | CGCACGAAAGCCGCCGTGGCGCAATGAA | 165.5 | -9.5 | 1.7 | -5.7 | 1.2E-08 | 3.0E-08 | BT20/MB468 | MITO |
| >trnaMT_SerGCT_MT_+_12207_12265@1.25.25\|GAGAAAGCTCACAAGAACTGCTAAC | >trnaMT_SerGCT_MT_+_12207_12265 | 1 | 25 | 25 | GAGAAAGCTCACAAGAACTGCTAAC | 107493.0 | 7.1 | 1.2 | 5.7 | 1.2E-08 | 3.0E-08 | BT20/MB468 | MITO |
| >hg38_21_+_8433172_8446622_RNA45SN1_45S_50ntFlanks@12927.12960.34\|CCTGGCGCTAAACCATTCGTAGACGACCTGCTTC | >hg38_21_+_8433172_8446622_RNA45SN1_45S_50ntFlanks | 12927 | 12960 | 34 | CCTGGCGCTAAACCATTCGTAGACGACCTGCTTC | 65.6 | -9.9 | 1.7 | -5.7 | 1.4E-08 | 3.0E-08 | MB231/MB468 | MP |
| >hg38_21_+_8433172_8446622_RNA45SN1_45S_50ntFlanks@5078.5098.21\|CGATAACGAACGAGACTCTGG | >hg38_21_+_8433172_8446622_RNA45SN1_45S_50ntFlanks | 5078 | 5098 | 21 | CGATAACGAACGAGACTCTGG | 813.8 | -8.2 | 1.5 | -5.6 | 1.6E-08 | 3.0E-08 | MB231/MB468 | MITO |
| >hg38_21_+_8433172_8446622_RNA45SN1_45S_50ntFlanks@10534.10563.30\|CCGCGAGGCGTCCAGTGCGGTAACGCGACC | >hg38_21_+_8433172_8446622_RNA45SN1_45S_50ntFlanks | 10534 | 10563 | 30 | CCGCGAGGCGTCCAGTGCGGTAACGCGACC | 113.6 | -9.9 | 1.7 | -6.0 | 2.2E-09 | 3.0E-08 | BT20/MB468 | MP |
| >trnaMT_SerGCT_MT_+_12207_12265@6.38.33\|AGCTCACAAGAACTGCTAACTCATGCCCCCATG | >trnaMT_SerGCT_MT_+_12207_12265 | 6 | 38 | 33 | AGCTCACAAGAACTGCTAACTCATGCCCCCATG | 318.0 | -7.7 | 1.4 | -5.7 | 1.5E-08 | 3.0E-08 | MB231/MB468 | NUC |
| >MI0000465\|hsa-mir-191&WithFlank&3\|-\|49020612\|49020715@22.42.21(+1C)&[MIMAT0000440&hsa-miR-191-5p&offsets\|0\|-2(+1C);m-48&3\|-\|49020672\|49020694&offsets\|0\|-2(+1C)]\|CAACGGAATCCCAAAAGCAGCC | >MI0000465\|hsa-mir-191&WithFlank&3\|-\|49020612\|49020715 | 22 | 42 | 21(+1C) | CAACGGAATCCCAAAAGCAGCC | 516.0 | 2.4 | 0.4 | 5.6 | 1.7E-08 | 3.0E-08 | BT20/MB468 | NUC |
| >trna20_ThrTGT_14_-_21099319_21099391@35.54.20,trna21_ThrTGT_14_-_21081949_21082021@35.54.20,trna4_ThrTGT_14_+_21149849_21149921@35.54.20,trna56_ThrTGT_1_+_222638347_222638419@35.54.20\|GTAAACCAGGGGTCGCGAGT | >trna20_ThrTGT_14_-_21099319_21099391 | 35 | 54 | 20 | GTAAACCAGGGGTCGCGAGT | 365.7 | 7.8 | 1.4 | 5.7 | 1.3E-08 | 3.0E-08 | BT20/MB468 | CYTO |
| >hg38_21_+_8433172_8446622_RNA45SN1_45S_50ntFlanks@10389.10430.42\|TGAACAGCAGTTGAACATGGGTCAGTCGGTCCTGAGAGATGG | >hg38_21_+_8433172_8446622_RNA45SN1_45S_50ntFlanks | 10389 | 10430 | 42 | TGAACAGCAGTTGAACATGGGTCAGTCGGTCCTGAGAGATGG | 295.6 | -5.5 | 1.0 | -5.7 | 1.5E-08 | 3.0E-08 | MB231/MB468 | CYTO |
| >hg38_21_+_8433172_8446622_RNA45SN1_45S_50ntFlanks@4007.4025.19\|CCTCGGGCCGATCGCACGC | >hg38_21_+_8433172_8446622_RNA45SN1_45S_50ntFlanks | 4007 | 4025 | 19 | CCTCGGGCCGATCGCACGC | 243.0 | -2.7 | 0.5 | -5.7 | 1.5E-08 | 3.0E-08 | MB231/MB468 | CYTO |
| >hg38_21_+_8433172_8446622_RNA45SN1_45S_50ntFlanks@6646.6664.19\|TCGTACGACTCTTAGCGGT | >hg38_21_+_8433172_8446622_RNA45SN1_45S_50ntFlanks | 6646 | 6664 | 19 | TCGTACGACTCTTAGCGGT | 11526.3 | 4.0 | 0.7 | 5.7 | 1.5E-08 | 3.0E-08 | MB231/MB468 | CYTO |
| >MI0000266\|hsa-mir-10a&WithFlank&17\|-\|48579832\|48579953@28.48.21(+1G)&[MIMAT0000253&hsa-miR-10a-5p&offsets\|0\|-2(+1G);m-18&17\|-\|48579905\|48579926&offsets\|0\|-1(+1G)]\|TACCCTGTAGATCCGAATTTGG | >MI0000266\|hsa-mir-10a&WithFlank&17\|-\|48579832\|48579953 | 28 | 48 | 21(+1G) | TACCCTGTAGATCCGAATTTGG | 228.6 | 11.6 | 1.9 | 6.0 | 2.2E-09 | 3.0E-08 | BT20/MB468 | MP |
| >hg38_21_+_8433172_8446622_RNA45SN1_45S_50ntFlanks@4297.4317.21\|CATTGGAGGGCAAGTCTGGTG | >hg38_21_+_8433172_8446622_RNA45SN1_45S_50ntFlanks | 4297 | 4317 | 21 | CATTGGAGGGCAAGTCTGGTG | 412.1 | -5.8 | 1.0 | -5.7 | 1.5E-08 | 3.0E-08 | MB231/MB468 | NUC |
| >hg38_21_+_8433172_8446622_RNA45SN1_45S_50ntFlanks@11876.11898.23\|AAGACCCTGTTGAGCTTGACTCT | >hg38_21_+_8433172_8446622_RNA45SN1_45S_50ntFlanks | 11876 | 11898 | 23 | AAGACCCTGTTGAGCTTGACTCT | 492.6 | -7.8 | 1.4 | -5.7 | 1.4E-08 | 3.0E-08 | MB231/MB468 | MP |
| >hg38_21_+_8433172_8446622_RNA45SN1_45S_50ntFlanks@12289.12320.32\|CCGTGAAAGCGGGGCCTCACGATCCTTCTGAC | >hg38_21_+_8433172_8446622_RNA45SN1_45S_50ntFlanks | 12289 | 12320 | 32 | CCGTGAAAGCGGGGCCTCACGATCCTTCTGAC | 530.9 | -8.6 | 1.5 | -5.6 | 1.7E-08 | 3.0E-08 | MB231/MB468 | MITO |
| >hg38_21_+_8433172_8446622_RNA45SN1_45S_50ntFlanks@6661.6688.28\|CGGTGGATCACTCGGCTCGTGCGTCGAT | >hg38_21_+_8433172_8446622_RNA45SN1_45S_50ntFlanks | 6661 | 6688 | 28 | CGGTGGATCACTCGGCTCGTGCGTCGAT | 8822.3 | 4.8 | 0.8 | 6.1 | 1.2E-09 | 3.0E-08 | BT20/MB231 | MITO |
| >hg38_21_+_8433172_8446622_RNA45SN1_45S_50ntFlanks@11778.11804.27\|ACGCGCATGAATGGATGAACGAGATTC | >hg38_21_+_8433172_8446622_RNA45SN1_45S_50ntFlanks | 11778 | 11804 | 27 | ACGCGCATGAATGGATGAACGAGATTC | 601.4 | -8.6 | 1.5 | -5.6 | 1.7E-08 | 3.1E-08 | BT20/MB468 | NUC |
| >MI0000255\|hsa-mir-30d&WithFlank&8\|-\|134804870\|134804951@12.35.24(+1A)&[MIMAT0000245&hsa-miR-30d-5p&offsets\|0\|+2(+1A);m-21&8\|-\|134804919\|134804940&offsets\|0\|+2(+1A)]\|TGTAAACATCCCCGACTGGAAGCTA | >MI0000255\|hsa-mir-30d&WithFlank&8\|-\|134804870\|134804951 | 12 | 35 | 24(+1A) | TGTAAACATCCCCGACTGGAAGCTA | 513.4 | 9.2 | 1.6 | 5.7 | 1.3E-08 | 3.1E-08 | BT20/MB468 | CYTO |
| >hg38_21_+_8433172_8446622_RNA45SN1_45S_50ntFlanks@7981.8012.32\|CTCAGATCAGACGTGGCGACCCGCTGAATTTA | >hg38_21_+_8433172_8446622_RNA45SN1_45S_50ntFlanks | 7981 | 8012 | 32 | CTCAGATCAGACGTGGCGACCCGCTGAATTTA | 371.9 | -7.6 | 1.3 | -5.7 | 1.4E-08 | 3.1E-08 | MB231/MB468 | MP |
| >hg38_21_+_8433172_8446622_RNA45SN1_45S_50ntFlanks@10786.10830.45\|AACAGCCTCTGGCATGTTGGAACAATGTAGGTAAGGGAAGTCGGC | >hg38_21_+_8433172_8446622_RNA45SN1_45S_50ntFlanks | 10786 | 10830 | 45 | AACAGCCTCTGGCATGTTGGAACAATGTAGGTAAGGGAAGTCGGC | 400.4 | -6.8 | 1.2 | -5.6 | 1.7E-08 | 3.1E-08 | MB231/MB468 | MITO |
| >hg38_21_+_8433172_8446622_RNA45SN1_45S_50ntFlanks@3838.3868.31\|CTCTCCTACTTGGATAACTGTGGTAATTCTA | >hg38_21_+_8433172_8446622_RNA45SN1_45S_50ntFlanks | 3838 | 3868 | 31 | CTCTCCTACTTGGATAACTGTGGTAATTCTA | 107.2 | -10.6 | 1.9 | -5.7 | 1.4E-08 | 3.1E-08 | MB231/MB468 | MP |
| >MI0000811\|hsa-mir-148b&WithFlank&12\|+\|54337210\|54337320@69.89.21(+1C)&[MIMAT0000759&hsa-miR-148b-3p&offsets\|0\|-1(+1C);m-77&12\|+\|54337278\|54337299&offsets\|0\|-1(+1C)]\|TCAGTGCATCACAGAACTTTGC | >MI0000811\|hsa-mir-148b&WithFlank&12\|+\|54337210\|54337320 | 69 | 89 | 21(+1C) | TCAGTGCATCACAGAACTTTGC | 854.9 | 6.5 | 1.1 | 5.6 | 1.7E-08 | 3.1E-08 | MB231/MB468 | MITO |
| >hg38_21_+_8433172_8446622_RNA45SN1_45S_50ntFlanks@11785.11806.22\|TGAATGGATGAACGAGATTCCC | >hg38_21_+_8433172_8446622_RNA45SN1_45S_50ntFlanks | 11785 | 11806 | 22 | TGAATGGATGAACGAGATTCCC | 582.8 | -6.4 | 1.1 | -5.6 | 1.7E-08 | 3.1E-08 | MB231/MB468 | MITO |
| >hg38_21_+_8433172_8446622_RNA45SN1_45S_50ntFlanks@11876.11906.31\|AAGACCCTGTTGAGCTTGACTCTAGTCTGGC | >hg38_21_+_8433172_8446622_RNA45SN1_45S_50ntFlanks | 11876 | 11906 | 31 | AAGACCCTGTTGAGCTTGACTCTAGTCTGGC | 608.2 | -3.8 | 0.7 | -5.7 | 1.2E-08 | 3.1E-08 | BT20/MB468 | MITO |
| >hg38_21_+_8433172_8446622_RNA45SN1_45S_50ntFlanks@3858.3898.41\|TGGTAATTCTAGAGCTAATACATGCCGACGGGCGCTGACCC | >hg38_21_+_8433172_8446622_RNA45SN1_45S_50ntFlanks | 3858 | 3898 | 41 | TGGTAATTCTAGAGCTAATACATGCCGACGGGCGCTGACCC | 308.9 | -7.5 | 1.3 | -5.6 | 1.7E-08 | 3.1E-08 | MB231/MB468 | MITO |
| >hg38_21_+_8433172_8446622_RNA45SN1_45S_50ntFlanks@5492.5513.22\|AGAAGACGGTCGAACTTGACTA | >hg38_21_+_8433172_8446622_RNA45SN1_45S_50ntFlanks | 5492 | 5513 | 22 | AGAAGACGGTCGAACTTGACTA | 410.4 | -7.7 | 1.4 | -5.6 | 1.7E-08 | 3.1E-08 | MB231/MB468 | MITO |
| >hg38_21_+_8433172_8446622_RNA45SN1_45S_50ntFlanks@10346.10366.21\|TCAAACGAGAACTTTGAAGGC | >hg38_21_+_8433172_8446622_RNA45SN1_45S_50ntFlanks | 10346 | 10366 | 21 | TCAAACGAGAACTTTGAAGGC | 141.2 | -10.2 | 1.7 | -6.0 | 2.2E-09 | 3.1E-08 | BT20/MB468 | MP |
| >hg38_21_+_8433172_8446622_RNA45SN1_45S_50ntFlanks@12674.12708.35\|TCAGAATCCCGCCCAGGCGGAACGATACGGCAGCG | >hg38_21_+_8433172_8446622_RNA45SN1_45S_50ntFlanks | 12674 | 12708 | 35 | TCAGAATCCCGCCCAGGCGGAACGATACGGCAGCG | 109.4 | -9.9 | 1.6 | -6.0 | 2.2E-09 | 3.1E-08 | BT20/MB468 | MP |
| >hg38_21_+_8433172_8446622_RNA45SN1_45S_50ntFlanks@8437.8459.23\|AGGATTCAACCCGGCGGCGGGTC | >hg38_21_+_8433172_8446622_RNA45SN1_45S_50ntFlanks | 8437 | 8459 | 23 | AGGATTCAACCCGGCGGCGGGTC | 199.5 | -10.7 | 1.8 | -6.0 | 2.2E-09 | 3.1E-08 | BT20/MB468 | MP |
| >hg38_21_+_8433172_8446622_RNA45SN1_45S_50ntFlanks@12071.12097.27\|CCCCGAGGGGCTCTCGCTTCTGGCGCC | >hg38_21_+_8433172_8446622_RNA45SN1_45S_50ntFlanks | 12071 | 12097 | 27 | CCCCGAGGGGCTCTCGCTTCTGGCGCC | 666.1 | -6.4 | 1.1 | -5.6 | 1.7E-08 | 3.1E-08 | MB231/MB468 | MITO |
| >hg38_21_+_8433172_8446622_RNA45SN1_45S_50ntFlanks@3843.3860.18\|CTACTTGGATAACTGTGG | >hg38_21_+_8433172_8446622_RNA45SN1_45S_50ntFlanks | 3843 | 3860 | 18 | CTACTTGGATAACTGTGG | 489.9 | -8.1 | 1.4 | -5.6 | 1.7E-08 | 3.1E-08 | MB231/MB468 | MITO |
| >hg38_21_+_8433172_8446622_RNA45SN1_45S_50ntFlanks@11781.11806.26\|CGCATGAATGGATGAACGAGATTCCC | >hg38_21_+_8433172_8446622_RNA45SN1_45S_50ntFlanks | 11781 | 11806 | 26 | CGCATGAATGGATGAACGAGATTCCC | 870.9 | -7.8 | 1.4 | -5.7 | 1.6E-08 | 3.1E-08 | MB231/MB468 | NUC |
| >MI0000342\|hsa-mir-200b&WithFlank&1\|+\|1167098\|1167204@63.85.23(+1C)&[MIMAT0000318&hsa-miR-200b-3p&offsets\|0\|+1(+1C);m-93&1\|+\|1167160\|1167182&offsets\|0\|0(+1C)]\|TAATACTGCCTGGTAATGATGACC | >MI0000342\|hsa-mir-200b&WithFlank&1\|+\|1167098\|1167204 | 63 | 85 | 23(+1C) | TAATACTGCCTGGTAATGATGACC | 162.4 | -5.5 | 1.0 | -5.7 | 1.5E-08 | 3.1E-08 | MB231/MB468 | CYTO |
| >hg38_21_+_8433172_8446622_RNA45SN1_45S_50ntFlanks@4364.4394.31\|CTGCAGTTAAAAAGCTCGTAGTTGGATCTTG | >hg38_21_+_8433172_8446622_RNA45SN1_45S_50ntFlanks | 4364 | 4394 | 31 | CTGCAGTTAAAAAGCTCGTAGTTGGATCTTG | 57.4 | -9.7 | 1.7 | -5.7 | 1.4E-08 | 3.1E-08 | MB231/MB468 | MP |
| >hg38_21_+_8433172_8446622_RNA45SN1_45S_50ntFlanks@3776.3801.26\|CCGGTACAGTGAAACTGCGAATGGCT | >hg38_21_+_8433172_8446622_RNA45SN1_45S_50ntFlanks | 3776 | 3801 | 26 | CCGGTACAGTGAAACTGCGAATGGCT | 368.5 | -6.9 | 1.2 | -5.7 | 1.2E-08 | 3.1E-08 | BT20/MB468 | MITO |
| >hg38_21_+_8433172_8446622_RNA45SN1_45S_50ntFlanks@475.504.30\|CTTCGTGATCGATGTGGTGACGTCGTGCTC | >hg38_21_+_8433172_8446622_RNA45SN1_45S_50ntFlanks | 475 | 504 | 30 | CTTCGTGATCGATGTGGTGACGTCGTGCTC | 28581.6 | 4.3 | 0.8 | 5.7 | 1.6E-08 | 3.1E-08 | MB231/MB468 | NUC |
| >MI0000078\|hsa-mir-22&WithFlank&17\|-\|1713897\|1713993@59.80.22&[MIMAT0000077&hsa-miR-22-3p&offsets\|0\|0;m-2&17\|-\|1713914\|1713935&offsets\|0\|0]\|AAGCTGCCAGTTGAAGAACTGT | >MI0000078\|hsa-mir-22&WithFlank&17\|-\|1713897\|1713993 | 59 | 80 | 22 | AAGCTGCCAGTTGAAGAACTGT | 34463.5 | 5.2 | 0.9 | 5.7 | 1.4E-08 | 3.1E-08 | MB231/MB468 | MP |
| >hg38_21_+_8433172_8446622_RNA45SN1_45S_50ntFlanks@5078.5102.25\|CGATAACGAACGAGACTCTGGCATG | >hg38_21_+_8433172_8446622_RNA45SN1_45S_50ntFlanks | 5078 | 5102 | 25 | CGATAACGAACGAGACTCTGGCATG | 769.0 | -5.4 | 1.0 | -5.7 | 1.6E-08 | 3.1E-08 | MB231/MB468 | NUC |
| >MI0000082\|hsa-mir-25&WithFlank&7\|-\|100093554\|100093649@58.79.22(+1U)&[MIMAT0000081&hsa-miR-25-3p&offsets\|0\|0(+1U);m-23&7\|-\|100093571\|100093592&offsets\|0\|0(+1U)]\|CATTGCACTTGTCTCGGTCTGAT | >MI0000082\|hsa-mir-25&WithFlank&7\|-\|100093554\|100093649 | 58 | 79 | 22(+1U) | CATTGCACTTGTCTCGGTCTGAT | 816.6 | 5.4 | 0.9 | 5.6 | 1.7E-08 | 3.1E-08 | MB231/MB468 | MITO |
| >hg38_21_+_8433172_8446622_RNA45SN1_45S_50ntFlanks@12416.12449.34\|TTCGATGTCGGCTCTTCCTATCATTGTGAAGCAG | >hg38_21_+_8433172_8446622_RNA45SN1_45S_50ntFlanks | 12416 | 12449 | 34 | TTCGATGTCGGCTCTTCCTATCATTGTGAAGCAG | 496.1 | 5.0 | 0.9 | 5.7 | 1.6E-08 | 3.1E-08 | MB231/MB468 | NUC |
| >hg38_21_+_8433172_8446622_RNA45SN1_45S_50ntFlanks@9822.9850.29\|CTAGTGGGCCACTTTTGGTAAGCAGAACT | >hg38_21_+_8433172_8446622_RNA45SN1_45S_50ntFlanks | 9822 | 9850 | 29 | CTAGTGGGCCACTTTTGGTAAGCAGAACT | 557.4 | -3.8 | 0.7 | -5.7 | 1.6E-08 | 3.1E-08 | MB231/MB468 | NUC |
| >hg38_21_+_8433172_8446622_RNA45SN1_45S_50ntFlanks@11768.11791.24\|CTAATTAGTGACGCGCATGAATGG | >hg38_21_+_8433172_8446622_RNA45SN1_45S_50ntFlanks | 11768 | 11791 | 24 | CTAATTAGTGACGCGCATGAATGG | 625.5 | -7.2 | 1.3 | -5.7 | 1.3E-08 | 3.1E-08 | BT20/MB468 | CYTO |
| >hg38_21_+_8433172_8446622_RNA45SN1_45S_50ntFlanks@10476.10497.22\|GGCCGATCGAAAGGGAGTCGGG | >hg38_21_+_8433172_8446622_RNA45SN1_45S_50ntFlanks | 10476 | 10497 | 22 | GGCCGATCGAAAGGGAGTCGGG | 78.7 | -10.2 | 1.8 | -5.7 | 1.5E-08 | 3.1E-08 | MB231/MB468 | MP |
| >hg38_21_+_8433172_8446622_RNA45SN1_45S_50ntFlanks@11918.11959.42\|ACATGAGAGGTGTAGAATAAGTGGGAGGCCCCCGGCGCCCCC | >hg38_21_+_8433172_8446622_RNA45SN1_45S_50ntFlanks | 11918 | 11959 | 42 | ACATGAGAGGTGTAGAATAAGTGGGAGGCCCCCGGCGCCCCC | 171.5 | -6.3 | 1.1 | -5.7 | 1.6E-08 | 3.1E-08 | MB231/MB468 | CYTO |
| >hg38_21_+_8433172_8446622_RNA45SN1_45S_50ntFlanks@11534.11557.24\|GTTCCGCGCGGCGCCTCGCCTCGG | >hg38_21_+_8433172_8446622_RNA45SN1_45S_50ntFlanks | 11534 | 11557 | 24 | GTTCCGCGCGGCGCCTCGCCTCGG | 314.3 | -7.7 | 1.3 | -5.7 | 1.3E-08 | 3.1E-08 | BT20/MB468 | CYTO |

<table>
<tr><td>>hg38_21_+_8433172_8446622_RNA45SN1_45S_50ntFlanks@5420.5466.47|CCGATTGGATGGTTTAGTGAGGCCCTCGGATCGGCCCCGCCGGGGTC</td><td>>hg38_21_+_8433172_8446622_RNA45SN1_45S_50ntFlanks</td><td>5420</td><td>5466</td><td>47</td><td>CCGATTGGATGGTTTAGTGAGGCCCTCGGATCGGCCCCGCCGGGGTC</td><td>131.7</td><td>-9.3</td><td>1.6</td><td>-5.7</td><td>1.3E-08</td><td>3.1E-08</td><td>BT20/MB468</td><td>CYTO</td></tr>
<tr><td>>am_trna133_GlyCCC_1_-_16872434_16872504@1.34.34,trna4_GlyCCC_1_+_17188416_17188486@1.34.34|GCATTGGTGGTTCAGTGGTAGAATTCTCGCCTCC</td><td>>am_trna133_GlyCCC_1_-_16872434_16872504</td><td>1</td><td>34</td><td>34</td><td>GCATTGGTGGTTCAGTGGTAGAATTCTCGCCTCC</td><td>259.5</td><td>11.8</td><td>2.0</td><td>6.0</td><td>2.3E-09</td><td>3.1E-08</td><td>BT20/MB468</td><td>MP</td></tr>
<tr><td>>hg38_21_+_8433172_8446622_RNA45SN1_45S_50ntFlanks@11580.11600.21|GAACTGGTGCGGACCAGGGGA</td><td>>hg38_21_+_8433172_8446622_RNA45SN1_45S_50ntFlanks</td><td>11580</td><td>11600</td><td>21</td><td>GAACTGGTGCGGACCAGGGGA</td><td>208.9</td><td>-7.3</td><td>1.3</td><td>-5.6</td><td>1.7E-08</td><td>3.1E-08</td><td>MB231/MB468</td><td>MITO</td></tr>
<tr><td>>hg38_21_+_8433172_8446622_RNA45SN1_45S_50ntFlanks@10457.10497.41|GATGGCCTCCGTTGCCCTCGGCCGATCGAAAGGGAGTCGGG</td><td>>hg38_21_+_8433172_8446622_RNA45SN1_45S_50ntFlanks</td><td>10457</td><td>10497</td><td>41</td><td>GATGGCCTCCGTTGCCCTCGGCCGATCGAAAGGGAGTCGGG</td><td>124.8</td><td>-10.0</td><td>1.7</td><td>-6.0</td><td>2.3E-09</td><td>3.2E-08</td><td>BT20/MB468</td><td>MP</td></tr>
<tr><td>>hg38_21_+_8433172_8446622_RNA45SN1_45S_50ntFlanks@11834.11872.39|CCACAGCCAAGGGAACGGGCTTGGCGGAATCAGCGGGGA</td><td>>hg38_21_+_8433172_8446622_RNA45SN1_45S_50ntFlanks</td><td>11834</td><td>11872</td><td>39</td><td>CCACAGCCAAGGGAACGGGCTTGGCGGAATCAGCGGGGA</td><td>155.0</td><td>-10.4</td><td>1.7</td><td>-6.0</td><td>2.3E-09</td><td>3.2E-08</td><td>BT20/MB468</td><td>MP</td></tr>
<tr><td>>hg38_21_+_8433172_8446622_RNA45SN1_45S_50ntFlanks@12076.12097.22|AGGGGCTCTCGCTTCTGGCGCC</td><td>>hg38_21_+_8433172_8446622_RNA45SN1_45S_50ntFlanks</td><td>12076</td><td>12097</td><td>22</td><td>AGGGGCTCTCGCTTCTGGCGCC</td><td>288.3</td><td>-5.8</td><td>1.0</td><td>-5.6</td><td>1.7E-08</td><td>3.2E-08</td><td>MB231/MB468</td><td>MITO</td></tr>
<tr><td>>hg38_21_+_8433172_8446622_RNA45SN1_45S_50ntFlanks@8182.8205.24|ATCGAGGCCCAGCCCGTGGACGGT</td><td>>hg38_21_+_8433172_8446622_RNA45SN1_45S_50ntFlanks</td><td>8182</td><td>8205</td><td>24</td><td>ATCGAGGCCCAGCCCGTGGACGGT</td><td>119.4</td><td>-10.0</td><td>1.7</td><td>-6.0</td><td>2.3E-09</td><td>3.2E-08</td><td>BT20/MB468</td><td>MP</td></tr>
<tr><td>>hg38_21_+_8433172_8446622_RNA45SN1_45S_50ntFlanks@11455.11474.20|CTCGGCGCGCGGCGGCGGCG</td><td>>hg38_21_+_8433172_8446622_RNA45SN1_45S_50ntFlanks</td><td>11455</td><td>11474</td><td>20</td><td>CTCGGCGCGCGGCGGCGGCG</td><td>132.8</td><td>-10.1</td><td>1.7</td><td>-6.0</td><td>2.3E-09</td><td>3.2E-08</td><td>BT20/MB468</td><td>MP</td></tr>
<tr><td>>am_trna111_HisGTG_1_-_147774845_147774916@3.29.27,trna118_HisGTG_1_-_145396881_145396952@3.29.27,trna16_HisGTG_1_+_146544773_146544844@3.29.27,trna1_HisGTG_15_+_45493349_45493420@3.29.27,trna21_HisGTG_1_+_147753471_147753542@3.29.27,trna33_HisGTG_6_+_27125906_27125977@3.29.27,trna7_HisGTG_9_-_14433938_14434009@3.29.27,trna8_HisGTG_15_-_45492611_45492682@3.29.27,trna9_HisGTG_15_-_45490804_45490875@3.29.27|CGTGATCGTATAGTGGTTAGTACTCTG</td><td>>am_trna111_HisGTG_1_-_147774845_147774916</td><td>3</td><td>29</td><td>27</td><td>CGTGATCGTATAGTGGTTAGTACTCTG</td><td>806.8</td><td>-6.9</td><td>1.2</td><td>-5.7</td><td>1.5E-08</td><td>3.2E-08</td><td>MB231/MB468</td><td>MP</td></tr>
<tr><td>>MI0003560|hsa-mir-92b&WithFlank&1|+|155195171|155195278@67.87.21(+1A)&[MIMAT0003218&hsa-miR-92b-3p&offsets|0|-1(+1A);m-115&1|+|155195237|155195258&offsets|0|-1(+1A)]|TATTGCACTCGTCCCGGCCTCA</td><td>>MI0003560|hsa-mir-92b&WithFlank&1|+|155195171|155195278</td><td>67</td><td>87</td><td>21(+1A)</td><td>TATTGCACTCGTCCCGGCCTCA</td><td>671.6</td><td>7.1</td><td>1.3</td><td>5.6</td><td>1.7E-08</td><td>3.2E-08</td><td>MB231/MB468</td><td>MITO</td></tr>
<tr><td>>hg38_21_+_8433172_8446622_RNA45SN1_45S_50ntFlanks@10689.10709.21|GCGTCCGGTGAGCTCTCGCTG</td><td>>hg38_21_+_8433172_8446622_RNA45SN1_45S_50ntFlanks</td><td>10689</td><td>10709</td><td>21</td><td>GCGTCCGGTGAGCTCTCGCTG</td><td>726.2</td><td>-5.5</td><td>1.0</td><td>-5.7</td><td>1.3E-08</td><td>3.2E-08</td><td>BT20/MB468</td><td>CYTO</td></tr>
<tr><td>>hg38_21_+_8433172_8446622_RNA45SN1_45S_50ntFlanks@9504.9529.26|GTGAACTATGCCTGGGCAGGGCGAAG</td><td>>hg38_21_+_8433172_8446622_RNA45SN1_45S_50ntFlanks</td><td>9504</td><td>9529</td><td>26</td><td>GTGAACTATGCCTGGGCAGGGCGAAG</td><td>372.4</td><td>-7.9</td><td>1.4</td><td>-5.7</td><td>1.3E-08</td><td>3.2E-08</td><td>BT20/MB468</td><td>CYTO</td></tr>
<tr><td>>hg38_21_+_8433172_8446622_RNA45SN1_45S_50ntFlanks@7988.8010.23|CAGACGTGGCGACCCGCTGAATT</td><td>>hg38_21_+_8433172_8446622_RNA45SN1_45S_50ntFlanks</td><td>7988</td><td>8010</td><td>23</td><td>CAGACGTGGCGACCCGCTGAATT</td><td>1033.8</td><td>-6.4</td><td>1.1</td><td>-5.7</td><td>1.5E-08</td><td>3.2E-08</td><td>MB231/MB468</td><td>MP</td></tr>
<tr><td>>hg38_21_+_8433172_8446622_RNA45SN1_45S_50ntFlanks@11886.11910.25|TGAGCTTGACTCTAGTCTGGCACGG</td><td>>hg38_21_+_8433172_8446622_RNA45SN1_45S_50ntFlanks</td><td>11886</td><td>11910</td><td>25</td><td>TGAGCTTGACTCTAGTCTGGCACGG</td><td>57.4</td><td>-9.7</td><td>1.7</td><td>-5.7</td><td>1.5E-08</td><td>3.2E-08</td><td>MB231/MB468</td><td>MP</td></tr>
<tr><td>>hg38_21_+_8433172_8446622_RNA45SN1_45S_50ntFlanks@5472.5502.31|ACGGCCCTGGCGGAGCGCTGAGAAGACGGTC</td><td>>hg38_21_+_8433172_8446622_RNA45SN1_45S_50ntFlanks</td><td>5472</td><td>5502</td><td>31</td><td>ACGGCCCTGGCGGAGCGCTGAGAAGACGGTC</td><td>238.0</td><td>3.3</td><td>0.6</td><td>5.6</td><td>1.6E-08</td><td>3.2E-08</td><td>MB231/MB468</td><td>NUC</td></tr>
<tr><td>>am_trna119_LysCTT_1_-_145395522_145395594@7.35.29,trna11_LysCTT_5_-_180648979_180649051@7.35.29,trna128_LysCTT_1_-_55423542_55423614@7.35.29,trna13_LysCTT_6_+_26556774_26556846@7.35.29,trna6_LysCTT_19_-_52425396_52425468@7.35.29,trna7_LysCTT_16_+_3225692_3225764@7.35.29,trna9_LysCTT_5_+_180634755_180634827@7.35.29|CTAGCTCAGTCGGTAGAGCATGAGACTCT</td><td>>am_trna119_LysCTT_1_-_145395522_145395594</td><td>7</td><td>35</td><td>29</td><td>CTAGCTCAGTCGGTAGAGCATGAGACTCT</td><td>876.9</td><td>7.2</td><td>1.3</td><td>5.6</td><td>1.7E-08</td><td>3.2E-08</td><td>MB231/MB468</td><td>MITO</td></tr>
<tr><td>>hg38_21_+_8433172_8446622_RNA45SN1_45S_50ntFlanks@9554.9596.43|CGTAGCGGTCCTGACGTGCAAATCGGTCGTCCGACCTGGGTAT</td><td>>hg38_21_+_8433172_8446622_RNA45SN1_45S_50ntFlanks</td><td>9554</td><td>9596</td><td>43</td><td>CGTAGCGGTCCTGACGTGCAAATCGGTCGTCCGACCTGGGTAT</td><td>286.4</td><td>-5.7</td><td>1.0</td><td>-5.7</td><td>1.6E-08</td><td>3.2E-08</td><td>MB231/MB468</td><td>CYTO</td></tr>
<tr><td>>hg38_21_+_8433172_8446622_RNA45SN1_45S_50ntFlanks@12581.12604.24|AACCGCAGGTTCAGACATTTGGTG</td><td>>hg38_21_+_8433172_8446622_RNA45SN1_45S_50ntFlanks</td><td>12581</td><td>12604</td><td>24</td><td>AACCGCAGGTTCAGACATTTGGTG</td><td>142.7</td><td>-10.2</td><td>1.7</td><td>-6.0</td><td>2.3E-09</td><td>3.2E-08</td><td>BT20/MB468</td><td>MP</td></tr>
<tr><td>>MI0000061|hsa-let-7a-2&WithFlank&11|-|122146516|122146599@11.28.18;MI0000064|hsa-let-7c&WithFlank&21|+|16539822|16539917@17.34.18;MI0000062|hsa-let-7a-3&WithFlank&22|+|46112743|46112828@10.27.18;MI0000060|hsa-let-7a-1&WithFlank&9|+|94175951|94176042@12.29.18&[MIMAT0000062&hsa-let-7a-5p&offsets|0|-4;m-6&11|-|122146568|122146589&offsets|0|-4];[MIMAT0000064&hsa-let-7c-5p&offsets|0|-4;m-37&21|+|16539838|16539859&offsets|0|-4];[MIMAT0000062_1&hsa-let-7a-5p&offsets|0|-4;m-5&22|+|46112752|46112773&offsets|0|-4];[MIMAT0000062_2&hsa-let-7a-5p&offsets|0|-4;m-4&9|+|94175962|94175983&offsets|0|-4]|TGAGGTAGTAGGTTGTAT</td><td>>MI0000061|hsa-let-7a-2&WithFlank&11|-|122146516|122146599</td><td>11</td><td>28</td><td>18</td><td>TGAGGTAGTAGGTTGTAT</td><td>3305.1</td><td>5.0</td><td>0.9</td><td>5.6</td><td>1.6E-08</td><td>3.2E-08</td><td>MB231/MB468</td><td>NUC</td></tr>
<tr><td>>hg38_MT_+_1621_3279_RNR2_16S_50ntFlanks@1367.1404.38|TCGATGTTGGATCAGGACATCCCGATGGTGCAGCCGCT</td><td>>hg38_MT_+_1621_3279_RNR2_16S_50ntFlanks</td><td>1367</td><td>1404</td><td>38</td><td>TCGATGTTGGATCAGGACATCCCGATGGTGCAGCCGCT</td><td>69.0</td><td>9.3</td><td>1.6</td><td>5.7</td><td>1.5E-08</td><td>3.2E-08</td><td>MB231/MB468</td><td>MP</td></tr>
<tr><td>>MI0000101|hsa-mir-99a&WithFlank&21|+|16539083|16539175@19.38.20(+1C)&[MIMAT0000097&hsa-miR-99a-5p&offsets|0|-2(+1C);m-64&21|+|16539101|16539121&offsets|0|-1(+1C)]|AACCCGTAGATCCGATCTTGC</td><td>>MI0000101|hsa-mir-99a&WithFlank&21|+|16539083|16539175</td><td>19</td><td>38</td><td>20(+1C)</td><td>AACCCGTAGATCCGATCTTGC</td><td>242.6</td><td>-11.3</td><td>1.8</td><td>-6.2</td><td>4.3E-10</td><td>3.2E-08</td><td>BT20/MB231</td><td>MP</td></tr>
<tr><td>>hg38_21_+_8433172_8446622_RNA45SN1_45S_50ntFlanks@12711.12745.35|GCGGAGCCTCGGTTGGCCTCGGATAGCCGGTCCCC</td><td>>hg38_21_+_8433172_8446622_RNA45SN1_45S_50ntFlanks</td><td>12711</td><td>12745</td><td>35</td><td>GCGGAGCCTCGGTTGGCCTCGGATAGCCGGTCCCC</td><td>136.6</td><td>-10.2</td><td>1.7</td><td>-6.0</td><td>2.3E-09</td><td>3.2E-08</td><td>BT20/MB468</td><td>MP</td></tr>
<tr><td>>hg38_21_+_8433172_8446622_RNA45SN1_45S_50ntFlanks@11670.11697.28|CCCAGTGCTCTGAATGTCAAAGTGAAGA</td><td>>hg38_21_+_8433172_8446622_RNA45SN1_45S_50ntFlanks</td><td>11670</td><td>11697</td><td>28</td><td>CCCAGTGCTCTGAATGTCAAAGTGAAGA</td><td>318.8</td><td>-2.7</td><td>0.5</td><td>-5.6</td><td>1.6E-08</td><td>3.2E-08</td><td>MB231/MB468</td><td>NUC</td></tr>
<tr><td>>hg38_21_+_8433172_8446622_RNA45SN1_45S_50ntFlanks@11580.11600.21|GAACTGGTGCGGACCAGGGGA</td><td>>hg38_21_+_8433172_8446622_RNA45SN1_45S_50ntFlanks</td><td>11580</td><td>11600</td><td>21</td><td>GAACTGGTGCGGACCAGGGGA</td><td>620.9</td><td>-4.4</td><td>0.8</td><td>-5.7</td><td>1.4E-08</td><td>3.2E-08</td><td>BT20/MB468</td><td>CYTO</td></tr>
<tr><td>>MI0000459|hsa-mir-143&WithFlank&5|+|149428912|149429029@67.87.21(+1U)&[MIMAT0000435&hsa-miR-143-3p&offsets|0|0(+1U);m-3&5|+|149428978|149428997&offsets|0|+1(+1U)]|TGAGATGAAGCACTGTAGCTCT</td><td>>MI0000459|hsa-mir-143&WithFlank&5|+|149428912|149429029</td><td>67</td><td>87</td><td>21(+1U)</td><td>TGAGATGAAGCACTGTAGCTCT</td><td>109.7</td><td>-9.9</td><td>1.7</td><td>-6.0</td><td>2.3E-09</td><td>3.2E-08</td><td>BT20/MB468</td><td>MP</td></tr>
<tr><td>>MI0000748|hsa-mir-130b&WithFlank&22|+|21653298|21653391@19.40.22&[MIMAT0004680&hsa-miR-130b-5p&offsets|0|+1;m-234&22|+|21653316|21653337&offsets|0|0]|ACTCTTTCCCTGTTGCACTACT</td><td>>MI0000748|hsa-mir-130b&WithFlank&22|+|21653298|21653391</td><td>19</td><td>40</td><td>22</td><td>ACTCTTTCCCTGTTGCACTACT</td><td>1211.7</td><td>5.1</td><td>0.9</td><td>5.6</td><td>1.6E-08</td><td>3.2E-08</td><td>MB231/MB468</td><td>NUC</td></tr>
<tr><td>>hg38_21_+_8433172_8446622_RNA45SN1_45S_50ntFlanks@10819.10837.19|AGGGAAGTCGGCAAGCCGG</td><td>>hg38_21_+_8433172_8446622_RNA45SN1_45S_50ntFlanks</td><td>10819</td><td>10837</td><td>19</td><td>AGGGAAGTCGGCAAGCCGG</td><td>1338.6</td><td>-6.7</td><td>1.2</td><td>-5.7</td><td>1.5E-08</td><td>3.2E-08</td><td>MB231/MB468</td><td>MP</td></tr>
<tr><td>>hg38_21_+_8433172_8446622_RNA45SN1_45S_50ntFlanks@12348.12365.18|GAAAAGTTACCACAGGGA</td><td>>hg38_21_+_8433172_8446622_RNA45SN1_45S_50ntFlanks</td><td>12348</td><td>12365</td><td>18</td><td>GAAAAGTTACCACAGGGA</td><td>63.1</td><td>-9.8</td><td>1.7</td><td>-5.7</td><td>1.5E-08</td><td>3.2E-08</td><td>MB231/MB468</td><td>MP</td></tr>
<tr><td>>hg38_21_+_8433172_8446622_RNA45SN1_45S_50ntFlanks@10534.10566.33|CCGCGAGGCGTCCAGTGCGGTAACGCGACCGAT</td><td>>hg38_21_+_8433172_8446622_RNA45SN1_45S_50ntFlanks</td><td>10534</td><td>10566</td><td>33</td><td>CCGCGAGGCGTCCAGTGCGGTAACGCGACCGAT</td><td>107.7</td><td>-9.8</td><td>1.6</td><td>-6.0</td><td>2.4E-09</td><td>3.2E-08</td><td>BT20/MB468</td><td>MP</td></tr>
<tr><td>>hg38_21_+_8433172_8446622_RNA45SN1_45S_50ntFlanks@12456.12481.26|CCAAGCGTTGGATTGTTCACCCACTA</td><td>>hg38_21_+_8433172_8446622_RNA45SN1_45S_50ntFlanks</td><td>12456</td><td>12481</td><td>26</td><td>CCAAGCGTTGGATTGTTCACCCACTA</td><td>1044.7</td><td>-7.8</td><td>1.4</td><td>-5.7</td><td>1.4E-08</td><td>3.2E-08</td><td>BT20/MB468</td><td>CYTO</td></tr>
<tr><td>>hg38_1_-_228634819_228635039_RNA5S12_5S_50ntFlanks@102.123.22|CTAAGCAGGGTCGGGCCTGGTT</td><td>>hg38_1_-_228634819_228635039_RNA5S12_5S_50ntFlanks</td><td>102</td><td>123</td><td>22</td><td>CTAAGCAGGGTCGGGCCTGGTT</td><td>78.6</td><td>-10.2</td><td>1.8</td><td>-5.7</td><td>1.5E-08</td><td>3.2E-08</td><td>MB231/MB468</td><td>MP</td></tr>
<tr><td>>hg38_21_+_8433172_8446622_RNA45SN1_45S_50ntFlanks@6753.6771.19|AACGCACTTGCGGCCCCGG</td><td>>hg38_21_+_8433172_8446622_RNA45SN1_45S_50ntFlanks</td><td>6753</td><td>6771</td><td>19</td><td>AACGCACTTGCGGCCCCGG</td><td>125.5</td><td>-10.1</td><td>1.7</td><td>-6.0</td><td>2.4E-09</td><td>3.2E-08</td><td>BT20/MB468</td><td>MP</td></tr>
<tr><td>>hg38_21_+_8433172_8446622_RNA45SN1_45S_50ntFlanks@7977.8010.34|CGACCTCAGATCAGACGTGGCGACCCGCTGAATT</td><td>>hg38_21_+_8433172_8446622_RNA45SN1_45S_50ntFlanks</td><td>7977</td><td>8010</td><td>34</td><td>CGACCTCAGATCAGACGTGGCGACCCGCTGAATT</td><td>452.5</td><td>3.6</td><td>0.6</td><td>5.6</td><td>1.6E-08</td><td>3.2E-08</td><td>MB231/MB468</td><td>NUC</td></tr>
<tr><td>>hg38_21_+_8433172_8446622_RNA45SN1_45S_50ntFlanks@11575.11592.18|ACTTAGAACTGGTGCGGA</td><td>>hg38_21_+_8433172_8446622_RNA45SN1_45S_50ntFlanks</td><td>11575</td><td>11592</td><td>18</td><td>ACTTAGAACTGGTGCGGA</td><td>818.2</td><td>-3.5</td><td>0.6</td><td>-5.7</td><td>1.2E-08</td><td>3.2E-08</td><td>BT20/MB468</td><td>MITO</td></tr>
<tr><td>>hg38_21_+_8433172_8446622_RNA45SN1_45S_50ntFlanks@11235.11255.21|CTTCCCGTGGATCGCCCCAGC</td><td>>hg38_21_+_8433172_8446622_RNA45SN1_45S_50ntFlanks</td><td>11235</td><td>11255</td><td>21</td><td>CTTCCCGTGGATCGCCCCAGC</td><td>438.8</td><td>-3.4</td><td>0.6</td><td>-5.6</td><td>1.6E-08</td><td>3.2E-08</td><td>MB231/MB468</td><td>NUC</td></tr>
<tr><td>>hg38_21_+_8433172_8446622_RNA45SN1_45S_50ntFlanks@6651.6677.27|CGACTCTTAGCGGTGGATCACTCGGCT</td><td>>hg38_21_+_8433172_8446622_RNA45SN1_45S_50ntFlanks</td><td>6651</td><td>6677</td><td>27</td><td>CGACTCTTAGCGGTGGATCACTCGGCT</td><td>5619.2</td><td>4.0</td><td>0.7</td><td>5.7</td><td>1.4E-08</td><td>3.2E-08</td><td>BT20/MB468</td><td>CYTO</td></tr>
<tr><td>>hg38_21_+_8433172_8446622_RNA45SN1_45S_50ntFlanks@12419.12436.18|GATGTCGGCTCTTCCTAT</td><td>>hg38_21_+_8433172_8446622_RNA45SN1_45S_50ntFlanks</td><td>12419</td><td>12436</td><td>18</td><td>GATGTCGGCTCTTCCTAT</td><td>1584.5</td><td>-3.9</td><td>0.7</td><td>-5.6</td><td>1.8E-08</td><td>3.2E-08</td><td>MB231/MB468</td><td>MITO</td></tr>
<tr><td>>trna119_LysCTT_1_-_145395522_145395594@1.36.36,trna11_LysCTT_5_-_180648979_180649051@1.36.36,trna13_LysCTT_6_+_26556774_26556846@1.36.36,trna7_LysCTT_16_+_3225692_3225764@1.36.36,trna9_LysCTT_5_+_180634755_180634827@1.36.36|GCCCGGCTAGCTCAGTCGGTAGAGCATGAGACTCTT</td><td>>trna119_LysCTT_1_-_145395522_145395594</td><td>1</td><td>36</td><td>36</td><td>GCCCGGCTAGCTCAGTCGGTAGAGCATGAGACTCTT</td><td>615.5</td><td>5.6</td><td>1.0</td><td>5.6</td><td>1.7E-08</td><td>3.2E-08</td><td>MB231/MB468</td><td>NUC</td></tr>
<tr><td>>hg38_21_+_8433172_8446622_RNA45SN1_45S_50ntFlanks@12960.12979.20|CTGGGTCGGGGTTTCGTACG</td><td>>hg38_21_+_8433172_8446622_RNA45SN1_45S_50ntFlanks</td><td>12960</td><td>12979</td><td>20</td><td>CTGGGTCGGGGTTTCGTACG</td><td>277.7</td><td>-6.0</td><td>1.1</td><td>-5.6</td><td>1.7E-08</td><td>3.2E-08</td><td>MB231/MB468</td><td>NUC</td></tr>
<tr><td>>hg38_21_+_8433172_8446622_RNA45SN1_45S_50ntFlanks@2959.2979.21|CGACGTGTGGCGTGGGTCGAC</td><td>>hg38_21_+_8433172_8446622_RNA45SN1_45S_50ntFlanks</td><td>2959</td><td>2979</td><td>21</td><td>CGACGTGTGGCGTGGGTCGAC</td><td>401.3</td><td>-7.3</td><td>1.3</td><td>-5.6</td><td>1.7E-08</td><td>3.2E-08</td><td>MB231/MB468</td><td>NUC</td></tr>
<tr><td>>trnaMT_PheGAA_MT_+_577_647@36.60.25|AAAATGTTTAGACGGGCTCACATCA</td><td>>trnaMT_PheGAA_MT_+_577_647</td><td>36</td><td>60</td><td>25</td><td>AAAATGTTTAGACGGGCTCACATCA</td><td>160.2</td><td>11.1</td><td>1.9</td><td>6.0</td><td>2.4E-09</td><td>3.3E-08</td><td>BT20/MB468</td><td>MP</td></tr>
<tr><td>>hg38_21_+_8433172_8446622_RNA45SN1_45S_50ntFlanks@10247.10284.38|GCGACGAGTAGGAGGGCCGCTGCGGTGAGCCTTGAAGC</td><td>>hg38_21_+_8433172_8446622_RNA45SN1_45S_50ntFlanks</td><td>10247</td><td>10284</td><td>38</td><td>GCGACGAGTAGGAGGGCCGCTGCGGTGAGCCTTGAAGC</td><td>88.6</td><td>10.2</td><td>1.7</td><td>6.0</td><td>2.4E-09</td><td>3.3E-08</td><td>BT20/MB468</td><td>MP</td></tr>
<tr><td>>hg38_21_+_8433172_8446622_RNA45SN1_45S_50ntFlanks@10818.10840.23|AAGGGAAGTCGGCAAGCCGGATC</td><td>>hg38_21_+_8433172_8446622_RNA45SN1_45S_50ntFlanks</td><td>10818</td><td>10840</td><td>23</td><td>AAGGGAAGTCGGCAAGCCGGATC</td><td>367.6</td><td>-6.0</td><td>1.1</td><td>-5.7</td><td>1.5E-08</td><td>3.3E-08</td><td>MB231/MB468</td><td>MP</td></tr>
</table>

| | | | | | | | | | | | | | |
|---|---|---|---|---|---|---|---|---|---|---|---|---|---|
| >hg38_21_+_8433172_8446622_RNA45SN1_45S_50ntFlanks@10513.10533.21\|CCGGAGTGGCGGAGATGGGCG | >hg38_21_+_8433172_8446622_RNA45SN1_45S_50ntFlanks | 10513 | 10533 | 21 | CCGGAGTGGCGGAGATGGGCG | 13326.9 | -6.3 | 1.1 | -6.0 | 2.4E-09 | 3.3E-08 | BT20/MB468 | MP |
| >hg38_21_+_8433172_8446622_RNA45SN1_45S_50ntFlanks@8171.8212.42\|AAGTCCTTCTGATCGAGGCCCAGCCCGTGGACGGTGTGAGGC | >hg38_21_+_8433172_8446622_RNA45SN1_45S_50ntFlanks | 8171 | 8212 | 42 | AAGTCCTTCTGATCGAGGCCCAGCCCGTGGACGGTGTGAGGC | 780.4 | -8.0 | 1.4 | -5.7 | 1.5E-08 | 3.3E-08 | MB231/MB468 | MP |
| >hg38_21_+_8433172_8446622_RNA45SN1_45S_50ntFlanks@8358.8379.22\|AAAGAACTTTGAAGAGAGAGTT | >hg38_21_+_8433172_8446622_RNA45SN1_45S_50ntFlanks | 8358 | 8379 | 22 | AAAGAACTTTGAAGAGAGAGTT | 876.1 | -4.6 | 0.8 | -5.6 | 1.7E-08 | 3.3E-08 | MB231/MB468 | NUC |
| >hg38_21_+_8433172_8446622_RNA45SN1_45S_50ntFlanks@6730.6756.27\|ACACATTGATCATCGACACTTCGAACG | >hg38_21_+_8433172_8446622_RNA45SN1_45S_50ntFlanks | 6730 | 6756 | 27 | ACACATTGATCATCGACACTTCGAACG | 175.7 | -2.2 | 0.4 | -5.7 | 1.4E-08 | 3.3E-08 | BT20/MB468 | CYTO |
| >hg38_21_+_8433172_8446622_RNA45SN1_45S_50ntFlanks@5186.5205.20\|CAATAACAGGTCTGTGATGC | >hg38_21_+_8433172_8446622_RNA45SN1_45S_50ntFlanks | 5186 | 5205 | 20 | CAATAACAGGTCTGTGATGC | 761.4 | -7.1 | 1.3 | -5.6 | 1.8E-08 | 3.3E-08 | MB231/MB468 | MITO |
| >MI0033673\|hsa-mir-10526&WithFlank&11\|-\|122152097\|122152162@41.61.21&[MIMAT0041996&hsa-miR-10526-3p&offsets\|0\|+1]\|AAAAGGGGGCTGAGGTGGAGG | >MI0033673\|hsa-mir-10526&WithFlank&11\|-\|122152097\|122152162 | 41 | 61 | 21 | AAAAGGGGGCTGAGGTGGAGG | 72.1 | 9.4 | 1.7 | 5.7 | 1.5E-08 | 3.3E-08 | MB231/MB468 | MP |
| >hg38_21_+_8433172_8446622_RNA45SN1_45S_50ntFlanks@8176.8203.28\|CTTCTGATCGAGGCCCAGCCCGTGGACG | >hg38_21_+_8433172_8446622_RNA45SN1_45S_50ntFlanks | 8176 | 8203 | 28 | CTTCTGATCGAGGCCCAGCCCGTGGACG | 4139.3 | -5.1 | 0.9 | -5.6 | 1.8E-08 | 3.3E-08 | MB231/MB468 | MITO |
| >hg38_21_+_8433172_8446622_RNA45SN1_45S_50ntFlanks@11323.11347.25\|CGTCTCGTCGCGCGCGCGTCCGCTG | >hg38_21_+_8433172_8446622_RNA45SN1_45S_50ntFlanks | 11323 | 11347 | 25 | CGTCTCGTCGCGCGCGCGTCCGCTG | 3965.2 | 7.0 | 1.2 | 6.1 | 1.3E-09 | 3.3E-08 | BT20/MB231 | MITO |
| >MI0000084\|hsa-mir-26b&WithFlank&2\|+\|218402640\|218402728@18.38.21(+1A)&[MIMAT0000083&hsa-miR-26b-5p&offsets\|0\|0(+1A);m-53&2\|+\|218402657\|218402678&offsets\|0\|-1(+1A)]\|TTCAAGTAATTCAGGATAGGTA | >MI0000084\|hsa-mir-26b&WithFlank&2\|+\|218402640\|218402728 | 18 | 38 | 21(+1A) | TTCAAGTAATTCAGGATAGGTA | 796.6 | 7.7 | 1.4 | 5.7 | 1.4E-08 | 3.3E-08 | BT20/MB468 | CYTO |
| >hg38_21_+_8433172_8446622_RNA45SN1_45S_50ntFlanks@12940.12972.33\|CATTCGTAGACGACCTGCTTCTGGGTCGGGGTT | >hg38_21_+_8433172_8446622_RNA45SN1_45S_50ntFlanks | 12940 | 12972 | 33 | CATTCGTAGACGACCTGCTTCTGGGTCGGGGTT | 575.6 | -7.4 | 1.3 | -5.7 | 1.4E-08 | 3.3E-08 | BT20/MB468 | CYTO |
| >hg38_21_+_8433172_8446622_RNA45SN1_45S_50ntFlanks@4370.4391.22\|TTAAAAAGCTCGTAGTTGGATC | >hg38_21_+_8433172_8446622_RNA45SN1_45S_50ntFlanks | 4370 | 4391 | 22 | TTAAAAAGCTCGTAGTTGGATC | 352.5 | -7.7 | 1.4 | -5.7 | 1.5E-08 | 3.3E-08 | MB231/MB468 | MP |
| >MI0000736\|hsa-mir-30c-1&WithFlank&1\|+\|40757278\|40757378@23.41.19(+1U);MI0000254\|hsa-mir-30c-2&WithFlank&6\|-\|71376954\|71377037@13.31.19(+1U)&[MIMAT0000244&hsa-miR-30c-5p&offsets\|0\|-4(+1U);m-57&1\|+\|40757300\|40757323&offsets\|0\|-5(+1U)];[MIMAT0000244_1&hsa-miR-30c-5p&offsets\|0\|-4(+1U);m-56&6\|-\|71377002\|71377025&offsets\|0\|-5(+1U)]\|TGTAAACATCCTACACTCTT | >MI0000736\|hsa-mir-30c-1&WithFlank&1\|+\|40757278\|40757378 | 23 | 41 | 19(+1U) | TGTAAACATCCTACACTCTT | 132.3 | 10.3 | 1.8 | 5.7 | 1.5E-08 | 3.3E-08 | MB231/MB468 | MP |
| >hg38_21_+_8433172_8446622_RNA45SN1_45S_50ntFlanks@11581.11607.27\|AACTGGTGCGGACCAGGGGAATCCGAC | >hg38_21_+_8433172_8446622_RNA45SN1_45S_50ntFlanks | 11581 | 11607 | 27 | AACTGGTGCGGACCAGGGGAATCCGAC | 353.9 | -3.9 | 0.7 | -5.6 | 1.9E-08 | 3.3E-08 | BT20/MB468 | NUC |
| >hg38_21_+_8433172_8446622_RNA45SN1_45S_50ntFlanks@4099.4121.23\|GTGACCACGGGTGACGGGGAATC | >hg38_21_+_8433172_8446622_RNA45SN1_45S_50ntFlanks | 4099 | 4121 | 23 | GTGACCACGGGTGACGGGGAATC | 126.2 | -9.1 | 1.6 | -5.7 | 1.3E-08 | 3.3E-08 | BT20/MB468 | MITO |
| >am_trnaMT_TyrGTA_MT_-_5826_5891@13.36.24\|GAGTGAAGCATTGGACTGTAAATC | >am_trnaMT_TyrGTA_MT_-_5826_5891 | 13 | 36 | 24 | GAGTGAAGCATTGGACTGTAAATC | 174.6 | 11.2 | 1.9 | 6.0 | 2.4E-09 | 3.3E-08 | BT20/MB468 | MP |
| >hg38_MT_+_598_1651_RNR1_12S_50ntFlanks@905.946.42\|AGTATACTTCAAAGGACATTTAACTAAAACCCCTACGCATTT | >hg38_MT_+_598_1651_RNR1_12S_50ntFlanks | 905 | 946 | 42 | AGTATACTTCAAAGGACATTTAACTAAAACCCCTACGCATTT | 355.0 | 12.2 | 2.0 | 6.0 | 2.4E-09 | 3.3E-08 | BT20/MB468 | MP |
| >hg38_21_+_8433172_8446622_RNA45SN1_45S_50ntFlanks@9204.9224.21\|TCGGGGCCACGCGCGCGTCCC | >hg38_21_+_8433172_8446622_RNA45SN1_45S_50ntFlanks | 9204 | 9224 | 21 | TCGGGGCCACGCGCGCGTCCC | 165.8 | -10.5 | 1.8 | -6.0 | 2.4E-09 | 3.3E-08 | BT20/MB468 | MP |
| >MI0000063\|hsa-let-7b&WithFlank&22\|+\|46113680\|46113774@12.33.22&[MIMAT0000063&hsa-let-7b-5p&offsets\|0\|0;m-14&22\|+\|46113691\|46113712&offsets\|0\|0]\|TGAGGTAGTAGGTTGTGTGGTT | >MI0000063\|hsa-let-7b&WithFlank&22\|+\|46113680\|46113774 | 12 | 33 | 22 | TGAGGTAGTAGGTTGTGTGGTT | 7144.2 | 4.7 | 0.8 | 5.6 | 1.8E-08 | 3.3E-08 | MB231/MB468 | MITO |
| >MI0000272\|hsa-mir-182&WithFlank&7\|-\|129770377\|129770498@29.53.25(+1A)&[MIMAT0000259&hsa-miR-182-5p&offsets\|0\|+1(+1A);m-22&7\|-\|129770449\|129770470&offsets\|0\|+3(+1A)]\|TTTGGCAATGGTAGAACTCACACTGA | >MI0000272\|hsa-mir-182&WithFlank&7\|-\|129770377\|129770498 | 29 | 53 | 25(+1A) | TTTGGCAATGGTAGAACTCACACTGA | 953.7 | 3.3 | 0.5 | 6.0 | 1.7E-09 | 3.3E-08 | BT20/MB231 | NUC |
| >hg38_21_+_8433172_8446622_RNA45SN1_45S_50ntFlanks@12676.12711.36\|AGAATCCCGCCCAGGCGGAACGATACGGCAGCGCCG | >hg38_21_+_8433172_8446622_RNA45SN1_45S_50ntFlanks | 12676 | 12711 | 36 | AGAATCCCGCCCAGGCGGAACGATACGGCAGCGCCG | 114.7 | -9.9 | 1.7 | -6.0 | 2.4E-09 | 3.3E-08 | BT20/MB468 | MP |
| >hg38_21_+_8433172_8446622_RNA45SN1_45S_50ntFlanks@11768.11803.36\|CTAATTAGTGACGCGCATGAATGGATGAACGAGATT | >hg38_21_+_8433172_8446622_RNA45SN1_45S_50ntFlanks | 11768 | 11803 | 36 | CTAATTAGTGACGCGCATGAATGGATGAACGAGATT | 1057.7 | -7.9 | 1.4 | -5.6 | 1.8E-08 | 3.3E-08 | MB231/MB468 | MITO |
| >am_trna106_HisGTG_1_-_149155828_149155899@11.29.19,trna111_HisGTG_1_-_147774845_147774916@11.29.19,trna118_HisGTG_1_-_145396881_145396952@11.29.19,trna16_HisGTG_1_+_146544773_146544844@11.29.19,trna1_HisGTG_15_+_45493349_45493420@11.29.19,trna21_HisGTG_1_+_147753471_147753542@11.29.19,trna33_HisGTG_6_+_27125906_27125977@11.29.19,trna7_HisGTG_9_-_14433938_14434009@11.29.19,trna8_HisGTG_15_-_45492611_45492682@11.29.19,trna9_HisGTG_15_-_45490804_45490875@11.29.19\|TATAGTGGTTAGTACTCTG | >am_trna106_HisGTG_1_-_149155828_149155899 | 11 | 29 | 19 | TATAGTGGTTAGTACTCTG | 103.2 | -9.8 | 1.6 | -6.0 | 2.4E-09 | 3.3E-08 | BT20/MB468 | MP |
| >MI0000077\|hsa-mir-21&WithFlank&17\|+\|59841260\|59841343@14.35.22(+2U)&[MIMAT0000076&hsa-miR-21-5p&offsets\|0\|0(+2U);m-1&17\|+\|59841273\|59841294&offsets\|0\|0(+2U)]\|TAGCTTATCAGACTGATGTTGATT | >MI0000077\|hsa-mir-21&WithFlank&17\|+\|59841260\|59841343 | 14 | 35 | 22(+2U) | TAGCTTATCAGACTGATGTTGATT | 131.1 | 4.1 | 0.7 | 5.6 | 1.9E-08 | 3.3E-08 | BT20/MB468 | NUC |
| >hg38_21_+_8433172_8446622_RNA45SN1_45S_50ntFlanks@10827.10864.38\|CGGCAAGCCGGATCCGTAACTTCGGGATAAGGATTGGC | >hg38_21_+_8433172_8446622_RNA45SN1_45S_50ntFlanks | 10827 | 10864 | 38 | CGGCAAGCCGGATCCGTAACTTCGGGATAAGGATTGGC | 122.8 | -7.0 | 1.2 | -5.7 | 1.4E-08 | 3.3E-08 | BT20/MB468 | CYTO |
| >MI0000061\|hsa-let-7a-2&WithFlank&11\|-\|122146516\|122146599@11.31.21(+1C);MI0000062\|hsa-let-7a-3&WithFlank&22\|+\|46112743\|46112828@10.30.21(+1C);MI0000060\|hsa-let-7a-1&WithFlank&9\|+\|94175951\|94176042@12.32.21(+1C)&[MIMAT0000062&hsa-let-7a-5p&offsets\|0\|-1(+1C);m-6&11\|-\|122146568\|122146589&offsets\|0\|-1(+1C)];[MIMAT0000062_1&hsa-let-7a-5p&offsets\|0\|-1(+1C);m-5&22\|+\|46112752\|46112773&offsets\|0\|-1(+1C)];[MIMAT0000062_2&hsa-let-7a-5p&offsets\|0\|-1(+1C);m-4&9\|+\|94175962\|94175983&offsets\|0\|-1(+1C)]\|TGAGGTAGTAGGTTGTATAGTC | >MI0000061\|hsa-let-7a-2&WithFlank&11\|-\|122146516\|122146599 | 11 | 31 | 21(+1C) | TGAGGTAGTAGGTTGTATAGTC | 4167.9 | 2.4 | 0.4 | 5.6 | 1.9E-08 | 3.3E-08 | BT20/MB468 | NUC |
| >MI0000062\|hsa-let-7a-3&WithFlank&22\|+\|46112743\|46112828@58.78.21(+1U);MI0000060\|hsa-let-7a-1&WithFlank&9\|+\|94175951\|94176042@63.83.21(+1U)&[MIMAT0004481&hsa-let-7a-3p&offsets\|0\|0(+1U);m-180&22\|+\|46112800\|46112821&offsets\|0\|-1(+1U)];[MIMAT0004481_1&hsa-let-7a-3p&offsets\|0\|0(+1U);m-181&9\|+\|94176013\|94176034&offsets\|0\|-1(+1U)]\|CTATACAATCTACTGTCTTTCT | >MI0000062\|hsa-let-7a-3&WithFlank&22\|+\|46112743\|46112828 | 58 | 78 | 21(+1U) | CTATACAATCTACTGTCTTTCT | 2071.2 | 5.5 | 1.0 | 5.6 | 1.8E-08 | 3.3E-08 | MB231/MB468 | MITO |
| >trnaMT_GlyTCC_MT_+_9991_10058@28.66.39\|ACTTCCAATTAACTAGTTTTGACAACATTCAAAAAAGAG | >trnaMT_GlyTCC_MT_+_9991_10058 | 28 | 66 | 39 | ACTTCCAATTAACTAGTTTTGACAACATTCAAAAAAGAG | 209.1 | -10.7 | 1.9 | -5.6 | 1.7E-08 | 3.3E-08 | MB231/MB468 | NUC |
| >trnaMT_HisGTG_MT_+_12138_12206@19.58.40\|AAACATCAGATTGTGAATCTGACAACAGAGGCTTACGACC | >trnaMT_HisGTG_MT_+_12138_12206 | 19 | 58 | 40 | AAACATCAGATTGTGAATCTGACAACAGAGGCTTACGACC | 261.4 | 11.8 | 2.0 | 6.0 | 2.5E-09 | 3.3E-08 | BT20/MB468 | MP |
| >hg38_21_+_8433172_8446622_RNA45SN1_45S_50ntFlanks@11209.11233.25\|CGGCGACTCTGGACGCGAGCCGGGC | >hg38_21_+_8433172_8446622_RNA45SN1_45S_50ntFlanks | 11209 | 11233 | 25 | CGGCGACTCTGGACGCGAGCCGGGC | 2530.8 | -3.8 | 0.7 | -5.7 | 1.4E-08 | 3.3E-08 | BT20/MB468 | CYTO |
| >hg38_21_+_8433172_8446622_RNA45SN1_45S_50ntFlanks@11593.11611.19\|CCAGGGGAATCCGACTGTT | >hg38_21_+_8433172_8446622_RNA45SN1_45S_50ntFlanks | 11593 | 11611 | 19 | CCAGGGGAATCCGACTGTT | 2940.7 | -4.2 | 0.7 | -5.7 | 1.4E-08 | 3.3E-08 | BT20/MB468 | CYTO |
| >hg38_21_+_8433172_8446622_RNA45SN1_45S_50ntFlanks@9534.9552.19\|AGGAAACTCTGGTGGAGGT | >hg38_21_+_8433172_8446622_RNA45SN1_45S_50ntFlanks | 9534 | 9552 | 19 | AGGAAACTCTGGTGGAGGT | 724.4 | -7.6 | 1.3 | -5.6 | 1.8E-08 | 3.3E-08 | MB231/MB468 | MITO |
| >hg38_21_+_8433172_8446622_RNA45SN1_45S_50ntFlanks@9906.9928.23\|AGACCCCAGAAAAGGTGTTGGTT | >hg38_21_+_8433172_8446622_RNA45SN1_45S_50ntFlanks | 9906 | 9928 | 23 | AGACCCCAGAAAAGGTGTTGGTT | 747.4 | 8.6 | 1.5 | 5.7 | 1.3E-08 | 3.3E-08 | BT20/MB468 | MITO |
| >hg38_21_+_8433172_8446622_RNA45SN1_45S_50ntFlanks@6656.6689.34\|CTTAGCGGTGGATCACTCGGCTCGTGCGTCGATG | >hg38_21_+_8433172_8446622_RNA45SN1_45S_50ntFlanks | 6656 | 6689 | 34 | CTTAGCGGTGGATCACTCGGCTCGTGCGTCGATG | 660.0 | -3.5 | 0.6 | -5.6 | 1.7E-08 | 3.3E-08 | MB231/MB468 | NUC |
| >hg38_21_+_8433172_8446622_RNA45SN1_45S_50ntFlanks@9495.9531.37\|CGAAAGATGGTGAACTATGCCTGGGCAGGGCGAAGCC | >hg38_21_+_8433172_8446622_RNA45SN1_45S_50ntFlanks | 9495 | 9531 | 37 | CGAAAGATGGTGAACTATGCCTGGGCAGGGCGAAGCC | 126.6 | 10.7 | 1.8 | 6.0 | 2.5E-09 | 3.3E-08 | BT20/MB468 | MP |
| >hg38_21_+_8433172_8446622_RNA45SN1_45S_50ntFlanks@12349.12376.28\|AAAAGTTACCACAGGGATAACTGGCTTG | >hg38_21_+_8433172_8446622_RNA45SN1_45S_50ntFlanks | 12349 | 12376 | 28 | AAAAGTTACCACAGGGATAACTGGCTTG | 135.8 | -10.2 | 1.7 | -6.0 | 2.5E-09 | 3.4E-08 | BT20/MB468 | MP |
| >MI0000255\|hsa-mir-30d&WithFlank&8\|-\|134804870\|134804951@12.33.22&[MIMAT0000245&hsa-miR-30d-5p&offsets\|0\|0;m-21&8\|-\|134804919\|134804940&offsets\|0\|0]\|TGTAAACATCCCCGACTGGAAG | >MI0000255\|hsa-mir-30d&WithFlank&8\|-\|134804870\|134804951 | 12 | 33 | 22 | TGTAAACATCCCCGACTGGAAG | 875.9 | 5.6 | 1.0 | 5.6 | 1.7E-08 | 3.4E-08 | MB231/MB468 | CYTO |
| >hg38_21_+_8433172_8446622_RNA45SN1_45S_50ntFlanks@12998.13039.42\|GCTGCGATCTATTGAAAGTCAGCCCTCGACACAAGGGTTTGT | >hg38_21_+_8433172_8446622_RNA45SN1_45S_50ntFlanks | 12998 | 13039 | 42 | GCTGCGATCTATTGAAAGTCAGCCCTCGACACAAGGGTTTGT | 7235.2 | 7.0 | 1.2 | 5.7 | 1.4E-08 | 3.4E-08 | BT20/MB468 | CYTO |
| >am_trnaMT_GluTTC_MT_-_14674_14742@27.52.26,trnalookalike8_GluTTC_5_-_93905172_93905240@27.52.26\|GTTTTTCATATCATTGGTCGTGGTTG | >am_trnaMT_GluTTC_MT_-_14674_14742 | 27 | 52 | 26 | GTTTTTCATATCATTGGTCGTGGTTG | 250.3 | -7.6 | 1.3 | -5.7 | 1.4E-08 | 3.4E-08 | BT20/MB468 | CYTO |
| >hg38_21_+_8433172_8446622_RNA45SN1_45S_50ntFlanks@7975.7998.24\|CGCGACCTCAGATCAGACGTGGCG | >hg38_21_+_8433172_8446622_RNA45SN1_45S_50ntFlanks | 7975 | 7998 | 24 | CGCGACCTCAGATCAGACGTGGCG | 1490.0 | 4.7 | 0.8 | 6.0 | 1.8E-09 | 3.4E-08 | BT20/MB231 | NUC |
| >hg38_21_+_8433172_8446622_RNA45SN1_45S_50ntFlanks@4735.4761.27\|ACGAAAGTCGGAGGTTCGAAGACGATC | >hg38_21_+_8433172_8446622_RNA45SN1_45S_50ntFlanks | 4735 | 4761 | 27 | ACGAAAGTCGGAGGTTCGAAGACGATC | 470.9 | 4.2 | 0.7 | 5.6 | 1.7E-08 | 3.4E-08 | MB231/MB468 | NUC |
| >hg38_21_+_8433172_8446622_RNA45SN1_45S_50ntFlanks@12479.12498.20\|CTAATAGGGAACGTGAGCTG | >hg38_21_+_8433172_8446622_RNA45SN1_45S_50ntFlanks | 12479 | 12498 | 20 | CTAATAGGGAACGTGAGCTG | 1121.9 | -6.3 | 1.1 | -5.6 | 1.7E-08 | 3.4E-08 | MB231/MB468 | NUC |
| >hg38_21_+_8433172_8446622_RNA45SN1_45S_50ntFlanks@9712.9737.26\|AGCGAATGATTAGAGGTCTTGGGGCC | >hg38_21_+_8433172_8446622_RNA45SN1_45S_50ntFlanks | 9712 | 9737 | 26 | AGCGAATGATTAGAGGTCTTGGGGCC | 389.6 | -8.1 | 1.4 | -5.7 | 1.6E-08 | 3.4E-08 | MB231/MB468 | MP |
| >hg38_21_+_8433172_8446622_RNA45SN1_45S_50ntFlanks@10116.10139.24\|TGCGTCGGAGGGCGGCGGCGGCGG | >hg38_21_+_8433172_8446622_RNA45SN1_45S_50ntFlanks | 10116 | 10139 | 24 | TGCGTCGGAGGGCGGCGGCGGCGG | 586.3 | -8.7 | 1.5 | -5.6 | 1.9E-08 | 3.4E-08 | BT20/MB468 | NUC |

<table>
<tr><td>>hg38_21_+_8433172_8446622_RNA45SN1_45S_50ntFlanks@11668.11688.21|TGCCCAGTGCTCTGAATGTCA</td><td>>hg38_21_+_8433172_8446622_RNA45SN1_45S_50ntFlanks</td><td>11668</td><td>11688</td><td>21</td><td>TGCCCAGTGCTCTGAATGTCA</td><td>383.7</td><td>4.3</td><td>0.8</td><td>5.6</td><td>1.7E-08</td><td>3.4E-08</td><td>MB231/MB468</td><td>NUC</td></tr>
<tr><td>>hg38_21_+_8433172_8446622_RNA45SN1_45S_50ntFlanks@4657.4689.33|CTAGAGGTGAAATTCTTGGACCGGCGCAAGACG</td><td>>hg38_21_+_8433172_8446622_RNA45SN1_45S_50ntFlanks</td><td>4657</td><td>4689</td><td>33</td><td>CTAGAGGTGAAATTCTTGGACCGGCGCAAGACG</td><td>3926.4</td><td>-5.3</td><td>0.9</td><td>-5.7</td><td>1.4E-08</td><td>3.4E-08</td><td>BT20/MB468</td><td>CYTO</td></tr>
<tr><td>>hg38_21_+_8433172_8446622_RNA45SN1_45S_50ntFlanks@10348.10376.29|AAACGAGAACTTTGAAGGCCGAAGTGGAG</td><td>>hg38_21_+_8433172_8446622_RNA45SN1_45S_50ntFlanks</td><td>10348</td><td>10376</td><td>29</td><td>AAACGAGAACTTTGAAGGCCGAAGTGGAG</td><td>596.3</td><td>-6.7</td><td>1.2</td><td>-5.6</td><td>1.9E-08</td><td>3.4E-08</td><td>BT20/MB468</td><td>NUC</td></tr>
<tr><td>>trna10_LysCTT_16_+_3241501_3241573@2.36.35,trna13_LysCTT_14_-_58706613_58706685@2.36.35,trna2_LysCTT_15_+_79152904_79152976@2.36.35|CCCGGCTAGCTCAGTCGGTAGAGCATGGGACTCTT</td><td>>trna10_LysCTT_16_+_3241501_3241573</td><td>2</td><td>36</td><td>35</td><td>CCCGGCTAGCTCAGTCGGTAGAGCATGGGACTCTT</td><td>74.0</td><td>9.4</td><td>1.7</td><td>5.7</td><td>1.6E-08</td><td>3.4E-08</td><td>MB231/MB468</td><td>MP</td></tr>
<tr><td>>hg38_21_+_8433172_8446622_RNA45SN1_45S_50ntFlanks@8196.8216.21|CGTGGACGGTGTGAGGCCGGT</td><td>>hg38_21_+_8433172_8446622_RNA45SN1_45S_50ntFlanks</td><td>8196</td><td>8216</td><td>21</td><td>CGTGGACGGTGTGAGGCCGGT</td><td>996.7</td><td>-4.2</td><td>0.7</td><td>-5.7</td><td>1.6E-08</td><td>3.4E-08</td><td>MB231/MB468</td><td>MP</td></tr>
<tr><td>>hg38_21_+_8433172_8446622_RNA45SN1_45S_50ntFlanks@12679.12703.25|ATCCCGCCCAGGCGGAACGATACGG</td><td>>hg38_21_+_8433172_8446622_RNA45SN1_45S_50ntFlanks</td><td>12679</td><td>12703</td><td>25</td><td>ATCCCGCCCAGGCGGAACGATACGG</td><td>139.3</td><td>-10.2</td><td>1.7</td><td>-6.0</td><td>2.5E-09</td><td>3.4E-08</td><td>BT20/MB468</td><td>MP</td></tr>
<tr><td>>hg38_21_+_8433172_8446622_RNA45SN1_45S_50ntFlanks@10534.10571.38|CCGCGAGGCGTCCAGTGCGGTAACGCGACCGATCCCGG</td><td>>hg38_21_+_8433172_8446622_RNA45SN1_45S_50ntFlanks</td><td>10534</td><td>10571</td><td>38</td><td>CCGCGAGGCGTCCAGTGCGGTAACGCGACCGATCCCGG</td><td>115.1</td><td>-9.9</td><td>1.7</td><td>-6.0</td><td>2.5E-09</td><td>3.4E-08</td><td>BT20/MB468</td><td>MP</td></tr>
<tr><td>>trna10_LysCTT_16_+_3241501_3241573@5.35.31,trna13_LysCTT_14_-_58706613_58706685@5.35.31,trna2_LysCTT_15_+_79152904_79152976@5.35.31|GGCTAGCTCAGTCGGTAGAGCATGGGACTCT</td><td>>trna10_LysCTT_16_+_3241501_3241573</td><td>5</td><td>35</td><td>31</td><td>GGCTAGCTCAGTCGGTAGAGCATGGGACTCT</td><td>91.8</td><td>9.7</td><td>1.7</td><td>5.7</td><td>1.6E-08</td><td>3.4E-08</td><td>MB231/MB468</td><td>MP</td></tr>
<tr><td>>hg38_21_+_8433172_8446622_RNA45SN1_45S_50ntFlanks@8472.8491.20|CGGCCCGGCGGATCTTTCCC</td><td>>hg38_21_+_8433172_8446622_RNA45SN1_45S_50ntFlanks</td><td>8472</td><td>8491</td><td>20</td><td>CGGCCCGGCGGATCTTTCCC</td><td>323.2</td><td>-6.4</td><td>1.1</td><td>-5.6</td><td>1.7E-08</td><td>3.4E-08</td><td>MB231/MB468</td><td>CYTO</td></tr>
<tr><td>>hg38_21_+_8433172_8446622_RNA45SN1_45S_50ntFlanks@6732.6774.43|ACATTGATCATCGACACTTCGAACGCACTTGCGGCCCCGGGTT</td><td>>hg38_21_+_8433172_8446622_RNA45SN1_45S_50ntFlanks</td><td>6732</td><td>6774</td><td>43</td><td>ACATTGATCATCGACACTTCGAACGCACTTGCGGCCCCGGGTT</td><td>414.9</td><td>9.5</td><td>1.7</td><td>5.7</td><td>1.4E-08</td><td>3.4E-08</td><td>BT20/MB468</td><td>CYTO</td></tr>
<tr><td>>MI0000446|hsa-mir-125b-1&WithFlank&11|-|122099751|122099850@21.42.22(+2A);MI0000470|hsa-mir-125b-2&WithFlank&21|+|16590231|16590331@23.44.22(+2A)&[MIMAT0000423&hsa-miR-125b-5p&offsets|0|0(+2A);m-67&11|-|122099809|122099830&offsets|0|0(+2A)];[MIMAT0000423_1&hsa-miR-125b-5p&offsets|0|0(+2A);m-68&21|+|16590253|16590274&offsets|0|0(+2A)]|TCCCTGAGACCCTAACTTGTGAAA</td><td>>MI0000446|hsa-mir-125b-1&WithFlank&11|-|122099751|122099850</td><td>21</td><td>42</td><td>22(+2A)</td><td>TCCCTGAGACCCTAACTTGTGAAA</td><td>993.6</td><td>8.3</td><td>1.5</td><td>5.6</td><td>1.9E-08</td><td>3.4E-08</td><td>MB231/MB468</td><td>MITO</td></tr>
<tr><td>>hg38_1_-_228634819_228635039_RNA5S12_5S_50ntFlanks@128.171.44|CTTGGATGGGAGACCGCCTGGGAATACCGGGTGCTGTAGGCTTT</td><td>>hg38_1_-_228634819_228635039_RNA5S12_5S_50ntFlanks</td><td>128</td><td>171</td><td>44</td><td>CTTGGATGGGAGACCGCCTGGGAATACCGGGTGCTGTAGGCTTT</td><td>713.1</td><td>6.0</td><td>1.1</td><td>5.7</td><td>1.3E-08</td><td>3.4E-08</td><td>BT20/MB468</td><td>MITO</td></tr>
<tr><td>>MI0000087|hsa-mir-29a&WithFlank&7|-|130876741|130876816@48.66.19&[MIMAT0000086&hsa-miR-29a-3p&offsets|0|-3;m-26&7|-|130876749|130876769&offsets|0|-2]|TAGCACCATCTGAAATCGG</td><td>>MI0000087|hsa-mir-29a&WithFlank&7|-|130876741|130876816</td><td>48</td><td>66</td><td>19</td><td>TAGCACCATCTGAAATCGG</td><td>209.0</td><td>11.5</td><td>1.9</td><td>6.0</td><td>2.5E-09</td><td>3.4E-08</td><td>BT20/MB468</td><td>MP</td></tr>
<tr><td>>hg38_21_+_8433172_8446622_RNA45SN1_45S_50ntFlanks@6652.6695.44|GACTCTTAGCGGTGGATCACTCGGCTCGTGCGTCGATGAAGAAC</td><td>>hg38_21_+_8433172_8446622_RNA45SN1_45S_50ntFlanks</td><td>6652</td><td>6695</td><td>44</td><td>GACTCTTAGCGGTGGATCACTCGGCTCGTGCGTCGATGAAGAAC</td><td>831.0</td><td>7.2</td><td>1.3</td><td>5.7</td><td>1.5E-08</td><td>3.4E-08</td><td>BT20/MB468</td><td>CYTO</td></tr>
<tr><td>>hg38_21_+_8433172_8446622_RNA45SN1_45S_50ntFlanks@12479.12497.19|CTAATAGGGAACGTGAGCT</td><td>>hg38_21_+_8433172_8446622_RNA45SN1_45S_50ntFlanks</td><td>12479</td><td>12497</td><td>19</td><td>CTAATAGGGAACGTGAGCT</td><td>462.8</td><td>2.3</td><td>0.4</td><td>5.7</td><td>1.3E-08</td><td>3.4E-08</td><td>BT20/MB468</td><td>MITO</td></tr>
<tr><td>>hg38_21_+_8433172_8446622_RNA45SN1_45S_50ntFlanks@10248.10284.37|CGACGAGTAGGAGGGCCGCTGCGGTGAGCCTTGAAGC</td><td>>hg38_21_+_8433172_8446622_RNA45SN1_45S_50ntFlanks</td><td>10248</td><td>10284</td><td>37</td><td>CGACGAGTAGGAGGGCCGCTGCGGTGAGCCTTGAAGC</td><td>97.7</td><td>10.4</td><td>1.7</td><td>6.0</td><td>2.6E-09</td><td>3.4E-08</td><td>BT20/MB468</td><td>MP</td></tr>
<tr><td>>MI0000737|hsa-mir-200a&WithFlank&1|+|1167857|1167958@22.43.22(+1A)&[MIMAT0001620&hsa-miR-200a-5p&offsets|0|0(+1A);m-154&1|+|1167878|1167899&offsets|0|0(+1A)]|CATCTTACCGGACAGTGCTGGAA</td><td>>MI0000737|hsa-mir-200a&WithFlank&1|+|1167857|1167958</td><td>22</td><td>43</td><td>22(+1A)</td><td>CATCTTACCGGACAGTGCTGGAA</td><td>98.2</td><td>-9.7</td><td>1.6</td><td>-6.0</td><td>2.6E-09</td><td>3.4E-08</td><td>BT20/MB468</td><td>MP</td></tr>
<tr><td>>am_trnaMT_ProTGG_MT_-_15956_16023@52.71.20|AAAGACTTTTTCTCTGACCA</td><td>>am_trnaMT_ProTGG_MT_-_15956_16023</td><td>52</td><td>71</td><td>20</td><td>AAAGACTTTTTCTCTGACCA</td><td>1932.1</td><td>8.7</td><td>1.5</td><td>5.7</td><td>1.5E-08</td><td>3.4E-08</td><td>BT20/MB468</td><td>CYTO</td></tr>
<tr><td>>hg38_21_+_8433172_8446622_RNA45SN1_45S_50ntFlanks@8105.8139.35|CGCGGCGGGGCGCGGGACATGTGGCGTACGGAAGA</td><td>>hg38_21_+_8433172_8446622_RNA45SN1_45S_50ntFlanks</td><td>8105</td><td>8139</td><td>35</td><td>CGCGGCGGGGCGCGGGACATGTGGCGTACGGAAGA</td><td>1340.9</td><td>-7.1</td><td>1.3</td><td>-5.6</td><td>1.9E-08</td><td>3.4E-08</td><td>MB231/MB468</td><td>MITO</td></tr>
<tr><td>>hg38_21_+_8433172_8446622_RNA45SN1_45S_50ntFlanks@9575.9594.20|ATCGGTCGTCCGACCTGGGT</td><td>>hg38_21_+_8433172_8446622_RNA45SN1_45S_50ntFlanks</td><td>9575</td><td>9594</td><td>20</td><td>ATCGGTCGTCCGACCTGGGT</td><td>2282.5</td><td>-8.9</td><td>1.6</td><td>-5.6</td><td>1.6E-08</td><td>3.4E-08</td><td>MB231/MB468</td><td>MP</td></tr>
<tr><td>>hg38_21_+_8433172_8446622_RNA45SN1_45S_50ntFlanks@12994.13013.20|CCTCGCTGCGATCTATTGAA</td><td>>hg38_21_+_8433172_8446622_RNA45SN1_45S_50ntFlanks</td><td>12994</td><td>13013</td><td>20</td><td>CCTCGCTGCGATCTATTGAA</td><td>335.6</td><td>-7.6</td><td>1.3</td><td>-5.6</td><td>1.6E-08</td><td>3.4E-08</td><td>MB231/MB468</td><td>MP</td></tr>
<tr><td>>hg38_21_+_8433172_8446622_RNA45SN1_45S_50ntFlanks@9578.9604.27|GGTCGTCCGACCTGGGTATAGGGGCGA</td><td>>hg38_21_+_8433172_8446622_RNA45SN1_45S_50ntFlanks</td><td>9578</td><td>9604</td><td>27</td><td>GGTCGTCCGACCTGGGTATAGGGGCGA</td><td>91.7</td><td>10.3</td><td>1.7</td><td>6.0</td><td>2.6E-09</td><td>3.4E-08</td><td>BT20/MB468</td><td>MP</td></tr>
<tr><td>>MI0000746|hsa-mir-99b&WithFlank&19|+|51692606|51692687@51.71.21&[MIMAT0004678&hsa-miR-99b-3p&offsets|0|-1;m-151&19|+|51692656|51692678&offsets|0|-2]|CAAGCTCGTGTCTGTGGGTCC</td><td>>MI0000746|hsa-mir-99b&WithFlank&19|+|51692606|51692687</td><td>51</td><td>71</td><td>21</td><td>CAAGCTCGTGTCTGTGGGTCC</td><td>109.3</td><td>2.5</td><td>0.4</td><td>5.6</td><td>1.9E-08</td><td>3.4E-08</td><td>BT20/MB468</td><td>NUC</td></tr>
<tr><td>>hg38_21_+_8433172_8446622_RNA45SN1_45S_50ntFlanks@3912.3937.26|GATGCGTGCATTTATCAGATCAAAAC</td><td>>hg38_21_+_8433172_8446622_RNA45SN1_45S_50ntFlanks</td><td>3912</td><td>3937</td><td>26</td><td>GATGCGTGCATTTATCAGATCAAAAC</td><td>84.3</td><td>-10.3</td><td>1.8</td><td>-5.6</td><td>1.6E-08</td><td>3.4E-08</td><td>MB231/MB468</td><td>MP</td></tr>
<tr><td>>hg38_21_+_8433172_8446622_RNA45SN1_45S_50ntFlanks@8380.8404.25|CAAGAGGGCGTGAAACCGTTAAGAG</td><td>>hg38_21_+_8433172_8446622_RNA45SN1_45S_50ntFlanks</td><td>8380</td><td>8404</td><td>25</td><td>CAAGAGGGCGTGAAACCGTTAAGAG</td><td>316.9</td><td>-7.1</td><td>1.3</td><td>-5.6</td><td>1.8E-08</td><td>3.5E-08</td><td>MB231/MB468</td><td>NUC</td></tr>
<tr><td>>hg38_21_+_8433172_8446622_RNA45SN1_45S_50ntFlanks@10443.10460.18|CCGAAGGGACGGGCGATG</td><td>>hg38_21_+_8433172_8446622_RNA45SN1_45S_50ntFlanks</td><td>10443</td><td>10460</td><td>18</td><td>CCGAAGGGACGGGCGATG</td><td>279.1</td><td>-6.1</td><td>1.1</td><td>-5.6</td><td>1.6E-08</td><td>3.5E-08</td><td>MB231/MB468</td><td>MP</td></tr>
<tr><td>>MI0000077|hsa-mir-21&WithFlank&17|+|59841260|59841343@14.35.22(+2A)&[MIMAT0000076&hsa-miR-21-5p&offsets|0|0(+2A);m-1&17|+|59841273|59841294&offsets|0|0(+2A)]|TAGCTTATCAGACTGATGTTGAAA</td><td>>MI0000077|hsa-mir-21&WithFlank&17|+|59841260|59841343</td><td>14</td><td>35</td><td>22(+2A)</td><td>TAGCTTATCAGACTGATGTTGAAA</td><td>275.7</td><td>2.6</td><td>0.5</td><td>5.6</td><td>1.9E-08</td><td>3.5E-08</td><td>BT20/MB468</td><td>NUC</td></tr>
<tr><td>>hg38_21_+_8433172_8446622_RNA45SN1_45S_50ntFlanks@10388.10424.37|GTGAACAGCAGTTGAACATGGGTCAGTCGGTCCTGAG</td><td>>hg38_21_+_8433172_8446622_RNA45SN1_45S_50ntFlanks</td><td>10388</td><td>10424</td><td>37</td><td>GTGAACAGCAGTTGAACATGGGTCAGTCGGTCCTGAG</td><td>6123.7</td><td>7.0</td><td>1.2</td><td>6.0</td><td>2.6E-09</td><td>3.5E-08</td><td>BT20/MB468</td><td>MP</td></tr>
<tr><td>>hg38_21_+_8433172_8446622_RNA45SN1_45S_50ntFlanks@10411.10433.23|CAGTCGGTCCTGAGAGATGGGCG</td><td>>hg38_21_+_8433172_8446622_RNA45SN1_45S_50ntFlanks</td><td>10411</td><td>10433</td><td>23</td><td>CAGTCGGTCCTGAGAGATGGGCG</td><td>2553.6</td><td>-7.5</td><td>1.3</td><td>-6.0</td><td>2.6E-09</td><td>3.5E-08</td><td>BT20/MB468</td><td>MP</td></tr>
<tr><td>>hg38_21_+_8433172_8446622_RNA45SN1_45S_50ntFlanks@9470.9506.37|GAGCACGAGCGCACGTGTTAGGACCCGAAAGATGGTG</td><td>>hg38_21_+_8433172_8446622_RNA45SN1_45S_50ntFlanks</td><td>9470</td><td>9506</td><td>37</td><td>GAGCACGAGCGCACGTGTTAGGACCCGAAAGATGGTG</td><td>55.7</td><td>-9.7</td><td>1.7</td><td>-5.6</td><td>1.6E-08</td><td>3.5E-08</td><td>MB231/MB468</td><td>MP</td></tr>
<tr><td>>hg38_21_+_8433172_8446622_RNA45SN1_45S_50ntFlanks@7983.8018.36|CAGATCAGACGTGGCGACCCGCTGAATTTAAGCATA</td><td>>hg38_21_+_8433172_8446622_RNA45SN1_45S_50ntFlanks</td><td>7983</td><td>8018</td><td>36</td><td>CAGATCAGACGTGGCGACCCGCTGAATTTAAGCATA</td><td>88.2</td><td>-10.3</td><td>1.8</td><td>-5.6</td><td>1.6E-08</td><td>3.5E-08</td><td>MB231/MB468</td><td>MP</td></tr>
<tr><td>>hg38_21_+_8433172_8446622_RNA45SN1_45S_50ntFlanks@11581.11606.26|AACTGGTGCGGACCAGGGGAATCCGA</td><td>>hg38_21_+_8433172_8446622_RNA45SN1_45S_50ntFlanks</td><td>11581</td><td>11606</td><td>26</td><td>AACTGGTGCGGACCAGGGGAATCCGA</td><td>421.6</td><td>-4.4</td><td>0.8</td><td>-5.6</td><td>1.9E-08</td><td>3.5E-08</td><td>MB231/MB468</td><td>MITO</td></tr>
<tr><td>>MI0000272|hsa-mir-182&WithFlank&7|-|129770377|129770498@29.54.26&[MIMAT0000259&hsa-miR-182-5p&offsets|0|+2;m-22&7|-|129770449|129770470&offsets|0|+4]|TTTGGCAATGGTAGAACTCACACTGG</td><td>>MI0000272|hsa-mir-182&WithFlank&7|-|129770377|129770498</td><td>29</td><td>54</td><td>26</td><td>TTTGGCAATGGTAGAACTCACACTGG</td><td>616.1</td><td>3.9</td><td>0.7</td><td>5.6</td><td>1.7E-08</td><td>3.5E-08</td><td>MB231/MB468</td><td>CYTO</td></tr>
<tr><td>>hg38_21_+_8433172_8446622_RNA45SN1_45S_50ntFlanks@12415.12441.27|CTTCGATGTCGGCTCTTCCTATCATTG</td><td>>hg38_21_+_8433172_8446622_RNA45SN1_45S_50ntFlanks</td><td>12415</td><td>12441</td><td>27</td><td>CTTCGATGTCGGCTCTTCCTATCATTG</td><td>5025.4</td><td>-5.2</td><td>0.9</td><td>-5.6</td><td>1.9E-08</td><td>3.5E-08</td><td>MB231/MB468</td><td>MITO</td></tr>
<tr><td>>trna5_AspGTC_12_+_98897281_98897352@35.75.41|TCACGCGGGAGACCGGGGTTCAATTCCCCGACGGGGAGCCA</td><td>>trna5_AspGTC_12_+_98897281_98897352</td><td>35</td><td>75</td><td>41</td><td>TCACGCGGGAGACCGGGGTTCAATTCCCCGACGGGGAGCCA</td><td>6553.9</td><td>-6.9</td><td>1.1</td><td>-6.1</td><td>1.0E-09</td><td>3.5E-08</td><td>BT20/MB231</td><td>CYTO</td></tr>
<tr><td>>hg38_21_+_8433172_8446622_RNA45SN1_45S_50ntFlanks@10232.10273.42|CGCGGACGCTACGCCGCGACGAGTAGGAGGGCCGCTGCGGTG</td><td>>hg38_21_+_8433172_8446622_RNA45SN1_45S_50ntFlanks</td><td>10232</td><td>10273</td><td>42</td><td>CGCGGACGCTACGCCGCGACGAGTAGGAGGGCCGCTGCGGTG</td><td>74.2</td><td>-10.1</td><td>1.8</td><td>-5.6</td><td>1.6E-08</td><td>3.5E-08</td><td>MB231/MB468</td><td>MP</td></tr>
<tr><td>>hg38_21_+_8433172_8446622_RNA45SN1_45S_50ntFlanks@6654.6681.28|CTCTTAGCGGTGGATCACTCGGCTCGTG</td><td>>hg38_21_+_8433172_8446622_RNA45SN1_45S_50ntFlanks</td><td>6654</td><td>6681</td><td>28</td><td>CTCTTAGCGGTGGATCACTCGGCTCGTG</td><td>2910.0</td><td>-4.7</td><td>0.8</td><td>-5.6</td><td>1.9E-08</td><td>3.5E-08</td><td>MB231/MB468</td><td>MITO</td></tr>
<tr><td>>MI0000102|hsa-mir-100&WithFlank&11|-|122152223|122152314@23.40.18&[MIMAT0000098&hsa-miR-100-5p&offsets|+4|0;m-27&11|-|122152276|122152296&offsets|+4|+1]|CGTAGATCCGAACTTGTG</td><td>>MI0000102|hsa-mir-100&WithFlank&11|-|122152223|122152314</td><td>23</td><td>40</td><td>18</td><td>CGTAGATCCGAACTTGTG</td><td>230.7</td><td>-11.2</td><td>1.8</td><td>-6.2</td><td>4.7E-10</td><td>3.5E-08</td><td>BT20/MB231</td><td>MP</td></tr>
<tr><td>>am_trna111_HisGTG_1_-_147774845_147774916@-1G.20.21,trna118_HisGTG_1_-_145396881_145396952@-1G.20.21,trna16_HisGTG_1_+_146544773_146544844@-1G.20.21,trna1_HisGTG_15_+_45493349_45493420@-1G.20.21,trna21_HisGTG_1_+_147753471_147753542@-1G.20.21,trna33_HisGTG_6_+_27125906_27125977@-1G.20.21,trna7_HisGTG_9_-_14433938_14434009@-1G.20.21,trna8_HisGTG_15_-_45492611_45492682@-1G.20.21,trna9_HisGTG_15_-_45490804_45490875@-1G.20.21|GGCCGTGATCGTATAGTGGTT</td><td>>am_trna111_HisGTG_1_-_147774845_147774916</td><td>-1G</td><td>20</td><td>21</td><td>GGCCGTGATCGTATAGTGGTT</td><td>260.4</td><td>-11.4</td><td>1.8</td><td>-6.2</td><td>4.7E-10</td><td>3.5E-08</td><td>BT20/MB231</td><td>MP</td></tr>
<tr><td>>hg38_21_+_8433172_8446622_RNA45SN1_45S_50ntFlanks@10366.10383.18|CCGAAGTGGAGAAGGGTT</td><td>>hg38_21_+_8433172_8446622_RNA45SN1_45S_50ntFlanks</td><td>10366</td><td>10383</td><td>18</td><td>CCGAAGTGGAGAAGGGTT</td><td>147.3</td><td>-4.1</td><td>0.7</td><td>-5.6</td><td>1.7E-08</td><td>3.5E-08</td><td>MB231/MB468</td><td>CYTO</td></tr>
<tr><td>>hg38_21_+_8433172_8446622_RNA45SN1_45S_50ntFlanks@3782.3810.29|CAGTGAAACTGCGAATGGCTCATTAAATC</td><td>>hg38_21_+_8433172_8446622_RNA45SN1_45S_50ntFlanks</td><td>3782</td><td>3810</td><td>29</td><td>CAGTGAAACTGCGAATGGCTCATTAAATC</td><td>315.6</td><td>-6.5</td><td>1.1</td><td>-5.6</td><td>1.6E-08</td><td>3.5E-08</td><td>MB231/MB468</td><td>MP</td></tr>
<tr><td>>hg38_1_-_228634819_228635039_RNA5S12_5S_50ntFlanks@126.170.45|TACTTGGATGGGAGACCGCCTGGGAATACCGGGTGCTGTAGGCTT</td><td>>hg38_1_-_228634819_228635039_RNA5S12_5S_50ntFlanks</td><td>126</td><td>170</td><td>45</td><td>TACTTGGATGGGAGACCGCCTGGGAATACCGGGTGCTGTAGGCTT</td><td>2292.0</td><td>7.6</td><td>1.3</td><td>5.7</td><td>1.5E-08</td><td>3.5E-08</td><td>BT20/MB468</td><td>CYTO</td></tr>
<tr><td>>hg38_21_+_8433172_8446622_RNA45SN1_45S_50ntFlanks@10771.10791.21|CAGGTCTCCAAGGTGAACAGC</td><td>>hg38_21_+_8433172_8446622_RNA45SN1_45S_50ntFlanks</td><td>10771</td><td>10791</td><td>21</td><td>CAGGTCTCCAAGGTGAACAGC</td><td>765.7</td><td>-8.7</td><td>1.5</td><td>-5.6</td><td>2.0E-08</td><td>3.5E-08</td><td>BT20/MB468</td><td>NUC</td></tr>
<tr><td>>hg38_21_+_8433172_8446622_RNA45SN1_45S_50ntFlanks@11918.11958.41|ACATGAGAGGTGTAGAATAAGTGGGAGGCCCCCGGCGCCCC</td><td>>hg38_21_+_8433172_8446622_RNA45SN1_45S_50ntFlanks</td><td>11918</td><td>11958</td><td>41</td><td>ACATGAGAGGTGTAGAATAAGTGGGAGGCCCCCGGCGCCCC</td><td>88.6</td><td>-10.3</td><td>1.8</td><td>-5.6</td><td>1.6E-08</td><td>3.5E-08</td><td>MB231/MB468</td><td>MP</td></tr>
<tr><td>>MI0000100|hsa-mir-98&WithFlank&X|-|53556217|53556347@28.48.21(+1A)&[MIMAT0000096&hsa-miR-98-5p&offsets|0|-1(+1A);m-143&X|-|53556299|53556320&offsets|0|-1(+1A)]|TGAGGTAGTAAGTTGTATTGTA</td><td>>MI0000100|hsa-mir-98&WithFlank&X|-|53556217|53556347</td><td>28</td><td>48</td><td>21(+1A)</td><td>TGAGGTAGTAAGTTGTATTGTA</td><td>421.4</td><td>5.7</td><td>1.0</td><td>5.7</td><td>1.5E-08</td><td>3.5E-08</td><td>BT20/MB468</td><td>CYTO</td></tr>
</table>

| | | | | | | | | | | | | | |
|---|---|---|---|---|---|---|---|---|---|---|---|---|---|
| >hg38_21_+_8433172_8446622_RNA45SN1_45S_50ntFlanks@10393.10422.30\|CAGCAGTTGAACATGGGTCAGTCGGTCCTG | >hg38_21_+_8433172_8446622_RNA45SN1_45S_50ntFlanks | 10393 | 10422 | 30 | CAGCAGTTGAACATGGGTCAGTCGGTCCTG | 496.5 | -6.3 | 1.1 | -5.6 | 1.6E-08 | 3.5E-08 | MB231/MB468 | MP |
| >hg38_21_+_8433172_8446622_RNA45SN1_45S_50ntFlanks@9489.9506.18\|AGGACCCGAAAGATGGTG | >hg38_21_+_8433172_8446622_RNA45SN1_45S_50ntFlanks | 9489 | 9506 | 18 | AGGACCCGAAAGATGGTG | 327.6 | -8.4 | 1.5 | -5.6 | 1.9E-08 | 3.5E-08 | MB231/MB468 | MITO |
| >hg38_21_+_8433172_8446622_RNA45SN1_45S_50ntFlanks@8921.8941.21\|CCGGGGCCGAGGGAGCGAGAC | >hg38_21_+_8433172_8446622_RNA45SN1_45S_50ntFlanks | 8921 | 8941 | 21 | CCGGGGCCGAGGGAGCGAGAC | 123.9 | 10.7 | 1.8 | 6.0 | 2.6E-09 | 3.5E-08 | BT20/MB468 | MP |
| >MI0000750\|hsa-mir-26a-2&WithFlank&12\|-\|57824603\|57824698@20.42.23;MI0000083\|hsa-mir-26a-1&WithFlank&3\|+\|37969398\|37969486@16.38.23&[MIMAT0000082&hsa-miR-26a-5p&offsets\|0\|+1;m-33&12\|-\|57824658\|57824679&offsets\|0\|+1];[MIMAT0000082_1&hsa-miR-26a-5p&offsets\|0\|+1;m-34&3\|+\|37969413\|37969434&offsets\|0\|+1]\|TTCAAGTAATCCAGGATAGGCTG | >MI0000750\|hsa-mir-26a-2&WithFlank&12\|-\|57824603\|57824698 | 20 | 42 | 23 | TTCAAGTAATCCAGGATAGGCTG | 216.7 | 4.4 | 0.8 | 5.6 | 1.8E-08 | 3.5E-08 | MB231/MB468 | NUC |
| >hg38_21_+_8433172_8446622_RNA45SN1_45S_50ntFlanks@5423.5466.44\|ATTGGATGGTTTAGTGAGGCCCTCGGATCGGCCCCGCCGGGGTC | >hg38_21_+_8433172_8446622_RNA45SN1_45S_50ntFlanks | 5423 | 5466 | 44 | ATTGGATGGTTTAGTGAGGCCCTCGGATCGGCCCCGCCGGGGTC | 1581.6 | -8.7 | 1.6 | -5.6 | 1.8E-08 | 3.5E-08 | MB231/MB468 | NUC |
| >hg38_21_+_8433172_8446622_RNA45SN1_45S_50ntFlanks@11642.11660.19\|GGCGGGTGTTGACGCGATG | >hg38_21_+_8433172_8446622_RNA45SN1_45S_50ntFlanks | 11642 | 11660 | 19 | GGCGGGTGTTGACGCGATG | 1462.6 | -7.3 | 1.3 | -5.6 | 1.6E-08 | 3.5E-08 | MB231/MB468 | MP |
| >hg38_21_+_8433172_8446622_RNA45SN1_45S_50ntFlanks@12241.12258.18\|CAGAAGGGCAAAAGCTCG | >hg38_21_+_8433172_8446622_RNA45SN1_45S_50ntFlanks | 12241 | 12258 | 18 | CAGAAGGGCAAAAGCTCG | 72.7 | -10.0 | 1.8 | -5.6 | 1.6E-08 | 3.5E-08 | MB231/MB468 | MP |
| >hg38_21_+_8433172_8446622_RNA45SN1_45S_50ntFlanks@11783.11810.28\|CATGAATGGATGAACGAGATTCCCACTG | >hg38_21_+_8433172_8446622_RNA45SN1_45S_50ntFlanks | 11783 | 11810 | 28 | CATGAATGGATGAACGAGATTCCCACTG | 209.5 | -10.8 | 1.8 | -6.0 | 2.6E-09 | 3.5E-08 | BT20/MB468 | MP |
| >hg38_21_+_8433172_8446622_RNA45SN1_45S_50ntFlanks@6719.6747.29\|TGAATTGCAGGACACATTGATCATCGACA | >hg38_21_+_8433172_8446622_RNA45SN1_45S_50ntFlanks | 6719 | 6747 | 29 | TGAATTGCAGGACACATTGATCATCGACA | 1823.5 | -9.3 | 1.7 | -5.6 | 2.0E-08 | 3.5E-08 | BT20/MB468 | NUC |
| >trna10_LysCTT_16_+_3241501_3241573@1.35.35,trna13_LysCTT_14_-_58706613_58706685@1.35.35,trna2_LysCTT_15_+_79152904_79152976@1.35.35\|GCCCGGCTAGCTCAGTCGGTAGAGCATGGGACTCT | >trna10_LysCTT_16_+_3241501_3241573 | 1 | 35 | 35 | GCCCGGCTAGCTCAGTCGGTAGAGCATGGGACTCT | 6800.3 | 6.3 | 1.1 | 5.6 | 1.9E-08 | 3.5E-08 | MB231/MB468 | MITO |
| >hg38_21_+_8433172_8446622_RNA45SN1_45S_50ntFlanks@12044.12072.29\|ACTGACCCGGTGAGGCGGGGGGGCGAGCC | >hg38_21_+_8433172_8446622_RNA45SN1_45S_50ntFlanks | 12044 | 12072 | 29 | ACTGACCCGGTGAGGCGGGGGGGCGAGCC | 235.9 | -6.2 | 1.1 | -5.6 | 1.7E-08 | 3.5E-08 | MB231/MB468 | CYTO |
| >hg38_21_+_8433172_8446622_RNA45SN1_45S_50ntFlanks@10845.10864.20\|ACTTCGGGATAAGGATTGGC | >hg38_21_+_8433172_8446622_RNA45SN1_45S_50ntFlanks | 10845 | 10864 | 20 | ACTTCGGGATAAGGATTGGC | 1130.2 | -5.4 | 1.0 | -5.6 | 1.7E-08 | 3.5E-08 | MB231/MB468 | CYTO |
| >hg38_21_+_8433172_8446622_RNA45SN1_45S_50ntFlanks@3838.3868.31\|CTCTCCTACTTGGATAACTGTGGTAATTCTA | >hg38_21_+_8433172_8446622_RNA45SN1_45S_50ntFlanks | 3838 | 3868 | 31 | CTCTCCTACTTGGATAACTGTGGTAATTCTA | 559.0 | -3.5 | 0.6 | -5.6 | 1.8E-08 | 3.5E-08 | MB231/MB468 | NUC |
| >hg38_21_+_8433172_8446622_RNA45SN1_45S_50ntFlanks@9564.9598.35\|CTGACGTGCAAATCGGTCGTCCGACCTGGGTATAG | >hg38_21_+_8433172_8446622_RNA45SN1_45S_50ntFlanks | 9564 | 9598 | 35 | CTGACGTGCAAATCGGTCGTCCGACCTGGGTATAG | 366.4 | -8.3 | 1.5 | -5.6 | 2.0E-08 | 3.5E-08 | BT20/MB468 | NUC |
| >hg38_21_+_8433172_8446622_RNA45SN1_45S_50ntFlanks@9539.9572.34\|ACTCTGGTGGAGGTCCGTAGCGGTCCTGACGTGC | >hg38_21_+_8433172_8446622_RNA45SN1_45S_50ntFlanks | 9539 | 9572 | 34 | ACTCTGGTGGAGGTCCGTAGCGGTCCTGACGTGC | 431.4 | 4.4 | 0.8 | 5.7 | 1.4E-08 | 3.5E-08 | BT20/MB468 | MITO |
| >hg38_21_+_8433172_8446622_RNA45SN1_45S_50ntFlanks@12084.12110.27\|TCGCTTCTGGCGCCAAGCGCCCGGCCG | >hg38_21_+_8433172_8446622_RNA45SN1_45S_50ntFlanks | 12084 | 12110 | 27 | TCGCTTCTGGCGCCAAGCGCCCGGCCG | 181.4 | -10.6 | 1.8 | -6.0 | 2.7E-09 | 3.5E-08 | BT20/MB468 | MP |
| >hg38_MT_+_1621_3279_RNR2_16S_50ntFlanks@51.73.23\|GCTAAACCTAGCCCCAAACCCAC | >hg38_MT_+_1621_3279_RNR2_16S_50ntFlanks | 51 | 73 | 23 | GCTAAACCTAGCCCCAAACCCAC | 3425.9 | 5.2 | 0.9 | 5.7 | 1.4E-08 | 3.6E-08 | BT20/MB468 | MITO |
| >hg38_21_+_8433172_8446622_RNA45SN1_45S_50ntFlanks@11209.11231.23\|CGGCGACTCTGGACGCGAGCCGG | >hg38_21_+_8433172_8446622_RNA45SN1_45S_50ntFlanks | 11209 | 11231 | 23 | CGGCGACTCTGGACGCGAGCCGG | 1848.2 | -6.4 | 1.1 | -5.6 | 1.8E-08 | 3.6E-08 | MB231/MB468 | CYTO |
| >hg38_21_+_8433172_8446622_RNA45SN1_45S_50ntFlanks@11885.11905.21\|TTGAGCTTGACTCTAGTCTGG | >hg38_21_+_8433172_8446622_RNA45SN1_45S_50ntFlanks | 11885 | 11905 | 21 | TTGAGCTTGACTCTAGTCTGG | 453.5 | -7.3 | 1.3 | -5.6 | 1.8E-08 | 3.6E-08 | MB231/MB468 | NUC |
| >hg38_21_+_8433172_8446622_RNA45SN1_45S_50ntFlanks@3930.3959.30\|ATCAAAACCAACCCGGTCAGCCCCTCTCCG | >hg38_21_+_8433172_8446622_RNA45SN1_45S_50ntFlanks | 3930 | 3959 | 30 | ATCAAAACCAACCCGGTCAGCCCCTCTCCG | 97.4 | -9.7 | 1.6 | -6.0 | 2.7E-09 | 3.6E-08 | BT20/MB468 | MP |
| >hg38_21_+_8433172_8446622_RNA45SN1_45S_50ntFlanks@10361.10388.28\|GAAGGCCGAAGTGGAGAAGGGTTCCATG | >hg38_21_+_8433172_8446622_RNA45SN1_45S_50ntFlanks | 10361 | 10388 | 28 | GAAGGCCGAAGTGGAGAAGGGTTCCATG | 680.0 | -8.2 | 1.5 | -5.6 | 1.8E-08 | 3.6E-08 | MB231/MB468 | NUC |
| >trna111_HisGTG_1_-_147774845_147774916@-1T.31.32,trna118_HisGTG_1_-_145396881_145396952@-1T.31.32,trna16_HisGTG_1_+_146544773_146544844@-1T.31.32,trna1_HisGTG_15_+_45493349_45493420@-1T.31.32,trna21_HisGTG_1_+_147753471_147753542@-1T.31.32,trna33_HisGTG_6_+_27125906_27125977@-1T.31.32,trna7_HisGTG_9_-_14433938_14434009@-1T.31.32,trna8_HisGTG_15_-_45492611_45492682@-1T.31.32,trna9_HisGTG_15_-_45490804_45490875@-1T.31.32\|TGCCGTGATCGTATAGTGGTTAGTACTCTGCG | >trna111_HisGTG_1_-_147774845_147774916 | -1T | 31 | 32 | TGCCGTGATCGTATAGTGGTTAGTACTCTGCG | 9105.0 | -4.3 | 0.8 | -5.6 | 1.7E-08 | 3.6E-08 | MB231/MB468 | MP |
| >MI0000434\|hsa-let-7i&WithFlank&12\|+\|62603680\|62603775@12.32.21(+1A)&[MIMAT0000415&hsa-let-7i-5p&offsets\|0\|-1(+1A);m-49&12\|+\|62603691\|62603712&offsets\|0\|-1(+1A)]\|TGAGGTAGTAGTTTGTGCTGTA | >MI0000434\|hsa-let-7i&WithFlank&12\|+\|62603680\|62603775 | 12 | 32 | 21(+1A) | TGAGGTAGTAGTTTGTGCTGTA | 1441.5 | 5.5 | 1.0 | 5.6 | 2.0E-08 | 3.6E-08 | MB231/MB468 | MITO |
| >hg38_MT_+_1621_3279_RNR2_16S_50ntFlanks@51.90.40\|GCTAAACCTAGCCCCAAACCCACTCCACCTTACTACCAGA | >hg38_MT_+_1621_3279_RNR2_16S_50ntFlanks | 51 | 90 | 40 | GCTAAACCTAGCCCCAAACCCACTCCACCTTACTACCAGA | 1487.7 | 5.0 | 0.9 | 5.7 | 1.4E-08 | 3.6E-08 | BT20/MB468 | MITO |
| >MI0000749\|hsa-mir-30e&WithFlank&1\|+\|40754349\|40754452@23.46.24&[MIMAT0000692&hsa-miR-30e-5p&offsets\|0\|+2;m-24&1\|+\|40754371\|40754394&offsets\|0\|0]\|TGTAAACATCCTTGACTGGAAGCT | >MI0000749\|hsa-mir-30e&WithFlank&1\|+\|40754349\|40754452 | 23 | 46 | 24 | TGTAAACATCCTTGACTGGAAGCT | 2692.2 | 3.7 | 0.6 | 5.7 | 1.4E-08 | 3.6E-08 | BT20/MB468 | MITO |
| >hg38_21_+_8433172_8446622_RNA45SN1_45S_50ntFlanks@10783.10818.36\|GTGAACAGCCTCTGGCATGTTGGAACAATGTAGGTA | >hg38_21_+_8433172_8446622_RNA45SN1_45S_50ntFlanks | 10783 | 10818 | 36 | GTGAACAGCCTCTGGCATGTTGGAACAATGTAGGTA | 713.6 | -3.8 | 0.7 | -5.7 | 1.4E-08 | 3.6E-08 | BT20/MB468 | MITO |
| >hg38_21_+_8433172_8446622_RNA45SN1_45S_50ntFlanks@3766.3794.29\|GTACGCACGGCCGGTACAGTGAAACTGCG | >hg38_21_+_8433172_8446622_RNA45SN1_45S_50ntFlanks | 3766 | 3794 | 29 | GTACGCACGGCCGGTACAGTGAAACTGCG | 303.8 | 5.8 | 1.0 | 5.6 | 1.8E-08 | 3.6E-08 | MB231/MB468 | CYTO |
| >hg38_21_+_8433172_8446622_RNA45SN1_45S_50ntFlanks@3765.3792.28\|AGTACGCACGGCCGGTACAGTGAAACTG | >hg38_21_+_8433172_8446622_RNA45SN1_45S_50ntFlanks | 3765 | 3792 | 28 | AGTACGCACGGCCGGTACAGTGAAACTG | 637.5 | -8.3 | 1.5 | -5.6 | 2.0E-08 | 3.6E-08 | MB231/MB468 | MITO |
| >hg38_1_-_228634819_228635039_RNA5S12_5S_50ntFlanks@51.78.28\|GTCTACGGCCATACCACCCTGAACGCGC | >hg38_1_-_228634819_228635039_RNA5S12_5S_50ntFlanks | 51 | 78 | 28 | GTCTACGGCCATACCACCCTGAACGCGC | 1106.9 | 8.6 | 1.5 | 5.7 | 1.5E-08 | 3.6E-08 | BT20/MB468 | CYTO |
| >hg38_21_+_8433172_8446622_RNA45SN1_45S_50ntFlanks@9545.9572.28\|GTGGAGGTCCGTAGCGGTCCTGACGTGC | >hg38_21_+_8433172_8446622_RNA45SN1_45S_50ntFlanks | 9545 | 9572 | 28 | GTGGAGGTCCGTAGCGGTCCTGACGTGC | 910.2 | -5.2 | 0.9 | -5.6 | 1.7E-08 | 3.6E-08 | MB231/MB468 | MP |
| >hg38_21_+_8433172_8446622_RNA45SN1_45S_50ntFlanks@5494.5532.39\|AAGACGGTCGAACTTGACTATCTAGAGGAAGTAAAAGTC | >hg38_21_+_8433172_8446622_RNA45SN1_45S_50ntFlanks | 5494 | 5532 | 39 | AAGACGGTCGAACTTGACTATCTAGAGGAAGTAAAAGTC | 524.7 | 6.5 | 1.2 | 5.6 | 1.8E-08 | 3.6E-08 | MB231/MB468 | CYTO |
| >hg38_21_+_8433172_8446622_RNA45SN1_45S_50ntFlanks@3785.3810.26\|TGAAACTGCGAATGGCTCATTAAATC | >hg38_21_+_8433172_8446622_RNA45SN1_45S_50ntFlanks | 3785 | 3810 | 26 | TGAAACTGCGAATGGCTCATTAAATC | 1119.9 | -8.9 | 1.6 | -5.6 | 2.0E-08 | 3.6E-08 | BT20/MB468 | NUC |
| >MI0000082\|hsa-mir-25&WithFlank&7\|-\|100093554\|100093649@58.78.21(+1C)&[MIMAT0000081&hsa-miR-25-3p&offsets\|0\|-1(+1C);m-23&7\|-\|100093571\|100093592&offsets\|0\|-1(+1C)]\|CATTGCACTTGTCTCGGTCTGC | >MI0000082\|hsa-mir-25&WithFlank&7\|-\|100093554\|100093649 | 58 | 78 | 21(+1C) | CATTGCACTTGTCTCGGTCTGC | 208.9 | 5.6 | 1.0 | 5.6 | 1.8E-08 | 3.6E-08 | MB231/MB468 | NUC |
| >hg38_21_+_8433172_8446622_RNA45SN1_45S_50ntFlanks@4668.4695.28\|ATTCTTGGACCGGCGCAAGACGGACCAG | >hg38_21_+_8433172_8446622_RNA45SN1_45S_50ntFlanks | 4668 | 4695 | 28 | ATTCTTGGACCGGCGCAAGACGGACCAG | 61.2 | -9.8 | 1.7 | -5.6 | 1.7E-08 | 3.6E-08 | MB231/MB468 | MP |
| >hg38_21_+_8433172_8446622_RNA45SN1_45S_50ntFlanks@12202.12235.34\|CTAAGGCGAGCTCAGGGAGGACAGAAACCTCCCG | >hg38_21_+_8433172_8446622_RNA45SN1_45S_50ntFlanks | 12202 | 12235 | 34 | CTAAGGCGAGCTCAGGGAGGACAGAAACCTCCCG | 88.7 | -10.3 | 1.8 | -5.6 | 1.7E-08 | 3.6E-08 | MB231/MB468 | MP |
| >hg38_21_+_8433172_8446622_RNA45SN1_45S_50ntFlanks@4727.4767.41\|AATCAAGAACGAAAGTCGGAGGTTCGAAGACGATCAGATAC | >hg38_21_+_8433172_8446622_RNA45SN1_45S_50ntFlanks | 4727 | 4767 | 41 | AATCAAGAACGAAAGTCGGAGGTTCGAAGACGATCAGATAC | 2147.5 | 10.5 | 1.9 | 5.7 | 1.5E-08 | 3.6E-08 | BT20/MB468 | CYTO |
| >hg38_21_+_8433172_8446622_RNA45SN1_45S_50ntFlanks@11641.11662.22\|CGGCGGGTGTTGACGCGATGTG | >hg38_21_+_8433172_8446622_RNA45SN1_45S_50ntFlanks | 11641 | 11662 | 22 | CGGCGGGTGTTGACGCGATGTG | 2854.8 | -9.7 | 1.7 | -5.6 | 2.0E-08 | 3.6E-08 | BT20/MB468 | NUC |
| >hg38_21_+_8433172_8446622_RNA45SN1_45S_50ntFlanks@3874.3898.25\|AATACATGCCGACGGGCGCTGACCC | >hg38_21_+_8433172_8446622_RNA45SN1_45S_50ntFlanks | 3874 | 3898 | 25 | AATACATGCCGACGGGCGCTGACCC | 4257.8 | -7.7 | 1.4 | -5.6 | 1.8E-08 | 3.6E-08 | MB231/MB468 | NUC |
| >hg38_21_+_8433172_8446622_RNA45SN1_45S_50ntFlanks@10355.10388.34\|AACTTTGAAGGCCGAAGTGGAGAAGGGTTCCATG | >hg38_21_+_8433172_8446622_RNA45SN1_45S_50ntFlanks | 10355 | 10388 | 34 | AACTTTGAAGGCCGAAGTGGAGAAGGGTTCCATG | 983.2 | -8.4 | 1.5 | -5.6 | 1.8E-08 | 3.6E-08 | MB231/MB468 | NUC |
| >hg38_21_+_8433172_8446622_RNA45SN1_45S_50ntFlanks@9536.9563.28\|GAAACTCTGGTGGAGGTCCGTAGCGGTC | >hg38_21_+_8433172_8446622_RNA45SN1_45S_50ntFlanks | 9536 | 9563 | 28 | GAAACTCTGGTGGAGGTCCGTAGCGGTC | 594.4 | -8.5 | 1.5 | -5.6 | 2.0E-08 | 3.6E-08 | BT20/MB468 | NUC |
| >hg38_21_+_8433172_8446622_RNA45SN1_45S_50ntFlanks@4047.4071.25\|ATTCGAACGTCTGCCCTATCAACTT | >hg38_21_+_8433172_8446622_RNA45SN1_45S_50ntFlanks | 4047 | 4071 | 25 | ATTCGAACGTCTGCCCTATCAACTT | 537.3 | -8.0 | 1.4 | -5.6 | 2.1E-08 | 3.6E-08 | BT20/MB468 | NUC |
| >hg38_1_-_228634819_228635039_RNA5S12_5S_50ntFlanks@147.171.25\|GGGAATACCGGGTGCTGTAGGCTTT | >hg38_1_-_228634819_228635039_RNA5S12_5S_50ntFlanks | 147 | 171 | 25 | GGGAATACCGGGTGCTGTAGGCTTT | 1896.6 | 3.2 | 0.6 | 5.7 | 1.4E-08 | 3.6E-08 | BT20/MB468 | MITO |
| >hg38_21_+_8433172_8446622_RNA45SN1_45S_50ntFlanks@12625.12657.33\|CAATGGGGCGAAGCTACCATCTGTGGGATTATG | >hg38_21_+_8433172_8446622_RNA45SN1_45S_50ntFlanks | 12625 | 12657 | 33 | CAATGGGGCGAAGCTACCATCTGTGGGATTATG | 546.3 | -8.4 | 1.5 | -5.6 | 2.1E-08 | 3.6E-08 | BT20/MB468 | NUC |
| >hg38_21_+_8433172_8446622_RNA45SN1_45S_50ntFlanks@11314.11342.29\|CCCACCCCACGTCTCGTCGCGCGCGCGTC | >hg38_21_+_8433172_8446622_RNA45SN1_45S_50ntFlanks | 11314 | 11342 | 29 | CCCACCCCACGTCTCGTCGCGCGCGCGTC | 73.6 | -10.1 | 1.8 | -5.6 | 1.7E-08 | 3.6E-08 | MB231/MB468 | MP |
| >hg38_21_+_8433172_8446622_RNA45SN1_45S_50ntFlanks@6651.6685.35\|CGACTCTTAGCGGTGGATCACTCGGCTCGTGCGTC | >hg38_21_+_8433172_8446622_RNA45SN1_45S_50ntFlanks | 6651 | 6685 | 35 | CGACTCTTAGCGGTGGATCACTCGGCTCGTGCGTC | 8615.5 | 3.0 | 0.5 | 5.7 | 1.4E-08 | 3.6E-08 | BT20/MB468 | MITO |
| >hg38_21_+_8433172_8446622_RNA45SN1_45S_50ntFlanks@6652.6683.32\|GACTCTTAGCGGTGGATCACTCGGCTCGTGCG | >hg38_21_+_8433172_8446622_RNA45SN1_45S_50ntFlanks | 6652 | 6683 | 32 | GACTCTTAGCGGTGGATCACTCGGCTCGTGCG | 23735.7 | 5.4 | 1.0 | 5.6 | 1.9E-08 | 3.6E-08 | MB231/MB468 | NUC |
| >hg38_21_+_8433172_8446622_RNA45SN1_45S_50ntFlanks@9623.9648.26\|TAGTAGCTGGTTCCCTCCGAAGTTTC | >hg38_21_+_8433172_8446622_RNA45SN1_45S_50ntFlanks | 9623 | 9648 | 26 | TAGTAGCTGGTTCCCTCCGAAGTTTC | 310.3 | -8.1 | 1.4 | -5.6 | 2.1E-08 | 3.6E-08 | BT20/MB468 | NUC |

| | | | | | | | | | | | | | |
|---|---|---|---|---|---|---|---|---|---|---|---|---|---|
| >hg38_21_+_8433172_8446622_RNA45SN1_45S_50ntFlanks@7984.8006.23\|AGATCAGACGTGGCGACCCGCTG | >hg38_21_+_8433172_8446622_RNA45SN1_45S_50ntFlanks | 7984 | 8006 | 23 | AGATCAGACGTGGCGACCCGCTG | 9342.8 | -6.2 | 1.0 | -5.9 | 2.7E-09 | 3.6E-08 | BT20/MB468 | MP |
| >hg38_21_+_8433172_8446622_RNA45SN1_45S_50ntFlanks@10317.10354.38\|GTGCAGATCTTGGTGGTAGTAGCAAATATTCAAACGAG | >hg38_21_+_8433172_8446622_RNA45SN1_45S_50ntFlanks | 10317 | 10354 | 38 | GTGCAGATCTTGGTGGTAGTAGCAAATATTCAAACGAG | 85.6 | 10.2 | 1.7 | 5.9 | 2.7E-09 | 3.6E-08 | BT20/MB468 | MP |
| >hg38_21_+_8433172_8446622_RNA45SN1_45S_50ntFlanks@9914.9940.27\|GAAAAGGTGTTGGTTGATATAGACAGC | >hg38_21_+_8433172_8446622_RNA45SN1_45S_50ntFlanks | 9914 | 9940 | 27 | GAAAAGGTGTTGGTTGATATAGACAGC | 103.8 | 10.4 | 1.8 | 5.9 | 2.7E-09 | 3.6E-08 | BT20/MB468 | MP |
| >hg38_21_+_8433172_8446622_RNA45SN1_45S_50ntFlanks@12909.12933.25\|CCGCACGTTCGTGGGGAACCTGGCG | >hg38_21_+_8433172_8446622_RNA45SN1_45S_50ntFlanks | 12909 | 12933 | 25 | CCGCACGTTCGTGGGGAACCTGGCG | 339.1 | -8.4 | 1.5 | -5.6 | 2.1E-08 | 3.6E-08 | BT20/MB468 | NUC |
| >hg38_21_+_8433172_8446622_RNA45SN1_45S_50ntFlanks@12957.12979.23\|CTTCTGGGTCGGGGTTTCGTACG | >hg38_21_+_8433172_8446622_RNA45SN1_45S_50ntFlanks | 12957 | 12979 | 23 | CTTCTGGGTCGGGGTTTCGTACG | 572.9 | -4.0 | 0.7 | -5.7 | 1.4E-08 | 3.7E-08 | BT20/MB468 | MITO |
| >MI0000301\|hsa-mir-224&WithFlank&X\|-\|151958572\|151958664@14.35.22&[MIMAT0000281&hsa-miR-224-5p&offsets\|+1\|-2;m-238&X\|-\|151958630\|151958651&offsets\|0\|0]\|CAAGTCACTAGTGGTTCCGTTT | >MI0000301\|hsa-mir-224&WithFlank&X\|-\|151958572\|151958664 | 14 | 35 | 22 | CAAGTCACTAGTGGTTCCGTTT | 7440.6 | 10.3 | 1.7 | 5.9 | 2.8E-09 | 3.7E-08 | BT20/MB468 | MP |
| >MI0000542\|hsa-mir-320a&WithFlank&8\|-\|22244960\|22245043@49.68.20(+3U)&[MIMAT0000510&hsa-miR-320a-3p&offsets\|+1\|-1(+3U);m-60&8\|-\|22244975\|22244996&offsets\|+1\|-1(+3U)]\|AAAGCTGGGTTGAGAGGGCGTTT | >MI0000542\|hsa-mir-320a&WithFlank&8\|-\|22244960\|22245043 | 49 | 68 | 20(+3U) | AAAGCTGGGTTGAGAGGGCGTTT | 223.8 | 11.6 | 1.9 | 5.9 | 2.8E-09 | 3.7E-08 | BT20/MB468 | MP |
| >MI0016746\|hsa-mir-548o-2&WithFlank&20\|+\|38516557\|38516638@51.72.22;MI0006402\|hsa-mir-548o&WithFlank&7\|-\|102405736\|102405861@93.114.22&[MIMAT0005919&hsa-miR-548o-3p&offsets\|0\|0;m-431&20\|+\|38516607\|38516628&offsets\|0\|0];[MIMAT0005919_1&hsa-miR-548o-3p&offsets\|0\|0;m-430&7\|-\|102405748\|102405769&offsets\|0\|0]\|CCAAAACTGCAGTTACTTTTGC | >MI0016746\|hsa-mir-548o-2&WithFlank&20\|+\|38516557\|38516638 | 51 | 72 | 22 | CCAAAACTGCAGTTACTTTTGC | 469.8 | 3.7 | 0.6 | 5.6 | 1.7E-08 | 3.7E-08 | MB231/MB468 | MP |
| >MI0000299\|hsa-mir-222&WithFlank&X\|-\|45747009\|45747130@75.97.23(+1U)&[MIMAT0000279&hsa-miR-222-3p&offsets\|0\|+2(+1U);m-140&X\|-\|45747033\|45747056&offsets\|0\|-1(+1U)]\|AGCTACATCTGGCTACTGGGTCTT | >MI0000299\|hsa-mir-222&WithFlank&X\|-\|45747009\|45747130 | 75 | 97 | 23(+1U) | AGCTACATCTGGCTACTGGGTCTT | 2062.0 | 4.9 | 0.9 | 5.6 | 1.7E-08 | 3.7E-08 | MB231/MB468 | MP |
| >hg38_21_+_8433172_8446622_RNA45SN1_45S_50ntFlanks@10873.10902.30\|CTGGGTCGGTCGGGCTGGGGCGCGAAGCGG | >hg38_21_+_8433172_8446622_RNA45SN1_45S_50ntFlanks | 10873 | 10902 | 30 | CTGGGTCGGTCGGGCTGGGGCGCGAAGCGG | 2584.7 | -9.6 | 1.7 | -5.6 | 2.1E-08 | 3.7E-08 | BT20/MB468 | NUC |
| >hg38_1_-_228634819_228635039_RNA5S12_5S_50ntFlanks@135.170.36\|GGGAGACCGCCTGGGAATACCGGGTGCTGTAGGCTT | >hg38_1_-_228634819_228635039_RNA5S12_5S_50ntFlanks | 135 | 170 | 36 | GGGAGACCGCCTGGGAATACCGGGTGCTGTAGGCTT | 39566.4 | 5.5 | 1.0 | 5.7 | 1.4E-08 | 3.7E-08 | BT20/MB468 | MITO |
| >hg38_21_+_8433172_8446622_RNA45SN1_45S_50ntFlanks@12678.12702.25\|AATCCCGCCCAGGCGGAACGATACG | >hg38_21_+_8433172_8446622_RNA45SN1_45S_50ntFlanks | 12678 | 12702 | 25 | AATCCCGCCCAGGCGGAACGATACG | 211.5 | -10.8 | 1.8 | -5.9 | 2.8E-09 | 3.7E-08 | BT20/MB468 | MP |
| >hg38_21_+_8433172_8446622_RNA45SN1_45S_50ntFlanks@11318.11344.27\|CCCCACGTCTCGTCGCGCGCGCGTCCG | >hg38_21_+_8433172_8446622_RNA45SN1_45S_50ntFlanks | 11318 | 11344 | 27 | CCCCACGTCTCGTCGCGCGCGCGTCCG | 33863.9 | -11.1 | 2.0 | -5.6 | 2.1E-08 | 3.7E-08 | BT20/MB468 | NUC |
| >MI0000299\|hsa-mir-222&WithFlank&X\|-\|45747009\|45747130@75.97.23(+3U)&[MIMAT0000279&hsa-miR-222-3p&offsets\|0\|+2(+3U);m-140&X\|-\|45747033\|45747056&offsets\|0\|-1(+3U)]\|AGCTACATCTGGCTACTGGGTCTTTT | >MI0000299\|hsa-mir-222&WithFlank&X\|-\|45747009\|45747130 | 75 | 97 | 23(+3U) | AGCTACATCTGGCTACTGGGTCTTTT | 186.7 | 7.0 | 1.2 | 5.6 | 1.9E-08 | 3.7E-08 | MB231/MB468 | NUC |
| >hg38_21_+_8433172_8446622_RNA45SN1_45S_50ntFlanks@4069.4094.26\|CTTTCGATGGTAGTCGCCGTGCCTAC | >hg38_21_+_8433172_8446622_RNA45SN1_45S_50ntFlanks | 4069 | 4094 | 26 | CTTTCGATGGTAGTCGCCGTGCCTAC | 194.9 | -3.6 | 0.6 | -5.6 | 1.9E-08 | 3.7E-08 | MB231/MB468 | NUC |
| >hg38_21_+_8433172_8446622_RNA45SN1_45S_50ntFlanks@4069.4094.26\|CTTTCGATGGTAGTCGCCGTGCCTAC | >hg38_21_+_8433172_8446622_RNA45SN1_45S_50ntFlanks | 4069 | 4094 | 26 | CTTTCGATGGTAGTCGCCGTGCCTAC | 234.7 | -2.0 | 0.4 | -5.7 | 1.6E-08 | 3.7E-08 | BT20/MB468 | CYTO |
| >hg38_21_+_8433172_8446622_RNA45SN1_45S_50ntFlanks@12635.12653.19\|AAGCTACCATCTGTGGGAT | >hg38_21_+_8433172_8446622_RNA45SN1_45S_50ntFlanks | 12635 | 12653 | 19 | AAGCTACCATCTGTGGGAT | 608.4 | -7.0 | 1.2 | -5.7 | 1.6E-08 | 3.7E-08 | BT20/MB468 | CYTO |
| >hg38_21_+_8433172_8446622_RNA45SN1_45S_50ntFlanks@12475.12498.24\|CCCACTAATAGGGAACGTGAGCTG | >hg38_21_+_8433172_8446622_RNA45SN1_45S_50ntFlanks | 12475 | 12498 | 24 | CCCACTAATAGGGAACGTGAGCTG | 285.8 | -7.5 | 1.3 | -5.6 | 1.7E-08 | 3.7E-08 | MB231/MB468 | MP |
| >hg38_21_+_8433172_8446622_RNA45SN1_45S_50ntFlanks@10684.10709.26\|CGGCGGCGTCCGGTGAGCTCTCGCTG | >hg38_21_+_8433172_8446622_RNA45SN1_45S_50ntFlanks | 10684 | 10709 | 26 | CGGCGGCGTCCGGTGAGCTCTCGCTG | 1749.5 | -8.3 | 1.5 | -5.6 | 1.7E-08 | 3.7E-08 | MB231/MB468 | MP |
| >hg38_21_+_8433172_8446622_RNA45SN1_45S_50ntFlanks@6771.6807.37\|GGTTCCTCCCGGGGCTACGCCTGTCTGAGCGTCGCTT | >hg38_21_+_8433172_8446622_RNA45SN1_45S_50ntFlanks | 6771 | 6807 | 37 | GGTTCCTCCCGGGGCTACGCCTGTCTGAGCGTCGCTT | 302.8 | -4.3 | 0.8 | -5.6 | 1.7E-08 | 3.7E-08 | MB231/MB468 | MP |
| >hg38_21_+_8433172_8446622_RNA45SN1_45S_50ntFlanks@9739.9762.24\|AAACGATCTCAACCTATTCTCAAA | >hg38_21_+_8433172_8446622_RNA45SN1_45S_50ntFlanks | 9739 | 9762 | 24 | AAACGATCTCAACCTATTCTCAAA | 724.2 | -6.8 | 1.2 | -5.6 | 2.0E-08 | 3.7E-08 | MB231/MB468 | MITO |
| >hg38_21_+_8433172_8446622_RNA45SN1_45S_50ntFlanks@11682.11699.18\|AATGTCAAAGTGAAGAAA | >hg38_21_+_8433172_8446622_RNA45SN1_45S_50ntFlanks | 11682 | 11699 | 18 | AATGTCAAAGTGAAGAAA | 61.7 | -9.8 | 1.7 | -5.6 | 1.7E-08 | 3.7E-08 | MB231/MB468 | MP |
| >hg38_21_+_8433172_8446622_RNA45SN1_45S_50ntFlanks@11200.11232.33\|CGGCGGCGGCGGCGACTCTGGACGCGAGCCGGG | >hg38_21_+_8433172_8446622_RNA45SN1_45S_50ntFlanks | 11200 | 11232 | 33 | CGGCGGCGGCGGCGACTCTGGACGCGAGCCGGG | 1587.2 | -4.7 | 0.8 | -5.9 | 2.8E-09 | 3.7E-08 | BT20/MB468 | MP |
| >MI0000087\|hsa-mir-29a&WithFlank&7\|-\|130876741\|130876816@47.67.21(+1A)&[MIMAT0000086&hsa-miR-29a-3p&offsets\|-1\|-2(+1A);m-26&7\|-\|130876749\|130876769&offsets\|-1\|-1(+1A)]\|CTAGCACCATCTGAAATCGGTA | >MI0000087\|hsa-mir-29a&WithFlank&7\|-\|130876741\|130876816 | 47 | 67 | 21(+1A) | CTAGCACCATCTGAAATCGGTA | 88.4 | 5.6 | 1.0 | 5.6 | 2.1E-08 | 3.7E-08 | BT20/MB468 | NUC |
| >hg38_21_+_8433172_8446622_RNA45SN1_45S_50ntFlanks@12941.12966.26\|ATTCGTAGACGACCTGCTTCTGGGTC | >hg38_21_+_8433172_8446622_RNA45SN1_45S_50ntFlanks | 12941 | 12966 | 26 | ATTCGTAGACGACCTGCTTCTGGGTC | 1244.6 | -7.9 | 1.4 | -5.7 | 1.6E-08 | 3.7E-08 | BT20/MB468 | CYTO |
| >hg38_21_+_8433172_8446622_RNA45SN1_45S_50ntFlanks@8386.8405.20\|GGCGTGAAACCGTTAAGAGG | >hg38_21_+_8433172_8446622_RNA45SN1_45S_50ntFlanks | 8386 | 8405 | 20 | GGCGTGAAACCGTTAAGAGG | 113.9 | -9.9 | 1.7 | -5.9 | 2.8E-09 | 3.7E-08 | BT20/MB468 | MP |
| >hg38_21_+_8433172_8446622_RNA45SN1_45S_50ntFlanks@9808.9838.31\|TGGAATGCGAGTGCCTAGTGGGCCACTTTTG | >hg38_21_+_8433172_8446622_RNA45SN1_45S_50ntFlanks | 9808 | 9838 | 31 | TGGAATGCGAGTGCCTAGTGGGCCACTTTTG | 90.2 | -10.4 | 1.8 | -5.6 | 1.8E-08 | 3.7E-08 | MB231/MB468 | MP |
| >hg38_21_+_8433172_8446622_RNA45SN1_45S_50ntFlanks@10801.10818.18\|GTTGGAACAATGTAGGTA | >hg38_21_+_8433172_8446622_RNA45SN1_45S_50ntFlanks | 10801 | 10818 | 18 | GTTGGAACAATGTAGGTA | 123.8 | -10.0 | 1.7 | -5.9 | 2.8E-09 | 3.7E-08 | BT20/MB468 | MP |
| >hg38_21_+_8433172_8446622_RNA45SN1_45S_50ntFlanks@4380.4408.29\|CGTAGTTGGATCTTGGGAGCGGGCGGGCG | >hg38_21_+_8433172_8446622_RNA45SN1_45S_50ntFlanks | 4380 | 4408 | 29 | CGTAGTTGGATCTTGGGAGCGGGCGGGCG | 339.2 | -7.6 | 1.4 | -5.6 | 1.8E-08 | 3.7E-08 | MB231/MB468 | MP |
| >hg38_21_+_8433172_8446622_RNA45SN1_45S_50ntFlanks@6648.6689.42\|GTACGACTCTTAGCGGTGGATCACTCGGCTCGTGCGTCGATG | >hg38_21_+_8433172_8446622_RNA45SN1_45S_50ntFlanks | 6648 | 6689 | 42 | GTACGACTCTTAGCGGTGGATCACTCGGCTCGTGCGTCGATG | 21240.2 | -4.3 | 0.8 | -5.6 | 2.0E-08 | 3.7E-08 | MB231/MB468 | MITO |
| >MI0000066\|hsa-let-7e&WithFlank&19\|+\|51692780\|51692870@14.35.22(+1U)&[MIMAT0000066&hsa-let-7e-5p&offsets\|0\|0(+1U);m-50&19\|+\|51692793\|51692814&offsets\|0\|0(+1U)]\|TGAGGTAGGAGGTTGTATAGTTT | >MI0000066\|hsa-let-7e&WithFlank&19\|+\|51692780\|51692870 | 14 | 35 | 22(+1U) | TGAGGTAGGAGGTTGTATAGTTT | 480.0 | 3.5 | 0.6 | 5.6 | 1.9E-08 | 3.7E-08 | MB231/MB468 | NUC |
| >hg38_21_+_8433172_8446622_RNA45SN1_45S_50ntFlanks@11200.11230.31\|CGGCGGCGGCGGCGACTCTGGACGCGAGCCG | >hg38_21_+_8433172_8446622_RNA45SN1_45S_50ntFlanks | 11200 | 11230 | 31 | CGGCGGCGGCGGCGACTCTGGACGCGAGCCG | 55.1 | -9.6 | 1.7 | -5.6 | 1.8E-08 | 3.7E-08 | MB231/MB468 | MP |
| >hg38_21_+_8433172_8446622_RNA45SN1_45S_50ntFlanks@11562.11588.27\|CGCCTAGCAGCCGACTTAGAACTGGTG | >hg38_21_+_8433172_8446622_RNA45SN1_45S_50ntFlanks | 11562 | 11588 | 27 | CGCCTAGCAGCCGACTTAGAACTGGTG | 426.9 | -3.4 | 0.6 | -5.6 | 1.9E-08 | 3.7E-08 | MB231/MB468 | NUC |
| >hg38_21_+_8433172_8446622_RNA45SN1_45S_50ntFlanks@12991.13029.39\|CTCCCTCGCTGCGATCTATTGAAAGTCAGCCCTCGACAC | >hg38_21_+_8433172_8446622_RNA45SN1_45S_50ntFlanks | 12991 | 13029 | 39 | CTCCCTCGCTGCGATCTATTGAAAGTCAGCCCTCGACAC | 106.5 | -8.8 | 1.6 | -5.7 | 1.4E-08 | 3.7E-08 | BT20/MB468 | MITO |
| >MI0000063\|hsa-let-7b&WithFlank&22\|+\|46113680\|46113774@66.86.21(+1U)&[MIMAT0004482&hsa-let-7b-3p&offsets\|0\|-1(+1U);m-171&22\|+\|46113745\|46113766&offsets\|0\|-1(+1U)]\|CTATACAACCTACTGCCTTCCT | >MI0000063\|hsa-let-7b&WithFlank&22\|+\|46113680\|46113774 | 66 | 86 | 21(+1U) | CTATACAACCTACTGCCTTCCT | 496.5 | 7.8 | 1.4 | 5.7 | 1.6E-08 | 3.7E-08 | BT20/MB468 | CYTO |
| >hg38_21_+_8433172_8446622_RNA45SN1_45S_50ntFlanks@10664.10711.48\|CTTGGAAAGCGTCGCGGTTCCGGCGGCGTCCGGTGAGCTCTCGCTGGC | >hg38_21_+_8433172_8446622_RNA45SN1_45S_50ntFlanks | 10664 | 10711 | 48 | CTTGGAAAGCGTCGCGGTTCCGGCGGCGTCCGGTGAGCTCTCGCTGGC | 170.0 | -9.7 | 1.7 | -5.7 | 1.6E-08 | 3.7E-08 | BT20/MB468 | CYTO |
| >hg38_21_+_8433172_8446622_RNA45SN1_45S_50ntFlanks@6646.6672.27\|TCGTACGACTCTTAGCGGTGGATCACT | >hg38_21_+_8433172_8446622_RNA45SN1_45S_50ntFlanks | 6646 | 6672 | 27 | TCGTACGACTCTTAGCGGTGGATCACT | 2408.7 | 4.4 | 0.8 | 5.6 | 1.9E-08 | 3.7E-08 | MB231/MB468 | CYTO |
| >hg38_21_+_8433172_8446622_RNA45SN1_45S_50ntFlanks@11881.11899.19\|CCTGTTGAGCTTGACTCTA | >hg38_21_+_8433172_8446622_RNA45SN1_45S_50ntFlanks | 11881 | 11899 | 19 | CCTGTTGAGCTTGACTCTA | 176.8 | -4.7 | 0.8 | -5.6 | 1.9E-08 | 3.7E-08 | MB231/MB468 | CYTO |
| >hg38_21_+_8433172_8446622_RNA45SN1_45S_50ntFlanks@8105.8138.34\|CGCGGCGGGGCGCGGGACATGTGGCGTACGGAAG | >hg38_21_+_8433172_8446622_RNA45SN1_45S_50ntFlanks | 8105 | 8138 | 34 | CGCGGCGGGGCGCGGGACATGTGGCGTACGGAAG | 562.5 | -6.1 | 1.1 | -5.6 | 1.9E-08 | 3.7E-08 | MB231/MB468 | NUC |
| >hg38_21_+_8433172_8446622_RNA45SN1_45S_50ntFlanks@12437.12473.37\|CATTGTGAAGCAGAATTCACCAAGCGTTGGATTGTTC | >hg38_21_+_8433172_8446622_RNA45SN1_45S_50ntFlanks | 12437 | 12473 | 37 | CATTGTGAAGCAGAATTCACCAAGCGTTGGATTGTTC | 2702.7 | -7.3 | 1.3 | -5.6 | 1.8E-08 | 3.8E-08 | MB231/MB468 | MP |
| >hg38_21_+_8433172_8446622_RNA45SN1_45S_50ntFlanks@10800.10818.19\|TGTTGGAACAATGTAGGTA | >hg38_21_+_8433172_8446622_RNA45SN1_45S_50ntFlanks | 10800 | 10818 | 19 | TGTTGGAACAATGTAGGTA | 424.5 | -5.9 | 1.1 | -5.6 | 1.9E-08 | 3.8E-08 | MB231/MB468 | CYTO |
| >MI0000102\|hsa-mir-100&WithFlank&11\|-\|122152223\|122152314@19.36.18(+1C)&[MIMAT0000098&hsa-miR-100-5p&offsets\|0\|-4(+1C);m-27&11\|-\|122152276\|122152296&offsets\|0\|-3(+1C)]\|AACCCGTAGATCCGAACTC | >MI0000102\|hsa-mir-100&WithFlank&11\|-\|122152223\|122152314 | 19 | 36 | 18(+1C) | AACCCGTAGATCCGAACTC | 84.1 | 9.6 | 1.7 | 5.6 | 1.8E-08 | 3.8E-08 | MB231/MB468 | MP |
| >hg38_21_+_8433172_8446622_RNA45SN1_45S_50ntFlanks@474.504.31\|TCTTCGTGATCGATGTGGTGACGTCGTGCTC | >hg38_21_+_8433172_8446622_RNA45SN1_45S_50ntFlanks | 474 | 504 | 31 | TCTTCGTGATCGATGTGGTGACGTCGTGCTC | 1159.3 | 4.5 | 0.8 | 5.6 | 1.9E-08 | 3.8E-08 | MB231/MB468 | NUC |
| >hg38_21_+_8433172_8446622_RNA45SN1_45S_50ntFlanks@8111.8136.26\|GGGGCGCGGGACATGTGGCGTACGGA | >hg38_21_+_8433172_8446622_RNA45SN1_45S_50ntFlanks | 8111 | 8136 | 26 | GGGGCGCGGGACATGTGGCGTACGGA | 1765.7 | 14.1 | 2.3 | 6.2 | 5.2E-10 | 3.8E-08 | BT20/MB231 | MP |
| >hg38_21_+_8433172_8446622_RNA45SN1_45S_50ntFlanks@10346.10367.22\|TCAAACGAGAACTTTGAAGGCC | >hg38_21_+_8433172_8446622_RNA45SN1_45S_50ntFlanks | 10346 | 10367 | 22 | TCAAACGAGAACTTTGAAGGCC | 338.8 | -4.1 | 0.7 | -5.6 | 2.1E-08 | 3.8E-08 | MB231/MB468 | MITO |
| >hg38_21_+_8433172_8446622_RNA45SN1_45S_50ntFlanks@12907.12930.24\|CACCGCACGTTCGTGGGGAACCTG | >hg38_21_+_8433172_8446622_RNA45SN1_45S_50ntFlanks | 12907 | 12930 | 24 | CACCGCACGTTCGTGGGGAACCTG | 389.7 | -8.0 | 1.4 | -5.6 | 1.8E-08 | 3.8E-08 | MB231/MB468 | MP |
| >MI0000255\|hsa-mir-30d&WithFlank&8\|-\|134804870\|134804951@12.31.20&[MIMAT0000245&hsa-miR-30d-5p&offsets\|0\|-2;m-21&8\|-\|134804919\|134804940&offsets\|0\|-2]\|TGTAAACATCCCCGACTGGA | >MI0000255\|hsa-mir-30d&WithFlank&8\|-\|134804870\|134804951 | 12 | 31 | 20 | TGTAAACATCCCCGACTGGA | 889.8 | 5.1 | 0.9 | 5.6 | 2.1E-08 | 3.8E-08 | MB231/MB468 | MITO |

| | | | | | | | | | | | | | |
|---|---|---|---|---|---|---|---|---|---|---|---|---|---|
| >MI0000070\|hsa-mir-16-1&WithFlank&13\|-\|50048967\|50049067@20.40.21(+1A);MI0000115\|hsa-mir-16-2&WithFlank&3\|+\|160404739\|160404831@16.36.21(+1A)&[MIMAT0000069&hsa-miR-16-5p&offsets\|0\|-1(+1A);m-62&13\|-\|50049027\|50049048&offsets\|0\|-1(+1A)];[MIMAT0000069_1&hsa-miR-16-5p&offsets\|0\|-1(+1A);m-61&3\|+\|160404754\|160404775&offsets\|0\|-1(+1A)]\|TAGCAGCACGTAAATATTGGCA | >MI0000070\|hsa-mir-16-1&WithFlank&13\|-\|50048967\|50049067 | 20 | 40 | 21(+1A) | TAGCAGCACGTAAATATTGGCA | 470.7 | 4.3 | 0.8 | 5.6 | 2.1E-08 | 3.8E-08 | BT20/MB468 | NUC |
| >hg38_21_+_8433172_8446622_RNA45SN1_45S_50ntFlanks@12949.12982.34\|ACGACCTGCTTCTGGGTCGGGGTTTCGTACGTAG | >hg38_21_+_8433172_8446622_RNA45SN1_45S_50ntFlanks | 12949 | 12982 | 34 | ACGACCTGCTTCTGGGTCGGGGTTTCGTACGTAG | 308.1 | -6.3 | 1.1 | -5.6 | 1.9E-08 | 3.8E-08 | MB231/MB468 | CYTO |
| >hg38_21_+_8433172_8446622_RNA45SN1_45S_50ntFlanks@4000.4024.25\|TAGATAACCTCGGGCCGATCGCACG | >hg38_21_+_8433172_8446622_RNA45SN1_45S_50ntFlanks | 4000 | 4024 | 25 | TAGATAACCTCGGGCCGATCGCACG | 655.6 | -2.9 | 0.5 | -5.6 | 1.9E-08 | 3.8E-08 | MB231/MB468 | CYTO |
| >hg38_21_+_8433172_8446622_RNA45SN1_45S_50ntFlanks@4002.4025.24\|GATAACCTCGGGCCGATCGCACGC | >hg38_21_+_8433172_8446622_RNA45SN1_45S_50ntFlanks | 4002 | 4025 | 24 | GATAACCTCGGGCCGATCGCACGC | 1163.1 | -8.3 | 1.5 | -5.6 | 1.9E-08 | 3.8E-08 | MB231/MB468 | NUC |
| >MI0001448\|hsa-mir-425&WithFlank&3\|-\|49020142\|49020240@64.81.18&[MIMAT0001343&hsa-miR-425-3p&offsets\|+3\|-1;m-258&3\|-\|49020160\|49020181&offsets\|+4\|0]\|GGGAATGTCGTGTCCGCC | >MI0001448\|hsa-mir-425&WithFlank&3\|-\|49020142\|49020240 | 64 | 81 | 18 | GGGAATGTCGTGTCCGCC | 239.0 | 11.6 | 2.0 | 5.9 | 2.9E-09 | 3.8E-08 | BT20/MB468 | MP |
| >hg38_21_+_8433172_8446622_RNA45SN1_45S_50ntFlanks@11579.11600.22\|AGAACTGGTGCGGACCAGGGGA | >hg38_21_+_8433172_8446622_RNA45SN1_45S_50ntFlanks | 11579 | 11600 | 22 | AGAACTGGTGCGGACCAGGGGA | 200.4 | -10.7 | 1.8 | -5.9 | 2.9E-09 | 3.8E-08 | BT20/MB468 | MP |
| >hg38_21_+_8433172_8446622_RNA45SN1_45S_50ntFlanks@3769.3792.24\|CGCACGGCCGGTACAGTGAAACTG | >hg38_21_+_8433172_8446622_RNA45SN1_45S_50ntFlanks | 3769 | 3792 | 24 | CGCACGGCCGGTACAGTGAAACTG | 812.2 | -7.5 | 1.3 | -5.6 | 1.9E-08 | 3.8E-08 | MB231/MB468 | NUC |
| >hg38_21_+_8433172_8446622_RNA45SN1_45S_50ntFlanks@11207.11233.27\|GGCGGCGACTCTGGACGCGAGCCGGGC | >hg38_21_+_8433172_8446622_RNA45SN1_45S_50ntFlanks | 11207 | 11233 | 27 | GGCGGCGACTCTGGACGCGAGCCGGGC | 197.5 | -6.7 | 1.2 | -5.6 | 1.9E-08 | 3.8E-08 | MB231/MB468 | NUC |
| >hg38_21_+_8433172_8446622_RNA45SN1_45S_50ntFlanks@11316.11334.19\|CACCCCACGTCTCGTCGCG | >hg38_21_+_8433172_8446622_RNA45SN1_45S_50ntFlanks | 11316 | 11334 | 19 | CACCCCACGTCTCGTCGCG | 1009.8 | -7.0 | 1.2 | -5.6 | 1.9E-08 | 3.8E-08 | MB231/MB468 | NUC |
| >hg38_21_+_8433172_8446622_RNA45SN1_45S_50ntFlanks@5471.5516.46\|CACGGCCCTGGCGGAGCGCTGAGAAGACGGTCGAACTTGACTATCT | >hg38_21_+_8433172_8446622_RNA45SN1_45S_50ntFlanks | 5471 | 5516 | 46 | CACGGCCCTGGCGGAGCGCTGAGAAGACGGTCGAACTTGACTATCT | 2416.1 | 7.3 | 1.3 | 5.7 | 1.5E-08 | 3.8E-08 | BT20/MB468 | MITO |
| >hg38_21_+_8433172_8446622_RNA45SN1_45S_50ntFlanks@5422.5450.29\|GATTGGATGGTTTAGTGAGGCCCTCGGAT | >hg38_21_+_8433172_8446622_RNA45SN1_45S_50ntFlanks | 5422 | 5450 | 29 | GATTGGATGGTTTAGTGAGGCCCTCGGAT | 70.6 | 9.4 | 1.7 | 5.6 | 1.8E-08 | 3.8E-08 | MB231/MB468 | MP |
| >hg38_21_+_8433172_8446622_RNA45SN1_45S_50ntFlanks@3855.3874.20\|CTGTGGTAATTCTAGAGCTA | >hg38_21_+_8433172_8446622_RNA45SN1_45S_50ntFlanks | 3855 | 3874 | 20 | CTGTGGTAATTCTAGAGCTA | 91.9 | -9.6 | 1.6 | -5.9 | 2.9E-09 | 3.8E-08 | BT20/MB468 | MP |
| >MI0000805\|hsa-mir-342&WithFlank&14\|+\|100109649\|100109759@67.89.23&[MIMAT0000753&hsa-miR-342-3p&offsets\|0\|0;m-128&14\|+\|100109715\|100109737&offsets\|0\|0]\|TCTCACACAGAAATCGCACCCGT | >MI0000805\|hsa-mir-342&WithFlank&14\|+\|100109649\|100109759 | 67 | 89 | 23 | TCTCACACAGAAATCGCACCCGT | 402.6 | 9.1 | 1.6 | 5.6 | 1.6E-08 | 3.8E-08 | BT20/MB468 | CYTO |
| >trnaMT_ValTAC_MT_+_1602_1670@54.72.19\|AACTTGACCGCTCTGACCA | >trnaMT_ValTAC_MT_+_1602_1670 | 54 | 72 | 19 | AACTTGACCGCTCTGACCA | 3213.0 | 5.0 | 0.9 | 5.6 | 2.0E-08 | 3.8E-08 | MB231/MB468 | NUC |
| >hg38_21_+_8433172_8446622_RNA45SN1_45S_50ntFlanks@8325.8354.30\|CGATAGTCAACAAGTACCGTAAGGGAAAGT | >hg38_21_+_8433172_8446622_RNA45SN1_45S_50ntFlanks | 8325 | 8354 | 30 | CGATAGTCAACAAGTACCGTAAGGGAAAGT | 60.1 | -9.8 | 1.7 | -5.6 | 1.8E-08 | 3.8E-08 | MB231/MB468 | MP |
| >hg38_21_+_8433172_8446622_RNA45SN1_45S_50ntFlanks@10539.10570.32\|AGGCGTCCAGTGCGGTAACGCGACCGATCCCG | >hg38_21_+_8433172_8446622_RNA45SN1_45S_50ntFlanks | 10539 | 10570 | 32 | AGGCGTCCAGTGCGGTAACGCGACCGATCCCG | 94.5 | -9.6 | 1.6 | -5.9 | 2.9E-09 | 3.8E-08 | BT20/MB468 | MP |
| >hg38_21_+_8433172_8446622_RNA45SN1_45S_50ntFlanks@10784.10822.39\|TGAACAGCCTCTGGCATGTTGGAACAATGTAGGTAAGGG | >hg38_21_+_8433172_8446622_RNA45SN1_45S_50ntFlanks | 10784 | 10822 | 39 | TGAACAGCCTCTGGCATGTTGGAACAATGTAGGTAAGGG | 66.6 | -9.9 | 1.8 | -5.6 | 1.8E-08 | 3.8E-08 | MB231/MB468 | MP |
| >MI0000101\|hsa-mir-99a&WithFlank&21\|+\|16539083\|16539175@19.40.22(+1U)&[MIMAT0000097&hsa-miR-99a-5p&offsets\|0\|0(+1U);m-64&21\|+\|16539101\|16539121&offsets\|0\|+1(+1U)]\|AACCCGTAGATCCGATCTTGTGT | >MI0000101\|hsa-mir-99a&WithFlank&21\|+\|16539083\|16539175 | 19 | 40 | 22(+1U) | AACCCGTAGATCCGATCTTGTGT | 213.9 | -11.1 | 1.8 | -6.2 | 5.3E-10 | 3.8E-08 | BT20/MB231 | MP |
| >hg38_21_+_8433172_8446622_RNA45SN1_45S_50ntFlanks@8118.8138.21\|GGGACATGTGGCGTACGGAAG | >hg38_21_+_8433172_8446622_RNA45SN1_45S_50ntFlanks | 8118 | 8138 | 21 | GGGACATGTGGCGTACGGAAG | 1521.7 | 13.9 | 2.2 | 6.2 | 5.3E-10 | 3.8E-08 | BT20/MB231 | MP |
| >hg38_21_+_8433172_8446622_RNA45SN1_45S_50ntFlanks@11566.11611.46\|TAGCAGCCGACTTAGAACTGGTGCGGACCAGGGGAATCCGACTGTT | >hg38_21_+_8433172_8446622_RNA45SN1_45S_50ntFlanks | 11566 | 11611 | 46 | TAGCAGCCGACTTAGAACTGGTGCGGACCAGGGGAATCCGACTGTT | 4991.6 | 6.3 | 1.1 | 5.7 | 1.5E-08 | 3.8E-08 | BT20/MB468 | MITO |
| >hg38_21_+_8433172_8446622_RNA45SN1_45S_50ntFlanks@6670.6699.30\|ACTCGGCTCGTGCGTCGATGAAGAACGCAG | >hg38_21_+_8433172_8446622_RNA45SN1_45S_50ntFlanks | 6670 | 6699 | 30 | ACTCGGCTCGTGCGTCGATGAAGAACGCAG | 254.8 | -3.2 | 0.6 | -5.6 | 1.6E-08 | 3.8E-08 | BT20/MB468 | CYTO |
| >hg38_21_+_8433172_8446622_RNA45SN1_45S_50ntFlanks@10384.10406.23\|CCATGTGAACAGCAGTTGAACAT | >hg38_21_+_8433172_8446622_RNA45SN1_45S_50ntFlanks | 10384 | 10406 | 23 | CCATGTGAACAGCAGTTGAACAT | 223.1 | -10.9 | 1.8 | -5.9 | 2.9E-09 | 3.8E-08 | BT20/MB468 | MP |
| >hg38_21_+_8433172_8446622_RNA45SN1_45S_50ntFlanks@3765.3798.34\|AGTACGCACGGCCGGTACAGTGAAACTGCGAATG | >hg38_21_+_8433172_8446622_RNA45SN1_45S_50ntFlanks | 3765 | 3798 | 34 | AGTACGCACGGCCGGTACAGTGAAACTGCGAATG | 244.6 | 3.6 | 0.6 | 5.6 | 1.9E-08 | 3.8E-08 | MB231/MB468 | CYTO |
| >hg38_21_+_8433172_8446622_RNA45SN1_45S_50ntFlanks@7991.8018.28\|ACGTGGCGACCCGCTGAATTTAAGCATA | >hg38_21_+_8433172_8446622_RNA45SN1_45S_50ntFlanks | 7991 | 8018 | 28 | ACGTGGCGACCCGCTGAATTTAAGCATA | 376.9 | -7.6 | 1.3 | -5.6 | 1.8E-08 | 3.8E-08 | MB231/MB468 | MP |
| >MI0000063\|hsa-let-7b&WithFlank&22\|+\|46113680\|46113774@12.33.22&[MIMAT0000063&hsa-let-7b-5p&offsets\|0\|0;m-14&22\|+\|46113691\|46113712&offsets\|0\|0]\|TGAGGTAGTAGGTTGTGTGGTT | >MI0000063\|hsa-let-7b&WithFlank&22\|+\|46113680\|46113774 | 12 | 33 | 22 | TGAGGTAGTAGGTTGTGTGGTT | 11576.2 | 3.4 | 0.6 | 5.6 | 2.0E-08 | 3.8E-08 | MB231/MB468 | NUC |
| >hg38_21_+_8433172_8446622_RNA45SN1_45S_50ntFlanks@12413.12432.20\|TCCTTCGATGTCGGCTCTTC | >hg38_21_+_8433172_8446622_RNA45SN1_45S_50ntFlanks | 12413 | 12432 | 20 | TCCTTCGATGTCGGCTCTTC | 389.0 | -5.8 | 1.0 | -5.6 | 2.1E-08 | 3.8E-08 | MB231/MB468 | MITO |
| >hg38_21_+_8433172_8446622_RNA45SN1_45S_50ntFlanks@6650.6667.18\|ACGACTCTTAGCGGTGGA | >hg38_21_+_8433172_8446622_RNA45SN1_45S_50ntFlanks | 6650 | 6667 | 18 | ACGACTCTTAGCGGTGGA | 537.2 | 6.5 | 1.1 | 5.6 | 1.6E-08 | 3.8E-08 | BT20/MB468 | CYTO |
| >hg38_MT_+_1621_3279_RNR2_16S_50ntFlanks@718.740.23\|AGCCTGCGTCAGATTAAAACACT | >hg38_MT_+_1621_3279_RNR2_16S_50ntFlanks | 718 | 740 | 23 | AGCCTGCGTCAGATTAAAACACT | 123.8 | 10.7 | 1.8 | 5.9 | 2.9E-09 | 3.8E-08 | BT20/MB468 | MP |
| >hg38_21_+_8433172_8446622_RNA45SN1_45S_50ntFlanks@12539.12559.21\|ATGATGTGTTGTTGCCATGGT | >hg38_21_+_8433172_8446622_RNA45SN1_45S_50ntFlanks | 12539 | 12559 | 21 | ATGATGTGTTGTTGCCATGGT | 123.5 | -10.0 | 1.7 | -5.9 | 2.9E-09 | 3.8E-08 | BT20/MB468 | MP |
| >hg38_21_+_8433172_8446622_RNA45SN1_45S_50ntFlanks@4473.4494.22\|TTAGCTGAGTGTCCCGCGGGGC | >hg38_21_+_8433172_8446622_RNA45SN1_45S_50ntFlanks | 4473 | 4494 | 22 | TTAGCTGAGTGTCCCGCGGGGC | 97.7 | -9.7 | 1.6 | -5.9 | 2.9E-09 | 3.8E-08 | BT20/MB468 | MP |
| >hg38_21_+_8433172_8446622_RNA45SN1_45S_50ntFlanks@12002.12030.29\|TGCGGGCCGCCGGTGAAATACCACTACTC | >hg38_21_+_8433172_8446622_RNA45SN1_45S_50ntFlanks | 12002 | 12030 | 29 | TGCGGGCCGCCGGTGAAATACCACTACTC | 649.7 | -7.5 | 1.3 | -5.6 | 2.1E-08 | 3.8E-08 | MB231/MB468 | MITO |
| >hg38_21_+_8433172_8446622_RNA45SN1_45S_50ntFlanks@10346.10373.28\|TCAAACGAGAACTTTGAAGGCCGAAGTG | >hg38_21_+_8433172_8446622_RNA45SN1_45S_50ntFlanks | 10346 | 10373 | 28 | TCAAACGAGAACTTTGAAGGCCGAAGTG | 558.1 | -4.0 | 0.7 | -5.6 | 1.8E-08 | 3.9E-08 | MB231/MB468 | MP |
| >MI0000255\|hsa-mir-30d&WithFlank&8\|-\|134804870\|134804951@12.34.23&[MIMAT0000245&hsa-miR-30d-5p&offsets\|0\|+1;m-21&8\|-\|134804919\|134804940&offsets\|0\|+1]\|TGTAAACATCCCCGACTGGAAGC | >MI0000255\|hsa-mir-30d&WithFlank&8\|-\|134804870\|134804951 | 12 | 34 | 23 | TGTAAACATCCCCGACTGGAAGC | 11969.8 | 4.9 | 0.8 | 5.9 | 2.9E-09 | 3.9E-08 | BT20/MB468 | MP |
| >MI0000542\|hsa-mir-320a&WithFlank&8\|-\|22244960\|22245043@49.69.21(+1U)&[MIMAT0000510&hsa-miR-320a-3p&offsets\|+1\|0(+1U);m-60&8\|-\|22244975\|22244996&offsets\|+1\|0(+1U)]\|AAAGCTGGGTTGAGAGGGCGAT | >MI0000542\|hsa-mir-320a&WithFlank&8\|-\|22244960\|22245043 | 49 | 69 | 21(+1U) | AAAGCTGGGTTGAGAGGGCGAT | 131.1 | 3.3 | 0.6 | 5.6 | 2.2E-08 | 3.9E-08 | BT20/MB468 | NUC |
| >MI0000101\|hsa-mir-99a&WithFlank&21\|+\|16539083\|16539175@19.38.20(+1C)&[MIMAT0000097&hsa-miR-99a-5p&offsets\|0\|-2(+1C);m-64&21\|+\|16539101\|16539121&offsets\|0\|-1(+1C)]\|AACCCGTAGATCCGATCTTGC | >MI0000101\|hsa-mir-99a&WithFlank&21\|+\|16539083\|16539175 | 19 | 38 | 20(+1C) | AACCCGTAGATCCGATCTTGC | 189.0 | -6.7 | 1.1 | -6.0 | 1.6E-09 | 3.9E-08 | BT20/MB231 | MITO |
| >hg38_21_+_8433172_8446622_RNA45SN1_45S_50ntFlanks@10735.10752.18\|TGTAAATCTCGCGCCGGG | >hg38_21_+_8433172_8446622_RNA45SN1_45S_50ntFlanks | 10735 | 10752 | 18 | TGTAAATCTCGCGCCGGG | 310.5 | -6.2 | 1.1 | -5.6 | 1.9E-08 | 3.9E-08 | MB231/MB468 | CYTO |
| >hg38_21_+_8433172_8446622_RNA45SN1_45S_50ntFlanks@9545.9570.26\|GTGGAGGTCCGTAGCGGTCCTGACGT | >hg38_21_+_8433172_8446622_RNA45SN1_45S_50ntFlanks | 9545 | 9570 | 26 | GTGGAGGTCCGTAGCGGTCCTGACGT | 2444.6 | -3.3 | 0.6 | -5.6 | 1.9E-08 | 3.9E-08 | MB231/MB468 | CYTO |
| >hg38_21_+_8433172_8446622_RNA45SN1_45S_50ntFlanks@9561.9596.36\|GTCCTGACGTGCAAATCGGTCGTCCGACCTGGGTAT | >hg38_21_+_8433172_8446622_RNA45SN1_45S_50ntFlanks | 9561 | 9596 | 36 | GTCCTGACGTGCAAATCGGTCGTCCGACCTGGGTAT | 589.1 | -8.7 | 1.5 | -5.6 | 2.2E-08 | 3.9E-08 | BT20/MB468 | NUC |
| >MI0000542\|hsa-mir-320a&WithFlank&8\|-\|22244960\|22245043@48.68.21(+2U)&[MIMAT0000510&hsa-miR-320a-3p&offsets\|0\|-1(+2U);m-60&8\|-\|22244975\|22244996&offsets\|0\|-1(+2U)]\|AAAAGCTGGGTTGAGAGGGCGTT | >MI0000542\|hsa-mir-320a&WithFlank&8\|-\|22244960\|22245043 | 48 | 68 | 21(+2U) | AAAAGCTGGGTTGAGAGGGCGTT | 5649.0 | 8.3 | 1.4 | 5.9 | 3.0E-09 | 3.9E-08 | BT20/MB468 | MP |
| >hg38_21_+_8433172_8446622_RNA45SN1_45S_50ntFlanks@10768.10785.18\|CAGCAGGTCTCCAAGGTG | >hg38_21_+_8433172_8446622_RNA45SN1_45S_50ntFlanks | 10768 | 10785 | 18 | CAGCAGGTCTCCAAGGTG | 576.5 | -5.5 | 1.0 | -5.6 | 1.9E-08 | 3.9E-08 | MB231/MB468 | CYTO |
| >hg38_21_+_8433172_8446622_RNA45SN1_45S_50ntFlanks@10737.10757.21\|TAAATCTCGCGCCGGGCCGTA | >hg38_21_+_8433172_8446622_RNA45SN1_45S_50ntFlanks | 10737 | 10757 | 21 | TAAATCTCGCGCCGGGCCGTA | 1707.5 | -9.3 | 1.7 | -5.6 | 2.2E-08 | 3.9E-08 | BT20/MB468 | NUC |
| >hg38_21_+_8433172_8446622_RNA45SN1_45S_50ntFlanks@8341.8379.39\|CCGTAAGGGAAAGTTGAAAAGAACTTTGAAGAGAGAGTT | >hg38_21_+_8433172_8446622_RNA45SN1_45S_50ntFlanks | 8341 | 8379 | 39 | CCGTAAGGGAAAGTTGAAAAGAACTTTGAAGAGAGAGTT | 62.2 | -9.8 | 1.7 | -5.6 | 1.8E-08 | 3.9E-08 | MB231/MB468 | MP |
| >hg38_21_+_8433172_8446622_RNA45SN1_45S_50ntFlanks@4982.5015.34\|ACGGACAGGATTGACAGATTGATAGCTCTTTCTC | >hg38_21_+_8433172_8446622_RNA45SN1_45S_50ntFlanks | 4982 | 5015 | 34 | ACGGACAGGATTGACAGATTGATAGCTCTTTCTC | 268.3 | -10.2 | 1.8 | -5.7 | 1.5E-08 | 3.9E-08 | BT20/MB468 | MITO |
| >MI0005566\|hsa-mir-374b&WithFlank&X\|-\|74218541\|74218624@17.37.21&[MIMAT0004955&hsa-miR-374b-5p&offsets\|0\|-1;m-153&X\|-\|74218587\|74218608&offsets\|0\|-1]\|ATATAATACAACCTGCTAAGT | >MI0005566\|hsa-mir-374b&WithFlank&X\|-\|74218541\|74218624 | 17 | 37 | 21 | ATATAATACAACCTGCTAAGT | 715.5 | 5.5 | 1.0 | 5.7 | 1.5E-08 | 3.9E-08 | BT20/MB468 | MITO |
| >hg38_21_+_8433172_8446622_RNA45SN1_45S_50ntFlanks@12423.12443.21\|TCGGCTCTTCCTATCATTGTG | >hg38_21_+_8433172_8446622_RNA45SN1_45S_50ntFlanks | 12423 | 12443 | 21 | TCGGCTCTTCCTATCATTGTG | 1262.8 | -8.9 | 1.6 | -5.6 | 2.2E-08 | 3.9E-08 | BT20/MB468 | NUC |
| >hg38_21_+_8433172_8446622_RNA45SN1_45S_50ntFlanks@9482.9513.32\|ACGTGTTAGGACCCGAAAGATGGTGAACTATG | >hg38_21_+_8433172_8446622_RNA45SN1_45S_50ntFlanks | 9482 | 9513 | 32 | ACGTGTTAGGACCCGAAAGATGGTGAACTATG | 119.9 | 10.7 | 1.8 | 5.9 | 3.0E-09 | 3.9E-08 | BT20/MB468 | MP |
| >hg38_21_+_8433172_8446622_RNA45SN1_45S_50ntFlanks@8158.8178.21\|TCGTGGGGGGCCCAAGTCCTT | >hg38_21_+_8433172_8446622_RNA45SN1_45S_50ntFlanks | 8158 | 8178 | 21 | TCGTGGGGGGCCCAAGTCCTT | 104.9 | -9.8 | 1.7 | -5.9 | 3.0E-09 | 3.9E-08 | BT20/MB468 | MP |
| >hg38_21_+_8433172_8446622_RNA45SN1_45S_50ntFlanks@11562.11588.27\|CGCCTAGCAGCCGACTTAGAACTGGTG | >hg38_21_+_8433172_8446622_RNA45SN1_45S_50ntFlanks | 11562 | 11588 | 27 | CGCCTAGCAGCCGACTTAGAACTGGTG | 503.7 | -5.8 | 1.0 | -5.6 | 2.1E-08 | 3.9E-08 | MB231/MB468 | MITO |
| >hg38_21_+_8433172_8446622_RNA45SN1_45S_50ntFlanks@12539.12563.25\|ATGATGTGTTGTTGCCATGGTAATC | >hg38_21_+_8433172_8446622_RNA45SN1_45S_50ntFlanks | 12539 | 12563 | 25 | ATGATGTGTTGTTGCCATGGTAATC | 735.5 | -7.7 | 1.4 | -5.6 | 2.0E-08 | 3.9E-08 | MB231/MB468 | NUC |
| >hg38_21_+_8433172_8446622_RNA45SN1_45S_50ntFlanks@7984.8008.25\|AGATCAGACGTGGCGACCCGCTGAA | >hg38_21_+_8433172_8446622_RNA45SN1_45S_50ntFlanks | 7984 | 8008 | 25 | AGATCAGACGTGGCGACCCGCTGAA | 265.8 | -3.6 | 0.6 | -5.6 | 1.9E-08 | 3.9E-08 | MB231/MB468 | CYTO |

| | | | | | | | | | | | | | |
|---|---|---|---|---|---|---|---|---|---|---|---|---|---|
| >MI0000471\|hsa-mir-126&WithFlank&9\|+\|136670596\|136670692@57.78.22(+1A)&[MIMAT0000445&hsa-miR-126-3p&offsets\|-1\|-1(+1A);m-40&9\|+\|136670653\|136670674&offsets\|-1\|-1(+1A)]\|CTCGTACCGTGAGTAATAATGCA | >MI0000471\|hsa-mir-126&WithFlank&9\|+\|136670596\|136670692 | 57 | 78 | 22(+1A) | CTCGTACCGTGAGTAATAATGCA | 73.7 | 9.4 | 1.7 | 5.6 | 1.8E-08 | 3.9E-08 | MB231/MB468 | MP |
| >hg38_21_+_8433172_8446622_RNA45SN1_45S_50ntFlanks@9554.9595.42\|CGTAGCGGTCCTGACGTGCAAATCGGTCGTCCGACCTGGGTA | >hg38_21_+_8433172_8446622_RNA45SN1_45S_50ntFlanks | 9554 | 9595 | 42 | CGTAGCGGTCCTGACGTGCAAATCGGTCGTCCGACCTGGGTA | 75.1 | -10.1 | 1.8 | -5.6 | 1.8E-08 | 3.9E-08 | MB231/MB468 | MP |
| >hg38_21_+_8433172_8446622_RNA45SN1_45S_50ntFlanks@11682.11701.20\|AATGTCAAAGTGAAGAAATT | >hg38_21_+_8433172_8446622_RNA45SN1_45S_50ntFlanks | 11682 | 11701 | 20 | AATGTCAAAGTGAAGAAATT | 55.3 | -9.7 | 1.7 | -5.6 | 1.8E-08 | 3.9E-08 | MB231/MB468 | MP |
| >hg38_21_+_8433172_8446622_RNA45SN1_45S_50ntFlanks@11808.11835.28\|CTGTCCCTACCTACTATCCAGCGAAACC | >hg38_21_+_8433172_8446622_RNA45SN1_45S_50ntFlanks | 11808 | 11835 | 28 | CTGTCCCTACCTACTATCCAGCGAAACC | 61.9 | -9.8 | 1.7 | -5.6 | 1.8E-08 | 3.9E-08 | MB231/MB468 | MP |
| >hg38_21_+_8433172_8446622_RNA45SN1_45S_50ntFlanks@6700.6719.20\|CTAGCTGCGAGAATTAATGT | >hg38_21_+_8433172_8446622_RNA45SN1_45S_50ntFlanks | 6700 | 6719 | 20 | CTAGCTGCGAGAATTAATGT | 779.8 | -6.5 | 1.2 | -5.6 | 1.9E-08 | 3.9E-08 | MB231/MB468 | MP |
| >hg38_21_+_8433172_8446622_RNA45SN1_45S_50ntFlanks@8179.8212.34\|CTGATCGAGGCCCAGCCCGTGGACGGTGTGAGGC | >hg38_21_+_8433172_8446622_RNA45SN1_45S_50ntFlanks | 8179 | 8212 | 34 | CTGATCGAGGCCCAGCCCGTGGACGGTGTGAGGC | 716.1 | -8.7 | 1.5 | -5.9 | 3.0E-09 | 3.9E-08 | BT20/MB468 | MP |
| >hg38_21_+_8433172_8446622_RNA45SN1_45S_50ntFlanks@12394.12419.26\|TAGCGACGTCGCTTTTTGATCCTTCG | >hg38_21_+_8433172_8446622_RNA45SN1_45S_50ntFlanks | 12394 | 12419 | 26 | TAGCGACGTCGCTTTTTGATCCTTCG | 2382.5 | -6.6 | 1.2 | -5.6 | 2.2E-08 | 3.9E-08 | MB231/MB468 | MITO |
| >MI0000239\|hsa-mir-197&WithFlank&1\|+\|109598887\|109598973@54.76.23&[MIMAT0000227&hsa-miR-197-3p&offsets\|0\|+1;m-99&1\|+\|109598940\|109598961&offsets\|0\|+1]\|TTCACCACCTTCTCCACCCAGCA | >MI0000239\|hsa-mir-197&WithFlank&1\|+\|109598887\|109598973 | 54 | 76 | 23 | TTCACCACCTTCTCCACCCAGCA | 238.1 | 3.9 | 0.6 | 6.0 | 2.1E-09 | 3.9E-08 | BT20/MB231 | NUC |
| >hg38_21_+_8433172_8446622_RNA45SN1_45S_50ntFlanks@5472.5516.45\|ACGGCCCTGGCGGAGCGCTGAGAAGACGGTCGAACTTGACTATCT | >hg38_21_+_8433172_8446622_RNA45SN1_45S_50ntFlanks | 5472 | 5516 | 45 | ACGGCCCTGGCGGAGCGCTGAGAAGACGGTCGAACTTGACTATCT | 79567.4 | -4.6 | 0.8 | -6.1 | 1.2E-09 | 3.9E-08 | BT20/MB231 | CYTO |
| >trna20_GluTTC_2_-_131094701_131094772@1.34.34,trna5_GluTTC_13_-_41634874_41634945@1.34.34\|TCCCATATGGTCTAGCGGTTAGGATTCCTGGTTT | >trna20_GluTTC_2_-_131094701_131094772 | 1 | 34 | 34 | TCCCATATGGTCTAGCGGTTAGGATTCCTGGTTT | 97.7 | 9.8 | 1.7 | 5.6 | 1.9E-08 | 3.9E-08 | MB231/MB468 | MP |
| >MI0000289\|hsa-mir-181a-1&WithFlank&1\|-\|198859038\|198859159@30.51.22(+1A);MI0000269\|hsa-mir-181a-2&WithFlank&9\|+\|124692436\|124692557@45.66.22(+1A)&[MIMAT0000256&hsa-miR-181a-5p&offsets\|0\|-1(+1A);m-32&1\|-\|198859108\|198859130&offsets\|0\|-1(+1A)];[MIMAT0000256_1&hsa-miR-181a-5p&offsets\|0\|-1(+1A);m-32&9\|+\|124692480\|124692502&offsets\|0\|-1(+1A)]\|AACATTCAACGCTGTCGGTGAGA | >MI0000289\|hsa-mir-181a-1&WithFlank&1\|-\|198859038\|198859159 | 30 | 51 | 22(+1A) | AACATTCAACGCTGTCGGTGAGA | 339.8 | 4.6 | 0.8 | 5.6 | 2.0E-08 | 3.9E-08 | MB231/MB468 | NUC |
| >hg38_21_+_8433172_8446622_RNA45SN1_45S_50ntFlanks@9738.9768.31\|GAAACGATCTCAACCTATTCTCAAACTTTAA | >hg38_21_+_8433172_8446622_RNA45SN1_45S_50ntFlanks | 9738 | 9768 | 31 | GAAACGATCTCAACCTATTCTCAAACTTTAA | 69.0 | -10.0 | 1.8 | -5.6 | 1.9E-08 | 3.9E-08 | MB231/MB468 | MP |
| >hg38_21_+_8433172_8446622_RNA45SN1_45S_50ntFlanks@4732.4764.33\|AGAACGAAAGTCGGAGGTTCGAAGACGATCAGA | >hg38_21_+_8433172_8446622_RNA45SN1_45S_50ntFlanks | 4732 | 4764 | 33 | AGAACGAAAGTCGGAGGTTCGAAGACGATCAGA | 513.4 | 8.3 | 1.5 | 5.6 | 1.7E-08 | 3.9E-08 | BT20/MB468 | CYTO |
| >hg38_21_+_8433172_8446622_RNA45SN1_45S_50ntFlanks@10116.10136.21\|TGCGTCGGAGGGCGGCGGCGG | >hg38_21_+_8433172_8446622_RNA45SN1_45S_50ntFlanks | 10116 | 10136 | 21 | TGCGTCGGAGGGCGGCGGCGG | 743.6 | -8.7 | 1.6 | -5.6 | 2.2E-08 | 4.0E-08 | BT20/MB468 | NUC |
| >hg38_21_+_8433172_8446622_RNA45SN1_45S_50ntFlanks@8116.8139.24\|GCGGGACATGTGGCGTACGGAAGA | >hg38_21_+_8433172_8446622_RNA45SN1_45S_50ntFlanks | 8116 | 8139 | 24 | GCGGGACATGTGGCGTACGGAAGA | 1195.2 | 13.6 | 2.2 | 6.2 | 5.6E-10 | 4.0E-08 | BT20/MB231 | MP |
| >hg38_21_+_8433172_8446622_RNA45SN1_45S_50ntFlanks@4759.4783.25\|ATCAGATACCGTCGTAGTTCCGACC | >hg38_21_+_8433172_8446622_RNA45SN1_45S_50ntFlanks | 4759 | 4783 | 25 | ATCAGATACCGTCGTAGTTCCGACC | 2419.8 | -8.8 | 1.6 | -5.6 | 1.9E-08 | 4.0E-08 | MB231/MB468 | MP |
| >hg38_21_+_8433172_8446622_RNA45SN1_45S_50ntFlanks@8839.8861.23\|CGGTGGCGGCGGCGGCGGCGGCG | >hg38_21_+_8433172_8446622_RNA45SN1_45S_50ntFlanks | 8839 | 8861 | 23 | CGGTGGCGGCGGCGGCGGCGGCG | 158.3 | -10.4 | 1.8 | -5.9 | 3.0E-09 | 4.0E-08 | BT20/MB468 | MP |
| >MI0000289\|hsa-mir-181a-1&WithFlank&1\|-\|198859038\|198859159@30.54.25;MI0000269\|hsa-mir-181a-2&WithFlank&9\|+\|124692436\|124692557@45.69.25&[MIMAT0000256&hsa-miR-181a-5p&offsets\|0\|+2;m-32&1\|-\|198859108\|198859130&offsets\|0\|+2];[MIMAT0000256_1&hsa-miR-181a-5p&offsets\|0\|+2;m-32&9\|+\|124692480\|124692502&offsets\|0\|+2]\|AACATTCAACGCTGTCGGTGAGTTT | >MI0000289\|hsa-mir-181a-1&WithFlank&1\|-\|198859038\|198859159 | 30 | 54 | 25 | AACATTCAACGCTGTCGGTGAGTTT | 540.2 | 5.6 | 1.0 | 5.6 | 2.0E-08 | 4.0E-08 | MB231/MB468 | CYTO |
| >hg38_21_+_8433172_8446622_RNA45SN1_45S_50ntFlanks@7985.8014.30\|GATCAGACGTGGCGACCCGCTGAATTTAAG | >hg38_21_+_8433172_8446622_RNA45SN1_45S_50ntFlanks | 7985 | 8014 | 30 | GATCAGACGTGGCGACCCGCTGAATTTAAG | 91.9 | -9.6 | 1.6 | -5.9 | 3.1E-09 | 4.0E-08 | BT20/MB468 | MP |
| >hg38_21_+_8433172_8446622_RNA45SN1_45S_50ntFlanks@8443.8464.22\|CAACCCGGCGGCGGGTCCGGCC | >hg38_21_+_8433172_8446622_RNA45SN1_45S_50ntFlanks | 8443 | 8464 | 22 | CAACCCGGCGGCGGGTCCGGCC | 130.3 | -10.1 | 1.7 | -5.9 | 3.1E-09 | 4.0E-08 | BT20/MB468 | MP |
| >hg38_21_+_8433172_8446622_RNA45SN1_45S_50ntFlanks@6737.6772.36\|GATCATCGACACTTCGAACGCACTTGCGGCCCCGGG | >hg38_21_+_8433172_8446622_RNA45SN1_45S_50ntFlanks | 6737 | 6772 | 36 | GATCATCGACACTTCGAACGCACTTGCGGCCCCGGG | 101.2 | -10.5 | 1.9 | -5.6 | 1.9E-08 | 4.0E-08 | MB231/MB468 | MP |
| >hg38_21_+_8433172_8446622_RNA45SN1_45S_50ntFlanks@10357.10374.18\|CTTTGAAGGCCGAAGTGG | >hg38_21_+_8433172_8446622_RNA45SN1_45S_50ntFlanks | 10357 | 10374 | 18 | CTTTGAAGGCCGAAGTGG | 128.4 | -10.1 | 1.7 | -5.9 | 3.1E-09 | 4.0E-08 | BT20/MB468 | MP |
| >hg38_1_-_228634819_228635039_RNA5S12_5S_50ntFlanks@131.169.39\|GGATGGGAGACCGCCTGGGAATACCGGGTGCTGTAGGCT | >hg38_1_-_228634819_228635039_RNA5S12_5S_50ntFlanks | 131 | 169 | 39 | GGATGGGAGACCGCCTGGGAATACCGGGTGCTGTAGGCT | 692.5 | 5.6 | 0.9 | 5.9 | 3.1E-09 | 4.0E-08 | BT20/MB468 | MP |
| >hg38_21_+_8433172_8446622_RNA45SN1_45S_50ntFlanks@8170.8212.43\|CAAGTCCTTCTGATCGAGGCCCAGCCCGTGGACGGTGTGAGGC | >hg38_21_+_8433172_8446622_RNA45SN1_45S_50ntFlanks | 8170 | 8212 | 43 | CAAGTCCTTCTGATCGAGGCCCAGCCCGTGGACGGTGTGAGGC | 849.2 | -4.6 | 0.8 | -5.6 | 1.7E-08 | 4.0E-08 | BT20/MB468 | CYTO |
| >trnaMT_HisGTG_MT_+_12138_12206@20.55.36\|AACATCAGATTGTGAATCTGACAACAGAGGCTTACG | >trnaMT_HisGTG_MT_+_12138_12206 | 20 | 55 | 36 | AACATCAGATTGTGAATCTGACAACAGAGGCTTACG | 231.4 | 11.6 | 2.0 | 5.9 | 3.1E-09 | 4.0E-08 | BT20/MB468 | MP |
| >hg38_21_+_8433172_8446622_RNA45SN1_45S_50ntFlanks@11703.11727.25\|AATGAAGCGCGGGTAAACGGCGGGA | >hg38_21_+_8433172_8446622_RNA45SN1_45S_50ntFlanks | 11703 | 11727 | 25 | AATGAAGCGCGGGTAAACGGCGGGA | 482.9 | -8.2 | 1.5 | -5.6 | 2.3E-08 | 4.0E-08 | BT20/MB468 | NUC |
| >hg38_21_+_8433172_8446622_RNA45SN1_45S_50ntFlanks@4375.4408.34\|AAGCTCGTAGTTGGATCTTGGGAGCGGGCGGGCG | >hg38_21_+_8433172_8446622_RNA45SN1_45S_50ntFlanks | 4375 | 4408 | 34 | AAGCTCGTAGTTGGATCTTGGGAGCGGGCGGGCG | 56.9 | -9.7 | 1.7 | -5.6 | 1.9E-08 | 4.0E-08 | MB231/MB468 | MP |
| >MI0000102\|hsa-mir-100&WithFlank&11\|-\|122152223\|122152314@19.39.21&[MIMAT0000098&hsa-miR-100-5p&offsets\|0\|-1;m-27&11\|-\|122152276\|122152296&offsets\|0\|0]\|AACCCGTAGATCCGAACTTGT | >MI0000102\|hsa-mir-100&WithFlank&11\|-\|122152223\|122152314 | 19 | 39 | 21 | AACCCGTAGATCCGAACTTGT | 294328.2 | -7.2 | 1.2 | -6.2 | 5.7E-10 | 4.0E-08 | BT20/MB231 | MP |
| >am_trna10_ValCAC_5_-_180649395_180649467@1.29.29,trna12_ValAAC_5_-_180645270_180645342@1.29.29,trna132_ValAAC_6_-_27721179_27721251@1.29.29,trna136_ValAAC_6_-_27648885_27648957@1.29.29,trna139_ValAAC_6_-_27618707_27618779@1.29.29,trna18_ValCAC_5_-_180529253_180529325@1.29.29,trna2_ValAAC_3_+_169490018_169490090@1.29.29,trna2_ValCAC_5_+_180524070_180524142@1.29.29,trna4_ValAAC_5_+_180591154_180591226@1.29.29,trna5_ValAAC_5_+_180596610_180596682@1.29.29,trna6_ValCAC_5_+_180600650_180600722@1.29.29,trna85_ValCAC_1_-_161369490_161369562@1.29.29,trna90_ValCAC_1_-_149684088_149684161@1.29.29,trna98_ValCAC_1_-_149298555_149298627@1.29.29,trna9_ValCAC_6_+_26538282_26538354@1.29.29\|GTTTCCGTAGTGTAGTGGTTATCACGTTC | >am_trna10_ValCAC_5_-_180649395_180649467 | 1 | 29 | 29 | GTTTCCGTAGTGTAGTGGTTATCACGTTC | 26446.6 | -5.5 | 1.0 | -5.6 | 1.9E-08 | 4.0E-08 | MB231/MB468 | MP |
| >hg38_21_+_8433172_8446622_RNA45SN1_45S_50ntFlanks@12457.12488.32\|CAAGCGTTGGATTGTTCACCCACTAATAGGGA | >hg38_21_+_8433172_8446622_RNA45SN1_45S_50ntFlanks | 12457 | 12488 | 32 | CAAGCGTTGGATTGTTCACCCACTAATAGGGA | 653.7 | -5.9 | 1.0 | -5.6 | 2.0E-08 | 4.0E-08 | MB231/MB468 | CYTO |
| >hg38_21_+_8433172_8446622_RNA45SN1_45S_50ntFlanks@10882.10902.21\|TCGGGCTGGGGCGCGAAGCGG | >hg38_21_+_8433172_8446622_RNA45SN1_45S_50ntFlanks | 10882 | 10902 | 21 | TCGGGCTGGGGCGCGAAGCGG | 356.8 | -5.9 | 1.0 | -5.6 | 2.1E-08 | 4.0E-08 | MB231/MB468 | NUC |
| >hg38_21_+_8433172_8446622_RNA45SN1_45S_50ntFlanks@10423.10456.34\|AGAGATGGGCGAGCGCCGTTCCGAAGGGACGGGC | >hg38_21_+_8433172_8446622_RNA45SN1_45S_50ntFlanks | 10423 | 10456 | 34 | AGAGATGGGCGAGCGCCGTTCCGAAGGGACGGGC | 2093.0 | -9.3 | 1.7 | -5.6 | 2.3E-08 | 4.0E-08 | BT20/MB468 | NUC |
| >hg38_21_+_8433172_8446622_RNA45SN1_45S_50ntFlanks@11891.11912.22\|TTGACTCTAGTCTGGCACGGTG | >hg38_21_+_8433172_8446622_RNA45SN1_45S_50ntFlanks | 11891 | 11912 | 22 | TTGACTCTAGTCTGGCACGGTG | 1334.8 | -6.5 | 1.2 | -5.6 | 2.1E-08 | 4.0E-08 | MB231/MB468 | NUC |
| >hg38_21_+_8433172_8446622_RNA45SN1_45S_50ntFlanks@4995.5028.34\|ACAGATTGATAGCTCTTTCTCGATTCCGTGGGTG | >hg38_21_+_8433172_8446622_RNA45SN1_45S_50ntFlanks | 4995 | 5028 | 34 | ACAGATTGATAGCTCTTTCTCGATTCCGTGGGTG | 96.5 | -10.5 | 1.9 | -5.6 | 1.9E-08 | 4.0E-08 | MB231/MB468 | MP |
| >hg38_21_+_8433172_8446622_RNA45SN1_45S_50ntFlanks@11561.11586.26\|GCGCCTAGCAGCCGACTTAGAACTGG | >hg38_21_+_8433172_8446622_RNA45SN1_45S_50ntFlanks | 11561 | 11586 | 26 | GCGCCTAGCAGCCGACTTAGAACTGG | 337.8 | -4.5 | 0.8 | -5.6 | 2.0E-08 | 4.0E-08 | MB231/MB468 | CYTO |
| >hg38_21_+_8433172_8446622_RNA45SN1_45S_50ntFlanks@4512.4537.26\|AAAAAATTAGAGTGTTCAAAGCAGGC | >hg38_21_+_8433172_8446622_RNA45SN1_45S_50ntFlanks | 4512 | 4537 | 26 | AAAAAATTAGAGTGTTCAAAGCAGGC | 110.1 | -9.9 | 1.7 | -5.9 | 3.1E-09 | 4.0E-08 | BT20/MB468 | MP |
| >hg38_21_+_8433172_8446622_RNA45SN1_45S_50ntFlanks@4445.4470.26\|CTTGCCTCTCGGCGCCCCCTCGATGC | >hg38_21_+_8433172_8446622_RNA45SN1_45S_50ntFlanks | 4445 | 4470 | 26 | CTTGCCTCTCGGCGCCCCCTCGATGC | 63.7 | -9.9 | 1.8 | -5.6 | 1.9E-08 | 4.0E-08 | MB231/MB468 | MP |
| >MI0016005\|hsa-mir-3615&WithFlank&17\|+\|74748607\|74748705@57.77.21(+1A)&[MIMAT0017994&hsa-miR-3615&offsets\|0\|0(+1A);m-271&17\|+\|74748663\|74748684&offsets\|0\|-1(+1A)]\|TCTCTCGGCTCCTCGCGGCTCA | >MI0016005\|hsa-mir-3615&WithFlank&17\|+\|74748607\|74748705 | 57 | 77 | 21(+1A) | TCTCTCGGCTCCTCGCGGCTCA | 315.1 | 6.0 | 1.1 | 5.6 | 2.0E-08 | 4.1E-08 | MB231/MB468 | CYTO |
| >MI0000301\|hsa-mir-224&WithFlank&X\|-\|151958572\|151958664@14.36.23&[MIMAT0000281&hsa-miR-224-5p&offsets\|+1\|-1;m-238&X\|-\|151958630\|151958651&offsets\|0\|+1]\|CAAGTCACTAGTGGTTCCGTTTA | >MI0000301\|hsa-mir-224&WithFlank&X\|-\|151958572\|151958664 | 14 | 36 | 23 | CAAGTCACTAGTGGTTCCGTTTA | 399.0 | 7.1 | 1.3 | 5.6 | 2.3E-08 | 4.1E-08 | BT20/MB468 | NUC |
| >MI0000077\|hsa-mir-21&WithFlank&17\|+\|59841260\|59841343@15.36.22&[MIMAT0000076&hsa-miR-21-5p&offsets\|+1\|+1;m-1&17\|+\|59841273\|59841294&offsets\|+1\|+1]\|AGCTTATCAGACTGATGTTGAC | >MI0000077\|hsa-mir-21&WithFlank&17\|+\|59841260\|59841343 | 15 | 36 | 22 | AGCTTATCAGACTGATGTTGAC | 335.1 | 5.4 | 1.0 | 5.6 | 2.0E-08 | 4.1E-08 | MB231/MB468 | CYTO |
| >hg38_21_+_8433172_8446622_RNA45SN1_45S_50ntFlanks@11663.11687.25\|ATTTCTGCCCAGTGCTCTGAATGTC | >hg38_21_+_8433172_8446622_RNA45SN1_45S_50ntFlanks | 11663 | 11687 | 25 | ATTTCTGCCCAGTGCTCTGAATGTC | 255.3 | 3.3 | 0.6 | 5.6 | 2.0E-08 | 4.1E-08 | MB231/MB468 | CYTO |
| >am_trnaMT_PheGAA_MT_+_577_647@40.57.18\|TGTTTAGACGGGCTCACA | >am_trnaMT_PheGAA_MT_+_577_647 | 40 | 57 | 18 | TGTTTAGACGGGCTCACA | 758.4 | 3.8 | 0.7 | 5.6 | 2.2E-08 | 4.1E-08 | MB231/MB468 | MITO |
| >hg38_21_+_8433172_8446622_RNA45SN1_45S_50ntFlanks@5082.5100.19\|AACGAACGAGACTCTGGCA | >hg38_21_+_8433172_8446622_RNA45SN1_45S_50ntFlanks | 5082 | 5100 | 19 | AACGAACGAGACTCTGGCA | 432.6 | 7.8 | 1.4 | 5.6 | 1.7E-08 | 4.1E-08 | BT20/MB468 | CYTO |

| | | | | | | | | | | | | | |
|---|---|---|---|---|---|---|---|---|---|---|---|---|---|
| >hg38_21_+_8433172_8446622_RNA45SN1_45S_50ntFlanks@12456.12493.38\|CCAAGCGTTGGATTGTTCACCCACTAATAGGGAACGTG | >hg38_21_+_8433172_8446622_RNA45SN1_45S_50ntFlanks | 12456 | 12493 | 38 | CCAAGCGTTGGATTGTTCACCCACTAATAGGGAACGTG | 528.1 | -3.3 | 0.6 | -5.6 | 2.0E-08 | 4.1E-08 | MB231/MB468 | CYTO |
| >hg38_21_+_8433172_8446622_RNA45SN1_45S_50ntFlanks@11784.11802.19\|ATGAATGGATGAACGAGAT | >hg38_21_+_8433172_8446622_RNA45SN1_45S_50ntFlanks | 11784 | 11802 | 19 | ATGAATGGATGAACGAGAT | 409.1 | -8.3 | 1.5 | -5.6 | 1.9E-08 | 4.1E-08 | MB231/MB468 | MP |
| >hg38_21_+_8433172_8446622_RNA45SN1_45S_50ntFlanks@4509.4528.20\|TTGAAAAAATTAGAGTGTTC | >hg38_21_+_8433172_8446622_RNA45SN1_45S_50ntFlanks | 4509 | 4528 | 20 | TTGAAAAAATTAGAGTGTTC | 96.7 | -9.7 | 1.6 | -5.9 | 3.1E-09 | 4.1E-08 | BT20/MB468 | MP |
| >hg38_21_+_8433172_8446622_RNA45SN1_45S_50ntFlanks@10788.10826.39\|CAGCCTCTGGCATGTTGGAACAATGTAGGTAAGGGAAGT | >hg38_21_+_8433172_8446622_RNA45SN1_45S_50ntFlanks | 10788 | 10826 | 39 | CAGCCTCTGGCATGTTGGAACAATGTAGGTAAGGGAAGT | 463.9 | -6.1 | 1.1 | -5.6 | 2.2E-08 | 4.1E-08 | MB231/MB468 | MITO |
| >hg38_21_+_8433172_8446622_RNA45SN1_45S_50ntFlanks@3986.4026.41\|GGCTTTGGTGACTCTAGATAACCTCGGGCCGATCGCACGCC | >hg38_21_+_8433172_8446622_RNA45SN1_45S_50ntFlanks | 3986 | 4026 | 41 | GGCTTTGGTGACTCTAGATAACCTCGGGCCGATCGCACGCC | 81.8 | -10.2 | 1.8 | -5.6 | 1.9E-08 | 4.1E-08 | MB231/MB468 | MP |
| >hg38_21_+_8433172_8446622_RNA45SN1_45S_50ntFlanks@8116.8134.19\|GCGGGACATGTGGCGTACG | >hg38_21_+_8433172_8446622_RNA45SN1_45S_50ntFlanks | 8116 | 8134 | 19 | GCGGGACATGTGGCGTACG | 712.0 | 12.8 | 2.1 | 6.2 | 5.9E-10 | 4.1E-08 | BT20/MB231 | MP |
| >MI0000077\|hsa-mir-21&WithFlank&17\|+\|59841260\|59841343@14.36.23(+1G)&[MIMAT0000076&hsa-miR-21-5p&offsets\|0\|+1(+1G);m-1&17\|+\|59841273\|59841294&offsets\|0\|+1(+1G)]\|TAGCTTATCAGACTGATGTTGACG | >MI0000077\|hsa-mir-21&WithFlank&17\|+\|59841260\|59841343 | 14 | 36 | 23(+1G) | TAGCTTATCAGACTGATGTTGACG | 5552.1 | 5.4 | 1.0 | 5.6 | 2.2E-08 | 4.1E-08 | MB231/MB468 | MITO |
| >hg38_MT_+_1621_3279_RNR2_16S_50ntFlanks@1348.1387.40\|AACAATAGGGTTTACGACCTCGATGTTGGATCAGGACATC | >hg38_MT_+_1621_3279_RNR2_16S_50ntFlanks | 1348 | 1387 | 40 | AACAATAGGGTTTACGACCTCGATGTTGGATCAGGACATC | 60.2 | 9.1 | 1.6 | 5.6 | 1.9E-08 | 4.1E-08 | MB231/MB468 | MP |
| >hg38_21_+_8433172_8446622_RNA45SN1_45S_50ntFlanks@12934.12969.36\|CTAAACCATTCGTAGACGACCTGCTTCTGGGTCGGG | >hg38_21_+_8433172_8446622_RNA45SN1_45S_50ntFlanks | 12934 | 12969 | 36 | CTAAACCATTCGTAGACGACCTGCTTCTGGGTCGGG | 580.6 | -7.5 | 1.3 | -5.6 | 1.8E-08 | 4.1E-08 | BT20/MB468 | CYTO |
| >hg38_21_+_8433172_8446622_RNA45SN1_45S_50ntFlanks@12456.12489.34\|CCAAGCGTTGGATTGTTCACCCACTAATAGGGAA | >hg38_21_+_8433172_8446622_RNA45SN1_45S_50ntFlanks | 12456 | 12489 | 34 | CCAAGCGTTGGATTGTTCACCCACTAATAGGGAA | 988.2 | -7.4 | 1.3 | -5.6 | 1.8E-08 | 4.1E-08 | BT20/MB468 | CYTO |
| >hg38_21_+_8433172_8446622_RNA45SN1_45S_50ntFlanks@3872.3898.27\|CTAATACATGCCGACGGGCGCTGACCC | >hg38_21_+_8433172_8446622_RNA45SN1_45S_50ntFlanks | 3872 | 3898 | 27 | CTAATACATGCCGACGGGCGCTGACCC | 1487.6 | -4.7 | 0.8 | -5.6 | 2.1E-08 | 4.1E-08 | MB231/MB468 | CYTO |
| >hg38_1_-_228634819_228635039_RNA5S12_5S_50ntFlanks@148.171.24\|GGAATACCGGGTGCTGTAGGCTTT | >hg38_1_-_228634819_228635039_RNA5S12_5S_50ntFlanks | 148 | 171 | 24 | GGAATACCGGGTGCTGTAGGCTTT | 5858.6 | 3.4 | 0.6 | 5.6 | 1.8E-08 | 4.1E-08 | BT20/MB468 | CYTO |
| >hg38_21_+_8433172_8446622_RNA45SN1_45S_50ntFlanks@12949.12973.25\|ACGACCTGCTTCTGGGTCGGGGTTT | >hg38_21_+_8433172_8446622_RNA45SN1_45S_50ntFlanks | 12949 | 12973 | 25 | ACGACCTGCTTCTGGGTCGGGGTTT | 83.1 | -10.2 | 1.8 | -5.6 | 1.9E-08 | 4.1E-08 | MB231/MB468 | MP |
| >hg38_21_+_8433172_8446622_RNA45SN1_45S_50ntFlanks@9572.9603.32\|CAAATCGGTCGTCCGACCTGGGTATAGGGGCG | >hg38_21_+_8433172_8446622_RNA45SN1_45S_50ntFlanks | 9572 | 9603 | 32 | CAAATCGGTCGTCCGACCTGGGTATAGGGGCG | 1159.4 | -3.4 | 0.6 | -5.6 | 1.8E-08 | 4.1E-08 | BT20/MB468 | CYTO |
| >MI0000253\|hsa-mir-148a&WithFlank&7\|-\|25949913\|25949992@50.71.22&[MIMAT0000243&hsa-miR-148a-3p&offsets\|0\|0;m-7&7\|-\|25949922\|25949943&offsets\|0\|0]\|TCAGTGCACTACAGAACTTTGT | >MI0000253\|hsa-mir-148a&WithFlank&7\|-\|25949913\|25949992 | 50 | 71 | 22 | TCAGTGCACTACAGAACTTTGT | 137345.2 | 4.9 | 0.8 | 6.1 | 1.2E-09 | 4.1E-08 | BT20/MB231 | CYTO |
| >hg38_21_+_8433172_8446622_RNA45SN1_45S_50ntFlanks@12676.12707.32\|AGAATCCCGCCCAGGCGGAACGATACGGCAGC | >hg38_21_+_8433172_8446622_RNA45SN1_45S_50ntFlanks | 12676 | 12707 | 32 | AGAATCCCGCCCAGGCGGAACGATACGGCAGC | 92.0 | -9.6 | 1.6 | -5.9 | 3.2E-09 | 4.1E-08 | BT20/MB468 | MP |
| >hg38_21_+_8433172_8446622_RNA45SN1_45S_50ntFlanks@11921.11956.36\|TGAGAGGTGTAGAATAAGTGGGAGGCCCCCGGCGCC | >hg38_21_+_8433172_8446622_RNA45SN1_45S_50ntFlanks | 11921 | 11956 | 36 | TGAGAGGTGTAGAATAAGTGGGAGGCCCCCGGCGCC | 68.6 | -10.0 | 1.8 | -5.6 | 1.9E-08 | 4.1E-08 | MB231/MB468 | MP |
| >hg38_21_+_8433172_8446622_RNA45SN1_45S_50ntFlanks@12935.12966.32\|TAAACCATTCGTAGACGACCTGCTTCTGGGTC | >hg38_21_+_8433172_8446622_RNA45SN1_45S_50ntFlanks | 12935 | 12966 | 32 | TAAACCATTCGTAGACGACCTGCTTCTGGGTC | 64.1 | -9.9 | 1.8 | -5.6 | 1.9E-08 | 4.1E-08 | MB231/MB468 | MP |
| >hg38_21_+_8433172_8446622_RNA45SN1_45S_50ntFlanks@11781.11820.40\|CGCATGAATGGATGAACGAGATTCCCACTGTCCCTACCTA | >hg38_21_+_8433172_8446622_RNA45SN1_45S_50ntFlanks | 11781 | 11820 | 40 | CGCATGAATGGATGAACGAGATTCCCACTGTCCCTACCTA | 131.8 | -9.1 | 1.6 | -5.7 | 1.6E-08 | 4.1E-08 | BT20/MB468 | MITO |
| >hg38_1_-_228634819_228635039_RNA5S12_5S_50ntFlanks@51.90.40\|GTCTACGGCCATACCACCCTGAACGCGCCCGATCTCGTCT | >hg38_1_-_228634819_228635039_RNA5S12_5S_50ntFlanks | 51 | 90 | 40 | GTCTACGGCCATACCACCCTGAACGCGCCCGATCTCGTCT | 2328.6 | 8.0 | 1.4 | 5.6 | 1.8E-08 | 4.1E-08 | BT20/MB468 | CYTO |
| >am_trnaMT_ProTGG_MT_-_15956_16023@54.71.18\|AGACTTTTTCTCTGACCA | >am_trnaMT_ProTGG_MT_-_15956_16023 | 54 | 71 | 18 | AGACTTTTTCTCTGACCA | 3646.7 | 5.0 | 0.9 | 5.6 | 1.8E-08 | 4.1E-08 | BT20/MB468 | CYTO |
| >hg38_21_+_8433172_8446622_RNA45SN1_45S_50ntFlanks@11515.11539.25\|GTCCGCCCCCGGGGCCGCGGTTCCG | >hg38_21_+_8433172_8446622_RNA45SN1_45S_50ntFlanks | 11515 | 11539 | 25 | GTCCGCCCCCGGGGCCGCGGTTCCG | 204.2 | -6.1 | 1.1 | -5.6 | 2.1E-08 | 4.1E-08 | MB231/MB468 | CYTO |
| >MI0000253\|hsa-mir-148a&WithFlank&7\|-\|25949913\|25949992@50.71.22(+1U)&[MIMAT0000243&hsa-miR-148a-3p&offsets\|0\|0(+1U);m-7&7\|-\|25949922\|25949943&offsets\|0\|0(+1U)]\|TCAGTGCACTACAGAACTTTGTT | >MI0000253\|hsa-mir-148a&WithFlank&7\|-\|25949913\|25949992 | 50 | 71 | 22(+1U) | TCAGTGCACTACAGAACTTTGTT | 234.4 | 3.7 | 0.7 | 5.6 | 2.3E-08 | 4.1E-08 | BT20/MB468 | NUC |
| >trna116_GluCTC_1_-_145399233_145399304@3.35.33,trna59_GluCTC_1_+_249168447_249168518@3.35.33,trna71_GluCTC_1_-_161439189_161439260@3.35.33,trna74_GluCTC_1_-_161431809_161431880@3.35.33,trna77_GluCTC_1_-_161424398_161424469@3.35.33,trna77_GluCTC_6_+_28949976_28950047@3.35.33,trna80_GluCTC_1_-_161417018_161417089@3.35.33,trna87_GluCTC_6_-_126101393_126101464@3.35.33\|CCTGGTGGTCTAGTGGTTAGGATTCGGCGCTCT | >trna116_GluCTC_1_-_145399233_145399304 | 3 | 35 | 33 | CCTGGTGGTCTAGTGGTTAGGATTCGGCGCTCT | 617.8 | 4.6 | 0.8 | 5.6 | 2.3E-08 | 4.1E-08 | MB231/MB468 | MITO |
| >MI0000433\|hsa-let-7g&WithFlank&3\|-\|52268272\|52268367@11.33.23&[MIMAT0000414&hsa-let-7g-5p&offsets\|0\|+1;m-58&3\|-\|52268336\|52268357&offsets\|0\|+1]\|TGAGGTAGTAGTTTGTACAGTTT | >MI0000433\|hsa-let-7g&WithFlank&3\|-\|52268272\|52268367 | 11 | 33 | 23 | TGAGGTAGTAGTTTGTACAGTTT | 610.1 | 5.0 | 0.9 | 5.6 | 2.3E-08 | 4.1E-08 | MB231/MB468 | MITO |
| >hg38_21_+_8433172_8446622_RNA45SN1_45S_50ntFlanks@5494.5526.33\|AAGACGGTCGAACTTGACTATCTAGAGGAAGTA | >hg38_21_+_8433172_8446622_RNA45SN1_45S_50ntFlanks | 5494 | 5526 | 33 | AAGACGGTCGAACTTGACTATCTAGAGGAAGTA | 526.5 | -4.1 | 0.7 | -5.7 | 1.6E-08 | 4.1E-08 | BT20/MB468 | MITO |
| >MI0000255\|hsa-mir-30d&WithFlank&8\|-\|134804870\|134804951@12.33.22(+1A)&[MIMAT0000245&hsa-miR-30d-5p&offsets\|0\|0(+1A);m-21&8\|-\|134804919\|134804940&offsets\|0\|0(+1A)]\|TGTAAACATCCCCGACTGGAAGA | >MI0000255\|hsa-mir-30d&WithFlank&8\|-\|134804870\|134804951 | 12 | 33 | 22(+1A) | TGTAAACATCCCCGACTGGAAGA | 664.3 | 5.1 | 0.9 | 5.6 | 2.3E-08 | 4.1E-08 | MB231/MB468 | MITO |
| >hg38_21_+_8433172_8446622_RNA45SN1_45S_50ntFlanks@10841.10861.21\|CGTAACTTCGGGATAAGGATT | >hg38_21_+_8433172_8446622_RNA45SN1_45S_50ntFlanks | 10841 | 10861 | 21 | CGTAACTTCGGGATAAGGATT | 979.1 | -5.1 | 0.9 | -5.6 | 2.3E-08 | 4.1E-08 | MB231/MB468 | MITO |
| >hg38_21_+_8433172_8446622_RNA45SN1_45S_50ntFlanks@10781.10799.19\|AGGTGAACAGCCTCTGGCA | >hg38_21_+_8433172_8446622_RNA45SN1_45S_50ntFlanks | 10781 | 10799 | 19 | AGGTGAACAGCCTCTGGCA | 362.4 | -7.5 | 1.3 | -5.6 | 2.0E-08 | 4.2E-08 | MB231/MB468 | MP |
| >hg38_21_+_8433172_8446622_RNA45SN1_45S_50ntFlanks@3979.4001.23\|CGCCGGCGGCTTTGGTGACTCTA | >hg38_21_+_8433172_8446622_RNA45SN1_45S_50ntFlanks | 3979 | 4001 | 23 | CGCCGGCGGCTTTGGTGACTCTA | 123.1 | -10.8 | 1.9 | -5.6 | 2.0E-08 | 4.2E-08 | MB231/MB468 | MP |
| >hg38_21_+_8433172_8446622_RNA45SN1_45S_50ntFlanks@9337.9370.34\|CTCGCACGAAAGCCGCCGTGGCGCAATGAAGGTG | >hg38_21_+_8433172_8446622_RNA45SN1_45S_50ntFlanks | 9337 | 9370 | 34 | CTCGCACGAAAGCCGCCGTGGCGCAATGAAGGTG | 129.0 | -9.1 | 1.6 | -5.6 | 1.6E-08 | 4.2E-08 | BT20/MB468 | MITO |
| >hg38_21_+_8433172_8446622_RNA45SN1_45S_50ntFlanks@7975.8005.31\|CGCGACCTCAGATCAGACGTGGCGACCCGCT | >hg38_21_+_8433172_8446622_RNA45SN1_45S_50ntFlanks | 7975 | 8005 | 31 | CGCGACCTCAGATCAGACGTGGCGACCCGCT | 13019.4 | 3.5 | 0.6 | 5.6 | 2.1E-08 | 4.2E-08 | MB231/MB468 | CYTO |
| >hg38_21_+_8433172_8446622_RNA45SN1_45S_50ntFlanks@6723.6744.22\|TTGCAGGACACATTGATCATCG | >hg38_21_+_8433172_8446622_RNA45SN1_45S_50ntFlanks | 6723 | 6744 | 22 | TTGCAGGACACATTGATCATCG | 272.5 | -5.7 | 1.0 | -5.6 | 2.3E-08 | 4.2E-08 | MB231/MB468 | MITO |
| >hg38_21_+_8433172_8446622_RNA45SN1_45S_50ntFlanks@12894.12924.31\|CTCGCCCGTCACGCACCGCACGTTCGTGGGG | >hg38_21_+_8433172_8446622_RNA45SN1_45S_50ntFlanks | 12894 | 12924 | 31 | CTCGCCCGTCACGCACCGCACGTTCGTGGGG | 100.8 | -9.7 | 1.6 | -5.9 | 3.2E-09 | 4.2E-08 | BT20/MB468 | MP |
| >MI0000285\|hsa-mir-205&WithFlank&1\|+\|209432127\|209432248@40.59.20(+1A)&[MIMAT0000266&hsa-miR-205-5p&offsets\|0\|-2(+1A);m-351&1\|+\|209432166\|209432187&offsets\|0\|-2(+1A)]\|TCCTTCATTCCACCGGAGTCA | >MI0000285\|hsa-mir-205&WithFlank&1\|+\|209432127\|209432248 | 40 | 59 | 20(+1A) | TCCTTCATTCCACCGGAGTCA | 353.6 | 11.8 | 1.9 | 6.2 | 6.1E-10 | 4.2E-08 | BT20/MB231 | MP |
| >MI0005544\|hsa-mir-147b&WithFlank&15\|+\|45433044\|45433135@55.73.19&[MIMAT0004928&hsa-miR-147b-3p&offsets\|0\|-2;m-575&15\|+\|45433098\|45433118&offsets\|0\|-2]\|GTGTGCGGAAATGCTTCTG | >MI0005544\|hsa-mir-147b&WithFlank&15\|+\|45433044\|45433135 | 55 | 73 | 19 | GTGTGCGGAAATGCTTCTG | 188.1 | -11.0 | 1.8 | -6.2 | 6.1E-10 | 4.2E-08 | BT20/MB231 | MP |
| >hg38_21_+_8433172_8446622_RNA45SN1_45S_50ntFlanks@11670.11701.32\|CCCAGTGCTCTGAATGTCAAAGTGAAGAAATT | >hg38_21_+_8433172_8446622_RNA45SN1_45S_50ntFlanks | 11670 | 11701 | 32 | CCCAGTGCTCTGAATGTCAAAGTGAAGAAATT | 6094.2 | -10.1 | 1.8 | -5.6 | 2.4E-08 | 4.2E-08 | BT20/MB468 | NUC |
| >hg38_21_+_8433172_8446622_RNA45SN1_45S_50ntFlanks@9545.9575.31\|GTGGAGGTCCGTAGCGGTCCTGACGTGCAAA | >hg38_21_+_8433172_8446622_RNA45SN1_45S_50ntFlanks | 9545 | 9575 | 31 | GTGGAGGTCCGTAGCGGTCCTGACGTGCAAA | 406.4 | -7.4 | 1.3 | -5.6 | 2.3E-08 | 4.2E-08 | MB231/MB468 | MITO |
| >hg38_21_+_8433172_8446622_RNA45SN1_45S_50ntFlanks@9862.9893.32\|ATGAACCGAACGCCGGGTTAAGGCGCCCGATG | >hg38_21_+_8433172_8446622_RNA45SN1_45S_50ntFlanks | 9862 | 9893 | 32 | ATGAACCGAACGCCGGGTTAAGGCGCCCGATG | 519.2 | -8.2 | 1.5 | -5.6 | 2.3E-08 | 4.2E-08 | MB231/MB468 | MITO |
| >hg38_21_+_8433172_8446622_RNA45SN1_45S_50ntFlanks@9534.9562.29\|AGGAAACTCTGGTGGAGGTCCGTAGCGGT | >hg38_21_+_8433172_8446622_RNA45SN1_45S_50ntFlanks | 9534 | 9562 | 29 | AGGAAACTCTGGTGGAGGTCCGTAGCGGT | 3366.1 | -8.8 | 1.6 | -5.6 | 2.0E-08 | 4.2E-08 | MB231/MB468 | MP |
| >hg38_21_+_8433172_8446622_RNA45SN1_45S_50ntFlanks@10381.10424.44\|GTTCCATGTGAACAGCAGTTGAACATGGGTCAGTCGGTCCTGAG | >hg38_21_+_8433172_8446622_RNA45SN1_45S_50ntFlanks | 10381 | 10424 | 44 | GTTCCATGTGAACAGCAGTTGAACATGGGTCAGTCGGTCCTGAG | 705.5 | 9.7 | 1.6 | 5.9 | 3.3E-09 | 4.2E-08 | BT20/MB468 | MP |
| >hg38_21_+_8433172_8446622_RNA45SN1_45S_50ntFlanks@4683.4707.25\|CAAGACGGACCAGAGCGAAAGCATT | >hg38_21_+_8433172_8446622_RNA45SN1_45S_50ntFlanks | 4683 | 4707 | 25 | CAAGACGGACCAGAGCGAAAGCATT | 500.8 | -4.4 | 0.8 | -5.6 | 2.2E-08 | 4.2E-08 | MB231/MB468 | NUC |
| >hg38_21_+_8433172_8446622_RNA45SN1_45S_50ntFlanks@12628.12654.27\|TGGGGCGAAGCTACCATCTGTGGGATT | >hg38_21_+_8433172_8446622_RNA45SN1_45S_50ntFlanks | 12628 | 12654 | 27 | TGGGGCGAAGCTACCATCTGTGGGATT | 1655.6 | -4.5 | 0.8 | -5.6 | 1.8E-08 | 4.2E-08 | BT20/MB468 | CYTO |
| >hg38_21_+_8433172_8446622_RNA45SN1_45S_50ntFlanks@11729.11747.19\|TAACTATGACTCTCTTAAG | >hg38_21_+_8433172_8446622_RNA45SN1_45S_50ntFlanks | 11729 | 11747 | 19 | TAACTATGACTCTCTTAAG | 5508.4 | -4.4 | 0.8 | -5.6 | 1.8E-08 | 4.2E-08 | BT20/MB468 | CYTO |
| >trnaMT_ValTAC_MT_+_1602_1670@49.72.24\|AACTTAACTTGACCGCTCTGACCA | >trnaMT_ValTAC_MT_+_1602_1670 | 49 | 72 | 24 | AACTTAACTTGACCGCTCTGACCA | 229.0 | 5.5 | 1.0 | 5.6 | 2.1E-08 | 4.2E-08 | MB231/MB468 | CYTO |
| >hg38_21_+_8433172_8446622_RNA45SN1_45S_50ntFlanks@4296.4317.22\|CCATTGGAGGGCAAGTCTGGTG | >hg38_21_+_8433172_8446622_RNA45SN1_45S_50ntFlanks | 4296 | 4317 | 22 | CCATTGGAGGGCAAGTCTGGTG | 104.5 | -9.8 | 1.7 | -5.9 | 3.3E-09 | 4.2E-08 | BT20/MB468 | MP |
| >hg38_21_+_8433172_8446622_RNA45SN1_45S_50ntFlanks@4367.4408.42\|CAGTTAAAAAGCTCGTAGTTGGATCTTGGGAGCGGGCGGGCG | >hg38_21_+_8433172_8446622_RNA45SN1_45S_50ntFlanks | 4367 | 4408 | 42 | CAGTTAAAAAGCTCGTAGTTGGATCTTGGGAGCGGGCGGGCG | 112.1 | -9.1 | 1.6 | -5.6 | 1.8E-08 | 4.2E-08 | BT20/MB468 | CYTO |

| | | | | | | | | | | | | | |
|---|---|---|---|---|---|---|---|---|---|---|---|---|---|
| >hg38_21_+_8433172_8446622_RNA45SN1_45S_50ntFlanks@12413.12441.29\|TCCTTCGATGTCGGCTCTTCCTATCATTG | >hg38_21_+_8433172_8446622_RNA45SN1_45S_50ntFlanks | 12413 | 12441 | 29 | TCCTTCGATGTCGGCTCTTCCTATCATTG | 812.1 | -8.7 | 1.6 | -5.6 | 2.4E-08 | 4.2E-08 | BT20/MB468 | NUC |
| >hg38_21_+_8433172_8446622_RNA45SN1_45S_50ntFlanks@12044.12065.22\|ACTGACCCGGTGAGGCGGGGGG | >hg38_21_+_8433172_8446622_RNA45SN1_45S_50ntFlanks | 12044 | 12065 | 22 | ACTGACCCGGTGAGGCGGGGGG | 88.6 | -10.3 | 1.8 | -5.6 | 2.0E-08 | 4.3E-08 | MB231/MB468 | MP |
| >trna112_GlnCTG_1_-_147737382_147737453@30.52.23,trna131_GlnCTG_6_-_27759135_27759206@30.52.23,trna146_GlnCTG_6_-_27515531_27515602@30.52.23,trna15_GlnCTG_1_+_145963304_145963375@30.52.23,trna19_GlnCTG_1_+_147505038_147505109@30.52.23,trna1_GlnCTG_6_+_18836402_18836473@30.52.23,trna28_GlnCTG_1_+_149186125_149186196@30.52.23,trna3_GlnCTG_17_+_8023070_8023141@30.52.23,trna49_GlnCTG_6_+_27487308_27487379@30.52.23,trna7_GlnCTG_15_-_66161400_66161471@30.52.23,trna99_GlnCTG_6_-_28909378_28909449@30.52.23\|GACTCTGAATCCAGCGATCCGAG | >trna112_GlnCTG_1_-_147737382_147737453 | 30 | 52 | 23 | GACTCTGAATCCAGCGATCCGAG | 204.3 | -10.8 | 1.8 | -5.9 | 3.3E-09 | 4.3E-08 | BT20/MB468 | MP |
| >hg38_21_+_8433172_8446622_RNA45SN1_45S_50ntFlanks@5077.5110.34\|CCGATAACGAACGAGACTCTGGCATGCTAACTAG | >hg38_21_+_8433172_8446622_RNA45SN1_45S_50ntFlanks | 5077 | 5110 | 34 | CCGATAACGAACGAGACTCTGGCATGCTAACTAG | 160.3 | -11.2 | 2.0 | -5.6 | 2.0E-08 | 4.3E-08 | MB231/MB468 | MP |
| >hg38_21_+_8433172_8446622_RNA45SN1_45S_50ntFlanks@8114.8132.19\|GCGCGGGACATGTGGCGTA | >hg38_21_+_8433172_8446622_RNA45SN1_45S_50ntFlanks | 8114 | 8132 | 19 | GCGCGGGACATGTGGCGTA | 367.3 | -8.0 | 1.4 | -5.6 | 2.0E-08 | 4.3E-08 | MB231/MB468 | MP |
| >MI0000078\|hsa-mir-22&WithFlank&17\|-\|1713897\|1713993@59.79.21(+1A)&[MIMAT0000077&hsa-miR-22-3p&offsets\|0\|-1(+1A);m-2&17\|-\|1713914\|1713935&offsets\|0\|-1(+1A)]\|AAGCTGCCAGTTGAAGAACTGA | >MI0000078\|hsa-mir-22&WithFlank&17\|-\|1713897\|1713993 | 59 | 79 | 21(+1A) | AAGCTGCCAGTTGAAGAACTGA | 89.7 | 6.5 | 1.2 | 5.6 | 2.4E-08 | 4.3E-08 | BT20/MB468 | NUC |
| >MI0000077\|hsa-mir-21&WithFlank&17\|+\|59841260\|59841343@13.35.23&[MIMAT0000076&hsa-miR-21-5p&offsets\|-1\|0;m-1&17\|+\|59841273\|59841294&offsets\|-1\|0]\|GTAGCTTATCAGACTGATGTTGA | >MI0000077\|hsa-mir-21&WithFlank&17\|+\|59841260\|59841343 | 13 | 35 | 23 | GTAGCTTATCAGACTGATGTTGA | 580.9 | 5.5 | 1.0 | 5.6 | 2.3E-08 | 4.3E-08 | MB231/MB468 | MITO |
| >hg38_21_+_8433172_8446622_RNA45SN1_45S_50ntFlanks@10346.10371.26\|TCAAACGAGAACTTTGAAGGCCGAAG | >hg38_21_+_8433172_8446622_RNA45SN1_45S_50ntFlanks | 10346 | 10371 | 26 | TCAAACGAGAACTTTGAAGGCCGAAG | 229.5 | -3.7 | 0.7 | -5.6 | 2.0E-08 | 4.3E-08 | MB231/MB468 | MP |
| >hg38_21_+_8433172_8446622_RNA45SN1_45S_50ntFlanks@5017.5034.18\|ATTCCGTGGGTGGTGGTG | >hg38_21_+_8433172_8446622_RNA45SN1_45S_50ntFlanks | 5017 | 5034 | 18 | ATTCCGTGGGTGGTGGTG | 738.0 | -11.6 | 2.1 | -5.6 | 1.6E-08 | 4.3E-08 | BT20/MB468 | MITO |
| >hg38_21_+_8433172_8446622_RNA45SN1_45S_50ntFlanks@8196.8225.30\|CGTGGACGGTGTGAGGCCGGTAGCGGCCCC | >hg38_21_+_8433172_8446622_RNA45SN1_45S_50ntFlanks | 8196 | 8225 | 30 | CGTGGACGGTGTGAGGCCGGTAGCGGCCCC | 332.2 | -7.6 | 1.4 | -5.6 | 2.0E-08 | 4.3E-08 | MB231/MB468 | MP |
| >hg38_21_+_8433172_8446622_RNA45SN1_45S_50ntFlanks@12627.12661.35\|ATGGGGCGAAGCTACCATCTGTGGGATTATGACTG | >hg38_21_+_8433172_8446622_RNA45SN1_45S_50ntFlanks | 12627 | 12661 | 35 | ATGGGGCGAAGCTACCATCTGTGGGATTATGACTG | 1702.5 | -9.2 | 1.6 | -5.6 | 2.4E-08 | 4.3E-08 | BT20/MB468 | NUC |
| >hg38_21_+_8433172_8446622_RNA45SN1_45S_50ntFlanks@10465.10497.33\|CCGTTGCCCTCGGCCGATCGAAAGGGAGTCGGG | >hg38_21_+_8433172_8446622_RNA45SN1_45S_50ntFlanks | 10465 | 10497 | 33 | CCGTTGCCCTCGGCCGATCGAAAGGGAGTCGGG | 100.9 | -9.7 | 1.6 | -5.9 | 3.3E-09 | 4.3E-08 | BT20/MB468 | MP |
| >MI0000294\|hsa-mir-218-1&WithFlank&4\|+\|20528269\|20528390@31.50.20(+1C);MI0000295\|hsa-mir-218-2&WithFlank&5\|-\|168768140\|168768261@31.50.20(+1C)&[MIMAT0000275&hsa-miR-218-5p&offsets\|0\|-1(+1C);m-185&4\|+\|20528299\|20528319&offsets\|0\|-1(+1C)];[MIMAT0000275_1&hsa-miR-218-5p&offsets\|0\|-1(+1C);m-187&5\|-\|168768211\|168768231&offsets\|0\|-1(+1C)]\|TTGTGCTTGATCTAACCATGC | >MI0000294\|hsa-mir-218-1&WithFlank&4\|+\|20528269\|20528390 | 31 | 50 | 20(+1C) | TTGTGCTTGATCTAACCATGC | 87.4 | 9.7 | 1.7 | 5.6 | 2.0E-08 | 4.3E-08 | MB231/MB468 | MP |
| >hg38_21_+_8433172_8446622_RNA45SN1_45S_50ntFlanks@11804.11821.18\|CCCACTGTCCCTACCTAC | >hg38_21_+_8433172_8446622_RNA45SN1_45S_50ntFlanks | 11804 | 11821 | 18 | CCCACTGTCCCTACCTAC | 232.4 | -4.8 | 0.8 | -5.6 | 2.1E-08 | 4.3E-08 | MB231/MB468 | CYTO |
| >hg38_21_+_8433172_8446622_RNA45SN1_45S_50ntFlanks@8176.8220.45\|CTTCTGATCGAGGCCCAGCCCGTGGACGGTGTGAGGCCGGTAGCG | >hg38_21_+_8433172_8446622_RNA45SN1_45S_50ntFlanks | 8176 | 8220 | 45 | CTTCTGATCGAGGCCCAGCCCGTGGACGGTGTGAGGCCGGTAGCG | 989.2 | -5.7 | 1.0 | -5.6 | 2.4E-08 | 4.3E-08 | MB231/MB468 | MITO |
| >hg38_21_+_8433172_8446622_RNA45SN1_45S_50ntFlanks@6650.6669.20\|ACGACTCTTAGCGGTGGATC | >hg38_21_+_8433172_8446622_RNA45SN1_45S_50ntFlanks | 6650 | 6669 | 20 | ACGACTCTTAGCGGTGGATC | 1430.1 | 3.3 | 0.6 | 5.6 | 1.8E-08 | 4.3E-08 | BT20/MB468 | CYTO |
| >hg38_21_+_8433172_8446622_RNA45SN1_45S_50ntFlanks@6733.6752.20\|CATTGATCATCGACACTTCG | >hg38_21_+_8433172_8446622_RNA45SN1_45S_50ntFlanks | 6733 | 6752 | 20 | CATTGATCATCGACACTTCG | 1350.5 | -6.5 | 1.2 | -5.6 | 2.0E-08 | 4.3E-08 | MB231/MB468 | MP |
| >hg38_21_+_8433172_8446622_RNA45SN1_45S_50ntFlanks@11573.11591.19\|CGACTTAGAACTGGTGCGG | >hg38_21_+_8433172_8446622_RNA45SN1_45S_50ntFlanks | 11573 | 11591 | 19 | CGACTTAGAACTGGTGCGG | 362.4 | -4.9 | 0.9 | -5.6 | 2.1E-08 | 4.3E-08 | MB231/MB468 | CYTO |
| >MI0000076\|hsa-mir-20a&WithFlank&13\|+\|91351059\|91351141@14.35.22&[MIMAT0000075&hsa-miR-20a-5p&offsets\|0\|-1;m-96&13\|+\|91351072\|91351094&offsets\|0\|-1]\|TAAAGTGCTTATAGTGCAGGTA | >MI0000076\|hsa-mir-20a&WithFlank&13\|+\|91351059\|91351141 | 14 | 35 | 22 | TAAAGTGCTTATAGTGCAGGTA | 289.7 | 6.0 | 1.1 | 5.6 | 2.2E-08 | 4.3E-08 | MB231/MB468 | NUC |
| >hg38_21_+_8433172_8446622_RNA45SN1_45S_50ntFlanks@7990.8010.21\|GACGTGGCGACCCGCTGAATT | >hg38_21_+_8433172_8446622_RNA45SN1_45S_50ntFlanks | 7990 | 8010 | 21 | GACGTGGCGACCCGCTGAATT | 245.4 | 5.7 | 1.0 | 5.6 | 1.8E-08 | 4.3E-08 | BT20/MB468 | CYTO |
| >hg38_21_+_8433172_8446622_RNA45SN1_45S_50ntFlanks@10726.10755.30\|GGGAGAGGGTGTAAATCTCGCGCCGGGCCG | >hg38_21_+_8433172_8446622_RNA45SN1_45S_50ntFlanks | 10726 | 10755 | 30 | GGGAGAGGGTGTAAATCTCGCGCCGGGCCG | 165.0 | -5.7 | 1.0 | -5.6 | 2.2E-08 | 4.3E-08 | MB231/MB468 | CYTO |
| >hg38_21_+_8433172_8446622_RNA45SN1_45S_50ntFlanks@10823.10865.43\|AAGTCGGCAAGCCGGATCCGTAACTTCGGGATAAGGATTGGCT | >hg38_21_+_8433172_8446622_RNA45SN1_45S_50ntFlanks | 10823 | 10865 | 43 | AAGTCGGCAAGCCGGATCCGTAACTTCGGGATAAGGATTGGCT | 681.6 | -8.0 | 1.4 | -5.6 | 1.8E-08 | 4.3E-08 | BT20/MB468 | CYTO |
| >hg38_21_+_8433172_8446622_RNA45SN1_45S_50ntFlanks@485.521.37\|GATGTGGTGACGTCGTGCTCTCCCGGGCCGGGTCCGA | >hg38_21_+_8433172_8446622_RNA45SN1_45S_50ntFlanks | 485 | 521 | 37 | GATGTGGTGACGTCGTGCTCTCCCGGGCCGGGTCCGA | 136.0 | 10.8 | 1.8 | 5.9 | 3.3E-09 | 4.3E-08 | BT20/MB468 | MP |
| >hg38_21_+_8433172_8446622_RNA45SN1_45S_50ntFlanks@11873.11902.30\|AAGAAGACCCTGTTGAGCTTGACTCTAGTC | >hg38_21_+_8433172_8446622_RNA45SN1_45S_50ntFlanks | 11873 | 11902 | 30 | AAGAAGACCCTGTTGAGCTTGACTCTAGTC | 204.6 | -7.2 | 1.3 | -5.6 | 2.4E-08 | 4.3E-08 | MB231/MB468 | MITO |
| >hg38_21_+_8433172_8446622_RNA45SN1_45S_50ntFlanks@3903.3932.30\|CGCGGGGGGGATGCGTGCATTTATCAGATC | >hg38_21_+_8433172_8446622_RNA45SN1_45S_50ntFlanks | 3903 | 3932 | 30 | CGCGGGGGGGATGCGTGCATTTATCAGATC | 106.9 | -10.6 | 1.9 | -5.6 | 2.0E-08 | 4.3E-08 | MB231/MB468 | MP |
| >hg38_21_+_8433172_8446622_RNA45SN1_45S_50ntFlanks@12008.12040.33\|CCGCCGGTGAAATACCACTACTCTGATCGTTTT | >hg38_21_+_8433172_8446622_RNA45SN1_45S_50ntFlanks | 12008 | 12040 | 33 | CCGCCGGTGAAATACCACTACTCTGATCGTTTT | 62.0 | -9.8 | 1.8 | -5.6 | 2.0E-08 | 4.3E-08 | MB231/MB468 | MP |
| >hg38_21_+_8433172_8446622_RNA45SN1_45S_50ntFlanks@12420.12443.24\|ATGTCGGCTCTTCCTATCATTGTG | >hg38_21_+_8433172_8446622_RNA45SN1_45S_50ntFlanks | 12420 | 12443 | 24 | ATGTCGGCTCTTCCTATCATTGTG | 1848.8 | -7.2 | 1.3 | -5.6 | 2.0E-08 | 4.3E-08 | MB231/MB468 | MP |
| >hg38_21_+_8433172_8446622_RNA45SN1_45S_50ntFlanks@5045.5065.21\|CTTAGTTGGTGGAGCGATTTG | >hg38_21_+_8433172_8446622_RNA45SN1_45S_50ntFlanks | 5045 | 5065 | 21 | CTTAGTTGGTGGAGCGATTTG | 122.2 | -10.8 | 1.9 | -5.6 | 2.0E-08 | 4.3E-08 | MB231/MB468 | MP |
| >am_trnaMT_PheGAA_MT_+_577_647@42.62.21\|TTTAGACGGGCTCACATCACC | >am_trnaMT_PheGAA_MT_+_577_647 | 42 | 62 | 21 | TTTAGACGGGCTCACATCACC | 237.3 | 6.6 | 1.2 | 5.6 | 2.2E-08 | 4.3E-08 | MB231/MB468 | CYTO |
| >hg38_21_+_8433172_8446622_RNA45SN1_45S_50ntFlanks@10795.10818.24\|TGGCATGTTGGAACAATGTAGGTA | >hg38_21_+_8433172_8446622_RNA45SN1_45S_50ntFlanks | 10795 | 10818 | 24 | TGGCATGTTGGAACAATGTAGGTA | 1171.8 | -5.6 | 1.0 | -5.6 | 2.2E-08 | 4.3E-08 | MB231/MB468 | CYTO |
| >hg38_21_+_8433172_8446622_RNA45SN1_45S_50ntFlanks@11318.11344.27\|CCCCACGTCTCGTCGCGCGCGCGTCCG | >hg38_21_+_8433172_8446622_RNA45SN1_45S_50ntFlanks | 11318 | 11344 | 27 | CCCCACGTCTCGTCGCGCGCGCGTCCG | 5237.6 | -6.7 | 1.2 | -5.6 | 1.9E-08 | 4.3E-08 | BT20/MB468 | CYTO |
| >hg38_21_+_8433172_8446622_RNA45SN1_45S_50ntFlanks@11593.11611.19\|CCAGGGGAATCCGACTGTT | >hg38_21_+_8433172_8446622_RNA45SN1_45S_50ntFlanks | 11593 | 11611 | 19 | CCAGGGGAATCCGACTGTT | 3430.2 | -4.6 | 0.8 | -5.6 | 2.2E-08 | 4.3E-08 | MB231/MB468 | CYTO |
| >hg38_21_+_8433172_8446622_RNA45SN1_45S_50ntFlanks@10766.10785.20\|CGCAGCAGGTCTCCAAGGTG | >hg38_21_+_8433172_8446622_RNA45SN1_45S_50ntFlanks | 10766 | 10785 | 20 | CGCAGCAGGTCTCCAAGGTG | 552.8 | -8.3 | 1.5 | -5.6 | 2.5E-08 | 4.3E-08 | BT20/MB468 | NUC |
| >hg38_21_+_8433172_8446622_RNA45SN1_45S_50ntFlanks@9554.9582.29\|CGTAGCGGTCCTGACGTGCAAATCGGTCG | >hg38_21_+_8433172_8446622_RNA45SN1_45S_50ntFlanks | 9554 | 9582 | 29 | CGTAGCGGTCCTGACGTGCAAATCGGTCG | 2510.9 | -9.7 | 1.7 | -5.6 | 2.5E-08 | 4.3E-08 | BT20/MB468 | NUC |
| >hg38_21_+_8433172_8446622_RNA45SN1_45S_50ntFlanks@7984.8007.24\|AGATCAGACGTGGCGACCCGCTGA | >hg38_21_+_8433172_8446622_RNA45SN1_45S_50ntFlanks | 7984 | 8007 | 24 | AGATCAGACGTGGCGACCCGCTGA | 496.7 | -4.1 | 0.7 | -5.6 | 1.7E-08 | 4.3E-08 | BT20/MB468 | MITO |
| >hg38_21_+_8433172_8446622_RNA45SN1_45S_50ntFlanks@5073.5093.21\|AATTCCGATAACGAACGAGAC | >hg38_21_+_8433172_8446622_RNA45SN1_45S_50ntFlanks | 5073 | 5093 | 21 | AATTCCGATAACGAACGAGAC | 855.5 | -9.2 | 1.6 | -5.6 | 2.0E-08 | 4.3E-08 | MB231/MB468 | MP |
| >hg38_21_+_8433172_8446622_RNA45SN1_45S_50ntFlanks@9573.9596.24\|AAATCGGTCGTCCGACCTGGGTAT | >hg38_21_+_8433172_8446622_RNA45SN1_45S_50ntFlanks | 9573 | 9596 | 24 | AAATCGGTCGTCCGACCTGGGTAT | 1105.4 | -9.0 | 1.6 | -5.6 | 2.5E-08 | 4.3E-08 | BT20/MB468 | NUC |
| >hg38_21_+_8433172_8446622_RNA45SN1_45S_50ntFlanks@12251.12271.21\|AAAGCTCGCTTGATCTTGATT | >hg38_21_+_8433172_8446622_RNA45SN1_45S_50ntFlanks | 12251 | 12271 | 21 | AAAGCTCGCTTGATCTTGATT | 598.7 | 5.1 | 0.9 | 5.6 | 2.2E-08 | 4.3E-08 | MB231/MB468 | CYTO |
| >hg38_21_+_8433172_8446622_RNA45SN1_45S_50ntFlanks@8116.8137.22\|GCGGGACATGTGGCGTACGGAA | >hg38_21_+_8433172_8446622_RNA45SN1_45S_50ntFlanks | 8116 | 8137 | 22 | GCGGGACATGTGGCGTACGGAA | 2258.1 | 14.5 | 2.3 | 6.2 | 6.4E-10 | 4.3E-08 | BT20/MB231 | MP |
| >hg38_21_+_8433172_8446622_RNA45SN1_45S_50ntFlanks@12418.12437.20\|CGATGTCGGCTCTTCCTATC | >hg38_21_+_8433172_8446622_RNA45SN1_45S_50ntFlanks | 12418 | 12437 | 20 | CGATGTCGGCTCTTCCTATC | 1141.7 | -6.7 | 1.2 | -5.6 | 2.1E-08 | 4.3E-08 | MB231/MB468 | MP |
| >hg38_21_+_8433172_8446622_RNA45SN1_45S_50ntFlanks@3782.3810.29\|CAGTGAAACTGCGAATGGCTCATTAAATC | >hg38_21_+_8433172_8446622_RNA45SN1_45S_50ntFlanks | 3782 | 3810 | 29 | CAGTGAAACTGCGAATGGCTCATTAAATC | 862.6 | -8.7 | 1.6 | -5.6 | 2.5E-08 | 4.3E-08 | BT20/MB468 | NUC |
| >hg38_21_+_8433172_8446622_RNA45SN1_45S_50ntFlanks@11564.11586.23\|CCTAGCAGCCGACTTAGAACTGG | >hg38_21_+_8433172_8446622_RNA45SN1_45S_50ntFlanks | 11564 | 11586 | 23 | CCTAGCAGCCGACTTAGAACTGG | 1606.9 | -6.4 | 1.1 | -5.6 | 2.4E-08 | 4.3E-08 | MB231/MB468 | MITO |
| >hg38_21_+_8433172_8446622_RNA45SN1_45S_50ntFlanks@11235.11255.21\|CTTCCCGTGGATCGCCCCAGC | >hg38_21_+_8433172_8446622_RNA45SN1_45S_50ntFlanks | 11235 | 11255 | 21 | CTTCCCGTGGATCGCCCCAGC | 467.6 | -3.9 | 0.7 | -5.6 | 2.4E-08 | 4.4E-08 | MB231/MB468 | MITO |
| >MI0000749\|hsa-mir-30e&WithFlank&1\|+\|40754349\|40754452@23.46.24&[MIMAT0000692&hsa-miR-30e-5p&offsets\|0\|+2;m-24&1\|+\|40754371\|40754394&offsets\|0\|0]\|TGTAAACATCCTTGACTGGAAGCT | >MI0000749\|hsa-mir-30e&WithFlank&1\|+\|40754349\|40754452 | 23 | 46 | 24 | TGTAAACATCCTTGACTGGAAGCT | 2525.1 | 4.5 | 0.8 | 5.6 | 1.9E-08 | 4.4E-08 | BT20/MB468 | CYTO |
| >hg38_21_+_8433172_8446622_RNA45SN1_45S_50ntFlanks@11602.11620.19\|TCCGACTGTTTAATTAAAA | >hg38_21_+_8433172_8446622_RNA45SN1_45S_50ntFlanks | 11602 | 11620 | 19 | TCCGACTGTTTAATTAAAA | 1023.9 | -4.6 | 0.8 | -5.6 | 1.9E-08 | 4.4E-08 | BT20/MB468 | CYTO |
| >hg38_21_+_8433172_8446622_RNA45SN1_45S_50ntFlanks@7988.8016.29\|CAGACGTGGCGACCCGCTGAATTTAAGCA | >hg38_21_+_8433172_8446622_RNA45SN1_45S_50ntFlanks | 7988 | 8016 | 29 | CAGACGTGGCGACCCGCTGAATTTAAGCA | 1541.7 | -3.5 | 0.6 | -5.6 | 2.4E-08 | 4.4E-08 | MB231/MB468 | MITO |
| >hg38_21_+_8433172_8446622_RNA45SN1_45S_50ntFlanks@5526.5550.25\|AAAAGTCGTAACAAGGTTTCCGTAG | >hg38_21_+_8433172_8446622_RNA45SN1_45S_50ntFlanks | 5526 | 5550 | 25 | AAAAGTCGTAACAAGGTTTCCGTAG | 1029.9 | -8.9 | 1.6 | -5.6 | 2.5E-08 | 4.4E-08 | BT20/MB468 | NUC |
| >hg38_21_+_8433172_8446622_RNA45SN1_45S_50ntFlanks@6651.6687.37\|CGACTCTTAGCGGTGGATCACTCGGCTCGTGCGTCGA | >hg38_21_+_8433172_8446622_RNA45SN1_45S_50ntFlanks | 6651 | 6687 | 37 | CGACTCTTAGCGGTGGATCACTCGGCTCGTGCGTCGA | 2753.6 | 4.6 | 0.8 | 5.6 | 2.2E-08 | 4.4E-08 | MB231/MB468 | NUC |
| >hg38_21_+_8433172_8446622_RNA45SN1_45S_50ntFlanks@4367.4398.32\|CAGTTAAAAAGCTCGTAGTTGGATCTTGGGAG | >hg38_21_+_8433172_8446622_RNA45SN1_45S_50ntFlanks | 4367 | 4398 | 32 | CAGTTAAAAAGCTCGTAGTTGGATCTTGGGAG | 398.5 | -5.2 | 0.9 | -5.6 | 2.2E-08 | 4.4E-08 | MB231/MB468 | CYTO |

<table>
<tr><td>>hg38_1_-_228634819_228635039_RNA5S12_5S_50ntFlanks@51.79.29|GTCTACGGCCATACCACCCTGAACGCGCC</td><td>>hg38_1_-_228634819_228635039_RNA5S12_5S_50ntFlanks</td><td>51</td><td>79</td><td>29</td><td>GTCTACGGCCATACCACCCTGAACGCGCC</td><td>1756.7</td><td>5.3</td><td>0.9</td><td>5.6</td><td>2.2E-08</td><td>4.4E-08</td><td>MB231/MB468</td><td>CYTO</td></tr>
<tr><td>>hg38_21_+_8433172_8446622_RNA45SN1_45S_50ntFlanks@6646.6664.19|TCGTACGACTCTTAGCGGT</td><td>>hg38_21_+_8433172_8446622_RNA45SN1_45S_50ntFlanks</td><td>6646</td><td>6664</td><td>19</td><td>TCGTACGACTCTTAGCGGT</td><td>32095.9</td><td>4.6</td><td>0.8</td><td>5.9</td><td>3.4E-09</td><td>4.4E-08</td><td>BT20/MB468</td><td>MP</td></tr>
<tr><td>>hg38_21_+_8433172_8446622_RNA45SN1_45S_50ntFlanks@3990.4026.37|TTGGTGACTCTAGATAACCTCGGGCCGATCGCACGCC</td><td>>hg38_21_+_8433172_8446622_RNA45SN1_45S_50ntFlanks</td><td>3990</td><td>4026</td><td>37</td><td>TTGGTGACTCTAGATAACCTCGGGCCGATCGCACGCC</td><td>895.6</td><td>-8.8</td><td>1.6</td><td>-5.6</td><td>2.5E-08</td><td>4.4E-08</td><td>BT20/MB468</td><td>NUC</td></tr>
<tr><td>>hg38_21_+_8433172_8446622_RNA45SN1_45S_50ntFlanks@12386.12414.29|AGCGTTCATAGCGACGTCGCTTTTTGATC</td><td>>hg38_21_+_8433172_8446622_RNA45SN1_45S_50ntFlanks</td><td>12386</td><td>12414</td><td>29</td><td>AGCGTTCATAGCGACGTCGCTTTTTGATC</td><td>1065.9</td><td>-5.2</td><td>0.9</td><td>-5.6</td><td>1.7E-08</td><td>4.4E-08</td><td>BT20/MB468</td><td>MITO</td></tr>
<tr><td>>hg38_21_+_8433172_8446622_RNA45SN1_45S_50ntFlanks@11876.11899.24|AAGACCCTGTTGAGCTTGACTCTA</td><td>>hg38_21_+_8433172_8446622_RNA45SN1_45S_50ntFlanks</td><td>11876</td><td>11899</td><td>24</td><td>AAGACCCTGTTGAGCTTGACTCTA</td><td>419.6</td><td>-3.6</td><td>0.6</td><td>-5.6</td><td>2.2E-08</td><td>4.4E-08</td><td>MB231/MB468</td><td>CYTO</td></tr>
<tr><td>>hg38_21_+_8433172_8446622_RNA45SN1_45S_50ntFlanks@12937.12955.19|AACCATTCGTAGACGACCT</td><td>>hg38_21_+_8433172_8446622_RNA45SN1_45S_50ntFlanks</td><td>12937</td><td>12955</td><td>19</td><td>AACCATTCGTAGACGACCT</td><td>519.5</td><td>-5.8</td><td>1.0</td><td>-5.6</td><td>2.3E-08</td><td>4.4E-08</td><td>MB231/MB468</td><td>NUC</td></tr>
<tr><td>>MI0003628|hsa-mir-615&WithFlank&12|+|54033944|54034051@67.85.19&[MIMAT0003283&hsa-miR-615-3p&offsets|0|-3;m-495&12|+|54034010|54034030&offsets|0|-2]|TCCGAGCCTGGGTCTCCCT</td><td>>MI0003628|hsa-mir-615&WithFlank&12|+|54033944|54034051</td><td>67</td><td>85</td><td>19</td><td>TCCGAGCCTGGGTCTCCCT</td><td>356.1</td><td>7.6</td><td>1.4</td><td>5.6</td><td>1.9E-08</td><td>4.4E-08</td><td>BT20/MB468</td><td>CYTO</td></tr>
<tr><td>>hg38_21_+_8433172_8446622_RNA45SN1_45S_50ntFlanks@12054.12071.18|TGAGGCGGGGGGGCGAGC</td><td>>hg38_21_+_8433172_8446622_RNA45SN1_45S_50ntFlanks</td><td>12054</td><td>12071</td><td>18</td><td>TGAGGCGGGGGGGCGAGC</td><td>110.4</td><td>-9.9</td><td>1.7</td><td>-5.9</td><td>3.4E-09</td><td>4.4E-08</td><td>BT20/MB468</td><td>MP</td></tr>
<tr><td>>hg38_MT_+_598_1651_RNR1_12S_50ntFlanks@811.840.30|AAGGGTCGAAGGTGGATTTAGCAGTAAACT</td><td>>hg38_MT_+_598_1651_RNR1_12S_50ntFlanks</td><td>811</td><td>840</td><td>30</td><td>AAGGGTCGAAGGTGGATTTAGCAGTAAACT</td><td>118.8</td><td>10.6</td><td>1.8</td><td>5.9</td><td>3.4E-09</td><td>4.4E-08</td><td>BT20/MB468</td><td>MP</td></tr>
<tr><td>>hg38_21_+_8433172_8446622_RNA45SN1_45S_50ntFlanks@11552.11574.23|CCTCGGCCGGCGCCTAGCAGCCG</td><td>>hg38_21_+_8433172_8446622_RNA45SN1_45S_50ntFlanks</td><td>11552</td><td>11574</td><td>23</td><td>CCTCGGCCGGCGCCTAGCAGCCG</td><td>109.5</td><td>-9.9</td><td>1.7</td><td>-5.9</td><td>3.4E-09</td><td>4.4E-08</td><td>BT20/MB468</td><td>MP</td></tr>
<tr><td>>hg38_21_+_8433172_8446622_RNA45SN1_45S_50ntFlanks@485.533.49|GATGTGGTGACGTCGTGCTCTCCCGGGCCGGGTCCGAGCCGCGACGGGC</td><td>>hg38_21_+_8433172_8446622_RNA45SN1_45S_50ntFlanks</td><td>485</td><td>533</td><td>49</td><td>GATGTGGTGACGTCGTGCTCTCCCGGGCCGGGTCCGAGCCGCGACGGGC</td><td>106.5</td><td>9.6</td><td>1.7</td><td>5.6</td><td>2.5E-08</td><td>4.4E-08</td><td>BT20/MB468</td><td>NUC</td></tr>
<tr><td>>hg38_21_+_8433172_8446622_RNA45SN1_45S_50ntFlanks@10444.10462.19|CGAAGGGACGGGCGATGGC</td><td>>hg38_21_+_8433172_8446622_RNA45SN1_45S_50ntFlanks</td><td>10444</td><td>10462</td><td>19</td><td>CGAAGGGACGGGCGATGGC</td><td>303.6</td><td>-7.6</td><td>1.3</td><td>-5.6</td><td>2.1E-08</td><td>4.4E-08</td><td>MB231/MB468</td><td>MP</td></tr>
<tr><td>>MI0000081|hsa-mir-24-2&WithFlank&19|-|13836281|13836365@56.77.22(+1A);MI0000080|hsa-mir-24-1&WithFlank&9|+|95086015|95086094@50.71.22(+1A)&[MIMAT0000080&hsa-miR-24-3p&offsets|0|0(+1A);m-36&19|-|13836288|13836310&offsets|0|-1(+1A)];[MIMAT0000080_1&hsa-miR-24-3p&offsets|0|0(+1A);m-35&9|+|95086064|95086086&offsets|0|-1(+1A)]|TGGCTCAGTTCAGCAGGAACAGA</td><td>>MI0000081|hsa-mir-24-2&WithFlank&19|-|13836281|13836365</td><td>56</td><td>77</td><td>22(+1A)</td><td>TGGCTCAGTTCAGCAGGAACAGA</td><td>304.5</td><td>3.7</td><td>0.7</td><td>5.6</td><td>2.3E-08</td><td>4.4E-08</td><td>MB231/MB468</td><td>NUC</td></tr>
<tr><td>>hg38_21_+_8433172_8446622_RNA45SN1_45S_50ntFlanks@5417.5445.29|CTACCGATTGGATGGTTTAGTGAGGCCCT</td><td>>hg38_21_+_8433172_8446622_RNA45SN1_45S_50ntFlanks</td><td>5417</td><td>5445</td><td>29</td><td>CTACCGATTGGATGGTTTAGTGAGGCCCT</td><td>475.2</td><td>-5.0</td><td>0.9</td><td>-5.6</td><td>1.9E-08</td><td>4.4E-08</td><td>BT20/MB468</td><td>CYTO</td></tr>
<tr><td>>trnaMT_GluTTC_MT_-_14674_14742@41.71.31,trnalookalike8_GluTTC_5_-_93905172_93905240@41.71.31|TGGTCGTGGTTGTAGTCCGTGCGAGAATACC</td><td>>trnaMT_GluTTC_MT_-_14674_14742</td><td>41</td><td>71</td><td>31</td><td>TGGTCGTGGTTGTAGTCCGTGCGAGAATACC</td><td>597.6</td><td>4.9</td><td>0.9</td><td>5.6</td><td>1.7E-08</td><td>4.4E-08</td><td>BT20/MB468</td><td>MITO</td></tr>
<tr><td>>hg38_21_+_8433172_8446622_RNA45SN1_45S_50ntFlanks@11324.11344.21|GTCTCGTCGCGCGCGCGTCCG</td><td>>hg38_21_+_8433172_8446622_RNA45SN1_45S_50ntFlanks</td><td>11324</td><td>11344</td><td>21</td><td>GTCTCGTCGCGCGCGCGTCCG</td><td>406.5</td><td>-4.1</td><td>0.7</td><td>-5.6</td><td>2.3E-08</td><td>4.4E-08</td><td>MB231/MB468</td><td>NUC</td></tr>
<tr><td>>hg38_21_+_8433172_8446622_RNA45SN1_45S_50ntFlanks@11602.11640.39|TCCGACTGTTTAATTAAAACAAAGCATCGCGAAGGCCCG</td><td>>hg38_21_+_8433172_8446622_RNA45SN1_45S_50ntFlanks</td><td>11602</td><td>11640</td><td>39</td><td>TCCGACTGTTTAATTAAAACAAAGCATCGCGAAGGCCCG</td><td>74.7</td><td>-10.1</td><td>1.8</td><td>-5.6</td><td>2.1E-08</td><td>4.4E-08</td><td>MB231/MB468</td><td>MP</td></tr>
<tr><td>>MI0000076|hsa-mir-20a&WithFlank&13|+|91351059|91351141@14.34.21&[MIMAT0000075&hsa-miR-20a-5p&offsets|0|-2;m-96&13|+|91351072|91351094&offsets|0|-2]|TAAAGTGCTTATAGTGCAGGT</td><td>>MI0000076|hsa-mir-20a&WithFlank&13|+|91351059|91351141</td><td>14</td><td>34</td><td>21</td><td>TAAAGTGCTTATAGTGCAGGT</td><td>2010.5</td><td>-5.7</td><td>0.9</td><td>-6.0</td><td>1.8E-09</td><td>4.4E-08</td><td>BT20/MB231</td><td>MITO</td></tr>
<tr><td>>hg38_MT_+_598_1651_RNR1_12S_50ntFlanks@843.872.30|GAGTAGAGTGCTTAGTTGAACAGGGCCCTG</td><td>>hg38_MT_+_598_1651_RNR1_12S_50ntFlanks</td><td>843</td><td>872</td><td>30</td><td>GAGTAGAGTGCTTAGTTGAACAGGGCCCTG</td><td>178.0</td><td>11.2</td><td>1.9</td><td>5.9</td><td>3.5E-09</td><td>4.4E-08</td><td>BT20/MB468</td><td>MP</td></tr>
<tr><td>>MI0000063|hsa-let-7b&WithFlank&22|+|46113680|46113774@66.86.21(+1U)&[MIMAT0004482&hsa-let-7b-3p&offsets|0|-1(+1U);m-171&22|+|46113745|46113766&offsets|0|-1(+1U)]|CTATACAACCTACTGCCTTCCT</td><td>>MI0000063|hsa-let-7b&WithFlank&22|+|46113680|46113774</td><td>66</td><td>86</td><td>21(+1U)</td><td>CTATACAACCTACTGCCTTCCT</td><td>252.1</td><td>6.6</td><td>1.2</td><td>5.6</td><td>2.2E-08</td><td>4.5E-08</td><td>MB231/MB468</td><td>CYTO</td></tr>
<tr><td>>hg38_21_+_8433172_8446622_RNA45SN1_45S_50ntFlanks@11572.11593.22|CCGACTTAGAACTGGTGCGGAC</td><td>>hg38_21_+_8433172_8446622_RNA45SN1_45S_50ntFlanks</td><td>11572</td><td>11593</td><td>22</td><td>CCGACTTAGAACTGGTGCGGAC</td><td>755.0</td><td>-5.1</td><td>0.9</td><td>-5.6</td><td>1.9E-08</td><td>4.5E-08</td><td>BT20/MB468</td><td>CYTO</td></tr>
<tr><td>>MI0000465|hsa-mir-191&WithFlank&3|-|49020612|49020715@22.43.22(+1A)&[MIMAT0000440&hsa-miR-191-5p&offsets|0|-1(+1A);m-48&3|-|49020672|49020694&offsets|0|-1(+1A)]|CAACGGAATCCCAAAAGCAGCTA</td><td>>MI0000465|hsa-mir-191&WithFlank&3|-|49020612|49020715</td><td>22</td><td>43</td><td>22(+1A)</td><td>CAACGGAATCCCAAAAGCAGCTA</td><td>1035.0</td><td>5.2</td><td>0.9</td><td>5.6</td><td>2.5E-08</td><td>4.5E-08</td><td>MB231/MB468</td><td>MITO</td></tr>
<tr><td>>hg38_21_+_8433172_8446622_RNA45SN1_45S_50ntFlanks@5152.5183.32|AGGGACAAGTGGCGTTCAGCCACCCGAGATTG</td><td>>hg38_21_+_8433172_8446622_RNA45SN1_45S_50ntFlanks</td><td>5152</td><td>5183</td><td>32</td><td>AGGGACAAGTGGCGTTCAGCCACCCGAGATTG</td><td>90.1</td><td>-10.4</td><td>1.8</td><td>-5.6</td><td>2.1E-08</td><td>4.5E-08</td><td>MB231/MB468</td><td>MP</td></tr>
<tr><td>>MI0000088|hsa-mir-30a&WithFlank&6|-|71403545|71403627@12.35.24(+1U)&[MIMAT0000087&hsa-miR-30a-5p&offsets|0|+2(+1U);m-15&6|-|71403595|71403616&offsets|0|+2(+1U)]|TGTAAACATCCTCGACTGGAAGCTT</td><td>>MI0000088|hsa-mir-30a&WithFlank&6|-|71403545|71403627</td><td>12</td><td>35</td><td>24(+1U)</td><td>TGTAAACATCCTCGACTGGAAGCTT</td><td>1328.4</td><td>6.6</td><td>1.2</td><td>5.6</td><td>2.2E-08</td><td>4.5E-08</td><td>MB231/MB468</td><td>CYTO</td></tr>
<tr><td>>hg38_21_+_8433172_8446622_RNA45SN1_45S_50ntFlanks@10798.10821.24|CATGTTGGAACAATGTAGGTAAGG</td><td>>hg38_21_+_8433172_8446622_RNA45SN1_45S_50ntFlanks</td><td>10798</td><td>10821</td><td>24</td><td>CATGTTGGAACAATGTAGGTAAGG</td><td>185.4</td><td>-5.9</td><td>1.1</td><td>-5.6</td><td>2.1E-08</td><td>4.5E-08</td><td>MB231/MB468</td><td>MP</td></tr>
<tr><td>>MI0000067|hsa-let-7f-1&WithFlank&9|+|94176341|94176439@13.33.21(+1G);MI0000068|hsa-let-7f-2&WithFlank&X|-|53557186|53557280@14.34.21(+1G)&[MIMAT0000067&hsa-let-7f-5p&offsets|0|-1(+1G);m-20&9|+|94176353|94176374&offsets|0|-1(+1G)];[MIMAT0000067_1&hsa-let-7f-5p&offsets|0|-1(+1G);m-19&X|-|53557246|53557267&offsets|0|-1(+1G)]|TGAGGTAGTAGATTGTATAGTG</td><td>>MI0000067|hsa-let-7f-1&WithFlank&9|+|94176341|94176439</td><td>13</td><td>33</td><td>21(+1G)</td><td>TGAGGTAGTAGATTGTATAGTG</td><td>1201.9</td><td>4.2</td><td>0.7</td><td>5.6</td><td>2.3E-08</td><td>4.5E-08</td><td>MB231/MB468</td><td>NUC</td></tr>
<tr><td>>hg38_21_+_8433172_8446622_RNA45SN1_45S_50ntFlanks@12709.12742.34|CCGCGGAGCCTCGGTTGGCCTCGGATAGCCGGTC</td><td>>hg38_21_+_8433172_8446622_RNA45SN1_45S_50ntFlanks</td><td>12709</td><td>12742</td><td>34</td><td>CCGCGGAGCCTCGGTTGGCCTCGGATAGCCGGTC</td><td>623.5</td><td>-7.0</td><td>1.2</td><td>-5.6</td><td>1.7E-08</td><td>4.5E-08</td><td>BT20/MB468</td><td>MITO</td></tr>
<tr><td>>hg38_21_+_8433172_8446622_RNA45SN1_45S_50ntFlanks@6771.6797.27|GGTTCCTCCCGGGGCTACGCCTGTCTG</td><td>>hg38_21_+_8433172_8446622_RNA45SN1_45S_50ntFlanks</td><td>6771</td><td>6797</td><td>27</td><td>GGTTCCTCCCGGGGCTACGCCTGTCTG</td><td>151.6</td><td>-10.3</td><td>1.7</td><td>-5.9</td><td>3.5E-09</td><td>4.5E-08</td><td>BT20/MB468</td><td>MP</td></tr>
<tr><td>>hg38_21_+_8433172_8446622_RNA45SN1_45S_50ntFlanks@11641.11664.24|CGGCGGGTGTTGACGCGATGTGAT</td><td>>hg38_21_+_8433172_8446622_RNA45SN1_45S_50ntFlanks</td><td>11641</td><td>11664</td><td>24</td><td>CGGCGGGTGTTGACGCGATGTGAT</td><td>1948.5</td><td>-9.4</td><td>1.7</td><td>-5.6</td><td>2.6E-08</td><td>4.5E-08</td><td>BT20/MB468</td><td>NUC</td></tr>
<tr><td>>hg38_21_+_8433172_8446622_RNA45SN1_45S_50ntFlanks@11769.11806.38|TAATTAGTGACGCGCATGAATGGATGAACGAGATTCCC</td><td>>hg38_21_+_8433172_8446622_RNA45SN1_45S_50ntFlanks</td><td>11769</td><td>11806</td><td>38</td><td>TAATTAGTGACGCGCATGAATGGATGAACGAGATTCCC</td><td>195.6</td><td>-3.0</td><td>0.5</td><td>-5.6</td><td>1.9E-08</td><td>4.5E-08</td><td>BT20/MB468</td><td>CYTO</td></tr>
<tr><td>>hg38_21_+_8433172_8446622_RNA45SN1_45S_50ntFlanks@12936.12958.23|AAACCATTCGTAGACGACCTGCT</td><td>>hg38_21_+_8433172_8446622_RNA45SN1_45S_50ntFlanks</td><td>12936</td><td>12958</td><td>23</td><td>AAACCATTCGTAGACGACCTGCT</td><td>296.4</td><td>-4.0</td><td>0.7</td><td>-5.6</td><td>2.5E-08</td><td>4.5E-08</td><td>MB231/MB468</td><td>MITO</td></tr>
<tr><td>>hg38_21_+_8433172_8446622_RNA45SN1_45S_50ntFlanks@8114.8139.26|GCGCGGGACATGTGGCGTACGGAAGA</td><td>>hg38_21_+_8433172_8446622_RNA45SN1_45S_50ntFlanks</td><td>8114</td><td>8139</td><td>26</td><td>GCGCGGGACATGTGGCGTACGGAAGA</td><td>1106.0</td><td>13.4</td><td>2.2</td><td>6.2</td><td>6.7E-10</td><td>4.5E-08</td><td>BT20/MB231</td><td>MP</td></tr>
<tr><td>>hg38_21_+_8433172_8446622_RNA45SN1_45S_50ntFlanks@11641.11682.42|CGGCGGGTGTTGACGCGATGTGATTTCTGCCCAGTGCTCTGA</td><td>>hg38_21_+_8433172_8446622_RNA45SN1_45S_50ntFlanks</td><td>11641</td><td>11682</td><td>42</td><td>CGGCGGGTGTTGACGCGATGTGATTTCTGCCCAGTGCTCTGA</td><td>79.6</td><td>-10.2</td><td>1.8</td><td>-5.6</td><td>2.1E-08</td><td>4.5E-08</td><td>MB231/MB468</td><td>MP</td></tr>
<tr><td>>hg38_21_+_8433172_8446622_RNA45SN1_45S_50ntFlanks@11781.11805.25|CGCATGAATGGATGAACGAGATTCC</td><td>>hg38_21_+_8433172_8446622_RNA45SN1_45S_50ntFlanks</td><td>11781</td><td>11805</td><td>25</td><td>CGCATGAATGGATGAACGAGATTCC</td><td>878.8</td><td>-6.2</td><td>1.1</td><td>-5.6</td><td>2.3E-08</td><td>4.5E-08</td><td>MB231/MB468</td><td>NUC</td></tr>
<tr><td>>MI0001445|hsa-mir-423&WithFlank&17|+|30117073|30117178@59.77.19(+1U)&[MIMAT0001340&hsa-miR-423-3p&offsets|0|-4(+1U);m-100&17|+|30117131|30117153&offsets|0|-4(+1U)]|AGCTCGGTCTGAGGCCCCTT</td><td>>MI0001445|hsa-mir-423&WithFlank&17|+|30117073|30117178</td><td>59</td><td>77</td><td>19(+1U)</td><td>AGCTCGGTCTGAGGCCCCTT</td><td>134.9</td><td>3.3</td><td>0.6</td><td>5.6</td><td>2.6E-08</td><td>4.5E-08</td><td>BT20/MB468</td><td>NUC</td></tr>
<tr><td>>hg38_21_+_8433172_8446622_RNA45SN1_45S_50ntFlanks@12954.12979.26|CTGCTTCTGGGTCGGGGTTTCGTACG</td><td>>hg38_21_+_8433172_8446622_RNA45SN1_45S_50ntFlanks</td><td>12954</td><td>12979</td><td>26</td><td>CTGCTTCTGGGTCGGGGTTTCGTACG</td><td>232.9</td><td>-7.6</td><td>1.4</td><td>-5.6</td><td>2.3E-08</td><td>4.5E-08</td><td>MB231/MB468</td><td>NUC</td></tr>
<tr><td>>hg38_21_+_8433172_8446622_RNA45SN1_45S_50ntFlanks@12126.12150.25|CGACCCGCTCCGGGGACAGTGCCAG</td><td>>hg38_21_+_8433172_8446622_RNA45SN1_45S_50ntFlanks</td><td>12126</td><td>12150</td><td>25</td><td>CGACCCGCTCCGGGGACAGTGCCAG</td><td>559.5</td><td>-8.2</td><td>1.4</td><td>-5.9</td><td>3.5E-09</td><td>4.5E-08</td><td>BT20/MB468</td><td>MP</td></tr>
<tr><td>>hg38_21_+_8433172_8446622_RNA45SN1_45S_50ntFlanks@8381.8399.19|AAGAGGGCGTGAAACCGTT</td><td>>hg38_21_+_8433172_8446622_RNA45SN1_45S_50ntFlanks</td><td>8381</td><td>8399</td><td>19</td><td>AAGAGGGCGTGAAACCGTT</td><td>1265.6</td><td>-8.5</td><td>1.5</td><td>-5.6</td><td>2.1E-08</td><td>4.5E-08</td><td>MB231/MB468</td><td>MP</td></tr>
<tr><td>>hg38_21_+_8433172_8446622_RNA45SN1_45S_50ntFlanks@10346.10390.45|TCAAACGAGAACTTTGAAGGCCGAAGTGGAGAAGGGTTCCATGTG</td><td>>hg38_21_+_8433172_8446622_RNA45SN1_45S_50ntFlanks</td><td>10346</td><td>10390</td><td>45</td><td>TCAAACGAGAACTTTGAAGGCCGAAGTGGAGAAGGGTTCCATGTG</td><td>1045.1</td><td>-5.2</td><td>0.9</td><td>-5.6</td><td>2.5E-08</td><td>4.6E-08</td><td>MB231/MB468</td><td>MITO</td></tr>
<tr><td>>hg38_21_+_8433172_8446622_RNA45SN1_45S_50ntFlanks@10392.10426.35|ACAGCAGTTGAACATGGGTCAGTCGGTCCTGAGAG</td><td>>hg38_21_+_8433172_8446622_RNA45SN1_45S_50ntFlanks</td><td>10392</td><td>10426</td><td>35</td><td>ACAGCAGTTGAACATGGGTCAGTCGGTCCTGAGAG</td><td>625.9</td><td>-6.0</td><td>1.1</td><td>-5.6</td><td>2.5E-08</td><td>4.6E-08</td><td>MB231/MB468</td><td>MITO</td></tr>
<tr><td>>hg38_21_+_8433172_8446622_RNA45SN1_45S_50ntFlanks@10869.10890.22|AGGGCTGGGTCGGTCGGGCTGG</td><td>>hg38_21_+_8433172_8446622_RNA45SN1_45S_50ntFlanks</td><td>10869</td><td>10890</td><td>22</td><td>AGGGCTGGGTCGGTCGGGCTGG</td><td>1146.2</td><td>-6.7</td><td>1.2</td><td>-5.6</td><td>2.3E-08</td><td>4.6E-08</td><td>MB231/MB468</td><td>NUC</td></tr>
<tr><td>>hg38_21_+_8433172_8446622_RNA45SN1_45S_50ntFlanks@8323.8355.33|ACCGATAGTCAACAAGTACCGTAAGGGAAAGTT</td><td>>hg38_21_+_8433172_8446622_RNA45SN1_45S_50ntFlanks</td><td>8323</td><td>8355</td><td>33</td><td>ACCGATAGTCAACAAGTACCGTAAGGGAAAGTT</td><td>98.0</td><td>-9.7</td><td>1.6</td><td>-5.9</td><td>3.6E-09</td><td>4.6E-08</td><td>BT20/MB468</td><td>MP</td></tr>
<tr><td>>hg38_21_+_8433172_8446622_RNA45SN1_45S_50ntFlanks@11565.11582.18|CTAGCAGCCGACTTAGAA</td><td>>hg38_21_+_8433172_8446622_RNA45SN1_45S_50ntFlanks</td><td>11565</td><td>11582</td><td>18</td><td>CTAGCAGCCGACTTAGAA</td><td>222.0</td><td>-3.3</td><td>0.6</td><td>-5.6</td><td>2.3E-08</td><td>4.6E-08</td><td>MB231/MB468</td><td>NUC</td></tr>
<tr><td>>hg38_21_+_8433172_8446622_RNA45SN1_45S_50ntFlanks@11836.11873.38|ACAGCCAAGGGAACGGGCTTGGCGGAATCAGCGGGGAA</td><td>>hg38_21_+_8433172_8446622_RNA45SN1_45S_50ntFlanks</td><td>11836</td><td>11873</td><td>38</td><td>ACAGCCAAGGGAACGGGCTTGGCGGAATCAGCGGGGAA</td><td>56.2</td><td>-9.7</td><td>1.7</td><td>-5.6</td><td>2.2E-08</td><td>4.6E-08</td><td>MB231/MB468</td><td>MP</td></tr>
<tr><td>>MI0000736|hsa-mir-30c-1&WithFlank&1|+|40757278|40757378@23.46.24;MI0000254|hsa-mir-30c-2&WithFlank&6|-|71376954|71377037@13.36.24&[MIMAT0000244&hsa-miR-30c-5p&offsets|0|+1;m-57&1|+|40757300|40757323&offsets|0|0];[MIMAT0000244_1&hsa-miR-30c-5p&offsets|0|+1;m-56&6|-|71377002|71377025&offsets|0|0]|TGTAAACATCCTACACTCTCAGCT</td><td>>MI0000736|hsa-mir-30c-1&WithFlank&1|+|40757278|40757378</td><td>23</td><td>46</td><td>24</td><td>TGTAAACATCCTACACTCTCAGCT</td><td>11087.6</td><td>4.8</td><td>0.9</td><td>5.6</td><td>2.3E-08</td><td>4.6E-08</td><td>MB231/MB468</td><td>CYTO</td></tr>
<tr><td>>MI0000478|hsa-mir-149&WithFlank&2|+|240455995|240456095@21.42.22(+1U)&[MIMAT0000450&hsa-miR-149-5p&offsets|0|-1(+1U);m-195&2|+|240456015|240456037&offsets|0|-1(+1U)]|TCTGGCTCCGTGTCTTCACTCCT</td><td>>MI0000478|hsa-mir-149&WithFlank&2|+|240455995|240456095</td><td>21</td><td>42</td><td>22(+1U)</td><td>TCTGGCTCCGTGTCTTCACTCCT</td><td>1173.9</td><td>4.6</td><td>0.8</td><td>6.0</td><td>2.5E-09</td><td>4.6E-08</td><td>BT20/MB231</td><td>NUC</td></tr>
<tr><td>>MI0000542|hsa-mir-320a&WithFlank&8|-|22244960|22245043@50.68.19&[MIMAT0000510&hsa-miR-320a-3p&offsets|+2|-1;m-60&8|-|22244975|22244996&offsets|+2|-1]|AAGCTGGGTTGAGAGGGCG</td><td>>MI0000542|hsa-mir-320a&WithFlank&8|-|22244960|22245043</td><td>50</td><td>68</td><td>19</td><td>AAGCTGGGTTGAGAGGGCG</td><td>124.7</td><td>10.7</td><td>1.8</td><td>5.9</td><td>3.6E-09</td><td>4.6E-08</td><td>BT20/MB468</td><td>MP</td></tr>
</table>

| | | | | | | | | | | | | | |
|---|---|---|---|---|---|---|---|---|---|---|---|---|---|
| >hg38_1_-_228634819_228635039_RNA5S12_5S_50ntFlanks@85.110.26\|TCGTCTGATCTCGGAAGCTAAGCAGG | >hg38_1_-_228634819_228635039_RNA5S12_5S_50ntFlanks | 85 | 110 | 26 | TCGTCTGATCTCGGAAGCTAAGCAGG | 156.0 | 11.0 | 1.9 | 5.9 | 3.6E-09 | 4.6E-08 | BT20/MB468 | MP |
| >hg38_21_+_8433172_8446622_RNA45SN1_45S_50ntFlanks@6780.6806.27\|CGGGGCTACGCCTGTCTGAGCGTCGCT | >hg38_21_+_8433172_8446622_RNA45SN1_45S_50ntFlanks | 6780 | 6806 | 27 | CGGGGCTACGCCTGTCTGAGCGTCGCT | 1695.9 | 5.0 | 0.9 | 5.6 | 1.8E-08 | 4.6E-08 | BT20/MB468 | MITO |
| >hg38_21_+_8433172_8446622_RNA45SN1_45S_50ntFlanks@12690.12711.22\|GCGGAACGATACGGCAGCGCCG | >hg38_21_+_8433172_8446622_RNA45SN1_45S_50ntFlanks | 12690 | 12711 | 22 | GCGGAACGATACGGCAGCGCCG | 381.9 | -6.3 | 1.1 | -5.6 | 2.5E-08 | 4.6E-08 | MB231/MB468 | MITO |
| >hg38_21_+_8433172_8446622_RNA45SN1_45S_50ntFlanks@11670.11687.18\|CCCAGTGCTCTGAATGTC | >hg38_21_+_8433172_8446622_RNA45SN1_45S_50ntFlanks | 11670 | 11687 | 18 | CCCAGTGCTCTGAATGTC | 552.0 | -4.2 | 0.7 | -5.6 | 2.5E-08 | 4.6E-08 | MB231/MB468 | MITO |
| >hg38_21_+_8433172_8446622_RNA45SN1_45S_50ntFlanks@11189.11225.37\|ACCCGGGGGGCCGGCGGCGGCGGCGACTCTGGACGCG | >hg38_21_+_8433172_8446622_RNA45SN1_45S_50ntFlanks | 11189 | 11225 | 37 | ACCCGGGGGGCCGGCGGCGGCGGCGACTCTGGACGCG | 1229.7 | -9.3 | 1.7 | -5.6 | 2.6E-08 | 4.6E-08 | BT20/MB468 | NUC |
| >hg38_21_+_8433172_8446622_RNA45SN1_45S_50ntFlanks@8105.8142.38\|CGCGGCGGGGCGCGGGACATGTGGCGTACGGAAGACCC | >hg38_21_+_8433172_8446622_RNA45SN1_45S_50ntFlanks | 8105 | 8142 | 38 | CGCGGCGGGGCGCGGGACATGTGGCGTACGGAAGACCC | 207.9 | -6.8 | 1.2 | -5.6 | 2.4E-08 | 4.6E-08 | MB231/MB468 | NUC |
| >hg38_21_+_8433172_8446622_RNA45SN1_45S_50ntFlanks@9718.9737.20\|TGATTAGAGGTCTTGGGGCC | >hg38_21_+_8433172_8446622_RNA45SN1_45S_50ntFlanks | 9718 | 9737 | 20 | TGATTAGAGGTCTTGGGGCC | 151.5 | -9.3 | 1.7 | -5.6 | 1.8E-08 | 4.6E-08 | BT20/MB468 | MITO |
| >hg38_21_+_8433172_8446622_RNA45SN1_45S_50ntFlanks@12988.13013.26\|CAGCTCCCTCGCTGCGATCTATTGAA | >hg38_21_+_8433172_8446622_RNA45SN1_45S_50ntFlanks | 12988 | 13013 | 26 | CAGCTCCCTCGCTGCGATCTATTGAA | 2576.2 | -4.9 | 0.9 | -5.6 | 2.5E-08 | 4.6E-08 | MB231/MB468 | MITO |
| >MI0000095\|hsa-mir-93&WithFlank&7\|-\|100093762\|100093853@17.37.21&[MIMAT0000093&hsa-miR-93-5p&offsets\|0\|-2;m-29&7\|-\|100093815\|100093837&offsets\|0\|-2]\|CAAAGTGCTGTTCGTGCAGGT | >MI0000095\|hsa-mir-93&WithFlank&7\|-\|100093762\|100093853 | 17 | 37 | 21 | CAAAGTGCTGTTCGTGCAGGT | 1436.7 | 5.5 | 1.0 | 5.6 | 2.5E-08 | 4.6E-08 | MB231/MB468 | MITO |
| >hg38_21_+_8433172_8446622_RNA45SN1_45S_50ntFlanks@8194.8211.18\|CCCGTGGACGGTGTGAGG | >hg38_21_+_8433172_8446622_RNA45SN1_45S_50ntFlanks | 8194 | 8211 | 18 | CCCGTGGACGGTGTGAGG | 781.9 | -3.5 | 0.6 | -5.6 | 2.3E-08 | 4.6E-08 | MB231/MB468 | CYTO |
| >hg38_21_+_8433172_8446622_RNA45SN1_45S_50ntFlanks@12414.12441.28\|CCTTCGATGTCGGCTCTTCCTATCATTG | >hg38_21_+_8433172_8446622_RNA45SN1_45S_50ntFlanks | 12414 | 12441 | 28 | CCTTCGATGTCGGCTCTTCCTATCATTG | 296.5 | 3.5 | 0.6 | 5.6 | 2.3E-08 | 4.6E-08 | MB231/MB468 | CYTO |
| >hg38_21_+_8433172_8446622_RNA45SN1_45S_50ntFlanks@10786.10823.38\|AACAGCCTCTGGCATGTTGGAACAATGTAGGTAAGGGA | >hg38_21_+_8433172_8446622_RNA45SN1_45S_50ntFlanks | 10786 | 10823 | 38 | AACAGCCTCTGGCATGTTGGAACAATGTAGGTAAGGGA | 824.1 | -8.7 | 1.6 | -5.6 | 2.6E-08 | 4.6E-08 | BT20/MB468 | NUC |
| >hg38_21_+_8433172_8446622_RNA45SN1_45S_50ntFlanks@3705.3739.35\|TACCTGGTTGATCCTGCCAGTAGCATATGCTTGTC | >hg38_21_+_8433172_8446622_RNA45SN1_45S_50ntFlanks | 3705 | 3739 | 35 | TACCTGGTTGATCCTGCCAGTAGCATATGCTTGTC | 255.8 | 2.3 | 0.4 | 5.6 | 2.4E-08 | 4.6E-08 | MB231/MB468 | NUC |
| >hg38_21_+_8433172_8446622_RNA45SN1_45S_50ntFlanks@12397.12417.21\|CGACGTCGCTTTTTGATCCTT | >hg38_21_+_8433172_8446622_RNA45SN1_45S_50ntFlanks | 12397 | 12417 | 21 | CGACGTCGCTTTTTGATCCTT | 1070.3 | -3.3 | 0.6 | -5.6 | 2.4E-08 | 4.6E-08 | MB231/MB468 | NUC |
| >hg38_21_+_8433172_8446622_RNA45SN1_45S_50ntFlanks@3813.3834.22\|TTATGGTTCCTTTGGTCGCTCG | >hg38_21_+_8433172_8446622_RNA45SN1_45S_50ntFlanks | 3813 | 3834 | 22 | TTATGGTTCCTTTGGTCGCTCG | 528.8 | -6.1 | 1.1 | -5.6 | 2.4E-08 | 4.6E-08 | MB231/MB468 | NUC |
| >hg38_21_+_8433172_8446622_RNA45SN1_45S_50ntFlanks@12992.13012.21\|TCCCTCGCTGCGATCTATTGA | >hg38_21_+_8433172_8446622_RNA45SN1_45S_50ntFlanks | 12992 | 13012 | 21 | TCCCTCGCTGCGATCTATTGA | 290.2 | -5.7 | 1.0 | -5.6 | 2.6E-08 | 4.6E-08 | MB231/MB468 | MITO |
| >hg38_21_+_8433172_8446622_RNA45SN1_45S_50ntFlanks@10737.10755.19\|TAAATCTCGCGCCGGGCCG | >hg38_21_+_8433172_8446622_RNA45SN1_45S_50ntFlanks | 10737 | 10755 | 19 | TAAATCTCGCGCCGGGCCG | 4244.7 | -8.8 | 1.6 | -5.6 | 2.4E-08 | 4.6E-08 | MB231/MB468 | NUC |
| >hg38_1_-_228634819_228635039_RNA5S12_5S_50ntFlanks@127.170.44\|ACTTGGATGGGAGACCGCCTGGGAATACCGGGTGCTGTAGGCTT | >hg38_1_-_228634819_228635039_RNA5S12_5S_50ntFlanks | 127 | 170 | 44 | ACTTGGATGGGAGACCGCCTGGGAATACCGGGTGCTGTAGGCTT | 1772.8 | 8.8 | 1.6 | 5.6 | 1.8E-08 | 4.6E-08 | BT20/MB468 | MITO |
| >hg38_21_+_8433172_8446622_RNA45SN1_45S_50ntFlanks@10389.10430.42\|TGAACAGCAGTTGAACATGGGTCAGTCGGTCCTGAGAGATGG | >hg38_21_+_8433172_8446622_RNA45SN1_45S_50ntFlanks | 10389 | 10430 | 42 | TGAACAGCAGTTGAACATGGGTCAGTCGGTCCTGAGAGATGG | 249.6 | -7.6 | 1.4 | -5.6 | 2.0E-08 | 4.6E-08 | BT20/MB468 | CYTO |
| >MI0000736\|hsa-mir-30c-1&WithFlank&1\|+\|40757278\|40757378@23.46.24;MI0000254\|hsa-mir-30c-2&WithFlank&6\|-\|71376954\|71377037@13.36.24&[MIMAT0000244&hsa-miR-30c-5p&offsets\|0\|+1;m-57&1\|+\|40757300\|40757323&offsets\|0\|0];[MIMAT0000244_1&hsa-miR-30c-5p&offsets\|0\|+1;m-56&6\|-\|71377002\|71377025&offsets\|0\|0]\|TGTAAACATCCTACACTCTCAGCT | >MI0000736\|hsa-mir-30c-1&WithFlank&1\|+\|40757278\|40757378 | 23 | 46 | 24 | TGTAAACATCCTACACTCTCAGCT | 44882.2 | 5.4 | 1.0 | 5.6 | 2.6E-08 | 4.6E-08 | MB231/MB468 | MITO |
| >hg38_21_+_8433172_8446622_RNA45SN1_45S_50ntFlanks@11769.11794.26\|TAATTAGTGACGCGCATGAATGGATG | >hg38_21_+_8433172_8446622_RNA45SN1_45S_50ntFlanks | 11769 | 11794 | 26 | TAATTAGTGACGCGCATGAATGGATG | 5681.4 | -10.0 | 1.8 | -5.6 | 2.7E-08 | 4.7E-08 | BT20/MB468 | NUC |
| >hg38_21_+_8433172_8446622_RNA45SN1_45S_50ntFlanks@12073.12108.36\|CCGAGGGGCTCTCGCTTCTGGCGCCAAGCGCCCGGC | >hg38_21_+_8433172_8446622_RNA45SN1_45S_50ntFlanks | 12073 | 12108 | 36 | CCGAGGGGCTCTCGCTTCTGGCGCCAAGCGCCCGGC | 356.3 | -7.4 | 1.3 | -5.6 | 2.2E-08 | 4.7E-08 | MB231/MB468 | MP |
| >hg38_21_+_8433172_8446622_RNA45SN1_45S_50ntFlanks@12007.12042.36\|GCCGCCGGTGAAATACCACTACTCTGATCGTTTTTT | >hg38_21_+_8433172_8446622_RNA45SN1_45S_50ntFlanks | 12007 | 12042 | 36 | GCCGCCGGTGAAATACCACTACTCTGATCGTTTTTT | 895.5 | -8.2 | 1.5 | -5.6 | 2.6E-08 | 4.7E-08 | MB231/MB468 | MITO |
| >hg38_21_+_8433172_8446622_RNA45SN1_45S_50ntFlanks@12785.12805.21\|CCCCGCGCGCGCGGGAGGGCG | >hg38_21_+_8433172_8446622_RNA45SN1_45S_50ntFlanks | 12785 | 12805 | 21 | CCCCGCGCGCGCGGGAGGGCG | 748.7 | -6.8 | 1.2 | -5.6 | 2.3E-08 | 4.7E-08 | MB231/MB468 | CYTO |
| >hg38_21_+_8433172_8446622_RNA45SN1_45S_50ntFlanks@7974.8007.34\|ACGCGACCTCAGATCAGACGTGGCGACCCGCTGA | >hg38_21_+_8433172_8446622_RNA45SN1_45S_50ntFlanks | 7974 | 8007 | 34 | ACGCGACCTCAGATCAGACGTGGCGACCCGCTGA | 11942.4 | 3.7 | 0.7 | 5.6 | 1.8E-08 | 4.7E-08 | BT20/MB468 | MITO |
| >trna111_HisGTG_1_-_147774845_147774916@-1T.25.26,trna118_HisGTG_1_-_145396881_145396952@-1T.25.26,trna16_HisGTG_1_+_146544773_146544844@-1T.25.26,trna1_HisGTG_15_+_45493349_45493420@-1T.25.26,trna21_HisGTG_1_+_147753471_147753542@-1T.25.26,trna33_HisGTG_6_+_27125906_27125977@-1T.25.26,trna7_HisGTG_9_-_14433938_14434009@-1T.25.26,trna8_HisGTG_15_-_45492611_45492682@-1T.25.26,trna9_HisGTG_15_-_45490804_45490875@-1T.25.26\|TGCCGTGATCGTATAGTGGTTAGTAC | >trna111_HisGTG_1_-_147774845_147774916 | -1T | 25 | 26 | TGCCGTGATCGTATAGTGGTTAGTAC | 609.2 | 4.4 | 0.8 | 5.6 | 2.4E-08 | 4.7E-08 | MB231/MB468 | NUC |
| >MI0000085\|hsa-mir-27a&WithFlank&19\|-\|13836434\|13836523@57.76.20(+3U)&[MIMAT0000084&hsa-miR-27a-3p&offsets\|0\|-1(+3U);m-43&19\|-\|13836447\|13836467&offsets\|0\|-1(+3U)]\|TTCACAGTGGCTAAGTTCCGTTT | >MI0000085\|hsa-mir-27a&WithFlank&19\|-\|13836434\|13836523 | 57 | 76 | 20(+3U) | TTCACAGTGGCTAAGTTCCGTTT | 1704.0 | 5.6 | 1.0 | 5.6 | 2.6E-08 | 4.7E-08 | MB231/MB468 | MITO |
| >hg38_21_+_8433172_8446622_RNA45SN1_45S_50ntFlanks@5035.5070.36\|CATGGCCGTTCTTAGTTGGTGGAGCGATTTGTCTGG | >hg38_21_+_8433172_8446622_RNA45SN1_45S_50ntFlanks | 5035 | 5070 | 36 | CATGGCCGTTCTTAGTTGGTGGAGCGATTTGTCTGG | 229.0 | -9.9 | 1.8 | -5.6 | 1.8E-08 | 4.7E-08 | BT20/MB468 | MITO |
| >MI0000446\|hsa-mir-125b-1&WithFlank&11\|-\|122099751\|122099850@21.42.22;MI0000470\|hsa-mir-125b-2&WithFlank&21\|+\|16590231\|16590331@23.44.22&[MIMAT0000423&hsa-miR-125b-5p&offsets\|0\|0;m-67&11\|-\|122099809\|122099830&offsets\|0\|0];[MIMAT0000423_1&hsa-miR-125b-5p&offsets\|0\|0;m-68&21\|+\|16590253\|16590274&offsets\|0\|0]\|TCCCTGAGACCCTAACTTGTGA | >MI0000446\|hsa-mir-125b-1&WithFlank&11\|-\|122099751\|122099850 | 21 | 42 | 22 | TCCCTGAGACCCTAACTTGTGA | 1597.1 | 5.8 | 1.0 | 5.6 | 2.0E-08 | 4.7E-08 | BT20/MB468 | CYTO |
| >hg38_1_-_228634819_228635039_RNA5S12_5S_50ntFlanks@125.169.45\|GTACTTGGATGGGAGACCGCCTGGGAATACCGGGTGCTGTAGGCT | >hg38_1_-_228634819_228635039_RNA5S12_5S_50ntFlanks | 125 | 169 | 45 | GTACTTGGATGGGAGACCGCCTGGGAATACCGGGTGCTGTAGGCT | 149.4 | 11.0 | 1.9 | 5.9 | 3.7E-09 | 4.7E-08 | BT20/MB468 | MP |
| >hg38_21_+_8433172_8446622_RNA45SN1_45S_50ntFlanks@3780.3801.22\|TACAGTGAAACTGCGAATGGCT | >hg38_21_+_8433172_8446622_RNA45SN1_45S_50ntFlanks | 3780 | 3801 | 22 | TACAGTGAAACTGCGAATGGCT | 748.3 | -9.1 | 1.6 | -5.6 | 2.7E-08 | 4.7E-08 | BT20/MB468 | NUC |
| >hg38_21_+_8433172_8446622_RNA45SN1_45S_50ntFlanks@6743.6770.28\|CGACACTTCGAACGCACTTGCGGCCCCG | >hg38_21_+_8433172_8446622_RNA45SN1_45S_50ntFlanks | 6743 | 6770 | 28 | CGACACTTCGAACGCACTTGCGGCCCCG | 2268.1 | -9.6 | 1.7 | -5.6 | 2.7E-08 | 4.7E-08 | BT20/MB468 | NUC |
| >hg38_21_+_8433172_8446622_RNA45SN1_45S_50ntFlanks@11208.11227.20\|GCGGCGACTCTGGACGCGAG | >hg38_21_+_8433172_8446622_RNA45SN1_45S_50ntFlanks | 11208 | 11227 | 20 | GCGGCGACTCTGGACGCGAG | 733.2 | -7.6 | 1.4 | -5.6 | 2.4E-08 | 4.7E-08 | MB231/MB468 | NUC |
| >hg38_21_+_8433172_8446622_RNA45SN1_45S_50ntFlanks@9573.9603.31\|AAATCGGTCGTCCGACCTGGGTATAGGGGCG | >hg38_21_+_8433172_8446622_RNA45SN1_45S_50ntFlanks | 9573 | 9603 | 31 | AAATCGGTCGTCCGACCTGGGTATAGGGGCG | 2760.2 | -5.3 | 0.9 | -5.6 | 2.2E-08 | 4.7E-08 | MB231/MB468 | MP |
| >hg38_21_+_8433172_8446622_RNA45SN1_45S_50ntFlanks@3802.3836.35\|CATTAAATCAGTTATGGTTCCTTTGGTCGCTCGCT | >hg38_21_+_8433172_8446622_RNA45SN1_45S_50ntFlanks | 3802 | 3836 | 35 | CATTAAATCAGTTATGGTTCCTTTGGTCGCTCGCT | 475.3 | -6.1 | 1.1 | -5.6 | 2.4E-08 | 4.7E-08 | MB231/MB468 | NUC |
| >MI0000760\|hsa-mir-361&WithFlank&X\|-\|85903630\|85903713@51.74.24&[MIMAT0004682&hsa-miR-361-3p&offsets\|0\|+1;m-124&X\|-\|85903642\|85903663&offsets\|0\|+2]\|TCCCCCAGGTGTGATTCTGATTTG | >MI0000760\|hsa-mir-361&WithFlank&X\|-\|85903630\|85903713 | 51 | 74 | 24 | TCCCCCAGGTGTGATTCTGATTTG | 550.1 | 6.5 | 1.2 | 5.6 | 2.4E-08 | 4.7E-08 | MB231/MB468 | CYTO |
| >MI0000811\|hsa-mir-148b&WithFlank&12\|+\|54337210\|54337320@69.89.21(+1C)&[MIMAT0000759&hsa-miR-148b-3p&offsets\|0\|-1(+1C);m-77&12\|+\|54337278\|54337299&offsets\|0\|-1(+1C)]\|TCAGTGCATCACAGAACTTTGC | >MI0000811\|hsa-mir-148b&WithFlank&12\|+\|54337210\|54337320 | 69 | 89 | 21(+1C) | TCAGTGCATCACAGAACTTTGC | 681.8 | 5.7 | 1.0 | 5.6 | 2.4E-08 | 4.7E-08 | MB231/MB468 | CYTO |
| >hg38_21_+_8433172_8446622_RNA45SN1_45S_50ntFlanks@13008.13025.18\|ATTGAAAGTCAGCCCTCG | >hg38_21_+_8433172_8446622_RNA45SN1_45S_50ntFlanks | 13008 | 13025 | 18 | ATTGAAAGTCAGCCCTCG | 1202.1 | -6.6 | 1.2 | -5.6 | 2.4E-08 | 4.7E-08 | MB231/MB468 | NUC |
| >hg38_21_+_8433172_8446622_RNA45SN1_45S_50ntFlanks@5485.5512.28\|AGCGCTGAGAAGACGGTCGAACTTGACT | >hg38_21_+_8433172_8446622_RNA45SN1_45S_50ntFlanks | 5485 | 5512 | 28 | AGCGCTGAGAAGACGGTCGAACTTGACT | 349.2 | -7.2 | 1.3 | -5.6 | 2.6E-08 | 4.7E-08 | MB231/MB468 | MITO |
| >hg38_21_+_8433172_8446622_RNA45SN1_45S_50ntFlanks@11641.11662.22\|CGGCGGGTGTTGACGCGATGTG | >hg38_21_+_8433172_8446622_RNA45SN1_45S_50ntFlanks | 11641 | 11662 | 22 | CGGCGGGTGTTGACGCGATGTG | 8823.8 | -7.9 | 1.3 | -5.9 | 3.7E-09 | 4.7E-08 | BT20/MB468 | MP |
| >hg38_21_+_8433172_8446622_RNA45SN1_45S_50ntFlanks@9572.9605.34\|CAAATCGGTCGTCCGACCTGGGTATAGGGGCGAA | >hg38_21_+_8433172_8446622_RNA45SN1_45S_50ntFlanks | 9572 | 9605 | 34 | CAAATCGGTCGTCCGACCTGGGTATAGGGGCGAA | 358.4 | -3.0 | 0.5 | -5.6 | 2.4E-08 | 4.7E-08 | MB231/MB468 | NUC |
| >hg38_21_+_8433172_8446622_RNA45SN1_45S_50ntFlanks@6649.6680.32\|TACGACTCTTAGCGGTGGATCACTCGGCTCGT | >hg38_21_+_8433172_8446622_RNA45SN1_45S_50ntFlanks | 6649 | 6680 | 32 | TACGACTCTTAGCGGTGGATCACTCGGCTCGT | 351.6 | -4.4 | 0.8 | -5.6 | 2.2E-08 | 4.7E-08 | MB231/MB468 | MP |
| >hg38_21_+_8433172_8446622_RNA45SN1_45S_50ntFlanks@3782.3805.24\|CAGTGAAACTGCGAATGGCTCATT | >hg38_21_+_8433172_8446622_RNA45SN1_45S_50ntFlanks | 3782 | 3805 | 24 | CAGTGAAACTGCGAATGGCTCATT | 835.8 | -6.8 | 1.2 | -5.6 | 2.7E-08 | 4.7E-08 | BT20/MB468 | NUC |
| >MI0000272\|hsa-mir-182&WithFlank&7\|-\|129770377\|129770498@29.52.24(+1U)&[MIMAT0000259&hsa-miR-182-5p&offsets\|0\|0(+1U);m-22&7\|-\|129770449\|129770470&offsets\|0\|+2(+1U)]\|TTTGGCAATGGTAGAACTCACACTT | >MI0000272\|hsa-mir-182&WithFlank&7\|-\|129770377\|129770498 | 29 | 52 | 24(+1U) | TTTGGCAATGGTAGAACTCACACTT | 403.4 | 4.4 | 0.7 | 6.1 | 1.4E-09 | 4.7E-08 | BT20/MB231 | CYTO |
| >MI0003137\|hsa-mir-193b&WithFlank&16\|+\|14303961\|14304055@57.77.21(+1A)&[MIMAT0002819&hsa-miR-193b-3p&offsets\|0\|-1(+1A);m-166&16\|+\|14304017\|14304038&offsets\|0\|-1(+1A)]\|AACTGGCCCTCAAAGTCCCGCA | >MI0003137\|hsa-mir-193b&WithFlank&16\|+\|14303961\|14304055 | 57 | 77 | 21(+1A) | AACTGGCCCTCAAAGTCCCGCA | 349.7 | 5.0 | 0.8 | 6.1 | 1.4E-09 | 4.7E-08 | BT20/MB231 | CYTO |

| | | | | | | | | | | | | | |
|---|---|---|---|---|---|---|---|---|---|---|---|---|---|
| >hg38_21_+_8433172_8446622_RNA45SN1_45S_50ntFlanks@11568.11588.21\|GCAGCCGACTTAGAACTGGTG | >hg38_21_+_8433172_8446622_RNA45SN1_45S_50ntFlanks | 11568 | 11588 | 21 | GCAGCCGACTTAGAACTGGTG | 461.4 | -2.5 | 0.4 | -5.6 | 2.0E-08 | 4.7E-08 | BT20/MB468 | CYTO |
| >hg38_21_+_8433172_8446622_RNA45SN1_45S_50ntFlanks@12004.12043.40\|CGGGCCGCCGGTGAAATACCACTACTCTGATCGTTTTTTC | >hg38_21_+_8433172_8446622_RNA45SN1_45S_50ntFlanks | 12004 | 12043 | 40 | CGGGCCGCCGGTGAAATACCACTACTCTGATCGTTTTTTC | 134.6 | -10.9 | 2.0 | -5.6 | 2.2E-08 | 4.7E-08 | MB231/MB468 | MP |
| >hg38_21_+_8433172_8446622_RNA45SN1_45S_50ntFlanks@10841.10862.22\|CGTAACTTCGGGATAAGGATTG | >hg38_21_+_8433172_8446622_RNA45SN1_45S_50ntFlanks | 10841 | 10862 | 22 | CGTAACTTCGGGATAAGGATTG | 618.2 | -4.0 | 0.7 | -5.6 | 1.8E-08 | 4.7E-08 | BT20/MB468 | MITO |
| >hg38_MT_+_598_1651_RNR1_12S_50ntFlanks@843.869.27\|GAGTAGAGTGCTTAGTTGAACAGGGCC | >hg38_MT_+_598_1651_RNR1_12S_50ntFlanks | 843 | 869 | 27 | GAGTAGAGTGCTTAGTTGAACAGGGCC | 117.3 | 10.6 | 1.8 | 5.9 | 3.7E-09 | 4.7E-08 | BT20/MB468 | MP |
| >hg38_21_+_8433172_8446622_RNA45SN1_45S_50ntFlanks@8194.8217.24\|CCCGTGGACGGTGTGAGGCCGGTA | >hg38_21_+_8433172_8446622_RNA45SN1_45S_50ntFlanks | 8194 | 8217 | 24 | CCCGTGGACGGTGTGAGGCCGGTA | 94.8 | -9.6 | 1.6 | -5.9 | 3.7E-09 | 4.7E-08 | BT20/MB468 | MP |
| >hg38_21_+_8433172_8446622_RNA45SN1_45S_50ntFlanks@9653.9682.30\|AGGATAGCTGGCGCTCTCGCAGACCCGACG | >hg38_21_+_8433172_8446622_RNA45SN1_45S_50ntFlanks | 9653 | 9682 | 30 | AGGATAGCTGGCGCTCTCGCAGACCCGACG | 286.8 | -4.2 | 0.8 | -5.6 | 2.6E-08 | 4.7E-08 | MB231/MB468 | MITO |
| >hg38_21_+_8433172_8446622_RNA45SN1_45S_50ntFlanks@8337.8373.37\|AGTACCGTAAGGGAAAGTTGAAAAGAACTTTGAAGAG | >hg38_21_+_8433172_8446622_RNA45SN1_45S_50ntFlanks | 8337 | 8373 | 37 | AGTACCGTAAGGGAAAGTTGAAAAGAACTTTGAAGAG | 103.0 | 10.4 | 1.8 | 5.9 | 3.7E-09 | 4.7E-08 | BT20/MB468 | MP |
| >hg38_21_+_8433172_8446622_RNA45SN1_45S_50ntFlanks@10402.10425.24\|AACATGGGTCAGTCGGTCCTGAGA | >hg38_21_+_8433172_8446622_RNA45SN1_45S_50ntFlanks | 10402 | 10425 | 24 | AACATGGGTCAGTCGGTCCTGAGA | 228.5 | -7.3 | 1.3 | -5.6 | 2.2E-08 | 4.7E-08 | MB231/MB468 | MP |
| >hg38_21_+_8433172_8446622_RNA45SN1_45S_50ntFlanks@10798.10822.25\|CATGTTGGAACAATGTAGGTAAGGG | >hg38_21_+_8433172_8446622_RNA45SN1_45S_50ntFlanks | 10798 | 10822 | 25 | CATGTTGGAACAATGTAGGTAAGGG | 255.8 | -3.3 | 0.6 | -5.6 | 2.4E-08 | 4.7E-08 | MB231/MB468 | CYTO |
| >trna152_ValCAC_6_-_27248049_27248121@1.33.33\|GCTTCTGTAGTGTAGTGGTTATCACGTTCGCCT | >trna152_ValCAC_6_-_27248049_27248121 | 1 | 33 | 33 | GCTTCTGTAGTGTAGTGGTTATCACGTTCGCCT | 924.6 | 8.1 | 1.4 | 5.6 | 2.6E-08 | 4.7E-08 | MB231/MB468 | MITO |
| >hg38_21_+_8433172_8446622_RNA45SN1_45S_50ntFlanks@11888.11912.25\|AGCTTGACTCTAGTCTGGCACGGTG | >hg38_21_+_8433172_8446622_RNA45SN1_45S_50ntFlanks | 11888 | 11912 | 25 | AGCTTGACTCTAGTCTGGCACGGTG | 117.9 | -10.0 | 1.7 | -5.9 | 3.7E-09 | 4.7E-08 | BT20/MB468 | MP |
| >hg38_21_+_8433172_8446622_RNA45SN1_45S_50ntFlanks@9792.9815.24\|GGCGTGGAGCCGGGCGTGGAATGC | >hg38_21_+_8433172_8446622_RNA45SN1_45S_50ntFlanks | 9792 | 9815 | 24 | GGCGTGGAGCCGGGCGTGGAATGC | 166.8 | 11.1 | 1.9 | 5.9 | 3.8E-09 | 4.7E-08 | BT20/MB468 | MP |
| >MI0000299\|hsa-mir-222&WithFlank&X\|-\|45747009\|45747130@75.94.20&[MIMAT0000279&hsa-miR-222-3p&offsets\|0\|-1;m-140&X\|-\|45747033\|45747056&offsets\|0\|-4]\|AGCTACATCTGGCTACTGGG | >MI0000299\|hsa-mir-222&WithFlank&X\|-\|45747009\|45747130 | 75 | 94 | 20 | AGCTACATCTGGCTACTGGG | 226.6 | 5.9 | 1.1 | 5.6 | 2.5E-08 | 4.7E-08 | MB231/MB468 | NUC |
| >hg38_21_+_8433172_8446622_RNA45SN1_45S_50ntFlanks@5074.5102.29\|ATTCCGATAACGAACGAGACTCTGGCATG | >hg38_21_+_8433172_8446622_RNA45SN1_45S_50ntFlanks | 5074 | 5102 | 29 | ATTCCGATAACGAACGAGACTCTGGCATG | 720.4 | -9.1 | 1.6 | -5.6 | 2.7E-08 | 4.7E-08 | BT20/MB468 | NUC |
| >MI0000749\|hsa-mir-30e&WithFlank&1\|+\|40754349\|40754452@65.85.21(+1U)&[MIMAT0000693&hsa-miR-30e-3p&offsets\|0\|-1(+1U);m-25&1\|+\|40754413\|40754434&offsets\|0\|-1(+1U)]\|CTTTCAGTCGGATGTTTACAGT | >MI0000749\|hsa-mir-30e&WithFlank&1\|+\|40754349\|40754452 | 65 | 85 | 21(+1U) | CTTTCAGTCGGATGTTTACAGT | 1756.6 | 5.0 | 0.9 | 5.6 | 2.4E-08 | 4.8E-08 | MB231/MB468 | CYTO |
| >hg38_21_+_8433172_8446622_RNA45SN1_45S_50ntFlanks@6653.6688.36\|ACTCTTAGCGGTGGATCACTCGGCTCGTGCGTCGAT | >hg38_21_+_8433172_8446622_RNA45SN1_45S_50ntFlanks | 6653 | 6688 | 36 | ACTCTTAGCGGTGGATCACTCGGCTCGTGCGTCGAT | 3889.6 | 3.4 | 0.6 | 5.6 | 1.8E-08 | 4.8E-08 | BT20/MB468 | MITO |
| >hg38_21_+_8433172_8446622_RNA45SN1_45S_50ntFlanks@10404.10426.23\|CATGGGTCAGTCGGTCCTGAGAG | >hg38_21_+_8433172_8446622_RNA45SN1_45S_50ntFlanks | 10404 | 10426 | 23 | CATGGGTCAGTCGGTCCTGAGAG | 191.6 | -7.2 | 1.3 | -5.6 | 2.3E-08 | 4.8E-08 | MB231/MB468 | MP |
| >trna10_AspGTC_12_-_125424193_125424264@32.71.40,trna12_AspGTC_12_-_125411891_125411962@32.71.40,trna144_AspGTC_6_-_27551236_27551307@32.71.40,trna38_AspGTC_17_-_8125556_8125627@32.71.40,trna45_AspGTC_6_+_27447453_27447524@32.71.40,trna48_AspGTC_6_+_27471523_27471594@32.71.40,trna4_AspGTC_12_+_96429799_96429870@32.71.40,trna69_AspGTC_1_-_161440205_161440276@32.71.40,trna72_AspGTC_1_-_161432824_161432895@32.71.40,trna75_AspGTC_1_-_161425414_161425485@32.71.40,trna78_AspGTC_1_-_161418033_161418104@32.71.40,trna81_AspGTC_1_-_161410615_161410686@32.71.40\|CTGTCACGCGGGAGACCGGGGTTCGATTCCCCGACGGGGA | >trna10_AspGTC_12_-_125424193_125424264 | 32 | 71 | 40 | CTGTCACGCGGGAGACCGGGGTTCGATTCCCCGACGGGGA | 222.1 | -6.5 | 1.2 | -5.6 | 2.1E-08 | 4.8E-08 | BT20/MB468 | CYTO |
| >hg38_21_+_8433172_8446622_RNA45SN1_45S_50ntFlanks@8401.8418.18\|AGAGGTAAACGGGTGGGG | >hg38_21_+_8433172_8446622_RNA45SN1_45S_50ntFlanks | 8401 | 8418 | 18 | AGAGGTAAACGGGTGGGG | 111.4 | -9.9 | 1.7 | -5.9 | 3.8E-09 | 4.8E-08 | BT20/MB468 | MP |
| >hg38_21_+_8433172_8446622_RNA45SN1_45S_50ntFlanks@11663.11685.23\|ATTTCTGCCCAGTGCTCTGAATG | >hg38_21_+_8433172_8446622_RNA45SN1_45S_50ntFlanks | 11663 | 11685 | 23 | ATTTCTGCCCAGTGCTCTGAATG | 822.2 | -5.0 | 0.9 | -5.6 | 2.7E-08 | 4.8E-08 | BT20/MB468 | NUC |
| >hg38_21_+_8433172_8446622_RNA45SN1_45S_50ntFlanks@10873.10901.29\|CTGGGTCGGTCGGGCTGGGGCGCGAAGCG | >hg38_21_+_8433172_8446622_RNA45SN1_45S_50ntFlanks | 10873 | 10901 | 29 | CTGGGTCGGTCGGGCTGGGGCGCGAAGCG | 405.0 | -7.3 | 1.3 | -5.6 | 2.5E-08 | 4.8E-08 | MB231/MB468 | NUC |
| >hg38_21_+_8433172_8446622_RNA45SN1_45S_50ntFlanks@4291.4317.27\|AGGATCCATTGGAGGGCAAGTCTGGTG | >hg38_21_+_8433172_8446622_RNA45SN1_45S_50ntFlanks | 4291 | 4317 | 27 | AGGATCCATTGGAGGGCAAGTCTGGTG | 389.6 | -8.2 | 1.5 | -5.6 | 2.7E-08 | 4.8E-08 | BT20/MB468 | NUC |
| >hg38_21_+_8433172_8446622_RNA45SN1_45S_50ntFlanks@10839.10865.27\|TCCGTAACTTCGGGATAAGGATTGGCT | >hg38_21_+_8433172_8446622_RNA45SN1_45S_50ntFlanks | 10839 | 10865 | 27 | TCCGTAACTTCGGGATAAGGATTGGCT | 1312.2 | -7.9 | 1.4 | -5.6 | 2.1E-08 | 4.8E-08 | BT20/MB468 | CYTO |
| >MI0001445\|hsa-mir-423&WithFlank&17\|+\|30117073\|30117178@23.45.23&[MIMAT0004748&hsa-miR-423-5p&offsets\|0\|0;m-141&17\|+\|30117095\|30117117&offsets\|0\|0]\|TGAGGGGCAGAGAGCGAGACTTT | >MI0001445\|hsa-mir-423&WithFlank&17\|+\|30117073\|30117178 | 23 | 45 | 23 | TGAGGGGCAGAGAGCGAGACTTT | 1129.7 | 2.3 | 0.4 | 5.6 | 2.7E-08 | 4.8E-08 | BT20/MB468 | NUC |
| >hg38_21_+_8433172_8446622_RNA45SN1_45S_50ntFlanks@12081.12110.30\|CTCTCGCTTCTGGCGCCAAGCGCCCGGCCG | >hg38_21_+_8433172_8446622_RNA45SN1_45S_50ntFlanks | 12081 | 12110 | 30 | CTCTCGCTTCTGGCGCCAAGCGCCCGGCCG | 1012.4 | -5.8 | 1.0 | -5.6 | 1.9E-08 | 4.8E-08 | BT20/MB468 | MITO |
| >MI0000270\|hsa-mir-181b-1&WithFlank&1\|-\|198858867\|198858988@42.63.22(+1A);MI0000683\|hsa-mir-181b-2&WithFlank&9\|+\|124693704\|124693804@22.43.22(+1A)&[MIMAT0000257&hsa-miR-181b-5p&offsets\|0\|-1(+1A);m-73&1\|-\|198858925\|198858947&offsets\|0\|-1(+1A)];[MIMAT0000257_1&hsa-miR-181b-5p&offsets\|0\|-1(+1A);m-72&9\|+\|124693725\|124693747&offsets\|0\|-1(+1A)]\|AACATTCATTGCTGTCGGTGGGA | >MI0000270\|hsa-mir-181b-1&WithFlank&1\|-\|198858867\|198858988 | 42 | 63 | 22(+1A) | AACATTCATTGCTGTCGGTGGGA | 301.0 | 4.9 | 0.9 | 5.6 | 2.4E-08 | 4.8E-08 | MB231/MB468 | CYTO |
| >hg38_21_+_8433172_8446622_RNA45SN1_45S_50ntFlanks@11586.11604.19\|GTGCGGACCAGGGGAATCC | >hg38_21_+_8433172_8446622_RNA45SN1_45S_50ntFlanks | 11586 | 11604 | 19 | GTGCGGACCAGGGGAATCC | 399.4 | -8.0 | 1.4 | -5.6 | 2.8E-08 | 4.8E-08 | BT20/MB468 | NUC |
| >MI0000462\|hsa-mir-152&WithFlank&17\|-\|48037155\|48037253@60.82.23&[MIMAT0000438&hsa-miR-152-3p&offsets\|0\|+2;m-97&17\|-\|48037174\|48037194&offsets\|0\|+2]\|TCAGTGCATGACAGAACTTGGGC | >MI0000462\|hsa-mir-152&WithFlank&17\|-\|48037155\|48037253 | 60 | 82 | 23 | TCAGTGCATGACAGAACTTGGGC | 76.0 | 9.5 | 1.7 | 5.6 | 2.3E-08 | 4.8E-08 | MB231/MB468 | MP |
| >hg38_21_+_8433172_8446622_RNA45SN1_45S_50ntFlanks@12159.12180.22\|TTTGACTGGGGCGGTACACCTG | >hg38_21_+_8433172_8446622_RNA45SN1_45S_50ntFlanks | 12159 | 12180 | 22 | TTTGACTGGGGCGGTACACCTG | 1669.5 | -6.1 | 1.1 | -5.6 | 1.9E-08 | 4.8E-08 | BT20/MB468 | MITO |
| >hg38_21_+_8433172_8446622_RNA45SN1_45S_50ntFlanks@12258.12287.30\|GCTTGATCTTGATTTTCAGTACGAATACAG | >hg38_21_+_8433172_8446622_RNA45SN1_45S_50ntFlanks | 12258 | 12287 | 30 | GCTTGATCTTGATTTTCAGTACGAATACAG | 65.5 | -9.9 | 1.8 | -5.6 | 2.3E-08 | 4.8E-08 | MB231/MB468 | MP |
| >MI0000809\|hsa-mir-151a&WithFlank&8\|-\|140732558\|140732659@53.74.22(+2A)&[MIMAT0000757&hsa-miR-151a-3p&offsets\|0\|+1(+2A);m-30&8\|-\|140732587\|140732607&offsets\|0\|+1(+2A)]\|CTAGACTGAAGCTCCTTGAGGAAA | >MI0000809\|hsa-mir-151a&WithFlank&8\|-\|140732558\|140732659 | 53 | 74 | 22(+2A) | CTAGACTGAAGCTCCTTGAGGAAA | 236.7 | 2.4 | 0.4 | 5.6 | 2.8E-08 | 4.8E-08 | BT20/MB468 | NUC |
| >hg38_21_+_8433172_8446622_RNA45SN1_45S_50ntFlanks@5485.5515.31\|AGCGCTGAGAAGACGGTCGAACTTGACTATC | >hg38_21_+_8433172_8446622_RNA45SN1_45S_50ntFlanks | 5485 | 5515 | 31 | AGCGCTGAGAAGACGGTCGAACTTGACTATC | 1090.0 | 13.4 | 2.2 | 6.2 | 7.2E-10 | 4.8E-08 | BT20/MB231 | MP |
| >MI0000102\|hsa-mir-100&WithFlank&11\|-\|122152223\|122152314@19.38.20(+2A)&[MIMAT0000098&hsa-miR-100-5p&offsets\|0\|-2(+2A);m-27&11\|-\|122152276\|122152296&offsets\|0\|-1(+2A)]\|AACCCGTAGATCCGAACTTGAA | >MI0000102\|hsa-mir-100&WithFlank&11\|-\|122152223\|122152314 | 19 | 38 | 20(+2A) | AACCCGTAGATCCGAACTTGAA | 206.3 | -11.1 | 1.8 | -6.2 | 7.3E-10 | 4.8E-08 | BT20/MB231 | MP |
| >hg38_21_+_8433172_8446622_RNA45SN1_45S_50ntFlanks@8114.8140.27\|GCGCGGGACATGTGGCGTACGGAAGAC | >hg38_21_+_8433172_8446622_RNA45SN1_45S_50ntFlanks | 8114 | 8140 | 27 | GCGCGGGACATGTGGCGTACGGAAGAC | 1605.9 | -4.2 | 0.7 | -5.6 | 1.9E-08 | 4.8E-08 | BT20/MB468 | MITO |
| >hg38_21_+_8433172_8446622_RNA45SN1_45S_50ntFlanks@4002.4023.22\|GATAACCTCGGGCCGATCGCAC | >hg38_21_+_8433172_8446622_RNA45SN1_45S_50ntFlanks | 4002 | 4023 | 22 | GATAACCTCGGGCCGATCGCAC | 680.1 | -4.3 | 0.8 | -5.6 | 2.7E-08 | 4.8E-08 | MB231/MB468 | MITO |
| >hg38_21_+_8433172_8446622_RNA45SN1_45S_50ntFlanks@10905.10932.28\|CTGGGCGCGCGCCGCGGCTGGACGAGGC | >hg38_21_+_8433172_8446622_RNA45SN1_45S_50ntFlanks | 10905 | 10932 | 28 | CTGGGCGCGCGCCGCGGCTGGACGAGGC | 1029.5 | -9.3 | 1.7 | -5.6 | 2.8E-08 | 4.8E-08 | BT20/MB468 | NUC |
| >hg38_21_+_8433172_8446622_RNA45SN1_45S_50ntFlanks@12404.12443.40\|GCTTTTTGATCCTTCGATGTCGGCTCTTCCTATCATTGTG | >hg38_21_+_8433172_8446622_RNA45SN1_45S_50ntFlanks | 12404 | 12443 | 40 | GCTTTTTGATCCTTCGATGTCGGCTCTTCCTATCATTGTG | 2200.6 | -10.0 | 1.8 | -5.6 | 2.8E-08 | 4.8E-08 | BT20/MB468 | NUC |
| >MI0000101\|hsa-mir-99a&WithFlank&21\|+\|16539083\|16539175@19.40.22&[MIMAT0000097&hsa-miR-99a-5p&offsets\|0\|0;m-64&21\|+\|16539101\|16539121&offsets\|0\|+1]\|AACCCGTAGATCCGATCTTGTG | >MI0000101\|hsa-mir-99a&WithFlank&21\|+\|16539083\|16539175 | 19 | 40 | 22 | AACCCGTAGATCCGATCTTGTG | 1515.2 | -8.7 | 1.5 | -5.9 | 3.8E-09 | 4.9E-08 | BT20/MB468 | MP |
| >hg38_21_+_8433172_8446622_RNA45SN1_45S_50ntFlanks@5473.5502.30\|CGGCCCTGGCGGAGCGCTGAGAAGACGGTC | >hg38_21_+_8433172_8446622_RNA45SN1_45S_50ntFlanks | 5473 | 5502 | 30 | CGGCCCTGGCGGAGCGCTGAGAAGACGGTC | 517.5 | 5.1 | 0.9 | 5.6 | 2.4E-08 | 4.9E-08 | MB231/MB468 | CYTO |
| >hg38_21_+_8433172_8446622_RNA45SN1_45S_50ntFlanks@6655.6681.27\|TCTTAGCGGTGGATCACTCGGCTCGTG | >hg38_21_+_8433172_8446622_RNA45SN1_45S_50ntFlanks | 6655 | 6681 | 27 | TCTTAGCGGTGGATCACTCGGCTCGTG | 1752.0 | -3.2 | 0.6 | -5.6 | 2.4E-08 | 4.9E-08 | MB231/MB468 | CYTO |
| >hg38_21_+_8433172_8446622_RNA45SN1_45S_50ntFlanks@11456.11473.18\|TCGGCGCGCGGCGGCGGC | >hg38_21_+_8433172_8446622_RNA45SN1_45S_50ntFlanks | 11456 | 11473 | 18 | TCGGCGCGCGGCGGCGGC | 624.0 | 6.3 | 1.1 | 5.6 | 2.5E-08 | 4.9E-08 | MB231/MB468 | CYTO |
| >hg38_21_+_8433172_8446622_RNA45SN1_45S_50ntFlanks@12420.12449.30\|ATGTCGGCTCTTCCTATCATTGTGAAGCAG | >hg38_21_+_8433172_8446622_RNA45SN1_45S_50ntFlanks | 12420 | 12449 | 30 | ATGTCGGCTCTTCCTATCATTGTGAAGCAG | 2153.1 | -7.5 | 1.3 | -5.6 | 2.8E-08 | 4.9E-08 | BT20/MB468 | NUC |
| >hg38_21_+_8433172_8446622_RNA45SN1_45S_50ntFlanks@11125.11145.21\|CGGCGGCGGCCCGCCGCGGGG | >hg38_21_+_8433172_8446622_RNA45SN1_45S_50ntFlanks | 11125 | 11145 | 21 | CGGCGGCGGCCCGCCGCGGGG | 109.6 | -10.6 | 1.9 | -5.6 | 2.3E-08 | 4.9E-08 | MB231/MB468 | MP |
| >hg38_21_+_8433172_8446622_RNA45SN1_45S_50ntFlanks@11588.11611.24\|GCGGACCAGGGGAATCCGACTGTT | >hg38_21_+_8433172_8446622_RNA45SN1_45S_50ntFlanks | 11588 | 11611 | 24 | GCGGACCAGGGGAATCCGACTGTT | 500.1 | 3.1 | 0.6 | 5.6 | 2.5E-08 | 4.9E-08 | MB231/MB468 | CYTO |
| >hg38_21_+_8433172_8446622_RNA45SN1_45S_50ntFlanks@4007.4024.18\|CCTCGGGCCGATCGCACG | >hg38_21_+_8433172_8446622_RNA45SN1_45S_50ntFlanks | 4007 | 4024 | 18 | CCTCGGGCCGATCGCACG | 1045.9 | -6.3 | 1.1 | -5.6 | 2.7E-08 | 4.9E-08 | MB231/MB468 | MITO |
| >hg38_21_+_8433172_8446622_RNA45SN1_45S_50ntFlanks@10794.10825.32\|CTGGCATGTTGGAACAATGTAGGTAAGGGAAG | >hg38_21_+_8433172_8446622_RNA45SN1_45S_50ntFlanks | 10794 | 10825 | 32 | CTGGCATGTTGGAACAATGTAGGTAAGGGAAG | 819.0 | -8.8 | 1.6 | -5.6 | 2.8E-08 | 4.9E-08 | BT20/MB468 | NUC |

| | | | | | | | | | | | | | |
|---|---|---|---|---|---|---|---|---|---|---|---|---|---|
| >hg38_21_+_8433172_8446622_RNA45SN1_45S_50ntFlanks@6741.6772.32\|ATCGACACTTCGAACGCACTTGCGGCCCCGGG | >hg38_21_+_8433172_8446622_RNA45SN1_45S_50ntFlanks | 6741 | 6772 | 32 | ATCGACACTTCGAACGCACTTGCGGCCCCGGG | 414.8 | -7.5 | 1.4 | -5.6 | 2.5E-08 | 4.9E-08 | MB231/MB468 | NUC |
| >hg38_21_+_8433172_8446622_RNA45SN1_45S_50ntFlanks@11572.11606.35\|CCGACTTAGAACTGGTGCGGACCAGGGGAATCCGA | >hg38_21_+_8433172_8446622_RNA45SN1_45S_50ntFlanks | 11572 | 11606 | 35 | CCGACTTAGAACTGGTGCGGACCAGGGGAATCCGA | 861.9 | -6.3 | 1.1 | -5.6 | 2.7E-08 | 4.9E-08 | MB231/MB468 | MITO |
| >hg38_21_+_8433172_8446622_RNA45SN1_45S_50ntFlanks@11783.11819.37\|CATGAATGGATGAACGAGATTCCCACTGTCCCTACCT | >hg38_21_+_8433172_8446622_RNA45SN1_45S_50ntFlanks | 11783 | 11819 | 37 | CATGAATGGATGAACGAGATTCCCACTGTCCCTACCT | 358.6 | -6.6 | 1.2 | -5.6 | 2.3E-08 | 4.9E-08 | MB231/MB468 | MP |
| >hg38_21_+_8433172_8446622_RNA45SN1_45S_50ntFlanks@9791.9815.25\|TGGCGTGGAGCCGGGCGTGGAATGC | >hg38_21_+_8433172_8446622_RNA45SN1_45S_50ntFlanks | 9791 | 9815 | 25 | TGGCGTGGAGCCGGGCGTGGAATGC | 263.3 | 11.8 | 2.0 | 5.9 | 3.9E-09 | 4.9E-08 | BT20/MB468 | MP |
| >hg38_21_+_8433172_8446622_RNA45SN1_45S_50ntFlanks@6651.6694.44\|CGACTCTTAGCGGTGGATCACTCGGCTCGTGCGTCGATGAAGAA | >hg38_21_+_8433172_8446622_RNA45SN1_45S_50ntFlanks | 6651 | 6694 | 44 | CGACTCTTAGCGGTGGATCACTCGGCTCGTGCGTCGATGAAGAA | 991.5 | 5.6 | 1.0 | 5.6 | 2.1E-08 | 4.9E-08 | BT20/MB468 | CYTO |
| >hg38_21_+_8433172_8446622_RNA45SN1_45S_50ntFlanks@6732.6760.29\|ACATTGATCATCGACACTTCGAACGCACT | >hg38_21_+_8433172_8446622_RNA45SN1_45S_50ntFlanks | 6732 | 6760 | 29 | ACATTGATCATCGACACTTCGAACGCACT | 354.9 | 3.3 | 0.6 | 5.6 | 2.1E-08 | 4.9E-08 | BT20/MB468 | CYTO |
| >MI0000542\|hsa-mir-320a&WithFlank&8\|-\|22244960\|22245043@48.69.22&[MIMAT0000510&hsa-miR-320a-3p&offsets\|0\|0;m-60&8\|-\|22244975\|22244996&offsets\|0\|0]\|AAAAGCTGGGTTGAGAGGGCGA | >MI0000542\|hsa-mir-320a&WithFlank&8\|-\|22244960\|22245043 | 48 | 69 | 22 | AAAAGCTGGGTTGAGAGGGCGA | 930.3 | 5.3 | 1.0 | 5.6 | 2.7E-08 | 4.9E-08 | MB231/MB468 | MITO |
| >hg38_21_+_8433172_8446622_RNA45SN1_45S_50ntFlanks@8171.8215.45\|AAGTCCTTCTGATCGAGGCCCAGCCCGTGGACGGTGTGAGGCCGG | >hg38_21_+_8433172_8446622_RNA45SN1_45S_50ntFlanks | 8171 | 8215 | 45 | AAGTCCTTCTGATCGAGGCCCAGCCCGTGGACGGTGTGAGGCCGG | 511.9 | -4.8 | 0.9 | -5.6 | 2.7E-08 | 4.9E-08 | MB231/MB468 | MITO |
| >hg38_21_+_8433172_8446622_RNA45SN1_45S_50ntFlanks@6721.6743.23\|AATTGCAGGACACATTGATCATC | >hg38_21_+_8433172_8446622_RNA45SN1_45S_50ntFlanks | 6721 | 6743 | 23 | AATTGCAGGACACATTGATCATC | 6739.7 | -4.6 | 0.8 | -5.6 | 2.5E-08 | 4.9E-08 | MB231/MB468 | CYTO |
| >hg38_21_+_8433172_8446622_RNA45SN1_45S_50ntFlanks@10410.10430.21\|TCAGTCGGTCCTGAGAGATGG | >hg38_21_+_8433172_8446622_RNA45SN1_45S_50ntFlanks | 10410 | 10430 | 21 | TCAGTCGGTCCTGAGAGATGG | 6983.9 | -7.0 | 1.3 | -5.6 | 2.5E-08 | 4.9E-08 | MB231/MB468 | CYTO |
| >hg38_21_+_8433172_8446622_RNA45SN1_45S_50ntFlanks@4729.4752.24\|TCAAGAACGAAAGTCGGAGGTTCG | >hg38_21_+_8433172_8446622_RNA45SN1_45S_50ntFlanks | 4729 | 4752 | 24 | TCAAGAACGAAAGTCGGAGGTTCG | 196.2 | -4.3 | 0.8 | -5.6 | 2.5E-08 | 4.9E-08 | MB231/MB468 | CYTO |
| >hg38_1_-_228634819_228635039_RNA5S12_5S_50ntFlanks@131.170.40\|GGATGGGAGACCGCCTGGGAATACCGGGTGCTGTAGGCTT | >hg38_1_-_228634819_228635039_RNA5S12_5S_50ntFlanks | 131 | 170 | 40 | GGATGGGAGACCGCCTGGGAATACCGGGTGCTGTAGGCTT | 3618.1 | 6.0 | 1.0 | 5.9 | 3.9E-09 | 4.9E-08 | BT20/MB468 | MP |
| >MI0006389\|hsa-mir-1255a&WithFlank&4\|-\|101330296\|101330420@34.56.23&[MIMAT0005906&hsa-miR-1255a&offsets\|0\|0;m-998&4\|-\|101330365\|101330387&offsets\|0\|0]\|AGGATGAGCAAAGAAAGTAGATT | >MI0006389\|hsa-mir-1255a&WithFlank&4\|-\|101330296\|101330420 | 34 | 56 | 23 | AGGATGAGCAAAGAAAGTAGATT | 65.2 | 9.2 | 1.7 | 5.6 | 2.3E-08 | 4.9E-08 | MB231/MB468 | MP |
| >trnaMT_GluTTC_MT_-_14674_14742@39.58.20,trnalookalike8_GluTTC_5_-_93905172_93905240@39.58.20\|ATTGGTCGTGGTTGTAGTCC | >trnaMT_GluTTC_MT_-_14674_14742 | 39 | 58 | 20 | ATTGGTCGTGGTTGTAGTCC | 1314.7 | 5.4 | 0.9 | 5.9 | 3.9E-09 | 4.9E-08 | BT20/MB468 | MP |
| >trna128_GlyGCC_6_-_27870686_27870756@19.38.20,trna19_GlyGCC_16_+_70823410_70823480@19.38.20,trna19_GlyGCC_2_-_157257659_157257729@19.38.20,trna24_GlyGCC_16_-_70812942_70813012@19.38.20,trna25_GlyGCC_16_-_70812114_70812184@19.38.20,trna2_GlyGCC_21_-_18827107_18827177@19.38.20,trna35_GlyGCC_1_+_161413094_161413164@19.38.20,trna37_GlyGCC_1_+_161420467_161420537@19.38.20,trna39_GlyGCC_1_+_161427898_161427968@19.38.20,trna41_GlyGCC_1_+_161435258_161435328@19.38.20,trna5_GlyGCC_17_+_8029064_8029134@19.38.20,trna68_GlyGCC_1_-_161493637_161493707@19.38.20\|TAGAATTCTCGCCTGCCACG | >trna128_GlyGCC_6_-_27870686_27870756 | 19 | 38 | 20 | TAGAATTCTCGCCTGCCACG | 4156.6 | 7.8 | 1.4 | 5.6 | 2.1E-08 | 4.9E-08 | BT20/MB468 | CYTO |
| >hg38_21_+_8433172_8446622_RNA45SN1_45S_50ntFlanks@8321.8356.36\|AGACCGATAGTCAACAAGTACCGTAAGGGAAAGTTG | >hg38_21_+_8433172_8446622_RNA45SN1_45S_50ntFlanks | 8321 | 8356 | 36 | AGACCGATAGTCAACAAGTACCGTAAGGGAAAGTTG | 110.0 | -9.9 | 1.7 | -5.9 | 3.9E-09 | 4.9E-08 | BT20/MB468 | MP |
| >MI0000108\|hsa-mir-103a-2&WithFlank&20\|+\|3917488\|3917577@54.76.23(+1C);MI0000109\|hsa-mir-103a-1&WithFlank&5\|-\|168560890\|168560979@54.76.23(+1C)&[MIMAT0000101&hsa-miR-103a-3p&offsets\|0\|0(+1C);m-12&20\|+\|3917541\|3917563&offsets\|0\|0(+1C)];[MIMAT0000101_1&hsa-miR-103a-3p&offsets\|0\|0(+1C);m-13&5\|-\|168560904\|168560926&offsets\|0\|0(+1C);m-44220&5\|-\|168560908\|168560926&offsets\|0\|+4(+1C)]\|AGCAGCATTGTACAGGGCTATGAC | >MI0000108\|hsa-mir-103a-2&WithFlank&20\|+\|3917488\|3917577 | 54 | 76 | 23(+1C) | AGCAGCATTGTACAGGGCTATGAC | 82.2 | 2.9 | 0.5 | 5.6 | 2.8E-08 | 4.9E-08 | BT20/MB468 | NUC |
| >hg38_21_+_8433172_8446622_RNA45SN1_45S_50ntFlanks@12907.12933.27\|CACCGCACGTTCGTGGGGAACCTGGCG | >hg38_21_+_8433172_8446622_RNA45SN1_45S_50ntFlanks | 12907 | 12933 | 27 | CACCGCACGTTCGTGGGGAACCTGGCG | 1240.0 | -8.3 | 1.5 | -5.6 | 2.3E-08 | 4.9E-08 | MB231/MB468 | MP |
| >hg38_21_+_8433172_8446622_RNA45SN1_45S_50ntFlanks@12027.12055.29\|ACTCTGATCGTTTTTTCACTGACCCGGTG | >hg38_21_+_8433172_8446622_RNA45SN1_45S_50ntFlanks | 12027 | 12055 | 29 | ACTCTGATCGTTTTTTCACTGACCCGGTG | 96.0 | 10.3 | 1.8 | 5.9 | 3.9E-09 | 4.9E-08 | BT20/MB468 | MP |
| >MI0005544\|hsa-mir-147b&WithFlank&15\|+\|45433044\|45433135@55.73.19(+1U)&[MIMAT0004928&hsa-miR-147b-3p&offsets\|0\|-2(+1U);m-575&15\|+\|45433098\|45433118&offsets\|0\|-2(+1U)]\|GTGTGCGGAAATGCTTCTGT | >MI0005544\|hsa-mir-147b&WithFlank&15\|+\|45433044\|45433135 | 55 | 73 | 19(+1U) | GTGTGCGGAAATGCTTCTGT | 70.0 | 9.4 | 1.7 | 5.6 | 2.3E-08 | 4.9E-08 | MB231/MB468 | MP |
| >MI0000446\|hsa-mir-125b-1&WithFlank&11\|-\|122099751\|122099850@21.40.20(+1A);MI0000470\|hsa-mir-125b-2&WithFlank&21\|+\|16590231\|16590331@23.42.20(+1A)&[MIMAT0000423&hsa-miR-125b-5p&offsets\|0\|-2(+1A);m-67&11\|-\|122099809\|122099830&offsets\|0\|-2(+1A)];[MIMAT0000423_1&hsa-miR-125b-5p&offsets\|0\|-2(+1A);m-68&21\|+\|16590253\|16590274&offsets\|0\|-2(+1A)]\|TCCCTGAGACCCTAACTTGTA | >MI0000446\|hsa-mir-125b-1&WithFlank&11\|-\|122099751\|122099850 | 21 | 40 | 20(+1A) | TCCCTGAGACCCTAACTTGTA | 1363.5 | 8.3 | 1.5 | 5.6 | 2.7E-08 | 4.9E-08 | MB231/MB468 | MITO |
| >hg38_21_+_8433172_8446622_RNA45SN1_45S_50ntFlanks@6742.6771.30\|TCGACACTTCGAACGCACTTGCGGCCCCGG | >hg38_21_+_8433172_8446622_RNA45SN1_45S_50ntFlanks | 6742 | 6771 | 30 | TCGACACTTCGAACGCACTTGCGGCCCCGG | 393.2 | -3.0 | 0.5 | -5.6 | 1.9E-08 | 4.9E-08 | BT20/MB468 | MITO |
| >hg38_21_+_8433172_8446622_RNA45SN1_45S_50ntFlanks@11718.11737.20\|AACGGCGGGAGTAACTATGA | >hg38_21_+_8433172_8446622_RNA45SN1_45S_50ntFlanks | 11718 | 11737 | 20 | AACGGCGGGAGTAACTATGA | 359.3 | -7.3 | 1.3 | -5.6 | 2.6E-08 | 4.9E-08 | MB231/MB468 | NUC |
| >hg38_21_+_8433172_8446622_RNA45SN1_45S_50ntFlanks@5189.5226.38\|TAACAGGTCTGTGATGCCCTTAGATGTCCGGGGCTGCA | >hg38_21_+_8433172_8446622_RNA45SN1_45S_50ntFlanks | 5189 | 5226 | 38 | TAACAGGTCTGTGATGCCCTTAGATGTCCGGGGCTGCA | 221.6 | -9.9 | 1.8 | -5.6 | 1.9E-08 | 4.9E-08 | BT20/MB468 | MITO |
| >trna10_LysCTT_16_+_3241501_3241573@1.35.35,trna13_LysCTT_14_-_58706613_58706685@1.35.35,trna2_LysCTT_15_+_79152904_79152976@1.35.35\|GCCCGGCTAGCTCAGTCGGTAGAGCATGGGACTCT | >trna10_LysCTT_16_+_3241501_3241573 | 1 | 35 | 35 | GCCCGGCTAGCTCAGTCGGTAGAGCATGGGACTCT | 1767.3 | 3.9 | 0.7 | 5.6 | 2.3E-08 | 4.9E-08 | MB231/MB468 | MP |
| >hg38_21_+_8433172_8446622_RNA45SN1_45S_50ntFlanks@12142.12179.38\|CAGTGCCAGGTGGGGAGTTTGACTGGGGCGGTACACCT | >hg38_21_+_8433172_8446622_RNA45SN1_45S_50ntFlanks | 12142 | 12179 | 38 | CAGTGCCAGGTGGGGAGTTTGACTGGGGCGGTACACCT | 86.3 | -10.3 | 1.8 | -5.6 | 2.4E-08 | 5.0E-08 | MB231/MB468 | MP |
| >hg38_21_+_8433172_8446622_RNA45SN1_45S_50ntFlanks@11999.12043.45\|CCCTGCGGGCCGCCGGTGAAATACCACTACTCTGATCGTTTTTTC | >hg38_21_+_8433172_8446622_RNA45SN1_45S_50ntFlanks | 11999 | 12043 | 45 | CCCTGCGGGCCGCCGGTGAAATACCACTACTCTGATCGTTTTTTC | 622.2 | -7.7 | 1.4 | -5.6 | 2.5E-08 | 5.0E-08 | MB231/MB468 | CYTO |
| >hg38_21_+_8433172_8446622_RNA45SN1_45S_50ntFlanks@475.502.28\|CTTCGTGATCGATGTGGTGACGTCGTGC | >hg38_21_+_8433172_8446622_RNA45SN1_45S_50ntFlanks | 475 | 502 | 28 | CTTCGTGATCGATGTGGTGACGTCGTGC | 1842.8 | 3.9 | 0.7 | 5.6 | 2.6E-08 | 5.0E-08 | MB231/MB468 | NUC |
| >hg38_21_+_8433172_8446622_RNA45SN1_45S_50ntFlanks@11670.11699.30\|CCCAGTGCTCTGAATGTCAAAGTGAAGAAA | >hg38_21_+_8433172_8446622_RNA45SN1_45S_50ntFlanks | 11670 | 11699 | 30 | CCCAGTGCTCTGAATGTCAAAGTGAAGAAA | 684.2 | -7.5 | 1.4 | -5.6 | 2.6E-08 | 5.0E-08 | MB231/MB468 | NUC |
| >hg38_21_+_8433172_8446622_RNA45SN1_45S_50ntFlanks@10384.10404.21\|CCATGTGAACAGCAGTTGAAC | >hg38_21_+_8433172_8446622_RNA45SN1_45S_50ntFlanks | 10384 | 10404 | 21 | CCATGTGAACAGCAGTTGAAC | 145.0 | -10.3 | 1.7 | -5.9 | 4.0E-09 | 5.0E-08 | BT20/MB468 | MP |
| >hg38_21_+_8433172_8446622_RNA45SN1_45S_50ntFlanks@11675.11709.35\|TGCTCTGAATGTCAAAGTGAAGAAATTCAATGAAG | >hg38_21_+_8433172_8446622_RNA45SN1_45S_50ntFlanks | 11675 | 11709 | 35 | TGCTCTGAATGTCAAAGTGAAGAAATTCAATGAAG | 445.9 | 3.3 | 0.6 | 5.6 | 2.6E-08 | 5.0E-08 | MB231/MB468 | NUC |
| >hg38_21_+_8433172_8446622_RNA45SN1_45S_50ntFlanks@3984.4024.41\|GCGGCTTTGGTGACTCTAGATAACCTCGGGCCGATCGCACG | >hg38_21_+_8433172_8446622_RNA45SN1_45S_50ntFlanks | 3984 | 4024 | 41 | GCGGCTTTGGTGACTCTAGATAACCTCGGGCCGATCGCACG | 1368.3 | -7.9 | 1.4 | -5.6 | 1.9E-08 | 5.0E-08 | BT20/MB468 | MITO |
| >MI0000272\|hsa-mir-182&WithFlank&7\|-\|129770377\|129770498@29.52.24&[MIMAT0000259&hsa-miR-182-5p&offsets\|0\|0;m-22&7\|-\|129770449\|129770470&offsets\|0\|+2]\|TTTGGCAATGGTAGAACTCACACT | >MI0000272\|hsa-mir-182&WithFlank&7\|-\|129770377\|129770498 | 29 | 52 | 24 | TTTGGCAATGGTAGAACTCACACT | 93303.5 | 4.7 | 0.8 | 6.0 | 1.5E-09 | 5.0E-08 | BT20/MB231 | CYTO |
| >MI0000070\|hsa-mir-16-1&WithFlank&13\|-\|50048967\|50049067@20.42.23;MI0000115\|hsa-mir-16-2&WithFlank&3\|+\|160404739\|160404831@16.38.23&[MIMAT0000069&hsa-miR-16-5p&offsets\|0\|+1;m-62&13\|-\|50049027\|50049048&offsets\|0\|+1];[MIMAT0000069_1&hsa-miR-16-5p&offsets\|0\|+1;m-61&3\|+\|160404754\|160404775&offsets\|0\|+1]\|TAGCAGCACGTAAATATTGGCGT | >MI0000070\|hsa-mir-16-1&WithFlank&13\|-\|50048967\|50049067 | 20 | 42 | 23 | TAGCAGCACGTAAATATTGGCGT | 2363.0 | 6.2 | 1.1 | 5.6 | 2.8E-08 | 5.0E-08 | MB231/MB468 | MITO |
| >hg38_21_+_8433172_8446622_RNA45SN1_45S_50ntFlanks@11538.11584.47\|CGCGCGGCGCCTCGCCTCGGCCGGCGCCTAGCAGCCGACTTAGAACT | >hg38_21_+_8433172_8446622_RNA45SN1_45S_50ntFlanks | 11538 | 11584 | 47 | CGCGCGGCGCCTCGCCTCGGCCGGCGCCTAGCAGCCGACTTAGAACT | 407.0 | 8.0 | 1.4 | 5.6 | 2.5E-08 | 5.0E-08 | MB231/MB468 | CYTO |
| >hg38_21_+_8433172_8446622_RNA45SN1_45S_50ntFlanks@5158.5185.28\|AAGTGGCGTTCAGCCACCCGAGATTGAG | >hg38_21_+_8433172_8446622_RNA45SN1_45S_50ntFlanks | 5158 | 5185 | 28 | AAGTGGCGTTCAGCCACCCGAGATTGAG | 109.7 | -10.6 | 1.9 | -5.6 | 2.4E-08 | 5.0E-08 | MB231/MB468 | MP |
| >hg38_21_+_8433172_8446622_RNA45SN1_45S_50ntFlanks@8170.8195.26\|CAAGTCCTTCTGATCGAGGCCCAGCC | >hg38_21_+_8433172_8446622_RNA45SN1_45S_50ntFlanks | 8170 | 8195 | 26 | CAAGTCCTTCTGATCGAGGCCCAGCC | 1252.8 | -5.7 | 1.0 | -5.6 | 2.5E-08 | 5.0E-08 | MB231/MB468 | CYTO |
| >hg38_21_+_8433172_8446622_RNA45SN1_45S_50ntFlanks@4006.4025.20\|ACCTCGGGCCGATCGCACGC | >hg38_21_+_8433172_8446622_RNA45SN1_45S_50ntFlanks | 4006 | 4025 | 20 | ACCTCGGGCCGATCGCACGC | 667.1 | -6.4 | 1.1 | -5.6 | 2.6E-08 | 5.0E-08 | MB231/MB468 | NUC |
| >hg38_21_+_8433172_8446622_RNA45SN1_45S_50ntFlanks@5473.5506.34\|CGGCCCTGGCGGAGCGCTGAGAAGACGGTCGAAC | >hg38_21_+_8433172_8446622_RNA45SN1_45S_50ntFlanks | 5473 | 5506 | 34 | CGGCCCTGGCGGAGCGCTGAGAAGACGGTCGAAC | 1149.2 | 7.4 | 1.3 | 5.6 | 2.2E-08 | 5.0E-08 | BT20/MB468 | CYTO |
| >hg38_21_+_8433172_8446622_RNA45SN1_45S_50ntFlanks@3843.3869.27\|CTACTTGGATAACTGTGGTAATTCTAG | >hg38_21_+_8433172_8446622_RNA45SN1_45S_50ntFlanks | 3843 | 3869 | 27 | CTACTTGGATAACTGTGGTAATTCTAG | 447.3 | -7.9 | 1.4 | -5.6 | 2.8E-08 | 5.0E-08 | MB231/MB468 | MITO |
| >hg38_21_+_8433172_8446622_RNA45SN1_45S_50ntFlanks@4293.4317.25\|GATCCATTGGAGGGCAAGTCTGGTG | >hg38_21_+_8433172_8446622_RNA45SN1_45S_50ntFlanks | 4293 | 4317 | 25 | GATCCATTGGAGGGCAAGTCTGGTG | 106.3 | -9.8 | 1.7 | -5.9 | 4.0E-09 | 5.0E-08 | BT20/MB468 | MP |
| >hg38_21_+_8433172_8446622_RNA45SN1_45S_50ntFlanks@9339.9366.28\|CGCACGAAAGCCGCCGTGGCGCAATGAA | >hg38_21_+_8433172_8446622_RNA45SN1_45S_50ntFlanks | 9339 | 9366 | 28 | CGCACGAAAGCCGCCGTGGCGCAATGAA | 152.7 | -10.3 | 1.8 | -5.9 | 4.0E-09 | 5.0E-08 | BT20/MB468 | MP |

<table>
<tr><td>>MI0006444|hsa-mir-1307&WithFlank&10|-|103394247|103394407@86.109.24&[MIMAT0005951&hsa-miR-1307-3p&offsets|0|+2;m-66&10|-|103394300|103394322&offsets|0|+1]|ACTCGGCGTGGCGTCGGTCGTGGT</td><td>>MI0006444|hsa-mir-1307&WithFlank&10|-|103394247|103394407</td><td>86</td><td>109</td><td>24</td><td>ACTCGGCGTGGCGTCGGTCGTGGT</td><td>412.7</td><td>5.6</td><td>1.0</td><td>5.6</td><td>2.8E-08</td><td>5.0E-08</td><td>MB231/MB468</td><td>MITO</td></tr>
<tr><td>>hg38_21_+_8433172_8446622_RNA45SN1_45S_50ntFlanks@9570.9597.28|TGCAAATCGGTCGTCCGACCTGGGTATA</td><td>>hg38_21_+_8433172_8446622_RNA45SN1_45S_50ntFlanks</td><td>9570</td><td>9597</td><td>28</td><td>TGCAAATCGGTCGTCCGACCTGGGTATA</td><td>649.6</td><td>-6.0</td><td>1.1</td><td>-5.6</td><td>2.6E-08</td><td>5.0E-08</td><td>MB231/MB468</td><td>NUC</td></tr>
<tr><td>>hg38_21_+_8433172_8446622_RNA45SN1_45S_50ntFlanks@12202.12219.18|CTAAGGCGAGCTCAGGGA</td><td>>hg38_21_+_8433172_8446622_RNA45SN1_45S_50ntFlanks</td><td>12202</td><td>12219</td><td>18</td><td>CTAAGGCGAGCTCAGGGA</td><td>218.1</td><td>-7.4</td><td>1.3</td><td>-5.6</td><td>2.4E-08</td><td>5.0E-08</td><td>MB231/MB468</td><td>MP</td></tr>
<tr><td>>hg38_21_+_8433172_8446622_RNA45SN1_45S_50ntFlanks@12635.12653.19|AAGCTACCATCTGTGGGAT</td><td>>hg38_21_+_8433172_8446622_RNA45SN1_45S_50ntFlanks</td><td>12635</td><td>12653</td><td>19</td><td>AAGCTACCATCTGTGGGAT</td><td>722.0</td><td>-6.8</td><td>1.2</td><td>-5.6</td><td>2.6E-08</td><td>5.0E-08</td><td>MB231/MB468</td><td>CYTO</td></tr>
<tr><td>>hg38_21_+_8433172_8446622_RNA45SN1_45S_50ntFlanks@8197.8215.19|GTGGACGGTGTGAGGCCGG</td><td>>hg38_21_+_8433172_8446622_RNA45SN1_45S_50ntFlanks</td><td>8197</td><td>8215</td><td>19</td><td>GTGGACGGTGTGAGGCCGG</td><td>569.5</td><td>-4.6</td><td>0.8</td><td>-5.6</td><td>2.4E-08</td><td>5.0E-08</td><td>MB231/MB468</td><td>MP</td></tr>
<tr><td>>MI0000815|hsa-mir-339&WithFlank&7|-|1022927|1023032@21.43.23&[MIMAT0000764&hsa-miR-339-5p&offsets|0|0;m-163&7|-|1022990|1023012&offsets|0|0]|TCCCTGTCCTCCAGGAGCTCACG</td><td>>MI0000815|hsa-mir-339&WithFlank&7|-|1022927|1023032</td><td>21</td><td>43</td><td>23</td><td>TCCCTGTCCTCCAGGAGCTCACG</td><td>963.0</td><td>5.8</td><td>1.0</td><td>5.6</td><td>2.6E-08</td><td>5.1E-08</td><td>MB231/MB468</td><td>CYTO</td></tr>
<tr><td>>hg38_21_+_8433172_8446622_RNA45SN1_45S_50ntFlanks@8799.8820.22|CCTCGGGAGGGCGCGCGGGTCG</td><td>>hg38_21_+_8433172_8446622_RNA45SN1_45S_50ntFlanks</td><td>8799</td><td>8820</td><td>22</td><td>CCTCGGGAGGGCGCGCGGGTCG</td><td>175.3</td><td>-10.5</td><td>1.8</td><td>-5.9</td><td>4.0E-09</td><td>5.1E-08</td><td>BT20/MB468</td><td>MP</td></tr>
<tr><td>>hg38_21_+_8433172_8446622_RNA45SN1_45S_50ntFlanks@9575.9604.30|ATCGGTCGTCCGACCTGGGTATAGGGGCGA</td><td>>hg38_21_+_8433172_8446622_RNA45SN1_45S_50ntFlanks</td><td>9575</td><td>9604</td><td>30</td><td>ATCGGTCGTCCGACCTGGGTATAGGGGCGA</td><td>114.6</td><td>-9.9</td><td>1.7</td><td>-5.9</td><td>4.0E-09</td><td>5.1E-08</td><td>BT20/MB468</td><td>MP</td></tr>
<tr><td>>hg38_21_+_8433172_8446622_RNA45SN1_45S_50ntFlanks@12072.12095.24|CCCGAGGGGCTCTCGCTTCTGGCG</td><td>>hg38_21_+_8433172_8446622_RNA45SN1_45S_50ntFlanks</td><td>12072</td><td>12095</td><td>24</td><td>CCCGAGGGGCTCTCGCTTCTGGCG</td><td>598.6</td><td>-5.7</td><td>1.0</td><td>-5.6</td><td>2.0E-08</td><td>5.1E-08</td><td>BT20/MB468</td><td>MITO</td></tr>
<tr><td>>hg38_21_+_8433172_8446622_RNA45SN1_45S_50ntFlanks@13022.13039.18|CTCGACACAAGGGTTTGT</td><td>>hg38_21_+_8433172_8446622_RNA45SN1_45S_50ntFlanks</td><td>13022</td><td>13039</td><td>18</td><td>CTCGACACAAGGGTTTGT</td><td>1343.2</td><td>4.1</td><td>0.7</td><td>5.6</td><td>2.0E-08</td><td>5.1E-08</td><td>BT20/MB468</td><td>MITO</td></tr>
<tr><td>>hg38_21_+_8433172_8446622_RNA45SN1_45S_50ntFlanks@12937.12954.18|AACCATTCGTAGACGACC</td><td>>hg38_21_+_8433172_8446622_RNA45SN1_45S_50ntFlanks</td><td>12937</td><td>12954</td><td>18</td><td>AACCATTCGTAGACGACC</td><td>356.6</td><td>-7.9</td><td>1.4</td><td>-5.5</td><td>2.9E-08</td><td>5.1E-08</td><td>BT20/MB468</td><td>NUC</td></tr>
<tr><td>>hg38_21_+_8433172_8446622_RNA45SN1_45S_50ntFlanks@12072.12091.20|CCCGAGGGGCTCTCGCTTCT</td><td>>hg38_21_+_8433172_8446622_RNA45SN1_45S_50ntFlanks</td><td>12072</td><td>12091</td><td>20</td><td>CCCGAGGGGCTCTCGCTTCT</td><td>130.3</td><td>-4.4</td><td>0.8</td><td>-5.6</td><td>2.6E-08</td><td>5.1E-08</td><td>MB231/MB468</td><td>CYTO</td></tr>
<tr><td>>hg38_21_+_8433172_8446622_RNA45SN1_45S_50ntFlanks@11663.11685.23|ATTTCTGCCCAGTGCTCTGAATG</td><td>>hg38_21_+_8433172_8446622_RNA45SN1_45S_50ntFlanks</td><td>11663</td><td>11685</td><td>23</td><td>ATTTCTGCCCAGTGCTCTGAATG</td><td>660.2</td><td>-6.2</td><td>1.1</td><td>-5.6</td><td>2.8E-08</td><td>5.1E-08</td><td>MB231/MB468</td><td>MITO</td></tr>
<tr><td>>MI0000746|hsa-mir-99b&WithFlank&19|+|51692606|51692687@13.33.21&[MIMAT0000689&hsa-miR-99b-5p&offsets|0|-1;m-10&19|+|51692618|51692639&offsets|0|-1]|CACCCGTAGAACCGACCTTGC</td><td>>MI0000746|hsa-mir-99b&WithFlank&19|+|51692606|51692687</td><td>13</td><td>33</td><td>21</td><td>CACCCGTAGAACCGACCTTGC</td><td>11118.3</td><td>4.0</td><td>0.7</td><td>5.6</td><td>2.6E-08</td><td>5.1E-08</td><td>MB231/MB468</td><td>NUC</td></tr>
<tr><td>>hg38_21_+_8433172_8446622_RNA45SN1_45S_50ntFlanks@4069.4094.26|CTTTCGATGGTAGTCGCCGTGCCTAC</td><td>>hg38_21_+_8433172_8446622_RNA45SN1_45S_50ntFlanks</td><td>4069</td><td>4094</td><td>26</td><td>CTTTCGATGGTAGTCGCCGTGCCTAC</td><td>373.0</td><td>-7.7</td><td>1.4</td><td>-5.5</td><td>2.9E-08</td><td>5.1E-08</td><td>BT20/MB468</td><td>NUC</td></tr>
<tr><td>>hg38_21_+_8433172_8446622_RNA45SN1_45S_50ntFlanks@3829.3860.32|CGCTCGCTCCTCTCCTACTTGGATAACTGTGG</td><td>>hg38_21_+_8433172_8446622_RNA45SN1_45S_50ntFlanks</td><td>3829</td><td>3860</td><td>32</td><td>CGCTCGCTCCTCTCCTACTTGGATAACTGTGG</td><td>273.7</td><td>-3.9</td><td>0.7</td><td>-5.6</td><td>2.6E-08</td><td>5.1E-08</td><td>MB231/MB468</td><td>NUC</td></tr>
<tr><td>>hg38_21_+_8433172_8446622_RNA45SN1_45S_50ntFlanks@11561.11593.33|GCGCCTAGCAGCCGACTTAGAACTGGTGCGGAC</td><td>>hg38_21_+_8433172_8446622_RNA45SN1_45S_50ntFlanks</td><td>11561</td><td>11593</td><td>33</td><td>GCGCCTAGCAGCCGACTTAGAACTGGTGCGGAC</td><td>399.5</td><td>-3.6</td><td>0.6</td><td>-5.6</td><td>2.0E-08</td><td>5.1E-08</td><td>BT20/MB468</td><td>MITO</td></tr>
<tr><td>>hg38_21_+_8433172_8446622_RNA45SN1_45S_50ntFlanks@11671.11700.30|CCAGTGCTCTGAATGTCAAAGTGAAGAAAT</td><td>>hg38_21_+_8433172_8446622_RNA45SN1_45S_50ntFlanks</td><td>11671</td><td>11700</td><td>30</td><td>CCAGTGCTCTGAATGTCAAAGTGAAGAAAT</td><td>1082.4</td><td>-8.8</td><td>1.6</td><td>-5.5</td><td>2.9E-08</td><td>5.1E-08</td><td>BT20/MB468</td><td>NUC</td></tr>
<tr><td>>MI0000462|hsa-mir-152&WithFlank&17|-|48037155|48037253@60.78.19&[MIMAT0000438&hsa-miR-152-3p&offsets|0|-2;m-97&17|-|48037174|48037194&offsets|0|-2]|TCAGTGCATGACAGAACTT</td><td>>MI0000462|hsa-mir-152&WithFlank&17|-|48037155|48037253</td><td>60</td><td>78</td><td>19</td><td>TCAGTGCATGACAGAACTT</td><td>144.2</td><td>10.9</td><td>1.9</td><td>5.9</td><td>4.1E-09</td><td>5.1E-08</td><td>BT20/MB468</td><td>MP</td></tr>
<tr><td>>MI0000261|hsa-mir-139&WithFlank&11|-|72615057|72615136@13.34.22&[MIMAT0000250&hsa-miR-139-5p&offsets|0|-1;m-186&11|-|72615102|72615124&offsets|0|-1]|TCTACAGTGCACGTGTCTCCAG</td><td>>MI0000261|hsa-mir-139&WithFlank&11|-|72615057|72615136</td><td>13</td><td>34</td><td>22</td><td>TCTACAGTGCACGTGTCTCCAG</td><td>72.6</td><td>9.4</td><td>1.7</td><td>5.6</td><td>2.4E-08</td><td>5.1E-08</td><td>MB231/MB468</td><td>MP</td></tr>
<tr><td>>hg38_21_+_8433172_8446622_RNA45SN1_45S_50ntFlanks@12405.12436.32|CTTTTTGATCCTTCGATGTCGGCTCTTCCTAT</td><td>>hg38_21_+_8433172_8446622_RNA45SN1_45S_50ntFlanks</td><td>12405</td><td>12436</td><td>32</td><td>CTTTTTGATCCTTCGATGTCGGCTCTTCCTAT</td><td>8238.0</td><td>-7.0</td><td>1.3</td><td>-5.6</td><td>2.6E-08</td><td>5.1E-08</td><td>MB231/MB468</td><td>CYTO</td></tr>
<tr><td>>hg38_21_+_8433172_8446622_RNA45SN1_45S_50ntFlanks@12936.12955.20|AAACCATTCGTAGACGACCT</td><td>>hg38_21_+_8433172_8446622_RNA45SN1_45S_50ntFlanks</td><td>12936</td><td>12955</td><td>20</td><td>AAACCATTCGTAGACGACCT</td><td>398.9</td><td>-2.9</td><td>0.5</td><td>-5.6</td><td>2.6E-08</td><td>5.1E-08</td><td>MB231/MB468</td><td>CYTO</td></tr>
<tr><td>>hg38_21_+_8433172_8446622_RNA45SN1_45S_50ntFlanks@6661.6689.29|CGGTGGATCACTCGGCTCGTGCGTCGATG</td><td>>hg38_21_+_8433172_8446622_RNA45SN1_45S_50ntFlanks</td><td>6661</td><td>6689</td><td>29</td><td>CGGTGGATCACTCGGCTCGTGCGTCGATG</td><td>10919.9</td><td>-5.4</td><td>1.0</td><td>-5.6</td><td>2.4E-08</td><td>5.1E-08</td><td>MB231/MB468</td><td>MP</td></tr>
<tr><td>>hg38_21_+_8433172_8446622_RNA45SN1_45S_50ntFlanks@6655.6700.46|TCTTAGCGGTGGATCACTCGGCTCGTGCGTCGATGAAGAACGCAGC</td><td>>hg38_21_+_8433172_8446622_RNA45SN1_45S_50ntFlanks</td><td>6655</td><td>6700</td><td>46</td><td>TCTTAGCGGTGGATCACTCGGCTCGTGCGTCGATGAAGAACGCAGC</td><td>212.6</td><td>11.5</td><td>2.0</td><td>5.9</td><td>4.1E-09</td><td>5.1E-08</td><td>BT20/MB468</td><td>MP</td></tr>
<tr><td>>hg38_21_+_8433172_8446622_RNA45SN1_45S_50ntFlanks@4129.4159.31|GATTCCGGAGAGGGAGCCTGAGAAACGGCTA</td><td>>hg38_21_+_8433172_8446622_RNA45SN1_45S_50ntFlanks</td><td>4129</td><td>4159</td><td>31</td><td>GATTCCGGAGAGGGAGCCTGAGAAACGGCTA</td><td>54.7</td><td>-9.6</td><td>1.7</td><td>-5.6</td><td>2.4E-08</td><td>5.1E-08</td><td>MB231/MB468</td><td>MP</td></tr>
<tr><td>>hg38_21_+_8433172_8446622_RNA45SN1_45S_50ntFlanks@6649.6669.21|TACGACTCTTAGCGGTGGATC</td><td>>hg38_21_+_8433172_8446622_RNA45SN1_45S_50ntFlanks</td><td>6649</td><td>6669</td><td>21</td><td>TACGACTCTTAGCGGTGGATC</td><td>7633.6</td><td>4.1</td><td>0.7</td><td>5.6</td><td>2.0E-08</td><td>5.1E-08</td><td>BT20/MB468</td><td>MITO</td></tr>
<tr><td>>hg38_21_+_8433172_8446622_RNA45SN1_45S_50ntFlanks@12996.13036.41|TCGCTGCGATCTATTGAAAGTCAGCCCTCGACACAAGGGTT</td><td>>hg38_21_+_8433172_8446622_RNA45SN1_45S_50ntFlanks</td><td>12996</td><td>13036</td><td>41</td><td>TCGCTGCGATCTATTGAAAGTCAGCCCTCGACACAAGGGTT</td><td>102.4</td><td>10.4</td><td>1.8</td><td>5.9</td><td>4.1E-09</td><td>5.1E-08</td><td>BT20/MB468</td><td>MP</td></tr>
<tr><td>>hg38_21_+_8433172_8446622_RNA45SN1_45S_50ntFlanks@4370.4389.20|TTAAAAAGCTCGTAGTTGGA</td><td>>hg38_21_+_8433172_8446622_RNA45SN1_45S_50ntFlanks</td><td>4370</td><td>4389</td><td>20</td><td>TTAAAAAGCTCGTAGTTGGA</td><td>152.1</td><td>-10.3</td><td>1.8</td><td>-5.9</td><td>4.1E-09</td><td>5.1E-08</td><td>BT20/MB468</td><td>MP</td></tr>
<tr><td>>hg38_21_+_8433172_8446622_RNA45SN1_45S_50ntFlanks@5491.5531.41|GAGAAGACGGTCGAACTTGACTATCTAGAGGAAGTAAAAGT</td><td>>hg38_21_+_8433172_8446622_RNA45SN1_45S_50ntFlanks</td><td>5491</td><td>5531</td><td>41</td><td>GAGAAGACGGTCGAACTTGACTATCTAGAGGAAGTAAAAGT</td><td>195.9</td><td>11.4</td><td>1.9</td><td>5.9</td><td>4.1E-09</td><td>5.1E-08</td><td>BT20/MB468</td><td>MP</td></tr>
<tr><td>>MI0003778|hsa-mir-320c-1&WithFlank&18|+|21683512|21683595@48.67.20;MI0008191|hsa-mir-320c-2&WithFlank&18|+|24321669|24321752@48.67.20&[MIMAT0005793&hsa-miR-320c&offsets|0|0;m-222&18|+|21683559|21683578&offsets|0|0];[MIMAT0005793_1&hsa-miR-320c&offsets|0|0;m-219&18|+|24321716|24321735&offsets|0|0]|AAAAGCTGGGTTGAGAGGGT</td><td>>MI0003778|hsa-mir-320c-1&WithFlank&18|+|21683512|21683595</td><td>48</td><td>67</td><td>20</td><td>AAAAGCTGGGTTGAGAGGGT</td><td>3983.7</td><td>8.2</td><td>1.4</td><td>5.9</td><td>4.2E-09</td><td>5.2E-08</td><td>BT20/MB468</td><td>MP</td></tr>
<tr><td>>hg38_21_+_8433172_8446622_RNA45SN1_45S_50ntFlanks@10131.10153.23|CGGCGGCGGCGGCGGCGGGGGTG</td><td>>hg38_21_+_8433172_8446622_RNA45SN1_45S_50ntFlanks</td><td>10131</td><td>10153</td><td>23</td><td>CGGCGGCGGCGGCGGCGGGGGTG</td><td>115.7</td><td>-9.9</td><td>1.7</td><td>-5.9</td><td>4.2E-09</td><td>5.2E-08</td><td>BT20/MB468</td><td>MP</td></tr>
<tr><td>>hg38_21_+_8433172_8446622_RNA45SN1_45S_50ntFlanks@9575.9598.24|ATCGGTCGTCCGACCTGGGTATAG</td><td>>hg38_21_+_8433172_8446622_RNA45SN1_45S_50ntFlanks</td><td>9575</td><td>9598</td><td>24</td><td>ATCGGTCGTCCGACCTGGGTATAG</td><td>99.8</td><td>-9.7</td><td>1.7</td><td>-5.9</td><td>4.2E-09</td><td>5.2E-08</td><td>BT20/MB468</td><td>MP</td></tr>
<tr><td>>hg38_21_+_8433172_8446622_RNA45SN1_45S_50ntFlanks@5189.5206.18|TAACAGGTCTGTGATGCC</td><td>>hg38_21_+_8433172_8446622_RNA45SN1_45S_50ntFlanks</td><td>5189</td><td>5206</td><td>18</td><td>TAACAGGTCTGTGATGCC</td><td>87.8</td><td>-10.3</td><td>1.9</td><td>-5.6</td><td>2.5E-08</td><td>5.2E-08</td><td>MB231/MB468</td><td>MP</td></tr>
<tr><td>>hg38_21_+_8433172_8446622_RNA45SN1_45S_50ntFlanks@11491.11516.26|CCGCGGGCCGGTCCCCCCCGCCGGGT</td><td>>hg38_21_+_8433172_8446622_RNA45SN1_45S_50ntFlanks</td><td>11491</td><td>11516</td><td>26</td><td>CCGCGGGCCGGTCCCCCCCGCCGGGT</td><td>119.9</td><td>-6.0</td><td>1.1</td><td>-5.6</td><td>2.2E-08</td><td>5.2E-08</td><td>BT20/MB468</td><td>CYTO</td></tr>
<tr><td>>hg38_21_+_8433172_8446622_RNA45SN1_45S_50ntFlanks@12456.12488.33|CCAAGCGTTGGATTGTTCACCCACTAATAGGGA</td><td>>hg38_21_+_8433172_8446622_RNA45SN1_45S_50ntFlanks</td><td>12456</td><td>12488</td><td>33</td><td>CCAAGCGTTGGATTGTTCACCCACTAATAGGGA</td><td>5520.3</td><td>-6.7</td><td>1.2</td><td>-5.6</td><td>2.2E-08</td><td>5.2E-08</td><td>BT20/MB468</td><td>CYTO</td></tr>
<tr><td>>hg38_21_+_8433172_8446622_RNA45SN1_45S_50ntFlanks@12073.12096.24|CCGAGGGGCTCTCGCTTCTGGCGC</td><td>>hg38_21_+_8433172_8446622_RNA45SN1_45S_50ntFlanks</td><td>12073</td><td>12096</td><td>24</td><td>CCGAGGGGCTCTCGCTTCTGGCGC</td><td>1822.9</td><td>-4.7</td><td>0.8</td><td>-5.6</td><td>2.2E-08</td><td>5.2E-08</td><td>BT20/MB468</td><td>CYTO</td></tr>
<tr><td>>hg38_21_+_8433172_8446622_RNA45SN1_45S_50ntFlanks@11641.11685.45|CGGCGGGTGTTGACGCGATGTGATTTCTGCCCAGTGCTCTGAATG</td><td>>hg38_21_+_8433172_8446622_RNA45SN1_45S_50ntFlanks</td><td>11641</td><td>11685</td><td>45</td><td>CGGCGGGTGTTGACGCGATGTGATTTCTGCCCAGTGCTCTGAATG</td><td>420.0</td><td>-7.4</td><td>1.3</td><td>-5.6</td><td>2.3E-08</td><td>5.2E-08</td><td>BT20/MB468</td><td>CYTO</td></tr>
<tr><td>>hg38_21_+_8433172_8446622_RNA45SN1_45S_50ntFlanks@9339.9364.26|CGCACGAAAGCCGCCGTGGCGCAATG</td><td>>hg38_21_+_8433172_8446622_RNA45SN1_45S_50ntFlanks</td><td>9339</td><td>9364</td><td>26</td><td>CGCACGAAAGCCGCCGTGGCGCAATG</td><td>235.0</td><td>-11.0</td><td>1.9</td><td>-5.9</td><td>4.2E-09</td><td>5.2E-08</td><td>BT20/MB468</td><td>MP</td></tr>
<tr><td>>hg38_21_+_8433172_8446622_RNA45SN1_45S_50ntFlanks@10871.10902.32|GGCTGGGTCGGTCGGGCTGGGGCGCGAAGCGG</td><td>>hg38_21_+_8433172_8446622_RNA45SN1_45S_50ntFlanks</td><td>10871</td><td>10902</td><td>32</td><td>GGCTGGGTCGGTCGGGCTGGGGCGCGAAGCGG</td><td>366.8</td><td>-8.0</td><td>1.4</td><td>-5.5</td><td>3.0E-08</td><td>5.2E-08</td><td>BT20/MB468</td><td>NUC</td></tr>
<tr><td>>hg38_21_+_8433172_8446622_RNA45SN1_45S_50ntFlanks@11671.11691.21|CCAGTGCTCTGAATGTCAAAG</td><td>>hg38_21_+_8433172_8446622_RNA45SN1_45S_50ntFlanks</td><td>11671</td><td>11691</td><td>21</td><td>CCAGTGCTCTGAATGTCAAAG</td><td>468.6</td><td>-4.7</td><td>0.8</td><td>-5.6</td><td>2.5E-08</td><td>5.2E-08</td><td>MB231/MB468</td><td>MP</td></tr>
<tr><td>>hg38_21_+_8433172_8446622_RNA45SN1_45S_50ntFlanks@3835.3872.38|CTCCTCTCCTACTTGGATAACTGTGGTAATTCTAGAGC</td><td>>hg38_21_+_8433172_8446622_RNA45SN1_45S_50ntFlanks</td><td>3835</td><td>3872</td><td>38</td><td>CTCCTCTCCTACTTGGATAACTGTGGTAATTCTAGAGC</td><td>521.7</td><td>-4.3</td><td>0.8</td><td>-5.6</td><td>2.7E-08</td><td>5.2E-08</td><td>MB231/MB468</td><td>NUC</td></tr>
<tr><td>>trnaMT_GluTTC_MT_-_14674_14742@38.71.34,trnalookalike8_GluTTC_5_-_93905172_93905240@38.71.34|CATTGGTCGTGGTTGTAGTCCGTGCGAGAATACC</td><td>>trnaMT_GluTTC_MT_-_14674_14742</td><td>38</td><td>71</td><td>34</td><td>CATTGGTCGTGGTTGTAGTCCGTGCGAGAATACC</td><td>329.6</td><td>3.9</td><td>0.7</td><td>5.5</td><td>2.9E-08</td><td>5.2E-08</td><td>MB231/MB468</td><td>MITO</td></tr>
<tr><td>>trnaMT_TyrGTA_MT_-_5826_5891@23.58.36|TTGGACTGTAAATCTAAAGACAGGGGTTAGGCCTCT</td><td>>trnaMT_TyrGTA_MT_-_5826_5891</td><td>23</td><td>58</td><td>36</td><td>TTGGACTGTAAATCTAAAGACAGGGGTTAGGCCTCT</td><td>141.2</td><td>10.9</td><td>1.9</td><td>5.9</td><td>4.2E-09</td><td>5.2E-08</td><td>BT20/MB468</td><td>MP</td></tr>
<tr><td>>am_trnaMT_ValTAC_MT_+_1602_1670@45.67.23|TTTCAACTTAACTTGACCGCTCT</td><td>>am_trnaMT_ValTAC_MT_+_1602_1670</td><td>45</td><td>67</td><td>23</td><td>TTTCAACTTAACTTGACCGCTCT</td><td>296.3</td><td>8.9</td><td>1.6</td><td>5.6</td><td>2.3E-08</td><td>5.2E-08</td><td>BT20/MB468</td><td>CYTO</td></tr>
<tr><td>>hg38_21_+_8433172_8446622_RNA45SN1_45S_50ntFlanks@11769.11790.22|TAATTAGTGACGCGCATGAATG</td><td>>hg38_21_+_8433172_8446622_RNA45SN1_45S_50ntFlanks</td><td>11769</td><td>11790</td><td>22</td><td>TAATTAGTGACGCGCATGAATG</td><td>728.5</td><td>-3.4</td><td>0.6</td><td>-5.6</td><td>2.7E-08</td><td>5.2E-08</td><td>MB231/MB468</td><td>NUC</td></tr>
<tr><td>>hg38_21_+_8433172_8446622_RNA45SN1_45S_50ntFlanks@12438.12455.18|ATTGTGAAGCAGAATTCA</td><td>>hg38_21_+_8433172_8446622_RNA45SN1_45S_50ntFlanks</td><td>12438</td><td>12455</td><td>18</td><td>ATTGTGAAGCAGAATTCA</td><td>1101.2</td><td>-6.2</td><td>1.1</td><td>-5.6</td><td>2.7E-08</td><td>5.2E-08</td><td>MB231/MB468</td><td>NUC</td></tr>
<tr><td>>hg38_21_+_8433172_8446622_RNA45SN1_45S_50ntFlanks@11933.11957.25|AATAAGTGGGAGGCCCCCGGCGCCC</td><td>>hg38_21_+_8433172_8446622_RNA45SN1_45S_50ntFlanks</td><td>11933</td><td>11957</td><td>25</td><td>AATAAGTGGGAGGCCCCCGGCGCCC</td><td>114.2</td><td>-9.9</td><td>1.7</td><td>-5.9</td><td>4.2E-09</td><td>5.3E-08</td><td>BT20/MB468</td><td>MP</td></tr>
<tr><td>>hg38_21_+_8433172_8446622_RNA45SN1_45S_50ntFlanks@9504.9528.25|GTGAACTATGCCTGGGCAGGGCGAA</td><td>>hg38_21_+_8433172_8446622_RNA45SN1_45S_50ntFlanks</td><td>9504</td><td>9528</td><td>25</td><td>GTGAACTATGCCTGGGCAGGGCGAA</td><td>131.7</td><td>-10.1</td><td>1.7</td><td>-5.9</td><td>4.2E-09</td><td>5.3E-08</td><td>BT20/MB468</td><td>MP</td></tr>
<tr><td>>MI0015866|hsa-mir-4326&WithFlank&20|+|63286802|63286872@17.39.23&[MIMAT0016888&hsa-miR-4326&offsets|0|+3;m-339&20|+|63286818|63286841&offsets|0|-1]|TGTTCCTCTGTCTCCCAGACTCT</td><td>>MI0015866|hsa-mir-4326&WithFlank&20|+|63286802|63286872</td><td>17</td><td>39</td><td>23</td><td>TGTTCCTCTGTCTCCCAGACTCT</td><td>73.8</td><td>9.4</td><td>1.7</td><td>5.6</td><td>2.5E-08</td><td>5.3E-08</td><td>MB231/MB468</td><td>MP</td></tr>
<tr><td>>hg38_21_+_8433172_8446622_RNA45SN1_45S_50ntFlanks@11198.11225.28|GCCGGCGGCGGCGGCGACTCTGGACGCG</td><td>>hg38_21_+_8433172_8446622_RNA45SN1_45S_50ntFlanks</td><td>11198</td><td>11225</td><td>28</td><td>GCCGGCGGCGGCGGCGACTCTGGACGCG</td><td>112.8</td><td>-6.1</td><td>1.1</td><td>-5.6</td><td>2.3E-08</td><td>5.3E-08</td><td>BT20/MB468</td><td>CYTO</td></tr>
</table>

<table>
<tr><td>>am_trna10_LysCTT_16_+_3241501_3241573@11.28.18,trna119_LysCTT_1_-_145395522_145395594@11.28.18,trna11_LysCTT_5_-_180648979_180649051@11.28.18,trna128_LysCTT_1_-_55423542_55423614@11.28.18,trna13_LysCTT_14_-_58706613_58706685@11.28.18,trna13_LysCTT_6_+_26556774_26556846@11.28.18,trna2_LysCTT_15_+_79152904_79152976@11.28.18,trna32_LysCTT_16_-_3207406_3207478@11.28.18,trna4_LysCTT_18_-_43669271_43669343@11.28.18,trna6_LysCTT_19_-_52425396_52425468@11.28.18,trna7_LysCTT_16_+_3225692_3225764@11.28.18,trna9_LysCTT_5_+_180634755_180634827@11.28.18|CTCAGTCGGTAGAGCATG</td><td>>am_trna10_LysCTT_16_+_3241501_3241573</td><td>11</td><td>28</td><td>18</td><td>CTCAGTCGGTAGAGCATG</td><td>847.3</td><td>-4.6</td><td>0.8</td><td>-5.6</td><td>2.7E-08</td><td>5.3E-08</td><td>MB231/MB468</td><td>CYTO</td></tr>
<tr><td>>MI0000111|hsa-mir-105-1&WithFlank&X|-|152392213|152392305@21.42.22;MI0000112|hsa-mir-105-2&WithFlank&X|-|152394406|152394498@21.42.22&[MIMAT0000102&hsa-miR-105-5p&offsets|+2|+1;m-735&X|-|152392266|152392287&offsets|+2|+2];[MIMAT0000102_1&hsa-miR-105-5p&offsets|+2|+1;m-735&X|-|152394459|152394480&offsets|+2|+2]|AAATGCTCAGACTCCTGTGGTG</td><td>>MI0000111|hsa-mir-105-1&WithFlank&X|-|152392213|152392305</td><td>21</td><td>42</td><td>22</td><td>AAATGCTCAGACTCCTGTGGTG</td><td>258.6</td><td>11.3</td><td>1.8</td><td>6.1</td><td>8.0E-10</td><td>5.3E-08</td><td>BT20/MB231</td><td>MP</td></tr>
<tr><td>>hg38_21_+_8433172_8446622_RNA45SN1_45S_50ntFlanks@11670.11692.23|CCCAGTGCTCTGAATGTCAAAGT</td><td>>hg38_21_+_8433172_8446622_RNA45SN1_45S_50ntFlanks</td><td>11670</td><td>11692</td><td>23</td><td>CCCAGTGCTCTGAATGTCAAAGT</td><td>598.7</td><td>-2.6</td><td>0.5</td><td>-5.6</td><td>2.7E-08</td><td>5.3E-08</td><td>MB231/MB468</td><td>NUC</td></tr>
<tr><td>>hg38_21_+_8433172_8446622_RNA45SN1_45S_50ntFlanks@3771.3799.29|CACGGCCGGTACAGTGAAACTGCGAATGG</td><td>>hg38_21_+_8433172_8446622_RNA45SN1_45S_50ntFlanks</td><td>3771</td><td>3799</td><td>29</td><td>CACGGCCGGTACAGTGAAACTGCGAATGG</td><td>249.1</td><td>-7.3</td><td>1.3</td><td>-5.5</td><td>2.9E-08</td><td>5.3E-08</td><td>MB231/MB468</td><td>MITO</td></tr>
<tr><td>>hg38_21_+_8433172_8446622_RNA45SN1_45S_50ntFlanks@8114.8135.22|GCGCGGGACATGTGGCGTACGG</td><td>>hg38_21_+_8433172_8446622_RNA45SN1_45S_50ntFlanks</td><td>8114</td><td>8135</td><td>22</td><td>GCGCGGGACATGTGGCGTACGG</td><td>688.1</td><td>12.8</td><td>2.1</td><td>6.1</td><td>8.1E-10</td><td>5.3E-08</td><td>BT20/MB231</td><td>MP</td></tr>
<tr><td>>am_MI0005763|hsa-mir-941-1&WithFlank&20|+|63919443|63919526@53.75.23(+1U);MI0005764|hsa-mir-941-2&WithFlank&20|+|63919499|63919582@53.75.23(+1U);MI0005765|hsa-mir-941-3&WithFlank&20|+|63919555|63919638@53.75.23(+1U);MI0005766|hsa-mir-941-4&WithFlank&20|+|63919750|63919833@53.75.23(+1U);MI0031520|hsa-mir-941-5&WithFlank&20|+|63919862|63919945@53.75.23(+1U)&[MIMAT0004984&hsa-miR-941&offsets|0|0(+1U);m-263&20|+|63919495|63919517&offsets|0|0(+1U)];[MIMAT0004984_1&hsa-miR-941&offsets|0|0(+1U);m-263&20|+|63919551|63919573&offsets|0|0(+1U)];[MIMAT0004984_2&hsa-miR-941&offsets|0|0(+1U);m-263&20|+|63919607|63919629&offsets|0|0(+1U)];[MIMAT0004984_3&hsa-miR-941&offsets|0|0(+1U);m-263&20|+|63919802|63919824&offsets|0|0(+1U)];[MIMAT0004984_4&hsa-miR-941&offsets|0|0(+1U);m-263&20|+|63919914|63919936&offsets|0|0(+1U)]|CACCCGGCTGTGTGCACATGTGCT</td><td>>am_MI0005763|hsa-mir-941-1&WithFlank&20|+|63919443|63919526</td><td>53</td><td>75</td><td>23(+1U)</td><td>CACCCGGCTGTGTGCACATGTGCT</td><td>466.3</td><td>7.2</td><td>1.3</td><td>5.6</td><td>2.7E-08</td><td>5.3E-08</td><td>MB231/MB468</td><td>CYTO</td></tr>
<tr><td>>hg38_21_+_8433172_8446622_RNA45SN1_45S_50ntFlanks@11670.11691.22|CCCAGTGCTCTGAATGTCAAAG</td><td>>hg38_21_+_8433172_8446622_RNA45SN1_45S_50ntFlanks</td><td>11670</td><td>11691</td><td>22</td><td>CCCAGTGCTCTGAATGTCAAAG</td><td>671.8</td><td>-6.2</td><td>1.1</td><td>-5.6</td><td>2.5E-08</td><td>5.3E-08</td><td>MB231/MB468</td><td>MP</td></tr>
<tr><td>>hg38_21_+_8433172_8446622_RNA45SN1_45S_50ntFlanks@6699.6717.19|GCTAGCTGCGAGAATTAAT</td><td>>hg38_21_+_8433172_8446622_RNA45SN1_45S_50ntFlanks</td><td>6699</td><td>6717</td><td>19</td><td>GCTAGCTGCGAGAATTAAT</td><td>1434.0</td><td>-8.6</td><td>1.5</td><td>-5.6</td><td>2.5E-08</td><td>5.3E-08</td><td>MB231/MB468</td><td>MP</td></tr>
<tr><td>>MI0000650|hsa-mir-200c&WithFlank&12|+|6963693|6963772@49.70.22&[MIMAT0000617&hsa-miR-200c-3p&offsets|-1|-2;m-54&12|+|6963742|6963764&offsets|-1|-2]|CTAATACTGCCGGGTAATGATG</td><td>>MI0000650|hsa-mir-200c&WithFlank&12|+|6963693|6963772</td><td>49</td><td>70</td><td>22</td><td>CTAATACTGCCGGGTAATGATG</td><td>191.1</td><td>10.9</td><td>1.8</td><td>6.1</td><td>8.2E-10</td><td>5.3E-08</td><td>BT20/MB231</td><td>MP</td></tr>
<tr><td>>hg38_21_+_8433172_8446622_RNA45SN1_45S_50ntFlanks@6664.6689.26|TGGATCACTCGGCTCGTGCGTCGATG</td><td>>hg38_21_+_8433172_8446622_RNA45SN1_45S_50ntFlanks</td><td>6664</td><td>6689</td><td>26</td><td>TGGATCACTCGGCTCGTGCGTCGATG</td><td>2084.9</td><td>-8.4</td><td>1.5</td><td>-5.6</td><td>2.5E-08</td><td>5.3E-08</td><td>MB231/MB468</td><td>MP</td></tr>
<tr><td>>hg38_21_+_8433172_8446622_RNA45SN1_45S_50ntFlanks@12351.12376.26|AAGTTACCACAGGGATAACTGGCTTG</td><td>>hg38_21_+_8433172_8446622_RNA45SN1_45S_50ntFlanks</td><td>12351</td><td>12376</td><td>26</td><td>AAGTTACCACAGGGATAACTGGCTTG</td><td>93.4</td><td>-9.6</td><td>1.6</td><td>-5.9</td><td>4.3E-09</td><td>5.3E-08</td><td>BT20/MB468</td><td>MP</td></tr>
<tr><td>>hg38_21_+_8433172_8446622_RNA45SN1_45S_50ntFlanks@12404.12431.28|GCTTTTTGATCCTTCGATGTCGGCTCTT</td><td>>hg38_21_+_8433172_8446622_RNA45SN1_45S_50ntFlanks</td><td>12404</td><td>12431</td><td>28</td><td>GCTTTTTGATCCTTCGATGTCGGCTCTT</td><td>504.4</td><td>-3.3</td><td>0.6</td><td>-5.6</td><td>2.7E-08</td><td>5.3E-08</td><td>MB231/MB468</td><td>CYTO</td></tr>
<tr><td>>hg38_21_+_8433172_8446622_RNA45SN1_45S_50ntFlanks@12892.12933.42|CCCTCGCCCGTCACGCACCGCACGTTCGTGGGGAACCTGGCG</td><td>>hg38_21_+_8433172_8446622_RNA45SN1_45S_50ntFlanks</td><td>12892</td><td>12933</td><td>42</td><td>CCCTCGCCCGTCACGCACCGCACGTTCGTGGGGAACCTGGCG</td><td>228.9</td><td>-7.2</td><td>1.3</td><td>-5.6</td><td>2.8E-08</td><td>5.3E-08</td><td>MB231/MB468</td><td>NUC</td></tr>
<tr><td>>hg38_21_+_8433172_8446622_RNA45SN1_45S_50ntFlanks@10401.10439.39|GAACATGGGTCAGTCGGTCCTGAGAGATGGGCGAGCGCC</td><td>>hg38_21_+_8433172_8446622_RNA45SN1_45S_50ntFlanks</td><td>10401</td><td>10439</td><td>39</td><td>GAACATGGGTCAGTCGGTCCTGAGAGATGGGCGAGCGCC</td><td>282.5</td><td>4.8</td><td>0.9</td><td>5.6</td><td>2.8E-08</td><td>5.3E-08</td><td>MB231/MB468</td><td>NUC</td></tr>
<tr><td>>trna10_AspGTC_12_-_125424193_125424264@38.64.27,trna12_AspGTC_12_-_125411891_125411962@38.64.27,trna144_AspGTC_6_-_27551236_27551307@38.64.27,trna38_AspGTC_17_-_8125556_8125627@38.64.27,trna45_AspGTC_6_+_27447453_27447524@38.64.27,trna48_AspGTC_6_+_27471523_27471594@38.64.27,trna4_AspGTC_12_+_96429799_96429870@38.64.27,trna69_AspGTC_1_-_161440205_161440276@38.64.27,trna72_AspGTC_1_-_161432824_161432895@38.64.27,trna75_AspGTC_1_-_161425414_161425485@38.64.27,trna78_AspGTC_1_-_161418033_161418104@38.64.27,trna81_AspGTC_1_-_161410615_161410686@38.64.27|CGCGGGAGACCGGGGTTCGATTCCCCG</td><td>>trna10_AspGTC_12_-_125424193_125424264</td><td>38</td><td>64</td><td>27</td><td>CGCGGGAGACCGGGGTTCGATTCCCCG</td><td>317.2</td><td>-7.8</td><td>1.4</td><td>-5.6</td><td>2.5E-08</td><td>5.3E-08</td><td>MB231/MB468</td><td>MP</td></tr>
<tr><td>>hg38_21_+_8433172_8446622_RNA45SN1_45S_50ntFlanks@8327.8356.30|ATAGTCAACAAGTACCGTAAGGGAAAGTTG</td><td>>hg38_21_+_8433172_8446622_RNA45SN1_45S_50ntFlanks</td><td>8327</td><td>8356</td><td>30</td><td>ATAGTCAACAAGTACCGTAAGGGAAAGTTG</td><td>93.6</td><td>-9.6</td><td>1.6</td><td>-5.9</td><td>4.3E-09</td><td>5.3E-08</td><td>BT20/MB468</td><td>MP</td></tr>
<tr><td>>hg38_21_+_8433172_8446622_RNA45SN1_45S_50ntFlanks@4759.4778.20|ATCAGATACCGTCGTAGTTC</td><td>>hg38_21_+_8433172_8446622_RNA45SN1_45S_50ntFlanks</td><td>4759</td><td>4778</td><td>20</td><td>ATCAGATACCGTCGTAGTTC</td><td>1382.0</td><td>-7.1</td><td>1.3</td><td>-5.6</td><td>2.7E-08</td><td>5.3E-08</td><td>MB231/MB468</td><td>CYTO</td></tr>
<tr><td>>hg38_21_+_8433172_8446622_RNA45SN1_45S_50ntFlanks@10351.10385.35|CGAGAACTTTGAAGGCCGAAGTGGAGAAGGGTTCC</td><td>>hg38_21_+_8433172_8446622_RNA45SN1_45S_50ntFlanks</td><td>10351</td><td>10385</td><td>35</td><td>CGAGAACTTTGAAGGCCGAAGTGGAGAAGGGTTCC</td><td>550.6</td><td>-8.2</td><td>1.5</td><td>-5.6</td><td>2.5E-08</td><td>5.3E-08</td><td>MB231/MB468</td><td>MP</td></tr>
<tr><td>>hg38_21_+_8433172_8446622_RNA45SN1_45S_50ntFlanks@11217.11234.18|CTGGACGCGAGCCGGGCC</td><td>>hg38_21_+_8433172_8446622_RNA45SN1_45S_50ntFlanks</td><td>11217</td><td>11234</td><td>18</td><td>CTGGACGCGAGCCGGGCC</td><td>3663.0</td><td>-5.1</td><td>0.9</td><td>-5.6</td><td>2.7E-08</td><td>5.3E-08</td><td>MB231/MB468</td><td>CYTO</td></tr>
<tr><td>>hg38_21_+_8433172_8446622_RNA45SN1_45S_50ntFlanks@12907.12934.28|CACCGCACGTTCGTGGGGAACCTGGCGC</td><td>>hg38_21_+_8433172_8446622_RNA45SN1_45S_50ntFlanks</td><td>12907</td><td>12934</td><td>28</td><td>CACCGCACGTTCGTGGGGAACCTGGCGC</td><td>207.3</td><td>-7.2</td><td>1.3</td><td>-5.5</td><td>3.0E-08</td><td>5.3E-08</td><td>MB231/MB468</td><td>MITO</td></tr>
<tr><td>>hg38_21_+_8433172_8446622_RNA45SN1_45S_50ntFlanks@10355.10383.29|AACTTTGAAGGCCGAAGTGGAGAAGGGTT</td><td>>hg38_21_+_8433172_8446622_RNA45SN1_45S_50ntFlanks</td><td>10355</td><td>10383</td><td>29</td><td>AACTTTGAAGGCCGAAGTGGAGAAGGGTT</td><td>180.3</td><td>-4.4</td><td>0.8</td><td>-5.6</td><td>2.7E-08</td><td>5.4E-08</td><td>MB231/MB468</td><td>CYTO</td></tr>
<tr><td>>hg38_21_+_8433172_8446622_RNA45SN1_45S_50ntFlanks@8184.8211.28|CGAGGCCCAGCCCGTGGACGGTGTGAGG</td><td>>hg38_21_+_8433172_8446622_RNA45SN1_45S_50ntFlanks</td><td>8184</td><td>8211</td><td>28</td><td>CGAGGCCCAGCCCGTGGACGGTGTGAGG</td><td>495.0</td><td>-8.3</td><td>1.5</td><td>-5.5</td><td>3.1E-08</td><td>5.4E-08</td><td>BT20/MB468</td><td>NUC</td></tr>
<tr><td>>hg38_21_+_8433172_8446622_RNA45SN1_45S_50ntFlanks@12044.12068.25|ACTGACCCGGTGAGGCGGGGGGGCG</td><td>>hg38_21_+_8433172_8446622_RNA45SN1_45S_50ntFlanks</td><td>12044</td><td>12068</td><td>25</td><td>ACTGACCCGGTGAGGCGGGGGGGCG</td><td>1199.7</td><td>-7.6</td><td>1.4</td><td>-5.6</td><td>2.1E-08</td><td>5.4E-08</td><td>BT20/MB468</td><td>MITO</td></tr>
<tr><td>>hg38_21_+_8433172_8446622_RNA45SN1_45S_50ntFlanks@4050.4081.32|CGAACGTCTGCCCTATCAACTTTCGATGGTAG</td><td>>hg38_21_+_8433172_8446622_RNA45SN1_45S_50ntFlanks</td><td>4050</td><td>4081</td><td>32</td><td>CGAACGTCTGCCCTATCAACTTTCGATGGTAG</td><td>141.8</td><td>-4.4</td><td>0.8</td><td>-5.6</td><td>2.3E-08</td><td>5.4E-08</td><td>BT20/MB468</td><td>CYTO</td></tr>
<tr><td>>hg38_21_+_8433172_8446622_RNA45SN1_45S_50ntFlanks@6670.6698.29|ACTCGGCTCGTGCGTCGATGAAGAACGCA</td><td>>hg38_21_+_8433172_8446622_RNA45SN1_45S_50ntFlanks</td><td>6670</td><td>6698</td><td>29</td><td>ACTCGGCTCGTGCGTCGATGAAGAACGCA</td><td>363.5</td><td>-6.2</td><td>1.1</td><td>-5.5</td><td>3.1E-08</td><td>5.4E-08</td><td>BT20/MB468</td><td>NUC</td></tr>
<tr><td>>hg38_21_+_8433172_8446622_RNA45SN1_45S_50ntFlanks@3985.4010.26|CGGCTTTGGTGACTCTAGATAACCTC</td><td>>hg38_21_+_8433172_8446622_RNA45SN1_45S_50ntFlanks</td><td>3985</td><td>4010</td><td>26</td><td>CGGCTTTGGTGACTCTAGATAACCTC</td><td>371.4</td><td>-6.0</td><td>1.1</td><td>-5.5</td><td>3.0E-08</td><td>5.4E-08</td><td>MB231/MB468</td><td>MITO</td></tr>
<tr><td>>hg38_MT_+_598_1651_RNR1_12S_50ntFlanks@366.404.39|CCCAATAAAGCTAAAACTCACCTGAGTTGTAAAAAACTC</td><td>>hg38_MT_+_598_1651_RNR1_12S_50ntFlanks</td><td>366</td><td>404</td><td>39</td><td>CCCAATAAAGCTAAAACTCACCTGAGTTGTAAAAAACTC</td><td>561.7</td><td>5.1</td><td>0.9</td><td>5.6</td><td>2.1E-08</td><td>5.4E-08</td><td>BT20/MB468</td><td>MITO</td></tr>
<tr><td>>hg38_21_+_8433172_8446622_RNA45SN1_45S_50ntFlanks@4723.4757.35|CATTAATCAAGAACGAAAGTCGGAGGTTCGAAGAC</td><td>>hg38_21_+_8433172_8446622_RNA45SN1_45S_50ntFlanks</td><td>4723</td><td>4757</td><td>35</td><td>CATTAATCAAGAACGAAAGTCGGAGGTTCGAAGAC</td><td>372.3</td><td>-7.6</td><td>1.4</td><td>-5.5</td><td>3.1E-08</td><td>5.4E-08</td><td>BT20/MB468</td><td>NUC</td></tr>
<tr><td>>hg38_21_+_8433172_8446622_RNA45SN1_45S_50ntFlanks@6660.6686.27|GCGGTGGATCACTCGGCTCGTGCGTCG</td><td>>hg38_21_+_8433172_8446622_RNA45SN1_45S_50ntFlanks</td><td>6660</td><td>6686</td><td>27</td><td>GCGGTGGATCACTCGGCTCGTGCGTCG</td><td>272.9</td><td>-6.2</td><td>1.1</td><td>-5.6</td><td>2.6E-08</td><td>5.4E-08</td><td>MB231/MB468</td><td>MP</td></tr>
<tr><td>>hg38_21_+_8433172_8446622_RNA45SN1_45S_50ntFlanks@8176.8223.48|CTTCTGATCGAGGCCCAGCCCGTGGACGGTGTGAGGCCGGTAGCGGCC</td><td>>hg38_21_+_8433172_8446622_RNA45SN1_45S_50ntFlanks</td><td>8176</td><td>8223</td><td>48</td><td>CTTCTGATCGAGGCCCAGCCCGTGGACGGTGTGAGGCCGGTAGCGGCC</td><td>184.5</td><td>-7.2</td><td>1.3</td><td>-5.6</td><td>2.3E-08</td><td>5.4E-08</td><td>BT20/MB468</td><td>CYTO</td></tr>
<tr><td>>hg38_21_+_8433172_8446622_RNA45SN1_45S_50ntFlanks@4733.4767.35|GAACGAAAGTCGGAGGTTCGAAGACGATCAGATAC</td><td>>hg38_21_+_8433172_8446622_RNA45SN1_45S_50ntFlanks</td><td>4733</td><td>4767</td><td>35</td><td>GAACGAAAGTCGGAGGTTCGAAGACGATCAGATAC</td><td>323.6</td><td>6.8</td><td>1.2</td><td>5.6</td><td>2.8E-08</td><td>5.4E-08</td><td>MB231/MB468</td><td>NUC</td></tr>
<tr><td>>hg38_MT_+_1621_3279_RNR2_16S_50ntFlanks@718.740.23|AGCCTGCGTCAGATTAAAACACT</td><td>>hg38_MT_+_1621_3279_RNR2_16S_50ntFlanks</td><td>718</td><td>740</td><td>23</td><td>AGCCTGCGTCAGATTAAAACACT</td><td>77.7</td><td>9.5</td><td>1.7</td><td>5.6</td><td>2.6E-08</td><td>5.4E-08</td><td>MB231/MB468</td><td>MP</td></tr>
<tr><td>>hg38_21_+_8433172_8446622_RNA45SN1_45S_50ntFlanks@11480.11502.23|CGGCGGAGGGGCCGCGGGCCGGT</td><td>>hg38_21_+_8433172_8446622_RNA45SN1_45S_50ntFlanks</td><td>11480</td><td>11502</td><td>23</td><td>CGGCGGAGGGGCCGCGGGCCGGT</td><td>160.0</td><td>-10.4</td><td>1.8</td><td>-5.9</td><td>4.4E-09</td><td>5.4E-08</td><td>BT20/MB468</td><td>MP</td></tr>
<tr><td>>hg38_21_+_8433172_8446622_RNA45SN1_45S_50ntFlanks@12407.12424.18|TTTTGATCCTTCGATGTC</td><td>>hg38_21_+_8433172_8446622_RNA45SN1_45S_50ntFlanks</td><td>12407</td><td>12424</td><td>18</td><td>TTTTGATCCTTCGATGTC</td><td>95.5</td><td>-9.7</td><td>1.6</td><td>-5.9</td><td>4.4E-09</td><td>5.4E-08</td><td>BT20/MB468</td><td>MP</td></tr>
<tr><td>>hg38_21_+_8433172_8446622_RNA45SN1_45S_50ntFlanks@11821.11845.25|CTATCCAGCGAAACCACAGCCAAGG</td><td>>hg38_21_+_8433172_8446622_RNA45SN1_45S_50ntFlanks</td><td>11821</td><td>11845</td><td>25</td><td>CTATCCAGCGAAACCACAGCCAAGG</td><td>103.4</td><td>-9.8</td><td>1.7</td><td>-5.9</td><td>4.4E-09</td><td>5.4E-08</td><td>BT20/MB468</td><td>MP</td></tr>
<tr><td>>hg38_21_+_8433172_8446622_RNA45SN1_45S_50ntFlanks@4557.4587.31|CAGCTAGGAATAATGGAATAGGACCGCGGTT</td><td>>hg38_21_+_8433172_8446622_RNA45SN1_45S_50ntFlanks</td><td>4557</td><td>4587</td><td>31</td><td>CAGCTAGGAATAATGGAATAGGACCGCGGTT</td><td>307.7</td><td>-2.7</td><td>0.5</td><td>-5.6</td><td>2.4E-08</td><td>5.4E-08</td><td>BT20/MB468</td><td>CYTO</td></tr>
<tr><td>>am_trnaMT_PheGAA_MT_+_577_647@40.59.20|TGTTTAGACGGGCTCACATC</td><td>>am_trnaMT_PheGAA_MT_+_577_647</td><td>40</td><td>59</td><td>20</td><td>TGTTTAGACGGGCTCACATC</td><td>461.8</td><td>9.2</td><td>1.6</td><td>5.6</td><td>2.4E-08</td><td>5.4E-08</td><td>BT20/MB468</td><td>CYTO</td></tr>
<tr><td>>hg38_21_+_8433172_8446622_RNA45SN1_45S_50ntFlanks@10234.10275.42|CGGACGCTACGCCGCGACGAGTAGGAGGGCCGCTGCGGTGAG</td><td>>hg38_21_+_8433172_8446622_RNA45SN1_45S_50ntFlanks</td><td>10234</td><td>10275</td><td>42</td><td>CGGACGCTACGCCGCGACGAGTAGGAGGGCCGCTGCGGTGAG</td><td>67.6</td><td>-9.9</td><td>1.8</td><td>-5.6</td><td>2.6E-08</td><td>5.4E-08</td><td>MB231/MB468</td><td>MP</td></tr>
<tr><td>>hg38_21_+_8433172_8446622_RNA45SN1_45S_50ntFlanks@12633.12657.25|CGAAGCTACCATCTGTGGGATTATG</td><td>>hg38_21_+_8433172_8446622_RNA45SN1_45S_50ntFlanks</td><td>12633</td><td>12657</td><td>25</td><td>CGAAGCTACCATCTGTGGGATTATG</td><td>1249.9</td><td>-4.2</td><td>0.7</td><td>-5.6</td><td>2.1E-08</td><td>5.4E-08</td><td>BT20/MB468</td><td>MITO</td></tr>
</table>

| | | | | | | | | | | | | | |
|---|---|---|---|---|---|---|---|---|---|---|---|---|---|
| >hg38_21_+_8433172_8446622_RNA45SN1_45S_50ntFlanks@12001.12029.29\|CTGCGGGCCGCCGGTGAAATACCACTACT | >hg38_21_+_8433172_8446622_RNA45SN1_45S_50ntFlanks | 12001 | 12029 | 29 | CTGCGGGCCGCCGGTGAAATACCACTACT | 308.4 | -7.7 | 1.4 | -5.5 | 3.1E-08 | 5.4E-08 | BT20/MB468 | NUC |
| >hg38_21_+_8433172_8446622_RNA45SN1_45S_50ntFlanks@11647.11687.41\|GTGTTGACGCGATGTGATTTCTGCCCAGTGCTCTGAATGTC | >hg38_21_+_8433172_8446622_RNA45SN1_45S_50ntFlanks | 11647 | 11687 | 41 | GTGTTGACGCGATGTGATTTCTGCCCAGTGCTCTGAATGTC | 2082.8 | 14.4 | 2.3 | 6.1 | 8.5E-10 | 5.4E-08 | BT20/MB231 | MP |
| >hg38_21_+_8433172_8446622_RNA45SN1_45S_50ntFlanks@5078.5110.33\|CGATAACGAACGAGACTCTGGCATGCTAACTAG | >hg38_21_+_8433172_8446622_RNA45SN1_45S_50ntFlanks | 5078 | 5110 | 33 | CGATAACGAACGAGACTCTGGCATGCTAACTAG | 739.3 | -6.9 | 1.3 | -5.5 | 3.0E-08 | 5.4E-08 | MB231/MB468 | MITO |
| >trna10_LysCTT_16_+_3241501_3241573@5.36.32,trna13_LysCTT_14_-_58706613_58706685@5.36.32,trna2_LysCTT_15_+_79152904_79152976@5.36.32\|GGCTAGCTCAGTCGGTAGAGCATGGGACTCTT | >trna10_LysCTT_16_+_3241501_3241573 | 5 | 36 | 32 | GGCTAGCTCAGTCGGTAGAGCATGGGACTCTT | 105.0 | 9.9 | 1.8 | 5.6 | 2.6E-08 | 5.5E-08 | MB231/MB468 | MP |
| >hg38_21_+_8433172_8446622_RNA45SN1_45S_50ntFlanks@12332.12366.35\|TAAGCAGGAGGTGTCAGAAAAGTTACCACAGGGAT | >hg38_21_+_8433172_8446622_RNA45SN1_45S_50ntFlanks | 12332 | 12366 | 35 | TAAGCAGGAGGTGTCAGAAAAGTTACCACAGGGAT | 64.4 | -9.9 | 1.8 | -5.6 | 2.6E-08 | 5.5E-08 | MB231/MB468 | MP |
| >hg38_21_+_8433172_8446622_RNA45SN1_45S_50ntFlanks@12934.12957.24\|CTAAACCATTCGTAGACGACCTGC | >hg38_21_+_8433172_8446622_RNA45SN1_45S_50ntFlanks | 12934 | 12957 | 24 | CTAAACCATTCGTAGACGACCTGC | 621.4 | -4.0 | 0.7 | -5.6 | 2.8E-08 | 5.5E-08 | MB231/MB468 | NUC |
| >hg38_21_+_8433172_8446622_RNA45SN1_45S_50ntFlanks@10680.10710.31\|GTTCCGGCGGCGTCCGGTGAGCTCTCGCTGG | >hg38_21_+_8433172_8446622_RNA45SN1_45S_50ntFlanks | 10680 | 10710 | 31 | GTTCCGGCGGCGTCCGGTGAGCTCTCGCTGG | 19191.7 | -5.6 | 1.0 | -5.6 | 2.4E-08 | 5.5E-08 | BT20/MB468 | CYTO |
| >trnaMT_ValTAC_MT_+_1602_1670@33.72.40\|ACACTTAGGAGATTTCAACTTAACTTGACCGCTCTGACCA | >trnaMT_ValTAC_MT_+_1602_1670 | 33 | 72 | 40 | ACACTTAGGAGATTTCAACTTAACTTGACCGCTCTGACCA | 204.8 | 7.2 | 1.3 | 5.6 | 2.8E-08 | 5.5E-08 | MB231/MB468 | NUC |
| >hg38_21_+_8433172_8446622_RNA45SN1_45S_50ntFlanks@12418.12444.27\|CGATGTCGGCTCTTCCTATCATTGTGA | >hg38_21_+_8433172_8446622_RNA45SN1_45S_50ntFlanks | 12418 | 12444 | 27 | CGATGTCGGCTCTTCCTATCATTGTGA | 407.2 | -6.0 | 1.1 | -5.6 | 2.6E-08 | 5.5E-08 | MB231/MB468 | MP |
| >hg38_21_+_8433172_8446622_RNA45SN1_45S_50ntFlanks@12218.12236.19\|GAGGACAGAAACCTCCCGT | >hg38_21_+_8433172_8446622_RNA45SN1_45S_50ntFlanks | 12218 | 12236 | 19 | GAGGACAGAAACCTCCCGT | 138.3 | 10.9 | 1.9 | 5.9 | 4.5E-09 | 5.5E-08 | BT20/MB468 | MP |
| >hg38_21_+_8433172_8446622_RNA45SN1_45S_50ntFlanks@9925.9949.25\|GGTTGATATAGACAGCAGGACGGTG | >hg38_21_+_8433172_8446622_RNA45SN1_45S_50ntFlanks | 9925 | 9949 | 25 | GGTTGATATAGACAGCAGGACGGTG | 92.2 | 10.3 | 1.8 | 5.9 | 4.4E-09 | 5.5E-08 | BT20/MB468 | MP |
| >hg38_21_+_8433172_8446622_RNA45SN1_45S_50ntFlanks@8169.8203.35\|CCAAGTCCTTCTGATCGAGGCCCAGCCCGTGGACG | >hg38_21_+_8433172_8446622_RNA45SN1_45S_50ntFlanks | 8169 | 8203 | 35 | CCAAGTCCTTCTGATCGAGGCCCAGCCCGTGGACG | 180.0 | -10.6 | 1.8 | -5.9 | 4.5E-09 | 5.5E-08 | BT20/MB468 | MP |
| >hg38_21_+_8433172_8446622_RNA45SN1_45S_50ntFlanks@11569.11606.38\|CAGCCGACTTAGAACTGGTGCGGACCAGGGGAATCCGA | >hg38_21_+_8433172_8446622_RNA45SN1_45S_50ntFlanks | 11569 | 11606 | 38 | CAGCCGACTTAGAACTGGTGCGGACCAGGGGAATCCGA | 514.0 | -8.5 | 1.5 | -5.5 | 3.2E-08 | 5.5E-08 | BT20/MB468 | NUC |
| >hg38_21_+_8433172_8446622_RNA45SN1_45S_50ntFlanks@12140.12179.40\|GACAGTGCCAGGTGGGGAGTTTGACTGGGGCGGTACACCT | >hg38_21_+_8433172_8446622_RNA45SN1_45S_50ntFlanks | 12140 | 12179 | 40 | GACAGTGCCAGGTGGGGAGTTTGACTGGGGCGGTACACCT | 86.7 | -10.3 | 1.9 | -5.6 | 2.6E-08 | 5.5E-08 | MB231/MB468 | MP |
| >hg38_21_+_8433172_8446622_RNA45SN1_45S_50ntFlanks@7975.8019.45\|CGCGACCTCAGATCAGACGTGGCGACCCGCTGAATTTAAGCATAT | >hg38_21_+_8433172_8446622_RNA45SN1_45S_50ntFlanks | 7975 | 8019 | 45 | CGCGACCTCAGATCAGACGTGGCGACCCGCTGAATTTAAGCATAT | 906.3 | 5.8 | 1.0 | 5.6 | 2.1E-08 | 5.5E-08 | BT20/MB468 | MITO |
| >hg38_21_+_8433172_8446622_RNA45SN1_45S_50ntFlanks@8357.8387.31\|AAAAGAACTTTGAAGAGAGAGTTCAAGAGGG | >hg38_21_+_8433172_8446622_RNA45SN1_45S_50ntFlanks | 8357 | 8387 | 31 | AAAAGAACTTTGAAGAGAGAGTTCAAGAGGG | 386.9 | -7.3 | 1.3 | -5.5 | 3.0E-08 | 5.5E-08 | MB231/MB468 | MITO |
| >hg38_21_+_8433172_8446622_RNA45SN1_45S_50ntFlanks@6719.6749.31\|TGAATTGCAGGACACATTGATCATCGACACT | >hg38_21_+_8433172_8446622_RNA45SN1_45S_50ntFlanks | 6719 | 6749 | 31 | TGAATTGCAGGACACATTGATCATCGACACT | 662.6 | -5.9 | 1.1 | -5.6 | 2.9E-08 | 5.5E-08 | MB231/MB468 | NUC |
| >hg38_21_+_8433172_8446622_RNA45SN1_45S_50ntFlanks@7984.8004.21\|AGATCAGACGTGGCGACCCGC | >hg38_21_+_8433172_8446622_RNA45SN1_45S_50ntFlanks | 7984 | 8004 | 21 | AGATCAGACGTGGCGACCCGC | 318.3 | -5.5 | 1.0 | -5.5 | 2.9E-08 | 5.5E-08 | MB231/MB468 | NUC |
| >hg38_21_+_8433172_8446622_RNA45SN1_45S_50ntFlanks@9554.9574.21\|CGTAGCGGTCCTGACGTGCAA | >hg38_21_+_8433172_8446622_RNA45SN1_45S_50ntFlanks | 9554 | 9574 | 21 | CGTAGCGGTCCTGACGTGCAA | 2493.4 | -3.7 | 0.7 | -5.5 | 2.9E-08 | 5.5E-08 | MB231/MB468 | NUC |
| >hg38_21_+_8433172_8446622_RNA45SN1_45S_50ntFlanks@6644.6663.20\|CCTCGTACGACTCTTAGCGG | >hg38_21_+_8433172_8446622_RNA45SN1_45S_50ntFlanks | 6644 | 6663 | 20 | CCTCGTACGACTCTTAGCGG | 621.2 | 3.8 | 0.7 | 5.6 | 2.1E-08 | 5.5E-08 | BT20/MB468 | MITO |
| >hg38_21_+_8433172_8446622_RNA45SN1_45S_50ntFlanks@5073.5109.37\|AATTCCGATAACGAACGAGACTCTGGCATGCTAACTA | >hg38_21_+_8433172_8446622_RNA45SN1_45S_50ntFlanks | 5073 | 5109 | 37 | AATTCCGATAACGAACGAGACTCTGGCATGCTAACTA | 148.4 | -9.3 | 1.7 | -5.6 | 2.1E-08 | 5.5E-08 | BT20/MB468 | MITO |
| >MI0000078\|hsa-mir-22&WithFlank&17\|-\|1713897\|1713993@59.81.23&[MIMAT0000077&hsa-miR-22-3p&offsets\|0\|+1;m-2&17\|-\|1713914\|1713935&offsets\|0\|+1]\|AAGCTGCCAGTTGAAGAACTGTT | >MI0000078\|hsa-mir-22&WithFlank&17\|-\|1713897\|1713993 | 59 | 81 | 23 | AAGCTGCCAGTTGAAGAACTGTT | 506.4 | 9.1 | 1.6 | 5.5 | 3.1E-08 | 5.5E-08 | MB231/MB468 | MITO |
| >MI0000299\|hsa-mir-222&WithFlank&X\|-\|45747009\|45747130@75.99.25(+4U)&[MIMAT0000279&hsa-miR-222-3p&offsets\|0\|+4(+4U);m-140&X\|-\|45747033\|45747056&offsets\|0\|+1(+4U)]\|AGCTACATCTGGCTACTGGGTCTCTTTTT | >MI0000299\|hsa-mir-222&WithFlank&X\|-\|45747009\|45747130 | 75 | 99 | 25(+4U) | AGCTACATCTGGCTACTGGGTCTCTTTTT | 502.7 | 8.8 | 1.6 | 5.5 | 3.1E-08 | 5.5E-08 | MB231/MB468 | MITO |
| >MI0000760\|hsa-mir-361&WithFlank&X\|-\|85903630\|85903713@52.74.23(+1U)&[MIMAT0004682&hsa-miR-361-3p&offsets\|+1\|+1(+1U);m-124&X\|-\|85903642\|85903663&offsets\|+1\|+2(+1U)]\|CCCCCAGGTGTGATTCTGATTTGT | >MI0000760\|hsa-mir-361&WithFlank&X\|-\|85903630\|85903713 | 52 | 74 | 23(+1U) | CCCCCAGGTGTGATTCTGATTTGT | 290.9 | 8.6 | 1.5 | 5.6 | 2.4E-08 | 5.5E-08 | BT20/MB468 | CYTO |
| >hg38_21_+_8433172_8446622_RNA45SN1_45S_50ntFlanks@7975.8011.37\|CGCGACCTCAGATCAGACGTGGCGACCCGCTGAATTT | >hg38_21_+_8433172_8446622_RNA45SN1_45S_50ntFlanks | 7975 | 8011 | 37 | CGCGACCTCAGATCAGACGTGGCGACCCGCTGAATTT | 137797.8 | 3.2 | 0.6 | 5.5 | 3.2E-08 | 5.5E-08 | BT20/MB468 | NUC |
| >hg38_21_+_8433172_8446622_RNA45SN1_45S_50ntFlanks@4760.4779.20\|TCAGATACCGTCGTAGTTCC | >hg38_21_+_8433172_8446622_RNA45SN1_45S_50ntFlanks | 4760 | 4779 | 20 | TCAGATACCGTCGTAGTTCC | 459.5 | -4.4 | 0.8 | -5.5 | 2.9E-08 | 5.5E-08 | MB231/MB468 | NUC |
| >hg38_21_+_8433172_8446622_RNA45SN1_45S_50ntFlanks@8157.8175.19\|CTCGTGGGGGGCCCAAGTC | >hg38_21_+_8433172_8446622_RNA45SN1_45S_50ntFlanks | 8157 | 8175 | 19 | CTCGTGGGGGGCCCAAGTC | 532.7 | -7.6 | 1.4 | -5.5 | 3.1E-08 | 5.5E-08 | MB231/MB468 | MITO |
| >hg38_21_+_8433172_8446622_RNA45SN1_45S_50ntFlanks@4738.4767.30\|AAAGTCGGAGGTTCGAAGACGATCAGATAC | >hg38_21_+_8433172_8446622_RNA45SN1_45S_50ntFlanks | 4738 | 4767 | 30 | AAAGTCGGAGGTTCGAAGACGATCAGATAC | 926.5 | 7.6 | 1.4 | 5.6 | 2.8E-08 | 5.5E-08 | MB231/MB468 | CYTO |
| >hg38_21_+_8433172_8446622_RNA45SN1_45S_50ntFlanks@9835.9875.41\|TTTGGTAAGCAGAACTGGCGCTGCGGGATGAACCGAACGCC | >hg38_21_+_8433172_8446622_RNA45SN1_45S_50ntFlanks | 9835 | 9875 | 41 | TTTGGTAAGCAGAACTGGCGCTGCGGGATGAACCGAACGCC | 102.6 | 10.4 | 1.8 | 5.9 | 4.5E-09 | 5.5E-08 | BT20/MB468 | MP |
| >hg38_21_+_8433172_8446622_RNA45SN1_45S_50ntFlanks@4728.4757.30\|ATCAAGAACGAAAGTCGGAGGTTCGAAGAC | >hg38_21_+_8433172_8446622_RNA45SN1_45S_50ntFlanks | 4728 | 4757 | 30 | ATCAAGAACGAAAGTCGGAGGTTCGAAGAC | 90.9 | -9.6 | 1.6 | -5.9 | 4.5E-09 | 5.5E-08 | BT20/MB468 | MP |
| >hg38_21_+_8433172_8446622_RNA45SN1_45S_50ntFlanks@9604.9622.19\|AAAGACTAATCGAACCATC | >hg38_21_+_8433172_8446622_RNA45SN1_45S_50ntFlanks | 9604 | 9622 | 19 | AAAGACTAATCGAACCATC | 1390.2 | -4.5 | 0.8 | -5.5 | 2.9E-08 | 5.5E-08 | MB231/MB468 | NUC |
| >hg38_21_+_8433172_8446622_RNA45SN1_45S_50ntFlanks@3881.3899.19\|GCCGACGGGCGCTGACCCC | >hg38_21_+_8433172_8446622_RNA45SN1_45S_50ntFlanks | 3881 | 3899 | 19 | GCCGACGGGCGCTGACCCC | 301.6 | -7.6 | 1.4 | -5.5 | 3.1E-08 | 5.5E-08 | MB231/MB468 | MITO |
| >hg38_21_+_8433172_8446622_RNA45SN1_45S_50ntFlanks@12681.12703.23\|CCCGCCCAGGCGGAACGATACGG | >hg38_21_+_8433172_8446622_RNA45SN1_45S_50ntFlanks | 12681 | 12703 | 23 | CCCGCCCAGGCGGAACGATACGG | 146.9 | -10.3 | 1.8 | -5.9 | 4.5E-09 | 5.5E-08 | BT20/MB468 | MP |
| >trnaMT_SerGCT_MT_+_12207_12265@8.30.23\|CTCACAAGAACTGCTAACTCATG | >trnaMT_SerGCT_MT_+_12207_12265 | 8 | 30 | 23 | CTCACAAGAACTGCTAACTCATG | 958.4 | -9.5 | 1.7 | -5.5 | 3.2E-08 | 5.5E-08 | BT20/MB468 | NUC |
| >hg38_21_+_8433172_8446622_RNA45SN1_45S_50ntFlanks@11579.11601.23\|AGAACTGGTGCGGACCAGGGGAA | >hg38_21_+_8433172_8446622_RNA45SN1_45S_50ntFlanks | 11579 | 11601 | 23 | AGAACTGGTGCGGACCAGGGGAA | 1188.3 | -8.9 | 1.6 | -5.5 | 3.2E-08 | 5.6E-08 | BT20/MB468 | NUC |
| >hg38_21_+_8433172_8446622_RNA45SN1_45S_50ntFlanks@8814.8844.31\|CGGGTCGGGGCGGCGGCGGCGGCGGCGGTGG | >hg38_21_+_8433172_8446622_RNA45SN1_45S_50ntFlanks | 8814 | 8844 | 31 | CGGGTCGGGGCGGCGGCGGCGGCGGCGGTGG | 68.9 | -10.0 | 1.8 | -5.6 | 2.7E-08 | 5.6E-08 | MB231/MB468 | MP |
| >hg38_21_+_8433172_8446622_RNA45SN1_45S_50ntFlanks@11215.11234.20\|CTCTGGACGCGAGCCGGGCC | >hg38_21_+_8433172_8446622_RNA45SN1_45S_50ntFlanks | 11215 | 11234 | 20 | CTCTGGACGCGAGCCGGGCC | 379.9 | -6.6 | 1.2 | -5.6 | 2.4E-08 | 5.6E-08 | BT20/MB468 | CYTO |
| >hg38_21_+_8433172_8446622_RNA45SN1_45S_50ntFlanks@11572.11591.20\|CCGACTTAGAACTGGTGCGG | >hg38_21_+_8433172_8446622_RNA45SN1_45S_50ntFlanks | 11572 | 11591 | 20 | CCGACTTAGAACTGGTGCGG | 5159.8 | -7.5 | 1.3 | -5.6 | 2.4E-08 | 5.6E-08 | BT20/MB468 | CYTO |
| >hg38_21_+_8433172_8446622_RNA45SN1_45S_50ntFlanks@10572.10591.20\|AGAAGCCGGCGGGAGCCCCG | >hg38_21_+_8433172_8446622_RNA45SN1_45S_50ntFlanks | 10572 | 10591 | 20 | AGAAGCCGGCGGGAGCCCCG | 941.1 | -7.0 | 1.3 | -5.6 | 2.8E-08 | 5.6E-08 | MB231/MB468 | CYTO |
| >hg38_21_+_8433172_8446622_RNA45SN1_45S_50ntFlanks@12685.12703.19\|CCCAGGCGGAACGATACGG | >hg38_21_+_8433172_8446622_RNA45SN1_45S_50ntFlanks | 12685 | 12703 | 19 | CCCAGGCGGAACGATACGG | 158.3 | -6.7 | 1.2 | -5.5 | 2.9E-08 | 5.6E-08 | MB231/MB468 | NUC |
| >am_MI0000809\|hsa-mir-151a&WithFlank&8\|-\|140732558\|140732659@53.72.20(+1A)&[MIMAT0000757&hsa-miR-151a-3p&offsets\|0\|-1(+1A);m-30&8\|-\|140732587\|140732607&offsets\|0\|-1(+1A)]\|CTAGACTGAAGCTCCTTGAGA | >am_MI0000809\|hsa-mir-151a&WithFlank&8\|-\|140732558\|140732659 | 53 | 72 | 20(+1A) | CTAGACTGAAGCTCCTTGAGA | 4812.1 | 3.9 | 0.7 | 5.6 | 2.7E-08 | 5.6E-08 | MB231/MB468 | MP |
| >MI0000746\|hsa-mir-99b&WithFlank&19\|+\|51692606\|51692687@13.32.20(+1U)&[MIMAT0000689&hsa-miR-99b-5p&offsets\|0\|-2(+1U);m-10&19\|+\|51692618\|51692639&offsets\|0\|-2(+1U)]\|CACCCGTAGAACCGACCTTGT | >MI0000746\|hsa-mir-99b&WithFlank&19\|+\|51692606\|51692687 | 13 | 32 | 20(+1U) | CACCCGTAGAACCGACCTTGT | 1149.7 | 4.7 | 0.9 | 5.5 | 3.1E-08 | 5.6E-08 | MB231/MB468 | MITO |
| >hg38_21_+_8433172_8446622_RNA45SN1_45S_50ntFlanks@12934.12968.35\|CTAAACCATTCGTAGACGACCTGCTTCTGGGTCGG | >hg38_21_+_8433172_8446622_RNA45SN1_45S_50ntFlanks | 12934 | 12968 | 35 | CTAAACCATTCGTAGACGACCTGCTTCTGGGTCGG | 81.1 | -10.2 | 1.8 | -5.6 | 2.7E-08 | 5.6E-08 | MB231/MB468 | MP |
| >hg38_21_+_8433172_8446622_RNA45SN1_45S_50ntFlanks@9554.9592.39\|CGTAGCGGTCCTGACGTGCAAATCGGTCGTCCGACCTGG | >hg38_21_+_8433172_8446622_RNA45SN1_45S_50ntFlanks | 9554 | 9592 | 39 | CGTAGCGGTCCTGACGTGCAAATCGGTCGTCCGACCTGG | 76.7 | -10.1 | 1.8 | -5.6 | 2.7E-08 | 5.6E-08 | MB231/MB468 | MP |
| >hg38_21_+_8433172_8446622_RNA45SN1_45S_50ntFlanks@10420.10453.34\|CTGAGAGATGGGCGAGCGCCGTTCCGAAGGGACG | >hg38_21_+_8433172_8446622_RNA45SN1_45S_50ntFlanks | 10420 | 10453 | 34 | CTGAGAGATGGGCGAGCGCCGTTCCGAAGGGACG | 514.1 | -7.5 | 1.3 | -5.5 | 3.1E-08 | 5.6E-08 | MB231/MB468 | MITO |
| >hg38_21_+_8433172_8446622_RNA45SN1_45S_50ntFlanks@10840.10865.26\|CCGTAACTTCGGGATAAGGATTGGCT | >hg38_21_+_8433172_8446622_RNA45SN1_45S_50ntFlanks | 10840 | 10865 | 26 | CCGTAACTTCGGGATAAGGATTGGCT | 2742.3 | -8.6 | 1.5 | -5.6 | 2.7E-08 | 5.6E-08 | MB231/MB468 | MP |
| >hg38_21_+_8433172_8446622_RNA45SN1_45S_50ntFlanks@9546.9563.18\|TGGAGGTCCGTAGCGGTC | >hg38_21_+_8433172_8446622_RNA45SN1_45S_50ntFlanks | 9546 | 9563 | 18 | TGGAGGTCCGTAGCGGTC | 319.5 | -3.1 | 0.6 | -5.5 | 2.9E-08 | 5.6E-08 | MB231/MB468 | NUC |
| >MI0000746\|hsa-mir-99b&WithFlank&19\|+\|51692606\|51692687@13.34.22(+1C)&[MIMAT0000689&hsa-miR-99b-5p&offsets\|0\|0(+1C);m-10&19\|+\|51692618\|51692639&offsets\|0\|0(+1C)]\|CACCCGTAGAACCGACCTTGCGC | >MI0000746\|hsa-mir-99b&WithFlank&19\|+\|51692606\|51692687 | 13 | 34 | 22(+1C) | CACCCGTAGAACCGACCTTGCGC | 376.6 | 7.3 | 1.3 | 5.6 | 2.9E-08 | 5.6E-08 | MB231/MB468 | CYTO |
| >hg38_21_+_8433172_8446622_RNA45SN1_45S_50ntFlanks@10771.10796.26\|CAGGTCTCCAAGGTGAACAGCCTCTG | >hg38_21_+_8433172_8446622_RNA45SN1_45S_50ntFlanks | 10771 | 10796 | 26 | CAGGTCTCCAAGGTGAACAGCCTCTG | 763.0 | -7.8 | 1.4 | -5.6 | 2.7E-08 | 5.6E-08 | MB231/MB468 | MP |
| >hg38_21_+_8433172_8446622_RNA45SN1_45S_50ntFlanks@12406.12441.36\|TTTTTGATCCTTCGATGTCGGCTCTTCCTATCATTG | >hg38_21_+_8433172_8446622_RNA45SN1_45S_50ntFlanks | 12406 | 12441 | 36 | TTTTTGATCCTTCGATGTCGGCTCTTCCTATCATTG | 1433.0 | -9.2 | 1.7 | -5.5 | 3.2E-08 | 5.6E-08 | BT20/MB468 | NUC |
| >hg38_MT_+_598_1651_RNR1_12S_50ntFlanks@843.877.35\|GAGTAGAGTGCTTAGTTGAACAGGGCCCTGAAGCG | >hg38_MT_+_598_1651_RNR1_12S_50ntFlanks | 843 | 877 | 35 | GAGTAGAGTGCTTAGTTGAACAGGGCCCTGAAGCG | 236.5 | 11.6 | 2.0 | 5.9 | 4.6E-09 | 5.6E-08 | BT20/MB468 | MP |

<table>
<tr><td>>hg38_21_+_8433172_8446622_RNA45SN1_45S_50ntFlanks@11694.11717.24|AAGAAATTCAATGAAGCGCGGGTA</td><td>>hg38_21_+_8433172_8446622_RNA45SN1_45S_50ntFlanks</td><td>11694</td><td>11717</td><td>24</td><td>AAGAAATTCAATGAAGCGCGGGTA</td><td>666.2</td><td>-8.6</td><td>1.6</td><td>-5.5</td><td>3.2E-08</td><td>5.6E-08</td><td>BT20/MB468</td><td>NUC</td></tr>
<tr><td>>MI0001445|hsa-mir-423&WithFlank&17|+|30117073|30117178@59.80.22(+1A)&[MIMAT0001340&hsa-miR-423-3p&offsets|0|-1(+1A);m-100&17|+|30117131|30117153&offsets|0|-1(+1A)]|AGCTCGGTCTGAGGCCCCTCAGA</td><td>>MI0001445|hsa-mir-423&WithFlank&17|+|30117073|30117178</td><td>59</td><td>80</td><td>22(+1A)</td><td>AGCTCGGTCTGAGGCCCCTCAGA</td><td>717.8</td><td>5.0</td><td>0.8</td><td>5.9</td><td>4.6E-09</td><td>5.6E-08</td><td>BT20/MB468</td><td>MP</td></tr>
<tr><td>>hg38_21_+_8433172_8446622_RNA45SN1_45S_50ntFlanks@5515.5533.19|CTAGAGGAAGTAAAAGTCG</td><td>>hg38_21_+_8433172_8446622_RNA45SN1_45S_50ntFlanks</td><td>5515</td><td>5533</td><td>19</td><td>CTAGAGGAAGTAAAAGTCG</td><td>223.3</td><td>-5.1</td><td>0.9</td><td>-5.6</td><td>2.9E-08</td><td>5.6E-08</td><td>MB231/MB468</td><td>CYTO</td></tr>
<tr><td>>MI0000433|hsa-let-7g&WithFlank&3|-|52268272|52268367@11.32.22&[MIMAT0000414&hsa-let-7g-5p&offsets|0|0;m-58&3|-|52268336|52268357&offsets|0|0]|TGAGGTAGTAGTTTGTACAGTT</td><td>>MI0000433|hsa-let-7g&WithFlank&3|-|52268272|52268367</td><td>11</td><td>32</td><td>22</td><td>TGAGGTAGTAGTTTGTACAGTT</td><td>31194.0</td><td>4.7</td><td>0.8</td><td>5.5</td><td>3.1E-08</td><td>5.6E-08</td><td>MB231/MB468</td><td>MITO</td></tr>
<tr><td>>MI0000284|hsa-mir-204&WithFlank&9|-|70809969|70810090@39.60.22&[MIMAT0000265&hsa-miR-204-5p&offsets|0|0;m-296&9|-|70810031|70810052&offsets|0|0]|TTCCCTTTGTCATCCTATGCCT</td><td>>MI0000284|hsa-mir-204&WithFlank&9|-|70809969|70810090</td><td>39</td><td>60</td><td>22</td><td>TTCCCTTTGTCATCCTATGCCT</td><td>184.0</td><td>-10.9</td><td>1.8</td><td>-6.1</td><td>8.8E-10</td><td>5.6E-08</td><td>BT20/MB231</td><td>MP</td></tr>
<tr><td>>hg38_21_+_8433172_8446622_RNA45SN1_45S_50ntFlanks@11891.11910.20|TTGACTCTAGTCTGGCACGG</td><td>>hg38_21_+_8433172_8446622_RNA45SN1_45S_50ntFlanks</td><td>11891</td><td>11910</td><td>20</td><td>TTGACTCTAGTCTGGCACGG</td><td>583.3</td><td>-7.7</td><td>1.4</td><td>-5.5</td><td>3.1E-08</td><td>5.6E-08</td><td>MB231/MB468</td><td>MITO</td></tr>
<tr><td>>hg38_21_+_8433172_8446622_RNA45SN1_45S_50ntFlanks@12405.12435.31|CTTTTTGATCCTTCGATGTCGGCTCTTCCTA</td><td>>hg38_21_+_8433172_8446622_RNA45SN1_45S_50ntFlanks</td><td>12405</td><td>12435</td><td>31</td><td>CTTTTTGATCCTTCGATGTCGGCTCTTCCTA</td><td>4072.3</td><td>-6.9</td><td>1.2</td><td>-5.6</td><td>2.5E-08</td><td>5.6E-08</td><td>BT20/MB468</td><td>CYTO</td></tr>
<tr><td>>hg38_21_+_8433172_8446622_RNA45SN1_45S_50ntFlanks@12357.12376.20|CCACAGGGATAACTGGCTTG</td><td>>hg38_21_+_8433172_8446622_RNA45SN1_45S_50ntFlanks</td><td>12357</td><td>12376</td><td>20</td><td>CCACAGGGATAACTGGCTTG</td><td>714.6</td><td>-8.5</td><td>1.5</td><td>-5.5</td><td>3.3E-08</td><td>5.6E-08</td><td>BT20/MB468</td><td>NUC</td></tr>
<tr><td>>hg38_21_+_8433172_8446622_RNA45SN1_45S_50ntFlanks@10846.10865.20|CTTCGGGATAAGGATTGGCT</td><td>>hg38_21_+_8433172_8446622_RNA45SN1_45S_50ntFlanks</td><td>10846</td><td>10865</td><td>20</td><td>CTTCGGGATAAGGATTGGCT</td><td>556.7</td><td>-3.3</td><td>0.6</td><td>-5.6</td><td>2.2E-08</td><td>5.6E-08</td><td>BT20/MB468</td><td>MITO</td></tr>
<tr><td>>hg38_21_+_8433172_8446622_RNA45SN1_45S_50ntFlanks@11694.11728.35|AAGAAATTCAATGAAGCGCGGGTAAACGGCGGGAG</td><td>>hg38_21_+_8433172_8446622_RNA45SN1_45S_50ntFlanks</td><td>11694</td><td>11728</td><td>35</td><td>AAGAAATTCAATGAAGCGCGGGTAAACGGCGGGAG</td><td>506.5</td><td>-6.3</td><td>1.1</td><td>-5.5</td><td>3.2E-08</td><td>5.7E-08</td><td>MB231/MB468</td><td>MITO</td></tr>
<tr><td>>hg38_21_+_8433172_8446622_RNA45SN1_45S_50ntFlanks@6663.6692.30|GTGGATCACTCGGCTCGTGCGTCGATGAAG</td><td>>hg38_21_+_8433172_8446622_RNA45SN1_45S_50ntFlanks</td><td>6663</td><td>6692</td><td>30</td><td>GTGGATCACTCGGCTCGTGCGTCGATGAAG</td><td>246.3</td><td>-7.3</td><td>1.3</td><td>-5.6</td><td>2.7E-08</td><td>5.7E-08</td><td>MB231/MB468</td><td>MP</td></tr>
<tr><td>>hg38_21_+_8433172_8446622_RNA45SN1_45S_50ntFlanks@10758.10785.28|CCCATATCCGCAGCAGGTCTCCAAGGTG</td><td>>hg38_21_+_8433172_8446622_RNA45SN1_45S_50ntFlanks</td><td>10758</td><td>10785</td><td>28</td><td>CCCATATCCGCAGCAGGTCTCCAAGGTG</td><td>97.2</td><td>-8.7</td><td>1.6</td><td>-5.6</td><td>2.2E-08</td><td>5.7E-08</td><td>BT20/MB468</td><td>MITO</td></tr>
<tr><td>>hg38_21_+_8433172_8446622_RNA45SN1_45S_50ntFlanks@10587.10607.21|CCCCGGGGAGAGTTCTCTTTT</td><td>>hg38_21_+_8433172_8446622_RNA45SN1_45S_50ntFlanks</td><td>10587</td><td>10607</td><td>21</td><td>CCCCGGGGAGAGTTCTCTTTT</td><td>305.7</td><td>-6.4</td><td>1.1</td><td>-5.5</td><td>2.9E-08</td><td>5.7E-08</td><td>MB231/MB468</td><td>CYTO</td></tr>
<tr><td>>hg38_21_+_8433172_8446622_RNA45SN1_45S_50ntFlanks@12442.12480.39|TGAAGCAGAATTCACCAAGCGTTGGATTGTTCACCCACT</td><td>>hg38_21_+_8433172_8446622_RNA45SN1_45S_50ntFlanks</td><td>12442</td><td>12480</td><td>39</td><td>TGAAGCAGAATTCACCAAGCGTTGGATTGTTCACCCACT</td><td>158.9</td><td>4.6</td><td>0.8</td><td>5.5</td><td>3.0E-08</td><td>5.7E-08</td><td>MB231/MB468</td><td>NUC</td></tr>
<tr><td>>hg38_21_+_8433172_8446622_RNA45SN1_45S_50ntFlanks@12073.12095.23|CCGAGGGGCTCTCGCTTCTGGCG</td><td>>hg38_21_+_8433172_8446622_RNA45SN1_45S_50ntFlanks</td><td>12073</td><td>12095</td><td>23</td><td>CCGAGGGGCTCTCGCTTCTGGCG</td><td>2556.5</td><td>-8.9</td><td>1.5</td><td>-5.9</td><td>4.6E-09</td><td>5.7E-08</td><td>BT20/MB468</td><td>MP</td></tr>
<tr><td>>hg38_21_+_8433172_8446622_RNA45SN1_45S_50ntFlanks@9567.9586.20|ACGTGCAAATCGGTCGTCCG</td><td>>hg38_21_+_8433172_8446622_RNA45SN1_45S_50ntFlanks</td><td>9567</td><td>9586</td><td>20</td><td>ACGTGCAAATCGGTCGTCCG</td><td>285.3</td><td>-7.2</td><td>1.3</td><td>-5.5</td><td>3.2E-08</td><td>5.7E-08</td><td>MB231/MB468</td><td>MITO</td></tr>
<tr><td>>hg38_21_+_8433172_8446622_RNA45SN1_45S_50ntFlanks@8107.8143.37|CGGCGGGGCGCGGGACATGTGGCGTACGGAAGACCCG</td><td>>hg38_21_+_8433172_8446622_RNA45SN1_45S_50ntFlanks</td><td>8107</td><td>8143</td><td>37</td><td>CGGCGGGGCGCGGGACATGTGGCGTACGGAAGACCCG</td><td>1489.1</td><td>-6.9</td><td>1.2</td><td>-5.9</td><td>4.7E-09</td><td>5.7E-08</td><td>BT20/MB468</td><td>MP</td></tr>
<tr><td>>hg38_1_-_228634819_228635039_RNA5S12_5S_50ntFlanks@129.170.42|TTGGATGGGAGACCGCCTGGGAATACCGGGTGCTGTAGGCTT</td><td>>hg38_1_-_228634819_228635039_RNA5S12_5S_50ntFlanks</td><td>129</td><td>170</td><td>42</td><td>TTGGATGGGAGACCGCCTGGGAATACCGGGTGCTGTAGGCTT</td><td>3017.9</td><td>7.1</td><td>1.2</td><td>5.9</td><td>4.7E-09</td><td>5.7E-08</td><td>BT20/MB468</td><td>MP</td></tr>
<tr><td>>hg38_21_+_8433172_8446622_RNA45SN1_45S_50ntFlanks@11874.11892.19|AGAAGACCCTGTTGAGCTT</td><td>>hg38_21_+_8433172_8446622_RNA45SN1_45S_50ntFlanks</td><td>11874</td><td>11892</td><td>19</td><td>AGAAGACCCTGTTGAGCTT</td><td>997.9</td><td>-8.6</td><td>1.6</td><td>-5.5</td><td>3.3E-08</td><td>5.7E-08</td><td>BT20/MB468</td><td>NUC</td></tr>
<tr><td>>trna152_ValCAC_6_-_27248049_27248121@1.36.36|GCTTCTGTAGTGTAGTGGTTATCACGTTCGCCTCAC</td><td>>trna152_ValCAC_6_-_27248049_27248121</td><td>1</td><td>36</td><td>36</td><td>GCTTCTGTAGTGTAGTGGTTATCACGTTCGCCTCAC</td><td>532.1</td><td>6.0</td><td>1.1</td><td>5.5</td><td>3.2E-08</td><td>5.7E-08</td><td>MB231/MB468</td><td>MITO</td></tr>
<tr><td>>MI0003591|hsa-mir-584&WithFlank&5|-|149062307|149062415@22.40.19&[MIMAT0003249&hsa-miR-584-5p&offsets|0|-3;m-167&5|-|149062374|149062394&offsets|0|-2]|TTATGGTTTGCCTGGGACT</td><td>>MI0003591|hsa-mir-584&WithFlank&5|-|149062307|149062415</td><td>22</td><td>40</td><td>19</td><td>TTATGGTTTGCCTGGGACT</td><td>528.2</td><td>7.8</td><td>1.4</td><td>5.5</td><td>3.2E-08</td><td>5.7E-08</td><td>MB231/MB468</td><td>MITO</td></tr>
<tr><td>>hg38_21_+_8433172_8446622_RNA45SN1_45S_50ntFlanks@11583.11609.27|CTGGTGCGGACCAGGGGAATCCGACTG</td><td>>hg38_21_+_8433172_8446622_RNA45SN1_45S_50ntFlanks</td><td>11583</td><td>11609</td><td>27</td><td>CTGGTGCGGACCAGGGGAATCCGACTG</td><td>1092.1</td><td>-6.3</td><td>1.1</td><td>-5.6</td><td>2.7E-08</td><td>5.7E-08</td><td>MB231/MB468</td><td>MP</td></tr>
<tr><td>>hg38_21_+_8433172_8446622_RNA45SN1_45S_50ntFlanks@11217.11234.18|CTGGACGCGAGCCGGGCC</td><td>>hg38_21_+_8433172_8446622_RNA45SN1_45S_50ntFlanks</td><td>11217</td><td>11234</td><td>18</td><td>CTGGACGCGAGCCGGGCC</td><td>1393.7</td><td>-4.4</td><td>0.8</td><td>-5.5</td><td>3.2E-08</td><td>5.7E-08</td><td>MB231/MB468</td><td>MITO</td></tr>
<tr><td>>hg38_21_+_8433172_8446622_RNA45SN1_45S_50ntFlanks@10689.10706.18|GCGTCCGGTGAGCTCTCG</td><td>>hg38_21_+_8433172_8446622_RNA45SN1_45S_50ntFlanks</td><td>10689</td><td>10706</td><td>18</td><td>GCGTCCGGTGAGCTCTCG</td><td>1668.7</td><td>-6.3</td><td>1.1</td><td>-5.6</td><td>2.5E-08</td><td>5.7E-08</td><td>BT20/MB468</td><td>CYTO</td></tr>
<tr><td>>hg38_21_+_8433172_8446622_RNA45SN1_45S_50ntFlanks@11766.11794.29|ATCTAATTAGTGACGCGCATGAATGGATG</td><td>>hg38_21_+_8433172_8446622_RNA45SN1_45S_50ntFlanks</td><td>11766</td><td>11794</td><td>29</td><td>ATCTAATTAGTGACGCGCATGAATGGATG</td><td>94.3</td><td>-9.6</td><td>1.6</td><td>-5.9</td><td>4.7E-09</td><td>5.7E-08</td><td>BT20/MB468</td><td>MP</td></tr>
<tr><td>>hg38_21_+_8433172_8446622_RNA45SN1_45S_50ntFlanks@4731.4767.37|AAGAACGAAAGTCGGAGGTTCGAAGACGATCAGATAC</td><td>>hg38_21_+_8433172_8446622_RNA45SN1_45S_50ntFlanks</td><td>4731</td><td>4767</td><td>37</td><td>AAGAACGAAAGTCGGAGGTTCGAAGACGATCAGATAC</td><td>331.9</td><td>5.5</td><td>1.0</td><td>5.5</td><td>3.0E-08</td><td>5.7E-08</td><td>MB231/MB468</td><td>NUC</td></tr>
<tr><td>>hg38_21_+_8433172_8446622_RNA45SN1_45S_50ntFlanks@9790.9831.42|CTGGCGTGGAGCCGGGCGTGGAATGCGAGTGCCTAGTGGGCC</td><td>>hg38_21_+_8433172_8446622_RNA45SN1_45S_50ntFlanks</td><td>9790</td><td>9831</td><td>42</td><td>CTGGCGTGGAGCCGGGCGTGGAATGCGAGTGCCTAGTGGGCC</td><td>299.1</td><td>-7.7</td><td>1.4</td><td>-5.5</td><td>3.2E-08</td><td>5.7E-08</td><td>MB231/MB468</td><td>MITO</td></tr>
<tr><td>>trnaMT_GluTTC_MT_-_14674_14742@37.72.36,trnalookalike8_GluTTC_5_-_93905172_93905240@37.72.36|TCATTGGTCGTGGTTGTAGTCCGTGCGAGAATACCA</td><td>>trnaMT_GluTTC_MT_-_14674_14742</td><td>37</td><td>72</td><td>36</td><td>TCATTGGTCGTGGTTGTAGTCCGTGCGAGAATACCA</td><td>2423.5</td><td>7.8</td><td>1.4</td><td>5.6</td><td>2.5E-08</td><td>5.7E-08</td><td>BT20/MB468</td><td>CYTO</td></tr>
<tr><td>>MI0000299|hsa-mir-222&WithFlank&X|-|45747009|45747130@75.98.24(+1C)&[MIMAT0000279&hsa-miR-222-3p&offsets|0|+3(+1C);m-140&X|-|45747033|45747056&offsets|0|0(+1C)]|AGCTACATCTGGCTACTGGGTCTCC</td><td>>MI0000299|hsa-mir-222&WithFlank&X|-|45747009|45747130</td><td>75</td><td>98</td><td>24(+1C)</td><td>AGCTACATCTGGCTACTGGGTCTCC</td><td>2144.5</td><td>4.5</td><td>0.8</td><td>5.6</td><td>2.7E-08</td><td>5.7E-08</td><td>MB231/MB468</td><td>MP</td></tr>
<tr><td>>hg38_21_+_8433172_8446622_RNA45SN1_45S_50ntFlanks@9573.9598.26|AAATCGGTCGTCCGACCTGGGTATAG</td><td>>hg38_21_+_8433172_8446622_RNA45SN1_45S_50ntFlanks</td><td>9573</td><td>9598</td><td>26</td><td>AAATCGGTCGTCCGACCTGGGTATAG</td><td>224.0</td><td>-7.2</td><td>1.3</td><td>-5.5</td><td>3.2E-08</td><td>5.7E-08</td><td>MB231/MB468</td><td>MITO</td></tr>
<tr><td>>hg38_21_+_8433172_8446622_RNA45SN1_45S_50ntFlanks@12676.12702.27|AGAATCCCGCCCAGGCGGAACGATACG</td><td>>hg38_21_+_8433172_8446622_RNA45SN1_45S_50ntFlanks</td><td>12676</td><td>12702</td><td>27</td><td>AGAATCCCGCCCAGGCGGAACGATACG</td><td>268.5</td><td>-11.1</td><td>1.9</td><td>-5.9</td><td>4.7E-09</td><td>5.7E-08</td><td>BT20/MB468</td><td>MP</td></tr>
<tr><td>>hg38_21_+_8433172_8446622_RNA45SN1_45S_50ntFlanks@10116.10134.19|TGCGTCGGAGGGCGGCGGC</td><td>>hg38_21_+_8433172_8446622_RNA45SN1_45S_50ntFlanks</td><td>10116</td><td>10134</td><td>19</td><td>TGCGTCGGAGGGCGGCGGC</td><td>182.2</td><td>-6.2</td><td>1.1</td><td>-5.6</td><td>2.5E-08</td><td>5.8E-08</td><td>BT20/MB468</td><td>CYTO</td></tr>
<tr><td>>hg38_21_+_8433172_8446622_RNA45SN1_45S_50ntFlanks@12633.12661.29|CGAAGCTACCATCTGTGGGATTATGACTG</td><td>>hg38_21_+_8433172_8446622_RNA45SN1_45S_50ntFlanks</td><td>12633</td><td>12661</td><td>29</td><td>CGAAGCTACCATCTGTGGGATTATGACTG</td><td>5816.7</td><td>-4.0</td><td>0.7</td><td>-5.6</td><td>2.2E-08</td><td>5.8E-08</td><td>BT20/MB468</td><td>MITO</td></tr>
<tr><td>>hg38_21_+_8433172_8446622_RNA45SN1_45S_50ntFlanks@9623.9656.34|TAGTAGCTGGTTCCCTCCGAAGTTTCCCTCAGGA</td><td>>hg38_21_+_8433172_8446622_RNA45SN1_45S_50ntFlanks</td><td>9623</td><td>9656</td><td>34</td><td>TAGTAGCTGGTTCCCTCCGAAGTTTCCCTCAGGA</td><td>59.1</td><td>-9.8</td><td>1.8</td><td>-5.6</td><td>2.8E-08</td><td>5.8E-08</td><td>MB231/MB468</td><td>MP</td></tr>
<tr><td>>hg38_21_+_8433172_8446622_RNA45SN1_45S_50ntFlanks@7978.8006.29|GACCTCAGATCAGACGTGGCGACCCGCTG</td><td>>hg38_21_+_8433172_8446622_RNA45SN1_45S_50ntFlanks</td><td>7978</td><td>8006</td><td>29</td><td>GACCTCAGATCAGACGTGGCGACCCGCTG</td><td>327.9</td><td>-7.5</td><td>1.4</td><td>-5.6</td><td>2.8E-08</td><td>5.8E-08</td><td>MB231/MB468</td><td>MP</td></tr>
<tr><td>>MI0000734|hsa-mir-106b&WithFlank&7|-|100093987|100094080@56.76.21(+1C)&[MIMAT0004672&hsa-miR-106b-3p&offsets|-2|-3(+1C);m-87&7|-|100094003|100094023&offsets|-2|-2(+1C)]|TACCGCACTGTGGGTACTTGCC</td><td>>MI0000734|hsa-mir-106b&WithFlank&7|-|100093987|100094080</td><td>56</td><td>76</td><td>21(+1C)</td><td>TACCGCACTGTGGGTACTTGCC</td><td>86.2</td><td>9.7</td><td>1.7</td><td>5.6</td><td>2.8E-08</td><td>5.8E-08</td><td>MB231/MB468</td><td>MP</td></tr>
<tr><td>>hg38_21_+_8433172_8446622_RNA45SN1_45S_50ntFlanks@10404.10424.21|CATGGGTCAGTCGGTCCTGAG</td><td>>hg38_21_+_8433172_8446622_RNA45SN1_45S_50ntFlanks</td><td>10404</td><td>10424</td><td>21</td><td>CATGGGTCAGTCGGTCCTGAG</td><td>1635.4</td><td>2.9</td><td>0.5</td><td>5.6</td><td>2.5E-08</td><td>5.8E-08</td><td>BT20/MB468</td><td>CYTO</td></tr>
<tr><td>>hg38_21_+_8433172_8446622_RNA45SN1_45S_50ntFlanks@10820.10840.21|GGGAAGTCGGCAAGCCGGATC</td><td>>hg38_21_+_8433172_8446622_RNA45SN1_45S_50ntFlanks</td><td>10820</td><td>10840</td><td>21</td><td>GGGAAGTCGGCAAGCCGGATC</td><td>1399.1</td><td>-3.4</td><td>0.6</td><td>-5.6</td><td>2.5E-08</td><td>5.8E-08</td><td>BT20/MB468</td><td>CYTO</td></tr>
<tr><td>>MI0000439|hsa-mir-23b&WithFlank&9|+|95085202|95085310@67.86.20(+1A)&[MIMAT0000418&hsa-miR-23b-3p&offsets|+3|0(+1A);m-42&9|+|95085265|95085285&offsets|+3|+2(+1A)]|ACATTGCCAGGGATTACCACA</td><td>>MI0000439|hsa-mir-23b&WithFlank&9|+|95085202|95085310</td><td>67</td><td>86</td><td>20(+1A)</td><td>ACATTGCCAGGGATTACCACA</td><td>114.5</td><td>10.6</td><td>1.8</td><td>5.9</td><td>4.8E-09</td><td>5.8E-08</td><td>BT20/MB468</td><td>MP</td></tr>
<tr><td>>MI0003628|hsa-mir-615&WithFlank&12|+|54033944|54034051@67.85.19&[MIMAT0003283&hsa-miR-615-3p&offsets|0|-3;m-495&12|+|54034010|54034030&offsets|0|-2]|TCCGAGCCTGGGTCTCCCT</td><td>>MI0003628|hsa-mir-615&WithFlank&12|+|54033944|54034051</td><td>67</td><td>85</td><td>19</td><td>TCCGAGCCTGGGTCTCCCT</td><td>236.9</td><td>4.2</td><td>0.7</td><td>5.9</td><td>3.2E-09</td><td>5.8E-08</td><td>BT20/MB231</td><td>NUC</td></tr>
<tr><td>>hg38_21_+_8433172_8446622_RNA45SN1_45S_50ntFlanks@11666.11697.32|TCTGCCCAGTGCTCTGAATGTCAAAGTGAAGA</td><td>>hg38_21_+_8433172_8446622_RNA45SN1_45S_50ntFlanks</td><td>11666</td><td>11697</td><td>32</td><td>TCTGCCCAGTGCTCTGAATGTCAAAGTGAAGA</td><td>210.6</td><td>3.6</td><td>0.6</td><td>5.5</td><td>2.9E-08</td><td>5.8E-08</td><td>MB231/MB468</td><td>CYTO</td></tr>
<tr><td>>hg38_21_+_8433172_8446622_RNA45SN1_45S_50ntFlanks@9535.9570.36|GGAAACTCTGGTGGAGGTCCGTAGCGGTCCTGACGT</td><td>>hg38_21_+_8433172_8446622_RNA45SN1_45S_50ntFlanks</td><td>9535</td><td>9570</td><td>36</td><td>GGAAACTCTGGTGGAGGTCCGTAGCGGTCCTGACGT</td><td>736.5</td><td>4.5</td><td>0.8</td><td>5.5</td><td>3.0E-08</td><td>5.8E-08</td><td>MB231/MB468</td><td>CYTO</td></tr>
<tr><td>>hg38_21_+_8433172_8446622_RNA45SN1_45S_50ntFlanks@10378.10401.24|AGGGTTCCATGTGAACAGCAGTTG</td><td>>hg38_21_+_8433172_8446622_RNA45SN1_45S_50ntFlanks</td><td>10378</td><td>10401</td><td>24</td><td>AGGGTTCCATGTGAACAGCAGTTG</td><td>72.5</td><td>-10.0</td><td>1.8</td><td>-5.6</td><td>2.8E-08</td><td>5.8E-08</td><td>MB231/MB468</td><td>MP</td></tr>
<tr><td>>hg38_21_+_8433172_8446622_RNA45SN1_45S_50ntFlanks@11621.11660.40|CAAAGCATCGCGAAGGCCCGCGGCGGGTGTTGACGCGATG</td><td>>hg38_21_+_8433172_8446622_RNA45SN1_45S_50ntFlanks</td><td>11621</td><td>11660</td><td>40</td><td>CAAAGCATCGCGAAGGCCCGCGGCGGGTGTTGACGCGATG</td><td>513.7</td><td>-7.4</td><td>1.3</td><td>-5.6</td><td>2.3E-08</td><td>5.8E-08</td><td>BT20/MB468</td><td>MITO</td></tr>
<tr><td>>trnaMT_ValTAC_MT_+_1602_1670@45.72.28|TTTCAACTTAACTTGACCGCTCTGACCA</td><td>>trnaMT_ValTAC_MT_+_1602_1670</td><td>45</td><td>72</td><td>28</td><td>TTTCAACTTAACTTGACCGCTCTGACCA</td><td>2044.2</td><td>5.8</td><td>1.0</td><td>5.6</td><td>2.3E-08</td><td>5.8E-08</td><td>BT20/MB468</td><td>MITO</td></tr>
<tr><td>>hg38_21_+_8433172_8446622_RNA45SN1_45S_50ntFlanks@9430.9453.24|CGCACCACCGGCCCGTCTCGCCCG</td><td>>hg38_21_+_8433172_8446622_RNA45SN1_45S_50ntFlanks</td><td>9430</td><td>9453</td><td>24</td><td>CGCACCACCGGCCCGTCTCGCCCG</td><td>173.8</td><td>-9.5</td><td>1.7</td><td>-5.6</td><td>2.3E-08</td><td>5.8E-08</td><td>BT20/MB468</td><td>MITO</td></tr>
<tr><td>>hg38_21_+_8433172_8446622_RNA45SN1_45S_50ntFlanks@4367.4389.23|CAGTTAAAAAGCTCGTAGTTGGA</td><td>>hg38_21_+_8433172_8446622_RNA45SN1_45S_50ntFlanks</td><td>4367</td><td>4389</td><td>23</td><td>CAGTTAAAAAGCTCGTAGTTGGA</td><td>821.3</td><td>-8.3</td><td>1.4</td><td>-5.9</td><td>4.8E-09</td><td>5.8E-08</td><td>BT20/MB468</td><td>MP</td></tr>
<tr><td>>hg38_21_+_8433172_8446622_RNA45SN1_45S_50ntFlanks@5485.5516.32|AGCGCTGAGAAGACGGTCGAACTTGACTATCT</td><td>>hg38_21_+_8433172_8446622_RNA45SN1_45S_50ntFlanks</td><td>5485</td><td>5516</td><td>32</td><td>AGCGCTGAGAAGACGGTCGAACTTGACTATCT</td><td>11934.7</td><td>6.8</td><td>1.1</td><td>6.0</td><td>2.4E-09</td><td>5.9E-08</td><td>BT20/MB231</td><td>MITO</td></tr>
<tr><td>>hg38_21_+_8433172_8446622_RNA45SN1_45S_50ntFlanks@4725.4761.37|TTAATCAAGAACGAAAGTCGGAGGTTCGAAGACGATC</td><td>>hg38_21_+_8433172_8446622_RNA45SN1_45S_50ntFlanks</td><td>4725</td><td>4761</td><td>37</td><td>TTAATCAAGAACGAAAGTCGGAGGTTCGAAGACGATC</td><td>349.1</td><td>5.2</td><td>0.9</td><td>5.5</td><td>3.0E-08</td><td>5.9E-08</td><td>MB231/MB468</td><td>CYTO</td></tr>
<tr><td>>hg38_21_+_8433172_8446622_RNA45SN1_45S_50ntFlanks@9914.9948.35|GAAAAGGTGTTGGTTGATATAGACAGCAGGACGGT</td><td>>hg38_21_+_8433172_8446622_RNA45SN1_45S_50ntFlanks</td><td>9914</td><td>9948</td><td>35</td><td>GAAAAGGTGTTGGTTGATATAGACAGCAGGACGGT</td><td>85.5</td><td>10.2</td><td>1.7</td><td>5.9</td><td>4.8E-09</td><td>5.9E-08</td><td>BT20/MB468</td><td>MP</td></tr>
<tr><td>>MI0001445|hsa-mir-423&WithFlank&17|+|30117073|30117178@23.45.23&[MIMAT0004748&hsa-miR-423-5p&offsets|0|0;m-141&17|+|30117095|30117117&offsets|0|0]|TGAGGGGCAGAGAGCGAGACTTT</td><td>>MI0001445|hsa-mir-423&WithFlank&17|+|30117073|30117178</td><td>23</td><td>45</td><td>23</td><td>TGAGGGGCAGAGAGCGAGACTTT</td><td>798.9</td><td>5.1</td><td>0.9</td><td>5.5</td><td>3.3E-08</td><td>5.9E-08</td><td>MB231/MB468</td><td>MITO</td></tr>
</table>

| | | | | | | | | | | | | | |
|---|---|---|---|---|---|---|---|---|---|---|---|---|---|
| >MI0003591\|hsa-mir-584&WithFlank&5\|-\|149062307\|149062415@22.41.20(+1U)&[MIMAT0003249&hsa-miR-584-5p&offsets\|0\|-2(+1U);m-167&5\|-\|149062374\|149062394&offsets\|0\|-1(+1U)]\|TTATGGTTTGCCTGGGACTGT | >MI0003591\|hsa-mir-584&WithFlank&5\|-\|149062307\|149062415 | 22 | 41 | 20(+1U) | TTATGGTTTGCCTGGGACTGT | 670.0 | 9.0 | 1.6 | 5.5 | 3.3E-08 | 5.9E-08 | MB231/MB468 | MITO |
| >hg38_21_+_8433172_8446622_RNA45SN1_45S_50ntFlanks@12405.12429.25\|CTTTTTGATCCTTCGATGTCGGCTC | >hg38_21_+_8433172_8446622_RNA45SN1_45S_50ntFlanks | 12405 | 12429 | 25 | CTTTTTGATCCTTCGATGTCGGCTC | 797.7 | -3.3 | 0.6 | -5.6 | 2.3E-08 | 5.9E-08 | BT20/MB468 | MITO |
| >hg38_21_+_8433172_8446622_RNA45SN1_45S_50ntFlanks@4894.4918.25\|ACTTAAAGGAATTGACGGAAGGGCA | >hg38_21_+_8433172_8446622_RNA45SN1_45S_50ntFlanks | 4894 | 4918 | 25 | ACTTAAAGGAATTGACGGAAGGGCA | 70.4 | -10.0 | 1.8 | -5.6 | 2.8E-08 | 5.9E-08 | MB231/MB468 | MP |
| >hg38_21_+_8433172_8446622_RNA45SN1_45S_50ntFlanks@9913.9948.36\|AGAAAAGGTGTTGGTTGATATAGACAGCAGGACGGT | >hg38_21_+_8433172_8446622_RNA45SN1_45S_50ntFlanks | 9913 | 9948 | 36 | AGAAAAGGTGTTGGTTGATATAGACAGCAGGACGGT | 81.4 | 10.1 | 1.7 | 5.9 | 4.8E-09 | 5.9E-08 | BT20/MB468 | MP |
| >am_trnaMT_SerGCT_MT_+_12207_12265@12.35.24\|CAAGAACTGCTAACTCATGCCCCC | >am_trnaMT_SerGCT_MT_+_12207_12265 | 12 | 35 | 24 | CAAGAACTGCTAACTCATGCCCCC | 223.1 | 11.5 | 2.0 | 5.9 | 4.9E-09 | 5.9E-08 | BT20/MB468 | MP |
| >trnaMT_ValTAC_MT_+_1602_1670@46.72.27\|TTCAACTTAACTTGACCGCTCTGACCA | >trnaMT_ValTAC_MT_+_1602_1670 | 46 | 72 | 27 | TTCAACTTAACTTGACCGCTCTGACCA | 3305.9 | 5.1 | 0.9 | 5.6 | 2.3E-08 | 5.9E-08 | BT20/MB468 | MITO |
| >hg38_21_+_8433172_8446622_RNA45SN1_45S_50ntFlanks@11208.11253.46\|GCGGCGACTCTGGACGCGAGCCGGGCCCTTCCCGTGGATCGCCCCA | >hg38_21_+_8433172_8446622_RNA45SN1_45S_50ntFlanks | 11208 | 11253 | 46 | GCGGCGACTCTGGACGCGAGCCGGGCCCTTCCCGTGGATCGCCCCA | 397.1 | -8.3 | 1.5 | -5.5 | 3.1E-08 | 5.9E-08 | MB231/MB468 | NUC |
| >hg38_21_+_8433172_8446622_RNA45SN1_45S_50ntFlanks@12936.12962.27\|AAACCATTCGTAGACGACCTGCTTCTG | >hg38_21_+_8433172_8446622_RNA45SN1_45S_50ntFlanks | 12936 | 12962 | 27 | AAACCATTCGTAGACGACCTGCTTCTG | 1157.0 | -8.8 | 1.6 | -5.5 | 3.4E-08 | 5.9E-08 | BT20/MB468 | NUC |
| >hg38_21_+_8433172_8446622_RNA45SN1_45S_50ntFlanks@11209.11254.46\|CGGCGACTCTGGACGCGAGCCGGGCCCTTCCCGTGGATCGCCCCAG | >hg38_21_+_8433172_8446622_RNA45SN1_45S_50ntFlanks | 11209 | 11254 | 46 | CGGCGACTCTGGACGCGAGCCGGGCCCTTCCCGTGGATCGCCCCAG | 10399.2 | -5.6 | 1.0 | -5.6 | 2.6E-08 | 5.9E-08 | BT20/MB468 | CYTO |
| >hg38_21_+_8433172_8446622_RNA45SN1_45S_50ntFlanks@11919.11951.33\|CATGAGAGGTGTAGAATAAGTGGGAGGCCCCCG | >hg38_21_+_8433172_8446622_RNA45SN1_45S_50ntFlanks | 11919 | 11951 | 33 | CATGAGAGGTGTAGAATAAGTGGGAGGCCCCCG | 273.3 | -6.5 | 1.2 | -5.6 | 2.6E-08 | 5.9E-08 | BT20/MB468 | CYTO |
| >MI0000272\|hsa-mir-182&WithFlank&7\|-\|129770377\|129770498@29.51.23(+1G)&[MIMAT0000259&hsa-miR-182-5p&offsets\|0\|-1(+1G);m-22&7\|-\|129770449\|129770470&offsets\|0\|+1(+1G)]\|TTTGGCAATGGTAGAACTCACACG | >MI0000272\|hsa-mir-182&WithFlank&7\|-\|129770377\|129770498 | 29 | 51 | 23(+1G) | TTTGGCAATGGTAGAACTCACACG | 3225.3 | 5.5 | 0.9 | 6.0 | 1.8E-09 | 5.9E-08 | BT20/MB231 | CYTO |
| >hg38_21_+_8433172_8446622_RNA45SN1_45S_50ntFlanks@3985.4024.40\|CGGCTTTGGTGACTCTAGATAACCTCGGGCCGATCGCACG | >hg38_21_+_8433172_8446622_RNA45SN1_45S_50ntFlanks | 3985 | 4024 | 40 | CGGCTTTGGTGACTCTAGATAACCTCGGGCCGATCGCACG | 3549.7 | -9.7 | 1.7 | -5.9 | 4.9E-09 | 5.9E-08 | BT20/MB468 | MP |
| >MI0000102\|hsa-mir-100&WithFlank&11\|-\|122152223\|122152314@19.40.22(+1A)&[MIMAT0000098&hsa-miR-100-5p&offsets\|0\|0(+1A);m-27&11\|-\|122152276\|122152296&offsets\|0\|+1(+1A)]\|AACCCGTAGATCCGAACTTGTGA | >MI0000102\|hsa-mir-100&WithFlank&11\|-\|122152223\|122152314 | 19 | 40 | 22(+1A) | AACCCGTAGATCCGAACTTGTGA | 182.7 | 7.2 | 1.3 | 5.5 | 3.4E-08 | 5.9E-08 | BT20/MB468 | NUC |
| >trnaMT_GlyTCC_MT_+_9991_10058@27.68.42\|AACTTCCAATTAACTAGTTTTGACAACATTCAAAAAAGAGTA | >trnaMT_GlyTCC_MT_+_9991_10058 | 27 | 68 | 42 | AACTTCCAATTAACTAGTTTTGACAACATTCAAAAAAGAGTA | 136.5 | -10.1 | 1.8 | -5.5 | 3.1E-08 | 5.9E-08 | MB231/MB468 | NUC |
| >hg38_21_+_8433172_8446622_RNA45SN1_45S_50ntFlanks@10385.10405.21\|CATGTGAACAGCAGTTGAACA | >hg38_21_+_8433172_8446622_RNA45SN1_45S_50ntFlanks | 10385 | 10405 | 21 | CATGTGAACAGCAGTTGAACA | 260.7 | -11.1 | 1.9 | -5.9 | 4.9E-09 | 5.9E-08 | BT20/MB468 | MP |
| >hg38_21_+_8433172_8446622_RNA45SN1_45S_50ntFlanks@4732.4759.28\|AGAACGAAAGTCGGAGGTTCGAAGACGA | >hg38_21_+_8433172_8446622_RNA45SN1_45S_50ntFlanks | 4732 | 4759 | 28 | AGAACGAAAGTCGGAGGTTCGAAGACGA | 1092.0 | -6.7 | 1.2 | -5.5 | 3.4E-08 | 5.9E-08 | BT20/MB468 | NUC |
| >hg38_21_+_8433172_8446622_RNA45SN1_45S_50ntFlanks@11729.11761.33\|TAACTATGACTCTCTTAAGGTAGCCAAATGCCT | >hg38_21_+_8433172_8446622_RNA45SN1_45S_50ntFlanks | 11729 | 11761 | 33 | TAACTATGACTCTCTTAAGGTAGCCAAATGCCT | 289.2 | -7.3 | 1.3 | -5.5 | 3.3E-08 | 5.9E-08 | MB231/MB468 | MITO |
| >hg38_21_+_8433172_8446622_RNA45SN1_45S_50ntFlanks@10473.10494.22\|CTCGGCCGATCGAAAGGGAGTC | >hg38_21_+_8433172_8446622_RNA45SN1_45S_50ntFlanks | 10473 | 10494 | 22 | CTCGGCCGATCGAAAGGGAGTC | 192.5 | -2.7 | 0.5 | -5.5 | 3.1E-08 | 5.9E-08 | MB231/MB468 | NUC |
| >hg38_21_+_8433172_8446622_RNA45SN1_45S_50ntFlanks@8472.8491.20\|CGGCCCGGCGGATCTTTCCC | >hg38_21_+_8433172_8446622_RNA45SN1_45S_50ntFlanks | 8472 | 8491 | 20 | CGGCCCGGCGGATCTTTCCC | 639.9 | -7.4 | 1.3 | -5.6 | 2.3E-08 | 5.9E-08 | BT20/MB468 | MITO |
| >MI0016746\|hsa-mir-548o-2&WithFlank&20\|+\|38516557\|38516638@51.72.22;MI0006402\|hsa-mir-548o&WithFlank&7\|-\|102405736\|102405861@93.114.22&[MIMAT0005919&hsa-miR-548o-3p&offsets\|0\|0;m-431&20\|+\|38516607\|38516628&offsets\|0\|0];[MIMAT0005919_1&hsa-miR-548o-3p&offsets\|0\|0;m-430&7\|-\|102405748\|102405769&offsets\|0\|0]\|CCAAAACTGCAGTTACTTTTGC | >MI0016746\|hsa-mir-548o-2&WithFlank&20\|+\|38516557\|38516638 | 51 | 72 | 22 | CCAAAACTGCAGTTACTTTTGC | 499.4 | 6.2 | 1.1 | 5.5 | 3.0E-08 | 5.9E-08 | MB231/MB468 | CYTO |
| >hg38_21_+_8433172_8446622_RNA45SN1_45S_50ntFlanks@12995.13035.41\|CTCGCTGCGATCTATTGAAAGTCAGCCCTCGACACAAGGGT | >hg38_21_+_8433172_8446622_RNA45SN1_45S_50ntFlanks | 12995 | 13035 | 41 | CTCGCTGCGATCTATTGAAAGTCAGCCCTCGACACAAGGGT | 369.7 | 9.5 | 1.7 | 5.6 | 2.6E-08 | 5.9E-08 | BT20/MB468 | CYTO |
| >hg38_21_+_8433172_8446622_RNA45SN1_45S_50ntFlanks@8081.8101.21\|GAGCCCAGCGCCGAATCCCCG | >hg38_21_+_8433172_8446622_RNA45SN1_45S_50ntFlanks | 8081 | 8101 | 21 | GAGCCCAGCGCCGAATCCCCG | 447.0 | 7.3 | 1.3 | 5.6 | 2.3E-08 | 5.9E-08 | BT20/MB468 | MITO |
| >hg38_21_+_8433172_8446622_RNA45SN1_45S_50ntFlanks@4983.5020.38\|CGGACAGGATTGACAGATTGATAGCTCTTTCTCGATTC | >hg38_21_+_8433172_8446622_RNA45SN1_45S_50ntFlanks | 4983 | 5020 | 38 | CGGACAGGATTGACAGATTGATAGCTCTTTCTCGATTC | 519.7 | -5.3 | 1.0 | -5.5 | 3.3E-08 | 5.9E-08 | MB231/MB468 | MITO |
| >hg38_21_+_8433172_8446622_RNA45SN1_45S_50ntFlanks@10823.10844.22\|AAGTCGGCAAGCCGGATCCGTA | >hg38_21_+_8433172_8446622_RNA45SN1_45S_50ntFlanks | 10823 | 10844 | 22 | AAGTCGGCAAGCCGGATCCGTA | 112.1 | -9.9 | 1.7 | -5.9 | 4.9E-09 | 5.9E-08 | BT20/MB468 | MP |
| >hg38_21_+_8433172_8446622_RNA45SN1_45S_50ntFlanks@6646.6666.21\|TCGTACGACTCTTAGCGGTGG | >hg38_21_+_8433172_8446622_RNA45SN1_45S_50ntFlanks | 6646 | 6666 | 21 | TCGTACGACTCTTAGCGGTGG | 77824.1 | 4.7 | 0.8 | 5.6 | 2.3E-08 | 5.9E-08 | BT20/MB468 | MITO |
| >hg38_21_+_8433172_8446622_RNA45SN1_45S_50ntFlanks@11581.11601.21\|AACTGGTGCGGACCAGGGGAA | >hg38_21_+_8433172_8446622_RNA45SN1_45S_50ntFlanks | 11581 | 11601 | 21 | AACTGGTGCGGACCAGGGGAA | 1749.3 | -4.6 | 0.8 | -5.6 | 2.6E-08 | 5.9E-08 | BT20/MB468 | CYTO |
| >am_trnaMT_PheGAA_MT_+_577_647@41.58.18\|GTTTAGACGGGCTCACAT | >am_trnaMT_PheGAA_MT_+_577_647 | 41 | 58 | 18 | GTTTAGACGGGCTCACAT | 1516.6 | 4.5 | 0.8 | 5.5 | 3.3E-08 | 5.9E-08 | MB231/MB468 | MITO |
| >hg38_21_+_8433172_8446622_RNA45SN1_45S_50ntFlanks@10686.10709.24\|GCGGCGTCCGGTGAGCTCTCGCTG | >hg38_21_+_8433172_8446622_RNA45SN1_45S_50ntFlanks | 10686 | 10709 | 24 | GCGGCGTCCGGTGAGCTCTCGCTG | 1661.2 | -4.0 | 0.7 | -5.5 | 3.3E-08 | 5.9E-08 | MB231/MB468 | MITO |
| >hg38_21_+_8433172_8446622_RNA45SN1_45S_50ntFlanks@12640.12669.30\|ACCATCTGTGGGATTATGACTGAACGCCTC | >hg38_21_+_8433172_8446622_RNA45SN1_45S_50ntFlanks | 12640 | 12669 | 30 | ACCATCTGTGGGATTATGACTGAACGCCTC | 249.9 | 4.1 | 0.7 | 5.5 | 3.1E-08 | 6.0E-08 | MB231/MB468 | NUC |
| >MI0000093\|hsa-mir-92a-1&WithFlank&13\|+\|91351308\|91351397@54.74.21(+1G);MI0000094\|hsa-mir-92a-2&WithFlank&X\|-\|134169532\|134169618@54.74.21(+1G)&[MIMAT0000092&hsa-miR-92a-3p&offsets\|0\|-1(+1G);m-9&13\|+\|91351361\|91351382&offsets\|0\|-1(+1G)];[MIMAT0000092_1&hsa-miR-92a-3p&offsets\|0\|-1(+1G);m-11&X\|-\|134169544\|134169565&offsets\|0\|-1(+1G)]\|TATTGCACTTGTCCCGGCCTGG | >MI0000093\|hsa-mir-92a-1&WithFlank&13\|+\|91351308\|91351397 | 54 | 74 | 21(+1G) | TATTGCACTTGTCCCGGCCTGG | 415.5 | 4.5 | 0.8 | 5.6 | 2.3E-08 | 6.0E-08 | BT20/MB468 | MITO |
| >hg38_21_+_8433172_8446622_RNA45SN1_45S_50ntFlanks@4297.4319.23\|CATTGGAGGGCAAGTCTGGTGCC | >hg38_21_+_8433172_8446622_RNA45SN1_45S_50ntFlanks | 4297 | 4319 | 23 | CATTGGAGGGCAAGTCTGGTGCC | 258.4 | -7.3 | 1.3 | -5.6 | 2.9E-08 | 6.0E-08 | MB231/MB468 | MP |
| >MI0000089\|hsa-mir-31&WithFlank&9\|-\|21512109\|21512191@14.35.22&[MIMAT0000089&hsa-miR-31-5p&offsets\|0\|+1;m-360&9\|-\|21512157\|21512178&offsets\|0\|0]\|AGGCAAGATGCTGGCATAGCTG | >MI0000089\|hsa-mir-31&WithFlank&9\|-\|21512109\|21512191 | 14 | 35 | 22 | AGGCAAGATGCTGGCATAGCTG | 404.0 | 12.0 | 2.0 | 6.1 | 9.5E-10 | 6.0E-08 | BT20/MB231 | MP |
| >hg38_21_+_8433172_8446622_RNA45SN1_45S_50ntFlanks@8117.8135.19\|CGGGACATGTGGCGTACGG | >hg38_21_+_8433172_8446622_RNA45SN1_45S_50ntFlanks | 8117 | 8135 | 19 | CGGGACATGTGGCGTACGG | 348.3 | 11.8 | 1.9 | 6.1 | 9.5E-10 | 6.0E-08 | BT20/MB231 | MP |
| >MI0000301\|hsa-mir-224&WithFlank&X\|-\|151958572\|151958664@14.34.21(+1A)&[MIMAT0000281&hsa-miR-224-5p&offsets\|+1\|-3(+1A);m-238&X\|-\|151958630\|151958651&offsets\|0\|-1(+1A)]\|CAAGTCACTAGTGGTTCCGTTA | >MI0000301\|hsa-mir-224&WithFlank&X\|-\|151958572\|151958664 | 14 | 34 | 21(+1A) | CAAGTCACTAGTGGTTCCGTTA | 145.8 | 10.9 | 1.9 | 5.8 | 4.9E-09 | 6.0E-08 | BT20/MB468 | MP |
| >hg38_21_+_8433172_8446622_RNA45SN1_45S_50ntFlanks@11602.11619.18\|TCCGACTGTTTAATTAAA | >hg38_21_+_8433172_8446622_RNA45SN1_45S_50ntFlanks | 11602 | 11619 | 18 | TCCGACTGTTTAATTAAA | 1319.5 | -4.3 | 0.8 | -5.6 | 2.6E-08 | 6.0E-08 | BT20/MB468 | CYTO |
| >hg38_21_+_8433172_8446622_RNA45SN1_45S_50ntFlanks@11741.11759.19\|TCTTAAGGTAGCCAAATGC | >hg38_21_+_8433172_8446622_RNA45SN1_45S_50ntFlanks | 11741 | 11759 | 19 | TCTTAAGGTAGCCAAATGC | 327.5 | -7.0 | 1.3 | -5.5 | 3.0E-08 | 6.0E-08 | MB231/MB468 | CYTO |
| >MI0000440\|hsa-mir-27b&WithFlank&9\|+\|95085439\|95085547@67.87.21(+2U)&[MIMAT0000419&hsa-miR-27b-3p&offsets\|0\|0(+2U);m-39&9\|+\|95085505\|95085525&offsets\|0\|0(+2U)]\|TTCACAGTGGCTAAGTTCTGCTT | >MI0000440\|hsa-mir-27b&WithFlank&9\|+\|95085439\|95085547 | 67 | 87 | 21(+2U) | TTCACAGTGGCTAAGTTCTGCTT | 470.8 | 4.5 | 0.8 | 5.5 | 2.9E-08 | 6.0E-08 | MB231/MB468 | MP |
| >hg38_21_+_8433172_8446622_RNA45SN1_45S_50ntFlanks@11572.11592.21\|CCGACTTAGAACTGGTGCGGA | >hg38_21_+_8433172_8446622_RNA45SN1_45S_50ntFlanks | 11572 | 11592 | 21 | CCGACTTAGAACTGGTGCGGA | 583.5 | -7.5 | 1.4 | -5.5 | 3.4E-08 | 6.0E-08 | MB231/MB468 | MITO |
| >hg38_21_+_8433172_8446622_RNA45SN1_45S_50ntFlanks@10905.10933.29\|CTGGGCGCGCGCCGCGGCTGGACGAGGCG | >hg38_21_+_8433172_8446622_RNA45SN1_45S_50ntFlanks | 10905 | 10933 | 29 | CTGGGCGCGCGCCGCGGCTGGACGAGGCG | 752.4 | -8.2 | 1.5 | -5.5 | 3.1E-08 | 6.0E-08 | MB231/MB468 | NUC |
| >MI0000264\|hsa-mir-7-2&WithFlank&15\|+\|88611819\|88611940@38.61.24;MI0000265\|hsa-mir-7-3&WithFlank&19\|+\|4770664\|4770785@37.60.24;MI0000263\|hsa-mir-7-1&WithFlank&9\|-\|83969742\|83969863@30.53.24&[MIMAT0000252&hsa-miR-7-5p&offsets\|0\|0;m-400&15\|+\|88611856\|88611879&offsets\|0\|0];[MIMAT0000252_1&hsa-miR-7-5p&offsets\|0\|0;m-402&19\|+\|4770700\|4770723&offsets\|0\|0];[MIMAT0000252_2&hsa-miR-7-5p&offsets\|0\|0;m-401&9\|-\|83969811\|83969834&offsets\|0\|0]\|TGGAAGACTAGTGATTTTGTTGTT | >MI0000264\|hsa-mir-7-2&WithFlank&15\|+\|88611819\|88611940 | 38 | 61 | 24 | TGGAAGACTAGTGATTTTGTTGTT | 39631.2 | 5.2 | 1.0 | 5.5 | 3.4E-08 | 6.0E-08 | MB231/MB468 | MITO |
| >hg38_21_+_8433172_8446622_RNA45SN1_45S_50ntFlanks@12539.12558.20\|ATGATGTGTTGTTGCCATGG | >hg38_21_+_8433172_8446622_RNA45SN1_45S_50ntFlanks | 12539 | 12558 | 20 | ATGATGTGTTGTTGCCATGG | 114.2 | -9.9 | 1.7 | -5.8 | 5.0E-09 | 6.0E-08 | BT20/MB468 | MP |
| >hg38_21_+_8433172_8446622_RNA45SN1_45S_50ntFlanks@11199.11227.29\|CCGGCGGCGGCGGCGACTCTGGACGCGAG | >hg38_21_+_8433172_8446622_RNA45SN1_45S_50ntFlanks | 11199 | 11227 | 29 | CCGGCGGCGGCGGCGACTCTGGACGCGAG | 916.1 | -5.5 | 1.0 | -5.6 | 2.6E-08 | 6.0E-08 | BT20/MB468 | CYTO |
| >trnaMT_PheGAA_MT_+_577_647@39.58.20\|ATGTTTAGACGGGCTCACAT | >trnaMT_PheGAA_MT_+_577_647 | 39 | 58 | 20 | ATGTTTAGACGGGCTCACAT | 440.0 | 4.7 | 0.8 | 5.5 | 3.4E-08 | 6.0E-08 | MB231/MB468 | MITO |
| >hg38_21_+_8433172_8446622_RNA45SN1_45S_50ntFlanks@7973.8010.38\|GACGCGACCTCAGATCAGACGTGGCGACCCGCTGAATT | >hg38_21_+_8433172_8446622_RNA45SN1_45S_50ntFlanks | 7973 | 8010 | 38 | GACGCGACCTCAGATCAGACGTGGCGACCCGCTGAATT | 4233.0 | 4.6 | 0.8 | 5.5 | 3.5E-08 | 6.0E-08 | BT20/MB468 | NUC |
| >hg38_21_+_8433172_8446622_RNA45SN1_45S_50ntFlanks@4732.4765.34\|AGAACGAAAGTCGGAGGTTCGAAGACGATCAGAT | >hg38_21_+_8433172_8446622_RNA45SN1_45S_50ntFlanks | 4732 | 4765 | 34 | AGAACGAAAGTCGGAGGTTCGAAGACGATCAGAT | 957.9 | 6.0 | 1.1 | 5.5 | 3.1E-08 | 6.0E-08 | MB231/MB468 | CYTO |
| >hg38_21_+_8433172_8446622_RNA45SN1_45S_50ntFlanks@10402.10432.31\|AACATGGGTCAGTCGGTCCTGAGAGATGGGC | >hg38_21_+_8433172_8446622_RNA45SN1_45S_50ntFlanks | 10402 | 10432 | 31 | AACATGGGTCAGTCGGTCCTGAGAGATGGGC | 266.9 | -2.5 | 0.5 | -5.5 | 3.1E-08 | 6.0E-08 | MB231/MB468 | CYTO |
| >hg38_21_+_8433172_8446622_RNA45SN1_45S_50ntFlanks@4003.4042.40\|ATAACCTCGGGCCGATCGCACGCCCCCCGTGGCGGCGACG | >hg38_21_+_8433172_8446622_RNA45SN1_45S_50ntFlanks | 4003 | 4042 | 40 | ATAACCTCGGGCCGATCGCACGCCCCCCGTGGCGGCGACG | 2735.3 | -4.9 | 0.9 | -5.5 | 3.1E-08 | 6.0E-08 | MB231/MB468 | CYTO |
| >hg38_21_+_8433172_8446622_RNA45SN1_45S_50ntFlanks@4594.4616.23\|TGTTGGTTTTCGGAACTGAGGCC | >hg38_21_+_8433172_8446622_RNA45SN1_45S_50ntFlanks | 4594 | 4616 | 23 | TGTTGGTTTTCGGAACTGAGGCC | 503.4 | -6.5 | 1.2 | -5.5 | 2.9E-08 | 6.0E-08 | MB231/MB468 | MP |

| | | | | | | | | | | | | | |
|---|---|---|---|---|---|---|---|---|---|---|---|---|---|
| >hg38_21_+_8433172_8446622_RNA45SN1_45S_50ntFlanks@11771.11792.22\|ATTAGTGACGCGCATGAATGGA | >hg38_21_+_8433172_8446622_RNA45SN1_45S_50ntFlanks | 11771 | 11792 | 22 | ATTAGTGACGCGCATGAATGGA | 267.4 | -5.6 | 1.0 | -5.5 | 3.1E-08 | 6.0E-08 | MB231/MB468 | NUC |
| >MI0003776\|hsa-mir-320b-1&WithFlank&1\|+\|116671740\|116671823@48.67.20;MI0003839\|hsa-mir-320b-2&WithFlank&1\|-\|224257034\|224257116@47.66.20;MI0000542\|hsa-mir-320a&WithFlank&8\|-\|22244960\|22245043@48.67.20&[MIMAT0005792&hsa-miR-320b&offsets\|0\|-2;m-183&1\|+\|116671787\|116671806&offsets\|0\|0];[MIMAT0005792_1&hsa-miR-320b&offsets\|0\|-2;m-183&1\|-\|224257051\|224257070&offsets\|0\|0];[MIMAT0000510&hsa-miR-320a-3p&offsets\|0\|-2;m-60&8\|-\|22244975\|22244996&offsets\|0\|-2]\|AAAAGCTGGGTTGAGAGGGC | >MI0003776\|hsa-mir-320b-1&WithFlank&1\|+\|116671740\|116671823 | 48 | 67 | 20 | AAAAGCTGGGTTGAGAGGGC | 399.4 | 2.9 | 0.5 | 5.5 | 3.5E-08 | 6.0E-08 | BT20/MB468 | NUC |
| >MI0000446\|hsa-mir-125b-1&WithFlank&11\|-\|122099751\|122099850@61.80.20(+3U)&[MIMAT0004592&hsa-miR-125b-1-3p&offsets\|0\|-2(+3U);m-485&11\|-\|122099769\|122099790&offsets\|0\|-2(+3U)]\|ACGGGTTAGGCTCTTGGGAGTTT | >MI0000446\|hsa-mir-125b-1&WithFlank&11\|-\|122099751\|122099850 | 61 | 80 | 20(+3U) | ACGGGTTAGGCTCTTGGGAGTTT | 79.7 | 9.5 | 1.7 | 5.5 | 2.9E-08 | 6.0E-08 | MB231/MB468 | MP |
| >hg38_21_+_8433172_8446622_RNA45SN1_45S_50ntFlanks@10411.10437.27\|CAGTCGGTCCTGAGAGATGGGCGAGCG | >hg38_21_+_8433172_8446622_RNA45SN1_45S_50ntFlanks | 10411 | 10437 | 27 | CAGTCGGTCCTGAGAGATGGGCGAGCG | 200.2 | -10.7 | 1.8 | -5.8 | 5.0E-09 | 6.0E-08 | BT20/MB468 | MP |
| >hg38_21_+_8433172_8446622_RNA45SN1_45S_50ntFlanks@11587.11623.37\|TGCGGACCAGGGGAATCCGACTGTTTAATTAAAACAA | >hg38_21_+_8433172_8446622_RNA45SN1_45S_50ntFlanks | 11587 | 11623 | 37 | TGCGGACCAGGGGAATCCGACTGTTTAATTAAAACAA | 646.1 | 7.9 | 1.4 | 5.6 | 2.4E-08 | 6.0E-08 | BT20/MB468 | MITO |
| >trnaMT_MetCAT_MT_+_4402_4469@53.71.19\|TACCCTTCCCGTACTACCA | >trnaMT_MetCAT_MT_+_4402_4469 | 53 | 71 | 19 | TACCCTTCCCGTACTACCA | 609.3 | 9.2 | 1.7 | 5.6 | 2.6E-08 | 6.0E-08 | BT20/MB468 | CYTO |
| >hg38_21_+_8433172_8446622_RNA45SN1_45S_50ntFlanks@10821.10838.18\|GGAAGTCGGCAAGCCGGA | >hg38_21_+_8433172_8446622_RNA45SN1_45S_50ntFlanks | 10821 | 10838 | 18 | GGAAGTCGGCAAGCCGGA | 665.1 | -4.0 | 0.7 | -5.6 | 2.6E-08 | 6.0E-08 | BT20/MB468 | CYTO |
| >hg38_21_+_8433172_8446622_RNA45SN1_45S_50ntFlanks@8115.8139.25\|CGCGGGACATGTGGCGTACGGAAGA | >hg38_21_+_8433172_8446622_RNA45SN1_45S_50ntFlanks | 8115 | 8139 | 25 | CGCGGGACATGTGGCGTACGGAAGA | 1166.6 | 13.5 | 2.2 | 6.1 | 9.7E-10 | 6.0E-08 | BT20/MB231 | MP |
| >am_MI0000786\|hsa-mir-378a&WithFlank&5\|+\|149732819\|149732896@50.70.21(+1U)&[MIMAT0000732&hsa-miR-378a-3p&offsets\|+1\|0(+1U);m-44&5\|+\|149732867\|149732888&offsets\|+1\|0(+1U)]\|CTGGACTTGGAGTCAGAAGGCT | >am_MI0000786\|hsa-mir-378a&WithFlank&5\|+\|149732819\|149732896 | 50 | 70 | 21(+1U) | CTGGACTTGGAGTCAGAAGGCT | 558.0 | 4.3 | 0.8 | 5.5 | 3.1E-08 | 6.1E-08 | MB231/MB468 | CYTO |
| >MI0003560\|hsa-mir-92b&WithFlank&1\|+\|155195171\|155195278@67.87.21&[MIMAT0003218&hsa-miR-92b-3p&offsets\|0\|-1;m-115&1\|+\|155195237\|155195258&offsets\|0\|-1]\|TATTGCACTCGTCCCGGCCTC | >MI0003560\|hsa-mir-92b&WithFlank&1\|+\|155195171\|155195278 | 67 | 87 | 21 | TATTGCACTCGTCCCGGCCTC | 649.4 | 5.5 | 1.0 | 5.5 | 2.9E-08 | 6.1E-08 | MB231/MB468 | MP |
| >hg38_21_+_8433172_8446622_RNA45SN1_45S_50ntFlanks@10411.10432.22\|CAGTCGGTCCTGAGAGATGGGC | >hg38_21_+_8433172_8446622_RNA45SN1_45S_50ntFlanks | 10411 | 10432 | 22 | CAGTCGGTCCTGAGAGATGGGC | 1572.0 | -4.6 | 0.8 | -5.8 | 5.0E-09 | 6.1E-08 | BT20/MB468 | MP |
| >hg38_21_+_8433172_8446622_RNA45SN1_45S_50ntFlanks@10843.10863.21\|TAACTTCGGGATAAGGATTGG | >hg38_21_+_8433172_8446622_RNA45SN1_45S_50ntFlanks | 10843 | 10863 | 21 | TAACTTCGGGATAAGGATTGG | 190.5 | -7.0 | 1.3 | -5.5 | 3.2E-08 | 6.1E-08 | MB231/MB468 | NUC |
| >hg38_21_+_8433172_8446622_RNA45SN1_45S_50ntFlanks@5016.5034.19\|GATTCCGTGGGTGGTGGTG | >hg38_21_+_8433172_8446622_RNA45SN1_45S_50ntFlanks | 5016 | 5034 | 19 | GATTCCGTGGGTGGTGGTG | 574.5 | -11.3 | 2.0 | -5.6 | 2.4E-08 | 6.1E-08 | BT20/MB468 | MITO |
| >hg38_21_+_8433172_8446622_RNA45SN1_45S_50ntFlanks@6671.6697.27\|CTCGGCTCGTGCGTCGATGAAGAACGC | >hg38_21_+_8433172_8446622_RNA45SN1_45S_50ntFlanks | 6671 | 6697 | 27 | CTCGGCTCGTGCGTCGATGAAGAACGC | 3055.1 | -5.3 | 0.9 | -5.6 | 2.4E-08 | 6.1E-08 | BT20/MB468 | MITO |
| >hg38_21_+_8433172_8446622_RNA45SN1_45S_50ntFlanks@6721.6751.31\|AATTGCAGGACACATTGATCATCGACACTTC | >hg38_21_+_8433172_8446622_RNA45SN1_45S_50ntFlanks | 6721 | 6751 | 31 | AATTGCAGGACACATTGATCATCGACACTTC | 342.4 | -7.9 | 1.4 | -5.5 | 3.5E-08 | 6.1E-08 | BT20/MB468 | NUC |
| >MI0000089\|hsa-mir-31&WithFlank&9\|-\|21512109\|21512191@14.35.22(+1A)&[MIMAT0000089&hsa-miR-31-5p&offsets\|0\|+1(+1A);m-360&9\|-\|21512157\|21512178&offsets\|0\|0(+1A)]\|AGGCAAGATGCTGGCATAGCTGA | >MI0000089\|hsa-mir-31&WithFlank&9\|-\|21512109\|21512191 | 14 | 35 | 22(+1A) | AGGCAAGATGCTGGCATAGCTGA | 102.8 | 3.3 | 0.6 | 5.5 | 3.5E-08 | 6.1E-08 | BT20/MB468 | NUC |
| >hg38_21_+_8433172_8446622_RNA45SN1_45S_50ntFlanks@4964.4994.31\|ACCTCACCCGGCCCGGACACGGACAGGATTG | >hg38_21_+_8433172_8446622_RNA45SN1_45S_50ntFlanks | 4964 | 4994 | 31 | ACCTCACCCGGCCCGGACACGGACAGGATTG | 64.1 | -9.9 | 1.8 | -5.5 | 2.9E-08 | 6.1E-08 | MB231/MB468 | MP |
| >hg38_21_+_8433172_8446622_RNA45SN1_45S_50ntFlanks@9571.9595.25\|GCAAATCGGTCGTCCGACCTGGGTA | >hg38_21_+_8433172_8446622_RNA45SN1_45S_50ntFlanks | 9571 | 9595 | 25 | GCAAATCGGTCGTCCGACCTGGGTA | 308.8 | -6.1 | 1.1 | -5.5 | 3.4E-08 | 6.1E-08 | MB231/MB468 | MITO |
| >hg38_21_+_8433172_8446622_RNA45SN1_45S_50ntFlanks@9623.9654.32\|TAGTAGCTGGTTCCCTCCGAAGTTTCCCTCAG | >hg38_21_+_8433172_8446622_RNA45SN1_45S_50ntFlanks | 9623 | 9654 | 32 | TAGTAGCTGGTTCCCTCCGAAGTTTCCCTCAG | 53.3 | -9.6 | 1.7 | -5.5 | 2.9E-08 | 6.1E-08 | MB231/MB468 | MP |
| >hg38_21_+_8433172_8446622_RNA45SN1_45S_50ntFlanks@3813.3841.29\|TTATGGTTCCTTTGGTCGCTCGCTCCTCT | >hg38_21_+_8433172_8446622_RNA45SN1_45S_50ntFlanks | 3813 | 3841 | 29 | TTATGGTTCCTTTGGTCGCTCGCTCCTCT | 521.9 | -8.4 | 1.5 | -5.5 | 3.5E-08 | 6.1E-08 | BT20/MB468 | NUC |
| >hg38_MT_+_1621_3279_RNR2_16S_50ntFlanks@51.74.24\|GCTAAACCTAGCCCCAAACCCACT | >hg38_MT_+_1621_3279_RNR2_16S_50ntFlanks | 51 | 74 | 24 | GCTAAACCTAGCCCCAAACCCACT | 13314.4 | 5.5 | 1.0 | 5.6 | 2.4E-08 | 6.1E-08 | BT20/MB468 | MITO |
| >hg38_21_+_8433172_8446622_RNA45SN1_45S_50ntFlanks@4003.4025.23\|ATAACCTCGGGCCGATCGCACGC | >hg38_21_+_8433172_8446622_RNA45SN1_45S_50ntFlanks | 4003 | 4025 | 23 | ATAACCTCGGGCCGATCGCACGC | 805.3 | -6.0 | 1.1 | -5.6 | 2.4E-08 | 6.1E-08 | BT20/MB468 | MITO |
| >hg38_21_+_8433172_8446622_RNA45SN1_45S_50ntFlanks@11589.11609.21\|CGGACCAGGGGAATCCGACTG | >hg38_21_+_8433172_8446622_RNA45SN1_45S_50ntFlanks | 11589 | 11609 | 21 | CGGACCAGGGGAATCCGACTG | 573.8 | -6.0 | 1.1 | -5.5 | 2.9E-08 | 6.1E-08 | MB231/MB468 | MP |
| >hg38_21_+_8433172_8446622_RNA45SN1_45S_50ntFlanks@4763.4780.18\|GATACCGTCGTAGTTCCG | >hg38_21_+_8433172_8446622_RNA45SN1_45S_50ntFlanks | 4763 | 4780 | 18 | GATACCGTCGTAGTTCCG | 1343.1 | -5.5 | 1.0 | -5.5 | 3.1E-08 | 6.1E-08 | MB231/MB468 | CYTO |
| >MI0000067\|hsa-let-7f-1&WithFlank&9\|+\|94176341\|94176439@13.35.23&[MIMAT0000067&hsa-let-7f-5p&offsets\|0\|+1;m-20&9\|+\|94176353\|94176374&offsets\|0\|+1]\|TGAGGTAGTAGATTGTATAGTTG | >MI0000067\|hsa-let-7f-1&WithFlank&9\|+\|94176341\|94176439 | 13 | 35 | 23 | TGAGGTAGTAGATTGTATAGTTG | 107.5 | 2.0 | 0.4 | 5.5 | 3.6E-08 | 6.1E-08 | BT20/MB468 | NUC |
| >hg38_21_+_8433172_8446622_RNA45SN1_45S_50ntFlanks@11906.11929.24\|CACGGTGAAGAGACATGAGAGGTG | >hg38_21_+_8433172_8446622_RNA45SN1_45S_50ntFlanks | 11906 | 11929 | 24 | CACGGTGAAGAGACATGAGAGGTG | 207.5 | -6.8 | 1.2 | -5.5 | 3.2E-08 | 6.1E-08 | MB231/MB468 | NUC |
| >MI0000481\|hsa-mir-184&WithFlank&15\|+\|79209782\|79209877@59.80.22&[MIMAT0000454&hsa-miR-184&offsets\|0\|0;m-460&15\|+\|79209840\|79209861&offsets\|0\|0]\|TGGACGGAGAACTGATAAGGGT | >MI0000481\|hsa-mir-184&WithFlank&15\|+\|79209782\|79209877 | 59 | 80 | 22 | TGGACGGAGAACTGATAAGGGT | 556.3 | 9.8 | 1.7 | 5.8 | 5.1E-09 | 6.1E-08 | BT20/MB468 | MP |
| >hg38_21_+_8433172_8446622_RNA45SN1_45S_50ntFlanks@7974.8007.34\|ACGCGACCTCAGATCAGACGTGGCGACCCGCTGA | >hg38_21_+_8433172_8446622_RNA45SN1_45S_50ntFlanks | 7974 | 8007 | 34 | ACGCGACCTCAGATCAGACGTGGCGACCCGCTGA | 1147.6 | 3.8 | 0.7 | 5.5 | 3.1E-08 | 6.1E-08 | MB231/MB468 | CYTO |
| >MI0000455\|hsa-mir-138-2&WithFlank&16\|+\|56858512\|56858607@16.37.22;MI0000476\|hsa-mir-138-1&WithFlank&3\|+\|44114206\|44114316@29.50.22&[MIMAT0000430&hsa-miR-138-5p&offsets\|0\|-1;m-509&16\|+\|56858527\|56858549&offsets\|0\|-1];[MIMAT0000430_1&hsa-miR-138-5p&offsets\|0\|-1;m-497&3\|+\|44114234\|44114256&offsets\|0\|-1]\|AGCTGGTGTTGTGAATCAGGCC | >MI0000455\|hsa-mir-138-2&WithFlank&16\|+\|56858512\|56858607 | 16 | 37 | 22 | AGCTGGTGTTGTGAATCAGGCC | 58.8 | 9.1 | 1.6 | 5.5 | 2.9E-08 | 6.2E-08 | MB231/MB468 | MP |
| >hg38_21_+_8433172_8446622_RNA45SN1_45S_50ntFlanks@11315.11336.22\|CCACCCCACGTCTCGTCGCGCG | >hg38_21_+_8433172_8446622_RNA45SN1_45S_50ntFlanks | 11315 | 11336 | 22 | CCACCCCACGTCTCGTCGCGCG | 108.7 | -9.8 | 1.7 | -5.8 | 5.1E-09 | 6.2E-08 | BT20/MB468 | MP |
| >hg38_1_-_228634819_228635039_RNA5S12_5S_50ntFlanks@123.170.48\|TAGTACTTGGATGGGAGACCGCCTGGGAATACCGGGTGCTGTAGGCTT | >hg38_1_-_228634819_228635039_RNA5S12_5S_50ntFlanks | 123 | 170 | 48 | TAGTACTTGGATGGGAGACCGCCTGGGAATACCGGGTGCTGTAGGCTT | 903.3 | 5.7 | 1.0 | 5.5 | 3.1E-08 | 6.2E-08 | MB231/MB468 | CYTO |
| >hg38_21_+_8433172_8446622_RNA45SN1_45S_50ntFlanks@4727.4765.39\|AATCAAGAACGAAAGTCGGAGGTTCGAAGACGATCAGAT | >hg38_21_+_8433172_8446622_RNA45SN1_45S_50ntFlanks | 4727 | 4765 | 39 | AATCAAGAACGAAAGTCGGAGGTTCGAAGACGATCAGAT | 2505.9 | 5.5 | 1.0 | 5.5 | 3.1E-08 | 6.2E-08 | MB231/MB468 | CYTO |
| >hg38_21_+_8433172_8446622_RNA45SN1_45S_50ntFlanks@12489.12509.21\|ACGTGAGCTGGGTTTAGACCG | >hg38_21_+_8433172_8446622_RNA45SN1_45S_50ntFlanks | 12489 | 12509 | 21 | ACGTGAGCTGGGTTTAGACCG | 735.2 | -6.7 | 1.2 | -5.5 | 3.1E-08 | 6.2E-08 | MB231/MB468 | CYTO |
| >MI0001733\|hsa-mir-452&WithFlank&X\|-\|151959622\|151959718@23.44.22&[MIMAT0001635&hsa-miR-452-5p&offsets\|+3\|+3;m-188&X\|-\|151959677\|151959699&offsets\|+3\|+2]\|TGTTTGCAGAGGAAACTGAGAC | >MI0001733\|hsa-mir-452&WithFlank&X\|-\|151959622\|151959718 | 23 | 44 | 22 | TGTTTGCAGAGGAAACTGAGAC | 1676.4 | 10.9 | 1.9 | 5.8 | 5.1E-09 | 6.2E-08 | BT20/MB468 | MP |
| >hg38_21_+_8433172_8446622_RNA45SN1_45S_50ntFlanks@7986.8003.18\|ATCAGACGTGGCGACCCG | >hg38_21_+_8433172_8446622_RNA45SN1_45S_50ntFlanks | 7986 | 8003 | 18 | ATCAGACGTGGCGACCCG | 2415.2 | -8.3 | 1.5 | -5.5 | 3.2E-08 | 6.2E-08 | MB231/MB468 | NUC |
| >hg38_21_+_8433172_8446622_RNA45SN1_45S_50ntFlanks@9534.9562.29\|AGGAAACTCTGGTGGAGGTCCGTAGCGGT | >hg38_21_+_8433172_8446622_RNA45SN1_45S_50ntFlanks | 9534 | 9562 | 29 | AGGAAACTCTGGTGGAGGTCCGTAGCGGT | 665.6 | -3.1 | 0.5 | -5.6 | 2.4E-08 | 6.2E-08 | BT20/MB468 | MITO |
| >MI0000289\|hsa-mir-181a-1&WithFlank&1\|-\|198859038\|198859159@30.51.22(+1C);MI0000269\|hsa-mir-181a-2&WithFlank&9\|+\|124692436\|124692557@45.66.22(+1C)&[MIMAT0000256&hsa-miR-181a-5p&offsets\|0\|-1(+1C);m-32&1\|-\|198859108\|198859130&offsets\|0\|-1(+1C)];[MIMAT0000256_1&hsa-miR-181a-5p&offsets\|0\|-1(+1C);m-32&9\|+\|124692480\|124692502&offsets\|0\|-1(+1C)]\|AACATTCAACGCTGTCGGTGAGC | >MI0000289\|hsa-mir-181a-1&WithFlank&1\|-\|198859038\|198859159 | 30 | 51 | 22(+1C) | AACATTCAACGCTGTCGGTGAGC | 958.1 | 6.0 | 1.1 | 5.5 | 3.5E-08 | 6.2E-08 | MB231/MB468 | MITO |
| >hg38_21_+_8433172_8446622_RNA45SN1_45S_50ntFlanks@10466.10497.32\|CGTTGCCCTCGGCCGATCGAAAGGGAGTCGGG | >hg38_21_+_8433172_8446622_RNA45SN1_45S_50ntFlanks | 10466 | 10497 | 32 | CGTTGCCCTCGGCCGATCGAAAGGGAGTCGGG | 764.0 | -7.1 | 1.3 | -5.6 | 2.7E-08 | 6.2E-08 | BT20/MB468 | CYTO |
| >MI0000446\|hsa-mir-125b-1&WithFlank&11\|-\|122099751\|122099850@60.80.21(+1U)&[MIMAT0004592&hsa-miR-125b-1-3p&offsets\|-1\|-2(+1U);m-485&11\|-\|122099769\|122099790&offsets\|-1\|-2(+1U)]\|CACGGGTTAGGCTCTTGGGAGT | >MI0000446\|hsa-mir-125b-1&WithFlank&11\|-\|122099751\|122099850 | 60 | 80 | 21(+1U) | CACGGGTTAGGCTCTTGGGAGT | 69.3 | 9.3 | 1.7 | 5.5 | 3.0E-08 | 6.2E-08 | MB231/MB468 | MP |
| >hg38_21_+_8433172_8446622_RNA45SN1_45S_50ntFlanks@3855.3872.18\|CTGTGGTAATTCTAGAGC | >hg38_21_+_8433172_8446622_RNA45SN1_45S_50ntFlanks | 3855 | 3872 | 18 | CTGTGGTAATTCTAGAGC | 219.7 | -7.3 | 1.3 | -5.5 | 3.0E-08 | 6.2E-08 | MB231/MB468 | MP |
| >hg38_21_+_8433172_8446622_RNA45SN1_45S_50ntFlanks@10344.10373.30\|ATTCAAACGAGAACTTTGAAGGCCGAAGTG | >hg38_21_+_8433172_8446622_RNA45SN1_45S_50ntFlanks | 10344 | 10373 | 30 | ATTCAAACGAGAACTTTGAAGGCCGAAGTG | 289.7 | 5.7 | 1.0 | 5.6 | 2.7E-08 | 6.2E-08 | BT20/MB468 | CYTO |
| >trna2_GlyGCC_21_-_18827107_18827177@1.28.28,trna35_GlyGCC_1_+_161413094_161413164@1.28.28,trna37_GlyGCC_1_+_161420467_161420537@1.28.28,trna39_GlyGCC_1_+_161427898_161427968@1.28.28,trna41_GlyGCC_1_+_161435258_161435328@1.28.28\|GCATGGGTGGTTCAGTGGTAGAATTCTC | >trna2_GlyGCC_21_-_18827107_18827177 | 1 | 28 | 28 | GCATGGGTGGTTCAGTGGTAGAATTCTC | 2151.3 | 14.7 | 2.5 | 6.0 | 2.6E-09 | 6.2E-08 | BT20/MB231 | MITO |
| >MI0000100\|hsa-mir-98&WithFlank&X\|-\|53556217\|53556347@28.48.21(+1C)&[MIMAT0000096&hsa-miR-98-5p&offsets\|0\|-1(+1C);m-143&X\|-\|53556299\|53556320&offsets\|0\|-1(+1C)]\|TGAGGTAGTAAGTTGTATTGTC | >MI0000100\|hsa-mir-98&WithFlank&X\|-\|53556217\|53556347 | 28 | 48 | 21(+1C) | TGAGGTAGTAAGTTGTATTGTC | 1072.4 | 4.9 | 0.9 | 5.6 | 2.7E-08 | 6.2E-08 | BT20/MB468 | CYTO |

<table>
<tr><td>>trna10_AlaCGC_6_+_26553731_26553802@1.24.24,trna119_AlaCGC_6_-_28641613_28641684@1.24.24,trna120_AlaAGC_6_-_28626014_28626085@1.24.24,trna13_AlaCGC_2_+_157257281_157257352@1.24.24,trna13_AlaTGC_12_-_125406301_125406372@1.24.24,trna66_AlaTGC_6_+_28611222_28611293@1.24.24,trna8_AlaTGC_12_+_125424512_125424583@1.24.24,trna8_AlaTGC_5_+_180633868_180633939@1.24.24|GGGGATGTAGCTCAGTGGTAGAGC</td><td>>trna10_AlaCGC_6_+_26553731_26553802</td><td>1</td><td>24</td><td>24</td><td>GGGGATGTAGCTCAGTGGTAGAGC</td><td>558.9</td><td>4.5</td><td>0.8</td><td>5.6</td><td>2.7E-08</td><td>6.2E-08</td><td>BT20/MB468</td><td>CYTO</td></tr>
<tr><td>>hg38_21_+_8433172_8446622_RNA45SN1_45S_50ntFlanks@11569.11601.33|CAGCCGACTTAGAACTGGTGCGGACCAGGGGAA</td><td>>hg38_21_+_8433172_8446622_RNA45SN1_45S_50ntFlanks</td><td>11569</td><td>11601</td><td>33</td><td>CAGCCGACTTAGAACTGGTGCGGACCAGGGGAA</td><td>6493.0</td><td>-5.1</td><td>0.9</td><td>-5.8</td><td>5.2E-09</td><td>6.2E-08</td><td>BT20/MB468</td><td>MP</td></tr>
<tr><td>>hg38_21_+_8433172_8446622_RNA45SN1_45S_50ntFlanks@12456.12485.30|CCAAGCGTTGGATTGTTCACCCACTAATAG</td><td>>hg38_21_+_8433172_8446622_RNA45SN1_45S_50ntFlanks</td><td>12456</td><td>12485</td><td>30</td><td>CCAAGCGTTGGATTGTTCACCCACTAATAG</td><td>2039.1</td><td>-6.5</td><td>1.2</td><td>-5.6</td><td>2.7E-08</td><td>6.2E-08</td><td>BT20/MB468</td><td>CYTO</td></tr>
<tr><td>>hg38_21_+_8433172_8446622_RNA45SN1_45S_50ntFlanks@12949.12982.34|ACGACCTGCTTCTGGGTCGGGGTTTCGTACGTAG</td><td>>hg38_21_+_8433172_8446622_RNA45SN1_45S_50ntFlanks</td><td>12949</td><td>12982</td><td>34</td><td>ACGACCTGCTTCTGGGTCGGGGTTTCGTACGTAG</td><td>262.0</td><td>-6.7</td><td>1.2</td><td>-5.6</td><td>2.7E-08</td><td>6.2E-08</td><td>BT20/MB468</td><td>CYTO</td></tr>
<tr><td>>hg38_21_+_8433172_8446622_RNA45SN1_45S_50ntFlanks@9554.9597.44|CGTAGCGGTCCTGACGTGCAAATCGGTCGTCCGACCTGGGTATA</td><td>>hg38_21_+_8433172_8446622_RNA45SN1_45S_50ntFlanks</td><td>9554</td><td>9597</td><td>44</td><td>CGTAGCGGTCCTGACGTGCAAATCGGTCGTCCGACCTGGGTATA</td><td>2773.2</td><td>-5.4</td><td>1.0</td><td>-5.6</td><td>2.4E-08</td><td>6.2E-08</td><td>BT20/MB468</td><td>MITO</td></tr>
<tr><td>>hg38_1_-_228634819_228635039_RNA5S12_5S_50ntFlanks@51.90.40|GTCTACGGCCATACCACCCTGAACGCGCCCGATCTCGTCT</td><td>>hg38_1_-_228634819_228635039_RNA5S12_5S_50ntFlanks</td><td>51</td><td>90</td><td>40</td><td>GTCTACGGCCATACCACCCTGAACGCGCCCGATCTCGTCT</td><td>810.2</td><td>6.3</td><td>1.1</td><td>5.5</td><td>3.2E-08</td><td>6.2E-08</td><td>MB231/MB468</td><td>CYTO</td></tr>
<tr><td>>hg38_21_+_8433172_8446622_RNA45SN1_45S_50ntFlanks@10347.10382.36|CAAACGAGAACTTTGAAGGCCGAAGTGGAGAAGGGT</td><td>>hg38_21_+_8433172_8446622_RNA45SN1_45S_50ntFlanks</td><td>10347</td><td>10382</td><td>36</td><td>CAAACGAGAACTTTGAAGGCCGAAGTGGAGAAGGGT</td><td>403.8</td><td>4.3</td><td>0.8</td><td>5.5</td><td>3.2E-08</td><td>6.2E-08</td><td>MB231/MB468</td><td>CYTO</td></tr>
<tr><td>>hg38_21_+_8433172_8446622_RNA45SN1_45S_50ntFlanks@12002.12038.37|TGCGGGCCGCCGGTGAAATACCACTACTCTGATCGTT</td><td>>hg38_21_+_8433172_8446622_RNA45SN1_45S_50ntFlanks</td><td>12002</td><td>12038</td><td>37</td><td>TGCGGGCCGCCGGTGAAATACCACTACTCTGATCGTT</td><td>72.5</td><td>-10.0</td><td>1.8</td><td>-5.5</td><td>3.0E-08</td><td>6.2E-08</td><td>MB231/MB468</td><td>MP</td></tr>
<tr><td>>hg38_21_+_8433172_8446622_RNA45SN1_45S_50ntFlanks@12934.12972.39|CTAAACCATTCGTAGACGACCTGCTTCTGGGTCGGGGTT</td><td>>hg38_21_+_8433172_8446622_RNA45SN1_45S_50ntFlanks</td><td>12934</td><td>12972</td><td>39</td><td>CTAAACCATTCGTAGACGACCTGCTTCTGGGTCGGGGTT</td><td>1319.0</td><td>-8.2</td><td>1.5</td><td>-5.5</td><td>3.3E-08</td><td>6.2E-08</td><td>MB231/MB468</td><td>NUC</td></tr>
<tr><td>>am_trna111_HisGTG_1_-_147774845_147774916@1.29.29,trna118_HisGTG_1_-_145396881_145396952@1.29.29,trna16_HisGTG_1_+_146544773_146544844@1.29.29,trna1_HisGTG_15_+_45493349_45493420@1.29.29,trna21_HisGTG_1_+_147753471_147753542@1.29.29,trna33_HisGTG_6_+_27125906_27125977@1.29.29,trna7_HisGTG_9_-_14433938_14434009@1.29.29,trna8_HisGTG_15_-_45492611_45492682@1.29.29,trna9_HisGTG_15_-_45490804_45490875@1.29.29|GCCGTGATCGTATAGTGGTTAGTACTCTG</td><td>>am_trna111_HisGTG_1_-_147774845_147774916</td><td>1</td><td>29</td><td>29</td><td>GCCGTGATCGTATAGTGGTTAGTACTCTG</td><td>22416.4</td><td>-5.0</td><td>0.9</td><td>-5.5</td><td>3.2E-08</td><td>6.2E-08</td><td>MB231/MB468</td><td>CYTO</td></tr>
<tr><td>>hg38_21_+_8433172_8446622_RNA45SN1_45S_50ntFlanks@10420.10457.38|CTGAGAGATGGGCGAGCGCCGTTCCGAAGGGACGGGCG</td><td>>hg38_21_+_8433172_8446622_RNA45SN1_45S_50ntFlanks</td><td>10420</td><td>10457</td><td>38</td><td>CTGAGAGATGGGCGAGCGCCGTTCCGAAGGGACGGGCG</td><td>1141.1</td><td>-5.6</td><td>1.0</td><td>-5.5</td><td>3.2E-08</td><td>6.2E-08</td><td>MB231/MB468</td><td>CYTO</td></tr>
<tr><td>>hg38_21_+_8433172_8446622_RNA45SN1_45S_50ntFlanks@10690.10708.19|CGTCCGGTGAGCTCTCGCT</td><td>>hg38_21_+_8433172_8446622_RNA45SN1_45S_50ntFlanks</td><td>10690</td><td>10708</td><td>19</td><td>CGTCCGGTGAGCTCTCGCT</td><td>3646.8</td><td>-4.0</td><td>0.7</td><td>-5.6</td><td>2.5E-08</td><td>6.2E-08</td><td>BT20/MB468</td><td>MITO</td></tr>
<tr><td>>MI0000071|hsa-mir-17&WithFlank&13|+|91350599|91350694@20.42.23(+1A)&[MIMAT0000070&hsa-miR-17-5p&offsets|0|0(+1A);m-88&13|+|91350618|91350640&offsets|0|0(+1A)]|CAAAGTGCTTACAGTGCAGGTAGA</td><td>>MI0000071|hsa-mir-17&WithFlank&13|+|91350599|91350694</td><td>20</td><td>42</td><td>23(+1A)</td><td>CAAAGTGCTTACAGTGCAGGTAGA</td><td>1200.3</td><td>4.4</td><td>0.8</td><td>5.6</td><td>2.5E-08</td><td>6.3E-08</td><td>BT20/MB468</td><td>MITO</td></tr>
<tr><td>>MI0000809|hsa-mir-151a&WithFlank&8|-|140732558|140732659@53.74.22(+3U)&[MIMAT0000757&hsa-miR-151a-3p&offsets|0|+1(+3U);m-30&8|-|140732587|140732607&offsets|0|+1(+3U)]|CTAGACTGAAGCTCCTTGAGGATTT</td><td>>MI0000809|hsa-mir-151a&WithFlank&8|-|140732558|140732659</td><td>53</td><td>74</td><td>22(+3U)</td><td>CTAGACTGAAGCTCCTTGAGGATTT</td><td>62.3</td><td>9.2</td><td>1.7</td><td>5.5</td><td>3.0E-08</td><td>6.3E-08</td><td>MB231/MB468</td><td>MP</td></tr>
<tr><td>>hg38_21_+_8433172_8446622_RNA45SN1_45S_50ntFlanks@12676.12709.34|AGAATCCCGCCCAGGCGGAACGATACGGCAGCGC</td><td>>hg38_21_+_8433172_8446622_RNA45SN1_45S_50ntFlanks</td><td>12676</td><td>12709</td><td>34</td><td>AGAATCCCGCCCAGGCGGAACGATACGGCAGCGC</td><td>85.2</td><td>-9.5</td><td>1.6</td><td>-5.8</td><td>5.2E-09</td><td>6.3E-08</td><td>BT20/MB468</td><td>MP</td></tr>
<tr><td>>hg38_21_+_8433172_8446622_RNA45SN1_45S_50ntFlanks@9956.9982.27|GAAGTCGGAATCCGCTAAGGAGTGTGT</td><td>>hg38_21_+_8433172_8446622_RNA45SN1_45S_50ntFlanks</td><td>9956</td><td>9982</td><td>27</td><td>GAAGTCGGAATCCGCTAAGGAGTGTGT</td><td>91.2</td><td>10.3</td><td>1.8</td><td>5.8</td><td>5.3E-09</td><td>6.3E-08</td><td>BT20/MB468</td><td>MP</td></tr>
<tr><td>>hg38_21_+_8433172_8446622_RNA45SN1_45S_50ntFlanks@12717.12751.35|CCTCGGTTGGCCTCGGATAGCCGGTCCCCCGCCTG</td><td>>hg38_21_+_8433172_8446622_RNA45SN1_45S_50ntFlanks</td><td>12717</td><td>12751</td><td>35</td><td>CCTCGGTTGGCCTCGGATAGCCGGTCCCCCGCCTG</td><td>230.6</td><td>-10.9</td><td>1.9</td><td>-5.8</td><td>5.3E-09</td><td>6.3E-08</td><td>BT20/MB468</td><td>MP</td></tr>
<tr><td>>hg38_21_+_8433172_8446622_RNA45SN1_45S_50ntFlanks@11209.11229.21|CGGCGACTCTGGACGCGAGCC</td><td>>hg38_21_+_8433172_8446622_RNA45SN1_45S_50ntFlanks</td><td>11209</td><td>11229</td><td>21</td><td>CGGCGACTCTGGACGCGAGCC</td><td>550.1</td><td>-3.9</td><td>0.7</td><td>-5.5</td><td>3.5E-08</td><td>6.3E-08</td><td>MB231/MB468</td><td>MITO</td></tr>
<tr><td>>hg38_21_+_8433172_8446622_RNA45SN1_45S_50ntFlanks@9534.9560.27|AGGAAACTCTGGTGGAGGTCCGTAGCG</td><td>>hg38_21_+_8433172_8446622_RNA45SN1_45S_50ntFlanks</td><td>9534</td><td>9560</td><td>27</td><td>AGGAAACTCTGGTGGAGGTCCGTAGCG</td><td>1534.6</td><td>-6.8</td><td>1.2</td><td>-5.5</td><td>3.0E-08</td><td>6.3E-08</td><td>MB231/MB468</td><td>MP</td></tr>
<tr><td>>hg38_21_+_8433172_8446622_RNA45SN1_45S_50ntFlanks@11594.11613.20|CAGGGGAATCCGACTGTTTA</td><td>>hg38_21_+_8433172_8446622_RNA45SN1_45S_50ntFlanks</td><td>11594</td><td>11613</td><td>20</td><td>CAGGGGAATCCGACTGTTTA</td><td>737.8</td><td>-6.6</td><td>1.2</td><td>-5.5</td><td>3.6E-08</td><td>6.3E-08</td><td>BT20/MB468</td><td>NUC</td></tr>
<tr><td>>hg38_21_+_8433172_8446622_RNA45SN1_45S_50ntFlanks@11873.11912.40|AAGAAGACCCTGTTGAGCTTGACTCTAGTCTGGCACGGTG</td><td>>hg38_21_+_8433172_8446622_RNA45SN1_45S_50ntFlanks</td><td>11873</td><td>11912</td><td>40</td><td>AAGAAGACCCTGTTGAGCTTGACTCTAGTCTGGCACGGTG</td><td>165.7</td><td>-7.0</td><td>1.3</td><td>-5.6</td><td>2.8E-08</td><td>6.3E-08</td><td>BT20/MB468</td><td>CYTO</td></tr>
<tr><td>>hg38_21_+_8433172_8446622_RNA45SN1_45S_50ntFlanks@11552.11591.40|CCTCGGCCGGCGCCTAGCAGCCGACTTAGAACTGGTGCGG</td><td>>hg38_21_+_8433172_8446622_RNA45SN1_45S_50ntFlanks</td><td>11552</td><td>11591</td><td>40</td><td>CCTCGGCCGGCGCCTAGCAGCCGACTTAGAACTGGTGCGG</td><td>973.4</td><td>-5.0</td><td>0.9</td><td>-5.5</td><td>3.3E-08</td><td>6.3E-08</td><td>MB231/MB468</td><td>NUC</td></tr>
<tr><td>>hg38_21_+_8433172_8446622_RNA45SN1_45S_50ntFlanks@11739.11768.30|TCTCTTAAGGTAGCCAAATGCCTCGTCATC</td><td>>hg38_21_+_8433172_8446622_RNA45SN1_45S_50ntFlanks</td><td>11739</td><td>11768</td><td>30</td><td>TCTCTTAAGGTAGCCAAATGCCTCGTCATC</td><td>99.7</td><td>-9.7</td><td>1.7</td><td>-5.8</td><td>5.3E-09</td><td>6.3E-08</td><td>BT20/MB468</td><td>MP</td></tr>
<tr><td>>hg38_21_+_8433172_8446622_RNA45SN1_45S_50ntFlanks@8380.8404.25|CAAGAGGGCGTGAAACCGTTAAGAG</td><td>>hg38_21_+_8433172_8446622_RNA45SN1_45S_50ntFlanks</td><td>8380</td><td>8404</td><td>25</td><td>CAAGAGGGCGTGAAACCGTTAAGAG</td><td>688.5</td><td>-4.6</td><td>0.8</td><td>-5.6</td><td>2.5E-08</td><td>6.3E-08</td><td>BT20/MB468</td><td>MITO</td></tr>
<tr><td>>hg38_21_+_8433172_8446622_RNA45SN1_45S_50ntFlanks@12288.12319.32|ACCGTGAAAGCGGGGCCTCACGATCCTTCTGA</td><td>>hg38_21_+_8433172_8446622_RNA45SN1_45S_50ntFlanks</td><td>12288</td><td>12319</td><td>32</td><td>ACCGTGAAAGCGGGGCCTCACGATCCTTCTGA</td><td>57.5</td><td>-9.7</td><td>1.8</td><td>-5.5</td><td>3.0E-08</td><td>6.3E-08</td><td>MB231/MB468</td><td>MP</td></tr>
<tr><td>>hg38_21_+_8433172_8446622_RNA45SN1_45S_50ntFlanks@11203.11225.23|CGGCGGCGGCGACTCTGGACGCG</td><td>>hg38_21_+_8433172_8446622_RNA45SN1_45S_50ntFlanks</td><td>11203</td><td>11225</td><td>23</td><td>CGGCGGCGGCGACTCTGGACGCG</td><td>289.6</td><td>-2.7</td><td>0.5</td><td>-5.6</td><td>2.8E-08</td><td>6.3E-08</td><td>BT20/MB468</td><td>CYTO</td></tr>
<tr><td>>hg38_21_+_8433172_8446622_RNA45SN1_45S_50ntFlanks@12822.12847.26|CCGGGACCGGGGTCCGGTGCGGAGTG</td><td>>hg38_21_+_8433172_8446622_RNA45SN1_45S_50ntFlanks</td><td>12822</td><td>12847</td><td>26</td><td>CCGGGACCGGGGTCCGGTGCGGAGTG</td><td>296.7</td><td>-11.3</td><td>1.9</td><td>-5.8</td><td>5.3E-09</td><td>6.3E-08</td><td>BT20/MB468</td><td>MP</td></tr>
<tr><td>>hg38_21_+_8433172_8446622_RNA45SN1_45S_50ntFlanks@4972.4994.23|CGGCCCGGACACGGACAGGATTG</td><td>>hg38_21_+_8433172_8446622_RNA45SN1_45S_50ntFlanks</td><td>4972</td><td>4994</td><td>23</td><td>CGGCCCGGACACGGACAGGATTG</td><td>272.5</td><td>-7.4</td><td>1.3</td><td>-5.5</td><td>3.0E-08</td><td>6.4E-08</td><td>MB231/MB468</td><td>MP</td></tr>
<tr><td>>hg38_21_+_8433172_8446622_RNA45SN1_45S_50ntFlanks@8176.8203.28|CTTCTGATCGAGGCCCAGCCCGTGGACG</td><td>>hg38_21_+_8433172_8446622_RNA45SN1_45S_50ntFlanks</td><td>8176</td><td>8203</td><td>28</td><td>CTTCTGATCGAGGCCCAGCCCGTGGACG</td><td>7935.3</td><td>-5.7</td><td>1.0</td><td>-5.8</td><td>5.3E-09</td><td>6.4E-08</td><td>BT20/MB468</td><td>MP</td></tr>
<tr><td>>hg38_21_+_8433172_8446622_RNA45SN1_45S_50ntFlanks@8850.8867.18|GCGGCGGCGGCGGCGGGA</td><td>>hg38_21_+_8433172_8446622_RNA45SN1_45S_50ntFlanks</td><td>8850</td><td>8867</td><td>18</td><td>GCGGCGGCGGCGGCGGGA</td><td>173.1</td><td>-10.5</td><td>1.8</td><td>-5.8</td><td>5.3E-09</td><td>6.4E-08</td><td>BT20/MB468</td><td>MP</td></tr>
<tr><td>>hg38_21_+_8433172_8446622_RNA45SN1_45S_50ntFlanks@4727.4765.39|AATCAAGAACGAAAGTCGGAGGTTCGAAGACGATCAGAT</td><td>>hg38_21_+_8433172_8446622_RNA45SN1_45S_50ntFlanks</td><td>4727</td><td>4765</td><td>39</td><td>AATCAAGAACGAAAGTCGGAGGTTCGAAGACGATCAGAT</td><td>371.2</td><td>3.5</td><td>0.6</td><td>5.5</td><td>3.3E-08</td><td>6.4E-08</td><td>MB231/MB468</td><td>NUC</td></tr>
<tr><td>>hg38_21_+_8433172_8446622_RNA45SN1_45S_50ntFlanks@4975.4999.25|CCCGGACACGGACAGGATTGACAGA</td><td>>hg38_21_+_8433172_8446622_RNA45SN1_45S_50ntFlanks</td><td>4975</td><td>4999</td><td>25</td><td>CCCGGACACGGACAGGATTGACAGA</td><td>881.0</td><td>-4.7</td><td>0.8</td><td>-5.8</td><td>5.3E-09</td><td>6.4E-08</td><td>BT20/MB468</td><td>MP</td></tr>
<tr><td>>hg38_21_+_8433172_8446622_RNA45SN1_45S_50ntFlanks@6722.6747.26|ATTGCAGGACACATTGATCATCGACA</td><td>>hg38_21_+_8433172_8446622_RNA45SN1_45S_50ntFlanks</td><td>6722</td><td>6747</td><td>26</td><td>ATTGCAGGACACATTGATCATCGACA</td><td>142.0</td><td>-5.5</td><td>1.0</td><td>-5.5</td><td>3.1E-08</td><td>6.4E-08</td><td>MB231/MB468</td><td>MP</td></tr>
<tr><td>>MI0000542|hsa-mir-320a&WithFlank&8|-|22244960|22245043@48.71.24(+1G)&[MIMAT0000510&hsa-miR-320a-3p&offsets|0|+2(+1G);m-60&8|-|22244975|22244996&offsets|0|+2(+1G)]|AAAAGCTGGGTTGAGAGGGCGAAAG</td><td>>MI0000542|hsa-mir-320a&WithFlank&8|-|22244960|22245043</td><td>48</td><td>71</td><td>24(+1G)</td><td>AAAAGCTGGGTTGAGAGGGCGAAAG</td><td>94.7</td><td>10.3</td><td>1.8</td><td>5.8</td><td>5.4E-09</td><td>6.4E-08</td><td>BT20/MB468</td><td>MP</td></tr>
<tr><td>>hg38_21_+_8433172_8446622_RNA45SN1_45S_50ntFlanks@5084.5112.29|CGAACGAGACTCTGGCATGCTAACTAGTT</td><td>>hg38_21_+_8433172_8446622_RNA45SN1_45S_50ntFlanks</td><td>5084</td><td>5112</td><td>29</td><td>CGAACGAGACTCTGGCATGCTAACTAGTT</td><td>1441.5</td><td>-8.7</td><td>1.6</td><td>-5.5</td><td>3.6E-08</td><td>6.4E-08</td><td>MB231/MB468</td><td>MITO</td></tr>
<tr><td>>hg38_21_+_8433172_8446622_RNA45SN1_45S_50ntFlanks@4918.4941.24|ACCACCAGGAGTGGAGCCTGCGGC</td><td>>hg38_21_+_8433172_8446622_RNA45SN1_45S_50ntFlanks</td><td>4918</td><td>4941</td><td>24</td><td>ACCACCAGGAGTGGAGCCTGCGGC</td><td>231.0</td><td>-9.9</td><td>1.8</td><td>-5.6</td><td>2.5E-08</td><td>6.4E-08</td><td>BT20/MB468</td><td>MITO</td></tr>
<tr><td>>hg38_21_+_8433172_8446622_RNA45SN1_45S_50ntFlanks@11667.11692.26|CTGCCCAGTGCTCTGAATGTCAAAGT</td><td>>hg38_21_+_8433172_8446622_RNA45SN1_45S_50ntFlanks</td><td>11667</td><td>11692</td><td>26</td><td>CTGCCCAGTGCTCTGAATGTCAAAGT</td><td>285.5</td><td>4.5</td><td>0.8</td><td>5.5</td><td>3.3E-08</td><td>6.4E-08</td><td>MB231/MB468</td><td>NUC</td></tr>
<tr><td>>hg38_21_+_8433172_8446622_RNA45SN1_45S_50ntFlanks@11778.11806.29|ACGCGCATGAATGGATGAACGAGATTCCC</td><td>>hg38_21_+_8433172_8446622_RNA45SN1_45S_50ntFlanks</td><td>11778</td><td>11806</td><td>29</td><td>ACGCGCATGAATGGATGAACGAGATTCCC</td><td>218.2</td><td>-3.4</td><td>0.6</td><td>-5.5</td><td>3.3E-08</td><td>6.4E-08</td><td>MB231/MB468</td><td>NUC</td></tr>
<tr><td>>MI0000809|hsa-mir-151a&WithFlank&8|-|140732558|140732659@53.74.22(+1U)&[MIMAT0000757&hsa-miR-151a-3p&offsets|0|+1(+1U);m-30&8|-|140732587|140732607&offsets|0|+1(+1U)]|CTAGACTGAAGCTCCTTGAGGAT</td><td>>MI0000809|hsa-mir-151a&WithFlank&8|-|140732558|140732659</td><td>53</td><td>74</td><td>22(+1U)</td><td>CTAGACTGAAGCTCCTTGAGGAT</td><td>1037.7</td><td>4.9</td><td>0.8</td><td>5.8</td><td>5.4E-09</td><td>6.4E-08</td><td>BT20/MB468</td><td>MP</td></tr>
<tr><td>>hg38_21_+_8433172_8446622_RNA45SN1_45S_50ntFlanks@11641.11681.41|CGGCGGGTGTTGACGCGATGTGATTTCTGCCCAGTGCTCTG</td><td>>hg38_21_+_8433172_8446622_RNA45SN1_45S_50ntFlanks</td><td>11641</td><td>11681</td><td>41</td><td>CGGCGGGTGTTGACGCGATGTGATTTCTGCCCAGTGCTCTG</td><td>837.4</td><td>-4.6</td><td>0.8</td><td>-5.6</td><td>2.5E-08</td><td>6.4E-08</td><td>BT20/MB468</td><td>MITO</td></tr>
<tr><td>>MI0000446|hsa-mir-125b-1&WithFlank&11|-|122099751|122099850@61.81.21(+1A)&[MIMAT0004592&hsa-miR-125b-1-3p&offsets|0|-1(+1A);m-485&11|-|122099769|122099790&offsets|0|-1(+1A)]|ACGGGTTAGGCTCTTGGGAGCA</td><td>>MI0000446|hsa-mir-125b-1&WithFlank&11|-|122099751|122099850</td><td>61</td><td>81</td><td>21(+1A)</td><td>ACGGGTTAGGCTCTTGGGAGCA</td><td>226.1</td><td>-11.2</td><td>1.8</td><td>-6.1</td><td>1.0E-09</td><td>6.4E-08</td><td>BT20/MB231</td><td>MP</td></tr>
<tr><td>>hg38_21_+_8433172_8446622_RNA45SN1_45S_50ntFlanks@11573.11593.21|CGACTTAGAACTGGTGCGGAC</td><td>>hg38_21_+_8433172_8446622_RNA45SN1_45S_50ntFlanks</td><td>11573</td><td>11593</td><td>21</td><td>CGACTTAGAACTGGTGCGGAC</td><td>430.0</td><td>-3.1</td><td>0.6</td><td>-5.5</td><td>3.4E-08</td><td>6.4E-08</td><td>MB231/MB468</td><td>NUC</td></tr>
<tr><td>>hg38_21_+_8433172_8446622_RNA45SN1_45S_50ntFlanks@8116.8135.20|GCGGGACATGTGGCGTACGG</td><td>>hg38_21_+_8433172_8446622_RNA45SN1_45S_50ntFlanks</td><td>8116</td><td>8135</td><td>20</td><td>GCGGGACATGTGGCGTACGG</td><td>791.4</td><td>13.0</td><td>2.1</td><td>6.1</td><td>1.0E-09</td><td>6.4E-08</td><td>BT20/MB231</td><td>MP</td></tr>
<tr><td>>MI0000065|hsa-let-7d&WithFlank&9|+|94178828|94178926@68.88.21&[MIMAT0004484&hsa-let-7d-3p&offsets|0|-1;m-98&9|+|94178895|94178916&offsets|0|-1]|CTATACGACCTGCTGCCTTTC</td><td>>MI0000065|hsa-let-7d&WithFlank&9|+|94178828|94178926</td><td>68</td><td>88</td><td>21</td><td>CTATACGACCTGCTGCCTTTC</td><td>689.1</td><td>4.5</td><td>0.8</td><td>5.5</td><td>3.4E-08</td><td>6.4E-08</td><td>MB231/MB468</td><td>NUC</td></tr>
</table>

<table>
<tr><th></th><th></th><th></th><th></th><th></th><th></th><th></th><th></th><th></th><th></th><th></th><th></th><th></th><th></th></tr>
<tr><td>>hg38_21_+_8433172_8446622_RNA45SN1_45S_50ntFlanks@12241.12273.33|CAGAAGGGCAAAAGCTCGCTTGATCTTGATTTT</td><td>>hg38_21_+_8433172_8446622_RNA45SN1_45S_50ntFlanks</td><td>12241</td><td>12273</td><td>33</td><td>CAGAAGGGCAAAAGCTCGCTTGATCTTGATTTT</td><td>347.6</td><td>-6.1</td><td>1.1</td><td>-5.5</td><td>3.7E-08</td><td>6.4E-08</td><td>BT20/MB468</td><td>NUC</td></tr>
<tr><td>>hg38_21_+_8433172_8446622_RNA45SN1_45S_50ntFlanks@3843.3871.29|CTACTTGGATAACTGTGGTAATTCTAGAG</td><td>>hg38_21_+_8433172_8446622_RNA45SN1_45S_50ntFlanks</td><td>3843</td><td>3871</td><td>29</td><td>CTACTTGGATAACTGTGGTAATTCTAGAG</td><td>398.0</td><td>-6.3</td><td>1.1</td><td>-5.5</td><td>3.7E-08</td><td>6.4E-08</td><td>BT20/MB468</td><td>NUC</td></tr>
<tr><td>>hg38_21_+_8433172_8446622_RNA45SN1_45S_50ntFlanks@3843.3860.18|CTACTTGGATAACTGTGG</td><td>>hg38_21_+_8433172_8446622_RNA45SN1_45S_50ntFlanks</td><td>3843</td><td>3860</td><td>18</td><td>CTACTTGGATAACTGTGG</td><td>306.4</td><td>-7.9</td><td>1.4</td><td>-5.5</td><td>3.7E-08</td><td>6.4E-08</td><td>BT20/MB468</td><td>NUC</td></tr>
<tr><td>>am_MI0000809|hsa-mir-151a&WithFlank&8|-|140732558|140732659@53.70.18&[MIMAT0000757&hsa-miR-151a-3p&offsets|0|-3;m-30&8|-|140732587|140732607&offsets|0|-3]|CTAGACTGAAGCTCCTTG</td><td>>am_MI0000809|hsa-mir-151a&WithFlank&8|-|140732558|140732659</td><td>53</td><td>70</td><td>18</td><td>CTAGACTGAAGCTCCTTG</td><td>970.0</td><td>4.2</td><td>0.8</td><td>5.5</td><td>3.1E-08</td><td>6.4E-08</td><td>MB231/MB468</td><td>MP</td></tr>
<tr><td>>hg38_21_+_8433172_8446622_RNA45SN1_45S_50ntFlanks@8154.8173.20|CCGCTCGTGGGGGGCCCAAG</td><td>>hg38_21_+_8433172_8446622_RNA45SN1_45S_50ntFlanks</td><td>8154</td><td>8173</td><td>20</td><td>CCGCTCGTGGGGGGCCCAAG</td><td>147.3</td><td>-9.3</td><td>1.7</td><td>-5.6</td><td>2.5E-08</td><td>6.4E-08</td><td>BT20/MB468</td><td>MITO</td></tr>
<tr><td>>hg38_1_-_228634819_228635039_RNA5S12_5S_50ntFlanks@140.169.30|ACCGCCTGGGAATACCGGGTGCTGTAGGCT</td><td>>hg38_1_-_228634819_228635039_RNA5S12_5S_50ntFlanks</td><td>140</td><td>169</td><td>30</td><td>ACCGCCTGGGAATACCGGGTGCTGTAGGCT</td><td>9910.8</td><td>4.6</td><td>0.8</td><td>5.6</td><td>2.8E-08</td><td>6.4E-08</td><td>BT20/MB468</td><td>CYTO</td></tr>
<tr><td>>hg38_21_+_8433172_8446622_RNA45SN1_45S_50ntFlanks@11592.11612.21|ACCAGGGGAATCCGACTGTTT</td><td>>hg38_21_+_8433172_8446622_RNA45SN1_45S_50ntFlanks</td><td>11592</td><td>11612</td><td>21</td><td>ACCAGGGGAATCCGACTGTTT</td><td>534.3</td><td>-6.1</td><td>1.1</td><td>-5.5</td><td>3.3E-08</td><td>6.4E-08</td><td>MB231/MB468</td><td>CYTO</td></tr>
<tr><td>>hg38_21_+_8433172_8446622_RNA45SN1_45S_50ntFlanks@12502.12520.19|TTAGACCGTCGTGAGACAG</td><td>>hg38_21_+_8433172_8446622_RNA45SN1_45S_50ntFlanks</td><td>12502</td><td>12520</td><td>19</td><td>TTAGACCGTCGTGAGACAG</td><td>76.9</td><td>-10.1</td><td>1.8</td><td>-5.5</td><td>3.1E-08</td><td>6.4E-08</td><td>MB231/MB468</td><td>MP</td></tr>
<tr><td>>hg38_21_+_8433172_8446622_RNA45SN1_45S_50ntFlanks@11924.11948.25|GAGGTGTAGAATAAGTGGGAGGCCC</td><td>>hg38_21_+_8433172_8446622_RNA45SN1_45S_50ntFlanks</td><td>11924</td><td>11948</td><td>25</td><td>GAGGTGTAGAATAAGTGGGAGGCCC</td><td>879.9</td><td>13.1</td><td>2.1</td><td>6.1</td><td>1.1E-09</td><td>6.4E-08</td><td>BT20/MB231</td><td>MP</td></tr>
<tr><td>>hg38_21_+_8433172_8446622_RNA45SN1_45S_50ntFlanks@10664.10711.48|CTTGGAAAGCGTCGCGGTTCCGGCGGCGTCCGGTGAGCTCTCGCTGGC</td><td>>hg38_21_+_8433172_8446622_RNA45SN1_45S_50ntFlanks</td><td>10664</td><td>10711</td><td>48</td><td>CTTGGAAAGCGTCGCGGTTCCGGCGGCGTCCGGTGAGCTCTCGCTGGC</td><td>199.6</td><td>-6.9</td><td>1.2</td><td>-5.5</td><td>3.3E-08</td><td>6.5E-08</td><td>MB231/MB468</td><td>CYTO</td></tr>
<tr><td>>hg38_21_+_8433172_8446622_RNA45SN1_45S_50ntFlanks@4128.4145.18|CGATTCCGGAGAGGGAGC</td><td>>hg38_21_+_8433172_8446622_RNA45SN1_45S_50ntFlanks</td><td>4128</td><td>4145</td><td>18</td><td>CGATTCCGGAGAGGGAGC</td><td>116.6</td><td>-9.9</td><td>1.7</td><td>-5.8</td><td>5.4E-09</td><td>6.5E-08</td><td>BT20/MB468</td><td>MP</td></tr>
<tr><td>>hg38_21_+_8433172_8446622_RNA45SN1_45S_50ntFlanks@12416.12440.25|TTCGATGTCGGCTCTTCCTATCATT</td><td>>hg38_21_+_8433172_8446622_RNA45SN1_45S_50ntFlanks</td><td>12416</td><td>12440</td><td>25</td><td>TTCGATGTCGGCTCTTCCTATCATT</td><td>855.3</td><td>4.2</td><td>0.8</td><td>5.5</td><td>3.4E-08</td><td>6.5E-08</td><td>MB231/MB468</td><td>NUC</td></tr>
<tr><td>>hg38_21_+_8433172_8446622_RNA45SN1_45S_50ntFlanks@10680.10710.31|GTTCCGGCGGCGTCCGGTGAGCTCTCGCTGG</td><td>>hg38_21_+_8433172_8446622_RNA45SN1_45S_50ntFlanks</td><td>10680</td><td>10710</td><td>31</td><td>GTTCCGGCGGCGTCCGGTGAGCTCTCGCTGG</td><td>33559.8</td><td>-9.3</td><td>1.7</td><td>-5.5</td><td>3.7E-08</td><td>6.5E-08</td><td>BT20/MB468</td><td>NUC</td></tr>
<tr><td>>hg38_MT_+_598_1651_RNR1_12S_50ntFlanks@52.77.26|ATAGGTTTGGTCCTAGCCTTTCTATT</td><td>>hg38_MT_+_598_1651_RNR1_12S_50ntFlanks</td><td>52</td><td>77</td><td>26</td><td>ATAGGTTTGGTCCTAGCCTTTCTATT</td><td>12264.8</td><td>6.4</td><td>1.1</td><td>5.6</td><td>2.6E-08</td><td>6.5E-08</td><td>BT20/MB468</td><td>MITO</td></tr>
<tr><td>>hg38_21_+_8433172_8446622_RNA45SN1_45S_50ntFlanks@12607.12626.20|TGTGCTTGGCTGAGGAGCCA</td><td>>hg38_21_+_8433172_8446622_RNA45SN1_45S_50ntFlanks</td><td>12607</td><td>12626</td><td>20</td><td>TGTGCTTGGCTGAGGAGCCA</td><td>331.7</td><td>-8.0</td><td>1.5</td><td>-5.5</td><td>3.8E-08</td><td>6.5E-08</td><td>BT20/MB468</td><td>NUC</td></tr>
<tr><td>>hg38_1_-_228634819_228635039_RNA5S12_5S_50ntFlanks@137.169.33|GAGACCGCCTGGGAATACCGGGTGCTGTAGGCT</td><td>>hg38_1_-_228634819_228635039_RNA5S12_5S_50ntFlanks</td><td>137</td><td>169</td><td>33</td><td>GAGACCGCCTGGGAATACCGGGTGCTGTAGGCT</td><td>585.6</td><td>8.0</td><td>1.4</td><td>5.6</td><td>2.8E-08</td><td>6.5E-08</td><td>BT20/MB468</td><td>CYTO</td></tr>
<tr><td>>hg38_21_+_8433172_8446622_RNA45SN1_45S_50ntFlanks@10781.10806.26|AGGTGAACAGCCTCTGGCATGTTGGA</td><td>>hg38_21_+_8433172_8446622_RNA45SN1_45S_50ntFlanks</td><td>10781</td><td>10806</td><td>26</td><td>AGGTGAACAGCCTCTGGCATGTTGGA</td><td>333.1</td><td>-7.1</td><td>1.3</td><td>-5.6</td><td>2.8E-08</td><td>6.5E-08</td><td>BT20/MB468</td><td>CYTO</td></tr>
<tr><td>>hg38_21_+_8433172_8446622_RNA45SN1_45S_50ntFlanks@485.522.38|GATGTGGTGACGTCGTGCTCTCCCGGGCCGGGTCCGAG</td><td>>hg38_21_+_8433172_8446622_RNA45SN1_45S_50ntFlanks</td><td>485</td><td>522</td><td>38</td><td>GATGTGGTGACGTCGTGCTCTCCCGGGCCGGGTCCGAG</td><td>138.4</td><td>10.9</td><td>1.9</td><td>5.8</td><td>5.5E-09</td><td>6.5E-08</td><td>BT20/MB468</td><td>MP</td></tr>
<tr><td>>hg38_21_+_8433172_8446622_RNA45SN1_45S_50ntFlanks@11660.11685.26|GTGATTTCTGCCCAGTGCTCTGAATG</td><td>>hg38_21_+_8433172_8446622_RNA45SN1_45S_50ntFlanks</td><td>11660</td><td>11685</td><td>26</td><td>GTGATTTCTGCCCAGTGCTCTGAATG</td><td>442.5</td><td>-3.3</td><td>0.6</td><td>-5.5</td><td>3.3E-08</td><td>6.5E-08</td><td>MB231/MB468</td><td>CYTO</td></tr>
<tr><td>>hg38_21_+_8433172_8446622_RNA45SN1_45S_50ntFlanks@9566.9596.31|GACGTGCAAATCGGTCGTCCGACCTGGGTAT</td><td>>hg38_21_+_8433172_8446622_RNA45SN1_45S_50ntFlanks</td><td>9566</td><td>9596</td><td>31</td><td>GACGTGCAAATCGGTCGTCCGACCTGGGTAT</td><td>1520.6</td><td>-8.3</td><td>1.5</td><td>-5.5</td><td>3.1E-08</td><td>6.5E-08</td><td>MB231/MB468</td><td>MP</td></tr>
<tr><td>>hg38_21_+_8433172_8446622_RNA45SN1_45S_50ntFlanks@10826.10848.23|TCGGCAAGCCGGATCCGTAACTT</td><td>>hg38_21_+_8433172_8446622_RNA45SN1_45S_50ntFlanks</td><td>10826</td><td>10848</td><td>23</td><td>TCGGCAAGCCGGATCCGTAACTT</td><td>274.0</td><td>-5.8</td><td>1.1</td><td>-5.5</td><td>3.3E-08</td><td>6.5E-08</td><td>MB231/MB468</td><td>CYTO</td></tr>
<tr><td>>hg38_21_+_8433172_8446622_RNA45SN1_45S_50ntFlanks@11575.11606.32|ACTTAGAACTGGTGCGGACCAGGGGAATCCGA</td><td>>hg38_21_+_8433172_8446622_RNA45SN1_45S_50ntFlanks</td><td>11575</td><td>11606</td><td>32</td><td>ACTTAGAACTGGTGCGGACCAGGGGAATCCGA</td><td>365.7</td><td>-4.9</td><td>0.9</td><td>-5.5</td><td>3.3E-08</td><td>6.5E-08</td><td>MB231/MB468</td><td>CYTO</td></tr>
<tr><td>>hg38_21_+_8433172_8446622_RNA45SN1_45S_50ntFlanks@12242.12268.27|AGAAGGGCAAAAGCTCGCTTGATCTTG</td><td>>hg38_21_+_8433172_8446622_RNA45SN1_45S_50ntFlanks</td><td>12242</td><td>12268</td><td>27</td><td>AGAAGGGCAAAAGCTCGCTTGATCTTG</td><td>565.1</td><td>-7.7</td><td>1.4</td><td>-5.5</td><td>3.1E-08</td><td>6.5E-08</td><td>MB231/MB468</td><td>MP</td></tr>
<tr><td>>hg38_21_+_8433172_8446622_RNA45SN1_45S_50ntFlanks@12320.12346.27|CCTTTTGGGTTTTAAGCAGGAGGTGTC</td><td>>hg38_21_+_8433172_8446622_RNA45SN1_45S_50ntFlanks</td><td>12320</td><td>12346</td><td>27</td><td>CCTTTTGGGTTTTAAGCAGGAGGTGTC</td><td>64.9</td><td>-9.9</td><td>1.8</td><td>-5.5</td><td>3.1E-08</td><td>6.5E-08</td><td>MB231/MB468</td><td>MP</td></tr>
<tr><td>>hg38_21_+_8433172_8446622_RNA45SN1_45S_50ntFlanks@12074.12095.22|CGAGGGGCTCTCGCTTCTGGCG</td><td>>hg38_21_+_8433172_8446622_RNA45SN1_45S_50ntFlanks</td><td>12074</td><td>12095</td><td>22</td><td>CGAGGGGCTCTCGCTTCTGGCG</td><td>1218.7</td><td>-8.9</td><td>1.6</td><td>-5.5</td><td>3.8E-08</td><td>6.5E-08</td><td>BT20/MB468</td><td>NUC</td></tr>
<tr><td>>hg38_21_+_8433172_8446622_RNA45SN1_45S_50ntFlanks@9925.9952.28|GGTTGATATAGACAGCAGGACGGTGGCC</td><td>>hg38_21_+_8433172_8446622_RNA45SN1_45S_50ntFlanks</td><td>9925</td><td>9952</td><td>28</td><td>GGTTGATATAGACAGCAGGACGGTGGCC</td><td>94.5</td><td>10.3</td><td>1.8</td><td>5.8</td><td>5.5E-09</td><td>6.5E-08</td><td>BT20/MB468</td><td>MP</td></tr>
<tr><td>>hg38_21_+_8433172_8446622_RNA45SN1_45S_50ntFlanks@10758.10785.28|CCCATATCCGCAGCAGGTCTCCAAGGTG</td><td>>hg38_21_+_8433172_8446622_RNA45SN1_45S_50ntFlanks</td><td>10758</td><td>10785</td><td>28</td><td>CCCATATCCGCAGCAGGTCTCCAAGGTG</td><td>129.1</td><td>-10.1</td><td>1.7</td><td>-5.8</td><td>5.5E-09</td><td>6.5E-08</td><td>BT20/MB468</td><td>MP</td></tr>
<tr><td>>MI0000234|hsa-mir-192&WithFlank&11|-|64891131|64891252@30.51.22(+2U)&[MIMAT0000222&hsa-miR-192-5p&offsets|0|+1(+2U);m-59&11|-|64891203|64891223&offsets|0|+1(+2U)]|CTGACCTATGAATTGACAGCCATT</td><td>>MI0000234|hsa-mir-192&WithFlank&11|-|64891131|64891252</td><td>30</td><td>51</td><td>22(+2U)</td><td>CTGACCTATGAATTGACAGCCATT</td><td>174.7</td><td>-10.8</td><td>1.8</td><td>-6.1</td><td>1.1E-09</td><td>6.5E-08</td><td>BT20/MB231</td><td>MP</td></tr>
<tr><td>>hg38_21_+_8433172_8446622_RNA45SN1_45S_50ntFlanks@5497.5526.30|ACGGTCGAACTTGACTATCTAGAGGAAGTA</td><td>>hg38_21_+_8433172_8446622_RNA45SN1_45S_50ntFlanks</td><td>5497</td><td>5526</td><td>30</td><td>ACGGTCGAACTTGACTATCTAGAGGAAGTA</td><td>458.8</td><td>-6.0</td><td>1.1</td><td>-5.5</td><td>3.7E-08</td><td>6.5E-08</td><td>MB231/MB468</td><td>MITO</td></tr>
<tr><td>>hg38_21_+_8433172_8446622_RNA45SN1_45S_50ntFlanks@11570.11612.43|AGCCGACTTAGAACTGGTGCGGACCAGGGGAATCCGACTGTTT</td><td>>hg38_21_+_8433172_8446622_RNA45SN1_45S_50ntFlanks</td><td>11570</td><td>11612</td><td>43</td><td>AGCCGACTTAGAACTGGTGCGGACCAGGGGAATCCGACTGTTT</td><td>1084.2</td><td>-5.4</td><td>1.0</td><td>-5.5</td><td>3.8E-08</td><td>6.5E-08</td><td>BT20/MB468</td><td>NUC</td></tr>
<tr><td>>hg38_21_+_8433172_8446622_RNA45SN1_45S_50ntFlanks@11572.11599.28|CCGACTTAGAACTGGTGCGGACCAGGGG</td><td>>hg38_21_+_8433172_8446622_RNA45SN1_45S_50ntFlanks</td><td>11572</td><td>11599</td><td>28</td><td>CCGACTTAGAACTGGTGCGGACCAGGGG</td><td>786.6</td><td>-3.7</td><td>0.7</td><td>-5.5</td><td>3.4E-08</td><td>6.5E-08</td><td>MB231/MB468</td><td>CYTO</td></tr>
<tr><td>>MI0003560|hsa-mir-92b&WithFlank&1|+|155195171|155195278@67.87.21&[MIMAT0003218&hsa-miR-92b-3p&offsets|0|-1;m-115&1|+|155195237|155195258&offsets|0|-1]|TATTGCACTCGTCCCGGCCTC</td><td>>MI0003560|hsa-mir-92b&WithFlank&1|+|155195171|155195278</td><td>67</td><td>87</td><td>21</td><td>TATTGCACTCGTCCCGGCCTC</td><td>1874.9</td><td>7.6</td><td>1.3</td><td>5.8</td><td>5.5E-09</td><td>6.6E-08</td><td>BT20/MB468</td><td>MP</td></tr>
<tr><td>>hg38_21_+_8433172_8446622_RNA45SN1_45S_50ntFlanks@3913.3938.26|ATGCGTGCATTTATCAGATCAAAACC</td><td>>hg38_21_+_8433172_8446622_RNA45SN1_45S_50ntFlanks</td><td>3913</td><td>3938</td><td>26</td><td>ATGCGTGCATTTATCAGATCAAAACC</td><td>129.1</td><td>-10.1</td><td>1.7</td><td>-5.8</td><td>5.5E-09</td><td>6.6E-08</td><td>BT20/MB468</td><td>MP</td></tr>
<tr><td>>hg38_21_+_8433172_8446622_RNA45SN1_45S_50ntFlanks@11564.11585.22|CCTAGCAGCCGACTTAGAACTG</td><td>>hg38_21_+_8433172_8446622_RNA45SN1_45S_50ntFlanks</td><td>11564</td><td>11585</td><td>22</td><td>CCTAGCAGCCGACTTAGAACTG</td><td>1636.5</td><td>-3.5</td><td>0.6</td><td>-5.6</td><td>2.6E-08</td><td>6.6E-08</td><td>BT20/MB468</td><td>MITO</td></tr>
<tr><td>>MI0003820|hsa-mir-454&WithFlank&17|-|59137752|59137878@70.92.23(+1C)&[MIMAT0003885&hsa-miR-454-3p&offsets|0|0(+1C);m-272&17|-|59137787|59137809&offsets|0|0(+1C)]|TAGTGCAATATTGCTTATAGGGTC</td><td>>MI0003820|hsa-mir-454&WithFlank&17|-|59137752|59137878</td><td>70</td><td>92</td><td>23(+1C)</td><td>TAGTGCAATATTGCTTATAGGGTC</td><td>201.3</td><td>-11.1</td><td>1.8</td><td>-6.1</td><td>1.1E-09</td><td>6.6E-08</td><td>BT20/MB231</td><td>MP</td></tr>
<tr><td>>hg38_21_+_8433172_8446622_RNA45SN1_45S_50ntFlanks@10781.10817.37|AGGTGAACAGCCTCTGGCATGTTGGAACAATGTAGGT</td><td>>hg38_21_+_8433172_8446622_RNA45SN1_45S_50ntFlanks</td><td>10781</td><td>10817</td><td>37</td><td>AGGTGAACAGCCTCTGGCATGTTGGAACAATGTAGGT</td><td>316.5</td><td>-3.8</td><td>0.7</td><td>-5.5</td><td>3.4E-08</td><td>6.6E-08</td><td>MB231/MB468</td><td>CYTO</td></tr>
<tr><td>>hg38_21_+_8433172_8446622_RNA45SN1_45S_50ntFlanks@10127.10145.19|GCGGCGGCGGCGGCGGCGG</td><td>>hg38_21_+_8433172_8446622_RNA45SN1_45S_50ntFlanks</td><td>10127</td><td>10145</td><td>19</td><td>GCGGCGGCGGCGGCGGCGG</td><td>100.5</td><td>-9.7</td><td>1.7</td><td>-5.8</td><td>5.6E-09</td><td>6.6E-08</td><td>BT20/MB468</td><td>MP</td></tr>
<tr><td>>hg38_21_+_8433172_8446622_RNA45SN1_45S_50ntFlanks@10130.10148.19|GCGGCGGCGGCGGCGGCGG</td><td>>hg38_21_+_8433172_8446622_RNA45SN1_45S_50ntFlanks</td><td>10130</td><td>10148</td><td>19</td><td>GCGGCGGCGGCGGCGGCGG</td><td>100.5</td><td>-9.7</td><td>1.7</td><td>-5.8</td><td>5.6E-09</td><td>6.6E-08</td><td>BT20/MB468</td><td>MP</td></tr>
<tr><td>>hg38_21_+_8433172_8446622_RNA45SN1_45S_50ntFlanks@8823.8841.19|GCGGCGGCGGCGGCGGCGG</td><td>>hg38_21_+_8433172_8446622_RNA45SN1_45S_50ntFlanks</td><td>8823</td><td>8841</td><td>19</td><td>GCGGCGGCGGCGGCGGCGG</td><td>100.5</td><td>-9.7</td><td>1.7</td><td>-5.8</td><td>5.6E-09</td><td>6.6E-08</td><td>BT20/MB468</td><td>MP</td></tr>
<tr><td>>hg38_21_+_8433172_8446622_RNA45SN1_45S_50ntFlanks@8844.8862.19|GCGGCGGCGGCGGCGGCGG</td><td>>hg38_21_+_8433172_8446622_RNA45SN1_45S_50ntFlanks</td><td>8844</td><td>8862</td><td>19</td><td>GCGGCGGCGGCGGCGGCGG</td><td>100.5</td><td>-9.7</td><td>1.7</td><td>-5.8</td><td>5.6E-09</td><td>6.6E-08</td><td>BT20/MB468</td><td>MP</td></tr>
<tr><td>>hg38_21_+_8433172_8446622_RNA45SN1_45S_50ntFlanks@8847.8865.19|GCGGCGGCGGCGGCGGCGG</td><td>>hg38_21_+_8433172_8446622_RNA45SN1_45S_50ntFlanks</td><td>8847</td><td>8865</td><td>19</td><td>GCGGCGGCGGCGGCGGCGG</td><td>100.5</td><td>-9.7</td><td>1.7</td><td>-5.8</td><td>5.6E-09</td><td>6.6E-08</td><td>BT20/MB468</td><td>MP</td></tr>
<tr><td>>hg38_21_+_8433172_8446622_RNA45SN1_45S_50ntFlanks@9545.9569.25|GTGGAGGTCCGTAGCGGTCCTGACG</td><td>>hg38_21_+_8433172_8446622_RNA45SN1_45S_50ntFlanks</td><td>9545</td><td>9569</td><td>25</td><td>GTGGAGGTCCGTAGCGGTCCTGACG</td><td>777.7</td><td>-6.2</td><td>1.1</td><td>-5.5</td><td>3.2E-08</td><td>6.6E-08</td><td>MB231/MB468</td><td>MP</td></tr>
<tr><td>>hg38_21_+_8433172_8446622_RNA45SN1_45S_50ntFlanks@10361.10390.30|GAAGGCCGAAGTGGAGAAGGGTTCCATGTG</td><td>>hg38_21_+_8433172_8446622_RNA45SN1_45S_50ntFlanks</td><td>10361</td><td>10390</td><td>30</td><td>GAAGGCCGAAGTGGAGAAGGGTTCCATGTG</td><td>425.4</td><td>-7.4</td><td>1.3</td><td>-5.5</td><td>3.5E-08</td><td>6.6E-08</td><td>MB231/MB468</td><td>NUC</td></tr>
<tr><td>>MI0000471|hsa-mir-126&WithFlank&9|+|136670596|136670692@59.78.20&[MIMAT0000445&hsa-miR-126-3p&offsets|+1|-1;m-40&9|+|136670653|136670674&offsets|+1|-1]|CGTACCGTGAGTAATAATGC</td><td>>MI0000471|hsa-mir-126&WithFlank&9|+|136670596|136670692</td><td>59</td><td>78</td><td>20</td><td>CGTACCGTGAGTAATAATGC</td><td>685.7</td><td>9.2</td><td>1.6</td><td>5.8</td><td>5.6E-09</td><td>6.6E-08</td><td>BT20/MB468</td><td>MP</td></tr>
<tr><td>>hg38_21_+_8433172_8446622_RNA45SN1_45S_50ntFlanks@4283.4302.20|CTTTAACGAGGATCCATTGG</td><td>>hg38_21_+_8433172_8446622_RNA45SN1_45S_50ntFlanks</td><td>4283</td><td>4302</td><td>20</td><td>CTTTAACGAGGATCCATTGG</td><td>89.2</td><td>-9.6</td><td>1.6</td><td>-5.8</td><td>5.6E-09</td><td>6.6E-08</td><td>BT20/MB468</td><td>MP</td></tr>
<tr><td>>hg38_21_+_8433172_8446622_RNA45SN1_45S_50ntFlanks@11902.11931.30|CTGGCACGGTGAAGAGACATGAGAGGTGTA</td><td>>hg38_21_+_8433172_8446622_RNA45SN1_45S_50ntFlanks</td><td>11902</td><td>11931</td><td>30</td><td>CTGGCACGGTGAAGAGACATGAGAGGTGTA</td><td>119.3</td><td>10.6</td><td>1.8</td><td>5.8</td><td>5.6E-09</td><td>6.6E-08</td><td>BT20/MB468</td><td>MP</td></tr>
<tr><td>>hg38_21_+_8433172_8446622_RNA45SN1_45S_50ntFlanks@9481.9513.33|CACGTGTTAGGACCCGAAAGATGGTGAACTATG</td><td>>hg38_21_+_8433172_8446622_RNA45SN1_45S_50ntFlanks</td><td>9481</td><td>9513</td><td>33</td><td>CACGTGTTAGGACCCGAAAGATGGTGAACTATG</td><td>95.8</td><td>10.3</td><td>1.8</td><td>5.8</td><td>5.7E-09</td><td>6.6E-08</td><td>BT20/MB468</td><td>MP</td></tr>
<tr><td>>hg38_21_+_8433172_8446622_RNA45SN1_45S_50ntFlanks@10912.10939.28|CGCGCCGCGGCTGGACGAGGCGCCGCCG</td><td>>hg38_21_+_8433172_8446622_RNA45SN1_45S_50ntFlanks</td><td>10912</td><td>10939</td><td>28</td><td>CGCGCCGCGGCTGGACGAGGCGCCGCCG</td><td>126.3</td><td>-10.1</td><td>1.7</td><td>-5.8</td><td>5.7E-09</td><td>6.6E-08</td><td>BT20/MB468</td><td>MP</td></tr>
<tr><td>>hg38_21_+_8433172_8446622_RNA45SN1_45S_50ntFlanks@4783.4803.21|CATAAACGATGCCGACCGGCG</td><td>>hg38_21_+_8433172_8446622_RNA45SN1_45S_50ntFlanks</td><td>4783</td><td>4803</td><td>21</td><td>CATAAACGATGCCGACCGGCG</td><td>101.0</td><td>-9.7</td><td>1.7</td><td>-5.8</td><td>5.7E-09</td><td>6.6E-08</td><td>BT20/MB468</td><td>MP</td></tr>
<tr><td>>hg38_21_+_8433172_8446622_RNA45SN1_45S_50ntFlanks@9534.9562.29|AGGAAACTCTGGTGGAGGTCCGTAGCGGT</td><td>>hg38_21_+_8433172_8446622_RNA45SN1_45S_50ntFlanks</td><td>9534</td><td>9562</td><td>29</td><td>AGGAAACTCTGGTGGAGGTCCGTAGCGGT</td><td>7489.5</td><td>-10.2</td><td>1.9</td><td>-5.5</td><td>3.9E-08</td><td>6.6E-08</td><td>BT20/MB468</td><td>NUC</td></tr>
<tr><td>>hg38_21_+_8433172_8446622_RNA45SN1_45S_50ntFlanks@11843.11860.18|AGGGAACGGGCTTGGCGG</td><td>>hg38_21_+_8433172_8446622_RNA45SN1_45S_50ntFlanks</td><td>11843</td><td>11860</td><td>18</td><td>AGGGAACGGGCTTGGCGG</td><td>149.1</td><td>-4.9</td><td>0.9</td><td>-5.5</td><td>3.4E-08</td><td>6.6E-08</td><td>MB231/MB468</td><td>CYTO</td></tr>
<tr><td>>hg38_21_+_8433172_8446622_RNA45SN1_45S_50ntFlanks@11580.11602.23|GAACTGGTGCGGACCAGGGGAAT</td><td>>hg38_21_+_8433172_8446622_RNA45SN1_45S_50ntFlanks</td><td>11580</td><td>11602</td><td>23</td><td>GAACTGGTGCGGACCAGGGGAAT</td><td>839.4</td><td>-5.7</td><td>1.0</td><td>-5.8</td><td>5.7E-09</td><td>6.7E-08</td><td>BT20/MB468</td><td>MP</td></tr>
</table>

<table>
<tr><th></th><th></th><th></th><th></th><th></th><th></th><th></th><th></th><th></th><th></th><th></th><th></th><th></th><th></th></tr>
<tr><td>>hg38_21_+_8433172_8446622_RNA45SN1_45S_50ntFlanks@4046.4081.36|CATTCGAACGTCTGCCCTATCAACTTTCGATGGTAG</td><td>>hg38_21_+_8433172_8446622_RNA45SN1_45S_50ntFlanks</td><td>4046</td><td>4081</td><td>36</td><td>CATTCGAACGTCTGCCCTATCAACTTTCGATGGTAG</td><td>65.2</td><td>-9.9</td><td>1.8</td><td>-5.5</td><td>3.2E-08</td><td>6.7E-08</td><td>MB231/MB468</td><td>MP</td></tr>
<tr><td>>hg38_21_+_8433172_8446622_RNA45SN1_45S_50ntFlanks@6741.6770.30|ATCGACACTTCGAACGCACTTGCGGCCCCG</td><td>>hg38_21_+_8433172_8446622_RNA45SN1_45S_50ntFlanks</td><td>6741</td><td>6770</td><td>30</td><td>ATCGACACTTCGAACGCACTTGCGGCCCCG</td><td>2204.6</td><td>-9.5</td><td>1.7</td><td>-5.5</td><td>3.9E-08</td><td>6.7E-08</td><td>BT20/MB468</td><td>NUC</td></tr>
<tr><td>>hg38_21_+_8433172_8446622_RNA45SN1_45S_50ntFlanks@6700.6717.18|CTAGCTGCGAGAATTAAT</td><td>>hg38_21_+_8433172_8446622_RNA45SN1_45S_50ntFlanks</td><td>6700</td><td>6717</td><td>18</td><td>CTAGCTGCGAGAATTAAT</td><td>423.9</td><td>-7.5</td><td>1.4</td><td>-5.5</td><td>3.7E-08</td><td>6.7E-08</td><td>MB231/MB468</td><td>MITO</td></tr>
<tr><td>>hg38_21_+_8433172_8446622_RNA45SN1_45S_50ntFlanks@9575.9593.19|ATCGGTCGTCCGACCTGGG</td><td>>hg38_21_+_8433172_8446622_RNA45SN1_45S_50ntFlanks</td><td>9575</td><td>9593</td><td>19</td><td>ATCGGTCGTCCGACCTGGG</td><td>125.5</td><td>-10.1</td><td>1.7</td><td>-5.8</td><td>5.7E-09</td><td>6.7E-08</td><td>BT20/MB468</td><td>MP</td></tr>
<tr><td>>trna2_GlyGCC_21_-_18827107_18827177@1.33.33,trna35_GlyGCC_1_+_161413094_161413164@1.33.33,trna37_GlyGCC_1_+_161420467_161420537@1.33.33,trna39_GlyGCC_1_+_161427898_161427968@1.33.33,trna41_GlyGCC_1_+_161435258_161435328@1.33.33|GCATGGGTGGTTCAGTGGTAGAATTCTCGCCTG</td><td>>trna2_GlyGCC_21_-_18827107_18827177</td><td>1</td><td>33</td><td>33</td><td>GCATGGGTGGTTCAGTGGTAGAATTCTCGCCTG</td><td>1413.9</td><td>4.9</td><td>0.8</td><td>6.0</td><td>2.1E-09</td><td>6.7E-08</td><td>BT20/MB231</td><td>CYTO</td></tr>
<tr><td>>trna5_AspGTC_12_+_98897281_98897352@37.73.37|ACGCGGGAGACCGGGGTTCAATTCCCCGACGGGGAGC</td><td>>trna5_AspGTC_12_+_98897281_98897352</td><td>37</td><td>73</td><td>37</td><td>ACGCGGGAGACCGGGGTTCAATTCCCCGACGGGGAGC</td><td>2912.7</td><td>-6.4</td><td>1.1</td><td>-6.0</td><td>2.1E-09</td><td>6.7E-08</td><td>BT20/MB231</td><td>CYTO</td></tr>
<tr><td>>trnaMT_GluTTC_MT_-_14674_14742@26.66.41,trnalookalike8_GluTTC_5_-_93905172_93905240@26.66.41|GGTTTTTCATATCATTGGTCGTGGTTGTAGTCCGTGCGAGA</td><td>>trnaMT_GluTTC_MT_-_14674_14742</td><td>26</td><td>66</td><td>41</td><td>GGTTTTTCATATCATTGGTCGTGGTTGTAGTCCGTGCGAGA</td><td>324.0</td><td>7.6</td><td>1.4</td><td>5.6</td><td>2.6E-08</td><td>6.7E-08</td><td>BT20/MB468</td><td>MITO</td></tr>
<tr><td>>MI0000285|hsa-mir-205&WithFlank&1|+|209432127|209432248@40.61.22(+1A)&[MIMAT0000266&hsa-miR-205-5p&offsets|0|0(+1A);m-351&1|+|209432166|209432187&offsets|0|0(+1A)]|TCCTTCATTCCACCGGAGTCTGA</td><td>>MI0000285|hsa-mir-205&WithFlank&1|+|209432127|209432248</td><td>40</td><td>61</td><td>22(+1A)</td><td>TCCTTCATTCCACCGGAGTCTGA</td><td>1465.7</td><td>-3.9</td><td>0.7</td><td>-5.5</td><td>3.4E-08</td><td>6.7E-08</td><td>MB231/MB468</td><td>CYTO</td></tr>
<tr><td>>hg38_21_+_8433172_8446622_RNA45SN1_45S_50ntFlanks@10404.10431.28|CATGGGTCAGTCGGTCCTGAGAGATGGG</td><td>>hg38_21_+_8433172_8446622_RNA45SN1_45S_50ntFlanks</td><td>10404</td><td>10431</td><td>28</td><td>CATGGGTCAGTCGGTCCTGAGAGATGGG</td><td>822.4</td><td>-8.7</td><td>1.6</td><td>-5.5</td><td>3.9E-08</td><td>6.7E-08</td><td>BT20/MB468</td><td>NUC</td></tr>
<tr><td>>MI0000102|hsa-mir-100&WithFlank&11|-|122152223|122152314@19.37.19(+1C)&[MIMAT0000098&hsa-miR-100-5p&offsets|0|-3(+1C);m-27&11|-|122152276|122152296&offsets|0|-2(+1C)]|AACCCGTAGATCCGAACTTC</td><td>>MI0000102|hsa-mir-100&WithFlank&11|-|122152223|122152314</td><td>19</td><td>37</td><td>19(+1C)</td><td>AACCCGTAGATCCGAACTTC</td><td>125.2</td><td>10.2</td><td>1.8</td><td>5.5</td><td>3.2E-08</td><td>6.7E-08</td><td>MB231/MB468</td><td>MP</td></tr>
<tr><td>>hg38_21_+_8433172_8446622_RNA45SN1_45S_50ntFlanks@9545.9563.19|GTGGAGGTCCGTAGCGGTC</td><td>>hg38_21_+_8433172_8446622_RNA45SN1_45S_50ntFlanks</td><td>9545</td><td>9563</td><td>19</td><td>GTGGAGGTCCGTAGCGGTC</td><td>5114.6</td><td>-4.8</td><td>0.9</td><td>-5.5</td><td>3.4E-08</td><td>6.7E-08</td><td>MB231/MB468</td><td>CYTO</td></tr>
<tr><td>>hg38_21_+_8433172_8446622_RNA45SN1_45S_50ntFlanks@11575.11611.37|ACTTAGAACTGGTGCGGACCAGGGGAATCCGACTGTT</td><td>>hg38_21_+_8433172_8446622_RNA45SN1_45S_50ntFlanks</td><td>11575</td><td>11611</td><td>37</td><td>ACTTAGAACTGGTGCGGACCAGGGGAATCCGACTGTT</td><td>486.4</td><td>-3.3</td><td>0.6</td><td>-5.5</td><td>3.5E-08</td><td>6.7E-08</td><td>MB231/MB468</td><td>NUC</td></tr>
<tr><td>>hg38_1_-_228634819_228635039_RNA5S12_5S_50ntFlanks@141.169.29|CCGCCTGGGAATACCGGGTGCTGTAGGCT</td><td>>hg38_1_-_228634819_228635039_RNA5S12_5S_50ntFlanks</td><td>141</td><td>169</td><td>29</td><td>CCGCCTGGGAATACCGGGTGCTGTAGGCT</td><td>949.1</td><td>3.8</td><td>0.7</td><td>5.5</td><td>3.5E-08</td><td>6.7E-08</td><td>MB231/MB468</td><td>NUC</td></tr>
<tr><td>>hg38_21_+_8433172_8446622_RNA45SN1_45S_50ntFlanks@8380.8403.24|CAAGAGGGCGTGAAACCGTTAAGA</td><td>>hg38_21_+_8433172_8446622_RNA45SN1_45S_50ntFlanks</td><td>8380</td><td>8403</td><td>24</td><td>CAAGAGGGCGTGAAACCGTTAAGA</td><td>650.9</td><td>-8.1</td><td>1.5</td><td>-5.5</td><td>3.9E-08</td><td>6.7E-08</td><td>BT20/MB468</td><td>NUC</td></tr>
<tr><td>>hg38_21_+_8433172_8446622_RNA45SN1_45S_50ntFlanks@12033.12052.20|ATCGTTTTTTCACTGACCCG</td><td>>hg38_21_+_8433172_8446622_RNA45SN1_45S_50ntFlanks</td><td>12033</td><td>12052</td><td>20</td><td>ATCGTTTTTTCACTGACCCG</td><td>97.4</td><td>-9.7</td><td>1.7</td><td>-5.8</td><td>5.8E-09</td><td>6.7E-08</td><td>BT20/MB468</td><td>MP</td></tr>
<tr><td>>MI0003643|hsa-mir-629&WithFlank&15|-|70079366|70079474@28.49.22(+1A)&[MIMAT0004810&hsa-miR-629-5p&offsets|0|+1(+1A);m-150&15|-|70079427|70079447&offsets|0|+1(+1A)]|TGGGTTTACGTTGGGAGAACTTA</td><td>>MI0003643|hsa-mir-629&WithFlank&15|-|70079366|70079474</td><td>28</td><td>49</td><td>22(+1A)</td><td>TGGGTTTACGTTGGGAGAACTTA</td><td>57.3</td><td>9.1</td><td>1.6</td><td>5.5</td><td>3.2E-08</td><td>6.7E-08</td><td>MB231/MB468</td><td>MP</td></tr>
<tr><td>>hg38_21_+_8433172_8446622_RNA45SN1_45S_50ntFlanks@11769.11791.23|TAATTAGTGACGCGCATGAATGG</td><td>>hg38_21_+_8433172_8446622_RNA45SN1_45S_50ntFlanks</td><td>11769</td><td>11791</td><td>23</td><td>TAATTAGTGACGCGCATGAATGG</td><td>1122.3</td><td>-3.8</td><td>0.7</td><td>-5.5</td><td>3.5E-08</td><td>6.7E-08</td><td>MB231/MB468</td><td>CYTO</td></tr>
<tr><td>>hg38_21_+_8433172_8446622_RNA45SN1_45S_50ntFlanks@12002.12030.29|TGCGGGCCGCCGGTGAAATACCACTACTC</td><td>>hg38_21_+_8433172_8446622_RNA45SN1_45S_50ntFlanks</td><td>12002</td><td>12030</td><td>29</td><td>TGCGGGCCGCCGGTGAAATACCACTACTC</td><td>491.5</td><td>-3.7</td><td>0.7</td><td>-5.5</td><td>3.5E-08</td><td>6.7E-08</td><td>MB231/MB468</td><td>CYTO</td></tr>
<tr><td>>hg38_21_+_8433172_8446622_RNA45SN1_45S_50ntFlanks@9533.9557.25|GAGGAAACTCTGGTGGAGGTCCGTA</td><td>>hg38_21_+_8433172_8446622_RNA45SN1_45S_50ntFlanks</td><td>9533</td><td>9557</td><td>25</td><td>GAGGAAACTCTGGTGGAGGTCCGTA</td><td>86.3</td><td>-10.3</td><td>1.9</td><td>-5.5</td><td>3.2E-08</td><td>6.7E-08</td><td>MB231/MB468</td><td>MP</td></tr>
<tr><td>>hg38_MT_+_598_1651_RNR1_12S_50ntFlanks@569.611.43|AAAGGACCTGGCGGTGCTTCATATCCCTCTAGAGGAGCCTGTT</td><td>>hg38_MT_+_598_1651_RNR1_12S_50ntFlanks</td><td>569</td><td>611</td><td>43</td><td>AAAGGACCTGGCGGTGCTTCATATCCCTCTAGAGGAGCCTGTT</td><td>54.8</td><td>9.0</td><td>1.6</td><td>5.5</td><td>3.2E-08</td><td>6.7E-08</td><td>MB231/MB468</td><td>MP</td></tr>
<tr><td>>MI0001445|hsa-mir-423&WithFlank&17|+|30117073|30117178@58.78.21&[MIMAT0001340&hsa-miR-423-3p&offsets|-1|-3;m-100&17|+|30117131|30117153&offsets|-1|-3]|AAGCTCGGTCTGAGGCCCCTC</td><td>>MI0001445|hsa-mir-423&WithFlank&17|+|30117073|30117178</td><td>58</td><td>78</td><td>21</td><td>AAGCTCGGTCTGAGGCCCCTC</td><td>1453.5</td><td>8.4</td><td>1.5</td><td>5.5</td><td>3.0E-08</td><td>6.8E-08</td><td>BT20/MB468</td><td>CYTO</td></tr>
<tr><td>>hg38_21_+_8433172_8446622_RNA45SN1_45S_50ntFlanks@12434.12455.22|TATCATTGTGAAGCAGAATTCA</td><td>>hg38_21_+_8433172_8446622_RNA45SN1_45S_50ntFlanks</td><td>12434</td><td>12455</td><td>22</td><td>TATCATTGTGAAGCAGAATTCA</td><td>375.9</td><td>-7.3</td><td>1.3</td><td>-5.5</td><td>3.8E-08</td><td>6.8E-08</td><td>MB231/MB468</td><td>MITO</td></tr>
<tr><td>>hg38_21_+_8433172_8446622_RNA45SN1_45S_50ntFlanks@12416.12437.22|TTCGATGTCGGCTCTTCCTATC</td><td>>hg38_21_+_8433172_8446622_RNA45SN1_45S_50ntFlanks</td><td>12416</td><td>12437</td><td>22</td><td>TTCGATGTCGGCTCTTCCTATC</td><td>1154.0</td><td>-3.3</td><td>0.6</td><td>-5.5</td><td>3.9E-08</td><td>6.8E-08</td><td>BT20/MB468</td><td>NUC</td></tr>
<tr><td>>hg38_21_+_8433172_8446622_RNA45SN1_45S_50ntFlanks@12936.12954.19|AAACCATTCGTAGACGACC</td><td>>hg38_21_+_8433172_8446622_RNA45SN1_45S_50ntFlanks</td><td>12936</td><td>12954</td><td>19</td><td>AAACCATTCGTAGACGACC</td><td>508.6</td><td>-7.9</td><td>1.4</td><td>-5.5</td><td>3.9E-08</td><td>6.8E-08</td><td>BT20/MB468</td><td>NUC</td></tr>
<tr><td>>hg38_21_+_8433172_8446622_RNA45SN1_45S_50ntFlanks@12836.12871.36|CGGTGCGGAGTGCCCTTCGTCCTGGGAAACGGGGCG</td><td>>hg38_21_+_8433172_8446622_RNA45SN1_45S_50ntFlanks</td><td>12836</td><td>12871</td><td>36</td><td>CGGTGCGGAGTGCCCTTCGTCCTGGGAAACGGGGCG</td><td>1019.2</td><td>-8.0</td><td>1.4</td><td>-5.5</td><td>3.8E-08</td><td>6.8E-08</td><td>MB231/MB468</td><td>MITO</td></tr>
<tr><td>>MI0000433|hsa-let-7g&WithFlank&3|-|52268272|52268367@11.33.23&[MIMAT0000414&hsa-let-7g-5p&offsets|0|+1;m-58&3|-|52268336|52268357&offsets|0|+1]|TGAGGTAGTAGTTTGTACAGTTT</td><td>>MI0000433|hsa-let-7g&WithFlank&3|-|52268272|52268367</td><td>11</td><td>33</td><td>23</td><td>TGAGGTAGTAGTTTGTACAGTTT</td><td>2226.3</td><td>5.4</td><td>1.0</td><td>5.5</td><td>3.3E-08</td><td>6.8E-08</td><td>MB231/MB468</td><td>MP</td></tr>
<tr><td>>hg38_21_+_8433172_8446622_RNA45SN1_45S_50ntFlanks@485.534.50|GATGTGGTGACGTCGTGCTCTCCCGGGCCGGGTCCGAGCCGCGACGGGCG</td><td>>hg38_21_+_8433172_8446622_RNA45SN1_45S_50ntFlanks</td><td>485</td><td>534</td><td>50</td><td>GATGTGGTGACGTCGTGCTCTCCCGGGCCGGGTCCGAGCCGCGACGGGCG</td><td>160.9</td><td>10.2</td><td>1.9</td><td>5.5</td><td>3.9E-08</td><td>6.8E-08</td><td>BT20/MB468</td><td>NUC</td></tr>
<tr><td>>hg38_21_+_8433172_8446622_RNA45SN1_45S_50ntFlanks@11562.11588.27|CGCCTAGCAGCCGACTTAGAACTGGTG</td><td>>hg38_21_+_8433172_8446622_RNA45SN1_45S_50ntFlanks</td><td>11562</td><td>11588</td><td>27</td><td>CGCCTAGCAGCCGACTTAGAACTGGTG</td><td>839.3</td><td>-8.6</td><td>1.6</td><td>-5.5</td><td>3.9E-08</td><td>6.8E-08</td><td>BT20/MB468</td><td>NUC</td></tr>
<tr><td>>MI0000734|hsa-mir-106b&WithFlank&7|-|100093987|100094080@56.77.22&[MIMAT0004672&hsa-miR-106b-3p&offsets|-2|-2;m-87&7|-|100094003|100094023&offsets|-2|-1]|TACCGCACTGTGGGTACTTGCT</td><td>>MI0000734|hsa-mir-106b&WithFlank&7|-|100093987|100094080</td><td>56</td><td>77</td><td>22</td><td>TACCGCACTGTGGGTACTTGCT</td><td>730.3</td><td>7.8</td><td>1.4</td><td>5.5</td><td>3.8E-08</td><td>6.8E-08</td><td>MB231/MB468</td><td>MITO</td></tr>
<tr><td>>hg38_21_+_8433172_8446622_RNA45SN1_45S_50ntFlanks@11805.11823.19|CCACTGTCCCTACCTACTA</td><td>>hg38_21_+_8433172_8446622_RNA45SN1_45S_50ntFlanks</td><td>11805</td><td>11823</td><td>19</td><td>CCACTGTCCCTACCTACTA</td><td>627.8</td><td>-8.5</td><td>1.5</td><td>-5.5</td><td>4.0E-08</td><td>6.8E-08</td><td>BT20/MB468</td><td>NUC</td></tr>
<tr><td>>hg38_21_+_8433172_8446622_RNA45SN1_45S_50ntFlanks@10805.10840.36|GAACAATGTAGGTAAGGGAAGTCGGCAAGCCGGATC</td><td>>hg38_21_+_8433172_8446622_RNA45SN1_45S_50ntFlanks</td><td>10805</td><td>10840</td><td>36</td><td>GAACAATGTAGGTAAGGGAAGTCGGCAAGCCGGATC</td><td>244.2</td><td>-5.5</td><td>1.0</td><td>-5.5</td><td>3.6E-08</td><td>6.8E-08</td><td>MB231/MB468</td><td>NUC</td></tr>
<tr><td>>hg38_21_+_8433172_8446622_RNA45SN1_45S_50ntFlanks@8357.8375.19|AAAAGAACTTTGAAGAGAG</td><td>>hg38_21_+_8433172_8446622_RNA45SN1_45S_50ntFlanks</td><td>8357</td><td>8375</td><td>19</td><td>AAAAGAACTTTGAAGAGAG</td><td>1209.7</td><td>5.6</td><td>1.0</td><td>5.9</td><td>2.8E-09</td><td>6.8E-08</td><td>BT20/MB231</td><td>MITO</td></tr>
<tr><td>>MI0003560|hsa-mir-92b&WithFlank&1|+|155195171|155195278@67.88.22&[MIMAT0003218&hsa-miR-92b-3p&offsets|0|0;m-115&1|+|155195237|155195258&offsets|0|0]|TATTGCACTCGTCCCGGCCTCC</td><td>>MI0003560|hsa-mir-92b&WithFlank&1|+|155195171|155195278</td><td>67</td><td>88</td><td>22</td><td>TATTGCACTCGTCCCGGCCTCC</td><td>2220.2</td><td>5.0</td><td>0.9</td><td>5.5</td><td>3.3E-08</td><td>6.8E-08</td><td>MB231/MB468</td><td>MP</td></tr>
<tr><td>>hg38_21_+_8433172_8446622_RNA45SN1_45S_50ntFlanks@10458.10492.35|ATGGCCTCCGTTGCCCTCGGCCGATCGAAAGGGAG</td><td>>hg38_21_+_8433172_8446622_RNA45SN1_45S_50ntFlanks</td><td>10458</td><td>10492</td><td>35</td><td>ATGGCCTCCGTTGCCCTCGGCCGATCGAAAGGGAG</td><td>130.1</td><td>-10.1</td><td>1.7</td><td>-5.8</td><td>5.8E-09</td><td>6.8E-08</td><td>BT20/MB468</td><td>MP</td></tr>
<tr><td>>hg38_21_+_8433172_8446622_RNA45SN1_45S_50ntFlanks@9545.9570.26|GTGGAGGTCCGTAGCGGTCCTGACGT</td><td>>hg38_21_+_8433172_8446622_RNA45SN1_45S_50ntFlanks</td><td>9545</td><td>9570</td><td>26</td><td>GTGGAGGTCCGTAGCGGTCCTGACGT</td><td>924.0</td><td>-4.4</td><td>0.8</td><td>-5.5</td><td>3.6E-08</td><td>6.8E-08</td><td>MB231/MB468</td><td>NUC</td></tr>
<tr><td>>MI0000102|hsa-mir-100&WithFlank&11|-|122152223|122152314@19.40.22(+1C)&[MIMAT0000098&hsa-miR-100-5p&offsets|0|0(+1C);m-27&11|-|122152276|122152296&offsets|0|+1(+1C)]|AACCCGTAGATCCGAACTTGTGC</td><td>>MI0000102|hsa-mir-100&WithFlank&11|-|122152223|122152314</td><td>19</td><td>40</td><td>22(+1C)</td><td>AACCCGTAGATCCGAACTTGTGC</td><td>10258.4</td><td>-9.0</td><td>1.5</td><td>-6.1</td><td>1.1E-09</td><td>6.8E-08</td><td>BT20/MB231</td><td>MP</td></tr>
<tr><td>>hg38_21_+_8433172_8446622_RNA45SN1_45S_50ntFlanks@12007.12029.23|GCCGCCGGTGAAATACCACTACT</td><td>>hg38_21_+_8433172_8446622_RNA45SN1_45S_50ntFlanks</td><td>12007</td><td>12029</td><td>23</td><td>GCCGCCGGTGAAATACCACTACT</td><td>3477.9</td><td>-5.1</td><td>0.9</td><td>-5.5</td><td>3.8E-08</td><td>6.8E-08</td><td>MB231/MB468</td><td>MITO</td></tr>
<tr><td>>hg38_21_+_8433172_8446622_RNA45SN1_45S_50ntFlanks@10771.10797.27|CAGGTCTCCAAGGTGAACAGCCTCTGG</td><td>>hg38_21_+_8433172_8446622_RNA45SN1_45S_50ntFlanks</td><td>10771</td><td>10797</td><td>27</td><td>CAGGTCTCCAAGGTGAACAGCCTCTGG</td><td>311.9</td><td>-7.1</td><td>1.3</td><td>-5.5</td><td>3.6E-08</td><td>6.8E-08</td><td>MB231/MB468</td><td>NUC</td></tr>
<tr><td>>hg38_21_+_8433172_8446622_RNA45SN1_45S_50ntFlanks@5084.5112.29|CGAACGAGACTCTGGCATGCTAACTAGTT</td><td>>hg38_21_+_8433172_8446622_RNA45SN1_45S_50ntFlanks</td><td>5084</td><td>5112</td><td>29</td><td>CGAACGAGACTCTGGCATGCTAACTAGTT</td><td>1495.7</td><td>-9.0</td><td>1.6</td><td>-5.5</td><td>3.3E-08</td><td>6.8E-08</td><td>MB231/MB468</td><td>MP</td></tr>
<tr><td>>hg38_21_+_8433172_8446622_RNA45SN1_45S_50ntFlanks@11930.11967.38|TAGAATAAGTGGGAGGCCCCCGGCGCCCCCCCGGTGTC</td><td>>hg38_21_+_8433172_8446622_RNA45SN1_45S_50ntFlanks</td><td>11930</td><td>11967</td><td>38</td><td>TAGAATAAGTGGGAGGCCCCCGGCGCCCCCCCGGTGTC</td><td>113.5</td><td>-9.9</td><td>1.7</td><td>-5.8</td><td>5.9E-09</td><td>6.8E-08</td><td>BT20/MB468</td><td>MP</td></tr>
<tr><td>>hg38_21_+_8433172_8446622_RNA45SN1_45S_50ntFlanks@9339.9374.36|CGCACGAAAGCCGCCGTGGCGCAATGAAGGTGAAGG</td><td>>hg38_21_+_8433172_8446622_RNA45SN1_45S_50ntFlanks</td><td>9339</td><td>9374</td><td>36</td><td>CGCACGAAAGCCGCCGTGGCGCAATGAAGGTGAAGG</td><td>131.6</td><td>-10.1</td><td>1.7</td><td>-5.8</td><td>5.9E-09</td><td>6.8E-08</td><td>BT20/MB468</td><td>MP</td></tr>
<tr><td>>hg38_21_+_8433172_8446622_RNA45SN1_45S_50ntFlanks@5073.5108.36|AATTCCGATAACGAACGAGACTCTGGCATGCTAACT</td><td>>hg38_21_+_8433172_8446622_RNA45SN1_45S_50ntFlanks</td><td>5073</td><td>5108</td><td>36</td><td>AATTCCGATAACGAACGAGACTCTGGCATGCTAACT</td><td>151.8</td><td>-11.1</td><td>2.0</td><td>-5.5</td><td>3.3E-08</td><td>6.8E-08</td><td>MB231/MB468</td><td>MP</td></tr>
<tr><td>>MI0000650|hsa-mir-200c&WithFlank&12|+|6963693|6963772@50.71.22(+1C)&[MIMAT0000617&hsa-miR-200c-3p&offsets|0|-1(+1C);m-54&12|+|6963742|6963764&offsets|0|-1(+1C)]|TAATACTGCCGGGTAATGATGGC</td><td>>MI0000650|hsa-mir-200c&WithFlank&12|+|6963693|6963772</td><td>50</td><td>71</td><td>22(+1C)</td><td>TAATACTGCCGGGTAATGATGGC</td><td>222.9</td><td>11.1</td><td>1.8</td><td>6.1</td><td>1.2E-09</td><td>6.8E-08</td><td>BT20/MB231</td><td>MP</td></tr>
<tr><td>>hg38_21_+_8433172_8446622_RNA45SN1_45S_50ntFlanks@8112.8136.25|GGGCGCGGGACATGTGGCGTACGGA</td><td>>hg38_21_+_8433172_8446622_RNA45SN1_45S_50ntFlanks</td><td>8112</td><td>8136</td><td>25</td><td>GGGCGCGGGACATGTGGCGTACGGA</td><td>1012.9</td><td>13.3</td><td>2.2</td><td>6.1</td><td>1.2E-09</td><td>6.8E-08</td><td>BT20/MB231</td><td>MP</td></tr>
<tr><td>>hg38_21_+_8433172_8446622_RNA45SN1_45S_50ntFlanks@8357.8387.31|AAAAGAACTTTGAAGAGAGAGTTCAAGAGGG</td><td>>hg38_21_+_8433172_8446622_RNA45SN1_45S_50ntFlanks</td><td>8357</td><td>8387</td><td>31</td><td>AAAAGAACTTTGAAGAGAGAGTTCAAGAGGG</td><td>684.2</td><td>-8.5</td><td>1.6</td><td>-5.5</td><td>4.0E-08</td><td>6.8E-08</td><td>BT20/MB468</td><td>NUC</td></tr>
<tr><td>>hg38_21_+_8433172_8446622_RNA45SN1_45S_50ntFlanks@10795.10818.24|TGGCATGTTGGAACAATGTAGGTA</td><td>>hg38_21_+_8433172_8446622_RNA45SN1_45S_50ntFlanks</td><td>10795</td><td>10818</td><td>24</td><td>TGGCATGTTGGAACAATGTAGGTA</td><td>532.9</td><td>-4.3</td><td>0.7</td><td>-5.8</td><td>5.9E-09</td><td>6.9E-08</td><td>BT20/MB468</td><td>MP</td></tr>
<tr><td>>hg38_21_+_8433172_8446622_RNA45SN1_45S_50ntFlanks@12072.12095.24|CCCGAGGGGCTCTCGCTTCTGGCG</td><td>>hg38_21_+_8433172_8446622_RNA45SN1_45S_50ntFlanks</td><td>12072</td><td>12095</td><td>24</td><td>CCCGAGGGGCTCTCGCTTCTGGCG</td><td>3054.9</td><td>-5.9</td><td>1.1</td><td>-5.5</td><td>3.5E-08</td><td>6.9E-08</td><td>MB231/MB468</td><td>CYTO</td></tr>
<tr><td>>MI0003643|hsa-mir-629&WithFlank&15|-|70079366|70079474@28.47.20(+1A)&[MIMAT0004810&hsa-miR-629-5p&offsets|0|-1(+1A);m-150&15|-|70079427|70079447&offsets|0|-1(+1A)]|TGGGTTTACGTTGGGAGAACA</td><td>>MI0003643|hsa-mir-629&WithFlank&15|-|70079366|70079474</td><td>28</td><td>47</td><td>20(+1A)</td><td>TGGGTTTACGTTGGGAGAACA</td><td>78.3</td><td>9.5</td><td>1.7</td><td>5.5</td><td>3.3E-08</td><td>6.9E-08</td><td>MB231/MB468</td><td>MP</td></tr>
<tr><td>>hg38_21_+_8433172_8446622_RNA45SN1_45S_50ntFlanks@4759.4781.23|ATCAGATACCGTCGTAGTTCCGA</td><td>>hg38_21_+_8433172_8446622_RNA45SN1_45S_50ntFlanks</td><td>4759</td><td>4781</td><td>23</td><td>ATCAGATACCGTCGTAGTTCCGA</td><td>410.7</td><td>-7.7</td><td>1.4</td><td>-5.5</td><td>3.3E-08</td><td>6.9E-08</td><td>MB231/MB468</td><td>MP</td></tr>
</table>

| | | | | | | | | | | | | | |
|---|---|---|---|---|---|---|---|---|---|---|---|---|---|
| >hg38_21_+_8433172_8446622_RNA45SN1_45S_50ntFlanks@11694.11728.35\|AAGAAATTCAATGAAGCGCGGGTAAACGGCGGGAG | >hg38_21_+_8433172_8446622_RNA45SN1_45S_50ntFlanks | 11694 | 11728 | 35 | AAGAAATTCAATGAAGCGCGGGTAAACGGCGGGAG | 184.1 | -4.4 | 0.8 | -5.5 | 3.5E-08 | 6.9E-08 | MB231/MB468 | CYTO |
| >hg38_21_+_8433172_8446622_RNA45SN1_45S_50ntFlanks@12152.12183.32\|TGGGGAGTTTGACTGGGGCGGTACACCTGTCA | >hg38_21_+_8433172_8446622_RNA45SN1_45S_50ntFlanks | 12152 | 12183 | 32 | TGGGGAGTTTGACTGGGGCGGTACACCTGTCA | 217.8 | -7.2 | 1.3 | -5.5 | 3.3E-08 | 6.9E-08 | MB231/MB468 | MP |
| >hg38_21_+_8433172_8446622_RNA45SN1_45S_50ntFlanks@11212.11253.42\|CGACTCTGGACGCGAGCCGGGCCCTTCCCGTGGATCGCCCCA | >hg38_21_+_8433172_8446622_RNA45SN1_45S_50ntFlanks | 11212 | 11253 | 42 | CGACTCTGGACGCGAGCCGGGCCCTTCCCGTGGATCGCCCCA | 1268.3 | -7.6 | 1.4 | -5.5 | 4.0E-08 | 6.9E-08 | BT20/MB468 | NUC |
| >hg38_21_+_8433172_8446622_RNA45SN1_45S_50ntFlanks@4202.4222.21\|CGACCCGGGGAGGTAGTGACG | >hg38_21_+_8433172_8446622_RNA45SN1_45S_50ntFlanks | 4202 | 4222 | 21 | CGACCCGGGGAGGTAGTGACG | 465.3 | -7.7 | 1.4 | -5.5 | 3.3E-08 | 6.9E-08 | MB231/MB468 | MP |
| >hg38_21_+_8433172_8446622_RNA45SN1_45S_50ntFlanks@11582.11623.42\|ACTGGTGCGGACCAGGGGAATCCGACTGTTTAATTAAAACAA | >hg38_21_+_8433172_8446622_RNA45SN1_45S_50ntFlanks | 11582 | 11623 | 42 | ACTGGTGCGGACCAGGGGAATCCGACTGTTTAATTAAAACAA | 517.8 | 9.2 | 1.7 | 5.6 | 2.7E-08 | 6.9E-08 | BT20/MB468 | MITO |
| >hg38_21_+_8433172_8446622_RNA45SN1_45S_50ntFlanks@11210.11231.22\|GGCGACTCTGGACGCGAGCCGG | >hg38_21_+_8433172_8446622_RNA45SN1_45S_50ntFlanks | 11210 | 11231 | 22 | GGCGACTCTGGACGCGAGCCGG | 211.7 | -5.2 | 0.9 | -5.5 | 3.6E-08 | 6.9E-08 | MB231/MB468 | CYTO |
| >hg38_21_+_8433172_8446622_RNA45SN1_45S_50ntFlanks@12935.12955.21\|TAAACCATTCGTAGACGACCT | >hg38_21_+_8433172_8446622_RNA45SN1_45S_50ntFlanks | 12935 | 12955 | 21 | TAAACCATTCGTAGACGACCT | 244.4 | -5.3 | 1.0 | -5.5 | 3.7E-08 | 6.9E-08 | MB231/MB468 | NUC |
| >hg38_21_+_8433172_8446622_RNA45SN1_45S_50ntFlanks@8115.8137.23\|CGCGGGACATGTGGCGTACGGAA | >hg38_21_+_8433172_8446622_RNA45SN1_45S_50ntFlanks | 8115 | 8137 | 23 | CGCGGGACATGTGGCGTACGGAA | 2098.3 | 14.4 | 2.4 | 6.1 | 1.2E-09 | 6.9E-08 | BT20/MB231 | MP |
| >MI0000255\|hsa-mir-30d&WithFlank&8\|-\|134804870\|134804951@12.35.24(+1C)&[MIMAT0000245&hsa-miR-30d-5p&offsets\|0\|+2(+1C);m-21&8\|-\|134804919\|134804940&offsets\|0\|+2(+1C)]\|TGTAAACATCCCCGACTGGAAGCTC | >MI0000255\|hsa-mir-30d&WithFlank&8\|-\|134804870\|134804951 | 12 | 35 | 24(+1C) | TGTAAACATCCCCGACTGGAAGCTC | 64.2 | 3.6 | 0.7 | 5.5 | 4.0E-08 | 6.9E-08 | BT20/MB468 | NUC |
| >trnaMT_GluTTC_MT_-_14674_14742@39.58.20,trnalookalike8_GluTTC_5_-_93905172_93905240@39.58.20\|ATTGGTCGTGGTTGTAGTCC | >trnaMT_GluTTC_MT_-_14674_14742 | 39 | 58 | 20 | ATTGGTCGTGGTTGTAGTCC | 547.3 | 9.1 | 1.6 | 5.5 | 3.1E-08 | 7.0E-08 | BT20/MB468 | CYTO |
| >hg38_21_+_8433172_8446622_RNA45SN1_45S_50ntFlanks@9537.9563.27\|AAACTCTGGTGGAGGTCCGTAGCGGTC | >hg38_21_+_8433172_8446622_RNA45SN1_45S_50ntFlanks | 9537 | 9563 | 27 | AAACTCTGGTGGAGGTCCGTAGCGGTC | 257.9 | -7.1 | 1.3 | -5.5 | 3.7E-08 | 7.0E-08 | MB231/MB468 | NUC |
| >hg38_21_+_8433172_8446622_RNA45SN1_45S_50ntFlanks@3991.4026.36\|TGGTGACTCTAGATAACCTCGGGCCGATCGCACGCC | >hg38_21_+_8433172_8446622_RNA45SN1_45S_50ntFlanks | 3991 | 4026 | 36 | TGGTGACTCTAGATAACCTCGGGCCGATCGCACGCC | 1202.1 | -8.9 | 1.6 | -5.5 | 4.1E-08 | 7.0E-08 | BT20/MB468 | NUC |
| >hg38_21_+_8433172_8446622_RNA45SN1_45S_50ntFlanks@10680.10711.32\|GTTCCGGCGGCGTCCGGTGAGCTCTCGCTGGC | >hg38_21_+_8433172_8446622_RNA45SN1_45S_50ntFlanks | 10680 | 10711 | 32 | GTTCCGGCGGCGTCCGGTGAGCTCTCGCTGGC | 10608.1 | -5.7 | 1.0 | -5.5 | 3.1E-08 | 7.0E-08 | BT20/MB468 | CYTO |
| >hg38_21_+_8433172_8446622_RNA45SN1_45S_50ntFlanks@6728.6756.29\|GGACACATTGATCATCGACACTTCGAACG | >hg38_21_+_8433172_8446622_RNA45SN1_45S_50ntFlanks | 6728 | 6756 | 29 | GGACACATTGATCATCGACACTTCGAACG | 334.0 | 3.8 | 0.7 | 5.5 | 3.6E-08 | 7.0E-08 | MB231/MB468 | CYTO |
| >hg38_21_+_8433172_8446622_RNA45SN1_45S_50ntFlanks@12406.12441.36\|TTTTTGATCCTTCGATGTCGGCTCTTCCTATCATTG | >hg38_21_+_8433172_8446622_RNA45SN1_45S_50ntFlanks | 12406 | 12441 | 36 | TTTTTGATCCTTCGATGTCGGCTCTTCCTATCATTG | 719.4 | -3.7 | 0.7 | -5.5 | 3.7E-08 | 7.0E-08 | MB231/MB468 | NUC |
| >hg38_21_+_8433172_8446622_RNA45SN1_45S_50ntFlanks@6648.6672.25\|GTACGACTCTTAGCGGTGGATCACT | >hg38_21_+_8433172_8446622_RNA45SN1_45S_50ntFlanks | 6648 | 6672 | 25 | GTACGACTCTTAGCGGTGGATCACT | 346.5 | 4.3 | 0.8 | 5.5 | 3.7E-08 | 7.0E-08 | MB231/MB468 | NUC |
| >hg38_21_+_8433172_8446622_RNA45SN1_45S_50ntFlanks@10781.10826.46\|AGGTGAACAGCCTCTGGCATGTTGGAACAATGTAGGTAAGGGAAGT | >hg38_21_+_8433172_8446622_RNA45SN1_45S_50ntFlanks | 10781 | 10826 | 46 | AGGTGAACAGCCTCTGGCATGTTGGAACAATGTAGGTAAGGGAAGT | 79.1 | -10.2 | 1.8 | -5.5 | 3.4E-08 | 7.0E-08 | MB231/MB468 | MP |
| >hg38_21_+_8433172_8446622_RNA45SN1_45S_50ntFlanks@8358.8395.38\|AAAGAACTTTGAAGAGAGAGTTCAAGAGGGCGTGAAAC | >hg38_21_+_8433172_8446622_RNA45SN1_45S_50ntFlanks | 8358 | 8395 | 38 | AAAGAACTTTGAAGAGAGAGTTCAAGAGGGCGTGAAAC | 79.8 | 10.1 | 1.7 | 5.8 | 6.0E-09 | 7.0E-08 | BT20/MB468 | MP |
| >hg38_21_+_8433172_8446622_RNA45SN1_45S_50ntFlanks@10866.10883.18\|CTAAGGGCTGGGTCGGTC | >hg38_21_+_8433172_8446622_RNA45SN1_45S_50ntFlanks | 10866 | 10883 | 18 | CTAAGGGCTGGGTCGGTC | 292.7 | -4.8 | 0.9 | -5.5 | 3.6E-08 | 7.0E-08 | MB231/MB468 | CYTO |
| >MI0000737\|hsa-mir-200a&WithFlank&1\|+\|1167857\|1167958@23.44.22&[MIMAT0001620&hsa-miR-200a-5p&offsets\|+1\|+1;m-154&1\|+\|1167878\|1167899&offsets\|+1\|+1]\|ATCTTACCGGACAGTGCTGGAT | >MI0000737\|hsa-mir-200a&WithFlank&1\|+\|1167857\|1167958 | 23 | 44 | 22 | ATCTTACCGGACAGTGCTGGAT | 89.5 | -9.6 | 1.6 | -5.8 | 6.0E-09 | 7.0E-08 | BT20/MB468 | MP |
| >hg38_21_+_8433172_8446622_RNA45SN1_45S_50ntFlanks@5086.5104.19\|AACGAGACTCTGGCATGCT | >hg38_21_+_8433172_8446622_RNA45SN1_45S_50ntFlanks | 5086 | 5104 | 19 | AACGAGACTCTGGCATGCT | 409.0 | 6.4 | 1.2 | 5.5 | 3.1E-08 | 7.0E-08 | BT20/MB468 | CYTO |
| >hg38_21_+_8433172_8446622_RNA45SN1_45S_50ntFlanks@4566.4587.22\|ATAATGGAATAGGACCGCGGTT | >hg38_21_+_8433172_8446622_RNA45SN1_45S_50ntFlanks | 4566 | 4587 | 22 | ATAATGGAATAGGACCGCGGTT | 299.0 | -3.0 | 0.5 | -5.5 | 3.1E-08 | 7.0E-08 | BT20/MB468 | CYTO |
| >trnaMT_HisGTG_MT_+_12138_12206@37.72.36\|CTGACAACAGAGGCTTACGACCCCTTATTTACCCCA | >trnaMT_HisGTG_MT_+_12138_12206 | 37 | 72 | 36 | CTGACAACAGAGGCTTACGACCCCTTATTTACCCCA | 118.0 | 10.6 | 1.8 | 5.8 | 6.0E-09 | 7.0E-08 | BT20/MB468 | MP |
| >hg38_21_+_8433172_8446622_RNA45SN1_45S_50ntFlanks@5421.5443.23\|CGATTGGATGGTTTAGTGAGGCC | >hg38_21_+_8433172_8446622_RNA45SN1_45S_50ntFlanks | 5421 | 5443 | 23 | CGATTGGATGGTTTAGTGAGGCC | 1173.5 | -6.2 | 1.1 | -5.5 | 3.1E-08 | 7.0E-08 | BT20/MB468 | CYTO |
| >hg38_21_+_8433172_8446622_RNA45SN1_45S_50ntFlanks@10406.10424.19\|TGGGTCAGTCGGTCCTGAG | >hg38_21_+_8433172_8446622_RNA45SN1_45S_50ntFlanks | 10406 | 10424 | 19 | TGGGTCAGTCGGTCCTGAG | 548.3 | 3.9 | 0.7 | 5.5 | 3.7E-08 | 7.0E-08 | MB231/MB468 | NUC |
| >hg38_21_+_8433172_8446622_RNA45SN1_45S_50ntFlanks@4161.4182.22\|CACATCCAAGGAAGGCAGCAGG | >hg38_21_+_8433172_8446622_RNA45SN1_45S_50ntFlanks | 4161 | 4182 | 22 | CACATCCAAGGAAGGCAGCAGG | 132.8 | -10.1 | 1.7 | -5.8 | 6.1E-09 | 7.0E-08 | BT20/MB468 | MP |
| >hg38_21_+_8433172_8446622_RNA45SN1_45S_50ntFlanks@9574.9605.32\|AATCGGTCGTCCGACCTGGGTATAGGGGCGAA | >hg38_21_+_8433172_8446622_RNA45SN1_45S_50ntFlanks | 9574 | 9605 | 32 | AATCGGTCGTCCGACCTGGGTATAGGGGCGAA | 634.7 | -3.9 | 0.7 | -5.5 | 4.1E-08 | 7.1E-08 | BT20/MB468 | NUC |
| >hg38_21_+_8433172_8446622_RNA45SN1_45S_50ntFlanks@11537.11560.24\|CCGCGCGGCGCCTCGCCTCGGCCG | >hg38_21_+_8433172_8446622_RNA45SN1_45S_50ntFlanks | 11537 | 11560 | 24 | CCGCGCGGCGCCTCGCCTCGGCCG | 108.8 | -9.8 | 1.7 | -5.8 | 6.1E-09 | 7.1E-08 | BT20/MB468 | MP |
| >MI0000446\|hsa-mir-125b-1&WithFlank&11\|-\|122099751\|122099850@21.41.21;MI0000470\|hsa-mir-125b-2&WithFlank&21\|+\|16590231\|16590331@23.43.21&[MIMAT0000423&hsa-miR-125b-5p&offsets\|0\|-1;m-67&11\|-\|122099809\|122099830&offsets\|0\|-1];[MIMAT0000423_1&hsa-miR-125b-5p&offsets\|0\|-1;m-68&21\|+\|16590253\|16590274&offsets\|0\|-1]\|TCCCTGAGACCCTAACTTGTG | >MI0000446\|hsa-mir-125b-1&WithFlank&11\|-\|122099751\|122099850 | 21 | 41 | 21 | TCCCTGAGACCCTAACTTGTG | 5300.6 | 4.7 | 0.9 | 5.5 | 3.4E-08 | 7.1E-08 | MB231/MB468 | MP |
| >hg38_21_+_8433172_8446622_RNA45SN1_45S_50ntFlanks@8114.8138.25\|GCGCGGGACATGTGGCGTACGGAAG | >hg38_21_+_8433172_8446622_RNA45SN1_45S_50ntFlanks | 8114 | 8138 | 25 | GCGCGGGACATGTGGCGTACGGAAG | 767.7 | 12.9 | 2.1 | 6.1 | 1.2E-09 | 7.1E-08 | BT20/MB231 | MP |
| >trna152_ValCAC_6_-_27248049_27248121@1.34.34\|GCTTCTGTAGTGTAGTGGTTATCACGTTCGCCTC | >trna152_ValCAC_6_-_27248049_27248121 | 1 | 34 | 34 | GCTTCTGTAGTGTAGTGGTTATCACGTTCGCCTC | 1920.8 | 8.3 | 1.4 | 5.8 | 6.1E-09 | 7.1E-08 | BT20/MB468 | MP |
| >hg38_21_+_8433172_8446622_RNA45SN1_45S_50ntFlanks@4734.4752.19\|AACGAAAGTCGGAGGTTCG | >hg38_21_+_8433172_8446622_RNA45SN1_45S_50ntFlanks | 4734 | 4752 | 19 | AACGAAAGTCGGAGGTTCG | 691.5 | -6.8 | 1.2 | -5.5 | 4.1E-08 | 7.1E-08 | BT20/MB468 | NUC |
| >hg38_21_+_8433172_8446622_RNA45SN1_45S_50ntFlanks@12170.12188.19\|CGGTACACCTGTCAAACGG | >hg38_21_+_8433172_8446622_RNA45SN1_45S_50ntFlanks | 12170 | 12188 | 19 | CGGTACACCTGTCAAACGG | 383.2 | -10.7 | 1.9 | -5.6 | 2.8E-08 | 7.1E-08 | BT20/MB468 | MITO |
| >hg38_21_+_8433172_8446622_RNA45SN1_45S_50ntFlanks@10682.10711.30\|TCCGGCGGCGTCCGGTGAGCTCTCGCTGGC | >hg38_21_+_8433172_8446622_RNA45SN1_45S_50ntFlanks | 10682 | 10711 | 30 | TCCGGCGGCGTCCGGTGAGCTCTCGCTGGC | 343.5 | -4.5 | 0.8 | -5.5 | 4.0E-08 | 7.1E-08 | MB231/MB468 | MITO |
| >hg38_21_+_8433172_8446622_RNA45SN1_45S_50ntFlanks@12324.12356.33\|TTGGGTTTTAAGCAGGAGGTGTCAGAAAAGTTA | >hg38_21_+_8433172_8446622_RNA45SN1_45S_50ntFlanks | 12324 | 12356 | 33 | TTGGGTTTTAAGCAGGAGGTGTCAGAAAAGTTA | 105.9 | -8.8 | 1.6 | -5.6 | 2.8E-08 | 7.1E-08 | BT20/MB468 | MITO |
| >hg38_21_+_8433172_8446622_RNA45SN1_45S_50ntFlanks@10788.10818.31\|CAGCCTCTGGCATGTTGGAACAATGTAGGTA | >hg38_21_+_8433172_8446622_RNA45SN1_45S_50ntFlanks | 10788 | 10818 | 31 | CAGCCTCTGGCATGTTGGAACAATGTAGGTA | 327.1 | -3.0 | 0.5 | -5.5 | 3.7E-08 | 7.1E-08 | MB231/MB468 | NUC |
| >hg38_21_+_8433172_8446622_RNA45SN1_45S_50ntFlanks@4297.4314.18\|CATTGGAGGGCAAGTCTG | >hg38_21_+_8433172_8446622_RNA45SN1_45S_50ntFlanks | 4297 | 4314 | 18 | CATTGGAGGGCAAGTCTG | 543.7 | -7.7 | 1.4 | -5.5 | 3.4E-08 | 7.1E-08 | MB231/MB468 | MP |
| >hg38_21_+_8433172_8446622_RNA45SN1_45S_50ntFlanks@4975.4999.25\|CCCGGACACGGACAGGATTGACAGA | >hg38_21_+_8433172_8446622_RNA45SN1_45S_50ntFlanks | 4975 | 4999 | 25 | CCCGGACACGGACAGGATTGACAGA | 1224.0 | -6.7 | 1.2 | -5.5 | 3.4E-08 | 7.1E-08 | MB231/MB468 | MP |
| >hg38_21_+_8433172_8446622_RNA45SN1_45S_50ntFlanks@6661.6698.38\|CGGTGGATCACTCGGCTCGTGCGTCGATGAAGAACGCA | >hg38_21_+_8433172_8446622_RNA45SN1_45S_50ntFlanks | 6661 | 6698 | 38 | CGGTGGATCACTCGGCTCGTGCGTCGATGAAGAACGCA | 444.6 | 2.9 | 0.5 | 5.5 | 3.1E-08 | 7.1E-08 | BT20/MB468 | CYTO |
| >hg38_21_+_8433172_8446622_RNA45SN1_45S_50ntFlanks@3837.3868.32\|CCTCTCCTACTTGGATAACTGTGGTAATTCTA | >hg38_21_+_8433172_8446622_RNA45SN1_45S_50ntFlanks | 3837 | 3868 | 32 | CCTCTCCTACTTGGATAACTGTGGTAATTCTA | 484.9 | -3.7 | 0.7 | -5.5 | 3.7E-08 | 7.1E-08 | MB231/MB468 | NUC |
| >hg38_21_+_8433172_8446622_RNA45SN1_45S_50ntFlanks@12489.12521.33\|ACGTGAGCTGGGTTTAGACCGTCGTGAGACAGG | >hg38_21_+_8433172_8446622_RNA45SN1_45S_50ntFlanks | 12489 | 12521 | 33 | ACGTGAGCTGGGTTTAGACCGTCGTGAGACAGG | 87.9 | -9.5 | 1.6 | -5.8 | 6.1E-09 | 7.1E-08 | BT20/MB468 | MP |
| >hg38_21_+_8433172_8446622_RNA45SN1_45S_50ntFlanks@4472.4496.25\|CTTAGCTGAGTGTCCCGCGGGGCCC | >hg38_21_+_8433172_8446622_RNA45SN1_45S_50ntFlanks | 4472 | 4496 | 25 | CTTAGCTGAGTGTCCCGCGGGGCCC | 234.9 | -7.2 | 1.3 | -5.5 | 3.8E-08 | 7.1E-08 | MB231/MB468 | NUC |
| >hg38_21_+_8433172_8446622_RNA45SN1_45S_50ntFlanks@12717.12753.37\|CCTCGGTTGGCCTCGGATAGCCGGTCCCCCGCCTGTC | >hg38_21_+_8433172_8446622_RNA45SN1_45S_50ntFlanks | 12717 | 12753 | 37 | CCTCGGTTGGCCTCGGATAGCCGGTCCCCCGCCTGTC | 113.9 | -10.7 | 1.9 | -5.5 | 3.4E-08 | 7.1E-08 | MB231/MB468 | MP |
| >hg38_21_+_8433172_8446622_RNA45SN1_45S_50ntFlanks@4145.4162.18\|CCTGAGAAACGGCTACCA | >hg38_21_+_8433172_8446622_RNA45SN1_45S_50ntFlanks | 4145 | 4162 | 18 | CCTGAGAAACGGCTACCA | 134.3 | -10.2 | 1.7 | -5.8 | 6.2E-09 | 7.1E-08 | BT20/MB468 | MP |
| >hg38_21_+_8433172_8446622_RNA45SN1_45S_50ntFlanks@4003.4044.42\|ATAACCTCGGGCCGATCGCACGCCCCCCGTGGCGGCGACGAC | >hg38_21_+_8433172_8446622_RNA45SN1_45S_50ntFlanks | 4003 | 4044 | 42 | ATAACCTCGGGCCGATCGCACGCCCCCCGTGGCGGCGACGAC | 117.7 | -9.2 | 1.7 | -5.5 | 3.2E-08 | 7.2E-08 | BT20/MB468 | CYTO |
| >hg38_21_+_8433172_8446622_RNA45SN1_45S_50ntFlanks@8354.8373.20\|TTGAAAAGAACTTTGAAGAG | >hg38_21_+_8433172_8446622_RNA45SN1_45S_50ntFlanks | 8354 | 8373 | 20 | TTGAAAAGAACTTTGAAGAG | 622.8 | 9.5 | 1.6 | 5.8 | 6.2E-09 | 7.2E-08 | BT20/MB468 | MP |
| >hg38_21_+_8433172_8446622_RNA45SN1_45S_50ntFlanks@10869.10901.33\|AGGGCTGGGTCGGTCGGGCTGGGGCGCGAAGCG | >hg38_21_+_8433172_8446622_RNA45SN1_45S_50ntFlanks | 10869 | 10901 | 33 | AGGGCTGGGTCGGTCGGGCTGGGGCGCGAAGCG | 3070.0 | -9.5 | 1.7 | -5.5 | 4.2E-08 | 7.2E-08 | BT20/MB468 | NUC |
| >hg38_21_+_8433172_8446622_RNA45SN1_45S_50ntFlanks@12456.12475.20\|CCAAGCGTTGGATTGTTCAC | >hg38_21_+_8433172_8446622_RNA45SN1_45S_50ntFlanks | 12456 | 12475 | 20 | CCAAGCGTTGGATTGTTCAC | 3871.6 | -8.2 | 1.5 | -5.6 | 2.8E-08 | 7.2E-08 | BT20/MB468 | MITO |
| >hg38_21_+_8433172_8446622_RNA45SN1_45S_50ntFlanks@5495.5514.20\|AGACGGTCGAACTTGACTAT | >hg38_21_+_8433172_8446622_RNA45SN1_45S_50ntFlanks | 5495 | 5514 | 20 | AGACGGTCGAACTTGACTAT | 431.1 | -7.9 | 1.4 | -5.5 | 3.5E-08 | 7.2E-08 | MB231/MB468 | MP |
| >hg38_21_+_8433172_8446622_RNA45SN1_45S_50ntFlanks@5423.5445.23\|ATTGGATGGTTTAGTGAGGCCCT | >hg38_21_+_8433172_8446622_RNA45SN1_45S_50ntFlanks | 5423 | 5445 | 23 | ATTGGATGGTTTAGTGAGGCCCT | 153.7 | -6.3 | 1.1 | -5.5 | 3.2E-08 | 7.2E-08 | BT20/MB468 | CYTO |
| >hg38_21_+_8433172_8446622_RNA45SN1_45S_50ntFlanks@4102.4119.18\|ACCACGGGTGACGGGGAA | >hg38_21_+_8433172_8446622_RNA45SN1_45S_50ntFlanks | 4102 | 4119 | 18 | ACCACGGGTGACGGGGAA | 92.3 | -9.6 | 1.7 | -5.8 | 6.2E-09 | 7.2E-08 | BT20/MB468 | MP |
| >hg38_21_+_8433172_8446622_RNA45SN1_45S_50ntFlanks@8144.8164.21\|CTCCCCGGCGCCGCTCGTGGG | >hg38_21_+_8433172_8446622_RNA45SN1_45S_50ntFlanks | 8144 | 8164 | 21 | CTCCCCGGCGCCGCTCGTGGG | 216.0 | -10.8 | 1.9 | -5.8 | 6.2E-09 | 7.2E-08 | BT20/MB468 | MP |
| >hg38_21_+_8433172_8446622_RNA45SN1_45S_50ntFlanks@8350.8390.41\|AAAGTTGAAAAGAACTTTGAAGAGAGAGTTCAAGAGGGCGT | >hg38_21_+_8433172_8446622_RNA45SN1_45S_50ntFlanks | 8350 | 8390 | 41 | AAAGTTGAAAAGAACTTTGAAGAGAGAGTTCAAGAGGGCGT | 81.0 | 10.1 | 1.7 | 5.8 | 6.2E-09 | 7.2E-08 | BT20/MB468 | MP |
| >MI0000433\|hsa-let-7g&WithFlank&3\|-\|52268272\|52268367@11.31.21&[MIMAT0000414&hsa-let-7g-5p&offsets\|0\|-1;m-58&3\|-\|52268336\|52268357&offsets\|0\|-1]\|TGAGGTAGTAGTTTGTACAGT | >MI0000433\|hsa-let-7g&WithFlank&3\|-\|52268272\|52268367 | 11 | 31 | 21 | TGAGGTAGTAGTTTGTACAGT | 1330.6 | 4.9 | 0.9 | 5.5 | 4.0E-08 | 7.2E-08 | MB231/MB468 | MITO |

<table>
<tr><td>>MI0000478|hsa-mir-149&WithFlank&2|+|240455995|240456095@21.41.21&[MIMAT0000450&hsa-miR-149-5p&offsets|0|-2;m-195&2|+|240456015|240456037&offsets|0|-2]|TCTGGCTCCGTGTCTTCACTC</td><td>>MI0000478|hsa-mir-149&WithFlank&2|+|240455995|240456095</td><td>21</td><td>41</td><td>21</td><td>TCTGGCTCCGTGTCTTCACTC</td><td>92.3</td><td>10.3</td><td>1.8</td><td>5.8</td><td>6.2E-09</td><td>7.2E-08</td><td>BT20/MB468</td><td>MP</td></tr>
<tr><td>>MI0000085|hsa-mir-27a&WithFlank&19|-|13836434|13836523@57.77.21&[MIMAT0000084&hsa-miR-27a-3p&offsets|0|0;m-43&19|-|13836447|13836467&offsets|0|0]|TTCACAGTGGCTAAGTTCCGC</td><td>>MI0000085|hsa-mir-27a&WithFlank&19|-|13836434|13836523</td><td>57</td><td>77</td><td>21</td><td>TTCACAGTGGCTAAGTTCCGC</td><td>716.5</td><td>4.0</td><td>0.7</td><td>5.6</td><td>2.8E-08</td><td>7.2E-08</td><td>BT20/MB468</td><td>MITO</td></tr>
<tr><td>>hg38_21_+_8433172_8446622_RNA45SN1_45S_50ntFlanks@5486.5516.31|GCGCTGAGAAGACGGTCGAACTTGACTATCT</td><td>>hg38_21_+_8433172_8446622_RNA45SN1_45S_50ntFlanks</td><td>5486</td><td>5516</td><td>31</td><td>GCGCTGAGAAGACGGTCGAACTTGACTATCT</td><td>1237.8</td><td>4.7</td><td>0.9</td><td>5.6</td><td>2.9E-08</td><td>7.2E-08</td><td>BT20/MB468</td><td>MITO</td></tr>
<tr><td>>hg38_21_+_8433172_8446622_RNA45SN1_45S_50ntFlanks@7975.7993.19|CGCGACCTCAGATCAGACG</td><td>>hg38_21_+_8433172_8446622_RNA45SN1_45S_50ntFlanks</td><td>7975</td><td>7993</td><td>19</td><td>CGCGACCTCAGATCAGACG</td><td>32126.6</td><td>4.8</td><td>0.9</td><td>5.5</td><td>3.2E-08</td><td>7.2E-08</td><td>BT20/MB468</td><td>CYTO</td></tr>
<tr><td>>hg38_21_+_8433172_8446622_RNA45SN1_45S_50ntFlanks@12488.12508.21|AACGTGAGCTGGGTTTAGACC</td><td>>hg38_21_+_8433172_8446622_RNA45SN1_45S_50ntFlanks</td><td>12488</td><td>12508</td><td>21</td><td>AACGTGAGCTGGGTTTAGACC</td><td>139.8</td><td>-5.2</td><td>0.9</td><td>-5.5</td><td>3.8E-08</td><td>7.2E-08</td><td>MB231/MB468</td><td>NUC</td></tr>
<tr><td>>hg38_21_+_8433172_8446622_RNA45SN1_45S_50ntFlanks@12998.13039.42|GCTGCGATCTATTGAAAGTCAGCCCTCGACACAAGGGTTTGT</td><td>>hg38_21_+_8433172_8446622_RNA45SN1_45S_50ntFlanks</td><td>12998</td><td>13039</td><td>42</td><td>GCTGCGATCTATTGAAAGTCAGCCCTCGACACAAGGGTTTGT</td><td>844.9</td><td>5.4</td><td>1.0</td><td>5.5</td><td>3.5E-08</td><td>7.2E-08</td><td>MB231/MB468</td><td>MP</td></tr>
<tr><td>>hg38_21_+_8433172_8446622_RNA45SN1_45S_50ntFlanks@11592.11611.20|ACCAGGGGAATCCGACTGTT</td><td>>hg38_21_+_8433172_8446622_RNA45SN1_45S_50ntFlanks</td><td>11592</td><td>11611</td><td>20</td><td>ACCAGGGGAATCCGACTGTT</td><td>965.6</td><td>-8.8</td><td>1.5</td><td>-5.8</td><td>6.3E-09</td><td>7.2E-08</td><td>BT20/MB468</td><td>MP</td></tr>
<tr><td>>hg38_21_+_8433172_8446622_RNA45SN1_45S_50ntFlanks@12628.12653.26|TGGGGCGAAGCTACCATCTGTGGGAT</td><td>>hg38_21_+_8433172_8446622_RNA45SN1_45S_50ntFlanks</td><td>12628</td><td>12653</td><td>26</td><td>TGGGGCGAAGCTACCATCTGTGGGAT</td><td>1647.8</td><td>-8.5</td><td>1.5</td><td>-5.8</td><td>6.3E-09</td><td>7.2E-08</td><td>BT20/MB468</td><td>MP</td></tr>
<tr><td>>hg38_21_+_8433172_8446622_RNA45SN1_45S_50ntFlanks@9204.9221.18|TCGGGGCCACGCGCGCGT</td><td>>hg38_21_+_8433172_8446622_RNA45SN1_45S_50ntFlanks</td><td>9204</td><td>9221</td><td>18</td><td>TCGGGGCCACGCGCGCGT</td><td>96.3</td><td>-9.7</td><td>1.7</td><td>-5.8</td><td>6.3E-09</td><td>7.2E-08</td><td>BT20/MB468</td><td>MP</td></tr>
<tr><td>>hg38_21_+_8433172_8446622_RNA45SN1_45S_50ntFlanks@11575.11599.25|ACTTAGAACTGGTGCGGACCAGGGG</td><td>>hg38_21_+_8433172_8446622_RNA45SN1_45S_50ntFlanks</td><td>11575</td><td>11599</td><td>25</td><td>ACTTAGAACTGGTGCGGACCAGGGG</td><td>155.2</td><td>-10.4</td><td>1.8</td><td>-5.8</td><td>6.3E-09</td><td>7.2E-08</td><td>BT20/MB468</td><td>MP</td></tr>
<tr><td>>hg38_21_+_8433172_8446622_RNA45SN1_45S_50ntFlanks@11921.11956.36|TGAGAGGTGTAGAATAAGTGGGAGGCCCCCGGCGCC</td><td>>hg38_21_+_8433172_8446622_RNA45SN1_45S_50ntFlanks</td><td>11921</td><td>11956</td><td>36</td><td>TGAGAGGTGTAGAATAAGTGGGAGGCCCCCGGCGCC</td><td>423.3</td><td>-8.3</td><td>1.5</td><td>-5.5</td><td>4.2E-08</td><td>7.2E-08</td><td>BT20/MB468</td><td>NUC</td></tr>
<tr><td>>hg38_21_+_8433172_8446622_RNA45SN1_45S_50ntFlanks@12456.12479.24|CCAAGCGTTGGATTGTTCACCCAC</td><td>>hg38_21_+_8433172_8446622_RNA45SN1_45S_50ntFlanks</td><td>12456</td><td>12479</td><td>24</td><td>CCAAGCGTTGGATTGTTCACCCAC</td><td>883.0</td><td>-8.5</td><td>1.5</td><td>-5.5</td><td>4.2E-08</td><td>7.2E-08</td><td>BT20/MB468</td><td>NUC</td></tr>
<tr><td>>MI0005559|hsa-mir-744&WithFlank&17|+|12081893|12082002@17.37.21&[MIMAT0004945&hsa-miR-744-5p&offsets|0|-1;m-172&17|+|12081909|12081929&offsets|0|0]|TGCGGGGCTAGGGCTAACAGC</td><td>>MI0005559|hsa-mir-744&WithFlank&17|+|12081893|12082002</td><td>17</td><td>37</td><td>21</td><td>TGCGGGGCTAGGGCTAACAGC</td><td>509.0</td><td>2.5</td><td>0.5</td><td>5.5</td><td>4.2E-08</td><td>7.2E-08</td><td>BT20/MB468</td><td>NUC</td></tr>
<tr><td>>trnaMT_PheGAA_MT_+_577_647@23.53.31|AAGCAATACACTGAAAATGTTTAGACGGGCT</td><td>>trnaMT_PheGAA_MT_+_577_647</td><td>23</td><td>53</td><td>31</td><td>AAGCAATACACTGAAAATGTTTAGACGGGCT</td><td>163.9</td><td>11.1</td><td>1.9</td><td>5.8</td><td>6.3E-09</td><td>7.2E-08</td><td>BT20/MB468</td><td>MP</td></tr>
<tr><td>>MI0000066|hsa-let-7e&WithFlank&19|+|51692780|51692870@14.35.22&[MIMAT0000066&hsa-let-7e-5p&offsets|0|0;m-50&19|+|51692793|51692814&offsets|0|0]|TGAGGTAGGAGGTTGTATAGTT</td><td>>MI0000066|hsa-let-7e&WithFlank&19|+|51692780|51692870</td><td>14</td><td>35</td><td>22</td><td>TGAGGTAGGAGGTTGTATAGTT</td><td>5229.9</td><td>4.7</td><td>0.9</td><td>5.5</td><td>4.1E-08</td><td>7.2E-08</td><td>MB231/MB468</td><td>MITO</td></tr>
<tr><td>>hg38_21_+_8433172_8446622_RNA45SN1_45S_50ntFlanks@10403.10429.27|ACATGGGTCAGTCGGTCCTGAGAGATG</td><td>>hg38_21_+_8433172_8446622_RNA45SN1_45S_50ntFlanks</td><td>10403</td><td>10429</td><td>27</td><td>ACATGGGTCAGTCGGTCCTGAGAGATG</td><td>413.6</td><td>-3.9</td><td>0.7</td><td>-5.5</td><td>4.1E-08</td><td>7.2E-08</td><td>MB231/MB468</td><td>MITO</td></tr>
<tr><td>>hg38_21_+_8433172_8446622_RNA45SN1_45S_50ntFlanks@6785.6806.22|CTACGCCTGTCTGAGCGTCGCT</td><td>>hg38_21_+_8433172_8446622_RNA45SN1_45S_50ntFlanks</td><td>6785</td><td>6806</td><td>22</td><td>CTACGCCTGTCTGAGCGTCGCT</td><td>1616.2</td><td>4.1</td><td>0.7</td><td>5.5</td><td>3.7E-08</td><td>7.3E-08</td><td>MB231/MB468</td><td>CYTO</td></tr>
<tr><td>>MI0000289|hsa-mir-181a-1&WithFlank&1|-|198859038|198859159@30.54.25(+1C);MI0000269|hsa-mir-181a-2&WithFlank&9|+|124692436|124692557@45.69.25(+1C)&[MIMAT0000256&hsa-miR-181a-5p&offsets|0|+2(+1C);m-32&1|-|198859108|198859130&offsets|0|+2(+1C)];[MIMAT0000256_1&hsa-miR-181a-5p&offsets|0|+2(+1C);m-32&9|+|124692480|124692502&offsets|0|+2(+1C)]|AACATTCAACGCTGTCGGTGAGTTTC</td><td>>MI0000289|hsa-mir-181a-1&WithFlank&1|-|198859038|198859159</td><td>30</td><td>54</td><td>25(+1C)</td><td>AACATTCAACGCTGTCGGTGAGTTTC</td><td>190.5</td><td>-11.0</td><td>1.8</td><td>-6.1</td><td>1.3E-09</td><td>7.3E-08</td><td>BT20/MB231</td><td>MP</td></tr>
<tr><td>>hg38_21_+_8433172_8446622_RNA45SN1_45S_50ntFlanks@5075.5102.28|TTCCGATAACGAACGAGACTCTGGCATG</td><td>>hg38_21_+_8433172_8446622_RNA45SN1_45S_50ntFlanks</td><td>5075</td><td>5102</td><td>28</td><td>TTCCGATAACGAACGAGACTCTGGCATG</td><td>112.5</td><td>-10.7</td><td>1.9</td><td>-5.5</td><td>3.5E-08</td><td>7.3E-08</td><td>MB231/MB468</td><td>MP</td></tr>
<tr><td>>hg38_21_+_8433172_8446622_RNA45SN1_45S_50ntFlanks@11710.11740.31|CGCGGGTAAACGGCGGGAGTAACTATGACTC</td><td>>hg38_21_+_8433172_8446622_RNA45SN1_45S_50ntFlanks</td><td>11710</td><td>11740</td><td>31</td><td>CGCGGGTAAACGGCGGGAGTAACTATGACTC</td><td>817.2</td><td>-8.8</td><td>1.6</td><td>-5.5</td><td>4.2E-08</td><td>7.3E-08</td><td>BT20/MB468</td><td>NUC</td></tr>
<tr><td>>hg38_21_+_8433172_8446622_RNA45SN1_45S_50ntFlanks@3855.3898.44|CTGTGGTAATTCTAGAGCTAATACATGCCGACGGGCGCTGACCC</td><td>>hg38_21_+_8433172_8446622_RNA45SN1_45S_50ntFlanks</td><td>3855</td><td>3898</td><td>44</td><td>CTGTGGTAATTCTAGAGCTAATACATGCCGACGGGCGCTGACCC</td><td>280.2</td><td>-8.3</td><td>1.5</td><td>-5.5</td><td>3.8E-08</td><td>7.3E-08</td><td>MB231/MB468</td><td>NUC</td></tr>
<tr><td>>hg38_21_+_8433172_8446622_RNA45SN1_45S_50ntFlanks@9833.9851.19|CTTTTGGTAAGCAGAACTG</td><td>>hg38_21_+_8433172_8446622_RNA45SN1_45S_50ntFlanks</td><td>9833</td><td>9851</td><td>19</td><td>CTTTTGGTAAGCAGAACTG</td><td>67.7</td><td>-9.9</td><td>1.8</td><td>-5.5</td><td>3.5E-08</td><td>7.3E-08</td><td>MB231/MB468</td><td>MP</td></tr>
<tr><td>>am_trna111_HisGTG_1_-_147774845_147774916@-1G.29.30,trna118_HisGTG_1_-_145396881_145396952@-1G.29.30,trna16_HisGTG_1_+_146544773_146544844@-1G.29.30,trna1_HisGTG_15_+_45493349_45493420@-1G.29.30,trna21_HisGTG_1_+_147753471_147753542@-1G.29.30,trna33_HisGTG_6_+_27125906_27125977@-1G.29.30,trna7_HisGTG_9_-_14433938_14434009@-1G.29.30,trna8_HisGTG_15_-_45492611_45492682@-1G.29.30,trna9_HisGTG_15_-_45490804_45490875@-1G.29.30|GGCCGTGATCGTATAGTGGTTAGTACTCTG</td><td>>am_trna111_HisGTG_1_-_147774845_147774916</td><td>-1G</td><td>29</td><td>30</td><td>GGCCGTGATCGTATAGTGGTTAGTACTCTG</td><td>10209.8</td><td>-5.6</td><td>1.0</td><td>-5.5</td><td>3.2E-08</td><td>7.3E-08</td><td>BT20/MB468</td><td>CYTO</td></tr>
<tr><td>>MI0000115|hsa-mir-16-2&WithFlank&3|+|160404739|160404831@58.79.22&[MIMAT0004518&hsa-miR-16-2-3p&offsets|-1|-1;m-252&3|+|160404796|160404816&offsets|0|+1]|ACCAATATTACTGTGCTGCTTT</td><td>>MI0000115|hsa-mir-16-2&WithFlank&3|+|160404739|160404831</td><td>58</td><td>79</td><td>22</td><td>ACCAATATTACTGTGCTGCTTT</td><td>1183.9</td><td>5.1</td><td>0.9</td><td>5.5</td><td>4.1E-08</td><td>7.3E-08</td><td>MB231/MB468</td><td>MITO</td></tr>
<tr><td>>MI0000283|hsa-mir-203a&WithFlank&14|+|104117399|104117520@72.92.21(+1U)&[MIMAT0000264&hsa-miR-203a-3p&offsets|+1|0(+1U);m-79&14|+|104117469|104117490&offsets|+1|0(+1U)]|TGAAATGTTTAGGACCACTAGT</td><td>>MI0000283|hsa-mir-203a&WithFlank&14|+|104117399|104117520</td><td>72</td><td>92</td><td>21(+1U)</td><td>TGAAATGTTTAGGACCACTAGT</td><td>181.5</td><td>10.8</td><td>1.8</td><td>6.1</td><td>1.3E-09</td><td>7.3E-08</td><td>BT20/MB231</td><td>MP</td></tr>
<tr><td>>hg38_21_+_8433172_8446622_RNA45SN1_45S_50ntFlanks@8423.8471.49|CGCAGTCCGCCCGGAGGATTCAACCCGGCGGCGGGTCCGGCCGTGTCGG</td><td>>hg38_21_+_8433172_8446622_RNA45SN1_45S_50ntFlanks</td><td>8423</td><td>8471</td><td>49</td><td>CGCAGTCCGCCCGGAGGATTCAACCCGGCGGCGGGTCCGGCCGTGTCGG</td><td>379.3</td><td>-11.6</td><td>2.0</td><td>-5.8</td><td>6.4E-09</td><td>7.3E-08</td><td>BT20/MB468</td><td>MP</td></tr>
<tr><td>>hg38_21_+_8433172_8446622_RNA45SN1_45S_50ntFlanks@4735.4761.27|ACGAAAGTCGGAGGTTCGAAGACGATC</td><td>>hg38_21_+_8433172_8446622_RNA45SN1_45S_50ntFlanks</td><td>4735</td><td>4761</td><td>27</td><td>ACGAAAGTCGGAGGTTCGAAGACGATC</td><td>834.1</td><td>6.3</td><td>1.1</td><td>5.5</td><td>3.2E-08</td><td>7.3E-08</td><td>BT20/MB468</td><td>CYTO</td></tr>
<tr><td>>hg38_21_+_8433172_8446622_RNA45SN1_45S_50ntFlanks@11873.11904.32|AAGAAGACCCTGTTGAGCTTGACTCTAGTCTG</td><td>>hg38_21_+_8433172_8446622_RNA45SN1_45S_50ntFlanks</td><td>11873</td><td>11904</td><td>32</td><td>AAGAAGACCCTGTTGAGCTTGACTCTAGTCTG</td><td>300.7</td><td>-6.6</td><td>1.2</td><td>-5.5</td><td>3.2E-08</td><td>7.3E-08</td><td>BT20/MB468</td><td>CYTO</td></tr>
<tr><td>>hg38_21_+_8433172_8446622_RNA45SN1_45S_50ntFlanks@9621.9647.27|TCTAGTAGCTGGTTCCCTCCGAAGTTT</td><td>>hg38_21_+_8433172_8446622_RNA45SN1_45S_50ntFlanks</td><td>9621</td><td>9647</td><td>27</td><td>TCTAGTAGCTGGTTCCCTCCGAAGTTT</td><td>219.7</td><td>-7.2</td><td>1.3</td><td>-5.5</td><td>4.1E-08</td><td>7.3E-08</td><td>MB231/MB468</td><td>MITO</td></tr>
<tr><td>>hg38_21_+_8433172_8446622_RNA45SN1_45S_50ntFlanks@7988.8014.27|CAGACGTGGCGACCCGCTGAATTTAAG</td><td>>hg38_21_+_8433172_8446622_RNA45SN1_45S_50ntFlanks</td><td>7988</td><td>8014</td><td>27</td><td>CAGACGTGGCGACCCGCTGAATTTAAG</td><td>1002.6</td><td>-3.6</td><td>0.7</td><td>-5.5</td><td>2.9E-08</td><td>7.3E-08</td><td>BT20/MB468</td><td>MITO</td></tr>
<tr><td>>am_trna11_ProAGG_16_+_3241989_3242060@1.28.28,trna12_ProAGG_6_+_26555498_26555569@1.28.28,trna12_ProTGG_11_-_75946869_75946940@1.28.28,trna14_ProTGG_5_-_180615854_180615925@1.28.28,trna22_ProAGG_14_-_21081560_21081631@1.28.28,trna23_ProAGG_14_-_21077495_21077566@1.28.28,trna28_ProTGG_16_-_3234133_3234204@1.28.28,trna29_ProAGG_16_-_3232635_3232706@1.28.28,trna2_ProAGG_7_+_128423504_128423575@1.28.28,trna30_ProCGG_6_+_27059521_27059592@1.28.28,trna37_ProCGG_17_-_8126151_8126222@1.28.28,trna3_ProTGG_16_+_3208923_3208994@1.28.28,trna52_ProCGG_1_+_167683962_167684033@1.28.28,trna65_ProAGG_1_-_167684725_167684796@1.28.28,trna6_ProCGG_16_+_3222049_3222120@1.28.28,trna6_ProTGG_14_+_21152175_21152246@1.28.28,trna8_ProTGG_16_+_3238094_3238165@1.28.28,trna9_ProAGG_11_+_75946557_75946628@1.28.28,trna9_ProAGG_16_+_3239634_3239705@1.28.28|GGCTCGTTGGTCTAGGGGTATGATTCTC</td><td>>am_trna11_ProAGG_16_+_3241989_3242060</td><td>1</td><td>28</td><td>28</td><td>GGCTCGTTGGTCTAGGGGTATGATTCTC</td><td>3031.3</td><td>-6.3</td><td>1.1</td><td>-5.5</td><td>3.8E-08</td><td>7.3E-08</td><td>MB231/MB468</td><td>CYTO</td></tr>
<tr><td>>hg38_21_+_8433172_8446622_RNA45SN1_45S_50ntFlanks@5484.5512.29|GAGCGCTGAGAAGACGGTCGAACTTGACT</td><td>>hg38_21_+_8433172_8446622_RNA45SN1_45S_50ntFlanks</td><td>5484</td><td>5512</td><td>29</td><td>GAGCGCTGAGAAGACGGTCGAACTTGACT</td><td>348.7</td><td>-5.9</td><td>1.1</td><td>-5.5</td><td>3.5E-08</td><td>7.3E-08</td><td>MB231/MB468</td><td>MP</td></tr>
<tr><td>>hg38_21_+_8433172_8446622_RNA45SN1_45S_50ntFlanks@5073.5113.41|AATTCCGATAACGAACGAGACTCTGGCATGCTAACTAGTTA</td><td>>hg38_21_+_8433172_8446622_RNA45SN1_45S_50ntFlanks</td><td>5073</td><td>5113</td><td>41</td><td>AATTCCGATAACGAACGAGACTCTGGCATGCTAACTAGTTA</td><td>609.3</td><td>-8.5</td><td>1.6</td><td>-5.5</td><td>4.3E-08</td><td>7.3E-08</td><td>BT20/MB468</td><td>NUC</td></tr>
<tr><td>>hg38_21_+_8433172_8446622_RNA45SN1_45S_50ntFlanks@7983.8003.21|CAGATCAGACGTGGCGACCCG</td><td>>hg38_21_+_8433172_8446622_RNA45SN1_45S_50ntFlanks</td><td>7983</td><td>8003</td><td>21</td><td>CAGATCAGACGTGGCGACCCG</td><td>819.7</td><td>-7.6</td><td>1.4</td><td>-5.5</td><td>4.1E-08</td><td>7.3E-08</td><td>MB231/MB468</td><td>MITO</td></tr>
<tr><td>>hg38_21_+_8433172_8446622_RNA45SN1_45S_50ntFlanks@10845.10866.22|ACTTCGGGATAAGGATTGGCTC</td><td>>hg38_21_+_8433172_8446622_RNA45SN1_45S_50ntFlanks</td><td>10845</td><td>10866</td><td>22</td><td>ACTTCGGGATAAGGATTGGCTC</td><td>468.2</td><td>-7.5</td><td>1.4</td><td>-5.5</td><td>3.9E-08</td><td>7.3E-08</td><td>MB231/MB468</td><td>NUC</td></tr>
<tr><td>>hg38_21_+_8433172_8446622_RNA45SN1_45S_50ntFlanks@12997.13014.18|CGCTGCGATCTATTGAAA</td><td>>hg38_21_+_8433172_8446622_RNA45SN1_45S_50ntFlanks</td><td>12997</td><td>13014</td><td>18</td><td>CGCTGCGATCTATTGAAA</td><td>3205.0</td><td>3.8</td><td>0.7</td><td>5.5</td><td>3.2E-08</td><td>7.3E-08</td><td>BT20/MB468</td><td>CYTO</td></tr>
<tr><td>>hg38_21_+_8433172_8446622_RNA45SN1_45S_50ntFlanks@12913.12933.21|ACGTTCGTGGGGAACCTGGCG</td><td>>hg38_21_+_8433172_8446622_RNA45SN1_45S_50ntFlanks</td><td>12913</td><td>12933</td><td>21</td><td>ACGTTCGTGGGGAACCTGGCG</td><td>678.4</td><td>-8.2</td><td>1.5</td><td>-5.5</td><td>3.5E-08</td><td>7.3E-08</td><td>MB231/MB468</td><td>MP</td></tr>
<tr><td>>hg38_21_+_8433172_8446622_RNA45SN1_45S_50ntFlanks@9240.9274.35|GCGGAGCGAGCGCACGGGGTCGGCGGCGACGTCGG</td><td>>hg38_21_+_8433172_8446622_RNA45SN1_45S_50ntFlanks</td><td>9240</td><td>9274</td><td>35</td><td>GCGGAGCGAGCGCACGGGGTCGGCGGCGACGTCGG</td><td>137.9</td><td>-9.2</td><td>1.7</td><td>-5.5</td><td>2.9E-08</td><td>7.3E-08</td><td>BT20/MB468</td><td>MITO</td></tr>
<tr><td>>hg38_21_+_8433172_8446622_RNA45SN1_45S_50ntFlanks@7981.7999.19|CTCAGATCAGACGTGGCGA</td><td>>hg38_21_+_8433172_8446622_RNA45SN1_45S_50ntFlanks</td><td>7981</td><td>7999</td><td>19</td><td>CTCAGATCAGACGTGGCGA</td><td>118.1</td><td>-6.2</td><td>1.1</td><td>-5.5</td><td>3.3E-08</td><td>7.3E-08</td><td>BT20/MB468</td><td>CYTO</td></tr>
<tr><td>>hg38_21_+_8433172_8446622_RNA45SN1_45S_50ntFlanks@9565.9595.31|TGACGTGCAAATCGGTCGTCCGACCTGGGTA</td><td>>hg38_21_+_8433172_8446622_RNA45SN1_45S_50ntFlanks</td><td>9565</td><td>9595</td><td>31</td><td>TGACGTGCAAATCGGTCGTCCGACCTGGGTA</td><td>932.1</td><td>-3.6</td><td>0.7</td><td>-5.5</td><td>3.3E-08</td><td>7.4E-08</td><td>BT20/MB468</td><td>CYTO</td></tr>
<tr><td>>MI0000471|hsa-mir-126&WithFlank&9|+|136670596|136670692@59.77.19&[MIMAT0000445&hsa-miR-126-3p&offsets|+1|-2;m-40&9|+|136670653|136670674&offsets|+1|-2]|CGTACCGTGAGTAATAATG</td><td>>MI0000471|hsa-mir-126&WithFlank&9|+|136670596|136670692</td><td>59</td><td>77</td><td>19</td><td>CGTACCGTGAGTAATAATG</td><td>127.4</td><td>10.7</td><td>1.9</td><td>5.8</td><td>6.4E-09</td><td>7.4E-08</td><td>BT20/MB468</td><td>MP</td></tr>
<tr><td>>hg38_21_+_8433172_8446622_RNA45SN1_45S_50ntFlanks@6719.6738.20|TGAATTGCAGGACACATTGA</td><td>>hg38_21_+_8433172_8446622_RNA45SN1_45S_50ntFlanks</td><td>6719</td><td>6738</td><td>20</td><td>TGAATTGCAGGACACATTGA</td><td>109.8</td><td>-9.9</td><td>1.7</td><td>-5.8</td><td>6.4E-09</td><td>7.4E-08</td><td>BT20/MB468</td><td>MP</td></tr>
<tr><td>>hg38_21_+_8433172_8446622_RNA45SN1_45S_50ntFlanks@12637.12659.23|GCTACCATCTGTGGGATTATGAC</td><td>>hg38_21_+_8433172_8446622_RNA45SN1_45S_50ntFlanks</td><td>12637</td><td>12659</td><td>23</td><td>GCTACCATCTGTGGGATTATGAC</td><td>558.3</td><td>-7.0</td><td>1.3</td><td>-5.5</td><td>4.3E-08</td><td>7.4E-08</td><td>BT20/MB468</td><td>NUC</td></tr>
</table>

| | | | | | | | | | | | | | |
|---|---|---|---|---|---|---|---|---|---|---|---|---|---|
| >hg38_21_+_8433172_8446622_RNA45SN1_45S_50ntFlanks@12419.12460.42\|GATGTCGGCTCTTCCTATCATTGTGAAGCAGAATTCACCAAG | >hg38_21_+_8433172_8446622_RNA45SN1_45S_50ntFlanks | 12419 | 12460 | 42 | GATGTCGGCTCTTCCTATCATTGTGAAGCAGAATTCACCAAG | 415.8 | 5.6 | 1.0 | 5.5 | 3.8E-08 | 7.4E-08 | MB231/MB468 | CYTO |
| >hg38_21_+_8433172_8446622_RNA45SN1_45S_50ntFlanks@12163.12182.20\|ACTGGGGCGGTACACCTGTC | >hg38_21_+_8433172_8446622_RNA45SN1_45S_50ntFlanks | 12163 | 12182 | 20 | ACTGGGGCGGTACACCTGTC | 1428.3 | -9.0 | 1.7 | -5.5 | 4.3E-08 | 7.4E-08 | BT20/MB468 | NUC |
| >hg38_21_+_8433172_8446622_RNA45SN1_45S_50ntFlanks@11923.11957.35\|AGAGGTGTAGAATAAGTGGGAGGCCCCCGGCGCCC | >hg38_21_+_8433172_8446622_RNA45SN1_45S_50ntFlanks | 11923 | 11957 | 35 | AGAGGTGTAGAATAAGTGGGAGGCCCCCGGCGCCC | 60.7 | -9.8 | 1.8 | -5.5 | 3.6E-08 | 7.4E-08 | MB231/MB468 | MP |
| >MI0003591\|hsa-mir-584&WithFlank&5\|-\|149062307\|149062415@22.42.21&[MIMAT0003249&hsa-miR-584-5p&offsets\|0\|-1;m-167&5\|-\|149062374\|149062394&offsets\|0\|0]\|TTATGGTTTGCCTGGGACTGA | >MI0003591\|hsa-mir-584&WithFlank&5\|-\|149062307\|149062415 | 22 | 42 | 21 | TTATGGTTTGCCTGGGACTGA | 6787.2 | 6.1 | 1.1 | 5.8 | 6.5E-09 | 7.4E-08 | BT20/MB468 | MP |
| >hg38_21_+_8433172_8446622_RNA45SN1_45S_50ntFlanks@8189.8212.24\|CCCAGCCCGTGGACGGTGTGAGGC | >hg38_21_+_8433172_8446622_RNA45SN1_45S_50ntFlanks | 8189 | 8212 | 24 | CCCAGCCCGTGGACGGTGTGAGGC | 387.5 | -7.4 | 1.4 | -5.5 | 4.2E-08 | 7.4E-08 | MB231/MB468 | MITO |
| >hg38_21_+_8433172_8446622_RNA45SN1_45S_50ntFlanks@10387.10410.24\|TGTGAACAGCAGTTGAACATGGGT | >hg38_21_+_8433172_8446622_RNA45SN1_45S_50ntFlanks | 10387 | 10410 | 24 | TGTGAACAGCAGTTGAACATGGGT | 440.2 | -7.9 | 1.4 | -5.5 | 3.6E-08 | 7.4E-08 | MB231/MB468 | MP |
| >MI0003560\|hsa-mir-92b&WithFlank&1\|+\|155195171\|155195278@67.87.21&[MIMAT0003218&hsa-miR-92b-3p&offsets\|0\|-1;m-115&1\|+\|155195237\|155195258&offsets\|0\|-1]\|TATTGCACTCGTCCCGGCCTC | >MI0003560\|hsa-mir-92b&WithFlank&1\|+\|155195171\|155195278 | 67 | 87 | 21 | TATTGCACTCGTCCCGGCCTC | 656.5 | 7.7 | 1.4 | 5.5 | 4.2E-08 | 7.4E-08 | MB231/MB468 | MITO |
| >hg38_21_+_8433172_8446622_RNA45SN1_45S_50ntFlanks@10420.10456.37\|CTGAGAGATGGGCGAGCGCCGTTCCGAAGGGACGGGC | >hg38_21_+_8433172_8446622_RNA45SN1_45S_50ntFlanks | 10420 | 10456 | 37 | CTGAGAGATGGGCGAGCGCCGTTCCGAAGGGACGGGC | 3564.8 | -3.8 | 0.7 | -5.5 | 3.3E-08 | 7.4E-08 | BT20/MB468 | CYTO |
| >MI0001445\|hsa-mir-423&WithFlank&17\|+\|30117073\|30117178@59.80.22&[MIMAT0001340&hsa-miR-423-3p&offsets\|0\|-1;m-100&17\|+\|30117131\|30117153&offsets\|0\|-1]\|AGCTCGGTCTGAGGCCCCTCAG | >MI0001445\|hsa-mir-423&WithFlank&17\|+\|30117073\|30117178 | 59 | 80 | 22 | AGCTCGGTCTGAGGCCCCTCAG | 1171.7 | 5.0 | 0.9 | 5.5 | 3.9E-08 | 7.4E-08 | MB231/MB468 | NUC |
| >MI0000273\|hsa-mir-183&WithFlank&7\|-\|129774899\|129775020@33.53.21(+1G)&[MIMAT0000261&hsa-miR-183-5p&offsets\|0\|-1(+1G);m-41&7\|-\|129774966\|129774987&offsets\|-1\|-2(+1G)]\|TATGGCACTGGTAGAATTCACG | >MI0000273\|hsa-mir-183&WithFlank&7\|-\|129774899\|129775020 | 33 | 53 | 21(+1G) | TATGGCACTGGTAGAATTCACG | 109.4 | 2.2 | 0.4 | 5.5 | 4.3E-08 | 7.4E-08 | BT20/MB468 | NUC |
| >am_trnaMT_ValTAC_MT_+_1602_1670@43.67.25\|GATTTCAACTTAACTTGACCGCTCT | >am_trnaMT_ValTAC_MT_+_1602_1670 | 43 | 67 | 25 | GATTTCAACTTAACTTGACCGCTCT | 569.0 | 8.0 | 1.4 | 5.5 | 3.3E-08 | 7.4E-08 | BT20/MB468 | CYTO |
| >hg38_21_+_8433172_8446622_RNA45SN1_45S_50ntFlanks@10838.10865.28\|ATCCGTAACTTCGGGATAAGGATTGGCT | >hg38_21_+_8433172_8446622_RNA45SN1_45S_50ntFlanks | 10838 | 10865 | 28 | ATCCGTAACTTCGGGATAAGGATTGGCT | 350.6 | -7.4 | 1.3 | -5.5 | 3.6E-08 | 7.4E-08 | MB231/MB468 | MP |
| >hg38_21_+_8433172_8446622_RNA45SN1_45S_50ntFlanks@11670.11703.34\|CCCAGTGCTCTGAATGTCAAAGTGAAGAAATTCA | >hg38_21_+_8433172_8446622_RNA45SN1_45S_50ntFlanks | 11670 | 11703 | 34 | CCCAGTGCTCTGAATGTCAAAGTGAAGAAATTCA | 491.1 | -3.0 | 0.5 | -5.5 | 3.3E-08 | 7.5E-08 | BT20/MB468 | CYTO |
| >hg38_21_+_8433172_8446622_RNA45SN1_45S_50ntFlanks@9568.9597.30\|CGTGCAAATCGGTCGTCCGACCTGGGTATA | >hg38_21_+_8433172_8446622_RNA45SN1_45S_50ntFlanks | 9568 | 9597 | 30 | CGTGCAAATCGGTCGTCCGACCTGGGTATA | 3134.4 | -6.4 | 1.2 | -5.5 | 3.3E-08 | 7.5E-08 | BT20/MB468 | CYTO |
| >hg38_21_+_8433172_8446622_RNA45SN1_45S_50ntFlanks@6690.6718.29\|AAGAACGCAGCTAGCTGCGAGAATTAATG | >hg38_21_+_8433172_8446622_RNA45SN1_45S_50ntFlanks | 6690 | 6718 | 29 | AAGAACGCAGCTAGCTGCGAGAATTAATG | 149.9 | -6.9 | 1.2 | -5.5 | 3.3E-08 | 7.5E-08 | BT20/MB468 | CYTO |
| >hg38_21_+_8433172_8446622_RNA45SN1_45S_50ntFlanks@11583.11601.19\|CTGGTGCGGACCAGGGGAA | >hg38_21_+_8433172_8446622_RNA45SN1_45S_50ntFlanks | 11583 | 11601 | 19 | CTGGTGCGGACCAGGGGAA | 804.6 | -8.3 | 1.4 | -5.8 | 6.5E-09 | 7.5E-08 | BT20/MB468 | MP |
| >hg38_21_+_8433172_8446622_RNA45SN1_45S_50ntFlanks@8168.8208.41\|CCCAAGTCCTTCTGATCGAGGCCCAGCCCGTGGACGGTGTG | >hg38_21_+_8433172_8446622_RNA45SN1_45S_50ntFlanks | 8168 | 8208 | 41 | CCCAAGTCCTTCTGATCGAGGCCCAGCCCGTGGACGGTGTG | 59.7 | -9.8 | 1.8 | -5.5 | 3.6E-08 | 7.5E-08 | MB231/MB468 | MP |
| >hg38_21_+_8433172_8446622_RNA45SN1_45S_50ntFlanks@12933.12972.40\|GCTAAACCATTCGTAGACGACCTGCTTCTGGGTCGGGGTT | >hg38_21_+_8433172_8446622_RNA45SN1_45S_50ntFlanks | 12933 | 12972 | 40 | GCTAAACCATTCGTAGACGACCTGCTTCTGGGTCGGGGTT | 233.9 | -7.5 | 1.4 | -5.5 | 3.3E-08 | 7.5E-08 | BT20/MB468 | CYTO |
| >MI0000440\|hsa-mir-27b&WithFlank&9\|+\|95085439\|95085547@25.46.22&[MIMAT0004588&hsa-miR-27b-5p&offsets\|0\|0;m-240&9\|+\|95085463\|95085484&offsets\|0\|0]\|AGAGCTTAGCTGATTGGTGAAC | >MI0000440\|hsa-mir-27b&WithFlank&9\|+\|95085439\|95085547 | 25 | 46 | 22 | AGAGCTTAGCTGATTGGTGAAC | 571.5 | 8.2 | 1.5 | 5.5 | 3.3E-08 | 7.5E-08 | BT20/MB468 | CYTO |
| >hg38_21_+_8433172_8446622_RNA45SN1_45S_50ntFlanks@10409.10431.23\|GTCAGTCGGTCCTGAGAGATGGG | >hg38_21_+_8433172_8446622_RNA45SN1_45S_50ntFlanks | 10409 | 10431 | 23 | GTCAGTCGGTCCTGAGAGATGGG | 1233.5 | -8.9 | 1.6 | -5.5 | 4.4E-08 | 7.5E-08 | BT20/MB468 | NUC |
| >MI0001445\|hsa-mir-423&WithFlank&17\|+\|30117073\|30117178@59.78.20&[MIMAT0001340&hsa-miR-423-3p&offsets\|0\|-3;m-100&17\|+\|30117131\|30117153&offsets\|0\|-3]\|AGCTCGGTCTGAGGCCCCTC | >MI0001445\|hsa-mir-423&WithFlank&17\|+\|30117073\|30117178 | 59 | 78 | 20 | AGCTCGGTCTGAGGCCCCTC | 2641.5 | 5.9 | 1.0 | 5.8 | 6.6E-09 | 7.5E-08 | BT20/MB468 | MP |
| >hg38_21_+_8433172_8446622_RNA45SN1_45S_50ntFlanks@12615.12634.20\|GCTGAGGAGCCAATGGGGCG | >hg38_21_+_8433172_8446622_RNA45SN1_45S_50ntFlanks | 12615 | 12634 | 20 | GCTGAGGAGCCAATGGGGCG | 91.9 | -9.6 | 1.7 | -5.8 | 6.6E-09 | 7.5E-08 | BT20/MB468 | MP |
| >hg38_21_+_8433172_8446622_RNA45SN1_45S_50ntFlanks@476.501.26\|TTCGTGATCGATGTGGTGACGTCGTG | >hg38_21_+_8433172_8446622_RNA45SN1_45S_50ntFlanks | 476 | 501 | 26 | TTCGTGATCGATGTGGTGACGTCGTG | 609.4 | 5.2 | 0.9 | 5.5 | 4.2E-08 | 7.5E-08 | MB231/MB468 | MITO |
| >MI0000749\|hsa-mir-30e&WithFlank&1\|+\|40754349\|40754452@65.85.21(+1A)&[MIMAT0000693&hsa-miR-30e-3p&offsets\|0\|-1(+1A);m-25&1\|+\|40754413\|40754434&offsets\|0\|-1(+1A)]\|CTTTCAGTCGGATGTTTACAGA | >MI0000749\|hsa-mir-30e&WithFlank&1\|+\|40754349\|40754452 | 65 | 85 | 21(+1A) | CTTTCAGTCGGATGTTTACAGA | 359.5 | 6.3 | 1.1 | 5.5 | 3.3E-08 | 7.5E-08 | BT20/MB468 | CYTO |
| >MI0000108\|hsa-mir-103a-2&WithFlank&20\|+\|3917488\|3917577@54.77.24;MI0000109\|hsa-mir-103a-1&WithFlank&5\|-\|168560890\|168560979@54.77.24&[MIMAT0000101&hsa-miR-103a-3p&offsets\|0\|+1;m-12&20\|+\|3917541\|3917563&offsets\|0\|+1];[MIMAT0000101_1&hsa-miR-103a-3p&offsets\|0\|+1;m-13&5\|-\|168560904\|168560926&offsets\|0\|+1;m-44220&5\|-\|168560908\|168560926&offsets\|0\|+5]\|AGCAGCATTGTACAGGGCTATGAA | >MI0000108\|hsa-mir-103a-2&WithFlank&20\|+\|3917488\|3917577 | 54 | 77 | 24 | AGCAGCATTGTACAGGGCTATGAA | 462.8 | 5.3 | 1.0 | 5.5 | 3.9E-08 | 7.5E-08 | MB231/MB468 | CYTO |
| >hg38_21_+_8433172_8446622_RNA45SN1_45S_50ntFlanks@12933.12956.24\|GCTAAACCATTCGTAGACGACCTG | >hg38_21_+_8433172_8446622_RNA45SN1_45S_50ntFlanks | 12933 | 12956 | 24 | GCTAAACCATTCGTAGACGACCTG | 1220.5 | -3.7 | 0.7 | -5.5 | 3.9E-08 | 7.5E-08 | MB231/MB468 | CYTO |
| >MI0000650\|hsa-mir-200c&WithFlank&12\|+\|6963693\|6963772@50.73.24&[MIMAT0000617&hsa-miR-200c-3p&offsets\|0\|+1;m-54&12\|+\|6963742\|6963764&offsets\|0\|+1]\|TAATACTGCCGGGTAATGATGGAG | >MI0000650\|hsa-mir-200c&WithFlank&12\|+\|6963693\|6963772 | 50 | 73 | 24 | TAATACTGCCGGGTAATGATGGAG | 163.9 | 3.7 | 0.7 | 5.5 | 4.4E-08 | 7.5E-08 | BT20/MB468 | NUC |
| >trnaMT_ValTAC_MT_+_1602_1670@42.72.31\|AGATTTCAACTTAACTTGACCGCTCTGACCA | >trnaMT_ValTAC_MT_+_1602_1670 | 42 | 72 | 31 | AGATTTCAACTTAACTTGACCGCTCTGACCA | 32811.8 | 6.1 | 1.1 | 5.5 | 3.0E-08 | 7.5E-08 | BT20/MB468 | MITO |
| >hg38_21_+_8433172_8446622_RNA45SN1_45S_50ntFlanks@12007.12042.36\|GCCGCCGGTGAAATACCACTACTCTGATCGTTTTTT | >hg38_21_+_8433172_8446622_RNA45SN1_45S_50ntFlanks | 12007 | 12042 | 36 | GCCGCCGGTGAAATACCACTACTCTGATCGTTTTTT | 98.2 | -10.5 | 1.9 | -5.5 | 3.6E-08 | 7.5E-08 | MB231/MB468 | MP |
| >MI0000079\|hsa-mir-23a&WithFlank&19\|-\|13836581\|13836665@51.70.20&[MIMAT0000078&hsa-miR-23a-3p&offsets\|0\|-1;m-31&19\|-\|13836595\|13836615&offsets\|0\|-1]\|ATCACATTGCCAGGGATTTC | >MI0000079\|hsa-mir-23a&WithFlank&19\|-\|13836581\|13836665 | 51 | 70 | 20 | ATCACATTGCCAGGGATTTC | 178.9 | -10.6 | 1.8 | -5.8 | 6.6E-09 | 7.5E-08 | BT20/MB468 | MP |
| >am_trnaMT_PheGAA_MT_+_577_647@43.62.20\|TTAGACGGGCTCACATCACC | >am_trnaMT_PheGAA_MT_+_577_647 | 43 | 62 | 20 | TTAGACGGGCTCACATCACC | 191.4 | 4.6 | 0.8 | 5.5 | 4.0E-08 | 7.5E-08 | MB231/MB468 | NUC |
| >MI0000285\|hsa-mir-205&WithFlank&1\|+\|209432127\|209432248@40.59.20(+1G)&[MIMAT0000266&hsa-miR-205-5p&offsets\|0\|-2(+1G);m-351&1\|+\|209432166\|209432187&offsets\|0\|-2(+1G)]\|TCCTTCATTCCACCGGAGTCG | >MI0000285\|hsa-mir-205&WithFlank&1\|+\|209432127\|209432248 | 40 | 59 | 20(+1G) | TCCTTCATTCCACCGGAGTCG | 64.8 | -9.9 | 1.8 | -5.5 | 3.6E-08 | 7.5E-08 | MB231/MB468 | MP |
| >hg38_21_+_8433172_8446622_RNA45SN1_45S_50ntFlanks@5084.5120.37\|CGAACGAGACTCTGGCATGCTAACTAGTTACGCGACC | >hg38_21_+_8433172_8446622_RNA45SN1_45S_50ntFlanks | 5084 | 5120 | 37 | CGAACGAGACTCTGGCATGCTAACTAGTTACGCGACC | 916.7 | -9.1 | 1.6 | -5.5 | 3.6E-08 | 7.6E-08 | MB231/MB468 | MP |
| >hg38_21_+_8433172_8446622_RNA45SN1_45S_50ntFlanks@11767.11803.37\|TCTAATTAGTGACGCGCATGAATGGATGAACGAGATT | >hg38_21_+_8433172_8446622_RNA45SN1_45S_50ntFlanks | 11767 | 11803 | 37 | TCTAATTAGTGACGCGCATGAATGGATGAACGAGATT | 1521.3 | -9.0 | 1.6 | -5.5 | 4.4E-08 | 7.6E-08 | BT20/MB468 | NUC |
| >MI0000746\|hsa-mir-99b&WithFlank&19\|+\|51692606\|51692687@13.32.20(+1A)&[MIMAT0000689&hsa-miR-99b-5p&offsets\|0\|-2(+1A);m-10&19\|+\|51692618\|51692639&offsets\|0\|-2(+1A)]\|CACCCGTAGAACCGACCTTGA | >MI0000746\|hsa-mir-99b&WithFlank&19\|+\|51692606\|51692687 | 13 | 32 | 20(+1A) | CACCCGTAGAACCGACCTTGA | 279.6 | 5.1 | 0.9 | 5.5 | 4.3E-08 | 7.6E-08 | MB231/MB468 | MITO |
| >MI0000465\|hsa-mir-191&WithFlank&3\|-\|49020612\|49020715@22.44.23(+1A)&[MIMAT0000440&hsa-miR-191-5p&offsets\|0\|0(+1A);m-48&3\|-\|49020672\|49020694&offsets\|0\|0(+1A)]\|CAACGGAATCCCAAAAGCAGCTGA | >MI0000465\|hsa-mir-191&WithFlank&3\|-\|49020612\|49020715 | 22 | 44 | 23(+1A) | CAACGGAATCCCAAAAGCAGCTGA | 524.4 | 5.4 | 1.0 | 5.5 | 4.3E-08 | 7.6E-08 | MB231/MB468 | MITO |
| >hg38_21_+_8433172_8446622_RNA45SN1_45S_50ntFlanks@12893.12932.40\|CCTCGCCCGTCACGCACCGCACGTTCGTGGGGAACCTGGC | >hg38_21_+_8433172_8446622_RNA45SN1_45S_50ntFlanks | 12893 | 12932 | 40 | CCTCGCCCGTCACGCACCGCACGTTCGTGGGGAACCTGGC | 2873.9 | -4.4 | 0.8 | -5.5 | 3.0E-08 | 7.6E-08 | BT20/MB468 | MITO |
| >hg38_21_+_8433172_8446622_RNA45SN1_45S_50ntFlanks@12932.12973.42\|CGCTAAACCATTCGTAGACGACCTGCTTCTGGGTCGGGGTTT | >hg38_21_+_8433172_8446622_RNA45SN1_45S_50ntFlanks | 12932 | 12973 | 42 | CGCTAAACCATTCGTAGACGACCTGCTTCTGGGTCGGGGTTT | 93.2 | -10.4 | 1.9 | -5.5 | 3.7E-08 | 7.6E-08 | MB231/MB468 | MP |
| >hg38_21_+_8433172_8446622_RNA45SN1_45S_50ntFlanks@4291.4317.27\|AGGATCCATTGGAGGGCAAGTCTGGTG | >hg38_21_+_8433172_8446622_RNA45SN1_45S_50ntFlanks | 4291 | 4317 | 27 | AGGATCCATTGGAGGGCAAGTCTGGTG | 454.5 | -6.4 | 1.2 | -5.5 | 3.7E-08 | 7.6E-08 | MB231/MB468 | MP |
| >hg38_21_+_8433172_8446622_RNA45SN1_45S_50ntFlanks@12394.12417.24\|TAGCGACGTCGCTTTTTGATCCTT | >hg38_21_+_8433172_8446622_RNA45SN1_45S_50ntFlanks | 12394 | 12417 | 24 | TAGCGACGTCGCTTTTTGATCCTT | 1833.5 | -8.4 | 1.5 | -5.5 | 4.3E-08 | 7.6E-08 | MB231/MB468 | MITO |
| >hg38_21_+_8433172_8446622_RNA45SN1_45S_50ntFlanks@9544.9572.29\|GGTGGAGGTCCGTAGCGGTCCTGACGTGC | >hg38_21_+_8433172_8446622_RNA45SN1_45S_50ntFlanks | 9544 | 9572 | 29 | GGTGGAGGTCCGTAGCGGTCCTGACGTGC | 911.9 | -3.4 | 0.6 | -5.5 | 4.0E-08 | 7.6E-08 | MB231/MB468 | NUC |
| >hg38_21_+_8433172_8446622_RNA45SN1_45S_50ntFlanks@8176.8199.24\|CTTCTGATCGAGGCCCAGCCCGTG | >hg38_21_+_8433172_8446622_RNA45SN1_45S_50ntFlanks | 8176 | 8199 | 24 | CTTCTGATCGAGGCCCAGCCCGTG | 367.2 | -4.1 | 0.7 | -5.5 | 4.3E-08 | 7.6E-08 | MB231/MB468 | MITO |
| >hg38_21_+_8433172_8446622_RNA45SN1_45S_50ntFlanks@12406.12436.31\|TTTTTGATCCTTCGATGTCGGCTCTTCCTAT | >hg38_21_+_8433172_8446622_RNA45SN1_45S_50ntFlanks | 12406 | 12436 | 31 | TTTTTGATCCTTCGATGTCGGCTCTTCCTAT | 433.5 | -7.2 | 1.3 | -5.5 | 4.3E-08 | 7.6E-08 | MB231/MB468 | MITO |

<table>
<tr><td>>am_MI0003776|hsa-mir-320b-1&WithFlank&1|+|116671740|116671823@48.65.18(+1U);MI0003839|hsa-mir-320b-2&WithFlank&1|-|224257034|224257116@47.64.18(+1U);MI0008190|hsa-mir-320d-1&WithFlank&13|-|40727810|40727893@48.65.18(+1U);MI0003778|hsa-mir-320c-1&WithFlank&18|+|21683512|21683595@48.65.18(+1U);MI0008191|hsa-mir-320c-2&WithFlank&18|+|24321669|24321752@48.65.18(+1U);MI0000542|hsa-mir-320a&WithFlank&8|-|22244960|22245043@48.65.18(+1U);MI0008192|hsa-mir-320d-2&WithFlank&X|-|140926154|140926237@48.65.18(+1U)&[MIMAT0005792&hsa-miR-320b&offsets|0|-4(+1U);m-183&1|+|116671787|116671806&offsets|0|-2(+1U)];[MIMAT0005792_1&hsa-miR-320b&offsets|0|-4(+1U);m-183&1|-|224257051|224257070&offsets|0|-2(+1U)];[MIMAT0006764&hsa-miR-320d&offsets|0|-1(+1U);m-246&13|-|40727829|40727846&offsets|0|0(+1U)];[MIMAT0005793&hsa-miR-320c&offsets|0|-2(+1U);m-222&18|+|21683559|21683578&offsets|0|-2(+1U)];[MIMAT0005793_1&hsa-miR-320c&offsets|0|-2(+1U);m-219&18|+|24321716|24321735&offsets|0|-2(+1U)];[MIMAT0000510&hsa-miR-320a-3p&offsets|0|-4(+1U);m-60&8|-|22244975|22244996&offsets|0|-4(+1U)];[MIMAT0006764_1&hsa-miR-320d&offsets|0|-1(+1U);m-253&X|-|140926173|140926190&offsets|0|0(+1U)]|AAAAGCTGGGTTGAGAGGT</td><td>>am_MI0003776|hsa-mir-320b-1&WithFlank&1|+|116671740|116671823</td><td>48</td><td>65</td><td>18(+1U)</td><td>AAAAGCTGGGTTGAGAGGT</td><td>136.2</td><td>10.8</td><td>1.9</td><td>5.8</td><td>6.7E-09</td><td>7.6E-08</td><td>BT20/MB468</td><td>MP</td></tr>
<tr><td>>MI0000063|hsa-let-7b&WithFlank&22|+|46113680|46113774@12.32.21(+1C)&[MIMAT0000063&hsa-let-7b-5p&offsets|0|-1(+1C);m-14&22|+|46113691|46113712&offsets|0|-1(+1C)]|TGAGGTAGTAGGTTGTGTGGTC</td><td>>MI0000063|hsa-let-7b&WithFlank&22|+|46113680|46113774</td><td>12</td><td>32</td><td>21(+1C)</td><td>TGAGGTAGTAGGTTGTGTGGTC</td><td>7377.1</td><td>3.1</td><td>0.5</td><td>5.9</td><td>4.2E-09</td><td>7.6E-08</td><td>BT20/MB231</td><td>NUC</td></tr>
<tr><td>>MI0000462|hsa-mir-152&WithFlank&17|-|48037155|48037253@60.81.22&[MIMAT0000438&hsa-miR-152-3p&offsets|0|+1;m-97&17|-|48037174|48037194&offsets|0|+1]|TCAGTGCATGACAGAACTTGGG</td><td>>MI0000462|hsa-mir-152&WithFlank&17|-|48037155|48037253</td><td>60</td><td>81</td><td>22</td><td>TCAGTGCATGACAGAACTTGGG</td><td>779.9</td><td>7.8</td><td>1.4</td><td>5.5</td><td>3.7E-08</td><td>7.6E-08</td><td>MB231/MB468</td><td>MP</td></tr>
<tr><td>>hg38_21_+_8433172_8446622_RNA45SN1_45S_50ntFlanks@12423.12456.34|TCGGCTCTTCCTATCATTGTGAAGCAGAATTCAC</td><td>>hg38_21_+_8433172_8446622_RNA45SN1_45S_50ntFlanks</td><td>12423</td><td>12456</td><td>34</td><td>TCGGCTCTTCCTATCATTGTGAAGCAGAATTCAC</td><td>622.4</td><td>-8.6</td><td>1.6</td><td>-5.5</td><td>4.5E-08</td><td>7.6E-08</td><td>BT20/MB468</td><td>NUC</td></tr>
<tr><td>>MI0000102|hsa-mir-100&WithFlank&11|-|122152223|122152314@20.40.21(+1U)&[MIMAT0000098&hsa-miR-100-5p&offsets|+1|0(+1U);m-27&11|-|122152276|122152296&offsets|+1|+1(+1U)]|ACCCGTAGATCCGAACTTGTGT</td><td>>MI0000102|hsa-mir-100&WithFlank&11|-|122152223|122152314</td><td>20</td><td>40</td><td>21(+1U)</td><td>ACCCGTAGATCCGAACTTGTGT</td><td>63.5</td><td>9.2</td><td>1.7</td><td>5.5</td><td>3.7E-08</td><td>7.6E-08</td><td>MB231/MB468</td><td>MP</td></tr>
<tr><td>>hg38_21_+_8433172_8446622_RNA45SN1_45S_50ntFlanks@5492.5531.40|AGAAGACGGTCGAACTTGACTATCTAGAGGAAGTAAAAGT</td><td>>hg38_21_+_8433172_8446622_RNA45SN1_45S_50ntFlanks</td><td>5492</td><td>5531</td><td>40</td><td>AGAAGACGGTCGAACTTGACTATCTAGAGGAAGTAAAAGT</td><td>380.9</td><td>6.4</td><td>1.2</td><td>5.5</td><td>3.4E-08</td><td>7.6E-08</td><td>BT20/MB468</td><td>CYTO</td></tr>
<tr><td>>hg38_21_+_8433172_8446622_RNA45SN1_45S_50ntFlanks@10408.10426.19|GGTCAGTCGGTCCTGAGAG</td><td>>hg38_21_+_8433172_8446622_RNA45SN1_45S_50ntFlanks</td><td>10408</td><td>10426</td><td>19</td><td>GGTCAGTCGGTCCTGAGAG</td><td>769.2</td><td>-5.4</td><td>1.0</td><td>-5.5</td><td>3.9E-08</td><td>7.6E-08</td><td>MB231/MB468</td><td>CYTO</td></tr>
<tr><td>>hg38_21_+_8433172_8446622_RNA45SN1_45S_50ntFlanks@8328.8375.48|TAGTCAACAAGTACCGTAAGGGAAAGTTGAAAAGAACTTTGAAGAGAG</td><td>>hg38_21_+_8433172_8446622_RNA45SN1_45S_50ntFlanks</td><td>8328</td><td>8375</td><td>48</td><td>TAGTCAACAAGTACCGTAAGGGAAAGTTGAAAAGAACTTTGAAGAGAG</td><td>102.9</td><td>10.4</td><td>1.8</td><td>5.8</td><td>6.7E-09</td><td>7.6E-08</td><td>BT20/MB468</td><td>MP</td></tr>
<tr><td>>hg38_21_+_8433172_8446622_RNA45SN1_45S_50ntFlanks@4723.4751.29|CATTAATCAAGAACGAAAGTCGGAGGTTC</td><td>>hg38_21_+_8433172_8446622_RNA45SN1_45S_50ntFlanks</td><td>4723</td><td>4751</td><td>29</td><td>CATTAATCAAGAACGAAAGTCGGAGGTTC</td><td>502.5</td><td>-4.2</td><td>0.8</td><td>-5.5</td><td>3.4E-08</td><td>7.6E-08</td><td>BT20/MB468</td><td>CYTO</td></tr>
<tr><td>>hg38_21_+_8433172_8446622_RNA45SN1_45S_50ntFlanks@10783.10816.34|GTGAACAGCCTCTGGCATGTTGGAACAATGTAGG</td><td>>hg38_21_+_8433172_8446622_RNA45SN1_45S_50ntFlanks</td><td>10783</td><td>10816</td><td>34</td><td>GTGAACAGCCTCTGGCATGTTGGAACAATGTAGG</td><td>326.8</td><td>-5.2</td><td>0.9</td><td>-5.5</td><td>3.9E-08</td><td>7.6E-08</td><td>MB231/MB468</td><td>CYTO</td></tr>
<tr><td>>hg38_21_+_8433172_8446622_RNA45SN1_45S_50ntFlanks@10873.10892.20|CTGGGTCGGTCGGGCTGGGG</td><td>>hg38_21_+_8433172_8446622_RNA45SN1_45S_50ntFlanks</td><td>10873</td><td>10892</td><td>20</td><td>CTGGGTCGGTCGGGCTGGGG</td><td>1108.8</td><td>-4.7</td><td>0.8</td><td>-5.8</td><td>6.7E-09</td><td>7.7E-08</td><td>BT20/MB468</td><td>MP</td></tr>
<tr><td>>hg38_21_+_8433172_8446622_RNA45SN1_45S_50ntFlanks@7988.8012.25|CAGACGTGGCGACCCGCTGAATTTA</td><td>>hg38_21_+_8433172_8446622_RNA45SN1_45S_50ntFlanks</td><td>7988</td><td>8012</td><td>25</td><td>CAGACGTGGCGACCCGCTGAATTTA</td><td>4663.1</td><td>-5.6</td><td>1.0</td><td>-5.8</td><td>6.7E-09</td><td>7.7E-08</td><td>BT20/MB468</td><td>MP</td></tr>
<tr><td>>hg38_21_+_8433172_8446622_RNA45SN1_45S_50ntFlanks@9141.9163.23|CGGGGCGGACTGTCCCCAGTGCG</td><td>>hg38_21_+_8433172_8446622_RNA45SN1_45S_50ntFlanks</td><td>9141</td><td>9163</td><td>23</td><td>CGGGGCGGACTGTCCCCAGTGCG</td><td>97.2</td><td>-9.7</td><td>1.7</td><td>-5.8</td><td>6.7E-09</td><td>7.7E-08</td><td>BT20/MB468</td><td>MP</td></tr>
<tr><td>>hg38_21_+_8433172_8446622_RNA45SN1_45S_50ntFlanks@5481.5516.36|GCGGAGCGCTGAGAAGACGGTCGAACTTGACTATCT</td><td>>hg38_21_+_8433172_8446622_RNA45SN1_45S_50ntFlanks</td><td>5481</td><td>5516</td><td>36</td><td>GCGGAGCGCTGAGAAGACGGTCGAACTTGACTATCT</td><td>122.5</td><td>10.7</td><td>1.8</td><td>5.8</td><td>6.8E-09</td><td>7.7E-08</td><td>BT20/MB468</td><td>MP</td></tr>
<tr><td>>hg38_21_+_8433172_8446622_RNA45SN1_45S_50ntFlanks@12622.12653.32|AGCCAATGGGGCGAAGCTACCATCTGTGGGAT</td><td>>hg38_21_+_8433172_8446622_RNA45SN1_45S_50ntFlanks</td><td>12622</td><td>12653</td><td>32</td><td>AGCCAATGGGGCGAAGCTACCATCTGTGGGAT</td><td>96.3</td><td>-9.7</td><td>1.7</td><td>-5.8</td><td>6.8E-09</td><td>7.7E-08</td><td>BT20/MB468</td><td>MP</td></tr>
<tr><td>>hg38_21_+_8433172_8446622_RNA45SN1_45S_50ntFlanks@12939.12966.28|CCATTCGTAGACGACCTGCTTCTGGGTC</td><td>>hg38_21_+_8433172_8446622_RNA45SN1_45S_50ntFlanks</td><td>12939</td><td>12966</td><td>28</td><td>CCATTCGTAGACGACCTGCTTCTGGGTC</td><td>127.4</td><td>-10.1</td><td>1.7</td><td>-5.8</td><td>6.8E-09</td><td>7.7E-08</td><td>BT20/MB468</td><td>MP</td></tr>
<tr><td>>hg38_21_+_8433172_8446622_RNA45SN1_45S_50ntFlanks@7974.8017.44|ACGCGACCTCAGATCAGACGTGGCGACCCGCTGAATTTAAGCAT</td><td>>hg38_21_+_8433172_8446622_RNA45SN1_45S_50ntFlanks</td><td>7974</td><td>8017</td><td>44</td><td>ACGCGACCTCAGATCAGACGTGGCGACCCGCTGAATTTAAGCAT</td><td>283.4</td><td>5.7</td><td>1.0</td><td>5.5</td><td>4.0E-08</td><td>7.7E-08</td><td>MB231/MB468</td><td>CYTO</td></tr>
<tr><td>>hg38_21_+_8433172_8446622_RNA45SN1_45S_50ntFlanks@12417.12452.36|TCGATGTCGGCTCTTCCTATCATTGTGAAGCAGAAT</td><td>>hg38_21_+_8433172_8446622_RNA45SN1_45S_50ntFlanks</td><td>12417</td><td>12452</td><td>36</td><td>TCGATGTCGGCTCTTCCTATCATTGTGAAGCAGAAT</td><td>393.2</td><td>3.8</td><td>0.7</td><td>5.5</td><td>4.0E-08</td><td>7.7E-08</td><td>MB231/MB468</td><td>CYTO</td></tr>
<tr><td>>hg38_21_+_8433172_8446622_RNA45SN1_45S_50ntFlanks@10794.10816.23|CTGGCATGTTGGAACAATGTAGG</td><td>>hg38_21_+_8433172_8446622_RNA45SN1_45S_50ntFlanks</td><td>10794</td><td>10816</td><td>23</td><td>CTGGCATGTTGGAACAATGTAGG</td><td>714.9</td><td>-4.1</td><td>0.8</td><td>-5.5</td><td>4.0E-08</td><td>7.7E-08</td><td>MB231/MB468</td><td>CYTO</td></tr>
<tr><td>>hg38_21_+_8433172_8446622_RNA45SN1_45S_50ntFlanks@5374.5395.22|CGTTGATTAAGTCCCTGCCCTT</td><td>>hg38_21_+_8433172_8446622_RNA45SN1_45S_50ntFlanks</td><td>5374</td><td>5395</td><td>22</td><td>CGTTGATTAAGTCCCTGCCCTT</td><td>252.6</td><td>-10.1</td><td>1.8</td><td>-5.5</td><td>3.0E-08</td><td>7.7E-08</td><td>BT20/MB468</td><td>MITO</td></tr>
<tr><td>>hg38_21_+_8433172_8446622_RNA45SN1_45S_50ntFlanks@10590.10608.19|CGGGGAGAGTTCTCTTTTC</td><td>>hg38_21_+_8433172_8446622_RNA45SN1_45S_50ntFlanks</td><td>10590</td><td>10608</td><td>19</td><td>CGGGGAGAGTTCTCTTTTC</td><td>99.9</td><td>-9.7</td><td>1.7</td><td>-5.8</td><td>6.8E-09</td><td>7.7E-08</td><td>BT20/MB468</td><td>MP</td></tr>
<tr><td>>am_trna128_GlyGCC_6_-_27870686_27870756@1.28.28,trna133_GlyCCC_1_-_16872434_16872504@1.28.28,trna18_GlyGCC_16_+_70822597_70822667@1.28.28,trna19_GlyGCC_16_+_70823410_70823480@1.28.28,trna19_GlyGCC_2_-_157257659_157257729@1.28.28,trna24_GlyGCC_16_-_70812942_70813012@1.28.28,trna25_GlyGCC_16_-_70812114_70812184@1.28.28,trna4_GlyCCC_1_+_17188416_17188486@1.28.28,trna5_GlyGCC_17_+_8029064_8029134@1.28.28,trna68_GlyGCC_1_-_161493637_161493707@1.28.28|GCATTGGTGGTTCAGTGGTAGAATTCTC</td><td>>am_trna128_GlyGCC_6_-_27870686_27870756</td><td>1</td><td>28</td><td>28</td><td>GCATTGGTGGTTCAGTGGTAGAATTCTC</td><td>14135.5</td><td>-4.6</td><td>0.8</td><td>-5.5</td><td>4.3E-08</td><td>7.7E-08</td><td>MB231/MB468</td><td>MITO</td></tr>
<tr><td>>hg38_21_+_8433172_8446622_RNA45SN1_45S_50ntFlanks@4763.4780.18|GATACCGTCGTAGTTCCG</td><td>>hg38_21_+_8433172_8446622_RNA45SN1_45S_50ntFlanks</td><td>4763</td><td>4780</td><td>18</td><td>GATACCGTCGTAGTTCCG</td><td>341.8</td><td>-6.5</td><td>1.2</td><td>-5.5</td><td>3.1E-08</td><td>7.7E-08</td><td>BT20/MB468</td><td>MITO</td></tr>
<tr><td>>hg38_21_+_8433172_8446622_RNA45SN1_45S_50ntFlanks@11199.11224.26|CCGGCGGCGGCGGCGACTCTGGACGC</td><td>>hg38_21_+_8433172_8446622_RNA45SN1_45S_50ntFlanks</td><td>11199</td><td>11224</td><td>26</td><td>CCGGCGGCGGCGGCGACTCTGGACGC</td><td>629.6</td><td>-8.6</td><td>1.6</td><td>-5.5</td><td>4.5E-08</td><td>7.7E-08</td><td>BT20/MB468</td><td>NUC</td></tr>
<tr><td>>hg38_21_+_8433172_8446622_RNA45SN1_45S_50ntFlanks@4046.4063.18|CATTCGAACGTCTGCCCT</td><td>>hg38_21_+_8433172_8446622_RNA45SN1_45S_50ntFlanks</td><td>4046</td><td>4063</td><td>18</td><td>CATTCGAACGTCTGCCCT</td><td>4084.5</td><td>-4.6</td><td>0.8</td><td>-5.5</td><td>4.0E-08</td><td>7.7E-08</td><td>MB231/MB468</td><td>CYTO</td></tr>
<tr><td>>hg38_21_+_8433172_8446622_RNA45SN1_45S_50ntFlanks@11767.11790.24|TCTAATTAGTGACGCGCATGAATG</td><td>>hg38_21_+_8433172_8446622_RNA45SN1_45S_50ntFlanks</td><td>11767</td><td>11790</td><td>24</td><td>TCTAATTAGTGACGCGCATGAATG</td><td>699.7</td><td>-8.4</td><td>1.5</td><td>-5.5</td><td>4.5E-08</td><td>7.7E-08</td><td>BT20/MB468</td><td>NUC</td></tr>
<tr><td>>hg38_21_+_8433172_8446622_RNA45SN1_45S_50ntFlanks@12488.12522.35|AACGTGAGCTGGGTTTAGACCGTCGTGAGACAGGT</td><td>>hg38_21_+_8433172_8446622_RNA45SN1_45S_50ntFlanks</td><td>12488</td><td>12522</td><td>35</td><td>AACGTGAGCTGGGTTTAGACCGTCGTGAGACAGGT</td><td>889.1</td><td>-8.8</td><td>1.6</td><td>-5.5</td><td>4.5E-08</td><td>7.7E-08</td><td>BT20/MB468</td><td>NUC</td></tr>
<tr><td>>hg38_21_+_8433172_8446622_RNA45SN1_45S_50ntFlanks@11778.11806.29|ACGCGCATGAATGGATGAACGAGATTCCC</td><td>>hg38_21_+_8433172_8446622_RNA45SN1_45S_50ntFlanks</td><td>11778</td><td>11806</td><td>29</td><td>ACGCGCATGAATGGATGAACGAGATTCCC</td><td>417.1</td><td>-8.0</td><td>1.5</td><td>-5.5</td><td>4.5E-08</td><td>7.7E-08</td><td>BT20/MB468</td><td>NUC</td></tr>
<tr><td>>hg38_21_+_8433172_8446622_RNA45SN1_45S_50ntFlanks@12288.12319.32|ACCGTGAAAGCGGGGCCTCACGATCCTTCTGA</td><td>>hg38_21_+_8433172_8446622_RNA45SN1_45S_50ntFlanks</td><td>12288</td><td>12319</td><td>32</td><td>ACCGTGAAAGCGGGGCCTCACGATCCTTCTGA</td><td>780.7</td><td>-6.4</td><td>1.2</td><td>-5.5</td><td>4.3E-08</td><td>7.7E-08</td><td>MB231/MB468</td><td>MITO</td></tr>
<tr><td>>hg38_21_+_8433172_8446622_RNA45SN1_45S_50ntFlanks@10728.10755.28|GAGAGGGTGTAAATCTCGCGCCGGGCCG</td><td>>hg38_21_+_8433172_8446622_RNA45SN1_45S_50ntFlanks</td><td>10728</td><td>10755</td><td>28</td><td>GAGAGGGTGTAAATCTCGCGCCGGGCCG</td><td>91.1</td><td>-9.6</td><td>1.7</td><td>-5.8</td><td>6.8E-09</td><td>7.7E-08</td><td>BT20/MB468</td><td>MP</td></tr>
<tr><td>>hg38_21_+_8433172_8446622_RNA45SN1_45S_50ntFlanks@12358.12375.18|CACAGGGATAACTGGCTT</td><td>>hg38_21_+_8433172_8446622_RNA45SN1_45S_50ntFlanks</td><td>12358</td><td>12375</td><td>18</td><td>CACAGGGATAACTGGCTT</td><td>86.3</td><td>-9.5</td><td>1.6</td><td>-5.8</td><td>6.8E-09</td><td>7.7E-08</td><td>BT20/MB468</td><td>MP</td></tr>
<tr><td>>hg38_21_+_8433172_8446622_RNA45SN1_45S_50ntFlanks@9577.9596.20|CGGTCGTCCGACCTGGGTAT</td><td>>hg38_21_+_8433172_8446622_RNA45SN1_45S_50ntFlanks</td><td>9577</td><td>9596</td><td>20</td><td>CGGTCGTCCGACCTGGGTAT</td><td>124.0</td><td>-10.0</td><td>1.7</td><td>-5.8</td><td>6.8E-09</td><td>7.7E-08</td><td>BT20/MB468</td><td>MP</td></tr>
<tr><td>>hg38_21_+_8433172_8446622_RNA45SN1_45S_50ntFlanks@9621.9648.28|TCTAGTAGCTGGTTCCCTCCGAAGTTTC</td><td>>hg38_21_+_8433172_8446622_RNA45SN1_45S_50ntFlanks</td><td>9621</td><td>9648</td><td>28</td><td>TCTAGTAGCTGGTTCCCTCCGAAGTTTC</td><td>562.3</td><td>-3.9</td><td>0.7</td><td>-5.5</td><td>3.1E-08</td><td>7.7E-08</td><td>BT20/MB468</td><td>MITO</td></tr>
<tr><td>>hg38_21_+_8433172_8446622_RNA45SN1_45S_50ntFlanks@7982.8003.22|TCAGATCAGACGTGGCGACCCG</td><td>>hg38_21_+_8433172_8446622_RNA45SN1_45S_50ntFlanks</td><td>7982</td><td>8003</td><td>22</td><td>TCAGATCAGACGTGGCGACCCG</td><td>329.8</td><td>-6.3</td><td>1.1</td><td>-5.5</td><td>4.5E-08</td><td>7.7E-08</td><td>BT20/MB468</td><td>NUC</td></tr>
<tr><td>>hg38_21_+_8433172_8446622_RNA45SN1_45S_50ntFlanks@5125.5158.34|AGCGGTCGGCGTCCCCCAACTTCTTAGAGGGACA</td><td>>hg38_21_+_8433172_8446622_RNA45SN1_45S_50ntFlanks</td><td>5125</td><td>5158</td><td>34</td><td>AGCGGTCGGCGTCCCCCAACTTCTTAGAGGGACA</td><td>506.5</td><td>6.0</td><td>1.1</td><td>5.5</td><td>4.0E-08</td><td>7.7E-08</td><td>MB231/MB468</td><td>CYTO</td></tr>
<tr><td>>hg38_21_+_8433172_8446622_RNA45SN1_45S_50ntFlanks@9736.9757.22|CCGAAACGATCTCAACCTATTC</td><td>>hg38_21_+_8433172_8446622_RNA45SN1_45S_50ntFlanks</td><td>9736</td><td>9757</td><td>22</td><td>CCGAAACGATCTCAACCTATTC</td><td>201.4</td><td>-5.8</td><td>1.1</td><td>-5.5</td><td>4.0E-08</td><td>7.7E-08</td><td>MB231/MB468</td><td>CYTO</td></tr>
<tr><td>>hg38_21_+_8433172_8446622_RNA45SN1_45S_50ntFlanks@3776.3799.24|CCGGTACAGTGAAACTGCGAATGG</td><td>>hg38_21_+_8433172_8446622_RNA45SN1_45S_50ntFlanks</td><td>3776</td><td>3799</td><td>24</td><td>CCGGTACAGTGAAACTGCGAATGG</td><td>91.4</td><td>-9.6</td><td>1.7</td><td>-5.8</td><td>6.9E-09</td><td>7.7E-08</td><td>BT20/MB468</td><td>MP</td></tr>
<tr><td>>hg38_21_+_8433172_8446622_RNA45SN1_45S_50ntFlanks@8317.8356.40|CACGAGACCGATAGTCAACAAGTACCGTAAGGGAAAGTTG</td><td>>hg38_21_+_8433172_8446622_RNA45SN1_45S_50ntFlanks</td><td>8317</td><td>8356</td><td>40</td><td>CACGAGACCGATAGTCAACAAGTACCGTAAGGGAAAGTTG</td><td>679.9</td><td>-4.1</td><td>0.7</td><td>-5.5</td><td>4.1E-08</td><td>7.7E-08</td><td>MB231/MB468</td><td>NUC</td></tr>
<tr><td>>MI0000286|hsa-mir-210&WithFlank&11|-|568083|568204@72.93.22&[MIMAT0000267&hsa-miR-210-3p&offsets|0|0;m-102&11|-|568112|568133&offsets|0|0]|CTGTGCGTGTGACAGCGGCTGA</td><td>>MI0000286|hsa-mir-210&WithFlank&11|-|568083|568204</td><td>72</td><td>93</td><td>22</td><td>CTGTGCGTGTGACAGCGGCTGA</td><td>195.0</td><td>7.3</td><td>1.3</td><td>5.5</td><td>4.5E-08</td><td>7.7E-08</td><td>BT20/MB468</td><td>NUC</td></tr>
<tr><td>>hg38_MT_+_1621_3279_RNR2_16S_50ntFlanks@729.759.31|GATTAAAACACTGAACTGACAATTAACAGCC</td><td>>hg38_MT_+_1621_3279_RNR2_16S_50ntFlanks</td><td>729</td><td>759</td><td>31</td><td>GATTAAAACACTGAACTGACAATTAACAGCC</td><td>106.9</td><td>10.5</td><td>1.8</td><td>5.8</td><td>6.9E-09</td><td>7.8E-08</td><td>BT20/MB468</td><td>MP</td></tr>
<tr><td>>hg38_21_+_8433172_8446622_RNA45SN1_45S_50ntFlanks@12539.12569.31|ATGATGTGTTGTTGCCATGGTAATCCTGCTC</td><td>>hg38_21_+_8433172_8446622_RNA45SN1_45S_50ntFlanks</td><td>12539</td><td>12569</td><td>31</td><td>ATGATGTGTTGTTGCCATGGTAATCCTGCTC</td><td>1769.3</td><td>-3.9</td><td>0.7</td><td>-5.5</td><td>3.1E-08</td><td>7.8E-08</td><td>BT20/MB468</td><td>MITO</td></tr>
</table>

<table>
<tr><td>>MI0000736|hsa-mir-30c-1&WithFlank&1|+|40757278|40757378@23.45.23(+1C);MI0000254|hsa-mir-30c-2&WithFlank&6|-|71376954|71377037@13.35.23(+1C)&[MIMAT0000244&hsa-miR-30c-5p&offsets|0|0(+1C);m-57&1|+|40757300|40757323&offsets|0|-1(+1C)];[MIMAT0000244_1&hsa-miR-30c-5p&offsets|0|0(+1C);m-56&6|-|71377002|71377025&offsets|0|-1(+1C)]|TGTAAACATCCTACACTCTCAGCC</td><td>>MI0000736|hsa-mir-30c-1&WithFlank&1|+|40757278|40757378</td><td>23</td><td>45</td><td>23(+1C)</td><td>TGTAAACATCCTACACTCTCAGCC</td><td>7594.6</td><td>5.7</td><td>1.0</td><td>5.5</td><td>4.4E-08</td><td>7.8E-08</td><td>MB231/MB468</td><td>MITO</td></tr>
<tr><td>>MI0000457|hsa-mir-141&WithFlank&12|+|6964091|6964197@23.44.22&[MIMAT0004598&hsa-miR-141-5p&offsets|0|0;m-221&12|+|6964113|6964133&offsets|0|+1]|CATCTTCCAGTACAGTGTTGGA</td><td>>MI0000457|hsa-mir-141&WithFlank&12|+|6964091|6964197</td><td>23</td><td>44</td><td>22</td><td>CATCTTCCAGTACAGTGTTGGA</td><td>274.8</td><td>11.4</td><td>1.9</td><td>6.1</td><td>1.4E-09</td><td>7.8E-08</td><td>BT20/MB231</td><td>MP</td></tr>
<tr><td>>hg38_21_+_8433172_8446622_RNA45SN1_45S_50ntFlanks@10344.10380.37|ATTCAAACGAGAACTTTGAAGGCCGAAGTGGAGAAGG</td><td>>hg38_21_+_8433172_8446622_RNA45SN1_45S_50ntFlanks</td><td>10344</td><td>10380</td><td>37</td><td>ATTCAAACGAGAACTTTGAAGGCCGAAGTGGAGAAGG</td><td>92.3</td><td>10.3</td><td>1.8</td><td>5.8</td><td>6.9E-09</td><td>7.8E-08</td><td>BT20/MB468</td><td>MP</td></tr>
<tr><td>>hg38_21_+_8433172_8446622_RNA45SN1_45S_50ntFlanks@10838.10868.31|ATCCGTAACTTCGGGATAAGGATTGGCTCTA</td><td>>hg38_21_+_8433172_8446622_RNA45SN1_45S_50ntFlanks</td><td>10838</td><td>10868</td><td>31</td><td>ATCCGTAACTTCGGGATAAGGATTGGCTCTA</td><td>105.1</td><td>-9.8</td><td>1.7</td><td>-5.8</td><td>6.9E-09</td><td>7.8E-08</td><td>BT20/MB468</td><td>MP</td></tr>
<tr><td>>hg38_21_+_8433172_8446622_RNA45SN1_45S_50ntFlanks@12944.12973.30|CGTAGACGACCTGCTTCTGGGTCGGGGTTT</td><td>>hg38_21_+_8433172_8446622_RNA45SN1_45S_50ntFlanks</td><td>12944</td><td>12973</td><td>30</td><td>CGTAGACGACCTGCTTCTGGGTCGGGGTTT</td><td>107.5</td><td>-9.8</td><td>1.7</td><td>-5.8</td><td>7.0E-09</td><td>7.8E-08</td><td>BT20/MB468</td><td>MP</td></tr>
<tr><td>>MI0000070|hsa-mir-16-1&WithFlank&13|-|50048967|50049067@20.41.22;MI0000115|hsa-mir-16-2&WithFlank&3|+|160404739|160404831@16.37.22&[MIMAT0000069&hsa-miR-16-5p&offsets|0|0;m-62&13|-|50049027|50049048&offsets|0|0];[MIMAT0000069_1&hsa-miR-16-5p&offsets|0|0;m-61&3|+|160404754|160404775&offsets|0|0]|TAGCAGCACGTAAATATTGGCG</td><td>>MI0000070|hsa-mir-16-1&WithFlank&13|-|50048967|50049067</td><td>20</td><td>41</td><td>22</td><td>TAGCAGCACGTAAATATTGGCG</td><td>1837.0</td><td>3.8</td><td>0.7</td><td>5.5</td><td>4.6E-08</td><td>7.8E-08</td><td>BT20/MB468</td><td>NUC</td></tr>
<tr><td>>hg38_21_+_8433172_8446622_RNA45SN1_45S_50ntFlanks@11795.11819.25|AACGAGATTCCCACTGTCCCTACCT</td><td>>hg38_21_+_8433172_8446622_RNA45SN1_45S_50ntFlanks</td><td>11795</td><td>11819</td><td>25</td><td>AACGAGATTCCCACTGTCCCTACCT</td><td>460.5</td><td>-8.2</td><td>1.5</td><td>-5.5</td><td>4.6E-08</td><td>7.8E-08</td><td>BT20/MB468</td><td>NUC</td></tr>
<tr><td>>hg38_21_+_8433172_8446622_RNA45SN1_45S_50ntFlanks@5487.5526.40|CGCTGAGAAGACGGTCGAACTTGACTATCTAGAGGAAGTA</td><td>>hg38_21_+_8433172_8446622_RNA45SN1_45S_50ntFlanks</td><td>5487</td><td>5526</td><td>40</td><td>CGCTGAGAAGACGGTCGAACTTGACTATCTAGAGGAAGTA</td><td>1031.6</td><td>-8.9</td><td>1.6</td><td>-5.5</td><td>4.6E-08</td><td>7.8E-08</td><td>BT20/MB468</td><td>NUC</td></tr>
<tr><td>>hg38_21_+_8433172_8446622_RNA45SN1_45S_50ntFlanks@8552.8577.26|CTCCCCGGAGGGGGCGGGCTCCGGCG</td><td>>hg38_21_+_8433172_8446622_RNA45SN1_45S_50ntFlanks</td><td>8552</td><td>8577</td><td>26</td><td>CTCCCCGGAGGGGGCGGGCTCCGGCG</td><td>900.8</td><td>-8.7</td><td>1.6</td><td>-5.5</td><td>4.6E-08</td><td>7.8E-08</td><td>BT20/MB468</td><td>NUC</td></tr>
<tr><td>>MI0000063|hsa-let-7b&WithFlank&22|+|46113680|46113774@12.34.23&[MIMAT0000063&hsa-let-7b-5p&offsets|0|+1;m-14&22|+|46113691|46113712&offsets|0|+1]|TGAGGTAGTAGGTTGTGTGGTTT</td><td>>MI0000063|hsa-let-7b&WithFlank&22|+|46113680|46113774</td><td>12</td><td>34</td><td>23</td><td>TGAGGTAGTAGGTTGTGTGGTTT</td><td>14378.7</td><td>5.7</td><td>1.0</td><td>5.8</td><td>7.0E-09</td><td>7.8E-08</td><td>BT20/MB468</td><td>MP</td></tr>
<tr><td>>hg38_21_+_8433172_8446622_RNA45SN1_45S_50ntFlanks@10821.10862.42|GGAAGTCGGCAAGCCGGATCCGTAACTTCGGGATAAGGATTG</td><td>>hg38_21_+_8433172_8446622_RNA45SN1_45S_50ntFlanks</td><td>10821</td><td>10862</td><td>42</td><td>GGAAGTCGGCAAGCCGGATCCGTAACTTCGGGATAAGGATTG</td><td>70.4</td><td>-10.0</td><td>1.8</td><td>-5.5</td><td>3.8E-08</td><td>7.8E-08</td><td>MB231/MB468</td><td>MP</td></tr>
<tr><td>>hg38_21_+_8433172_8446622_RNA45SN1_45S_50ntFlanks@10821.10865.45|GGAAGTCGGCAAGCCGGATCCGTAACTTCGGGATAAGGATTGGCT</td><td>>hg38_21_+_8433172_8446622_RNA45SN1_45S_50ntFlanks</td><td>10821</td><td>10865</td><td>45</td><td>GGAAGTCGGCAAGCCGGATCCGTAACTTCGGGATAAGGATTGGCT</td><td>596.6</td><td>-7.9</td><td>1.4</td><td>-5.5</td><td>4.4E-08</td><td>7.8E-08</td><td>MB231/MB468</td><td>MITO</td></tr>
<tr><td>>hg38_21_+_8433172_8446622_RNA45SN1_45S_50ntFlanks@12605.12622.18|TATGTGCTTGGCTGAGGA</td><td>>hg38_21_+_8433172_8446622_RNA45SN1_45S_50ntFlanks</td><td>12605</td><td>12622</td><td>18</td><td>TATGTGCTTGGCTGAGGA</td><td>1006.0</td><td>-8.7</td><td>1.6</td><td>-5.5</td><td>4.6E-08</td><td>7.9E-08</td><td>BT20/MB468</td><td>NUC</td></tr>
<tr><td>>hg38_21_+_8433172_8446622_RNA45SN1_45S_50ntFlanks@9534.9556.23|AGGAAACTCTGGTGGAGGTCCGT</td><td>>hg38_21_+_8433172_8446622_RNA45SN1_45S_50ntFlanks</td><td>9534</td><td>9556</td><td>23</td><td>AGGAAACTCTGGTGGAGGTCCGT</td><td>204.7</td><td>-10.8</td><td>1.9</td><td>-5.8</td><td>7.0E-09</td><td>7.9E-08</td><td>BT20/MB468</td><td>MP</td></tr>
<tr><td>>hg38_21_+_8433172_8446622_RNA45SN1_45S_50ntFlanks@12394.12419.26|TAGCGACGTCGCTTTTTGATCCTTCG</td><td>>hg38_21_+_8433172_8446622_RNA45SN1_45S_50ntFlanks</td><td>12394</td><td>12419</td><td>26</td><td>TAGCGACGTCGCTTTTTGATCCTTCG</td><td>1040.2</td><td>-3.6</td><td>0.6</td><td>-5.5</td><td>3.1E-08</td><td>7.9E-08</td><td>BT20/MB468</td><td>MITO</td></tr>
<tr><td>>MI0000239|hsa-mir-197&WithFlank&1|+|109598887|109598973@54.75.22&[MIMAT0000227&hsa-miR-197-3p&offsets|0|0;m-99&1|+|109598940|109598961&offsets|0|0]|TTCACCACCTTCTCCACCCAGC</td><td>>MI0000239|hsa-mir-197&WithFlank&1|+|109598887|109598973</td><td>54</td><td>75</td><td>22</td><td>TTCACCACCTTCTCCACCCAGC</td><td>3227.2</td><td>2.9</td><td>0.5</td><td>5.5</td><td>4.6E-08</td><td>7.9E-08</td><td>BT20/MB468</td><td>NUC</td></tr>
<tr><td>>hg38_21_+_8433172_8446622_RNA45SN1_45S_50ntFlanks@11575.11612.38|ACTTAGAACTGGTGCGGACCAGGGGAATCCGACTGTTT</td><td>>hg38_21_+_8433172_8446622_RNA45SN1_45S_50ntFlanks</td><td>11575</td><td>11612</td><td>38</td><td>ACTTAGAACTGGTGCGGACCAGGGGAATCCGACTGTTT</td><td>465.4</td><td>-4.4</td><td>0.8</td><td>-5.5</td><td>4.6E-08</td><td>7.9E-08</td><td>BT20/MB468</td><td>NUC</td></tr>
<tr><td>>hg38_21_+_8433172_8446622_RNA45SN1_45S_50ntFlanks@11212.11229.18|CGACTCTGGACGCGAGCC</td><td>>hg38_21_+_8433172_8446622_RNA45SN1_45S_50ntFlanks</td><td>11212</td><td>11229</td><td>18</td><td>CGACTCTGGACGCGAGCC</td><td>418.5</td><td>-3.6</td><td>0.6</td><td>-5.5</td><td>3.5E-08</td><td>7.9E-08</td><td>BT20/MB468</td><td>CYTO</td></tr>
<tr><td>>hg38_21_+_8433172_8446622_RNA45SN1_45S_50ntFlanks@11587.11611.25|TGCGGACCAGGGGAATCCGACTGTT</td><td>>hg38_21_+_8433172_8446622_RNA45SN1_45S_50ntFlanks</td><td>11587</td><td>11611</td><td>25</td><td>TGCGGACCAGGGGAATCCGACTGTT</td><td>963.6</td><td>-3.4</td><td>0.6</td><td>-5.5</td><td>4.4E-08</td><td>7.9E-08</td><td>MB231/MB468</td><td>MITO</td></tr>
<tr><td>>hg38_21_+_8433172_8446622_RNA45SN1_45S_50ntFlanks@11190.11225.36|CCCGGGGGGCCGGCGGCGGCGGCGACTCTGGACGCG</td><td>>hg38_21_+_8433172_8446622_RNA45SN1_45S_50ntFlanks</td><td>11190</td><td>11225</td><td>36</td><td>CCCGGGGGGCCGGCGGCGGCGGCGACTCTGGACGCG</td><td>520.7</td><td>-6.8</td><td>1.2</td><td>-5.5</td><td>4.1E-08</td><td>7.9E-08</td><td>MB231/MB468</td><td>CYTO</td></tr>
<tr><td>>hg38_21_+_8433172_8446622_RNA45SN1_45S_50ntFlanks@9565.9597.33|TGACGTGCAAATCGGTCGTCCGACCTGGGTATA</td><td>>hg38_21_+_8433172_8446622_RNA45SN1_45S_50ntFlanks</td><td>9565</td><td>9597</td><td>33</td><td>TGACGTGCAAATCGGTCGTCCGACCTGGGTATA</td><td>2809.4</td><td>-3.8</td><td>0.7</td><td>-5.5</td><td>4.1E-08</td><td>7.9E-08</td><td>MB231/MB468</td><td>CYTO</td></tr>
<tr><td>>hg38_1_-_228634819_228635039_RNA5S12_5S_50ntFlanks@129.171.43|TTGGATGGGAGACCGCCTGGGAATACCGGGTGCTGTAGGCTTT</td><td>>hg38_1_-_228634819_228635039_RNA5S12_5S_50ntFlanks</td><td>129</td><td>171</td><td>43</td><td>TTGGATGGGAGACCGCCTGGGAATACCGGGTGCTGTAGGCTTT</td><td>2964.8</td><td>10.1</td><td>1.8</td><td>5.5</td><td>3.5E-08</td><td>7.9E-08</td><td>BT20/MB468</td><td>CYTO</td></tr>
<tr><td>>trnaMT_GluTTC_MT_-_14674_14742@39.63.25,trnalookalike8_GluTTC_5_-_93905172_93905240@39.63.25|ATTGGTCGTGGTTGTAGTCCGTGCG</td><td>>trnaMT_GluTTC_MT_-_14674_14742</td><td>39</td><td>63</td><td>25</td><td>ATTGGTCGTGGTTGTAGTCCGTGCG</td><td>237.3</td><td>4.4</td><td>0.8</td><td>5.5</td><td>4.2E-08</td><td>7.9E-08</td><td>MB231/MB468</td><td>NUC</td></tr>
<tr><td>>hg38_21_+_8433172_8446622_RNA45SN1_45S_50ntFlanks@10362.10379.18|AAGGCCGAAGTGGAGAAG</td><td>>hg38_21_+_8433172_8446622_RNA45SN1_45S_50ntFlanks</td><td>10362</td><td>10379</td><td>18</td><td>AAGGCCGAAGTGGAGAAG</td><td>284.1</td><td>-7.3</td><td>1.3</td><td>-5.5</td><td>3.8E-08</td><td>7.9E-08</td><td>MB231/MB468</td><td>MP</td></tr>
<tr><td>>MI0000085|hsa-mir-27a&WithFlank&19|-|13836434|13836523@57.76.20(+1A)&[MIMAT0000084&hsa-miR-27a-3p&offsets|0|-1(+1A);m-43&19|-|13836447|13836467&offsets|0|-1(+1A)]|TTCACAGTGGCTAAGTTCCGA</td><td>>MI0000085|hsa-mir-27a&WithFlank&19|-|13836434|13836523</td><td>57</td><td>76</td><td>20(+1A)</td><td>TTCACAGTGGCTAAGTTCCGA</td><td>1429.1</td><td>4.2</td><td>0.8</td><td>5.5</td><td>3.1E-08</td><td>7.9E-08</td><td>BT20/MB468</td><td>MITO</td></tr>
<tr><td>>hg38_21_+_8433172_8446622_RNA45SN1_45S_50ntFlanks@3811.3834.24|AGTTATGGTTCCTTTGGTCGCTCG</td><td>>hg38_21_+_8433172_8446622_RNA45SN1_45S_50ntFlanks</td><td>3811</td><td>3834</td><td>24</td><td>AGTTATGGTTCCTTTGGTCGCTCG</td><td>954.9</td><td>-8.7</td><td>1.6</td><td>-5.5</td><td>4.7E-08</td><td>7.9E-08</td><td>BT20/MB468</td><td>NUC</td></tr>
<tr><td>>hg38_21_+_8433172_8446622_RNA45SN1_45S_50ntFlanks@6733.6753.21|CATTGATCATCGACACTTCGA</td><td>>hg38_21_+_8433172_8446622_RNA45SN1_45S_50ntFlanks</td><td>6733</td><td>6753</td><td>21</td><td>CATTGATCATCGACACTTCGA</td><td>369.9</td><td>-3.9</td><td>0.7</td><td>-5.5</td><td>4.7E-08</td><td>7.9E-08</td><td>BT20/MB468</td><td>NUC</td></tr>
<tr><td>>hg38_21_+_8433172_8446622_RNA45SN1_45S_50ntFlanks@12636.12654.19|AGCTACCATCTGTGGGATT</td><td>>hg38_21_+_8433172_8446622_RNA45SN1_45S_50ntFlanks</td><td>12636</td><td>12654</td><td>19</td><td>AGCTACCATCTGTGGGATT</td><td>541.2</td><td>-8.3</td><td>1.5</td><td>-5.5</td><td>4.7E-08</td><td>7.9E-08</td><td>BT20/MB468</td><td>NUC</td></tr>
<tr><td>>MI0000455|hsa-mir-138-2&WithFlank&16|+|56858512|56858607@16.39.24&[MIMAT0000430&hsa-miR-138-5p&offsets|0|+1;m-509&16|+|56858527|56858549&offsets|0|+1]|AGCTGGTGTTGTGAATCAGGCCGA</td><td>>MI0000455|hsa-mir-138-2&WithFlank&16|+|56858512|56858607</td><td>16</td><td>39</td><td>24</td><td>AGCTGGTGTTGTGAATCAGGCCGA</td><td>58.3</td><td>9.1</td><td>1.7</td><td>5.5</td><td>3.8E-08</td><td>7.9E-08</td><td>MB231/MB468</td><td>MP</td></tr>
<tr><td>>am_trnaMT_ValTAC_MT_+_1602_1670@29.66.38|ACTTACACTTAGGAGATTTCAACTTAACTTGACCGCTC</td><td>>am_trnaMT_ValTAC_MT_+_1602_1670</td><td>29</td><td>66</td><td>38</td><td>ACTTACACTTAGGAGATTTCAACTTAACTTGACCGCTC</td><td>53.9</td><td>9.0</td><td>1.6</td><td>5.5</td><td>3.8E-08</td><td>8.0E-08</td><td>MB231/MB468</td><td>MP</td></tr>
<tr><td>>hg38_21_+_8433172_8446622_RNA45SN1_45S_50ntFlanks@11581.11605.25|AACTGGTGCGGACCAGGGGAATCCG</td><td>>hg38_21_+_8433172_8446622_RNA45SN1_45S_50ntFlanks</td><td>11581</td><td>11605</td><td>25</td><td>AACTGGTGCGGACCAGGGGAATCCG</td><td>2498.1</td><td>-4.8</td><td>0.8</td><td>-5.8</td><td>7.1E-09</td><td>8.0E-08</td><td>BT20/MB468</td><td>MP</td></tr>
<tr><td>>hg38_21_+_8433172_8446622_RNA45SN1_45S_50ntFlanks@6659.6700.42|AGCGGTGGATCACTCGGCTCGTGCGTCGATGAAGAACGCAGC</td><td>>hg38_21_+_8433172_8446622_RNA45SN1_45S_50ntFlanks</td><td>6659</td><td>6700</td><td>42</td><td>AGCGGTGGATCACTCGGCTCGTGCGTCGATGAAGAACGCAGC</td><td>148.7</td><td>11.0</td><td>1.9</td><td>5.8</td><td>7.1E-09</td><td>8.0E-08</td><td>BT20/MB468</td><td>MP</td></tr>
<tr><td>>hg38_21_+_8433172_8446622_RNA45SN1_45S_50ntFlanks@10794.10818.25|CTGGCATGTTGGAACAATGTAGGTA</td><td>>hg38_21_+_8433172_8446622_RNA45SN1_45S_50ntFlanks</td><td>10794</td><td>10818</td><td>25</td><td>CTGGCATGTTGGAACAATGTAGGTA</td><td>2145.9</td><td>-8.9</td><td>1.5</td><td>-5.8</td><td>7.1E-09</td><td>8.0E-08</td><td>BT20/MB468</td><td>MP</td></tr>
<tr><td>>hg38_21_+_8433172_8446622_RNA45SN1_45S_50ntFlanks@11498.11516.19|CCGGTCCCCCCCGCCGGGT</td><td>>hg38_21_+_8433172_8446622_RNA45SN1_45S_50ntFlanks</td><td>11498</td><td>11516</td><td>19</td><td>CCGGTCCCCCCCGCCGGGT</td><td>143.4</td><td>-10.2</td><td>1.8</td><td>-5.8</td><td>7.1E-09</td><td>8.0E-08</td><td>BT20/MB468</td><td>MP</td></tr>
<tr><td>>hg38_21_+_8433172_8446622_RNA45SN1_45S_50ntFlanks@4367.4393.27|CAGTTAAAAAGCTCGTAGTTGGATCTT</td><td>>hg38_21_+_8433172_8446622_RNA45SN1_45S_50ntFlanks</td><td>4367</td><td>4393</td><td>27</td><td>CAGTTAAAAAGCTCGTAGTTGGATCTT</td><td>93.6</td><td>-9.6</td><td>1.7</td><td>-5.8</td><td>7.1E-09</td><td>8.0E-08</td><td>BT20/MB468</td><td>MP</td></tr>
<tr><td>>MI0000087|hsa-mir-29a&WithFlank&7|-|130876741|130876816@47.68.22(+1U)&[MIMAT0000086&hsa-miR-29a-3p&offsets|-1|-1(+1U);m-26&7|-|130876749|130876769&offsets|-1|0(+1U)]|CTAGCACCATCTGAAATCGGTTT</td><td>>MI0000087|hsa-mir-29a&WithFlank&7|-|130876741|130876816</td><td>47</td><td>68</td><td>22(+1U)</td><td>CTAGCACCATCTGAAATCGGTTT</td><td>2850.3</td><td>6.9</td><td>1.2</td><td>5.8</td><td>7.1E-09</td><td>8.0E-08</td><td>BT20/MB468</td><td>MP</td></tr>
<tr><td>>hg38_21_+_8433172_8446622_RNA45SN1_45S_50ntFlanks@12418.12445.28|CGATGTCGGCTCTTCCTATCATTGTGAA</td><td>>hg38_21_+_8433172_8446622_RNA45SN1_45S_50ntFlanks</td><td>12418</td><td>12445</td><td>28</td><td>CGATGTCGGCTCTTCCTATCATTGTGAA</td><td>1221.4</td><td>-4.1</td><td>0.7</td><td>-5.5</td><td>4.5E-08</td><td>8.0E-08</td><td>MB231/MB468</td><td>MITO</td></tr>
<tr><td>>hg38_21_+_8433172_8446622_RNA45SN1_45S_50ntFlanks@4722.4751.30|TCATTAATCAAGAACGAAAGTCGGAGGTTC</td><td>>hg38_21_+_8433172_8446622_RNA45SN1_45S_50ntFlanks</td><td>4722</td><td>4751</td><td>30</td><td>TCATTAATCAAGAACGAAAGTCGGAGGTTC</td><td>535.4</td><td>-5.9</td><td>1.1</td><td>-5.5</td><td>3.6E-08</td><td>8.0E-08</td><td>BT20/MB468</td><td>CYTO</td></tr>
<tr><td>>hg38_21_+_8433172_8446622_RNA45SN1_45S_50ntFlanks@4615.4632.18|CCATGATTAAGAGGGACG</td><td>>hg38_21_+_8433172_8446622_RNA45SN1_45S_50ntFlanks</td><td>4615</td><td>4632</td><td>18</td><td>CCATGATTAAGAGGGACG</td><td>109.7</td><td>-9.9</td><td>1.7</td><td>-5.8</td><td>7.2E-09</td><td>8.0E-08</td><td>BT20/MB468</td><td>MP</td></tr>
<tr><td>>MI0000298|hsa-mir-221&WithFlank&X|-|45746151|45746272@71.93.23(+2U)&[MIMAT0000278&hsa-miR-221-3p&offsets|0|0(+2U);m-91&X|-|45746181|45746202&offsets|0|+1(+2U)]|AGCTACATTGTCTGCTGGGTTTCTT</td><td>>MI0000298|hsa-mir-221&WithFlank&X|-|45746151|45746272</td><td>71</td><td>93</td><td>23(+2U)</td><td>AGCTACATTGTCTGCTGGGTTTCTT</td><td>365.7</td><td>6.0</td><td>1.1</td><td>5.5</td><td>4.1E-08</td><td>8.0E-08</td><td>MB231/MB468</td><td>CYTO</td></tr>
<tr><td>>hg38_21_+_8433172_8446622_RNA45SN1_45S_50ntFlanks@6646.6672.27|TCGTACGACTCTTAGCGGTGGATCACT</td><td>>hg38_21_+_8433172_8446622_RNA45SN1_45S_50ntFlanks</td><td>6646</td><td>6672</td><td>27</td><td>TCGTACGACTCTTAGCGGTGGATCACT</td><td>3263.6</td><td>3.7</td><td>0.7</td><td>5.5</td><td>4.2E-08</td><td>8.0E-08</td><td>MB231/MB468</td><td>NUC</td></tr>
<tr><td>>hg38_21_+_8433172_8446622_RNA45SN1_45S_50ntFlanks@3840.3872.33|CTCCTACTTGGATAACTGTGGTAATTCTAGAGC</td><td>>hg38_21_+_8433172_8446622_RNA45SN1_45S_50ntFlanks</td><td>3840</td><td>3872</td><td>33</td><td>CTCCTACTTGGATAACTGTGGTAATTCTAGAGC</td><td>104.4</td><td>-10.6</td><td>1.9</td><td>-5.5</td><td>3.9E-08</td><td>8.0E-08</td><td>MB231/MB468</td><td>MP</td></tr>
<tr><td>>hg38_21_+_8433172_8446622_RNA45SN1_45S_50ntFlanks@10834.10865.32|CCGGATCCGTAACTTCGGGATAAGGATTGGCT</td><td>>hg38_21_+_8433172_8446622_RNA45SN1_45S_50ntFlanks</td><td>10834</td><td>10865</td><td>32</td><td>CCGGATCCGTAACTTCGGGATAAGGATTGGCT</td><td>337.8</td><td>-11.5</td><td>2.0</td><td>-5.8</td><td>7.2E-09</td><td>8.0E-08</td><td>BT20/MB468</td><td>MP</td></tr>
<tr><td>>hg38_21_+_8433172_8446622_RNA45SN1_45S_50ntFlanks@8170.8193.24|CAAGTCCTTCTGATCGAGGCCCAG</td><td>>hg38_21_+_8433172_8446622_RNA45SN1_45S_50ntFlanks</td><td>8170</td><td>8193</td><td>24</td><td>CAAGTCCTTCTGATCGAGGCCCAG</td><td>113.0</td><td>-9.9</td><td>1.7</td><td>-5.8</td><td>7.2E-09</td><td>8.0E-08</td><td>BT20/MB468</td><td>MP</td></tr>
<tr><td>>hg38_21_+_8433172_8446622_RNA45SN1_45S_50ntFlanks@8405.8422.18|GTAAACGGGTGGGGTCCG</td><td>>hg38_21_+_8433172_8446622_RNA45SN1_45S_50ntFlanks</td><td>8405</td><td>8422</td><td>18</td><td>GTAAACGGGTGGGGTCCG</td><td>92.8</td><td>-9.6</td><td>1.7</td><td>-5.8</td><td>7.2E-09</td><td>8.0E-08</td><td>BT20/MB468</td><td>MP</td></tr>
<tr><td>>hg38_21_+_8433172_8446622_RNA45SN1_45S_50ntFlanks@11582.11621.40|ACTGGTGCGGACCAGGGGAATCCGACTGTTTAATTAAAAC</td><td>>hg38_21_+_8433172_8446622_RNA45SN1_45S_50ntFlanks</td><td>11582</td><td>11621</td><td>40</td><td>ACTGGTGCGGACCAGGGGAATCCGACTGTTTAATTAAAAC</td><td>89.7</td><td>10.2</td><td>1.8</td><td>5.8</td><td>7.2E-09</td><td>8.0E-08</td><td>BT20/MB468</td><td>MP</td></tr>
<tr><td>>hg38_21_+_8433172_8446622_RNA45SN1_45S_50ntFlanks@3839.3869.31|TCTCCTACTTGGATAACTGTGGTAATTCTAG</td><td>>hg38_21_+_8433172_8446622_RNA45SN1_45S_50ntFlanks</td><td>3839</td><td>3869</td><td>31</td><td>TCTCCTACTTGGATAACTGTGGTAATTCTAG</td><td>424.7</td><td>5.4</td><td>1.0</td><td>5.5</td><td>4.1E-08</td><td>8.0E-08</td><td>MB231/MB468</td><td>CYTO</td></tr>
<tr><td>>hg38_21_+_8433172_8446622_RNA45SN1_45S_50ntFlanks@9549.9571.23|AGGTCCGTAGCGGTCCTGACGTG</td><td>>hg38_21_+_8433172_8446622_RNA45SN1_45S_50ntFlanks</td><td>9549</td><td>9571</td><td>23</td><td>AGGTCCGTAGCGGTCCTGACGTG</td><td>178.9</td><td>-3.8</td><td>0.7</td><td>-5.5</td><td>3.9E-08</td><td>8.0E-08</td><td>MB231/MB468</td><td>MP</td></tr>
<tr><td>>hg38_21_+_8433172_8446622_RNA45SN1_45S_50ntFlanks@11885.11903.19|TTGAGCTTGACTCTAGTCT</td><td>>hg38_21_+_8433172_8446622_RNA45SN1_45S_50ntFlanks</td><td>11885</td><td>11903</td><td>19</td><td>TTGAGCTTGACTCTAGTCT</td><td>620.3</td><td>-7.6</td><td>1.4</td><td>-5.5</td><td>3.6E-08</td><td>8.0E-08</td><td>BT20/MB468</td><td>CYTO</td></tr>
<tr><td>>hg38_21_+_8433172_8446622_RNA45SN1_45S_50ntFlanks@8194.8216.23|CCCGTGGACGGTGTGAGGCCGGT</td><td>>hg38_21_+_8433172_8446622_RNA45SN1_45S_50ntFlanks</td><td>8194</td><td>8216</td><td>23</td><td>CCCGTGGACGGTGTGAGGCCGGT</td><td>428.0</td><td>-6.3</td><td>1.1</td><td>-5.5</td><td>4.7E-08</td><td>8.0E-08</td><td>BT20/MB468</td><td>NUC</td></tr>
</table>

| | | | | | | | | | | | | | |
|---|---|---|---|---|---|---|---|---|---|---|---|---|---|
| >hg38_21_+_8433172_8446622_RNA45SN1_45S_50ntFlanks@10514.10533.20\|CGGAGTGGCGGAGATGGGCG | >hg38_21_+_8433172_8446622_RNA45SN1_45S_50ntFlanks | 10514 | 10533 | 20 | CGGAGTGGCGGAGATGGGCG | 514.7 | -7.8 | 1.4 | -5.5 | 4.3E-08 | 8.1E-08 | MB231/MB468 | NUC |
| >hg38_21_+_8433172_8446622_RNA45SN1_45S_50ntFlanks@12008.12039.32\|CCGCCGGTGAAATACCACTACTCTGATCGTTT | >hg38_21_+_8433172_8446622_RNA45SN1_45S_50ntFlanks | 12008 | 12039 | 32 | CCGCCGGTGAAATACCACTACTCTGATCGTTT | 412.0 | -3.0 | 0.5 | -5.5 | 3.2E-08 | 8.1E-08 | BT20/MB468 | MITO |
| >hg38_21_+_8433172_8446622_RNA45SN1_45S_50ntFlanks@11594.11611.18\|CAGGGGAATCCGACTGTT | >hg38_21_+_8433172_8446622_RNA45SN1_45S_50ntFlanks | 11594 | 11611 | 18 | CAGGGGAATCCGACTGTT | 1974.6 | -3.1 | 0.6 | -5.5 | 3.6E-08 | 8.1E-08 | BT20/MB468 | CYTO |
| >MI0016005\|hsa-mir-3615&WithFlank&17\|+\|74748607\|74748705@57.78.22(+2U)&[MIMAT0017994&hsa-miR-3615&offsets\|0\|+1(+2U);m-271&17\|+\|74748663\|74748684&offsets\|0\|0(+2U)]\|TCTCTCGGCTCCTCGCGGCTCGTT | >MI0016005\|hsa-mir-3615&WithFlank&17\|+\|74748607\|74748705 | 57 | 78 | 22(+2U) | TCTCTCGGCTCCTCGCGGCTCGTT | 342.5 | 7.4 | 1.3 | 5.5 | 3.6E-08 | 8.1E-08 | BT20/MB468 | CYTO |
| >MI0000462\|hsa-mir-152&WithFlank&17\|-\|48037155\|48037253@60.80.21(+1U)&[MIMAT0000438&hsa-miR-152-3p&offsets\|0\|0(+1U);m-97&17\|-\|48037174\|48037194&offsets\|0\|0(+1U)]\|TCAGTGCATGACAGAACTTGGT | >MI0000462\|hsa-mir-152&WithFlank&17\|-\|48037155\|48037253 | 60 | 80 | 21(+1U) | TCAGTGCATGACAGAACTTGGT | 558.8 | 7.5 | 1.4 | 5.5 | 3.9E-08 | 8.1E-08 | MB231/MB468 | MP |
| >hg38_21_+_8433172_8446622_RNA45SN1_45S_50ntFlanks@11586.11604.19\|GTGCGGACCAGGGGAATCC | >hg38_21_+_8433172_8446622_RNA45SN1_45S_50ntFlanks | 11586 | 11604 | 19 | GTGCGGACCAGGGGAATCC | 193.9 | -5.4 | 1.0 | -5.5 | 4.3E-08 | 8.1E-08 | MB231/MB468 | NUC |
| >hg38_21_+_8433172_8446622_RNA45SN1_45S_50ntFlanks@8358.8379.22\|AAAGAACTTTGAAGAGAGAGTT | >hg38_21_+_8433172_8446622_RNA45SN1_45S_50ntFlanks | 8358 | 8379 | 22 | AAAGAACTTTGAAGAGAGAGTT | 773.8 | -5.0 | 0.9 | -5.5 | 4.6E-08 | 8.1E-08 | MB231/MB468 | MITO |
| >hg38_21_+_8433172_8446622_RNA45SN1_45S_50ntFlanks@10684.10710.27\|CGGCGGCGTCCGGTGAGCTCTCGCTGG | >hg38_21_+_8433172_8446622_RNA45SN1_45S_50ntFlanks | 10684 | 10710 | 27 | CGGCGGCGTCCGGTGAGCTCTCGCTGG | 7986.7 | -5.1 | 0.9 | -5.5 | 3.6E-08 | 8.1E-08 | BT20/MB468 | CYTO |
| >hg38_21_+_8433172_8446622_RNA45SN1_45S_50ntFlanks@12418.12445.28\|CGATGTCGGCTCTTCCTATCATTGTGAA | >hg38_21_+_8433172_8446622_RNA45SN1_45S_50ntFlanks | 12418 | 12445 | 28 | CGATGTCGGCTCTTCCTATCATTGTGAA | 3604.4 | -7.7 | 1.4 | -5.5 | 4.8E-08 | 8.1E-08 | BT20/MB468 | NUC |
| >hg38_21_+_8433172_8446622_RNA45SN1_45S_50ntFlanks@3855.3873.19\|CTGTGGTAATTCTAGAGCT | >hg38_21_+_8433172_8446622_RNA45SN1_45S_50ntFlanks | 3855 | 3873 | 19 | CTGTGGTAATTCTAGAGCT | 205.1 | -7.1 | 1.3 | -5.5 | 3.9E-08 | 8.1E-08 | MB231/MB468 | MP |
| >hg38_21_+_8433172_8446622_RNA45SN1_45S_50ntFlanks@10401.10429.29\|GAACATGGGTCAGTCGGTCCTGAGAGATG | >hg38_21_+_8433172_8446622_RNA45SN1_45S_50ntFlanks | 10401 | 10429 | 29 | GAACATGGGTCAGTCGGTCCTGAGAGATG | 1601.3 | -5.4 | 1.0 | -5.5 | 3.9E-08 | 8.1E-08 | MB231/MB468 | MP |
| >hg38_21_+_8433172_8446622_RNA45SN1_45S_50ntFlanks@10684.10705.22\|CGGCGGCGTCCGGTGAGCTCTC | >hg38_21_+_8433172_8446622_RNA45SN1_45S_50ntFlanks | 10684 | 10705 | 22 | CGGCGGCGTCCGGTGAGCTCTC | 261.7 | -4.3 | 0.8 | -5.5 | 4.2E-08 | 8.1E-08 | MB231/MB468 | CYTO |
| >hg38_21_+_8433172_8446622_RNA45SN1_45S_50ntFlanks@9533.9558.26\|GAGGAAACTCTGGTGGAGGTCCGTAG | >hg38_21_+_8433172_8446622_RNA45SN1_45S_50ntFlanks | 9533 | 9558 | 26 | GAGGAAACTCTGGTGGAGGTCCGTAG | 99.9 | -9.7 | 1.7 | -5.8 | 7.3E-09 | 8.1E-08 | BT20/MB468 | MP |
| >hg38_MT_+_598_1651_RNR1_12S_50ntFlanks@428.468.41\|AAAGTGGCTTTAACATATCTGAACACACAATAGCTAAGACC | >hg38_MT_+_598_1651_RNR1_12S_50ntFlanks | 428 | 468 | 41 | AAAGTGGCTTTAACATATCTGAACACACAATAGCTAAGACC | 867.3 | 6.1 | 1.1 | 5.5 | 4.6E-08 | 8.1E-08 | MB231/MB468 | MITO |
| >hg38_21_+_8433172_8446622_RNA45SN1_45S_50ntFlanks@7967.8005.39\|CTCCGAGACGCGACCTCAGATCAGACGTGGCGACCCGCT | >hg38_21_+_8433172_8446622_RNA45SN1_45S_50ntFlanks | 7967 | 8005 | 39 | CTCCGAGACGCGACCTCAGATCAGACGTGGCGACCCGCT | 663.1 | 5.6 | 1.0 | 5.5 | 4.8E-08 | 8.1E-08 | BT20/MB468 | NUC |
| >am_trna116_GluCTC_1_-_145399233_145399304@12.29.18,trna25_GluCTC_2_-_71273488_71273560@12.29.18,trna59_GluCTC_1_+_249168447_249168518@12.29.18,trna71_GluCTC_1_-_161439189_161439260@12.29.18,trna74_GluCTC_1_-_161431809_161431880@12.29.18,trna77_GluCTC_1_-_161424398_161424469@12.29.18,trna77_GluCTC_6_+_28949976_28950047@12.29.18,trna80_GluCTC_1_-_161417018_161417089@12.29.18,trna87_GluCTC_6_-_126101393_126101464@12.29.18\|CTAGTGGTTAGGATTCGG | >am_trna116_GluCTC_1_-_145399233_145399304 | 12 | 29 | 18 | CTAGTGGTTAGGATTCGG | 102.7 | -9.8 | 1.7 | -5.8 | 7.3E-09 | 8.2E-08 | BT20/MB468 | MP |
| >hg38_21_+_8433172_8446622_RNA45SN1_45S_50ntFlanks@11567.11591.25\|AGCAGCCGACTTAGAACTGGTGCGG | >hg38_21_+_8433172_8446622_RNA45SN1_45S_50ntFlanks | 11567 | 11591 | 25 | AGCAGCCGACTTAGAACTGGTGCGG | 472.0 | -6.9 | 1.3 | -5.5 | 4.0E-08 | 8.2E-08 | MB231/MB468 | MP |
| >hg38_1_-_228634819_228635039_RNA5S12_5S_50ntFlanks@51.89.39\|GTCTACGGCCATACCACCCTGAACGCGCCCGATCTCGTC | >hg38_1_-_228634819_228635039_RNA5S12_5S_50ntFlanks | 51 | 89 | 39 | GTCTACGGCCATACCACCCTGAACGCGCCCGATCTCGTC | 29377.6 | 5.7 | 1.0 | 5.5 | 4.3E-08 | 8.2E-08 | MB231/MB468 | NUC |
| >hg38_21_+_8433172_8446622_RNA45SN1_45S_50ntFlanks@4367.4405.39\|CAGTTAAAAAGCTCGTAGTTGGATCTTGGGAGCGGGCGG | >hg38_21_+_8433172_8446622_RNA45SN1_45S_50ntFlanks | 4367 | 4405 | 39 | CAGTTAAAAAGCTCGTAGTTGGATCTTGGGAGCGGGCGG | 86.2 | -10.3 | 1.9 | -5.5 | 4.0E-08 | 8.2E-08 | MB231/MB468 | MP |
| >hg38_21_+_8433172_8446622_RNA45SN1_45S_50ntFlanks@4760.4784.25\|TCAGATACCGTCGTAGTTCCGACCA | >hg38_21_+_8433172_8446622_RNA45SN1_45S_50ntFlanks | 4760 | 4784 | 25 | TCAGATACCGTCGTAGTTCCGACCA | 89.8 | -9.6 | 1.7 | -5.8 | 7.4E-09 | 8.2E-08 | BT20/MB468 | MP |
| >hg38_21_+_8433172_8446622_RNA45SN1_45S_50ntFlanks@6645.6681.37\|CTCGTACGACTCTTAGCGGTGGATCACTCGGCTCGTG | >hg38_21_+_8433172_8446622_RNA45SN1_45S_50ntFlanks | 6645 | 6681 | 37 | CTCGTACGACTCTTAGCGGTGGATCACTCGGCTCGTG | 4863.6 | -5.1 | 0.9 | -5.5 | 4.0E-08 | 8.2E-08 | MB231/MB468 | MP |
| >hg38_21_+_8433172_8446622_RNA45SN1_45S_50ntFlanks@12395.12414.20\|AGCGACGTCGCTTTTTGATC | >hg38_21_+_8433172_8446622_RNA45SN1_45S_50ntFlanks | 12395 | 12414 | 20 | AGCGACGTCGCTTTTTGATC | 528.2 | 4.0 | 0.7 | 5.5 | 3.3E-08 | 8.2E-08 | BT20/MB468 | MITO |
| >hg38_21_+_8433172_8446622_RNA45SN1_45S_50ntFlanks@10693.10711.19\|CCGGTGAGCTCTCGCTGGC | >hg38_21_+_8433172_8446622_RNA45SN1_45S_50ntFlanks | 10693 | 10711 | 19 | CCGGTGAGCTCTCGCTGGC | 181.1 | -10.6 | 1.8 | -5.8 | 7.4E-09 | 8.2E-08 | BT20/MB468 | MP |
| >MI0000063\|hsa-let-7b&WithFlank&22\|+\|46113680\|46113774@12.32.21(+1C)&[MIMAT0000063&hsa-let-7b-5p&offsets\|0\|-1(+1C);m-14&22\|+\|46113691\|46113712&offsets\|0\|-1(+1C)]\|TGAGGTAGTAGGTTGTGTGGTC | >MI0000063\|hsa-let-7b&WithFlank&22\|+\|46113680\|46113774 | 12 | 32 | 21(+1C) | TGAGGTAGTAGGTTGTGTGGTC | 1082.9 | 5.1 | 0.9 | 5.5 | 4.6E-08 | 8.2E-08 | MB231/MB468 | MITO |
| >trnaMT_LeuTAA_MT_+_3230_3304@1.30.30\|GTTAAGATGGCAGAGCCCGGTAATCGCATA | >trnaMT_LeuTAA_MT_+_3230_3304 | 1 | 30 | 30 | GTTAAGATGGCAGAGCCCGGTAATCGCATA | 3132.6 | 6.3 | 1.1 | 5.5 | 3.3E-08 | 8.2E-08 | BT20/MB468 | MITO |
| >hg38_21_+_8433172_8446622_RNA45SN1_45S_50ntFlanks@3861.3898.38\|TAATTCTAGAGCTAATACATGCCGACGGGCGCTGACCC | >hg38_21_+_8433172_8446622_RNA45SN1_45S_50ntFlanks | 3861 | 3898 | 38 | TAATTCTAGAGCTAATACATGCCGACGGGCGCTGACCC | 244.1 | -6.3 | 1.1 | -5.5 | 3.3E-08 | 8.2E-08 | BT20/MB468 | MITO |
| >hg38_21_+_8433172_8446622_RNA45SN1_45S_50ntFlanks@12457.12474.18\|CAAGCGTTGGATTGTTCA | >hg38_21_+_8433172_8446622_RNA45SN1_45S_50ntFlanks | 12457 | 12474 | 18 | CAAGCGTTGGATTGTTCA | 1764.9 | -5.5 | 1.0 | -5.5 | 3.7E-08 | 8.3E-08 | BT20/MB468 | CYTO |
| >hg38_21_+_8433172_8446622_RNA45SN1_45S_50ntFlanks@12457.12477.21\|CAAGCGTTGGATTGTTCACCC | >hg38_21_+_8433172_8446622_RNA45SN1_45S_50ntFlanks | 12457 | 12477 | 21 | CAAGCGTTGGATTGTTCACCC | 969.5 | -3.3 | 0.6 | -5.5 | 3.3E-08 | 8.3E-08 | BT20/MB468 | MITO |
| >hg38_21_+_8433172_8446622_RNA45SN1_45S_50ntFlanks@12631.12654.24\|GGCGAAGCTACCATCTGTGGGATT | >hg38_21_+_8433172_8446622_RNA45SN1_45S_50ntFlanks | 12631 | 12654 | 24 | GGCGAAGCTACCATCTGTGGGATT | 546.0 | -3.6 | 0.7 | -5.5 | 3.3E-08 | 8.3E-08 | BT20/MB468 | MITO |
| >hg38_MT_+_1621_3279_RNR2_16S_50ntFlanks@286.310.25\|GAAACCAGACGAGCTACCTAAGAAC | >hg38_MT_+_1621_3279_RNR2_16S_50ntFlanks | 286 | 310 | 25 | GAAACCAGACGAGCTACCTAAGAAC | 144.5 | 10.9 | 1.9 | 5.8 | 7.5E-09 | 8.3E-08 | BT20/MB468 | MP |
| >hg38_21_+_8433172_8446622_RNA45SN1_45S_50ntFlanks@10713.10736.24\|CTTGAAAATCCGGGGGAGAGGGTG | >hg38_21_+_8433172_8446622_RNA45SN1_45S_50ntFlanks | 10713 | 10736 | 24 | CTTGAAAATCCGGGGGAGAGGGTG | 370.0 | -3.5 | 0.6 | -5.5 | 4.3E-08 | 8.3E-08 | MB231/MB468 | CYTO |
| >hg38_21_+_8433172_8446622_RNA45SN1_45S_50ntFlanks@12996.13035.40\|TCGCTGCGATCTATTGAAAGTCAGCCCTCGACACAAGGGT | >hg38_21_+_8433172_8446622_RNA45SN1_45S_50ntFlanks | 12996 | 13035 | 40 | TCGCTGCGATCTATTGAAAGTCAGCCCTCGACACAAGGGT | 337.3 | 9.3 | 1.7 | 5.5 | 3.7E-08 | 8.3E-08 | BT20/MB468 | CYTO |
| >hg38_21_+_8433172_8446622_RNA45SN1_45S_50ntFlanks@12008.12031.24\|CCGCCGGTGAAATACCACTACTCT | >hg38_21_+_8433172_8446622_RNA45SN1_45S_50ntFlanks | 12008 | 12031 | 24 | CCGCCGGTGAAATACCACTACTCT | 2208.1 | -4.3 | 0.8 | -5.5 | 3.7E-08 | 8.3E-08 | BT20/MB468 | CYTO |
| >hg38_21_+_8433172_8446622_RNA45SN1_45S_50ntFlanks@10355.10383.29\|AACTTTGAAGGCCGAAGTGGAGAAGGGTT | >hg38_21_+_8433172_8446622_RNA45SN1_45S_50ntFlanks | 10355 | 10383 | 29 | AACTTTGAAGGCCGAAGTGGAGAAGGGTT | 479.8 | -5.0 | 0.9 | -5.5 | 4.0E-08 | 8.3E-08 | MB231/MB468 | MP |
| >trna119_LysCTT_1_-_145395522_145395594@1.35.35,trna11_LysCTT_5_-_180648979_180649051@1.35.35,trna13_LysCTT_6_+_26556774_26556846@1.35.35,trna7_LysCTT_16_+_3225692_3225764@1.35.35,trna9_LysCTT_5_+_180634755_180634827@1.35.35\|GCCCGGCTAGCTCAGTCGGTAGAGCATGAGACTCT | >trna119_LysCTT_1_-_145395522_145395594 | 1 | 35 | 35 | GCCCGGCTAGCTCAGTCGGTAGAGCATGAGACTCT | 1250.6 | 4.4 | 0.8 | 5.5 | 4.0E-08 | 8.3E-08 | MB231/MB468 | MP |
| >hg38_21_+_8433172_8446622_RNA45SN1_45S_50ntFlanks@10727.10747.21\|GGAGAGGGTGTAAATCTCGCG | >hg38_21_+_8433172_8446622_RNA45SN1_45S_50ntFlanks | 10727 | 10747 | 21 | GGAGAGGGTGTAAATCTCGCG | 124.9 | -10.0 | 1.7 | -5.8 | 7.5E-09 | 8.3E-08 | BT20/MB468 | MP |
| >hg38_21_+_8433172_8446622_RNA45SN1_45S_50ntFlanks@10350.10377.28\|ACGAGAACTTTGAAGGCCGAAGTGGAGA | >hg38_21_+_8433172_8446622_RNA45SN1_45S_50ntFlanks | 10350 | 10377 | 28 | ACGAGAACTTTGAAGGCCGAAGTGGAGA | 1336.4 | -2.7 | 0.5 | -5.5 | 3.7E-08 | 8.3E-08 | BT20/MB468 | CYTO |
| >hg38_21_+_8433172_8446622_RNA45SN1_45S_50ntFlanks@12983.13001.19\|CAGAGCAGCTCCCTCGCTG | >hg38_21_+_8433172_8446622_RNA45SN1_45S_50ntFlanks | 12983 | 13001 | 19 | CAGAGCAGCTCCCTCGCTG | 102.0 | -9.8 | 1.7 | -5.8 | 7.5E-09 | 8.3E-08 | BT20/MB468 | MP |
| >trnaMT_HisGTG_MT_+_12138_12206@38.72.35\|TGACAACAGAGGCTTACGACCCCTTATTTACCCCA | >trnaMT_HisGTG_MT_+_12138_12206 | 38 | 72 | 35 | TGACAACAGAGGCTTACGACCCCTTATTTACCCCA | 141.1 | 10.9 | 1.9 | 5.8 | 7.6E-09 | 8.4E-08 | BT20/MB468 | MP |
| >MI0003513\|hsa-mir-455&WithFlank&9\|+\|114209428\|114209535@60.81.22&[MIMAT0004784&hsa-miR-455-3p&offsets\|0\|+1;m-135&9\|+\|114209487\|114209508&offsets\|0\|0]\|GCAGTCCATGGGCATATACACT | >MI0003513\|hsa-mir-455&WithFlank&9\|+\|114209428\|114209535 | 60 | 81 | 22 | GCAGTCCATGGGCATATACACT | 314.4 | -3.5 | 0.6 | -5.9 | 4.6E-09 | 8.4E-08 | BT20/MB231 | NUC |
| >hg38_21_+_8433172_8446622_RNA45SN1_45S_50ntFlanks@10797.10818.22\|GCATGTTGGAACAATGTAGGTA | >hg38_21_+_8433172_8446622_RNA45SN1_45S_50ntFlanks | 10797 | 10818 | 22 | GCATGTTGGAACAATGTAGGTA | 3012.0 | -3.9 | 0.7 | -5.5 | 3.7E-08 | 8.4E-08 | BT20/MB468 | CYTO |
| >hg38_21_+_8433172_8446622_RNA45SN1_45S_50ntFlanks@12633.12656.24\|CGAAGCTACCATCTGTGGGATTAT | >hg38_21_+_8433172_8446622_RNA45SN1_45S_50ntFlanks | 12633 | 12656 | 24 | CGAAGCTACCATCTGTGGGATTAT | 690.0 | -7.9 | 1.4 | -5.5 | 4.7E-08 | 8.4E-08 | MB231/MB468 | MITO |
| >hg38_21_+_8433172_8446622_RNA45SN1_45S_50ntFlanks@10796.10818.23\|GGCATGTTGGAACAATGTAGGTA | >hg38_21_+_8433172_8446622_RNA45SN1_45S_50ntFlanks | 10796 | 10818 | 23 | GGCATGTTGGAACAATGTAGGTA | 817.1 | -7.8 | 1.4 | -5.5 | 4.1E-08 | 8.4E-08 | MB231/MB468 | MP |
| >hg38_21_+_8433172_8446622_RNA45SN1_45S_50ntFlanks@11919.11948.30\|CATGAGAGGTGTAGAATAAGTGGGAGGCCC | >hg38_21_+_8433172_8446622_RNA45SN1_45S_50ntFlanks | 11919 | 11948 | 30 | CATGAGAGGTGTAGAATAAGTGGGAGGCCC | 244.7 | -5.3 | 1.0 | -5.5 | 4.3E-08 | 8.4E-08 | MB231/MB468 | CYTO |
| >hg38_21_+_8433172_8446622_RNA45SN1_45S_50ntFlanks@8327.8375.49\|ATAGTCAACAAGTACCGTAAGGGAAAGTTGAAAAGAACTTTGAAGAGAG | >hg38_21_+_8433172_8446622_RNA45SN1_45S_50ntFlanks | 8327 | 8375 | 49 | ATAGTCAACAAGTACCGTAAGGGAAAGTTGAAAAGAACTTTGAAGAGAG | 124.0 | 10.7 | 1.9 | 5.8 | 7.6E-09 | 8.4E-08 | BT20/MB468 | MP |
| >trnaMT_TyrGTA_MT_-_5826_5891@24.58.35\|TGGACTGTAAATCTAAAGACAGGGGTTAGGCCTCT | >trnaMT_TyrGTA_MT_-_5826_5891 | 24 | 58 | 35 | TGGACTGTAAATCTAAAGACAGGGGTTAGGCCTCT | 108.8 | 10.5 | 1.8 | 5.8 | 7.6E-09 | 8.4E-08 | BT20/MB468 | MP |
| >hg38_21_+_8433172_8446622_RNA45SN1_45S_50ntFlanks@12936.12966.31\|AAACCATTCGTAGACGACCTGCTTCTGGGTC | >hg38_21_+_8433172_8446622_RNA45SN1_45S_50ntFlanks | 12936 | 12966 | 31 | AAACCATTCGTAGACGACCTGCTTCTGGGTC | 125.4 | -10.1 | 1.7 | -5.8 | 7.6E-09 | 8.4E-08 | BT20/MB468 | MP |
| >MI0000542\|hsa-mir-320a&WithFlank&8\|-\|22244960\|22245043@48.68.21(+4U)&[MIMAT0000510&hsa-miR-320a-3p&offsets\|0\|-1(+4U);m-60&8\|-\|22244975\|22244996&offsets\|0\|-1(+4U)]\|AAAAGCTGGGTTGAGAGGGCGTTTT | >MI0000542\|hsa-mir-320a&WithFlank&8\|-\|22244960\|22245043 | 48 | 68 | 21(+4U) | AAAAGCTGGGTTGAGAGGGCGTTTT | 142.1 | 10.9 | 1.9 | 5.8 | 7.6E-09 | 8.4E-08 | BT20/MB468 | MP |
| >hg38_21_+_8433172_8446622_RNA45SN1_45S_50ntFlanks@10865.10902.38\|TCTAAGGGCTGGGTCGGTCGGGCTGGGGCGCGAAGCGG | >hg38_21_+_8433172_8446622_RNA45SN1_45S_50ntFlanks | 10865 | 10902 | 38 | TCTAAGGGCTGGGTCGGTCGGGCTGGGGCGCGAAGCGG | 232.4 | -7.2 | 1.3 | -5.5 | 4.5E-08 | 8.4E-08 | MB231/MB468 | NUC |

<table>
<tr><td>>hg38_21_+_8433172_8446622_RNA45SN1_45S_50ntFlanks@11896.11931.36|TCTAGTCTGGCACGGTGAAGAGACATGAGAGGTGTA</td><td>>hg38_21_+_8433172_8446622_RNA45SN1_45S_50ntFlanks</td><td>11896</td><td>11931</td><td>36</td><td>TCTAGTCTGGCACGGTGAAGAGACATGAGAGGTGTA</td><td>95.2</td><td>10.3</td><td>1.8</td><td>5.8</td><td>7.7E-09</td><td>8.4E-08</td><td>BT20/MB468</td><td>MP</td></tr>
<tr><td>>hg38_21_+_8433172_8446622_RNA45SN1_45S_50ntFlanks@9835.9868.34|TTTGGTAAGCAGAACTGGCGCTGCGGGATGAACC</td><td>>hg38_21_+_8433172_8446622_RNA45SN1_45S_50ntFlanks</td><td>9835</td><td>9868</td><td>34</td><td>TTTGGTAAGCAGAACTGGCGCTGCGGGATGAACC</td><td>85.2</td><td>10.2</td><td>1.8</td><td>5.8</td><td>7.7E-09</td><td>8.4E-08</td><td>BT20/MB468</td><td>MP</td></tr>
<tr><td>>trnaMT_ValTAC_MT_+_1602_1670@50.71.22|ACTTAACTTGACCGCTCTGACC</td><td>>trnaMT_ValTAC_MT_+_1602_1670</td><td>50</td><td>71</td><td>22</td><td>ACTTAACTTGACCGCTCTGACC</td><td>1397.7</td><td>7.3</td><td>1.3</td><td>5.5</td><td>3.8E-08</td><td>8.4E-08</td><td>BT20/MB468</td><td>CYTO</td></tr>
<tr><td>>hg38_21_+_8433172_8446622_RNA45SN1_45S_50ntFlanks@483.518.36|TCGATGTGGTGACGTCGTGCTCTCCCGGGCCGGGTC</td><td>>hg38_21_+_8433172_8446622_RNA45SN1_45S_50ntFlanks</td><td>483</td><td>518</td><td>36</td><td>TCGATGTGGTGACGTCGTGCTCTCCCGGGCCGGGTC</td><td>293.2</td><td>4.3</td><td>0.8</td><td>5.5</td><td>4.5E-08</td><td>8.5E-08</td><td>MB231/MB468</td><td>NUC</td></tr>
<tr><td>>hg38_21_+_8433172_8446622_RNA45SN1_45S_50ntFlanks@4722.4751.30|TCATTAATCAAGAACGAAAGTCGGAGGTTC</td><td>>hg38_21_+_8433172_8446622_RNA45SN1_45S_50ntFlanks</td><td>4722</td><td>4751</td><td>30</td><td>TCATTAATCAAGAACGAAAGTCGGAGGTTC</td><td>660.9</td><td>-4.4</td><td>0.8</td><td>-5.5</td><td>4.4E-08</td><td>8.5E-08</td><td>MB231/MB468</td><td>CYTO</td></tr>
<tr><td>>hg38_21_+_8433172_8446622_RNA45SN1_45S_50ntFlanks@5496.5516.21|GACGGTCGAACTTGACTATCT</td><td>>hg38_21_+_8433172_8446622_RNA45SN1_45S_50ntFlanks</td><td>5496</td><td>5516</td><td>21</td><td>GACGGTCGAACTTGACTATCT</td><td>854.0</td><td>8.6</td><td>1.6</td><td>5.5</td><td>3.4E-08</td><td>8.5E-08</td><td>BT20/MB468</td><td>MITO</td></tr>
<tr><td>>MI0003137|hsa-mir-193b&WithFlank&16|+|14303961|14304055@57.77.21&[MIMAT0002819&hsa-miR-193b-3p&offsets|0|-1;m-166&16|+|14304017|14304038&offsets|0|-1]|AACTGGCCCTCAAAGTCCCGC</td><td>>MI0003137|hsa-mir-193b&WithFlank&16|+|14303961|14304055</td><td>57</td><td>77</td><td>21</td><td>AACTGGCCCTCAAAGTCCCGC</td><td>204.5</td><td>7.8</td><td>1.3</td><td>5.9</td><td>4.7E-09</td><td>8.5E-08</td><td>BT20/MB231</td><td>NUC</td></tr>
<tr><td>>hg38_21_+_8433172_8446622_RNA45SN1_45S_50ntFlanks@12940.12973.34|CATTCGTAGACGACCTGCTTCTGGGTCGGGGTTT</td><td>>hg38_21_+_8433172_8446622_RNA45SN1_45S_50ntFlanks</td><td>12940</td><td>12973</td><td>34</td><td>CATTCGTAGACGACCTGCTTCTGGGTCGGGGTTT</td><td>116.6</td><td>-9.9</td><td>1.7</td><td>-5.8</td><td>7.7E-09</td><td>8.5E-08</td><td>BT20/MB468</td><td>MP</td></tr>
<tr><td>>hg38_21_+_8433172_8446622_RNA45SN1_45S_50ntFlanks@12457.12490.34|CAAGCGTTGGATTGTTCACCCACTAATAGGGAAC</td><td>>hg38_21_+_8433172_8446622_RNA45SN1_45S_50ntFlanks</td><td>12457</td><td>12490</td><td>34</td><td>CAAGCGTTGGATTGTTCACCCACTAATAGGGAAC</td><td>382.2</td><td>-6.3</td><td>1.2</td><td>-5.5</td><td>5.0E-08</td><td>8.5E-08</td><td>BT20/MB468</td><td>NUC</td></tr>
<tr><td>>hg38_21_+_8433172_8446622_RNA45SN1_45S_50ntFlanks@11918.11951.34|ACATGAGAGGTGTAGAATAAGTGGGAGGCCCCCG</td><td>>hg38_21_+_8433172_8446622_RNA45SN1_45S_50ntFlanks</td><td>11918</td><td>11951</td><td>34</td><td>ACATGAGAGGTGTAGAATAAGTGGGAGGCCCCCG</td><td>1403.7</td><td>-7.6</td><td>1.4</td><td>-5.5</td><td>3.8E-08</td><td>8.5E-08</td><td>BT20/MB468</td><td>CYTO</td></tr>
<tr><td>>hg38_21_+_8433172_8446622_RNA45SN1_45S_50ntFlanks@12406.12431.26|TTTTTGATCCTTCGATGTCGGCTCTT</td><td>>hg38_21_+_8433172_8446622_RNA45SN1_45S_50ntFlanks</td><td>12406</td><td>12431</td><td>26</td><td>TTTTTGATCCTTCGATGTCGGCTCTT</td><td>679.8</td><td>-5.7</td><td>1.0</td><td>-5.5</td><td>3.8E-08</td><td>8.5E-08</td><td>BT20/MB468</td><td>CYTO</td></tr>
<tr><td>>hg38_21_+_8433172_8446622_RNA45SN1_45S_50ntFlanks@9540.9571.32|CTCTGGTGGAGGTCCGTAGCGGTCCTGACGTG</td><td>>hg38_21_+_8433172_8446622_RNA45SN1_45S_50ntFlanks</td><td>9540</td><td>9571</td><td>32</td><td>CTCTGGTGGAGGTCCGTAGCGGTCCTGACGTG</td><td>687.0</td><td>-2.8</td><td>0.5</td><td>-5.5</td><td>3.8E-08</td><td>8.5E-08</td><td>BT20/MB468</td><td>CYTO</td></tr>
<tr><td>>hg38_21_+_8433172_8446622_RNA45SN1_45S_50ntFlanks@12474.12496.23|ACCCACTAATAGGGAACGTGAGC</td><td>>hg38_21_+_8433172_8446622_RNA45SN1_45S_50ntFlanks</td><td>12474</td><td>12496</td><td>23</td><td>ACCCACTAATAGGGAACGTGAGC</td><td>538.9</td><td>-4.3</td><td>0.8</td><td>-5.5</td><td>5.0E-08</td><td>8.5E-08</td><td>BT20/MB468</td><td>NUC</td></tr>
<tr><td>>trnaMT_HisGTG_MT_+_12138_12206@27.63.37|GATTGTGAATCTGACAACAGAGGCTTACGACCCCTTA</td><td>>trnaMT_HisGTG_MT_+_12138_12206</td><td>27</td><td>63</td><td>37</td><td>GATTGTGAATCTGACAACAGAGGCTTACGACCCCTTA</td><td>178.1</td><td>11.2</td><td>1.9</td><td>5.8</td><td>7.7E-09</td><td>8.5E-08</td><td>BT20/MB468</td><td>MP</td></tr>
<tr><td>>MI0000462|hsa-mir-152&WithFlank&17|-|48037155|48037253@60.81.22&[MIMAT0000438&hsa-miR-152-3p&offsets|0|+1;m-97&17|-|48037174|48037194&offsets|0|+1]|TCAGTGCATGACAGAACTTGGG</td><td>>MI0000462|hsa-mir-152&WithFlank&17|-|48037155|48037253</td><td>60</td><td>81</td><td>22</td><td>TCAGTGCATGACAGAACTTGGG</td><td>964.0</td><td>8.8</td><td>1.5</td><td>5.8</td><td>7.8E-09</td><td>8.5E-08</td><td>BT20/MB468</td><td>MP</td></tr>
<tr><td>>hg38_21_+_8433172_8446622_RNA45SN1_45S_50ntFlanks@9005.9026.22|CTCCCCCGCGGGGGCGCGCCGG</td><td>>hg38_21_+_8433172_8446622_RNA45SN1_45S_50ntFlanks</td><td>9005</td><td>9026</td><td>22</td><td>CTCCCCCGCGGGGGCGCGCCGG</td><td>359.0</td><td>-8.0</td><td>1.5</td><td>-5.5</td><td>5.0E-08</td><td>8.5E-08</td><td>BT20/MB468</td><td>NUC</td></tr>
<tr><td>>hg38_21_+_8433172_8446622_RNA45SN1_45S_50ntFlanks@9546.9573.28|TGGAGGTCCGTAGCGGTCCTGACGTGCA</td><td>>hg38_21_+_8433172_8446622_RNA45SN1_45S_50ntFlanks</td><td>9546</td><td>9573</td><td>28</td><td>TGGAGGTCCGTAGCGGTCCTGACGTGCA</td><td>392.5</td><td>2.9</td><td>0.5</td><td>5.5</td><td>3.8E-08</td><td>8.5E-08</td><td>BT20/MB468</td><td>CYTO</td></tr>
<tr><td>>trnaMT_GluTTC_MT_-_14674_14742@40.58.19,trnalookalike8_GluTTC_5_-_93905172_93905240@40.58.19|TTGGTCGTGGTTGTAGTCC</td><td>>trnaMT_GluTTC_MT_-_14674_14742</td><td>40</td><td>58</td><td>19</td><td>TTGGTCGTGGTTGTAGTCC</td><td>861.5</td><td>5.1</td><td>0.9</td><td>5.8</td><td>7.8E-09</td><td>8.5E-08</td><td>BT20/MB468</td><td>MP</td></tr>
<tr><td>>hg38_21_+_8433172_8446622_RNA45SN1_45S_50ntFlanks@11924.11949.26|GAGGTGTAGAATAAGTGGGAGGCCCC</td><td>>hg38_21_+_8433172_8446622_RNA45SN1_45S_50ntFlanks</td><td>11924</td><td>11949</td><td>26</td><td>GAGGTGTAGAATAAGTGGGAGGCCCC</td><td>1631.1</td><td>14.0</td><td>2.3</td><td>6.0</td><td>1.5E-09</td><td>8.5E-08</td><td>BT20/MB231</td><td>MP</td></tr>
<tr><td>>hg38_21_+_8433172_8446622_RNA45SN1_45S_50ntFlanks@12417.12446.30|TCGATGTCGGCTCTTCCTATCATTGTGAAG</td><td>>hg38_21_+_8433172_8446622_RNA45SN1_45S_50ntFlanks</td><td>12417</td><td>12446</td><td>30</td><td>TCGATGTCGGCTCTTCCTATCATTGTGAAG</td><td>1580.0</td><td>-3.1</td><td>0.6</td><td>-5.5</td><td>4.5E-08</td><td>8.5E-08</td><td>MB231/MB468</td><td>NUC</td></tr>
<tr><td>>hg38_21_+_8433172_8446622_RNA45SN1_45S_50ntFlanks@12214.12232.19|CAGGGAGGACAGAAACCTC</td><td>>hg38_21_+_8433172_8446622_RNA45SN1_45S_50ntFlanks</td><td>12214</td><td>12232</td><td>19</td><td>CAGGGAGGACAGAAACCTC</td><td>83.5</td><td>-9.5</td><td>1.6</td><td>-5.8</td><td>7.8E-09</td><td>8.5E-08</td><td>BT20/MB468</td><td>MP</td></tr>
<tr><td>>hg38_21_+_8433172_8446622_RNA45SN1_45S_50ntFlanks@6784.6802.19|GCTACGCCTGTCTGAGCGT</td><td>>hg38_21_+_8433172_8446622_RNA45SN1_45S_50ntFlanks</td><td>6784</td><td>6802</td><td>19</td><td>GCTACGCCTGTCTGAGCGT</td><td>58.4</td><td>-9.7</td><td>1.8</td><td>-5.5</td><td>4.1E-08</td><td>8.5E-08</td><td>MB231/MB468</td><td>MP</td></tr>
<tr><td>>hg38_21_+_8433172_8446622_RNA45SN1_45S_50ntFlanks@10663.10706.44|CCTTGGAAAGCGTCGCGGTTCCGGCGGCGTCCGGTGAGCTCTCG</td><td>>hg38_21_+_8433172_8446622_RNA45SN1_45S_50ntFlanks</td><td>10663</td><td>10706</td><td>44</td><td>CCTTGGAAAGCGTCGCGGTTCCGGCGGCGTCCGGTGAGCTCTCG</td><td>289.4</td><td>-5.4</td><td>1.0</td><td>-5.5</td><td>4.4E-08</td><td>8.6E-08</td><td>MB231/MB468</td><td>CYTO</td></tr>
<tr><td>>hg38_21_+_8433172_8446622_RNA45SN1_45S_50ntFlanks@12004.12035.32|CGGGCCGCCGGTGAAATACCACTACTCTGATC</td><td>>hg38_21_+_8433172_8446622_RNA45SN1_45S_50ntFlanks</td><td>12004</td><td>12035</td><td>32</td><td>CGGGCCGCCGGTGAAATACCACTACTCTGATC</td><td>884.6</td><td>5.4</td><td>0.9</td><td>5.8</td><td>7.8E-09</td><td>8.6E-08</td><td>BT20/MB468</td><td>MP</td></tr>
<tr><td>>hg38_21_+_8433172_8446622_RNA45SN1_45S_50ntFlanks@5083.5110.28|ACGAACGAGACTCTGGCATGCTAACTAG</td><td>>hg38_21_+_8433172_8446622_RNA45SN1_45S_50ntFlanks</td><td>5083</td><td>5110</td><td>28</td><td>ACGAACGAGACTCTGGCATGCTAACTAG</td><td>1218.5</td><td>-5.9</td><td>1.1</td><td>-5.5</td><td>4.8E-08</td><td>8.6E-08</td><td>MB231/MB468</td><td>MITO</td></tr>
<tr><td>>hg38_21_+_8433172_8446622_RNA45SN1_45S_50ntFlanks@12624.12660.37|CCAATGGGGCGAAGCTACCATCTGTGGGATTATGACT</td><td>>hg38_21_+_8433172_8446622_RNA45SN1_45S_50ntFlanks</td><td>12624</td><td>12660</td><td>37</td><td>CCAATGGGGCGAAGCTACCATCTGTGGGATTATGACT</td><td>68.9</td><td>-10.0</td><td>1.8</td><td>-5.5</td><td>4.1E-08</td><td>8.6E-08</td><td>MB231/MB468</td><td>MP</td></tr>
<tr><td>>hg38_21_+_8433172_8446622_RNA45SN1_45S_50ntFlanks@9570.9596.27|TGCAAATCGGTCGTCCGACCTGGGTAT</td><td>>hg38_21_+_8433172_8446622_RNA45SN1_45S_50ntFlanks</td><td>9570</td><td>9596</td><td>27</td><td>TGCAAATCGGTCGTCCGACCTGGGTAT</td><td>822.7</td><td>-7.4</td><td>1.3</td><td>-5.5</td><td>4.5E-08</td><td>8.6E-08</td><td>MB231/MB468</td><td>NUC</td></tr>
<tr><td>>MI0000289|hsa-mir-181a-1&WithFlank&1|-|198859038|198859159@70.90.21(+1A)&[MIMAT0000270&hsa-miR-181a-3p&offsets|0|-1(+1A);m-125&1|-|198859069|198859090&offsets|0|-1(+1A)]|ACCATCGACCGTTGATTGTACA</td><td>>MI0000289|hsa-mir-181a-1&WithFlank&1|-|198859038|198859159</td><td>70</td><td>90</td><td>21(+1A)</td><td>ACCATCGACCGTTGATTGTACA</td><td>178.7</td><td>-10.9</td><td>1.8</td><td>-6.0</td><td>1.5E-09</td><td>8.6E-08</td><td>BT20/MB231</td><td>MP</td></tr>
<tr><td>>hg38_21_+_8433172_8446622_RNA45SN1_45S_50ntFlanks@12415.12435.21|CTTCGATGTCGGCTCTTCCTA</td><td>>hg38_21_+_8433172_8446622_RNA45SN1_45S_50ntFlanks</td><td>12415</td><td>12435</td><td>21</td><td>CTTCGATGTCGGCTCTTCCTA</td><td>2148.5</td><td>-3.6</td><td>0.7</td><td>-5.5</td><td>3.4E-08</td><td>8.6E-08</td><td>BT20/MB468</td><td>MITO</td></tr>
<tr><td>>hg38_21_+_8433172_8446622_RNA45SN1_45S_50ntFlanks@5071.5098.28|TTAATTCCGATAACGAACGAGACTCTGG</td><td>>hg38_21_+_8433172_8446622_RNA45SN1_45S_50ntFlanks</td><td>5071</td><td>5098</td><td>28</td><td>TTAATTCCGATAACGAACGAGACTCTGG</td><td>1519.2</td><td>-5.7</td><td>1.0</td><td>-5.5</td><td>4.6E-08</td><td>8.6E-08</td><td>MB231/MB468</td><td>NUC</td></tr>
<tr><td>>hg38_21_+_8433172_8446622_RNA45SN1_45S_50ntFlanks@4160.4186.27|CCACATCCAAGGAAGGCAGCAGGCGCG</td><td>>hg38_21_+_8433172_8446622_RNA45SN1_45S_50ntFlanks</td><td>4160</td><td>4186</td><td>27</td><td>CCACATCCAAGGAAGGCAGCAGGCGCG</td><td>260.1</td><td>-7.6</td><td>1.4</td><td>-5.5</td><td>4.2E-08</td><td>8.6E-08</td><td>MB231/MB468</td><td>MP</td></tr>
<tr><td>>hg38_21_+_8433172_8446622_RNA45SN1_45S_50ntFlanks@10788.10815.28|CAGCCTCTGGCATGTTGGAACAATGTAG</td><td>>hg38_21_+_8433172_8446622_RNA45SN1_45S_50ntFlanks</td><td>10788</td><td>10815</td><td>28</td><td>CAGCCTCTGGCATGTTGGAACAATGTAG</td><td>515.3</td><td>-3.9</td><td>0.7</td><td>-5.5</td><td>4.5E-08</td><td>8.6E-08</td><td>MB231/MB468</td><td>CYTO</td></tr>
<tr><td>>hg38_21_+_8433172_8446622_RNA45SN1_45S_50ntFlanks@12406.12444.39|TTTTTGATCCTTCGATGTCGGCTCTTCCTATCATTGTGA</td><td>>hg38_21_+_8433172_8446622_RNA45SN1_45S_50ntFlanks</td><td>12406</td><td>12444</td><td>39</td><td>TTTTTGATCCTTCGATGTCGGCTCTTCCTATCATTGTGA</td><td>668.7</td><td>-8.8</td><td>1.6</td><td>-5.4</td><td>5.1E-08</td><td>8.6E-08</td><td>BT20/MB468</td><td>NUC</td></tr>
<tr><td>>hg38_21_+_8433172_8446622_RNA45SN1_45S_50ntFlanks@3840.3872.33|CTCCTACTTGGATAACTGTGGTAATTCTAGAGC</td><td>>hg38_21_+_8433172_8446622_RNA45SN1_45S_50ntFlanks</td><td>3840</td><td>3872</td><td>33</td><td>CTCCTACTTGGATAACTGTGGTAATTCTAGAGC</td><td>749.7</td><td>3.5</td><td>0.6</td><td>5.5</td><td>3.9E-08</td><td>8.6E-08</td><td>BT20/MB468</td><td>CYTO</td></tr>
<tr><td>>hg38_21_+_8433172_8446622_RNA45SN1_45S_50ntFlanks@11774.11794.21|AGTGACGCGCATGAATGGATG</td><td>>hg38_21_+_8433172_8446622_RNA45SN1_45S_50ntFlanks</td><td>11774</td><td>11794</td><td>21</td><td>AGTGACGCGCATGAATGGATG</td><td>659.8</td><td>-4.9</td><td>0.9</td><td>-5.4</td><td>5.1E-08</td><td>8.6E-08</td><td>BT20/MB468</td><td>NUC</td></tr>
<tr><td>>am_trna130_GlnTTG_6_-_27763640_27763711@13.30.18,trna146_GlnCTG_6_-_27515531_27515602@13.30.18,trna16_GlnTTG_17_+_47269890_47269961@13.30.18,trna173_GlnTTG_6_-_26311975_26312046@13.30.18,trna174_GlnTTG_6_-_26311424_26311495@13.30.18,trna1_GlnCTG_6_+_18836402_18836473@13.30.18,trna3_GlnCTG_17_+_8023070_8023141@13.30.18,trna42_GlnCTG_6_+_27263212_27263283@13.30.18,trna49_GlnCTG_6_+_27487308_27487379@13.30.18,trna64_GlnTTG_6_+_28557156_28557227@13.30.18,trna7_GlnCTG_15_-_66161400_66161471@13.30.18,trna84_GlnTTG_6_+_145503859_145503930@13.30.18,trna99_GlnCTG_6_-_28909378_28909449@13.30.18|TAATGGTTAGCACTCTGG</td><td>>am_trna130_GlnTTG_6_-_27763640_27763711</td><td>13</td><td>30</td><td>18</td><td>TAATGGTTAGCACTCTGG</td><td>228.8</td><td>-7.2</td><td>1.3</td><td>-5.5</td><td>4.6E-08</td><td>8.6E-08</td><td>MB231/MB468</td><td>NUC</td></tr>
<tr><td>>hg38_21_+_8433172_8446622_RNA45SN1_45S_50ntFlanks@5490.5532.43|TGAGAAGACGGTCGAACTTGACTATCTAGAGGAAGTAAAAGTC</td><td>>hg38_21_+_8433172_8446622_RNA45SN1_45S_50ntFlanks</td><td>5490</td><td>5532</td><td>43</td><td>TGAGAAGACGGTCGAACTTGACTATCTAGAGGAAGTAAAAGTC</td><td>95.6</td><td>10.3</td><td>1.8</td><td>5.8</td><td>7.9E-09</td><td>8.6E-08</td><td>BT20/MB468</td><td>MP</td></tr>
<tr><td>>MI0000283|hsa-mir-203a&WithFlank&14|+|104117399|104117520@71.93.23&[MIMAT0000264&hsa-miR-203a-3p&offsets|0|+1;m-79&14|+|104117469|104117490&offsets|0|+1]|GTGAAATGTTTAGGACCACTAGA</td><td>>MI0000283|hsa-mir-203a&WithFlank&14|+|104117399|104117520</td><td>71</td><td>93</td><td>23</td><td>GTGAAATGTTTAGGACCACTAGA</td><td>88.3</td><td>3.8</td><td>0.7</td><td>5.4</td><td>5.1E-08</td><td>8.6E-08</td><td>BT20/MB468</td><td>NUC</td></tr>
<tr><td>>hg38_21_+_8433172_8446622_RNA45SN1_45S_50ntFlanks@10727.10745.19|GGAGAGGGTGTAAATCTCG</td><td>>hg38_21_+_8433172_8446622_RNA45SN1_45S_50ntFlanks</td><td>10727</td><td>10745</td><td>19</td><td>GGAGAGGGTGTAAATCTCG</td><td>147.5</td><td>-10.3</td><td>1.8</td><td>-5.8</td><td>7.9E-09</td><td>8.7E-08</td><td>BT20/MB468</td><td>MP</td></tr>
<tr><td>>hg38_21_+_8433172_8446622_RNA45SN1_45S_50ntFlanks@10797.10829.33|GCATGTTGGAACAATGTAGGTAAGGGAAGTCGG</td><td>>hg38_21_+_8433172_8446622_RNA45SN1_45S_50ntFlanks</td><td>10797</td><td>10829</td><td>33</td><td>GCATGTTGGAACAATGTAGGTAAGGGAAGTCGG</td><td>241.4</td><td>-7.2</td><td>1.3</td><td>-5.5</td><td>4.2E-08</td><td>8.7E-08</td><td>MB231/MB468</td><td>MP</td></tr>
<tr><td>>hg38_21_+_8433172_8446622_RNA45SN1_45S_50ntFlanks@7975.7992.18|CGCGACCTCAGATCAGAC</td><td>>hg38_21_+_8433172_8446622_RNA45SN1_45S_50ntFlanks</td><td>7975</td><td>7992</td><td>18</td><td>CGCGACCTCAGATCAGAC</td><td>8542.2</td><td>3.7</td><td>0.7</td><td>5.5</td><td>3.9E-08</td><td>8.7E-08</td><td>BT20/MB468</td><td>CYTO</td></tr>
<tr><td>>MI0000114|hsa-mir-107&WithFlank&10|-|89592741|89592833@56.76.21;MI0000108|hsa-mir-103a-2&WithFlank&20|+|3917488|3917577@54.74.21;MI0000109|hsa-mir-103a-1&WithFlank&5|-|168560890|168560979@54.74.21&[MIMAT0000104&hsa-miR-107&offsets|0|-2;m-78&10|-|89592758|89592778&offsets|0|0];[MIMAT0000101&hsa-miR-103a-3p&offsets|0|-2;m-12&20|+|3917541|3917563&offsets|0|-2];[MIMAT0000101_1&hsa-miR-103a-3p&offsets|0|-2;m-13&5|-|168560904|168560926&offsets|0|-2;m-44220&5|-|168560908|168560926&offsets|0|+2]|AGCAGCATTGTACAGGGCTAT</td><td>>MI0000114|hsa-mir-107&WithFlank&10|-|89592741|89592833</td><td>56</td><td>76</td><td>21</td><td>AGCAGCATTGTACAGGGCTAT</td><td>242.2</td><td>4.1</td><td>0.7</td><td>5.4</td><td>5.1E-08</td><td>8.7E-08</td><td>BT20/MB468</td><td>NUC</td></tr>
<tr><td>>hg38_21_+_8433172_8446622_RNA45SN1_45S_50ntFlanks@4687.4707.21|ACGGACCAGAGCGAAAGCATT</td><td>>hg38_21_+_8433172_8446622_RNA45SN1_45S_50ntFlanks</td><td>4687</td><td>4707</td><td>21</td><td>ACGGACCAGAGCGAAAGCATT</td><td>306.8</td><td>-4.0</td><td>0.7</td><td>-5.5</td><td>4.6E-08</td><td>8.7E-08</td><td>MB231/MB468</td><td>NUC</td></tr>
<tr><td>>MI0000469|hsa-mir-125a&WithFlank&19|+|51693248|51693345@21.44.24&[MIMAT0000443&hsa-miR-125a-5p&offsets|0|0;m-63&19|+|51693268|51693289&offsets|0|+2]|TCCCTGAGACCCTTTAACCTGTGA</td><td>>MI0000469|hsa-mir-125a&WithFlank&19|+|51693248|51693345</td><td>21</td><td>44</td><td>24</td><td>TCCCTGAGACCCTTTAACCTGTGA</td><td>2809.6</td><td>3.8</td><td>0.7</td><td>5.5</td><td>4.6E-08</td><td>8.7E-08</td><td>MB231/MB468</td><td>NUC</td></tr>
<tr><td>>hg38_21_+_8433172_8446622_RNA45SN1_45S_50ntFlanks@5487.5522.36|CGCTGAGAAGACGGTCGAACTTGACTATCTAGAGGA</td><td>>hg38_21_+_8433172_8446622_RNA45SN1_45S_50ntFlanks</td><td>5487</td><td>5522</td><td>36</td><td>CGCTGAGAAGACGGTCGAACTTGACTATCTAGAGGA</td><td>273.2</td><td>-7.6</td><td>1.4</td><td>-5.5</td><td>4.2E-08</td><td>8.7E-08</td><td>MB231/MB468</td><td>MP</td></tr>
<tr><td>>hg38_21_+_8433172_8446622_RNA45SN1_45S_50ntFlanks@10791.10826.36|CCTCTGGCATGTTGGAACAATGTAGGTAAGGGAAGT</td><td>>hg38_21_+_8433172_8446622_RNA45SN1_45S_50ntFlanks</td><td>10791</td><td>10826</td><td>36</td><td>CCTCTGGCATGTTGGAACAATGTAGGTAAGGGAAGT</td><td>46.6</td><td>-9.4</td><td>1.7</td><td>-5.5</td><td>4.2E-08</td><td>8.7E-08</td><td>MB231/MB468</td><td>MP</td></tr>
<tr><td>>hg38_21_+_8433172_8446622_RNA45SN1_45S_50ntFlanks@51.71.21|GCTGACACGCTGTCCTCTGGC</td><td>>hg38_21_+_8433172_8446622_RNA45SN1_45S_50ntFlanks</td><td>51</td><td>71</td><td>21</td><td>GCTGACACGCTGTCCTCTGGC</td><td>1014.1</td><td>-3.5</td><td>0.6</td><td>-5.4</td><td>5.1E-08</td><td>8.7E-08</td><td>BT20/MB468</td><td>NUC</td></tr>
</table>

| | | | | | | | | | | | | | |
|---|---|---|---|---|---|---|---|---|---|---|---|---|---|
| >hg38_21_+_8433172_8446622_RNA45SN1_45S_50ntFlanks@6698.6721.24\|AGCTAGCTGCGAGAATTAATGTGA | >hg38_21_+_8433172_8446622_RNA45SN1_45S_50ntFlanks | 6698 | 6721 | 24 | AGCTAGCTGCGAGAATTAATGTGA | 706.3 | -6.6 | 1.2 | -5.4 | 5.1E-08 | 8.7E-08 | BT20/MB468 | NUC |
| >hg38_21_+_8433172_8446622_RNA45SN1_45S_50ntFlanks@8176.8197.22\|CTTCTGATCGAGGCCCAGCCCG | >hg38_21_+_8433172_8446622_RNA45SN1_45S_50ntFlanks | 8176 | 8197 | 22 | CTTCTGATCGAGGCCCAGCCCG | 1601.6 | -9.0 | 1.7 | -5.4 | 5.2E-08 | 8.7E-08 | BT20/MB468 | NUC |
| >hg38_21_+_8433172_8446622_RNA45SN1_45S_50ntFlanks@10805.10823.19\|GAACAATGTAGGTAAGGGA | >hg38_21_+_8433172_8446622_RNA45SN1_45S_50ntFlanks | 10805 | 10823 | 19 | GAACAATGTAGGTAAGGGA | 222.8 | -6.2 | 1.1 | -5.5 | 3.5E-08 | 8.7E-08 | BT20/MB468 | MITO |
| >hg38_21_+_8433172_8446622_RNA45SN1_45S_50ntFlanks@12456.12479.24\|CCAAGCGTTGGATTGTTCACCCAC | >hg38_21_+_8433172_8446622_RNA45SN1_45S_50ntFlanks | 12456 | 12479 | 24 | CCAAGCGTTGGATTGTTCACCCAC | 413.4 | -5.9 | 1.1 | -5.5 | 4.6E-08 | 8.7E-08 | MB231/MB468 | NUC |
| >MI0000077\|hsa-mir-21&WithFlank&17\|+\|59841260\|59841343@14.33.20(+1A)&[MIMAT0000076&hsa-miR-21-5p&offsets\|0\|-2(+1A);m-1&17\|+\|59841273\|59841294&offsets\|0\|-2(+1A)]\|TAGCTTATCAGACTGATGTTA | >MI0000077\|hsa-mir-21&WithFlank&17\|+\|59841260\|59841343 | 14 | 33 | 20(+1A) | TAGCTTATCAGACTGATGTTA | 1247.8 | 5.8 | 1.1 | 5.5 | 4.9E-08 | 8.7E-08 | MB231/MB468 | MITO |
| >MI0001445\|hsa-mir-423&WithFlank&17\|+\|30117073\|30117178@59.80.22&[MIMAT0001340&hsa-miR-423-3p&offsets\|0\|-1;m-100&17\|+\|30117131\|30117153&offsets\|0\|-1]\|AGCTCGGTCTGAGGCCCCTCAG | >MI0001445\|hsa-mir-423&WithFlank&17\|+\|30117073\|30117178 | 59 | 80 | 22 | AGCTCGGTCTGAGGCCCCTCAG | 984.0 | 4.7 | 0.9 | 5.5 | 4.9E-08 | 8.7E-08 | MB231/MB468 | MITO |
| >hg38_21_+_8433172_8446622_RNA45SN1_45S_50ntFlanks@475.503.29\|CTTCGTGATCGATGTGGTGACGTCGTGCT | >hg38_21_+_8433172_8446622_RNA45SN1_45S_50ntFlanks | 475 | 503 | 29 | CTTCGTGATCGATGTGGTGACGTCGTGCT | 1590.0 | 5.3 | 1.0 | 5.5 | 4.9E-08 | 8.7E-08 | MB231/MB468 | MITO |
| >hg38_21_+_8433172_8446622_RNA45SN1_45S_50ntFlanks@6653.6700.48\|ACTCTTAGCGGTGGATCACTCGGCTCGTGCGTCGATGAAGAACGCAGC | >hg38_21_+_8433172_8446622_RNA45SN1_45S_50ntFlanks | 6653 | 6700 | 48 | ACTCTTAGCGGTGGATCACTCGGCTCGTGCGTCGATGAAGAACGCAGC | 508.9 | 7.9 | 1.4 | 5.5 | 3.5E-08 | 8.8E-08 | BT20/MB468 | MITO |
| >hg38_21_+_8433172_8446622_RNA45SN1_45S_50ntFlanks@4048.4066.19\|TTCGAACGTCTGCCCTATC | >hg38_21_+_8433172_8446622_RNA45SN1_45S_50ntFlanks | 4048 | 4066 | 19 | TTCGAACGTCTGCCCTATC | 1033.7 | -5.2 | 0.9 | -5.5 | 5.0E-08 | 8.8E-08 | MB231/MB468 | MITO |
| >hg38_21_+_8433172_8446622_RNA45SN1_45S_50ntFlanks@12997.13038.42\|CGCTGCGATCTATTGAAAGTCAGCCCTCGACACAAGGGTTTG | >hg38_21_+_8433172_8446622_RNA45SN1_45S_50ntFlanks | 12997 | 13038 | 42 | CGCTGCGATCTATTGAAAGTCAGCCCTCGACACAAGGGTTTG | 1293.0 | 8.1 | 1.5 | 5.5 | 3.9E-08 | 8.8E-08 | BT20/MB468 | CYTO |
| >hg38_21_+_8433172_8446622_RNA45SN1_45S_50ntFlanks@11682.11706.25\|AATGTCAAAGTGAAGAAATTCAATG | >hg38_21_+_8433172_8446622_RNA45SN1_45S_50ntFlanks | 11682 | 11706 | 25 | AATGTCAAAGTGAAGAAATTCAATG | 287.9 | -7.5 | 1.4 | -5.5 | 4.3E-08 | 8.8E-08 | MB231/MB468 | MP |
| >hg38_21_+_8433172_8446622_RNA45SN1_45S_50ntFlanks@12241.12268.28\|CAGAAGGGCAAAAGCTCGCTTGATCTTG | >hg38_21_+_8433172_8446622_RNA45SN1_45S_50ntFlanks | 12241 | 12268 | 28 | CAGAAGGGCAAAAGCTCGCTTGATCTTG | 428.7 | -7.4 | 1.4 | -5.5 | 5.0E-08 | 8.8E-08 | MB231/MB468 | MITO |
| >hg38_21_+_8433172_8446622_RNA45SN1_45S_50ntFlanks@11770.11794.25\|AATTAGTGACGCGCATGAATGGATG | >hg38_21_+_8433172_8446622_RNA45SN1_45S_50ntFlanks | 11770 | 11794 | 25 | AATTAGTGACGCGCATGAATGGATG | 1020.9 | -4.7 | 0.9 | -5.4 | 5.2E-08 | 8.8E-08 | BT20/MB468 | NUC |
| >hg38_21_+_8433172_8446622_RNA45SN1_45S_50ntFlanks@10781.10825.45\|AGGTGAACAGCCTCTGGCATGTTGGAACAATGTAGGTAAGGGAAG | >hg38_21_+_8433172_8446622_RNA45SN1_45S_50ntFlanks | 10781 | 10825 | 45 | AGGTGAACAGCCTCTGGCATGTTGGAACAATGTAGGTAAGGGAAG | 76.7 | -10.1 | 1.8 | -5.5 | 4.3E-08 | 8.8E-08 | MB231/MB468 | MP |
| >MI0000068\|hsa-let-7f-2&WithFlank&X\|-\|53557186\|53557280@14.36.23&[MIMAT0000067_1&hsa-let-7f-5p&offsets\|0\|+1;m-19&X\|-\|53557246\|53557267&offsets\|0\|+1]\|TGAGGTAGTAGATTGTATAGTTT | >MI0000068\|hsa-let-7f-2&WithFlank&X\|-\|53557186\|53557280 | 14 | 36 | 23 | TGAGGTAGTAGATTGTATAGTTT | 4640.5 | 4.7 | 0.9 | 5.5 | 4.3E-08 | 8.8E-08 | MB231/MB468 | MP |
| >hg38_21_+_8433172_8446622_RNA45SN1_45S_50ntFlanks@11217.11250.34\|CTGGACGCGAGCCGGGCCCTTCCCGTGGATCGCC | >hg38_21_+_8433172_8446622_RNA45SN1_45S_50ntFlanks | 11217 | 11250 | 34 | CTGGACGCGAGCCGGGCCCTTCCCGTGGATCGCC | 239.8 | -7.2 | 1.3 | -5.5 | 4.3E-08 | 8.8E-08 | MB231/MB468 | MP |
| >hg38_21_+_8433172_8446622_RNA45SN1_45S_50ntFlanks@476.506.31\|TTCGTGATCGATGTGGTGACGTCGTGCTCTC | >hg38_21_+_8433172_8446622_RNA45SN1_45S_50ntFlanks | 476 | 506 | 31 | TTCGTGATCGATGTGGTGACGTCGTGCTCTC | 966.7 | 5.0 | 0.9 | 5.5 | 5.0E-08 | 8.8E-08 | MB231/MB468 | MITO |
| >hg38_21_+_8433172_8446622_RNA45SN1_45S_50ntFlanks@5084.5117.34\|CGAACGAGACTCTGGCATGCTAACTAGTTACGCG | >hg38_21_+_8433172_8446622_RNA45SN1_45S_50ntFlanks | 5084 | 5117 | 34 | CGAACGAGACTCTGGCATGCTAACTAGTTACGCG | 418.6 | -7.8 | 1.4 | -5.5 | 5.0E-08 | 8.8E-08 | MB231/MB468 | MITO |
| >hg38_21_+_8433172_8446622_RNA45SN1_45S_50ntFlanks@3776.3805.30\|CCGGTACAGTGAAACTGCGAATGGCTCATT | >hg38_21_+_8433172_8446622_RNA45SN1_45S_50ntFlanks | 3776 | 3805 | 30 | CCGGTACAGTGAAACTGCGAATGGCTCATT | 715.4 | -3.2 | 0.6 | -5.5 | 4.7E-08 | 8.8E-08 | MB231/MB468 | NUC |
| >MI0000108\|hsa-mir-103a-2&WithFlank&20\|+\|3917488\|3917577@54.77.24(+1U);MI0000109\|hsa-mir-103a-1&WithFlank&5\|-\|168560890\|168560979@54.77.24(+1U)&[MIMAT0000101&hsa-miR-103a-3p&offsets\|0\|+1(+1U);m-12&20\|+\|3917541\|3917563&offsets\|0\|+1(+1U)];[MIMAT0000101_1&hsa-miR-103a-3p&offsets\|0\|+1(+1U);m-13&5\|-\|168560904\|168560926&offsets\|0\|+1(+1U);m-44220&5\|-\|168560908\|168560926&offsets\|0\|+5(+1U)]\|AGCAGCATTGTACAGGGCTATGAAT | >MI0000108\|hsa-mir-103a-2&WithFlank&20\|+\|3917488\|3917577 | 54 | 77 | 24(+1U) | AGCAGCATTGTACAGGGCTATGAAT | 296.1 | 7.4 | 1.4 | 5.5 | 4.0E-08 | 8.8E-08 | BT20/MB468 | CYTO |
| >hg38_21_+_8433172_8446622_RNA45SN1_45S_50ntFlanks@10351.10385.35\|CGAGAACTTTGAAGGCCGAAGTGGAGAAGGGTTCC | >hg38_21_+_8433172_8446622_RNA45SN1_45S_50ntFlanks | 10351 | 10385 | 35 | CGAGAACTTTGAAGGCCGAAGTGGAGAAGGGTTCC | 332.1 | -7.3 | 1.3 | -5.5 | 5.0E-08 | 8.8E-08 | MB231/MB468 | MITO |
| >hg38_21_+_8433172_8446622_RNA45SN1_45S_50ntFlanks@6721.6750.30\|AATTGCAGGACACATTGATCATCGACACTT | >hg38_21_+_8433172_8446622_RNA45SN1_45S_50ntFlanks | 6721 | 6750 | 30 | AATTGCAGGACACATTGATCATCGACACTT | 640.3 | -5.8 | 1.1 | -5.5 | 5.0E-08 | 8.8E-08 | MB231/MB468 | MITO |
| >hg38_21_+_8433172_8446622_RNA45SN1_45S_50ntFlanks@9629.9648.20\|CTGGTTCCCTCCGAAGTTTC | >hg38_21_+_8433172_8446622_RNA45SN1_45S_50ntFlanks | 9629 | 9648 | 20 | CTGGTTCCCTCCGAAGTTTC | 112.7 | -10.7 | 1.9 | -5.5 | 4.3E-08 | 8.9E-08 | MB231/MB468 | MP |
| >hg38_21_+_8433172_8446622_RNA45SN1_45S_50ntFlanks@12934.12970.37\|CTAAACCATTCGTAGACGACCTGCTTCTGGGTCGGGG | >hg38_21_+_8433172_8446622_RNA45SN1_45S_50ntFlanks | 12934 | 12970 | 37 | CTAAACCATTCGTAGACGACCTGCTTCTGGGTCGGGG | 169.5 | -10.5 | 1.8 | -5.8 | 8.1E-09 | 8.9E-08 | BT20/MB468 | MP |
| >hg38_21_+_8433172_8446622_RNA45SN1_45S_50ntFlanks@4279.4314.36\|AATCCTTTAACGAGGATCCATTGGAGGGCAAGTCTG | >hg38_21_+_8433172_8446622_RNA45SN1_45S_50ntFlanks | 4279 | 4314 | 36 | AATCCTTTAACGAGGATCCATTGGAGGGCAAGTCTG | 409.8 | -5.8 | 1.1 | -5.5 | 4.7E-08 | 8.9E-08 | MB231/MB468 | NUC |
| >hg38_21_+_8433172_8446622_RNA45SN1_45S_50ntFlanks@3783.3801.19\|AGTGAAACTGCGAATGGCT | >hg38_21_+_8433172_8446622_RNA45SN1_45S_50ntFlanks | 3783 | 3801 | 19 | AGTGAAACTGCGAATGGCT | 857.3 | -5.5 | 1.0 | -5.5 | 4.3E-08 | 8.9E-08 | MB231/MB468 | MP |
| >hg38_21_+_8433172_8446622_RNA45SN1_45S_50ntFlanks@12243.12271.29\|GAAGGGCAAAAGCTCGCTTGATCTTGATT | >hg38_21_+_8433172_8446622_RNA45SN1_45S_50ntFlanks | 12243 | 12271 | 29 | GAAGGGCAAAAGCTCGCTTGATCTTGATT | 257.1 | 5.4 | 1.0 | 5.5 | 4.6E-08 | 8.9E-08 | MB231/MB468 | CYTO |
| >MI0001445\|hsa-mir-423&WithFlank&17\|+\|30117073\|30117178@59.79.21&[MIMAT0001340&hsa-miR-423-3p&offsets\|0\|-2;m-100&17\|+\|30117131\|30117153&offsets\|0\|-2]\|AGCTCGGTCTGAGGCCCCTCA | >MI0001445\|hsa-mir-423&WithFlank&17\|+\|30117073\|30117178 | 59 | 79 | 21 | AGCTCGGTCTGAGGCCCCTCA | 546.4 | 5.4 | 0.9 | 5.8 | 8.2E-09 | 8.9E-08 | BT20/MB468 | MP |
| >hg38_21_+_8433172_8446622_RNA45SN1_45S_50ntFlanks@12434.12455.22\|TATCATTGTGAAGCAGAATTCA | >hg38_21_+_8433172_8446622_RNA45SN1_45S_50ntFlanks | 12434 | 12455 | 22 | TATCATTGTGAAGCAGAATTCA | 249.2 | -5.3 | 1.0 | -5.5 | 4.6E-08 | 8.9E-08 | MB231/MB468 | CYTO |
| >hg38_21_+_8433172_8446622_RNA45SN1_45S_50ntFlanks@5084.5110.27\|CGAACGAGACTCTGGCATGCTAACTAG | >hg38_21_+_8433172_8446622_RNA45SN1_45S_50ntFlanks | 5084 | 5110 | 27 | CGAACGAGACTCTGGCATGCTAACTAG | 2098.8 | -9.2 | 1.7 | -5.5 | 4.3E-08 | 9.0E-08 | MB231/MB468 | MP |
| >hg38_21_+_8433172_8446622_RNA45SN1_45S_50ntFlanks@7973.7991.19\|GACGCGACCTCAGATCAGA | >hg38_21_+_8433172_8446622_RNA45SN1_45S_50ntFlanks | 7973 | 7991 | 19 | GACGCGACCTCAGATCAGA | 1169.1 | 5.1 | 0.9 | 5.5 | 4.6E-08 | 9.0E-08 | MB231/MB468 | CYTO |
| >MI0000750\|hsa-mir-26a-2&WithFlank&12\|-\|57824603\|57824698@20.41.22(+3U);MI0000083\|hsa-mir-26a-1&WithFlank&3\|+\|37969398\|37969486@16.37.22(+3U)&[MIMAT0000082&hsa-miR-26a-5p&offsets\|0\|0(+3U);m-33&12\|-\|57824658\|57824679&offsets\|0\|0(+3U)];[MIMAT0000082_1&hsa-miR-26a-5p&offsets\|0\|0(+3U);m-34&3\|+\|37969413\|37969434&offsets\|0\|0(+3U)]\|TTCAAGTAATCCAGGATAGGCTTTT | >MI0000750\|hsa-mir-26a-2&WithFlank&12\|-\|57824603\|57824698 | 20 | 41 | 22(+3U) | TTCAAGTAATCCAGGATAGGCTTTT | 370.1 | 8.8 | 1.6 | 5.4 | 5.1E-08 | 9.0E-08 | MB231/MB468 | MITO |
| >hg38_21_+_8433172_8446622_RNA45SN1_45S_50ntFlanks@9545.9572.28\|GTGGAGGTCCGTAGCGGTCCTGACGTGC | >hg38_21_+_8433172_8446622_RNA45SN1_45S_50ntFlanks | 9545 | 9572 | 28 | GTGGAGGTCCGTAGCGGTCCTGACGTGC | 2670.8 | -2.9 | 0.5 | -5.5 | 4.0E-08 | 9.0E-08 | BT20/MB468 | CYTO |
| >hg38_21_+_8433172_8446622_RNA45SN1_45S_50ntFlanks@475.497.23\|CTTCGTGATCGATGTGGTGACGT | >hg38_21_+_8433172_8446622_RNA45SN1_45S_50ntFlanks | 475 | 497 | 23 | CTTCGTGATCGATGTGGTGACGT | 792.4 | 8.5 | 1.5 | 5.5 | 4.0E-08 | 9.0E-08 | BT20/MB468 | CYTO |
| >hg38_MT_+_1621_3279_RNR2_16S_50ntFlanks@51.71.21\|GCTAAACCTAGCCCCAAACCC | >hg38_MT_+_1621_3279_RNR2_16S_50ntFlanks | 51 | 71 | 21 | GCTAAACCTAGCCCCAAACCC | 5223.9 | 5.7 | 1.0 | 5.5 | 3.6E-08 | 9.0E-08 | BT20/MB468 | MITO |
| >hg38_21_+_8433172_8446622_RNA45SN1_45S_50ntFlanks@12081.12108.28\|CTCTCGCTTCTGGCGCCAAGCGCCCGGC | >hg38_21_+_8433172_8446622_RNA45SN1_45S_50ntFlanks | 12081 | 12108 | 28 | CTCTCGCTTCTGGCGCCAAGCGCCCGGC | 218.3 | -6.6 | 1.2 | -5.5 | 4.8E-08 | 9.0E-08 | MB231/MB468 | NUC |
| >hg38_21_+_8433172_8446622_RNA45SN1_45S_50ntFlanks@10375.10406.32\|AGAAGGGTTCCATGTGAACAGCAGTTGAACAT | >hg38_21_+_8433172_8446622_RNA45SN1_45S_50ntFlanks | 10375 | 10406 | 32 | AGAAGGGTTCCATGTGAACAGCAGTTGAACAT | 247.4 | -6.6 | 1.2 | -5.5 | 4.0E-08 | 9.0E-08 | BT20/MB468 | CYTO |
| >hg38_21_+_8433172_8446622_RNA45SN1_45S_50ntFlanks@11561.11600.40\|GCGCCTAGCAGCCGACTTAGAACTGGTGCGGACCAGGGGA | >hg38_21_+_8433172_8446622_RNA45SN1_45S_50ntFlanks | 11561 | 11600 | 40 | GCGCCTAGCAGCCGACTTAGAACTGGTGCGGACCAGGGGA | 1731.0 | -6.5 | 1.2 | -5.5 | 4.8E-08 | 9.0E-08 | MB231/MB468 | NUC |
| >hg38_21_+_8433172_8446622_RNA45SN1_45S_50ntFlanks@10873.10892.20\|CTGGGTCGGTCGGGCTGGGG | >hg38_21_+_8433172_8446622_RNA45SN1_45S_50ntFlanks | 10873 | 10892 | 20 | CTGGGTCGGTCGGGCTGGGG | 844.8 | -3.4 | 0.6 | -5.5 | 4.0E-08 | 9.0E-08 | BT20/MB468 | CYTO |
| >hg38_21_+_8433172_8446622_RNA45SN1_45S_50ntFlanks@12589.12614.26\|GTTCAGACATTTGGTGTATGTGCTTG | >hg38_21_+_8433172_8446622_RNA45SN1_45S_50ntFlanks | 12589 | 12614 | 26 | GTTCAGACATTTGGTGTATGTGCTTG | 277.7 | -7.5 | 1.4 | -5.5 | 4.1E-08 | 9.0E-08 | BT20/MB468 | CYTO |
| >hg38_21_+_8433172_8446622_RNA45SN1_45S_50ntFlanks@12416.12439.24\|TTCGATGTCGGCTCTTCCTATCAT | >hg38_21_+_8433172_8446622_RNA45SN1_45S_50ntFlanks | 12416 | 12439 | 24 | TTCGATGTCGGCTCTTCCTATCAT | 613.2 | 3.9 | 0.7 | 5.5 | 4.8E-08 | 9.0E-08 | MB231/MB468 | NUC |
| >hg38_21_+_8433172_8446622_RNA45SN1_45S_50ntFlanks@11577.11602.26\|TTAGAACTGGTGCGGACCAGGGGAAT | >hg38_21_+_8433172_8446622_RNA45SN1_45S_50ntFlanks | 11577 | 11602 | 26 | TTAGAACTGGTGCGGACCAGGGGAAT | 185.0 | -5.2 | 1.0 | -5.5 | 4.8E-08 | 9.0E-08 | MB231/MB468 | NUC |
| >hg38_21_+_8433172_8446622_RNA45SN1_45S_50ntFlanks@9545.9573.29\|GTGGAGGTCCGTAGCGGTCCTGACGTGCA | >hg38_21_+_8433172_8446622_RNA45SN1_45S_50ntFlanks | 9545 | 9573 | 29 | GTGGAGGTCCGTAGCGGTCCTGACGTGCA | 715.8 | -2.9 | 0.5 | -5.5 | 3.6E-08 | 9.0E-08 | BT20/MB468 | MITO |
| >hg38_21_+_8433172_8446622_RNA45SN1_45S_50ntFlanks@11562.11592.31\|CGCCTAGCAGCCGACTTAGAACTGGTGCGGA | >hg38_21_+_8433172_8446622_RNA45SN1_45S_50ntFlanks | 11562 | 11592 | 31 | CGCCTAGCAGCCGACTTAGAACTGGTGCGGA | 880.1 | -4.4 | 0.8 | -5.5 | 4.7E-08 | 9.0E-08 | MB231/MB468 | CYTO |
| >MI0000542\|hsa-mir-320a&WithFlank&8\|-\|22244960\|22245043@49.69.21(+1C)&[MIMAT0000510&hsa-miR-320a-3p&offsets\|+1\|0(+1C);m-60&8\|-\|22244975\|22244996&offsets\|+1\|0(+1C)]\|AAAGCTGGGTTGAGAGGGCGAC | >MI0000542\|hsa-mir-320a&WithFlank&8\|-\|22244960\|22245043 | 49 | 69 | 21(+1C) | AAAGCTGGGTTGAGAGGGCGAC | 124.2 | 10.7 | 1.9 | 5.8 | 8.3E-09 | 9.1E-08 | BT20/MB468 | MP |
| >hg38_21_+_8433172_8446622_RNA45SN1_45S_50ntFlanks@9563.9592.30\|CCTGACGTGCAAATCGGTCGTCCGACCTGG | >hg38_21_+_8433172_8446622_RNA45SN1_45S_50ntFlanks | 9563 | 9592 | 30 | CCTGACGTGCAAATCGGTCGTCCGACCTGG | 192.4 | -4.4 | 0.8 | -5.5 | 4.7E-08 | 9.1E-08 | MB231/MB468 | CYTO |
| >hg38_21_+_8433172_8446622_RNA45SN1_45S_50ntFlanks@12617.12634.18\|TGAGGAGCCAATGGGGCG | >hg38_21_+_8433172_8446622_RNA45SN1_45S_50ntFlanks | 12617 | 12634 | 18 | TGAGGAGCCAATGGGGCG | 53.3 | -9.6 | 1.8 | -5.5 | 4.4E-08 | 9.1E-08 | MB231/MB468 | MP |
| >MI0000289\|hsa-mir-181a-1&WithFlank&1\|-\|198859038\|198859159@70.90.21&[MIMAT0000270&hsa-miR-181a-3p&offsets\|0\|-1;m-125&1\|-\|198859069\|198859090&offsets\|0\|-1]\|ACCATCGACCGTTGATTGTAC | >MI0000289\|hsa-mir-181a-1&WithFlank&1\|-\|198859038\|198859159 | 70 | 90 | 21 | ACCATCGACCGTTGATTGTAC | 172.0 | -10.8 | 1.8 | -6.0 | 1.6E-09 | 9.1E-08 | BT20/MB231 | MP |
| >hg38_21_+_8433172_8446622_RNA45SN1_45S_50ntFlanks@7973.7992.20\|GACGCGACCTCAGATCAGAC | >hg38_21_+_8433172_8446622_RNA45SN1_45S_50ntFlanks | 7973 | 7992 | 20 | GACGCGACCTCAGATCAGAC | 1351.6 | 5.1 | 0.9 | 5.5 | 4.7E-08 | 9.1E-08 | MB231/MB468 | CYTO |

<table>
<tr><th></th><th></th><th></th><th></th><th></th><th></th><th></th><th></th><th></th><th></th><th></th><th></th><th></th><th></th></tr>
<tr><td>>hg38_21_+_8433172_8446622_RNA45SN1_45S_50ntFlanks@8104.8143.40|CCGCGGCGGGGCGCGGGACATGTGGCGTACGGAAGACCCG</td><td>>hg38_21_+_8433172_8446622_RNA45SN1_45S_50ntFlanks</td><td>8104</td><td>8143</td><td>40</td><td>CCGCGGCGGGGCGCGGGACATGTGGCGTACGGAAGACCCG</td><td>15883.0</td><td>-5.7</td><td>1.0</td><td>-5.5</td><td>3.7E-08</td><td>9.1E-08</td><td>BT20/MB468</td><td>MITO</td></tr>
<tr><td>>hg38_21_+_8433172_8446622_RNA45SN1_45S_50ntFlanks@10783.10818.36|GTGAACAGCCTCTGGCATGTTGGAACAATGTAGGTA</td><td>>hg38_21_+_8433172_8446622_RNA45SN1_45S_50ntFlanks</td><td>10783</td><td>10818</td><td>36</td><td>GTGAACAGCCTCTGGCATGTTGGAACAATGTAGGTA</td><td>1794.0</td><td>-5.0</td><td>0.9</td><td>-5.5</td><td>4.1E-08</td><td>9.2E-08</td><td>BT20/MB468</td><td>CYTO</td></tr>
<tr><td>>hg38_21_+_8433172_8446622_RNA45SN1_45S_50ntFlanks@9340.9370.31|GCACGAAAGCCGCCGTGGCGCAATGAAGGTG</td><td>>hg38_21_+_8433172_8446622_RNA45SN1_45S_50ntFlanks</td><td>9340</td><td>9370</td><td>31</td><td>GCACGAAAGCCGCCGTGGCGCAATGAAGGTG</td><td>60.5</td><td>-9.8</td><td>1.8</td><td>-5.5</td><td>4.4E-08</td><td>9.2E-08</td><td>MB231/MB468</td><td>MP</td></tr>
<tr><td>>hg38_21_+_8433172_8446622_RNA45SN1_45S_50ntFlanks@5527.5550.24|AAAGTCGTAACAAGGTTTCCGTAG</td><td>>hg38_21_+_8433172_8446622_RNA45SN1_45S_50ntFlanks</td><td>5527</td><td>5550</td><td>24</td><td>AAAGTCGTAACAAGGTTTCCGTAG</td><td>644.0</td><td>-7.1</td><td>1.3</td><td>-5.5</td><td>4.1E-08</td><td>9.2E-08</td><td>BT20/MB468</td><td>CYTO</td></tr>
<tr><td>>hg38_21_+_8433172_8446622_RNA45SN1_45S_50ntFlanks@4729.4759.31|TCAAGAACGAAAGTCGGAGGTTCGAAGACGA</td><td>>hg38_21_+_8433172_8446622_RNA45SN1_45S_50ntFlanks</td><td>4729</td><td>4759</td><td>31</td><td>TCAAGAACGAAAGTCGGAGGTTCGAAGACGA</td><td>594.7</td><td>-4.2</td><td>0.8</td><td>-5.5</td><td>4.9E-08</td><td>9.2E-08</td><td>MB231/MB468</td><td>NUC</td></tr>
<tr><td>>hg38_21_+_8433172_8446622_RNA45SN1_45S_50ntFlanks@3990.4024.35|TTGGTGACTCTAGATAACCTCGGGCCGATCGCACG</td><td>>hg38_21_+_8433172_8446622_RNA45SN1_45S_50ntFlanks</td><td>3990</td><td>4024</td><td>35</td><td>TTGGTGACTCTAGATAACCTCGGGCCGATCGCACG</td><td>379.7</td><td>-4.5</td><td>0.8</td><td>-5.5</td><td>4.5E-08</td><td>9.2E-08</td><td>MB231/MB468</td><td>MP</td></tr>
<tr><td>>hg38_21_+_8433172_8446622_RNA45SN1_45S_50ntFlanks@8836.8858.23|CGGCGGTGGCGGCGGCGGCGGCG</td><td>>hg38_21_+_8433172_8446622_RNA45SN1_45S_50ntFlanks</td><td>8836</td><td>8858</td><td>23</td><td>CGGCGGTGGCGGCGGCGGCGGCG</td><td>103.1</td><td>-9.8</td><td>1.7</td><td>-5.8</td><td>8.4E-09</td><td>9.2E-08</td><td>BT20/MB468</td><td>MP</td></tr>
<tr><td>>hg38_1_-_228634819_228635039_RNA5S12_5S_50ntFlanks@51.82.32|GTCTACGGCCATACCACCCTGAACGCGCCCGA</td><td>>hg38_1_-_228634819_228635039_RNA5S12_5S_50ntFlanks</td><td>51</td><td>82</td><td>32</td><td>GTCTACGGCCATACCACCCTGAACGCGCCCGA</td><td>2105.0</td><td>4.8</td><td>0.9</td><td>5.5</td><td>4.8E-08</td><td>9.2E-08</td><td>MB231/MB468</td><td>CYTO</td></tr>
<tr><td>>hg38_21_+_8433172_8446622_RNA45SN1_45S_50ntFlanks@4585.4621.37|GTTCTATTTTGTTGGTTTTCGGAACTGAGGCCATGAT</td><td>>hg38_21_+_8433172_8446622_RNA45SN1_45S_50ntFlanks</td><td>4585</td><td>4621</td><td>37</td><td>GTTCTATTTTGTTGGTTTTCGGAACTGAGGCCATGAT</td><td>522.2</td><td>6.1</td><td>1.1</td><td>5.5</td><td>4.8E-08</td><td>9.2E-08</td><td>MB231/MB468</td><td>CYTO</td></tr>
<tr><td>>hg38_21_+_8433172_8446622_RNA45SN1_45S_50ntFlanks@9545.9575.31|GTGGAGGTCCGTAGCGGTCCTGACGTGCAAA</td><td>>hg38_21_+_8433172_8446622_RNA45SN1_45S_50ntFlanks</td><td>9545</td><td>9575</td><td>31</td><td>GTGGAGGTCCGTAGCGGTCCTGACGTGCAAA</td><td>455.1</td><td>-3.8</td><td>0.7</td><td>-5.5</td><td>4.9E-08</td><td>9.2E-08</td><td>MB231/MB468</td><td>NUC</td></tr>
<tr><td>>hg38_21_+_8433172_8446622_RNA45SN1_45S_50ntFlanks@12991.13011.21|CTCCCTCGCTGCGATCTATTG</td><td>>hg38_21_+_8433172_8446622_RNA45SN1_45S_50ntFlanks</td><td>12991</td><td>13011</td><td>21</td><td>CTCCCTCGCTGCGATCTATTG</td><td>1707.0</td><td>-3.8</td><td>0.7</td><td>-5.5</td><td>4.8E-08</td><td>9.2E-08</td><td>MB231/MB468</td><td>CYTO</td></tr>
<tr><td>>MI0000433|hsa-let-7g&WithFlank&3|-|52268272|52268367@11.31.21(+1A)&[MIMAT0000414&hsa-let-7g-5p&offsets|0|-1(+1A);m-58&3|-|52268336|52268357&offsets|0|-1(+1A)]|TGAGGTAGTAGTTTGTACAGTA</td><td>>MI0000433|hsa-let-7g&WithFlank&3|-|52268272|52268367</td><td>11</td><td>31</td><td>21(+1A)</td><td>TGAGGTAGTAGTTTGTACAGTA</td><td>933.4</td><td>4.9</td><td>0.9</td><td>5.4</td><td>5.2E-08</td><td>9.2E-08</td><td>MB231/MB468</td><td>MITO</td></tr>
<tr><td>>hg38_21_+_8433172_8446622_RNA45SN1_45S_50ntFlanks@11579.11601.23|AGAACTGGTGCGGACCAGGGGAA</td><td>>hg38_21_+_8433172_8446622_RNA45SN1_45S_50ntFlanks</td><td>11579</td><td>11601</td><td>23</td><td>AGAACTGGTGCGGACCAGGGGAA</td><td>949.8</td><td>-4.6</td><td>0.8</td><td>-5.5</td><td>4.2E-08</td><td>9.3E-08</td><td>BT20/MB468</td><td>CYTO</td></tr>
<tr><td>>hg38_21_+_8433172_8446622_RNA45SN1_45S_50ntFlanks@9554.9574.21|CGTAGCGGTCCTGACGTGCAA</td><td>>hg38_21_+_8433172_8446622_RNA45SN1_45S_50ntFlanks</td><td>9554</td><td>9574</td><td>21</td><td>CGTAGCGGTCCTGACGTGCAA</td><td>2770.2</td><td>-2.5</td><td>0.4</td><td>-5.5</td><td>4.2E-08</td><td>9.3E-08</td><td>BT20/MB468</td><td>CYTO</td></tr>
<tr><td>>hg38_21_+_8433172_8446622_RNA45SN1_45S_50ntFlanks@11718.11739.22|AACGGCGGGAGTAACTATGACT</td><td>>hg38_21_+_8433172_8446622_RNA45SN1_45S_50ntFlanks</td><td>11718</td><td>11739</td><td>22</td><td>AACGGCGGGAGTAACTATGACT</td><td>1110.4</td><td>-8.9</td><td>1.6</td><td>-5.4</td><td>5.5E-08</td><td>9.3E-08</td><td>BT20/MB468</td><td>NUC</td></tr>
<tr><td>>trna134_GluTTC_1_-_16861774_16861845@1.31.31,trna5_GluTTC_1_+_17199078_17199149@1.31.31,trna84_GluTTC_1_-_161391883_161391954@1.31.31,trna94_GluTTC_1_-_149664355_149664427@1.31.31|TCCCTGGTGGTCTAGTGGCTAGGATTCGGCG</td><td>>trna134_GluTTC_1_-_16861774_16861845</td><td>1</td><td>31</td><td>31</td><td>TCCCTGGTGGTCTAGTGGCTAGGATTCGGCG</td><td>572.4</td><td>-6.7</td><td>1.2</td><td>-5.5</td><td>4.5E-08</td><td>9.3E-08</td><td>MB231/MB468</td><td>MP</td></tr>
<tr><td>>hg38_21_+_8433172_8446622_RNA45SN1_45S_50ntFlanks@10668.10706.39|GAAAGCGTCGCGGTTCCGGCGGCGTCCGGTGAGCTCTCG</td><td>>hg38_21_+_8433172_8446622_RNA45SN1_45S_50ntFlanks</td><td>10668</td><td>10706</td><td>39</td><td>GAAAGCGTCGCGGTTCCGGCGGCGTCCGGTGAGCTCTCG</td><td>1346.8</td><td>-7.0</td><td>1.3</td><td>-5.5</td><td>4.9E-08</td><td>9.3E-08</td><td>MB231/MB468</td><td>NUC</td></tr>
<tr><td>>hg38_21_+_8433172_8446622_RNA45SN1_45S_50ntFlanks@4006.4024.19|ACCTCGGGCCGATCGCACG</td><td>>hg38_21_+_8433172_8446622_RNA45SN1_45S_50ntFlanks</td><td>4006</td><td>4024</td><td>19</td><td>ACCTCGGGCCGATCGCACG</td><td>1224.3</td><td>-5.6</td><td>1.0</td><td>-5.5</td><td>4.2E-08</td><td>9.3E-08</td><td>BT20/MB468</td><td>CYTO</td></tr>
<tr><td>>trnaMT_PheGAA_MT_+_577_647@27.57.31|AATACACTGAAAATGTTTAGACGGGCTCACA</td><td>>trnaMT_PheGAA_MT_+_577_647</td><td>27</td><td>57</td><td>31</td><td>AATACACTGAAAATGTTTAGACGGGCTCACA</td><td>162.0</td><td>11.1</td><td>1.9</td><td>5.8</td><td>8.5E-09</td><td>9.3E-08</td><td>BT20/MB468</td><td>MP</td></tr>
<tr><td>>hg38_21_+_8433172_8446622_RNA45SN1_45S_50ntFlanks@12250.12271.22|AAAAGCTCGCTTGATCTTGATT</td><td>>hg38_21_+_8433172_8446622_RNA45SN1_45S_50ntFlanks</td><td>12250</td><td>12271</td><td>22</td><td>AAAAGCTCGCTTGATCTTGATT</td><td>293.2</td><td>5.3</td><td>1.0</td><td>5.5</td><td>4.8E-08</td><td>9.3E-08</td><td>MB231/MB468</td><td>CYTO</td></tr>
<tr><td>>hg38_1_-_228634819_228635039_RNA5S12_5S_50ntFlanks@140.169.30|ACCGCCTGGGAATACCGGGTGCTGTAGGCT</td><td>>hg38_1_-_228634819_228635039_RNA5S12_5S_50ntFlanks</td><td>140</td><td>169</td><td>30</td><td>ACCGCCTGGGAATACCGGGTGCTGTAGGCT</td><td>8835.0</td><td>-4.0</td><td>0.7</td><td>-5.5</td><td>4.5E-08</td><td>9.3E-08</td><td>MB231/MB468</td><td>MP</td></tr>
<tr><td>>MI0005756|hsa-mir-934&WithFlank&X|+|136550872|136550966@21.42.22(+1C)&[MIMAT0004977&hsa-miR-934&offsets|0|0(+1C);m-943&X|+|136550892|136550913&offsets|0|0(+1C)]|TGTCTACTACTGGAGACACTGGC</td><td>>MI0005756|hsa-mir-934&WithFlank&X|+|136550872|136550966</td><td>21</td><td>42</td><td>22(+1C)</td><td>TGTCTACTACTGGAGACACTGGC</td><td>375.5</td><td>7.9</td><td>1.4</td><td>5.5</td><td>4.2E-08</td><td>9.3E-08</td><td>BT20/MB468</td><td>CYTO</td></tr>
<tr><td>>trnaMT_PheGAA_MT_+_577_647@27.55.29|AATACACTGAAAATGTTTAGACGGGCTCA</td><td>>trnaMT_PheGAA_MT_+_577_647</td><td>27</td><td>55</td><td>29</td><td>AATACACTGAAAATGTTTAGACGGGCTCA</td><td>156.8</td><td>11.0</td><td>1.9</td><td>5.8</td><td>8.6E-09</td><td>9.3E-08</td><td>BT20/MB468</td><td>MP</td></tr>
<tr><td>>hg38_21_+_8433172_8446622_RNA45SN1_45S_50ntFlanks@4735.4767.33|ACGAAAGTCGGAGGTTCGAAGACGATCAGATAC</td><td>>hg38_21_+_8433172_8446622_RNA45SN1_45S_50ntFlanks</td><td>4735</td><td>4767</td><td>33</td><td>ACGAAAGTCGGAGGTTCGAAGACGATCAGATAC</td><td>225.2</td><td>8.7</td><td>1.6</td><td>5.5</td><td>5.0E-08</td><td>9.3E-08</td><td>MB231/MB468</td><td>NUC</td></tr>
<tr><td>>hg38_21_+_8433172_8446622_RNA45SN1_45S_50ntFlanks@10458.10487.30|ATGGCCTCCGTTGCCCTCGGCCGATCGAAA</td><td>>hg38_21_+_8433172_8446622_RNA45SN1_45S_50ntFlanks</td><td>10458</td><td>10487</td><td>30</td><td>ATGGCCTCCGTTGCCCTCGGCCGATCGAAA</td><td>752.2</td><td>-4.2</td><td>0.8</td><td>-5.5</td><td>5.0E-08</td><td>9.3E-08</td><td>MB231/MB468</td><td>NUC</td></tr>
<tr><td>>MI0000088|hsa-mir-30a&WithFlank&6|-|71403545|71403627@12.33.22&[MIMAT0000087&hsa-miR-30a-5p&offsets|0|0;m-15&6|-|71403595|71403616&offsets|0|0]|TGTAAACATCCTCGACTGGAAG</td><td>>MI0000088|hsa-mir-30a&WithFlank&6|-|71403545|71403627</td><td>12</td><td>33</td><td>22</td><td>TGTAAACATCCTCGACTGGAAG</td><td>922.2</td><td>5.3</td><td>1.0</td><td>5.5</td><td>4.9E-08</td><td>9.3E-08</td><td>MB231/MB468</td><td>CYTO</td></tr>
<tr><td>>hg38_21_+_8433172_8446622_RNA45SN1_45S_50ntFlanks@12007.12032.26|GCCGCCGGTGAAATACCACTACTCTG</td><td>>hg38_21_+_8433172_8446622_RNA45SN1_45S_50ntFlanks</td><td>12007</td><td>12032</td><td>26</td><td>GCCGCCGGTGAAATACCACTACTCTG</td><td>6845.3</td><td>-5.6</td><td>1.0</td><td>-5.5</td><td>5.0E-08</td><td>9.4E-08</td><td>MB231/MB468</td><td>NUC</td></tr>
<tr><td>>hg38_21_+_8433172_8446622_RNA45SN1_45S_50ntFlanks@12589.12615.27|GTTCAGACATTTGGTGTATGTGCTTGG</td><td>>hg38_21_+_8433172_8446622_RNA45SN1_45S_50ntFlanks</td><td>12589</td><td>12615</td><td>27</td><td>GTTCAGACATTTGGTGTATGTGCTTGG</td><td>93.7</td><td>-9.6</td><td>1.7</td><td>-5.8</td><td>8.6E-09</td><td>9.4E-08</td><td>BT20/MB468</td><td>MP</td></tr>
<tr><td>>hg38_21_+_8433172_8446622_RNA45SN1_45S_50ntFlanks@6745.6767.23|ACACTTCGAACGCACTTGCGGCC</td><td>>hg38_21_+_8433172_8446622_RNA45SN1_45S_50ntFlanks</td><td>6745</td><td>6767</td><td>23</td><td>ACACTTCGAACGCACTTGCGGCC</td><td>96.4</td><td>-9.7</td><td>1.7</td><td>-5.8</td><td>8.6E-09</td><td>9.4E-08</td><td>BT20/MB468</td><td>MP</td></tr>
<tr><td>>MI0000086|hsa-mir-28&WithFlank&3|+|188688775|188688872@61.81.21&[MIMAT0004502&hsa-miR-28-3p&offsets|+1|0;m-28&3|+|188688834|188688855&offsets|+1|0]|ACTAGATTGTGAGCTCCTGGA</td><td>>MI0000086|hsa-mir-28&WithFlank&3|+|188688775|188688872</td><td>61</td><td>81</td><td>21</td><td>ACTAGATTGTGAGCTCCTGGA</td><td>80.5</td><td>10.1</td><td>1.8</td><td>5.8</td><td>8.7E-09</td><td>9.4E-08</td><td>BT20/MB468</td><td>MP</td></tr>
<tr><td>>MI0001733|hsa-mir-452&WithFlank&X|-|151959622|151959718@23.43.21(+1U)&[MIMAT0001635&hsa-miR-452-5p&offsets|+3|+2(+1U);m-188&X|-|151959677|151959699&offsets|+3|+1(+1U)]|TGTTTGCAGAGGAAACTGAGAT</td><td>>MI0001733|hsa-mir-452&WithFlank&X|-|151959622|151959718</td><td>23</td><td>43</td><td>21(+1U)</td><td>TGTTTGCAGAGGAAACTGAGAT</td><td>218.0</td><td>11.5</td><td>2.0</td><td>5.8</td><td>8.7E-09</td><td>9.4E-08</td><td>BT20/MB468</td><td>MP</td></tr>
<tr><td>>MI0000441|hsa-mir-30b&WithFlank&8|-|134800514|134800613@23.43.21(+1A)&[MIMAT0000420&hsa-miR-30b-5p&offsets|0|-1(+1A);m-76&8|-|134800570|134800591&offsets|0|-1(+1A)]|TGTAAACATCCTACACTCAGCA</td><td>>MI0000441|hsa-mir-30b&WithFlank&8|-|134800514|134800613</td><td>23</td><td>43</td><td>21(+1A)</td><td>TGTAAACATCCTACACTCAGCA</td><td>688.7</td><td>9.6</td><td>1.7</td><td>5.5</td><td>4.2E-08</td><td>9.4E-08</td><td>BT20/MB468</td><td>CYTO</td></tr>
<tr><td>>hg38_21_+_8433172_8446622_RNA45SN1_45S_50ntFlanks@10389.10419.31|TGAACAGCAGTTGAACATGGGTCAGTCGGTC</td><td>>hg38_21_+_8433172_8446622_RNA45SN1_45S_50ntFlanks</td><td>10389</td><td>10419</td><td>31</td><td>TGAACAGCAGTTGAACATGGGTCAGTCGGTC</td><td>169.8</td><td>-6.8</td><td>1.2</td><td>-5.5</td><td>5.0E-08</td><td>9.4E-08</td><td>MB231/MB468</td><td>NUC</td></tr>
<tr><td>>hg38_21_+_8433172_8446622_RNA45SN1_45S_50ntFlanks@12008.12037.30|CCGCCGGTGAAATACCACTACTCTGATCGT</td><td>>hg38_21_+_8433172_8446622_RNA45SN1_45S_50ntFlanks</td><td>12008</td><td>12037</td><td>30</td><td>CCGCCGGTGAAATACCACTACTCTGATCGT</td><td>333.8</td><td>-5.6</td><td>1.0</td><td>-5.5</td><td>5.0E-08</td><td>9.4E-08</td><td>MB231/MB468</td><td>NUC</td></tr>
<tr><td>>hg38_21_+_8433172_8446622_RNA45SN1_45S_50ntFlanks@10410.10442.33|TCAGTCGGTCCTGAGAGATGGGCGAGCGCCGTT</td><td>>hg38_21_+_8433172_8446622_RNA45SN1_45S_50ntFlanks</td><td>10410</td><td>10442</td><td>33</td><td>TCAGTCGGTCCTGAGAGATGGGCGAGCGCCGTT</td><td>362.2</td><td>-3.3</td><td>0.6</td><td>-5.5</td><td>4.2E-08</td><td>9.4E-08</td><td>BT20/MB468</td><td>CYTO</td></tr>
<tr><td>>hg38_21_+_8433172_8446622_RNA45SN1_45S_50ntFlanks@11365.11387.23|GGCGGCGGCGGTCGGCGGGCGGC</td><td>>hg38_21_+_8433172_8446622_RNA45SN1_45S_50ntFlanks</td><td>11365</td><td>11387</td><td>23</td><td>GGCGGCGGCGGTCGGCGGGCGGC</td><td>168.1</td><td>11.1</td><td>1.9</td><td>5.8</td><td>8.7E-09</td><td>9.4E-08</td><td>BT20/MB468</td><td>MP</td></tr>
<tr><td>>hg38_21_+_8433172_8446622_RNA45SN1_45S_50ntFlanks@5485.5518.34|AGCGCTGAGAAGACGGTCGAACTTGACTATCTAG</td><td>>hg38_21_+_8433172_8446622_RNA45SN1_45S_50ntFlanks</td><td>5485</td><td>5518</td><td>34</td><td>AGCGCTGAGAAGACGGTCGAACTTGACTATCTAG</td><td>1731.2</td><td>-5.5</td><td>1.0</td><td>-5.5</td><td>4.9E-08</td><td>9.4E-08</td><td>MB231/MB468</td><td>CYTO</td></tr>
<tr><td>>hg38_21_+_8433172_8446622_RNA45SN1_45S_50ntFlanks@7970.8010.41|CGAGACGCGACCTCAGATCAGACGTGGCGACCCGCTGAATT</td><td>>hg38_21_+_8433172_8446622_RNA45SN1_45S_50ntFlanks</td><td>7970</td><td>8010</td><td>41</td><td>CGAGACGCGACCTCAGATCAGACGTGGCGACCCGCTGAATT</td><td>298.1</td><td>4.5</td><td>0.8</td><td>5.4</td><td>5.0E-08</td><td>9.5E-08</td><td>MB231/MB468</td><td>NUC</td></tr>
<tr><td>>hg38_21_+_8433172_8446622_RNA45SN1_45S_50ntFlanks@4364.4398.35|CTGCAGTTAAAAAGCTCGTAGTTGGATCTTGGGAG</td><td>>hg38_21_+_8433172_8446622_RNA45SN1_45S_50ntFlanks</td><td>4364</td><td>4398</td><td>35</td><td>CTGCAGTTAAAAAGCTCGTAGTTGGATCTTGGGAG</td><td>72.7</td><td>-10.0</td><td>1.8</td><td>-5.5</td><td>4.6E-08</td><td>9.5E-08</td><td>MB231/MB468</td><td>MP</td></tr>
<tr><td>>hg38_21_+_8433172_8446622_RNA45SN1_45S_50ntFlanks@11455.11478.24|CTCGGCGCGCGGCGGCGGCGGCGG</td><td>>hg38_21_+_8433172_8446622_RNA45SN1_45S_50ntFlanks</td><td>11455</td><td>11478</td><td>24</td><td>CTCGGCGCGCGGCGGCGGCGGCGG</td><td>184.8</td><td>-6.2</td><td>1.1</td><td>-5.5</td><td>4.3E-08</td><td>9.5E-08</td><td>BT20/MB468</td><td>CYTO</td></tr>
<tr><td>>hg38_21_+_8433172_8446622_RNA45SN1_45S_50ntFlanks@4297.4317.21|CATTGGAGGGCAAGTCTGGTG</td><td>>hg38_21_+_8433172_8446622_RNA45SN1_45S_50ntFlanks</td><td>4297</td><td>4317</td><td>21</td><td>CATTGGAGGGCAAGTCTGGTG</td><td>603.6</td><td>-3.8</td><td>0.7</td><td>-5.5</td><td>4.3E-08</td><td>9.5E-08</td><td>BT20/MB468</td><td>CYTO</td></tr>
<tr><td>>hg38_21_+_8433172_8446622_RNA45SN1_45S_50ntFlanks@12971.13010.40|TTTCGTACGTAGCAGAGCAGCTCCCTCGCTGCGATCTATT</td><td>>hg38_21_+_8433172_8446622_RNA45SN1_45S_50ntFlanks</td><td>12971</td><td>13010</td><td>40</td><td>TTTCGTACGTAGCAGAGCAGCTCCCTCGCTGCGATCTATT</td><td>426.2</td><td>4.7</td><td>0.9</td><td>5.5</td><td>3.8E-08</td><td>9.5E-08</td><td>BT20/MB468</td><td>MITO</td></tr>
<tr><td>>hg38_21_+_8433172_8446622_RNA45SN1_45S_50ntFlanks@9033.9061.29|CTCGTGGGGGGGCCGGGCCACCCCTCCCA</td><td>>hg38_21_+_8433172_8446622_RNA45SN1_45S_50ntFlanks</td><td>9033</td><td>9061</td><td>29</td><td>CTCGTGGGGGGGCCGGGCCACCCCTCCCA</td><td>130.9</td><td>-10.1</td><td>1.8</td><td>-5.8</td><td>8.8E-09</td><td>9.5E-08</td><td>BT20/MB468</td><td>MP</td></tr>
<tr><td>>hg38_21_+_8433172_8446622_RNA45SN1_45S_50ntFlanks@3984.4010.27|GCGGCTTTGGTGACTCTAGATAACCTC</td><td>>hg38_21_+_8433172_8446622_RNA45SN1_45S_50ntFlanks</td><td>3984</td><td>4010</td><td>27</td><td>GCGGCTTTGGTGACTCTAGATAACCTC</td><td>319.6</td><td>-3.9</td><td>0.7</td><td>-5.4</td><td>5.1E-08</td><td>9.5E-08</td><td>MB231/MB468</td><td>NUC</td></tr>
<tr><td>>MI0000094|hsa-mir-92a-2&WithFlank&X|-|134169532|134169618@54.76.23&[MIMAT0000092_1&hsa-miR-92a-3p&offsets|0|+1;m-11&X|-|134169544|134169565&offsets|0|+1]|TATTGCACTTGTCCCGGCCTGTG</td><td>>MI0000094|hsa-mir-92a-2&WithFlank&X|-|134169532|134169618</td><td>54</td><td>76</td><td>23</td><td>TATTGCACTTGTCCCGGCCTGTG</td><td>234.6</td><td>3.6</td><td>0.7</td><td>5.4</td><td>5.1E-08</td><td>9.5E-08</td><td>MB231/MB468</td><td>NUC</td></tr>
<tr><td>>hg38_21_+_8433172_8446622_RNA45SN1_45S_50ntFlanks@4139.4157.19|AGGGAGCCTGAGAAACGGC</td><td>>hg38_21_+_8433172_8446622_RNA45SN1_45S_50ntFlanks</td><td>4139</td><td>4157</td><td>19</td><td>AGGGAGCCTGAGAAACGGC</td><td>184.2</td><td>-7.2</td><td>1.3</td><td>-5.5</td><td>4.6E-08</td><td>9.5E-08</td><td>MB231/MB468</td><td>MP</td></tr>
<tr><td>>hg38_21_+_8433172_8446622_RNA45SN1_45S_50ntFlanks@8114.8138.25|GCGCGGGACATGTGGCGTACGGAAG</td><td>>hg38_21_+_8433172_8446622_RNA45SN1_45S_50ntFlanks</td><td>8114</td><td>8138</td><td>25</td><td>GCGCGGGACATGTGGCGTACGGAAG</td><td>511.0</td><td>-7.6</td><td>1.4</td><td>-5.4</td><td>5.1E-08</td><td>9.5E-08</td><td>MB231/MB468</td><td>NUC</td></tr>
<tr><td>>hg38_21_+_8433172_8446622_RNA45SN1_45S_50ntFlanks@9534.9560.27|AGGAAACTCTGGTGGAGGTCCGTAGCG</td><td>>hg38_21_+_8433172_8446622_RNA45SN1_45S_50ntFlanks</td><td>9534</td><td>9560</td><td>27</td><td>AGGAAACTCTGGTGGAGGTCCGTAGCG</td><td>1361.1</td><td>-9.0</td><td>1.7</td><td>-5.4</td><td>5.6E-08</td><td>9.5E-08</td><td>BT20/MB468</td><td>NUC</td></tr>
</table>

| | | | | | | | | | | | | | |
|---|---|---|---|---|---|---|---|---|---|---|---|---|---|
| >hg38_21_+_8433172_8446622_RNA45SN1_45S_50ntFlanks@11671.11693.23\|CCAGTGCTCTGAATGTCAAAGTG | >hg38_21_+_8433172_8446622_RNA45SN1_45S_50ntFlanks | 11671 | 11693 | 23 | CCAGTGCTCTGAATGTCAAAGTG | 749.6 | -6.9 | 1.3 | -5.5 | 4.6E-08 | 9.5E-08 | MB231/MB468 | MP |
| >hg38_21_+_8433172_8446622_RNA45SN1_45S_50ntFlanks@11601.11619.19\|ATCCGACTGTTTAATTAAA | >hg38_21_+_8433172_8446622_RNA45SN1_45S_50ntFlanks | 11601 | 11619 | 19 | ATCCGACTGTTTAATTAAA | 84.4 | -9.5 | 1.6 | -5.8 | 8.8E-09 | 9.5E-08 | BT20/MB468 | MP |
| >hg38_21_+_8433172_8446622_RNA45SN1_45S_50ntFlanks@9925.9948.24\|GGTTGATATAGACAGCAGGACGGT | >hg38_21_+_8433172_8446622_RNA45SN1_45S_50ntFlanks | 9925 | 9948 | 24 | GGTTGATATAGACAGCAGGACGGT | 91.3 | 10.3 | 1.8 | 5.8 | 8.8E-09 | 9.5E-08 | BT20/MB468 | MP |
| >MI0000101\|hsa-mir-99a&WithFlank&21\|+\|16539083\|16539175@19.38.20(+1A)&[MIMAT0000097&hsa-miR-99a-5p&offsets\|0\|-2(+1A);m-64&21\|+\|16539101\|16539121&offsets\|0\|-1(+1A)]\|AACCCGTAGATCCGATCTTGA | >MI0000101\|hsa-mir-99a&WithFlank&21\|+\|16539083\|16539175 | 19 | 38 | 20(+1A) | AACCCGTAGATCCGATCTTGA | 80.8 | -9.4 | 1.6 | -5.8 | 8.8E-09 | 9.5E-08 | BT20/MB468 | MP |
| >hg38_21_+_8433172_8446622_RNA45SN1_45S_50ntFlanks@11189.11225.37\|ACCCGGGGGGCCGGCGGCGGCGGCGACTCTGGACGCG | >hg38_21_+_8433172_8446622_RNA45SN1_45S_50ntFlanks | 11189 | 11225 | 37 | ACCCGGGGGGCCGGCGGCGGCGGCGACTCTGGACGCG | 375.7 | -6.4 | 1.2 | -5.5 | 5.0E-08 | 9.5E-08 | MB231/MB468 | CYTO |
| >hg38_21_+_8433172_8446622_RNA45SN1_45S_50ntFlanks@8198.8224.27\|TGGACGGTGTGAGGCCGGTAGCGGCCC | >hg38_21_+_8433172_8446622_RNA45SN1_45S_50ntFlanks | 8198 | 8224 | 27 | TGGACGGTGTGAGGCCGGTAGCGGCCC | 405.0 | -5.5 | 1.0 | -5.4 | 5.1E-08 | 9.5E-08 | MB231/MB468 | NUC |
| >hg38_21_+_8433172_8446622_RNA45SN1_45S_50ntFlanks@6719.6737.19\|TGAATTGCAGGACACATTG | >hg38_21_+_8433172_8446622_RNA45SN1_45S_50ntFlanks | 6719 | 6737 | 19 | TGAATTGCAGGACACATTG | 645.4 | -3.4 | 0.6 | -5.5 | 3.8E-08 | 9.5E-08 | BT20/MB468 | MITO |
| >hg38_21_+_8433172_8446622_RNA45SN1_45S_50ntFlanks@11315.11342.28\|CCACCCCACGTCTCGTCGCGCGCGCGTC | >hg38_21_+_8433172_8446622_RNA45SN1_45S_50ntFlanks | 11315 | 11342 | 28 | CCACCCCACGTCTCGTCGCGCGCGCGTC | 554.2 | -5.2 | 1.0 | -5.5 | 4.3E-08 | 9.6E-08 | BT20/MB468 | CYTO |
| >hg38_21_+_8433172_8446622_RNA45SN1_45S_50ntFlanks@11731.11751.21\|ACTATGACTCTCTTAAGGTAG | >hg38_21_+_8433172_8446622_RNA45SN1_45S_50ntFlanks | 11731 | 11751 | 21 | ACTATGACTCTCTTAAGGTAG | 452.4 | -3.7 | 0.7 | -5.4 | 5.1E-08 | 9.6E-08 | MB231/MB468 | NUC |
| >hg38_21_+_8433172_8446622_RNA45SN1_45S_50ntFlanks@11760.11777.18\|CTCGTCATCTAATTAGTG | >hg38_21_+_8433172_8446622_RNA45SN1_45S_50ntFlanks | 11760 | 11777 | 18 | CTCGTCATCTAATTAGTG | 92.9 | -9.6 | 1.7 | -5.8 | 8.9E-09 | 9.6E-08 | BT20/MB468 | MP |
| >hg38_21_+_8433172_8446622_RNA45SN1_45S_50ntFlanks@9736.9754.19\|CCGAAACGATCTCAACCTA | >hg38_21_+_8433172_8446622_RNA45SN1_45S_50ntFlanks | 9736 | 9754 | 19 | CCGAAACGATCTCAACCTA | 432.1 | -7.9 | 1.4 | -5.8 | 8.9E-09 | 9.6E-08 | BT20/MB468 | MP |
| >hg38_21_+_8433172_8446622_RNA45SN1_45S_50ntFlanks@12419.12447.29\|GATGTCGGCTCTTCCTATCATTGTGAAGC | >hg38_21_+_8433172_8446622_RNA45SN1_45S_50ntFlanks | 12419 | 12447 | 29 | GATGTCGGCTCTTCCTATCATTGTGAAGC | 516.8 | 3.8 | 0.7 | 5.4 | 5.1E-08 | 9.6E-08 | MB231/MB468 | NUC |
| >MI0000434\|hsa-let-7i&WithFlank&12\|+\|62603680\|62603775@12.33.22(+1U)&[MIMAT0000415&hsa-let-7i-5p&offsets\|0\|0(+1U);m-49&12\|+\|62603691\|62603712&offsets\|0\|0(+1U)]\|TGAGGTAGTAGTTTGTGCTGTTT | >MI0000434\|hsa-let-7i&WithFlank&12\|+\|62603680\|62603775 | 12 | 33 | 22(+1U) | TGAGGTAGTAGTTTGTGCTGTTT | 1960.2 | 5.4 | 1.0 | 5.4 | 5.4E-08 | 9.6E-08 | MB231/MB468 | MITO |
| >hg38_21_+_8433172_8446622_RNA45SN1_45S_50ntFlanks@10846.10863.18\|CTTCGGGATAAGGATTGG | >hg38_21_+_8433172_8446622_RNA45SN1_45S_50ntFlanks | 10846 | 10863 | 18 | CTTCGGGATAAGGATTGG | 634.1 | -8.5 | 1.6 | -5.4 | 5.7E-08 | 9.6E-08 | BT20/MB468 | NUC |
| >hg38_21_+_8433172_8446622_RNA45SN1_45S_50ntFlanks@12634.12659.26\|GAAGCTACCATCTGTGGGATTATGAC | >hg38_21_+_8433172_8446622_RNA45SN1_45S_50ntFlanks | 12634 | 12659 | 26 | GAAGCTACCATCTGTGGGATTATGAC | 862.5 | -6.7 | 1.2 | -5.4 | 5.5E-08 | 9.6E-08 | MB231/MB468 | MITO |
| >hg38_21_+_8433172_8446622_RNA45SN1_45S_50ntFlanks@9032.9059.28\|CCTCGTGGGGGGGCCGGGCCACCCCTCC | >hg38_21_+_8433172_8446622_RNA45SN1_45S_50ntFlanks | 9032 | 9059 | 28 | CCTCGTGGGGGGGCCGGGCCACCCCTCC | 516.6 | -7.1 | 1.3 | -5.5 | 3.9E-08 | 9.6E-08 | BT20/MB468 | MITO |
| >hg38_21_+_8433172_8446622_RNA45SN1_45S_50ntFlanks@12633.12653.21\|CGAAGCTACCATCTGTGGGAT | >hg38_21_+_8433172_8446622_RNA45SN1_45S_50ntFlanks | 12633 | 12653 | 21 | CGAAGCTACCATCTGTGGGAT | 2009.7 | -7.7 | 1.4 | -5.5 | 4.3E-08 | 9.6E-08 | BT20/MB468 | CYTO |
| >hg38_21_+_8433172_8446622_RNA45SN1_45S_50ntFlanks@7990.8017.28\|GACGTGGCGACCCGCTGAATTTAAGCAT | >hg38_21_+_8433172_8446622_RNA45SN1_45S_50ntFlanks | 7990 | 8017 | 28 | GACGTGGCGACCCGCTGAATTTAAGCAT | 492.0 | 8.4 | 1.5 | 5.5 | 5.0E-08 | 9.6E-08 | MB231/MB468 | CYTO |
| >hg38_21_+_8433172_8446622_RNA45SN1_45S_50ntFlanks@3705.3748.44\|TACCTGGTTGATCCTGCCAGTAGCATATGCTTGTCTCAAAGATT | >hg38_21_+_8433172_8446622_RNA45SN1_45S_50ntFlanks | 3705 | 3748 | 44 | TACCTGGTTGATCCTGCCAGTAGCATATGCTTGTCTCAAAGATT | 781.1 | 4.8 | 0.9 | 5.5 | 3.9E-08 | 9.7E-08 | BT20/MB468 | MITO |
| >hg38_21_+_8433172_8446622_RNA45SN1_45S_50ntFlanks@6654.6688.35\|CTCTTAGCGGTGGATCACTCGGCTCGTGCGTCGAT | >hg38_21_+_8433172_8446622_RNA45SN1_45S_50ntFlanks | 6654 | 6688 | 35 | CTCTTAGCGGTGGATCACTCGGCTCGTGCGTCGAT | 53.2 | -9.6 | 1.8 | -5.5 | 4.7E-08 | 9.7E-08 | MB231/MB468 | MP |
| >hg38_21_+_8433172_8446622_RNA45SN1_45S_50ntFlanks@12993.13039.47\|CCCTCGCTGCGATCTATTGAAAGTCAGCCCTCGACACAAGGGTTTGT | >hg38_21_+_8433172_8446622_RNA45SN1_45S_50ntFlanks | 12993 | 13039 | 47 | CCCTCGCTGCGATCTATTGAAAGTCAGCCCTCGACACAAGGGTTTGT | 5423.9 | 4.6 | 0.9 | 5.5 | 5.0E-08 | 9.7E-08 | MB231/MB468 | CYTO |
| >hg38_21_+_8433172_8446622_RNA45SN1_45S_50ntFlanks@10733.10755.23\|GGTGTAAATCTCGCGCCGGGCCG | >hg38_21_+_8433172_8446622_RNA45SN1_45S_50ntFlanks | 10733 | 10755 | 23 | GGTGTAAATCTCGCGCCGGGCCG | 91.4 | -9.6 | 1.7 | -5.7 | 9.0E-09 | 9.7E-08 | BT20/MB468 | MP |
| >hg38_21_+_8433172_8446622_RNA45SN1_45S_50ntFlanks@9539.9572.34\|ACTCTGGTGGAGGTCCGTAGCGGTCCTGACGTGC | >hg38_21_+_8433172_8446622_RNA45SN1_45S_50ntFlanks | 9539 | 9572 | 34 | ACTCTGGTGGAGGTCCGTAGCGGTCCTGACGTGC | 322.2 | -3.5 | 0.6 | -5.4 | 5.2E-08 | 9.7E-08 | MB231/MB468 | NUC |
| >MI0000736\|hsa-mir-30c-1&WithFlank&1\|+\|40757278\|40757378@23.44.22(+1U);MI0000254\|hsa-mir-30c-2&WithFlank&6\|-\|71376954\|71377037@13.34.22(+1U)&[MIMAT0000244&hsa-miR-30c-5p&offsets\|0\|-1(+1U);m-57&1\|+\|40757300\|40757323&offsets\|0\|-2(+1U)];[MIMAT0000244_1&hsa-miR-30c-5p&offsets\|0\|-1(+1U);m-56&6\|-\|71377002\|71377025&offsets\|0\|-2(+1U)]\|TGTAAACATCCTACACTCTCAGT | >MI0000736\|hsa-mir-30c-1&WithFlank&1\|+\|40757278\|40757378 | 23 | 44 | 22(+1U) | TGTAAACATCCTACACTCTCAGT | 1950.1 | 4.9 | 0.9 | 5.5 | 3.9E-08 | 9.7E-08 | BT20/MB468 | MITO |
| >hg38_21_+_8433172_8446622_RNA45SN1_45S_50ntFlanks@12404.12431.28\|GCTTTTTGATCCTTCGATGTCGGCTCTT | >hg38_21_+_8433172_8446622_RNA45SN1_45S_50ntFlanks | 12404 | 12431 | 28 | GCTTTTTGATCCTTCGATGTCGGCTCTT | 519.0 | -6.5 | 1.2 | -5.4 | 5.7E-08 | 9.7E-08 | BT20/MB468 | NUC |
| >hg38_21_+_8433172_8446622_RNA45SN1_45S_50ntFlanks@8388.8406.19\|CGTGAAACCGTTAAGAGGT | >hg38_21_+_8433172_8446622_RNA45SN1_45S_50ntFlanks | 8388 | 8406 | 19 | CGTGAAACCGTTAAGAGGT | 103.4 | -9.8 | 1.7 | -5.7 | 9.0E-09 | 9.7E-08 | BT20/MB468 | MP |
| >MI0005566\|hsa-mir-374b&WithFlank&X\|-\|74218541\|74218624@17.39.23&[MIMAT0004955&hsa-miR-374b-5p&offsets\|0\|+1;m-153&X\|-\|74218587\|74218608&offsets\|0\|+1]\|ATATAATACAACCTGCTAAGTGT | >MI0005566\|hsa-mir-374b&WithFlank&X\|-\|74218541\|74218624 | 17 | 39 | 23 | ATATAATACAACCTGCTAAGTGT | 517.3 | 6.0 | 1.1 | 5.4 | 5.5E-08 | 9.7E-08 | MB231/MB468 | MITO |
| >hg38_21_+_8433172_8446622_RNA45SN1_45S_50ntFlanks@12619.12653.35\|AGGAGCCAATGGGGCGAAGCTACCATCTGTGGGAT | >hg38_21_+_8433172_8446622_RNA45SN1_45S_50ntFlanks | 12619 | 12653 | 35 | AGGAGCCAATGGGGCGAAGCTACCATCTGTGGGAT | 584.7 | -5.4 | 1.0 | -5.4 | 5.1E-08 | 9.7E-08 | MB231/MB468 | CYTO |
| >hg38_21_+_8433172_8446622_RNA45SN1_45S_50ntFlanks@11572.11605.34\|CCGACTTAGAACTGGTGCGGACCAGGGGAATCCG | >hg38_21_+_8433172_8446622_RNA45SN1_45S_50ntFlanks | 11572 | 11605 | 34 | CCGACTTAGAACTGGTGCGGACCAGGGGAATCCG | 4316.9 | -4.2 | 0.8 | -5.5 | 3.9E-08 | 9.7E-08 | BT20/MB468 | MITO |
| >hg38_21_+_8433172_8446622_RNA45SN1_45S_50ntFlanks@10479.10512.34\|CGATCGAAAGGGAGTCGGGTTCAGATCCCCGAAT | >hg38_21_+_8433172_8446622_RNA45SN1_45S_50ntFlanks | 10479 | 10512 | 34 | CGATCGAAAGGGAGTCGGGTTCAGATCCCCGAAT | 100.0 | -9.7 | 1.7 | -5.7 | 9.0E-09 | 9.7E-08 | BT20/MB468 | MP |
| >MI0003674\|hsa-mir-653&WithFlank&7\|-\|93482754\|93482861@22.43.22&[MIMAT0003328&hsa-miR-653-5p&offsets\|+3\|+4;m-368&7\|-\|93482819\|93482840&offsets\|0\|0]\|TTGAAACAATCTCTACTGAACC | >MI0003674\|hsa-mir-653&WithFlank&7\|-\|93482754\|93482861 | 22 | 43 | 22 | TTGAAACAATCTCTACTGAACC | 189.1 | -11.0 | 1.8 | -6.0 | 1.8E-09 | 9.7E-08 | BT20/MB231 | MP |
| >trnaMT_SerGCT_MT_+_12207_12265@1.41.41\|GAGAAAGCTCACAAGAACTGCTAACTCATGCCCCCATGTCT | >trnaMT_SerGCT_MT_+_12207_12265 | 1 | 41 | 41 | GAGAAAGCTCACAAGAACTGCTAACTCATGCCCCCATGTCT | 679.7 | 5.3 | 1.0 | 5.5 | 3.9E-08 | 9.7E-08 | BT20/MB468 | MITO |
| >trna14_LysTTT_11_-_59327808_59327880@1.33.33,trna2_LysTTT_17_+_8022473_8022545@1.33.33,trna54_LysTTT_1_+_204475655_204475727@1.33.33,trna5_LysTTT_11_+_59323902_59323974@1.33.33,trna62_LysTTT_1_-_204476158_204476230@1.33.33,trna76_LysTTT_6_+_28918806_28918878@1.33.33\|GCCCGGATAGCTCAGTCGGTAGAGCATCAGACT | >trna14_LysTTT_11_-_59327808_59327880 | 1 | 33 | 33 | GCCCGGATAGCTCAGTCGGTAGAGCATCAGACT | 221.7 | 3.7 | 0.7 | 5.4 | 5.8E-08 | 9.7E-08 | BT20/MB468 | NUC |
| >hg38_21_+_8433172_8446622_RNA45SN1_45S_50ntFlanks@4724.4757.34\|ATTAATCAAGAACGAAAGTCGGAGGTTCGAAGAC | >hg38_21_+_8433172_8446622_RNA45SN1_45S_50ntFlanks | 4724 | 4757 | 34 | ATTAATCAAGAACGAAAGTCGGAGGTTCGAAGAC | 287.2 | -5.3 | 1.0 | -5.4 | 5.2E-08 | 9.8E-08 | MB231/MB468 | NUC |
| >hg38_21_+_8433172_8446622_RNA45SN1_45S_50ntFlanks@3837.3862.26\|CCTCTCCTACTTGGATAACTGTGGTA | >hg38_21_+_8433172_8446622_RNA45SN1_45S_50ntFlanks | 3837 | 3862 | 26 | CCTCTCCTACTTGGATAACTGTGGTA | 347.8 | -7.9 | 1.5 | -5.4 | 5.5E-08 | 9.8E-08 | MB231/MB468 | MITO |
| >MI0000087\|hsa-mir-29a&WithFlank&7\|-\|130876741\|130876816@47.70.24&[MIMAT0000086&hsa-miR-29a-3p&offsets\|-1\|+1;m-26&7\|-\|130876749\|130876769&offsets\|-1\|+2]\|CTAGCACCATCTGAAATCGGTTAT | >MI0000087\|hsa-mir-29a&WithFlank&7\|-\|130876741\|130876816 | 47 | 70 | 24 | CTAGCACCATCTGAAATCGGTTAT | 556.9 | 8.8 | 1.6 | 5.5 | 4.7E-08 | 9.8E-08 | MB231/MB468 | MP |
| >hg38_21_+_8433172_8446622_RNA45SN1_45S_50ntFlanks@8472.8490.19\|CGGCCCGGCGGATCTTTCC | >hg38_21_+_8433172_8446622_RNA45SN1_45S_50ntFlanks | 8472 | 8490 | 19 | CGGCCCGGCGGATCTTTCC | 137.9 | -6.4 | 1.2 | -5.5 | 4.4E-08 | 9.8E-08 | BT20/MB468 | CYTO |
| >hg38_21_+_8433172_8446622_RNA45SN1_45S_50ntFlanks@12934.12951.18\|CTAAACCATTCGTAGACG | >hg38_21_+_8433172_8446622_RNA45SN1_45S_50ntFlanks | 12934 | 12951 | 18 | CTAAACCATTCGTAGACG | 434.2 | -2.7 | 0.5 | -5.5 | 3.9E-08 | 9.8E-08 | BT20/MB468 | MITO |
| >trnaMT_ValTAC_MT_+_1602_1670@52.72.21\|TTAACTTGACCGCTCTGACCA | >trnaMT_ValTAC_MT_+_1602_1670 | 52 | 72 | 21 | TTAACTTGACCGCTCTGACCA | 3716.8 | 5.8 | 1.1 | 5.5 | 3.9E-08 | 9.8E-08 | BT20/MB468 | MITO |
| >hg38_21_+_8433172_8446622_RNA45SN1_45S_50ntFlanks@9572.9593.22\|CAAATCGGTCGTCCGACCTGGG | >hg38_21_+_8433172_8446622_RNA45SN1_45S_50ntFlanks | 9572 | 9593 | 22 | CAAATCGGTCGTCCGACCTGGG | 123.4 | -6.3 | 1.1 | -5.5 | 4.4E-08 | 9.8E-08 | BT20/MB468 | CYTO |
| >MI0000283\|hsa-mir-203a&WithFlank&14\|+\|104117399\|104117520@71.90.20&[MIMAT0000264&hsa-miR-203a-3p&offsets\|0\|-2;m-79&14\|+\|104117469\|104117490&offsets\|0\|-2]\|GTGAAATGTTTAGGACCACT | >MI0000283\|hsa-mir-203a&WithFlank&14\|+\|104117399\|104117520 | 71 | 90 | 20 | GTGAAATGTTTAGGACCACT | 404.6 | 7.9 | 1.4 | 5.5 | 3.9E-08 | 9.8E-08 | BT20/MB468 | MITO |
| >hg38_21_+_8433172_8446622_RNA45SN1_45S_50ntFlanks@11567.11600.34\|AGCAGCCGACTTAGAACTGGTGCGGACCAGGGGA | >hg38_21_+_8433172_8446622_RNA45SN1_45S_50ntFlanks | 11567 | 11600 | 34 | AGCAGCCGACTTAGAACTGGTGCGGACCAGGGGA | 743.9 | -2.7 | 0.5 | -5.4 | 5.3E-08 | 9.8E-08 | MB231/MB468 | NUC |
| >trnaMT_ValTAC_MT_+_1602_1670@28.72.45\|AACTTACACTTAGGAGATTTCAACTTAACTTGACCGCTCTGACCA | >trnaMT_ValTAC_MT_+_1602_1670 | 28 | 72 | 45 | AACTTACACTTAGGAGATTTCAACTTAACTTGACCGCTCTGACCA | 1533.1 | 5.1 | 0.9 | 5.5 | 4.0E-08 | 9.8E-08 | BT20/MB468 | MITO |
| >MI0000465\|hsa-mir-191&WithFlank&3\|-\|49020612\|49020715@22.44.23&[MIMAT0000440&hsa-miR-191-5p&offsets\|0\|0;m-48&3\|-\|49020672\|49020694&offsets\|0\|0]\|CAACGGAATCCCAAAAGCAGCTG | >MI0000465\|hsa-mir-191&WithFlank&3\|-\|49020612\|49020715 | 22 | 44 | 23 | CAACGGAATCCCAAAAGCAGCTG | 11130.8 | 4.3 | 0.8 | 5.4 | 5.3E-08 | 9.9E-08 | MB231/MB468 | NUC |
| >am_trna128_GlyGCC_6_-_27870686_27870756@7.28.22,trna133_GlyCCC_1_-_16872434_16872504@7.28.22,trna17_GluTTC_2_-_204229131_204229201@7.28.22,trna18_GlyGCC_16_+_70822597_70822667@7.28.22,trna19_GlyGCC_16_+_70823410_70823480@7.28.22,trna19_GlyGCC_2_-_157257659_157257729@7.28.22,trna24_GlyGCC_16_-_70812942_70813012@7.28.22,trna25_GlyGCC_16_-_70812114_70812184@7.28.22,trna2_GlyGCC_21_-_18827107_18827177@7.28.22,trna35_GlyGCC_1_+_161413094_161413164@7.28.22,trna37_GlyGCC_1_+_161420467_161420537@7.28.22,trna39_GlyGCC_1_+_161427898_161427968@7.28.22,trna41_GlyGCC_1_+_161435258_161435328@7.28.22,trna4_GlyCCC_1_+_17188416_17188486@7.28.22,trna5_GlyGCC_17_+_8029064_8029134@7.28.22,trna68_GlyGCC_1_-_161493637_161493707@7.28.22,trna91_GlyCCC_1_-_149680210_149680280@7.28.22,trna99_ValCAC_1_-_149294666_149294736@7.28.22\|GTGGTTCAGTGGTAGAATTCTC | >am_trna128_GlyGCC_6_-_27870686_27870756 | 7 | 28 | 22 | GTGGTTCAGTGGTAGAATTCTC | 313.3 | -6.3 | 1.1 | -5.5 | 4.8E-08 | 9.9E-08 | MB231/MB468 | MP |
| >trnaMT_ValTAC_MT_+_1602_1670@54.72.19\|AACTTGACCGCTCTGACCA | >trnaMT_ValTAC_MT_+_1602_1670 | 54 | 72 | 19 | AACTTGACCGCTCTGACCA | 524.3 | 8.3 | 1.5 | 5.4 | 5.2E-08 | 9.9E-08 | MB231/MB468 | CYTO |

| | | | | | | | | | | | | | |
|---|---|---|---|---|---|---|---|---|---|---|---|---|---|
| >hg38_21_+_8433172_8446622_RNA45SN1_45S_50ntFlanks@10392.10419.28\|ACAGCAGTTGAACATGGGTCAGTCGGTC | >hg38_21_+_8433172_8446622_RNA45SN1_45S_50ntFlanks | 10392 | 10419 | 28 | ACAGCAGTTGAACATGGGTCAGTCGGTC | 515.9 | -5.0 | 0.9 | -5.4 | 5.2E-08 | 9.9E-08 | MB231/MB468 | CYTO |
| >MI0000086\|hsa-mir-28&WithFlank&3\|+\|188688775\|188688872@62.82.21(+1C)&[MIMAT0004502&hsa-miR-28-3p&offsets\|+2\|+1(+1C);m-28&3\|+\|188688834\|188688855&offsets\|+2\|+1(+1C)]\|CTAGATTGTGAGCTCCTGGAGC | >MI0000086\|hsa-mir-28&WithFlank&3\|+\|188688775\|188688872 | 62 | 82 | 21(+1C) | CTAGATTGTGAGCTCCTGGAGC | 121.7 | 10.7 | 1.9 | 5.7 | 9.2E-09 | 9.9E-08 | BT20/MB468 | MP |
| >trnaMT_GluTTC_MT_-_14674_14742@21.63.43,trnalookalike8_GluTTC_5_-_93905172_93905240@21.63.43\|ACGATGGTTTTTCATATCATTGGTCGTGGTTGTAGTCCGTGCG | >trnaMT_GluTTC_MT_-_14674_14742 | 21 | 63 | 43 | ACGATGGTTTTTCATATCATTGGTCGTGGTTGTAGTCCGTGCG | 338.8 | 12.2 | 2.1 | 5.7 | 9.2E-09 | 9.9E-08 | BT20/MB468 | MP |
| >hg38_21_+_8433172_8446622_RNA45SN1_45S_50ntFlanks@9544.9576.33\|GGTGGAGGTCCGTAGCGGTCCTGACGTGCAAAT | >hg38_21_+_8433172_8446622_RNA45SN1_45S_50ntFlanks | 9544 | 9576 | 33 | GGTGGAGGTCCGTAGCGGTCCTGACGTGCAAAT | 190.7 | 6.3 | 1.1 | 5.4 | 5.2E-08 | 9.9E-08 | MB231/MB468 | CYTO |
| >hg38_21_+_8433172_8446622_RNA45SN1_45S_50ntFlanks@10384.10430.47\|CCATGTGAACAGCAGTTGAACATGGGTCAGTCGGTCCTGAGAGATGG | >hg38_21_+_8433172_8446622_RNA45SN1_45S_50ntFlanks | 10384 | 10430 | 47 | CCATGTGAACAGCAGTTGAACATGGGTCAGTCGGTCCTGAGAGATGG | 121.8 | -9.2 | 1.7 | -5.5 | 4.5E-08 | 9.9E-08 | BT20/MB468 | CYTO |
| >hg38_21_+_8433172_8446622_RNA45SN1_45S_50ntFlanks@6699.6719.21\|GCTAGCTGCGAGAATTAATGT | >hg38_21_+_8433172_8446622_RNA45SN1_45S_50ntFlanks | 6699 | 6719 | 21 | GCTAGCTGCGAGAATTAATGT | 1877.3 | -3.5 | 0.6 | -5.4 | 5.2E-08 | 9.9E-08 | MB231/MB468 | CYTO |
| >hg38_21_+_8433172_8446622_RNA45SN1_45S_50ntFlanks@10411.10456.46\|CAGTCGGTCCTGAGAGATGGGCGAGCGCCGTTCCGAAGGGACGGGC | >hg38_21_+_8433172_8446622_RNA45SN1_45S_50ntFlanks | 10411 | 10456 | 46 | CAGTCGGTCCTGAGAGATGGGCGAGCGCCGTTCCGAAGGGACGGGC | 426.1 | -7.6 | 1.4 | -5.5 | 4.5E-08 | 9.9E-08 | BT20/MB468 | CYTO |
| >hg38_21_+_8433172_8446622_RNA45SN1_45S_50ntFlanks@10992.11010.19\|CGCGCGCCGCTCGCTCCCT | >hg38_21_+_8433172_8446622_RNA45SN1_45S_50ntFlanks | 10992 | 11010 | 19 | CGCGCGCCGCTCGCTCCCT | 196.8 | -4.5 | 0.8 | -5.5 | 4.5E-08 | 1.0E-07 | BT20/MB468 | CYTO |
| >hg38_21_+_8433172_8446622_RNA45SN1_45S_50ntFlanks@3835.3868.34\|CTCCTCTCCTACTTGGATAACTGTGGTAATTCTA | >hg38_21_+_8433172_8446622_RNA45SN1_45S_50ntFlanks | 3835 | 3868 | 34 | CTCCTCTCCTACTTGGATAACTGTGGTAATTCTA | 458.9 | -6.1 | 1.1 | -5.4 | 5.7E-08 | 1.0E-07 | MB231/MB468 | MITO |
| >hg38_21_+_8433172_8446622_RNA45SN1_45S_50ntFlanks@3782.3801.20\|CAGTGAAACTGCGAATGGCT | >hg38_21_+_8433172_8446622_RNA45SN1_45S_50ntFlanks | 3782 | 3801 | 20 | CAGTGAAACTGCGAATGGCT | 1996.8 | -7.2 | 1.3 | -5.4 | 5.3E-08 | 1.0E-07 | MB231/MB468 | NUC |
| >hg38_21_+_8433172_8446622_RNA45SN1_45S_50ntFlanks@8115.8135.21\|CGCGGGACATGTGGCGTACGG | >hg38_21_+_8433172_8446622_RNA45SN1_45S_50ntFlanks | 8115 | 8135 | 21 | CGCGGGACATGTGGCGTACGG | 696.7 | 12.8 | 2.1 | 6.0 | 1.8E-09 | 1.0E-07 | BT20/MB231 | MP |
| >hg38_21_+_8433172_8446622_RNA45SN1_45S_50ntFlanks@8357.8402.46\|AAAAGAACTTTGAAGAGAGAGTTCAAGAGGGCGTGAAACCGTTAAG | >hg38_21_+_8433172_8446622_RNA45SN1_45S_50ntFlanks | 8357 | 8402 | 46 | AAAAGAACTTTGAAGAGAGAGTTCAAGAGGGCGTGAAACCGTTAAG | 109.4 | -5.1 | 0.9 | -5.4 | 5.2E-08 | 1.0E-07 | MB231/MB468 | CYTO |
| >hg38_21_+_8433172_8446622_RNA45SN1_45S_50ntFlanks@11569.11609.41\|CAGCCGACTTAGAACTGGTGCGGACCAGGGGAATCCGACTG | >hg38_21_+_8433172_8446622_RNA45SN1_45S_50ntFlanks | 11569 | 11609 | 41 | CAGCCGACTTAGAACTGGTGCGGACCAGGGGAATCCGACTG | 306.9 | -6.6 | 1.2 | -5.5 | 4.9E-08 | 1.0E-07 | MB231/MB468 | MP |
| >trna116_GluCTC_1_-_145399233_145399304@1.33.33,trna59_GluCTC_1_+_249168447_249168518@1.33.33,trna71_GluCTC_1_-_161439189_161439260@1.33.33,trna74_GluCTC_1_-_161431809_161431880@1.33.33,trna77_GluCTC_1_-_161424398_161424469@1.33.33,trna77_GluCTC_6_+_28949976_28950047@1.33.33,trna80_GluCTC_1_-_161417018_161417089@1.33.33,trna87_GluCTC_6_-_126101393_126101464@1.33.33\|TCCCTGGTGGTCTAGTGGTTAGGATTCGGCGCT | >trna116_GluCTC_1_-_145399233_145399304 | 1 | 33 | 33 | TCCCTGGTGGTCTAGTGGTTAGGATTCGGCGCT | 57494.8 | 6.1 | 1.1 | 5.4 | 5.4E-08 | 1.0E-07 | MB231/MB468 | NUC |
| >hg38_21_+_8433172_8446622_RNA45SN1_45S_50ntFlanks@12416.12458.43\|TTCGATGTCGGCTCTTCCTATCATTGTGAAGCAGAATTCACCA | >hg38_21_+_8433172_8446622_RNA45SN1_45S_50ntFlanks | 12416 | 12458 | 43 | TTCGATGTCGGCTCTTCCTATCATTGTGAAGCAGAATTCACCA | 760.1 | -5.0 | 0.8 | -5.9 | 3.1E-09 | 1.0E-07 | BT20/MB231 | CYTO |
| >hg38_21_+_8433172_8446622_RNA45SN1_45S_50ntFlanks@9740.9765.26\|AACGATCTCAACCTATTCTCAAACTT | >hg38_21_+_8433172_8446622_RNA45SN1_45S_50ntFlanks | 9740 | 9765 | 26 | AACGATCTCAACCTATTCTCAAACTT | 170.5 | -7.1 | 1.3 | -5.5 | 4.9E-08 | 1.0E-07 | MB231/MB468 | MP |
| >hg38_21_+_8433172_8446622_RNA45SN1_45S_50ntFlanks@11873.11902.30\|AAGAAGACCCTGTTGAGCTTGACTCTAGTC | >hg38_21_+_8433172_8446622_RNA45SN1_45S_50ntFlanks | 11873 | 11902 | 30 | AAGAAGACCCTGTTGAGCTTGACTCTAGTC | 588.0 | -8.3 | 1.5 | -5.4 | 5.9E-08 | 1.0E-07 | BT20/MB468 | NUC |
| >hg38_21_+_8433172_8446622_RNA45SN1_45S_50ntFlanks@12072.12110.39\|CCCGAGGGGCTCTCGCTTCTGGCGCCAAGCGCCCGGCCG | >hg38_21_+_8433172_8446622_RNA45SN1_45S_50ntFlanks | 12072 | 12110 | 39 | CCCGAGGGGCTCTCGCTTCTGGCGCCAAGCGCCCGGCCG | 58.1 | -9.7 | 1.8 | -5.5 | 4.9E-08 | 1.0E-07 | MB231/MB468 | MP |
| >hg38_21_+_8433172_8446622_RNA45SN1_45S_50ntFlanks@7979.8012.34\|ACCTCAGATCAGACGTGGCGACCCGCTGAATTTA | >hg38_21_+_8433172_8446622_RNA45SN1_45S_50ntFlanks | 7979 | 8012 | 34 | ACCTCAGATCAGACGTGGCGACCCGCTGAATTTA | 2157.6 | -4.5 | 0.8 | -5.4 | 5.7E-08 | 1.0E-07 | MB231/MB468 | MITO |
| >hg38_21_+_8433172_8446622_RNA45SN1_45S_50ntFlanks@9299.9330.32\|CACGGACCAAGGAGTCTAACACGTGCGCGAGT | >hg38_21_+_8433172_8446622_RNA45SN1_45S_50ntFlanks | 9299 | 9330 | 32 | CACGGACCAAGGAGTCTAACACGTGCGCGAGT | 82.1 | 10.1 | 1.8 | 5.7 | 9.4E-09 | 1.0E-07 | BT20/MB468 | MP |
| >hg38_MT_+_598_1651_RNR1_12S_50ntFlanks@428.455.28\|AAAGTGGCTTTAACATATCTGAACACAC | >hg38_MT_+_598_1651_RNR1_12S_50ntFlanks | 428 | 455 | 28 | AAAGTGGCTTTAACATATCTGAACACAC | 273.7 | 8.5 | 1.6 | 5.4 | 5.4E-08 | 1.0E-07 | MB231/MB468 | NUC |
| >hg38_21_+_8433172_8446622_RNA45SN1_45S_50ntFlanks@10824.10865.42\|AGTCGGCAAGCCGGATCCGTAACTTCGGGATAAGGATTGGCT | >hg38_21_+_8433172_8446622_RNA45SN1_45S_50ntFlanks | 10824 | 10865 | 42 | AGTCGGCAAGCCGGATCCGTAACTTCGGGATAAGGATTGGCT | 107.3 | -9.1 | 1.7 | -5.5 | 4.6E-08 | 1.0E-07 | BT20/MB468 | CYTO |
| >hg38_21_+_8433172_8446622_RNA45SN1_45S_50ntFlanks@5067.5098.32\|CTGGTTAATTCCGATAACGAACGAGACTCTGG | >hg38_21_+_8433172_8446622_RNA45SN1_45S_50ntFlanks | 5067 | 5098 | 32 | CTGGTTAATTCCGATAACGAACGAGACTCTGG | 91.0 | -10.4 | 1.9 | -5.5 | 4.9E-08 | 1.0E-07 | MB231/MB468 | MP |
| >hg38_21_+_8433172_8446622_RNA45SN1_45S_50ntFlanks@10792.10818.27\|CTCTGGCATGTTGGAACAATGTAGGTA | >hg38_21_+_8433172_8446622_RNA45SN1_45S_50ntFlanks | 10792 | 10818 | 27 | CTCTGGCATGTTGGAACAATGTAGGTA | 492.1 | -5.8 | 1.1 | -5.4 | 5.7E-08 | 1.0E-07 | MB231/MB468 | MITO |
| >hg38_21_+_8433172_8446622_RNA45SN1_45S_50ntFlanks@5186.5205.20\|CAATAACAGGTCTGTGATGC | >hg38_21_+_8433172_8446622_RNA45SN1_45S_50ntFlanks | 5186 | 5205 | 20 | CAATAACAGGTCTGTGATGC | 817.7 | -8.9 | 1.6 | -5.4 | 6.0E-08 | 1.0E-07 | BT20/MB468 | NUC |
| >hg38_21_+_8433172_8446622_RNA45SN1_45S_50ntFlanks@11569.11600.32\|CAGCCGACTTAGAACTGGTGCGGACCAGGGGA | >hg38_21_+_8433172_8446622_RNA45SN1_45S_50ntFlanks | 11569 | 11600 | 32 | CAGCCGACTTAGAACTGGTGCGGACCAGGGGA | 1608.2 | -3.6 | 0.7 | -5.5 | 4.6E-08 | 1.0E-07 | BT20/MB468 | CYTO |
| >hg38_21_+_8433172_8446622_RNA45SN1_45S_50ntFlanks@11875.11893.19\|GAAGACCCTGTTGAGCTTG | >hg38_21_+_8433172_8446622_RNA45SN1_45S_50ntFlanks | 11875 | 11893 | 19 | GAAGACCCTGTTGAGCTTG | 506.7 | -6.3 | 1.2 | -5.4 | 5.7E-08 | 1.0E-07 | MB231/MB468 | MITO |
| >hg38_21_+_8433172_8446622_RNA45SN1_45S_50ntFlanks@9565.9592.28\|TGACGTGCAAATCGGTCGTCCGACCTGG | >hg38_21_+_8433172_8446622_RNA45SN1_45S_50ntFlanks | 9565 | 9592 | 28 | TGACGTGCAAATCGGTCGTCCGACCTGG | 1030.3 | -6.8 | 1.3 | -5.4 | 6.0E-08 | 1.0E-07 | BT20/MB468 | NUC |
| >MI0000469\|hsa-mir-125a&WithFlank&19\|+\|51693248\|51693345@59.79.21&[MIMAT0004602&hsa-miR-125a-3p&offsets\|0\|-1;m-333&19\|+\|51693306\|51693326&offsets\|0\|0]\|ACAGGTGAGGTTCTTGGGAGC | >MI0000469\|hsa-mir-125a&WithFlank&19\|+\|51693248\|51693345 | 59 | 79 | 21 | ACAGGTGAGGTTCTTGGGAGC | 320.2 | 6.9 | 1.3 | 5.4 | 5.3E-08 | 1.0E-07 | MB231/MB468 | CYTO |
| >hg38_21_+_8433172_8446622_RNA45SN1_45S_50ntFlanks@9740.9758.19\|AACGATCTCAACCTATTCT | >hg38_21_+_8433172_8446622_RNA45SN1_45S_50ntFlanks | 9740 | 9758 | 19 | AACGATCTCAACCTATTCT | 381.3 | -3.5 | 0.6 | -5.4 | 5.3E-08 | 1.0E-07 | MB231/MB468 | CYTO |
| >hg38_21_+_8433172_8446622_RNA45SN1_45S_50ntFlanks@4367.4408.42\|CAGTTAAAAAGCTCGTAGTTGGATCTTGGGAGCGGGCGGGCG | >hg38_21_+_8433172_8446622_RNA45SN1_45S_50ntFlanks | 4367 | 4408 | 42 | CAGTTAAAAAGCTCGTAGTTGGATCTTGGGAGCGGGCGGGCG | 102.7 | -8.8 | 1.6 | -5.5 | 4.1E-08 | 1.0E-07 | BT20/MB468 | MITO |
| >hg38_21_+_8433172_8446622_RNA45SN1_45S_50ntFlanks@4367.4406.40\|CAGTTAAAAAGCTCGTAGTTGGATCTTGGGAGCGGGCGGG | >hg38_21_+_8433172_8446622_RNA45SN1_45S_50ntFlanks | 4367 | 4406 | 40 | CAGTTAAAAAGCTCGTAGTTGGATCTTGGGAGCGGGCGGG | 154.5 | -4.7 | 0.9 | -5.4 | 5.3E-08 | 1.0E-07 | MB231/MB468 | CYTO |
| >hg38_21_+_8433172_8446622_RNA45SN1_45S_50ntFlanks@11569.11590.22\|CAGCCGACTTAGAACTGGTGCG | >hg38_21_+_8433172_8446622_RNA45SN1_45S_50ntFlanks | 11569 | 11590 | 22 | CAGCCGACTTAGAACTGGTGCG | 1128.9 | -7.1 | 1.3 | -5.5 | 4.9E-08 | 1.0E-07 | MB231/MB468 | MP |
| >trnaMT_PheGAA_MT_+_577_647@36.67.32\|AAAATGTTTAGACGGGCTCACATCACCCCATA | >trnaMT_PheGAA_MT_+_577_647 | 36 | 67 | 32 | AAAATGTTTAGACGGGCTCACATCACCCCATA | 61.1 | 9.2 | 1.7 | 5.5 | 4.9E-08 | 1.0E-07 | MB231/MB468 | MP |
| >hg38_21_+_8433172_8446622_RNA45SN1_45S_50ntFlanks@3992.4023.32\|GGTGACTCTAGATAACCTCGGGCCGATCGCAC | >hg38_21_+_8433172_8446622_RNA45SN1_45S_50ntFlanks | 3992 | 4023 | 32 | GGTGACTCTAGATAACCTCGGGCCGATCGCAC | 428.6 | 4.0 | 0.7 | 5.4 | 5.5E-08 | 1.0E-07 | MB231/MB468 | NUC |
| >hg38_21_+_8433172_8446622_RNA45SN1_45S_50ntFlanks@4064.4101.38\|ATCAACTTTCGATGGTAGTCGCCGTGCCTACCATGGTG | >hg38_21_+_8433172_8446622_RNA45SN1_45S_50ntFlanks | 4064 | 4101 | 38 | ATCAACTTTCGATGGTAGTCGCCGTGCCTACCATGGTG | 118.1 | -10.0 | 1.7 | -5.7 | 9.5E-09 | 1.0E-07 | BT20/MB468 | MP |
| >trnaMT_HisGTG_MT_+_12138_12206@32.69.38\|TGAATCTGACAACAGAGGCTTACGACCCCTTATTTACC | >trnaMT_HisGTG_MT_+_12138_12206 | 32 | 69 | 38 | TGAATCTGACAACAGAGGCTTACGACCCCTTATTTACC | 562.5 | 5.2 | 1.0 | 5.4 | 5.5E-08 | 1.0E-07 | MB231/MB468 | NUC |
| >hg38_21_+_8433172_8446622_RNA45SN1_45S_50ntFlanks@12324.12365.42\|TTGGGTTTTAAGCAGGAGGTGTCAGAAAAGTTACCACAGGGA | >hg38_21_+_8433172_8446622_RNA45SN1_45S_50ntFlanks | 12324 | 12365 | 42 | TTGGGTTTTAAGCAGGAGGTGTCAGAAAAGTTACCACAGGGA | 135.6 | -9.2 | 1.7 | -5.5 | 4.1E-08 | 1.0E-07 | BT20/MB468 | MITO |
| >trnaMT_GluTTC_MT_-_14674_14742@40.70.31,trnalookalike8_GluTTC_5_-_93905172_93905240@40.70.31\|TTGGTCGTGGTTGTAGTCCGTGCGAGAATAC | >trnaMT_GluTTC_MT_-_14674_14742 | 40 | 70 | 31 | TTGGTCGTGGTTGTAGTCCGTGCGAGAATAC | 580.4 | 9.3 | 1.7 | 5.5 | 4.1E-08 | 1.0E-07 | BT20/MB468 | MITO |
| >hg38_MT_+_1621_3279_RNR2_16S_50ntFlanks@51.96.46\|GCTAAACCTAGCCCCAAACCCACTCCACCTTACTACCAGACAACCT | >hg38_MT_+_1621_3279_RNR2_16S_50ntFlanks | 51 | 96 | 46 | GCTAAACCTAGCCCCAAACCCACTCCACCTTACTACCAGACAACCT | 1006.8 | 5.4 | 1.0 | 5.5 | 4.1E-08 | 1.0E-07 | BT20/MB468 | MITO |
| >MI0000101\|hsa-mir-99a&WithFlank&21\|+\|16539083\|16539175@19.40.22&[MIMAT0000097&hsa-miR-99a-5p&offsets\|0\|0;m-64&21\|+\|16539101\|16539121&offsets\|0\|+1]\|AACCCGTAGATCCGATCTTGTG | >MI0000101\|hsa-mir-99a&WithFlank&21\|+\|16539083\|16539175 | 19 | 40 | 22 | AACCCGTAGATCCGATCTTGTG | 1942.4 | -8.8 | 1.6 | -5.4 | 6.1E-08 | 1.0E-07 | BT20/MB468 | NUC |
| >hg38_21_+_8433172_8446622_RNA45SN1_45S_50ntFlanks@12412.12455.44\|ATCCTTCGATGTCGGCTCTTCCTATCATTGTGAAGCAGAATTCA | >hg38_21_+_8433172_8446622_RNA45SN1_45S_50ntFlanks | 12412 | 12455 | 44 | ATCCTTCGATGTCGGCTCTTCCTATCATTGTGAAGCAGAATTCA | 153.0 | -9.3 | 1.7 | -5.5 | 4.1E-08 | 1.0E-07 | BT20/MB468 | MITO |
| >hg38_21_+_8433172_8446622_RNA45SN1_45S_50ntFlanks@9534.9561.28\|AGGAAACTCTGGTGGAGGTCCGTAGCGG | >hg38_21_+_8433172_8446622_RNA45SN1_45S_50ntFlanks | 9534 | 9561 | 28 | AGGAAACTCTGGTGGAGGTCCGTAGCGG | 904.2 | -4.0 | 0.7 | -5.5 | 4.1E-08 | 1.0E-07 | BT20/MB468 | MITO |
| >hg38_21_+_8433172_8446622_RNA45SN1_45S_50ntFlanks@10458.10487.30\|ATGGCCTCCGTTGCCCTCGGCCGATCGAAA | >hg38_21_+_8433172_8446622_RNA45SN1_45S_50ntFlanks | 10458 | 10487 | 30 | ATGGCCTCCGTTGCCCTCGGCCGATCGAAA | 1427.9 | -9.1 | 1.7 | -5.4 | 6.1E-08 | 1.0E-07 | BT20/MB468 | NUC |
| >hg38_21_+_8433172_8446622_RNA45SN1_45S_50ntFlanks@475.502.28\|CTTCGTGATCGATGTGGTGACGTCGTGC | >hg38_21_+_8433172_8446622_RNA45SN1_45S_50ntFlanks | 475 | 502 | 28 | CTTCGTGATCGATGTGGTGACGTCGTGC | 543.8 | 5.4 | 1.0 | 5.4 | 5.8E-08 | 1.0E-07 | MB231/MB468 | MITO |
| >hg38_21_+_8433172_8446622_RNA45SN1_45S_50ntFlanks@3705.3725.21\|TACCTGGTTGATCCTGCCAGT | >hg38_21_+_8433172_8446622_RNA45SN1_45S_50ntFlanks | 3705 | 3725 | 21 | TACCTGGTTGATCCTGCCAGT | 3910.9 | 5.4 | 1.0 | 5.5 | 4.1E-08 | 1.0E-07 | BT20/MB468 | MITO |
| >hg38_21_+_8433172_8446622_RNA45SN1_45S_50ntFlanks@12995.13036.42\|CTCGCTGCGATCTATTGAAAGTCAGCCCTCGACACAAGGGTT | >hg38_21_+_8433172_8446622_RNA45SN1_45S_50ntFlanks | 12995 | 13036 | 42 | CTCGCTGCGATCTATTGAAAGTCAGCCCTCGACACAAGGGTT | 92.8 | 10.3 | 1.8 | 5.7 | 9.6E-09 | 1.0E-07 | BT20/MB468 | MP |
| >hg38_21_+_8433172_8446622_RNA45SN1_45S_50ntFlanks@9929.9961.33\|GATATAGACAGCAGGACGGTGGCCATGGAAGTC | >hg38_21_+_8433172_8446622_RNA45SN1_45S_50ntFlanks | 9929 | 9961 | 33 | GATATAGACAGCAGGACGGTGGCCATGGAAGTC | 222.6 | -6.2 | 1.1 | -5.5 | 5.0E-08 | 1.0E-07 | MB231/MB468 | MP |
| >hg38_21_+_8433172_8446622_RNA45SN1_45S_50ntFlanks@12156.12183.28\|GAGTTTGACTGGGGCGGTACACCTGTCA | >hg38_21_+_8433172_8446622_RNA45SN1_45S_50ntFlanks | 12156 | 12183 | 28 | GAGTTTGACTGGGGCGGTACACCTGTCA | 101.4 | -9.7 | 1.7 | -5.7 | 9.6E-09 | 1.0E-07 | BT20/MB468 | MP |
| >hg38_21_+_8433172_8446622_RNA45SN1_45S_50ntFlanks@12946.12990.45\|TAGACGACCTGCTTCTGGGTCGGGGTTTCGTACGTAGCAGAGCAG | >hg38_21_+_8433172_8446622_RNA45SN1_45S_50ntFlanks | 12946 | 12990 | 45 | TAGACGACCTGCTTCTGGGTCGGGGTTTCGTACGTAGCAGAGCAG | 88.3 | -8.8 | 1.6 | -5.5 | 4.7E-08 | 1.0E-07 | BT20/MB468 | CYTO |
| >hg38_21_+_8433172_8446622_RNA45SN1_45S_50ntFlanks@11589.11612.24\|CGGACCAGGGGAATCCGACTGTTT | >hg38_21_+_8433172_8446622_RNA45SN1_45S_50ntFlanks | 11589 | 11612 | 24 | CGGACCAGGGGAATCCGACTGTTT | 602.0 | -6.2 | 1.1 | -5.5 | 5.0E-08 | 1.0E-07 | MB231/MB468 | MP |

| | | | | | | | | | | | | | |
|---|---|---|---|---|---|---|---|---|---|---|---|---|---|
| >MI0000441\|hsa-mir-30b&WithFlank&8\|-\|134800514\|134800613@23.43.21(+1C)&[MIMAT0000420&hsa-miR-30b-5p&offsets\|0\|-1(+1C);m-76&8\|-\|134800570\|134800591&offsets\|0\|-1(+1C)]\|TGTAAACATCCTACACTCAGCC | >MI0000441\|hsa-mir-30b&WithFlank&8\|-\|134800514\|134800613 | 23 | 43 | 21(+1C) | TGTAAACATCCTACACTCAGCC | 1731.5 | 8.9 | 1.6 | 5.4 | 5.9E-08 | 1.0E-07 | MB231/MB468 | MITO |
| >hg38_21_+_8433172_8446622_RNA45SN1_45S_50ntFlanks@6662.6700.39\|GGTGGATCACTCGGCTCGTGCGTCGATGAAGAACGCAGC | >hg38_21_+_8433172_8446622_RNA45SN1_45S_50ntFlanks | 6662 | 6700 | 39 | GGTGGATCACTCGGCTCGTGCGTCGATGAAGAACGCAGC | 459.2 | 9.4 | 1.6 | 5.7 | 9.7E-09 | 1.0E-07 | BT20/MB468 | MP |
| >hg38_21_+_8433172_8446622_RNA45SN1_45S_50ntFlanks@10539.10564.26\|AGGCGTCCAGTGCGGTAACGCGACCG | >hg38_21_+_8433172_8446622_RNA45SN1_45S_50ntFlanks | 10539 | 10564 | 26 | AGGCGTCCAGTGCGGTAACGCGACCG | 99.2 | -9.7 | 1.7 | -5.7 | 9.7E-09 | 1.0E-07 | BT20/MB468 | MP |
| >hg38_21_+_8433172_8446622_RNA45SN1_45S_50ntFlanks@10876.10894.19\|GGTCGGTCGGGCTGGGGCG | >hg38_21_+_8433172_8446622_RNA45SN1_45S_50ntFlanks | 10876 | 10894 | 19 | GGTCGGTCGGGCTGGGGCG | 101.8 | -9.8 | 1.7 | -5.7 | 9.7E-09 | 1.0E-07 | BT20/MB468 | MP |
| >hg38_21_+_8433172_8446622_RNA45SN1_45S_50ntFlanks@11783.11805.23\|CATGAATGGATGAACGAGATTCC | >hg38_21_+_8433172_8446622_RNA45SN1_45S_50ntFlanks | 11783 | 11805 | 23 | CATGAATGGATGAACGAGATTCC | 1618.1 | -4.6 | 0.8 | -5.7 | 9.7E-09 | 1.0E-07 | BT20/MB468 | MP |
| >hg38_21_+_8433172_8446622_RNA45SN1_45S_50ntFlanks@12420.12437.18\|ATGTCGGCTCTTCCTATC | >hg38_21_+_8433172_8446622_RNA45SN1_45S_50ntFlanks | 12420 | 12437 | 18 | ATGTCGGCTCTTCCTATC | 12389.8 | -3.3 | 0.6 | -5.4 | 5.5E-08 | 1.0E-07 | MB231/MB468 | NUC |
| >MI0000298\|hsa-mir-221&WithFlank&X\|-\|45746151\|45746272@71.91.21(+1A)&[MIMAT0000278&hsa-miR-221-3p&offsets\|0\|-2(+1A);m-91&X\|-\|45746181\|45746202&offsets\|0\|-1(+1A)]\|AGCTACATTGTCTGCTGGGTTA | >MI0000298\|hsa-mir-221&WithFlank&X\|-\|45746151\|45746272 | 71 | 91 | 21(+1A) | AGCTACATTGTCTGCTGGGTTA | 476.6 | 7.0 | 1.3 | 5.4 | 5.4E-08 | 1.0E-07 | MB231/MB468 | CYTO |
| >hg38_1_-_228634819_228635039_RNA5S12_5S_50ntFlanks@90.123.34\|TGATCTCGGAAGCTAAGCAGGGTCGGGCCTGGTT | >hg38_1_-_228634819_228635039_RNA5S12_5S_50ntFlanks | 90 | 123 | 34 | TGATCTCGGAAGCTAAGCAGGGTCGGGCCTGGTT | 500.5 | -4.9 | 0.8 | -5.9 | 3.3E-09 | 1.0E-07 | BT20/MB231 | CYTO |
| >hg38_21_+_8433172_8446622_RNA45SN1_45S_50ntFlanks@12999.13027.29\|CTGCGATCTATTGAAAGTCAGCCCTCGAC | >hg38_21_+_8433172_8446622_RNA45SN1_45S_50ntFlanks | 12999 | 13027 | 29 | CTGCGATCTATTGAAAGTCAGCCCTCGAC | 888.9 | -3.8 | 0.7 | -5.4 | 5.9E-08 | 1.0E-07 | MB231/MB468 | MITO |
| >hg38_21_+_8433172_8446622_RNA45SN1_45S_50ntFlanks@10516.10533.18\|GAGTGGCGGAGATGGGCG | >hg38_21_+_8433172_8446622_RNA45SN1_45S_50ntFlanks | 10516 | 10533 | 18 | GAGTGGCGGAGATGGGCG | 594.0 | -7.0 | 1.3 | -5.5 | 4.2E-08 | 1.0E-07 | BT20/MB468 | MITO |
| >hg38_21_+_8433172_8446622_RNA45SN1_45S_50ntFlanks@4372.4398.27\|AAAAAGCTCGTAGTTGGATCTTGGGAG | >hg38_21_+_8433172_8446622_RNA45SN1_45S_50ntFlanks | 4372 | 4398 | 27 | AAAAAGCTCGTAGTTGGATCTTGGGAG | 538.6 | -8.4 | 1.6 | -5.4 | 6.1E-08 | 1.0E-07 | BT20/MB468 | NUC |
| >hg38_21_+_8433172_8446622_RNA45SN1_45S_50ntFlanks@11668.11685.18\|TGCCCAGTGCTCTGAATG | >hg38_21_+_8433172_8446622_RNA45SN1_45S_50ntFlanks | 11668 | 11685 | 18 | TGCCCAGTGCTCTGAATG | 584.3 | -3.5 | 0.6 | -5.4 | 5.0E-08 | 1.0E-07 | MB231/MB468 | MP |
| >hg38_21_+_8433172_8446622_RNA45SN1_45S_50ntFlanks@11784.11804.21\|ATGAATGGATGAACGAGATTC | >hg38_21_+_8433172_8446622_RNA45SN1_45S_50ntFlanks | 11784 | 11804 | 21 | ATGAATGGATGAACGAGATTC | 6202.7 | -5.2 | 0.9 | -5.7 | 9.8E-09 | 1.0E-07 | BT20/MB468 | MP |
| >trnaMT_ProTGG_MT_-_15956_16023@42.67.26\|TGGTGGAGTTAAAGACTTTTTCTCTG | >trnaMT_ProTGG_MT_-_15956_16023 | 42 | 67 | 26 | TGGTGGAGTTAAAGACTTTTTCTCTG | 86.3 | -9.5 | 1.7 | -5.7 | 9.8E-09 | 1.0E-07 | BT20/MB468 | MP |
| >hg38_21_+_8433172_8446622_RNA45SN1_45S_50ntFlanks@9027.9068.42\|CGTCTCCTCGTGGGGGGGCCGGGCCACCCCTCCCACGGCGCG | >hg38_21_+_8433172_8446622_RNA45SN1_45S_50ntFlanks | 9027 | 9068 | 42 | CGTCTCCTCGTGGGGGGGCCGGGCCACCCCTCCCACGGCGCG | 95.4 | -8.7 | 1.6 | -5.5 | 4.2E-08 | 1.0E-07 | BT20/MB468 | MITO |
| >MI0000746\|hsa-mir-99b&WithFlank&19\|+\|51692606\|51692687@13.34.22(+1C)&[MIMAT0000689&hsa-miR-99b-5p&offsets\|0\|0(+1C);m-10&19\|+\|51692618\|51692639&offsets\|0\|0(+1C)]\|CACCCGTAGAACCGACCTTGCGC | >MI0000746\|hsa-mir-99b&WithFlank&19\|+\|51692606\|51692687 | 13 | 34 | 22(+1C) | CACCCGTAGAACCGACCTTGCGC | 91.0 | 2.3 | 0.4 | 5.4 | 6.2E-08 | 1.0E-07 | BT20/MB468 | NUC |
| >hg38_21_+_8433172_8446622_RNA45SN1_45S_50ntFlanks@12044.12064.21\|ACTGACCCGGTGAGGCGGGGG | >hg38_21_+_8433172_8446622_RNA45SN1_45S_50ntFlanks | 12044 | 12064 | 21 | ACTGACCCGGTGAGGCGGGGG | 170.1 | -10.5 | 1.8 | -5.7 | 9.8E-09 | 1.0E-07 | BT20/MB468 | MP |
| >MI0015866\|hsa-mir-4326&WithFlank&20\|+\|63286802\|63286872@17.39.23&[MIMAT0016888&hsa-miR-4326&offsets\|0\|+3;m-339&20\|+\|63286818\|63286841&offsets\|0\|-1]\|TGTTCCTCTGTCTCCCAGACTCT | >MI0015866\|hsa-mir-4326&WithFlank&20\|+\|63286802\|63286872 | 17 | 39 | 23 | TGTTCCTCTGTCTCCCAGACTCT | 100.3 | 10.4 | 1.8 | 5.7 | 9.8E-09 | 1.0E-07 | BT20/MB468 | MP |
| >hg38_21_+_8433172_8446622_RNA45SN1_45S_50ntFlanks@4555.4587.33\|CGCAGCTAGGAATAATGGAATAGGACCGCGGTT | >hg38_21_+_8433172_8446622_RNA45SN1_45S_50ntFlanks | 4555 | 4587 | 33 | CGCAGCTAGGAATAATGGAATAGGACCGCGGTT | 114.1 | -9.9 | 1.7 | -5.7 | 9.8E-09 | 1.0E-07 | BT20/MB468 | MP |
| >hg38_21_+_8433172_8446622_RNA45SN1_45S_50ntFlanks@10664.10710.47\|CTTGGAAAGCGTCGCGGTTCCGGCGGCGTCCGGTGAGCTCTCGCTGG | >hg38_21_+_8433172_8446622_RNA45SN1_45S_50ntFlanks | 10664 | 10710 | 47 | CTTGGAAAGCGTCGCGGTTCCGGCGGCGTCCGGTGAGCTCTCGCTGG | 74.6 | -10.1 | 1.9 | -5.4 | 5.1E-08 | 1.0E-07 | MB231/MB468 | MP |
| >hg38_21_+_8433172_8446622_RNA45SN1_45S_50ntFlanks@11570.11591.22\|AGCCGACTTAGAACTGGTGCGG | >hg38_21_+_8433172_8446622_RNA45SN1_45S_50ntFlanks | 11570 | 11591 | 22 | AGCCGACTTAGAACTGGTGCGG | 623.5 | -8.3 | 1.5 | -5.4 | 6.2E-08 | 1.0E-07 | BT20/MB468 | NUC |
| >MI0000086\|hsa-mir-28&WithFlank&3\|+\|188688775\|188688872@62.80.19&[MIMAT0004502&hsa-miR-28-3p&offsets\|+2\|-1;m-28&3\|+\|188688834\|188688855&offsets\|+2\|-1]\|CTAGATTGTGAGCTCCTGG | >MI0000086\|hsa-mir-28&WithFlank&3\|+\|188688775\|188688872 | 62 | 80 | 19 | CTAGATTGTGAGCTCCTGG | 132.9 | 10.8 | 1.9 | 5.7 | 9.8E-09 | 1.0E-07 | BT20/MB468 | MP |
| >am_trna111_HisGTG_1_-_147774845_147774916@1.29.29,trna118_HisGTG_1_-_145396881_145396952@1.29.29,trna16_HisGTG_1_+_146544773_146544844@1.29.29,trna1_HisGTG_15_+_45493349_45493420@1.29.29,trna21_HisGTG_1_+_147753471_147753542@1.29.29,trna33_HisGTG_6_+_27125906_27125977@1.29.29,trna7_HisGTG_9_-_14433938_14434009@1.29.29,trna8_HisGTG_15_-_45492611_45492682@1.29.29,trna9_HisGTG_15_-_45490804_45490875@1.29.29\|GCCGTGATCGTATAGTGGTTAGTACTCTG | >am_trna111_HisGTG_1_-_147774845_147774916 | 1 | 29 | 29 | GCCGTGATCGTATAGTGGTTAGTACTCTG | 3210.4 | -5.1 | 0.9 | -5.7 | 9.8E-09 | 1.0E-07 | BT20/MB468 | MP |
| >hg38_21_+_8433172_8446622_RNA45SN1_45S_50ntFlanks@9554.9571.18\|CGTAGCGGTCCTGACGTG | >hg38_21_+_8433172_8446622_RNA45SN1_45S_50ntFlanks | 9554 | 9571 | 18 | CGTAGCGGTCCTGACGTG | 203034.9 | -4.0 | 0.7 | -5.4 | 5.5E-08 | 1.0E-07 | MB231/MB468 | CYTO |
| >hg38_21_+_8433172_8446622_RNA45SN1_45S_50ntFlanks@10351.10384.34\|CGAGAACTTTGAAGGCCGAAGTGGAGAAGGGTTC | >hg38_21_+_8433172_8446622_RNA45SN1_45S_50ntFlanks | 10351 | 10384 | 34 | CGAGAACTTTGAAGGCCGAAGTGGAGAAGGGTTC | 7673.2 | -7.0 | 1.2 | -5.7 | 9.9E-09 | 1.0E-07 | BT20/MB468 | MP |
| >hg38_21_+_8433172_8446622_RNA45SN1_45S_50ntFlanks@4722.4747.26\|TCATTAATCAAGAACGAAAGTCGGAG | >hg38_21_+_8433172_8446622_RNA45SN1_45S_50ntFlanks | 4722 | 4747 | 26 | TCATTAATCAAGAACGAAAGTCGGAG | 92.2 | -9.6 | 1.7 | -5.7 | 9.9E-09 | 1.0E-07 | BT20/MB468 | MP |
| >hg38_21_+_8433172_8446622_RNA45SN1_45S_50ntFlanks@12405.12422.18\|CTTTTTGATCCTTCGATG | >hg38_21_+_8433172_8446622_RNA45SN1_45S_50ntFlanks | 12405 | 12422 | 18 | CTTTTTGATCCTTCGATG | 278.8 | -6.5 | 1.2 | -5.5 | 4.2E-08 | 1.0E-07 | BT20/MB468 | MITO |
| >hg38_21_+_8433172_8446622_RNA45SN1_45S_50ntFlanks@12622.12654.33\|AGCCAATGGGGCGAAGCTACCATCTGTGGGATT | >hg38_21_+_8433172_8446622_RNA45SN1_45S_50ntFlanks | 12622 | 12654 | 33 | AGCCAATGGGGCGAAGCTACCATCTGTGGGATT | 536.2 | -3.2 | 0.6 | -5.5 | 4.2E-08 | 1.0E-07 | BT20/MB468 | MITO |
| >hg38_21_+_8433172_8446622_RNA45SN1_45S_50ntFlanks@12991.13019.29\|CTCCCTCGCTGCGATCTATTGAAAGTCAG | >hg38_21_+_8433172_8446622_RNA45SN1_45S_50ntFlanks | 12991 | 13019 | 29 | CTCCCTCGCTGCGATCTATTGAAAGTCAG | 369.2 | -6.2 | 1.1 | -5.4 | 5.5E-08 | 1.1E-07 | MB231/MB468 | CYTO |
| >hg38_21_+_8433172_8446622_RNA45SN1_45S_50ntFlanks@10678.10707.30\|CGGTTCCGGCGGCGTCCGGTGAGCTCTCGC | >hg38_21_+_8433172_8446622_RNA45SN1_45S_50ntFlanks | 10678 | 10707 | 30 | CGGTTCCGGCGGCGTCCGGTGAGCTCTCGC | 548.7 | -7.9 | 1.4 | -5.4 | 5.1E-08 | 1.1E-07 | MB231/MB468 | MP |
| >MI0000294\|hsa-mir-218-1&WithFlank&4\|+\|20528269\|20528390@31.52.22(+1A);MI0000295\|hsa-mir-218-2&WithFlank&5\|-\|168768140\|168768261@31.52.22(+1A)&[MIMAT0000275&hsa-miR-218-5p&offsets\|0\|+1(+1A);m-185&4\|+\|20528299\|20528319&offsets\|0\|+1(+1A)];[MIMAT0000275_1&hsa-miR-218-5p&offsets\|0\|+1(+1A);m-187&5\|-\|168768211\|168768231&offsets\|0\|+1(+1A)]\|TTGTGCTTGATCTAACCATGTGA | >MI0000294\|hsa-mir-218-1&WithFlank&4\|+\|20528269\|20528390 | 31 | 52 | 22(+1A) | TTGTGCTTGATCTAACCATGTGA | 81.4 | 9.6 | 1.8 | 5.4 | 5.1E-08 | 1.1E-07 | MB231/MB468 | MP |
| >MI0000108\|hsa-mir-103a-2&WithFlank&20\|+\|3917488\|3917577@54.77.24;MI0000109\|hsa-mir-103a-1&WithFlank&5\|-\|168560890\|168560979@54.77.24&[MIMAT0000101&hsa-miR-103a-3p&offsets\|0\|+1;m-12&20\|+\|3917541\|3917563&offsets\|0\|+1];[MIMAT0000101_1&hsa-miR-103a-3p&offsets\|0\|+1;m-13&5\|-\|168560904\|168560926&offsets\|0\|+1;m-44220&5\|-\|168560908\|168560926&offsets\|0\|+5]\|AGCAGCATTGTACAGGGCTATGAA | >MI0000108\|hsa-mir-103a-2&WithFlank&20\|+\|3917488\|3917577 | 54 | 77 | 24 | AGCAGCATTGTACAGGGCTATGAA | 714.8 | 5.7 | 1.0 | 5.4 | 6.0E-08 | 1.1E-07 | MB231/MB468 | MITO |
| >hg38_1_-_228634819_228635039_RNA5S12_5S_50ntFlanks@85.109.25\|TCGTCTGATCTCGGAAGCTAAGCAG | >hg38_1_-_228634819_228635039_RNA5S12_5S_50ntFlanks | 85 | 109 | 25 | TCGTCTGATCTCGGAAGCTAAGCAG | 803.1 | 13.0 | 2.2 | 6.0 | 2.0E-09 | 1.1E-07 | BT20/MB231 | MP |
| >MI0000061\|hsa-let-7a-2&WithFlank&11\|-\|122146516\|122146599@56.77.22&[MIMAT0010195&hsa-let-7a-2-3p&offsets\|0\|0;m-426&11\|-\|122146523\|122146544&offsets\|0\|0]\|CTGTACAGCCTCCTAGCTTTCC | >MI0000061\|hsa-let-7a-2&WithFlank&11\|-\|122146516\|122146599 | 56 | 77 | 22 | CTGTACAGCCTCCTAGCTTTCC | 183.4 | -10.9 | 1.8 | -6.0 | 1.9E-09 | 1.1E-07 | BT20/MB231 | MP |
| >am_trnaMT_ValTAC_MT_+_1602_1670@28.65.38\|AACTTACACTTAGGAGATTTCAACTTAACTTGACCGCT | >am_trnaMT_ValTAC_MT_+_1602_1670 | 28 | 65 | 38 | AACTTACACTTAGGAGATTTCAACTTAACTTGACCGCT | 266.5 | 11.8 | 2.1 | 5.7 | 1.0E-08 | 1.1E-07 | BT20/MB468 | MP |
| >trna5_AspGTC_12_+_98897281_98897352@32.72.41\|CTGTCACGCGGGAGACCGGGGTTCAATTCCCCGACGGGGAG | >trna5_AspGTC_12_+_98897281_98897352 | 32 | 72 | 41 | CTGTCACGCGGGAGACCGGGGTTCAATTCCCCGACGGGGAG | 143.6 | -6.0 | 1.1 | -5.5 | 4.8E-08 | 1.1E-07 | BT20/MB468 | CYTO |
| >hg38_21_+_8433172_8446622_RNA45SN1_45S_50ntFlanks@6700.6719.20\|CTAGCTGCGAGAATTAATGT | >hg38_21_+_8433172_8446622_RNA45SN1_45S_50ntFlanks | 6700 | 6719 | 20 | CTAGCTGCGAGAATTAATGT | 510.9 | -4.9 | 0.9 | -5.5 | 4.8E-08 | 1.1E-07 | BT20/MB468 | CYTO |
| >hg38_21_+_8433172_8446622_RNA45SN1_45S_50ntFlanks@6762.6806.45\|GCGGCCCCGGGTTCCTCCCGGGGCTACGCCTGTCTGAGCGTCGCT | >hg38_21_+_8433172_8446622_RNA45SN1_45S_50ntFlanks | 6762 | 6806 | 45 | GCGGCCCCGGGTTCCTCCCGGGGCTACGCCTGTCTGAGCGTCGCT | 555.8 | 7.7 | 1.4 | 5.5 | 4.3E-08 | 1.1E-07 | BT20/MB468 | MITO |
| >hg38_21_+_8433172_8446622_RNA45SN1_45S_50ntFlanks@9554.9597.44\|CGTAGCGGTCCTGACGTGCAAATCGGTCGTCCGACCTGGGTATA | >hg38_21_+_8433172_8446622_RNA45SN1_45S_50ntFlanks | 9554 | 9597 | 44 | CGTAGCGGTCCTGACGTGCAAATCGGTCGTCCGACCTGGGTATA | 461.1 | -7.6 | 1.4 | -5.5 | 4.8E-08 | 1.1E-07 | BT20/MB468 | CYTO |
| >hg38_21_+_8433172_8446622_RNA45SN1_45S_50ntFlanks@13005.13027.23\|TCTATTGAAAGTCAGCCCTCGAC | >hg38_21_+_8433172_8446622_RNA45SN1_45S_50ntFlanks | 13005 | 13027 | 23 | TCTATTGAAAGTCAGCCCTCGAC | 436.8 | -7.3 | 1.3 | -5.5 | 4.8E-08 | 1.1E-07 | BT20/MB468 | CYTO |
| >hg38_21_+_8433172_8446622_RNA45SN1_45S_50ntFlanks@10914.10931.18\|CGCCGCGGCTGGACGAGG | >hg38_21_+_8433172_8446622_RNA45SN1_45S_50ntFlanks | 10914 | 10931 | 18 | CGCCGCGGCTGGACGAGG | 1162.7 | -4.4 | 0.8 | -5.7 | 1.0E-08 | 1.1E-07 | BT20/MB468 | MP |
| >hg38_21_+_8433172_8446622_RNA45SN1_45S_50ntFlanks@5522.5550.29\|AAGTAAAAGTCGTAACAAGGTTTCCGTAG | >hg38_21_+_8433172_8446622_RNA45SN1_45S_50ntFlanks | 5522 | 5550 | 29 | AAGTAAAAGTCGTAACAAGGTTTCCGTAG | 229.0 | -6.6 | 1.2 | -5.4 | 5.7E-08 | 1.1E-07 | MB231/MB468 | NUC |
| >hg38_21_+_8433172_8446622_RNA45SN1_45S_50ntFlanks@11314.11343.30\|CCCACCCCACGTCTCGTCGCGCGCGCGTCC | >hg38_21_+_8433172_8446622_RNA45SN1_45S_50ntFlanks | 11314 | 11343 | 30 | CCCACCCCACGTCTCGTCGCGCGCGCGTCC | 470.8 | -8.3 | 1.5 | -5.4 | 6.3E-08 | 1.1E-07 | BT20/MB468 | NUC |
| >hg38_21_+_8433172_8446622_RNA45SN1_45S_50ntFlanks@5471.5491.21\|CACGGCCCTGGCGGAGCGCTG | >hg38_21_+_8433172_8446622_RNA45SN1_45S_50ntFlanks | 5471 | 5491 | 21 | CACGGCCCTGGCGGAGCGCTG | 194.1 | -7.2 | 1.3 | -5.4 | 5.2E-08 | 1.1E-07 | MB231/MB468 | MP |
| >MI0000077\|hsa-mir-21&WithFlank&17\|+\|59841260\|59841343@14.35.22(+2A)&[MIMAT0000076&hsa-miR-21-5p&offsets\|0\|0(+2A);m-1&17\|+\|59841273\|59841294&offsets\|0\|0(+2A)]\|TAGCTTATCAGACTGATGTTGAAA | >MI0000077\|hsa-mir-21&WithFlank&17\|+\|59841260\|59841343 | 14 | 35 | 22(+2A) | TAGCTTATCAGACTGATGTTGAAA | 560.9 | 5.0 | 0.9 | 5.4 | 6.1E-08 | 1.1E-07 | MB231/MB468 | MITO |
| >hg38_21_+_8433172_8446622_RNA45SN1_45S_50ntFlanks@12071.12092.22\|CCCCGAGGGGCTCTCGCTTCTG | >hg38_21_+_8433172_8446622_RNA45SN1_45S_50ntFlanks | 12071 | 12092 | 22 | CCCCGAGGGGCTCTCGCTTCTG | 279.9 | -7.2 | 1.3 | -5.4 | 6.1E-08 | 1.1E-07 | MB231/MB468 | MITO |

<table>
<tr><th></th><th></th><th></th><th></th><th></th><th></th><th></th><th></th><th></th><th></th><th></th><th></th><th></th><th></th></tr>
<tr><td>>MI0000264|hsa-mir-7-2&WithFlank&15|+|88611819|88611940@38.61.24;MI0000265|hsa-mir-7-3&WithFlank&19|+|4770664|4770785@37.60.24;MI0000263|hsa-mir-7-1&WithFlank&9|-|83969742|83969863@30.53.24&[MIMAT0000252&hsa-miR-7-5p&offsets|0|0;m-400&15|+|88611856|88611879&offsets|0|0];[MIMAT0000252_1&hsa-miR-7-5p&offsets|0|0;m-402&19|+|4770700|4770723&offsets|0|0];[MIMAT0000252_2&hsa-miR-7-5p&offsets|0|0;m-401&9|-|83969811|83969834&offsets|0|0]|TGGAAGACTAGTGATTTTGTTGTT</td><td>>MI0000264|hsa-mir-7-2&WithFlank&15|+|88611819|88611940</td><td>38</td><td>61</td><td>24</td><td>TGGAAGACTAGTGATTTTGTTGTT</td><td>14846.2</td><td>3.6</td><td>0.7</td><td>5.4</td><td>5.6E-08</td><td>1.1E-07</td><td>MB231/MB468</td><td>CYTO</td></tr>
<tr><td>>hg38_21_+_8433172_8446622_RNA45SN1_45S_50ntFlanks@11999.12033.35|CCCTGCGGGCCGCCGGTGAAATACCACTACTCTGA</td><td>>hg38_21_+_8433172_8446622_RNA45SN1_45S_50ntFlanks</td><td>11999</td><td>12033</td><td>35</td><td>CCCTGCGGGCCGCCGGTGAAATACCACTACTCTGA</td><td>397.8</td><td>-4.6</td><td>0.8</td><td>-5.4</td><td>5.6E-08</td><td>1.1E-07</td><td>MB231/MB468</td><td>CYTO</td></tr>
<tr><td>>hg38_21_+_8433172_8446622_RNA45SN1_45S_50ntFlanks@476.498.23|TTCGTGATCGATGTGGTGACGTC</td><td>>hg38_21_+_8433172_8446622_RNA45SN1_45S_50ntFlanks</td><td>476</td><td>498</td><td>23</td><td>TTCGTGATCGATGTGGTGACGTC</td><td>2354.3</td><td>7.3</td><td>1.3</td><td>5.4</td><td>6.1E-08</td><td>1.1E-07</td><td>MB231/MB468</td><td>MITO</td></tr>
<tr><td>>hg38_21_+_8433172_8446622_RNA45SN1_45S_50ntFlanks@10409.10426.18|GTCAGTCGGTCCTGAGAG</td><td>>hg38_21_+_8433172_8446622_RNA45SN1_45S_50ntFlanks</td><td>10409</td><td>10426</td><td>18</td><td>GTCAGTCGGTCCTGAGAG</td><td>355.2</td><td>-7.9</td><td>1.5</td><td>-5.4</td><td>6.3E-08</td><td>1.1E-07</td><td>BT20/MB468</td><td>NUC</td></tr>
<tr><td>>hg38_21_+_8433172_8446622_RNA45SN1_45S_50ntFlanks@9766.9795.30|TAAATGGGTAAGAAGCCCGGCTCGCTGGCG</td><td>>hg38_21_+_8433172_8446622_RNA45SN1_45S_50ntFlanks</td><td>9766</td><td>9795</td><td>30</td><td>TAAATGGGTAAGAAGCCCGGCTCGCTGGCG</td><td>483.1</td><td>-8.0</td><td>1.5</td><td>-5.4</td><td>6.1E-08</td><td>1.1E-07</td><td>MB231/MB468</td><td>MITO</td></tr>
<tr><td>>hg38_21_+_8433172_8446622_RNA45SN1_45S_50ntFlanks@6734.6771.38|ATTGATCATCGACACTTCGAACGCACTTGCGGCCCCGG</td><td>>hg38_21_+_8433172_8446622_RNA45SN1_45S_50ntFlanks</td><td>6734</td><td>6771</td><td>38</td><td>ATTGATCATCGACACTTCGAACGCACTTGCGGCCCCGG</td><td>497.9</td><td>-8.0</td><td>1.5</td><td>-5.4</td><td>5.3E-08</td><td>1.1E-07</td><td>MB231/MB468</td><td>MP</td></tr>
<tr><td>>hg38_21_+_8433172_8446622_RNA45SN1_45S_50ntFlanks@10829.10852.24|GCAAGCCGGATCCGTAACTTCGGG</td><td>>hg38_21_+_8433172_8446622_RNA45SN1_45S_50ntFlanks</td><td>10829</td><td>10852</td><td>24</td><td>GCAAGCCGGATCCGTAACTTCGGG</td><td>407.0</td><td>-8.0</td><td>1.5</td><td>-5.4</td><td>6.4E-08</td><td>1.1E-07</td><td>BT20/MB468</td><td>NUC</td></tr>
<tr><td>>hg38_21_+_8433172_8446622_RNA45SN1_45S_50ntFlanks@12942.12972.31|TTCGTAGACGACCTGCTTCTGGGTCGGGGTT</td><td>>hg38_21_+_8433172_8446622_RNA45SN1_45S_50ntFlanks</td><td>12942</td><td>12972</td><td>31</td><td>TTCGTAGACGACCTGCTTCTGGGTCGGGGTT</td><td>915.1</td><td>-7.5</td><td>1.4</td><td>-5.5</td><td>4.9E-08</td><td>1.1E-07</td><td>BT20/MB468</td><td>CYTO</td></tr>
<tr><td>>hg38_21_+_8433172_8446622_RNA45SN1_45S_50ntFlanks@10353.10383.31|AGAACTTTGAAGGCCGAAGTGGAGAAGGGTT</td><td>>hg38_21_+_8433172_8446622_RNA45SN1_45S_50ntFlanks</td><td>10353</td><td>10383</td><td>31</td><td>AGAACTTTGAAGGCCGAAGTGGAGAAGGGTT</td><td>595.6</td><td>-4.1</td><td>0.7</td><td>-5.4</td><td>5.3E-08</td><td>1.1E-07</td><td>MB231/MB468</td><td>MP</td></tr>
<tr><td>>trna27_GlyCCC_2_-_70476123_70476193@1.29.29,trna34_GlyCCC_16_-_686736_686806@1.29.29|GCGCCGCTGGTGTAGTGGTATCATGCAAG</td><td>>trna27_GlyCCC_2_-_70476123_70476193</td><td>1</td><td>29</td><td>29</td><td>GCGCCGCTGGTGTAGTGGTATCATGCAAG</td><td>426.4</td><td>-11.8</td><td>2.1</td><td>-5.7</td><td>1.0E-08</td><td>1.1E-07</td><td>BT20/MB468</td><td>MP</td></tr>
<tr><td>>hg38_21_+_8433172_8446622_RNA45SN1_45S_50ntFlanks@12993.13016.24|CCCTCGCTGCGATCTATTGAAAGT</td><td>>hg38_21_+_8433172_8446622_RNA45SN1_45S_50ntFlanks</td><td>12993</td><td>13016</td><td>24</td><td>CCCTCGCTGCGATCTATTGAAAGT</td><td>185.2</td><td>3.1</td><td>0.6</td><td>5.4</td><td>5.6E-08</td><td>1.1E-07</td><td>MB231/MB468</td><td>CYTO</td></tr>
<tr><td>>hg38_21_+_8433172_8446622_RNA45SN1_45S_50ntFlanks@12627.12660.34|ATGGGGCGAAGCTACCATCTGTGGGATTATGACT</td><td>>hg38_21_+_8433172_8446622_RNA45SN1_45S_50ntFlanks</td><td>12627</td><td>12660</td><td>34</td><td>ATGGGGCGAAGCTACCATCTGTGGGATTATGACT</td><td>427.2</td><td>-6.0</td><td>1.1</td><td>-5.4</td><td>6.2E-08</td><td>1.1E-07</td><td>MB231/MB468</td><td>MITO</td></tr>
<tr><td>>MI0003129|hsa-mir-146b&WithFlank&10|+|102436506|102436590@15.34.20(+2U)&[MIMAT0002809&hsa-miR-146b-5p&offsets|0|-3(+2U);m-70&10|+|102436520|102436543&offsets|0|-4(+2U)]|TGAGAACTGAATTCCATAGGTT</td><td>>MI0003129|hsa-mir-146b&WithFlank&10|+|102436506|102436590</td><td>15</td><td>34</td><td>20(+2U)</td><td>TGAGAACTGAATTCCATAGGTT</td><td>87.7</td><td>9.7</td><td>1.8</td><td>5.4</td><td>5.3E-08</td><td>1.1E-07</td><td>MB231/MB468</td><td>MP</td></tr>
<tr><td>>hg38_21_+_8433172_8446622_RNA45SN1_45S_50ntFlanks@11199.11225.27|CCGGCGGCGGCGGCGACTCTGGACGCG</td><td>>hg38_21_+_8433172_8446622_RNA45SN1_45S_50ntFlanks</td><td>11199</td><td>11225</td><td>27</td><td>CCGGCGGCGGCGGCGACTCTGGACGCG</td><td>1077.0</td><td>-3.4</td><td>0.6</td><td>-5.5</td><td>4.4E-08</td><td>1.1E-07</td><td>BT20/MB468</td><td>MITO</td></tr>
<tr><td>>hg38_21_+_8433172_8446622_RNA45SN1_45S_50ntFlanks@10756.10785.30|TACCCATATCCGCAGCAGGTCTCCAAGGTG</td><td>>hg38_21_+_8433172_8446622_RNA45SN1_45S_50ntFlanks</td><td>10756</td><td>10785</td><td>30</td><td>TACCCATATCCGCAGCAGGTCTCCAAGGTG</td><td>98.5</td><td>-8.7</td><td>1.6</td><td>-5.5</td><td>4.4E-08</td><td>1.1E-07</td><td>BT20/MB468</td><td>MITO</td></tr>
<tr><td>>hg38_21_+_8433172_8446622_RNA45SN1_45S_50ntFlanks@4727.4768.42|AATCAAGAACGAAAGTCGGAGGTTCGAAGACGATCAGATACC</td><td>>hg38_21_+_8433172_8446622_RNA45SN1_45S_50ntFlanks</td><td>4727</td><td>4768</td><td>42</td><td>AATCAAGAACGAAAGTCGGAGGTTCGAAGACGATCAGATACC</td><td>1055.0</td><td>9.9</td><td>1.8</td><td>5.5</td><td>4.9E-08</td><td>1.1E-07</td><td>BT20/MB468</td><td>CYTO</td></tr>
<tr><td>>hg38_21_+_8433172_8446622_RNA45SN1_45S_50ntFlanks@11211.11232.22|GCGACTCTGGACGCGAGCCGGG</td><td>>hg38_21_+_8433172_8446622_RNA45SN1_45S_50ntFlanks</td><td>11211</td><td>11232</td><td>22</td><td>GCGACTCTGGACGCGAGCCGGG</td><td>514.8</td><td>-5.9</td><td>1.1</td><td>-5.4</td><td>5.8E-08</td><td>1.1E-07</td><td>MB231/MB468</td><td>NUC</td></tr>
<tr><td>>trna14_LysTTT_11_-_59327808_59327880@1.24.24,trna2_LysTTT_17_+_8022473_8022545@1.24.24,trna54_LysTTT_1_+_204475655_204475727@1.24.24,trna5_LysTTT_11_+_59323902_59323974@1.24.24,trna62_LysTTT_1_-_204476158_204476230@1.24.24,trna76_LysTTT_6_+_28918806_28918878@1.24.24|GCCCGGATAGCTCAGTCGGTAGAG</td><td>>trna14_LysTTT_11_-_59327808_59327880</td><td>1</td><td>24</td><td>24</td><td>GCCCGGATAGCTCAGTCGGTAGAG</td><td>253.2</td><td>-4.7</td><td>0.9</td><td>-5.4</td><td>5.7E-08</td><td>1.1E-07</td><td>MB231/MB468</td><td>CYTO</td></tr>
<tr><td>>hg38_21_+_8433172_8446622_RNA45SN1_45S_50ntFlanks@3985.4024.40|CGGCTTTGGTGACTCTAGATAACCTCGGGCCGATCGCACG</td><td>>hg38_21_+_8433172_8446622_RNA45SN1_45S_50ntFlanks</td><td>3985</td><td>4024</td><td>40</td><td>CGGCTTTGGTGACTCTAGATAACCTCGGGCCGATCGCACG</td><td>1243.0</td><td>-6.4</td><td>1.2</td><td>-5.4</td><td>5.7E-08</td><td>1.1E-07</td><td>MB231/MB468</td><td>CYTO</td></tr>
<tr><td>>MI0000098|hsa-mir-96&WithFlank&7|-|129774686|129774775@15.37.23&[MIMAT0000095&hsa-miR-96-5p&offsets|0|0;m-248&7|-|129774739|129774761&offsets|0|0]|TTTGGCACTAGCACATTTTTGCT</td><td>>MI0000098|hsa-mir-96&WithFlank&7|-|129774686|129774775</td><td>15</td><td>37</td><td>23</td><td>TTTGGCACTAGCACATTTTTGCT</td><td>1899.2</td><td>4.0</td><td>0.7</td><td>5.4</td><td>6.4E-08</td><td>1.1E-07</td><td>BT20/MB468</td><td>NUC</td></tr>
<tr><td>>MI0000746|hsa-mir-99b&WithFlank&19|+|51692606|51692687@13.32.20(+1G)&[MIMAT0000689&hsa-miR-99b-5p&offsets|0|-2(+1G);m-10&19|+|51692618|51692639&offsets|0|-2(+1G)]|CACCCGTAGAACCGACCTTGG</td><td>>MI0000746|hsa-mir-99b&WithFlank&19|+|51692606|51692687</td><td>13</td><td>32</td><td>20(+1G)</td><td>CACCCGTAGAACCGACCTTGG</td><td>833.3</td><td>5.3</td><td>0.9</td><td>5.9</td><td>3.5E-09</td><td>1.1E-07</td><td>BT20/MB231</td><td>CYTO</td></tr>
<tr><td>>hg38_21_+_8433172_8446622_RNA45SN1_45S_50ntFlanks@6661.6695.35|CGGTGGATCACTCGGCTCGTGCGTCGATGAAGAAC</td><td>>hg38_21_+_8433172_8446622_RNA45SN1_45S_50ntFlanks</td><td>6661</td><td>6695</td><td>35</td><td>CGGTGGATCACTCGGCTCGTGCGTCGATGAAGAAC</td><td>2961.1</td><td>4.4</td><td>0.8</td><td>5.7</td><td>1.0E-08</td><td>1.1E-07</td><td>BT20/MB468</td><td>MP</td></tr>
<tr><td>>hg38_21_+_8433172_8446622_RNA45SN1_45S_50ntFlanks@4558.4587.30|AGCTAGGAATAATGGAATAGGACCGCGGTT</td><td>>hg38_21_+_8433172_8446622_RNA45SN1_45S_50ntFlanks</td><td>4558</td><td>4587</td><td>30</td><td>AGCTAGGAATAATGGAATAGGACCGCGGTT</td><td>281.9</td><td>-7.0</td><td>1.3</td><td>-5.4</td><td>5.9E-08</td><td>1.1E-07</td><td>MB231/MB468</td><td>NUC</td></tr>
<tr><td>>hg38_21_+_8433172_8446622_RNA45SN1_45S_50ntFlanks@6790.6807.18|CCTGTCTGAGCGTCGCTT</td><td>>hg38_21_+_8433172_8446622_RNA45SN1_45S_50ntFlanks</td><td>6790</td><td>6807</td><td>18</td><td>CCTGTCTGAGCGTCGCTT</td><td>6186.6</td><td>4.1</td><td>0.7</td><td>5.5</td><td>4.4E-08</td><td>1.1E-07</td><td>BT20/MB468</td><td>MITO</td></tr>
<tr><td>>hg38_21_+_8433172_8446622_RNA45SN1_45S_50ntFlanks@9534.9559.26|AGGAAACTCTGGTGGAGGTCCGTAGC</td><td>>hg38_21_+_8433172_8446622_RNA45SN1_45S_50ntFlanks</td><td>9534</td><td>9559</td><td>26</td><td>AGGAAACTCTGGTGGAGGTCCGTAGC</td><td>1112.7</td><td>-7.1</td><td>1.2</td><td>-5.7</td><td>1.0E-08</td><td>1.1E-07</td><td>BT20/MB468</td><td>MP</td></tr>
<tr><td>>MI0000102|hsa-mir-100&WithFlank&11|-|122152223|122152314@19.42.24&[MIMAT0000098&hsa-miR-100-5p&offsets|0|+2;m-27&11|-|122152276|122152296&offsets|0|+3]|AACCCGTAGATCCGAACTTGTGGT</td><td>>MI0000102|hsa-mir-100&WithFlank&11|-|122152223|122152314</td><td>19</td><td>42</td><td>24</td><td>AACCCGTAGATCCGAACTTGTGGT</td><td>187.5</td><td>-10.9</td><td>1.8</td><td>-6.0</td><td>2.0E-09</td><td>1.1E-07</td><td>BT20/MB231</td><td>MP</td></tr>
<tr><td>>hg38_21_+_8433172_8446622_RNA45SN1_45S_50ntFlanks@11657.11685.29|GATGTGATTTCTGCCCAGTGCTCTGAATG</td><td>>hg38_21_+_8433172_8446622_RNA45SN1_45S_50ntFlanks</td><td>11657</td><td>11685</td><td>29</td><td>GATGTGATTTCTGCCCAGTGCTCTGAATG</td><td>1660.4</td><td>14.0</td><td>2.3</td><td>6.0</td><td>2.1E-09</td><td>1.1E-07</td><td>BT20/MB231</td><td>MP</td></tr>
<tr><td>>hg38_21_+_8433172_8446622_RNA45SN1_45S_50ntFlanks@10663.10710.48|CCTTGGAAAGCGTCGCGGTTCCGGCGGCGTCCGGTGAGCTCTCGCTGG</td><td>>hg38_21_+_8433172_8446622_RNA45SN1_45S_50ntFlanks</td><td>10663</td><td>10710</td><td>48</td><td>CCTTGGAAAGCGTCGCGGTTCCGGCGGCGTCCGGTGAGCTCTCGCTGG</td><td>336.0</td><td>-8.1</td><td>1.5</td><td>-5.5</td><td>5.0E-08</td><td>1.1E-07</td><td>BT20/MB468</td><td>CYTO</td></tr>
<tr><td>>hg38_21_+_8433172_8446622_RNA45SN1_45S_50ntFlanks@11918.11954.37|ACATGAGAGGTGTAGAATAAGTGGGAGGCCCCCGGCG</td><td>>hg38_21_+_8433172_8446622_RNA45SN1_45S_50ntFlanks</td><td>11918</td><td>11954</td><td>37</td><td>ACATGAGAGGTGTAGAATAAGTGGGAGGCCCCCGGCG</td><td>289.4</td><td>-7.1</td><td>1.3</td><td>-5.5</td><td>5.0E-08</td><td>1.1E-07</td><td>BT20/MB468</td><td>CYTO</td></tr>
<tr><td>>hg38_21_+_8433172_8446622_RNA45SN1_45S_50ntFlanks@10346.10371.26|TCAAACGAGAACTTTGAAGGCCGAAG</td><td>>hg38_21_+_8433172_8446622_RNA45SN1_45S_50ntFlanks</td><td>10346</td><td>10371</td><td>26</td><td>TCAAACGAGAACTTTGAAGGCCGAAG</td><td>249.0</td><td>2.7</td><td>0.5</td><td>5.4</td><td>5.7E-08</td><td>1.1E-07</td><td>MB231/MB468</td><td>CYTO</td></tr>
<tr><td>>hg38_21_+_8433172_8446622_RNA45SN1_45S_50ntFlanks@6780.6802.23|CGGGGCTACGCCTGTCTGAGCGT</td><td>>hg38_21_+_8433172_8446622_RNA45SN1_45S_50ntFlanks</td><td>6780</td><td>6802</td><td>23</td><td>CGGGGCTACGCCTGTCTGAGCGT</td><td>511.2</td><td>3.9</td><td>0.7</td><td>5.4</td><td>5.9E-08</td><td>1.1E-07</td><td>MB231/MB468</td><td>NUC</td></tr>
<tr><td>>hg38_21_+_8433172_8446622_RNA45SN1_45S_50ntFlanks@7972.8009.38|AGACGCGACCTCAGATCAGACGTGGCGACCCGCTGAAT</td><td>>hg38_21_+_8433172_8446622_RNA45SN1_45S_50ntFlanks</td><td>7972</td><td>8009</td><td>38</td><td>AGACGCGACCTCAGATCAGACGTGGCGACCCGCTGAAT</td><td>189.8</td><td>3.5</td><td>0.6</td><td>5.4</td><td>6.5E-08</td><td>1.1E-07</td><td>BT20/MB468</td><td>NUC</td></tr>
<tr><td>>MI0000081|hsa-mir-24-2&WithFlank&19|-|13836281|13836365@56.75.20;MI0000080|hsa-mir-24-1&WithFlank&9|+|95086015|95086094@50.69.20&[MIMAT0000080&hsa-miR-24-3p&offsets|0|-2;m-36&19|-|13836288|13836310&offsets|0|-3];[MIMAT0000080_1&hsa-miR-24-3p&offsets|0|-2;m-35&9|+|95086064|95086086&offsets|0|-3]|TGGCTCAGTTCAGCAGGAAC</td><td>>MI0000081|hsa-mir-24-2&WithFlank&19|-|13836281|13836365</td><td>56</td><td>75</td><td>20</td><td>TGGCTCAGTTCAGCAGGAAC</td><td>5302.5</td><td>4.5</td><td>0.8</td><td>5.4</td><td>5.8E-08</td><td>1.1E-07</td><td>MB231/MB468</td><td>CYTO</td></tr>
<tr><td>>hg38_21_+_8433172_8446622_RNA45SN1_45S_50ntFlanks@8381.8405.25|AAGAGGGCGTGAAACCGTTAAGAGG</td><td>>hg38_21_+_8433172_8446622_RNA45SN1_45S_50ntFlanks</td><td>8381</td><td>8405</td><td>25</td><td>AAGAGGGCGTGAAACCGTTAAGAGG</td><td>864.5</td><td>-8.6</td><td>1.6</td><td>-5.4</td><td>6.6E-08</td><td>1.1E-07</td><td>BT20/MB468</td><td>NUC</td></tr>
<tr><td>>hg38_21_+_8433172_8446622_RNA45SN1_45S_50ntFlanks@3838.3874.37|CTCTCCTACTTGGATAACTGTGGTAATTCTAGAGCTA</td><td>>hg38_21_+_8433172_8446622_RNA45SN1_45S_50ntFlanks</td><td>3838</td><td>3874</td><td>37</td><td>CTCTCCTACTTGGATAACTGTGGTAATTCTAGAGCTA</td><td>372.4</td><td>-7.9</td><td>1.5</td><td>-5.4</td><td>6.6E-08</td><td>1.1E-07</td><td>BT20/MB468</td><td>NUC</td></tr>
<tr><td>>trna116_GluCTC_1_-_145399233_145399304@36.75.40,trna134_GluTTC_1_-_16861774_16861845@36.75.40,trna71_GluCTC_1_-_161439189_161439260@36.75.40,trna74_GluCTC_1_-_161431809_161431880@36.75.40,trna77_GluCTC_1_-_161424398_161424469@36.75.40,trna77_GluCTC_6_+_28949976_28950047@36.75.40,trna80_GluCTC_1_-_161417018_161417089@36.75.40,trna84_GluTTC_1_-_161391883_161391954@36.75.40,trna87_GluCTC_6_-_126101393_126101464@36.75.40|CACCGCCGCGGCCCGGGTTCGATTCCCGGTCAGGGAACCA</td><td>>trna116_GluCTC_1_-_145399233_145399304</td><td>36</td><td>75</td><td>40</td><td>CACCGCCGCGGCCCGGGTTCGATTCCCGGTCAGGGAACCA</td><td>5932.3</td><td>6.4</td><td>1.2</td><td>5.4</td><td>5.8E-08</td><td>1.1E-07</td><td>MB231/MB468</td><td>CYTO</td></tr>
<tr><td>>hg38_21_+_8433172_8446622_RNA45SN1_45S_50ntFlanks@8182.8210.29|ATCGAGGCCCAGCCCGTGGACGGTGTGAG</td><td>>hg38_21_+_8433172_8446622_RNA45SN1_45S_50ntFlanks</td><td>8182</td><td>8210</td><td>29</td><td>ATCGAGGCCCAGCCCGTGGACGGTGTGAG</td><td>615.9</td><td>-7.9</td><td>1.5</td><td>-5.4</td><td>5.4E-08</td><td>1.1E-07</td><td>MB231/MB468</td><td>MP</td></tr>
<tr><td>>hg38_21_+_8433172_8446622_RNA45SN1_45S_50ntFlanks@8380.8399.20|CAAGAGGGCGTGAAACCGTT</td><td>>hg38_21_+_8433172_8446622_RNA45SN1_45S_50ntFlanks</td><td>8380</td><td>8399</td><td>20</td><td>CAAGAGGGCGTGAAACCGTT</td><td>1509.6</td><td>-5.2</td><td>0.9</td><td>-5.7</td><td>1.0E-08</td><td>1.1E-07</td><td>BT20/MB468</td><td>MP</td></tr>
<tr><td>>trna119_LysCTT_1_-_145395522_145395594@1.35.35,trna11_LysCTT_5_-_180648979_180649051@1.35.35,trna13_LysCTT_6_+_26556774_26556846@1.35.35,trna7_LysCTT_16_+_3225692_3225764@1.35.35,trna9_LysCTT_5_+_180634755_180634827@1.35.35|GCCCGGCTAGCTCAGTCGGTAGAGCATGAGACTCT</td><td>>trna119_LysCTT_1_-_145395522_145395594</td><td>1</td><td>35</td><td>35</td><td>GCCCGGCTAGCTCAGTCGGTAGAGCATGAGACTCT</td><td>3625.8</td><td>6.7</td><td>1.2</td><td>5.5</td><td>4.5E-08</td><td>1.1E-07</td><td>BT20/MB468</td><td>MITO</td></tr>
<tr><td>>hg38_21_+_8433172_8446622_RNA45SN1_45S_50ntFlanks@6652.6678.27|GACTCTTAGCGGTGGATCACTCGGCTC</td><td>>hg38_21_+_8433172_8446622_RNA45SN1_45S_50ntFlanks</td><td>6652</td><td>6678</td><td>27</td><td>GACTCTTAGCGGTGGATCACTCGGCTC</td><td>13732.6</td><td>4.2</td><td>0.8</td><td>5.5</td><td>4.5E-08</td><td>1.1E-07</td><td>BT20/MB468</td><td>MITO</td></tr>
</table>

<table>
<tr><td>>MI0000108|hsa-mir-103a-2&WithFlank&20|+|3917488|3917577@54.75.22;MI0000109|hsa-mir-103a-1&WithFlank&5|-|168560890|168560979@54.75.22&[MIMAT0000101&hsa-miR-103a-3p&offsets|0|-1;m-12&20|+|3917541|3917563&offsets|0|-1];[MIMAT0000101_1&hsa-miR-103a-3p&offsets|0|-1;m-13&5|-|168560904|168560926&offsets|0|-1;m-44220&5|-|168560908|168560926&offsets|0|+3]|AGCAGCATTGTACAGGGCTATG</td><td>>MI0000108|hsa-mir-103a-2&WithFlank&20|+|3917488|3917577</td><td>54</td><td>75</td><td>22</td><td>AGCAGCATTGTACAGGGCTATG</td><td>1701.5</td><td>2.6</td><td>0.5</td><td>5.4</td><td>6.6E-08</td><td>1.1E-07</td><td>BT20/MB468</td><td>NUC</td></tr>
<tr><td>>hg38_21_+_8433172_8446622_RNA45SN1_45S_50ntFlanks@12045.12066.22|CTGACCCGGTGAGGCGGGGGGG</td><td>>hg38_21_+_8433172_8446622_RNA45SN1_45S_50ntFlanks</td><td>12045</td><td>12066</td><td>22</td><td>CTGACCCGGTGAGGCGGGGGGG</td><td>63.9</td><td>-9.9</td><td>1.8</td><td>-5.4</td><td>5.4E-08</td><td>1.1E-07</td><td>MB231/MB468</td><td>MP</td></tr>
<tr><td>>hg38_21_+_8433172_8446622_RNA45SN1_45S_50ntFlanks@12394.12416.23|TAGCGACGTCGCTTTTTGATCCT</td><td>>hg38_21_+_8433172_8446622_RNA45SN1_45S_50ntFlanks</td><td>12394</td><td>12416</td><td>23</td><td>TAGCGACGTCGCTTTTTGATCCT</td><td>911.4</td><td>-6.7</td><td>1.2</td><td>-5.4</td><td>6.3E-08</td><td>1.1E-07</td><td>MB231/MB468</td><td>MITO</td></tr>
<tr><td>>am_trna128_GlyGCC_6_-_27870686_27870756@1.28.28,trna133_GlyCCC_1_-_16872434_16872504@1.28.28,trna18_GlyGCC_16_+_70822597_70822667@1.28.28,trna19_GlyGCC_16_+_70823410_70823480@1.28.28,trna19_GlyGCC_2_-_157257659_157257729@1.28.28,trna24_GlyGCC_16_-_70812942_70813012@1.28.28,trna25_GlyGCC_16_-_70812114_70812184@1.28.28,trna4_GlyCCC_1_+_17188416_17188486@1.28.28,trna5_GlyGCC_17_+_8029064_8029134@1.28.28,trna68_GlyGCC_1_-_161493637_161493707@1.28.28|GCATTGGTGGTTCAGTGGTAGAATTCTC</td><td>>am_trna128_GlyGCC_6_-_27870686_27870756</td><td>1</td><td>28</td><td>28</td><td>GCATTGGTGGTTCAGTGGTAGAATTCTC</td><td>68695.7</td><td>-3.6</td><td>0.7</td><td>-5.4</td><td>5.8E-08</td><td>1.1E-07</td><td>MB231/MB468</td><td>CYTO</td></tr>
<tr><td>>hg38_21_+_8433172_8446622_RNA45SN1_45S_50ntFlanks@7993.8012.20|GTGGCGACCCGCTGAATTTA</td><td>>hg38_21_+_8433172_8446622_RNA45SN1_45S_50ntFlanks</td><td>7993</td><td>8012</td><td>20</td><td>GTGGCGACCCGCTGAATTTA</td><td>530.6</td><td>-5.9</td><td>1.1</td><td>-5.4</td><td>6.3E-08</td><td>1.1E-07</td><td>MB231/MB468</td><td>MITO</td></tr>
<tr><td>>hg38_21_+_8433172_8446622_RNA45SN1_45S_50ntFlanks@4497.4524.28|GAAGCGTTTACTTTGAAAAAATTAGAGT</td><td>>hg38_21_+_8433172_8446622_RNA45SN1_45S_50ntFlanks</td><td>4497</td><td>4524</td><td>28</td><td>GAAGCGTTTACTTTGAAAAAATTAGAGT</td><td>319.8</td><td>8.8</td><td>1.6</td><td>5.4</td><td>5.0E-08</td><td>1.1E-07</td><td>BT20/MB468</td><td>CYTO</td></tr>
<tr><td>>hg38_21_+_8433172_8446622_RNA45SN1_45S_50ntFlanks@12399.12419.21|ACGTCGCTTTTTGATCCTTCG</td><td>>hg38_21_+_8433172_8446622_RNA45SN1_45S_50ntFlanks</td><td>12399</td><td>12419</td><td>21</td><td>ACGTCGCTTTTTGATCCTTCG</td><td>539.4</td><td>-7.5</td><td>1.4</td><td>-5.4</td><td>6.3E-08</td><td>1.1E-07</td><td>MB231/MB468</td><td>MITO</td></tr>
<tr><td>>hg38_21_+_8433172_8446622_RNA45SN1_45S_50ntFlanks@8171.8195.25|AAGTCCTTCTGATCGAGGCCCAGCC</td><td>>hg38_21_+_8433172_8446622_RNA45SN1_45S_50ntFlanks</td><td>8171</td><td>8195</td><td>25</td><td>AAGTCCTTCTGATCGAGGCCCAGCC</td><td>550.3</td><td>-6.6</td><td>1.2</td><td>-5.4</td><td>5.4E-08</td><td>1.1E-07</td><td>MB231/MB468</td><td>MP</td></tr>
<tr><td>>hg38_21_+_8433172_8446622_RNA45SN1_45S_50ntFlanks@3782.3810.29|CAGTGAAACTGCGAATGGCTCATTAAATC</td><td>>hg38_21_+_8433172_8446622_RNA45SN1_45S_50ntFlanks</td><td>3782</td><td>3810</td><td>29</td><td>CAGTGAAACTGCGAATGGCTCATTAAATC</td><td>446.1</td><td>-3.9</td><td>0.7</td><td>-5.4</td><td>6.0E-08</td><td>1.1E-07</td><td>MB231/MB468</td><td>NUC</td></tr>
<tr><td>>hg38_21_+_8433172_8446622_RNA45SN1_45S_50ntFlanks@5424.5466.43|TTGGATGGTTTAGTGAGGCCCTCGGATCGGCCCCGCCGGGGTC</td><td>>hg38_21_+_8433172_8446622_RNA45SN1_45S_50ntFlanks</td><td>5424</td><td>5466</td><td>43</td><td>TTGGATGGTTTAGTGAGGCCCTCGGATCGGCCCCGCCGGGGTC</td><td>787.0</td><td>-8.8</td><td>1.6</td><td>-5.4</td><td>6.6E-08</td><td>1.1E-07</td><td>BT20/MB468</td><td>NUC</td></tr>
<tr><td>>MI0000061|hsa-let-7a-2&WithFlank&11|-|122146516|122146599@11.31.21;MI0000062|hsa-let-7a-3&WithFlank&22|+|46112743|46112828@10.30.21;MI0000060|hsa-let-7a-1&WithFlank&9|+|94175951|94176042@12.32.21&[MIMAT0000062&hsa-let-7a-5p&offsets|0|-1;m-6&11|-|122146568|122146589&offsets|0|-1];[MIMAT0000062_1&hsa-let-7a-5p&offsets|0|-1;m-5&22|+|46112752|46112773&offsets|0|-1];[MIMAT0000062_2&hsa-let-7a-5p&offsets|0|-1;m-4&9|+|94175962|94175983&offsets|0|-1]|TGAGGTAGTAGGTTGTATAGT</td><td>>MI0000061|hsa-let-7a-2&WithFlank&11|-|122146516|122146599</td><td>11</td><td>31</td><td>21</td><td>TGAGGTAGTAGGTTGTATAGT</td><td>2769.0</td><td>2.1</td><td>0.4</td><td>5.4</td><td>6.6E-08</td><td>1.1E-07</td><td>BT20/MB468</td><td>NUC</td></tr>
<tr><td>>MI0000253|hsa-mir-148a&WithFlank&7|-|25949913|25949992@50.72.23&[MIMAT0000243&hsa-miR-148a-3p&offsets|0|+1;m-7&7|-|25949922|25949943&offsets|0|+1]|TCAGTGCACTACAGAACTTTGTC</td><td>>MI0000253|hsa-mir-148a&WithFlank&7|-|25949913|25949992</td><td>50</td><td>72</td><td>23</td><td>TCAGTGCACTACAGAACTTTGTC</td><td>2134.4</td><td>5.4</td><td>0.9</td><td>5.9</td><td>3.6E-09</td><td>1.1E-07</td><td>BT20/MB231</td><td>CYTO</td></tr>
<tr><td>>MI0000478|hsa-mir-149&WithFlank&2|+|240455995|240456095@21.42.22(+1U)&[MIMAT0000450&hsa-miR-149-5p&offsets|0|-1(+1U);m-195&2|+|240456015|240456037&offsets|0|-1(+1U)]|TCTGGCTCCGTGTCTTCACTCCT</td><td>>MI0000478|hsa-mir-149&WithFlank&2|+|240455995|240456095</td><td>21</td><td>42</td><td>22(+1U)</td><td>TCTGGCTCCGTGTCTTCACTCCT</td><td>1531.2</td><td>4.7</td><td>0.8</td><td>5.9</td><td>3.6E-09</td><td>1.1E-07</td><td>BT20/MB231</td><td>CYTO</td></tr>
<tr><td>>hg38_21_+_8433172_8446622_RNA45SN1_45S_50ntFlanks@9485.9509.25|TGTTAGGACCCGAAAGATGGTGAAC</td><td>>hg38_21_+_8433172_8446622_RNA45SN1_45S_50ntFlanks</td><td>9485</td><td>9509</td><td>25</td><td>TGTTAGGACCCGAAAGATGGTGAAC</td><td>84.3</td><td>10.1</td><td>1.8</td><td>5.7</td><td>1.1E-08</td><td>1.1E-07</td><td>BT20/MB468</td><td>MP</td></tr>
<tr><td>>hg38_21_+_8433172_8446622_RNA45SN1_45S_50ntFlanks@10401.10430.30|GAACATGGGTCAGTCGGTCCTGAGAGATGG</td><td>>hg38_21_+_8433172_8446622_RNA45SN1_45S_50ntFlanks</td><td>10401</td><td>10430</td><td>30</td><td>GAACATGGGTCAGTCGGTCCTGAGAGATGG</td><td>356.8</td><td>-6.2</td><td>1.1</td><td>-5.4</td><td>5.4E-08</td><td>1.1E-07</td><td>MB231/MB468</td><td>MP</td></tr>
<tr><td>>trnaMT_LeuTAA_MT_+_3230_3304@1.35.35|GTTAAGATGGCAGAGCCCGGTAATCGCATAAAACT</td><td>>trnaMT_LeuTAA_MT_+_3230_3304</td><td>1</td><td>35</td><td>35</td><td>GTTAAGATGGCAGAGCCCGGTAATCGCATAAAACT</td><td>59.4</td><td>5.9</td><td>1.1</td><td>5.4</td><td>6.7E-08</td><td>1.1E-07</td><td>BT20/MB468</td><td>NUC</td></tr>
<tr><td>>hg38_21_+_8433172_8446622_RNA45SN1_45S_50ntFlanks@6722.6747.26|ATTGCAGGACACATTGATCATCGACA</td><td>>hg38_21_+_8433172_8446622_RNA45SN1_45S_50ntFlanks</td><td>6722</td><td>6747</td><td>26</td><td>ATTGCAGGACACATTGATCATCGACA</td><td>337.4</td><td>-6.1</td><td>1.1</td><td>-5.4</td><td>6.7E-08</td><td>1.1E-07</td><td>BT20/MB468</td><td>NUC</td></tr>
<tr><td>>hg38_21_+_8433172_8446622_RNA45SN1_45S_50ntFlanks@6723.6744.22|TTGCAGGACACATTGATCATCG</td><td>>hg38_21_+_8433172_8446622_RNA45SN1_45S_50ntFlanks</td><td>6723</td><td>6744</td><td>22</td><td>TTGCAGGACACATTGATCATCG</td><td>1454.5</td><td>-4.9</td><td>0.9</td><td>-5.4</td><td>5.1E-08</td><td>1.1E-07</td><td>BT20/MB468</td><td>CYTO</td></tr>
<tr><td>>hg38_21_+_8433172_8446622_RNA45SN1_45S_50ntFlanks@7983.8007.25|CAGATCAGACGTGGCGACCCGCTGA</td><td>>hg38_21_+_8433172_8446622_RNA45SN1_45S_50ntFlanks</td><td>7983</td><td>8007</td><td>25</td><td>CAGATCAGACGTGGCGACCCGCTGA</td><td>933.6</td><td>-4.2</td><td>0.8</td><td>-5.5</td><td>4.5E-08</td><td>1.1E-07</td><td>BT20/MB468</td><td>MITO</td></tr>
<tr><td>>hg38_21_+_8433172_8446622_RNA45SN1_45S_50ntFlanks@7989.8014.26|AGACGTGGCGACCCGCTGAATTTAAG</td><td>>hg38_21_+_8433172_8446622_RNA45SN1_45S_50ntFlanks</td><td>7989</td><td>8014</td><td>26</td><td>AGACGTGGCGACCCGCTGAATTTAAG</td><td>1321.0</td><td>-4.1</td><td>0.8</td><td>-5.4</td><td>6.4E-08</td><td>1.1E-07</td><td>MB231/MB468</td><td>MITO</td></tr>
<tr><td>>hg38_21_+_8433172_8446622_RNA45SN1_45S_50ntFlanks@3899.3934.36|CCTTCGCGGGGGGGATGCGTGCATTTATCAGATCAA</td><td>>hg38_21_+_8433172_8446622_RNA45SN1_45S_50ntFlanks</td><td>3899</td><td>3934</td><td>36</td><td>CCTTCGCGGGGGGGATGCGTGCATTTATCAGATCAA</td><td>66.5</td><td>-9.9</td><td>1.8</td><td>-5.4</td><td>5.5E-08</td><td>1.1E-07</td><td>MB231/MB468</td><td>MP</td></tr>
<tr><td>>hg38_21_+_8433172_8446622_RNA45SN1_45S_50ntFlanks@12998.13039.42|GCTGCGATCTATTGAAAGTCAGCCCTCGACACAAGGGTTTGT</td><td>>hg38_21_+_8433172_8446622_RNA45SN1_45S_50ntFlanks</td><td>12998</td><td>13039</td><td>42</td><td>GCTGCGATCTATTGAAAGTCAGCCCTCGACACAAGGGTTTGT</td><td>8511.6</td><td>5.9</td><td>1.1</td><td>5.5</td><td>4.5E-08</td><td>1.1E-07</td><td>BT20/MB468</td><td>MITO</td></tr>
<tr><td>>hg38_21_+_8433172_8446622_RNA45SN1_45S_50ntFlanks@8351.8373.23|AAGTTGAAAAGAACTTTGAAGAG</td><td>>hg38_21_+_8433172_8446622_RNA45SN1_45S_50ntFlanks</td><td>8351</td><td>8373</td><td>23</td><td>AAGTTGAAAAGAACTTTGAAGAG</td><td>1363.6</td><td>9.9</td><td>1.7</td><td>5.7</td><td>1.1E-08</td><td>1.1E-07</td><td>BT20/MB468</td><td>MP</td></tr>
<tr><td>>hg38_21_+_8433172_8446622_RNA45SN1_45S_50ntFlanks@8836.8859.24|CGGCGGTGGCGGCGGCGGCGGCGG</td><td>>hg38_21_+_8433172_8446622_RNA45SN1_45S_50ntFlanks</td><td>8836</td><td>8859</td><td>24</td><td>CGGCGGTGGCGGCGGCGGCGGCGG</td><td>113.2</td><td>-9.9</td><td>1.7</td><td>-5.7</td><td>1.1E-08</td><td>1.1E-07</td><td>BT20/MB468</td><td>MP</td></tr>
<tr><td>>trnaMT_ValTAC_MT_+_1602_1670@53.71.19|TAACTTGACCGCTCTGACC</td><td>>trnaMT_ValTAC_MT_+_1602_1670</td><td>53</td><td>71</td><td>19</td><td>TAACTTGACCGCTCTGACC</td><td>436.2</td><td>5.4</td><td>1.0</td><td>5.4</td><td>6.0E-08</td><td>1.1E-07</td><td>MB231/MB468</td><td>NUC</td></tr>
<tr><td>>MI0000115|hsa-mir-16-2&WithFlank&3|+|160404739|160404831@58.79.22&[MIMAT0004518&hsa-miR-16-2-3p&offsets|-1|-1;m-252&3|+|160404796|160404816&offsets|0|+1]|ACCAATATTACTGTGCTGCTTT</td><td>>MI0000115|hsa-mir-16-2&WithFlank&3|+|160404739|160404831</td><td>58</td><td>79</td><td>22</td><td>ACCAATATTACTGTGCTGCTTT</td><td>705.4</td><td>4.8</td><td>0.9</td><td>5.4</td><td>5.9E-08</td><td>1.1E-07</td><td>MB231/MB468</td><td>CYTO</td></tr>
<tr><td>>hg38_21_+_8433172_8446622_RNA45SN1_45S_50ntFlanks@11769.11792.24|TAATTAGTGACGCGCATGAATGGA</td><td>>hg38_21_+_8433172_8446622_RNA45SN1_45S_50ntFlanks</td><td>11769</td><td>11792</td><td>24</td><td>TAATTAGTGACGCGCATGAATGGA</td><td>676.8</td><td>-5.5</td><td>1.0</td><td>-5.4</td><td>5.1E-08</td><td>1.1E-07</td><td>BT20/MB468</td><td>CYTO</td></tr>
<tr><td>>hg38_21_+_8433172_8446622_RNA45SN1_45S_50ntFlanks@5071.5097.27|TTAATTCCGATAACGAACGAGACTCTG</td><td>>hg38_21_+_8433172_8446622_RNA45SN1_45S_50ntFlanks</td><td>5071</td><td>5097</td><td>27</td><td>TTAATTCCGATAACGAACGAGACTCTG</td><td>486.7</td><td>-6.8</td><td>1.3</td><td>-5.4</td><td>6.4E-08</td><td>1.1E-07</td><td>MB231/MB468</td><td>MITO</td></tr>
<tr><td>>hg38_21_+_8433172_8446622_RNA45SN1_45S_50ntFlanks@9534.9558.25|AGGAAACTCTGGTGGAGGTCCGTAG</td><td>>hg38_21_+_8433172_8446622_RNA45SN1_45S_50ntFlanks</td><td>9534</td><td>9558</td><td>25</td><td>AGGAAACTCTGGTGGAGGTCCGTAG</td><td>526.7</td><td>-6.5</td><td>1.2</td><td>-5.4</td><td>5.9E-08</td><td>1.1E-07</td><td>MB231/MB468</td><td>CYTO</td></tr>
<tr><td>>hg38_21_+_8433172_8446622_RNA45SN1_45S_50ntFlanks@10690.10708.19|CGTCCGGTGAGCTCTCGCT</td><td>>hg38_21_+_8433172_8446622_RNA45SN1_45S_50ntFlanks</td><td>10690</td><td>10708</td><td>19</td><td>CGTCCGGTGAGCTCTCGCT</td><td>4988.7</td><td>-5.4</td><td>0.9</td><td>-5.7</td><td>1.1E-08</td><td>1.1E-07</td><td>BT20/MB468</td><td>MP</td></tr>
<tr><td>>hg38_21_+_8433172_8446622_RNA45SN1_45S_50ntFlanks@3838.3862.25|CTCTCCTACTTGGATAACTGTGGTA</td><td>>hg38_21_+_8433172_8446622_RNA45SN1_45S_50ntFlanks</td><td>3838</td><td>3862</td><td>25</td><td>CTCTCCTACTTGGATAACTGTGGTA</td><td>173.5</td><td>-3.5</td><td>0.6</td><td>-5.4</td><td>6.1E-08</td><td>1.1E-07</td><td>MB231/MB468</td><td>NUC</td></tr>
<tr><td>>hg38_21_+_8433172_8446622_RNA45SN1_45S_50ntFlanks@12415.12446.32|CTTCGATGTCGGCTCTTCCTATCATTGTGAAG</td><td>>hg38_21_+_8433172_8446622_RNA45SN1_45S_50ntFlanks</td><td>12415</td><td>12446</td><td>32</td><td>CTTCGATGTCGGCTCTTCCTATCATTGTGAAG</td><td>1238.2</td><td>-5.0</td><td>0.9</td><td>-5.4</td><td>6.7E-08</td><td>1.1E-07</td><td>BT20/MB468</td><td>NUC</td></tr>
<tr><td>>hg38_21_+_8433172_8446622_RNA45SN1_45S_50ntFlanks@10101.10127.27|CGCGCGCGTGTGGTGTGCGTCGGAGGG</td><td>>hg38_21_+_8433172_8446622_RNA45SN1_45S_50ntFlanks</td><td>10101</td><td>10127</td><td>27</td><td>CGCGCGCGTGTGGTGTGCGTCGGAGGG</td><td>608.4</td><td>-7.1</td><td>1.3</td><td>-5.4</td><td>5.1E-08</td><td>1.1E-07</td><td>BT20/MB468</td><td>CYTO</td></tr>
<tr><td>>hg38_21_+_8433172_8446622_RNA45SN1_45S_50ntFlanks@10510.10533.24|AATCCGGAGTGGCGGAGATGGGCG</td><td>>hg38_21_+_8433172_8446622_RNA45SN1_45S_50ntFlanks</td><td>10510</td><td>10533</td><td>24</td><td>AATCCGGAGTGGCGGAGATGGGCG</td><td>4010.1</td><td>-9.9</td><td>1.8</td><td>-5.4</td><td>6.7E-08</td><td>1.1E-07</td><td>BT20/MB468</td><td>NUC</td></tr>
<tr><td>>hg38_21_+_8433172_8446622_RNA45SN1_45S_50ntFlanks@11188.11208.21|CACCCGGGGGGCCGGCGGCGG</td><td>>hg38_21_+_8433172_8446622_RNA45SN1_45S_50ntFlanks</td><td>11188</td><td>11208</td><td>21</td><td>CACCCGGGGGGCCGGCGGCGG</td><td>127.1</td><td>-10.1</td><td>1.8</td><td>-5.7</td><td>1.1E-08</td><td>1.1E-07</td><td>BT20/MB468</td><td>MP</td></tr>
<tr><td>>hg38_21_+_8433172_8446622_RNA45SN1_45S_50ntFlanks@6721.6741.21|AATTGCAGGACACATTGATCA</td><td>>hg38_21_+_8433172_8446622_RNA45SN1_45S_50ntFlanks</td><td>6721</td><td>6741</td><td>21</td><td>AATTGCAGGACACATTGATCA</td><td>542.6</td><td>-5.3</td><td>1.0</td><td>-5.4</td><td>5.9E-08</td><td>1.1E-07</td><td>MB231/MB468</td><td>CYTO</td></tr>
<tr><td>>hg38_21_+_8433172_8446622_RNA45SN1_45S_50ntFlanks@4734.4759.26|AACGAAAGTCGGAGGTTCGAAGACGA</td><td>>hg38_21_+_8433172_8446622_RNA45SN1_45S_50ntFlanks</td><td>4734</td><td>4759</td><td>26</td><td>AACGAAAGTCGGAGGTTCGAAGACGA</td><td>783.4</td><td>-4.6</td><td>0.8</td><td>-5.4</td><td>6.5E-08</td><td>1.1E-07</td><td>MB231/MB468</td><td>MITO</td></tr>
<tr><td>>hg38_21_+_8433172_8446622_RNA45SN1_45S_50ntFlanks@3839.3868.30|TCTCCTACTTGGATAACTGTGGTAATTCTA</td><td>>hg38_21_+_8433172_8446622_RNA45SN1_45S_50ntFlanks</td><td>3839</td><td>3868</td><td>30</td><td>TCTCCTACTTGGATAACTGTGGTAATTCTA</td><td>1030.4</td><td>5.0</td><td>0.9</td><td>5.4</td><td>5.1E-08</td><td>1.1E-07</td><td>BT20/MB468</td><td>CYTO</td></tr>
<tr><td>>hg38_21_+_8433172_8446622_RNA45SN1_45S_50ntFlanks@10826.10858.33|TCGGCAAGCCGGATCCGTAACTTCGGGATAAGG</td><td>>hg38_21_+_8433172_8446622_RNA45SN1_45S_50ntFlanks</td><td>10826</td><td>10858</td><td>33</td><td>TCGGCAAGCCGGATCCGTAACTTCGGGATAAGG</td><td>450.0</td><td>-7.9</td><td>1.5</td><td>-5.4</td><td>5.2E-08</td><td>1.1E-07</td><td>BT20/MB468</td><td>CYTO</td></tr>
<tr><td>>hg38_21_+_8433172_8446622_RNA45SN1_45S_50ntFlanks@8194.8212.19|CCCGTGGACGGTGTGAGGC</td><td>>hg38_21_+_8433172_8446622_RNA45SN1_45S_50ntFlanks</td><td>8194</td><td>8212</td><td>19</td><td>CCCGTGGACGGTGTGAGGC</td><td>820.0</td><td>-7.6</td><td>1.4</td><td>-5.4</td><td>6.5E-08</td><td>1.1E-07</td><td>MB231/MB468</td><td>MITO</td></tr>
<tr><td>>hg38_21_+_8433172_8446622_RNA45SN1_45S_50ntFlanks@10393.10428.36|CAGCAGTTGAACATGGGTCAGTCGGTCCTGAGAGAT</td><td>>hg38_21_+_8433172_8446622_RNA45SN1_45S_50ntFlanks</td><td>10393</td><td>10428</td><td>36</td><td>CAGCAGTTGAACATGGGTCAGTCGGTCCTGAGAGAT</td><td>446.4</td><td>-6.3</td><td>1.2</td><td>-5.4</td><td>6.8E-08</td><td>1.1E-07</td><td>BT20/MB468</td><td>NUC</td></tr>
<tr><td>>MI0000101|hsa-mir-99a&WithFlank&21|+|16539083|16539175@19.39.21(+1A)&[MIMAT0000097&hsa-miR-99a-5p&offsets|0|-1(+1A);m-64&21|+|16539101|16539121&offsets|0|0(+1A)]|AACCCGTAGATCCGATCTTGTA</td><td>>MI0000101|hsa-mir-99a&WithFlank&21|+|16539083|16539175</td><td>19</td><td>39</td><td>21(+1A)</td><td>AACCCGTAGATCCGATCTTGTA</td><td>206.0</td><td>-11.1</td><td>1.9</td><td>-6.0</td><td>2.1E-09</td><td>1.1E-07</td><td>BT20/MB231</td><td>MP</td></tr>
<tr><td>>hg38_21_+_8433172_8446622_RNA45SN1_45S_50ntFlanks@3989.4026.38|TTTGGTGACTCTAGATAACCTCGGGCCGATCGCACGCC</td><td>>hg38_21_+_8433172_8446622_RNA45SN1_45S_50ntFlanks</td><td>3989</td><td>4026</td><td>38</td><td>TTTGGTGACTCTAGATAACCTCGGGCCGATCGCACGCC</td><td>56.0</td><td>-9.7</td><td>1.8</td><td>-5.4</td><td>5.5E-08</td><td>1.1E-07</td><td>MB231/MB468</td><td>MP</td></tr>
<tr><td>>hg38_21_+_8433172_8446622_RNA45SN1_45S_50ntFlanks@12626.12659.34|AATGGGGCGAAGCTACCATCTGTGGGATTATGAC</td><td>>hg38_21_+_8433172_8446622_RNA45SN1_45S_50ntFlanks</td><td>12626</td><td>12659</td><td>34</td><td>AATGGGGCGAAGCTACCATCTGTGGGATTATGAC</td><td>2586.6</td><td>-5.2</td><td>1.0</td><td>-5.5</td><td>4.6E-08</td><td>1.1E-07</td><td>BT20/MB468</td><td>MITO</td></tr>
<tr><td>>hg38_21_+_8433172_8446622_RNA45SN1_45S_50ntFlanks@4547.4564.18|CTGGATACCGCAGCTAGG</td><td>>hg38_21_+_8433172_8446622_RNA45SN1_45S_50ntFlanks</td><td>4547</td><td>4564</td><td>18</td><td>CTGGATACCGCAGCTAGG</td><td>108.3</td><td>-9.8</td><td>1.7</td><td>-5.7</td><td>1.1E-08</td><td>1.1E-07</td><td>BT20/MB468</td><td>MP</td></tr>
</table>

| | | | | | | | | | | | | | |
|---|---|---|---|---|---|---|---|---|---|---|---|---|---|
| >hg38_1_-_228634819_228635039_RNA5S12_5S_50ntFlanks@104.121.18\|AAGCAGGGTCGGGCCTGG | >hg38_1_-_228634819_228635039_RNA5S12_5S_50ntFlanks | 104 | 121 | 18 | AAGCAGGGTCGGGCCTGG | 58.7 | -9.7 | 1.8 | -5.4 | 5.5E-08 | 1.1E-07 | MB231/MB468 | MP |
| >hg38_21_+_8433172_8446622_RNA45SN1_45S_50ntFlanks@11576.11599.24\|CTTAGAACTGGTGCGGACCAGGGG | >hg38_21_+_8433172_8446622_RNA45SN1_45S_50ntFlanks | 11576 | 11599 | 24 | CTTAGAACTGGTGCGGACCAGGGG | 149.8 | -6.9 | 1.3 | -5.4 | 5.2E-08 | 1.1E-07 | BT20/MB468 | CYTO |
| >hg38_21_+_8433172_8446622_RNA45SN1_45S_50ntFlanks@8342.8379.38\|CGTAAGGGAAAGTTGAAAAGAACTTTGAAGAGAGAGTT | >hg38_21_+_8433172_8446622_RNA45SN1_45S_50ntFlanks | 8342 | 8379 | 38 | CGTAAGGGAAAGTTGAAAAGAACTTTGAAGAGAGAGTT | 1166.6 | 6.9 | 1.3 | 5.4 | 5.2E-08 | 1.1E-07 | BT20/MB468 | CYTO |
| >hg38_MT_+_1621_3279_RNR2_16S_50ntFlanks@1209.1231.23\|CAGAACCCAACCTCCGAGCAGTA | >hg38_MT_+_1621_3279_RNR2_16S_50ntFlanks | 1209 | 1231 | 23 | CAGAACCCAACCTCCGAGCAGTA | 457.0 | -6.6 | 1.2 | -5.4 | 6.5E-08 | 1.1E-07 | MB231/MB468 | MITO |
| >hg38_21_+_8433172_8446622_RNA45SN1_45S_50ntFlanks@8184.8222.39\|CGAGGCCCAGCCCGTGGACGGTGTGAGGCCGGTAGCGGC | >hg38_21_+_8433172_8446622_RNA45SN1_45S_50ntFlanks | 8184 | 8222 | 39 | CGAGGCCCAGCCCGTGGACGGTGTGAGGCCGGTAGCGGC | 55.8 | -9.7 | 1.8 | -5.4 | 5.6E-08 | 1.1E-07 | MB231/MB468 | MP |
| >hg38_21_+_8433172_8446622_RNA45SN1_45S_50ntFlanks@12619.12661.43\|AGGAGCCAATGGGGCGAAGCTACCATCTGTGGGATTATGACTG | >hg38_21_+_8433172_8446622_RNA45SN1_45S_50ntFlanks | 12619 | 12661 | 43 | AGGAGCCAATGGGGCGAAGCTACCATCTGTGGGATTATGACTG | 82.6 | -10.2 | 1.9 | -5.4 | 5.6E-08 | 1.1E-07 | MB231/MB468 | MP |
| >hg38_21_+_8433172_8446622_RNA45SN1_45S_50ntFlanks@4007.4026.20\|CCTCGGGCCGATCGCACGCC | >hg38_21_+_8433172_8446622_RNA45SN1_45S_50ntFlanks | 4007 | 4026 | 20 | CCTCGGGCCGATCGCACGCC | 220.0 | -4.7 | 0.9 | -5.4 | 6.0E-08 | 1.1E-07 | MB231/MB468 | CYTO |
| >hg38_21_+_8433172_8446622_RNA45SN1_45S_50ntFlanks@9931.9955.25\|TATAGACAGCAGGACGGTGGCCATG | >hg38_21_+_8433172_8446622_RNA45SN1_45S_50ntFlanks | 9931 | 9955 | 25 | TATAGACAGCAGGACGGTGGCCATG | 84.8 | 10.2 | 1.8 | 5.7 | 1.1E-08 | 1.1E-07 | BT20/MB468 | MP |
| >hg38_21_+_8433172_8446622_RNA45SN1_45S_50ntFlanks@7974.8016.43\|ACGCGACCTCAGATCAGACGTGGCGACCCGCTGAATTTAAGCA | >hg38_21_+_8433172_8446622_RNA45SN1_45S_50ntFlanks | 7974 | 8016 | 43 | ACGCGACCTCAGATCAGACGTGGCGACCCGCTGAATTTAAGCA | 613.6 | 5.8 | 1.1 | 5.4 | 5.2E-08 | 1.1E-07 | BT20/MB468 | CYTO |
| >trna10_LysCTT_16_+_3241501_3241573@1.34.34,trna13_LysCTT_14_-_58706613_58706685@1.34.34,trna2_LysCTT_15_+_79152904_79152976@1.34.34\|GCCCGGCTAGCTCAGTCGGTAGAGCATGGGACTC | >trna10_LysCTT_16_+_3241501_3241573 | 1 | 34 | 34 | GCCCGGCTAGCTCAGTCGGTAGAGCATGGGACTC | 7040.3 | 4.7 | 0.9 | 5.4 | 5.6E-08 | 1.1E-07 | MB231/MB468 | MP |
| >hg38_21_+_8433172_8446622_RNA45SN1_45S_50ntFlanks@12045.12072.28\|CTGACCCGGTGAGGCGGGGGGGCGAGCC | >hg38_21_+_8433172_8446622_RNA45SN1_45S_50ntFlanks | 12045 | 12072 | 28 | CTGACCCGGTGAGGCGGGGGGGCGAGCC | 276.1 | -7.2 | 1.3 | -5.4 | 6.2E-08 | 1.1E-07 | MB231/MB468 | NUC |
| >hg38_1_-_228634819_228635039_RNA5S12_5S_50ntFlanks@124.147.24\|AGTACTTGGATGGGAGACCGCCTG | >hg38_1_-_228634819_228635039_RNA5S12_5S_50ntFlanks | 124 | 147 | 24 | AGTACTTGGATGGGAGACCGCCTG | 468.8 | -7.7 | 1.4 | -5.4 | 6.2E-08 | 1.1E-07 | MB231/MB468 | NUC |
| >hg38_21_+_8433172_8446622_RNA45SN1_45S_50ntFlanks@11712.11742.31\|CGGGTAAACGGCGGGAGTAACTATGACTCTC | >hg38_21_+_8433172_8446622_RNA45SN1_45S_50ntFlanks | 11712 | 11742 | 31 | CGGGTAAACGGCGGGAGTAACTATGACTCTC | 164.1 | -3.5 | 0.6 | -5.4 | 6.2E-08 | 1.1E-07 | MB231/MB468 | NUC |
| >hg38_21_+_8433172_8446622_RNA45SN1_45S_50ntFlanks@12385.12414.30\|AAGCGTTCATAGCGACGTCGCTTTTTGATC | >hg38_21_+_8433172_8446622_RNA45SN1_45S_50ntFlanks | 12385 | 12414 | 30 | AAGCGTTCATAGCGACGTCGCTTTTTGATC | 92.1 | -9.6 | 1.7 | -5.7 | 1.1E-08 | 1.2E-07 | BT20/MB468 | MP |
| >hg38_21_+_8433172_8446622_RNA45SN1_45S_50ntFlanks@3899.3919.21\|CCTTCGCGGGGGGGATGCGTG | >hg38_21_+_8433172_8446622_RNA45SN1_45S_50ntFlanks | 3899 | 3919 | 21 | CCTTCGCGGGGGGGATGCGTG | 3479.0 | -8.2 | 1.5 | -5.5 | 4.7E-08 | 1.2E-07 | BT20/MB468 | MITO |
| >hg38_21_+_8433172_8446622_RNA45SN1_45S_50ntFlanks@12459.12477.19\|AGCGTTGGATTGTTCACCC | >hg38_21_+_8433172_8446622_RNA45SN1_45S_50ntFlanks | 12459 | 12477 | 19 | AGCGTTGGATTGTTCACCC | 822.9 | -3.5 | 0.6 | -5.4 | 5.3E-08 | 1.2E-07 | BT20/MB468 | CYTO |
| >hg38_21_+_8433172_8446622_RNA45SN1_45S_50ntFlanks@7989.8018.30\|AGACGTGGCGACCCGCTGAATTTAAGCATA | >hg38_21_+_8433172_8446622_RNA45SN1_45S_50ntFlanks | 7989 | 8018 | 30 | AGACGTGGCGACCCGCTGAATTTAAGCATA | 311.3 | 5.9 | 1.1 | 5.4 | 6.1E-08 | 1.2E-07 | MB231/MB468 | CYTO |
| >hg38_21_+_8433172_8446622_RNA45SN1_45S_50ntFlanks@6664.6681.18\|TGGATCACTCGGCTCGTG | >hg38_21_+_8433172_8446622_RNA45SN1_45S_50ntFlanks | 6664 | 6681 | 18 | TGGATCACTCGGCTCGTG | 1652.6 | -2.2 | 0.4 | -5.5 | 4.7E-08 | 1.2E-07 | BT20/MB468 | MITO |
| >hg38_21_+_8433172_8446622_RNA45SN1_45S_50ntFlanks@10679.10705.27\|GGTTCCGGCGGCGTCCGGTGAGCTCTC | >hg38_21_+_8433172_8446622_RNA45SN1_45S_50ntFlanks | 10679 | 10705 | 27 | GGTTCCGGCGGCGTCCGGTGAGCTCTC | 250.2 | -6.8 | 1.3 | -5.4 | 6.2E-08 | 1.2E-07 | MB231/MB468 | NUC |
| >hg38_21_+_8433172_8446622_RNA45SN1_45S_50ntFlanks@12481.12498.18\|AATAGGGAACGTGAGCTG | >hg38_21_+_8433172_8446622_RNA45SN1_45S_50ntFlanks | 12481 | 12498 | 18 | AATAGGGAACGTGAGCTG | 432.2 | -3.3 | 0.6 | -5.4 | 6.2E-08 | 1.2E-07 | MB231/MB468 | NUC |
| >MI0000542\|hsa-mir-320a&WithFlank&8\|-\|22244960\|22245043@48.68.21(+3U)&[MIMAT0000510&hsa-miR-320a-3p&offsets\|0\|-1(+3U);m-60&8\|-\|22244975\|22244996&offsets\|0\|-1(+3U)]\|AAAAGCTGGGTTGAGAGGGCGTTT | >MI0000542\|hsa-mir-320a&WithFlank&8\|-\|22244960\|22245043 | 48 | 68 | 21(+3U) | AAAAGCTGGGTTGAGAGGGCGTTT | 4883.3 | 5.2 | 0.9 | 5.9 | 3.8E-09 | 1.2E-07 | BT20/MB231 | CYTO |
| >hg38_21_+_8433172_8446622_RNA45SN1_45S_50ntFlanks@7979.8012.34\|ACCTCAGATCAGACGTGGCGACCCGCTGAATTTA | >hg38_21_+_8433172_8446622_RNA45SN1_45S_50ntFlanks | 7979 | 8012 | 34 | ACCTCAGATCAGACGTGGCGACCCGCTGAATTTA | 545.3 | -6.4 | 1.2 | -5.4 | 5.6E-08 | 1.2E-07 | MB231/MB468 | MP |
| >hg38_21_+_8433172_8446622_RNA45SN1_45S_50ntFlanks@12636.12653.18\|AGCTACCATCTGTGGGAT | >hg38_21_+_8433172_8446622_RNA45SN1_45S_50ntFlanks | 12636 | 12653 | 18 | AGCTACCATCTGTGGGAT | 681.6 | -7.6 | 1.4 | -5.5 | 4.7E-08 | 1.2E-07 | BT20/MB468 | MITO |
| >hg38_21_+_8433172_8446622_RNA45SN1_45S_50ntFlanks@8381.8400.20\|AAGAGGGCGTGAAACCGTTA | >hg38_21_+_8433172_8446622_RNA45SN1_45S_50ntFlanks | 8381 | 8400 | 20 | AAGAGGGCGTGAAACCGTTA | 740.8 | -7.1 | 1.3 | -5.4 | 5.3E-08 | 1.2E-07 | BT20/MB468 | CYTO |
| >hg38_21_+_8433172_8446622_RNA45SN1_45S_50ntFlanks@11872.11893.22\|AAAGAAGACCCTGTTGAGCTTG | >hg38_21_+_8433172_8446622_RNA45SN1_45S_50ntFlanks | 11872 | 11893 | 22 | AAAGAAGACCCTGTTGAGCTTG | 514.8 | -8.2 | 1.5 | -5.4 | 6.9E-08 | 1.2E-07 | BT20/MB468 | NUC |
| >hg38_21_+_8433172_8446622_RNA45SN1_45S_50ntFlanks@11537.11560.24\|CCGCGCGGCGCCTCGCCTCGGCCG | >hg38_21_+_8433172_8446622_RNA45SN1_45S_50ntFlanks | 11537 | 11560 | 24 | CCGCGCGGCGCCTCGCCTCGGCCG | 344.1 | -7.5 | 1.4 | -5.4 | 6.2E-08 | 1.2E-07 | MB231/MB468 | NUC |
| >MI0000301\|hsa-mir-224&WithFlank&X\|-\|151958572\|151958664@14.36.23&[MIMAT0000281&hsa-miR-224-5p&offsets\|+1\|-1;m-238&X\|-\|151958630\|151958651&offsets\|0\|+1]\|CAAGTCACTAGTGGTTCCGTTTA | >MI0000301\|hsa-mir-224&WithFlank&X\|-\|151958572\|151958664 | 14 | 36 | 23 | CAAGTCACTAGTGGTTCCGTTTA | 79.6 | 9.5 | 1.8 | 5.4 | 5.7E-08 | 1.2E-07 | MB231/MB468 | MP |
| >am_trnaMT_ValTAC_MT_+_1602_1670@30.68.39\|CTTACACTTAGGAGATTTCAACTTAACTTGACCGCTCTG | >am_trnaMT_ValTAC_MT_+_1602_1670 | 30 | 68 | 39 | CTTACACTTAGGAGATTTCAACTTAACTTGACCGCTCTG | 1771.5 | -10.1 | 1.9 | -5.4 | 6.9E-08 | 1.2E-07 | BT20/MB468 | NUC |
| >MI0005559\|hsa-mir-744&WithFlank&17\|+\|12081893\|12082002@17.38.22&[MIMAT0004945&hsa-miR-744-5p&offsets\|0\|0;m-172&17\|+\|12081909\|12081929&offsets\|0\|+1]\|TGCGGGGCTAGGGCTAACAGCA | >MI0005559\|hsa-mir-744&WithFlank&17\|+\|12081893\|12082002 | 17 | 38 | 22 | TGCGGGGCTAGGGCTAACAGCA | 193.4 | 2.7 | 0.5 | 5.4 | 6.9E-08 | 1.2E-07 | BT20/MB468 | NUC |
| >hg38_21_+_8433172_8446622_RNA45SN1_45S_50ntFlanks@4732.4767.36\|AGAACGAAAGTCGGAGGTTCGAAGACGATCAGATAC | >hg38_21_+_8433172_8446622_RNA45SN1_45S_50ntFlanks | 4732 | 4767 | 36 | AGAACGAAAGTCGGAGGTTCGAAGACGATCAGATAC | 273.7 | 5.6 | 1.0 | 5.4 | 6.3E-08 | 1.2E-07 | MB231/MB468 | NUC |
| >MI0000077\|hsa-mir-21&WithFlank&17\|+\|59841260\|59841343@14.36.23(+1A)&[MIMAT0000076&hsa-miR-21-5p&offsets\|0\|+1(+1A);m-1&17\|+\|59841273\|59841294&offsets\|0\|+1(+1A)]\|TAGCTTATCAGACTGATGTTGACA | >MI0000077\|hsa-mir-21&WithFlank&17\|+\|59841260\|59841343 | 14 | 36 | 23(+1A) | TAGCTTATCAGACTGATGTTGACA | 55072.7 | 4.7 | 0.9 | 5.4 | 6.1E-08 | 1.2E-07 | MB231/MB468 | CYTO |
| >hg38_21_+_8433172_8446622_RNA45SN1_45S_50ntFlanks@4046.4067.22\|CATTCGAACGTCTGCCCTATCA | >hg38_21_+_8433172_8446622_RNA45SN1_45S_50ntFlanks | 4046 | 4067 | 22 | CATTCGAACGTCTGCCCTATCA | 420.4 | -5.7 | 1.1 | -5.4 | 6.3E-08 | 1.2E-07 | MB231/MB468 | NUC |
| >hg38_21_+_8433172_8446622_RNA45SN1_45S_50ntFlanks@10389.10426.38\|TGAACAGCAGTTGAACATGGGTCAGTCGGTCCTGAGAG | >hg38_21_+_8433172_8446622_RNA45SN1_45S_50ntFlanks | 10389 | 10426 | 38 | TGAACAGCAGTTGAACATGGGTCAGTCGGTCCTGAGAG | 299.5 | -6.1 | 1.1 | -5.4 | 6.3E-08 | 1.2E-07 | MB231/MB468 | NUC |
| >hg38_21_+_8433172_8446622_RNA45SN1_45S_50ntFlanks@5075.5094.20\|TTCCGATAACGAACGAGACT | >hg38_21_+_8433172_8446622_RNA45SN1_45S_50ntFlanks | 5075 | 5094 | 20 | TTCCGATAACGAACGAGACT | 134.5 | -10.9 | 2.0 | -5.4 | 5.7E-08 | 1.2E-07 | MB231/MB468 | MP |
| >hg38_21_+_8433172_8446622_RNA45SN1_45S_50ntFlanks@11552.11591.40\|CCTCGGCCGGCGCCTAGCAGCCGACTTAGAACTGGTGCGG | >hg38_21_+_8433172_8446622_RNA45SN1_45S_50ntFlanks | 11552 | 11591 | 40 | CCTCGGCCGGCGCCTAGCAGCCGACTTAGAACTGGTGCGG | 1986.4 | -9.7 | 1.8 | -5.4 | 7.0E-08 | 1.2E-07 | BT20/MB468 | NUC |
| >MI0000108\|hsa-mir-103a-2&WithFlank&20\|+\|3917488\|3917577@54.76.23;MI0000109\|hsa-mir-103a-1&WithFlank&5\|-\|168560890\|168560979@54.76.23&[MIMAT0000101&hsa-miR-103a-3p&offsets\|0\|0;m-12&20\|+\|3917541\|3917563&offsets\|0\|0];[MIMAT0000101_1&hsa-miR-103a-3p&offsets\|0\|0;m-13&5\|-\|168560904\|168560926&offsets\|0\|0;m-44220&5\|-\|168560908\|168560926&offsets\|0\|+4]\|AGCAGCATTGTACAGGGCTATGA | >MI0000108\|hsa-mir-103a-2&WithFlank&20\|+\|3917488\|3917577 | 54 | 76 | 23 | AGCAGCATTGTACAGGGCTATGA | 8065.1 | 2.3 | 0.4 | 5.4 | 7.0E-08 | 1.2E-07 | BT20/MB468 | NUC |
| >MI0000750\|hsa-mir-26a-2&WithFlank&12\|-\|57824603\|57824698@20.41.22(+1C);MI0000083\|hsa-mir-26a-1&WithFlank&3\|+\|37969398\|37969486@16.37.22(+1C)&[MIMAT0000082&hsa-miR-26a-5p&offsets\|0\|0(+1C);m-33&12\|-\|57824658\|57824679&offsets\|0\|0(+1C)];[MIMAT0000082_1&hsa-miR-26a-5p&offsets\|0\|0(+1C);m-34&3\|+\|37969413\|37969434&offsets\|0\|0(+1C)]\|TTCAAGTAATCCAGGATAGGCTC | >MI0000750\|hsa-mir-26a-2&WithFlank&12\|-\|57824603\|57824698 | 20 | 41 | 22(+1C) | TTCAAGTAATCCAGGATAGGCTC | 787.9 | 6.9 | 1.3 | 5.4 | 6.7E-08 | 1.2E-07 | MB231/MB468 | MITO |
| >am_trna128_GlyGCC_6_-_27870686_27870756@1.34.34,trna18_GlyGCC_16_+_70822597_70822667@1.34.34,trna19_GlyGCC_16_+_70823410_70823480@1.34.34,trna19_GlyGCC_2_-_157257659_157257729@1.34.34,trna24_GlyGCC_16_-_70812942_70813012@1.34.34,trna25_GlyGCC_16_-_70812114_70812184@1.34.34,trna5_GlyGCC_17_+_8029064_8029134@1.34.34,trna68_GlyGCC_1_-_161493637_161493707@1.34.34\|GCATTGGTGGTTCAGTGGTAGAATTCTCGCCTGC | >am_trna128_GlyGCC_6_-_27870686_27870756 | 1 | 34 | 34 | GCATTGGTGGTTCAGTGGTAGAATTCTCGCCTGC | 1641.3 | 5.6 | 1.0 | 5.4 | 5.7E-08 | 1.2E-07 | MB231/MB468 | MP |
| >hg38_21_+_8433172_8446622_RNA45SN1_45S_50ntFlanks@11873.11893.21\|AAGAAGACCCTGTTGAGCTTG | >hg38_21_+_8433172_8446622_RNA45SN1_45S_50ntFlanks | 11873 | 11893 | 21 | AAGAAGACCCTGTTGAGCTTG | 6020.2 | -5.4 | 0.9 | -5.7 | 1.1E-08 | 1.2E-07 | BT20/MB468 | MP |
| >hg38_21_+_8433172_8446622_RNA45SN1_45S_50ntFlanks@10806.10830.25\|AACAATGTAGGTAAGGGAAGTCGGC | >hg38_21_+_8433172_8446622_RNA45SN1_45S_50ntFlanks | 10806 | 10830 | 25 | AACAATGTAGGTAAGGGAAGTCGGC | 199.8 | -5.3 | 1.0 | -5.4 | 6.3E-08 | 1.2E-07 | MB231/MB468 | NUC |
| >am_trna128_GlyGCC_6_-_27870686_27870756@1.27.27,trna133_GlyCCC_1_-_16872434_16872504@1.27.27,trna18_GlyGCC_16_+_70822597_70822667@1.27.27,trna19_GlyGCC_16_+_70823410_70823480@1.27.27,trna19_GlyGCC_2_-_157257659_157257729@1.27.27,trna24_GlyGCC_16_-_70812942_70813012@1.27.27,trna25_GlyGCC_16_-_70812114_70812184@1.27.27,trna4_GlyCCC_1_+_17188416_17188486@1.27.27,trna5_GlyGCC_17_+_8029064_8029134@1.27.27,trna68_GlyGCC_1_-_161493637_161493707@1.27.27\|GCATTGGTGGTTCAGTGGTAGAATTCT | >am_trna128_GlyGCC_6_-_27870686_27870756 | 1 | 27 | 27 | GCATTGGTGGTTCAGTGGTAGAATTCT | 4306.1 | -5.6 | 1.0 | -5.4 | 7.0E-08 | 1.2E-07 | BT20/MB468 | NUC |
| >hg38_21_+_8433172_8446622_RNA45SN1_45S_50ntFlanks@9739.9769.31\|AAACGATCTCAACCTATTCTCAAACTTTAAA | >hg38_21_+_8433172_8446622_RNA45SN1_45S_50ntFlanks | 9739 | 9769 | 31 | AAACGATCTCAACCTATTCTCAAACTTTAAA | 1234.2 | -2.7 | 0.5 | -5.5 | 4.8E-08 | 1.2E-07 | BT20/MB468 | MITO |
| >hg38_21_+_8433172_8446622_RNA45SN1_45S_50ntFlanks@10259.10290.32\|AGGGCCGCTGCGGTGAGCCTTGAAGCCTAGGG | >hg38_21_+_8433172_8446622_RNA45SN1_45S_50ntFlanks | 10259 | 10290 | 32 | AGGGCCGCTGCGGTGAGCCTTGAAGCCTAGGG | 434.9 | 4.1 | 0.8 | 5.4 | 6.3E-08 | 1.2E-07 | MB231/MB468 | NUC |
| >hg38_21_+_8433172_8446622_RNA45SN1_45S_50ntFlanks@4048.4069.22\|TTCGAACGTCTGCCCTATCAAC | >hg38_21_+_8433172_8446622_RNA45SN1_45S_50ntFlanks | 4048 | 4069 | 22 | TTCGAACGTCTGCCCTATCAAC | 235.7 | 5.4 | 1.0 | 5.4 | 6.2E-08 | 1.2E-07 | MB231/MB468 | CYTO |
| >hg38_21_+_8433172_8446622_RNA45SN1_45S_50ntFlanks@4027.4046.20\|CCCCGTGGCGGCGACGACCC | >hg38_21_+_8433172_8446622_RNA45SN1_45S_50ntFlanks | 4027 | 4046 | 20 | CCCCGTGGCGGCGACGACCC | 133.7 | -9.2 | 1.7 | -5.5 | 4.8E-08 | 1.2E-07 | BT20/MB468 | MITO |
| >hg38_21_+_8433172_8446622_RNA45SN1_45S_50ntFlanks@5484.5524.41\|GAGCGCTGAGAAGACGGTCGAACTTGACTATCTAGAGGAAG | >hg38_21_+_8433172_8446622_RNA45SN1_45S_50ntFlanks | 5484 | 5524 | 41 | GAGCGCTGAGAAGACGGTCGAACTTGACTATCTAGAGGAAG | 53.9 | -9.6 | 1.8 | -5.4 | 5.8E-08 | 1.2E-07 | MB231/MB468 | MP |

<table>
<tr><td></td><td></td><td></td><td></td><td></td><td></td><td></td><td></td><td></td><td></td><td></td><td></td><td></td><td></td></tr>
<tr><td>>hg38_21_+_8433172_8446622_RNA45SN1_45S_50ntFlanks@6646.6685.40|TCGTACGACTCTTAGCGGTGGATCACTCGGCTCGTGCGTC</td><td>>hg38_21_+_8433172_8446622_RNA45SN1_45S_50ntFlanks</td><td>6646</td><td>6685</td><td>40</td><td>TCGTACGACTCTTAGCGGTGGATCACTCGGCTCGTGCGTC</td><td>15908.5</td><td>5.4</td><td>0.9</td><td>5.7</td><td>1.1E-08</td><td>1.2E-07</td><td>BT20/MB468</td><td>MP</td></tr>
<tr><td>>hg38_21_+_8433172_8446622_RNA45SN1_45S_50ntFlanks@9339.9373.35|CGCACGAAAGCCGCCGTGGCGCAATGAAGGTGAAG</td><td>>hg38_21_+_8433172_8446622_RNA45SN1_45S_50ntFlanks</td><td>9339</td><td>9373</td><td>35</td><td>CGCACGAAAGCCGCCGTGGCGCAATGAAGGTGAAG</td><td>120.9</td><td>-10.0</td><td>1.8</td><td>-5.7</td><td>1.1E-08</td><td>1.2E-07</td><td>BT20/MB468</td><td>MP</td></tr>
<tr><td>>hg38_21_+_8433172_8446622_RNA45SN1_45S_50ntFlanks@3802.3837.36|CATTAAATCAGTTATGGTTCCTTTGGTCGCTCGCTC</td><td>>hg38_21_+_8433172_8446622_RNA45SN1_45S_50ntFlanks</td><td>3802</td><td>3837</td><td>36</td><td>CATTAAATCAGTTATGGTTCCTTTGGTCGCTCGCTC</td><td>68.0</td><td>-10.0</td><td>1.8</td><td>-5.4</td><td>5.8E-08</td><td>1.2E-07</td><td>MB231/MB468</td><td>MP</td></tr>
<tr><td>>hg38_21_+_8433172_8446622_RNA45SN1_45S_50ntFlanks@5086.5119.34|AACGAGACTCTGGCATGCTAACTAGTTACGCGAC</td><td>>hg38_21_+_8433172_8446622_RNA45SN1_45S_50ntFlanks</td><td>5086</td><td>5119</td><td>34</td><td>AACGAGACTCTGGCATGCTAACTAGTTACGCGAC</td><td>86.7</td><td>-10.3</td><td>1.9</td><td>-5.4</td><td>5.8E-08</td><td>1.2E-07</td><td>MB231/MB468</td><td>MP</td></tr>
<tr><td>>trna116_GluCTC_1_-_145399233_145399304@1.34.34,trna59_GluCTC_1_+_249168447_249168518@1.34.34,trna71_GluCTC_1_-_161439189_161439260@1.34.34,trna74_GluCTC_1_-_161431809_161431880@1.34.34,trna77_GluCTC_1_-_161424398_161424469@1.34.34,trna77_GluCTC_6_+_28949976_28950047@1.34.34,trna80_GluCTC_1_-_161417018_161417089@1.34.34,trna87_GluCTC_6_-_126101393_126101464@1.34.34|TCCCTGGTGGTCTAGTGGTTAGGATTCGGCGCTC</td><td>>trna116_GluCTC_1_-_145399233_145399304</td><td>1</td><td>34</td><td>34</td><td>TCCCTGGTGGTCTAGTGGTTAGGATTCGGCGCTC</td><td>3023.9</td><td>4.4</td><td>0.8</td><td>5.4</td><td>6.8E-08</td><td>1.2E-07</td><td>MB231/MB468</td><td>MITO</td></tr>
<tr><td>>hg38_21_+_8433172_8446622_RNA45SN1_45S_50ntFlanks@11561.11580.20|GCGCCTAGCAGCCGACTTAG</td><td>>hg38_21_+_8433172_8446622_RNA45SN1_45S_50ntFlanks</td><td>11561</td><td>11580</td><td>20</td><td>GCGCCTAGCAGCCGACTTAG</td><td>118.6</td><td>-10.0</td><td>1.7</td><td>-5.7</td><td>1.1E-08</td><td>1.2E-07</td><td>BT20/MB468</td><td>MP</td></tr>
<tr><td>>hg38_21_+_8433172_8446622_RNA45SN1_45S_50ntFlanks@12243.12268.26|GAAGGGCAAAAGCTCGCTTGATCTTG</td><td>>hg38_21_+_8433172_8446622_RNA45SN1_45S_50ntFlanks</td><td>12243</td><td>12268</td><td>26</td><td>GAAGGGCAAAAGCTCGCTTGATCTTG</td><td>554.1</td><td>-7.6</td><td>1.4</td><td>-5.4</td><td>5.8E-08</td><td>1.2E-07</td><td>MB231/MB468</td><td>MP</td></tr>
<tr><td>>hg38_21_+_8433172_8446622_RNA45SN1_45S_50ntFlanks@10547.10581.35|AGTGCGGTAACGCGACCGATCCCGGAGAAGCCGGC</td><td>>hg38_21_+_8433172_8446622_RNA45SN1_45S_50ntFlanks</td><td>10547</td><td>10581</td><td>35</td><td>AGTGCGGTAACGCGACCGATCCCGGAGAAGCCGGC</td><td>88.9</td><td>10.2</td><td>1.8</td><td>5.7</td><td>1.1E-08</td><td>1.2E-07</td><td>BT20/MB468</td><td>MP</td></tr>
<tr><td>>hg38_21_+_8433172_8446622_RNA45SN1_45S_50ntFlanks@485.527.43|GATGTGGTGACGTCGTGCTCTCCCGGGCCGGGTCCGAGCCGCG</td><td>>hg38_21_+_8433172_8446622_RNA45SN1_45S_50ntFlanks</td><td>485</td><td>527</td><td>43</td><td>GATGTGGTGACGTCGTGCTCTCCCGGGCCGGGTCCGAGCCGCG</td><td>163.3</td><td>11.1</td><td>1.9</td><td>5.7</td><td>1.1E-08</td><td>1.2E-07</td><td>BT20/MB468</td><td>MP</td></tr>
<tr><td>>hg38_21_+_8433172_8446622_RNA45SN1_45S_50ntFlanks@6671.6698.28|CTCGGCTCGTGCGTCGATGAAGAACGCA</td><td>>hg38_21_+_8433172_8446622_RNA45SN1_45S_50ntFlanks</td><td>6671</td><td>6698</td><td>28</td><td>CTCGGCTCGTGCGTCGATGAAGAACGCA</td><td>810.7</td><td>-5.3</td><td>1.0</td><td>-5.4</td><td>6.2E-08</td><td>1.2E-07</td><td>MB231/MB468</td><td>CYTO</td></tr>
<tr><td>>trnaMT_SerGCT_MT_+_12207_12265@9.38.30|TCACAAGAACTGCTAACTCATGCCCCCATG</td><td>>trnaMT_SerGCT_MT_+_12207_12265</td><td>9</td><td>38</td><td>30</td><td>TCACAAGAACTGCTAACTCATGCCCCCATG</td><td>200.3</td><td>-7.4</td><td>1.4</td><td>-5.4</td><td>6.4E-08</td><td>1.2E-07</td><td>MB231/MB468</td><td>NUC</td></tr>
<tr><td>>hg38_21_+_8433172_8446622_RNA45SN1_45S_50ntFlanks@11769.11804.36|TAATTAGTGACGCGCATGAATGGATGAACGAGATTC</td><td>>hg38_21_+_8433172_8446622_RNA45SN1_45S_50ntFlanks</td><td>11769</td><td>11804</td><td>36</td><td>TAATTAGTGACGCGCATGAATGGATGAACGAGATTC</td><td>1036.0</td><td>-6.1</td><td>1.1</td><td>-5.4</td><td>6.4E-08</td><td>1.2E-07</td><td>MB231/MB468</td><td>NUC</td></tr>
<tr><td>>hg38_21_+_8433172_8446622_RNA45SN1_45S_50ntFlanks@11212.11232.21|CGACTCTGGACGCGAGCCGGG</td><td>>hg38_21_+_8433172_8446622_RNA45SN1_45S_50ntFlanks</td><td>11212</td><td>11232</td><td>21</td><td>CGACTCTGGACGCGAGCCGGG</td><td>1757.7</td><td>-6.7</td><td>1.2</td><td>-5.7</td><td>1.1E-08</td><td>1.2E-07</td><td>BT20/MB468</td><td>MP</td></tr>
<tr><td>>hg38_21_+_8433172_8446622_RNA45SN1_45S_50ntFlanks@9554.9574.21|CGTAGCGGTCCTGACGTGCAA</td><td>>hg38_21_+_8433172_8446622_RNA45SN1_45S_50ntFlanks</td><td>9554</td><td>9574</td><td>21</td><td>CGTAGCGGTCCTGACGTGCAA</td><td>907.5</td><td>-2.9</td><td>0.5</td><td>-5.5</td><td>4.8E-08</td><td>1.2E-07</td><td>BT20/MB468</td><td>MITO</td></tr>
<tr><td>>hg38_21_+_8433172_8446622_RNA45SN1_45S_50ntFlanks@10683.10705.23|CCGGCGGCGTCCGGTGAGCTCTC</td><td>>hg38_21_+_8433172_8446622_RNA45SN1_45S_50ntFlanks</td><td>10683</td><td>10705</td><td>23</td><td>CCGGCGGCGTCCGGTGAGCTCTC</td><td>2150.1</td><td>-7.2</td><td>1.3</td><td>-5.4</td><td>7.1E-08</td><td>1.2E-07</td><td>BT20/MB468</td><td>NUC</td></tr>
<tr><td>>trna116_GluCTC_1_-_145399233_145399304@1.38.38,trna59_GluCTC_1_+_249168447_249168518@1.38.38,trna71_GluCTC_1_-_161439189_161439260@1.38.38,trna74_GluCTC_1_-_161431809_161431880@1.38.38,trna77_GluCTC_1_-_161424398_161424469@1.38.38,trna77_GluCTC_6_+_28949976_28950047@1.38.38,trna80_GluCTC_1_-_161417018_161417089@1.38.38,trna87_GluCTC_6_-_126101393_126101464@1.38.38|TCCCTGGTGGTCTAGTGGTTAGGATTCGGCGCTCTCAC</td><td>>trna116_GluCTC_1_-_145399233_145399304</td><td>1</td><td>38</td><td>38</td><td>TCCCTGGTGGTCTAGTGGTTAGGATTCGGCGCTCTCAC</td><td>374.6</td><td>6.9</td><td>1.3</td><td>5.4</td><td>6.2E-08</td><td>1.2E-07</td><td>MB231/MB468</td><td>CYTO</td></tr>
<tr><td>>hg38_21_+_8433172_8446622_RNA45SN1_45S_50ntFlanks@6700.6720.21|CTAGCTGCGAGAATTAATGTG</td><td>>hg38_21_+_8433172_8446622_RNA45SN1_45S_50ntFlanks</td><td>6700</td><td>6720</td><td>21</td><td>CTAGCTGCGAGAATTAATGTG</td><td>18532.6</td><td>-3.8</td><td>0.7</td><td>-5.4</td><td>6.3E-08</td><td>1.2E-07</td><td>MB231/MB468</td><td>CYTO</td></tr>
<tr><td>>hg38_21_+_8433172_8446622_RNA45SN1_45S_50ntFlanks@11561.11588.28|GCGCCTAGCAGCCGACTTAGAACTGGTG</td><td>>hg38_21_+_8433172_8446622_RNA45SN1_45S_50ntFlanks</td><td>11561</td><td>11588</td><td>28</td><td>GCGCCTAGCAGCCGACTTAGAACTGGTG</td><td>496.6</td><td>-3.3</td><td>0.6</td><td>-5.5</td><td>4.8E-08</td><td>1.2E-07</td><td>BT20/MB468</td><td>MITO</td></tr>
<tr><td>>hg38_21_+_8433172_8446622_RNA45SN1_45S_50ntFlanks@6775.6806.32|CCTCCCGGGGCTACGCCTGTCTGAGCGTCGCT</td><td>>hg38_21_+_8433172_8446622_RNA45SN1_45S_50ntFlanks</td><td>6775</td><td>6806</td><td>32</td><td>CCTCCCGGGGCTACGCCTGTCTGAGCGTCGCT</td><td>1910.9</td><td>4.5</td><td>0.8</td><td>5.5</td><td>4.8E-08</td><td>1.2E-07</td><td>BT20/MB468</td><td>MITO</td></tr>
<tr><td>>hg38_21_+_8433172_8446622_RNA45SN1_45S_50ntFlanks@8115.8140.26|CGCGGGACATGTGGCGTACGGAAGAC</td><td>>hg38_21_+_8433172_8446622_RNA45SN1_45S_50ntFlanks</td><td>8115</td><td>8140</td><td>26</td><td>CGCGGGACATGTGGCGTACGGAAGAC</td><td>690.7</td><td>12.8</td><td>2.1</td><td>6.0</td><td>2.3E-09</td><td>1.2E-07</td><td>BT20/MB231</td><td>MP</td></tr>
<tr><td>>hg38_21_+_8433172_8446622_RNA45SN1_45S_50ntFlanks@8168.8195.28|CCCAAGTCCTTCTGATCGAGGCCCAGCC</td><td>>hg38_21_+_8433172_8446622_RNA45SN1_45S_50ntFlanks</td><td>8168</td><td>8195</td><td>28</td><td>CCCAAGTCCTTCTGATCGAGGCCCAGCC</td><td>346.9</td><td>-4.2</td><td>0.8</td><td>-5.4</td><td>6.5E-08</td><td>1.2E-07</td><td>MB231/MB468</td><td>NUC</td></tr>
<tr><td>>hg38_21_+_8433172_8446622_RNA45SN1_45S_50ntFlanks@9759.9789.31|CAAACTTTAAATGGGTAAGAAGCCCGGCTCG</td><td>>hg38_21_+_8433172_8446622_RNA45SN1_45S_50ntFlanks</td><td>9759</td><td>9789</td><td>31</td><td>CAAACTTTAAATGGGTAAGAAGCCCGGCTCG</td><td>120.3</td><td>-10.0</td><td>1.8</td><td>-5.7</td><td>1.1E-08</td><td>1.2E-07</td><td>BT20/MB468</td><td>MP</td></tr>
<tr><td>>hg38_21_+_8433172_8446622_RNA45SN1_45S_50ntFlanks@476.510.35|TTCGTGATCGATGTGGTGACGTCGTGCTCTCCCGG</td><td>>hg38_21_+_8433172_8446622_RNA45SN1_45S_50ntFlanks</td><td>476</td><td>510</td><td>35</td><td>TTCGTGATCGATGTGGTGACGTCGTGCTCTCCCGG</td><td>6584.2</td><td>3.8</td><td>0.7</td><td>5.4</td><td>6.5E-08</td><td>1.2E-07</td><td>MB231/MB468</td><td>NUC</td></tr>
<tr><td>>MI0000299|hsa-mir-222&WithFlank&X|-|45747009|45747130@75.97.23(+3U)&[MIMAT0000279&hsa-miR-222-3p&offsets|0|+2(+3U);m-140&X|-|45747033|45747056&offsets|0|-1(+3U)]|AGCTACATCTGGCTACTGGGTCTTTT</td><td>>MI0000299|hsa-mir-222&WithFlank&X|-|45747009|45747130</td><td>75</td><td>97</td><td>23(+3U)</td><td>AGCTACATCTGGCTACTGGGTCTTTT</td><td>362.7</td><td>8.5</td><td>1.6</td><td>5.4</td><td>6.8E-08</td><td>1.2E-07</td><td>MB231/MB468</td><td>MITO</td></tr>
<tr><td>>MI0000062|hsa-let-7a-3&WithFlank&22|+|46112743|46112828@58.79.22;MI0000060|hsa-let-7a-1&WithFlank&9|+|94175951|94176042@63.84.22&[MIMAT0004481&hsa-let-7a-3p&offsets|0|+1;m-180&22|+|46112800|46112821&offsets|0|0];[MIMAT0004481_1&hsa-let-7a-3p&offsets|0|+1;m-181&9|+|94176013|94176034&offsets|0|0]|CTATACAATCTACTGTCTTTCC</td><td>>MI0000062|hsa-let-7a-3&WithFlank&22|+|46112743|46112828</td><td>58</td><td>79</td><td>22</td><td>CTATACAATCTACTGTCTTTCC</td><td>409.7</td><td>5.6</td><td>1.0</td><td>5.4</td><td>6.8E-08</td><td>1.2E-07</td><td>MB231/MB468</td><td>MITO</td></tr>
<tr><td>>hg38_21_+_8433172_8446622_RNA45SN1_45S_50ntFlanks@11660.11681.22|GTGATTTCTGCCCAGTGCTCTG</td><td>>hg38_21_+_8433172_8446622_RNA45SN1_45S_50ntFlanks</td><td>11660</td><td>11681</td><td>22</td><td>GTGATTTCTGCCCAGTGCTCTG</td><td>438.8</td><td>-7.9</td><td>1.5</td><td>-5.4</td><td>6.8E-08</td><td>1.2E-07</td><td>MB231/MB468</td><td>MITO</td></tr>
<tr><td>>MI0000750|hsa-mir-26a-2&WithFlank&12|-|57824603|57824698@20.41.22(+1U);MI0000083|hsa-mir-26a-1&WithFlank&3|+|37969398|37969486@16.37.22(+1U)&[MIMAT0000082&hsa-miR-26a-5p&offsets|0|0(+1U);m-33&12|-|57824658|57824679&offsets|0|0(+1U)];[MIMAT0000082_1&hsa-miR-26a-5p&offsets|0|0(+1U);m-34&3|+|37969413|37969434&offsets|0|0(+1U)]|TTCAAGTAATCCAGGATAGGCTT</td><td>>MI0000750|hsa-mir-26a-2&WithFlank&12|-|57824603|57824698</td><td>20</td><td>41</td><td>22(+1U)</td><td>TTCAAGTAATCCAGGATAGGCTT</td><td>4610.8</td><td>4.7</td><td>0.9</td><td>5.4</td><td>5.8E-08</td><td>1.2E-07</td><td>MB231/MB468</td><td>MP</td></tr>
<tr><td>>MI0000273|hsa-mir-183&WithFlank&7|-|129774899|129775020@34.55.22&[MIMAT0000261&hsa-miR-183-5p&offsets|+1|+1;m-41&7|-|129774966|129774987&offsets|0|0]|ATGGCACTGGTAGAATTCACTG</td><td>>MI0000273|hsa-mir-183&WithFlank&7|-|129774899|129775020</td><td>34</td><td>55</td><td>22</td><td>ATGGCACTGGTAGAATTCACTG</td><td>3389.9</td><td>3.2</td><td>0.6</td><td>5.8</td><td>6.7E-09</td><td>1.2E-07</td><td>BT20/MB231</td><td>NUC</td></tr>
<tr><td>>hg38_21_+_8433172_8446622_RNA45SN1_45S_50ntFlanks@10350.10381.32|ACGAGAACTTTGAAGGCCGAAGTGGAGAAGGG</td><td>>hg38_21_+_8433172_8446622_RNA45SN1_45S_50ntFlanks</td><td>10350</td><td>10381</td><td>32</td><td>ACGAGAACTTTGAAGGCCGAAGTGGAGAAGGG</td><td>711.4</td><td>-5.8</td><td>1.1</td><td>-5.4</td><td>6.5E-08</td><td>1.2E-07</td><td>MB231/MB468</td><td>NUC</td></tr>
<tr><td>>MI0000073|hsa-mir-19a&WithFlank&13|+|91350885|91350978@55.78.24&[MIMAT0000073&hsa-miR-19a-3p&offsets|0|+1;m-201&13|+|91350939|91350961&offsets|0|+1]|TGTGCAAATCTATGCAAAACTGAT</td><td>>MI0000073|hsa-mir-19a&WithFlank&13|+|91350885|91350978</td><td>55</td><td>78</td><td>24</td><td>TGTGCAAATCTATGCAAAACTGAT</td><td>418.1</td><td>6.1</td><td>1.1</td><td>5.4</td><td>6.9E-08</td><td>1.2E-07</td><td>MB231/MB468</td><td>MITO</td></tr>
<tr><td>>hg38_21_+_8433172_8446622_RNA45SN1_45S_50ntFlanks@12615.12652.38|GCTGAGGAGCCAATGGGGCGAAGCTACCATCTGTGGGA</td><td>>hg38_21_+_8433172_8446622_RNA45SN1_45S_50ntFlanks</td><td>12615</td><td>12652</td><td>38</td><td>GCTGAGGAGCCAATGGGGCGAAGCTACCATCTGTGGGA</td><td>104.5</td><td>-8.8</td><td>1.6</td><td>-5.5</td><td>4.9E-08</td><td>1.2E-07</td><td>BT20/MB468</td><td>MITO</td></tr>
<tr><td>>hg38_21_+_8433172_8446622_RNA45SN1_45S_50ntFlanks@10873.10890.18|CTGGGTCGGTCGGGCTGG</td><td>>hg38_21_+_8433172_8446622_RNA45SN1_45S_50ntFlanks</td><td>10873</td><td>10890</td><td>18</td><td>CTGGGTCGGTCGGGCTGG</td><td>446.6</td><td>-5.9</td><td>1.1</td><td>-5.4</td><td>6.5E-08</td><td>1.2E-07</td><td>MB231/MB468</td><td>NUC</td></tr>
<tr><td>>MI0000803|hsa-mir-330&WithFlank&19|-|45638988|45639093@63.86.24(+1U)&[MIMAT0000751&hsa-miR-330-3p&offsets|0|+1(+1U);m-347&19|-|45639009|45639030&offsets|-1|+1(+1U)]|GCAAAGCACACGGCCTGCAGAGAGT</td><td>>MI0000803|hsa-mir-330&WithFlank&19|-|45638988|45639093</td><td>63</td><td>86</td><td>24(+1U)</td><td>GCAAAGCACACGGCCTGCAGAGAGT</td><td>408.4</td><td>5.0</td><td>0.9</td><td>5.4</td><td>6.9E-08</td><td>1.2E-07</td><td>MB231/MB468</td><td>MITO</td></tr>
<tr><td>>am_trna2_GlyGCC_21_-_18827107_18827177@1.25.25,trna35_GlyGCC_1_+_161413094_161413164@1.25.25,trna37_GlyGCC_1_+_161420467_161420537@1.25.25,trna39_GlyGCC_1_+_161427898_161427968@1.25.25,trna41_GlyGCC_1_+_161435258_161435328@1.25.25|GCATGGGTGGTTCAGTGGTAGAATT</td><td>>am_trna2_GlyGCC_21_-_18827107_18827177</td><td>1</td><td>25</td><td>25</td><td>GCATGGGTGGTTCAGTGGTAGAATT</td><td>489.2</td><td>4.9</td><td>0.9</td><td>5.5</td><td>4.9E-08</td><td>1.2E-07</td><td>BT20/MB468</td><td>MITO</td></tr>
<tr><td>>trna10_LysCTT_16_+_3241501_3241573@1.28.28,trna119_LysCTT_1_-_145395522_145395594@1.28.28,trna11_LysCTT_5_-_180648979_180649051@1.28.28,trna13_LysCTT_14_-_58706613_58706685@1.28.28,trna13_LysCTT_6_+_26556774_26556846@1.28.28,trna2_LysCTT_15_+_79152904_79152976@1.28.28,trna32_LysCTT_16_-_3207406_3207478@1.28.28,trna7_LysCTT_16_+_3225692_3225764@1.28.28,trna9_LysCTT_5_+_180634755_180634827@1.28.28|GCCCGGCTAGCTCAGTCGGTAGAGCATG</td><td>>trna10_LysCTT_16_+_3241501_3241573</td><td>1</td><td>28</td><td>28</td><td>GCCCGGCTAGCTCAGTCGGTAGAGCATG</td><td>1443.0</td><td>-4.7</td><td>0.9</td><td>-5.4</td><td>5.9E-08</td><td>1.2E-07</td><td>MB231/MB468</td><td>MP</td></tr>
<tr><td>>am_trnaMT_ValTAC_MT_+_1602_1670@51.69.19|CTTAACTTGACCGCTCTGA</td><td>>am_trnaMT_ValTAC_MT_+_1602_1670</td><td>51</td><td>69</td><td>19</td><td>CTTAACTTGACCGCTCTGA</td><td>1010.0</td><td>-9.4</td><td>1.8</td><td>-5.4</td><td>7.2E-08</td><td>1.2E-07</td><td>BT20/MB468</td><td>NUC</td></tr>
<tr><td>>hg38_21_+_8433172_8446622_RNA45SN1_45S_50ntFlanks@12416.12447.32|TTCGATGTCGGCTCTTCCTATCATTGTGAAGC</td><td>>hg38_21_+_8433172_8446622_RNA45SN1_45S_50ntFlanks</td><td>12416</td><td>12447</td><td>32</td><td>TTCGATGTCGGCTCTTCCTATCATTGTGAAGC</td><td>284.8</td><td>4.2</td><td>0.8</td><td>5.4</td><td>6.9E-08</td><td>1.2E-07</td><td>MB231/MB468</td><td>MITO</td></tr>
<tr><td>>hg38_21_+_8433172_8446622_RNA45SN1_45S_50ntFlanks@10678.10706.29|CGGTTCCGGCGGCGTCCGGTGAGCTCTCG</td><td>>hg38_21_+_8433172_8446622_RNA45SN1_45S_50ntFlanks</td><td>10678</td><td>10706</td><td>29</td><td>CGGTTCCGGCGGCGTCCGGTGAGCTCTCG</td><td>19216.0</td><td>-5.2</td><td>1.0</td><td>-5.4</td><td>5.5E-08</td><td>1.2E-07</td><td>BT20/MB468</td><td>CYTO</td></tr>
<tr><td>>MI0000542|hsa-mir-320a&WithFlank&8|-|22244960|22245043@50.69.20(+2U)&[MIMAT0000510&hsa-miR-320a-3p&offsets|+2|0(+2U);m-60&8|-|22244975|22244996&offsets|+2|0(+2U)]|AAGCTGGGTTGAGAGGGCGATT</td><td>>MI0000542|hsa-mir-320a&WithFlank&8|-|22244960|22245043</td><td>50</td><td>69</td><td>20(+2U)</td><td>AAGCTGGGTTGAGAGGGCGATT</td><td>113.5</td><td>10.6</td><td>1.9</td><td>5.7</td><td>1.2E-08</td><td>1.2E-07</td><td>BT20/MB468</td><td>MP</td></tr>
</table>

| | | | | | | | | | | | | | |
|---|---|---|---|---|---|---|---|---|---|---|---|---|---|
| >hg38_21_+_8433172_8446622_RNA45SN1_45S_50ntFlanks@12996.13038.43\|TCGCTGCGATCTATTGAAAGTCAGCCCTCGACACAAGGGTTTG | >hg38_21_+_8433172_8446622_RNA45SN1_45S_50ntFlanks | 12996 | 13038 | 43 | TCGCTGCGATCTATTGAAAGTCAGCCCTCGACACAAGGGTTTG | 1460.6 | 8.3 | 1.5 | 5.4 | 5.5E-08 | 1.2E-07 | BT20/MB468 | CYTO |
| >am_trna128_GlyGCC_6_-_27870686_27870756@3.34.32,trna18_GlyGCC_16_+_70822597_70822667@3.34.32,trna19_GlyGCC_16_+_70823410_70823480@3.34.32,trna19_GlyGCC_2_-_157257659_157257729@3.34.32,trna24_GlyGCC_16_-_70812942_70813012@3.34.32,trna25_GlyGCC_16_-_70812114_70812184@3.34.32,trna5_GlyGCC_17_+_8029064_8029134@3.34.32,trna68_GlyGCC_1_-_161493637_161493707@3.34.32\|ATTGGTGGTTCAGTGGTAGAATTCTCGCCTGC | >am_trna128_GlyGCC_6_-_27870686_27870756 | 3 | 34 | 32 | ATTGGTGGTTCAGTGGTAGAATTCTCGCCTGC | 80.0 | 9.5 | 1.8 | 5.4 | 5.9E-08 | 1.2E-07 | MB231/MB468 | MP |
| >hg38_21_+_8433172_8446622_RNA45SN1_45S_50ntFlanks@8356.8391.36\|GAAAAGAACTTTGAAGAGAGAGTTCAAGAGGGCGTG | >hg38_21_+_8433172_8446622_RNA45SN1_45S_50ntFlanks | 8356 | 8391 | 36 | GAAAAGAACTTTGAAGAGAGAGTTCAAGAGGGCGTG | 230.6 | -4.1 | 0.8 | -5.4 | 5.9E-08 | 1.2E-07 | MB231/MB468 | MP |
| >MI0000077\|hsa-mir-21&WithFlank&17\|+\|59841260\|59841343@14.35.22(+1G)&[MIMAT0000076&hsa-miR-21-5p&offsets\|0\|0(+1G);m-1&17\|+\|59841273\|59841294&offsets\|0\|0(+1G)]\|TAGCTTATCAGACTGATGTTGAG | >MI0000077\|hsa-mir-21&WithFlank&17\|+\|59841260\|59841343 | 14 | 35 | 22(+1G) | TAGCTTATCAGACTGATGTTGAG | 373.3 | 3.9 | 0.7 | 5.4 | 7.2E-08 | 1.2E-07 | BT20/MB468 | NUC |
| >MI0000065\|hsa-let-7d&WithFlank&9\|+\|94178828\|94178926@68.88.21&[MIMAT0004484&hsa-let-7d-3p&offsets\|0\|-1;m-98&9\|+\|94178895\|94178916&offsets\|0\|-1]\|CTATACGACCTGCTGCCTTTC | >MI0000065\|hsa-let-7d&WithFlank&9\|+\|94178828\|94178926 | 68 | 88 | 21 | CTATACGACCTGCTGCCTTTC | 368.8 | 5.1 | 0.9 | 5.4 | 6.9E-08 | 1.2E-07 | MB231/MB468 | MITO |
| >hg38_21_+_8433172_8446622_RNA45SN1_45S_50ntFlanks@12420.12444.25\|ATGTCGGCTCTTCCTATCATTGTGA | >hg38_21_+_8433172_8446622_RNA45SN1_45S_50ntFlanks | 12420 | 12444 | 25 | ATGTCGGCTCTTCCTATCATTGTGA | 4725.6 | -3.6 | 0.7 | -5.4 | 6.6E-08 | 1.2E-07 | MB231/MB468 | NUC |
| >MI0000063\|hsa-let-7b&WithFlank&22\|+\|46113680\|46113774@12.32.21&[MIMAT0000063&hsa-let-7b-5p&offsets\|0\|-1;m-14&22\|+\|46113691\|46113712&offsets\|0\|-1]\|TGAGGTAGTAGGTTGTGTGGT | >MI0000063\|hsa-let-7b&WithFlank&22\|+\|46113680\|46113774 | 12 | 32 | 21 | TGAGGTAGTAGGTTGTGTGGT | 863.1 | 4.7 | 0.9 | 5.4 | 6.9E-08 | 1.2E-07 | MB231/MB468 | MITO |
| >hg38_21_+_8433172_8446622_RNA45SN1_45S_50ntFlanks@5070.5102.33\|GTTAATTCCGATAACGAACGAGACTCTGGCATG | >hg38_21_+_8433172_8446622_RNA45SN1_45S_50ntFlanks | 5070 | 5102 | 33 | GTTAATTCCGATAACGAACGAGACTCTGGCATG | 96.3 | -10.5 | 1.9 | -5.4 | 5.9E-08 | 1.2E-07 | MB231/MB468 | MP |
| >hg38_21_+_8433172_8446622_RNA45SN1_45S_50ntFlanks@475.496.22\|CTTCGTGATCGATGTGGTGACG | >hg38_21_+_8433172_8446622_RNA45SN1_45S_50ntFlanks | 475 | 496 | 22 | CTTCGTGATCGATGTGGTGACG | 567.4 | 8.1 | 1.5 | 5.4 | 5.5E-08 | 1.2E-07 | BT20/MB468 | CYTO |
| >hg38_21_+_8433172_8446622_RNA45SN1_45S_50ntFlanks@12569.12587.19\|CAGTACGAGAGGAACCGCA | >hg38_21_+_8433172_8446622_RNA45SN1_45S_50ntFlanks | 12569 | 12587 | 19 | CAGTACGAGAGGAACCGCA | 349.3 | -7.1 | 1.3 | -5.4 | 7.0E-08 | 1.2E-07 | MB231/MB468 | MITO |
| >hg38_21_+_8433172_8446622_RNA45SN1_45S_50ntFlanks@11876.11897.22\|AAGACCCTGTTGAGCTTGACTC | >hg38_21_+_8433172_8446622_RNA45SN1_45S_50ntFlanks | 11876 | 11897 | 22 | AAGACCCTGTTGAGCTTGACTC | 572.9 | -4.4 | 0.8 | -5.4 | 7.0E-08 | 1.2E-07 | MB231/MB468 | MITO |
| >hg38_21_+_8433172_8446622_RNA45SN1_45S_50ntFlanks@4047.4073.27\|ATTCGAACGTCTGCCCTATCAACTTTC | >hg38_21_+_8433172_8446622_RNA45SN1_45S_50ntFlanks | 4047 | 4073 | 27 | ATTCGAACGTCTGCCCTATCAACTTTC | 440.6 | -7.7 | 1.4 | -5.4 | 7.0E-08 | 1.2E-07 | MB231/MB468 | MITO |
| >hg38_MT_+_598_1651_RNR1_12S_50ntFlanks@843.879.37\|GAGTAGAGTGCTTAGTTGAACAGGGCCCTGAAGCGCG | >hg38_MT_+_598_1651_RNR1_12S_50ntFlanks | 843 | 879 | 37 | GAGTAGAGTGCTTAGTTGAACAGGGCCCTGAAGCGCG | 153.8 | 11.0 | 1.9 | 5.7 | 1.2E-08 | 1.2E-07 | BT20/MB468 | MP |
| >hg38_21_+_8433172_8446622_RNA45SN1_45S_50ntFlanks@12255.12284.30\|CTCGCTTGATCTTGATTTTCAGTACGAATA | >hg38_21_+_8433172_8446622_RNA45SN1_45S_50ntFlanks | 12255 | 12284 | 30 | CTCGCTTGATCTTGATTTTCAGTACGAATA | 415.3 | -6.7 | 1.2 | -5.4 | 7.0E-08 | 1.2E-07 | MB231/MB468 | MITO |
| >MI0000446\|hsa-mir-125b-1&WithFlank&11\|-\|122099751\|122099850@21.42.22(+1C);MI0000470\|hsa-mir-125b-2&WithFlank&21\|+\|16590231\|16590331@23.44.22(+1C)&[MIMAT0000423&hsa-miR-125b-5p&offsets\|0\|0(+1C);m-67&11\|-\|122099809\|122099830&offsets\|0\|0(+1C)];[MIMAT0000423_1&hsa-miR-125b-5p&offsets\|0\|0(+1C);m-68&21\|+\|16590253\|16590274&offsets\|0\|0(+1C)]\|TCCCTGAGACCCTAACTTGTGAC | >MI0000446\|hsa-mir-125b-1&WithFlank&11\|-\|122099751\|122099850 | 21 | 42 | 22(+1C) | TCCCTGAGACCCTAACTTGTGAC | 68.2 | 9.3 | 1.7 | 5.4 | 6.0E-08 | 1.2E-07 | MB231/MB468 | MP |
| >hg38_21_+_8433172_8446622_RNA45SN1_45S_50ntFlanks@10313.10340.28\|GCAGGTGCAGATCTTGGTGGTAGTAGCA | >hg38_21_+_8433172_8446622_RNA45SN1_45S_50ntFlanks | 10313 | 10340 | 28 | GCAGGTGCAGATCTTGGTGGTAGTAGCA | 170.7 | 11.2 | 2.0 | 5.7 | 1.2E-08 | 1.2E-07 | BT20/MB468 | MP |
| >trna10_LysCTT_16_+_3241501_3241573@1.28.28,trna119_LysCTT_1_-_145395522_145395594@1.28.28,trna11_LysCTT_5_-_180648979_180649051@1.28.28,trna13_LysCTT_14_-_58706613_58706685@1.28.28,trna13_LysCTT_6_+_26556774_26556846@1.28.28,trna2_LysCTT_15_+_79152904_79152976@1.28.28,trna32_LysCTT_16_-_3207406_3207478@1.28.28,trna7_LysCTT_16_+_3225692_3225764@1.28.28,trna9_LysCTT_5_+_180634755_180634827@1.28.28\|GCCCGGCTAGCTCAGTCGGTAGAGCATG | >trna10_LysCTT_16_+_3241501_3241573 | 1 | 28 | 28 | GCCCGGCTAGCTCAGTCGGTAGAGCATG | 4868.7 | -4.4 | 0.8 | -5.4 | 5.6E-08 | 1.2E-07 | BT20/MB468 | CYTO |
| >hg38_1_-_228634819_228635039_RNA5S12_5S_50ntFlanks@127.170.44\|ACTTGGATGGGAGACCGCCTGGGAATACCGGGTGCTGTAGGCTT | >hg38_1_-_228634819_228635039_RNA5S12_5S_50ntFlanks | 127 | 170 | 44 | ACTTGGATGGGAGACCGCCTGGGAATACCGGGTGCTGTAGGCTT | 2210.1 | 9.7 | 1.8 | 5.4 | 5.6E-08 | 1.2E-07 | BT20/MB468 | CYTO |
| >hg38_21_+_8433172_8446622_RNA45SN1_45S_50ntFlanks@10663.10706.44\|CCTTGGAAAGCGTCGCGGTTCCGGCGGCGTCCGGTGAGCTCTCG | >hg38_21_+_8433172_8446622_RNA45SN1_45S_50ntFlanks | 10663 | 10706 | 44 | CCTTGGAAAGCGTCGCGGTTCCGGCGGCGTCCGGTGAGCTCTCG | 97.7 | -10.5 | 1.9 | -5.4 | 6.0E-08 | 1.2E-07 | MB231/MB468 | MP |
| >am_MI0003581\|hsa-mir-574&WithFlank&4\|+\|38868026\|38868133@31.55.25&[MIMAT0004795&hsa-miR-574-5p&offsets\|0\|+2;m-213&4\|+\|38868056\|38868078&offsets\|0\|+2]\|TGAGTGTGTGTGTGTGAGTGTGTGT | >am_MI0003581\|hsa-mir-574&WithFlank&4\|+\|38868026\|38868133 | 31 | 55 | 25 | TGAGTGTGTGTGTGTGAGTGTGTGT | 818.3 | 8.0 | 1.5 | 5.4 | 7.0E-08 | 1.2E-07 | MB231/MB468 | MITO |
| >hg38_21_+_8433172_8446622_RNA45SN1_45S_50ntFlanks@10679.10709.31\|GGTTCCGGCGGCGTCCGGTGAGCTCTCGCTG | >hg38_21_+_8433172_8446622_RNA45SN1_45S_50ntFlanks | 10679 | 10709 | 31 | GGTTCCGGCGGCGTCCGGTGAGCTCTCGCTG | 5315.7 | -6.8 | 1.3 | -5.4 | 6.5E-08 | 1.2E-07 | MB231/MB468 | CYTO |
| >hg38_21_+_8433172_8446622_RNA45SN1_45S_50ntFlanks@9496.9531.36\|GAAAGATGGTGAACTATGCCTGGGCAGGGCGAAGCC | >hg38_21_+_8433172_8446622_RNA45SN1_45S_50ntFlanks | 9496 | 9531 | 36 | GAAAGATGGTGAACTATGCCTGGGCAGGGCGAAGCC | 85.3 | 10.2 | 1.8 | 5.7 | 1.2E-08 | 1.2E-07 | BT20/MB468 | MP |
| >hg38_21_+_8433172_8446622_RNA45SN1_45S_50ntFlanks@11604.11640.37\|CGACTGTTTAATTAAAACAAAGCATCGCGAAGGCCCG | >hg38_21_+_8433172_8446622_RNA45SN1_45S_50ntFlanks | 11604 | 11640 | 37 | CGACTGTTTAATTAAAACAAAGCATCGCGAAGGCCCG | 172.8 | -9.5 | 1.7 | -5.5 | 5.0E-08 | 1.2E-07 | BT20/MB468 | MITO |
| >MI0000650\|hsa-mir-200c&WithFlank&12\|+\|6963693\|6963772@50.72.23(+2A)&[MIMAT0000617&hsa-miR-200c-3p&offsets\|0\|0(+2A);m-54&12\|+\|6963742\|6963764&offsets\|0\|0(+2A)]\|TAATACTGCCGGGTAATGATGGAAA | >MI0000650\|hsa-mir-200c&WithFlank&12\|+\|6963693\|6963772 | 50 | 72 | 23(+2A) | TAATACTGCCGGGTAATGATGGAAA | 294.6 | 2.1 | 0.4 | 5.4 | 7.4E-08 | 1.2E-07 | BT20/MB468 | NUC |
| >hg38_21_+_8433172_8446622_RNA45SN1_45S_50ntFlanks@7975.8007.33\|CGCGACCTCAGATCAGACGTGGCGACCCGCTGA | >hg38_21_+_8433172_8446622_RNA45SN1_45S_50ntFlanks | 7975 | 8007 | 33 | CGCGACCTCAGATCAGACGTGGCGACCCGCTGA | 8958.9 | 4.0 | 0.7 | 5.4 | 6.5E-08 | 1.2E-07 | MB231/MB468 | CYTO |
| >hg38_21_+_8433172_8446622_RNA45SN1_45S_50ntFlanks@12910.12933.24\|CGCACGTTCGTGGGGAACCTGGCG | >hg38_21_+_8433172_8446622_RNA45SN1_45S_50ntFlanks | 12910 | 12933 | 24 | CGCACGTTCGTGGGGAACCTGGCG | 408.1 | -7.9 | 1.5 | -5.4 | 6.0E-08 | 1.2E-07 | MB231/MB468 | MP |
| >hg38_21_+_8433172_8446622_RNA45SN1_45S_50ntFlanks@4884.4917.34\|CAAAGCTGAAACTTAAAGGAATTGACGGAAGGGC | >hg38_21_+_8433172_8446622_RNA45SN1_45S_50ntFlanks | 4884 | 4917 | 34 | CAAAGCTGAAACTTAAAGGAATTGACGGAAGGGC | 377.2 | -7.5 | 1.4 | -5.4 | 6.0E-08 | 1.2E-07 | MB231/MB468 | MP |
| >hg38_21_+_8433172_8446622_RNA45SN1_45S_50ntFlanks@9534.9560.27\|AGGAAACTCTGGTGGAGGTCCGTAGCG | >hg38_21_+_8433172_8446622_RNA45SN1_45S_50ntFlanks | 9534 | 9560 | 27 | AGGAAACTCTGGTGGAGGTCCGTAGCG | 238.7 | -2.3 | 0.4 | -5.5 | 5.0E-08 | 1.2E-07 | BT20/MB468 | MITO |
| >MI0000063\|hsa-let-7b&WithFlank&22\|+\|46113680\|46113774@12.34.23(+1U)&[MIMAT0000063&hsa-let-7b-5p&offsets\|0\|+1(+1U);m-14&22\|+\|46113691\|46113712&offsets\|0\|+1(+1U)]\|TGAGGTAGTAGGTTGTGTGGTTTT | >MI0000063\|hsa-let-7b&WithFlank&22\|+\|46113680\|46113774 | 12 | 34 | 23(+1U) | TGAGGTAGTAGGTTGTGTGGTTTT | 231.3 | 3.9 | 0.7 | 5.4 | 6.7E-08 | 1.2E-07 | MB231/MB468 | NUC |
| >hg38_21_+_8433172_8446622_RNA45SN1_45S_50ntFlanks@11104.11132.29\|AAGGGTCGGGGCGGCAGGGGCCGGCGGCG | >hg38_21_+_8433172_8446622_RNA45SN1_45S_50ntFlanks | 11104 | 11132 | 29 | AAGGGTCGGGGCGGCAGGGGCCGGCGGCG | 432.4 | -8.0 | 1.5 | -5.4 | 7.4E-08 | 1.2E-07 | BT20/MB468 | NUC |
| >hg38_21_+_8433172_8446622_RNA45SN1_45S_50ntFlanks@6731.6752.22\|CACATTGATCATCGACACTTCG | >hg38_21_+_8433172_8446622_RNA45SN1_45S_50ntFlanks | 6731 | 6752 | 22 | CACATTGATCATCGACACTTCG | 815.4 | -8.5 | 1.6 | -5.4 | 7.4E-08 | 1.2E-07 | BT20/MB468 | NUC |
| >hg38_21_+_8433172_8446622_RNA45SN1_45S_50ntFlanks@3843.3860.18\|CTACTTGGATAACTGTGG | >hg38_21_+_8433172_8446622_RNA45SN1_45S_50ntFlanks | 3843 | 3860 | 18 | CTACTTGGATAACTGTGG | 387.1 | -3.7 | 0.7 | -5.4 | 6.5E-08 | 1.2E-07 | MB231/MB468 | CYTO |
| >hg38_21_+_8433172_8446622_RNA45SN1_45S_50ntFlanks@9498.9520.23\|AAGATGGTGAACTATGCCTGGGC | >hg38_21_+_8433172_8446622_RNA45SN1_45S_50ntFlanks | 9498 | 9520 | 23 | AAGATGGTGAACTATGCCTGGGC | 75.7 | 10.0 | 1.8 | 5.7 | 1.2E-08 | 1.2E-07 | BT20/MB468 | MP |
| >hg38_21_+_8433172_8446622_RNA45SN1_45S_50ntFlanks@6732.6771.40\|ACATTGATCATCGACACTTCGAACGCACTTGCGGCCCCGG | >hg38_21_+_8433172_8446622_RNA45SN1_45S_50ntFlanks | 6732 | 6771 | 40 | ACATTGATCATCGACACTTCGAACGCACTTGCGGCCCCGG | 2095.0 | -3.2 | 0.6 | -5.4 | 5.0E-08 | 1.2E-07 | BT20/MB468 | MITO |
| >MI0000298\|hsa-mir-221&WithFlank&X\|-\|45746151\|45746272@71.92.22(+2U)&[MIMAT0000278&hsa-miR-221-3p&offsets\|0\|-1(+2U);m-91&X\|-\|45746181\|45746202&offsets\|0\|0(+2U)]\|AGCTACATTGTCTGCTGGGTTTTT | >MI0000298\|hsa-mir-221&WithFlank&X\|-\|45746151\|45746272 | 71 | 92 | 22(+2U) | AGCTACATTGTCTGCTGGGTTTTT | 984.0 | -5.0 | 0.8 | -6.0 | 2.4E-09 | 1.2E-07 | BT20/MB231 | MP |
| >am_MI0000786\|hsa-mir-378a&WithFlank&5\|+\|149732819\|149732896@50.70.21&[MIMAT0000732&hsa-miR-378a-3p&offsets\|+1\|0;m-44&5\|+\|149732867\|149732888&offsets\|+1\|0]\|CTGGACTTGGAGTCAGAAGGC | >am_MI0000786\|hsa-mir-378a&WithFlank&5\|+\|149732819\|149732896 | 50 | 70 | 21 | CTGGACTTGGAGTCAGAAGGC | 1327.3 | 4.4 | 0.8 | 5.4 | 6.5E-08 | 1.2E-07 | MB231/MB468 | CYTO |
| >hg38_21_+_8433172_8446622_RNA45SN1_45S_50ntFlanks@10392.10430.39\|ACAGCAGTTGAACATGGGTCAGTCGGTCCTGAGAGATGG | >hg38_21_+_8433172_8446622_RNA45SN1_45S_50ntFlanks | 10392 | 10430 | 39 | ACAGCAGTTGAACATGGGTCAGTCGGTCCTGAGAGATGG | 922.4 | -5.7 | 1.1 | -5.4 | 6.5E-08 | 1.2E-07 | MB231/MB468 | CYTO |
| >trnaMT_PheGAA_MT_+_577_647@36.57.22\|AAAATGTTTAGACGGGCTCACA | >trnaMT_PheGAA_MT_+_577_647 | 36 | 57 | 22 | AAAATGTTTAGACGGGCTCACA | 196.4 | 3.3 | 0.6 | 5.4 | 6.1E-08 | 1.2E-07 | MB231/MB468 | MP |
| >hg38_21_+_8433172_8446622_RNA45SN1_45S_50ntFlanks@11884.11903.20\|GTTGAGCTTGACTCTAGTCT | >hg38_21_+_8433172_8446622_RNA45SN1_45S_50ntFlanks | 11884 | 11903 | 20 | GTTGAGCTTGACTCTAGTCT | 1073.5 | 6.6 | 1.1 | 5.8 | 5.2E-09 | 1.2E-07 | BT20/MB231 | MITO |
| >MI0000471\|hsa-mir-126&WithFlank&9\|+\|136670596\|136670692@58.78.21(+1A)&[MIMAT0000445&hsa-miR-126-3p&offsets\|0\|-1(+1A);m-40&9\|+\|136670653\|136670674&offsets\|0\|-1(+1A)]\|TCGTACCGTGAGTAATAATGCA | >MI0000471\|hsa-mir-126&WithFlank&9\|+\|136670596\|136670692 | 58 | 78 | 21(+1A) | TCGTACCGTGAGTAATAATGCA | 1354.1 | 8.3 | 1.5 | 5.4 | 6.1E-08 | 1.2E-07 | MB231/MB468 | MP |
| >hg38_21_+_8433172_8446622_RNA45SN1_45S_50ntFlanks@4773.4793.21\|TAGTTCCGACCATAAACGATG | >hg38_21_+_8433172_8446622_RNA45SN1_45S_50ntFlanks | 4773 | 4793 | 21 | TAGTTCCGACCATAAACGATG | 2230.3 | -3.5 | 0.7 | -5.4 | 6.5E-08 | 1.2E-07 | MB231/MB468 | CYTO |
| >MI0000087\|hsa-mir-29a&WithFlank&7\|-\|130876741\|130876816@47.68.22&[MIMAT0000086&hsa-miR-29a-3p&offsets\|-1\|-1;m-26&7\|-\|130876749\|130876769&offsets\|-1\|0]\|CTAGCACCATCTGAAATCGGTT | >MI0000087\|hsa-mir-29a&WithFlank&7\|-\|130876741\|130876816 | 47 | 68 | 22 | CTAGCACCATCTGAAATCGGTT | 15954.1 | 6.0 | 1.0 | 5.7 | 1.2E-08 | 1.2E-07 | BT20/MB468 | MP |

| | | | | | | | | | | | | | |
|---|---|---|---|---|---|---|---|---|---|---|---|---|---|
| >MI0000471\|hsa-mir-126&WithFlank&9\|+\|136670596\|136670692@58.78.21&[MIMAT0000445&hsa-miR-126-3p&offsets\|0\|-1;m-40&9\|+\|136670653\|136670674&offsets\|0\|-1]\|TCGTACCGTGAGTAATAATGC | >MI0000471\|hsa-mir-126&WithFlank&9\|+\|136670596\|136670692 | 58 | 78 | 21 | TCGTACCGTGAGTAATAATGC | 3668.0 | 7.4 | 1.4 | 5.4 | 6.1E-08 | 1.2E-07 | MB231/MB468 | MP |
| >MI0000093\|hsa-mir-92a-1&WithFlank&13\|+\|91351308\|91351397@54.76.23(+1U)&[MIMAT0000092&hsa-miR-92a-3p&offsets\|0\|+1(+1U);m-9&13\|+\|91351361\|91351382&offsets\|0\|+1(+1U)]\|TATTGCACTTGTCCCGGCCTGTTT | >MI0000093\|hsa-mir-92a-1&WithFlank&13\|+\|91351308\|91351397 | 54 | 76 | 23(+1U) | TATTGCACTTGTCCCGGCCTGTTT | 490.5 | 5.1 | 0.9 | 5.4 | 7.1E-08 | 1.2E-07 | MB231/MB468 | MITO |
| >MI0000076\|hsa-mir-20a&WithFlank&13\|+\|91351059\|91351141@14.36.23&[MIMAT0000075&hsa-miR-20a-5p&offsets\|0\|0;m-96&13\|+\|91351072\|91351094&offsets\|0\|0]\|TAAAGTGCTTATAGTGCAGGTAG | >MI0000076\|hsa-mir-20a&WithFlank&13\|+\|91351059\|91351141 | 14 | 36 | 23 | TAAAGTGCTTATAGTGCAGGTAG | 24724.7 | 5.5 | 1.0 | 5.4 | 7.1E-08 | 1.2E-07 | MB231/MB468 | MITO |
| >MI0000342\|hsa-mir-200b&WithFlank&1\|+\|1167098\|1167204@63.85.23(+1U)&[MIMAT0000318&hsa-miR-200b-3p&offsets\|0\|+1(+1U);m-93&1\|+\|1167160\|1167182&offsets\|0\|0(+1U)]\|TAATACTGCCTGGTAATGATGACT | >MI0000342\|hsa-mir-200b&WithFlank&1\|+\|1167098\|1167204 | 63 | 85 | 23(+1U) | TAATACTGCCTGGTAATGATGACT | 1767.0 | -4.0 | 0.7 | -5.4 | 6.7E-08 | 1.2E-07 | MB231/MB468 | NUC |
| >hg38_21_+_8433172_8446622_RNA45SN1_45S_50ntFlanks@12934.12964.31\|CTAAACCATTCGTAGACGACCTGCTTCTGGG | >hg38_21_+_8433172_8446622_RNA45SN1_45S_50ntFlanks | 12934 | 12964 | 31 | CTAAACCATTCGTAGACGACCTGCTTCTGGG | 86.8 | -9.5 | 1.7 | -5.7 | 1.2E-08 | 1.2E-07 | BT20/MB468 | MP |
| >hg38_21_+_8433172_8446622_RNA45SN1_45S_50ntFlanks@12957.12981.25\|CTTCTGGGTCGGGGTTTCGTACGTA | >hg38_21_+_8433172_8446622_RNA45SN1_45S_50ntFlanks | 12957 | 12981 | 25 | CTTCTGGGTCGGGGTTTCGTACGTA | 210.9 | -4.6 | 0.8 | -5.4 | 6.6E-08 | 1.2E-07 | MB231/MB468 | CYTO |
| >hg38_21_+_8433172_8446622_RNA45SN1_45S_50ntFlanks@10402.10419.18\|AACATGGGTCAGTCGGTC | >hg38_21_+_8433172_8446622_RNA45SN1_45S_50ntFlanks | 10402 | 10419 | 18 | AACATGGGTCAGTCGGTC | 463.0 | -7.2 | 1.3 | -5.4 | 7.1E-08 | 1.2E-07 | MB231/MB468 | MITO |
| >hg38_21_+_8433172_8446622_RNA45SN1_45S_50ntFlanks@12957.12982.26\|CTTCTGGGTCGGGGTTTCGTACGTAG | >hg38_21_+_8433172_8446622_RNA45SN1_45S_50ntFlanks | 12957 | 12982 | 26 | CTTCTGGGTCGGGGTTTCGTACGTAG | 1313.4 | -5.7 | 1.0 | -5.4 | 5.1E-08 | 1.2E-07 | BT20/MB468 | MITO |
| >hg38_21_+_8433172_8446622_RNA45SN1_45S_50ntFlanks@3851.3874.24\|ATAACTGTGGTAATTCTAGAGCTA | >hg38_21_+_8433172_8446622_RNA45SN1_45S_50ntFlanks | 3851 | 3874 | 24 | ATAACTGTGGTAATTCTAGAGCTA | 305.2 | -7.3 | 1.3 | -5.4 | 6.1E-08 | 1.2E-07 | MB231/MB468 | MP |
| >MI0000066\|hsa-let-7e&WithFlank&19\|+\|51692780\|51692870@14.34.21(+1C)&[MIMAT0000066&hsa-let-7e-5p&offsets\|0\|-1(+1C);m-50&19\|+\|51692793\|51692814&offsets\|0\|-1(+1C)]\|TGAGGTAGGAGGTTGTATAGTC | >MI0000066\|hsa-let-7e&WithFlank&19\|+\|51692780\|51692870 | 14 | 34 | 21(+1C) | TGAGGTAGGAGGTTGTATAGTC | 529.9 | 1.8 | 0.3 | 5.4 | 7.5E-08 | 1.2E-07 | BT20/MB468 | NUC |
| >hg38_1_-_228634819_228635039_RNA5S12_5S_50ntFlanks@129.169.41\|TTGGATGGGAGACCGCCTGGGAATACCGGGTGCTGTAGGCT | >hg38_1_-_228634819_228635039_RNA5S12_5S_50ntFlanks | 129 | 169 | 41 | TTGGATGGGAGACCGCCTGGGAATACCGGGTGCTGTAGGCT | 487.3 | 9.0 | 1.6 | 5.4 | 5.1E-08 | 1.2E-07 | BT20/MB468 | MITO |
| >hg38_21_+_8433172_8446622_RNA45SN1_45S_50ntFlanks@10683.10712.30\|CCGGCGGCGTCCGGTGAGCTCTCGCTGGCC | >hg38_21_+_8433172_8446622_RNA45SN1_45S_50ntFlanks | 10683 | 10712 | 30 | CCGGCGGCGTCCGGTGAGCTCTCGCTGGCC | 1793.0 | -3.8 | 0.7 | -5.4 | 5.1E-08 | 1.2E-07 | BT20/MB468 | MITO |
| >hg38_21_+_8433172_8446622_RNA45SN1_45S_50ntFlanks@12405.12445.41\|CTTTTTGATCCTTCGATGTCGGCTCTTCCTATCATTGTGAA | >hg38_21_+_8433172_8446622_RNA45SN1_45S_50ntFlanks | 12405 | 12445 | 41 | CTTTTTGATCCTTCGATGTCGGCTCTTCCTATCATTGTGAA | 820.5 | -4.9 | 0.9 | -5.4 | 6.8E-08 | 1.3E-07 | MB231/MB468 | NUC |
| >hg38_21_+_8433172_8446622_RNA45SN1_45S_50ntFlanks@12076.12095.20\|AGGGGCTCTCGCTTCTGGCG | >hg38_21_+_8433172_8446622_RNA45SN1_45S_50ntFlanks | 12076 | 12095 | 20 | AGGGGCTCTCGCTTCTGGCG | 8147.3 | -4.3 | 0.8 | -5.4 | 5.7E-08 | 1.3E-07 | BT20/MB468 | CYTO |
| >hg38_21_+_8433172_8446622_RNA45SN1_45S_50ntFlanks@3882.3899.18\|CCGACGGGCGCTGACCCC | >hg38_21_+_8433172_8446622_RNA45SN1_45S_50ntFlanks | 3882 | 3899 | 18 | CCGACGGGCGCTGACCCC | 1926.6 | -4.1 | 0.8 | -5.4 | 5.7E-08 | 1.3E-07 | BT20/MB468 | CYTO |
| >trnaMT_PheGAA_MT_+_577_647@36.57.22\|AAAATGTTTAGACGGGCTCACA | >trnaMT_PheGAA_MT_+_577_647 | 36 | 57 | 22 | AAAATGTTTAGACGGGCTCACA | 184.0 | 8.1 | 1.5 | 5.4 | 7.5E-08 | 1.3E-07 | BT20/MB468 | NUC |
| >hg38_21_+_8433172_8446622_RNA45SN1_45S_50ntFlanks@10391.10432.42\|AACAGCAGTTGAACATGGGTCAGTCGGTCCTGAGAGATGGGC | >hg38_21_+_8433172_8446622_RNA45SN1_45S_50ntFlanks | 10391 | 10432 | 42 | AACAGCAGTTGAACATGGGTCAGTCGGTCCTGAGAGATGGGC | 63.6 | -9.9 | 1.8 | -5.4 | 6.1E-08 | 1.3E-07 | MB231/MB468 | MP |
| >hg38_21_+_8433172_8446622_RNA45SN1_45S_50ntFlanks@5222.5244.23\|CTGCACGCGCGCTACACTGACTG | >hg38_21_+_8433172_8446622_RNA45SN1_45S_50ntFlanks | 5222 | 5244 | 23 | CTGCACGCGCGCTACACTGACTG | 446.3 | -10.9 | 2.0 | -5.4 | 5.1E-08 | 1.3E-07 | BT20/MB468 | MITO |
| >hg38_21_+_8433172_8446622_RNA45SN1_45S_50ntFlanks@12457.12475.19\|CAAGCGTTGGATTGTTCAC | >hg38_21_+_8433172_8446622_RNA45SN1_45S_50ntFlanks | 12457 | 12475 | 19 | CAAGCGTTGGATTGTTCAC | 401.7 | -5.4 | 1.0 | -5.4 | 5.1E-08 | 1.3E-07 | BT20/MB468 | MITO |
| >MI0000102\|hsa-mir-100&WithFlank&11\|-\|122152223\|122152314@19.37.19(+1U)&[MIMAT0000098&hsa-miR-100-5p&offsets\|0\|-3(+1U);m-27&11\|-\|122152276\|122152296&offsets\|0\|-2(+1U)]\|AACCCGTAGATCCGAACTTT | >MI0000102\|hsa-mir-100&WithFlank&11\|-\|122152223\|122152314 | 19 | 37 | 19(+1U) | AACCCGTAGATCCGAACTTT | 121.5 | -10.0 | 1.7 | -5.8 | 7.1E-09 | 1.3E-07 | BT20/MB231 | NUC |
| >MI0000087\|hsa-mir-29a&WithFlank&7\|-\|130876741\|130876816@47.69.23(+1C)&[MIMAT0000086&hsa-miR-29a-3p&offsets\|-1\|0(+1C);m-26&7\|-\|130876749\|130876769&offsets\|-1\|+1(+1C)]\|CTAGCACCATCTGAAATCGGTTAC | >MI0000087\|hsa-mir-29a&WithFlank&7\|-\|130876741\|130876816 | 47 | 69 | 23(+1C) | CTAGCACCATCTGAAATCGGTTAC | 81.9 | 9.6 | 1.8 | 5.4 | 6.2E-08 | 1.3E-07 | MB231/MB468 | MP |
| >hg38_21_+_8433172_8446622_RNA45SN1_45S_50ntFlanks@12715.12747.33\|AGCCTCGGTTGGCCTCGGATAGCCGGTCCCCCG | >hg38_21_+_8433172_8446622_RNA45SN1_45S_50ntFlanks | 12715 | 12747 | 33 | AGCCTCGGTTGGCCTCGGATAGCCGGTCCCCCG | 598.2 | -8.2 | 1.5 | -5.4 | 6.2E-08 | 1.3E-07 | MB231/MB468 | MP |
| >hg38_21_+_8433172_8446622_RNA45SN1_45S_50ntFlanks@10346.10376.31\|TCAAACGAGAACTTTGAAGGCCGAAGTGGAG | >hg38_21_+_8433172_8446622_RNA45SN1_45S_50ntFlanks | 10346 | 10376 | 31 | TCAAACGAGAACTTTGAAGGCCGAAGTGGAG | 234.5 | -5.5 | 1.0 | -5.4 | 7.2E-08 | 1.3E-07 | MB231/MB468 | MITO |
| >hg38_21_+_8433172_8446622_RNA45SN1_45S_50ntFlanks@11208.11232.25\|GCGGCGACTCTGGACGCGAGCCGGG | >hg38_21_+_8433172_8446622_RNA45SN1_45S_50ntFlanks | 11208 | 11232 | 25 | GCGGCGACTCTGGACGCGAGCCGGG | 8742.5 | -6.0 | 1.0 | -5.7 | 1.2E-08 | 1.3E-07 | BT20/MB468 | MP |
| >hg38_21_+_8433172_8446622_RNA45SN1_45S_50ntFlanks@9129.9148.20\|CGGGTCGGGGGGCGGGGCGG | >hg38_21_+_8433172_8446622_RNA45SN1_45S_50ntFlanks | 9129 | 9148 | 20 | CGGGTCGGGGGGCGGGGCGG | 150.3 | -10.3 | 1.8 | -5.7 | 1.2E-08 | 1.3E-07 | BT20/MB468 | MP |
| >hg38_21_+_8433172_8446622_RNA45SN1_45S_50ntFlanks@10802.10822.21\|TTGGAACAATGTAGGTAAGGG | >hg38_21_+_8433172_8446622_RNA45SN1_45S_50ntFlanks | 10802 | 10822 | 21 | TTGGAACAATGTAGGTAAGGG | 384.8 | -4.1 | 0.7 | -5.7 | 1.2E-08 | 1.3E-07 | BT20/MB468 | MP |
| >MI0000433\|hsa-let-7g&WithFlank&3\|-\|52268272\|52268367@11.29.19&[MIMAT0000414&hsa-let-7g-5p&offsets\|0\|-3;m-58&3\|-\|52268336\|52268357&offsets\|0\|-3]\|TGAGGTAGTAGTTTGTACA | >MI0000433\|hsa-let-7g&WithFlank&3\|-\|52268272\|52268367 | 11 | 29 | 19 | TGAGGTAGTAGTTTGTACA | 433.3 | 4.2 | 0.7 | 5.9 | 4.2E-09 | 1.3E-07 | BT20/MB231 | CYTO |
| >hg38_21_+_8433172_8446622_RNA45SN1_45S_50ntFlanks@10421.10456.36\|TGAGAGATGGGCGAGCGCCGTTCCGAAGGGACGGGC | >hg38_21_+_8433172_8446622_RNA45SN1_45S_50ntFlanks | 10421 | 10456 | 36 | TGAGAGATGGGCGAGCGCCGTTCCGAAGGGACGGGC | 198.5 | -7.3 | 1.4 | -5.4 | 6.2E-08 | 1.3E-07 | MB231/MB468 | MP |
| >hg38_21_+_8433172_8446622_RNA45SN1_45S_50ntFlanks@11573.11591.19\|CGACTTAGAACTGGTGCGG | >hg38_21_+_8433172_8446622_RNA45SN1_45S_50ntFlanks | 11573 | 11591 | 19 | CGACTTAGAACTGGTGCGG | 569.3 | -8.3 | 1.5 | -5.4 | 7.6E-08 | 1.3E-07 | BT20/MB468 | NUC |
| >MI0000760\|hsa-mir-361&WithFlank&X\|-\|85903630\|85903713@51.73.23&[MIMAT0004682&hsa-miR-361-3p&offsets\|0\|0;m-124&X\|-\|85903642\|85903663&offsets\|0\|+1]\|TCCCCCAGGTGTGATTCTGATTT | >MI0000760\|hsa-mir-361&WithFlank&X\|-\|85903630\|85903713 | 51 | 73 | 23 | TCCCCCAGGTGTGATTCTGATTT | 426.2 | 6.4 | 1.2 | 5.4 | 6.7E-08 | 1.3E-07 | MB231/MB468 | CYTO |
| >trnaMT_SerGCT_MT_+_12207_12265@8.31.24\|CTCACAAGAACTGCTAACTCATGC | >trnaMT_SerGCT_MT_+_12207_12265 | 8 | 31 | 24 | CTCACAAGAACTGCTAACTCATGC | 425.3 | -7.0 | 1.3 | -5.4 | 6.9E-08 | 1.3E-07 | MB231/MB468 | NUC |
| >hg38_21_+_8433172_8446622_RNA45SN1_45S_50ntFlanks@10677.10710.34\|GCGGTTCCGGCGGCGTCCGGTGAGCTCTCGCTGG | >hg38_21_+_8433172_8446622_RNA45SN1_45S_50ntFlanks | 10677 | 10710 | 34 | GCGGTTCCGGCGGCGTCCGGTGAGCTCTCGCTGG | 3678.1 | -4.2 | 0.8 | -5.4 | 7.3E-08 | 1.3E-07 | MB231/MB468 | MITO |
| >MI0000650\|hsa-mir-200c&WithFlank&12\|+\|6963693\|6963772@50.72.23(+2U)&[MIMAT0000617&hsa-miR-200c-3p&offsets\|0\|0(+2U);m-54&12\|+\|6963742\|6963764&offsets\|0\|0(+2U)]\|TAATACTGCCGGGTAATGATGGATT | >MI0000650\|hsa-mir-200c&WithFlank&12\|+\|6963693\|6963772 | 50 | 72 | 23(+2U) | TAATACTGCCGGGTAATGATGGATT | 589.7 | 8.0 | 1.5 | 5.4 | 5.2E-08 | 1.3E-07 | BT20/MB468 | MITO |
| >am_trna119_LysCTT_1_-_145395522_145395594@13.35.23,trna11_LysCTT_5_-_180648979_180649051@13.35.23,trna128_LysCTT_1_-_55423542_55423614@13.35.23,trna13_LysCTT_6_+_26556774_26556846@13.35.23,trna6_LysCTT_19_-_52425396_52425468@13.35.23,trna7_LysCTT_16_+_3225692_3225764@13.35.23,trna9_LysCTT_5_+_180634755_180634827@13.35.23\|CAGTCGGTAGAGCATGAGACTCT | >am_trna119_LysCTT_1_-_145395522_145395594 | 13 | 35 | 23 | CAGTCGGTAGAGCATGAGACTCT | 654.2 | 6.9 | 1.3 | 5.4 | 7.3E-08 | 1.3E-07 | MB231/MB468 | MITO |
| >trna106_HisGTG_1_-_149155828_149155899@13.31.19,trna111_HisGTG_1_-_147774845_147774916@13.31.19,trna118_HisGTG_1_-_145396881_145396952@13.31.19,trna16_HisGTG_1_+_146544773_146544844@13.31.19,trna1_HisGTG_15_+_45493349_45493420@13.31.19,trna21_HisGTG_1_+_147753471_147753542@13.31.19,trna33_HisGTG_6_+_27125906_27125977@13.31.19,trna7_HisGTG_9_-_14433938_14434009@13.31.19,trna8_HisGTG_15_-_45492611_45492682@13.31.19,trna9_HisGTG_15_-_45490804_45490875@13.31.19\|TAGTGGTTAGTACTCTGCG | >trna106_HisGTG_1_-_149155828_149155899 | 13 | 31 | 19 | TAGTGGTTAGTACTCTGCG | 187.4 | -4.4 | 0.8 | -5.4 | 6.7E-08 | 1.3E-07 | MB231/MB468 | CYTO |
| >hg38_21_+_8433172_8446622_RNA45SN1_45S_50ntFlanks@10349.10383.35\|AACGAGAACTTTGAAGGCCGAAGTGGAGAAGGGTT | >hg38_21_+_8433172_8446622_RNA45SN1_45S_50ntFlanks | 10349 | 10383 | 35 | AACGAGAACTTTGAAGGCCGAAGTGGAGAAGGGTT | 4593.0 | -4.5 | 0.8 | -5.4 | 7.3E-08 | 1.3E-07 | MB231/MB468 | MITO |
| >hg38_21_+_8433172_8446622_RNA45SN1_45S_50ntFlanks@4449.4471.23\|CCTCTCGGCGCCCCCTCGATGCT | >hg38_21_+_8433172_8446622_RNA45SN1_45S_50ntFlanks | 4449 | 4471 | 23 | CCTCTCGGCGCCCCCTCGATGCT | 1013.8 | -6.0 | 1.1 | -5.4 | 6.7E-08 | 1.3E-07 | MB231/MB468 | CYTO |
| >hg38_21_+_8433172_8446622_RNA45SN1_45S_50ntFlanks@12404.12435.32\|GCTTTTTGATCCTTCGATGTCGGCTCTTCCTA | >hg38_21_+_8433172_8446622_RNA45SN1_45S_50ntFlanks | 12404 | 12435 | 32 | GCTTTTTGATCCTTCGATGTCGGCTCTTCCTA | 1183.3 | -9.1 | 1.7 | -5.4 | 7.6E-08 | 1.3E-07 | BT20/MB468 | NUC |
| >hg38_21_+_8433172_8446622_RNA45SN1_45S_50ntFlanks@4917.4941.25\|CACCACCAGGAGTGGAGCCTGCGGC | >hg38_21_+_8433172_8446622_RNA45SN1_45S_50ntFlanks | 4917 | 4941 | 25 | CACCACCAGGAGTGGAGCCTGCGGC | 362.4 | -7.4 | 1.4 | -5.4 | 6.2E-08 | 1.3E-07 | MB231/MB468 | MP |
| >hg38_21_+_8433172_8446622_RNA45SN1_45S_50ntFlanks@9029.9064.36\|TCTCCTCGTGGGGGGGCCGGGCCACCCCTCCCACGG | >hg38_21_+_8433172_8446622_RNA45SN1_45S_50ntFlanks | 9029 | 9064 | 36 | TCTCCTCGTGGGGGGGCCGGGCCACCCCTCCCACGG | 88.6 | -9.6 | 1.7 | -5.7 | 1.2E-08 | 1.3E-07 | BT20/MB468 | MP |
| >hg38_21_+_8433172_8446622_RNA45SN1_45S_50ntFlanks@7973.7995.23\|GACGCGACCTCAGATCAGACGTG | >hg38_21_+_8433172_8446622_RNA45SN1_45S_50ntFlanks | 7973 | 7995 | 23 | GACGCGACCTCAGATCAGACGTG | 559.4 | 2.5 | 0.5 | 5.4 | 6.9E-08 | 1.3E-07 | MB231/MB468 | NUC |
| >hg38_21_+_8433172_8446622_RNA45SN1_45S_50ntFlanks@12633.12659.27\|CGAAGCTACCATCTGTGGGATTATGAC | >hg38_21_+_8433172_8446622_RNA45SN1_45S_50ntFlanks | 12633 | 12659 | 27 | CGAAGCTACCATCTGTGGGATTATGAC | 3976.6 | -4.5 | 0.8 | -5.4 | 5.2E-08 | 1.3E-07 | BT20/MB468 | MITO |
| >hg38_21_+_8433172_8446622_RNA45SN1_45S_50ntFlanks@12081.12102.22\|CTCTCGCTTCTGGCGCCAAGCG | >hg38_21_+_8433172_8446622_RNA45SN1_45S_50ntFlanks | 12081 | 12102 | 22 | CTCTCGCTTCTGGCGCCAAGCG | 199.3 | -6.7 | 1.2 | -5.4 | 5.8E-08 | 1.3E-07 | BT20/MB468 | CYTO |
| >hg38_21_+_8433172_8446622_RNA45SN1_45S_50ntFlanks@6652.6688.37\|GACTCTTAGCGGTGGATCACTCGGCTCGTGCGTCGAT | >hg38_21_+_8433172_8446622_RNA45SN1_45S_50ntFlanks | 6652 | 6688 | 37 | GACTCTTAGCGGTGGATCACTCGGCTCGTGCGTCGAT | 1250.3 | 4.3 | 0.8 | 5.4 | 5.8E-08 | 1.3E-07 | BT20/MB468 | CYTO |
| >hg38_21_+_8433172_8446622_RNA45SN1_45S_50ntFlanks@12072.12096.25\|CCCGAGGGGCTCTCGCTTCTGGCGC | >hg38_21_+_8433172_8446622_RNA45SN1_45S_50ntFlanks | 12072 | 12096 | 25 | CCCGAGGGGCTCTCGCTTCTGGCGC | 836.0 | -4.2 | 0.8 | -5.4 | 6.8E-08 | 1.3E-07 | MB231/MB468 | CYTO |

<table>
<tr><td>>hg38_21_+_8433172_8446622_RNA45SN1_45S_50ntFlanks@4069.4095.27|CTTTCGATGGTAGTCGCCGTGCCTACC</td><td>>hg38_21_+_8433172_8446622_RNA45SN1_45S_50ntFlanks</td><td>4069</td><td>4095</td><td>27</td><td>CTTTCGATGGTAGTCGCCGTGCCTACC</td><td>228.5</td><td>-4.6</td><td>0.8</td><td>-5.4</td><td>5.9E-08</td><td>1.3E-07</td><td>BT20/MB468</td><td>CYTO</td></tr>
<tr><td>>hg38_21_+_8433172_8446622_RNA45SN1_45S_50ntFlanks@10805.10840.36|GAACAATGTAGGTAAGGGAAGTCGGCAAGCCGGATC</td><td>>hg38_21_+_8433172_8446622_RNA45SN1_45S_50ntFlanks</td><td>10805</td><td>10840</td><td>36</td><td>GAACAATGTAGGTAAGGGAAGTCGGCAAGCCGGATC</td><td>207.6</td><td>-4.3</td><td>0.8</td><td>-5.4</td><td>5.9E-08</td><td>1.3E-07</td><td>BT20/MB468</td><td>CYTO</td></tr>
<tr><td>>hg38_21_+_8433172_8446622_RNA45SN1_45S_50ntFlanks@9574.9610.37|AATCGGTCGTCCGACCTGGGTATAGGGGCGAAAGACT</td><td>>hg38_21_+_8433172_8446622_RNA45SN1_45S_50ntFlanks</td><td>9574</td><td>9610</td><td>37</td><td>AATCGGTCGTCCGACCTGGGTATAGGGGCGAAAGACT</td><td>260.2</td><td>5.1</td><td>1.0</td><td>5.4</td><td>7.0E-08</td><td>1.3E-07</td><td>MB231/MB468</td><td>NUC</td></tr>
<tr><td>>MI0000746|hsa-mir-99b&WithFlank&19|+|51692606|51692687@13.34.22(+1U)&[MIMAT0000689&hsa-miR-99b-5p&offsets|0|0(+1U);m-10&19|+|51692618|51692639&offsets|0|0(+1U)]|CACCCGTAGAACCGACCTTGCGT</td><td>>MI0000746|hsa-mir-99b&WithFlank&19|+|51692606|51692687</td><td>13</td><td>34</td><td>22(+1U)</td><td>CACCCGTAGAACCGACCTTGCGT</td><td>653.7</td><td>2.0</td><td>0.4</td><td>5.4</td><td>7.7E-08</td><td>1.3E-07</td><td>BT20/MB468</td><td>NUC</td></tr>
<tr><td>>MI0000095|hsa-mir-93&WithFlank&7|-|100093762|100093853@17.39.23&[MIMAT0000093&hsa-miR-93-5p&offsets|0|0;m-29&7|-|100093815|100093837&offsets|0|0]|CAAAGTGCTGTTCGTGCAGGTAG</td><td>>MI0000095|hsa-mir-93&WithFlank&7|-|100093762|100093853</td><td>17</td><td>39</td><td>23</td><td>CAAAGTGCTGTTCGTGCAGGTAG</td><td>8321.2</td><td>4.7</td><td>0.9</td><td>5.4</td><td>7.4E-08</td><td>1.3E-07</td><td>MB231/MB468</td><td>MITO</td></tr>
<tr><td>>hg38_21_+_8433172_8446622_RNA45SN1_45S_50ntFlanks@12444.12485.42|AAGCAGAATTCACCAAGCGTTGGATTGTTCACCCACTAATAG</td><td>>hg38_21_+_8433172_8446622_RNA45SN1_45S_50ntFlanks</td><td>12444</td><td>12485</td><td>42</td><td>AAGCAGAATTCACCAAGCGTTGGATTGTTCACCCACTAATAG</td><td>127.8</td><td>-9.1</td><td>1.7</td><td>-5.4</td><td>5.3E-08</td><td>1.3E-07</td><td>BT20/MB468</td><td>MITO</td></tr>
<tr><td>>MI0005757|hsa-mir-935&WithFlank&19|+|53982301|53982403@63.84.22&[MIMAT0004978&hsa-miR-935&offsets|+1|0;m-549&19|+|53982363|53982384&offsets|0|0]|CAGTTACCGCTTCCGCTACCGC</td><td>>MI0005757|hsa-mir-935&WithFlank&19|+|53982301|53982403</td><td>63</td><td>84</td><td>22</td><td>CAGTTACCGCTTCCGCTACCGC</td><td>119.2</td><td>10.6</td><td>1.9</td><td>5.7</td><td>1.2E-08</td><td>1.3E-07</td><td>BT20/MB468</td><td>MP</td></tr>
<tr><td>>hg38_21_+_8433172_8446622_RNA45SN1_45S_50ntFlanks@5186.5207.22|CAATAACAGGTCTGTGATGCCC</td><td>>hg38_21_+_8433172_8446622_RNA45SN1_45S_50ntFlanks</td><td>5186</td><td>5207</td><td>22</td><td>CAATAACAGGTCTGTGATGCCC</td><td>143.2</td><td>-10.2</td><td>1.8</td><td>-5.7</td><td>1.2E-08</td><td>1.3E-07</td><td>BT20/MB468</td><td>MP</td></tr>
<tr><td>>hg38_21_+_8433172_8446622_RNA45SN1_45S_50ntFlanks@6648.6693.46|GTACGACTCTTAGCGGTGGATCACTCGGCTCGTGCGTCGATGAAGA</td><td>>hg38_21_+_8433172_8446622_RNA45SN1_45S_50ntFlanks</td><td>6648</td><td>6693</td><td>46</td><td>GTACGACTCTTAGCGGTGGATCACTCGGCTCGTGCGTCGATGAAGA</td><td>1101.9</td><td>4.3</td><td>0.8</td><td>5.4</td><td>5.3E-08</td><td>1.3E-07</td><td>BT20/MB468</td><td>MITO</td></tr>
<tr><td>>hg38_21_+_8433172_8446622_RNA45SN1_45S_50ntFlanks@11316.11336.21|CACCCCACGTCTCGTCGCGCG</td><td>>hg38_21_+_8433172_8446622_RNA45SN1_45S_50ntFlanks</td><td>11316</td><td>11336</td><td>21</td><td>CACCCCACGTCTCGTCGCGCG</td><td>2034.5</td><td>-5.9</td><td>1.1</td><td>-5.4</td><td>5.9E-08</td><td>1.3E-07</td><td>BT20/MB468</td><td>CYTO</td></tr>
<tr><td>>hg38_21_+_8433172_8446622_RNA45SN1_45S_50ntFlanks@11598.11623.26|GGAATCCGACTGTTTAATTAAAACAA</td><td>>hg38_21_+_8433172_8446622_RNA45SN1_45S_50ntFlanks</td><td>11598</td><td>11623</td><td>26</td><td>GGAATCCGACTGTTTAATTAAAACAA</td><td>105.6</td><td>10.5</td><td>1.8</td><td>5.7</td><td>1.3E-08</td><td>1.3E-07</td><td>BT20/MB468</td><td>MP</td></tr>
<tr><td>>trnaMT_GluTTC_MT_-_14674_14742@40.72.33,trnalookalike8_GluTTC_5_-_93905172_93905240@40.72.33|TTGGTCGTGGTTGTAGTCCGTGCGAGAATACCA</td><td>>trnaMT_GluTTC_MT_-_14674_14742</td><td>40</td><td>72</td><td>33</td><td>TTGGTCGTGGTTGTAGTCCGTGCGAGAATACCA</td><td>2509.6</td><td>5.1</td><td>0.9</td><td>5.4</td><td>5.9E-08</td><td>1.3E-07</td><td>BT20/MB468</td><td>CYTO</td></tr>
<tr><td>>MI0000076|hsa-mir-20a&WithFlank&13|+|91351059|91351141@14.37.24&[MIMAT0000075&hsa-miR-20a-5p&offsets|0|+1;m-96&13|+|91351072|91351094&offsets|0|+1]|TAAAGTGCTTATAGTGCAGGTAGT</td><td>>MI0000076|hsa-mir-20a&WithFlank&13|+|91351059|91351141</td><td>14</td><td>37</td><td>24</td><td>TAAAGTGCTTATAGTGCAGGTAGT</td><td>2825.2</td><td>-5.8</td><td>1.0</td><td>-5.8</td><td>5.5E-09</td><td>1.3E-07</td><td>BT20/MB231</td><td>MITO</td></tr>
<tr><td>>hg38_21_+_8433172_8446622_RNA45SN1_45S_50ntFlanks@4045.4067.23|CCATTCGAACGTCTGCCCTATCA</td><td>>hg38_21_+_8433172_8446622_RNA45SN1_45S_50ntFlanks</td><td>4045</td><td>4067</td><td>23</td><td>CCATTCGAACGTCTGCCCTATCA</td><td>356.4</td><td>-6.3</td><td>1.2</td><td>-5.4</td><td>7.4E-08</td><td>1.3E-07</td><td>MB231/MB468</td><td>MITO</td></tr>
<tr><td>>hg38_21_+_8433172_8446622_RNA45SN1_45S_50ntFlanks@12937.12955.19|AACCATTCGTAGACGACCT</td><td>>hg38_21_+_8433172_8446622_RNA45SN1_45S_50ntFlanks</td><td>12937</td><td>12955</td><td>19</td><td>AACCATTCGTAGACGACCT</td><td>258.5</td><td>-7.1</td><td>1.3</td><td>-5.4</td><td>7.5E-08</td><td>1.3E-07</td><td>MB231/MB468</td><td>MITO</td></tr>
<tr><td>>hg38_21_+_8433172_8446622_RNA45SN1_45S_50ntFlanks@9562.9594.33|TCCTGACGTGCAAATCGGTCGTCCGACCTGGGT</td><td>>hg38_21_+_8433172_8446622_RNA45SN1_45S_50ntFlanks</td><td>9562</td><td>9594</td><td>33</td><td>TCCTGACGTGCAAATCGGTCGTCCGACCTGGGT</td><td>267.6</td><td>-7.1</td><td>1.3</td><td>-5.4</td><td>7.5E-08</td><td>1.3E-07</td><td>MB231/MB468</td><td>MITO</td></tr>
<tr><td>>hg38_21_+_8433172_8446622_RNA45SN1_45S_50ntFlanks@11662.11685.24|GATTTCTGCCCAGTGCTCTGAATG</td><td>>hg38_21_+_8433172_8446622_RNA45SN1_45S_50ntFlanks</td><td>11662</td><td>11685</td><td>24</td><td>GATTTCTGCCCAGTGCTCTGAATG</td><td>175.5</td><td>-2.5</td><td>0.5</td><td>-5.4</td><td>7.0E-08</td><td>1.3E-07</td><td>MB231/MB468</td><td>NUC</td></tr>
<tr><td>>hg38_21_+_8433172_8446622_RNA45SN1_45S_50ntFlanks@12960.12982.23|CTGGGTCGGGGTTTCGTACGTAG</td><td>>hg38_21_+_8433172_8446622_RNA45SN1_45S_50ntFlanks</td><td>12960</td><td>12982</td><td>23</td><td>CTGGGTCGGGGTTTCGTACGTAG</td><td>241.2</td><td>-4.2</td><td>0.8</td><td>-5.4</td><td>6.9E-08</td><td>1.3E-07</td><td>MB231/MB468</td><td>CYTO</td></tr>
<tr><td>>hg38_21_+_8433172_8446622_RNA45SN1_45S_50ntFlanks@7985.8012.28|GATCAGACGTGGCGACCCGCTGAATTTA</td><td>>hg38_21_+_8433172_8446622_RNA45SN1_45S_50ntFlanks</td><td>7985</td><td>8012</td><td>28</td><td>GATCAGACGTGGCGACCCGCTGAATTTA</td><td>1158.4</td><td>-7.8</td><td>1.5</td><td>-5.4</td><td>7.5E-08</td><td>1.3E-07</td><td>MB231/MB468</td><td>MITO</td></tr>
<tr><td>>hg38_21_+_8433172_8446622_RNA45SN1_45S_50ntFlanks@7975.8007.33|CGCGACCTCAGATCAGACGTGGCGACCCGCTGA</td><td>>hg38_21_+_8433172_8446622_RNA45SN1_45S_50ntFlanks</td><td>7975</td><td>8007</td><td>33</td><td>CGCGACCTCAGATCAGACGTGGCGACCCGCTGA</td><td>24586.0</td><td>5.8</td><td>1.1</td><td>5.4</td><td>6.0E-08</td><td>1.3E-07</td><td>BT20/MB468</td><td>CYTO</td></tr>
<tr><td>>hg38_21_+_8433172_8446622_RNA45SN1_45S_50ntFlanks@11570.11603.34|AGCCGACTTAGAACTGGTGCGGACCAGGGGAATC</td><td>>hg38_21_+_8433172_8446622_RNA45SN1_45S_50ntFlanks</td><td>11570</td><td>11603</td><td>34</td><td>AGCCGACTTAGAACTGGTGCGGACCAGGGGAATC</td><td>394.6</td><td>-2.2</td><td>0.4</td><td>-5.4</td><td>6.0E-08</td><td>1.3E-07</td><td>BT20/MB468</td><td>CYTO</td></tr>
<tr><td>>hg38_21_+_8433172_8446622_RNA45SN1_45S_50ntFlanks@10926.10946.21|ACGAGGCGCCGCCGCCCCCCC</td><td>>hg38_21_+_8433172_8446622_RNA45SN1_45S_50ntFlanks</td><td>10926</td><td>10946</td><td>21</td><td>ACGAGGCGCCGCCGCCCCCCC</td><td>80.3</td><td>-8.4</td><td>1.5</td><td>-5.4</td><td>5.3E-08</td><td>1.3E-07</td><td>BT20/MB468</td><td>MITO</td></tr>
<tr><td>>hg38_21_+_8433172_8446622_RNA45SN1_45S_50ntFlanks@12366.12404.39|TAACTGGCTTGTGGCGGCCAAGCGTTCATAGCGACGTCG</td><td>>hg38_21_+_8433172_8446622_RNA45SN1_45S_50ntFlanks</td><td>12366</td><td>12404</td><td>39</td><td>TAACTGGCTTGTGGCGGCCAAGCGTTCATAGCGACGTCG</td><td>54.6</td><td>-9.6</td><td>1.8</td><td>-5.4</td><td>6.4E-08</td><td>1.3E-07</td><td>MB231/MB468</td><td>MP</td></tr>
<tr><td>>hg38_21_+_8433172_8446622_RNA45SN1_45S_50ntFlanks@9955.9982.28|GGAAGTCGGAATCCGCTAAGGAGTGTGT</td><td>>hg38_21_+_8433172_8446622_RNA45SN1_45S_50ntFlanks</td><td>9955</td><td>9982</td><td>28</td><td>GGAAGTCGGAATCCGCTAAGGAGTGTGT</td><td>93.6</td><td>10.3</td><td>1.8</td><td>5.7</td><td>1.3E-08</td><td>1.3E-07</td><td>BT20/MB468</td><td>MP</td></tr>
<tr><td>>hg38_21_+_8433172_8446622_RNA45SN1_45S_50ntFlanks@11683.11706.24|ATGTCAAAGTGAAGAAATTCAATG</td><td>>hg38_21_+_8433172_8446622_RNA45SN1_45S_50ntFlanks</td><td>11683</td><td>11706</td><td>24</td><td>ATGTCAAAGTGAAGAAATTCAATG</td><td>95.1</td><td>-9.7</td><td>1.7</td><td>-5.7</td><td>1.3E-08</td><td>1.3E-07</td><td>BT20/MB468</td><td>MP</td></tr>
<tr><td>>hg38_21_+_8433172_8446622_RNA45SN1_45S_50ntFlanks@4002.4020.19|GATAACCTCGGGCCGATCG</td><td>>hg38_21_+_8433172_8446622_RNA45SN1_45S_50ntFlanks</td><td>4002</td><td>4020</td><td>19</td><td>GATAACCTCGGGCCGATCG</td><td>348.2</td><td>-4.8</td><td>0.9</td><td>-5.4</td><td>6.0E-08</td><td>1.3E-07</td><td>BT20/MB468</td><td>CYTO</td></tr>
<tr><td>>hg38_21_+_8433172_8446622_RNA45SN1_45S_50ntFlanks@6725.6744.20|GCAGGACACATTGATCATCG</td><td>>hg38_21_+_8433172_8446622_RNA45SN1_45S_50ntFlanks</td><td>6725</td><td>6744</td><td>20</td><td>GCAGGACACATTGATCATCG</td><td>915.4</td><td>-8.8</td><td>1.6</td><td>-5.4</td><td>7.9E-08</td><td>1.3E-07</td><td>BT20/MB468</td><td>NUC</td></tr>
<tr><td>>MI0000102|hsa-mir-100&WithFlank&11|-|122152223|122152314@19.39.21(+1A)&[MIMAT0000098&hsa-miR-100-5p&offsets|0|-1(+1A);m-27&11|-|122152276|122152296&offsets|0|0(+1A)]|AACCCGTAGATCCGAACTTGTA</td><td>>MI0000102|hsa-mir-100&WithFlank&11|-|122152223|122152314</td><td>19</td><td>39</td><td>21(+1A)</td><td>AACCCGTAGATCCGAACTTGTA</td><td>48707.8</td><td>-5.5</td><td>0.9</td><td>-5.8</td><td>7.4E-09</td><td>1.3E-07</td><td>BT20/MB231</td><td>NUC</td></tr>
<tr><td>>hg38_21_+_8433172_8446622_RNA45SN1_45S_50ntFlanks@11561.11602.42|GCGCCTAGCAGCCGACTTAGAACTGGTGCGGACCAGGGGAAT</td><td>>hg38_21_+_8433172_8446622_RNA45SN1_45S_50ntFlanks</td><td>11561</td><td>11602</td><td>42</td><td>GCGCCTAGCAGCCGACTTAGAACTGGTGCGGACCAGGGGAAT</td><td>2266.9</td><td>-6.7</td><td>1.3</td><td>-5.4</td><td>7.1E-08</td><td>1.3E-07</td><td>MB231/MB468</td><td>NUC</td></tr>
<tr><td>>hg38_21_+_8433172_8446622_RNA45SN1_45S_50ntFlanks@12478.12497.20|ACTAATAGGGAACGTGAGCT</td><td>>hg38_21_+_8433172_8446622_RNA45SN1_45S_50ntFlanks</td><td>12478</td><td>12497</td><td>20</td><td>ACTAATAGGGAACGTGAGCT</td><td>2383.8</td><td>3.0</td><td>0.6</td><td>5.4</td><td>6.9E-08</td><td>1.3E-07</td><td>MB231/MB468</td><td>CYTO</td></tr>
<tr><td>>hg38_21_+_8433172_8446622_RNA45SN1_45S_50ntFlanks@12502.12521.20|TTAGACCGTCGTGAGACAGG</td><td>>hg38_21_+_8433172_8446622_RNA45SN1_45S_50ntFlanks</td><td>12502</td><td>12521</td><td>20</td><td>TTAGACCGTCGTGAGACAGG</td><td>56.9</td><td>-9.7</td><td>1.8</td><td>-5.4</td><td>6.4E-08</td><td>1.3E-07</td><td>MB231/MB468</td><td>MP</td></tr>
<tr><td>>hg38_21_+_8433172_8446622_RNA45SN1_45S_50ntFlanks@4722.4748.27|TCATTAATCAAGAACGAAAGTCGGAGG</td><td>>hg38_21_+_8433172_8446622_RNA45SN1_45S_50ntFlanks</td><td>4722</td><td>4748</td><td>27</td><td>TCATTAATCAAGAACGAAAGTCGGAGG</td><td>400.7</td><td>-6.7</td><td>1.2</td><td>-5.4</td><td>5.4E-08</td><td>1.3E-07</td><td>BT20/MB468</td><td>MITO</td></tr>
<tr><td>>hg38_21_+_8433172_8446622_RNA45SN1_45S_50ntFlanks@12415.12435.21|CTTCGATGTCGGCTCTTCCTA</td><td>>hg38_21_+_8433172_8446622_RNA45SN1_45S_50ntFlanks</td><td>12415</td><td>12435</td><td>21</td><td>CTTCGATGTCGGCTCTTCCTA</td><td>11566.4</td><td>-6.6</td><td>1.2</td><td>-5.4</td><td>7.9E-08</td><td>1.3E-07</td><td>BT20/MB468</td><td>NUC</td></tr>
<tr><td>>hg38_21_+_8433172_8446622_RNA45SN1_45S_50ntFlanks@12412.12431.20|ATCCTTCGATGTCGGCTCTT</td><td>>hg38_21_+_8433172_8446622_RNA45SN1_45S_50ntFlanks</td><td>12412</td><td>12431</td><td>20</td><td>ATCCTTCGATGTCGGCTCTT</td><td>78.4</td><td>-9.4</td><td>1.6</td><td>-5.7</td><td>1.3E-08</td><td>1.3E-07</td><td>BT20/MB468</td><td>MP</td></tr>
<tr><td>>hg38_21_+_8433172_8446622_RNA45SN1_45S_50ntFlanks@11729.11758.30|TAACTATGACTCTCTTAAGGTAGCCAAATG</td><td>>hg38_21_+_8433172_8446622_RNA45SN1_45S_50ntFlanks</td><td>11729</td><td>11758</td><td>30</td><td>TAACTATGACTCTCTTAAGGTAGCCAAATG</td><td>144.5</td><td>-10.3</td><td>1.8</td><td>-5.7</td><td>1.3E-08</td><td>1.3E-07</td><td>BT20/MB468</td><td>MP</td></tr>
<tr><td>>hg38_21_+_8433172_8446622_RNA45SN1_45S_50ntFlanks@6690.6720.31|AAGAACGCAGCTAGCTGCGAGAATTAATGTG</td><td>>hg38_21_+_8433172_8446622_RNA45SN1_45S_50ntFlanks</td><td>6690</td><td>6720</td><td>31</td><td>AAGAACGCAGCTAGCTGCGAGAATTAATGTG</td><td>2453.5</td><td>-7.9</td><td>1.5</td><td>-5.4</td><td>6.0E-08</td><td>1.3E-07</td><td>BT20/MB468</td><td>CYTO</td></tr>
<tr><td>>hg38_21_+_8433172_8446622_RNA45SN1_45S_50ntFlanks@6646.6673.28|TCGTACGACTCTTAGCGGTGGATCACTC</td><td>>hg38_21_+_8433172_8446622_RNA45SN1_45S_50ntFlanks</td><td>6646</td><td>6673</td><td>28</td><td>TCGTACGACTCTTAGCGGTGGATCACTC</td><td>796.4</td><td>4.3</td><td>0.8</td><td>5.4</td><td>7.0E-08</td><td>1.3E-07</td><td>MB231/MB468</td><td>CYTO</td></tr>
<tr><td>>hg38_21_+_8433172_8446622_RNA45SN1_45S_50ntFlanks@11661.11681.21|TGATTTCTGCCCAGTGCTCTG</td><td>>hg38_21_+_8433172_8446622_RNA45SN1_45S_50ntFlanks</td><td>11661</td><td>11681</td><td>21</td><td>TGATTTCTGCCCAGTGCTCTG</td><td>5971.5</td><td>-6.2</td><td>1.2</td><td>-5.4</td><td>7.0E-08</td><td>1.3E-07</td><td>MB231/MB468</td><td>CYTO</td></tr>
<tr><td>>hg38_21_+_8433172_8446622_RNA45SN1_45S_50ntFlanks@6663.6686.24|GTGGATCACTCGGCTCGTGCGTCG</td><td>>hg38_21_+_8433172_8446622_RNA45SN1_45S_50ntFlanks</td><td>6663</td><td>6686</td><td>24</td><td>GTGGATCACTCGGCTCGTGCGTCG</td><td>560.0</td><td>-3.4</td><td>0.6</td><td>-5.4</td><td>7.2E-08</td><td>1.3E-07</td><td>MB231/MB468</td><td>NUC</td></tr>
<tr><td>>MI0003188|hsa-mir-503&WithFlank&X|-|134546322|134546404@12.32.21&[MIMAT0002874&hsa-miR-503-5p&offsets|0|-2;m-297&X|-|134546371|134546393&offsets|0|-2]|TAGCAGCGGGAACAGTTCTGC</td><td>>MI0003188|hsa-mir-503&WithFlank&X|-|134546322|134546404</td><td>12</td><td>32</td><td>21</td><td>TAGCAGCGGGAACAGTTCTGC</td><td>62.5</td><td>9.2</td><td>1.7</td><td>5.4</td><td>6.5E-08</td><td>1.3E-07</td><td>MB231/MB468</td><td>MP</td></tr>
<tr><td>>MI0000081|hsa-mir-24-2&WithFlank&19|-|13836281|13836365@56.75.20;MI0000080|hsa-mir-24-1&WithFlank&9|+|95086015|95086094@50.69.20&[MIMAT0000080&hsa-miR-24-3p&offsets|0|-2;m-36&19|-|13836288|13836310&offsets|0|-3];[MIMAT0000080_1&hsa-miR-24-3p&offsets|0|-2;m-35&9|+|95086064|95086086&offsets|0|-3]|TGGCTCAGTTCAGCAGGAAC</td><td>>MI0000081|hsa-mir-24-2&WithFlank&19|-|13836281|13836365</td><td>56</td><td>75</td><td>20</td><td>TGGCTCAGTTCAGCAGGAAC</td><td>12457.9</td><td>4.7</td><td>0.9</td><td>5.4</td><td>7.6E-08</td><td>1.3E-07</td><td>MB231/MB468</td><td>MITO</td></tr>
<tr><td>>hg38_21_+_8433172_8446622_RNA45SN1_45S_50ntFlanks@12008.12033.26|CCGCCGGTGAAATACCACTACTCTGA</td><td>>hg38_21_+_8433172_8446622_RNA45SN1_45S_50ntFlanks</td><td>12008</td><td>12033</td><td>26</td><td>CCGCCGGTGAAATACCACTACTCTGA</td><td>688.7</td><td>-8.1</td><td>1.4</td><td>-5.7</td><td>1.3E-08</td><td>1.3E-07</td><td>BT20/MB468</td><td>MP</td></tr>
<tr><td>>MI0000070|hsa-mir-16-1&WithFlank&13|-|50048967|50049067@20.40.21;MI0000115|hsa-mir-16-2&WithFlank&3|+|160404739|160404831@16.36.21&[MIMAT0000069&hsa-miR-16-5p&offsets|0|-1;m-62&13|-|50049027|50049048&offsets|0|-1];[MIMAT0000069_1&hsa-miR-16-5p&offsets|0|-1;m-61&3|+|160404754|160404775&offsets|0|-1]|TAGCAGCACGTAAATATTGGC</td><td>>MI0000070|hsa-mir-16-1&WithFlank&13|-|50048967|50049067</td><td>20</td><td>40</td><td>21</td><td>TAGCAGCACGTAAATATTGGC</td><td>716.2</td><td>5.0</td><td>0.9</td><td>5.4</td><td>7.0E-08</td><td>1.3E-07</td><td>MB231/MB468</td><td>CYTO</td></tr>
<tr><td>>MI0000264|hsa-mir-7-2&WithFlank&15|+|88611819|88611940@38.59.22;MI0000265|hsa-mir-7-3&WithFlank&19|+|4770664|4770785@37.58.22;MI0000263|hsa-mir-7-1&WithFlank&9|-|83969742|83969863@30.51.22&[MIMAT0000252&hsa-miR-7-5p&offsets|0|-2;m-400&15|+|88611856|88611879&offsets|0|-2];[MIMAT0000252_1&hsa-miR-7-5p&offsets|0|-2;m-402&19|+|4770700|4770723&offsets|0|-2];[MIMAT0000252_2&hsa-miR-7-5p&offsets|0|-2;m-401&9|-|83969811|83969834&offsets|0|-2]|TGGAAGACTAGTGATTTTGTTG</td><td>>MI0000264|hsa-mir-7-2&WithFlank&15|+|88611819|88611940</td><td>38</td><td>59</td><td>22</td><td>TGGAAGACTAGTGATTTTGTTG</td><td>328.3</td><td>3.3</td><td>0.6</td><td>5.4</td><td>8.0E-08</td><td>1.3E-07</td><td>BT20/MB468</td><td>NUC</td></tr>
<tr><td>>am_MI0000809|hsa-mir-151a&WithFlank&8|-|140732558|140732659@53.71.19(+1A)&[MIMAT0000757&hsa-miR-151a-3p&offsets|0|-2(+1A);m-30&8|-|140732587|140732607&offsets|0|-2(+1A)]|CTAGACTGAAGCTCCTTGAA</td><td>>am_MI0000809|hsa-mir-151a&WithFlank&8|-|140732558|140732659</td><td>53</td><td>71</td><td>19(+1A)</td><td>CTAGACTGAAGCTCCTTGAA</td><td>86.0</td><td>5.0</td><td>0.9</td><td>5.4</td><td>8.0E-08</td><td>1.3E-07</td><td>BT20/MB468</td><td>NUC</td></tr>
<tr><td>>hg38_21_+_8433172_8446622_RNA45SN1_45S_50ntFlanks@4640.4662.23|GCATTCGTATTGCGCCGCTAGAG</td><td>>hg38_21_+_8433172_8446622_RNA45SN1_45S_50ntFlanks</td><td>4640</td><td>4662</td><td>23</td><td>GCATTCGTATTGCGCCGCTAGAG</td><td>79.9</td><td>-9.4</td><td>1.7</td><td>-5.7</td><td>1.3E-08</td><td>1.3E-07</td><td>BT20/MB468</td><td>MP</td></tr>
<tr><td>>hg38_21_+_8433172_8446622_RNA45SN1_45S_50ntFlanks@6647.6666.20|CGTACGACTCTTAGCGGTGG</td><td>>hg38_21_+_8433172_8446622_RNA45SN1_45S_50ntFlanks</td><td>6647</td><td>6666</td><td>20</td><td>CGTACGACTCTTAGCGGTGG</td><td>9415.9</td><td>4.2</td><td>0.8</td><td>5.4</td><td>5.4E-08</td><td>1.3E-07</td><td>BT20/MB468</td><td>MITO</td></tr>
</table>

<table>
<tr><td></td><td></td><td></td><td></td><td></td><td></td><td></td><td></td><td></td><td></td><td></td><td></td><td></td><td></td></tr>
<tr><td>>hg38_21_+_8433172_8446622_RNA45SN1_45S_50ntFlanks@10410.10433.24|TCAGTCGGTCCTGAGAGATGGGCG</td><td>>hg38_21_+_8433172_8446622_RNA45SN1_45S_50ntFlanks</td><td>10410</td><td>10433</td><td>24</td><td>TCAGTCGGTCCTGAGAGATGGGCG</td><td>1037.6</td><td>-7.3</td><td>1.4</td><td>-5.4</td><td>6.1E-08</td><td>1.3E-07</td><td>BT20/MB468</td><td>CYTO</td></tr>
<tr><td>>hg38_1_-_228634819_228635039_RNA5S12_5S_50ntFlanks@146.169.24|TGGGAATACCGGGTGCTGTAGGCT</td><td>>hg38_1_-_228634819_228635039_RNA5S12_5S_50ntFlanks</td><td>146</td><td>169</td><td>24</td><td>TGGGAATACCGGGTGCTGTAGGCT</td><td>235.6</td><td>3.8</td><td>0.7</td><td>5.4</td><td>7.2E-08</td><td>1.3E-07</td><td>MB231/MB468</td><td>NUC</td></tr>
<tr><td>>hg38_21_+_8433172_8446622_RNA45SN1_45S_50ntFlanks@6744.6770.27|GACACTTCGAACGCACTTGCGGCCCCG</td><td>>hg38_21_+_8433172_8446622_RNA45SN1_45S_50ntFlanks</td><td>6744</td><td>6770</td><td>27</td><td>GACACTTCGAACGCACTTGCGGCCCCG</td><td>2427.7</td><td>-6.5</td><td>1.2</td><td>-5.4</td><td>6.1E-08</td><td>1.3E-07</td><td>BT20/MB468</td><td>CYTO</td></tr>
<tr><td>>hg38_21_+_8433172_8446622_RNA45SN1_45S_50ntFlanks@11315.11332.18|CCACCCCACGTCTCGTCG</td><td>>hg38_21_+_8433172_8446622_RNA45SN1_45S_50ntFlanks</td><td>11315</td><td>11332</td><td>18</td><td>CCACCCCACGTCTCGTCG</td><td>84.6</td><td>-9.5</td><td>1.7</td><td>-5.7</td><td>1.3E-08</td><td>1.3E-07</td><td>BT20/MB468</td><td>MP</td></tr>
<tr><td>>hg38_21_+_8433172_8446622_RNA45SN1_45S_50ntFlanks@7981.8012.32|CTCAGATCAGACGTGGCGACCCGCTGAATTTA</td><td>>hg38_21_+_8433172_8446622_RNA45SN1_45S_50ntFlanks</td><td>7981</td><td>8012</td><td>32</td><td>CTCAGATCAGACGTGGCGACCCGCTGAATTTA</td><td>934.7</td><td>-5.9</td><td>1.1</td><td>-5.4</td><td>7.7E-08</td><td>1.3E-07</td><td>MB231/MB468</td><td>MITO</td></tr>
<tr><td>>trna143_LysTTT_6_-_27559593_27559665@13.30.18,trna149_LysTTT_6_-_27302769_27302841@13.30.18,trna14_LysTTT_11_-_59327808_59327880@13.30.18,trna2_LysTTT_17_+_8022473_8022545@13.30.18,trna54_LysTTT_1_+_204475655_204475727@13.30.18,trna5_LysTTT_11_+_59323902_59323974@13.30.18,trna62_LysTTT_1_-_204476158_204476230@13.30.18,trna76_LysTTT_6_+_28918806_28918878@13.30.18|CAGTCGGTAGAGCATCAG</td><td>>trna143_LysTTT_6_-_27559593_27559665</td><td>13</td><td>30</td><td>18</td><td>CAGTCGGTAGAGCATCAG</td><td>963.9</td><td>-4.9</td><td>0.9</td><td>-5.4</td><td>7.0E-08</td><td>1.3E-07</td><td>MB231/MB468</td><td>CYTO</td></tr>
<tr><td>>hg38_21_+_8433172_8446622_RNA45SN1_45S_50ntFlanks@11641.11664.24|CGGCGGGTGTTGACGCGATGTGAT</td><td>>hg38_21_+_8433172_8446622_RNA45SN1_45S_50ntFlanks</td><td>11641</td><td>11664</td><td>24</td><td>CGGCGGGTGTTGACGCGATGTGAT</td><td>323.6</td><td>-4.8</td><td>0.9</td><td>-5.4</td><td>7.0E-08</td><td>1.3E-07</td><td>MB231/MB468</td><td>CYTO</td></tr>
<tr><td>>hg38_21_+_8433172_8446622_RNA45SN1_45S_50ntFlanks@10771.10805.35|CAGGTCTCCAAGGTGAACAGCCTCTGGCATGTTGG</td><td>>hg38_21_+_8433172_8446622_RNA45SN1_45S_50ntFlanks</td><td>10771</td><td>10805</td><td>35</td><td>CAGGTCTCCAAGGTGAACAGCCTCTGGCATGTTGG</td><td>610.6</td><td>-5.0</td><td>0.9</td><td>-5.7</td><td>1.3E-08</td><td>1.3E-07</td><td>BT20/MB468</td><td>MP</td></tr>
<tr><td>>hg38_21_+_8433172_8446622_RNA45SN1_45S_50ntFlanks@9569.9596.28|GTGCAAATCGGTCGTCCGACCTGGGTAT</td><td>>hg38_21_+_8433172_8446622_RNA45SN1_45S_50ntFlanks</td><td>9569</td><td>9596</td><td>28</td><td>GTGCAAATCGGTCGTCCGACCTGGGTAT</td><td>387.6</td><td>-5.2</td><td>1.0</td><td>-5.4</td><td>7.1E-08</td><td>1.3E-07</td><td>MB231/MB468</td><td>CYTO</td></tr>
<tr><td>>hg38_21_+_8433172_8446622_RNA45SN1_45S_50ntFlanks@4892.4917.26|AAACTTAAAGGAATTGACGGAAGGGC</td><td>>hg38_21_+_8433172_8446622_RNA45SN1_45S_50ntFlanks</td><td>4892</td><td>4917</td><td>26</td><td>AAACTTAAAGGAATTGACGGAAGGGC</td><td>169.6</td><td>-10.5</td><td>1.8</td><td>-5.7</td><td>1.3E-08</td><td>1.3E-07</td><td>BT20/MB468</td><td>MP</td></tr>
<tr><td>>hg38_21_+_8433172_8446622_RNA45SN1_45S_50ntFlanks@4722.4761.40|TCATTAATCAAGAACGAAAGTCGGAGGTTCGAAGACGATC</td><td>>hg38_21_+_8433172_8446622_RNA45SN1_45S_50ntFlanks</td><td>4722</td><td>4761</td><td>40</td><td>TCATTAATCAAGAACGAAAGTCGGAGGTTCGAAGACGATC</td><td>233.5</td><td>-4.4</td><td>0.8</td><td>-5.4</td><td>7.7E-08</td><td>1.3E-07</td><td>MB231/MB468</td><td>MITO</td></tr>
<tr><td>>hg38_21_+_8433172_8446622_RNA45SN1_45S_50ntFlanks@11212.11253.42|CGACTCTGGACGCGAGCCGGGCCCTTCCCGTGGATCGCCCCA</td><td>>hg38_21_+_8433172_8446622_RNA45SN1_45S_50ntFlanks</td><td>11212</td><td>11253</td><td>42</td><td>CGACTCTGGACGCGAGCCGGGCCCTTCCCGTGGATCGCCCCA</td><td>334.7</td><td>-3.7</td><td>0.7</td><td>-5.4</td><td>7.1E-08</td><td>1.3E-07</td><td>MB231/MB468</td><td>CYTO</td></tr>
<tr><td>>hg38_21_+_8433172_8446622_RNA45SN1_45S_50ntFlanks@11582.11603.22|ACTGGTGCGGACCAGGGGAATC</td><td>>hg38_21_+_8433172_8446622_RNA45SN1_45S_50ntFlanks</td><td>11582</td><td>11603</td><td>22</td><td>ACTGGTGCGGACCAGGGGAATC</td><td>717.4</td><td>-3.1</td><td>0.6</td><td>-5.4</td><td>7.3E-08</td><td>1.3E-07</td><td>MB231/MB468</td><td>NUC</td></tr>
<tr><td>>hg38_MT_+_1621_3279_RNR2_16S_50ntFlanks@382.416.35|GAGCCTGGTGATAGCTGGTTGTCCAAGATAGAATC</td><td>>hg38_MT_+_1621_3279_RNR2_16S_50ntFlanks</td><td>382</td><td>416</td><td>35</td><td>GAGCCTGGTGATAGCTGGTTGTCCAAGATAGAATC</td><td>62.3</td><td>9.2</td><td>1.7</td><td>5.4</td><td>6.6E-08</td><td>1.3E-07</td><td>MB231/MB468</td><td>MP</td></tr>
<tr><td>>hg38_21_+_8433172_8446622_RNA45SN1_45S_50ntFlanks@10783.10818.36|GTGAACAGCCTCTGGCATGTTGGAACAATGTAGGTA</td><td>>hg38_21_+_8433172_8446622_RNA45SN1_45S_50ntFlanks</td><td>10783</td><td>10818</td><td>36</td><td>GTGAACAGCCTCTGGCATGTTGGAACAATGTAGGTA</td><td>768.6</td><td>-6.1</td><td>1.1</td><td>-5.4</td><td>7.3E-08</td><td>1.3E-07</td><td>MB231/MB468</td><td>NUC</td></tr>
<tr><td>>MI0001446|hsa-mir-424&WithFlank&X|-|134546608|134546717@54.72.19&[MIMAT0004749&hsa-miR-424-3p&offsets|0|-2;m-305&X|-|134546644|134546664&offsets|0|-2]|CAAAACGTGAGGCGCTGCT</td><td>>MI0001446|hsa-mir-424&WithFlank&X|-|134546608|134546717</td><td>54</td><td>72</td><td>19</td><td>CAAAACGTGAGGCGCTGCT</td><td>129.7</td><td>10.8</td><td>1.9</td><td>5.7</td><td>1.3E-08</td><td>1.3E-07</td><td>BT20/MB468</td><td>MP</td></tr>
<tr><td>>hg38_21_+_8433172_8446622_RNA45SN1_45S_50ntFlanks@3846.3874.29|CTTGGATAACTGTGGTAATTCTAGAGCTA</td><td>>hg38_21_+_8433172_8446622_RNA45SN1_45S_50ntFlanks</td><td>3846</td><td>3874</td><td>29</td><td>CTTGGATAACTGTGGTAATTCTAGAGCTA</td><td>255.9</td><td>-7.5</td><td>1.4</td><td>-5.4</td><td>6.6E-08</td><td>1.3E-07</td><td>MB231/MB468</td><td>MP</td></tr>
<tr><td>>MI0000076|hsa-mir-20a&WithFlank&13|+|91351059|91351141@14.36.23(+1A)&[MIMAT0000075&hsa-miR-20a-5p&offsets|0|0(+1A);m-96&13|+|91351072|91351094&offsets|0|0(+1A)]|TAAAGTGCTTATAGTGCAGGTAGA</td><td>>MI0000076|hsa-mir-20a&WithFlank&13|+|91351059|91351141</td><td>14</td><td>36</td><td>23(+1A)</td><td>TAAAGTGCTTATAGTGCAGGTAGA</td><td>5327.6</td><td>5.7</td><td>1.1</td><td>5.4</td><td>7.7E-08</td><td>1.3E-07</td><td>MB231/MB468</td><td>MITO</td></tr>
<tr><td>>hg38_21_+_8433172_8446622_RNA45SN1_45S_50ntFlanks@4685.4723.39|AGACGGACCAGAGCGAAAGCATTTGCCAAGAATGTTTTC</td><td>>hg38_21_+_8433172_8446622_RNA45SN1_45S_50ntFlanks</td><td>4685</td><td>4723</td><td>39</td><td>AGACGGACCAGAGCGAAAGCATTTGCCAAGAATGTTTTC</td><td>916.8</td><td>-6.5</td><td>1.2</td><td>-5.4</td><td>7.8E-08</td><td>1.3E-07</td><td>MB231/MB468</td><td>MITO</td></tr>
<tr><td>>hg38_21_+_8433172_8446622_RNA45SN1_45S_50ntFlanks@11207.11254.48|GGCGGCGACTCTGGACGCGAGCCGGGCCCTTCCCGTGGATCGCCCCAG</td><td>>hg38_21_+_8433172_8446622_RNA45SN1_45S_50ntFlanks</td><td>11207</td><td>11254</td><td>48</td><td>GGCGGCGACTCTGGACGCGAGCCGGGCCCTTCCCGTGGATCGCCCCAG</td><td>3112.8</td><td>-10.0</td><td>1.9</td><td>-5.4</td><td>8.1E-08</td><td>1.3E-07</td><td>BT20/MB468</td><td>NUC</td></tr>
<tr><td>>hg38_21_+_8433172_8446622_RNA45SN1_45S_50ntFlanks@12416.12442.27|TTCGATGTCGGCTCTTCCTATCATTGT</td><td>>hg38_21_+_8433172_8446622_RNA45SN1_45S_50ntFlanks</td><td>12416</td><td>12442</td><td>27</td><td>TTCGATGTCGGCTCTTCCTATCATTGT</td><td>573.1</td><td>5.0</td><td>0.9</td><td>5.4</td><td>7.3E-08</td><td>1.3E-07</td><td>MB231/MB468</td><td>NUC</td></tr>
<tr><td>>hg38_21_+_8433172_8446622_RNA45SN1_45S_50ntFlanks@10116.10135.20|TGCGTCGGAGGGCGGCGGCG</td><td>>hg38_21_+_8433172_8446622_RNA45SN1_45S_50ntFlanks</td><td>10116</td><td>10135</td><td>20</td><td>TGCGTCGGAGGGCGGCGGCG</td><td>423.1</td><td>-8.1</td><td>1.5</td><td>-5.4</td><td>8.1E-08</td><td>1.3E-07</td><td>BT20/MB468</td><td>NUC</td></tr>
<tr><td>>hg38_21_+_8433172_8446622_RNA45SN1_45S_50ntFlanks@12399.12434.36|ACGTCGCTTTTTGATCCTTCGATGTCGGCTCTTCCT</td><td>>hg38_21_+_8433172_8446622_RNA45SN1_45S_50ntFlanks</td><td>12399</td><td>12434</td><td>36</td><td>ACGTCGCTTTTTGATCCTTCGATGTCGGCTCTTCCT</td><td>548.2</td><td>-4.7</td><td>0.9</td><td>-5.4</td><td>8.1E-08</td><td>1.3E-07</td><td>BT20/MB468</td><td>NUC</td></tr>
<tr><td>>hg38_21_+_8433172_8446622_RNA45SN1_45S_50ntFlanks@8168.8203.36|CCCAAGTCCTTCTGATCGAGGCCCAGCCCGTGGACG</td><td>>hg38_21_+_8433172_8446622_RNA45SN1_45S_50ntFlanks</td><td>8168</td><td>8203</td><td>36</td><td>CCCAAGTCCTTCTGATCGAGGCCCAGCCCGTGGACG</td><td>827.6</td><td>-8.0</td><td>1.5</td><td>-5.4</td><td>6.6E-08</td><td>1.3E-07</td><td>MB231/MB468</td><td>MP</td></tr>
<tr><td>>hg38_21_+_8433172_8446622_RNA45SN1_45S_50ntFlanks@10873.10894.22|CTGGGTCGGTCGGGCTGGGGCG</td><td>>hg38_21_+_8433172_8446622_RNA45SN1_45S_50ntFlanks</td><td>10873</td><td>10894</td><td>22</td><td>CTGGGTCGGTCGGGCTGGGGCG</td><td>387.2</td><td>-2.7</td><td>0.5</td><td>-5.4</td><td>6.2E-08</td><td>1.4E-07</td><td>BT20/MB468</td><td>CYTO</td></tr>
<tr><td>>hg38_21_+_8433172_8446622_RNA45SN1_45S_50ntFlanks@11804.11821.18|CCCACTGTCCCTACCTAC</td><td>>hg38_21_+_8433172_8446622_RNA45SN1_45S_50ntFlanks</td><td>11804</td><td>11821</td><td>18</td><td>CCCACTGTCCCTACCTAC</td><td>558.9</td><td>-7.7</td><td>1.4</td><td>-5.4</td><td>7.3E-08</td><td>1.4E-07</td><td>MB231/MB468</td><td>NUC</td></tr>
<tr><td>>MI0000809|hsa-mir-151a&WithFlank&8|-|140732558|140732659@51.71.21(+1A)&[MIMAT0000757&hsa-miR-151a-3p&offsets|-2|-2(+1A);m-30&8|-|140732587|140732607&offsets|-2|-2(+1A)]|TACTAGACTGAAGCTCCTTGAA</td><td>>MI0000809|hsa-mir-151a&WithFlank&8|-|140732558|140732659</td><td>51</td><td>71</td><td>21(+1A)</td><td>TACTAGACTGAAGCTCCTTGAA</td><td>67.6</td><td>9.3</td><td>1.7</td><td>5.4</td><td>6.6E-08</td><td>1.4E-07</td><td>MB231/MB468</td><td>MP</td></tr>
<tr><td>>trna10_AspGTC_12_-_125424193_125424264@37.74.38,trna12_AspGTC_12_-_125411891_125411962@37.74.38,trna144_AspGTC_6_-_27551236_27551307@37.74.38,trna38_AspGTC_17_-_8125556_8125627@37.74.38,trna45_AspGTC_6_+_27447453_27447524@37.74.38,trna48_AspGTC_6_+_27471523_27471594@37.74.38,trna4_AspGTC_12_+_96429799_96429870@37.74.38,trna69_AspGTC_1_-_161440205_161440276@37.74.38,trna72_AspGTC_1_-_161432824_161432895@37.74.38,trna75_AspGTC_1_-_161425414_161425485@37.74.38,trna78_AspGTC_1_-_161418033_161418104@37.74.38,trna81_AspGTC_1_-_161410615_161410686@37.74.38|ACGCGGGAGACCGGGGTTCGATTCCCCGACGGGGAGCC</td><td>>trna10_AspGTC_12_-_125424193_125424264</td><td>37</td><td>74</td><td>38</td><td>ACGCGGGAGACCGGGGTTCGATTCCCCGACGGGGAGCC</td><td>24115.3</td><td>-6.7</td><td>1.1</td><td>-5.9</td><td>4.5E-09</td><td>1.4E-07</td><td>BT20/MB231</td><td>CYTO</td></tr>
<tr><td>>hg38_21_+_8433172_8446622_RNA45SN1_45S_50ntFlanks@10846.10868.23|CTTCGGGATAAGGATTGGCTCTA</td><td>>hg38_21_+_8433172_8446622_RNA45SN1_45S_50ntFlanks</td><td>10846</td><td>10868</td><td>23</td><td>CTTCGGGATAAGGATTGGCTCTA</td><td>875.9</td><td>-5.2</td><td>1.0</td><td>-5.4</td><td>6.2E-08</td><td>1.4E-07</td><td>BT20/MB468</td><td>CYTO</td></tr>
<tr><td>>hg38_21_+_8433172_8446622_RNA45SN1_45S_50ntFlanks@8114.8137.24|GCGCGGGACATGTGGCGTACGGAA</td><td>>hg38_21_+_8433172_8446622_RNA45SN1_45S_50ntFlanks</td><td>8114</td><td>8137</td><td>24</td><td>GCGCGGGACATGTGGCGTACGGAA</td><td>1652.4</td><td>14.0</td><td>2.4</td><td>6.0</td><td>2.6E-09</td><td>1.4E-07</td><td>BT20/MB231</td><td>MP</td></tr>
<tr><td>>MI0003820|hsa-mir-454&WithFlank&17|-|59137752|59137878@70.95.26&[MIMAT0003885&hsa-miR-454-3p&offsets|0|+3;m-272&17|-|59137787|59137809&offsets|0|+3]|TAGTGCAATATTGCTTATAGGGTTTT</td><td>>MI0003820|hsa-mir-454&WithFlank&17|-|59137752|59137878</td><td>70</td><td>95</td><td>26</td><td>TAGTGCAATATTGCTTATAGGGTTTT</td><td>168.7</td><td>-10.8</td><td>1.8</td><td>-6.0</td><td>2.6E-09</td><td>1.4E-07</td><td>BT20/MB231</td><td>MP</td></tr>
<tr><td>>hg38_21_+_8433172_8446622_RNA45SN1_45S_50ntFlanks@8379.8397.19|TCAAGAGGGCGTGAAACCG</td><td>>hg38_21_+_8433172_8446622_RNA45SN1_45S_50ntFlanks</td><td>8379</td><td>8397</td><td>19</td><td>TCAAGAGGGCGTGAAACCG</td><td>158.7</td><td>-7.0</td><td>1.3</td><td>-5.4</td><td>6.7E-08</td><td>1.4E-07</td><td>MB231/MB468</td><td>MP</td></tr>
<tr><td>>hg38_21_+_8433172_8446622_RNA45SN1_45S_50ntFlanks@4730.4752.23|CAAGAACGAAAGTCGGAGGTTCG</td><td>>hg38_21_+_8433172_8446622_RNA45SN1_45S_50ntFlanks</td><td>4730</td><td>4752</td><td>23</td><td>CAAGAACGAAAGTCGGAGGTTCG</td><td>175.1</td><td>-5.1</td><td>1.0</td><td>-5.4</td><td>7.4E-08</td><td>1.4E-07</td><td>MB231/MB468</td><td>NUC</td></tr>
<tr><td>>MI0000746|hsa-mir-99b&WithFlank&19|+|51692606|51692687@13.32.20(+2G)&[MIMAT0000689&hsa-miR-99b-5p&offsets|0|-2(+2G);m-10&19|+|51692618|51692639&offsets|0|-2(+2G)]|CACCCGTAGAACCGACCTTGGG</td><td>>MI0000746|hsa-mir-99b&WithFlank&19|+|51692606|51692687</td><td>13</td><td>32</td><td>20(+2G)</td><td>CACCCGTAGAACCGACCTTGGG</td><td>576.9</td><td>9.4</td><td>1.7</td><td>5.4</td><td>6.2E-08</td><td>1.4E-07</td><td>BT20/MB468</td><td>CYTO</td></tr>
<tr><td>>hg38_21_+_8433172_8446622_RNA45SN1_45S_50ntFlanks@6662.6689.28|GGTGGATCACTCGGCTCGTGCGTCGATG</td><td>>hg38_21_+_8433172_8446622_RNA45SN1_45S_50ntFlanks</td><td>6662</td><td>6689</td><td>28</td><td>GGTGGATCACTCGGCTCGTGCGTCGATG</td><td>304.7</td><td>4.1</td><td>0.7</td><td>5.8</td><td>5.8E-09</td><td>1.4E-07</td><td>BT20/MB231</td><td>MITO</td></tr>
<tr><td>>MI0000299|hsa-mir-222&WithFlank&X|-|45747009|45747130@75.99.25(+1C)&[MIMAT0000279&hsa-miR-222-3p&offsets|0|+4(+1C);m-140&X|-|45747033|45747056&offsets|0|+1(+1C)]|AGCTACATCTGGCTACTGGGTCTCTC</td><td>>MI0000299|hsa-mir-222&WithFlank&X|-|45747009|45747130</td><td>75</td><td>99</td><td>25(+1C)</td><td>AGCTACATCTGGCTACTGGGTCTCTC</td><td>628.6</td><td>8.7</td><td>1.6</td><td>5.4</td><td>7.8E-08</td><td>1.4E-07</td><td>MB231/MB468</td><td>MITO</td></tr>
<tr><td>>hg38_MT_+_1621_3279_RNR2_16S_50ntFlanks@388.416.29|GGTGATAGCTGGTTGTCCAAGATAGAATC</td><td>>hg38_MT_+_1621_3279_RNR2_16S_50ntFlanks</td><td>388</td><td>416</td><td>29</td><td>GGTGATAGCTGGTTGTCCAAGATAGAATC</td><td>131.6</td><td>10.8</td><td>1.9</td><td>5.7</td><td>1.3E-08</td><td>1.4E-07</td><td>BT20/MB468</td><td>MP</td></tr>
<tr><td>>hg38_21_+_8433172_8446622_RNA45SN1_45S_50ntFlanks@4763.4780.18|GATACCGTCGTAGTTCCG</td><td>>hg38_21_+_8433172_8446622_RNA45SN1_45S_50ntFlanks</td><td>4763</td><td>4780</td><td>18</td><td>GATACCGTCGTAGTTCCG</td><td>1140.0</td><td>-5.8</td><td>1.1</td><td>-5.4</td><td>6.2E-08</td><td>1.4E-07</td><td>BT20/MB468</td><td>CYTO</td></tr>
<tr><td>>hg38_21_+_8433172_8446622_RNA45SN1_45S_50ntFlanks@11669.11706.38|GCCCAGTGCTCTGAATGTCAAAGTGAAGAAATTCAATG</td><td>>hg38_21_+_8433172_8446622_RNA45SN1_45S_50ntFlanks</td><td>11669</td><td>11706</td><td>38</td><td>GCCCAGTGCTCTGAATGTCAAAGTGAAGAAATTCAATG</td><td>315.9</td><td>7.4</td><td>1.4</td><td>5.4</td><td>5.6E-08</td><td>1.4E-07</td><td>BT20/MB468</td><td>MITO</td></tr>
<tr><td>>hg38_MT_+_598_1651_RNR1_12S_50ntFlanks@848.877.30|GAGTGCTTAGTTGAACAGGGCCCTGAAGCG</td><td>>hg38_MT_+_598_1651_RNR1_12S_50ntFlanks</td><td>848</td><td>877</td><td>30</td><td>GAGTGCTTAGTTGAACAGGGCCCTGAAGCG</td><td>151.0</td><td>11.0</td><td>1.9</td><td>5.7</td><td>1.3E-08</td><td>1.4E-07</td><td>BT20/MB468</td><td>MP</td></tr>
<tr><td>>hg38_21_+_8433172_8446622_RNA45SN1_45S_50ntFlanks@8114.8140.27|GCGCGGGACATGTGGCGTACGGAAGAC</td><td>>hg38_21_+_8433172_8446622_RNA45SN1_45S_50ntFlanks</td><td>8114</td><td>8140</td><td>27</td><td>GCGCGGGACATGTGGCGTACGGAAGAC</td><td>532.4</td><td>12.4</td><td>2.1</td><td>6.0</td><td>2.7E-09</td><td>1.4E-07</td><td>BT20/MB231</td><td>MP</td></tr>
<tr><td>>hg38_21_+_8433172_8446622_RNA45SN1_45S_50ntFlanks@8172.8212.41|AGTCCTTCTGATCGAGGCCCAGCCCGTGGACGGTGTGAGGC</td><td>>hg38_21_+_8433172_8446622_RNA45SN1_45S_50ntFlanks</td><td>8172</td><td>8212</td><td>41</td><td>AGTCCTTCTGATCGAGGCCCAGCCCGTGGACGGTGTGAGGC</td><td>1150.6</td><td>-4.1</td><td>0.8</td><td>-5.4</td><td>7.4E-08</td><td>1.4E-07</td><td>MB231/MB468</td><td>NUC</td></tr>
<tr><td>>hg38_21_+_8433172_8446622_RNA45SN1_45S_50ntFlanks@8347.8373.27|GGGAAAGTTGAAAAGAACTTTGAAGAG</td><td>>hg38_21_+_8433172_8446622_RNA45SN1_45S_50ntFlanks</td><td>8347</td><td>8373</td><td>27</td><td>GGGAAAGTTGAAAAGAACTTTGAAGAG</td><td>769.2</td><td>9.6</td><td>1.7</td><td>5.7</td><td>1.3E-08</td><td>1.4E-07</td><td>BT20/MB468</td><td>MP</td></tr>
<tr><td>>hg38_21_+_8433172_8446622_RNA45SN1_45S_50ntFlanks@12822.12849.28|CCGGGACCGGGGTCCGGTGCGGAGTGCC</td><td>>hg38_21_+_8433172_8446622_RNA45SN1_45S_50ntFlanks</td><td>12822</td><td>12849</td><td>28</td><td>CCGGGACCGGGGTCCGGTGCGGAGTGCC</td><td>213.7</td><td>-10.8</td><td>1.9</td><td>-5.7</td><td>1.3E-08</td><td>1.4E-07</td><td>BT20/MB468</td><td>MP</td></tr>
<tr><td>>trnaMT_LeuTAA_MT_+_3230_3304@18.44.27|CGGTAATCGCATAAAACTTAAAACTTT</td><td>>trnaMT_LeuTAA_MT_+_3230_3304</td><td>18</td><td>44</td><td>27</td><td>CGGTAATCGCATAAAACTTAAAACTTT</td><td>1313.3</td><td>-5.4</td><td>1.0</td><td>-5.4</td><td>8.2E-08</td><td>1.4E-07</td><td>BT20/MB468</td><td>NUC</td></tr>
<tr><td>>hg38_21_+_8433172_8446622_RNA45SN1_45S_50ntFlanks@4005.4025.21|AACCTCGGGCCGATCGCACGC</td><td>>hg38_21_+_8433172_8446622_RNA45SN1_45S_50ntFlanks</td><td>4005</td><td>4025</td><td>21</td><td>AACCTCGGGCCGATCGCACGC</td><td>1239.3</td><td>-7.2</td><td>1.3</td><td>-5.4</td><td>8.2E-08</td><td>1.4E-07</td><td>BT20/MB468</td><td>NUC</td></tr>
</table>

<table>
<tr><td>>hg38_21_+_8433172_8446622_RNA45SN1_45S_50ntFlanks@11592.11612.21|ACCAGGGGAATCCGACTGTTT</td><td>>hg38_21_+_8433172_8446622_RNA45SN1_45S_50ntFlanks</td><td>11592</td><td>11612</td><td>21</td><td>ACCAGGGGAATCCGACTGTTT</td><td>381.1</td><td>-3.9</td><td>0.7</td><td>-5.4</td><td>5.6E-08</td><td>1.4E-07</td><td>BT20/MB468</td><td>MITO</td></tr>
<tr><td>>hg38_21_+_8433172_8446622_RNA45SN1_45S_50ntFlanks@4045.4065.21|CCATTCGAACGTCTGCCCTAT</td><td>>hg38_21_+_8433172_8446622_RNA45SN1_45S_50ntFlanks</td><td>4045</td><td>4065</td><td>21</td><td>CCATTCGAACGTCTGCCCTAT</td><td>151.3</td><td>-2.4</td><td>0.4</td><td>-5.4</td><td>6.3E-08</td><td>1.4E-07</td><td>BT20/MB468</td><td>CYTO</td></tr>
<tr><td>>MI0000108|hsa-mir-103a-2&WithFlank&20|+|3917488|3917577@54.75.22(+1G);MI0000109|hsa-mir-103a-1&WithFlank&5|-|168560890|168560979@54.75.22(+1G)&[MIMAT0000101&hsa-miR-103a-3p&offsets|0|-1(+1G);m-12&20|+|3917541|3917563&offsets|0|-1(+1G)];[MIMAT0000101_1&hsa-miR-103a-3p&offsets|0|-1(+1G);m-13&5|-|168560904|168560926&offsets|0|-1(+1G);m-44220&5|-|168560908|168560926&offsets|0|+3(+1G)]|AGCAGCATTGTACAGGGCTATGG</td><td>>MI0000108|hsa-mir-103a-2&WithFlank&20|+|3917488|3917577</td><td>54</td><td>75</td><td>22(+1G)</td><td>AGCAGCATTGTACAGGGCTATGG</td><td>208.1</td><td>3.0</td><td>0.6</td><td>5.4</td><td>8.2E-08</td><td>1.4E-07</td><td>BT20/MB468</td><td>NUC</td></tr>
<tr><td>>hg38_21_+_8433172_8446622_RNA45SN1_45S_50ntFlanks@11200.11233.34|CGGCGGCGGCGGCGACTCTGGACGCGAGCCGGGC</td><td>>hg38_21_+_8433172_8446622_RNA45SN1_45S_50ntFlanks</td><td>11200</td><td>11233</td><td>34</td><td>CGGCGGCGGCGGCGACTCTGGACGCGAGCCGGGC</td><td>336.8</td><td>-7.7</td><td>1.4</td><td>-5.4</td><td>8.2E-08</td><td>1.4E-07</td><td>BT20/MB468</td><td>NUC</td></tr>
<tr><td>>MI0000736|hsa-mir-30c-1&WithFlank&1|+|40757278|40757378@23.42.20;MI0000254|hsa-mir-30c-2&WithFlank&6|-|71376954|71377037@13.32.20&[MIMAT0000244&hsa-miR-30c-5p&offsets|0|-3;m-57&1|+|40757300|40757323&offsets|0|-4];[MIMAT0000244_1&hsa-miR-30c-5p&offsets|0|-3;m-56&6|-|71377002|71377025&offsets|0|-4]|TGTAAACATCCTACACTCTC</td><td>>MI0000736|hsa-mir-30c-1&WithFlank&1|+|40757278|40757378</td><td>23</td><td>42</td><td>20</td><td>TGTAAACATCCTACACTCTC</td><td>816.5</td><td>4.8</td><td>0.9</td><td>5.4</td><td>6.7E-08</td><td>1.4E-07</td><td>MB231/MB468</td><td>MP</td></tr>
<tr><td>>hg38_21_+_8433172_8446622_RNA45SN1_45S_50ntFlanks@9533.9563.31|GAGGAAACTCTGGTGGAGGTCCGTAGCGGTC</td><td>>hg38_21_+_8433172_8446622_RNA45SN1_45S_50ntFlanks</td><td>9533</td><td>9563</td><td>31</td><td>GAGGAAACTCTGGTGGAGGTCCGTAGCGGTC</td><td>2959.8</td><td>-4.3</td><td>0.8</td><td>-5.4</td><td>6.3E-08</td><td>1.4E-07</td><td>BT20/MB468</td><td>CYTO</td></tr>
<tr><td>>hg38_21_+_8433172_8446622_RNA45SN1_45S_50ntFlanks@8104.8143.40|CCGCGGCGGGGCGCGGGACATGTGGCGTACGGAAGACCCG</td><td>>hg38_21_+_8433172_8446622_RNA45SN1_45S_50ntFlanks</td><td>8104</td><td>8143</td><td>40</td><td>CCGCGGCGGGGCGCGGGACATGTGGCGTACGGAAGACCCG</td><td>13552.6</td><td>-10.5</td><td>2.0</td><td>-5.4</td><td>8.3E-08</td><td>1.4E-07</td><td>BT20/MB468</td><td>NUC</td></tr>
<tr><td>>hg38_21_+_8433172_8446622_RNA45SN1_45S_50ntFlanks@9572.9602.31|CAAATCGGTCGTCCGACCTGGGTATAGGGGC</td><td>>hg38_21_+_8433172_8446622_RNA45SN1_45S_50ntFlanks</td><td>9572</td><td>9602</td><td>31</td><td>CAAATCGGTCGTCCGACCTGGGTATAGGGGC</td><td>664.2</td><td>-4.9</td><td>0.9</td><td>-5.4</td><td>6.8E-08</td><td>1.4E-07</td><td>MB231/MB468</td><td>MP</td></tr>
<tr><td>>MI0000088|hsa-mir-30a&WithFlank&6|-|71403545|71403627@12.34.23(+1G)&[MIMAT0000087&hsa-miR-30a-5p&offsets|0|+1(+1G);m-15&6|-|71403595|71403616&offsets|0|+1(+1G)]|TGTAAACATCCTCGACTGGAAGCG</td><td>>MI0000088|hsa-mir-30a&WithFlank&6|-|71403545|71403627</td><td>12</td><td>34</td><td>23(+1G)</td><td>TGTAAACATCCTCGACTGGAAGCG</td><td>1050.8</td><td>5.1</td><td>0.9</td><td>5.4</td><td>6.3E-08</td><td>1.4E-07</td><td>BT20/MB468</td><td>CYTO</td></tr>
<tr><td>>hg38_21_+_8433172_8446622_RNA45SN1_45S_50ntFlanks@9120.9138.19|TGCCGCGCGCGGGTCGGGG</td><td>>hg38_21_+_8433172_8446622_RNA45SN1_45S_50ntFlanks</td><td>9120</td><td>9138</td><td>19</td><td>TGCCGCGCGCGGGTCGGGG</td><td>102.4</td><td>-9.8</td><td>1.7</td><td>-5.7</td><td>1.4E-08</td><td>1.4E-07</td><td>BT20/MB468</td><td>MP</td></tr>
<tr><td>>hg38_21_+_8433172_8446622_RNA45SN1_45S_50ntFlanks@6742.6770.29|TCGACACTTCGAACGCACTTGCGGCCCCG</td><td>>hg38_21_+_8433172_8446622_RNA45SN1_45S_50ntFlanks</td><td>6742</td><td>6770</td><td>29</td><td>TCGACACTTCGAACGCACTTGCGGCCCCG</td><td>539.3</td><td>-7.8</td><td>1.4</td><td>-5.7</td><td>1.4E-08</td><td>1.4E-07</td><td>BT20/MB468</td><td>MP</td></tr>
<tr><td>>hg38_21_+_8433172_8446622_RNA45SN1_45S_50ntFlanks@10412.10429.18|AGTCGGTCCTGAGAGATG</td><td>>hg38_21_+_8433172_8446622_RNA45SN1_45S_50ntFlanks</td><td>10412</td><td>10429</td><td>18</td><td>AGTCGGTCCTGAGAGATG</td><td>170.8</td><td>-5.9</td><td>1.1</td><td>-5.4</td><td>6.4E-08</td><td>1.4E-07</td><td>BT20/MB468</td><td>CYTO</td></tr>
<tr><td>>hg38_21_+_8433172_8446622_RNA45SN1_45S_50ntFlanks@11641.11679.39|CGGCGGGTGTTGACGCGATGTGATTTCTGCCCAGTGCTC</td><td>>hg38_21_+_8433172_8446622_RNA45SN1_45S_50ntFlanks</td><td>11641</td><td>11679</td><td>39</td><td>CGGCGGGTGTTGACGCGATGTGATTTCTGCCCAGTGCTC</td><td>788.7</td><td>-7.1</td><td>1.3</td><td>-5.4</td><td>5.7E-08</td><td>1.4E-07</td><td>BT20/MB468</td><td>MITO</td></tr>
<tr><td>>hg38_21_+_8433172_8446622_RNA45SN1_45S_50ntFlanks@4738.4758.21|AAAGTCGGAGGTTCGAAGACG</td><td>>hg38_21_+_8433172_8446622_RNA45SN1_45S_50ntFlanks</td><td>4738</td><td>4758</td><td>21</td><td>AAAGTCGGAGGTTCGAAGACG</td><td>3635.4</td><td>-2.4</td><td>0.4</td><td>-5.4</td><td>7.6E-08</td><td>1.4E-07</td><td>MB231/MB468</td><td>NUC</td></tr>
<tr><td>>trnaMT_ValTAC_MT_+_1602_1670@27.72.46|CAACTTACACTTAGGAGATTTCAACTTAACTTGACCGCTCTGACCA</td><td>>trnaMT_ValTAC_MT_+_1602_1670</td><td>27</td><td>72</td><td>46</td><td>CAACTTACACTTAGGAGATTTCAACTTAACTTGACCGCTCTGACCA</td><td>554.5</td><td>5.4</td><td>1.0</td><td>5.4</td><td>5.7E-08</td><td>1.4E-07</td><td>BT20/MB468</td><td>MITO</td></tr>
<tr><td>>hg38_21_+_8433172_8446622_RNA45SN1_45S_50ntFlanks@7968.8005.38|TCCGAGACGCGACCTCAGATCAGACGTGGCGACCCGCT</td><td>>hg38_21_+_8433172_8446622_RNA45SN1_45S_50ntFlanks</td><td>7968</td><td>8005</td><td>38</td><td>TCCGAGACGCGACCTCAGATCAGACGTGGCGACCCGCT</td><td>505.9</td><td>5.1</td><td>1.0</td><td>5.4</td><td>8.4E-08</td><td>1.4E-07</td><td>BT20/MB468</td><td>NUC</td></tr>
<tr><td>>MI0000089|hsa-mir-31&WithFlank&9|-|21512109|21512191@14.35.22(+1A)&[MIMAT0000089&hsa-miR-31-5p&offsets|0|+1(+1A);m-360&9|-|21512157|21512178&offsets|0|0(+1A)]|AGGCAAGATGCTGGCATAGCTGA</td><td>>MI0000089|hsa-mir-31&WithFlank&9|-|21512109|21512191</td><td>14</td><td>35</td><td>22(+1A)</td><td>AGGCAAGATGCTGGCATAGCTGA</td><td>195.5</td><td>10.9</td><td>1.8</td><td>5.9</td><td>2.7E-09</td><td>1.4E-07</td><td>BT20/MB231</td><td>MP</td></tr>
<tr><td>>hg38_21_+_8433172_8446622_RNA45SN1_45S_50ntFlanks@11564.11600.37|CCTAGCAGCCGACTTAGAACTGGTGCGGACCAGGGGA</td><td>>hg38_21_+_8433172_8446622_RNA45SN1_45S_50ntFlanks</td><td>11564</td><td>11600</td><td>37</td><td>CCTAGCAGCCGACTTAGAACTGGTGCGGACCAGGGGA</td><td>566.2</td><td>-3.7</td><td>0.7</td><td>-5.4</td><td>6.4E-08</td><td>1.4E-07</td><td>BT20/MB468</td><td>CYTO</td></tr>
<tr><td>>trnaMT_PheGAA_MT_+_577_647@39.56.18|ATGTTTAGACGGGCTCAC</td><td>>trnaMT_PheGAA_MT_+_577_647</td><td>39</td><td>56</td><td>18</td><td>ATGTTTAGACGGGCTCAC</td><td>1663.8</td><td>9.8</td><td>1.8</td><td>5.4</td><td>6.4E-08</td><td>1.4E-07</td><td>BT20/MB468</td><td>CYTO</td></tr>
<tr><td>>hg38_1_-_228634819_228635039_RNA5S12_5S_50ntFlanks@140.171.32|ACCGCCTGGGAATACCGGGTGCTGTAGGCTTT</td><td>>hg38_1_-_228634819_228635039_RNA5S12_5S_50ntFlanks</td><td>140</td><td>171</td><td>32</td><td>ACCGCCTGGGAATACCGGGTGCTGTAGGCTTT</td><td>7909.5</td><td>4.9</td><td>0.9</td><td>5.4</td><td>6.4E-08</td><td>1.4E-07</td><td>BT20/MB468</td><td>CYTO</td></tr>
<tr><td>>hg38_21_+_8433172_8446622_RNA45SN1_45S_50ntFlanks@10411.10453.43|CAGTCGGTCCTGAGAGATGGGCGAGCGCCGTTCCGAAGGGACG</td><td>>hg38_21_+_8433172_8446622_RNA45SN1_45S_50ntFlanks</td><td>10411</td><td>10453</td><td>43</td><td>CAGTCGGTCCTGAGAGATGGGCGAGCGCCGTTCCGAAGGGACG</td><td>1355.0</td><td>-5.6</td><td>1.0</td><td>-5.4</td><td>7.4E-08</td><td>1.4E-07</td><td>MB231/MB468</td><td>CYTO</td></tr>
<tr><td>>hg38_21_+_8433172_8446622_RNA45SN1_45S_50ntFlanks@11781.11798.18|CGCATGAATGGATGAACG</td><td>>hg38_21_+_8433172_8446622_RNA45SN1_45S_50ntFlanks</td><td>11781</td><td>11798</td><td>18</td><td>CGCATGAATGGATGAACG</td><td>398.2</td><td>-7.2</td><td>1.3</td><td>-5.4</td><td>8.0E-08</td><td>1.4E-07</td><td>MB231/MB468</td><td>MITO</td></tr>
<tr><td>>hg38_21_+_8433172_8446622_RNA45SN1_45S_50ntFlanks@8381.8399.19|AAGAGGGCGTGAAACCGTT</td><td>>hg38_21_+_8433172_8446622_RNA45SN1_45S_50ntFlanks</td><td>8381</td><td>8399</td><td>19</td><td>AAGAGGGCGTGAAACCGTT</td><td>163.6</td><td>-6.0</td><td>1.1</td><td>-5.4</td><td>6.4E-08</td><td>1.4E-07</td><td>BT20/MB468</td><td>CYTO</td></tr>
<tr><td>>hg38_21_+_8433172_8446622_RNA45SN1_45S_50ntFlanks@11885.11906.22|TTGAGCTTGACTCTAGTCTGGC</td><td>>hg38_21_+_8433172_8446622_RNA45SN1_45S_50ntFlanks</td><td>11885</td><td>11906</td><td>22</td><td>TTGAGCTTGACTCTAGTCTGGC</td><td>108.2</td><td>-9.8</td><td>1.7</td><td>-5.7</td><td>1.4E-08</td><td>1.4E-07</td><td>BT20/MB468</td><td>MP</td></tr>
<tr><td>>MI0003513|hsa-mir-455&WithFlank&9|+|114209428|114209535@22.43.22(+1A)&[MIMAT0003150&hsa-miR-455-5p&offsets|0|0(+1A);m-232&9|+|114209449|114209470&offsets|0|0(+1A)]|TATGTGCCTTTGGACTACATCGA</td><td>>MI0003513|hsa-mir-455&WithFlank&9|+|114209428|114209535</td><td>22</td><td>43</td><td>22(+1A)</td><td>TATGTGCCTTTGGACTACATCGA</td><td>55.9</td><td>9.0</td><td>1.7</td><td>5.4</td><td>6.9E-08</td><td>1.4E-07</td><td>MB231/MB468</td><td>MP</td></tr>
<tr><td>>hg38_21_+_8433172_8446622_RNA45SN1_45S_50ntFlanks@12405.12432.28|CTTTTTGATCCTTCGATGTCGGCTCTTC</td><td>>hg38_21_+_8433172_8446622_RNA45SN1_45S_50ntFlanks</td><td>12405</td><td>12432</td><td>28</td><td>CTTTTTGATCCTTCGATGTCGGCTCTTC</td><td>12984.6</td><td>-7.4</td><td>1.4</td><td>-5.4</td><td>6.5E-08</td><td>1.4E-07</td><td>BT20/MB468</td><td>CYTO</td></tr>
<tr><td>>hg38_21_+_8433172_8446622_RNA45SN1_45S_50ntFlanks@10350.10382.33|ACGAGAACTTTGAAGGCCGAAGTGGAGAAGGGT</td><td>>hg38_21_+_8433172_8446622_RNA45SN1_45S_50ntFlanks</td><td>10350</td><td>10382</td><td>33</td><td>ACGAGAACTTTGAAGGCCGAAGTGGAGAAGGGT</td><td>1983.0</td><td>-4.6</td><td>0.9</td><td>-5.4</td><td>6.9E-08</td><td>1.4E-07</td><td>MB231/MB468</td><td>MP</td></tr>
<tr><td>>hg38_21_+_8433172_8446622_RNA45SN1_45S_50ntFlanks@10396.10422.27|CAGTTGAACATGGGTCAGTCGGTCCTG</td><td>>hg38_21_+_8433172_8446622_RNA45SN1_45S_50ntFlanks</td><td>10396</td><td>10422</td><td>27</td><td>CAGTTGAACATGGGTCAGTCGGTCCTG</td><td>267.0</td><td>-4.8</td><td>0.9</td><td>-5.4</td><td>6.9E-08</td><td>1.4E-07</td><td>MB231/MB468</td><td>MP</td></tr>
<tr><td>>hg38_21_+_8433172_8446622_RNA45SN1_45S_50ntFlanks@12712.12749.38|CGGAGCCTCGGTTGGCCTCGGATAGCCGGTCCCCCGCC</td><td>>hg38_21_+_8433172_8446622_RNA45SN1_45S_50ntFlanks</td><td>12712</td><td>12749</td><td>38</td><td>CGGAGCCTCGGTTGGCCTCGGATAGCCGGTCCCCCGCC</td><td>81.2</td><td>-10.2</td><td>1.9</td><td>-5.4</td><td>6.9E-08</td><td>1.4E-07</td><td>MB231/MB468</td><td>MP</td></tr>
<tr><td>>hg38_21_+_8433172_8446622_RNA45SN1_45S_50ntFlanks@9566.9597.32|GACGTGCAAATCGGTCGTCCGACCTGGGTATA</td><td>>hg38_21_+_8433172_8446622_RNA45SN1_45S_50ntFlanks</td><td>9566</td><td>9597</td><td>32</td><td>GACGTGCAAATCGGTCGTCCGACCTGGGTATA</td><td>2747.4</td><td>-9.3</td><td>1.7</td><td>-5.4</td><td>8.5E-08</td><td>1.4E-07</td><td>BT20/MB468</td><td>NUC</td></tr>
<tr><td>>hg38_21_+_8433172_8446622_RNA45SN1_45S_50ntFlanks@11576.11609.34|CTTAGAACTGGTGCGGACCAGGGGAATCCGACTG</td><td>>hg38_21_+_8433172_8446622_RNA45SN1_45S_50ntFlanks</td><td>11576</td><td>11609</td><td>34</td><td>CTTAGAACTGGTGCGGACCAGGGGAATCCGACTG</td><td>672.1</td><td>-5.1</td><td>0.9</td><td>-5.4</td><td>7.4E-08</td><td>1.4E-07</td><td>MB231/MB468</td><td>CYTO</td></tr>
<tr><td>>MI0003834|hsa-mir-769&WithFlank&19|+|46018926|46019055@36.57.22&[MIMAT0003886&hsa-miR-769-5p&offsets|0|0;m-216&19|+|46018961|46018982&offsets|0|0]|TGAGACCTCTGGGTTCTGAGCT</td><td>>MI0003834|hsa-mir-769&WithFlank&19|+|46018926|46019055</td><td>36</td><td>57</td><td>22</td><td>TGAGACCTCTGGGTTCTGAGCT</td><td>1106.3</td><td>5.4</td><td>1.0</td><td>5.4</td><td>8.1E-08</td><td>1.4E-07</td><td>MB231/MB468</td><td>MITO</td></tr>
<tr><td>>hg38_21_+_8433172_8446622_RNA45SN1_45S_50ntFlanks@12942.12979.38|TTCGTAGACGACCTGCTTCTGGGTCGGGGTTTCGTACG</td><td>>hg38_21_+_8433172_8446622_RNA45SN1_45S_50ntFlanks</td><td>12942</td><td>12979</td><td>38</td><td>TTCGTAGACGACCTGCTTCTGGGTCGGGGTTTCGTACG</td><td>51.9</td><td>-9.6</td><td>1.8</td><td>-5.4</td><td>6.9E-08</td><td>1.4E-07</td><td>MB231/MB468</td><td>MP</td></tr>
<tr><td>>hg38_21_+_8433172_8446622_RNA45SN1_45S_50ntFlanks@11999.12034.36|CCCTGCGGGCCGCCGGTGAAATACCACTACTCTGAT</td><td>>hg38_21_+_8433172_8446622_RNA45SN1_45S_50ntFlanks</td><td>11999</td><td>12034</td><td>36</td><td>CCCTGCGGGCCGCCGGTGAAATACCACTACTCTGAT</td><td>50.2</td><td>-9.5</td><td>1.8</td><td>-5.4</td><td>6.9E-08</td><td>1.4E-07</td><td>MB231/MB468</td><td>MP</td></tr>
<tr><td>>hg38_21_+_8433172_8446622_RNA45SN1_45S_50ntFlanks@10431.10456.26|GCGAGCGCCGTTCCGAAGGGACGGGC</td><td>>hg38_21_+_8433172_8446622_RNA45SN1_45S_50ntFlanks</td><td>10431</td><td>10456</td><td>26</td><td>GCGAGCGCCGTTCCGAAGGGACGGGC</td><td>200.9</td><td>-5.8</td><td>1.1</td><td>-5.4</td><td>7.5E-08</td><td>1.4E-07</td><td>MB231/MB468</td><td>CYTO</td></tr>
<tr><td>>hg38_21_+_8433172_8446622_RNA45SN1_45S_50ntFlanks@4760.4777.18|TCAGATACCGTCGTAGTT</td><td>>hg38_21_+_8433172_8446622_RNA45SN1_45S_50ntFlanks</td><td>4760</td><td>4777</td><td>18</td><td>TCAGATACCGTCGTAGTT</td><td>244.7</td><td>-7.3</td><td>1.4</td><td>-5.4</td><td>6.9E-08</td><td>1.4E-07</td><td>MB231/MB468</td><td>MP</td></tr>
<tr><td>>hg38_21_+_8433172_8446622_RNA45SN1_45S_50ntFlanks@11574.11603.30|GACTTAGAACTGGTGCGGACCAGGGGAATC</td><td>>hg38_21_+_8433172_8446622_RNA45SN1_45S_50ntFlanks</td><td>11574</td><td>11603</td><td>30</td><td>GACTTAGAACTGGTGCGGACCAGGGGAATC</td><td>524.7</td><td>-8.1</td><td>1.5</td><td>-5.4</td><td>8.5E-08</td><td>1.4E-07</td><td>BT20/MB468</td><td>NUC</td></tr>
<tr><td>>hg38_21_+_8433172_8446622_RNA45SN1_45S_50ntFlanks@10408.10430.23|GGTCAGTCGGTCCTGAGAGATGG</td><td>>hg38_21_+_8433172_8446622_RNA45SN1_45S_50ntFlanks</td><td>10408</td><td>10430</td><td>23</td><td>GGTCAGTCGGTCCTGAGAGATGG</td><td>1667.9</td><td>-7.6</td><td>1.4</td><td>-5.4</td><td>6.5E-08</td><td>1.4E-07</td><td>BT20/MB468</td><td>CYTO</td></tr>
<tr><td>>hg38_21_+_8433172_8446622_RNA45SN1_45S_50ntFlanks@5084.5111.28|CGAACGAGACTCTGGCATGCTAACTAGT</td><td>>hg38_21_+_8433172_8446622_RNA45SN1_45S_50ntFlanks</td><td>5084</td><td>5111</td><td>28</td><td>CGAACGAGACTCTGGCATGCTAACTAGT</td><td>87.4</td><td>-10.3</td><td>1.9</td><td>-5.4</td><td>6.9E-08</td><td>1.4E-07</td><td>MB231/MB468</td><td>MP</td></tr>
<tr><td>>hg38_1_-_228634819_228635039_RNA5S12_5S_50ntFlanks@51.79.29|GTCTACGGCCATACCACCCTGAACGCGCC</td><td>>hg38_1_-_228634819_228635039_RNA5S12_5S_50ntFlanks</td><td>51</td><td>79</td><td>29</td><td>GTCTACGGCCATACCACCCTGAACGCGCC</td><td>1998.9</td><td>5.7</td><td>1.1</td><td>5.4</td><td>6.5E-08</td><td>1.4E-07</td><td>BT20/MB468</td><td>CYTO</td></tr>
<tr><td>>hg38_21_+_8433172_8446622_RNA45SN1_45S_50ntFlanks@10349.10378.30|AACGAGAACTTTGAAGGCCGAAGTGGAGAA</td><td>>hg38_21_+_8433172_8446622_RNA45SN1_45S_50ntFlanks</td><td>10349</td><td>10378</td><td>30</td><td>AACGAGAACTTTGAAGGCCGAAGTGGAGAA</td><td>576.0</td><td>-3.2</td><td>0.6</td><td>-5.4</td><td>7.7E-08</td><td>1.4E-07</td><td>MB231/MB468</td><td>NUC</td></tr>
<tr><td>>trna111_HisGTG_1_-_147774845_147774916@-1A.20.21,trna118_HisGTG_1_-_145396881_145396952@-1A.20.21,trna16_HisGTG_1_+_146544773_146544844@-1A.20.21,trna1_HisGTG_15_+_45493349_45493420@-1A.20.21,trna21_HisGTG_1_+_147753471_147753542@-1A.20.21,trna33_HisGTG_6_+_27125906_27125977@-1A.20.21,trna7_HisGTG_9_-_14433938_14434009@-1A.20.21,trna8_HisGTG_15_-_45492611_45492682@-1A.20.21,trna9_HisGTG_15_-_45490804_45490875@-1A.20.21|AGCCGTGATCGTATAGTGGTT</td><td>>trna111_HisGTG_1_-_147774845_147774916</td><td>-1A</td><td>20</td><td>21</td><td>AGCCGTGATCGTATAGTGGTT</td><td>79.7</td><td>9.5</td><td>1.8</td><td>5.4</td><td>7.0E-08</td><td>1.4E-07</td><td>MB231/MB468</td><td>MP</td></tr>
<tr><td>>hg38_21_+_8433172_8446622_RNA45SN1_45S_50ntFlanks@6654.6686.33|CTCTTAGCGGTGGATCACTCGGCTCGTGCGTCG</td><td>>hg38_21_+_8433172_8446622_RNA45SN1_45S_50ntFlanks</td><td>6654</td><td>6686</td><td>33</td><td>CTCTTAGCGGTGGATCACTCGGCTCGTGCGTCG</td><td>487.1</td><td>-4.1</td><td>0.8</td><td>-5.4</td><td>8.2E-08</td><td>1.4E-07</td><td>MB231/MB468</td><td>MITO</td></tr>
<tr><td>>hg38_21_+_8433172_8446622_RNA45SN1_45S_50ntFlanks@3776.3794.19|CCGGTACAGTGAAACTGCG</td><td>>hg38_21_+_8433172_8446622_RNA45SN1_45S_50ntFlanks</td><td>3776</td><td>3794</td><td>19</td><td>CCGGTACAGTGAAACTGCG</td><td>82.5</td><td>-9.4</td><td>1.7</td><td>-5.7</td><td>1.4E-08</td><td>1.4E-07</td><td>BT20/MB468</td><td>MP</td></tr>
<tr><td>>hg38_21_+_8433172_8446622_RNA45SN1_45S_50ntFlanks@3802.3829.28|CATTAAATCAGTTATGGTTCCTTTGGTC</td><td>>hg38_21_+_8433172_8446622_RNA45SN1_45S_50ntFlanks</td><td>3802</td><td>3829</td><td>28</td><td>CATTAAATCAGTTATGGTTCCTTTGGTC</td><td>310.9</td><td>-5.3</td><td>1.0</td><td>-5.4</td><td>7.5E-08</td><td>1.4E-07</td><td>MB231/MB468</td><td>CYTO</td></tr>
<tr><td>>hg38_21_+_8433172_8446622_RNA45SN1_45S_50ntFlanks@12008.12033.26|CCGCCGGTGAAATACCACTACTCTGA</td><td>>hg38_21_+_8433172_8446622_RNA45SN1_45S_50ntFlanks</td><td>12008</td><td>12033</td><td>26</td><td>CCGCCGGTGAAATACCACTACTCTGA</td><td>444.8</td><td>-7.2</td><td>1.3</td><td>-5.4</td><td>7.7E-08</td><td>1.4E-07</td><td>MB231/MB468</td><td>NUC</td></tr>
<tr><td>>hg38_21_+_8433172_8446622_RNA45SN1_45S_50ntFlanks@12003.12032.30|GCGGGCCGCCGGTGAAATACCACTACTCTG</td><td>>hg38_21_+_8433172_8446622_RNA45SN1_45S_50ntFlanks</td><td>12003</td><td>12032</td><td>30</td><td>GCGGGCCGCCGGTGAAATACCACTACTCTG</td><td>2794.2</td><td>-4.1</td><td>0.8</td><td>-5.4</td><td>6.5E-08</td><td>1.4E-07</td><td>BT20/MB468</td><td>CYTO</td></tr>
</table>

| | | | | | | | | | | | | | |
|---|---|---|---|---|---|---|---|---|---|---|---|---|---|
| >hg38_21_+_8433172_8446622_RNA45SN1_45S_50ntFlanks@10797.10823.27\|GCATGTTGGAACAATGTAGGTAAGGGA | >hg38_21_+_8433172_8446622_RNA45SN1_45S_50ntFlanks | 10797 | 10823 | 27 | GCATGTTGGAACAATGTAGGTAAGGGA | 317.1 | -7.0 | 1.3 | -5.4 | 8.2E-08 | 1.4E-07 | MB231/MB468 | MITO |
| >hg38_21_+_8433172_8446622_RNA45SN1_45S_50ntFlanks@475.501.27\|CTTCGTGATCGATGTGGTGACGTCGTG | >hg38_21_+_8433172_8446622_RNA45SN1_45S_50ntFlanks | 475 | 501 | 27 | CTTCGTGATCGATGTGGTGACGTCGTG | 1240.3 | 4.2 | 0.8 | 5.4 | 7.7E-08 | 1.4E-07 | MB231/MB468 | NUC |
| >hg38_21_+_8433172_8446622_RNA45SN1_45S_50ntFlanks@9565.9594.30\|TGACGTGCAAATCGGTCGTCCGACCTGGGT | >hg38_21_+_8433172_8446622_RNA45SN1_45S_50ntFlanks | 9565 | 9594 | 30 | TGACGTGCAAATCGGTCGTCCGACCTGGGT | 397.0 | -4.0 | 0.8 | -5.4 | 8.2E-08 | 1.4E-07 | MB231/MB468 | MITO |
| >hg38_21_+_8433172_8446622_RNA45SN1_45S_50ntFlanks@10873.10894.22\|CTGGGTCGGTCGGGCTGGGGCG | >hg38_21_+_8433172_8446622_RNA45SN1_45S_50ntFlanks | 10873 | 10894 | 22 | CTGGGTCGGTCGGGCTGGGGCG | 1342.5 | -8.9 | 1.7 | -5.4 | 8.6E-08 | 1.4E-07 | BT20/MB468 | NUC |
| >hg38_21_+_8433172_8446622_RNA45SN1_45S_50ntFlanks@9034.9056.23\|TCGTGGGGGGGCCGGGCCACCCC | >hg38_21_+_8433172_8446622_RNA45SN1_45S_50ntFlanks | 9034 | 9056 | 23 | TCGTGGGGGGGCCGGGCCACCCC | 111.0 | -9.9 | 1.7 | -5.7 | 1.4E-08 | 1.4E-07 | BT20/MB468 | MP |
| >hg38_21_+_8433172_8446622_RNA45SN1_45S_50ntFlanks@12202.12238.37\|CTAAGGCGAGCTCAGGGAGGACAGAAACCTCCCGTGG | >hg38_21_+_8433172_8446622_RNA45SN1_45S_50ntFlanks | 12202 | 12238 | 37 | CTAAGGCGAGCTCAGGGAGGACAGAAACCTCCCGTGG | 92.2 | -10.4 | 1.9 | -5.4 | 7.0E-08 | 1.4E-07 | MB231/MB468 | MP |
| >hg38_21_+_8433172_8446622_RNA45SN1_45S_50ntFlanks@5334.5352.19\|ATGAACGAGGAATTCCCAG | >hg38_21_+_8433172_8446622_RNA45SN1_45S_50ntFlanks | 5334 | 5352 | 19 | ATGAACGAGGAATTCCCAG | 85.6 | -9.5 | 1.7 | -5.7 | 1.4E-08 | 1.4E-07 | BT20/MB468 | MP |
| >hg38_21_+_8433172_8446622_RNA45SN1_45S_50ntFlanks@5421.5443.23\|CGATTGGATGGTTTAGTGAGGCC | >hg38_21_+_8433172_8446622_RNA45SN1_45S_50ntFlanks | 5421 | 5443 | 23 | CGATTGGATGGTTTAGTGAGGCC | 363.4 | -6.7 | 1.2 | -5.4 | 5.9E-08 | 1.4E-07 | BT20/MB468 | MITO |
| >hg38_21_+_8433172_8446622_RNA45SN1_45S_50ntFlanks@12932.12956.25\|CGCTAAACCATTCGTAGACGACCTG | >hg38_21_+_8433172_8446622_RNA45SN1_45S_50ntFlanks | 12932 | 12956 | 25 | CGCTAAACCATTCGTAGACGACCTG | 684.6 | -3.3 | 0.6 | -5.4 | 7.6E-08 | 1.4E-07 | MB231/MB468 | CYTO |
| >hg38_MT_+_1621_3279_RNR2_16S_50ntFlanks@714.740.27\|CATAAGCCTGCGTCAGATTAAAACACT | >hg38_MT_+_1621_3279_RNR2_16S_50ntFlanks | 714 | 740 | 27 | CATAAGCCTGCGTCAGATTAAAACACT | 65.9 | 9.3 | 1.7 | 5.4 | 7.0E-08 | 1.4E-07 | MB231/MB468 | MP |
| >am_trna111_HisGTG_1_-_147774845_147774916@-1G.29.30,trna118_HisGTG_1_-_145396881_145396952@-1G.29.30,trna16_HisGTG_1_+_146544773_146544844@-1G.29.30,trna1_HisGTG_15_+_45493349_45493420@-1G.29.30,trna21_HisGTG_1_+_147753471_147753542@-1G.29.30,trna33_HisGTG_6_+_27125906_27125977@-1G.29.30,trna7_HisGTG_9_-_14433938_14434009@-1G.29.30,trna8_HisGTG_15_-_45492611_45492682@-1G.29.30,trna9_HisGTG_15_-_45490804_45490875@-1G.29.30\|GGCCGTGATCGTATAGTGGTTAGTACTCTG | >am_trna111_HisGTG_1_-_147774845_147774916 | -1G | 29 | 30 | GGCCGTGATCGTATAGTGGTTAGTACTCTG | 4252.2 | -4.0 | 0.8 | -5.4 | 8.3E-08 | 1.4E-07 | MB231/MB468 | MITO |
| >hg38_21_+_8433172_8446622_RNA45SN1_45S_50ntFlanks@11325.11344.20\|TCTCGTCGCGCGCGCGTCCG | >hg38_21_+_8433172_8446622_RNA45SN1_45S_50ntFlanks | 11325 | 11344 | 20 | TCTCGTCGCGCGCGCGTCCG | 1063.1 | -6.6 | 1.2 | -5.4 | 8.3E-08 | 1.4E-07 | MB231/MB468 | MITO |
| >hg38_21_+_8433172_8446622_RNA45SN1_45S_50ntFlanks@11654.11685.32\|CGCGATGTGATTTCTGCCCAGTGCTCTGAATG | >hg38_21_+_8433172_8446622_RNA45SN1_45S_50ntFlanks | 11654 | 11685 | 32 | CGCGATGTGATTTCTGCCCAGTGCTCTGAATG | 367.1 | -6.4 | 1.2 | -5.4 | 8.3E-08 | 1.4E-07 | MB231/MB468 | MITO |
| >hg38_21_+_8433172_8446622_RNA45SN1_45S_50ntFlanks@9468.9503.36\|TGGAGCACGAGCGCACGTGTTAGGACCCGAAAGATG | >hg38_21_+_8433172_8446622_RNA45SN1_45S_50ntFlanks | 9468 | 9503 | 36 | TGGAGCACGAGCGCACGTGTTAGGACCCGAAAGATG | 329.9 | -7.3 | 1.4 | -5.4 | 8.2E-08 | 1.4E-07 | MB231/MB468 | MITO |
| >hg38_21_+_8433172_8446622_RNA45SN1_45S_50ntFlanks@12324.12348.25\|TTGGGTTTTAAGCAGGAGGTGTCAG | >hg38_21_+_8433172_8446622_RNA45SN1_45S_50ntFlanks | 12324 | 12348 | 25 | TTGGGTTTTAAGCAGGAGGTGTCAG | 798.6 | -8.0 | 1.4 | -5.7 | 1.4E-08 | 1.4E-07 | BT20/MB468 | MP |
| >hg38_21_+_8433172_8446622_RNA45SN1_45S_50ntFlanks@9554.9595.42\|CGTAGCGGTCCTGACGTGCAAATCGGTCGTCCGACCTGGGTA | >hg38_21_+_8433172_8446622_RNA45SN1_45S_50ntFlanks | 9554 | 9595 | 42 | CGTAGCGGTCCTGACGTGCAAATCGGTCGTCCGACCTGGGTA | 1759.1 | -10.4 | 1.9 | -5.4 | 8.6E-08 | 1.4E-07 | BT20/MB468 | NUC |
| >MI0000446\|hsa-mir-125b-1&WithFlank&11\|-\|122099751\|122099850@21.42.22(+1A);MI0000470\|hsa-mir-125b-2&WithFlank&21\|+\|16590231\|16590331@23.44.22(+1A)&[MIMAT0000423&hsa-miR-125b-5p&offsets\|0\|0(+1A);m-67&11\|-\|122099809\|122099830&offsets\|0\|0(+1A)];[MIMAT0000423_1&hsa-miR-125b-5p&offsets\|0\|0(+1A);m-68&21\|+\|16590253\|16590274&offsets\|0\|0(+1A)]\|TCCCTGAGACCCTAACTTGTGAA | >MI0000446\|hsa-mir-125b-1&WithFlank&11\|-\|122099751\|122099850 | 21 | 42 | 22(+1A) | TCCCTGAGACCCTAACTTGTGAA | 865.4 | 4.4 | 0.8 | 5.4 | 7.0E-08 | 1.4E-07 | MB231/MB468 | MP |
| >hg38_21_+_8433172_8446622_RNA45SN1_45S_50ntFlanks@8439.8469.31\|GATTCAACCCGGCGGCGGGTCCGGCCGTGTC | >hg38_21_+_8433172_8446622_RNA45SN1_45S_50ntFlanks | 8439 | 8469 | 31 | GATTCAACCCGGCGGCGGGTCCGGCCGTGTC | 99.5 | -9.7 | 1.7 | -5.7 | 1.4E-08 | 1.4E-07 | BT20/MB468 | MP |
| >hg38_21_+_8433172_8446622_RNA45SN1_45S_50ntFlanks@8155.8175.21\|CGCTCGTGGGGGGCCCAAGTC | >hg38_21_+_8433172_8446622_RNA45SN1_45S_50ntFlanks | 8155 | 8175 | 21 | CGCTCGTGGGGGGCCCAAGTC | 91.6 | -9.6 | 1.7 | -5.7 | 1.4E-08 | 1.4E-07 | BT20/MB468 | MP |
| >am_trnaMT_ProTGG_MT_-_15956_16023@52.71.20\|AAAGACTTTTTCTCTGACCA | >am_trnaMT_ProTGG_MT_-_15956_16023 | 52 | 71 | 20 | AAAGACTTTTTCTCTGACCA | 4736.4 | 5.6 | 1.0 | 5.4 | 5.9E-08 | 1.4E-07 | BT20/MB468 | MITO |
| >hg38_21_+_8433172_8446622_RNA45SN1_45S_50ntFlanks@12044.12070.27\|ACTGACCCGGTGAGGCGGGGGGGCGAG | >hg38_21_+_8433172_8446622_RNA45SN1_45S_50ntFlanks | 12044 | 12070 | 27 | ACTGACCCGGTGAGGCGGGGGGGCGAG | 1645.5 | -7.7 | 1.4 | -5.4 | 5.9E-08 | 1.4E-07 | BT20/MB468 | MITO |
| >hg38_21_+_8433172_8446622_RNA45SN1_45S_50ntFlanks@8381.8402.22\|AAGAGGGCGTGAAACCGTTAAG | >hg38_21_+_8433172_8446622_RNA45SN1_45S_50ntFlanks | 8381 | 8402 | 22 | AAGAGGGCGTGAAACCGTTAAG | 5948.4 | -6.0 | 1.1 | -5.7 | 1.4E-08 | 1.4E-07 | BT20/MB468 | MP |
| >hg38_21_+_8433172_8446622_RNA45SN1_45S_50ntFlanks@5423.5466.44\|ATTGGATGGTTTAGTGAGGCCCTCGGATCGGCCCCGCCGGGGTC | >hg38_21_+_8433172_8446622_RNA45SN1_45S_50ntFlanks | 5423 | 5466 | 44 | ATTGGATGGTTTAGTGAGGCCCTCGGATCGGCCCCGCCGGGGTC | 3144.2 | -9.8 | 1.8 | -5.4 | 8.7E-08 | 1.4E-07 | BT20/MB468 | NUC |
| >hg38_21_+_8433172_8446622_RNA45SN1_45S_50ntFlanks@6699.6720.22\|GCTAGCTGCGAGAATTAATGTG | >hg38_21_+_8433172_8446622_RNA45SN1_45S_50ntFlanks | 6699 | 6720 | 22 | GCTAGCTGCGAGAATTAATGTG | 38781.7 | -4.0 | 0.7 | -5.4 | 7.6E-08 | 1.4E-07 | MB231/MB468 | CYTO |
| >hg38_21_+_8433172_8446622_RNA45SN1_45S_50ntFlanks@9929.9957.29\|GATATAGACAGCAGGACGGTGGCCATGGA | >hg38_21_+_8433172_8446622_RNA45SN1_45S_50ntFlanks | 9929 | 9957 | 29 | GATATAGACAGCAGGACGGTGGCCATGGA | 593.9 | -6.4 | 1.2 | -5.4 | 7.1E-08 | 1.4E-07 | MB231/MB468 | MP |
| >hg38_21_+_8433172_8446622_RNA45SN1_45S_50ntFlanks@12709.12747.39\|CCGCGGAGCCTCGGTTGGCCTCGGATAGCCGGTCCCCCG | >hg38_21_+_8433172_8446622_RNA45SN1_45S_50ntFlanks | 12709 | 12747 | 39 | CCGCGGAGCCTCGGTTGGCCTCGGATAGCCGGTCCCCCG | 127.5 | -10.1 | 1.8 | -5.7 | 1.4E-08 | 1.4E-07 | BT20/MB468 | MP |
| >am_trna128_GlyGCC_6_-_27870686_27870756@10.28.19,trna133_GlyCCC_1_-_16872434_16872504@10.28.19,trna17_GluTTC_2_-_204229131_204229201@10.28.19,trna18_GlyGCC_16_+_70822597_70822667@10.28.19,trna19_GlyGCC_16_+_70823410_70823480@10.28.19,trna19_GlyGCC_2_-_157257659_157257729@10.28.19,trna24_GlyGCC_16_-_70812942_70813012@10.28.19,trna25_GlyGCC_16_-_70812114_70812184@10.28.19,trna2_GlyGCC_21_-_18827107_18827177@10.28.19,trna35_GlyGCC_1_+_161413094_161413164@10.28.19,trna37_GlyGCC_1_+_161420467_161420537@10.28.19,trna39_GlyGCC_1_+_161427898_161427968@10.28.19,trna41_GlyGCC_1_+_161435258_161435328@10.28.19,trna4_GlyCCC_1_+_17188416_17188486@10.28.19,trna5_GlyGCC_17_+_8029064_8029134@10.28.19,trna68_GlyGCC_1_-_161493637_161493707@10.28.19,trna91_GlyCCC_1_-_149680210_149680280@10.28.19,trna99_ValCAC_1_-_149294666_149294736@10.28.19\|GTTCAGTGGTAGAATTCTC | >am_trna128_GlyGCC_6_-_27870686_27870756 | 10 | 28 | 19 | GTTCAGTGGTAGAATTCTC | 163.9 | -10.4 | 1.8 | -5.7 | 1.4E-08 | 1.4E-07 | BT20/MB468 | MP |
| >hg38_21_+_8433172_8446622_RNA45SN1_45S_50ntFlanks@12716.12742.27\|GCCTCGGTTGGCCTCGGATAGCCGGTC | >hg38_21_+_8433172_8446622_RNA45SN1_45S_50ntFlanks | 12716 | 12742 | 27 | GCCTCGGTTGGCCTCGGATAGCCGGTC | 95.8 | -9.7 | 1.7 | -5.7 | 1.4E-08 | 1.4E-07 | BT20/MB468 | MP |
| >trnaMT_PheGAA_MT_+_577_647@36.55.20\|AAAATGTTTAGACGGGCTCA | >trnaMT_PheGAA_MT_+_577_647 | 36 | 55 | 20 | AAAATGTTTAGACGGGCTCA | 842.9 | 7.6 | 1.3 | 5.7 | 1.4E-08 | 1.4E-07 | BT20/MB468 | MP |
| >hg38_21_+_8433172_8446622_RNA45SN1_45S_50ntFlanks@10461.10497.37\|GCCTCCGTTGCCCTCGGCCGATCGAAAGGGAGTCGGG | >hg38_21_+_8433172_8446622_RNA45SN1_45S_50ntFlanks | 10461 | 10497 | 37 | GCCTCCGTTGCCCTCGGCCGATCGAAAGGGAGTCGGG | 100.2 | -9.7 | 1.7 | -5.7 | 1.4E-08 | 1.4E-07 | BT20/MB468 | MP |
| >hg38_21_+_8433172_8446622_RNA45SN1_45S_50ntFlanks@11319.11342.24\|CCCACGTCTCGTCGCGCGCGCGTC | >hg38_21_+_8433172_8446622_RNA45SN1_45S_50ntFlanks | 11319 | 11342 | 24 | CCCACGTCTCGTCGCGCGCGCGTC | 96.2 | -9.7 | 1.7 | -5.7 | 1.4E-08 | 1.4E-07 | BT20/MB468 | MP |
| >hg38_21_+_8433172_8446622_RNA45SN1_45S_50ntFlanks@11933.11958.26\|AATAAGTGGGAGGCCCCCGGCGCCCC | >hg38_21_+_8433172_8446622_RNA45SN1_45S_50ntFlanks | 11933 | 11958 | 26 | AATAAGTGGGAGGCCCCCGGCGCCCC | 92.4 | -9.6 | 1.7 | -5.7 | 1.4E-08 | 1.4E-07 | BT20/MB468 | MP |
| >hg38_21_+_8433172_8446622_RNA45SN1_45S_50ntFlanks@5492.5532.41\|AGAAGACGGTCGAACTTGACTATCTAGAGGAAGTAAAAGTC | >hg38_21_+_8433172_8446622_RNA45SN1_45S_50ntFlanks | 5492 | 5532 | 41 | AGAAGACGGTCGAACTTGACTATCTAGAGGAAGTAAAAGTC | 664.8 | 5.5 | 1.0 | 5.4 | 6.7E-08 | 1.4E-07 | BT20/MB468 | CYTO |
| >MI0003581\|hsa-mir-574&WithFlank&4\|+\|38868026\|38868133@67.88.22&[MIMAT0003239&hsa-miR-574-3p&offsets\|0\|0;m-149&4\|+\|38868092\|38868113&offsets\|0\|0]\|CACGCTCATGCACACACCCACA | >MI0003581\|hsa-mir-574&WithFlank&4\|+\|38868026\|38868133 | 67 | 88 | 22 | CACGCTCATGCACACACCCACA | 1049.1 | 5.1 | 0.9 | 5.4 | 8.3E-08 | 1.4E-07 | MB231/MB468 | MITO |
| >hg38_21_+_8433172_8446622_RNA45SN1_45S_50ntFlanks@12607.12625.19\|TGTGCTTGGCTGAGGAGCC | >hg38_21_+_8433172_8446622_RNA45SN1_45S_50ntFlanks | 12607 | 12625 | 19 | TGTGCTTGGCTGAGGAGCC | 331.0 | -3.6 | 0.7 | -5.4 | 7.1E-08 | 1.4E-07 | MB231/MB468 | MP |
| >hg38_21_+_8433172_8446622_RNA45SN1_45S_50ntFlanks@12461.12487.27\|CGTTGGATTGTTCACCCACTAATAGGG | >hg38_21_+_8433172_8446622_RNA45SN1_45S_50ntFlanks | 12461 | 12487 | 27 | CGTTGGATTGTTCACCCACTAATAGGG | 97.0 | -9.7 | 1.7 | -5.7 | 1.4E-08 | 1.4E-07 | BT20/MB468 | MP |
| >hg38_21_+_8433172_8446622_RNA45SN1_45S_50ntFlanks@11235.11252.18\|CTTCCCGTGGATCGCCCC | >hg38_21_+_8433172_8446622_RNA45SN1_45S_50ntFlanks | 11235 | 11252 | 18 | CTTCCCGTGGATCGCCCC | 1992.0 | -6.1 | 1.1 | -5.4 | 6.7E-08 | 1.4E-07 | BT20/MB468 | CYTO |
| >hg38_21_+_8433172_8446622_RNA45SN1_45S_50ntFlanks@3705.3747.43\|TACCTGGTTGATCCTGCCAGTAGCATATGCTTGTCTCAAAGAT | >hg38_21_+_8433172_8446622_RNA45SN1_45S_50ntFlanks | 3705 | 3747 | 43 | TACCTGGTTGATCCTGCCAGTAGCATATGCTTGTCTCAAAGAT | 3018.2 | 10.0 | 1.8 | 5.4 | 6.7E-08 | 1.4E-07 | BT20/MB468 | CYTO |
| >hg38_MT_+_598_1651_RNR1_12S_50ntFlanks@570.612.43\|AAGGACCTGGCGGTGCTTCATATCCCTCTAGAGGAGCCTGTTC | >hg38_MT_+_598_1651_RNR1_12S_50ntFlanks | 570 | 612 | 43 | AAGGACCTGGCGGTGCTTCATATCCCTCTAGAGGAGCCTGTTC | 52.4 | 8.9 | 1.7 | 5.4 | 7.1E-08 | 1.4E-07 | MB231/MB468 | MP |
| >hg38_21_+_8433172_8446622_RNA45SN1_45S_50ntFlanks@4006.4042.37\|ACCTCGGGCCGATCGCACGCCCCCCGTGGCGGCGACG | >hg38_21_+_8433172_8446622_RNA45SN1_45S_50ntFlanks | 4006 | 4042 | 37 | ACCTCGGGCCGATCGCACGCCCCCCGTGGCGGCGACG | 51.6 | -9.6 | 1.8 | -5.4 | 7.1E-08 | 1.4E-07 | MB231/MB468 | MP |
| >hg38_21_+_8433172_8446622_RNA45SN1_45S_50ntFlanks@9345.9368.24\|AAAGCCGCCGTGGCGCAATGAAGG | >hg38_21_+_8433172_8446622_RNA45SN1_45S_50ntFlanks | 9345 | 9368 | 24 | AAAGCCGCCGTGGCGCAATGAAGG | 54.1 | -9.6 | 1.8 | -5.4 | 7.1E-08 | 1.4E-07 | MB231/MB468 | MP |
| >hg38_21_+_8433172_8446622_RNA45SN1_45S_50ntFlanks@11323.11345.23\|CGTCTCGTCGCGCGCGCGTCCGC | >hg38_21_+_8433172_8446622_RNA45SN1_45S_50ntFlanks | 11323 | 11345 | 23 | CGTCTCGTCGCGCGCGCGTCCGC | 1326.7 | -6.8 | 1.3 | -5.4 | 8.4E-08 | 1.4E-07 | MB231/MB468 | MITO |
| >hg38_21_+_8433172_8446622_RNA45SN1_45S_50ntFlanks@12998.13039.42\|GCTGCGATCTATTGAAAGTCAGCCCTCGACACAAGGGTTTGT | >hg38_21_+_8433172_8446622_RNA45SN1_45S_50ntFlanks | 12998 | 13039 | 42 | GCTGCGATCTATTGAAAGTCAGCCCTCGACACAAGGGTTTGT | 5598.2 | 5.8 | 1.1 | 5.4 | 7.9E-08 | 1.4E-07 | MB231/MB468 | NUC |

<table>
<tr><th></th><th></th><th></th><th></th><th></th><th></th><th></th><th></th><th></th><th></th><th></th><th></th><th></th><th></th></tr>
<tr><td>>hg38_21_+_8433172_8446622_RNA45SN1_45S_50ntFlanks@11363.11381.19|CGGGCGGCGGCGGTCGGCG</td><td>>hg38_21_+_8433172_8446622_RNA45SN1_45S_50ntFlanks</td><td>11363</td><td>11381</td><td>19</td><td>CGGGCGGCGGCGGTCGGCG</td><td>114.9</td><td>-9.9</td><td>1.8</td><td>-5.7</td><td>1.4E-08</td><td>1.5E-07</td><td>BT20/MB468</td><td>MP</td></tr>
<tr><td>>hg38_21_+_8433172_8446622_RNA45SN1_45S_50ntFlanks@11718.11740.23|AACGGCGGGAGTAACTATGACTC</td><td>>hg38_21_+_8433172_8446622_RNA45SN1_45S_50ntFlanks</td><td>11718</td><td>11740</td><td>23</td><td>AACGGCGGGAGTAACTATGACTC</td><td>1931.2</td><td>-5.4</td><td>1.0</td><td>-5.4</td><td>6.7E-08</td><td>1.5E-07</td><td>BT20/MB468</td><td>CYTO</td></tr>
<tr><td>>hg38_21_+_8433172_8446622_RNA45SN1_45S_50ntFlanks@8359.8379.21|AAGAACTTTGAAGAGAGAGTT</td><td>>hg38_21_+_8433172_8446622_RNA45SN1_45S_50ntFlanks</td><td>8359</td><td>8379</td><td>21</td><td>AAGAACTTTGAAGAGAGAGTT</td><td>505.5</td><td>-4.8</td><td>0.9</td><td>-5.4</td><td>8.4E-08</td><td>1.5E-07</td><td>MB231/MB468</td><td>MITO</td></tr>
<tr><td>>hg38_21_+_8433172_8446622_RNA45SN1_45S_50ntFlanks@8381.8405.25|AAGAGGGCGTGAAACCGTTAAGAGG</td><td>>hg38_21_+_8433172_8446622_RNA45SN1_45S_50ntFlanks</td><td>8381</td><td>8405</td><td>25</td><td>AAGAGGGCGTGAAACCGTTAAGAGG</td><td>1063.0</td><td>-6.8</td><td>1.3</td><td>-5.4</td><td>8.4E-08</td><td>1.5E-07</td><td>MB231/MB468</td><td>MITO</td></tr>
<tr><td>>hg38_MT_+_598_1651_RNR1_12S_50ntFlanks@843.870.28|GAGTAGAGTGCTTAGTTGAACAGGGCCC</td><td>>hg38_MT_+_598_1651_RNR1_12S_50ntFlanks</td><td>843</td><td>870</td><td>28</td><td>GAGTAGAGTGCTTAGTTGAACAGGGCCC</td><td>144.3</td><td>10.9</td><td>1.9</td><td>5.7</td><td>1.4E-08</td><td>1.5E-07</td><td>BT20/MB468</td><td>MP</td></tr>
<tr><td>>hg38_21_+_8433172_8446622_RNA45SN1_45S_50ntFlanks@9930.9968.39|ATATAGACAGCAGGACGGTGGCCATGGAAGTCGGAATCC</td><td>>hg38_21_+_8433172_8446622_RNA45SN1_45S_50ntFlanks</td><td>9930</td><td>9968</td><td>39</td><td>ATATAGACAGCAGGACGGTGGCCATGGAAGTCGGAATCC</td><td>567.3</td><td>12.5</td><td>2.1</td><td>5.9</td><td>2.9E-09</td><td>1.5E-07</td><td>BT20/MB231</td><td>MP</td></tr>
<tr><td>>hg38_21_+_8433172_8446622_RNA45SN1_45S_50ntFlanks@10687.10705.19|CGGCGTCCGGTGAGCTCTC</td><td>>hg38_21_+_8433172_8446622_RNA45SN1_45S_50ntFlanks</td><td>10687</td><td>10705</td><td>19</td><td>CGGCGTCCGGTGAGCTCTC</td><td>433.8</td><td>-7.2</td><td>1.3</td><td>-5.4</td><td>7.9E-08</td><td>1.5E-07</td><td>MB231/MB468</td><td>NUC</td></tr>
<tr><td>>hg38_21_+_8433172_8446622_RNA45SN1_45S_50ntFlanks@8836.8861.26|CGGCGGTGGCGGCGGCGGCGGCGGCG</td><td>>hg38_21_+_8433172_8446622_RNA45SN1_45S_50ntFlanks</td><td>8836</td><td>8861</td><td>26</td><td>CGGCGGTGGCGGCGGCGGCGGCGGCG</td><td>104.7</td><td>-9.8</td><td>1.7</td><td>-5.7</td><td>1.4E-08</td><td>1.5E-07</td><td>BT20/MB468</td><td>MP</td></tr>
<tr><td>>hg38_21_+_8433172_8446622_RNA45SN1_45S_50ntFlanks@12628.12654.27|TGGGGCGAAGCTACCATCTGTGGGATT</td><td>>hg38_21_+_8433172_8446622_RNA45SN1_45S_50ntFlanks</td><td>12628</td><td>12654</td><td>27</td><td>TGGGGCGAAGCTACCATCTGTGGGATT</td><td>1983.1</td><td>-4.6</td><td>0.9</td><td>-5.4</td><td>7.7E-08</td><td>1.5E-07</td><td>MB231/MB468</td><td>CYTO</td></tr>
<tr><td>>hg38_21_+_8433172_8446622_RNA45SN1_45S_50ntFlanks@9554.9596.43|CGTAGCGGTCCTGACGTGCAAATCGGTCGTCCGACCTGGGTAT</td><td>>hg38_21_+_8433172_8446622_RNA45SN1_45S_50ntFlanks</td><td>9554</td><td>9596</td><td>43</td><td>CGTAGCGGTCCTGACGTGCAAATCGGTCGTCCGACCTGGGTAT</td><td>239.6</td><td>-7.6</td><td>1.4</td><td>-5.4</td><td>6.7E-08</td><td>1.5E-07</td><td>BT20/MB468</td><td>CYTO</td></tr>
<tr><td>>hg38_21_+_8433172_8446622_RNA45SN1_45S_50ntFlanks@11593.11613.21|CCAGGGGAATCCGACTGTTTA</td><td>>hg38_21_+_8433172_8446622_RNA45SN1_45S_50ntFlanks</td><td>11593</td><td>11613</td><td>21</td><td>CCAGGGGAATCCGACTGTTTA</td><td>428.1</td><td>-4.4</td><td>0.8</td><td>-5.4</td><td>7.8E-08</td><td>1.5E-07</td><td>MB231/MB468</td><td>CYTO</td></tr>
<tr><td>>hg38_21_+_8433172_8446622_RNA45SN1_45S_50ntFlanks@8213.8237.25|CGGTAGCGGCCCCCGGCGCGCCGGG</td><td>>hg38_21_+_8433172_8446622_RNA45SN1_45S_50ntFlanks</td><td>8213</td><td>8237</td><td>25</td><td>CGGTAGCGGCCCCCGGCGCGCCGGG</td><td>65.7</td><td>-9.9</td><td>1.8</td><td>-5.4</td><td>7.2E-08</td><td>1.5E-07</td><td>MB231/MB468</td><td>MP</td></tr>
<tr><td>>trnaMT_PheGAA_MT_+_577_647@35.62.28|GAAAATGTTTAGACGGGCTCACATCACC</td><td>>trnaMT_PheGAA_MT_+_577_647</td><td>35</td><td>62</td><td>28</td><td>GAAAATGTTTAGACGGGCTCACATCACC</td><td>1754.1</td><td>8.9</td><td>1.6</td><td>5.4</td><td>6.0E-08</td><td>1.5E-07</td><td>BT20/MB468</td><td>MITO</td></tr>
<tr><td>>hg38_21_+_8433172_8446622_RNA45SN1_45S_50ntFlanks@10402.10422.21|AACATGGGTCAGTCGGTCCTG</td><td>>hg38_21_+_8433172_8446622_RNA45SN1_45S_50ntFlanks</td><td>10402</td><td>10422</td><td>21</td><td>AACATGGGTCAGTCGGTCCTG</td><td>275.3</td><td>-4.7</td><td>0.9</td><td>-5.4</td><td>6.8E-08</td><td>1.5E-07</td><td>BT20/MB468</td><td>CYTO</td></tr>
<tr><td>>hg38_21_+_8433172_8446622_RNA45SN1_45S_50ntFlanks@6745.6769.25|ACACTTCGAACGCACTTGCGGCCCC</td><td>>hg38_21_+_8433172_8446622_RNA45SN1_45S_50ntFlanks</td><td>6745</td><td>6769</td><td>25</td><td>ACACTTCGAACGCACTTGCGGCCCC</td><td>317.9</td><td>-6.5</td><td>1.2</td><td>-5.4</td><td>6.8E-08</td><td>1.5E-07</td><td>BT20/MB468</td><td>CYTO</td></tr>
<tr><td>>hg38_21_+_8433172_8446622_RNA45SN1_45S_50ntFlanks@10786.10824.39|AACAGCCTCTGGCATGTTGGAACAATGTAGGTAAGGGAA</td><td>>hg38_21_+_8433172_8446622_RNA45SN1_45S_50ntFlanks</td><td>10786</td><td>10824</td><td>39</td><td>AACAGCCTCTGGCATGTTGGAACAATGTAGGTAAGGGAA</td><td>59.9</td><td>-9.8</td><td>1.8</td><td>-5.4</td><td>7.2E-08</td><td>1.5E-07</td><td>MB231/MB468</td><td>MP</td></tr>
<tr><td>>hg38_21_+_8433172_8446622_RNA45SN1_45S_50ntFlanks@10866.10891.26|CTAAGGGCTGGGTCGGTCGGGCTGGG</td><td>>hg38_21_+_8433172_8446622_RNA45SN1_45S_50ntFlanks</td><td>10866</td><td>10891</td><td>26</td><td>CTAAGGGCTGGGTCGGTCGGGCTGGG</td><td>1200.4</td><td>-8.7</td><td>1.6</td><td>-5.3</td><td>8.8E-08</td><td>1.5E-07</td><td>BT20/MB468</td><td>NUC</td></tr>
<tr><td>>MI0000447|hsa-mir-128-1&WithFlank&2|+|135665391|135665484@56.77.22&[MIMAT0000424&hsa-miR-128-3p&offsets|0|+1;m-108&2|+|135665446|135665466&offsets|0|+1]|TCACAGTGAACCGGTCTCTTTT</td><td>>MI0000447|hsa-mir-128-1&WithFlank&2|+|135665391|135665484</td><td>56</td><td>77</td><td>22</td><td>TCACAGTGAACCGGTCTCTTTT</td><td>1202.9</td><td>3.7</td><td>0.7</td><td>5.4</td><td>7.8E-08</td><td>1.5E-07</td><td>MB231/MB468</td><td>CYTO</td></tr>
<tr><td>>hg38_21_+_8433172_8446622_RNA45SN1_45S_50ntFlanks@10248.10286.39|CGACGAGTAGGAGGGCCGCTGCGGTGAGCCTTGAAGCCT</td><td>>hg38_21_+_8433172_8446622_RNA45SN1_45S_50ntFlanks</td><td>10248</td><td>10286</td><td>39</td><td>CGACGAGTAGGAGGGCCGCTGCGGTGAGCCTTGAAGCCT</td><td>399.8</td><td>4.8</td><td>0.9</td><td>5.4</td><td>6.0E-08</td><td>1.5E-07</td><td>BT20/MB468</td><td>MITO</td></tr>
<tr><td>>hg38_21_+_8433172_8446622_RNA45SN1_45S_50ntFlanks@10680.10712.33|GTTCCGGCGGCGTCCGGTGAGCTCTCGCTGGCC</td><td>>hg38_21_+_8433172_8446622_RNA45SN1_45S_50ntFlanks</td><td>10680</td><td>10712</td><td>33</td><td>GTTCCGGCGGCGTCCGGTGAGCTCTCGCTGGCC</td><td>4294.1</td><td>-4.3</td><td>0.8</td><td>-5.4</td><td>6.0E-08</td><td>1.5E-07</td><td>BT20/MB468</td><td>MITO</td></tr>
<tr><td>>hg38_21_+_8433172_8446622_RNA45SN1_45S_50ntFlanks@8118.8135.18|GGGACATGTGGCGTACGG</td><td>>hg38_21_+_8433172_8446622_RNA45SN1_45S_50ntFlanks</td><td>8118</td><td>8135</td><td>18</td><td>GGGACATGTGGCGTACGG</td><td>771.0</td><td>12.9</td><td>2.2</td><td>5.9</td><td>2.9E-09</td><td>1.5E-07</td><td>BT20/MB231</td><td>MP</td></tr>
<tr><td>>hg38_1_-_228634819_228635039_RNA5S12_5S_50ntFlanks@51.94.44|GTCTACGGCCATACCACCCTGAACGCGCCCGATCTCGTCTGATC</td><td>>hg38_1_-_228634819_228635039_RNA5S12_5S_50ntFlanks</td><td>51</td><td>94</td><td>44</td><td>GTCTACGGCCATACCACCCTGAACGCGCCCGATCTCGTCTGATC</td><td>1313.8</td><td>8.0</td><td>1.5</td><td>5.4</td><td>7.8E-08</td><td>1.5E-07</td><td>MB231/MB468</td><td>CYTO</td></tr>
<tr><td>>MI0000086|hsa-mir-28&WithFlank&3|+|188688775|188688872@60.80.21(+1G)&[MIMAT0004502&hsa-miR-28-3p&offsets|0|-1(+1G);m-28&3|+|188688834|188688855&offsets|0|-1(+1G)]|CACTAGATTGTGAGCTCCTGGG</td><td>>MI0000086|hsa-mir-28&WithFlank&3|+|188688775|188688872</td><td>60</td><td>80</td><td>21(+1G)</td><td>CACTAGATTGTGAGCTCCTGGG</td><td>123.4</td><td>10.7</td><td>1.9</td><td>5.7</td><td>1.5E-08</td><td>1.5E-07</td><td>BT20/MB468</td><td>MP</td></tr>
<tr><td>>hg38_21_+_8433172_8446622_RNA45SN1_45S_50ntFlanks@8847.8867.21|GCGGCGGCGGCGGCGGCGGGA</td><td>>hg38_21_+_8433172_8446622_RNA45SN1_45S_50ntFlanks</td><td>8847</td><td>8867</td><td>21</td><td>GCGGCGGCGGCGGCGGCGGGA</td><td>97.8</td><td>-9.7</td><td>1.7</td><td>-5.7</td><td>1.5E-08</td><td>1.5E-07</td><td>BT20/MB468</td><td>MP</td></tr>
<tr><td>>MI0003137|hsa-mir-193b&WithFlank&16|+|14303961|14304055@19.40.22&[MIMAT0004767&hsa-miR-193b-5p&offsets|-1|-1;m-343&16|+|14303980|14304001&offsets|-1|-1]|TCGGGGTTTTGAGGGCGAGATG</td><td>>MI0003137|hsa-mir-193b&WithFlank&16|+|14303961|14304055</td><td>19</td><td>40</td><td>22</td><td>TCGGGGTTTTGAGGGCGAGATG</td><td>160.0</td><td>10.7</td><td>1.8</td><td>5.9</td><td>2.9E-09</td><td>1.5E-07</td><td>BT20/MB231</td><td>MP</td></tr>
<tr><td>>hg38_21_+_8433172_8446622_RNA45SN1_45S_50ntFlanks@8168.8185.18|CCCAAGTCCTTCTGATCG</td><td>>hg38_21_+_8433172_8446622_RNA45SN1_45S_50ntFlanks</td><td>8168</td><td>8185</td><td>18</td><td>CCCAAGTCCTTCTGATCG</td><td>640.3</td><td>-4.0</td><td>0.7</td><td>-5.4</td><td>8.0E-08</td><td>1.5E-07</td><td>MB231/MB468</td><td>NUC</td></tr>
<tr><td>>hg38_21_+_8433172_8446622_RNA45SN1_45S_50ntFlanks@6653.6696.44|ACTCTTAGCGGTGGATCACTCGGCTCGTGCGTCGATGAAGAACG</td><td>>hg38_21_+_8433172_8446622_RNA45SN1_45S_50ntFlanks</td><td>6653</td><td>6696</td><td>44</td><td>ACTCTTAGCGGTGGATCACTCGGCTCGTGCGTCGATGAAGAACG</td><td>521.5</td><td>4.4</td><td>0.8</td><td>5.4</td><td>8.0E-08</td><td>1.5E-07</td><td>MB231/MB468</td><td>NUC</td></tr>
<tr><td>>hg38_21_+_8433172_8446622_RNA45SN1_45S_50ntFlanks@10317.10352.36|GTGCAGATCTTGGTGGTAGTAGCAAATATTCAAACG</td><td>>hg38_21_+_8433172_8446622_RNA45SN1_45S_50ntFlanks</td><td>10317</td><td>10352</td><td>36</td><td>GTGCAGATCTTGGTGGTAGTAGCAAATATTCAAACG</td><td>74.4</td><td>10.0</td><td>1.8</td><td>5.7</td><td>1.5E-08</td><td>1.5E-07</td><td>BT20/MB468</td><td>MP</td></tr>
<tr><td>>hg38_MT_+_1621_3279_RNR2_16S_50ntFlanks@341.373.33|AAATAGTGGGAAGATTTATAGGTAGAGGCGACA</td><td>>hg38_MT_+_1621_3279_RNR2_16S_50ntFlanks</td><td>341</td><td>373</td><td>33</td><td>AAATAGTGGGAAGATTTATAGGTAGAGGCGACA</td><td>116.2</td><td>10.6</td><td>1.9</td><td>5.7</td><td>1.5E-08</td><td>1.5E-07</td><td>BT20/MB468</td><td>MP</td></tr>
<tr><td>>hg38_21_+_8433172_8446622_RNA45SN1_45S_50ntFlanks@10324.10352.29|TCTTGGTGGTAGTAGCAAATATTCAAACG</td><td>>hg38_21_+_8433172_8446622_RNA45SN1_45S_50ntFlanks</td><td>10324</td><td>10352</td><td>29</td><td>TCTTGGTGGTAGTAGCAAATATTCAAACG</td><td>74.8</td><td>10.0</td><td>1.8</td><td>5.7</td><td>1.5E-08</td><td>1.5E-07</td><td>BT20/MB468</td><td>MP</td></tr>
<tr><td>>hg38_21_+_8433172_8446622_RNA45SN1_45S_50ntFlanks@11565.11601.37|CTAGCAGCCGACTTAGAACTGGTGCGGACCAGGGGAA</td><td>>hg38_21_+_8433172_8446622_RNA45SN1_45S_50ntFlanks</td><td>11565</td><td>11601</td><td>37</td><td>CTAGCAGCCGACTTAGAACTGGTGCGGACCAGGGGAA</td><td>239.7</td><td>-7.6</td><td>1.4</td><td>-5.4</td><td>7.3E-08</td><td>1.5E-07</td><td>MB231/MB468</td><td>MP</td></tr>
<tr><td>>hg38_21_+_8433172_8446622_RNA45SN1_45S_50ntFlanks@8776.8796.21|CCGCCCCCCGGCCCCGCGTCC</td><td>>hg38_21_+_8433172_8446622_RNA45SN1_45S_50ntFlanks</td><td>8776</td><td>8796</td><td>21</td><td>CCGCCCCCCGGCCCCGCGTCC</td><td>327.0</td><td>-11.4</td><td>2.0</td><td>-5.7</td><td>1.5E-08</td><td>1.5E-07</td><td>BT20/MB468</td><td>MP</td></tr>
<tr><td>>hg38_21_+_8433172_8446622_RNA45SN1_45S_50ntFlanks@6659.6689.31|AGCGGTGGATCACTCGGCTCGTGCGTCGATG</td><td>>hg38_21_+_8433172_8446622_RNA45SN1_45S_50ntFlanks</td><td>6659</td><td>6689</td><td>31</td><td>AGCGGTGGATCACTCGGCTCGTGCGTCGATG</td><td>811.3</td><td>-6.4</td><td>1.2</td><td>-5.4</td><td>7.3E-08</td><td>1.5E-07</td><td>MB231/MB468</td><td>MP</td></tr>
<tr><td>>hg38_21_+_8433172_8446622_RNA45SN1_45S_50ntFlanks@5215.5233.19|TCCGGGGCTGCACGCGCGC</td><td>>hg38_21_+_8433172_8446622_RNA45SN1_45S_50ntFlanks</td><td>5215</td><td>5233</td><td>19</td><td>TCCGGGGCTGCACGCGCGC</td><td>333.7</td><td>8.6</td><td>1.6</td><td>5.4</td><td>8.1E-08</td><td>1.5E-07</td><td>MB231/MB468</td><td>NUC</td></tr>
<tr><td>>hg38_21_+_8433172_8446622_RNA45SN1_45S_50ntFlanks@8184.8208.25|CGAGGCCCAGCCCGTGGACGGTGTG</td><td>>hg38_21_+_8433172_8446622_RNA45SN1_45S_50ntFlanks</td><td>8184</td><td>8208</td><td>25</td><td>CGAGGCCCAGCCCGTGGACGGTGTG</td><td>295.8</td><td>-3.8</td><td>0.7</td><td>-5.4</td><td>8.1E-08</td><td>1.5E-07</td><td>MB231/MB468</td><td>NUC</td></tr>
<tr><td>>hg38_21_+_8433172_8446622_RNA45SN1_45S_50ntFlanks@12087.12106.20|CTTCTGGCGCCAAGCGCCCG</td><td>>hg38_21_+_8433172_8446622_RNA45SN1_45S_50ntFlanks</td><td>12087</td><td>12106</td><td>20</td><td>CTTCTGGCGCCAAGCGCCCG</td><td>85.0</td><td>-9.5</td><td>1.7</td><td>-5.7</td><td>1.5E-08</td><td>1.5E-07</td><td>BT20/MB468</td><td>MP</td></tr>
<tr><td>>hg38_21_+_8433172_8446622_RNA45SN1_45S_50ntFlanks@9759.9788.30|CAAACTTTAAATGGGTAAGAAGCCCGGCTC</td><td>>hg38_21_+_8433172_8446622_RNA45SN1_45S_50ntFlanks</td><td>9759</td><td>9788</td><td>30</td><td>CAAACTTTAAATGGGTAAGAAGCCCGGCTC</td><td>119.6</td><td>-10.0</td><td>1.8</td><td>-5.7</td><td>1.5E-08</td><td>1.5E-07</td><td>BT20/MB468</td><td>MP</td></tr>
<tr><td>>hg38_21_+_8433172_8446622_RNA45SN1_45S_50ntFlanks@476.507.32|TTCGTGATCGATGTGGTGACGTCGTGCTCTCC</td><td>>hg38_21_+_8433172_8446622_RNA45SN1_45S_50ntFlanks</td><td>476</td><td>507</td><td>32</td><td>TTCGTGATCGATGTGGTGACGTCGTGCTCTCC</td><td>456.4</td><td>4.9</td><td>0.9</td><td>5.4</td><td>8.6E-08</td><td>1.5E-07</td><td>MB231/MB468</td><td>MITO</td></tr>
<tr><td>>hg38_21_+_8433172_8446622_RNA45SN1_45S_50ntFlanks@10781.10808.28|AGGTGAACAGCCTCTGGCATGTTGGAAC</td><td>>hg38_21_+_8433172_8446622_RNA45SN1_45S_50ntFlanks</td><td>10781</td><td>10808</td><td>28</td><td>AGGTGAACAGCCTCTGGCATGTTGGAAC</td><td>78.4</td><td>-9.4</td><td>1.7</td><td>-5.7</td><td>1.5E-08</td><td>1.5E-07</td><td>BT20/MB468</td><td>MP</td></tr>
<tr><td>>hg38_21_+_8433172_8446622_RNA45SN1_45S_50ntFlanks@12712.12746.35|CGGAGCCTCGGTTGGCCTCGGATAGCCGGTCCCCC</td><td>>hg38_21_+_8433172_8446622_RNA45SN1_45S_50ntFlanks</td><td>12712</td><td>12746</td><td>35</td><td>CGGAGCCTCGGTTGGCCTCGGATAGCCGGTCCCCC</td><td>95.0</td><td>-9.7</td><td>1.7</td><td>-5.7</td><td>1.5E-08</td><td>1.5E-07</td><td>BT20/MB468</td><td>MP</td></tr>
<tr><td>>hg38_21_+_8433172_8446622_RNA45SN1_45S_50ntFlanks@9928.9952.25|TGATATAGACAGCAGGACGGTGGCC</td><td>>hg38_21_+_8433172_8446622_RNA45SN1_45S_50ntFlanks</td><td>9928</td><td>9952</td><td>25</td><td>TGATATAGACAGCAGGACGGTGGCC</td><td>108.0</td><td>10.5</td><td>1.9</td><td>5.7</td><td>1.5E-08</td><td>1.5E-07</td><td>BT20/MB468</td><td>MP</td></tr>
<tr><td>>hg38_21_+_8433172_8446622_RNA45SN1_45S_50ntFlanks@10186.10209.24|TCCTCCCCTCCTCCCGCCCACGCC</td><td>>hg38_21_+_8433172_8446622_RNA45SN1_45S_50ntFlanks</td><td>10186</td><td>10209</td><td>24</td><td>TCCTCCCCTCCTCCCGCCCACGCC</td><td>797.5</td><td>-8.5</td><td>1.6</td><td>-5.3</td><td>8.9E-08</td><td>1.5E-07</td><td>BT20/MB468</td><td>NUC</td></tr>
<tr><td>>hg38_21_+_8433172_8446622_RNA45SN1_45S_50ntFlanks@12991.13013.23|CTCCCTCGCTGCGATCTATTGAA</td><td>>hg38_21_+_8433172_8446622_RNA45SN1_45S_50ntFlanks</td><td>12991</td><td>13013</td><td>23</td><td>CTCCCTCGCTGCGATCTATTGAA</td><td>7562.3</td><td>-4.8</td><td>0.9</td><td>-5.4</td><td>6.9E-08</td><td>1.5E-07</td><td>BT20/MB468</td><td>CYTO</td></tr>
<tr><td>>hg38_21_+_8433172_8446622_RNA45SN1_45S_50ntFlanks@3705.3725.21|TACCTGGTTGATCCTGCCAGT</td><td>>hg38_21_+_8433172_8446622_RNA45SN1_45S_50ntFlanks</td><td>3705</td><td>3725</td><td>21</td><td>TACCTGGTTGATCCTGCCAGT</td><td>20885.8</td><td>4.9</td><td>0.8</td><td>5.8</td><td>5.0E-09</td><td>1.5E-07</td><td>BT20/MB231</td><td>CYTO</td></tr>
<tr><td>>trnaMT_ValTAC_MT_+_1602_1670@52.72.21|TTAACTTGACCGCTCTGACCA</td><td>>trnaMT_ValTAC_MT_+_1602_1670</td><td>52</td><td>72</td><td>21</td><td>TTAACTTGACCGCTCTGACCA</td><td>963.9</td><td>4.7</td><td>0.9</td><td>5.4</td><td>8.1E-08</td><td>1.5E-07</td><td>MB231/MB468</td><td>NUC</td></tr>
<tr><td>>hg38_21_+_8433172_8446622_RNA45SN1_45S_50ntFlanks@10376.10424.49|GAAGGGTTCCATGTGAACAGCAGTTGAACATGGGTCAGTCGGTCCTGAG</td><td>>hg38_21_+_8433172_8446622_RNA45SN1_45S_50ntFlanks</td><td>10376</td><td>10424</td><td>49</td><td>GAAGGGTTCCATGTGAACAGCAGTTGAACATGGGTCAGTCGGTCCTGAG</td><td>103.0</td><td>10.4</td><td>1.8</td><td>5.7</td><td>1.5E-08</td><td>1.5E-07</td><td>BT20/MB468</td><td>MP</td></tr>
<tr><td>>MI0000434|hsa-let-7i&WithFlank&12|+|62603680|62603775@12.32.21(+1C)&[MIMAT0000415&hsa-let-7i-5p&offsets|0|-1(+1C);m-49&12|+|62603691|62603712&offsets|0|-1(+1C)]|TGAGGTAGTAGTTTGTGCTGTC</td><td>>MI0000434|hsa-let-7i&WithFlank&12|+|62603680|62603775</td><td>12</td><td>32</td><td>21(+1C)</td><td>TGAGGTAGTAGTTTGTGCTGTC</td><td>3755.8</td><td>2.1</td><td>0.4</td><td>5.3</td><td>9.0E-08</td><td>1.5E-07</td><td>BT20/MB468</td><td>NUC</td></tr>
<tr><td>>hg38_21_+_8433172_8446622_RNA45SN1_45S_50ntFlanks@6650.6693.44|ACGACTCTTAGCGGTGGATCACTCGGCTCGTGCGTCGATGAAGA</td><td>>hg38_21_+_8433172_8446622_RNA45SN1_45S_50ntFlanks</td><td>6650</td><td>6693</td><td>44</td><td>ACGACTCTTAGCGGTGGATCACTCGGCTCGTGCGTCGATGAAGA</td><td>1249.9</td><td>4.7</td><td>0.9</td><td>5.4</td><td>6.1E-08</td><td>1.5E-07</td><td>BT20/MB468</td><td>MITO</td></tr>
<tr><td>>hg38_21_+_8433172_8446622_RNA45SN1_45S_50ntFlanks@12219.12236.18|AGGACAGAAACCTCCCGT</td><td>>hg38_21_+_8433172_8446622_RNA45SN1_45S_50ntFlanks</td><td>12219</td><td>12236</td><td>18</td><td>AGGACAGAAACCTCCCGT</td><td>2036.1</td><td>5.5</td><td>0.9</td><td>5.8</td><td>6.5E-09</td><td>1.5E-07</td><td>BT20/MB231</td><td>MITO</td></tr>
<tr><td>>hg38_21_+_8433172_8446622_RNA45SN1_45S_50ntFlanks@3900.3934.35|CTTCGCGGGGGGGATGCGTGCATTTATCAGATCAA</td><td>>hg38_21_+_8433172_8446622_RNA45SN1_45S_50ntFlanks</td><td>3900</td><td>3934</td><td>35</td><td>CTTCGCGGGGGGGATGCGTGCATTTATCAGATCAA</td><td>60.8</td><td>-9.8</td><td>1.8</td><td>-5.4</td><td>7.4E-08</td><td>1.5E-07</td><td>MB231/MB468</td><td>MP</td></tr>
<tr><td>>hg38_21_+_8433172_8446622_RNA45SN1_45S_50ntFlanks@12714.12747.34|GAGCCTCGGTTGGCCTCGGATAGCCGGTCCCCCG</td><td>>hg38_21_+_8433172_8446622_RNA45SN1_45S_50ntFlanks</td><td>12714</td><td>12747</td><td>34</td><td>GAGCCTCGGTTGGCCTCGGATAGCCGGTCCCCCG</td><td>1200.2</td><td>-5.3</td><td>1.0</td><td>-5.4</td><td>7.4E-08</td><td>1.5E-07</td><td>MB231/MB468</td><td>MP</td></tr>
<tr><td>>hg38_21_+_8433172_8446622_RNA45SN1_45S_50ntFlanks@10411.10453.43|CAGTCGGTCCTGAGAGATGGGCGAGCGCCGTTCCGAAGGGACG</td><td>>hg38_21_+_8433172_8446622_RNA45SN1_45S_50ntFlanks</td><td>10411</td><td>10453</td><td>43</td><td>CAGTCGGTCCTGAGAGATGGGCGAGCGCCGTTCCGAAGGGACG</td><td>1562.0</td><td>-4.6</td><td>0.8</td><td>-5.4</td><td>6.2E-08</td><td>1.5E-07</td><td>BT20/MB468</td><td>MITO</td></tr>
</table>

<table>
<tr><td>>am_trna111_HisGTG_1_-_147774845_147774916@-1G.23.24,trna118_HisGTG_1_-_145396881_145396952@-1G.23.24,trna16_HisGTG_1_+_146544773_146544844@-1G.23.24,trna1_HisGTG_15_+_45493349_45493420@-1G.23.24,trna21_HisGTG_1_+_147753471_147753542@-1G.23.24,trna33_HisGTG_6_+_27125906_27125977@-1G.23.24,trna7_HisGTG_9_-_14433938_14434009@-1G.23.24,trna8_HisGTG_15_-_45492611_45492682@-1G.23.24,trna9_HisGTG_15_-_45490804_45490875@-1G.23.24|GGCCGTGATCGTATAGTGGTTAGT</td><td>>am_trna111_HisGTG_1_-_147774845_147774916</td><td>-1G</td><td>23</td><td>24</td><td>GGCCGTGATCGTATAGTGGTTAGT</td><td>487.4</td><td>8.8</td><td>1.6</td><td>5.4</td><td>7.4E-08</td><td>1.5E-07</td><td>MB231/MB468</td><td>MP</td></tr>
<tr><td>>hg38_21_+_8433172_8446622_RNA45SN1_45S_50ntFlanks@10869.10903.35|AGGGCTGGGTCGGTCGGGCTGGGGCGCGAAGCGGG</td><td>>hg38_21_+_8433172_8446622_RNA45SN1_45S_50ntFlanks</td><td>10869</td><td>10903</td><td>35</td><td>AGGGCTGGGTCGGTCGGGCTGGGGCGCGAAGCGGG</td><td>599.9</td><td>-8.3</td><td>1.6</td><td>-5.3</td><td>9.1E-08</td><td>1.5E-07</td><td>BT20/MB468</td><td>NUC</td></tr>
<tr><td>>hg38_21_+_8433172_8446622_RNA45SN1_45S_50ntFlanks@11848.11872.25|ACGGGCTTGGCGGAATCAGCGGGGA</td><td>>hg38_21_+_8433172_8446622_RNA45SN1_45S_50ntFlanks</td><td>11848</td><td>11872</td><td>25</td><td>ACGGGCTTGGCGGAATCAGCGGGGA</td><td>968.0</td><td>-4.6</td><td>0.8</td><td>-5.7</td><td>1.5E-08</td><td>1.5E-07</td><td>BT20/MB468</td><td>MP</td></tr>
<tr><td>>trnaMT_HisGTG_MT_+_12138_12206@26.62.37|AGATTGTGAATCTGACAACAGAGGCTTACGACCCCTT</td><td>>trnaMT_HisGTG_MT_+_12138_12206</td><td>26</td><td>62</td><td>37</td><td>AGATTGTGAATCTGACAACAGAGGCTTACGACCCCTT</td><td>102.3</td><td>10.4</td><td>1.8</td><td>5.7</td><td>1.5E-08</td><td>1.5E-07</td><td>BT20/MB468</td><td>MP</td></tr>
<tr><td>>hg38_21_+_8433172_8446622_RNA45SN1_45S_50ntFlanks@3920.3959.40|CATTTATCAGATCAAAACCAACCCGGTCAGCCCCTCTCCG</td><td>>hg38_21_+_8433172_8446622_RNA45SN1_45S_50ntFlanks</td><td>3920</td><td>3959</td><td>40</td><td>CATTTATCAGATCAAAACCAACCCGGTCAGCCCCTCTCCG</td><td>316.9</td><td>-7.4</td><td>1.4</td><td>-5.4</td><td>7.0E-08</td><td>1.5E-07</td><td>BT20/MB468</td><td>CYTO</td></tr>
<tr><td>>hg38_21_+_8433172_8446622_RNA45SN1_45S_50ntFlanks@4557.4574.18|CAGCTAGGAATAATGGAA</td><td>>hg38_21_+_8433172_8446622_RNA45SN1_45S_50ntFlanks</td><td>4557</td><td>4574</td><td>18</td><td>CAGCTAGGAATAATGGAA</td><td>282.1</td><td>-6.4</td><td>1.2</td><td>-5.3</td><td>9.1E-08</td><td>1.5E-07</td><td>BT20/MB468</td><td>NUC</td></tr>
<tr><td>>MI0000736|hsa-mir-30c-1&WithFlank&1|+|40757278|40757378@23.45.23(+1A);MI0000254|hsa-mir-30c-2&WithFlank&6|-|71376954|71377037@13.35.23(+1A)&[MIMAT0000244&hsa-miR-30c-5p&offsets|0|0(+1A);m-57&1|+|40757300|40757323&offsets|0|-1(+1A)];[MIMAT0000244_1&hsa-miR-30c-5p&offsets|0|0(+1A);m-56&6|-|71377002|71377025&offsets|0|-1(+1A)]|TGTAAACATCCTACACTCTCAGCA</td><td>>MI0000736|hsa-mir-30c-1&WithFlank&1|+|40757278|40757378</td><td>23</td><td>45</td><td>23(+1A)</td><td>TGTAAACATCCTACACTCTCAGCA</td><td>5289.4</td><td>5.6</td><td>1.0</td><td>5.4</td><td>8.7E-08</td><td>1.5E-07</td><td>MB231/MB468</td><td>MITO</td></tr>
<tr><td>>hg38_21_+_8433172_8446622_RNA45SN1_45S_50ntFlanks@8117.8139.23|CGGGACATGTGGCGTACGGAAGA</td><td>>hg38_21_+_8433172_8446622_RNA45SN1_45S_50ntFlanks</td><td>8117</td><td>8139</td><td>23</td><td>CGGGACATGTGGCGTACGGAAGA</td><td>459.6</td><td>12.2</td><td>2.1</td><td>5.9</td><td>3.0E-09</td><td>1.5E-07</td><td>BT20/MB231</td><td>MP</td></tr>
<tr><td>>hg38_21_+_8433172_8446622_RNA45SN1_45S_50ntFlanks@9941.9979.39|AGGACGGTGGCCATGGAAGTCGGAATCCGCTAAGGAGTG</td><td>>hg38_21_+_8433172_8446622_RNA45SN1_45S_50ntFlanks</td><td>9941</td><td>9979</td><td>39</td><td>AGGACGGTGGCCATGGAAGTCGGAATCCGCTAAGGAGTG</td><td>150.5</td><td>11.0</td><td>1.9</td><td>5.7</td><td>1.5E-08</td><td>1.5E-07</td><td>BT20/MB468</td><td>MP</td></tr>
<tr><td>>hg38_21_+_8433172_8446622_RNA45SN1_45S_50ntFlanks@12033.12068.36|ATCGTTTTTTCACTGACCCGGTGAGGCGGGGGGGCG</td><td>>hg38_21_+_8433172_8446622_RNA45SN1_45S_50ntFlanks</td><td>12033</td><td>12068</td><td>36</td><td>ATCGTTTTTTCACTGACCCGGTGAGGCGGGGGGGCG</td><td>107.5</td><td>-9.8</td><td>1.7</td><td>-5.7</td><td>1.5E-08</td><td>1.5E-07</td><td>BT20/MB468</td><td>MP</td></tr>
<tr><td>>am_trnaMT_IleGAT_MT_+_4263_4331@1.33.33|AGAAATATGTCTGATAAAAGAGTTACTTTGATA</td><td>>am_trnaMT_IleGAT_MT_+_4263_4331</td><td>1</td><td>33</td><td>33</td><td>AGAAATATGTCTGATAAAAGAGTTACTTTGATA</td><td>61.4</td><td>9.2</td><td>1.7</td><td>5.4</td><td>7.5E-08</td><td>1.5E-07</td><td>MB231/MB468</td><td>MP</td></tr>
<tr><td>>hg38_21_+_8433172_8446622_RNA45SN1_45S_50ntFlanks@7975.8018.44|CGCGACCTCAGATCAGACGTGGCGACCCGCTGAATTTAAGCATA</td><td>>hg38_21_+_8433172_8446622_RNA45SN1_45S_50ntFlanks</td><td>7975</td><td>8018</td><td>44</td><td>CGCGACCTCAGATCAGACGTGGCGACCCGCTGAATTTAAGCATA</td><td>40934.1</td><td>5.2</td><td>1.0</td><td>5.4</td><td>6.3E-08</td><td>1.5E-07</td><td>BT20/MB468</td><td>MITO</td></tr>
<tr><td>>hg38_21_+_8433172_8446622_RNA45SN1_45S_50ntFlanks@3931.3959.29|TCAAAACCAACCCGGTCAGCCCCTCTCCG</td><td>>hg38_21_+_8433172_8446622_RNA45SN1_45S_50ntFlanks</td><td>3931</td><td>3959</td><td>29</td><td>TCAAAACCAACCCGGTCAGCCCCTCTCCG</td><td>1072.6</td><td>-8.6</td><td>1.6</td><td>-5.3</td><td>9.2E-08</td><td>1.5E-07</td><td>BT20/MB468</td><td>NUC</td></tr>
<tr><td>>hg38_21_+_8433172_8446622_RNA45SN1_45S_50ntFlanks@10387.10410.24|TGTGAACAGCAGTTGAACATGGGT</td><td>>hg38_21_+_8433172_8446622_RNA45SN1_45S_50ntFlanks</td><td>10387</td><td>10410</td><td>24</td><td>TGTGAACAGCAGTTGAACATGGGT</td><td>220.2</td><td>-2.7</td><td>0.5</td><td>-5.4</td><td>7.0E-08</td><td>1.5E-07</td><td>BT20/MB468</td><td>CYTO</td></tr>
<tr><td>>hg38_21_+_8433172_8446622_RNA45SN1_45S_50ntFlanks@10409.10433.25|GTCAGTCGGTCCTGAGAGATGGGCG</td><td>>hg38_21_+_8433172_8446622_RNA45SN1_45S_50ntFlanks</td><td>10409</td><td>10433</td><td>25</td><td>GTCAGTCGGTCCTGAGAGATGGGCG</td><td>275.7</td><td>-7.2</td><td>1.3</td><td>-5.4</td><td>7.5E-08</td><td>1.5E-07</td><td>MB231/MB468</td><td>MP</td></tr>
<tr><td>>hg38_21_+_8433172_8446622_RNA45SN1_45S_50ntFlanks@9566.9596.31|GACGTGCAAATCGGTCGTCCGACCTGGGTAT</td><td>>hg38_21_+_8433172_8446622_RNA45SN1_45S_50ntFlanks</td><td>9566</td><td>9596</td><td>31</td><td>GACGTGCAAATCGGTCGTCCGACCTGGGTAT</td><td>7122.3</td><td>-9.9</td><td>1.9</td><td>-5.3</td><td>9.2E-08</td><td>1.5E-07</td><td>BT20/MB468</td><td>NUC</td></tr>
<tr><td>>am_trna128_GlyGCC_6_-_27870686_27870756@4.31.28,trna133_GlyCCC_1_-_16872434_16872504@4.31.28,trna18_GlyGCC_16_+_70822597_70822667@4.31.28,trna19_GlyGCC_16_+_70823410_70823480@4.31.28,trna19_GlyGCC_2_-_157257659_157257729@4.31.28,trna24_GlyGCC_16_-_70812942_70813012@4.31.28,trna25_GlyGCC_16_-_70812114_70812184@4.31.28,trna4_GlyCCC_1_+_17188416_17188486@4.31.28,trna5_GlyGCC_17_+_8029064_8029134@4.31.28,trna68_GlyGCC_1_-_161493637_161493707@4.31.28|TTGGTGGTTCAGTGGTAGAATTCTCGCC</td><td>>am_trna128_GlyGCC_6_-_27870686_27870756</td><td>4</td><td>31</td><td>28</td><td>TTGGTGGTTCAGTGGTAGAATTCTCGCC</td><td>61.3</td><td>9.2</td><td>1.7</td><td>5.4</td><td>7.5E-08</td><td>1.5E-07</td><td>MB231/MB468</td><td>MP</td></tr>
<tr><td>>hg38_21_+_8433172_8446622_RNA45SN1_45S_50ntFlanks@10411.10432.22|CAGTCGGTCCTGAGAGATGGGC</td><td>>hg38_21_+_8433172_8446622_RNA45SN1_45S_50ntFlanks</td><td>10411</td><td>10432</td><td>22</td><td>CAGTCGGTCCTGAGAGATGGGC</td><td>503.5</td><td>-5.6</td><td>1.1</td><td>-5.4</td><td>8.3E-08</td><td>1.5E-07</td><td>MB231/MB468</td><td>NUC</td></tr>
<tr><td>>hg38_21_+_8433172_8446622_RNA45SN1_45S_50ntFlanks@4727.4764.38|AATCAAGAACGAAAGTCGGAGGTTCGAAGACGATCAGA</td><td>>hg38_21_+_8433172_8446622_RNA45SN1_45S_50ntFlanks</td><td>4727</td><td>4764</td><td>38</td><td>AATCAAGAACGAAAGTCGGAGGTTCGAAGACGATCAGA</td><td>446.4</td><td>7.2</td><td>1.3</td><td>5.4</td><td>8.1E-08</td><td>1.5E-07</td><td>MB231/MB468</td><td>CYTO</td></tr>
<tr><td>>hg38_21_+_8433172_8446622_RNA45SN1_45S_50ntFlanks@11321.11343.23|CACGTCTCGTCGCGCGCGCGTCC</td><td>>hg38_21_+_8433172_8446622_RNA45SN1_45S_50ntFlanks</td><td>11321</td><td>11343</td><td>23</td><td>CACGTCTCGTCGCGCGCGCGTCC</td><td>766.1</td><td>-4.4</td><td>0.8</td><td>-5.4</td><td>8.1E-08</td><td>1.5E-07</td><td>MB231/MB468</td><td>CYTO</td></tr>
<tr><td>>hg38_21_+_8433172_8446622_RNA45SN1_45S_50ntFlanks@12394.12414.21|TAGCGACGTCGCTTTTTGATC</td><td>>hg38_21_+_8433172_8446622_RNA45SN1_45S_50ntFlanks</td><td>12394</td><td>12414</td><td>21</td><td>TAGCGACGTCGCTTTTTGATC</td><td>3605.0</td><td>-5.2</td><td>1.0</td><td>-5.3</td><td>8.8E-08</td><td>1.5E-07</td><td>MB231/MB468</td><td>MITO</td></tr>
<tr><td>>hg38_21_+_8433172_8446622_RNA45SN1_45S_50ntFlanks@10907.10931.25|GGGCGCGCGCCGCGGCTGGACGAGG</td><td>>hg38_21_+_8433172_8446622_RNA45SN1_45S_50ntFlanks</td><td>10907</td><td>10931</td><td>25</td><td>GGGCGCGCGCCGCGGCTGGACGAGG</td><td>119.3</td><td>10.6</td><td>1.9</td><td>5.7</td><td>1.5E-08</td><td>1.5E-07</td><td>BT20/MB468</td><td>MP</td></tr>
<tr><td>>MI0001445|hsa-mir-423&WithFlank&17|+|30117073|30117178@59.80.22&[MIMAT0001340&hsa-miR-423-3p&offsets|0|-1;m-100&17|+|30117131|30117153&offsets|0|-1]|AGCTCGGTCTGAGGCCCCTCAG</td><td>>MI0001445|hsa-mir-423&WithFlank&17|+|30117073|30117178</td><td>59</td><td>80</td><td>22</td><td>AGCTCGGTCTGAGGCCCCTCAG</td><td>904.1</td><td>5.4</td><td>0.9</td><td>5.7</td><td>1.5E-08</td><td>1.5E-07</td><td>BT20/MB468</td><td>MP</td></tr>
<tr><td>>hg38_21_+_8433172_8446622_RNA45SN1_45S_50ntFlanks@11666.11687.22|TCTGCCCAGTGCTCTGAATGTC</td><td>>hg38_21_+_8433172_8446622_RNA45SN1_45S_50ntFlanks</td><td>11666</td><td>11687</td><td>22</td><td>TCTGCCCAGTGCTCTGAATGTC</td><td>1848.7</td><td>3.8</td><td>0.7</td><td>5.4</td><td>8.1E-08</td><td>1.5E-07</td><td>MB231/MB468</td><td>CYTO</td></tr>
<tr><td>>MI0003560|hsa-mir-92b&WithFlank&1|+|155195171|155195278@67.86.20&[MIMAT0003218&hsa-miR-92b-3p&offsets|0|-2;m-115&1|+|155195237|155195258&offsets|0|-2]|TATTGCACTCGTCCCGGCCT</td><td>>MI0003560|hsa-mir-92b&WithFlank&1|+|155195171|155195278</td><td>67</td><td>86</td><td>20</td><td>TATTGCACTCGTCCCGGCCT</td><td>325.1</td><td>8.4</td><td>1.6</td><td>5.3</td><td>8.9E-08</td><td>1.5E-07</td><td>MB231/MB468</td><td>MITO</td></tr>
<tr><td>>hg38_21_+_8433172_8446622_RNA45SN1_45S_50ntFlanks@12407.12445.39|TTTTGATCCTTCGATGTCGGCTCTTCCTATCATTGTGAA</td><td>>hg38_21_+_8433172_8446622_RNA45SN1_45S_50ntFlanks</td><td>12407</td><td>12445</td><td>39</td><td>TTTTGATCCTTCGATGTCGGCTCTTCCTATCATTGTGAA</td><td>520.1</td><td>-6.6</td><td>1.2</td><td>-5.3</td><td>9.3E-08</td><td>1.5E-07</td><td>BT20/MB468</td><td>NUC</td></tr>
<tr><td>>hg38_21_+_8433172_8446622_RNA45SN1_45S_50ntFlanks@11641.11658.18|CGGCGGGTGTTGACGCGA</td><td>>hg38_21_+_8433172_8446622_RNA45SN1_45S_50ntFlanks</td><td>11641</td><td>11658</td><td>18</td><td>CGGCGGGTGTTGACGCGA</td><td>1071.9</td><td>-8.7</td><td>1.5</td><td>-5.7</td><td>1.6E-08</td><td>1.5E-07</td><td>BT20/MB468</td><td>MP</td></tr>
<tr><td>>hg38_21_+_8433172_8446622_RNA45SN1_45S_50ntFlanks@8158.8181.24|TCGTGGGGGGCCCAAGTCCTTCTG</td><td>>hg38_21_+_8433172_8446622_RNA45SN1_45S_50ntFlanks</td><td>8158</td><td>8181</td><td>24</td><td>TCGTGGGGGGCCCAAGTCCTTCTG</td><td>203.3</td><td>-9.8</td><td>1.8</td><td>-5.4</td><td>6.3E-08</td><td>1.5E-07</td><td>BT20/MB468</td><td>MITO</td></tr>
<tr><td>>hg38_21_+_8433172_8446622_RNA45SN1_45S_50ntFlanks@8737.8756.20|CCCGTCCGTCCGTCCGTCCG</td><td>>hg38_21_+_8433172_8446622_RNA45SN1_45S_50ntFlanks</td><td>8737</td><td>8756</td><td>20</td><td>CCCGTCCGTCCGTCCGTCCG</td><td>202.7</td><td>-6.4</td><td>1.2</td><td>-5.4</td><td>7.1E-08</td><td>1.5E-07</td><td>BT20/MB468</td><td>CYTO</td></tr>
<tr><td>>hg38_21_+_8433172_8446622_RNA45SN1_45S_50ntFlanks@10240.10259.20|CTACGCCGCGACGAGTAGGA</td><td>>hg38_21_+_8433172_8446622_RNA45SN1_45S_50ntFlanks</td><td>10240</td><td>10259</td><td>20</td><td>CTACGCCGCGACGAGTAGGA</td><td>4094.2</td><td>4.9</td><td>0.9</td><td>5.7</td><td>1.6E-08</td><td>1.5E-07</td><td>BT20/MB468</td><td>MP</td></tr>
<tr><td>>hg38_21_+_8433172_8446622_RNA45SN1_45S_50ntFlanks@10247.10278.32|GCGACGAGTAGGAGGGCCGCTGCGGTGAGCCT</td><td>>hg38_21_+_8433172_8446622_RNA45SN1_45S_50ntFlanks</td><td>10247</td><td>10278</td><td>32</td><td>GCGACGAGTAGGAGGGCCGCTGCGGTGAGCCT</td><td>79.0</td><td>10.1</td><td>1.8</td><td>5.7</td><td>1.6E-08</td><td>1.5E-07</td><td>BT20/MB468</td><td>MP</td></tr>
<tr><td>>hg38_21_+_8433172_8446622_RNA45SN1_45S_50ntFlanks@11792.11816.25|ATGAACGAGATTCCCACTGTCCCTA</td><td>>hg38_21_+_8433172_8446622_RNA45SN1_45S_50ntFlanks</td><td>11792</td><td>11816</td><td>25</td><td>ATGAACGAGATTCCCACTGTCCCTA</td><td>91.5</td><td>-9.6</td><td>1.7</td><td>-5.7</td><td>1.6E-08</td><td>1.5E-07</td><td>BT20/MB468</td><td>MP</td></tr>
<tr><td>>hg38_21_+_8433172_8446622_RNA45SN1_45S_50ntFlanks@12324.12349.26|TTGGGTTTTAAGCAGGAGGTGTCAGA</td><td>>hg38_21_+_8433172_8446622_RNA45SN1_45S_50ntFlanks</td><td>12324</td><td>12349</td><td>26</td><td>TTGGGTTTTAAGCAGGAGGTGTCAGA</td><td>178.2</td><td>-6.0</td><td>1.1</td><td>-5.4</td><td>7.2E-08</td><td>1.5E-07</td><td>BT20/MB468</td><td>CYTO</td></tr>
<tr><td>>hg38_21_+_8433172_8446622_RNA45SN1_45S_50ntFlanks@10362.10384.23|AAGGCCGAAGTGGAGAAGGGTTC</td><td>>hg38_21_+_8433172_8446622_RNA45SN1_45S_50ntFlanks</td><td>10362</td><td>10384</td><td>23</td><td>AAGGCCGAAGTGGAGAAGGGTTC</td><td>682.9</td><td>-5.1</td><td>0.9</td><td>-5.4</td><td>8.2E-08</td><td>1.5E-07</td><td>MB231/MB468</td><td>CYTO</td></tr>
<tr><td>>hg38_21_+_8433172_8446622_RNA45SN1_45S_50ntFlanks@12991.13012.22|CTCCCTCGCTGCGATCTATTGA</td><td>>hg38_21_+_8433172_8446622_RNA45SN1_45S_50ntFlanks</td><td>12991</td><td>13012</td><td>22</td><td>CTCCCTCGCTGCGATCTATTGA</td><td>2615.9</td><td>-3.4</td><td>0.6</td><td>-5.4</td><td>8.2E-08</td><td>1.5E-07</td><td>MB231/MB468</td><td>CYTO</td></tr>
<tr><td>>MI0000266|hsa-mir-10a&WithFlank&17|-|48579832|48579953@27.49.23&[MIMAT0000253&hsa-miR-10a-5p&offsets|-1|-1;m-18&17|-|48579905|48579926&offsets|-1|0]|ATACCCTGTAGATCCGAATTTGT</td><td>>MI0000266|hsa-mir-10a&WithFlank&17|-|48579832|48579953</td><td>27</td><td>49</td><td>23</td><td>ATACCCTGTAGATCCGAATTTGT</td><td>52.0</td><td>8.9</td><td>1.7</td><td>5.4</td><td>7.6E-08</td><td>1.5E-07</td><td>MB231/MB468</td><td>MP</td></tr>
<tr><td>>MI0000114|hsa-mir-107&WithFlank&10|-|89592741|89592833@56.76.21(+1U);MI0000108|hsa-mir-103a-2&WithFlank&20|+|3917488|3917577@54.74.21(+1U);MI0000109|hsa-mir-103a-1&WithFlank&5|-|168560890|168560979@54.74.21(+1U)&[MIMAT0000104&hsa-miR-107&offsets|0|-2(+1U);m-78&10|-|89592758|89592778&offsets|0|0(+1U)];[MIMAT0000101&hsa-miR-103a-3p&offsets|0|-2(+1U);m-12&20|+|3917541|3917563&offsets|0|-2(+1U)];[MIMAT0000101_1&hsa-miR-103a-3p&offsets|0|-2(+1U);m-13&5|-|168560904|168560926&offsets|0|-2(+1U);m-44220&5|-|168560908|168560926&offsets|0|+2(+1U)]|AGCAGCATTGTACAGGGCTATT</td><td>>MI0000114|hsa-mir-107&WithFlank&10|-|89592741|89592833</td><td>56</td><td>76</td><td>21(+1U)</td><td>AGCAGCATTGTACAGGGCTATT</td><td>186.3</td><td>7.0</td><td>1.3</td><td>5.4</td><td>8.5E-08</td><td>1.5E-07</td><td>MB231/MB468</td><td>NUC</td></tr>
<tr><td>>hg38_21_+_8433172_8446622_RNA45SN1_45S_50ntFlanks@12073.12091.19|CCGAGGGGCTCTCGCTTCT</td><td>>hg38_21_+_8433172_8446622_RNA45SN1_45S_50ntFlanks</td><td>12073</td><td>12091</td><td>19</td><td>CCGAGGGGCTCTCGCTTCT</td><td>92.6</td><td>-8.6</td><td>1.6</td><td>-5.4</td><td>6.4E-08</td><td>1.6E-07</td><td>BT20/MB468</td><td>MITO</td></tr>
<tr><td>>hg38_1_-_228634819_228635039_RNA5S12_5S_50ntFlanks@130.171.42|TGGATGGGAGACCGCCTGGGAATACCGGGTGCTGTAGGCTTT</td><td>>hg38_1_-_228634819_228635039_RNA5S12_5S_50ntFlanks</td><td>130</td><td>171</td><td>42</td><td>TGGATGGGAGACCGCCTGGGAATACCGGGTGCTGTAGGCTTT</td><td>1012.8</td><td>6.1</td><td>1.1</td><td>5.4</td><td>8.5E-08</td><td>1.6E-07</td><td>MB231/MB468</td><td>NUC</td></tr>
<tr><td>>hg38_21_+_8433172_8446622_RNA45SN1_45S_50ntFlanks@8357.8402.46|AAAAGAACTTTGAAGAGAGAGTTCAAGAGGGCGTGAAACCGTTAAG</td><td>>hg38_21_+_8433172_8446622_RNA45SN1_45S_50ntFlanks</td><td>8357</td><td>8402</td><td>46</td><td>AAAAGAACTTTGAAGAGAGAGTTCAAGAGGGCGTGAAACCGTTAAG</td><td>91.2</td><td>-8.8</td><td>1.6</td><td>-5.4</td><td>7.2E-08</td><td>1.6E-07</td><td>BT20/MB468</td><td>CYTO</td></tr>
<tr><td>>hg38_21_+_8433172_8446622_RNA45SN1_45S_50ntFlanks@4731.4764.34|AAGAACGAAAGTCGGAGGTTCGAAGACGATCAGA</td><td>>hg38_21_+_8433172_8446622_RNA45SN1_45S_50ntFlanks</td><td>4731</td><td>4764</td><td>34</td><td>AAGAACGAAAGTCGGAGGTTCGAAGACGATCAGA</td><td>835.0</td><td>7.4</td><td>1.4</td><td>5.4</td><td>8.3E-08</td><td>1.6E-07</td><td>MB231/MB468</td><td>CYTO</td></tr>
<tr><td>>MI0000746|hsa-mir-99b&WithFlank&19|+|51692606|51692687@13.30.18&[MIMAT0000689&hsa-miR-99b-5p&offsets|0|-4;m-10&19|+|51692618|51692639&offsets|0|-4]|CACCCGTAGAACCGACCT</td><td>>MI0000746|hsa-mir-99b&WithFlank&19|+|51692606|51692687</td><td>13</td><td>30</td><td>18</td><td>CACCCGTAGAACCGACCT</td><td>126.6</td><td>10.7</td><td>1.9</td><td>5.7</td><td>1.6E-08</td><td>1.6E-07</td><td>BT20/MB468</td><td>MP</td></tr>
<tr><td>>hg38_21_+_8433172_8446622_RNA45SN1_45S_50ntFlanks@8170.8216.47|CAAGTCCTTCTGATCGAGGCCCAGCCCGTGGACGGTGTGAGGCCGGT</td><td>>hg38_21_+_8433172_8446622_RNA45SN1_45S_50ntFlanks</td><td>8170</td><td>8216</td><td>47</td><td>CAAGTCCTTCTGATCGAGGCCCAGCCCGTGGACGGTGTGAGGCCGGT</td><td>567.3</td><td>-5.6</td><td>1.0</td><td>-5.3</td><td>9.0E-08</td><td>1.6E-07</td><td>MB231/MB468</td><td>MITO</td></tr>
</table>

<table>
<tr><td>>hg38_21_+_8433172_8446622_RNA45SN1_45S_50ntFlanks@9566.9596.31|GACGTGCAAATCGGTCGTCCGACCTGGGTAT</td><td>>hg38_21_+_8433172_8446622_RNA45SN1_45S_50ntFlanks</td><td>9566</td><td>9596</td><td>31</td><td>GACGTGCAAATCGGTCGTCCGACCTGGGTAT</td><td>2339.4</td><td>-5.9</td><td>1.1</td><td>-5.4</td><td>7.2E-08</td><td>1.6E-07</td><td>BT20/MB468</td><td>CYTO</td></tr>
<tr><td>>hg38_21_+_8433172_8446622_RNA45SN1_45S_50ntFlanks@11579.11604.26|AGAACTGGTGCGGACCAGGGGAATCC</td><td>>hg38_21_+_8433172_8446622_RNA45SN1_45S_50ntFlanks</td><td>11579</td><td>11604</td><td>26</td><td>AGAACTGGTGCGGACCAGGGGAATCC</td><td>436.8</td><td>-6.4</td><td>1.2</td><td>-5.3</td><td>9.4E-08</td><td>1.6E-07</td><td>BT20/MB468</td><td>NUC</td></tr>
<tr><td>>hg38_21_+_8433172_8446622_RNA45SN1_45S_50ntFlanks@5472.5490.19|ACGGCCCTGGCGGAGCGCT</td><td>>hg38_21_+_8433172_8446622_RNA45SN1_45S_50ntFlanks</td><td>5472</td><td>5490</td><td>19</td><td>ACGGCCCTGGCGGAGCGCT</td><td>276.8</td><td>5.5</td><td>1.0</td><td>5.4</td><td>8.5E-08</td><td>1.6E-07</td><td>MB231/MB468</td><td>NUC</td></tr>
<tr><td>>hg38_21_+_8433172_8446622_RNA45SN1_45S_50ntFlanks@11572.11593.22|CCGACTTAGAACTGGTGCGGAC</td><td>>hg38_21_+_8433172_8446622_RNA45SN1_45S_50ntFlanks</td><td>11572</td><td>11593</td><td>22</td><td>CCGACTTAGAACTGGTGCGGAC</td><td>889.5</td><td>-4.7</td><td>0.9</td><td>-5.4</td><td>8.3E-08</td><td>1.6E-07</td><td>MB231/MB468</td><td>CYTO</td></tr>
<tr><td>>hg38_21_+_8433172_8446622_RNA45SN1_45S_50ntFlanks@11874.11917.44|AGAAGACCCTGTTGAGCTTGACTCTAGTCTGGCACGGTGAAGAG</td><td>>hg38_21_+_8433172_8446622_RNA45SN1_45S_50ntFlanks</td><td>11874</td><td>11917</td><td>44</td><td>AGAAGACCCTGTTGAGCTTGACTCTAGTCTGGCACGGTGAAGAG</td><td>65.5</td><td>-9.9</td><td>1.8</td><td>-5.4</td><td>7.7E-08</td><td>1.6E-07</td><td>MB231/MB468</td><td>MP</td></tr>
<tr><td>>hg38_21_+_8433172_8446622_RNA45SN1_45S_50ntFlanks@10687.10710.24|CGGCGTCCGGTGAGCTCTCGCTGG</td><td>>hg38_21_+_8433172_8446622_RNA45SN1_45S_50ntFlanks</td><td>10687</td><td>10710</td><td>24</td><td>CGGCGTCCGGTGAGCTCTCGCTGG</td><td>3512.5</td><td>-5.6</td><td>1.0</td><td>-5.7</td><td>1.6E-08</td><td>1.6E-07</td><td>BT20/MB468</td><td>MP</td></tr>
<tr><td>>trnaMT_GluTTC_MT_-_14674_14742@40.71.32,trnalookalike8_GluTTC_5_-_93905172_93905240@40.71.32|TTGGTCGTGGTTGTAGTCCGTGCGAGAATACC</td><td>>trnaMT_GluTTC_MT_-_14674_14742</td><td>40</td><td>71</td><td>32</td><td>TTGGTCGTGGTTGTAGTCCGTGCGAGAATACC</td><td>850.2</td><td>4.0</td><td>0.8</td><td>5.3</td><td>9.0E-08</td><td>1.6E-07</td><td>MB231/MB468</td><td>MITO</td></tr>
<tr><td>>hg38_21_+_8433172_8446622_RNA45SN1_45S_50ntFlanks@6771.6806.36|GGTTCCTCCCGGGGCTACGCCTGTCTGAGCGTCGCT</td><td>>hg38_21_+_8433172_8446622_RNA45SN1_45S_50ntFlanks</td><td>6771</td><td>6806</td><td>36</td><td>GGTTCCTCCCGGGGCTACGCCTGTCTGAGCGTCGCT</td><td>599.0</td><td>-3.4</td><td>0.6</td><td>-5.3</td><td>9.4E-08</td><td>1.6E-07</td><td>BT20/MB468</td><td>NUC</td></tr>
<tr><td>>hg38_21_+_8433172_8446622_RNA45SN1_45S_50ntFlanks@12642.12664.23|CATCTGTGGGATTATGACTGAAC</td><td>>hg38_21_+_8433172_8446622_RNA45SN1_45S_50ntFlanks</td><td>12642</td><td>12664</td><td>23</td><td>CATCTGTGGGATTATGACTGAAC</td><td>87.7</td><td>10.2</td><td>1.8</td><td>5.7</td><td>1.6E-08</td><td>1.6E-07</td><td>BT20/MB468</td><td>MP</td></tr>
<tr><td>>hg38_21_+_8433172_8446622_RNA45SN1_45S_50ntFlanks@12722.12742.21|GTTGGCCTCGGATAGCCGGTC</td><td>>hg38_21_+_8433172_8446622_RNA45SN1_45S_50ntFlanks</td><td>12722</td><td>12742</td><td>21</td><td>GTTGGCCTCGGATAGCCGGTC</td><td>101.1</td><td>-9.7</td><td>1.7</td><td>-5.7</td><td>1.6E-08</td><td>1.6E-07</td><td>BT20/MB468</td><td>MP</td></tr>
<tr><td>>MI0000270|hsa-mir-181b-1&WithFlank&1|-|198858867|198858988@42.64.23(+1C);MI0000683|hsa-mir-181b-2&WithFlank&9|+|124693704|124693804@22.44.23(+1C)&[MIMAT0000257&hsa-miR-181b-5p&offsets|0|0(+1C);m-73&1|-|198858925|198858947&offsets|0|0(+1C)];[MIMAT0000257_1&hsa-miR-181b-5p&offsets|0|0(+1C);m-72&9|+|124693725|124693747&offsets|0|0(+1C)]|AACATTCATTGCTGTCGGTGGGTC</td><td>>MI0000270|hsa-mir-181b-1&WithFlank&1|-|198858867|198858988</td><td>42</td><td>64</td><td>23(+1C)</td><td>AACATTCATTGCTGTCGGTGGGTC</td><td>432.8</td><td>8.8</td><td>1.6</td><td>5.3</td><td>9.1E-08</td><td>1.6E-07</td><td>MB231/MB468</td><td>MITO</td></tr>
<tr><td>>hg38_21_+_8433172_8446622_RNA45SN1_45S_50ntFlanks@3835.3860.26|CTCCTCTCCTACTTGGATAACTGTGG</td><td>>hg38_21_+_8433172_8446622_RNA45SN1_45S_50ntFlanks</td><td>3835</td><td>3860</td><td>26</td><td>CTCCTCTCCTACTTGGATAACTGTGG</td><td>2112.8</td><td>-4.4</td><td>0.8</td><td>-5.4</td><td>6.5E-08</td><td>1.6E-07</td><td>BT20/MB468</td><td>MITO</td></tr>
<tr><td>>hg38_21_+_8433172_8446622_RNA45SN1_45S_50ntFlanks@10463.10481.19|CTCCGTTGCCCTCGGCCGA</td><td>>hg38_21_+_8433172_8446622_RNA45SN1_45S_50ntFlanks</td><td>10463</td><td>10481</td><td>19</td><td>CTCCGTTGCCCTCGGCCGA</td><td>90.6</td><td>-9.6</td><td>1.7</td><td>-5.7</td><td>1.6E-08</td><td>1.6E-07</td><td>BT20/MB468</td><td>MP</td></tr>
<tr><td>>hg38_21_+_8433172_8446622_RNA45SN1_45S_50ntFlanks@5419.5445.27|ACCGATTGGATGGTTTAGTGAGGCCCT</td><td>>hg38_21_+_8433172_8446622_RNA45SN1_45S_50ntFlanks</td><td>5419</td><td>5445</td><td>27</td><td>ACCGATTGGATGGTTTAGTGAGGCCCT</td><td>287.4</td><td>-7.2</td><td>1.4</td><td>-5.4</td><td>8.6E-08</td><td>1.6E-07</td><td>MB231/MB468</td><td>NUC</td></tr>
<tr><td>>hg38_21_+_8433172_8446622_RNA45SN1_45S_50ntFlanks@4729.4751.23|TCAAGAACGAAAGTCGGAGGTTC</td><td>>hg38_21_+_8433172_8446622_RNA45SN1_45S_50ntFlanks</td><td>4729</td><td>4751</td><td>23</td><td>TCAAGAACGAAAGTCGGAGGTTC</td><td>913.9</td><td>-6.1</td><td>1.1</td><td>-5.4</td><td>7.7E-08</td><td>1.6E-07</td><td>MB231/MB468</td><td>MP</td></tr>
<tr><td>>hg38_21_+_8433172_8446622_RNA45SN1_45S_50ntFlanks@5473.5512.40|CGGCCCTGGCGGAGCGCTGAGAAGACGGTCGAACTTGACT</td><td>>hg38_21_+_8433172_8446622_RNA45SN1_45S_50ntFlanks</td><td>5473</td><td>5512</td><td>40</td><td>CGGCCCTGGCGGAGCGCTGAGAAGACGGTCGAACTTGACT</td><td>203.9</td><td>3.9</td><td>0.7</td><td>5.4</td><td>8.6E-08</td><td>1.6E-07</td><td>MB231/MB468</td><td>NUC</td></tr>
<tr><td>>hg38_21_+_8433172_8446622_RNA45SN1_45S_50ntFlanks@11573.11602.30|CGACTTAGAACTGGTGCGGACCAGGGGAAT</td><td>>hg38_21_+_8433172_8446622_RNA45SN1_45S_50ntFlanks</td><td>11573</td><td>11602</td><td>30</td><td>CGACTTAGAACTGGTGCGGACCAGGGGAAT</td><td>1060.5</td><td>-6.3</td><td>1.2</td><td>-5.4</td><td>7.7E-08</td><td>1.6E-07</td><td>MB231/MB468</td><td>MP</td></tr>
<tr><td>>hg38_21_+_8433172_8446622_RNA45SN1_45S_50ntFlanks@12072.12093.22|CCCGAGGGGCTCTCGCTTCTGG</td><td>>hg38_21_+_8433172_8446622_RNA45SN1_45S_50ntFlanks</td><td>12072</td><td>12093</td><td>22</td><td>CCCGAGGGGCTCTCGCTTCTGG</td><td>78.6</td><td>-9.4</td><td>1.7</td><td>-5.7</td><td>1.6E-08</td><td>1.6E-07</td><td>BT20/MB468</td><td>MP</td></tr>
<tr><td>>hg38_21_+_8433172_8446622_RNA45SN1_45S_50ntFlanks@9198.9222.25|GTTCTCTCGGGGCCACGCGCGCGTC</td><td>>hg38_21_+_8433172_8446622_RNA45SN1_45S_50ntFlanks</td><td>9198</td><td>9222</td><td>25</td><td>GTTCTCTCGGGGCCACGCGCGCGTC</td><td>663.1</td><td>-7.8</td><td>1.5</td><td>-5.4</td><td>8.6E-08</td><td>1.6E-07</td><td>MB231/MB468</td><td>NUC</td></tr>
<tr><td>>hg38_21_+_8433172_8446622_RNA45SN1_45S_50ntFlanks@11873.11892.20|AAGAAGACCCTGTTGAGCTT</td><td>>hg38_21_+_8433172_8446622_RNA45SN1_45S_50ntFlanks</td><td>11873</td><td>11892</td><td>20</td><td>AAGAAGACCCTGTTGAGCTT</td><td>705.6</td><td>-5.8</td><td>1.1</td><td>-5.4</td><td>8.6E-08</td><td>1.6E-07</td><td>MB231/MB468</td><td>NUC</td></tr>
<tr><td>>hg38_21_+_8433172_8446622_RNA45SN1_45S_50ntFlanks@11620.11660.41|ACAAAGCATCGCGAAGGCCCGCGGCGGGTGTTGACGCGATG</td><td>>hg38_21_+_8433172_8446622_RNA45SN1_45S_50ntFlanks</td><td>11620</td><td>11660</td><td>41</td><td>ACAAAGCATCGCGAAGGCCCGCGGCGGGTGTTGACGCGATG</td><td>101.6</td><td>-9.7</td><td>1.7</td><td>-5.7</td><td>1.6E-08</td><td>1.6E-07</td><td>BT20/MB468</td><td>MP</td></tr>
<tr><td>>am_trna119_LysCTT_1_-_145395522_145395594@16.33.18,trna11_LysCTT_5_-_180648979_180649051@16.33.18,trna128_LysCTT_1_-_55423542_55423614@16.33.18,trna13_LysCTT_6_+_26556774_26556846@16.33.18,trna6_LysCTT_19_-_52425396_52425468@16.33.18,trna7_LysCTT_16_+_3225692_3225764@16.33.18,trna9_LysCTT_5_+_180634755_180634827@16.33.18|TCGGTAGAGCATGAGACT</td><td>>am_trna119_LysCTT_1_-_145395522_145395594</td><td>16</td><td>33</td><td>18</td><td>TCGGTAGAGCATGAGACT</td><td>412.2</td><td>8.0</td><td>1.5</td><td>5.4</td><td>8.4E-08</td><td>1.6E-07</td><td>MB231/MB468</td><td>CYTO</td></tr>
<tr><td>>hg38_21_+_8433172_8446622_RNA45SN1_45S_50ntFlanks@10846.10866.21|CTTCGGGATAAGGATTGGCTC</td><td>>hg38_21_+_8433172_8446622_RNA45SN1_45S_50ntFlanks</td><td>10846</td><td>10866</td><td>21</td><td>CTTCGGGATAAGGATTGGCTC</td><td>270.3</td><td>-5.6</td><td>1.0</td><td>-5.4</td><td>8.6E-08</td><td>1.6E-07</td><td>MB231/MB468</td><td>NUC</td></tr>
<tr><td>>am_MI0000086|hsa-mir-28&WithFlank&3|+|188688775|188688872@20.39.20&[MIMAT0000085&hsa-miR-28-5p&offsets|0|-2;m-133&3|+|188688794|188688815&offsets|0|-2]|AAGGAGCTCACAGTCTATTG</td><td>>am_MI0000086|hsa-mir-28&WithFlank&3|+|188688775|188688872</td><td>20</td><td>39</td><td>20</td><td>AAGGAGCTCACAGTCTATTG</td><td>62.4</td><td>3.4</td><td>0.6</td><td>5.3</td><td>9.5E-08</td><td>1.6E-07</td><td>BT20/MB468</td><td>NUC</td></tr>
<tr><td>>am_trnaMT_PheGAA_MT_+_577_647@41.59.19|GTTTAGACGGGCTCACATC</td><td>>am_trnaMT_PheGAA_MT_+_577_647</td><td>41</td><td>59</td><td>19</td><td>GTTTAGACGGGCTCACATC</td><td>175.7</td><td>5.6</td><td>1.1</td><td>5.4</td><td>8.6E-08</td><td>1.6E-07</td><td>MB231/MB468</td><td>NUC</td></tr>
<tr><td>>hg38_21_+_8433172_8446622_RNA45SN1_45S_50ntFlanks@10788.10816.29|CAGCCTCTGGCATGTTGGAACAATGTAGG</td><td>>hg38_21_+_8433172_8446622_RNA45SN1_45S_50ntFlanks</td><td>10788</td><td>10816</td><td>29</td><td>CAGCCTCTGGCATGTTGGAACAATGTAGG</td><td>638.7</td><td>-4.0</td><td>0.8</td><td>-5.4</td><td>8.6E-08</td><td>1.6E-07</td><td>MB231/MB468</td><td>NUC</td></tr>
<tr><td>>hg38_21_+_8433172_8446622_RNA45SN1_45S_50ntFlanks@4733.4759.27|GAACGAAAGTCGGAGGTTCGAAGACGA</td><td>>hg38_21_+_8433172_8446622_RNA45SN1_45S_50ntFlanks</td><td>4733</td><td>4759</td><td>27</td><td>GAACGAAAGTCGGAGGTTCGAAGACGA</td><td>379.2</td><td>-4.6</td><td>0.9</td><td>-5.3</td><td>9.1E-08</td><td>1.6E-07</td><td>MB231/MB468</td><td>MITO</td></tr>
<tr><td>>hg38_21_+_8433172_8446622_RNA45SN1_45S_50ntFlanks@4047.4069.23|ATTCGAACGTCTGCCCTATCAAC</td><td>>hg38_21_+_8433172_8446622_RNA45SN1_45S_50ntFlanks</td><td>4047</td><td>4069</td><td>23</td><td>ATTCGAACGTCTGCCCTATCAAC</td><td>1195.1</td><td>-6.9</td><td>1.3</td><td>-5.3</td><td>9.2E-08</td><td>1.6E-07</td><td>MB231/MB468</td><td>MITO</td></tr>
<tr><td>>hg38_21_+_8433172_8446622_RNA45SN1_45S_50ntFlanks@3840.3879.40|CTCCTACTTGGATAACTGTGGTAATTCTAGAGCTAATACA</td><td>>hg38_21_+_8433172_8446622_RNA45SN1_45S_50ntFlanks</td><td>3840</td><td>3879</td><td>40</td><td>CTCCTACTTGGATAACTGTGGTAATTCTAGAGCTAATACA</td><td>574.2</td><td>5.9</td><td>1.1</td><td>5.4</td><td>8.4E-08</td><td>1.6E-07</td><td>MB231/MB468</td><td>CYTO</td></tr>
<tr><td>>hg38_21_+_8433172_8446622_RNA45SN1_45S_50ntFlanks@12075.12095.21|GAGGGGCTCTCGCTTCTGGCG</td><td>>hg38_21_+_8433172_8446622_RNA45SN1_45S_50ntFlanks</td><td>12075</td><td>12095</td><td>21</td><td>GAGGGGCTCTCGCTTCTGGCG</td><td>334.5</td><td>-3.0</td><td>0.6</td><td>-5.4</td><td>8.7E-08</td><td>1.6E-07</td><td>MB231/MB468</td><td>NUC</td></tr>
<tr><td>>MI0000102|hsa-mir-100&WithFlank&11|-|122152223|122152314@19.40.22(+1A)&[MIMAT0000098&hsa-miR-100-5p&offsets|0|0(+1A);m-27&11|-|122152276|122152296&offsets|0|+1(+1A)]|AACCCGTAGATCCGAACTTGTGA</td><td>>MI0000102|hsa-mir-100&WithFlank&11|-|122152223|122152314</td><td>19</td><td>40</td><td>22(+1A)</td><td>AACCCGTAGATCCGAACTTGTGA</td><td>46255.4</td><td>-7.0</td><td>1.2</td><td>-5.9</td><td>3.2E-09</td><td>1.6E-07</td><td>BT20/MB231</td><td>MP</td></tr>
<tr><td>>MI0000650|hsa-mir-200c&WithFlank&12|+|6963693|6963772@50.73.24&[MIMAT0000617&hsa-miR-200c-3p&offsets|0|+1;m-54&12|+|6963742|6963764&offsets|0|+1]|TAATACTGCCGGGTAATGATGGAG</td><td>>MI0000650|hsa-mir-200c&WithFlank&12|+|6963693|6963772</td><td>50</td><td>73</td><td>24</td><td>TAATACTGCCGGGTAATGATGGAG</td><td>191.5</td><td>10.9</td><td>1.8</td><td>5.9</td><td>3.2E-09</td><td>1.6E-07</td><td>BT20/MB231</td><td>MP</td></tr>
<tr><td>>trna111_HisGTG_1_-_147774845_147774916@-1T.23.24,trna118_HisGTG_1_-_145396881_145396952@-1T.23.24,trna16_HisGTG_1_+_146544773_146544844@-1T.23.24,trna1_HisGTG_15_+_45493349_45493420@-1T.23.24,trna21_HisGTG_1_+_147753471_147753542@-1T.23.24,trna33_HisGTG_6_+_27125906_27125977@-1T.23.24,trna7_HisGTG_9_-_14433938_14434009@-1T.23.24,trna8_HisGTG_15_-_45492611_45492682@-1T.23.24,trna9_HisGTG_15_-_45490804_45490875@-1T.23.24|TGCCGTGATCGTATAGTGGTTAGT</td><td>>trna111_HisGTG_1_-_147774845_147774916</td><td>-1T</td><td>23</td><td>24</td><td>TGCCGTGATCGTATAGTGGTTAGT</td><td>1661.1</td><td>-6.1</td><td>1.0</td><td>-5.9</td><td>3.2E-09</td><td>1.6E-07</td><td>BT20/MB231</td><td>MP</td></tr>
<tr><td>>hg38_21_+_8433172_8446622_RNA45SN1_45S_50ntFlanks@12510.12546.37|TCGTGAGACAGGTTAGTTTTACCCTACTGATGATGTG</td><td>>hg38_21_+_8433172_8446622_RNA45SN1_45S_50ntFlanks</td><td>12510</td><td>12546</td><td>37</td><td>TCGTGAGACAGGTTAGTTTTACCCTACTGATGATGTG</td><td>83.3</td><td>-8.5</td><td>1.6</td><td>-5.4</td><td>6.5E-08</td><td>1.6E-07</td><td>BT20/MB468</td><td>MITO</td></tr>
<tr><td>>hg38_21_+_8433172_8446622_RNA45SN1_45S_50ntFlanks@6661.6708.48|CGGTGGATCACTCGGCTCGTGCGTCGATGAAGAACGCAGCTAGCTGCG</td><td>>hg38_21_+_8433172_8446622_RNA45SN1_45S_50ntFlanks</td><td>6661</td><td>6708</td><td>48</td><td>CGGTGGATCACTCGGCTCGTGCGTCGATGAAGAACGCAGCTAGCTGCG</td><td>12002.2</td><td>5.0</td><td>0.9</td><td>5.4</td><td>6.5E-08</td><td>1.6E-07</td><td>BT20/MB468</td><td>MITO</td></tr>
<tr><td>>hg38_21_+_8433172_8446622_RNA45SN1_45S_50ntFlanks@10786.10828.43|AACAGCCTCTGGCATGTTGGAACAATGTAGGTAAGGGAAGTCG</td><td>>hg38_21_+_8433172_8446622_RNA45SN1_45S_50ntFlanks</td><td>10786</td><td>10828</td><td>43</td><td>AACAGCCTCTGGCATGTTGGAACAATGTAGGTAAGGGAAGTCG</td><td>195.4</td><td>-7.0</td><td>1.3</td><td>-5.4</td><td>7.3E-08</td><td>1.6E-07</td><td>BT20/MB468</td><td>CYTO</td></tr>
<tr><td>>trnaMT_HisGTG_MT_+_12138_12206@39.72.34|GACAACAGAGGCTTACGACCCCTTATTTACCCCA</td><td>>trnaMT_HisGTG_MT_+_12138_12206</td><td>39</td><td>72</td><td>34</td><td>GACAACAGAGGCTTACGACCCCTTATTTACCCCA</td><td>212.7</td><td>11.5</td><td>2.0</td><td>5.6</td><td>1.6E-08</td><td>1.6E-07</td><td>BT20/MB468</td><td>MP</td></tr>
<tr><td>>hg38_21_+_8433172_8446622_RNA45SN1_45S_50ntFlanks@3842.3879.38|CCTACTTGGATAACTGTGGTAATTCTAGAGCTAATACA</td><td>>hg38_21_+_8433172_8446622_RNA45SN1_45S_50ntFlanks</td><td>3842</td><td>3879</td><td>38</td><td>CCTACTTGGATAACTGTGGTAATTCTAGAGCTAATACA</td><td>579.0</td><td>5.4</td><td>1.0</td><td>5.4</td><td>7.4E-08</td><td>1.6E-07</td><td>BT20/MB468</td><td>CYTO</td></tr>
<tr><td>>hg38_21_+_8433172_8446622_RNA45SN1_45S_50ntFlanks@9503.9520.18|GGTGAACTATGCCTGGGC</td><td>>hg38_21_+_8433172_8446622_RNA45SN1_45S_50ntFlanks</td><td>9503</td><td>9520</td><td>18</td><td>GGTGAACTATGCCTGGGC</td><td>301.6</td><td>-6.0</td><td>1.1</td><td>-5.4</td><td>7.8E-08</td><td>1.6E-07</td><td>MB231/MB468</td><td>MP</td></tr>
<tr><td>>hg38_21_+_8433172_8446622_RNA45SN1_45S_50ntFlanks@12488.12524.37|AACGTGAGCTGGGTTTAGACCGTCGTGAGACAGGTTA</td><td>>hg38_21_+_8433172_8446622_RNA45SN1_45S_50ntFlanks</td><td>12488</td><td>12524</td><td>37</td><td>AACGTGAGCTGGGTTTAGACCGTCGTGAGACAGGTTA</td><td>197.5</td><td>-4.0</td><td>0.8</td><td>-5.4</td><td>8.4E-08</td><td>1.6E-07</td><td>MB231/MB468</td><td>CYTO</td></tr>
<tr><td>>hg38_21_+_8433172_8446622_RNA45SN1_45S_50ntFlanks@12432.12454.23|CCTATCATTGTGAAGCAGAATTC</td><td>>hg38_21_+_8433172_8446622_RNA45SN1_45S_50ntFlanks</td><td>12432</td><td>12454</td><td>23</td><td>CCTATCATTGTGAAGCAGAATTC</td><td>313.5</td><td>-6.1</td><td>1.1</td><td>-5.4</td><td>7.8E-08</td><td>1.6E-07</td><td>MB231/MB468</td><td>MP</td></tr>
<tr><td>>hg38_MT_+_598_1651_RNR1_12S_50ntFlanks@427.468.42|GAAAGTGGCTTTAACATATCTGAACACACAATAGCTAAGACC</td><td>>hg38_MT_+_598_1651_RNR1_12S_50ntFlanks</td><td>427</td><td>468</td><td>42</td><td>GAAAGTGGCTTTAACATATCTGAACACACAATAGCTAAGACC</td><td>190.6</td><td>11.3</td><td>2.0</td><td>5.6</td><td>1.6E-08</td><td>1.6E-07</td><td>BT20/MB468</td><td>MP</td></tr>
<tr><td>>hg38_21_+_8433172_8446622_RNA45SN1_45S_50ntFlanks@3991.4024.34|TGGTGACTCTAGATAACCTCGGGCCGATCGCACG</td><td>>hg38_21_+_8433172_8446622_RNA45SN1_45S_50ntFlanks</td><td>3991</td><td>4024</td><td>34</td><td>TGGTGACTCTAGATAACCTCGGGCCGATCGCACG</td><td>412.3</td><td>-6.0</td><td>1.1</td><td>-5.4</td><td>7.9E-08</td><td>1.6E-07</td><td>MB231/MB468</td><td>MP</td></tr>
<tr><td>>hg38_21_+_8433172_8446622_RNA45SN1_45S_50ntFlanks@10916.10936.21|CCGCGGCTGGACGAGGCGCCG</td><td>>hg38_21_+_8433172_8446622_RNA45SN1_45S_50ntFlanks</td><td>10916</td><td>10936</td><td>21</td><td>CCGCGGCTGGACGAGGCGCCG</td><td>477.1</td><td>-8.4</td><td>1.6</td><td>-5.3</td><td>9.6E-08</td><td>1.6E-07</td><td>BT20/MB468</td><td>NUC</td></tr>
<tr><td>>hg38_21_+_8433172_8446622_RNA45SN1_45S_50ntFlanks@475.494.20|CTTCGTGATCGATGTGGTGA</td><td>>hg38_21_+_8433172_8446622_RNA45SN1_45S_50ntFlanks</td><td>475</td><td>494</td><td>20</td><td>CTTCGTGATCGATGTGGTGA</td><td>1259.0</td><td>5.2</td><td>1.0</td><td>5.4</td><td>8.7E-08</td><td>1.6E-07</td><td>MB231/MB468</td><td>NUC</td></tr>
<tr><td>>hg38_1_-_228634819_228635039_RNA5S12_5S_50ntFlanks@140.170.31|ACCGCCTGGGAATACCGGGTGCTGTAGGCTT</td><td>>hg38_1_-_228634819_228635039_RNA5S12_5S_50ntFlanks</td><td>140</td><td>170</td><td>31</td><td>ACCGCCTGGGAATACCGGGTGCTGTAGGCTT</td><td>20994.9</td><td>-3.2</td><td>0.6</td><td>-5.4</td><td>7.9E-08</td><td>1.6E-07</td><td>MB231/MB468</td><td>MP</td></tr>
<tr><td>>MI0015866|hsa-mir-4326&WithFlank&20|+|63286802|63286872@17.39.23&[MIMAT0016888&hsa-miR-4326&offsets|0|+3;m-339&20|+|63286818|63286841&offsets|0|-1]|TGTTCCTCTGTCTCCCAGACTCT</td><td>>MI0015866|hsa-mir-4326&WithFlank&20|+|63286802|63286872</td><td>17</td><td>39</td><td>23</td><td>TGTTCCTCTGTCTCCCAGACTCT</td><td>210.3</td><td>6.9</td><td>1.3</td><td>5.4</td><td>8.7E-08</td><td>1.6E-07</td><td>MB231/MB468</td><td>NUC</td></tr>
</table>

<table>
<tr><td>>am_trna146_GlnCTG_6_-_27515531_27515602@1.29.29,trna1_GlnCTG_6_+_18836402_18836473@1.29.29,trna3_GlnCTG_17_+_8023070_8023141@1.29.29,trna42_GlnCTG_6_+_27263212_27263283@1.29.29,trna49_GlnCTG_6_+_27487308_27487379@1.29.29,trna7_GlnCTG_15_-_66161400_66161471@1.29.29,trna99_GlnCTG_6_-_28909378_28909449@1.29.29|GGTTCCATGGTGTAATGGTTAGCACTCTG</td><td>>am_trna146_GlnCTG_6_-_27515531_27515602</td><td>1</td><td>29</td><td>29</td><td>GGTTCCATGGTGTAATGGTTAGCACTCTG</td><td>3142.4</td><td>-6.5</td><td>1.2</td><td>-5.4</td><td>8.7E-08</td><td>1.6E-07</td><td>MB231/MB468</td><td>NUC</td></tr>
<tr><td>>hg38_21_+_8433172_8446622_RNA45SN1_45S_50ntFlanks@11885.11903.19|TTGAGCTTGACTCTAGTCT</td><td>>hg38_21_+_8433172_8446622_RNA45SN1_45S_50ntFlanks</td><td>11885</td><td>11903</td><td>19</td><td>TTGAGCTTGACTCTAGTCT</td><td>160.7</td><td>-5.0</td><td>0.9</td><td>-5.4</td><td>8.7E-08</td><td>1.6E-07</td><td>MB231/MB468</td><td>NUC</td></tr>
<tr><td>>hg38_21_+_8433172_8446622_RNA45SN1_45S_50ntFlanks@6733.6758.26|CATTGATCATCGACACTTCGAACGCA</td><td>>hg38_21_+_8433172_8446622_RNA45SN1_45S_50ntFlanks</td><td>6733</td><td>6758</td><td>26</td><td>CATTGATCATCGACACTTCGAACGCA</td><td>324.4</td><td>-3.0</td><td>0.6</td><td>-5.4</td><td>8.8E-08</td><td>1.6E-07</td><td>MB231/MB468</td><td>NUC</td></tr>
<tr><td>>hg38_21_+_8433172_8446622_RNA45SN1_45S_50ntFlanks@5472.5507.36|ACGGCCCTGGCGGAGCGCTGAGAAGACGGTCGAACT</td><td>>hg38_21_+_8433172_8446622_RNA45SN1_45S_50ntFlanks</td><td>5472</td><td>5507</td><td>36</td><td>ACGGCCCTGGCGGAGCGCTGAGAAGACGGTCGAACT</td><td>327.2</td><td>6.3</td><td>1.2</td><td>5.4</td><td>8.8E-08</td><td>1.6E-07</td><td>MB231/MB468</td><td>NUC</td></tr>
<tr><td>>hg38_21_+_8433172_8446622_RNA45SN1_45S_50ntFlanks@3875.3899.25|ATACATGCCGACGGGCGCTGACCCC</td><td>>hg38_21_+_8433172_8446622_RNA45SN1_45S_50ntFlanks</td><td>3875</td><td>3899</td><td>25</td><td>ATACATGCCGACGGGCGCTGACCCC</td><td>920.4</td><td>-3.5</td><td>0.7</td><td>-5.4</td><td>8.5E-08</td><td>1.6E-07</td><td>MB231/MB468</td><td>CYTO</td></tr>
<tr><td>>hg38_21_+_8433172_8446622_RNA45SN1_45S_50ntFlanks@6780.6803.24|CGGGGCTACGCCTGTCTGAGCGTC</td><td>>hg38_21_+_8433172_8446622_RNA45SN1_45S_50ntFlanks</td><td>6780</td><td>6803</td><td>24</td><td>CGGGGCTACGCCTGTCTGAGCGTC</td><td>529.7</td><td>-6.0</td><td>1.1</td><td>-5.4</td><td>7.9E-08</td><td>1.6E-07</td><td>MB231/MB468</td><td>MP</td></tr>
<tr><td>>MI0000066|hsa-let-7e&WithFlank&19|+|51692780|51692870@14.34.21(+1C)&[MIMAT0000066&hsa-let-7e-5p&offsets|0|-1(+1C);m-50&19|+|51692793|51692814&offsets|0|-1(+1C)]|TGAGGTAGGAGGTTGTATAGTC</td><td>>MI0000066|hsa-let-7e&WithFlank&19|+|51692780|51692870</td><td>14</td><td>34</td><td>21(+1C)</td><td>TGAGGTAGGAGGTTGTATAGTC</td><td>735.2</td><td>5.0</td><td>0.9</td><td>5.3</td><td>9.3E-08</td><td>1.6E-07</td><td>MB231/MB468</td><td>MITO</td></tr>
<tr><td>>hg38_21_+_8433172_8446622_RNA45SN1_45S_50ntFlanks@12676.12705.30|AGAATCCCGCCCAGGCGGAACGATACGGCA</td><td>>hg38_21_+_8433172_8446622_RNA45SN1_45S_50ntFlanks</td><td>12676</td><td>12705</td><td>30</td><td>AGAATCCCGCCCAGGCGGAACGATACGGCA</td><td>100.5</td><td>-9.7</td><td>1.7</td><td>-5.6</td><td>1.6E-08</td><td>1.6E-07</td><td>BT20/MB468</td><td>MP</td></tr>
<tr><td>>hg38_21_+_8433172_8446622_RNA45SN1_45S_50ntFlanks@10361.10388.28|GAAGGCCGAAGTGGAGAAGGGTTCCATG</td><td>>hg38_21_+_8433172_8446622_RNA45SN1_45S_50ntFlanks</td><td>10361</td><td>10388</td><td>28</td><td>GAAGGCCGAAGTGGAGAAGGGTTCCATG</td><td>891.2</td><td>-6.2</td><td>1.2</td><td>-5.4</td><td>8.5E-08</td><td>1.6E-07</td><td>MB231/MB468</td><td>CYTO</td></tr>
<tr><td>>hg38_21_+_8433172_8446622_RNA45SN1_45S_50ntFlanks@5485.5514.30|AGCGCTGAGAAGACGGTCGAACTTGACTAT</td><td>>hg38_21_+_8433172_8446622_RNA45SN1_45S_50ntFlanks</td><td>5485</td><td>5514</td><td>30</td><td>AGCGCTGAGAAGACGGTCGAACTTGACTAT</td><td>2640.3</td><td>-8.3</td><td>1.5</td><td>-5.4</td><td>8.8E-08</td><td>1.6E-07</td><td>MB231/MB468</td><td>NUC</td></tr>
<tr><td>>hg38_21_+_8433172_8446622_RNA45SN1_45S_50ntFlanks@12396.12416.21|GCGACGTCGCTTTTTGATCCT</td><td>>hg38_21_+_8433172_8446622_RNA45SN1_45S_50ntFlanks</td><td>12396</td><td>12416</td><td>21</td><td>GCGACGTCGCTTTTTGATCCT</td><td>81.7</td><td>-10.2</td><td>1.9</td><td>-5.4</td><td>7.9E-08</td><td>1.6E-07</td><td>MB231/MB468</td><td>MP</td></tr>
<tr><td>>hg38_21_+_8433172_8446622_RNA45SN1_45S_50ntFlanks@11315.11334.20|CCACCCCACGTCTCGTCGCG</td><td>>hg38_21_+_8433172_8446622_RNA45SN1_45S_50ntFlanks</td><td>11315</td><td>11334</td><td>20</td><td>CCACCCCACGTCTCGTCGCG</td><td>517.0</td><td>-8.1</td><td>1.5</td><td>-5.3</td><td>8.8E-08</td><td>1.6E-07</td><td>MB231/MB468</td><td>NUC</td></tr>
<tr><td>>hg38_21_+_8433172_8446622_RNA45SN1_45S_50ntFlanks@10687.10712.26|CGGCGTCCGGTGAGCTCTCGCTGGCC</td><td>>hg38_21_+_8433172_8446622_RNA45SN1_45S_50ntFlanks</td><td>10687</td><td>10712</td><td>26</td><td>CGGCGTCCGGTGAGCTCTCGCTGGCC</td><td>986.9</td><td>-6.3</td><td>1.2</td><td>-5.3</td><td>8.8E-08</td><td>1.6E-07</td><td>MB231/MB468</td><td>NUC</td></tr>
<tr><td>>hg38_21_+_8433172_8446622_RNA45SN1_45S_50ntFlanks@4003.4044.42|ATAACCTCGGGCCGATCGCACGCCCCCCGTGGCGGCGACGAC</td><td>>hg38_21_+_8433172_8446622_RNA45SN1_45S_50ntFlanks</td><td>4003</td><td>4044</td><td>42</td><td>ATAACCTCGGGCCGATCGCACGCCCCCCGTGGCGGCGACGAC</td><td>142.5</td><td>-5.3</td><td>1.0</td><td>-5.4</td><td>8.6E-08</td><td>1.6E-07</td><td>MB231/MB468</td><td>CYTO</td></tr>
<tr><td>>trna116_GluCTC_1_-_145399233_145399304@37.75.39,trna134_GluTTC_1_-_16861774_16861845@37.75.39,trna71_GluCTC_1_-_161439189_161439260@37.75.39,trna74_GluCTC_1_-_161431809_161431880@37.75.39,trna77_GluCTC_1_-_161424398_161424469@37.75.39,trna77_GluCTC_6_+_28949976_28950047@37.75.39,trna80_GluCTC_1_-_161417018_161417089@37.75.39,trna84_GluTTC_1_-_161391883_161391954@37.75.39,trna87_GluCTC_6_-_126101393_126101464@37.75.39|ACCGCCGCGGCCCGGGTTCGATTCCCGGTCAGGGAACCA</td><td>>trna116_GluCTC_1_-_145399233_145399304</td><td>37</td><td>75</td><td>39</td><td>ACCGCCGCGGCCCGGGTTCGATTCCCGGTCAGGGAACCA</td><td>1686.5</td><td>4.4</td><td>0.8</td><td>5.4</td><td>7.5E-08</td><td>1.6E-07</td><td>BT20/MB468</td><td>CYTO</td></tr>
<tr><td>>hg38_21_+_8433172_8446622_RNA45SN1_45S_50ntFlanks@3920.3959.40|CATTTATCAGATCAAAACCAACCCGGTCAGCCCCTCTCCG</td><td>>hg38_21_+_8433172_8446622_RNA45SN1_45S_50ntFlanks</td><td>3920</td><td>3959</td><td>40</td><td>CATTTATCAGATCAAAACCAACCCGGTCAGCCCCTCTCCG</td><td>386.3</td><td>-7.0</td><td>1.3</td><td>-5.4</td><td>6.7E-08</td><td>1.6E-07</td><td>BT20/MB468</td><td>MITO</td></tr>
<tr><td>>MI0000063|hsa-let-7b&WithFlank&22|+|46113680|46113774@12.31.20(+1A)&[MIMAT0000063&hsa-let-7b-5p&offsets|0|-2(+1A);m-14&22|+|46113691|46113712&offsets|0|-2(+1A)]|TGAGGTAGTAGGTTGTGTGGA</td><td>>MI0000063|hsa-let-7b&WithFlank&22|+|46113680|46113774</td><td>12</td><td>31</td><td>20(+1A)</td><td>TGAGGTAGTAGGTTGTGTGGA</td><td>237.7</td><td>3.4</td><td>0.6</td><td>5.7</td><td>9.2E-09</td><td>1.6E-07</td><td>BT20/MB231</td><td>NUC</td></tr>
<tr><td>>hg38_1_-_228634819_228635039_RNA5S12_5S_50ntFlanks@134.171.38|TGGGAGACCGCCTGGGAATACCGGGTGCTGTAGGCTTT</td><td>>hg38_1_-_228634819_228635039_RNA5S12_5S_50ntFlanks</td><td>134</td><td>171</td><td>38</td><td>TGGGAGACCGCCTGGGAATACCGGGTGCTGTAGGCTTT</td><td>12242.8</td><td>5.0</td><td>0.9</td><td>5.3</td><td>9.3E-08</td><td>1.6E-07</td><td>MB231/MB468</td><td>MITO</td></tr>
<tr><td>>hg38_21_+_8433172_8446622_RNA45SN1_45S_50ntFlanks@11671.11702.32|CCAGTGCTCTGAATGTCAAAGTGAAGAAATTC</td><td>>hg38_21_+_8433172_8446622_RNA45SN1_45S_50ntFlanks</td><td>11671</td><td>11702</td><td>32</td><td>CCAGTGCTCTGAATGTCAAAGTGAAGAAATTC</td><td>1073.3</td><td>-6.5</td><td>1.2</td><td>-5.4</td><td>8.0E-08</td><td>1.6E-07</td><td>MB231/MB468</td><td>MP</td></tr>
<tr><td>>hg38_21_+_8433172_8446622_RNA45SN1_45S_50ntFlanks@476.509.34|TTCGTGATCGATGTGGTGACGTCGTGCTCTCCCG</td><td>>hg38_21_+_8433172_8446622_RNA45SN1_45S_50ntFlanks</td><td>476</td><td>509</td><td>34</td><td>TTCGTGATCGATGTGGTGACGTCGTGCTCTCCCG</td><td>941.3</td><td>6.9</td><td>1.3</td><td>5.4</td><td>6.7E-08</td><td>1.6E-07</td><td>BT20/MB468</td><td>MITO</td></tr>
<tr><td>>hg38_21_+_8433172_8446622_RNA45SN1_45S_50ntFlanks@12457.12484.28|CAAGCGTTGGATTGTTCACCCACTAATA</td><td>>hg38_21_+_8433172_8446622_RNA45SN1_45S_50ntFlanks</td><td>12457</td><td>12484</td><td>28</td><td>CAAGCGTTGGATTGTTCACCCACTAATA</td><td>806.5</td><td>-6.6</td><td>1.2</td><td>-5.4</td><td>8.6E-08</td><td>1.6E-07</td><td>MB231/MB468</td><td>CYTO</td></tr>
<tr><td>>MI0000743|hsa-mir-34c&WithFlank&11|+|111513433|111513521@19.39.21&[MIMAT0000686&hsa-miR-34c-5p&offsets|0|-2;m-336&11|+|111513451|111513473&offsets|0|-2]|AGGCAGTGTAGTTAGCTGATT</td><td>>MI0000743|hsa-mir-34c&WithFlank&11|+|111513433|111513521</td><td>19</td><td>39</td><td>21</td><td>AGGCAGTGTAGTTAGCTGATT</td><td>56.0</td><td>9.0</td><td>1.7</td><td>5.4</td><td>8.0E-08</td><td>1.6E-07</td><td>MB231/MB468</td><td>MP</td></tr>
<tr><td>>hg38_21_+_8433172_8446622_RNA45SN1_45S_50ntFlanks@9567.9586.20|ACGTGCAAATCGGTCGTCCG</td><td>>hg38_21_+_8433172_8446622_RNA45SN1_45S_50ntFlanks</td><td>9567</td><td>9586</td><td>20</td><td>ACGTGCAAATCGGTCGTCCG</td><td>212.2</td><td>-3.6</td><td>0.7</td><td>-5.3</td><td>8.9E-08</td><td>1.6E-07</td><td>MB231/MB468</td><td>NUC</td></tr>
<tr><td>>hg38_21_+_8433172_8446622_RNA45SN1_45S_50ntFlanks@12936.12966.31|AAACCATTCGTAGACGACCTGCTTCTGGGTC</td><td>>hg38_21_+_8433172_8446622_RNA45SN1_45S_50ntFlanks</td><td>12936</td><td>12966</td><td>31</td><td>AAACCATTCGTAGACGACCTGCTTCTGGGTC</td><td>194.4</td><td>-7.0</td><td>1.3</td><td>-5.3</td><td>8.9E-08</td><td>1.6E-07</td><td>MB231/MB468</td><td>NUC</td></tr>
<tr><td>>hg38_21_+_8433172_8446622_RNA45SN1_45S_50ntFlanks@9712.9729.18|AGCGAATGATTAGAGGTC</td><td>>hg38_21_+_8433172_8446622_RNA45SN1_45S_50ntFlanks</td><td>9712</td><td>9729</td><td>18</td><td>AGCGAATGATTAGAGGTC</td><td>213.5</td><td>-7.5</td><td>1.4</td><td>-5.4</td><td>8.0E-08</td><td>1.6E-07</td><td>MB231/MB468</td><td>MP</td></tr>
<tr><td>>hg38_21_+_8433172_8446622_RNA45SN1_45S_50ntFlanks@10841.10868.28|CGTAACTTCGGGATAAGGATTGGCTCTA</td><td>>hg38_21_+_8433172_8446622_RNA45SN1_45S_50ntFlanks</td><td>10841</td><td>10868</td><td>28</td><td>CGTAACTTCGGGATAAGGATTGGCTCTA</td><td>4728.5</td><td>-7.3</td><td>1.4</td><td>-5.4</td><td>8.0E-08</td><td>1.6E-07</td><td>MB231/MB468</td><td>MP</td></tr>
<tr><td>>MI0000077|hsa-mir-21&WithFlank&17|+|59841260|59841343@14.36.23(+1C)&[MIMAT0000076&hsa-miR-21-5p&offsets|0|+1(+1C);m-1&17|+|59841273|59841294&offsets|0|+1(+1C)]|TAGCTTATCAGACTGATGTTGACC</td><td>>MI0000077|hsa-mir-21&WithFlank&17|+|59841260|59841343</td><td>14</td><td>36</td><td>23(+1C)</td><td>TAGCTTATCAGACTGATGTTGACC</td><td>7961.1</td><td>4.1</td><td>0.8</td><td>5.4</td><td>8.0E-08</td><td>1.6E-07</td><td>MB231/MB468</td><td>MP</td></tr>
<tr><td>>hg38_21_+_8433172_8446622_RNA45SN1_45S_50ntFlanks@4753.4783.31|AAGACGATCAGATACCGTCGTAGTTCCGACC</td><td>>hg38_21_+_8433172_8446622_RNA45SN1_45S_50ntFlanks</td><td>4753</td><td>4783</td><td>31</td><td>AAGACGATCAGATACCGTCGTAGTTCCGACC</td><td>1677.0</td><td>-6.8</td><td>1.3</td><td>-5.3</td><td>9.4E-08</td><td>1.6E-07</td><td>MB231/MB468</td><td>MITO</td></tr>
<tr><td>>hg38_21_+_8433172_8446622_RNA45SN1_45S_50ntFlanks@3874.3899.26|AATACATGCCGACGGGCGCTGACCCC</td><td>>hg38_21_+_8433172_8446622_RNA45SN1_45S_50ntFlanks</td><td>3874</td><td>3899</td><td>26</td><td>AATACATGCCGACGGGCGCTGACCCC</td><td>451.6</td><td>-3.1</td><td>0.6</td><td>-5.4</td><td>6.7E-08</td><td>1.6E-07</td><td>BT20/MB468</td><td>MITO</td></tr>
<tr><td>>am_trna116_GluCTC_1_-_145399233_145399304@1.28.28,trna59_GluCTC_1_+_249168447_249168518@1.28.28,trna71_GluCTC_1_-_161439189_161439260@1.28.28,trna74_GluCTC_1_-_161431809_161431880@1.28.28,trna77_GluCTC_1_-_161424398_161424469@1.28.28,trna77_GluCTC_6_+_28949976_28950047@1.28.28,trna80_GluCTC_1_-_161417018_161417089@1.28.28,trna87_GluCTC_6_-_126101393_126101464@1.28.28|TCCCTGGTGGTCTAGTGGTTAGGATTCG</td><td>>am_trna116_GluCTC_1_-_145399233_145399304</td><td>1</td><td>28</td><td>28</td><td>TCCCTGGTGGTCTAGTGGTTAGGATTCG</td><td>20356.8</td><td>-2.7</td><td>0.5</td><td>-5.4</td><td>7.6E-08</td><td>1.6E-07</td><td>BT20/MB468</td><td>CYTO</td></tr>
<tr><td>>MI0000465|hsa-mir-191&WithFlank&3|-|49020612|49020715@22.42.21(+1A)&[MIMAT0000440&hsa-miR-191-5p&offsets|0|-2(+1A);m-48&3|-|49020672|49020694&offsets|0|-2(+1A)]|CAACGGAATCCCAAAAGCAGCA</td><td>>MI0000465|hsa-mir-191&WithFlank&3|-|49020612|49020715</td><td>22</td><td>42</td><td>21(+1A)</td><td>CAACGGAATCCCAAAAGCAGCA</td><td>285.8</td><td>2.5</td><td>0.5</td><td>5.3</td><td>9.9E-08</td><td>1.6E-07</td><td>BT20/MB468</td><td>NUC</td></tr>
<tr><td>>hg38_21_+_8433172_8446622_RNA45SN1_45S_50ntFlanks@4588.4611.24|CTATTTTGTTGGTTTTCGGAACTG</td><td>>hg38_21_+_8433172_8446622_RNA45SN1_45S_50ntFlanks</td><td>4588</td><td>4611</td><td>24</td><td>CTATTTTGTTGGTTTTCGGAACTG</td><td>248.3</td><td>-4.8</td><td>0.9</td><td>-5.4</td><td>8.7E-08</td><td>1.6E-07</td><td>MB231/MB468</td><td>CYTO</td></tr>
<tr><td>>hg38_21_+_8433172_8446622_RNA45SN1_45S_50ntFlanks@12674.12703.30|TCAGAATCCCGCCCAGGCGGAACGATACGG</td><td>>hg38_21_+_8433172_8446622_RNA45SN1_45S_50ntFlanks</td><td>12674</td><td>12703</td><td>30</td><td>TCAGAATCCCGCCCAGGCGGAACGATACGG</td><td>138.6</td><td>-10.2</td><td>1.8</td><td>-5.6</td><td>1.7E-08</td><td>1.6E-07</td><td>BT20/MB468</td><td>MP</td></tr>
<tr><td>>hg38_21_+_8433172_8446622_RNA45SN1_45S_50ntFlanks@6669.6689.21|CACTCGGCTCGTGCGTCGATG</td><td>>hg38_21_+_8433172_8446622_RNA45SN1_45S_50ntFlanks</td><td>6669</td><td>6689</td><td>21</td><td>CACTCGGCTCGTGCGTCGATG</td><td>163.5</td><td>-10.4</td><td>1.8</td><td>-5.6</td><td>1.7E-08</td><td>1.6E-07</td><td>BT20/MB468</td><td>MP</td></tr>
<tr><td>>hg38_21_+_8433172_8446622_RNA45SN1_45S_50ntFlanks@8191.8213.23|CAGCCCGTGGACGGTGTGAGGCC</td><td>>hg38_21_+_8433172_8446622_RNA45SN1_45S_50ntFlanks</td><td>8191</td><td>8213</td><td>23</td><td>CAGCCCGTGGACGGTGTGAGGCC</td><td>1023.5</td><td>-6.0</td><td>1.1</td><td>-5.3</td><td>9.5E-08</td><td>1.6E-07</td><td>MB231/MB468</td><td>MITO</td></tr>
<tr><td>>hg38_21_+_8433172_8446622_RNA45SN1_45S_50ntFlanks@7979.8012.34|ACCTCAGATCAGACGTGGCGACCCGCTGAATTTA</td><td>>hg38_21_+_8433172_8446622_RNA45SN1_45S_50ntFlanks</td><td>7979</td><td>8012</td><td>34</td><td>ACCTCAGATCAGACGTGGCGACCCGCTGAATTTA</td><td>849.3</td><td>-6.7</td><td>1.3</td><td>-5.3</td><td>9.9E-08</td><td>1.6E-07</td><td>BT20/MB468</td><td>NUC</td></tr>
<tr><td>>hg38_21_+_8433172_8446622_RNA45SN1_45S_50ntFlanks@8179.8205.27|CTGATCGAGGCCCAGCCCGTGGACGGT</td><td>>hg38_21_+_8433172_8446622_RNA45SN1_45S_50ntFlanks</td><td>8179</td><td>8205</td><td>27</td><td>CTGATCGAGGCCCAGCCCGTGGACGGT</td><td>199.2</td><td>-7.4</td><td>1.4</td><td>-5.4</td><td>8.1E-08</td><td>1.6E-07</td><td>MB231/MB468</td><td>MP</td></tr>
<tr><td>>hg38_21_+_8433172_8446622_RNA45SN1_45S_50ntFlanks@11581.11602.22|AACTGGTGCGGACCAGGGGAAT</td><td>>hg38_21_+_8433172_8446622_RNA45SN1_45S_50ntFlanks</td><td>11581</td><td>11602</td><td>22</td><td>AACTGGTGCGGACCAGGGGAAT</td><td>1705.5</td><td>-4.3</td><td>0.8</td><td>-5.4</td><td>8.7E-08</td><td>1.6E-07</td><td>MB231/MB468</td><td>CYTO</td></tr>
<tr><td>>MI0001733|hsa-mir-452&WithFlank&X|-|151959622|151959718@20.43.24&[MIMAT0001635&hsa-miR-452-5p&offsets|0|+2;m-188&X|-|151959677|151959699&offsets|0|+1]|AACTGTTTGCAGAGGAAACTGAGA</td><td>>MI0001733|hsa-mir-452&WithFlank&X|-|151959622|151959718</td><td>20</td><td>43</td><td>24</td><td>AACTGTTTGCAGAGGAAACTGAGA</td><td>1144.8</td><td>3.0</td><td>0.5</td><td>5.7</td><td>9.4E-09</td><td>1.6E-07</td><td>BT20/MB231</td><td>NUC</td></tr>
<tr><td>>hg38_21_+_8433172_8446622_RNA45SN1_45S_50ntFlanks@10868.10902.35|AAGGGCTGGGTCGGTCGGGCTGGGGCGCGAAGCGG</td><td>>hg38_21_+_8433172_8446622_RNA45SN1_45S_50ntFlanks</td><td>10868</td><td>10902</td><td>35</td><td>AAGGGCTGGGTCGGTCGGGCTGGGGCGCGAAGCGG</td><td>185.2</td><td>-7.1</td><td>1.3</td><td>-5.4</td><td>8.1E-08</td><td>1.6E-07</td><td>MB231/MB468</td><td>MP</td></tr>
<tr><td>>hg38_21_+_8433172_8446622_RNA45SN1_45S_50ntFlanks@3835.3872.38|CTCCTCTCCTACTTGGATAACTGTGGTAATTCTAGAGC</td><td>>hg38_21_+_8433172_8446622_RNA45SN1_45S_50ntFlanks</td><td>3835</td><td>3872</td><td>38</td><td>CTCCTCTCCTACTTGGATAACTGTGGTAATTCTAGAGC</td><td>445.6</td><td>-5.7</td><td>1.1</td><td>-5.3</td><td>9.5E-08</td><td>1.6E-07</td><td>MB231/MB468</td><td>MITO</td></tr>
<tr><td>>hg38_21_+_8433172_8446622_RNA45SN1_45S_50ntFlanks@10664.10712.49|CTTGGAAAGCGTCGCGGTTCCGGCGGCGTCCGGTGAGCTCTCGCTGGCC</td><td>>hg38_21_+_8433172_8446622_RNA45SN1_45S_50ntFlanks</td><td>10664</td><td>10712</td><td>49</td><td>CTTGGAAAGCGTCGCGGTTCCGGCGGCGTCCGGTGAGCTCTCGCTGGCC</td><td>132.9</td><td>-9.4</td><td>1.7</td><td>-5.4</td><td>7.6E-08</td><td>1.6E-07</td><td>BT20/MB468</td><td>CYTO</td></tr>
</table>

<table>
<tr><th></th><th></th><th></th><th></th><th></th><th></th><th></th><th></th><th></th><th></th><th></th><th></th><th></th><th></th><th></th></tr>
<tr><td>>hg38_21_+_8433172_8446622_RNA45SN1_45S_50ntFlanks@10843.10863.21|TAACTTCGGGATAAGGATTGG</td><td>>hg38_21_+_8433172_8446622_RNA45SN1_45S_50ntFlanks</td><td>10843</td><td>10863</td><td>21</td><td>TAACTTCGGGATAAGGATTGG</td><td>609.7</td><td>-7.0</td><td>1.3</td><td>-5.4</td><td>7.6E-08</td><td>1.6E-07</td><td>BT20/MB468</td><td>CYTO</td></tr>
<tr><td>>hg38_21_+_8433172_8446622_RNA45SN1_45S_50ntFlanks@10819.10839.21|AGGGAAGTCGGCAAGCCGGAT</td><td>>hg38_21_+_8433172_8446622_RNA45SN1_45S_50ntFlanks</td><td>10819</td><td>10839</td><td>21</td><td>AGGGAAGTCGGCAAGCCGGAT</td><td>191.6</td><td>-3.9</td><td>0.7</td><td>-5.4</td><td>8.8E-08</td><td>1.6E-07</td><td>MB231/MB468</td><td>CYTO</td></tr>
<tr><td>>hg38_21_+_8433172_8446622_RNA45SN1_45S_50ntFlanks@7979.7999.21|ACCTCAGATCAGACGTGGCGA</td><td>>hg38_21_+_8433172_8446622_RNA45SN1_45S_50ntFlanks</td><td>7979</td><td>7999</td><td>21</td><td>ACCTCAGATCAGACGTGGCGA</td><td>205.7</td><td>-7.0</td><td>1.3</td><td>-5.4</td><td>8.1E-08</td><td>1.6E-07</td><td>MB231/MB468</td><td>MP</td></tr>
<tr><td>>hg38_21_+_8433172_8446622_RNA45SN1_45S_50ntFlanks@8117.8137.21|CGGGACATGTGGCGTACGGAA</td><td>>hg38_21_+_8433172_8446622_RNA45SN1_45S_50ntFlanks</td><td>8117</td><td>8137</td><td>21</td><td>CGGGACATGTGGCGTACGGAA</td><td>691.4</td><td>12.8</td><td>2.2</td><td>5.9</td><td>3.4E-09</td><td>1.6E-07</td><td>BT20/MB231</td><td>MP</td></tr>
<tr><td>>hg38_21_+_8433172_8446622_RNA45SN1_45S_50ntFlanks@9792.9816.25|GGCGTGGAGCCGGGCGTGGAATGCG</td><td>>hg38_21_+_8433172_8446622_RNA45SN1_45S_50ntFlanks</td><td>9792</td><td>9816</td><td>25</td><td>GGCGTGGAGCCGGGCGTGGAATGCG</td><td>356.3</td><td>11.8</td><td>2.0</td><td>5.9</td><td>3.4E-09</td><td>1.6E-07</td><td>BT20/MB231</td><td>MP</td></tr>
<tr><td>>hg38_21_+_8433172_8446622_RNA45SN1_45S_50ntFlanks@10788.10817.30|CAGCCTCTGGCATGTTGGAACAATGTAGGT</td><td>>hg38_21_+_8433172_8446622_RNA45SN1_45S_50ntFlanks</td><td>10788</td><td>10817</td><td>30</td><td>CAGCCTCTGGCATGTTGGAACAATGTAGGT</td><td>372.3</td><td>-5.7</td><td>1.1</td><td>-5.3</td><td>9.6E-08</td><td>1.6E-07</td><td>MB231/MB468</td><td>MITO</td></tr>
<tr><td>>MI0000478|hsa-mir-149&WithFlank&2|+|240455995|240456095@21.42.22(+1A)&[MIMAT0000450&hsa-miR-149-5p&offsets|0|-1(+1A);m-195&2|+|240456015|240456037&offsets|0|-1(+1A)]|TCTGGCTCCGTGTCTTCACTCCA</td><td>>MI0000478|hsa-mir-149&WithFlank&2|+|240455995|240456095</td><td>21</td><td>42</td><td>22(+1A)</td><td>TCTGGCTCCGTGTCTTCACTCCA</td><td>778.0</td><td>5.0</td><td>0.9</td><td>5.8</td><td>5.6E-09</td><td>1.6E-07</td><td>BT20/MB231</td><td>CYTO</td></tr>
<tr><td>>MI0000459|hsa-mir-143&WithFlank&5|+|149428912|149429029@67.86.20(+1U)&[MIMAT0000435&hsa-miR-143-3p&offsets|0|-1(+1U);m-3&5|+|149428978|149428997&offsets|0|0(+1U)]|TGAGATGAAGCACTGTAGCTT</td><td>>MI0000459|hsa-mir-143&WithFlank&5|+|149428912|149429029</td><td>67</td><td>86</td><td>20(+1U)</td><td>TGAGATGAAGCACTGTAGCTT</td><td>81.0</td><td>-9.4</td><td>1.7</td><td>-5.6</td><td>1.7E-08</td><td>1.7E-07</td><td>BT20/MB468</td><td>MP</td></tr>
<tr><td>>hg38_21_+_8433172_8446622_RNA45SN1_45S_50ntFlanks@11571.11601.31|GCCGACTTAGAACTGGTGCGGACCAGGGGAA</td><td>>hg38_21_+_8433172_8446622_RNA45SN1_45S_50ntFlanks</td><td>11571</td><td>11601</td><td>31</td><td>GCCGACTTAGAACTGGTGCGGACCAGGGGAA</td><td>2620.7</td><td>-5.8</td><td>1.1</td><td>-5.3</td><td>1.0E-07</td><td>1.7E-07</td><td>BT20/MB468</td><td>NUC</td></tr>
<tr><td>>hg38_21_+_8433172_8446622_RNA45SN1_45S_50ntFlanks@8170.8195.26|CAAGTCCTTCTGATCGAGGCCCAGCC</td><td>>hg38_21_+_8433172_8446622_RNA45SN1_45S_50ntFlanks</td><td>8170</td><td>8195</td><td>26</td><td>CAAGTCCTTCTGATCGAGGCCCAGCC</td><td>871.5</td><td>-5.9</td><td>1.1</td><td>-5.4</td><td>6.8E-08</td><td>1.7E-07</td><td>BT20/MB468</td><td>MITO</td></tr>
<tr><td>>hg38_21_+_8433172_8446622_RNA45SN1_45S_50ntFlanks@9564.9593.30|CTGACGTGCAAATCGGTCGTCCGACCTGGG</td><td>>hg38_21_+_8433172_8446622_RNA45SN1_45S_50ntFlanks</td><td>9564</td><td>9593</td><td>30</td><td>CTGACGTGCAAATCGGTCGTCCGACCTGGG</td><td>167.7</td><td>-2.6</td><td>0.5</td><td>-5.3</td><td>9.1E-08</td><td>1.7E-07</td><td>MB231/MB468</td><td>NUC</td></tr>
<tr><td>>hg38_21_+_8433172_8446622_RNA45SN1_45S_50ntFlanks@12073.12095.23|CCGAGGGGCTCTCGCTTCTGGCG</td><td>>hg38_21_+_8433172_8446622_RNA45SN1_45S_50ntFlanks</td><td>12073</td><td>12095</td><td>23</td><td>CCGAGGGGCTCTCGCTTCTGGCG</td><td>4901.9</td><td>-6.5</td><td>1.2</td><td>-5.4</td><td>7.7E-08</td><td>1.7E-07</td><td>BT20/MB468</td><td>CYTO</td></tr>
<tr><td>>hg38_MT_+_1621_3279_RNR2_16S_50ntFlanks@114.146.33|CAAATAAAGTATAGGCGATAGAAATTGAAACCT</td><td>>hg38_MT_+_1621_3279_RNR2_16S_50ntFlanks</td><td>114</td><td>146</td><td>33</td><td>CAAATAAAGTATAGGCGATAGAAATTGAAACCT</td><td>78.3</td><td>9.5</td><td>1.8</td><td>5.4</td><td>8.2E-08</td><td>1.7E-07</td><td>MB231/MB468</td><td>MP</td></tr>
<tr><td>>hg38_21_+_8433172_8446622_RNA45SN1_45S_50ntFlanks@6660.6683.24|GCGGTGGATCACTCGGCTCGTGCG</td><td>>hg38_21_+_8433172_8446622_RNA45SN1_45S_50ntFlanks</td><td>6660</td><td>6683</td><td>24</td><td>GCGGTGGATCACTCGGCTCGTGCG</td><td>1192.9</td><td>4.3</td><td>0.8</td><td>5.3</td><td>9.1E-08</td><td>1.7E-07</td><td>MB231/MB468</td><td>NUC</td></tr>
<tr><td>>hg38_21_+_8433172_8446622_RNA45SN1_45S_50ntFlanks@12416.12456.41|TTCGATGTCGGCTCTTCCTATCATTGTGAAGCAGAATTCAC</td><td>>hg38_21_+_8433172_8446622_RNA45SN1_45S_50ntFlanks</td><td>12416</td><td>12456</td><td>41</td><td>TTCGATGTCGGCTCTTCCTATCATTGTGAAGCAGAATTCAC</td><td>391.3</td><td>4.3</td><td>0.8</td><td>5.3</td><td>8.8E-08</td><td>1.7E-07</td><td>MB231/MB468</td><td>CYTO</td></tr>
<tr><td>>MI0000066|hsa-let-7e&WithFlank&19|+|51692780|51692870@14.34.21&[MIMAT0000066&hsa-let-7e-5p&offsets|0|-1;m-50&19|+|51692793|51692814&offsets|0|-1]|TGAGGTAGGAGGTTGTATAGT</td><td>>MI0000066|hsa-let-7e&WithFlank&19|+|51692780|51692870</td><td>14</td><td>34</td><td>21</td><td>TGAGGTAGGAGGTTGTATAGT</td><td>753.8</td><td>4.8</td><td>0.9</td><td>5.3</td><td>9.6E-08</td><td>1.7E-07</td><td>MB231/MB468</td><td>MITO</td></tr>
<tr><td>>MI0000061|hsa-let-7a-2&WithFlank&11|-|122146516|122146599@11.28.18;MI0000064|hsa-let-7c&WithFlank&21|+|16539822|16539917@17.34.18;MI0000062|hsa-let-7a-3&WithFlank&22|+|46112743|46112828@10.27.18;MI0000060|hsa-let-7a-1&WithFlank&9|+|94175951|94176042@12.29.18&[MIMAT0000062&hsa-let-7a-5p&offsets|0|-4;m-6&11|-|122146568|122146589&offsets|0|-4];[MIMAT0000064&hsa-let-7c-5p&offsets|0|-4;m-37&21|+|16539838|16539859&offsets|0|-4];[MIMAT0000062_1&hsa-let-7a-5p&offsets|0|-4;m-5&22|+|46112752|46112773&offsets|0|-4];[MIMAT0000062_2&hsa-let-7a-5p&offsets|0|-4;m-4&9|+|94175962|94175983&offsets|0|-4]|TGAGGTAGTAGGTTGTAT</td><td>>MI0000061|hsa-let-7a-2&WithFlank&11|-|122146516|122146599</td><td>11</td><td>28</td><td>18</td><td>TGAGGTAGTAGGTTGTAT</td><td>788.4</td><td>4.6</td><td>0.8</td><td>5.6</td><td>1.7E-08</td><td>1.7E-07</td><td>BT20/MB468</td><td>MP</td></tr>
<tr><td>>hg38_21_+_8433172_8446622_RNA45SN1_45S_50ntFlanks@11671.11688.18|CCAGTGCTCTGAATGTCA</td><td>>hg38_21_+_8433172_8446622_RNA45SN1_45S_50ntFlanks</td><td>11671</td><td>11688</td><td>18</td><td>CCAGTGCTCTGAATGTCA</td><td>883.3</td><td>-8.8</td><td>1.6</td><td>-5.3</td><td>1.0E-07</td><td>1.7E-07</td><td>BT20/MB468</td><td>NUC</td></tr>
<tr><td>>hg38_21_+_8433172_8446622_RNA45SN1_45S_50ntFlanks@9821.9850.30|CCTAGTGGGCCACTTTTGGTAAGCAGAACT</td><td>>hg38_21_+_8433172_8446622_RNA45SN1_45S_50ntFlanks</td><td>9821</td><td>9850</td><td>30</td><td>CCTAGTGGGCCACTTTTGGTAAGCAGAACT</td><td>1248.2</td><td>-8.8</td><td>1.7</td><td>-5.3</td><td>1.0E-07</td><td>1.7E-07</td><td>BT20/MB468</td><td>NUC</td></tr>
<tr><td>>hg38_21_+_8433172_8446622_RNA45SN1_45S_50ntFlanks@7977.8019.43|CGACCTCAGATCAGACGTGGCGACCCGCTGAATTTAAGCATAT</td><td>>hg38_21_+_8433172_8446622_RNA45SN1_45S_50ntFlanks</td><td>7977</td><td>8019</td><td>43</td><td>CGACCTCAGATCAGACGTGGCGACCCGCTGAATTTAAGCATAT</td><td>737.2</td><td>-6.9</td><td>1.2</td><td>-5.8</td><td>5.7E-09</td><td>1.7E-07</td><td>BT20/MB231</td><td>CYTO</td></tr>
<tr><td>>MI0000650|hsa-mir-200c&WithFlank&12|+|6963693|6963772@50.69.20&[MIMAT0000617&hsa-miR-200c-3p&offsets|0|-3;m-54&12|+|6963742|6963764&offsets|0|-3]|TAATACTGCCGGGTAATGAT</td><td>>MI0000650|hsa-mir-200c&WithFlank&12|+|6963693|6963772</td><td>50</td><td>69</td><td>20</td><td>TAATACTGCCGGGTAATGAT</td><td>605.9</td><td>3.9</td><td>0.7</td><td>5.3</td><td>1.0E-07</td><td>1.7E-07</td><td>BT20/MB468</td><td>NUC</td></tr>
<tr><td>>hg38_21_+_8433172_8446622_RNA45SN1_45S_50ntFlanks@11999.12030.32|CCCTGCGGGCCGCCGGTGAAATACCACTACTC</td><td>>hg38_21_+_8433172_8446622_RNA45SN1_45S_50ntFlanks</td><td>11999</td><td>12030</td><td>32</td><td>CCCTGCGGGCCGCCGGTGAAATACCACTACTC</td><td>283.8</td><td>-6.4</td><td>1.2</td><td>-5.4</td><td>6.9E-08</td><td>1.7E-07</td><td>BT20/MB468</td><td>MITO</td></tr>
<tr><td>>hg38_21_+_8433172_8446622_RNA45SN1_45S_50ntFlanks@11212.11232.21|CGACTCTGGACGCGAGCCGGG</td><td>>hg38_21_+_8433172_8446622_RNA45SN1_45S_50ntFlanks</td><td>11212</td><td>11232</td><td>21</td><td>CGACTCTGGACGCGAGCCGGG</td><td>452.0</td><td>-7.1</td><td>1.3</td><td>-5.3</td><td>9.2E-08</td><td>1.7E-07</td><td>MB231/MB468</td><td>NUC</td></tr>
<tr><td>>hg38_21_+_8433172_8446622_RNA45SN1_45S_50ntFlanks@10794.10817.24|CTGGCATGTTGGAACAATGTAGGT</td><td>>hg38_21_+_8433172_8446622_RNA45SN1_45S_50ntFlanks</td><td>10794</td><td>10817</td><td>24</td><td>CTGGCATGTTGGAACAATGTAGGT</td><td>284.1</td><td>-4.7</td><td>0.9</td><td>-5.3</td><td>8.9E-08</td><td>1.7E-07</td><td>MB231/MB468</td><td>CYTO</td></tr>
<tr><td>>hg38_21_+_8433172_8446622_RNA45SN1_45S_50ntFlanks@12259.12288.30|CTTGATCTTGATTTTCAGTACGAATACAGA</td><td>>hg38_21_+_8433172_8446622_RNA45SN1_45S_50ntFlanks</td><td>12259</td><td>12288</td><td>30</td><td>CTTGATCTTGATTTTCAGTACGAATACAGA</td><td>671.3</td><td>-8.3</td><td>1.6</td><td>-5.3</td><td>9.7E-08</td><td>1.7E-07</td><td>MB231/MB468</td><td>MITO</td></tr>
<tr><td>>hg38_21_+_8433172_8446622_RNA45SN1_45S_50ntFlanks@7976.8009.34|GCGACCTCAGATCAGACGTGGCGACCCGCTGAAT</td><td>>hg38_21_+_8433172_8446622_RNA45SN1_45S_50ntFlanks</td><td>7976</td><td>8009</td><td>34</td><td>GCGACCTCAGATCAGACGTGGCGACCCGCTGAAT</td><td>784.5</td><td>5.1</td><td>1.0</td><td>5.3</td><td>9.2E-08</td><td>1.7E-07</td><td>MB231/MB468</td><td>NUC</td></tr>
<tr><td>>hg38_1_-_228634819_228635039_RNA5S12_5S_50ntFlanks@150.169.20|AATACCGGGTGCTGTAGGCT</td><td>>hg38_1_-_228634819_228635039_RNA5S12_5S_50ntFlanks</td><td>150</td><td>169</td><td>20</td><td>AATACCGGGTGCTGTAGGCT</td><td>931.5</td><td>-4.2</td><td>0.8</td><td>-5.3</td><td>9.7E-08</td><td>1.7E-07</td><td>MB231/MB468</td><td>MITO</td></tr>
<tr><td>>hg38_21_+_8433172_8446622_RNA45SN1_45S_50ntFlanks@10318.10336.19|TGCAGATCTTGGTGGTAGT</td><td>>hg38_21_+_8433172_8446622_RNA45SN1_45S_50ntFlanks</td><td>10318</td><td>10336</td><td>19</td><td>TGCAGATCTTGGTGGTAGT</td><td>85.4</td><td>10.2</td><td>1.8</td><td>5.6</td><td>1.7E-08</td><td>1.7E-07</td><td>BT20/MB468</td><td>MP</td></tr>
<tr><td>>hg38_21_+_8433172_8446622_RNA45SN1_45S_50ntFlanks@11600.11629.30|AATCCGACTGTTTAATTAAAACAAAGCATC</td><td>>hg38_21_+_8433172_8446622_RNA45SN1_45S_50ntFlanks</td><td>11600</td><td>11629</td><td>30</td><td>AATCCGACTGTTTAATTAAAACAAAGCATC</td><td>79.0</td><td>10.1</td><td>1.8</td><td>5.6</td><td>1.7E-08</td><td>1.7E-07</td><td>BT20/MB468</td><td>MP</td></tr>
<tr><td>>hg38_21_+_8433172_8446622_RNA45SN1_45S_50ntFlanks@11778.11803.26|ACGCGCATGAATGGATGAACGAGATT</td><td>>hg38_21_+_8433172_8446622_RNA45SN1_45S_50ntFlanks</td><td>11778</td><td>11803</td><td>26</td><td>ACGCGCATGAATGGATGAACGAGATT</td><td>300.9</td><td>-3.6</td><td>0.7</td><td>-5.4</td><td>7.8E-08</td><td>1.7E-07</td><td>BT20/MB468</td><td>CYTO</td></tr>
<tr><td>>MI0000734|hsa-mir-106b&WithFlank&7|-|100093987|100094080@18.39.22&[MIMAT0000680&hsa-miR-106b-5p&offsets|0|+1;m-103&7|-|100094043|100094063&offsets|0|+1]|TAAAGTGCTGACAGTGCAGATA</td><td>>MI0000734|hsa-mir-106b&WithFlank&7|-|100093987|100094080</td><td>18</td><td>39</td><td>22</td><td>TAAAGTGCTGACAGTGCAGATA</td><td>681.7</td><td>3.6</td><td>0.7</td><td>5.3</td><td>1.0E-07</td><td>1.7E-07</td><td>BT20/MB468</td><td>NUC</td></tr>
<tr><td>>hg38_21_+_8433172_8446622_RNA45SN1_45S_50ntFlanks@9580.9597.18|TCGTCCGACCTGGGTATA</td><td>>hg38_21_+_8433172_8446622_RNA45SN1_45S_50ntFlanks</td><td>9580</td><td>9597</td><td>18</td><td>TCGTCCGACCTGGGTATA</td><td>91.0</td><td>-9.6</td><td>1.7</td><td>-5.6</td><td>1.7E-08</td><td>1.7E-07</td><td>BT20/MB468</td><td>MP</td></tr>
<tr><td>>trna2_GlyGCC_21_-_18827107_18827177@1.30.30,trna35_GlyGCC_1_+_161413094_161413164@1.30.30,trna37_GlyGCC_1_+_161420467_161420537@1.30.30,trna39_GlyGCC_1_+_161427898_161427968@1.30.30,trna41_GlyGCC_1_+_161435258_161435328@1.30.30|GCATGGGTGGTTCAGTGGTAGAATTCTCGC</td><td>>trna2_GlyGCC_21_-_18827107_18827177</td><td>1</td><td>30</td><td>30</td><td>GCATGGGTGGTTCAGTGGTAGAATTCTCGC</td><td>16635.1</td><td>-5.0</td><td>0.9</td><td>-5.3</td><td>9.0E-08</td><td>1.7E-07</td><td>MB231/MB468</td><td>CYTO</td></tr>
<tr><td>>hg38_21_+_8433172_8446622_RNA45SN1_45S_50ntFlanks@11569.11605.37|CAGCCGACTTAGAACTGGTGCGGACCAGGGGAATCCG</td><td>>hg38_21_+_8433172_8446622_RNA45SN1_45S_50ntFlanks</td><td>11569</td><td>11605</td><td>37</td><td>CAGCCGACTTAGAACTGGTGCGGACCAGGGGAATCCG</td><td>1626.5</td><td>-4.2</td><td>0.8</td><td>-5.4</td><td>7.0E-08</td><td>1.7E-07</td><td>BT20/MB468</td><td>MITO</td></tr>
<tr><td>>hg38_21_+_8433172_8446622_RNA45SN1_45S_50ntFlanks@12233.12258.26|CCGTGGAGCAGAAGGGCAAAAGCTCG</td><td>>hg38_21_+_8433172_8446622_RNA45SN1_45S_50ntFlanks</td><td>12233</td><td>12258</td><td>26</td><td>CCGTGGAGCAGAAGGGCAAAAGCTCG</td><td>124.4</td><td>-9.0</td><td>1.7</td><td>-5.4</td><td>7.0E-08</td><td>1.7E-07</td><td>BT20/MB468</td><td>MITO</td></tr>
<tr><td>>MI0000465|hsa-mir-191&WithFlank&3|-|49020612|49020715@22.42.21(+1A)&[MIMAT0000440&hsa-miR-191-5p&offsets|0|-2(+1A);m-48&3|-|49020672|49020694&offsets|0|-2(+1A)]|CAACGGAATCCCAAAAGCAGCA</td><td>>MI0000465|hsa-mir-191&WithFlank&3|-|49020612|49020715</td><td>22</td><td>42</td><td>21(+1A)</td><td>CAACGGAATCCCAAAAGCAGCA</td><td>529.4</td><td>5.3</td><td>1.0</td><td>5.3</td><td>9.8E-08</td><td>1.7E-07</td><td>MB231/MB468</td><td>MITO</td></tr>
<tr><td>>hg38_MT_+_598_1651_RNR1_12S_50ntFlanks@416.455.40|AAATAGACTACGAAAGTGGCTTTAACATATCTGAACACAC</td><td>>hg38_MT_+_598_1651_RNR1_12S_50ntFlanks</td><td>416</td><td>455</td><td>40</td><td>AAATAGACTACGAAAGTGGCTTTAACATATCTGAACACAC</td><td>59.4</td><td>9.1</td><td>1.7</td><td>5.4</td><td>8.3E-08</td><td>1.7E-07</td><td>MB231/MB468</td><td>MP</td></tr>
<tr><td>>hg38_21_+_8433172_8446622_RNA45SN1_45S_50ntFlanks@9482.9509.28|ACGTGTTAGGACCCGAAAGATGGTGAAC</td><td>>hg38_21_+_8433172_8446622_RNA45SN1_45S_50ntFlanks</td><td>9482</td><td>9509</td><td>28</td><td>ACGTGTTAGGACCCGAAAGATGGTGAAC</td><td>107.5</td><td>10.5</td><td>1.9</td><td>5.6</td><td>1.7E-08</td><td>1.7E-07</td><td>BT20/MB468</td><td>MP</td></tr>
<tr><td>>hg38_21_+_8433172_8446622_RNA45SN1_45S_50ntFlanks@5067.5097.31|CTGGTTAATTCCGATAACGAACGAGACTCTG</td><td>>hg38_21_+_8433172_8446622_RNA45SN1_45S_50ntFlanks</td><td>5067</td><td>5097</td><td>31</td><td>CTGGTTAATTCCGATAACGAACGAGACTCTG</td><td>75.0</td><td>-10.1</td><td>1.9</td><td>-5.4</td><td>8.4E-08</td><td>1.7E-07</td><td>MB231/MB468</td><td>MP</td></tr>
<tr><td>>hg38_21_+_8433172_8446622_RNA45SN1_45S_50ntFlanks@12914.12933.20|CGTTCGTGGGGAACCTGGCG</td><td>>hg38_21_+_8433172_8446622_RNA45SN1_45S_50ntFlanks</td><td>12914</td><td>12933</td><td>20</td><td>CGTTCGTGGGGAACCTGGCG</td><td>332.3</td><td>-6.3</td><td>1.2</td><td>-5.3</td><td>1.0E-07</td><td>1.7E-07</td><td>BT20/MB468</td><td>NUC</td></tr>
<tr><td>>hg38_21_+_8433172_8446622_RNA45SN1_45S_50ntFlanks@11581.11606.26|AACTGGTGCGGACCAGGGGAATCCGA</td><td>>hg38_21_+_8433172_8446622_RNA45SN1_45S_50ntFlanks</td><td>11581</td><td>11606</td><td>26</td><td>AACTGGTGCGGACCAGGGGAATCCGA</td><td>302.5</td><td>-4.8</td><td>0.9</td><td>-5.3</td><td>9.0E-08</td><td>1.7E-07</td><td>MB231/MB468</td><td>CYTO</td></tr>
<tr><td>>hg38_21_+_8433172_8446622_RNA45SN1_45S_50ntFlanks@6778.6802.25|CCCGGGGCTACGCCTGTCTGAGCGT</td><td>>hg38_21_+_8433172_8446622_RNA45SN1_45S_50ntFlanks</td><td>6778</td><td>6802</td><td>25</td><td>CCCGGGGCTACGCCTGTCTGAGCGT</td><td>967.7</td><td>4.8</td><td>0.9</td><td>5.4</td><td>7.9E-08</td><td>1.7E-07</td><td>BT20/MB468</td><td>CYTO</td></tr>
<tr><td>>hg38_21_+_8433172_8446622_RNA45SN1_45S_50ntFlanks@9574.9597.24|AATCGGTCGTCCGACCTGGGTATA</td><td>>hg38_21_+_8433172_8446622_RNA45SN1_45S_50ntFlanks</td><td>9574</td><td>9597</td><td>24</td><td>AATCGGTCGTCCGACCTGGGTATA</td><td>10743.1</td><td>-8.2</td><td>1.5</td><td>-5.6</td><td>1.7E-08</td><td>1.7E-07</td><td>BT20/MB468</td><td>MP</td></tr>
<tr><td>>hg38_21_+_8433172_8446622_RNA45SN1_45S_50ntFlanks@12405.12446.42|CTTTTTGATCCTTCGATGTCGGCTCTTCCTATCATTGTGAAG</td><td>>hg38_21_+_8433172_8446622_RNA45SN1_45S_50ntFlanks</td><td>12405</td><td>12446</td><td>42</td><td>CTTTTTGATCCTTCGATGTCGGCTCTTCCTATCATTGTGAAG</td><td>1285.5</td><td>-7.6</td><td>1.4</td><td>-5.4</td><td>7.9E-08</td><td>1.7E-07</td><td>BT20/MB468</td><td>CYTO</td></tr>
<tr><td>>hg38_21_+_8433172_8446622_RNA45SN1_45S_50ntFlanks@9545.9569.25|GTGGAGGTCCGTAGCGGTCCTGACG</td><td>>hg38_21_+_8433172_8446622_RNA45SN1_45S_50ntFlanks</td><td>9545</td><td>9569</td><td>25</td><td>GTGGAGGTCCGTAGCGGTCCTGACG</td><td>1024.0</td><td>-3.8</td><td>0.7</td><td>-5.4</td><td>7.9E-08</td><td>1.7E-07</td><td>BT20/MB468</td><td>CYTO</td></tr>
</table>

<table>
<tr><td>>hg38_21_+_8433172_8446622_RNA45SN1_45S_50ntFlanks@4367.4391.25|CAGTTAAAAAGCTCGTAGTTGGATC</td><td>>hg38_21_+_8433172_8446622_RNA45SN1_45S_50ntFlanks</td><td>4367</td><td>4391</td><td>25</td><td>CAGTTAAAAAGCTCGTAGTTGGATC</td><td>844.6</td><td>-6.5</td><td>1.1</td><td>-5.6</td><td>1.7E-08</td><td>1.7E-07</td><td>BT20/MB468</td><td>MP</td></tr>
<tr><td>>hg38_21_+_8433172_8446622_RNA45SN1_45S_50ntFlanks@11561.11584.24|GCGCCTAGCAGCCGACTTAGAACT</td><td>>hg38_21_+_8433172_8446622_RNA45SN1_45S_50ntFlanks</td><td>11561</td><td>11584</td><td>24</td><td>GCGCCTAGCAGCCGACTTAGAACT</td><td>490.2</td><td>-6.3</td><td>1.2</td><td>-5.3</td><td>1.0E-07</td><td>1.7E-07</td><td>BT20/MB468</td><td>NUC</td></tr>
<tr><td>>hg38_21_+_8433172_8446622_RNA45SN1_45S_50ntFlanks@7992.8015.24|CGTGGCGACCCGCTGAATTTAAGC</td><td>>hg38_21_+_8433172_8446622_RNA45SN1_45S_50ntFlanks</td><td>7992</td><td>8015</td><td>24</td><td>CGTGGCGACCCGCTGAATTTAAGC</td><td>205.4</td><td>6.1</td><td>1.1</td><td>5.3</td><td>9.1E-08</td><td>1.7E-07</td><td>MB231/MB468</td><td>CYTO</td></tr>
<tr><td>>hg38_21_+_8433172_8446622_RNA45SN1_45S_50ntFlanks@12686.12703.18|CCAGGCGGAACGATACGG</td><td>>hg38_21_+_8433172_8446622_RNA45SN1_45S_50ntFlanks</td><td>12686</td><td>12703</td><td>18</td><td>CCAGGCGGAACGATACGG</td><td>305.5</td><td>-6.0</td><td>1.1</td><td>-5.3</td><td>9.9E-08</td><td>1.7E-07</td><td>MB231/MB468</td><td>MITO</td></tr>
<tr><td>>MI0003776|hsa-mir-320b-1&WithFlank&1|+|116671740|116671823@49.66.18(+1A);MI0003839|hsa-mir-320b-2&WithFlank&1|-|224257034|224257116@48.65.18(+1A);MI0003778|hsa-mir-320c-1&WithFlank&18|+|21683512|21683595@49.66.18(+1A);MI0008191|hsa-mir-320c-2&WithFlank&18|+|24321669|24321752@49.66.18(+1A);MI0000542|hsa-mir-320a&WithFlank&8|-|22244960|22245043@49.66.18(+1A)&[MIMAT0005792&hsa-miR-320b&offsets|+1|-3(+1A);m-183&1|+|116671787|116671806&offsets|+1|-1(+1A)];[MIMAT0005792_1&hsa-miR-320b&offsets|+1|-3(+1A);m-183&1|-|224257051|224257070&offsets|+1|-1(+1A)];[MIMAT0005793&hsa-miR-320c&offsets|+1|-1(+1A);m-222&18|+|21683559|21683578&offsets|+1|-1(+1A)];[MIMAT0005793_1&hsa-miR-320c&offsets|+1|-1(+1A);m-219&18|+|24321716|24321735&offsets|+1|-1(+1A)];[MIMAT0000510&hsa-miR-320a-3p&offsets|+1|-3(+1A);m-60&8|-|22244975|22244996&offsets|+1|-3(+1A)]|AAAGCTGGGTTGAGAGGGA</td><td>>MI0003776|hsa-mir-320b-1&WithFlank&1|+|116671740|116671823</td><td>49</td><td>66</td><td>18(+1A)</td><td>AAAGCTGGGTTGAGAGGGA</td><td>109.5</td><td>10.5</td><td>1.9</td><td>5.6</td><td>1.7E-08</td><td>1.7E-07</td><td>BT20/MB468</td><td>MP</td></tr>
<tr><td>>hg38_21_+_8433172_8446622_RNA45SN1_45S_50ntFlanks@10819.10837.19|AGGGAAGTCGGCAAGCCGG</td><td>>hg38_21_+_8433172_8446622_RNA45SN1_45S_50ntFlanks</td><td>10819</td><td>10837</td><td>19</td><td>AGGGAAGTCGGCAAGCCGG</td><td>886.6</td><td>-4.4</td><td>0.8</td><td>-5.6</td><td>1.7E-08</td><td>1.7E-07</td><td>BT20/MB468</td><td>MP</td></tr>
<tr><td>>MI0000650|hsa-mir-200c&WithFlank&12|+|6963693|6963772@50.68.19(+1C)&[MIMAT0000617&hsa-miR-200c-3p&offsets|0|-4(+1C);m-54&12|+|6963742|6963764&offsets|0|-4(+1C)]|TAATACTGCCGGGTAATGAC</td><td>>MI0000650|hsa-mir-200c&WithFlank&12|+|6963693|6963772</td><td>50</td><td>68</td><td>19(+1C)</td><td>TAATACTGCCGGGTAATGAC</td><td>151.0</td><td>10.6</td><td>1.8</td><td>5.9</td><td>3.5E-09</td><td>1.7E-07</td><td>BT20/MB231</td><td>MP</td></tr>
<tr><td>>hg38_21_+_8433172_8446622_RNA45SN1_45S_50ntFlanks@4738.4765.28|AAAGTCGGAGGTTCGAAGACGATCAGAT</td><td>>hg38_21_+_8433172_8446622_RNA45SN1_45S_50ntFlanks</td><td>4738</td><td>4765</td><td>28</td><td>AAAGTCGGAGGTTCGAAGACGATCAGAT</td><td>450.5</td><td>5.6</td><td>1.0</td><td>5.3</td><td>9.1E-08</td><td>1.7E-07</td><td>MB231/MB468</td><td>CYTO</td></tr>
<tr><td>>MI0006444|hsa-mir-1307&WithFlank&10|-|103394247|103394407@86.107.22(+1A)&[MIMAT0005951&hsa-miR-1307-3p&offsets|0|0(+1A);m-66&10|-|103394300|103394322&offsets|0|-1(+1A)]|ACTCGGCGTGGCGTCGGTCGTGA</td><td>>MI0006444|hsa-mir-1307&WithFlank&10|-|103394247|103394407</td><td>86</td><td>107</td><td>22(+1A)</td><td>ACTCGGCGTGGCGTCGGTCGTGA</td><td>406.0</td><td>4.9</td><td>0.9</td><td>5.3</td><td>9.9E-08</td><td>1.7E-07</td><td>MB231/MB468</td><td>MITO</td></tr>
<tr><td>>hg38_21_+_8433172_8446622_RNA45SN1_45S_50ntFlanks@12397.12416.20|CGACGTCGCTTTTTGATCCT</td><td>>hg38_21_+_8433172_8446622_RNA45SN1_45S_50ntFlanks</td><td>12397</td><td>12416</td><td>20</td><td>CGACGTCGCTTTTTGATCCT</td><td>623.0</td><td>-4.8</td><td>0.9</td><td>-5.3</td><td>9.9E-08</td><td>1.7E-07</td><td>MB231/MB468</td><td>MITO</td></tr>
<tr><td>>hg38_21_+_8433172_8446622_RNA45SN1_45S_50ntFlanks@3793.3812.20|CGAATGGCTCATTAAATCAG</td><td>>hg38_21_+_8433172_8446622_RNA45SN1_45S_50ntFlanks</td><td>3793</td><td>3812</td><td>20</td><td>CGAATGGCTCATTAAATCAG</td><td>77.7</td><td>-9.4</td><td>1.7</td><td>-5.6</td><td>1.8E-08</td><td>1.7E-07</td><td>BT20/MB468</td><td>MP</td></tr>
<tr><td>>hg38_21_+_8433172_8446622_RNA45SN1_45S_50ntFlanks@5447.5466.20|GGATCGGCCCCGCCGGGGTC</td><td>>hg38_21_+_8433172_8446622_RNA45SN1_45S_50ntFlanks</td><td>5447</td><td>5466</td><td>20</td><td>GGATCGGCCCCGCCGGGGTC</td><td>130.5</td><td>-10.1</td><td>1.8</td><td>-5.6</td><td>1.8E-08</td><td>1.7E-07</td><td>BT20/MB468</td><td>MP</td></tr>
<tr><td>>hg38_21_+_8433172_8446622_RNA45SN1_45S_50ntFlanks@11583.11603.21|CTGGTGCGGACCAGGGGAATC</td><td>>hg38_21_+_8433172_8446622_RNA45SN1_45S_50ntFlanks</td><td>11583</td><td>11603</td><td>21</td><td>CTGGTGCGGACCAGGGGAATC</td><td>2765.2</td><td>-3.7</td><td>0.7</td><td>-5.3</td><td>9.1E-08</td><td>1.7E-07</td><td>MB231/MB468</td><td>CYTO</td></tr>
<tr><td>>MI0000782|hsa-mir-374a&WithFlank&X|-|74287280|74287363@18.40.23&[MIMAT0000727&hsa-miR-374a-5p&offsets|0|+1;m-184&X|-|74287325|74287346&offsets|0|+1]|TTATAATACAACCTGATAAGTGT</td><td>>MI0000782|hsa-mir-374a&WithFlank&X|-|74287280|74287363</td><td>18</td><td>40</td><td>23</td><td>TTATAATACAACCTGATAAGTGT</td><td>181.6</td><td>-10.9</td><td>1.8</td><td>-5.9</td><td>3.6E-09</td><td>1.7E-07</td><td>BT20/MB231</td><td>MP</td></tr>
<tr><td>>MI0000063|hsa-let-7b&WithFlank&22|+|46113680|46113774@12.33.22(+1A)&[MIMAT0000063&hsa-let-7b-5p&offsets|0|0(+1A);m-14&22|+|46113691|46113712&offsets|0|0(+1A)]|TGAGGTAGTAGGTTGTGTGGTTA</td><td>>MI0000063|hsa-let-7b&WithFlank&22|+|46113680|46113774</td><td>12</td><td>33</td><td>22(+1A)</td><td>TGAGGTAGTAGGTTGTGTGGTTA</td><td>818.9</td><td>2.5</td><td>0.5</td><td>5.3</td><td>1.0E-07</td><td>1.7E-07</td><td>BT20/MB468</td><td>NUC</td></tr>
<tr><td>>hg38_21_+_8433172_8446622_RNA45SN1_45S_50ntFlanks@10846.10864.19|CTTCGGGATAAGGATTGGC</td><td>>hg38_21_+_8433172_8446622_RNA45SN1_45S_50ntFlanks</td><td>10846</td><td>10864</td><td>19</td><td>CTTCGGGATAAGGATTGGC</td><td>768.7</td><td>-5.6</td><td>1.0</td><td>-5.4</td><td>8.0E-08</td><td>1.7E-07</td><td>BT20/MB468</td><td>CYTO</td></tr>
<tr><td>>hg38_21_+_8433172_8446622_RNA45SN1_45S_50ntFlanks@12432.12455.24|CCTATCATTGTGAAGCAGAATTCA</td><td>>hg38_21_+_8433172_8446622_RNA45SN1_45S_50ntFlanks</td><td>12432</td><td>12455</td><td>24</td><td>CCTATCATTGTGAAGCAGAATTCA</td><td>211.8</td><td>-6.4</td><td>1.2</td><td>-5.4</td><td>8.0E-08</td><td>1.7E-07</td><td>BT20/MB468</td><td>CYTO</td></tr>
<tr><td>>hg38_21_+_8433172_8446622_RNA45SN1_45S_50ntFlanks@8168.8195.28|CCCAAGTCCTTCTGATCGAGGCCCAGCC</td><td>>hg38_21_+_8433172_8446622_RNA45SN1_45S_50ntFlanks</td><td>8168</td><td>8195</td><td>28</td><td>CCCAAGTCCTTCTGATCGAGGCCCAGCC</td><td>448.9</td><td>-5.7</td><td>1.1</td><td>-5.3</td><td>1.0E-07</td><td>1.7E-07</td><td>MB231/MB468</td><td>MITO</td></tr>
<tr><td>>hg38_21_+_8433172_8446622_RNA45SN1_45S_50ntFlanks@7975.7993.19|CGCGACCTCAGATCAGACG</td><td>>hg38_21_+_8433172_8446622_RNA45SN1_45S_50ntFlanks</td><td>7975</td><td>7993</td><td>19</td><td>CGCGACCTCAGATCAGACG</td><td>154507.6</td><td>4.8</td><td>0.9</td><td>5.4</td><td>7.1E-08</td><td>1.7E-07</td><td>BT20/MB468</td><td>MITO</td></tr>
<tr><td>>hg38_21_+_8433172_8446622_RNA45SN1_45S_50ntFlanks@10781.10807.27|AGGTGAACAGCCTCTGGCATGTTGGAA</td><td>>hg38_21_+_8433172_8446622_RNA45SN1_45S_50ntFlanks</td><td>10781</td><td>10807</td><td>27</td><td>AGGTGAACAGCCTCTGGCATGTTGGAA</td><td>385.6</td><td>-7.3</td><td>1.4</td><td>-5.4</td><td>8.0E-08</td><td>1.7E-07</td><td>BT20/MB468</td><td>CYTO</td></tr>
<tr><td>>hg38_21_+_8433172_8446622_RNA45SN1_45S_50ntFlanks@4723.4765.43|CATTAATCAAGAACGAAAGTCGGAGGTTCGAAGACGATCAGAT</td><td>>hg38_21_+_8433172_8446622_RNA45SN1_45S_50ntFlanks</td><td>4723</td><td>4765</td><td>43</td><td>CATTAATCAAGAACGAAAGTCGGAGGTTCGAAGACGATCAGAT</td><td>425.3</td><td>5.8</td><td>1.1</td><td>5.3</td><td>9.2E-08</td><td>1.7E-07</td><td>MB231/MB468</td><td>CYTO</td></tr>
<tr><td>>hg38_21_+_8433172_8446622_RNA45SN1_45S_50ntFlanks@11568.11592.25|GCAGCCGACTTAGAACTGGTGCGGA</td><td>>hg38_21_+_8433172_8446622_RNA45SN1_45S_50ntFlanks</td><td>11568</td><td>11592</td><td>25</td><td>GCAGCCGACTTAGAACTGGTGCGGA</td><td>354.6</td><td>-4.6</td><td>0.9</td><td>-5.3</td><td>9.2E-08</td><td>1.7E-07</td><td>MB231/MB468</td><td>CYTO</td></tr>
<tr><td>>hg38_21_+_8433172_8446622_RNA45SN1_45S_50ntFlanks@12413.12433.21|TCCTTCGATGTCGGCTCTTCC</td><td>>hg38_21_+_8433172_8446622_RNA45SN1_45S_50ntFlanks</td><td>12413</td><td>12433</td><td>21</td><td>TCCTTCGATGTCGGCTCTTCC</td><td>535.0</td><td>-4.6</td><td>0.9</td><td>-5.3</td><td>1.0E-07</td><td>1.7E-07</td><td>BT20/MB468</td><td>NUC</td></tr>
<tr><td>>hg38_21_+_8433172_8446622_RNA45SN1_45S_50ntFlanks@4640.4663.24|GCATTCGTATTGCGCCGCTAGAGG</td><td>>hg38_21_+_8433172_8446622_RNA45SN1_45S_50ntFlanks</td><td>4640</td><td>4663</td><td>24</td><td>GCATTCGTATTGCGCCGCTAGAGG</td><td>76.2</td><td>-9.3</td><td>1.7</td><td>-5.6</td><td>1.8E-08</td><td>1.7E-07</td><td>BT20/MB468</td><td>MP</td></tr>
<tr><td>>MI0005531|hsa-mir-450b&WithFlank&X|-|134540179|134540268@17.37.21&[MIMAT0004909&hsa-miR-450b-5p&offsets|0|-1;m-266&X|-|134540232|134540252&offsets|0|0]|TTTTGCAATATGTTCCTGAAT</td><td>>MI0005531|hsa-mir-450b&WithFlank&X|-|134540179|134540268</td><td>17</td><td>37</td><td>21</td><td>TTTTGCAATATGTTCCTGAAT</td><td>116.4</td><td>5.2</td><td>1.0</td><td>5.3</td><td>1.0E-07</td><td>1.7E-07</td><td>BT20/MB468</td><td>NUC</td></tr>
<tr><td>>hg38_21_+_8433172_8446622_RNA45SN1_45S_50ntFlanks@9858.9900.43|CGGGATGAACCGAACGCCGGGTTAAGGCGCCCGATGCCGACGC</td><td>>hg38_21_+_8433172_8446622_RNA45SN1_45S_50ntFlanks</td><td>9858</td><td>9900</td><td>43</td><td>CGGGATGAACCGAACGCCGGGTTAAGGCGCCCGATGCCGACGC</td><td>46.6</td><td>-9.4</td><td>1.8</td><td>-5.4</td><td>8.5E-08</td><td>1.7E-07</td><td>MB231/MB468</td><td>MP</td></tr>
<tr><td>>hg38_21_+_8433172_8446622_RNA45SN1_45S_50ntFlanks@5491.5516.26|GAGAAGACGGTCGAACTTGACTATCT</td><td>>hg38_21_+_8433172_8446622_RNA45SN1_45S_50ntFlanks</td><td>5491</td><td>5516</td><td>26</td><td>GAGAAGACGGTCGAACTTGACTATCT</td><td>4372.3</td><td>8.5</td><td>1.5</td><td>5.6</td><td>1.8E-08</td><td>1.7E-07</td><td>BT20/MB468</td><td>MP</td></tr>
<tr><td>>MI0000743|hsa-mir-34c&WithFlank&11|+|111513433|111513521@19.40.22&[MIMAT0000686&hsa-miR-34c-5p&offsets|0|-1;m-336&11|+|111513451|111513473&offsets|0|-1]|AGGCAGTGTAGTTAGCTGATTG</td><td>>MI0000743|hsa-mir-34c&WithFlank&11|+|111513433|111513521</td><td>19</td><td>40</td><td>22</td><td>AGGCAGTGTAGTTAGCTGATTG</td><td>53.5</td><td>9.0</td><td>1.7</td><td>5.4</td><td>8.5E-08</td><td>1.7E-07</td><td>MB231/MB468</td><td>MP</td></tr>
<tr><td>>MI0000087|hsa-mir-29a&WithFlank&7|-|130876741|130876816@48.69.22&[MIMAT0000086&hsa-miR-29a-3p&offsets|0|0;m-26&7|-|130876749|130876769&offsets|0|+1]|TAGCACCATCTGAAATCGGTTA</td><td>>MI0000087|hsa-mir-29a&WithFlank&7|-|130876741|130876816</td><td>48</td><td>69</td><td>22</td><td>TAGCACCATCTGAAATCGGTTA</td><td>31080.9</td><td>5.8</td><td>1.1</td><td>5.4</td><td>8.5E-08</td><td>1.7E-07</td><td>MB231/MB468</td><td>MP</td></tr>
<tr><td>>hg38_21_+_8433172_8446622_RNA45SN1_45S_50ntFlanks@10393.10435.43|CAGCAGTTGAACATGGGTCAGTCGGTCCTGAGAGATGGGCGAG</td><td>>hg38_21_+_8433172_8446622_RNA45SN1_45S_50ntFlanks</td><td>10393</td><td>10435</td><td>43</td><td>CAGCAGTTGAACATGGGTCAGTCGGTCCTGAGAGATGGGCGAG</td><td>459.8</td><td>-5.7</td><td>1.1</td><td>-5.3</td><td>9.2E-08</td><td>1.7E-07</td><td>MB231/MB468</td><td>CYTO</td></tr>
<tr><td>>hg38_21_+_8433172_8446622_RNA45SN1_45S_50ntFlanks@6645.6680.36|CTCGTACGACTCTTAGCGGTGGATCACTCGGCTCGT</td><td>>hg38_21_+_8433172_8446622_RNA45SN1_45S_50ntFlanks</td><td>6645</td><td>6680</td><td>36</td><td>CTCGTACGACTCTTAGCGGTGGATCACTCGGCTCGT</td><td>945.8</td><td>3.9</td><td>0.7</td><td>5.4</td><td>8.0E-08</td><td>1.7E-07</td><td>BT20/MB468</td><td>CYTO</td></tr>
<tr><td>>hg38_21_+_8433172_8446622_RNA45SN1_45S_50ntFlanks@9623.9649.27|TAGTAGCTGGTTCCCTCCGAAGTTTCC</td><td>>hg38_21_+_8433172_8446622_RNA45SN1_45S_50ntFlanks</td><td>9623</td><td>9649</td><td>27</td><td>TAGTAGCTGGTTCCCTCCGAAGTTTCC</td><td>540.5</td><td>-2.5</td><td>0.5</td><td>-5.4</td><td>7.1E-08</td><td>1.7E-07</td><td>BT20/MB468</td><td>MITO</td></tr>
<tr><td>>MI0000272|hsa-mir-182&WithFlank&7|-|129770377|129770498@29.54.26&[MIMAT0000259&hsa-miR-182-5p&offsets|0|+2;m-22&7|-|129770449|129770470&offsets|0|+4]|TTTGGCAATGGTAGAACTCACACTGG</td><td>>MI0000272|hsa-mir-182&WithFlank&7|-|129770377|129770498</td><td>29</td><td>54</td><td>26</td><td>TTTGGCAATGGTAGAACTCACACTGG</td><td>4963.8</td><td>3.0</td><td>0.5</td><td>5.7</td><td>1.0E-08</td><td>1.7E-07</td><td>BT20/MB231</td><td>NUC</td></tr>
<tr><td>>hg38_21_+_8433172_8446622_RNA45SN1_45S_50ntFlanks@10845.10867.23|ACTTCGGGATAAGGATTGGCTCT</td><td>>hg38_21_+_8433172_8446622_RNA45SN1_45S_50ntFlanks</td><td>10845</td><td>10867</td><td>23</td><td>ACTTCGGGATAAGGATTGGCTCT</td><td>1152.8</td><td>-8.7</td><td>1.6</td><td>-5.3</td><td>1.0E-07</td><td>1.7E-07</td><td>BT20/MB468</td><td>NUC</td></tr>
<tr><td>>hg38_21_+_8433172_8446622_RNA45SN1_45S_50ntFlanks@11665.11693.29|TTCTGCCCAGTGCTCTGAATGTCAAAGTG</td><td>>hg38_21_+_8433172_8446622_RNA45SN1_45S_50ntFlanks</td><td>11665</td><td>11693</td><td>29</td><td>TTCTGCCCAGTGCTCTGAATGTCAAAGTG</td><td>254.8</td><td>5.7</td><td>1.1</td><td>5.3</td><td>9.2E-08</td><td>1.7E-07</td><td>MB231/MB468</td><td>CYTO</td></tr>
<tr><td>>hg38_1_-_228634819_228635039_RNA5S12_5S_50ntFlanks@51.75.25|GTCTACGGCCATACCACCCTGAACG</td><td>>hg38_1_-_228634819_228635039_RNA5S12_5S_50ntFlanks</td><td>51</td><td>75</td><td>25</td><td>GTCTACGGCCATACCACCCTGAACG</td><td>1375.7</td><td>4.6</td><td>0.9</td><td>5.3</td><td>9.2E-08</td><td>1.7E-07</td><td>MB231/MB468</td><td>CYTO</td></tr>
<tr><td>>hg38_21_+_8433172_8446622_RNA45SN1_45S_50ntFlanks@12419.12436.18|GATGTCGGCTCTTCCTAT</td><td>>hg38_21_+_8433172_8446622_RNA45SN1_45S_50ntFlanks</td><td>12419</td><td>12436</td><td>18</td><td>GATGTCGGCTCTTCCTAT</td><td>739.1</td><td>-2.2</td><td>0.4</td><td>-5.4</td><td>7.2E-08</td><td>1.7E-07</td><td>BT20/MB468</td><td>MITO</td></tr>
<tr><td>>hg38_21_+_8433172_8446622_RNA45SN1_45S_50ntFlanks@12957.12990.34|CTTCTGGGTCGGGGTTTCGTACGTAGCAGAGCAG</td><td>>hg38_21_+_8433172_8446622_RNA45SN1_45S_50ntFlanks</td><td>12957</td><td>12990</td><td>34</td><td>CTTCTGGGTCGGGGTTTCGTACGTAGCAGAGCAG</td><td>469.8</td><td>-5.2</td><td>1.0</td><td>-5.3</td><td>9.3E-08</td><td>1.7E-07</td><td>MB231/MB468</td><td>CYTO</td></tr>
<tr><td>>hg38_21_+_8433172_8446622_RNA45SN1_45S_50ntFlanks@4006.4026.21|ACCTCGGGCCGATCGCACGCC</td><td>>hg38_21_+_8433172_8446622_RNA45SN1_45S_50ntFlanks</td><td>4006</td><td>4026</td><td>21</td><td>ACCTCGGGCCGATCGCACGCC</td><td>347.0</td><td>-6.2</td><td>1.2</td><td>-5.3</td><td>9.3E-08</td><td>1.7E-07</td><td>MB231/MB468</td><td>CYTO</td></tr>
<tr><td>>hg38_21_+_8433172_8446622_RNA45SN1_45S_50ntFlanks@12477.12497.21|CACTAATAGGGAACGTGAGCT</td><td>>hg38_21_+_8433172_8446622_RNA45SN1_45S_50ntFlanks</td><td>12477</td><td>12497</td><td>21</td><td>CACTAATAGGGAACGTGAGCT</td><td>2481.5</td><td>2.9</td><td>0.5</td><td>5.4</td><td>8.1E-08</td><td>1.7E-07</td><td>BT20/MB468</td><td>CYTO</td></tr>
<tr><td>>hg38_21_+_8433172_8446622_RNA45SN1_45S_50ntFlanks@12327.12346.20|GGTTTTAAGCAGGAGGTGTC</td><td>>hg38_21_+_8433172_8446622_RNA45SN1_45S_50ntFlanks</td><td>12327</td><td>12346</td><td>20</td><td>GGTTTTAAGCAGGAGGTGTC</td><td>133.5</td><td>-5.9</td><td>1.1</td><td>-5.4</td><td>8.1E-08</td><td>1.7E-07</td><td>BT20/MB468</td><td>CYTO</td></tr>
<tr><td>>hg38_21_+_8433172_8446622_RNA45SN1_45S_50ntFlanks@13022.13039.18|CTCGACACAAGGGTTTGT</td><td>>hg38_21_+_8433172_8446622_RNA45SN1_45S_50ntFlanks</td><td>13022</td><td>13039</td><td>18</td><td>CTCGACACAAGGGTTTGT</td><td>883.8</td><td>3.3</td><td>0.6</td><td>5.3</td><td>9.3E-08</td><td>1.7E-07</td><td>MB231/MB468</td><td>CYTO</td></tr>
<tr><td>>hg38_21_+_8433172_8446622_RNA45SN1_45S_50ntFlanks@11573.11591.19|CGACTTAGAACTGGTGCGG</td><td>>hg38_21_+_8433172_8446622_RNA45SN1_45S_50ntFlanks</td><td>11573</td><td>11591</td><td>19</td><td>CGACTTAGAACTGGTGCGG</td><td>273.3</td><td>-5.3</td><td>1.0</td><td>-5.3</td><td>9.5E-08</td><td>1.7E-07</td><td>MB231/MB468</td><td>NUC</td></tr>
<tr><td>>hg38_21_+_8433172_8446622_RNA45SN1_45S_50ntFlanks@12928.12973.46|CTGGCGCTAAACCATTCGTAGACGACCTGCTTCTGGGTCGGGGTTT</td><td>>hg38_21_+_8433172_8446622_RNA45SN1_45S_50ntFlanks</td><td>12928</td><td>12973</td><td>46</td><td>CTGGCGCTAAACCATTCGTAGACGACCTGCTTCTGGGTCGGGGTTT</td><td>450.2</td><td>-7.7</td><td>1.5</td><td>-5.3</td><td>1.0E-07</td><td>1.7E-07</td><td>MB231/MB468</td><td>MITO</td></tr>
<tr><td>>hg38_21_+_8433172_8446622_RNA45SN1_45S_50ntFlanks@9556.9574.19|TAGCGGTCCTGACGTGCAA</td><td>>hg38_21_+_8433172_8446622_RNA45SN1_45S_50ntFlanks</td><td>9556</td><td>9574</td><td>19</td><td>TAGCGGTCCTGACGTGCAA</td><td>105.4</td><td>-9.8</td><td>1.7</td><td>-5.6</td><td>1.8E-08</td><td>1.7E-07</td><td>BT20/MB468</td><td>MP</td></tr>
<tr><td>>hg38_21_+_8433172_8446622_RNA45SN1_45S_50ntFlanks@6778.6807.30|CCCGGGGCTACGCCTGTCTGAGCGTCGCTT</td><td>>hg38_21_+_8433172_8446622_RNA45SN1_45S_50ntFlanks</td><td>6778</td><td>6807</td><td>30</td><td>CCCGGGGCTACGCCTGTCTGAGCGTCGCTT</td><td>1510.6</td><td>4.4</td><td>0.8</td><td>5.4</td><td>7.2E-08</td><td>1.7E-07</td><td>BT20/MB468</td><td>MITO</td></tr>
</table>

<table>
<tr><th></th><th></th><th></th><th></th><th></th><th></th><th></th><th></th><th></th><th></th><th></th><th></th><th></th><th></th></tr>
<tr><td>>trna116_GluCTC_1_-_145399233_145399304@32.72.41,trna71_GluCTC_1_-_161439189_161439260@32.72.41,trna74_GluCTC_1_-_161431809_161431880@32.72.41,trna77_GluCTC_1_-_161424398_161424469@32.72.41,trna77_GluCTC_6_+_28949976_28950047@32.72.41,trna80_GluCTC_1_-_161417018_161417089@32.72.41,trna87_GluCTC_6_-_126101393_126101464@32.72.41|CTCTCACCGCCGCGGCCCGGGTTCGATTCCCGGTCAGGGAA</td><td>>trna116_GluCTC_1_-_145399233_145399304</td><td>32</td><td>72</td><td>41</td><td>CTCTCACCGCCGCGGCCCGGGTTCGATTCCCGGTCAGGGAA</td><td>654.5</td><td>-7.0</td><td>1.3</td><td>-5.4</td><td>8.1E-08</td><td>1.7E-07</td><td>BT20/MB468</td><td>CYTO</td></tr>
<tr><td>>hg38_1_-_228634819_228635039_RNA5S12_5S_50ntFlanks@51.88.38|GTCTACGGCCATACCACCCTGAACGCGCCCGATCTCGT</td><td>>hg38_1_-_228634819_228635039_RNA5S12_5S_50ntFlanks</td><td>51</td><td>88</td><td>38</td><td>GTCTACGGCCATACCACCCTGAACGCGCCCGATCTCGT</td><td>4260.3</td><td>5.4</td><td>1.0</td><td>5.3</td><td>9.6E-08</td><td>1.7E-07</td><td>MB231/MB468</td><td>NUC</td></tr>
<tr><td>>hg38_21_+_8433172_8446622_RNA45SN1_45S_50ntFlanks@10403.10430.28|ACATGGGTCAGTCGGTCCTGAGAGATGG</td><td>>hg38_21_+_8433172_8446622_RNA45SN1_45S_50ntFlanks</td><td>10403</td><td>10430</td><td>28</td><td>ACATGGGTCAGTCGGTCCTGAGAGATGG</td><td>207.7</td><td>-3.0</td><td>0.6</td><td>-5.3</td><td>9.3E-08</td><td>1.7E-07</td><td>MB231/MB468</td><td>CYTO</td></tr>
<tr><td>>am_trna128_GlyGCC_6_-_27870686_27870756@3.34.32,trna18_GlyGCC_16_+_70822597_70822667@3.34.32,trna19_GlyGCC_16_+_70823410_70823480@3.34.32,trna19_GlyGCC_2_-_157257659_157257729@3.34.32,trna24_GlyGCC_16_-_70812942_70813012@3.34.32,trna25_GlyGCC_16_-_70812114_70812184@3.34.32,trna5_GlyGCC_17_+_8029064_8029134@3.34.32,trna68_GlyGCC_1_-_161493637_161493707@3.34.32|ATTGGTGGTTCAGTGGTAGAATTCTCGCCTGC</td><td>>am_trna128_GlyGCC_6_-_27870686_27870756</td><td>3</td><td>34</td><td>32</td><td>ATTGGTGGTTCAGTGGTAGAATTCTCGCCTGC</td><td>475.2</td><td>6.2</td><td>1.2</td><td>5.3</td><td>1.0E-07</td><td>1.7E-07</td><td>MB231/MB468</td><td>MITO</td></tr>
<tr><td>>hg38_21_+_8433172_8446622_RNA45SN1_45S_50ntFlanks@11781.11806.26|CGCATGAATGGATGAACGAGATTCCC</td><td>>hg38_21_+_8433172_8446622_RNA45SN1_45S_50ntFlanks</td><td>11781</td><td>11806</td><td>26</td><td>CGCATGAATGGATGAACGAGATTCCC</td><td>514.1</td><td>-6.8</td><td>1.3</td><td>-5.4</td><td>8.2E-08</td><td>1.8E-07</td><td>BT20/MB468</td><td>CYTO</td></tr>
<tr><td>>hg38_21_+_8433172_8446622_RNA45SN1_45S_50ntFlanks@6647.6680.34|CGTACGACTCTTAGCGGTGGATCACTCGGCTCGT</td><td>>hg38_21_+_8433172_8446622_RNA45SN1_45S_50ntFlanks</td><td>6647</td><td>6680</td><td>34</td><td>CGTACGACTCTTAGCGGTGGATCACTCGGCTCGT</td><td>840.4</td><td>3.9</td><td>0.7</td><td>5.4</td><td>8.2E-08</td><td>1.8E-07</td><td>BT20/MB468</td><td>CYTO</td></tr>
<tr><td>>hg38_21_+_8433172_8446622_RNA45SN1_45S_50ntFlanks@8170.8187.18|CAAGTCCTTCTGATCGAG</td><td>>hg38_21_+_8433172_8446622_RNA45SN1_45S_50ntFlanks</td><td>8170</td><td>8187</td><td>18</td><td>CAAGTCCTTCTGATCGAG</td><td>647.0</td><td>-5.1</td><td>1.0</td><td>-5.3</td><td>9.4E-08</td><td>1.8E-07</td><td>MB231/MB468</td><td>CYTO</td></tr>
<tr><td>>hg38_21_+_8433172_8446622_RNA45SN1_45S_50ntFlanks@11453.11478.26|TCCTCGGCGCGCGGCGGCGGCGGCGG</td><td>>hg38_21_+_8433172_8446622_RNA45SN1_45S_50ntFlanks</td><td>11453</td><td>11478</td><td>26</td><td>TCCTCGGCGCGCGGCGGCGGCGGCGG</td><td>89.5</td><td>-9.6</td><td>1.7</td><td>-5.6</td><td>1.8E-08</td><td>1.8E-07</td><td>BT20/MB468</td><td>MP</td></tr>
<tr><td>>hg38_21_+_8433172_8446622_RNA45SN1_45S_50ntFlanks@12912.12932.21|CACGTTCGTGGGGAACCTGGC</td><td>>hg38_21_+_8433172_8446622_RNA45SN1_45S_50ntFlanks</td><td>12912</td><td>12932</td><td>21</td><td>CACGTTCGTGGGGAACCTGGC</td><td>328.8</td><td>-6.2</td><td>1.2</td><td>-5.4</td><td>8.7E-08</td><td>1.8E-07</td><td>MB231/MB468</td><td>MP</td></tr>
<tr><td>>hg38_21_+_8433172_8446622_RNA45SN1_45S_50ntFlanks@3834.3860.27|GCTCCTCTCCTACTTGGATAACTGTGG</td><td>>hg38_21_+_8433172_8446622_RNA45SN1_45S_50ntFlanks</td><td>3834</td><td>3860</td><td>27</td><td>GCTCCTCTCCTACTTGGATAACTGTGG</td><td>300.6</td><td>-3.0</td><td>0.6</td><td>-5.3</td><td>9.7E-08</td><td>1.8E-07</td><td>MB231/MB468</td><td>NUC</td></tr>
<tr><td>>hg38_21_+_8433172_8446622_RNA45SN1_45S_50ntFlanks@12159.12181.23|TTTGACTGGGGCGGTACACCTGT</td><td>>hg38_21_+_8433172_8446622_RNA45SN1_45S_50ntFlanks</td><td>12159</td><td>12181</td><td>23</td><td>TTTGACTGGGGCGGTACACCTGT</td><td>310.3</td><td>-6.5</td><td>1.2</td><td>-5.4</td><td>8.2E-08</td><td>1.8E-07</td><td>BT20/MB468</td><td>CYTO</td></tr>
<tr><td>>MI0000272|hsa-mir-182&WithFlank&7|-|129770377|129770498@29.51.23(+1A)&[MIMAT0000259&hsa-miR-182-5p&offsets|0|-1(+1A);m-22&7|-|129770449|129770470&offsets|0|+1(+1A)]|TTTGGCAATGGTAGAACTCACACA</td><td>>MI0000272|hsa-mir-182&WithFlank&7|-|129770377|129770498</td><td>29</td><td>51</td><td>23(+1A)</td><td>TTTGGCAATGGTAGAACTCACACA</td><td>300.3</td><td>3.0</td><td>0.6</td><td>5.3</td><td>9.7E-08</td><td>1.8E-07</td><td>MB231/MB468</td><td>NUC</td></tr>
<tr><td>>MI0000285|hsa-mir-205&WithFlank&1|+|209432127|209432248@40.59.20&[MIMAT0000266&hsa-miR-205-5p&offsets|0|-2;m-351&1|+|209432166|209432187&offsets|0|-2]|TCCTTCATTCCACCGGAGTC</td><td>>MI0000285|hsa-mir-205&WithFlank&1|+|209432127|209432248</td><td>40</td><td>59</td><td>20</td><td>TCCTTCATTCCACCGGAGTC</td><td>293.4</td><td>11.5</td><td>2.0</td><td>5.9</td><td>3.7E-09</td><td>1.8E-07</td><td>BT20/MB231</td><td>MP</td></tr>
<tr><td>>hg38_21_+_8433172_8446622_RNA45SN1_45S_50ntFlanks@10384.10403.20|CCATGTGAACAGCAGTTGAA</td><td>>hg38_21_+_8433172_8446622_RNA45SN1_45S_50ntFlanks</td><td>10384</td><td>10403</td><td>20</td><td>CCATGTGAACAGCAGTTGAA</td><td>127.1</td><td>-10.1</td><td>1.8</td><td>-5.6</td><td>1.8E-08</td><td>1.8E-07</td><td>BT20/MB468</td><td>MP</td></tr>
<tr><td>>hg38_21_+_8433172_8446622_RNA45SN1_45S_50ntFlanks@11576.11601.26|CTTAGAACTGGTGCGGACCAGGGGAA</td><td>>hg38_21_+_8433172_8446622_RNA45SN1_45S_50ntFlanks</td><td>11576</td><td>11601</td><td>26</td><td>CTTAGAACTGGTGCGGACCAGGGGAA</td><td>711.8</td><td>-6.7</td><td>1.2</td><td>-5.6</td><td>1.8E-08</td><td>1.8E-07</td><td>BT20/MB468</td><td>MP</td></tr>
<tr><td>>hg38_21_+_8433172_8446622_RNA45SN1_45S_50ntFlanks@12615.12654.40|GCTGAGGAGCCAATGGGGCGAAGCTACCATCTGTGGGATT</td><td>>hg38_21_+_8433172_8446622_RNA45SN1_45S_50ntFlanks</td><td>12615</td><td>12654</td><td>40</td><td>GCTGAGGAGCCAATGGGGCGAAGCTACCATCTGTGGGATT</td><td>1904.4</td><td>-5.4</td><td>1.0</td><td>-5.6</td><td>1.8E-08</td><td>1.8E-07</td><td>BT20/MB468</td><td>MP</td></tr>
<tr><td>>hg38_21_+_8433172_8446622_RNA45SN1_45S_50ntFlanks@9202.9219.18|TCTCGGGGCCACGCGCGC</td><td>>hg38_21_+_8433172_8446622_RNA45SN1_45S_50ntFlanks</td><td>9202</td><td>9219</td><td>18</td><td>TCTCGGGGCCACGCGCGC</td><td>80.9</td><td>-9.4</td><td>1.7</td><td>-5.6</td><td>1.8E-08</td><td>1.8E-07</td><td>BT20/MB468</td><td>MP</td></tr>
<tr><td>>hg38_21_+_8433172_8446622_RNA45SN1_45S_50ntFlanks@12126.12149.24|CGACCCGCTCCGGGGACAGTGCCA</td><td>>hg38_21_+_8433172_8446622_RNA45SN1_45S_50ntFlanks</td><td>12126</td><td>12149</td><td>24</td><td>CGACCCGCTCCGGGGACAGTGCCA</td><td>84.9</td><td>-9.5</td><td>1.7</td><td>-5.6</td><td>1.8E-08</td><td>1.8E-07</td><td>BT20/MB468</td><td>MP</td></tr>
<tr><td>>MI0000255|hsa-mir-30d&WithFlank&8|-|134804870|134804951@12.32.21&[MIMAT0000245&hsa-miR-30d-5p&offsets|0|-1;m-21&8|-|134804919|134804940&offsets|0|-1]|TGTAAACATCCCCGACTGGAA</td><td>>MI0000255|hsa-mir-30d&WithFlank&8|-|134804870|134804951</td><td>12</td><td>32</td><td>21</td><td>TGTAAACATCCCCGACTGGAA</td><td>297.2</td><td>5.0</td><td>0.9</td><td>5.3</td><td>1.0E-07</td><td>1.8E-07</td><td>MB231/MB468</td><td>MITO</td></tr>
<tr><td>>hg38_21_+_8433172_8446622_RNA45SN1_45S_50ntFlanks@12974.12992.19|CGTACGTAGCAGAGCAGCT</td><td>>hg38_21_+_8433172_8446622_RNA45SN1_45S_50ntFlanks</td><td>12974</td><td>12992</td><td>19</td><td>CGTACGTAGCAGAGCAGCT</td><td>262.5</td><td>-5.5</td><td>1.0</td><td>-5.3</td><td>1.0E-07</td><td>1.8E-07</td><td>MB231/MB468</td><td>MITO</td></tr>
<tr><td>>hg38_21_+_8433172_8446622_RNA45SN1_45S_50ntFlanks@5236.5260.25|CACTGACTGGCTCAGCGTGTGCCTA</td><td>>hg38_21_+_8433172_8446622_RNA45SN1_45S_50ntFlanks</td><td>5236</td><td>5260</td><td>25</td><td>CACTGACTGGCTCAGCGTGTGCCTA</td><td>366.3</td><td>-10.6</td><td>2.0</td><td>-5.4</td><td>7.3E-08</td><td>1.8E-07</td><td>BT20/MB468</td><td>MITO</td></tr>
<tr><td>>am_trna10_ValCAC_5_-_180649395_180649467@1.32.32,trna12_ValAAC_5_-_180645270_180645342@1.32.32,trna132_ValAAC_6_-_27721179_27721251@1.32.32,trna136_ValAAC_6_-_27648885_27648957@1.32.32,trna139_ValAAC_6_-_27618707_27618779@1.32.32,trna18_ValCAC_5_-_180529253_180529325@1.32.32,trna2_ValAAC_3_+_169490018_169490090@1.32.32,trna2_ValCAC_5_+_180524070_180524142@1.32.32,trna4_ValAAC_5_+_180591154_180591226@1.32.32,trna5_ValAAC_5_+_180596610_180596682@1.32.32,trna6_ValCAC_5_+_180600650_180600722@1.32.32,trna85_ValCAC_1_-_161369490_161369562@1.32.32,trna90_ValCAC_1_-_149684088_149684161@1.32.32,trna98_ValCAC_1_-_149298555_149298627@1.32.32,trna9_ValCAC_6_+_26538282_26538354@1.32.32|GTTTCCGTAGTGTAGTGGTTATCACGTTCGCC</td><td>>am_trna10_ValCAC_5_-_180649395_180649467</td><td>1</td><td>32</td><td>32</td><td>GTTTCCGTAGTGTAGTGGTTATCACGTTCGCC</td><td>4824.1</td><td>6.1</td><td>1.2</td><td>5.3</td><td>9.8E-08</td><td>1.8E-07</td><td>MB231/MB468</td><td>NUC</td></tr>
<tr><td>>hg38_21_+_8433172_8446622_RNA45SN1_45S_50ntFlanks@12912.12933.22|CACGTTCGTGGGGAACCTGGCG</td><td>>hg38_21_+_8433172_8446622_RNA45SN1_45S_50ntFlanks</td><td>12912</td><td>12933</td><td>22</td><td>CACGTTCGTGGGGAACCTGGCG</td><td>1265.0</td><td>-9.6</td><td>1.8</td><td>-5.3</td><td>1.1E-07</td><td>1.8E-07</td><td>BT20/MB468</td><td>NUC</td></tr>
<tr><td>>hg38_21_+_8433172_8446622_RNA45SN1_45S_50ntFlanks@10241.10259.19|TACGCCGCGACGAGTAGGA</td><td>>hg38_21_+_8433172_8446622_RNA45SN1_45S_50ntFlanks</td><td>10241</td><td>10259</td><td>19</td><td>TACGCCGCGACGAGTAGGA</td><td>83.3</td><td>10.1</td><td>1.8</td><td>5.6</td><td>1.8E-08</td><td>1.8E-07</td><td>BT20/MB468</td><td>MP</td></tr>
<tr><td>>hg38_21_+_8433172_8446622_RNA45SN1_45S_50ntFlanks@12407.12443.37|TTTTGATCCTTCGATGTCGGCTCTTCCTATCATTGTG</td><td>>hg38_21_+_8433172_8446622_RNA45SN1_45S_50ntFlanks</td><td>12407</td><td>12443</td><td>37</td><td>TTTTGATCCTTCGATGTCGGCTCTTCCTATCATTGTG</td><td>2472.5</td><td>-4.4</td><td>0.8</td><td>-5.3</td><td>1.0E-07</td><td>1.8E-07</td><td>MB231/MB468</td><td>MITO</td></tr>
<tr><td>>hg38_21_+_8433172_8446622_RNA45SN1_45S_50ntFlanks@12415.12440.26|CTTCGATGTCGGCTCTTCCTATCATT</td><td>>hg38_21_+_8433172_8446622_RNA45SN1_45S_50ntFlanks</td><td>12415</td><td>12440</td><td>26</td><td>CTTCGATGTCGGCTCTTCCTATCATT</td><td>748.8</td><td>-3.2</td><td>0.6</td><td>-5.3</td><td>1.1E-07</td><td>1.8E-07</td><td>BT20/MB468</td><td>NUC</td></tr>
<tr><td>>hg38_21_+_8433172_8446622_RNA45SN1_45S_50ntFlanks@8737.8758.22|CCCGTCCGTCCGTCCGTCCGTC</td><td>>hg38_21_+_8433172_8446622_RNA45SN1_45S_50ntFlanks</td><td>8737</td><td>8758</td><td>22</td><td>CCCGTCCGTCCGTCCGTCCGTC</td><td>288.8</td><td>-6.4</td><td>1.2</td><td>-5.4</td><td>8.3E-08</td><td>1.8E-07</td><td>BT20/MB468</td><td>CYTO</td></tr>
<tr><td>>hg38_21_+_8433172_8446622_RNA45SN1_45S_50ntFlanks@11918.11948.31|ACATGAGAGGTGTAGAATAAGTGGGAGGCCC</td><td>>hg38_21_+_8433172_8446622_RNA45SN1_45S_50ntFlanks</td><td>11918</td><td>11948</td><td>31</td><td>ACATGAGAGGTGTAGAATAAGTGGGAGGCCC</td><td>838.2</td><td>-4.8</td><td>0.9</td><td>-5.4</td><td>8.3E-08</td><td>1.8E-07</td><td>BT20/MB468</td><td>CYTO</td></tr>
<tr><td>>hg38_21_+_8433172_8446622_RNA45SN1_45S_50ntFlanks@11573.11603.31|CGACTTAGAACTGGTGCGGACCAGGGGAATC</td><td>>hg38_21_+_8433172_8446622_RNA45SN1_45S_50ntFlanks</td><td>11573</td><td>11603</td><td>31</td><td>CGACTTAGAACTGGTGCGGACCAGGGGAATC</td><td>739.6</td><td>-2.8</td><td>0.5</td><td>-5.3</td><td>9.6E-08</td><td>1.8E-07</td><td>MB231/MB468</td><td>CYTO</td></tr>
<tr><td>>hg38_21_+_8433172_8446622_RNA45SN1_45S_50ntFlanks@10679.10712.34|GGTTCCGGCGGCGTCCGGTGAGCTCTCGCTGGCC</td><td>>hg38_21_+_8433172_8446622_RNA45SN1_45S_50ntFlanks</td><td>10679</td><td>10712</td><td>34</td><td>GGTTCCGGCGGCGTCCGGTGAGCTCTCGCTGGCC</td><td>1180.9</td><td>-9.1</td><td>1.7</td><td>-5.3</td><td>1.1E-07</td><td>1.8E-07</td><td>BT20/MB468</td><td>NUC</td></tr>
<tr><td>>MI0000088|hsa-mir-30a&WithFlank&6|-|71403545|71403627@13.35.23(+1U)&[MIMAT0000087&hsa-miR-30a-5p&offsets|+1|+2(+1U);m-15&6|-|71403595|71403616&offsets|+1|+2(+1U)]|GTAAACATCCTCGACTGGAAGCTT</td><td>>MI0000088|hsa-mir-30a&WithFlank&6|-|71403545|71403627</td><td>13</td><td>35</td><td>23(+1U)</td><td>GTAAACATCCTCGACTGGAAGCTT</td><td>744.8</td><td>6.0</td><td>1.1</td><td>5.3</td><td>8.9E-08</td><td>1.8E-07</td><td>MB231/MB468</td><td>MP</td></tr>
<tr><td>>hg38_21_+_8433172_8446622_RNA45SN1_45S_50ntFlanks@13011.13039.29|GAAAGTCAGCCCTCGACACAAGGGTTTGT</td><td>>hg38_21_+_8433172_8446622_RNA45SN1_45S_50ntFlanks</td><td>13011</td><td>13039</td><td>29</td><td>GAAAGTCAGCCCTCGACACAAGGGTTTGT</td><td>678.2</td><td>8.4</td><td>1.6</td><td>5.4</td><td>7.4E-08</td><td>1.8E-07</td><td>BT20/MB468</td><td>MITO</td></tr>
<tr><td>>hg38_21_+_8433172_8446622_RNA45SN1_45S_50ntFlanks@4002.4025.24|GATAACCTCGGGCCGATCGCACGC</td><td>>hg38_21_+_8433172_8446622_RNA45SN1_45S_50ntFlanks</td><td>4002</td><td>4025</td><td>24</td><td>GATAACCTCGGGCCGATCGCACGC</td><td>470.0</td><td>-4.8</td><td>0.9</td><td>-5.3</td><td>9.6E-08</td><td>1.8E-07</td><td>MB231/MB468</td><td>CYTO</td></tr>
<tr><td>>hg38_21_+_8433172_8446622_RNA45SN1_45S_50ntFlanks@10791.10823.33|CCTCTGGCATGTTGGAACAATGTAGGTAAGGGA</td><td>>hg38_21_+_8433172_8446622_RNA45SN1_45S_50ntFlanks</td><td>10791</td><td>10823</td><td>33</td><td>CCTCTGGCATGTTGGAACAATGTAGGTAAGGGA</td><td>424.1</td><td>-5.6</td><td>1.1</td><td>-5.3</td><td>1.0E-07</td><td>1.8E-07</td><td>MB231/MB468</td><td>MITO</td></tr>
<tr><td>>hg38_21_+_8433172_8446622_RNA45SN1_45S_50ntFlanks@10786.10820.35|AACAGCCTCTGGCATGTTGGAACAATGTAGGTAAG</td><td>>hg38_21_+_8433172_8446622_RNA45SN1_45S_50ntFlanks</td><td>10786</td><td>10820</td><td>35</td><td>AACAGCCTCTGGCATGTTGGAACAATGTAGGTAAG</td><td>1061.9</td><td>-6.0</td><td>1.1</td><td>-5.3</td><td>1.0E-07</td><td>1.8E-07</td><td>MB231/MB468</td><td>MITO</td></tr>
<tr><td>>hg38_21_+_8433172_8446622_RNA45SN1_45S_50ntFlanks@12003.12029.27|GCGGGCCGCCGGTGAAATACCACTACT</td><td>>hg38_21_+_8433172_8446622_RNA45SN1_45S_50ntFlanks</td><td>12003</td><td>12029</td><td>27</td><td>GCGGGCCGCCGGTGAAATACCACTACT</td><td>1771.8</td><td>4.7</td><td>0.9</td><td>5.3</td><td>9.7E-08</td><td>1.8E-07</td><td>MB231/MB468</td><td>CYTO</td></tr>
<tr><td>>MI0000279|hsa-mir-196a-2&WithFlank&12|+|53991732|53991853@31.50.20;MI0000238|hsa-mir-196a-1&WithFlank&17|-|48632484|48632565@13.32.20&[MIMAT0000226&hsa-miR-196a-5p&offsets|0|-2;m-261&12|+|53991762|53991783&offsets|0|-2];[MIMAT0000226_1&hsa-miR-196a-5p&offsets|0|-2;m-268&17|-|48632532|48632553&offsets|0|-2]|TAGGTAGTTTCATGTTGTTG</td><td>>MI0000279|hsa-mir-196a-2&WithFlank&12|+|53991732|53991853</td><td>31</td><td>50</td><td>20</td><td>TAGGTAGTTTCATGTTGTTG</td><td>155.9</td><td>10.6</td><td>1.8</td><td>5.9</td><td>3.8E-09</td><td>1.8E-07</td><td>BT20/MB231</td><td>MP</td></tr>
<tr><td>>hg38_21_+_8433172_8446622_RNA45SN1_45S_50ntFlanks@9922.9940.19|GTTGGTTGATATAGACAGC</td><td>>hg38_21_+_8433172_8446622_RNA45SN1_45S_50ntFlanks</td><td>9922</td><td>9940</td><td>19</td><td>GTTGGTTGATATAGACAGC</td><td>101.6</td><td>10.4</td><td>1.9</td><td>5.6</td><td>1.9E-08</td><td>1.8E-07</td><td>BT20/MB468</td><td>MP</td></tr>
<tr><td>>hg38_21_+_8433172_8446622_RNA45SN1_45S_50ntFlanks@11849.11872.24|CGGGCTTGGCGGAATCAGCGGGGA</td><td>>hg38_21_+_8433172_8446622_RNA45SN1_45S_50ntFlanks</td><td>11849</td><td>11872</td><td>24</td><td>CGGGCTTGGCGGAATCAGCGGGGA</td><td>1344.8</td><td>-8.7</td><td>1.6</td><td>-5.3</td><td>1.1E-07</td><td>1.8E-07</td><td>BT20/MB468</td><td>NUC</td></tr>
<tr><td>>hg38_21_+_8433172_8446622_RNA45SN1_45S_50ntFlanks@13009.13027.19|TTGAAAGTCAGCCCTCGAC</td><td>>hg38_21_+_8433172_8446622_RNA45SN1_45S_50ntFlanks</td><td>13009</td><td>13027</td><td>19</td><td>TTGAAAGTCAGCCCTCGAC</td><td>587.6</td><td>-8.1</td><td>1.5</td><td>-5.4</td><td>8.5E-08</td><td>1.8E-07</td><td>BT20/MB468</td><td>CYTO</td></tr>
<tr><td>>MI0000255|hsa-mir-30d&WithFlank&8|-|134804870|134804951@12.33.22(+1U)&[MIMAT0000245&hsa-miR-30d-5p&offsets|0|0(+1U);m-21&8|-|134804919|134804940&offsets|0|0(+1U)]|TGTAAACATCCCCGACTGGAAGT</td><td>>MI0000255|hsa-mir-30d&WithFlank&8|-|134804870|134804951</td><td>12</td><td>33</td><td>22(+1U)</td><td>TGTAAACATCCCCGACTGGAAGT</td><td>3542.7</td><td>5.4</td><td>1.0</td><td>5.6</td><td>1.9E-08</td><td>1.8E-07</td><td>BT20/MB468</td><td>MP</td></tr>
</table>

| | | | | | | | | | | | | | |
|---|---|---|---|---|---|---|---|---|---|---|---|---|---|
| >hg38_21_+_8433172_8446622_RNA45SN1_45S_50ntFlanks@11664.11685.22\|TTTCTGCCCAGTGCTCTGAATG | >hg38_21_+_8433172_8446622_RNA45SN1_45S_50ntFlanks | 11664 | 11685 | 22 | TTTCTGCCCAGTGCTCTGAATG | 274.1 | -5.6 | 1.1 | -5.3 | 1.1E-07 | 1.8E-07 | MB231/MB468 | MITO |
| >hg38_21_+_8433172_8446622_RNA45SN1_45S_50ntFlanks@10463.10489.27\|CTCCGTTGCCCTCGGCCGATCGAAAGG | >hg38_21_+_8433172_8446622_RNA45SN1_45S_50ntFlanks | 10463 | 10489 | 27 | CTCCGTTGCCCTCGGCCGATCGAAAGG | 224.0 | -6.9 | 1.3 | -5.3 | 1.0E-07 | 1.8E-07 | MB231/MB468 | NUC |
| >hg38_21_+_8433172_8446622_RNA45SN1_45S_50ntFlanks@11572.11609.38\|CCGACTTAGAACTGGTGCGGACCAGGGGAATCCGACTG | >hg38_21_+_8433172_8446622_RNA45SN1_45S_50ntFlanks | 11572 | 11609 | 38 | CCGACTTAGAACTGGTGCGGACCAGGGGAATCCGACTG | 648.6 | -6.5 | 1.2 | -5.3 | 9.0E-08 | 1.8E-07 | MB231/MB468 | MP |
| >hg38_21_+_8433172_8446622_RNA45SN1_45S_50ntFlanks@11881.11899.19\|CCTGTTGAGCTTGACTCTA | >hg38_21_+_8433172_8446622_RNA45SN1_45S_50ntFlanks | 11881 | 11899 | 19 | CCTGTTGAGCTTGACTCTA | 146.2 | -6.3 | 1.2 | -5.4 | 8.5E-08 | 1.8E-07 | BT20/MB468 | CYTO |
| >hg38_21_+_8433172_8446622_RNA45SN1_45S_50ntFlanks@3840.3873.34\|CTCCTACTTGGATAACTGTGGTAATTCTAGAGCT | >hg38_21_+_8433172_8446622_RNA45SN1_45S_50ntFlanks | 3840 | 3873 | 34 | CTCCTACTTGGATAACTGTGGTAATTCTAGAGCT | 1188.0 | 4.3 | 0.8 | 5.4 | 8.5E-08 | 1.8E-07 | BT20/MB468 | CYTO |
| >hg38_21_+_8433172_8446622_RNA45SN1_45S_50ntFlanks@7974.7991.18\|ACGCGACCTCAGATCAGA | >hg38_21_+_8433172_8446622_RNA45SN1_45S_50ntFlanks | 7974 | 7991 | 18 | ACGCGACCTCAGATCAGA | 5836.1 | 5.4 | 1.0 | 5.3 | 9.8E-08 | 1.8E-07 | MB231/MB468 | CYTO |
| >hg38_21_+_8433172_8446622_RNA45SN1_45S_50ntFlanks@10357.10384.28\|CTTTGAAGGCCGAAGTGGAGAAGGGTTC | >hg38_21_+_8433172_8446622_RNA45SN1_45S_50ntFlanks | 10357 | 10384 | 28 | CTTTGAAGGCCGAAGTGGAGAAGGGTTC | 251.8 | -6.0 | 1.1 | -5.3 | 9.0E-08 | 1.8E-07 | MB231/MB468 | MP |
| >hg38_21_+_8433172_8446622_RNA45SN1_45S_50ntFlanks@476.503.28\|TTCGTGATCGATGTGGTGACGTCGTGCT | >hg38_21_+_8433172_8446622_RNA45SN1_45S_50ntFlanks | 476 | 503 | 28 | TTCGTGATCGATGTGGTGACGTCGTGCT | 628.2 | 5.2 | 1.0 | 5.3 | 1.1E-07 | 1.8E-07 | MB231/MB468 | MITO |
| >MI0000286\|hsa-mir-210&WithFlank&11\|-\|568083\|568204@72.93.22(+1A)&[MIMAT0000267&hsa-miR-210-3p&offsets\|0\|0(+1A);m-102&11\|-\|568112\|568133&offsets\|0\|0(+1A)]\|CTGTGCGTGTGACAGCGGCTGAA | >MI0000286\|hsa-mir-210&WithFlank&11\|-\|568083\|568204 | 72 | 93 | 22(+1A) | CTGTGCGTGTGACAGCGGCTGAA | 209.0 | 7.6 | 1.4 | 5.3 | 1.1E-07 | 1.8E-07 | BT20/MB468 | NUC |
| >hg38_21_+_8433172_8446622_RNA45SN1_45S_50ntFlanks@4759.4776.18\|ATCAGATACCGTCGTAGT | >hg38_21_+_8433172_8446622_RNA45SN1_45S_50ntFlanks | 4759 | 4776 | 18 | ATCAGATACCGTCGTAGT | 266.5 | -6.4 | 1.2 | -5.3 | 9.8E-08 | 1.8E-07 | MB231/MB468 | CYTO |
| >hg38_21_+_8433172_8446622_RNA45SN1_45S_50ntFlanks@483.522.40\|TCGATGTGGTGACGTCGTGCTCTCCCGGGCCGGGTCCGAG | >hg38_21_+_8433172_8446622_RNA45SN1_45S_50ntFlanks | 483 | 522 | 40 | TCGATGTGGTGACGTCGTGCTCTCCCGGGCCGGGTCCGAG | 780.8 | 4.0 | 0.8 | 5.3 | 1.0E-07 | 1.8E-07 | MB231/MB468 | NUC |
| >hg38_21_+_8433172_8446622_RNA45SN1_45S_50ntFlanks@7973.8005.33\|GACGCGACCTCAGATCAGACGTGGCGACCCGCT | >hg38_21_+_8433172_8446622_RNA45SN1_45S_50ntFlanks | 7973 | 8005 | 33 | GACGCGACCTCAGATCAGACGTGGCGACCCGCT | 3367.6 | 4.9 | 0.9 | 5.3 | 1.1E-07 | 1.8E-07 | BT20/MB468 | NUC |
| >hg38_21_+_8433172_8446622_RNA45SN1_45S_50ntFlanks@5446.5467.22\|CGGATCGGCCCCGCCGGGGTCG | >hg38_21_+_8433172_8446622_RNA45SN1_45S_50ntFlanks | 5446 | 5467 | 22 | CGGATCGGCCCCGCCGGGGTCG | 309.4 | -6.5 | 1.2 | -5.4 | 8.6E-08 | 1.8E-07 | BT20/MB468 | CYTO |
| >MI0000077\|hsa-mir-21&WithFlank&17\|+\|59841260\|59841343@14.34.21(+2C)&[MIMAT0000076&hsa-miR-21-5p&offsets\|0\|-1(+2C);m-1&17\|+\|59841273\|59841294&offsets\|0\|-1(+2C)]\|TAGCTTATCAGACTGATGTTGCC | >MI0000077\|hsa-mir-21&WithFlank&17\|+\|59841260\|59841343 | 14 | 34 | 21(+2C) | TAGCTTATCAGACTGATGTTGCC | 2590.9 | -4.9 | 0.9 | -5.8 | 8.0E-09 | 1.8E-07 | BT20/MB231 | MITO |
| >hg38_21_+_8433172_8446622_RNA45SN1_45S_50ntFlanks@12329.12365.37\|TTTTAAGCAGGAGGTGTCAGAAAAGTTACCACAGGGA | >hg38_21_+_8433172_8446622_RNA45SN1_45S_50ntFlanks | 12329 | 12365 | 37 | TTTTAAGCAGGAGGTGTCAGAAAAGTTACCACAGGGA | 49.9 | -9.5 | 1.8 | -5.3 | 9.1E-08 | 1.8E-07 | MB231/MB468 | MP |
| >hg38_21_+_8433172_8446622_RNA45SN1_45S_50ntFlanks@4753.4784.32\|AAGACGATCAGATACCGTCGTAGTTCCGACCA | >hg38_21_+_8433172_8446622_RNA45SN1_45S_50ntFlanks | 4753 | 4784 | 32 | AAGACGATCAGATACCGTCGTAGTTCCGACCA | 140.0 | -9.2 | 1.7 | -5.4 | 7.6E-08 | 1.8E-07 | BT20/MB468 | MITO |
| >hg38_21_+_8433172_8446622_RNA45SN1_45S_50ntFlanks@11211.11251.41\|GCGACTCTGGACGCGAGCCGGGCCCTTCCCGTGGATCGCCC | >hg38_21_+_8433172_8446622_RNA45SN1_45S_50ntFlanks | 11211 | 11251 | 41 | GCGACTCTGGACGCGAGCCGGGCCCTTCCCGTGGATCGCCC | 372.2 | 4.8 | 0.9 | 5.4 | 7.6E-08 | 1.8E-07 | BT20/MB468 | MITO |
| >hg38_21_+_8433172_8446622_RNA45SN1_45S_50ntFlanks@11876.11899.24\|AAGACCCTGTTGAGCTTGACTCTA | >hg38_21_+_8433172_8446622_RNA45SN1_45S_50ntFlanks | 11876 | 11899 | 24 | AAGACCCTGTTGAGCTTGACTCTA | 537.3 | -3.9 | 0.7 | -5.3 | 1.0E-07 | 1.8E-07 | MB231/MB468 | NUC |
| >hg38_21_+_8433172_8446622_RNA45SN1_45S_50ntFlanks@12996.13039.44\|TCGCTGCGATCTATTGAAAGTCAGCCCTCGACACAAGGGTTTGT | >hg38_21_+_8433172_8446622_RNA45SN1_45S_50ntFlanks | 12996 | 13039 | 44 | TCGCTGCGATCTATTGAAAGTCAGCCCTCGACACAAGGGTTTGT | 151022.8 | 7.0 | 1.3 | 5.4 | 8.6E-08 | 1.8E-07 | BT20/MB468 | CYTO |
| >hg38_21_+_8433172_8446622_RNA45SN1_45S_50ntFlanks@6726.6744.19\|CAGGACACATTGATCATCG | >hg38_21_+_8433172_8446622_RNA45SN1_45S_50ntFlanks | 6726 | 6744 | 19 | CAGGACACATTGATCATCG | 147.1 | -10.3 | 1.8 | -5.6 | 1.9E-08 | 1.8E-07 | BT20/MB468 | MP |
| >hg38_21_+_8433172_8446622_RNA45SN1_45S_50ntFlanks@9461.9503.43\|GGGGAGGTGGAGCACGAGCGCACGTGTTAGGACCCGAAAGATG | >hg38_21_+_8433172_8446622_RNA45SN1_45S_50ntFlanks | 9461 | 9503 | 43 | GGGGAGGTGGAGCACGAGCGCACGTGTTAGGACCCGAAAGATG | 90.4 | -10.4 | 1.9 | -5.3 | 9.1E-08 | 1.8E-07 | MB231/MB468 | MP |
| >hg38_21_+_8433172_8446622_RNA45SN1_45S_50ntFlanks@10797.10828.32\|GCATGTTGGAACAATGTAGGTAAGGGAAGTCG | >hg38_21_+_8433172_8446622_RNA45SN1_45S_50ntFlanks | 10797 | 10828 | 32 | GCATGTTGGAACAATGTAGGTAAGGGAAGTCG | 358.6 | -7.4 | 1.4 | -5.3 | 9.1E-08 | 1.8E-07 | MB231/MB468 | MP |
| >MI0003685\|hsa-mir-421&WithFlank&X\|-\|74218371\|74218467@57.76.20&[MIMAT0003339&hsa-miR-421&offsets\|+3\|0;m-309&X\|-\|74218392\|74218414&offsets\|+3\|0]\|AACAGACATTAATTGGGCGC | >MI0003685\|hsa-mir-421&WithFlank&X\|-\|74218371\|74218467 | 57 | 76 | 20 | AACAGACATTAATTGGGCGC | 145.0 | 10.9 | 1.9 | 5.6 | 1.9E-08 | 1.8E-07 | BT20/MB468 | MP |
| >MI0001733\|hsa-mir-452&WithFlank&X\|-\|151959622\|151959718@20.42.23&[MIMAT0001635&hsa-miR-452-5p&offsets\|0\|+1;m-188&X\|-\|151959677\|151959699&offsets\|0\|0]\|AACTGTTTGCAGAGGAAACTGAG | >MI0001733\|hsa-mir-452&WithFlank&X\|-\|151959622\|151959718 | 20 | 42 | 23 | AACTGTTTGCAGAGGAAACTGAG | 660.0 | 10.1 | 1.8 | 5.6 | 1.9E-08 | 1.8E-07 | BT20/MB468 | MP |
| >hg38_21_+_8433172_8446622_RNA45SN1_45S_50ntFlanks@6730.6771.42\|ACACATTGATCATCGACACTTCGAACGCACTTGCGGCCCCGG | >hg38_21_+_8433172_8446622_RNA45SN1_45S_50ntFlanks | 6730 | 6771 | 42 | ACACATTGATCATCGACACTTCGAACGCACTTGCGGCCCCGG | 108.4 | -6.1 | 1.1 | -5.4 | 8.6E-08 | 1.8E-07 | BT20/MB468 | CYTO |
| >hg38_21_+_8433172_8446622_RNA45SN1_45S_50ntFlanks@11208.11231.24\|GCGGCGACTCTGGACGCGAGCCGG | >hg38_21_+_8433172_8446622_RNA45SN1_45S_50ntFlanks | 11208 | 11231 | 24 | GCGGCGACTCTGGACGCGAGCCGG | 2150.7 | -6.6 | 1.2 | -5.3 | 1.0E-07 | 1.8E-07 | MB231/MB468 | NUC |
| >hg38_21_+_8433172_8446622_RNA45SN1_45S_50ntFlanks@9736.9757.22\|CCGAAACGATCTCAACCTATTC | >hg38_21_+_8433172_8446622_RNA45SN1_45S_50ntFlanks | 9736 | 9757 | 22 | CCGAAACGATCTCAACCTATTC | 595.7 | -7.6 | 1.4 | -5.4 | 7.7E-08 | 1.8E-07 | BT20/MB468 | MITO |
| >hg38_21_+_8433172_8446622_RNA45SN1_45S_50ntFlanks@10369.10390.22\|AAGTGGAGAAGGGTTCCATGTG | >hg38_21_+_8433172_8446622_RNA45SN1_45S_50ntFlanks | 10369 | 10390 | 22 | AAGTGGAGAAGGGTTCCATGTG | 86.1 | -9.5 | 1.7 | -5.6 | 1.9E-08 | 1.9E-07 | BT20/MB468 | MP |
| >hg38_21_+_8433172_8446622_RNA45SN1_45S_50ntFlanks@10786.10830.45\|AACAGCCTCTGGCATGTTGGAACAATGTAGGTAAGGGAAGTCGGC | >hg38_21_+_8433172_8446622_RNA45SN1_45S_50ntFlanks | 10786 | 10830 | 45 | AACAGCCTCTGGCATGTTGGAACAATGTAGGTAAGGGAAGTCGGC | 60.2 | -9.8 | 1.8 | -5.3 | 9.2E-08 | 1.9E-07 | MB231/MB468 | MP |
| >hg38_21_+_8433172_8446622_RNA45SN1_45S_50ntFlanks@10807.10828.22\|ACAATGTAGGTAAGGGAAGTCG | >hg38_21_+_8433172_8446622_RNA45SN1_45S_50ntFlanks | 10807 | 10828 | 22 | ACAATGTAGGTAAGGGAAGTCG | 261.6 | -6.2 | 1.2 | -5.4 | 7.7E-08 | 1.9E-07 | BT20/MB468 | MITO |
| >hg38_21_+_8433172_8446622_RNA45SN1_45S_50ntFlanks@11661.11679.19\|TGATTTCTGCCCAGTGCTC | >hg38_21_+_8433172_8446622_RNA45SN1_45S_50ntFlanks | 11661 | 11679 | 19 | TGATTTCTGCCCAGTGCTC | 3070.8 | -8.6 | 1.6 | -5.4 | 8.7E-08 | 1.9E-07 | BT20/MB468 | CYTO |
| >hg38_21_+_8433172_8446622_RNA45SN1_45S_50ntFlanks@10468.10488.21\|TTGCCCTCGGCCGATCGAAAG | >hg38_21_+_8433172_8446622_RNA45SN1_45S_50ntFlanks | 10468 | 10488 | 21 | TTGCCCTCGGCCGATCGAAAG | 1569.0 | -7.4 | 1.4 | -5.3 | 1.1E-07 | 1.9E-07 | BT20/MB468 | NUC |
| >hg38_21_+_8433172_8446622_RNA45SN1_45S_50ntFlanks@12632.12659.28\|GCGAAGCTACCATCTGTGGGATTATGAC | >hg38_21_+_8433172_8446622_RNA45SN1_45S_50ntFlanks | 12632 | 12659 | 28 | GCGAAGCTACCATCTGTGGGATTATGAC | 415.4 | -3.6 | 0.7 | -5.3 | 1.0E-07 | 1.9E-07 | MB231/MB468 | NUC |
| >MI0001448\|hsa-mir-425&WithFlank&3\|-\|49020142\|49020240@64.82.19&[MIMAT0001343&hsa-miR-425-3p&offsets\|+3\|0;m-258&3\|-\|49020160\|49020181&offsets\|+4\|+1]\|GGGAATGTCGTGTCCGCCC | >MI0001448\|hsa-mir-425&WithFlank&3\|-\|49020142\|49020240 | 64 | 82 | 19 | GGGAATGTCGTGTCCGCCC | 113.2 | 10.6 | 1.9 | 5.6 | 1.9E-08 | 1.9E-07 | BT20/MB468 | MP |
| >hg38_MT_+_598_1651_RNR1_12S_50ntFlanks@906.946.41\|GTATACTTCAAAGGACATTTAACTAAAACCCCTACGCATTT | >hg38_MT_+_598_1651_RNR1_12S_50ntFlanks | 906 | 946 | 41 | GTATACTTCAAAGGACATTTAACTAAAACCCCTACGCATTT | 178.7 | 11.2 | 2.0 | 5.6 | 1.9E-08 | 1.9E-07 | BT20/MB468 | MP |
| >hg38_21_+_8433172_8446622_RNA45SN1_45S_50ntFlanks@5470.5499.30\|CCACGGCCCTGGCGGAGCGCTGAGAAGACG | >hg38_21_+_8433172_8446622_RNA45SN1_45S_50ntFlanks | 5470 | 5499 | 30 | CCACGGCCCTGGCGGAGCGCTGAGAAGACG | 365.7 | -3.2 | 0.6 | -5.4 | 7.7E-08 | 1.9E-07 | BT20/MB468 | MITO |
| >MI0000478\|hsa-mir-149&WithFlank&2\|+\|240455995\|240456095@21.42.22(+1A)&[MIMAT0000450&hsa-miR-149-5p&offsets\|0\|-1(+1A);m-195&2\|+\|240456015\|240456037&offsets\|0\|-1(+1A)]\|TCTGGCTCCGTGTCTTCACTCCA | >MI0000478\|hsa-mir-149&WithFlank&2\|+\|240455995\|240456095 | 21 | 42 | 22(+1A) | TCTGGCTCCGTGTCTTCACTCCA | 416.2 | 4.6 | 0.8 | 5.7 | 1.1E-08 | 1.9E-07 | BT20/MB231 | NUC |
| >MI0000084\|hsa-mir-26b&WithFlank&2\|+\|218402640\|218402728@18.39.22(+1U)&[MIMAT0000083&hsa-miR-26b-5p&offsets\|0\|+1(+1U);m-53&2\|+\|218402657\|218402678&offsets\|0\|0(+1U)]\|TTCAAGTAATTCAGGATAGGTTT | >MI0000084\|hsa-mir-26b&WithFlank&2\|+\|218402640\|218402728 | 18 | 39 | 22(+1U) | TTCAAGTAATTCAGGATAGGTTT | 732.7 | 5.4 | 1.0 | 5.3 | 1.1E-07 | 1.9E-07 | MB231/MB468 | MITO |
| >hg38_21_+_8433172_8446622_RNA45SN1_45S_50ntFlanks@12499.12516.18\|GGTTTAGACCGTCGTGAG | >hg38_21_+_8433172_8446622_RNA45SN1_45S_50ntFlanks | 12499 | 12516 | 18 | GGTTTAGACCGTCGTGAG | 143.7 | -5.8 | 1.1 | -5.4 | 8.7E-08 | 1.9E-07 | BT20/MB468 | CYTO |
| >MI0000650\|hsa-mir-200c&WithFlank&12\|+\|6963693\|6963772@50.71.22&[MIMAT0000617&hsa-miR-200c-3p&offsets\|0\|-1;m-54&12\|+\|6963742\|6963764&offsets\|0\|-1]\|TAATACTGCCGGGTAATGATGG | >MI0000650\|hsa-mir-200c&WithFlank&12\|+\|6963693\|6963772 | 50 | 71 | 22 | TAATACTGCCGGGTAATGATGG | 24962.4 | 5.8 | 1.0 | 5.6 | 1.9E-08 | 1.9E-07 | BT20/MB468 | MP |
| >hg38_21_+_8433172_8446622_RNA45SN1_45S_50ntFlanks@12396.12416.21\|GCGACGTCGCTTTTTGATCCT | >hg38_21_+_8433172_8446622_RNA45SN1_45S_50ntFlanks | 12396 | 12416 | 21 | GCGACGTCGCTTTTTGATCCT | 697.0 | -4.8 | 0.9 | -5.3 | 1.1E-07 | 1.9E-07 | MB231/MB468 | MITO |
| >hg38_21_+_8433172_8446622_RNA45SN1_45S_50ntFlanks@4004.4023.20\|TAACCTCGGGCCGATCGCAC | >hg38_21_+_8433172_8446622_RNA45SN1_45S_50ntFlanks | 4004 | 4023 | 20 | TAACCTCGGGCCGATCGCAC | 444.1 | 4.4 | 0.8 | 5.3 | 1.0E-07 | 1.9E-07 | MB231/MB468 | CYTO |
| >hg38_21_+_8433172_8446622_RNA45SN1_45S_50ntFlanks@12635.12660.26\|AAGCTACCATCTGTGGGATTATGACT | >hg38_21_+_8433172_8446622_RNA45SN1_45S_50ntFlanks | 12635 | 12660 | 26 | AAGCTACCATCTGTGGGATTATGACT | 925.8 | -6.1 | 1.1 | -5.3 | 1.1E-07 | 1.9E-07 | MB231/MB468 | MITO |
| >MI0000683\|hsa-mir-181b-2&WithFlank&9\|+\|124693704\|124693804@22.46.25(+1C)&[MIMAT0000257_1&hsa-miR-181b-5p&offsets\|0\|+2(+1C);m-72&9\|+\|124693725\|124693747&offsets\|0\|+2(+1C)]\|AACATTCATTGCTGTCGGTGGGTTTC | >MI0000683\|hsa-mir-181b-2&WithFlank&9\|+\|124693704\|124693804 | 22 | 46 | 25(+1C) | AACATTCATTGCTGTCGGTGGGTTTC | 65.8 | 9.3 | 1.7 | 5.3 | 9.3E-08 | 1.9E-07 | MB231/MB468 | MP |
| >hg38_21_+_8433172_8446622_RNA45SN1_45S_50ntFlanks@10229.10252.24\|CCCCGCGGACGCTACGCCGCGACG | >hg38_21_+_8433172_8446622_RNA45SN1_45S_50ntFlanks | 10229 | 10252 | 24 | CCCCGCGGACGCTACGCCGCGACG | 777.1 | 10.3 | 1.8 | 5.6 | 2.0E-08 | 1.9E-07 | BT20/MB468 | MP |
| >hg38_21_+_8433172_8446622_RNA45SN1_45S_50ntFlanks@482.506.25\|ATCGATGTGGTGACGTCGTGCTCTC | >hg38_21_+_8433172_8446622_RNA45SN1_45S_50ntFlanks | 482 | 506 | 25 | ATCGATGTGGTGACGTCGTGCTCTC | 99.4 | -9.7 | 1.7 | -5.6 | 2.0E-08 | 1.9E-07 | BT20/MB468 | MP |
| >MI0000782\|hsa-mir-374a&WithFlank&X\|-\|74287280\|74287363@18.39.22&[MIMAT0000727&hsa-miR-374a-5p&offsets\|0\|0;m-184&X\|-\|74287325\|74287346&offsets\|0\|0]\|TTATAATACAACCTGATAAGTG | >MI0000782\|hsa-mir-374a&WithFlank&X\|-\|74287280\|74287363 | 18 | 39 | 22 | TTATAATACAACCTGATAAGTG | 568.6 | 4.8 | 0.9 | 5.4 | 8.8E-08 | 1.9E-07 | BT20/MB468 | CYTO |
| >hg38_21_+_8433172_8446622_RNA45SN1_45S_50ntFlanks@10411.10430.20\|CAGTCGGTCCTGAGAGATGG | >hg38_21_+_8433172_8446622_RNA45SN1_45S_50ntFlanks | 10411 | 10430 | 20 | CAGTCGGTCCTGAGAGATGG | 12810.5 | -7.2 | 1.3 | -5.3 | 1.0E-07 | 1.9E-07 | MB231/MB468 | CYTO |
| >hg38_21_+_8433172_8446622_RNA45SN1_45S_50ntFlanks@8386.8405.20\|GGCGTGAAACCGTTAAGAGG | >hg38_21_+_8433172_8446622_RNA45SN1_45S_50ntFlanks | 8386 | 8405 | 20 | GGCGTGAAACCGTTAAGAGG | 161.3 | -7.0 | 1.3 | -5.3 | 9.3E-08 | 1.9E-07 | MB231/MB468 | MP |
| >hg38_21_+_8433172_8446622_RNA45SN1_45S_50ntFlanks@12417.12446.30\|TCGATGTCGGCTCTTCCTATCATTGTGAAG | >hg38_21_+_8433172_8446622_RNA45SN1_45S_50ntFlanks | 12417 | 12446 | 30 | TCGATGTCGGCTCTTCCTATCATTGTGAAG | 539.0 | -5.9 | 1.1 | -5.3 | 9.3E-08 | 1.9E-07 | MB231/MB468 | MP |

<table>
<tr><td>>MI0000456|hsa-mir-140&WithFlank&16|+|69933075|69933186@69.90.22(+1G)&[MIMAT0004597&hsa-miR-140-3p&offsets|+1|+2(+1G);m-45&16|+|69933143|69933164&offsets|0|0(+1G)]|ACCACAGGGTAGAACCACGGACG</td><td>>MI0000456|hsa-mir-140&WithFlank&16|+|69933075|69933186</td><td>69</td><td>90</td><td>22(+1G)</td><td>ACCACAGGGTAGAACCACGGACG</td><td>134.0</td><td>10.8</td><td>1.9</td><td>5.6</td><td>2.0E-08</td><td>1.9E-07</td><td>BT20/MB468</td><td>MP</td></tr>
<tr><td>>MI0000086|hsa-mir-28&WithFlank&3|+|188688775|188688872@20.41.22&[MIMAT0000085&hsa-miR-28-5p&offsets|0|0;m-133&3|+|188688794|188688815&offsets|0|0]|AAGGAGCTCACAGTCTATTGAG</td><td>>MI0000086|hsa-mir-28&WithFlank&3|+|188688775|188688872</td><td>20</td><td>41</td><td>22</td><td>AAGGAGCTCACAGTCTATTGAG</td><td>202.0</td><td>4.7</td><td>0.9</td><td>5.3</td><td>1.1E-07</td><td>1.9E-07</td><td>BT20/MB468</td><td>NUC</td></tr>
<tr><td>>hg38_21_+_8433172_8446622_RNA45SN1_45S_50ntFlanks@12141.12181.41|ACAGTGCCAGGTGGGGAGTTTGACTGGGGCGGTACACCTGT</td><td>>hg38_21_+_8433172_8446622_RNA45SN1_45S_50ntFlanks</td><td>12141</td><td>12181</td><td>41</td><td>ACAGTGCCAGGTGGGGAGTTTGACTGGGGCGGTACACCTGT</td><td>81.9</td><td>-8.4</td><td>1.6</td><td>-5.4</td><td>7.8E-08</td><td>1.9E-07</td><td>BT20/MB468</td><td>MITO</td></tr>
<tr><td>>hg38_21_+_8433172_8446622_RNA45SN1_45S_50ntFlanks@12636.12661.26|AGCTACCATCTGTGGGATTATGACTG</td><td>>hg38_21_+_8433172_8446622_RNA45SN1_45S_50ntFlanks</td><td>12636</td><td>12661</td><td>26</td><td>AGCTACCATCTGTGGGATTATGACTG</td><td>3534.0</td><td>-4.0</td><td>0.7</td><td>-5.4</td><td>7.8E-08</td><td>1.9E-07</td><td>BT20/MB468</td><td>MITO</td></tr>
<tr><td>>hg38_21_+_8433172_8446622_RNA45SN1_45S_50ntFlanks@4242.4264.23|TCTTTCGAGGCCCTGTAATTGGA</td><td>>hg38_21_+_8433172_8446622_RNA45SN1_45S_50ntFlanks</td><td>4242</td><td>4264</td><td>23</td><td>TCTTTCGAGGCCCTGTAATTGGA</td><td>57.1</td><td>-9.7</td><td>1.8</td><td>-5.3</td><td>9.4E-08</td><td>1.9E-07</td><td>MB231/MB468</td><td>MP</td></tr>
<tr><td>>hg38_21_+_8433172_8446622_RNA45SN1_45S_50ntFlanks@10792.10818.27|CTCTGGCATGTTGGAACAATGTAGGTA</td><td>>hg38_21_+_8433172_8446622_RNA45SN1_45S_50ntFlanks</td><td>10792</td><td>10818</td><td>27</td><td>CTCTGGCATGTTGGAACAATGTAGGTA</td><td>243.8</td><td>-2.9</td><td>0.5</td><td>-5.3</td><td>1.0E-07</td><td>1.9E-07</td><td>MB231/MB468</td><td>CYTO</td></tr>
<tr><td>>hg38_21_+_8433172_8446622_RNA45SN1_45S_50ntFlanks@11315.11344.30|CCACCCCACGTCTCGTCGCGCGCGCGTCCG</td><td>>hg38_21_+_8433172_8446622_RNA45SN1_45S_50ntFlanks</td><td>11315</td><td>11344</td><td>30</td><td>CCACCCCACGTCTCGTCGCGCGCGCGTCCG</td><td>2699.3</td><td>-7.4</td><td>1.4</td><td>-5.3</td><td>1.0E-07</td><td>1.9E-07</td><td>MB231/MB468</td><td>CYTO</td></tr>
<tr><td>>hg38_21_+_8433172_8446622_RNA45SN1_45S_50ntFlanks@10393.10430.38|CAGCAGTTGAACATGGGTCAGTCGGTCCTGAGAGATGG</td><td>>hg38_21_+_8433172_8446622_RNA45SN1_45S_50ntFlanks</td><td>10393</td><td>10430</td><td>38</td><td>CAGCAGTTGAACATGGGTCAGTCGGTCCTGAGAGATGG</td><td>492.1</td><td>-7.8</td><td>1.5</td><td>-5.3</td><td>9.4E-08</td><td>1.9E-07</td><td>MB231/MB468</td><td>MP</td></tr>
<tr><td>>hg38_21_+_8433172_8446622_RNA45SN1_45S_50ntFlanks@7992.8010.19|CGTGGCGACCCGCTGAATT</td><td>>hg38_21_+_8433172_8446622_RNA45SN1_45S_50ntFlanks</td><td>7992</td><td>8010</td><td>19</td><td>CGTGGCGACCCGCTGAATT</td><td>355.5</td><td>4.5</td><td>0.8</td><td>5.3</td><td>1.0E-07</td><td>1.9E-07</td><td>MB231/MB468</td><td>NUC</td></tr>
<tr><td>>hg38_21_+_8433172_8446622_RNA45SN1_45S_50ntFlanks@8194.8212.19|CCCGTGGACGGTGTGAGGC</td><td>>hg38_21_+_8433172_8446622_RNA45SN1_45S_50ntFlanks</td><td>8194</td><td>8212</td><td>19</td><td>CCCGTGGACGGTGTGAGGC</td><td>3632.7</td><td>-9.5</td><td>1.8</td><td>-5.3</td><td>1.2E-07</td><td>1.9E-07</td><td>BT20/MB468</td><td>NUC</td></tr>
<tr><td>>MI0003513|hsa-mir-455&WithFlank&9|+|114209428|114209535@22.44.23&[MIMAT0003150&hsa-miR-455-5p&offsets|0|+1;m-232&9|+|114209449|114209470&offsets|0|+1]|TATGTGCCTTTGGACTACATCGT</td><td>>MI0003513|hsa-mir-455&WithFlank&9|+|114209428|114209535</td><td>22</td><td>44</td><td>23</td><td>TATGTGCCTTTGGACTACATCGT</td><td>1299.1</td><td>-4.3</td><td>0.7</td><td>-5.8</td><td>8.3E-09</td><td>1.9E-07</td><td>BT20/MB231</td><td>MITO</td></tr>
<tr><td>>hg38_21_+_8433172_8446622_RNA45SN1_45S_50ntFlanks@6698.6720.23|AGCTAGCTGCGAGAATTAATGTG</td><td>>hg38_21_+_8433172_8446622_RNA45SN1_45S_50ntFlanks</td><td>6698</td><td>6720</td><td>23</td><td>AGCTAGCTGCGAGAATTAATGTG</td><td>6652.9</td><td>-3.4</td><td>0.6</td><td>-5.4</td><td>7.9E-08</td><td>1.9E-07</td><td>BT20/MB468</td><td>MITO</td></tr>
<tr><td>>hg38_21_+_8433172_8446622_RNA45SN1_45S_50ntFlanks@9554.9595.42|CGTAGCGGTCCTGACGTGCAAATCGGTCGTCCGACCTGGGTA</td><td>>hg38_21_+_8433172_8446622_RNA45SN1_45S_50ntFlanks</td><td>9554</td><td>9595</td><td>42</td><td>CGTAGCGGTCCTGACGTGCAAATCGGTCGTCCGACCTGGGTA</td><td>2011.8</td><td>-7.2</td><td>1.4</td><td>-5.3</td><td>1.1E-07</td><td>1.9E-07</td><td>MB231/MB468</td><td>MITO</td></tr>
<tr><td>>hg38_21_+_8433172_8446622_RNA45SN1_45S_50ntFlanks@12046.12070.25|TGACCCGGTGAGGCGGGGGGGCGAG</td><td>>hg38_21_+_8433172_8446622_RNA45SN1_45S_50ntFlanks</td><td>12046</td><td>12070</td><td>25</td><td>TGACCCGGTGAGGCGGGGGGGCGAG</td><td>83.3</td><td>-9.5</td><td>1.7</td><td>-5.6</td><td>2.0E-08</td><td>1.9E-07</td><td>BT20/MB468</td><td>MP</td></tr>
<tr><td>>MI0000342|hsa-mir-200b&WithFlank&1|+|1167098|1167204@63.85.23&[MIMAT0000318&hsa-miR-200b-3p&offsets|0|+1;m-93&1|+|1167160|1167182&offsets|0|0]|TAATACTGCCTGGTAATGATGAC</td><td>>MI0000342|hsa-mir-200b&WithFlank&1|+|1167098|1167204</td><td>63</td><td>85</td><td>23</td><td>TAATACTGCCTGGTAATGATGAC</td><td>1949.2</td><td>-3.3</td><td>0.6</td><td>-5.3</td><td>1.0E-07</td><td>1.9E-07</td><td>MB231/MB468</td><td>CYTO</td></tr>
<tr><td>>hg38_21_+_8433172_8446622_RNA45SN1_45S_50ntFlanks@7972.8005.34|AGACGCGACCTCAGATCAGACGTGGCGACCCGCT</td><td>>hg38_21_+_8433172_8446622_RNA45SN1_45S_50ntFlanks</td><td>7972</td><td>8005</td><td>34</td><td>AGACGCGACCTCAGATCAGACGTGGCGACCCGCT</td><td>283.9</td><td>5.8</td><td>1.1</td><td>5.3</td><td>1.2E-07</td><td>1.9E-07</td><td>BT20/MB468</td><td>NUC</td></tr>
<tr><td>>hg38_21_+_8433172_8446622_RNA45SN1_45S_50ntFlanks@10914.10946.33|CGCCGCGGCTGGACGAGGCGCCGCCGCCCCCCC</td><td>>hg38_21_+_8433172_8446622_RNA45SN1_45S_50ntFlanks</td><td>10914</td><td>10946</td><td>33</td><td>CGCCGCGGCTGGACGAGGCGCCGCCGCCCCCCC</td><td>457.3</td><td>-7.2</td><td>1.3</td><td>-5.4</td><td>7.9E-08</td><td>1.9E-07</td><td>BT20/MB468</td><td>MITO</td></tr>
<tr><td>>MI0000263|hsa-mir-7-1&WithFlank&9|-|83969742|83969863@30.54.25&[MIMAT0000252_2&hsa-miR-7-5p&offsets|0|+1;m-401&9|-|83969811|83969834&offsets|0|+1]|TGGAAGACTAGTGATTTTGTTGTTT</td><td>>MI0000263|hsa-mir-7-1&WithFlank&9|-|83969742|83969863</td><td>30</td><td>54</td><td>25</td><td>TGGAAGACTAGTGATTTTGTTGTTT</td><td>1903.8</td><td>4.2</td><td>0.8</td><td>5.3</td><td>9.4E-08</td><td>1.9E-07</td><td>MB231/MB468</td><td>MP</td></tr>
<tr><td>>hg38_21_+_8433172_8446622_RNA45SN1_45S_50ntFlanks@5078.5115.38|CGATAACGAACGAGACTCTGGCATGCTAACTAGTTACG</td><td>>hg38_21_+_8433172_8446622_RNA45SN1_45S_50ntFlanks</td><td>5078</td><td>5115</td><td>38</td><td>CGATAACGAACGAGACTCTGGCATGCTAACTAGTTACG</td><td>321.4</td><td>-3.8</td><td>0.7</td><td>-5.3</td><td>1.1E-07</td><td>1.9E-07</td><td>MB231/MB468</td><td>MITO</td></tr>
<tr><td>>hg38_21_+_8433172_8446622_RNA45SN1_45S_50ntFlanks@8359.8379.21|AAGAACTTTGAAGAGAGAGTT</td><td>>hg38_21_+_8433172_8446622_RNA45SN1_45S_50ntFlanks</td><td>8359</td><td>8379</td><td>21</td><td>AAGAACTTTGAAGAGAGAGTT</td><td>507.4</td><td>-4.3</td><td>0.8</td><td>-5.3</td><td>1.1E-07</td><td>1.9E-07</td><td>MB231/MB468</td><td>NUC</td></tr>
<tr><td>>hg38_21_+_8433172_8446622_RNA45SN1_45S_50ntFlanks@10361.10384.24|GAAGGCCGAAGTGGAGAAGGGTTC</td><td>>hg38_21_+_8433172_8446622_RNA45SN1_45S_50ntFlanks</td><td>10361</td><td>10384</td><td>24</td><td>GAAGGCCGAAGTGGAGAAGGGTTC</td><td>152.2</td><td>-6.1</td><td>1.1</td><td>-5.3</td><td>1.1E-07</td><td>1.9E-07</td><td>MB231/MB468</td><td>NUC</td></tr>
<tr><td>>hg38_21_+_8433172_8446622_RNA45SN1_45S_50ntFlanks@10807.10840.34|ACAATGTAGGTAAGGGAAGTCGGCAAGCCGGATC</td><td>>hg38_21_+_8433172_8446622_RNA45SN1_45S_50ntFlanks</td><td>10807</td><td>10840</td><td>34</td><td>ACAATGTAGGTAAGGGAAGTCGGCAAGCCGGATC</td><td>642.9</td><td>-8.3</td><td>1.6</td><td>-5.3</td><td>1.2E-07</td><td>1.9E-07</td><td>BT20/MB468</td><td>NUC</td></tr>
<tr><td>>hg38_21_+_8433172_8446622_RNA45SN1_45S_50ntFlanks@6773.6793.21|TTCCTCCCGGGGCTACGCCTG</td><td>>hg38_21_+_8433172_8446622_RNA45SN1_45S_50ntFlanks</td><td>6773</td><td>6793</td><td>21</td><td>TTCCTCCCGGGGCTACGCCTG</td><td>390.2</td><td>-6.3</td><td>1.2</td><td>-5.3</td><td>1.0E-07</td><td>1.9E-07</td><td>MB231/MB468</td><td>CYTO</td></tr>
<tr><td>>hg38_21_+_8433172_8446622_RNA45SN1_45S_50ntFlanks@11564.11587.24|CCTAGCAGCCGACTTAGAACTGGT</td><td>>hg38_21_+_8433172_8446622_RNA45SN1_45S_50ntFlanks</td><td>11564</td><td>11587</td><td>24</td><td>CCTAGCAGCCGACTTAGAACTGGT</td><td>304.9</td><td>-3.9</td><td>0.7</td><td>-5.3</td><td>1.1E-07</td><td>1.9E-07</td><td>MB231/MB468</td><td>MITO</td></tr>
<tr><td>>hg38_21_+_8433172_8446622_RNA45SN1_45S_50ntFlanks@12420.12455.36|ATGTCGGCTCTTCCTATCATTGTGAAGCAGAATTCA</td><td>>hg38_21_+_8433172_8446622_RNA45SN1_45S_50ntFlanks</td><td>12420</td><td>12455</td><td>36</td><td>ATGTCGGCTCTTCCTATCATTGTGAAGCAGAATTCA</td><td>3516.0</td><td>-4.1</td><td>0.8</td><td>-5.3</td><td>1.0E-07</td><td>1.9E-07</td><td>MB231/MB468</td><td>CYTO</td></tr>
<tr><td>>MI0000478|hsa-mir-149&WithFlank&2|+|240455995|240456095@21.42.22(+1U)&[MIMAT0000450&hsa-miR-149-5p&offsets|0|-1(+1U);m-195&2|+|240456015|240456037&offsets|0|-1(+1U)]|TCTGGCTCCGTGTCTTCACTCCT</td><td>>MI0000478|hsa-mir-149&WithFlank&2|+|240455995|240456095</td><td>21</td><td>42</td><td>22(+1U)</td><td>TCTGGCTCCGTGTCTTCACTCCT</td><td>347.3</td><td>9.1</td><td>1.6</td><td>5.6</td><td>2.0E-08</td><td>1.9E-07</td><td>BT20/MB468</td><td>MP</td></tr>
<tr><td>>hg38_21_+_8433172_8446622_RNA45SN1_45S_50ntFlanks@12354.12376.23|TTACCACAGGGATAACTGGCTTG</td><td>>hg38_21_+_8433172_8446622_RNA45SN1_45S_50ntFlanks</td><td>12354</td><td>12376</td><td>23</td><td>TTACCACAGGGATAACTGGCTTG</td><td>75.5</td><td>-9.3</td><td>1.7</td><td>-5.6</td><td>2.0E-08</td><td>1.9E-07</td><td>BT20/MB468</td><td>MP</td></tr>
<tr><td>>hg38_21_+_8433172_8446622_RNA45SN1_45S_50ntFlanks@3876.3898.23|TACATGCCGACGGGCGCTGACCC</td><td>>hg38_21_+_8433172_8446622_RNA45SN1_45S_50ntFlanks</td><td>3876</td><td>3898</td><td>23</td><td>TACATGCCGACGGGCGCTGACCC</td><td>1695.6</td><td>-3.6</td><td>0.7</td><td>-5.3</td><td>1.0E-07</td><td>1.9E-07</td><td>MB231/MB468</td><td>CYTO</td></tr>
<tr><td>>hg38_1_-_228634819_228635039_RNA5S12_5S_50ntFlanks@140.171.32|ACCGCCTGGGAATACCGGGTGCTGTAGGCTTT</td><td>>hg38_1_-_228634819_228635039_RNA5S12_5S_50ntFlanks</td><td>140</td><td>171</td><td>32</td><td>ACCGCCTGGGAATACCGGGTGCTGTAGGCTTT</td><td>7718.2</td><td>4.6</td><td>0.9</td><td>5.3</td><td>1.0E-07</td><td>1.9E-07</td><td>MB231/MB468</td><td>CYTO</td></tr>
<tr><td>>hg38_21_+_8433172_8446622_RNA45SN1_45S_50ntFlanks@10686.10706.21|GCGGCGTCCGGTGAGCTCTCG</td><td>>hg38_21_+_8433172_8446622_RNA45SN1_45S_50ntFlanks</td><td>10686</td><td>10706</td><td>21</td><td>GCGGCGTCCGGTGAGCTCTCG</td><td>1666.3</td><td>-6.7</td><td>1.3</td><td>-5.3</td><td>1.1E-07</td><td>1.9E-07</td><td>MB231/MB468</td><td>NUC</td></tr>
<tr><td>>MI0000433|hsa-let-7g&WithFlank&3|-|52268272|52268367@11.31.21(+1A)&[MIMAT0000414&hsa-let-7g-5p&offsets|0|-1(+1A);m-58&3|-|52268336|52268357&offsets|0|-1(+1A)]|TGAGGTAGTAGTTTGTACAGTA</td><td>>MI0000433|hsa-let-7g&WithFlank&3|-|52268272|52268367</td><td>11</td><td>31</td><td>21(+1A)</td><td>TGAGGTAGTAGTTTGTACAGTA</td><td>1003.9</td><td>5.0</td><td>0.9</td><td>5.3</td><td>1.0E-07</td><td>1.9E-07</td><td>MB231/MB468</td><td>CYTO</td></tr>
<tr><td>>hg38_21_+_8433172_8446622_RNA45SN1_45S_50ntFlanks@4453.4473.21|TCGGCGCCCCCTCGATGCTCT</td><td>>hg38_21_+_8433172_8446622_RNA45SN1_45S_50ntFlanks</td><td>4453</td><td>4473</td><td>21</td><td>TCGGCGCCCCCTCGATGCTCT</td><td>50.5</td><td>8.9</td><td>1.7</td><td>5.3</td><td>9.5E-08</td><td>1.9E-07</td><td>MB231/MB468</td><td>MP</td></tr>
<tr><td>>hg38_21_+_8433172_8446622_RNA45SN1_45S_50ntFlanks@12635.12661.27|AAGCTACCATCTGTGGGATTATGACTG</td><td>>hg38_21_+_8433172_8446622_RNA45SN1_45S_50ntFlanks</td><td>12635</td><td>12661</td><td>27</td><td>AAGCTACCATCTGTGGGATTATGACTG</td><td>911.3</td><td>-3.2</td><td>0.6</td><td>-5.3</td><td>1.0E-07</td><td>1.9E-07</td><td>MB231/MB468</td><td>CYTO</td></tr>
<tr><td>>hg38_21_+_8433172_8446622_RNA45SN1_45S_50ntFlanks@9534.9551.18|AGGAAACTCTGGTGGAGG</td><td>>hg38_21_+_8433172_8446622_RNA45SN1_45S_50ntFlanks</td><td>9534</td><td>9551</td><td>18</td><td>AGGAAACTCTGGTGGAGG</td><td>5568.3</td><td>-5.3</td><td>1.0</td><td>-5.3</td><td>9.0E-08</td><td>1.9E-07</td><td>BT20/MB468</td><td>CYTO</td></tr>
<tr><td>>hg38_21_+_8433172_8446622_RNA45SN1_45S_50ntFlanks@3837.3871.35|CCTCTCCTACTTGGATAACTGTGGTAATTCTAGAG</td><td>>hg38_21_+_8433172_8446622_RNA45SN1_45S_50ntFlanks</td><td>3837</td><td>3871</td><td>35</td><td>CCTCTCCTACTTGGATAACTGTGGTAATTCTAGAG</td><td>952.8</td><td>-8.5</td><td>1.6</td><td>-5.3</td><td>1.2E-07</td><td>1.9E-07</td><td>BT20/MB468</td><td>NUC</td></tr>
<tr><td>>hg38_21_+_8433172_8446622_RNA45SN1_45S_50ntFlanks@11235.11268.34|CTTCCCGTGGATCGCCCCAGCTGCGGCGGGCGTC</td><td>>hg38_21_+_8433172_8446622_RNA45SN1_45S_50ntFlanks</td><td>11235</td><td>11268</td><td>34</td><td>CTTCCCGTGGATCGCCCCAGCTGCGGCGGGCGTC</td><td>457.2</td><td>-7.4</td><td>1.4</td><td>-5.3</td><td>1.1E-07</td><td>1.9E-07</td><td>MB231/MB468</td><td>NUC</td></tr>
<tr><td>>hg38_21_+_8433172_8446622_RNA45SN1_45S_50ntFlanks@9574.9601.28|AATCGGTCGTCCGACCTGGGTATAGGGG</td><td>>hg38_21_+_8433172_8446622_RNA45SN1_45S_50ntFlanks</td><td>9574</td><td>9601</td><td>28</td><td>AATCGGTCGTCCGACCTGGGTATAGGGG</td><td>1114.6</td><td>-3.3</td><td>0.6</td><td>-5.3</td><td>9.6E-08</td><td>1.9E-07</td><td>MB231/MB468</td><td>MP</td></tr>
<tr><td>>MI0003560|hsa-mir-92b&WithFlank&1|+|155195171|155195278@67.87.21(+1A)&[MIMAT0003218&hsa-miR-92b-3p&offsets|0|-1(+1A);m-115&1|+|155195237|155195258&offsets|0|-1(+1A)]|TATTGCACTCGTCCCGGCCTCA</td><td>>MI0003560|hsa-mir-92b&WithFlank&1|+|155195171|155195278</td><td>67</td><td>87</td><td>21(+1A)</td><td>TATTGCACTCGTCCCGGCCTCA</td><td>286.7</td><td>3.4</td><td>0.6</td><td>5.3</td><td>1.2E-07</td><td>1.9E-07</td><td>BT20/MB468</td><td>NUC</td></tr>
<tr><td>>hg38_21_+_8433172_8446622_RNA45SN1_45S_50ntFlanks@5084.5112.29|CGAACGAGACTCTGGCATGCTAACTAGTT</td><td>>hg38_21_+_8433172_8446622_RNA45SN1_45S_50ntFlanks</td><td>5084</td><td>5112</td><td>29</td><td>CGAACGAGACTCTGGCATGCTAACTAGTT</td><td>1963.5</td><td>-8.3</td><td>1.6</td><td>-5.3</td><td>1.2E-07</td><td>1.9E-07</td><td>BT20/MB468</td><td>NUC</td></tr>
<tr><td>>MI0000102|hsa-mir-100&WithFlank&11|-|122152223|122152314@20.40.21&[MIMAT0000098&hsa-miR-100-5p&offsets|+1|0;m-27&11|-|122152276|122152296&offsets|+1|+1]|ACCCGTAGATCCGAACTTGTG</td><td>>MI0000102|hsa-mir-100&WithFlank&11|-|122152223|122152314</td><td>20</td><td>40</td><td>21</td><td>ACCCGTAGATCCGAACTTGTG</td><td>1152.2</td><td>-7.9</td><td>1.4</td><td>-5.7</td><td>1.1E-08</td><td>1.9E-07</td><td>BT20/MB231</td><td>NUC</td></tr>
<tr><td>>hg38_21_+_8433172_8446622_RNA45SN1_45S_50ntFlanks@5487.5521.35|CGCTGAGAAGACGGTCGAACTTGACTATCTAGAGG</td><td>>hg38_21_+_8433172_8446622_RNA45SN1_45S_50ntFlanks</td><td>5487</td><td>5521</td><td>35</td><td>CGCTGAGAAGACGGTCGAACTTGACTATCTAGAGG</td><td>371.6</td><td>-6.0</td><td>1.1</td><td>-5.3</td><td>9.6E-08</td><td>1.9E-07</td><td>MB231/MB468</td><td>MP</td></tr>
<tr><td>>hg38_21_+_8433172_8446622_RNA45SN1_45S_50ntFlanks@3845.3879.35|ACTTGGATAACTGTGGTAATTCTAGAGCTAATACA</td><td>>hg38_21_+_8433172_8446622_RNA45SN1_45S_50ntFlanks</td><td>3845</td><td>3879</td><td>35</td><td>ACTTGGATAACTGTGGTAATTCTAGAGCTAATACA</td><td>282.5</td><td>3.8</td><td>0.7</td><td>5.3</td><td>1.1E-07</td><td>1.9E-07</td><td>MB231/MB468</td><td>NUC</td></tr>
<tr><td>>MI0000078|hsa-mir-22&WithFlank&17|-|1713897|1713993@21.41.21&[MIMAT0004495&hsa-miR-22-5p&offsets|0|-1;m-217&17|-|1713953|1713973&offsets|0|0]|AGTTCTTCAGTGGCAAGCTTT</td><td>>MI0000078|hsa-mir-22&WithFlank&17|-|1713897|1713993</td><td>21</td><td>41</td><td>21</td><td>AGTTCTTCAGTGGCAAGCTTT</td><td>597.4</td><td>6.6</td><td>1.2</td><td>5.3</td><td>1.1E-07</td><td>1.9E-07</td><td>MB231/MB468</td><td>MITO</td></tr>
<tr><td>>hg38_21_+_8433172_8446622_RNA45SN1_45S_50ntFlanks@8425.8469.45|CAGTCCGCCCGGAGGATTCAACCCGGCGGCGGGTCCGGCCGTGTC</td><td>>hg38_21_+_8433172_8446622_RNA45SN1_45S_50ntFlanks</td><td>8425</td><td>8469</td><td>45</td><td>CAGTCCGCCCGGAGGATTCAACCCGGCGGCGGGTCCGGCCGTGTC</td><td>91.6</td><td>-10.4</td><td>1.9</td><td>-5.3</td><td>9.6E-08</td><td>1.9E-07</td><td>MB231/MB468</td><td>MP</td></tr>
<tr><td>>MI0000071|hsa-mir-17&WithFlank&13|+|91350599|91350694@20.40.21&[MIMAT0000070&hsa-miR-17-5p&offsets|0|-2;m-88&13|+|91350618|91350640&offsets|0|-2]|CAAAGTGCTTACAGTGCAGGT</td><td>>MI0000071|hsa-mir-17&WithFlank&13|+|91350599|91350694</td><td>20</td><td>40</td><td>21</td><td>CAAAGTGCTTACAGTGCAGGT</td><td>1298.0</td><td>5.5</td><td>1.0</td><td>5.3</td><td>1.1E-07</td><td>1.9E-07</td><td>MB231/MB468</td><td>MITO</td></tr>
<tr><td>>hg38_21_+_8433172_8446622_RNA45SN1_45S_50ntFlanks@11875.11897.23|GAAGACCCTGTTGAGCTTGACTC</td><td>>hg38_21_+_8433172_8446622_RNA45SN1_45S_50ntFlanks</td><td>11875</td><td>11897</td><td>23</td><td>GAAGACCCTGTTGAGCTTGACTC</td><td>486.5</td><td>-6.1</td><td>1.2</td><td>-5.3</td><td>1.2E-07</td><td>1.9E-07</td><td>BT20/MB468</td><td>NUC</td></tr>
<tr><td>>hg38_21_+_8433172_8446622_RNA45SN1_45S_50ntFlanks@9574.9596.23|AATCGGTCGTCCGACCTGGGTAT</td><td>>hg38_21_+_8433172_8446622_RNA45SN1_45S_50ntFlanks</td><td>9574</td><td>9596</td><td>23</td><td>AATCGGTCGTCCGACCTGGGTAT</td><td>1838.6</td><td>-9.0</td><td>1.7</td><td>-5.3</td><td>1.2E-07</td><td>1.9E-07</td><td>BT20/MB468</td><td>NUC</td></tr>
<tr><td>>MI0000482|hsa-mir-185&WithFlank&22|+|20033133|20033226@21.43.23&[MIMAT0000455&hsa-miR-185-5p&offsets|0|+1;m-145&22|+|20033153|20033174&offsets|0|+1]|TGGAGAGAAAGGCAGTTCCTGAT</td><td>>MI0000482|hsa-mir-185&WithFlank&22|+|20033133|20033226</td><td>21</td><td>43</td><td>23</td><td>TGGAGAGAAAGGCAGTTCCTGAT</td><td>120.9</td><td>1.9</td><td>0.4</td><td>5.3</td><td>1.2E-07</td><td>1.9E-07</td><td>BT20/MB468</td><td>NUC</td></tr>
<tr><td>>hg38_21_+_8433172_8446622_RNA45SN1_45S_50ntFlanks@11670.11706.37|CCCAGTGCTCTGAATGTCAAAGTGAAGAAATTCAATG</td><td>>hg38_21_+_8433172_8446622_RNA45SN1_45S_50ntFlanks</td><td>11670</td><td>11706</td><td>37</td><td>CCCAGTGCTCTGAATGTCAAAGTGAAGAAATTCAATG</td><td>1127.6</td><td>-7.0</td><td>1.3</td><td>-5.3</td><td>1.2E-07</td><td>1.9E-07</td><td>BT20/MB468</td><td>NUC</td></tr>
</table>

<table>
<tr><td>>hg38_21_+_8433172_8446622_RNA45SN1_45S_50ntFlanks@7982.8005.24|TCAGATCAGACGTGGCGACCCGCT</td><td>>hg38_21_+_8433172_8446622_RNA45SN1_45S_50ntFlanks</td><td>7982</td><td>8005</td><td>24</td><td>TCAGATCAGACGTGGCGACCCGCT</td><td>213.8</td><td>-3.1</td><td>0.6</td><td>-5.3</td><td>1.1E-07</td><td>1.9E-07</td><td>MB231/MB468</td><td>NUC</td></tr>
<tr><td>>hg38_21_+_8433172_8446622_RNA45SN1_45S_50ntFlanks@11613.11660.48|AATTAAAACAAAGCATCGCGAAGGCCCGCGGCGGGTGTTGACGCGATG</td><td>>hg38_21_+_8433172_8446622_RNA45SN1_45S_50ntFlanks</td><td>11613</td><td>11660</td><td>48</td><td>AATTAAAACAAAGCATCGCGAAGGCCCGCGGCGGGTGTTGACGCGATG</td><td>151.2</td><td>-9.3</td><td>1.7</td><td>-5.4</td><td>8.1E-08</td><td>1.9E-07</td><td>BT20/MB468</td><td>MITO</td></tr>
<tr><td>>MI0000272|hsa-mir-182&WithFlank&7|-|129770377|129770498@29.53.25&[MIMAT0000259&hsa-miR-182-5p&offsets|0|+1;m-22&7|-|129770449|129770470&offsets|0|+3]|TTTGGCAATGGTAGAACTCACACTG</td><td>>MI0000272|hsa-mir-182&WithFlank&7|-|129770377|129770498</td><td>29</td><td>53</td><td>25</td><td>TTTGGCAATGGTAGAACTCACACTG</td><td>9630.8</td><td>2.8</td><td>0.5</td><td>5.7</td><td>1.1E-08</td><td>1.9E-07</td><td>BT20/MB231</td><td>NUC</td></tr>
<tr><td>>MI0000810|hsa-mir-135b&WithFlank&1|-|205448296|205448404@22.44.23&[MIMAT0000758&hsa-miR-135b-5p&offsets|0|0;m-321&1|-|205448361|205448383&offsets|0|0]|TATGGCTTTTCATTCCTATGTGA</td><td>>MI0000810|hsa-mir-135b&WithFlank&1|-|205448296|205448404</td><td>22</td><td>44</td><td>23</td><td>TATGGCTTTTCATTCCTATGTGA</td><td>1212.2</td><td>5.3</td><td>1.0</td><td>5.4</td><td>8.1E-08</td><td>1.9E-07</td><td>BT20/MB468</td><td>MITO</td></tr>
<tr><td>>hg38_21_+_8433172_8446622_RNA45SN1_45S_50ntFlanks@10468.10497.30|TTGCCCTCGGCCGATCGAAAGGGAGTCGGG</td><td>>hg38_21_+_8433172_8446622_RNA45SN1_45S_50ntFlanks</td><td>10468</td><td>10497</td><td>30</td><td>TTGCCCTCGGCCGATCGAAAGGGAGTCGGG</td><td>264.3</td><td>-5.3</td><td>1.0</td><td>-5.3</td><td>1.1E-07</td><td>1.9E-07</td><td>MB231/MB468</td><td>NUC</td></tr>
<tr><td>>hg38_21_+_8433172_8446622_RNA45SN1_45S_50ntFlanks@9533.9567.35|GAGGAAACTCTGGTGGAGGTCCGTAGCGGTCCTGA</td><td>>hg38_21_+_8433172_8446622_RNA45SN1_45S_50ntFlanks</td><td>9533</td><td>9567</td><td>35</td><td>GAGGAAACTCTGGTGGAGGTCCGTAGCGGTCCTGA</td><td>584.6</td><td>-7.6</td><td>1.4</td><td>-5.3</td><td>1.1E-07</td><td>1.9E-07</td><td>MB231/MB468</td><td>NUC</td></tr>
<tr><td>>hg38_21_+_8433172_8446622_RNA45SN1_45S_50ntFlanks@12994.13013.20|CCTCGCTGCGATCTATTGAA</td><td>>hg38_21_+_8433172_8446622_RNA45SN1_45S_50ntFlanks</td><td>12994</td><td>13013</td><td>20</td><td>CCTCGCTGCGATCTATTGAA</td><td>582.7</td><td>-4.0</td><td>0.7</td><td>-5.4</td><td>8.1E-08</td><td>1.9E-07</td><td>BT20/MB468</td><td>MITO</td></tr>
<tr><td>>MI0000272|hsa-mir-182&WithFlank&7|-|129770377|129770498@29.52.24(+1U)&[MIMAT0000259&hsa-miR-182-5p&offsets|0|0(+1U);m-22&7|-|129770449|129770470&offsets|0|+2(+1U)]|TTTGGCAATGGTAGAACTCACACTT</td><td>>MI0000272|hsa-mir-182&WithFlank&7|-|129770377|129770498</td><td>29</td><td>52</td><td>24(+1U)</td><td>TTTGGCAATGGTAGAACTCACACTT</td><td>286.0</td><td>3.5</td><td>0.6</td><td>5.7</td><td>1.2E-08</td><td>1.9E-07</td><td>BT20/MB231</td><td>NUC</td></tr>
<tr><td>>hg38_21_+_8433172_8446622_RNA45SN1_45S_50ntFlanks@9622.9656.35|CTAGTAGCTGGTTCCCTCCGAAGTTTCCCTCAGGA</td><td>>hg38_21_+_8433172_8446622_RNA45SN1_45S_50ntFlanks</td><td>9622</td><td>9656</td><td>35</td><td>CTAGTAGCTGGTTCCCTCCGAAGTTTCCCTCAGGA</td><td>51.9</td><td>-9.6</td><td>1.8</td><td>-5.3</td><td>9.6E-08</td><td>1.9E-07</td><td>MB231/MB468</td><td>MP</td></tr>
<tr><td>>hg38_21_+_8433172_8446622_RNA45SN1_45S_50ntFlanks@8359.8379.21|AAGAACTTTGAAGAGAGAGTT</td><td>>hg38_21_+_8433172_8446622_RNA45SN1_45S_50ntFlanks</td><td>8359</td><td>8379</td><td>21</td><td>AAGAACTTTGAAGAGAGAGTT</td><td>505.8</td><td>-5.5</td><td>1.0</td><td>-5.3</td><td>1.0E-07</td><td>1.9E-07</td><td>MB231/MB468</td><td>CYTO</td></tr>
<tr><td>>hg38_21_+_8433172_8446622_RNA45SN1_45S_50ntFlanks@4773.4793.21|TAGTTCCGACCATAAACGATG</td><td>>hg38_21_+_8433172_8446622_RNA45SN1_45S_50ntFlanks</td><td>4773</td><td>4793</td><td>21</td><td>TAGTTCCGACCATAAACGATG</td><td>563.7</td><td>-3.6</td><td>0.7</td><td>-5.4</td><td>8.1E-08</td><td>1.9E-07</td><td>BT20/MB468</td><td>MITO</td></tr>
<tr><td>>hg38_21_+_8433172_8446622_RNA45SN1_45S_50ntFlanks@3811.3829.19|AGTTATGGTTCCTTTGGTC</td><td>>hg38_21_+_8433172_8446622_RNA45SN1_45S_50ntFlanks</td><td>3811</td><td>3829</td><td>19</td><td>AGTTATGGTTCCTTTGGTC</td><td>451.2</td><td>-6.4</td><td>1.2</td><td>-5.3</td><td>1.2E-07</td><td>2.0E-07</td><td>BT20/MB468</td><td>NUC</td></tr>
<tr><td>>hg38_21_+_8433172_8446622_RNA45SN1_45S_50ntFlanks@476.507.32|TTCGTGATCGATGTGGTGACGTCGTGCTCTCC</td><td>>hg38_21_+_8433172_8446622_RNA45SN1_45S_50ntFlanks</td><td>476</td><td>507</td><td>32</td><td>TTCGTGATCGATGTGGTGACGTCGTGCTCTCC</td><td>7145.3</td><td>3.7</td><td>0.7</td><td>5.3</td><td>1.1E-07</td><td>2.0E-07</td><td>MB231/MB468</td><td>NUC</td></tr>
<tr><td>>hg38_21_+_8433172_8446622_RNA45SN1_45S_50ntFlanks@8182.8209.28|ATCGAGGCCCAGCCCGTGGACGGTGTGA</td><td>>hg38_21_+_8433172_8446622_RNA45SN1_45S_50ntFlanks</td><td>8182</td><td>8209</td><td>28</td><td>ATCGAGGCCCAGCCCGTGGACGGTGTGA</td><td>102.1</td><td>-9.8</td><td>1.7</td><td>-5.6</td><td>2.0E-08</td><td>2.0E-07</td><td>BT20/MB468</td><td>MP</td></tr>
<tr><td>>MI0000088|hsa-mir-30a&WithFlank&6|-|71403545|71403627@12.35.24(+1A)&[MIMAT0000087&hsa-miR-30a-5p&offsets|0|+2(+1A);m-15&6|-|71403595|71403616&offsets|0|+2(+1A)]|TGTAAACATCCTCGACTGGAAGCTA</td><td>>MI0000088|hsa-mir-30a&WithFlank&6|-|71403545|71403627</td><td>12</td><td>35</td><td>24(+1A)</td><td>TGTAAACATCCTCGACTGGAAGCTA</td><td>1061.1</td><td>8.2</td><td>1.5</td><td>5.3</td><td>1.1E-07</td><td>2.0E-07</td><td>MB231/MB468</td><td>MITO</td></tr>
<tr><td>>hg38_21_+_8433172_8446622_RNA45SN1_45S_50ntFlanks@12957.12988.32|CTTCTGGGTCGGGGTTTCGTACGTAGCAGAGC</td><td>>hg38_21_+_8433172_8446622_RNA45SN1_45S_50ntFlanks</td><td>12957</td><td>12988</td><td>32</td><td>CTTCTGGGTCGGGGTTTCGTACGTAGCAGAGC</td><td>441.4</td><td>-5.6</td><td>1.0</td><td>-5.3</td><td>1.1E-07</td><td>2.0E-07</td><td>MB231/MB468</td><td>NUC</td></tr>
<tr><td>>hg38_21_+_8433172_8446622_RNA45SN1_45S_50ntFlanks@3876.3899.24|TACATGCCGACGGGCGCTGACCCC</td><td>>hg38_21_+_8433172_8446622_RNA45SN1_45S_50ntFlanks</td><td>3876</td><td>3899</td><td>24</td><td>TACATGCCGACGGGCGCTGACCCC</td><td>372.7</td><td>-6.1</td><td>1.1</td><td>-5.3</td><td>1.2E-07</td><td>2.0E-07</td><td>BT20/MB468</td><td>NUC</td></tr>
<tr><td>>hg38_21_+_8433172_8446622_RNA45SN1_45S_50ntFlanks@3854.3881.28|ACTGTGGTAATTCTAGAGCTAATACATG</td><td>>hg38_21_+_8433172_8446622_RNA45SN1_45S_50ntFlanks</td><td>3854</td><td>3881</td><td>28</td><td>ACTGTGGTAATTCTAGAGCTAATACATG</td><td>818.1</td><td>-7.3</td><td>1.4</td><td>-5.4</td><td>8.2E-08</td><td>2.0E-07</td><td>BT20/MB468</td><td>MITO</td></tr>
<tr><td>>hg38_21_+_8433172_8446622_RNA45SN1_45S_50ntFlanks@10510.10533.24|AATCCGGAGTGGCGGAGATGGGCG</td><td>>hg38_21_+_8433172_8446622_RNA45SN1_45S_50ntFlanks</td><td>10510</td><td>10533</td><td>24</td><td>AATCCGGAGTGGCGGAGATGGGCG</td><td>811.5</td><td>-5.9</td><td>1.1</td><td>-5.3</td><td>1.1E-07</td><td>2.0E-07</td><td>MB231/MB468</td><td>CYTO</td></tr>
<tr><td>>MI0000088|hsa-mir-30a&WithFlank&6|-|71403545|71403627@13.31.19&[MIMAT0000087&hsa-miR-30a-5p&offsets|+1|-2;m-15&6|-|71403595|71403616&offsets|+1|-2]|GTAAACATCCTCGACTGGA</td><td>>MI0000088|hsa-mir-30a&WithFlank&6|-|71403545|71403627</td><td>13</td><td>31</td><td>19</td><td>GTAAACATCCTCGACTGGA</td><td>62.1</td><td>9.2</td><td>1.7</td><td>5.3</td><td>9.7E-08</td><td>2.0E-07</td><td>MB231/MB468</td><td>MP</td></tr>
<tr><td>>hg38_21_+_8433172_8446622_RNA45SN1_45S_50ntFlanks@12912.12951.40|CACGTTCGTGGGGAACCTGGCGCTAAACCATTCGTAGACG</td><td>>hg38_21_+_8433172_8446622_RNA45SN1_45S_50ntFlanks</td><td>12912</td><td>12951</td><td>40</td><td>CACGTTCGTGGGGAACCTGGCGCTAAACCATTCGTAGACG</td><td>68.0</td><td>-10.0</td><td>1.9</td><td>-5.3</td><td>9.8E-08</td><td>2.0E-07</td><td>MB231/MB468</td><td>MP</td></tr>
<tr><td>>hg38_21_+_8433172_8446622_RNA45SN1_45S_50ntFlanks@8355.8392.38|TGAAAAGAACTTTGAAGAGAGAGTTCAAGAGGGCGTGA</td><td>>hg38_21_+_8433172_8446622_RNA45SN1_45S_50ntFlanks</td><td>8355</td><td>8392</td><td>38</td><td>TGAAAAGAACTTTGAAGAGAGAGTTCAAGAGGGCGTGA</td><td>69.0</td><td>9.9</td><td>1.8</td><td>5.6</td><td>2.1E-08</td><td>2.0E-07</td><td>BT20/MB468</td><td>MP</td></tr>
<tr><td>>MI0000088|hsa-mir-30a&WithFlank&6|-|71403545|71403627@12.34.23(+1A)&[MIMAT0000087&hsa-miR-30a-5p&offsets|0|+1(+1A);m-15&6|-|71403595|71403616&offsets|0|+1(+1A)]|TGTAAACATCCTCGACTGGAAGCA</td><td>>MI0000088|hsa-mir-30a&WithFlank&6|-|71403545|71403627</td><td>12</td><td>34</td><td>23(+1A)</td><td>TGTAAACATCCTCGACTGGAAGCA</td><td>26741.4</td><td>4.3</td><td>0.8</td><td>5.3</td><td>9.8E-08</td><td>2.0E-07</td><td>MB231/MB468</td><td>MP</td></tr>
<tr><td>>hg38_21_+_8433172_8446622_RNA45SN1_45S_50ntFlanks@11572.11597.26|CCGACTTAGAACTGGTGCGGACCAGG</td><td>>hg38_21_+_8433172_8446622_RNA45SN1_45S_50ntFlanks</td><td>11572</td><td>11597</td><td>26</td><td>CCGACTTAGAACTGGTGCGGACCAGG</td><td>292.5</td><td>11.5</td><td>2.0</td><td>5.9</td><td>4.2E-09</td><td>2.0E-07</td><td>BT20/MB231</td><td>MP</td></tr>
<tr><td>>hg38_21_+_8433172_8446622_RNA45SN1_45S_50ntFlanks@11739.11758.20|TCTCTTAAGGTAGCCAAATG</td><td>>hg38_21_+_8433172_8446622_RNA45SN1_45S_50ntFlanks</td><td>11739</td><td>11758</td><td>20</td><td>TCTCTTAAGGTAGCCAAATG</td><td>389.6</td><td>-7.2</td><td>1.3</td><td>-5.3</td><td>9.3E-08</td><td>2.0E-07</td><td>BT20/MB468</td><td>CYTO</td></tr>
<tr><td>>hg38_1_-_228634819_228635039_RNA5S12_5S_50ntFlanks@86.103.18|CGTCTGATCTCGGAAGCT</td><td>>hg38_1_-_228634819_228635039_RNA5S12_5S_50ntFlanks</td><td>86</td><td>103</td><td>18</td><td>CGTCTGATCTCGGAAGCT</td><td>252.5</td><td>-4.8</td><td>0.9</td><td>-5.3</td><td>1.1E-07</td><td>2.0E-07</td><td>MB231/MB468</td><td>CYTO</td></tr>
<tr><td>>hg38_1_-_228634819_228635039_RNA5S12_5S_50ntFlanks@152.169.18|TACCGGGTGCTGTAGGCT</td><td>>hg38_1_-_228634819_228635039_RNA5S12_5S_50ntFlanks</td><td>152</td><td>169</td><td>18</td><td>TACCGGGTGCTGTAGGCT</td><td>1558.3</td><td>-3.7</td><td>0.7</td><td>-5.3</td><td>9.8E-08</td><td>2.0E-07</td><td>MB231/MB468</td><td>MP</td></tr>
<tr><td>>hg38_21_+_8433172_8446622_RNA45SN1_45S_50ntFlanks@5494.5524.31|AAGACGGTCGAACTTGACTATCTAGAGGAAG</td><td>>hg38_21_+_8433172_8446622_RNA45SN1_45S_50ntFlanks</td><td>5494</td><td>5524</td><td>31</td><td>AAGACGGTCGAACTTGACTATCTAGAGGAAG</td><td>184.8</td><td>-5.4</td><td>1.0</td><td>-5.3</td><td>9.8E-08</td><td>2.0E-07</td><td>MB231/MB468</td><td>MP</td></tr>
<tr><td>>hg38_21_+_8433172_8446622_RNA45SN1_45S_50ntFlanks@11580.11609.30|GAACTGGTGCGGACCAGGGGAATCCGACTG</td><td>>hg38_21_+_8433172_8446622_RNA45SN1_45S_50ntFlanks</td><td>11580</td><td>11609</td><td>30</td><td>GAACTGGTGCGGACCAGGGGAATCCGACTG</td><td>626.8</td><td>-2.3</td><td>0.4</td><td>-5.3</td><td>9.3E-08</td><td>2.0E-07</td><td>BT20/MB468</td><td>CYTO</td></tr>
<tr><td>>am_trna10_ValCAC_5_-_180649395_180649467@1.34.34,trna18_ValCAC_5_-_180529253_180529325@1.34.34,trna2_ValCAC_5_+_180524070_180524142@1.34.34,trna6_ValCAC_5_+_180600650_180600722@1.34.34,trna85_ValCAC_1_-_161369490_161369562@1.34.34,trna90_ValCAC_1_-_149684088_149684161@1.34.34,trna98_ValCAC_1_-_149298555_149298627@1.34.34,trna9_ValCAC_6_+_26538282_26538354@1.34.34|GTTTCCGTAGTGTAGTGGTTATCACGTTCGCCTC</td><td>>am_trna10_ValCAC_5_-_180649395_180649467</td><td>1</td><td>34</td><td>34</td><td>GTTTCCGTAGTGTAGTGGTTATCACGTTCGCCTC</td><td>644.0</td><td>8.0</td><td>1.5</td><td>5.3</td><td>9.3E-08</td><td>2.0E-07</td><td>BT20/MB468</td><td>CYTO</td></tr>
<tr><td>>hg38_21_+_8433172_8446622_RNA45SN1_45S_50ntFlanks@9941.9971.31|AGGACGGTGGCCATGGAAGTCGGAATCCGCT</td><td>>hg38_21_+_8433172_8446622_RNA45SN1_45S_50ntFlanks</td><td>9941</td><td>9971</td><td>31</td><td>AGGACGGTGGCCATGGAAGTCGGAATCCGCT</td><td>70.8</td><td>9.9</td><td>1.8</td><td>5.6</td><td>2.1E-08</td><td>2.0E-07</td><td>BT20/MB468</td><td>MP</td></tr>
<tr><td>>trna32_LysCTT_16_-_3207406_3207478@4.33.30|CGGCTAGCTCAGTCGGTAGAGCATGAGACC</td><td>>trna32_LysCTT_16_-_3207406_3207478</td><td>4</td><td>33</td><td>30</td><td>CGGCTAGCTCAGTCGGTAGAGCATGAGACC</td><td>445.3</td><td>-6.0</td><td>1.0</td><td>-5.8</td><td>8.8E-09</td><td>2.0E-07</td><td>BT20/MB231</td><td>MITO</td></tr>
<tr><td>>hg38_21_+_8433172_8446622_RNA45SN1_45S_50ntFlanks@6652.6693.42|GACTCTTAGCGGTGGATCACTCGGCTCGTGCGTCGATGAAGA</td><td>>hg38_21_+_8433172_8446622_RNA45SN1_45S_50ntFlanks</td><td>6652</td><td>6693</td><td>42</td><td>GACTCTTAGCGGTGGATCACTCGGCTCGTGCGTCGATGAAGA</td><td>1380.0</td><td>4.3</td><td>0.8</td><td>5.3</td><td>1.1E-07</td><td>2.0E-07</td><td>MB231/MB468</td><td>NUC</td></tr>
<tr><td>>hg38_21_+_8433172_8446622_RNA45SN1_45S_50ntFlanks@9339.9368.30|CGCACGAAAGCCGCCGTGGCGCAATGAAGG</td><td>>hg38_21_+_8433172_8446622_RNA45SN1_45S_50ntFlanks</td><td>9339</td><td>9368</td><td>30</td><td>CGCACGAAAGCCGCCGTGGCGCAATGAAGG</td><td>89.6</td><td>-8.6</td><td>1.6</td><td>-5.4</td><td>8.3E-08</td><td>2.0E-07</td><td>BT20/MB468</td><td>MITO</td></tr>
<tr><td>>hg38_21_+_8433172_8446622_RNA45SN1_45S_50ntFlanks@12999.13027.29|CTGCGATCTATTGAAAGTCAGCCCTCGAC</td><td>>hg38_21_+_8433172_8446622_RNA45SN1_45S_50ntFlanks</td><td>12999</td><td>13027</td><td>29</td><td>CTGCGATCTATTGAAAGTCAGCCCTCGAC</td><td>212.9</td><td>-5.5</td><td>1.0</td><td>-5.3</td><td>9.9E-08</td><td>2.0E-07</td><td>MB231/MB468</td><td>MP</td></tr>
<tr><td>>MI0001445|hsa-mir-423&WithFlank&17|+|30117073|30117178@59.78.20&[MIMAT0001340&hsa-miR-423-3p&offsets|0|-3;m-100&17|+|30117131|30117153&offsets|0|-3]|AGCTCGGTCTGAGGCCCCTC</td><td>>MI0001445|hsa-mir-423&WithFlank&17|+|30117073|30117178</td><td>59</td><td>78</td><td>20</td><td>AGCTCGGTCTGAGGCCCCTC</td><td>3665.4</td><td>4.8</td><td>0.9</td><td>5.3</td><td>1.1E-07</td><td>2.0E-07</td><td>MB231/MB468</td><td>NUC</td></tr>
<tr><td>>hg38_21_+_8433172_8446622_RNA45SN1_45S_50ntFlanks@7972.8009.38|AGACGCGACCTCAGATCAGACGTGGCGACCCGCTGAAT</td><td>>hg38_21_+_8433172_8446622_RNA45SN1_45S_50ntFlanks</td><td>7972</td><td>8009</td><td>38</td><td>AGACGCGACCTCAGATCAGACGTGGCGACCCGCTGAAT</td><td>317.0</td><td>5.7</td><td>1.1</td><td>5.4</td><td>8.3E-08</td><td>2.0E-07</td><td>BT20/MB468</td><td>MITO</td></tr>
<tr><td>>hg38_21_+_8433172_8446622_RNA45SN1_45S_50ntFlanks@10409.10430.22|GTCAGTCGGTCCTGAGAGATGG</td><td>>hg38_21_+_8433172_8446622_RNA45SN1_45S_50ntFlanks</td><td>10409</td><td>10430</td><td>22</td><td>GTCAGTCGGTCCTGAGAGATGG</td><td>696.7</td><td>-5.3</td><td>1.0</td><td>-5.3</td><td>9.4E-08</td><td>2.0E-07</td><td>BT20/MB468</td><td>CYTO</td></tr>
<tr><td>>hg38_21_+_8433172_8446622_RNA45SN1_45S_50ntFlanks@8113.8137.25|GGCGCGGGACATGTGGCGTACGGAA</td><td>>hg38_21_+_8433172_8446622_RNA45SN1_45S_50ntFlanks</td><td>8113</td><td>8137</td><td>25</td><td>GGCGCGGGACATGTGGCGTACGGAA</td><td>925.0</td><td>13.2</td><td>2.2</td><td>5.9</td><td>4.3E-09</td><td>2.0E-07</td><td>BT20/MB231</td><td>MP</td></tr>
<tr><td>>hg38_21_+_8433172_8446622_RNA45SN1_45S_50ntFlanks@8113.8139.27|GGCGCGGGACATGTGGCGTACGGAAGA</td><td>>hg38_21_+_8433172_8446622_RNA45SN1_45S_50ntFlanks</td><td>8113</td><td>8139</td><td>27</td><td>GGCGCGGGACATGTGGCGTACGGAAGA</td><td>715.4</td><td>12.8</td><td>2.2</td><td>5.9</td><td>4.3E-09</td><td>2.0E-07</td><td>BT20/MB231</td><td>MP</td></tr>
<tr><td>>hg38_21_+_8433172_8446622_RNA45SN1_45S_50ntFlanks@10820.10840.21|GGGAAGTCGGCAAGCCGGATC</td><td>>hg38_21_+_8433172_8446622_RNA45SN1_45S_50ntFlanks</td><td>10820</td><td>10840</td><td>21</td><td>GGGAAGTCGGCAAGCCGGATC</td><td>1647.4</td><td>-3.4</td><td>0.6</td><td>-5.3</td><td>1.1E-07</td><td>2.0E-07</td><td>MB231/MB468</td><td>CYTO</td></tr>
<tr><td>>hg38_21_+_8433172_8446622_RNA45SN1_45S_50ntFlanks@5472.5510.39|ACGGCCCTGGCGGAGCGCTGAGAAGACGGTCGAACTTGA</td><td>>hg38_21_+_8433172_8446622_RNA45SN1_45S_50ntFlanks</td><td>5472</td><td>5510</td><td>39</td><td>ACGGCCCTGGCGGAGCGCTGAGAAGACGGTCGAACTTGA</td><td>430.3</td><td>4.4</td><td>0.8</td><td>5.3</td><td>1.1E-07</td><td>2.0E-07</td><td>MB231/MB468</td><td>NUC</td></tr>
<tr><td>>MI0000433|hsa-let-7g&WithFlank&3|-|52268272|52268367@11.31.21(+1C)&[MIMAT0000414&hsa-let-7g-5p&offsets|0|-1(+1C);m-58&3|-|52268336|52268357&offsets|0|-1(+1C)]|TGAGGTAGTAGTTTGTACAGTC</td><td>>MI0000433|hsa-let-7g&WithFlank&3|-|52268272|52268367</td><td>11</td><td>31</td><td>21(+1C)</td><td>TGAGGTAGTAGTTTGTACAGTC</td><td>4492.1</td><td>2.3</td><td>0.4</td><td>5.3</td><td>1.2E-07</td><td>2.0E-07</td><td>BT20/MB468</td><td>NUC</td></tr>
<tr><td>>hg38_21_+_8433172_8446622_RNA45SN1_45S_50ntFlanks@10498.10533.36|TTCAGATCCCCGAATCCGGAGTGGCGGAGATGGGCG</td><td>>hg38_21_+_8433172_8446622_RNA45SN1_45S_50ntFlanks</td><td>10498</td><td>10533</td><td>36</td><td>TTCAGATCCCCGAATCCGGAGTGGCGGAGATGGGCG</td><td>84.8</td><td>-8.5</td><td>1.6</td><td>-5.4</td><td>8.4E-08</td><td>2.0E-07</td><td>BT20/MB468</td><td>MITO</td></tr>
<tr><td>>hg38_21_+_8433172_8446622_RNA45SN1_45S_50ntFlanks@6698.6719.22|AGCTAGCTGCGAGAATTAATGT</td><td>>hg38_21_+_8433172_8446622_RNA45SN1_45S_50ntFlanks</td><td>6698</td><td>6719</td><td>22</td><td>AGCTAGCTGCGAGAATTAATGT</td><td>750.3</td><td>-6.0</td><td>1.1</td><td>-5.3</td><td>1.2E-07</td><td>2.0E-07</td><td>MB231/MB468</td><td>MITO</td></tr>
<tr><td>>hg38_21_+_8433172_8446622_RNA45SN1_45S_50ntFlanks@12993.13013.21|CCCTCGCTGCGATCTATTGAA</td><td>>hg38_21_+_8433172_8446622_RNA45SN1_45S_50ntFlanks</td><td>12993</td><td>13013</td><td>21</td><td>CCCTCGCTGCGATCTATTGAA</td><td>560.7</td><td>-5.8</td><td>1.1</td><td>-5.3</td><td>1.2E-07</td><td>2.0E-07</td><td>MB231/MB468</td><td>MITO</td></tr>
<tr><td>>hg38_21_+_8433172_8446622_RNA45SN1_45S_50ntFlanks@10786.10828.43|AACAGCCTCTGGCATGTTGGAACAATGTAGGTAAGGGAAGTCG</td><td>>hg38_21_+_8433172_8446622_RNA45SN1_45S_50ntFlanks</td><td>10786</td><td>10828</td><td>43</td><td>AACAGCCTCTGGCATGTTGGAACAATGTAGGTAAGGGAAGTCG</td><td>138.5</td><td>-10.2</td><td>1.8</td><td>-5.6</td><td>2.1E-08</td><td>2.0E-07</td><td>BT20/MB468</td><td>MP</td></tr>
<tr><td>>hg38_21_+_8433172_8446622_RNA45SN1_45S_50ntFlanks@8170.8187.18|CAAGTCCTTCTGATCGAG</td><td>>hg38_21_+_8433172_8446622_RNA45SN1_45S_50ntFlanks</td><td>8170</td><td>8187</td><td>18</td><td>CAAGTCCTTCTGATCGAG</td><td>542.6</td><td>-7.0</td><td>1.3</td><td>-5.3</td><td>9.5E-08</td><td>2.0E-07</td><td>BT20/MB468</td><td>CYTO</td></tr>
</table>

| | | | | | | | | | | | | | |
|---|---|---|---|---|---|---|---|---|---|---|---|---|---|
| >hg38_21_+_8433172_8446622_RNA45SN1_45S_50ntFlanks@11575.11606.32\|ACTTAGAACTGGTGCGGACCAGGGGAATCCGA | >hg38_21_+_8433172_8446622_RNA45SN1_45S_50ntFlanks | 11575 | 11606 | 32 | ACTTAGAACTGGTGCGGACCAGGGGAATCCGA | 441.1 | -8.0 | 1.5 | -5.3 | 1.2E-07 | 2.0E-07 | BT20/MB468 | NUC |
| >hg38_21_+_8433172_8446622_RNA45SN1_45S_50ntFlanks@10389.10429.41\|TGAACAGCAGTTGAACATGGGTCAGTCGGTCCTGAGAGATG | >hg38_21_+_8433172_8446622_RNA45SN1_45S_50ntFlanks | 10389 | 10429 | 41 | TGAACAGCAGTTGAACATGGGTCAGTCGGTCCTGAGAGATG | 2212.7 | -9.5 | 1.8 | -5.3 | 1.2E-07 | 2.0E-07 | BT20/MB468 | NUC |
| >hg38_21_+_8433172_8446622_RNA45SN1_45S_50ntFlanks@6652.6691.40\|GACTCTTAGCGGTGGATCACTCGGCTCGTGCGTCGATGAA | >hg38_21_+_8433172_8446622_RNA45SN1_45S_50ntFlanks | 6652 | 6691 | 40 | GACTCTTAGCGGTGGATCACTCGGCTCGTGCGTCGATGAA | 417.9 | 4.3 | 0.8 | 5.3 | 1.1E-07 | 2.0E-07 | MB231/MB468 | NUC |
| >hg38_21_+_8433172_8446622_RNA45SN1_45S_50ntFlanks@10388.10410.23\|GTGAACAGCAGTTGAACATGGGT | >hg38_21_+_8433172_8446622_RNA45SN1_45S_50ntFlanks | 10388 | 10410 | 23 | GTGAACAGCAGTTGAACATGGGT | 900.7 | -7.6 | 1.4 | -5.3 | 1.1E-07 | 2.0E-07 | MB231/MB468 | NUC |
| >hg38_21_+_8433172_8446622_RNA45SN1_45S_50ntFlanks@12437.12454.18\|CATTGTGAAGCAGAATTC | >hg38_21_+_8433172_8446622_RNA45SN1_45S_50ntFlanks | 12437 | 12454 | 18 | CATTGTGAAGCAGAATTC | 463.2 | -3.5 | 0.7 | -5.4 | 8.5E-08 | 2.0E-07 | BT20/MB468 | MITO |
| >am_trnaMT_HisGTG_MT_+_12138_12206@17.51.35\|CAAAACATCAGATTGTGAATCTGACAACAGAGGCT | >am_trnaMT_HisGTG_MT_+_12138_12206 | 17 | 51 | 35 | CAAAACATCAGATTGTGAATCTGACAACAGAGGCT | 112.6 | 10.6 | 1.9 | 5.6 | 2.1E-08 | 2.0E-07 | BT20/MB468 | MP |
| >hg38_21_+_8433172_8446622_RNA45SN1_45S_50ntFlanks@7988.8006.19\|CAGACGTGGCGACCCGCTG | >hg38_21_+_8433172_8446622_RNA45SN1_45S_50ntFlanks | 7988 | 8006 | 19 | CAGACGTGGCGACCCGCTG | 1791.5 | -7.9 | 1.4 | -5.6 | 2.1E-08 | 2.0E-07 | BT20/MB468 | MP |
| >hg38_21_+_8433172_8446622_RNA45SN1_45S_50ntFlanks@8168.8185.18\|CCCAAGTCCTTCTGATCG | >hg38_21_+_8433172_8446622_RNA45SN1_45S_50ntFlanks | 8168 | 8185 | 18 | CCCAAGTCCTTCTGATCG | 644.3 | -5.8 | 1.1 | -5.3 | 1.2E-07 | 2.0E-07 | MB231/MB468 | MITO |
| >hg38_21_+_8433172_8446622_RNA45SN1_45S_50ntFlanks@475.492.18\|CTTCGTGATCGATGTGGT | >hg38_21_+_8433172_8446622_RNA45SN1_45S_50ntFlanks | 475 | 492 | 18 | CTTCGTGATCGATGTGGT | 746.0 | 7.2 | 1.3 | 5.4 | 8.5E-08 | 2.0E-07 | BT20/MB468 | MITO |
| >hg38_21_+_8433172_8446622_RNA45SN1_45S_50ntFlanks@11235.11268.34\|CTTCCCGTGGATCGCCCCAGCTGCGGCGGGCGTC | >hg38_21_+_8433172_8446622_RNA45SN1_45S_50ntFlanks | 11235 | 11268 | 34 | CTTCCCGTGGATCGCCCCAGCTGCGGCGGGCGTC | 155.0 | -4.1 | 0.8 | -5.3 | 9.5E-08 | 2.0E-07 | BT20/MB468 | CYTO |
| >hg38_21_+_8433172_8446622_RNA45SN1_45S_50ntFlanks@9033.9068.36\|CTCGTGGGGGGGCCGGGCCACCCCTCCCACGGCGCG | >hg38_21_+_8433172_8446622_RNA45SN1_45S_50ntFlanks | 9033 | 9068 | 36 | CTCGTGGGGGGGCCGGGCCACCCCTCCCACGGCGCG | 77.9 | -9.4 | 1.7 | -5.6 | 2.1E-08 | 2.0E-07 | BT20/MB468 | MP |
| >hg38_21_+_8433172_8446622_RNA45SN1_45S_50ntFlanks@10534.10562.29\|CCGCGAGGCGTCCAGTGCGGTAACGCGAC | >hg38_21_+_8433172_8446622_RNA45SN1_45S_50ntFlanks | 10534 | 10562 | 29 | CCGCGAGGCGTCCAGTGCGGTAACGCGAC | 83.1 | -9.5 | 1.7 | -5.6 | 2.1E-08 | 2.0E-07 | BT20/MB468 | MP |
| >hg38_21_+_8433172_8446622_RNA45SN1_45S_50ntFlanks@11999.12029.31\|CCCTGCGGGCCGCCGGTGAAATACCACTACT | >hg38_21_+_8433172_8446622_RNA45SN1_45S_50ntFlanks | 11999 | 12029 | 31 | CCCTGCGGGCCGCCGGTGAAATACCACTACT | 650.5 | -7.3 | 1.4 | -5.3 | 1.1E-07 | 2.0E-07 | MB231/MB468 | NUC |
| >hg38_21_+_8433172_8446622_RNA45SN1_45S_50ntFlanks@4726.4752.27\|TAATCAAGAACGAAAGTCGGAGGTTCG | >hg38_21_+_8433172_8446622_RNA45SN1_45S_50ntFlanks | 4726 | 4752 | 27 | TAATCAAGAACGAAAGTCGGAGGTTCG | 387.8 | -5.0 | 0.9 | -5.3 | 1.1E-07 | 2.0E-07 | MB231/MB468 | CYTO |
| >hg38_21_+_8433172_8446622_RNA45SN1_45S_50ntFlanks@4046.4065.20\|CATTCGAACGTCTGCCCTAT | >hg38_21_+_8433172_8446622_RNA45SN1_45S_50ntFlanks | 4046 | 4065 | 20 | CATTCGAACGTCTGCCCTAT | 301.9 | -7.4 | 1.4 | -5.3 | 1.2E-07 | 2.0E-07 | MB231/MB468 | MITO |
| >hg38_21_+_8433172_8446622_RNA45SN1_45S_50ntFlanks@5066.5097.32\|TCTGGTTAATTCCGATAACGAACGAGACTCTG | >hg38_21_+_8433172_8446622_RNA45SN1_45S_50ntFlanks | 5066 | 5097 | 32 | TCTGGTTAATTCCGATAACGAACGAGACTCTG | 106.2 | -10.6 | 2.0 | -5.3 | 1.0E-07 | 2.0E-07 | MB231/MB468 | MP |
| >hg38_21_+_8433172_8446622_RNA45SN1_45S_50ntFlanks@10389.10419.31\|TGAACAGCAGTTGAACATGGGTCAGTCGGTC | >hg38_21_+_8433172_8446622_RNA45SN1_45S_50ntFlanks | 10389 | 10419 | 31 | TGAACAGCAGTTGAACATGGGTCAGTCGGTC | 79.4 | -9.4 | 1.7 | -5.6 | 2.1E-08 | 2.0E-07 | BT20/MB468 | MP |
| >hg38_21_+_8433172_8446622_RNA45SN1_45S_50ntFlanks@10420.10462.43\|CTGAGAGATGGGCGAGCGCCGTTCCGAAGGGACGGGCGATGGC | >hg38_21_+_8433172_8446622_RNA45SN1_45S_50ntFlanks | 10420 | 10462 | 43 | CTGAGAGATGGGCGAGCGCCGTTCCGAAGGGACGGGCGATGGC | 113.6 | -4.3 | 0.8 | -5.3 | 1.1E-07 | 2.0E-07 | MB231/MB468 | CYTO |
| >MI0003820\|hsa-mir-454&WithFlank&17\|-\|59137752\|59137878@70.93.24(+1C)&[MIMAT0003885&hsa-miR-454-3p&offsets\|0\|+1(+1C);m-272&17\|-\|59137787\|59137809&offsets\|0\|+1(+1C)]\|TAGTGCAATATTGCTTATAGGGTTC | >MI0003820\|hsa-mir-454&WithFlank&17\|-\|59137752\|59137878 | 70 | 93 | 24(+1C) | TAGTGCAATATTGCTTATAGGGTTC | 175.4 | -10.9 | 1.8 | -5.9 | 4.4E-09 | 2.0E-07 | BT20/MB231 | MP |
| >MI0000734\|hsa-mir-106b&WithFlank&7\|-\|100093987\|100094080@58.78.21(+1U)&[MIMAT0004672&hsa-miR-106b-3p&offsets\|0\|-1(+1U);m-87&7\|-\|100094003\|100094023&offsets\|0\|0(+1U)]\|CCGCACTGTGGGTACTTGCTGT | >MI0000734\|hsa-mir-106b&WithFlank&7\|-\|100093987\|100094080 | 58 | 78 | 21(+1U) | CCGCACTGTGGGTACTTGCTGT | 487.4 | 1.6 | 0.3 | 5.3 | 1.2E-07 | 2.0E-07 | BT20/MB468 | NUC |
| >hg38_21_+_8433172_8446622_RNA45SN1_45S_50ntFlanks@10384.10401.18\|CCATGTGAACAGCAGTTG | >hg38_21_+_8433172_8446622_RNA45SN1_45S_50ntFlanks | 10384 | 10401 | 18 | CCATGTGAACAGCAGTTG | 159.5 | -10.4 | 1.9 | -5.6 | 2.2E-08 | 2.0E-07 | BT20/MB468 | MP |
| >hg38_21_+_8433172_8446622_RNA45SN1_45S_50ntFlanks@6731.6756.26\|CACATTGATCATCGACACTTCGAACG | >hg38_21_+_8433172_8446622_RNA45SN1_45S_50ntFlanks | 6731 | 6756 | 26 | CACATTGATCATCGACACTTCGAACG | 767.7 | -3.4 | 0.6 | -5.3 | 1.1E-07 | 2.0E-07 | MB231/MB468 | CYTO |
| >hg38_21_+_8433172_8446622_RNA45SN1_45S_50ntFlanks@12149.12179.31\|AGGTGGGGAGTTTGACTGGGGCGGTACACCT | >hg38_21_+_8433172_8446622_RNA45SN1_45S_50ntFlanks | 12149 | 12179 | 31 | AGGTGGGGAGTTTGACTGGGGCGGTACACCT | 73.7 | -9.3 | 1.7 | -5.6 | 2.2E-08 | 2.0E-07 | BT20/MB468 | MP |
| >hg38_21_+_8433172_8446622_RNA45SN1_45S_50ntFlanks@12008.12029.22\|CCGCCGGTGAAATACCACTACT | >hg38_21_+_8433172_8446622_RNA45SN1_45S_50ntFlanks | 12008 | 12029 | 22 | CCGCCGGTGAAATACCACTACT | 919.8 | -5.6 | 1.0 | -5.4 | 8.6E-08 | 2.0E-07 | BT20/MB468 | MITO |
| >hg38_21_+_8433172_8446622_RNA45SN1_45S_50ntFlanks@6680.6718.39\|TGCGTCGATGAAGAACGCAGCTAGCTGCGAGAATTAATG | >hg38_21_+_8433172_8446622_RNA45SN1_45S_50ntFlanks | 6680 | 6718 | 39 | TGCGTCGATGAAGAACGCAGCTAGCTGCGAGAATTAATG | 189.1 | -6.9 | 1.3 | -5.3 | 1.1E-07 | 2.0E-07 | MB231/MB468 | NUC |
| >hg38_21_+_8433172_8446622_RNA45SN1_45S_50ntFlanks@5084.5106.23\|CGAACGAGACTCTGGCATGCTAA | >hg38_21_+_8433172_8446622_RNA45SN1_45S_50ntFlanks | 5084 | 5106 | 23 | CGAACGAGACTCTGGCATGCTAA | 266.1 | -10.1 | 1.9 | -5.4 | 8.6E-08 | 2.1E-07 | BT20/MB468 | MITO |
| >hg38_21_+_8433172_8446622_RNA45SN1_45S_50ntFlanks@12488.12525.38\|AACGTGAGCTGGGTTTAGACCGTCGTGAGACAGGTTAG | >hg38_21_+_8433172_8446622_RNA45SN1_45S_50ntFlanks | 12488 | 12525 | 38 | AACGTGAGCTGGGTTTAGACCGTCGTGAGACAGGTTAG | 1133.6 | -9.2 | 1.7 | -5.3 | 1.3E-07 | 2.1E-07 | BT20/MB468 | NUC |
| >hg38_1_-_228634819_228635039_RNA5S12_5S_50ntFlanks@138.169.32\|AGACCGCCTGGGAATACCGGGTGCTGTAGGCT | >hg38_1_-_228634819_228635039_RNA5S12_5S_50ntFlanks | 138 | 169 | 32 | AGACCGCCTGGGAATACCGGGTGCTGTAGGCT | 439.5 | 4.4 | 0.8 | 5.3 | 1.1E-07 | 2.1E-07 | MB231/MB468 | NUC |
| >MI0000749\|hsa-mir-30e&WithFlank&1\|+\|40754349\|40754452@65.85.21&[MIMAT0000693&hsa-miR-30e-3p&offsets\|0\|-1;m-25&1\|+\|40754413\|40754434&offsets\|0\|-1]\|CTTTCAGTCGGATGTTTACAG | >MI0000749\|hsa-mir-30e&WithFlank&1\|+\|40754349\|40754452 | 65 | 85 | 21 | CTTTCAGTCGGATGTTTACAG | 358.2 | 3.7 | 0.7 | 5.3 | 1.1E-07 | 2.1E-07 | MB231/MB468 | NUC |
| >hg38_21_+_8433172_8446622_RNA45SN1_45S_50ntFlanks@11561.11588.28\|GCGCCTAGCAGCCGACTTAGAACTGGTG | >hg38_21_+_8433172_8446622_RNA45SN1_45S_50ntFlanks | 11561 | 11588 | 28 | GCGCCTAGCAGCCGACTTAGAACTGGTG | 889.9 | -4.8 | 0.9 | -5.6 | 2.2E-08 | 2.1E-07 | BT20/MB468 | MP |
| >MI0000107\|hsa-mir-29b-2&WithFlank&1\|-\|207802437\|207802529@58.80.23;MI0000105\|hsa-mir-29b-1&WithFlank&7\|-\|130877453\|130877545@57.79.23&[MIMAT0000100&hsa-miR-29b-3p&offsets\|0\|0;m-81&1\|-\|207802450\|207802472&offsets\|0\|0];[MIMAT0000100_1&hsa-miR-29b-3p&offsets\|0\|0;m-82&7\|-\|130877467\|130877489&offsets\|0\|0]\|TAGCACCATTTGAAATCAGTGTT | >MI0000107\|hsa-mir-29b-2&WithFlank&1\|-\|207802437\|207802529 | 58 | 80 | 23 | TAGCACCATTTGAAATCAGTGTT | 486.4 | 8.1 | 1.5 | 5.3 | 1.3E-07 | 2.1E-07 | BT20/MB468 | NUC |
| >MI0000446\|hsa-mir-125b-1&WithFlank&11\|-\|122099751\|122099850@21.39.19;MI0000470\|hsa-mir-125b-2&WithFlank&21\|+\|16590231\|16590331@23.41.19&[MIMAT0000423&hsa-miR-125b-5p&offsets\|0\|-3;m-67&11\|-\|122099809\|122099830&offsets\|0\|-3];[MIMAT0000423_1&hsa-miR-125b-5p&offsets\|0\|-3;m-68&21\|+\|16590253\|16590274&offsets\|0\|-3]\|TCCCTGAGACCCTAACTTG | >MI0000446\|hsa-mir-125b-1&WithFlank&11\|-\|122099751\|122099850 | 21 | 39 | 19 | TCCCTGAGACCCTAACTTG | 452.1 | 9.0 | 1.7 | 5.3 | 1.0E-07 | 2.1E-07 | MB231/MB468 | MP |
| >MI0000542\|hsa-mir-320a&WithFlank&8\|-\|22244960\|22245043@48.70.23(+1G)&[MIMAT0000510&hsa-miR-320a-3p&offsets\|0\|+1(+1G);m-60&8\|-\|22244975\|22244996&offsets\|0\|+1(+1G)]\|AAAAGCTGGGTTGAGAGGGCGAAG | >MI0000542\|hsa-mir-320a&WithFlank&8\|-\|22244960\|22245043 | 48 | 70 | 23(+1G) | AAAAGCTGGGTTGAGAGGGCGAAG | 1834.4 | 4.5 | 0.8 | 5.8 | 7.1E-09 | 2.1E-07 | BT20/MB231 | CYTO |
| >MI0003628\|hsa-mir-615&WithFlank&12\|+\|54033944\|54034051@67.86.20&[MIMAT0003283&hsa-miR-615-3p&offsets\|0\|-2;m-495&12\|+\|54034010\|54034030&offsets\|0\|-1]\|TCCGAGCCTGGGTCTCCCTC | >MI0003628\|hsa-mir-615&WithFlank&12\|+\|54033944\|54034051 | 67 | 86 | 20 | TCCGAGCCTGGGTCTCCCTC | 585.9 | 4.8 | 0.8 | 5.8 | 7.2E-09 | 2.1E-07 | BT20/MB231 | CYTO |
| >hg38_21_+_8433172_8446622_RNA45SN1_45S_50ntFlanks@11999.12042.44\|CCCTGCGGGCCGCCGGTGAAATACCACTACTCTGATCGTTTTTT | >hg38_21_+_8433172_8446622_RNA45SN1_45S_50ntFlanks | 11999 | 12042 | 44 | CCCTGCGGGCCGCCGGTGAAATACCACTACTCTGATCGTTTTTT | 169.8 | -9.5 | 1.8 | -5.4 | 8.7E-08 | 2.1E-07 | BT20/MB468 | MITO |
| >hg38_21_+_8433172_8446622_RNA45SN1_45S_50ntFlanks@5071.5097.27\|TTAATTCCGATAACGAACGAGACTCTG | >hg38_21_+_8433172_8446622_RNA45SN1_45S_50ntFlanks | 5071 | 5097 | 27 | TTAATTCCGATAACGAACGAGACTCTG | 915.1 | -7.2 | 1.4 | -5.3 | 1.3E-07 | 2.1E-07 | BT20/MB468 | NUC |
| >hg38_21_+_8433172_8446622_RNA45SN1_45S_50ntFlanks@9604.9622.19\|AAAGACTAATCGAACCATC | >hg38_21_+_8433172_8446622_RNA45SN1_45S_50ntFlanks | 9604 | 9622 | 19 | AAAGACTAATCGAACCATC | 2523.9 | -5.4 | 1.0 | -5.6 | 2.2E-08 | 2.1E-07 | BT20/MB468 | MP |
| >hg38_21_+_8433172_8446622_RNA45SN1_45S_50ntFlanks@10420.10455.36\|CTGAGAGATGGGCGAGCGCCGTTCCGAAGGGACGGG | >hg38_21_+_8433172_8446622_RNA45SN1_45S_50ntFlanks | 10420 | 10455 | 36 | CTGAGAGATGGGCGAGCGCCGTTCCGAAGGGACGGG | 393.6 | -3.8 | 0.7 | -5.3 | 9.8E-08 | 2.1E-07 | BT20/MB468 | CYTO |
| >hg38_1_-_228634819_228635039_RNA5S12_5S_50ntFlanks@51.88.38\|GTCTACGGCCATACCACCCTGAACGCGCCCGATCTCGT | >hg38_1_-_228634819_228635039_RNA5S12_5S_50ntFlanks | 51 | 88 | 38 | GTCTACGGCCATACCACCCTGAACGCGCCCGATCTCGT | 1437.4 | 4.2 | 0.8 | 5.3 | 1.2E-07 | 2.1E-07 | MB231/MB468 | MITO |
| >hg38_21_+_8433172_8446622_RNA45SN1_45S_50ntFlanks@485.532.48\|GATGTGGTGACGTCGTGCTCTCCCGGGCCGGGTCCGAGCCGCGACGGG | >hg38_21_+_8433172_8446622_RNA45SN1_45S_50ntFlanks | 485 | 532 | 48 | GATGTGGTGACGTCGTGCTCTCCCGGGCCGGGTCCGAGCCGCGACGGG | 188.4 | 11.3 | 2.0 | 5.6 | 2.2E-08 | 2.1E-07 | BT20/MB468 | MP |
| >hg38_21_+_8433172_8446622_RNA45SN1_45S_50ntFlanks@11829.11872.44\|CGAAACCACAGCCAAGGGAACGGGCTTGGCGGAATCAGCGGGGA | >hg38_21_+_8433172_8446622_RNA45SN1_45S_50ntFlanks | 11829 | 11872 | 44 | CGAAACCACAGCCAAGGGAACGGGCTTGGCGGAATCAGCGGGGA | 93.4 | -8.6 | 1.6 | -5.4 | 8.7E-08 | 2.1E-07 | BT20/MB468 | MITO |
| >hg38_21_+_8433172_8446622_RNA45SN1_45S_50ntFlanks@11741.11758.18\|TCTTAAGGTAGCCAAATG | >hg38_21_+_8433172_8446622_RNA45SN1_45S_50ntFlanks | 11741 | 11758 | 18 | TCTTAAGGTAGCCAAATG | 558.2 | -7.5 | 1.4 | -5.3 | 1.1E-07 | 2.1E-07 | MB231/MB468 | CYTO |
| >hg38_21_+_8433172_8446622_RNA45SN1_45S_50ntFlanks@10766.10797.32\|CGCAGCAGGTCTCCAAGGTGAACAGCCTCTGG | >hg38_21_+_8433172_8446622_RNA45SN1_45S_50ntFlanks | 10766 | 10797 | 32 | CGCAGCAGGTCTCCAAGGTGAACAGCCTCTGG | 178.4 | -7.5 | 1.4 | -5.3 | 1.2E-07 | 2.1E-07 | MB231/MB468 | MITO |
| >hg38_21_+_8433172_8446622_RNA45SN1_45S_50ntFlanks@12202.12221.20\|CTAAGGCGAGCTCAGGGAGG | >hg38_21_+_8433172_8446622_RNA45SN1_45S_50ntFlanks | 12202 | 12221 | 20 | CTAAGGCGAGCTCAGGGAGG | 397.8 | -5.6 | 1.1 | -5.3 | 1.2E-07 | 2.1E-07 | MB231/MB468 | NUC |
| >MI0000811\|hsa-mir-148b&WithFlank&12\|+\|54337210\|54337320@69.89.21&[MIMAT0000759&hsa-miR-148b-3p&offsets\|0\|-1;m-77&12\|+\|54337278\|54337299&offsets\|0\|-1]\|TCAGTGCATCACAGAACTTTG | >MI0000811\|hsa-mir-148b&WithFlank&12\|+\|54337210\|54337320 | 69 | 89 | 21 | TCAGTGCATCACAGAACTTTG | 472.8 | 5.5 | 1.0 | 5.3 | 1.2E-07 | 2.1E-07 | MB231/MB468 | MITO |
| >hg38_21_+_8433172_8446622_RNA45SN1_45S_50ntFlanks@10348.10388.41\|AAACGAGAACTTTGAAGGCCGAAGTGGAGAAGGGTTCCATG | >hg38_21_+_8433172_8446622_RNA45SN1_45S_50ntFlanks | 10348 | 10388 | 41 | AAACGAGAACTTTGAAGGCCGAAGTGGAGAAGGGTTCCATG | 689.4 | -4.4 | 0.8 | -5.3 | 1.2E-07 | 2.1E-07 | MB231/MB468 | MITO |
| >hg38_21_+_8433172_8446622_RNA45SN1_45S_50ntFlanks@11629.11660.32\|CGCGAAGGCCCGCGGCGGGTGTTGACGCGATG | >hg38_21_+_8433172_8446622_RNA45SN1_45S_50ntFlanks | 11629 | 11660 | 32 | CGCGAAGGCCCGCGGCGGGTGTTGACGCGATG | 407.6 | -7.8 | 1.5 | -5.3 | 1.0E-07 | 2.1E-07 | MB231/MB468 | MP |
| >MI0000542\|hsa-mir-320a&WithFlank&8\|-\|22244960\|22245043@48.69.22(+2U)&[MIMAT0000510&hsa-miR-320a-3p&offsets\|0\|0(+2U);m-60&8\|-\|22244975\|22244996&offsets\|0\|0(+2U)]\|AAAAGCTGGGTTGAGAGGGCGATT | >MI0000542\|hsa-mir-320a&WithFlank&8\|-\|22244960\|22245043 | 48 | 69 | 22(+2U) | AAAAGCTGGGTTGAGAGGGCGATT | 722.9 | 3.4 | 0.6 | 5.3 | 1.3E-07 | 2.1E-07 | BT20/MB468 | NUC |

<table>
<tr><th></th><th></th><th></th><th></th><th></th><th></th><th></th><th></th><th></th><th></th><th></th><th></th><th></th><th></th></tr>
<tr><td>>MI0003776|hsa-mir-320b-1&WithFlank&1|+|116671740|116671823@48.67.20;MI0003839|hsa-mir-320b-2&WithFlank&1|-|224257034|224257116@47.66.20;MI0000542|hsa-mir-320a&WithFlank&8|-|22244960|22245043@48.67.20&[MIMAT0005792&hsa-miR-320b&offsets|0|-2;m-183&1|+|116671787|116671806&offsets|0|0];[MIMAT0005792_1&hsa-miR-320b&offsets|0|-2;m-183&1|-|224257051|224257070&offsets|0|0];[MIMAT0000510&hsa-miR-320a-3p&offsets|0|-2;m-60&8|-|22244975|22244996&offsets|0|-2]|AAAAGCTGGGTTGAGAGGGC</td><td>>MI0003776|hsa-mir-320b-1&WithFlank&1|+|116671740|116671823</td><td>48</td><td>67</td><td>20</td><td>AAAAGCTGGGTTGAGAGGGC</td><td>406.3</td><td>7.0</td><td>1.3</td><td>5.3</td><td>1.1E-07</td><td>2.1E-07</td><td>MB231/MB468</td><td>CYTO</td></tr>
<tr><td>>hg38_21_+_8433172_8446622_RNA45SN1_45S_50ntFlanks@3990.4026.37|TTGGTGACTCTAGATAACCTCGGGCCGATCGCACGCC</td><td>>hg38_21_+_8433172_8446622_RNA45SN1_45S_50ntFlanks</td><td>3990</td><td>4026</td><td>37</td><td>TTGGTGACTCTAGATAACCTCGGGCCGATCGCACGCC</td><td>295.5</td><td>-5.5</td><td>1.0</td><td>-5.3</td><td>1.2E-07</td><td>2.1E-07</td><td>MB231/MB468</td><td>MITO</td></tr>
<tr><td>>hg38_21_+_8433172_8446622_RNA45SN1_45S_50ntFlanks@6732.6774.43|ACATTGATCATCGACACTTCGAACGCACTTGCGGCCCCGGGTT</td><td>>hg38_21_+_8433172_8446622_RNA45SN1_45S_50ntFlanks</td><td>6732</td><td>6774</td><td>43</td><td>ACATTGATCATCGACACTTCGAACGCACTTGCGGCCCCGGGTT</td><td>95.6</td><td>10.3</td><td>1.8</td><td>5.6</td><td>2.2E-08</td><td>2.1E-07</td><td>BT20/MB468</td><td>MP</td></tr>
<tr><td>>trna10_LysCTT_16_+_3241501_3241573@1.36.36,trna13_LysCTT_14_-_58706613_58706685@1.36.36,trna2_LysCTT_15_+_79152904_79152976@1.36.36|GCCCGGCTAGCTCAGTCGGTAGAGCATGGGACTCTT</td><td>>trna10_LysCTT_16_+_3241501_3241573</td><td>1</td><td>36</td><td>36</td><td>GCCCGGCTAGCTCAGTCGGTAGAGCATGGGACTCTT</td><td>2867.4</td><td>4.5</td><td>0.8</td><td>5.3</td><td>1.0E-07</td><td>2.1E-07</td><td>MB231/MB468</td><td>MP</td></tr>
<tr><td>>trna119_LysCTT_1_-_145395522_145395594@5.36.32,trna11_LysCTT_5_-_180648979_180649051@5.36.32,trna13_LysCTT_6_+_26556774_26556846@5.36.32,trna7_LysCTT_16_+_3225692_3225764@5.36.32,trna9_LysCTT_5_+_180634755_180634827@5.36.32|GGCTAGCTCAGTCGGTAGAGCATGAGACTCTT</td><td>>trna119_LysCTT_1_-_145395522_145395594</td><td>5</td><td>36</td><td>32</td><td>GGCTAGCTCAGTCGGTAGAGCATGAGACTCTT</td><td>1392.5</td><td>-6.2</td><td>1.1</td><td>-5.7</td><td>9.3E-09</td><td>2.1E-07</td><td>BT20/MB231</td><td>MITO</td></tr>
<tr><td>>hg38_21_+_8433172_8446622_RNA45SN1_45S_50ntFlanks@4919.4940.22|CCACCAGGAGTGGAGCCTGCGG</td><td>>hg38_21_+_8433172_8446622_RNA45SN1_45S_50ntFlanks</td><td>4919</td><td>4940</td><td>22</td><td>CCACCAGGAGTGGAGCCTGCGG</td><td>256.0</td><td>-10.1</td><td>1.9</td><td>-5.4</td><td>8.8E-08</td><td>2.1E-07</td><td>BT20/MB468</td><td>MITO</td></tr>
<tr><td>>hg38_21_+_8433172_8446622_RNA45SN1_45S_50ntFlanks@12627.12654.28|ATGGGGCGAAGCTACCATCTGTGGGATT</td><td>>hg38_21_+_8433172_8446622_RNA45SN1_45S_50ntFlanks</td><td>12627</td><td>12654</td><td>28</td><td>ATGGGGCGAAGCTACCATCTGTGGGATT</td><td>10235.3</td><td>-9.4</td><td>1.7</td><td>-5.6</td><td>2.2E-08</td><td>2.1E-07</td><td>BT20/MB468</td><td>MP</td></tr>
<tr><td>>hg38_21_+_8433172_8446622_RNA45SN1_45S_50ntFlanks@9241.9273.33|CGGAGCGAGCGCACGGGGTCGGCGGCGACGTCG</td><td>>hg38_21_+_8433172_8446622_RNA45SN1_45S_50ntFlanks</td><td>9241</td><td>9273</td><td>33</td><td>CGGAGCGAGCGCACGGGGTCGGCGGCGACGTCG</td><td>72.4</td><td>-9.3</td><td>1.7</td><td>-5.6</td><td>2.2E-08</td><td>2.1E-07</td><td>BT20/MB468</td><td>MP</td></tr>
<tr><td>>hg38_21_+_8433172_8446622_RNA45SN1_45S_50ntFlanks@12356.12375.20|ACCACAGGGATAACTGGCTT</td><td>>hg38_21_+_8433172_8446622_RNA45SN1_45S_50ntFlanks</td><td>12356</td><td>12375</td><td>20</td><td>ACCACAGGGATAACTGGCTT</td><td>77.4</td><td>-9.4</td><td>1.7</td><td>-5.6</td><td>2.2E-08</td><td>2.1E-07</td><td>BT20/MB468</td><td>MP</td></tr>
<tr><td>>hg38_21_+_8433172_8446622_RNA45SN1_45S_50ntFlanks@11671.11702.32|CCAGTGCTCTGAATGTCAAAGTGAAGAAATTC</td><td>>hg38_21_+_8433172_8446622_RNA45SN1_45S_50ntFlanks</td><td>11671</td><td>11702</td><td>32</td><td>CCAGTGCTCTGAATGTCAAAGTGAAGAAATTC</td><td>1096.1</td><td>-3.1</td><td>0.6</td><td>-5.3</td><td>9.9E-08</td><td>2.1E-07</td><td>BT20/MB468</td><td>CYTO</td></tr>
<tr><td>>hg38_21_+_8433172_8446622_RNA45SN1_45S_50ntFlanks@10841.10872.32|CGTAACTTCGGGATAAGGATTGGCTCTAAGGG</td><td>>hg38_21_+_8433172_8446622_RNA45SN1_45S_50ntFlanks</td><td>10841</td><td>10872</td><td>32</td><td>CGTAACTTCGGGATAAGGATTGGCTCTAAGGG</td><td>950.5</td><td>-3.0</td><td>0.6</td><td>-5.3</td><td>9.9E-08</td><td>2.1E-07</td><td>BT20/MB468</td><td>CYTO</td></tr>
<tr><td>>hg38_21_+_8433172_8446622_RNA45SN1_45S_50ntFlanks@8196.8224.29|CGTGGACGGTGTGAGGCCGGTAGCGGCCC</td><td>>hg38_21_+_8433172_8446622_RNA45SN1_45S_50ntFlanks</td><td>8196</td><td>8224</td><td>29</td><td>CGTGGACGGTGTGAGGCCGGTAGCGGCCC</td><td>1280.5</td><td>-3.4</td><td>0.6</td><td>-5.3</td><td>9.9E-08</td><td>2.1E-07</td><td>BT20/MB468</td><td>CYTO</td></tr>
<tr><td>>hg38_21_+_8433172_8446622_RNA45SN1_45S_50ntFlanks@5500.5526.27|GTCGAACTTGACTATCTAGAGGAAGTA</td><td>>hg38_21_+_8433172_8446622_RNA45SN1_45S_50ntFlanks</td><td>5500</td><td>5526</td><td>27</td><td>GTCGAACTTGACTATCTAGAGGAAGTA</td><td>133.6</td><td>-9.2</td><td>1.7</td><td>-5.3</td><td>8.8E-08</td><td>2.1E-07</td><td>BT20/MB468</td><td>MITO</td></tr>
<tr><td>>hg38_21_+_8433172_8446622_RNA45SN1_45S_50ntFlanks@10794.10816.23|CTGGCATGTTGGAACAATGTAGG</td><td>>hg38_21_+_8433172_8446622_RNA45SN1_45S_50ntFlanks</td><td>10794</td><td>10816</td><td>23</td><td>CTGGCATGTTGGAACAATGTAGG</td><td>1581.2</td><td>-4.4</td><td>0.8</td><td>-5.6</td><td>2.2E-08</td><td>2.1E-07</td><td>BT20/MB468</td><td>MP</td></tr>
<tr><td>>hg38_21_+_8433172_8446622_RNA45SN1_45S_50ntFlanks@11593.11613.21|CCAGGGGAATCCGACTGTTTA</td><td>>hg38_21_+_8433172_8446622_RNA45SN1_45S_50ntFlanks</td><td>11593</td><td>11613</td><td>21</td><td>CCAGGGGAATCCGACTGTTTA</td><td>538.3</td><td>-5.9</td><td>1.1</td><td>-5.3</td><td>1.2E-07</td><td>2.1E-07</td><td>MB231/MB468</td><td>NUC</td></tr>
<tr><td>>am_MI0015825|hsa-mir-378c&WithFlank&10|-|130962582|130962674@17.35.19;MI0016749|hsa-mir-378d-1&WithFlank&4|-|5923269|5923334@37.55.19;MI0000786|hsa-mir-378a&WithFlank&5|+|149732819|149732896@49.67.19;MI0003840|hsa-mir-378d-2&WithFlank&8|-|93916016|93916125@28.46.19&[MIMAT0016847&hsa-miR-378c&offsets|0|-6;m-130&10|-|130962639|130962658&offsets|0|-1];[MIMAT0018926&hsa-miR-378d&offsets|0|-1;m-228&4|-|5923281|5923298&offsets|0|+1];[MIMAT0000732&hsa-miR-378a-3p&offsets|0|-3;m-44&5|+|149732867|149732888&offsets|0|-3];[MIMAT0018926_1&hsa-miR-378d&offsets|0|-1;m-227&8|-|93916081|93916098&offsets|0|+1]|ACTGGACTTGGAGTCAGAA</td><td>>am_MI0015825|hsa-mir-378c&WithFlank&10|-|130962582|130962674</td><td>17</td><td>35</td><td>19</td><td>ACTGGACTTGGAGTCAGAA</td><td>227.3</td><td>3.9</td><td>0.7</td><td>5.3</td><td>1.2E-07</td><td>2.1E-07</td><td>MB231/MB468</td><td>NUC</td></tr>
<tr><td>>hg38_21_+_8433172_8446622_RNA45SN1_45S_50ntFlanks@9567.9594.28|ACGTGCAAATCGGTCGTCCGACCTGGGT</td><td>>hg38_21_+_8433172_8446622_RNA45SN1_45S_50ntFlanks</td><td>9567</td><td>9594</td><td>28</td><td>ACGTGCAAATCGGTCGTCCGACCTGGGT</td><td>726.5</td><td>-8.4</td><td>1.6</td><td>-5.3</td><td>1.3E-07</td><td>2.1E-07</td><td>BT20/MB468</td><td>NUC</td></tr>
<tr><td>>MI0000734|hsa-mir-106b&WithFlank&7|-|100093987|100094080@58.80.23&[MIMAT0004672&hsa-miR-106b-3p&offsets|0|+1;m-87&7|-|100094003|100094023&offsets|0|+2]|CCGCACTGTGGGTACTTGCTGCT</td><td>>MI0000734|hsa-mir-106b&WithFlank&7|-|100093987|100094080</td><td>58</td><td>80</td><td>23</td><td>CCGCACTGTGGGTACTTGCTGCT</td><td>426.3</td><td>7.4</td><td>1.4</td><td>5.3</td><td>9.9E-08</td><td>2.1E-07</td><td>BT20/MB468</td><td>CYTO</td></tr>
<tr><td>>hg38_21_+_8433172_8446622_RNA45SN1_45S_50ntFlanks@5090.5120.31|AGACTCTGGCATGCTAACTAGTTACGCGACC</td><td>>hg38_21_+_8433172_8446622_RNA45SN1_45S_50ntFlanks</td><td>5090</td><td>5120</td><td>31</td><td>AGACTCTGGCATGCTAACTAGTTACGCGACC</td><td>83.3</td><td>-10.2</td><td>1.9</td><td>-5.3</td><td>1.0E-07</td><td>2.1E-07</td><td>MB231/MB468</td><td>MP</td></tr>
<tr><td>>hg38_21_+_8433172_8446622_RNA45SN1_45S_50ntFlanks@9835.9864.30|TTTGGTAAGCAGAACTGGCGCTGCGGGATG</td><td>>hg38_21_+_8433172_8446622_RNA45SN1_45S_50ntFlanks</td><td>9835</td><td>9864</td><td>30</td><td>TTTGGTAAGCAGAACTGGCGCTGCGGGATG</td><td>84.4</td><td>10.1</td><td>1.8</td><td>5.6</td><td>2.2E-08</td><td>2.1E-07</td><td>BT20/MB468</td><td>MP</td></tr>
<tr><td>>hg38_21_+_8433172_8446622_RNA45SN1_45S_50ntFlanks@10348.10374.27|AAACGAGAACTTTGAAGGCCGAAGTGG</td><td>>hg38_21_+_8433172_8446622_RNA45SN1_45S_50ntFlanks</td><td>10348</td><td>10374</td><td>27</td><td>AAACGAGAACTTTGAAGGCCGAAGTGG</td><td>2605.6</td><td>-7.1</td><td>1.3</td><td>-5.3</td><td>1.1E-07</td><td>2.1E-07</td><td>MB231/MB468</td><td>MP</td></tr>
<tr><td>>hg38_21_+_8433172_8446622_RNA45SN1_45S_50ntFlanks@10350.10379.30|ACGAGAACTTTGAAGGCCGAAGTGGAGAAG</td><td>>hg38_21_+_8433172_8446622_RNA45SN1_45S_50ntFlanks</td><td>10350</td><td>10379</td><td>30</td><td>ACGAGAACTTTGAAGGCCGAAGTGGAGAAG</td><td>1237.2</td><td>-6.5</td><td>1.2</td><td>-5.3</td><td>1.1E-07</td><td>2.1E-07</td><td>MB231/MB468</td><td>MP</td></tr>
<tr><td>>trna11_GluTTC_15_-_26327381_26327452@41.75.35,trna3_GluTTC_13_-_45492062_45492133@41.75.35|AGGCGGCCCGGGTTCGACTCCCGGTGTGGGAACCA</td><td>>trna11_GluTTC_15_-_26327381_26327452</td><td>41</td><td>75</td><td>35</td><td>AGGCGGCCCGGGTTCGACTCCCGGTGTGGGAACCA</td><td>82.6</td><td>10.1</td><td>1.8</td><td>5.6</td><td>2.3E-08</td><td>2.1E-07</td><td>BT20/MB468</td><td>MP</td></tr>
<tr><td>>hg38_1_-_228634819_228635039_RNA5S12_5S_50ntFlanks@148.171.24|GGAATACCGGGTGCTGTAGGCTTT</td><td>>hg38_1_-_228634819_228635039_RNA5S12_5S_50ntFlanks</td><td>148</td><td>171</td><td>24</td><td>GGAATACCGGGTGCTGTAGGCTTT</td><td>3938.4</td><td>-3.6</td><td>0.7</td><td>-5.3</td><td>1.2E-07</td><td>2.1E-07</td><td>MB231/MB468</td><td>MITO</td></tr>
<tr><td>>hg38_21_+_8433172_8446622_RNA45SN1_45S_50ntFlanks@8321.8356.36|AGACCGATAGTCAACAAGTACCGTAAGGGAAAGTTG</td><td>>hg38_21_+_8433172_8446622_RNA45SN1_45S_50ntFlanks</td><td>8321</td><td>8356</td><td>36</td><td>AGACCGATAGTCAACAAGTACCGTAAGGGAAAGTTG</td><td>180.3</td><td>-6.0</td><td>1.1</td><td>-5.3</td><td>1.0E-07</td><td>2.1E-07</td><td>BT20/MB468</td><td>CYTO</td></tr>
<tr><td>>hg38_21_+_8433172_8446622_RNA45SN1_45S_50ntFlanks@12932.12972.41|CGCTAAACCATTCGTAGACGACCTGCTTCTGGGTCGGGGTT</td><td>>hg38_21_+_8433172_8446622_RNA45SN1_45S_50ntFlanks</td><td>12932</td><td>12972</td><td>41</td><td>CGCTAAACCATTCGTAGACGACCTGCTTCTGGGTCGGGGTT</td><td>1123.1</td><td>-5.6</td><td>1.1</td><td>-5.3</td><td>1.1E-07</td><td>2.1E-07</td><td>MB231/MB468</td><td>CYTO</td></tr>
<tr><td>>hg38_21_+_8433172_8446622_RNA45SN1_45S_50ntFlanks@12459.12477.19|AGCGTTGGATTGTTCACCC</td><td>>hg38_21_+_8433172_8446622_RNA45SN1_45S_50ntFlanks</td><td>12459</td><td>12477</td><td>19</td><td>AGCGTTGGATTGTTCACCC</td><td>449.6</td><td>-5.8</td><td>1.1</td><td>-5.3</td><td>1.2E-07</td><td>2.1E-07</td><td>MB231/MB468</td><td>MITO</td></tr>
<tr><td>>hg38_21_+_8433172_8446622_RNA45SN1_45S_50ntFlanks@4721.4758.38|TTCATTAATCAAGAACGAAAGTCGGAGGTTCGAAGACG</td><td>>hg38_21_+_8433172_8446622_RNA45SN1_45S_50ntFlanks</td><td>4721</td><td>4758</td><td>38</td><td>TTCATTAATCAAGAACGAAAGTCGGAGGTTCGAAGACG</td><td>489.5</td><td>-8.1</td><td>1.5</td><td>-5.3</td><td>1.3E-07</td><td>2.1E-07</td><td>BT20/MB468</td><td>NUC</td></tr>
<tr><td>>hg38_21_+_8433172_8446622_RNA45SN1_45S_50ntFlanks@11767.11794.28|TCTAATTAGTGACGCGCATGAATGGATG</td><td>>hg38_21_+_8433172_8446622_RNA45SN1_45S_50ntFlanks</td><td>11767</td><td>11794</td><td>28</td><td>TCTAATTAGTGACGCGCATGAATGGATG</td><td>5919.9</td><td>-9.7</td><td>1.8</td><td>-5.3</td><td>1.3E-07</td><td>2.1E-07</td><td>BT20/MB468</td><td>NUC</td></tr>
<tr><td>>hg38_21_+_8433172_8446622_RNA45SN1_45S_50ntFlanks@8116.8140.25|GCGGGACATGTGGCGTACGGAAGAC</td><td>>hg38_21_+_8433172_8446622_RNA45SN1_45S_50ntFlanks</td><td>8116</td><td>8140</td><td>25</td><td>GCGGGACATGTGGCGTACGGAAGAC</td><td>692.7</td><td>12.8</td><td>2.2</td><td>5.9</td><td>4.6E-09</td><td>2.1E-07</td><td>BT20/MB231</td><td>MP</td></tr>
<tr><td>>hg38_21_+_8433172_8446622_RNA45SN1_45S_50ntFlanks@8335.8375.41|CAAGTACCGTAAGGGAAAGTTGAAAAGAACTTTGAAGAGAG</td><td>>hg38_21_+_8433172_8446622_RNA45SN1_45S_50ntFlanks</td><td>8335</td><td>8375</td><td>41</td><td>CAAGTACCGTAAGGGAAAGTTGAAAAGAACTTTGAAGAGAG</td><td>646.2</td><td>9.3</td><td>1.7</td><td>5.6</td><td>2.3E-08</td><td>2.1E-07</td><td>BT20/MB468</td><td>MP</td></tr>
<tr><td>>hg38_21_+_8433172_8446622_RNA45SN1_45S_50ntFlanks@9923.9949.27|TTGGTTGATATAGACAGCAGGACGGTG</td><td>>hg38_21_+_8433172_8446622_RNA45SN1_45S_50ntFlanks</td><td>9923</td><td>9949</td><td>27</td><td>TTGGTTGATATAGACAGCAGGACGGTG</td><td>68.3</td><td>9.8</td><td>1.8</td><td>5.6</td><td>2.3E-08</td><td>2.1E-07</td><td>BT20/MB468</td><td>MP</td></tr>
<tr><td>>hg38_21_+_8433172_8446622_RNA45SN1_45S_50ntFlanks@8329.8375.47|AGTCAACAAGTACCGTAAGGGAAAGTTGAAAAGAACTTTGAAGAGAG</td><td>>hg38_21_+_8433172_8446622_RNA45SN1_45S_50ntFlanks</td><td>8329</td><td>8375</td><td>47</td><td>AGTCAACAAGTACCGTAAGGGAAAGTTGAAAAGAACTTTGAAGAGAG</td><td>81.8</td><td>10.1</td><td>1.8</td><td>5.6</td><td>2.3E-08</td><td>2.1E-07</td><td>BT20/MB468</td><td>MP</td></tr>
<tr><td>>MI0000283|hsa-mir-203a&WithFlank&14|+|104117399|104117520@71.92.22&[MIMAT0000264&hsa-miR-203a-3p&offsets|0|0;m-79&14|+|104117469|104117490&offsets|0|0]|GTGAAATGTTTAGGACCACTAG</td><td>>MI0000283|hsa-mir-203a&WithFlank&14|+|104117399|104117520</td><td>71</td><td>92</td><td>22</td><td>GTGAAATGTTTAGGACCACTAG</td><td>680.7</td><td>5.7</td><td>1.0</td><td>5.6</td><td>2.3E-08</td><td>2.1E-07</td><td>BT20/MB468</td><td>MP</td></tr>
<tr><td>>hg38_21_+_8433172_8446622_RNA45SN1_45S_50ntFlanks@5019.5050.32|TCCGTGGGTGGTGGTGCATGGCCGTTCTTAGT</td><td>>hg38_21_+_8433172_8446622_RNA45SN1_45S_50ntFlanks</td><td>5019</td><td>5050</td><td>32</td><td>TCCGTGGGTGGTGGTGCATGGCCGTTCTTAGT</td><td>163.0</td><td>11.1</td><td>2.0</td><td>5.6</td><td>2.3E-08</td><td>2.1E-07</td><td>BT20/MB468</td><td>MP</td></tr>
<tr><td>>hg38_21_+_8433172_8446622_RNA45SN1_45S_50ntFlanks@12709.12743.35|CCGCGGAGCCTCGGTTGGCCTCGGATAGCCGGTCC</td><td>>hg38_21_+_8433172_8446622_RNA45SN1_45S_50ntFlanks</td><td>12709</td><td>12743</td><td>35</td><td>CCGCGGAGCCTCGGTTGGCCTCGGATAGCCGGTCC</td><td>85.1</td><td>-9.5</td><td>1.7</td><td>-5.6</td><td>2.3E-08</td><td>2.1E-07</td><td>BT20/MB468</td><td>MP</td></tr>
<tr><td>>hg38_21_+_8433172_8446622_RNA45SN1_45S_50ntFlanks@12934.12959.26|CTAAACCATTCGTAGACGACCTGCTT</td><td>>hg38_21_+_8433172_8446622_RNA45SN1_45S_50ntFlanks</td><td>12934</td><td>12959</td><td>26</td><td>CTAAACCATTCGTAGACGACCTGCTT</td><td>381.3</td><td>-2.9</td><td>0.5</td><td>-5.3</td><td>9.0E-08</td><td>2.1E-07</td><td>BT20/MB468</td><td>MITO</td></tr>
<tr><td>>hg38_21_+_8433172_8446622_RNA45SN1_45S_50ntFlanks@12942.12959.18|TTCGTAGACGACCTGCTT</td><td>>hg38_21_+_8433172_8446622_RNA45SN1_45S_50ntFlanks</td><td>12942</td><td>12959</td><td>18</td><td>TTCGTAGACGACCTGCTT</td><td>294.4</td><td>-4.2</td><td>0.8</td><td>-5.3</td><td>1.3E-07</td><td>2.1E-07</td><td>MB231/MB468</td><td>MITO</td></tr>
<tr><td>>MI0003137|hsa-mir-193b&WithFlank&16|+|14303961|14304055@20.40.21&[MIMAT0004767&hsa-miR-193b-5p&offsets|0|-1;m-343&16|+|14303980|14304001&offsets|0|-1]|CGGGGTTTTGAGGGCGAGATG</td><td>>MI0003137|hsa-mir-193b&WithFlank&16|+|14303961|14304055</td><td>20</td><td>40</td><td>21</td><td>CGGGGTTTTGAGGGCGAGATG</td><td>385.7</td><td>9.4</td><td>1.7</td><td>5.6</td><td>2.3E-08</td><td>2.1E-07</td><td>BT20/MB468</td><td>MP</td></tr>
<tr><td>>hg38_21_+_8433172_8446622_RNA45SN1_45S_50ntFlanks@12675.12708.34|CAGAATCCCGCCCAGGCGGAACGATACGGCAGCG</td><td>>hg38_21_+_8433172_8446622_RNA45SN1_45S_50ntFlanks</td><td>12675</td><td>12708</td><td>34</td><td>CAGAATCCCGCCCAGGCGGAACGATACGGCAGCG</td><td>77.8</td><td>-9.4</td><td>1.7</td><td>-5.6</td><td>2.3E-08</td><td>2.1E-07</td><td>BT20/MB468</td><td>MP</td></tr>
<tr><td>>hg38_21_+_8433172_8446622_RNA45SN1_45S_50ntFlanks@11322.11344.23|ACGTCTCGTCGCGCGCGCGTCCG</td><td>>hg38_21_+_8433172_8446622_RNA45SN1_45S_50ntFlanks</td><td>11322</td><td>11344</td><td>23</td><td>ACGTCTCGTCGCGCGCGCGTCCG</td><td>29117.5</td><td>-7.1</td><td>1.3</td><td>-5.3</td><td>1.3E-07</td><td>2.1E-07</td><td>BT20/MB468</td><td>NUC</td></tr>
<tr><td>>hg38_21_+_8433172_8446622_RNA45SN1_45S_50ntFlanks@12407.12431.25|TTTTGATCCTTCGATGTCGGCTCTT</td><td>>hg38_21_+_8433172_8446622_RNA45SN1_45S_50ntFlanks</td><td>12407</td><td>12431</td><td>25</td><td>TTTTGATCCTTCGATGTCGGCTCTT</td><td>2781.8</td><td>-3.7</td><td>0.7</td><td>-5.3</td><td>1.2E-07</td><td>2.1E-07</td><td>MB231/MB468</td><td>CYTO</td></tr>
<tr><td>>hg38_21_+_8433172_8446622_RNA45SN1_45S_50ntFlanks@10319.10352.34|GCAGATCTTGGTGGTAGTAGCAAATATTCAAACG</td><td>>hg38_21_+_8433172_8446622_RNA45SN1_45S_50ntFlanks</td><td>10319</td><td>10352</td><td>34</td><td>GCAGATCTTGGTGGTAGTAGCAAATATTCAAACG</td><td>69.2</td><td>9.9</td><td>1.8</td><td>5.6</td><td>2.3E-08</td><td>2.1E-07</td><td>BT20/MB468</td><td>MP</td></tr>
<tr><td>>hg38_21_+_8433172_8446622_RNA45SN1_45S_50ntFlanks@10402.10432.31|AACATGGGTCAGTCGGTCCTGAGAGATGGGC</td><td>>hg38_21_+_8433172_8446622_RNA45SN1_45S_50ntFlanks</td><td>10402</td><td>10432</td><td>31</td><td>AACATGGGTCAGTCGGTCCTGAGAGATGGGC</td><td>1151.4</td><td>-8.7</td><td>1.6</td><td>-5.3</td><td>1.3E-07</td><td>2.1E-07</td><td>BT20/MB468</td><td>NUC</td></tr>
</table>

<table>
<tr><th></th><th></th><th></th><th></th><th></th><th></th><th></th><th></th><th></th><th></th><th></th><th></th><th></th><th></th></tr>
<tr><td>>hg38_21_+_8433172_8446622_RNA45SN1_45S_50ntFlanks@10404.10433.30|CATGGGTCAGTCGGTCCTGAGAGATGGGCG</td><td>>hg38_21_+_8433172_8446622_RNA45SN1_45S_50ntFlanks</td><td>10404</td><td>10433</td><td>30</td><td>CATGGGTCAGTCGGTCCTGAGAGATGGGCG</td><td>130.6</td><td>-5.7</td><td>1.1</td><td>-5.3</td><td>1.0E-07</td><td>2.1E-07</td><td>BT20/MB468</td><td>CYTO</td></tr>
<tr><td>>MI0000068|hsa-let-7f-2&WithFlank&X|-|53557186|53557280@14.36.23&[MIMAT0000067_1&hsa-let-7f-5p&offsets|0|+1;m-19&X|-|53557246|53557267&offsets|0|+1]|TGAGGTAGTAGATTGTATAGTTT</td><td>>MI0000068|hsa-let-7f-2&WithFlank&X|-|53557186|53557280</td><td>14</td><td>36</td><td>23</td><td>TGAGGTAGTAGATTGTATAGTTT</td><td>1321.3</td><td>4.9</td><td>0.9</td><td>5.3</td><td>1.3E-07</td><td>2.1E-07</td><td>MB231/MB468</td><td>MITO</td></tr>
<tr><td>>hg38_21_+_8433172_8446622_RNA45SN1_45S_50ntFlanks@10234.10260.27|CGGACGCTACGCCGCGACGAGTAGGAG</td><td>>hg38_21_+_8433172_8446622_RNA45SN1_45S_50ntFlanks</td><td>10234</td><td>10260</td><td>27</td><td>CGGACGCTACGCCGCGACGAGTAGGAG</td><td>316.4</td><td>-7.1</td><td>1.3</td><td>-5.3</td><td>1.3E-07</td><td>2.1E-07</td><td>MB231/MB468</td><td>MITO</td></tr>
<tr><td>>MI0025242|hsa-mir-7706&WithFlank&15|+|85380590|85380668@50.72.23(+1U)&[MIMAT0030021&hsa-miR-7706&offsets|0|-1(+1U);m-390&15|+|85380639|85380662&offsets|0|-1(+1U)]|TGAAGCGCCTGTGCTCTGCCGAGT</td><td>>MI0025242|hsa-mir-7706&WithFlank&15|+|85380590|85380668</td><td>50</td><td>72</td><td>23(+1U)</td><td>TGAAGCGCCTGTGCTCTGCCGAGT</td><td>388.4</td><td>7.5</td><td>1.4</td><td>5.3</td><td>1.0E-07</td><td>2.1E-07</td><td>BT20/MB468</td><td>CYTO</td></tr>
<tr><td>>MI0001448|hsa-mir-425&WithFlank&3|-|49020142|49020240@21.44.24&[MIMAT0003393&hsa-miR-425-5p&offsets|+1|+2;m-95&3|-|49020200|49020221&offsets|+1|+3]|ATGACACGATCACTCCCGTTGAGT</td><td>>MI0001448|hsa-mir-425&WithFlank&3|-|49020142|49020240</td><td>21</td><td>44</td><td>24</td><td>ATGACACGATCACTCCCGTTGAGT</td><td>94.7</td><td>2.6</td><td>0.5</td><td>5.3</td><td>1.3E-07</td><td>2.1E-07</td><td>BT20/MB468</td><td>NUC</td></tr>
<tr><td>>MI0000061|hsa-let-7a-2&WithFlank&11|-|122146516|122146599@11.28.18;MI0000064|hsa-let-7c&WithFlank&21|+|16539822|16539917@17.34.18;MI0000062|hsa-let-7a-3&WithFlank&22|+|46112743|46112828@10.27.18;MI0000060|hsa-let-7a-1&WithFlank&9|+|94175951|94176042@12.29.18&[MIMAT0000062&hsa-let-7a-5p&offsets|0|-4;m-6&11|-|122146568|122146589&offsets|0|-4];[MIMAT0000064&hsa-let-7c-5p&offsets|0|-4;m-37&21|+|16539838|16539859&offsets|0|-4];[MIMAT0000062_1&hsa-let-7a-5p&offsets|0|-4;m-5&22|+|46112752|46112773&offsets|0|-4];[MIMAT0000062_2&hsa-let-7a-5p&offsets|0|-4;m-4&9|+|94175962|94175983&offsets|0|-4]|TGAGGTAGTAGGTTGTAT</td><td>>MI0000061|hsa-let-7a-2&WithFlank&11|-|122146516|122146599</td><td>11</td><td>28</td><td>18</td><td>TGAGGTAGTAGGTTGTAT</td><td>2254.4</td><td>5.5</td><td>1.0</td><td>5.3</td><td>1.1E-07</td><td>2.2E-07</td><td>MB231/MB468</td><td>MP</td></tr>
<tr><td>>hg38_21_+_8433172_8446622_RNA45SN1_45S_50ntFlanks@4380.4406.27|CGTAGTTGGATCTTGGGAGCGGGCGGG</td><td>>hg38_21_+_8433172_8446622_RNA45SN1_45S_50ntFlanks</td><td>4380</td><td>4406</td><td>27</td><td>CGTAGTTGGATCTTGGGAGCGGGCGGG</td><td>206.5</td><td>-5.3</td><td>1.0</td><td>-5.3</td><td>1.2E-07</td><td>2.2E-07</td><td>MB231/MB468</td><td>NUC</td></tr>
<tr><td>>hg38_21_+_8433172_8446622_RNA45SN1_45S_50ntFlanks@12141.12182.42|ACAGTGCCAGGTGGGGAGTTTGACTGGGGCGGTACACCTGTC</td><td>>hg38_21_+_8433172_8446622_RNA45SN1_45S_50ntFlanks</td><td>12141</td><td>12182</td><td>42</td><td>ACAGTGCCAGGTGGGGAGTTTGACTGGGGCGGTACACCTGTC</td><td>111.4</td><td>-8.9</td><td>1.7</td><td>-5.3</td><td>9.0E-08</td><td>2.2E-07</td><td>BT20/MB468</td><td>MITO</td></tr>
<tr><td>>MI0000081|hsa-mir-24-2&WithFlank&19|-|13836281|13836365@56.77.22(+1C);MI0000080|hsa-mir-24-1&WithFlank&9|+|95086015|95086094@50.71.22(+1C)&[MIMAT0000080&hsa-miR-24-3p&offsets|0|0(+1C);m-36&19|-|13836288|13836310&offsets|0|-1(+1C)];[MIMAT0000080_1&hsa-miR-24-3p&offsets|0|0(+1C);m-35&9|+|95086064|95086086&offsets|0|-1(+1C)]|TGGCTCAGTTCAGCAGGAACAGC</td><td>>MI0000081|hsa-mir-24-2&WithFlank&19|-|13836281|13836365</td><td>56</td><td>77</td><td>22(+1C)</td><td>TGGCTCAGTTCAGCAGGAACAGC</td><td>292.9</td><td>3.1</td><td>0.6</td><td>5.3</td><td>1.3E-07</td><td>2.2E-07</td><td>BT20/MB468</td><td>NUC</td></tr>
<tr><td>>MI0000063|hsa-let-7b&WithFlank&22|+|46113680|46113774@12.32.21&[MIMAT0000063&hsa-let-7b-5p&offsets|0|-1;m-14&22|+|46113691|46113712&offsets|0|-1]|TGAGGTAGTAGGTTGTGTGGT</td><td>>MI0000063|hsa-let-7b&WithFlank&22|+|46113680|46113774</td><td>12</td><td>32</td><td>21</td><td>TGAGGTAGTAGGTTGTGTGGT</td><td>10145.9</td><td>5.2</td><td>0.9</td><td>5.6</td><td>2.3E-08</td><td>2.2E-07</td><td>BT20/MB468</td><td>MP</td></tr>
<tr><td>>MI0000650|hsa-mir-200c&WithFlank&12|+|6963693|6963772@49.71.23&[MIMAT0000617&hsa-miR-200c-3p&offsets|-1|-1;m-54&12|+|6963742|6963764&offsets|-1|-1]|CTAATACTGCCGGGTAATGATGG</td><td>>MI0000650|hsa-mir-200c&WithFlank&12|+|6963693|6963772</td><td>49</td><td>71</td><td>23</td><td>CTAATACTGCCGGGTAATGATGG</td><td>151.5</td><td>10.6</td><td>1.8</td><td>5.9</td><td>4.7E-09</td><td>2.2E-07</td><td>BT20/MB231</td><td>MP</td></tr>
<tr><td>>hg38_21_+_8433172_8446622_RNA45SN1_45S_50ntFlanks@10715.10736.22|TGAAAATCCGGGGGAGAGGGTG</td><td>>hg38_21_+_8433172_8446622_RNA45SN1_45S_50ntFlanks</td><td>10715</td><td>10736</td><td>22</td><td>TGAAAATCCGGGGGAGAGGGTG</td><td>198.1</td><td>-5.7</td><td>1.1</td><td>-5.3</td><td>1.1E-07</td><td>2.2E-07</td><td>MB231/MB468</td><td>MP</td></tr>
<tr><td>>hg38_21_+_8433172_8446622_RNA45SN1_45S_50ntFlanks@11572.11602.31|CCGACTTAGAACTGGTGCGGACCAGGGGAAT</td><td>>hg38_21_+_8433172_8446622_RNA45SN1_45S_50ntFlanks</td><td>11572</td><td>11602</td><td>31</td><td>CCGACTTAGAACTGGTGCGGACCAGGGGAAT</td><td>6320.9</td><td>-4.1</td><td>0.8</td><td>-5.3</td><td>1.2E-07</td><td>2.2E-07</td><td>MB231/MB468</td><td>CYTO</td></tr>
<tr><td>>hg38_21_+_8433172_8446622_RNA45SN1_45S_50ntFlanks@11561.11594.34|GCGCCTAGCAGCCGACTTAGAACTGGTGCGGACC</td><td>>hg38_21_+_8433172_8446622_RNA45SN1_45S_50ntFlanks</td><td>11561</td><td>11594</td><td>34</td><td>GCGCCTAGCAGCCGACTTAGAACTGGTGCGGACC</td><td>544.7</td><td>-8.2</td><td>1.6</td><td>-5.3</td><td>1.3E-07</td><td>2.2E-07</td><td>BT20/MB468</td><td>NUC</td></tr>
<tr><td>>hg38_MT_+_598_1651_RNR1_12S_50ntFlanks@427.468.42|GAAAGTGGCTTTAACATATCTGAACACACAATAGCTAAGACC</td><td>>hg38_MT_+_598_1651_RNR1_12S_50ntFlanks</td><td>427</td><td>468</td><td>42</td><td>GAAAGTGGCTTTAACATATCTGAACACACAATAGCTAAGACC</td><td>60.6</td><td>9.1</td><td>1.7</td><td>5.3</td><td>1.1E-07</td><td>2.2E-07</td><td>MB231/MB468</td><td>MP</td></tr>
<tr><td>>hg38_21_+_8433172_8446622_RNA45SN1_45S_50ntFlanks@6645.6672.28|CTCGTACGACTCTTAGCGGTGGATCACT</td><td>>hg38_21_+_8433172_8446622_RNA45SN1_45S_50ntFlanks</td><td>6645</td><td>6672</td><td>28</td><td>CTCGTACGACTCTTAGCGGTGGATCACT</td><td>1603.7</td><td>6.5</td><td>1.2</td><td>5.3</td><td>1.0E-07</td><td>2.2E-07</td><td>BT20/MB468</td><td>CYTO</td></tr>
<tr><td>>hg38_21_+_8433172_8446622_RNA45SN1_45S_50ntFlanks@9739.9759.21|AAACGATCTCAACCTATTCTC</td><td>>hg38_21_+_8433172_8446622_RNA45SN1_45S_50ntFlanks</td><td>9739</td><td>9759</td><td>21</td><td>AAACGATCTCAACCTATTCTC</td><td>459.5</td><td>-3.2</td><td>0.6</td><td>-5.3</td><td>1.0E-07</td><td>2.2E-07</td><td>BT20/MB468</td><td>CYTO</td></tr>
<tr><td>>hg38_21_+_8433172_8446622_RNA45SN1_45S_50ntFlanks@9929.9947.19|GATATAGACAGCAGGACGG</td><td>>hg38_21_+_8433172_8446622_RNA45SN1_45S_50ntFlanks</td><td>9929</td><td>9947</td><td>19</td><td>GATATAGACAGCAGGACGG</td><td>335.2</td><td>-6.2</td><td>1.2</td><td>-5.3</td><td>1.1E-07</td><td>2.2E-07</td><td>MB231/MB468</td><td>MP</td></tr>
<tr><td>>hg38_21_+_8433172_8446622_RNA45SN1_45S_50ntFlanks@7974.7992.19|ACGCGACCTCAGATCAGAC</td><td>>hg38_21_+_8433172_8446622_RNA45SN1_45S_50ntFlanks</td><td>7974</td><td>7992</td><td>19</td><td>ACGCGACCTCAGATCAGAC</td><td>2255.9</td><td>4.4</td><td>0.8</td><td>5.3</td><td>1.2E-07</td><td>2.2E-07</td><td>MB231/MB468</td><td>CYTO</td></tr>
<tr><td>>trna111_HisGTG_1_-_147774845_147774916@-1G.33.34,trna118_HisGTG_1_-_145396881_145396952@-1G.33.34,trna16_HisGTG_1_+_146544773_146544844@-1G.33.34,trna1_HisGTG_15_+_45493349_45493420@-1G.33.34,trna21_HisGTG_1_+_147753471_147753542@-1G.33.34,trna33_HisGTG_6_+_27125906_27125977@-1G.33.34,trna7_HisGTG_9_-_14433938_14434009@-1G.33.34,trna8_HisGTG_15_-_45492611_45492682@-1G.33.34,trna9_HisGTG_15_-_45490804_45490875@-1G.33.34|GGCCGTGATCGTATAGTGGTTAGTACTCTGCGTT</td><td>>trna111_HisGTG_1_-_147774845_147774916</td><td>-1G</td><td>33</td><td>34</td><td>GGCCGTGATCGTATAGTGGTTAGTACTCTGCGTT</td><td>12277.7</td><td>8.6</td><td>1.5</td><td>5.6</td><td>2.3E-08</td><td>2.2E-07</td><td>BT20/MB468</td><td>MP</td></tr>
<tr><td>>hg38_21_+_8433172_8446622_RNA45SN1_45S_50ntFlanks@12004.12032.29|CGGGCCGCCGGTGAAATACCACTACTCTG</td><td>>hg38_21_+_8433172_8446622_RNA45SN1_45S_50ntFlanks</td><td>12004</td><td>12032</td><td>29</td><td>CGGGCCGCCGGTGAAATACCACTACTCTG</td><td>1136.6</td><td>-3.7</td><td>0.7</td><td>-5.6</td><td>2.3E-08</td><td>2.2E-07</td><td>BT20/MB468</td><td>MP</td></tr>
<tr><td>>hg38_21_+_8433172_8446622_RNA45SN1_45S_50ntFlanks@12126.12147.22|CGACCCGCTCCGGGGACAGTGC</td><td>>hg38_21_+_8433172_8446622_RNA45SN1_45S_50ntFlanks</td><td>12126</td><td>12147</td><td>22</td><td>CGACCCGCTCCGGGGACAGTGC</td><td>87.1</td><td>-9.5</td><td>1.7</td><td>-5.6</td><td>2.3E-08</td><td>2.2E-07</td><td>BT20/MB468</td><td>MP</td></tr>
<tr><td>>hg38_21_+_8433172_8446622_RNA45SN1_45S_50ntFlanks@12785.12805.21|CCCCGCGCGCGCGGGAGGGCG</td><td>>hg38_21_+_8433172_8446622_RNA45SN1_45S_50ntFlanks</td><td>12785</td><td>12805</td><td>21</td><td>CCCCGCGCGCGCGGGAGGGCG</td><td>1114.4</td><td>-8.3</td><td>1.5</td><td>-5.6</td><td>2.3E-08</td><td>2.2E-07</td><td>BT20/MB468</td><td>MP</td></tr>
<tr><td>>hg38_21_+_8433172_8446622_RNA45SN1_45S_50ntFlanks@12008.12036.29|CCGCCGGTGAAATACCACTACTCTGATCG</td><td>>hg38_21_+_8433172_8446622_RNA45SN1_45S_50ntFlanks</td><td>12008</td><td>12036</td><td>29</td><td>CCGCCGGTGAAATACCACTACTCTGATCG</td><td>326.2</td><td>-5.7</td><td>1.1</td><td>-5.3</td><td>1.2E-07</td><td>2.2E-07</td><td>MB231/MB468</td><td>NUC</td></tr>
<tr><td>>hg38_21_+_8433172_8446622_RNA45SN1_45S_50ntFlanks@9505.9525.21|TGAACTATGCCTGGGCAGGGC</td><td>>hg38_21_+_8433172_8446622_RNA45SN1_45S_50ntFlanks</td><td>9505</td><td>9525</td><td>21</td><td>TGAACTATGCCTGGGCAGGGC</td><td>137.6</td><td>-10.2</td><td>1.8</td><td>-5.6</td><td>2.3E-08</td><td>2.2E-07</td><td>BT20/MB468</td><td>MP</td></tr>
<tr><td>>MI0000273|hsa-mir-183&WithFlank&7|-|129774899|129775020@34.54.21(+1A)&[MIMAT0000261&hsa-miR-183-5p&offsets|+1|0(+1A);m-41&7|-|129774966|129774987&offsets|0|-1(+1A)]|ATGGCACTGGTAGAATTCACTA</td><td>>MI0000273|hsa-mir-183&WithFlank&7|-|129774899|129775020</td><td>34</td><td>54</td><td>21(+1A)</td><td>ATGGCACTGGTAGAATTCACTA</td><td>717.0</td><td>4.7</td><td>0.8</td><td>5.8</td><td>7.6E-09</td><td>2.2E-07</td><td>BT20/MB231</td><td>CYTO</td></tr>
<tr><td>>MI0005756|hsa-mir-934&WithFlank&X|+|136550872|136550966@21.41.21(+1A)&[MIMAT0004977&hsa-miR-934&offsets|0|-1(+1A);m-943&X|+|136550892|136550913&offsets|0|-1(+1A)]|TGTCTACTACTGGAGACACTGA</td><td>>MI0005756|hsa-mir-934&WithFlank&X|+|136550872|136550966</td><td>21</td><td>41</td><td>21(+1A)</td><td>TGTCTACTACTGGAGACACTGA</td><td>377.0</td><td>8.8</td><td>1.6</td><td>5.3</td><td>1.0E-07</td><td>2.2E-07</td><td>BT20/MB468</td><td>CYTO</td></tr>
<tr><td>>MI0000286|hsa-mir-210&WithFlank&11|-|568083|568204@72.93.22(+2A)&[MIMAT0000267&hsa-miR-210-3p&offsets|0|0(+2A);m-102&11|-|568112|568133&offsets|0|0(+2A)]|CTGTGCGTGTGACAGCGGCTGAAA</td><td>>MI0000286|hsa-mir-210&WithFlank&11|-|568083|568204</td><td>72</td><td>93</td><td>22(+2A)</td><td>CTGTGCGTGTGACAGCGGCTGAAA</td><td>2552.7</td><td>5.6</td><td>1.0</td><td>5.6</td><td>2.3E-08</td><td>2.2E-07</td><td>BT20/MB468</td><td>MP</td></tr>
<tr><td>>hg38_21_+_8433172_8446622_RNA45SN1_45S_50ntFlanks@3920.3938.19|CATTTATCAGATCAAAACC</td><td>>hg38_21_+_8433172_8446622_RNA45SN1_45S_50ntFlanks</td><td>3920</td><td>3938</td><td>19</td><td>CATTTATCAGATCAAAACC</td><td>81.9</td><td>-9.4</td><td>1.7</td><td>-5.6</td><td>2.4E-08</td><td>2.2E-07</td><td>BT20/MB468</td><td>MP</td></tr>
<tr><td>>MI0000542|hsa-mir-320a&WithFlank&8|-|22244960|22245043@48.72.25&[MIMAT0000510&hsa-miR-320a-3p&offsets|0|+3;m-60&8|-|22244975|22244996&offsets|0|+3]|AAAAGCTGGGTTGAGAGGGCGAAAA</td><td>>MI0000542|hsa-mir-320a&WithFlank&8|-|22244960|22245043</td><td>48</td><td>72</td><td>25</td><td>AAAAGCTGGGTTGAGAGGGCGAAAA</td><td>717.7</td><td>4.4</td><td>0.8</td><td>5.8</td><td>7.7E-09</td><td>2.2E-07</td><td>BT20/MB231</td><td>CYTO</td></tr>
<tr><td>>hg38_21_+_8433172_8446622_RNA45SN1_45S_50ntFlanks@8195.8215.21|CCGTGGACGGTGTGAGGCCGG</td><td>>hg38_21_+_8433172_8446622_RNA45SN1_45S_50ntFlanks</td><td>8195</td><td>8215</td><td>21</td><td>CCGTGGACGGTGTGAGGCCGG</td><td>358.0</td><td>-4.5</td><td>0.8</td><td>-5.3</td><td>1.1E-07</td><td>2.2E-07</td><td>MB231/MB468</td><td>MP</td></tr>
<tr><td>>hg38_21_+_8433172_8446622_RNA45SN1_45S_50ntFlanks@10362.10388.27|AAGGCCGAAGTGGAGAAGGGTTCCATG</td><td>>hg38_21_+_8433172_8446622_RNA45SN1_45S_50ntFlanks</td><td>10362</td><td>10388</td><td>27</td><td>AAGGCCGAAGTGGAGAAGGGTTCCATG</td><td>4073.0</td><td>-7.4</td><td>1.4</td><td>-5.3</td><td>1.0E-07</td><td>2.2E-07</td><td>BT20/MB468</td><td>CYTO</td></tr>
<tr><td>>hg38_21_+_8433172_8446622_RNA45SN1_45S_50ntFlanks@9534.9573.40|AGGAAACTCTGGTGGAGGTCCGTAGCGGTCCTGACGTGCA</td><td>>hg38_21_+_8433172_8446622_RNA45SN1_45S_50ntFlanks</td><td>9534</td><td>9573</td><td>40</td><td>AGGAAACTCTGGTGGAGGTCCGTAGCGGTCCTGACGTGCA</td><td>562.9</td><td>-6.4</td><td>1.2</td><td>-5.3</td><td>1.1E-07</td><td>2.2E-07</td><td>MB231/MB468</td><td>MP</td></tr>
<tr><td>>am_trna10_ValCAC_5_-_180649395_180649467@6.29.24,trna12_ValAAC_5_-_180645270_180645342@6.29.24,trna132_ValAAC_6_-_27721179_27721251@6.29.24,trna136_ValAAC_6_-_27648885_27648957@6.29.24,trna139_ValAAC_6_-_27618707_27618779@6.29.24,trna18_ValCAC_5_-_180529253_180529325@6.29.24,trna2_ValAAC_3_+_169490018_169490090@6.29.24,trna2_ValCAC_5_+_180524070_180524142@6.29.24,trna4_ValAAC_5_+_180591154_180591226@6.29.24,trna5_ValAAC_5_+_180596610_180596682@6.29.24,trna6_ValCAC_5_+_180600650_180600722@6.29.24,trna85_ValCAC_1_-_161369490_161369562@6.29.24,trna90_ValCAC_1_-_149684088_149684161@6.29.24,trna98_ValCAC_1_-_149298555_149298627@6.29.24,trna9_ValCAC_6_+_26538282_26538354@6.29.24|CGTAGTGTAGTGGTTATCACGTTC</td><td>>am_trna10_ValCAC_5_-_180649395_180649467</td><td>6</td><td>29</td><td>24</td><td>CGTAGTGTAGTGGTTATCACGTTC</td><td>127.5</td><td>-10.1</td><td>1.8</td><td>-5.6</td><td>2.4E-08</td><td>2.2E-07</td><td>BT20/MB468</td><td>MP</td></tr>
<tr><td>>MI0000487|hsa-mir-193a&WithFlank&17|+|31559990|31560089@27.48.22&[MIMAT0004614&hsa-miR-193a-5p&offsets|0|0;m-123&17|+|31560016|31560037&offsets|0|0]|TGGGTCTTTGCGGGCGAGATGA</td><td>>MI0000487|hsa-mir-193a&WithFlank&17|+|31559990|31560089</td><td>27</td><td>48</td><td>22</td><td>TGGGTCTTTGCGGGCGAGATGA</td><td>314.7</td><td>4.1</td><td>0.7</td><td>5.7</td><td>9.9E-09</td><td>2.2E-07</td><td>BT20/MB231</td><td>MITO</td></tr>
<tr><td>>hg38_21_+_8433172_8446622_RNA45SN1_45S_50ntFlanks@12273.12290.18|TCAGTACGAATACAGACC</td><td>>hg38_21_+_8433172_8446622_RNA45SN1_45S_50ntFlanks</td><td>12273</td><td>12290</td><td>18</td><td>TCAGTACGAATACAGACC</td><td>192.0</td><td>-5.8</td><td>1.1</td><td>-5.3</td><td>1.1E-07</td><td>2.2E-07</td><td>MB231/MB468</td><td>MP</td></tr>
<tr><td>>hg38_21_+_8433172_8446622_RNA45SN1_45S_50ntFlanks@4770.4793.24|TCGTAGTTCCGACCATAAACGATG</td><td>>hg38_21_+_8433172_8446622_RNA45SN1_45S_50ntFlanks</td><td>4770</td><td>4793</td><td>24</td><td>TCGTAGTTCCGACCATAAACGATG</td><td>478.6</td><td>-2.7</td><td>0.5</td><td>-5.3</td><td>1.0E-07</td><td>2.2E-07</td><td>BT20/MB468</td><td>CYTO</td></tr>
<tr><td>>hg38_21_+_8433172_8446622_RNA45SN1_45S_50ntFlanks@6652.6688.37|GACTCTTAGCGGTGGATCACTCGGCTCGTGCGTCGAT</td><td>>hg38_21_+_8433172_8446622_RNA45SN1_45S_50ntFlanks</td><td>6652</td><td>6688</td><td>37</td><td>GACTCTTAGCGGTGGATCACTCGGCTCGTGCGTCGAT</td><td>13411.4</td><td>2.9</td><td>0.5</td><td>5.3</td><td>9.2E-08</td><td>2.2E-07</td><td>BT20/MB468</td><td>MITO</td></tr>
</table>

<table>
<tr><td>>hg38_21_+_8433172_8446622_RNA45SN1_45S_50ntFlanks@11873.11893.21|AAGAAGACCCTGTTGAGCTTG</td><td>>hg38_21_+_8433172_8446622_RNA45SN1_45S_50ntFlanks</td><td>11873</td><td>11893</td><td>21</td><td>AAGAAGACCCTGTTGAGCTTG</td><td>1359.2</td><td>-3.6</td><td>0.7</td><td>-5.3</td><td>9.2E-08</td><td>2.2E-07</td><td>BT20/MB468</td><td>MITO</td></tr>
<tr><td>>hg38_21_+_8433172_8446622_RNA45SN1_45S_50ntFlanks@11873.11904.32|AAGAAGACCCTGTTGAGCTTGACTCTAGTCTG</td><td>>hg38_21_+_8433172_8446622_RNA45SN1_45S_50ntFlanks</td><td>11873</td><td>11904</td><td>32</td><td>AAGAAGACCCTGTTGAGCTTGACTCTAGTCTG</td><td>2251.9</td><td>-7.3</td><td>1.3</td><td>-5.6</td><td>2.4E-08</td><td>2.2E-07</td><td>BT20/MB468</td><td>MP</td></tr>
<tr><td>>hg38_21_+_8433172_8446622_RNA45SN1_45S_50ntFlanks@12455.12473.19|ACCAAGCGTTGGATTGTTC</td><td>>hg38_21_+_8433172_8446622_RNA45SN1_45S_50ntFlanks</td><td>12455</td><td>12473</td><td>19</td><td>ACCAAGCGTTGGATTGTTC</td><td>1227.5</td><td>-4.0</td><td>0.8</td><td>-5.3</td><td>1.1E-07</td><td>2.2E-07</td><td>MB231/MB468</td><td>MP</td></tr>
<tr><td>>hg38_21_+_8433172_8446622_RNA45SN1_45S_50ntFlanks@12640.12667.28|ACCATCTGTGGGATTATGACTGAACGCC</td><td>>hg38_21_+_8433172_8446622_RNA45SN1_45S_50ntFlanks</td><td>12640</td><td>12667</td><td>28</td><td>ACCATCTGTGGGATTATGACTGAACGCC</td><td>218.3</td><td>5.7</td><td>1.1</td><td>5.3</td><td>1.2E-07</td><td>2.2E-07</td><td>MB231/MB468</td><td>NUC</td></tr>
<tr><td>>hg38_21_+_8433172_8446622_RNA45SN1_45S_50ntFlanks@9527.9571.45|AAGCCAGAGGAAACTCTGGTGGAGGTCCGTAGCGGTCCTGACGTG</td><td>>hg38_21_+_8433172_8446622_RNA45SN1_45S_50ntFlanks</td><td>9527</td><td>9571</td><td>45</td><td>AAGCCAGAGGAAACTCTGGTGGAGGTCCGTAGCGGTCCTGACGTG</td><td>828.7</td><td>-9.1</td><td>1.7</td><td>-5.3</td><td>1.4E-07</td><td>2.2E-07</td><td>BT20/MB468</td><td>NUC</td></tr>
<tr><td>>MI0003628|hsa-mir-615&WithFlank&12|+|54033944|54034051@67.86.20(+1C)&[MIMAT0003283&hsa-miR-615-3p&offsets|0|-2(+1C);m-495&12|+|54034010|54034030&offsets|0|-1(+1C)]|TCCGAGCCTGGGTCTCCCTCC</td><td>>MI0003628|hsa-mir-615&WithFlank&12|+|54033944|54034051</td><td>67</td><td>86</td><td>20(+1C)</td><td>TCCGAGCCTGGGTCTCCCTCC</td><td>265.4</td><td>6.1</td><td>1.1</td><td>5.7</td><td>1.3E-08</td><td>2.2E-07</td><td>BT20/MB231</td><td>NUC</td></tr>
<tr><td>>hg38_21_+_8433172_8446622_RNA45SN1_45S_50ntFlanks@11322.11340.19|ACGTCTCGTCGCGCGCGCG</td><td>>hg38_21_+_8433172_8446622_RNA45SN1_45S_50ntFlanks</td><td>11322</td><td>11340</td><td>19</td><td>ACGTCTCGTCGCGCGCGCG</td><td>767.6</td><td>-6.6</td><td>1.3</td><td>-5.3</td><td>1.3E-07</td><td>2.2E-07</td><td>MB231/MB468</td><td>MITO</td></tr>
<tr><td>>hg38_21_+_8433172_8446622_RNA45SN1_45S_50ntFlanks@12073.12094.22|CCGAGGGGCTCTCGCTTCTGGC</td><td>>hg38_21_+_8433172_8446622_RNA45SN1_45S_50ntFlanks</td><td>12073</td><td>12094</td><td>22</td><td>CCGAGGGGCTCTCGCTTCTGGC</td><td>325.7</td><td>-7.2</td><td>1.4</td><td>-5.3</td><td>1.3E-07</td><td>2.2E-07</td><td>MB231/MB468</td><td>MITO</td></tr>
<tr><td>>hg38_21_+_8433172_8446622_RNA45SN1_45S_50ntFlanks@12357.12376.20|CCACAGGGATAACTGGCTTG</td><td>>hg38_21_+_8433172_8446622_RNA45SN1_45S_50ntFlanks</td><td>12357</td><td>12376</td><td>20</td><td>CCACAGGGATAACTGGCTTG</td><td>516.1</td><td>-6.9</td><td>1.3</td><td>-5.3</td><td>1.0E-07</td><td>2.2E-07</td><td>BT20/MB468</td><td>CYTO</td></tr>
<tr><td>>MI0000114|hsa-mir-107&WithFlank&10|-|89592741|89592833@56.77.22&[MIMAT0000104&hsa-miR-107&offsets|0|-1;m-78&10|-|89592758|89592778&offsets|0|+1]|AGCAGCATTGTACAGGGCTATC</td><td>>MI0000114|hsa-mir-107&WithFlank&10|-|89592741|89592833</td><td>56</td><td>77</td><td>22</td><td>AGCAGCATTGTACAGGGCTATC</td><td>901.7</td><td>5.4</td><td>1.0</td><td>5.3</td><td>1.3E-07</td><td>2.2E-07</td><td>MB231/MB468</td><td>MITO</td></tr>
<tr><td>>hg38_21_+_8433172_8446622_RNA45SN1_45S_50ntFlanks@8384.8402.19|AGGGCGTGAAACCGTTAAG</td><td>>hg38_21_+_8433172_8446622_RNA45SN1_45S_50ntFlanks</td><td>8384</td><td>8402</td><td>19</td><td>AGGGCGTGAAACCGTTAAG</td><td>299.0</td><td>-7.7</td><td>1.5</td><td>-5.3</td><td>1.3E-07</td><td>2.2E-07</td><td>MB231/MB468</td><td>MITO</td></tr>
<tr><td>>hg38_21_+_8433172_8446622_RNA45SN1_45S_50ntFlanks@6645.6667.23|CTCGTACGACTCTTAGCGGTGGA</td><td>>hg38_21_+_8433172_8446622_RNA45SN1_45S_50ntFlanks</td><td>6645</td><td>6667</td><td>23</td><td>CTCGTACGACTCTTAGCGGTGGA</td><td>1194.5</td><td>4.3</td><td>0.8</td><td>5.3</td><td>1.2E-07</td><td>2.2E-07</td><td>MB231/MB468</td><td>CYTO</td></tr>
<tr><td>>hg38_21_+_8433172_8446622_RNA45SN1_45S_50ntFlanks@12992.13013.22|TCCCTCGCTGCGATCTATTGAA</td><td>>hg38_21_+_8433172_8446622_RNA45SN1_45S_50ntFlanks</td><td>12992</td><td>13013</td><td>22</td><td>TCCCTCGCTGCGATCTATTGAA</td><td>508.7</td><td>-5.5</td><td>1.0</td><td>-5.3</td><td>9.3E-08</td><td>2.2E-07</td><td>BT20/MB468</td><td>MITO</td></tr>
<tr><td>>MI0000095|hsa-mir-93&WithFlank&7|-|100093762|100093853@17.40.24&[MIMAT0000093&hsa-miR-93-5p&offsets|0|+1;m-29&7|-|100093815|100093837&offsets|0|+1]|CAAAGTGCTGTTCGTGCAGGTAGT</td><td>>MI0000095|hsa-mir-93&WithFlank&7|-|100093762|100093853</td><td>17</td><td>40</td><td>24</td><td>CAAAGTGCTGTTCGTGCAGGTAGT</td><td>1907.7</td><td>5.2</td><td>1.0</td><td>5.3</td><td>1.2E-07</td><td>2.2E-07</td><td>MB231/MB468</td><td>CYTO</td></tr>
<tr><td>>hg38_21_+_8433172_8446622_RNA45SN1_45S_50ntFlanks@10346.10374.29|TCAAACGAGAACTTTGAAGGCCGAAGTGG</td><td>>hg38_21_+_8433172_8446622_RNA45SN1_45S_50ntFlanks</td><td>10346</td><td>10374</td><td>29</td><td>TCAAACGAGAACTTTGAAGGCCGAAGTGG</td><td>810.9</td><td>-2.7</td><td>0.5</td><td>-5.3</td><td>1.2E-07</td><td>2.2E-07</td><td>MB231/MB468</td><td>CYTO</td></tr>
<tr><td>>MI0003560|hsa-mir-92b&WithFlank&1|+|155195171|155195278@67.87.21&[MIMAT0003218&hsa-miR-92b-3p&offsets|0|-1;m-115&1|+|155195237|155195258&offsets|0|-1]|TATTGCACTCGTCCCGGCCTC</td><td>>MI0003560|hsa-mir-92b&WithFlank&1|+|155195171|155195278</td><td>67</td><td>87</td><td>21</td><td>TATTGCACTCGTCCCGGCCTC</td><td>262.8</td><td>3.0</td><td>0.6</td><td>5.3</td><td>1.4E-07</td><td>2.2E-07</td><td>BT20/MB468</td><td>NUC</td></tr>
<tr><td>>hg38_21_+_8433172_8446622_RNA45SN1_45S_50ntFlanks@10409.10431.23|GTCAGTCGGTCCTGAGAGATGGG</td><td>>hg38_21_+_8433172_8446622_RNA45SN1_45S_50ntFlanks</td><td>10409</td><td>10431</td><td>23</td><td>GTCAGTCGGTCCTGAGAGATGGG</td><td>288.7</td><td>-4.5</td><td>0.8</td><td>-5.3</td><td>1.0E-07</td><td>2.2E-07</td><td>BT20/MB468</td><td>CYTO</td></tr>
<tr><td>>hg38_21_+_8433172_8446622_RNA45SN1_45S_50ntFlanks@11570.11600.31|AGCCGACTTAGAACTGGTGCGGACCAGGGGA</td><td>>hg38_21_+_8433172_8446622_RNA45SN1_45S_50ntFlanks</td><td>11570</td><td>11600</td><td>31</td><td>AGCCGACTTAGAACTGGTGCGGACCAGGGGA</td><td>2361.1</td><td>-7.2</td><td>1.4</td><td>-5.3</td><td>1.4E-07</td><td>2.2E-07</td><td>BT20/MB468</td><td>NUC</td></tr>
<tr><td>>hg38_21_+_8433172_8446622_RNA45SN1_45S_50ntFlanks@3920.3958.39|CATTTATCAGATCAAAACCAACCCGGTCAGCCCCTCTCC</td><td>>hg38_21_+_8433172_8446622_RNA45SN1_45S_50ntFlanks</td><td>3920</td><td>3958</td><td>39</td><td>CATTTATCAGATCAAAACCAACCCGGTCAGCCCCTCTCC</td><td>89.0</td><td>-8.6</td><td>1.6</td><td>-5.3</td><td>9.3E-08</td><td>2.2E-07</td><td>BT20/MB468</td><td>MITO</td></tr>
<tr><td>>am_trnaMT_SerGCT_MT_+_12207_12265@13.32.20|AAGAACTGCTAACTCATGCC</td><td>>am_trnaMT_SerGCT_MT_+_12207_12265</td><td>13</td><td>32</td><td>20</td><td>AAGAACTGCTAACTCATGCC</td><td>409.7</td><td>5.9</td><td>1.1</td><td>5.3</td><td>1.1E-07</td><td>2.2E-07</td><td>BT20/MB468</td><td>CYTO</td></tr>
<tr><td>>MI0000272|hsa-mir-182&WithFlank&7|-|129770377|129770498@29.53.25&[MIMAT0000259&hsa-miR-182-5p&offsets|0|+1;m-22&7|-|129770449|129770470&offsets|0|+3]|TTTGGCAATGGTAGAACTCACACTG</td><td>>MI0000272|hsa-mir-182&WithFlank&7|-|129770377|129770498</td><td>29</td><td>53</td><td>25</td><td>TTTGGCAATGGTAGAACTCACACTG</td><td>1681.4</td><td>3.9</td><td>0.7</td><td>5.3</td><td>1.2E-07</td><td>2.2E-07</td><td>MB231/MB468</td><td>CYTO</td></tr>
<tr><td>>hg38_21_+_8433172_8446622_RNA45SN1_45S_50ntFlanks@5059.5098.40|CGATTTGTCTGGTTAATTCCGATAACGAACGAGACTCTGG</td><td>>hg38_21_+_8433172_8446622_RNA45SN1_45S_50ntFlanks</td><td>5059</td><td>5098</td><td>40</td><td>CGATTTGTCTGGTTAATTCCGATAACGAACGAGACTCTGG</td><td>1826.5</td><td>-8.7</td><td>1.7</td><td>-5.3</td><td>1.4E-07</td><td>2.2E-07</td><td>BT20/MB468</td><td>NUC</td></tr>
<tr><td>>hg38_21_+_8433172_8446622_RNA45SN1_45S_50ntFlanks@5488.5516.29|GCTGAGAAGACGGTCGAACTTGACTATCT</td><td>>hg38_21_+_8433172_8446622_RNA45SN1_45S_50ntFlanks</td><td>5488</td><td>5516</td><td>29</td><td>GCTGAGAAGACGGTCGAACTTGACTATCT</td><td>451.8</td><td>5.5</td><td>1.0</td><td>5.6</td><td>2.4E-08</td><td>2.2E-07</td><td>BT20/MB468</td><td>MP</td></tr>
<tr><td>>trnaMT_PheGAA_MT_+_577_647@39.62.24|ATGTTTAGACGGGCTCACATCACC</td><td>>trnaMT_PheGAA_MT_+_577_647</td><td>39</td><td>62</td><td>24</td><td>ATGTTTAGACGGGCTCACATCACC</td><td>302.1</td><td>8.6</td><td>1.6</td><td>5.3</td><td>1.1E-07</td><td>2.2E-07</td><td>BT20/MB468</td><td>CYTO</td></tr>
<tr><td>>hg38_21_+_8433172_8446622_RNA45SN1_45S_50ntFlanks@6732.6753.22|ACATTGATCATCGACACTTCGA</td><td>>hg38_21_+_8433172_8446622_RNA45SN1_45S_50ntFlanks</td><td>6732</td><td>6753</td><td>22</td><td>ACATTGATCATCGACACTTCGA</td><td>359.0</td><td>-5.6</td><td>1.1</td><td>-5.3</td><td>1.1E-07</td><td>2.2E-07</td><td>MB231/MB468</td><td>MP</td></tr>
<tr><td>>hg38_21_+_8433172_8446622_RNA45SN1_45S_50ntFlanks@12418.12444.27|CGATGTCGGCTCTTCCTATCATTGTGA</td><td>>hg38_21_+_8433172_8446622_RNA45SN1_45S_50ntFlanks</td><td>12418</td><td>12444</td><td>27</td><td>CGATGTCGGCTCTTCCTATCATTGTGA</td><td>2593.9</td><td>-4.3</td><td>0.8</td><td>-5.3</td><td>1.3E-07</td><td>2.2E-07</td><td>MB231/MB468</td><td>MITO</td></tr>
<tr><td>>trna119_LysCTT_1_-_145395522_145395594@2.33.32,trna11_LysCTT_5_-_180648979_180649051@2.33.32,trna13_LysCTT_6_+_26556774_26556846@2.33.32,trna7_LysCTT_16_+_3225692_3225764@2.33.32,trna9_LysCTT_5_+_180634755_180634827@2.33.32|CCCGGCTAGCTCAGTCGGTAGAGCATGAGACT</td><td>>trna119_LysCTT_1_-_145395522_145395594</td><td>2</td><td>33</td><td>32</td><td>CCCGGCTAGCTCAGTCGGTAGAGCATGAGACT</td><td>60.2</td><td>9.1</td><td>1.7</td><td>5.3</td><td>1.1E-07</td><td>2.2E-07</td><td>MB231/MB468</td><td>MP</td></tr>
<tr><td>>MI0016887|hsa-mir-4521&WithFlank&17|+|8186939|8187010@10.35.26&[MIMAT0019058&hsa-miR-4521&offsets|0|+4;m-765&17|+|8186948|8186969&offsets|0|+4]|GCTAAGGAAGTCCTGTGCTCAGTTTT</td><td>>MI0016887|hsa-mir-4521&WithFlank&17|+|8186939|8187010</td><td>10</td><td>35</td><td>26</td><td>GCTAAGGAAGTCCTGTGCTCAGTTTT</td><td>2546.5</td><td>5.6</td><td>1.1</td><td>5.3</td><td>1.1E-07</td><td>2.2E-07</td><td>BT20/MB468</td><td>CYTO</td></tr>
<tr><td>>hg38_21_+_8433172_8446622_RNA45SN1_45S_50ntFlanks@8184.8212.29|CGAGGCCCAGCCCGTGGACGGTGTGAGGC</td><td>>hg38_21_+_8433172_8446622_RNA45SN1_45S_50ntFlanks</td><td>8184</td><td>8212</td><td>29</td><td>CGAGGCCCAGCCCGTGGACGGTGTGAGGC</td><td>541.4</td><td>-8.2</td><td>1.5</td><td>-5.3</td><td>1.1E-07</td><td>2.2E-07</td><td>MB231/MB468</td><td>MP</td></tr>
<tr><td>>hg38_21_+_8433172_8446622_RNA45SN1_45S_50ntFlanks@12539.12579.41|ATGATGTGTTGTTGCCATGGTAATCCTGCTCAGTACGAGAG</td><td>>hg38_21_+_8433172_8446622_RNA45SN1_45S_50ntFlanks</td><td>12539</td><td>12579</td><td>41</td><td>ATGATGTGTTGTTGCCATGGTAATCCTGCTCAGTACGAGAG</td><td>52.0</td><td>-9.6</td><td>1.8</td><td>-5.3</td><td>1.1E-07</td><td>2.2E-07</td><td>MB231/MB468</td><td>MP</td></tr>
<tr><td>>hg38_21_+_8433172_8446622_RNA45SN1_45S_50ntFlanks@6671.6692.22|CTCGGCTCGTGCGTCGATGAAG</td><td>>hg38_21_+_8433172_8446622_RNA45SN1_45S_50ntFlanks</td><td>6671</td><td>6692</td><td>22</td><td>CTCGGCTCGTGCGTCGATGAAG</td><td>295.0</td><td>-7.2</td><td>1.4</td><td>-5.3</td><td>1.1E-07</td><td>2.2E-07</td><td>MB231/MB468</td><td>MP</td></tr>
<tr><td>>hg38_21_+_8433172_8446622_RNA45SN1_45S_50ntFlanks@12444.12475.32|AAGCAGAATTCACCAAGCGTTGGATTGTTCAC</td><td>>hg38_21_+_8433172_8446622_RNA45SN1_45S_50ntFlanks</td><td>12444</td><td>12475</td><td>32</td><td>AAGCAGAATTCACCAAGCGTTGGATTGTTCAC</td><td>138.9</td><td>-10.2</td><td>1.8</td><td>-5.6</td><td>2.4E-08</td><td>2.2E-07</td><td>BT20/MB468</td><td>MP</td></tr>
<tr><td>>hg38_21_+_8433172_8446622_RNA45SN1_45S_50ntFlanks@10347.10382.36|CAAACGAGAACTTTGAAGGCCGAAGTGGAGAAGGGT</td><td>>hg38_21_+_8433172_8446622_RNA45SN1_45S_50ntFlanks</td><td>10347</td><td>10382</td><td>36</td><td>CAAACGAGAACTTTGAAGGCCGAAGTGGAGAAGGGT</td><td>293.5</td><td>4.0</td><td>0.8</td><td>5.3</td><td>1.1E-07</td><td>2.2E-07</td><td>BT20/MB468</td><td>CYTO</td></tr>
<tr><td>>trnaMT_GluTTC_MT_-_14674_14742@37.72.36,trnalookalike8_GluTTC_5_-_93905172_93905240@37.72.36|TCATTGGTCGTGGTTGTAGTCCGTGCGAGAATACCA</td><td>>trnaMT_GluTTC_MT_-_14674_14742</td><td>37</td><td>72</td><td>36</td><td>TCATTGGTCGTGGTTGTAGTCCGTGCGAGAATACCA</td><td>3475.5</td><td>4.6</td><td>0.9</td><td>5.3</td><td>9.4E-08</td><td>2.2E-07</td><td>BT20/MB468</td><td>MITO</td></tr>
<tr><td>>hg38_21_+_8433172_8446622_RNA45SN1_45S_50ntFlanks@12033.12072.40|ATCGTTTTTTCACTGACCCGGTGAGGCGGGGGGGCGAGCC</td><td>>hg38_21_+_8433172_8446622_RNA45SN1_45S_50ntFlanks</td><td>12033</td><td>12072</td><td>40</td><td>ATCGTTTTTTCACTGACCCGGTGAGGCGGGGGGGCGAGCC</td><td>98.2</td><td>-8.7</td><td>1.6</td><td>-5.3</td><td>9.4E-08</td><td>2.2E-07</td><td>BT20/MB468</td><td>MITO</td></tr>
<tr><td>>MI0000063|hsa-let-7b&WithFlank&22|+|46113680|46113774@12.33.22(+1C)&[MIMAT0000063&hsa-let-7b-5p&offsets|0|0(+1C);m-14&22|+|46113691|46113712&offsets|0|0(+1C)]|TGAGGTAGTAGGTTGTGTGGTTC</td><td>>MI0000063|hsa-let-7b&WithFlank&22|+|46113680|46113774</td><td>12</td><td>33</td><td>22(+1C)</td><td>TGAGGTAGTAGGTTGTGTGGTTC</td><td>1931.2</td><td>5.8</td><td>1.0</td><td>5.6</td><td>2.4E-08</td><td>2.2E-07</td><td>BT20/MB468</td><td>MP</td></tr>
<tr><td>>hg38_21_+_8433172_8446622_RNA45SN1_45S_50ntFlanks@6645.6676.32|CTCGTACGACTCTTAGCGGTGGATCACTCGGC</td><td>>hg38_21_+_8433172_8446622_RNA45SN1_45S_50ntFlanks</td><td>6645</td><td>6676</td><td>32</td><td>CTCGTACGACTCTTAGCGGTGGATCACTCGGC</td><td>786.9</td><td>4.4</td><td>0.8</td><td>5.6</td><td>2.4E-08</td><td>2.2E-07</td><td>BT20/MB468</td><td>MP</td></tr>
<tr><td>>hg38_21_+_8433172_8446622_RNA45SN1_45S_50ntFlanks@8168.8206.39|CCCAAGTCCTTCTGATCGAGGCCCAGCCCGTGGACGGTG</td><td>>hg38_21_+_8433172_8446622_RNA45SN1_45S_50ntFlanks</td><td>8168</td><td>8206</td><td>39</td><td>CCCAAGTCCTTCTGATCGAGGCCCAGCCCGTGGACGGTG</td><td>140.1</td><td>-10.2</td><td>1.8</td><td>-5.6</td><td>2.4E-08</td><td>2.2E-07</td><td>BT20/MB468</td><td>MP</td></tr>
<tr><td>>hg38_21_+_8433172_8446622_RNA45SN1_45S_50ntFlanks@6645.6672.28|CTCGTACGACTCTTAGCGGTGGATCACT</td><td>>hg38_21_+_8433172_8446622_RNA45SN1_45S_50ntFlanks</td><td>6645</td><td>6672</td><td>28</td><td>CTCGTACGACTCTTAGCGGTGGATCACT</td><td>743.3</td><td>3.9</td><td>0.7</td><td>5.3</td><td>1.2E-07</td><td>2.2E-07</td><td>MB231/MB468</td><td>NUC</td></tr>
<tr><td>>hg38_21_+_8433172_8446622_RNA45SN1_45S_50ntFlanks@9504.9526.23|GTGAACTATGCCTGGGCAGGGCG</td><td>>hg38_21_+_8433172_8446622_RNA45SN1_45S_50ntFlanks</td><td>9504</td><td>9526</td><td>23</td><td>GTGAACTATGCCTGGGCAGGGCG</td><td>4325.9</td><td>-8.1</td><td>1.5</td><td>-5.3</td><td>1.1E-07</td><td>2.2E-07</td><td>BT20/MB468</td><td>CYTO</td></tr>
<tr><td>>MI0000093|hsa-mir-92a-1&WithFlank&13|+|91351308|91351397@54.73.20;MI0000094|hsa-mir-92a-2&WithFlank&X|-|134169532|134169618@54.73.20&[MIMAT0000092&hsa-miR-92a-3p&offsets|0|-2;m-9&13|+|91351361|91351382&offsets|0|-2];[MIMAT0000092_1&hsa-miR-92a-3p&offsets|0|-2;m-11&X|-|134169544|134169565&offsets|0|-2]|TATTGCACTTGTCCCGGCCT</td><td>>MI0000093|hsa-mir-92a-1&WithFlank&13|+|91351308|91351397</td><td>54</td><td>73</td><td>20</td><td>TATTGCACTTGTCCCGGCCT</td><td>282.9</td><td>7.3</td><td>1.4</td><td>5.3</td><td>1.1E-07</td><td>2.2E-07</td><td>BT20/MB468</td><td>CYTO</td></tr>
<tr><td>>hg38_21_+_8433172_8446622_RNA45SN1_45S_50ntFlanks@9538.9571.34|AACTCTGGTGGAGGTCCGTAGCGGTCCTGACGTG</td><td>>hg38_21_+_8433172_8446622_RNA45SN1_45S_50ntFlanks</td><td>9538</td><td>9571</td><td>34</td><td>AACTCTGGTGGAGGTCCGTAGCGGTCCTGACGTG</td><td>328.4</td><td>-5.5</td><td>1.0</td><td>-5.3</td><td>1.3E-07</td><td>2.3E-07</td><td>MB231/MB468</td><td>MITO</td></tr>
<tr><td>>hg38_21_+_8433172_8446622_RNA45SN1_45S_50ntFlanks@5436.5466.31|GTGAGGCCCTCGGATCGGCCCCGCCGGGGTC</td><td>>hg38_21_+_8433172_8446622_RNA45SN1_45S_50ntFlanks</td><td>5436</td><td>5466</td><td>31</td><td>GTGAGGCCCTCGGATCGGCCCCGCCGGGGTC</td><td>490.0</td><td>-7.2</td><td>1.4</td><td>-5.3</td><td>1.3E-07</td><td>2.3E-07</td><td>MB231/MB468</td><td>NUC</td></tr>
<tr><td>>hg38_21_+_8433172_8446622_RNA45SN1_45S_50ntFlanks@10389.10407.19|TGAACAGCAGTTGAACATG</td><td>>hg38_21_+_8433172_8446622_RNA45SN1_45S_50ntFlanks</td><td>10389</td><td>10407</td><td>19</td><td>TGAACAGCAGTTGAACATG</td><td>1293.0</td><td>-6.5</td><td>1.2</td><td>-5.6</td><td>2.4E-08</td><td>2.3E-07</td><td>BT20/MB468</td><td>MP</td></tr>
<tr><td>>hg38_21_+_8433172_8446622_RNA45SN1_45S_50ntFlanks@11770.11794.25|AATTAGTGACGCGCATGAATGGATG</td><td>>hg38_21_+_8433172_8446622_RNA45SN1_45S_50ntFlanks</td><td>11770</td><td>11794</td><td>25</td><td>AATTAGTGACGCGCATGAATGGATG</td><td>672.6</td><td>2.7</td><td>0.5</td><td>5.3</td><td>1.1E-07</td><td>2.3E-07</td><td>BT20/MB468</td><td>CYTO</td></tr>
<tr><td>>hg38_21_+_8433172_8446622_RNA45SN1_45S_50ntFlanks@9786.9819.34|CTCGCTGGCGTGGAGCCGGGCGTGGAATGCGAGT</td><td>>hg38_21_+_8433172_8446622_RNA45SN1_45S_50ntFlanks</td><td>9786</td><td>9819</td><td>34</td><td>CTCGCTGGCGTGGAGCCGGGCGTGGAATGCGAGT</td><td>310.1</td><td>11.6</td><td>2.0</td><td>5.8</td><td>5.0E-09</td><td>2.3E-07</td><td>BT20/MB231</td><td>MP</td></tr>
<tr><td>>MI0000086|hsa-mir-28&WithFlank&3|+|188688775|188688872@62.82.21&[MIMAT0004502&hsa-miR-28-3p&offsets|+2|+1;m-28&3|+|188688834|188688855&offsets|+2|+1]|CTAGATTGTGAGCTCCTGGAG</td><td>>MI0000086|hsa-mir-28&WithFlank&3|+|188688775|188688872</td><td>62</td><td>82</td><td>21</td><td>CTAGATTGTGAGCTCCTGGAG</td><td>380.3</td><td>9.4</td><td>1.7</td><td>5.6</td><td>2.4E-08</td><td>2.3E-07</td><td>BT20/MB468</td><td>MP</td></tr>
<tr><td>>hg38_21_+_8433172_8446622_RNA45SN1_45S_50ntFlanks@11786.11804.19|GAATGGATGAACGAGATTC</td><td>>hg38_21_+_8433172_8446622_RNA45SN1_45S_50ntFlanks</td><td>11786</td><td>11804</td><td>19</td><td>GAATGGATGAACGAGATTC</td><td>503.6</td><td>-7.2</td><td>1.4</td><td>-5.3</td><td>1.3E-07</td><td>2.3E-07</td><td>MB231/MB468</td><td>MITO</td></tr>
<tr><td>>hg38_21_+_8433172_8446622_RNA45SN1_45S_50ntFlanks@6737.6770.34|GATCATCGACACTTCGAACGCACTTGCGGCCCCG</td><td>>hg38_21_+_8433172_8446622_RNA45SN1_45S_50ntFlanks</td><td>6737</td><td>6770</td><td>34</td><td>GATCATCGACACTTCGAACGCACTTGCGGCCCCG</td><td>326.4</td><td>-6.1</td><td>1.2</td><td>-5.3</td><td>1.2E-07</td><td>2.3E-07</td><td>MB231/MB468</td><td>CYTO</td></tr>
<tr><td>>hg38_21_+_8433172_8446622_RNA45SN1_45S_50ntFlanks@5078.5097.20|CGATAACGAACGAGACTCTG</td><td>>hg38_21_+_8433172_8446622_RNA45SN1_45S_50ntFlanks</td><td>5078</td><td>5097</td><td>20</td><td>CGATAACGAACGAGACTCTG</td><td>1759.0</td><td>-8.7</td><td>1.6</td><td>-5.3</td><td>1.1E-07</td><td>2.3E-07</td><td>MB231/MB468</td><td>MP</td></tr>
</table>

<table>
<tr><td>>hg38_21_+_8433172_8446622_RNA45SN1_45S_50ntFlanks@10409.10428.20|GTCAGTCGGTCCTGAGAGAT</td><td>>hg38_21_+_8433172_8446622_RNA45SN1_45S_50ntFlanks</td><td>10409</td><td>10428</td><td>20</td><td>GTCAGTCGGTCCTGAGAGAT</td><td>290.9</td><td>-3.3</td><td>0.6</td><td>-5.3</td><td>1.3E-07</td><td>2.3E-07</td><td>MB231/MB468</td><td>NUC</td></tr>
<tr><td>>trna111_HisGTG_1_-_147774845_147774916@1.35.35,trna118_HisGTG_1_-_145396881_145396952@1.35.35,trna16_HisGTG_1_+_146544773_146544844@1.35.35,trna1_HisGTG_15_+_45493349_45493420@1.35.35,trna21_HisGTG_1_+_147753471_147753542@1.35.35,trna33_HisGTG_6_+_27125906_27125977@1.35.35,trna7_HisGTG_9_-_14433938_14434009@1.35.35,trna8_HisGTG_15_-_45492611_45492682@1.35.35,trna9_HisGTG_15_-_45490804_45490875@1.35.35|GCCGTGATCGTATAGTGGTTAGTACTCTGCGTTGT</td><td>>trna111_HisGTG_1_-_147774845_147774916</td><td>1</td><td>35</td><td>35</td><td>GCCGTGATCGTATAGTGGTTAGTACTCTGCGTTGT</td><td>303.3</td><td>6.8</td><td>1.3</td><td>5.3</td><td>1.4E-07</td><td>2.3E-07</td><td>BT20/MB468</td><td>NUC</td></tr>
<tr><td>>hg38_21_+_8433172_8446622_RNA45SN1_45S_50ntFlanks@11769.11792.24|TAATTAGTGACGCGCATGAATGGA</td><td>>hg38_21_+_8433172_8446622_RNA45SN1_45S_50ntFlanks</td><td>11769</td><td>11792</td><td>24</td><td>TAATTAGTGACGCGCATGAATGGA</td><td>217.5</td><td>-6.9</td><td>1.3</td><td>-5.3</td><td>1.3E-07</td><td>2.3E-07</td><td>MB231/MB468</td><td>MITO</td></tr>
<tr><td>>trna111_HisGTG_1_-_147774845_147774916@-1G.34.35,trna118_HisGTG_1_-_145396881_145396952@-1G.34.35,trna16_HisGTG_1_+_146544773_146544844@-1G.34.35,trna1_HisGTG_15_+_45493349_45493420@-1G.34.35,trna21_HisGTG_1_+_147753471_147753542@-1G.34.35,trna33_HisGTG_6_+_27125906_27125977@-1G.34.35,trna7_HisGTG_9_-_14433938_14434009@-1G.34.35,trna8_HisGTG_15_-_45492611_45492682@-1G.34.35,trna9_HisGTG_15_-_45490804_45490875@-1G.34.35|GGCCGTGATCGTATAGTGGTTAGTACTCTGCGTTG</td><td>>trna111_HisGTG_1_-_147774845_147774916</td><td>-1G</td><td>34</td><td>35</td><td>GGCCGTGATCGTATAGTGGTTAGTACTCTGCGTTG</td><td>3957.5</td><td>6.0</td><td>1.1</td><td>5.3</td><td>1.1E-07</td><td>2.3E-07</td><td>MB231/MB468</td><td>MP</td></tr>
<tr><td>>hg38_21_+_8433172_8446622_RNA45SN1_45S_50ntFlanks@9903.9928.26|ATCAGACCCCAGAAAAGGTGTTGGTT</td><td>>hg38_21_+_8433172_8446622_RNA45SN1_45S_50ntFlanks</td><td>9903</td><td>9928</td><td>26</td><td>ATCAGACCCCAGAAAAGGTGTTGGTT</td><td>90.7</td><td>-9.6</td><td>1.7</td><td>-5.6</td><td>2.5E-08</td><td>2.3E-07</td><td>BT20/MB468</td><td>MP</td></tr>
<tr><td>>hg38_21_+_8433172_8446622_RNA45SN1_45S_50ntFlanks@4924.4943.20|AGGAGTGGAGCCTGCGGCTT</td><td>>hg38_21_+_8433172_8446622_RNA45SN1_45S_50ntFlanks</td><td>4924</td><td>4943</td><td>20</td><td>AGGAGTGGAGCCTGCGGCTT</td><td>463.8</td><td>6.3</td><td>1.2</td><td>5.3</td><td>1.2E-07</td><td>2.3E-07</td><td>MB231/MB468</td><td>CYTO</td></tr>
<tr><td>>hg38_21_+_8433172_8446622_RNA45SN1_45S_50ntFlanks@4498.4517.20|AAGCGTTTACTTTGAAAAAA</td><td>>hg38_21_+_8433172_8446622_RNA45SN1_45S_50ntFlanks</td><td>4498</td><td>4517</td><td>20</td><td>AAGCGTTTACTTTGAAAAAA</td><td>792.8</td><td>-8.4</td><td>1.6</td><td>-5.3</td><td>1.3E-07</td><td>2.3E-07</td><td>MB231/MB468</td><td>MITO</td></tr>
<tr><td>>hg38_21_+_8433172_8446622_RNA45SN1_45S_50ntFlanks@11575.11599.25|ACTTAGAACTGGTGCGGACCAGGGG</td><td>>hg38_21_+_8433172_8446622_RNA45SN1_45S_50ntFlanks</td><td>11575</td><td>11599</td><td>25</td><td>ACTTAGAACTGGTGCGGACCAGGGG</td><td>300.6</td><td>-6.6</td><td>1.2</td><td>-5.3</td><td>1.1E-07</td><td>2.3E-07</td><td>BT20/MB468</td><td>CYTO</td></tr>
<tr><td>>hg38_21_+_8433172_8446622_RNA45SN1_45S_50ntFlanks@12397.12414.18|CGACGTCGCTTTTTGATC</td><td>>hg38_21_+_8433172_8446622_RNA45SN1_45S_50ntFlanks</td><td>12397</td><td>12414</td><td>18</td><td>CGACGTCGCTTTTTGATC</td><td>1606.8</td><td>-4.8</td><td>0.9</td><td>-5.3</td><td>1.4E-07</td><td>2.3E-07</td><td>BT20/MB468</td><td>NUC</td></tr>
<tr><td>>hg38_21_+_8433172_8446622_RNA45SN1_45S_50ntFlanks@4548.4568.21|TGGATACCGCAGCTAGGAATA</td><td>>hg38_21_+_8433172_8446622_RNA45SN1_45S_50ntFlanks</td><td>4548</td><td>4568</td><td>21</td><td>TGGATACCGCAGCTAGGAATA</td><td>207.8</td><td>3.7</td><td>0.7</td><td>5.3</td><td>1.2E-07</td><td>2.3E-07</td><td>MB231/MB468</td><td>CYTO</td></tr>
<tr><td>>hg38_21_+_8433172_8446622_RNA45SN1_45S_50ntFlanks@12412.12432.21|ATCCTTCGATGTCGGCTCTTC</td><td>>hg38_21_+_8433172_8446622_RNA45SN1_45S_50ntFlanks</td><td>12412</td><td>12432</td><td>21</td><td>ATCCTTCGATGTCGGCTCTTC</td><td>851.9</td><td>-7.5</td><td>1.4</td><td>-5.3</td><td>1.3E-07</td><td>2.3E-07</td><td>MB231/MB468</td><td>MITO</td></tr>
<tr><td>>hg38_21_+_8433172_8446622_RNA45SN1_45S_50ntFlanks@11325.11342.18|TCTCGTCGCGCGCGCGTC</td><td>>hg38_21_+_8433172_8446622_RNA45SN1_45S_50ntFlanks</td><td>11325</td><td>11342</td><td>18</td><td>TCTCGTCGCGCGCGCGTC</td><td>169.6</td><td>-5.7</td><td>1.1</td><td>-5.3</td><td>1.2E-07</td><td>2.3E-07</td><td>MB231/MB468</td><td>CYTO</td></tr>
<tr><td>>hg38_21_+_8433172_8446622_RNA45SN1_45S_50ntFlanks@9565.9592.28|TGACGTGCAAATCGGTCGTCCGACCTGG</td><td>>hg38_21_+_8433172_8446622_RNA45SN1_45S_50ntFlanks</td><td>9565</td><td>9592</td><td>28</td><td>TGACGTGCAAATCGGTCGTCCGACCTGG</td><td>1488.6</td><td>-3.8</td><td>0.7</td><td>-5.3</td><td>1.2E-07</td><td>2.3E-07</td><td>MB231/MB468</td><td>CYTO</td></tr>
<tr><td>>hg38_21_+_8433172_8446622_RNA45SN1_45S_50ntFlanks@12029.12050.22|TCTGATCGTTTTTTCACTGACC</td><td>>hg38_21_+_8433172_8446622_RNA45SN1_45S_50ntFlanks</td><td>12029</td><td>12050</td><td>22</td><td>TCTGATCGTTTTTTCACTGACC</td><td>106.4</td><td>10.5</td><td>1.9</td><td>5.6</td><td>2.5E-08</td><td>2.3E-07</td><td>BT20/MB468</td><td>MP</td></tr>
<tr><td>>hg38_21_+_8433172_8446622_RNA45SN1_45S_50ntFlanks@5443.5467.25|CCTCGGATCGGCCCCGCCGGGGTCG</td><td>>hg38_21_+_8433172_8446622_RNA45SN1_45S_50ntFlanks</td><td>5443</td><td>5467</td><td>25</td><td>CCTCGGATCGGCCCCGCCGGGGTCG</td><td>251.0</td><td>-4.7</td><td>0.9</td><td>-5.3</td><td>1.3E-07</td><td>2.3E-07</td><td>MB231/MB468</td><td>MITO</td></tr>
<tr><td>>trnaMT_PheGAA_MT_+_577_647@23.55.33|AAGCAATACACTGAAAATGTTTAGACGGGCTCA</td><td>>trnaMT_PheGAA_MT_+_577_647</td><td>23</td><td>55</td><td>33</td><td>AAGCAATACACTGAAAATGTTTAGACGGGCTCA</td><td>128.1</td><td>10.7</td><td>1.9</td><td>5.6</td><td>2.5E-08</td><td>2.3E-07</td><td>BT20/MB468</td><td>MP</td></tr>
<tr><td>>MI0000542|hsa-mir-320a&WithFlank&8|-|22244960|22245043@48.70.23(+1C)&[MIMAT0000510&hsa-miR-320a-3p&offsets|0|+1(+1C);m-60&8|-|22244975|22244996&offsets|0|+1(+1C)]|AAAAGCTGGGTTGAGAGGGCGAAC</td><td>>MI0000542|hsa-mir-320a&WithFlank&8|-|22244960|22245043</td><td>48</td><td>70</td><td>23(+1C)</td><td>AAAAGCTGGGTTGAGAGGGCGAAC</td><td>1266.9</td><td>7.2</td><td>1.3</td><td>5.6</td><td>2.5E-08</td><td>2.3E-07</td><td>BT20/MB468</td><td>MP</td></tr>
<tr><td>>hg38_21_+_8433172_8446622_RNA45SN1_45S_50ntFlanks@11671.11706.36|CCAGTGCTCTGAATGTCAAAGTGAAGAAATTCAATG</td><td>>hg38_21_+_8433172_8446622_RNA45SN1_45S_50ntFlanks</td><td>11671</td><td>11706</td><td>36</td><td>CCAGTGCTCTGAATGTCAAAGTGAAGAAATTCAATG</td><td>1072.9</td><td>-3.2</td><td>0.6</td><td>-5.3</td><td>1.3E-07</td><td>2.3E-07</td><td>MB231/MB468</td><td>NUC</td></tr>
<tr><td>>hg38_21_+_8433172_8446622_RNA45SN1_45S_50ntFlanks@12995.13037.43|CTCGCTGCGATCTATTGAAAGTCAGCCCTCGACACAAGGGTTT</td><td>>hg38_21_+_8433172_8446622_RNA45SN1_45S_50ntFlanks</td><td>12995</td><td>13037</td><td>43</td><td>CTCGCTGCGATCTATTGAAAGTCAGCCCTCGACACAAGGGTTT</td><td>318.3</td><td>7.8</td><td>1.5</td><td>5.3</td><td>1.3E-07</td><td>2.3E-07</td><td>MB231/MB468</td><td>MITO</td></tr>
<tr><td>>hg38_21_+_8433172_8446622_RNA45SN1_45S_50ntFlanks@10016.10033.18|AATGGATGGCGCTGGAGC</td><td>>hg38_21_+_8433172_8446622_RNA45SN1_45S_50ntFlanks</td><td>10016</td><td>10033</td><td>18</td><td>AATGGATGGCGCTGGAGC</td><td>179.6</td><td>11.2</td><td>2.0</td><td>5.6</td><td>2.5E-08</td><td>2.3E-07</td><td>BT20/MB468</td><td>MP</td></tr>
<tr><td>>hg38_21_+_8433172_8446622_RNA45SN1_45S_50ntFlanks@10683.10708.26|CCGGCGGCGTCCGGTGAGCTCTCGCT</td><td>>hg38_21_+_8433172_8446622_RNA45SN1_45S_50ntFlanks</td><td>10683</td><td>10708</td><td>26</td><td>CCGGCGGCGTCCGGTGAGCTCTCGCT</td><td>261.6</td><td>-5.7</td><td>1.1</td><td>-5.3</td><td>1.1E-07</td><td>2.3E-07</td><td>MB231/MB468</td><td>MP</td></tr>
<tr><td>>hg38_21_+_8433172_8446622_RNA45SN1_45S_50ntFlanks@12990.13013.24|GCTCCCTCGCTGCGATCTATTGAA</td><td>>hg38_21_+_8433172_8446622_RNA45SN1_45S_50ntFlanks</td><td>12990</td><td>13013</td><td>24</td><td>GCTCCCTCGCTGCGATCTATTGAA</td><td>1054.0</td><td>-3.2</td><td>0.6</td><td>-5.3</td><td>9.6E-08</td><td>2.3E-07</td><td>BT20/MB468</td><td>MITO</td></tr>
<tr><td>>hg38_21_+_8433172_8446622_RNA45SN1_45S_50ntFlanks@11912.11940.29|GAAGAGACATGAGAGGTGTAGAATAAGTG</td><td>>hg38_21_+_8433172_8446622_RNA45SN1_45S_50ntFlanks</td><td>11912</td><td>11940</td><td>29</td><td>GAAGAGACATGAGAGGTGTAGAATAAGTG</td><td>87.9</td><td>10.2</td><td>1.8</td><td>5.6</td><td>2.5E-08</td><td>2.3E-07</td><td>BT20/MB468</td><td>MP</td></tr>
<tr><td>>hg38_21_+_8433172_8446622_RNA45SN1_45S_50ntFlanks@12936.12959.24|AAACCATTCGTAGACGACCTGCTT</td><td>>hg38_21_+_8433172_8446622_RNA45SN1_45S_50ntFlanks</td><td>12936</td><td>12959</td><td>24</td><td>AAACCATTCGTAGACGACCTGCTT</td><td>73.1</td><td>-9.3</td><td>1.7</td><td>-5.6</td><td>2.5E-08</td><td>2.3E-07</td><td>BT20/MB468</td><td>MP</td></tr>
<tr><td>>hg38_21_+_8433172_8446622_RNA45SN1_45S_50ntFlanks@9738.9756.19|GAAACGATCTCAACCTATT</td><td>>hg38_21_+_8433172_8446622_RNA45SN1_45S_50ntFlanks</td><td>9738</td><td>9756</td><td>19</td><td>GAAACGATCTCAACCTATT</td><td>91.4</td><td>-9.6</td><td>1.7</td><td>-5.6</td><td>2.5E-08</td><td>2.3E-07</td><td>BT20/MB468</td><td>MP</td></tr>
<tr><td>>trna101_AlaAGC_6_-_28831462_28831533@58.75.18,trna102_AlaAGC_6_-_28806221_28806292@58.75.18,trna110_AlaTGC_6_-_28757547_28757618@58.75.18,trna113_AlaTGC_6_-_28726141_28726212@58.75.18,trna65_AlaAGC_6_+_28574933_28575004@58.75.18|TCCCCGGCACCTCCACCA</td><td>>trna101_AlaAGC_6_-_28831462_28831533</td><td>58</td><td>75</td><td>18</td><td>TCCCCGGCACCTCCACCA</td><td>875.9</td><td>4.2</td><td>0.8</td><td>5.3</td><td>1.3E-07</td><td>2.3E-07</td><td>MB231/MB468</td><td>MITO</td></tr>
<tr><td>>MI0000270|hsa-mir-181b-1&WithFlank&1|-|198858867|198858988@42.62.21;MI0000683|hsa-mir-181b-2&WithFlank&9|+|124693704|124693804@22.42.21&[MIMAT0000257&hsa-miR-181b-5p&offsets|0|-2;m-73&1|-|198858925|198858947&offsets|0|-2];[MIMAT0000257_1&hsa-miR-181b-5p&offsets|0|-2;m-72&9|+|124693725|124693747&offsets|0|-2]|AACATTCATTGCTGTCGGTGG</td><td>>MI0000270|hsa-mir-181b-1&WithFlank&1|-|198858867|198858988</td><td>42</td><td>62</td><td>21</td><td>AACATTCATTGCTGTCGGTGG</td><td>366.2</td><td>4.9</td><td>0.9</td><td>5.3</td><td>1.3E-07</td><td>2.3E-07</td><td>MB231/MB468</td><td>MITO</td></tr>
<tr><td>>hg38_MT_+_1621_3279_RNR2_16S_50ntFlanks@488.521.34|GAACAGCTCTTTGGACACTAGGAAAAAACCTTGT</td><td>>hg38_MT_+_1621_3279_RNR2_16S_50ntFlanks</td><td>488</td><td>521</td><td>34</td><td>GAACAGCTCTTTGGACACTAGGAAAAAACCTTGT</td><td>69.8</td><td>9.9</td><td>1.8</td><td>5.6</td><td>2.5E-08</td><td>2.3E-07</td><td>BT20/MB468</td><td>MP</td></tr>
<tr><td>>hg38_21_+_8433172_8446622_RNA45SN1_45S_50ntFlanks@11999.12035.37|CCCTGCGGGCCGCCGGTGAAATACCACTACTCTGATC</td><td>>hg38_21_+_8433172_8446622_RNA45SN1_45S_50ntFlanks</td><td>11999</td><td>12035</td><td>37</td><td>CCCTGCGGGCCGCCGGTGAAATACCACTACTCTGATC</td><td>454.1</td><td>-7.3</td><td>1.4</td><td>-5.3</td><td>1.3E-07</td><td>2.3E-07</td><td>MB231/MB468</td><td>NUC</td></tr>
<tr><td>>hg38_21_+_8433172_8446622_RNA45SN1_45S_50ntFlanks@3774.3792.19|GGCCGGTACAGTGAAACTG</td><td>>hg38_21_+_8433172_8446622_RNA45SN1_45S_50ntFlanks</td><td>3774</td><td>3792</td><td>19</td><td>GGCCGGTACAGTGAAACTG</td><td>180.6</td><td>-5.5</td><td>1.0</td><td>-5.3</td><td>1.3E-07</td><td>2.3E-07</td><td>MB231/MB468</td><td>NUC</td></tr>
<tr><td>>hg38_21_+_8433172_8446622_RNA45SN1_45S_50ntFlanks@8168.8204.37|CCCAAGTCCTTCTGATCGAGGCCCAGCCCGTGGACGG</td><td>>hg38_21_+_8433172_8446622_RNA45SN1_45S_50ntFlanks</td><td>8168</td><td>8204</td><td>37</td><td>CCCAAGTCCTTCTGATCGAGGCCCAGCCCGTGGACGG</td><td>152.0</td><td>-10.3</td><td>1.9</td><td>-5.6</td><td>2.5E-08</td><td>2.3E-07</td><td>BT20/MB468</td><td>MP</td></tr>
<tr><td>>hg38_21_+_8433172_8446622_RNA45SN1_45S_50ntFlanks@11876.11905.30|AAGACCCTGTTGAGCTTGACTCTAGTCTGG</td><td>>hg38_21_+_8433172_8446622_RNA45SN1_45S_50ntFlanks</td><td>11876</td><td>11905</td><td>30</td><td>AAGACCCTGTTGAGCTTGACTCTAGTCTGG</td><td>443.4</td><td>-4.1</td><td>0.8</td><td>-5.3</td><td>9.7E-08</td><td>2.3E-07</td><td>BT20/MB468</td><td>MITO</td></tr>
<tr><td>>hg38_21_+_8433172_8446622_RNA45SN1_45S_50ntFlanks@4687.4707.21|ACGGACCAGAGCGAAAGCATT</td><td>>hg38_21_+_8433172_8446622_RNA45SN1_45S_50ntFlanks</td><td>4687</td><td>4707</td><td>21</td><td>ACGGACCAGAGCGAAAGCATT</td><td>653.1</td><td>-6.7</td><td>1.3</td><td>-5.3</td><td>1.4E-07</td><td>2.3E-07</td><td>BT20/MB468</td><td>NUC</td></tr>
<tr><td>>hg38_21_+_8433172_8446622_RNA45SN1_45S_50ntFlanks@11037.11060.24|CTCTCCCCCGCTCCCCGTCCTCCC</td><td>>hg38_21_+_8433172_8446622_RNA45SN1_45S_50ntFlanks</td><td>11037</td><td>11060</td><td>24</td><td>CTCTCCCCCGCTCCCCGTCCTCCC</td><td>217.2</td><td>-6.0</td><td>1.1</td><td>-5.3</td><td>1.3E-07</td><td>2.3E-07</td><td>MB231/MB468</td><td>CYTO</td></tr>
<tr><td>>hg38_21_+_8433172_8446622_RNA45SN1_45S_50ntFlanks@10845.10866.22|ACTTCGGGATAAGGATTGGCTC</td><td>>hg38_21_+_8433172_8446622_RNA45SN1_45S_50ntFlanks</td><td>10845</td><td>10866</td><td>22</td><td>ACTTCGGGATAAGGATTGGCTC</td><td>417.5</td><td>-3.7</td><td>0.7</td><td>-5.3</td><td>1.3E-07</td><td>2.3E-07</td><td>MB231/MB468</td><td>CYTO</td></tr>
<tr><td>>hg38_21_+_8433172_8446622_RNA45SN1_45S_50ntFlanks@11999.12038.40|CCCTGCGGGCCGCCGGTGAAATACCACTACTCTGATCGTT</td><td>>hg38_21_+_8433172_8446622_RNA45SN1_45S_50ntFlanks</td><td>11999</td><td>12038</td><td>40</td><td>CCCTGCGGGCCGCCGGTGAAATACCACTACTCTGATCGTT</td><td>55.3</td><td>-9.7</td><td>1.8</td><td>-5.3</td><td>1.2E-07</td><td>2.3E-07</td><td>MB231/MB468</td><td>MP</td></tr>
<tr><td>>am_trnaMT_SerGCT_MT_+_12207_12265@14.32.19|AGAACTGCTAACTCATGCC</td><td>>am_trnaMT_SerGCT_MT_+_12207_12265</td><td>14</td><td>32</td><td>19</td><td>AGAACTGCTAACTCATGCC</td><td>884.3</td><td>9.3</td><td>1.8</td><td>5.3</td><td>1.1E-07</td><td>2.3E-07</td><td>BT20/MB468</td><td>CYTO</td></tr>
<tr><td>>hg38_21_+_8433172_8446622_RNA45SN1_45S_50ntFlanks@9565.9594.30|TGACGTGCAAATCGGTCGTCCGACCTGGGT</td><td>>hg38_21_+_8433172_8446622_RNA45SN1_45S_50ntFlanks</td><td>9565</td><td>9594</td><td>30</td><td>TGACGTGCAAATCGGTCGTCCGACCTGGGT</td><td>145.9</td><td>-5.4</td><td>1.0</td><td>-5.3</td><td>1.2E-07</td><td>2.3E-07</td><td>MB231/MB468</td><td>MP</td></tr>
<tr><td>>hg38_21_+_8433172_8446622_RNA45SN1_45S_50ntFlanks@11517.11539.23|CCGCCCCCGGGGCCGCGGTTCCG</td><td>>hg38_21_+_8433172_8446622_RNA45SN1_45S_50ntFlanks</td><td>11517</td><td>11539</td><td>23</td><td>CCGCCCCCGGGGCCGCGGTTCCG</td><td>299.2</td><td>-5.2</td><td>1.0</td><td>-5.3</td><td>1.3E-07</td><td>2.3E-07</td><td>MB231/MB468</td><td>CYTO</td></tr>
<tr><td>>hg38_21_+_8433172_8446622_RNA45SN1_45S_50ntFlanks@4976.4993.18|CCGGACACGGACAGGATT</td><td>>hg38_21_+_8433172_8446622_RNA45SN1_45S_50ntFlanks</td><td>4976</td><td>4993</td><td>18</td><td>CCGGACACGGACAGGATT</td><td>93.9</td><td>2.9</td><td>0.6</td><td>5.3</td><td>1.4E-07</td><td>2.3E-07</td><td>BT20/MB468</td><td>NUC</td></tr>
<tr><td>>hg38_21_+_8433172_8446622_RNA45SN1_45S_50ntFlanks@6670.6698.29|ACTCGGCTCGTGCGTCGATGAAGAACGCA</td><td>>hg38_21_+_8433172_8446622_RNA45SN1_45S_50ntFlanks</td><td>6670</td><td>6698</td><td>29</td><td>ACTCGGCTCGTGCGTCGATGAAGAACGCA</td><td>152.7</td><td>-2.3</td><td>0.4</td><td>-5.3</td><td>1.1E-07</td><td>2.3E-07</td><td>BT20/MB468</td><td>CYTO</td></tr>
<tr><td>>hg38_21_+_8433172_8446622_RNA45SN1_45S_50ntFlanks@12033.12071.39|ATCGTTTTTTCACTGACCCGGTGAGGCGGGGGGGCGAGC</td><td>>hg38_21_+_8433172_8446622_RNA45SN1_45S_50ntFlanks</td><td>12033</td><td>12071</td><td>39</td><td>ATCGTTTTTTCACTGACCCGGTGAGGCGGGGGGGCGAGC</td><td>116.6</td><td>-9.0</td><td>1.7</td><td>-5.3</td><td>9.8E-08</td><td>2.3E-07</td><td>BT20/MB468</td><td>MITO</td></tr>
<tr><td>>hg38_21_+_8433172_8446622_RNA45SN1_45S_50ntFlanks@10402.10426.25|AACATGGGTCAGTCGGTCCTGAGAG</td><td>>hg38_21_+_8433172_8446622_RNA45SN1_45S_50ntFlanks</td><td>10402</td><td>10426</td><td>25</td><td>AACATGGGTCAGTCGGTCCTGAGAG</td><td>514.1</td><td>-5.2</td><td>1.0</td><td>-5.3</td><td>1.3E-07</td><td>2.3E-07</td><td>MB231/MB468</td><td>CYTO</td></tr>
<tr><td>>hg38_21_+_8433172_8446622_RNA45SN1_45S_50ntFlanks@10408.10435.28|GGTCAGTCGGTCCTGAGAGATGGGCGAG</td><td>>hg38_21_+_8433172_8446622_RNA45SN1_45S_50ntFlanks</td><td>10408</td><td>10435</td><td>28</td><td>GGTCAGTCGGTCCTGAGAGATGGGCGAG</td><td>839.6</td><td>-7.5</td><td>1.4</td><td>-5.3</td><td>9.8E-08</td><td>2.3E-07</td><td>BT20/MB468</td><td>MITO</td></tr>
<tr><td>>hg38_21_+_8433172_8446622_RNA45SN1_45S_50ntFlanks@10907.10929.23|GGGCGCGCGCCGCGGCTGGACGA</td><td>>hg38_21_+_8433172_8446622_RNA45SN1_45S_50ntFlanks</td><td>10907</td><td>10929</td><td>23</td><td>GGGCGCGCGCCGCGGCTGGACGA</td><td>95.3</td><td>10.3</td><td>1.9</td><td>5.6</td><td>2.5E-08</td><td>2.3E-07</td><td>BT20/MB468</td><td>MP</td></tr>
<tr><td>>hg38_21_+_8433172_8446622_RNA45SN1_45S_50ntFlanks@12407.12432.26|TTTTGATCCTTCGATGTCGGCTCTTC</td><td>>hg38_21_+_8433172_8446622_RNA45SN1_45S_50ntFlanks</td><td>12407</td><td>12432</td><td>26</td><td>TTTTGATCCTTCGATGTCGGCTCTTC</td><td>456.7</td><td>-2.8</td><td>0.5</td><td>-5.3</td><td>1.3E-07</td><td>2.3E-07</td><td>MB231/MB468</td><td>NUC</td></tr>
<tr><td>>hg38_21_+_8433172_8446622_RNA45SN1_45S_50ntFlanks@11575.11605.31|ACTTAGAACTGGTGCGGACCAGGGGAATCCG</td><td>>hg38_21_+_8433172_8446622_RNA45SN1_45S_50ntFlanks</td><td>11575</td><td>11605</td><td>31</td><td>ACTTAGAACTGGTGCGGACCAGGGGAATCCG</td><td>842.6</td><td>-3.9</td><td>0.7</td><td>-5.3</td><td>9.8E-08</td><td>2.3E-07</td><td>BT20/MB468</td><td>MITO</td></tr>
</table>

<table>
<tr><td>>hg38_21_+_8433172_8446622_RNA45SN1_45S_50ntFlanks@12973.12994.22|TCGTACGTAGCAGAGCAGCTCC</td><td>>hg38_21_+_8433172_8446622_RNA45SN1_45S_50ntFlanks</td><td>12973</td><td>12994</td><td>22</td><td>TCGTACGTAGCAGAGCAGCTCC</td><td>268.9</td><td>-7.4</td><td>1.4</td><td>-5.3</td><td>1.4E-07</td><td>2.3E-07</td><td>BT20/MB468</td><td>NUC</td></tr>
<tr><td>>hg38_21_+_8433172_8446622_RNA45SN1_45S_50ntFlanks@6646.6663.18|TCGTACGACTCTTAGCGG</td><td>>hg38_21_+_8433172_8446622_RNA45SN1_45S_50ntFlanks</td><td>6646</td><td>6663</td><td>18</td><td>TCGTACGACTCTTAGCGG</td><td>20138.8</td><td>3.4</td><td>0.6</td><td>5.3</td><td>1.1E-07</td><td>2.3E-07</td><td>BT20/MB468</td><td>CYTO</td></tr>
<tr><td>>hg38_21_+_8433172_8446622_RNA45SN1_45S_50ntFlanks@9544.9577.34|GGTGGAGGTCCGTAGCGGTCCTGACGTGCAAATC</td><td>>hg38_21_+_8433172_8446622_RNA45SN1_45S_50ntFlanks</td><td>9544</td><td>9577</td><td>34</td><td>GGTGGAGGTCCGTAGCGGTCCTGACGTGCAAATC</td><td>206.5</td><td>6.5</td><td>1.2</td><td>5.3</td><td>1.3E-07</td><td>2.3E-07</td><td>MB231/MB468</td><td>CYTO</td></tr>
<tr><td>>hg38_21_+_8433172_8446622_RNA45SN1_45S_50ntFlanks@12838.12871.34|GTGCGGAGTGCCCTTCGTCCTGGGAAACGGGGCG</td><td>>hg38_21_+_8433172_8446622_RNA45SN1_45S_50ntFlanks</td><td>12838</td><td>12871</td><td>34</td><td>GTGCGGAGTGCCCTTCGTCCTGGGAAACGGGGCG</td><td>225.7</td><td>-7.6</td><td>1.4</td><td>-5.3</td><td>1.4E-07</td><td>2.3E-07</td><td>MB231/MB468</td><td>MITO</td></tr>
<tr><td>>hg38_21_+_8433172_8446622_RNA45SN1_45S_50ntFlanks@3704.3725.22|CTACCTGGTTGATCCTGCCAGT</td><td>>hg38_21_+_8433172_8446622_RNA45SN1_45S_50ntFlanks</td><td>3704</td><td>3725</td><td>22</td><td>CTACCTGGTTGATCCTGCCAGT</td><td>60.9</td><td>5.0</td><td>0.9</td><td>5.3</td><td>1.4E-07</td><td>2.3E-07</td><td>BT20/MB468</td><td>NUC</td></tr>
<tr><td>>hg38_21_+_8433172_8446622_RNA45SN1_45S_50ntFlanks@12997.13039.43|CGCTGCGATCTATTGAAAGTCAGCCCTCGACACAAGGGTTTGT</td><td>>hg38_21_+_8433172_8446622_RNA45SN1_45S_50ntFlanks</td><td>12997</td><td>13039</td><td>43</td><td>CGCTGCGATCTATTGAAAGTCAGCCCTCGACACAAGGGTTTGT</td><td>172385.8</td><td>6.3</td><td>1.2</td><td>5.3</td><td>1.3E-07</td><td>2.3E-07</td><td>MB231/MB468</td><td>NUC</td></tr>
<tr><td>>hg38_21_+_8433172_8446622_RNA45SN1_45S_50ntFlanks@8357.8389.33|AAAAGAACTTTGAAGAGAGAGTTCAAGAGGGCG</td><td>>hg38_21_+_8433172_8446622_RNA45SN1_45S_50ntFlanks</td><td>8357</td><td>8389</td><td>33</td><td>AAAAGAACTTTGAAGAGAGAGTTCAAGAGGGCG</td><td>235.3</td><td>-5.6</td><td>1.1</td><td>-5.3</td><td>1.3E-07</td><td>2.3E-07</td><td>MB231/MB468</td><td>NUC</td></tr>
<tr><td>>MI0000486|hsa-mir-190a&WithFlank&15|+|62823951|62824047@21.44.24&[MIMAT0000458&hsa-miR-190a-5p&offsets|0|+2;m-349&15|+|62823971|62823993&offsets|0|+1]|TGATATGTTTGATATATTAGGTTG</td><td>>MI0000486|hsa-mir-190a&WithFlank&15|+|62823951|62824047</td><td>21</td><td>44</td><td>24</td><td>TGATATGTTTGATATATTAGGTTG</td><td>153.4</td><td>-10.7</td><td>1.8</td><td>-5.8</td><td>5.2E-09</td><td>2.3E-07</td><td>BT20/MB231</td><td>MP</td></tr>
<tr><td>>hg38_21_+_8433172_8446622_RNA45SN1_45S_50ntFlanks@3842.3859.18|CCTACTTGGATAACTGTG</td><td>>hg38_21_+_8433172_8446622_RNA45SN1_45S_50ntFlanks</td><td>3842</td><td>3859</td><td>18</td><td>CCTACTTGGATAACTGTG</td><td>389.8</td><td>-4.2</td><td>0.8</td><td>-5.3</td><td>1.4E-07</td><td>2.3E-07</td><td>MB231/MB468</td><td>MITO</td></tr>
<tr><td>>hg38_21_+_8433172_8446622_RNA45SN1_45S_50ntFlanks@10409.10435.27|GTCAGTCGGTCCTGAGAGATGGGCGAG</td><td>>hg38_21_+_8433172_8446622_RNA45SN1_45S_50ntFlanks</td><td>10409</td><td>10435</td><td>27</td><td>GTCAGTCGGTCCTGAGAGATGGGCGAG</td><td>184.9</td><td>-6.1</td><td>1.1</td><td>-5.3</td><td>1.1E-07</td><td>2.3E-07</td><td>BT20/MB468</td><td>CYTO</td></tr>
<tr><td>>hg38_21_+_8433172_8446622_RNA45SN1_45S_50ntFlanks@11540.11562.23|CGCGGCGCCTCGCCTCGGCCGGC</td><td>>hg38_21_+_8433172_8446622_RNA45SN1_45S_50ntFlanks</td><td>11540</td><td>11562</td><td>23</td><td>CGCGGCGCCTCGCCTCGGCCGGC</td><td>306.6</td><td>6.4</td><td>1.2</td><td>5.3</td><td>1.3E-07</td><td>2.3E-07</td><td>MB231/MB468</td><td>CYTO</td></tr>
<tr><td>>hg38_21_+_8433172_8446622_RNA45SN1_45S_50ntFlanks@4067.4089.23|AACTTTCGATGGTAGTCGCCGTG</td><td>>hg38_21_+_8433172_8446622_RNA45SN1_45S_50ntFlanks</td><td>4067</td><td>4089</td><td>23</td><td>AACTTTCGATGGTAGTCGCCGTG</td><td>1232.6</td><td>-6.3</td><td>1.2</td><td>-5.3</td><td>1.4E-07</td><td>2.3E-07</td><td>MB231/MB468</td><td>MITO</td></tr>
<tr><td>>hg38_21_+_8433172_8446622_RNA45SN1_45S_50ntFlanks@11876.11904.29|AAGACCCTGTTGAGCTTGACTCTAGTCTG</td><td>>hg38_21_+_8433172_8446622_RNA45SN1_45S_50ntFlanks</td><td>11876</td><td>11904</td><td>29</td><td>AAGACCCTGTTGAGCTTGACTCTAGTCTG</td><td>669.5</td><td>-8.2</td><td>1.5</td><td>-5.6</td><td>2.6E-08</td><td>2.3E-07</td><td>BT20/MB468</td><td>MP</td></tr>
<tr><td>>hg38_21_+_8433172_8446622_RNA45SN1_45S_50ntFlanks@5227.5245.19|CGCGCGCTACACTGACTGG</td><td>>hg38_21_+_8433172_8446622_RNA45SN1_45S_50ntFlanks</td><td>5227</td><td>5245</td><td>19</td><td>CGCGCGCTACACTGACTGG</td><td>476.6</td><td>-11.0</td><td>2.1</td><td>-5.3</td><td>9.9E-08</td><td>2.4E-07</td><td>BT20/MB468</td><td>MITO</td></tr>
<tr><td>>hg38_21_+_8433172_8446622_RNA45SN1_45S_50ntFlanks@12007.12032.26|GCCGCCGGTGAAATACCACTACTCTG</td><td>>hg38_21_+_8433172_8446622_RNA45SN1_45S_50ntFlanks</td><td>12007</td><td>12032</td><td>26</td><td>GCCGCCGGTGAAATACCACTACTCTG</td><td>3299.0</td><td>-4.8</td><td>0.9</td><td>-5.3</td><td>1.3E-07</td><td>2.4E-07</td><td>MB231/MB468</td><td>CYTO</td></tr>
<tr><td>>hg38_21_+_8433172_8446622_RNA45SN1_45S_50ntFlanks@12008.12030.23|CCGCCGGTGAAATACCACTACTC</td><td>>hg38_21_+_8433172_8446622_RNA45SN1_45S_50ntFlanks</td><td>12008</td><td>12030</td><td>23</td><td>CCGCCGGTGAAATACCACTACTC</td><td>4677.5</td><td>-9.0</td><td>1.6</td><td>-5.6</td><td>2.6E-08</td><td>2.4E-07</td><td>BT20/MB468</td><td>MP</td></tr>
<tr><td>>hg38_21_+_8433172_8446622_RNA45SN1_45S_50ntFlanks@11667.11696.30|CTGCCCAGTGCTCTGAATGTCAAAGTGAAG</td><td>>hg38_21_+_8433172_8446622_RNA45SN1_45S_50ntFlanks</td><td>11667</td><td>11696</td><td>30</td><td>CTGCCCAGTGCTCTGAATGTCAAAGTGAAG</td><td>101.7</td><td>10.4</td><td>1.9</td><td>5.6</td><td>2.6E-08</td><td>2.4E-07</td><td>BT20/MB468</td><td>MP</td></tr>
<tr><td>>hg38_21_+_8433172_8446622_RNA45SN1_45S_50ntFlanks@4726.4751.26|TAATCAAGAACGAAAGTCGGAGGTTC</td><td>>hg38_21_+_8433172_8446622_RNA45SN1_45S_50ntFlanks</td><td>4726</td><td>4751</td><td>26</td><td>TAATCAAGAACGAAAGTCGGAGGTTC</td><td>228.9</td><td>-4.4</td><td>0.8</td><td>-5.3</td><td>1.3E-07</td><td>2.4E-07</td><td>MB231/MB468</td><td>CYTO</td></tr>
<tr><td>>hg38_1_-_228634819_228635039_RNA5S12_5S_50ntFlanks@152.171.20|TACCGGGTGCTGTAGGCTTT</td><td>>hg38_1_-_228634819_228635039_RNA5S12_5S_50ntFlanks</td><td>152</td><td>171</td><td>20</td><td>TACCGGGTGCTGTAGGCTTT</td><td>7461.4</td><td>-4.0</td><td>0.8</td><td>-5.3</td><td>1.2E-07</td><td>2.4E-07</td><td>MB231/MB468</td><td>MP</td></tr>
<tr><td>>hg38_21_+_8433172_8446622_RNA45SN1_45S_50ntFlanks@11572.11599.28|CCGACTTAGAACTGGTGCGGACCAGGGG</td><td>>hg38_21_+_8433172_8446622_RNA45SN1_45S_50ntFlanks</td><td>11572</td><td>11599</td><td>28</td><td>CCGACTTAGAACTGGTGCGGACCAGGGG</td><td>405.5</td><td>-3.0</td><td>0.6</td><td>-5.3</td><td>1.3E-07</td><td>2.4E-07</td><td>MB231/MB468</td><td>NUC</td></tr>
<tr><td>>hg38_21_+_8433172_8446622_RNA45SN1_45S_50ntFlanks@6721.6770.50|AATTGCAGGACACATTGATCATCGACACTTCGAACGCACTTGCGGCCCCG</td><td>>hg38_21_+_8433172_8446622_RNA45SN1_45S_50ntFlanks</td><td>6721</td><td>6770</td><td>50</td><td>AATTGCAGGACACATTGATCATCGACACTTCGAACGCACTTGCGGCCCCG</td><td>353.7</td><td>-11.5</td><td>2.1</td><td>-5.6</td><td>2.6E-08</td><td>2.4E-07</td><td>BT20/MB468</td><td>MP</td></tr>
<tr><td>>hg38_21_+_8433172_8446622_RNA45SN1_45S_50ntFlanks@12417.12455.39|TCGATGTCGGCTCTTCCTATCATTGTGAAGCAGAATTCA</td><td>>hg38_21_+_8433172_8446622_RNA45SN1_45S_50ntFlanks</td><td>12417</td><td>12455</td><td>39</td><td>TCGATGTCGGCTCTTCCTATCATTGTGAAGCAGAATTCA</td><td>364.6</td><td>-6.6</td><td>1.3</td><td>-5.3</td><td>1.2E-07</td><td>2.4E-07</td><td>MB231/MB468</td><td>MP</td></tr>
<tr><td>>hg38_21_+_8433172_8446622_RNA45SN1_45S_50ntFlanks@11918.11956.39|ACATGAGAGGTGTAGAATAAGTGGGAGGCCCCCGGCGCC</td><td>>hg38_21_+_8433172_8446622_RNA45SN1_45S_50ntFlanks</td><td>11918</td><td>11956</td><td>39</td><td>ACATGAGAGGTGTAGAATAAGTGGGAGGCCCCCGGCGCC</td><td>171.6</td><td>-5.8</td><td>1.1</td><td>-5.3</td><td>1.3E-07</td><td>2.4E-07</td><td>MB231/MB468</td><td>CYTO</td></tr>
<tr><td>>MI0000477|hsa-mir-146a&WithFlank&5|+|160485346|160485456@27.47.21&[MIMAT0000449&hsa-miR-146a-5p&offsets|0|-1;m-129&5|+|160485372|160485393&offsets|0|-1]|TGAGAACTGAATTCCATGGGT</td><td>>MI0000477|hsa-mir-146a&WithFlank&5|+|160485346|160485456</td><td>27</td><td>47</td><td>21</td><td>TGAGAACTGAATTCCATGGGT</td><td>15080.8</td><td>-9.3</td><td>1.6</td><td>-5.8</td><td>5.3E-09</td><td>2.4E-07</td><td>BT20/MB231</td><td>MP</td></tr>
<tr><td>>hg38_21_+_8433172_8446622_RNA45SN1_45S_50ntFlanks@10784.10817.34|TGAACAGCCTCTGGCATGTTGGAACAATGTAGGT</td><td>>hg38_21_+_8433172_8446622_RNA45SN1_45S_50ntFlanks</td><td>10784</td><td>10817</td><td>34</td><td>TGAACAGCCTCTGGCATGTTGGAACAATGTAGGT</td><td>158.5</td><td>-5.5</td><td>1.0</td><td>-5.3</td><td>1.2E-07</td><td>2.4E-07</td><td>MB231/MB468</td><td>MP</td></tr>
<tr><td>>hg38_21_+_8433172_8446622_RNA45SN1_45S_50ntFlanks@12457.12474.18|CAAGCGTTGGATTGTTCA</td><td>>hg38_21_+_8433172_8446622_RNA45SN1_45S_50ntFlanks</td><td>12457</td><td>12474</td><td>18</td><td>CAAGCGTTGGATTGTTCA</td><td>258.1</td><td>-7.0</td><td>1.3</td><td>-5.3</td><td>1.4E-07</td><td>2.4E-07</td><td>MB231/MB468</td><td>MITO</td></tr>
<tr><td>>hg38_21_+_8433172_8446622_RNA45SN1_45S_50ntFlanks@10458.10477.20|ATGGCCTCCGTTGCCCTCGG</td><td>>hg38_21_+_8433172_8446622_RNA45SN1_45S_50ntFlanks</td><td>10458</td><td>10477</td><td>20</td><td>ATGGCCTCCGTTGCCCTCGG</td><td>1651.3</td><td>-8.4</td><td>1.5</td><td>-5.6</td><td>2.6E-08</td><td>2.4E-07</td><td>BT20/MB468</td><td>MP</td></tr>
<tr><td>>hg38_21_+_8433172_8446622_RNA45SN1_45S_50ntFlanks@3769.3786.18|CGCACGGCCGGTACAGTG</td><td>>hg38_21_+_8433172_8446622_RNA45SN1_45S_50ntFlanks</td><td>3769</td><td>3786</td><td>18</td><td>CGCACGGCCGGTACAGTG</td><td>509.0</td><td>-6.4</td><td>1.2</td><td>-5.3</td><td>1.4E-07</td><td>2.4E-07</td><td>MB231/MB468</td><td>MITO</td></tr>
<tr><td>>hg38_21_+_8433172_8446622_RNA45SN1_45S_50ntFlanks@10498.10533.36|TTCAGATCCCCGAATCCGGAGTGGCGGAGATGGGCG</td><td>>hg38_21_+_8433172_8446622_RNA45SN1_45S_50ntFlanks</td><td>10498</td><td>10533</td><td>36</td><td>TTCAGATCCCCGAATCCGGAGTGGCGGAGATGGGCG</td><td>86.7</td><td>-9.5</td><td>1.7</td><td>-5.6</td><td>2.6E-08</td><td>2.4E-07</td><td>BT20/MB468</td><td>MP</td></tr>
<tr><td>>hg38_21_+_8433172_8446622_RNA45SN1_45S_50ntFlanks@6729.6770.42|GACACATTGATCATCGACACTTCGAACGCACTTGCGGCCCCG</td><td>>hg38_21_+_8433172_8446622_RNA45SN1_45S_50ntFlanks</td><td>6729</td><td>6770</td><td>42</td><td>GACACATTGATCATCGACACTTCGAACGCACTTGCGGCCCCG</td><td>991.6</td><td>-8.8</td><td>1.7</td><td>-5.3</td><td>1.5E-07</td><td>2.4E-07</td><td>BT20/MB468</td><td>NUC</td></tr>
<tr><td>>hg38_21_+_8433172_8446622_RNA45SN1_45S_50ntFlanks@5084.5115.32|CGAACGAGACTCTGGCATGCTAACTAGTTACG</td><td>>hg38_21_+_8433172_8446622_RNA45SN1_45S_50ntFlanks</td><td>5084</td><td>5115</td><td>32</td><td>CGAACGAGACTCTGGCATGCTAACTAGTTACG</td><td>573.3</td><td>-8.1</td><td>1.5</td><td>-5.3</td><td>1.2E-07</td><td>2.4E-07</td><td>MB231/MB468</td><td>MP</td></tr>
<tr><td>>hg38_21_+_8433172_8446622_RNA45SN1_45S_50ntFlanks@11641.11668.28|CGGCGGGTGTTGACGCGATGTGATTTCT</td><td>>hg38_21_+_8433172_8446622_RNA45SN1_45S_50ntFlanks</td><td>11641</td><td>11668</td><td>28</td><td>CGGCGGGTGTTGACGCGATGTGATTTCT</td><td>1080.2</td><td>-8.6</td><td>1.6</td><td>-5.3</td><td>1.5E-07</td><td>2.4E-07</td><td>BT20/MB468</td><td>NUC</td></tr>
<tr><td>>hg38_21_+_8433172_8446622_RNA45SN1_45S_50ntFlanks@11774.11794.21|AGTGACGCGCATGAATGGATG</td><td>>hg38_21_+_8433172_8446622_RNA45SN1_45S_50ntFlanks</td><td>11774</td><td>11794</td><td>21</td><td>AGTGACGCGCATGAATGGATG</td><td>782.3</td><td>3.5</td><td>0.7</td><td>5.3</td><td>1.3E-07</td><td>2.4E-07</td><td>MB231/MB468</td><td>CYTO</td></tr>
<tr><td>>hg38_21_+_8433172_8446622_RNA45SN1_45S_50ntFlanks@3705.3725.21|TACCTGGTTGATCCTGCCAGT</td><td>>hg38_21_+_8433172_8446622_RNA45SN1_45S_50ntFlanks</td><td>3705</td><td>3725</td><td>21</td><td>TACCTGGTTGATCCTGCCAGT</td><td>4380.6</td><td>4.8</td><td>0.9</td><td>5.7</td><td>1.4E-08</td><td>2.4E-07</td><td>BT20/MB231</td><td>NUC</td></tr>
<tr><td>>hg38_21_+_8433172_8446622_RNA45SN1_45S_50ntFlanks@10687.10710.24|CGGCGTCCGGTGAGCTCTCGCTGG</td><td>>hg38_21_+_8433172_8446622_RNA45SN1_45S_50ntFlanks</td><td>10687</td><td>10710</td><td>24</td><td>CGGCGTCCGGTGAGCTCTCGCTGG</td><td>10668.5</td><td>-10.0</td><td>1.9</td><td>-5.3</td><td>1.5E-07</td><td>2.4E-07</td><td>BT20/MB468</td><td>NUC</td></tr>
<tr><td>>hg38_21_+_8433172_8446622_RNA45SN1_45S_50ntFlanks@9565.9592.28|TGACGTGCAAATCGGTCGTCCGACCTGG</td><td>>hg38_21_+_8433172_8446622_RNA45SN1_45S_50ntFlanks</td><td>9565</td><td>9592</td><td>28</td><td>TGACGTGCAAATCGGTCGTCCGACCTGG</td><td>517.4</td><td>-3.2</td><td>0.6</td><td>-5.3</td><td>1.3E-07</td><td>2.4E-07</td><td>MB231/MB468</td><td>NUC</td></tr>
<tr><td>>MI0000455|hsa-mir-138-2&WithFlank&16|+|56858512|56858607@16.36.21;MI0000476|hsa-mir-138-1&WithFlank&3|+|44114206|44114316@29.49.21&[MIMAT0000430&hsa-miR-138-5p&offsets|0|-2;m-509&16|+|56858527|56858549&offsets|0|-2];[MIMAT0000430_1&hsa-miR-138-5p&offsets|0|-2;m-497&3|+|44114234|44114256&offsets|0|-2]|AGCTGGTGTTGTGAATCAGGC</td><td>>MI0000455|hsa-mir-138-2&WithFlank&16|+|56858512|56858607</td><td>16</td><td>36</td><td>21</td><td>AGCTGGTGTTGTGAATCAGGC</td><td>150.3</td><td>-10.6</td><td>1.8</td><td>-5.8</td><td>5.3E-09</td><td>2.4E-07</td><td>BT20/MB231</td><td>MP</td></tr>
<tr><td>>hg38_21_+_8433172_8446622_RNA45SN1_45S_50ntFlanks@11601.11620.20|ATCCGACTGTTTAATTAAAA</td><td>>hg38_21_+_8433172_8446622_RNA45SN1_45S_50ntFlanks</td><td>11601</td><td>11620</td><td>20</td><td>ATCCGACTGTTTAATTAAAA</td><td>72.3</td><td>-9.3</td><td>1.7</td><td>-5.6</td><td>2.6E-08</td><td>2.4E-07</td><td>BT20/MB468</td><td>MP</td></tr>
<tr><td>>hg38_21_+_8433172_8446622_RNA45SN1_45S_50ntFlanks@6657.6683.27|TTAGCGGTGGATCACTCGGCTCGTGCG</td><td>>hg38_21_+_8433172_8446622_RNA45SN1_45S_50ntFlanks</td><td>6657</td><td>6683</td><td>27</td><td>TTAGCGGTGGATCACTCGGCTCGTGCG</td><td>224.2</td><td>3.5</td><td>0.7</td><td>5.3</td><td>1.3E-07</td><td>2.4E-07</td><td>MB231/MB468</td><td>NUC</td></tr>
<tr><td>>hg38_21_+_8433172_8446622_RNA45SN1_45S_50ntFlanks@3815.3842.28|ATGGTTCCTTTGGTCGCTCGCTCCTCTC</td><td>>hg38_21_+_8433172_8446622_RNA45SN1_45S_50ntFlanks</td><td>3815</td><td>3842</td><td>28</td><td>ATGGTTCCTTTGGTCGCTCGCTCCTCTC</td><td>289.1</td><td>-7.1</td><td>1.4</td><td>-5.3</td><td>1.4E-07</td><td>2.4E-07</td><td>MB231/MB468</td><td>MITO</td></tr>
<tr><td>>MI0000086|hsa-mir-28&WithFlank&3|+|188688775|188688872@60.81.22(+1A)&[MIMAT0004502&hsa-miR-28-3p&offsets|0|0(+1A);m-28&3|+|188688834|188688855&offsets|0|0(+1A)]|CACTAGATTGTGAGCTCCTGGAA</td><td>>MI0000086|hsa-mir-28&WithFlank&3|+|188688775|188688872</td><td>60</td><td>81</td><td>22(+1A)</td><td>CACTAGATTGTGAGCTCCTGGAA</td><td>384.5</td><td>7.4</td><td>1.4</td><td>5.3</td><td>1.4E-07</td><td>2.4E-07</td><td>MB231/MB468</td><td>MITO</td></tr>
<tr><td>>hg38_21_+_8433172_8446622_RNA45SN1_45S_50ntFlanks@8170.8199.30|CAAGTCCTTCTGATCGAGGCCCAGCCCGTG</td><td>>hg38_21_+_8433172_8446622_RNA45SN1_45S_50ntFlanks</td><td>8170</td><td>8199</td><td>30</td><td>CAAGTCCTTCTGATCGAGGCCCAGCCCGTG</td><td>577.3</td><td>-8.5</td><td>1.6</td><td>-5.3</td><td>1.5E-07</td><td>2.4E-07</td><td>BT20/MB468</td><td>NUC</td></tr>
<tr><td>>hg38_21_+_8433172_8446622_RNA45SN1_45S_50ntFlanks@4007.4042.36|CCTCGGGCCGATCGCACGCCCCCCGTGGCGGCGACG</td><td>>hg38_21_+_8433172_8446622_RNA45SN1_45S_50ntFlanks</td><td>4007</td><td>4042</td><td>36</td><td>CCTCGGGCCGATCGCACGCCCCCCGTGGCGGCGACG</td><td>574.5</td><td>-6.3</td><td>1.2</td><td>-5.3</td><td>1.5E-07</td><td>2.4E-07</td><td>BT20/MB468</td><td>NUC</td></tr>
<tr><td>>hg38_21_+_8433172_8446622_RNA45SN1_45S_50ntFlanks@8168.8203.36|CCCAAGTCCTTCTGATCGAGGCCCAGCCCGTGGACG</td><td>>hg38_21_+_8433172_8446622_RNA45SN1_45S_50ntFlanks</td><td>8168</td><td>8203</td><td>36</td><td>CCCAAGTCCTTCTGATCGAGGCCCAGCCCGTGGACG</td><td>332.1</td><td>-6.6</td><td>1.2</td><td>-5.3</td><td>1.3E-07</td><td>2.4E-07</td><td>MB231/MB468</td><td>CYTO</td></tr>
<tr><td>>hg38_21_+_8433172_8446622_RNA45SN1_45S_50ntFlanks@10808.10837.30|CAATGTAGGTAAGGGAAGTCGGCAAGCCGG</td><td>>hg38_21_+_8433172_8446622_RNA45SN1_45S_50ntFlanks</td><td>10808</td><td>10837</td><td>30</td><td>CAATGTAGGTAAGGGAAGTCGGCAAGCCGG</td><td>672.1</td><td>-7.4</td><td>1.4</td><td>-5.3</td><td>1.4E-07</td><td>2.4E-07</td><td>MB231/MB468</td><td>MITO</td></tr>
<tr><td>>hg38_21_+_8433172_8446622_RNA45SN1_45S_50ntFlanks@6653.6684.32|ACTCTTAGCGGTGGATCACTCGGCTCGTGCGT</td><td>>hg38_21_+_8433172_8446622_RNA45SN1_45S_50ntFlanks</td><td>6653</td><td>6684</td><td>32</td><td>ACTCTTAGCGGTGGATCACTCGGCTCGTGCGT</td><td>373.4</td><td>3.9</td><td>0.7</td><td>5.3</td><td>1.0E-07</td><td>2.4E-07</td><td>BT20/MB468</td><td>MITO</td></tr>
<tr><td>>hg38_21_+_8433172_8446622_RNA45SN1_45S_50ntFlanks@11716.11740.25|TAAACGGCGGGAGTAACTATGACTC</td><td>>hg38_21_+_8433172_8446622_RNA45SN1_45S_50ntFlanks</td><td>11716</td><td>11740</td><td>25</td><td>TAAACGGCGGGAGTAACTATGACTC</td><td>254.4</td><td>-6.8</td><td>1.3</td><td>-5.3</td><td>1.3E-07</td><td>2.4E-07</td><td>MB231/MB468</td><td>NUC</td></tr>
<tr><td>>hg38_MT_+_598_1651_RNR1_12S_50ntFlanks@425.469.45|ACGAAAGTGGCTTTAACATATCTGAACACACAATAGCTAAGACCC</td><td>>hg38_MT_+_598_1651_RNR1_12S_50ntFlanks</td><td>425</td><td>469</td><td>45</td><td>ACGAAAGTGGCTTTAACATATCTGAACACACAATAGCTAAGACCC</td><td>227.4</td><td>11.6</td><td>2.1</td><td>5.6</td><td>2.6E-08</td><td>2.4E-07</td><td>BT20/MB468</td><td>MP</td></tr>
<tr><td>>hg38_21_+_8433172_8446622_RNA45SN1_45S_50ntFlanks@10677.10706.30|GCGGTTCCGGCGGCGTCCGGTGAGCTCTCG</td><td>>hg38_21_+_8433172_8446622_RNA45SN1_45S_50ntFlanks</td><td>10677</td><td>10706</td><td>30</td><td>GCGGTTCCGGCGGCGTCCGGTGAGCTCTCG</td><td>202.5</td><td>11.0</td><td>1.9</td><td>5.8</td><td>5.4E-09</td><td>2.4E-07</td><td>BT20/MB231</td><td>MP</td></tr>
<tr><td>>MI0000102|hsa-mir-100&WithFlank&11|-|122152223|122152314@19.37.19(+2U)&[MIMAT0000098&hsa-miR-100-5p&offsets|0|-3(+2U);m-27&11|-|122152276|122152296&offsets|0|-2(+2U)]|AACCCGTAGATCCGAACTTTT</td><td>>MI0000102|hsa-mir-100&WithFlank&11|-|122152223|122152314</td><td>19</td><td>37</td><td>19(+2U)</td><td>AACCCGTAGATCCGAACTTTT</td><td>168.5</td><td>-10.8</td><td>1.9</td><td>-5.8</td><td>5.4E-09</td><td>2.4E-07</td><td>BT20/MB231</td><td>MP</td></tr>
<tr><td>>hg38_21_+_8433172_8446622_RNA45SN1_45S_50ntFlanks@12654.12675.22|TATGACTGAACGCCTCTAAGTC</td><td>>hg38_21_+_8433172_8446622_RNA45SN1_45S_50ntFlanks</td><td>12654</td><td>12675</td><td>22</td><td>TATGACTGAACGCCTCTAAGTC</td><td>1560.8</td><td>-4.0</td><td>0.8</td><td>-5.3</td><td>1.0E-07</td><td>2.4E-07</td><td>BT20/MB468</td><td>MITO</td></tr>
</table>

<table>
<tr><th></th><th></th><th></th><th></th><th></th><th></th><th></th><th></th><th></th><th></th><th></th><th></th><th></th><th></th></tr>
<tr><td>>hg38_21_+_8433172_8446622_RNA45SN1_45S_50ntFlanks@6773.6797.25|TTCCTCCCGGGGCTACGCCTGTCTG</td><td>>hg38_21_+_8433172_8446622_RNA45SN1_45S_50ntFlanks</td><td>6773</td><td>6797</td><td>25</td><td>TTCCTCCCGGGGCTACGCCTGTCTG</td><td>213.9</td><td>-6.3</td><td>1.2</td><td>-5.3</td><td>1.1E-07</td><td>2.4E-07</td><td>BT20/MB468</td><td>CYTO</td></tr>
<tr><td>>hg38_21_+_8433172_8446622_RNA45SN1_45S_50ntFlanks@10351.10378.28|CGAGAACTTTGAAGGCCGAAGTGGAGAA</td><td>>hg38_21_+_8433172_8446622_RNA45SN1_45S_50ntFlanks</td><td>10351</td><td>10378</td><td>28</td><td>CGAGAACTTTGAAGGCCGAAGTGGAGAA</td><td>576.6</td><td>-3.2</td><td>0.6</td><td>-5.3</td><td>1.1E-07</td><td>2.4E-07</td><td>BT20/MB468</td><td>CYTO</td></tr>
<tr><td>>am_trna2_GlyGCC_21_-_18827107_18827177@1.25.25,trna35_GlyGCC_1_+_161413094_161413164@1.25.25,trna37_GlyGCC_1_+_161420467_161420537@1.25.25,trna39_GlyGCC_1_+_161427898_161427968@1.25.25,trna41_GlyGCC_1_+_161435258_161435328@1.25.25|GCATGGGTGGTTCAGTGGTAGAATT</td><td>>am_trna2_GlyGCC_21_-_18827107_18827177</td><td>1</td><td>25</td><td>25</td><td>GCATGGGTGGTTCAGTGGTAGAATT</td><td>2761.1</td><td>9.7</td><td>1.7</td><td>5.8</td><td>5.5E-09</td><td>2.4E-07</td><td>BT20/MB231</td><td>MP</td></tr>
<tr><td>>hg38_21_+_8433172_8446622_RNA45SN1_45S_50ntFlanks@12459.12478.20|AGCGTTGGATTGTTCACCCA</td><td>>hg38_21_+_8433172_8446622_RNA45SN1_45S_50ntFlanks</td><td>12459</td><td>12478</td><td>20</td><td>AGCGTTGGATTGTTCACCCA</td><td>730.0</td><td>-4.3</td><td>0.8</td><td>-5.3</td><td>1.1E-07</td><td>2.4E-07</td><td>BT20/MB468</td><td>CYTO</td></tr>
<tr><td>>MI0000272|hsa-mir-182&WithFlank&7|-|129770377|129770498@29.50.22&[MIMAT0000259&hsa-miR-182-5p&offsets|0|-2;m-22&7|-|129770449|129770470&offsets|0|0]|TTTGGCAATGGTAGAACTCACA</td><td>>MI0000272|hsa-mir-182&WithFlank&7|-|129770377|129770498</td><td>29</td><td>50</td><td>22</td><td>TTTGGCAATGGTAGAACTCACA</td><td>458.8</td><td>2.7</td><td>0.5</td><td>5.3</td><td>1.3E-07</td><td>2.4E-07</td><td>MB231/MB468</td><td>NUC</td></tr>
<tr><td>>hg38_21_+_8433172_8446622_RNA45SN1_45S_50ntFlanks@12416.12453.38|TTCGATGTCGGCTCTTCCTATCATTGTGAAGCAGAATT</td><td>>hg38_21_+_8433172_8446622_RNA45SN1_45S_50ntFlanks</td><td>12416</td><td>12453</td><td>38</td><td>TTCGATGTCGGCTCTTCCTATCATTGTGAAGCAGAATT</td><td>495.0</td><td>6.0</td><td>1.1</td><td>5.3</td><td>1.3E-07</td><td>2.4E-07</td><td>MB231/MB468</td><td>CYTO</td></tr>
<tr><td>>hg38_21_+_8433172_8446622_RNA45SN1_45S_50ntFlanks@8168.8193.26|CCCAAGTCCTTCTGATCGAGGCCCAG</td><td>>hg38_21_+_8433172_8446622_RNA45SN1_45S_50ntFlanks</td><td>8168</td><td>8193</td><td>26</td><td>CCCAAGTCCTTCTGATCGAGGCCCAG</td><td>76.7</td><td>-9.3</td><td>1.7</td><td>-5.6</td><td>2.6E-08</td><td>2.4E-07</td><td>BT20/MB468</td><td>MP</td></tr>
<tr><td>>hg38_21_+_8433172_8446622_RNA45SN1_45S_50ntFlanks@12499.12521.23|GGTTTAGACCGTCGTGAGACAGG</td><td>>hg38_21_+_8433172_8446622_RNA45SN1_45S_50ntFlanks</td><td>12499</td><td>12521</td><td>23</td><td>GGTTTAGACCGTCGTGAGACAGG</td><td>145.8</td><td>-10.3</td><td>1.8</td><td>-5.6</td><td>2.6E-08</td><td>2.4E-07</td><td>BT20/MB468</td><td>MP</td></tr>
<tr><td>>hg38_21_+_8433172_8446622_RNA45SN1_45S_50ntFlanks@6663.6698.36|GTGGATCACTCGGCTCGTGCGTCGATGAAGAACGCA</td><td>>hg38_21_+_8433172_8446622_RNA45SN1_45S_50ntFlanks</td><td>6663</td><td>6698</td><td>36</td><td>GTGGATCACTCGGCTCGTGCGTCGATGAAGAACGCA</td><td>372.0</td><td>-5.6</td><td>1.1</td><td>-5.3</td><td>1.4E-07</td><td>2.4E-07</td><td>MB231/MB468</td><td>MITO</td></tr>
<tr><td>>hg38_21_+_8433172_8446622_RNA45SN1_45S_50ntFlanks@9534.9575.42|AGGAAACTCTGGTGGAGGTCCGTAGCGGTCCTGACGTGCAAA</td><td>>hg38_21_+_8433172_8446622_RNA45SN1_45S_50ntFlanks</td><td>9534</td><td>9575</td><td>42</td><td>AGGAAACTCTGGTGGAGGTCCGTAGCGGTCCTGACGTGCAAA</td><td>70.9</td><td>-10.0</td><td>1.9</td><td>-5.3</td><td>1.2E-07</td><td>2.4E-07</td><td>MB231/MB468</td><td>MP</td></tr>
<tr><td>>hg38_21_+_8433172_8446622_RNA45SN1_45S_50ntFlanks@8839.8856.18|CGGTGGCGGCGGCGGCGG</td><td>>hg38_21_+_8433172_8446622_RNA45SN1_45S_50ntFlanks</td><td>8839</td><td>8856</td><td>18</td><td>CGGTGGCGGCGGCGGCGG</td><td>86.2</td><td>-9.5</td><td>1.7</td><td>-5.6</td><td>2.7E-08</td><td>2.4E-07</td><td>BT20/MB468</td><td>MP</td></tr>
<tr><td>>hg38_21_+_8433172_8446622_RNA45SN1_45S_50ntFlanks@9922.9949.28|GTTGGTTGATATAGACAGCAGGACGGTG</td><td>>hg38_21_+_8433172_8446622_RNA45SN1_45S_50ntFlanks</td><td>9922</td><td>9949</td><td>28</td><td>GTTGGTTGATATAGACAGCAGGACGGTG</td><td>71.5</td><td>9.9</td><td>1.8</td><td>5.6</td><td>2.7E-08</td><td>2.4E-07</td><td>BT20/MB468</td><td>MP</td></tr>
<tr><td>>hg38_21_+_8433172_8446622_RNA45SN1_45S_50ntFlanks@9339.9365.27|CGCACGAAAGCCGCCGTGGCGCAATGA</td><td>>hg38_21_+_8433172_8446622_RNA45SN1_45S_50ntFlanks</td><td>9339</td><td>9365</td><td>27</td><td>CGCACGAAAGCCGCCGTGGCGCAATGA</td><td>91.6</td><td>-8.6</td><td>1.6</td><td>-5.3</td><td>1.0E-07</td><td>2.4E-07</td><td>BT20/MB468</td><td>MITO</td></tr>
<tr><td>>hg38_21_+_8433172_8446622_RNA45SN1_45S_50ntFlanks@5035.5060.26|CATGGCCGTTCTTAGTTGGTGGAGCG</td><td>>hg38_21_+_8433172_8446622_RNA45SN1_45S_50ntFlanks</td><td>5035</td><td>5060</td><td>26</td><td>CATGGCCGTTCTTAGTTGGTGGAGCG</td><td>399.4</td><td>-10.7</td><td>2.0</td><td>-5.3</td><td>1.0E-07</td><td>2.4E-07</td><td>BT20/MB468</td><td>MITO</td></tr>
<tr><td>>hg38_21_+_8433172_8446622_RNA45SN1_45S_50ntFlanks@10827.10853.27|CGGCAAGCCGGATCCGTAACTTCGGGA</td><td>>hg38_21_+_8433172_8446622_RNA45SN1_45S_50ntFlanks</td><td>10827</td><td>10853</td><td>27</td><td>CGGCAAGCCGGATCCGTAACTTCGGGA</td><td>296.2</td><td>-7.5</td><td>1.4</td><td>-5.3</td><td>1.2E-07</td><td>2.4E-07</td><td>MB231/MB468</td><td>MP</td></tr>
<tr><td>>MI0000272|hsa-mir-182&WithFlank&7|-|129770377|129770498@29.53.25(+1U)&[MIMAT0000259&hsa-miR-182-5p&offsets|0|+1(+1U);m-22&7|-|129770449|129770470&offsets|0|+3(+1U)]|TTTGGCAATGGTAGAACTCACACTGT</td><td>>MI0000272|hsa-mir-182&WithFlank&7|-|129770377|129770498</td><td>29</td><td>53</td><td>25(+1U)</td><td>TTTGGCAATGGTAGAACTCACACTGT</td><td>598.4</td><td>4.2</td><td>0.8</td><td>5.3</td><td>1.3E-07</td><td>2.4E-07</td><td>MB231/MB468</td><td>CYTO</td></tr>
<tr><td>>MI0001445|hsa-mir-423&WithFlank&17|+|30117073|30117178@58.81.24(+1U)&[MIMAT0001340&hsa-miR-423-3p&offsets|-1|0(+1U);m-100&17|+|30117131|30117153&offsets|-1|0(+1U)]|AAGCTCGGTCTGAGGCCCCTCAGTT</td><td>>MI0001445|hsa-mir-423&WithFlank&17|+|30117073|30117178</td><td>58</td><td>81</td><td>24(+1U)</td><td>AAGCTCGGTCTGAGGCCCCTCAGTT</td><td>296.8</td><td>6.6</td><td>1.2</td><td>5.3</td><td>1.4E-07</td><td>2.4E-07</td><td>MB231/MB468</td><td>MITO</td></tr>
<tr><td>>hg38_21_+_8433172_8446622_RNA45SN1_45S_50ntFlanks@6661.6692.32|CGGTGGATCACTCGGCTCGTGCGTCGATGAAG</td><td>>hg38_21_+_8433172_8446622_RNA45SN1_45S_50ntFlanks</td><td>6661</td><td>6692</td><td>32</td><td>CGGTGGATCACTCGGCTCGTGCGTCGATGAAG</td><td>555.0</td><td>-3.8</td><td>0.7</td><td>-5.3</td><td>1.2E-07</td><td>2.4E-07</td><td>MB231/MB468</td><td>MP</td></tr>
<tr><td>>hg38_21_+_8433172_8446622_RNA45SN1_45S_50ntFlanks@8190.8211.22|CCAGCCCGTGGACGGTGTGAGG</td><td>>hg38_21_+_8433172_8446622_RNA45SN1_45S_50ntFlanks</td><td>8190</td><td>8211</td><td>22</td><td>CCAGCCCGTGGACGGTGTGAGG</td><td>126.9</td><td>-10.1</td><td>1.8</td><td>-5.6</td><td>2.7E-08</td><td>2.4E-07</td><td>BT20/MB468</td><td>MP</td></tr>
<tr><td>>hg38_21_+_8433172_8446622_RNA45SN1_45S_50ntFlanks@9837.9875.39|TGGTAAGCAGAACTGGCGCTGCGGGATGAACCGAACGCC</td><td>>hg38_21_+_8433172_8446622_RNA45SN1_45S_50ntFlanks</td><td>9837</td><td>9875</td><td>39</td><td>TGGTAAGCAGAACTGGCGCTGCGGGATGAACCGAACGCC</td><td>76.4</td><td>10.0</td><td>1.8</td><td>5.6</td><td>2.7E-08</td><td>2.4E-07</td><td>BT20/MB468</td><td>MP</td></tr>
<tr><td>>hg38_21_+_8433172_8446622_RNA45SN1_45S_50ntFlanks@4588.4614.27|CTATTTTGTTGGTTTTCGGAACTGAGG</td><td>>hg38_21_+_8433172_8446622_RNA45SN1_45S_50ntFlanks</td><td>4588</td><td>4614</td><td>27</td><td>CTATTTTGTTGGTTTTCGGAACTGAGG</td><td>645.3</td><td>-7.3</td><td>1.4</td><td>-5.3</td><td>1.4E-07</td><td>2.4E-07</td><td>MB231/MB468</td><td>NUC</td></tr>
<tr><td>>MI0001445|hsa-mir-423&WithFlank&17|+|30117073|30117178@59.80.22(+1A)&[MIMAT0001340&hsa-miR-423-3p&offsets|0|-1(+1A);m-100&17|+|30117131|30117153&offsets|0|-1(+1A)]|AGCTCGGTCTGAGGCCCCTCAGA</td><td>>MI0001445|hsa-mir-423&WithFlank&17|+|30117073|30117178</td><td>59</td><td>80</td><td>22(+1A)</td><td>AGCTCGGTCTGAGGCCCCTCAGA</td><td>493.3</td><td>6.0</td><td>1.1</td><td>5.3</td><td>1.3E-07</td><td>2.4E-07</td><td>MB231/MB468</td><td>CYTO</td></tr>
<tr><td>>hg38_21_+_8433172_8446622_RNA45SN1_45S_50ntFlanks@10727.10748.22|GGAGAGGGTGTAAATCTCGCGC</td><td>>hg38_21_+_8433172_8446622_RNA45SN1_45S_50ntFlanks</td><td>10727</td><td>10748</td><td>22</td><td>GGAGAGGGTGTAAATCTCGCGC</td><td>124.8</td><td>-10.0</td><td>1.8</td><td>-5.6</td><td>2.7E-08</td><td>2.4E-07</td><td>BT20/MB468</td><td>MP</td></tr>
<tr><td>>hg38_21_+_8433172_8446622_RNA45SN1_45S_50ntFlanks@10786.10827.42|AACAGCCTCTGGCATGTTGGAACAATGTAGGTAAGGGAAGTC</td><td>>hg38_21_+_8433172_8446622_RNA45SN1_45S_50ntFlanks</td><td>10786</td><td>10827</td><td>42</td><td>AACAGCCTCTGGCATGTTGGAACAATGTAGGTAAGGGAAGTC</td><td>315.2</td><td>-7.9</td><td>1.5</td><td>-5.3</td><td>1.5E-07</td><td>2.4E-07</td><td>BT20/MB468</td><td>NUC</td></tr>
<tr><td>>hg38_21_+_8433172_8446622_RNA45SN1_45S_50ntFlanks@6729.6770.42|GACACATTGATCATCGACACTTCGAACGCACTTGCGGCCCCG</td><td>>hg38_21_+_8433172_8446622_RNA45SN1_45S_50ntFlanks</td><td>6729</td><td>6770</td><td>42</td><td>GACACATTGATCATCGACACTTCGAACGCACTTGCGGCCCCG</td><td>241.3</td><td>-11.8</td><td>2.2</td><td>-5.3</td><td>1.2E-07</td><td>2.4E-07</td><td>MB231/MB468</td><td>MP</td></tr>
<tr><td>>hg38_21_+_8433172_8446622_RNA45SN1_45S_50ntFlanks@11534.11555.22|GTTCCGCGCGGCGCCTCGCCTC</td><td>>hg38_21_+_8433172_8446622_RNA45SN1_45S_50ntFlanks</td><td>11534</td><td>11555</td><td>22</td><td>GTTCCGCGCGGCGCCTCGCCTC</td><td>380.9</td><td>-6.7</td><td>1.3</td><td>-5.3</td><td>1.2E-07</td><td>2.4E-07</td><td>BT20/MB468</td><td>CYTO</td></tr>
<tr><td>>hg38_21_+_8433172_8446622_RNA45SN1_45S_50ntFlanks@12002.12043.42|TGCGGGCCGCCGGTGAAATACCACTACTCTGATCGTTTTTTC</td><td>>hg38_21_+_8433172_8446622_RNA45SN1_45S_50ntFlanks</td><td>12002</td><td>12043</td><td>42</td><td>TGCGGGCCGCCGGTGAAATACCACTACTCTGATCGTTTTTTC</td><td>289.0</td><td>-11.3</td><td>2.0</td><td>-5.6</td><td>2.7E-08</td><td>2.4E-07</td><td>BT20/MB468</td><td>MP</td></tr>
<tr><td>>hg38_21_+_8433172_8446622_RNA45SN1_45S_50ntFlanks@9556.9596.41|TAGCGGTCCTGACGTGCAAATCGGTCGTCCGACCTGGGTAT</td><td>>hg38_21_+_8433172_8446622_RNA45SN1_45S_50ntFlanks</td><td>9556</td><td>9596</td><td>41</td><td>TAGCGGTCCTGACGTGCAAATCGGTCGTCCGACCTGGGTAT</td><td>184.0</td><td>-9.6</td><td>1.8</td><td>-5.3</td><td>1.0E-07</td><td>2.4E-07</td><td>BT20/MB468</td><td>MITO</td></tr>
<tr><td>>MI0000078|hsa-mir-22&WithFlank&17|-|1713897|1713993@59.77.19(+1C)&[MIMAT0000077&hsa-miR-22-3p&offsets|0|-3(+1C);m-2&17|-|1713914|1713935&offsets|0|-3(+1C)]|AAGCTGCCAGTTGAAGAACC</td><td>>MI0000078|hsa-mir-22&WithFlank&17|-|1713897|1713993</td><td>59</td><td>77</td><td>19(+1C)</td><td>AAGCTGCCAGTTGAAGAACC</td><td>70.2</td><td>9.4</td><td>1.8</td><td>5.3</td><td>1.2E-07</td><td>2.4E-07</td><td>MB231/MB468</td><td>MP</td></tr>
<tr><td>>MI0000270|hsa-mir-181b-1&WithFlank&1|-|198858867|198858988@42.65.24;MI0000683|hsa-mir-181b-2&WithFlank&9|+|124693704|124693804@22.45.24&[MIMAT0000257&hsa-miR-181b-5p&offsets|0|+1;m-73&1|-|198858925|198858947&offsets|0|+1];[MIMAT0000257_1&hsa-miR-181b-5p&offsets|0|+1;m-72&9|+|124693725|124693747&offsets|0|+1]|AACATTCATTGCTGTCGGTGGGTT</td><td>>MI0000270|hsa-mir-181b-1&WithFlank&1|-|198858867|198858988</td><td>42</td><td>65</td><td>24</td><td>AACATTCATTGCTGTCGGTGGGTT</td><td>928.5</td><td>5.0</td><td>1.0</td><td>5.3</td><td>1.3E-07</td><td>2.5E-07</td><td>MB231/MB468</td><td>CYTO</td></tr>
<tr><td>>trnaMT_GluTTC_MT_-_14674_14742@39.66.28,trnalookalike8_GluTTC_5_-_93905172_93905240@39.66.28|ATTGGTCGTGGTTGTAGTCCGTGCGAGA</td><td>>trnaMT_GluTTC_MT_-_14674_14742</td><td>39</td><td>66</td><td>28</td><td>ATTGGTCGTGGTTGTAGTCCGTGCGAGA</td><td>358.1</td><td>4.5</td><td>0.9</td><td>5.3</td><td>1.4E-07</td><td>2.5E-07</td><td>MB231/MB468</td><td>NUC</td></tr>
<tr><td>>hg38_21_+_8433172_8446622_RNA45SN1_45S_50ntFlanks@12631.12654.24|GGCGAAGCTACCATCTGTGGGATT</td><td>>hg38_21_+_8433172_8446622_RNA45SN1_45S_50ntFlanks</td><td>12631</td><td>12654</td><td>24</td><td>GGCGAAGCTACCATCTGTGGGATT</td><td>202.3</td><td>-4.2</td><td>0.8</td><td>-5.3</td><td>1.2E-07</td><td>2.5E-07</td><td>BT20/MB468</td><td>CYTO</td></tr>
<tr><td>>hg38_21_+_8433172_8446622_RNA45SN1_45S_50ntFlanks@11888.11905.18|AGCTTGACTCTAGTCTGG</td><td>>hg38_21_+_8433172_8446622_RNA45SN1_45S_50ntFlanks</td><td>11888</td><td>11905</td><td>18</td><td>AGCTTGACTCTAGTCTGG</td><td>188.9</td><td>-6.7</td><td>1.3</td><td>-5.3</td><td>1.2E-07</td><td>2.5E-07</td><td>BT20/MB468</td><td>CYTO</td></tr>
<tr><td>>hg38_21_+_8433172_8446622_RNA45SN1_45S_50ntFlanks@9564.9594.31|CTGACGTGCAAATCGGTCGTCCGACCTGGGT</td><td>>hg38_21_+_8433172_8446622_RNA45SN1_45S_50ntFlanks</td><td>9564</td><td>9594</td><td>31</td><td>CTGACGTGCAAATCGGTCGTCCGACCTGGGT</td><td>595.3</td><td>-3.0</td><td>0.6</td><td>-5.3</td><td>1.3E-07</td><td>2.5E-07</td><td>MB231/MB468</td><td>CYTO</td></tr>
<tr><td>>hg38_21_+_8433172_8446622_RNA45SN1_45S_50ntFlanks@10806.10828.23|AACAATGTAGGTAAGGGAAGTCG</td><td>>hg38_21_+_8433172_8446622_RNA45SN1_45S_50ntFlanks</td><td>10806</td><td>10828</td><td>23</td><td>AACAATGTAGGTAAGGGAAGTCG</td><td>412.2</td><td>-6.9</td><td>1.3</td><td>-5.3</td><td>1.4E-07</td><td>2.5E-07</td><td>MB231/MB468</td><td>NUC</td></tr>
<tr><td>>hg38_21_+_8433172_8446622_RNA45SN1_45S_50ntFlanks@11564.11601.38|CCTAGCAGCCGACTTAGAACTGGTGCGGACCAGGGGAA</td><td>>hg38_21_+_8433172_8446622_RNA45SN1_45S_50ntFlanks</td><td>11564</td><td>11601</td><td>38</td><td>CCTAGCAGCCGACTTAGAACTGGTGCGGACCAGGGGAA</td><td>4700.3</td><td>-3.7</td><td>0.7</td><td>-5.3</td><td>1.0E-07</td><td>2.5E-07</td><td>BT20/MB468</td><td>MITO</td></tr>
<tr><td>>hg38_21_+_8433172_8446622_RNA45SN1_45S_50ntFlanks@11567.11602.36|AGCAGCCGACTTAGAACTGGTGCGGACCAGGGGAAT</td><td>>hg38_21_+_8433172_8446622_RNA45SN1_45S_50ntFlanks</td><td>11567</td><td>11602</td><td>36</td><td>AGCAGCCGACTTAGAACTGGTGCGGACCAGGGGAAT</td><td>644.6</td><td>-2.7</td><td>0.5</td><td>-5.3</td><td>1.4E-07</td><td>2.5E-07</td><td>MB231/MB468</td><td>NUC</td></tr>
<tr><td>>am_trnaMT_LeuTAG_MT_+_12266_12336@42.61.20|AAAATTTTGGTGCAACTCCA</td><td>>am_trnaMT_LeuTAG_MT_+_12266_12336</td><td>42</td><td>61</td><td>20</td><td>AAAATTTTGGTGCAACTCCA</td><td>85.2</td><td>-9.5</td><td>1.7</td><td>-5.6</td><td>2.7E-08</td><td>2.5E-07</td><td>BT20/MB468</td><td>MP</td></tr>
<tr><td>>hg38_21_+_8433172_8446622_RNA45SN1_45S_50ntFlanks@6661.6688.28|CGGTGGATCACTCGGCTCGTGCGTCGAT</td><td>>hg38_21_+_8433172_8446622_RNA45SN1_45S_50ntFlanks</td><td>6661</td><td>6688</td><td>28</td><td>CGGTGGATCACTCGGCTCGTGCGTCGAT</td><td>1010.3</td><td>-3.7</td><td>0.7</td><td>-5.3</td><td>1.2E-07</td><td>2.5E-07</td><td>MB231/MB468</td><td>MP</td></tr>
<tr><td>>am_trnaMT_PheGAA_MT_+_577_647@41.58.18|GTTTAGACGGGCTCACAT</td><td>>am_trnaMT_PheGAA_MT_+_577_647</td><td>41</td><td>58</td><td>18</td><td>GTTTAGACGGGCTCACAT</td><td>689.3</td><td>4.9</td><td>0.9</td><td>5.6</td><td>2.7E-08</td><td>2.5E-07</td><td>BT20/MB468</td><td>MP</td></tr>
<tr><td>>hg38_21_+_8433172_8446622_RNA45SN1_45S_50ntFlanks@10458.10495.38|ATGGCCTCCGTTGCCCTCGGCCGATCGAAAGGGAGTCG</td><td>>hg38_21_+_8433172_8446622_RNA45SN1_45S_50ntFlanks</td><td>10458</td><td>10495</td><td>38</td><td>ATGGCCTCCGTTGCCCTCGGCCGATCGAAAGGGAGTCG</td><td>550.8</td><td>-7.6</td><td>1.4</td><td>-5.3</td><td>1.2E-07</td><td>2.5E-07</td><td>MB231/MB468</td><td>MP</td></tr>
<tr><td>>hg38_21_+_8433172_8446622_RNA45SN1_45S_50ntFlanks@12159.12181.23|TTTGACTGGGGCGGTACACCTGT</td><td>>hg38_21_+_8433172_8446622_RNA45SN1_45S_50ntFlanks</td><td>12159</td><td>12181</td><td>23</td><td>TTTGACTGGGGCGGTACACCTGT</td><td>4058.2</td><td>-8.9</td><td>1.6</td><td>-5.6</td><td>2.7E-08</td><td>2.5E-07</td><td>BT20/MB468</td><td>MP</td></tr>
<tr><td>>trnaMT_SerGCT_MT_+_12207_12265@8.30.23|CTCACAAGAACTGCTAACTCATG</td><td>>trnaMT_SerGCT_MT_+_12207_12265</td><td>8</td><td>30</td><td>23</td><td>CTCACAAGAACTGCTAACTCATG</td><td>423.5</td><td>-7.8</td><td>1.5</td><td>-5.3</td><td>1.4E-07</td><td>2.5E-07</td><td>MB231/MB468</td><td>NUC</td></tr>
<tr><td>>hg38_21_+_8433172_8446622_RNA45SN1_45S_50ntFlanks@10357.10390.34|CTTTGAAGGCCGAAGTGGAGAAGGGTTCCATGTG</td><td>>hg38_21_+_8433172_8446622_RNA45SN1_45S_50ntFlanks</td><td>10357</td><td>10390</td><td>34</td><td>CTTTGAAGGCCGAAGTGGAGAAGGGTTCCATGTG</td><td>1267.4</td><td>-8.9</td><td>1.7</td><td>-5.2</td><td>1.5E-07</td><td>2.5E-07</td><td>BT20/MB468</td><td>NUC</td></tr>
<tr><td>>hg38_21_+_8433172_8446622_RNA45SN1_45S_50ntFlanks@6646.6674.29|TCGTACGACTCTTAGCGGTGGATCACTCG</td><td>>hg38_21_+_8433172_8446622_RNA45SN1_45S_50ntFlanks</td><td>6646</td><td>6674</td><td>29</td><td>TCGTACGACTCTTAGCGGTGGATCACTCG</td><td>1779.7</td><td>4.4</td><td>0.8</td><td>5.3</td><td>1.2E-07</td><td>2.5E-07</td><td>BT20/MB468</td><td>CYTO</td></tr>
<tr><td>>hg38_21_+_8433172_8446622_RNA45SN1_45S_50ntFlanks@475.505.31|CTTCGTGATCGATGTGGTGACGTCGTGCTCT</td><td>>hg38_21_+_8433172_8446622_RNA45SN1_45S_50ntFlanks</td><td>475</td><td>505</td><td>31</td><td>CTTCGTGATCGATGTGGTGACGTCGTGCTCT</td><td>1959.6</td><td>5.5</td><td>1.0</td><td>5.6</td><td>2.7E-08</td><td>2.5E-07</td><td>BT20/MB468</td><td>MP</td></tr>
<tr><td>>hg38_21_+_8433172_8446622_RNA45SN1_45S_50ntFlanks@5150.5186.37|AGAGGGACAAGTGGCGTTCAGCCACCCGAGATTGAGC</td><td>>hg38_21_+_8433172_8446622_RNA45SN1_45S_50ntFlanks</td><td>5150</td><td>5186</td><td>37</td><td>AGAGGGACAAGTGGCGTTCAGCCACCCGAGATTGAGC</td><td>76.8</td><td>10.0</td><td>1.8</td><td>5.6</td><td>2.7E-08</td><td>2.5E-07</td><td>BT20/MB468</td><td>MP</td></tr>
<tr><td>>hg38_21_+_8433172_8446622_RNA45SN1_45S_50ntFlanks@10690.10707.18|CGTCCGGTGAGCTCTCGC</td><td>>hg38_21_+_8433172_8446622_RNA45SN1_45S_50ntFlanks</td><td>10690</td><td>10707</td><td>18</td><td>CGTCCGGTGAGCTCTCGC</td><td>405.3</td><td>-6.2</td><td>1.2</td><td>-5.2</td><td>1.5E-07</td><td>2.5E-07</td><td>BT20/MB468</td><td>NUC</td></tr>
<tr><td>>hg38_21_+_8433172_8446622_RNA45SN1_45S_50ntFlanks@6730.6756.27|ACACATTGATCATCGACACTTCGAACG</td><td>>hg38_21_+_8433172_8446622_RNA45SN1_45S_50ntFlanks</td><td>6730</td><td>6756</td><td>27</td><td>ACACATTGATCATCGACACTTCGAACG</td><td>544.4</td><td>-8.1</td><td>1.5</td><td>-5.2</td><td>1.5E-07</td><td>2.5E-07</td><td>BT20/MB468</td><td>NUC</td></tr>
<tr><td>>hg38_21_+_8433172_8446622_RNA45SN1_45S_50ntFlanks@11587.11612.26|TGCGGACCAGGGGAATCCGACTGTTT</td><td>>hg38_21_+_8433172_8446622_RNA45SN1_45S_50ntFlanks</td><td>11587</td><td>11612</td><td>26</td><td>TGCGGACCAGGGGAATCCGACTGTTT</td><td>386.4</td><td>-3.0</td><td>0.6</td><td>-5.2</td><td>1.5E-07</td><td>2.5E-07</td><td>BT20/MB468</td><td>NUC</td></tr>
</table>

| | | | | | | | | | | | | | |
|---|---|---|---|---|---|---|---|---|---|---|---|---|---|
| >hg38_21_+_8433172_8446622_RNA45SN1_45S_50ntFlanks@10806.10826.21\|AACAATGTAGGTAAGGGAAGT | >hg38_21_+_8433172_8446622_RNA45SN1_45S_50ntFlanks | 10806 | 10826 | 21 | AACAATGTAGGTAAGGGAAGT | 478.0 | -5.1 | 1.0 | -5.3 | 1.0E-07 | 2.5E-07 | BT20/MB468 | MITO |
| >hg38_21_+_8433172_8446622_RNA45SN1_45S_50ntFlanks@12940.12973.34\|CATTCGTAGACGACCTGCTTCTGGGTCGGGGTTT | >hg38_21_+_8433172_8446622_RNA45SN1_45S_50ntFlanks | 12940 | 12973 | 34 | CATTCGTAGACGACCTGCTTCTGGGTCGGGGTTT | 135.6 | -6.7 | 1.3 | -5.3 | 1.2E-07 | 2.5E-07 | BT20/MB468 | CYTO |
| >hg38_21_+_8433172_8446622_RNA45SN1_45S_50ntFlanks@12407.12448.42\|TTTTGATCCTTCGATGTCGGCTCTTCCTATCATTGTGAAGCA | >hg38_21_+_8433172_8446622_RNA45SN1_45S_50ntFlanks | 12407 | 12448 | 42 | TTTTGATCCTTCGATGTCGGCTCTTCCTATCATTGTGAAGCA | 657.6 | 6.7 | 1.3 | 5.3 | 1.4E-07 | 2.5E-07 | MB231/MB468 | CYTO |
| >hg38_21_+_8433172_8446622_RNA45SN1_45S_50ntFlanks@7977.8010.34\|CGACCTCAGATCAGACGTGGCGACCCGCTGAATT | >hg38_21_+_8433172_8446622_RNA45SN1_45S_50ntFlanks | 7977 | 8010 | 34 | CGACCTCAGATCAGACGTGGCGACCCGCTGAATT | 333.9 | 7.3 | 1.4 | 5.3 | 1.2E-07 | 2.5E-07 | BT20/MB468 | CYTO |
| >hg38_21_+_8433172_8446622_RNA45SN1_45S_50ntFlanks@10680.10706.27\|GTTCCGGCGGCGTCCGGTGAGCTCTCG | >hg38_21_+_8433172_8446622_RNA45SN1_45S_50ntFlanks | 10680 | 10706 | 27 | GTTCCGGCGGCGTCCGGTGAGCTCTCG | 4943.8 | -4.1 | 0.8 | -5.3 | 1.1E-07 | 2.5E-07 | BT20/MB468 | MITO |
| >hg38_21_+_8433172_8446622_RNA45SN1_45S_50ntFlanks@12635.12657.23\|AAGCTACCATCTGTGGGATTATG | >hg38_21_+_8433172_8446622_RNA45SN1_45S_50ntFlanks | 12635 | 12657 | 23 | AAGCTACCATCTGTGGGATTATG | 210.6 | -5.2 | 1.0 | -5.3 | 1.4E-07 | 2.5E-07 | MB231/MB468 | CYTO |
| >hg38_21_+_8433172_8446622_RNA45SN1_45S_50ntFlanks@12616.12657.42\|CTGAGGAGCCAATGGGGCGAAGCTACCATCTGTGGGATTATG | >hg38_21_+_8433172_8446622_RNA45SN1_45S_50ntFlanks | 12616 | 12657 | 42 | CTGAGGAGCCAATGGGGCGAAGCTACCATCTGTGGGATTATG | 68.7 | -10.0 | 1.9 | -5.3 | 1.2E-07 | 2.5E-07 | MB231/MB468 | MP |
| >hg38_21_+_8433172_8446622_RNA45SN1_45S_50ntFlanks@12407.12435.29\|TTTTGATCCTTCGATGTCGGCTCTTCCTA | >hg38_21_+_8433172_8446622_RNA45SN1_45S_50ntFlanks | 12407 | 12435 | 29 | TTTTGATCCTTCGATGTCGGCTCTTCCTA | 3634.6 | -9.6 | 1.8 | -5.2 | 1.5E-07 | 2.5E-07 | BT20/MB468 | NUC |
| >hg38_21_+_8433172_8446622_RNA45SN1_45S_50ntFlanks@10845.10874.30\|ACTTCGGGATAAGGATTGGCTCTAAGGGCT | >hg38_21_+_8433172_8446622_RNA45SN1_45S_50ntFlanks | 10845 | 10874 | 30 | ACTTCGGGATAAGGATTGGCTCTAAGGGCT | 375.9 | -7.8 | 1.5 | -5.2 | 1.5E-07 | 2.5E-07 | BT20/MB468 | NUC |
| >hg38_21_+_8433172_8446622_RNA45SN1_45S_50ntFlanks@4403.4423.21\|CGGGCGGTCCGCCGCGAGGCG | >hg38_21_+_8433172_8446622_RNA45SN1_45S_50ntFlanks | 4403 | 4423 | 21 | CGGGCGGTCCGCCGCGAGGCG | 244.6 | -10.0 | 1.9 | -5.3 | 1.1E-07 | 2.5E-07 | BT20/MB468 | MITO |
| >hg38_21_+_8433172_8446622_RNA45SN1_45S_50ntFlanks@3872.3898.27\|CTAATACATGCCGACGGGCGCTGACCC | >hg38_21_+_8433172_8446622_RNA45SN1_45S_50ntFlanks | 3872 | 3898 | 27 | CTAATACATGCCGACGGGCGCTGACCC | 1309.7 | -7.9 | 1.5 | -5.3 | 1.4E-07 | 2.5E-07 | MB231/MB468 | NUC |
| >hg38_21_+_8433172_8446622_RNA45SN1_45S_50ntFlanks@11534.11560.27\|GTTCCGCGCGGCGCCTCGCCTCGGCCG | >hg38_21_+_8433172_8446622_RNA45SN1_45S_50ntFlanks | 11534 | 11560 | 27 | GTTCCGCGCGGCGCCTCGCCTCGGCCG | 608.5 | -7.1 | 1.3 | -5.3 | 1.2E-07 | 2.5E-07 | BT20/MB468 | CYTO |
| >hg38_21_+_8433172_8446622_RNA45SN1_45S_50ntFlanks@10684.10712.29\|CGGCGGCGTCCGGTGAGCTCTCGCTGGCC | >hg38_21_+_8433172_8446622_RNA45SN1_45S_50ntFlanks | 10684 | 10712 | 29 | CGGCGGCGTCCGGTGAGCTCTCGCTGGCC | 501.0 | -5.9 | 1.1 | -5.3 | 1.3E-07 | 2.5E-07 | MB231/MB468 | MP |
| >hg38_21_+_8433172_8446622_RNA45SN1_45S_50ntFlanks@11583.11602.20\|CTGGTGCGGACCAGGGGAAT | >hg38_21_+_8433172_8446622_RNA45SN1_45S_50ntFlanks | 11583 | 11602 | 20 | CTGGTGCGGACCAGGGGAAT | 1005.7 | -6.6 | 1.3 | -5.3 | 1.5E-07 | 2.5E-07 | MB231/MB468 | MITO |
| >hg38_21_+_8433172_8446622_RNA45SN1_45S_50ntFlanks@8116.8138.23\|GCGGGACATGTGGCGTACGGAAG | >hg38_21_+_8433172_8446622_RNA45SN1_45S_50ntFlanks | 8116 | 8138 | 23 | GCGGGACATGTGGCGTACGGAAG | 555.0 | 12.4 | 2.1 | 5.8 | 5.7E-09 | 2.5E-07 | BT20/MB231 | MP |
| >hg38_21_+_8433172_8446622_RNA45SN1_45S_50ntFlanks@9460.9503.44\|CGGGGAGGTGGAGCACGAGCGCACGTGTTAGGACCCGAAAGATG | >hg38_21_+_8433172_8446622_RNA45SN1_45S_50ntFlanks | 9460 | 9503 | 44 | CGGGGAGGTGGAGCACGAGCGCACGTGTTAGGACCCGAAAGATG | 69.0 | -10.0 | 1.9 | -5.3 | 1.3E-07 | 2.5E-07 | MB231/MB468 | MP |
| >MI0000065\|hsa-let-7d&WithFlank&9\|+\|94178828\|94178926@68.88.21&[MIMAT0004484&hsa-let-7d-3p&offsets\|0\|-1;m-98&9\|+\|94178895\|94178916&offsets\|0\|-1]\|CTATACGACCTGCTGCCTTTC | >MI0000065\|hsa-let-7d&WithFlank&9\|+\|94178828\|94178926 | 68 | 88 | 21 | CTATACGACCTGCTGCCTTTC | 462.1 | 3.9 | 0.7 | 5.3 | 1.3E-07 | 2.5E-07 | MB231/MB468 | MP |
| >hg38_21_+_8433172_8446622_RNA45SN1_45S_50ntFlanks@5078.5102.25\|CGATAACGAACGAGACTCTGGCATG | >hg38_21_+_8433172_8446622_RNA45SN1_45S_50ntFlanks | 5078 | 5102 | 25 | CGATAACGAACGAGACTCTGGCATG | 1456.0 | -9.3 | 1.8 | -5.2 | 1.6E-07 | 2.5E-07 | BT20/MB468 | NUC |
| >hg38_21_+_8433172_8446622_RNA45SN1_45S_50ntFlanks@10349.10368.20\|AACGAGAACTTTGAAGGCCG | >hg38_21_+_8433172_8446622_RNA45SN1_45S_50ntFlanks | 10349 | 10368 | 20 | AACGAGAACTTTGAAGGCCG | 4731.5 | -7.0 | 1.3 | -5.3 | 1.3E-07 | 2.5E-07 | MB231/MB468 | MP |
| >trna116_GluCTC_1_-_145399233_145399304@1.31.31,trna59_GluCTC_1_+_249168447_249168518@1.31.31,trna71_GluCTC_1_-_161439189_161439260@1.31.31,trna74_GluCTC_1_-_161431809_161431880@1.31.31,trna77_GluCTC_1_-_161424398_161424469@1.31.31,trna77_GluCTC_6_+_28949976_28950047@1.31.31,trna80_GluCTC_1_-_161417018_161417089@1.31.31,trna87_GluCTC_6_-_126101393_126101464@1.31.31\|TCCCTGGTGGTCTAGTGGTTAGGATTCGGCG | >trna116_GluCTC_1_-_145399233_145399304 | 1 | 31 | 31 | TCCCTGGTGGTCTAGTGGTTAGGATTCGGCG | 139786.9 | -5.1 | 1.0 | -5.3 | 1.4E-07 | 2.5E-07 | MB231/MB468 | NUC |
| >hg38_21_+_8433172_8446622_RNA45SN1_45S_50ntFlanks@8169.8187.19\|CCAAGTCCTTCTGATCGAG | >hg38_21_+_8433172_8446622_RNA45SN1_45S_50ntFlanks | 8169 | 8187 | 19 | CCAAGTCCTTCTGATCGAG | 278.3 | -5.5 | 1.1 | -5.3 | 1.4E-07 | 2.5E-07 | MB231/MB468 | NUC |
| >hg38_21_+_8433172_8446622_RNA45SN1_45S_50ntFlanks@3843.3861.19\|CTACTTGGATAACTGTGGT | >hg38_21_+_8433172_8446622_RNA45SN1_45S_50ntFlanks | 3843 | 3861 | 19 | CTACTTGGATAACTGTGGT | 1017.3 | -8.6 | 1.6 | -5.3 | 1.3E-07 | 2.5E-07 | MB231/MB468 | MP |
| >hg38_1_-_228634819_228635039_RNA5S12_5S_50ntFlanks@127.171.45\|ACTTGGATGGGAGACCGCCTGGGAATACCGGGTGCTGTAGGCTTT | >hg38_1_-_228634819_228635039_RNA5S12_5S_50ntFlanks | 127 | 171 | 45 | ACTTGGATGGGAGACCGCCTGGGAATACCGGGTGCTGTAGGCTTT | 778.1 | 8.2 | 1.5 | 5.3 | 1.1E-07 | 2.5E-07 | BT20/MB468 | MITO |
| >hg38_21_+_8433172_8446622_RNA45SN1_45S_50ntFlanks@3771.3792.22\|CACGGCCGGTACAGTGAAACTG | >hg38_21_+_8433172_8446622_RNA45SN1_45S_50ntFlanks | 3771 | 3792 | 22 | CACGGCCGGTACAGTGAAACTG | 707.7 | -7.5 | 1.4 | -5.3 | 1.1E-07 | 2.5E-07 | BT20/MB468 | MITO |
| >hg38_21_+_8433172_8446622_RNA45SN1_45S_50ntFlanks@11567.11592.26\|AGCAGCCGACTTAGAACTGGTGCGGA | >hg38_21_+_8433172_8446622_RNA45SN1_45S_50ntFlanks | 11567 | 11592 | 26 | AGCAGCCGACTTAGAACTGGTGCGGA | 1086.5 | -7.1 | 1.3 | -5.3 | 1.3E-07 | 2.5E-07 | MB231/MB468 | MP |
| >hg38_21_+_8433172_8446622_RNA45SN1_45S_50ntFlanks@11233.11269.37\|CCCTTCCCGTGGATCGCCCCAGCTGCGGCGGGCGTCG | >hg38_21_+_8433172_8446622_RNA45SN1_45S_50ntFlanks | 11233 | 11269 | 37 | CCCTTCCCGTGGATCGCCCCAGCTGCGGCGGGCGTCG | 218.0 | -6.2 | 1.2 | -5.3 | 1.1E-07 | 2.5E-07 | BT20/MB468 | MITO |
| >hg38_21_+_8433172_8446622_RNA45SN1_45S_50ntFlanks@4731.4765.35\|AAGAACGAAAGTCGGAGGTTCGAAGACGATCAGAT | >hg38_21_+_8433172_8446622_RNA45SN1_45S_50ntFlanks | 4731 | 4765 | 35 | AAGAACGAAAGTCGGAGGTTCGAAGACGATCAGAT | 975.0 | 5.0 | 0.9 | 5.6 | 2.8E-08 | 2.5E-07 | BT20/MB468 | MP |
| >hg38_21_+_8433172_8446622_RNA45SN1_45S_50ntFlanks@11602.11621.20\|TCCGACTGTTTAATTAAAAC | >hg38_21_+_8433172_8446622_RNA45SN1_45S_50ntFlanks | 11602 | 11621 | 20 | TCCGACTGTTTAATTAAAAC | 566.4 | -6.5 | 1.2 | -5.2 | 1.6E-07 | 2.5E-07 | BT20/MB468 | NUC |
| >hg38_21_+_8433172_8446622_RNA45SN1_45S_50ntFlanks@12646.12667.22\|TGTGGGATTATGACTGAACGCC | >hg38_21_+_8433172_8446622_RNA45SN1_45S_50ntFlanks | 12646 | 12667 | 22 | TGTGGGATTATGACTGAACGCC | 75.5 | 10.0 | 1.8 | 5.6 | 2.8E-08 | 2.5E-07 | BT20/MB468 | MP |
| >hg38_21_+_8433172_8446622_RNA45SN1_45S_50ntFlanks@8170.8208.39\|CAAGTCCTTCTGATCGAGGCCCAGCCCGTGGACGGTGTG | >hg38_21_+_8433172_8446622_RNA45SN1_45S_50ntFlanks | 8170 | 8208 | 39 | CAAGTCCTTCTGATCGAGGCCCAGCCCGTGGACGGTGTG | 145.7 | -10.3 | 1.8 | -5.6 | 2.8E-08 | 2.5E-07 | BT20/MB468 | MP |
| >hg38_21_+_8433172_8446622_RNA45SN1_45S_50ntFlanks@3900.3919.20\|CTTCGCGGGGGGGATGCGTG | >hg38_21_+_8433172_8446622_RNA45SN1_45S_50ntFlanks | 3900 | 3919 | 20 | CTTCGCGGGGGGGATGCGTG | 331.5 | -4.6 | 0.9 | -5.3 | 1.4E-07 | 2.5E-07 | MB231/MB468 | CYTO |
| >hg38_21_+_8433172_8446622_RNA45SN1_45S_50ntFlanks@12420.12450.31\|ATGTCGGCTCTTCCTATCATTGTGAAGCAGA | >hg38_21_+_8433172_8446622_RNA45SN1_45S_50ntFlanks | 12420 | 12450 | 31 | ATGTCGGCTCTTCCTATCATTGTGAAGCAGA | 616.4 | -6.6 | 1.3 | -5.2 | 1.6E-07 | 2.5E-07 | BT20/MB468 | NUC |
| >hg38_21_+_8433172_8446622_RNA45SN1_45S_50ntFlanks@10781.10800.20\|AGGTGAACAGCCTCTGGCAT | >hg38_21_+_8433172_8446622_RNA45SN1_45S_50ntFlanks | 10781 | 10800 | 20 | AGGTGAACAGCCTCTGGCAT | 196.3 | -7.0 | 1.3 | -5.3 | 1.3E-07 | 2.5E-07 | MB231/MB468 | MP |
| >hg38_21_+_8433172_8446622_RNA45SN1_45S_50ntFlanks@10832.10862.31\|AGCCGGATCCGTAACTTCGGGATAAGGATTG | >hg38_21_+_8433172_8446622_RNA45SN1_45S_50ntFlanks | 10832 | 10862 | 31 | AGCCGGATCCGTAACTTCGGGATAAGGATTG | 436.6 | -7.3 | 1.4 | -5.3 | 1.2E-07 | 2.5E-07 | BT20/MB468 | CYTO |
| >hg38_21_+_8433172_8446622_RNA45SN1_45S_50ntFlanks@7975.8001.27\|CGCGACCTCAGATCAGACGTGGCGACC | >hg38_21_+_8433172_8446622_RNA45SN1_45S_50ntFlanks | 7975 | 8001 | 27 | CGCGACCTCAGATCAGACGTGGCGACC | 467.7 | 4.3 | 0.8 | 5.3 | 1.4E-07 | 2.5E-07 | MB231/MB468 | CYTO |
| >MI0000456\|hsa-mir-140&WithFlank&16\|+\|69933075\|69933186@70.90.21&[MIMAT0004597&hsa-miR-140-3p&offsets\|+2\|+2;m-45&16\|+\|69933143\|69933164&offsets\|+1\|0]\|CCACAGGGTAGAACCACGGAC | >MI0000456\|hsa-mir-140&WithFlank&16\|+\|69933075\|69933186 | 70 | 90 | 21 | CCACAGGGTAGAACCACGGAC | 57.5 | 9.1 | 1.7 | 5.3 | 1.3E-07 | 2.5E-07 | MB231/MB468 | MP |
| >hg38_21_+_8433172_8446622_RNA45SN1_45S_50ntFlanks@3840.3860.21\|CTCCTACTTGGATAACTGTGG | >hg38_21_+_8433172_8446622_RNA45SN1_45S_50ntFlanks | 3840 | 3860 | 21 | CTCCTACTTGGATAACTGTGG | 958.7 | -9.2 | 1.7 | -5.3 | 1.3E-07 | 2.5E-07 | MB231/MB468 | MP |
| >hg38_21_+_8433172_8446622_RNA45SN1_45S_50ntFlanks@11670.11702.33\|CCCAGTGCTCTGAATGTCAAAGTGAAGAAATTC | >hg38_21_+_8433172_8446622_RNA45SN1_45S_50ntFlanks | 11670 | 11702 | 33 | CCCAGTGCTCTGAATGTCAAAGTGAAGAAATTC | 1282.8 | -2.3 | 0.4 | -5.3 | 1.1E-07 | 2.5E-07 | BT20/MB468 | MITO |
| >hg38_21_+_8433172_8446622_RNA45SN1_45S_50ntFlanks@4068.4089.22\|ACTTTCGATGGTAGTCGCCGTG | >hg38_21_+_8433172_8446622_RNA45SN1_45S_50ntFlanks | 4068 | 4089 | 22 | ACTTTCGATGGTAGTCGCCGTG | 168.1 | -4.4 | 0.8 | -5.3 | 1.4E-07 | 2.5E-07 | MB231/MB468 | CYTO |
| >trnaMT_ProTGG_MT_-_15956_16023@31.70.40\|TTGGGTGCTAATGGTGGAGTTAAAGACTTTTTCTCTGACC | >trnaMT_ProTGG_MT_-_15956_16023 | 31 | 70 | 40 | TTGGGTGCTAATGGTGGAGTTAAAGACTTTTTCTCTGACC | 159.0 | 11.1 | 2.0 | 5.6 | 2.8E-08 | 2.5E-07 | BT20/MB468 | MP |
| >hg38_21_+_8433172_8446622_RNA45SN1_45S_50ntFlanks@5472.5510.39\|ACGGCCCTGGCGGAGCGCTGAGAAGACGGTCGAACTTGA | >hg38_21_+_8433172_8446622_RNA45SN1_45S_50ntFlanks | 5472 | 5510 | 39 | ACGGCCCTGGCGGAGCGCTGAGAAGACGGTCGAACTTGA | 1336.0 | 4.8 | 0.9 | 5.3 | 1.4E-07 | 2.6E-07 | MB231/MB468 | CYTO |
| >hg38_21_+_8433172_8446622_RNA45SN1_45S_50ntFlanks@12488.12508.21\|AACGTGAGCTGGGTTTAGACC | >hg38_21_+_8433172_8446622_RNA45SN1_45S_50ntFlanks | 12488 | 12508 | 21 | AACGTGAGCTGGGTTTAGACC | 293.3 | -7.5 | 1.4 | -5.2 | 1.6E-07 | 2.6E-07 | BT20/MB468 | NUC |
| >hg38_21_+_8433172_8446622_RNA45SN1_45S_50ntFlanks@10347.10368.22\|CAAACGAGAACTTTGAAGGCCG | >hg38_21_+_8433172_8446622_RNA45SN1_45S_50ntFlanks | 10347 | 10368 | 22 | CAAACGAGAACTTTGAAGGCCG | 475.6 | -5.6 | 1.1 | -5.3 | 1.5E-07 | 2.6E-07 | MB231/MB468 | MITO |
| >hg38_21_+_8433172_8446622_RNA45SN1_45S_50ntFlanks@10808.10826.19\|CAATGTAGGTAAGGGAAGT | >hg38_21_+_8433172_8446622_RNA45SN1_45S_50ntFlanks | 10808 | 10826 | 19 | CAATGTAGGTAAGGGAAGT | 558.7 | -7.8 | 1.4 | -5.6 | 2.8E-08 | 2.6E-07 | BT20/MB468 | MP |
| >hg38_21_+_8433172_8446622_RNA45SN1_45S_50ntFlanks@11766.11803.38\|ATCTAATTAGTGACGCGCATGAATGGATGAACGAGATT | >hg38_21_+_8433172_8446622_RNA45SN1_45S_50ntFlanks | 11766 | 11803 | 38 | ATCTAATTAGTGACGCGCATGAATGGATGAACGAGATT | 128.9 | -10.1 | 1.8 | -5.6 | 2.8E-08 | 2.6E-07 | BT20/MB468 | MP |
| >hg38_21_+_8433172_8446622_RNA45SN1_45S_50ntFlanks@10510.10533.24\|AATCCGGAGTGGCGGAGATGGGCG | >hg38_21_+_8433172_8446622_RNA45SN1_45S_50ntFlanks | 10510 | 10533 | 24 | AATCCGGAGTGGCGGAGATGGGCG | 688.0 | -5.4 | 1.0 | -5.3 | 1.2E-07 | 2.6E-07 | BT20/MB468 | CYTO |
| >hg38_21_+_8433172_8446622_RNA45SN1_45S_50ntFlanks@10362.10381.20\|AAGGCCGAAGTGGAGAAGGG | >hg38_21_+_8433172_8446622_RNA45SN1_45S_50ntFlanks | 10362 | 10381 | 20 | AAGGCCGAAGTGGAGAAGGG | 293.6 | -7.2 | 1.4 | -5.3 | 1.3E-07 | 2.6E-07 | MB231/MB468 | MP |
| >hg38_21_+_8433172_8446622_RNA45SN1_45S_50ntFlanks@12937.12959.23\|AACCATTCGTAGACGACCTGCTT | >hg38_21_+_8433172_8446622_RNA45SN1_45S_50ntFlanks | 12937 | 12959 | 23 | AACCATTCGTAGACGACCTGCTT | 262.9 | -3.9 | 0.7 | -5.3 | 1.5E-07 | 2.6E-07 | MB231/MB468 | MITO |
| >hg38_21_+_8433172_8446622_RNA45SN1_45S_50ntFlanks@8176.8202.27\|CTTCTGATCGAGGCCCAGCCCGTGGAC | >hg38_21_+_8433172_8446622_RNA45SN1_45S_50ntFlanks | 8176 | 8202 | 27 | CTTCTGATCGAGGCCCAGCCCGTGGAC | 2484.8 | 2.8 | 0.5 | 5.3 | 1.2E-07 | 2.6E-07 | BT20/MB468 | CYTO |

<table>
<tr><td>>hg38_21_+_8433172_8446622_RNA45SN1_45S_50ntFlanks@7975.8018.44|CGCGACCTCAGATCAGACGTGGCGACCCGCTGAATTTAAGCATA</td><td>>hg38_21_+_8433172_8446622_RNA45SN1_45S_50ntFlanks</td><td>7975</td><td>8018</td><td>44</td><td>CGCGACCTCAGATCAGACGTGGCGACCCGCTGAATTTAAGCATA</td><td>1281.1</td><td>6.0</td><td>1.1</td><td>5.3</td><td>1.4E-07</td><td>2.6E-07</td><td>MB231/MB468</td><td>CYTO</td></tr>
<tr><td>>hg38_1_-_228634819_228635039_RNA5S12_5S_50ntFlanks@134.169.36|TGGGAGACCGCCTGGGAATACCGGGTGCTGTAGGCT</td><td>>hg38_1_-_228634819_228635039_RNA5S12_5S_50ntFlanks</td><td>134</td><td>169</td><td>36</td><td>TGGGAGACCGCCTGGGAATACCGGGTGCTGTAGGCT</td><td>5191.0</td><td>4.4</td><td>0.8</td><td>5.3</td><td>1.4E-07</td><td>2.6E-07</td><td>MB231/MB468</td><td>NUC</td></tr>
<tr><td>>hg38_21_+_8433172_8446622_RNA45SN1_45S_50ntFlanks@11571.11594.24|GCCGACTTAGAACTGGTGCGGACC</td><td>>hg38_21_+_8433172_8446622_RNA45SN1_45S_50ntFlanks</td><td>11571</td><td>11594</td><td>24</td><td>GCCGACTTAGAACTGGTGCGGACC</td><td>306.6</td><td>3.2</td><td>0.6</td><td>5.3</td><td>1.4E-07</td><td>2.6E-07</td><td>MB231/MB468</td><td>NUC</td></tr>
<tr><td>>hg38_21_+_8433172_8446622_RNA45SN1_45S_50ntFlanks@4353.4391.39|TATTAAAGTTGCTGCAGTTAAAAAGCTCGTAGTTGGATC</td><td>>hg38_21_+_8433172_8446622_RNA45SN1_45S_50ntFlanks</td><td>4353</td><td>4391</td><td>39</td><td>TATTAAAGTTGCTGCAGTTAAAAAGCTCGTAGTTGGATC</td><td>228.7</td><td>-3.5</td><td>0.7</td><td>-5.3</td><td>1.4E-07</td><td>2.6E-07</td><td>MB231/MB468</td><td>NUC</td></tr>
<tr><td>>hg38_21_+_8433172_8446622_RNA45SN1_45S_50ntFlanks@11315.11344.30|CCACCCCACGTCTCGTCGCGCGCGCGTCCG</td><td>>hg38_21_+_8433172_8446622_RNA45SN1_45S_50ntFlanks</td><td>11315</td><td>11344</td><td>30</td><td>CCACCCCACGTCTCGTCGCGCGCGCGTCCG</td><td>18861.2</td><td>-10.4</td><td>2.0</td><td>-5.2</td><td>1.6E-07</td><td>2.6E-07</td><td>BT20/MB468</td><td>NUC</td></tr>
<tr><td>>hg38_21_+_8433172_8446622_RNA45SN1_45S_50ntFlanks@4976.5000.25|CCGGACACGGACAGGATTGACAGAT</td><td>>hg38_21_+_8433172_8446622_RNA45SN1_45S_50ntFlanks</td><td>4976</td><td>5000</td><td>25</td><td>CCGGACACGGACAGGATTGACAGAT</td><td>135.6</td><td>-5.3</td><td>1.0</td><td>-5.3</td><td>1.3E-07</td><td>2.6E-07</td><td>MB231/MB468</td><td>MP</td></tr>
<tr><td>>MI0000087|hsa-mir-29a&WithFlank&7|-|130876741|130876816@47.67.21(+1C)&[MIMAT0000086&hsa-miR-29a-3p&offsets|-1|-2(+1C);m-26&7|-|130876749|130876769&offsets|-1|-1(+1C)]|CTAGCACCATCTGAAATCGGTC</td><td>>MI0000087|hsa-mir-29a&WithFlank&7|-|130876741|130876816</td><td>47</td><td>67</td><td>21(+1C)</td><td>CTAGCACCATCTGAAATCGGTC</td><td>2016.2</td><td>6.0</td><td>1.1</td><td>5.6</td><td>2.9E-08</td><td>2.6E-07</td><td>BT20/MB468</td><td>MP</td></tr>
<tr><td>>hg38_21_+_8433172_8446622_RNA45SN1_45S_50ntFlanks@11829.11855.27|CGAAACCACAGCCAAGGGAACGGGCTT</td><td>>hg38_21_+_8433172_8446622_RNA45SN1_45S_50ntFlanks</td><td>11829</td><td>11855</td><td>27</td><td>CGAAACCACAGCCAAGGGAACGGGCTT</td><td>86.3</td><td>-9.5</td><td>1.7</td><td>-5.6</td><td>2.9E-08</td><td>2.6E-07</td><td>BT20/MB468</td><td>MP</td></tr>
<tr><td>>hg38_21_+_8433172_8446622_RNA45SN1_45S_50ntFlanks@8357.8379.23|AAAAGAACTTTGAAGAGAGAGTT</td><td>>hg38_21_+_8433172_8446622_RNA45SN1_45S_50ntFlanks</td><td>8357</td><td>8379</td><td>23</td><td>AAAAGAACTTTGAAGAGAGAGTT</td><td>64550.2</td><td>-10.4</td><td>1.9</td><td>-5.5</td><td>2.9E-08</td><td>2.6E-07</td><td>BT20/MB468</td><td>MP</td></tr>
<tr><td>>hg38_21_+_8433172_8446622_RNA45SN1_45S_50ntFlanks@12183.12211.29|AAACGGTAACGCAGGTGTCCTAAGGCGAG</td><td>>hg38_21_+_8433172_8446622_RNA45SN1_45S_50ntFlanks</td><td>12183</td><td>12211</td><td>29</td><td>AAACGGTAACGCAGGTGTCCTAAGGCGAG</td><td>84.9</td><td>-9.5</td><td>1.7</td><td>-5.5</td><td>2.9E-08</td><td>2.6E-07</td><td>BT20/MB468</td><td>MP</td></tr>
<tr><td>>hg38_21_+_8433172_8446622_RNA45SN1_45S_50ntFlanks@9925.9955.31|GGTTGATATAGACAGCAGGACGGTGGCCATG</td><td>>hg38_21_+_8433172_8446622_RNA45SN1_45S_50ntFlanks</td><td>9925</td><td>9955</td><td>31</td><td>GGTTGATATAGACAGCAGGACGGTGGCCATG</td><td>86.0</td><td>10.2</td><td>1.8</td><td>5.5</td><td>2.9E-08</td><td>2.6E-07</td><td>BT20/MB468</td><td>MP</td></tr>
<tr><td>>hg38_21_+_8433172_8446622_RNA45SN1_45S_50ntFlanks@8200.8223.24|GACGGTGTGAGGCCGGTAGCGGCC</td><td>>hg38_21_+_8433172_8446622_RNA45SN1_45S_50ntFlanks</td><td>8200</td><td>8223</td><td>24</td><td>GACGGTGTGAGGCCGGTAGCGGCC</td><td>84.6</td><td>-9.5</td><td>1.7</td><td>-5.5</td><td>2.9E-08</td><td>2.6E-07</td><td>BT20/MB468</td><td>MP</td></tr>
<tr><td>>MI0001733|hsa-mir-452&WithFlank&X|-|151959622|151959718@23.43.21&[MIMAT0001635&hsa-miR-452-5p&offsets|+3|+2;m-188&X|-|151959677|151959699&offsets|+3|+1]|TGTTTGCAGAGGAAACTGAGA</td><td>>MI0001733|hsa-mir-452&WithFlank&X|-|151959622|151959718</td><td>23</td><td>43</td><td>21</td><td>TGTTTGCAGAGGAAACTGAGA</td><td>680.0</td><td>3.1</td><td>0.6</td><td>5.7</td><td>1.6E-08</td><td>2.6E-07</td><td>BT20/MB231</td><td>NUC</td></tr>
<tr><td>>hg38_21_+_8433172_8446622_RNA45SN1_45S_50ntFlanks@10408.10443.36|GGTCAGTCGGTCCTGAGAGATGGGCGAGCGCCGTTC</td><td>>hg38_21_+_8433172_8446622_RNA45SN1_45S_50ntFlanks</td><td>10408</td><td>10443</td><td>36</td><td>GGTCAGTCGGTCCTGAGAGATGGGCGAGCGCCGTTC</td><td>1309.8</td><td>-6.7</td><td>1.3</td><td>-5.3</td><td>1.3E-07</td><td>2.6E-07</td><td>MB231/MB468</td><td>MP</td></tr>
<tr><td>>hg38_21_+_8433172_8446622_RNA45SN1_45S_50ntFlanks@6744.6770.27|GACACTTCGAACGCACTTGCGGCCCCG</td><td>>hg38_21_+_8433172_8446622_RNA45SN1_45S_50ntFlanks</td><td>6744</td><td>6770</td><td>27</td><td>GACACTTCGAACGCACTTGCGGCCCCG</td><td>2392.7</td><td>-9.3</td><td>1.8</td><td>-5.2</td><td>1.6E-07</td><td>2.6E-07</td><td>BT20/MB468</td><td>NUC</td></tr>
<tr><td>>hg38_21_+_8433172_8446622_RNA45SN1_45S_50ntFlanks@8379.8400.22|TCAAGAGGGCGTGAAACCGTTA</td><td>>hg38_21_+_8433172_8446622_RNA45SN1_45S_50ntFlanks</td><td>8379</td><td>8400</td><td>22</td><td>TCAAGAGGGCGTGAAACCGTTA</td><td>130.4</td><td>-5.9</td><td>1.1</td><td>-5.3</td><td>1.2E-07</td><td>2.6E-07</td><td>BT20/MB468</td><td>CYTO</td></tr>
<tr><td>>hg38_21_+_8433172_8446622_RNA45SN1_45S_50ntFlanks@10687.10712.26|CGGCGTCCGGTGAGCTCTCGCTGGCC</td><td>>hg38_21_+_8433172_8446622_RNA45SN1_45S_50ntFlanks</td><td>10687</td><td>10712</td><td>26</td><td>CGGCGTCCGGTGAGCTCTCGCTGGCC</td><td>1647.5</td><td>-5.7</td><td>1.1</td><td>-5.3</td><td>1.2E-07</td><td>2.6E-07</td><td>BT20/MB468</td><td>CYTO</td></tr>
<tr><td>>hg38_21_+_8433172_8446622_RNA45SN1_45S_50ntFlanks@4730.4759.30|CAAGAACGAAAGTCGGAGGTTCGAAGACGA</td><td>>hg38_21_+_8433172_8446622_RNA45SN1_45S_50ntFlanks</td><td>4730</td><td>4759</td><td>30</td><td>CAAGAACGAAAGTCGGAGGTTCGAAGACGA</td><td>670.8</td><td>-4.0</td><td>0.8</td><td>-5.3</td><td>1.5E-07</td><td>2.6E-07</td><td>MB231/MB468</td><td>NUC</td></tr>
<tr><td>>hg38_21_+_8433172_8446622_RNA45SN1_45S_50ntFlanks@11569.11601.33|CAGCCGACTTAGAACTGGTGCGGACCAGGGGAA</td><td>>hg38_21_+_8433172_8446622_RNA45SN1_45S_50ntFlanks</td><td>11569</td><td>11601</td><td>33</td><td>CAGCCGACTTAGAACTGGTGCGGACCAGGGGAA</td><td>3123.9</td><td>-4.4</td><td>0.8</td><td>-5.3</td><td>1.2E-07</td><td>2.6E-07</td><td>BT20/MB468</td><td>CYTO</td></tr>
<tr><td>>am_trnaMT_HisGTG_MT_+_12138_12206@21.51.31|ACATCAGATTGTGAATCTGACAACAGAGGCT</td><td>>am_trnaMT_HisGTG_MT_+_12138_12206</td><td>21</td><td>51</td><td>31</td><td>ACATCAGATTGTGAATCTGACAACAGAGGCT</td><td>110.8</td><td>10.5</td><td>1.9</td><td>5.5</td><td>2.9E-08</td><td>2.6E-07</td><td>BT20/MB468</td><td>MP</td></tr>
<tr><td>>hg38_21_+_8433172_8446622_RNA45SN1_45S_50ntFlanks@485.523.39|GATGTGGTGACGTCGTGCTCTCCCGGGCCGGGTCCGAGC</td><td>>hg38_21_+_8433172_8446622_RNA45SN1_45S_50ntFlanks</td><td>485</td><td>523</td><td>39</td><td>GATGTGGTGACGTCGTGCTCTCCCGGGCCGGGTCCGAGC</td><td>215.4</td><td>11.5</td><td>2.1</td><td>5.5</td><td>2.9E-08</td><td>2.6E-07</td><td>BT20/MB468</td><td>MP</td></tr>
<tr><td>>hg38_21_+_8433172_8446622_RNA45SN1_45S_50ntFlanks@3746.3763.18|ATTAAGCCATGCATGTCT</td><td>>hg38_21_+_8433172_8446622_RNA45SN1_45S_50ntFlanks</td><td>3746</td><td>3763</td><td>18</td><td>ATTAAGCCATGCATGTCT</td><td>81.5</td><td>-9.4</td><td>1.7</td><td>-5.5</td><td>2.9E-08</td><td>2.6E-07</td><td>BT20/MB468</td><td>MP</td></tr>
<tr><td>>hg38_21_+_8433172_8446622_RNA45SN1_45S_50ntFlanks@12054.12072.19|TGAGGCGGGGGGGCGAGCC</td><td>>hg38_21_+_8433172_8446622_RNA45SN1_45S_50ntFlanks</td><td>12054</td><td>12072</td><td>19</td><td>TGAGGCGGGGGGGCGAGCC</td><td>92.8</td><td>-9.6</td><td>1.7</td><td>-5.5</td><td>2.9E-08</td><td>2.6E-07</td><td>BT20/MB468</td><td>MP</td></tr>
<tr><td>>hg38_21_+_8433172_8446622_RNA45SN1_45S_50ntFlanks@11641.11680.40|CGGCGGGTGTTGACGCGATGTGATTTCTGCCCAGTGCTCT</td><td>>hg38_21_+_8433172_8446622_RNA45SN1_45S_50ntFlanks</td><td>11641</td><td>11680</td><td>40</td><td>CGGCGGGTGTTGACGCGATGTGATTTCTGCCCAGTGCTCT</td><td>3204.3</td><td>-6.9</td><td>1.3</td><td>-5.2</td><td>1.5E-07</td><td>2.6E-07</td><td>MB231/MB468</td><td>MITO</td></tr>
<tr><td>>hg38_21_+_8433172_8446622_RNA45SN1_45S_50ntFlanks@11670.11703.34|CCCAGTGCTCTGAATGTCAAAGTGAAGAAATTCA</td><td>>hg38_21_+_8433172_8446622_RNA45SN1_45S_50ntFlanks</td><td>11670</td><td>11703</td><td>34</td><td>CCCAGTGCTCTGAATGTCAAAGTGAAGAAATTCA</td><td>3507.4</td><td>-9.4</td><td>1.8</td><td>-5.2</td><td>1.6E-07</td><td>2.6E-07</td><td>BT20/MB468</td><td>NUC</td></tr>
<tr><td>>hg38_21_+_8433172_8446622_RNA45SN1_45S_50ntFlanks@4297.4318.22|CATTGGAGGGCAAGTCTGGTGC</td><td>>hg38_21_+_8433172_8446622_RNA45SN1_45S_50ntFlanks</td><td>4297</td><td>4318</td><td>22</td><td>CATTGGAGGGCAAGTCTGGTGC</td><td>166.6</td><td>-7.0</td><td>1.3</td><td>-5.3</td><td>1.3E-07</td><td>2.6E-07</td><td>MB231/MB468</td><td>MP</td></tr>
<tr><td>>MI0000102|hsa-mir-100&WithFlank&11|-|122152223|122152314@20.40.21(+1A)&[MIMAT0000098&hsa-miR-100-5p&offsets|+1|0(+1A);m-27&11|-|122152276|122152296&offsets|+1|+1(+1A)]|ACCCGTAGATCCGAACTTGTGA</td><td>>MI0000102|hsa-mir-100&WithFlank&11|-|122152223|122152314</td><td>20</td><td>40</td><td>21(+1A)</td><td>ACCCGTAGATCCGAACTTGTGA</td><td>56.6</td><td>9.0</td><td>1.7</td><td>5.3</td><td>1.3E-07</td><td>2.6E-07</td><td>MB231/MB468</td><td>MP</td></tr>
<tr><td>>hg38_21_+_8433172_8446622_RNA45SN1_45S_50ntFlanks@11881.11898.18|CCTGTTGAGCTTGACTCT</td><td>>hg38_21_+_8433172_8446622_RNA45SN1_45S_50ntFlanks</td><td>11881</td><td>11898</td><td>18</td><td>CCTGTTGAGCTTGACTCT</td><td>212.6</td><td>-7.0</td><td>1.3</td><td>-5.3</td><td>1.3E-07</td><td>2.6E-07</td><td>MB231/MB468</td><td>MP</td></tr>
<tr><td>>hg38_21_+_8433172_8446622_RNA45SN1_45S_50ntFlanks@9939.9975.37|GCAGGACGGTGGCCATGGAAGTCGGAATCCGCTAAGG</td><td>>hg38_21_+_8433172_8446622_RNA45SN1_45S_50ntFlanks</td><td>9939</td><td>9975</td><td>37</td><td>GCAGGACGGTGGCCATGGAAGTCGGAATCCGCTAAGG</td><td>67.9</td><td>9.8</td><td>1.8</td><td>5.5</td><td>2.9E-08</td><td>2.6E-07</td><td>BT20/MB468</td><td>MP</td></tr>
<tr><td>>hg38_21_+_8433172_8446622_RNA45SN1_45S_50ntFlanks@9533.9553.21|GAGGAAACTCTGGTGGAGGTC</td><td>>hg38_21_+_8433172_8446622_RNA45SN1_45S_50ntFlanks</td><td>9533</td><td>9553</td><td>21</td><td>GAGGAAACTCTGGTGGAGGTC</td><td>16932.5</td><td>-4.2</td><td>0.8</td><td>-5.3</td><td>1.3E-07</td><td>2.6E-07</td><td>BT20/MB468</td><td>CYTO</td></tr>
<tr><td>>hg38_21_+_8433172_8446622_RNA45SN1_45S_50ntFlanks@9198.9220.23|GTTCTCTCGGGGCCACGCGCGCG</td><td>>hg38_21_+_8433172_8446622_RNA45SN1_45S_50ntFlanks</td><td>9198</td><td>9220</td><td>23</td><td>GTTCTCTCGGGGCCACGCGCGCG</td><td>256.5</td><td>-7.7</td><td>1.5</td><td>-5.3</td><td>1.5E-07</td><td>2.6E-07</td><td>MB231/MB468</td><td>NUC</td></tr>
<tr><td>>MI0000102|hsa-mir-100&WithFlank&11|-|122152223|122152314@19.38.20(+1C)&[MIMAT0000098&hsa-miR-100-5p&offsets|0|-2(+1C);m-27&11|-|122152276|122152296&offsets|0|-1(+1C)]|AACCCGTAGATCCGAACTTGC</td><td>>MI0000102|hsa-mir-100&WithFlank&11|-|122152223|122152314</td><td>19</td><td>38</td><td>20(+1C)</td><td>AACCCGTAGATCCGAACTTGC</td><td>16602.4</td><td>-5.9</td><td>1.0</td><td>-5.7</td><td>1.6E-08</td><td>2.6E-07</td><td>BT20/MB231</td><td>NUC</td></tr>
<tr><td>>hg38_21_+_8433172_8446622_RNA45SN1_45S_50ntFlanks@4100.4117.18|TGACCACGGGTGACGGGG</td><td>>hg38_21_+_8433172_8446622_RNA45SN1_45S_50ntFlanks</td><td>4100</td><td>4117</td><td>18</td><td>TGACCACGGGTGACGGGG</td><td>89.1</td><td>-9.6</td><td>1.7</td><td>-5.5</td><td>2.9E-08</td><td>2.6E-07</td><td>BT20/MB468</td><td>MP</td></tr>
<tr><td>>hg38_21_+_8433172_8446622_RNA45SN1_45S_50ntFlanks@11319.11338.20|CCCACGTCTCGTCGCGCGCG</td><td>>hg38_21_+_8433172_8446622_RNA45SN1_45S_50ntFlanks</td><td>11319</td><td>11338</td><td>20</td><td>CCCACGTCTCGTCGCGCGCG</td><td>235.3</td><td>-5.2</td><td>1.0</td><td>-5.3</td><td>1.3E-07</td><td>2.6E-07</td><td>BT20/MB468</td><td>CYTO</td></tr>
<tr><td>>hg38_21_+_8433172_8446622_RNA45SN1_45S_50ntFlanks@11515.11539.25|GTCCGCCCCCGGGGCCGCGGTTCCG</td><td>>hg38_21_+_8433172_8446622_RNA45SN1_45S_50ntFlanks</td><td>11515</td><td>11539</td><td>25</td><td>GTCCGCCCCCGGGGCCGCGGTTCCG</td><td>171.9</td><td>-6.3</td><td>1.2</td><td>-5.3</td><td>1.3E-07</td><td>2.6E-07</td><td>BT20/MB468</td><td>CYTO</td></tr>
<tr><td>>am_MI0000809|hsa-mir-151a&WithFlank&8|-|140732558|140732659@53.73.21(+3U)&[MIMAT0000757&hsa-miR-151a-3p&offsets|0|0(+3U);m-30&8|-|140732587|140732607&offsets|0|0(+3U)]|CTAGACTGAAGCTCCTTGAGGTTT</td><td>>am_MI0000809|hsa-mir-151a&WithFlank&8|-|140732558|140732659</td><td>53</td><td>73</td><td>21(+3U)</td><td>CTAGACTGAAGCTCCTTGAGGTTT</td><td>81.2</td><td>3.1</td><td>0.6</td><td>5.2</td><td>1.6E-07</td><td>2.6E-07</td><td>BT20/MB468</td><td>NUC</td></tr>
<tr><td>>hg38_21_+_8433172_8446622_RNA45SN1_45S_50ntFlanks@7975.7995.21|CGCGACCTCAGATCAGACGTG</td><td>>hg38_21_+_8433172_8446622_RNA45SN1_45S_50ntFlanks</td><td>7975</td><td>7995</td><td>21</td><td>CGCGACCTCAGATCAGACGTG</td><td>51393.6</td><td>4.4</td><td>0.8</td><td>5.3</td><td>1.1E-07</td><td>2.6E-07</td><td>BT20/MB468</td><td>MITO</td></tr>
<tr><td>>am_trnaMT_SerGCT_MT_+_12207_12265@15.32.18|GAACTGCTAACTCATGCC</td><td>>am_trnaMT_SerGCT_MT_+_12207_12265</td><td>15</td><td>32</td><td>18</td><td>GAACTGCTAACTCATGCC</td><td>241.9</td><td>8.4</td><td>1.6</td><td>5.3</td><td>1.3E-07</td><td>2.6E-07</td><td>BT20/MB468</td><td>CYTO</td></tr>
<tr><td>>hg38_21_+_8433172_8446622_RNA45SN1_45S_50ntFlanks@12347.12364.18|AGAAAAGTTACCACAGGG</td><td>>hg38_21_+_8433172_8446622_RNA45SN1_45S_50ntFlanks</td><td>12347</td><td>12364</td><td>18</td><td>AGAAAAGTTACCACAGGG</td><td>115.5</td><td>-9.9</td><td>1.8</td><td>-5.5</td><td>3.0E-08</td><td>2.6E-07</td><td>BT20/MB468</td><td>MP</td></tr>
<tr><td>>hg38_21_+_8433172_8446622_RNA45SN1_45S_50ntFlanks@7992.8011.20|CGTGGCGACCCGCTGAATTT</td><td>>hg38_21_+_8433172_8446622_RNA45SN1_45S_50ntFlanks</td><td>7992</td><td>8011</td><td>20</td><td>CGTGGCGACCCGCTGAATTT</td><td>751.2</td><td>4.1</td><td>0.8</td><td>5.3</td><td>1.5E-07</td><td>2.6E-07</td><td>MB231/MB468</td><td>NUC</td></tr>
<tr><td>>hg38_21_+_8433172_8446622_RNA45SN1_45S_50ntFlanks@9507.9525.19|AACTATGCCTGGGCAGGGC</td><td>>hg38_21_+_8433172_8446622_RNA45SN1_45S_50ntFlanks</td><td>9507</td><td>9525</td><td>19</td><td>AACTATGCCTGGGCAGGGC</td><td>144.5</td><td>-10.3</td><td>1.8</td><td>-5.5</td><td>3.0E-08</td><td>2.6E-07</td><td>BT20/MB468</td><td>MP</td></tr>
<tr><td>>MI0000736|hsa-mir-30c-1&WithFlank&1|+|40757278|40757378@23.46.24;MI0000254|hsa-mir-30c-2&WithFlank&6|-|71376954|71377037@13.36.24&[MIMAT0000244&hsa-miR-30c-5p&offsets|0|+1;m-57&1|+|40757300|40757323&offsets|0|0];[MIMAT0000244_1&hsa-miR-30c-5p&offsets|0|+1;m-56&6|-|71377002|71377025&offsets|0|0]|TGTAAACATCCTACACTCTCAGCT</td><td>>MI0000736|hsa-mir-30c-1&WithFlank&1|+|40757278|40757378</td><td>23</td><td>46</td><td>24</td><td>TGTAAACATCCTACACTCTCAGCT</td><td>12592.9</td><td>4.8</td><td>0.9</td><td>5.3</td><td>1.1E-07</td><td>2.6E-07</td><td>BT20/MB468</td><td>MITO</td></tr>
<tr><td>>trnaMT_MetCAT_MT_+_4402_4469@40.71.32|AAAATGTTGGTTATACCCTTCCCGTACTACCA</td><td>>trnaMT_MetCAT_MT_+_4402_4469</td><td>40</td><td>71</td><td>32</td><td>AAAATGTTGGTTATACCCTTCCCGTACTACCA</td><td>607.8</td><td>-3.7</td><td>0.7</td><td>-5.2</td><td>1.6E-07</td><td>2.6E-07</td><td>MB231/MB468</td><td>MITO</td></tr>
<tr><td>>hg38_21_+_8433172_8446622_RNA45SN1_45S_50ntFlanks@11170.11189.20|CCGCGAGGGGGGCCCGGGCA</td><td>>hg38_21_+_8433172_8446622_RNA45SN1_45S_50ntFlanks</td><td>11170</td><td>11189</td><td>20</td><td>CCGCGAGGGGGGCCCGGGCA</td><td>411.8</td><td>6.9</td><td>1.3</td><td>5.3</td><td>1.5E-07</td><td>2.6E-07</td><td>MB231/MB468</td><td>NUC</td></tr>
<tr><td>>hg38_21_+_8433172_8446622_RNA45SN1_45S_50ntFlanks@4730.4764.35|CAAGAACGAAAGTCGGAGGTTCGAAGACGATCAGA</td><td>>hg38_21_+_8433172_8446622_RNA45SN1_45S_50ntFlanks</td><td>4730</td><td>4764</td><td>35</td><td>CAAGAACGAAAGTCGGAGGTTCGAAGACGATCAGA</td><td>403.3</td><td>7.9</td><td>1.5</td><td>5.3</td><td>1.3E-07</td><td>2.6E-07</td><td>BT20/MB468</td><td>CYTO</td></tr>
<tr><td>>hg38_21_+_8433172_8446622_RNA45SN1_45S_50ntFlanks@10351.10391.41|CGAGAACTTTGAAGGCCGAAGTGGAGAAGGGTTCCATGTGA</td><td>>hg38_21_+_8433172_8446622_RNA45SN1_45S_50ntFlanks</td><td>10351</td><td>10391</td><td>41</td><td>CGAGAACTTTGAAGGCCGAAGTGGAGAAGGGTTCCATGTGA</td><td>652.7</td><td>-8.0</td><td>1.5</td><td>-5.3</td><td>1.3E-07</td><td>2.6E-07</td><td>MB231/MB468</td><td>MP</td></tr>
<tr><td>>hg38_21_+_8433172_8446622_RNA45SN1_45S_50ntFlanks@10384.10429.46|CCATGTGAACAGCAGTTGAACATGGGTCAGTCGGTCCTGAGAGATG</td><td>>hg38_21_+_8433172_8446622_RNA45SN1_45S_50ntFlanks</td><td>10384</td><td>10429</td><td>46</td><td>CCATGTGAACAGCAGTTGAACATGGGTCAGTCGGTCCTGAGAGATG</td><td>215.6</td><td>-6.1</td><td>1.2</td><td>-5.2</td><td>1.6E-07</td><td>2.6E-07</td><td>MB231/MB468</td><td>MITO</td></tr>
<tr><td>>hg38_21_+_8433172_8446622_RNA45SN1_45S_50ntFlanks@10683.10702.20|CCGGCGGCGTCCGGTGAGCT</td><td>>hg38_21_+_8433172_8446622_RNA45SN1_45S_50ntFlanks</td><td>10683</td><td>10702</td><td>20</td><td>CCGGCGGCGTCCGGTGAGCT</td><td>76.7</td><td>-9.3</td><td>1.7</td><td>-5.5</td><td>3.0E-08</td><td>2.6E-07</td><td>BT20/MB468</td><td>MP</td></tr>
<tr><td>>hg38_21_+_8433172_8446622_RNA45SN1_45S_50ntFlanks@12157.12180.24|AGTTTGACTGGGGCGGTACACCTG</td><td>>hg38_21_+_8433172_8446622_RNA45SN1_45S_50ntFlanks</td><td>12157</td><td>12180</td><td>24</td><td>AGTTTGACTGGGGCGGTACACCTG</td><td>389.0</td><td>-6.4</td><td>1.2</td><td>-5.3</td><td>1.3E-07</td><td>2.7E-07</td><td>BT20/MB468</td><td>CYTO</td></tr>
<tr><td>>hg38_21_+_8433172_8446622_RNA45SN1_45S_50ntFlanks@7990.8019.30|GACGTGGCGACCCGCTGAATTTAAGCATAT</td><td>>hg38_21_+_8433172_8446622_RNA45SN1_45S_50ntFlanks</td><td>7990</td><td>8019</td><td>30</td><td>GACGTGGCGACCCGCTGAATTTAAGCATAT</td><td>216.9</td><td>8.5</td><td>1.6</td><td>5.3</td><td>1.3E-07</td><td>2.7E-07</td><td>BT20/MB468</td><td>CYTO</td></tr>
<tr><td>>hg38_21_+_8433172_8446622_RNA45SN1_45S_50ntFlanks@8383.8402.20|GAGGGCGTGAAACCGTTAAG</td><td>>hg38_21_+_8433172_8446622_RNA45SN1_45S_50ntFlanks</td><td>8383</td><td>8402</td><td>20</td><td>GAGGGCGTGAAACCGTTAAG</td><td>262.5</td><td>-5.7</td><td>1.1</td><td>-5.3</td><td>1.3E-07</td><td>2.7E-07</td><td>MB231/MB468</td><td>MP</td></tr>
</table>

<table>
<tr><td>>hg38_21_+_8433172_8446622_RNA45SN1_45S_50ntFlanks@11669.11707.39|GCCCAGTGCTCTGAATGTCAAAGTGAAGAAATTCAATGA</td><td>>hg38_21_+_8433172_8446622_RNA45SN1_45S_50ntFlanks</td><td>11669</td><td>11707</td><td>39</td><td>GCCCAGTGCTCTGAATGTCAAAGTGAAGAAATTCAATGA</td><td>256.0</td><td>6.0</td><td>1.1</td><td>5.3</td><td>1.5E-07</td><td>2.7E-07</td><td>MB231/MB468</td><td>NUC</td></tr>
<tr><td>>hg38_21_+_8433172_8446622_RNA45SN1_45S_50ntFlanks@10684.10712.29|CGGCGGCGTCCGGTGAGCTCTCGCTGGCC</td><td>>hg38_21_+_8433172_8446622_RNA45SN1_45S_50ntFlanks</td><td>10684</td><td>10712</td><td>29</td><td>CGGCGGCGTCCGGTGAGCTCTCGCTGGCC</td><td>639.6</td><td>-4.1</td><td>0.8</td><td>-5.3</td><td>1.5E-07</td><td>2.7E-07</td><td>MB231/MB468</td><td>CYTO</td></tr>
<tr><td>>hg38_21_+_8433172_8446622_RNA45SN1_45S_50ntFlanks@10737.10757.21|TAAATCTCGCGCCGGGCCGTA</td><td>>hg38_21_+_8433172_8446622_RNA45SN1_45S_50ntFlanks</td><td>10737</td><td>10757</td><td>21</td><td>TAAATCTCGCGCCGGGCCGTA</td><td>853.2</td><td>-7.6</td><td>1.4</td><td>-5.3</td><td>1.5E-07</td><td>2.7E-07</td><td>MB231/MB468</td><td>NUC</td></tr>
<tr><td>>MI0003513|hsa-mir-455&WithFlank&9|+|114209428|114209535@22.44.23&[MIMAT0003150&hsa-miR-455-5p&offsets|0|+1;m-232&9|+|114209449|114209470&offsets|0|+1]|TATGTGCCTTTGGACTACATCGT</td><td>>MI0003513|hsa-mir-455&WithFlank&9|+|114209428|114209535</td><td>22</td><td>44</td><td>23</td><td>TATGTGCCTTTGGACTACATCGT</td><td>91.0</td><td>10.3</td><td>1.9</td><td>5.5</td><td>3.0E-08</td><td>2.7E-07</td><td>BT20/MB468</td><td>MP</td></tr>
<tr><td>>hg38_21_+_8433172_8446622_RNA45SN1_45S_50ntFlanks@8475.8497.23|CCCGGCGGATCTTTCCCGCCCCC</td><td>>hg38_21_+_8433172_8446622_RNA45SN1_45S_50ntFlanks</td><td>8475</td><td>8497</td><td>23</td><td>CCCGGCGGATCTTTCCCGCCCCC</td><td>464.2</td><td>-8.0</td><td>1.4</td><td>-5.5</td><td>3.0E-08</td><td>2.7E-07</td><td>BT20/MB468</td><td>MP</td></tr>
<tr><td>>trna119_LysCTT_1_-_145395522_145395594@1.34.34,trna11_LysCTT_5_-_180648979_180649051@1.34.34,trna13_LysCTT_6_+_26556774_26556846@1.34.34,trna7_LysCTT_16_+_3225692_3225764@1.34.34,trna9_LysCTT_5_+_180634755_180634827@1.34.34|GCCCGGCTAGCTCAGTCGGTAGAGCATGAGACTC</td><td>>trna119_LysCTT_1_-_145395522_145395594</td><td>1</td><td>34</td><td>34</td><td>GCCCGGCTAGCTCAGTCGGTAGAGCATGAGACTC</td><td>1287.8</td><td>4.0</td><td>0.8</td><td>5.3</td><td>1.3E-07</td><td>2.7E-07</td><td>MB231/MB468</td><td>MP</td></tr>
<tr><td>>hg38_21_+_8433172_8446622_RNA45SN1_45S_50ntFlanks@7990.8012.23|GACGTGGCGACCCGCTGAATTTA</td><td>>hg38_21_+_8433172_8446622_RNA45SN1_45S_50ntFlanks</td><td>7990</td><td>8012</td><td>23</td><td>GACGTGGCGACCCGCTGAATTTA</td><td>1875.7</td><td>-3.6</td><td>0.7</td><td>-5.2</td><td>1.6E-07</td><td>2.7E-07</td><td>MB231/MB468</td><td>MITO</td></tr>
<tr><td>>am_trnaMT_ProTGG_MT_-_15956_16023@52.71.20|AAAGACTTTTTCTCTGACCA</td><td>>am_trnaMT_ProTGG_MT_-_15956_16023</td><td>52</td><td>71</td><td>20</td><td>AAAGACTTTTTCTCTGACCA</td><td>1162.0</td><td>7.7</td><td>1.5</td><td>5.3</td><td>1.5E-07</td><td>2.7E-07</td><td>MB231/MB468</td><td>CYTO</td></tr>
<tr><td>>hg38_21_+_8433172_8446622_RNA45SN1_45S_50ntFlanks@7983.8012.30|CAGATCAGACGTGGCGACCCGCTGAATTTA</td><td>>hg38_21_+_8433172_8446622_RNA45SN1_45S_50ntFlanks</td><td>7983</td><td>8012</td><td>30</td><td>CAGATCAGACGTGGCGACCCGCTGAATTTA</td><td>247.0</td><td>-7.6</td><td>1.4</td><td>-5.2</td><td>1.7E-07</td><td>2.7E-07</td><td>BT20/MB468</td><td>NUC</td></tr>
<tr><td>>hg38_21_+_8433172_8446622_RNA45SN1_45S_50ntFlanks@11562.11585.24|CGCCTAGCAGCCGACTTAGAACTG</td><td>>hg38_21_+_8433172_8446622_RNA45SN1_45S_50ntFlanks</td><td>11562</td><td>11585</td><td>24</td><td>CGCCTAGCAGCCGACTTAGAACTG</td><td>370.9</td><td>-7.3</td><td>1.4</td><td>-5.2</td><td>1.6E-07</td><td>2.7E-07</td><td>MB231/MB468</td><td>MITO</td></tr>
<tr><td>>hg38_MT_+_1621_3279_RNR2_16S_50ntFlanks@493.517.25|GCTCTTTGGACACTAGGAAAAAACC</td><td>>hg38_MT_+_1621_3279_RNR2_16S_50ntFlanks</td><td>493</td><td>517</td><td>25</td><td>GCTCTTTGGACACTAGGAAAAAACC</td><td>657.9</td><td>3.9</td><td>0.7</td><td>5.2</td><td>1.6E-07</td><td>2.7E-07</td><td>MB231/MB468</td><td>MITO</td></tr>
<tr><td>>hg38_21_+_8433172_8446622_RNA45SN1_45S_50ntFlanks@10317.10336.20|GTGCAGATCTTGGTGGTAGT</td><td>>hg38_21_+_8433172_8446622_RNA45SN1_45S_50ntFlanks</td><td>10317</td><td>10336</td><td>20</td><td>GTGCAGATCTTGGTGGTAGT</td><td>330.2</td><td>11.7</td><td>2.0</td><td>5.8</td><td>6.1E-09</td><td>2.7E-07</td><td>BT20/MB231</td><td>MP</td></tr>
<tr><td>>trnaMT_LeuTAA_MT_+_3230_3304@16.44.29|CCCGGTAATCGCATAAAACTTAAAACTTT</td><td>>trnaMT_LeuTAA_MT_+_3230_3304</td><td>16</td><td>44</td><td>29</td><td>CCCGGTAATCGCATAAAACTTAAAACTTT</td><td>1235.1</td><td>-4.4</td><td>0.8</td><td>-5.3</td><td>1.5E-07</td><td>2.7E-07</td><td>MB231/MB468</td><td>CYTO</td></tr>
<tr><td>>MI0008329|hsa-mir-1908&WithFlank&11|-|61815155|61815246@18.38.21&[MIMAT0007881&hsa-miR-1908-5p&offsets|0|0;m-915&11|-|61815209|61815229&offsets|0|0]|CGGCGGGGACGGCGATTGGTC</td><td>>MI0008329|hsa-mir-1908&WithFlank&11|-|61815155|61815246</td><td>18</td><td>38</td><td>21</td><td>CGGCGGGGACGGCGATTGGTC</td><td>64.0</td><td>9.2</td><td>1.7</td><td>5.3</td><td>1.3E-07</td><td>2.7E-07</td><td>MB231/MB468</td><td>MP</td></tr>
<tr><td>>hg38_21_+_8433172_8446622_RNA45SN1_45S_50ntFlanks@7988.8008.21|CAGACGTGGCGACCCGCTGAA</td><td>>hg38_21_+_8433172_8446622_RNA45SN1_45S_50ntFlanks</td><td>7988</td><td>8008</td><td>21</td><td>CAGACGTGGCGACCCGCTGAA</td><td>3016.0</td><td>-5.0</td><td>1.0</td><td>-5.2</td><td>1.7E-07</td><td>2.7E-07</td><td>BT20/MB468</td><td>NUC</td></tr>
<tr><td>>hg38_21_+_8433172_8446622_RNA45SN1_45S_50ntFlanks@5487.5526.40|CGCTGAGAAGACGGTCGAACTTGACTATCTAGAGGAAGTA</td><td>>hg38_21_+_8433172_8446622_RNA45SN1_45S_50ntFlanks</td><td>5487</td><td>5526</td><td>40</td><td>CGCTGAGAAGACGGTCGAACTTGACTATCTAGAGGAAGTA</td><td>1193.0</td><td>-5.5</td><td>1.0</td><td>-5.3</td><td>1.1E-07</td><td>2.7E-07</td><td>BT20/MB468</td><td>MITO</td></tr>
<tr><td>>hg38_21_+_8433172_8446622_RNA45SN1_45S_50ntFlanks@9483.9510.28|CGTGTTAGGACCCGAAAGATGGTGAACT</td><td>>hg38_21_+_8433172_8446622_RNA45SN1_45S_50ntFlanks</td><td>9483</td><td>9510</td><td>28</td><td>CGTGTTAGGACCCGAAAGATGGTGAACT</td><td>370.3</td><td>11.9</td><td>2.0</td><td>5.8</td><td>6.2E-09</td><td>2.7E-07</td><td>BT20/MB231</td><td>MP</td></tr>
<tr><td>>hg38_21_+_8433172_8446622_RNA45SN1_45S_50ntFlanks@9534.9564.31|AGGAAACTCTGGTGGAGGTCCGTAGCGGTCC</td><td>>hg38_21_+_8433172_8446622_RNA45SN1_45S_50ntFlanks</td><td>9534</td><td>9564</td><td>31</td><td>AGGAAACTCTGGTGGAGGTCCGTAGCGGTCC</td><td>1686.3</td><td>-8.2</td><td>1.6</td><td>-5.3</td><td>1.3E-07</td><td>2.7E-07</td><td>MB231/MB468</td><td>MP</td></tr>
<tr><td>>hg38_21_+_8433172_8446622_RNA45SN1_45S_50ntFlanks@10792.10815.24|CTCTGGCATGTTGGAACAATGTAG</td><td>>hg38_21_+_8433172_8446622_RNA45SN1_45S_50ntFlanks</td><td>10792</td><td>10815</td><td>24</td><td>CTCTGGCATGTTGGAACAATGTAG</td><td>446.6</td><td>-6.4</td><td>1.2</td><td>-5.2</td><td>1.7E-07</td><td>2.7E-07</td><td>BT20/MB468</td><td>NUC</td></tr>
<tr><td>>hg38_21_+_8433172_8446622_RNA45SN1_45S_50ntFlanks@5084.5113.30|CGAACGAGACTCTGGCATGCTAACTAGTTA</td><td>>hg38_21_+_8433172_8446622_RNA45SN1_45S_50ntFlanks</td><td>5084</td><td>5113</td><td>30</td><td>CGAACGAGACTCTGGCATGCTAACTAGTTA</td><td>162.4</td><td>-9.4</td><td>1.8</td><td>-5.3</td><td>1.1E-07</td><td>2.7E-07</td><td>BT20/MB468</td><td>MITO</td></tr>
<tr><td>>hg38_21_+_8433172_8446622_RNA45SN1_45S_50ntFlanks@12525.12546.22|GTTTTACCCTACTGATGATGTG</td><td>>hg38_21_+_8433172_8446622_RNA45SN1_45S_50ntFlanks</td><td>12525</td><td>12546</td><td>22</td><td>GTTTTACCCTACTGATGATGTG</td><td>107.9</td><td>-9.8</td><td>1.8</td><td>-5.5</td><td>3.0E-08</td><td>2.7E-07</td><td>BT20/MB468</td><td>MP</td></tr>
<tr><td>>hg38_21_+_8433172_8446622_RNA45SN1_45S_50ntFlanks@10350.10390.41|ACGAGAACTTTGAAGGCCGAAGTGGAGAAGGGTTCCATGTG</td><td>>hg38_21_+_8433172_8446622_RNA45SN1_45S_50ntFlanks</td><td>10350</td><td>10390</td><td>41</td><td>ACGAGAACTTTGAAGGCCGAAGTGGAGAAGGGTTCCATGTG</td><td>3196.7</td><td>-3.5</td><td>0.7</td><td>-5.3</td><td>1.5E-07</td><td>2.7E-07</td><td>MB231/MB468</td><td>CYTO</td></tr>
<tr><td>>hg38_MT_+_598_1651_RNR1_12S_50ntFlanks@430.469.40|AGTGGCTTTAACATATCTGAACACACAATAGCTAAGACCC</td><td>>hg38_MT_+_598_1651_RNR1_12S_50ntFlanks</td><td>430</td><td>469</td><td>40</td><td>AGTGGCTTTAACATATCTGAACACACAATAGCTAAGACCC</td><td>52.0</td><td>8.9</td><td>1.7</td><td>5.3</td><td>1.3E-07</td><td>2.7E-07</td><td>MB231/MB468</td><td>MP</td></tr>
<tr><td>>hg38_21_+_8433172_8446622_RNA45SN1_45S_50ntFlanks@7975.8014.40|CGCGACCTCAGATCAGACGTGGCGACCCGCTGAATTTAAG</td><td>>hg38_21_+_8433172_8446622_RNA45SN1_45S_50ntFlanks</td><td>7975</td><td>8014</td><td>40</td><td>CGCGACCTCAGATCAGACGTGGCGACCCGCTGAATTTAAG</td><td>4171.8</td><td>4.9</td><td>0.9</td><td>5.3</td><td>1.3E-07</td><td>2.7E-07</td><td>BT20/MB468</td><td>CYTO</td></tr>
<tr><td>>hg38_21_+_8433172_8446622_RNA45SN1_45S_50ntFlanks@8114.8138.25|GCGCGGGACATGTGGCGTACGGAAG</td><td>>hg38_21_+_8433172_8446622_RNA45SN1_45S_50ntFlanks</td><td>8114</td><td>8138</td><td>25</td><td>GCGCGGGACATGTGGCGTACGGAAG</td><td>1675.8</td><td>-4.1</td><td>0.8</td><td>-5.3</td><td>1.1E-07</td><td>2.7E-07</td><td>BT20/MB468</td><td>MITO</td></tr>
<tr><td>>hg38_21_+_8433172_8446622_RNA45SN1_45S_50ntFlanks@12676.12710.35|AGAATCCCGCCCAGGCGGAACGATACGGCAGCGCC</td><td>>hg38_21_+_8433172_8446622_RNA45SN1_45S_50ntFlanks</td><td>12676</td><td>12710</td><td>35</td><td>AGAATCCCGCCCAGGCGGAACGATACGGCAGCGCC</td><td>74.3</td><td>-9.3</td><td>1.7</td><td>-5.5</td><td>3.0E-08</td><td>2.7E-07</td><td>BT20/MB468</td><td>MP</td></tr>
<tr><td>>MI0000477|hsa-mir-146a&WithFlank&5|+|160485346|160485456@27.46.20(+1G)&[MIMAT0000449&hsa-miR-146a-5p&offsets|0|-2(+1G);m-129&5|+|160485372|160485393&offsets|0|-2(+1G)]|TGAGAACTGAATTCCATGGGG</td><td>>MI0000477|hsa-mir-146a&WithFlank&5|+|160485346|160485456</td><td>27</td><td>46</td><td>20(+1G)</td><td>TGAGAACTGAATTCCATGGGG</td><td>256.1</td><td>-11.4</td><td>2.0</td><td>-5.8</td><td>6.3E-09</td><td>2.7E-07</td><td>BT20/MB231</td><td>MP</td></tr>
<tr><td>>hg38_21_+_8433172_8446622_RNA45SN1_45S_50ntFlanks@10463.10497.35|CTCCGTTGCCCTCGGCCGATCGAAAGGGAGTCGGG</td><td>>hg38_21_+_8433172_8446622_RNA45SN1_45S_50ntFlanks</td><td>10463</td><td>10497</td><td>35</td><td>CTCCGTTGCCCTCGGCCGATCGAAAGGGAGTCGGG</td><td>246.1</td><td>-6.2</td><td>1.2</td><td>-5.3</td><td>1.3E-07</td><td>2.7E-07</td><td>BT20/MB468</td><td>CYTO</td></tr>
<tr><td>>hg38_21_+_8433172_8446622_RNA45SN1_45S_50ntFlanks@12413.12436.24|TCCTTCGATGTCGGCTCTTCCTAT</td><td>>hg38_21_+_8433172_8446622_RNA45SN1_45S_50ntFlanks</td><td>12413</td><td>12436</td><td>24</td><td>TCCTTCGATGTCGGCTCTTCCTAT</td><td>219.0</td><td>-4.8</td><td>0.9</td><td>-5.3</td><td>1.5E-07</td><td>2.7E-07</td><td>MB231/MB468</td><td>CYTO</td></tr>
<tr><td>>hg38_21_+_8433172_8446622_RNA45SN1_45S_50ntFlanks@7991.8017.27|ACGTGGCGACCCGCTGAATTTAAGCAT</td><td>>hg38_21_+_8433172_8446622_RNA45SN1_45S_50ntFlanks</td><td>7991</td><td>8017</td><td>27</td><td>ACGTGGCGACCCGCTGAATTTAAGCAT</td><td>388.7</td><td>-4.0</td><td>0.8</td><td>-5.2</td><td>1.7E-07</td><td>2.7E-07</td><td>BT20/MB468</td><td>NUC</td></tr>
<tr><td>>hg38_21_+_8433172_8446622_RNA45SN1_45S_50ntFlanks@7974.7991.18|ACGCGACCTCAGATCAGA</td><td>>hg38_21_+_8433172_8446622_RNA45SN1_45S_50ntFlanks</td><td>7974</td><td>7991</td><td>18</td><td>ACGCGACCTCAGATCAGA</td><td>2143.8</td><td>4.2</td><td>0.8</td><td>5.3</td><td>1.3E-07</td><td>2.7E-07</td><td>BT20/MB468</td><td>CYTO</td></tr>
<tr><td>>hg38_21_+_8433172_8446622_RNA45SN1_45S_50ntFlanks@10789.10818.30|AGCCTCTGGCATGTTGGAACAATGTAGGTA</td><td>>hg38_21_+_8433172_8446622_RNA45SN1_45S_50ntFlanks</td><td>10789</td><td>10818</td><td>30</td><td>AGCCTCTGGCATGTTGGAACAATGTAGGTA</td><td>849.1</td><td>-6.1</td><td>1.2</td><td>-5.2</td><td>1.6E-07</td><td>2.7E-07</td><td>MB231/MB468</td><td>MITO</td></tr>
<tr><td>>hg38_21_+_8433172_8446622_RNA45SN1_45S_50ntFlanks@11316.11343.28|CACCCCACGTCTCGTCGCGCGCGCGTCC</td><td>>hg38_21_+_8433172_8446622_RNA45SN1_45S_50ntFlanks</td><td>11316</td><td>11343</td><td>28</td><td>CACCCCACGTCTCGTCGCGCGCGCGTCC</td><td>94.0</td><td>-9.6</td><td>1.7</td><td>-5.5</td><td>3.1E-08</td><td>2.7E-07</td><td>BT20/MB468</td><td>MP</td></tr>
<tr><td>>hg38_21_+_8433172_8446622_RNA45SN1_45S_50ntFlanks@11785.11803.19|TGAATGGATGAACGAGATT</td><td>>hg38_21_+_8433172_8446622_RNA45SN1_45S_50ntFlanks</td><td>11785</td><td>11803</td><td>19</td><td>TGAATGGATGAACGAGATT</td><td>1711.0</td><td>-4.9</td><td>0.9</td><td>-5.5</td><td>3.1E-08</td><td>2.7E-07</td><td>BT20/MB468</td><td>MP</td></tr>
<tr><td>>hg38_21_+_8433172_8446622_RNA45SN1_45S_50ntFlanks@7975.8005.31|CGCGACCTCAGATCAGACGTGGCGACCCGCT</td><td>>hg38_21_+_8433172_8446622_RNA45SN1_45S_50ntFlanks</td><td>7975</td><td>8005</td><td>31</td><td>CGCGACCTCAGATCAGACGTGGCGACCCGCT</td><td>42549.2</td><td>4.8</td><td>0.9</td><td>5.5</td><td>3.1E-08</td><td>2.7E-07</td><td>BT20/MB468</td><td>MP</td></tr>
<tr><td>>hg38_21_+_8433172_8446622_RNA45SN1_45S_50ntFlanks@9741.9758.18|ACGATCTCAACCTATTCT</td><td>>hg38_21_+_8433172_8446622_RNA45SN1_45S_50ntFlanks</td><td>9741</td><td>9758</td><td>18</td><td>ACGATCTCAACCTATTCT</td><td>72.2</td><td>-9.3</td><td>1.7</td><td>-5.5</td><td>3.1E-08</td><td>2.7E-07</td><td>BT20/MB468</td><td>MP</td></tr>
<tr><td>>hg38_21_+_8433172_8446622_RNA45SN1_45S_50ntFlanks@10534.10564.31|CCGCGAGGCGTCCAGTGCGGTAACGCGACCG</td><td>>hg38_21_+_8433172_8446622_RNA45SN1_45S_50ntFlanks</td><td>10534</td><td>10564</td><td>31</td><td>CCGCGAGGCGTCCAGTGCGGTAACGCGACCG</td><td>106.2</td><td>-8.8</td><td>1.7</td><td>-5.3</td><td>1.2E-07</td><td>2.7E-07</td><td>BT20/MB468</td><td>MITO</td></tr>
<tr><td>>hg38_21_+_8433172_8446622_RNA45SN1_45S_50ntFlanks@8380.8397.18|CAAGAGGGCGTGAAACCG</td><td>>hg38_21_+_8433172_8446622_RNA45SN1_45S_50ntFlanks</td><td>8380</td><td>8397</td><td>18</td><td>CAAGAGGGCGTGAAACCG</td><td>541.9</td><td>-7.2</td><td>1.4</td><td>-5.2</td><td>1.5E-07</td><td>2.7E-07</td><td>MB231/MB468</td><td>NUC</td></tr>
<tr><td>>hg38_21_+_8433172_8446622_RNA45SN1_45S_50ntFlanks@10767.10785.19|GCAGCAGGTCTCCAAGGTG</td><td>>hg38_21_+_8433172_8446622_RNA45SN1_45S_50ntFlanks</td><td>10767</td><td>10785</td><td>19</td><td>GCAGCAGGTCTCCAAGGTG</td><td>248.5</td><td>-3.9</td><td>0.7</td><td>-5.2</td><td>1.5E-07</td><td>2.7E-07</td><td>MB231/MB468</td><td>NUC</td></tr>
<tr><td>>MI0000736|hsa-mir-30c-1&WithFlank&1|+|40757278|40757378@23.45.23(+1A);MI0000254|hsa-mir-30c-2&WithFlank&6|-|71376954|71377037@13.35.23(+1A)&[MIMAT0000244&hsa-miR-30c-5p&offsets|0|0(+1A);m-57&1|+|40757300|40757323&offsets|0|-1(+1A)];[MIMAT0000244_1&hsa-miR-30c-5p&offsets|0|0(+1A);m-56&6|-|71377002|71377025&offsets|0|-1(+1A)]|TGTAAACATCCTACACTCTCAGCA</td><td>>MI0000736|hsa-mir-30c-1&WithFlank&1|+|40757278|40757378</td><td>23</td><td>45</td><td>23(+1A)</td><td>TGTAAACATCCTACACTCTCAGCA</td><td>1266.0</td><td>5.0</td><td>1.0</td><td>5.3</td><td>1.5E-07</td><td>2.7E-07</td><td>MB231/MB468</td><td>CYTO</td></tr>
<tr><td>>MI0001445|hsa-mir-423&WithFlank&17|+|30117073|30117178@59.79.21&[MIMAT0001340&hsa-miR-423-3p&offsets|0|-2;m-100&17|+|30117131|30117153&offsets|0|-2]|AGCTCGGTCTGAGGCCCCTCA</td><td>>MI0001445|hsa-mir-423&WithFlank&17|+|30117073|30117178</td><td>59</td><td>79</td><td>21</td><td>AGCTCGGTCTGAGGCCCCTCA</td><td>573.9</td><td>7.3</td><td>1.4</td><td>5.3</td><td>1.5E-07</td><td>2.7E-07</td><td>MB231/MB468</td><td>CYTO</td></tr>
<tr><td>>hg38_21_+_8433172_8446622_RNA45SN1_45S_50ntFlanks@4128.4162.35|CGATTCCGGAGAGGGAGCCTGAGAAACGGCTACCA</td><td>>hg38_21_+_8433172_8446622_RNA45SN1_45S_50ntFlanks</td><td>4128</td><td>4162</td><td>35</td><td>CGATTCCGGAGAGGGAGCCTGAGAAACGGCTACCA</td><td>341.4</td><td>-6.6</td><td>1.3</td><td>-5.3</td><td>1.2E-07</td><td>2.7E-07</td><td>BT20/MB468</td><td>MITO</td></tr>
<tr><td>>hg38_21_+_8433172_8446622_RNA45SN1_45S_50ntFlanks@9555.9597.43|GTAGCGGTCCTGACGTGCAAATCGGTCGTCCGACCTGGGTATA</td><td>>hg38_21_+_8433172_8446622_RNA45SN1_45S_50ntFlanks</td><td>9555</td><td>9597</td><td>43</td><td>GTAGCGGTCCTGACGTGCAAATCGGTCGTCCGACCTGGGTATA</td><td>568.0</td><td>-6.3</td><td>1.2</td><td>-5.2</td><td>1.6E-07</td><td>2.7E-07</td><td>MB231/MB468</td><td>MITO</td></tr>
<tr><td>>hg38_21_+_8433172_8446622_RNA45SN1_45S_50ntFlanks@10458.10477.20|ATGGCCTCCGTTGCCCTCGG</td><td>>hg38_21_+_8433172_8446622_RNA45SN1_45S_50ntFlanks</td><td>10458</td><td>10477</td><td>20</td><td>ATGGCCTCCGTTGCCCTCGG</td><td>4902.3</td><td>-8.6</td><td>1.6</td><td>-5.3</td><td>1.3E-07</td><td>2.7E-07</td><td>BT20/MB468</td><td>CYTO</td></tr>
<tr><td>>MI0000746|hsa-mir-99b&WithFlank&19|+|51692606|51692687@13.32.20(+1G)&[MIMAT0000689&hsa-miR-99b-5p&offsets|0|-2(+1G);m-10&19|+|51692618|51692639&offsets|0|-2(+1G)]|CACCCGTAGAACCGACCTTGG</td><td>>MI0000746|hsa-mir-99b&WithFlank&19|+|51692606|51692687</td><td>13</td><td>32</td><td>20(+1G)</td><td>CACCCGTAGAACCGACCTTGG</td><td>103.4</td><td>10.4</td><td>1.9</td><td>5.5</td><td>3.1E-08</td><td>2.7E-07</td><td>BT20/MB468</td><td>MP</td></tr>
<tr><td>>MI0000477|hsa-mir-146a&WithFlank&5|+|160485346|160485456@28.48.21&[MIMAT0000449&hsa-miR-146a-5p&offsets|+1|0;m-129&5|+|160485372|160485393&offsets|+1|0]|GAGAACTGAATTCCATGGGTT</td><td>>MI0000477|hsa-mir-146a&WithFlank&5|+|160485346|160485456</td><td>28</td><td>48</td><td>21</td><td>GAGAACTGAATTCCATGGGTT</td><td>198.5</td><td>-11.0</td><td>1.9</td><td>-5.8</td><td>6.4E-09</td><td>2.7E-07</td><td>BT20/MB231</td><td>MP</td></tr>
<tr><td>>MI0005544|hsa-mir-147b&WithFlank&15|+|45433044|45433135@55.74.20&[MIMAT0004928&hsa-miR-147b-3p&offsets|0|-1;m-575&15|+|45433098|45433118&offsets|0|-1]|GTGTGCGGAAATGCTTCTGC</td><td>>MI0005544|hsa-mir-147b&WithFlank&15|+|45433044|45433135</td><td>55</td><td>74</td><td>20</td><td>GTGTGCGGAAATGCTTCTGC</td><td>144.6</td><td>-10.6</td><td>1.8</td><td>-5.8</td><td>6.4E-09</td><td>2.7E-07</td><td>BT20/MB231</td><td>MP</td></tr>
<tr><td>>hg38_21_+_8433172_8446622_RNA45SN1_45S_50ntFlanks@10346.10368.23|TCAAACGAGAACTTTGAAGGCCG</td><td>>hg38_21_+_8433172_8446622_RNA45SN1_45S_50ntFlanks</td><td>10346</td><td>10368</td><td>23</td><td>TCAAACGAGAACTTTGAAGGCCG</td><td>3167.2</td><td>-5.8</td><td>1.0</td><td>-5.5</td><td>3.1E-08</td><td>2.7E-07</td><td>BT20/MB468</td><td>MP</td></tr>
<tr><td>>hg38_21_+_8433172_8446622_RNA45SN1_45S_50ntFlanks@11255.11283.29|CTGCGGCGGGCGTCGCGGCCGCCCCCGGG</td><td>>hg38_21_+_8433172_8446622_RNA45SN1_45S_50ntFlanks</td><td>11255</td><td>11283</td><td>29</td><td>CTGCGGCGGGCGTCGCGGCCGCCCCCGGG</td><td>80.2</td><td>-10.2</td><td>1.9</td><td>-5.3</td><td>1.4E-07</td><td>2.7E-07</td><td>MB231/MB468</td><td>MP</td></tr>
</table>

<table>
<tr><th></th><th></th><th></th><th></th><th></th><th></th><th></th><th></th><th></th><th></th><th></th><th></th><th></th><th></th></tr>
<tr><td>>MI0000746|hsa-mir-99b&WithFlank&19|+|51692606|51692687@13.32.20(+1U)&[MIMAT0000689&hsa-miR-99b-5p&offsets|0|-2(+1U);m-10&19|+|51692618|51692639&offsets|0|-2(+1U)]|CACCCGTAGAACCGACCTTGT</td><td>>MI0000746|hsa-mir-99b&WithFlank&19|+|51692606|51692687</td><td>13</td><td>32</td><td>20(+1U)</td><td>CACCCGTAGAACCGACCTTGT</td><td>28733.7</td><td>4.4</td><td>0.8</td><td>5.7</td><td>9.8E-09</td><td>2.7E-07</td><td>BT20/MB231</td><td>CYTO</td></tr>
<tr><td>>hg38_21_+_8433172_8446622_RNA45SN1_45S_50ntFlanks@8168.8212.45|CCCAAGTCCTTCTGATCGAGGCCCAGCCCGTGGACGGTGTGAGGC</td><td>>hg38_21_+_8433172_8446622_RNA45SN1_45S_50ntFlanks</td><td>8168</td><td>8212</td><td>45</td><td>CCCAAGTCCTTCTGATCGAGGCCCAGCCCGTGGACGGTGTGAGGC</td><td>246.2</td><td>-11.8</td><td>2.2</td><td>-5.3</td><td>1.4E-07</td><td>2.7E-07</td><td>MB231/MB468</td><td>MP</td></tr>
<tr><td>>am_trnaMT_ValTAC_MT_+_1602_1670@49.66.18|AACTTAACTTGACCGCTC</td><td>>am_trnaMT_ValTAC_MT_+_1602_1670</td><td>49</td><td>66</td><td>18</td><td>AACTTAACTTGACCGCTC</td><td>599.8</td><td>8.0</td><td>1.5</td><td>5.3</td><td>1.3E-07</td><td>2.7E-07</td><td>BT20/MB468</td><td>CYTO</td></tr>
<tr><td>>MI0000683|hsa-mir-181b-2&WithFlank&9|+|124693704|124693804@22.46.25&[MIMAT0000257_1&hsa-miR-181b-5p&offsets|0|+2;m-72&9|+|124693725|124693747&offsets|0|+2]|AACATTCATTGCTGTCGGTGGGTTT</td><td>>MI0000683|hsa-mir-181b-2&WithFlank&9|+|124693704|124693804</td><td>22</td><td>46</td><td>25</td><td>AACATTCATTGCTGTCGGTGGGTTT</td><td>302.7</td><td>5.2</td><td>1.0</td><td>5.3</td><td>1.5E-07</td><td>2.8E-07</td><td>MB231/MB468</td><td>CYTO</td></tr>
<tr><td>>hg38_21_+_8433172_8446622_RNA45SN1_45S_50ntFlanks@4007.4042.36|CCTCGGGCCGATCGCACGCCCCCCGTGGCGGCGACG</td><td>>hg38_21_+_8433172_8446622_RNA45SN1_45S_50ntFlanks</td><td>4007</td><td>4042</td><td>36</td><td>CCTCGGGCCGATCGCACGCCCCCCGTGGCGGCGACG</td><td>366.8</td><td>-3.8</td><td>0.7</td><td>-5.2</td><td>1.6E-07</td><td>2.8E-07</td><td>MB231/MB468</td><td>MITO</td></tr>
<tr><td>>MI0000264|hsa-mir-7-2&WithFlank&15|+|88611819|88611940@38.59.22;MI0000265|hsa-mir-7-3&WithFlank&19|+|4770664|4770785@37.58.22;MI0000263|hsa-mir-7-1&WithFlank&9|-|83969742|83969863@30.51.22&[MIMAT0000252&hsa-miR-7-5p&offsets|0|-2;m-400&15|+|88611856|88611879&offsets|0|-2];[MIMAT0000252_1&hsa-miR-7-5p&offsets|0|-2;m-402&19|+|4770700|4770723&offsets|0|-2];[MIMAT0000252_2&hsa-miR-7-5p&offsets|0|-2;m-401&9|-|83969811|83969834&offsets|0|-2]|TGGAAGACTAGTGATTTTGTTG</td><td>>MI0000264|hsa-mir-7-2&WithFlank&15|+|88611819|88611940</td><td>38</td><td>59</td><td>22</td><td>TGGAAGACTAGTGATTTTGTTG</td><td>923.4</td><td>6.6</td><td>1.3</td><td>5.2</td><td>1.6E-07</td><td>2.8E-07</td><td>MB231/MB468</td><td>MITO</td></tr>
<tr><td>>hg38_21_+_8433172_8446622_RNA45SN1_45S_50ntFlanks@10346.10372.27|TCAAACGAGAACTTTGAAGGCCGAAGT</td><td>>hg38_21_+_8433172_8446622_RNA45SN1_45S_50ntFlanks</td><td>10346</td><td>10372</td><td>27</td><td>TCAAACGAGAACTTTGAAGGCCGAAGT</td><td>437.4</td><td>-3.9</td><td>0.7</td><td>-5.2</td><td>1.6E-07</td><td>2.8E-07</td><td>MB231/MB468</td><td>MITO</td></tr>
<tr><td>>hg38_21_+_8433172_8446622_RNA45SN1_45S_50ntFlanks@10386.10410.25|ATGTGAACAGCAGTTGAACATGGGT</td><td>>hg38_21_+_8433172_8446622_RNA45SN1_45S_50ntFlanks</td><td>10386</td><td>10410</td><td>25</td><td>ATGTGAACAGCAGTTGAACATGGGT</td><td>276.2</td><td>-4.1</td><td>0.8</td><td>-5.3</td><td>1.5E-07</td><td>2.8E-07</td><td>MB231/MB468</td><td>CYTO</td></tr>
<tr><td>>MI0000077|hsa-mir-21&WithFlank&17|+|59841260|59841343@14.33.20&[MIMAT0000076&hsa-miR-21-5p&offsets|0|-2;m-1&17|+|59841273|59841294&offsets|0|-2]|TAGCTTATCAGACTGATGTT</td><td>>MI0000077|hsa-mir-21&WithFlank&17|+|59841260|59841343</td><td>14</td><td>33</td><td>20</td><td>TAGCTTATCAGACTGATGTT</td><td>634.6</td><td>5.1</td><td>1.0</td><td>5.2</td><td>1.6E-07</td><td>2.8E-07</td><td>MB231/MB468</td><td>MITO</td></tr>
<tr><td>>hg38_21_+_8433172_8446622_RNA45SN1_45S_50ntFlanks@12478.12497.20|ACTAATAGGGAACGTGAGCT</td><td>>hg38_21_+_8433172_8446622_RNA45SN1_45S_50ntFlanks</td><td>12478</td><td>12497</td><td>20</td><td>ACTAATAGGGAACGTGAGCT</td><td>2822.6</td><td>3.6</td><td>0.7</td><td>5.3</td><td>1.3E-07</td><td>2.8E-07</td><td>BT20/MB468</td><td>CYTO</td></tr>
<tr><td>>hg38_21_+_8433172_8446622_RNA45SN1_45S_50ntFlanks@12640.12668.29|ACCATCTGTGGGATTATGACTGAACGCCT</td><td>>hg38_21_+_8433172_8446622_RNA45SN1_45S_50ntFlanks</td><td>12640</td><td>12668</td><td>29</td><td>ACCATCTGTGGGATTATGACTGAACGCCT</td><td>219.3</td><td>7.1</td><td>1.3</td><td>5.3</td><td>1.3E-07</td><td>2.8E-07</td><td>BT20/MB468</td><td>CYTO</td></tr>
<tr><td>>hg38_21_+_8433172_8446622_RNA45SN1_45S_50ntFlanks@4892.4916.25|AAACTTAAAGGAATTGACGGAAGGG</td><td>>hg38_21_+_8433172_8446622_RNA45SN1_45S_50ntFlanks</td><td>4892</td><td>4916</td><td>25</td><td>AAACTTAAAGGAATTGACGGAAGGG</td><td>98.8</td><td>-9.7</td><td>1.8</td><td>-5.5</td><td>3.1E-08</td><td>2.8E-07</td><td>BT20/MB468</td><td>MP</td></tr>
<tr><td>>MI0000253|hsa-mir-148a&WithFlank&7|-|25949913|25949992@50.72.23&[MIMAT0000243&hsa-miR-148a-3p&offsets|0|+1;m-7&7|-|25949922|25949943&offsets|0|+1]|TCAGTGCACTACAGAACTTTGTC</td><td>>MI0000253|hsa-mir-148a&WithFlank&7|-|25949913|25949992</td><td>50</td><td>72</td><td>23</td><td>TCAGTGCACTACAGAACTTTGTC</td><td>725.0</td><td>3.0</td><td>0.6</td><td>5.2</td><td>1.7E-07</td><td>2.8E-07</td><td>BT20/MB468</td><td>NUC</td></tr>
<tr><td>>MI0000440|hsa-mir-27b&WithFlank&9|+|95085439|95085547@67.85.19&[MIMAT0000419&hsa-miR-27b-3p&offsets|0|-2;m-39&9|+|95085505|95085525&offsets|0|-2]|TTCACAGTGGCTAAGTTCT</td><td>>MI0000440|hsa-mir-27b&WithFlank&9|+|95085439|95085547</td><td>67</td><td>85</td><td>19</td><td>TTCACAGTGGCTAAGTTCT</td><td>1346.3</td><td>4.8</td><td>0.9</td><td>5.2</td><td>1.6E-07</td><td>2.8E-07</td><td>MB231/MB468</td><td>MITO</td></tr>
<tr><td>>hg38_21_+_8433172_8446622_RNA45SN1_45S_50ntFlanks@3782.3802.21|CAGTGAAACTGCGAATGGCTC</td><td>>hg38_21_+_8433172_8446622_RNA45SN1_45S_50ntFlanks</td><td>3782</td><td>3802</td><td>21</td><td>CAGTGAAACTGCGAATGGCTC</td><td>405.7</td><td>-5.9</td><td>1.1</td><td>-5.2</td><td>1.6E-07</td><td>2.8E-07</td><td>MB231/MB468</td><td>NUC</td></tr>
<tr><td>>hg38_21_+_8433172_8446622_RNA45SN1_45S_50ntFlanks@10348.10383.36|AAACGAGAACTTTGAAGGCCGAAGTGGAGAAGGGTT</td><td>>hg38_21_+_8433172_8446622_RNA45SN1_45S_50ntFlanks</td><td>10348</td><td>10383</td><td>36</td><td>AAACGAGAACTTTGAAGGCCGAAGTGGAGAAGGGTT</td><td>1803.9</td><td>-3.7</td><td>0.7</td><td>-5.2</td><td>1.6E-07</td><td>2.8E-07</td><td>MB231/MB468</td><td>NUC</td></tr>
<tr><td>>hg38_21_+_8433172_8446622_RNA45SN1_45S_50ntFlanks@10767.10785.19|GCAGCAGGTCTCCAAGGTG</td><td>>hg38_21_+_8433172_8446622_RNA45SN1_45S_50ntFlanks</td><td>10767</td><td>10785</td><td>19</td><td>GCAGCAGGTCTCCAAGGTG</td><td>481.4</td><td>-8.0</td><td>1.5</td><td>-5.2</td><td>1.7E-07</td><td>2.8E-07</td><td>BT20/MB468</td><td>NUC</td></tr>
<tr><td>>hg38_21_+_8433172_8446622_RNA45SN1_45S_50ntFlanks@9568.9603.36|CGTGCAAATCGGTCGTCCGACCTGGGTATAGGGGCG</td><td>>hg38_21_+_8433172_8446622_RNA45SN1_45S_50ntFlanks</td><td>9568</td><td>9603</td><td>36</td><td>CGTGCAAATCGGTCGTCCGACCTGGGTATAGGGGCG</td><td>3063.0</td><td>-6.5</td><td>1.2</td><td>-5.2</td><td>1.6E-07</td><td>2.8E-07</td><td>MB231/MB468</td><td>MITO</td></tr>
<tr><td>>hg38_21_+_8433172_8446622_RNA45SN1_45S_50ntFlanks@6763.6806.44|CGGCCCCGGGTTCCTCCCGGGGCTACGCCTGTCTGAGCGTCGCT</td><td>>hg38_21_+_8433172_8446622_RNA45SN1_45S_50ntFlanks</td><td>6763</td><td>6806</td><td>44</td><td>CGGCCCCGGGTTCCTCCCGGGGCTACGCCTGTCTGAGCGTCGCT</td><td>506.4</td><td>7.6</td><td>1.4</td><td>5.3</td><td>1.2E-07</td><td>2.8E-07</td><td>BT20/MB468</td><td>MITO</td></tr>
<tr><td>>MI0000749|hsa-mir-30e&WithFlank&1|+|40754349|40754452@65.86.22&[MIMAT0000693&hsa-miR-30e-3p&offsets|0|0;m-25&1|+|40754413|40754434&offsets|0|0]|CTTTCAGTCGGATGTTTACAGC</td><td>>MI0000749|hsa-mir-30e&WithFlank&1|+|40754349|40754452</td><td>65</td><td>86</td><td>22</td><td>CTTTCAGTCGGATGTTTACAGC</td><td>954.8</td><td>3.6</td><td>0.7</td><td>5.3</td><td>1.2E-07</td><td>2.8E-07</td><td>BT20/MB468</td><td>MITO</td></tr>
<tr><td>>hg38_21_+_8433172_8446622_RNA45SN1_45S_50ntFlanks@9545.9562.18|GTGGAGGTCCGTAGCGGT</td><td>>hg38_21_+_8433172_8446622_RNA45SN1_45S_50ntFlanks</td><td>9545</td><td>9562</td><td>18</td><td>GTGGAGGTCCGTAGCGGT</td><td>1071.9</td><td>-8.6</td><td>1.6</td><td>-5.2</td><td>1.7E-07</td><td>2.8E-07</td><td>BT20/MB468</td><td>NUC</td></tr>
<tr><td>>hg38_21_+_8433172_8446622_RNA45SN1_45S_50ntFlanks@12786.12805.20|CCCGCGCGCGCGGGAGGGCG</td><td>>hg38_21_+_8433172_8446622_RNA45SN1_45S_50ntFlanks</td><td>12786</td><td>12805</td><td>20</td><td>CCCGCGCGCGCGGGAGGGCG</td><td>87.5</td><td>-9.5</td><td>1.7</td><td>-5.5</td><td>3.1E-08</td><td>2.8E-07</td><td>BT20/MB468</td><td>MP</td></tr>
<tr><td>>hg38_21_+_8433172_8446622_RNA45SN1_45S_50ntFlanks@3899.3935.37|CCTTCGCGGGGGGGATGCGTGCATTTATCAGATCAAA</td><td>>hg38_21_+_8433172_8446622_RNA45SN1_45S_50ntFlanks</td><td>3899</td><td>3935</td><td>37</td><td>CCTTCGCGGGGGGGATGCGTGCATTTATCAGATCAAA</td><td>227.3</td><td>-10.9</td><td>2.0</td><td>-5.5</td><td>3.1E-08</td><td>2.8E-07</td><td>BT20/MB468</td><td>MP</td></tr>
<tr><td>>hg38_MT_+_1621_3279_RNR2_16S_50ntFlanks@1380.1415.36|AGGACATCCCGATGGTGCAGCCGCTATTAAAGGTTC</td><td>>hg38_MT_+_1621_3279_RNR2_16S_50ntFlanks</td><td>1380</td><td>1415</td><td>36</td><td>AGGACATCCCGATGGTGCAGCCGCTATTAAAGGTTC</td><td>50.3</td><td>8.9</td><td>1.7</td><td>5.3</td><td>1.4E-07</td><td>2.8E-07</td><td>MB231/MB468</td><td>MP</td></tr>
<tr><td>>hg38_21_+_8433172_8446622_RNA45SN1_45S_50ntFlanks@4731.4768.38|AAGAACGAAAGTCGGAGGTTCGAAGACGATCAGATACC</td><td>>hg38_21_+_8433172_8446622_RNA45SN1_45S_50ntFlanks</td><td>4731</td><td>4768</td><td>38</td><td>AAGAACGAAAGTCGGAGGTTCGAAGACGATCAGATACC</td><td>258.5</td><td>5.2</td><td>1.0</td><td>5.2</td><td>1.6E-07</td><td>2.8E-07</td><td>MB231/MB468</td><td>NUC</td></tr>
<tr><td>>hg38_21_+_8433172_8446622_RNA45SN1_45S_50ntFlanks@9623.9650.28|TAGTAGCTGGTTCCCTCCGAAGTTTCCC</td><td>>hg38_21_+_8433172_8446622_RNA45SN1_45S_50ntFlanks</td><td>9623</td><td>9650</td><td>28</td><td>TAGTAGCTGGTTCCCTCCGAAGTTTCCC</td><td>211.5</td><td>-5.1</td><td>1.0</td><td>-5.2</td><td>1.6E-07</td><td>2.8E-07</td><td>MB231/MB468</td><td>NUC</td></tr>
<tr><td>>hg38_21_+_8433172_8446622_RNA45SN1_45S_50ntFlanks@12399.12419.21|ACGTCGCTTTTTGATCCTTCG</td><td>>hg38_21_+_8433172_8446622_RNA45SN1_45S_50ntFlanks</td><td>12399</td><td>12419</td><td>21</td><td>ACGTCGCTTTTTGATCCTTCG</td><td>71.9</td><td>-9.2</td><td>1.7</td><td>-5.5</td><td>3.2E-08</td><td>2.8E-07</td><td>BT20/MB468</td><td>MP</td></tr>
<tr><td>>hg38_21_+_8433172_8446622_RNA45SN1_45S_50ntFlanks@8423.8469.47|CGCAGTCCGCCCGGAGGATTCAACCCGGCGGCGGGTCCGGCCGTGTC</td><td>>hg38_21_+_8433172_8446622_RNA45SN1_45S_50ntFlanks</td><td>8423</td><td>8469</td><td>47</td><td>CGCAGTCCGCCCGGAGGATTCAACCCGGCGGCGGGTCCGGCCGTGTC</td><td>240.4</td><td>-11.0</td><td>2.0</td><td>-5.5</td><td>3.2E-08</td><td>2.8E-07</td><td>BT20/MB468</td><td>MP</td></tr>
<tr><td>>hg38_21_+_8433172_8446622_RNA45SN1_45S_50ntFlanks@3785.3802.18|TGAAACTGCGAATGGCTC</td><td>>hg38_21_+_8433172_8446622_RNA45SN1_45S_50ntFlanks</td><td>3785</td><td>3802</td><td>18</td><td>TGAAACTGCGAATGGCTC</td><td>175.9</td><td>-5.3</td><td>1.0</td><td>-5.2</td><td>1.6E-07</td><td>2.8E-07</td><td>MB231/MB468</td><td>NUC</td></tr>
<tr><td>>hg38_21_+_8433172_8446622_RNA45SN1_45S_50ntFlanks@10239.10259.21|GCTACGCCGCGACGAGTAGGA</td><td>>hg38_21_+_8433172_8446622_RNA45SN1_45S_50ntFlanks</td><td>10239</td><td>10259</td><td>21</td><td>GCTACGCCGCGACGAGTAGGA</td><td>497.1</td><td>5.2</td><td>0.9</td><td>5.5</td><td>3.2E-08</td><td>2.8E-07</td><td>BT20/MB468</td><td>MP</td></tr>
<tr><td>>hg38_21_+_8433172_8446622_RNA45SN1_45S_50ntFlanks@8176.8197.22|CTTCTGATCGAGGCCCAGCCCG</td><td>>hg38_21_+_8433172_8446622_RNA45SN1_45S_50ntFlanks</td><td>8176</td><td>8197</td><td>22</td><td>CTTCTGATCGAGGCCCAGCCCG</td><td>545.3</td><td>-3.1</td><td>0.6</td><td>-5.3</td><td>1.2E-07</td><td>2.8E-07</td><td>BT20/MB468</td><td>MITO</td></tr>
<tr><td>>hg38_21_+_8433172_8446622_RNA45SN1_45S_50ntFlanks@6650.6672.23|ACGACTCTTAGCGGTGGATCACT</td><td>>hg38_21_+_8433172_8446622_RNA45SN1_45S_50ntFlanks</td><td>6650</td><td>6672</td><td>23</td><td>ACGACTCTTAGCGGTGGATCACT</td><td>457.6</td><td>6.4</td><td>1.2</td><td>5.3</td><td>1.3E-07</td><td>2.8E-07</td><td>BT20/MB468</td><td>CYTO</td></tr>
<tr><td>>MI0000103|hsa-mir-101-1&WithFlank&1|-|65058428|65058514@53.74.22;MI0000739|hsa-mir-101-2&WithFlank&9|+|4850291|4850381@55.76.22&[MIMAT0000099&hsa-miR-101-3p&offsets|0|+1;m-17&1|-|65058443|65058462&offsets|0|+2];[MIMAT0000099_1&hsa-miR-101-3p&offsets|0|+1;m-16&9|+|4850345|4850364&offsets|0|+2]|TACAGTACTGTGATAACTGAAG</td><td>>MI0000103|hsa-mir-101-1&WithFlank&1|-|65058428|65058514</td><td>53</td><td>74</td><td>22</td><td>TACAGTACTGTGATAACTGAAG</td><td>1386.0</td><td>5.8</td><td>1.1</td><td>5.2</td><td>1.7E-07</td><td>2.8E-07</td><td>MB231/MB468</td><td>MITO</td></tr>
<tr><td>>hg38_21_+_8433172_8446622_RNA45SN1_45S_50ntFlanks@6650.6676.27|ACGACTCTTAGCGGTGGATCACTCGGC</td><td>>hg38_21_+_8433172_8446622_RNA45SN1_45S_50ntFlanks</td><td>6650</td><td>6676</td><td>27</td><td>ACGACTCTTAGCGGTGGATCACTCGGC</td><td>1198.1</td><td>5.4</td><td>1.0</td><td>5.5</td><td>3.2E-08</td><td>2.8E-07</td><td>BT20/MB468</td><td>MP</td></tr>
<tr><td>>hg38_21_+_8433172_8446622_RNA45SN1_45S_50ntFlanks@7975.7997.23|CGCGACCTCAGATCAGACGTGGC</td><td>>hg38_21_+_8433172_8446622_RNA45SN1_45S_50ntFlanks</td><td>7975</td><td>7997</td><td>23</td><td>CGCGACCTCAGATCAGACGTGGC</td><td>9169.5</td><td>3.5</td><td>0.7</td><td>5.2</td><td>1.6E-07</td><td>2.8E-07</td><td>MB231/MB468</td><td>NUC</td></tr>
<tr><td>>MI0000471|hsa-mir-126&WithFlank&9|+|136670596|136670692@58.77.20(+1A)&[MIMAT0000445&hsa-miR-126-3p&offsets|0|-2(+1A);m-40&9|+|136670653|136670674&offsets|0|-2(+1A)]|TCGTACCGTGAGTAATAATGA</td><td>>MI0000471|hsa-mir-126&WithFlank&9|+|136670596|136670692</td><td>58</td><td>77</td><td>20(+1A)</td><td>TCGTACCGTGAGTAATAATGA</td><td>91.7</td><td>10.3</td><td>1.9</td><td>5.5</td><td>3.2E-08</td><td>2.8E-07</td><td>BT20/MB468</td><td>MP</td></tr>
<tr><td>>hg38_21_+_8433172_8446622_RNA45SN1_45S_50ntFlanks@10808.10830.23|CAATGTAGGTAAGGGAAGTCGGC</td><td>>hg38_21_+_8433172_8446622_RNA45SN1_45S_50ntFlanks</td><td>10808</td><td>10830</td><td>23</td><td>CAATGTAGGTAAGGGAAGTCGGC</td><td>436.2</td><td>-7.2</td><td>1.4</td><td>-5.2</td><td>1.6E-07</td><td>2.8E-07</td><td>MB231/MB468</td><td>NUC</td></tr>
<tr><td>>hg38_MT_+_1621_3279_RNR2_16S_50ntFlanks@51.97.47|GCTAAACCTAGCCCCAAACCCACTCCACCTTACTACCAGACAACCTT</td><td>>hg38_MT_+_1621_3279_RNR2_16S_50ntFlanks</td><td>51</td><td>97</td><td>47</td><td>GCTAAACCTAGCCCCAAACCCACTCCACCTTACTACCAGACAACCTT</td><td>522.4</td><td>5.4</td><td>1.0</td><td>5.3</td><td>1.2E-07</td><td>2.8E-07</td><td>BT20/MB468</td><td>MITO</td></tr>
<tr><td>>MI0000076|hsa-mir-20a&WithFlank&13|+|91351059|91351141@14.35.22(+1A)&[MIMAT0000075&hsa-miR-20a-5p&offsets|0|-1(+1A);m-96&13|+|91351072|91351094&offsets|0|-1(+1A)]|TAAAGTGCTTATAGTGCAGGTAA</td><td>>MI0000076|hsa-mir-20a&WithFlank&13|+|91351059|91351141</td><td>14</td><td>35</td><td>22(+1A)</td><td>TAAAGTGCTTATAGTGCAGGTAA</td><td>3124.5</td><td>6.1</td><td>1.2</td><td>5.2</td><td>1.7E-07</td><td>2.8E-07</td><td>MB231/MB468</td><td>MITO</td></tr>
<tr><td>>hg38_21_+_8433172_8446622_RNA45SN1_45S_50ntFlanks@5086.5120.35|AACGAGACTCTGGCATGCTAACTAGTTACGCGACC</td><td>>hg38_21_+_8433172_8446622_RNA45SN1_45S_50ntFlanks</td><td>5086</td><td>5120</td><td>35</td><td>AACGAGACTCTGGCATGCTAACTAGTTACGCGACC</td><td>544.8</td><td>-5.0</td><td>1.0</td><td>-5.2</td><td>1.7E-07</td><td>2.8E-07</td><td>BT20/MB468</td><td>NUC</td></tr>
<tr><td>>hg38_21_+_8433172_8446622_RNA45SN1_45S_50ntFlanks@12944.12972.29|CGTAGACGACCTGCTTCTGGGTCGGGGTT</td><td>>hg38_21_+_8433172_8446622_RNA45SN1_45S_50ntFlanks</td><td>12944</td><td>12972</td><td>29</td><td>CGTAGACGACCTGCTTCTGGGTCGGGGTT</td><td>319.9</td><td>-4.2</td><td>0.8</td><td>-5.3</td><td>1.3E-07</td><td>2.8E-07</td><td>BT20/MB468</td><td>CYTO</td></tr>
<tr><td>>MI0000434|hsa-let-7i&WithFlank&12|+|62603680|62603775@12.32.21(+1C)&[MIMAT0000415&hsa-let-7i-5p&offsets|0|-1(+1C);m-49&12|+|62603691|62603712&offsets|0|-1(+1C)]|TGAGGTAGTAGTTTGTGCTGTC</td><td>>MI0000434|hsa-let-7i&WithFlank&12|+|62603680|62603775</td><td>12</td><td>32</td><td>21(+1C)</td><td>TGAGGTAGTAGTTTGTGCTGTC</td><td>22430.6</td><td>4.7</td><td>0.9</td><td>5.3</td><td>1.4E-07</td><td>2.8E-07</td><td>MB231/MB468</td><td>MP</td></tr>
<tr><td>>hg38_21_+_8433172_8446622_RNA45SN1_45S_50ntFlanks@11575.11592.18|ACTTAGAACTGGTGCGGA</td><td>>hg38_21_+_8433172_8446622_RNA45SN1_45S_50ntFlanks</td><td>11575</td><td>11592</td><td>18</td><td>ACTTAGAACTGGTGCGGA</td><td>3330.7</td><td>-7.3</td><td>1.3</td><td>-5.5</td><td>3.2E-08</td><td>2.8E-07</td><td>BT20/MB468</td><td>MP</td></tr>
<tr><td>>hg38_21_+_8433172_8446622_RNA45SN1_45S_50ntFlanks@6738.6760.23|ATCATCGACACTTCGAACGCACT</td><td>>hg38_21_+_8433172_8446622_RNA45SN1_45S_50ntFlanks</td><td>6738</td><td>6760</td><td>23</td><td>ATCATCGACACTTCGAACGCACT</td><td>68.6</td><td>-9.2</td><td>1.7</td><td>-5.5</td><td>3.2E-08</td><td>2.8E-07</td><td>BT20/MB468</td><td>MP</td></tr>
<tr><td>>hg38_21_+_8433172_8446622_RNA45SN1_45S_50ntFlanks@10058.10081.24|CGGCAGTCGAGAGTGGACGGGAGC</td><td>>hg38_21_+_8433172_8446622_RNA45SN1_45S_50ntFlanks</td><td>10058</td><td>10081</td><td>24</td><td>CGGCAGTCGAGAGTGGACGGGAGC</td><td>78.7</td><td>10.0</td><td>1.8</td><td>5.5</td><td>3.2E-08</td><td>2.8E-07</td><td>BT20/MB468</td><td>MP</td></tr>
<tr><td>>hg38_21_+_8433172_8446622_RNA45SN1_45S_50ntFlanks@6721.6743.23|AATTGCAGGACACATTGATCATC</td><td>>hg38_21_+_8433172_8446622_RNA45SN1_45S_50ntFlanks</td><td>6721</td><td>6743</td><td>23</td><td>AATTGCAGGACACATTGATCATC</td><td>1548.3</td><td>-3.1</td><td>0.6</td><td>-5.3</td><td>1.2E-07</td><td>2.8E-07</td><td>BT20/MB468</td><td>MITO</td></tr>
</table>

| | | | | | | | | | | | | | |
|---|---|---|---|---|---|---|---|---|---|---|---|---|---|
| >MI0000255\|hsa-mir-30d&WithFlank&8\|-\|134804870\|134804951@12.33.22(+1A)&[MIMAT0000245&hsa-miR-30d-5p&offsets\|0\|0(+1A);m-21&8\|-\|134804919\|134804940&offsets\|0\|0(+1A)]\|TGTAAACATCCCCGACTGGAAGA | >MI0000255\|hsa-mir-30d&WithFlank&8\|-\|134804870\|134804951 | 12 | 33 | 22(+1A) | TGTAAACATCCCCGACTGGAAGA | 503.4 | 6.3 | 1.2 | 5.2 | 1.5E-07 | 2.8E-07 | MB231/MB468 | CYTO |
| >hg38_21_+_8433172_8446622_RNA45SN1_45S_50ntFlanks@11648.11687.40\|TGTTGACGCGATGTGATTTCTGCCCAGTGCTCTGAATGTC | >hg38_21_+_8433172_8446622_RNA45SN1_45S_50ntFlanks | 11648 | 11687 | 40 | TGTTGACGCGATGTGATTTCTGCCCAGTGCTCTGAATGTC | 153.4 | 11.0 | 2.0 | 5.5 | 3.2E-08 | 2.8E-07 | BT20/MB468 | MP |
| >hg38_21_+_8433172_8446622_RNA45SN1_45S_50ntFlanks@10347.10374.28\|CAAACGAGAACTTTGAAGGCCGAAGTGG | >hg38_21_+_8433172_8446622_RNA45SN1_45S_50ntFlanks | 10347 | 10374 | 28 | CAAACGAGAACTTTGAAGGCCGAAGTGG | 313.4 | -3.3 | 0.6 | -5.2 | 1.6E-07 | 2.8E-07 | MB231/MB468 | NUC |
| >hg38_21_+_8433172_8446622_RNA45SN1_45S_50ntFlanks@9534.9560.27\|AGGAAACTCTGGTGGAGGTCCGTAGCG | >hg38_21_+_8433172_8446622_RNA45SN1_45S_50ntFlanks | 9534 | 9560 | 27 | AGGAAACTCTGGTGGAGGTCCGTAGCG | 498.3 | -7.1 | 1.4 | -5.2 | 1.7E-07 | 2.8E-07 | MB231/MB468 | MITO |
| >MI0000471\|hsa-mir-126&WithFlank&9\|+\|136670596\|136670692@59.79.21(+1U)&[MIMAT0000445&hsa-miR-126-3p&offsets\|+1\|0(+1U);m-40&9\|+\|136670653\|136670674&offsets\|+1\|0(+1U)]\|CGTACCGTGAGTAATAATGCGT | >MI0000471\|hsa-mir-126&WithFlank&9\|+\|136670596\|136670692 | 59 | 79 | 21(+1U) | CGTACCGTGAGTAATAATGCGT | 141.4 | 10.9 | 2.0 | 5.5 | 3.2E-08 | 2.8E-07 | BT20/MB468 | MP |
| >MI0000299\|hsa-mir-222&WithFlank&X\|-\|45747009\|45747130@75.97.23&[MIMAT0000279&hsa-miR-222-3p&offsets\|0\|+2;m-140&X\|-\|45747033\|45747056&offsets\|0\|-1]\|AGCTACATCTGGCTACTGGGTCT | >MI0000299\|hsa-mir-222&WithFlank&X\|-\|45747009\|45747130 | 75 | 97 | 23 | AGCTACATCTGGCTACTGGGTCT | 6693.1 | 4.6 | 0.9 | 5.3 | 1.4E-07 | 2.8E-07 | MB231/MB468 | MP |
| >hg38_21_+_8433172_8446622_RNA45SN1_45S_50ntFlanks@12970.12990.21\|GTTTCGTACGTAGCAGAGCAG | >hg38_21_+_8433172_8446622_RNA45SN1_45S_50ntFlanks | 12970 | 12990 | 21 | GTTTCGTACGTAGCAGAGCAG | 141.5 | -6.2 | 1.2 | -5.3 | 1.4E-07 | 2.8E-07 | BT20/MB468 | CYTO |
| >hg38_21_+_8433172_8446622_RNA45SN1_45S_50ntFlanks@8356.8387.32\|GAAAAGAACTTTGAAGAGAGAGTTCAAGAGGG | >hg38_21_+_8433172_8446622_RNA45SN1_45S_50ntFlanks | 8356 | 8387 | 32 | GAAAAGAACTTTGAAGAGAGAGTTCAAGAGGG | 264.2 | -4.6 | 0.9 | -5.3 | 1.4E-07 | 2.8E-07 | MB231/MB468 | MP |
| >hg38_21_+_8433172_8446622_RNA45SN1_45S_50ntFlanks@8115.8138.24\|CGCGGGACATGTGGCGTACGGAAG | >hg38_21_+_8433172_8446622_RNA45SN1_45S_50ntFlanks | 8115 | 8138 | 24 | CGCGGGACATGTGGCGTACGGAAG | 565.5 | 12.5 | 2.2 | 5.8 | 6.7E-09 | 2.8E-07 | BT20/MB231 | MP |
| >MI0003513\|hsa-mir-455&WithFlank&9\|+\|114209428\|114209535@22.42.21(+1A)&[MIMAT0003150&hsa-miR-455-5p&offsets\|0\|-1(+1A);m-232&9\|+\|114209449\|114209470&offsets\|0\|-1(+1A)]\|TATGTGCCTTTGGACTACATCA | >MI0003513\|hsa-mir-455&WithFlank&9\|+\|114209428\|114209535 | 22 | 42 | 21(+1A) | TATGTGCCTTTGGACTACATCA | 153.1 | -10.7 | 1.8 | -5.8 | 6.7E-09 | 2.8E-07 | BT20/MB231 | MP |
| >hg38_21_+_8433172_8446622_RNA45SN1_45S_50ntFlanks@11675.11707.33\|TGCTCTGAATGTCAAAGTGAAGAAATTCAATGA | >hg38_21_+_8433172_8446622_RNA45SN1_45S_50ntFlanks | 11675 | 11707 | 33 | TGCTCTGAATGTCAAAGTGAAGAAATTCAATGA | 562.6 | 4.4 | 0.8 | 5.2 | 1.6E-07 | 2.8E-07 | MB231/MB468 | CYTO |
| >hg38_21_+_8433172_8446622_RNA45SN1_45S_50ntFlanks@9567.9586.20\|ACGTGCAAATCGGTCGTCCG | >hg38_21_+_8433172_8446622_RNA45SN1_45S_50ntFlanks | 9567 | 9586 | 20 | ACGTGCAAATCGGTCGTCCG | 664.9 | -5.1 | 1.0 | -5.2 | 1.6E-07 | 2.8E-07 | MB231/MB468 | CYTO |
| >hg38_21_+_8433172_8446622_RNA45SN1_45S_50ntFlanks@4760.4779.20\|TCAGATACCGTCGTAGTTCC | >hg38_21_+_8433172_8446622_RNA45SN1_45S_50ntFlanks | 4760 | 4779 | 20 | TCAGATACCGTCGTAGTTCC | 396.7 | -4.7 | 0.9 | -5.3 | 1.4E-07 | 2.8E-07 | BT20/MB468 | CYTO |
| >hg38_21_+_8433172_8446622_RNA45SN1_45S_50ntFlanks@8111.8141.31\|GGGGCGCGGGACATGTGGCGTACGGAAGACC | >hg38_21_+_8433172_8446622_RNA45SN1_45S_50ntFlanks | 8111 | 8141 | 31 | GGGGCGCGGGACATGTGGCGTACGGAAGACC | 536.9 | 12.4 | 2.1 | 5.8 | 6.8E-09 | 2.8E-07 | BT20/MB231 | MP |
| >hg38_21_+_8433172_8446622_RNA45SN1_45S_50ntFlanks@11270.11295.26\|CGGCCGCCCCCGGGGAGCCCGGCGGG | >hg38_21_+_8433172_8446622_RNA45SN1_45S_50ntFlanks | 11270 | 11295 | 26 | CGGCCGCCCCCGGGGAGCCCGGCGGG | 1762.9 | -7.6 | 1.4 | -5.3 | 1.2E-07 | 2.8E-07 | BT20/MB468 | MITO |
| >hg38_21_+_8433172_8446622_RNA45SN1_45S_50ntFlanks@5444.5467.24\|CTCGGATCGGCCCCGCCGGGGTCG | >hg38_21_+_8433172_8446622_RNA45SN1_45S_50ntFlanks | 5444 | 5467 | 24 | CTCGGATCGGCCCCGCCGGGGTCG | 1199.6 | -4.4 | 0.8 | -5.2 | 1.6E-07 | 2.8E-07 | MB231/MB468 | CYTO |
| >hg38_21_+_8433172_8446622_RNA45SN1_45S_50ntFlanks@474.498.25\|TCTTCGTGATCGATGTGGTGACGTC | >hg38_21_+_8433172_8446622_RNA45SN1_45S_50ntFlanks | 474 | 498 | 25 | TCTTCGTGATCGATGTGGTGACGTC | 905.2 | 5.1 | 1.0 | 5.2 | 1.6E-07 | 2.8E-07 | MB231/MB468 | NUC |
| >MI0000737\|hsa-mir-200a&WithFlank&1\|+\|1167857\|1167958@60.81.22(+1G)&[MIMAT0000682&hsa-miR-200a-3p&offsets\|0\|0(+1G);m-142&1\|+\|1167916\|1167938&offsets\|0\|-1(+1G)]\|TAACACTGTCTGGTAACGATGTG | >MI0000737\|hsa-mir-200a&WithFlank&1\|+\|1167857\|1167958 | 60 | 81 | 22(+1G) | TAACACTGTCTGGTAACGATGTG | 129.4 | -10.1 | 1.8 | -5.5 | 3.2E-08 | 2.8E-07 | BT20/MB468 | MP |
| >hg38_21_+_8433172_8446622_RNA45SN1_45S_50ntFlanks@10392.10410.19\|ACAGCAGTTGAACATGGGT | >hg38_21_+_8433172_8446622_RNA45SN1_45S_50ntFlanks | 10392 | 10410 | 19 | ACAGCAGTTGAACATGGGT | 3591.3 | -5.2 | 1.0 | -5.3 | 1.4E-07 | 2.8E-07 | BT20/MB468 | CYTO |
| >hg38_21_+_8433172_8446622_RNA45SN1_45S_50ntFlanks@11598.11626.29\|GGAATCCGACTGTTTAATTAAAACAAAGC | >hg38_21_+_8433172_8446622_RNA45SN1_45S_50ntFlanks | 11598 | 11626 | 29 | GGAATCCGACTGTTTAATTAAAACAAAGC | 76.1 | 10.0 | 1.8 | 5.5 | 3.2E-08 | 2.9E-07 | BT20/MB468 | MP |
| >hg38_21_+_8433172_8446622_RNA45SN1_45S_50ntFlanks@4223.4242.20\|AAAAATAACAATACAGGACT | >hg38_21_+_8433172_8446622_RNA45SN1_45S_50ntFlanks | 4223 | 4242 | 20 | AAAAATAACAATACAGGACT | 332.6 | -6.0 | 1.1 | -5.2 | 1.8E-07 | 2.9E-07 | BT20/MB468 | NUC |
| >hg38_21_+_8433172_8446622_RNA45SN1_45S_50ntFlanks@4003.4026.24\|ATAACCTCGGGCCGATCGCACGCC | >hg38_21_+_8433172_8446622_RNA45SN1_45S_50ntFlanks | 4003 | 4026 | 24 | ATAACCTCGGGCCGATCGCACGCC | 703.9 | -6.8 | 1.3 | -5.3 | 1.4E-07 | 2.9E-07 | BT20/MB468 | CYTO |
| >hg38_21_+_8433172_8446622_RNA45SN1_45S_50ntFlanks@5186.5207.22\|CAATAACAGGTCTGTGATGCCC | >hg38_21_+_8433172_8446622_RNA45SN1_45S_50ntFlanks | 5186 | 5207 | 22 | CAATAACAGGTCTGTGATGCCC | 373.5 | -10.6 | 2.0 | -5.3 | 1.2E-07 | 2.9E-07 | BT20/MB468 | MITO |
| >hg38_21_+_8433172_8446622_RNA45SN1_45S_50ntFlanks@10134.10158.25\|CGGCGGCGGCGGCGGGGGTGTGGGG | >hg38_21_+_8433172_8446622_RNA45SN1_45S_50ntFlanks | 10134 | 10158 | 25 | CGGCGGCGGCGGCGGGGGTGTGGGG | 84.9 | -9.5 | 1.7 | -5.5 | 3.3E-08 | 2.9E-07 | BT20/MB468 | MP |
| >hg38_21_+_8433172_8446622_RNA45SN1_45S_50ntFlanks@12413.12432.20\|TCCTTCGATGTCGGCTCTTC | >hg38_21_+_8433172_8446622_RNA45SN1_45S_50ntFlanks | 12413 | 12432 | 20 | TCCTTCGATGTCGGCTCTTC | 2099.1 | -9.0 | 1.7 | -5.2 | 1.8E-07 | 2.9E-07 | BT20/MB468 | NUC |
| >trna129_MetCAT_6_-_27870271_27870342@29.51.23,trna142_MetCAT_6_-_27560600_27560671@29.51.23,trna150_MetCAT_6_-_27300764_27300835@29.51.23,trna169_MetCAT_6_-_26330529_26330600@29.51.23,trna171_MetCAT_6_-_26313352_26313423@29.51.23,trna20_MetCAT_17_-_80452597_80452668@29.51.23,trna2_MetCAT_6_+_26286754_26286825@29.51.23,trna32_MetCAT_1_+_153643726_153643797@29.51.23,trna61_MetCAT_6_+_27745664_27745735@29.51.23\|GGCCCATAACCCAGAGGTCGATG | >trna129_MetCAT_6_-_27870271_27870342 | 29 | 51 | 23 | GGCCCATAACCCAGAGGTCGATG | 111.9 | -9.9 | 1.8 | -5.5 | 3.3E-08 | 2.9E-07 | BT20/MB468 | MP |
| >hg38_21_+_8433172_8446622_RNA45SN1_45S_50ntFlanks@10367.10388.22\|CGAAGTGGAGAAGGGTTCCATG | >hg38_21_+_8433172_8446622_RNA45SN1_45S_50ntFlanks | 10367 | 10388 | 22 | CGAAGTGGAGAAGGGTTCCATG | 80.8 | -9.4 | 1.7 | -5.5 | 3.3E-08 | 2.9E-07 | BT20/MB468 | MP |
| >hg38_21_+_8433172_8446622_RNA45SN1_45S_50ntFlanks@8192.8211.20\|AGCCCGTGGACGGTGTGAGG | >hg38_21_+_8433172_8446622_RNA45SN1_45S_50ntFlanks | 8192 | 8211 | 20 | AGCCCGTGGACGGTGTGAGG | 562.1 | -8.5 | 1.6 | -5.2 | 1.8E-07 | 2.9E-07 | BT20/MB468 | NUC |
| >hg38_21_+_8433172_8446622_RNA45SN1_45S_50ntFlanks@10393.10430.38\|CAGCAGTTGAACATGGGTCAGTCGGTCCTGAGAGATGG | >hg38_21_+_8433172_8446622_RNA45SN1_45S_50ntFlanks | 10393 | 10430 | 38 | CAGCAGTTGAACATGGGTCAGTCGGTCCTGAGAGATGG | 429.3 | -3.8 | 0.7 | -5.3 | 1.2E-07 | 2.9E-07 | BT20/MB468 | MITO |
| >hg38_21_+_8433172_8446622_RNA45SN1_45S_50ntFlanks@7982.8012.31\|TCAGATCAGACGTGGCGACCCGCTGAATTTA | >hg38_21_+_8433172_8446622_RNA45SN1_45S_50ntFlanks | 7982 | 8012 | 31 | TCAGATCAGACGTGGCGACCCGCTGAATTTA | 584.9 | -5.9 | 1.1 | -5.2 | 1.7E-07 | 2.9E-07 | MB231/MB468 | MITO |
| >hg38_21_+_8433172_8446622_RNA45SN1_45S_50ntFlanks@10669.10706.38\|AAAGCGTCGCGGTTCCGGCGGCGTCCGGTGAGCTCTCG | >hg38_21_+_8433172_8446622_RNA45SN1_45S_50ntFlanks | 10669 | 10706 | 38 | AAAGCGTCGCGGTTCCGGCGGCGTCCGGTGAGCTCTCG | 609.4 | -6.1 | 1.2 | -5.2 | 1.6E-07 | 2.9E-07 | MB231/MB468 | CYTO |
| >hg38_21_+_8433172_8446622_RNA45SN1_45S_50ntFlanks@4760.4783.24\|TCAGATACCGTCGTAGTTCCGACC | >hg38_21_+_8433172_8446622_RNA45SN1_45S_50ntFlanks | 4760 | 4783 | 24 | TCAGATACCGTCGTAGTTCCGACC | 723.7 | -2.9 | 0.5 | -5.2 | 1.6E-07 | 2.9E-07 | MB231/MB468 | CYTO |
| >hg38_21_+_8433172_8446622_RNA45SN1_45S_50ntFlanks@4045.4065.21\|CCATTCGAACGTCTGCCCTAT | >hg38_21_+_8433172_8446622_RNA45SN1_45S_50ntFlanks | 4045 | 4065 | 21 | CCATTCGAACGTCTGCCCTAT | 213.9 | -5.4 | 1.0 | -5.2 | 1.6E-07 | 2.9E-07 | MB231/MB468 | NUC |
| >hg38_21_+_8433172_8446622_RNA45SN1_45S_50ntFlanks@12905.12933.29\|CGCACCGCACGTTCGTGGGGAACCTGGCG | >hg38_21_+_8433172_8446622_RNA45SN1_45S_50ntFlanks | 12905 | 12933 | 29 | CGCACCGCACGTTCGTGGGGAACCTGGCG | 146.1 | -10.3 | 1.9 | -5.5 | 3.3E-08 | 2.9E-07 | BT20/MB468 | MP |
| >hg38_21_+_8433172_8446622_RNA45SN1_45S_50ntFlanks@6661.6706.46\|CGGTGGATCACTCGGCTCGTGCGTCGATGAAGAACGCAGCTAGCTG | >hg38_21_+_8433172_8446622_RNA45SN1_45S_50ntFlanks | 6661 | 6706 | 46 | CGGTGGATCACTCGGCTCGTGCGTCGATGAAGAACGCAGCTAGCTG | 1303.7 | 5.2 | 1.0 | 5.3 | 1.4E-07 | 2.9E-07 | BT20/MB468 | CYTO |
| >hg38_21_+_8433172_8446622_RNA45SN1_45S_50ntFlanks@12407.12434.28\|TTTTGATCCTTCGATGTCGGCTCTTCCT | >hg38_21_+_8433172_8446622_RNA45SN1_45S_50ntFlanks | 12407 | 12434 | 28 | TTTTGATCCTTCGATGTCGGCTCTTCCT | 4544.8 | -3.1 | 0.6 | -5.2 | 1.6E-07 | 2.9E-07 | MB231/MB468 | NUC |
| >hg38_21_+_8433172_8446622_RNA45SN1_45S_50ntFlanks@10509.10533.25\|GAATCCGGAGTGGCGGAGATGGGCG | >hg38_21_+_8433172_8446622_RNA45SN1_45S_50ntFlanks | 10509 | 10533 | 25 | GAATCCGGAGTGGCGGAGATGGGCG | 295.6 | -7.5 | 1.4 | -5.2 | 1.6E-07 | 2.9E-07 | MB231/MB468 | NUC |
| >hg38_21_+_8433172_8446622_RNA45SN1_45S_50ntFlanks@4664.4695.32\|TGAAATTCTTGGACCGGCGCAAGACGGACCAG | >hg38_21_+_8433172_8446622_RNA45SN1_45S_50ntFlanks | 4664 | 4695 | 32 | TGAAATTCTTGGACCGGCGCAAGACGGACCAG | 70.5 | -9.2 | 1.7 | -5.5 | 3.3E-08 | 2.9E-07 | BT20/MB468 | MP |
| >hg38_21_+_8433172_8446622_RNA45SN1_45S_50ntFlanks@4090.4121.32\|CCTACCATGGTGACCACGGGTGACGGGGAATC | >hg38_21_+_8433172_8446622_RNA45SN1_45S_50ntFlanks | 4090 | 4121 | 32 | CCTACCATGGTGACCACGGGTGACGGGGAATC | 195.7 | -9.7 | 1.8 | -5.3 | 1.2E-07 | 2.9E-07 | BT20/MB468 | MITO |
| >MI0000809\|hsa-mir-151a&WithFlank&8\|-\|140732558\|140732659@51.73.23&[MIMAT0000757&hsa-miR-151a-3p&offsets\|-2\|0;m-30&8\|-\|140732587\|140732607&offsets\|-2\|0]\|TACTAGACTGAAGCTCCTTGAGG | >MI0000809\|hsa-mir-151a&WithFlank&8\|-\|140732558\|140732659 | 51 | 73 | 23 | TACTAGACTGAAGCTCCTTGAGG | 2267.8 | 3.8 | 0.7 | 5.3 | 1.4E-07 | 2.9E-07 | MB231/MB468 | MP |
| >hg38_21_+_8433172_8446622_RNA45SN1_45S_50ntFlanks@8202.8223.22\|CGGTGTGAGGCCGGTAGCGGCC | >hg38_21_+_8433172_8446622_RNA45SN1_45S_50ntFlanks | 8202 | 8223 | 22 | CGGTGTGAGGCCGGTAGCGGCC | 69.4 | -9.2 | 1.7 | -5.5 | 3.3E-08 | 2.9E-07 | BT20/MB468 | MP |
| >hg38_21_+_8433172_8446622_RNA45SN1_45S_50ntFlanks@11589.11623.35\|CGGACCAGGGGAATCCGACTGTTTAATTAAAACAA | >hg38_21_+_8433172_8446622_RNA45SN1_45S_50ntFlanks | 11589 | 11623 | 35 | CGGACCAGGGGAATCCGACTGTTTAATTAAAACAA | 530.4 | 9.5 | 1.7 | 5.5 | 3.3E-08 | 2.9E-07 | BT20/MB468 | MP |
| >hg38_21_+_8433172_8446622_RNA45SN1_45S_50ntFlanks@10410.10437.28\|TCAGTCGGTCCTGAGAGATGGGCGAGCG | >hg38_21_+_8433172_8446622_RNA45SN1_45S_50ntFlanks | 10410 | 10437 | 28 | TCAGTCGGTCCTGAGAGATGGGCGAGCG | 115.8 | -9.9 | 1.8 | -5.5 | 3.3E-08 | 2.9E-07 | BT20/MB468 | MP |
| >hg38_21_+_8433172_8446622_RNA45SN1_45S_50ntFlanks@4683.4707.25\|CAAGACGGACCAGAGCGAAAGCATT | >hg38_21_+_8433172_8446622_RNA45SN1_45S_50ntFlanks | 4683 | 4707 | 25 | CAAGACGGACCAGAGCGAAAGCATT | 565.5 | -3.3 | 0.6 | -5.3 | 1.2E-07 | 2.9E-07 | BT20/MB468 | MITO |
| >hg38_21_+_8433172_8446622_RNA45SN1_45S_50ntFlanks@12447.12484.38\|CAGAATTCACCAAGCGTTGGATTGTTCACCCACTAATA | >hg38_21_+_8433172_8446622_RNA45SN1_45S_50ntFlanks | 12447 | 12484 | 38 | CAGAATTCACCAAGCGTTGGATTGTTCACCCACTAATA | 256.8 | -3.6 | 0.7 | -5.2 | 1.6E-07 | 2.9E-07 | MB231/MB468 | NUC |
| >hg38_21_+_8433172_8446622_RNA45SN1_45S_50ntFlanks@12914.12935.22\|CGTTCGTGGGGAACCTGGCGCT | >hg38_21_+_8433172_8446622_RNA45SN1_45S_50ntFlanks | 12914 | 12935 | 22 | CGTTCGTGGGGAACCTGGCGCT | 213.9 | -7.1 | 1.4 | -5.2 | 1.7E-07 | 2.9E-07 | MB231/MB468 | MITO |

<table>
<tr><td></td><td></td><td></td><td></td><td></td><td></td><td></td><td></td><td></td><td></td><td></td><td></td><td></td><td></td><td></td></tr>
<tr><td>>MI0000449|hsa-mir-132&WithFlank&17|-|2049902|2050014@65.85.21(+1A)&[MIMAT0000426&hsa-miR-132-3p&offsets|0|-1(+1A);m-138&17|-|2049929|2049950&offsets|0|-1(+1A)]|TAACAGTCTACAGCCATGGTCA</td><td>>MI0000449|hsa-mir-132&WithFlank&17|-|2049902|2050014</td><td>65</td><td>85</td><td>21(+1A)</td><td>TAACAGTCTACAGCCATGGTCA</td><td>174.7</td><td>11.2</td><td>2.0</td><td>5.5</td><td>3.3E-08</td><td>2.9E-07</td><td>BT20/MB468</td><td>MP</td></tr>
<tr><td>>hg38_21_+_8433172_8446622_RNA45SN1_45S_50ntFlanks@12021.12055.35|ACCACTACTCTGATCGTTTTTTCACTGACCCGGTG</td><td>>hg38_21_+_8433172_8446622_RNA45SN1_45S_50ntFlanks</td><td>12021</td><td>12055</td><td>35</td><td>ACCACTACTCTGATCGTTTTTTCACTGACCCGGTG</td><td>70.3</td><td>9.9</td><td>1.8</td><td>5.5</td><td>3.3E-08</td><td>2.9E-07</td><td>BT20/MB468</td><td>MP</td></tr>
<tr><td>>hg38_21_+_8433172_8446622_RNA45SN1_45S_50ntFlanks@10680.10705.26|GTTCCGGCGGCGTCCGGTGAGCTCTC</td><td>>hg38_21_+_8433172_8446622_RNA45SN1_45S_50ntFlanks</td><td>10680</td><td>10705</td><td>26</td><td>GTTCCGGCGGCGTCCGGTGAGCTCTC</td><td>823.5</td><td>-3.4</td><td>0.6</td><td>-5.3</td><td>1.4E-07</td><td>2.9E-07</td><td>BT20/MB468</td><td>CYTO</td></tr>
<tr><td>>hg38_21_+_8433172_8446622_RNA45SN1_45S_50ntFlanks@4567.4587.21|TAATGGAATAGGACCGCGGTT</td><td>>hg38_21_+_8433172_8446622_RNA45SN1_45S_50ntFlanks</td><td>4567</td><td>4587</td><td>21</td><td>TAATGGAATAGGACCGCGGTT</td><td>585.4</td><td>-5.2</td><td>1.0</td><td>-5.2</td><td>1.6E-07</td><td>2.9E-07</td><td>MB231/MB468</td><td>CYTO</td></tr>
<tr><td>>hg38_21_+_8433172_8446622_RNA45SN1_45S_50ntFlanks@8380.8406.27|CAAGAGGGCGTGAAACCGTTAAGAGGT</td><td>>hg38_21_+_8433172_8446622_RNA45SN1_45S_50ntFlanks</td><td>8380</td><td>8406</td><td>27</td><td>CAAGAGGGCGTGAAACCGTTAAGAGGT</td><td>630.4</td><td>-4.1</td><td>0.8</td><td>-5.2</td><td>1.6E-07</td><td>2.9E-07</td><td>MB231/MB468</td><td>NUC</td></tr>
<tr><td>>hg38_21_+_8433172_8446622_RNA45SN1_45S_50ntFlanks@11718.11736.19|AACGGCGGGAGTAACTATG</td><td>>hg38_21_+_8433172_8446622_RNA45SN1_45S_50ntFlanks</td><td>11718</td><td>11736</td><td>19</td><td>AACGGCGGGAGTAACTATG</td><td>885.1</td><td>-8.1</td><td>1.5</td><td>-5.3</td><td>1.5E-07</td><td>2.9E-07</td><td>MB231/MB468</td><td>MP</td></tr>
<tr><td>>hg38_21_+_8433172_8446622_RNA45SN1_45S_50ntFlanks@10822.10840.19|GAAGTCGGCAAGCCGGATC</td><td>>hg38_21_+_8433172_8446622_RNA45SN1_45S_50ntFlanks</td><td>10822</td><td>10840</td><td>19</td><td>GAAGTCGGCAAGCCGGATC</td><td>596.2</td><td>-6.3</td><td>1.2</td><td>-5.3</td><td>1.5E-07</td><td>2.9E-07</td><td>MB231/MB468</td><td>MP</td></tr>
<tr><td>>hg38_21_+_8433172_8446622_RNA45SN1_45S_50ntFlanks@12674.12704.31|TCAGAATCCCGCCCAGGCGGAACGATACGGC</td><td>>hg38_21_+_8433172_8446622_RNA45SN1_45S_50ntFlanks</td><td>12674</td><td>12704</td><td>31</td><td>TCAGAATCCCGCCCAGGCGGAACGATACGGC</td><td>79.1</td><td>-9.4</td><td>1.7</td><td>-5.5</td><td>3.3E-08</td><td>2.9E-07</td><td>BT20/MB468</td><td>MP</td></tr>
<tr><td>>hg38_21_+_8433172_8446622_RNA45SN1_45S_50ntFlanks@4380.4406.27|CGTAGTTGGATCTTGGGAGCGGGCGGG</td><td>>hg38_21_+_8433172_8446622_RNA45SN1_45S_50ntFlanks</td><td>4380</td><td>4406</td><td>27</td><td>CGTAGTTGGATCTTGGGAGCGGGCGGG</td><td>235.4</td><td>-7.3</td><td>1.4</td><td>-5.3</td><td>1.5E-07</td><td>2.9E-07</td><td>MB231/MB468</td><td>MP</td></tr>
<tr><td>>hg38_21_+_8433172_8446622_RNA45SN1_45S_50ntFlanks@9156.9188.33|CCAGTGCGCCCCGGGCGGGTCGCGCCGTCGGGC</td><td>>hg38_21_+_8433172_8446622_RNA45SN1_45S_50ntFlanks</td><td>9156</td><td>9188</td><td>33</td><td>CCAGTGCGCCCCGGGCGGGTCGCGCCGTCGGGC</td><td>1526.6</td><td>14.6</td><td>2.8</td><td>5.2</td><td>1.6E-07</td><td>2.9E-07</td><td>MB231/MB468</td><td>CYTO</td></tr>
<tr><td>>hg38_21_+_8433172_8446622_RNA45SN1_45S_50ntFlanks@12936.12953.18|AAACCATTCGTAGACGAC</td><td>>hg38_21_+_8433172_8446622_RNA45SN1_45S_50ntFlanks</td><td>12936</td><td>12953</td><td>18</td><td>AAACCATTCGTAGACGAC</td><td>781.2</td><td>-5.2</td><td>1.0</td><td>-5.2</td><td>1.6E-07</td><td>2.9E-07</td><td>MB231/MB468</td><td>CYTO</td></tr>
<tr><td>>hg38_21_+_8433172_8446622_RNA45SN1_45S_50ntFlanks@4892.4918.27|AAACTTAAAGGAATTGACGGAAGGGCA</td><td>>hg38_21_+_8433172_8446622_RNA45SN1_45S_50ntFlanks</td><td>4892</td><td>4918</td><td>27</td><td>AAACTTAAAGGAATTGACGGAAGGGCA</td><td>102.2</td><td>-9.8</td><td>1.8</td><td>-5.5</td><td>3.3E-08</td><td>2.9E-07</td><td>BT20/MB468</td><td>MP</td></tr>
<tr><td>>hg38_21_+_8433172_8446622_RNA45SN1_45S_50ntFlanks@12457.12477.21|CAAGCGTTGGATTGTTCACCC</td><td>>hg38_21_+_8433172_8446622_RNA45SN1_45S_50ntFlanks</td><td>12457</td><td>12477</td><td>21</td><td>CAAGCGTTGGATTGTTCACCC</td><td>4613.7</td><td>-4.4</td><td>0.8</td><td>-5.3</td><td>1.4E-07</td><td>2.9E-07</td><td>BT20/MB468</td><td>CYTO</td></tr>
<tr><td>>hg38_21_+_8433172_8446622_RNA45SN1_45S_50ntFlanks@3783.3800.18|AGTGAAACTGCGAATGGC</td><td>>hg38_21_+_8433172_8446622_RNA45SN1_45S_50ntFlanks</td><td>3783</td><td>3800</td><td>18</td><td>AGTGAAACTGCGAATGGC</td><td>492.3</td><td>-6.5</td><td>1.2</td><td>-5.2</td><td>1.8E-07</td><td>2.9E-07</td><td>BT20/MB468</td><td>NUC</td></tr>
<tr><td>>hg38_21_+_8433172_8446622_RNA45SN1_45S_50ntFlanks@9570.9603.34|TGCAAATCGGTCGTCCGACCTGGGTATAGGGGCG</td><td>>hg38_21_+_8433172_8446622_RNA45SN1_45S_50ntFlanks</td><td>9570</td><td>9603</td><td>34</td><td>TGCAAATCGGTCGTCCGACCTGGGTATAGGGGCG</td><td>790.6</td><td>-5.0</td><td>0.9</td><td>-5.3</td><td>1.5E-07</td><td>2.9E-07</td><td>MB231/MB468</td><td>MP</td></tr>
<tr><td>>am_trnaMT_LysTTT_MT_+_8295_8364@51.69.19,trnalookalike7_LysTTT_1_+_568844_568913@51.69.19|CCAACACCTCTTTACAGTG</td><td>>am_trnaMT_LysTTT_MT_+_8295_8364</td><td>51</td><td>69</td><td>19</td><td>CCAACACCTCTTTACAGTG</td><td>95.3</td><td>-9.7</td><td>1.7</td><td>-5.5</td><td>3.4E-08</td><td>2.9E-07</td><td>BT20/MB468</td><td>MP</td></tr>
<tr><td>>hg38_MT_+_598_1651_RNR1_12S_50ntFlanks@746.766.21|ATGAGGTGGCAAGAAATGGGC</td><td>>hg38_MT_+_598_1651_RNR1_12S_50ntFlanks</td><td>746</td><td>766</td><td>21</td><td>ATGAGGTGGCAAGAAATGGGC</td><td>66.8</td><td>9.8</td><td>1.8</td><td>5.5</td><td>3.4E-08</td><td>2.9E-07</td><td>BT20/MB468</td><td>MP</td></tr>
<tr><td>>hg38_21_+_8433172_8446622_RNA45SN1_45S_50ntFlanks@11272.11295.24|GCCGCCCCCGGGGAGCCCGGCGGG</td><td>>hg38_21_+_8433172_8446622_RNA45SN1_45S_50ntFlanks</td><td>11272</td><td>11295</td><td>24</td><td>GCCGCCCCCGGGGAGCCCGGCGGG</td><td>103.7</td><td>-9.8</td><td>1.8</td><td>-5.5</td><td>3.4E-08</td><td>2.9E-07</td><td>BT20/MB468</td><td>MP</td></tr>
<tr><td>>hg38_21_+_8433172_8446622_RNA45SN1_45S_50ntFlanks@10461.10484.24|GCCTCCGTTGCCCTCGGCCGATCG</td><td>>hg38_21_+_8433172_8446622_RNA45SN1_45S_50ntFlanks</td><td>10461</td><td>10484</td><td>24</td><td>GCCTCCGTTGCCCTCGGCCGATCG</td><td>241.4</td><td>-5.9</td><td>1.1</td><td>-5.2</td><td>1.6E-07</td><td>2.9E-07</td><td>MB231/MB468</td><td>NUC</td></tr>
<tr><td>>am_trna130_GlnTTG_6_-_27763640_27763711@13.30.18,trna146_GlnCTG_6_-_27515531_27515602@13.30.18,trna16_GlnTTG_17_+_47269890_47269961@13.30.18,trna173_GlnTTG_6_-_26311975_26312046@13.30.18,trna174_GlnTTG_6_-_26311424_26311495@13.30.18,trna1_GlnCTG_6_+_18836402_18836473@13.30.18,trna3_GlnCTG_17_+_8023070_8023141@13.30.18,trna42_GlnCTG_6_+_27263212_27263283@13.30.18,trna49_GlnCTG_6_+_27487308_27487379@13.30.18,trna64_GlnTTG_6_+_28557156_28557227@13.30.18,trna7_GlnCTG_15_-_66161400_66161471@13.30.18,trna84_GlnTTG_6_+_145503859_145503930@13.30.18,trna99_GlnCTG_6_-_28909378_28909449@13.30.18|TAATGGTTAGCACTCTGG</td><td>>am_trna130_GlnTTG_6_-_27763640_27763711</td><td>13</td><td>30</td><td>18</td><td>TAATGGTTAGCACTCTGG</td><td>485.5</td><td>-8.2</td><td>1.6</td><td>-5.2</td><td>1.8E-07</td><td>2.9E-07</td><td>BT20/MB468</td><td>NUC</td></tr>
<tr><td>>hg38_21_+_8433172_8446622_RNA45SN1_45S_50ntFlanks@12417.12457.41|TCGATGTCGGCTCTTCCTATCATTGTGAAGCAGAATTCACC</td><td>>hg38_21_+_8433172_8446622_RNA45SN1_45S_50ntFlanks</td><td>12417</td><td>12457</td><td>41</td><td>TCGATGTCGGCTCTTCCTATCATTGTGAAGCAGAATTCACC</td><td>649.5</td><td>5.7</td><td>1.1</td><td>5.2</td><td>1.6E-07</td><td>2.9E-07</td><td>MB231/MB468</td><td>CYTO</td></tr>
<tr><td>>MI0000082|hsa-mir-25&WithFlank&7|-|100093554|100093649@58.78.21&[MIMAT0000081&hsa-miR-25-3p&offsets|0|-1;m-23&7|-|100093571|100093592&offsets|0|-1]|CATTGCACTTGTCTCGGTCTG</td><td>>MI0000082|hsa-mir-25&WithFlank&7|-|100093554|100093649</td><td>58</td><td>78</td><td>21</td><td>CATTGCACTTGTCTCGGTCTG</td><td>5280.0</td><td>3.7</td><td>0.7</td><td>5.3</td><td>1.5E-07</td><td>2.9E-07</td><td>MB231/MB468</td><td>MP</td></tr>
<tr><td>>hg38_21_+_8433172_8446622_RNA45SN1_45S_50ntFlanks@10402.10427.26|AACATGGGTCAGTCGGTCCTGAGAGA</td><td>>hg38_21_+_8433172_8446622_RNA45SN1_45S_50ntFlanks</td><td>10402</td><td>10427</td><td>26</td><td>AACATGGGTCAGTCGGTCCTGAGAGA</td><td>206.8</td><td>-7.0</td><td>1.3</td><td>-5.3</td><td>1.5E-07</td><td>2.9E-07</td><td>MB231/MB468</td><td>MP</td></tr>
<tr><td>>hg38_21_+_8433172_8446622_RNA45SN1_45S_50ntFlanks@11874.11899.26|AGAAGACCCTGTTGAGCTTGACTCTA</td><td>>hg38_21_+_8433172_8446622_RNA45SN1_45S_50ntFlanks</td><td>11874</td><td>11899</td><td>26</td><td>AGAAGACCCTGTTGAGCTTGACTCTA</td><td>457.6</td><td>-4.7</td><td>0.9</td><td>-5.3</td><td>1.4E-07</td><td>2.9E-07</td><td>BT20/MB468</td><td>CYTO</td></tr>
<tr><td>>hg38_21_+_8433172_8446622_RNA45SN1_45S_50ntFlanks@12994.13013.20|CCTCGCTGCGATCTATTGAA</td><td>>hg38_21_+_8433172_8446622_RNA45SN1_45S_50ntFlanks</td><td>12994</td><td>13013</td><td>20</td><td>CCTCGCTGCGATCTATTGAA</td><td>1055.7</td><td>-4.3</td><td>0.8</td><td>-5.3</td><td>1.4E-07</td><td>2.9E-07</td><td>BT20/MB468</td><td>CYTO</td></tr>
<tr><td>>hg38_21_+_8433172_8446622_RNA45SN1_45S_50ntFlanks@9941.9968.28|AGGACGGTGGCCATGGAAGTCGGAATCC</td><td>>hg38_21_+_8433172_8446622_RNA45SN1_45S_50ntFlanks</td><td>9941</td><td>9968</td><td>28</td><td>AGGACGGTGGCCATGGAAGTCGGAATCC</td><td>874.8</td><td>7.5</td><td>1.4</td><td>5.5</td><td>3.4E-08</td><td>2.9E-07</td><td>BT20/MB468</td><td>MP</td></tr>
<tr><td>>hg38_21_+_8433172_8446622_RNA45SN1_45S_50ntFlanks@10808.10844.37|CAATGTAGGTAAGGGAAGTCGGCAAGCCGGATCCGTA</td><td>>hg38_21_+_8433172_8446622_RNA45SN1_45S_50ntFlanks</td><td>10808</td><td>10844</td><td>37</td><td>CAATGTAGGTAAGGGAAGTCGGCAAGCCGGATCCGTA</td><td>367.2</td><td>-7.9</td><td>1.5</td><td>-5.2</td><td>1.8E-07</td><td>2.9E-07</td><td>BT20/MB468</td><td>NUC</td></tr>
<tr><td>>hg38_21_+_8433172_8446622_RNA45SN1_45S_50ntFlanks@10663.10709.47|CCTTGGAAAGCGTCGCGGTTCCGGCGGCGTCCGGTGAGCTCTCGCTG</td><td>>hg38_21_+_8433172_8446622_RNA45SN1_45S_50ntFlanks</td><td>10663</td><td>10709</td><td>47</td><td>CCTTGGAAAGCGTCGCGGTTCCGGCGGCGTCCGGTGAGCTCTCGCTG</td><td>92.8</td><td>-8.8</td><td>1.7</td><td>-5.3</td><td>1.4E-07</td><td>2.9E-07</td><td>BT20/MB468</td><td>CYTO</td></tr>
<tr><td>>hg38_21_+_8433172_8446622_RNA45SN1_45S_50ntFlanks@11208.11232.25|GCGGCGACTCTGGACGCGAGCCGGG</td><td>>hg38_21_+_8433172_8446622_RNA45SN1_45S_50ntFlanks</td><td>11208</td><td>11232</td><td>25</td><td>GCGGCGACTCTGGACGCGAGCCGGG</td><td>18028.7</td><td>-7.4</td><td>1.4</td><td>-5.2</td><td>1.6E-07</td><td>2.9E-07</td><td>MB231/MB468</td><td>CYTO</td></tr>
<tr><td>>hg38_21_+_8433172_8446622_RNA45SN1_45S_50ntFlanks@9572.9604.33|CAAATCGGTCGTCCGACCTGGGTATAGGGGCGA</td><td>>hg38_21_+_8433172_8446622_RNA45SN1_45S_50ntFlanks</td><td>9572</td><td>9604</td><td>33</td><td>CAAATCGGTCGTCCGACCTGGGTATAGGGGCGA</td><td>273.7</td><td>-6.1</td><td>1.2</td><td>-5.3</td><td>1.5E-07</td><td>2.9E-07</td><td>MB231/MB468</td><td>MP</td></tr>
<tr><td>>hg38_21_+_8433172_8446622_RNA45SN1_45S_50ntFlanks@10571.10589.19|GAGAAGCCGGCGGGAGCCC</td><td>>hg38_21_+_8433172_8446622_RNA45SN1_45S_50ntFlanks</td><td>10571</td><td>10589</td><td>19</td><td>GAGAAGCCGGCGGGAGCCC</td><td>204.3</td><td>-5.8</td><td>1.1</td><td>-5.2</td><td>1.6E-07</td><td>2.9E-07</td><td>MB231/MB468</td><td>CYTO</td></tr>
<tr><td>>hg38_21_+_8433172_8446622_RNA45SN1_45S_50ntFlanks@8839.8862.24|CGGTGGCGGCGGCGGCGGCGGCGG</td><td>>hg38_21_+_8433172_8446622_RNA45SN1_45S_50ntFlanks</td><td>8839</td><td>8862</td><td>24</td><td>CGGTGGCGGCGGCGGCGGCGGCGG</td><td>108.5</td><td>-9.8</td><td>1.8</td><td>-5.5</td><td>3.4E-08</td><td>2.9E-07</td><td>BT20/MB468</td><td>MP</td></tr>
<tr><td>>hg38_21_+_8433172_8446622_RNA45SN1_45S_50ntFlanks@12997.13039.43|CGCTGCGATCTATTGAAAGTCAGCCCTCGACACAAGGGTTTGT</td><td>>hg38_21_+_8433172_8446622_RNA45SN1_45S_50ntFlanks</td><td>12997</td><td>13039</td><td>43</td><td>CGCTGCGATCTATTGAAAGTCAGCCCTCGACACAAGGGTTTGT</td><td>154581.8</td><td>5.6</td><td>1.1</td><td>5.2</td><td>1.7E-07</td><td>2.9E-07</td><td>MB231/MB468</td><td>MITO</td></tr>
<tr><td>>hg38_21_+_8433172_8446622_RNA45SN1_45S_50ntFlanks@4498.4520.23|AAGCGTTTACTTTGAAAAAATTA</td><td>>hg38_21_+_8433172_8446622_RNA45SN1_45S_50ntFlanks</td><td>4498</td><td>4520</td><td>23</td><td>AAGCGTTTACTTTGAAAAAATTA</td><td>1351.0</td><td>4.7</td><td>0.9</td><td>5.3</td><td>1.4E-07</td><td>2.9E-07</td><td>BT20/MB468</td><td>CYTO</td></tr>
<tr><td>>hg38_21_+_8433172_8446622_RNA45SN1_45S_50ntFlanks@10128.10147.20|CGGCGGCGGCGGCGGCGGCG</td><td>>hg38_21_+_8433172_8446622_RNA45SN1_45S_50ntFlanks</td><td>10128</td><td>10147</td><td>20</td><td>CGGCGGCGGCGGCGGCGGCG</td><td>410.8</td><td>-7.8</td><td>1.5</td><td>-5.2</td><td>1.8E-07</td><td>2.9E-07</td><td>BT20/MB468</td><td>NUC</td></tr>
<tr><td>>hg38_21_+_8433172_8446622_RNA45SN1_45S_50ntFlanks@8845.8864.20|CGGCGGCGGCGGCGGCGGCG</td><td>>hg38_21_+_8433172_8446622_RNA45SN1_45S_50ntFlanks</td><td>8845</td><td>8864</td><td>20</td><td>CGGCGGCGGCGGCGGCGGCG</td><td>410.8</td><td>-7.8</td><td>1.5</td><td>-5.2</td><td>1.8E-07</td><td>2.9E-07</td><td>BT20/MB468</td><td>NUC</td></tr>
<tr><td>>hg38_21_+_8433172_8446622_RNA45SN1_45S_50ntFlanks@9540.9571.32|CTCTGGTGGAGGTCCGTAGCGGTCCTGACGTG</td><td>>hg38_21_+_8433172_8446622_RNA45SN1_45S_50ntFlanks</td><td>9540</td><td>9571</td><td>32</td><td>CTCTGGTGGAGGTCCGTAGCGGTCCTGACGTG</td><td>3886.7</td><td>-9.4</td><td>1.8</td><td>-5.2</td><td>1.8E-07</td><td>2.9E-07</td><td>BT20/MB468</td><td>NUC</td></tr>
<tr><td>>MI0000095|hsa-mir-93&WithFlank&7|-|100093762|100093853@17.39.23(+1C)&[MIMAT0000093&hsa-miR-93-5p&offsets|0|0(+1C);m-29&7|-|100093815|100093837&offsets|0|0(+1C)]|CAAAGTGCTGTTCGTGCAGGTAGC</td><td>>MI0000095|hsa-mir-93&WithFlank&7|-|100093762|100093853</td><td>17</td><td>39</td><td>23(+1C)</td><td>CAAAGTGCTGTTCGTGCAGGTAGC</td><td>439.3</td><td>5.6</td><td>1.1</td><td>5.2</td><td>1.8E-07</td><td>3.0E-07</td><td>MB231/MB468</td><td>MITO</td></tr>
<tr><td>>am_trna117_GlyTCC_1_-_145397864_145397935@1.32.32,trna45_GlyTCC_1_+_161500903_161500974@1.32.32,trna70_GlyTCC_1_-_161439547_161439618@1.32.32,trna73_GlyTCC_1_-_161432166_161432237@1.32.32,trna76_GlyTCC_1_-_161424756_161424827@1.32.32,trna79_GlyTCC_1_-_161417375_161417446@1.32.32|GCGTTGGTGGTATAGTGGTGAGCATAGCTGCC</td><td>>am_trna117_GlyTCC_1_-_145397864_145397935</td><td>1</td><td>32</td><td>32</td><td>GCGTTGGTGGTATAGTGGTGAGCATAGCTGCC</td><td>492.0</td><td>5.8</td><td>1.1</td><td>5.3</td><td>1.4E-07</td><td>3.0E-07</td><td>BT20/MB468</td><td>CYTO</td></tr>
<tr><td>>hg38_21_+_8433172_8446622_RNA45SN1_45S_50ntFlanks@11675.11707.33|TGCTCTGAATGTCAAAGTGAAGAAATTCAATGA</td><td>>hg38_21_+_8433172_8446622_RNA45SN1_45S_50ntFlanks</td><td>11675</td><td>11707</td><td>33</td><td>TGCTCTGAATGTCAAAGTGAAGAAATTCAATGA</td><td>483.7</td><td>4.5</td><td>0.8</td><td>5.3</td><td>1.4E-07</td><td>3.0E-07</td><td>BT20/MB468</td><td>CYTO</td></tr>
<tr><td>>MI0000542|hsa-mir-320a&WithFlank&8|-|22244960|22245043@50.72.23&[MIMAT0000510&hsa-miR-320a-3p&offsets|+2|+3;m-60&8|-|22244975|22244996&offsets|+2|+3]|AAGCTGGGTTGAGAGGGCGAAAA</td><td>>MI0000542|hsa-mir-320a&WithFlank&8|-|22244960|22245043</td><td>50</td><td>72</td><td>23</td><td>AAGCTGGGTTGAGAGGGCGAAAA</td><td>75.1</td><td>10.0</td><td>1.8</td><td>5.5</td><td>3.4E-08</td><td>3.0E-07</td><td>BT20/MB468</td><td>MP</td></tr>
<tr><td>>hg38_21_+_8433172_8446622_RNA45SN1_45S_50ntFlanks@6721.6747.27|AATTGCAGGACACATTGATCATCGACA</td><td>>hg38_21_+_8433172_8446622_RNA45SN1_45S_50ntFlanks</td><td>6721</td><td>6747</td><td>27</td><td>AATTGCAGGACACATTGATCATCGACA</td><td>1775.0</td><td>-3.2</td><td>0.6</td><td>-5.3</td><td>1.4E-07</td><td>3.0E-07</td><td>BT20/MB468</td><td>CYTO</td></tr>
<tr><td>>hg38_21_+_8433172_8446622_RNA45SN1_45S_50ntFlanks@4972.4994.23|CGGCCCGGACACGGACAGGATTG</td><td>>hg38_21_+_8433172_8446622_RNA45SN1_45S_50ntFlanks</td><td>4972</td><td>4994</td><td>23</td><td>CGGCCCGGACACGGACAGGATTG</td><td>338.8</td><td>-8.1</td><td>1.5</td><td>-5.2</td><td>1.8E-07</td><td>3.0E-07</td><td>MB231/MB468</td><td>MITO</td></tr>
<tr><td>>hg38_21_+_8433172_8446622_RNA45SN1_45S_50ntFlanks@9033.9059.27|CTCGTGGGGGGGCCGGGCCACCCCTCC</td><td>>hg38_21_+_8433172_8446622_RNA45SN1_45S_50ntFlanks</td><td>9033</td><td>9059</td><td>27</td><td>CTCGTGGGGGGGCCGGGCCACCCCTCC</td><td>1084.7</td><td>-7.4</td><td>1.4</td><td>-5.3</td><td>1.3E-07</td><td>3.0E-07</td><td>BT20/MB468</td><td>MITO</td></tr>
<tr><td>>trnaMT_GluTTC_MT_-_14674_14742@38.70.33,trnalookalike8_GluTTC_5_-_93905172_93905240@38.70.33|CATTGGTCGTGGTTGTAGTCCGTGCGAGAATAC</td><td>>trnaMT_GluTTC_MT_-_14674_14742</td><td>38</td><td>70</td><td>33</td><td>CATTGGTCGTGGTTGTAGTCCGTGCGAGAATAC</td><td>156.1</td><td>11.0</td><td>2.0</td><td>5.5</td><td>3.4E-08</td><td>3.0E-07</td><td>BT20/MB468</td><td>MP</td></tr>
</table>

<table>
<tr><td>>hg38_21_+_8433172_8446622_RNA45SN1_45S_50ntFlanks@8191.8213.23|CAGCCCGTGGACGGTGTGAGGCC</td><td>>hg38_21_+_8433172_8446622_RNA45SN1_45S_50ntFlanks</td><td>8191</td><td>8213</td><td>23</td><td>CAGCCCGTGGACGGTGTGAGGCC</td><td>1188.8</td><td>-6.2</td><td>1.2</td><td>-5.3</td><td>1.5E-07</td><td>3.0E-07</td><td>MB231/MB468</td><td>MP</td></tr>
<tr><td>>hg38_21_+_8433172_8446622_RNA45SN1_45S_50ntFlanks@10736.10755.20|GTAAATCTCGCGCCGGGCCG</td><td>>hg38_21_+_8433172_8446622_RNA45SN1_45S_50ntFlanks</td><td>10736</td><td>10755</td><td>20</td><td>GTAAATCTCGCGCCGGGCCG</td><td>668.1</td><td>-7.5</td><td>1.4</td><td>-5.2</td><td>1.7E-07</td><td>3.0E-07</td><td>MB231/MB468</td><td>NUC</td></tr>
<tr><td>>hg38_21_+_8433172_8446622_RNA45SN1_45S_50ntFlanks@11572.11592.21|CCGACTTAGAACTGGTGCGGA</td><td>>hg38_21_+_8433172_8446622_RNA45SN1_45S_50ntFlanks</td><td>11572</td><td>11592</td><td>21</td><td>CCGACTTAGAACTGGTGCGGA</td><td>4327.7</td><td>-7.7</td><td>1.4</td><td>-5.5</td><td>3.4E-08</td><td>3.0E-07</td><td>BT20/MB468</td><td>MP</td></tr>
<tr><td>>hg38_21_+_8433172_8446622_RNA45SN1_45S_50ntFlanks@8381.8404.24|AAGAGGGCGTGAAACCGTTAAGAG</td><td>>hg38_21_+_8433172_8446622_RNA45SN1_45S_50ntFlanks</td><td>8381</td><td>8404</td><td>24</td><td>AAGAGGGCGTGAAACCGTTAAGAG</td><td>259.2</td><td>-3.7</td><td>0.7</td><td>-5.2</td><td>1.6E-07</td><td>3.0E-07</td><td>MB231/MB468</td><td>CYTO</td></tr>
<tr><td>>hg38_21_+_8433172_8446622_RNA45SN1_45S_50ntFlanks@10239.10261.23|GCTACGCCGCGACGAGTAGGAGG</td><td>>hg38_21_+_8433172_8446622_RNA45SN1_45S_50ntFlanks</td><td>10239</td><td>10261</td><td>23</td><td>GCTACGCCGCGACGAGTAGGAGG</td><td>227.8</td><td>11.2</td><td>1.9</td><td>5.8</td><td>7.1E-09</td><td>3.0E-07</td><td>BT20/MB231</td><td>MP</td></tr>
<tr><td>>hg38_21_+_8433172_8446622_RNA45SN1_45S_50ntFlanks@12407.12426.20|TTTTGATCCTTCGATGTCGG</td><td>>hg38_21_+_8433172_8446622_RNA45SN1_45S_50ntFlanks</td><td>12407</td><td>12426</td><td>20</td><td>TTTTGATCCTTCGATGTCGG</td><td>490.4</td><td>-6.6</td><td>1.3</td><td>-5.3</td><td>1.4E-07</td><td>3.0E-07</td><td>BT20/MB468</td><td>CYTO</td></tr>
<tr><td>>hg38_21_+_8433172_8446622_RNA45SN1_45S_50ntFlanks@10393.10419.27|CAGCAGTTGAACATGGGTCAGTCGGTC</td><td>>hg38_21_+_8433172_8446622_RNA45SN1_45S_50ntFlanks</td><td>10393</td><td>10419</td><td>27</td><td>CAGCAGTTGAACATGGGTCAGTCGGTC</td><td>258.2</td><td>-7.0</td><td>1.3</td><td>-5.2</td><td>1.8E-07</td><td>3.0E-07</td><td>MB231/MB468</td><td>MITO</td></tr>
<tr><td>>trnaMT_SerGCT_MT_+_12207_12265@1.18.18|GAGAAAGCTCACAAGAAC</td><td>>trnaMT_SerGCT_MT_+_12207_12265</td><td>1</td><td>18</td><td>18</td><td>GAGAAAGCTCACAAGAAC</td><td>1276.0</td><td>7.6</td><td>1.5</td><td>5.2</td><td>1.6E-07</td><td>3.0E-07</td><td>MB231/MB468</td><td>CYTO</td></tr>
<tr><td>>hg38_21_+_8433172_8446622_RNA45SN1_45S_50ntFlanks@11876.11897.22|AAGACCCTGTTGAGCTTGACTC</td><td>>hg38_21_+_8433172_8446622_RNA45SN1_45S_50ntFlanks</td><td>11876</td><td>11897</td><td>22</td><td>AAGACCCTGTTGAGCTTGACTC</td><td>1128.3</td><td>-6.6</td><td>1.3</td><td>-5.2</td><td>1.9E-07</td><td>3.0E-07</td><td>BT20/MB468</td><td>NUC</td></tr>
<tr><td>>hg38_21_+_8433172_8446622_RNA45SN1_45S_50ntFlanks@4725.4759.35|TTAATCAAGAACGAAAGTCGGAGGTTCGAAGACGA</td><td>>hg38_21_+_8433172_8446622_RNA45SN1_45S_50ntFlanks</td><td>4725</td><td>4759</td><td>35</td><td>TTAATCAAGAACGAAAGTCGGAGGTTCGAAGACGA</td><td>407.5</td><td>-7.5</td><td>1.4</td><td>-5.3</td><td>1.5E-07</td><td>3.0E-07</td><td>MB231/MB468</td><td>MP</td></tr>
<tr><td>>hg38_MT_+_598_1651_RNR1_12S_50ntFlanks@52.88.37|ATAGGTTTGGTCCTAGCCTTTCTATTAGCTCTTAGTA</td><td>>hg38_MT_+_598_1651_RNR1_12S_50ntFlanks</td><td>52</td><td>88</td><td>37</td><td>ATAGGTTTGGTCCTAGCCTTTCTATTAGCTCTTAGTA</td><td>790.5</td><td>3.7</td><td>0.7</td><td>5.2</td><td>1.7E-07</td><td>3.0E-07</td><td>MB231/MB468</td><td>NUC</td></tr>
<tr><td>>MI0000746|hsa-mir-99b&WithFlank&19|+|51692606|51692687@13.32.20&[MIMAT0000689&hsa-miR-99b-5p&offsets|0|-2;m-10&19|+|51692618|51692639&offsets|0|-2]|CACCCGTAGAACCGACCTTG</td><td>>MI0000746|hsa-mir-99b&WithFlank&19|+|51692606|51692687</td><td>13</td><td>32</td><td>20</td><td>CACCCGTAGAACCGACCTTG</td><td>694.1</td><td>4.6</td><td>0.9</td><td>5.2</td><td>1.7E-07</td><td>3.0E-07</td><td>MB231/MB468</td><td>NUC</td></tr>
<tr><td>>hg38_21_+_8433172_8446622_RNA45SN1_45S_50ntFlanks@9807.9830.24|GTGGAATGCGAGTGCCTAGTGGGC</td><td>>hg38_21_+_8433172_8446622_RNA45SN1_45S_50ntFlanks</td><td>9807</td><td>9830</td><td>24</td><td>GTGGAATGCGAGTGCCTAGTGGGC</td><td>402.4</td><td>12.0</td><td>2.1</td><td>5.8</td><td>7.2E-09</td><td>3.0E-07</td><td>BT20/MB231</td><td>MP</td></tr>
<tr><td>>MI0000101|hsa-mir-99a&WithFlank&21|+|16539083|16539175@19.39.21&[MIMAT0000097&hsa-miR-99a-5p&offsets|0|-1;m-64&21|+|16539101|16539121&offsets|0|0]|AACCCGTAGATCCGATCTTGT</td><td>>MI0000101|hsa-mir-99a&WithFlank&21|+|16539083|16539175</td><td>19</td><td>39</td><td>21</td><td>AACCCGTAGATCCGATCTTGT</td><td>1527.8</td><td>-5.6</td><td>1.0</td><td>-5.8</td><td>7.2E-09</td><td>3.0E-07</td><td>BT20/MB231</td><td>MP</td></tr>
<tr><td>>hg38_21_+_8433172_8446622_RNA45SN1_45S_50ntFlanks@11212.11254.43|CGACTCTGGACGCGAGCCGGGCCCTTCCCGTGGATCGCCCCAG</td><td>>hg38_21_+_8433172_8446622_RNA45SN1_45S_50ntFlanks</td><td>11212</td><td>11254</td><td>43</td><td>CGACTCTGGACGCGAGCCGGGCCCTTCCCGTGGATCGCCCCAG</td><td>75.8</td><td>-9.3</td><td>1.7</td><td>-5.5</td><td>3.5E-08</td><td>3.0E-07</td><td>BT20/MB468</td><td>MP</td></tr>
<tr><td>>hg38_MT_+_598_1651_RNR1_12S_50ntFlanks@850.872.23|GTGCTTAGTTGAACAGGGCCCTG</td><td>>hg38_MT_+_598_1651_RNR1_12S_50ntFlanks</td><td>850</td><td>872</td><td>23</td><td>GTGCTTAGTTGAACAGGGCCCTG</td><td>92.4</td><td>10.3</td><td>1.9</td><td>5.5</td><td>3.5E-08</td><td>3.0E-07</td><td>BT20/MB468</td><td>MP</td></tr>
<tr><td>>hg38_21_+_8433172_8446622_RNA45SN1_45S_50ntFlanks@12896.12932.37|CGCCCGTCACGCACCGCACGTTCGTGGGGAACCTGGC</td><td>>hg38_21_+_8433172_8446622_RNA45SN1_45S_50ntFlanks</td><td>12896</td><td>12932</td><td>37</td><td>CGCCCGTCACGCACCGCACGTTCGTGGGGAACCTGGC</td><td>467.9</td><td>-4.4</td><td>0.8</td><td>-5.2</td><td>1.8E-07</td><td>3.0E-07</td><td>MB231/MB468</td><td>MITO</td></tr>
<tr><td>>trna116_GluCTC_1_-_145399233_145399304@32.50.19,trna59_GluCTC_1_+_249168447_249168518@32.50.19,trna71_GluCTC_1_-_161439189_161439260@32.50.19,trna74_GluCTC_1_-_161431809_161431880@32.50.19,trna77_GluCTC_1_-_161424398_161424469@32.50.19,trna77_GluCTC_6_+_28949976_28950047@32.50.19,trna80_GluCTC_1_-_161417018_161417089@32.50.19,trna87_GluCTC_6_-_126101393_126101464@32.50.19|CTCTCACCGCCGCGGCCCG</td><td>>trna116_GluCTC_1_-_145399233_145399304</td><td>32</td><td>50</td><td>19</td><td>CTCTCACCGCCGCGGCCCG</td><td>613.6</td><td>-9.4</td><td>1.8</td><td>-5.2</td><td>1.9E-07</td><td>3.0E-07</td><td>BT20/MB468</td><td>NUC</td></tr>
<tr><td>>hg38_21_+_8433172_8446622_RNA45SN1_45S_50ntFlanks@11217.11246.30|CTGGACGCGAGCCGGGCCCTTCCCGTGGAT</td><td>>hg38_21_+_8433172_8446622_RNA45SN1_45S_50ntFlanks</td><td>11217</td><td>11246</td><td>30</td><td>CTGGACGCGAGCCGGGCCCTTCCCGTGGAT</td><td>254.5</td><td>3.3</td><td>0.6</td><td>5.2</td><td>1.7E-07</td><td>3.0E-07</td><td>MB231/MB468</td><td>NUC</td></tr>
<tr><td>>trnaMT_GluTTC_MT_-_14674_14742@26.67.42,trnalookalike8_GluTTC_5_-_93905172_93905240@26.67.42|GGTTTTTCATATCATTGGTCGTGGTTGTAGTCCGTGCGAGAA</td><td>>trnaMT_GluTTC_MT_-_14674_14742</td><td>26</td><td>67</td><td>42</td><td>GGTTTTTCATATCATTGGTCGTGGTTGTAGTCCGTGCGAGAA</td><td>127.3</td><td>10.7</td><td>1.9</td><td>5.5</td><td>3.5E-08</td><td>3.0E-07</td><td>BT20/MB468</td><td>MP</td></tr>
<tr><td>>trna134_GluTTC_1_-_16861774_16861845@1.33.33,trna5_GluTTC_1_+_17199078_17199149@1.33.33,trna84_GluTTC_1_-_161391883_161391954@1.33.33,trna94_GluTTC_1_-_149664355_149664427@1.33.33|TCCCTGGTGGTCTAGTGGCTAGGATTCGGCGCT</td><td>>trna134_GluTTC_1_-_16861774_16861845</td><td>1</td><td>33</td><td>33</td><td>TCCCTGGTGGTCTAGTGGCTAGGATTCGGCGCT</td><td>1031.8</td><td>4.5</td><td>0.8</td><td>5.7</td><td>1.4E-08</td><td>3.0E-07</td><td>BT20/MB231</td><td>MITO</td></tr>
<tr><td>>hg38_21_+_8433172_8446622_RNA45SN1_45S_50ntFlanks@8175.8216.42|CCTTCTGATCGAGGCCCAGCCCGTGGACGGTGTGAGGCCGGT</td><td>>hg38_21_+_8433172_8446622_RNA45SN1_45S_50ntFlanks</td><td>8175</td><td>8216</td><td>42</td><td>CCTTCTGATCGAGGCCCAGCCCGTGGACGGTGTGAGGCCGGT</td><td>976.8</td><td>6.2</td><td>1.2</td><td>5.3</td><td>1.3E-07</td><td>3.0E-07</td><td>BT20/MB468</td><td>MITO</td></tr>
<tr><td>>hg38_21_+_8433172_8446622_RNA45SN1_45S_50ntFlanks@11569.11599.31|CAGCCGACTTAGAACTGGTGCGGACCAGGGG</td><td>>hg38_21_+_8433172_8446622_RNA45SN1_45S_50ntFlanks</td><td>11569</td><td>11599</td><td>31</td><td>CAGCCGACTTAGAACTGGTGCGGACCAGGGG</td><td>409.8</td><td>-2.9</td><td>0.5</td><td>-5.3</td><td>1.5E-07</td><td>3.0E-07</td><td>BT20/MB468</td><td>CYTO</td></tr>
<tr><td>>MI0000760|hsa-mir-361&WithFlank&X|-|85903630|85903713@52.75.24&[MIMAT0004682&hsa-miR-361-3p&offsets|+1|+2;m-124&X|-|85903642|85903663&offsets|+1|+3]|CCCCCAGGTGTGATTCTGATTTGC</td><td>>MI0000760|hsa-mir-361&WithFlank&X|-|85903630|85903713</td><td>52</td><td>75</td><td>24</td><td>CCCCCAGGTGTGATTCTGATTTGC</td><td>201.4</td><td>4.6</td><td>0.9</td><td>5.2</td><td>1.8E-07</td><td>3.0E-07</td><td>MB231/MB468</td><td>MITO</td></tr>
<tr><td>>hg38_21_+_8433172_8446622_RNA45SN1_45S_50ntFlanks@7983.8008.26|CAGATCAGACGTGGCGACCCGCTGAA</td><td>>hg38_21_+_8433172_8446622_RNA45SN1_45S_50ntFlanks</td><td>7983</td><td>8008</td><td>26</td><td>CAGATCAGACGTGGCGACCCGCTGAA</td><td>480.7</td><td>-3.4</td><td>0.7</td><td>-5.2</td><td>1.7E-07</td><td>3.0E-07</td><td>MB231/MB468</td><td>CYTO</td></tr>
<tr><td>>MI0000081|hsa-mir-24-2&WithFlank&19|-|13836281|13836365@56.77.22(+1A);MI0000080|hsa-mir-24-1&WithFlank&9|+|95086015|95086094@50.71.22(+1A)&[MIMAT0000080&hsa-miR-24-3p&offsets|0|0(+1A);m-36&19|-|13836288|13836310&offsets|0|-1(+1A)];[MIMAT0000080_1&hsa-miR-24-3p&offsets|0|0(+1A);m-35&9|+|95086064|95086086&offsets|0|-1(+1A)]|TGGCTCAGTTCAGCAGGAACAGA</td><td>>MI0000081|hsa-mir-24-2&WithFlank&19|-|13836281|13836365</td><td>56</td><td>77</td><td>22(+1A)</td><td>TGGCTCAGTTCAGCAGGAACAGA</td><td>255.5</td><td>2.2</td><td>0.4</td><td>5.2</td><td>1.9E-07</td><td>3.0E-07</td><td>BT20/MB468</td><td>NUC</td></tr>
<tr><td>>hg38_21_+_8433172_8446622_RNA45SN1_45S_50ntFlanks@3910.3935.26|GGGATGCGTGCATTTATCAGATCAAA</td><td>>hg38_21_+_8433172_8446622_RNA45SN1_45S_50ntFlanks</td><td>3910</td><td>3935</td><td>26</td><td>GGGATGCGTGCATTTATCAGATCAAA</td><td>66.7</td><td>-9.9</td><td>1.9</td><td>-5.3</td><td>1.5E-07</td><td>3.0E-07</td><td>MB231/MB468</td><td>MP</td></tr>
<tr><td>>hg38_21_+_8433172_8446622_RNA45SN1_45S_50ntFlanks@6653.6686.34|ACTCTTAGCGGTGGATCACTCGGCTCGTGCGTCG</td><td>>hg38_21_+_8433172_8446622_RNA45SN1_45S_50ntFlanks</td><td>6653</td><td>6686</td><td>34</td><td>ACTCTTAGCGGTGGATCACTCGGCTCGTGCGTCG</td><td>699.1</td><td>-3.0</td><td>0.6</td><td>-5.2</td><td>1.9E-07</td><td>3.0E-07</td><td>BT20/MB468</td><td>NUC</td></tr>
<tr><td>>hg38_21_+_8433172_8446622_RNA45SN1_45S_50ntFlanks@11564.11603.40|CCTAGCAGCCGACTTAGAACTGGTGCGGACCAGGGGAATC</td><td>>hg38_21_+_8433172_8446622_RNA45SN1_45S_50ntFlanks</td><td>11564</td><td>11603</td><td>40</td><td>CCTAGCAGCCGACTTAGAACTGGTGCGGACCAGGGGAATC</td><td>887.3</td><td>-4.2</td><td>0.8</td><td>-5.2</td><td>1.7E-07</td><td>3.0E-07</td><td>MB231/MB468</td><td>CYTO</td></tr>
<tr><td>>hg38_21_+_8433172_8446622_RNA45SN1_45S_50ntFlanks@12627.12652.26|ATGGGGCGAAGCTACCATCTGTGGGA</td><td>>hg38_21_+_8433172_8446622_RNA45SN1_45S_50ntFlanks</td><td>12627</td><td>12652</td><td>26</td><td>ATGGGGCGAAGCTACCATCTGTGGGA</td><td>371.5</td><td>-5.2</td><td>1.0</td><td>-5.3</td><td>1.3E-07</td><td>3.0E-07</td><td>BT20/MB468</td><td>MITO</td></tr>
<tr><td>>hg38_21_+_8433172_8446622_RNA45SN1_45S_50ntFlanks@8176.8200.25|CTTCTGATCGAGGCCCAGCCCGTGG</td><td>>hg38_21_+_8433172_8446622_RNA45SN1_45S_50ntFlanks</td><td>8176</td><td>8200</td><td>25</td><td>CTTCTGATCGAGGCCCAGCCCGTGG</td><td>110.9</td><td>-9.9</td><td>1.8</td><td>-5.5</td><td>3.5E-08</td><td>3.0E-07</td><td>BT20/MB468</td><td>MP</td></tr>
<tr><td>>hg38_21_+_8433172_8446622_RNA45SN1_45S_50ntFlanks@6719.6758.40|TGAATTGCAGGACACATTGATCATCGACACTTCGAACGCA</td><td>>hg38_21_+_8433172_8446622_RNA45SN1_45S_50ntFlanks</td><td>6719</td><td>6758</td><td>40</td><td>TGAATTGCAGGACACATTGATCATCGACACTTCGAACGCA</td><td>365.0</td><td>-5.1</td><td>1.0</td><td>-5.2</td><td>1.7E-07</td><td>3.0E-07</td><td>MB231/MB468</td><td>NUC</td></tr>
<tr><td>>MI0000477|hsa-mir-146a&WithFlank&5|+|160485346|160485456@27.45.19(+3U)&[MIMAT0000449&hsa-miR-146a-5p&offsets|0|-3(+3U);m-129&5|+|160485372|160485393&offsets|0|-3(+3U)]|TGAGAACTGAATTCCATGGTTT</td><td>>MI0000477|hsa-mir-146a&WithFlank&5|+|160485346|160485456</td><td>27</td><td>45</td><td>19(+3U)</td><td>TGAGAACTGAATTCCATGGTTT</td><td>96.8</td><td>9.8</td><td>1.9</td><td>5.2</td><td>1.5E-07</td><td>3.0E-07</td><td>MB231/MB468</td><td>MP</td></tr>
<tr><td>>hg38_21_+_8433172_8446622_RNA45SN1_45S_50ntFlanks@10840.10868.29|CCGTAACTTCGGGATAAGGATTGGCTCTA</td><td>>hg38_21_+_8433172_8446622_RNA45SN1_45S_50ntFlanks</td><td>10840</td><td>10868</td><td>29</td><td>CCGTAACTTCGGGATAAGGATTGGCTCTA</td><td>667.0</td><td>-4.1</td><td>0.8</td><td>-5.2</td><td>1.7E-07</td><td>3.0E-07</td><td>MB231/MB468</td><td>CYTO</td></tr>
<tr><td>>hg38_21_+_8433172_8446622_RNA45SN1_45S_50ntFlanks@10681.10710.30|TTCCGGCGGCGTCCGGTGAGCTCTCGCTGG</td><td>>hg38_21_+_8433172_8446622_RNA45SN1_45S_50ntFlanks</td><td>10681</td><td>10710</td><td>30</td><td>TTCCGGCGGCGTCCGGTGAGCTCTCGCTGG</td><td>2589.0</td><td>-9.3</td><td>1.8</td><td>-5.2</td><td>1.9E-07</td><td>3.0E-07</td><td>BT20/MB468</td><td>NUC</td></tr>
<tr><td>>hg38_21_+_8433172_8446622_RNA45SN1_45S_50ntFlanks@6651.6690.40|CGACTCTTAGCGGTGGATCACTCGGCTCGTGCGTCGATGA</td><td>>hg38_21_+_8433172_8446622_RNA45SN1_45S_50ntFlanks</td><td>6651</td><td>6690</td><td>40</td><td>CGACTCTTAGCGGTGGATCACTCGGCTCGTGCGTCGATGA</td><td>638.0</td><td>3.3</td><td>0.6</td><td>5.2</td><td>1.7E-07</td><td>3.0E-07</td><td>MB231/MB468</td><td>CYTO</td></tr>
<tr><td>>hg38_21_+_8433172_8446622_RNA45SN1_45S_50ntFlanks@6697.6720.24|CAGCTAGCTGCGAGAATTAATGTG</td><td>>hg38_21_+_8433172_8446622_RNA45SN1_45S_50ntFlanks</td><td>6697</td><td>6720</td><td>24</td><td>CAGCTAGCTGCGAGAATTAATGTG</td><td>1526.1</td><td>-4.3</td><td>0.8</td><td>-5.2</td><td>1.8E-07</td><td>3.0E-07</td><td>MB231/MB468</td><td>MITO</td></tr>
<tr><td>>hg38_21_+_8433172_8446622_RNA45SN1_45S_50ntFlanks@11564.11588.25|CCTAGCAGCCGACTTAGAACTGGTG</td><td>>hg38_21_+_8433172_8446622_RNA45SN1_45S_50ntFlanks</td><td>11564</td><td>11588</td><td>25</td><td>CCTAGCAGCCGACTTAGAACTGGTG</td><td>1502.9</td><td>-4.0</td><td>0.7</td><td>-5.5</td><td>3.5E-08</td><td>3.0E-07</td><td>BT20/MB468</td><td>MP</td></tr>
<tr><td>>hg38_21_+_8433172_8446622_RNA45SN1_45S_50ntFlanks@12489.12509.21|ACGTGAGCTGGGTTTAGACCG</td><td>>hg38_21_+_8433172_8446622_RNA45SN1_45S_50ntFlanks</td><td>12489</td><td>12509</td><td>21</td><td>ACGTGAGCTGGGTTTAGACCG</td><td>669.4</td><td>-5.9</td><td>1.1</td><td>-5.2</td><td>1.7E-07</td><td>3.1E-07</td><td>MB231/MB468</td><td>NUC</td></tr>
<tr><td>>MI0000085|hsa-mir-27a&WithFlank&19|-|13836434|13836523@57.75.19(+1A)&[MIMAT0000084&hsa-miR-27a-3p&offsets|0|-2(+1A);m-43&19|-|13836447|13836467&offsets|0|-2(+1A)]|TTCACAGTGGCTAAGTTCCA</td><td>>MI0000085|hsa-mir-27a&WithFlank&19|-|13836434|13836523</td><td>57</td><td>75</td><td>19(+1A)</td><td>TTCACAGTGGCTAAGTTCCA</td><td>880.7</td><td>4.3</td><td>0.8</td><td>5.3</td><td>1.3E-07</td><td>3.1E-07</td><td>BT20/MB468</td><td>MITO</td></tr>
<tr><td>>hg38_21_+_8433172_8446622_RNA45SN1_45S_50ntFlanks@6773.6799.27|TTCCTCCCGGGGCTACGCCTGTCTGAG</td><td>>hg38_21_+_8433172_8446622_RNA45SN1_45S_50ntFlanks</td><td>6773</td><td>6799</td><td>27</td><td>TTCCTCCCGGGGCTACGCCTGTCTGAG</td><td>680.1</td><td>-5.5</td><td>1.0</td><td>-5.3</td><td>1.3E-07</td><td>3.1E-07</td><td>BT20/MB468</td><td>MITO</td></tr>
<tr><td>>hg38_21_+_8433172_8446622_RNA45SN1_45S_50ntFlanks@10351.10381.31|CGAGAACTTTGAAGGCCGAAGTGGAGAAGGG</td><td>>hg38_21_+_8433172_8446622_RNA45SN1_45S_50ntFlanks</td><td>10351</td><td>10381</td><td>31</td><td>CGAGAACTTTGAAGGCCGAAGTGGAGAAGGG</td><td>668.2</td><td>-4.9</td><td>0.9</td><td>-5.2</td><td>1.7E-07</td><td>3.1E-07</td><td>MB231/MB468</td><td>CYTO</td></tr>
<tr><td>>hg38_21_+_8433172_8446622_RNA45SN1_45S_50ntFlanks@12407.12438.32|TTTTGATCCTTCGATGTCGGCTCTTCCTATCA</td><td>>hg38_21_+_8433172_8446622_RNA45SN1_45S_50ntFlanks</td><td>12407</td><td>12438</td><td>32</td><td>TTTTGATCCTTCGATGTCGGCTCTTCCTATCA</td><td>713.9</td><td>-4.2</td><td>0.8</td><td>-5.2</td><td>1.9E-07</td><td>3.1E-07</td><td>BT20/MB468</td><td>NUC</td></tr>
<tr><td>>hg38_21_+_8433172_8446622_RNA45SN1_45S_50ntFlanks@11552.11574.23|CCTCGGCCGGCGCCTAGCAGCCG</td><td>>hg38_21_+_8433172_8446622_RNA45SN1_45S_50ntFlanks</td><td>11552</td><td>11574</td><td>23</td><td>CCTCGGCCGGCGCCTAGCAGCCG</td><td>467.8</td><td>-5.9</td><td>1.1</td><td>-5.2</td><td>1.8E-07</td><td>3.1E-07</td><td>MB231/MB468</td><td>MITO</td></tr>
</table>

<table>
<tr><th></th><th></th><th></th><th></th><th></th><th></th><th></th><th></th><th></th><th></th><th></th><th></th><th></th><th></th></tr>
<tr><td>>hg38_21_+_8433172_8446622_RNA45SN1_45S_50ntFlanks@3802.3834.33|CATTAAATCAGTTATGGTTCCTTTGGTCGCTCG</td><td>>hg38_21_+_8433172_8446622_RNA45SN1_45S_50ntFlanks</td><td>3802</td><td>3834</td><td>33</td><td>CATTAAATCAGTTATGGTTCCTTTGGTCGCTCG</td><td>369.4</td><td>-6.9</td><td>1.3</td><td>-5.3</td><td>1.3E-07</td><td>3.1E-07</td><td>BT20/MB468</td><td>MITO</td></tr>
<tr><td>>hg38_21_+_8433172_8446622_RNA45SN1_45S_50ntFlanks@3780.3801.22|TACAGTGAAACTGCGAATGGCT</td><td>>hg38_21_+_8433172_8446622_RNA45SN1_45S_50ntFlanks</td><td>3780</td><td>3801</td><td>22</td><td>TACAGTGAAACTGCGAATGGCT</td><td>211.0</td><td>-3.1</td><td>0.6</td><td>-5.2</td><td>1.7E-07</td><td>3.1E-07</td><td>MB231/MB468</td><td>CYTO</td></tr>
<tr><td>>hg38_21_+_8433172_8446622_RNA45SN1_45S_50ntFlanks@11562.11602.41|CGCCTAGCAGCCGACTTAGAACTGGTGCGGACCAGGGGAAT</td><td>>hg38_21_+_8433172_8446622_RNA45SN1_45S_50ntFlanks</td><td>11562</td><td>11602</td><td>41</td><td>CGCCTAGCAGCCGACTTAGAACTGGTGCGGACCAGGGGAAT</td><td>459.2</td><td>-8.0</td><td>1.5</td><td>-5.2</td><td>1.9E-07</td><td>3.1E-07</td><td>BT20/MB468</td><td>NUC</td></tr>
<tr><td>>hg38_21_+_8433172_8446622_RNA45SN1_45S_50ntFlanks@479.505.27|GTGATCGATGTGGTGACGTCGTGCTCT</td><td>>hg38_21_+_8433172_8446622_RNA45SN1_45S_50ntFlanks</td><td>479</td><td>505</td><td>27</td><td>GTGATCGATGTGGTGACGTCGTGCTCT</td><td>220.5</td><td>7.6</td><td>1.4</td><td>5.2</td><td>1.7E-07</td><td>3.1E-07</td><td>MB231/MB468</td><td>NUC</td></tr>
<tr><td>>hg38_21_+_8433172_8446622_RNA45SN1_45S_50ntFlanks@10590.10607.18|CGGGGAGAGTTCTCTTTT</td><td>>hg38_21_+_8433172_8446622_RNA45SN1_45S_50ntFlanks</td><td>10590</td><td>10607</td><td>18</td><td>CGGGGAGAGTTCTCTTTT</td><td>109.6</td><td>-9.9</td><td>1.8</td><td>-5.5</td><td>3.6E-08</td><td>3.1E-07</td><td>BT20/MB468</td><td>MP</td></tr>
<tr><td>>hg38_21_+_8433172_8446622_RNA45SN1_45S_50ntFlanks@12578.12601.24|AGGAACCGCAGGTTCAGACATTTG</td><td>>hg38_21_+_8433172_8446622_RNA45SN1_45S_50ntFlanks</td><td>12578</td><td>12601</td><td>24</td><td>AGGAACCGCAGGTTCAGACATTTG</td><td>71.2</td><td>-9.2</td><td>1.7</td><td>-5.5</td><td>3.6E-08</td><td>3.1E-07</td><td>BT20/MB468</td><td>MP</td></tr>
<tr><td>>MI0000298|hsa-mir-221&WithFlank&X|-|45746151|45746272@71.93.23(+1C)&[MIMAT0000278&hsa-miR-221-3p&offsets|0|0(+1C);m-91&X|-|45746181|45746202&offsets|0|+1(+1C)]|AGCTACATTGTCTGCTGGGTTTCC</td><td>>MI0000298|hsa-mir-221&WithFlank&X|-|45746151|45746272</td><td>71</td><td>93</td><td>23(+1C)</td><td>AGCTACATTGTCTGCTGGGTTTCC</td><td>1172.5</td><td>4.2</td><td>0.8</td><td>5.2</td><td>1.5E-07</td><td>3.1E-07</td><td>MB231/MB468</td><td>MP</td></tr>
<tr><td>>hg38_21_+_8433172_8446622_RNA45SN1_45S_50ntFlanks@11670.11701.32|CCCAGTGCTCTGAATGTCAAAGTGAAGAAATT</td><td>>hg38_21_+_8433172_8446622_RNA45SN1_45S_50ntFlanks</td><td>11670</td><td>11701</td><td>32</td><td>CCCAGTGCTCTGAATGTCAAAGTGAAGAAATT</td><td>430.4</td><td>-2.9</td><td>0.5</td><td>-5.3</td><td>1.5E-07</td><td>3.1E-07</td><td>BT20/MB468</td><td>CYTO</td></tr>
<tr><td>>hg38_21_+_8433172_8446622_RNA45SN1_45S_50ntFlanks@10351.10376.26|CGAGAACTTTGAAGGCCGAAGTGGAG</td><td>>hg38_21_+_8433172_8446622_RNA45SN1_45S_50ntFlanks</td><td>10351</td><td>10376</td><td>26</td><td>CGAGAACTTTGAAGGCCGAAGTGGAG</td><td>845.2</td><td>-4.4</td><td>0.8</td><td>-5.3</td><td>1.3E-07</td><td>3.1E-07</td><td>BT20/MB468</td><td>MITO</td></tr>
<tr><td>>hg38_21_+_8433172_8446622_RNA45SN1_45S_50ntFlanks@5485.5514.30|AGCGCTGAGAAGACGGTCGAACTTGACTAT</td><td>>hg38_21_+_8433172_8446622_RNA45SN1_45S_50ntFlanks</td><td>5485</td><td>5514</td><td>30</td><td>AGCGCTGAGAAGACGGTCGAACTTGACTAT</td><td>984.5</td><td>13.3</td><td>2.3</td><td>5.8</td><td>7.5E-09</td><td>3.1E-07</td><td>BT20/MB231</td><td>MP</td></tr>
<tr><td>>MI0000070|hsa-mir-16-1&WithFlank&13|-|50048967|50049067@20.40.21;MI0000115|hsa-mir-16-2&WithFlank&3|+|160404739|160404831@16.36.21&[MIMAT0000069&hsa-miR-16-5p&offsets|0|-1;m-62&13|-|50049027|50049048&offsets|0|-1];[MIMAT0000069_1&hsa-miR-16-5p&offsets|0|-1;m-61&3|+|160404754|160404775&offsets|0|-1]|TAGCAGCACGTAAATATTGGC</td><td>>MI0000070|hsa-mir-16-1&WithFlank&13|-|50048967|50049067</td><td>20</td><td>40</td><td>21</td><td>TAGCAGCACGTAAATATTGGC</td><td>1011.5</td><td>5.0</td><td>1.0</td><td>5.2</td><td>1.8E-07</td><td>3.1E-07</td><td>MB231/MB468</td><td>MITO</td></tr>
<tr><td>>hg38_21_+_8433172_8446622_RNA45SN1_45S_50ntFlanks@6730.6771.42|ACACATTGATCATCGACACTTCGAACGCACTTGCGGCCCCGG</td><td>>hg38_21_+_8433172_8446622_RNA45SN1_45S_50ntFlanks</td><td>6730</td><td>6771</td><td>42</td><td>ACACATTGATCATCGACACTTCGAACGCACTTGCGGCCCCGG</td><td>720.2</td><td>-3.7</td><td>0.7</td><td>-5.3</td><td>1.3E-07</td><td>3.1E-07</td><td>BT20/MB468</td><td>MITO</td></tr>
<tr><td>>hg38_21_+_8433172_8446622_RNA45SN1_45S_50ntFlanks@10805.10825.21|GAACAATGTAGGTAAGGGAAG</td><td>>hg38_21_+_8433172_8446622_RNA45SN1_45S_50ntFlanks</td><td>10805</td><td>10825</td><td>21</td><td>GAACAATGTAGGTAAGGGAAG</td><td>181.0</td><td>-6.2</td><td>1.2</td><td>-5.3</td><td>1.5E-07</td><td>3.1E-07</td><td>BT20/MB468</td><td>CYTO</td></tr>
<tr><td>>hg38_21_+_8433172_8446622_RNA45SN1_45S_50ntFlanks@11759.11780.22|CCTCGTCATCTAATTAGTGACG</td><td>>hg38_21_+_8433172_8446622_RNA45SN1_45S_50ntFlanks</td><td>11759</td><td>11780</td><td>22</td><td>CCTCGTCATCTAATTAGTGACG</td><td>108.6</td><td>-9.8</td><td>1.8</td><td>-5.5</td><td>3.6E-08</td><td>3.1E-07</td><td>BT20/MB468</td><td>MP</td></tr>
<tr><td>>hg38_21_+_8433172_8446622_RNA45SN1_45S_50ntFlanks@6655.6685.31|TCTTAGCGGTGGATCACTCGGCTCGTGCGTC</td><td>>hg38_21_+_8433172_8446622_RNA45SN1_45S_50ntFlanks</td><td>6655</td><td>6685</td><td>31</td><td>TCTTAGCGGTGGATCACTCGGCTCGTGCGTC</td><td>221.6</td><td>4.0</td><td>0.7</td><td>5.5</td><td>3.6E-08</td><td>3.1E-07</td><td>BT20/MB468</td><td>MP</td></tr>
<tr><td>>hg38_21_+_8433172_8446622_RNA45SN1_45S_50ntFlanks@5174.5197.24|CCCGAGATTGAGCAATAACAGGTC</td><td>>hg38_21_+_8433172_8446622_RNA45SN1_45S_50ntFlanks</td><td>5174</td><td>5197</td><td>24</td><td>CCCGAGATTGAGCAATAACAGGTC</td><td>60.6</td><td>-9.8</td><td>1.9</td><td>-5.2</td><td>1.6E-07</td><td>3.1E-07</td><td>MB231/MB468</td><td>MP</td></tr>
<tr><td>>hg38_21_+_8433172_8446622_RNA45SN1_45S_50ntFlanks@11325.11344.20|TCTCGTCGCGCGCGCGTCCG</td><td>>hg38_21_+_8433172_8446622_RNA45SN1_45S_50ntFlanks</td><td>11325</td><td>11344</td><td>20</td><td>TCTCGTCGCGCGCGCGTCCG</td><td>854.2</td><td>-4.3</td><td>0.8</td><td>-5.2</td><td>1.7E-07</td><td>3.1E-07</td><td>MB231/MB468</td><td>NUC</td></tr>
<tr><td>>hg38_21_+_8433172_8446622_RNA45SN1_45S_50ntFlanks@10393.10433.41|CAGCAGTTGAACATGGGTCAGTCGGTCCTGAGAGATGGGCG</td><td>>hg38_21_+_8433172_8446622_RNA45SN1_45S_50ntFlanks</td><td>10393</td><td>10433</td><td>41</td><td>CAGCAGTTGAACATGGGTCAGTCGGTCCTGAGAGATGGGCG</td><td>513.8</td><td>-4.5</td><td>0.9</td><td>-5.2</td><td>1.7E-07</td><td>3.1E-07</td><td>MB231/MB468</td><td>NUC</td></tr>
<tr><td>>hg38_21_+_8433172_8446622_RNA45SN1_45S_50ntFlanks@7989.8016.28|AGACGTGGCGACCCGCTGAATTTAAGCA</td><td>>hg38_21_+_8433172_8446622_RNA45SN1_45S_50ntFlanks</td><td>7989</td><td>8016</td><td>28</td><td>AGACGTGGCGACCCGCTGAATTTAAGCA</td><td>568.2</td><td>-3.5</td><td>0.7</td><td>-5.2</td><td>1.6E-07</td><td>3.1E-07</td><td>MB231/MB468</td><td>MP</td></tr>
<tr><td>>hg38_21_+_8433172_8446622_RNA45SN1_45S_50ntFlanks@11873.11905.33|AAGAAGACCCTGTTGAGCTTGACTCTAGTCTGG</td><td>>hg38_21_+_8433172_8446622_RNA45SN1_45S_50ntFlanks</td><td>11873</td><td>11905</td><td>33</td><td>AAGAAGACCCTGTTGAGCTTGACTCTAGTCTGG</td><td>218.1</td><td>-6.6</td><td>1.3</td><td>-5.2</td><td>1.8E-07</td><td>3.1E-07</td><td>MB231/MB468</td><td>NUC</td></tr>
<tr><td>>MI0000076|hsa-mir-20a&WithFlank&13|+|91351059|91351141@14.36.23&[MIMAT0000075&hsa-miR-20a-5p&offsets|0|0;m-96&13|+|91351072|91351094&offsets|0|0]|TAAAGTGCTTATAGTGCAGGTAG</td><td>>MI0000076|hsa-mir-20a&WithFlank&13|+|91351059|91351141</td><td>14</td><td>36</td><td>23</td><td>TAAAGTGCTTATAGTGCAGGTAG</td><td>4188.1</td><td>4.4</td><td>0.8</td><td>5.3</td><td>1.5E-07</td><td>3.1E-07</td><td>BT20/MB468</td><td>CYTO</td></tr>
<tr><td>>hg38_21_+_8433172_8446622_RNA45SN1_45S_50ntFlanks@10727.10752.26|GGAGAGGGTGTAAATCTCGCGCCGGG</td><td>>hg38_21_+_8433172_8446622_RNA45SN1_45S_50ntFlanks</td><td>10727</td><td>10752</td><td>26</td><td>GGAGAGGGTGTAAATCTCGCGCCGGG</td><td>157.4</td><td>-5.8</td><td>1.1</td><td>-5.3</td><td>1.5E-07</td><td>3.1E-07</td><td>BT20/MB468</td><td>CYTO</td></tr>
<tr><td>>hg38_21_+_8433172_8446622_RNA45SN1_45S_50ntFlanks@10574.10591.18|AAGCCGGCGGGAGCCCCG</td><td>>hg38_21_+_8433172_8446622_RNA45SN1_45S_50ntFlanks</td><td>10574</td><td>10591</td><td>18</td><td>AAGCCGGCGGGAGCCCCG</td><td>414.6</td><td>-6.7</td><td>1.3</td><td>-5.3</td><td>1.5E-07</td><td>3.1E-07</td><td>BT20/MB468</td><td>CYTO</td></tr>
<tr><td>>hg38_21_+_8433172_8446622_RNA45SN1_45S_50ntFlanks@7977.7995.19|CGACCTCAGATCAGACGTG</td><td>>hg38_21_+_8433172_8446622_RNA45SN1_45S_50ntFlanks</td><td>7977</td><td>7995</td><td>19</td><td>CGACCTCAGATCAGACGTG</td><td>250.0</td><td>-3.0</td><td>0.6</td><td>-5.2</td><td>1.8E-07</td><td>3.1E-07</td><td>MB231/MB468</td><td>NUC</td></tr>
<tr><td>>hg38_21_+_8433172_8446622_RNA45SN1_45S_50ntFlanks@10794.10823.30|CTGGCATGTTGGAACAATGTAGGTAAGGGA</td><td>>hg38_21_+_8433172_8446622_RNA45SN1_45S_50ntFlanks</td><td>10794</td><td>10823</td><td>30</td><td>CTGGCATGTTGGAACAATGTAGGTAAGGGA</td><td>279.9</td><td>-2.4</td><td>0.4</td><td>-5.3</td><td>1.3E-07</td><td>3.1E-07</td><td>BT20/MB468</td><td>MITO</td></tr>
<tr><td>>hg38_21_+_8433172_8446622_RNA45SN1_45S_50ntFlanks@10806.10823.18|AACAATGTAGGTAAGGGA</td><td>>hg38_21_+_8433172_8446622_RNA45SN1_45S_50ntFlanks</td><td>10806</td><td>10823</td><td>18</td><td>AACAATGTAGGTAAGGGA</td><td>482.8</td><td>-6.8</td><td>1.3</td><td>-5.3</td><td>1.5E-07</td><td>3.1E-07</td><td>BT20/MB468</td><td>CYTO</td></tr>
<tr><td>>MI0000434|hsa-let-7i&WithFlank&12|+|62603680|62603775@12.31.20(+1C)&[MIMAT0000415&hsa-let-7i-5p&offsets|0|-2(+1C);m-49&12|+|62603691|62603712&offsets|0|-2(+1C)]|TGAGGTAGTAGTTTGTGCTGC</td><td>>MI0000434|hsa-let-7i&WithFlank&12|+|62603680|62603775</td><td>12</td><td>31</td><td>20(+1C)</td><td>TGAGGTAGTAGTTTGTGCTGC</td><td>365.3</td><td>8.6</td><td>1.6</td><td>5.3</td><td>1.3E-07</td><td>3.1E-07</td><td>BT20/MB468</td><td>MITO</td></tr>
<tr><td>>hg38_21_+_8433172_8446622_RNA45SN1_45S_50ntFlanks@8172.8212.41|AGTCCTTCTGATCGAGGCCCAGCCCGTGGACGGTGTGAGGC</td><td>>hg38_21_+_8433172_8446622_RNA45SN1_45S_50ntFlanks</td><td>8172</td><td>8212</td><td>41</td><td>AGTCCTTCTGATCGAGGCCCAGCCCGTGGACGGTGTGAGGC</td><td>323.1</td><td>-5.6</td><td>1.1</td><td>-5.2</td><td>1.9E-07</td><td>3.1E-07</td><td>MB231/MB468</td><td>MITO</td></tr>
<tr><td>>MI0000746|hsa-mir-99b&WithFlank&19|+|51692606|51692687@13.33.21(+1U)&[MIMAT0000689&hsa-miR-99b-5p&offsets|0|-1(+1U);m-10&19|+|51692618|51692639&offsets|0|-1(+1U)]|CACCCGTAGAACCGACCTTGCT</td><td>>MI0000746|hsa-mir-99b&WithFlank&19|+|51692606|51692687</td><td>13</td><td>33</td><td>21(+1U)</td><td>CACCCGTAGAACCGACCTTGCT</td><td>514.5</td><td>5.3</td><td>1.0</td><td>5.2</td><td>1.9E-07</td><td>3.1E-07</td><td>MB231/MB468</td><td>MITO</td></tr>
<tr><td>>hg38_21_+_8433172_8446622_RNA45SN1_45S_50ntFlanks@9534.9565.32|AGGAAACTCTGGTGGAGGTCCGTAGCGGTCCT</td><td>>hg38_21_+_8433172_8446622_RNA45SN1_45S_50ntFlanks</td><td>9534</td><td>9565</td><td>32</td><td>AGGAAACTCTGGTGGAGGTCCGTAGCGGTCCT</td><td>872.2</td><td>-6.2</td><td>1.2</td><td>-5.2</td><td>1.9E-07</td><td>3.1E-07</td><td>MB231/MB468</td><td>MITO</td></tr>
<tr><td>>MI0000070|hsa-mir-16-1&WithFlank&13|-|50048967|50049067@20.41.22;MI0000115|hsa-mir-16-2&WithFlank&3|+|160404739|160404831@16.37.22&[MIMAT0000069&hsa-miR-16-5p&offsets|0|0;m-62&13|-|50049027|50049048&offsets|0|0];[MIMAT0000069_1&hsa-miR-16-5p&offsets|0|0;m-61&3|+|160404754|160404775&offsets|0|0]|TAGCAGCACGTAAATATTGGCG</td><td>>MI0000070|hsa-mir-16-1&WithFlank&13|-|50048967|50049067</td><td>20</td><td>41</td><td>22</td><td>TAGCAGCACGTAAATATTGGCG</td><td>1390.3</td><td>5.2</td><td>1.0</td><td>5.2</td><td>1.7E-07</td><td>3.1E-07</td><td>MB231/MB468</td><td>CYTO</td></tr>
<tr><td>>hg38_21_+_8433172_8446622_RNA45SN1_45S_50ntFlanks@12072.12092.21|CCCGAGGGGCTCTCGCTTCTG</td><td>>hg38_21_+_8433172_8446622_RNA45SN1_45S_50ntFlanks</td><td>12072</td><td>12092</td><td>21</td><td>CCCGAGGGGCTCTCGCTTCTG</td><td>662.2</td><td>-3.3</td><td>0.6</td><td>-5.2</td><td>1.7E-07</td><td>3.1E-07</td><td>MB231/MB468</td><td>CYTO</td></tr>
<tr><td>>am_trnaMT_ArgTCG_MT_+_10405_10469@31.48.18|TCGACTCATTAAATTATG</td><td>>am_trnaMT_ArgTCG_MT_+_10405_10469</td><td>31</td><td>48</td><td>18</td><td>TCGACTCATTAAATTATG</td><td>348.9</td><td>-8.4</td><td>1.6</td><td>-5.2</td><td>1.9E-07</td><td>3.1E-07</td><td>BT20/MB468</td><td>NUC</td></tr>
<tr><td>>hg38_21_+_8433172_8446622_RNA45SN1_45S_50ntFlanks@11572.11606.35|CCGACTTAGAACTGGTGCGGACCAGGGGAATCCGA</td><td>>hg38_21_+_8433172_8446622_RNA45SN1_45S_50ntFlanks</td><td>11572</td><td>11606</td><td>35</td><td>CCGACTTAGAACTGGTGCGGACCAGGGGAATCCGA</td><td>391.6</td><td>-7.6</td><td>1.4</td><td>-5.2</td><td>1.6E-07</td><td>3.1E-07</td><td>MB231/MB468</td><td>MP</td></tr>
<tr><td>>MI0000342|hsa-mir-200b&WithFlank&1|+|1167098|1167204@63.86.24&[MIMAT0000318&hsa-miR-200b-3p&offsets|0|+2;m-93&1|+|1167160|1167182&offsets|0|+1]|TAATACTGCCTGGTAATGATGACG</td><td>>MI0000342|hsa-mir-200b&WithFlank&1|+|1167098|1167204</td><td>63</td><td>86</td><td>24</td><td>TAATACTGCCTGGTAATGATGACG</td><td>68.7</td><td>-9.2</td><td>1.7</td><td>-5.5</td><td>3.6E-08</td><td>3.1E-07</td><td>BT20/MB468</td><td>MP</td></tr>
<tr><td>>hg38_21_+_8433172_8446622_RNA45SN1_45S_50ntFlanks@12456.12498.43|CCAAGCGTTGGATTGTTCACCCACTAATAGGGAACGTGAGCTG</td><td>>hg38_21_+_8433172_8446622_RNA45SN1_45S_50ntFlanks</td><td>12456</td><td>12498</td><td>43</td><td>CCAAGCGTTGGATTGTTCACCCACTAATAGGGAACGTGAGCTG</td><td>52.4</td><td>-9.6</td><td>1.8</td><td>-5.2</td><td>1.6E-07</td><td>3.1E-07</td><td>MB231/MB468</td><td>MP</td></tr>
<tr><td>>am_trnaMT_SerGCT_MT_+_12207_12265@14.31.18|AGAACTGCTAACTCATGC</td><td>>am_trnaMT_SerGCT_MT_+_12207_12265</td><td>14</td><td>31</td><td>18</td><td>AGAACTGCTAACTCATGC</td><td>735.8</td><td>6.3</td><td>1.2</td><td>5.3</td><td>1.5E-07</td><td>3.1E-07</td><td>BT20/MB468</td><td>CYTO</td></tr>
<tr><td>>am_trna10_LysCTT_16_+_3241501_3241573@31.51.21,trna119_LysCTT_1_-_145395522_145395594@31.51.21,trna11_LysCTT_5_-_180648979_180649051@31.51.21,trna13_LysCTT_16_+_3246153_3246225@31.51.21,trna13_LysCTT_6_+_26556774_26556846@31.51.21,trna30_LysCTT_16_-_3230555_3230627@31.51.21,trna5_LysCTT_16_+_3214939_3215011@31.51.21,trna7_LysCTT_16_+_3225692_3225764@31.51.21,trna8_LysCTT_1_+_39970195_39970267@31.51.21,trna9_LysCTT_5_+_180634755_180634827@31.51.21|ACTCTTAATCTCAGGGTCGTG</td><td>>am_trna10_LysCTT_16_+_3241501_3241573</td><td>31</td><td>51</td><td>21</td><td>ACTCTTAATCTCAGGGTCGTG</td><td>795.5</td><td>-5.9</td><td>1.1</td><td>-5.3</td><td>1.5E-07</td><td>3.1E-07</td><td>BT20/MB468</td><td>CYTO</td></tr>
<tr><td>>hg38_21_+_8433172_8446622_RNA45SN1_45S_50ntFlanks@12241.12265.25|CAGAAGGGCAAAAGCTCGCTTGATC</td><td>>hg38_21_+_8433172_8446622_RNA45SN1_45S_50ntFlanks</td><td>12241</td><td>12265</td><td>25</td><td>CAGAAGGGCAAAAGCTCGCTTGATC</td><td>385.9</td><td>-5.7</td><td>1.1</td><td>-5.2</td><td>1.6E-07</td><td>3.1E-07</td><td>MB231/MB468</td><td>MP</td></tr>
<tr><td>>hg38_21_+_8433172_8446622_RNA45SN1_45S_50ntFlanks@5476.5516.41|CCCTGGCGGAGCGCTGAGAAGACGGTCGAACTTGACTATCT</td><td>>hg38_21_+_8433172_8446622_RNA45SN1_45S_50ntFlanks</td><td>5476</td><td>5516</td><td>41</td><td>CCCTGGCGGAGCGCTGAGAAGACGGTCGAACTTGACTATCT</td><td>2260.1</td><td>5.6</td><td>1.0</td><td>5.7</td><td>1.4E-08</td><td>3.1E-07</td><td>BT20/MB231</td><td>MITO</td></tr>
<tr><td>>hg38_MT_+_1621_3279_RNR2_16S_50ntFlanks@729.752.24|GATTAAAACACTGAACTGACAATT</td><td>>hg38_MT_+_1621_3279_RNR2_16S_50ntFlanks</td><td>729</td><td>752</td><td>24</td><td>GATTAAAACACTGAACTGACAATT</td><td>84.6</td><td>9.6</td><td>1.8</td><td>5.2</td><td>1.6E-07</td><td>3.1E-07</td><td>MB231/MB468</td><td>MP</td></tr>
<tr><td>>hg38_21_+_8433172_8446622_RNA45SN1_45S_50ntFlanks@7975.8012.38|CGCGACCTCAGATCAGACGTGGCGACCCGCTGAATTTA</td><td>>hg38_21_+_8433172_8446622_RNA45SN1_45S_50ntFlanks</td><td>7975</td><td>8012</td><td>38</td><td>CGCGACCTCAGATCAGACGTGGCGACCCGCTGAATTTA</td><td>9383.1</td><td>3.8</td><td>0.7</td><td>5.3</td><td>1.5E-07</td><td>3.1E-07</td><td>BT20/MB468</td><td>CYTO</td></tr>
<tr><td>>hg38_21_+_8433172_8446622_RNA45SN1_45S_50ntFlanks@4128.4148.21|CGATTCCGGAGAGGGAGCCTG</td><td>>hg38_21_+_8433172_8446622_RNA45SN1_45S_50ntFlanks</td><td>4128</td><td>4148</td><td>21</td><td>CGATTCCGGAGAGGGAGCCTG</td><td>286.1</td><td>-4.0</td><td>0.8</td><td>-5.2</td><td>1.6E-07</td><td>3.1E-07</td><td>MB231/MB468</td><td>MP</td></tr>
<tr><td>>hg38_21_+_8433172_8446622_RNA45SN1_45S_50ntFlanks@3823.3852.30|TTTGGTCGCTCGCTCCTCTCCTACTTGGAT</td><td>>hg38_21_+_8433172_8446622_RNA45SN1_45S_50ntFlanks</td><td>3823</td><td>3852</td><td>30</td><td>TTTGGTCGCTCGCTCCTCTCCTACTTGGAT</td><td>1163.4</td><td>-8.9</td><td>1.7</td><td>-5.2</td><td>2.0E-07</td><td>3.1E-07</td><td>BT20/MB468</td><td>NUC</td></tr>
<tr><td>>hg38_21_+_8433172_8446622_RNA45SN1_45S_50ntFlanks@11208.11227.20|GCGGCGACTCTGGACGCGAG</td><td>>hg38_21_+_8433172_8446622_RNA45SN1_45S_50ntFlanks</td><td>11208</td><td>11227</td><td>20</td><td>GCGGCGACTCTGGACGCGAG</td><td>2579.0</td><td>-6.1</td><td>1.2</td><td>-5.3</td><td>1.5E-07</td><td>3.1E-07</td><td>BT20/MB468</td><td>CYTO</td></tr>
<tr><td>>hg38_21_+_8433172_8446622_RNA45SN1_45S_50ntFlanks@12437.12473.37|CATTGTGAAGCAGAATTCACCAAGCGTTGGATTGTTC</td><td>>hg38_21_+_8433172_8446622_RNA45SN1_45S_50ntFlanks</td><td>12437</td><td>12473</td><td>37</td><td>CATTGTGAAGCAGAATTCACCAAGCGTTGGATTGTTC</td><td>1985.6</td><td>-5.6</td><td>1.0</td><td>-5.5</td><td>3.7E-08</td><td>3.1E-07</td><td>BT20/MB468</td><td>MP</td></tr>
<tr><td>>hg38_21_+_8433172_8446622_RNA45SN1_45S_50ntFlanks@11312.11344.33|CCCCCACCCCACGTCTCGTCGCGCGCGCGTCCG</td><td>>hg38_21_+_8433172_8446622_RNA45SN1_45S_50ntFlanks</td><td>11312</td><td>11344</td><td>33</td><td>CCCCCACCCCACGTCTCGTCGCGCGCGCGTCCG</td><td>1520.3</td><td>-8.7</td><td>1.7</td><td>-5.2</td><td>2.0E-07</td><td>3.1E-07</td><td>BT20/MB468</td><td>NUC</td></tr>
<tr><td>>hg38_21_+_8433172_8446622_RNA45SN1_45S_50ntFlanks@6657.6681.25|TTAGCGGTGGATCACTCGGCTCGTG</td><td>>hg38_21_+_8433172_8446622_RNA45SN1_45S_50ntFlanks</td><td>6657</td><td>6681</td><td>25</td><td>TTAGCGGTGGATCACTCGGCTCGTG</td><td>1456.0</td><td>-4.9</td><td>0.9</td><td>-5.2</td><td>1.9E-07</td><td>3.2E-07</td><td>MB231/MB468</td><td>MITO</td></tr>
</table>

| | | | | | | | | | | | | | |
|---|---|---|---|---|---|---|---|---|---|---|---|---|---|
| >hg38_21_+_8433172_8446622_RNA45SN1_45S_50ntFlanks@475.521.47\|CTTCGTGATCGATGTGGTGACGTCGTGCTCTCCCGGGCCGGGTCCGA | >hg38_21_+_8433172_8446622_RNA45SN1_45S_50ntFlanks | 475 | 521 | 47 | CTTCGTGATCGATGTGGTGACGTCGTGCTCTCCCGGGCCGGGTCCGA | 91.4 | 10.3 | 1.9 | 5.5 | 3.7E-08 | 3.2E-07 | BT20/MB468 | MP |
| >hg38_21_+_8433172_8446622_RNA45SN1_45S_50ntFlanks@9923.9948.26\|TTGGTTGATATAGACAGCAGGACGGT | >hg38_21_+_8433172_8446622_RNA45SN1_45S_50ntFlanks | 9923 | 9948 | 26 | TTGGTTGATATAGACAGCAGGACGGT | 68.1 | 9.8 | 1.8 | 5.5 | 3.7E-08 | 3.2E-07 | BT20/MB468 | MP |
| >hg38_21_+_8433172_8446622_RNA45SN1_45S_50ntFlanks@12162.12180.19\|GACTGGGGCGGTACACCTG | >hg38_21_+_8433172_8446622_RNA45SN1_45S_50ntFlanks | 12162 | 12180 | 19 | GACTGGGGCGGTACACCTG | 1528.9 | -8.2 | 1.5 | -5.5 | 3.7E-08 | 3.2E-07 | BT20/MB468 | MP |
| >hg38_21_+_8433172_8446622_RNA45SN1_45S_50ntFlanks@6721.6758.38\|AATTGCAGGACACATTGATCATCGACACTTCGAACGCA | >hg38_21_+_8433172_8446622_RNA45SN1_45S_50ntFlanks | 6721 | 6758 | 38 | AATTGCAGGACACATTGATCATCGACACTTCGAACGCA | 2069.8 | -3.1 | 0.6 | -5.2 | 1.7E-07 | 3.2E-07 | MB231/MB468 | CYTO |
| >hg38_21_+_8433172_8446622_RNA45SN1_45S_50ntFlanks@10393.10431.39\|CAGCAGTTGAACATGGGTCAGTCGGTCCTGAGAGATGGG | >hg38_21_+_8433172_8446622_RNA45SN1_45S_50ntFlanks | 10393 | 10431 | 39 | CAGCAGTTGAACATGGGTCAGTCGGTCCTGAGAGATGGG | 347.8 | -7.3 | 1.4 | -5.2 | 1.8E-07 | 3.2E-07 | MB231/MB468 | NUC |
| >MI0000066\|hsa-let-7e&WithFlank&19\|+\|51692780\|51692870@14.34.21(+1A)&[MIMAT0000066&hsa-let-7e-5p&offsets\|0\|-1(+1A);m-50&19\|+\|51692793\|51692814&offsets\|0\|-1(+1A)]\|TGAGGTAGGAGGTTGTATAGTA | >MI0000066\|hsa-let-7e&WithFlank&19\|+\|51692780\|51692870 | 14 | 34 | 21(+1A) | TGAGGTAGGAGGTTGTATAGTA | 386.7 | 6.6 | 1.3 | 5.2 | 1.7E-07 | 3.2E-07 | MB231/MB468 | CYTO |
| >hg38_21_+_8433172_8446622_RNA45SN1_45S_50ntFlanks@12957.12975.19\|CTTCTGGGTCGGGGTTTCG | >hg38_21_+_8433172_8446622_RNA45SN1_45S_50ntFlanks | 12957 | 12975 | 19 | CTTCTGGGTCGGGGTTTCG | 691.5 | -8.2 | 1.6 | -5.2 | 2.0E-07 | 3.2E-07 | BT20/MB468 | NUC |
| >MI0000108\|hsa-mir-103a-2&WithFlank&20\|+\|3917488\|3917577@54.76.23(+2U);MI0000109\|hsa-mir-103a-1&WithFlank&5\|-\|168560890\|168560979@54.76.23(+2U)&[MIMAT0000101&hsa-miR-103a-3p&offsets\|0\|0(+2U);m-12&20\|+\|3917541\|3917563&offsets\|0\|0(+2U)];[MIMAT0000101_1&hsa-miR-103a-3p&offsets\|0\|0(+2U);m-13&5\|-\|168560904\|168560926&offsets\|0\|0(+2U);m-44220&5\|-\|168560908\|168560926&offsets\|0\|+4(+2U)]\|AGCAGCATTGTACAGGGCTATGATT | >MI0000108\|hsa-mir-103a-2&WithFlank&20\|+\|3917488\|3917577 | 54 | 76 | 23(+2U) | AGCAGCATTGTACAGGGCTATGATT | 1043.6 | 6.6 | 1.3 | 5.2 | 1.9E-07 | 3.2E-07 | MB231/MB468 | MITO |
| >MI0000743\|hsa-mir-34c&WithFlank&11\|+\|111513433\|111513521@19.40.22(+1A)&[MIMAT0000686&hsa-miR-34c-5p&offsets\|0\|-1(+1A);m-336&11\|+\|111513451\|111513473&offsets\|0\|-1(+1A)]\|AGGCAGTGTAGTTAGCTGATTGA | >MI0000743\|hsa-mir-34c&WithFlank&11\|+\|111513433\|111513521 | 19 | 40 | 22(+1A) | AGGCAGTGTAGTTAGCTGATTGA | 143.3 | -10.6 | 1.8 | -5.8 | 7.8E-09 | 3.2E-07 | BT20/MB231 | MP |
| >hg38_21_+_8433172_8446622_RNA45SN1_45S_50ntFlanks@10392.10430.39\|ACAGCAGTTGAACATGGGTCAGTCGGTCCTGAGAGATGG | >hg38_21_+_8433172_8446622_RNA45SN1_45S_50ntFlanks | 10392 | 10430 | 39 | ACAGCAGTTGAACATGGGTCAGTCGGTCCTGAGAGATGG | 585.1 | -3.9 | 0.7 | -5.3 | 1.4E-07 | 3.2E-07 | BT20/MB468 | MITO |
| >hg38_21_+_8433172_8446622_RNA45SN1_45S_50ntFlanks@10878.10902.25\|TCGGTCGGGCTGGGGCGCGAAGCGG | >hg38_21_+_8433172_8446622_RNA45SN1_45S_50ntFlanks | 10878 | 10902 | 25 | TCGGTCGGGCTGGGGCGCGAAGCGG | 1237.4 | -8.7 | 1.7 | -5.2 | 2.0E-07 | 3.2E-07 | BT20/MB468 | NUC |
| >hg38_21_+_8433172_8446622_RNA45SN1_45S_50ntFlanks@3992.4026.35\|GGTGACTCTAGATAACCTCGGGCCGATCGCACGCC | >hg38_21_+_8433172_8446622_RNA45SN1_45S_50ntFlanks | 3992 | 4026 | 35 | GGTGACTCTAGATAACCTCGGGCCGATCGCACGCC | 426.8 | -5.6 | 1.1 | -5.2 | 1.9E-07 | 3.2E-07 | MB231/MB468 | MITO |
| >hg38_21_+_8433172_8446622_RNA45SN1_45S_50ntFlanks@11565.11585.21\|CTAGCAGCCGACTTAGAACTG | >hg38_21_+_8433172_8446622_RNA45SN1_45S_50ntFlanks | 11565 | 11585 | 21 | CTAGCAGCCGACTTAGAACTG | 251.7 | -2.5 | 0.5 | -5.2 | 1.8E-07 | 3.2E-07 | MB231/MB468 | CYTO |
| >hg38_21_+_8433172_8446622_RNA45SN1_45S_50ntFlanks@8350.8375.26\|AAAGTTGAAAAGAACTTTGAAGAGAG | >hg38_21_+_8433172_8446622_RNA45SN1_45S_50ntFlanks | 8350 | 8375 | 26 | AAAGTTGAAAAGAACTTTGAAGAGAG | 2665.9 | 10.2 | 1.8 | 5.5 | 3.7E-08 | 3.2E-07 | BT20/MB468 | MP |
| >hg38_21_+_8433172_8446622_RNA45SN1_45S_50ntFlanks@10682.10711.30\|TCCGGCGGCGTCCGGTGAGCTCTCGCTGGC | >hg38_21_+_8433172_8446622_RNA45SN1_45S_50ntFlanks | 10682 | 10711 | 30 | TCCGGCGGCGTCCGGTGAGCTCTCGCTGGC | 69.9 | -9.2 | 1.7 | -5.5 | 3.7E-08 | 3.2E-07 | BT20/MB468 | MP |
| >hg38_21_+_8433172_8446622_RNA45SN1_45S_50ntFlanks@9536.9553.18\|GAAACTCTGGTGGAGGTC | >hg38_21_+_8433172_8446622_RNA45SN1_45S_50ntFlanks | 9536 | 9553 | 18 | GAAACTCTGGTGGAGGTC | 143.8 | -10.2 | 1.9 | -5.5 | 3.7E-08 | 3.2E-07 | BT20/MB468 | MP |
| >hg38_21_+_8433172_8446622_RNA45SN1_45S_50ntFlanks@6738.6772.35\|ATCATCGACACTTCGAACGCACTTGCGGCCCCGGG | >hg38_21_+_8433172_8446622_RNA45SN1_45S_50ntFlanks | 6738 | 6772 | 35 | ATCATCGACACTTCGAACGCACTTGCGGCCCCGGG | 245.3 | -5.8 | 1.1 | -5.2 | 1.8E-07 | 3.2E-07 | MB231/MB468 | CYTO |
| >hg38_21_+_8433172_8446622_RNA45SN1_45S_50ntFlanks@8176.8196.21\|CTTCTGATCGAGGCCCAGCCC | >hg38_21_+_8433172_8446622_RNA45SN1_45S_50ntFlanks | 8176 | 8196 | 21 | CTTCTGATCGAGGCCCAGCCC | 429.8 | -3.6 | 0.7 | -5.2 | 1.9E-07 | 3.2E-07 | MB231/MB468 | MITO |
| >hg38_21_+_8433172_8446622_RNA45SN1_45S_50ntFlanks@5486.5519.34\|GCGCTGAGAAGACGGTCGAACTTGACTATCTAGA | >hg38_21_+_8433172_8446622_RNA45SN1_45S_50ntFlanks | 5486 | 5519 | 34 | GCGCTGAGAAGACGGTCGAACTTGACTATCTAGA | 389.8 | 6.3 | 1.2 | 5.2 | 1.5E-07 | 3.2E-07 | BT20/MB468 | CYTO |
| >hg38_21_+_8433172_8446622_RNA45SN1_45S_50ntFlanks@12960.12979.20\|CTGGGTCGGGGTTTCGTACG | >hg38_21_+_8433172_8446622_RNA45SN1_45S_50ntFlanks | 12960 | 12979 | 20 | CTGGGTCGGGGTTTCGTACG | 545.6 | -8.2 | 1.6 | -5.2 | 2.0E-07 | 3.2E-07 | BT20/MB468 | NUC |
| >hg38_21_+_8433172_8446622_RNA45SN1_45S_50ntFlanks@12619.12654.36\|AGGAGCCAATGGGGCGAAGCTACCATCTGTGGGATT | >hg38_21_+_8433172_8446622_RNA45SN1_45S_50ntFlanks | 12619 | 12654 | 36 | AGGAGCCAATGGGGCGAAGCTACCATCTGTGGGATT | 4500.4 | -9.6 | 1.8 | -5.2 | 2.0E-07 | 3.2E-07 | BT20/MB468 | NUC |
| >hg38_21_+_8433172_8446622_RNA45SN1_45S_50ntFlanks@7988.8018.31\|CAGACGTGGCGACCCGCTGAATTTAAGCATA | >hg38_21_+_8433172_8446622_RNA45SN1_45S_50ntFlanks | 7988 | 8018 | 31 | CAGACGTGGCGACCCGCTGAATTTAAGCATA | 489.8 | -3.4 | 0.6 | -5.3 | 1.4E-07 | 3.2E-07 | BT20/MB468 | MITO |
| >hg38_21_+_8433172_8446622_RNA45SN1_45S_50ntFlanks@4753.4782.30\|AAGACGATCAGATACCGTCGTAGTTCCGAC | >hg38_21_+_8433172_8446622_RNA45SN1_45S_50ntFlanks | 4753 | 4782 | 30 | AAGACGATCAGATACCGTCGTAGTTCCGAC | 540.9 | -3.3 | 0.6 | -5.2 | 1.8E-07 | 3.2E-07 | MB231/MB468 | CYTO |
| >hg38_21_+_8433172_8446622_RNA45SN1_45S_50ntFlanks@8182.8208.27\|ATCGAGGCCCAGCCCGTGGACGGTGTG | >hg38_21_+_8433172_8446622_RNA45SN1_45S_50ntFlanks | 8182 | 8208 | 27 | ATCGAGGCCCAGCCCGTGGACGGTGTG | 292.2 | -7.0 | 1.3 | -5.2 | 1.9E-07 | 3.2E-07 | MB231/MB468 | MITO |
| >MI0000289\|hsa-mir-181a-1&WithFlank&1\|-\|198859038\|198859159@30.49.20;MI0000269\|hsa-mir-181a-2&WithFlank&9\|+\|124692436\|124692557@45.64.20&[MIMAT0000256&hsa-miR-181a-5p&offsets\|0\|-3;m-32&1\|-\|198859108\|198859130&offsets\|0\|-3];[MIMAT0000256_1&hsa-miR-181a-5p&offsets\|0\|-3;m-32&9\|+\|124692480\|124692502&offsets\|0\|-3]\|AACATTCAACGCTGTCGGTG | >MI0000289\|hsa-mir-181a-1&WithFlank&1\|-\|198859038\|198859159 | 30 | 49 | 20 | AACATTCAACGCTGTCGGTG | 591.0 | 4.7 | 0.9 | 5.2 | 1.9E-07 | 3.2E-07 | MB231/MB468 | MITO |
| >hg38_21_+_8433172_8446622_RNA45SN1_45S_50ntFlanks@9544.9563.20\|GGTGGAGGTCCGTAGCGGTC | >hg38_21_+_8433172_8446622_RNA45SN1_45S_50ntFlanks | 9544 | 9563 | 20 | GGTGGAGGTCCGTAGCGGTC | 528.1 | -6.1 | 1.2 | -5.2 | 1.8E-07 | 3.2E-07 | MB231/MB468 | NUC |
| >hg38_21_+_8433172_8446622_RNA45SN1_45S_50ntFlanks@11729.11752.24\|TAACTATGACTCTCTTAAGGTAGC | >hg38_21_+_8433172_8446622_RNA45SN1_45S_50ntFlanks | 11729 | 11752 | 24 | TAACTATGACTCTCTTAAGGTAGC | 803.4 | -5.3 | 1.0 | -5.2 | 1.5E-07 | 3.2E-07 | BT20/MB468 | CYTO |
| >MI0000098\|hsa-mir-96&WithFlank&7\|-\|129774686\|129774775@15.37.23&[MIMAT0000095&hsa-miR-96-5p&offsets\|0\|0;m-248&7\|-\|129774739\|129774761&offsets\|0\|0]\|TTTGGCACTAGCACATTTTTGCT | >MI0000098\|hsa-mir-96&WithFlank&7\|-\|129774686\|129774775 | 15 | 37 | 23 | TTTGGCACTAGCACATTTTTGCT | 2776.1 | 4.5 | 0.8 | 5.7 | 1.1E-08 | 3.2E-07 | BT20/MB231 | CYTO |
| >hg38_21_+_8433172_8446622_RNA45SN1_45S_50ntFlanks@5236.5261.26\|CACTGACTGGCTCAGCGTGTGCCTAC | >hg38_21_+_8433172_8446622_RNA45SN1_45S_50ntFlanks | 5236 | 5261 | 26 | CACTGACTGGCTCAGCGTGTGCCTAC | 357.5 | -8.3 | 1.6 | -5.2 | 1.9E-07 | 3.2E-07 | MB231/MB468 | MITO |
| >MI0000734\|hsa-mir-106b&WithFlank&7\|-\|100093987\|100094080@58.75.18(+1U)&[MIMAT0004672&hsa-miR-106b-3p&offsets\|0\|-4(+1U);m-87&7\|-\|100094003\|100094023&offsets\|0\|-3(+1U)]\|CCGCACTGTGGGTACTTGT | >MI0000734\|hsa-mir-106b&WithFlank&7\|-\|100093987\|100094080 | 58 | 75 | 18(+1U) | CCGCACTGTGGGTACTTGT | 89.8 | 10.2 | 1.9 | 5.5 | 3.7E-08 | 3.2E-07 | BT20/MB468 | MP |
| >hg38_21_+_8433172_8446622_RNA45SN1_45S_50ntFlanks@9554.9594.41\|CGTAGCGGTCCTGACGTGCAAATCGGTCGTCCGACCTGGGT | >hg38_21_+_8433172_8446622_RNA45SN1_45S_50ntFlanks | 9554 | 9594 | 41 | CGTAGCGGTCCTGACGTGCAAATCGGTCGTCCGACCTGGGT | 612.1 | -6.5 | 1.2 | -5.2 | 1.9E-07 | 3.2E-07 | MB231/MB468 | MITO |
| >hg38_21_+_8433172_8446622_RNA45SN1_45S_50ntFlanks@11671.11698.28\|CCAGTGCTCTGAATGTCAAAGTGAAGAA | >hg38_21_+_8433172_8446622_RNA45SN1_45S_50ntFlanks | 11671 | 11698 | 28 | CCAGTGCTCTGAATGTCAAAGTGAAGAA | 232.4 | -2.8 | 0.5 | -5.2 | 1.8E-07 | 3.2E-07 | MB231/MB468 | NUC |
| >hg38_21_+_8433172_8446622_RNA45SN1_45S_50ntFlanks@12249.12266.18\|CAAAAGCTCGCTTGATCT | >hg38_21_+_8433172_8446622_RNA45SN1_45S_50ntFlanks | 12249 | 12266 | 18 | CAAAAGCTCGCTTGATCT | 71.0 | -9.2 | 1.7 | -5.5 | 3.7E-08 | 3.2E-07 | BT20/MB468 | MP |
| >hg38_21_+_8433172_8446622_RNA45SN1_45S_50ntFlanks@4160.4181.22\|CCACATCCAAGGAAGGCAGCAG | >hg38_21_+_8433172_8446622_RNA45SN1_45S_50ntFlanks | 4160 | 4181 | 22 | CCACATCCAAGGAAGGCAGCAG | 99.1 | -9.7 | 1.8 | -5.5 | 3.8E-08 | 3.2E-07 | BT20/MB468 | MP |
| >hg38_21_+_8433172_8446622_RNA45SN1_45S_50ntFlanks@6652.6692.41\|GACTCTTAGCGGTGGATCACTCGGCTCGTGCGTCGATGAAG | >hg38_21_+_8433172_8446622_RNA45SN1_45S_50ntFlanks | 6652 | 6692 | 41 | GACTCTTAGCGGTGGATCACTCGGCTCGTGCGTCGATGAAG | 707.8 | 3.4 | 0.6 | 5.2 | 1.8E-07 | 3.2E-07 | MB231/MB468 | CYTO |
| >hg38_21_+_8433172_8446622_RNA45SN1_45S_50ntFlanks@11785.11802.18\|TGAATGGATGAACGAGAT | >hg38_21_+_8433172_8446622_RNA45SN1_45S_50ntFlanks | 11785 | 11802 | 18 | TGAATGGATGAACGAGAT | 405.3 | -7.9 | 1.5 | -5.2 | 2.0E-07 | 3.2E-07 | BT20/MB468 | NUC |
| >hg38_21_+_8433172_8446622_RNA45SN1_45S_50ntFlanks@9129.9152.24\|CGGGTCGGGGGGCGGGGCGGACTG | >hg38_21_+_8433172_8446622_RNA45SN1_45S_50ntFlanks | 9129 | 9152 | 24 | CGGGTCGGGGGGCGGGGCGGACTG | 1514.1 | -7.5 | 1.4 | -5.3 | 1.4E-07 | 3.2E-07 | BT20/MB468 | MITO |
| >hg38_21_+_8433172_8446622_RNA45SN1_45S_50ntFlanks@10794.10826.33\|CTGGCATGTTGGAACAATGTAGGTAAGGGAAGT | >hg38_21_+_8433172_8446622_RNA45SN1_45S_50ntFlanks | 10794 | 10826 | 33 | CTGGCATGTTGGAACAATGTAGGTAAGGGAAGT | 623.6 | -8.2 | 1.6 | -5.2 | 2.0E-07 | 3.2E-07 | BT20/MB468 | NUC |
| >hg38_21_+_8433172_8446622_RNA45SN1_45S_50ntFlanks@6726.6744.19\|CAGGACACATTGATCATCG | >hg38_21_+_8433172_8446622_RNA45SN1_45S_50ntFlanks | 6726 | 6744 | 19 | CAGGACACATTGATCATCG | 536.4 | -5.9 | 1.1 | -5.2 | 1.8E-07 | 3.2E-07 | MB231/MB468 | CYTO |
| >hg38_21_+_8433172_8446622_RNA45SN1_45S_50ntFlanks@11454.11471.18\|CCTCGGCGCGCGGCGGCG | >hg38_21_+_8433172_8446622_RNA45SN1_45S_50ntFlanks | 11454 | 11471 | 18 | CCTCGGCGCGCGGCGGCG | 163.6 | -5.9 | 1.1 | -5.2 | 1.6E-07 | 3.2E-07 | BT20/MB468 | CYTO |
| >hg38_21_+_8433172_8446622_RNA45SN1_45S_50ntFlanks@10476.10495.20\|GGCCGATCGAAAGGGAGTCG | >hg38_21_+_8433172_8446622_RNA45SN1_45S_50ntFlanks | 10476 | 10495 | 20 | GGCCGATCGAAAGGGAGTCG | 190.4 | -7.2 | 1.4 | -5.2 | 1.6E-07 | 3.2E-07 | MB231/MB468 | MP |
| >MI0000285\|hsa-mir-205&WithFlank&1\|+\|209432127\|209432248@40.62.23(+1U)&[MIMAT0000266&hsa-miR-205-5p&offsets\|0\|+1(+1U);m-351&1\|+\|209432166\|209432187&offsets\|0\|+1(+1U)]\|TCCTTCATTCCACCGGAGTCTGTT | >MI0000285\|hsa-mir-205&WithFlank&1\|+\|209432127\|209432248 | 40 | 62 | 23(+1U) | TCCTTCATTCCACCGGAGTCTGTT | 1919.2 | 10.3 | 1.8 | 5.7 | 1.2E-08 | 3.2E-07 | BT20/MB231 | CYTO |
| >hg38_21_+_8433172_8446622_RNA45SN1_45S_50ntFlanks@6759.6806.48\|CTTGCGGCCCCGGGTTCCTCCCGGGGCTACGCCTGTCTGAGCGTCGCT | >hg38_21_+_8433172_8446622_RNA45SN1_45S_50ntFlanks | 6759 | 6806 | 48 | CTTGCGGCCCCGGGTTCCTCCCGGGGCTACGCCTGTCTGAGCGTCGCT | 1957.0 | 5.6 | 1.1 | 5.3 | 1.4E-07 | 3.2E-07 | BT20/MB468 | MITO |
| >hg38_21_+_8433172_8446622_RNA45SN1_45S_50ntFlanks@11452.11478.27\|CTCCTCGGCGCGCGGCGGCGGCGG | >hg38_21_+_8433172_8446622_RNA45SN1_45S_50ntFlanks | 11452 | 11478 | 27 | CTCCTCGGCGCGCGGCGGCGGCGG | 1973.0 | -9.0 | 1.7 | -5.2 | 2.0E-07 | 3.2E-07 | BT20/MB468 | NUC |
| >hg38_21_+_8433172_8446622_RNA45SN1_45S_50ntFlanks@6654.6683.30\|CTCTTAGCGGTGGATCACTCGGCTCGTGCG | >hg38_21_+_8433172_8446622_RNA45SN1_45S_50ntFlanks | 6654 | 6683 | 30 | CTCTTAGCGGTGGATCACTCGGCTCGTGCG | 1732.4 | 4.3 | 0.8 | 5.2 | 1.8E-07 | 3.2E-07 | MB231/MB468 | NUC |
| >hg38_21_+_8433172_8446622_RNA45SN1_45S_50ntFlanks@10786.10822.37\|AACAGCCTCTGGCATGTTGGAACAATGTAGGTAAGGG | >hg38_21_+_8433172_8446622_RNA45SN1_45S_50ntFlanks | 10786 | 10822 | 37 | AACAGCCTCTGGCATGTTGGAACAATGTAGGTAAGGG | 451.5 | -5.7 | 1.1 | -5.2 | 1.8E-07 | 3.2E-07 | MB231/MB468 | NUC |
| >hg38_21_+_8433172_8446622_RNA45SN1_45S_50ntFlanks@3997.4024.28\|CTCTAGATAACCTCGGGCCGATCGCACG | >hg38_21_+_8433172_8446622_RNA45SN1_45S_50ntFlanks | 3997 | 4024 | 28 | CTCTAGATAACCTCGGGCCGATCGCACG | 709.3 | -8.3 | 1.6 | -5.2 | 2.0E-07 | 3.2E-07 | BT20/MB468 | NUC |
| >hg38_21_+_8433172_8446622_RNA45SN1_45S_50ntFlanks@10819.10836.18\|AGGGAAGTCGGCAAGCCG | >hg38_21_+_8433172_8446622_RNA45SN1_45S_50ntFlanks | 10819 | 10836 | 18 | AGGGAAGTCGGCAAGCCG | 534.2 | -7.4 | 1.4 | -5.2 | 1.6E-07 | 3.2E-07 | MB231/MB468 | MP |

| | | | | | | | | | | | | | |
|---|---|---|---|---|---|---|---|---|---|---|---|---|---|
| >hg38_21_+_8433172_8446622_RNA45SN1_45S_50ntFlanks@7988.8013.26\|CAGACGTGGCGACCCGCTGAATTTAA | >hg38_21_+_8433172_8446622_RNA45SN1_45S_50ntFlanks | 7988 | 8013 | 26 | CAGACGTGGCGACCCGCTGAATTTAA | 440.6 | -7.2 | 1.4 | -5.2 | 1.6E-07 | 3.3E-07 | MB231/MB468 | MP |
| >trna128_GlyGCC_6_-_27870686_27870756@1.42.42,trna19_GlyGCC_16_+_70823410_70823480@1.42.42,trna19_GlyGCC_2_-_157257659_157257729@1.42.42,trna24_GlyGCC_16_-_70812942_70813012@1.42.42,trna25_GlyGCC_16_-_70812114_70812184@1.42.42,trna5_GlyGCC_17_+_8029064_8029134@1.42.42,trna68_GlyGCC_1_-_161493637_161493707@1.42.42\|GCATTGGTGGTTCAGTGGTAGAATTCTCGCCTGCCACGCGGG | >trna128_GlyGCC_6_-_27870686_27870756 | 1 | 42 | 42 | GCATTGGTGGTTCAGTGGTAGAATTCTCGCCTGCCACGCGGG | 252.7 | 6.5 | 1.3 | 5.2 | 1.8E-07 | 3.3E-07 | MB231/MB468 | CYTO |
| >hg38_21_+_8433172_8446622_RNA45SN1_45S_50ntFlanks@9947.9978.32\|GTGGCCATGGAAGTCGGAATCCGCTAAGGAGT | >hg38_21_+_8433172_8446622_RNA45SN1_45S_50ntFlanks | 9947 | 9978 | 32 | GTGGCCATGGAAGTCGGAATCCGCTAAGGAGT | 65.0 | 9.8 | 1.8 | 5.5 | 3.8E-08 | 3.3E-07 | BT20/MB468 | MP |
| >am_trna15_CysGCA_17_+_37023898_37023969@14.32.19,trna1_CysACA_5_+_151988596_151988771@14.32.19,trna26_CysGCA_17_-_37310744_37310815@14.32.19,trna27_CysGCA_17_-_37309987_37310058@14.32.19,trna28_CysGCA_17_-_37025545_37025616@14.32.19,trna3_CysGCA_4_-_124430005_124430076@14.32.19,trna5_CysGCA_7_+_149007281_149007352@14.32.19,trna7_CysGCA_3_-_131947944_131948015@14.32.19\|AGTGGTAGAGCATTTGACT | >am_trna15_CysGCA_17_+_37023898_37023969 | 14 | 32 | 19 | AGTGGTAGAGCATTTGACT | 47.8 | 8.8 | 1.7 | 5.2 | 1.6E-07 | 3.3E-07 | MB231/MB468 | MP |
| >MI0000471\|hsa-mir-126&WithFlank&9\|+\|136670596\|136670692@21.40.20(+1A)&[MIMAT0000444&hsa-miR-126-5p&offsets\|0\|-1(+1A);m-114&9\|+\|136670616\|136670636&offsets\|0\|-1(+1A)]\|CATTATTACTTTTGGTACGCA | >MI0000471\|hsa-mir-126&WithFlank&9\|+\|136670596\|136670692 | 21 | 40 | 20(+1A) | CATTATTACTTTTGGTACGCA | 72.7 | 9.4 | 1.8 | 5.2 | 1.6E-07 | 3.3E-07 | MB231/MB468 | MP |
| >hg38_21_+_8433172_8446622_RNA45SN1_45S_50ntFlanks@8839.8867.29\|CGGTGGCGGCGGCGGCGGCGGCGGCGGGA | >hg38_21_+_8433172_8446622_RNA45SN1_45S_50ntFlanks | 8839 | 8867 | 29 | CGGTGGCGGCGGCGGCGGCGGCGGCGGGA | 116.8 | -9.9 | 1.8 | -5.5 | 3.8E-08 | 3.3E-07 | BT20/MB468 | MP |
| >hg38_21_+_8433172_8446622_RNA45SN1_45S_50ntFlanks@11786.11804.19\|GAATGGATGAACGAGATTC | >hg38_21_+_8433172_8446622_RNA45SN1_45S_50ntFlanks | 11786 | 11804 | 19 | GAATGGATGAACGAGATTC | 1123.2 | -5.6 | 1.1 | -5.2 | 1.8E-07 | 3.3E-07 | MB231/MB468 | CYTO |
| >hg38_21_+_8433172_8446622_RNA45SN1_45S_50ntFlanks@9540.9571.32\|CTCTGGTGGAGGTCCGTAGCGGTCCTGACGTG | >hg38_21_+_8433172_8446622_RNA45SN1_45S_50ntFlanks | 9540 | 9571 | 32 | CTCTGGTGGAGGTCCGTAGCGGTCCTGACGTG | 280.5 | -7.3 | 1.4 | -5.2 | 1.6E-07 | 3.3E-07 | MB231/MB468 | MP |
| >hg38_21_+_8433172_8446622_RNA45SN1_45S_50ntFlanks@10412.10429.18\|AGTCGGTCCTGAGAGATG | >hg38_21_+_8433172_8446622_RNA45SN1_45S_50ntFlanks | 10412 | 10429 | 18 | AGTCGGTCCTGAGAGATG | 203.8 | -5.4 | 1.0 | -5.2 | 1.8E-07 | 3.3E-07 | MB231/MB468 | CYTO |
| >hg38_21_+_8433172_8446622_RNA45SN1_45S_50ntFlanks@8344.8373.30\|TAAGGGAAAGTTGAAAAGAACTTTGAAGAG | >hg38_21_+_8433172_8446622_RNA45SN1_45S_50ntFlanks | 8344 | 8373 | 30 | TAAGGGAAAGTTGAAAAGAACTTTGAAGAG | 428.0 | 9.3 | 1.7 | 5.5 | 3.8E-08 | 3.3E-07 | BT20/MB468 | MP |
| >hg38_MT_+_598_1651_RNR1_12S_50ntFlanks@913.950.38\|TCAAAGGACATTTAACTAAAACCCCTACGCATTTATAT | >hg38_MT_+_598_1651_RNR1_12S_50ntFlanks | 913 | 950 | 38 | TCAAAGGACATTTAACTAAAACCCCTACGCATTTATAT | 139.7 | 10.9 | 2.0 | 5.5 | 3.8E-08 | 3.3E-07 | BT20/MB468 | MP |
| >hg38_21_+_8433172_8446622_RNA45SN1_45S_50ntFlanks@11586.11603.18\|GTGCGGACCAGGGGAATC | >hg38_21_+_8433172_8446622_RNA45SN1_45S_50ntFlanks | 11586 | 11603 | 18 | GTGCGGACCAGGGGAATC | 521.4 | -5.2 | 1.0 | -5.3 | 1.4E-07 | 3.3E-07 | BT20/MB468 | MITO |
| >trna116_GluCTC_1_-_145399233_145399304@1.35.35,trna59_GluCTC_1_+_249168447_249168518@1.35.35,trna71_GluCTC_1_-_161439189_161439260@1.35.35,trna74_GluCTC_1_-_161431809_161431880@1.35.35,trna77_GluCTC_1_-_161424398_161424469@1.35.35,trna77_GluCTC_6_+_28949976_28950047@1.35.35,trna80_GluCTC_1_-_161417018_161417089@1.35.35,trna87_GluCTC_6_-_126101393_126101464@1.35.35\|TCCCTGGTGGTCTAGTGGTTAGGATTCGGCGCTCT | >trna116_GluCTC_1_-_145399233_145399304 | 1 | 35 | 35 | TCCCTGGTGGTCTAGTGGTTAGGATTCGGCGCTCT | 2970.6 | 5.3 | 1.0 | 5.2 | 1.9E-07 | 3.3E-07 | MB231/MB468 | NUC |
| >hg38_21_+_8433172_8446622_RNA45SN1_45S_50ntFlanks@3813.3837.25\|TTATGGTTCCTTTGGTCGCTCGCTC | >hg38_21_+_8433172_8446622_RNA45SN1_45S_50ntFlanks | 3813 | 3837 | 25 | TTATGGTTCCTTTGGTCGCTCGCTC | 1350.4 | -8.8 | 1.7 | -5.2 | 2.1E-07 | 3.3E-07 | BT20/MB468 | NUC |
| >trnaMT_ValTAC_MT_+_1602_1670@52.71.20\|TTAACTTGACCGCTCTGACC | >trnaMT_ValTAC_MT_+_1602_1670 | 52 | 71 | 20 | TTAACTTGACCGCTCTGACC | 542.1 | 5.1 | 1.0 | 5.3 | 1.4E-07 | 3.3E-07 | BT20/MB468 | MITO |
| >hg38_21_+_8433172_8446622_RNA45SN1_45S_50ntFlanks@13005.13039.35\|TCTATTGAAAGTCAGCCCTCGACACAAGGGTTTGT | >hg38_21_+_8433172_8446622_RNA45SN1_45S_50ntFlanks | 13005 | 13039 | 35 | TCTATTGAAAGTCAGCCCTCGACACAAGGGTTTGT | 852.8 | 5.2 | 1.0 | 5.3 | 1.4E-07 | 3.3E-07 | BT20/MB468 | MITO |
| >hg38_21_+_8433172_8446622_RNA45SN1_45S_50ntFlanks@5379.5399.21\|ATTAAGTCCCTGCCCTTTGTA | >hg38_21_+_8433172_8446622_RNA45SN1_45S_50ntFlanks | 5379 | 5399 | 21 | ATTAAGTCCCTGCCCTTTGTA | 1477.2 | -8.3 | 1.6 | -5.3 | 1.4E-07 | 3.3E-07 | BT20/MB468 | MITO |
| >hg38_21_+_8433172_8446622_RNA45SN1_45S_50ntFlanks@12007.12032.26\|GCCGCCGGTGAAATACCACTACTCTG | >hg38_21_+_8433172_8446622_RNA45SN1_45S_50ntFlanks | 12007 | 12032 | 26 | GCCGCCGGTGAAATACCACTACTCTG | 13383.8 | -8.5 | 1.6 | -5.2 | 2.1E-07 | 3.3E-07 | BT20/MB468 | NUC |
| >MI0000736\|hsa-mir-30c-1&WithFlank&1\|+\|40757278\|40757378@23.44.22;MI0000254\|hsa-mir-30c-2&WithFlank&6\|-\|71376954\|71377037@13.34.22&[MIMAT0000244&hsa-miR-30c-5p&offsets\|0\|-1;m-57&1\|+\|40757300\|40757323&offsets\|0\|-2];[MIMAT0000244_1&hsa-miR-30c-5p&offsets\|0\|-1;m-56&6\|-\|71377002\|71377025&offsets\|0\|-2]\|TGTAAACATCCTACACTCTCAG | >MI0000736\|hsa-mir-30c-1&WithFlank&1\|+\|40757278\|40757378 | 23 | 44 | 22 | TGTAAACATCCTACACTCTCAG | 282.5 | 5.7 | 1.1 | 5.2 | 1.9E-07 | 3.3E-07 | MB231/MB468 | NUC |
| >hg38_21_+_8433172_8446622_RNA45SN1_45S_50ntFlanks@11572.11606.35\|CCGACTTAGAACTGGTGCGGACCAGGGGAATCCGA | >hg38_21_+_8433172_8446622_RNA45SN1_45S_50ntFlanks | 11572 | 11606 | 35 | CCGACTTAGAACTGGTGCGGACCAGGGGAATCCGA | 313.1 | -2.7 | 0.5 | -5.2 | 1.8E-07 | 3.3E-07 | MB231/MB468 | CYTO |
| >hg38_21_+_8433172_8446622_RNA45SN1_45S_50ntFlanks@8190.8208.19\|CCAGCCCGTGGACGGTGTG | >hg38_21_+_8433172_8446622_RNA45SN1_45S_50ntFlanks | 8190 | 8208 | 19 | CCAGCCCGTGGACGGTGTG | 92.6 | -9.6 | 1.7 | -5.5 | 3.9E-08 | 3.3E-07 | BT20/MB468 | MP |
| >hg38_21_+_8433172_8446622_RNA45SN1_45S_50ntFlanks@10792.10822.31\|CTCTGGCATGTTGGAACAATGTAGGTAAGGG | >hg38_21_+_8433172_8446622_RNA45SN1_45S_50ntFlanks | 10792 | 10822 | 31 | CTCTGGCATGTTGGAACAATGTAGGTAAGGG | 385.7 | -6.2 | 1.2 | -5.2 | 1.7E-07 | 3.3E-07 | MB231/MB468 | MP |
| >MI0000095\|hsa-mir-93&WithFlank&7\|-\|100093762\|100093853@17.38.22&[MIMAT0000093&hsa-miR-93-5p&offsets\|0\|-1;m-29&7\|-\|100093815\|100093837&offsets\|0\|-1]\|CAAAGTGCTGTTCGTGCAGGTA | >MI0000095\|hsa-mir-93&WithFlank&7\|-\|100093762\|100093853 | 17 | 38 | 22 | CAAAGTGCTGTTCGTGCAGGTA | 869.9 | 5.5 | 1.1 | 5.2 | 2.0E-07 | 3.3E-07 | MB231/MB468 | MITO |
| >MI0000255\|hsa-mir-30d&WithFlank&8\|-\|134804870\|134804951@12.35.24(+3U)&[MIMAT0000245&hsa-miR-30d-5p&offsets\|0\|+2(+3U);m-21&8\|-\|134804919\|134804940&offsets\|0\|+2(+3U)]\|TGTAAACATCCCCGACTGGAAGCTTTT | >MI0000255\|hsa-mir-30d&WithFlank&8\|-\|134804870\|134804951 | 12 | 35 | 24(+3U) | TGTAAACATCCCCGACTGGAAGCTTTT | 402.2 | 8.3 | 1.6 | 5.2 | 2.0E-07 | 3.3E-07 | MB231/MB468 | MITO |
| >hg38_21_+_8433172_8446622_RNA45SN1_45S_50ntFlanks@8839.8866.28\|CGGTGGCGGCGGCGGCGGCGGCGGCGGG | >hg38_21_+_8433172_8446622_RNA45SN1_45S_50ntFlanks | 8839 | 8866 | 28 | CGGTGGCGGCGGCGGCGGCGGCGGCGGG | 113.6 | -9.9 | 1.8 | -5.5 | 3.9E-08 | 3.3E-07 | BT20/MB468 | MP |
| >hg38_21_+_8433172_8446622_RNA45SN1_45S_50ntFlanks@10391.10426.36\|AACAGCAGTTGAACATGGGTCAGTCGGTCCTGAGAG | >hg38_21_+_8433172_8446622_RNA45SN1_45S_50ntFlanks | 10391 | 10426 | 36 | AACAGCAGTTGAACATGGGTCAGTCGGTCCTGAGAG | 464.6 | -7.3 | 1.4 | -5.2 | 2.0E-07 | 3.3E-07 | MB231/MB468 | MITO |
| >hg38_21_+_8433172_8446622_RNA45SN1_45S_50ntFlanks@12396.12417.22\|GCGACGTCGCTTTTTGATCCTT | >hg38_21_+_8433172_8446622_RNA45SN1_45S_50ntFlanks | 12396 | 12417 | 22 | GCGACGTCGCTTTTTGATCCTT | 3045.1 | -9.3 | 1.8 | -5.2 | 2.1E-07 | 3.3E-07 | BT20/MB468 | NUC |
| >hg38_21_+_8433172_8446622_RNA45SN1_45S_50ntFlanks@8799.8827.29\|CCTCGGGAGGGCGCGCGGGTCGGGGCGGC | >hg38_21_+_8433172_8446622_RNA45SN1_45S_50ntFlanks | 8799 | 8827 | 29 | CCTCGGGAGGGCGCGCGGGTCGGGGCGGC | 104.4 | -9.8 | 1.8 | -5.5 | 3.9E-08 | 3.3E-07 | BT20/MB468 | MP |
| >hg38_21_+_8433172_8446622_RNA45SN1_45S_50ntFlanks@10794.10826.33\|CTGGCATGTTGGAACAATGTAGGTAAGGGAAGT | >hg38_21_+_8433172_8446622_RNA45SN1_45S_50ntFlanks | 10794 | 10826 | 33 | CTGGCATGTTGGAACAATGTAGGTAAGGGAAGT | 547.0 | -4.3 | 0.8 | -5.5 | 3.9E-08 | 3.3E-07 | BT20/MB468 | MP |
| >trna116_GluCTC_1_-_145399233_145399304@3.33.31,trna59_GluCTC_1_+_249168447_249168518@3.33.31,trna71_GluCTC_1_-_161439189_161439260@3.33.31,trna74_GluCTC_1_-_161431809_161431880@3.33.31,trna77_GluCTC_1_-_161424398_161424469@3.33.31,trna77_GluCTC_6_+_28949976_28950047@3.33.31,trna80_GluCTC_1_-_161417018_161417089@3.33.31,trna87_GluCTC_6_-_126101393_126101464@3.33.31\|CCTGGTGGTCTAGTGGTTAGGATTCGGCGCT | >trna116_GluCTC_1_-_145399233_145399304 | 3 | 33 | 31 | CCTGGTGGTCTAGTGGTTAGGATTCGGCGCT | 520.1 | 4.3 | 0.8 | 5.2 | 2.0E-07 | 3.3E-07 | MB231/MB468 | MITO |
| >hg38_21_+_8433172_8446622_RNA45SN1_45S_50ntFlanks@9300.9324.25\|ACGGACCAAGGAGTCTAACACGTGC | >hg38_21_+_8433172_8446622_RNA45SN1_45S_50ntFlanks | 9300 | 9324 | 25 | ACGGACCAAGGAGTCTAACACGTGC | 66.7 | 9.8 | 1.8 | 5.5 | 3.9E-08 | 3.3E-07 | BT20/MB468 | MP |
| >MI0000750\|hsa-mir-26a-2&WithFlank&12\|-\|57824603\|57824698@20.39.20(+1U);MI0000083\|hsa-mir-26a-1&WithFlank&3\|+\|37969398\|37969486@16.35.20(+1U)&[MIMAT0000082&hsa-miR-26a-5p&offsets\|0\|-2(+1U);m-33&12\|-\|57824658\|57824679&offsets\|0\|-2(+1U)];[MIMAT0000082_1&hsa-miR-26a-5p&offsets\|0\|-2(+1U);m-34&3\|+\|37969413\|37969434&offsets\|0\|-2(+1U)]\|TTCAAGTAATCCAGGATAGGT | >MI0000750\|hsa-mir-26a-2&WithFlank&12\|-\|57824603\|57824698 | 20 | 39 | 20(+1U) | TTCAAGTAATCCAGGATAGGT | 111.4 | 3.0 | 0.6 | 5.2 | 2.1E-07 | 3.3E-07 | BT20/MB468 | NUC |
| >MI0003560\|hsa-mir-92b&WithFlank&1\|+\|155195171\|155195278@67.87.21(+1U)&[MIMAT0003218&hsa-miR-92b-3p&offsets\|0\|-1(+1U);m-115&1\|+\|155195237\|155195258&offsets\|0\|-1(+1U)]\|TATTGCACTCGTCCCGGCCTCT | >MI0003560\|hsa-mir-92b&WithFlank&1\|+\|155195171\|155195278 | 67 | 87 | 21(+1U) | TATTGCACTCGTCCCGGCCTCT | 720.9 | 3.5 | 0.7 | 5.2 | 2.1E-07 | 3.3E-07 | BT20/MB468 | NUC |
| >trna119_LysCTT_1_-_145395522_145395594@4.36.33,trna11_LysCTT_5_-_180648979_180649051@4.36.33,trna13_LysCTT_6_+_26556774_26556846@4.36.33,trna7_LysCTT_16_+_3225692_3225764@4.36.33,trna9_LysCTT_5_+_180634755_180634827@4.36.33\|CGGCTAGCTCAGTCGGTAGAGCATGAGACTCTT | >trna119_LysCTT_1_-_145395522_145395594 | 4 | 36 | 33 | CGGCTAGCTCAGTCGGTAGAGCATGAGACTCTT | 2226.5 | 7.3 | 1.4 | 5.2 | 2.0E-07 | 3.3E-07 | MB231/MB468 | MITO |
| >hg38_21_+_8433172_8446622_RNA45SN1_45S_50ntFlanks@10766.10785.20\|CGCAGCAGGTCTCCAAGGTG | >hg38_21_+_8433172_8446622_RNA45SN1_45S_50ntFlanks | 10766 | 10785 | 20 | CGCAGCAGGTCTCCAAGGTG | 320.2 | -7.3 | 1.4 | -5.2 | 1.7E-07 | 3.3E-07 | MB231/MB468 | MP |
| >hg38_21_+_8433172_8446622_RNA45SN1_45S_50ntFlanks@11671.11692.22\|CCAGTGCTCTGAATGTCAAAGT | >hg38_21_+_8433172_8446622_RNA45SN1_45S_50ntFlanks | 11671 | 11692 | 22 | CCAGTGCTCTGAATGTCAAAGT | 835.8 | -4.1 | 0.8 | -5.2 | 2.1E-07 | 3.3E-07 | BT20/MB468 | NUC |
| >hg38_21_+_8433172_8446622_RNA45SN1_45S_50ntFlanks@10805.10823.19\|GAACAATGTAGGTAAGGGA | >hg38_21_+_8433172_8446622_RNA45SN1_45S_50ntFlanks | 10805 | 10823 | 19 | GAACAATGTAGGTAAGGGA | 240.0 | -6.2 | 1.2 | -5.2 | 1.6E-07 | 3.4E-07 | BT20/MB468 | CYTO |
| >hg38_21_+_8433172_8446622_RNA45SN1_45S_50ntFlanks@11686.11717.32\|TCAAAGTGAAGAAATTCAATGAAGCGCGGGTA | >hg38_21_+_8433172_8446622_RNA45SN1_45S_50ntFlanks | 11686 | 11717 | 32 | TCAAAGTGAAGAAATTCAATGAAGCGCGGGTA | 212.6 | -7.1 | 1.4 | -5.2 | 1.6E-07 | 3.4E-07 | BT20/MB468 | CYTO |
| >hg38_21_+_8433172_8446622_RNA45SN1_45S_50ntFlanks@12418.12453.36\|CGATGTCGGCTCTTCCTATCATTGTGAAGCAGAATT | >hg38_21_+_8433172_8446622_RNA45SN1_45S_50ntFlanks | 12418 | 12453 | 36 | CGATGTCGGCTCTTCCTATCATTGTGAAGCAGAATT | 295.0 | -4.9 | 0.9 | -5.2 | 2.1E-07 | 3.4E-07 | BT20/MB468 | NUC |

<table>
<tr><th></th><th></th><th></th><th></th><th></th><th></th><th></th><th></th><th></th><th></th><th></th><th></th><th></th><th></th></tr>
<tr><td>>MI0000433|hsa-let-7g&WithFlank&3|-|52268272|52268367@11.31.21&[MIMAT0000414&hsa-let-7g-5p&offsets|0|-1;m-58&3|-|52268336|52268357&offsets|0|-1]|TGAGGTAGTAGTTTGTACAGT</td><td>>MI0000433|hsa-let-7g&WithFlank&3|-|52268272|52268367</td><td>11</td><td>31</td><td>21</td><td>TGAGGTAGTAGTTTGTACAGT</td><td>1732.7</td><td>2.2</td><td>0.4</td><td>5.2</td><td>2.1E-07</td><td>3.4E-07</td><td>BT20/MB468</td><td>NUC</td></tr>
<tr><td>>MI0000289|hsa-mir-181a-1&WithFlank&1|-|198859038|198859159@30.51.22;MI0000269|hsa-mir-181a-2&WithFlank&9|+|124692436|124692557@45.66.22&[MIMAT0000256&hsa-miR-181a-5p&offsets|0|-1;m-32&1|-|198859108|198859130&offsets|0|-1];[MIMAT0000256_1&hsa-miR-181a-5p&offsets|0|-1;m-32&9|+|124692480|124692502&offsets|0|-1]|AACATTCAACGCTGTCGGTGAG</td><td>>MI0000289|hsa-mir-181a-1&WithFlank&1|-|198859038|198859159</td><td>30</td><td>51</td><td>22</td><td>AACATTCAACGCTGTCGGTGAG</td><td>447.7</td><td>4.7</td><td>0.9</td><td>5.2</td><td>2.0E-07</td><td>3.4E-07</td><td>MB231/MB468</td><td>MITO</td></tr>
<tr><td>>hg38_21_+_8433172_8446622_RNA45SN1_45S_50ntFlanks@12617.12653.37|TGAGGAGCCAATGGGGCGAAGCTACCATCTGTGGGAT</td><td>>hg38_21_+_8433172_8446622_RNA45SN1_45S_50ntFlanks</td><td>12617</td><td>12653</td><td>37</td><td>TGAGGAGCCAATGGGGCGAAGCTACCATCTGTGGGAT</td><td>569.5</td><td>-3.9</td><td>0.7</td><td>-5.3</td><td>1.4E-07</td><td>3.4E-07</td><td>BT20/MB468</td><td>MITO</td></tr>
<tr><td>>MI0003591|hsa-mir-584&WithFlank&5|-|149062307|149062415@22.41.20&[MIMAT0003249&hsa-miR-584-5p&offsets|0|-2;m-167&5|-|149062374|149062394&offsets|0|-1]|TTATGGTTTGCCTGGGACTG</td><td>>MI0003591|hsa-mir-584&WithFlank&5|-|149062307|149062415</td><td>22</td><td>41</td><td>20</td><td>TTATGGTTTGCCTGGGACTG</td><td>249.8</td><td>7.0</td><td>1.3</td><td>5.2</td><td>2.0E-07</td><td>3.4E-07</td><td>MB231/MB468</td><td>MITO</td></tr>
<tr><td>>trnaMT_HisGTG_MT_+_12138_12206@20.58.39|AACATCAGATTGTGAATCTGACAACAGAGGCTTACGACC</td><td>>trnaMT_HisGTG_MT_+_12138_12206</td><td>20</td><td>58</td><td>39</td><td>AACATCAGATTGTGAATCTGACAACAGAGGCTTACGACC</td><td>137.6</td><td>10.9</td><td>2.0</td><td>5.5</td><td>4.0E-08</td><td>3.4E-07</td><td>BT20/MB468</td><td>MP</td></tr>
<tr><td>>hg38_21_+_8433172_8446622_RNA45SN1_45S_50ntFlanks@10408.10426.19|GGTCAGTCGGTCCTGAGAG</td><td>>hg38_21_+_8433172_8446622_RNA45SN1_45S_50ntFlanks</td><td>10408</td><td>10426</td><td>19</td><td>GGTCAGTCGGTCCTGAGAG</td><td>646.1</td><td>-7.0</td><td>1.3</td><td>-5.2</td><td>1.6E-07</td><td>3.4E-07</td><td>BT20/MB468</td><td>CYTO</td></tr>
<tr><td>>hg38_21_+_8433172_8446622_RNA45SN1_45S_50ntFlanks@11589.11609.21|CGGACCAGGGGAATCCGACTG</td><td>>hg38_21_+_8433172_8446622_RNA45SN1_45S_50ntFlanks</td><td>11589</td><td>11609</td><td>21</td><td>CGGACCAGGGGAATCCGACTG</td><td>513.9</td><td>-3.8</td><td>0.7</td><td>-5.2</td><td>2.0E-07</td><td>3.4E-07</td><td>MB231/MB468</td><td>MITO</td></tr>
<tr><td>>am_MI0003776|hsa-mir-320b-1&WithFlank&1|+|116671740|116671823@48.65.18;MI0003839|hsa-mir-320b-2&WithFlank&1|-|224257034|224257116@47.64.18;MI0008190|hsa-mir-320d-1&WithFlank&13|-|40727810|40727893@48.65.18;MI0003778|hsa-mir-320c-1&WithFlank&18|+|21683512|21683595@48.65.18;MI0008191|hsa-mir-320c-2&WithFlank&18|+|24321669|24321752@48.65.18;MI0000542|hsa-mir-320a&WithFlank&8|-|22244960|22245043@48.65.18;MI0008192|hsa-mir-320d-2&WithFlank&X|-|140926154|140926237@48.65.18&[MIMAT0005792&hsa-miR-320b&offsets|0|-4;m-183&1|+|116671787|116671806&offsets|0|-2];[MIMAT0005792_1&hsa-miR-320b&offsets|0|-4;m-183&1|-|224257051|224257070&offsets|0|-2];[MIMAT0006764&hsa-miR-320d&offsets|0|-1;m-246&13|-|40727829|40727846&offsets|0|0];[MIMAT0005793&hsa-miR-320c&offsets|0|-2;m-222&18|+|21683559|21683578&offsets|0|-2];[MIMAT0005793_1&hsa-miR-320c&offsets|0|-2;m-219&18|+|24321716|24321735&offsets|0|-2];[MIMAT0000510&hsa-miR-320a-3p&offsets|0|-4;m-60&8|-|22244975|22244996&offsets|0|-4];[MIMAT0006764_1&hsa-miR-320d&offsets|0|-1;m-253&X|-|140926173|140926190&offsets|0|0]|AAAAGCTGGGTTGAGAGG</td><td>>am_MI0003776|hsa-mir-320b-1&WithFlank&1|+|116671740|116671823</td><td>48</td><td>65</td><td>18</td><td>AAAAGCTGGGTTGAGAGG</td><td>2592.1</td><td>5.9</td><td>1.1</td><td>5.5</td><td>4.0E-08</td><td>3.4E-07</td><td>BT20/MB468</td><td>MP</td></tr>
<tr><td>>hg38_21_+_8433172_8446622_RNA45SN1_45S_50ntFlanks@5492.5516.25|AGAAGACGGTCGAACTTGACTATCT</td><td>>hg38_21_+_8433172_8446622_RNA45SN1_45S_50ntFlanks</td><td>5492</td><td>5516</td><td>25</td><td>AGAAGACGGTCGAACTTGACTATCT</td><td>3153.5</td><td>5.3</td><td>1.0</td><td>5.5</td><td>4.0E-08</td><td>3.4E-07</td><td>BT20/MB468</td><td>MP</td></tr>
<tr><td>>hg38_21_+_8433172_8446622_RNA45SN1_45S_50ntFlanks@476.513.38|TTCGTGATCGATGTGGTGACGTCGTGCTCTCCCGGGCC</td><td>>hg38_21_+_8433172_8446622_RNA45SN1_45S_50ntFlanks</td><td>476</td><td>513</td><td>38</td><td>TTCGTGATCGATGTGGTGACGTCGTGCTCTCCCGGGCC</td><td>1567.1</td><td>4.0</td><td>0.8</td><td>5.2</td><td>1.9E-07</td><td>3.4E-07</td><td>MB231/MB468</td><td>NUC</td></tr>
<tr><td>>hg38_21_+_8433172_8446622_RNA45SN1_45S_50ntFlanks@10873.10890.18|CTGGGTCGGTCGGGCTGG</td><td>>hg38_21_+_8433172_8446622_RNA45SN1_45S_50ntFlanks</td><td>10873</td><td>10890</td><td>18</td><td>CTGGGTCGGTCGGGCTGG</td><td>891.3</td><td>-6.7</td><td>1.3</td><td>-5.2</td><td>2.1E-07</td><td>3.4E-07</td><td>BT20/MB468</td><td>NUC</td></tr>
<tr><td>>hg38_21_+_8433172_8446622_RNA45SN1_45S_50ntFlanks@10678.10709.32|CGGTTCCGGCGGCGTCCGGTGAGCTCTCGCTG</td><td>>hg38_21_+_8433172_8446622_RNA45SN1_45S_50ntFlanks</td><td>10678</td><td>10709</td><td>32</td><td>CGGTTCCGGCGGCGTCCGGTGAGCTCTCGCTG</td><td>12500.5</td><td>-5.9</td><td>1.1</td><td>-5.5</td><td>4.0E-08</td><td>3.4E-07</td><td>BT20/MB468</td><td>MP</td></tr>
<tr><td>>hg38_21_+_8433172_8446622_RNA45SN1_45S_50ntFlanks@11658.11685.28|ATGTGATTTCTGCCCAGTGCTCTGAATG</td><td>>hg38_21_+_8433172_8446622_RNA45SN1_45S_50ntFlanks</td><td>11658</td><td>11685</td><td>28</td><td>ATGTGATTTCTGCCCAGTGCTCTGAATG</td><td>832.7</td><td>13.0</td><td>2.3</td><td>5.8</td><td>8.4E-09</td><td>3.4E-07</td><td>BT20/MB231</td><td>MP</td></tr>
<tr><td>>hg38_21_+_8433172_8446622_RNA45SN1_45S_50ntFlanks@9791.9819.29|TGGCGTGGAGCCGGGCGTGGAATGCGAGT</td><td>>hg38_21_+_8433172_8446622_RNA45SN1_45S_50ntFlanks</td><td>9791</td><td>9819</td><td>29</td><td>TGGCGTGGAGCCGGGCGTGGAATGCGAGT</td><td>515.3</td><td>12.3</td><td>2.1</td><td>5.8</td><td>8.4E-09</td><td>3.4E-07</td><td>BT20/MB231</td><td>MP</td></tr>
<tr><td>>hg38_21_+_8433172_8446622_RNA45SN1_45S_50ntFlanks@11564.11602.39|CCTAGCAGCCGACTTAGAACTGGTGCGGACCAGGGGAAT</td><td>>hg38_21_+_8433172_8446622_RNA45SN1_45S_50ntFlanks</td><td>11564</td><td>11602</td><td>39</td><td>CCTAGCAGCCGACTTAGAACTGGTGCGGACCAGGGGAAT</td><td>799.8</td><td>-3.9</td><td>0.8</td><td>-5.2</td><td>1.6E-07</td><td>3.4E-07</td><td>BT20/MB468</td><td>CYTO</td></tr>
<tr><td>>hg38_21_+_8433172_8446622_RNA45SN1_45S_50ntFlanks@8380.8400.21|CAAGAGGGCGTGAAACCGTTA</td><td>>hg38_21_+_8433172_8446622_RNA45SN1_45S_50ntFlanks</td><td>8380</td><td>8400</td><td>21</td><td>CAAGAGGGCGTGAAACCGTTA</td><td>2254.4</td><td>-7.7</td><td>1.5</td><td>-5.2</td><td>1.6E-07</td><td>3.4E-07</td><td>BT20/MB468</td><td>CYTO</td></tr>
<tr><td>>hg38_21_+_8433172_8446622_RNA45SN1_45S_50ntFlanks@11567.11584.18|AGCAGCCGACTTAGAACT</td><td>>hg38_21_+_8433172_8446622_RNA45SN1_45S_50ntFlanks</td><td>11567</td><td>11584</td><td>18</td><td>AGCAGCCGACTTAGAACT</td><td>1070.3</td><td>4.0</td><td>0.8</td><td>5.2</td><td>2.0E-07</td><td>3.4E-07</td><td>MB231/MB468</td><td>MITO</td></tr>
<tr><td>>hg38_21_+_8433172_8446622_RNA45SN1_45S_50ntFlanks@5526.5550.25|AAAAGTCGTAACAAGGTTTCCGTAG</td><td>>hg38_21_+_8433172_8446622_RNA45SN1_45S_50ntFlanks</td><td>5526</td><td>5550</td><td>25</td><td>AAAAGTCGTAACAAGGTTTCCGTAG</td><td>495.7</td><td>-6.9</td><td>1.3</td><td>-5.2</td><td>1.9E-07</td><td>3.4E-07</td><td>MB231/MB468</td><td>NUC</td></tr>
<tr><td>>hg38_21_+_8433172_8446622_RNA45SN1_45S_50ntFlanks@12418.12441.24|CGATGTCGGCTCTTCCTATCATTG</td><td>>hg38_21_+_8433172_8446622_RNA45SN1_45S_50ntFlanks</td><td>12418</td><td>12441</td><td>24</td><td>CGATGTCGGCTCTTCCTATCATTG</td><td>1848.5</td><td>-3.4</td><td>0.7</td><td>-5.2</td><td>1.9E-07</td><td>3.4E-07</td><td>MB231/MB468</td><td>CYTO</td></tr>
<tr><td>>MI0000736|hsa-mir-30c-1&WithFlank&1|+|40757278|40757378@23.45.23(+1C);MI0000254|hsa-mir-30c-2&WithFlank&6|-|71376954|71377037@13.35.23(+1C)&[MIMAT0000244&hsa-miR-30c-5p&offsets|0|0(+1C);m-57&1|+|40757300|40757323&offsets|0|-1(+1C)];[MIMAT0000244_1&hsa-miR-30c-5p&offsets|0|0(+1C);m-56&6|-|71377002|71377025&offsets|0|-1(+1C)]|TGTAAACATCCTACACTCTCAGCC</td><td>>MI0000736|hsa-mir-30c-1&WithFlank&1|+|40757278|40757378</td><td>23</td><td>45</td><td>23(+1C)</td><td>TGTAAACATCCTACACTCTCAGCC</td><td>1391.2</td><td>4.5</td><td>0.9</td><td>5.2</td><td>1.9E-07</td><td>3.4E-07</td><td>MB231/MB468</td><td>CYTO</td></tr>
<tr><td>>hg38_21_+_8433172_8446622_RNA45SN1_45S_50ntFlanks@9604.9622.19|AAAGACTAATCGAACCATC</td><td>>hg38_21_+_8433172_8446622_RNA45SN1_45S_50ntFlanks</td><td>9604</td><td>9622</td><td>19</td><td>AAAGACTAATCGAACCATC</td><td>3017.3</td><td>-9.6</td><td>1.8</td><td>-5.2</td><td>2.1E-07</td><td>3.4E-07</td><td>BT20/MB468</td><td>NUC</td></tr>
<tr><td>>hg38_21_+_8433172_8446622_RNA45SN1_45S_50ntFlanks@11217.11250.34|CTGGACGCGAGCCGGGCCCTTCCCGTGGATCGCC</td><td>>hg38_21_+_8433172_8446622_RNA45SN1_45S_50ntFlanks</td><td>11217</td><td>11250</td><td>34</td><td>CTGGACGCGAGCCGGGCCCTTCCCGTGGATCGCC</td><td>305.9</td><td>-4.5</td><td>0.9</td><td>-5.2</td><td>2.0E-07</td><td>3.4E-07</td><td>MB231/MB468</td><td>MITO</td></tr>
<tr><td>>hg38_21_+_8433172_8446622_RNA45SN1_45S_50ntFlanks@10386.10409.24|ATGTGAACAGCAGTTGAACATGGG</td><td>>hg38_21_+_8433172_8446622_RNA45SN1_45S_50ntFlanks</td><td>10386</td><td>10409</td><td>24</td><td>ATGTGAACAGCAGTTGAACATGGG</td><td>309.6</td><td>-11.4</td><td>2.1</td><td>-5.5</td><td>4.0E-08</td><td>3.4E-07</td><td>BT20/MB468</td><td>MP</td></tr>
<tr><td>>hg38_21_+_8433172_8446622_RNA45SN1_45S_50ntFlanks@9360.9380.21|CAATGAAGGTGAAGGCCGGCG</td><td>>hg38_21_+_8433172_8446622_RNA45SN1_45S_50ntFlanks</td><td>9360</td><td>9380</td><td>21</td><td>CAATGAAGGTGAAGGCCGGCG</td><td>135.6</td><td>-10.2</td><td>1.9</td><td>-5.5</td><td>4.0E-08</td><td>3.4E-07</td><td>BT20/MB468</td><td>MP</td></tr>
<tr><td>>hg38_21_+_8433172_8446622_RNA45SN1_45S_50ntFlanks@11694.11717.24|AAGAAATTCAATGAAGCGCGGGTA</td><td>>hg38_21_+_8433172_8446622_RNA45SN1_45S_50ntFlanks</td><td>11694</td><td>11717</td><td>24</td><td>AAGAAATTCAATGAAGCGCGGGTA</td><td>555.5</td><td>-6.3</td><td>1.2</td><td>-5.2</td><td>1.7E-07</td><td>3.4E-07</td><td>MB231/MB468</td><td>MP</td></tr>
<tr><td>>hg38_21_+_8433172_8446622_RNA45SN1_45S_50ntFlanks@9561.9596.36|GTCCTGACGTGCAAATCGGTCGTCCGACCTGGGTAT</td><td>>hg38_21_+_8433172_8446622_RNA45SN1_45S_50ntFlanks</td><td>9561</td><td>9596</td><td>36</td><td>GTCCTGACGTGCAAATCGGTCGTCCGACCTGGGTAT</td><td>807.4</td><td>-5.9</td><td>1.1</td><td>-5.2</td><td>2.0E-07</td><td>3.4E-07</td><td>MB231/MB468</td><td>MITO</td></tr>
<tr><td>>hg38_21_+_8433172_8446622_RNA45SN1_45S_50ntFlanks@10319.10336.18|GCAGATCTTGGTGGTAGT</td><td>>hg38_21_+_8433172_8446622_RNA45SN1_45S_50ntFlanks</td><td>10319</td><td>10336</td><td>18</td><td>GCAGATCTTGGTGGTAGT</td><td>85.2</td><td>10.2</td><td>1.9</td><td>5.5</td><td>4.0E-08</td><td>3.4E-07</td><td>BT20/MB468</td><td>MP</td></tr>
<tr><td>>hg38_21_+_8433172_8446622_RNA45SN1_45S_50ntFlanks@7975.7998.24|CGCGACCTCAGATCAGACGTGGCG</td><td>>hg38_21_+_8433172_8446622_RNA45SN1_45S_50ntFlanks</td><td>7975</td><td>7998</td><td>24</td><td>CGCGACCTCAGATCAGACGTGGCG</td><td>2258.2</td><td>6.1</td><td>1.2</td><td>5.3</td><td>1.5E-07</td><td>3.4E-07</td><td>BT20/MB468</td><td>MITO</td></tr>
<tr><td>>hg38_21_+_8433172_8446622_RNA45SN1_45S_50ntFlanks@9572.9596.25|CAAATCGGTCGTCCGACCTGGGTAT</td><td>>hg38_21_+_8433172_8446622_RNA45SN1_45S_50ntFlanks</td><td>9572</td><td>9596</td><td>25</td><td>CAAATCGGTCGTCCGACCTGGGTAT</td><td>1248.9</td><td>-6.3</td><td>1.2</td><td>-5.2</td><td>1.9E-07</td><td>3.4E-07</td><td>MB231/MB468</td><td>NUC</td></tr>
<tr><td>>hg38_21_+_8433172_8446622_RNA45SN1_45S_50ntFlanks@12418.12455.38|CGATGTCGGCTCTTCCTATCATTGTGAAGCAGAATTCA</td><td>>hg38_21_+_8433172_8446622_RNA45SN1_45S_50ntFlanks</td><td>12418</td><td>12455</td><td>38</td><td>CGATGTCGGCTCTTCCTATCATTGTGAAGCAGAATTCA</td><td>3917.1</td><td>-4.6</td><td>0.9</td><td>-5.2</td><td>1.9E-07</td><td>3.4E-07</td><td>MB231/MB468</td><td>NUC</td></tr>
<tr><td>>hg38_21_+_8433172_8446622_RNA45SN1_45S_50ntFlanks@5488.5516.29|GCTGAGAAGACGGTCGAACTTGACTATCT</td><td>>hg38_21_+_8433172_8446622_RNA45SN1_45S_50ntFlanks</td><td>5488</td><td>5516</td><td>29</td><td>GCTGAGAAGACGGTCGAACTTGACTATCT</td><td>123.5</td><td>2.6</td><td>0.5</td><td>5.2</td><td>2.2E-07</td><td>3.4E-07</td><td>BT20/MB468</td><td>NUC</td></tr>
<tr><td>>hg38_1_-_228634819_228635039_RNA5S12_5S_50ntFlanks@90.127.38|TGATCTCGGAAGCTAAGCAGGGTCGGGCCTGGTTAGTA</td><td>>hg38_1_-_228634819_228635039_RNA5S12_5S_50ntFlanks</td><td>90</td><td>127</td><td>38</td><td>TGATCTCGGAAGCTAAGCAGGGTCGGGCCTGGTTAGTA</td><td>67.3</td><td>9.8</td><td>1.8</td><td>5.5</td><td>4.1E-08</td><td>3.4E-07</td><td>BT20/MB468</td><td>MP</td></tr>
<tr><td>>hg38_21_+_8433172_8446622_RNA45SN1_45S_50ntFlanks@11769.11786.18|TAATTAGTGACGCGCATG</td><td>>hg38_21_+_8433172_8446622_RNA45SN1_45S_50ntFlanks</td><td>11769</td><td>11786</td><td>18</td><td>TAATTAGTGACGCGCATG</td><td>320.1</td><td>-5.9</td><td>1.1</td><td>-5.2</td><td>2.0E-07</td><td>3.4E-07</td><td>MB231/MB468</td><td>NUC</td></tr>
<tr><td>>hg38_21_+_8433172_8446622_RNA45SN1_45S_50ntFlanks@11714.11740.27|GGTAAACGGCGGGAGTAACTATGACTC</td><td>>hg38_21_+_8433172_8446622_RNA45SN1_45S_50ntFlanks</td><td>11714</td><td>11740</td><td>27</td><td>GGTAAACGGCGGGAGTAACTATGACTC</td><td>304.1</td><td>-4.2</td><td>0.8</td><td>-5.2</td><td>2.0E-07</td><td>3.4E-07</td><td>MB231/MB468</td><td>NUC</td></tr>
<tr><td>>hg38_21_+_8433172_8446622_RNA45SN1_45S_50ntFlanks@12907.12931.25|CACCGCACGTTCGTGGGGAACCTGG</td><td>>hg38_21_+_8433172_8446622_RNA45SN1_45S_50ntFlanks</td><td>12907</td><td>12931</td><td>25</td><td>CACCGCACGTTCGTGGGGAACCTGG</td><td>304.6</td><td>-7.2</td><td>1.4</td><td>-5.2</td><td>2.1E-07</td><td>3.4E-07</td><td>MB231/MB468</td><td>MITO</td></tr>
<tr><td>>trna10_AspGTC_12_-_125424193_125424264@35.72.38,trna12_AspGTC_12_-_125411891_125411962@35.72.38,trna144_AspGTC_6_-_27551236_27551307@35.72.38,trna38_AspGTC_17_-_8125556_8125627@35.72.38,trna45_AspGTC_6_+_27447453_27447524@35.72.38,trna48_AspGTC_6_+_27471523_27471594@35.72.38,trna4_AspGTC_12_+_96429799_96429870@35.72.38,trna69_AspGTC_1_-_161440205_161440276@35.72.38,trna72_AspGTC_1_-_161432824_161432895@35.72.38,trna75_AspGTC_1_-_161425414_161425485@35.72.38,trna78_AspGTC_1_-_161418033_161418104@35.72.38,trna81_AspGTC_1_-_161410615_161410686@35.72.38|TCACGCGGGAGACCGGGGTTCGATTCCCCGACGGGGAG</td><td>>trna10_AspGTC_12_-_125424193_125424264</td><td>35</td><td>72</td><td>38</td><td>TCACGCGGGAGACCGGGGTTCGATTCCCCGACGGGGAG</td><td>895.1</td><td>-5.0</td><td>0.9</td><td>-5.2</td><td>1.7E-07</td><td>3.4E-07</td><td>MB231/MB468</td><td>MP</td></tr>
<tr><td>>MI0000088|hsa-mir-30a&WithFlank&6|-|71403545|71403627@54.74.21&[MIMAT0000088&hsa-miR-30a-3p&offsets|+1|0;m-38&6|-|71403554|71403575&offsets|+1|0]|TTTCAGTCGGATGTTTGCAGC</td><td>>MI0000088|hsa-mir-30a&WithFlank&6|-|71403545|71403627</td><td>54</td><td>74</td><td>21</td><td>TTTCAGTCGGATGTTTGCAGC</td><td>2655.4</td><td>4.3</td><td>0.8</td><td>5.2</td><td>1.7E-07</td><td>3.4E-07</td><td>MB231/MB468</td><td>MP</td></tr>
<tr><td>>trna10_LysCTT_16_+_3241501_3241573@3.34.32,trna13_LysCTT_14_-_58706613_58706685@3.34.32,trna2_LysCTT_15_+_79152904_79152976@3.34.32|CCGGCTAGCTCAGTCGGTAGAGCATGGGACTC</td><td>>trna10_LysCTT_16_+_3241501_3241573</td><td>3</td><td>34</td><td>32</td><td>CCGGCTAGCTCAGTCGGTAGAGCATGGGACTC</td><td>472.5</td><td>7.2</td><td>1.4</td><td>5.2</td><td>2.1E-07</td><td>3.4E-07</td><td>MB231/MB468</td><td>MITO</td></tr>
<tr><td>>hg38_21_+_8433172_8446622_RNA45SN1_45S_50ntFlanks@12456.12493.38|CCAAGCGTTGGATTGTTCACCCACTAATAGGGAACGTG</td><td>>hg38_21_+_8433172_8446622_RNA45SN1_45S_50ntFlanks</td><td>12456</td><td>12493</td><td>38</td><td>CCAAGCGTTGGATTGTTCACCCACTAATAGGGAACGTG</td><td>610.8</td><td>-7.7</td><td>1.5</td><td>-5.2</td><td>1.7E-07</td><td>3.4E-07</td><td>MB231/MB468</td><td>MP</td></tr>
<tr><td>>hg38_21_+_8433172_8446622_RNA45SN1_45S_50ntFlanks@12444.12481.38|AAGCAGAATTCACCAAGCGTTGGATTGTTCACCCACTA</td><td>>hg38_21_+_8433172_8446622_RNA45SN1_45S_50ntFlanks</td><td>12444</td><td>12481</td><td>38</td><td>AAGCAGAATTCACCAAGCGTTGGATTGTTCACCCACTA</td><td>307.7</td><td>-7.6</td><td>1.5</td><td>-5.2</td><td>2.2E-07</td><td>3.4E-07</td><td>BT20/MB468</td><td>NUC</td></tr>
</table>

| | | | | | | | | | | | | | |
|---|---|---|---|---|---|---|---|---|---|---|---|---|---|
| >hg38_21_+_8433172_8446622_RNA45SN1_45S_50ntFlanks@4602.4622.21\|TTCGGAACTGAGGCCATGATT | >hg38_21_+_8433172_8446622_RNA45SN1_45S_50ntFlanks | 4602 | 4622 | 21 | TTCGGAACTGAGGCCATGATT | 135.3 | -6.8 | 1.3 | -5.2 | 1.7E-07 | 3.4E-07 | MB231/MB468 | MP |
| >hg38_21_+_8433172_8446622_RNA45SN1_45S_50ntFlanks@10806.10828.23\|AACAATGTAGGTAAGGGAAGTCG | >hg38_21_+_8433172_8446622_RNA45SN1_45S_50ntFlanks | 10806 | 10828 | 23 | AACAATGTAGGTAAGGGAAGTCG | 823.0 | -8.4 | 1.6 | -5.2 | 2.2E-07 | 3.4E-07 | BT20/MB468 | NUC |
| >hg38_21_+_8433172_8446622_RNA45SN1_45S_50ntFlanks@11200.11223.24\|CGGCGGCGGCGGCGACTCTGGACG | >hg38_21_+_8433172_8446622_RNA45SN1_45S_50ntFlanks | 11200 | 11223 | 24 | CGGCGGCGGCGGCGACTCTGGACG | 1134.2 | -8.9 | 1.7 | -5.2 | 2.2E-07 | 3.5E-07 | BT20/MB468 | NUC |
| >hg38_21_+_8433172_8446622_RNA45SN1_45S_50ntFlanks@11728.11749.22\|GTAACTATGACTCTCTTAAGGT | >hg38_21_+_8433172_8446622_RNA45SN1_45S_50ntFlanks | 11728 | 11749 | 22 | GTAACTATGACTCTCTTAAGGT | 516.1 | -8.1 | 1.6 | -5.2 | 2.2E-07 | 3.5E-07 | BT20/MB468 | NUC |
| >trna10_AspGTC_12_-_125424193_125424264@1.30.30,trna12_AspGTC_12_-_125411891_125411962@1.30.30,trna38_AspGTC_17_-_8125556_8125627@1.30.30,trna45_AspGTC_6_+_27447453_27447524@1.30.30,trna48_AspGTC_6_+_27471523_27471594@1.30.30,trna4_AspGTC_12_+_96429799_96429870@1.30.30,trna69_AspGTC_1_-_161440205_161440276@1.30.30,trna72_AspGTC_1_-_161432824_161432895@1.30.30,trna75_AspGTC_1_-_161425414_161425485@1.30.30,trna78_AspGTC_1_-_161418033_161418104@1.30.30,trna81_AspGTC_1_-_161410615_161410686@1.30.30\|TCCTCGTTAGTATAGTGGTGAGTATCCCCG | >trna10_AspGTC_12_-_125424193_125424264 | 1 | 30 | 30 | TCCTCGTTAGTATAGTGGTGAGTATCCCCG | 342.0 | -4.9 | 0.9 | -5.2 | 2.1E-07 | 3.5E-07 | MB231/MB468 | MITO |
| >hg38_21_+_8433172_8446622_RNA45SN1_45S_50ntFlanks@12043.12070.28\|CACTGACCCGGTGAGGCGGGGGGGCGAG | >hg38_21_+_8433172_8446622_RNA45SN1_45S_50ntFlanks | 12043 | 12070 | 28 | CACTGACCCGGTGAGGCGGGGGGGCGAG | 86.9 | -9.5 | 1.7 | -5.5 | 4.1E-08 | 3.5E-07 | BT20/MB468 | MP |
| >hg38_21_+_8433172_8446622_RNA45SN1_45S_50ntFlanks@11575.11606.32\|ACTTAGAACTGGTGCGGACCAGGGGAATCCGA | >hg38_21_+_8433172_8446622_RNA45SN1_45S_50ntFlanks | 11575 | 11606 | 32 | ACTTAGAACTGGTGCGGACCAGGGGAATCCGA | 216.3 | -5.3 | 1.0 | -5.2 | 2.0E-07 | 3.5E-07 | MB231/MB468 | NUC |
| >hg38_21_+_8433172_8446622_RNA45SN1_45S_50ntFlanks@10403.10424.22\|ACATGGGTCAGTCGGTCCTGAG | >hg38_21_+_8433172_8446622_RNA45SN1_45S_50ntFlanks | 10403 | 10424 | 22 | ACATGGGTCAGTCGGTCCTGAG | 1224.7 | 3.7 | 0.7 | 5.2 | 1.9E-07 | 3.5E-07 | MB231/MB468 | CYTO |
| >hg38_21_+_8433172_8446622_RNA45SN1_45S_50ntFlanks@8472.8491.20\|CGGCCCGGCGGATCTTTCCC | >hg38_21_+_8433172_8446622_RNA45SN1_45S_50ntFlanks | 8472 | 8491 | 20 | CGGCCCGGCGGATCTTTCCC | 73.6 | -9.3 | 1.7 | -5.5 | 4.1E-08 | 3.5E-07 | BT20/MB468 | MP |
| >hg38_21_+_8433172_8446622_RNA45SN1_45S_50ntFlanks@11115.11133.19\|CGGCAGGGGCCGGCGGCGG | >hg38_21_+_8433172_8446622_RNA45SN1_45S_50ntFlanks | 11115 | 11133 | 19 | CGGCAGGGGCCGGCGGCGG | 322.3 | -6.4 | 1.2 | -5.2 | 1.7E-07 | 3.5E-07 | BT20/MB468 | CYTO |
| >hg38_21_+_8433172_8446622_RNA45SN1_45S_50ntFlanks@8354.8389.36\|TTGAAAAGAACTTTGAAGAGAGAGTTCAAGAGGGCG | >hg38_21_+_8433172_8446622_RNA45SN1_45S_50ntFlanks | 8354 | 8389 | 36 | TTGAAAAGAACTTTGAAGAGAGAGTTCAAGAGGGCG | 68.9 | 9.9 | 1.8 | 5.5 | 4.1E-08 | 3.5E-07 | BT20/MB468 | MP |
| >hg38_21_+_8433172_8446622_RNA45SN1_45S_50ntFlanks@4758.4780.23\|GATCAGATACCGTCGTAGTTCCG | >hg38_21_+_8433172_8446622_RNA45SN1_45S_50ntFlanks | 4758 | 4780 | 23 | GATCAGATACCGTCGTAGTTCCG | 93.5 | -9.6 | 1.8 | -5.5 | 4.1E-08 | 3.5E-07 | BT20/MB468 | MP |
| >hg38_21_+_8433172_8446622_RNA45SN1_45S_50ntFlanks@5074.5097.24\|ATTCCGATAACGAACGAGACTCTG | >hg38_21_+_8433172_8446622_RNA45SN1_45S_50ntFlanks | 5074 | 5097 | 24 | ATTCCGATAACGAACGAGACTCTG | 530.1 | -7.8 | 1.5 | -5.2 | 2.1E-07 | 3.5E-07 | MB231/MB468 | MITO |
| >hg38_21_+_8433172_8446622_RNA45SN1_45S_50ntFlanks@3815.3850.36\|ATGGTTCCTTTGGTCGCTCGCTCCTCTCCTACTTGG | >hg38_21_+_8433172_8446622_RNA45SN1_45S_50ntFlanks | 3815 | 3850 | 36 | ATGGTTCCTTTGGTCGCTCGCTCCTCTCCTACTTGG | 335.2 | -3.6 | 0.7 | -5.2 | 2.0E-07 | 3.5E-07 | MB231/MB468 | NUC |
| >hg38_21_+_8433172_8446622_RNA45SN1_45S_50ntFlanks@4573.4616.44\|AATAGGACCGCGGTTCTATTTTGTTGGTTTTCGGAACTGAGGCC | >hg38_21_+_8433172_8446622_RNA45SN1_45S_50ntFlanks | 4573 | 4616 | 44 | AATAGGACCGCGGTTCTATTTTGTTGGTTTTCGGAACTGAGGCC | 127.6 | -9.3 | 1.8 | -5.2 | 1.7E-07 | 3.5E-07 | BT20/MB468 | CYTO |
| >hg38_21_+_8433172_8446622_RNA45SN1_45S_50ntFlanks@8972.8990.19\|CCACCCCCGCGGGGAATCC | >hg38_21_+_8433172_8446622_RNA45SN1_45S_50ntFlanks | 8972 | 8990 | 19 | CCACCCCCGCGGGGAATCC | 162.1 | -9.4 | 1.8 | -5.3 | 1.5E-07 | 3.5E-07 | BT20/MB468 | MITO |
| >MI0006383\|hsa-mir-1248&WithFlank&3\|+\|186786666\|186786783@10.32.23&[MIMAT0005900&hsa-miR-1248&offsets\|0\|-4;m-827&3\|+\|186786675\|186786701&offsets\|0\|-4]\|ACCTTCTTGTATAAGCACTGTGC | >MI0006383\|hsa-mir-1248&WithFlank&3\|+\|186786666\|186786783 | 10 | 32 | 23 | ACCTTCTTGTATAAGCACTGTGC | 267.9 | 3.3 | 0.6 | 5.2 | 2.0E-07 | 3.5E-07 | MB231/MB468 | NUC |
| >hg38_21_+_8433172_8446622_RNA45SN1_45S_50ntFlanks@9758.9786.29\|TCAAACTTTAAATGGGTAAGAAGCCCGGC | >hg38_21_+_8433172_8446622_RNA45SN1_45S_50ntFlanks | 9758 | 9786 | 29 | TCAAACTTTAAATGGGTAAGAAGCCCGGC | 222.7 | -10.9 | 2.0 | -5.5 | 4.1E-08 | 3.5E-07 | BT20/MB468 | MP |
| >hg38_21_+_8433172_8446622_RNA45SN1_45S_50ntFlanks@8552.8581.30\|CTCCCCGGAGGGGGCGGGCTCCGGCGGGTG | >hg38_21_+_8433172_8446622_RNA45SN1_45S_50ntFlanks | 8552 | 8581 | 30 | CTCCCCGGAGGGGGCGGGCTCCGGCGGGTG | 142.8 | -10.2 | 1.9 | -5.5 | 4.1E-08 | 3.5E-07 | BT20/MB468 | MP |
| >hg38_21_+_8433172_8446622_RNA45SN1_45S_50ntFlanks@12008.12038.31\|CCGCCGGTGAAATACCACTACTCTGATCGTT | >hg38_21_+_8433172_8446622_RNA45SN1_45S_50ntFlanks | 12008 | 12038 | 31 | CCGCCGGTGAAATACCACTACTCTGATCGTT | 324.4 | -3.5 | 0.7 | -5.3 | 1.5E-07 | 3.5E-07 | BT20/MB468 | MITO |
| >hg38_21_+_8433172_8446622_RNA45SN1_45S_50ntFlanks@5484.5514.31\|GAGCGCTGAGAAGACGGTCGAACTTGACTAT | >hg38_21_+_8433172_8446622_RNA45SN1_45S_50ntFlanks | 5484 | 5514 | 31 | GAGCGCTGAGAAGACGGTCGAACTTGACTAT | 9537.6 | -8.3 | 1.6 | -5.2 | 2.2E-07 | 3.5E-07 | BT20/MB468 | NUC |
| >hg38_1_-_228634819_228635039_RNA5S12_5S_50ntFlanks@121.170.50\|GTTAGTACTTGGATGGGAGACCGCCTGGGAATACCGGGTGCTGTAGGCTT | >hg38_1_-_228634819_228635039_RNA5S12_5S_50ntFlanks | 121 | 170 | 50 | GTTAGTACTTGGATGGGAGACCGCCTGGGAATACCGGGTGCTGTAGGCTT | 593.9 | 5.1 | 1.0 | 5.2 | 1.9E-07 | 3.5E-07 | MB231/MB468 | CYTO |
| >hg38_21_+_8433172_8446622_RNA45SN1_45S_50ntFlanks@11561.11587.27\|GCGCCTAGCAGCCGACTTAGAACTGGT | >hg38_21_+_8433172_8446622_RNA45SN1_45S_50ntFlanks | 11561 | 11587 | 27 | GCGCCTAGCAGCCGACTTAGAACTGGT | 307.5 | -7.6 | 1.5 | -5.2 | 2.2E-07 | 3.5E-07 | BT20/MB468 | NUC |
| >hg38_21_+_8433172_8446622_RNA45SN1_45S_50ntFlanks@7986.8016.31\|ATCAGACGTGGCGACCCGCTGAATTTAAGCA | >hg38_21_+_8433172_8446622_RNA45SN1_45S_50ntFlanks | 7986 | 8016 | 31 | ATCAGACGTGGCGACCCGCTGAATTTAAGCA | 947.1 | -7.6 | 1.5 | -5.2 | 1.8E-07 | 3.5E-07 | MB231/MB468 | MP |
| >hg38_21_+_8433172_8446622_RNA45SN1_45S_50ntFlanks@4672.4689.18\|TTGGACCGGCGCAAGACG | >hg38_21_+_8433172_8446622_RNA45SN1_45S_50ntFlanks | 4672 | 4689 | 18 | TTGGACCGGCGCAAGACG | 199.4 | -5.6 | 1.1 | -5.2 | 1.8E-07 | 3.5E-07 | MB231/MB468 | MP |
| >hg38_21_+_8433172_8446622_RNA45SN1_45S_50ntFlanks@6651.6685.35\|CGACTCTTAGCGGTGGATCACTCGGCTCGTGCGTC | >hg38_21_+_8433172_8446622_RNA45SN1_45S_50ntFlanks | 6651 | 6685 | 35 | CGACTCTTAGCGGTGGATCACTCGGCTCGTGCGTC | 1699.9 | 3.3 | 0.6 | 5.2 | 2.0E-07 | 3.5E-07 | MB231/MB468 | NUC |
| >MI0000079\|hsa-mir-23a&WithFlank&19\|-\|13836581\|13836665@54.73.20&[MIMAT0000078&hsa-miR-23a-3p&offsets\|+3\|+2;m-31&19\|-\|13836595\|13836615&offsets\|+3\|+2]\|ACATTGCCAGGGATTTCCAA | >MI0000079\|hsa-mir-23a&WithFlank&19\|-\|13836581\|13836665 | 54 | 73 | 20 | ACATTGCCAGGGATTTCCAA | 98.5 | 10.4 | 1.9 | 5.5 | 4.1E-08 | 3.5E-07 | BT20/MB468 | MP |
| >hg38_21_+_8433172_8446622_RNA45SN1_45S_50ntFlanks@5485.5515.31\|AGCGCTGAGAAGACGGTCGAACTTGACTATC | >hg38_21_+_8433172_8446622_RNA45SN1_45S_50ntFlanks | 5485 | 5515 | 31 | AGCGCTGAGAAGACGGTCGAACTTGACTATC | 361.3 | -4.2 | 0.8 | -5.2 | 2.0E-07 | 3.5E-07 | MB231/MB468 | NUC |
| >hg38_21_+_8433172_8446622_RNA45SN1_45S_50ntFlanks@12995.13036.42\|CTCGCTGCGATCTATTGAAAGTCAGCCCTCGACACAAGGGTT | >hg38_21_+_8433172_8446622_RNA45SN1_45S_50ntFlanks | 12995 | 13036 | 42 | CTCGCTGCGATCTATTGAAAGTCAGCCCTCGACACAAGGGTT | 1414.5 | 5.0 | 1.0 | 5.2 | 2.0E-07 | 3.5E-07 | MB231/MB468 | NUC |
| >hg38_1_-_228634819_228635039_RNA5S12_5S_50ntFlanks@145.171.27\|CTGGGAATACCGGGTGCTGTAGGCTTT | >hg38_1_-_228634819_228635039_RNA5S12_5S_50ntFlanks | 145 | 171 | 27 | CTGGGAATACCGGGTGCTGTAGGCTTT | 1168.7 | 4.7 | 0.9 | 5.2 | 2.0E-07 | 3.5E-07 | MB231/MB468 | NUC |
| >hg38_MT_+_1621_3279_RNR2_16S_50ntFlanks@1367.1404.38\|TCGATGTTGGATCAGGACATCCCGATGGTGCAGCCGCT | >hg38_MT_+_1621_3279_RNR2_16S_50ntFlanks | 1367 | 1404 | 38 | TCGATGTTGGATCAGGACATCCCGATGGTGCAGCCGCT | 105.4 | 10.5 | 1.9 | 5.5 | 4.2E-08 | 3.5E-07 | BT20/MB468 | MP |
| >hg38_21_+_8433172_8446622_RNA45SN1_45S_50ntFlanks@10694.10712.19\|CGGTGAGCTCTCGCTGGCC | >hg38_21_+_8433172_8446622_RNA45SN1_45S_50ntFlanks | 10694 | 10712 | 19 | CGGTGAGCTCTCGCTGGCC | 123.8 | -10.0 | 1.8 | -5.5 | 4.2E-08 | 3.5E-07 | BT20/MB468 | MP |
| >hg38_21_+_8433172_8446622_RNA45SN1_45S_50ntFlanks@5423.5445.23\|ATTGGATGGTTTAGTGAGGCCCT | >hg38_21_+_8433172_8446622_RNA45SN1_45S_50ntFlanks | 5423 | 5445 | 23 | ATTGGATGGTTTAGTGAGGCCCT | 182.6 | -5.6 | 1.1 | -5.2 | 1.9E-07 | 3.5E-07 | MB231/MB468 | CYTO |
| >hg38_21_+_8433172_8446622_RNA45SN1_45S_50ntFlanks@6784.6801.18\|GCTACGCCTGTCTGAGCG | >hg38_21_+_8433172_8446622_RNA45SN1_45S_50ntFlanks | 6784 | 6801 | 18 | GCTACGCCTGTCTGAGCG | 369.1 | -6.2 | 1.2 | -5.2 | 2.1E-07 | 3.5E-07 | MB231/MB468 | MITO |
| >MI0000283\|hsa-mir-203a&WithFlank&14\|+\|104117399\|104117520@72.93.22&[MIMAT0000264&hsa-miR-203a-3p&offsets\|+1\|+1;m-79&14\|+\|104117469\|104117490&offsets\|+1\|+1]\|TGAAATGTTTAGGACCACTAGA | >MI0000283\|hsa-mir-203a&WithFlank&14\|+\|104117399\|104117520 | 72 | 93 | 22 | TGAAATGTTTAGGACCACTAGA | 161.1 | 10.7 | 1.9 | 5.8 | 8.7E-09 | 3.5E-07 | BT20/MB231 | MP |
| >MI0000102\|hsa-mir-100&WithFlank&11\|-\|122152223\|122152314@19.39.21(+1C)&[MIMAT0000098&hsa-miR-100-5p&offsets\|0\|-1(+1C);m-27&11\|-\|122152276\|122152296&offsets\|0\|0(+1C)]\|AACCCGTAGATCCGAACTTGTC | >MI0000102\|hsa-mir-100&WithFlank&11\|-\|122152223\|122152314 | 19 | 39 | 21(+1C) | AACCCGTAGATCCGAACTTGTC | 3551.8 | -8.4 | 1.5 | -5.8 | 8.8E-09 | 3.5E-07 | BT20/MB231 | MP |
| >hg38_21_+_8433172_8446622_RNA45SN1_45S_50ntFlanks@8168.8197.30\|CCCAAGTCCTTCTGATCGAGGCCCAGCCCG | >hg38_21_+_8433172_8446622_RNA45SN1_45S_50ntFlanks | 8168 | 8197 | 30 | CCCAAGTCCTTCTGATCGAGGCCCAGCCCG | 81.2 | -9.4 | 1.7 | -5.5 | 4.2E-08 | 3.5E-07 | BT20/MB468 | MP |
| >hg38_21_+_8433172_8446622_RNA45SN1_45S_50ntFlanks@12233.12265.33\|CCGTGGAGCAGAAGGGCAAAAGCTCGCTTGATC | >hg38_21_+_8433172_8446622_RNA45SN1_45S_50ntFlanks | 12233 | 12265 | 33 | CCGTGGAGCAGAAGGGCAAAAGCTCGCTTGATC | 282.3 | -7.3 | 1.4 | -5.2 | 1.8E-07 | 3.5E-07 | MB231/MB468 | MP |
| >hg38_21_+_8433172_8446622_RNA45SN1_45S_50ntFlanks@10771.10816.46\|CAGGTCTCCAAGGTGAACAGCCTCTGGCATGTTGGAACAATGTAGG | >hg38_21_+_8433172_8446622_RNA45SN1_45S_50ntFlanks | 10771 | 10816 | 46 | CAGGTCTCCAAGGTGAACAGCCTCTGGCATGTTGGAACAATGTAGG | 311.2 | -6.3 | 1.2 | -5.2 | 2.1E-07 | 3.5E-07 | MB231/MB468 | MITO |
| >hg38_21_+_8433172_8446622_RNA45SN1_45S_50ntFlanks@4982.4999.18\|ACGGACAGGATTGACAGA | >hg38_21_+_8433172_8446622_RNA45SN1_45S_50ntFlanks | 4982 | 4999 | 18 | ACGGACAGGATTGACAGA | 387.2 | -5.9 | 1.1 | -5.2 | 1.8E-07 | 3.5E-07 | MB231/MB468 | MP |
| >hg38_1_-_228634819_228635039_RNA5S12_5S_50ntFlanks@149.169.21\|GAATACCGGGTGCTGTAGGCT | >hg38_1_-_228634819_228635039_RNA5S12_5S_50ntFlanks | 149 | 169 | 21 | GAATACCGGGTGCTGTAGGCT | 1233.3 | -4.1 | 0.8 | -5.2 | 1.8E-07 | 3.5E-07 | MB231/MB468 | MP |
| >hg38_21_+_8433172_8446622_RNA45SN1_45S_50ntFlanks@11217.11241.25\|CTGGACGCGAGCCGGGCCCTTCCCG | >hg38_21_+_8433172_8446622_RNA45SN1_45S_50ntFlanks | 11217 | 11241 | 25 | CTGGACGCGAGCCGGGCCCTTCCCG | 468.7 | -7.3 | 1.4 | -5.2 | 2.1E-07 | 3.5E-07 | MB231/MB468 | MITO |
| >hg38_21_+_8433172_8446622_RNA45SN1_45S_50ntFlanks@11703.11738.36\|AATGAAGCGCGGGTAAACGGCGGGAGTAACTATGAC | >hg38_21_+_8433172_8446622_RNA45SN1_45S_50ntFlanks | 11703 | 11738 | 36 | AATGAAGCGCGGGTAAACGGCGGGAGTAACTATGAC | 408.2 | -7.9 | 1.5 | -5.2 | 2.2E-07 | 3.5E-07 | BT20/MB468 | NUC |
| >hg38_21_+_8433172_8446622_RNA45SN1_45S_50ntFlanks@11845.11884.40\|GGAACGGGCTTGGCGGAATCAGCGGGGAAAGAAGACCCTG | >hg38_21_+_8433172_8446622_RNA45SN1_45S_50ntFlanks | 11845 | 11884 | 40 | GGAACGGGCTTGGCGGAATCAGCGGGGAAAGAAGACCCTG | 72.0 | -10.0 | 1.9 | -5.2 | 1.8E-07 | 3.5E-07 | MB231/MB468 | MP |
| >hg38_21_+_8433172_8446622_RNA45SN1_45S_50ntFlanks@9534.9569.36\|AGGAAACTCTGGTGGAGGTCCGTAGCGGTCCTGACG | >hg38_21_+_8433172_8446622_RNA45SN1_45S_50ntFlanks | 9534 | 9569 | 36 | AGGAAACTCTGGTGGAGGTCCGTAGCGGTCCTGACG | 2398.5 | -8.4 | 1.6 | -5.2 | 1.8E-07 | 3.5E-07 | MB231/MB468 | MP |
| >MI0000734\|hsa-mir-106b&WithFlank&7\|-\|100093987\|100094080@58.80.23&[MIMAT0004672&hsa-miR-106b-3p&offsets\|0\|+1;m-87&7\|-\|100094003\|100094023&offsets\|0\|+2]\|CCGCACTGTGGGTACTTGCTGCT | >MI0000734\|hsa-mir-106b&WithFlank&7\|-\|100093987\|100094080 | 58 | 80 | 23 | CCGCACTGTGGGTACTTGCTGCT | 241.7 | 6.3 | 1.2 | 5.2 | 2.0E-07 | 3.5E-07 | MB231/MB468 | CYTO |
| >trnaMT_HisGTG_MT_+_12138_12206@45.72.28\|AGAGGCTTACGACCCCTTATTTACCCCA | >trnaMT_HisGTG_MT_+_12138_12206 | 45 | 72 | 28 | AGAGGCTTACGACCCCTTATTTACCCCA | 134.4 | 10.8 | 2.0 | 5.5 | 4.2E-08 | 3.5E-07 | BT20/MB468 | MP |
| >hg38_21_+_8433172_8446622_RNA45SN1_45S_50ntFlanks@11208.11243.36\|GCGGCGACTCTGGACGCGAGCCGGGCCCTTCCCGTG | >hg38_21_+_8433172_8446622_RNA45SN1_45S_50ntFlanks | 11208 | 11243 | 36 | GCGGCGACTCTGGACGCGAGCCGGGCCCTTCCCGTG | 55.0 | -9.6 | 1.8 | -5.2 | 1.8E-07 | 3.5E-07 | MB231/MB468 | MP |

<table>
<tr><th></th><th></th><th></th><th></th><th></th><th></th><th></th><th></th><th></th><th></th><th></th><th></th><th></th><th></th></tr>
<tr><td>>hg38_21_+_8433172_8446622_RNA45SN1_45S_50ntFlanks@10841.10868.28|CGTAACTTCGGGATAAGGATTGGCTCTA</td><td>>hg38_21_+_8433172_8446622_RNA45SN1_45S_50ntFlanks</td><td>10841</td><td>10868</td><td>28</td><td>CGTAACTTCGGGATAAGGATTGGCTCTA</td><td>2235.7</td><td>-6.4</td><td>1.2</td><td>-5.2</td><td>2.1E-07</td><td>3.5E-07</td><td>MB231/MB468</td><td>MITO</td></tr>
<tr><td>>hg38_21_+_8433172_8446622_RNA45SN1_45S_50ntFlanks@6650.6685.36|ACGACTCTTAGCGGTGGATCACTCGGCTCGTGCGTC</td><td>>hg38_21_+_8433172_8446622_RNA45SN1_45S_50ntFlanks</td><td>6650</td><td>6685</td><td>36</td><td>ACGACTCTTAGCGGTGGATCACTCGGCTCGTGCGTC</td><td>514.6</td><td>3.6</td><td>0.7</td><td>5.3</td><td>1.5E-07</td><td>3.5E-07</td><td>BT20/MB468</td><td>MITO</td></tr>
<tr><td>>hg38_21_+_8433172_8446622_RNA45SN1_45S_50ntFlanks@12416.12433.18|TTCGATGTCGGCTCTTCC</td><td>>hg38_21_+_8433172_8446622_RNA45SN1_45S_50ntFlanks</td><td>12416</td><td>12433</td><td>18</td><td>TTCGATGTCGGCTCTTCC</td><td>2862.0</td><td>4.0</td><td>0.8</td><td>5.2</td><td>1.7E-07</td><td>3.5E-07</td><td>BT20/MB468</td><td>CYTO</td></tr>
<tr><td>>hg38_21_+_8433172_8446622_RNA45SN1_45S_50ntFlanks@12366.12404.39|TAACTGGCTTGTGGCGGCCAAGCGTTCATAGCGACGTCG</td><td>>hg38_21_+_8433172_8446622_RNA45SN1_45S_50ntFlanks</td><td>12366</td><td>12404</td><td>39</td><td>TAACTGGCTTGTGGCGGCCAAGCGTTCATAGCGACGTCG</td><td>297.0</td><td>-8.3</td><td>1.6</td><td>-5.2</td><td>1.7E-07</td><td>3.5E-07</td><td>BT20/MB468</td><td>CYTO</td></tr>
<tr><td>>MI0000683|hsa-mir-181b-2&WithFlank&9|+|124693704|124693804@22.46.25(+2U)&[MIMAT0000257_1&hsa-miR-181b-5p&offsets|0|+2(+2U);m-72&9|+|124693725|124693747&offsets|0|+2(+2U)]|AACATTCATTGCTGTCGGTGGGTTTTT</td><td>>MI0000683|hsa-mir-181b-2&WithFlank&9|+|124693704|124693804</td><td>22</td><td>46</td><td>25(+2U)</td><td>AACATTCATTGCTGTCGGTGGGTTTTT</td><td>62.1</td><td>9.2</td><td>1.8</td><td>5.2</td><td>1.8E-07</td><td>3.5E-07</td><td>MB231/MB468</td><td>MP</td></tr>
<tr><td>>hg38_21_+_8433172_8446622_RNA45SN1_45S_50ntFlanks@8845.8867.23|CGGCGGCGGCGGCGGCGGCGGGA</td><td>>hg38_21_+_8433172_8446622_RNA45SN1_45S_50ntFlanks</td><td>8845</td><td>8867</td><td>23</td><td>CGGCGGCGGCGGCGGCGGCGGGA</td><td>649.0</td><td>-8.0</td><td>1.5</td><td>-5.5</td><td>4.2E-08</td><td>3.5E-07</td><td>BT20/MB468</td><td>MP</td></tr>
<tr><td>>hg38_21_+_8433172_8446622_RNA45SN1_45S_50ntFlanks@11731.11751.21|ACTATGACTCTCTTAAGGTAG</td><td>>hg38_21_+_8433172_8446622_RNA45SN1_45S_50ntFlanks</td><td>11731</td><td>11751</td><td>21</td><td>ACTATGACTCTCTTAAGGTAG</td><td>260.7</td><td>-4.2</td><td>0.8</td><td>-5.2</td><td>2.0E-07</td><td>3.5E-07</td><td>MB231/MB468</td><td>CYTO</td></tr>
<tr><td>>hg38_21_+_8433172_8446622_RNA45SN1_45S_50ntFlanks@4726.4748.23|TAATCAAGAACGAAAGTCGGAGG</td><td>>hg38_21_+_8433172_8446622_RNA45SN1_45S_50ntFlanks</td><td>4726</td><td>4748</td><td>23</td><td>TAATCAAGAACGAAAGTCGGAGG</td><td>276.0</td><td>-7.4</td><td>1.4</td><td>-5.2</td><td>2.2E-07</td><td>3.5E-07</td><td>BT20/MB468</td><td>NUC</td></tr>
<tr><td>>MI0000736|hsa-mir-30c-1&WithFlank&1|+|40757278|40757378@23.45.23(+1C);MI0000254|hsa-mir-30c-2&WithFlank&6|-|71376954|71377037@13.35.23(+1C)&[MIMAT0000244&hsa-miR-30c-5p&offsets|0|0(+1C);m-57&1|+|40757300|40757323&offsets|0|-1(+1C)];[MIMAT0000244_1&hsa-miR-30c-5p&offsets|0|0(+1C);m-56&6|-|71377002|71377025&offsets|0|-1(+1C)]|TGTAAACATCCTACACTCTCAGCC</td><td>>MI0000736|hsa-mir-30c-1&WithFlank&1|+|40757278|40757378</td><td>23</td><td>45</td><td>23(+1C)</td><td>TGTAAACATCCTACACTCTCAGCC</td><td>2078.4</td><td>5.0</td><td>1.0</td><td>5.2</td><td>1.5E-07</td><td>3.5E-07</td><td>BT20/MB468</td><td>MITO</td></tr>
<tr><td>>hg38_21_+_8433172_8446622_RNA45SN1_45S_50ntFlanks@12615.12653.39|GCTGAGGAGCCAATGGGGCGAAGCTACCATCTGTGGGAT</td><td>>hg38_21_+_8433172_8446622_RNA45SN1_45S_50ntFlanks</td><td>12615</td><td>12653</td><td>39</td><td>GCTGAGGAGCCAATGGGGCGAAGCTACCATCTGTGGGAT</td><td>2215.4</td><td>-6.1</td><td>1.2</td><td>-5.2</td><td>1.5E-07</td><td>3.5E-07</td><td>BT20/MB468</td><td>MITO</td></tr>
<tr><td>>hg38_21_+_8433172_8446622_RNA45SN1_45S_50ntFlanks@3789.3812.24|ACTGCGAATGGCTCATTAAATCAG</td><td>>hg38_21_+_8433172_8446622_RNA45SN1_45S_50ntFlanks</td><td>3789</td><td>3812</td><td>24</td><td>ACTGCGAATGGCTCATTAAATCAG</td><td>103.9</td><td>-9.8</td><td>1.8</td><td>-5.5</td><td>4.2E-08</td><td>3.6E-07</td><td>BT20/MB468</td><td>MP</td></tr>
<tr><td>>trna116_GluCTC_1_-_145399233_145399304@32.71.40,trna71_GluCTC_1_-_161439189_161439260@32.71.40,trna74_GluCTC_1_-_161431809_161431880@32.71.40,trna77_GluCTC_1_-_161424398_161424469@32.71.40,trna77_GluCTC_6_+_28949976_28950047@32.71.40,trna80_GluCTC_1_-_161417018_161417089@32.71.40,trna87_GluCTC_6_-_126101393_126101464@32.71.40|CTCTCACCGCCGCGGCCCGGGTTCGATTCCCGGTCAGGGA</td><td>>trna116_GluCTC_1_-_145399233_145399304</td><td>32</td><td>71</td><td>40</td><td>CTCTCACCGCCGCGGCCCGGGTTCGATTCCCGGTCAGGGA</td><td>1637.9</td><td>-7.5</td><td>1.4</td><td>-5.2</td><td>1.7E-07</td><td>3.6E-07</td><td>BT20/MB468</td><td>CYTO</td></tr>
<tr><td>>hg38_21_+_8433172_8446622_RNA45SN1_45S_50ntFlanks@7976.8015.40|GCGACCTCAGATCAGACGTGGCGACCCGCTGAATTTAAGC</td><td>>hg38_21_+_8433172_8446622_RNA45SN1_45S_50ntFlanks</td><td>7976</td><td>8015</td><td>40</td><td>GCGACCTCAGATCAGACGTGGCGACCCGCTGAATTTAAGC</td><td>400.5</td><td>7.4</td><td>1.4</td><td>5.2</td><td>1.5E-07</td><td>3.6E-07</td><td>BT20/MB468</td><td>MITO</td></tr>
<tr><td>>hg38_21_+_8433172_8446622_RNA45SN1_45S_50ntFlanks@6659.6681.23|AGCGGTGGATCACTCGGCTCGTG</td><td>>hg38_21_+_8433172_8446622_RNA45SN1_45S_50ntFlanks</td><td>6659</td><td>6681</td><td>23</td><td>AGCGGTGGATCACTCGGCTCGTG</td><td>2837.0</td><td>-5.2</td><td>1.0</td><td>-5.2</td><td>2.1E-07</td><td>3.6E-07</td><td>MB231/MB468</td><td>MITO</td></tr>
<tr><td>>hg38_21_+_8433172_8446622_RNA45SN1_45S_50ntFlanks@3811.3836.26|AGTTATGGTTCCTTTGGTCGCTCGCT</td><td>>hg38_21_+_8433172_8446622_RNA45SN1_45S_50ntFlanks</td><td>3811</td><td>3836</td><td>26</td><td>AGTTATGGTTCCTTTGGTCGCTCGCT</td><td>567.8</td><td>-8.1</td><td>1.6</td><td>-5.2</td><td>2.2E-07</td><td>3.6E-07</td><td>BT20/MB468</td><td>NUC</td></tr>
<tr><td>>hg38_21_+_8433172_8446622_RNA45SN1_45S_50ntFlanks@12444.12484.41|AAGCAGAATTCACCAAGCGTTGGATTGTTCACCCACTAATA</td><td>>hg38_21_+_8433172_8446622_RNA45SN1_45S_50ntFlanks</td><td>12444</td><td>12484</td><td>41</td><td>AAGCAGAATTCACCAAGCGTTGGATTGTTCACCCACTAATA</td><td>1056.3</td><td>-4.8</td><td>0.9</td><td>-5.2</td><td>2.0E-07</td><td>3.6E-07</td><td>MB231/MB468</td><td>NUC</td></tr>
<tr><td>>hg38_21_+_8433172_8446622_RNA45SN1_45S_50ntFlanks@9554.9572.19|CGTAGCGGTCCTGACGTGC</td><td>>hg38_21_+_8433172_8446622_RNA45SN1_45S_50ntFlanks</td><td>9554</td><td>9572</td><td>19</td><td>CGTAGCGGTCCTGACGTGC</td><td>3798.7</td><td>-2.2</td><td>0.4</td><td>-5.2</td><td>1.5E-07</td><td>3.6E-07</td><td>BT20/MB468</td><td>MITO</td></tr>
<tr><td>>MI0000264|hsa-mir-7-2&WithFlank&15|+|88611819|88611940@38.60.23(+1C);MI0000265|hsa-mir-7-3&WithFlank&19|+|4770664|4770785@37.59.23(+1C);MI0000263|hsa-mir-7-1&WithFlank&9|-|83969742|83969863@30.52.23(+1C)&[MIMAT0000252&hsa-miR-7-5p&offsets|0|-1(+1C);m-400&15|+|88611856|88611879&offsets|0|-1(+1C)];[MIMAT0000252_1&hsa-miR-7-5p&offsets|0|-1(+1C);m-402&19|+|4770700|4770723&offsets|0|-1(+1C)];[MIMAT0000252_2&hsa-miR-7-5p&offsets|0|-1(+1C);m-401&9|-|83969811|83969834&offsets|0|-1(+1C)]|TGGAAGACTAGTGATTTTGTTGTC</td><td>>MI0000264|hsa-mir-7-2&WithFlank&15|+|88611819|88611940</td><td>38</td><td>60</td><td>23(+1C)</td><td>TGGAAGACTAGTGATTTTGTTGTC</td><td>1628.5</td><td>3.6</td><td>0.7</td><td>5.2</td><td>2.0E-07</td><td>3.6E-07</td><td>MB231/MB468</td><td>CYTO</td></tr>
<tr><td>>MI0000273|hsa-mir-183&WithFlank&7|-|129774899|129775020@33.54.22(+1A)&[MIMAT0000261&hsa-miR-183-5p&offsets|0|0(+1A);m-41&7|-|129774966|129774987&offsets|-1|-1(+1A)]|TATGGCACTGGTAGAATTCACTA</td><td>>MI0000273|hsa-mir-183&WithFlank&7|-|129774899|129775020</td><td>33</td><td>54</td><td>22(+1A)</td><td>TATGGCACTGGTAGAATTCACTA</td><td>783.0</td><td>4.4</td><td>0.8</td><td>5.7</td><td>1.3E-08</td><td>3.6E-07</td><td>BT20/MB231</td><td>CYTO</td></tr>
<tr><td>>hg38_21_+_8433172_8446622_RNA45SN1_45S_50ntFlanks@7973.8008.36|GACGCGACCTCAGATCAGACGTGGCGACCCGCTGAA</td><td>>hg38_21_+_8433172_8446622_RNA45SN1_45S_50ntFlanks</td><td>7973</td><td>8008</td><td>36</td><td>GACGCGACCTCAGATCAGACGTGGCGACCCGCTGAA</td><td>6494.0</td><td>3.4</td><td>0.6</td><td>5.2</td><td>1.5E-07</td><td>3.6E-07</td><td>BT20/MB468</td><td>MITO</td></tr>
<tr><td>>hg38_21_+_8433172_8446622_RNA45SN1_45S_50ntFlanks@11881.11904.24|CCTGTTGAGCTTGACTCTAGTCTG</td><td>>hg38_21_+_8433172_8446622_RNA45SN1_45S_50ntFlanks</td><td>11881</td><td>11904</td><td>24</td><td>CCTGTTGAGCTTGACTCTAGTCTG</td><td>1041.5</td><td>-7.5</td><td>1.4</td><td>-5.2</td><td>2.0E-07</td><td>3.6E-07</td><td>MB231/MB468</td><td>NUC</td></tr>
<tr><td>>MI0000061|hsa-let-7a-2&WithFlank&11|-|122146516|122146599@11.31.21(+1G);MI0000062|hsa-let-7a-3&WithFlank&22|+|46112743|46112828@10.30.21(+1G);MI0000060|hsa-let-7a-1&WithFlank&9|+|94175951|94176042@12.32.21(+1G)&[MIMAT0000062&hsa-let-7a-5p&offsets|0|-1(+1G);m-6&11|-|122146568|122146589&offsets|0|-1(+1G)];[MIMAT0000062_1&hsa-let-7a-5p&offsets|0|-1(+1G);m-5&22|+|46112752|46112773&offsets|0|-1(+1G)];[MIMAT0000062_2&hsa-let-7a-5p&offsets|0|-1(+1G);m-4&9|+|94175962|94175983&offsets|0|-1(+1G)]|TGAGGTAGTAGGTTGTATAGTG</td><td>>MI0000061|hsa-let-7a-2&WithFlank&11|-|122146516|122146599</td><td>11</td><td>31</td><td>21(+1G)</td><td>TGAGGTAGTAGGTTGTATAGTG</td><td>516.6</td><td>5.3</td><td>1.0</td><td>5.2</td><td>2.2E-07</td><td>3.6E-07</td><td>MB231/MB468</td><td>MITO</td></tr>
<tr><td>>hg38_21_+_8433172_8446622_RNA45SN1_45S_50ntFlanks@7973.7995.23|GACGCGACCTCAGATCAGACGTG</td><td>>hg38_21_+_8433172_8446622_RNA45SN1_45S_50ntFlanks</td><td>7973</td><td>7995</td><td>23</td><td>GACGCGACCTCAGATCAGACGTG</td><td>1604.4</td><td>4.1</td><td>0.8</td><td>5.2</td><td>1.6E-07</td><td>3.6E-07</td><td>BT20/MB468</td><td>MITO</td></tr>
<tr><td>>hg38_21_+_8433172_8446622_RNA45SN1_45S_50ntFlanks@11581.11599.19|AACTGGTGCGGACCAGGGG</td><td>>hg38_21_+_8433172_8446622_RNA45SN1_45S_50ntFlanks</td><td>11581</td><td>11599</td><td>19</td><td>AACTGGTGCGGACCAGGGG</td><td>131.3</td><td>-10.1</td><td>1.8</td><td>-5.5</td><td>4.3E-08</td><td>3.6E-07</td><td>BT20/MB468</td><td>MP</td></tr>
<tr><td>>hg38_21_+_8433172_8446622_RNA45SN1_45S_50ntFlanks@9564.9597.34|CTGACGTGCAAATCGGTCGTCCGACCTGGGTATA</td><td>>hg38_21_+_8433172_8446622_RNA45SN1_45S_50ntFlanks</td><td>9564</td><td>9597</td><td>34</td><td>CTGACGTGCAAATCGGTCGTCCGACCTGGGTATA</td><td>6174.1</td><td>-6.4</td><td>1.2</td><td>-5.2</td><td>1.8E-07</td><td>3.6E-07</td><td>BT20/MB468</td><td>CYTO</td></tr>
<tr><td>>hg38_1_-_228634819_228635039_RNA5S12_5S_50ntFlanks@153.170.18|ACCGGGTGCTGTAGGCTT</td><td>>hg38_1_-_228634819_228635039_RNA5S12_5S_50ntFlanks</td><td>153</td><td>170</td><td>18</td><td>ACCGGGTGCTGTAGGCTT</td><td>8441.2</td><td>4.6</td><td>0.9</td><td>5.2</td><td>2.1E-07</td><td>3.6E-07</td><td>MB231/MB468</td><td>NUC</td></tr>
<tr><td>>hg38_21_+_8433172_8446622_RNA45SN1_45S_50ntFlanks@4732.4758.27|AGAACGAAAGTCGGAGGTTCGAAGACG</td><td>>hg38_21_+_8433172_8446622_RNA45SN1_45S_50ntFlanks</td><td>4732</td><td>4758</td><td>27</td><td>AGAACGAAAGTCGGAGGTTCGAAGACG</td><td>1982.6</td><td>-3.2</td><td>0.6</td><td>-5.2</td><td>2.1E-07</td><td>3.6E-07</td><td>MB231/MB468</td><td>NUC</td></tr>
<tr><td>>hg38_21_+_8433172_8446622_RNA45SN1_45S_50ntFlanks@476.512.37|TTCGTGATCGATGTGGTGACGTCGTGCTCTCCCGGGC</td><td>>hg38_21_+_8433172_8446622_RNA45SN1_45S_50ntFlanks</td><td>476</td><td>512</td><td>37</td><td>TTCGTGATCGATGTGGTGACGTCGTGCTCTCCCGGGC</td><td>2167.8</td><td>3.6</td><td>0.7</td><td>5.2</td><td>2.1E-07</td><td>3.6E-07</td><td>MB231/MB468</td><td>NUC</td></tr>
<tr><td>>hg38_21_+_8433172_8446622_RNA45SN1_45S_50ntFlanks@9572.9605.34|CAAATCGGTCGTCCGACCTGGGTATAGGGGCGAA</td><td>>hg38_21_+_8433172_8446622_RNA45SN1_45S_50ntFlanks</td><td>9572</td><td>9605</td><td>34</td><td>CAAATCGGTCGTCCGACCTGGGTATAGGGGCGAA</td><td>952.0</td><td>-5.5</td><td>1.1</td><td>-5.2</td><td>1.8E-07</td><td>3.6E-07</td><td>MB231/MB468</td><td>MP</td></tr>
<tr><td>>hg38_21_+_8433172_8446622_RNA45SN1_45S_50ntFlanks@12950.12990.41|CGACCTGCTTCTGGGTCGGGGTTTCGTACGTAGCAGAGCAG</td><td>>hg38_21_+_8433172_8446622_RNA45SN1_45S_50ntFlanks</td><td>12950</td><td>12990</td><td>41</td><td>CGACCTGCTTCTGGGTCGGGGTTTCGTACGTAGCAGAGCAG</td><td>166.5</td><td>-7.1</td><td>1.4</td><td>-5.2</td><td>2.1E-07</td><td>3.6E-07</td><td>MB231/MB468</td><td>NUC</td></tr>
<tr><td>>MI0000279|hsa-mir-196a-2&WithFlank&12|+|53991732|53991853@31.52.22(+1U);MI0000238|hsa-mir-196a-1&WithFlank&17|-|48632484|48632565@13.34.22(+1U)&[MIMAT0000226&hsa-miR-196a-5p&offsets|0|0(+1U);m-261&12|+|53991762|53991783&offsets|0|0(+1U)];[MIMAT0000226_1&hsa-miR-196a-5p&offsets|0|0(+1U);m-268&17|-|48632532|48632553&offsets|0|0(+1U)]|TAGGTAGTTTCATGTTGTTGGGT</td><td>>MI0000279|hsa-mir-196a-2&WithFlank&12|+|53991732|53991853</td><td>31</td><td>52</td><td>22(+1U)</td><td>TAGGTAGTTTCATGTTGTTGGGT</td><td>178.1</td><td>2.6</td><td>0.5</td><td>5.2</td><td>2.3E-07</td><td>3.6E-07</td><td>BT20/MB468</td><td>NUC</td></tr>
<tr><td>>hg38_21_+_8433172_8446622_RNA45SN1_45S_50ntFlanks@12607.12624.18|TGTGCTTGGCTGAGGAGC</td><td>>hg38_21_+_8433172_8446622_RNA45SN1_45S_50ntFlanks</td><td>12607</td><td>12624</td><td>18</td><td>TGTGCTTGGCTGAGGAGC</td><td>612.6</td><td>3.4</td><td>0.7</td><td>5.2</td><td>2.0E-07</td><td>3.6E-07</td><td>MB231/MB468</td><td>CYTO</td></tr>
<tr><td>>hg38_21_+_8433172_8446622_RNA45SN1_45S_50ntFlanks@12360.12384.25|CAGGGATAACTGGCTTGTGGCGGCC</td><td>>hg38_21_+_8433172_8446622_RNA45SN1_45S_50ntFlanks</td><td>12360</td><td>12384</td><td>25</td><td>CAGGGATAACTGGCTTGTGGCGGCC</td><td>528.1</td><td>-7.2</td><td>1.4</td><td>-5.2</td><td>2.2E-07</td><td>3.6E-07</td><td>MB231/MB468</td><td>MITO</td></tr>
<tr><td>>hg38_21_+_8433172_8446622_RNA45SN1_45S_50ntFlanks@11668.11693.26|TGCCCAGTGCTCTGAATGTCAAAGTG</td><td>>hg38_21_+_8433172_8446622_RNA45SN1_45S_50ntFlanks</td><td>11668</td><td>11693</td><td>26</td><td>TGCCCAGTGCTCTGAATGTCAAAGTG</td><td>376.3</td><td>3.8</td><td>0.7</td><td>5.2</td><td>2.1E-07</td><td>3.6E-07</td><td>MB231/MB468</td><td>NUC</td></tr>
<tr><td>>trnaMT_LeuTAG_MT_+_12266_12336@39.74.36|CCAAAAATTTTGGTGCAACTCCAAATAAAAGTACCA</td><td>>trnaMT_LeuTAG_MT_+_12266_12336</td><td>39</td><td>74</td><td>36</td><td>CCAAAAATTTTGGTGCAACTCCAAATAAAAGTACCA</td><td>185.9</td><td>-10.9</td><td>1.9</td><td>-5.7</td><td>9.1E-09</td><td>3.6E-07</td><td>BT20/MB231</td><td>MP</td></tr>
<tr><td>>hg38_21_+_8433172_8446622_RNA45SN1_45S_50ntFlanks@9568.9603.36|CGTGCAAATCGGTCGTCCGACCTGGGTATAGGGGCG</td><td>>hg38_21_+_8433172_8446622_RNA45SN1_45S_50ntFlanks</td><td>9568</td><td>9603</td><td>36</td><td>CGTGCAAATCGGTCGTCCGACCTGGGTATAGGGGCG</td><td>833.5</td><td>-7.0</td><td>1.3</td><td>-5.2</td><td>1.8E-07</td><td>3.6E-07</td><td>MB231/MB468</td><td>MP</td></tr>
<tr><td>>MI0000071|hsa-mir-17&WithFlank&13|+|91350599|91350694@20.42.23(+1A)&[MIMAT0000070&hsa-miR-17-5p&offsets|0|0(+1A);m-88&13|+|91350618|91350640&offsets|0|0(+1A)]|CAAAGTGCTTACAGTGCAGGTAGA</td><td>>MI0000071|hsa-mir-17&WithFlank&13|+|91350599|91350694</td><td>20</td><td>42</td><td>23(+1A)</td><td>CAAAGTGCTTACAGTGCAGGTAGA</td><td>4891.8</td><td>5.3</td><td>1.0</td><td>5.2</td><td>2.2E-07</td><td>3.6E-07</td><td>MB231/MB468</td><td>MITO</td></tr>
<tr><td>>MI0000085|hsa-mir-27a&WithFlank&19|-|13836434|13836523@57.75.19(+1U)&[MIMAT0000084&hsa-miR-27a-3p&offsets|0|-2(+1U);m-43&19|-|13836447|13836467&offsets|0|-2(+1U)]|TTCACAGTGGCTAAGTTCCT</td><td>>MI0000085|hsa-mir-27a&WithFlank&19|-|13836434|13836523</td><td>57</td><td>75</td><td>19(+1U)</td><td>TTCACAGTGGCTAAGTTCCT</td><td>232.8</td><td>6.0</td><td>1.2</td><td>5.2</td><td>2.1E-07</td><td>3.6E-07</td><td>MB231/MB468</td><td>NUC</td></tr>
<tr><td>>hg38_21_+_8433172_8446622_RNA45SN1_45S_50ntFlanks@11923.11950.28|AGAGGTGTAGAATAAGTGGGAGGCCCCC</td><td>>hg38_21_+_8433172_8446622_RNA45SN1_45S_50ntFlanks</td><td>11923</td><td>11950</td><td>28</td><td>AGAGGTGTAGAATAAGTGGGAGGCCCCC</td><td>266.3</td><td>-7.4</td><td>1.4</td><td>-5.2</td><td>1.8E-07</td><td>3.6E-07</td><td>MB231/MB468</td><td>MP</td></tr>
<tr><td>>hg38_21_+_8433172_8446622_RNA45SN1_45S_50ntFlanks@12071.12095.25|CCCCGAGGGGCTCTCGCTTCTGGCG</td><td>>hg38_21_+_8433172_8446622_RNA45SN1_45S_50ntFlanks</td><td>12071</td><td>12095</td><td>25</td><td>CCCCGAGGGGCTCTCGCTTCTGGCG</td><td>914.1</td><td>-5.7</td><td>1.1</td><td>-5.2</td><td>1.6E-07</td><td>3.6E-07</td><td>BT20/MB468</td><td>MITO</td></tr>
<tr><td>>hg38_21_+_8433172_8446622_RNA45SN1_45S_50ntFlanks@4732.4759.28|AGAACGAAAGTCGGAGGTTCGAAGACGA</td><td>>hg38_21_+_8433172_8446622_RNA45SN1_45S_50ntFlanks</td><td>4732</td><td>4759</td><td>28</td><td>AGAACGAAAGTCGGAGGTTCGAAGACGA</td><td>568.3</td><td>-3.0</td><td>0.6</td><td>-5.2</td><td>2.1E-07</td><td>3.6E-07</td><td>MB231/MB468</td><td>NUC</td></tr>
</table>

| | | | | | | | | | | | | | |
|---|---|---|---|---|---|---|---|---|---|---|---|---|---|
| >hg38_MT_+_1621_3279_RNR2_16S_50ntFlanks@485.521.37\|GAGGAACAGCTCTTTGGACACTAGGAAAAAACCTTGT | >hg38_MT_+_1621_3279_RNR2_16S_50ntFlanks | 485 | 521 | 37 | GAGGAACAGCTCTTTGGACACTAGGAAAAAACCTTGT | 69.9 | 9.9 | 1.8 | 5.5 | 4.3E-08 | 3.6E-07 | BT20/MB468 | MP |
| >hg38_21_+_8433172_8446622_RNA45SN1_45S_50ntFlanks@12251.12273.23\|AAAGCTCGCTTGATCTTGATTTT | >hg38_21_+_8433172_8446622_RNA45SN1_45S_50ntFlanks | 12251 | 12273 | 23 | AAAGCTCGCTTGATCTTGATTTT | 820.4 | -6.8 | 1.3 | -5.2 | 2.2E-07 | 3.6E-07 | MB231/MB468 | MITO |
| >trnaMT_ValTAC_MT_+_1602_1670@48.71.24\|CAACTTAACTTGACCGCTCTGACC | >trnaMT_ValTAC_MT_+_1602_1670 | 48 | 71 | 24 | CAACTTAACTTGACCGCTCTGACC | 535.0 | 4.5 | 0.9 | 5.2 | 2.1E-07 | 3.6E-07 | MB231/MB468 | NUC |
| >hg38_21_+_8433172_8446622_RNA45SN1_45S_50ntFlanks@11580.11600.21\|GAACTGGTGCGGACCAGGGGA | >hg38_21_+_8433172_8446622_RNA45SN1_45S_50ntFlanks | 11580 | 11600 | 21 | GAACTGGTGCGGACCAGGGGA | 720.4 | -4.8 | 0.9 | -5.2 | 2.0E-07 | 3.6E-07 | MB231/MB468 | CYTO |
| >MI0000085\|hsa-mir-27a&WithFlank&19\|-\|13836434\|13836523@16.35.20(+1U)&[MIMAT0004501&hsa-miR-27a-5p&offsets\|0\|-2(+1U);m-270&19\|-\|13836487\|13836508&offsets\|0\|-2(+1U)]\|AGGGCTTAGCTGCTTGTGAGT | >MI0000085\|hsa-mir-27a&WithFlank&19\|-\|13836434\|13836523 | 16 | 35 | 20(+1U) | AGGGCTTAGCTGCTTGTGAGT | 66.9 | 9.3 | 1.8 | 5.2 | 1.8E-07 | 3.6E-07 | MB231/MB468 | MP |
| >MI0000471\|hsa-mir-126&WithFlank&9\|+\|136670596\|136670692@58.79.22&[MIMAT0000445&hsa-miR-126-3p&offsets\|0\|0;m-40&9\|+\|136670653\|136670674&offsets\|0\|0]\|TCGTACCGTGAGTAATAATGCG | >MI0000471\|hsa-mir-126&WithFlank&9\|+\|136670596\|136670692 | 58 | 79 | 22 | TCGTACCGTGAGTAATAATGCG | 5311.4 | 7.3 | 1.4 | 5.2 | 1.8E-07 | 3.6E-07 | MB231/MB468 | MP |
| >hg38_21_+_8433172_8446622_RNA45SN1_45S_50ntFlanks@11570.11609.40\|AGCCGACTTAGAACTGGTGCGGACCAGGGGAATCCGACTG | >hg38_21_+_8433172_8446622_RNA45SN1_45S_50ntFlanks | 11570 | 11609 | 40 | AGCCGACTTAGAACTGGTGCGGACCAGGGGAATCCGACTG | 59.3 | -9.8 | 1.9 | -5.2 | 1.8E-07 | 3.6E-07 | MB231/MB468 | MP |
| >hg38_21_+_8433172_8446622_RNA45SN1_45S_50ntFlanks@8345.8373.29\|AAGGGAAAGTTGAAAAGAACTTTGAAGAG | >hg38_21_+_8433172_8446622_RNA45SN1_45S_50ntFlanks | 8345 | 8373 | 29 | AAGGGAAAGTTGAAAAGAACTTTGAAGAG | 721.0 | 9.5 | 1.7 | 5.5 | 4.4E-08 | 3.6E-07 | BT20/MB468 | MP |
| >hg38_21_+_8433172_8446622_RNA45SN1_45S_50ntFlanks@12490.12509.20\|CGTGAGCTGGGTTTAGACCG | >hg38_21_+_8433172_8446622_RNA45SN1_45S_50ntFlanks | 12490 | 12509 | 20 | CGTGAGCTGGGTTTAGACCG | 87.3 | -9.5 | 1.7 | -5.5 | 4.4E-08 | 3.6E-07 | BT20/MB468 | MP |
| >hg38_21_+_8433172_8446622_RNA45SN1_45S_50ntFlanks@12454.12480.27\|CACCAAGCGTTGGATTGTTCACCCACT | >hg38_21_+_8433172_8446622_RNA45SN1_45S_50ntFlanks | 12454 | 12480 | 27 | CACCAAGCGTTGGATTGTTCACCCACT | 191.0 | 3.7 | 0.7 | 5.2 | 2.0E-07 | 3.7E-07 | MB231/MB468 | CYTO |
| >hg38_21_+_8433172_8446622_RNA45SN1_45S_50ntFlanks@10348.10368.21\|AAACGAGAACTTTGAAGGCCG | >hg38_21_+_8433172_8446622_RNA45SN1_45S_50ntFlanks | 10348 | 10368 | 21 | AAACGAGAACTTTGAAGGCCG | 2366.6 | -5.9 | 1.1 | -5.5 | 4.4E-08 | 3.7E-07 | BT20/MB468 | MP |
| >hg38_21_+_8433172_8446622_RNA45SN1_45S_50ntFlanks@11930.11948.19\|TAGAATAAGTGGGAGGCCC | >hg38_21_+_8433172_8446622_RNA45SN1_45S_50ntFlanks | 11930 | 11948 | 19 | TAGAATAAGTGGGAGGCCC | 537.2 | -7.7 | 1.4 | -5.5 | 4.4E-08 | 3.7E-07 | BT20/MB468 | MP |
| >hg38_21_+_8433172_8446622_RNA45SN1_45S_50ntFlanks@3843.3861.19\|CTACTTGGATAACTGTGGT | >hg38_21_+_8433172_8446622_RNA45SN1_45S_50ntFlanks | 3843 | 3861 | 19 | CTACTTGGATAACTGTGGT | 1220.9 | -6.5 | 1.3 | -5.2 | 2.2E-07 | 3.7E-07 | MB231/MB468 | MITO |
| >hg38_21_+_8433172_8446622_RNA45SN1_45S_50ntFlanks@12934.12953.20\|CTAAACCATTCGTAGACGAC | >hg38_21_+_8433172_8446622_RNA45SN1_45S_50ntFlanks | 12934 | 12953 | 20 | CTAAACCATTCGTAGACGAC | 310.1 | -3.1 | 0.6 | -5.2 | 1.6E-07 | 3.7E-07 | BT20/MB468 | MITO |
| >hg38_21_+_8433172_8446622_RNA45SN1_45S_50ntFlanks@13004.13039.36\|ATCTATTGAAAGTCAGCCCTCGACACAAGGGTTTGT | >hg38_21_+_8433172_8446622_RNA45SN1_45S_50ntFlanks | 13004 | 13039 | 36 | ATCTATTGAAAGTCAGCCCTCGACACAAGGGTTTGT | 98.1 | -9.7 | 1.8 | -5.5 | 4.4E-08 | 3.7E-07 | BT20/MB468 | MP |
| >MI0000301\|hsa-mir-224&WithFlank&X\|-\|151958572\|151958664@14.38.25&[MIMAT0000281&hsa-miR-224-5p&offsets\|+1\|+1;m-238&X\|-\|151958630\|151958651&offsets\|0\|+3]\|CAAGTCACTAGTGGTTCCGTTTAGT | >MI0000301\|hsa-mir-224&WithFlank&X\|-\|151958572\|151958664 | 14 | 38 | 25 | CAAGTCACTAGTGGTTCCGTTTAGT | 124.0 | 5.7 | 1.1 | 5.2 | 2.3E-07 | 3.7E-07 | BT20/MB468 | NUC |
| >hg38_21_+_8433172_8446622_RNA45SN1_45S_50ntFlanks@4763.4780.18\|GATACCGTCGTAGTTCCG | >hg38_21_+_8433172_8446622_RNA45SN1_45S_50ntFlanks | 4763 | 4780 | 18 | GATACCGTCGTAGTTCCG | 501.1 | -5.8 | 1.1 | -5.2 | 2.1E-07 | 3.7E-07 | MB231/MB468 | NUC |
| >MI0000101\|hsa-mir-99a&WithFlank&21\|+\|16539083\|16539175@19.38.20(+1G)&[MIMAT0000097&hsa-miR-99a-5p&offsets\|0\|-2(+1G);m-64&21\|+\|16539101\|16539121&offsets\|0\|-1(+1G)]\|AACCCGTAGATCCGATCTTGG | >MI0000101\|hsa-mir-99a&WithFlank&21\|+\|16539083\|16539175 | 19 | 38 | 20(+1G) | AACCCGTAGATCCGATCTTGG | 70.8 | -9.2 | 1.7 | -5.5 | 4.4E-08 | 3.7E-07 | BT20/MB468 | MP |
| >hg38_21_+_8433172_8446622_RNA45SN1_45S_50ntFlanks@11729.11748.20\|TAACTATGACTCTCTTAAGG | >hg38_21_+_8433172_8446622_RNA45SN1_45S_50ntFlanks | 11729 | 11748 | 20 | TAACTATGACTCTCTTAAGG | 1323.5 | -2.5 | 0.5 | -5.2 | 1.6E-07 | 3.7E-07 | BT20/MB468 | MITO |
| >hg38_21_+_8433172_8446622_RNA45SN1_45S_50ntFlanks@11668.11707.40\|TGCCCAGTGCTCTGAATGTCAAAGTGAAGAAATTCAATGA | >hg38_21_+_8433172_8446622_RNA45SN1_45S_50ntFlanks | 11668 | 11707 | 40 | TGCCCAGTGCTCTGAATGTCAAAGTGAAGAAATTCAATGA | 497.9 | 5.5 | 1.1 | 5.2 | 2.1E-07 | 3.7E-07 | MB231/MB468 | NUC |
| >MI0000440\|hsa-mir-27b&WithFlank&9\|+\|95085439\|95085547@67.86.20(+1U)&[MIMAT0000419&hsa-miR-27b-3p&offsets\|0\|-1(+1U);m-39&9\|+\|95085505\|95085525&offsets\|0\|-1(+1U)]\|TTCACAGTGGCTAAGTTCTGT | >MI0000440\|hsa-mir-27b&WithFlank&9\|+\|95085439\|95085547 | 67 | 86 | 20(+1U) | TTCACAGTGGCTAAGTTCTGT | 6927.6 | 4.2 | 0.8 | 5.2 | 1.9E-07 | 3.7E-07 | MB231/MB468 | MP |
| >hg38_21_+_8433172_8446622_RNA45SN1_45S_50ntFlanks@4203.4223.21\|GACCCGGGGAGGTAGTGACGA | >hg38_21_+_8433172_8446622_RNA45SN1_45S_50ntFlanks | 4203 | 4223 | 21 | GACCCGGGGAGGTAGTGACGA | 254.6 | -4.3 | 0.8 | -5.2 | 2.1E-07 | 3.7E-07 | MB231/MB468 | NUC |
| >hg38_21_+_8433172_8446622_RNA45SN1_45S_50ntFlanks@11930.11951.22\|TAGAATAAGTGGGAGGCCCCCG | >hg38_21_+_8433172_8446622_RNA45SN1_45S_50ntFlanks | 11930 | 11951 | 22 | TAGAATAAGTGGGAGGCCCCCG | 492.3 | -8.1 | 1.5 | -5.5 | 4.4E-08 | 3.7E-07 | BT20/MB468 | MP |
| >hg38_21_+_8433172_8446622_RNA45SN1_45S_50ntFlanks@11581.11600.20\|AACTGGTGCGGACCAGGGGA | >hg38_21_+_8433172_8446622_RNA45SN1_45S_50ntFlanks | 11581 | 11600 | 20 | AACTGGTGCGGACCAGGGGA | 145.6 | -6.3 | 1.2 | -5.2 | 2.1E-07 | 3.7E-07 | MB231/MB468 | NUC |
| >hg38_21_+_8433172_8446622_RNA45SN1_45S_50ntFlanks@7989.8015.27\|AGACGTGGCGACCCGCTGAATTTAAGC | >hg38_21_+_8433172_8446622_RNA45SN1_45S_50ntFlanks | 7989 | 8015 | 27 | AGACGTGGCGACCCGCTGAATTTAAGC | 536.9 | 3.8 | 0.7 | 5.2 | 1.6E-07 | 3.7E-07 | BT20/MB468 | MITO |
| >hg38_21_+_8433172_8446622_RNA45SN1_45S_50ntFlanks@8176.8204.29\|CTTCTGATCGAGGCCCAGCCCGTGGACGG | >hg38_21_+_8433172_8446622_RNA45SN1_45S_50ntFlanks | 8176 | 8204 | 29 | CTTCTGATCGAGGCCCAGCCCGTGGACGG | 2423.6 | -5.1 | 0.9 | -5.5 | 4.4E-08 | 3.7E-07 | BT20/MB468 | MP |
| >hg38_21_+_8433172_8446622_RNA45SN1_45S_50ntFlanks@9483.9513.31\|CGTGTTAGGACCCGAAAGATGGTGAACTATG | >hg38_21_+_8433172_8446622_RNA45SN1_45S_50ntFlanks | 9483 | 9513 | 31 | CGTGTTAGGACCCGAAAGATGGTGAACTATG | 76.3 | 10.0 | 1.8 | 5.5 | 4.4E-08 | 3.7E-07 | BT20/MB468 | MP |
| >hg38_21_+_8433172_8446622_RNA45SN1_45S_50ntFlanks@12288.12318.31\|ACCGTGAAAGCGGGGCCTCACGATCCTTCTG | >hg38_21_+_8433172_8446622_RNA45SN1_45S_50ntFlanks | 12288 | 12318 | 31 | ACCGTGAAAGCGGGGCCTCACGATCCTTCTG | 1105.4 | -4.5 | 0.9 | -5.2 | 1.6E-07 | 3.7E-07 | BT20/MB468 | MITO |
| >hg38_21_+_8433172_8446622_RNA45SN1_45S_50ntFlanks@12074.12108.35\|CGAGGGGCTCTCGCTTCTGGCGCCAAGCGCCCGGC | >hg38_21_+_8433172_8446622_RNA45SN1_45S_50ntFlanks | 12074 | 12108 | 35 | CGAGGGGCTCTCGCTTCTGGCGCCAAGCGCCCGGC | 352.0 | -5.4 | 1.0 | -5.2 | 2.2E-07 | 3.7E-07 | MB231/MB468 | MITO |
| >hg38_21_+_8433172_8446622_RNA45SN1_45S_50ntFlanks@10794.10823.30\|CTGGCATGTTGGAACAATGTAGGTAAGGGA | >hg38_21_+_8433172_8446622_RNA45SN1_45S_50ntFlanks | 10794 | 10823 | 30 | CTGGCATGTTGGAACAATGTAGGTAAGGGA | 251.8 | -6.8 | 1.3 | -5.2 | 2.1E-07 | 3.7E-07 | MB231/MB468 | NUC |
| >hg38_21_+_8433172_8446622_RNA45SN1_45S_50ntFlanks@4399.4423.25\|CGGGCGGGCGGTCCGCCGCGAGGCG | >hg38_21_+_8433172_8446622_RNA45SN1_45S_50ntFlanks | 4399 | 4423 | 25 | CGGGCGGGCGGTCCGCCGCGAGGCG | 380.7 | -3.8 | 0.7 | -5.2 | 2.1E-07 | 3.7E-07 | MB231/MB468 | CYTO |
| >hg38_21_+_8433172_8446622_RNA45SN1_45S_50ntFlanks@9791.9815.25\|TGGCGTGGAGCCGGGCGTGGAATGC | >hg38_21_+_8433172_8446622_RNA45SN1_45S_50ntFlanks | 9791 | 9815 | 25 | TGGCGTGGAGCCGGGCGTGGAATGC | 745.2 | 12.9 | 2.2 | 5.7 | 9.4E-09 | 3.7E-07 | BT20/MB231 | MP |
| >MI0000446\|hsa-mir-125b-1&WithFlank&11\|-\|122099751\|122099850@21.42.22(+2A);MI0000470\|hsa-mir-125b-2&WithFlank&21\|+\|16590231\|16590331@23.44.22(+2A)&[MIMAT0000423&hsa-miR-125b-5p&offsets\|0\|0(+2A);m-67&11\|-\|122099809\|122099830&offsets\|0\|0(+2A)];[MIMAT0000423_1&hsa-miR-125b-5p&offsets\|0\|0(+2A);m-68&21\|+\|16590253\|16590274&offsets\|0\|0(+2A)]\|TCCCTGAGACCCTAACTTGTGAAA | >MI0000446\|hsa-mir-125b-1&WithFlank&11\|-\|122099751\|122099850 | 21 | 42 | 22(+2A) | TCCCTGAGACCCTAACTTGTGAAA | 139.3 | -10.5 | 1.8 | -5.7 | 9.4E-09 | 3.7E-07 | BT20/MB231 | MP |
| >hg38_21_+_8433172_8446622_RNA45SN1_45S_50ntFlanks@4047.4067.21\|ATTCGAACGTCTGCCCTATCA | >hg38_21_+_8433172_8446622_RNA45SN1_45S_50ntFlanks | 4047 | 4067 | 21 | ATTCGAACGTCTGCCCTATCA | 324.3 | -2.2 | 0.4 | -5.2 | 1.8E-07 | 3.7E-07 | BT20/MB468 | CYTO |
| >am_trnaMT_ValTAC_MT_+_1602_1670@33.62.30\|ACACTTAGGAGATTTCAACTTAACTTGACC | >am_trnaMT_ValTAC_MT_+_1602_1670 | 33 | 62 | 30 | ACACTTAGGAGATTTCAACTTAACTTGACC | 243.4 | 11.7 | 2.1 | 5.5 | 4.5E-08 | 3.7E-07 | BT20/MB468 | MP |
| >hg38_21_+_8433172_8446622_RNA45SN1_45S_50ntFlanks@11564.11609.46\|CCTAGCAGCCGACTTAGAACTGGTGCGGACCAGGGGAATCCGACTG | >hg38_21_+_8433172_8446622_RNA45SN1_45S_50ntFlanks | 11564 | 11609 | 46 | CCTAGCAGCCGACTTAGAACTGGTGCGGACCAGGGGAATCCGACTG | 394.0 | -7.7 | 1.5 | -5.2 | 2.2E-07 | 3.7E-07 | MB231/MB468 | MITO |
| >MI0000440\|hsa-mir-27b&WithFlank&9\|+\|95085439\|95085547@67.85.19(+1A)&[MIMAT0000419&hsa-miR-27b-3p&offsets\|0\|-2(+1A);m-39&9\|+\|95085505\|95085525&offsets\|0\|-2(+1A)]\|TTCACAGTGGCTAAGTTCTA | >MI0000440\|hsa-mir-27b&WithFlank&9\|+\|95085439\|95085547 | 67 | 85 | 19(+1A) | TTCACAGTGGCTAAGTTCTA | 648.9 | 4.3 | 0.8 | 5.2 | 1.9E-07 | 3.7E-07 | MB231/MB468 | MP |
| >hg38_21_+_8433172_8446622_RNA45SN1_45S_50ntFlanks@12971.12988.18\|TTTCGTACGTAGCAGAGC | >hg38_21_+_8433172_8446622_RNA45SN1_45S_50ntFlanks | 12971 | 12988 | 18 | TTTCGTACGTAGCAGAGC | 763.9 | -8.0 | 1.5 | -5.2 | 2.3E-07 | 3.7E-07 | BT20/MB468 | NUC |
| >hg38_21_+_8433172_8446622_RNA45SN1_45S_50ntFlanks@12990.13014.25\|GCTCCCTCGCTGCGATCTATTGAAA | >hg38_21_+_8433172_8446622_RNA45SN1_45S_50ntFlanks | 12990 | 13014 | 25 | GCTCCCTCGCTGCGATCTATTGAAA | 3937.6 | -9.5 | 1.8 | -5.2 | 2.3E-07 | 3.7E-07 | BT20/MB468 | NUC |
| >hg38_21_+_8433172_8446622_RNA45SN1_45S_50ntFlanks@10351.10373.23\|CGAGAACTTTGAAGGCCGAAGTG | >hg38_21_+_8433172_8446622_RNA45SN1_45S_50ntFlanks | 10351 | 10373 | 23 | CGAGAACTTTGAAGGCCGAAGTG | 712.8 | -5.6 | 1.1 | -5.2 | 2.2E-07 | 3.7E-07 | MB231/MB468 | MITO |
| >hg38_21_+_8433172_8446622_RNA45SN1_45S_50ntFlanks@10802.10826.25\|TTGGAACAATGTAGGTAAGGGAAGT | >hg38_21_+_8433172_8446622_RNA45SN1_45S_50ntFlanks | 10802 | 10826 | 25 | TTGGAACAATGTAGGTAAGGGAAGT | 239.1 | -6.9 | 1.3 | -5.2 | 2.2E-07 | 3.7E-07 | MB231/MB468 | MITO |
| >hg38_21_+_8433172_8446622_RNA45SN1_45S_50ntFlanks@10793.10818.26\|TCTGGCATGTTGGAACAATGTAGGTA | >hg38_21_+_8433172_8446622_RNA45SN1_45S_50ntFlanks | 10793 | 10818 | 26 | TCTGGCATGTTGGAACAATGTAGGTA | 482.0 | -5.7 | 1.1 | -5.2 | 2.2E-07 | 3.7E-07 | MB231/MB468 | MITO |
| >hg38_1_-_228634819_228635039_RNA5S12_5S_50ntFlanks@85.108.24\|TCGTCTGATCTCGGAAGCTAAGCA | >hg38_1_-_228634819_228635039_RNA5S12_5S_50ntFlanks | 85 | 108 | 24 | TCGTCTGATCTCGGAAGCTAAGCA | 112.1 | 10.6 | 1.9 | 5.5 | 4.5E-08 | 3.7E-07 | BT20/MB468 | MP |
| >MI0000088\|hsa-mir-30a&WithFlank&6\|-\|71403545\|71403627@53.73.21&[MIMAT0000088&hsa-miR-30a-3p&offsets\|0\|-1;m-38&6\|-\|71403554\|71403575&offsets\|0\|-1]\|CTTTCAGTCGGATGTTTGCAG | >MI0000088\|hsa-mir-30a&WithFlank&6\|-\|71403545\|71403627 | 53 | 73 | 21 | CTTTCAGTCGGATGTTTGCAG | 1700.2 | 3.9 | 0.8 | 5.2 | 1.9E-07 | 3.7E-07 | MB231/MB468 | MP |
| >hg38_21_+_8433172_8446622_RNA45SN1_45S_50ntFlanks@11670.11688.19\|CCCAGTGCTCTGAATGTCA | >hg38_21_+_8433172_8446622_RNA45SN1_45S_50ntFlanks | 11670 | 11688 | 19 | CCCAGTGCTCTGAATGTCA | 310.2 | -5.7 | 1.1 | -5.2 | 1.9E-07 | 3.7E-07 | MB231/MB468 | MP |
| >hg38_21_+_8433172_8446622_RNA45SN1_45S_50ntFlanks@12073.12090.18\|CCGAGGGGCTCTCGCTTC | >hg38_21_+_8433172_8446622_RNA45SN1_45S_50ntFlanks | 12073 | 12090 | 18 | CCGAGGGGCTCTCGCTTC | 339.4 | -7.6 | 1.5 | -5.2 | 2.4E-07 | 3.7E-07 | BT20/MB468 | NUC |
| >hg38_21_+_8433172_8446622_RNA45SN1_45S_50ntFlanks@11576.11594.19\|CTTAGAACTGGTGCGGACC | >hg38_21_+_8433172_8446622_RNA45SN1_45S_50ntFlanks | 11576 | 11594 | 19 | CTTAGAACTGGTGCGGACC | 1314.5 | -3.0 | 0.6 | -5.2 | 1.8E-07 | 3.7E-07 | BT20/MB468 | CYTO |
| >hg38_21_+_8433172_8446622_RNA45SN1_45S_50ntFlanks@10737.10767.31\|TAAATCTCGCGCCGGGCCGTACCCATATCCG | >hg38_21_+_8433172_8446622_RNA45SN1_45S_50ntFlanks | 10737 | 10767 | 31 | TAAATCTCGCGCCGGGCCGTACCCATATCCG | 164.4 | -6.2 | 1.2 | -5.2 | 1.8E-07 | 3.7E-07 | BT20/MB468 | CYTO |
| >hg38_21_+_8433172_8446622_RNA45SN1_45S_50ntFlanks@12645.12669.25\|CTGTGGGATTATGACTGAACGCCTC | >hg38_21_+_8433172_8446622_RNA45SN1_45S_50ntFlanks | 12645 | 12669 | 25 | CTGTGGGATTATGACTGAACGCCTC | 180.8 | 4.6 | 0.9 | 5.2 | 2.1E-07 | 3.7E-07 | MB231/MB468 | CYTO |
| >hg38_21_+_8433172_8446622_RNA45SN1_45S_50ntFlanks@11582.11623.42\|ACTGGTGCGGACCAGGGGAATCCGACTGTTTAATTAAAACAA | >hg38_21_+_8433172_8446622_RNA45SN1_45S_50ntFlanks | 11582 | 11623 | 42 | ACTGGTGCGGACCAGGGGAATCCGACTGTTTAATTAAAACAA | 87.4 | 10.2 | 1.9 | 5.5 | 4.5E-08 | 3.7E-07 | BT20/MB468 | MP |
| >trnaMT_PheGAA_MT_+_577_647@32.56.25\|ACTGAAAATGTTTAGACGGGCTCAC | >trnaMT_PheGAA_MT_+_577_647 | 32 | 56 | 25 | ACTGAAAATGTTTAGACGGGCTCAC | 98.6 | 10.4 | 1.9 | 5.5 | 4.5E-08 | 3.7E-07 | BT20/MB468 | MP |

<table>
<tr><td>>hg38_21_+_8433172_8446622_RNA45SN1_45S_50ntFlanks@10735.10757.23|TGTAAATCTCGCGCCGGGCCGTA</td><td>>hg38_21_+_8433172_8446622_RNA45SN1_45S_50ntFlanks</td><td>10735</td><td>10757</td><td>23</td><td>TGTAAATCTCGCGCCGGGCCGTA</td><td>699.7</td><td>-7.4</td><td>1.4</td><td>-5.2</td><td>2.1E-07</td><td>3.7E-07</td><td>MB231/MB468</td><td>NUC</td></tr>
<tr><td>>hg38_21_+_8433172_8446622_RNA45SN1_45S_50ntFlanks@3847.3868.22|TTGGATAACTGTGGTAATTCTA</td><td>>hg38_21_+_8433172_8446622_RNA45SN1_45S_50ntFlanks</td><td>3847</td><td>3868</td><td>22</td><td>TTGGATAACTGTGGTAATTCTA</td><td>240.1</td><td>2.8</td><td>0.5</td><td>5.2</td><td>2.1E-07</td><td>3.8E-07</td><td>MB231/MB468</td><td>CYTO</td></tr>
<tr><td>>hg38_21_+_8433172_8446622_RNA45SN1_45S_50ntFlanks@4758.4780.23|GATCAGATACCGTCGTAGTTCCG</td><td>>hg38_21_+_8433172_8446622_RNA45SN1_45S_50ntFlanks</td><td>4758</td><td>4780</td><td>23</td><td>GATCAGATACCGTCGTAGTTCCG</td><td>125.4</td><td>-5.1</td><td>1.0</td><td>-5.2</td><td>2.1E-07</td><td>3.8E-07</td><td>MB231/MB468</td><td>CYTO</td></tr>
<tr><td>>hg38_21_+_8433172_8446622_RNA45SN1_45S_50ntFlanks@12605.12625.21|TATGTGCTTGGCTGAGGAGCC</td><td>>hg38_21_+_8433172_8446622_RNA45SN1_45S_50ntFlanks</td><td>12605</td><td>12625</td><td>21</td><td>TATGTGCTTGGCTGAGGAGCC</td><td>1233.6</td><td>-3.3</td><td>0.6</td><td>-5.2</td><td>1.6E-07</td><td>3.8E-07</td><td>BT20/MB468</td><td>MITO</td></tr>
<tr><td>>am_trnaMT_ArgTCG_MT_+_10405_10469@31.48.18|TCGACTCATTAAATTATG</td><td>>am_trnaMT_ArgTCG_MT_+_10405_10469</td><td>31</td><td>48</td><td>18</td><td>TCGACTCATTAAATTATG</td><td>213.5</td><td>-6.4</td><td>1.2</td><td>-5.2</td><td>2.1E-07</td><td>3.8E-07</td><td>MB231/MB468</td><td>CYTO</td></tr>
<tr><td>>trna10_AlaCGC_6_+_26553731_26553802@1.24.24,trna119_AlaCGC_6_-_28641613_28641684@1.24.24,trna120_AlaAGC_6_-_28626014_28626085@1.24.24,trna13_AlaCGC_2_+_157257281_157257352@1.24.24,trna13_AlaTGC_12_-_125406301_125406372@1.24.24,trna66_AlaTGC_6_+_28611222_28611293@1.24.24,trna8_AlaTGC_12_+_125424512_125424583@1.24.24,trna8_AlaTGC_5_+_180633868_180633939@1.24.24|GGGGATGTAGCTCAGTGGTAGAGC</td><td>>trna10_AlaCGC_6_+_26553731_26553802</td><td>1</td><td>24</td><td>24</td><td>GGGGATGTAGCTCAGTGGTAGAGC</td><td>642.0</td><td>4.5</td><td>0.9</td><td>5.2</td><td>2.1E-07</td><td>3.8E-07</td><td>MB231/MB468</td><td>CYTO</td></tr>
<tr><td>>hg38_21_+_8433172_8446622_RNA45SN1_45S_50ntFlanks@10792.10816.25|CTCTGGCATGTTGGAACAATGTAGG</td><td>>hg38_21_+_8433172_8446622_RNA45SN1_45S_50ntFlanks</td><td>10792</td><td>10816</td><td>25</td><td>CTCTGGCATGTTGGAACAATGTAGG</td><td>438.4</td><td>-6.3</td><td>1.2</td><td>-5.2</td><td>2.4E-07</td><td>3.8E-07</td><td>BT20/MB468</td><td>NUC</td></tr>
<tr><td>>MI0000816|hsa-mir-335&WithFlank&7|+|130496105|130496210@57.79.23&[MIMAT0004703&hsa-miR-335-3p&offsets|-1|0;m-159&7|+|130496162|130496183&offsets|-1|0]|GTTTTTCATTATTGCTCCTGACC</td><td>>MI0000816|hsa-mir-335&WithFlank&7|+|130496105|130496210</td><td>57</td><td>79</td><td>23</td><td>GTTTTTCATTATTGCTCCTGACC</td><td>236.0</td><td>5.2</td><td>1.0</td><td>5.2</td><td>2.3E-07</td><td>3.8E-07</td><td>MB231/MB468</td><td>MITO</td></tr>
<tr><td>>hg38_21_+_8433172_8446622_RNA45SN1_45S_50ntFlanks@11313.11344.32|CCCCACCCCACGTCTCGTCGCGCGCGCGTCCG</td><td>>hg38_21_+_8433172_8446622_RNA45SN1_45S_50ntFlanks</td><td>11313</td><td>11344</td><td>32</td><td>CCCCACCCCACGTCTCGTCGCGCGCGCGTCCG</td><td>1188.9</td><td>-8.6</td><td>1.7</td><td>-5.2</td><td>2.4E-07</td><td>3.8E-07</td><td>BT20/MB468</td><td>NUC</td></tr>
<tr><td>>hg38_21_+_8433172_8446622_RNA45SN1_45S_50ntFlanks@12635.12652.18|AAGCTACCATCTGTGGGA</td><td>>hg38_21_+_8433172_8446622_RNA45SN1_45S_50ntFlanks</td><td>12635</td><td>12652</td><td>18</td><td>AAGCTACCATCTGTGGGA</td><td>1203.5</td><td>-8.4</td><td>1.5</td><td>-5.5</td><td>4.5E-08</td><td>3.8E-07</td><td>BT20/MB468</td><td>MP</td></tr>
<tr><td>>hg38_21_+_8433172_8446622_RNA45SN1_45S_50ntFlanks@12028.12058.31|CTCTGATCGTTTTTTCACTGACCCGGTGAGG</td><td>>hg38_21_+_8433172_8446622_RNA45SN1_45S_50ntFlanks</td><td>12028</td><td>12058</td><td>31</td><td>CTCTGATCGTTTTTTCACTGACCCGGTGAGG</td><td>91.9</td><td>10.3</td><td>1.9</td><td>5.5</td><td>4.5E-08</td><td>3.8E-07</td><td>BT20/MB468</td><td>MP</td></tr>
<tr><td>>MI0000477|hsa-mir-146a&WithFlank&5|+|160485346|160485456@27.45.19(+1A)&[MIMAT0000449&hsa-miR-146a-5p&offsets|0|-3(+1A);m-129&5|+|160485372|160485393&offsets|0|-3(+1A)]|TGAGAACTGAATTCCATGGA</td><td>>MI0000477|hsa-mir-146a&WithFlank&5|+|160485346|160485456</td><td>27</td><td>45</td><td>19(+1A)</td><td>TGAGAACTGAATTCCATGGA</td><td>75.0</td><td>9.5</td><td>1.8</td><td>5.2</td><td>1.9E-07</td><td>3.8E-07</td><td>MB231/MB468</td><td>MP</td></tr>
<tr><td>>hg38_21_+_8433172_8446622_RNA45SN1_45S_50ntFlanks@12893.12933.41|CCTCGCCCGTCACGCACCGCACGTTCGTGGGGAACCTGGCG</td><td>>hg38_21_+_8433172_8446622_RNA45SN1_45S_50ntFlanks</td><td>12893</td><td>12933</td><td>41</td><td>CCTCGCCCGTCACGCACCGCACGTTCGTGGGGAACCTGGCG</td><td>181.3</td><td>-5.5</td><td>1.1</td><td>-5.2</td><td>2.1E-07</td><td>3.8E-07</td><td>MB231/MB468</td><td>CYTO</td></tr>
<tr><td>>MI0003628|hsa-mir-615&WithFlank&12|+|54033944|54034051@67.86.20&[MIMAT0003283&hsa-miR-615-3p&offsets|0|-2;m-495&12|+|54034010|54034030&offsets|0|-1]|TCCGAGCCTGGGTCTCCCTC</td><td>>MI0003628|hsa-mir-615&WithFlank&12|+|54033944|54034051</td><td>67</td><td>86</td><td>20</td><td>TCCGAGCCTGGGTCTCCCTC</td><td>545.5</td><td>4.4</td><td>0.8</td><td>5.6</td><td>2.3E-08</td><td>3.8E-07</td><td>BT20/MB231</td><td>NUC</td></tr>
<tr><td>>hg38_21_+_8433172_8446622_RNA45SN1_45S_50ntFlanks@10782.10801.20|GGTGAACAGCCTCTGGCATG</td><td>>hg38_21_+_8433172_8446622_RNA45SN1_45S_50ntFlanks</td><td>10782</td><td>10801</td><td>20</td><td>GGTGAACAGCCTCTGGCATG</td><td>601.6</td><td>-3.8</td><td>0.7</td><td>-5.2</td><td>1.8E-07</td><td>3.8E-07</td><td>BT20/MB468</td><td>CYTO</td></tr>
<tr><td>>hg38_21_+_8433172_8446622_RNA45SN1_45S_50ntFlanks@9534.9576.43|AGGAAACTCTGGTGGAGGTCCGTAGCGGTCCTGACGTGCAAAT</td><td>>hg38_21_+_8433172_8446622_RNA45SN1_45S_50ntFlanks</td><td>9534</td><td>9576</td><td>43</td><td>AGGAAACTCTGGTGGAGGTCCGTAGCGGTCCTGACGTGCAAAT</td><td>360.4</td><td>4.0</td><td>0.8</td><td>5.2</td><td>2.1E-07</td><td>3.8E-07</td><td>MB231/MB468</td><td>CYTO</td></tr>
<tr><td>>MI0001445|hsa-mir-423&WithFlank&17|+|30117073|30117178@59.77.19(+1U)&[MIMAT0001340&hsa-miR-423-3p&offsets|0|-4(+1U);m-100&17|+|30117131|30117153&offsets|0|-4(+1U)]|AGCTCGGTCTGAGGCCCCTT</td><td>>MI0001445|hsa-mir-423&WithFlank&17|+|30117073|30117178</td><td>59</td><td>77</td><td>19(+1U)</td><td>AGCTCGGTCTGAGGCCCCTT</td><td>279.7</td><td>4.8</td><td>0.9</td><td>5.2</td><td>2.3E-07</td><td>3.8E-07</td><td>MB231/MB468</td><td>MITO</td></tr>
<tr><td>>hg38_21_+_8433172_8446622_RNA45SN1_45S_50ntFlanks@8198.8215.18|TGGACGGTGTGAGGCCGG</td><td>>hg38_21_+_8433172_8446622_RNA45SN1_45S_50ntFlanks</td><td>8198</td><td>8215</td><td>18</td><td>TGGACGGTGTGAGGCCGG</td><td>314.8</td><td>-6.7</td><td>1.3</td><td>-5.2</td><td>1.6E-07</td><td>3.8E-07</td><td>BT20/MB468</td><td>MITO</td></tr>
<tr><td>>hg38_21_+_8433172_8446622_RNA45SN1_45S_50ntFlanks@4110.4151.42|TGACGGGGAATCAGGGTTCGATTCCGGAGAGGGAGCCTGAGA</td><td>>hg38_21_+_8433172_8446622_RNA45SN1_45S_50ntFlanks</td><td>4110</td><td>4151</td><td>42</td><td>TGACGGGGAATCAGGGTTCGATTCCGGAGAGGGAGCCTGAGA</td><td>132.7</td><td>10.8</td><td>2.0</td><td>5.5</td><td>4.6E-08</td><td>3.8E-07</td><td>BT20/MB468</td><td>MP</td></tr>
<tr><td>>MI0000087|hsa-mir-29a&WithFlank&7|-|130876741|130876816@48.68.21(+1U)&[MIMAT0000086&hsa-miR-29a-3p&offsets|0|-1(+1U);m-26&7|-|130876749|130876769&offsets|0|0(+1U)]|TAGCACCATCTGAAATCGGTTT</td><td>>MI0000087|hsa-mir-29a&WithFlank&7|-|130876741|130876816</td><td>48</td><td>68</td><td>21(+1U)</td><td>TAGCACCATCTGAAATCGGTTT</td><td>1821.8</td><td>6.7</td><td>1.3</td><td>5.2</td><td>1.9E-07</td><td>3.8E-07</td><td>MB231/MB468</td><td>MP</td></tr>
<tr><td>>hg38_21_+_8433172_8446622_RNA45SN1_45S_50ntFlanks@10842.10865.24|GTAACTTCGGGATAAGGATTGGCT</td><td>>hg38_21_+_8433172_8446622_RNA45SN1_45S_50ntFlanks</td><td>10842</td><td>10865</td><td>24</td><td>GTAACTTCGGGATAAGGATTGGCT</td><td>630.6</td><td>-8.2</td><td>1.6</td><td>-5.2</td><td>2.4E-07</td><td>3.8E-07</td><td>BT20/MB468</td><td>NUC</td></tr>
<tr><td>>trnaMT_PheGAA_MT_+_577_647@35.59.25|GAAAATGTTTAGACGGGCTCACATC</td><td>>trnaMT_PheGAA_MT_+_577_647</td><td>35</td><td>59</td><td>25</td><td>GAAAATGTTTAGACGGGCTCACATC</td><td>279.5</td><td>4.1</td><td>0.8</td><td>5.2</td><td>2.3E-07</td><td>3.8E-07</td><td>MB231/MB468</td><td>MITO</td></tr>
<tr><td>>hg38_21_+_8433172_8446622_RNA45SN1_45S_50ntFlanks@10866.10903.38|CTAAGGGCTGGGTCGGTCGGGCTGGGGCGCGAAGCGGG</td><td>>hg38_21_+_8433172_8446622_RNA45SN1_45S_50ntFlanks</td><td>10866</td><td>10903</td><td>38</td><td>CTAAGGGCTGGGTCGGTCGGGCTGGGGCGCGAAGCGGG</td><td>477.6</td><td>-8.1</td><td>1.6</td><td>-5.2</td><td>2.4E-07</td><td>3.8E-07</td><td>BT20/MB468</td><td>NUC</td></tr>
<tr><td>>hg38_21_+_8433172_8446622_RNA45SN1_45S_50ntFlanks@9339.9378.40|CGCACGAAAGCCGCCGTGGCGCAATGAAGGTGAAGGCCGG</td><td>>hg38_21_+_8433172_8446622_RNA45SN1_45S_50ntFlanks</td><td>9339</td><td>9378</td><td>40</td><td>CGCACGAAAGCCGCCGTGGCGCAATGAAGGTGAAGGCCGG</td><td>105.8</td><td>-9.8</td><td>1.8</td><td>-5.5</td><td>4.6E-08</td><td>3.8E-07</td><td>BT20/MB468</td><td>MP</td></tr>
<tr><td>>hg38_21_+_8433172_8446622_RNA45SN1_45S_50ntFlanks@12002.12031.30|TGCGGGCCGCCGGTGAAATACCACTACTCT</td><td>>hg38_21_+_8433172_8446622_RNA45SN1_45S_50ntFlanks</td><td>12002</td><td>12031</td><td>30</td><td>TGCGGGCCGCCGGTGAAATACCACTACTCT</td><td>1967.9</td><td>-8.9</td><td>1.7</td><td>-5.2</td><td>2.4E-07</td><td>3.8E-07</td><td>BT20/MB468</td><td>NUC</td></tr>
<tr><td>>hg38_21_+_8433172_8446622_RNA45SN1_45S_50ntFlanks@9546.9573.28|TGGAGGTCCGTAGCGGTCCTGACGTGCA</td><td>>hg38_21_+_8433172_8446622_RNA45SN1_45S_50ntFlanks</td><td>9546</td><td>9573</td><td>28</td><td>TGGAGGTCCGTAGCGGTCCTGACGTGCA</td><td>590.2</td><td>3.3</td><td>0.6</td><td>5.2</td><td>2.1E-07</td><td>3.8E-07</td><td>MB231/MB468</td><td>CYTO</td></tr>
<tr><td>>hg38_21_+_8433172_8446622_RNA45SN1_45S_50ntFlanks@3866.3899.34|CTAGAGCTAATACATGCCGACGGGCGCTGACCCC</td><td>>hg38_21_+_8433172_8446622_RNA45SN1_45S_50ntFlanks</td><td>3866</td><td>3899</td><td>34</td><td>CTAGAGCTAATACATGCCGACGGGCGCTGACCCC</td><td>189.4</td><td>-4.1</td><td>0.8</td><td>-5.2</td><td>2.1E-07</td><td>3.8E-07</td><td>MB231/MB468</td><td>CYTO</td></tr>
<tr><td>>hg38_21_+_8433172_8446622_RNA45SN1_45S_50ntFlanks@11561.11591.31|GCGCCTAGCAGCCGACTTAGAACTGGTGCGG</td><td>>hg38_21_+_8433172_8446622_RNA45SN1_45S_50ntFlanks</td><td>11561</td><td>11591</td><td>31</td><td>GCGCCTAGCAGCCGACTTAGAACTGGTGCGG</td><td>281.0</td><td>-6.8</td><td>1.3</td><td>-5.2</td><td>2.2E-07</td><td>3.8E-07</td><td>MB231/MB468</td><td>NUC</td></tr>
<tr><td>>hg38_21_+_8433172_8446622_RNA45SN1_45S_50ntFlanks@9853.9899.47|CGCTGCGGGATGAACCGAACGCCGGGTTAAGGCGCCCGATGCCGACG</td><td>>hg38_21_+_8433172_8446622_RNA45SN1_45S_50ntFlanks</td><td>9853</td><td>9899</td><td>47</td><td>CGCTGCGGGATGAACCGAACGCCGGGTTAAGGCGCCCGATGCCGACG</td><td>231.5</td><td>-11.6</td><td>2.3</td><td>-5.2</td><td>2.3E-07</td><td>3.8E-07</td><td>MB231/MB468</td><td>MITO</td></tr>
<tr><td>>hg38_21_+_8433172_8446622_RNA45SN1_45S_50ntFlanks@9739.9759.21|AAACGATCTCAACCTATTCTC</td><td>>hg38_21_+_8433172_8446622_RNA45SN1_45S_50ntFlanks</td><td>9739</td><td>9759</td><td>21</td><td>AAACGATCTCAACCTATTCTC</td><td>2800.5</td><td>-4.1</td><td>0.8</td><td>-5.2</td><td>1.7E-07</td><td>3.8E-07</td><td>BT20/MB468</td><td>MITO</td></tr>
<tr><td>>hg38_21_+_8433172_8446622_RNA45SN1_45S_50ntFlanks@9534.9557.24|AGGAAACTCTGGTGGAGGTCCGTA</td><td>>hg38_21_+_8433172_8446622_RNA45SN1_45S_50ntFlanks</td><td>9534</td><td>9557</td><td>24</td><td>AGGAAACTCTGGTGGAGGTCCGTA</td><td>195.6</td><td>-10.7</td><td>2.0</td><td>-5.5</td><td>4.6E-08</td><td>3.8E-07</td><td>BT20/MB468</td><td>MP</td></tr>
<tr><td>>hg38_21_+_8433172_8446622_RNA45SN1_45S_50ntFlanks@11576.11594.19|CTTAGAACTGGTGCGGACC</td><td>>hg38_21_+_8433172_8446622_RNA45SN1_45S_50ntFlanks</td><td>11576</td><td>11594</td><td>19</td><td>CTTAGAACTGGTGCGGACC</td><td>704.1</td><td>-6.0</td><td>1.1</td><td>-5.2</td><td>1.9E-07</td><td>3.8E-07</td><td>MB231/MB468</td><td>MP</td></tr>
<tr><td>>trna5_AspGTC_12_+_98897281_98897352@35.75.41|TCACGCGGGAGACCGGGGTTCAATTCCCCGACGGGGAGCCA</td><td>>trna5_AspGTC_12_+_98897281_98897352</td><td>35</td><td>75</td><td>41</td><td>TCACGCGGGAGACCGGGGTTCAATTCCCCGACGGGGAGCCA</td><td>10811.7</td><td>13.6</td><td>2.6</td><td>5.2</td><td>2.1E-07</td><td>3.8E-07</td><td>MB231/MB468</td><td>CYTO</td></tr>
<tr><td>>hg38_21_+_8433172_8446622_RNA45SN1_45S_50ntFlanks@12626.12667.42|AATGGGGCGAAGCTACCATCTGTGGGATTATGACTGAACGCC</td><td>>hg38_21_+_8433172_8446622_RNA45SN1_45S_50ntFlanks</td><td>12626</td><td>12667</td><td>42</td><td>AATGGGGCGAAGCTACCATCTGTGGGATTATGACTGAACGCC</td><td>648.7</td><td>4.7</td><td>0.9</td><td>5.2</td><td>1.7E-07</td><td>3.9E-07</td><td>BT20/MB468</td><td>MITO</td></tr>
<tr><td>>trna16_ValTAC_11_-_59318460_59318532@35.76.42,trna17_ValTAC_11_-_59318102_59318174@35.76.42,trna4_ValTAC_X_-_18693029_18693101@35.76.42|ACACGCAGAAGGTCCTGGGTTCGAGCCCCAGTGGAACCACCA</td><td>>trna16_ValTAC_11_-_59318460_59318532</td><td>35</td><td>76</td><td>42</td><td>ACACGCAGAAGGTCCTGGGTTCGAGCCCCAGTGGAACCACCA</td><td>1140.3</td><td>14.2</td><td>2.7</td><td>5.2</td><td>2.1E-07</td><td>3.9E-07</td><td>MB231/MB468</td><td>CYTO</td></tr>
<tr><td>>hg38_21_+_8433172_8446622_RNA45SN1_45S_50ntFlanks@3705.3726.22|TACCTGGTTGATCCTGCCAGTA</td><td>>hg38_21_+_8433172_8446622_RNA45SN1_45S_50ntFlanks</td><td>3705</td><td>3726</td><td>22</td><td>TACCTGGTTGATCCTGCCAGTA</td><td>802.9</td><td>4.6</td><td>0.8</td><td>5.7</td><td>1.4E-08</td><td>3.9E-07</td><td>BT20/MB231</td><td>CYTO</td></tr>
<tr><td>>hg38_21_+_8433172_8446622_RNA45SN1_45S_50ntFlanks@11873.11900.28|AAGAAGACCCTGTTGAGCTTGACTCTAG</td><td>>hg38_21_+_8433172_8446622_RNA45SN1_45S_50ntFlanks</td><td>11873</td><td>11900</td><td>28</td><td>AAGAAGACCCTGTTGAGCTTGACTCTAG</td><td>899.9</td><td>-5.6</td><td>1.1</td><td>-5.2</td><td>1.9E-07</td><td>3.9E-07</td><td>BT20/MB468</td><td>CYTO</td></tr>
<tr><td>>MI0000273|hsa-mir-183&WithFlank&7|-|129774899|129775020@34.54.21&[MIMAT0000261&hsa-miR-183-5p&offsets|+1|0;m-41&7|-|129774966|129774987&offsets|0|-1]|ATGGCACTGGTAGAATTCACT</td><td>>MI0000273|hsa-mir-183&WithFlank&7|-|129774899|129775020</td><td>34</td><td>54</td><td>21</td><td>ATGGCACTGGTAGAATTCACT</td><td>3492.5</td><td>2.9</td><td>0.5</td><td>5.6</td><td>2.4E-08</td><td>3.9E-07</td><td>BT20/MB231</td><td>NUC</td></tr>
<tr><td>>hg38_21_+_8433172_8446622_RNA45SN1_45S_50ntFlanks@9204.9225.22|TCGGGGCCACGCGCGCGTCCCC</td><td>>hg38_21_+_8433172_8446622_RNA45SN1_45S_50ntFlanks</td><td>9204</td><td>9225</td><td>22</td><td>TCGGGGCCACGCGCGCGTCCCC</td><td>143.5</td><td>-10.2</td><td>1.9</td><td>-5.5</td><td>4.7E-08</td><td>3.9E-07</td><td>BT20/MB468</td><td>MP</td></tr>
<tr><td>>hg38_21_+_8433172_8446622_RNA45SN1_45S_50ntFlanks@12628.12647.20|TGGGGCGAAGCTACCATCTG</td><td>>hg38_21_+_8433172_8446622_RNA45SN1_45S_50ntFlanks</td><td>12628</td><td>12647</td><td>20</td><td>TGGGGCGAAGCTACCATCTG</td><td>86.6</td><td>-9.5</td><td>1.7</td><td>-5.5</td><td>4.7E-08</td><td>3.9E-07</td><td>BT20/MB468</td><td>MP</td></tr>
<tr><td>>hg38_21_+_8433172_8446622_RNA45SN1_45S_50ntFlanks@12992.13014.23|TCCCTCGCTGCGATCTATTGAAA</td><td>>hg38_21_+_8433172_8446622_RNA45SN1_45S_50ntFlanks</td><td>12992</td><td>13014</td><td>23</td><td>TCCCTCGCTGCGATCTATTGAAA</td><td>325.9</td><td>-5.5</td><td>1.1</td><td>-5.2</td><td>2.3E-07</td><td>3.9E-07</td><td>MB231/MB468</td><td>MITO</td></tr>
<tr><td>>hg38_21_+_8433172_8446622_RNA45SN1_45S_50ntFlanks@11575.11606.32|ACTTAGAACTGGTGCGGACCAGGGGAATCCGA</td><td>>hg38_21_+_8433172_8446622_RNA45SN1_45S_50ntFlanks</td><td>11575</td><td>11606</td><td>32</td><td>ACTTAGAACTGGTGCGGACCAGGGGAATCCGA</td><td>245.8</td><td>-7.1</td><td>1.4</td><td>-5.2</td><td>2.3E-07</td><td>3.9E-07</td><td>MB231/MB468</td><td>MITO</td></tr>
<tr><td>>MI0000487|hsa-mir-193a&WithFlank&17|+|31559990|31560089@27.46.20(+1U)&[MIMAT0004614&hsa-miR-193a-5p&offsets|0|-2(+1U);m-123&17|+|31560016|31560037&offsets|0|-2(+1U)]|TGGGTCTTTGCGGGCGAGATT</td><td>>MI0000487|hsa-mir-193a&WithFlank&17|+|31559990|31560089</td><td>27</td><td>46</td><td>20(+1U)</td><td>TGGGTCTTTGCGGGCGAGATT</td><td>144.8</td><td>10.9</td><td>2.0</td><td>5.5</td><td>4.7E-08</td><td>3.9E-07</td><td>BT20/MB468</td><td>MP</td></tr>
<tr><td>>hg38_21_+_8433172_8446622_RNA45SN1_45S_50ntFlanks@11189.11207.19|ACCCGGGGGGCCGGCGGCG</td><td>>hg38_21_+_8433172_8446622_RNA45SN1_45S_50ntFlanks</td><td>11189</td><td>11207</td><td>19</td><td>ACCCGGGGGGCCGGCGGCG</td><td>92.0</td><td>-9.6</td><td>1.8</td><td>-5.5</td><td>4.7E-08</td><td>3.9E-07</td><td>BT20/MB468</td><td>MP</td></tr>
<tr><td>>hg38_21_+_8433172_8446622_RNA45SN1_45S_50ntFlanks@9760.9786.27|AAACTTTAAATGGGTAAGAAGCCCGGC</td><td>>hg38_21_+_8433172_8446622_RNA45SN1_45S_50ntFlanks</td><td>9760</td><td>9786</td><td>27</td><td>AAACTTTAAATGGGTAAGAAGCCCGGC</td><td>268.1</td><td>-11.1</td><td>2.0</td><td>-5.5</td><td>4.7E-08</td><td>3.9E-07</td><td>BT20/MB468</td><td>MP</td></tr>
<tr><td>>am_trnaMT_ValTAC_MT_+_1602_1670@51.69.19|CTTAACTTGACCGCTCTGA</td><td>>am_trnaMT_ValTAC_MT_+_1602_1670</td><td>51</td><td>69</td><td>19</td><td>CTTAACTTGACCGCTCTGA</td><td>123.0</td><td>-10.0</td><td>1.8</td><td>-5.5</td><td>4.7E-08</td><td>3.9E-07</td><td>BT20/MB468</td><td>MP</td></tr>
<tr><td>>hg38_21_+_8433172_8446622_RNA45SN1_45S_50ntFlanks@12637.12654.18|GCTACCATCTGTGGGATT</td><td>>hg38_21_+_8433172_8446622_RNA45SN1_45S_50ntFlanks</td><td>12637</td><td>12654</td><td>18</td><td>GCTACCATCTGTGGGATT</td><td>983.0</td><td>-4.4</td><td>0.8</td><td>-5.2</td><td>1.7E-07</td><td>3.9E-07</td><td>BT20/MB468</td><td>MITO</td></tr>
<tr><td>>MI0003513|hsa-mir-455&WithFlank&9|+|114209428|114209535@60.81.22&[MIMAT0004784&hsa-miR-455-3p&offsets|0|+1;m-135&9|+|114209487|114209508&offsets|0|0]|GCAGTCCATGGGCATATACACT</td><td>>MI0003513|hsa-mir-455&WithFlank&9|+|114209428|114209535</td><td>60</td><td>81</td><td>22</td><td>GCAGTCCATGGGCATATACACT</td><td>400.5</td><td>-4.2</td><td>0.7</td><td>-5.6</td><td>1.8E-08</td><td>3.9E-07</td><td>BT20/MB231</td><td>MITO</td></tr>
</table>

| | | | | | | | | | | | | | |
|---|---|---|---|---|---|---|---|---|---|---|---|---|---|
| >MI0000542\|hsa-mir-320a&WithFlank&8\|-\|22244960\|22245043@48.71.24(+1U)&[MIMAT0000510&hsa-miR-320a-3p&offsets\|0\|+2(+1U);m-60&8\|-\|22244975\|22244996&offsets\|0\|+2(+1U)]\|AAAAGCTGGGTTGAGAGGGCGAAAT | >MI0000542\|hsa-mir-320a&WithFlank&8\|-\|22244960\|22245043 | 48 | 71 | 24(+1U) | AAAAGCTGGGTTGAGAGGGCGAAAT | 1109.0 | 7.1 | 1.3 | 5.5 | 4.7E-08 | 3.9E-07 | BT20/MB468 | MP |
| >hg38_21_+_8433172_8446622_RNA45SN1_45S_50ntFlanks@9421.9453.33\|CGCCGAGGGCGCACCACCGGCCCGTCTCGCCCG | >hg38_21_+_8433172_8446622_RNA45SN1_45S_50ntFlanks | 9421 | 9453 | 33 | CGCCGAGGGCGCACCACCGGCCCGTCTCGCCCG | 80.4 | -9.4 | 1.7 | -5.5 | 4.7E-08 | 3.9E-07 | BT20/MB468 | MP |
| >hg38_21_+_8433172_8446622_RNA45SN1_45S_50ntFlanks@12926.12956.31\|ACCTGGCGCTAAACCATTCGTAGACGACCTG | >hg38_21_+_8433172_8446622_RNA45SN1_45S_50ntFlanks | 12926 | 12956 | 31 | ACCTGGCGCTAAACCATTCGTAGACGACCTG | 118.1 | -4.4 | 0.8 | -5.2 | 2.2E-07 | 3.9E-07 | MB231/MB468 | CYTO |
| >hg38_21_+_8433172_8446622_RNA45SN1_45S_50ntFlanks@10731.10755.25\|AGGGTGTAAATCTCGCGCCGGGCCG | >hg38_21_+_8433172_8446622_RNA45SN1_45S_50ntFlanks | 10731 | 10755 | 25 | AGGGTGTAAATCTCGCGCCGGGCCG | 234.2 | -6.3 | 1.2 | -5.2 | 1.7E-07 | 3.9E-07 | BT20/MB468 | MITO |
| >hg38_21_+_8433172_8446622_RNA45SN1_45S_50ntFlanks@12007.12031.25\|GCCGCCGGTGAAATACCACTACTCT | >hg38_21_+_8433172_8446622_RNA45SN1_45S_50ntFlanks | 12007 | 12031 | 25 | GCCGCCGGTGAAATACCACTACTCT | 2683.8 | -5.3 | 1.0 | -5.2 | 2.5E-07 | 3.9E-07 | BT20/MB468 | NUC |
| >am_trnaMT_ValTAC_MT_+_1602_1670@50.67.18\|ACTTAACTTGACCGCTCT | >am_trnaMT_ValTAC_MT_+_1602_1670 | 50 | 67 | 18 | ACTTAACTTGACCGCTCT | 3028.2 | 5.5 | 1.1 | 5.2 | 1.7E-07 | 3.9E-07 | BT20/MB468 | MITO |
| >hg38_21_+_8433172_8446622_RNA45SN1_45S_50ntFlanks@8170.8197.28\|CAAGTCCTTCTGATCGAGGCCCAGCCCG | >hg38_21_+_8433172_8446622_RNA45SN1_45S_50ntFlanks | 8170 | 8197 | 28 | CAAGTCCTTCTGATCGAGGCCCAGCCCG | 960.8 | -5.5 | 1.1 | -5.2 | 2.2E-07 | 3.9E-07 | MB231/MB468 | CYTO |
| >hg38_21_+_8433172_8446622_RNA45SN1_45S_50ntFlanks@12678.12703.26\|AATCCCGCCCAGGCGGAACGATACGG | >hg38_21_+_8433172_8446622_RNA45SN1_45S_50ntFlanks | 12678 | 12703 | 26 | AATCCCGCCCAGGCGGAACGATACGG | 291.0 | -6.7 | 1.3 | -5.2 | 1.9E-07 | 3.9E-07 | BT20/MB468 | CYTO |
| >hg38_21_+_8433172_8446622_RNA45SN1_45S_50ntFlanks@6735.6770.36\|TTGATCATCGACACTTCGAACGCACTTGCGGCCCCG | >hg38_21_+_8433172_8446622_RNA45SN1_45S_50ntFlanks | 6735 | 6770 | 36 | TTGATCATCGACACTTCGAACGCACTTGCGGCCCCG | 189.2 | -5.9 | 1.1 | -5.2 | 1.9E-07 | 3.9E-07 | BT20/MB468 | CYTO |
| >MI0000342\|hsa-mir-200b&WithFlank&1\|+\|1167098\|1167204@63.85.23(+1A)&[MIMAT0000318&hsa-miR-200b-3p&offsets\|0\|+1(+1A);m-93&1\|+\|1167160\|1167182&offsets\|0\|0(+1A)]\|TAATACTGCCTGGTAATGATGACA | >MI0000342\|hsa-mir-200b&WithFlank&1\|+\|1167098\|1167204 | 63 | 85 | 23(+1A) | TAATACTGCCTGGTAATGATGACA | 502.5 | -4.3 | 0.8 | -5.2 | 2.2E-07 | 3.9E-07 | MB231/MB468 | CYTO |
| >trna116_GluCTC_1_-_145399233_145399304@3.33.31,trna59_GluCTC_1_+_249168447_249168518@3.33.31,trna71_GluCTC_1_-_161439189_161439260@3.33.31,trna74_GluCTC_1_-_161431809_161431880@3.33.31,trna77_GluCTC_1_-_161424398_161424469@3.33.31,trna77_GluCTC_6_+_28949976_28950047@3.33.31,trna80_GluCTC_1_-_161417018_161417089@3.33.31,trna87_GluCTC_6_-_126101393_126101464@3.33.31\|CCTGGTGGTCTAGTGGTTAGGATTCGGCGCT | >trna116_GluCTC_1_-_145399233_145399304 | 3 | 33 | 31 | CCTGGTGGTCTAGTGGTTAGGATTCGGCGCT | 598.0 | 7.0 | 1.3 | 5.2 | 2.2E-07 | 3.9E-07 | MB231/MB468 | CYTO |
| >hg38_21_+_8433172_8446622_RNA45SN1_45S_50ntFlanks@4738.4766.29\|AAAGTCGGAGGTTCGAAGACGATCAGATA | >hg38_21_+_8433172_8446622_RNA45SN1_45S_50ntFlanks | 4738 | 4766 | 29 | AAAGTCGGAGGTTCGAAGACGATCAGATA | 1170.4 | 6.5 | 1.3 | 5.2 | 1.9E-07 | 3.9E-07 | BT20/MB468 | CYTO |
| >hg38_21_+_8433172_8446622_RNA45SN1_45S_50ntFlanks@12624.12659.36\|CCAATGGGGCGAAGCTACCATCTGTGGGATTATGAC | >hg38_21_+_8433172_8446622_RNA45SN1_45S_50ntFlanks | 12624 | 12659 | 36 | CCAATGGGGCGAAGCTACCATCTGTGGGATTATGAC | 144.5 | -3.9 | 0.7 | -5.2 | 2.2E-07 | 3.9E-07 | MB231/MB468 | CYTO |
| >hg38_21_+_8433172_8446622_RNA45SN1_45S_50ntFlanks@8115.8134.20\|CGCGGGACATGTGGCGTACG | >hg38_21_+_8433172_8446622_RNA45SN1_45S_50ntFlanks | 8115 | 8134 | 20 | CGCGGGACATGTGGCGTACG | 349.6 | 11.8 | 2.1 | 5.7 | 1.0E-08 | 3.9E-07 | BT20/MB231 | MP |
| >hg38_21_+_8433172_8446622_RNA45SN1_45S_50ntFlanks@11209.11229.21\|CGGCGACTCTGGACGCGAGCC | >hg38_21_+_8433172_8446622_RNA45SN1_45S_50ntFlanks | 11209 | 11229 | 21 | CGGCGACTCTGGACGCGAGCC | 559.4 | -3.2 | 0.6 | -5.2 | 2.2E-07 | 3.9E-07 | MB231/MB468 | NUC |
| >hg38_21_+_8433172_8446622_RNA45SN1_45S_50ntFlanks@483.526.44\|TCGATGTGGTGACGTCGTGCTCTCCCGGGCCGGGTCCGAGCCGC | >hg38_21_+_8433172_8446622_RNA45SN1_45S_50ntFlanks | 483 | 526 | 44 | TCGATGTGGTGACGTCGTGCTCTCCCGGGCCGGGTCCGAGCCGC | 315.3 | 5.4 | 1.0 | 5.2 | 2.2E-07 | 3.9E-07 | MB231/MB468 | NUC |
| >hg38_21_+_8433172_8446622_RNA45SN1_45S_50ntFlanks@8379.8402.24\|TCAAGAGGGCGTGAAACCGTTAAG | >hg38_21_+_8433172_8446622_RNA45SN1_45S_50ntFlanks | 8379 | 8402 | 24 | TCAAGAGGGCGTGAAACCGTTAAG | 614.4 | -4.5 | 0.8 | -5.5 | 4.8E-08 | 3.9E-07 | BT20/MB468 | MP |
| >MI0000081\|hsa-mir-24-2&WithFlank&19\|-\|13836281\|13836365@56.77.22(+1U);MI0000080\|hsa-mir-24-1&WithFlank&9\|+\|95086015\|95086094@50.71.22(+1U)&[MIMAT0000080&hsa-miR-24-3p&offsets\|0\|0(+1U);m-36&19\|-\|13836288\|13836310&offsets\|0\|-1(+1U)];[MIMAT0000080_1&hsa-miR-24-3p&offsets\|0\|0(+1U);m-35&9\|+\|95086064\|95086086&offsets\|0\|-1(+1U)]\|TGGCTCAGTTCAGCAGGAACAGT | >MI0000081\|hsa-mir-24-2&WithFlank&19\|-\|13836281\|13836365 | 56 | 77 | 22(+1U) | TGGCTCAGTTCAGCAGGAACAGT | 2189.7 | 2.6 | 0.5 | 5.2 | 2.5E-07 | 4.0E-07 | BT20/MB468 | NUC |
| >hg38_21_+_8433172_8446622_RNA45SN1_45S_50ntFlanks@11570.11591.22\|AGCCGACTTAGAACTGGTGCGG | >hg38_21_+_8433172_8446622_RNA45SN1_45S_50ntFlanks | 11570 | 11591 | 22 | AGCCGACTTAGAACTGGTGCGG | 934.0 | -4.4 | 0.8 | -5.5 | 4.8E-08 | 4.0E-07 | BT20/MB468 | MP |
| >hg38_21_+_8433172_8446622_RNA45SN1_45S_50ntFlanks@12084.12106.23\|TCGCTTCTGGCGCCAAGCGCCCG | >hg38_21_+_8433172_8446622_RNA45SN1_45S_50ntFlanks | 12084 | 12106 | 23 | TCGCTTCTGGCGCCAAGCGCCCG | 75.9 | -9.3 | 1.7 | -5.5 | 4.8E-08 | 4.0E-07 | BT20/MB468 | MP |
| >hg38_21_+_8433172_8446622_RNA45SN1_45S_50ntFlanks@12415.12447.33\|CTTCGATGTCGGCTCTTCCTATCATTGTGAAGC | >hg38_21_+_8433172_8446622_RNA45SN1_45S_50ntFlanks | 12415 | 12447 | 33 | CTTCGATGTCGGCTCTTCCTATCATTGTGAAGC | 384.1 | 3.7 | 0.7 | 5.2 | 2.3E-07 | 4.0E-07 | MB231/MB468 | NUC |
| >hg38_21_+_8433172_8446622_RNA45SN1_45S_50ntFlanks@7971.8005.35\|GAGACGCGACCTCAGATCAGACGTGGCGACCCGCT | >hg38_21_+_8433172_8446622_RNA45SN1_45S_50ntFlanks | 7971 | 8005 | 35 | GAGACGCGACCTCAGATCAGACGTGGCGACCCGCT | 279.7 | 5.1 | 1.0 | 5.2 | 2.5E-07 | 4.0E-07 | BT20/MB468 | NUC |
| >am_trna128_GlyGCC_6_-_27870686_27870756@1.36.36,trna18_GlyGCC_16_+_70822597_70822667@1.36.36,trna19_GlyGCC_16_+_70823410_70823480@1.36.36,trna19_GlyGCC_2_-_157257659_157257729@1.36.36,trna24_GlyGCC_16_-_70812942_70813012@1.36.36,trna25_GlyGCC_16_-_70812114_70812184@1.36.36,trna5_GlyGCC_17_+_8029064_8029134@1.36.36,trna68_GlyGCC_1_-_161493637_161493707@1.36.36\|GCATTGGTGGTTCAGTGGTAGAATTCTCGCCTGCCA | >am_trna128_GlyGCC_6_-_27870686_27870756 | 1 | 36 | 36 | GCATTGGTGGTTCAGTGGTAGAATTCTCGCCTGCCA | 425.9 | 3.9 | 0.8 | 5.2 | 2.4E-07 | 4.0E-07 | MB231/MB468 | MITO |
| >hg38_21_+_8433172_8446622_RNA45SN1_45S_50ntFlanks@12996.13014.19\|TCGCTGCGATCTATTGAAA | >hg38_21_+_8433172_8446622_RNA45SN1_45S_50ntFlanks | 12996 | 13014 | 19 | TCGCTGCGATCTATTGAAA | 671.9 | -3.0 | 0.6 | -5.2 | 2.5E-07 | 4.0E-07 | BT20/MB468 | NUC |
| >MI0000102\|hsa-mir-100&WithFlank&11\|-\|122152223\|122152314@19.40.22(+2A)&[MIMAT0000098&hsa-miR-100-5p&offsets\|0\|0(+2A);m-27&11\|-\|122152276\|122152296&offsets\|0\|+1(+2A)]\|AACCCGTAGATCCGAACTTGTGAA | >MI0000102\|hsa-mir-100&WithFlank&11\|-\|122152223\|122152314 | 19 | 40 | 22(+2A) | AACCCGTAGATCCGAACTTGTGAA | 1550.6 | -7.9 | 1.4 | -5.7 | 1.0E-08 | 4.0E-07 | BT20/MB231 | MP |
| >hg38_21_+_8433172_8446622_RNA45SN1_45S_50ntFlanks@6777.6802.26\|TCCCGGGGCTACGCCTGTCTGAGCGT | >hg38_21_+_8433172_8446622_RNA45SN1_45S_50ntFlanks | 6777 | 6802 | 26 | TCCCGGGGCTACGCCTGTCTGAGCGT | 222.2 | 3.8 | 0.7 | 5.2 | 2.3E-07 | 4.0E-07 | MB231/MB468 | NUC |
| >hg38_21_+_8433172_8446622_RNA45SN1_45S_50ntFlanks@9501.9520.20\|ATGGTGAACTATGCCTGGGC | >hg38_21_+_8433172_8446622_RNA45SN1_45S_50ntFlanks | 9501 | 9520 | 20 | ATGGTGAACTATGCCTGGGC | 241.2 | -5.8 | 1.1 | -5.2 | 2.0E-07 | 4.0E-07 | MB231/MB468 | MP |
| >hg38_21_+_8433172_8446622_RNA45SN1_45S_50ntFlanks@485.525.41\|GATGTGGTGACGTCGTGCTCTCCCGGGCCGGGTCCGAGCCG | >hg38_21_+_8433172_8446622_RNA45SN1_45S_50ntFlanks | 485 | 525 | 41 | GATGTGGTGACGTCGTGCTCTCCCGGGCCGGGTCCGAGCCG | 100.5 | 10.4 | 1.9 | 5.5 | 4.8E-08 | 4.0E-07 | BT20/MB468 | MP |
| >hg38_21_+_8433172_8446622_RNA45SN1_45S_50ntFlanks@6734.6774.41\|ATTGATCATCGACACTTCGAACGCACTTGCGGCCCCGGGTT | >hg38_21_+_8433172_8446622_RNA45SN1_45S_50ntFlanks | 6734 | 6774 | 41 | ATTGATCATCGACACTTCGAACGCACTTGCGGCCCCGGGTT | 83.8 | 9.6 | 1.8 | 5.2 | 2.0E-07 | 4.0E-07 | MB231/MB468 | MP |
| >MI0000077\|hsa-mir-21&WithFlank&17\|+\|59841260\|59841343@14.32.19&[MIMAT0000076&hsa-miR-21-5p&offsets\|0\|-3;m-1&17\|+\|59841273\|59841294&offsets\|0\|-3]\|TAGCTTATCAGACTGATGT | >MI0000077\|hsa-mir-21&WithFlank&17\|+\|59841260\|59841343 | 14 | 32 | 19 | TAGCTTATCAGACTGATGT | 2981.2 | 4.2 | 0.8 | 5.2 | 2.0E-07 | 4.0E-07 | MB231/MB468 | MP |
| >hg38_21_+_8433172_8446622_RNA45SN1_45S_50ntFlanks@11572.11611.40\|CCGACTTAGAACTGGTGCGGACCAGGGGAATCCGACTGTT | >hg38_21_+_8433172_8446622_RNA45SN1_45S_50ntFlanks | 11572 | 11611 | 40 | CCGACTTAGAACTGGTGCGGACCAGGGGAATCCGACTGTT | 1354.2 | -4.2 | 0.8 | -5.2 | 2.3E-07 | 4.0E-07 | MB231/MB468 | NUC |
| >am_trnaMT_SerGCT_MT_+_12207_12265@16.38.23\|AACTGCTAACTCATGCCCCCATG | >am_trnaMT_SerGCT_MT_+_12207_12265 | 16 | 38 | 23 | AACTGCTAACTCATGCCCCCATG | 488.2 | -5.8 | 1.1 | -5.2 | 2.4E-07 | 4.0E-07 | MB231/MB468 | MITO |
| >hg38_21_+_8433172_8446622_RNA45SN1_45S_50ntFlanks@9939.9979.41\|GCAGGACGGTGGCCATGGAAGTCGGAATCCGCTAAGGAGTG | >hg38_21_+_8433172_8446622_RNA45SN1_45S_50ntFlanks | 9939 | 9979 | 41 | GCAGGACGGTGGCCATGGAAGTCGGAATCCGCTAAGGAGTG | 70.7 | 9.9 | 1.8 | 5.5 | 4.9E-08 | 4.0E-07 | BT20/MB468 | MP |
| >hg38_21_+_8433172_8446622_RNA45SN1_45S_50ntFlanks@9714.9732.19\|CGAATGATTAGAGGTCTTG | >hg38_21_+_8433172_8446622_RNA45SN1_45S_50ntFlanks | 9714 | 9732 | 19 | CGAATGATTAGAGGTCTTG | 82.1 | -9.4 | 1.7 | -5.5 | 4.9E-08 | 4.0E-07 | BT20/MB468 | MP |
| >hg38_21_+_8433172_8446622_RNA45SN1_45S_50ntFlanks@12996.13015.20\|TCGCTGCGATCTATTGAAAG | >hg38_21_+_8433172_8446622_RNA45SN1_45S_50ntFlanks | 12996 | 13015 | 20 | TCGCTGCGATCTATTGAAAG | 912.3 | 3.4 | 0.7 | 5.2 | 2.2E-07 | 4.0E-07 | MB231/MB468 | CYTO |
| >trnaMT_PheGAA_MT_+_577_647@33.59.27\|CTGAAAATGTTTAGACGGGCTCACATC | >trnaMT_PheGAA_MT_+_577_647 | 33 | 59 | 27 | CTGAAAATGTTTAGACGGGCTCACATC | 193.3 | 11.3 | 2.1 | 5.5 | 4.9E-08 | 4.0E-07 | BT20/MB468 | MP |
| >hg38_21_+_8433172_8446622_RNA45SN1_45S_50ntFlanks@8170.8191.22\|CAAGTCCTTCTGATCGAGGCCC | >hg38_21_+_8433172_8446622_RNA45SN1_45S_50ntFlanks | 8170 | 8191 | 22 | CAAGTCCTTCTGATCGAGGCCC | 83.9 | -9.5 | 1.7 | -5.5 | 4.9E-08 | 4.0E-07 | BT20/MB468 | MP |
| >hg38_21_+_8433172_8446622_RNA45SN1_45S_50ntFlanks@11161.11188.28\|CACGGTCCCCCGCGAGGGGGGCCCGGGC | >hg38_21_+_8433172_8446622_RNA45SN1_45S_50ntFlanks | 11161 | 11188 | 28 | CACGGTCCCCCGCGAGGGGGGCCCGGGC | 548.4 | -7.1 | 1.4 | -5.2 | 2.0E-07 | 4.0E-07 | BT20/MB468 | CYTO |
| >hg38_21_+_8433172_8446622_RNA45SN1_45S_50ntFlanks@3854.3872.19\|ACTGTGGTAATTCTAGAGC | >hg38_21_+_8433172_8446622_RNA45SN1_45S_50ntFlanks | 3854 | 3872 | 19 | ACTGTGGTAATTCTAGAGC | 427.2 | -7.8 | 1.5 | -5.2 | 2.5E-07 | 4.0E-07 | BT20/MB468 | NUC |
| >hg38_MT_+_1621_3279_RNR2_16S_50ntFlanks@51.73.23\|GCTAAACCTAGCCCCAAACCCAC | >hg38_MT_+_1621_3279_RNR2_16S_50ntFlanks | 51 | 73 | 23 | GCTAAACCTAGCCCCAAACCCAC | 97.0 | 2.1 | 0.4 | 5.2 | 2.5E-07 | 4.0E-07 | BT20/MB468 | NUC |
| >hg38_21_+_8433172_8446622_RNA45SN1_45S_50ntFlanks@10571.10598.28\|GAGAAGCCGGCGGGAGCCCCGGGGAGAG | >hg38_21_+_8433172_8446622_RNA45SN1_45S_50ntFlanks | 10571 | 10598 | 28 | GAGAAGCCGGCGGGAGCCCCGGGGAGAG | 87.5 | -9.5 | 1.7 | -5.5 | 4.9E-08 | 4.0E-07 | BT20/MB468 | MP |
| >hg38_21_+_8433172_8446622_RNA45SN1_45S_50ntFlanks@6652.6686.35\|GACTCTTAGCGGTGGATCACTCGGCTCGTGCGTCG | >hg38_21_+_8433172_8446622_RNA45SN1_45S_50ntFlanks | 6652 | 6686 | 35 | GACTCTTAGCGGTGGATCACTCGGCTCGTGCGTCG | 8289.8 | -3.3 | 0.6 | -5.2 | 2.4E-07 | 4.0E-07 | MB231/MB468 | MITO |
| >hg38_21_+_8433172_8446622_RNA45SN1_45S_50ntFlanks@3984.4008.25\|GCGGCTTTGGTGACTCTAGATAACC | >hg38_21_+_8433172_8446622_RNA45SN1_45S_50ntFlanks | 3984 | 4008 | 25 | GCGGCTTTGGTGACTCTAGATAACC | 63.4 | -9.9 | 1.9 | -5.2 | 2.0E-07 | 4.0E-07 | MB231/MB468 | MP |
| >hg38_MT_+_1621_3279_RNR2_16S_50ntFlanks@722.740.19\|TGCGTCAGATTAAAACACT | >hg38_MT_+_1621_3279_RNR2_16S_50ntFlanks | 722 | 740 | 19 | TGCGTCAGATTAAAACACT | 56.4 | 9.0 | 1.7 | 5.2 | 2.0E-07 | 4.0E-07 | MB231/MB468 | MP |
| >hg38_21_+_8433172_8446622_RNA45SN1_45S_50ntFlanks@10767.10785.19\|GCAGCAGGTCTCCAAGGTG | >hg38_21_+_8433172_8446622_RNA45SN1_45S_50ntFlanks | 10767 | 10785 | 19 | GCAGCAGGTCTCCAAGGTG | 283.2 | -6.9 | 1.3 | -5.2 | 2.4E-07 | 4.0E-07 | MB231/MB468 | MITO |
| >MI0000085\|hsa-mir-27a&WithFlank&19\|-\|13836434\|13836523@57.76.20(+3U)&[MIMAT0000084&hsa-miR-27a-3p&offsets\|0\|-1(+3U);m-43&19\|-\|13836447\|13836467&offsets\|0\|-1(+3U)]\|TTCACAGTGGCTAAGTTCCGTTT | >MI0000085\|hsa-mir-27a&WithFlank&19\|-\|13836434\|13836523 | 57 | 76 | 20(+3U) | TTCACAGTGGCTAAGTTCCGTTT | 1803.4 | -4.6 | 0.8 | -5.6 | 1.9E-08 | 4.0E-07 | BT20/MB231 | MITO |

<table>
<tr><td>>trna111_HisGTG_1_-_147774845_147774916@-1A.23.24,trna118_HisGTG_1_-_145396881_145396952@-1A.23.24,trna16_HisGTG_1_+_146544773_146544844@-1A.23.24,trna1_HisGTG_15_+_45493349_45493420@-1A.23.24,trna21_HisGTG_1_+_147753471_147753542@-1A.23.24,trna33_HisGTG_6_+_27125906_27125977@-1A.23.24,trna7_HisGTG_9_-_14433938_14434009@-1A.23.24,trna8_HisGTG_15_-_45492611_45492682@-1A.23.24,trna9_HisGTG_15_-_45490804_45490875@-1A.23.24|AGCCGTGATCGTATAGTGGTTAGT</td><td>>trna111_HisGTG_1_-_147774845_147774916</td><td>-1A</td><td>23</td><td>24</td><td>AGCCGTGATCGTATAGTGGTTAGT</td><td>691.3</td><td>-5.7</td><td>1.0</td><td>-5.6</td><td>1.9E-08</td><td>4.0E-07</td><td>BT20/MB231</td><td>MITO</td></tr>
<tr><td>>MI0000746|hsa-mir-99b&WithFlank&19|+|51692606|51692687@13.32.20&[MIMAT0000689&hsa-miR-99b-5p&offsets|0|-2;m-10&19|+|51692618|51692639&offsets|0|-2]|CACCCGTAGAACCGACCTTG</td><td>>MI0000746|hsa-mir-99b&WithFlank&19|+|51692606|51692687</td><td>13</td><td>32</td><td>20</td><td>CACCCGTAGAACCGACCTTG</td><td>555.8</td><td>3.2</td><td>0.6</td><td>5.2</td><td>2.6E-07</td><td>4.0E-07</td><td>BT20/MB468</td><td>NUC</td></tr>
<tr><td>>hg38_21_+_8433172_8446622_RNA45SN1_45S_50ntFlanks@12714.12753.40|GAGCCTCGGTTGGCCTCGGATAGCCGGTCCCCCGCCTGTC</td><td>>hg38_21_+_8433172_8446622_RNA45SN1_45S_50ntFlanks</td><td>12714</td><td>12753</td><td>40</td><td>GAGCCTCGGTTGGCCTCGGATAGCCGGTCCCCCGCCTGTC</td><td>67.2</td><td>-9.9</td><td>1.9</td><td>-5.2</td><td>2.0E-07</td><td>4.0E-07</td><td>MB231/MB468</td><td>MP</td></tr>
<tr><td>>hg38_21_+_8433172_8446622_RNA45SN1_45S_50ntFlanks@9481.9510.30|CACGTGTTAGGACCCGAAAGATGGTGAACT</td><td>>hg38_21_+_8433172_8446622_RNA45SN1_45S_50ntFlanks</td><td>9481</td><td>9510</td><td>30</td><td>CACGTGTTAGGACCCGAAAGATGGTGAACT</td><td>100.7</td><td>10.4</td><td>1.9</td><td>5.5</td><td>4.9E-08</td><td>4.0E-07</td><td>BT20/MB468</td><td>MP</td></tr>
<tr><td>>hg38_21_+_8433172_8446622_RNA45SN1_45S_50ntFlanks@10384.10405.22|CCATGTGAACAGCAGTTGAACA</td><td>>hg38_21_+_8433172_8446622_RNA45SN1_45S_50ntFlanks</td><td>10384</td><td>10405</td><td>22</td><td>CCATGTGAACAGCAGTTGAACA</td><td>512.5</td><td>-4.6</td><td>0.9</td><td>-5.2</td><td>2.3E-07</td><td>4.1E-07</td><td>MB231/MB468</td><td>NUC</td></tr>
<tr><td>>hg38_21_+_8433172_8446622_RNA45SN1_45S_50ntFlanks@12406.12434.29|TTTTTGATCCTTCGATGTCGGCTCTTCCT</td><td>>hg38_21_+_8433172_8446622_RNA45SN1_45S_50ntFlanks</td><td>12406</td><td>12434</td><td>29</td><td>TTTTTGATCCTTCGATGTCGGCTCTTCCT</td><td>439.8</td><td>-3.4</td><td>0.7</td><td>-5.2</td><td>2.4E-07</td><td>4.1E-07</td><td>MB231/MB468</td><td>MITO</td></tr>
<tr><td>>hg38_21_+_8433172_8446622_RNA45SN1_45S_50ntFlanks@3774.3791.18|GGCCGGTACAGTGAAACT</td><td>>hg38_21_+_8433172_8446622_RNA45SN1_45S_50ntFlanks</td><td>3774</td><td>3791</td><td>18</td><td>GGCCGGTACAGTGAAACT</td><td>187.8</td><td>3.4</td><td>0.6</td><td>5.2</td><td>2.3E-07</td><td>4.1E-07</td><td>MB231/MB468</td><td>CYTO</td></tr>
<tr><td>>hg38_21_+_8433172_8446622_RNA45SN1_45S_50ntFlanks@12419.12452.34|GATGTCGGCTCTTCCTATCATTGTGAAGCAGAAT</td><td>>hg38_21_+_8433172_8446622_RNA45SN1_45S_50ntFlanks</td><td>12419</td><td>12452</td><td>34</td><td>GATGTCGGCTCTTCCTATCATTGTGAAGCAGAAT</td><td>515.2</td><td>5.0</td><td>1.0</td><td>5.2</td><td>2.3E-07</td><td>4.1E-07</td><td>MB231/MB468</td><td>NUC</td></tr>
<tr><td>>hg38_21_+_8433172_8446622_RNA45SN1_45S_50ntFlanks@9104.9128.25|CCGGCGACGGGGGGGGTGCCGCGCG</td><td>>hg38_21_+_8433172_8446622_RNA45SN1_45S_50ntFlanks</td><td>9104</td><td>9128</td><td>25</td><td>CCGGCGACGGGGGGGGTGCCGCGCG</td><td>80.7</td><td>-9.4</td><td>1.7</td><td>-5.5</td><td>4.9E-08</td><td>4.1E-07</td><td>BT20/MB468</td><td>MP</td></tr>
<tr><td>>hg38_21_+_8433172_8446622_RNA45SN1_45S_50ntFlanks@3705.3744.40|TACCTGGTTGATCCTGCCAGTAGCATATGCTTGTCTCAAA</td><td>>hg38_21_+_8433172_8446622_RNA45SN1_45S_50ntFlanks</td><td>3705</td><td>3744</td><td>40</td><td>TACCTGGTTGATCCTGCCAGTAGCATATGCTTGTCTCAAA</td><td>503.9</td><td>4.0</td><td>0.8</td><td>5.2</td><td>1.8E-07</td><td>4.1E-07</td><td>BT20/MB468</td><td>MITO</td></tr>
<tr><td>>hg38_21_+_8433172_8446622_RNA45SN1_45S_50ntFlanks@11677.11702.26|CTCTGAATGTCAAAGTGAAGAAATTC</td><td>>hg38_21_+_8433172_8446622_RNA45SN1_45S_50ntFlanks</td><td>11677</td><td>11702</td><td>26</td><td>CTCTGAATGTCAAAGTGAAGAAATTC</td><td>610.2</td><td>-6.2</td><td>1.2</td><td>-5.2</td><td>2.4E-07</td><td>4.1E-07</td><td>MB231/MB468</td><td>MITO</td></tr>
<tr><td>>hg38_21_+_8433172_8446622_RNA45SN1_45S_50ntFlanks@12415.12457.43|CTTCGATGTCGGCTCTTCCTATCATTGTGAAGCAGAATTCACC</td><td>>hg38_21_+_8433172_8446622_RNA45SN1_45S_50ntFlanks</td><td>12415</td><td>12457</td><td>43</td><td>CTTCGATGTCGGCTCTTCCTATCATTGTGAAGCAGAATTCACC</td><td>323.5</td><td>6.7</td><td>1.3</td><td>5.2</td><td>2.3E-07</td><td>4.1E-07</td><td>MB231/MB468</td><td>CYTO</td></tr>
<tr><td>>MI0000746|hsa-mir-99b&WithFlank&19|+|51692606|51692687@51.73.23&[MIMAT0004678&hsa-miR-99b-3p&offsets|0|+1;m-151&19|+|51692656|51692678&offsets|0|0]|CAAGCTCGTGTCTGTGGGTCCGT</td><td>>MI0000746|hsa-mir-99b&WithFlank&19|+|51692606|51692687</td><td>51</td><td>73</td><td>23</td><td>CAAGCTCGTGTCTGTGGGTCCGT</td><td>522.3</td><td>4.6</td><td>0.8</td><td>5.5</td><td>5.0E-08</td><td>4.1E-07</td><td>BT20/MB468</td><td>MP</td></tr>
<tr><td>>hg38_21_+_8433172_8446622_RNA45SN1_45S_50ntFlanks@10794.10815.22|CTGGCATGTTGGAACAATGTAG</td><td>>hg38_21_+_8433172_8446622_RNA45SN1_45S_50ntFlanks</td><td>10794</td><td>10815</td><td>22</td><td>CTGGCATGTTGGAACAATGTAG</td><td>133.5</td><td>-5.0</td><td>1.0</td><td>-5.2</td><td>2.3E-07</td><td>4.1E-07</td><td>MB231/MB468</td><td>NUC</td></tr>
<tr><td>>hg38_21_+_8433172_8446622_RNA45SN1_45S_50ntFlanks@7984.8017.34|AGATCAGACGTGGCGACCCGCTGAATTTAAGCAT</td><td>>hg38_21_+_8433172_8446622_RNA45SN1_45S_50ntFlanks</td><td>7984</td><td>8017</td><td>34</td><td>AGATCAGACGTGGCGACCCGCTGAATTTAAGCAT</td><td>92.5</td><td>-9.6</td><td>1.8</td><td>-5.5</td><td>5.0E-08</td><td>4.1E-07</td><td>BT20/MB468</td><td>MP</td></tr>
<tr><td>>hg38_21_+_8433172_8446622_RNA45SN1_45S_50ntFlanks@5473.5506.34|CGGCCCTGGCGGAGCGCTGAGAAGACGGTCGAAC</td><td>>hg38_21_+_8433172_8446622_RNA45SN1_45S_50ntFlanks</td><td>5473</td><td>5506</td><td>34</td><td>CGGCCCTGGCGGAGCGCTGAGAAGACGGTCGAAC</td><td>206.2</td><td>4.5</td><td>0.9</td><td>5.2</td><td>2.3E-07</td><td>4.1E-07</td><td>MB231/MB468</td><td>CYTO</td></tr>
<tr><td>>hg38_21_+_8433172_8446622_RNA45SN1_45S_50ntFlanks@12439.12480.42|TTGTGAAGCAGAATTCACCAAGCGTTGGATTGTTCACCCACT</td><td>>hg38_21_+_8433172_8446622_RNA45SN1_45S_50ntFlanks</td><td>12439</td><td>12480</td><td>42</td><td>TTGTGAAGCAGAATTCACCAAGCGTTGGATTGTTCACCCACT</td><td>50.6</td><td>8.9</td><td>1.7</td><td>5.2</td><td>2.1E-07</td><td>4.1E-07</td><td>MB231/MB468</td><td>MP</td></tr>
<tr><td>>hg38_21_+_8433172_8446622_RNA45SN1_45S_50ntFlanks@8350.8375.26|AAAGTTGAAAAGAACTTTGAAGAGAG</td><td>>hg38_21_+_8433172_8446622_RNA45SN1_45S_50ntFlanks</td><td>8350</td><td>8375</td><td>26</td><td>AAAGTTGAAAAGAACTTTGAAGAGAG</td><td>484.8</td><td>6.1</td><td>1.1</td><td>5.6</td><td>1.9E-08</td><td>4.1E-07</td><td>BT20/MB231</td><td>MITO</td></tr>
<tr><td>>hg38_21_+_8433172_8446622_RNA45SN1_45S_50ntFlanks@13007.13025.19|TATTGAAAGTCAGCCCTCG</td><td>>hg38_21_+_8433172_8446622_RNA45SN1_45S_50ntFlanks</td><td>13007</td><td>13025</td><td>19</td><td>TATTGAAAGTCAGCCCTCG</td><td>1246.3</td><td>-4.2</td><td>0.8</td><td>-5.2</td><td>2.3E-07</td><td>4.1E-07</td><td>MB231/MB468</td><td>CYTO</td></tr>
<tr><td>>hg38_21_+_8433172_8446622_RNA45SN1_45S_50ntFlanks@12071.12106.36|CCCCGAGGGGCTCTCGCTTCTGGCGCCAAGCGCCCG</td><td>>hg38_21_+_8433172_8446622_RNA45SN1_45S_50ntFlanks</td><td>12071</td><td>12106</td><td>36</td><td>CCCCGAGGGGCTCTCGCTTCTGGCGCCAAGCGCCCG</td><td>1390.2</td><td>-5.9</td><td>1.1</td><td>-5.2</td><td>2.5E-07</td><td>4.1E-07</td><td>MB231/MB468</td><td>MITO</td></tr>
<tr><td>>hg38_21_+_8433172_8446622_RNA45SN1_45S_50ntFlanks@12938.12961.24|ACCATTCGTAGACGACCTGCTTCT</td><td>>hg38_21_+_8433172_8446622_RNA45SN1_45S_50ntFlanks</td><td>12938</td><td>12961</td><td>24</td><td>ACCATTCGTAGACGACCTGCTTCT</td><td>522.5</td><td>-7.9</td><td>1.5</td><td>-5.2</td><td>2.6E-07</td><td>4.1E-07</td><td>BT20/MB468</td><td>NUC</td></tr>
<tr><td>>MI0000301|hsa-mir-224&WithFlank&X|-|151958572|151958664@14.34.21(+1C)&[MIMAT0000281&hsa-miR-224-5p&offsets|+1|-3(+1C);m-238&X|-|151958630|151958651&offsets|0|-1(+1C)]|CAAGTCACTAGTGGTTCCGTTC</td><td>>MI0000301|hsa-mir-224&WithFlank&X|-|151958572|151958664</td><td>14</td><td>34</td><td>21(+1C)</td><td>CAAGTCACTAGTGGTTCCGTTC</td><td>810.5</td><td>9.8</td><td>1.8</td><td>5.5</td><td>5.0E-08</td><td>4.1E-07</td><td>BT20/MB468</td><td>MP</td></tr>
<tr><td>>hg38_21_+_8433172_8446622_RNA45SN1_45S_50ntFlanks@12640.12659.20|ACCATCTGTGGGATTATGAC</td><td>>hg38_21_+_8433172_8446622_RNA45SN1_45S_50ntFlanks</td><td>12640</td><td>12659</td><td>20</td><td>ACCATCTGTGGGATTATGAC</td><td>450.7</td><td>-6.7</td><td>1.3</td><td>-5.2</td><td>2.6E-07</td><td>4.1E-07</td><td>BT20/MB468</td><td>NUC</td></tr>
<tr><td>>MI0000060|hsa-let-7a-1&WithFlank&9|+|94175951|94176042@12.35.24&[MIMAT0000062_2&hsa-let-7a-5p&offsets|0|+2;m-4&9|+|94175962|94175983&offsets|0|+2]|TGAGGTAGTAGGTTGTATAGTTTT</td><td>>MI0000060|hsa-let-7a-1&WithFlank&9|+|94175951|94176042</td><td>12</td><td>35</td><td>24</td><td>TGAGGTAGTAGGTTGTATAGTTTT</td><td>64.2</td><td>9.2</td><td>1.8</td><td>5.2</td><td>2.1E-07</td><td>4.1E-07</td><td>MB231/MB468</td><td>MP</td></tr>
<tr><td>>hg38_1_-_228634819_228635039_RNA5S12_5S_50ntFlanks@51.94.44|GTCTACGGCCATACCACCCTGAACGCGCCCGATCTCGTCTGATC</td><td>>hg38_1_-_228634819_228635039_RNA5S12_5S_50ntFlanks</td><td>51</td><td>94</td><td>44</td><td>GTCTACGGCCATACCACCCTGAACGCGCCCGATCTCGTCTGATC</td><td>11119.2</td><td>11.3</td><td>2.2</td><td>5.2</td><td>2.0E-07</td><td>4.1E-07</td><td>BT20/MB468</td><td>CYTO</td></tr>
<tr><td>>MI0000066|hsa-let-7e&WithFlank&19|+|51692780|51692870@14.35.22(+1A)&[MIMAT0000066&hsa-let-7e-5p&offsets|0|0(+1A);m-50&19|+|51692793|51692814&offsets|0|0(+1A)]|TGAGGTAGGAGGTTGTATAGTTA</td><td>>MI0000066|hsa-let-7e&WithFlank&19|+|51692780|51692870</td><td>14</td><td>35</td><td>22(+1A)</td><td>TGAGGTAGGAGGTTGTATAGTTA</td><td>335.6</td><td>6.1</td><td>1.2</td><td>5.2</td><td>2.3E-07</td><td>4.1E-07</td><td>MB231/MB468</td><td>CYTO</td></tr>
<tr><td>>hg38_21_+_8433172_8446622_RNA45SN1_45S_50ntFlanks@4735.4768.34|ACGAAAGTCGGAGGTTCGAAGACGATCAGATACC</td><td>>hg38_21_+_8433172_8446622_RNA45SN1_45S_50ntFlanks</td><td>4735</td><td>4768</td><td>34</td><td>ACGAAAGTCGGAGGTTCGAAGACGATCAGATACC</td><td>211.8</td><td>7.4</td><td>1.4</td><td>5.2</td><td>2.4E-07</td><td>4.1E-07</td><td>MB231/MB468</td><td>NUC</td></tr>
<tr><td>>hg38_21_+_8433172_8446622_RNA45SN1_45S_50ntFlanks@12456.12479.24|CCAAGCGTTGGATTGTTCACCCAC</td><td>>hg38_21_+_8433172_8446622_RNA45SN1_45S_50ntFlanks</td><td>12456</td><td>12479</td><td>24</td><td>CCAAGCGTTGGATTGTTCACCCAC</td><td>583.0</td><td>-5.1</td><td>1.0</td><td>-5.2</td><td>2.3E-07</td><td>4.1E-07</td><td>MB231/MB468</td><td>CYTO</td></tr>
<tr><td>>hg38_21_+_8433172_8446622_RNA45SN1_45S_50ntFlanks@12638.12661.24|CTACCATCTGTGGGATTATGACTG</td><td>>hg38_21_+_8433172_8446622_RNA45SN1_45S_50ntFlanks</td><td>12638</td><td>12661</td><td>24</td><td>CTACCATCTGTGGGATTATGACTG</td><td>525.0</td><td>-3.4</td><td>0.7</td><td>-5.2</td><td>2.4E-07</td><td>4.1E-07</td><td>MB231/MB468</td><td>NUC</td></tr>
<tr><td>>hg38_21_+_8433172_8446622_RNA45SN1_45S_50ntFlanks@4722.4748.27|TCATTAATCAAGAACGAAAGTCGGAGG</td><td>>hg38_21_+_8433172_8446622_RNA45SN1_45S_50ntFlanks</td><td>4722</td><td>4748</td><td>27</td><td>TCATTAATCAAGAACGAAAGTCGGAGG</td><td>384.4</td><td>-6.9</td><td>1.3</td><td>-5.2</td><td>2.4E-07</td><td>4.1E-07</td><td>MB231/MB468</td><td>NUC</td></tr>
<tr><td>>hg38_21_+_8433172_8446622_RNA45SN1_45S_50ntFlanks@11317.11344.28|ACCCCACGTCTCGTCGCGCGCGCGTCCG</td><td>>hg38_21_+_8433172_8446622_RNA45SN1_45S_50ntFlanks</td><td>11317</td><td>11344</td><td>28</td><td>ACCCCACGTCTCGTCGCGCGCGCGTCCG</td><td>6982.3</td><td>-7.8</td><td>1.5</td><td>-5.2</td><td>2.3E-07</td><td>4.1E-07</td><td>MB231/MB468</td><td>CYTO</td></tr>
<tr><td>>hg38_21_+_8433172_8446622_RNA45SN1_45S_50ntFlanks@4971.4999.29|CCGGCCCGGACACGGACAGGATTGACAGA</td><td>>hg38_21_+_8433172_8446622_RNA45SN1_45S_50ntFlanks</td><td>4971</td><td>4999</td><td>29</td><td>CCGGCCCGGACACGGACAGGATTGACAGA</td><td>1067.3</td><td>-6.5</td><td>1.3</td><td>-5.2</td><td>2.1E-07</td><td>4.1E-07</td><td>MB231/MB468</td><td>MP</td></tr>
<tr><td>>hg38_21_+_8433172_8446622_RNA45SN1_45S_50ntFlanks@9544.9574.31|GGTGGAGGTCCGTAGCGGTCCTGACGTGCAA</td><td>>hg38_21_+_8433172_8446622_RNA45SN1_45S_50ntFlanks</td><td>9544</td><td>9574</td><td>31</td><td>GGTGGAGGTCCGTAGCGGTCCTGACGTGCAA</td><td>616.1</td><td>-8.1</td><td>1.6</td><td>-5.1</td><td>2.6E-07</td><td>4.1E-07</td><td>BT20/MB468</td><td>NUC</td></tr>
<tr><td>>hg38_21_+_8433172_8446622_RNA45SN1_45S_50ntFlanks@12073.12094.22|CCGAGGGGCTCTCGCTTCTGGC</td><td>>hg38_21_+_8433172_8446622_RNA45SN1_45S_50ntFlanks</td><td>12073</td><td>12094</td><td>22</td><td>CCGAGGGGCTCTCGCTTCTGGC</td><td>717.7</td><td>-3.5</td><td>0.7</td><td>-5.2</td><td>2.0E-07</td><td>4.1E-07</td><td>BT20/MB468</td><td>CYTO</td></tr>
<tr><td>>hg38_21_+_8433172_8446622_RNA45SN1_45S_50ntFlanks@12991.13012.22|CTCCCTCGCTGCGATCTATTGA</td><td>>hg38_21_+_8433172_8446622_RNA45SN1_45S_50ntFlanks</td><td>12991</td><td>13012</td><td>22</td><td>CTCCCTCGCTGCGATCTATTGA</td><td>1101.7</td><td>-7.1</td><td>1.4</td><td>-5.2</td><td>1.8E-07</td><td>4.1E-07</td><td>BT20/MB468</td><td>MITO</td></tr>
<tr><td>>hg38_21_+_8433172_8446622_RNA45SN1_45S_50ntFlanks@6703.6720.18|GCTGCGAGAATTAATGTG</td><td>>hg38_21_+_8433172_8446622_RNA45SN1_45S_50ntFlanks</td><td>6703</td><td>6720</td><td>18</td><td>GCTGCGAGAATTAATGTG</td><td>777.0</td><td>-6.1</td><td>1.2</td><td>-5.2</td><td>2.1E-07</td><td>4.1E-07</td><td>MB231/MB468</td><td>MP</td></tr>
<tr><td>>MI0000289|hsa-mir-181a-1&WithFlank&1|-|198859038|198859159@70.91.22&[MIMAT0000270&hsa-miR-181a-3p&offsets|0|0;m-125&1|-|198859069|198859090&offsets|0|0]|ACCATCGACCGTTGATTGTACC</td><td>>MI0000289|hsa-mir-181a-1&WithFlank&1|-|198859038|198859159</td><td>70</td><td>91</td><td>22</td><td>ACCATCGACCGTTGATTGTACC</td><td>891.8</td><td>-8.8</td><td>1.5</td><td>-5.7</td><td>1.1E-08</td><td>4.1E-07</td><td>BT20/MB231</td><td>MP</td></tr>
<tr><td>>hg38_21_+_8433172_8446622_RNA45SN1_45S_50ntFlanks@5471.5493.23|CACGGCCCTGGCGGAGCGCTGAG</td><td>>hg38_21_+_8433172_8446622_RNA45SN1_45S_50ntFlanks</td><td>5471</td><td>5493</td><td>23</td><td>CACGGCCCTGGCGGAGCGCTGAG</td><td>200.4</td><td>-7.0</td><td>1.4</td><td>-5.2</td><td>2.1E-07</td><td>4.1E-07</td><td>MB231/MB468</td><td>MP</td></tr>
<tr><td>>hg38_21_+_8433172_8446622_RNA45SN1_45S_50ntFlanks@10391.10411.21|AACAGCAGTTGAACATGGGTC</td><td>>hg38_21_+_8433172_8446622_RNA45SN1_45S_50ntFlanks</td><td>10391</td><td>10411</td><td>21</td><td>AACAGCAGTTGAACATGGGTC</td><td>519.6</td><td>-7.9</td><td>1.5</td><td>-5.1</td><td>2.6E-07</td><td>4.1E-07</td><td>BT20/MB468</td><td>NUC</td></tr>
<tr><td>>hg38_21_+_8433172_8446622_RNA45SN1_45S_50ntFlanks@6729.6771.43|GACACATTGATCATCGACACTTCGAACGCACTTGCGGCCCCGG</td><td>>hg38_21_+_8433172_8446622_RNA45SN1_45S_50ntFlanks</td><td>6729</td><td>6771</td><td>43</td><td>GACACATTGATCATCGACACTTCGAACGCACTTGCGGCCCCGG</td><td>937.7</td><td>-8.9</td><td>1.7</td><td>-5.1</td><td>2.6E-07</td><td>4.1E-07</td><td>BT20/MB468</td><td>NUC</td></tr>
<tr><td>>hg38_21_+_8433172_8446622_RNA45SN1_45S_50ntFlanks@4265.4282.18|ATGAGTCCACTTTAAATC</td><td>>hg38_21_+_8433172_8446622_RNA45SN1_45S_50ntFlanks</td><td>4265</td><td>4282</td><td>18</td><td>ATGAGTCCACTTTAAATC</td><td>65.5</td><td>-9.1</td><td>1.7</td><td>-5.4</td><td>5.1E-08</td><td>4.2E-07</td><td>BT20/MB468</td><td>MP</td></tr>
<tr><td>>hg38_21_+_8433172_8446622_RNA45SN1_45S_50ntFlanks@4444.4462.19|CCTTGCCTCTCGGCGCCCC</td><td>>hg38_21_+_8433172_8446622_RNA45SN1_45S_50ntFlanks</td><td>4444</td><td>4462</td><td>19</td><td>CCTTGCCTCTCGGCGCCCC</td><td>92.1</td><td>-9.6</td><td>1.8</td><td>-5.4</td><td>5.1E-08</td><td>4.2E-07</td><td>BT20/MB468</td><td>MP</td></tr>
<tr><td>>hg38_21_+_8433172_8446622_RNA45SN1_45S_50ntFlanks@6733.6758.26|CATTGATCATCGACACTTCGAACGCA</td><td>>hg38_21_+_8433172_8446622_RNA45SN1_45S_50ntFlanks</td><td>6733</td><td>6758</td><td>26</td><td>CATTGATCATCGACACTTCGAACGCA</td><td>803.6</td><td>-4.0</td><td>0.8</td><td>-5.2</td><td>2.5E-07</td><td>4.2E-07</td><td>MB231/MB468</td><td>MITO</td></tr>
<tr><td>>hg38_21_+_8433172_8446622_RNA45SN1_45S_50ntFlanks@8359.8395.37|AAGAACTTTGAAGAGAGAGTTCAAGAGGGCGTGAAAC</td><td>>hg38_21_+_8433172_8446622_RNA45SN1_45S_50ntFlanks</td><td>8359</td><td>8395</td><td>37</td><td>AAGAACTTTGAAGAGAGAGTTCAAGAGGGCGTGAAAC</td><td>73.0</td><td>9.9</td><td>1.8</td><td>5.4</td><td>5.1E-08</td><td>4.2E-07</td><td>BT20/MB468</td><td>MP</td></tr>
<tr><td>>hg38_21_+_8433172_8446622_RNA45SN1_45S_50ntFlanks@3841.3873.33|TCCTACTTGGATAACTGTGGTAATTCTAGAGCT</td><td>>hg38_21_+_8433172_8446622_RNA45SN1_45S_50ntFlanks</td><td>3841</td><td>3873</td><td>33</td><td>TCCTACTTGGATAACTGTGGTAATTCTAGAGCT</td><td>631.0</td><td>4.7</td><td>0.9</td><td>5.2</td><td>2.0E-07</td><td>4.2E-07</td><td>BT20/MB468</td><td>CYTO</td></tr>
<tr><td>>hg38_21_+_8433172_8446622_RNA45SN1_45S_50ntFlanks@9938.9978.41|AGCAGGACGGTGGCCATGGAAGTCGGAATCCGCTAAGGAGT</td><td>>hg38_21_+_8433172_8446622_RNA45SN1_45S_50ntFlanks</td><td>9938</td><td>9978</td><td>41</td><td>AGCAGGACGGTGGCCATGGAAGTCGGAATCCGCTAAGGAGT</td><td>540.7</td><td>9.5</td><td>1.7</td><td>5.4</td><td>5.1E-08</td><td>4.2E-07</td><td>BT20/MB468</td><td>MP</td></tr>
<tr><td>>hg38_21_+_8433172_8446622_RNA45SN1_45S_50ntFlanks@10350.10390.41|ACGAGAACTTTGAAGGCCGAAGTGGAGAAGGGTTCCATGTG</td><td>>hg38_21_+_8433172_8446622_RNA45SN1_45S_50ntFlanks</td><td>10350</td><td>10390</td><td>41</td><td>ACGAGAACTTTGAAGGCCGAAGTGGAGAAGGGTTCCATGTG</td><td>2658.4</td><td>-4.1</td><td>0.8</td><td>-5.2</td><td>2.0E-07</td><td>4.2E-07</td><td>BT20/MB468</td><td>CYTO</td></tr>
<tr><td>>hg38_21_+_8433172_8446622_RNA45SN1_45S_50ntFlanks@12950.12988.39|CGACCTGCTTCTGGGTCGGGGTTTCGTACGTAGCAGAGC</td><td>>hg38_21_+_8433172_8446622_RNA45SN1_45S_50ntFlanks</td><td>12950</td><td>12988</td><td>39</td><td>CGACCTGCTTCTGGGTCGGGGTTTCGTACGTAGCAGAGC</td><td>565.5</td><td>-8.1</td><td>1.6</td><td>-5.1</td><td>2.6E-07</td><td>4.2E-07</td><td>BT20/MB468</td><td>NUC</td></tr>
</table>

<table>
<tr><td>>hg38_21_+_8433172_8446622_RNA45SN1_45S_50ntFlanks@475.516.42|CTTCGTGATCGATGTGGTGACGTCGTGCTCTCCCGGGCCGGG</td><td>>hg38_21_+_8433172_8446622_RNA45SN1_45S_50ntFlanks</td><td>475</td><td>516</td><td>42</td><td>CTTCGTGATCGATGTGGTGACGTCGTGCTCTCCCGGGCCGGG</td><td>70.2</td><td>9.4</td><td>1.8</td><td>5.2</td><td>2.1E-07</td><td>4.2E-07</td><td>MB231/MB468</td><td>MP</td></tr>
<tr><td>>hg38_21_+_8433172_8446622_RNA45SN1_45S_50ntFlanks@11641.11661.21|CGGCGGGTGTTGACGCGATGT</td><td>>hg38_21_+_8433172_8446622_RNA45SN1_45S_50ntFlanks</td><td>11641</td><td>11661</td><td>21</td><td>CGGCGGGTGTTGACGCGATGT</td><td>436.7</td><td>-6.7</td><td>1.3</td><td>-5.2</td><td>1.8E-07</td><td>4.2E-07</td><td>BT20/MB468</td><td>MITO</td></tr>
<tr><td>>hg38_21_+_8433172_8446622_RNA45SN1_45S_50ntFlanks@4243.4262.20|CTTTCGAGGCCCTGTAATTG</td><td>>hg38_21_+_8433172_8446622_RNA45SN1_45S_50ntFlanks</td><td>4243</td><td>4262</td><td>20</td><td>CTTTCGAGGCCCTGTAATTG</td><td>244.5</td><td>-5.4</td><td>1.0</td><td>-5.2</td><td>2.4E-07</td><td>4.2E-07</td><td>MB231/MB468</td><td>NUC</td></tr>
<tr><td>>hg38_21_+_8433172_8446622_RNA45SN1_45S_50ntFlanks@7988.8018.31|CAGACGTGGCGACCCGCTGAATTTAAGCATA</td><td>>hg38_21_+_8433172_8446622_RNA45SN1_45S_50ntFlanks</td><td>7988</td><td>8018</td><td>31</td><td>CAGACGTGGCGACCCGCTGAATTTAAGCATA</td><td>423.7</td><td>-7.8</td><td>1.4</td><td>-5.4</td><td>5.1E-08</td><td>4.2E-07</td><td>BT20/MB468</td><td>MP</td></tr>
<tr><td>>hg38_MT_+_1621_3279_RNR2_16S_50ntFlanks@729.749.21|GATTAAAACACTGAACTGACA</td><td>>hg38_MT_+_1621_3279_RNR2_16S_50ntFlanks</td><td>729</td><td>749</td><td>21</td><td>GATTAAAACACTGAACTGACA</td><td>55.8</td><td>9.0</td><td>1.7</td><td>5.2</td><td>2.1E-07</td><td>4.2E-07</td><td>MB231/MB468</td><td>MP</td></tr>
<tr><td>>MI0000446|hsa-mir-125b-1&WithFlank&11|-|122099751|122099850@21.41.21(+1C);MI0000470|hsa-mir-125b-2&WithFlank&21|+|16590231|16590331@23.43.21(+1C)&[MIMAT0000423&hsa-miR-125b-5p&offsets|0|-1(+1C);m-67&11|-|122099809|122099830&offsets|0|-1(+1C)];[MIMAT0000423_1&hsa-miR-125b-5p&offsets|0|-1(+1C);m-68&21|+|16590253|16590274&offsets|0|-1(+1C)]|TCCCTGAGACCCTAACTTGTGC</td><td>>MI0000446|hsa-mir-125b-1&WithFlank&11|-|122099751|122099850</td><td>21</td><td>41</td><td>21(+1C)</td><td>TCCCTGAGACCCTAACTTGTGC</td><td>70.3</td><td>9.4</td><td>1.8</td><td>5.2</td><td>2.1E-07</td><td>4.2E-07</td><td>MB231/MB468</td><td>MP</td></tr>
<tr><td>>hg38_21_+_8433172_8446622_RNA45SN1_45S_50ntFlanks@8169.8195.27|CCAAGTCCTTCTGATCGAGGCCCAGCC</td><td>>hg38_21_+_8433172_8446622_RNA45SN1_45S_50ntFlanks</td><td>8169</td><td>8195</td><td>27</td><td>CCAAGTCCTTCTGATCGAGGCCCAGCC</td><td>253.6</td><td>-5.5</td><td>1.1</td><td>-5.2</td><td>2.5E-07</td><td>4.2E-07</td><td>MB231/MB468</td><td>MITO</td></tr>
<tr><td>>MI0000095|hsa-mir-93&WithFlank&7|-|100093762|100093853@17.38.22(+1A)&[MIMAT0000093&hsa-miR-93-5p&offsets|0|-1(+1A);m-29&7|-|100093815|100093837&offsets|0|-1(+1A)]|CAAAGTGCTGTTCGTGCAGGTAA</td><td>>MI0000095|hsa-mir-93&WithFlank&7|-|100093762|100093853</td><td>17</td><td>38</td><td>22(+1A)</td><td>CAAAGTGCTGTTCGTGCAGGTAA</td><td>1042.9</td><td>5.4</td><td>1.0</td><td>5.2</td><td>2.5E-07</td><td>4.2E-07</td><td>MB231/MB468</td><td>MITO</td></tr>
<tr><td>>hg38_21_+_8433172_8446622_RNA45SN1_45S_50ntFlanks@3840.3882.43|CTCCTACTTGGATAACTGTGGTAATTCTAGAGCTAATACATGC</td><td>>hg38_21_+_8433172_8446622_RNA45SN1_45S_50ntFlanks</td><td>3840</td><td>3882</td><td>43</td><td>CTCCTACTTGGATAACTGTGGTAATTCTAGAGCTAATACATGC</td><td>242.9</td><td>5.3</td><td>1.0</td><td>5.2</td><td>2.4E-07</td><td>4.2E-07</td><td>MB231/MB468</td><td>CYTO</td></tr>
<tr><td>>MI0000289|hsa-mir-181a-1&WithFlank&1|-|198859038|198859159@70.90.21(+1U)&[MIMAT0000270&hsa-miR-181a-3p&offsets|0|-1(+1U);m-125&1|-|198859069|198859090&offsets|0|-1(+1U)]|ACCATCGACCGTTGATTGTACT</td><td>>MI0000289|hsa-mir-181a-1&WithFlank&1|-|198859038|198859159</td><td>70</td><td>90</td><td>21(+1U)</td><td>ACCATCGACCGTTGATTGTACT</td><td>462.3</td><td>8.5</td><td>1.7</td><td>5.2</td><td>2.5E-07</td><td>4.2E-07</td><td>MB231/MB468</td><td>MITO</td></tr>
<tr><td>>MI0000433|hsa-let-7g&WithFlank&3|-|52268272|52268367@11.32.22(+1A)&[MIMAT0000414&hsa-let-7g-5p&offsets|0|0(+1A);m-58&3|-|52268336|52268357&offsets|0|0(+1A)]|TGAGGTAGTAGTTTGTACAGTTA</td><td>>MI0000433|hsa-let-7g&WithFlank&3|-|52268272|52268367</td><td>11</td><td>32</td><td>22(+1A)</td><td>TGAGGTAGTAGTTTGTACAGTTA</td><td>2055.5</td><td>5.6</td><td>1.0</td><td>5.4</td><td>5.2E-08</td><td>4.2E-07</td><td>BT20/MB468</td><td>MP</td></tr>
<tr><td>>hg38_21_+_8433172_8446622_RNA45SN1_45S_50ntFlanks@12003.12032.30|GCGGGCCGCCGGTGAAATACCACTACTCTG</td><td>>hg38_21_+_8433172_8446622_RNA45SN1_45S_50ntFlanks</td><td>12003</td><td>12032</td><td>30</td><td>GCGGGCCGCCGGTGAAATACCACTACTCTG</td><td>4693.2</td><td>-7.2</td><td>1.3</td><td>-5.4</td><td>5.2E-08</td><td>4.2E-07</td><td>BT20/MB468</td><td>MP</td></tr>
<tr><td>>hg38_21_+_8433172_8446622_RNA45SN1_45S_50ntFlanks@12719.12742.24|TCGGTTGGCCTCGGATAGCCGGTC</td><td>>hg38_21_+_8433172_8446622_RNA45SN1_45S_50ntFlanks</td><td>12719</td><td>12742</td><td>24</td><td>TCGGTTGGCCTCGGATAGCCGGTC</td><td>97.7</td><td>-9.7</td><td>1.8</td><td>-5.4</td><td>5.2E-08</td><td>4.2E-07</td><td>BT20/MB468</td><td>MP</td></tr>
<tr><td>>hg38_21_+_8433172_8446622_RNA45SN1_45S_50ntFlanks@9125.9142.18|CGCGCGGGTCGGGGGGCG</td><td>>hg38_21_+_8433172_8446622_RNA45SN1_45S_50ntFlanks</td><td>9125</td><td>9142</td><td>18</td><td>CGCGCGGGTCGGGGGGCG</td><td>93.1</td><td>-9.6</td><td>1.8</td><td>-5.4</td><td>5.2E-08</td><td>4.2E-07</td><td>BT20/MB468</td><td>MP</td></tr>
<tr><td>>hg38_21_+_8433172_8446622_RNA45SN1_45S_50ntFlanks@10411.10442.32|CAGTCGGTCCTGAGAGATGGGCGAGCGCCGTT</td><td>>hg38_21_+_8433172_8446622_RNA45SN1_45S_50ntFlanks</td><td>10411</td><td>10442</td><td>32</td><td>CAGTCGGTCCTGAGAGATGGGCGAGCGCCGTT</td><td>924.4</td><td>-8.5</td><td>1.7</td><td>-5.1</td><td>2.7E-07</td><td>4.2E-07</td><td>BT20/MB468</td><td>NUC</td></tr>
<tr><td>>hg38_21_+_8433172_8446622_RNA45SN1_45S_50ntFlanks@3782.3814.33|CAGTGAAACTGCGAATGGCTCATTAAATCAGTT</td><td>>hg38_21_+_8433172_8446622_RNA45SN1_45S_50ntFlanks</td><td>3782</td><td>3814</td><td>33</td><td>CAGTGAAACTGCGAATGGCTCATTAAATCAGTT</td><td>138.0</td><td>-10.2</td><td>1.9</td><td>-5.4</td><td>5.2E-08</td><td>4.2E-07</td><td>BT20/MB468</td><td>MP</td></tr>
<tr><td>>MI0003513|hsa-mir-455&WithFlank&9|+|114209428|114209535@60.80.21(+1C)&[MIMAT0004784&hsa-miR-455-3p&offsets|0|0(+1C);m-135&9|+|114209487|114209508&offsets|0|-1(+1C)]|GCAGTCCATGGGCATATACACC</td><td>>MI0003513|hsa-mir-455&WithFlank&9|+|114209428|114209535</td><td>60</td><td>80</td><td>21(+1C)</td><td>GCAGTCCATGGGCATATACACC</td><td>268.8</td><td>-3.3</td><td>0.6</td><td>-5.6</td><td>2.6E-08</td><td>4.2E-07</td><td>BT20/MB231</td><td>NUC</td></tr>
<tr><td>>MI0001445|hsa-mir-423&WithFlank&17|+|30117073|30117178@23.44.22(+1A)&[MIMAT0004748&hsa-miR-423-5p&offsets|0|-1(+1A);m-141&17|+|30117095|30117117&offsets|0|-1(+1A)]|TGAGGGGCAGAGAGCGAGACTTA</td><td>>MI0001445|hsa-mir-423&WithFlank&17|+|30117073|30117178</td><td>23</td><td>44</td><td>22(+1A)</td><td>TGAGGGGCAGAGAGCGAGACTTA</td><td>1403.3</td><td>7.4</td><td>1.4</td><td>5.4</td><td>5.2E-08</td><td>4.2E-07</td><td>BT20/MB468</td><td>MP</td></tr>
<tr><td>>hg38_21_+_8433172_8446622_RNA45SN1_45S_50ntFlanks@9955.9983.29|GGAAGTCGGAATCCGCTAAGGAGTGTGTA</td><td>>hg38_21_+_8433172_8446622_RNA45SN1_45S_50ntFlanks</td><td>9955</td><td>9983</td><td>29</td><td>GGAAGTCGGAATCCGCTAAGGAGTGTGTA</td><td>73.2</td><td>9.9</td><td>1.8</td><td>5.4</td><td>5.2E-08</td><td>4.2E-07</td><td>BT20/MB468</td><td>MP</td></tr>
<tr><td>>hg38_21_+_8433172_8446622_RNA45SN1_45S_50ntFlanks@10756.10785.30|TACCCATATCCGCAGCAGGTCTCCAAGGTG</td><td>>hg38_21_+_8433172_8446622_RNA45SN1_45S_50ntFlanks</td><td>10756</td><td>10785</td><td>30</td><td>TACCCATATCCGCAGCAGGTCTCCAAGGTG</td><td>76.7</td><td>-9.3</td><td>1.7</td><td>-5.4</td><td>5.2E-08</td><td>4.3E-07</td><td>BT20/MB468</td><td>MP</td></tr>
<tr><td>>hg38_21_+_8433172_8446622_RNA45SN1_45S_50ntFlanks@11668.11691.24|TGCCCAGTGCTCTGAATGTCAAAG</td><td>>hg38_21_+_8433172_8446622_RNA45SN1_45S_50ntFlanks</td><td>11668</td><td>11691</td><td>24</td><td>TGCCCAGTGCTCTGAATGTCAAAG</td><td>239.6</td><td>5.4</td><td>1.0</td><td>5.2</td><td>2.1E-07</td><td>4.3E-07</td><td>BT20/MB468</td><td>CYTO</td></tr>
<tr><td>>hg38_21_+_8433172_8446622_RNA45SN1_45S_50ntFlanks@11322.11343.22|ACGTCTCGTCGCGCGCGCGTCC</td><td>>hg38_21_+_8433172_8446622_RNA45SN1_45S_50ntFlanks</td><td>11322</td><td>11343</td><td>22</td><td>ACGTCTCGTCGCGCGCGCGTCC</td><td>1874.4</td><td>-4.4</td><td>0.9</td><td>-5.2</td><td>2.6E-07</td><td>4.3E-07</td><td>MB231/MB468</td><td>MITO</td></tr>
<tr><td>>hg38_21_+_8433172_8446622_RNA45SN1_45S_50ntFlanks@12458.12476.19|AAGCGTTGGATTGTTCACC</td><td>>hg38_21_+_8433172_8446622_RNA45SN1_45S_50ntFlanks</td><td>12458</td><td>12476</td><td>19</td><td>AAGCGTTGGATTGTTCACC</td><td>2012.4</td><td>-4.1</td><td>0.8</td><td>-5.2</td><td>2.2E-07</td><td>4.3E-07</td><td>MB231/MB468</td><td>MP</td></tr>
<tr><td>>hg38_21_+_8433172_8446622_RNA45SN1_45S_50ntFlanks@10845.10864.20|ACTTCGGGATAAGGATTGGC</td><td>>hg38_21_+_8433172_8446622_RNA45SN1_45S_50ntFlanks</td><td>10845</td><td>10864</td><td>20</td><td>ACTTCGGGATAAGGATTGGC</td><td>948.4</td><td>-5.4</td><td>1.0</td><td>-5.2</td><td>2.1E-07</td><td>4.3E-07</td><td>BT20/MB468</td><td>CYTO</td></tr>
<tr><td>>hg38_21_+_8433172_8446622_RNA45SN1_45S_50ntFlanks@8184.8204.21|CGAGGCCCAGCCCGTGGACGG</td><td>>hg38_21_+_8433172_8446622_RNA45SN1_45S_50ntFlanks</td><td>8184</td><td>8204</td><td>21</td><td>CGAGGCCCAGCCCGTGGACGG</td><td>326.3</td><td>-5.9</td><td>1.1</td><td>-5.2</td><td>2.2E-07</td><td>4.3E-07</td><td>MB231/MB468</td><td>MP</td></tr>
<tr><td>>hg38_21_+_8433172_8446622_RNA45SN1_45S_50ntFlanks@10766.10785.20|CGCAGCAGGTCTCCAAGGTG</td><td>>hg38_21_+_8433172_8446622_RNA45SN1_45S_50ntFlanks</td><td>10766</td><td>10785</td><td>20</td><td>CGCAGCAGGTCTCCAAGGTG</td><td>318.3</td><td>-6.9</td><td>1.3</td><td>-5.2</td><td>2.6E-07</td><td>4.3E-07</td><td>MB231/MB468</td><td>MITO</td></tr>
<tr><td>>hg38_21_+_8433172_8446622_RNA45SN1_45S_50ntFlanks@10351.10370.20|CGAGAACTTTGAAGGCCGAA</td><td>>hg38_21_+_8433172_8446622_RNA45SN1_45S_50ntFlanks</td><td>10351</td><td>10370</td><td>20</td><td>CGAGAACTTTGAAGGCCGAA</td><td>1474.7</td><td>-6.5</td><td>1.3</td><td>-5.2</td><td>2.2E-07</td><td>4.3E-07</td><td>MB231/MB468</td><td>MP</td></tr>
<tr><td>>hg38_21_+_8433172_8446622_RNA45SN1_45S_50ntFlanks@11163.11189.27|CGGTCCCCCGCGAGGGGGGCCCGGGCA</td><td>>hg38_21_+_8433172_8446622_RNA45SN1_45S_50ntFlanks</td><td>11163</td><td>11189</td><td>27</td><td>CGGTCCCCCGCGAGGGGGGCCCGGGCA</td><td>176.2</td><td>-10.5</td><td>1.9</td><td>-5.4</td><td>5.3E-08</td><td>4.3E-07</td><td>BT20/MB468</td><td>MP</td></tr>
<tr><td>>hg38_21_+_8433172_8446622_RNA45SN1_45S_50ntFlanks@12627.12656.30|ATGGGGCGAAGCTACCATCTGTGGGATTAT</td><td>>hg38_21_+_8433172_8446622_RNA45SN1_45S_50ntFlanks</td><td>12627</td><td>12656</td><td>30</td><td>ATGGGGCGAAGCTACCATCTGTGGGATTAT</td><td>79.9</td><td>-9.4</td><td>1.7</td><td>-5.4</td><td>5.3E-08</td><td>4.3E-07</td><td>BT20/MB468</td><td>MP</td></tr>
<tr><td>>hg38_21_+_8433172_8446622_RNA45SN1_45S_50ntFlanks@10869.10894.26|AGGGCTGGGTCGGTCGGGCTGGGGCG</td><td>>hg38_21_+_8433172_8446622_RNA45SN1_45S_50ntFlanks</td><td>10869</td><td>10894</td><td>26</td><td>AGGGCTGGGTCGGTCGGGCTGGGGCG</td><td>711.1</td><td>-7.5</td><td>1.4</td><td>-5.2</td><td>2.5E-07</td><td>4.3E-07</td><td>MB231/MB468</td><td>NUC</td></tr>
<tr><td>>trnaMT_HisGTG_MT_+_12138_12206@19.58.40|AAACATCAGATTGTGAATCTGACAACAGAGGCTTACGACC</td><td>>trnaMT_HisGTG_MT_+_12138_12206</td><td>19</td><td>58</td><td>40</td><td>AAACATCAGATTGTGAATCTGACAACAGAGGCTTACGACC</td><td>47.3</td><td>8.8</td><td>1.7</td><td>5.2</td><td>2.2E-07</td><td>4.3E-07</td><td>MB231/MB468</td><td>MP</td></tr>
<tr><td>>hg38_21_+_8433172_8446622_RNA45SN1_45S_50ntFlanks@476.508.33|TTCGTGATCGATGTGGTGACGTCGTGCTCTCCC</td><td>>hg38_21_+_8433172_8446622_RNA45SN1_45S_50ntFlanks</td><td>476</td><td>508</td><td>33</td><td>TTCGTGATCGATGTGGTGACGTCGTGCTCTCCC</td><td>1713.0</td><td>3.4</td><td>0.7</td><td>5.2</td><td>2.5E-07</td><td>4.3E-07</td><td>MB231/MB468</td><td>NUC</td></tr>
<tr><td>>hg38_21_+_8433172_8446622_RNA45SN1_45S_50ntFlanks@9534.9566.33|AGGAAACTCTGGTGGAGGTCCGTAGCGGTCCTG</td><td>>hg38_21_+_8433172_8446622_RNA45SN1_45S_50ntFlanks</td><td>9534</td><td>9566</td><td>33</td><td>AGGAAACTCTGGTGGAGGTCCGTAGCGGTCCTG</td><td>2172.9</td><td>-5.9</td><td>1.1</td><td>-5.2</td><td>2.1E-07</td><td>4.3E-07</td><td>BT20/MB468</td><td>CYTO</td></tr>
<tr><td>>hg38_21_+_8433172_8446622_RNA45SN1_45S_50ntFlanks@4204.4222.19|ACCCGGGGAGGTAGTGACG</td><td>>hg38_21_+_8433172_8446622_RNA45SN1_45S_50ntFlanks</td><td>4204</td><td>4222</td><td>19</td><td>ACCCGGGGAGGTAGTGACG</td><td>296.9</td><td>-7.5</td><td>1.5</td><td>-5.2</td><td>2.2E-07</td><td>4.3E-07</td><td>MB231/MB468</td><td>MP</td></tr>
<tr><td>>am_MI0039740|hsa-mir-12136&WithFlank&1|-|632609|632691@7.32.26&[MIMAT0049032&hsa-miR-12136&offsets|0|+8]|GAAAAAGTCATGGAGGCCATGGGGTT</td><td>>am_MI0039740|hsa-mir-12136&WithFlank&1|-|632609|632691</td><td>7</td><td>32</td><td>26</td><td>GAAAAAGTCATGGAGGCCATGGGGTT</td><td>248.2</td><td>5.6</td><td>1.1</td><td>5.2</td><td>2.1E-07</td><td>4.3E-07</td><td>BT20/MB468</td><td>CYTO</td></tr>
<tr><td>>trnaMT_SerTGA_MT_-_7446_7514@1.26.26,trnalookalike6_SerTGA_1_-_567997_568065@1.26.26|GAAAAAGTCATGGAGGCCATGGGGTT</td><td>>trnaMT_SerTGA_MT_-_7446_7514</td><td>1</td><td>26</td><td>26</td><td>GAAAAAGTCATGGAGGCCATGGGGTT</td><td>248.2</td><td>5.6</td><td>1.1</td><td>5.2</td><td>2.1E-07</td><td>4.3E-07</td><td>BT20/MB468</td><td>CYTO</td></tr>
<tr><td>>hg38_21_+_8433172_8446622_RNA45SN1_45S_50ntFlanks@9760.9787.28|AAACTTTAAATGGGTAAGAAGCCCGGCT</td><td>>hg38_21_+_8433172_8446622_RNA45SN1_45S_50ntFlanks</td><td>9760</td><td>9787</td><td>28</td><td>AAACTTTAAATGGGTAAGAAGCCCGGCT</td><td>198.1</td><td>-10.7</td><td>2.0</td><td>-5.4</td><td>5.3E-08</td><td>4.3E-07</td><td>BT20/MB468</td><td>MP</td></tr>
<tr><td>>hg38_21_+_8433172_8446622_RNA45SN1_45S_50ntFlanks@9922.9951.30|GTTGGTTGATATAGACAGCAGGACGGTGGC</td><td>>hg38_21_+_8433172_8446622_RNA45SN1_45S_50ntFlanks</td><td>9922</td><td>9951</td><td>30</td><td>GTTGGTTGATATAGACAGCAGGACGGTGGC</td><td>79.2</td><td>10.1</td><td>1.8</td><td>5.4</td><td>5.3E-08</td><td>4.3E-07</td><td>BT20/MB468</td><td>MP</td></tr>
<tr><td>>hg38_21_+_8433172_8446622_RNA45SN1_45S_50ntFlanks@8357.8376.20|AAAAGAACTTTGAAGAGAGA</td><td>>hg38_21_+_8433172_8446622_RNA45SN1_45S_50ntFlanks</td><td>8357</td><td>8376</td><td>20</td><td>AAAAGAACTTTGAAGAGAGA</td><td>323.0</td><td>-5.7</td><td>1.1</td><td>-5.2</td><td>2.6E-07</td><td>4.3E-07</td><td>MB231/MB468</td><td>MITO</td></tr>
<tr><td>>hg38_21_+_8433172_8446622_RNA45SN1_45S_50ntFlanks@4367.4397.31|CAGTTAAAAAGCTCGTAGTTGGATCTTGGGA</td><td>>hg38_21_+_8433172_8446622_RNA45SN1_45S_50ntFlanks</td><td>4367</td><td>4397</td><td>31</td><td>CAGTTAAAAAGCTCGTAGTTGGATCTTGGGA</td><td>246.6</td><td>-6.6</td><td>1.3</td><td>-5.2</td><td>2.5E-07</td><td>4.3E-07</td><td>MB231/MB468</td><td>NUC</td></tr>
<tr><td>>hg38_21_+_8433172_8446622_RNA45SN1_45S_50ntFlanks@5472.5512.41|ACGGCCCTGGCGGAGCGCTGAGAAGACGGTCGAACTTGACT</td><td>>hg38_21_+_8433172_8446622_RNA45SN1_45S_50ntFlanks</td><td>5472</td><td>5512</td><td>41</td><td>ACGGCCCTGGCGGAGCGCTGAGAAGACGGTCGAACTTGACT</td><td>1858.6</td><td>7.0</td><td>1.3</td><td>5.2</td><td>1.9E-07</td><td>4.3E-07</td><td>BT20/MB468</td><td>MITO</td></tr>
<tr><td>>hg38_21_+_8433172_8446622_RNA45SN1_45S_50ntFlanks@7983.8004.22|CAGATCAGACGTGGCGACCCGC</td><td>>hg38_21_+_8433172_8446622_RNA45SN1_45S_50ntFlanks</td><td>7983</td><td>8004</td><td>22</td><td>CAGATCAGACGTGGCGACCCGC</td><td>853.5</td><td>-5.9</td><td>1.1</td><td>-5.2</td><td>2.5E-07</td><td>4.3E-07</td><td>MB231/MB468</td><td>NUC</td></tr>
<tr><td>>hg38_21_+_8433172_8446622_RNA45SN1_45S_50ntFlanks@8105.8140.36|CGCGGCGGGGCGCGGGACATGTGGCGTACGGAAGAC</td><td>>hg38_21_+_8433172_8446622_RNA45SN1_45S_50ntFlanks</td><td>8105</td><td>8140</td><td>36</td><td>CGCGGCGGGGCGCGGGACATGTGGCGTACGGAAGAC</td><td>741.8</td><td>-7.3</td><td>1.4</td><td>-5.2</td><td>2.5E-07</td><td>4.3E-07</td><td>MB231/MB468</td><td>NUC</td></tr>
<tr><td>>hg38_21_+_8433172_8446622_RNA45SN1_45S_50ntFlanks@9906.9928.23|AGACCCCAGAAAAGGTGTTGGTT</td><td>>hg38_21_+_8433172_8446622_RNA45SN1_45S_50ntFlanks</td><td>9906</td><td>9928</td><td>23</td><td>AGACCCCAGAAAAGGTGTTGGTT</td><td>373.3</td><td>6.7</td><td>1.2</td><td>5.6</td><td>2.7E-08</td><td>4.3E-07</td><td>BT20/MB231</td><td>NUC</td></tr>
<tr><td>>hg38_21_+_8433172_8446622_RNA45SN1_45S_50ntFlanks@10786.10812.27|AACAGCCTCTGGCATGTTGGAACAATG</td><td>>hg38_21_+_8433172_8446622_RNA45SN1_45S_50ntFlanks</td><td>10786</td><td>10812</td><td>27</td><td>AACAGCCTCTGGCATGTTGGAACAATG</td><td>321.7</td><td>-7.3</td><td>1.4</td><td>-5.2</td><td>2.2E-07</td><td>4.3E-07</td><td>MB231/MB468</td><td>MP</td></tr>
<tr><td>>hg38_21_+_8433172_8446622_RNA45SN1_45S_50ntFlanks@10906.10925.20|TGGGCGCGCGCCGCGGCTGG</td><td>>hg38_21_+_8433172_8446622_RNA45SN1_45S_50ntFlanks</td><td>10906</td><td>10925</td><td>20</td><td>TGGGCGCGCGCCGCGGCTGG</td><td>74.8</td><td>-9.3</td><td>1.7</td><td>-5.4</td><td>5.4E-08</td><td>4.3E-07</td><td>BT20/MB468</td><td>MP</td></tr>
<tr><td>>MI0000239|hsa-mir-197&WithFlank&1|+|109598887|109598973@54.75.22&[MIMAT0000227&hsa-miR-197-3p&offsets|0|0;m-99&1|+|109598940|109598961&offsets|0|0]|TTCACCACCTTCTCCACCCAGC</td><td>>MI0000239|hsa-mir-197&WithFlank&1|+|109598887|109598973</td><td>54</td><td>75</td><td>22</td><td>TTCACCACCTTCTCCACCCAGC</td><td>10199.4</td><td>3.4</td><td>0.6</td><td>5.6</td><td>2.7E-08</td><td>4.3E-07</td><td>BT20/MB231</td><td>NUC</td></tr>
<tr><td>>hg38_21_+_8433172_8446622_RNA45SN1_45S_50ntFlanks@6650.6672.23|ACGACTCTTAGCGGTGGATCACT</td><td>>hg38_21_+_8433172_8446622_RNA45SN1_45S_50ntFlanks</td><td>6650</td><td>6672</td><td>23</td><td>ACGACTCTTAGCGGTGGATCACT</td><td>401.2</td><td>4.5</td><td>0.9</td><td>5.2</td><td>2.5E-07</td><td>4.3E-07</td><td>MB231/MB468</td><td>NUC</td></tr>
<tr><td>>MI0000809|hsa-mir-151a&WithFlank&8|-|140732558|140732659@53.74.22(+2A)&[MIMAT0000757&hsa-miR-151a-3p&offsets|0|+1(+2A);m-30&8|-|140732587|140732607&offsets|0|+1(+2A)]|CTAGACTGAAGCTCCTTGAGGAAA</td><td>>MI0000809|hsa-mir-151a&WithFlank&8|-|140732558|140732659</td><td>53</td><td>74</td><td>22(+2A)</td><td>CTAGACTGAAGCTCCTTGAGGAAA</td><td>265.2</td><td>5.1</td><td>1.0</td><td>5.2</td><td>2.4E-07</td><td>4.4E-07</td><td>MB231/MB468</td><td>CYTO</td></tr>
</table>

| | | | | | | | | | | | | | |
|---|---|---|---|---|---|---|---|---|---|---|---|---|---|
| >hg38_21_+_8433172_8446622_RNA45SN1_45S_50ntFlanks@9621.9650.30\|TCTAGTAGCTGGTTCCCTCCGAAGTTTCCC | >hg38_21_+_8433172_8446622_RNA45SN1_45S_50ntFlanks | 9621 | 9650 | 30 | TCTAGTAGCTGGTTCCCTCCGAAGTTTCCC | 391.7 | -6.4 | 1.2 | -5.2 | 1.9E-07 | 4.4E-07 | BT20/MB468 | MITO |
| >hg38_21_+_8433172_8446622_RNA45SN1_45S_50ntFlanks@9338.9370.33\|TCGCACGAAAGCCGCCGTGGCGCAATGAAGGTG | >hg38_21_+_8433172_8446622_RNA45SN1_45S_50ntFlanks | 9338 | 9370 | 33 | TCGCACGAAAGCCGCCGTGGCGCAATGAAGGTG | 510.8 | -8.0 | 1.5 | -5.4 | 5.4E-08 | 4.4E-07 | BT20/MB468 | MP |
| >hg38_21_+_8433172_8446622_RNA45SN1_45S_50ntFlanks@11783.11805.23\|CATGAATGGATGAACGAGATTCC | >hg38_21_+_8433172_8446622_RNA45SN1_45S_50ntFlanks | 11783 | 11805 | 23 | CATGAATGGATGAACGAGATTCC | 1965.1 | -4.9 | 0.9 | -5.2 | 2.1E-07 | 4.4E-07 | BT20/MB468 | CYTO |
| >MI0000261\|hsa-mir-139&WithFlank&11\|-\|72615057\|72615136@13.34.22(+1A)&[MIMAT0000250&hsa-miR-139-5p&offsets\|0\|-1(+1A);m-186&11\|-\|72615102\|72615124&offsets\|0\|-1(+1A)]\|TCTACAGTGCACGTGTCTCCAGA | >MI0000261\|hsa-mir-139&WithFlank&11\|-\|72615057\|72615136 | 13 | 34 | 22(+1A) | TCTACAGTGCACGTGTCTCCAGA | 58.5 | 9.1 | 1.8 | 5.2 | 2.2E-07 | 4.4E-07 | MB231/MB468 | MP |
| >hg38_21_+_8433172_8446622_RNA45SN1_45S_50ntFlanks@12072.12089.18\|CCCGAGGGGCTCTCGCTT | >hg38_21_+_8433172_8446622_RNA45SN1_45S_50ntFlanks | 12072 | 12089 | 18 | CCCGAGGGGCTCTCGCTT | 132.8 | -5.7 | 1.1 | -5.2 | 2.1E-07 | 4.4E-07 | BT20/MB468 | CYTO |
| >hg38_21_+_8433172_8446622_RNA45SN1_45S_50ntFlanks@8354.8392.39\|TTGAAAAGAACTTTGAAGAGAGAGTTCAAGAGGGCGTGA | >hg38_21_+_8433172_8446622_RNA45SN1_45S_50ntFlanks | 8354 | 8392 | 39 | TTGAAAAGAACTTTGAAGAGAGAGTTCAAGAGGGCGTGA | 65.6 | 9.8 | 1.8 | 5.4 | 5.4E-08 | 4.4E-07 | BT20/MB468 | MP |
| >MI0022688\|hsa-mir-6842&WithFlank&8\|+\|27433364\|27433440@50.71.22&[MIMAT0027587&hsa-miR-6842-3p&offsets\|0\|0;m-370&8\|+\|27433413\|27433435&offsets\|0\|-1]\|TTGGCTGGTCTCTGCTCCGCAG | >MI0022688\|hsa-mir-6842&WithFlank&8\|+\|27433364\|27433440 | 50 | 71 | 22 | TTGGCTGGTCTCTGCTCCGCAG | 77.6 | 10.0 | 1.8 | 5.4 | 5.4E-08 | 4.4E-07 | BT20/MB468 | MP |
| >hg38_21_+_8433172_8446622_RNA45SN1_45S_50ntFlanks@12941.12973.33\|ATTCGTAGACGACCTGCTTCTGGGTCGGGGTTT | >hg38_21_+_8433172_8446622_RNA45SN1_45S_50ntFlanks | 12941 | 12973 | 33 | ATTCGTAGACGACCTGCTTCTGGGTCGGGGTTT | 76.4 | -9.3 | 1.7 | -5.4 | 5.4E-08 | 4.4E-07 | BT20/MB468 | MP |
| >hg38_21_+_8433172_8446622_RNA45SN1_45S_50ntFlanks@12204.12235.32\|AAGGCGAGCTCAGGGAGGACAGAAACCTCCCG | >hg38_21_+_8433172_8446622_RNA45SN1_45S_50ntFlanks | 12204 | 12235 | 32 | AAGGCGAGCTCAGGGAGGACAGAAACCTCCCG | 335.8 | 5.2 | 1.0 | 5.2 | 2.4E-07 | 4.4E-07 | MB231/MB468 | CYTO |
| >hg38_21_+_8433172_8446622_RNA45SN1_45S_50ntFlanks@12456.12498.43\|CCAAGCGTTGGATTGTTCACCCACTAATAGGGAACGTGAGCTG | >hg38_21_+_8433172_8446622_RNA45SN1_45S_50ntFlanks | 12456 | 12498 | 43 | CCAAGCGTTGGATTGTTCACCCACTAATAGGGAACGTGAGCTG | 288.9 | -5.4 | 1.1 | -5.2 | 2.4E-07 | 4.4E-07 | MB231/MB468 | CYTO |
| >hg38_21_+_8433172_8446622_RNA45SN1_45S_50ntFlanks@12941.12960.20\|ATTCGTAGACGACCTGCTTC | >hg38_21_+_8433172_8446622_RNA45SN1_45S_50ntFlanks | 12941 | 12960 | 20 | ATTCGTAGACGACCTGCTTC | 724.8 | 3.0 | 0.6 | 5.2 | 2.4E-07 | 4.4E-07 | MB231/MB468 | CYTO |
| >hg38_21_+_8433172_8446622_RNA45SN1_45S_50ntFlanks@10819.10838.20\|AGGGAAGTCGGCAAGCCGGA | >hg38_21_+_8433172_8446622_RNA45SN1_45S_50ntFlanks | 10819 | 10838 | 20 | AGGGAAGTCGGCAAGCCGGA | 265.7 | -4.1 | 0.8 | -5.2 | 2.4E-07 | 4.4E-07 | MB231/MB468 | CYTO |
| >hg38_21_+_8433172_8446622_RNA45SN1_45S_50ntFlanks@12539.12557.19\|ATGATGTGTTGTTGCCATG | >hg38_21_+_8433172_8446622_RNA45SN1_45S_50ntFlanks | 12539 | 12557 | 19 | ATGATGTGTTGTTGCCATG | 111.5 | -6.1 | 1.2 | -5.2 | 2.1E-07 | 4.4E-07 | BT20/MB468 | CYTO |
| >MI0000542\|hsa-mir-320a&WithFlank&8\|-\|22244960\|22245043@48.71.24&[MIMAT0000510&hsa-miR-320a-3p&offsets\|0\|+2;m-60&8\|-\|22244975\|22244996&offsets\|0\|+2]\|AAAAGCTGGGTTGAGAGGGCGAAA | >MI0000542\|hsa-mir-320a&WithFlank&8\|-\|22244960\|22245043 | 48 | 71 | 24 | AAAAGCTGGGTTGAGAGGGCGAAA | 2572.1 | 7.2 | 1.3 | 5.4 | 5.4E-08 | 4.4E-07 | BT20/MB468 | MP |
| >hg38_21_+_8433172_8446622_RNA45SN1_45S_50ntFlanks@4983.5003.21\|CGGACAGGATTGACAGATTGA | >hg38_21_+_8433172_8446622_RNA45SN1_45S_50ntFlanks | 4983 | 5003 | 21 | CGGACAGGATTGACAGATTGA | 429.4 | -10.8 | 2.1 | -5.2 | 1.9E-07 | 4.4E-07 | BT20/MB468 | MITO |
| >am_trna128_GlyGCC_6_-_27870686_27870756@7.28.22,trna133_GlyCCC_1_-_16872434_16872504@7.28.22,trna17_GluTTC_2_-_204229131_204229201@7.28.22,trna18_GlyGCC_16_+_70822597_70822667@7.28.22,trna19_GlyGCC_16_+_70823410_70823480@7.28.22,trna19_GlyGCC_2_-_157257659_157257729@7.28.22,trna24_GlyGCC_16_-_70812942_70813012@7.28.22,trna25_GlyGCC_16_-_70812114_70812184@7.28.22,trna2_GlyGCC_21_-_18827107_18827177@7.28.22,trna35_GlyGCC_1_+_161413094_161413164@7.28.22,trna37_GlyGCC_1_+_161420467_161420537@7.28.22,trna39_GlyGCC_1_+_161427898_161427968@7.28.22,trna41_GlyGCC_1_+_161435258_161435328@7.28.22,trna4_GlyCCC_1_+_17188416_17188486@7.28.22,trna5_GlyGCC_17_+_8029064_8029134@7.28.22,trna68_GlyGCC_1_-_161493637_161493707@7.28.22,trna91_GlyCCC_1_-_149680210_149680280@7.28.22,trna99_ValCAC_1_-_149294666_149294736@7.28.22\|GTGGTTCAGTGGTAGAATTCTC | >am_trna128_GlyGCC_6_-_27870686_27870756 | 7 | 28 | 22 | GTGGTTCAGTGGTAGAATTCTC | 230.1 | -5.1 | 1.0 | -5.2 | 2.5E-07 | 4.4E-07 | MB231/MB468 | CYTO |
| >trna152_ValCAC_6_-_27248049_27248121@1.33.33\|GCTTCTGTAGTGTAGTGGTTATCACGTTCGCCT | >trna152_ValCAC_6_-_27248049_27248121 | 1 | 33 | 33 | GCTTCTGTAGTGTAGTGGTTATCACGTTCGCCT | 760.8 | 8.1 | 1.6 | 5.2 | 2.1E-07 | 4.4E-07 | BT20/MB468 | CYTO |
| >hg38_21_+_8433172_8446622_RNA45SN1_45S_50ntFlanks@10770.10805.36\|GCAGGTCTCCAAGGTGAACAGCCTCTGGCATGTTGG | >hg38_21_+_8433172_8446622_RNA45SN1_45S_50ntFlanks | 10770 | 10805 | 36 | GCAGGTCTCCAAGGTGAACAGCCTCTGGCATGTTGG | 340.2 | -6.9 | 1.3 | -5.1 | 2.6E-07 | 4.4E-07 | MB231/MB468 | MITO |
| >hg38_21_+_8433172_8446622_RNA45SN1_45S_50ntFlanks@10458.10478.21\|ATGGCCTCCGTTGCCCTCGGC | >hg38_21_+_8433172_8446622_RNA45SN1_45S_50ntFlanks | 10458 | 10478 | 21 | ATGGCCTCCGTTGCCCTCGGC | 3025.5 | -6.5 | 1.3 | -5.2 | 2.2E-07 | 4.4E-07 | BT20/MB468 | CYTO |
| >hg38_21_+_8433172_8446622_RNA45SN1_45S_50ntFlanks@476.517.42\|TTCGTGATCGATGTGGTGACGTCGTGCTCTCCCGGGCCGGGT | >hg38_21_+_8433172_8446622_RNA45SN1_45S_50ntFlanks | 476 | 517 | 42 | TTCGTGATCGATGTGGTGACGTCGTGCTCTCCCGGGCCGGGT | 1472.5 | 8.0 | 1.5 | 5.2 | 1.9E-07 | 4.4E-07 | BT20/MB468 | MITO |
| >hg38_21_+_8433172_8446622_RNA45SN1_45S_50ntFlanks@10355.10390.36\|AACTTTGAAGGCCGAAGTGGAGAAGGGTTCCATGTG | >hg38_21_+_8433172_8446622_RNA45SN1_45S_50ntFlanks | 10355 | 10390 | 36 | AACTTTGAAGGCCGAAGTGGAGAAGGGTTCCATGTG | 241.2 | -4.6 | 0.9 | -5.2 | 2.2E-07 | 4.4E-07 | MB231/MB468 | MP |
| >hg38_21_+_8433172_8446622_RNA45SN1_45S_50ntFlanks@12912.12933.22\|CACGTTCGTGGGGAACCTGGCG | >hg38_21_+_8433172_8446622_RNA45SN1_45S_50ntFlanks | 12912 | 12933 | 22 | CACGTTCGTGGGGAACCTGGCG | 560.5 | -6.7 | 1.3 | -5.2 | 2.5E-07 | 4.4E-07 | MB231/MB468 | NUC |
| >hg38_21_+_8433172_8446622_RNA45SN1_45S_50ntFlanks@10116.10136.21\|TGCGTCGGAGGGCGGCGGCGG | >hg38_21_+_8433172_8446622_RNA45SN1_45S_50ntFlanks | 10116 | 10136 | 21 | TGCGTCGGAGGGCGGCGGCGG | 391.2 | -6.4 | 1.2 | -5.2 | 2.2E-07 | 4.4E-07 | BT20/MB468 | CYTO |
| >hg38_MT_+_598_1651_RNR1_12S_50ntFlanks@853.871.19\|CTTAGTTGAACAGGGCCCT | >hg38_MT_+_598_1651_RNR1_12S_50ntFlanks | 853 | 871 | 19 | CTTAGTTGAACAGGGCCCT | 95.7 | 10.3 | 1.9 | 5.4 | 5.5E-08 | 4.4E-07 | BT20/MB468 | MP |
| >MI0000077\|hsa-mir-21&WithFlank&17\|+\|59841260\|59841343@14.35.22(+1U)&[MIMAT0000076&hsa-miR-21-5p&offsets\|0\|0(+1U);m-1&17\|+\|59841273\|59841294&offsets\|0\|0(+1U)]\|TAGCTTATCAGACTGATGTTGAT | >MI0000077\|hsa-mir-21&WithFlank&17\|+\|59841260\|59841343 | 14 | 35 | 22(+1U) | TAGCTTATCAGACTGATGTTGAT | 43282.8 | 5.7 | 1.1 | 5.2 | 2.5E-07 | 4.4E-07 | MB231/MB468 | CYTO |
| >hg38_21_+_8433172_8446622_RNA45SN1_45S_50ntFlanks@8105.8142.38\|CGCGGCGGGGCGCGGGACATGTGGCGTACGGAAGACCC | >hg38_21_+_8433172_8446622_RNA45SN1_45S_50ntFlanks | 8105 | 8142 | 38 | CGCGGCGGGGCGCGGGACATGTGGCGTACGGAAGACCC | 2587.0 | -8.8 | 1.7 | -5.2 | 2.2E-07 | 4.4E-07 | MB231/MB468 | MP |
| >hg38_21_+_8433172_8446622_RNA45SN1_45S_50ntFlanks@11665.11691.27\|TTCTGCCCAGTGCTCTGAATGTCAAAG | >hg38_21_+_8433172_8446622_RNA45SN1_45S_50ntFlanks | 11665 | 11691 | 27 | TTCTGCCCAGTGCTCTGAATGTCAAAG | 297.3 | 4.6 | 0.9 | 5.2 | 1.9E-07 | 4.4E-07 | BT20/MB468 | MITO |
| >trnaMT_GluTTC_MT_-_14674_14742@26.63.38,trnalookalike8_GluTTC_5_-_93905172_93905240@26.63.38\|GGTTTTTCATATCATTGGTCGTGGTTGTAGTCCGTGCG | >trnaMT_GluTTC_MT_-_14674_14742 | 26 | 63 | 38 | GGTTTTTCATATCATTGGTCGTGGTTGTAGTCCGTGCG | 141.0 | 10.9 | 2.0 | 5.4 | 5.5E-08 | 4.4E-07 | BT20/MB468 | MP |
| >hg38_1_-_228634819_228635039_RNA5S12_5S_50ntFlanks@135.169.35\|GGGAGACCGCCTGGGAATACCGGGTGCTGTAGGCT | >hg38_1_-_228634819_228635039_RNA5S12_5S_50ntFlanks | 135 | 169 | 35 | GGGAGACCGCCTGGGAATACCGGGTGCTGTAGGCT | 734.0 | 3.1 | 0.6 | 5.2 | 2.5E-07 | 4.4E-07 | MB231/MB468 | NUC |
| >trnaMT_PheGAA_MT_+_577_647@35.54.20\|GAAAATGTTTAGACGGGCTC | >trnaMT_PheGAA_MT_+_577_647 | 35 | 54 | 20 | GAAAATGTTTAGACGGGCTC | 107.9 | 4.8 | 0.9 | 5.1 | 2.8E-07 | 4.4E-07 | BT20/MB468 | NUC |
| >hg38_MT_+_598_1651_RNR1_12S_50ntFlanks@376.404.29\|CTAAAACTCACCTGAGTTGTAAAAAACTC | >hg38_MT_+_598_1651_RNR1_12S_50ntFlanks | 376 | 404 | 29 | CTAAAACTCACCTGAGTTGTAAAAAACTC | 483.4 | 4.5 | 0.9 | 5.2 | 1.9E-07 | 4.4E-07 | BT20/MB468 | MITO |
| >hg38_21_+_8433172_8446622_RNA45SN1_45S_50ntFlanks@6741.6771.31\|ATCGACACTTCGAACGCACTTGCGGCCCCGG | >hg38_21_+_8433172_8446622_RNA45SN1_45S_50ntFlanks | 6741 | 6771 | 31 | ATCGACACTTCGAACGCACTTGCGGCCCCGG | 2015.5 | -8.9 | 1.7 | -5.1 | 2.8E-07 | 4.4E-07 | BT20/MB468 | NUC |
| >hg38_21_+_8433172_8446622_RNA45SN1_45S_50ntFlanks@9454.9493.40\|CCGCGCCGGGGAGGTGGAGCACGAGCGCACGTGTTAGGAC | >hg38_21_+_8433172_8446622_RNA45SN1_45S_50ntFlanks | 9454 | 9493 | 40 | CCGCGCCGGGGAGGTGGAGCACGAGCGCACGTGTTAGGAC | 284.2 | -6.4 | 1.2 | -5.2 | 1.9E-07 | 4.4E-07 | BT20/MB468 | MITO |
| >hg38_21_+_8433172_8446622_RNA45SN1_45S_50ntFlanks@5486.5509.24\|GCGCTGAGAAGACGGTCGAACTTG | >hg38_21_+_8433172_8446622_RNA45SN1_45S_50ntFlanks | 5486 | 5509 | 24 | GCGCTGAGAAGACGGTCGAACTTG | 712.8 | -7.5 | 1.4 | -5.2 | 2.3E-07 | 4.4E-07 | MB231/MB468 | MP |
| >hg38_21_+_8433172_8446622_RNA45SN1_45S_50ntFlanks@11562.11592.31\|CGCCTAGCAGCCGACTTAGAACTGGTGCGGA | >hg38_21_+_8433172_8446622_RNA45SN1_45S_50ntFlanks | 11562 | 11592 | 31 | CGCCTAGCAGCCGACTTAGAACTGGTGCGGA | 607.1 | -8.1 | 1.6 | -5.1 | 2.8E-07 | 4.4E-07 | BT20/MB468 | NUC |
| >hg38_21_+_8433172_8446622_RNA45SN1_45S_50ntFlanks@4002.4025.24\|GATAACCTCGGGCCGATCGCACGC | >hg38_21_+_8433172_8446622_RNA45SN1_45S_50ntFlanks | 4002 | 4025 | 24 | GATAACCTCGGGCCGATCGCACGC | 2276.3 | -9.0 | 1.7 | -5.1 | 2.8E-07 | 4.4E-07 | BT20/MB468 | NUC |
| >hg38_21_+_8433172_8446622_RNA45SN1_45S_50ntFlanks@11568.11602.35\|GCAGCCGACTTAGAACTGGTGCGGACCAGGGGAAT | >hg38_21_+_8433172_8446622_RNA45SN1_45S_50ntFlanks | 11568 | 11602 | 35 | GCAGCCGACTTAGAACTGGTGCGGACCAGGGGAAT | 832.1 | -4.2 | 0.8 | -5.1 | 2.7E-07 | 4.4E-07 | MB231/MB468 | MITO |
| >hg38_21_+_8433172_8446622_RNA45SN1_45S_50ntFlanks@6670.6696.27\|ACTCGGCTCGTGCGTCGATGAAGAACG | >hg38_21_+_8433172_8446622_RNA45SN1_45S_50ntFlanks | 6670 | 6696 | 27 | ACTCGGCTCGTGCGTCGATGAAGAACG | 1057.1 | -3.5 | 0.7 | -5.1 | 2.7E-07 | 4.4E-07 | MB231/MB468 | MITO |
| >hg38_21_+_8433172_8446622_RNA45SN1_45S_50ntFlanks@8556.8577.22\|CCGGAGGGGGCGGGCTCCGGCG | >hg38_21_+_8433172_8446622_RNA45SN1_45S_50ntFlanks | 8556 | 8577 | 22 | CCGGAGGGGGCGGGCTCCGGCG | 87.9 | -9.5 | 1.8 | -5.4 | 5.5E-08 | 4.4E-07 | BT20/MB468 | MP |
| >hg38_21_+_8433172_8446622_RNA45SN1_45S_50ntFlanks@12182.12200.19\|CAAACGGTAACGCAGGTGT | >hg38_21_+_8433172_8446622_RNA45SN1_45S_50ntFlanks | 12182 | 12200 | 19 | CAAACGGTAACGCAGGTGT | 105.0 | -9.8 | 1.8 | -5.4 | 5.5E-08 | 4.5E-07 | BT20/MB468 | MP |
| >hg38_21_+_8433172_8446622_RNA45SN1_45S_50ntFlanks@4759.4787.29\|ATCAGATACCGTCGTAGTTCCGACCATAA | >hg38_21_+_8433172_8446622_RNA45SN1_45S_50ntFlanks | 4759 | 4787 | 29 | ATCAGATACCGTCGTAGTTCCGACCATAA | 75.5 | -9.3 | 1.7 | -5.4 | 5.5E-08 | 4.5E-07 | BT20/MB468 | MP |
| >hg38_21_+_8433172_8446622_RNA45SN1_45S_50ntFlanks@7980.8012.33\|CCTCAGATCAGACGTGGCGACCCGCTGAATTTA | >hg38_21_+_8433172_8446622_RNA45SN1_45S_50ntFlanks | 7980 | 8012 | 33 | CCTCAGATCAGACGTGGCGACCCGCTGAATTTA | 249.7 | -5.3 | 1.0 | -5.2 | 2.6E-07 | 4.5E-07 | MB231/MB468 | NUC |
| >hg38_21_+_8433172_8446622_RNA45SN1_45S_50ntFlanks@6652.6678.27\|GACTCTTAGCGGTGGATCACTCGGCTC | >hg38_21_+_8433172_8446622_RNA45SN1_45S_50ntFlanks | 6652 | 6678 | 27 | GACTCTTAGCGGTGGATCACTCGGCTC | 1673.7 | 3.7 | 0.7 | 5.2 | 2.2E-07 | 4.5E-07 | BT20/MB468 | CYTO |
| >hg38_21_+_8433172_8446622_RNA45SN1_45S_50ntFlanks@12618.12654.37\|GAGGAGCCAATGGGGCGAAGCTACCATCTGTGGGATT | >hg38_21_+_8433172_8446622_RNA45SN1_45S_50ntFlanks | 12618 | 12654 | 37 | GAGGAGCCAATGGGGCGAAGCTACCATCTGTGGGATT | 244.6 | -3.4 | 0.7 | -5.2 | 2.6E-07 | 4.5E-07 | MB231/MB468 | NUC |
| >hg38_21_+_8433172_8446622_RNA45SN1_45S_50ntFlanks@12631.12654.24\|GGCGAAGCTACCATCTGTGGGATT | >hg38_21_+_8433172_8446622_RNA45SN1_45S_50ntFlanks | 12631 | 12654 | 24 | GGCGAAGCTACCATCTGTGGGATT | 261.1 | -6.7 | 1.3 | -5.2 | 2.6E-07 | 4.5E-07 | MB231/MB468 | NUC |
| >hg38_21_+_8433172_8446622_RNA45SN1_45S_50ntFlanks@10689.10707.19\|GCGTCCGGTGAGCTCTCGC | >hg38_21_+_8433172_8446622_RNA45SN1_45S_50ntFlanks | 10689 | 10707 | 19 | GCGTCCGGTGAGCTCTCGC | 533.2 | -6.3 | 1.2 | -5.1 | 2.8E-07 | 4.5E-07 | BT20/MB468 | NUC |
| >am_trnaMT_PheGAA_MT_+_577_647@41.62.22\|GTTTAGACGGGCTCACATCACC | >am_trnaMT_PheGAA_MT_+_577_647 | 41 | 62 | 22 | GTTTAGACGGGCTCACATCACC | 1315.5 | 8.2 | 1.6 | 5.2 | 2.2E-07 | 4.5E-07 | BT20/MB468 | CYTO |

<table>
<tr><td>>hg38_21_+_8433172_8446622_RNA45SN1_45S_50ntFlanks@11667.11697.31|CTGCCCAGTGCTCTGAATGTCAAAGTGAAGA</td><td>>hg38_21_+_8433172_8446622_RNA45SN1_45S_50ntFlanks</td><td>11667</td><td>11697</td><td>31</td><td>CTGCCCAGTGCTCTGAATGTCAAAGTGAAGA</td><td>70.2</td><td>9.9</td><td>1.8</td><td>5.4</td><td>5.6E-08</td><td>4.5E-07</td><td>BT20/MB468</td><td>MP</td></tr>
<tr><td>>hg38_21_+_8433172_8446622_RNA45SN1_45S_50ntFlanks@8168.8189.22|CCCAAGTCCTTCTGATCGAGGC</td><td>>hg38_21_+_8433172_8446622_RNA45SN1_45S_50ntFlanks</td><td>8168</td><td>8189</td><td>22</td><td>CCCAAGTCCTTCTGATCGAGGC</td><td>69.7</td><td>-9.2</td><td>1.7</td><td>-5.4</td><td>5.6E-08</td><td>4.5E-07</td><td>BT20/MB468</td><td>MP</td></tr>
<tr><td>>hg38_21_+_8433172_8446622_RNA45SN1_45S_50ntFlanks@8168.8188.21|CCCAAGTCCTTCTGATCGAGG</td><td>>hg38_21_+_8433172_8446622_RNA45SN1_45S_50ntFlanks</td><td>8168</td><td>8188</td><td>21</td><td>CCCAAGTCCTTCTGATCGAGG</td><td>426.7</td><td>-7.9</td><td>1.5</td><td>-5.2</td><td>2.2E-07</td><td>4.5E-07</td><td>BT20/MB468</td><td>CYTO</td></tr>
<tr><td>>hg38_21_+_8433172_8446622_RNA45SN1_45S_50ntFlanks@4069.4089.21|CTTTCGATGGTAGTCGCCGTG</td><td>>hg38_21_+_8433172_8446622_RNA45SN1_45S_50ntFlanks</td><td>4069</td><td>4089</td><td>21</td><td>CTTTCGATGGTAGTCGCCGTG</td><td>436.3</td><td>-4.0</td><td>0.7</td><td>-5.4</td><td>5.6E-08</td><td>4.5E-07</td><td>BT20/MB468</td><td>MP</td></tr>
<tr><td>>MI0000465|hsa-mir-191&WithFlank&3|-|49020612|49020715@22.44.23(+1C)&[MIMAT0000440&hsa-miR-191-5p&offsets|0|0(+1C);m-48&3|-|49020672|49020694&offsets|0|0(+1C)]|CAACGGAATCCCAAAAGCAGCTGC</td><td>>MI0000465|hsa-mir-191&WithFlank&3|-|49020612|49020715</td><td>22</td><td>44</td><td>23(+1C)</td><td>CAACGGAATCCCAAAAGCAGCTGC</td><td>855.2</td><td>7.9</td><td>1.5</td><td>5.1</td><td>2.7E-07</td><td>4.5E-07</td><td>MB231/MB468</td><td>MITO</td></tr>
<tr><td>>hg38_21_+_8433172_8446622_RNA45SN1_45S_50ntFlanks@10786.10816.31|AACAGCCTCTGGCATGTTGGAACAATGTAGG</td><td>>hg38_21_+_8433172_8446622_RNA45SN1_45S_50ntFlanks</td><td>10786</td><td>10816</td><td>31</td><td>AACAGCCTCTGGCATGTTGGAACAATGTAGG</td><td>1265.0</td><td>-6.0</td><td>1.2</td><td>-5.2</td><td>2.6E-07</td><td>4.5E-07</td><td>MB231/MB468</td><td>NUC</td></tr>
<tr><td>>hg38_21_+_8433172_8446622_RNA45SN1_45S_50ntFlanks@10355.10390.36|AACTTTGAAGGCCGAAGTGGAGAAGGGTTCCATGTG</td><td>>hg38_21_+_8433172_8446622_RNA45SN1_45S_50ntFlanks</td><td>10355</td><td>10390</td><td>36</td><td>AACTTTGAAGGCCGAAGTGGAGAAGGGTTCCATGTG</td><td>1029.5</td><td>-8.6</td><td>1.7</td><td>-5.1</td><td>2.8E-07</td><td>4.5E-07</td><td>BT20/MB468</td><td>NUC</td></tr>
<tr><td>>hg38_21_+_8433172_8446622_RNA45SN1_45S_50ntFlanks@4297.4317.21|CATTGGAGGGCAAGTCTGGTG</td><td>>hg38_21_+_8433172_8446622_RNA45SN1_45S_50ntFlanks</td><td>4297</td><td>4317</td><td>21</td><td>CATTGGAGGGCAAGTCTGGTG</td><td>1894.3</td><td>-6.6</td><td>1.3</td><td>-5.2</td><td>2.3E-07</td><td>4.5E-07</td><td>MB231/MB468</td><td>MP</td></tr>
<tr><td>>MI0001445|hsa-mir-423&WithFlank&17|+|30117073|30117178@58.78.21&[MIMAT0001340&hsa-miR-423-3p&offsets|-1|-3;m-100&17|+|30117131|30117153&offsets|-1|-3]|AAGCTCGGTCTGAGGCCCCTC</td><td>>MI0001445|hsa-mir-423&WithFlank&17|+|30117073|30117178</td><td>58</td><td>78</td><td>21</td><td>AAGCTCGGTCTGAGGCCCCTC</td><td>454.3</td><td>4.4</td><td>0.9</td><td>5.1</td><td>2.7E-07</td><td>4.5E-07</td><td>MB231/MB468</td><td>MITO</td></tr>
<tr><td>>hg38_21_+_8433172_8446622_RNA45SN1_45S_50ntFlanks@5073.5094.22|AATTCCGATAACGAACGAGACT</td><td>>hg38_21_+_8433172_8446622_RNA45SN1_45S_50ntFlanks</td><td>5073</td><td>5094</td><td>22</td><td>AATTCCGATAACGAACGAGACT</td><td>1366.4</td><td>-9.0</td><td>1.7</td><td>-5.2</td><td>2.3E-07</td><td>4.5E-07</td><td>MB231/MB468</td><td>MP</td></tr>
<tr><td>>hg38_21_+_8433172_8446622_RNA45SN1_45S_50ntFlanks@13000.13039.40|TGCGATCTATTGAAAGTCAGCCCTCGACACAAGGGTTTGT</td><td>>hg38_21_+_8433172_8446622_RNA45SN1_45S_50ntFlanks</td><td>13000</td><td>13039</td><td>40</td><td>TGCGATCTATTGAAAGTCAGCCCTCGACACAAGGGTTTGT</td><td>669.3</td><td>4.9</td><td>1.0</td><td>5.2</td><td>2.6E-07</td><td>4.5E-07</td><td>MB231/MB468</td><td>NUC</td></tr>
<tr><td>>trna111_HisGTG_1_-_147774845_147774916@-1A.26.27,trna118_HisGTG_1_-_145396881_145396952@-1A.26.27,trna16_HisGTG_1_+_146544773_146544844@-1A.26.27,trna1_HisGTG_15_+_45493349_45493420@-1A.26.27,trna21_HisGTG_1_+_147753471_147753542@-1A.26.27,trna33_HisGTG_6_+_27125906_27125977@-1A.26.27,trna7_HisGTG_9_-_14433938_14434009@-1A.26.27,trna8_HisGTG_15_-_45492611_45492682@-1A.26.27,trna9_HisGTG_15_-_45490804_45490875@-1A.26.27|AGCCGTGATCGTATAGTGGTTAGTACT</td><td>>trna111_HisGTG_1_-_147774845_147774916</td><td>-1A</td><td>26</td><td>27</td><td>AGCCGTGATCGTATAGTGGTTAGTACT</td><td>429.0</td><td>7.1</td><td>1.4</td><td>5.1</td><td>2.7E-07</td><td>4.5E-07</td><td>MB231/MB468</td><td>MITO</td></tr>
<tr><td>>hg38_21_+_8433172_8446622_RNA45SN1_45S_50ntFlanks@7978.8016.39|GACCTCAGATCAGACGTGGCGACCCGCTGAATTTAAGCA</td><td>>hg38_21_+_8433172_8446622_RNA45SN1_45S_50ntFlanks</td><td>7978</td><td>8016</td><td>39</td><td>GACCTCAGATCAGACGTGGCGACCCGCTGAATTTAAGCA</td><td>307.7</td><td>5.1</td><td>1.0</td><td>5.2</td><td>1.9E-07</td><td>4.5E-07</td><td>BT20/MB468</td><td>MITO</td></tr>
<tr><td>>MI0000095|hsa-mir-93&WithFlank&7|-|100093762|100093853@17.39.23(+1A)&[MIMAT0000093&hsa-miR-93-5p&offsets|0|0(+1A);m-29&7|-|100093815|100093837&offsets|0|0(+1A)]|CAAAGTGCTGTTCGTGCAGGTAGA</td><td>>MI0000095|hsa-mir-93&WithFlank&7|-|100093762|100093853</td><td>17</td><td>39</td><td>23(+1A)</td><td>CAAAGTGCTGTTCGTGCAGGTAGA</td><td>1618.1</td><td>5.1</td><td>1.0</td><td>5.2</td><td>2.5E-07</td><td>4.5E-07</td><td>MB231/MB468</td><td>CYTO</td></tr>
<tr><td>>hg38_21_+_8433172_8446622_RNA45SN1_45S_50ntFlanks@9561.9597.37|GTCCTGACGTGCAAATCGGTCGTCCGACCTGGGTATA</td><td>>hg38_21_+_8433172_8446622_RNA45SN1_45S_50ntFlanks</td><td>9561</td><td>9597</td><td>37</td><td>GTCCTGACGTGCAAATCGGTCGTCCGACCTGGGTATA</td><td>681.2</td><td>-8.3</td><td>1.6</td><td>-5.1</td><td>2.9E-07</td><td>4.5E-07</td><td>BT20/MB468</td><td>NUC</td></tr>
<tr><td>>hg38_21_+_8433172_8446622_RNA45SN1_45S_50ntFlanks@11641.11666.26|CGGCGGGTGTTGACGCGATGTGATTT</td><td>>hg38_21_+_8433172_8446622_RNA45SN1_45S_50ntFlanks</td><td>11641</td><td>11666</td><td>26</td><td>CGGCGGGTGTTGACGCGATGTGATTT</td><td>292.0</td><td>-6.6</td><td>1.3</td><td>-5.2</td><td>2.0E-07</td><td>4.5E-07</td><td>BT20/MB468</td><td>MITO</td></tr>
<tr><td>>hg38_21_+_8433172_8446622_RNA45SN1_45S_50ntFlanks@10402.10430.29|AACATGGGTCAGTCGGTCCTGAGAGATGG</td><td>>hg38_21_+_8433172_8446622_RNA45SN1_45S_50ntFlanks</td><td>10402</td><td>10430</td><td>29</td><td>AACATGGGTCAGTCGGTCCTGAGAGATGG</td><td>1741.2</td><td>-5.6</td><td>1.1</td><td>-5.2</td><td>2.2E-07</td><td>4.5E-07</td><td>BT20/MB468</td><td>CYTO</td></tr>
<tr><td>>hg38_21_+_8433172_8446622_RNA45SN1_45S_50ntFlanks@11578.11609.32|TAGAACTGGTGCGGACCAGGGGAATCCGACTG</td><td>>hg38_21_+_8433172_8446622_RNA45SN1_45S_50ntFlanks</td><td>11578</td><td>11609</td><td>32</td><td>TAGAACTGGTGCGGACCAGGGGAATCCGACTG</td><td>217.1</td><td>-5.8</td><td>1.1</td><td>-5.2</td><td>2.3E-07</td><td>4.5E-07</td><td>MB231/MB468</td><td>MP</td></tr>
<tr><td>>MI0000809|hsa-mir-151a&WithFlank&8|-|140732558|140732659@51.72.22(+1U)&[MIMAT0000757&hsa-miR-151a-3p&offsets|-2|-1(+1U);m-30&8|-|140732587|140732607&offsets|-2|-1(+1U)]|TACTAGACTGAAGCTCCTTGAGT</td><td>>MI0000809|hsa-mir-151a&WithFlank&8|-|140732558|140732659</td><td>51</td><td>72</td><td>22(+1U)</td><td>TACTAGACTGAAGCTCCTTGAGT</td><td>642.3</td><td>7.4</td><td>1.4</td><td>5.1</td><td>2.7E-07</td><td>4.5E-07</td><td>MB231/MB468</td><td>MITO</td></tr>
<tr><td>>MI0000067|hsa-let-7f-1&WithFlank&9|+|94176341|94176439@13.34.22(+1A);MI0000068|hsa-let-7f-2&WithFlank&X|-|53557186|53557280@14.35.22(+1A)&[MIMAT0000067&hsa-let-7f-5p&offsets|0|0(+1A);m-20&9|+|94176353|94176374&offsets|0|0(+1A)];[MIMAT0000067_1&hsa-let-7f-5p&offsets|0|0(+1A);m-19&X|-|53557246|53557267&offsets|0|0(+1A)]|TGAGGTAGTAGATTGTATAGTTA</td><td>>MI0000067|hsa-let-7f-1&WithFlank&9|+|94176341|94176439</td><td>13</td><td>34</td><td>22(+1A)</td><td>TGAGGTAGTAGATTGTATAGTTA</td><td>636.6</td><td>3.6</td><td>0.7</td><td>5.2</td><td>2.5E-07</td><td>4.5E-07</td><td>MB231/MB468</td><td>CYTO</td></tr>
<tr><td>>hg38_21_+_8433172_8446622_RNA45SN1_45S_50ntFlanks@12944.12988.45|CGTAGACGACCTGCTTCTGGGTCGGGGTTTCGTACGTAGCAGAGC</td><td>>hg38_21_+_8433172_8446622_RNA45SN1_45S_50ntFlanks</td><td>12944</td><td>12988</td><td>45</td><td>CGTAGACGACCTGCTTCTGGGTCGGGGTTTCGTACGTAGCAGAGC</td><td>198.4</td><td>3.6</td><td>0.7</td><td>5.2</td><td>2.5E-07</td><td>4.5E-07</td><td>MB231/MB468</td><td>CYTO</td></tr>
<tr><td>>hg38_21_+_8433172_8446622_RNA45SN1_45S_50ntFlanks@10347.10368.22|CAAACGAGAACTTTGAAGGCCG</td><td>>hg38_21_+_8433172_8446622_RNA45SN1_45S_50ntFlanks</td><td>10347</td><td>10368</td><td>22</td><td>CAAACGAGAACTTTGAAGGCCG</td><td>765.4</td><td>-6.5</td><td>1.3</td><td>-5.2</td><td>2.3E-07</td><td>4.5E-07</td><td>MB231/MB468</td><td>MP</td></tr>
<tr><td>>hg38_21_+_8433172_8446622_RNA45SN1_45S_50ntFlanks@12632.12661.30|GCGAAGCTACCATCTGTGGGATTATGACTG</td><td>>hg38_21_+_8433172_8446622_RNA45SN1_45S_50ntFlanks</td><td>12632</td><td>12661</td><td>30</td><td>GCGAAGCTACCATCTGTGGGATTATGACTG</td><td>3385.0</td><td>-6.6</td><td>1.3</td><td>-5.1</td><td>2.7E-07</td><td>4.5E-07</td><td>MB231/MB468</td><td>MITO</td></tr>
<tr><td>>hg38_21_+_8433172_8446622_RNA45SN1_45S_50ntFlanks@7993.8014.22|GTGGCGACCCGCTGAATTTAAG</td><td>>hg38_21_+_8433172_8446622_RNA45SN1_45S_50ntFlanks</td><td>7993</td><td>8014</td><td>22</td><td>GTGGCGACCCGCTGAATTTAAG</td><td>342.7</td><td>-4.2</td><td>0.8</td><td>-5.1</td><td>2.7E-07</td><td>4.5E-07</td><td>MB231/MB468</td><td>MITO</td></tr>
<tr><td>>hg38_21_+_8433172_8446622_RNA45SN1_45S_50ntFlanks@11769.11804.36|TAATTAGTGACGCGCATGAATGGATGAACGAGATTC</td><td>>hg38_21_+_8433172_8446622_RNA45SN1_45S_50ntFlanks</td><td>11769</td><td>11804</td><td>36</td><td>TAATTAGTGACGCGCATGAATGGATGAACGAGATTC</td><td>343.9</td><td>-5.7</td><td>1.1</td><td>-5.1</td><td>2.7E-07</td><td>4.5E-07</td><td>MB231/MB468</td><td>MITO</td></tr>
<tr><td>>hg38_21_+_8433172_8446622_RNA45SN1_45S_50ntFlanks@4919.4939.21|CCACCAGGAGTGGAGCCTGCG</td><td>>hg38_21_+_8433172_8446622_RNA45SN1_45S_50ntFlanks</td><td>4919</td><td>4939</td><td>21</td><td>CCACCAGGAGTGGAGCCTGCG</td><td>231.4</td><td>-9.9</td><td>1.9</td><td>-5.2</td><td>2.0E-07</td><td>4.5E-07</td><td>BT20/MB468</td><td>MITO</td></tr>
<tr><td>>hg38_21_+_8433172_8446622_RNA45SN1_45S_50ntFlanks@9574.9595.22|AATCGGTCGTCCGACCTGGGTA</td><td>>hg38_21_+_8433172_8446622_RNA45SN1_45S_50ntFlanks</td><td>9574</td><td>9595</td><td>22</td><td>AATCGGTCGTCCGACCTGGGTA</td><td>1954.5</td><td>-4.7</td><td>0.9</td><td>-5.2</td><td>2.0E-07</td><td>4.5E-07</td><td>BT20/MB468</td><td>MITO</td></tr>
<tr><td>>hg38_21_+_8433172_8446622_RNA45SN1_45S_50ntFlanks@8196.8214.19|CGTGGACGGTGTGAGGCCG</td><td>>hg38_21_+_8433172_8446622_RNA45SN1_45S_50ntFlanks</td><td>8196</td><td>8214</td><td>19</td><td>CGTGGACGGTGTGAGGCCG</td><td>2383.6</td><td>-3.3</td><td>0.6</td><td>-5.2</td><td>2.2E-07</td><td>4.5E-07</td><td>BT20/MB468</td><td>CYTO</td></tr>
<tr><td>>hg38_21_+_8433172_8446622_RNA45SN1_45S_50ntFlanks@12420.12441.22|ATGTCGGCTCTTCCTATCATTG</td><td>>hg38_21_+_8433172_8446622_RNA45SN1_45S_50ntFlanks</td><td>12420</td><td>12441</td><td>22</td><td>ATGTCGGCTCTTCCTATCATTG</td><td>971.1</td><td>-3.1</td><td>0.6</td><td>-5.2</td><td>2.0E-07</td><td>4.5E-07</td><td>BT20/MB468</td><td>MITO</td></tr>
<tr><td>>hg38_21_+_8433172_8446622_RNA45SN1_45S_50ntFlanks@10240.10279.40|CTACGCCGCGACGAGTAGGAGGGCCGCTGCGGTGAGCCTT</td><td>>hg38_21_+_8433172_8446622_RNA45SN1_45S_50ntFlanks</td><td>10240</td><td>10279</td><td>40</td><td>CTACGCCGCGACGAGTAGGAGGGCCGCTGCGGTGAGCCTT</td><td>117.2</td><td>10.6</td><td>2.0</td><td>5.4</td><td>5.7E-08</td><td>4.6E-07</td><td>BT20/MB468</td><td>MP</td></tr>
<tr><td>>hg38_21_+_8433172_8446622_RNA45SN1_45S_50ntFlanks@9720.9748.29|ATTAGAGGTCTTGGGGCCGAAACGATCTC</td><td>>hg38_21_+_8433172_8446622_RNA45SN1_45S_50ntFlanks</td><td>9720</td><td>9748</td><td>29</td><td>ATTAGAGGTCTTGGGGCCGAAACGATCTC</td><td>86.9</td><td>-9.5</td><td>1.8</td><td>-5.4</td><td>5.7E-08</td><td>4.6E-07</td><td>BT20/MB468</td><td>MP</td></tr>
<tr><td>>MI0001446|hsa-mir-424&WithFlank&X|-|134546608|134546717@54.74.21&[MIMAT0004749&hsa-miR-424-3p&offsets|0|0;m-305&X|-|134546644|134546664&offsets|0|0]|CAAAACGTGAGGCGCTGCTAT</td><td>>MI0001446|hsa-mir-424&WithFlank&X|-|134546608|134546717</td><td>54</td><td>74</td><td>21</td><td>CAAAACGTGAGGCGCTGCTAT</td><td>428.0</td><td>8.2</td><td>1.5</td><td>5.4</td><td>5.7E-08</td><td>4.6E-07</td><td>BT20/MB468</td><td>MP</td></tr>
<tr><td>>hg38_21_+_8433172_8446622_RNA45SN1_45S_50ntFlanks@11786.11806.21|GAATGGATGAACGAGATTCCC</td><td>>hg38_21_+_8433172_8446622_RNA45SN1_45S_50ntFlanks</td><td>11786</td><td>11806</td><td>21</td><td>GAATGGATGAACGAGATTCCC</td><td>99.1</td><td>-9.7</td><td>1.8</td><td>-5.4</td><td>5.7E-08</td><td>4.6E-07</td><td>BT20/MB468</td><td>MP</td></tr>
<tr><td>>hg38_21_+_8433172_8446622_RNA45SN1_45S_50ntFlanks@11789.11806.18|TGGATGAACGAGATTCCC</td><td>>hg38_21_+_8433172_8446622_RNA45SN1_45S_50ntFlanks</td><td>11789</td><td>11806</td><td>18</td><td>TGGATGAACGAGATTCCC</td><td>105.2</td><td>-9.8</td><td>1.8</td><td>-5.4</td><td>5.7E-08</td><td>4.6E-07</td><td>BT20/MB468</td><td>MP</td></tr>
<tr><td>>hg38_21_+_8433172_8446622_RNA45SN1_45S_50ntFlanks@10922.10947.26|CTGGACGAGGCGCCGCCGCCCCCCCC</td><td>>hg38_21_+_8433172_8446622_RNA45SN1_45S_50ntFlanks</td><td>10922</td><td>10947</td><td>26</td><td>CTGGACGAGGCGCCGCCGCCCCCCCC</td><td>244.8</td><td>-7.0</td><td>1.4</td><td>-5.1</td><td>2.6E-07</td><td>4.6E-07</td><td>MB231/MB468</td><td>NUC</td></tr>
<tr><td>>hg38_21_+_8433172_8446622_RNA45SN1_45S_50ntFlanks@6759.6806.48|CTTGCGGCCCCGGGTTCCTCCCGGGGCTACGCCTGTCTGAGCGTCGCT</td><td>>hg38_21_+_8433172_8446622_RNA45SN1_45S_50ntFlanks</td><td>6759</td><td>6806</td><td>48</td><td>CTTGCGGCCCCGGGTTCCTCCCGGGGCTACGCCTGTCTGAGCGTCGCT</td><td>816.9</td><td>6.5</td><td>1.3</td><td>5.2</td><td>2.6E-07</td><td>4.6E-07</td><td>MB231/MB468</td><td>CYTO</td></tr>
<tr><td>>hg38_21_+_8433172_8446622_RNA45SN1_45S_50ntFlanks@10677.10711.35|GCGGTTCCGGCGGCGTCCGGTGAGCTCTCGCTGGC</td><td>>hg38_21_+_8433172_8446622_RNA45SN1_45S_50ntFlanks</td><td>10677</td><td>10711</td><td>35</td><td>GCGGTTCCGGCGGCGTCCGGTGAGCTCTCGCTGGC</td><td>511.6</td><td>-2.9</td><td>0.6</td><td>-5.2</td><td>2.6E-07</td><td>4.6E-07</td><td>MB231/MB468</td><td>CYTO</td></tr>
<tr><td>>hg38_21_+_8433172_8446622_RNA45SN1_45S_50ntFlanks@10408.10427.20|GGTCAGTCGGTCCTGAGAGA</td><td>>hg38_21_+_8433172_8446622_RNA45SN1_45S_50ntFlanks</td><td>10408</td><td>10427</td><td>20</td><td>GGTCAGTCGGTCCTGAGAGA</td><td>169.5</td><td>-4.2</td><td>0.8</td><td>-5.2</td><td>2.6E-07</td><td>4.6E-07</td><td>MB231/MB468</td><td>CYTO</td></tr>
<tr><td>>hg38_21_+_8433172_8446622_RNA45SN1_45S_50ntFlanks@11569.11607.39|CAGCCGACTTAGAACTGGTGCGGACCAGGGGAATCCGAC</td><td>>hg38_21_+_8433172_8446622_RNA45SN1_45S_50ntFlanks</td><td>11569</td><td>11607</td><td>39</td><td>CAGCCGACTTAGAACTGGTGCGGACCAGGGGAATCCGAC</td><td>463.1</td><td>-5.8</td><td>1.1</td><td>-5.1</td><td>2.8E-07</td><td>4.6E-07</td><td>MB231/MB468</td><td>MITO</td></tr>
<tr><td>>hg38_21_+_8433172_8446622_RNA45SN1_45S_50ntFlanks@3854.3873.20|ACTGTGGTAATTCTAGAGCT</td><td>>hg38_21_+_8433172_8446622_RNA45SN1_45S_50ntFlanks</td><td>3854</td><td>3873</td><td>20</td><td>ACTGTGGTAATTCTAGAGCT</td><td>677.7</td><td>-5.2</td><td>1.0</td><td>-5.1</td><td>2.8E-07</td><td>4.6E-07</td><td>MB231/MB468</td><td>MITO</td></tr>
<tr><td>>MI0003776|hsa-mir-320b-1&WithFlank&1|+|116671740|116671823@49.66.18;MI0003839|hsa-mir-320b-2&WithFlank&1|-|224257034|224257116@48.65.18;MI0003778|hsa-mir-320c-1&WithFlank&18|+|21683512|21683595@49.66.18;MI0008191|hsa-mir-320c-2&WithFlank&18|+|24321669|24321752@49.66.18;MI0000542|hsa-mir-320a&WithFlank&8|-|22244960|22245043@49.66.18&[MIMAT0005792&hsa-miR-320b&offsets|+1|-3;m-183&1|+|116671787|116671806&offsets|+1|-1];[MIMAT0005792_1&hsa-miR-320b&offsets|+1|-3;m-183&1|-|224257051|224257070&offsets|+1|-1];[MIMAT0005793&hsa-miR-320c&offsets|+1|-1;m-222&18|+|21683559|21683578&offsets|+1|-1];[MIMAT0005793_1&hsa-miR-320c&offsets|+1|-1;m-219&18|+|24321716|24321735&offsets|+1|-1];[MIMAT0000510&hsa-miR-320a-3p&offsets|+1|-3;m-60&8|-|22244975|22244996&offsets|+1|-3]|AAAGCTGGGTTGAGAGGG</td><td>>MI0003776|hsa-mir-320b-1&WithFlank&1|+|116671740|116671823</td><td>49</td><td>66</td><td>18</td><td>AAAGCTGGGTTGAGAGGG</td><td>94.2</td><td>10.3</td><td>1.9</td><td>5.4</td><td>5.7E-08</td><td>4.6E-07</td><td>BT20/MB468</td><td>MP</td></tr>
<tr><td>>trnaMT_HisGTG_MT_+_12138_12206@26.60.35|AGATTGTGAATCTGACAACAGAGGCTTACGACCCC</td><td>>trnaMT_HisGTG_MT_+_12138_12206</td><td>26</td><td>60</td><td>35</td><td>AGATTGTGAATCTGACAACAGAGGCTTACGACCCC</td><td>99.0</td><td>10.4</td><td>1.9</td><td>5.4</td><td>5.7E-08</td><td>4.6E-07</td><td>BT20/MB468</td><td>MP</td></tr>
<tr><td>>hg38_21_+_8433172_8446622_RNA45SN1_45S_50ntFlanks@10905.10941.37|CTGGGCGCGCGCCGCGGCTGGACGAGGCGCCGCCGCC</td><td>>hg38_21_+_8433172_8446622_RNA45SN1_45S_50ntFlanks</td><td>10905</td><td>10941</td><td>37</td><td>CTGGGCGCGCGCCGCGGCTGGACGAGGCGCCGCCGCC</td><td>86.0</td><td>-9.5</td><td>1.8</td><td>-5.4</td><td>5.7E-08</td><td>4.6E-07</td><td>BT20/MB468</td><td>MP</td></tr>
</table>

| | | | | | | | | | | | | | |
|---|---|---|---|---|---|---|---|---|---|---|---|---|---|
| >hg38_21_+_8433172_8446622_RNA45SN1_45S_50ntFlanks@7979.8009.31\|ACCTCAGATCAGACGTGGCGACCCGCTGAAT | >hg38_21_+_8433172_8446622_RNA45SN1_45S_50ntFlanks | 7979 | 8009 | 31 | ACCTCAGATCAGACGTGGCGACCCGCTGAAT | 757.1 | -2.0 | 0.4 | -5.1 | 2.9E-07 | 4.6E-07 | BT20/MB468 | NUC |
| >MI0000301\|hsa-mir-224&WithFlank&X\|-\|151958572\|151958664@14.33.20(+1C)&[MIMAT0000281&hsa-miR-224-5p&offsets\|+1\|-4(+1C);m-238&X\|-\|151958630\|151958651&offsets\|0\|-2(+1C)]\|CAAGTCACTAGTGGTTCCGTC | >MI0000301\|hsa-mir-224&WithFlank&X\|-\|151958572\|151958664 | 14 | 33 | 20(+1C) | CAAGTCACTAGTGGTTCCGTC | 86.2 | 10.2 | 1.9 | 5.4 | 5.7E-08 | 4.6E-07 | BT20/MB468 | MP |
| >hg38_21_+_8433172_8446622_RNA45SN1_45S_50ntFlanks@9535.9553.19\|GGAAACTCTGGTGGAGGTC | >hg38_21_+_8433172_8446622_RNA45SN1_45S_50ntFlanks | 9535 | 9553 | 19 | GGAAACTCTGGTGGAGGTC | 424.5 | -6.3 | 1.2 | -5.2 | 2.6E-07 | 4.6E-07 | MB231/MB468 | CYTO |
| >trnaMT_GluTTC_MT_-_14674_14742@39.63.25,trnalookalike8_GluTTC_5_-_93905172_93905240@39.63.25\|ATTGGTCGTGGTTGTAGTCCGTGCG | >trnaMT_GluTTC_MT_-_14674_14742 | 39 | 63 | 25 | ATTGGTCGTGGTTGTAGTCCGTGCG | 1069.4 | 5.8 | 1.1 | 5.4 | 5.7E-08 | 4.6E-07 | BT20/MB468 | MP |
| >hg38_21_+_8433172_8446622_RNA45SN1_45S_50ntFlanks@12607.12624.18\|TGTGCTTGGCTGAGGAGC | >hg38_21_+_8433172_8446622_RNA45SN1_45S_50ntFlanks | 12607 | 12624 | 18 | TGTGCTTGGCTGAGGAGC | 427.1 | 3.1 | 0.6 | 5.2 | 2.3E-07 | 4.6E-07 | BT20/MB468 | CYTO |
| >MI0000542\|hsa-mir-320a&WithFlank&8\|-\|22244960\|22245043@48.69.22(+1G)&[MIMAT0000510&hsa-miR-320a-3p&offsets\|0\|0(+1G);m-60&8\|-\|22244975\|22244996&offsets\|0\|0(+1G)]\|AAAAGCTGGGTTGAGAGGGCGAG | >MI0000542\|hsa-mir-320a&WithFlank&8\|-\|22244960\|22245043 | 48 | 69 | 22(+1G) | AAAAGCTGGGTTGAGAGGGCGAG | 1912.7 | 4.7 | 0.8 | 5.6 | 1.7E-08 | 4.6E-07 | BT20/MB231 | CYTO |
| >MI0000102\|hsa-mir-100&WithFlank&11\|-\|122152223\|122152314@21.40.20&[MIMAT0000098&hsa-miR-100-5p&offsets\|+2\|0;m-27&11\|-\|122152276\|122152296&offsets\|+2\|+1]\|CCCGTAGATCCGAACTTGTG | >MI0000102\|hsa-mir-100&WithFlank&11\|-\|122152223\|122152314 | 21 | 40 | 20 | CCCGTAGATCCGAACTTGTG | 247.0 | -5.8 | 1.0 | -5.6 | 1.7E-08 | 4.6E-07 | BT20/MB231 | CYTO |
| >MI0000102\|hsa-mir-100&WithFlank&11\|-\|122152223\|122152314@20.41.22&[MIMAT0000098&hsa-miR-100-5p&offsets\|+1\|+1;m-27&11\|-\|122152276\|122152296&offsets\|+1\|+2]\|ACCCGTAGATCCGAACTTGTGG | >MI0000102\|hsa-mir-100&WithFlank&11\|-\|122152223\|122152314 | 20 | 41 | 22 | ACCCGTAGATCCGAACTTGTGG | 136.6 | -10.5 | 1.8 | -5.7 | 1.2E-08 | 4.6E-07 | BT20/MB231 | MP |
| >MI0000289\|hsa-mir-181a-1&WithFlank&1\|-\|198859038\|198859159@30.54.25(+1U);MI0000269\|hsa-mir-181a-2&WithFlank&9\|+\|124692436\|124692557@45.69.25(+1U)&[MIMAT0000256&hsa-miR-181a-5p&offsets\|0\|+2(+1U);m-32&1\|-\|198859108\|198859130&offsets\|0\|+2(+1U)];[MIMAT0000256_1&hsa-miR-181a-5p&offsets\|0\|+2(+1U);m-32&9\|+\|124692480\|124692502&offsets\|0\|+2(+1U)]\|AACATTCAACGCTGTCGGTGAGTTTT | >MI0000289\|hsa-mir-181a-1&WithFlank&1\|-\|198859038\|198859159 | 30 | 54 | 25(+1U) | AACATTCAACGCTGTCGGTGAGTTTT | 1423.5 | -7.3 | 1.3 | -5.7 | 1.2E-08 | 4.6E-07 | BT20/MB231 | MP |
| >hg38_21_+_8433172_8446622_RNA45SN1_45S_50ntFlanks@12434.12455.22\|TATCATTGTGAAGCAGAATTCA | >hg38_21_+_8433172_8446622_RNA45SN1_45S_50ntFlanks | 12434 | 12455 | 22 | TATCATTGTGAAGCAGAATTCA | 333.9 | -5.5 | 1.1 | -5.1 | 2.7E-07 | 4.6E-07 | MB231/MB468 | NUC |
| >hg38_21_+_8433172_8446622_RNA45SN1_45S_50ntFlanks@8176.8204.29\|CTTCTGATCGAGGCCCAGCCCGTGGACGG | >hg38_21_+_8433172_8446622_RNA45SN1_45S_50ntFlanks | 8176 | 8204 | 29 | CTTCTGATCGAGGCCCAGCCCGTGGACGG | 1100.4 | -2.9 | 0.6 | -5.2 | 2.6E-07 | 4.6E-07 | MB231/MB468 | CYTO |
| >hg38_21_+_8433172_8446622_RNA45SN1_45S_50ntFlanks@7991.8014.24\|ACGTGGCGACCCGCTGAATTTAAG | >hg38_21_+_8433172_8446622_RNA45SN1_45S_50ntFlanks | 7991 | 8014 | 24 | ACGTGGCGACCCGCTGAATTTAAG | 831.0 | -5.1 | 1.0 | -5.1 | 2.9E-07 | 4.6E-07 | BT20/MB468 | NUC |
| >hg38_21_+_8433172_8446622_RNA45SN1_45S_50ntFlanks@476.504.29\|TTCGTGATCGATGTGGTGACGTCGTGCTC | >hg38_21_+_8433172_8446622_RNA45SN1_45S_50ntFlanks | 476 | 504 | 29 | TTCGTGATCGATGTGGTGACGTCGTGCTC | 872.3 | 5.5 | 1.1 | 5.1 | 2.8E-07 | 4.6E-07 | MB231/MB468 | MITO |
| >MI0000066\|hsa-let-7e&WithFlank&19\|+\|51692780\|51692870@14.31.18&[MIMAT0000066&hsa-let-7e-5p&offsets\|0\|-4;m-50&19\|+\|51692793\|51692814&offsets\|0\|-4]\|TGAGGTAGGAGGTTGTAT | >MI0000066\|hsa-let-7e&WithFlank&19\|+\|51692780\|51692870 | 14 | 31 | 18 | TGAGGTAGGAGGTTGTAT | 562.5 | 7.5 | 1.5 | 5.2 | 2.6E-07 | 4.6E-07 | MB231/MB468 | CYTO |
| >hg38_21_+_8433172_8446622_RNA45SN1_45S_50ntFlanks@10814.10840.27\|AGGTAAGGGAAGTCGGCAAGCCGGATC | >hg38_21_+_8433172_8446622_RNA45SN1_45S_50ntFlanks | 10814 | 10840 | 27 | AGGTAAGGGAAGTCGGCAAGCCGGATC | 671.0 | 12.7 | 2.2 | 5.7 | 1.2E-08 | 4.6E-07 | BT20/MB231 | MP |
| >hg38_MT_+_598_1651_RNR1_12S_50ntFlanks@570.611.42\|AAGGACCTGGCGGTGCTTCATATCCCTCTAGAGGAGCCTGTT | >hg38_MT_+_598_1651_RNR1_12S_50ntFlanks | 570 | 611 | 42 | AAGGACCTGGCGGTGCTTCATATCCCTCTAGAGGAGCCTGTT | 131.0 | -10.4 | 1.8 | -5.7 | 1.2E-08 | 4.6E-07 | BT20/MB231 | MP |
| >hg38_21_+_8433172_8446622_RNA45SN1_45S_50ntFlanks@6732.6753.22\|ACATTGATCATCGACACTTCGA | >hg38_21_+_8433172_8446622_RNA45SN1_45S_50ntFlanks | 6732 | 6753 | 22 | ACATTGATCATCGACACTTCGA | 392.3 | -3.1 | 0.6 | -5.1 | 2.9E-07 | 4.6E-07 | BT20/MB468 | NUC |
| >hg38_21_+_8433172_8446622_RNA45SN1_45S_50ntFlanks@5417.5445.29\|CTACCGATTGGATGGTTTAGTGAGGCCCT | >hg38_21_+_8433172_8446622_RNA45SN1_45S_50ntFlanks | 5417 | 5445 | 29 | CTACCGATTGGATGGTTTAGTGAGGCCCT | 580.4 | -4.3 | 0.8 | -5.1 | 2.6E-07 | 4.6E-07 | MB231/MB468 | CYTO |
| >hg38_21_+_8433172_8446622_RNA45SN1_45S_50ntFlanks@12949.12990.42\|ACGACCTGCTTCTGGGTCGGGGTTTCGTACGTAGCAGAGCAG | >hg38_21_+_8433172_8446622_RNA45SN1_45S_50ntFlanks | 12949 | 12990 | 42 | ACGACCTGCTTCTGGGTCGGGGTTTCGTACGTAGCAGAGCAG | 167.7 | -7.0 | 1.4 | -5.2 | 2.3E-07 | 4.6E-07 | BT20/MB468 | CYTO |
| >hg38_21_+_8433172_8446622_RNA45SN1_45S_50ntFlanks@11821.11843.23\|CTATCCAGCGAAACCACAGCCAA | >hg38_21_+_8433172_8446622_RNA45SN1_45S_50ntFlanks | 11821 | 11843 | 23 | CTATCCAGCGAAACCACAGCCAA | 83.0 | -9.5 | 1.7 | -5.4 | 5.8E-08 | 4.7E-07 | BT20/MB468 | MP |
| >hg38_21_+_8433172_8446622_RNA45SN1_45S_50ntFlanks@5168.5185.18\|CAGCCACCCGAGATTGAG | >hg38_21_+_8433172_8446622_RNA45SN1_45S_50ntFlanks | 5168 | 5185 | 18 | CAGCCACCCGAGATTGAG | 242.6 | -10.0 | 1.9 | -5.2 | 2.0E-07 | 4.7E-07 | BT20/MB468 | MITO |
| >hg38_21_+_8433172_8446622_RNA45SN1_45S_50ntFlanks@11316.11344.29\|CACCCCACGTCTCGTCGCGCGCGCGTCCG | >hg38_21_+_8433172_8446622_RNA45SN1_45S_50ntFlanks | 11316 | 11344 | 29 | CACCCCACGTCTCGTCGCGCGCGCGTCCG | 10281.9 | -6.8 | 1.3 | -5.2 | 2.3E-07 | 4.7E-07 | BT20/MB468 | CYTO |
| >trna2_GlyGCC_21_-_18827107_18827177@1.28.28,trna35_GlyGCC_1_+_161413094_161413164@1.28.28,trna37_GlyGCC_1_+_161420467_161420537@1.28.28,trna39_GlyGCC_1_+_161427898_161427968@1.28.28,trna41_GlyGCC_1_+_161435258_161435328@1.28.28\|GCATGGGTGGTTCAGTGGTAGAATTCTC | >trna2_GlyGCC_21_-_18827107_18827177 | 1 | 28 | 28 | GCATGGGTGGTTCAGTGGTAGAATTCTC | 48727.4 | -6.6 | 1.3 | -5.1 | 3.0E-07 | 4.7E-07 | BT20/MB468 | NUC |
| >trnaMT_MetCAT_MT_+_4402_4469@38.70.33\|CGAAAATGTTGGTTATACCCTTCCCGTACTACC | >trnaMT_MetCAT_MT_+_4402_4469 | 38 | 70 | 33 | CGAAAATGTTGGTTATACCCTTCCCGTACTACC | 47.9 | 8.8 | 1.7 | 5.2 | 2.4E-07 | 4.7E-07 | MB231/MB468 | MP |
| >hg38_21_+_8433172_8446622_RNA45SN1_45S_50ntFlanks@12893.12931.39\|CCTCGCCCGTCACGCACCGCACGTTCGTGGGGAACCTGG | >hg38_21_+_8433172_8446622_RNA45SN1_45S_50ntFlanks | 12893 | 12931 | 39 | CCTCGCCCGTCACGCACCGCACGTTCGTGGGGAACCTGG | 92.5 | -9.6 | 1.8 | -5.4 | 5.9E-08 | 4.7E-07 | BT20/MB468 | MP |
| >hg38_21_+_8433172_8446622_RNA45SN1_45S_50ntFlanks@9738.9766.29\|GAAACGATCTCAACCTATTCTCAAACTTT | >hg38_21_+_8433172_8446622_RNA45SN1_45S_50ntFlanks | 9738 | 9766 | 29 | GAAACGATCTCAACCTATTCTCAAACTTT | 513.2 | -6.1 | 1.2 | -5.1 | 3.0E-07 | 4.7E-07 | BT20/MB468 | NUC |
| >hg38_21_+_8433172_8446622_RNA45SN1_45S_50ntFlanks@3853.3887.35\|AACTGTGGTAATTCTAGAGCTAATACATGCCGACG | >hg38_21_+_8433172_8446622_RNA45SN1_45S_50ntFlanks | 3853 | 3887 | 35 | AACTGTGGTAATTCTAGAGCTAATACATGCCGACG | 267.7 | 3.7 | 0.7 | 5.1 | 2.6E-07 | 4.7E-07 | MB231/MB468 | CYTO |
| >hg38_21_+_8433172_8446622_RNA45SN1_45S_50ntFlanks@11209.11253.45\|CGGCGACTCTGGACGCGAGCCGGGCCCTTCCCGTGGATCGCCCCA | >hg38_21_+_8433172_8446622_RNA45SN1_45S_50ntFlanks | 11209 | 11253 | 45 | CGGCGACTCTGGACGCGAGCCGGGCCCTTCCCGTGGATCGCCCCA | 189.7 | -6.1 | 1.2 | -5.1 | 2.6E-07 | 4.7E-07 | MB231/MB468 | CYTO |
| >hg38_21_+_8433172_8446622_RNA45SN1_45S_50ntFlanks@12259.12285.27\|CTTGATCTTGATTTTCAGTACGAATAC | >hg38_21_+_8433172_8446622_RNA45SN1_45S_50ntFlanks | 12259 | 12285 | 27 | CTTGATCTTGATTTTCAGTACGAATAC | 229.9 | -7.5 | 1.4 | -5.2 | 2.4E-07 | 4.7E-07 | MB231/MB468 | MP |
| >MI0000481\|hsa-mir-184&WithFlank&15\|+\|79209782\|79209877@59.80.22(+1U)&[MIMAT0000454&hsa-miR-184&offsets\|0\|0(+1U);m-460&15\|+\|79209840\|79209861&offsets\|0\|0(+1U)]\|TGGACGGAGAACTGATAAGGGTT | >MI0000481\|hsa-mir-184&WithFlank&15\|+\|79209782\|79209877 | 59 | 80 | 22(+1U) | TGGACGGAGAACTGATAAGGGTT | 98.7 | 10.4 | 1.9 | 5.4 | 5.9E-08 | 4.7E-07 | BT20/MB468 | MP |
| >hg38_21_+_8433172_8446622_RNA45SN1_45S_50ntFlanks@9766.9789.24\|TAAATGGGTAAGAAGCCCGGCTCG | >hg38_21_+_8433172_8446622_RNA45SN1_45S_50ntFlanks | 9766 | 9789 | 24 | TAAATGGGTAAGAAGCCCGGCTCG | 91.2 | -9.6 | 1.8 | -5.4 | 5.9E-08 | 4.7E-07 | BT20/MB468 | MP |
| >MI0003599\|hsa-mir-589&WithFlank&7\|-\|5495813\|5495923@30.51.22(+1U)&[MIMAT0004799&hsa-miR-589-5p&offsets\|0\|0(+1U);m-162&7\|-\|5495874\|5495894&offsets\|0\|+1(+1U)]\|TGAGAACCACGTCTGCTCTGAGT | >MI0003599\|hsa-mir-589&WithFlank&7\|-\|5495813\|5495923 | 30 | 51 | 22(+1U) | TGAGAACCACGTCTGCTCTGAGT | 82.3 | 10.1 | 1.9 | 5.4 | 5.9E-08 | 4.7E-07 | BT20/MB468 | MP |
| >hg38_21_+_8433172_8446622_RNA45SN1_45S_50ntFlanks@6671.6696.26\|CTCGGCTCGTGCGTCGATGAAGAACG | >hg38_21_+_8433172_8446622_RNA45SN1_45S_50ntFlanks | 6671 | 6696 | 26 | CTCGGCTCGTGCGTCGATGAAGAACG | 2544.6 | -4.6 | 0.8 | -5.4 | 5.9E-08 | 4.7E-07 | BT20/MB468 | MP |
| >hg38_21_+_8433172_8446622_RNA45SN1_45S_50ntFlanks@6733.6752.20\|CATTGATCATCGACACTTCG | >hg38_21_+_8433172_8446622_RNA45SN1_45S_50ntFlanks | 6733 | 6752 | 20 | CATTGATCATCGACACTTCG | 687.1 | -3.3 | 0.6 | -5.2 | 2.3E-07 | 4.7E-07 | BT20/MB468 | CYTO |
| >hg38_21_+_8433172_8446622_RNA45SN1_45S_50ntFlanks@4557.4574.18\|CAGCTAGGAATAATGGAA | >hg38_21_+_8433172_8446622_RNA45SN1_45S_50ntFlanks | 4557 | 4574 | 18 | CAGCTAGGAATAATGGAA | 212.3 | -5.4 | 1.0 | -5.2 | 2.4E-07 | 4.7E-07 | MB231/MB468 | MP |
| >hg38_21_+_8433172_8446622_RNA45SN1_45S_50ntFlanks@9910.9928.19\|CCCAGAAAAGGTGTTGGTT | >hg38_21_+_8433172_8446622_RNA45SN1_45S_50ntFlanks | 9910 | 9928 | 19 | CCCAGAAAAGGTGTTGGTT | 96.8 | -9.7 | 1.8 | -5.4 | 5.9E-08 | 4.7E-07 | BT20/MB468 | MP |
| >MI0000077\|hsa-mir-21&WithFlank&17\|+\|59841260\|59841343@14.37.24&[MIMAT0000076&hsa-miR-21-5p&offsets\|0\|+2;m-1&17\|+\|59841273\|59841294&offsets\|0\|+2]\|TAGCTTATCAGACTGATGTTGACT | >MI0000077\|hsa-mir-21&WithFlank&17\|+\|59841260\|59841343 | 14 | 37 | 24 | TAGCTTATCAGACTGATGTTGACT | 40592.9 | 4.0 | 0.8 | 5.2 | 2.4E-07 | 4.7E-07 | MB231/MB468 | MP |
| >hg38_21_+_8433172_8446622_RNA45SN1_45S_50ntFlanks@11200.11225.26\|CGGCGGCGGCGGCGACTCTGGACGCG | >hg38_21_+_8433172_8446622_RNA45SN1_45S_50ntFlanks | 11200 | 11225 | 26 | CGGCGGCGGCGGCGACTCTGGACGCG | 1604.6 | -8.7 | 1.7 | -5.1 | 3.0E-07 | 4.7E-07 | BT20/MB468 | NUC |
| >hg38_21_+_8433172_8446622_RNA45SN1_45S_50ntFlanks@9855.9885.31\|CTGCGGGATGAACCGAACGCCGGGTTAAGGC | >hg38_21_+_8433172_8446622_RNA45SN1_45S_50ntFlanks | 9855 | 9885 | 31 | CTGCGGGATGAACCGAACGCCGGGTTAAGGC | 68.9 | 9.9 | 1.8 | 5.4 | 5.9E-08 | 4.7E-07 | BT20/MB468 | MP |
| >hg38_21_+_8433172_8446622_RNA45SN1_45S_50ntFlanks@5486.5526.41\|GCGCTGAGAAGACGGTCGAACTTGACTATCTAGAGGAAGTA | >hg38_21_+_8433172_8446622_RNA45SN1_45S_50ntFlanks | 5486 | 5526 | 41 | GCGCTGAGAAGACGGTCGAACTTGACTATCTAGAGGAAGTA | 262.3 | -7.4 | 1.4 | -5.1 | 2.7E-07 | 4.7E-07 | MB231/MB468 | NUC |
| >MI0003560\|hsa-mir-92b&WithFlank&1\|+\|155195171\|155195278@67.86.20&[MIMAT0003218&hsa-miR-92b-3p&offsets\|0\|-2;m-115&1\|+\|155195237\|155195258&offsets\|0\|-2]\|TATTGCACTCGTCCCGGCCT | >MI0003560\|hsa-mir-92b&WithFlank&1\|+\|155195171\|155195278 | 67 | 86 | 20 | TATTGCACTCGTCCCGGCCT | 321.3 | 8.3 | 1.6 | 5.2 | 2.4E-07 | 4.7E-07 | MB231/MB468 | MP |
| >hg38_21_+_8433172_8446622_RNA45SN1_45S_50ntFlanks@11553.11591.39\|CTCGGCCGGCGCCTAGCAGCCGACTTAGAACTGGTGCGG | >hg38_21_+_8433172_8446622_RNA45SN1_45S_50ntFlanks | 11553 | 11591 | 39 | CTCGGCCGGCGCCTAGCAGCCGACTTAGAACTGGTGCGG | 313.4 | -5.4 | 1.1 | -5.1 | 2.9E-07 | 4.7E-07 | MB231/MB468 | MITO |
| >hg38_21_+_8433172_8446622_RNA45SN1_45S_50ntFlanks@8176.8208.33\|CTTCTGATCGAGGCCCAGCCCGTGGACGGTGTG | >hg38_21_+_8433172_8446622_RNA45SN1_45S_50ntFlanks | 8176 | 8208 | 33 | CTTCTGATCGAGGCCCAGCCCGTGGACGGTGTG | 2192.5 | -7.2 | 1.4 | -5.1 | 3.0E-07 | 4.7E-07 | BT20/MB468 | NUC |
| >hg38_21_+_8433172_8446622_RNA45SN1_45S_50ntFlanks@9569.9595.27\|GTGCAAATCGGTCGTCCGACCTGGGTA | >hg38_21_+_8433172_8446622_RNA45SN1_45S_50ntFlanks | 9569 | 9595 | 27 | GTGCAAATCGGTCGTCCGACCTGGGTA | 412.9 | -2.7 | 0.5 | -5.1 | 2.7E-07 | 4.7E-07 | MB231/MB468 | CYTO |
| >hg38_21_+_8433172_8446622_RNA45SN1_45S_50ntFlanks@9505.9533.29\|TGAACTATGCCTGGGCAGGGCGAAGCCAG | >hg38_21_+_8433172_8446622_RNA45SN1_45S_50ntFlanks | 9505 | 9533 | 29 | TGAACTATGCCTGGGCAGGGCGAAGCCAG | 108.3 | -9.8 | 1.8 | -5.4 | 6.0E-08 | 4.7E-07 | BT20/MB468 | MP |
| >MI0000760\|hsa-mir-361&WithFlank&X\|-\|85903630\|85903713@12.33.22&[MIMAT0000703&hsa-miR-361-5p&offsets\|0\|0;m-89&X\|-\|85903681\|85903702&offsets\|0\|0]\|TTATCAGAATCTCCAGGGGTAC | >MI0000760\|hsa-mir-361&WithFlank&X\|-\|85903630\|85903713 | 12 | 33 | 22 | TTATCAGAATCTCCAGGGGTAC | 404.5 | 5.3 | 1.0 | 5.1 | 2.9E-07 | 4.7E-07 | MB231/MB468 | MITO |
| >hg38_21_+_8433172_8446622_RNA45SN1_45S_50ntFlanks@11273.11295.23\|CCGCCCCCGGGGAGCCCGGCGGG | >hg38_21_+_8433172_8446622_RNA45SN1_45S_50ntFlanks | 11273 | 11295 | 23 | CCGCCCCCGGGGAGCCCGGCGGG | 113.2 | -9.9 | 1.8 | -5.4 | 6.0E-08 | 4.7E-07 | BT20/MB468 | MP |
| >hg38_21_+_8433172_8446622_RNA45SN1_45S_50ntFlanks@8356.8377.22\|GAAAAGAACTTTGAAGAGAGAG | >hg38_21_+_8433172_8446622_RNA45SN1_45S_50ntFlanks | 8356 | 8377 | 22 | GAAAAGAACTTTGAAGAGAGAG | 642.1 | -4.3 | 0.8 | -5.1 | 2.7E-07 | 4.7E-07 | MB231/MB468 | NUC |
| >hg38_21_+_8433172_8446622_RNA45SN1_45S_50ntFlanks@12002.12031.30\|TGCGGGCCGCCGGTGAAATACCACTACTCT | >hg38_21_+_8433172_8446622_RNA45SN1_45S_50ntFlanks | 12002 | 12031 | 30 | TGCGGGCCGCCGGTGAAATACCACTACTCT | 1082.1 | -4.0 | 0.8 | -5.1 | 2.7E-07 | 4.8E-07 | MB231/MB468 | CYTO |

<table>
<tr><td>>hg38_21_+_8433172_8446622_RNA45SN1_45S_50ntFlanks@5143.5162.20|ACTTCTTAGAGGGACAAGTG</td><td>>hg38_21_+_8433172_8446622_RNA45SN1_45S_50ntFlanks</td><td>5143</td><td>5162</td><td>20</td><td>ACTTCTTAGAGGGACAAGTG</td><td>291.7</td><td>-10.3</td><td>2.0</td><td>-5.2</td><td>2.1E-07</td><td>4.8E-07</td><td>BT20/MB468</td><td>MITO</td></tr>
<tr><td>>trna10_AspGTC_12_-_125424193_125424264@36.75.40,trna12_AspGTC_12_-_125411891_125411962@36.75.40,trna144_AspGTC_6_-_27551236_27551307@36.75.40,trna38_AspGTC_17_-_8125556_8125627@36.75.40,trna45_AspGTC_6_+_27447453_27447524@36.75.40,trna48_AspGTC_6_+_27471523_27471594@36.75.40,trna4_AspGTC_12_+_96429799_96429870@36.75.40,trna69_AspGTC_1_-_161440205_161440276@36.75.40,trna72_AspGTC_1_-_161432824_161432895@36.75.40,trna75_AspGTC_1_-_161425414_161425485@36.75.40,trna78_AspGTC_1_-_161418033_161418104@36.75.40,trna81_AspGTC_1_-_161410615_161410686@36.75.40|CACGCGGGAGACCGGGGTTCGATTCCCCGACGGGGAGCCA</td><td>>trna10_AspGTC_12_-_125424193_125424264</td><td>36</td><td>75</td><td>40</td><td>CACGCGGGAGACCGGGGTTCGATTCCCCGACGGGGAGCCA</td><td>571.1</td><td>5.2</td><td>1.0</td><td>5.1</td><td>2.7E-07</td><td>4.8E-07</td><td>MB231/MB468</td><td>NUC</td></tr>
<tr><td>>hg38_21_+_8433172_8446622_RNA45SN1_45S_50ntFlanks@4005.4042.38|AACCTCGGGCCGATCGCACGCCCCCCGTGGCGGCGACG</td><td>>hg38_21_+_8433172_8446622_RNA45SN1_45S_50ntFlanks</td><td>4005</td><td>4042</td><td>38</td><td>AACCTCGGGCCGATCGCACGCCCCCCGTGGCGGCGACG</td><td>719.8</td><td>-5.8</td><td>1.1</td><td>-5.1</td><td>2.7E-07</td><td>4.8E-07</td><td>MB231/MB468</td><td>NUC</td></tr>
<tr><td>>hg38_21_+_8433172_8446622_RNA45SN1_45S_50ntFlanks@10764.10785.22|TCCGCAGCAGGTCTCCAAGGTG</td><td>>hg38_21_+_8433172_8446622_RNA45SN1_45S_50ntFlanks</td><td>10764</td><td>10785</td><td>22</td><td>TCCGCAGCAGGTCTCCAAGGTG</td><td>68.5</td><td>-9.2</td><td>1.7</td><td>-5.4</td><td>6.0E-08</td><td>4.8E-07</td><td>BT20/MB468</td><td>MP</td></tr>
<tr><td>>hg38_21_+_8433172_8446622_RNA45SN1_45S_50ntFlanks@10423.10457.35|AGAGATGGGCGAGCGCCGTTCCGAAGGGACGGGCG</td><td>>hg38_21_+_8433172_8446622_RNA45SN1_45S_50ntFlanks</td><td>10423</td><td>10457</td><td>35</td><td>AGAGATGGGCGAGCGCCGTTCCGAAGGGACGGGCG</td><td>623.0</td><td>-6.7</td><td>1.3</td><td>-5.2</td><td>2.3E-07</td><td>4.8E-07</td><td>BT20/MB468</td><td>CYTO</td></tr>
<tr><td>>hg38_21_+_8433172_8446622_RNA45SN1_45S_50ntFlanks@11704.11726.23|ATGAAGCGCGGGTAAACGGCGGG</td><td>>hg38_21_+_8433172_8446622_RNA45SN1_45S_50ntFlanks</td><td>11704</td><td>11726</td><td>23</td><td>ATGAAGCGCGGGTAAACGGCGGG</td><td>356.6</td><td>-7.6</td><td>1.5</td><td>-5.1</td><td>3.0E-07</td><td>4.8E-07</td><td>BT20/MB468</td><td>NUC</td></tr>
<tr><td>>hg38_21_+_8433172_8446622_RNA45SN1_45S_50ntFlanks@8351.8375.25|AAGTTGAAAAGAACTTTGAAGAGAG</td><td>>hg38_21_+_8433172_8446622_RNA45SN1_45S_50ntFlanks</td><td>8351</td><td>8375</td><td>25</td><td>AAGTTGAAAAGAACTTTGAAGAGAG</td><td>829.3</td><td>6.4</td><td>1.2</td><td>5.6</td><td>2.2E-08</td><td>4.8E-07</td><td>BT20/MB231</td><td>MITO</td></tr>
<tr><td>>hg38_MT_+_598_1651_RNR1_12S_50ntFlanks@569.612.44|AAAGGACCTGGCGGTGCTTCATATCCCTCTAGAGGAGCCTGTTC</td><td>>hg38_MT_+_598_1651_RNR1_12S_50ntFlanks</td><td>569</td><td>612</td><td>44</td><td>AAAGGACCTGGCGGTGCTTCATATCCCTCTAGAGGAGCCTGTTC</td><td>46.1</td><td>8.7</td><td>1.7</td><td>5.2</td><td>2.4E-07</td><td>4.8E-07</td><td>MB231/MB468</td><td>MP</td></tr>
<tr><td>>hg38_21_+_8433172_8446622_RNA45SN1_45S_50ntFlanks@10247.10281.35|GCGACGAGTAGGAGGGCCGCTGCGGTGAGCCTTGA</td><td>>hg38_21_+_8433172_8446622_RNA45SN1_45S_50ntFlanks</td><td>10247</td><td>10281</td><td>35</td><td>GCGACGAGTAGGAGGGCCGCTGCGGTGAGCCTTGA</td><td>67.6</td><td>9.8</td><td>1.8</td><td>5.4</td><td>6.0E-08</td><td>4.8E-07</td><td>BT20/MB468</td><td>MP</td></tr>
<tr><td>>hg38_21_+_8433172_8446622_RNA45SN1_45S_50ntFlanks@11540.11583.44|CGCGGCGCCTCGCCTCGGCCGGCGCCTAGCAGCCGACTTAGAAC</td><td>>hg38_21_+_8433172_8446622_RNA45SN1_45S_50ntFlanks</td><td>11540</td><td>11583</td><td>44</td><td>CGCGGCGCCTCGCCTCGGCCGGCGCCTAGCAGCCGACTTAGAAC</td><td>596.8</td><td>4.3</td><td>0.8</td><td>5.2</td><td>2.1E-07</td><td>4.8E-07</td><td>BT20/MB468</td><td>MITO</td></tr>
<tr><td>>trna111_HisGTG_1_-_147774845_147774916@-1A.31.32,trna118_HisGTG_1_-_145396881_145396952@-1A.31.32,trna16_HisGTG_1_+_146544773_146544844@-1A.31.32,trna1_HisGTG_15_+_45493349_45493420@-1A.31.32,trna21_HisGTG_1_+_147753471_147753542@-1A.31.32,trna33_HisGTG_6_+_27125906_27125977@-1A.31.32,trna7_HisGTG_9_-_14433938_14434009@-1A.31.32,trna8_HisGTG_15_-_45492611_45492682@-1A.31.32,trna9_HisGTG_15_-_45490804_45490875@-1A.31.32|AGCCGTGATCGTATAGTGGTTAGTACTCTGCG</td><td>>trna111_HisGTG_1_-_147774845_147774916</td><td>-1A</td><td>31</td><td>32</td><td>AGCCGTGATCGTATAGTGGTTAGTACTCTGCG</td><td>1813.2</td><td>-4.3</td><td>0.8</td><td>-5.1</td><td>2.8E-07</td><td>4.8E-07</td><td>MB231/MB468</td><td>NUC</td></tr>
<tr><td>>hg38_21_+_8433172_8446622_RNA45SN1_45S_50ntFlanks@12457.12489.33|CAAGCGTTGGATTGTTCACCCACTAATAGGGAA</td><td>>hg38_21_+_8433172_8446622_RNA45SN1_45S_50ntFlanks</td><td>12457</td><td>12489</td><td>33</td><td>CAAGCGTTGGATTGTTCACCCACTAATAGGGAA</td><td>172.4</td><td>-6.3</td><td>1.2</td><td>-5.1</td><td>2.8E-07</td><td>4.8E-07</td><td>MB231/MB468</td><td>NUC</td></tr>
<tr><td>>trnaMT_SerGCT_MT_+_12207_12265@8.30.23|CTCACAAGAACTGCTAACTCATG</td><td>>trnaMT_SerGCT_MT_+_12207_12265</td><td>8</td><td>30</td><td>23</td><td>CTCACAAGAACTGCTAACTCATG</td><td>314.9</td><td>-7.4</td><td>1.4</td><td>-5.1</td><td>2.9E-07</td><td>4.8E-07</td><td>MB231/MB468</td><td>MITO</td></tr>
<tr><td>>MI0000061|hsa-let-7a-2&WithFlank&11|-|122146516|122146599@11.32.22(+1A);MI0000062|hsa-let-7a-3&WithFlank&22|+|46112743|46112828@10.31.22(+1A);MI0000060|hsa-let-7a-1&WithFlank&9|+|94175951|94176042@12.33.22(+1A)&[MIMAT0000062&hsa-let-7a-5p&offsets|0|0(+1A);m-6&11|-|122146568|122146589&offsets|0|0(+1A)];[MIMAT0000062_1&hsa-let-7a-5p&offsets|0|0(+1A);m-5&22|+|46112752|46112773&offsets|0|0(+1A)];[MIMAT0000062_2&hsa-let-7a-5p&offsets|0|0(+1A);m-4&9|+|94175962|94175983&offsets|0|0(+1A)]|TGAGGTAGTAGGTTGTATAGTTA</td><td>>MI0000061|hsa-let-7a-2&WithFlank&11|-|122146516|122146599</td><td>11</td><td>32</td><td>22(+1A)</td><td>TGAGGTAGTAGGTTGTATAGTTA</td><td>1368.3</td><td>4.2</td><td>0.8</td><td>5.2</td><td>2.4E-07</td><td>4.8E-07</td><td>MB231/MB468</td><td>MP</td></tr>
<tr><td>>hg38_21_+_8433172_8446622_RNA45SN1_45S_50ntFlanks@7988.8008.21|CAGACGTGGCGACCCGCTGAA</td><td>>hg38_21_+_8433172_8446622_RNA45SN1_45S_50ntFlanks</td><td>7988</td><td>8008</td><td>21</td><td>CAGACGTGGCGACCCGCTGAA</td><td>11297.0</td><td>-5.5</td><td>1.0</td><td>-5.4</td><td>6.1E-08</td><td>4.8E-07</td><td>BT20/MB468</td><td>MP</td></tr>
<tr><td>>hg38_21_+_8433172_8446622_RNA45SN1_45S_50ntFlanks@8527.8551.25|CCTTCCCCCGCCGCCCCTCCTCCTC</td><td>>hg38_21_+_8433172_8446622_RNA45SN1_45S_50ntFlanks</td><td>8527</td><td>8551</td><td>25</td><td>CCTTCCCCCGCCGCCCCTCCTCCTC</td><td>96.1</td><td>-9.7</td><td>1.8</td><td>-5.4</td><td>6.1E-08</td><td>4.8E-07</td><td>BT20/MB468</td><td>MP</td></tr>
<tr><td>>hg38_21_+_8433172_8446622_RNA45SN1_45S_50ntFlanks@10348.10382.35|AAACGAGAACTTTGAAGGCCGAAGTGGAGAAGGGT</td><td>>hg38_21_+_8433172_8446622_RNA45SN1_45S_50ntFlanks</td><td>10348</td><td>10382</td><td>35</td><td>AAACGAGAACTTTGAAGGCCGAAGTGGAGAAGGGT</td><td>1210.7</td><td>3.7</td><td>0.7</td><td>5.1</td><td>2.7E-07</td><td>4.8E-07</td><td>MB231/MB468</td><td>CYTO</td></tr>
<tr><td>>hg38_21_+_8433172_8446622_RNA45SN1_45S_50ntFlanks@12419.12451.33|GATGTCGGCTCTTCCTATCATTGTGAAGCAGAA</td><td>>hg38_21_+_8433172_8446622_RNA45SN1_45S_50ntFlanks</td><td>12419</td><td>12451</td><td>33</td><td>GATGTCGGCTCTTCCTATCATTGTGAAGCAGAA</td><td>250.7</td><td>3.2</td><td>0.6</td><td>5.1</td><td>2.7E-07</td><td>4.8E-07</td><td>MB231/MB468</td><td>CYTO</td></tr>
<tr><td>>hg38_21_+_8433172_8446622_RNA45SN1_45S_50ntFlanks@10679.10708.30|GGTTCCGGCGGCGTCCGGTGAGCTCTCGCT</td><td>>hg38_21_+_8433172_8446622_RNA45SN1_45S_50ntFlanks</td><td>10679</td><td>10708</td><td>30</td><td>GGTTCCGGCGGCGTCCGGTGAGCTCTCGCT</td><td>19252.0</td><td>-8.5</td><td>1.7</td><td>-5.1</td><td>3.1E-07</td><td>4.8E-07</td><td>BT20/MB468</td><td>NUC</td></tr>
<tr><td>>hg38_21_+_8433172_8446622_RNA45SN1_45S_50ntFlanks@12933.12965.33|GCTAAACCATTCGTAGACGACCTGCTTCTGGGT</td><td>>hg38_21_+_8433172_8446622_RNA45SN1_45S_50ntFlanks</td><td>12933</td><td>12965</td><td>33</td><td>GCTAAACCATTCGTAGACGACCTGCTTCTGGGT</td><td>659.3</td><td>-8.2</td><td>1.6</td><td>-5.1</td><td>3.1E-07</td><td>4.8E-07</td><td>BT20/MB468</td><td>NUC</td></tr>
<tr><td>>hg38_21_+_8433172_8446622_RNA45SN1_45S_50ntFlanks@7979.8008.30|ACCTCAGATCAGACGTGGCGACCCGCTGAA</td><td>>hg38_21_+_8433172_8446622_RNA45SN1_45S_50ntFlanks</td><td>7979</td><td>8008</td><td>30</td><td>ACCTCAGATCAGACGTGGCGACCCGCTGAA</td><td>1755.5</td><td>-4.4</td><td>0.8</td><td>-5.4</td><td>6.1E-08</td><td>4.8E-07</td><td>BT20/MB468</td><td>MP</td></tr>
<tr><td>>hg38_21_+_8433172_8446622_RNA45SN1_45S_50ntFlanks@8824.8844.21|CGGCGGCGGCGGCGGCGGTGG</td><td>>hg38_21_+_8433172_8446622_RNA45SN1_45S_50ntFlanks</td><td>8824</td><td>8844</td><td>21</td><td>CGGCGGCGGCGGCGGCGGTGG</td><td>92.2</td><td>-9.6</td><td>1.8</td><td>-5.4</td><td>6.1E-08</td><td>4.8E-07</td><td>BT20/MB468</td><td>MP</td></tr>
<tr><td>>MI0000272|hsa-mir-182&WithFlank&7|-|129770377|129770498@29.49.21&[MIMAT0000259&hsa-miR-182-5p&offsets|0|-3;m-22&7|-|129770449|129770470&offsets|0|-1]|TTTGGCAATGGTAGAACTCAC</td><td>>MI0000272|hsa-mir-182&WithFlank&7|-|129770377|129770498</td><td>29</td><td>49</td><td>21</td><td>TTTGGCAATGGTAGAACTCAC</td><td>934.0</td><td>4.9</td><td>0.9</td><td>5.6</td><td>1.8E-08</td><td>4.8E-07</td><td>BT20/MB231</td><td>CYTO</td></tr>
<tr><td>>MI0000483|hsa-mir-186&WithFlank&1|-|71067625|71067722@21.44.24&[MIMAT0000456&hsa-miR-186-5p&offsets|0|+2;m-84&1|-|71067681|71067702&offsets|0|+2]|CAAAGAATTCTCCTTTTGGGCTTT</td><td>>MI0000483|hsa-mir-186&WithFlank&1|-|71067625|71067722</td><td>21</td><td>44</td><td>24</td><td>CAAAGAATTCTCCTTTTGGGCTTT</td><td>549.2</td><td>5.8</td><td>1.1</td><td>5.1</td><td>2.9E-07</td><td>4.8E-07</td><td>MB231/MB468</td><td>MITO</td></tr>
<tr><td>>MI0000065|hsa-let-7d&WithFlank&9|+|94178828|94178926@68.88.21&[MIMAT0004484&hsa-let-7d-3p&offsets|0|-1;m-98&9|+|94178895|94178916&offsets|0|-1]|CTATACGACCTGCTGCCTTTC</td><td>>MI0000065|hsa-let-7d&WithFlank&9|+|94178828|94178926</td><td>68</td><td>88</td><td>21</td><td>CTATACGACCTGCTGCCTTTC</td><td>1130.9</td><td>5.8</td><td>1.1</td><td>5.4</td><td>6.1E-08</td><td>4.8E-07</td><td>BT20/MB468</td><td>MP</td></tr>
<tr><td>>hg38_21_+_8433172_8446622_RNA45SN1_45S_50ntFlanks@12623.12653.31|GCCAATGGGGCGAAGCTACCATCTGTGGGAT</td><td>>hg38_21_+_8433172_8446622_RNA45SN1_45S_50ntFlanks</td><td>12623</td><td>12653</td><td>31</td><td>GCCAATGGGGCGAAGCTACCATCTGTGGGAT</td><td>73.4</td><td>-9.3</td><td>1.7</td><td>-5.4</td><td>6.1E-08</td><td>4.8E-07</td><td>BT20/MB468</td><td>MP</td></tr>
<tr><td>>hg38_21_+_8433172_8446622_RNA45SN1_45S_50ntFlanks@4283.4310.28|CTTTAACGAGGATCCATTGGAGGGCAAG</td><td>>hg38_21_+_8433172_8446622_RNA45SN1_45S_50ntFlanks</td><td>4283</td><td>4310</td><td>28</td><td>CTTTAACGAGGATCCATTGGAGGGCAAG</td><td>70.3</td><td>-9.2</td><td>1.7</td><td>-5.4</td><td>6.1E-08</td><td>4.8E-07</td><td>BT20/MB468</td><td>MP</td></tr>
<tr><td>>MI0003832|hsa-mir-1283-1&WithFlank&19|+|53688475|53688573@21.43.23;MI0006430|hsa-mir-1283-2&WithFlank&19|+|53758226|53758324@21.43.23&[MIMAT0005799&hsa-miR-1283&offsets|+1|+2;m-905&19|+|53688495|53688516&offsets|0|+1];[MIMAT0005799_1&hsa-miR-1283&offsets|+1|+2;m-905&19|+|53758246|53758267&offsets|0|+1]|CTACAAAGGAAAGCGCTTTCTGT</td><td>>MI0003832|hsa-mir-1283-1&WithFlank&19|+|53688475|53688573</td><td>21</td><td>43</td><td>23</td><td>CTACAAAGGAAAGCGCTTTCTGT</td><td>152.6</td><td>-10.7</td><td>1.9</td><td>-5.7</td><td>1.3E-08</td><td>4.8E-07</td><td>BT20/MB231</td><td>MP</td></tr>
<tr><td>>MI0001445|hsa-mir-423&WithFlank&17|+|30117073|30117178@23.44.22(+1C)&[MIMAT0004748&hsa-miR-423-5p&offsets|0|-1(+1C);m-141&17|+|30117095|30117117&offsets|0|-1(+1C)]|TGAGGGGCAGAGAGCGAGACTTC</td><td>>MI0001445|hsa-mir-423&WithFlank&17|+|30117073|30117178</td><td>23</td><td>44</td><td>22(+1C)</td><td>TGAGGGGCAGAGAGCGAGACTTC</td><td>1914.3</td><td>4.8</td><td>0.9</td><td>5.2</td><td>2.5E-07</td><td>4.8E-07</td><td>MB231/MB468</td><td>MP</td></tr>
<tr><td>>hg38_21_+_8433172_8446622_RNA45SN1_45S_50ntFlanks@12463.12487.25|TTGGATTGTTCACCCACTAATAGGG</td><td>>hg38_21_+_8433172_8446622_RNA45SN1_45S_50ntFlanks</td><td>12463</td><td>12487</td><td>25</td><td>TTGGATTGTTCACCCACTAATAGGG</td><td>69.3</td><td>-9.2</td><td>1.7</td><td>-5.4</td><td>6.1E-08</td><td>4.8E-07</td><td>BT20/MB468</td><td>MP</td></tr>
<tr><td>>MI0005566|hsa-mir-374b&WithFlank&X|-|74218541|74218624@17.38.22&[MIMAT0004955&hsa-miR-374b-5p&offsets|0|0;m-153&X|-|74218587|74218608&offsets|0|0]|ATATAATACAACCTGCTAAGTG</td><td>>MI0005566|hsa-mir-374b&WithFlank&X|-|74218541|74218624</td><td>17</td><td>38</td><td>22</td><td>ATATAATACAACCTGCTAAGTG</td><td>2109.3</td><td>5.4</td><td>1.0</td><td>5.1</td><td>2.9E-07</td><td>4.8E-07</td><td>MB231/MB468</td><td>MITO</td></tr>
<tr><td>>hg38_21_+_8433172_8446622_RNA45SN1_45S_50ntFlanks@9556.9573.18|TAGCGGTCCTGACGTGCA</td><td>>hg38_21_+_8433172_8446622_RNA45SN1_45S_50ntFlanks</td><td>9556</td><td>9573</td><td>18</td><td>TAGCGGTCCTGACGTGCA</td><td>360.9</td><td>-3.0</td><td>0.6</td><td>-5.2</td><td>2.4E-07</td><td>4.8E-07</td><td>BT20/MB468</td><td>CYTO</td></tr>
<tr><td>>hg38_21_+_8433172_8446622_RNA45SN1_45S_50ntFlanks@9504.9531.28|GTGAACTATGCCTGGGCAGGGCGAAGCC</td><td>>hg38_21_+_8433172_8446622_RNA45SN1_45S_50ntFlanks</td><td>9504</td><td>9531</td><td>28</td><td>GTGAACTATGCCTGGGCAGGGCGAAGCC</td><td>379.7</td><td>-7.4</td><td>1.4</td><td>-5.2</td><td>2.5E-07</td><td>4.8E-07</td><td>MB231/MB468</td><td>MP</td></tr>
<tr><td>>hg38_21_+_8433172_8446622_RNA45SN1_45S_50ntFlanks@10392.10432.41|ACAGCAGTTGAACATGGGTCAGTCGGTCCTGAGAGATGGGC</td><td>>hg38_21_+_8433172_8446622_RNA45SN1_45S_50ntFlanks</td><td>10392</td><td>10432</td><td>41</td><td>ACAGCAGTTGAACATGGGTCAGTCGGTCCTGAGAGATGGGC</td><td>897.0</td><td>-8.7</td><td>1.7</td><td>-5.1</td><td>3.1E-07</td><td>4.8E-07</td><td>BT20/MB468</td><td>NUC</td></tr>
<tr><td>>hg38_21_+_8433172_8446622_RNA45SN1_45S_50ntFlanks@7985.8005.21|GATCAGACGTGGCGACCCGCT</td><td>>hg38_21_+_8433172_8446622_RNA45SN1_45S_50ntFlanks</td><td>7985</td><td>8005</td><td>21</td><td>GATCAGACGTGGCGACCCGCT</td><td>782.4</td><td>-7.5</td><td>1.5</td><td>-5.2</td><td>2.5E-07</td><td>4.8E-07</td><td>MB231/MB468</td><td>MP</td></tr>
<tr><td>>MI0000088|hsa-mir-30a&WithFlank&6|-|71403545|71403627@53.71.19&[MIMAT0000088&hsa-miR-30a-3p&offsets|0|-3;m-38&6|-|71403554|71403575&offsets|0|-3]|CTTTCAGTCGGATGTTTGC</td><td>>MI0000088|hsa-mir-30a&WithFlank&6|-|71403545|71403627</td><td>53</td><td>71</td><td>19</td><td>CTTTCAGTCGGATGTTTGC</td><td>1107.9</td><td>5.0</td><td>1.0</td><td>5.1</td><td>2.9E-07</td><td>4.9E-07</td><td>MB231/MB468</td><td>MITO</td></tr>
<tr><td>>hg38_21_+_8433172_8446622_RNA45SN1_45S_50ntFlanks@6745.6762.18|ACACTTCGAACGCACTTG</td><td>>hg38_21_+_8433172_8446622_RNA45SN1_45S_50ntFlanks</td><td>6745</td><td>6762</td><td>18</td><td>ACACTTCGAACGCACTTG</td><td>226.3</td><td>-7.0</td><td>1.4</td><td>-5.1</td><td>2.9E-07</td><td>4.9E-07</td><td>MB231/MB468</td><td>MITO</td></tr>
<tr><td>>hg38_21_+_8433172_8446622_RNA45SN1_45S_50ntFlanks@10344.10385.42|ATTCAAACGAGAACTTTGAAGGCCGAAGTGGAGAAGGGTTCC</td><td>>hg38_21_+_8433172_8446622_RNA45SN1_45S_50ntFlanks</td><td>10344</td><td>10385</td><td>42</td><td>ATTCAAACGAGAACTTTGAAGGCCGAAGTGGAGAAGGGTTCC</td><td>69.3</td><td>9.9</td><td>1.8</td><td>5.4</td><td>6.1E-08</td><td>4.9E-07</td><td>BT20/MB468</td><td>MP</td></tr>
<tr><td>>hg38_21_+_8433172_8446622_RNA45SN1_45S_50ntFlanks@12347.12366.20|AGAAAAGTTACCACAGGGAT</td><td>>hg38_21_+_8433172_8446622_RNA45SN1_45S_50ntFlanks</td><td>12347</td><td>12366</td><td>20</td><td>AGAAAAGTTACCACAGGGAT</td><td>90.3</td><td>-9.6</td><td>1.8</td><td>-5.4</td><td>6.1E-08</td><td>4.9E-07</td><td>BT20/MB468</td><td>MP</td></tr>
<tr><td>>hg38_21_+_8433172_8446622_RNA45SN1_45S_50ntFlanks@4733.4757.25|GAACGAAAGTCGGAGGTTCGAAGAC</td><td>>hg38_21_+_8433172_8446622_RNA45SN1_45S_50ntFlanks</td><td>4733</td><td>4757</td><td>25</td><td>GAACGAAAGTCGGAGGTTCGAAGAC</td><td>186.6</td><td>4.8</td><td>0.9</td><td>5.1</td><td>2.7E-07</td><td>4.9E-07</td><td>MB231/MB468</td><td>CYTO</td></tr>
<tr><td>>hg38_21_+_8433172_8446622_RNA45SN1_45S_50ntFlanks@12182.12201.20|CAAACGGTAACGCAGGTGTC</td><td>>hg38_21_+_8433172_8446622_RNA45SN1_45S_50ntFlanks</td><td>12182</td><td>12201</td><td>20</td><td>CAAACGGTAACGCAGGTGTC</td><td>102.9</td><td>-9.8</td><td>1.8</td><td>-5.4</td><td>6.2E-08</td><td>4.9E-07</td><td>BT20/MB468</td><td>MP</td></tr>
</table>

<table>
<tr><td></td><td></td><td></td><td></td><td></td><td></td><td></td><td></td><td></td><td></td><td></td><td></td><td></td><td></td></tr>
<tr><td>>am_MI0000809|hsa-mir-151a&WithFlank&8|-|140732558|140732659@17.38.22&[MIMAT0004697&hsa-miR-151a-5p&offsets|0|+1;m-75&8|-|140732623|140732643&offsets|0|+1]|TCGAGGAGCTCACAGTCTAGTA</td><td>>am_MI0000809|hsa-mir-151a&WithFlank&8|-|140732558|140732659</td><td>17</td><td>38</td><td>22</td><td>TCGAGGAGCTCACAGTCTAGTA</td><td>286.9</td><td>5.1</td><td>1.0</td><td>5.1</td><td>2.9E-07</td><td>4.9E-07</td><td>MB231/MB468</td><td>MITO</td></tr>
<tr><td>>hg38_21_+_8433172_8446622_RNA45SN1_45S_50ntFlanks@5492.5532.41|AGAAGACGGTCGAACTTGACTATCTAGAGGAAGTAAAAGTC</td><td>>hg38_21_+_8433172_8446622_RNA45SN1_45S_50ntFlanks</td><td>5492</td><td>5532</td><td>41</td><td>AGAAGACGGTCGAACTTGACTATCTAGAGGAAGTAAAAGTC</td><td>671.5</td><td>5.2</td><td>1.0</td><td>5.1</td><td>2.7E-07</td><td>4.9E-07</td><td>MB231/MB468</td><td>CYTO</td></tr>
<tr><td>>hg38_21_+_8433172_8446622_RNA45SN1_45S_50ntFlanks@9598.9618.21|GGGGCGAAAGACTAATCGAAC</td><td>>hg38_21_+_8433172_8446622_RNA45SN1_45S_50ntFlanks</td><td>9598</td><td>9618</td><td>21</td><td>GGGGCGAAAGACTAATCGAAC</td><td>189.8</td><td>-7.0</td><td>1.4</td><td>-5.2</td><td>2.5E-07</td><td>4.9E-07</td><td>MB231/MB468</td><td>MP</td></tr>
<tr><td>>hg38_21_+_8433172_8446622_RNA45SN1_45S_50ntFlanks@6733.6758.26|CATTGATCATCGACACTTCGAACGCA</td><td>>hg38_21_+_8433172_8446622_RNA45SN1_45S_50ntFlanks</td><td>6733</td><td>6758</td><td>26</td><td>CATTGATCATCGACACTTCGAACGCA</td><td>646.3</td><td>-6.3</td><td>1.2</td><td>-5.1</td><td>3.1E-07</td><td>4.9E-07</td><td>BT20/MB468</td><td>NUC</td></tr>
<tr><td>>hg38_21_+_8433172_8446622_RNA45SN1_45S_50ntFlanks@8921.8942.22|CCGGGGCCGAGGGAGCGAGACC</td><td>>hg38_21_+_8433172_8446622_RNA45SN1_45S_50ntFlanks</td><td>8921</td><td>8942</td><td>22</td><td>CCGGGGCCGAGGGAGCGAGACC</td><td>76.1</td><td>10.0</td><td>1.8</td><td>5.4</td><td>6.2E-08</td><td>4.9E-07</td><td>BT20/MB468</td><td>MP</td></tr>
<tr><td>>hg38_21_+_8433172_8446622_RNA45SN1_45S_50ntFlanks@10101.10127.27|CGCGCGCGTGTGGTGTGCGTCGGAGGG</td><td>>hg38_21_+_8433172_8446622_RNA45SN1_45S_50ntFlanks</td><td>10101</td><td>10127</td><td>27</td><td>CGCGCGCGTGTGGTGTGCGTCGGAGGG</td><td>694.9</td><td>-7.9</td><td>1.5</td><td>-5.4</td><td>6.2E-08</td><td>4.9E-07</td><td>BT20/MB468</td><td>MP</td></tr>
<tr><td>>MI0000070|hsa-mir-16-1&WithFlank&13|-|50048967|50049067@20.39.20;MI0000115|hsa-mir-16-2&WithFlank&3|+|160404739|160404831@16.35.20&[MIMAT0000069&hsa-miR-16-5p&offsets|0|-2;m-62&13|-|50049027|50049048&offsets|0|-2];[MIMAT0000069_1&hsa-miR-16-5p&offsets|0|-2;m-61&3|+|160404754|160404775&offsets|0|-2]|TAGCAGCACGTAAATATTGG</td><td>>MI0000070|hsa-mir-16-1&WithFlank&13|-|50048967|50049067</td><td>20</td><td>39</td><td>20</td><td>TAGCAGCACGTAAATATTGG</td><td>140.1</td><td>3.4</td><td>0.7</td><td>5.1</td><td>3.1E-07</td><td>4.9E-07</td><td>BT20/MB468</td><td>NUC</td></tr>
<tr><td>>hg38_21_+_8433172_8446622_RNA45SN1_45S_50ntFlanks@9482.9510.29|ACGTGTTAGGACCCGAAAGATGGTGAACT</td><td>>hg38_21_+_8433172_8446622_RNA45SN1_45S_50ntFlanks</td><td>9482</td><td>9510</td><td>29</td><td>ACGTGTTAGGACCCGAAAGATGGTGAACT</td><td>402.1</td><td>12.0</td><td>2.1</td><td>5.7</td><td>1.3E-08</td><td>4.9E-07</td><td>BT20/MB231</td><td>MP</td></tr>
<tr><td>>hg38_21_+_8433172_8446622_RNA45SN1_45S_50ntFlanks@6655.6672.18|TCTTAGCGGTGGATCACT</td><td>>hg38_21_+_8433172_8446622_RNA45SN1_45S_50ntFlanks</td><td>6655</td><td>6672</td><td>18</td><td>TCTTAGCGGTGGATCACT</td><td>883.6</td><td>8.2</td><td>1.6</td><td>5.2</td><td>2.1E-07</td><td>4.9E-07</td><td>BT20/MB468</td><td>MITO</td></tr>
<tr><td>>hg38_21_+_8433172_8446622_RNA45SN1_45S_50ntFlanks@10396.10429.34|CAGTTGAACATGGGTCAGTCGGTCCTGAGAGATG</td><td>>hg38_21_+_8433172_8446622_RNA45SN1_45S_50ntFlanks</td><td>10396</td><td>10429</td><td>34</td><td>CAGTTGAACATGGGTCAGTCGGTCCTGAGAGATG</td><td>385.8</td><td>-5.7</td><td>1.1</td><td>-5.2</td><td>2.5E-07</td><td>4.9E-07</td><td>MB231/MB468</td><td>MP</td></tr>
<tr><td>>hg38_21_+_8433172_8446622_RNA45SN1_45S_50ntFlanks@476.511.36|TTCGTGATCGATGTGGTGACGTCGTGCTCTCCCGGG</td><td>>hg38_21_+_8433172_8446622_RNA45SN1_45S_50ntFlanks</td><td>476</td><td>511</td><td>36</td><td>TTCGTGATCGATGTGGTGACGTCGTGCTCTCCCGGG</td><td>741.5</td><td>7.7</td><td>1.5</td><td>5.2</td><td>2.1E-07</td><td>4.9E-07</td><td>BT20/MB468</td><td>MITO</td></tr>
<tr><td>>hg38_21_+_8433172_8446622_RNA45SN1_45S_50ntFlanks@12081.12106.26|CTCTCGCTTCTGGCGCCAAGCGCCCG</td><td>>hg38_21_+_8433172_8446622_RNA45SN1_45S_50ntFlanks</td><td>12081</td><td>12106</td><td>26</td><td>CTCTCGCTTCTGGCGCCAAGCGCCCG</td><td>242.4</td><td>-5.3</td><td>1.0</td><td>-5.1</td><td>2.8E-07</td><td>4.9E-07</td><td>MB231/MB468</td><td>NUC</td></tr>
<tr><td>>hg38_21_+_8433172_8446622_RNA45SN1_45S_50ntFlanks@5084.5115.32|CGAACGAGACTCTGGCATGCTAACTAGTTACG</td><td>>hg38_21_+_8433172_8446622_RNA45SN1_45S_50ntFlanks</td><td>5084</td><td>5115</td><td>32</td><td>CGAACGAGACTCTGGCATGCTAACTAGTTACG</td><td>785.1</td><td>-7.0</td><td>1.4</td><td>-5.1</td><td>3.1E-07</td><td>4.9E-07</td><td>BT20/MB468</td><td>NUC</td></tr>
<tr><td>>hg38_21_+_8433172_8446622_RNA45SN1_45S_50ntFlanks@11208.11252.45|GCGGCGACTCTGGACGCGAGCCGGGCCCTTCCCGTGGATCGCCCC</td><td>>hg38_21_+_8433172_8446622_RNA45SN1_45S_50ntFlanks</td><td>11208</td><td>11252</td><td>45</td><td>GCGGCGACTCTGGACGCGAGCCGGGCCCTTCCCGTGGATCGCCCC</td><td>4359.2</td><td>-7.7</td><td>1.5</td><td>-5.1</td><td>3.1E-07</td><td>4.9E-07</td><td>BT20/MB468</td><td>NUC</td></tr>
<tr><td>>hg38_21_+_8433172_8446622_RNA45SN1_45S_50ntFlanks@11571.11591.21|GCCGACTTAGAACTGGTGCGG</td><td>>hg38_21_+_8433172_8446622_RNA45SN1_45S_50ntFlanks</td><td>11571</td><td>11591</td><td>21</td><td>GCCGACTTAGAACTGGTGCGG</td><td>635.1</td><td>-6.1</td><td>1.2</td><td>-5.2</td><td>2.5E-07</td><td>4.9E-07</td><td>MB231/MB468</td><td>MP</td></tr>
<tr><td>>hg38_21_+_8433172_8446622_RNA45SN1_45S_50ntFlanks@12478.12496.19|ACTAATAGGGAACGTGAGC</td><td>>hg38_21_+_8433172_8446622_RNA45SN1_45S_50ntFlanks</td><td>12478</td><td>12496</td><td>19</td><td>ACTAATAGGGAACGTGAGC</td><td>2095.1</td><td>3.3</td><td>0.6</td><td>5.2</td><td>2.4E-07</td><td>4.9E-07</td><td>BT20/MB468</td><td>CYTO</td></tr>
<tr><td>>am_trna128_GlyGCC_6_-_27870686_27870756@1.33.33,trna18_GlyGCC_16_+_70822597_70822667@1.33.33,trna19_GlyGCC_16_+_70823410_70823480@1.33.33,trna19_GlyGCC_2_-_157257659_157257729@1.33.33,trna24_GlyGCC_16_-_70812942_70813012@1.33.33,trna25_GlyGCC_16_-_70812114_70812184@1.33.33,trna5_GlyGCC_17_+_8029064_8029134@1.33.33,trna68_GlyGCC_1_-_161493637_161493707@1.33.33|GCATTGGTGGTTCAGTGGTAGAATTCTCGCCTG</td><td>>am_trna128_GlyGCC_6_-_27870686_27870756</td><td>1</td><td>33</td><td>33</td><td>GCATTGGTGGTTCAGTGGTAGAATTCTCGCCTG</td><td>3612.5</td><td>4.6</td><td>0.9</td><td>5.1</td><td>3.0E-07</td><td>4.9E-07</td><td>MB231/MB468</td><td>MITO</td></tr>
<tr><td>>hg38_21_+_8433172_8446622_RNA45SN1_45S_50ntFlanks@12076.12095.20|AGGGGCTCTCGCTTCTGGCG</td><td>>hg38_21_+_8433172_8446622_RNA45SN1_45S_50ntFlanks</td><td>12076</td><td>12095</td><td>20</td><td>AGGGGCTCTCGCTTCTGGCG</td><td>9741.3</td><td>-4.0</td><td>0.8</td><td>-5.1</td><td>2.8E-07</td><td>4.9E-07</td><td>MB231/MB468</td><td>CYTO</td></tr>
<tr><td>>MI0000272|hsa-mir-182&WithFlank&7|-|129770377|129770498@29.50.22(+1U)&[MIMAT0000259&hsa-miR-182-5p&offsets|0|-2(+1U);m-22&7|-|129770449|129770470&offsets|0|0(+1U)]|TTTGGCAATGGTAGAACTCACAT</td><td>>MI0000272|hsa-mir-182&WithFlank&7|-|129770377|129770498</td><td>29</td><td>50</td><td>22(+1U)</td><td>TTTGGCAATGGTAGAACTCACAT</td><td>816.9</td><td>5.2</td><td>0.9</td><td>5.6</td><td>1.8E-08</td><td>4.9E-07</td><td>BT20/MB231</td><td>CYTO</td></tr>
<tr><td>>MI0001445|hsa-mir-423&WithFlank&17|+|30117073|30117178@59.80.22(+1A)&[MIMAT0001340&hsa-miR-423-3p&offsets|0|-1(+1A);m-100&17|+|30117131|30117153&offsets|0|-1(+1A)]|AGCTCGGTCTGAGGCCCCTCAGA</td><td>>MI0001445|hsa-mir-423&WithFlank&17|+|30117073|30117178</td><td>59</td><td>80</td><td>22(+1A)</td><td>AGCTCGGTCTGAGGCCCCTCAGA</td><td>445.7</td><td>4.3</td><td>0.8</td><td>5.1</td><td>2.8E-07</td><td>4.9E-07</td><td>MB231/MB468</td><td>NUC</td></tr>
<tr><td>>hg38_21_+_8433172_8446622_RNA45SN1_45S_50ntFlanks@11821.11852.32|CTATCCAGCGAAACCACAGCCAAGGGAACGGG</td><td>>hg38_21_+_8433172_8446622_RNA45SN1_45S_50ntFlanks</td><td>11821</td><td>11852</td><td>32</td><td>CTATCCAGCGAAACCACAGCCAAGGGAACGGG</td><td>66.7</td><td>-9.1</td><td>1.7</td><td>-5.4</td><td>6.2E-08</td><td>4.9E-07</td><td>BT20/MB468</td><td>MP</td></tr>
<tr><td>>hg38_21_+_8433172_8446622_RNA45SN1_45S_50ntFlanks@8353.8373.21|GTTGAAAAGAACTTTGAAGAG</td><td>>hg38_21_+_8433172_8446622_RNA45SN1_45S_50ntFlanks</td><td>8353</td><td>8373</td><td>21</td><td>GTTGAAAAGAACTTTGAAGAG</td><td>1708.0</td><td>9.8</td><td>1.8</td><td>5.4</td><td>6.2E-08</td><td>4.9E-07</td><td>BT20/MB468</td><td>MP</td></tr>
<tr><td>>MI0006273|hsa-mir-1180&WithFlank&17|-|19344500|19344580@47.68.22&[MIMAT0005825&hsa-miR-1180-3p&offsets|0|0;m-214&17|-|19344513|19344534&offsets|0|0]|TTTCCGGCTCGCGTGGGTGTGT</td><td>>MI0006273|hsa-mir-1180&WithFlank&17|-|19344500|19344580</td><td>47</td><td>68</td><td>22</td><td>TTTCCGGCTCGCGTGGGTGTGT</td><td>1754.5</td><td>4.5</td><td>0.8</td><td>5.4</td><td>6.3E-08</td><td>4.9E-07</td><td>BT20/MB468</td><td>MP</td></tr>
<tr><td>>hg38_21_+_8433172_8446622_RNA45SN1_45S_50ntFlanks@10482.10512.31|TCGAAAGGGAGTCGGGTTCAGATCCCCGAAT</td><td>>hg38_21_+_8433172_8446622_RNA45SN1_45S_50ntFlanks</td><td>10482</td><td>10512</td><td>31</td><td>TCGAAAGGGAGTCGGGTTCAGATCCCCGAAT</td><td>74.4</td><td>-9.3</td><td>1.7</td><td>-5.4</td><td>6.3E-08</td><td>4.9E-07</td><td>BT20/MB468</td><td>MP</td></tr>
<tr><td>>hg38_21_+_8433172_8446622_RNA45SN1_45S_50ntFlanks@4977.5000.24|CGGACACGGACAGGATTGACAGAT</td><td>>hg38_21_+_8433172_8446622_RNA45SN1_45S_50ntFlanks</td><td>4977</td><td>5000</td><td>24</td><td>CGGACACGGACAGGATTGACAGAT</td><td>70.2</td><td>-9.2</td><td>1.7</td><td>-5.4</td><td>6.3E-08</td><td>4.9E-07</td><td>BT20/MB468</td><td>MP</td></tr>
<tr><td>>hg38_21_+_8433172_8446622_RNA45SN1_45S_50ntFlanks@8839.8858.20|CGGTGGCGGCGGCGGCGGCG</td><td>>hg38_21_+_8433172_8446622_RNA45SN1_45S_50ntFlanks</td><td>8839</td><td>8858</td><td>20</td><td>CGGTGGCGGCGGCGGCGGCG</td><td>91.4</td><td>-9.6</td><td>1.8</td><td>-5.4</td><td>6.3E-08</td><td>4.9E-07</td><td>BT20/MB468</td><td>MP</td></tr>
<tr><td>>hg38_21_+_8433172_8446622_RNA45SN1_45S_50ntFlanks@6722.6746.25|ATTGCAGGACACATTGATCATCGAC</td><td>>hg38_21_+_8433172_8446622_RNA45SN1_45S_50ntFlanks</td><td>6722</td><td>6746</td><td>25</td><td>ATTGCAGGACACATTGATCATCGAC</td><td>234.4</td><td>-6.9</td><td>1.4</td><td>-5.1</td><td>3.0E-07</td><td>4.9E-07</td><td>MB231/MB468</td><td>MITO</td></tr>
<tr><td>>trna2_GlyGCC_21_-_18827107_18827177@1.31.31,trna35_GlyGCC_1_+_161413094_161413164@1.31.31,trna37_GlyGCC_1_+_161420467_161420537@1.31.31,trna39_GlyGCC_1_+_161427898_161427968@1.31.31,trna41_GlyGCC_1_+_161435258_161435328@1.31.31|GCATGGGTGGTTCAGTGGTAGAATTCTCGCC</td><td>>trna2_GlyGCC_21_-_18827107_18827177</td><td>1</td><td>31</td><td>31</td><td>GCATGGGTGGTTCAGTGGTAGAATTCTCGCC</td><td>4755.2</td><td>5.2</td><td>0.9</td><td>5.5</td><td>3.1E-08</td><td>4.9E-07</td><td>BT20/MB231</td><td>NUC</td></tr>
<tr><td>>hg38_21_+_8433172_8446622_RNA45SN1_45S_50ntFlanks@6661.6699.39|CGGTGGATCACTCGGCTCGTGCGTCGATGAAGAACGCAG</td><td>>hg38_21_+_8433172_8446622_RNA45SN1_45S_50ntFlanks</td><td>6661</td><td>6699</td><td>39</td><td>CGGTGGATCACTCGGCTCGTGCGTCGATGAAGAACGCAG</td><td>2347.6</td><td>-4.5</td><td>0.9</td><td>-5.2</td><td>2.5E-07</td><td>4.9E-07</td><td>MB231/MB468</td><td>MP</td></tr>
<tr><td>>MI0000434|hsa-let-7i&WithFlank&12|+|62603680|62603775@12.32.21&[MIMAT0000415&hsa-let-7i-5p&offsets|0|-1;m-49&12|+|62603691|62603712&offsets|0|-1]|TGAGGTAGTAGTTTGTGCTGT</td><td>>MI0000434|hsa-let-7i&WithFlank&12|+|62603680|62603775</td><td>12</td><td>32</td><td>21</td><td>TGAGGTAGTAGTTTGTGCTGT</td><td>6371.2</td><td>4.8</td><td>0.9</td><td>5.2</td><td>2.5E-07</td><td>4.9E-07</td><td>MB231/MB468</td><td>MP</td></tr>
<tr><td>>trnaMT_PheGAA_MT_+_577_647@22.57.36|AAAGCAATACACTGAAAATGTTTAGACGGGCTCACA</td><td>>trnaMT_PheGAA_MT_+_577_647</td><td>22</td><td>57</td><td>36</td><td>AAAGCAATACACTGAAAATGTTTAGACGGGCTCACA</td><td>158.4</td><td>11.1</td><td>2.0</td><td>5.4</td><td>6.3E-08</td><td>4.9E-07</td><td>BT20/MB468</td><td>MP</td></tr>
<tr><td>>hg38_21_+_8433172_8446622_RNA45SN1_45S_50ntFlanks@9937.9978.42|CAGCAGGACGGTGGCCATGGAAGTCGGAATCCGCTAAGGAGT</td><td>>hg38_21_+_8433172_8446622_RNA45SN1_45S_50ntFlanks</td><td>9937</td><td>9978</td><td>42</td><td>CAGCAGGACGGTGGCCATGGAAGTCGGAATCCGCTAAGGAGT</td><td>64.4</td><td>9.8</td><td>1.8</td><td>5.4</td><td>6.3E-08</td><td>4.9E-07</td><td>BT20/MB468</td><td>MP</td></tr>
<tr><td>>hg38_21_+_8433172_8446622_RNA45SN1_45S_50ntFlanks@12415.12433.19|CTTCGATGTCGGCTCTTCC</td><td>>hg38_21_+_8433172_8446622_RNA45SN1_45S_50ntFlanks</td><td>12415</td><td>12433</td><td>19</td><td>CTTCGATGTCGGCTCTTCC</td><td>415.4</td><td>-4.1</td><td>0.8</td><td>-5.2</td><td>2.5E-07</td><td>4.9E-07</td><td>MB231/MB468</td><td>MP</td></tr>
<tr><td>>hg38_21_+_8433172_8446622_RNA45SN1_45S_50ntFlanks@12249.12272.24|CAAAAGCTCGCTTGATCTTGATTT</td><td>>hg38_21_+_8433172_8446622_RNA45SN1_45S_50ntFlanks</td><td>12249</td><td>12272</td><td>24</td><td>CAAAAGCTCGCTTGATCTTGATTT</td><td>609.5</td><td>-8.1</td><td>1.6</td><td>-5.1</td><td>3.2E-07</td><td>4.9E-07</td><td>BT20/MB468</td><td>NUC</td></tr>
<tr><td>>trna116_GluCTC_1_-_145399233_145399304@35.70.36,trna134_GluTTC_1_-_16861774_16861845@35.70.36,trna71_GluCTC_1_-_161439189_161439260@35.70.36,trna74_GluCTC_1_-_161431809_161431880@35.70.36,trna77_GluCTC_1_-_161424398_161424469@35.70.36,trna77_GluCTC_6_+_28949976_28950047@35.70.36,trna80_GluCTC_1_-_161417018_161417089@35.70.36,trna84_GluTTC_1_-_161391883_161391954@35.70.36,trna87_GluCTC_6_-_126101393_126101464@35.70.36|TCACCGCCGCGGCCCGGGTTCGATTCCCGGTCAGGG</td><td>>trna116_GluCTC_1_-_145399233_145399304</td><td>35</td><td>70</td><td>36</td><td>TCACCGCCGCGGCCCGGGTTCGATTCCCGGTCAGGG</td><td>180.5</td><td>-6.0</td><td>1.2</td><td>-5.2</td><td>2.4E-07</td><td>5.0E-07</td><td>BT20/MB468</td><td>CYTO</td></tr>
<tr><td>>hg38_21_+_8433172_8446622_RNA45SN1_45S_50ntFlanks@7978.8011.34|GACCTCAGATCAGACGTGGCGACCCGCTGAATTT</td><td>>hg38_21_+_8433172_8446622_RNA45SN1_45S_50ntFlanks</td><td>7978</td><td>8011</td><td>34</td><td>GACCTCAGATCAGACGTGGCGACCCGCTGAATTT</td><td>586.8</td><td>4.2</td><td>0.8</td><td>5.1</td><td>2.9E-07</td><td>5.0E-07</td><td>MB231/MB468</td><td>NUC</td></tr>
<tr><td>>MI0000542|hsa-mir-320a&WithFlank&8|-|22244960|22245043@48.68.21(+1C)&[MIMAT0000510&hsa-miR-320a-3p&offsets|0|-1(+1C);m-60&8|-|22244975|22244996&offsets|0|-1(+1C)]|AAAAGCTGGGTTGAGAGGGCGC</td><td>>MI0000542|hsa-mir-320a&WithFlank&8|-|22244960|22245043</td><td>48</td><td>68</td><td>21(+1C)</td><td>AAAAGCTGGGTTGAGAGGGCGC</td><td>3486.5</td><td>4.4</td><td>0.8</td><td>5.6</td><td>1.9E-08</td><td>5.0E-07</td><td>BT20/MB231</td><td>CYTO</td></tr>
<tr><td>>MI0003628|hsa-mir-615&WithFlank&12|+|54033944|54034051@67.85.19&[MIMAT0003283&hsa-miR-615-3p&offsets|0|-3;m-495&12|+|54034010|54034030&offsets|0|-2]|TCCGAGCCTGGGTCTCCCT</td><td>>MI0003628|hsa-mir-615&WithFlank&12|+|54033944|54034051</td><td>67</td><td>85</td><td>19</td><td>TCCGAGCCTGGGTCTCCCT</td><td>212.3</td><td>5.6</td><td>1.0</td><td>5.6</td><td>1.9E-08</td><td>5.0E-07</td><td>BT20/MB231</td><td>CYTO</td></tr>
<tr><td>>hg38_21_+_8433172_8446622_RNA45SN1_45S_50ntFlanks@4045.4066.22|CCATTCGAACGTCTGCCCTATC</td><td>>hg38_21_+_8433172_8446622_RNA45SN1_45S_50ntFlanks</td><td>4045</td><td>4066</td><td>22</td><td>CCATTCGAACGTCTGCCCTATC</td><td>1420.4</td><td>-4.1</td><td>0.8</td><td>-5.1</td><td>2.9E-07</td><td>5.0E-07</td><td>MB231/MB468</td><td>NUC</td></tr>
<tr><td>>MI0000070|hsa-mir-16-1&WithFlank&13|-|50048967|50049067@20.41.22;MI0000115|hsa-mir-16-2&WithFlank&3|+|160404739|160404831@16.37.22&[MIMAT0000069&hsa-miR-16-5p&offsets|0|0;m-62&13|-|50049027|50049048&offsets|0|0];[MIMAT0000069_1&hsa-miR-16-5p&offsets|0|0;m-61&3|+|160404754|160404775&offsets|0|0]|TAGCAGCACGTAAATATTGGCG</td><td>>MI0000070|hsa-mir-16-1&WithFlank&13|-|50048967|50049067</td><td>20</td><td>41</td><td>22</td><td>TAGCAGCACGTAAATATTGGCG</td><td>2262.5</td><td>5.4</td><td>1.1</td><td>5.1</td><td>3.0E-07</td><td>5.0E-07</td><td>MB231/MB468</td><td>MITO</td></tr>
<tr><td>>hg38_21_+_8433172_8446622_RNA45SN1_45S_50ntFlanks@4730.4748.19|CAAGAACGAAAGTCGGAGG</td><td>>hg38_21_+_8433172_8446622_RNA45SN1_45S_50ntFlanks</td><td>4730</td><td>4748</td><td>19</td><td>CAAGAACGAAAGTCGGAGG</td><td>610.3</td><td>-3.6</td><td>0.7</td><td>-5.2</td><td>2.5E-07</td><td>5.0E-07</td><td>MB231/MB468</td><td>MP</td></tr>
<tr><td>>MI0000749|hsa-mir-30e&WithFlank&1|+|40754349|40754452@65.86.22&[MIMAT0000693&hsa-miR-30e-3p&offsets|0|0;m-25&1|+|40754413|40754434&offsets|0|0]|CTTTCAGTCGGATGTTTACAGC</td><td>>MI0000749|hsa-mir-30e&WithFlank&1|+|40754349|40754452</td><td>65</td><td>86</td><td>22</td><td>CTTTCAGTCGGATGTTTACAGC</td><td>2105.6</td><td>4.1</td><td>0.8</td><td>5.1</td><td>2.8E-07</td><td>5.0E-07</td><td>MB231/MB468</td><td>CYTO</td></tr>
</table>

<table>
<tr><td></td><td></td><td></td><td></td><td></td><td></td><td></td><td></td><td></td><td></td><td></td><td></td><td></td><td></td><td></td></tr>
<tr><td>>hg38_21_+_8433172_8446622_RNA45SN1_45S_50ntFlanks@11574.11603.30|GACTTAGAACTGGTGCGGACCAGGGGAATC</td><td>>hg38_21_+_8433172_8446622_RNA45SN1_45S_50ntFlanks</td><td>11574</td><td>11603</td><td>30</td><td>GACTTAGAACTGGTGCGGACCAGGGGAATC</td><td>229.6</td><td>-3.4</td><td>0.7</td><td>-5.1</td><td>2.8E-07</td><td>5.0E-07</td><td>MB231/MB468</td><td>CYTO</td></tr>
<tr><td>>hg38_21_+_8433172_8446622_RNA45SN1_45S_50ntFlanks@10508.10533.26|CGAATCCGGAGTGGCGGAGATGGGCG</td><td>>hg38_21_+_8433172_8446622_RNA45SN1_45S_50ntFlanks</td><td>10508</td><td>10533</td><td>26</td><td>CGAATCCGGAGTGGCGGAGATGGGCG</td><td>629.0</td><td>-8.3</td><td>1.6</td><td>-5.1</td><td>3.2E-07</td><td>5.0E-07</td><td>BT20/MB468</td><td>NUC</td></tr>
<tr><td>>hg38_21_+_8433172_8446622_RNA45SN1_45S_50ntFlanks@11675.11706.32|TGCTCTGAATGTCAAAGTGAAGAAATTCAATG</td><td>>hg38_21_+_8433172_8446622_RNA45SN1_45S_50ntFlanks</td><td>11675</td><td>11706</td><td>32</td><td>TGCTCTGAATGTCAAAGTGAAGAAATTCAATG</td><td>479.5</td><td>3.2</td><td>0.6</td><td>5.1</td><td>2.9E-07</td><td>5.0E-07</td><td>MB231/MB468</td><td>NUC</td></tr>
<tr><td>>hg38_21_+_8433172_8446622_RNA45SN1_45S_50ntFlanks@11769.11791.23|TAATTAGTGACGCGCATGAATGG</td><td>>hg38_21_+_8433172_8446622_RNA45SN1_45S_50ntFlanks</td><td>11769</td><td>11791</td><td>23</td><td>TAATTAGTGACGCGCATGAATGG</td><td>334.3</td><td>-7.2</td><td>1.4</td><td>-5.2</td><td>2.5E-07</td><td>5.0E-07</td><td>MB231/MB468</td><td>MP</td></tr>
<tr><td>>hg38_21_+_8433172_8446622_RNA45SN1_45S_50ntFlanks@9791.9816.26|TGGCGTGGAGCCGGGCGTGGAATGCG</td><td>>hg38_21_+_8433172_8446622_RNA45SN1_45S_50ntFlanks</td><td>9791</td><td>9816</td><td>26</td><td>TGGCGTGGAGCCGGGCGTGGAATGCG</td><td>507.8</td><td>12.3</td><td>2.2</td><td>5.7</td><td>1.3E-08</td><td>5.0E-07</td><td>BT20/MB231</td><td>MP</td></tr>
<tr><td>>hg38_21_+_8433172_8446622_RNA45SN1_45S_50ntFlanks@12053.12071.19|GTGAGGCGGGGGGGCGAGC</td><td>>hg38_21_+_8433172_8446622_RNA45SN1_45S_50ntFlanks</td><td>12053</td><td>12071</td><td>19</td><td>GTGAGGCGGGGGGGCGAGC</td><td>76.9</td><td>-9.3</td><td>1.7</td><td>-5.4</td><td>6.3E-08</td><td>5.0E-07</td><td>BT20/MB468</td><td>MP</td></tr>
<tr><td>>hg38_21_+_8433172_8446622_RNA45SN1_45S_50ntFlanks@11209.11227.19|CGGCGACTCTGGACGCGAG</td><td>>hg38_21_+_8433172_8446622_RNA45SN1_45S_50ntFlanks</td><td>11209</td><td>11227</td><td>19</td><td>CGGCGACTCTGGACGCGAG</td><td>1819.4</td><td>-5.9</td><td>1.2</td><td>-5.2</td><td>2.5E-07</td><td>5.0E-07</td><td>BT20/MB468</td><td>CYTO</td></tr>
<tr><td>>hg38_21_+_8433172_8446622_RNA45SN1_45S_50ntFlanks@12786.12805.20|CCCGCGCGCGCGGGAGGGCG</td><td>>hg38_21_+_8433172_8446622_RNA45SN1_45S_50ntFlanks</td><td>12786</td><td>12805</td><td>20</td><td>CCCGCGCGCGCGGGAGGGCG</td><td>250.2</td><td>-6.2</td><td>1.2</td><td>-5.2</td><td>2.2E-07</td><td>5.0E-07</td><td>BT20/MB468</td><td>MITO</td></tr>
<tr><td>>hg38_21_+_8433172_8446622_RNA45SN1_45S_50ntFlanks@12540.12569.30|TGATGTGTTGTTGCCATGGTAATCCTGCTC</td><td>>hg38_21_+_8433172_8446622_RNA45SN1_45S_50ntFlanks</td><td>12540</td><td>12569</td><td>30</td><td>TGATGTGTTGTTGCCATGGTAATCCTGCTC</td><td>149.4</td><td>-2.6</td><td>0.5</td><td>-5.2</td><td>2.5E-07</td><td>5.0E-07</td><td>BT20/MB468</td><td>CYTO</td></tr>
<tr><td>>hg38_21_+_8433172_8446622_RNA45SN1_45S_50ntFlanks@10771.10791.21|CAGGTCTCCAAGGTGAACAGC</td><td>>hg38_21_+_8433172_8446622_RNA45SN1_45S_50ntFlanks</td><td>10771</td><td>10791</td><td>21</td><td>CAGGTCTCCAAGGTGAACAGC</td><td>402.0</td><td>-3.3</td><td>0.6</td><td>-5.1</td><td>2.9E-07</td><td>5.0E-07</td><td>MB231/MB468</td><td>NUC</td></tr>
<tr><td>>hg38_21_+_8433172_8446622_RNA45SN1_45S_50ntFlanks@4583.4601.19|CGGTTCTATTTTGTTGGTT</td><td>>hg38_21_+_8433172_8446622_RNA45SN1_45S_50ntFlanks</td><td>4583</td><td>4601</td><td>19</td><td>CGGTTCTATTTTGTTGGTT</td><td>77.1</td><td>-9.3</td><td>1.7</td><td>-5.4</td><td>6.4E-08</td><td>5.0E-07</td><td>BT20/MB468</td><td>MP</td></tr>
<tr><td>>hg38_21_+_8433172_8446622_RNA45SN1_45S_50ntFlanks@3842.3860.19|CCTACTTGGATAACTGTGG</td><td>>hg38_21_+_8433172_8446622_RNA45SN1_45S_50ntFlanks</td><td>3842</td><td>3860</td><td>19</td><td>CCTACTTGGATAACTGTGG</td><td>265.6</td><td>-7.6</td><td>1.5</td><td>-5.2</td><td>2.5E-07</td><td>5.0E-07</td><td>MB231/MB468</td><td>MP</td></tr>
<tr><td>>hg38_21_+_8433172_8446622_RNA45SN1_45S_50ntFlanks@12406.12432.27|TTTTTGATCCTTCGATGTCGGCTCTTC</td><td>>hg38_21_+_8433172_8446622_RNA45SN1_45S_50ntFlanks</td><td>12406</td><td>12432</td><td>27</td><td>TTTTTGATCCTTCGATGTCGGCTCTTC</td><td>173.5</td><td>-7.2</td><td>1.4</td><td>-5.2</td><td>2.6E-07</td><td>5.0E-07</td><td>MB231/MB468</td><td>MP</td></tr>
<tr><td>>hg38_21_+_8433172_8446622_RNA45SN1_45S_50ntFlanks@5471.5514.44|CACGGCCCTGGCGGAGCGCTGAGAAGACGGTCGAACTTGACTAT</td><td>>hg38_21_+_8433172_8446622_RNA45SN1_45S_50ntFlanks</td><td>5471</td><td>5514</td><td>44</td><td>CACGGCCCTGGCGGAGCGCTGAGAAGACGGTCGAACTTGACTAT</td><td>279.2</td><td>-4.1</td><td>0.7</td><td>-5.6</td><td>1.9E-08</td><td>5.0E-07</td><td>BT20/MB231</td><td>CYTO</td></tr>
<tr><td>>hg38_21_+_8433172_8446622_RNA45SN1_45S_50ntFlanks@9493.9515.23|CCCGAAAGATGGTGAACTATGCC</td><td>>hg38_21_+_8433172_8446622_RNA45SN1_45S_50ntFlanks</td><td>9493</td><td>9515</td><td>23</td><td>CCCGAAAGATGGTGAACTATGCC</td><td>285.7</td><td>11.5</td><td>2.0</td><td>5.7</td><td>1.3E-08</td><td>5.0E-07</td><td>BT20/MB231</td><td>MP</td></tr>
<tr><td>>MI0000102|hsa-mir-100&WithFlank&11|-|122152223|122152314@19.39.21(+1U)&[MIMAT0000098&hsa-miR-100-5p&offsets|0|-1(+1U);m-27&11|-|122152276|122152296&offsets|0|0(+1U)]|AACCCGTAGATCCGAACTTGTT</td><td>>MI0000102|hsa-mir-100&WithFlank&11|-|122152223|122152314</td><td>19</td><td>39</td><td>21(+1U)</td><td>AACCCGTAGATCCGAACTTGTT</td><td>4501.2</td><td>-5.4</td><td>1.0</td><td>-5.5</td><td>3.2E-08</td><td>5.0E-07</td><td>BT20/MB231</td><td>NUC</td></tr>
<tr><td>>hg38_21_+_8433172_8446622_RNA45SN1_45S_50ntFlanks@10410.10453.44|TCAGTCGGTCCTGAGAGATGGGCGAGCGCCGTTCCGAAGGGACG</td><td>>hg38_21_+_8433172_8446622_RNA45SN1_45S_50ntFlanks</td><td>10410</td><td>10453</td><td>44</td><td>TCAGTCGGTCCTGAGAGATGGGCGAGCGCCGTTCCGAAGGGACG</td><td>566.7</td><td>-7.1</td><td>1.4</td><td>-5.2</td><td>2.5E-07</td><td>5.0E-07</td><td>BT20/MB468</td><td>CYTO</td></tr>
<tr><td>>hg38_21_+_8433172_8446622_RNA45SN1_45S_50ntFlanks@11932.11951.20|GAATAAGTGGGAGGCCCCCG</td><td>>hg38_21_+_8433172_8446622_RNA45SN1_45S_50ntFlanks</td><td>11932</td><td>11951</td><td>20</td><td>GAATAAGTGGGAGGCCCCCG</td><td>85.5</td><td>-9.5</td><td>1.8</td><td>-5.4</td><td>6.4E-08</td><td>5.0E-07</td><td>BT20/MB468</td><td>MP</td></tr>
<tr><td>>hg38_21_+_8433172_8446622_RNA45SN1_45S_50ntFlanks@11665.11692.28|TTCTGCCCAGTGCTCTGAATGTCAAAGT</td><td>>hg38_21_+_8433172_8446622_RNA45SN1_45S_50ntFlanks</td><td>11665</td><td>11692</td><td>28</td><td>TTCTGCCCAGTGCTCTGAATGTCAAAGT</td><td>67.6</td><td>9.8</td><td>1.8</td><td>5.4</td><td>6.4E-08</td><td>5.0E-07</td><td>BT20/MB468</td><td>MP</td></tr>
<tr><td>>hg38_21_+_8433172_8446622_RNA45SN1_45S_50ntFlanks@6752.6770.19|GAACGCACTTGCGGCCCCG</td><td>>hg38_21_+_8433172_8446622_RNA45SN1_45S_50ntFlanks</td><td>6752</td><td>6770</td><td>19</td><td>GAACGCACTTGCGGCCCCG</td><td>73.5</td><td>-9.3</td><td>1.7</td><td>-5.4</td><td>6.4E-08</td><td>5.0E-07</td><td>BT20/MB468</td><td>MP</td></tr>
<tr><td>>hg38_21_+_8433172_8446622_RNA45SN1_45S_50ntFlanks@12971.12992.22|TTTCGTACGTAGCAGAGCAGCT</td><td>>hg38_21_+_8433172_8446622_RNA45SN1_45S_50ntFlanks</td><td>12971</td><td>12992</td><td>22</td><td>TTTCGTACGTAGCAGAGCAGCT</td><td>561.4</td><td>-7.9</td><td>1.6</td><td>-5.1</td><td>3.2E-07</td><td>5.1E-07</td><td>BT20/MB468</td><td>NUC</td></tr>
<tr><td>>hg38_21_+_8433172_8446622_RNA45SN1_45S_50ntFlanks@12437.12474.38|CATTGTGAAGCAGAATTCACCAAGCGTTGGATTGTTCA</td><td>>hg38_21_+_8433172_8446622_RNA45SN1_45S_50ntFlanks</td><td>12437</td><td>12474</td><td>38</td><td>CATTGTGAAGCAGAATTCACCAAGCGTTGGATTGTTCA</td><td>626.6</td><td>-5.3</td><td>1.0</td><td>-5.1</td><td>2.8E-07</td><td>5.1E-07</td><td>MB231/MB468</td><td>CYTO</td></tr>
<tr><td>>trnaMT_HisGTG_MT_+_12138_12206@20.55.36|AACATCAGATTGTGAATCTGACAACAGAGGCTTACG</td><td>>trnaMT_HisGTG_MT_+_12138_12206</td><td>20</td><td>55</td><td>36</td><td>AACATCAGATTGTGAATCTGACAACAGAGGCTTACG</td><td>45.3</td><td>8.7</td><td>1.7</td><td>5.2</td><td>2.6E-07</td><td>5.1E-07</td><td>MB231/MB468</td><td>MP</td></tr>
<tr><td>>hg38_21_+_8433172_8446622_RNA45SN1_45S_50ntFlanks@11051.11075.25|CCGTCCTCCCCCCTCCCCGGGGGAG</td><td>>hg38_21_+_8433172_8446622_RNA45SN1_45S_50ntFlanks</td><td>11051</td><td>11075</td><td>25</td><td>CCGTCCTCCCCCCTCCCCGGGGGAG</td><td>108.2</td><td>-9.8</td><td>1.8</td><td>-5.4</td><td>6.5E-08</td><td>5.1E-07</td><td>BT20/MB468</td><td>MP</td></tr>
<tr><td>>hg38_21_+_8433172_8446622_RNA45SN1_45S_50ntFlanks@11670.11700.31|CCCAGTGCTCTGAATGTCAAAGTGAAGAAAT</td><td>>hg38_21_+_8433172_8446622_RNA45SN1_45S_50ntFlanks</td><td>11670</td><td>11700</td><td>31</td><td>CCCAGTGCTCTGAATGTCAAAGTGAAGAAAT</td><td>159.5</td><td>-6.9</td><td>1.3</td><td>-5.2</td><td>2.6E-07</td><td>5.1E-07</td><td>MB231/MB468</td><td>MP</td></tr>
<tr><td>>MI0000650|hsa-mir-200c&WithFlank&12|+|6963693|6963772@50.70.21(+1A)&[MIMAT0000617&hsa-miR-200c-3p&offsets|0|-2(+1A);m-54&12|+|6963742|6963764&offsets|0|-2(+1A)]|TAATACTGCCGGGTAATGATGA</td><td>>MI0000650|hsa-mir-200c&WithFlank&12|+|6963693|6963772</td><td>50</td><td>70</td><td>21(+1A)</td><td>TAATACTGCCGGGTAATGATGA</td><td>3141.9</td><td>5.8</td><td>1.1</td><td>5.4</td><td>6.5E-08</td><td>5.1E-07</td><td>BT20/MB468</td><td>MP</td></tr>
<tr><td>>hg38_21_+_8433172_8446622_RNA45SN1_45S_50ntFlanks@6648.6673.26|GTACGACTCTTAGCGGTGGATCACTC</td><td>>hg38_21_+_8433172_8446622_RNA45SN1_45S_50ntFlanks</td><td>6648</td><td>6673</td><td>26</td><td>GTACGACTCTTAGCGGTGGATCACTC</td><td>320.3</td><td>4.9</td><td>1.0</td><td>5.1</td><td>2.9E-07</td><td>5.1E-07</td><td>MB231/MB468</td><td>NUC</td></tr>
<tr><td>>hg38_21_+_8433172_8446622_RNA45SN1_45S_50ntFlanks@7980.8012.33|CCTCAGATCAGACGTGGCGACCCGCTGAATTTA</td><td>>hg38_21_+_8433172_8446622_RNA45SN1_45S_50ntFlanks</td><td>7980</td><td>8012</td><td>33</td><td>CCTCAGATCAGACGTGGCGACCCGCTGAATTTA</td><td>527.0</td><td>-8.0</td><td>1.6</td><td>-5.1</td><td>3.2E-07</td><td>5.1E-07</td><td>BT20/MB468</td><td>NUC</td></tr>
<tr><td>>hg38_21_+_8433172_8446622_RNA45SN1_45S_50ntFlanks@3912.3938.27|GATGCGTGCATTTATCAGATCAAAACC</td><td>>hg38_21_+_8433172_8446622_RNA45SN1_45S_50ntFlanks</td><td>3912</td><td>3938</td><td>27</td><td>GATGCGTGCATTTATCAGATCAAAACC</td><td>98.1</td><td>-9.7</td><td>1.8</td><td>-5.4</td><td>6.5E-08</td><td>5.1E-07</td><td>BT20/MB468</td><td>MP</td></tr>
<tr><td>>hg38_21_+_8433172_8446622_RNA45SN1_45S_50ntFlanks@4297.4317.21|CATTGGAGGGCAAGTCTGGTG</td><td>>hg38_21_+_8433172_8446622_RNA45SN1_45S_50ntFlanks</td><td>4297</td><td>4317</td><td>21</td><td>CATTGGAGGGCAAGTCTGGTG</td><td>828.1</td><td>-8.3</td><td>1.6</td><td>-5.1</td><td>3.2E-07</td><td>5.1E-07</td><td>BT20/MB468</td><td>NUC</td></tr>
<tr><td>>hg38_21_+_8433172_8446622_RNA45SN1_45S_50ntFlanks@4684.4707.24|AAGACGGACCAGAGCGAAAGCATT</td><td>>hg38_21_+_8433172_8446622_RNA45SN1_45S_50ntFlanks</td><td>4684</td><td>4707</td><td>24</td><td>AAGACGGACCAGAGCGAAAGCATT</td><td>205.4</td><td>-7.0</td><td>1.4</td><td>-5.2</td><td>2.6E-07</td><td>5.1E-07</td><td>MB231/MB468</td><td>MP</td></tr>
<tr><td>>hg38_21_+_8433172_8446622_RNA45SN1_45S_50ntFlanks@11577.11611.35|TTAGAACTGGTGCGGACCAGGGGAATCCGACTGTT</td><td>>hg38_21_+_8433172_8446622_RNA45SN1_45S_50ntFlanks</td><td>11577</td><td>11611</td><td>35</td><td>TTAGAACTGGTGCGGACCAGGGGAATCCGACTGTT</td><td>364.3</td><td>-7.6</td><td>1.5</td><td>-5.2</td><td>2.6E-07</td><td>5.1E-07</td><td>MB231/MB468</td><td>MP</td></tr>
<tr><td>>hg38_21_+_8433172_8446622_RNA45SN1_45S_50ntFlanks@6652.6678.27|GACTCTTAGCGGTGGATCACTCGGCTC</td><td>>hg38_21_+_8433172_8446622_RNA45SN1_45S_50ntFlanks</td><td>6652</td><td>6678</td><td>27</td><td>GACTCTTAGCGGTGGATCACTCGGCTC</td><td>1260.5</td><td>3.0</td><td>0.6</td><td>5.1</td><td>2.9E-07</td><td>5.1E-07</td><td>MB231/MB468</td><td>CYTO</td></tr>
<tr><td>>hg38_21_+_8433172_8446622_RNA45SN1_45S_50ntFlanks@6721.6750.30|AATTGCAGGACACATTGATCATCGACACTT</td><td>>hg38_21_+_8433172_8446622_RNA45SN1_45S_50ntFlanks</td><td>6721</td><td>6750</td><td>30</td><td>AATTGCAGGACACATTGATCATCGACACTT</td><td>394.5</td><td>-7.3</td><td>1.4</td><td>-5.2</td><td>2.6E-07</td><td>5.1E-07</td><td>MB231/MB468</td><td>MP</td></tr>
<tr><td>>MI0000482|hsa-mir-185&WithFlank&22|+|20033133|20033226@21.42.22&[MIMAT0000455&hsa-miR-185-5p&offsets|0|0;m-145&22|+|20033153|20033174&offsets|0|0]|TGGAGAGAAAGGCAGTTCCTGA</td><td>>MI0000482|hsa-mir-185&WithFlank&22|+|20033133|20033226</td><td>21</td><td>42</td><td>22</td><td>TGGAGAGAAAGGCAGTTCCTGA</td><td>572.7</td><td>1.7</td><td>0.3</td><td>5.1</td><td>3.3E-07</td><td>5.1E-07</td><td>BT20/MB468</td><td>NUC</td></tr>
<tr><td>>trnaMT_ValTAC_MT_+_1602_1670@54.72.19|AACTTGACCGCTCTGACCA</td><td>>trnaMT_ValTAC_MT_+_1602_1670</td><td>54</td><td>72</td><td>19</td><td>AACTTGACCGCTCTGACCA</td><td>7139.2</td><td>4.9</td><td>0.9</td><td>5.2</td><td>2.2E-07</td><td>5.1E-07</td><td>BT20/MB468</td><td>MITO</td></tr>
<tr><td>>hg38_21_+_8433172_8446622_RNA45SN1_45S_50ntFlanks@11619.11660.42|AACAAAGCATCGCGAAGGCCCGCGGCGGGTGTTGACGCGATG</td><td>>hg38_21_+_8433172_8446622_RNA45SN1_45S_50ntFlanks</td><td>11619</td><td>11660</td><td>42</td><td>AACAAAGCATCGCGAAGGCCCGCGGCGGGTGTTGACGCGATG</td><td>95.0</td><td>-8.7</td><td>1.7</td><td>-5.2</td><td>2.2E-07</td><td>5.1E-07</td><td>BT20/MB468</td><td>MITO</td></tr>
<tr><td>>hg38_21_+_8433172_8446622_RNA45SN1_45S_50ntFlanks@8176.8204.29|CTTCTGATCGAGGCCCAGCCCGTGGACGG</td><td>>hg38_21_+_8433172_8446622_RNA45SN1_45S_50ntFlanks</td><td>8176</td><td>8204</td><td>29</td><td>CTTCTGATCGAGGCCCAGCCCGTGGACGG</td><td>1665.3</td><td>-5.9</td><td>1.2</td><td>-5.1</td><td>3.1E-07</td><td>5.1E-07</td><td>MB231/MB468</td><td>MITO</td></tr>
<tr><td>>hg38_21_+_8433172_8446622_RNA45SN1_45S_50ntFlanks@8107.8143.37|CGGCGGGGCGCGGGACATGTGGCGTACGGAAGACCCG</td><td>>hg38_21_+_8433172_8446622_RNA45SN1_45S_50ntFlanks</td><td>8107</td><td>8143</td><td>37</td><td>CGGCGGGGCGCGGGACATGTGGCGTACGGAAGACCCG</td><td>482.3</td><td>-7.8</td><td>1.5</td><td>-5.1</td><td>3.3E-07</td><td>5.1E-07</td><td>BT20/MB468</td><td>NUC</td></tr>
<tr><td>>MI0000087|hsa-mir-29a&WithFlank&7|-|130876741|130876816@48.69.22(+1A)&[MIMAT0000086&hsa-miR-29a-3p&offsets|0|0(+1A);m-26&7|-|130876749|130876769&offsets|0|+1(+1A)]|TAGCACCATCTGAAATCGGTTAA</td><td>>MI0000087|hsa-mir-29a&WithFlank&7|-|130876741|130876816</td><td>48</td><td>69</td><td>22(+1A)</td><td>TAGCACCATCTGAAATCGGTTAA</td><td>319.5</td><td>8.6</td><td>1.7</td><td>5.1</td><td>3.1E-07</td><td>5.1E-07</td><td>MB231/MB468</td><td>MITO</td></tr>
<tr><td>>hg38_21_+_8433172_8446622_RNA45SN1_45S_50ntFlanks@9858.9893.36|CGGGATGAACCGAACGCCGGGTTAAGGCGCCCGATG</td><td>>hg38_21_+_8433172_8446622_RNA45SN1_45S_50ntFlanks</td><td>9858</td><td>9893</td><td>36</td><td>CGGGATGAACCGAACGCCGGGTTAAGGCGCCCGATG</td><td>240.0</td><td>-11.7</td><td>2.3</td><td>-5.1</td><td>3.1E-07</td><td>5.1E-07</td><td>MB231/MB468</td><td>MITO</td></tr>
<tr><td>>hg38_21_+_8433172_8446622_RNA45SN1_45S_50ntFlanks@10574.10596.23|AAGCCGGCGGGAGCCCCGGGGAG</td><td>>hg38_21_+_8433172_8446622_RNA45SN1_45S_50ntFlanks</td><td>10574</td><td>10596</td><td>23</td><td>AAGCCGGCGGGAGCCCCGGGGAG</td><td>256.0</td><td>-6.6</td><td>1.3</td><td>-5.1</td><td>3.0E-07</td><td>5.1E-07</td><td>MB231/MB468</td><td>NUC</td></tr>
<tr><td>>hg38_21_+_8433172_8446622_RNA45SN1_45S_50ntFlanks@8356.8378.23|GAAAAGAACTTTGAAGAGAGAGT</td><td>>hg38_21_+_8433172_8446622_RNA45SN1_45S_50ntFlanks</td><td>8356</td><td>8378</td><td>23</td><td>GAAAAGAACTTTGAAGAGAGAGT</td><td>809.9</td><td>-3.3</td><td>0.7</td><td>-5.1</td><td>2.9E-07</td><td>5.1E-07</td><td>MB231/MB468</td><td>CYTO</td></tr>
<tr><td>>hg38_21_+_8433172_8446622_RNA45SN1_45S_50ntFlanks@5482.5499.18|CGGAGCGCTGAGAAGACG</td><td>>hg38_21_+_8433172_8446622_RNA45SN1_45S_50ntFlanks</td><td>5482</td><td>5499</td><td>18</td><td>CGGAGCGCTGAGAAGACG</td><td>228.1</td><td>-4.0</td><td>0.8</td><td>-5.1</td><td>3.0E-07</td><td>5.1E-07</td><td>MB231/MB468</td><td>NUC</td></tr>
<tr><td>>hg38_21_+_8433172_8446622_RNA45SN1_45S_50ntFlanks@11209.11230.22|CGGCGACTCTGGACGCGAGCCG</td><td>>hg38_21_+_8433172_8446622_RNA45SN1_45S_50ntFlanks</td><td>11209</td><td>11230</td><td>22</td><td>CGGCGACTCTGGACGCGAGCCG</td><td>713.8</td><td>-5.7</td><td>1.1</td><td>-5.1</td><td>3.0E-07</td><td>5.1E-07</td><td>MB231/MB468</td><td>NUC</td></tr>
<tr><td>>MI0000746|hsa-mir-99b&WithFlank&19|+|51692606|51692687@13.32.20(+1U)&[MIMAT0000689&hsa-miR-99b-5p&offsets|0|-2(+1U);m-10&19|+|51692618|51692639&offsets|0|-2(+1U)]|CACCCGTAGAACCGACCTTGT</td><td>>MI0000746|hsa-mir-99b&WithFlank&19|+|51692606|51692687</td><td>13</td><td>32</td><td>20(+1U)</td><td>CACCCGTAGAACCGACCTTGT</td><td>9417.7</td><td>3.0</td><td>0.5</td><td>5.5</td><td>3.3E-08</td><td>5.1E-07</td><td>BT20/MB231</td><td>NUC</td></tr>
<tr><td>>hg38_21_+_8433172_8446622_RNA45SN1_45S_50ntFlanks@11584.11611.28|TGGTGCGGACCAGGGGAATCCGACTGTT</td><td>>hg38_21_+_8433172_8446622_RNA45SN1_45S_50ntFlanks</td><td>11584</td><td>11611</td><td>28</td><td>TGGTGCGGACCAGGGGAATCCGACTGTT</td><td>924.6</td><td>4.2</td><td>0.8</td><td>5.5</td><td>3.3E-08</td><td>5.1E-07</td><td>BT20/MB231</td><td>NUC</td></tr>
<tr><td>>hg38_21_+_8433172_8446622_RNA45SN1_45S_50ntFlanks@10947.10967.21|CACGCCCGGGGCACCCCCCTC</td><td>>hg38_21_+_8433172_8446622_RNA45SN1_45S_50ntFlanks</td><td>10947</td><td>10967</td><td>21</td><td>CACGCCCGGGGCACCCCCCTC</td><td>125.7</td><td>-6.0</td><td>1.2</td><td>-5.2</td><td>2.5E-07</td><td>5.2E-07</td><td>BT20/MB468</td><td>CYTO</td></tr>
<tr><td>>MI0000471|hsa-mir-126&WithFlank&9|+|136670596|136670692@59.77.19(+1A)&[MIMAT0000445&hsa-miR-126-3p&offsets|+1|-2(+1A);m-40&9|+|136670653|136670674&offsets|+1|-2(+1A)]|CGTACCGTGAGTAATAATGA</td><td>>MI0000471|hsa-mir-126&WithFlank&9|+|136670596|136670692</td><td>59</td><td>77</td><td>19(+1A)</td><td>CGTACCGTGAGTAATAATGA</td><td>71.1</td><td>9.4</td><td>1.8</td><td>5.1</td><td>2.6E-07</td><td>5.2E-07</td><td>MB231/MB468</td><td>MP</td></tr>
<tr><td>>hg38_21_+_8433172_8446622_RNA45SN1_45S_50ntFlanks@6651.6695.45|CGACTCTTAGCGGTGGATCACTCGGCTCGTGCGTCGATGAAGAAC</td><td>>hg38_21_+_8433172_8446622_RNA45SN1_45S_50ntFlanks</td><td>6651</td><td>6695</td><td>45</td><td>CGACTCTTAGCGGTGGATCACTCGGCTCGTGCGTCGATGAAGAAC</td><td>5209.1</td><td>5.1</td><td>1.0</td><td>5.1</td><td>3.0E-07</td><td>5.2E-07</td><td>MB231/MB468</td><td>NUC</td></tr>
</table>

<table>
<tr><td>>hg38_21_+_8433172_8446622_RNA45SN1_45S_50ntFlanks@10441.10460.20|TTCCGAAGGGACGGGCGATG</td><td>>hg38_21_+_8433172_8446622_RNA45SN1_45S_50ntFlanks</td><td>10441</td><td>10460</td><td>20</td><td>TTCCGAAGGGACGGGCGATG</td><td>608.4</td><td>-4.2</td><td>0.8</td><td>-5.1</td><td>3.0E-07</td><td>5.2E-07</td><td>MB231/MB468</td><td>NUC</td></tr>
<tr><td>>hg38_21_+_8433172_8446622_RNA45SN1_45S_50ntFlanks@7974.8003.30|ACGCGACCTCAGATCAGACGTGGCGACCCG</td><td>>hg38_21_+_8433172_8446622_RNA45SN1_45S_50ntFlanks</td><td>7974</td><td>8003</td><td>30</td><td>ACGCGACCTCAGATCAGACGTGGCGACCCG</td><td>311.3</td><td>-2.8</td><td>0.5</td><td>-5.1</td><td>2.6E-07</td><td>5.2E-07</td><td>MB231/MB468</td><td>MP</td></tr>
<tr><td>>am_MI0005763|hsa-mir-941-1&WithFlank&20|+|63919443|63919526@53.74.22(+1A);MI0005764|hsa-mir-941-2&WithFlank&20|+|63919499|63919582@53.74.22(+1A);MI0005765|hsa-mir-941-3&WithFlank&20|+|63919555|63919638@53.74.22(+1A);MI0005766|hsa-mir-941-4&WithFlank&20|+|63919750|63919833@53.74.22(+1A);MI0031520|hsa-mir-941-5&WithFlank&20|+|63919862|63919945@53.74.22(+1A)&[MIMAT0004984&hsa-miR-941&offsets|0|-1(+1A);m-263&20|+|63919495|63919517&offsets|0|-1(+1A)];[MIMAT0004984_1&hsa-miR-941&offsets|0|-1(+1A);m-263&20|+|63919551|63919573&offsets|0|-1(+1A)];[MIMAT0004984_2&hsa-miR-941&offsets|0|-1(+1A);m-263&20|+|63919607|63919629&offsets|0|-1(+1A)];[MIMAT0004984_3&hsa-miR-941&offsets|0|-1(+1A);m-263&20|+|63919802|63919824&offsets|0|-1(+1A)];[MIMAT0004984_4&hsa-miR-941&offsets|0|-1(+1A);m-263&20|+|63919914|63919936&offsets|0|-1(+1A)]|CACCCGGCTGTGTGCACATGTGA</td><td>>am_MI0005763|hsa-mir-941-1&WithFlank&20|+|63919443|63919526</td><td>53</td><td>74</td><td>22(+1A)</td><td>CACCCGGCTGTGTGCACATGTGA</td><td>427.0</td><td>6.2</td><td>1.2</td><td>5.1</td><td>3.1E-07</td><td>5.2E-07</td><td>MB231/MB468</td><td>MITO</td></tr>
<tr><td>>MI0000456|hsa-mir-140&WithFlank&16|+|69933075|69933186@69.89.21(+1U)&[MIMAT0004597&hsa-miR-140-3p&offsets|+1|+1(+1U);m-45&16|+|69933143|69933164&offsets|0|-1(+1U)]|ACCACAGGGTAGAACCACGGAT</td><td>>MI0000456|hsa-mir-140&WithFlank&16|+|69933075|69933186</td><td>69</td><td>89</td><td>21(+1U)</td><td>ACCACAGGGTAGAACCACGGAT</td><td>349.5</td><td>8.1</td><td>1.6</td><td>5.1</td><td>2.6E-07</td><td>5.2E-07</td><td>MB231/MB468</td><td>MP</td></tr>
<tr><td>>trnaMT_GluTTC_MT_-_14674_14742@40.66.27,trnalookalike8_GluTTC_5_-_93905172_93905240@40.66.27|TTGGTCGTGGTTGTAGTCCGTGCGAGA</td><td>>trnaMT_GluTTC_MT_-_14674_14742</td><td>40</td><td>66</td><td>27</td><td>TTGGTCGTGGTTGTAGTCCGTGCGAGA</td><td>789.0</td><td>5.3</td><td>1.0</td><td>5.4</td><td>6.6E-08</td><td>5.2E-07</td><td>BT20/MB468</td><td>MP</td></tr>
<tr><td>>hg38_21_+_8433172_8446622_RNA45SN1_45S_50ntFlanks@10808.10837.30|CAATGTAGGTAAGGGAAGTCGGCAAGCCGG</td><td>>hg38_21_+_8433172_8446622_RNA45SN1_45S_50ntFlanks</td><td>10808</td><td>10837</td><td>30</td><td>CAATGTAGGTAAGGGAAGTCGGCAAGCCGG</td><td>558.0</td><td>-8.1</td><td>1.6</td><td>-5.1</td><td>3.3E-07</td><td>5.2E-07</td><td>BT20/MB468</td><td>NUC</td></tr>
<tr><td>>hg38_21_+_8433172_8446622_RNA45SN1_45S_50ntFlanks@9575.9594.20|ATCGGTCGTCCGACCTGGGT</td><td>>hg38_21_+_8433172_8446622_RNA45SN1_45S_50ntFlanks</td><td>9575</td><td>9594</td><td>20</td><td>ATCGGTCGTCCGACCTGGGT</td><td>411.8</td><td>-7.3</td><td>1.4</td><td>-5.1</td><td>3.2E-07</td><td>5.2E-07</td><td>MB231/MB468</td><td>MITO</td></tr>
<tr><td>>am_trna10_ValCAC_5_-_180649395_180649467@1.19.19,trna12_ValAAC_5_-_180645270_180645342@1.19.19,trna132_ValAAC_6_-_27721179_27721251@1.19.19,trna133_ValCAC_6_-_27696327_27696399@1.19.19,trna136_ValAAC_6_-_27648885_27648957@1.19.19,trna139_ValAAC_6_-_27618707_27618779@1.19.19,trna15_ValAAC_5_-_180615416_180615488@1.19.19,trna18_ValCAC_5_-_180529253_180529325@1.19.19,trna2_ValAAC_3_+_169490018_169490090@1.19.19,trna2_ValCAC_5_+_180524070_180524142@1.19.19,trna37_ValAAC_6_+_27203288_27203360@1.19.19,trna4_ValAAC_5_+_180591154_180591226@1.19.19,trna5_ValAAC_5_+_180596610_180596682@1.19.19,trna6_ValCAC_5_+_180600650_180600722@1.19.19,trna85_ValCAC_1_-_161369490_161369562@1.19.19,trna90_ValCAC_1_-_149684088_149684161@1.19.19,trna98_ValCAC_1_-_149298555_149298627@1.19.19,trna9_ValCAC_6_+_26538282_26538354@1.19.19|GTTTCCGTAGTGTAGTGGT</td><td>>am_trna10_ValCAC_5_-_180649395_180649467</td><td>1</td><td>19</td><td>19</td><td>GTTTCCGTAGTGTAGTGGT</td><td>740.1</td><td>6.2</td><td>1.2</td><td>5.2</td><td>2.6E-07</td><td>5.2E-07</td><td>BT20/MB468</td><td>CYTO</td></tr>
<tr><td>>hg38_21_+_8433172_8446622_RNA45SN1_45S_50ntFlanks@8170.8203.34|CAAGTCCTTCTGATCGAGGCCCAGCCCGTGGACG</td><td>>hg38_21_+_8433172_8446622_RNA45SN1_45S_50ntFlanks</td><td>8170</td><td>8203</td><td>34</td><td>CAAGTCCTTCTGATCGAGGCCCAGCCCGTGGACG</td><td>556.4</td><td>-7.9</td><td>1.5</td><td>-5.1</td><td>2.6E-07</td><td>5.2E-07</td><td>MB231/MB468</td><td>MP</td></tr>
<tr><td>>hg38_21_+_8433172_8446622_RNA45SN1_45S_50ntFlanks@475.506.32|CTTCGTGATCGATGTGGTGACGTCGTGCTCTC</td><td>>hg38_21_+_8433172_8446622_RNA45SN1_45S_50ntFlanks</td><td>475</td><td>506</td><td>32</td><td>CTTCGTGATCGATGTGGTGACGTCGTGCTCTC</td><td>2423.4</td><td>5.1</td><td>1.0</td><td>5.1</td><td>3.2E-07</td><td>5.2E-07</td><td>MB231/MB468</td><td>MITO</td></tr>
<tr><td>>hg38_21_+_8433172_8446622_RNA45SN1_45S_50ntFlanks@6649.6670.22|TACGACTCTTAGCGGTGGATCA</td><td>>hg38_21_+_8433172_8446622_RNA45SN1_45S_50ntFlanks</td><td>6649</td><td>6670</td><td>22</td><td>TACGACTCTTAGCGGTGGATCA</td><td>689.8</td><td>-4.0</td><td>0.8</td><td>-5.1</td><td>3.2E-07</td><td>5.2E-07</td><td>MB231/MB468</td><td>MITO</td></tr>
<tr><td>>trna119_LysCTT_1_-_145395522_145395594@1.29.29,trna11_LysCTT_5_-_180648979_180649051@1.29.29,trna13_LysCTT_6_+_26556774_26556846@1.29.29,trna32_LysCTT_16_-_3207406_3207478@1.29.29,trna7_LysCTT_16_+_3225692_3225764@1.29.29,trna9_LysCTT_5_+_180634755_180634827@1.29.29|GCCCGGCTAGCTCAGTCGGTAGAGCATGA</td><td>>trna119_LysCTT_1_-_145395522_145395594</td><td>1</td><td>29</td><td>29</td><td>GCCCGGCTAGCTCAGTCGGTAGAGCATGA</td><td>701.8</td><td>-6.1</td><td>1.2</td><td>-5.1</td><td>2.7E-07</td><td>5.2E-07</td><td>MB231/MB468</td><td>MP</td></tr>
<tr><td>>hg38_21_+_8433172_8446622_RNA45SN1_45S_50ntFlanks@10361.10383.23|GAAGGCCGAAGTGGAGAAGGGTT</td><td>>hg38_21_+_8433172_8446622_RNA45SN1_45S_50ntFlanks</td><td>10361</td><td>10383</td><td>23</td><td>GAAGGCCGAAGTGGAGAAGGGTT</td><td>423.5</td><td>-2.9</td><td>0.6</td><td>-5.1</td><td>3.0E-07</td><td>5.2E-07</td><td>MB231/MB468</td><td>CYTO</td></tr>
<tr><td>>hg38_21_+_8433172_8446622_RNA45SN1_45S_50ntFlanks@12638.12661.24|CTACCATCTGTGGGATTATGACTG</td><td>>hg38_21_+_8433172_8446622_RNA45SN1_45S_50ntFlanks</td><td>12638</td><td>12661</td><td>24</td><td>CTACCATCTGTGGGATTATGACTG</td><td>256.8</td><td>-2.6</td><td>0.5</td><td>-5.2</td><td>2.6E-07</td><td>5.2E-07</td><td>BT20/MB468</td><td>CYTO</td></tr>
<tr><td>>hg38_21_+_8433172_8446622_RNA45SN1_45S_50ntFlanks@10393.10426.34|CAGCAGTTGAACATGGGTCAGTCGGTCCTGAGAG</td><td>>hg38_21_+_8433172_8446622_RNA45SN1_45S_50ntFlanks</td><td>10393</td><td>10426</td><td>34</td><td>CAGCAGTTGAACATGGGTCAGTCGGTCCTGAGAG</td><td>370.9</td><td>-7.4</td><td>1.4</td><td>-5.1</td><td>2.7E-07</td><td>5.2E-07</td><td>MB231/MB468</td><td>MP</td></tr>
<tr><td>>hg38_21_+_8433172_8446622_RNA45SN1_45S_50ntFlanks@476.511.36|TTCGTGATCGATGTGGTGACGTCGTGCTCTCCCGGG</td><td>>hg38_21_+_8433172_8446622_RNA45SN1_45S_50ntFlanks</td><td>476</td><td>511</td><td>36</td><td>TTCGTGATCGATGTGGTGACGTCGTGCTCTCCCGGG</td><td>4069.5</td><td>3.7</td><td>0.7</td><td>5.1</td><td>3.0E-07</td><td>5.2E-07</td><td>MB231/MB468</td><td>NUC</td></tr>
<tr><td>>hg38_21_+_8433172_8446622_RNA45SN1_45S_50ntFlanks@12417.12456.40|TCGATGTCGGCTCTTCCTATCATTGTGAAGCAGAATTCAC</td><td>>hg38_21_+_8433172_8446622_RNA45SN1_45S_50ntFlanks</td><td>12417</td><td>12456</td><td>40</td><td>TCGATGTCGGCTCTTCCTATCATTGTGAAGCAGAATTCAC</td><td>667.8</td><td>-8.4</td><td>1.6</td><td>-5.1</td><td>3.3E-07</td><td>5.2E-07</td><td>BT20/MB468</td><td>NUC</td></tr>
<tr><td>>hg38_21_+_8433172_8446622_RNA45SN1_45S_50ntFlanks@4727.4768.42|AATCAAGAACGAAAGTCGGAGGTTCGAAGACGATCAGATACC</td><td>>hg38_21_+_8433172_8446622_RNA45SN1_45S_50ntFlanks</td><td>4727</td><td>4768</td><td>42</td><td>AATCAAGAACGAAAGTCGGAGGTTCGAAGACGATCAGATACC</td><td>85.6</td><td>10.2</td><td>1.9</td><td>5.4</td><td>6.7E-08</td><td>5.2E-07</td><td>BT20/MB468</td><td>MP</td></tr>
<tr><td>>hg38_21_+_8433172_8446622_RNA45SN1_45S_50ntFlanks@8189.8208.20|CCCAGCCCGTGGACGGTGTG</td><td>>hg38_21_+_8433172_8446622_RNA45SN1_45S_50ntFlanks</td><td>8189</td><td>8208</td><td>20</td><td>CCCAGCCCGTGGACGGTGTG</td><td>290.8</td><td>-7.4</td><td>1.4</td><td>-5.4</td><td>6.7E-08</td><td>5.2E-07</td><td>BT20/MB468</td><td>MP</td></tr>
<tr><td>>hg38_21_+_8433172_8446622_RNA45SN1_45S_50ntFlanks@9504.9525.22|GTGAACTATGCCTGGGCAGGGC</td><td>>hg38_21_+_8433172_8446622_RNA45SN1_45S_50ntFlanks</td><td>9504</td><td>9525</td><td>22</td><td>GTGAACTATGCCTGGGCAGGGC</td><td>982.3</td><td>-7.0</td><td>1.4</td><td>-5.2</td><td>2.6E-07</td><td>5.2E-07</td><td>BT20/MB468</td><td>CYTO</td></tr>
<tr><td>>hg38_21_+_8433172_8446622_RNA45SN1_45S_50ntFlanks@12748.12774.27|CCTGTCCCCGCCGGCGGGCCGCCCCCC</td><td>>hg38_21_+_8433172_8446622_RNA45SN1_45S_50ntFlanks</td><td>12748</td><td>12774</td><td>27</td><td>CCTGTCCCCGCCGGCGGGCCGCCCCCC</td><td>262.0</td><td>-7.4</td><td>1.4</td><td>-5.1</td><td>3.0E-07</td><td>5.2E-07</td><td>MB231/MB468</td><td>NUC</td></tr>
<tr><td>>am_MI0000809|hsa-mir-151a&WithFlank&8|-|140732558|140732659@53.70.18(+3G)&[MIMAT0000757&hsa-miR-151a-3p&offsets|0|-3(+3G);m-30&8|-|140732587|140732607&offsets|0|-3(+3G)]|CTAGACTGAAGCTCCTTGGGG</td><td>>am_MI0000809|hsa-mir-151a&WithFlank&8|-|140732558|140732659</td><td>53</td><td>70</td><td>18(+3G)</td><td>CTAGACTGAAGCTCCTTGGGG</td><td>192.2</td><td>5.5</td><td>1.1</td><td>5.1</td><td>3.0E-07</td><td>5.3E-07</td><td>MB231/MB468</td><td>NUC</td></tr>
<tr><td>>hg38_21_+_8433172_8446622_RNA45SN1_45S_50ntFlanks@11576.11605.30|CTTAGAACTGGTGCGGACCAGGGGAATCCG</td><td>>hg38_21_+_8433172_8446622_RNA45SN1_45S_50ntFlanks</td><td>11576</td><td>11605</td><td>30</td><td>CTTAGAACTGGTGCGGACCAGGGGAATCCG</td><td>481.8</td><td>-3.4</td><td>0.7</td><td>-5.2</td><td>2.3E-07</td><td>5.3E-07</td><td>BT20/MB468</td><td>MITO</td></tr>
<tr><td>>hg38_21_+_8433172_8446622_RNA45SN1_45S_50ntFlanks@12008.12043.36|CCGCCGGTGAAATACCACTACTCTGATCGTTTTTTC</td><td>>hg38_21_+_8433172_8446622_RNA45SN1_45S_50ntFlanks</td><td>12008</td><td>12043</td><td>36</td><td>CCGCCGGTGAAATACCACTACTCTGATCGTTTTTTC</td><td>3864.3</td><td>-9.7</td><td>1.8</td><td>-5.4</td><td>6.8E-08</td><td>5.3E-07</td><td>BT20/MB468</td><td>MP</td></tr>
<tr><td>>hg38_21_+_8433172_8446622_RNA45SN1_45S_50ntFlanks@8117.8141.25|CGGGACATGTGGCGTACGGAAGACC</td><td>>hg38_21_+_8433172_8446622_RNA45SN1_45S_50ntFlanks</td><td>8117</td><td>8141</td><td>25</td><td>CGGGACATGTGGCGTACGGAAGACC</td><td>647.5</td><td>-7.5</td><td>1.5</td><td>-5.1</td><td>2.7E-07</td><td>5.3E-07</td><td>MB231/MB468</td><td>MP</td></tr>
<tr><td>>trnaMT_PheGAA_MT_+_577_647@36.56.21|AAAATGTTTAGACGGGCTCAC</td><td>>trnaMT_PheGAA_MT_+_577_647</td><td>36</td><td>56</td><td>21</td><td>AAAATGTTTAGACGGGCTCAC</td><td>688.8</td><td>6.2</td><td>1.1</td><td>5.4</td><td>6.8E-08</td><td>5.3E-07</td><td>BT20/MB468</td><td>MP</td></tr>
<tr><td>>hg38_21_+_8433172_8446622_RNA45SN1_45S_50ntFlanks@12709.12751.43|CCGCGGAGCCTCGGTTGGCCTCGGATAGCCGGTCCCCCGCCTG</td><td>>hg38_21_+_8433172_8446622_RNA45SN1_45S_50ntFlanks</td><td>12709</td><td>12751</td><td>43</td><td>CCGCGGAGCCTCGGTTGGCCTCGGATAGCCGGTCCCCCGCCTG</td><td>62.2</td><td>-9.8</td><td>1.9</td><td>-5.1</td><td>2.7E-07</td><td>5.3E-07</td><td>MB231/MB468</td><td>MP</td></tr>
<tr><td>>hg38_21_+_8433172_8446622_RNA45SN1_45S_50ntFlanks@9953.9983.31|ATGGAAGTCGGAATCCGCTAAGGAGTGTGTA</td><td>>hg38_21_+_8433172_8446622_RNA45SN1_45S_50ntFlanks</td><td>9953</td><td>9983</td><td>31</td><td>ATGGAAGTCGGAATCCGCTAAGGAGTGTGTA</td><td>68.1</td><td>9.8</td><td>1.8</td><td>5.4</td><td>6.8E-08</td><td>5.3E-07</td><td>BT20/MB468</td><td>MP</td></tr>
<tr><td>>MI0000650|hsa-mir-200c&WithFlank&12|+|6963693|6963772@50.72.23(+1U)&[MIMAT0000617&hsa-miR-200c-3p&offsets|0|0(+1U);m-54&12|+|6963742|6963764&offsets|0|0(+1U)]|TAATACTGCCGGGTAATGATGGAT</td><td>>MI0000650|hsa-mir-200c&WithFlank&12|+|6963693|6963772</td><td>50</td><td>72</td><td>23(+1U)</td><td>TAATACTGCCGGGTAATGATGGAT</td><td>1811.4</td><td>7.6</td><td>1.3</td><td>5.7</td><td>1.4E-08</td><td>5.3E-07</td><td>BT20/MB231</td><td>MP</td></tr>
<tr><td>>hg38_21_+_8433172_8446622_RNA45SN1_45S_50ntFlanks@10731.10757.27|AGGGTGTAAATCTCGCGCCGGGCCGTA</td><td>>hg38_21_+_8433172_8446622_RNA45SN1_45S_50ntFlanks</td><td>10731</td><td>10757</td><td>27</td><td>AGGGTGTAAATCTCGCGCCGGGCCGTA</td><td>143.1</td><td>-5.2</td><td>1.0</td><td>-5.1</td><td>3.0E-07</td><td>5.3E-07</td><td>MB231/MB468</td><td>CYTO</td></tr>
<tr><td>>hg38_21_+_8433172_8446622_RNA45SN1_45S_50ntFlanks@12201.12219.19|CCTAAGGCGAGCTCAGGGA</td><td>>hg38_21_+_8433172_8446622_RNA45SN1_45S_50ntFlanks</td><td>12201</td><td>12219</td><td>19</td><td>CCTAAGGCGAGCTCAGGGA</td><td>265.4</td><td>-5.4</td><td>1.0</td><td>-5.1</td><td>2.7E-07</td><td>5.3E-07</td><td>MB231/MB468</td><td>MP</td></tr>
<tr><td>>hg38_21_+_8433172_8446622_RNA45SN1_45S_50ntFlanks@10843.10864.22|TAACTTCGGGATAAGGATTGGC</td><td>>hg38_21_+_8433172_8446622_RNA45SN1_45S_50ntFlanks</td><td>10843</td><td>10864</td><td>22</td><td>TAACTTCGGGATAAGGATTGGC</td><td>377.1</td><td>-3.5</td><td>0.7</td><td>-5.1</td><td>3.0E-07</td><td>5.3E-07</td><td>MB231/MB468</td><td>CYTO</td></tr>
<tr><td>>trna10_AspGTC_12_-_125424193_125424264@35.74.40,trna12_AspGTC_12_-_125411891_125411962@35.74.40,trna144_AspGTC_6_-_27551236_27551307@35.74.40,trna38_AspGTC_17_-_8125556_8125627@35.74.40,trna45_AspGTC_6_+_27447453_27447524@35.74.40,trna48_AspGTC_6_+_27471523_27471594@35.74.40,trna4_AspGTC_12_+_96429799_96429870@35.74.40,trna69_AspGTC_1_-_161440205_161440276@35.74.40,trna72_AspGTC_1_-_161432824_161432895@35.74.40,trna75_AspGTC_1_-_161425414_161425485@35.74.40,trna78_AspGTC_1_-_161418033_161418104@35.74.40,trna81_AspGTC_1_-_161410615_161410686@35.74.40|TCACGCGGGAGACCGGGGTTCGATTCCCCGACGGGGAGCC</td><td>>trna10_AspGTC_12_-_125424193_125424264</td><td>35</td><td>74</td><td>40</td><td>TCACGCGGGAGACCGGGGTTCGATTCCCCGACGGGGAGCC</td><td>11711.1</td><td>-6.0</td><td>1.1</td><td>-5.6</td><td>2.1E-08</td><td>5.3E-07</td><td>BT20/MB231</td><td>CYTO</td></tr>
<tr><td>>am_trna2_GlyGCC_21_-_18827107_18827177@1.25.25,trna35_GlyGCC_1_+_161413094_161413164@1.25.25,trna37_GlyGCC_1_+_161420467_161420537@1.25.25,trna39_GlyGCC_1_+_161427898_161427968@1.25.25,trna41_GlyGCC_1_+_161435258_161435328@1.25.25|GCATGGGTGGTTCAGTGGTAGAATT</td><td>>am_trna2_GlyGCC_21_-_18827107_18827177</td><td>1</td><td>25</td><td>25</td><td>GCATGGGTGGTTCAGTGGTAGAATT</td><td>275.5</td><td>-7.2</td><td>1.4</td><td>-5.1</td><td>2.7E-07</td><td>5.3E-07</td><td>MB231/MB468</td><td>MP</td></tr>
<tr><td>>hg38_21_+_8433172_8446622_RNA45SN1_45S_50ntFlanks@6671.6694.24|CTCGGCTCGTGCGTCGATGAAGAA</td><td>>hg38_21_+_8433172_8446622_RNA45SN1_45S_50ntFlanks</td><td>6671</td><td>6694</td><td>24</td><td>CTCGGCTCGTGCGTCGATGAAGAA</td><td>76.1</td><td>-9.3</td><td>1.7</td><td>-5.4</td><td>6.8E-08</td><td>5.3E-07</td><td>BT20/MB468</td><td>MP</td></tr>
</table>

| | | | | | | | | | | | | | |
|---|---|---|---|---|---|---|---|---|---|---|---|---|---|
| >hg38_21_+_8433172_8446622_RNA45SN1_45S_50ntFlanks@4727.4765.39\|AATCAAGAACGAAAGTCGGAGGTTCGAAGACGATCAGAT | >hg38_21_+_8433172_8446622_RNA45SN1_45S_50ntFlanks | 4727 | 4765 | 39 | AATCAAGAACGAAAGTCGGAGGTTCGAAGACGATCAGAT | 903.0 | 8.4 | 1.6 | 5.4 | 6.9E-08 | 5.3E-07 | BT20/MB468 | MP |
| >hg38_21_+_8433172_8446622_RNA45SN1_45S_50ntFlanks@11317.11344.28\|ACCCCACGTCTCGTCGCGCGCGCGTCCG | >hg38_21_+_8433172_8446622_RNA45SN1_45S_50ntFlanks | 11317 | 11344 | 28 | ACCCCACGTCTCGTCGCGCGCGCGTCCG | 5886.8 | -6.4 | 1.2 | -5.1 | 2.6E-07 | 5.3E-07 | BT20/MB468 | CYTO |
| >hg38_21_+_8433172_8446622_RNA45SN1_45S_50ntFlanks@12995.13039.45\|CTCGCTGCGATCTATTGAAAGTCAGCCCTCGACACAAGGGTTTGT | >hg38_21_+_8433172_8446622_RNA45SN1_45S_50ntFlanks | 12995 | 13039 | 45 | CTCGCTGCGATCTATTGAAAGTCAGCCCTCGACACAAGGGTTTGT | 166557.0 | 5.5 | 1.1 | 5.1 | 3.3E-07 | 5.4E-07 | MB231/MB468 | MITO |
| >hg38_21_+_8433172_8446622_RNA45SN1_45S_50ntFlanks@10879.10902.24\|CGGTCGGGCTGGGGCGCGAAGCGG | >hg38_21_+_8433172_8446622_RNA45SN1_45S_50ntFlanks | 10879 | 10902 | 24 | CGGTCGGGCTGGGGCGCGAAGCGG | 230.1 | -4.1 | 0.8 | -5.1 | 3.0E-07 | 5.4E-07 | MB231/MB468 | CYTO |
| >hg38_21_+_8433172_8446622_RNA45SN1_45S_50ntFlanks@12073.12104.32\|CCGAGGGGCTCTCGCTTCTGGCGCCAAGCGCC | >hg38_21_+_8433172_8446622_RNA45SN1_45S_50ntFlanks | 12073 | 12104 | 32 | CCGAGGGGCTCTCGCTTCTGGCGCCAAGCGCC | 203.9 | -4.2 | 0.8 | -5.1 | 3.0E-07 | 5.4E-07 | MB231/MB468 | CYTO |
| >MI0000478\|hsa-mir-149&WithFlank&2\|+\|240455995\|240456095@21.41.21(+1U)&[MIMAT0000450&hsa-miR-149-5p&offsets\|0\|-2(+1U);m-195&2\|+\|240456015\|240456037&offsets\|0\|-2(+1U)]\|TCTGGCTCCGTGTCTTCACTCT | >MI0000478\|hsa-mir-149&WithFlank&2\|+\|240455995\|240456095 | 21 | 41 | 21(+1U) | TCTGGCTCCGTGTCTTCACTCT | 301.8 | 8.9 | 1.7 | 5.4 | 6.9E-08 | 5.4E-07 | BT20/MB468 | MP |
| >hg38_21_+_8433172_8446622_RNA45SN1_45S_50ntFlanks@5184.5201.18\|AGCAATAACAGGTCTGTG | >hg38_21_+_8433172_8446622_RNA45SN1_45S_50ntFlanks | 5184 | 5201 | 18 | AGCAATAACAGGTCTGTG | 91.0 | -9.6 | 1.8 | -5.4 | 6.9E-08 | 5.4E-07 | BT20/MB468 | MP |
| >am_MI0003581\|hsa-mir-574&WithFlank&4\|+\|38868026\|38868133@31.53.23&[MIMAT0004795&hsa-miR-574-5p&offsets\|0\|0;m-213&4\|+\|38868056\|38868078&offsets\|0\|0]\|TGAGTGTGTGTGTGTGAGTGTGT | >am_MI0003581\|hsa-mir-574&WithFlank&4\|+\|38868026\|38868133 | 31 | 53 | 23 | TGAGTGTGTGTGTGTGAGTGTGT | 1472.2 | 3.7 | 0.7 | 5.1 | 2.8E-07 | 5.4E-07 | MB231/MB468 | MP |
| >hg38_21_+_8433172_8446622_RNA45SN1_45S_50ntFlanks@11781.11804.24\|CGCATGAATGGATGAACGAGATTC | >hg38_21_+_8433172_8446622_RNA45SN1_45S_50ntFlanks | 11781 | 11804 | 24 | CGCATGAATGGATGAACGAGATTC | 4754.8 | -5.5 | 1.0 | -5.4 | 7.0E-08 | 5.4E-07 | BT20/MB468 | MP |
| >hg38_21_+_8433172_8446622_RNA45SN1_45S_50ntFlanks@9504.9533.30\|GTGAACTATGCCTGGGCAGGGCGAAGCCAG | >hg38_21_+_8433172_8446622_RNA45SN1_45S_50ntFlanks | 9504 | 9533 | 30 | GTGAACTATGCCTGGGCAGGGCGAAGCCAG | 1069.5 | -7.5 | 1.4 | -5.1 | 2.7E-07 | 5.4E-07 | BT20/MB468 | CYTO |
| >hg38_21_+_8433172_8446622_RNA45SN1_45S_50ntFlanks@6732.6771.40\|ACATTGATCATCGACACTTCGAACGCACTTGCGGCCCCGG | >hg38_21_+_8433172_8446622_RNA45SN1_45S_50ntFlanks | 6732 | 6771 | 40 | ACATTGATCATCGACACTTCGAACGCACTTGCGGCCCCGG | 288.8 | -3.0 | 0.6 | -5.1 | 2.7E-07 | 5.4E-07 | BT20/MB468 | CYTO |
| >hg38_21_+_8433172_8446622_RNA45SN1_45S_50ntFlanks@12090.12110.21\|CTGGCGCCAAGCGCCCGGCCG | >hg38_21_+_8433172_8446622_RNA45SN1_45S_50ntFlanks | 12090 | 12110 | 21 | CTGGCGCCAAGCGCCCGGCCG | 121.2 | -10.0 | 1.9 | -5.4 | 7.0E-08 | 5.4E-07 | BT20/MB468 | MP |
| >hg38_21_+_8433172_8446622_RNA45SN1_45S_50ntFlanks@9573.9601.29\|AAATCGGTCGTCCGACCTGGGTATAGGGG | >hg38_21_+_8433172_8446622_RNA45SN1_45S_50ntFlanks | 9573 | 9601 | 29 | AAATCGGTCGTCCGACCTGGGTATAGGGG | 523.7 | -3.6 | 0.7 | -5.1 | 3.3E-07 | 5.4E-07 | MB231/MB468 | MITO |
| >hg38_21_+_8433172_8446622_RNA45SN1_45S_50ntFlanks@3840.3874.35\|CTCCTACTTGGATAACTGTGGTAATTCTAGAGCTA | >hg38_21_+_8433172_8446622_RNA45SN1_45S_50ntFlanks | 3840 | 3874 | 35 | CTCCTACTTGGATAACTGTGGTAATTCTAGAGCTA | 384.5 | 3.9 | 0.8 | 5.1 | 2.7E-07 | 5.4E-07 | BT20/MB468 | CYTO |
| >hg38_21_+_8433172_8446622_RNA45SN1_45S_50ntFlanks@10781.10803.23\|AGGTGAACAGCCTCTGGCATGTT | >hg38_21_+_8433172_8446622_RNA45SN1_45S_50ntFlanks | 10781 | 10803 | 23 | AGGTGAACAGCCTCTGGCATGTT | 436.5 | -4.2 | 0.8 | -5.1 | 2.7E-07 | 5.4E-07 | BT20/MB468 | CYTO |
| >hg38_21_+_8433172_8446622_RNA45SN1_45S_50ntFlanks@4895.4918.24\|CTTAAAGGAATTGACGGAAGGGCA | >hg38_21_+_8433172_8446622_RNA45SN1_45S_50ntFlanks | 4895 | 4918 | 24 | CTTAAAGGAATTGACGGAAGGGCA | 86.8 | -9.5 | 1.8 | -5.4 | 7.0E-08 | 5.4E-07 | BT20/MB468 | MP |
| >am_trna10_LysCTT_16_+_3241501_3241573@11.28.18,trna119_LysCTT_1_-_145395522_145395594@11.28.18,trna11_LysCTT_5_-_180648979_180649051@11.28.18,trna128_LysCTT_1_-_55423542_55423614@11.28.18,trna13_LysCTT_14_-_58706613_58706685@11.28.18,trna13_LysCTT_6_+_26556774_26556846@11.28.18,trna2_LysCTT_15_+_79152904_79152976@11.28.18,trna32_LysCTT_16_-_3207406_3207478@11.28.18,trna4_LysCTT_18_-_43669271_43669343@11.28.18,trna6_LysCTT_19_-_52425396_52425468@11.28.18,trna7_LysCTT_16_+_3225692_3225764@11.28.18,trna9_LysCTT_5_+_180634755_180634827@11.28.18\|CTCAGTCGGTAGAGCATG | >am_trna10_LysCTT_16_+_3241501_3241573 | 11 | 28 | 18 | CTCAGTCGGTAGAGCATG | 729.3 | -4.3 | 0.8 | -5.1 | 2.7E-07 | 5.4E-07 | BT20/MB468 | CYTO |
| >hg38_21_+_8433172_8446622_RNA45SN1_45S_50ntFlanks@3704.3725.22\|CTACCTGGTTGATCCTGCCAGT | >hg38_21_+_8433172_8446622_RNA45SN1_45S_50ntFlanks | 3704 | 3725 | 22 | CTACCTGGTTGATCCTGCCAGT | 666.5 | 9.1 | 1.8 | 5.1 | 2.7E-07 | 5.4E-07 | BT20/MB468 | CYTO |
| >hg38_21_+_8433172_8446622_RNA45SN1_45S_50ntFlanks@5056.5082.27\|GAGCGATTTGTCTGGTTAATTCCGATA | >hg38_21_+_8433172_8446622_RNA45SN1_45S_50ntFlanks | 5056 | 5082 | 27 | GAGCGATTTGTCTGGTTAATTCCGATA | 1963.6 | -9.4 | 1.8 | -5.1 | 2.8E-07 | 5.4E-07 | MB231/MB468 | MP |
| >hg38_21_+_8433172_8446622_RNA45SN1_45S_50ntFlanks@12029.12055.27\|TCTGATCGTTTTTTCACTGACCCGGTG | >hg38_21_+_8433172_8446622_RNA45SN1_45S_50ntFlanks | 12029 | 12055 | 27 | TCTGATCGTTTTTTCACTGACCCGGTG | 220.0 | 8.6 | 1.6 | 5.4 | 7.0E-08 | 5.4E-07 | BT20/MB468 | MP |
| >hg38_21_+_8433172_8446622_RNA45SN1_45S_50ntFlanks@10905.10939.35\|CTGGGCGCGCGCCGCGGCTGGACGAGGCGCCGCCG | >hg38_21_+_8433172_8446622_RNA45SN1_45S_50ntFlanks | 10905 | 10939 | 35 | CTGGGCGCGCGCCGCGGCTGGACGAGGCGCCGCCG | 125.9 | -10.1 | 1.9 | -5.4 | 7.0E-08 | 5.4E-07 | BT20/MB468 | MP |
| >hg38_21_+_8433172_8446622_RNA45SN1_45S_50ntFlanks@6738.6762.25\|ATCATCGACACTTCGAACGCACTTG | >hg38_21_+_8433172_8446622_RNA45SN1_45S_50ntFlanks | 6738 | 6762 | 25 | ATCATCGACACTTCGAACGCACTTG | 64.5 | -9.1 | 1.7 | -5.4 | 7.0E-08 | 5.4E-07 | BT20/MB468 | MP |
| >hg38_21_+_8433172_8446622_RNA45SN1_45S_50ntFlanks@11273.11293.21\|CCGCCCCCGGGGAGCCCGGCG | >hg38_21_+_8433172_8446622_RNA45SN1_45S_50ntFlanks | 11273 | 11293 | 21 | CCGCCCCCGGGGAGCCCGGCG | 84.6 | -9.5 | 1.8 | -5.4 | 7.0E-08 | 5.4E-07 | BT20/MB468 | MP |
| >hg38_21_+_8433172_8446622_RNA45SN1_45S_50ntFlanks@12357.12375.19\|CCACAGGGATAACTGGCTT | >hg38_21_+_8433172_8446622_RNA45SN1_45S_50ntFlanks | 12357 | 12375 | 19 | CCACAGGGATAACTGGCTT | 276.4 | -6.1 | 1.2 | -5.1 | 2.7E-07 | 5.4E-07 | BT20/MB468 | CYTO |
| >hg38_21_+_8433172_8446622_RNA45SN1_45S_50ntFlanks@3985.4005.21\|CGGCTTTGGTGACTCTAGATA | >hg38_21_+_8433172_8446622_RNA45SN1_45S_50ntFlanks | 3985 | 4005 | 21 | CGGCTTTGGTGACTCTAGATA | 593.3 | -4.0 | 0.8 | -5.2 | 2.4E-07 | 5.4E-07 | BT20/MB468 | MITO |
| >hg38_21_+_8433172_8446622_RNA45SN1_45S_50ntFlanks@12007.12043.37\|GCCGCCGGTGAAATACCACTACTCTGATCGTTTTTTC | >hg38_21_+_8433172_8446622_RNA45SN1_45S_50ntFlanks | 12007 | 12043 | 37 | GCCGCCGGTGAAATACCACTACTCTGATCGTTTTTTC | 903.7 | -5.7 | 1.1 | -5.1 | 3.1E-07 | 5.4E-07 | MB231/MB468 | CYTO |
| >hg38_21_+_8433172_8446622_RNA45SN1_45S_50ntFlanks@9527.9571.45\|AAGCCAGAGGAAACTCTGGTGGAGGTCCGTAGCGGTCCTGACGTG | >hg38_21_+_8433172_8446622_RNA45SN1_45S_50ntFlanks | 9527 | 9571 | 45 | AAGCCAGAGGAAACTCTGGTGGAGGTCCGTAGCGGTCCTGACGTG | 451.0 | -4.6 | 0.9 | -5.1 | 3.3E-07 | 5.4E-07 | MB231/MB468 | MITO |
| >hg38_21_+_8433172_8446622_RNA45SN1_45S_50ntFlanks@6719.6752.34\|TGAATTGCAGGACACATTGATCATCGACACTTCG | >hg38_21_+_8433172_8446622_RNA45SN1_45S_50ntFlanks | 6719 | 6752 | 34 | TGAATTGCAGGACACATTGATCATCGACACTTCG | 330.6 | -6.0 | 1.2 | -5.1 | 3.2E-07 | 5.4E-07 | MB231/MB468 | NUC |
| >am_trna60_LeuTAA_1_-_238106956_238107030@27.44.18,trnaMT_LeuTAA_MT_+_3230_3304@27.44.18\|CATAAAACTTAAAACTTT | >am_trna60_LeuTAA_1_-_238106956_238107030 | 27 | 44 | 18 | CATAAAACTTAAAACTTT | 190.7 | -6.9 | 1.3 | -5.1 | 2.8E-07 | 5.5E-07 | MB231/MB468 | MP |
| >am_trna128_GlyGCC_6_-_27870686_27870756@1.30.30,trna133_GlyCCC_1_-_16872434_16872504@1.30.30,trna18_GlyGCC_16_+_70822597_70822667@1.30.30,trna19_GlyGCC_16_+_70823410_70823480@1.30.30,trna19_GlyGCC_2_-_157257659_157257729@1.30.30,trna24_GlyGCC_16_-_70812942_70813012@1.30.30,trna25_GlyGCC_16_-_70812114_70812184@1.30.30,trna4_GlyCCC_1_+_17188416_17188486@1.30.30,trna5_GlyGCC_17_+_8029064_8029134@1.30.30,trna68_GlyGCC_1_-_161493637_161493707@1.30.30\|GCATTGGTGGTTCAGTGGTAGAATTCTCGC | >am_trna128_GlyGCC_6_-_27870686_27870756 | 1 | 30 | 30 | GCATTGGTGGTTCAGTGGTAGAATTCTCGC | 27718.0 | -3.2 | 0.6 | -5.1 | 2.7E-07 | 5.5E-07 | BT20/MB468 | CYTO |
| >hg38_21_+_8433172_8446622_RNA45SN1_45S_50ntFlanks@4002.4022.21\|GATAACCTCGGGCCGATCGCA | >hg38_21_+_8433172_8446622_RNA45SN1_45S_50ntFlanks | 4002 | 4022 | 21 | GATAACCTCGGGCCGATCGCA | 2484.9 | -9.1 | 1.8 | -5.1 | 3.5E-07 | 5.5E-07 | BT20/MB468 | NUC |
| >hg38_21_+_8433172_8446622_RNA45SN1_45S_50ntFlanks@4726.4748.23\|TAATCAAGAACGAAAGTCGGAGG | >hg38_21_+_8433172_8446622_RNA45SN1_45S_50ntFlanks | 4726 | 4748 | 23 | TAATCAAGAACGAAAGTCGGAGG | 251.5 | -5.7 | 1.1 | -5.1 | 2.8E-07 | 5.5E-07 | MB231/MB468 | MP |
| >hg38_21_+_8433172_8446622_RNA45SN1_45S_50ntFlanks@12159.12181.23\|TTTGACTGGGGCGGTACACCTGT | >hg38_21_+_8433172_8446622_RNA45SN1_45S_50ntFlanks | 12159 | 12181 | 23 | TTTGACTGGGGCGGTACACCTGT | 372.7 | -6.2 | 1.2 | -5.1 | 3.1E-07 | 5.5E-07 | MB231/MB468 | CYTO |
| >hg38_21_+_8433172_8446622_RNA45SN1_45S_50ntFlanks@10349.10368.20\|AACGAGAACTTTGAAGGCCG | >hg38_21_+_8433172_8446622_RNA45SN1_45S_50ntFlanks | 10349 | 10368 | 20 | AACGAGAACTTTGAAGGCCG | 3328.8 | -5.3 | 1.0 | -5.4 | 7.1E-08 | 5.5E-07 | BT20/MB468 | MP |
| >MI0000087\|hsa-mir-29a&WithFlank&7\|-\|130876741\|130876816@47.67.21(+1A)&[MIMAT0000086&hsa-miR-29a-3p&offsets\|-1\|-2(+1A);m-26&7\|-\|130876749\|130876769&offsets\|-1\|-1(+1A)]\|CTAGCACCATCTGAAATCGGTA | >MI0000087\|hsa-mir-29a&WithFlank&7\|-\|130876741\|130876816 | 47 | 67 | 21(+1A) | CTAGCACCATCTGAAATCGGTA | 817.0 | 6.6 | 1.3 | 5.1 | 2.8E-07 | 5.5E-07 | MB231/MB468 | MP |
| >hg38_21_+_8433172_8446622_RNA45SN1_45S_50ntFlanks@4444.4471.28\|CCTTGCCTCTCGGCGCCCCCTCGATGCT | >hg38_21_+_8433172_8446622_RNA45SN1_45S_50ntFlanks | 4444 | 4471 | 28 | CCTTGCCTCTCGGCGCCCCCTCGATGCT | 290.7 | -4.7 | 0.9 | -5.1 | 2.7E-07 | 5.5E-07 | BT20/MB468 | CYTO |
| >hg38_21_+_8433172_8446622_RNA45SN1_45S_50ntFlanks@10391.10433.43\|AACAGCAGTTGAACATGGGTCAGTCGGTCCTGAGAGATGGGCG | >hg38_21_+_8433172_8446622_RNA45SN1_45S_50ntFlanks | 10391 | 10433 | 43 | AACAGCAGTTGAACATGGGTCAGTCGGTCCTGAGAGATGGGCG | 485.3 | -4.0 | 0.8 | -5.1 | 3.1E-07 | 5.5E-07 | MB231/MB468 | CYTO |
| >hg38_21_+_8433172_8446622_RNA45SN1_45S_50ntFlanks@10350.10388.39\|ACGAGAACTTTGAAGGCCGAAGTGGAGAAGGGTTCCATG | >hg38_21_+_8433172_8446622_RNA45SN1_45S_50ntFlanks | 10350 | 10388 | 39 | ACGAGAACTTTGAAGGCCGAAGTGGAGAAGGGTTCCATG | 2249.1 | -3.4 | 0.7 | -5.2 | 2.4E-07 | 5.5E-07 | BT20/MB468 | MITO |
| >hg38_21_+_8433172_8446622_RNA45SN1_45S_50ntFlanks@12991.13014.24\|CTCCCTCGCTGCGATCTATTGAAA | >hg38_21_+_8433172_8446622_RNA45SN1_45S_50ntFlanks | 12991 | 13014 | 24 | CTCCCTCGCTGCGATCTATTGAAA | 1434.9 | -7.3 | 1.4 | -5.2 | 2.4E-07 | 5.5E-07 | BT20/MB468 | MITO |
| >hg38_21_+_8433172_8446622_RNA45SN1_45S_50ntFlanks@8673.8706.34\|CGGTGCGCCGCGACCGGCTCCGGGACGGCTGGGA | >hg38_21_+_8433172_8446622_RNA45SN1_45S_50ntFlanks | 8673 | 8706 | 34 | CGGTGCGCCGCGACCGGCTCCGGGACGGCTGGGA | 597.5 | 12.6 | 2.2 | 5.7 | 1.5E-08 | 5.5E-07 | BT20/MB231 | MP |
| >am_trnaMT_IleGAT_MT_+_4263_4331@1.32.32\|AGAAATATGTCTGATAAAAGAGTTACTTTGAT | >am_trnaMT_IleGAT_MT_+_4263_4331 | 1 | 32 | 32 | AGAAATATGTCTGATAAAAGAGTTACTTTGAT | 46.0 | 8.7 | 1.7 | 5.1 | 2.8E-07 | 5.5E-07 | MB231/MB468 | MP |
| >hg38_21_+_8433172_8446622_RNA45SN1_45S_50ntFlanks@8184.8223.40\|CGAGGCCCAGCCCGTGGACGGTGTGAGGCCGGTAGCGGCC | >hg38_21_+_8433172_8446622_RNA45SN1_45S_50ntFlanks | 8184 | 8223 | 40 | CGAGGCCCAGCCCGTGGACGGTGTGAGGCCGGTAGCGGCC | 258.9 | -4.8 | 0.9 | -5.1 | 3.1E-07 | 5.5E-07 | MB231/MB468 | CYTO |
| >hg38_21_+_8433172_8446622_RNA45SN1_45S_50ntFlanks@9956.9980.25\|GAAGTCGGAATCCGCTAAGGAGTGT | >hg38_21_+_8433172_8446622_RNA45SN1_45S_50ntFlanks | 9956 | 9980 | 25 | GAAGTCGGAATCCGCTAAGGAGTGT | 64.4 | 9.8 | 1.8 | 5.4 | 7.1E-08 | 5.5E-07 | BT20/MB468 | MP |

<table>
<tr><th></th><th></th><th></th><th></th><th></th><th></th><th></th><th></th><th></th><th></th><th></th><th></th><th></th><th></th></tr>
<tr><td>>MI0000650|hsa-mir-200c&WithFlank&12|+|6963693|6963772@50.70.21&[MIMAT0000617&hsa-miR-200c-3p&offsets|0|-2;m-54&12|+|6963742|6963764&offsets|0|-2]|TAATACTGCCGGGTAATGATG</td><td>>MI0000650|hsa-mir-200c&WithFlank&12|+|6963693|6963772</td><td>50</td><td>70</td><td>21</td><td>TAATACTGCCGGGTAATGATG</td><td>940.2</td><td>3.5</td><td>0.7</td><td>5.1</td><td>3.5E-07</td><td>5.5E-07</td><td>BT20/MB468</td><td>NUC</td></tr>
<tr><td>>hg38_21_+_8433172_8446622_RNA45SN1_45S_50ntFlanks@11576.11607.32|CTTAGAACTGGTGCGGACCAGGGGAATCCGAC</td><td>>hg38_21_+_8433172_8446622_RNA45SN1_45S_50ntFlanks</td><td>11576</td><td>11607</td><td>32</td><td>CTTAGAACTGGTGCGGACCAGGGGAATCCGAC</td><td>761.5</td><td>-6.4</td><td>1.3</td><td>-5.1</td><td>3.5E-07</td><td>5.5E-07</td><td>BT20/MB468</td><td>NUC</td></tr>
<tr><td>>hg38_21_+_8433172_8446622_RNA45SN1_45S_50ntFlanks@8198.8221.24|TGGACGGTGTGAGGCCGGTAGCGG</td><td>>hg38_21_+_8433172_8446622_RNA45SN1_45S_50ntFlanks</td><td>8198</td><td>8221</td><td>24</td><td>TGGACGGTGTGAGGCCGGTAGCGG</td><td>190.0</td><td>-7.2</td><td>1.4</td><td>-5.1</td><td>2.8E-07</td><td>5.5E-07</td><td>MB231/MB468</td><td>MP</td></tr>
<tr><td>>hg38_21_+_8433172_8446622_RNA45SN1_45S_50ntFlanks@11665.11688.24|TTCTGCCCAGTGCTCTGAATGTCA</td><td>>hg38_21_+_8433172_8446622_RNA45SN1_45S_50ntFlanks</td><td>11665</td><td>11688</td><td>24</td><td>TTCTGCCCAGTGCTCTGAATGTCA</td><td>221.2</td><td>6.5</td><td>1.3</td><td>5.1</td><td>3.1E-07</td><td>5.5E-07</td><td>MB231/MB468</td><td>CYTO</td></tr>
<tr><td>>hg38_21_+_8433172_8446622_RNA45SN1_45S_50ntFlanks@4732.4751.20|AGAACGAAAGTCGGAGGTTC</td><td>>hg38_21_+_8433172_8446622_RNA45SN1_45S_50ntFlanks</td><td>4732</td><td>4751</td><td>20</td><td>AGAACGAAAGTCGGAGGTTC</td><td>939.3</td><td>-3.8</td><td>0.7</td><td>-5.1</td><td>2.8E-07</td><td>5.5E-07</td><td>MB231/MB468</td><td>MP</td></tr>
<tr><td>>hg38_21_+_8433172_8446622_RNA45SN1_45S_50ntFlanks@6722.6758.37|ATTGCAGGACACATTGATCATCGACACTTCGAACGCA</td><td>>hg38_21_+_8433172_8446622_RNA45SN1_45S_50ntFlanks</td><td>6722</td><td>6758</td><td>37</td><td>ATTGCAGGACACATTGATCATCGACACTTCGAACGCA</td><td>584.3</td><td>-8.2</td><td>1.6</td><td>-5.1</td><td>3.5E-07</td><td>5.5E-07</td><td>BT20/MB468</td><td>NUC</td></tr>
<tr><td>>hg38_21_+_8433172_8446622_RNA45SN1_45S_50ntFlanks@4733.4758.26|GAACGAAAGTCGGAGGTTCGAAGACG</td><td>>hg38_21_+_8433172_8446622_RNA45SN1_45S_50ntFlanks</td><td>4733</td><td>4758</td><td>26</td><td>GAACGAAAGTCGGAGGTTCGAAGACG</td><td>1366.4</td><td>-4.5</td><td>0.9</td><td>-5.1</td><td>3.4E-07</td><td>5.5E-07</td><td>MB231/MB468</td><td>MITO</td></tr>
<tr><td>>hg38_21_+_8433172_8446622_RNA45SN1_45S_50ntFlanks@6743.6772.30|CGACACTTCGAACGCACTTGCGGCCCCGGG</td><td>>hg38_21_+_8433172_8446622_RNA45SN1_45S_50ntFlanks</td><td>6743</td><td>6772</td><td>30</td><td>CGACACTTCGAACGCACTTGCGGCCCCGGG</td><td>741.9</td><td>-4.0</td><td>0.8</td><td>-5.2</td><td>2.4E-07</td><td>5.5E-07</td><td>BT20/MB468</td><td>MITO</td></tr>
<tr><td>>MI0000071|hsa-mir-17&WithFlank&13|+|91350599|91350694@20.42.23&[MIMAT0000070&hsa-miR-17-5p&offsets|0|0;m-88&13|+|91350618|91350640&offsets|0|0]|CAAAGTGCTTACAGTGCAGGTAG</td><td>>MI0000071|hsa-mir-17&WithFlank&13|+|91350599|91350694</td><td>20</td><td>42</td><td>23</td><td>CAAAGTGCTTACAGTGCAGGTAG</td><td>11344.0</td><td>5.0</td><td>1.0</td><td>5.1</td><td>3.4E-07</td><td>5.5E-07</td><td>MB231/MB468</td><td>MITO</td></tr>
<tr><td>>hg38_21_+_8433172_8446622_RNA45SN1_45S_50ntFlanks@3997.4024.28|CTCTAGATAACCTCGGGCCGATCGCACG</td><td>>hg38_21_+_8433172_8446622_RNA45SN1_45S_50ntFlanks</td><td>3997</td><td>4024</td><td>28</td><td>CTCTAGATAACCTCGGGCCGATCGCACG</td><td>357.6</td><td>-5.5</td><td>1.1</td><td>-5.1</td><td>3.2E-07</td><td>5.5E-07</td><td>MB231/MB468</td><td>NUC</td></tr>
<tr><td>>MI0003513|hsa-mir-455&WithFlank&9|+|114209428|114209535@24.45.22(+1U)&[MIMAT0003150&hsa-miR-455-5p&offsets|+2|+2(+1U);m-232&9|+|114209449|114209470&offsets|+2|+2(+1U)]|TGTGCCTTTGGACTACATCGTGT</td><td>>MI0003513|hsa-mir-455&WithFlank&9|+|114209428|114209535</td><td>24</td><td>45</td><td>22(+1U)</td><td>TGTGCCTTTGGACTACATCGTGT</td><td>49.2</td><td>8.8</td><td>1.7</td><td>5.1</td><td>2.8E-07</td><td>5.5E-07</td><td>MB231/MB468</td><td>MP</td></tr>
<tr><td>>hg38_21_+_8433172_8446622_RNA45SN1_45S_50ntFlanks@11572.11611.40|CCGACTTAGAACTGGTGCGGACCAGGGGAATCCGACTGTT</td><td>>hg38_21_+_8433172_8446622_RNA45SN1_45S_50ntFlanks</td><td>11572</td><td>11611</td><td>40</td><td>CCGACTTAGAACTGGTGCGGACCAGGGGAATCCGACTGTT</td><td>1082.0</td><td>-6.9</td><td>1.3</td><td>-5.1</td><td>2.8E-07</td><td>5.6E-07</td><td>MB231/MB468</td><td>MP</td></tr>
<tr><td>>hg38_21_+_8433172_8446622_RNA45SN1_45S_50ntFlanks@11581.11623.43|AACTGGTGCGGACCAGGGGAATCCGACTGTTTAATTAAAACAA</td><td>>hg38_21_+_8433172_8446622_RNA45SN1_45S_50ntFlanks</td><td>11581</td><td>11623</td><td>43</td><td>AACTGGTGCGGACCAGGGGAATCCGACTGTTTAATTAAAACAA</td><td>92.5</td><td>10.3</td><td>1.9</td><td>5.4</td><td>7.2E-08</td><td>5.6E-07</td><td>BT20/MB468</td><td>MP</td></tr>
<tr><td>>hg38_21_+_8433172_8446622_RNA45SN1_45S_50ntFlanks@6650.6690.41|ACGACTCTTAGCGGTGGATCACTCGGCTCGTGCGTCGATGA</td><td>>hg38_21_+_8433172_8446622_RNA45SN1_45S_50ntFlanks</td><td>6650</td><td>6690</td><td>41</td><td>ACGACTCTTAGCGGTGGATCACTCGGCTCGTGCGTCGATGA</td><td>282.2</td><td>3.5</td><td>0.7</td><td>5.1</td><td>3.2E-07</td><td>5.6E-07</td><td>MB231/MB468</td><td>NUC</td></tr>
<tr><td>>hg38_21_+_8433172_8446622_RNA45SN1_45S_50ntFlanks@485.532.48|GATGTGGTGACGTCGTGCTCTCCCGGGCCGGGTCCGAGCCGCGACGGG</td><td>>hg38_21_+_8433172_8446622_RNA45SN1_45S_50ntFlanks</td><td>485</td><td>532</td><td>48</td><td>GATGTGGTGACGTCGTGCTCTCCCGGGCCGGGTCCGAGCCGCGACGGG</td><td>1251.1</td><td>8.7</td><td>1.7</td><td>5.1</td><td>3.6E-07</td><td>5.6E-07</td><td>BT20/MB468</td><td>NUC</td></tr>
<tr><td>>hg38_21_+_8433172_8446622_RNA45SN1_45S_50ntFlanks@8122.8141.20|CATGTGGCGTACGGAAGACC</td><td>>hg38_21_+_8433172_8446622_RNA45SN1_45S_50ntFlanks</td><td>8122</td><td>8141</td><td>20</td><td>CATGTGGCGTACGGAAGACC</td><td>154.6</td><td>10.6</td><td>1.9</td><td>5.7</td><td>1.5E-08</td><td>5.6E-07</td><td>BT20/MB231</td><td>MP</td></tr>
<tr><td>>hg38_21_+_8433172_8446622_RNA45SN1_45S_50ntFlanks@4367.4406.40|CAGTTAAAAAGCTCGTAGTTGGATCTTGGGAGCGGGCGGG</td><td>>hg38_21_+_8433172_8446622_RNA45SN1_45S_50ntFlanks</td><td>4367</td><td>4406</td><td>40</td><td>CAGTTAAAAAGCTCGTAGTTGGATCTTGGGAGCGGGCGGG</td><td>128.7</td><td>-6.3</td><td>1.2</td><td>-5.1</td><td>2.8E-07</td><td>5.6E-07</td><td>BT20/MB468</td><td>CYTO</td></tr>
<tr><td>>hg38_21_+_8433172_8446622_RNA45SN1_45S_50ntFlanks@8192.8210.19|AGCCCGTGGACGGTGTGAG</td><td>>hg38_21_+_8433172_8446622_RNA45SN1_45S_50ntFlanks</td><td>8192</td><td>8210</td><td>19</td><td>AGCCCGTGGACGGTGTGAG</td><td>85.7</td><td>-9.5</td><td>1.8</td><td>-5.4</td><td>7.2E-08</td><td>5.6E-07</td><td>BT20/MB468</td><td>MP</td></tr>
<tr><td>>hg38_1_-_228634819_228635039_RNA5S12_5S_50ntFlanks@145.170.26|CTGGGAATACCGGGTGCTGTAGGCTT</td><td>>hg38_1_-_228634819_228635039_RNA5S12_5S_50ntFlanks</td><td>145</td><td>170</td><td>26</td><td>CTGGGAATACCGGGTGCTGTAGGCTT</td><td>1051.3</td><td>4.3</td><td>0.9</td><td>5.1</td><td>3.2E-07</td><td>5.6E-07</td><td>MB231/MB468</td><td>NUC</td></tr>
<tr><td>>hg38_21_+_8433172_8446622_RNA45SN1_45S_50ntFlanks@3843.3871.29|CTACTTGGATAACTGTGGTAATTCTAGAG</td><td>>hg38_21_+_8433172_8446622_RNA45SN1_45S_50ntFlanks</td><td>3843</td><td>3871</td><td>29</td><td>CTACTTGGATAACTGTGGTAATTCTAGAG</td><td>200.9</td><td>-3.0</td><td>0.6</td><td>-5.1</td><td>3.2E-07</td><td>5.6E-07</td><td>MB231/MB468</td><td>NUC</td></tr>
<tr><td>>hg38_21_+_8433172_8446622_RNA45SN1_45S_50ntFlanks@9572.9594.23|CAAATCGGTCGTCCGACCTGGGT</td><td>>hg38_21_+_8433172_8446622_RNA45SN1_45S_50ntFlanks</td><td>9572</td><td>9594</td><td>23</td><td>CAAATCGGTCGTCCGACCTGGGT</td><td>293.5</td><td>-6.0</td><td>1.2</td><td>-5.1</td><td>3.6E-07</td><td>5.6E-07</td><td>BT20/MB468</td><td>NUC</td></tr>
<tr><td>>hg38_21_+_8433172_8446622_RNA45SN1_45S_50ntFlanks@5471.5494.24|CACGGCCCTGGCGGAGCGCTGAGA</td><td>>hg38_21_+_8433172_8446622_RNA45SN1_45S_50ntFlanks</td><td>5471</td><td>5494</td><td>24</td><td>CACGGCCCTGGCGGAGCGCTGAGA</td><td>813.4</td><td>4.4</td><td>0.9</td><td>5.1</td><td>3.2E-07</td><td>5.6E-07</td><td>MB231/MB468</td><td>CYTO</td></tr>
<tr><td>>am_trna2_GlyGCC_21_-_18827107_18827177@1.22.22,trna35_GlyGCC_1_+_161413094_161413164@1.22.22,trna37_GlyGCC_1_+_161420467_161420537@1.22.22,trna39_GlyGCC_1_+_161427898_161427968@1.22.22,trna41_GlyGCC_1_+_161435258_161435328@1.22.22|GCATGGGTGGTTCAGTGGTAGA</td><td>>am_trna2_GlyGCC_21_-_18827107_18827177</td><td>1</td><td>22</td><td>22</td><td>GCATGGGTGGTTCAGTGGTAGA</td><td>147.2</td><td>10.5</td><td>1.9</td><td>5.7</td><td>1.5E-08</td><td>5.6E-07</td><td>BT20/MB231</td><td>MP</td></tr>
<tr><td>>hg38_21_+_8433172_8446622_RNA45SN1_45S_50ntFlanks@6661.6678.18|CGGTGGATCACTCGGCTC</td><td>>hg38_21_+_8433172_8446622_RNA45SN1_45S_50ntFlanks</td><td>6661</td><td>6678</td><td>18</td><td>CGGTGGATCACTCGGCTC</td><td>2032.3</td><td>5.6</td><td>1.0</td><td>5.4</td><td>7.2E-08</td><td>5.6E-07</td><td>BT20/MB468</td><td>MP</td></tr>
<tr><td>>MI0000093|hsa-mir-92a-1&WithFlank&13|+|91351308|91351397@54.74.21(+1G);MI0000094|hsa-mir-92a-2&WithFlank&X|-|134169532|134169618@54.74.21(+1G)&[MIMAT0000092&hsa-miR-92a-3p&offsets|0|-1(+1G);m-9&13|+|91351361|91351382&offsets|0|-1(+1G)];[MIMAT0000092_1&hsa-miR-92a-3p&offsets|0|-1(+1G);m-11&X|-|134169544|134169565&offsets|0|-1(+1G)]|TATTGCACTTGTCCCGGCCTGG</td><td>>MI0000093|hsa-mir-92a-1&WithFlank&13|+|91351308|91351397</td><td>54</td><td>74</td><td>21(+1G)</td><td>TATTGCACTTGTCCCGGCCTGG</td><td>1517.9</td><td>4.9</td><td>1.0</td><td>5.1</td><td>3.2E-07</td><td>5.6E-07</td><td>MB231/MB468</td><td>CYTO</td></tr>
<tr><td>>hg38_21_+_8433172_8446622_RNA45SN1_45S_50ntFlanks@11873.11900.28|AAGAAGACCCTGTTGAGCTTGACTCTAG</td><td>>hg38_21_+_8433172_8446622_RNA45SN1_45S_50ntFlanks</td><td>11873</td><td>11900</td><td>28</td><td>AAGAAGACCCTGTTGAGCTTGACTCTAG</td><td>427.7</td><td>-3.0</td><td>0.6</td><td>-5.2</td><td>2.4E-07</td><td>5.6E-07</td><td>BT20/MB468</td><td>MITO</td></tr>
<tr><td>>hg38_21_+_8433172_8446622_RNA45SN1_45S_50ntFlanks@3705.3733.29|TACCTGGTTGATCCTGCCAGTAGCATATG</td><td>>hg38_21_+_8433172_8446622_RNA45SN1_45S_50ntFlanks</td><td>3705</td><td>3733</td><td>29</td><td>TACCTGGTTGATCCTGCCAGTAGCATATG</td><td>371.0</td><td>5.2</td><td>1.0</td><td>5.1</td><td>3.2E-07</td><td>5.6E-07</td><td>MB231/MB468</td><td>CYTO</td></tr>
<tr><td>>hg38_21_+_8433172_8446622_RNA45SN1_45S_50ntFlanks@4555.4574.20|CGCAGCTAGGAATAATGGAA</td><td>>hg38_21_+_8433172_8446622_RNA45SN1_45S_50ntFlanks</td><td>4555</td><td>4574</td><td>20</td><td>CGCAGCTAGGAATAATGGAA</td><td>258.1</td><td>-5.2</td><td>1.0</td><td>-5.1</td><td>3.3E-07</td><td>5.6E-07</td><td>MB231/MB468</td><td>NUC</td></tr>
<tr><td>>am_trna128_GlyGCC_6_-_27870686_27870756@1.32.32,trna133_GlyCCC_1_-_16872434_16872504@1.32.32,trna18_GlyGCC_16_+_70822597_70822667@1.32.32,trna19_GlyGCC_16_+_70823410_70823480@1.32.32,trna19_GlyGCC_2_-_157257659_157257729@1.32.32,trna24_GlyGCC_16_-_70812942_70813012@1.32.32,trna25_GlyGCC_16_-_70812114_70812184@1.32.32,trna4_GlyCCC_1_+_17188416_17188486@1.32.32,trna5_GlyGCC_17_+_8029064_8029134@1.32.32,trna68_GlyGCC_1_-_161493637_161493707@1.32.32|GCATTGGTGGTTCAGTGGTAGAATTCTCGCCT</td><td>>am_trna128_GlyGCC_6_-_27870686_27870756</td><td>1</td><td>32</td><td>32</td><td>GCATTGGTGGTTCAGTGGTAGAATTCTCGCCT</td><td>408.2</td><td>3.0</td><td>0.6</td><td>5.1</td><td>3.6E-07</td><td>5.6E-07</td><td>BT20/MB468</td><td>NUC</td></tr>
<tr><td>>hg38_21_+_8433172_8446622_RNA45SN1_45S_50ntFlanks@6648.6668.21|GTACGACTCTTAGCGGTGGAT</td><td>>hg38_21_+_8433172_8446622_RNA45SN1_45S_50ntFlanks</td><td>6648</td><td>6668</td><td>21</td><td>GTACGACTCTTAGCGGTGGAT</td><td>444.1</td><td>3.2</td><td>0.6</td><td>5.1</td><td>3.3E-07</td><td>5.6E-07</td><td>MB231/MB468</td><td>NUC</td></tr>
<tr><td>>hg38_21_+_8433172_8446622_RNA45SN1_45S_50ntFlanks@6661.6690.30|CGGTGGATCACTCGGCTCGTGCGTCGATGA</td><td>>hg38_21_+_8433172_8446622_RNA45SN1_45S_50ntFlanks</td><td>6661</td><td>6690</td><td>30</td><td>CGGTGGATCACTCGGCTCGTGCGTCGATGA</td><td>978.7</td><td>4.1</td><td>0.8</td><td>5.2</td><td>2.5E-07</td><td>5.6E-07</td><td>BT20/MB468</td><td>MITO</td></tr>
<tr><td>>hg38_21_+_8433172_8446622_RNA45SN1_45S_50ntFlanks@10694.10711.18|CGGTGAGCTCTCGCTGGC</td><td>>hg38_21_+_8433172_8446622_RNA45SN1_45S_50ntFlanks</td><td>10694</td><td>10711</td><td>18</td><td>CGGTGAGCTCTCGCTGGC</td><td>347.1</td><td>-7.5</td><td>1.5</td><td>-5.1</td><td>3.6E-07</td><td>5.6E-07</td><td>BT20/MB468</td><td>NUC</td></tr>
<tr><td>>hg38_21_+_8433172_8446622_RNA45SN1_45S_50ntFlanks@10546.10581.36|CAGTGCGGTAACGCGACCGATCCCGGAGAAGCCGGC</td><td>>hg38_21_+_8433172_8446622_RNA45SN1_45S_50ntFlanks</td><td>10546</td><td>10581</td><td>36</td><td>CAGTGCGGTAACGCGACCGATCCCGGAGAAGCCGGC</td><td>61.4</td><td>9.7</td><td>1.8</td><td>5.4</td><td>7.3E-08</td><td>5.6E-07</td><td>BT20/MB468</td><td>MP</td></tr>
<tr><td>>hg38_21_+_8433172_8446622_RNA45SN1_45S_50ntFlanks@12008.12033.26|CCGCCGGTGAAATACCACTACTCTGA</td><td>>hg38_21_+_8433172_8446622_RNA45SN1_45S_50ntFlanks</td><td>12008</td><td>12033</td><td>26</td><td>CCGCCGGTGAAATACCACTACTCTGA</td><td>546.3</td><td>-6.5</td><td>1.3</td><td>-5.2</td><td>2.5E-07</td><td>5.6E-07</td><td>BT20/MB468</td><td>MITO</td></tr>
<tr><td>>MI0000734|hsa-mir-106b&WithFlank&7|-|100093987|100094080@58.78.21(+1U)&[MIMAT0004672&hsa-miR-106b-3p&offsets|0|-1(+1U);m-87&7|-|100094003|100094023&offsets|0|0(+1U)]|CCGCACTGTGGGTACTTGCTGT</td><td>>MI0000734|hsa-mir-106b&WithFlank&7|-|100093987|100094080</td><td>58</td><td>78</td><td>21(+1U)</td><td>CCGCACTGTGGGTACTTGCTGT</td><td>1743.5</td><td>5.6</td><td>1.0</td><td>5.4</td><td>7.3E-08</td><td>5.6E-07</td><td>BT20/MB468</td><td>MP</td></tr>
<tr><td>>hg38_21_+_8433172_8446622_RNA45SN1_45S_50ntFlanks@9570.9595.26|TGCAAATCGGTCGTCCGACCTGGGTA</td><td>>hg38_21_+_8433172_8446622_RNA45SN1_45S_50ntFlanks</td><td>9570</td><td>9595</td><td>26</td><td>TGCAAATCGGTCGTCCGACCTGGGTA</td><td>2397.8</td><td>-5.5</td><td>1.0</td><td>-5.4</td><td>7.3E-08</td><td>5.6E-07</td><td>BT20/MB468</td><td>MP</td></tr>
<tr><td>>MI0000101|hsa-mir-99a&WithFlank&21|+|16539083|16539175@19.39.21&[MIMAT0000097&hsa-miR-99a-5p&offsets|0|-1;m-64&21|+|16539101|16539121&offsets|0|0]|AACCCGTAGATCCGATCTTGT</td><td>>MI0000101|hsa-mir-99a&WithFlank&21|+|16539083|16539175</td><td>19</td><td>39</td><td>21</td><td>AACCCGTAGATCCGATCTTGT</td><td>1332.7</td><td>-5.7</td><td>1.0</td><td>-5.6</td><td>2.7E-08</td><td>5.6E-07</td><td>BT20/MB231</td><td>MITO</td></tr>
<tr><td>>hg38_21_+_8433172_8446622_RNA45SN1_45S_50ntFlanks@10684.10706.23|CGGCGGCGTCCGGTGAGCTCTCG</td><td>>hg38_21_+_8433172_8446622_RNA45SN1_45S_50ntFlanks</td><td>10684</td><td>10706</td><td>23</td><td>CGGCGGCGTCCGGTGAGCTCTCG</td><td>22500.1</td><td>-10.3</td><td>2.0</td><td>-5.1</td><td>3.6E-07</td><td>5.6E-07</td><td>BT20/MB468</td><td>NUC</td></tr>
<tr><td>>hg38_21_+_8433172_8446622_RNA45SN1_45S_50ntFlanks@8183.8208.26|TCGAGGCCCAGCCCGTGGACGGTGTG</td><td>>hg38_21_+_8433172_8446622_RNA45SN1_45S_50ntFlanks</td><td>8183</td><td>8208</td><td>26</td><td>TCGAGGCCCAGCCCGTGGACGGTGTG</td><td>211.5</td><td>-3.7</td><td>0.7</td><td>-5.1</td><td>3.3E-07</td><td>5.6E-07</td><td>MB231/MB468</td><td>NUC</td></tr>
<tr><td>>hg38_21_+_8433172_8446622_RNA45SN1_45S_50ntFlanks@4760.4778.19|TCAGATACCGTCGTAGTTC</td><td>>hg38_21_+_8433172_8446622_RNA45SN1_45S_50ntFlanks</td><td>4760</td><td>4778</td><td>19</td><td>TCAGATACCGTCGTAGTTC</td><td>544.1</td><td>-6.6</td><td>1.3</td><td>-5.1</td><td>2.8E-07</td><td>5.6E-07</td><td>BT20/MB468</td><td>CYTO</td></tr>
<tr><td>>hg38_21_+_8433172_8446622_RNA45SN1_45S_50ntFlanks@5487.5526.40|CGCTGAGAAGACGGTCGAACTTGACTATCTAGAGGAAGTA</td><td>>hg38_21_+_8433172_8446622_RNA45SN1_45S_50ntFlanks</td><td>5487</td><td>5526</td><td>40</td><td>CGCTGAGAAGACGGTCGAACTTGACTATCTAGAGGAAGTA</td><td>517.0</td><td>-7.3</td><td>1.4</td><td>-5.1</td><td>3.3E-07</td><td>5.6E-07</td><td>MB231/MB468</td><td>NUC</td></tr>
<tr><td>>hg38_21_+_8433172_8446622_RNA45SN1_45S_50ntFlanks@11208.11234.27|GCGGCGACTCTGGACGCGAGCCGGGCC</td><td>>hg38_21_+_8433172_8446622_RNA45SN1_45S_50ntFlanks</td><td>11208</td><td>11234</td><td>27</td><td>GCGGCGACTCTGGACGCGAGCCGGGCC</td><td>6829.2</td><td>-5.3</td><td>1.0</td><td>-5.1</td><td>2.8E-07</td><td>5.6E-07</td><td>BT20/MB468</td><td>CYTO</td></tr>
<tr><td>>MI0005561|hsa-mir-877&WithFlank&6|+|30584326|30584423@8.29.22&[MIMAT0004949&hsa-miR-877-5p&offsets|+1|+3;m-382&6|+|30584332|30584354&offsets|+1|0]|TAGAGGAGATGGCGCAGGGGAC</td><td>>MI0005561|hsa-mir-877&WithFlank&6|+|30584326|30584423</td><td>8</td><td>29</td><td>22</td><td>TAGAGGAGATGGCGCAGGGGAC</td><td>87.1</td><td>10.2</td><td>1.9</td><td>5.4</td><td>7.3E-08</td><td>5.6E-07</td><td>BT20/MB468</td><td>MP</td></tr>
<tr><td>>hg38_21_+_8433172_8446622_RNA45SN1_45S_50ntFlanks@3840.3881.42|CTCCTACTTGGATAACTGTGGTAATTCTAGAGCTAATACATG</td><td>>hg38_21_+_8433172_8446622_RNA45SN1_45S_50ntFlanks</td><td>3840</td><td>3881</td><td>42</td><td>CTCCTACTTGGATAACTGTGGTAATTCTAGAGCTAATACATG</td><td>427.9</td><td>-6.5</td><td>1.3</td><td>-5.1</td><td>3.6E-07</td><td>5.6E-07</td><td>BT20/MB468</td><td>NUC</td></tr>
<tr><td>>hg38_21_+_8433172_8446622_RNA45SN1_45S_50ntFlanks@11672.11704.33|CAGTGCTCTGAATGTCAAAGTGAAGAAATTCAA</td><td>>hg38_21_+_8433172_8446622_RNA45SN1_45S_50ntFlanks</td><td>11672</td><td>11704</td><td>33</td><td>CAGTGCTCTGAATGTCAAAGTGAAGAAATTCAA</td><td>1060.4</td><td>-6.7</td><td>1.3</td><td>-5.1</td><td>3.6E-07</td><td>5.6E-07</td><td>BT20/MB468</td><td>NUC</td></tr>
<tr><td>>am_MI0000786|hsa-mir-378a&WithFlank&5|+|149732819|149732896@49.69.21(+1A)&[MIMAT0000732&hsa-miR-378a-3p&offsets|0|-1(+1A);m-44&5|+|149732867|149732888&offsets|0|-1(+1A)]|ACTGGACTTGGAGTCAGAAGGA</td><td>>am_MI0000786|hsa-mir-378a&WithFlank&5|+|149732819|149732896</td><td>49</td><td>69</td><td>21(+1A)</td><td>ACTGGACTTGGAGTCAGAAGGA</td><td>711.6</td><td>2.9</td><td>0.6</td><td>5.1</td><td>3.3E-07</td><td>5.7E-07</td><td>MB231/MB468</td><td>NUC</td></tr>
</table>

| | | | | | | | | | | | | | |
|---|---|---|---|---|---|---|---|---|---|---|---|---|---|
| >MI0001733\|hsa-mir-452&WithFlank&X\|-\|151959622\|151959718@23.44.22&[MIMAT0001635&hsa-miR-452-5p&offsets\|+3\|+3;m-188&X\|-\|151959677\|151959699&offsets\|+3\|+2]\|TGTTTGCAGAGGAAACTGAGAC | >MI0001733\|hsa-mir-452&WithFlank&X\|-\|151959622\|151959718 | 23 | 44 | 22 | TGTTTGCAGAGGAAACTGAGAC | 333.9 | 8.0 | 1.6 | 5.1 | 2.9E-07 | 5.7E-07 | MB231/MB468 | MP |
| >MI0000079\|hsa-mir-23a&WithFlank&19\|-\|13836581\|13836665@51.73.23&[MIMAT0000078&hsa-miR-23a-3p&offsets\|0\|+2;m-31&19\|-\|13836595\|13836615&offsets\|0\|+2]\|ATCACATTGCCAGGGATTTCCAA | >MI0000079\|hsa-mir-23a&WithFlank&19\|-\|13836581\|13836665 | 51 | 73 | 23 | ATCACATTGCCAGGGATTTCCAA | 123.1 | -10.0 | 1.9 | -5.4 | 7.4E-08 | 5.7E-07 | BT20/MB468 | MP |
| >hg38_21_+_8433172_8446622_RNA45SN1_45S_50ntFlanks@9196.9218.23\|AGGTTCTCTCGGGGCCACGCGCG | >hg38_21_+_8433172_8446622_RNA45SN1_45S_50ntFlanks | 9196 | 9218 | 23 | AGGTTCTCTCGGGGCCACGCGCG | 90.2 | -9.6 | 1.8 | -5.4 | 7.4E-08 | 5.7E-07 | BT20/MB468 | MP |
| >hg38_21_+_8433172_8446622_RNA45SN1_45S_50ntFlanks@11567.11603.37\|AGCAGCCGACTTAGAACTGGTGCGGACCAGGGGAATC | >hg38_21_+_8433172_8446622_RNA45SN1_45S_50ntFlanks | 11567 | 11603 | 37 | AGCAGCCGACTTAGAACTGGTGCGGACCAGGGGAATC | 1971.5 | -7.1 | 1.4 | -5.1 | 3.6E-07 | 5.7E-07 | BT20/MB468 | NUC |
| >MI0000102\|hsa-mir-100&WithFlank&11\|-\|122152223\|122152314@19.37.19(+1A)&[MIMAT0000098&hsa-miR-100-5p&offsets\|0\|-3(+1A);m-27&11\|-\|122152276\|122152296&offsets\|0\|-2(+1A)]\|AACCCGTAGATCCGAACTTA | >MI0000102\|hsa-mir-100&WithFlank&11\|-\|122152223\|122152314 | 19 | 37 | 19(+1A) | AACCCGTAGATCCGAACTTA | 550.5 | 13.4 | 2.6 | 5.1 | 2.8E-07 | 5.7E-07 | BT20/MB468 | CYTO |
| >trna14_LysTTT_11_-_59327808_59327880@3.36.34,trna2_LysTTT_17_+_8022473_8022545@3.36.34,trna54_LysTTT_1_+_204475655_204475727@3.36.34,trna5_LysTTT_11_+_59323902_59323974@3.36.34,trna62_LysTTT_1_-_204476158_204476230@3.36.34,trna76_LysTTT_6_+_28918806_28918878@3.36.34\|CCGGATAGCTCAGTCGGTAGAGCATCAGACTTTT | >trna14_LysTTT_11_-_59327808_59327880 | 3 | 36 | 34 | CCGGATAGCTCAGTCGGTAGAGCATCAGACTTTT | 405.1 | 8.7 | 1.7 | 5.1 | 2.8E-07 | 5.7E-07 | BT20/MB468 | CYTO |
| >hg38_21_+_8433172_8446622_RNA45SN1_45S_50ntFlanks@11921.11959.39\|TGAGAGGTGTAGAATAAGTGGGAGGCCCCCGGCGCCCCC | >hg38_21_+_8433172_8446622_RNA45SN1_45S_50ntFlanks | 11921 | 11959 | 39 | TGAGAGGTGTAGAATAAGTGGGAGGCCCCCGGCGCCCCC | 71.2 | -10.0 | 2.0 | -5.1 | 2.9E-07 | 5.7E-07 | MB231/MB468 | MP |
| >hg38_21_+_8433172_8446622_RNA45SN1_45S_50ntFlanks@11604.11640.37\|CGACTGTTTAATTAAAACAAAGCATCGCGAAGGCCCG | >hg38_21_+_8433172_8446622_RNA45SN1_45S_50ntFlanks | 11604 | 11640 | 37 | CGACTGTTTAATTAAAACAAAGCATCGCGAAGGCCCG | 110.4 | -9.9 | 1.8 | -5.4 | 7.4E-08 | 5.7E-07 | BT20/MB468 | MP |
| >hg38_21_+_8433172_8446622_RNA45SN1_45S_50ntFlanks@11875.11904.30\|GAAGACCCTGTTGAGCTTGACTCTAGTCTG | >hg38_21_+_8433172_8446622_RNA45SN1_45S_50ntFlanks | 11875 | 11904 | 30 | GAAGACCCTGTTGAGCTTGACTCTAGTCTG | 116.0 | -9.9 | 1.8 | -5.4 | 7.4E-08 | 5.7E-07 | BT20/MB468 | MP |
| >hg38_21_+_8433172_8446622_RNA45SN1_45S_50ntFlanks@11890.11908.19\|CTTGACTCTAGTCTGGCAC | >hg38_21_+_8433172_8446622_RNA45SN1_45S_50ntFlanks | 11890 | 11908 | 19 | CTTGACTCTAGTCTGGCAC | 72.9 | -9.3 | 1.7 | -5.4 | 7.4E-08 | 5.7E-07 | BT20/MB468 | MP |
| >hg38_21_+_8433172_8446622_RNA45SN1_45S_50ntFlanks@6671.6697.27\|CTCGGCTCGTGCGTCGATGAAGAACGC | >hg38_21_+_8433172_8446622_RNA45SN1_45S_50ntFlanks | 6671 | 6697 | 27 | CTCGGCTCGTGCGTCGATGAAGAACGC | 434.5 | -4.4 | 0.9 | -5.1 | 3.2E-07 | 5.7E-07 | MB231/MB468 | CYTO |
| >hg38_21_+_8433172_8446622_RNA45SN1_45S_50ntFlanks@10680.10706.27\|GTTCCGGCGGCGTCCGGTGAGCTCTCG | >hg38_21_+_8433172_8446622_RNA45SN1_45S_50ntFlanks | 10680 | 10706 | 27 | GTTCCGGCGGCGTCCGGTGAGCTCTCG | 31437.1 | -8.7 | 1.7 | -5.1 | 3.6E-07 | 5.7E-07 | BT20/MB468 | NUC |
| >hg38_21_+_8433172_8446622_RNA45SN1_45S_50ntFlanks@11315.11338.24\|CCACCCCACGTCTCGTCGCGCGCG | >hg38_21_+_8433172_8446622_RNA45SN1_45S_50ntFlanks | 11315 | 11338 | 24 | CCACCCCACGTCTCGTCGCGCGCG | 551.4 | -5.2 | 1.0 | -5.1 | 2.8E-07 | 5.7E-07 | BT20/MB468 | CYTO |
| >hg38_21_+_8433172_8446622_RNA45SN1_45S_50ntFlanks@12417.12443.27\|TCGATGTCGGCTCTTCCTATCATTGTG | >hg38_21_+_8433172_8446622_RNA45SN1_45S_50ntFlanks | 12417 | 12443 | 27 | TCGATGTCGGCTCTTCCTATCATTGTG | 3265.6 | -3.3 | 0.6 | -5.2 | 2.5E-07 | 5.7E-07 | BT20/MB468 | MITO |
| >hg38_21_+_8433172_8446622_RNA45SN1_45S_50ntFlanks@9598.9622.25\|GGGGCGAAAGACTAATCGAACCATC | >hg38_21_+_8433172_8446622_RNA45SN1_45S_50ntFlanks | 9598 | 9622 | 25 | GGGGCGAAAGACTAATCGAACCATC | 146.9 | -10.3 | 1.9 | -5.4 | 7.4E-08 | 5.7E-07 | BT20/MB468 | MP |
| >hg38_21_+_8433172_8446622_RNA45SN1_45S_50ntFlanks@10357.10390.34\|CTTTGAAGGCCGAAGTGGAGAAGGGTTCCATGTG | >hg38_21_+_8433172_8446622_RNA45SN1_45S_50ntFlanks | 10357 | 10390 | 34 | CTTTGAAGGCCGAAGTGGAGAAGGGTTCCATGTG | 398.7 | -3.8 | 0.7 | -5.1 | 2.8E-07 | 5.7E-07 | BT20/MB468 | CYTO |
| >MI0000746\|hsa-mir-99b&WithFlank&19\|+\|51692606\|51692687@51.71.21&[MIMAT0004678&hsa-miR-99b-3p&offsets\|0\|-1;m-151&19\|+\|51692656\|51692678&offsets\|0\|-2]\|CAAGCTCGTGTCTGTGGGTCC | >MI0000746\|hsa-mir-99b&WithFlank&19\|+\|51692606\|51692687 | 51 | 71 | 21 | CAAGCTCGTGTCTGTGGGTCC | 657.2 | 5.6 | 1.0 | 5.4 | 7.4E-08 | 5.7E-07 | BT20/MB468 | MP |
| >hg38_21_+_8433172_8446622_RNA45SN1_45S_50ntFlanks@11538.11560.23\|CGCGCGGCGCCTCGCCTCGGCCG | >hg38_21_+_8433172_8446622_RNA45SN1_45S_50ntFlanks | 11538 | 11560 | 23 | CGCGCGGCGCCTCGCCTCGGCCG | 81.8 | -9.4 | 1.8 | -5.4 | 7.4E-08 | 5.7E-07 | BT20/MB468 | MP |
| >hg38_1_-_228634819_228635039_RNA5S12_5S_50ntFlanks@130.170.41\|TGGATGGGAGACCGCCTGGGAATACCGGGTGCTGTAGGCTT | >hg38_1_-_228634819_228635039_RNA5S12_5S_50ntFlanks | 130 | 170 | 41 | TGGATGGGAGACCGCCTGGGAATACCGGGTGCTGTAGGCTT | 1377.6 | 5.4 | 1.0 | 5.1 | 3.3E-07 | 5.7E-07 | MB231/MB468 | NUC |
| >hg38_21_+_8433172_8446622_RNA45SN1_45S_50ntFlanks@12948.12988.41\|GACGACCTGCTTCTGGGTCGGGGTTTCGTACGTAGCAGAGC | >hg38_21_+_8433172_8446622_RNA45SN1_45S_50ntFlanks | 12948 | 12988 | 41 | GACGACCTGCTTCTGGGTCGGGGTTTCGTACGTAGCAGAGC | 685.2 | 3.2 | 0.6 | 5.1 | 3.2E-07 | 5.7E-07 | MB231/MB468 | CYTO |
| >hg38_21_+_8433172_8446622_RNA45SN1_45S_50ntFlanks@12457.12475.19\|CAAGCGTTGGATTGTTCAC | >hg38_21_+_8433172_8446622_RNA45SN1_45S_50ntFlanks | 12457 | 12475 | 19 | CAAGCGTTGGATTGTTCAC | 1449.8 | -7.8 | 1.5 | -5.1 | 2.8E-07 | 5.7E-07 | BT20/MB468 | CYTO |
| >hg38_21_+_8433172_8446622_RNA45SN1_45S_50ntFlanks@12936.12954.19\|AAACCATTCGTAGACGACC | >hg38_21_+_8433172_8446622_RNA45SN1_45S_50ntFlanks | 12936 | 12954 | 19 | AAACCATTCGTAGACGACC | 274.5 | -2.8 | 0.6 | -5.1 | 3.3E-07 | 5.7E-07 | MB231/MB468 | NUC |
| >hg38_21_+_8433172_8446622_RNA45SN1_45S_50ntFlanks@8342.8365.24\|CGTAAGGGAAAGTTGAAAAGAACT | >hg38_21_+_8433172_8446622_RNA45SN1_45S_50ntFlanks | 8342 | 8365 | 24 | CGTAAGGGAAAGTTGAAAAGAACT | 442.5 | 8.7 | 1.7 | 5.2 | 2.5E-07 | 5.7E-07 | BT20/MB468 | MITO |
| >MI0015866\|hsa-mir-4326&WithFlank&20\|+\|63286802\|63286872@17.39.23&[MIMAT0016888&hsa-miR-4326&offsets\|0\|+3;m-339&20\|+\|63286818\|63286841&offsets\|0\|-1]\|TGTTCCTCTGTCTCCCAGACTCT | >MI0015866\|hsa-mir-4326&WithFlank&20\|+\|63286802\|63286872 | 17 | 39 | 23 | TGTTCCTCTGTCTCCCAGACTCT | 220.3 | 8.0 | 1.6 | 5.1 | 3.5E-07 | 5.7E-07 | MB231/MB468 | MITO |
| >MI0000085\|hsa-mir-27a&WithFlank&19\|-\|13836434\|13836523@57.75.19(+1A)&[MIMAT0000084&hsa-miR-27a-3p&offsets\|0\|-2(+1A);m-43&19\|-\|13836447\|13836467&offsets\|0\|-2(+1A)]\|TTCACAGTGGCTAAGTTCCA | >MI0000085\|hsa-mir-27a&WithFlank&19\|-\|13836434\|13836523 | 57 | 75 | 19(+1A) | TTCACAGTGGCTAAGTTCCA | 3044.0 | 4.9 | 1.0 | 5.1 | 3.5E-07 | 5.7E-07 | MB231/MB468 | MITO |
| >hg38_21_+_8433172_8446622_RNA45SN1_45S_50ntFlanks@3788.3810.23\|AACTGCGAATGGCTCATTAAATC | >hg38_21_+_8433172_8446622_RNA45SN1_45S_50ntFlanks | 3788 | 3810 | 23 | AACTGCGAATGGCTCATTAAATC | 365.5 | -5.7 | 1.1 | -5.1 | 3.5E-07 | 5.7E-07 | MB231/MB468 | MITO |
| >hg38_21_+_8433172_8446622_RNA45SN1_45S_50ntFlanks@12405.12422.18\|CTTTTTGATCCTTCGATG | >hg38_21_+_8433172_8446622_RNA45SN1_45S_50ntFlanks | 12405 | 12422 | 18 | CTTTTTGATCCTTCGATG | 538.5 | -7.7 | 1.4 | -5.4 | 7.5E-08 | 5.7E-07 | BT20/MB468 | MP |
| >MI0000542\|hsa-mir-320a&WithFlank&8\|-\|22244960\|22245043@48.68.21&[MIMAT0000510&hsa-miR-320a-3p&offsets\|0\|-1;m-60&8\|-\|22244975\|22244996&offsets\|0\|-1]\|AAAAGCTGGGTTGAGAGGGCG | >MI0000542\|hsa-mir-320a&WithFlank&8\|-\|22244960\|22245043 | 48 | 68 | 21 | AAAAGCTGGGTTGAGAGGGCG | 577.6 | 5.0 | 1.0 | 5.1 | 3.5E-07 | 5.7E-07 | MB231/MB468 | MITO |
| >hg38_21_+_8433172_8446622_RNA45SN1_45S_50ntFlanks@11561.11584.24\|GCGCCTAGCAGCCGACTTAGAACT | >hg38_21_+_8433172_8446622_RNA45SN1_45S_50ntFlanks | 11561 | 11584 | 24 | GCGCCTAGCAGCCGACTTAGAACT | 655.7 | -4.0 | 0.8 | -5.1 | 3.5E-07 | 5.7E-07 | MB231/MB468 | MITO |
| >MI0015866\|hsa-mir-4326&WithFlank&20\|+\|63286802\|63286872@17.39.23(+1A)&[MIMAT0016888&hsa-miR-4326&offsets\|0\|+3(+1A);m-339&20\|+\|63286818\|63286841&offsets\|0\|-1(+1A)]\|TGTTCCTCTGTCTCCCAGACTCTA | >MI0015866\|hsa-mir-4326&WithFlank&20\|+\|63286802\|63286872 | 17 | 39 | 23(+1A) | TGTTCCTCTGTCTCCCAGACTCTA | 46.5 | 8.8 | 1.7 | 5.1 | 2.9E-07 | 5.7E-07 | MB231/MB468 | MP |
| >trnaMT_HisGTG_MT_+_12138_12206@26.53.28\|AGATTGTGAATCTGACAACAGAGGCTTA | >trnaMT_HisGTG_MT_+_12138_12206 | 26 | 53 | 28 | AGATTGTGAATCTGACAACAGAGGCTTA | 114.0 | 10.6 | 2.0 | 5.4 | 7.5E-08 | 5.7E-07 | BT20/MB468 | MP |
| >hg38_21_+_8433172_8446622_RNA45SN1_45S_50ntFlanks@12367.12384.18\|AACTGGCTTGTGGCGGCC | >hg38_21_+_8433172_8446622_RNA45SN1_45S_50ntFlanks | 12367 | 12384 | 18 | AACTGGCTTGTGGCGGCC | 222.5 | -6.2 | 1.2 | -5.1 | 2.9E-07 | 5.7E-07 | BT20/MB468 | CYTO |
| >am_trna111_HisGTG_1_-_147774845_147774916@2.29.28,trna118_HisGTG_1_-_145396881_145396952@2.29.28,trna16_HisGTG_1_+_146544773_146544844@2.29.28,trna1_HisGTG_15_+_45493349_45493420@2.29.28,trna21_HisGTG_1_+_147753471_147753542@2.29.28,trna33_HisGTG_6_+_27125906_27125977@2.29.28,trna7_HisGTG_9_-_14433938_14434009@2.29.28,trna8_HisGTG_15_-_45492611_45492682@2.29.28,trna9_HisGTG_15_-_45490804_45490875@2.29.28\|CCGTGATCGTATAGTGGTTAGTACTCTG | >am_trna111_HisGTG_1_-_147774845_147774916 | 2 | 29 | 28 | CCGTGATCGTATAGTGGTTAGTACTCTG | 298.0 | -3.3 | 0.6 | -5.1 | 3.3E-07 | 5.7E-07 | MB231/MB468 | CYTO |
| >trnaMT_TrpTCA_MT_+_5512_5579@53.71.19,trnalookalike3_TrpTCA_1_+_566062_566129@53.71.19\|CAATACTTAATTTCTGCCA | >trnaMT_TrpTCA_MT_+_5512_5579 | 53 | 71 | 19 | CAATACTTAATTTCTGCCA | 941.7 | -3.4 | 0.7 | -5.1 | 2.9E-07 | 5.7E-07 | MB231/MB468 | MP |
| >MI0000542\|hsa-mir-320a&WithFlank&8\|-\|22244960\|22245043@48.69.22(+1G)&[MIMAT0000510&hsa-miR-320a-3p&offsets\|0\|0(+1G);m-60&8\|-\|22244975\|22244996&offsets\|0\|0(+1G)]\|AAAAGCTGGGTTGAGAGGGCGAG | >MI0000542\|hsa-mir-320a&WithFlank&8\|-\|22244960\|22245043 | 48 | 69 | 22(+1G) | AAAAGCTGGGTTGAGAGGGCGAG | 662.3 | 6.9 | 1.3 | 5.4 | 7.5E-08 | 5.8E-07 | BT20/MB468 | MP |
| >hg38_21_+_8433172_8446622_RNA45SN1_45S_50ntFlanks@10346.10383.38\|TCAAACGAGAACTTTGAAGGCCGAAGTGGAGAAGGGTT | >hg38_21_+_8433172_8446622_RNA45SN1_45S_50ntFlanks | 10346 | 10383 | 38 | TCAAACGAGAACTTTGAAGGCCGAAGTGGAGAAGGGTT | 544.9 | -3.3 | 0.7 | -5.1 | 3.4E-07 | 5.8E-07 | MB231/MB468 | NUC |
| >hg38_21_+_8433172_8446622_RNA45SN1_45S_50ntFlanks@8357.8375.19\|AAAAGAACTTTGAAGAGAG | >hg38_21_+_8433172_8446622_RNA45SN1_45S_50ntFlanks | 8357 | 8375 | 19 | AAAAGAACTTTGAAGAGAG | 20198.0 | -5.4 | 1.1 | -5.1 | 2.9E-07 | 5.8E-07 | MB231/MB468 | MP |
| >hg38_21_+_8433172_8446622_RNA45SN1_45S_50ntFlanks@6660.6696.37\|GCGGTGGATCACTCGGCTCGTGCGTCGATGAAGAACG | >hg38_21_+_8433172_8446622_RNA45SN1_45S_50ntFlanks | 6660 | 6696 | 37 | GCGGTGGATCACTCGGCTCGTGCGTCGATGAAGAACG | 310.6 | 2.7 | 0.5 | 5.1 | 3.4E-07 | 5.8E-07 | MB231/MB468 | NUC |
| >hg38_21_+_8433172_8446622_RNA45SN1_45S_50ntFlanks@10683.10705.23\|CCGGCGGCGTCCGGTGAGCTCTC | >hg38_21_+_8433172_8446622_RNA45SN1_45S_50ntFlanks | 10683 | 10705 | 23 | CCGGCGGCGTCCGGTGAGCTCTC | 807.4 | -3.0 | 0.6 | -5.1 | 3.3E-07 | 5.8E-07 | MB231/MB468 | CYTO |
| >hg38_21_+_8433172_8446622_RNA45SN1_45S_50ntFlanks@8198.8224.27\|TGGACGGTGTGAGGCCGGTAGCGGCCC | >hg38_21_+_8433172_8446622_RNA45SN1_45S_50ntFlanks | 8198 | 8224 | 27 | TGGACGGTGTGAGGCCGGTAGCGGCCC | 389.4 | -5.8 | 1.1 | -5.1 | 3.0E-07 | 5.8E-07 | MB231/MB468 | MP |
| >hg38_21_+_8433172_8446622_RNA45SN1_45S_50ntFlanks@9381.9409.29\|CGCTCGCCGGCCGAGGTGGGATCCCGAGG | >hg38_21_+_8433172_8446622_RNA45SN1_45S_50ntFlanks | 9381 | 9409 | 29 | CGCTCGCCGGCCGAGGTGGGATCCCGAGG | 567.0 | -7.3 | 1.4 | -5.1 | 3.4E-07 | 5.8E-07 | MB231/MB468 | NUC |
| >hg38_21_+_8433172_8446622_RNA45SN1_45S_50ntFlanks@9573.9596.24\|AAATCGGTCGTCCGACCTGGGTAT | >hg38_21_+_8433172_8446622_RNA45SN1_45S_50ntFlanks | 9573 | 9596 | 24 | AAATCGGTCGTCCGACCTGGGTAT | 539.6 | -3.7 | 0.7 | -5.1 | 3.4E-07 | 5.8E-07 | MB231/MB468 | NUC |
| >hg38_21_+_8433172_8446622_RNA45SN1_45S_50ntFlanks@4726.4751.26\|TAATCAAGAACGAAAGTCGGAGGTTC | >hg38_21_+_8433172_8446622_RNA45SN1_45S_50ntFlanks | 4726 | 4751 | 26 | TAATCAAGAACGAAAGTCGGAGGTTC | 387.5 | -7.5 | 1.4 | -5.4 | 7.6E-08 | 5.8E-07 | BT20/MB468 | MP |
| >hg38_21_+_8433172_8446622_RNA45SN1_45S_50ntFlanks@10348.10374.27\|AAACGAGAACTTTGAAGGCCGAAGTGG | >hg38_21_+_8433172_8446622_RNA45SN1_45S_50ntFlanks | 10348 | 10374 | 27 | AAACGAGAACTTTGAAGGCCGAAGTGG | 978.4 | -5.6 | 1.1 | -5.1 | 3.5E-07 | 5.8E-07 | MB231/MB468 | MITO |
| >MI0000750\|hsa-mir-26a-2&WithFlank&12\|-\|57824603\|57824698@20.41.22(+1A);MI0000083\|hsa-mir-26a-1&WithFlank&3\|+\|37969398\|37969486@16.37.22(+1A)&[MIMAT0000082&hsa-miR-26a-5p&offsets\|0\|0(+1A);m-33&12\|-\|57824658\|57824679&offsets\|0\|0(+1A)];[MIMAT0000082_1&hsa-miR-26a-5p&offsets\|0\|0(+1A);m-34&3\|+\|37969413\|37969434&offsets\|0\|0(+1A)]\|TTCAAGTAATCCAGGATAGGCTA | >MI0000750\|hsa-mir-26a-2&WithFlank&12\|-\|57824603\|57824698 | 20 | 41 | 22(+1A) | TTCAAGTAATCCAGGATAGGCTA | 1068.8 | 4.8 | 0.9 | 5.1 | 3.5E-07 | 5.8E-07 | MB231/MB468 | MITO |

<table>
<tr><td>>MI0006444|hsa-mir-1307&WithFlank&10|-|103394247|103394407@86.107.22&[MIMAT0005951&hsa-miR-1307-3p&offsets|0|0;m-66&10|-|103394300|103394322&offsets|0|-1]|ACTCGGCGTGGCGTCGGTCGTG</td><td>>MI0006444|hsa-mir-1307&WithFlank&10|-|103394247|103394407</td><td>86</td><td>107</td><td>22</td><td>ACTCGGCGTGGCGTCGGTCGTG</td><td>568.1</td><td>4.5</td><td>0.9</td><td>5.1</td><td>3.5E-07</td><td>5.8E-07</td><td>MB231/MB468</td><td>MITO</td></tr>
<tr><td>>hg38_21_+_8433172_8446622_RNA45SN1_45S_50ntFlanks@4724.4751.28|ATTAATCAAGAACGAAAGTCGGAGGTTC</td><td>>hg38_21_+_8433172_8446622_RNA45SN1_45S_50ntFlanks</td><td>4724</td><td>4751</td><td>28</td><td>ATTAATCAAGAACGAAAGTCGGAGGTTC</td><td>680.9</td><td>-3.7</td><td>0.7</td><td>-5.1</td><td>3.3E-07</td><td>5.8E-07</td><td>MB231/MB468</td><td>CYTO</td></tr>
<tr><td>>hg38_1_-_228634819_228635039_RNA5S12_5S_50ntFlanks@51.77.27|GTCTACGGCCATACCACCCTGAACGCG</td><td>>hg38_1_-_228634819_228635039_RNA5S12_5S_50ntFlanks</td><td>51</td><td>77</td><td>27</td><td>GTCTACGGCCATACCACCCTGAACGCG</td><td>810.6</td><td>3.7</td><td>0.7</td><td>5.1</td><td>2.9E-07</td><td>5.8E-07</td><td>BT20/MB468</td><td>CYTO</td></tr>
<tr><td>>hg38_21_+_8433172_8446622_RNA45SN1_45S_50ntFlanks@12477.12496.20|CACTAATAGGGAACGTGAGC</td><td>>hg38_21_+_8433172_8446622_RNA45SN1_45S_50ntFlanks</td><td>12477</td><td>12496</td><td>20</td><td>CACTAATAGGGAACGTGAGC</td><td>2537.8</td><td>2.6</td><td>0.5</td><td>5.1</td><td>2.9E-07</td><td>5.8E-07</td><td>BT20/MB468</td><td>CYTO</td></tr>
<tr><td>>hg38_21_+_8433172_8446622_RNA45SN1_45S_50ntFlanks@12129.12146.18|CCCGCTCCGGGGACAGTG</td><td>>hg38_21_+_8433172_8446622_RNA45SN1_45S_50ntFlanks</td><td>12129</td><td>12146</td><td>18</td><td>CCCGCTCCGGGGACAGTG</td><td>65.8</td><td>-9.1</td><td>1.7</td><td>-5.4</td><td>7.6E-08</td><td>5.8E-07</td><td>BT20/MB468</td><td>MP</td></tr>
<tr><td>>hg38_21_+_8433172_8446622_RNA45SN1_45S_50ntFlanks@12241.12273.33|CAGAAGGGCAAAAGCTCGCTTGATCTTGATTTT</td><td>>hg38_21_+_8433172_8446622_RNA45SN1_45S_50ntFlanks</td><td>12241</td><td>12273</td><td>33</td><td>CAGAAGGGCAAAAGCTCGCTTGATCTTGATTTT</td><td>294.8</td><td>-6.3</td><td>1.2</td><td>-5.1</td><td>3.0E-07</td><td>5.8E-07</td><td>MB231/MB468</td><td>MP</td></tr>
<tr><td>>am_MI0005763|hsa-mir-941-1&WithFlank&20|+|63919443|63919526@53.75.23(+1U);MI0005764|hsa-mir-941-2&WithFlank&20|+|63919499|63919582@53.75.23(+1U);MI0005765|hsa-mir-941-3&WithFlank&20|+|63919555|63919638@53.75.23(+1U);MI0005766|hsa-mir-941-4&WithFlank&20|+|63919750|63919833@53.75.23(+1U);MI0031520|hsa-mir-941-5&WithFlank&20|+|63919862|63919945@53.75.23(+1U)&[MIMAT0004984&hsa-miR-941&offsets|0|0(+1U);m-263&20|+|63919495|63919517&offsets|0|0(+1U)];[MIMAT0004984_1&hsa-miR-941&offsets|0|0(+1U);m-263&20|+|63919551|63919573&offsets|0|0(+1U)];[MIMAT0004984_2&hsa-miR-941&offsets|0|0(+1U);m-263&20|+|63919607|63919629&offsets|0|0(+1U)];[MIMAT0004984_3&hsa-miR-941&offsets|0|0(+1U);m-263&20|+|63919802|63919824&offsets|0|0(+1U)];[MIMAT0004984_4&hsa-miR-941&offsets|0|0(+1U);m-263&20|+|63919914|63919936&offsets|0|0(+1U)]|CACCCGGCTGTGTGCACATGTGCT</td><td>>am_MI0005763|hsa-mir-941-1&WithFlank&20|+|63919443|63919526</td><td>53</td><td>75</td><td>23(+1U)</td><td>CACCCGGCTGTGTGCACATGTGCT</td><td>374.4</td><td>7.1</td><td>1.4</td><td>5.1</td><td>3.6E-07</td><td>5.8E-07</td><td>MB231/MB468</td><td>MITO</td></tr>
<tr><td>>trnaMT_MetCAT_MT_+_4402_4469@52.71.20|ATACCCTTCCCGTACTACCA</td><td>>trnaMT_MetCAT_MT_+_4402_4469</td><td>52</td><td>71</td><td>20</td><td>ATACCCTTCCCGTACTACCA</td><td>907.8</td><td>9.4</td><td>1.8</td><td>5.1</td><td>2.9E-07</td><td>5.8E-07</td><td>BT20/MB468</td><td>CYTO</td></tr>
<tr><td>>hg38_21_+_8433172_8446622_RNA45SN1_45S_50ntFlanks@11575.11606.32|ACTTAGAACTGGTGCGGACCAGGGGAATCCGA</td><td>>hg38_21_+_8433172_8446622_RNA45SN1_45S_50ntFlanks</td><td>11575</td><td>11606</td><td>32</td><td>ACTTAGAACTGGTGCGGACCAGGGGAATCCGA</td><td>326.3</td><td>-3.2</td><td>0.6</td><td>-5.1</td><td>2.9E-07</td><td>5.8E-07</td><td>BT20/MB468</td><td>CYTO</td></tr>
<tr><td>>trna10_AspGTC_12_-_125424193_125424264@37.75.39,trna12_AspGTC_12_-_125411891_125411962@37.75.39,trna144_AspGTC_6_-_27551236_27551307@37.75.39,trna38_AspGTC_17_-_8125556_8125627@37.75.39,trna45_AspGTC_6_+_27447453_27447524@37.75.39,trna48_AspGTC_6_+_27471523_27471594@37.75.39,trna4_AspGTC_12_+_96429799_96429870@37.75.39,trna69_AspGTC_1_-_161440205_161440276@37.75.39,trna72_AspGTC_1_-_161432824_161432895@37.75.39,trna75_AspGTC_1_-_161425414_161425485@37.75.39,trna78_AspGTC_1_-_161418033_161418104@37.75.39,trna81_AspGTC_1_-_161410615_161410686@37.75.39|ACGCGGGAGACCGGGGTTCGATTCCCCGACGGGGAGCCA</td><td>>trna10_AspGTC_12_-_125424193_125424264</td><td>37</td><td>75</td><td>39</td><td>ACGCGGGAGACCGGGGTTCGATTCCCCGACGGGGAGCCA</td><td>1342.1</td><td>6.0</td><td>1.2</td><td>5.1</td><td>3.4E-07</td><td>5.8E-07</td><td>MB231/MB468</td><td>NUC</td></tr>
<tr><td>>hg38_21_+_8433172_8446622_RNA45SN1_45S_50ntFlanks@11704.11726.23|ATGAAGCGCGGGTAAACGGCGGG</td><td>>hg38_21_+_8433172_8446622_RNA45SN1_45S_50ntFlanks</td><td>11704</td><td>11726</td><td>23</td><td>ATGAAGCGCGGGTAAACGGCGGG</td><td>176.8</td><td>-4.9</td><td>1.0</td><td>-5.1</td><td>3.4E-07</td><td>5.8E-07</td><td>MB231/MB468</td><td>NUC</td></tr>
<tr><td>>hg38_21_+_8433172_8446622_RNA45SN1_45S_50ntFlanks@11594.11613.20|CAGGGGAATCCGACTGTTTA</td><td>>hg38_21_+_8433172_8446622_RNA45SN1_45S_50ntFlanks</td><td>11594</td><td>11613</td><td>20</td><td>CAGGGGAATCCGACTGTTTA</td><td>387.0</td><td>-3.5</td><td>0.7</td><td>-5.1</td><td>3.4E-07</td><td>5.8E-07</td><td>MB231/MB468</td><td>NUC</td></tr>
<tr><td>>trna2_GlyGCC_21_-_18827107_18827177@1.32.32,trna35_GlyGCC_1_+_161413094_161413164@1.32.32,trna37_GlyGCC_1_+_161420467_161420537@1.32.32,trna39_GlyGCC_1_+_161427898_161427968@1.32.32,trna41_GlyGCC_1_+_161435258_161435328@1.32.32|GCATGGGTGGTTCAGTGGTAGAATTCTCGCCT</td><td>>trna2_GlyGCC_21_-_18827107_18827177</td><td>1</td><td>32</td><td>32</td><td>GCATGGGTGGTTCAGTGGTAGAATTCTCGCCT</td><td>39806.3</td><td>8.3</td><td>1.5</td><td>5.4</td><td>7.6E-08</td><td>5.8E-07</td><td>BT20/MB468</td><td>MP</td></tr>
<tr><td>>hg38_21_+_8433172_8446622_RNA45SN1_45S_50ntFlanks@10095.10115.21|CGCGCGCGCGCGCGTGTGGTG</td><td>>hg38_21_+_8433172_8446622_RNA45SN1_45S_50ntFlanks</td><td>10095</td><td>10115</td><td>21</td><td>CGCGCGCGCGCGCGTGTGGTG</td><td>76.7</td><td>-9.3</td><td>1.7</td><td>-5.4</td><td>7.7E-08</td><td>5.8E-07</td><td>BT20/MB468</td><td>MP</td></tr>
<tr><td>>hg38_21_+_8433172_8446622_RNA45SN1_45S_50ntFlanks@12474.12498.25|ACCCACTAATAGGGAACGTGAGCTG</td><td>>hg38_21_+_8433172_8446622_RNA45SN1_45S_50ntFlanks</td><td>12474</td><td>12498</td><td>25</td><td>ACCCACTAATAGGGAACGTGAGCTG</td><td>340.0</td><td>-5.5</td><td>1.1</td><td>-5.1</td><td>3.6E-07</td><td>5.8E-07</td><td>MB231/MB468</td><td>MITO</td></tr>
<tr><td>>hg38_21_+_8433172_8446622_RNA45SN1_45S_50ntFlanks@10808.10827.20|CAATGTAGGTAAGGGAAGTC</td><td>>hg38_21_+_8433172_8446622_RNA45SN1_45S_50ntFlanks</td><td>10808</td><td>10827</td><td>20</td><td>CAATGTAGGTAAGGGAAGTC</td><td>525.5</td><td>-5.4</td><td>1.0</td><td>-5.1</td><td>2.9E-07</td><td>5.8E-07</td><td>BT20/MB468</td><td>CYTO</td></tr>
<tr><td>>hg38_21_+_8433172_8446622_RNA45SN1_45S_50ntFlanks@12633.12652.20|CGAAGCTACCATCTGTGGGA</td><td>>hg38_21_+_8433172_8446622_RNA45SN1_45S_50ntFlanks</td><td>12633</td><td>12652</td><td>20</td><td>CGAAGCTACCATCTGTGGGA</td><td>192.8</td><td>-6.4</td><td>1.3</td><td>-5.1</td><td>3.4E-07</td><td>5.9E-07</td><td>MB231/MB468</td><td>NUC</td></tr>
<tr><td>>MI0000289|hsa-mir-181a-1&WithFlank&1|-|198859038|198859159@30.54.25;MI0000269|hsa-mir-181a-2&WithFlank&9|+|124692436|124692557@45.69.25&[MIMAT0000256&hsa-miR-181a-5p&offsets|0|+2;m-32&1|-|198859108|198859130&offsets|0|+2];[MIMAT0000256_1&hsa-miR-181a-5p&offsets|0|+2;m-32&9|+|124692480|124692502&offsets|0|+2]|AACATTCAACGCTGTCGGTGAGTTT</td><td>>MI0000289|hsa-mir-181a-1&WithFlank&1|-|198859038|198859159</td><td>30</td><td>54</td><td>25</td><td>AACATTCAACGCTGTCGGTGAGTTT</td><td>3777.5</td><td>-5.1</td><td>0.9</td><td>-5.6</td><td>2.8E-08</td><td>5.9E-07</td><td>BT20/MB231</td><td>MITO</td></tr>
<tr><td>>hg38_21_+_8433172_8446622_RNA45SN1_45S_50ntFlanks@11656.11685.30|CGATGTGATTTCTGCCCAGTGCTCTGAATG</td><td>>hg38_21_+_8433172_8446622_RNA45SN1_45S_50ntFlanks</td><td>11656</td><td>11685</td><td>30</td><td>CGATGTGATTTCTGCCCAGTGCTCTGAATG</td><td>548.1</td><td>12.4</td><td>2.2</td><td>5.7</td><td>1.6E-08</td><td>5.9E-07</td><td>BT20/MB231</td><td>MP</td></tr>
<tr><td>>hg38_21_+_8433172_8446622_RNA45SN1_45S_50ntFlanks@6748.6770.23|CTTCGAACGCACTTGCGGCCCCG</td><td>>hg38_21_+_8433172_8446622_RNA45SN1_45S_50ntFlanks</td><td>6748</td><td>6770</td><td>23</td><td>CTTCGAACGCACTTGCGGCCCCG</td><td>446.2</td><td>-5.1</td><td>1.0</td><td>-5.2</td><td>2.6E-07</td><td>5.9E-07</td><td>BT20/MB468</td><td>MITO</td></tr>
<tr><td>>MI0000465|hsa-mir-191&WithFlank&3|-|49020612|49020715@22.43.22(+1U)&[MIMAT0000440&hsa-miR-191-5p&offsets|0|-1(+1U);m-48&3|-|49020672|49020694&offsets|0|-1(+1U)]|CAACGGAATCCCAAAAGCAGCTT</td><td>>MI0000465|hsa-mir-191&WithFlank&3|-|49020612|49020715</td><td>22</td><td>43</td><td>22(+1U)</td><td>CAACGGAATCCCAAAAGCAGCTT</td><td>148.7</td><td>2.0</td><td>0.4</td><td>5.1</td><td>3.8E-07</td><td>5.9E-07</td><td>BT20/MB468</td><td>NUC</td></tr>
<tr><td>>hg38_21_+_8433172_8446622_RNA45SN1_45S_50ntFlanks@12420.12462.43|ATGTCGGCTCTTCCTATCATTGTGAAGCAGAATTCACCAAGCG</td><td>>hg38_21_+_8433172_8446622_RNA45SN1_45S_50ntFlanks</td><td>12420</td><td>12462</td><td>43</td><td>ATGTCGGCTCTTCCTATCATTGTGAAGCAGAATTCACCAAGCG</td><td>1235.6</td><td>-5.0</td><td>0.9</td><td>-5.6</td><td>2.3E-08</td><td>5.9E-07</td><td>BT20/MB231</td><td>CYTO</td></tr>
<tr><td>>hg38_21_+_8433172_8446622_RNA45SN1_45S_50ntFlanks@6731.6758.28|CACATTGATCATCGACACTTCGAACGCA</td><td>>hg38_21_+_8433172_8446622_RNA45SN1_45S_50ntFlanks</td><td>6731</td><td>6758</td><td>28</td><td>CACATTGATCATCGACACTTCGAACGCA</td><td>295.6</td><td>-2.8</td><td>0.6</td><td>-5.1</td><td>3.3E-07</td><td>5.9E-07</td><td>MB231/MB468</td><td>CYTO</td></tr>
<tr><td>>trna119_LysCTT_1_-_145395522_145395594@3.35.33,trna11_LysCTT_5_-_180648979_180649051@3.35.33,trna13_LysCTT_6_+_26556774_26556846@3.35.33,trna7_LysCTT_16_+_3225692_3225764@3.35.33,trna9_LysCTT_5_+_180634755_180634827@3.35.33|CCGGCTAGCTCAGTCGGTAGAGCATGAGACTCT</td><td>>trna119_LysCTT_1_-_145395522_145395594</td><td>3</td><td>35</td><td>33</td><td>CCGGCTAGCTCAGTCGGTAGAGCATGAGACTCT</td><td>790.2</td><td>7.4</td><td>1.5</td><td>5.1</td><td>3.6E-07</td><td>5.9E-07</td><td>MB231/MB468</td><td>MITO</td></tr>
<tr><td>>hg38_21_+_8433172_8446622_RNA45SN1_45S_50ntFlanks@7974.7995.22|ACGCGACCTCAGATCAGACGTG</td><td>>hg38_21_+_8433172_8446622_RNA45SN1_45S_50ntFlanks</td><td>7974</td><td>7995</td><td>22</td><td>ACGCGACCTCAGATCAGACGTG</td><td>11784.2</td><td>3.9</td><td>0.8</td><td>5.2</td><td>2.6E-07</td><td>5.9E-07</td><td>BT20/MB468</td><td>MITO</td></tr>
<tr><td>>hg38_21_+_8433172_8446622_RNA45SN1_45S_50ntFlanks@12352.12376.25|AGTTACCACAGGGATAACTGGCTTG</td><td>>hg38_21_+_8433172_8446622_RNA45SN1_45S_50ntFlanks</td><td>12352</td><td>12376</td><td>25</td><td>AGTTACCACAGGGATAACTGGCTTG</td><td>62.2</td><td>-9.0</td><td>1.7</td><td>-5.4</td><td>7.7E-08</td><td>5.9E-07</td><td>BT20/MB468</td><td>MP</td></tr>
<tr><td>>hg38_21_+_8433172_8446622_RNA45SN1_45S_50ntFlanks@8171.8213.43|AAGTCCTTCTGATCGAGGCCCAGCCCGTGGACGGTGTGAGGCC</td><td>>hg38_21_+_8433172_8446622_RNA45SN1_45S_50ntFlanks</td><td>8171</td><td>8213</td><td>43</td><td>AAGTCCTTCTGATCGAGGCCCAGCCCGTGGACGGTGTGAGGCC</td><td>863.1</td><td>-6.5</td><td>1.3</td><td>-5.1</td><td>3.8E-07</td><td>5.9E-07</td><td>BT20/MB468</td><td>NUC</td></tr>
<tr><td>>hg38_21_+_8433172_8446622_RNA45SN1_45S_50ntFlanks@8356.8377.22|GAAAAGAACTTTGAAGAGAGAG</td><td>>hg38_21_+_8433172_8446622_RNA45SN1_45S_50ntFlanks</td><td>8356</td><td>8377</td><td>22</td><td>GAAAAGAACTTTGAAGAGAGAG</td><td>253.8</td><td>-4.1</td><td>0.8</td><td>-5.1</td><td>3.3E-07</td><td>5.9E-07</td><td>MB231/MB468</td><td>CYTO</td></tr>
<tr><td>>hg38_21_+_8433172_8446622_RNA45SN1_45S_50ntFlanks@10312.10340.29|CGCAGGTGCAGATCTTGGTGGTAGTAGCA</td><td>>hg38_21_+_8433172_8446622_RNA45SN1_45S_50ntFlanks</td><td>10312</td><td>10340</td><td>29</td><td>CGCAGGTGCAGATCTTGGTGGTAGTAGCA</td><td>91.0</td><td>10.3</td><td>1.9</td><td>5.4</td><td>7.8E-08</td><td>5.9E-07</td><td>BT20/MB468</td><td>MP</td></tr>
<tr><td>>hg38_21_+_8433172_8446622_RNA45SN1_45S_50ntFlanks@8349.8391.43|GAAAGTTGAAAAGAACTTTGAAGAGAGAGTTCAAGAGGGCGTG</td><td>>hg38_21_+_8433172_8446622_RNA45SN1_45S_50ntFlanks</td><td>8349</td><td>8391</td><td>43</td><td>GAAAGTTGAAAAGAACTTTGAAGAGAGAGTTCAAGAGGGCGTG</td><td>66.8</td><td>9.8</td><td>1.8</td><td>5.4</td><td>7.8E-08</td><td>5.9E-07</td><td>BT20/MB468</td><td>MP</td></tr>
<tr><td>>hg38_21_+_8433172_8446622_RNA45SN1_45S_50ntFlanks@10367.10390.24|CGAAGTGGAGAAGGGTTCCATGTG</td><td>>hg38_21_+_8433172_8446622_RNA45SN1_45S_50ntFlanks</td><td>10367</td><td>10390</td><td>24</td><td>CGAAGTGGAGAAGGGTTCCATGTG</td><td>79.1</td><td>-9.4</td><td>1.7</td><td>-5.4</td><td>7.8E-08</td><td>5.9E-07</td><td>BT20/MB468</td><td>MP</td></tr>
<tr><td>>MI0000342|hsa-mir-200b&WithFlank&1|+|1167098|1167204@63.83.21(+1U)&[MIMAT0000318&hsa-miR-200b-3p&offsets|0|-1(+1U);m-93&1|+|1167160|1167182&offsets|0|-2(+1U)]|TAATACTGCCTGGTAATGATGT</td><td>>MI0000342|hsa-mir-200b&WithFlank&1|+|1167098|1167204</td><td>63</td><td>83</td><td>21(+1U)</td><td>TAATACTGCCTGGTAATGATGT</td><td>71.3</td><td>-9.2</td><td>1.7</td><td>-5.4</td><td>7.8E-08</td><td>5.9E-07</td><td>BT20/MB468</td><td>MP</td></tr>
<tr><td>>trnaMT_GluTTC_MT_-_14674_14742@31.72.42,trnalookalike8_GluTTC_5_-_93905172_93905240@31.72.42|TTCATATCATTGGTCGTGGTTGTAGTCCGTGCGAGAATACCA</td><td>>trnaMT_GluTTC_MT_-_14674_14742</td><td>31</td><td>72</td><td>42</td><td>TTCATATCATTGGTCGTGGTTGTAGTCCGTGCGAGAATACCA</td><td>1296.2</td><td>4.7</td><td>0.9</td><td>5.1</td><td>3.6E-07</td><td>5.9E-07</td><td>MB231/MB468</td><td>MITO</td></tr>
<tr><td>>hg38_21_+_8433172_8446622_RNA45SN1_45S_50ntFlanks@6735.6770.36|TTGATCATCGACACTTCGAACGCACTTGCGGCCCCG</td><td>>hg38_21_+_8433172_8446622_RNA45SN1_45S_50ntFlanks</td><td>6735</td><td>6770</td><td>36</td><td>TTGATCATCGACACTTCGAACGCACTTGCGGCCCCG</td><td>945.8</td><td>-8.4</td><td>1.7</td><td>-5.1</td><td>3.8E-07</td><td>5.9E-07</td><td>BT20/MB468</td><td>NUC</td></tr>
<tr><td>>hg38_21_+_8433172_8446622_RNA45SN1_45S_50ntFlanks@6719.6758.40|TGAATTGCAGGACACATTGATCATCGACACTTCGAACGCA</td><td>>hg38_21_+_8433172_8446622_RNA45SN1_45S_50ntFlanks</td><td>6719</td><td>6758</td><td>40</td><td>TGAATTGCAGGACACATTGATCATCGACACTTCGAACGCA</td><td>760.4</td><td>-8.8</td><td>1.7</td><td>-5.1</td><td>3.8E-07</td><td>5.9E-07</td><td>BT20/MB468</td><td>NUC</td></tr>
<tr><td>>MI0000812|hsa-mir-331&WithFlank&12|+|95308414|95308519@30.52.23&[MIMAT0004700&hsa-miR-331-5p&offsets|-2|-1;TJU_CMC_MD2.ID00734.5p-miR&offsets|+18|+21;m-2747&12|+|95308425|95308444&offsets|+18|+21;m-377&12|+|95308445|95308465&offsets|-2|0]|TTCTAGGTATGGTCCCAGGGATC</td><td>>MI0000812|hsa-mir-331&WithFlank&12|+|95308414|95308519</td><td>30</td><td>52</td><td>23</td><td>TTCTAGGTATGGTCCCAGGGATC</td><td>57.8</td><td>9.1</td><td>1.8</td><td>5.1</td><td>3.0E-07</td><td>5.9E-07</td><td>MB231/MB468</td><td>MP</td></tr>
<tr><td>>hg38_21_+_8433172_8446622_RNA45SN1_45S_50ntFlanks@12416.12448.33|TTCGATGTCGGCTCTTCCTATCATTGTGAAGCA</td><td>>hg38_21_+_8433172_8446622_RNA45SN1_45S_50ntFlanks</td><td>12416</td><td>12448</td><td>33</td><td>TTCGATGTCGGCTCTTCCTATCATTGTGAAGCA</td><td>1148.3</td><td>5.1</td><td>1.0</td><td>5.1</td><td>3.5E-07</td><td>5.9E-07</td><td>MB231/MB468</td><td>NUC</td></tr>
</table>

| | | | | | | | | | | | | | |
|---|---|---|---|---|---|---|---|---|---|---|---|---|---|
| >trnaMT_HisGTG_MT_+_12138_12206@44.72.29\|CAGAGGCTTACGACCCCTTATTTACCCCA | >trnaMT_HisGTG_MT_+_12138_12206 | 44 | 72 | 29 | CAGAGGCTTACGACCCCTTATTTACCCCA | 150.1 | 11.0 | 2.0 | 5.4 | 7.8E-08 | 5.9E-07 | BT20/MB468 | MP |
| >MI0000815\|hsa-mir-339&WithFlank&7\|-\|1022927\|1023032@21.43.23&[MIMAT0000764&hsa-miR-339-5p&offsets\|0\|0;m-163&7\|-\|1022990\|1023012&offsets\|0\|0]\|TCCCTGTCCTCCAGGAGCTCACG | >MI0000815\|hsa-mir-339&WithFlank&7\|-\|1022927\|1023032 | 21 | 43 | 23 | TCCCTGTCCTCCAGGAGCTCACG | 389.6 | 4.4 | 0.9 | 5.1 | 3.5E-07 | 5.9E-07 | MB231/MB468 | NUC |
| >MI0000447\|hsa-mir-128-1&WithFlank&2\|+\|135665391\|135665484@56.76.21;MI0000727\|hsa-mir-128-2&WithFlank&3\|+\|35744470\|35744565@58.78.21&[MIMAT0000424&hsa-miR-128-3p&offsets\|0\|0;m-108&2\|+\|135665446\|135665466&offsets\|0\|0];[MIMAT0000424_1&hsa-miR-128-3p&offsets\|0\|0;m-117&3\|+\|35744527\|35744547&offsets\|0\|0]\|TCACAGTGAACCGGTCTCTTT | >MI0000447\|hsa-mir-128-1&WithFlank&2\|+\|135665391\|135665484 | 56 | 76 | 21 | TCACAGTGAACCGGTCTCTTT | 2830.9 | 3.5 | 0.7 | 5.1 | 3.4E-07 | 6.0E-07 | MB231/MB468 | CYTO |
| >hg38_21_+_8433172_8446622_RNA45SN1_45S_50ntFlanks@11729.11746.18\|TAACTATGACTCTCTTAA | >hg38_21_+_8433172_8446622_RNA45SN1_45S_50ntFlanks | 11729 | 11746 | 18 | TAACTATGACTCTCTTAA | 294.5 | -2.6 | 0.5 | -5.1 | 3.4E-07 | 6.0E-07 | MB231/MB468 | CYTO |
| >hg38_21_+_8433172_8446622_RNA45SN1_45S_50ntFlanks@11315.11344.30\|CCACCCCACGTCTCGTCGCGCGCGCGTCCG | >hg38_21_+_8433172_8446622_RNA45SN1_45S_50ntFlanks | 11315 | 11344 | 30 | CCACCCCACGTCTCGTCGCGCGCGCGTCCG | 2288.5 | -6.0 | 1.2 | -5.1 | 3.0E-07 | 6.0E-07 | BT20/MB468 | CYTO |
| >hg38_21_+_8433172_8446622_RNA45SN1_45S_50ntFlanks@12624.12656.33\|CCAATGGGGCGAAGCTACCATCTGTGGGATTAT | >hg38_21_+_8433172_8446622_RNA45SN1_45S_50ntFlanks | 12624 | 12656 | 33 | CCAATGGGGCGAAGCTACCATCTGTGGGATTAT | 239.5 | -6.1 | 1.2 | -5.1 | 2.6E-07 | 6.0E-07 | BT20/MB468 | MITO |
| >hg38_21_+_8433172_8446622_RNA45SN1_45S_50ntFlanks@12259.12278.20\|CTTGATCTTGATTTTCAGTA | >hg38_21_+_8433172_8446622_RNA45SN1_45S_50ntFlanks | 12259 | 12278 | 20 | CTTGATCTTGATTTTCAGTA | 64.5 | -9.1 | 1.7 | -5.4 | 7.8E-08 | 6.0E-07 | BT20/MB468 | MP |
| >hg38_21_+_8433172_8446622_RNA45SN1_45S_50ntFlanks@10668.10706.39\|GAAAGCGTCGCGGTTCCGGCGGCGTCCGGTGAGCTCTCG | >hg38_21_+_8433172_8446622_RNA45SN1_45S_50ntFlanks | 10668 | 10706 | 39 | GAAAGCGTCGCGGTTCCGGCGGCGTCCGGTGAGCTCTCG | 654.9 | -5.1 | 1.0 | -5.1 | 3.4E-07 | 6.0E-07 | MB231/MB468 | CYTO |
| >hg38_21_+_8433172_8446622_RNA45SN1_45S_50ntFlanks@11572.11592.21\|CCGACTTAGAACTGGTGCGGA | >hg38_21_+_8433172_8446622_RNA45SN1_45S_50ntFlanks | 11572 | 11592 | 21 | CCGACTTAGAACTGGTGCGGA | 758.6 | -5.5 | 1.1 | -5.1 | 3.0E-07 | 6.0E-07 | BT20/MB468 | CYTO |
| >MI0000446\|hsa-mir-125b-1&WithFlank&11\|-\|122099751\|122099850@61.80.20&[MIMAT0004592&hsa-miR-125b-1-3p&offsets\|0\|-2;m-485&11\|-\|122099769\|122099790&offsets\|0\|-2]\|ACGGGTTAGGCTCTTGGGAG | >MI0000446\|hsa-mir-125b-1&WithFlank&11\|-\|122099751\|122099850 | 61 | 80 | 20 | ACGGGTTAGGCTCTTGGGAG | 58.0 | 9.1 | 1.8 | 5.1 | 3.1E-07 | 6.0E-07 | MB231/MB468 | MP |
| >hg38_21_+_8433172_8446622_RNA45SN1_45S_50ntFlanks@4002.4023.22\|GATAACCTCGGGCCGATCGCAC | >hg38_21_+_8433172_8446622_RNA45SN1_45S_50ntFlanks | 4002 | 4023 | 22 | GATAACCTCGGGCCGATCGCAC | 1809.2 | -3.7 | 0.7 | -5.1 | 3.5E-07 | 6.0E-07 | MB231/MB468 | NUC |
| >hg38_21_+_8433172_8446622_RNA45SN1_45S_50ntFlanks@11362.11383.22\|TCGGGCGGCGGCGGTCGGCGGG | >hg38_21_+_8433172_8446622_RNA45SN1_45S_50ntFlanks | 11362 | 11383 | 22 | TCGGGCGGCGGCGGTCGGCGGG | 89.2 | -9.6 | 1.8 | -5.4 | 7.9E-08 | 6.0E-07 | BT20/MB468 | MP |
| >hg38_21_+_8433172_8446622_RNA45SN1_45S_50ntFlanks@7974.8007.34\|ACGCGACCTCAGATCAGACGTGGCGACCCGCTGA | >hg38_21_+_8433172_8446622_RNA45SN1_45S_50ntFlanks | 7974 | 8007 | 34 | ACGCGACCTCAGATCAGACGTGGCGACCCGCTGA | 3512.8 | 5.8 | 1.1 | 5.1 | 3.0E-07 | 6.0E-07 | BT20/MB468 | CYTO |
| >hg38_21_+_8433172_8446622_RNA45SN1_45S_50ntFlanks@3991.4024.34\|TGGTGACTCTAGATAACCTCGGGCCGATCGCACG | >hg38_21_+_8433172_8446622_RNA45SN1_45S_50ntFlanks | 3991 | 4024 | 34 | TGGTGACTCTAGATAACCTCGGGCCGATCGCACG | 673.4 | -2.9 | 0.6 | -5.1 | 3.5E-07 | 6.0E-07 | MB231/MB468 | NUC |
| >MI0000077\|hsa-mir-21&WithFlank&17\|+\|59841260\|59841343@14.35.22(+2U)&[MIMAT0000076&hsa-miR-21-5p&offsets\|0\|0(+2U);m-1&17\|+\|59841273\|59841294&offsets\|0\|0(+2U)]\|TAGCTTATCAGACTGATGTTGATT | >MI0000077\|hsa-mir-21&WithFlank&17\|+\|59841260\|59841343 | 14 | 35 | 22(+2U) | TAGCTTATCAGACTGATGTTGATT | 964.3 | 8.3 | 1.6 | 5.1 | 3.0E-07 | 6.0E-07 | BT20/MB468 | CYTO |
| >hg38_21_+_8433172_8446622_RNA45SN1_45S_50ntFlanks@4740.4758.19\|AGTCGGAGGTTCGAAGACG | >hg38_21_+_8433172_8446622_RNA45SN1_45S_50ntFlanks | 4740 | 4758 | 19 | AGTCGGAGGTTCGAAGACG | 391.1 | -3.1 | 0.6 | -5.1 | 3.1E-07 | 6.0E-07 | MB231/MB468 | MP |
| >hg38_21_+_8433172_8446622_RNA45SN1_45S_50ntFlanks@12044.12062.19\|ACTGACCCGGTGAGGCGGG | >hg38_21_+_8433172_8446622_RNA45SN1_45S_50ntFlanks | 12044 | 12062 | 19 | ACTGACCCGGTGAGGCGGG | 118.2 | -10.0 | 1.9 | -5.4 | 7.9E-08 | 6.0E-07 | BT20/MB468 | MP |
| >am_trnaMT_HisGTG_MT_+_12138_12206@18.49.32\|AAAACATCAGATTGTGAATCTGACAACAGAGG | >am_trnaMT_HisGTG_MT_+_12138_12206 | 18 | 49 | 32 | AAAACATCAGATTGTGAATCTGACAACAGAGG | 108.8 | 10.5 | 2.0 | 5.4 | 7.9E-08 | 6.0E-07 | BT20/MB468 | MP |
| >hg38_21_+_8433172_8446622_RNA45SN1_45S_50ntFlanks@10768.10785.18\|CAGCAGGTCTCCAAGGTG | >hg38_21_+_8433172_8446622_RNA45SN1_45S_50ntFlanks | 10768 | 10785 | 18 | CAGCAGGTCTCCAAGGTG | 761.0 | -7.9 | 1.5 | -5.4 | 7.9E-08 | 6.0E-07 | BT20/MB468 | MP |
| >hg38_21_+_8433172_8446622_RNA45SN1_45S_50ntFlanks@12126.12144.19\|CGACCCGCTCCGGGGACAG | >hg38_21_+_8433172_8446622_RNA45SN1_45S_50ntFlanks | 12126 | 12144 | 19 | CGACCCGCTCCGGGGACAG | 87.5 | -9.5 | 1.8 | -5.4 | 7.9E-08 | 6.0E-07 | BT20/MB468 | MP |
| >hg38_21_+_8433172_8446622_RNA45SN1_45S_50ntFlanks@10351.10385.35\|CGAGAACTTTGAAGGCCGAAGTGGAGAAGGGTTCC | >hg38_21_+_8433172_8446622_RNA45SN1_45S_50ntFlanks | 10351 | 10385 | 35 | CGAGAACTTTGAAGGCCGAAGTGGAGAAGGGTTCC | 237.9 | -3.1 | 0.6 | -5.1 | 3.4E-07 | 6.0E-07 | MB231/MB468 | CYTO |
| >hg38_21_+_8433172_8446622_RNA45SN1_45S_50ntFlanks@4738.4768.31\|AAAGTCGGAGGTTCGAAGACGATCAGATACC | >hg38_21_+_8433172_8446622_RNA45SN1_45S_50ntFlanks | 4738 | 4768 | 31 | AAAGTCGGAGGTTCGAAGACGATCAGATACC | 382.5 | 5.2 | 1.0 | 5.1 | 3.5E-07 | 6.0E-07 | MB231/MB468 | NUC |
| >hg38_21_+_8433172_8446622_RNA45SN1_45S_50ntFlanks@5074.5098.25\|ATTCCGATAACGAACGAGACTCTGG | >hg38_21_+_8433172_8446622_RNA45SN1_45S_50ntFlanks | 5074 | 5098 | 25 | ATTCCGATAACGAACGAGACTCTGG | 1142.2 | -7.6 | 1.5 | -5.1 | 3.9E-07 | 6.0E-07 | BT20/MB468 | NUC |
| >hg38_21_+_8433172_8446622_RNA45SN1_45S_50ntFlanks@5084.5109.26\|CGAACGAGACTCTGGCATGCTAACTA | >hg38_21_+_8433172_8446622_RNA45SN1_45S_50ntFlanks | 5084 | 5109 | 26 | CGAACGAGACTCTGGCATGCTAACTA | 210.8 | -9.8 | 1.9 | -5.1 | 2.6E-07 | 6.0E-07 | BT20/MB468 | MITO |
| >MI0003643\|hsa-mir-629&WithFlank&15\|-\|70079366\|70079474@28.48.21(+1G)&[MIMAT0004810&hsa-miR-629-5p&offsets\|0\|0(+1G);m-150&15\|-\|70079427\|70079447&offsets\|0\|0(+1G)]\|TGGGTTTACGTTGGGAGAACTG | >MI0003643\|hsa-mir-629&WithFlank&15\|-\|70079366\|70079474 | 28 | 48 | 21(+1G) | TGGGTTTACGTTGGGAGAACTG | 139.8 | 10.9 | 2.0 | 5.4 | 8.0E-08 | 6.0E-07 | BT20/MB468 | MP |
| >hg38_21_+_8433172_8446622_RNA45SN1_45S_50ntFlanks@11575.11607.33\|ACTTAGAACTGGTGCGGACCAGGGGAATCCGAC | >hg38_21_+_8433172_8446622_RNA45SN1_45S_50ntFlanks | 11575 | 11607 | 33 | ACTTAGAACTGGTGCGGACCAGGGGAATCCGAC | 383.2 | -7.5 | 1.5 | -5.1 | 3.1E-07 | 6.0E-07 | MB231/MB468 | MP |
| >MI0002468\|hsa-mir-484&WithFlank&16\|+\|15643288\|15643378@14.35.22&[MIMAT0002174&hsa-miR-484&offsets\|0\|0;m-127&16\|+\|15643301\|15643321&offsets\|0\|+1]\|TCAGGCTCAGTCCCCTCCCGAT | >MI0002468\|hsa-mir-484&WithFlank&16\|+\|15643288\|15643378 | 14 | 35 | 22 | TCAGGCTCAGTCCCCTCCCGAT | 876.4 | 4.0 | 0.8 | 5.1 | 3.5E-07 | 6.1E-07 | MB231/MB468 | NUC |
| >hg38_21_+_8433172_8446622_RNA45SN1_45S_50ntFlanks@10342.10361.20\|ATATTCAAACGAGAACTTTG | >hg38_21_+_8433172_8446622_RNA45SN1_45S_50ntFlanks | 10342 | 10361 | 20 | ATATTCAAACGAGAACTTTG | 495.0 | -6.4 | 1.3 | -5.1 | 3.9E-07 | 6.1E-07 | BT20/MB468 | NUC |
| >hg38_21_+_8433172_8446622_RNA45SN1_45S_50ntFlanks@10393.10419.27\|CAGCAGTTGAACATGGGTCAGTCGGTC | >hg38_21_+_8433172_8446622_RNA45SN1_45S_50ntFlanks | 10393 | 10419 | 27 | CAGCAGTTGAACATGGGTCAGTCGGTC | 400.4 | -7.3 | 1.4 | -5.1 | 3.1E-07 | 6.1E-07 | MB231/MB468 | MP |
| >hg38_21_+_8433172_8446622_RNA45SN1_45S_50ntFlanks@4641.4665.25\|CATTCGTATTGCGCCGCTAGAGGTG | >hg38_21_+_8433172_8446622_RNA45SN1_45S_50ntFlanks | 4641 | 4665 | 25 | CATTCGTATTGCGCCGCTAGAGGTG | 341.2 | -3.7 | 0.7 | -5.1 | 3.1E-07 | 6.1E-07 | MB231/MB468 | MP |
| >hg38_21_+_8433172_8446622_RNA45SN1_45S_50ntFlanks@11879.11898.20\|ACCCTGTTGAGCTTGACTCT | >hg38_21_+_8433172_8446622_RNA45SN1_45S_50ntFlanks | 11879 | 11898 | 20 | ACCCTGTTGAGCTTGACTCT | 331.2 | 3.2 | 0.6 | 5.1 | 2.7E-07 | 6.1E-07 | BT20/MB468 | MITO |
| >hg38_21_+_8433172_8446622_RNA45SN1_45S_50ntFlanks@11670.11691.22\|CCCAGTGCTCTGAATGTCAAAG | >hg38_21_+_8433172_8446622_RNA45SN1_45S_50ntFlanks | 11670 | 11691 | 22 | CCCAGTGCTCTGAATGTCAAAG | 1318.4 | -3.3 | 0.6 | -5.1 | 3.6E-07 | 6.1E-07 | MB231/MB468 | NUC |
| >hg38_21_+_8433172_8446622_RNA45SN1_45S_50ntFlanks@9570.9594.25\|TGCAAATCGGTCGTCCGACCTGGGT | >hg38_21_+_8433172_8446622_RNA45SN1_45S_50ntFlanks | 9570 | 9594 | 25 | TGCAAATCGGTCGTCCGACCTGGGT | 369.0 | -3.3 | 0.6 | -5.1 | 3.0E-07 | 6.1E-07 | BT20/MB468 | CYTO |
| >hg38_21_+_8433172_8446622_RNA45SN1_45S_50ntFlanks@5311.5330.20\|ATCGGGGATTGCAATTATTC | >hg38_21_+_8433172_8446622_RNA45SN1_45S_50ntFlanks | 5311 | 5330 | 20 | ATCGGGGATTGCAATTATTC | 98.1 | -9.7 | 1.8 | -5.4 | 8.0E-08 | 6.1E-07 | BT20/MB468 | MP |
| >hg38_21_+_8433172_8446622_RNA45SN1_45S_50ntFlanks@10765.10785.21\|CCGCAGCAGGTCTCCAAGGTG | >hg38_21_+_8433172_8446622_RNA45SN1_45S_50ntFlanks | 10765 | 10785 | 21 | CCGCAGCAGGTCTCCAAGGTG | 73.5 | -9.3 | 1.7 | -5.4 | 8.0E-08 | 6.1E-07 | BT20/MB468 | MP |
| >MI0002470\|hsa-mir-486-1&WithFlank&8\|-\|41660435\|41660514@10.31.22(+1U);MI0023622\|hsa-mir-486-2&WithFlank&8\|+\|41660438\|41660513@7.28.22(+1U)&[MIMAT0002177&hsa-miR-486-5p&offsets\|0\|0(+1U);m-107&8\|-\|41660484\|41660505&offsets\|0\|0(+1U)];[MIMAT0002177_1&hsa-miR-486-5p&offsets\|0\|0(+1U);TJU_CMC_MD2.ID03138.5p-miR&offsets\|0\|-2(+1U);m-106&8\|+\|41660444\|41660465&offsets\|0\|0(+1U)]\|TCCTGTACTGAGCTGCCCCGAGT | >MI0002470\|hsa-mir-486-1&WithFlank&8\|-\|41660435\|41660514 | 10 | 31 | 22(+1U) | TCCTGTACTGAGCTGCCCCGAGT | 314.9 | 8.1 | 1.6 | 5.1 | 3.1E-07 | 6.1E-07 | MB231/MB468 | MP |
| >MI0000298\|hsa-mir-221&WithFlank&X\|-\|45746151\|45746272@31.53.23&[MIMAT0004568&hsa-miR-221-5p&offsets\|0\|+1;m-405&X\|-\|45746220\|45746242&offsets\|0\|0]\|ACCTGGCATACAATGTAGATTTC | >MI0000298\|hsa-mir-221&WithFlank&X\|-\|45746151\|45746272 | 31 | 53 | 23 | ACCTGGCATACAATGTAGATTTC | 253.9 | 3.5 | 0.7 | 5.1 | 3.1E-07 | 6.1E-07 | MB231/MB468 | MP |
| >hg38_21_+_8433172_8446622_RNA45SN1_45S_50ntFlanks@4000.4026.27\|TAGATAACCTCGGGCCGATCGCACGCC | >hg38_21_+_8433172_8446622_RNA45SN1_45S_50ntFlanks | 4000 | 4026 | 27 | TAGATAACCTCGGGCCGATCGCACGCC | 67.0 | -9.1 | 1.7 | -5.4 | 8.1E-08 | 6.1E-07 | BT20/MB468 | MP |
| >trna119_LysCTT_1_-_145395522_145395594@1.33.33,trna11_LysCTT_5_-_180648979_180649051@1.33.33,trna13_LysCTT_6_+_26556774_26556846@1.33.33,trna7_LysCTT_16_+_3225692_3225764@1.33.33,trna9_LysCTT_5_+_180634755_180634827@1.33.33\|GCCCGGCTAGCTCAGTCGGTAGAGCATGAGACT | >trna119_LysCTT_1_-_145395522_145395594 | 1 | 33 | 33 | GCCCGGCTAGCTCAGTCGGTAGAGCATGAGACT | 4662.4 | 6.5 | 1.3 | 5.1 | 2.7E-07 | 6.1E-07 | BT20/MB468 | MITO |
| >hg38_21_+_8433172_8446622_RNA45SN1_45S_50ntFlanks@3993.4025.33\|GTGACTCTAGATAACCTCGGGCCGATCGCACGC | >hg38_21_+_8433172_8446622_RNA45SN1_45S_50ntFlanks | 3993 | 4025 | 33 | GTGACTCTAGATAACCTCGGGCCGATCGCACGC | 634.9 | -6.1 | 1.2 | -5.1 | 3.7E-07 | 6.1E-07 | MB231/MB468 | MITO |
| >hg38_21_+_8433172_8446622_RNA45SN1_45S_50ntFlanks@5496.5516.21\|GACGGTCGAACTTGACTATCT | >hg38_21_+_8433172_8446622_RNA45SN1_45S_50ntFlanks | 5496 | 5516 | 21 | GACGGTCGAACTTGACTATCT | 604.3 | 5.1 | 0.9 | 5.5 | 2.9E-08 | 6.1E-07 | BT20/MB231 | MITO |
| >hg38_21_+_8433172_8446622_RNA45SN1_45S_50ntFlanks@5056.5082.27\|GAGCGATTTGTCTGGTTAATTCCGATA | >hg38_21_+_8433172_8446622_RNA45SN1_45S_50ntFlanks | 5056 | 5082 | 27 | GAGCGATTTGTCTGGTTAATTCCGATA | 569.8 | -6.6 | 1.3 | -5.1 | 3.7E-07 | 6.1E-07 | MB231/MB468 | MITO |
| >hg38_21_+_8433172_8446622_RNA45SN1_45S_50ntFlanks@12454.12477.24\|CACCAAGCGTTGGATTGTTCACCC | >hg38_21_+_8433172_8446622_RNA45SN1_45S_50ntFlanks | 12454 | 12477 | 24 | CACCAAGCGTTGGATTGTTCACCC | 408.8 | 3.2 | 0.6 | 5.1 | 3.5E-07 | 6.1E-07 | MB231/MB468 | CYTO |
| >hg38_21_+_8433172_8446622_RNA45SN1_45S_50ntFlanks@10866.10894.29\|CTAAGGGCTGGGTCGGTCGGGCTGGGGCG | >hg38_21_+_8433172_8446622_RNA45SN1_45S_50ntFlanks | 10866 | 10894 | 29 | CTAAGGGCTGGGTCGGTCGGGCTGGGGCG | 765.1 | -7.9 | 1.5 | -5.4 | 8.1E-08 | 6.1E-07 | BT20/MB468 | MP |
| >hg38_21_+_8433172_8446622_RNA45SN1_45S_50ntFlanks@10350.10384.35\|ACGAGAACTTTGAAGGCCGAAGTGGAGAAGGGTTC | >hg38_21_+_8433172_8446622_RNA45SN1_45S_50ntFlanks | 10350 | 10384 | 35 | ACGAGAACTTTGAAGGCCGAAGTGGAGAAGGGTTC | 1496.4 | -3.4 | 0.7 | -5.1 | 3.5E-07 | 6.1E-07 | MB231/MB468 | CYTO |
| >hg38_21_+_8433172_8446622_RNA45SN1_45S_50ntFlanks@12002.12030.29\|TGCGGGCCGCCGGTGAAATACCACTACTC | >hg38_21_+_8433172_8446622_RNA45SN1_45S_50ntFlanks | 12002 | 12030 | 29 | TGCGGGCCGCCGGTGAAATACCACTACTC | 1871.0 | -8.8 | 1.7 | -5.1 | 3.9E-07 | 6.1E-07 | BT20/MB468 | NUC |
| >hg38_21_+_8433172_8446622_RNA45SN1_45S_50ntFlanks@12916.12933.18\|TTCGTGGGGAACCTGGCG | >hg38_21_+_8433172_8446622_RNA45SN1_45S_50ntFlanks | 12916 | 12933 | 18 | TTCGTGGGGAACCTGGCG | 214.7 | -7.4 | 1.4 | -5.1 | 3.1E-07 | 6.1E-07 | MB231/MB468 | MP |
| >hg38_21_+_8433172_8446622_RNA45SN1_45S_50ntFlanks@10680.10710.31\|GTTCCGGCGGCGTCCGGTGAGCTCTCGCTGG | >hg38_21_+_8433172_8446622_RNA45SN1_45S_50ntFlanks | 10680 | 10710 | 31 | GTTCCGGCGGCGTCCGGTGAGCTCTCGCTGG | 11319.8 | -5.9 | 1.1 | -5.4 | 8.1E-08 | 6.1E-07 | BT20/MB468 | MP |
| >MI0000342\|hsa-mir-200b&WithFlank&1\|+\|1167098\|1167204@64.85.22(+1A)&[MIMAT0000318&hsa-miR-200b-3p&offsets\|+1\|+1(+1A);m-93&1\|+\|1167160\|1167182&offsets\|+1\|0(+1A)]\|AATACTGCCTGGTAATGATGACA | >MI0000342\|hsa-mir-200b&WithFlank&1\|+\|1167098\|1167204 | 64 | 85 | 22(+1A) | AATACTGCCTGGTAATGATGACA | 63.2 | -9.1 | 1.7 | -5.4 | 8.1E-08 | 6.1E-07 | BT20/MB468 | MP |

<table>
<tr><th></th><th></th><th></th><th></th><th></th><th></th><th></th><th></th><th></th><th></th><th></th><th></th><th></th><th></th></tr>
<tr><td>>hg38_21_+_8433172_8446622_RNA45SN1_45S_50ntFlanks@12212.12232.21|CTCAGGGAGGACAGAAACCTC</td><td>>hg38_21_+_8433172_8446622_RNA45SN1_45S_50ntFlanks</td><td>12212</td><td>12232</td><td>21</td><td>CTCAGGGAGGACAGAAACCTC</td><td>79.8</td><td>-9.4</td><td>1.8</td><td>-5.4</td><td>8.1E-08</td><td>6.1E-07</td><td>BT20/MB468</td><td>MP</td></tr>
<tr><td>>hg38_21_+_8433172_8446622_RNA45SN1_45S_50ntFlanks@6736.6770.35|TGATCATCGACACTTCGAACGCACTTGCGGCCCCG</td><td>>hg38_21_+_8433172_8446622_RNA45SN1_45S_50ntFlanks</td><td>6736</td><td>6770</td><td>35</td><td>TGATCATCGACACTTCGAACGCACTTGCGGCCCCG</td><td>89.5</td><td>-9.6</td><td>1.8</td><td>-5.4</td><td>8.1E-08</td><td>6.1E-07</td><td>BT20/MB468</td><td>MP</td></tr>
<tr><td>>hg38_21_+_8433172_8446622_RNA45SN1_45S_50ntFlanks@10914.10941.28|CGCCGCGGCTGGACGAGGCGCCGCCGCC</td><td>>hg38_21_+_8433172_8446622_RNA45SN1_45S_50ntFlanks</td><td>10914</td><td>10941</td><td>28</td><td>CGCCGCGGCTGGACGAGGCGCCGCCGCC</td><td>91.0</td><td>-9.6</td><td>1.8</td><td>-5.4</td><td>8.1E-08</td><td>6.1E-07</td><td>BT20/MB468</td><td>MP</td></tr>
<tr><td>>hg38_21_+_8433172_8446622_RNA45SN1_45S_50ntFlanks@11641.11667.27|CGGCGGGTGTTGACGCGATGTGATTTC</td><td>>hg38_21_+_8433172_8446622_RNA45SN1_45S_50ntFlanks</td><td>11641</td><td>11667</td><td>27</td><td>CGGCGGGTGTTGACGCGATGTGATTTC</td><td>697.0</td><td>-6.0</td><td>1.2</td><td>-5.1</td><td>3.6E-07</td><td>6.1E-07</td><td>MB231/MB468</td><td>NUC</td></tr>
<tr><td>>hg38_21_+_8433172_8446622_RNA45SN1_45S_50ntFlanks@11767.11791.25|TCTAATTAGTGACGCGCATGAATGG</td><td>>hg38_21_+_8433172_8446622_RNA45SN1_45S_50ntFlanks</td><td>11767</td><td>11791</td><td>25</td><td>TCTAATTAGTGACGCGCATGAATGG</td><td>228.7</td><td>-5.6</td><td>1.1</td><td>-5.1</td><td>3.5E-07</td><td>6.1E-07</td><td>MB231/MB468</td><td>CYTO</td></tr>
<tr><td>>hg38_1_-_228634819_228635039_RNA5S12_5S_50ntFlanks@135.169.35|GGGAGACCGCCTGGGAATACCGGGTGCTGTAGGCT</td><td>>hg38_1_-_228634819_228635039_RNA5S12_5S_50ntFlanks</td><td>135</td><td>169</td><td>35</td><td>GGGAGACCGCCTGGGAATACCGGGTGCTGTAGGCT</td><td>2098.3</td><td>5.0</td><td>1.0</td><td>5.1</td><td>2.7E-07</td><td>6.2E-07</td><td>BT20/MB468</td><td>MITO</td></tr>
<tr><td>>am_trna10_ValCAC_5_-_180649395_180649467@1.34.34,trna18_ValCAC_5_-_180529253_180529325@1.34.34,trna2_ValCAC_5_+_180524070_180524142@1.34.34,trna6_ValCAC_5_+_180600650_180600722@1.34.34,trna85_ValCAC_1_-_161369490_161369562@1.34.34,trna90_ValCAC_1_-_149684088_149684161@1.34.34,trna98_ValCAC_1_-_149298555_149298627@1.34.34,trna9_ValCAC_6_+_26538282_26538354@1.34.34|GTTTCCGTAGTGTAGTGGTTATCACGTTCGCCTC</td><td>>am_trna10_ValCAC_5_-_180649395_180649467</td><td>1</td><td>34</td><td>34</td><td>GTTTCCGTAGTGTAGTGGTTATCACGTTCGCCTC</td><td>2157.4</td><td>11.5</td><td>2.1</td><td>5.4</td><td>8.2E-08</td><td>6.2E-07</td><td>BT20/MB468</td><td>MP</td></tr>
<tr><td>>hg38_21_+_8433172_8446622_RNA45SN1_45S_50ntFlanks@11932.11950.19|GAATAAGTGGGAGGCCCCC</td><td>>hg38_21_+_8433172_8446622_RNA45SN1_45S_50ntFlanks</td><td>11932</td><td>11950</td><td>19</td><td>GAATAAGTGGGAGGCCCCC</td><td>79.3</td><td>-9.4</td><td>1.8</td><td>-5.4</td><td>8.2E-08</td><td>6.2E-07</td><td>BT20/MB468</td><td>MP</td></tr>
<tr><td>>hg38_21_+_8433172_8446622_RNA45SN1_45S_50ntFlanks@12621.12653.33|GAGCCAATGGGGCGAAGCTACCATCTGTGGGAT</td><td>>hg38_21_+_8433172_8446622_RNA45SN1_45S_50ntFlanks</td><td>12621</td><td>12653</td><td>33</td><td>GAGCCAATGGGGCGAAGCTACCATCTGTGGGAT</td><td>74.5</td><td>-9.3</td><td>1.7</td><td>-5.4</td><td>8.2E-08</td><td>6.2E-07</td><td>BT20/MB468</td><td>MP</td></tr>
<tr><td>>MI0003581|hsa-mir-574&WithFlank&4|+|38868026|38868133@67.88.22&[MIMAT0003239&hsa-miR-574-3p&offsets|0|0;m-149&4|+|38868092|38868113&offsets|0|0]|CACGCTCATGCACACACCCACA</td><td>>MI0003581|hsa-mir-574&WithFlank&4|+|38868026|38868133</td><td>67</td><td>88</td><td>22</td><td>CACGCTCATGCACACACCCACA</td><td>908.5</td><td>4.2</td><td>0.8</td><td>5.1</td><td>3.6E-07</td><td>6.2E-07</td><td>MB231/MB468</td><td>NUC</td></tr>
<tr><td>>hg38_21_+_8433172_8446622_RNA45SN1_45S_50ntFlanks@10734.10755.22|GTGTAAATCTCGCGCCGGGCCG</td><td>>hg38_21_+_8433172_8446622_RNA45SN1_45S_50ntFlanks</td><td>10734</td><td>10755</td><td>22</td><td>GTGTAAATCTCGCGCCGGGCCG</td><td>519.6</td><td>-7.1</td><td>1.4</td><td>-5.1</td><td>3.6E-07</td><td>6.2E-07</td><td>MB231/MB468</td><td>NUC</td></tr>
<tr><td>>hg38_21_+_8433172_8446622_RNA45SN1_45S_50ntFlanks@483.517.35|TCGATGTGGTGACGTCGTGCTCTCCCGGGCCGGGT</td><td>>hg38_21_+_8433172_8446622_RNA45SN1_45S_50ntFlanks</td><td>483</td><td>517</td><td>35</td><td>TCGATGTGGTGACGTCGTGCTCTCCCGGGCCGGGT</td><td>393.7</td><td>4.3</td><td>0.8</td><td>5.1</td><td>3.6E-07</td><td>6.2E-07</td><td>MB231/MB468</td><td>NUC</td></tr>
<tr><td>>hg38_21_+_8433172_8446622_RNA45SN1_45S_50ntFlanks@3986.4024.39|GGCTTTGGTGACTCTAGATAACCTCGGGCCGATCGCACG</td><td>>hg38_21_+_8433172_8446622_RNA45SN1_45S_50ntFlanks</td><td>3986</td><td>4024</td><td>39</td><td>GGCTTTGGTGACTCTAGATAACCTCGGGCCGATCGCACG</td><td>382.5</td><td>-4.0</td><td>0.8</td><td>-5.1</td><td>3.8E-07</td><td>6.2E-07</td><td>MB231/MB468</td><td>MITO</td></tr>
<tr><td>>hg38_21_+_8433172_8446622_RNA45SN1_45S_50ntFlanks@11704.11726.23|ATGAAGCGCGGGTAAACGGCGGG</td><td>>hg38_21_+_8433172_8446622_RNA45SN1_45S_50ntFlanks</td><td>11704</td><td>11726</td><td>23</td><td>ATGAAGCGCGGGTAAACGGCGGG</td><td>226.9</td><td>-7.1</td><td>1.4</td><td>-5.1</td><td>3.2E-07</td><td>6.2E-07</td><td>MB231/MB468</td><td>MP</td></tr>
<tr><td>>hg38_21_+_8433172_8446622_RNA45SN1_45S_50ntFlanks@5423.5441.19|ATTGGATGGTTTAGTGAGG</td><td>>hg38_21_+_8433172_8446622_RNA45SN1_45S_50ntFlanks</td><td>5423</td><td>5441</td><td>19</td><td>ATTGGATGGTTTAGTGAGG</td><td>91.6</td><td>-9.6</td><td>1.8</td><td>-5.4</td><td>8.3E-08</td><td>6.2E-07</td><td>BT20/MB468</td><td>MP</td></tr>
<tr><td>>hg38_21_+_8433172_8446622_RNA45SN1_45S_50ntFlanks@10421.10455.35|TGAGAGATGGGCGAGCGCCGTTCCGAAGGGACGGG</td><td>>hg38_21_+_8433172_8446622_RNA45SN1_45S_50ntFlanks</td><td>10421</td><td>10455</td><td>35</td><td>TGAGAGATGGGCGAGCGCCGTTCCGAAGGGACGGG</td><td>250.5</td><td>-3.6</td><td>0.7</td><td>-5.1</td><td>3.1E-07</td><td>6.2E-07</td><td>BT20/MB468</td><td>CYTO</td></tr>
<tr><td>>hg38_21_+_8433172_8446622_RNA45SN1_45S_50ntFlanks@8403.8442.40|AGGTAAACGGGTGGGGTCCGCGCAGTCCGCCCGGAGGATT</td><td>>hg38_21_+_8433172_8446622_RNA45SN1_45S_50ntFlanks</td><td>8403</td><td>8442</td><td>40</td><td>AGGTAAACGGGTGGGGTCCGCGCAGTCCGCCCGGAGGATT</td><td>82.2</td><td>-9.4</td><td>1.8</td><td>-5.4</td><td>8.3E-08</td><td>6.2E-07</td><td>BT20/MB468</td><td>MP</td></tr>
<tr><td>>hg38_21_+_8433172_8446622_RNA45SN1_45S_50ntFlanks@12988.13013.26|CAGCTCCCTCGCTGCGATCTATTGAA</td><td>>hg38_21_+_8433172_8446622_RNA45SN1_45S_50ntFlanks</td><td>12988</td><td>13013</td><td>26</td><td>CAGCTCCCTCGCTGCGATCTATTGAA</td><td>1024.9</td><td>2.8</td><td>0.5</td><td>5.1</td><td>3.5E-07</td><td>6.2E-07</td><td>MB231/MB468</td><td>CYTO</td></tr>
<tr><td>>hg38_21_+_8433172_8446622_RNA45SN1_45S_50ntFlanks@485.506.22|GATGTGGTGACGTCGTGCTCTC</td><td>>hg38_21_+_8433172_8446622_RNA45SN1_45S_50ntFlanks</td><td>485</td><td>506</td><td>22</td><td>GATGTGGTGACGTCGTGCTCTC</td><td>798.3</td><td>13.6</td><td>2.7</td><td>5.1</td><td>3.5E-07</td><td>6.2E-07</td><td>MB231/MB468</td><td>CYTO</td></tr>
<tr><td>>hg38_21_+_8433172_8446622_RNA45SN1_45S_50ntFlanks@12939.12972.34|CCATTCGTAGACGACCTGCTTCTGGGTCGGGGTT</td><td>>hg38_21_+_8433172_8446622_RNA45SN1_45S_50ntFlanks</td><td>12939</td><td>12972</td><td>34</td><td>CCATTCGTAGACGACCTGCTTCTGGGTCGGGGTT</td><td>220.5</td><td>-5.7</td><td>1.1</td><td>-5.1</td><td>3.5E-07</td><td>6.2E-07</td><td>MB231/MB468</td><td>CYTO</td></tr>
<tr><td>>hg38_21_+_8433172_8446622_RNA45SN1_45S_50ntFlanks@476.516.41|TTCGTGATCGATGTGGTGACGTCGTGCTCTCCCGGGCCGGG</td><td>>hg38_21_+_8433172_8446622_RNA45SN1_45S_50ntFlanks</td><td>476</td><td>516</td><td>41</td><td>TTCGTGATCGATGTGGTGACGTCGTGCTCTCCCGGGCCGGG</td><td>16930.8</td><td>4.3</td><td>0.8</td><td>5.1</td><td>3.6E-07</td><td>6.2E-07</td><td>MB231/MB468</td><td>NUC</td></tr>
<tr><td>>hg38_21_+_8433172_8446622_RNA45SN1_45S_50ntFlanks@11569.11599.31|CAGCCGACTTAGAACTGGTGCGGACCAGGGG</td><td>>hg38_21_+_8433172_8446622_RNA45SN1_45S_50ntFlanks</td><td>11569</td><td>11599</td><td>31</td><td>CAGCCGACTTAGAACTGGTGCGGACCAGGGG</td><td>281.5</td><td>-5.2</td><td>1.0</td><td>-5.1</td><td>3.8E-07</td><td>6.2E-07</td><td>MB231/MB468</td><td>MITO</td></tr>
<tr><td>>MI0000342|hsa-mir-200b&WithFlank&1|+|1167098|1167204@64.85.22(+1C)&[MIMAT0000318&hsa-miR-200b-3p&offsets|+1|+1(+1C);m-93&1|+|1167160|1167182&offsets|+1|0(+1C)]|AATACTGCCTGGTAATGATGACC</td><td>>MI0000342|hsa-mir-200b&WithFlank&1|+|1167098|1167204</td><td>64</td><td>85</td><td>22(+1C)</td><td>AATACTGCCTGGTAATGATGACC</td><td>64.0</td><td>-9.1</td><td>1.7</td><td>-5.4</td><td>8.3E-08</td><td>6.2E-07</td><td>BT20/MB468</td><td>MP</td></tr>
<tr><td>>MI0000085|hsa-mir-27a&WithFlank&19|-|13836434|13836523@57.75.19(+1U)&[MIMAT0000084&hsa-miR-27a-3p&offsets|0|-2(+1U);m-43&19|-|13836447|13836467&offsets|0|-2(+1U)]|TTCACAGTGGCTAAGTTCCT</td><td>>MI0000085|hsa-mir-27a&WithFlank&19|-|13836434|13836523</td><td>57</td><td>75</td><td>19(+1U)</td><td>TTCACAGTGGCTAAGTTCCT</td><td>481.3</td><td>5.3</td><td>1.1</td><td>5.1</td><td>3.8E-07</td><td>6.2E-07</td><td>MB231/MB468</td><td>MITO</td></tr>
<tr><td>>hg38_21_+_8433172_8446622_RNA45SN1_45S_50ntFlanks@4585.4619.35|GTTCTATTTTGTTGGTTTTCGGAACTGAGGCCATG</td><td>>hg38_21_+_8433172_8446622_RNA45SN1_45S_50ntFlanks</td><td>4585</td><td>4619</td><td>35</td><td>GTTCTATTTTGTTGGTTTTCGGAACTGAGGCCATG</td><td>194.4</td><td>4.8</td><td>1.0</td><td>5.1</td><td>3.6E-07</td><td>6.2E-07</td><td>MB231/MB468</td><td>CYTO</td></tr>
<tr><td>>hg38_21_+_8433172_8446622_RNA45SN1_45S_50ntFlanks@12971.12991.21|TTTCGTACGTAGCAGAGCAGC</td><td>>hg38_21_+_8433172_8446622_RNA45SN1_45S_50ntFlanks</td><td>12971</td><td>12991</td><td>21</td><td>TTTCGTACGTAGCAGAGCAGC</td><td>1029.1</td><td>-7.4</td><td>1.5</td><td>-5.1</td><td>3.7E-07</td><td>6.3E-07</td><td>MB231/MB468</td><td>NUC</td></tr>
<tr><td>>hg38_21_+_8433172_8446622_RNA45SN1_45S_50ntFlanks@5471.5514.44|CACGGCCCTGGCGGAGCGCTGAGAAGACGGTCGAACTTGACTAT</td><td>>hg38_21_+_8433172_8446622_RNA45SN1_45S_50ntFlanks</td><td>5471</td><td>5514</td><td>44</td><td>CACGGCCCTGGCGGAGCGCTGAGAAGACGGTCGAACTTGACTAT</td><td>1589.1</td><td>-5.3</td><td>1.0</td><td>-5.1</td><td>3.8E-07</td><td>6.3E-07</td><td>MB231/MB468</td><td>MITO</td></tr>
<tr><td>>trna116_GluCTC_1_-_145399233_145399304@37.73.37,trna134_GluTTC_1_-_16861774_16861845@37.73.37,trna71_GluCTC_1_-_161439189_161439260@37.73.37,trna74_GluCTC_1_-_161431809_161431880@37.73.37,trna77_GluCTC_1_-_161424398_161424469@37.73.37,trna77_GluCTC_6_+_28949976_28950047@37.73.37,trna80_GluCTC_1_-_161417018_161417089@37.73.37,trna84_GluTTC_1_-_161391883_161391954@37.73.37,trna87_GluCTC_6_-_126101393_126101464@37.73.37|ACCGCCGCGGCCCGGGTTCGATTCCCGGTCAGGGAAC</td><td>>trna116_GluCTC_1_-_145399233_145399304</td><td>37</td><td>73</td><td>37</td><td>ACCGCCGCGGCCCGGGTTCGATTCCCGGTCAGGGAAC</td><td>1186.4</td><td>6.9</td><td>1.4</td><td>5.1</td><td>3.7E-07</td><td>6.3E-07</td><td>MB231/MB468</td><td>NUC</td></tr>
<tr><td>>hg38_MT_+_598_1651_RNR1_12S_50ntFlanks@430.469.40|AGTGGCTTTAACATATCTGAACACACAATAGCTAAGACCC</td><td>>hg38_MT_+_598_1651_RNR1_12S_50ntFlanks</td><td>430</td><td>469</td><td>40</td><td>AGTGGCTTTAACATATCTGAACACACAATAGCTAAGACCC</td><td>132.4</td><td>10.8</td><td>2.0</td><td>5.4</td><td>8.4E-08</td><td>6.3E-07</td><td>BT20/MB468</td><td>MP</td></tr>
<tr><td>>hg38_21_+_8433172_8446622_RNA45SN1_45S_50ntFlanks@12539.12569.31|ATGATGTGTTGTTGCCATGGTAATCCTGCTC</td><td>>hg38_21_+_8433172_8446622_RNA45SN1_45S_50ntFlanks</td><td>12539</td><td>12569</td><td>31</td><td>ATGATGTGTTGTTGCCATGGTAATCCTGCTC</td><td>4030.5</td><td>-8.3</td><td>1.6</td><td>-5.1</td><td>3.7E-07</td><td>6.3E-07</td><td>MB231/MB468</td><td>NUC</td></tr>
<tr><td>>hg38_21_+_8433172_8446622_RNA45SN1_45S_50ntFlanks@6722.6745.24|ATTGCAGGACACATTGATCATCGA</td><td>>hg38_21_+_8433172_8446622_RNA45SN1_45S_50ntFlanks</td><td>6722</td><td>6745</td><td>24</td><td>ATTGCAGGACACATTGATCATCGA</td><td>182.5</td><td>-6.4</td><td>1.2</td><td>-5.1</td><td>3.7E-07</td><td>6.3E-07</td><td>MB231/MB468</td><td>NUC</td></tr>
<tr><td>>MI0000750|hsa-mir-26a-2&WithFlank&12|-|57824603|57824698@20.41.22(+1A);MI0000083|hsa-mir-26a-1&WithFlank&3|+|37969398|37969486@16.37.22(+1A)&[MIMAT0000082&hsa-miR-26a-5p&offsets|0|0(+1A);m-33&12|-|57824658|57824679&offsets|0|0(+1A)];[MIMAT0000082_1&hsa-miR-26a-5p&offsets|0|0(+1A);m-34&3|+|37969413|37969434&offsets|0|0(+1A)]|TTCAAGTAATCCAGGATAGGCTA</td><td>>MI0000750|hsa-mir-26a-2&WithFlank&12|-|57824603|57824698</td><td>20</td><td>41</td><td>22(+1A)</td><td>TTCAAGTAATCCAGGATAGGCTA</td><td>517.7</td><td>5.0</td><td>1.0</td><td>5.1</td><td>3.6E-07</td><td>6.3E-07</td><td>MB231/MB468</td><td>CYTO</td></tr>
<tr><td>>MI0000483|hsa-mir-186&WithFlank&1|-|71067625|71067722@21.43.23&[MIMAT0000456&hsa-miR-186-5p&offsets|0|+1;m-84&1|-|71067681|71067702&offsets|0|+1]|CAAAGAATTCTCCTTTTGGGCTT</td><td>>MI0000483|hsa-mir-186&WithFlank&1|-|71067625|71067722</td><td>21</td><td>43</td><td>23</td><td>CAAAGAATTCTCCTTTTGGGCTT</td><td>808.9</td><td>5.2</td><td>1.0</td><td>5.1</td><td>3.9E-07</td><td>6.3E-07</td><td>MB231/MB468</td><td>MITO</td></tr>
<tr><td>>hg38_21_+_8433172_8446622_RNA45SN1_45S_50ntFlanks@11564.11602.39|CCTAGCAGCCGACTTAGAACTGGTGCGGACCAGGGGAAT</td><td>>hg38_21_+_8433172_8446622_RNA45SN1_45S_50ntFlanks</td><td>11564</td><td>11602</td><td>39</td><td>CCTAGCAGCCGACTTAGAACTGGTGCGGACCAGGGGAAT</td><td>957.0</td><td>-3.6</td><td>0.7</td><td>-5.1</td><td>3.6E-07</td><td>6.3E-07</td><td>MB231/MB468</td><td>CYTO</td></tr>
<tr><td>>MI0000542|hsa-mir-320a&WithFlank&8|-|22244960|22245043@49.71.23&[MIMAT0000510&hsa-miR-320a-3p&offsets|+1|+2;m-60&8|-|22244975|22244996&offsets|+1|+2]|AAAGCTGGGTTGAGAGGGCGAAA</td><td>>MI0000542|hsa-mir-320a&WithFlank&8|-|22244960|22245043</td><td>49</td><td>71</td><td>23</td><td>AAAGCTGGGTTGAGAGGGCGAAA</td><td>550.8</td><td>4.3</td><td>0.8</td><td>5.6</td><td>2.5E-08</td><td>6.3E-07</td><td>BT20/MB231</td><td>CYTO</td></tr>
<tr><td>>hg38_21_+_8433172_8446622_RNA45SN1_45S_50ntFlanks@6668.6689.22|TCACTCGGCTCGTGCGTCGATG</td><td>>hg38_21_+_8433172_8446622_RNA45SN1_45S_50ntFlanks</td><td>6668</td><td>6689</td><td>22</td><td>TCACTCGGCTCGTGCGTCGATG</td><td>86.4</td><td>-9.5</td><td>1.8</td><td>-5.4</td><td>8.4E-08</td><td>6.3E-07</td><td>BT20/MB468</td><td>MP</td></tr>
<tr><td>>MI0000063|hsa-let-7b&WithFlank&22|+|46113680|46113774@12.31.20(+1C)&[MIMAT0000063&hsa-let-7b-5p&offsets|0|-2(+1C);m-14&22|+|46113691|46113712&offsets|0|-2(+1C)]|TGAGGTAGTAGGTTGTGTGGC</td><td>>MI0000063|hsa-let-7b&WithFlank&22|+|46113680|46113774</td><td>12</td><td>31</td><td>20(+1C)</td><td>TGAGGTAGTAGGTTGTGTGGC</td><td>1429.0</td><td>5.3</td><td>1.0</td><td>5.4</td><td>8.4E-08</td><td>6.3E-07</td><td>BT20/MB468</td><td>MP</td></tr>
<tr><td>>hg38_21_+_8433172_8446622_RNA45SN1_45S_50ntFlanks@12714.12742.29|GAGCCTCGGTTGGCCTCGGATAGCCGGTC</td><td>>hg38_21_+_8433172_8446622_RNA45SN1_45S_50ntFlanks</td><td>12714</td><td>12742</td><td>29</td><td>GAGCCTCGGTTGGCCTCGGATAGCCGGTC</td><td>633.5</td><td>-5.1</td><td>1.0</td><td>-5.4</td><td>8.4E-08</td><td>6.3E-07</td><td>BT20/MB468</td><td>MP</td></tr>
<tr><td>>hg38_21_+_8433172_8446622_RNA45SN1_45S_50ntFlanks@10808.10830.23|CAATGTAGGTAAGGGAAGTCGGC</td><td>>hg38_21_+_8433172_8446622_RNA45SN1_45S_50ntFlanks</td><td>10808</td><td>10830</td><td>23</td><td>CAATGTAGGTAAGGGAAGTCGGC</td><td>445.3</td><td>-4.7</td><td>0.9</td><td>-5.1</td><td>3.2E-07</td><td>6.3E-07</td><td>BT20/MB468</td><td>CYTO</td></tr>
<tr><td>>hg38_21_+_8433172_8446622_RNA45SN1_45S_50ntFlanks@12907.12931.25|CACCGCACGTTCGTGGGGAACCTGG</td><td>>hg38_21_+_8433172_8446622_RNA45SN1_45S_50ntFlanks</td><td>12907</td><td>12931</td><td>25</td><td>CACCGCACGTTCGTGGGGAACCTGG</td><td>271.7</td><td>-7.3</td><td>1.4</td><td>-5.1</td><td>3.2E-07</td><td>6.3E-07</td><td>MB231/MB468</td><td>MP</td></tr>
<tr><td>>hg38_21_+_8433172_8446622_RNA45SN1_45S_50ntFlanks@9505.9523.19|TGAACTATGCCTGGGCAGG</td><td>>hg38_21_+_8433172_8446622_RNA45SN1_45S_50ntFlanks</td><td>9505</td><td>9523</td><td>19</td><td>TGAACTATGCCTGGGCAGG</td><td>188.0</td><td>-6.1</td><td>1.2</td><td>-5.1</td><td>3.2E-07</td><td>6.3E-07</td><td>BT20/MB468</td><td>CYTO</td></tr>
<tr><td>>trna11_GluTTC_15_-_26327381_26327452@1.33.33,trna3_GluTTC_13_-_45492062_45492133@1.33.33|TCCCACATGGTCTAGCGGTTAGGATTCCTGGTT</td><td>>trna11_GluTTC_15_-_26327381_26327452</td><td>1</td><td>33</td><td>33</td><td>TCCCACATGGTCTAGCGGTTAGGATTCCTGGTT</td><td>1117.6</td><td>6.1</td><td>1.2</td><td>5.1</td><td>3.2E-07</td><td>6.3E-07</td><td>MB231/MB468</td><td>MP</td></tr>
</table>

<table>
<tr><td>>MI0000298|hsa-mir-221&WithFlank&X|-|45746151|45746272@71.93.23(+1G)&[MIMAT0000278&hsa-miR-221-3p&offsets|0|0(+1G);m-91&X|-|45746181|45746202&offsets|0|+1(+1G)]|AGCTACATTGTCTGCTGGGTTTCG</td><td>>MI0000298|hsa-mir-221&WithFlank&X|-|45746151|45746272</td><td>71</td><td>93</td><td>23(+1G)</td><td>AGCTACATTGTCTGCTGGGTTTCG</td><td>253.5</td><td>4.8</td><td>1.0</td><td>5.1</td><td>3.9E-07</td><td>6.3E-07</td><td>MB231/MB468</td><td>MITO</td></tr>
<tr><td>>am_trnaMT_PheGAA_MT_+_577_647@41.59.19|GTTTAGACGGGCTCACATC</td><td>>am_trnaMT_PheGAA_MT_+_577_647</td><td>41</td><td>59</td><td>19</td><td>GTTTAGACGGGCTCACATC</td><td>235.8</td><td>3.8</td><td>0.7</td><td>5.1</td><td>3.9E-07</td><td>6.3E-07</td><td>MB231/MB468</td><td>MITO</td></tr>
<tr><td>>hg38_21_+_8433172_8446622_RNA45SN1_45S_50ntFlanks@3738.3763.26|TCTCAAAGATTAAGCCATGCATGTCT</td><td>>hg38_21_+_8433172_8446622_RNA45SN1_45S_50ntFlanks</td><td>3738</td><td>3763</td><td>26</td><td>TCTCAAAGATTAAGCCATGCATGTCT</td><td>65.4</td><td>-9.1</td><td>1.7</td><td>-5.4</td><td>8.5E-08</td><td>6.3E-07</td><td>BT20/MB468</td><td>MP</td></tr>
<tr><td>>trna116_GluCTC_1_-_145399233_145399304@1.35.35,trna59_GluCTC_1_+_249168447_249168518@1.35.35,trna71_GluCTC_1_-_161439189_161439260@1.35.35,trna74_GluCTC_1_-_161431809_161431880@1.35.35,trna77_GluCTC_1_-_161424398_161424469@1.35.35,trna77_GluCTC_6_+_28949976_28950047@1.35.35,trna80_GluCTC_1_-_161417018_161417089@1.35.35,trna87_GluCTC_6_-_126101393_126101464@1.35.35|TCCCTGGTGGTCTAGTGGTTAGGATTCGGCGCTCT</td><td>>trna116_GluCTC_1_-_145399233_145399304</td><td>1</td><td>35</td><td>35</td><td>TCCCTGGTGGTCTAGTGGTTAGGATTCGGCGCTCT</td><td>1914.6</td><td>4.3</td><td>0.9</td><td>5.1</td><td>3.9E-07</td><td>6.3E-07</td><td>MB231/MB468</td><td>MITO</td></tr>
<tr><td>>hg38_MT_+_598_1651_RNR1_12S_50ntFlanks@416.452.37|AAATAGACTACGAAAGTGGCTTTAACATATCTGAACA</td><td>>hg38_MT_+_598_1651_RNR1_12S_50ntFlanks</td><td>416</td><td>452</td><td>37</td><td>AAATAGACTACGAAAGTGGCTTTAACATATCTGAACA</td><td>56.2</td><td>9.0</td><td>1.8</td><td>5.1</td><td>3.2E-07</td><td>6.3E-07</td><td>MB231/MB468</td><td>MP</td></tr>
<tr><td>>trna111_HisGTG_1_-_147774845_147774916@-1A.29.30,trna118_HisGTG_1_-_145396881_145396952@-1A.29.30,trna16_HisGTG_1_+_146544773_146544844@-1A.29.30,trna1_HisGTG_15_+_45493349_45493420@-1A.29.30,trna21_HisGTG_1_+_147753471_147753542@-1A.29.30,trna33_HisGTG_6_+_27125906_27125977@-1A.29.30,trna7_HisGTG_9_-_14433938_14434009@-1A.29.30,trna8_HisGTG_15_-_45492611_45492682@-1A.29.30,trna9_HisGTG_15_-_45490804_45490875@-1A.29.30|AGCCGTGATCGTATAGTGGTTAGTACTCTG</td><td>>trna111_HisGTG_1_-_147774845_147774916</td><td>-1A</td><td>29</td><td>30</td><td>AGCCGTGATCGTATAGTGGTTAGTACTCTG</td><td>869.0</td><td>-6.0</td><td>1.2</td><td>-5.1</td><td>3.7E-07</td><td>6.3E-07</td><td>MB231/MB468</td><td>NUC</td></tr>
<tr><td>>hg38_21_+_8433172_8446622_RNA45SN1_45S_50ntFlanks@11199.11223.25|CCGGCGGCGGCGGCGACTCTGGACG</td><td>>hg38_21_+_8433172_8446622_RNA45SN1_45S_50ntFlanks</td><td>11199</td><td>11223</td><td>25</td><td>CCGGCGGCGGCGGCGACTCTGGACG</td><td>869.7</td><td>-7.7</td><td>1.5</td><td>-5.1</td><td>3.7E-07</td><td>6.3E-07</td><td>MB231/MB468</td><td>NUC</td></tr>
<tr><td>>hg38_21_+_8433172_8446622_RNA45SN1_45S_50ntFlanks@10797.10818.22|GCATGTTGGAACAATGTAGGTA</td><td>>hg38_21_+_8433172_8446622_RNA45SN1_45S_50ntFlanks</td><td>10797</td><td>10818</td><td>22</td><td>GCATGTTGGAACAATGTAGGTA</td><td>946.2</td><td>-3.4</td><td>0.7</td><td>-5.1</td><td>2.8E-07</td><td>6.4E-07</td><td>BT20/MB468</td><td>MITO</td></tr>
<tr><td>>hg38_21_+_8433172_8446622_RNA45SN1_45S_50ntFlanks@10408.10428.21|GGTCAGTCGGTCCTGAGAGAT</td><td>>hg38_21_+_8433172_8446622_RNA45SN1_45S_50ntFlanks</td><td>10408</td><td>10428</td><td>21</td><td>GGTCAGTCGGTCCTGAGAGAT</td><td>313.4</td><td>-4.4</td><td>0.9</td><td>-5.1</td><td>3.6E-07</td><td>6.4E-07</td><td>MB231/MB468</td><td>CYTO</td></tr>
<tr><td>>hg38_21_+_8433172_8446622_RNA45SN1_45S_50ntFlanks@6652.6669.18|GACTCTTAGCGGTGGATC</td><td>>hg38_21_+_8433172_8446622_RNA45SN1_45S_50ntFlanks</td><td>6652</td><td>6669</td><td>18</td><td>GACTCTTAGCGGTGGATC</td><td>163.9</td><td>2.3</td><td>0.4</td><td>5.1</td><td>4.1E-07</td><td>6.4E-07</td><td>BT20/MB468</td><td>NUC</td></tr>
<tr><td>>MI0000264|hsa-mir-7-2&WithFlank&15|+|88611819|88611940@38.60.23;MI0000265|hsa-mir-7-3&WithFlank&19|+|4770664|4770785@37.59.23;MI0000263|hsa-mir-7-1&WithFlank&9|-|83969742|83969863@30.52.23&[MIMAT0000252&hsa-miR-7-5p&offsets|0|-1;m-400&15|+|88611856|88611879&offsets|0|-1];[MIMAT0000252_1&hsa-miR-7-5p&offsets|0|-1;m-402&19|+|4770700|4770723&offsets|0|-1];[MIMAT0000252_2&hsa-miR-7-5p&offsets|0|-1;m-401&9|-|83969811|83969834&offsets|0|-1]|TGGAAGACTAGTGATTTTGTTGT</td><td>>MI0000264|hsa-mir-7-2&WithFlank&15|+|88611819|88611940</td><td>38</td><td>60</td><td>23</td><td>TGGAAGACTAGTGATTTTGTTGT</td><td>1673.3</td><td>4.1</td><td>0.8</td><td>5.1</td><td>3.6E-07</td><td>6.4E-07</td><td>MB231/MB468</td><td>CYTO</td></tr>
<tr><td>>hg38_21_+_8433172_8446622_RNA45SN1_45S_50ntFlanks@10846.10868.23|CTTCGGGATAAGGATTGGCTCTA</td><td>>hg38_21_+_8433172_8446622_RNA45SN1_45S_50ntFlanks</td><td>10846</td><td>10868</td><td>23</td><td>CTTCGGGATAAGGATTGGCTCTA</td><td>1015.4</td><td>-5.8</td><td>1.1</td><td>-5.1</td><td>3.6E-07</td><td>6.4E-07</td><td>MB231/MB468</td><td>CYTO</td></tr>
<tr><td>>hg38_21_+_8433172_8446622_RNA45SN1_45S_50ntFlanks@8673.8700.28|CGGTGCGCCGCGACCGGCTCCGGGACGG</td><td>>hg38_21_+_8433172_8446622_RNA45SN1_45S_50ntFlanks</td><td>8673</td><td>8700</td><td>28</td><td>CGGTGCGCCGCGACCGGCTCCGGGACGG</td><td>225.5</td><td>11.1</td><td>2.0</td><td>5.6</td><td>1.7E-08</td><td>6.4E-07</td><td>BT20/MB231</td><td>MP</td></tr>
<tr><td>>hg38_21_+_8433172_8446622_RNA45SN1_45S_50ntFlanks@9947.9968.22|GTGGCCATGGAAGTCGGAATCC</td><td>>hg38_21_+_8433172_8446622_RNA45SN1_45S_50ntFlanks</td><td>9947</td><td>9968</td><td>22</td><td>GTGGCCATGGAAGTCGGAATCC</td><td>442.7</td><td>12.1</td><td>2.2</td><td>5.6</td><td>1.7E-08</td><td>6.4E-07</td><td>BT20/MB231</td><td>MP</td></tr>
<tr><td>>hg38_21_+_8433172_8446622_RNA45SN1_45S_50ntFlanks@10921.10946.26|GCTGGACGAGGCGCCGCCGCCCCCCC</td><td>>hg38_21_+_8433172_8446622_RNA45SN1_45S_50ntFlanks</td><td>10921</td><td>10946</td><td>26</td><td>GCTGGACGAGGCGCCGCCGCCCCCCC</td><td>287.1</td><td>-6.8</td><td>1.3</td><td>-5.1</td><td>3.7E-07</td><td>6.4E-07</td><td>MB231/MB468</td><td>NUC</td></tr>
<tr><td>>MI0000272|hsa-mir-182&WithFlank&7|-|129770377|129770498@29.53.25(+1U)&[MIMAT0000259&hsa-miR-182-5p&offsets|0|+1(+1U);m-22&7|-|129770449|129770470&offsets|0|+3(+1U)]|TTTGGCAATGGTAGAACTCACACTGT</td><td>>MI0000272|hsa-mir-182&WithFlank&7|-|129770377|129770498</td><td>29</td><td>53</td><td>25(+1U)</td><td>TTTGGCAATGGTAGAACTCACACTGT</td><td>697.9</td><td>5.3</td><td>1.0</td><td>5.1</td><td>3.9E-07</td><td>6.4E-07</td><td>MB231/MB468</td><td>MITO</td></tr>
<tr><td>>hg38_21_+_8433172_8446622_RNA45SN1_45S_50ntFlanks@12626.12661.36|AATGGGGCGAAGCTACCATCTGTGGGATTATGACTG</td><td>>hg38_21_+_8433172_8446622_RNA45SN1_45S_50ntFlanks</td><td>12626</td><td>12661</td><td>36</td><td>AATGGGGCGAAGCTACCATCTGTGGGATTATGACTG</td><td>481.2</td><td>-2.1</td><td>0.4</td><td>-5.1</td><td>3.2E-07</td><td>6.4E-07</td><td>BT20/MB468</td><td>CYTO</td></tr>
<tr><td>>hg38_MT_+_1621_3279_RNR2_16S_50ntFlanks@497.526.30|TTTGGACACTAGGAAAAAACCTTGTAGAGA</td><td>>hg38_MT_+_1621_3279_RNR2_16S_50ntFlanks</td><td>497</td><td>526</td><td>30</td><td>TTTGGACACTAGGAAAAAACCTTGTAGAGA</td><td>79.8</td><td>10.1</td><td>1.9</td><td>5.4</td><td>8.5E-08</td><td>6.4E-07</td><td>BT20/MB468</td><td>MP</td></tr>
<tr><td>>hg38_21_+_8433172_8446622_RNA45SN1_45S_50ntFlanks@11228.11254.27|CCGGGCCCTTCCCGTGGATCGCCCCAG</td><td>>hg38_21_+_8433172_8446622_RNA45SN1_45S_50ntFlanks</td><td>11228</td><td>11254</td><td>27</td><td>CCGGGCCCTTCCCGTGGATCGCCCCAG</td><td>103.2</td><td>-9.8</td><td>1.8</td><td>-5.4</td><td>8.5E-08</td><td>6.4E-07</td><td>BT20/MB468</td><td>MP</td></tr>
<tr><td>>hg38_21_+_8433172_8446622_RNA45SN1_45S_50ntFlanks@8196.8214.19|CGTGGACGGTGTGAGGCCG</td><td>>hg38_21_+_8433172_8446622_RNA45SN1_45S_50ntFlanks</td><td>8196</td><td>8214</td><td>19</td><td>CGTGGACGGTGTGAGGCCG</td><td>3058.5</td><td>-4.4</td><td>0.8</td><td>-5.4</td><td>8.6E-08</td><td>6.4E-07</td><td>BT20/MB468</td><td>MP</td></tr>
<tr><td>>hg38_21_+_8433172_8446622_RNA45SN1_45S_50ntFlanks@10807.10825.19|ACAATGTAGGTAAGGGAAG</td><td>>hg38_21_+_8433172_8446622_RNA45SN1_45S_50ntFlanks</td><td>10807</td><td>10825</td><td>19</td><td>ACAATGTAGGTAAGGGAAG</td><td>280.4</td><td>-6.2</td><td>1.2</td><td>-5.1</td><td>2.8E-07</td><td>6.4E-07</td><td>BT20/MB468</td><td>MITO</td></tr>
<tr><td>>hg38_21_+_8433172_8446622_RNA45SN1_45S_50ntFlanks@4734.4764.31|AACGAAAGTCGGAGGTTCGAAGACGATCAGA</td><td>>hg38_21_+_8433172_8446622_RNA45SN1_45S_50ntFlanks</td><td>4734</td><td>4764</td><td>31</td><td>AACGAAAGTCGGAGGTTCGAAGACGATCAGA</td><td>301.9</td><td>7.6</td><td>1.5</td><td>5.1</td><td>3.2E-07</td><td>6.4E-07</td><td>BT20/MB468</td><td>CYTO</td></tr>
<tr><td>>hg38_21_+_8433172_8446622_RNA45SN1_45S_50ntFlanks@3811.3829.19|AGTTATGGTTCCTTTGGTC</td><td>>hg38_21_+_8433172_8446622_RNA45SN1_45S_50ntFlanks</td><td>3811</td><td>3829</td><td>19</td><td>AGTTATGGTTCCTTTGGTC</td><td>225.4</td><td>-5.2</td><td>1.0</td><td>-5.1</td><td>3.8E-07</td><td>6.4E-07</td><td>MB231/MB468</td><td>NUC</td></tr>
<tr><td>>MI0000438|hsa-mir-15b&WithFlank&3|+|160404582|160404691@26.47.22&[MIMAT0000417&hsa-miR-15b-5p&offsets|0|0;m-110&3|+|160404607|160404628&offsets|0|0]|TAGCAGCACATCATGGTTTACA</td><td>>MI0000438|hsa-mir-15b&WithFlank&3|+|160404582|160404691</td><td>26</td><td>47</td><td>22</td><td>TAGCAGCACATCATGGTTTACA</td><td>576.7</td><td>4.8</td><td>0.9</td><td>5.1</td><td>3.6E-07</td><td>6.4E-07</td><td>MB231/MB468</td><td>CYTO</td></tr>
<tr><td>>hg38_21_+_8433172_8446622_RNA45SN1_45S_50ntFlanks@8439.8467.29|GATTCAACCCGGCGGCGGGTCCGGCCGTG</td><td>>hg38_21_+_8433172_8446622_RNA45SN1_45S_50ntFlanks</td><td>8439</td><td>8467</td><td>29</td><td>GATTCAACCCGGCGGCGGGTCCGGCCGTG</td><td>72.4</td><td>-9.3</td><td>1.7</td><td>-5.4</td><td>8.6E-08</td><td>6.4E-07</td><td>BT20/MB468</td><td>MP</td></tr>
<tr><td>>hg38_21_+_8433172_8446622_RNA45SN1_45S_50ntFlanks@476.518.43|TTCGTGATCGATGTGGTGACGTCGTGCTCTCCCGGGCCGGGTC</td><td>>hg38_21_+_8433172_8446622_RNA45SN1_45S_50ntFlanks</td><td>476</td><td>518</td><td>43</td><td>TTCGTGATCGATGTGGTGACGTCGTGCTCTCCCGGGCCGGGTC</td><td>743.7</td><td>7.6</td><td>1.5</td><td>5.1</td><td>2.8E-07</td><td>6.4E-07</td><td>BT20/MB468</td><td>MITO</td></tr>
<tr><td>>MI0000815|hsa-mir-339&WithFlank&7|-|1022927|1023032@21.43.23&[MIMAT0000764&hsa-miR-339-5p&offsets|0|0;m-163&7|-|1022990|1023012&offsets|0|0]|TCCCTGTCCTCCAGGAGCTCACG</td><td>>MI0000815|hsa-mir-339&WithFlank&7|-|1022927|1023032</td><td>21</td><td>43</td><td>23</td><td>TCCCTGTCCTCCAGGAGCTCACG</td><td>479.0</td><td>5.0</td><td>1.0</td><td>5.1</td><td>3.9E-07</td><td>6.4E-07</td><td>MB231/MB468</td><td>MITO</td></tr>
<tr><td>>hg38_21_+_8433172_8446622_RNA45SN1_45S_50ntFlanks@8242.8269.28|GGTCTTCCCGGAGTCGGGTTGCTTGGGA</td><td>>hg38_21_+_8433172_8446622_RNA45SN1_45S_50ntFlanks</td><td>8242</td><td>8269</td><td>28</td><td>GGTCTTCCCGGAGTCGGGTTGCTTGGGA</td><td>251.8</td><td>-7.3</td><td>1.4</td><td>-5.1</td><td>3.3E-07</td><td>6.4E-07</td><td>MB231/MB468</td><td>MP</td></tr>
<tr><td>>hg38_21_+_8433172_8446622_RNA45SN1_45S_50ntFlanks@11323.11346.24|CGTCTCGTCGCGCGCGCGTCCGCT</td><td>>hg38_21_+_8433172_8446622_RNA45SN1_45S_50ntFlanks</td><td>11323</td><td>11346</td><td>24</td><td>CGTCTCGTCGCGCGCGCGTCCGCT</td><td>2908.7</td><td>6.0</td><td>1.2</td><td>5.1</td><td>2.8E-07</td><td>6.4E-07</td><td>BT20/MB468</td><td>MITO</td></tr>
<tr><td>>MI0000760|hsa-mir-361&WithFlank&X|-|85903630|85903713@51.75.25&[MIMAT0004682&hsa-miR-361-3p&offsets|0|+2;m-124&X|-|85903642|85903663&offsets|0|+3]|TCCCCCAGGTGTGATTCTGATTTGC</td><td>>MI0000760|hsa-mir-361&WithFlank&X|-|85903630|85903713</td><td>51</td><td>75</td><td>25</td><td>TCCCCCAGGTGTGATTCTGATTTGC</td><td>205.3</td><td>4.2</td><td>0.8</td><td>5.1</td><td>3.8E-07</td><td>6.4E-07</td><td>MB231/MB468</td><td>NUC</td></tr>
<tr><td>>hg38_21_+_8433172_8446622_RNA45SN1_45S_50ntFlanks@12444.12473.30|AAGCAGAATTCACCAAGCGTTGGATTGTTC</td><td>>hg38_21_+_8433172_8446622_RNA45SN1_45S_50ntFlanks</td><td>12444</td><td>12473</td><td>30</td><td>AAGCAGAATTCACCAAGCGTTGGATTGTTC</td><td>542.5</td><td>-5.9</td><td>1.2</td><td>-5.1</td><td>3.3E-07</td><td>6.4E-07</td><td>MB231/MB468</td><td>MP</td></tr>
<tr><td>>hg38_21_+_8433172_8446622_RNA45SN1_45S_50ntFlanks@475.517.43|CTTCGTGATCGATGTGGTGACGTCGTGCTCTCCCGGGCCGGGT</td><td>>hg38_21_+_8433172_8446622_RNA45SN1_45S_50ntFlanks</td><td>475</td><td>517</td><td>43</td><td>CTTCGTGATCGATGTGGTGACGTCGTGCTCTCCCGGGCCGGGT</td><td>4591.2</td><td>8.6</td><td>1.7</td><td>5.1</td><td>2.8E-07</td><td>6.4E-07</td><td>BT20/MB468</td><td>MITO</td></tr>
<tr><td>>MI0000067|hsa-let-7f-1&WithFlank&9|+|94176341|94176439@13.30.18;MI0000068|hsa-let-7f-2&WithFlank&X|-|53557186|53557280@14.31.18&[MIMAT0000067&hsa-let-7f-5p&offsets|0|-4;m-20&9|+|94176353|94176374&offsets|0|-4];[MIMAT0000067_1&hsa-let-7f-5p&offsets|0|-4;m-19&X|-|53557246|53557267&offsets|0|-4]|TGAGGTAGTAGATTGTAT</td><td>>MI0000067|hsa-let-7f-1&WithFlank&9|+|94176341|94176439</td><td>13</td><td>30</td><td>18</td><td>TGAGGTAGTAGATTGTAT</td><td>1613.5</td><td>4.9</td><td>1.0</td><td>5.1</td><td>3.9E-07</td><td>6.4E-07</td><td>MB231/MB468</td><td>MITO</td></tr>
<tr><td>>hg38_21_+_8433172_8446622_RNA45SN1_45S_50ntFlanks@10781.10822.42|AGGTGAACAGCCTCTGGCATGTTGGAACAATGTAGGTAAGGG</td><td>>hg38_21_+_8433172_8446622_RNA45SN1_45S_50ntFlanks</td><td>10781</td><td>10822</td><td>42</td><td>AGGTGAACAGCCTCTGGCATGTTGGAACAATGTAGGTAAGGG</td><td>152.7</td><td>-5.6</td><td>1.1</td><td>-5.1</td><td>3.7E-07</td><td>6.4E-07</td><td>MB231/MB468</td><td>CYTO</td></tr>
<tr><td>>hg38_21_+_8433172_8446622_RNA45SN1_45S_50ntFlanks@7992.8014.23|CGTGGCGACCCGCTGAATTTAAG</td><td>>hg38_21_+_8433172_8446622_RNA45SN1_45S_50ntFlanks</td><td>7992</td><td>8014</td><td>23</td><td>CGTGGCGACCCGCTGAATTTAAG</td><td>987.8</td><td>-3.9</td><td>0.8</td><td>-5.1</td><td>4.0E-07</td><td>6.4E-07</td><td>MB231/MB468</td><td>MITO</td></tr>
<tr><td>>am_trna128_GlyGCC_6_-_27870686_27870756@7.35.29,trna18_GlyGCC_16_+_70822597_70822667@7.35.29,trna19_GlyGCC_16_+_70823410_70823480@7.35.29,trna19_GlyGCC_2_-_157257659_157257729@7.35.29,trna24_GlyGCC_16_-_70812942_70813012@7.35.29,trna25_GlyGCC_16_-_70812114_70812184@7.35.29,trna2_GlyGCC_21_-_18827107_18827177@7.35.29,trna35_GlyGCC_1_+_161413094_161413164@7.35.29,trna37_GlyGCC_1_+_161420467_161420537@7.35.29,trna39_GlyGCC_1_+_161427898_161427968@7.35.29,trna41_GlyGCC_1_+_161435258_161435328@7.35.29,trna5_GlyGCC_17_+_8029064_8029134@7.35.29,trna68_GlyGCC_1_-_161493637_161493707@7.35.29|GTGGTTCAGTGGTAGAATTCTCGCCTGCC</td><td>>am_trna128_GlyGCC_6_-_27870686_27870756</td><td>7</td><td>35</td><td>29</td><td>GTGGTTCAGTGGTAGAATTCTCGCCTGCC</td><td>419.8</td><td>4.1</td><td>0.8</td><td>5.1</td><td>4.0E-07</td><td>6.5E-07</td><td>MB231/MB468</td><td>MITO</td></tr>
<tr><td>>trnaMT_PheGAA_MT_+_577_647@20.53.34|TCAAAGCAATACACTGAAAATGTTTAGACGGGCT</td><td>>trnaMT_PheGAA_MT_+_577_647</td><td>20</td><td>53</td><td>34</td><td>TCAAAGCAATACACTGAAAATGTTTAGACGGGCT</td><td>107.1</td><td>10.5</td><td>2.0</td><td>5.4</td><td>8.6E-08</td><td>6.5E-07</td><td>BT20/MB468</td><td>MP</td></tr>
<tr><td>>hg38_21_+_8433172_8446622_RNA45SN1_45S_50ntFlanks@5156.5185.30|ACAAGTGGCGTTCAGCCACCCGAGATTGAG</td><td>>hg38_21_+_8433172_8446622_RNA45SN1_45S_50ntFlanks</td><td>5156</td><td>5185</td><td>30</td><td>ACAAGTGGCGTTCAGCCACCCGAGATTGAG</td><td>234.8</td><td>-10.0</td><td>1.9</td><td>-5.1</td><td>2.8E-07</td><td>6.5E-07</td><td>BT20/MB468</td><td>MITO</td></tr>
<tr><td>>hg38_21_+_8433172_8446622_RNA45SN1_45S_50ntFlanks@11316.11342.27|CACCCCACGTCTCGTCGCGCGCGCGTC</td><td>>hg38_21_+_8433172_8446622_RNA45SN1_45S_50ntFlanks</td><td>11316</td><td>11342</td><td>27</td><td>CACCCCACGTCTCGTCGCGCGCGCGTC</td><td>3587.2</td><td>-6.2</td><td>1.2</td><td>-5.1</td><td>3.2E-07</td><td>6.5E-07</td><td>BT20/MB468</td><td>CYTO</td></tr>
<tr><td>>hg38_21_+_8433172_8446622_RNA45SN1_45S_50ntFlanks@6662.6688.27|GGTGGATCACTCGGCTCGTGCGTCGAT</td><td>>hg38_21_+_8433172_8446622_RNA45SN1_45S_50ntFlanks</td><td>6662</td><td>6688</td><td>27</td><td>GGTGGATCACTCGGCTCGTGCGTCGAT</td><td>371.4</td><td>11.9</td><td>2.1</td><td>5.6</td><td>1.8E-08</td><td>6.5E-07</td><td>BT20/MB231</td><td>MP</td></tr>
<tr><td>>hg38_21_+_8433172_8446622_RNA45SN1_45S_50ntFlanks@10316.10341.26|GGTGCAGATCTTGGTGGTAGTAGCAA</td><td>>hg38_21_+_8433172_8446622_RNA45SN1_45S_50ntFlanks</td><td>10316</td><td>10341</td><td>26</td><td>GGTGCAGATCTTGGTGGTAGTAGCAA</td><td>519.2</td><td>12.4</td><td>2.2</td><td>5.6</td><td>1.8E-08</td><td>6.5E-07</td><td>BT20/MB231</td><td>MP</td></tr>
</table>

<table>
<tr><td>>hg38_21_+_8433172_8446622_RNA45SN1_45S_50ntFlanks@8117.8138.22|CGGGACATGTGGCGTACGGAAG</td><td>>hg38_21_+_8433172_8446622_RNA45SN1_45S_50ntFlanks</td><td>8117</td><td>8138</td><td>22</td><td>CGGGACATGTGGCGTACGGAAG</td><td>236.1</td><td>11.2</td><td>2.0</td><td>5.6</td><td>1.8E-08</td><td>6.5E-07</td><td>BT20/MB231</td><td>MP</td></tr>
<tr><td>>MI0003513|hsa-mir-455&WithFlank&9|+|114209428|114209535@24.45.22&[MIMAT0003150&hsa-miR-455-5p&offsets|+2|+2;m-232&9|+|114209449|114209470&offsets|+2|+2]|TGTGCCTTTGGACTACATCGTG</td><td>>MI0003513|hsa-mir-455&WithFlank&9|+|114209428|114209535</td><td>24</td><td>45</td><td>22</td><td>TGTGCCTTTGGACTACATCGTG</td><td>129.6</td><td>-10.4</td><td>1.9</td><td>-5.6</td><td>1.8E-08</td><td>6.5E-07</td><td>BT20/MB231</td><td>MP</td></tr>
<tr><td>>trnaMT_SerGCT_MT_+_12207_12265@1.18.18|GAGAAAGCTCACAAGAAC</td><td>>trnaMT_SerGCT_MT_+_12207_12265</td><td>1</td><td>18</td><td>18</td><td>GAGAAAGCTCACAAGAAC</td><td>80162.3</td><td>5.8</td><td>1.1</td><td>5.1</td><td>4.0E-07</td><td>6.5E-07</td><td>MB231/MB468</td><td>MITO</td></tr>
<tr><td>>MI0001641|hsa-mir-429&WithFlank&1|+|1168999|1169093@57.77.21&[MIMAT0001536&hsa-miR-429&offsets|0|-1;m-207&1|+|1169055|1169076&offsets|0|-1]|TAATACTGTCTGGTAAAACCG</td><td>>MI0001641|hsa-mir-429&WithFlank&1|+|1168999|1169093</td><td>57</td><td>77</td><td>21</td><td>TAATACTGTCTGGTAAAACCG</td><td>64.0</td><td>-9.1</td><td>1.7</td><td>-5.4</td><td>8.7E-08</td><td>6.5E-07</td><td>BT20/MB468</td><td>MP</td></tr>
<tr><td>>trna111_HisGTG_1_-_147774845_147774916@-1C.31.32,trna118_HisGTG_1_-_145396881_145396952@-1C.31.32,trna16_HisGTG_1_+_146544773_146544844@-1C.31.32,trna1_HisGTG_15_+_45493349_45493420@-1C.31.32,trna21_HisGTG_1_+_147753471_147753542@-1C.31.32,trna33_HisGTG_6_+_27125906_27125977@-1C.31.32,trna7_HisGTG_9_-_14433938_14434009@-1C.31.32,trna8_HisGTG_15_-_45492611_45492682@-1C.31.32,trna9_HisGTG_15_-_45490804_45490875@-1C.31.32|CGCCGTGATCGTATAGTGGTTAGTACTCTGCG</td><td>>trna111_HisGTG_1_-_147774845_147774916</td><td>-1C</td><td>31</td><td>32</td><td>CGCCGTGATCGTATAGTGGTTAGTACTCTGCG</td><td>61.4</td><td>-9.0</td><td>1.7</td><td>-5.4</td><td>8.7E-08</td><td>6.5E-07</td><td>BT20/MB468</td><td>MP</td></tr>
<tr><td>>hg38_21_+_8433172_8446622_RNA45SN1_45S_50ntFlanks@11552.11574.23|CCTCGGCCGGCGCCTAGCAGCCG</td><td>>hg38_21_+_8433172_8446622_RNA45SN1_45S_50ntFlanks</td><td>11552</td><td>11574</td><td>23</td><td>CCTCGGCCGGCGCCTAGCAGCCG</td><td>204.2</td><td>-6.1</td><td>1.2</td><td>-5.1</td><td>3.2E-07</td><td>6.5E-07</td><td>BT20/MB468</td><td>CYTO</td></tr>
<tr><td>>hg38_21_+_8433172_8446622_RNA45SN1_45S_50ntFlanks@12054.12072.19|TGAGGCGGGGGGGCGAGCC</td><td>>hg38_21_+_8433172_8446622_RNA45SN1_45S_50ntFlanks</td><td>12054</td><td>12072</td><td>19</td><td>TGAGGCGGGGGGGCGAGCC</td><td>287.0</td><td>-5.2</td><td>1.0</td><td>-5.1</td><td>3.7E-07</td><td>6.5E-07</td><td>MB231/MB468</td><td>CYTO</td></tr>
<tr><td>>hg38_21_+_8433172_8446622_RNA45SN1_45S_50ntFlanks@12073.12094.22|CCGAGGGGCTCTCGCTTCTGGC</td><td>>hg38_21_+_8433172_8446622_RNA45SN1_45S_50ntFlanks</td><td>12073</td><td>12094</td><td>22</td><td>CCGAGGGGCTCTCGCTTCTGGC</td><td>497.2</td><td>-7.4</td><td>1.5</td><td>-5.1</td><td>3.3E-07</td><td>6.5E-07</td><td>MB231/MB468</td><td>MP</td></tr>
<tr><td>>hg38_21_+_8433172_8446622_RNA45SN1_45S_50ntFlanks@10868.10906.39|AAGGGCTGGGTCGGTCGGGCTGGGGCGCGAAGCGGGGCT</td><td>>hg38_21_+_8433172_8446622_RNA45SN1_45S_50ntFlanks</td><td>10868</td><td>10906</td><td>39</td><td>AAGGGCTGGGTCGGTCGGGCTGGGGCGCGAAGCGGGGCT</td><td>69.1</td><td>9.9</td><td>1.8</td><td>5.4</td><td>8.7E-08</td><td>6.5E-07</td><td>BT20/MB468</td><td>MP</td></tr>
<tr><td>>trnaMT_HisGTG_MT_+_12138_12206@25.55.31|CAGATTGTGAATCTGACAACAGAGGCTTACG</td><td>>trnaMT_HisGTG_MT_+_12138_12206</td><td>25</td><td>55</td><td>31</td><td>CAGATTGTGAATCTGACAACAGAGGCTTACG</td><td>96.0</td><td>10.3</td><td>1.9</td><td>5.4</td><td>8.7E-08</td><td>6.5E-07</td><td>BT20/MB468</td><td>MP</td></tr>
<tr><td>>hg38_21_+_8433172_8446622_RNA45SN1_45S_50ntFlanks@7973.8011.39|GACGCGACCTCAGATCAGACGTGGCGACCCGCTGAATTT</td><td>>hg38_21_+_8433172_8446622_RNA45SN1_45S_50ntFlanks</td><td>7973</td><td>8011</td><td>39</td><td>GACGCGACCTCAGATCAGACGTGGCGACCCGCTGAATTT</td><td>2556.2</td><td>3.2</td><td>0.6</td><td>5.1</td><td>4.2E-07</td><td>6.5E-07</td><td>BT20/MB468</td><td>NUC</td></tr>
<tr><td>>hg38_1_-_228634819_228635039_RNA5S12_5S_50ntFlanks@141.169.29|CCGCCTGGGAATACCGGGTGCTGTAGGCT</td><td>>hg38_1_-_228634819_228635039_RNA5S12_5S_50ntFlanks</td><td>141</td><td>169</td><td>29</td><td>CCGCCTGGGAATACCGGGTGCTGTAGGCT</td><td>1196.7</td><td>-2.8</td><td>0.6</td><td>-5.1</td><td>3.3E-07</td><td>6.5E-07</td><td>MB231/MB468</td><td>MP</td></tr>
<tr><td>>hg38_21_+_8433172_8446622_RNA45SN1_45S_50ntFlanks@12393.12414.22|ATAGCGACGTCGCTTTTTGATC</td><td>>hg38_21_+_8433172_8446622_RNA45SN1_45S_50ntFlanks</td><td>12393</td><td>12414</td><td>22</td><td>ATAGCGACGTCGCTTTTTGATC</td><td>309.9</td><td>-4.1</td><td>0.8</td><td>-5.1</td><td>3.3E-07</td><td>6.5E-07</td><td>MB231/MB468</td><td>MP</td></tr>
<tr><td>>hg38_21_+_8433172_8446622_RNA45SN1_45S_50ntFlanks@11209.11253.45|CGGCGACTCTGGACGCGAGCCGGGCCCTTCCCGTGGATCGCCCCA</td><td>>hg38_21_+_8433172_8446622_RNA45SN1_45S_50ntFlanks</td><td>11209</td><td>11253</td><td>45</td><td>CGGCGACTCTGGACGCGAGCCGGGCCCTTCCCGTGGATCGCCCCA</td><td>1214.1</td><td>-9.2</td><td>1.8</td><td>-5.1</td><td>4.2E-07</td><td>6.5E-07</td><td>BT20/MB468</td><td>NUC</td></tr>
<tr><td>>MI0000462|hsa-mir-152&WithFlank&17|-|48037155|48037253@60.80.21&[MIMAT0000438&hsa-miR-152-3p&offsets|0|0;m-97&17|-|48037174|48037194&offsets|0|0]|TCAGTGCATGACAGAACTTGG</td><td>>MI0000462|hsa-mir-152&WithFlank&17|-|48037155|48037253</td><td>60</td><td>80</td><td>21</td><td>TCAGTGCATGACAGAACTTGG</td><td>1020.3</td><td>7.3</td><td>1.4</td><td>5.1</td><td>3.4E-07</td><td>6.5E-07</td><td>MB231/MB468</td><td>MP</td></tr>
<tr><td>>hg38_21_+_8433172_8446622_RNA45SN1_45S_50ntFlanks@3899.3938.40|CCTTCGCGGGGGGGATGCGTGCATTTATCAGATCAAAACC</td><td>>hg38_21_+_8433172_8446622_RNA45SN1_45S_50ntFlanks</td><td>3899</td><td>3938</td><td>40</td><td>CCTTCGCGGGGGGGATGCGTGCATTTATCAGATCAAAACC</td><td>481.8</td><td>-7.5</td><td>1.5</td><td>-5.1</td><td>3.3E-07</td><td>6.5E-07</td><td>BT20/MB468</td><td>CYTO</td></tr>
<tr><td>>MI0000746|hsa-mir-99b&WithFlank&19|+|51692606|51692687@51.69.19(+1U)&[MIMAT0004678&hsa-miR-99b-3p&offsets|0|-3(+1U);m-151&19|+|51692656|51692678&offsets|0|-4(+1U)]|CAAGCTCGTGTCTGTGGGTT</td><td>>MI0000746|hsa-mir-99b&WithFlank&19|+|51692606|51692687</td><td>51</td><td>69</td><td>19(+1U)</td><td>CAAGCTCGTGTCTGTGGGTT</td><td>94.9</td><td>10.3</td><td>1.9</td><td>5.4</td><td>8.8E-08</td><td>6.5E-07</td><td>BT20/MB468</td><td>MP</td></tr>
<tr><td>>hg38_21_+_8433172_8446622_RNA45SN1_45S_50ntFlanks@10829.10858.30|GCAAGCCGGATCCGTAACTTCGGGATAAGG</td><td>>hg38_21_+_8433172_8446622_RNA45SN1_45S_50ntFlanks</td><td>10829</td><td>10858</td><td>30</td><td>GCAAGCCGGATCCGTAACTTCGGGATAAGG</td><td>520.0</td><td>-7.8</td><td>1.5</td><td>-5.1</td><td>3.4E-07</td><td>6.5E-07</td><td>MB231/MB468</td><td>MP</td></tr>
<tr><td>>hg38_1_-_228634819_228635039_RNA5S12_5S_50ntFlanks@153.172.20|ACCGGGTGCTGTAGGCTTTT</td><td>>hg38_1_-_228634819_228635039_RNA5S12_5S_50ntFlanks</td><td>153</td><td>172</td><td>20</td><td>ACCGGGTGCTGTAGGCTTTT</td><td>380.2</td><td>8.6</td><td>1.7</td><td>5.1</td><td>3.3E-07</td><td>6.5E-07</td><td>BT20/MB468</td><td>CYTO</td></tr>
<tr><td>>hg38_21_+_8433172_8446622_RNA45SN1_45S_50ntFlanks@10236.10261.26|GACGCTACGCCGCGACGAGTAGGAGG</td><td>>hg38_21_+_8433172_8446622_RNA45SN1_45S_50ntFlanks</td><td>10236</td><td>10261</td><td>26</td><td>GACGCTACGCCGCGACGAGTAGGAGG</td><td>272.4</td><td>11.4</td><td>2.0</td><td>5.6</td><td>1.8E-08</td><td>6.6E-07</td><td>BT20/MB231</td><td>MP</td></tr>
<tr><td>>hg38_21_+_8433172_8446622_RNA45SN1_45S_50ntFlanks@10349.10374.26|AACGAGAACTTTGAAGGCCGAAGTGG</td><td>>hg38_21_+_8433172_8446622_RNA45SN1_45S_50ntFlanks</td><td>10349</td><td>10374</td><td>26</td><td>AACGAGAACTTTGAAGGCCGAAGTGG</td><td>3049.6</td><td>-6.9</td><td>1.3</td><td>-5.1</td><td>3.4E-07</td><td>6.6E-07</td><td>MB231/MB468</td><td>MP</td></tr>
<tr><td>>hg38_21_+_8433172_8446622_RNA45SN1_45S_50ntFlanks@4008.4025.18|CTCGGGCCGATCGCACGC</td><td>>hg38_21_+_8433172_8446622_RNA45SN1_45S_50ntFlanks</td><td>4008</td><td>4025</td><td>18</td><td>CTCGGGCCGATCGCACGC</td><td>307.5</td><td>-2.3</td><td>0.5</td><td>-5.1</td><td>3.7E-07</td><td>6.6E-07</td><td>MB231/MB468</td><td>CYTO</td></tr>
<tr><td>>hg38_MT_+_1621_3279_RNR2_16S_50ntFlanks@714.745.32|CATAAGCCTGCGTCAGATTAAAACACTGAACT</td><td>>hg38_MT_+_1621_3279_RNR2_16S_50ntFlanks</td><td>714</td><td>745</td><td>32</td><td>CATAAGCCTGCGTCAGATTAAAACACTGAACT</td><td>54.3</td><td>9.0</td><td>1.8</td><td>5.1</td><td>3.4E-07</td><td>6.6E-07</td><td>MB231/MB468</td><td>MP</td></tr>
<tr><td>>trnaMT_ValTAC_MT_+_1602_1670@43.71.29|GATTTCAACTTAACTTGACCGCTCTGACC</td><td>>trnaMT_ValTAC_MT_+_1602_1670</td><td>43</td><td>71</td><td>29</td><td>GATTTCAACTTAACTTGACCGCTCTGACC</td><td>1562.0</td><td>5.4</td><td>1.0</td><td>5.1</td><td>2.9E-07</td><td>6.6E-07</td><td>BT20/MB468</td><td>MITO</td></tr>
<tr><td>>hg38_21_+_8433172_8446622_RNA45SN1_45S_50ntFlanks@5489.5518.30|CTGAGAAGACGGTCGAACTTGACTATCTAG</td><td>>hg38_21_+_8433172_8446622_RNA45SN1_45S_50ntFlanks</td><td>5489</td><td>5518</td><td>30</td><td>CTGAGAAGACGGTCGAACTTGACTATCTAG</td><td>567.4</td><td>-7.9</td><td>1.5</td><td>-5.3</td><td>8.9E-08</td><td>6.6E-07</td><td>BT20/MB468</td><td>MP</td></tr>
<tr><td>>MI0000076|hsa-mir-20a&WithFlank&13|+|91351059|91351141@14.35.22&[MIMAT0000075&hsa-miR-20a-5p&offsets|0|-1;m-96&13|+|91351072|91351094&offsets|0|-1]|TAAAGTGCTTATAGTGCAGGTA</td><td>>MI0000076|hsa-mir-20a&WithFlank&13|+|91351059|91351141</td><td>14</td><td>35</td><td>22</td><td>TAAAGTGCTTATAGTGCAGGTA</td><td>570.4</td><td>5.8</td><td>1.1</td><td>5.1</td><td>4.1E-07</td><td>6.6E-07</td><td>MB231/MB468</td><td>MITO</td></tr>
<tr><td>>trnaMT_GluTTC_MT_-_14674_14742@35.72.38,trnalookalike8_GluTTC_5_-_93905172_93905240@35.72.38|TATCATTGGTCGTGGTTGTAGTCCGTGCGAGAATACCA</td><td>>trnaMT_GluTTC_MT_-_14674_14742</td><td>35</td><td>72</td><td>38</td><td>TATCATTGGTCGTGGTTGTAGTCCGTGCGAGAATACCA</td><td>2694.6</td><td>3.7</td><td>0.7</td><td>5.1</td><td>4.1E-07</td><td>6.6E-07</td><td>MB231/MB468</td><td>MITO</td></tr>
<tr><td>>trnaMT_LeuTAG_MT_+_12266_12336@54.74.21|CAACTCCAAATAAAAGTACCA</td><td>>trnaMT_LeuTAG_MT_+_12266_12336</td><td>54</td><td>74</td><td>21</td><td>CAACTCCAAATAAAAGTACCA</td><td>311.1</td><td>-7.3</td><td>1.4</td><td>-5.1</td><td>4.1E-07</td><td>6.6E-07</td><td>MB231/MB468</td><td>MITO</td></tr>
<tr><td>>hg38_21_+_8433172_8446622_RNA45SN1_45S_50ntFlanks@10679.10709.31|GGTTCCGGCGGCGTCCGGTGAGCTCTCGCTG</td><td>>hg38_21_+_8433172_8446622_RNA45SN1_45S_50ntFlanks</td><td>10679</td><td>10709</td><td>31</td><td>GGTTCCGGCGGCGTCCGGTGAGCTCTCGCTG</td><td>11352.0</td><td>-4.9</td><td>1.0</td><td>-5.1</td><td>2.9E-07</td><td>6.6E-07</td><td>BT20/MB468</td><td>MITO</td></tr>
<tr><td>>hg38_21_+_8433172_8446622_RNA45SN1_45S_50ntFlanks@7973.8009.37|GACGCGACCTCAGATCAGACGTGGCGACCCGCTGAAT</td><td>>hg38_21_+_8433172_8446622_RNA45SN1_45S_50ntFlanks</td><td>7973</td><td>8009</td><td>37</td><td>GACGCGACCTCAGATCAGACGTGGCGACCCGCTGAAT</td><td>2301.6</td><td>3.0</td><td>0.6</td><td>5.1</td><td>4.3E-07</td><td>6.6E-07</td><td>BT20/MB468</td><td>NUC</td></tr>
<tr><td>>MI0000746|hsa-mir-99b&WithFlank&19|+|51692606|51692687@51.72.22(+1A)&[MIMAT0004678&hsa-miR-99b-3p&offsets|0|0(+1A);m-151&19|+|51692656|51692678&offsets|0|-1(+1A)]|CAAGCTCGTGTCTGTGGGTCCGA</td><td>>MI0000746|hsa-mir-99b&WithFlank&19|+|51692606|51692687</td><td>51</td><td>72</td><td>22(+1A)</td><td>CAAGCTCGTGTCTGTGGGTCCGA</td><td>618.6</td><td>4.8</td><td>0.9</td><td>5.3</td><td>8.9E-08</td><td>6.7E-07</td><td>BT20/MB468</td><td>MP</td></tr>
<tr><td>>hg38_21_+_8433172_8446622_RNA45SN1_45S_50ntFlanks@4557.4575.19|CAGCTAGGAATAATGGAAT</td><td>>hg38_21_+_8433172_8446622_RNA45SN1_45S_50ntFlanks</td><td>4557</td><td>4575</td><td>19</td><td>CAGCTAGGAATAATGGAAT</td><td>68.8</td><td>-9.2</td><td>1.7</td><td>-5.3</td><td>8.9E-08</td><td>6.7E-07</td><td>BT20/MB468</td><td>MP</td></tr>
<tr><td>>hg38_21_+_8433172_8446622_RNA45SN1_45S_50ntFlanks@10392.10433.42|ACAGCAGTTGAACATGGGTCAGTCGGTCCTGAGAGATGGGCG</td><td>>hg38_21_+_8433172_8446622_RNA45SN1_45S_50ntFlanks</td><td>10392</td><td>10433</td><td>42</td><td>ACAGCAGTTGAACATGGGTCAGTCGGTCCTGAGAGATGGGCG</td><td>676.2</td><td>-4.1</td><td>0.8</td><td>-5.1</td><td>3.8E-07</td><td>6.7E-07</td><td>MB231/MB468</td><td>CYTO</td></tr>
<tr><td>>hg38_21_+_8433172_8446622_RNA45SN1_45S_50ntFlanks@9341.9367.27|CACGAAAGCCGCCGTGGCGCAATGAAG</td><td>>hg38_21_+_8433172_8446622_RNA45SN1_45S_50ntFlanks</td><td>9341</td><td>9367</td><td>27</td><td>CACGAAAGCCGCCGTGGCGCAATGAAG</td><td>73.5</td><td>-9.3</td><td>1.7</td><td>-5.3</td><td>9.0E-08</td><td>6.7E-07</td><td>BT20/MB468</td><td>MP</td></tr>
<tr><td>>MI0000736|hsa-mir-30c-1&WithFlank&1|+|40757278|40757378@23.46.24(+1A);MI0000254|hsa-mir-30c-2&WithFlank&6|-|71376954|71377037@13.36.24(+1A)&[MIMAT0000244&hsa-miR-30c-5p&offsets|0|+1(+1A);m-57&1|+|40757300|40757323&offsets|0|0(+1A)];[MIMAT0000244_1&hsa-miR-30c-5p&offsets|0|+1(+1A);m-56&6|-|71377002|71377025&offsets|0|0(+1A)]|TGTAAACATCCTACACTCTCAGCTA</td><td>>MI0000736|hsa-mir-30c-1&WithFlank&1|+|40757278|40757378</td><td>23</td><td>46</td><td>24(+1A)</td><td>TGTAAACATCCTACACTCTCAGCTA</td><td>440.9</td><td>5.2</td><td>1.0</td><td>5.1</td><td>4.1E-07</td><td>6.7E-07</td><td>MB231/MB468</td><td>MITO</td></tr>
<tr><td>>MI0000299|hsa-mir-222&WithFlank&X|-|45747009|45747130@75.95.21&[MIMAT0000279&hsa-miR-222-3p&offsets|0|0;m-140&X|-|45747033|45747056&offsets|0|-3]|AGCTACATCTGGCTACTGGGT</td><td>>MI0000299|hsa-mir-222&WithFlank&X|-|45747009|45747130</td><td>75</td><td>95</td><td>21</td><td>AGCTACATCTGGCTACTGGGT</td><td>6599.8</td><td>4.0</td><td>0.8</td><td>5.1</td><td>3.4E-07</td><td>6.7E-07</td><td>MB231/MB468</td><td>MP</td></tr>
<tr><td>>hg38_MT_+_1621_3279_RNR2_16S_50ntFlanks@730.759.30|ATTAAAACACTGAACTGACAATTAACAGCC</td><td>>hg38_MT_+_1621_3279_RNR2_16S_50ntFlanks</td><td>730</td><td>759</td><td>30</td><td>ATTAAAACACTGAACTGACAATTAACAGCC</td><td>61.1</td><td>9.2</td><td>1.8</td><td>5.1</td><td>3.4E-07</td><td>6.7E-07</td><td>MB231/MB468</td><td>MP</td></tr>
<tr><td>>hg38_21_+_8433172_8446622_RNA45SN1_45S_50ntFlanks@3848.3866.19|TGGATAACTGTGGTAATTC</td><td>>hg38_21_+_8433172_8446622_RNA45SN1_45S_50ntFlanks</td><td>3848</td><td>3866</td><td>19</td><td>TGGATAACTGTGGTAATTC</td><td>136.0</td><td>5.5</td><td>1.1</td><td>5.1</td><td>3.9E-07</td><td>6.7E-07</td><td>MB231/MB468</td><td>NUC</td></tr>
<tr><td>>hg38_21_+_8433172_8446622_RNA45SN1_45S_50ntFlanks@8472.8489.18|CGGCCCGGCGGATCTTTC</td><td>>hg38_21_+_8433172_8446622_RNA45SN1_45S_50ntFlanks</td><td>8472</td><td>8489</td><td>18</td><td>CGGCCCGGCGGATCTTTC</td><td>68.1</td><td>-9.2</td><td>1.7</td><td>-5.3</td><td>9.0E-08</td><td>6.7E-07</td><td>BT20/MB468</td><td>MP</td></tr>
<tr><td>>hg38_21_+_8433172_8446622_RNA45SN1_45S_50ntFlanks@12714.12743.30|GAGCCTCGGTTGGCCTCGGATAGCCGGTCC</td><td>>hg38_21_+_8433172_8446622_RNA45SN1_45S_50ntFlanks</td><td>12714</td><td>12743</td><td>30</td><td>GAGCCTCGGTTGGCCTCGGATAGCCGGTCC</td><td>63.0</td><td>-9.1</td><td>1.7</td><td>-5.3</td><td>9.0E-08</td><td>6.7E-07</td><td>BT20/MB468</td><td>MP</td></tr>
<tr><td>>hg38_21_+_8433172_8446622_RNA45SN1_45S_50ntFlanks@10678.10709.32|CGGTTCCGGCGGCGTCCGGTGAGCTCTCGCTG</td><td>>hg38_21_+_8433172_8446622_RNA45SN1_45S_50ntFlanks</td><td>10678</td><td>10709</td><td>32</td><td>CGGTTCCGGCGGCGTCCGGTGAGCTCTCGCTG</td><td>133880.8</td><td>-9.5</td><td>1.9</td><td>-5.1</td><td>4.3E-07</td><td>6.7E-07</td><td>BT20/MB468</td><td>NUC</td></tr>
<tr><td>>hg38_21_+_8433172_8446622_RNA45SN1_45S_50ntFlanks@4367.4394.28|CAGTTAAAAAGCTCGTAGTTGGATCTTG</td><td>>hg38_21_+_8433172_8446622_RNA45SN1_45S_50ntFlanks</td><td>4367</td><td>4394</td><td>28</td><td>CAGTTAAAAAGCTCGTAGTTGGATCTTG</td><td>237.6</td><td>-6.7</td><td>1.3</td><td>-5.1</td><td>4.0E-07</td><td>6.7E-07</td><td>MB231/MB468</td><td>NUC</td></tr>
<tr><td>>trnaMT_PheGAA_MT_+_577_647@38.62.25|AATGTTTAGACGGGCTCACATCACC</td><td>>trnaMT_PheGAA_MT_+_577_647</td><td>38</td><td>62</td><td>25</td><td>AATGTTTAGACGGGCTCACATCACC</td><td>394.0</td><td>6.0</td><td>1.2</td><td>5.1</td><td>4.1E-07</td><td>6.7E-07</td><td>MB231/MB468</td><td>MITO</td></tr>
<tr><td>>hg38_21_+_8433172_8446622_RNA45SN1_45S_50ntFlanks@11202.11225.24|GCGGCGGCGGCGACTCTGGACGCG</td><td>>hg38_21_+_8433172_8446622_RNA45SN1_45S_50ntFlanks</td><td>11202</td><td>11225</td><td>24</td><td>GCGGCGGCGGCGACTCTGGACGCG</td><td>146.4</td><td>-6.9</td><td>1.4</td><td>-5.1</td><td>3.5E-07</td><td>6.7E-07</td><td>MB231/MB468</td><td>MP</td></tr>
<tr><td>>hg38_21_+_8433172_8446622_RNA45SN1_45S_50ntFlanks@9602.9621.20|CGAAAGACTAATCGAACCAT</td><td>>hg38_21_+_8433172_8446622_RNA45SN1_45S_50ntFlanks</td><td>9602</td><td>9621</td><td>20</td><td>CGAAAGACTAATCGAACCAT</td><td>73.0</td><td>-9.3</td><td>1.7</td><td>-5.3</td><td>9.1E-08</td><td>6.7E-07</td><td>BT20/MB468</td><td>MP</td></tr>
<tr><td>>trna10_AspGTC_12_-_125424193_125424264@37.64.28,trna12_AspGTC_12_-_125411891_125411962@37.64.28,trna144_AspGTC_6_-_27551236_27551307@37.64.28,trna38_AspGTC_17_-_8125556_8125627@37.64.28,trna45_AspGTC_6_+_27447453_27447524@37.64.28,trna48_AspGTC_6_+_27471523_27471594@37.64.28,trna4_AspGTC_12_+_96429799_96429870@37.64.28,trna69_AspGTC_1_-_161440205_161440276@37.64.28,trna72_AspGTC_1_-_161432824_161432895@37.64.28,trna75_AspGTC_1_-_161425414_161425485@37.64.28,trna78_AspGTC_1_-_161418033_161418104@37.64.28,trna81_AspGTC_1_-_161410615_161410686@37.64.28|ACGCGGGAGACCGGGGTTCGATTCCCCG</td><td>>trna10_AspGTC_12_-_125424193_125424264</td><td>37</td><td>64</td><td>28</td><td>ACGCGGGAGACCGGGGTTCGATTCCCCG</td><td>114.0</td><td>-9.9</td><td>1.9</td><td>-5.3</td><td>9.1E-08</td><td>6.7E-07</td><td>BT20/MB468</td><td>MP</td></tr>
<tr><td>>hg38_21_+_8433172_8446622_RNA45SN1_45S_50ntFlanks@3838.3871.34|CTCTCCTACTTGGATAACTGTGGTAATTCTAGAG</td><td>>hg38_21_+_8433172_8446622_RNA45SN1_45S_50ntFlanks</td><td>3838</td><td>3871</td><td>34</td><td>CTCTCCTACTTGGATAACTGTGGTAATTCTAGAG</td><td>409.6</td><td>-5.5</td><td>1.1</td><td>-5.1</td><td>4.1E-07</td><td>6.7E-07</td><td>MB231/MB468</td><td>MITO</td></tr>
</table>

<table>
<tr><td>>hg38_21_+_8433172_8446622_RNA45SN1_45S_50ntFlanks@6661.6697.37|CGGTGGATCACTCGGCTCGTGCGTCGATGAAGAACGC</td><td>>hg38_21_+_8433172_8446622_RNA45SN1_45S_50ntFlanks</td><td>6661</td><td>6697</td><td>37</td><td>CGGTGGATCACTCGGCTCGTGCGTCGATGAAGAACGC</td><td>960.1</td><td>-1.5</td><td>0.3</td><td>-5.1</td><td>4.3E-07</td><td>6.7E-07</td><td>BT20/MB468</td><td>NUC</td></tr>
<tr><td>>hg38_21_+_8433172_8446622_RNA45SN1_45S_50ntFlanks@11884.11904.21|GTTGAGCTTGACTCTAGTCTG</td><td>>hg38_21_+_8433172_8446622_RNA45SN1_45S_50ntFlanks</td><td>11884</td><td>11904</td><td>21</td><td>GTTGAGCTTGACTCTAGTCTG</td><td>188.1</td><td>-6.7</td><td>1.3</td><td>-5.1</td><td>3.4E-07</td><td>6.7E-07</td><td>BT20/MB468</td><td>CYTO</td></tr>
<tr><td>>hg38_21_+_8433172_8446622_RNA45SN1_45S_50ntFlanks@12973.12997.25|TCGTACGTAGCAGAGCAGCTCCCTC</td><td>>hg38_21_+_8433172_8446622_RNA45SN1_45S_50ntFlanks</td><td>12973</td><td>12997</td><td>25</td><td>TCGTACGTAGCAGAGCAGCTCCCTC</td><td>282.0</td><td>2.7</td><td>0.5</td><td>5.1</td><td>3.8E-07</td><td>6.7E-07</td><td>MB231/MB468</td><td>CYTO</td></tr>
<tr><td>>hg38_21_+_8433172_8446622_RNA45SN1_45S_50ntFlanks@6699.6717.19|GCTAGCTGCGAGAATTAAT</td><td>>hg38_21_+_8433172_8446622_RNA45SN1_45S_50ntFlanks</td><td>6699</td><td>6717</td><td>19</td><td>GCTAGCTGCGAGAATTAAT</td><td>871.4</td><td>-8.1</td><td>1.5</td><td>-5.3</td><td>9.1E-08</td><td>6.8E-07</td><td>BT20/MB468</td><td>MP</td></tr>
<tr><td>>hg38_21_+_8433172_8446622_RNA45SN1_45S_50ntFlanks@12007.12030.24|GCCGCCGGTGAAATACCACTACTC</td><td>>hg38_21_+_8433172_8446622_RNA45SN1_45S_50ntFlanks</td><td>12007</td><td>12030</td><td>24</td><td>GCCGCCGGTGAAATACCACTACTC</td><td>439.7</td><td>-7.5</td><td>1.4</td><td>-5.3</td><td>9.1E-08</td><td>6.8E-07</td><td>BT20/MB468</td><td>MP</td></tr>
<tr><td>>hg38_21_+_8433172_8446622_RNA45SN1_45S_50ntFlanks@12893.12924.32|CCTCGCCCGTCACGCACCGCACGTTCGTGGGG</td><td>>hg38_21_+_8433172_8446622_RNA45SN1_45S_50ntFlanks</td><td>12893</td><td>12924</td><td>32</td><td>CCTCGCCCGTCACGCACCGCACGTTCGTGGGG</td><td>80.9</td><td>-9.4</td><td>1.8</td><td>-5.3</td><td>9.1E-08</td><td>6.8E-07</td><td>BT20/MB468</td><td>MP</td></tr>
<tr><td>>hg38_21_+_8433172_8446622_RNA45SN1_45S_50ntFlanks@8184.8211.28|CGAGGCCCAGCCCGTGGACGGTGTGAGG</td><td>>hg38_21_+_8433172_8446622_RNA45SN1_45S_50ntFlanks</td><td>8184</td><td>8211</td><td>28</td><td>CGAGGCCCAGCCCGTGGACGGTGTGAGG</td><td>251.2</td><td>-5.3</td><td>1.1</td><td>-5.1</td><td>4.0E-07</td><td>6.8E-07</td><td>MB231/MB468</td><td>NUC</td></tr>
<tr><td>>hg38_21_+_8433172_8446622_RNA45SN1_45S_50ntFlanks@8112.8143.32|GGGCGCGGGACATGTGGCGTACGGAAGACCCG</td><td>>hg38_21_+_8433172_8446622_RNA45SN1_45S_50ntFlanks</td><td>8112</td><td>8143</td><td>32</td><td>GGGCGCGGGACATGTGGCGTACGGAAGACCCG</td><td>722.2</td><td>-7.8</td><td>1.5</td><td>-5.3</td><td>9.2E-08</td><td>6.8E-07</td><td>BT20/MB468</td><td>MP</td></tr>
<tr><td>>hg38_21_+_8433172_8446622_RNA45SN1_45S_50ntFlanks@11623.11660.38|AAGCATCGCGAAGGCCCGCGGCGGGTGTTGACGCGATG</td><td>>hg38_21_+_8433172_8446622_RNA45SN1_45S_50ntFlanks</td><td>11623</td><td>11660</td><td>38</td><td>AAGCATCGCGAAGGCCCGCGGCGGGTGTTGACGCGATG</td><td>75.5</td><td>-9.3</td><td>1.7</td><td>-5.3</td><td>9.2E-08</td><td>6.8E-07</td><td>BT20/MB468</td><td>MP</td></tr>
<tr><td>>hg38_21_+_8433172_8446622_RNA45SN1_45S_50ntFlanks@8337.8375.39|AGTACCGTAAGGGAAAGTTGAAAAGAACTTTGAAGAGAG</td><td>>hg38_21_+_8433172_8446622_RNA45SN1_45S_50ntFlanks</td><td>8337</td><td>8375</td><td>39</td><td>AGTACCGTAAGGGAAAGTTGAAAAGAACTTTGAAGAGAG</td><td>261.5</td><td>8.6</td><td>1.6</td><td>5.3</td><td>9.2E-08</td><td>6.8E-07</td><td>BT20/MB468</td><td>MP</td></tr>
<tr><td>>hg38_21_+_8433172_8446622_RNA45SN1_45S_50ntFlanks@6644.6662.19|CCTCGTACGACTCTTAGCG</td><td>>hg38_21_+_8433172_8446622_RNA45SN1_45S_50ntFlanks</td><td>6644</td><td>6662</td><td>19</td><td>CCTCGTACGACTCTTAGCG</td><td>723.6</td><td>3.9</td><td>0.8</td><td>5.1</td><td>4.0E-07</td><td>6.8E-07</td><td>MB231/MB468</td><td>NUC</td></tr>
<tr><td>>hg38_21_+_8433172_8446622_RNA45SN1_45S_50ntFlanks@11568.11585.18|GCAGCCGACTTAGAACTG</td><td>>hg38_21_+_8433172_8446622_RNA45SN1_45S_50ntFlanks</td><td>11568</td><td>11585</td><td>18</td><td>GCAGCCGACTTAGAACTG</td><td>2537.1</td><td>-4.3</td><td>0.9</td><td>-5.1</td><td>3.5E-07</td><td>6.8E-07</td><td>MB231/MB468</td><td>MP</td></tr>
<tr><td>>am_MI0000748|hsa-mir-130b&WithFlank&22|+|21653298|21653391@19.39.21&[MIMAT0004680&hsa-miR-130b-5p&offsets|0|0;m-234&22|+|21653316|21653337&offsets|0|-1]|ACTCTTTCCCTGTTGCACTAC</td><td>>am_MI0000748|hsa-mir-130b&WithFlank&22|+|21653298|21653391</td><td>19</td><td>39</td><td>21</td><td>ACTCTTTCCCTGTTGCACTAC</td><td>679.5</td><td>7.6</td><td>1.5</td><td>5.1</td><td>3.4E-07</td><td>6.8E-07</td><td>BT20/MB468</td><td>CYTO</td></tr>
<tr><td>>hg38_21_+_8433172_8446622_RNA45SN1_45S_50ntFlanks@6644.6665.22|CCTCGTACGACTCTTAGCGGTG</td><td>>hg38_21_+_8433172_8446622_RNA45SN1_45S_50ntFlanks</td><td>6644</td><td>6665</td><td>22</td><td>CCTCGTACGACTCTTAGCGGTG</td><td>641.0</td><td>4.6</td><td>0.9</td><td>5.3</td><td>9.2E-08</td><td>6.8E-07</td><td>BT20/MB468</td><td>MP</td></tr>
<tr><td>>MI0000082|hsa-mir-25&WithFlank&7|-|100093554|100093649@58.78.21(+1U)&[MIMAT0000081&hsa-miR-25-3p&offsets|0|-1(+1U);m-23&7|-|100093571|100093592&offsets|0|-1(+1U)]|CATTGCACTTGTCTCGGTCTGT</td><td>>MI0000082|hsa-mir-25&WithFlank&7|-|100093554|100093649</td><td>58</td><td>78</td><td>21(+1U)</td><td>CATTGCACTTGTCTCGGTCTGT</td><td>738.0</td><td>2.4</td><td>0.5</td><td>5.1</td><td>4.4E-07</td><td>6.8E-07</td><td>BT20/MB468</td><td>NUC</td></tr>
<tr><td>>hg38_21_+_8433172_8446622_RNA45SN1_45S_50ntFlanks@4243.4263.21|CTTTCGAGGCCCTGTAATTGG</td><td>>hg38_21_+_8433172_8446622_RNA45SN1_45S_50ntFlanks</td><td>4243</td><td>4263</td><td>21</td><td>CTTTCGAGGCCCTGTAATTGG</td><td>676.2</td><td>-7.2</td><td>1.4</td><td>-5.1</td><td>4.2E-07</td><td>6.8E-07</td><td>MB231/MB468</td><td>MITO</td></tr>
<tr><td>>hg38_21_+_8433172_8446622_RNA45SN1_45S_50ntFlanks@11604.11622.19|CGACTGTTTAATTAAAACA</td><td>>hg38_21_+_8433172_8446622_RNA45SN1_45S_50ntFlanks</td><td>11604</td><td>11622</td><td>19</td><td>CGACTGTTTAATTAAAACA</td><td>263.3</td><td>11.4</td><td>2.0</td><td>5.6</td><td>1.9E-08</td><td>6.8E-07</td><td>BT20/MB231</td><td>MP</td></tr>
<tr><td>>MI0000067|hsa-let-7f-1&WithFlank&9|+|94176341|94176439@13.33.21(+1A);MI0000068|hsa-let-7f-2&WithFlank&X|-|53557186|53557280@14.34.21(+1A)&[MIMAT0000067&hsa-let-7f-5p&offsets|0|-1(+1A);m-20&9|+|94176353|94176374&offsets|0|-1(+1A)];[MIMAT0000067_1&hsa-let-7f-5p&offsets|0|-1(+1A);m-19&X|-|53557246|53557267&offsets|0|-1(+1A)]|TGAGGTAGTAGATTGTATAGTA</td><td>>MI0000067|hsa-let-7f-1&WithFlank&9|+|94176341|94176439</td><td>13</td><td>33</td><td>21(+1A)</td><td>TGAGGTAGTAGATTGTATAGTA</td><td>1174.4</td><td>5.2</td><td>1.0</td><td>5.1</td><td>4.2E-07</td><td>6.8E-07</td><td>MB231/MB468</td><td>MITO</td></tr>
<tr><td>>hg38_21_+_8433172_8446622_RNA45SN1_45S_50ntFlanks@9483.9509.27|CGTGTTAGGACCCGAAAGATGGTGAAC</td><td>>hg38_21_+_8433172_8446622_RNA45SN1_45S_50ntFlanks</td><td>9483</td><td>9509</td><td>27</td><td>CGTGTTAGGACCCGAAAGATGGTGAAC</td><td>98.3</td><td>10.4</td><td>1.9</td><td>5.3</td><td>9.2E-08</td><td>6.8E-07</td><td>BT20/MB468</td><td>MP</td></tr>
<tr><td>>MI0003591|hsa-mir-584&WithFlank&5|-|149062307|149062415@22.44.23(+1U)&[MIMAT0003249&hsa-miR-584-5p&offsets|0|+1(+1U);m-167&5|-|149062374|149062394&offsets|0|+2(+1U)]|TTATGGTTTGCCTGGGACTGAGGT</td><td>>MI0003591|hsa-mir-584&WithFlank&5|-|149062307|149062415</td><td>22</td><td>44</td><td>23(+1U)</td><td>TTATGGTTTGCCTGGGACTGAGGT</td><td>72.1</td><td>9.9</td><td>1.9</td><td>5.3</td><td>9.3E-08</td><td>6.8E-07</td><td>BT20/MB468</td><td>MP</td></tr>
<tr><td>>MI0002470|hsa-mir-486-1&WithFlank&8|-|41660435|41660514@10.30.21;MI0023622|hsa-mir-486-2&WithFlank&8|+|41660438|41660513@7.27.21&[MIMAT0002177&hsa-miR-486-5p&offsets|0|-1;m-107&8|-|41660484|41660505&offsets|0|-1];[MIMAT0002177_1&hsa-miR-486-5p&offsets|0|-1;TJU_CMC_MD2.ID03138.5p-miR&offsets|0|-3;m-106&8|+|41660444|41660465&offsets|0|-1]|TCCTGTACTGAGCTGCCCCGA</td><td>>MI0002470|hsa-mir-486-1&WithFlank&8|-|41660435|41660514</td><td>10</td><td>30</td><td>21</td><td>TCCTGTACTGAGCTGCCCCGA</td><td>51.3</td><td>8.9</td><td>1.7</td><td>5.1</td><td>3.5E-07</td><td>6.8E-07</td><td>MB231/MB468</td><td>MP</td></tr>
<tr><td>>hg38_21_+_8433172_8446622_RNA45SN1_45S_50ntFlanks@11729.11747.19|TAACTATGACTCTCTTAAG</td><td>>hg38_21_+_8433172_8446622_RNA45SN1_45S_50ntFlanks</td><td>11729</td><td>11747</td><td>19</td><td>TAACTATGACTCTCTTAAG</td><td>6644.3</td><td>-3.6</td><td>0.7</td><td>-5.1</td><td>3.9E-07</td><td>6.8E-07</td><td>MB231/MB468</td><td>CYTO</td></tr>
<tr><td>>hg38_21_+_8433172_8446622_RNA45SN1_45S_50ntFlanks@11576.11604.29|CTTAGAACTGGTGCGGACCAGGGGAATCC</td><td>>hg38_21_+_8433172_8446622_RNA45SN1_45S_50ntFlanks</td><td>11576</td><td>11604</td><td>29</td><td>CTTAGAACTGGTGCGGACCAGGGGAATCC</td><td>185.1</td><td>-6.6</td><td>1.3</td><td>-5.1</td><td>4.0E-07</td><td>6.8E-07</td><td>MB231/MB468</td><td>NUC</td></tr>
<tr><td>>hg38_21_+_8433172_8446622_RNA45SN1_45S_50ntFlanks@12822.12840.19|CCGGGACCGGGGTCCGGTG</td><td>>hg38_21_+_8433172_8446622_RNA45SN1_45S_50ntFlanks</td><td>12822</td><td>12840</td><td>19</td><td>CCGGGACCGGGGTCCGGTG</td><td>224.7</td><td>-10.9</td><td>2.0</td><td>-5.3</td><td>9.3E-08</td><td>6.8E-07</td><td>BT20/MB468</td><td>MP</td></tr>
<tr><td>>trna16_ValTAC_11_-_59318460_59318532@37.76.40,trna17_ValTAC_11_-_59318102_59318174@37.76.40,trna4_ValTAC_X_-_18693029_18693101@37.76.40|ACGCAGAAGGTCCTGGGTTCGAGCCCCAGTGGAACCACCA</td><td>>trna16_ValTAC_11_-_59318460_59318532</td><td>37</td><td>76</td><td>40</td><td>ACGCAGAAGGTCCTGGGTTCGAGCCCCAGTGGAACCACCA</td><td>554.3</td><td>7.1</td><td>1.4</td><td>5.1</td><td>3.9E-07</td><td>6.8E-07</td><td>MB231/MB468</td><td>CYTO</td></tr>
<tr><td>>hg38_21_+_8433172_8446622_RNA45SN1_45S_50ntFlanks@12626.12661.36|AATGGGGCGAAGCTACCATCTGTGGGATTATGACTG</td><td>>hg38_21_+_8433172_8446622_RNA45SN1_45S_50ntFlanks</td><td>12626</td><td>12661</td><td>36</td><td>AATGGGGCGAAGCTACCATCTGTGGGATTATGACTG</td><td>2801.9</td><td>-7.2</td><td>1.4</td><td>-5.1</td><td>4.4E-07</td><td>6.8E-07</td><td>BT20/MB468</td><td>NUC</td></tr>
<tr><td>>hg38_21_+_8433172_8446622_RNA45SN1_45S_50ntFlanks@11565.11601.37|CTAGCAGCCGACTTAGAACTGGTGCGGACCAGGGGAA</td><td>>hg38_21_+_8433172_8446622_RNA45SN1_45S_50ntFlanks</td><td>11565</td><td>11601</td><td>37</td><td>CTAGCAGCCGACTTAGAACTGGTGCGGACCAGGGGAA</td><td>335.6</td><td>-5.5</td><td>1.1</td><td>-5.1</td><td>4.2E-07</td><td>6.8E-07</td><td>MB231/MB468</td><td>MITO</td></tr>
<tr><td>>hg38_21_+_8433172_8446622_RNA45SN1_45S_50ntFlanks@10362.10382.21|AAGGCCGAAGTGGAGAAGGGT</td><td>>hg38_21_+_8433172_8446622_RNA45SN1_45S_50ntFlanks</td><td>10362</td><td>10382</td><td>21</td><td>AAGGCCGAAGTGGAGAAGGGT</td><td>367.9</td><td>-3.7</td><td>0.7</td><td>-5.1</td><td>4.0E-07</td><td>6.9E-07</td><td>MB231/MB468</td><td>NUC</td></tr>
<tr><td>>hg38_21_+_8433172_8446622_RNA45SN1_45S_50ntFlanks@11755.11777.23|AATGCCTCGTCATCTAATTAGTG</td><td>>hg38_21_+_8433172_8446622_RNA45SN1_45S_50ntFlanks</td><td>11755</td><td>11777</td><td>23</td><td>AATGCCTCGTCATCTAATTAGTG</td><td>363.8</td><td>-7.0</td><td>1.4</td><td>-5.1</td><td>4.2E-07</td><td>6.9E-07</td><td>MB231/MB468</td><td>MITO</td></tr>
<tr><td>>hg38_21_+_8433172_8446622_RNA45SN1_45S_50ntFlanks@9736.9754.19|CCGAAACGATCTCAACCTA</td><td>>hg38_21_+_8433172_8446622_RNA45SN1_45S_50ntFlanks</td><td>9736</td><td>9754</td><td>19</td><td>CCGAAACGATCTCAACCTA</td><td>450.8</td><td>-6.9</td><td>1.4</td><td>-5.1</td><td>4.0E-07</td><td>6.9E-07</td><td>MB231/MB468</td><td>NUC</td></tr>
<tr><td>>hg38_21_+_8433172_8446622_RNA45SN1_45S_50ntFlanks@12453.12476.24|TCACCAAGCGTTGGATTGTTCACC</td><td>>hg38_21_+_8433172_8446622_RNA45SN1_45S_50ntFlanks</td><td>12453</td><td>12476</td><td>24</td><td>TCACCAAGCGTTGGATTGTTCACC</td><td>204.8</td><td>3.9</td><td>0.8</td><td>5.1</td><td>4.0E-07</td><td>6.9E-07</td><td>MB231/MB468</td><td>NUC</td></tr>
<tr><td>>hg38_21_+_8433172_8446622_RNA45SN1_45S_50ntFlanks@12003.12033.31|GCGGGCCGCCGGTGAAATACCACTACTCTGA</td><td>>hg38_21_+_8433172_8446622_RNA45SN1_45S_50ntFlanks</td><td>12003</td><td>12033</td><td>31</td><td>GCGGGCCGCCGGTGAAATACCACTACTCTGA</td><td>2487.7</td><td>5.4</td><td>1.1</td><td>5.1</td><td>3.9E-07</td><td>6.9E-07</td><td>MB231/MB468</td><td>CYTO</td></tr>
<tr><td>>hg38_21_+_8433172_8446622_RNA45SN1_45S_50ntFlanks@3775.3792.18|GCCGGTACAGTGAAACTG</td><td>>hg38_21_+_8433172_8446622_RNA45SN1_45S_50ntFlanks</td><td>3775</td><td>3792</td><td>18</td><td>GCCGGTACAGTGAAACTG</td><td>1908.5</td><td>-6.2</td><td>1.2</td><td>-5.1</td><td>3.0E-07</td><td>6.9E-07</td><td>BT20/MB468</td><td>MITO</td></tr>
<tr><td>>hg38_21_+_8433172_8446622_RNA45SN1_45S_50ntFlanks@9337.9370.34|CTCGCACGAAAGCCGCCGTGGCGCAATGAAGGTG</td><td>>hg38_21_+_8433172_8446622_RNA45SN1_45S_50ntFlanks</td><td>9337</td><td>9370</td><td>34</td><td>CTCGCACGAAAGCCGCCGTGGCGCAATGAAGGTG</td><td>343.1</td><td>-7.7</td><td>1.4</td><td>-5.3</td><td>9.4E-08</td><td>6.9E-07</td><td>BT20/MB468</td><td>MP</td></tr>
<tr><td>>MI0000088|hsa-mir-30a&WithFlank&6|-|71403545|71403627@12.33.22(+1U)&[MIMAT0000087&hsa-miR-30a-5p&offsets|0|0(+1U);m-15&6|-|71403595|71403616&offsets|0|0(+1U)]|TGTAAACATCCTCGACTGGAAGT</td><td>>MI0000088|hsa-mir-30a&WithFlank&6|-|71403545|71403627</td><td>12</td><td>33</td><td>22(+1U)</td><td>TGTAAACATCCTCGACTGGAAGT</td><td>1380.4</td><td>5.6</td><td>1.1</td><td>5.1</td><td>3.9E-07</td><td>6.9E-07</td><td>MB231/MB468</td><td>CYTO</td></tr>
<tr><td>>hg38_21_+_8433172_8446622_RNA45SN1_45S_50ntFlanks@10391.10431.41|AACAGCAGTTGAACATGGGTCAGTCGGTCCTGAGAGATGGG</td><td>>hg38_21_+_8433172_8446622_RNA45SN1_45S_50ntFlanks</td><td>10391</td><td>10431</td><td>41</td><td>AACAGCAGTTGAACATGGGTCAGTCGGTCCTGAGAGATGGG</td><td>305.3</td><td>-7.3</td><td>1.4</td><td>-5.1</td><td>4.1E-07</td><td>6.9E-07</td><td>MB231/MB468</td><td>NUC</td></tr>
<tr><td>>hg38_21_+_8433172_8446622_RNA45SN1_45S_50ntFlanks@10411.10431.21|CAGTCGGTCCTGAGAGATGGG</td><td>>hg38_21_+_8433172_8446622_RNA45SN1_45S_50ntFlanks</td><td>10411</td><td>10431</td><td>21</td><td>CAGTCGGTCCTGAGAGATGGG</td><td>5866.3</td><td>-6.7</td><td>1.3</td><td>-5.1</td><td>3.9E-07</td><td>6.9E-07</td><td>MB231/MB468</td><td>CYTO</td></tr>
<tr><td>>hg38_21_+_8433172_8446622_RNA45SN1_45S_50ntFlanks@11454.11478.25|CCTCGGCGCGCGGCGGCGGCGGCGG</td><td>>hg38_21_+_8433172_8446622_RNA45SN1_45S_50ntFlanks</td><td>11454</td><td>11478</td><td>25</td><td>CCTCGGCGCGCGGCGGCGGCGGCGG</td><td>79.2</td><td>-9.4</td><td>1.8</td><td>-5.3</td><td>9.4E-08</td><td>6.9E-07</td><td>BT20/MB468</td><td>MP</td></tr>
<tr><td>>hg38_1_-_228634819_228635039_RNA5S12_5S_50ntFlanks@149.169.21|GAATACCGGGTGCTGTAGGCT</td><td>>hg38_1_-_228634819_228635039_RNA5S12_5S_50ntFlanks</td><td>149</td><td>169</td><td>21</td><td>GAATACCGGGTGCTGTAGGCT</td><td>558.0</td><td>2.7</td><td>0.5</td><td>5.1</td><td>3.1E-07</td><td>6.9E-07</td><td>BT20/MB468</td><td>MITO</td></tr>
<tr><td>>hg38_21_+_8433172_8446622_RNA45SN1_45S_50ntFlanks@5237.5267.31|ACTGACTGGCTCAGCGTGTGCCTACCCTACG</td><td>>hg38_21_+_8433172_8446622_RNA45SN1_45S_50ntFlanks</td><td>5237</td><td>5267</td><td>31</td><td>ACTGACTGGCTCAGCGTGTGCCTACCCTACG</td><td>298.6</td><td>-7.5</td><td>1.5</td><td>-5.1</td><td>4.3E-07</td><td>6.9E-07</td><td>MB231/MB468</td><td>MITO</td></tr>
<tr><td>>hg38_21_+_8433172_8446622_RNA45SN1_45S_50ntFlanks@6646.6664.19|TCGTACGACTCTTAGCGGT</td><td>>hg38_21_+_8433172_8446622_RNA45SN1_45S_50ntFlanks</td><td>6646</td><td>6664</td><td>19</td><td>TCGTACGACTCTTAGCGGT</td><td>15647.3</td><td>3.1</td><td>0.6</td><td>5.1</td><td>4.1E-07</td><td>6.9E-07</td><td>MB231/MB468</td><td>NUC</td></tr>
<tr><td>>trnaMT_HisGTG_MT_+_12138_12206@19.52.34|AAACATCAGATTGTGAATCTGACAACAGAGGCTT</td><td>>trnaMT_HisGTG_MT_+_12138_12206</td><td>19</td><td>52</td><td>34</td><td>AAACATCAGATTGTGAATCTGACAACAGAGGCTT</td><td>101.3</td><td>10.4</td><td>1.9</td><td>5.3</td><td>9.4E-08</td><td>6.9E-07</td><td>BT20/MB468</td><td>MP</td></tr>
<tr><td>>hg38_21_+_8433172_8446622_RNA45SN1_45S_50ntFlanks@10679.10707.29|GGTTCCGGCGGCGTCCGGTGAGCTCTCGC</td><td>>hg38_21_+_8433172_8446622_RNA45SN1_45S_50ntFlanks</td><td>10679</td><td>10707</td><td>29</td><td>GGTTCCGGCGGCGTCCGGTGAGCTCTCGC</td><td>192.3</td><td>-6.9</td><td>1.4</td><td>-5.1</td><td>3.6E-07</td><td>6.9E-07</td><td>MB231/MB468</td><td>MP</td></tr>
<tr><td>>hg38_21_+_8433172_8446622_RNA45SN1_45S_50ntFlanks@11571.11612.42|GCCGACTTAGAACTGGTGCGGACCAGGGGAATCCGACTGTTT</td><td>>hg38_21_+_8433172_8446622_RNA45SN1_45S_50ntFlanks</td><td>11571</td><td>11612</td><td>42</td><td>GCCGACTTAGAACTGGTGCGGACCAGGGGAATCCGACTGTTT</td><td>380.6</td><td>4.2</td><td>0.8</td><td>5.1</td><td>4.0E-07</td><td>6.9E-07</td><td>MB231/MB468</td><td>CYTO</td></tr>
<tr><td>>hg38_21_+_8433172_8446622_RNA45SN1_45S_50ntFlanks@10116.10135.20|TGCGTCGGAGGGCGGCGGCG</td><td>>hg38_21_+_8433172_8446622_RNA45SN1_45S_50ntFlanks</td><td>10116</td><td>10135</td><td>20</td><td>TGCGTCGGAGGGCGGCGGCG</td><td>338.2</td><td>-6.2</td><td>1.2</td><td>-5.1</td><td>3.5E-07</td><td>6.9E-07</td><td>BT20/MB468</td><td>CYTO</td></tr>
<tr><td>>hg38_21_+_8433172_8446622_RNA45SN1_45S_50ntFlanks@11027.11061.35|CTCTCTCTCTCTCTCCCCCGCTCCCCGTCCTCCCC</td><td>>hg38_21_+_8433172_8446622_RNA45SN1_45S_50ntFlanks</td><td>11027</td><td>11061</td><td>35</td><td>CTCTCTCTCTCTCTCCCCCGCTCCCCGTCCTCCCC</td><td>1585.2</td><td>-9.0</td><td>1.8</td><td>-5.0</td><td>4.5E-07</td><td>6.9E-07</td><td>BT20/MB468</td><td>NUC</td></tr>
<tr><td>>hg38_21_+_8433172_8446622_RNA45SN1_45S_50ntFlanks@6652.6672.21|GACTCTTAGCGGTGGATCACT</td><td>>hg38_21_+_8433172_8446622_RNA45SN1_45S_50ntFlanks</td><td>6652</td><td>6672</td><td>21</td><td>GACTCTTAGCGGTGGATCACT</td><td>107.1</td><td>2.5</td><td>0.5</td><td>5.0</td><td>4.5E-07</td><td>6.9E-07</td><td>BT20/MB468</td><td>NUC</td></tr>
<tr><td>>hg38_21_+_8433172_8446622_RNA45SN1_45S_50ntFlanks@5078.5112.35|CGATAACGAACGAGACTCTGGCATGCTAACTAGTT</td><td>>hg38_21_+_8433172_8446622_RNA45SN1_45S_50ntFlanks</td><td>5078</td><td>5112</td><td>35</td><td>CGATAACGAACGAGACTCTGGCATGCTAACTAGTT</td><td>310.8</td><td>-6.9</td><td>1.3</td><td>-5.1</td><td>3.6E-07</td><td>6.9E-07</td><td>MB231/MB468</td><td>MP</td></tr>
<tr><td>>trna17_ValTAC_11_-_59318102_59318174@1.34.34,trna4_ValTAC_X_-_18693029_18693101@1.34.34|GGTTCCATAGTGTAGTGGTTATCACGTCTGCTTT</td><td>>trna17_ValTAC_11_-_59318102_59318174</td><td>1</td><td>34</td><td>34</td><td>GGTTCCATAGTGTAGTGGTTATCACGTCTGCTTT</td><td>386.6</td><td>8.4</td><td>1.7</td><td>5.1</td><td>4.3E-07</td><td>7.0E-07</td><td>MB231/MB468</td><td>MITO</td></tr>
<tr><td>>hg38_21_+_8433172_8446622_RNA45SN1_45S_50ntFlanks@13008.13025.18|ATTGAAAGTCAGCCCTCG</td><td>>hg38_21_+_8433172_8446622_RNA45SN1_45S_50ntFlanks</td><td>13008</td><td>13025</td><td>18</td><td>ATTGAAAGTCAGCCCTCG</td><td>1730.3</td><td>-4.5</td><td>0.9</td><td>-5.1</td><td>4.0E-07</td><td>7.0E-07</td><td>MB231/MB468</td><td>CYTO</td></tr>
<tr><td>>hg38_21_+_8433172_8446622_RNA45SN1_45S_50ntFlanks@10873.10894.22|CTGGGTCGGTCGGGCTGGGGCG</td><td>>hg38_21_+_8433172_8446622_RNA45SN1_45S_50ntFlanks</td><td>10873</td><td>10894</td><td>22</td><td>CTGGGTCGGTCGGGCTGGGGCG</td><td>661.1</td><td>-7.2</td><td>1.4</td><td>-5.1</td><td>4.1E-07</td><td>7.0E-07</td><td>MB231/MB468</td><td>NUC</td></tr>
</table>

<table>
<tr><td>>hg38_21_+_8433172_8446622_RNA45SN1_45S_50ntFlanks@9739.9763.25|AAACGATCTCAACCTATTCTCAAAC</td><td>>hg38_21_+_8433172_8446622_RNA45SN1_45S_50ntFlanks</td><td>9739</td><td>9763</td><td>25</td><td>AAACGATCTCAACCTATTCTCAAAC</td><td>209.6</td><td>-7.2</td><td>1.4</td><td>-5.1</td><td>3.6E-07</td><td>7.0E-07</td><td>MB231/MB468</td><td>MP</td></tr>
<tr><td>>hg38_21_+_8433172_8446622_RNA45SN1_45S_50ntFlanks@11768.11803.36|CTAATTAGTGACGCGCATGAATGGATGAACGAGATT</td><td>>hg38_21_+_8433172_8446622_RNA45SN1_45S_50ntFlanks</td><td>11768</td><td>11803</td><td>36</td><td>CTAATTAGTGACGCGCATGAATGGATGAACGAGATT</td><td>1462.9</td><td>-7.1</td><td>1.4</td><td>-5.1</td><td>3.6E-07</td><td>7.0E-07</td><td>MB231/MB468</td><td>MP</td></tr>
<tr><td>>trnaMT_HisGTG_MT_+_12138_12206@47.72.26|AGGCTTACGACCCCTTATTTACCCCA</td><td>>trnaMT_HisGTG_MT_+_12138_12206</td><td>47</td><td>72</td><td>26</td><td>AGGCTTACGACCCCTTATTTACCCCA</td><td>82.4</td><td>10.1</td><td>1.9</td><td>5.3</td><td>9.5E-08</td><td>7.0E-07</td><td>BT20/MB468</td><td>MP</td></tr>
<tr><td>>hg38_21_+_8433172_8446622_RNA45SN1_45S_50ntFlanks@6721.6753.33|AATTGCAGGACACATTGATCATCGACACTTCGA</td><td>>hg38_21_+_8433172_8446622_RNA45SN1_45S_50ntFlanks</td><td>6721</td><td>6753</td><td>33</td><td>AATTGCAGGACACATTGATCATCGACACTTCGA</td><td>239.4</td><td>-5.2</td><td>1.0</td><td>-5.1</td><td>4.1E-07</td><td>7.0E-07</td><td>MB231/MB468</td><td>NUC</td></tr>
<tr><td>>hg38_21_+_8433172_8446622_RNA45SN1_45S_50ntFlanks@10399.10424.26|TTGAACATGGGTCAGTCGGTCCTGAG</td><td>>hg38_21_+_8433172_8446622_RNA45SN1_45S_50ntFlanks</td><td>10399</td><td>10424</td><td>26</td><td>TTGAACATGGGTCAGTCGGTCCTGAG</td><td>1017.7</td><td>5.4</td><td>1.0</td><td>5.3</td><td>9.5E-08</td><td>7.0E-07</td><td>BT20/MB468</td><td>MP</td></tr>
<tr><td>>hg38_21_+_8433172_8446622_RNA45SN1_45S_50ntFlanks@13006.13039.34|CTATTGAAAGTCAGCCCTCGACACAAGGGTTTGT</td><td>>hg38_21_+_8433172_8446622_RNA45SN1_45S_50ntFlanks</td><td>13006</td><td>13039</td><td>34</td><td>CTATTGAAAGTCAGCCCTCGACACAAGGGTTTGT</td><td>1389.9</td><td>4.7</td><td>0.9</td><td>5.1</td><td>3.5E-07</td><td>7.0E-07</td><td>BT20/MB468</td><td>CYTO</td></tr>
<tr><td>>hg38_21_+_8433172_8446622_RNA45SN1_45S_50ntFlanks@3991.4024.34|TGGTGACTCTAGATAACCTCGGGCCGATCGCACG</td><td>>hg38_21_+_8433172_8446622_RNA45SN1_45S_50ntFlanks</td><td>3991</td><td>4024</td><td>34</td><td>TGGTGACTCTAGATAACCTCGGGCCGATCGCACG</td><td>1304.2</td><td>-8.6</td><td>1.7</td><td>-5.0</td><td>4.5E-07</td><td>7.0E-07</td><td>BT20/MB468</td><td>NUC</td></tr>
<tr><td>>hg38_21_+_8433172_8446622_RNA45SN1_45S_50ntFlanks@12499.12517.19|GGTTTAGACCGTCGTGAGA</td><td>>hg38_21_+_8433172_8446622_RNA45SN1_45S_50ntFlanks</td><td>12499</td><td>12517</td><td>19</td><td>GGTTTAGACCGTCGTGAGA</td><td>168.6</td><td>-6.1</td><td>1.2</td><td>-5.1</td><td>3.5E-07</td><td>7.0E-07</td><td>BT20/MB468</td><td>CYTO</td></tr>
<tr><td>>hg38_21_+_8433172_8446622_RNA45SN1_45S_50ntFlanks@10789.10816.28|AGCCTCTGGCATGTTGGAACAATGTAGG</td><td>>hg38_21_+_8433172_8446622_RNA45SN1_45S_50ntFlanks</td><td>10789</td><td>10816</td><td>28</td><td>AGCCTCTGGCATGTTGGAACAATGTAGG</td><td>229.3</td><td>11.2</td><td>2.0</td><td>5.6</td><td>2.0E-08</td><td>7.0E-07</td><td>BT20/MB231</td><td>MP</td></tr>
<tr><td>>hg38_21_+_8433172_8446622_RNA45SN1_45S_50ntFlanks@6689.6720.32|GAAGAACGCAGCTAGCTGCGAGAATTAATGTG</td><td>>hg38_21_+_8433172_8446622_RNA45SN1_45S_50ntFlanks</td><td>6689</td><td>6720</td><td>32</td><td>GAAGAACGCAGCTAGCTGCGAGAATTAATGTG</td><td>393.6</td><td>-4.8</td><td>0.9</td><td>-5.1</td><td>4.0E-07</td><td>7.0E-07</td><td>MB231/MB468</td><td>CYTO</td></tr>
<tr><td>>hg38_21_+_8433172_8446622_RNA45SN1_45S_50ntFlanks@11656.11685.30|CGATGTGATTTCTGCCCAGTGCTCTGAATG</td><td>>hg38_21_+_8433172_8446622_RNA45SN1_45S_50ntFlanks</td><td>11656</td><td>11685</td><td>30</td><td>CGATGTGATTTCTGCCCAGTGCTCTGAATG</td><td>342.7</td><td>-5.4</td><td>1.1</td><td>-5.1</td><td>4.3E-07</td><td>7.0E-07</td><td>MB231/MB468</td><td>MITO</td></tr>
<tr><td>>hg38_21_+_8433172_8446622_RNA45SN1_45S_50ntFlanks@9105.9128.24|CGGCGACGGGGGGGGTGCCGCGCG</td><td>>hg38_21_+_8433172_8446622_RNA45SN1_45S_50ntFlanks</td><td>9105</td><td>9128</td><td>24</td><td>CGGCGACGGGGGGGGTGCCGCGCG</td><td>87.0</td><td>-9.5</td><td>1.8</td><td>-5.3</td><td>9.6E-08</td><td>7.0E-07</td><td>BT20/MB468</td><td>MP</td></tr>
<tr><td>>hg38_21_+_8433172_8446622_RNA45SN1_45S_50ntFlanks@9391.9410.20|CCGAGGTGGGATCCCGAGGC</td><td>>hg38_21_+_8433172_8446622_RNA45SN1_45S_50ntFlanks</td><td>9391</td><td>9410</td><td>20</td><td>CCGAGGTGGGATCCCGAGGC</td><td>99.8</td><td>-9.7</td><td>1.8</td><td>-5.3</td><td>9.6E-08</td><td>7.0E-07</td><td>BT20/MB468</td><td>MP</td></tr>
<tr><td>>am_MI0001445|hsa-mir-423&WithFlank&17|+|30117073|30117178@23.40.18&[MIMAT0004748&hsa-miR-423-5p&offsets|0|-5;m-141&17|+|30117095|30117117&offsets|0|-5]|TGAGGGGCAGAGAGCGAG</td><td>>am_MI0001445|hsa-mir-423&WithFlank&17|+|30117073|30117178</td><td>23</td><td>40</td><td>18</td><td>TGAGGGGCAGAGAGCGAG</td><td>102.8</td><td>10.4</td><td>2.0</td><td>5.3</td><td>9.6E-08</td><td>7.0E-07</td><td>BT20/MB468</td><td>MP</td></tr>
<tr><td>>hg38_21_+_8433172_8446622_RNA45SN1_45S_50ntFlanks@5334.5362.29|ATGAACGAGGAATTCCCAGTAAGTGCGGG</td><td>>hg38_21_+_8433172_8446622_RNA45SN1_45S_50ntFlanks</td><td>5334</td><td>5362</td><td>29</td><td>ATGAACGAGGAATTCCCAGTAAGTGCGGG</td><td>173.6</td><td>-9.5</td><td>1.9</td><td>-5.1</td><td>3.1E-07</td><td>7.0E-07</td><td>BT20/MB468</td><td>MITO</td></tr>
<tr><td>>hg38_21_+_8433172_8446622_RNA45SN1_45S_50ntFlanks@9496.9523.28|GAAAGATGGTGAACTATGCCTGGGCAGG</td><td>>hg38_21_+_8433172_8446622_RNA45SN1_45S_50ntFlanks</td><td>9496</td><td>9523</td><td>28</td><td>GAAAGATGGTGAACTATGCCTGGGCAGG</td><td>527.9</td><td>12.4</td><td>2.2</td><td>5.6</td><td>2.0E-08</td><td>7.0E-07</td><td>BT20/MB231</td><td>MP</td></tr>
<tr><td>>hg38_21_+_8433172_8446622_RNA45SN1_45S_50ntFlanks@6745.6769.25|ACACTTCGAACGCACTTGCGGCCCC</td><td>>hg38_21_+_8433172_8446622_RNA45SN1_45S_50ntFlanks</td><td>6745</td><td>6769</td><td>25</td><td>ACACTTCGAACGCACTTGCGGCCCC</td><td>178.2</td><td>-5.1</td><td>1.0</td><td>-5.1</td><td>4.2E-07</td><td>7.0E-07</td><td>MB231/MB468</td><td>NUC</td></tr>
<tr><td>>hg38_21_+_8433172_8446622_RNA45SN1_45S_50ntFlanks@5492.5514.23|AGAAGACGGTCGAACTTGACTAT</td><td>>hg38_21_+_8433172_8446622_RNA45SN1_45S_50ntFlanks</td><td>5492</td><td>5514</td><td>23</td><td>AGAAGACGGTCGAACTTGACTAT</td><td>7907.6</td><td>-7.4</td><td>1.4</td><td>-5.3</td><td>9.6E-08</td><td>7.1E-07</td><td>BT20/MB468</td><td>MP</td></tr>
<tr><td>>hg38_21_+_8433172_8446622_RNA45SN1_45S_50ntFlanks@6657.6689.33|TTAGCGGTGGATCACTCGGCTCGTGCGTCGATG</td><td>>hg38_21_+_8433172_8446622_RNA45SN1_45S_50ntFlanks</td><td>6657</td><td>6689</td><td>33</td><td>TTAGCGGTGGATCACTCGGCTCGTGCGTCGATG</td><td>423.7</td><td>-5.4</td><td>1.1</td><td>-5.1</td><td>4.3E-07</td><td>7.1E-07</td><td>MB231/MB468</td><td>MITO</td></tr>
<tr><td>>hg38_21_+_8433172_8446622_RNA45SN1_45S_50ntFlanks@6731.6758.28|CACATTGATCATCGACACTTCGAACGCA</td><td>>hg38_21_+_8433172_8446622_RNA45SN1_45S_50ntFlanks</td><td>6731</td><td>6758</td><td>28</td><td>CACATTGATCATCGACACTTCGAACGCA</td><td>462.2</td><td>-5.5</td><td>1.1</td><td>-5.1</td><td>4.3E-07</td><td>7.1E-07</td><td>MB231/MB468</td><td>MITO</td></tr>
<tr><td>>hg38_21_+_8433172_8446622_RNA45SN1_45S_50ntFlanks@11569.11603.35|CAGCCGACTTAGAACTGGTGCGGACCAGGGGAATC</td><td>>hg38_21_+_8433172_8446622_RNA45SN1_45S_50ntFlanks</td><td>11569</td><td>11603</td><td>35</td><td>CAGCCGACTTAGAACTGGTGCGGACCAGGGGAATC</td><td>2259.5</td><td>-4.0</td><td>0.8</td><td>-5.1</td><td>4.0E-07</td><td>7.1E-07</td><td>MB231/MB468</td><td>CYTO</td></tr>
<tr><td>>hg38_MT_+_1621_3279_RNR2_16S_50ntFlanks@717.745.29|AAGCCTGCGTCAGATTAAAACACTGAACT</td><td>>hg38_MT_+_1621_3279_RNR2_16S_50ntFlanks</td><td>717</td><td>745</td><td>29</td><td>AAGCCTGCGTCAGATTAAAACACTGAACT</td><td>52.3</td><td>8.9</td><td>1.8</td><td>5.1</td><td>3.6E-07</td><td>7.1E-07</td><td>MB231/MB468</td><td>MP</td></tr>
<tr><td>>MI0000255|hsa-mir-30d&WithFlank&8|-|134804870|134804951@12.31.20&[MIMAT0000245&hsa-miR-30d-5p&offsets|0|-2;m-21&8|-|134804919|134804940&offsets|0|-2]|TGTAAACATCCCCGACTGGA</td><td>>MI0000255|hsa-mir-30d&WithFlank&8|-|134804870|134804951</td><td>12</td><td>31</td><td>20</td><td>TGTAAACATCCCCGACTGGA</td><td>968.3</td><td>4.3</td><td>0.8</td><td>5.1</td><td>3.6E-07</td><td>7.1E-07</td><td>MB231/MB468</td><td>MP</td></tr>
<tr><td>>hg38_21_+_8433172_8446622_RNA45SN1_45S_50ntFlanks@5420.5438.19|CCGATTGGATGGTTTAGTG</td><td>>hg38_21_+_8433172_8446622_RNA45SN1_45S_50ntFlanks</td><td>5420</td><td>5438</td><td>19</td><td>CCGATTGGATGGTTTAGTG</td><td>229.2</td><td>-6.4</td><td>1.3</td><td>-5.1</td><td>3.5E-07</td><td>7.1E-07</td><td>BT20/MB468</td><td>CYTO</td></tr>
<tr><td>>hg38_21_+_8433172_8446622_RNA45SN1_45S_50ntFlanks@12944.12962.19|CGTAGACGACCTGCTTCTG</td><td>>hg38_21_+_8433172_8446622_RNA45SN1_45S_50ntFlanks</td><td>12944</td><td>12962</td><td>19</td><td>CGTAGACGACCTGCTTCTG</td><td>316.0</td><td>-7.1</td><td>1.4</td><td>-5.1</td><td>3.6E-07</td><td>7.1E-07</td><td>MB231/MB468</td><td>MP</td></tr>
<tr><td>>trnaMT_PheGAA_MT_+_577_647@36.56.21|AAAATGTTTAGACGGGCTCAC</td><td>>trnaMT_PheGAA_MT_+_577_647</td><td>36</td><td>56</td><td>21</td><td>AAAATGTTTAGACGGGCTCAC</td><td>259.8</td><td>4.5</td><td>0.9</td><td>5.1</td><td>4.4E-07</td><td>7.1E-07</td><td>MB231/MB468</td><td>MITO</td></tr>
<tr><td>>hg38_21_+_8433172_8446622_RNA45SN1_45S_50ntFlanks@6645.6674.30|CTCGTACGACTCTTAGCGGTGGATCACTCG</td><td>>hg38_21_+_8433172_8446622_RNA45SN1_45S_50ntFlanks</td><td>6645</td><td>6674</td><td>30</td><td>CTCGTACGACTCTTAGCGGTGGATCACTCG</td><td>770.2</td><td>3.9</td><td>0.8</td><td>5.1</td><td>4.1E-07</td><td>7.1E-07</td><td>MB231/MB468</td><td>CYTO</td></tr>
<tr><td>>hg38_21_+_8433172_8446622_RNA45SN1_45S_50ntFlanks@5073.5098.26|AATTCCGATAACGAACGAGACTCTGG</td><td>>hg38_21_+_8433172_8446622_RNA45SN1_45S_50ntFlanks</td><td>5073</td><td>5098</td><td>26</td><td>AATTCCGATAACGAACGAGACTCTGG</td><td>5214.2</td><td>-8.0</td><td>1.6</td><td>-5.0</td><td>4.6E-07</td><td>7.1E-07</td><td>BT20/MB468</td><td>NUC</td></tr>
<tr><td>>hg38_MT_+_1621_3279_RNR2_16S_50ntFlanks@459.491.33|CTTGTAAATTTAACTGTTAGTCCAAAGAGGAAC</td><td>>hg38_MT_+_1621_3279_RNR2_16S_50ntFlanks</td><td>459</td><td>491</td><td>33</td><td>CTTGTAAATTTAACTGTTAGTCCAAAGAGGAAC</td><td>46.5</td><td>8.8</td><td>1.7</td><td>5.1</td><td>3.6E-07</td><td>7.1E-07</td><td>MB231/MB468</td><td>MP</td></tr>
<tr><td>>hg38_21_+_8433172_8446622_RNA45SN1_45S_50ntFlanks@12436.12473.38|TCATTGTGAAGCAGAATTCACCAAGCGTTGGATTGTTC</td><td>>hg38_21_+_8433172_8446622_RNA45SN1_45S_50ntFlanks</td><td>12436</td><td>12473</td><td>38</td><td>TCATTGTGAAGCAGAATTCACCAAGCGTTGGATTGTTC</td><td>1212.3</td><td>-3.5</td><td>0.7</td><td>-5.1</td><td>4.1E-07</td><td>7.1E-07</td><td>MB231/MB468</td><td>CYTO</td></tr>
<tr><td>>MI0000471|hsa-mir-126&WithFlank&9|+|136670596|136670692@21.41.21&[MIMAT0000444&hsa-miR-126-5p&offsets|0|0;m-114&9|+|136670616|136670636&offsets|0|0]|CATTATTACTTTTGGTACGCG</td><td>>MI0000471|hsa-mir-126&WithFlank&9|+|136670596|136670692</td><td>21</td><td>41</td><td>21</td><td>CATTATTACTTTTGGTACGCG</td><td>257.8</td><td>8.0</td><td>1.6</td><td>5.1</td><td>3.7E-07</td><td>7.1E-07</td><td>MB231/MB468</td><td>MP</td></tr>
<tr><td>>MI0000746|hsa-mir-99b&WithFlank&19|+|51692606|51692687@13.34.22(+1A)&[MIMAT0000689&hsa-miR-99b-5p&offsets|0|0(+1A);m-10&19|+|51692618|51692639&offsets|0|0(+1A)]|CACCCGTAGAACCGACCTTGCGA</td><td>>MI0000746|hsa-mir-99b&WithFlank&19|+|51692606|51692687</td><td>13</td><td>34</td><td>22(+1A)</td><td>CACCCGTAGAACCGACCTTGCGA</td><td>544.3</td><td>4.6</td><td>0.9</td><td>5.1</td><td>4.4E-07</td><td>7.1E-07</td><td>MB231/MB468</td><td>MITO</td></tr>
<tr><td>>hg38_21_+_8433172_8446622_RNA45SN1_45S_50ntFlanks@7980.8003.24|CCTCAGATCAGACGTGGCGACCCG</td><td>>hg38_21_+_8433172_8446622_RNA45SN1_45S_50ntFlanks</td><td>7980</td><td>8003</td><td>24</td><td>CCTCAGATCAGACGTGGCGACCCG</td><td>571.8</td><td>-7.1</td><td>1.4</td><td>-5.1</td><td>4.4E-07</td><td>7.1E-07</td><td>MB231/MB468</td><td>MITO</td></tr>
<tr><td>>hg38_21_+_8433172_8446622_RNA45SN1_45S_50ntFlanks@12639.12659.21|TACCATCTGTGGGATTATGAC</td><td>>hg38_21_+_8433172_8446622_RNA45SN1_45S_50ntFlanks</td><td>12639</td><td>12659</td><td>21</td><td>TACCATCTGTGGGATTATGAC</td><td>587.1</td><td>-4.8</td><td>1.0</td><td>-5.1</td><td>4.4E-07</td><td>7.1E-07</td><td>MB231/MB468</td><td>MITO</td></tr>
<tr><td>>hg38_21_+_8433172_8446622_RNA45SN1_45S_50ntFlanks@7969.8009.41|CCGAGACGCGACCTCAGATCAGACGTGGCGACCCGCTGAAT</td><td>>hg38_21_+_8433172_8446622_RNA45SN1_45S_50ntFlanks</td><td>7969</td><td>8009</td><td>41</td><td>CCGAGACGCGACCTCAGATCAGACGTGGCGACCCGCTGAAT</td><td>185.0</td><td>4.2</td><td>0.8</td><td>5.0</td><td>4.6E-07</td><td>7.1E-07</td><td>BT20/MB468</td><td>NUC</td></tr>
<tr><td>>MI0000285|hsa-mir-205&WithFlank&1|+|209432127|209432248@40.59.20(+1C)&[MIMAT0000266&hsa-miR-205-5p&offsets|0|-2(+1C);m-351&1|+|209432166|209432187&offsets|0|-2(+1C)]|TCCTTCATTCCACCGGAGTCC</td><td>>MI0000285|hsa-mir-205&WithFlank&1|+|209432127|209432248</td><td>40</td><td>59</td><td>20(+1C)</td><td>TCCTTCATTCCACCGGAGTCC</td><td>350.9</td><td>3.6</td><td>0.7</td><td>5.0</td><td>4.6E-07</td><td>7.1E-07</td><td>BT20/MB468</td><td>NUC</td></tr>
<tr><td>>hg38_21_+_8433172_8446622_RNA45SN1_45S_50ntFlanks@9792.9815.24|GGCGTGGAGCCGGGCGTGGAATGC</td><td>>hg38_21_+_8433172_8446622_RNA45SN1_45S_50ntFlanks</td><td>9792</td><td>9815</td><td>24</td><td>GGCGTGGAGCCGGGCGTGGAATGC</td><td>427.9</td><td>12.1</td><td>2.2</td><td>5.6</td><td>2.0E-08</td><td>7.1E-07</td><td>BT20/MB231</td><td>MP</td></tr>
<tr><td>>hg38_21_+_8433172_8446622_RNA45SN1_45S_50ntFlanks@3802.3830.29|CATTAAATCAGTTATGGTTCCTTTGGTCG</td><td>>hg38_21_+_8433172_8446622_RNA45SN1_45S_50ntFlanks</td><td>3802</td><td>3830</td><td>29</td><td>CATTAAATCAGTTATGGTTCCTTTGGTCG</td><td>70.5</td><td>-9.2</td><td>1.7</td><td>-5.3</td><td>9.7E-08</td><td>7.1E-07</td><td>BT20/MB468</td><td>MP</td></tr>
<tr><td>>trna20_GluTTC_2_-_131094701_131094772@1.31.31,trna5_GluTTC_13_-_41634874_41634945@1.31.31|TCCCATATGGTCTAGCGGTTAGGATTCCTGG</td><td>>trna20_GluTTC_2_-_131094701_131094772</td><td>1</td><td>31</td><td>31</td><td>TCCCATATGGTCTAGCGGTTAGGATTCCTGG</td><td>330.5</td><td>-5.6</td><td>1.0</td><td>-5.5</td><td>3.5E-08</td><td>7.1E-07</td><td>BT20/MB231</td><td>MITO</td></tr>
<tr><td>>hg38_21_+_8433172_8446622_RNA45SN1_45S_50ntFlanks@10387.10407.21|TGTGAACAGCAGTTGAACATG</td><td>>hg38_21_+_8433172_8446622_RNA45SN1_45S_50ntFlanks</td><td>10387</td><td>10407</td><td>21</td><td>TGTGAACAGCAGTTGAACATG</td><td>307.2</td><td>-5.6</td><td>1.1</td><td>-5.1</td><td>4.4E-07</td><td>7.1E-07</td><td>MB231/MB468</td><td>MITO</td></tr>
<tr><td>>hg38_21_+_8433172_8446622_RNA45SN1_45S_50ntFlanks@10796.10818.23|GGCATGTTGGAACAATGTAGGTA</td><td>>hg38_21_+_8433172_8446622_RNA45SN1_45S_50ntFlanks</td><td>10796</td><td>10818</td><td>23</td><td>GGCATGTTGGAACAATGTAGGTA</td><td>633.1</td><td>-5.8</td><td>1.2</td><td>-5.1</td><td>4.4E-07</td><td>7.1E-07</td><td>MB231/MB468</td><td>MITO</td></tr>
<tr><td>>MI0000767|hsa-mir-365a&WithFlank&16|+|14309279|14309377@62.81.20(+2U);MI0000769|hsa-mir-365b&WithFlank&17|+|31575405|31575527@74.93.20(+2U)&[MIMAT0000710&hsa-miR-365a-3p&offsets|0|-2(+2U);m-176&16|+|14309340|14309361&offsets|0|-2(+2U)];[MIMAT0022834&hsa-miR-365b-3p&offsets|0|-2(+2U);m-175&17|+|31575478|31575499&offsets|0|-2(+2U)]|TAATGCCCCTAAAAATCCTTTT</td><td>>MI0000767|hsa-mir-365a&WithFlank&16|+|14309279|14309377</td><td>62</td><td>81</td><td>20(+2U)</td><td>TAATGCCCCTAAAAATCCTTTT</td><td>289.3</td><td>3.7</td><td>0.7</td><td>5.5</td><td>4.6E-08</td><td>7.2E-07</td><td>BT20/MB231</td><td>NUC</td></tr>
<tr><td>>MI0003628|hsa-mir-615&WithFlank&12|+|54033944|54034051@67.86.20(+1A)&[MIMAT0003283&hsa-miR-615-3p&offsets|0|-2(+1A);m-495&12|+|54034010|54034030&offsets|0|-1(+1A)]|TCCGAGCCTGGGTCTCCCTCA</td><td>>MI0003628|hsa-mir-615&WithFlank&12|+|54033944|54034051</td><td>67</td><td>86</td><td>20(+1A)</td><td>TCCGAGCCTGGGTCTCCCTCA</td><td>214.7</td><td>5.9</td><td>1.1</td><td>5.5</td><td>4.7E-08</td><td>7.2E-07</td><td>BT20/MB231</td><td>NUC</td></tr>
<tr><td>>hg38_21_+_8433172_8446622_RNA45SN1_45S_50ntFlanks@9566.9597.32|GACGTGCAAATCGGTCGTCCGACCTGGGTATA</td><td>>hg38_21_+_8433172_8446622_RNA45SN1_45S_50ntFlanks</td><td>9566</td><td>9597</td><td>32</td><td>GACGTGCAAATCGGTCGTCCGACCTGGGTATA</td><td>1269.6</td><td>-3.3</td><td>0.7</td><td>-5.1</td><td>4.1E-07</td><td>7.2E-07</td><td>MB231/MB468</td><td>CYTO</td></tr>
<tr><td>>hg38_21_+_8433172_8446622_RNA45SN1_45S_50ntFlanks@12456.12500.45|CCAAGCGTTGGATTGTTCACCCACTAATAGGGAACGTGAGCTGGG</td><td>>hg38_21_+_8433172_8446622_RNA45SN1_45S_50ntFlanks</td><td>12456</td><td>12500</td><td>45</td><td>CCAAGCGTTGGATTGTTCACCCACTAATAGGGAACGTGAGCTGGG</td><td>302.7</td><td>-7.5</td><td>1.5</td><td>-5.0</td><td>4.4E-07</td><td>7.2E-07</td><td>MB231/MB468</td><td>MITO</td></tr>
<tr><td>>hg38_21_+_8433172_8446622_RNA45SN1_45S_50ntFlanks@13001.13027.27|GCGATCTATTGAAAGTCAGCCCTCGAC</td><td>>hg38_21_+_8433172_8446622_RNA45SN1_45S_50ntFlanks</td><td>13001</td><td>13027</td><td>27</td><td>GCGATCTATTGAAAGTCAGCCCTCGAC</td><td>382.4</td><td>-5.3</td><td>1.0</td><td>-5.0</td><td>4.4E-07</td><td>7.2E-07</td><td>MB231/MB468</td><td>MITO</td></tr>
<tr><td>>hg38_21_+_8433172_8446622_RNA45SN1_45S_50ntFlanks@5470.5514.45|CCACGGCCCTGGCGGAGCGCTGAGAAGACGGTCGAACTTGACTAT</td><td>>hg38_21_+_8433172_8446622_RNA45SN1_45S_50ntFlanks</td><td>5470</td><td>5514</td><td>45</td><td>CCACGGCCCTGGCGGAGCGCTGAGAAGACGGTCGAACTTGACTAT</td><td>244.3</td><td>-11.8</td><td>2.3</td><td>-5.1</td><td>3.7E-07</td><td>7.2E-07</td><td>MB231/MB468</td><td>MP</td></tr>
<tr><td>>hg38_21_+_8433172_8446622_RNA45SN1_45S_50ntFlanks@5515.5535.21|CTAGAGGAAGTAAAAGTCGTA</td><td>>hg38_21_+_8433172_8446622_RNA45SN1_45S_50ntFlanks</td><td>5515</td><td>5535</td><td>21</td><td>CTAGAGGAAGTAAAAGTCGTA</td><td>630.9</td><td>-7.6</td><td>1.4</td><td>-5.3</td><td>9.8E-08</td><td>7.2E-07</td><td>BT20/MB468</td><td>MP</td></tr>
<tr><td>>MI0000471|hsa-mir-126&WithFlank&9|+|136670596|136670692@59.77.19&[MIMAT0000445&hsa-miR-126-3p&offsets|+1|-2;m-40&9|+|136670653|136670674&offsets|+1|-2]|CGTACCGTGAGTAATAATG</td><td>>MI0000471|hsa-mir-126&WithFlank&9|+|136670596|136670692</td><td>59</td><td>77</td><td>19</td><td>CGTACCGTGAGTAATAATG</td><td>82.4</td><td>9.6</td><td>1.9</td><td>5.1</td><td>3.7E-07</td><td>7.2E-07</td><td>MB231/MB468</td><td>MP</td></tr>
<tr><td>>hg38_21_+_8433172_8446622_RNA45SN1_45S_50ntFlanks@8114.8139.26|GCGCGGGACATGTGGCGTACGGAAGA</td><td>>hg38_21_+_8433172_8446622_RNA45SN1_45S_50ntFlanks</td><td>8114</td><td>8139</td><td>26</td><td>GCGCGGGACATGTGGCGTACGGAAGA</td><td>128.1</td><td>-6.4</td><td>1.3</td><td>-5.1</td><td>4.2E-07</td><td>7.2E-07</td><td>MB231/MB468</td><td>NUC</td></tr>
</table>

| | | | | | | | | | | | | | |
|---|---|---|---|---|---|---|---|---|---|---|---|---|---|
| >hg38_21_+_8433172_8446622_RNA45SN1_45S_50ntFlanks@6748.6770.23\|CTTCGAACGCACTTGCGGCCCCG | >hg38_21_+_8433172_8446622_RNA45SN1_45S_50ntFlanks | 6748 | 6770 | 23 | CTTCGAACGCACTTGCGGCCCCG | 152.7 | -6.3 | 1.3 | -5.1 | 4.2E-07 | 7.2E-07 | MB231/MB468 | NUC |
| >am_trna128_GlyGCC_6_-_27870686_27870756@1.31.31,trna133_GlyCCC_1_-_16872434_16872504@1.31.31,trna18_GlyGCC_16_+_70822597_70822667@1.31.31,trna19_GlyGCC_16_+_70823410_70823480@1.31.31,trna19_GlyGCC_2_-_157257659_157257729@1.31.31,trna24_GlyGCC_16_-_70812942_70813012@1.31.31,trna25_GlyGCC_16_-_70812114_70812184@1.31.31,trna4_GlyCCC_1_+_17188416_17188486@1.31.31,trna5_GlyGCC_17_+_8029064_8029134@1.31.31,trna68_GlyGCC_1_-_161493637_161493707@1.31.31\|GCATTGGTGGTTCAGTGGTAGAATTCTCGCC | >am_trna128_GlyGCC_6_-_27870686_27870756 | 1 | 31 | 31 | GCATTGGTGGTTCAGTGGTAGAATTCTCGCC | 16603.7 | 5.4 | 1.1 | 5.1 | 4.2E-07 | 7.2E-07 | MB231/MB468 | NUC |
| >hg38_21_+_8433172_8446622_RNA45SN1_45S_50ntFlanks@3785.3802.18\|TGAAACTGCGAATGGCTC | >hg38_21_+_8433172_8446622_RNA45SN1_45S_50ntFlanks | 3785 | 3802 | 18 | TGAAACTGCGAATGGCTC | 265.7 | -6.8 | 1.4 | -5.0 | 4.4E-07 | 7.2E-07 | MB231/MB468 | MITO |
| >hg38_21_+_8433172_8446622_RNA45SN1_45S_50ntFlanks@5084.5107.24\|CGAACGAGACTCTGGCATGCTAAC | >hg38_21_+_8433172_8446622_RNA45SN1_45S_50ntFlanks | 5084 | 5107 | 24 | CGAACGAGACTCTGGCATGCTAAC | 544.2 | -8.2 | 1.6 | -5.0 | 4.4E-07 | 7.2E-07 | MB231/MB468 | MITO |
| >hg38_21_+_8433172_8446622_RNA45SN1_45S_50ntFlanks@6651.6688.38\|CGACTCTTAGCGGTGGATCACTCGGCTCGTGCGTCGAT | >hg38_21_+_8433172_8446622_RNA45SN1_45S_50ntFlanks | 6651 | 6688 | 38 | CGACTCTTAGCGGTGGATCACTCGGCTCGTGCGTCGAT | 3160.9 | 5.1 | 1.0 | 5.1 | 3.6E-07 | 7.2E-07 | BT20/MB468 | CYTO |
| >hg38_21_+_8433172_8446622_RNA45SN1_45S_50ntFlanks@9308.9338.31\|AGGAGTCTAACACGTGCGCGAGTCGGGGGCT | >hg38_21_+_8433172_8446622_RNA45SN1_45S_50ntFlanks | 9308 | 9338 | 31 | AGGAGTCTAACACGTGCGCGAGTCGGGGGCT | 3287.2 | -6.9 | 1.3 | -5.3 | 9.8E-08 | 7.2E-07 | BT20/MB468 | MP |
| >hg38_21_+_8433172_8446622_RNA45SN1_45S_50ntFlanks@9929.9956.28\|GATATAGACAGCAGGACGGTGGCCATGG | >hg38_21_+_8433172_8446622_RNA45SN1_45S_50ntFlanks | 9929 | 9956 | 28 | GATATAGACAGCAGGACGGTGGCCATGG | 1156.5 | -5.6 | 1.1 | -5.1 | 3.7E-07 | 7.2E-07 | MB231/MB468 | MP |
| >hg38_21_+_8433172_8446622_RNA45SN1_45S_50ntFlanks@11092.11114.23\|CGGCGGGGGGAGAAGGGTCGGGG | >hg38_21_+_8433172_8446622_RNA45SN1_45S_50ntFlanks | 11092 | 11114 | 23 | CGGCGGGGGGAGAAGGGTCGGGG | 86.4 | -9.5 | 1.8 | -5.3 | 9.9E-08 | 7.2E-07 | BT20/MB468 | MP |
| >hg38_21_+_8433172_8446622_RNA45SN1_45S_50ntFlanks@11568.11592.25\|GCAGCCGACTTAGAACTGGTGCGGA | >hg38_21_+_8433172_8446622_RNA45SN1_45S_50ntFlanks | 11568 | 11592 | 25 | GCAGCCGACTTAGAACTGGTGCGGA | 320.4 | -2.9 | 0.6 | -5.1 | 3.6E-07 | 7.2E-07 | BT20/MB468 | CYTO |
| >hg38_21_+_8433172_8446622_RNA45SN1_45S_50ntFlanks@12407.12439.33\|TTTTGATCCTTCGATGTCGGCTCTTCCTATCAT | >hg38_21_+_8433172_8446622_RNA45SN1_45S_50ntFlanks | 12407 | 12439 | 33 | TTTTGATCCTTCGATGTCGGCTCTTCCTATCAT | 240.5 | 4.0 | 0.8 | 5.1 | 4.1E-07 | 7.2E-07 | MB231/MB468 | CYTO |
| >hg38_21_+_8433172_8446622_RNA45SN1_45S_50ntFlanks@11774.11809.36\|AGTGACGCGCATGAATGGATGAACGAGATTCCCACT | >hg38_21_+_8433172_8446622_RNA45SN1_45S_50ntFlanks | 11774 | 11809 | 36 | AGTGACGCGCATGAATGGATGAACGAGATTCCCACT | 59.1 | 9.6 | 1.8 | 5.3 | 9.9E-08 | 7.2E-07 | BT20/MB468 | MP |
| >hg38_21_+_8433172_8446622_RNA45SN1_45S_50ntFlanks@11641.11685.45\|CGGCGGGTGTTGACGCGATGTGATTTCTGCCCAGTGCTCTGAATG | >hg38_21_+_8433172_8446622_RNA45SN1_45S_50ntFlanks | 11641 | 11685 | 45 | CGGCGGGTGTTGACGCGATGTGATTTCTGCCCAGTGCTCTGAATG | 519.0 | -4.0 | 0.8 | -5.1 | 4.1E-07 | 7.2E-07 | MB231/MB468 | CYTO |
| >hg38_21_+_8433172_8446622_RNA45SN1_45S_50ntFlanks@11670.11693.24\|CCCAGTGCTCTGAATGTCAAAGTG | >hg38_21_+_8433172_8446622_RNA45SN1_45S_50ntFlanks | 11670 | 11693 | 24 | CCCAGTGCTCTGAATGTCAAAGTG | 384.8 | -4.3 | 0.9 | -5.0 | 4.5E-07 | 7.3E-07 | MB231/MB468 | MITO |
| >hg38_21_+_8433172_8446622_RNA45SN1_45S_50ntFlanks@11582.11602.21\|ACTGGTGCGGACCAGGGGAAT | >hg38_21_+_8433172_8446622_RNA45SN1_45S_50ntFlanks | 11582 | 11602 | 21 | ACTGGTGCGGACCAGGGGAAT | 606.9 | -3.5 | 0.7 | -5.1 | 3.7E-07 | 7.3E-07 | MB231/MB468 | MP |
| >hg38_21_+_8433172_8446622_RNA45SN1_45S_50ntFlanks@12458.12487.30\|AAGCGTTGGATTGTTCACCCACTAATAGGG | >hg38_21_+_8433172_8446622_RNA45SN1_45S_50ntFlanks | 12458 | 12487 | 30 | AAGCGTTGGATTGTTCACCCACTAATAGGG | 1228.3 | -5.8 | 1.2 | -5.1 | 4.3E-07 | 7.3E-07 | MB231/MB468 | NUC |
| >hg38_21_+_8433172_8446622_RNA45SN1_45S_50ntFlanks@4727.4767.41\|AATCAAGAACGAAAGTCGGAGGTTCGAAGACGATCAGATAC | >hg38_21_+_8433172_8446622_RNA45SN1_45S_50ntFlanks | 4727 | 4767 | 41 | AATCAAGAACGAAAGTCGGAGGTTCGAAGACGATCAGATAC | 59.8 | 9.1 | 1.8 | 5.1 | 3.7E-07 | 7.3E-07 | MB231/MB468 | MP |
| >hg38_21_+_8433172_8446622_RNA45SN1_45S_50ntFlanks@12660.12681.22\|TGAACGCCTCTAAGTCAGAATC | >hg38_21_+_8433172_8446622_RNA45SN1_45S_50ntFlanks | 12660 | 12681 | 22 | TGAACGCCTCTAAGTCAGAATC | 94.7 | -9.6 | 1.8 | -5.3 | 9.9E-08 | 7.3E-07 | BT20/MB468 | MP |
| >MI0005559\|hsa-mir-744&WithFlank&17\|+\|12081893\|12082002@17.36.20&[MIMAT0004945&hsa-miR-744-5p&offsets\|0\|-2;m-172&17\|+\|12081909\|12081929&offsets\|0\|-1]\|TGCGGGGCTAGGGCTAACAG | >MI0005559\|hsa-mir-744&WithFlank&17\|+\|12081893\|12082002 | 17 | 36 | 20 | TGCGGGGCTAGGGCTAACAG | 58.7 | 9.6 | 1.8 | 5.3 | 1.0E-07 | 7.3E-07 | BT20/MB468 | MP |
| >hg38_21_+_8433172_8446622_RNA45SN1_45S_50ntFlanks@11569.11587.19\|CAGCCGACTTAGAACTGGT | >hg38_21_+_8433172_8446622_RNA45SN1_45S_50ntFlanks | 11569 | 11587 | 19 | CAGCCGACTTAGAACTGGT | 254.6 | -5.4 | 1.1 | -5.1 | 3.7E-07 | 7.3E-07 | MB231/MB468 | MP |
| >hg38_21_+_8433172_8446622_RNA45SN1_45S_50ntFlanks@9739.9763.25\|AAACGATCTCAACCTATTCTCAAAC | >hg38_21_+_8433172_8446622_RNA45SN1_45S_50ntFlanks | 9739 | 9763 | 25 | AAACGATCTCAACCTATTCTCAAAC | 420.7 | -6.3 | 1.2 | -5.0 | 4.7E-07 | 7.3E-07 | BT20/MB468 | NUC |
| >hg38_21_+_8433172_8446622_RNA45SN1_45S_50ntFlanks@12619.12653.35\|AGGAGCCAATGGGGCGAAGCTACCATCTGTGGGAT | >hg38_21_+_8433172_8446622_RNA45SN1_45S_50ntFlanks | 12619 | 12653 | 35 | AGGAGCCAATGGGGCGAAGCTACCATCTGTGGGAT | 701.3 | -3.3 | 0.6 | -5.1 | 3.2E-07 | 7.3E-07 | BT20/MB468 | MITO |
| >am_trnaMT_GluTTC_MT_-_14674_14742@27.55.29,trnalookalike8_GluTTC_5_-_93905172_93905240@27.55.29\|GTTTTTCATATCATTGGTCGTGGTTGTAG | >am_trnaMT_GluTTC_MT_-_14674_14742 | 27 | 55 | 29 | GTTTTTCATATCATTGGTCGTGGTTGTAG | 195.5 | -7.0 | 1.4 | -5.1 | 3.8E-07 | 7.3E-07 | MB231/MB468 | MP |
| >hg38_21_+_8433172_8446622_RNA45SN1_45S_50ntFlanks@12991.13011.21\|CTCCCTCGCTGCGATCTATTG | >hg38_21_+_8433172_8446622_RNA45SN1_45S_50ntFlanks | 12991 | 13011 | 21 | CTCCCTCGCTGCGATCTATTG | 62.3 | -9.0 | 1.7 | -5.3 | 1.0E-07 | 7.3E-07 | BT20/MB468 | MP |
| >hg38_21_+_8433172_8446622_RNA45SN1_45S_50ntFlanks@12641.12659.19\|CCATCTGTGGGATTATGAC | >hg38_21_+_8433172_8446622_RNA45SN1_45S_50ntFlanks | 12641 | 12659 | 19 | CCATCTGTGGGATTATGAC | 640.1 | -5.3 | 1.0 | -5.1 | 3.7E-07 | 7.3E-07 | BT20/MB468 | CYTO |
| >hg38_21_+_8433172_8446622_RNA45SN1_45S_50ntFlanks@6779.6803.25\|CCGGGGCTACGCCTGTCTGAGCGTC | >hg38_21_+_8433172_8446622_RNA45SN1_45S_50ntFlanks | 6779 | 6803 | 25 | CCGGGGCTACGCCTGTCTGAGCGTC | 2157.2 | -6.1 | 1.2 | -5.0 | 4.5E-07 | 7.3E-07 | MB231/MB468 | MITO |
| >hg38_21_+_8433172_8446622_RNA45SN1_45S_50ntFlanks@6645.6673.29\|CTCGTACGACTCTTAGCGGTGGATCACTC | >hg38_21_+_8433172_8446622_RNA45SN1_45S_50ntFlanks | 6645 | 6673 | 29 | CTCGTACGACTCTTAGCGGTGGATCACTC | 352.8 | 4.1 | 0.8 | 5.3 | 1.0E-07 | 7.3E-07 | BT20/MB468 | MP |
| >MI0000273\|hsa-mir-183&WithFlank&7\|-\|129774899\|129775020@34.55.22(+1A)&[MIMAT0000261&hsa-miR-183-5p&offsets\|+1\|+1(+1A);m-41&7\|-\|129774966\|129774987&offsets\|0\|0(+1A)]\|ATGGCACTGGTAGAATTCACTGA | >MI0000273\|hsa-mir-183&WithFlank&7\|-\|129774899\|129775020 | 34 | 55 | 22(+1A) | ATGGCACTGGTAGAATTCACTGA | 208.9 | 4.6 | 0.8 | 5.5 | 2.9E-08 | 7.3E-07 | BT20/MB231 | CYTO |
| >MI0000542\|hsa-mir-320a&WithFlank&8\|-\|22244960\|22245043@48.68.21(+1G)&[MIMAT0000510&hsa-miR-320a-3p&offsets\|0\|-1(+1G);m-60&8\|-\|22244975\|22244996&offsets\|0\|-1(+1G)]\|AAAAGCTGGGTTGAGAGGGCGG | >MI0000542\|hsa-mir-320a&WithFlank&8\|-\|22244960\|22245043 | 48 | 68 | 21(+1G) | AAAAGCTGGGTTGAGAGGGCGG | 1981.7 | 5.1 | 0.9 | 5.5 | 2.9E-08 | 7.3E-07 | BT20/MB231 | CYTO |
| >MI0003137\|hsa-mir-193b&WithFlank&16\|+\|14303961\|14304055@57.78.22&[MIMAT0002819&hsa-miR-193b-3p&offsets\|0\|0;m-166&16\|+\|14304017\|14304038&offsets\|0\|0]\|AACTGGCCCTCAAAGTCCCGCT | >MI0003137\|hsa-mir-193b&WithFlank&16\|+\|14303961\|14304055 | 57 | 78 | 22 | AACTGGCCCTCAAAGTCCCGCT | 2633.7 | 4.5 | 0.8 | 5.5 | 2.9E-08 | 7.3E-07 | BT20/MB231 | CYTO |
| >hg38_21_+_8433172_8446622_RNA45SN1_45S_50ntFlanks@10423.10453.31\|AGAGATGGGCGAGCGCCGTTCCGAAGGGACG | >hg38_21_+_8433172_8446622_RNA45SN1_45S_50ntFlanks | 10423 | 10453 | 31 | AGAGATGGGCGAGCGCCGTTCCGAAGGGACG | 491.5 | -3.2 | 0.6 | -5.1 | 3.7E-07 | 7.3E-07 | BT20/MB468 | CYTO |
| >hg38_21_+_8433172_8446622_RNA45SN1_45S_50ntFlanks@8463.8497.35\|CCGTGTCGGCGGCCCGGCGGATCTTTCCCGCCCCC | >hg38_21_+_8433172_8446622_RNA45SN1_45S_50ntFlanks | 8463 | 8497 | 35 | CCGTGTCGGCGGCCCGGCGGATCTTTCCCGCCCCC | 381.8 | -7.7 | 1.5 | -5.0 | 4.7E-07 | 7.3E-07 | BT20/MB468 | NUC |
| >MI0000815\|hsa-mir-339&WithFlank&7\|-\|1022927\|1023032@56.77.22(+1A)&[MIMAT0004702&hsa-miR-339-3p&offsets\|0\|-1(+1A);m-235&7\|-\|1022957\|1022977&offsets\|0\|+1(+1A)]\|TGAGCGCCTCGACGACAGAGCCA | >MI0000815\|hsa-mir-339&WithFlank&7\|-\|1022927\|1023032 | 56 | 77 | 22(+1A) | TGAGCGCCTCGACGACAGAGCCA | 128.3 | 10.8 | 2.0 | 5.3 | 1.0E-07 | 7.3E-07 | BT20/MB468 | MP |
| >hg38_21_+_8433172_8446622_RNA45SN1_45S_50ntFlanks@8380.8407.28\|CAAGAGGGCGTGAAACCGTTAAGAGGTA | >hg38_21_+_8433172_8446622_RNA45SN1_45S_50ntFlanks | 8380 | 8407 | 28 | CAAGAGGGCGTGAAACCGTTAAGAGGTA | 339.9 | -7.5 | 1.5 | -5.0 | 4.7E-07 | 7.4E-07 | BT20/MB468 | NUC |
| >hg38_21_+_8433172_8446622_RNA45SN1_45S_50ntFlanks@12616.12653.38\|CTGAGGAGCCAATGGGGCGAAGCTACCATCTGTGGGAT | >hg38_21_+_8433172_8446622_RNA45SN1_45S_50ntFlanks | 12616 | 12653 | 38 | CTGAGGAGCCAATGGGGCGAAGCTACCATCTGTGGGAT | 297.2 | -4.5 | 0.9 | -5.1 | 4.2E-07 | 7.4E-07 | MB231/MB468 | CYTO |
| >hg38_21_+_8433172_8446622_RNA45SN1_45S_50ntFlanks@10344.10381.38\|ATTCAAACGAGAACTTTGAAGGCCGAAGTGGAGAAGGG | >hg38_21_+_8433172_8446622_RNA45SN1_45S_50ntFlanks | 10344 | 10381 | 38 | ATTCAAACGAGAACTTTGAAGGCCGAAGTGGAGAAGGG | 65.0 | 9.8 | 1.8 | 5.3 | 1.0E-07 | 7.4E-07 | BT20/MB468 | MP |
| >hg38_21_+_8433172_8446622_RNA45SN1_45S_50ntFlanks@10685.10706.22\|GGCGGCGTCCGGTGAGCTCTCG | >hg38_21_+_8433172_8446622_RNA45SN1_45S_50ntFlanks | 10685 | 10706 | 22 | GGCGGCGTCCGGTGAGCTCTCG | 357.0 | -7.0 | 1.4 | -5.0 | 4.6E-07 | 7.4E-07 | MB231/MB468 | MITO |
| >am_trna119_LysCTT_1_-_145395522_145395594@13.34.22,trna11_LysCTT_5_-_180648979_180649051@13.34.22,trna128_LysCTT_1_-_55423542_55423614@13.34.22,trna13_LysCTT_6_+_26556774_26556846@13.34.22,trna6_LysCTT_19_-_52425396_52425468@13.34.22,trna7_LysCTT_16_+_3225692_3225764@13.34.22,trna9_LysCTT_5_+_180634755_180634827@13.34.22\|CAGTCGGTAGAGCATGAGACTC | >am_trna119_LysCTT_1_-_145395522_145395594 | 13 | 34 | 22 | CAGTCGGTAGAGCATGAGACTC | 332.5 | 8.1 | 1.6 | 5.0 | 4.6E-07 | 7.4E-07 | MB231/MB468 | MITO |
| >MI0000809\|hsa-mir-151a&WithFlank&8\|-\|140732558\|140732659@51.71.21(+1A)&[MIMAT0000757&hsa-miR-151a-3p&offsets\|-2\|-2(+1A);m-30&8\|-\|140732587\|140732607&offsets\|-2\|-2(+1A)]\|TACTAGACTGAAGCTCCTTGAA | >MI0000809\|hsa-mir-151a&WithFlank&8\|-\|140732558\|140732659 | 51 | 71 | 21(+1A) | TACTAGACTGAAGCTCCTTGAA | 98.7 | 10.4 | 1.9 | 5.3 | 1.0E-07 | 7.4E-07 | BT20/MB468 | MP |
| >hg38_21_+_8433172_8446622_RNA45SN1_45S_50ntFlanks@9032.9055.24\|CCTCGTGGGGGGGCCGGGCCACCC | >hg38_21_+_8433172_8446622_RNA45SN1_45S_50ntFlanks | 9032 | 9055 | 24 | CCTCGTGGGGGGGCCGGGCCACCC | 91.3 | -9.6 | 1.8 | -5.3 | 1.0E-07 | 7.4E-07 | BT20/MB468 | MP |
| >hg38_21_+_8433172_8446622_RNA45SN1_45S_50ntFlanks@4753.4786.34\|AAGACGATCAGATACCGTCGTAGTTCCGACCATA | >hg38_21_+_8433172_8446622_RNA45SN1_45S_50ntFlanks | 4753 | 4786 | 34 | AAGACGATCAGATACCGTCGTAGTTCCGACCATA | 1082.1 | -6.2 | 1.2 | -5.0 | 4.6E-07 | 7.4E-07 | MB231/MB468 | MITO |
| >hg38_21_+_8433172_8446622_RNA45SN1_45S_50ntFlanks@11564.11583.20\|CCTAGCAGCCGACTTAGAAC | >hg38_21_+_8433172_8446622_RNA45SN1_45S_50ntFlanks | 11564 | 11583 | 20 | CCTAGCAGCCGACTTAGAAC | 389.6 | -4.2 | 0.8 | -5.0 | 4.6E-07 | 7.4E-07 | MB231/MB468 | MITO |
| >hg38_21_+_8433172_8446622_RNA45SN1_45S_50ntFlanks@9758.9787.30\|TCAAACTTTAAATGGGTAAGAAGCCCGGCT | >hg38_21_+_8433172_8446622_RNA45SN1_45S_50ntFlanks | 9758 | 9787 | 30 | TCAAACTTTAAATGGGTAAGAAGCCCGGCT | 159.5 | -10.4 | 2.0 | -5.3 | 1.0E-07 | 7.4E-07 | BT20/MB468 | MP |
| >hg38_21_+_8433172_8446622_RNA45SN1_45S_50ntFlanks@11663.11685.23\|ATTTCTGCCCAGTGCTCTGAATG | >hg38_21_+_8433172_8446622_RNA45SN1_45S_50ntFlanks | 11663 | 11685 | 23 | ATTTCTGCCCAGTGCTCTGAATG | 387.1 | -3.9 | 0.8 | -5.1 | 4.4E-07 | 7.4E-07 | MB231/MB468 | NUC |
| >hg38_21_+_8433172_8446622_RNA45SN1_45S_50ntFlanks@10346.10377.32\|TCAAACGAGAACTTTGAAGGCCGAAGTGGAGA | >hg38_21_+_8433172_8446622_RNA45SN1_45S_50ntFlanks | 10346 | 10377 | 32 | TCAAACGAGAACTTTGAAGGCCGAAGTGGAGA | 240.6 | -3.6 | 0.7 | -5.0 | 4.6E-07 | 7.4E-07 | MB231/MB468 | MITO |
| >MI0003776\|hsa-mir-320b-1&WithFlank&1\|+\|116671740\|116671823@48.68.21;MI0003839\|hsa-mir-320b-2&WithFlank&1\|-\|224257034\|224257116@47.67.21&[MIMAT0005792&hsa-miR-320b&offsets\|0\|-1;m-183&1\|+\|116671787\|116671806&offsets\|0\|+1];[MIMAT0005792_1&hsa-miR-320b&offsets\|0\|-1;m-183&1\|-\|224257051\|224257070&offsets\|0\|+1]\|AAAAGCTGGGTTGAGAGGGCA | >MI0003776\|hsa-mir-320b-1&WithFlank&1\|+\|116671740\|116671823 | 48 | 68 | 21 | AAAAGCTGGGTTGAGAGGGCA | 4495.2 | 7.7 | 1.4 | 5.3 | 1.0E-07 | 7.4E-07 | BT20/MB468 | MP |

<table>
<tr><th></th><th></th><th></th><th></th><th></th><th></th><th></th><th></th><th></th><th></th><th></th><th></th><th></th><th></th></tr>
<tr><td>>hg38_MT_+_598_1651_RNR1_12S_50ntFlanks@429.469.41|AAGTGGCTTTAACATATCTGAACACACAATAGCTAAGACCC</td><td>>hg38_MT_+_598_1651_RNR1_12S_50ntFlanks</td><td>429</td><td>469</td><td>41</td><td>AAGTGGCTTTAACATATCTGAACACACAATAGCTAAGACCC</td><td>108.2</td><td>10.5</td><td>2.0</td><td>5.3</td><td>1.0E-07</td><td>7.4E-07</td><td>BT20/MB468</td><td>MP</td></tr>
<tr><td>>hg38_21_+_8433172_8446622_RNA45SN1_45S_50ntFlanks@9201.9220.20|CTCTCGGGGCCACGCGCGCG</td><td>>hg38_21_+_8433172_8446622_RNA45SN1_45S_50ntFlanks</td><td>9201</td><td>9220</td><td>20</td><td>CTCTCGGGGCCACGCGCGCG</td><td>457.5</td><td>-7.6</td><td>1.4</td><td>-5.3</td><td>1.0E-07</td><td>7.4E-07</td><td>BT20/MB468</td><td>MP</td></tr>
<tr><td>>hg38_21_+_8433172_8446622_RNA45SN1_45S_50ntFlanks@9559.9597.39|CGGTCCTGACGTGCAAATCGGTCGTCCGACCTGGGTATA</td><td>>hg38_21_+_8433172_8446622_RNA45SN1_45S_50ntFlanks</td><td>9559</td><td>9597</td><td>39</td><td>CGGTCCTGACGTGCAAATCGGTCGTCCGACCTGGGTATA</td><td>116.0</td><td>-8.9</td><td>1.8</td><td>-5.1</td><td>3.3E-07</td><td>7.4E-07</td><td>BT20/MB468</td><td>MITO</td></tr>
<tr><td>>hg38_21_+_8433172_8446622_RNA45SN1_45S_50ntFlanks@5485.5516.32|AGCGCTGAGAAGACGGTCGAACTTGACTATCT</td><td>>hg38_21_+_8433172_8446622_RNA45SN1_45S_50ntFlanks</td><td>5485</td><td>5516</td><td>32</td><td>AGCGCTGAGAAGACGGTCGAACTTGACTATCT</td><td>1552.2</td><td>-4.1</td><td>0.8</td><td>-5.0</td><td>4.6E-07</td><td>7.4E-07</td><td>MB231/MB468</td><td>MITO</td></tr>
<tr><td>>hg38_21_+_8433172_8446622_RNA45SN1_45S_50ntFlanks@5207.5235.29|CTTAGATGTCCGGGGCTGCACGCGCGCTA</td><td>>hg38_21_+_8433172_8446622_RNA45SN1_45S_50ntFlanks</td><td>5207</td><td>5235</td><td>29</td><td>CTTAGATGTCCGGGGCTGCACGCGCGCTA</td><td>68.5</td><td>-10.0</td><td>2.0</td><td>-5.1</td><td>3.8E-07</td><td>7.4E-07</td><td>MB231/MB468</td><td>MP</td></tr>
<tr><td>>hg38_21_+_8433172_8446622_RNA45SN1_45S_50ntFlanks@11572.11597.26|CCGACTTAGAACTGGTGCGGACCAGG</td><td>>hg38_21_+_8433172_8446622_RNA45SN1_45S_50ntFlanks</td><td>11572</td><td>11597</td><td>26</td><td>CCGACTTAGAACTGGTGCGGACCAGG</td><td>288.9</td><td>-5.7</td><td>1.1</td><td>-5.0</td><td>4.8E-07</td><td>7.4E-07</td><td>BT20/MB468</td><td>NUC</td></tr>
<tr><td>>am_trna116_GluCTC_1_-_145399233_145399304@1.27.27,trna59_GluCTC_1_+_249168447_249168518@1.27.27,trna71_GluCTC_1_-_161439189_161439260@1.27.27,trna74_GluCTC_1_-_161431809_161431880@1.27.27,trna77_GluCTC_1_-_161424398_161424469@1.27.27,trna77_GluCTC_6_+_28949976_28950047@1.27.27,trna80_GluCTC_1_-_161417018_161417089@1.27.27,trna87_GluCTC_6_-_126101393_126101464@1.27.27|TCCCTGGTGGTCTAGTGGTTAGGATTC</td><td>>am_trna116_GluCTC_1_-_145399233_145399304</td><td>1</td><td>27</td><td>27</td><td>TCCCTGGTGGTCTAGTGGTTAGGATTC</td><td>3144.6</td><td>-4.0</td><td>0.8</td><td>-5.1</td><td>4.4E-07</td><td>7.4E-07</td><td>MB231/MB468</td><td>NUC</td></tr>
<tr><td>>hg38_21_+_8433172_8446622_RNA45SN1_45S_50ntFlanks@12419.12457.39|GATGTCGGCTCTTCCTATCATTGTGAAGCAGAATTCACC</td><td>>hg38_21_+_8433172_8446622_RNA45SN1_45S_50ntFlanks</td><td>12419</td><td>12457</td><td>39</td><td>GATGTCGGCTCTTCCTATCATTGTGAAGCAGAATTCACC</td><td>733.7</td><td>5.6</td><td>1.1</td><td>5.1</td><td>4.4E-07</td><td>7.4E-07</td><td>MB231/MB468</td><td>NUC</td></tr>
<tr><td>>hg38_21_+_8433172_8446622_RNA45SN1_45S_50ntFlanks@10796.10818.23|GGCATGTTGGAACAATGTAGGTA</td><td>>hg38_21_+_8433172_8446622_RNA45SN1_45S_50ntFlanks</td><td>10796</td><td>10818</td><td>23</td><td>GGCATGTTGGAACAATGTAGGTA</td><td>577.7</td><td>-3.0</td><td>0.6</td><td>-5.1</td><td>3.7E-07</td><td>7.4E-07</td><td>BT20/MB468</td><td>CYTO</td></tr>
<tr><td>>MI0000102|hsa-mir-100&WithFlank&11|-|122152223|122152314@19.36.18(+1C)&[MIMAT0000098&hsa-miR-100-5p&offsets|0|-4(+1C);m-27&11|-|122152276|122152296&offsets|0|-3(+1C)]|AACCCGTAGATCCGAACTC</td><td>>MI0000102|hsa-mir-100&WithFlank&11|-|122152223|122152314</td><td>19</td><td>36</td><td>18(+1C)</td><td>AACCCGTAGATCCGAACTC</td><td>148.7</td><td>-10.6</td><td>1.9</td><td>-5.6</td><td>2.1E-08</td><td>7.5E-07</td><td>BT20/MB231</td><td>MP</td></tr>
<tr><td>>hg38_21_+_8433172_8446622_RNA45SN1_45S_50ntFlanks@12360.12382.23|CAGGGATAACTGGCTTGTGGCGG</td><td>>hg38_21_+_8433172_8446622_RNA45SN1_45S_50ntFlanks</td><td>12360</td><td>12382</td><td>23</td><td>CAGGGATAACTGGCTTGTGGCGG</td><td>186.0</td><td>-5.9</td><td>1.2</td><td>-5.1</td><td>3.7E-07</td><td>7.5E-07</td><td>BT20/MB468</td><td>CYTO</td></tr>
<tr><td>>hg38_21_+_8433172_8446622_RNA45SN1_45S_50ntFlanks@10349.10388.40|AACGAGAACTTTGAAGGCCGAAGTGGAGAAGGGTTCCATG</td><td>>hg38_21_+_8433172_8446622_RNA45SN1_45S_50ntFlanks</td><td>10349</td><td>10388</td><td>40</td><td>AACGAGAACTTTGAAGGCCGAAGTGGAGAAGGGTTCCATG</td><td>1428.9</td><td>-4.0</td><td>0.8</td><td>-5.1</td><td>4.4E-07</td><td>7.5E-07</td><td>MB231/MB468</td><td>NUC</td></tr>
<tr><td>>hg38_MT_+_598_1651_RNR1_12S_50ntFlanks@428.469.42|AAAGTGGCTTTAACATATCTGAACACACAATAGCTAAGACCC</td><td>>hg38_MT_+_598_1651_RNR1_12S_50ntFlanks</td><td>428</td><td>469</td><td>42</td><td>AAAGTGGCTTTAACATATCTGAACACACAATAGCTAAGACCC</td><td>989.4</td><td>9.1</td><td>1.8</td><td>5.1</td><td>3.3E-07</td><td>7.5E-07</td><td>BT20/MB468</td><td>MITO</td></tr>
<tr><td>>hg38_21_+_8433172_8446622_RNA45SN1_45S_50ntFlanks@12218.12241.24|GAGGACAGAAACCTCCCGTGGAGC</td><td>>hg38_21_+_8433172_8446622_RNA45SN1_45S_50ntFlanks</td><td>12218</td><td>12241</td><td>24</td><td>GAGGACAGAAACCTCCCGTGGAGC</td><td>59.3</td><td>9.6</td><td>1.8</td><td>5.3</td><td>1.0E-07</td><td>7.5E-07</td><td>BT20/MB468</td><td>MP</td></tr>
<tr><td>>hg38_21_+_8433172_8446622_RNA45SN1_45S_50ntFlanks@9305.9338.34|CCAAGGAGTCTAACACGTGCGCGAGTCGGGGGCT</td><td>>hg38_21_+_8433172_8446622_RNA45SN1_45S_50ntFlanks</td><td>9305</td><td>9338</td><td>34</td><td>CCAAGGAGTCTAACACGTGCGCGAGTCGGGGGCT</td><td>352.1</td><td>-6.7</td><td>1.3</td><td>-5.0</td><td>4.4E-07</td><td>7.5E-07</td><td>MB231/MB468</td><td>NUC</td></tr>
<tr><td>>hg38_21_+_8433172_8446622_RNA45SN1_45S_50ntFlanks@8443.8471.29|CAACCCGGCGGCGGGTCCGGCCGTGTCGG</td><td>>hg38_21_+_8433172_8446622_RNA45SN1_45S_50ntFlanks</td><td>8443</td><td>8471</td><td>29</td><td>CAACCCGGCGGCGGGTCCGGCCGTGTCGG</td><td>76.8</td><td>-9.3</td><td>1.8</td><td>-5.3</td><td>1.0E-07</td><td>7.5E-07</td><td>BT20/MB468</td><td>MP</td></tr>
<tr><td>>hg38_21_+_8433172_8446622_RNA45SN1_45S_50ntFlanks@12159.12182.24|TTTGACTGGGGCGGTACACCTGTC</td><td>>hg38_21_+_8433172_8446622_RNA45SN1_45S_50ntFlanks</td><td>12159</td><td>12182</td><td>24</td><td>TTTGACTGGGGCGGTACACCTGTC</td><td>220.6</td><td>-6.5</td><td>1.3</td><td>-5.1</td><td>3.8E-07</td><td>7.5E-07</td><td>BT20/MB468</td><td>CYTO</td></tr>
<tr><td>>hg38_21_+_8433172_8446622_RNA45SN1_45S_50ntFlanks@9133.9152.20|TCGGGGGGCGGGGCGGACTG</td><td>>hg38_21_+_8433172_8446622_RNA45SN1_45S_50ntFlanks</td><td>9133</td><td>9152</td><td>20</td><td>TCGGGGGGCGGGGCGGACTG</td><td>1215.7</td><td>-8.3</td><td>1.6</td><td>-5.3</td><td>1.0E-07</td><td>7.5E-07</td><td>BT20/MB468</td><td>MP</td></tr>
<tr><td>>hg38_21_+_8433172_8446622_RNA45SN1_45S_50ntFlanks@11572.11604.33|CCGACTTAGAACTGGTGCGGACCAGGGGAATCC</td><td>>hg38_21_+_8433172_8446622_RNA45SN1_45S_50ntFlanks</td><td>11572</td><td>11604</td><td>33</td><td>CCGACTTAGAACTGGTGCGGACCAGGGGAATCC</td><td>768.4</td><td>-6.4</td><td>1.3</td><td>-5.0</td><td>4.8E-07</td><td>7.5E-07</td><td>BT20/MB468</td><td>NUC</td></tr>
<tr><td>>MI0000542|hsa-mir-320a&WithFlank&8|-|22244960|22245043@49.68.20(+1G)&[MIMAT0000510&hsa-miR-320a-3p&offsets|+1|-1(+1G);m-60&8|-|22244975|22244996&offsets|+1|-1(+1G)]|AAAGCTGGGTTGAGAGGGCGG</td><td>>MI0000542|hsa-mir-320a&WithFlank&8|-|22244960|22245043</td><td>49</td><td>68</td><td>20(+1G)</td><td>AAAGCTGGGTTGAGAGGGCGG</td><td>87.2</td><td>10.2</td><td>1.9</td><td>5.3</td><td>1.0E-07</td><td>7.5E-07</td><td>BT20/MB468</td><td>MP</td></tr>
<tr><td>>hg38_21_+_8433172_8446622_RNA45SN1_45S_50ntFlanks@11569.11586.18|CAGCCGACTTAGAACTGG</td><td>>hg38_21_+_8433172_8446622_RNA45SN1_45S_50ntFlanks</td><td>11569</td><td>11586</td><td>18</td><td>CAGCCGACTTAGAACTGG</td><td>1427.8</td><td>-4.5</td><td>0.9</td><td>-5.0</td><td>4.6E-07</td><td>7.5E-07</td><td>MB231/MB468</td><td>MITO</td></tr>
<tr><td>>hg38_21_+_8433172_8446622_RNA45SN1_45S_50ntFlanks@12712.12755.44|CGGAGCCTCGGTTGGCCTCGGATAGCCGGTCCCCCGCCTGTCCC</td><td>>hg38_21_+_8433172_8446622_RNA45SN1_45S_50ntFlanks</td><td>12712</td><td>12755</td><td>44</td><td>CGGAGCCTCGGTTGGCCTCGGATAGCCGGTCCCCCGCCTGTCCC</td><td>65.2</td><td>-9.9</td><td>1.9</td><td>-5.1</td><td>3.9E-07</td><td>7.5E-07</td><td>MB231/MB468</td><td>MP</td></tr>
<tr><td>>hg38_21_+_8433172_8446622_RNA45SN1_45S_50ntFlanks@10788.10816.29|CAGCCTCTGGCATGTTGGAACAATGTAGG</td><td>>hg38_21_+_8433172_8446622_RNA45SN1_45S_50ntFlanks</td><td>10788</td><td>10816</td><td>29</td><td>CAGCCTCTGGCATGTTGGAACAATGTAGG</td><td>509.3</td><td>-5.7</td><td>1.1</td><td>-5.0</td><td>4.7E-07</td><td>7.5E-07</td><td>MB231/MB468</td><td>MITO</td></tr>
<tr><td>>hg38_21_+_8433172_8446622_RNA45SN1_45S_50ntFlanks@10683.10709.27|CCGGCGGCGTCCGGTGAGCTCTCGCTG</td><td>>hg38_21_+_8433172_8446622_RNA45SN1_45S_50ntFlanks</td><td>10683</td><td>10709</td><td>27</td><td>CCGGCGGCGTCCGGTGAGCTCTCGCTG</td><td>318.1</td><td>-5.6</td><td>1.1</td><td>-5.0</td><td>4.5E-07</td><td>7.5E-07</td><td>MB231/MB468</td><td>NUC</td></tr>
<tr><td>>MI0000103|hsa-mir-101-1&WithFlank&1|-|65058428|65058514@53.73.21(+1U);MI0000739|hsa-mir-101-2&WithFlank&9|+|4850291|4850381@55.75.21(+1U)&[MIMAT0000099&hsa-miR-101-3p&offsets|0|0(+1U);m-17&1|-|65058443|65058462&offsets|0|+1(+1U)];[MIMAT0000099_1&hsa-miR-101-3p&offsets|0|0(+1U);m-16&9|+|4850345|4850364&offsets|0|+1(+1U)]|TACAGTACTGTGATAACTGAAT</td><td>>MI0000103|hsa-mir-101-1&WithFlank&1|-|65058428|65058514</td><td>53</td><td>73</td><td>21(+1U)</td><td>TACAGTACTGTGATAACTGAAT</td><td>575.3</td><td>8.2</td><td>1.6</td><td>5.0</td><td>4.7E-07</td><td>7.5E-07</td><td>MB231/MB468</td><td>MITO</td></tr>
<tr><td>>hg38_21_+_8433172_8446622_RNA45SN1_45S_50ntFlanks@10407.10429.23|GGGTCAGTCGGTCCTGAGAGATG</td><td>>hg38_21_+_8433172_8446622_RNA45SN1_45S_50ntFlanks</td><td>10407</td><td>10429</td><td>23</td><td>GGGTCAGTCGGTCCTGAGAGATG</td><td>254.0</td><td>-4.4</td><td>0.9</td><td>-5.1</td><td>4.3E-07</td><td>7.6E-07</td><td>MB231/MB468</td><td>CYTO</td></tr>
<tr><td>>hg38_21_+_8433172_8446622_RNA45SN1_45S_50ntFlanks@4449.4466.18|CCTCTCGGCGCCCCCTCG</td><td>>hg38_21_+_8433172_8446622_RNA45SN1_45S_50ntFlanks</td><td>4449</td><td>4466</td><td>18</td><td>CCTCTCGGCGCCCCCTCG</td><td>912.3</td><td>-7.1</td><td>1.4</td><td>-5.0</td><td>4.9E-07</td><td>7.6E-07</td><td>BT20/MB468</td><td>NUC</td></tr>
<tr><td>>hg38_21_+_8433172_8446622_RNA45SN1_45S_50ntFlanks@6645.6668.24|CTCGTACGACTCTTAGCGGTGGAT</td><td>>hg38_21_+_8433172_8446622_RNA45SN1_45S_50ntFlanks</td><td>6645</td><td>6668</td><td>24</td><td>CTCGTACGACTCTTAGCGGTGGAT</td><td>936.9</td><td>3.1</td><td>0.6</td><td>5.0</td><td>4.5E-07</td><td>7.6E-07</td><td>MB231/MB468</td><td>NUC</td></tr>
<tr><td>>hg38_21_+_8433172_8446622_RNA45SN1_45S_50ntFlanks@9855.9878.24|CTGCGGGATGAACCGAACGCCGGG</td><td>>hg38_21_+_8433172_8446622_RNA45SN1_45S_50ntFlanks</td><td>9855</td><td>9878</td><td>24</td><td>CTGCGGGATGAACCGAACGCCGGG</td><td>147.9</td><td>-4.0</td><td>0.8</td><td>-5.1</td><td>3.9E-07</td><td>7.6E-07</td><td>MB231/MB468</td><td>MP</td></tr>
<tr><td>>hg38_21_+_8433172_8446622_RNA45SN1_45S_50ntFlanks@12436.12454.19|TCATTGTGAAGCAGAATTC</td><td>>hg38_21_+_8433172_8446622_RNA45SN1_45S_50ntFlanks</td><td>12436</td><td>12454</td><td>19</td><td>TCATTGTGAAGCAGAATTC</td><td>386.7</td><td>-5.9</td><td>1.2</td><td>-5.0</td><td>4.7E-07</td><td>7.6E-07</td><td>MB231/MB468</td><td>MITO</td></tr>
<tr><td>>MI0000101|hsa-mir-99a&WithFlank&21|+|16539083|16539175@19.39.21&[MIMAT0000097&hsa-miR-99a-5p&offsets|0|-1;m-64&21|+|16539101|16539121&offsets|0|0]|AACCCGTAGATCCGATCTTGT</td><td>>MI0000101|hsa-mir-99a&WithFlank&21|+|16539083|16539175</td><td>19</td><td>39</td><td>21</td><td>AACCCGTAGATCCGATCTTGT</td><td>3031.3</td><td>-5.6</td><td>1.1</td><td>-5.3</td><td>1.0E-07</td><td>7.6E-07</td><td>BT20/MB468</td><td>MP</td></tr>
<tr><td>>hg38_21_+_8433172_8446622_RNA45SN1_45S_50ntFlanks@12141.12180.40|ACAGTGCCAGGTGGGGAGTTTGACTGGGGCGGTACACCTG</td><td>>hg38_21_+_8433172_8446622_RNA45SN1_45S_50ntFlanks</td><td>12141</td><td>12180</td><td>40</td><td>ACAGTGCCAGGTGGGGAGTTTGACTGGGGCGGTACACCTG</td><td>99.3</td><td>-9.7</td><td>1.8</td><td>-5.3</td><td>1.0E-07</td><td>7.6E-07</td><td>BT20/MB468</td><td>MP</td></tr>
<tr><td>>hg38_MT_+_1621_3279_RNR2_16S_50ntFlanks@499.530.32|TGGACACTAGGAAAAAACCTTGTAGAGAGAGT</td><td>>hg38_MT_+_1621_3279_RNR2_16S_50ntFlanks</td><td>499</td><td>530</td><td>32</td><td>TGGACACTAGGAAAAAACCTTGTAGAGAGAGT</td><td>58.8</td><td>9.6</td><td>1.8</td><td>5.3</td><td>1.0E-07</td><td>7.6E-07</td><td>BT20/MB468</td><td>MP</td></tr>
<tr><td>>hg38_21_+_8433172_8446622_RNA45SN1_45S_50ntFlanks@5489.5514.26|CTGAGAAGACGGTCGAACTTGACTAT</td><td>>hg38_21_+_8433172_8446622_RNA45SN1_45S_50ntFlanks</td><td>5489</td><td>5514</td><td>26</td><td>CTGAGAAGACGGTCGAACTTGACTAT</td><td>916.1</td><td>-3.5</td><td>0.7</td><td>-5.1</td><td>4.4E-07</td><td>7.6E-07</td><td>MB231/MB468</td><td>CYTO</td></tr>
<tr><td>>hg38_21_+_8433172_8446622_RNA45SN1_45S_50ntFlanks@8157.8185.29|CTCGTGGGGGGCCCAAGTCCTTCTGATCG</td><td>>hg38_21_+_8433172_8446622_RNA45SN1_45S_50ntFlanks</td><td>8157</td><td>8185</td><td>29</td><td>CTCGTGGGGGGCCCAAGTCCTTCTGATCG</td><td>384.4</td><td>-3.5</td><td>0.7</td><td>-5.0</td><td>4.7E-07</td><td>7.6E-07</td><td>MB231/MB468</td><td>MITO</td></tr>
<tr><td>>hg38_21_+_8433172_8446622_RNA45SN1_45S_50ntFlanks@8190.8212.23|CCAGCCCGTGGACGGTGTGAGGC</td><td>>hg38_21_+_8433172_8446622_RNA45SN1_45S_50ntFlanks</td><td>8190</td><td>8212</td><td>23</td><td>CCAGCCCGTGGACGGTGTGAGGC</td><td>1869.4</td><td>-6.6</td><td>1.3</td><td>-5.1</td><td>3.9E-07</td><td>7.6E-07</td><td>MB231/MB468</td><td>MP</td></tr>
<tr><td>>hg38_21_+_8433172_8446622_RNA45SN1_45S_50ntFlanks@6699.6719.21|GCTAGCTGCGAGAATTAATGT</td><td>>hg38_21_+_8433172_8446622_RNA45SN1_45S_50ntFlanks</td><td>6699</td><td>6719</td><td>21</td><td>GCTAGCTGCGAGAATTAATGT</td><td>3293.9</td><td>-6.9</td><td>1.4</td><td>-5.1</td><td>3.9E-07</td><td>7.6E-07</td><td>MB231/MB468</td><td>MP</td></tr>
<tr><td>>hg38_21_+_8433172_8446622_RNA45SN1_45S_50ntFlanks@12626.12653.28|AATGGGGCGAAGCTACCATCTGTGGGAT</td><td>>hg38_21_+_8433172_8446622_RNA45SN1_45S_50ntFlanks</td><td>12626</td><td>12653</td><td>28</td><td>AATGGGGCGAAGCTACCATCTGTGGGAT</td><td>7772.0</td><td>-9.0</td><td>1.7</td><td>-5.3</td><td>1.1E-07</td><td>7.6E-07</td><td>BT20/MB468</td><td>MP</td></tr>
<tr><td>>hg38_21_+_8433172_8446622_RNA45SN1_45S_50ntFlanks@12029.12058.30|TCTGATCGTTTTTTCACTGACCCGGTGAGG</td><td>>hg38_21_+_8433172_8446622_RNA45SN1_45S_50ntFlanks</td><td>12029</td><td>12058</td><td>30</td><td>TCTGATCGTTTTTTCACTGACCCGGTGAGG</td><td>82.7</td><td>10.1</td><td>1.9</td><td>5.3</td><td>1.1E-07</td><td>7.6E-07</td><td>BT20/MB468</td><td>MP</td></tr>
<tr><td>>hg38_21_+_8433172_8446622_RNA45SN1_45S_50ntFlanks@6661.6678.18|CGGTGGATCACTCGGCTC</td><td>>hg38_21_+_8433172_8446622_RNA45SN1_45S_50ntFlanks</td><td>6661</td><td>6678</td><td>18</td><td>CGGTGGATCACTCGGCTC</td><td>878.1</td><td>3.2</td><td>0.6</td><td>5.5</td><td>5.0E-08</td><td>7.6E-07</td><td>BT20/MB231</td><td>NUC</td></tr>
<tr><td>>hg38_21_+_8433172_8446622_RNA45SN1_45S_50ntFlanks@11767.11790.24|TCTAATTAGTGACGCGCATGAATG</td><td>>hg38_21_+_8433172_8446622_RNA45SN1_45S_50ntFlanks</td><td>11767</td><td>11790</td><td>24</td><td>TCTAATTAGTGACGCGCATGAATG</td><td>366.7</td><td>-3.0</td><td>0.6</td><td>-5.1</td><td>4.4E-07</td><td>7.6E-07</td><td>MB231/MB468</td><td>CYTO</td></tr>
<tr><td>>MI0000255|hsa-mir-30d&WithFlank&8|-|134804870|134804951@12.35.24(+1C)&[MIMAT0000245&hsa-miR-30d-5p&offsets|0|+2(+1C);m-21&8|-|134804919|134804940&offsets|0|+2(+1C)]|TGTAAACATCCCCGACTGGAAGCTC</td><td>>MI0000255|hsa-mir-30d&WithFlank&8|-|134804870|134804951</td><td>12</td><td>35</td><td>24(+1C)</td><td>TGTAAACATCCCCGACTGGAAGCTC</td><td>472.2</td><td>8.4</td><td>1.7</td><td>5.0</td><td>4.7E-07</td><td>7.6E-07</td><td>MB231/MB468</td><td>MITO</td></tr>
<tr><td>>hg38_21_+_8433172_8446622_RNA45SN1_45S_50ntFlanks@4730.4767.38|CAAGAACGAAAGTCGGAGGTTCGAAGACGATCAGATAC</td><td>>hg38_21_+_8433172_8446622_RNA45SN1_45S_50ntFlanks</td><td>4730</td><td>4767</td><td>38</td><td>CAAGAACGAAAGTCGGAGGTTCGAAGACGATCAGATAC</td><td>48.3</td><td>8.8</td><td>1.7</td><td>5.1</td><td>3.9E-07</td><td>7.6E-07</td><td>MB231/MB468</td><td>MP</td></tr>
<tr><td>>hg38_21_+_8433172_8446622_RNA45SN1_45S_50ntFlanks@10689.10707.19|GCGTCCGGTGAGCTCTCGC</td><td>>hg38_21_+_8433172_8446622_RNA45SN1_45S_50ntFlanks</td><td>10689</td><td>10707</td><td>19</td><td>GCGTCCGGTGAGCTCTCGC</td><td>123.7</td><td>-5.2</td><td>1.0</td><td>-5.1</td><td>4.4E-07</td><td>7.6E-07</td><td>MB231/MB468</td><td>CYTO</td></tr>
<tr><td>>MI0000469|hsa-mir-125a&WithFlank&19|+|51693248|51693345@21.43.23(+1U)&[MIMAT0000443&hsa-miR-125a-5p&offsets|0|-1(+1U);m-63&19|+|51693268|51693289&offsets|0|+1(+1U)]|TCCCTGAGACCCTTTAACCTGTGT</td><td>>MI0000469|hsa-mir-125a&WithFlank&19|+|51693248|51693345</td><td>21</td><td>43</td><td>23(+1U)</td><td>TCCCTGAGACCCTTTAACCTGTGT</td><td>956.9</td><td>4.7</td><td>0.9</td><td>5.0</td><td>4.7E-07</td><td>7.6E-07</td><td>MB231/MB468</td><td>MITO</td></tr>
<tr><td>>hg38_21_+_8433172_8446622_RNA45SN1_45S_50ntFlanks@10377.10424.48|AAGGGTTCCATGTGAACAGCAGTTGAACATGGGTCAGTCGGTCCTGAG</td><td>>hg38_21_+_8433172_8446622_RNA45SN1_45S_50ntFlanks</td><td>10377</td><td>10424</td><td>48</td><td>AAGGGTTCCATGTGAACAGCAGTTGAACATGGGTCAGTCGGTCCTGAG</td><td>334.7</td><td>9.0</td><td>1.7</td><td>5.3</td><td>1.1E-07</td><td>7.6E-07</td><td>BT20/MB468</td><td>MP</td></tr>
<tr><td>>hg38_21_+_8433172_8446622_RNA45SN1_45S_50ntFlanks@10389.10420.32|TGAACAGCAGTTGAACATGGGTCAGTCGGTCC</td><td>>hg38_21_+_8433172_8446622_RNA45SN1_45S_50ntFlanks</td><td>10389</td><td>10420</td><td>32</td><td>TGAACAGCAGTTGAACATGGGTCAGTCGGTCC</td><td>67.1</td><td>-9.1</td><td>1.7</td><td>-5.3</td><td>1.1E-07</td><td>7.6E-07</td><td>BT20/MB468</td><td>MP</td></tr>
</table>

| | | | | | | | | | | | | | |
|---|---|---|---|---|---|---|---|---|---|---|---|---|---|
| >hg38_21_+_8433172_8446622_RNA45SN1_45S_50ntFlanks@11314.11344.31\|CCCACCCCACGTCTCGTCGCGCGCGCGTCCG | >hg38_21_+_8433172_8446622_RNA45SN1_45S_50ntFlanks | 11314 | 11344 | 31 | CCCACCCCACGTCTCGTCGCGCGCGCGTCCG | 67.0 | -9.1 | 1.7 | -5.3 | 1.1E-07 | 7.6E-07 | BT20/MB468 | MP |
| >hg38_21_+_8433172_8446622_RNA45SN1_45S_50ntFlanks@7982.8006.25\|TCAGATCAGACGTGGCGACCCGCTG | >hg38_21_+_8433172_8446622_RNA45SN1_45S_50ntFlanks | 7982 | 8006 | 25 | TCAGATCAGACGTGGCGACCCGCTG | 841.9 | -5.0 | 1.0 | -5.1 | 4.4E-07 | 7.6E-07 | MB231/MB468 | CYTO |
| >hg38_21_+_8433172_8446622_RNA45SN1_45S_50ntFlanks@10355.10390.36\|AACTTTGAAGGCCGAAGTGGAGAAGGGTTCCATGTG | >hg38_21_+_8433172_8446622_RNA45SN1_45S_50ntFlanks | 10355 | 10390 | 36 | AACTTTGAAGGCCGAAGTGGAGAAGGGTTCCATGTG | 521.6 | -5.9 | 1.2 | -5.0 | 4.5E-07 | 7.6E-07 | MB231/MB468 | NUC |
| >hg38_21_+_8433172_8446622_RNA45SN1_45S_50ntFlanks@10357.10383.27\|CTTTGAAGGCCGAAGTGGAGAAGGGTT | >hg38_21_+_8433172_8446622_RNA45SN1_45S_50ntFlanks | 10357 | 10383 | 27 | CTTTGAAGGCCGAAGTGGAGAAGGGTT | 341.8 | -3.0 | 0.6 | -5.1 | 4.4E-07 | 7.6E-07 | MB231/MB468 | CYTO |
| >hg38_21_+_8433172_8446622_RNA45SN1_45S_50ntFlanks@11567.11601.35\|AGCAGCCGACTTAGAACTGGTGCGGACCAGGGGAA | >hg38_21_+_8433172_8446622_RNA45SN1_45S_50ntFlanks | 11567 | 11601 | 35 | AGCAGCCGACTTAGAACTGGTGCGGACCAGGGGAA | 2770.3 | -7.3 | 1.5 | -5.0 | 4.9E-07 | 7.7E-07 | BT20/MB468 | NUC |
| >hg38_21_+_8433172_8446622_RNA45SN1_45S_50ntFlanks@10771.10792.22\|CAGGTCTCCAAGGTGAACAGCC | >hg38_21_+_8433172_8446622_RNA45SN1_45S_50ntFlanks | 10771 | 10792 | 22 | CAGGTCTCCAAGGTGAACAGCC | 61.3 | -9.0 | 1.7 | -5.3 | 1.1E-07 | 7.7E-07 | BT20/MB468 | MP |
| >hg38_21_+_8433172_8446622_RNA45SN1_45S_50ntFlanks@9577.9597.21\|CGGTCGTCCGACCTGGGTATA | >hg38_21_+_8433172_8446622_RNA45SN1_45S_50ntFlanks | 9577 | 9597 | 21 | CGGTCGTCCGACCTGGGTATA | 77.5 | -9.4 | 1.8 | -5.3 | 1.1E-07 | 7.7E-07 | BT20/MB468 | MP |
| >hg38_21_+_8433172_8446622_RNA45SN1_45S_50ntFlanks@9164.9183.20\|CCCCGGGCGGGTCGCGCCGT | >hg38_21_+_8433172_8446622_RNA45SN1_45S_50ntFlanks | 9164 | 9183 | 20 | CCCCGGGCGGGTCGCGCCGT | 63.9 | -9.1 | 1.7 | -5.3 | 1.1E-07 | 7.7E-07 | BT20/MB468 | MP |
| >hg38_21_+_8433172_8446622_RNA45SN1_45S_50ntFlanks@13007.13039.33\|TATTGAAAGTCAGCCCTCGACACAAGGGTTTGT | >hg38_21_+_8433172_8446622_RNA45SN1_45S_50ntFlanks | 13007 | 13039 | 33 | TATTGAAAGTCAGCCCTCGACACAAGGGTTTGT | 1338.8 | 4.1 | 0.8 | 5.0 | 4.5E-07 | 7.7E-07 | MB231/MB468 | NUC |
| >hg38_21_+_8433172_8446622_RNA45SN1_45S_50ntFlanks@10545.10567.23\|CCAGTGCGGTAACGCGACCGATC | >hg38_21_+_8433172_8446622_RNA45SN1_45S_50ntFlanks | 10545 | 10567 | 23 | CCAGTGCGGTAACGCGACCGATC | 185.6 | -6.8 | 1.4 | -5.1 | 4.0E-07 | 7.7E-07 | MB231/MB468 | MP |
| >hg38_21_+_8433172_8446622_RNA45SN1_45S_50ntFlanks@11031.11060.30\|CTCTCTCTCTCCCCCGCTCCCCGTCCTCCC | >hg38_21_+_8433172_8446622_RNA45SN1_45S_50ntFlanks | 11031 | 11060 | 30 | CTCTCTCTCTCCCCCGCTCCCCGTCCTCCC | 393.4 | -7.7 | 1.5 | -5.0 | 4.6E-07 | 7.7E-07 | MB231/MB468 | NUC |
| >hg38_21_+_8433172_8446622_RNA45SN1_45S_50ntFlanks@4981.5018.38\|CACGGACAGGATTGACAGATTGATAGCTCTTTCTCGAT | >hg38_21_+_8433172_8446622_RNA45SN1_45S_50ntFlanks | 4981 | 5018 | 38 | CACGGACAGGATTGACAGATTGATAGCTCTTTCTCGAT | 94.3 | 10.3 | 1.9 | 5.3 | 1.1E-07 | 7.7E-07 | BT20/MB468 | MP |
| >hg38_21_+_8433172_8446622_RNA45SN1_45S_50ntFlanks@10796.10818.23\|GGCATGTTGGAACAATGTAGGTA | >hg38_21_+_8433172_8446622_RNA45SN1_45S_50ntFlanks | 10796 | 10818 | 23 | GGCATGTTGGAACAATGTAGGTA | 272.7 | -3.6 | 0.7 | -5.0 | 4.6E-07 | 7.7E-07 | MB231/MB468 | NUC |
| >hg38_21_+_8433172_8446622_RNA45SN1_45S_50ntFlanks@12588.12605.18\|GGTTCAGACATTTGGTGT | >hg38_21_+_8433172_8446622_RNA45SN1_45S_50ntFlanks | 12588 | 12605 | 18 | GGTTCAGACATTTGGTGT | 61.9 | -9.0 | 1.7 | -5.3 | 1.1E-07 | 7.7E-07 | BT20/MB468 | MP |
| >hg38_21_+_8433172_8446622_RNA45SN1_45S_50ntFlanks@4163.4180.18\|CATCCAAGGAAGGCAGCA | >hg38_21_+_8433172_8446622_RNA45SN1_45S_50ntFlanks | 4163 | 4180 | 18 | CATCCAAGGAAGGCAGCA | 92.3 | -9.6 | 1.8 | -5.3 | 1.1E-07 | 7.7E-07 | BT20/MB468 | MP |
| >hg38_MT_+_1621_3279_RNR2_16S_50ntFlanks@243.276.34\|AACTAGAAATAACTTTGCAAGGAGAGCCAAAGCT | >hg38_MT_+_1621_3279_RNR2_16S_50ntFlanks | 243 | 276 | 34 | AACTAGAAATAACTTTGCAAGGAGAGCCAAAGCT | 112.0 | 10.6 | 2.0 | 5.3 | 1.1E-07 | 7.7E-07 | BT20/MB468 | MP |
| >hg38_21_+_8433172_8446622_RNA45SN1_45S_50ntFlanks@4449.4471.23\|CCTCTCGGCGCCCCCTCGATGCT | >hg38_21_+_8433172_8446622_RNA45SN1_45S_50ntFlanks | 4449 | 4471 | 23 | CCTCTCGGCGCCCCCTCGATGCT | 844.7 | -5.6 | 1.1 | -5.1 | 3.9E-07 | 7.7E-07 | BT20/MB468 | CYTO |
| >hg38_21_+_8433172_8446622_RNA45SN1_45S_50ntFlanks@10882.10902.21\|TCGGGCTGGGGCGCGAAGCGG | >hg38_21_+_8433172_8446622_RNA45SN1_45S_50ntFlanks | 10882 | 10902 | 21 | TCGGGCTGGGGCGCGAAGCGG | 326.2 | -4.8 | 0.9 | -5.0 | 4.4E-07 | 7.7E-07 | MB231/MB468 | CYTO |
| >MI0000255\|hsa-mir-30d&WithFlank&8\|-\|134804870\|134804951@12.34.23(+1G)&[MIMAT0000245&hsa-miR-30d-5p&offsets\|0\|+1(+1G);m-21&8\|-\|134804919\|134804940&offsets\|0\|+1(+1G)]\|TGTAAACATCCCCGACTGGAAGCG | >MI0000255\|hsa-mir-30d&WithFlank&8\|-\|134804870\|134804951 | 12 | 34 | 23(+1G) | TGTAAACATCCCCGACTGGAAGCG | 941.9 | 5.5 | 1.1 | 5.0 | 4.8E-07 | 7.7E-07 | MB231/MB468 | MITO |
| >hg38_21_+_8433172_8446622_RNA45SN1_45S_50ntFlanks@6651.6673.23\|CGACTCTTAGCGGTGGATCACTC | >hg38_21_+_8433172_8446622_RNA45SN1_45S_50ntFlanks | 6651 | 6673 | 23 | CGACTCTTAGCGGTGGATCACTC | 2875.8 | 4.5 | 0.9 | 5.0 | 4.6E-07 | 7.7E-07 | MB231/MB468 | NUC |
| >hg38_21_+_8433172_8446622_RNA45SN1_45S_50ntFlanks@11739.11758.20\|TCTCTTAAGGTAGCCAAATG | >hg38_21_+_8433172_8446622_RNA45SN1_45S_50ntFlanks | 11739 | 11758 | 20 | TCTCTTAAGGTAGCCAAATG | 223.7 | -5.8 | 1.1 | -5.0 | 4.6E-07 | 7.7E-07 | MB231/MB468 | NUC |
| >hg38_21_+_8433172_8446622_RNA45SN1_45S_50ntFlanks@12788.12805.18\|CGCGCGCGCGGGAGGGCG | >hg38_21_+_8433172_8446622_RNA45SN1_45S_50ntFlanks | 12788 | 12805 | 18 | CGCGCGCGCGGGAGGGCG | 77.6 | -9.4 | 1.8 | -5.3 | 1.1E-07 | 7.7E-07 | BT20/MB468 | MP |
| >hg38_21_+_8433172_8446622_RNA45SN1_45S_50ntFlanks@5078.5104.27\|CGATAACGAACGAGACTCTGGCATGCT | >hg38_21_+_8433172_8446622_RNA45SN1_45S_50ntFlanks | 5078 | 5104 | 27 | CGATAACGAACGAGACTCTGGCATGCT | 331.9 | -7.6 | 1.5 | -5.0 | 4.8E-07 | 7.7E-07 | MB231/MB468 | MITO |
| >hg38_21_+_8433172_8446622_RNA45SN1_45S_50ntFlanks@9553.9571.19\|CCGTAGCGGTCCTGACGTG | >hg38_21_+_8433172_8446622_RNA45SN1_45S_50ntFlanks | 9553 | 9571 | 19 | CCGTAGCGGTCCTGACGTG | 177.0 | -5.3 | 1.0 | -5.1 | 4.0E-07 | 7.7E-07 | MB231/MB468 | MP |
| >MI0000234\|hsa-mir-192&WithFlank&11\|-\|64891131\|64891252@30.51.22(+1U)&[MIMAT0000222&hsa-miR-192-5p&offsets\|0\|+1(+1U);m-59&11\|-\|64891203\|64891223&offsets\|0\|+1(+1U)]\|CTGACCTATGAATTGACAGCCAT | >MI0000234\|hsa-mir-192&WithFlank&11\|-\|64891131\|64891252 | 30 | 51 | 22(+1U) | CTGACCTATGAATTGACAGCCAT | 824.3 | 4.7 | 0.9 | 5.0 | 4.8E-07 | 7.7E-07 | MB231/MB468 | MITO |
| >MI0000294\|hsa-mir-218-1&WithFlank&4\|+\|20528269\|20528390@31.50.20(+1C);MI0000295\|hsa-mir-218-2&WithFlank&5\|-\|168768140\|168768261@31.50.20(+1C)&[MIMAT0000275&hsa-miR-218-5p&offsets\|0\|-1(+1C);m-185&4\|+\|20528299\|20528319&offsets\|0\|-1(+1C)];[MIMAT0000275_1&hsa-miR-218-5p&offsets\|0\|-1(+1C);m-187&5\|-\|168768211\|168768231&offsets\|0\|-1(+1C)]\|TTGTGCTTGATCTAACCATGC | >MI0000294\|hsa-mir-218-1&WithFlank&4\|+\|20528269\|20528390 | 31 | 50 | 20(+1C) | TTGTGCTTGATCTAACCATGC | 154.5 | -10.7 | 1.9 | -5.6 | 2.2E-08 | 7.7E-07 | BT20/MB231 | MP |
| >hg38_21_+_8433172_8446622_RNA45SN1_45S_50ntFlanks@6652.6684.33\|GACTCTTAGCGGTGGATCACTCGGCTCGTGCGT | >hg38_21_+_8433172_8446622_RNA45SN1_45S_50ntFlanks | 6652 | 6684 | 33 | GACTCTTAGCGGTGGATCACTCGGCTCGTGCGT | 2428.1 | 3.7 | 0.7 | 5.1 | 3.4E-07 | 7.7E-07 | BT20/MB468 | MITO |
| >hg38_21_+_8433172_8446622_RNA45SN1_45S_50ntFlanks@10348.10369.22\|AAACGAGAACTTTGAAGGCCGA | >hg38_21_+_8433172_8446622_RNA45SN1_45S_50ntFlanks | 10348 | 10369 | 22 | AAACGAGAACTTTGAAGGCCGA | 539.9 | 2.8 | 0.6 | 5.0 | 4.5E-07 | 7.8E-07 | MB231/MB468 | CYTO |
| >trna152_ValCAC_6_-_27248049_27248121@1.33.33\|GCTTCTGTAGTGTAGTGGTTATCACGTTCGCCT | >trna152_ValCAC_6_-_27248049_27248121 | 1 | 33 | 33 | GCTTCTGTAGTGTAGTGGTTATCACGTTCGCCT | 966.1 | 7.9 | 1.6 | 5.1 | 4.0E-07 | 7.8E-07 | MB231/MB468 | MP |
| >MI0000650\|hsa-mir-200c&WithFlank&12\|+\|6963693\|6963772@50.71.22(+2U)&[MIMAT0000617&hsa-miR-200c-3p&offsets\|0\|-1(+2U);m-54&12\|+\|6963742\|6963764&offsets\|0\|-1(+2U)]\|TAATACTGCCGGGTAATGATGGTT | >MI0000650\|hsa-mir-200c&WithFlank&12\|+\|6963693\|6963772 | 50 | 71 | 22(+2U) | TAATACTGCCGGGTAATGATGGTT | 207.4 | 11.0 | 2.0 | 5.6 | 2.2E-08 | 7.8E-07 | BT20/MB231 | MP |
| >hg38_21_+_8433172_8446622_RNA45SN1_45S_50ntFlanks@4722.4759.38\|TCATTAATCAAGAACGAAAGTCGGAGGTTCGAAGACGA | >hg38_21_+_8433172_8446622_RNA45SN1_45S_50ntFlanks | 4722 | 4759 | 38 | TCATTAATCAAGAACGAAAGTCGGAGGTTCGAAGACGA | 2380.2 | -6.9 | 1.4 | -5.1 | 4.0E-07 | 7.8E-07 | MB231/MB468 | MP |
| >hg38_21_+_8433172_8446622_RNA45SN1_45S_50ntFlanks@12241.12272.32\|CAGAAGGGCAAAAGCTCGCTTGATCTTGATTT | >hg38_21_+_8433172_8446622_RNA45SN1_45S_50ntFlanks | 12241 | 12272 | 32 | CAGAAGGGCAAAAGCTCGCTTGATCTTGATTT | 1068.8 | -3.5 | 0.7 | -5.1 | 3.4E-07 | 7.8E-07 | BT20/MB468 | MITO |
| >hg38_21_+_8433172_8446622_RNA45SN1_45S_50ntFlanks@9567.9588.22\|ACGTGCAAATCGGTCGTCCGAC | >hg38_21_+_8433172_8446622_RNA45SN1_45S_50ntFlanks | 9567 | 9588 | 22 | ACGTGCAAATCGGTCGTCCGAC | 195.4 | -6.3 | 1.2 | -5.1 | 3.9E-07 | 7.8E-07 | BT20/MB468 | CYTO |
| >trnaMT_PheGAA_MT_+_577_647@33.57.25\|CTGAAAATGTTTAGACGGGCTCACA | >trnaMT_PheGAA_MT_+_577_647 | 33 | 57 | 25 | CTGAAAATGTTTAGACGGGCTCACA | 113.3 | 10.6 | 2.0 | 5.3 | 1.1E-07 | 7.8E-07 | BT20/MB468 | MP |
| >hg38_21_+_8433172_8446622_RNA45SN1_45S_50ntFlanks@12366.12383.18\|TAACTGGCTTGTGGCGGC | >hg38_21_+_8433172_8446622_RNA45SN1_45S_50ntFlanks | 12366 | 12383 | 18 | TAACTGGCTTGTGGCGGC | 688.7 | -7.6 | 1.4 | -5.3 | 1.1E-07 | 7.8E-07 | BT20/MB468 | MP |
| >hg38_21_+_8433172_8446622_RNA45SN1_45S_50ntFlanks@12624.12652.29\|CCAATGGGGCGAAGCTACCATCTGTGGGA | >hg38_21_+_8433172_8446622_RNA45SN1_45S_50ntFlanks | 12624 | 12652 | 29 | CCAATGGGGCGAAGCTACCATCTGTGGGA | 530.9 | -8.0 | 1.5 | -5.3 | 1.1E-07 | 7.8E-07 | BT20/MB468 | MP |
| >hg38_21_+_8433172_8446622_RNA45SN1_45S_50ntFlanks@3782.3814.33\|CAGTGAAACTGCGAATGGCTCATTAAATCAGTT | >hg38_21_+_8433172_8446622_RNA45SN1_45S_50ntFlanks | 3782 | 3814 | 33 | CAGTGAAACTGCGAATGGCTCATTAAATCAGTT | 446.6 | -5.6 | 1.1 | -5.0 | 4.6E-07 | 7.8E-07 | MB231/MB468 | NUC |
| >hg38_21_+_8433172_8446622_RNA45SN1_45S_50ntFlanks@12071.12095.25\|CCCCGAGGGGCTCTCGCTTCTGGCG | >hg38_21_+_8433172_8446622_RNA45SN1_45S_50ntFlanks | 12071 | 12095 | 25 | CCCCGAGGGGCTCTCGCTTCTGGCG | 3380.4 | -5.9 | 1.2 | -5.0 | 4.5E-07 | 7.8E-07 | MB231/MB468 | CYTO |
| >hg38_21_+_8433172_8446622_RNA45SN1_45S_50ntFlanks@7975.8005.31\|CGCGACCTCAGATCAGACGTGGCGACCCGCT | >hg38_21_+_8433172_8446622_RNA45SN1_45S_50ntFlanks | 7975 | 8005 | 31 | CGCGACCTCAGATCAGACGTGGCGACCCGCT | 234003.7 | 4.4 | 0.8 | 5.4 | 5.1E-08 | 7.8E-07 | BT20/MB231 | NUC |
| >hg38_21_+_8433172_8446622_RNA45SN1_45S_50ntFlanks@4729.4751.23\|TCAAGAACGAAAGTCGGAGGTTC | >hg38_21_+_8433172_8446622_RNA45SN1_45S_50ntFlanks | 4729 | 4751 | 23 | TCAAGAACGAAAGTCGGAGGTTC | 629.4 | -4.1 | 0.8 | -5.3 | 1.1E-07 | 7.8E-07 | BT20/MB468 | MP |
| >MI0000102\|hsa-mir-100&WithFlank&11\|-\|122152223\|122152314@19.37.19(+1C)&[MIMAT0000098&hsa-miR-100-5p&offsets\|0\|-3(+1C);m-27&11\|-\|122152276\|122152296&offsets\|0\|-2(+1C)]\|AACCCGTAGATCCGAACTTC | >MI0000102\|hsa-mir-100&WithFlank&11\|-\|122152223\|122152314 | 19 | 37 | 19(+1C) | AACCCGTAGATCCGAACTTC | 216.9 | -11.2 | 2.0 | -5.6 | 2.3E-08 | 7.8E-07 | BT20/MB231 | MP |
| >MI0000079\|hsa-mir-23a&WithFlank&19\|-\|13836581\|13836665@54.72.19(+1U)&[MIMAT0000078&hsa-miR-23a-3p&offsets\|+3\|+1(+1U);m-31&19\|-\|13836595\|13836615&offsets\|+3\|+1(+1U)]\|ACATTGCCAGGGATTTCCAT | >MI0000079\|hsa-mir-23a&WithFlank&19\|-\|13836581\|13836665 | 54 | 72 | 19(+1U) | ACATTGCCAGGGATTTCCAT | 82.6 | 10.1 | 1.9 | 5.3 | 1.1E-07 | 7.8E-07 | BT20/MB468 | MP |
| >hg38_21_+_8433172_8446622_RNA45SN1_45S_50ntFlanks@6651.6669.19\|CGACTCTTAGCGGTGGATC | >hg38_21_+_8433172_8446622_RNA45SN1_45S_50ntFlanks | 6651 | 6669 | 19 | CGACTCTTAGCGGTGGATC | 2408.5 | 2.3 | 0.4 | 5.0 | 4.7E-07 | 7.8E-07 | MB231/MB468 | NUC |
| >MI0000272\|hsa-mir-182&WithFlank&7\|-\|129770377\|129770498@29.51.23&[MIMAT0000259&hsa-miR-182-5p&offsets\|0\|-1;m-22&7\|-\|129770449\|129770470&offsets\|0\|+1]\|TTTGGCAATGGTAGAACTCACAC | >MI0000272\|hsa-mir-182&WithFlank&7\|-\|129770377\|129770498 | 29 | 51 | 23 | TTTGGCAATGGTAGAACTCACAC | 3661.3 | 4.6 | 0.8 | 5.5 | 3.2E-08 | 7.9E-07 | BT20/MB231 | CYTO |
| >MI0000289\|hsa-mir-181a-1&WithFlank&1\|-\|198859038\|198859159@30.52.23;MI0000269\|hsa-mir-181a-2&WithFlank&9\|+\|124692436\|124692557@45.67.23&[MIMAT0000256&hsa-miR-181a-5p&offsets\|0\|0;m-32&1\|-\|198859108\|198859130&offsets\|0\|0];[MIMAT0000256_1&hsa-miR-181a-5p&offsets\|0\|0;m-32&9\|+\|124692480\|124692502&offsets\|0\|0]\|AACATTCAACGCTGTCGGTGAGT | >MI0000289\|hsa-mir-181a-1&WithFlank&1\|-\|198859038\|198859159 | 30 | 52 | 23 | AACATTCAACGCTGTCGGTGAGT | 2333.1 | 4.7 | 0.9 | 5.0 | 4.5E-07 | 7.9E-07 | MB231/MB468 | CYTO |
| >hg38_21_+_8433172_8446622_RNA45SN1_45S_50ntFlanks@11208.11252.45\|GCGGCGACTCTGGACGCGAGCCGGGCCCTTCCCGTGGATCGCCCC | >hg38_21_+_8433172_8446622_RNA45SN1_45S_50ntFlanks | 11208 | 11252 | 45 | GCGGCGACTCTGGACGCGAGCCGGGCCCTTCCCGTGGATCGCCCC | 2089.9 | -6.6 | 1.3 | -5.0 | 4.7E-07 | 7.9E-07 | MB231/MB468 | NUC |
| >trna116_GluCTC_1_-_145399233_145399304@39.74.36,trna134_GluTTC_1_-_16861774_16861845@39.74.36,trna71_GluCTC_1_-_161439189_161439260@39.74.36,trna74_GluCTC_1_-_161431809_161431880@39.74.36,trna77_GluCTC_1_-_161424398_161424469@39.74.36,trna77_GluCTC_6_+_28949976_28950047@39.74.36,trna80_GluCTC_1_-_161417018_161417089@39.74.36,trna84_GluTTC_1_-_161391883_161391954@39.74.36,trna87_GluCTC_6_-_126101393_126101464@39.74.36\|CGCCGCGGCCCGGGTTCGATTCCCGGTCAGGGAACC | >trna116_GluCTC_1_-_145399233_145399304 | 39 | 74 | 36 | CGCCGCGGCCCGGGTTCGATTCCCGGTCAGGGAACC | 1511.7 | 6.9 | 1.4 | 5.0 | 4.5E-07 | 7.9E-07 | MB231/MB468 | CYTO |

<table>
<tr><td>>hg38_21_+_8433172_8446622_RNA45SN1_45S_50ntFlanks@5237.5254.18|ACTGACTGGCTCAGCGTG</td><td>>hg38_21_+_8433172_8446622_RNA45SN1_45S_50ntFlanks</td><td>5237</td><td>5254</td><td>18</td><td>ACTGACTGGCTCAGCGTG</td><td>1044.9</td><td>-3.8</td><td>0.8</td><td>-5.1</td><td>4.1E-07</td><td>7.9E-07</td><td>MB231/MB468</td><td>MP</td></tr>
<tr><td>>hg38_21_+_8433172_8446622_RNA45SN1_45S_50ntFlanks@9622.9650.29|CTAGTAGCTGGTTCCCTCCGAAGTTTCCC</td><td>>hg38_21_+_8433172_8446622_RNA45SN1_45S_50ntFlanks</td><td>9622</td><td>9650</td><td>29</td><td>CTAGTAGCTGGTTCCCTCCGAAGTTTCCC</td><td>1016.8</td><td>-7.6</td><td>1.5</td><td>-5.0</td><td>4.9E-07</td><td>7.9E-07</td><td>MB231/MB468</td><td>MITO</td></tr>
<tr><td>>hg38_21_+_8433172_8446622_RNA45SN1_45S_50ntFlanks@10849.10868.20|CGGGATAAGGATTGGCTCTA</td><td>>hg38_21_+_8433172_8446622_RNA45SN1_45S_50ntFlanks</td><td>10849</td><td>10868</td><td>20</td><td>CGGGATAAGGATTGGCTCTA</td><td>754.5</td><td>-6.1</td><td>1.2</td><td>-5.0</td><td>4.5E-07</td><td>7.9E-07</td><td>MB231/MB468</td><td>CYTO</td></tr>
<tr><td>>hg38_21_+_8433172_8446622_RNA45SN1_45S_50ntFlanks@4200.4219.20|CCCGACCCGGGGAGGTAGTG</td><td>>hg38_21_+_8433172_8446622_RNA45SN1_45S_50ntFlanks</td><td>4200</td><td>4219</td><td>20</td><td>CCCGACCCGGGGAGGTAGTG</td><td>166.5</td><td>-4.0</td><td>0.8</td><td>-5.1</td><td>4.1E-07</td><td>7.9E-07</td><td>MB231/MB468</td><td>MP</td></tr>
<tr><td>>hg38_21_+_8433172_8446622_RNA45SN1_45S_50ntFlanks@7983.8002.20|CAGATCAGACGTGGCGACCC</td><td>>hg38_21_+_8433172_8446622_RNA45SN1_45S_50ntFlanks</td><td>7983</td><td>8002</td><td>20</td><td>CAGATCAGACGTGGCGACCC</td><td>341.9</td><td>11.8</td><td>2.1</td><td>5.6</td><td>2.3E-08</td><td>7.9E-07</td><td>BT20/MB231</td><td>MP</td></tr>
<tr><td>>hg38_21_+_8433172_8446622_RNA45SN1_45S_50ntFlanks@9739.9761.23|AAACGATCTCAACCTATTCTCAA</td><td>>hg38_21_+_8433172_8446622_RNA45SN1_45S_50ntFlanks</td><td>9739</td><td>9761</td><td>23</td><td>AAACGATCTCAACCTATTCTCAA</td><td>539.1</td><td>-7.6</td><td>1.5</td><td>-5.0</td><td>4.9E-07</td><td>7.9E-07</td><td>MB231/MB468</td><td>MITO</td></tr>
<tr><td>>hg38_21_+_8433172_8446622_RNA45SN1_45S_50ntFlanks@12623.12661.39|GCCAATGGGGCGAAGCTACCATCTGTGGGATTATGACTG</td><td>>hg38_21_+_8433172_8446622_RNA45SN1_45S_50ntFlanks</td><td>12623</td><td>12661</td><td>39</td><td>GCCAATGGGGCGAAGCTACCATCTGTGGGATTATGACTG</td><td>587.4</td><td>-7.4</td><td>1.5</td><td>-5.0</td><td>4.9E-07</td><td>7.9E-07</td><td>MB231/MB468</td><td>MITO</td></tr>
<tr><td>>hg38_21_+_8433172_8446622_RNA45SN1_45S_50ntFlanks@6652.6682.31|GACTCTTAGCGGTGGATCACTCGGCTCGTGC</td><td>>hg38_21_+_8433172_8446622_RNA45SN1_45S_50ntFlanks</td><td>6652</td><td>6682</td><td>31</td><td>GACTCTTAGCGGTGGATCACTCGGCTCGTGC</td><td>992.4</td><td>4.9</td><td>0.9</td><td>5.3</td><td>1.1E-07</td><td>7.9E-07</td><td>BT20/MB468</td><td>MP</td></tr>
<tr><td>>trna111_HisGTG_1_-_147774845_147774916@-1T.28.29,trna118_HisGTG_1_-_145396881_145396952@-1T.28.29,trna16_HisGTG_1_+_146544773_146544844@-1T.28.29,trna1_HisGTG_15_+_45493349_45493420@-1T.28.29,trna21_HisGTG_1_+_147753471_147753542@-1T.28.29,trna33_HisGTG_6_+_27125906_27125977@-1T.28.29,trna7_HisGTG_9_-_14433938_14434009@-1T.28.29,trna8_HisGTG_15_-_45492611_45492682@-1T.28.29,trna9_HisGTG_15_-_45490804_45490875@-1T.28.29|TGCCGTGATCGTATAGTGGTTAGTACTCT</td><td>>trna111_HisGTG_1_-_147774845_147774916</td><td>-1T</td><td>28</td><td>29</td><td>TGCCGTGATCGTATAGTGGTTAGTACTCT</td><td>2171.9</td><td>-4.9</td><td>1.0</td><td>-5.0</td><td>5.1E-07</td><td>7.9E-07</td><td>BT20/MB468</td><td>NUC</td></tr>
<tr><td>>hg38_21_+_8433172_8446622_RNA45SN1_45S_50ntFlanks@10783.10815.33|GTGAACAGCCTCTGGCATGTTGGAACAATGTAG</td><td>>hg38_21_+_8433172_8446622_RNA45SN1_45S_50ntFlanks</td><td>10783</td><td>10815</td><td>33</td><td>GTGAACAGCCTCTGGCATGTTGGAACAATGTAG</td><td>68.5</td><td>-9.2</td><td>1.7</td><td>-5.3</td><td>1.1E-07</td><td>7.9E-07</td><td>BT20/MB468</td><td>MP</td></tr>
<tr><td>>MI0000734|hsa-mir-106b&WithFlank&7|-|100093987|100094080@56.76.21(+1C)&[MIMAT0004672&hsa-miR-106b-3p&offsets|-2|-3(+1C);m-87&7|-|100094003|100094023&offsets|-2|-2(+1C)]|TACCGCACTGTGGGTACTTGCC</td><td>>MI0000734|hsa-mir-106b&WithFlank&7|-|100093987|100094080</td><td>56</td><td>76</td><td>21(+1C)</td><td>TACCGCACTGTGGGTACTTGCC</td><td>85.7</td><td>10.2</td><td>1.9</td><td>5.3</td><td>1.1E-07</td><td>7.9E-07</td><td>BT20/MB468</td><td>MP</td></tr>
<tr><td>>MI0000273|hsa-mir-183&WithFlank&7|-|129774899|129775020@33.55.23&[MIMAT0000261&hsa-miR-183-5p&offsets|0|+1;m-41&7|-|129774966|129774987&offsets|-1|0]|TATGGCACTGGTAGAATTCACTG</td><td>>MI0000273|hsa-mir-183&WithFlank&7|-|129774899|129775020</td><td>33</td><td>55</td><td>23</td><td>TATGGCACTGGTAGAATTCACTG</td><td>1321.4</td><td>1.9</td><td>0.4</td><td>5.0</td><td>5.1E-07</td><td>7.9E-07</td><td>BT20/MB468</td><td>NUC</td></tr>
<tr><td>>hg38_21_+_8433172_8446622_RNA45SN1_45S_50ntFlanks@4760.4781.22|TCAGATACCGTCGTAGTTCCGA</td><td>>hg38_21_+_8433172_8446622_RNA45SN1_45S_50ntFlanks</td><td>4760</td><td>4781</td><td>22</td><td>TCAGATACCGTCGTAGTTCCGA</td><td>65.5</td><td>-9.1</td><td>1.7</td><td>-5.3</td><td>1.1E-07</td><td>7.9E-07</td><td>BT20/MB468</td><td>MP</td></tr>
<tr><td>>hg38_21_+_8433172_8446622_RNA45SN1_45S_50ntFlanks@5219.5244.26|GGGCTGCACGCGCGCTACACTGACTG</td><td>>hg38_21_+_8433172_8446622_RNA45SN1_45S_50ntFlanks</td><td>5219</td><td>5244</td><td>26</td><td>GGGCTGCACGCGCGCTACACTGACTG</td><td>265.5</td><td>-10.1</td><td>2.0</td><td>-5.1</td><td>3.5E-07</td><td>7.9E-07</td><td>BT20/MB468</td><td>MITO</td></tr>
<tr><td>>hg38_21_+_8433172_8446622_RNA45SN1_45S_50ntFlanks@8079.8104.26|AAGAGCCCAGCGCCGAATCCCCGCCC</td><td>>hg38_21_+_8433172_8446622_RNA45SN1_45S_50ntFlanks</td><td>8079</td><td>8104</td><td>26</td><td>AAGAGCCCAGCGCCGAATCCCCGCCC</td><td>147.9</td><td>-6.3</td><td>1.2</td><td>-5.0</td><td>4.7E-07</td><td>7.9E-07</td><td>MB231/MB468</td><td>NUC</td></tr>
<tr><td>>hg38_21_+_8433172_8446622_RNA45SN1_45S_50ntFlanks@11564.11589.26|CCTAGCAGCCGACTTAGAACTGGTGC</td><td>>hg38_21_+_8433172_8446622_RNA45SN1_45S_50ntFlanks</td><td>11564</td><td>11589</td><td>26</td><td>CCTAGCAGCCGACTTAGAACTGGTGC</td><td>298.9</td><td>-7.1</td><td>1.4</td><td>-5.1</td><td>4.1E-07</td><td>8.0E-07</td><td>MB231/MB468</td><td>MP</td></tr>
<tr><td>>MI0000081|hsa-mir-24-2&WithFlank&19|-|13836281|13836365@56.76.21(+1A);MI0000080|hsa-mir-24-1&WithFlank&9|+|95086015|95086094@50.70.21(+1A)&[MIMAT0000080&hsa-miR-24-3p&offsets|0|-1(+1A);m-36&19|-|13836288|13836310&offsets|0|-2(+1A)];[MIMAT0000080_1&hsa-miR-24-3p&offsets|0|-1(+1A);m-35&9|+|95086064|95086086&offsets|0|-2(+1A)]|TGGCTCAGTTCAGCAGGAACAA</td><td>>MI0000081|hsa-mir-24-2&WithFlank&19|-|13836281|13836365</td><td>56</td><td>76</td><td>21(+1A)</td><td>TGGCTCAGTTCAGCAGGAACAA</td><td>392.5</td><td>2.0</td><td>0.4</td><td>5.0</td><td>5.2E-07</td><td>8.0E-07</td><td>BT20/MB468</td><td>NUC</td></tr>
<tr><td>>hg38_21_+_8433172_8446622_RNA45SN1_45S_50ntFlanks@12639.12657.19|TACCATCTGTGGGATTATG</td><td>>hg38_21_+_8433172_8446622_RNA45SN1_45S_50ntFlanks</td><td>12639</td><td>12657</td><td>19</td><td>TACCATCTGTGGGATTATG</td><td>65.3</td><td>-9.1</td><td>1.7</td><td>-5.3</td><td>1.1E-07</td><td>8.0E-07</td><td>BT20/MB468</td><td>MP</td></tr>
<tr><td>>hg38_21_+_8433172_8446622_RNA45SN1_45S_50ntFlanks@10786.10817.32|AACAGCCTCTGGCATGTTGGAACAATGTAGGT</td><td>>hg38_21_+_8433172_8446622_RNA45SN1_45S_50ntFlanks</td><td>10786</td><td>10817</td><td>32</td><td>AACAGCCTCTGGCATGTTGGAACAATGTAGGT</td><td>801.9</td><td>-3.8</td><td>0.7</td><td>-5.1</td><td>4.0E-07</td><td>8.0E-07</td><td>BT20/MB468</td><td>CYTO</td></tr>
<tr><td>>hg38_21_+_8433172_8446622_RNA45SN1_45S_50ntFlanks@9928.9956.29|TGATATAGACAGCAGGACGGTGGCCATGG</td><td>>hg38_21_+_8433172_8446622_RNA45SN1_45S_50ntFlanks</td><td>9928</td><td>9956</td><td>29</td><td>TGATATAGACAGCAGGACGGTGGCCATGG</td><td>65.8</td><td>9.8</td><td>1.8</td><td>5.3</td><td>1.1E-07</td><td>8.0E-07</td><td>BT20/MB468</td><td>MP</td></tr>
<tr><td>>hg38_21_+_8433172_8446622_RNA45SN1_45S_50ntFlanks@11671.11688.18|CCAGTGCTCTGAATGTCA</td><td>>hg38_21_+_8433172_8446622_RNA45SN1_45S_50ntFlanks</td><td>11671</td><td>11688</td><td>18</td><td>CCAGTGCTCTGAATGTCA</td><td>283.6</td><td>-5.7</td><td>1.1</td><td>-5.1</td><td>4.1E-07</td><td>8.0E-07</td><td>MB231/MB468</td><td>MP</td></tr>
<tr><td>>MI0000456|hsa-mir-140&WithFlank&16|+|69933075|69933186@69.89.21(+2U)&[MIMAT0004597&hsa-miR-140-3p&offsets|+1|+1(+2U);m-45&16|+|69933143|69933164&offsets|0|-1(+2U)]|ACCACAGGGTAGAACCACGGATT</td><td>>MI0000456|hsa-mir-140&WithFlank&16|+|69933075|69933186</td><td>69</td><td>89</td><td>21(+2U)</td><td>ACCACAGGGTAGAACCACGGATT</td><td>45.0</td><td>8.7</td><td>1.7</td><td>5.1</td><td>4.1E-07</td><td>8.0E-07</td><td>MB231/MB468</td><td>MP</td></tr>
<tr><td>>hg38_21_+_8433172_8446622_RNA45SN1_45S_50ntFlanks@8112.8137.26|GGGCGCGGGACATGTGGCGTACGGAA</td><td>>hg38_21_+_8433172_8446622_RNA45SN1_45S_50ntFlanks</td><td>8112</td><td>8137</td><td>26</td><td>GGGCGCGGGACATGTGGCGTACGGAA</td><td>557.6</td><td>12.5</td><td>2.2</td><td>5.6</td><td>2.3E-08</td><td>8.0E-07</td><td>BT20/MB231</td><td>MP</td></tr>
<tr><td>>hg38_21_+_8433172_8446622_RNA45SN1_45S_50ntFlanks@7973.8007.35|GACGCGACCTCAGATCAGACGTGGCGACCCGCTGA</td><td>>hg38_21_+_8433172_8446622_RNA45SN1_45S_50ntFlanks</td><td>7973</td><td>8007</td><td>35</td><td>GACGCGACCTCAGATCAGACGTGGCGACCCGCTGA</td><td>1164.1</td><td>3.4</td><td>0.7</td><td>5.1</td><td>3.5E-07</td><td>8.0E-07</td><td>BT20/MB468</td><td>MITO</td></tr>
<tr><td>>hg38_21_+_8433172_8446622_RNA45SN1_45S_50ntFlanks@7992.8009.18|CGTGGCGACCCGCTGAAT</td><td>>hg38_21_+_8433172_8446622_RNA45SN1_45S_50ntFlanks</td><td>7992</td><td>8009</td><td>18</td><td>CGTGGCGACCCGCTGAAT</td><td>1408.6</td><td>-4.5</td><td>0.9</td><td>-5.1</td><td>4.1E-07</td><td>8.0E-07</td><td>MB231/MB468</td><td>MP</td></tr>
<tr><td>>hg38_21_+_8433172_8446622_RNA45SN1_45S_50ntFlanks@6675.6700.26|GCTCGTGCGTCGATGAAGAACGCAGC</td><td>>hg38_21_+_8433172_8446622_RNA45SN1_45S_50ntFlanks</td><td>6675</td><td>6700</td><td>26</td><td>GCTCGTGCGTCGATGAAGAACGCAGC</td><td>74.1</td><td>10.0</td><td>1.9</td><td>5.3</td><td>1.1E-07</td><td>8.0E-07</td><td>BT20/MB468</td><td>MP</td></tr>
<tr><td>>am_trnaMT_LeuTAG_MT_+_12266_12336@42.59.18|AAAATTTTGGTGCAACTC</td><td>>am_trnaMT_LeuTAG_MT_+_12266_12336</td><td>42</td><td>59</td><td>18</td><td>AAAATTTTGGTGCAACTC</td><td>67.4</td><td>-9.2</td><td>1.7</td><td>-5.3</td><td>1.1E-07</td><td>8.0E-07</td><td>BT20/MB468</td><td>MP</td></tr>
<tr><td>>hg38_21_+_8433172_8446622_RNA45SN1_45S_50ntFlanks@11670.11687.18|CCCAGTGCTCTGAATGTC</td><td>>hg38_21_+_8433172_8446622_RNA45SN1_45S_50ntFlanks</td><td>11670</td><td>11687</td><td>18</td><td>CCCAGTGCTCTGAATGTC</td><td>1117.4</td><td>-3.1</td><td>0.6</td><td>-5.0</td><td>4.8E-07</td><td>8.0E-07</td><td>MB231/MB468</td><td>NUC</td></tr>
<tr><td>>hg38_21_+_8433172_8446622_RNA45SN1_45S_50ntFlanks@10841.10874.34|CGTAACTTCGGGATAAGGATTGGCTCTAAGGGCT</td><td>>hg38_21_+_8433172_8446622_RNA45SN1_45S_50ntFlanks</td><td>10841</td><td>10874</td><td>34</td><td>CGTAACTTCGGGATAAGGATTGGCTCTAAGGGCT</td><td>2633.4</td><td>-8.9</td><td>1.8</td><td>-5.0</td><td>5.2E-07</td><td>8.0E-07</td><td>BT20/MB468</td><td>NUC</td></tr>
<tr><td>>hg38_21_+_8433172_8446622_RNA45SN1_45S_50ntFlanks@12818.12840.23|CGCGCCGGGACCGGGGTCCGGTG</td><td>>hg38_21_+_8433172_8446622_RNA45SN1_45S_50ntFlanks</td><td>12818</td><td>12840</td><td>23</td><td>CGCGCCGGGACCGGGGTCCGGTG</td><td>326.2</td><td>-12.2</td><td>2.4</td><td>-5.1</td><td>4.1E-07</td><td>8.0E-07</td><td>MB231/MB468</td><td>MP</td></tr>
<tr><td>>hg38_21_+_8433172_8446622_RNA45SN1_45S_50ntFlanks@4730.4751.22|CAAGAACGAAAGTCGGAGGTTC</td><td>>hg38_21_+_8433172_8446622_RNA45SN1_45S_50ntFlanks</td><td>4730</td><td>4751</td><td>22</td><td>CAAGAACGAAAGTCGGAGGTTC</td><td>463.3</td><td>-3.8</td><td>0.8</td><td>-5.0</td><td>4.8E-07</td><td>8.0E-07</td><td>MB231/MB468</td><td>NUC</td></tr>
<tr><td>>hg38_21_+_8433172_8446622_RNA45SN1_45S_50ntFlanks@8472.8496.25|CGGCCCGGCGGATCTTTCCCGCCCC</td><td>>hg38_21_+_8433172_8446622_RNA45SN1_45S_50ntFlanks</td><td>8472</td><td>8496</td><td>25</td><td>CGGCCCGGCGGATCTTTCCCGCCCC</td><td>101.3</td><td>-9.7</td><td>1.8</td><td>-5.3</td><td>1.1E-07</td><td>8.0E-07</td><td>BT20/MB468</td><td>MP</td></tr>
<tr><td>>hg38_21_+_8433172_8446622_RNA45SN1_45S_50ntFlanks@11663.11681.19|ATTTCTGCCCAGTGCTCTG</td><td>>hg38_21_+_8433172_8446622_RNA45SN1_45S_50ntFlanks</td><td>11663</td><td>11681</td><td>19</td><td>ATTTCTGCCCAGTGCTCTG</td><td>247.8</td><td>-4.4</td><td>0.9</td><td>-5.0</td><td>4.6E-07</td><td>8.1E-07</td><td>MB231/MB468</td><td>CYTO</td></tr>
<tr><td>>MI0000446|hsa-mir-125b-1&WithFlank&11|-|122099751|122099850@22.42.21;MI0000470|hsa-mir-125b-2&WithFlank&21|+|16590231|16590331@24.44.21&[MIMAT0000423&hsa-miR-125b-5p&offsets|+1|0;m-67&11|-|122099809|122099830&offsets|+1|0];[MIMAT0000423_1&hsa-miR-125b-5p&offsets|+1|0;m-68&21|+|16590253|16590274&offsets|+1|0]|CCCTGAGACCCTAACTTGTGA</td><td>>MI0000446|hsa-mir-125b-1&WithFlank&11|-|122099751|122099850</td><td>22</td><td>42</td><td>21</td><td>CCCTGAGACCCTAACTTGTGA</td><td>125.9</td><td>-10.4</td><td>1.9</td><td>-5.6</td><td>2.4E-08</td><td>8.1E-07</td><td>BT20/MB231</td><td>MP</td></tr>
<tr><td>>hg38_21_+_8433172_8446622_RNA45SN1_45S_50ntFlanks@9483.9503.21|CGTGTTAGGACCCGAAAGATG</td><td>>hg38_21_+_8433172_8446622_RNA45SN1_45S_50ntFlanks</td><td>9483</td><td>9503</td><td>21</td><td>CGTGTTAGGACCCGAAAGATG</td><td>262.2</td><td>11.4</td><td>2.0</td><td>5.6</td><td>2.4E-08</td><td>8.1E-07</td><td>BT20/MB231</td><td>MP</td></tr>
<tr><td>>hg38_21_+_8433172_8446622_RNA45SN1_45S_50ntFlanks@10351.10388.38|CGAGAACTTTGAAGGCCGAAGTGGAGAAGGGTTCCATG</td><td>>hg38_21_+_8433172_8446622_RNA45SN1_45S_50ntFlanks</td><td>10351</td><td>10388</td><td>38</td><td>CGAGAACTTTGAAGGCCGAAGTGGAGAAGGGTTCCATG</td><td>5734.3</td><td>-5.1</td><td>1.0</td><td>-5.3</td><td>1.1E-07</td><td>8.1E-07</td><td>BT20/MB468</td><td>MP</td></tr>
<tr><td>>hg38_21_+_8433172_8446622_RNA45SN1_45S_50ntFlanks@9573.9597.25|AAATCGGTCGTCCGACCTGGGTATA</td><td>>hg38_21_+_8433172_8446622_RNA45SN1_45S_50ntFlanks</td><td>9573</td><td>9597</td><td>25</td><td>AAATCGGTCGTCCGACCTGGGTATA</td><td>4853.6</td><td>-8.9</td><td>1.7</td><td>-5.3</td><td>1.1E-07</td><td>8.1E-07</td><td>BT20/MB468</td><td>MP</td></tr>
<tr><td>>hg38_21_+_8433172_8446622_RNA45SN1_45S_50ntFlanks@10922.10947.26|CTGGACGAGGCGCCGCCGCCCCCCCC</td><td>>hg38_21_+_8433172_8446622_RNA45SN1_45S_50ntFlanks</td><td>10922</td><td>10947</td><td>26</td><td>CTGGACGAGGCGCCGCCGCCCCCCCC</td><td>491.5</td><td>-8.0</td><td>1.6</td><td>-5.0</td><td>5.2E-07</td><td>8.1E-07</td><td>BT20/MB468</td><td>NUC</td></tr>
<tr><td>>hg38_21_+_8433172_8446622_RNA45SN1_45S_50ntFlanks@3838.3861.24|CTCTCCTACTTGGATAACTGTGGT</td><td>>hg38_21_+_8433172_8446622_RNA45SN1_45S_50ntFlanks</td><td>3838</td><td>3861</td><td>24</td><td>CTCTCCTACTTGGATAACTGTGGT</td><td>361.4</td><td>-3.3</td><td>0.7</td><td>-5.0</td><td>4.8E-07</td><td>8.1E-07</td><td>MB231/MB468</td><td>NUC</td></tr>
<tr><td>>hg38_21_+_8433172_8446622_RNA45SN1_45S_50ntFlanks@10681.10706.26|TTCCGGCGGCGTCCGGTGAGCTCTCG</td><td>>hg38_21_+_8433172_8446622_RNA45SN1_45S_50ntFlanks</td><td>10681</td><td>10706</td><td>26</td><td>TTCCGGCGGCGTCCGGTGAGCTCTCG</td><td>875.9</td><td>-2.7</td><td>0.5</td><td>-5.1</td><td>3.6E-07</td><td>8.1E-07</td><td>BT20/MB468</td><td>MITO</td></tr>
<tr><td>>hg38_21_+_8433172_8446622_RNA45SN1_45S_50ntFlanks@10787.10821.35|ACAGCCTCTGGCATGTTGGAACAATGTAGGTAAGG</td><td>>hg38_21_+_8433172_8446622_RNA45SN1_45S_50ntFlanks</td><td>10787</td><td>10821</td><td>35</td><td>ACAGCCTCTGGCATGTTGGAACAATGTAGGTAAGG</td><td>143.8</td><td>-5.4</td><td>1.1</td><td>-5.1</td><td>4.2E-07</td><td>8.1E-07</td><td>MB231/MB468</td><td>MP</td></tr>
<tr><td>>hg38_21_+_8433172_8446622_RNA45SN1_45S_50ntFlanks@6722.6739.18|ATTGCAGGACACATTGAT</td><td>>hg38_21_+_8433172_8446622_RNA45SN1_45S_50ntFlanks</td><td>6722</td><td>6739</td><td>18</td><td>ATTGCAGGACACATTGAT</td><td>188.4</td><td>-5.2</td><td>1.0</td><td>-5.0</td><td>4.8E-07</td><td>8.1E-07</td><td>MB231/MB468</td><td>NUC</td></tr>
<tr><td>>trnaMT_PheGAA_MT_+_577_647@1.44.44|GTTTATGTAGCTTACCTCCTCAAAGCAATACACTGAAAATGTTT</td><td>>trnaMT_PheGAA_MT_+_577_647</td><td>1</td><td>44</td><td>44</td><td>GTTTATGTAGCTTACCTCCTCAAAGCAATACACTGAAAATGTTT</td><td>592.0</td><td>7.8</td><td>1.5</td><td>5.1</td><td>3.6E-07</td><td>8.1E-07</td><td>BT20/MB468</td><td>MITO</td></tr>
<tr><td>>MI0000065|hsa-let-7d&WithFlank&9|+|94178828|94178926@14.35.22&[MIMAT0000065&hsa-let-7d-5p&offsets|0|0;m-90&9|+|94178841|94178862&offsets|0|0]|AGAGGTAGTAGGTTGCATAGTT</td><td>>MI0000065|hsa-let-7d&WithFlank&9|+|94178828|94178926</td><td>14</td><td>35</td><td>22</td><td>AGAGGTAGTAGGTTGCATAGTT</td><td>4265.2</td><td>2.4</td><td>0.5</td><td>5.0</td><td>5.2E-07</td><td>8.1E-07</td><td>BT20/MB468</td><td>NUC</td></tr>
<tr><td>>hg38_21_+_8433172_8446622_RNA45SN1_45S_50ntFlanks@4130.4148.19|ATTCCGGAGAGGGAGCCTG</td><td>>hg38_21_+_8433172_8446622_RNA45SN1_45S_50ntFlanks</td><td>4130</td><td>4148</td><td>19</td><td>ATTCCGGAGAGGGAGCCTG</td><td>243.1</td><td>-6.9</td><td>1.4</td><td>-5.1</td><td>4.2E-07</td><td>8.1E-07</td><td>MB231/MB468</td><td>MP</td></tr>
<tr><td>>hg38_1_-_228634819_228635039_RNA5S12_5S_50ntFlanks@85.110.26|TCGTCTGATCTCGGAAGCTAAGCAGG</td><td>>hg38_1_-_228634819_228635039_RNA5S12_5S_50ntFlanks</td><td>85</td><td>110</td><td>26</td><td>TCGTCTGATCTCGGAAGCTAAGCAGG</td><td>379.4</td><td>11.9</td><td>2.1</td><td>5.6</td><td>2.4E-08</td><td>8.1E-07</td><td>BT20/MB231</td><td>MP</td></tr>
<tr><td>>hg38_21_+_8433172_8446622_RNA45SN1_45S_50ntFlanks@11907.11929.23|ACGGTGAAGAGACATGAGAGGTG</td><td>>hg38_21_+_8433172_8446622_RNA45SN1_45S_50ntFlanks</td><td>11907</td><td>11929</td><td>23</td><td>ACGGTGAAGAGACATGAGAGGTG</td><td>309.4</td><td>11.6</td><td>2.1</td><td>5.6</td><td>2.4E-08</td><td>8.1E-07</td><td>BT20/MB231</td><td>MP</td></tr>
<tr><td>>hg38_21_+_8433172_8446622_RNA45SN1_45S_50ntFlanks@3769.3786.18|CGCACGGCCGGTACAGTG</td><td>>hg38_21_+_8433172_8446622_RNA45SN1_45S_50ntFlanks</td><td>3769</td><td>3786</td><td>18</td><td>CGCACGGCCGGTACAGTG</td><td>291.3</td><td>-3.7</td><td>0.7</td><td>-5.1</td><td>4.2E-07</td><td>8.1E-07</td><td>MB231/MB468</td><td>MP</td></tr>
<tr><td>>MI0001445|hsa-mir-423&WithFlank&17|+|30117073|30117178@23.42.20(+1C)&[MIMAT0004748&hsa-miR-423-5p&offsets|0|-3(+1C);m-141&17|+|30117095|30117117&offsets|0|-3(+1C)]|TGAGGGGCAGAGAGCGAGACC</td><td>>MI0001445|hsa-mir-423&WithFlank&17|+|30117073|30117178</td><td>23</td><td>42</td><td>20(+1C)</td><td>TGAGGGGCAGAGAGCGAGACC</td><td>1446.0</td><td>7.4</td><td>1.4</td><td>5.3</td><td>1.1E-07</td><td>8.1E-07</td><td>BT20/MB468</td><td>MP</td></tr>
<tr><td>>hg38_21_+_8433172_8446622_RNA45SN1_45S_50ntFlanks@11575.11602.28|ACTTAGAACTGGTGCGGACCAGGGGAAT</td><td>>hg38_21_+_8433172_8446622_RNA45SN1_45S_50ntFlanks</td><td>11575</td><td>11602</td><td>28</td><td>ACTTAGAACTGGTGCGGACCAGGGGAAT</td><td>639.8</td><td>-4.7</td><td>0.9</td><td>-5.3</td><td>1.1E-07</td><td>8.1E-07</td><td>BT20/MB468</td><td>MP</td></tr>
</table>

<table>
<thead><tr><th></th><th></th><th></th><th></th><th></th><th></th><th></th><th></th><th></th><th></th><th></th><th></th><th></th><th></th></tr></thead>
<tbody>
<tr><td>>hg38_21_+_8433172_8446622_RNA45SN1_45S_50ntFlanks@4759.4783.25|ATCAGATACCGTCGTAGTTCCGACC</td><td>>hg38_21_+_8433172_8446622_RNA45SN1_45S_50ntFlanks</td><td>4759</td><td>4783</td><td>25</td><td>ATCAGATACCGTCGTAGTTCCGACC</td><td>4104.8</td><td>-8.4</td><td>1.7</td><td>-5.0</td><td>5.0E-07</td><td>8.1E-07</td><td>MB231/MB468</td><td>MITO</td></tr>
<tr><td>>hg38_21_+_8433172_8446622_RNA45SN1_45S_50ntFlanks@11821.11842.22|CTATCCAGCGAAACCACAGCCA</td><td>>hg38_21_+_8433172_8446622_RNA45SN1_45S_50ntFlanks</td><td>11821</td><td>11842</td><td>22</td><td>CTATCCAGCGAAACCACAGCCA</td><td>147.0</td><td>-5.9</td><td>1.2</td><td>-5.1</td><td>4.1E-07</td><td>8.1E-07</td><td>BT20/MB468</td><td>CYTO</td></tr>
<tr><td>>hg38_21_+_8433172_8446622_RNA45SN1_45S_50ntFlanks@10679.10708.30|GGTTCCGGCGGCGTCCGGTGAGCTCTCGCT</td><td>>hg38_21_+_8433172_8446622_RNA45SN1_45S_50ntFlanks</td><td>10679</td><td>10708</td><td>30</td><td>GGTTCCGGCGGCGTCCGGTGAGCTCTCGCT</td><td>7419.2</td><td>-4.2</td><td>0.8</td><td>-5.1</td><td>3.6E-07</td><td>8.1E-07</td><td>BT20/MB468</td><td>MITO</td></tr>
<tr><td>>hg38_MT_+_1621_3279_RNR2_16S_50ntFlanks@718.745.28|AGCCTGCGTCAGATTAAAACACTGAACT</td><td>>hg38_MT_+_1621_3279_RNR2_16S_50ntFlanks</td><td>718</td><td>745</td><td>28</td><td>AGCCTGCGTCAGATTAAAACACTGAACT</td><td>59.9</td><td>9.1</td><td>1.8</td><td>5.1</td><td>4.2E-07</td><td>8.1E-07</td><td>MB231/MB468</td><td>MP</td></tr>
<tr><td>>MI0000456|hsa-mir-140&WithFlank&16|+|69933075|69933186@69.90.22(+1G)&[MIMAT0004597&hsa-miR-140-3p&offsets|+1|+2(+1G);m-45&16|+|69933143|69933164&offsets|0|0(+1G)]|ACCACAGGGTAGAACCACGGACG</td><td>>MI0000456|hsa-mir-140&WithFlank&16|+|69933075|69933186</td><td>69</td><td>90</td><td>22(+1G)</td><td>ACCACAGGGTAGAACCACGGACG</td><td>46.9</td><td>8.8</td><td>1.7</td><td>5.1</td><td>4.2E-07</td><td>8.1E-07</td><td>MB231/MB468</td><td>MP</td></tr>
<tr><td>>am_trnaMT_ValTAC_MT_+_1602_1670@49.67.19|AACTTAACTTGACCGCTCT</td><td>>am_trnaMT_ValTAC_MT_+_1602_1670</td><td>49</td><td>67</td><td>19</td><td>AACTTAACTTGACCGCTCT</td><td>8353.7</td><td>7.8</td><td>1.5</td><td>5.1</td><td>4.1E-07</td><td>8.1E-07</td><td>BT20/MB468</td><td>CYTO</td></tr>
<tr><td>>trnaMT_SerGCT_MT_+_12207_12265@13.62.50|AAGAACTGCTAACTCATGCCCCCATGTCTAACAACATGGCTTTCTCACCA</td><td>>trnaMT_SerGCT_MT_+_12207_12265</td><td>13</td><td>62</td><td>50</td><td>AAGAACTGCTAACTCATGCCCCCATGTCTAACAACATGGCTTTCTCACCA</td><td>1223.8</td><td>6.9</td><td>1.4</td><td>5.1</td><td>3.6E-07</td><td>8.1E-07</td><td>BT20/MB468</td><td>MITO</td></tr>
<tr><td>>hg38_21_+_8433172_8446622_RNA45SN1_45S_50ntFlanks@11718.11743.26|AACGGCGGGAGTAACTATGACTCTCT</td><td>>hg38_21_+_8433172_8446622_RNA45SN1_45S_50ntFlanks</td><td>11718</td><td>11743</td><td>26</td><td>AACGGCGGGAGTAACTATGACTCTCT</td><td>854.3</td><td>-6.0</td><td>1.2</td><td>-5.0</td><td>4.7E-07</td><td>8.1E-07</td><td>MB231/MB468</td><td>CYTO</td></tr>
<tr><td>>hg38_21_+_8433172_8446622_RNA45SN1_45S_50ntFlanks@12475.12498.24|CCCACTAATAGGGAACGTGAGCTG</td><td>>hg38_21_+_8433172_8446622_RNA45SN1_45S_50ntFlanks</td><td>12475</td><td>12498</td><td>24</td><td>CCCACTAATAGGGAACGTGAGCTG</td><td>1455.6</td><td>-8.6</td><td>1.7</td><td>-5.0</td><td>5.3E-07</td><td>8.1E-07</td><td>BT20/MB468</td><td>NUC</td></tr>
<tr><td>>hg38_21_+_8433172_8446622_RNA45SN1_45S_50ntFlanks@8192.8215.24|AGCCCGTGGACGGTGTGAGGCCGG</td><td>>hg38_21_+_8433172_8446622_RNA45SN1_45S_50ntFlanks</td><td>8192</td><td>8215</td><td>24</td><td>AGCCCGTGGACGGTGTGAGGCCGG</td><td>224.8</td><td>-7.0</td><td>1.4</td><td>-5.1</td><td>4.2E-07</td><td>8.1E-07</td><td>MB231/MB468</td><td>MP</td></tr>
<tr><td>>am_trna128_GlyGCC_6_-_27870686_27870756@1.35.35,trna18_GlyGCC_16_+_70822597_70822667@1.35.35,trna19_GlyGCC_16_+_70823410_70823480@1.35.35,trna19_GlyGCC_2_-_157257659_157257729@1.35.35,trna24_GlyGCC_16_-_70812942_70813012@1.35.35,trna25_GlyGCC_16_-_70812114_70812184@1.35.35,trna5_GlyGCC_17_+_8029064_8029134@1.35.35,trna68_GlyGCC_1_-_161493637_161493707@1.35.35|GCATTGGTGGTTCAGTGGTAGAATTCTCGCCTGCC</td><td>>am_trna128_GlyGCC_6_-_27870686_27870756</td><td>1</td><td>35</td><td>35</td><td>GCATTGGTGGTTCAGTGGTAGAATTCTCGCCTGCC</td><td>1099.6</td><td>5.3</td><td>1.0</td><td>5.1</td><td>3.6E-07</td><td>8.1E-07</td><td>BT20/MB468</td><td>MITO</td></tr>
<tr><td>>trnaMT_GluTTC_MT_-_14674_14742@44.72.29,trnalookalike8_GluTTC_5_-_93905172_93905240@44.72.29|TCGTGGTTGTAGTCCGTGCGAGAATACCA</td><td>>trnaMT_GluTTC_MT_-_14674_14742</td><td>44</td><td>72</td><td>29</td><td>TCGTGGTTGTAGTCCGTGCGAGAATACCA</td><td>2926.4</td><td>5.1</td><td>1.0</td><td>5.1</td><td>3.6E-07</td><td>8.1E-07</td><td>BT20/MB468</td><td>MITO</td></tr>
<tr><td>>hg38_21_+_8433172_8446622_RNA45SN1_45S_50ntFlanks@7991.8009.19|ACGTGGCGACCCGCTGAAT</td><td>>hg38_21_+_8433172_8446622_RNA45SN1_45S_50ntFlanks</td><td>7991</td><td>8009</td><td>19</td><td>ACGTGGCGACCCGCTGAAT</td><td>4132.5</td><td>-2.5</td><td>0.5</td><td>-5.1</td><td>3.6E-07</td><td>8.1E-07</td><td>BT20/MB468</td><td>MITO</td></tr>
<tr><td>>hg38_21_+_8433172_8446622_RNA45SN1_45S_50ntFlanks@12606.12626.21|ATGTGCTTGGCTGAGGAGCCA</td><td>>hg38_21_+_8433172_8446622_RNA45SN1_45S_50ntFlanks</td><td>12606</td><td>12626</td><td>21</td><td>ATGTGCTTGGCTGAGGAGCCA</td><td>357.8</td><td>-4.8</td><td>1.0</td><td>-5.0</td><td>4.7E-07</td><td>8.2E-07</td><td>MB231/MB468</td><td>CYTO</td></tr>
<tr><td>>hg38_21_+_8433172_8446622_RNA45SN1_45S_50ntFlanks@11537.11580.44|CCGCGCGGCGCCTCGCCTCGGCCGGCGCCTAGCAGCCGACTTAG</td><td>>hg38_21_+_8433172_8446622_RNA45SN1_45S_50ntFlanks</td><td>11537</td><td>11580</td><td>44</td><td>CCGCGCGGCGCCTCGCCTCGGCCGGCGCCTAGCAGCCGACTTAG</td><td>964.0</td><td>-6.1</td><td>1.2</td><td>-5.0</td><td>5.1E-07</td><td>8.2E-07</td><td>MB231/MB468</td><td>MITO</td></tr>
<tr><td>>hg38_21_+_8433172_8446622_RNA45SN1_45S_50ntFlanks@4640.4665.26|GCATTCGTATTGCGCCGCTAGAGGTG</td><td>>hg38_21_+_8433172_8446622_RNA45SN1_45S_50ntFlanks</td><td>4640</td><td>4665</td><td>26</td><td>GCATTCGTATTGCGCCGCTAGAGGTG</td><td>58.9</td><td>-9.0</td><td>1.7</td><td>-5.3</td><td>1.1E-07</td><td>8.2E-07</td><td>BT20/MB468</td><td>MP</td></tr>
<tr><td>>hg38_21_+_8433172_8446622_RNA45SN1_45S_50ntFlanks@8079.8106.28|AAGAGCCCAGCGCCGAATCCCCGCCCCG</td><td>>hg38_21_+_8433172_8446622_RNA45SN1_45S_50ntFlanks</td><td>8079</td><td>8106</td><td>28</td><td>AAGAGCCCAGCGCCGAATCCCCGCCCCG</td><td>99.9</td><td>-9.7</td><td>1.8</td><td>-5.3</td><td>1.1E-07</td><td>8.2E-07</td><td>BT20/MB468</td><td>MP</td></tr>
<tr><td>>MI0000746|hsa-mir-99b&WithFlank&19|+|51692606|51692687@13.31.19(+1A)&[MIMAT0000689&hsa-miR-99b-5p&offsets|0|-3(+1A);m-10&19|+|51692618|51692639&offsets|0|-3(+1A)]|CACCCGTAGAACCGACCTTA</td><td>>MI0000746|hsa-mir-99b&WithFlank&19|+|51692606|51692687</td><td>13</td><td>31</td><td>19(+1A)</td><td>CACCCGTAGAACCGACCTTA</td><td>54.4</td><td>4.7</td><td>0.9</td><td>5.0</td><td>5.3E-07</td><td>8.2E-07</td><td>BT20/MB468</td><td>NUC</td></tr>
<tr><td>>hg38_21_+_8433172_8446622_RNA45SN1_45S_50ntFlanks@9200.9218.19|TCTCTCGGGGCCACGCGCG</td><td>>hg38_21_+_8433172_8446622_RNA45SN1_45S_50ntFlanks</td><td>9200</td><td>9218</td><td>19</td><td>TCTCTCGGGGCCACGCGCG</td><td>73.1</td><td>-9.3</td><td>1.7</td><td>-5.3</td><td>1.2E-07</td><td>8.2E-07</td><td>BT20/MB468</td><td>MP</td></tr>
<tr><td>>hg38_21_+_8433172_8446622_RNA45SN1_45S_50ntFlanks@10690.10707.18|CGTCCGGTGAGCTCTCGC</td><td>>hg38_21_+_8433172_8446622_RNA45SN1_45S_50ntFlanks</td><td>10690</td><td>10707</td><td>18</td><td>CGTCCGGTGAGCTCTCGC</td><td>204.9</td><td>-5.4</td><td>1.1</td><td>-5.0</td><td>4.9E-07</td><td>8.2E-07</td><td>MB231/MB468</td><td>NUC</td></tr>
<tr><td>>MI0000736|hsa-mir-30c-1&WithFlank&1|+|40757278|40757378@23.46.24(+2U);MI0000254|hsa-mir-30c-2&WithFlank&6|-|71376954|71377037@13.36.24(+2U)&[MIMAT0000244&hsa-miR-30c-5p&offsets|0|+1(+2U);m-57&1|+|40757300|40757323&offsets|0|0(+2U)];[MIMAT0000244_1&hsa-miR-30c-5p&offsets|0|+1(+2U);m-56&6|-|71377002|71377025&offsets|0|0(+2U)]|TGTAAACATCCTACACTCTCAGCTTT</td><td>>MI0000736|hsa-mir-30c-1&WithFlank&1|+|40757278|40757378</td><td>23</td><td>46</td><td>24(+2U)</td><td>TGTAAACATCCTACACTCTCAGCTTT</td><td>707.9</td><td>8.6</td><td>1.7</td><td>5.0</td><td>5.1E-07</td><td>8.2E-07</td><td>MB231/MB468</td><td>MITO</td></tr>
<tr><td>>hg38_21_+_8433172_8446622_RNA45SN1_45S_50ntFlanks@12994.13016.23|CCTCGCTGCGATCTATTGAAAGT</td><td>>hg38_21_+_8433172_8446622_RNA45SN1_45S_50ntFlanks</td><td>12994</td><td>13016</td><td>23</td><td>CCTCGCTGCGATCTATTGAAAGT</td><td>318.2</td><td>3.2</td><td>0.6</td><td>5.0</td><td>4.7E-07</td><td>8.2E-07</td><td>MB231/MB468</td><td>CYTO</td></tr>
<tr><td>>MI0000234|hsa-mir-192&WithFlank&11|-|64891131|64891252@30.51.22(+1U)&[MIMAT0000222&hsa-miR-192-5p&offsets|0|+1(+1U);m-59&11|-|64891203|64891223&offsets|0|+1(+1U)]|CTGACCTATGAATTGACAGCCAT</td><td>>MI0000234|hsa-mir-192&WithFlank&11|-|64891131|64891252</td><td>30</td><td>51</td><td>22(+1U)</td><td>CTGACCTATGAATTGACAGCCAT</td><td>324.8</td><td>4.3</td><td>0.8</td><td>5.0</td><td>4.7E-07</td><td>8.2E-07</td><td>MB231/MB468</td><td>CYTO</td></tr>
<tr><td>>hg38_21_+_8433172_8446622_RNA45SN1_45S_50ntFlanks@8179.8214.36|CTGATCGAGGCCCAGCCCGTGGACGGTGTGAGGCCG</td><td>>hg38_21_+_8433172_8446622_RNA45SN1_45S_50ntFlanks</td><td>8179</td><td>8214</td><td>36</td><td>CTGATCGAGGCCCAGCCCGTGGACGGTGTGAGGCCG</td><td>89.0</td><td>-9.6</td><td>1.8</td><td>-5.3</td><td>1.2E-07</td><td>8.2E-07</td><td>BT20/MB468</td><td>MP</td></tr>
<tr><td>>hg38_21_+_8433172_8446622_RNA45SN1_45S_50ntFlanks@11675.11702.28|TGCTCTGAATGTCAAAGTGAAGAAATTC</td><td>>hg38_21_+_8433172_8446622_RNA45SN1_45S_50ntFlanks</td><td>11675</td><td>11702</td><td>28</td><td>TGCTCTGAATGTCAAAGTGAAGAAATTC</td><td>1050.2</td><td>3.6</td><td>0.7</td><td>5.0</td><td>4.9E-07</td><td>8.3E-07</td><td>MB231/MB468</td><td>NUC</td></tr>
<tr><td>>trna103_AsnGTT_1_-_149230570_149230643@33.53.21,trna107_AsnGTT_1_-_148760356_148760429@33.53.21,trna108_AsnGTT_1_-_148598314_148598387@33.53.21,trna1_AsnGTT_19_+_1383562_1383635@33.53.21,trna26_AsnGTT_1_+_148248115_148248188@33.53.21,trna31_AsnGTT_17_-_36908034_36908107@33.53.21,trna47_AsnGTT_1_+_161510031_161510104@33.53.21,trna4_AsnGTT_10_-_22518438_22518511@33.53.21,trna7_AsnGTT_13_-_31248101_31248174@33.53.21,trna83_AsnGTT_1_-_161397867_161397940@33.53.21|CTGTTAACCGAAAGGTTGGTG</td><td>>trna103_AsnGTT_1_-_149230570_149230643</td><td>33</td><td>53</td><td>21</td><td>CTGTTAACCGAAAGGTTGGTG</td><td>100.9</td><td>-9.7</td><td>1.8</td><td>-5.3</td><td>1.2E-07</td><td>8.3E-07</td><td>BT20/MB468</td><td>MP</td></tr>
<tr><td>>hg38_21_+_8433172_8446622_RNA45SN1_45S_50ntFlanks@10841.10858.18|CGTAACTTCGGGATAAGG</td><td>>hg38_21_+_8433172_8446622_RNA45SN1_45S_50ntFlanks</td><td>10841</td><td>10858</td><td>18</td><td>CGTAACTTCGGGATAAGG</td><td>1634.1</td><td>-7.9</td><td>1.6</td><td>-5.1</td><td>4.2E-07</td><td>8.3E-07</td><td>BT20/MB468</td><td>CYTO</td></tr>
<tr><td>>hg38_21_+_8433172_8446622_RNA45SN1_45S_50ntFlanks@9788.9821.34|CGCTGGCGTGGAGCCGGGCGTGGAATGCGAGTGC</td><td>>hg38_21_+_8433172_8446622_RNA45SN1_45S_50ntFlanks</td><td>9788</td><td>9821</td><td>34</td><td>CGCTGGCGTGGAGCCGGGCGTGGAATGCGAGTGC</td><td>181.4</td><td>-6.7</td><td>1.3</td><td>-5.1</td><td>4.3E-07</td><td>8.3E-07</td><td>MB231/MB468</td><td>MP</td></tr>
<tr><td>>hg38_21_+_8433172_8446622_RNA45SN1_45S_50ntFlanks@10846.10867.22|CTTCGGGATAAGGATTGGCTCT</td><td>>hg38_21_+_8433172_8446622_RNA45SN1_45S_50ntFlanks</td><td>10846</td><td>10867</td><td>22</td><td>CTTCGGGATAAGGATTGGCTCT</td><td>129.5</td><td>-5.5</td><td>1.1</td><td>-5.1</td><td>4.3E-07</td><td>8.3E-07</td><td>MB231/MB468</td><td>MP</td></tr>
<tr><td>>MI0000281|hsa-mir-199a-2&WithFlank&1|-|172144529|172144650@76.97.22;MI0000242|hsa-mir-199a-1&WithFlank&19|-|10817420|10817502@53.74.22;MI0000282|hsa-mir-199b&WithFlank&9|-|128244715|128244836@71.92.22&[MIMAT0000232&hsa-miR-199a-3p&offsets|0|0;m-46&1|-|172144555|172144575&offsets|0|+1];[MIMAT0000232_1&hsa-miR-199a-3p&offsets|0|0;m-47&19|-|10817430|10817450&offsets|0|+1];[MIMAT0004563&hsa-miR-199b-3p&offsets|0|0;m-47&9|-|128244746|128244766&offsets|0|+1]|ACAGTAGTCTGCACATTGGTTA</td><td>>MI0000281|hsa-mir-199a-2&WithFlank&1|-|172144529|172144650</td><td>76</td><td>97</td><td>22</td><td>ACAGTAGTCTGCACATTGGTTA</td><td>106.1</td><td>10.5</td><td>2.0</td><td>5.3</td><td>1.2E-07</td><td>8.3E-07</td><td>BT20/MB468</td><td>MP</td></tr>
<tr><td>>trna119_LysCTT_1_-_145395522_145395594@3.34.32,trna11_LysCTT_5_-_180648979_180649051@3.34.32,trna13_LysCTT_6_+_26556774_26556846@3.34.32,trna7_LysCTT_16_+_3225692_3225764@3.34.32,trna9_LysCTT_5_+_180634755_180634827@3.34.32|CCGGCTAGCTCAGTCGGTAGAGCATGAGACTC</td><td>>trna119_LysCTT_1_-_145395522_145395594</td><td>3</td><td>34</td><td>32</td><td>CCGGCTAGCTCAGTCGGTAGAGCATGAGACTC</td><td>526.0</td><td>8.1</td><td>1.6</td><td>5.0</td><td>5.1E-07</td><td>8.3E-07</td><td>MB231/MB468</td><td>MITO</td></tr>
<tr><td>>hg38_21_+_8433172_8446622_RNA45SN1_45S_50ntFlanks@7973.8006.34|GACGCGACCTCAGATCAGACGTGGCGACCCGCTG</td><td>>hg38_21_+_8433172_8446622_RNA45SN1_45S_50ntFlanks</td><td>7973</td><td>8006</td><td>34</td><td>GACGCGACCTCAGATCAGACGTGGCGACCCGCTG</td><td>624.4</td><td>-4.2</td><td>0.8</td><td>-5.1</td><td>4.3E-07</td><td>8.3E-07</td><td>MB231/MB468</td><td>MP</td></tr>
<tr><td>>hg38_21_+_8433172_8446622_RNA45SN1_45S_50ntFlanks@12073.12094.22|CCGAGGGGCTCTCGCTTCTGGC</td><td>>hg38_21_+_8433172_8446622_RNA45SN1_45S_50ntFlanks</td><td>12073</td><td>12094</td><td>22</td><td>CCGAGGGGCTCTCGCTTCTGGC</td><td>845.5</td><td>-3.6</td><td>0.7</td><td>-5.0</td><td>4.8E-07</td><td>8.3E-07</td><td>MB231/MB468</td><td>CYTO</td></tr>
<tr><td>>MI0000455|hsa-mir-138-2&WithFlank&16|+|56858512|56858607@16.37.22(+1A);MI0000476|hsa-mir-138-1&WithFlank&3|+|44114206|44114316@29.50.22(+1A)&[MIMAT0000430&hsa-miR-138-5p&offsets|0|-1(+1A);m-509&16|+|56858527|56858549&offsets|0|-1(+1A)];[MIMAT0000430_1&hsa-miR-138-5p&offsets|0|-1(+1A);m-497&3|+|44114234|44114256&offsets|0|-1(+1A)]|AGCTGGTGTTGTGAATCAGGCCA</td><td>>MI0000455|hsa-mir-138-2&WithFlank&16|+|56858512|56858607</td><td>16</td><td>37</td><td>22(+1A)</td><td>AGCTGGTGTTGTGAATCAGGCCA</td><td>123.6</td><td>-10.3</td><td>1.9</td><td>-5.6</td><td>2.5E-08</td><td>8.3E-07</td><td>BT20/MB231</td><td>MP</td></tr>
<tr><td>>hg38_21_+_8433172_8446622_RNA45SN1_45S_50ntFlanks@10786.10816.31|AACAGCCTCTGGCATGTTGGAACAATGTAGG</td><td>>hg38_21_+_8433172_8446622_RNA45SN1_45S_50ntFlanks</td><td>10786</td><td>10816</td><td>31</td><td>AACAGCCTCTGGCATGTTGGAACAATGTAGG</td><td>712.7</td><td>-3.6</td><td>0.7</td><td>-5.3</td><td>1.2E-07</td><td>8.3E-07</td><td>BT20/MB468</td><td>MP</td></tr>
<tr><td>>hg38_21_+_8433172_8446622_RNA45SN1_45S_50ntFlanks@12437.12454.18|CATTGTGAAGCAGAATTC</td><td>>hg38_21_+_8433172_8446622_RNA45SN1_45S_50ntFlanks</td><td>12437</td><td>12454</td><td>18</td><td>CATTGTGAAGCAGAATTC</td><td>4483.6</td><td>-8.9</td><td>1.7</td><td>-5.3</td><td>1.2E-07</td><td>8.3E-07</td><td>BT20/MB468</td><td>MP</td></tr>
<tr><td>>hg38_21_+_8433172_8446622_RNA45SN1_45S_50ntFlanks@10679.10706.28|GGTTCCGGCGGCGTCCGGTGAGCTCTCG</td><td>>hg38_21_+_8433172_8446622_RNA45SN1_45S_50ntFlanks</td><td>10679</td><td>10706</td><td>28</td><td>GGTTCCGGCGGCGTCCGGTGAGCTCTCG</td><td>458.4</td><td>12.2</td><td>2.2</td><td>5.6</td><td>2.5E-08</td><td>8.3E-07</td><td>BT20/MB231</td><td>MP</td></tr>
<tr><td>>MI0033673|hsa-mir-10526&WithFlank&11|-|122152097|122152162@41.61.21&[MIMAT0041996&hsa-miR-10526-3p&offsets|0|+1]|AAAAGGGGGCTGAGGTGGAGG</td><td>>MI0033673|hsa-mir-10526&WithFlank&11|-|122152097|122152162</td><td>41</td><td>61</td><td>21</td><td>AAAAGGGGGCTGAGGTGGAGG</td><td>130.6</td><td>-10.4</td><td>1.9</td><td>-5.6</td><td>2.5E-08</td><td>8.3E-07</td><td>BT20/MB231</td><td>MP</td></tr>
<tr><td>>hg38_21_+_8433172_8446622_RNA45SN1_45S_50ntFlanks@7981.8012.32|CTCAGATCAGACGTGGCGACCCGCTGAATTTA</td><td>>hg38_21_+_8433172_8446622_RNA45SN1_45S_50ntFlanks</td><td>7981</td><td>8012</td><td>32</td><td>CTCAGATCAGACGTGGCGACCCGCTGAATTTA</td><td>436.7</td><td>-3.2</td><td>0.6</td><td>-5.1</td><td>3.7E-07</td><td>8.3E-07</td><td>BT20/MB468</td><td>MITO</td></tr>
<tr><td>>hg38_21_+_8433172_8446622_RNA45SN1_45S_50ntFlanks@10351.10368.18|CGAGAACTTTGAAGGCCG</td><td>>hg38_21_+_8433172_8446622_RNA45SN1_45S_50ntFlanks</td><td>10351</td><td>10368</td><td>18</td><td>CGAGAACTTTGAAGGCCG</td><td>19685.2</td><td>-6.2</td><td>1.2</td><td>-5.3</td><td>1.2E-07</td><td>8.3E-07</td><td>BT20/MB468</td><td>MP</td></tr>
<tr><td>>hg38_21_+_8433172_8446622_RNA45SN1_45S_50ntFlanks@11999.12032.34|CCCTGCGGGCCGCCGGTGAAATACCACTACTCTG</td><td>>hg38_21_+_8433172_8446622_RNA45SN1_45S_50ntFlanks</td><td>11999</td><td>12032</td><td>34</td><td>CCCTGCGGGCCGCCGGTGAAATACCACTACTCTG</td><td>13117.3</td><td>-8.0</td><td>1.6</td><td>-5.0</td><td>4.8E-07</td><td>8.3E-07</td><td>MB231/MB468</td><td>CYTO</td></tr>
<tr><td>>hg38_21_+_8433172_8446622_RNA45SN1_45S_50ntFlanks@9574.9603.30|AATCGGTCGTCCGACCTGGGTATAGGGGCG</td><td>>hg38_21_+_8433172_8446622_RNA45SN1_45S_50ntFlanks</td><td>9574</td><td>9603</td><td>30</td><td>AATCGGTCGTCCGACCTGGGTATAGGGGCG</td><td>2851.7</td><td>-4.2</td><td>0.8</td><td>-5.0</td><td>5.4E-07</td><td>8.3E-07</td><td>BT20/MB468</td><td>NUC</td></tr>
</tbody>
</table>

<table>
<tr><td>>hg38_21_+_8433172_8446622_RNA45SN1_45S_50ntFlanks@7991.8010.20|ACGTGGCGACCCGCTGAATT</td><td>>hg38_21_+_8433172_8446622_RNA45SN1_45S_50ntFlanks</td><td>7991</td><td>8010</td><td>20</td><td>ACGTGGCGACCCGCTGAATT</td><td>1452.4</td><td>-4.2</td><td>0.8</td><td>-5.1</td><td>4.3E-07</td><td>8.3E-07</td><td>MB231/MB468</td><td>MP</td></tr>
<tr><td>>hg38_21_+_8433172_8446622_RNA45SN1_45S_50ntFlanks@11884.11903.20|GTTGAGCTTGACTCTAGTCT</td><td>>hg38_21_+_8433172_8446622_RNA45SN1_45S_50ntFlanks</td><td>11884</td><td>11903</td><td>20</td><td>GTTGAGCTTGACTCTAGTCT</td><td>767.2</td><td>5.2</td><td>1.0</td><td>5.3</td><td>1.2E-07</td><td>8.3E-07</td><td>BT20/MB468</td><td>MP</td></tr>
<tr><td>>hg38_21_+_8433172_8446622_RNA45SN1_45S_50ntFlanks@9351.9370.20|GCCGTGGCGCAATGAAGGTG</td><td>>hg38_21_+_8433172_8446622_RNA45SN1_45S_50ntFlanks</td><td>9351</td><td>9370</td><td>20</td><td>GCCGTGGCGCAATGAAGGTG</td><td>73.5</td><td>-9.3</td><td>1.8</td><td>-5.3</td><td>1.2E-07</td><td>8.3E-07</td><td>BT20/MB468</td><td>MP</td></tr>
<tr><td>>MI0000285|hsa-mir-205&WithFlank&1|+|209432127|209432248@40.59.20(+1A)&[MIMAT0000266&hsa-miR-205-5p&offsets|0|-2(+1A);m-351&1|+|209432166|209432187&offsets|0|-2(+1A)]|TCCTTCATTCCACCGGAGTCA</td><td>>MI0000285|hsa-mir-205&WithFlank&1|+|209432127|209432248</td><td>40</td><td>59</td><td>20(+1A)</td><td>TCCTTCATTCCACCGGAGTCA</td><td>227.4</td><td>3.8</td><td>0.8</td><td>5.0</td><td>5.4E-07</td><td>8.4E-07</td><td>BT20/MB468</td><td>NUC</td></tr>
<tr><td>>hg38_21_+_8433172_8446622_RNA45SN1_45S_50ntFlanks@6647.6673.27|CGTACGACTCTTAGCGGTGGATCACTC</td><td>>hg38_21_+_8433172_8446622_RNA45SN1_45S_50ntFlanks</td><td>6647</td><td>6673</td><td>27</td><td>CGTACGACTCTTAGCGGTGGATCACTC</td><td>272.2</td><td>4.2</td><td>0.8</td><td>5.0</td><td>5.0E-07</td><td>8.4E-07</td><td>MB231/MB468</td><td>NUC</td></tr>
<tr><td>>hg38_21_+_8433172_8446622_RNA45SN1_45S_50ntFlanks@9346.9370.25|AAGCCGCCGTGGCGCAATGAAGGTG</td><td>>hg38_21_+_8433172_8446622_RNA45SN1_45S_50ntFlanks</td><td>9346</td><td>9370</td><td>25</td><td>AAGCCGCCGTGGCGCAATGAAGGTG</td><td>82.8</td><td>-9.5</td><td>1.8</td><td>-5.3</td><td>1.2E-07</td><td>8.4E-07</td><td>BT20/MB468</td><td>MP</td></tr>
<tr><td>>hg38_21_+_8433172_8446622_RNA45SN1_45S_50ntFlanks@12959.12982.24|TCTGGGTCGGGGTTTCGTACGTAG</td><td>>hg38_21_+_8433172_8446622_RNA45SN1_45S_50ntFlanks</td><td>12959</td><td>12982</td><td>24</td><td>TCTGGGTCGGGGTTTCGTACGTAG</td><td>239.9</td><td>-2.3</td><td>0.5</td><td>-5.0</td><td>4.8E-07</td><td>8.4E-07</td><td>MB231/MB468</td><td>CYTO</td></tr>
<tr><td>>trna9_ArgTCT_1_+_94313129_94313213@1.29.29|GGCTCCGTGGCGCAATGGATAGCGCATTG</td><td>>trna9_ArgTCT_1_+_94313129_94313213</td><td>1</td><td>29</td><td>29</td><td>GGCTCCGTGGCGCAATGGATAGCGCATTG</td><td>64.3</td><td>-9.1</td><td>1.7</td><td>-5.3</td><td>1.2E-07</td><td>8.4E-07</td><td>BT20/MB468</td><td>MP</td></tr>
<tr><td>>hg38_21_+_8433172_8446622_RNA45SN1_45S_50ntFlanks@8191.8210.20|CAGCCCGTGGACGGTGTGAG</td><td>>hg38_21_+_8433172_8446622_RNA45SN1_45S_50ntFlanks</td><td>8191</td><td>8210</td><td>20</td><td>CAGCCCGTGGACGGTGTGAG</td><td>81.4</td><td>-9.4</td><td>1.8</td><td>-5.3</td><td>1.2E-07</td><td>8.4E-07</td><td>BT20/MB468</td><td>MP</td></tr>
<tr><td>>hg38_21_+_8433172_8446622_RNA45SN1_45S_50ntFlanks@10404.10431.28|CATGGGTCAGTCGGTCCTGAGAGATGGG</td><td>>hg38_21_+_8433172_8446622_RNA45SN1_45S_50ntFlanks</td><td>10404</td><td>10431</td><td>28</td><td>CATGGGTCAGTCGGTCCTGAGAGATGGG</td><td>207.6</td><td>-4.6</td><td>0.9</td><td>-5.0</td><td>4.9E-07</td><td>8.4E-07</td><td>MB231/MB468</td><td>CYTO</td></tr>
<tr><td>>hg38_21_+_8433172_8446622_RNA45SN1_45S_50ntFlanks@11209.11254.46|CGGCGACTCTGGACGCGAGCCGGGCCCTTCCCGTGGATCGCCCCAG</td><td>>hg38_21_+_8433172_8446622_RNA45SN1_45S_50ntFlanks</td><td>11209</td><td>11254</td><td>46</td><td>CGGCGACTCTGGACGCGAGCCGGGCCCTTCCCGTGGATCGCCCCAG</td><td>12199.2</td><td>-5.2</td><td>1.0</td><td>-5.0</td><td>4.9E-07</td><td>8.4E-07</td><td>MB231/MB468</td><td>CYTO</td></tr>
<tr><td>>hg38_21_+_8433172_8446622_RNA45SN1_45S_50ntFlanks@6647.6690.44|CGTACGACTCTTAGCGGTGGATCACTCGGCTCGTGCGTCGATGA</td><td>>hg38_21_+_8433172_8446622_RNA45SN1_45S_50ntFlanks</td><td>6647</td><td>6690</td><td>44</td><td>CGTACGACTCTTAGCGGTGGATCACTCGGCTCGTGCGTCGATGA</td><td>402.4</td><td>3.0</td><td>0.6</td><td>5.1</td><td>3.7E-07</td><td>8.4E-07</td><td>BT20/MB468</td><td>MITO</td></tr>
<tr><td>>hg38_21_+_8433172_8446622_RNA45SN1_45S_50ntFlanks@10401.10424.24|GAACATGGGTCAGTCGGTCCTGAG</td><td>>hg38_21_+_8433172_8446622_RNA45SN1_45S_50ntFlanks</td><td>10401</td><td>10424</td><td>24</td><td>GAACATGGGTCAGTCGGTCCTGAG</td><td>2794.2</td><td>3.5</td><td>0.7</td><td>5.0</td><td>5.0E-07</td><td>8.4E-07</td><td>MB231/MB468</td><td>NUC</td></tr>
<tr><td>>hg38_21_+_8433172_8446622_RNA45SN1_45S_50ntFlanks@10781.10809.29|AGGTGAACAGCCTCTGGCATGTTGGAACA</td><td>>hg38_21_+_8433172_8446622_RNA45SN1_45S_50ntFlanks</td><td>10781</td><td>10809</td><td>29</td><td>AGGTGAACAGCCTCTGGCATGTTGGAACA</td><td>370.9</td><td>-7.6</td><td>1.5</td><td>-5.0</td><td>5.5E-07</td><td>8.4E-07</td><td>BT20/MB468</td><td>NUC</td></tr>
<tr><td>>MI0025750|hsa-mir-7974&WithFlank&19|-|11495538|11495628@62.83.22&[MIMAT0031177&hsa-miR-7974&offsets|0|-2;m-867&19|-|11495544|11495567&offsets|0|-2]|AGGCTGTGATGCTCTCCTGAGC</td><td>>MI0025750|hsa-mir-7974&WithFlank&19|-|11495538|11495628</td><td>62</td><td>83</td><td>22</td><td>AGGCTGTGATGCTCTCCTGAGC</td><td>325.6</td><td>-5.6</td><td>1.1</td><td>-5.0</td><td>5.5E-07</td><td>8.5E-07</td><td>BT20/MB468</td><td>NUC</td></tr>
<tr><td>>trna119_LysCTT_1_-_145395522_145395594@1.40.40,trna11_LysCTT_5_-_180648979_180649051@1.40.40,trna13_LysCTT_6_+_26556774_26556846@1.40.40,trna7_LysCTT_16_+_3225692_3225764@1.40.40,trna9_LysCTT_5_+_180634755_180634827@1.40.40|GCCCGGCTAGCTCAGTCGGTAGAGCATGAGACTCTTAATC</td><td>>trna119_LysCTT_1_-_145395522_145395594</td><td>1</td><td>40</td><td>40</td><td>GCCCGGCTAGCTCAGTCGGTAGAGCATGAGACTCTTAATC</td><td>159.3</td><td>4.3</td><td>0.9</td><td>5.0</td><td>5.3E-07</td><td>8.5E-07</td><td>MB231/MB468</td><td>MITO</td></tr>
<tr><td>>hg38_MT_+_1621_3279_RNR2_16S_50ntFlanks@622.663.42|TATAACTGAACTCCTCACACCCAATTGGACCAATCTATCACC</td><td>>hg38_MT_+_1621_3279_RNR2_16S_50ntFlanks</td><td>622</td><td>663</td><td>42</td><td>TATAACTGAACTCCTCACACCCAATTGGACCAATCTATCACC</td><td>42.7</td><td>8.6</td><td>1.7</td><td>5.1</td><td>4.4E-07</td><td>8.5E-07</td><td>MB231/MB468</td><td>MP</td></tr>
<tr><td>>hg38_21_+_8433172_8446622_RNA45SN1_45S_50ntFlanks@12525.12549.25|GTTTTACCCTACTGATGATGTGTTG</td><td>>hg38_21_+_8433172_8446622_RNA45SN1_45S_50ntFlanks</td><td>12525</td><td>12549</td><td>25</td><td>GTTTTACCCTACTGATGATGTGTTG</td><td>73.9</td><td>-9.3</td><td>1.8</td><td>-5.3</td><td>1.2E-07</td><td>8.5E-07</td><td>BT20/MB468</td><td>MP</td></tr>
<tr><td>>hg38_21_+_8433172_8446622_RNA45SN1_45S_50ntFlanks@11553.11592.40|CTCGGCCGGCGCCTAGCAGCCGACTTAGAACTGGTGCGGA</td><td>>hg38_21_+_8433172_8446622_RNA45SN1_45S_50ntFlanks</td><td>11553</td><td>11592</td><td>40</td><td>CTCGGCCGGCGCCTAGCAGCCGACTTAGAACTGGTGCGGA</td><td>232.3</td><td>-5.3</td><td>1.1</td><td>-5.0</td><td>5.3E-07</td><td>8.5E-07</td><td>MB231/MB468</td><td>MITO</td></tr>
<tr><td>>hg38_21_+_8433172_8446622_RNA45SN1_45S_50ntFlanks@11569.11591.23|CAGCCGACTTAGAACTGGTGCGG</td><td>>hg38_21_+_8433172_8446622_RNA45SN1_45S_50ntFlanks</td><td>11569</td><td>11591</td><td>23</td><td>CAGCCGACTTAGAACTGGTGCGG</td><td>910.9</td><td>-8.6</td><td>1.6</td><td>-5.3</td><td>1.2E-07</td><td>8.5E-07</td><td>BT20/MB468</td><td>MP</td></tr>
<tr><td>>hg38_21_+_8433172_8446622_RNA45SN1_45S_50ntFlanks@10348.10368.21|AAACGAGAACTTTGAAGGCCG</td><td>>hg38_21_+_8433172_8446622_RNA45SN1_45S_50ntFlanks</td><td>10348</td><td>10368</td><td>21</td><td>AAACGAGAACTTTGAAGGCCG</td><td>3357.9</td><td>-6.9</td><td>1.4</td><td>-5.1</td><td>4.4E-07</td><td>8.5E-07</td><td>MB231/MB468</td><td>MP</td></tr>
<tr><td>>hg38_21_+_8433172_8446622_RNA45SN1_45S_50ntFlanks@6661.6701.41|CGGTGGATCACTCGGCTCGTGCGTCGATGAAGAACGCAGCT</td><td>>hg38_21_+_8433172_8446622_RNA45SN1_45S_50ntFlanks</td><td>6661</td><td>6701</td><td>41</td><td>CGGTGGATCACTCGGCTCGTGCGTCGATGAAGAACGCAGCT</td><td>371.6</td><td>4.7</td><td>0.9</td><td>5.0</td><td>4.9E-07</td><td>8.5E-07</td><td>MB231/MB468</td><td>CYTO</td></tr>
<tr><td>>hg38_21_+_8433172_8446622_RNA45SN1_45S_50ntFlanks@8118.8136.19|GGGACATGTGGCGTACGGA</td><td>>hg38_21_+_8433172_8446622_RNA45SN1_45S_50ntFlanks</td><td>8118</td><td>8136</td><td>19</td><td>GGGACATGTGGCGTACGGA</td><td>1182.1</td><td>9.4</td><td>1.8</td><td>5.3</td><td>1.2E-07</td><td>8.5E-07</td><td>BT20/MB468</td><td>MP</td></tr>
<tr><td>>hg38_21_+_8433172_8446622_RNA45SN1_45S_50ntFlanks@8921.8939.19|CCGGGGCCGAGGGAGCGAG</td><td>>hg38_21_+_8433172_8446622_RNA45SN1_45S_50ntFlanks</td><td>8921</td><td>8939</td><td>19</td><td>CCGGGGCCGAGGGAGCGAG</td><td>67.6</td><td>9.8</td><td>1.9</td><td>5.3</td><td>1.2E-07</td><td>8.5E-07</td><td>BT20/MB468</td><td>MP</td></tr>
<tr><td>>hg38_21_+_8433172_8446622_RNA45SN1_45S_50ntFlanks@12894.12925.32|CTCGCCCGTCACGCACCGCACGTTCGTGGGGA</td><td>>hg38_21_+_8433172_8446622_RNA45SN1_45S_50ntFlanks</td><td>12894</td><td>12925</td><td>32</td><td>CTCGCCCGTCACGCACCGCACGTTCGTGGGGA</td><td>94.1</td><td>-9.6</td><td>1.8</td><td>-5.3</td><td>1.2E-07</td><td>8.5E-07</td><td>BT20/MB468</td><td>MP</td></tr>
<tr><td>>hg38_21_+_8433172_8446622_RNA45SN1_45S_50ntFlanks@11576.11599.24|CTTAGAACTGGTGCGGACCAGGGG</td><td>>hg38_21_+_8433172_8446622_RNA45SN1_45S_50ntFlanks</td><td>11576</td><td>11599</td><td>24</td><td>CTTAGAACTGGTGCGGACCAGGGG</td><td>84.8</td><td>-9.5</td><td>1.8</td><td>-5.3</td><td>1.2E-07</td><td>8.5E-07</td><td>BT20/MB468</td><td>MP</td></tr>
<tr><td>>trna116_GluCTC_1_-_145399233_145399304@32.74.43,trna71_GluCTC_1_-_161439189_161439260@32.74.43,trna74_GluCTC_1_-_161431809_161431880@32.74.43,trna77_GluCTC_1_-_161424398_161424469@32.74.43,trna77_GluCTC_6_+_28949976_28950047@32.74.43,trna80_GluCTC_1_-_161417018_161417089@32.74.43,trna87_GluCTC_6_-_126101393_126101464@32.74.43|CTCTCACCGCCGCGGCCCGGGTTCGATTCCCGGTCAGGGAACC</td><td>>trna116_GluCTC_1_-_145399233_145399304</td><td>32</td><td>74</td><td>43</td><td>CTCTCACCGCCGCGGCCCGGGTTCGATTCCCGGTCAGGGAACC</td><td>212.1</td><td>-4.7</td><td>0.9</td><td>-5.0</td><td>4.9E-07</td><td>8.5E-07</td><td>MB231/MB468</td><td>CYTO</td></tr>
<tr><td>>MI0000542|hsa-mir-320a&WithFlank&8|-|22244960|22245043@47.69.23&[MIMAT0000510&hsa-miR-320a-3p&offsets|-1|0;m-60&8|-|22244975|22244996&offsets|-1|0]|GAAAAGCTGGGTTGAGAGGGCGA</td><td>>MI0000542|hsa-mir-320a&WithFlank&8|-|22244960|22245043</td><td>47</td><td>69</td><td>23</td><td>GAAAAGCTGGGTTGAGAGGGCGA</td><td>545.9</td><td>8.2</td><td>1.6</td><td>5.3</td><td>1.2E-07</td><td>8.5E-07</td><td>BT20/MB468</td><td>MP</td></tr>
<tr><td>>trna116_GluCTC_1_-_145399233_145399304@1.43.43,trna59_GluCTC_1_+_249168447_249168518@1.43.43,trna71_GluCTC_1_-_161439189_161439260@1.43.43,trna74_GluCTC_1_-_161431809_161431880@1.43.43,trna77_GluCTC_1_-_161424398_161424469@1.43.43,trna77_GluCTC_6_+_28949976_28950047@1.43.43,trna80_GluCTC_1_-_161417018_161417089@1.43.43,trna87_GluCTC_6_-_126101393_126101464@1.43.43|TCCCTGGTGGTCTAGTGGTTAGGATTCGGCGCTCTCACCGCCG</td><td>>trna116_GluCTC_1_-_145399233_145399304</td><td>1</td><td>43</td><td>43</td><td>TCCCTGGTGGTCTAGTGGTTAGGATTCGGCGCTCTCACCGCCG</td><td>982.7</td><td>5.7</td><td>1.1</td><td>5.1</td><td>4.3E-07</td><td>8.5E-07</td><td>BT20/MB468</td><td>CYTO</td></tr>
<tr><td>>hg38_21_+_8433172_8446622_RNA45SN1_45S_50ntFlanks@6650.6695.46|ACGACTCTTAGCGGTGGATCACTCGGCTCGTGCGTCGATGAAGAAC</td><td>>hg38_21_+_8433172_8446622_RNA45SN1_45S_50ntFlanks</td><td>6650</td><td>6695</td><td>46</td><td>ACGACTCTTAGCGGTGGATCACTCGGCTCGTGCGTCGATGAAGAAC</td><td>344.1</td><td>7.8</td><td>1.5</td><td>5.1</td><td>4.3E-07</td><td>8.5E-07</td><td>BT20/MB468</td><td>CYTO</td></tr>
<tr><td>>hg38_21_+_8433172_8446622_RNA45SN1_45S_50ntFlanks@11212.11252.41|CGACTCTGGACGCGAGCCGGGCCCTTCCCGTGGATCGCCCC</td><td>>hg38_21_+_8433172_8446622_RNA45SN1_45S_50ntFlanks</td><td>11212</td><td>11252</td><td>41</td><td>CGACTCTGGACGCGAGCCGGGCCCTTCCCGTGGATCGCCCC</td><td>54.4</td><td>-9.6</td><td>1.9</td><td>-5.1</td><td>4.4E-07</td><td>8.5E-07</td><td>MB231/MB468</td><td>MP</td></tr>
<tr><td>>hg38_21_+_8433172_8446622_RNA45SN1_45S_50ntFlanks@6732.6773.42|ACATTGATCATCGACACTTCGAACGCACTTGCGGCCCCGGGT</td><td>>hg38_21_+_8433172_8446622_RNA45SN1_45S_50ntFlanks</td><td>6732</td><td>6773</td><td>42</td><td>ACATTGATCATCGACACTTCGAACGCACTTGCGGCCCCGGGT</td><td>926.3</td><td>4.3</td><td>0.9</td><td>5.0</td><td>5.3E-07</td><td>8.5E-07</td><td>MB231/MB468</td><td>MITO</td></tr>
<tr><td>>hg38_21_+_8433172_8446622_RNA45SN1_45S_50ntFlanks@10458.10484.27|ATGGCCTCCGTTGCCCTCGGCCGATCG</td><td>>hg38_21_+_8433172_8446622_RNA45SN1_45S_50ntFlanks</td><td>10458</td><td>10484</td><td>27</td><td>ATGGCCTCCGTTGCCCTCGGCCGATCG</td><td>520.5</td><td>-5.1</td><td>1.0</td><td>-5.1</td><td>4.3E-07</td><td>8.5E-07</td><td>BT20/MB468</td><td>CYTO</td></tr>
<tr><td>>hg38_21_+_8433172_8446622_RNA45SN1_45S_50ntFlanks@6744.6771.28|GACACTTCGAACGCACTTGCGGCCCCGG</td><td>>hg38_21_+_8433172_8446622_RNA45SN1_45S_50ntFlanks</td><td>6744</td><td>6771</td><td>28</td><td>GACACTTCGAACGCACTTGCGGCCCCGG</td><td>1044.9</td><td>-6.5</td><td>1.3</td><td>-5.0</td><td>5.5E-07</td><td>8.6E-07</td><td>BT20/MB468</td><td>NUC</td></tr>
<tr><td>>hg38_21_+_8433172_8446622_RNA45SN1_45S_50ntFlanks@5238.5282.45|CTGACTGGCTCAGCGTGTGCCTACCCTACGCCGGCAGGCGCGGGT</td><td>>hg38_21_+_8433172_8446622_RNA45SN1_45S_50ntFlanks</td><td>5238</td><td>5282</td><td>45</td><td>CTGACTGGCTCAGCGTGTGCCTACCCTACGCCGGCAGGCGCGGGT</td><td>94.0</td><td>10.3</td><td>1.9</td><td>5.3</td><td>1.2E-07</td><td>8.6E-07</td><td>BT20/MB468</td><td>MP</td></tr>
<tr><td>>hg38_21_+_8433172_8446622_RNA45SN1_45S_50ntFlanks@12447.12467.21|CAGAATTCACCAAGCGTTGGA</td><td>>hg38_21_+_8433172_8446622_RNA45SN1_45S_50ntFlanks</td><td>12447</td><td>12467</td><td>21</td><td>CAGAATTCACCAAGCGTTGGA</td><td>84.8</td><td>-9.5</td><td>1.8</td><td>-5.3</td><td>1.2E-07</td><td>8.6E-07</td><td>BT20/MB468</td><td>MP</td></tr>
<tr><td>>hg38_21_+_8433172_8446622_RNA45SN1_45S_50ntFlanks@11314.11344.31|CCCACCCCACGTCTCGTCGCGCGCGCGTCCG</td><td>>hg38_21_+_8433172_8446622_RNA45SN1_45S_50ntFlanks</td><td>11314</td><td>11344</td><td>31</td><td>CCCACCCCACGTCTCGTCGCGCGCGCGTCCG</td><td>570.2</td><td>-6.7</td><td>1.3</td><td>-5.1</td><td>4.3E-07</td><td>8.6E-07</td><td>BT20/MB468</td><td>CYTO</td></tr>
<tr><td>>hg38_21_+_8433172_8446622_RNA45SN1_45S_50ntFlanks@12415.12435.21|CTTCGATGTCGGCTCTTCCTA</td><td>>hg38_21_+_8433172_8446622_RNA45SN1_45S_50ntFlanks</td><td>12415</td><td>12435</td><td>21</td><td>CTTCGATGTCGGCTCTTCCTA</td><td>2224.0</td><td>-3.5</td><td>0.7</td><td>-5.1</td><td>4.3E-07</td><td>8.6E-07</td><td>BT20/MB468</td><td>CYTO</td></tr>
<tr><td>>hg38_21_+_8433172_8446622_RNA45SN1_45S_50ntFlanks@9791.9812.22|TGGCGTGGAGCCGGGCGTGGAA</td><td>>hg38_21_+_8433172_8446622_RNA45SN1_45S_50ntFlanks</td><td>9791</td><td>9812</td><td>22</td><td>TGGCGTGGAGCCGGGCGTGGAA</td><td>64.8</td><td>9.8</td><td>1.8</td><td>5.3</td><td>1.2E-07</td><td>8.6E-07</td><td>BT20/MB468</td><td>MP</td></tr>
<tr><td>>hg38_21_+_8433172_8446622_RNA45SN1_45S_50ntFlanks@11572.11589.18|CCGACTTAGAACTGGTGC</td><td>>hg38_21_+_8433172_8446622_RNA45SN1_45S_50ntFlanks</td><td>11572</td><td>11589</td><td>18</td><td>CCGACTTAGAACTGGTGC</td><td>724.6</td><td>-5.8</td><td>1.1</td><td>-5.0</td><td>4.4E-07</td><td>8.6E-07</td><td>MB231/MB468</td><td>MP</td></tr>
<tr><td>>hg38_21_+_8433172_8446622_RNA45SN1_45S_50ntFlanks@4560.4582.23|CTAGGAATAATGGAATAGGACCG</td><td>>hg38_21_+_8433172_8446622_RNA45SN1_45S_50ntFlanks</td><td>4560</td><td>4582</td><td>23</td><td>CTAGGAATAATGGAATAGGACCG</td><td>62.1</td><td>-9.0</td><td>1.7</td><td>-5.3</td><td>1.2E-07</td><td>8.6E-07</td><td>BT20/MB468</td><td>MP</td></tr>
<tr><td>>trna152_ValCAC_6_-_27248049_27248121@1.41.41|GCTTCTGTAGTGTAGTGGTTATCACGTTCGCCTCACACGCG</td><td>>trna152_ValCAC_6_-_27248049_27248121</td><td>1</td><td>41</td><td>41</td><td>GCTTCTGTAGTGTAGTGGTTATCACGTTCGCCTCACACGCG</td><td>366.4</td><td>4.9</td><td>1.0</td><td>5.0</td><td>5.0E-07</td><td>8.6E-07</td><td>MB231/MB468</td><td>CYTO</td></tr>
<tr><td>>hg38_21_+_8433172_8446622_RNA45SN1_45S_50ntFlanks@6738.6771.34|ATCATCGACACTTCGAACGCACTTGCGGCCCCGG</td><td>>hg38_21_+_8433172_8446622_RNA45SN1_45S_50ntFlanks</td><td>6738</td><td>6771</td><td>34</td><td>ATCATCGACACTTCGAACGCACTTGCGGCCCCGG</td><td>1219.5</td><td>-6.6</td><td>1.3</td><td>-5.0</td><td>5.0E-07</td><td>8.6E-07</td><td>MB231/MB468</td><td>CYTO</td></tr>
<tr><td>>hg38_MT_+_598_1651_RNR1_12S_50ntFlanks@428.465.38|AAAGTGGCTTTAACATATCTGAACACACAATAGCTAAG</td><td>>hg38_MT_+_598_1651_RNR1_12S_50ntFlanks</td><td>428</td><td>465</td><td>38</td><td>AAAGTGGCTTTAACATATCTGAACACACAATAGCTAAG</td><td>184.6</td><td>4.2</td><td>0.8</td><td>5.0</td><td>5.3E-07</td><td>8.6E-07</td><td>MB231/MB468</td><td>MITO</td></tr>
<tr><td>>hg38_21_+_8433172_8446622_RNA45SN1_45S_50ntFlanks@3853.3881.29|AACTGTGGTAATTCTAGAGCTAATACATG</td><td>>hg38_21_+_8433172_8446622_RNA45SN1_45S_50ntFlanks</td><td>3853</td><td>3881</td><td>29</td><td>AACTGTGGTAATTCTAGAGCTAATACATG</td><td>601.4</td><td>-4.1</td><td>0.8</td><td>-5.0</td><td>5.0E-07</td><td>8.6E-07</td><td>MB231/MB468</td><td>CYTO</td></tr>
</table>

<table>
<tr><td>>MI0000108|hsa-mir-103a-2&WithFlank&20|+|3917488|3917577@54.77.24(+1U);MI0000109|hsa-mir-103a-1&WithFlank&5|-|168560890|168560979@54.77.24(+1U)&[MIMAT0000101&hsa-miR-103a-3p&offsets|0|+1(+1U);m-12&20|+|3917541|3917563&offsets|0|+1(+1U)];[MIMAT0000101_1&hsa-miR-103a-3p&offsets|0|+1(+1U);m-13&5|-|168560904|168560926&offsets|0|+1(+1U);m-44220&5|-|168560908|168560926&offsets|0|+5(+1U)]|AGCAGCATTGTACAGGGCTATGAAT</td><td>>MI0000108|hsa-mir-103a-2&WithFlank&20|+|3917488|3917577</td><td>54</td><td>77</td><td>24(+1U)</td><td>AGCAGCATTGTACAGGGCTATGAAT</td><td>652.5</td><td>7.5</td><td>1.5</td><td>5.0</td><td>5.4E-07</td><td>8.6E-07</td><td>MB231/MB468</td><td>MITO</td></tr>
<tr><td>>hg38_21_+_8433172_8446622_RNA45SN1_45S_50ntFlanks@9507.9533.27|AACTATGCCTGGGCAGGGCGAAGCCAG</td><td>>hg38_21_+_8433172_8446622_RNA45SN1_45S_50ntFlanks</td><td>9507</td><td>9533</td><td>27</td><td>AACTATGCCTGGGCAGGGCGAAGCCAG</td><td>105.5</td><td>-9.8</td><td>1.9</td><td>-5.3</td><td>1.2E-07</td><td>8.6E-07</td><td>BT20/MB468</td><td>MP</td></tr>
<tr><td>>trna3_ArgTCT_11_+_59318767_59318852@1.29.29,trna4_ArgTCT_17_+_8024243_8024330@1.29.29,trna4_ArgTCT_9_-_131102355_131102445@1.29.29,trna52_ArgTCT_6_+_27529963_27530049@1.29.29|GGCTCTGTGGCGCAATGGATAGCGCATTG</td><td>>trna3_ArgTCT_11_+_59318767_59318852</td><td>1</td><td>29</td><td>29</td><td>GGCTCTGTGGCGCAATGGATAGCGCATTG</td><td>141.1</td><td>-5.8</td><td>1.2</td><td>-5.1</td><td>4.3E-07</td><td>8.6E-07</td><td>BT20/MB468</td><td>CYTO</td></tr>
<tr><td>>hg38_21_+_8433172_8446622_RNA45SN1_45S_50ntFlanks@12957.12987.31|CTTCTGGGTCGGGGTTTCGTACGTAGCAGAG</td><td>>hg38_21_+_8433172_8446622_RNA45SN1_45S_50ntFlanks</td><td>12957</td><td>12987</td><td>31</td><td>CTTCTGGGTCGGGGTTTCGTACGTAGCAGAG</td><td>245.8</td><td>-6.0</td><td>1.2</td><td>-5.1</td><td>4.3E-07</td><td>8.6E-07</td><td>BT20/MB468</td><td>CYTO</td></tr>
<tr><td>>hg38_21_+_8433172_8446622_RNA45SN1_45S_50ntFlanks@5236.5263.28|CACTGACTGGCTCAGCGTGTGCCTACCC</td><td>>hg38_21_+_8433172_8446622_RNA45SN1_45S_50ntFlanks</td><td>5236</td><td>5263</td><td>28</td><td>CACTGACTGGCTCAGCGTGTGCCTACCC</td><td>319.1</td><td>-7.9</td><td>1.6</td><td>-5.0</td><td>5.4E-07</td><td>8.6E-07</td><td>MB231/MB468</td><td>MITO</td></tr>
<tr><td>>hg38_21_+_8433172_8446622_RNA45SN1_45S_50ntFlanks@11669.11688.20|GCCCAGTGCTCTGAATGTCA</td><td>>hg38_21_+_8433172_8446622_RNA45SN1_45S_50ntFlanks</td><td>11669</td><td>11688</td><td>20</td><td>GCCCAGTGCTCTGAATGTCA</td><td>301.5</td><td>4.0</td><td>0.8</td><td>5.0</td><td>5.1E-07</td><td>8.6E-07</td><td>MB231/MB468</td><td>NUC</td></tr>
<tr><td>>hg38_21_+_8433172_8446622_RNA45SN1_45S_50ntFlanks@11217.11254.38|CTGGACGCGAGCCGGGCCCTTCCCGTGGATCGCCCCAG</td><td>>hg38_21_+_8433172_8446622_RNA45SN1_45S_50ntFlanks</td><td>11217</td><td>11254</td><td>38</td><td>CTGGACGCGAGCCGGGCCCTTCCCGTGGATCGCCCCAG</td><td>67.0</td><td>-9.1</td><td>1.7</td><td>-5.3</td><td>1.2E-07</td><td>8.6E-07</td><td>BT20/MB468</td><td>MP</td></tr>
<tr><td>>hg38_21_+_8433172_8446622_RNA45SN1_45S_50ntFlanks@11592.11612.21|ACCAGGGGAATCCGACTGTTT</td><td>>hg38_21_+_8433172_8446622_RNA45SN1_45S_50ntFlanks</td><td>11592</td><td>11612</td><td>21</td><td>ACCAGGGGAATCCGACTGTTT</td><td>464.8</td><td>-6.0</td><td>1.2</td><td>-5.0</td><td>5.6E-07</td><td>8.6E-07</td><td>BT20/MB468</td><td>NUC</td></tr>
<tr><td>>MI0000811|hsa-mir-148b&WithFlank&12|+|54337210|54337320@69.89.21&[MIMAT0000759&hsa-miR-148b-3p&offsets|0|-1;m-77&12|+|54337278|54337299&offsets|0|-1]|TCAGTGCATCACAGAACTTTG</td><td>>MI0000811|hsa-mir-148b&WithFlank&12|+|54337210|54337320</td><td>69</td><td>89</td><td>21</td><td>TCAGTGCATCACAGAACTTTG</td><td>113.6</td><td>2.6</td><td>0.5</td><td>5.0</td><td>5.6E-07</td><td>8.6E-07</td><td>BT20/MB468</td><td>NUC</td></tr>
<tr><td>>trnaMT_GluTTC_MT_-_14674_14742@39.61.23,trnalookalike8_GluTTC_5_-_93905172_93905240@39.61.23|ATTGGTCGTGGTTGTAGTCCGTG</td><td>>trnaMT_GluTTC_MT_-_14674_14742</td><td>39</td><td>61</td><td>23</td><td>ATTGGTCGTGGTTGTAGTCCGTG</td><td>1022.8</td><td>5.1</td><td>1.0</td><td>5.3</td><td>1.2E-07</td><td>8.6E-07</td><td>BT20/MB468</td><td>MP</td></tr>
<tr><td>>MI0000088|hsa-mir-30a&WithFlank&6|-|71403545|71403627@54.72.19(+1A)&[MIMAT0000088&hsa-miR-30a-3p&offsets|+1|-2(+1A);m-38&6|-|71403554|71403575&offsets|+1|-2(+1A)]|TTTCAGTCGGATGTTTGCAA</td><td>>MI0000088|hsa-mir-30a&WithFlank&6|-|71403545|71403627</td><td>54</td><td>72</td><td>19(+1A)</td><td>TTTCAGTCGGATGTTTGCAA</td><td>58.6</td><td>9.1</td><td>1.8</td><td>5.0</td><td>4.5E-07</td><td>8.6E-07</td><td>MB231/MB468</td><td>MP</td></tr>
<tr><td>>hg38_21_+_8433172_8446622_RNA45SN1_45S_50ntFlanks@9027.9066.40|CGTCTCCTCGTGGGGGGGCCGGGCCACCCCTCCCACGGCG</td><td>>hg38_21_+_8433172_8446622_RNA45SN1_45S_50ntFlanks</td><td>9027</td><td>9066</td><td>40</td><td>CGTCTCCTCGTGGGGGGGCCGGGCCACCCCTCCCACGGCG</td><td>594.7</td><td>-7.0</td><td>1.4</td><td>-5.1</td><td>3.8E-07</td><td>8.7E-07</td><td>BT20/MB468</td><td>MITO</td></tr>
<tr><td>>hg38_21_+_8433172_8446622_RNA45SN1_45S_50ntFlanks@3844.3879.36|TACTTGGATAACTGTGGTAATTCTAGAGCTAATACA</td><td>>hg38_21_+_8433172_8446622_RNA45SN1_45S_50ntFlanks</td><td>3844</td><td>3879</td><td>36</td><td>TACTTGGATAACTGTGGTAATTCTAGAGCTAATACA</td><td>312.8</td><td>4.3</td><td>0.8</td><td>5.0</td><td>5.0E-07</td><td>8.7E-07</td><td>MB231/MB468</td><td>CYTO</td></tr>
<tr><td>>hg38_21_+_8433172_8446622_RNA45SN1_45S_50ntFlanks@12893.12933.41|CCTCGCCCGTCACGCACCGCACGTTCGTGGGGAACCTGGCG</td><td>>hg38_21_+_8433172_8446622_RNA45SN1_45S_50ntFlanks</td><td>12893</td><td>12933</td><td>41</td><td>CCTCGCCCGTCACGCACCGCACGTTCGTGGGGAACCTGGCG</td><td>2983.1</td><td>-7.5</td><td>1.5</td><td>-5.1</td><td>3.9E-07</td><td>8.7E-07</td><td>BT20/MB468</td><td>MITO</td></tr>
<tr><td>>MI0003820|hsa-mir-454&WithFlank&17|-|59137752|59137878@70.94.25&[MIMAT0003885&hsa-miR-454-3p&offsets|0|+2;m-272&17|-|59137787|59137809&offsets|0|+2]|TAGTGCAATATTGCTTATAGGGTTT</td><td>>MI0003820|hsa-mir-454&WithFlank&17|-|59137752|59137878</td><td>70</td><td>94</td><td>25</td><td>TAGTGCAATATTGCTTATAGGGTTT</td><td>1388.0</td><td>-4.8</td><td>0.9</td><td>-5.5</td><td>4.2E-08</td><td>8.7E-07</td><td>BT20/MB231</td><td>MITO</td></tr>
<tr><td>>hg38_21_+_8433172_8446622_RNA45SN1_45S_50ntFlanks@12417.12443.27|TCGATGTCGGCTCTTCCTATCATTGTG</td><td>>hg38_21_+_8433172_8446622_RNA45SN1_45S_50ntFlanks</td><td>12417</td><td>12443</td><td>27</td><td>TCGATGTCGGCTCTTCCTATCATTGTG</td><td>3467.0</td><td>-3.4</td><td>0.7</td><td>-5.0</td><td>5.2E-07</td><td>8.7E-07</td><td>MB231/MB468</td><td>NUC</td></tr>
<tr><td>>hg38_21_+_8433172_8446622_RNA45SN1_45S_50ntFlanks@10770.10797.28|GCAGGTCTCCAAGGTGAACAGCCTCTGG</td><td>>hg38_21_+_8433172_8446622_RNA45SN1_45S_50ntFlanks</td><td>10770</td><td>10797</td><td>28</td><td>GCAGGTCTCCAAGGTGAACAGCCTCTGG</td><td>61.9</td><td>-9.0</td><td>1.7</td><td>-5.3</td><td>1.2E-07</td><td>8.7E-07</td><td>BT20/MB468</td><td>MP</td></tr>
<tr><td>>hg38_21_+_8433172_8446622_RNA45SN1_45S_50ntFlanks@12624.12657.34|CCAATGGGGCGAAGCTACCATCTGTGGGATTATG</td><td>>hg38_21_+_8433172_8446622_RNA45SN1_45S_50ntFlanks</td><td>12624</td><td>12657</td><td>34</td><td>CCAATGGGGCGAAGCTACCATCTGTGGGATTATG</td><td>279.0</td><td>-5.3</td><td>1.0</td><td>-5.1</td><td>4.4E-07</td><td>8.7E-07</td><td>BT20/MB468</td><td>CYTO</td></tr>
<tr><td>>hg38_21_+_8433172_8446622_RNA45SN1_45S_50ntFlanks@7970.8011.42|CGAGACGCGACCTCAGATCAGACGTGGCGACCCGCTGAATTT</td><td>>hg38_21_+_8433172_8446622_RNA45SN1_45S_50ntFlanks</td><td>7970</td><td>8011</td><td>42</td><td>CGAGACGCGACCTCAGATCAGACGTGGCGACCCGCTGAATTT</td><td>319.8</td><td>4.8</td><td>1.0</td><td>5.0</td><td>5.2E-07</td><td>8.7E-07</td><td>MB231/MB468</td><td>NUC</td></tr>
<tr><td>>hg38_21_+_8433172_8446622_RNA45SN1_45S_50ntFlanks@10677.10706.30|GCGGTTCCGGCGGCGTCCGGTGAGCTCTCG</td><td>>hg38_21_+_8433172_8446622_RNA45SN1_45S_50ntFlanks</td><td>10677</td><td>10706</td><td>30</td><td>GCGGTTCCGGCGGCGTCCGGTGAGCTCTCG</td><td>302.3</td><td>-3.3</td><td>0.6</td><td>-5.0</td><td>5.0E-07</td><td>8.7E-07</td><td>MB231/MB468</td><td>CYTO</td></tr>
<tr><td>>hg38_21_+_8433172_8446622_RNA45SN1_45S_50ntFlanks@9562.9597.36|TCCTGACGTGCAAATCGGTCGTCCGACCTGGGTATA</td><td>>hg38_21_+_8433172_8446622_RNA45SN1_45S_50ntFlanks</td><td>9562</td><td>9597</td><td>36</td><td>TCCTGACGTGCAAATCGGTCGTCCGACCTGGGTATA</td><td>565.3</td><td>-7.0</td><td>1.4</td><td>-5.1</td><td>4.4E-07</td><td>8.7E-07</td><td>BT20/MB468</td><td>CYTO</td></tr>
<tr><td>>hg38_21_+_8433172_8446622_RNA45SN1_45S_50ntFlanks@11482.11502.21|GCGGAGGGGCCGCGGGCCGGT</td><td>>hg38_21_+_8433172_8446622_RNA45SN1_45S_50ntFlanks</td><td>11482</td><td>11502</td><td>21</td><td>GCGGAGGGGCCGCGGGCCGGT</td><td>121.4</td><td>-5.6</td><td>1.1</td><td>-5.1</td><td>4.4E-07</td><td>8.7E-07</td><td>BT20/MB468</td><td>CYTO</td></tr>
<tr><td>>MI0003560|hsa-mir-92b&WithFlank&1|+|155195171|155195278@67.86.20&[MIMAT0003218&hsa-miR-92b-3p&offsets|0|-2;m-115&1|+|155195237|155195258&offsets|0|-2]|TATTGCACTCGTCCCGGCCT</td><td>>MI0003560|hsa-mir-92b&WithFlank&1|+|155195171|155195278</td><td>67</td><td>86</td><td>20</td><td>TATTGCACTCGTCCCGGCCT</td><td>441.5</td><td>9.6</td><td>1.8</td><td>5.3</td><td>1.2E-07</td><td>8.7E-07</td><td>BT20/MB468</td><td>MP</td></tr>
<tr><td>>hg38_21_+_8433172_8446622_RNA45SN1_45S_50ntFlanks@485.507.23|GATGTGGTGACGTCGTGCTCTCC</td><td>>hg38_21_+_8433172_8446622_RNA45SN1_45S_50ntFlanks</td><td>485</td><td>507</td><td>23</td><td>GATGTGGTGACGTCGTGCTCTCC</td><td>82.1</td><td>10.1</td><td>1.9</td><td>5.3</td><td>1.2E-07</td><td>8.7E-07</td><td>BT20/MB468</td><td>MP</td></tr>
<tr><td>>hg38_21_+_8433172_8446622_RNA45SN1_45S_50ntFlanks@12076.12097.22|AGGGGCTCTCGCTTCTGGCGCC</td><td>>hg38_21_+_8433172_8446622_RNA45SN1_45S_50ntFlanks</td><td>12076</td><td>12097</td><td>22</td><td>AGGGGCTCTCGCTTCTGGCGCC</td><td>445.0</td><td>-3.0</td><td>0.6</td><td>-5.0</td><td>5.0E-07</td><td>8.7E-07</td><td>MB231/MB468</td><td>CYTO</td></tr>
<tr><td>>hg38_21_+_8433172_8446622_RNA45SN1_45S_50ntFlanks@5490.5531.42|TGAGAAGACGGTCGAACTTGACTATCTAGAGGAAGTAAAAGT</td><td>>hg38_21_+_8433172_8446622_RNA45SN1_45S_50ntFlanks</td><td>5490</td><td>5531</td><td>42</td><td>TGAGAAGACGGTCGAACTTGACTATCTAGAGGAAGTAAAAGT</td><td>58.7</td><td>9.6</td><td>1.8</td><td>5.3</td><td>1.2E-07</td><td>8.7E-07</td><td>BT20/MB468</td><td>MP</td></tr>
<tr><td>>hg38_21_+_8433172_8446622_RNA45SN1_45S_50ntFlanks@10869.10886.18|AGGGCTGGGTCGGTCGGG</td><td>>hg38_21_+_8433172_8446622_RNA45SN1_45S_50ntFlanks</td><td>10869</td><td>10886</td><td>18</td><td>AGGGCTGGGTCGGTCGGG</td><td>1199.2</td><td>-8.7</td><td>1.7</td><td>-5.0</td><td>5.7E-07</td><td>8.7E-07</td><td>BT20/MB468</td><td>NUC</td></tr>
<tr><td>>hg38_21_+_8433172_8446622_RNA45SN1_45S_50ntFlanks@10401.10422.22|GAACATGGGTCAGTCGGTCCTG</td><td>>hg38_21_+_8433172_8446622_RNA45SN1_45S_50ntFlanks</td><td>10401</td><td>10422</td><td>22</td><td>GAACATGGGTCAGTCGGTCCTG</td><td>290.5</td><td>-3.4</td><td>0.7</td><td>-5.0</td><td>4.5E-07</td><td>8.8E-07</td><td>MB231/MB468</td><td>MP</td></tr>
<tr><td>>hg38_21_+_8433172_8446622_RNA45SN1_45S_50ntFlanks@9504.9522.19|GTGAACTATGCCTGGGCAG</td><td>>hg38_21_+_8433172_8446622_RNA45SN1_45S_50ntFlanks</td><td>9504</td><td>9522</td><td>19</td><td>GTGAACTATGCCTGGGCAG</td><td>1111.4</td><td>-7.3</td><td>1.4</td><td>-5.0</td><td>4.4E-07</td><td>8.8E-07</td><td>BT20/MB468</td><td>CYTO</td></tr>
<tr><td>>hg38_21_+_8433172_8446622_RNA45SN1_45S_50ntFlanks@10393.10426.34|CAGCAGTTGAACATGGGTCAGTCGGTCCTGAGAG</td><td>>hg38_21_+_8433172_8446622_RNA45SN1_45S_50ntFlanks</td><td>10393</td><td>10426</td><td>34</td><td>CAGCAGTTGAACATGGGTCAGTCGGTCCTGAGAG</td><td>1159.4</td><td>-8.5</td><td>1.7</td><td>-5.0</td><td>5.7E-07</td><td>8.8E-07</td><td>BT20/MB468</td><td>NUC</td></tr>
<tr><td>>hg38_21_+_8433172_8446622_RNA45SN1_45S_50ntFlanks@6669.6686.18|CACTCGGCTCGTGCGTCG</td><td>>hg38_21_+_8433172_8446622_RNA45SN1_45S_50ntFlanks</td><td>6669</td><td>6686</td><td>18</td><td>CACTCGGCTCGTGCGTCG</td><td>406.8</td><td>-7.3</td><td>1.4</td><td>-5.3</td><td>1.3E-07</td><td>8.8E-07</td><td>BT20/MB468</td><td>MP</td></tr>
<tr><td>>hg38_21_+_8433172_8446622_RNA45SN1_45S_50ntFlanks@10404.10430.27|CATGGGTCAGTCGGTCCTGAGAGATGG</td><td>>hg38_21_+_8433172_8446622_RNA45SN1_45S_50ntFlanks</td><td>10404</td><td>10430</td><td>27</td><td>CATGGGTCAGTCGGTCCTGAGAGATGG</td><td>441.8</td><td>-4.6</td><td>0.9</td><td>-5.0</td><td>5.1E-07</td><td>8.8E-07</td><td>MB231/MB468</td><td>CYTO</td></tr>
<tr><td>>MI0000462|hsa-mir-152&WithFlank&17|-|48037155|48037253@60.79.20(+1A)&[MIMAT0000438&hsa-miR-152-3p&offsets|0|-1(+1A);m-97&17|-|48037174|48037194&offsets|0|-1(+1A)]|TCAGTGCATGACAGAACTTGA</td><td>>MI0000462|hsa-mir-152&WithFlank&17|-|48037155|48037253</td><td>60</td><td>79</td><td>20(+1A)</td><td>TCAGTGCATGACAGAACTTGA</td><td>115.3</td><td>10.6</td><td>2.0</td><td>5.3</td><td>1.3E-07</td><td>8.8E-07</td><td>BT20/MB468</td><td>MP</td></tr>
<tr><td>>hg38_21_+_8433172_8446622_RNA45SN1_45S_50ntFlanks@9484.9509.26|GTGTTAGGACCCGAAAGATGGTGAAC</td><td>>hg38_21_+_8433172_8446622_RNA45SN1_45S_50ntFlanks</td><td>9484</td><td>9509</td><td>26</td><td>GTGTTAGGACCCGAAAGATGGTGAAC</td><td>65.6</td><td>9.8</td><td>1.9</td><td>5.3</td><td>1.3E-07</td><td>8.8E-07</td><td>BT20/MB468</td><td>MP</td></tr>
<tr><td>>hg38_21_+_8433172_8446622_RNA45SN1_45S_50ntFlanks@11808.11828.21|CTGTCCCTACCTACTATCCAG</td><td>>hg38_21_+_8433172_8446622_RNA45SN1_45S_50ntFlanks</td><td>11808</td><td>11828</td><td>21</td><td>CTGTCCCTACCTACTATCCAG</td><td>93.1</td><td>-9.6</td><td>1.8</td><td>-5.3</td><td>1.3E-07</td><td>8.8E-07</td><td>BT20/MB468</td><td>MP</td></tr>
<tr><td>>hg38_21_+_8433172_8446622_RNA45SN1_45S_50ntFlanks@12907.12933.27|CACCGCACGTTCGTGGGGAACCTGGCG</td><td>>hg38_21_+_8433172_8446622_RNA45SN1_45S_50ntFlanks</td><td>12907</td><td>12933</td><td>27</td><td>CACCGCACGTTCGTGGGGAACCTGGCG</td><td>2483.7</td><td>-9.4</td><td>1.9</td><td>-5.0</td><td>5.7E-07</td><td>8.8E-07</td><td>BT20/MB468</td><td>NUC</td></tr>
<tr><td>>hg38_21_+_8433172_8446622_RNA45SN1_45S_50ntFlanks@10849.10867.19|CGGGATAAGGATTGGCTCT</td><td>>hg38_21_+_8433172_8446622_RNA45SN1_45S_50ntFlanks</td><td>10849</td><td>10867</td><td>19</td><td>CGGGATAAGGATTGGCTCT</td><td>238.5</td><td>-3.2</td><td>0.6</td><td>-5.0</td><td>5.2E-07</td><td>8.8E-07</td><td>MB231/MB468</td><td>NUC</td></tr>
<tr><td>>hg38_21_+_8433172_8446622_RNA45SN1_45S_50ntFlanks@12405.12437.33|CTTTTTGATCCTTCGATGTCGGCTCTTCCTATC</td><td>>hg38_21_+_8433172_8446622_RNA45SN1_45S_50ntFlanks</td><td>12405</td><td>12437</td><td>33</td><td>CTTTTTGATCCTTCGATGTCGGCTCTTCCTATC</td><td>3464.0</td><td>-4.3</td><td>0.9</td><td>-5.0</td><td>5.1E-07</td><td>8.8E-07</td><td>MB231/MB468</td><td>CYTO</td></tr>
<tr><td>>hg38_21_+_8433172_8446622_RNA45SN1_45S_50ntFlanks@8336.8378.43|AAGTACCGTAAGGGAAAGTTGAAAAGAACTTTGAAGAGAGAGT</td><td>>hg38_21_+_8433172_8446622_RNA45SN1_45S_50ntFlanks</td><td>8336</td><td>8378</td><td>43</td><td>AAGTACCGTAAGGGAAAGTTGAAAAGAACTTTGAAGAGAGAGT</td><td>73.1</td><td>9.9</td><td>1.9</td><td>5.3</td><td>1.3E-07</td><td>8.8E-07</td><td>BT20/MB468</td><td>MP</td></tr>
<tr><td>>hg38_21_+_8433172_8446622_RNA45SN1_45S_50ntFlanks@10350.10384.35|ACGAGAACTTTGAAGGCCGAAGTGGAGAAGGGTTC</td><td>>hg38_21_+_8433172_8446622_RNA45SN1_45S_50ntFlanks</td><td>10350</td><td>10384</td><td>35</td><td>ACGAGAACTTTGAAGGCCGAAGTGGAGAAGGGTTC</td><td>902.9</td><td>-3.5</td><td>0.7</td><td>-5.1</td><td>3.9E-07</td><td>8.8E-07</td><td>BT20/MB468</td><td>MITO</td></tr>
<tr><td>>hg38_21_+_8433172_8446622_RNA45SN1_45S_50ntFlanks@11569.11592.24|CAGCCGACTTAGAACTGGTGCGGA</td><td>>hg38_21_+_8433172_8446622_RNA45SN1_45S_50ntFlanks</td><td>11569</td><td>11592</td><td>24</td><td>CAGCCGACTTAGAACTGGTGCGGA</td><td>3774.4</td><td>-9.5</td><td>1.8</td><td>-5.3</td><td>1.3E-07</td><td>8.8E-07</td><td>BT20/MB468</td><td>MP</td></tr>
<tr><td>>hg38_21_+_8433172_8446622_RNA45SN1_45S_50ntFlanks@11767.11794.28|TCTAATTAGTGACGCGCATGAATGGATG</td><td>>hg38_21_+_8433172_8446622_RNA45SN1_45S_50ntFlanks</td><td>11767</td><td>11794</td><td>28</td><td>TCTAATTAGTGACGCGCATGAATGGATG</td><td>783.8</td><td>-5.8</td><td>1.1</td><td>-5.0</td><td>4.6E-07</td><td>8.8E-07</td><td>MB231/MB468</td><td>MP</td></tr>
<tr><td>>hg38_21_+_8433172_8446622_RNA45SN1_45S_50ntFlanks@10735.10755.21|TGTAAATCTCGCGCCGGGCCG</td><td>>hg38_21_+_8433172_8446622_RNA45SN1_45S_50ntFlanks</td><td>10735</td><td>10755</td><td>21</td><td>TGTAAATCTCGCGCCGGGCCG</td><td>864.6</td><td>-6.9</td><td>1.4</td><td>-5.0</td><td>4.5E-07</td><td>8.8E-07</td><td>BT20/MB468</td><td>CYTO</td></tr>
<tr><td>>MI0000114|hsa-mir-107&WithFlank&10|-|89592741|89592833@56.76.21(+1A);MI0000108|hsa-mir-103a-2&WithFlank&20|+|3917488|3917577@54.74.21(+1A);MI0000109|hsa-mir-103a-1&WithFlank&5|-|168560890|168560979@54.74.21(+1A)&[MIMAT0000104&hsa-miR-107&offsets|0|-2(+1A);m-78&10|-|89592758|89592778&offsets|0|0(+1A)];[MIMAT0000101&hsa-miR-103a-3p&offsets|0|-2(+1A);m-12&20|+|3917541|3917563&offsets|0|-2(+1A)];[MIMAT0000101_1&hsa-miR-103a-3p&offsets|0|-2(+1A);m-13&5|-|168560904|168560926&offsets|0|-2(+1A);m-44220&5|-|168560908|168560926&offsets|0|+2(+1A)]|AGCAGCATTGTACAGGGCTATA</td><td>>MI0000114|hsa-mir-107&WithFlank&10|-|89592741|89592833</td><td>56</td><td>76</td><td>21(+1A)</td><td>AGCAGCATTGTACAGGGCTATA</td><td>324.7</td><td>5.5</td><td>1.1</td><td>5.0</td><td>5.1E-07</td><td>8.8E-07</td><td>MB231/MB468</td><td>CYTO</td></tr>
<tr><td>>hg38_21_+_8433172_8446622_RNA45SN1_45S_50ntFlanks@10841.10861.21|CGTAACTTCGGGATAAGGATT</td><td>>hg38_21_+_8433172_8446622_RNA45SN1_45S_50ntFlanks</td><td>10841</td><td>10861</td><td>21</td><td>CGTAACTTCGGGATAAGGATT</td><td>3441.9</td><td>-3.7</td><td>0.7</td><td>-5.0</td><td>5.1E-07</td><td>8.8E-07</td><td>MB231/MB468</td><td>CYTO</td></tr>
</table>

| | | | | | | | | | | | | | |
|---|---|---|---|---|---|---|---|---|---|---|---|---|---|
| >hg38_21_+_8433172_8446622_RNA45SN1_45S_50ntFlanks@9567.9597.31\|ACGTGCAAATCGGTCGTCCGACCTGGGTATA | >hg38_21_+_8433172_8446622_RNA45SN1_45S_50ntFlanks | 9567 | 9597 | 31 | ACGTGCAAATCGGTCGTCCGACCTGGGTATA | 3763.6 | -6.2 | 1.2 | -5.0 | 4.5E-07 | 8.8E-07 | BT20/MB468 | CYTO |
| >hg38_21_+_8433172_8446622_RNA45SN1_45S_50ntFlanks@3802.3832.31\|CATTAAATCAGTTATGGTTCCTTTGGTCGCT | >hg38_21_+_8433172_8446622_RNA45SN1_45S_50ntFlanks | 3802 | 3832 | 31 | CATTAAATCAGTTATGGTTCCTTTGGTCGCT | 156.0 | -5.9 | 1.2 | -5.0 | 5.3E-07 | 8.9E-07 | MB231/MB468 | NUC |
| >hg38_21_+_8433172_8446622_RNA45SN1_45S_50ntFlanks@485.529.45\|GATGTGGTGACGTCGTGCTCTCCCGGGCCGGGTCCGAGCCGCGAC | >hg38_21_+_8433172_8446622_RNA45SN1_45S_50ntFlanks | 485 | 529 | 45 | GATGTGGTGACGTCGTGCTCTCCCGGGCCGGGTCCGAGCCGCGAC | 114.6 | 10.6 | 2.0 | 5.3 | 1.3E-07 | 8.9E-07 | BT20/MB468 | MP |
| >hg38_21_+_8433172_8446622_RNA45SN1_45S_50ntFlanks@12457.12475.19\|CAAGCGTTGGATTGTTCAC | >hg38_21_+_8433172_8446622_RNA45SN1_45S_50ntFlanks | 12457 | 12475 | 19 | CAAGCGTTGGATTGTTCAC | 386.7 | -5.5 | 1.1 | -5.0 | 5.3E-07 | 8.9E-07 | MB231/MB468 | NUC |
| >hg38_21_+_8433172_8446622_RNA45SN1_45S_50ntFlanks@5157.5185.29\|CAAGTGGCGTTCAGCCACCCGAGATTGAG | >hg38_21_+_8433172_8446622_RNA45SN1_45S_50ntFlanks | 5157 | 5185 | 29 | CAAGTGGCGTTCAGCCACCCGAGATTGAG | 303.5 | -10.3 | 2.0 | -5.1 | 3.9E-07 | 8.9E-07 | BT20/MB468 | MITO |
| >MI0000253\|hsa-mir-148a&WithFlank&7\|-\|25949913\|25949992@12.33.22&[MIMAT0004549&hsa-miR-148a-5p&offsets\|0\|0;m-189&7\|-\|25949960\|25949981&offsets\|0\|0]\|AAAGTTCTGAGACACTCCGACT | >MI0000253\|hsa-mir-148a&WithFlank&7\|-\|25949913\|25949992 | 12 | 33 | 22 | AAAGTTCTGAGACACTCCGACT | 680.4 | 4.8 | 0.9 | 5.5 | 3.6E-08 | 8.9E-07 | BT20/MB231 | CYTO |
| >MI0000650\|hsa-mir-200c&WithFlank&12\|+\|6963693\|6963772@50.71.22(+1G)&[MIMAT0000617&hsa-miR-200c-3p&offsets\|0\|-1(+1G);m-54&12\|+\|6963742\|6963764&offsets\|0\|-1(+1G)]\|TAATACTGCCGGGTAATGATGGG | >MI0000650\|hsa-mir-200c&WithFlank&12\|+\|6963693\|6963772 | 50 | 71 | 22(+1G) | TAATACTGCCGGGTAATGATGGG | 3277.0 | 10.3 | 1.9 | 5.5 | 3.6E-08 | 8.9E-07 | BT20/MB231 | CYTO |
| >am_trna116_GluCTC_1_-_145399233_145399304@1.28.28,trna59_GluCTC_1_+_249168447_249168518@1.28.28,trna71_GluCTC_1_-_161439189_161439260@1.28.28,trna74_GluCTC_1_-_161431809_161431880@1.28.28,trna77_GluCTC_1_-_161424398_161424469@1.28.28,trna77_GluCTC_6_+_28949976_28950047@1.28.28,trna80_GluCTC_1_-_161417018_161417089@1.28.28,trna87_GluCTC_6_-_126101393_126101464@1.28.28\|TCCCTGGTGGTCTAGTGGTTAGGATTCG | >am_trna116_GluCTC_1_-_145399233_145399304 | 1 | 28 | 28 | TCCCTGGTGGTCTAGTGGTTAGGATTCG | 5039.7 | -3.9 | 0.8 | -5.0 | 5.5E-07 | 8.9E-07 | MB231/MB468 | MITO |
| >hg38_21_+_8433172_8446622_RNA45SN1_45S_50ntFlanks@4372.4391.20\|AAAAAGCTCGTAGTTGGATC | >hg38_21_+_8433172_8446622_RNA45SN1_45S_50ntFlanks | 4372 | 4391 | 20 | AAAAAGCTCGTAGTTGGATC | 429.3 | -5.7 | 1.1 | -5.0 | 4.6E-07 | 8.9E-07 | MB231/MB468 | MP |
| >hg38_21_+_8433172_8446622_RNA45SN1_45S_50ntFlanks@8344.8375.32\|TAAGGGAAAGTTGAAAAGAACTTTGAAGAGAG | >hg38_21_+_8433172_8446622_RNA45SN1_45S_50ntFlanks | 8344 | 8375 | 32 | TAAGGGAAAGTTGAAAAGAACTTTGAAGAGAG | 1092.0 | 9.6 | 1.8 | 5.3 | 1.3E-07 | 8.9E-07 | BT20/MB468 | MP |
| >hg38_21_+_8433172_8446622_RNA45SN1_45S_50ntFlanks@7974.8008.35\|ACGCGACCTCAGATCAGACGTGGCGACCCGCTGAA | >hg38_21_+_8433172_8446622_RNA45SN1_45S_50ntFlanks | 7974 | 8008 | 35 | ACGCGACCTCAGATCAGACGTGGCGACCCGCTGAA | 52431.7 | 3.2 | 0.6 | 5.1 | 4.0E-07 | 8.9E-07 | BT20/MB468 | MITO |
| >hg38_21_+_8433172_8446622_RNA45SN1_45S_50ntFlanks@11600.11623.24\|AATCCGACTGTTTAATTAAAACAA | >hg38_21_+_8433172_8446622_RNA45SN1_45S_50ntFlanks | 11600 | 11623 | 24 | AATCCGACTGTTTAATTAAAACAA | 1863.3 | 4.5 | 0.8 | 5.5 | 4.4E-08 | 8.9E-07 | BT20/MB231 | MITO |
| >trnaMT_GluTTC_MT_-_14674_14742@19.58.40,trnalookalike8_GluTTC_5_-_93905172_93905240@19.58.40\|CAACGATGGTTTTTCATATCATTGGTCGTGGTTGTAGTCC | >trnaMT_GluTTC_MT_-_14674_14742 | 19 | 58 | 40 | CAACGATGGTTTTTCATATCATTGGTCGTGGTTGTAGTCC | 112.9 | 10.6 | 2.0 | 5.3 | 1.3E-07 | 8.9E-07 | BT20/MB468 | MP |
| >hg38_21_+_8433172_8446622_RNA45SN1_45S_50ntFlanks@9952.9980.29\|CATGGAAGTCGGAATCCGCTAAGGAGTGT | >hg38_21_+_8433172_8446622_RNA45SN1_45S_50ntFlanks | 9952 | 9980 | 29 | CATGGAAGTCGGAATCCGCTAAGGAGTGT | 71.4 | 9.9 | 1.9 | 5.3 | 1.3E-07 | 8.9E-07 | BT20/MB468 | MP |
| >hg38_21_+_8433172_8446622_RNA45SN1_45S_50ntFlanks@12027.12049.23\|ACTCTGATCGTTTTTTCACTGAC | >hg38_21_+_8433172_8446622_RNA45SN1_45S_50ntFlanks | 12027 | 12049 | 23 | ACTCTGATCGTTTTTTCACTGAC | 82.5 | 10.1 | 1.9 | 5.3 | 1.3E-07 | 8.9E-07 | BT20/MB468 | MP |
| >hg38_21_+_8433172_8446622_RNA45SN1_45S_50ntFlanks@11561.11583.23\|GCGCCTAGCAGCCGACTTAGAAC | >hg38_21_+_8433172_8446622_RNA45SN1_45S_50ntFlanks | 11561 | 11583 | 23 | GCGCCTAGCAGCCGACTTAGAAC | 337.0 | -7.6 | 1.5 | -5.0 | 5.8E-07 | 8.9E-07 | BT20/MB468 | NUC |
| >trnaMT_HisGTG_MT_+_12138_12206@18.53.36\|AAAACATCAGATTGTGAATCTGACAACAGAGGCTTA | >trnaMT_HisGTG_MT_+_12138_12206 | 18 | 53 | 36 | AAAACATCAGATTGTGAATCTGACAACAGAGGCTTA | 46.2 | 8.8 | 1.7 | 5.0 | 4.6E-07 | 8.9E-07 | MB231/MB468 | MP |
| >hg38_21_+_8433172_8446622_RNA45SN1_45S_50ntFlanks@4770.4793.24\|TCGTAGTTCCGACCATAAACGATG | >hg38_21_+_8433172_8446622_RNA45SN1_45S_50ntFlanks | 4770 | 4793 | 24 | TCGTAGTTCCGACCATAAACGATG | 550.7 | -8.0 | 1.6 | -5.0 | 5.8E-07 | 8.9E-07 | BT20/MB468 | NUC |
| >hg38_21_+_8433172_8446622_RNA45SN1_45S_50ntFlanks@4045.4069.25\|CCATTCGAACGTCTGCCCTATCAAC | >hg38_21_+_8433172_8446622_RNA45SN1_45S_50ntFlanks | 4045 | 4069 | 25 | CCATTCGAACGTCTGCCCTATCAAC | 132.3 | -9.1 | 1.8 | -5.1 | 4.0E-07 | 8.9E-07 | BT20/MB468 | MITO |
| >hg38_21_+_8433172_8446622_RNA45SN1_45S_50ntFlanks@4243.4264.22\|CTTTCGAGGCCCTGTAATTGGA | >hg38_21_+_8433172_8446622_RNA45SN1_45S_50ntFlanks | 4243 | 4264 | 22 | CTTTCGAGGCCCTGTAATTGGA | 597.5 | -6.3 | 1.3 | -5.0 | 5.8E-07 | 9.0E-07 | BT20/MB468 | NUC |
| >hg38_21_+_8433172_8446622_RNA45SN1_45S_50ntFlanks@4730.4759.30\|CAAGAACGAAAGTCGGAGGTTCGAAGACGA | >hg38_21_+_8433172_8446622_RNA45SN1_45S_50ntFlanks | 4730 | 4759 | 30 | CAAGAACGAAAGTCGGAGGTTCGAAGACGA | 509.7 | -3.3 | 0.6 | -5.1 | 4.0E-07 | 9.0E-07 | BT20/MB468 | MITO |
| >hg38_21_+_8433172_8446622_RNA45SN1_45S_50ntFlanks@7983.8006.24\|CAGATCAGACGTGGCGACCCGCTG | >hg38_21_+_8433172_8446622_RNA45SN1_45S_50ntFlanks | 7983 | 8006 | 24 | CAGATCAGACGTGGCGACCCGCTG | 2391.3 | -4.5 | 0.9 | -5.0 | 4.5E-07 | 9.0E-07 | BT20/MB468 | CYTO |
| >hg38_21_+_8433172_8446622_RNA45SN1_45S_50ntFlanks@6646.6677.32\|TCGTACGACTCTTAGCGGTGGATCACTCGGCT | >hg38_21_+_8433172_8446622_RNA45SN1_45S_50ntFlanks | 6646 | 6677 | 32 | TCGTACGACTCTTAGCGGTGGATCACTCGGCT | 1032.9 | 3.1 | 0.6 | 5.0 | 5.2E-07 | 9.0E-07 | MB231/MB468 | CYTO |
| >hg38_21_+_8433172_8446622_RNA45SN1_45S_50ntFlanks@6732.6758.27\|ACATTGATCATCGACACTTCGAACGCA | >hg38_21_+_8433172_8446622_RNA45SN1_45S_50ntFlanks | 6732 | 6758 | 27 | ACATTGATCATCGACACTTCGAACGCA | 279.2 | -5.3 | 1.1 | -5.0 | 4.7E-07 | 9.0E-07 | MB231/MB468 | MP |
| >hg38_21_+_8433172_8446622_RNA45SN1_45S_50ntFlanks@11846.11886.41\|GAACGGGCTTGGCGGAATCAGCGGGGAAAGAAGACCCTGTT | >hg38_21_+_8433172_8446622_RNA45SN1_45S_50ntFlanks | 11846 | 11886 | 41 | GAACGGGCTTGGCGGAATCAGCGGGGAAAGAAGACCCTGTT | 64.1 | 9.8 | 1.8 | 5.3 | 1.3E-07 | 9.0E-07 | BT20/MB468 | MP |
| >MI0003686\|hsa-mir-542&WithFlank&X\|-\|134541335\|134541443@59.80.22&[MIMAT0003389&hsa-miR-542-3p&offsets\|0\|0;m-136&X\|-\|134541364\|134541385&offsets\|0\|0]\|TGTGACAGATTGATAACTGAAA | >MI0003686\|hsa-mir-542&WithFlank&X\|-\|134541335\|134541443 | 59 | 80 | 22 | TGTGACAGATTGATAACTGAAA | 44.6 | 8.7 | 1.7 | 5.0 | 4.7E-07 | 9.0E-07 | MB231/MB468 | MP |
| >hg38_21_+_8433172_8446622_RNA45SN1_45S_50ntFlanks@3854.3884.31\|ACTGTGGTAATTCTAGAGCTAATACATGCCG | >hg38_21_+_8433172_8446622_RNA45SN1_45S_50ntFlanks | 3854 | 3884 | 31 | ACTGTGGTAATTCTAGAGCTAATACATGCCG | 521.5 | -7.8 | 1.6 | -5.0 | 5.9E-07 | 9.0E-07 | BT20/MB468 | NUC |
| >hg38_21_+_8433172_8446622_RNA45SN1_45S_50ntFlanks@5158.5185.28\|AAGTGGCGTTCAGCCACCCGAGATTGAG | >hg38_21_+_8433172_8446622_RNA45SN1_45S_50ntFlanks | 5158 | 5185 | 28 | AAGTGGCGTTCAGCCACCCGAGATTGAG | 402.0 | -10.7 | 2.1 | -5.1 | 4.0E-07 | 9.0E-07 | BT20/MB468 | MITO |
| >hg38_21_+_8433172_8446622_RNA45SN1_45S_50ntFlanks@10783.10817.35\|GTGAACAGCCTCTGGCATGTTGGAACAATGTAGGT | >hg38_21_+_8433172_8446622_RNA45SN1_45S_50ntFlanks | 10783 | 10817 | 35 | GTGAACAGCCTCTGGCATGTTGGAACAATGTAGGT | 392.8 | -6.1 | 1.2 | -5.0 | 4.7E-07 | 9.0E-07 | MB231/MB468 | MP |
| >hg38_21_+_8433172_8446622_RNA45SN1_45S_50ntFlanks@12082.12108.27\|TCTCGCTTCTGGCGCCAAGCGCCCGGC | >hg38_21_+_8433172_8446622_RNA45SN1_45S_50ntFlanks | 12082 | 12108 | 27 | TCTCGCTTCTGGCGCCAAGCGCCCGGC | 64.7 | -9.1 | 1.7 | -5.3 | 1.3E-07 | 9.0E-07 | BT20/MB468 | MP |
| >am_trnaMT_PheGAA_MT_+_577_647@44.61.18\|TAGACGGGCTCACATCAC | >am_trnaMT_PheGAA_MT_+_577_647 | 44 | 61 | 18 | TAGACGGGCTCACATCAC | 205.2 | 5.8 | 1.1 | 5.0 | 4.6E-07 | 9.0E-07 | BT20/MB468 | CYTO |
| >hg38_21_+_8433172_8446622_RNA45SN1_45S_50ntFlanks@6661.6688.28\|CGGTGGATCACTCGGCTCGTGCGTCGAT | >hg38_21_+_8433172_8446622_RNA45SN1_45S_50ntFlanks | 6661 | 6688 | 28 | CGGTGGATCACTCGGCTCGTGCGTCGAT | 1207.6 | 4.3 | 0.8 | 5.0 | 4.6E-07 | 9.0E-07 | BT20/MB468 | CYTO |
| >MI0000071\|hsa-mir-17&WithFlank&13\|+\|91350599\|91350694@20.39.20(+1A)&[MIMAT0000070&hsa-miR-17-5p&offsets\|0\|-3(+1A);m-88&13\|+\|91350618\|91350640&offsets\|0\|-3(+1A)]\|CAAAGTGCTTACAGTGCAGGA | >MI0000071\|hsa-mir-17&WithFlank&13\|+\|91350599\|91350694 | 20 | 39 | 20(+1A) | CAAAGTGCTTACAGTGCAGGA | 146.2 | -10.6 | 1.9 | -5.6 | 2.7E-08 | 9.0E-07 | BT20/MB231 | MP |
| >hg38_21_+_8433172_8446622_RNA45SN1_45S_50ntFlanks@4738.4767.30\|AAAGTCGGAGGTTCGAAGACGATCAGATAC | >hg38_21_+_8433172_8446622_RNA45SN1_45S_50ntFlanks | 4738 | 4767 | 30 | AAAGTCGGAGGTTCGAAGACGATCAGATAC | 331.2 | 4.8 | 1.0 | 5.0 | 5.4E-07 | 9.0E-07 | MB231/MB468 | NUC |
| >hg38_21_+_8433172_8446622_RNA45SN1_45S_50ntFlanks@7985.8018.34\|GATCAGACGTGGCGACCCGCTGAATTTAAGCATA | >hg38_21_+_8433172_8446622_RNA45SN1_45S_50ntFlanks | 7985 | 8018 | 34 | GATCAGACGTGGCGACCCGCTGAATTTAAGCATA | 98.3 | -9.7 | 1.8 | -5.3 | 1.3E-07 | 9.0E-07 | BT20/MB468 | MP |
| >hg38_21_+_8433172_8446622_RNA45SN1_45S_50ntFlanks@9198.9220.23\|GTTCTCTCGGGGCCACGCGCGCG | >hg38_21_+_8433172_8446622_RNA45SN1_45S_50ntFlanks | 9198 | 9220 | 23 | GTTCTCTCGGGGCCACGCGCGCG | 385.8 | -7.5 | 1.4 | -5.3 | 1.3E-07 | 9.0E-07 | BT20/MB468 | MP |
| >hg38_21_+_8433172_8446622_RNA45SN1_45S_50ntFlanks@8356.8377.22\|GAAAAGAACTTTGAAGAGAGAG | >hg38_21_+_8433172_8446622_RNA45SN1_45S_50ntFlanks | 8356 | 8377 | 22 | GAAAAGAACTTTGAAGAGAGAG | 463.8 | -5.5 | 1.1 | -5.0 | 5.6E-07 | 9.1E-07 | MB231/MB468 | MITO |
| >hg38_21_+_8433172_8446622_RNA45SN1_45S_50ntFlanks@12181.12201.21\|TCAAACGGTAACGCAGGTGTC | >hg38_21_+_8433172_8446622_RNA45SN1_45S_50ntFlanks | 12181 | 12201 | 21 | TCAAACGGTAACGCAGGTGTC | 87.0 | -9.5 | 1.8 | -5.3 | 1.3E-07 | 9.1E-07 | BT20/MB468 | MP |
| >hg38_21_+_8433172_8446622_RNA45SN1_45S_50ntFlanks@10687.10708.22\|CGGCGTCCGGTGAGCTCTCGCT | >hg38_21_+_8433172_8446622_RNA45SN1_45S_50ntFlanks | 10687 | 10708 | 22 | CGGCGTCCGGTGAGCTCTCGCT | 2061.6 | -3.4 | 0.7 | -5.0 | 4.6E-07 | 9.1E-07 | BT20/MB468 | CYTO |
| >MI0000542\|hsa-mir-320a&WithFlank&8\|-\|22244960\|22245043@48.68.21(+1G)&[MIMAT0000510&hsa-miR-320a-3p&offsets\|0\|-1(+1G);m-60&8\|-\|22244975\|22244996&offsets\|0\|-1(+1G)]\|AAAAGCTGGGTTGAGAGGGCGG | >MI0000542\|hsa-mir-320a&WithFlank&8\|-\|22244960\|22245043 | 48 | 68 | 21(+1G) | AAAAGCTGGGTTGAGAGGGCGG | 1933.8 | 7.4 | 1.4 | 5.3 | 1.3E-07 | 9.1E-07 | BT20/MB468 | MP |
| >MI0000809\|hsa-mir-151a&WithFlank&8\|-\|140732558\|140732659@51.73.23&[MIMAT0000757&hsa-miR-151a-3p&offsets\|-2\|0;m-30&8\|-\|140732587\|140732607&offsets\|-2\|0]\|TACTAGACTGAAGCTCCTTGAGG | >MI0000809\|hsa-mir-151a&WithFlank&8\|-\|140732558\|140732659 | 51 | 73 | 23 | TACTAGACTGAAGCTCCTTGAGG | 2612.8 | 4.6 | 0.9 | 5.3 | 1.3E-07 | 9.1E-07 | BT20/MB468 | MP |
| >hg38_21_+_8433172_8446622_RNA45SN1_45S_50ntFlanks@12457.12476.20\|CAAGCGTTGGATTGTTCACC | >hg38_21_+_8433172_8446622_RNA45SN1_45S_50ntFlanks | 12457 | 12476 | 20 | CAAGCGTTGGATTGTTCACC | 4213.1 | -3.4 | 0.7 | -5.0 | 5.4E-07 | 9.1E-07 | MB231/MB468 | NUC |
| >hg38_21_+_8433172_8446622_RNA45SN1_45S_50ntFlanks@10466.10487.22\|CGTTGCCCTCGGCCGATCGAAA | >hg38_21_+_8433172_8446622_RNA45SN1_45S_50ntFlanks | 10466 | 10487 | 22 | CGTTGCCCTCGGCCGATCGAAA | 398.6 | -5.8 | 1.2 | -5.0 | 5.6E-07 | 9.1E-07 | MB231/MB468 | MITO |
| >hg38_21_+_8433172_8446622_RNA45SN1_45S_50ntFlanks@10439.10460.22\|CGTTCCGAAGGGACGGGCGATG | >hg38_21_+_8433172_8446622_RNA45SN1_45S_50ntFlanks | 10439 | 10460 | 22 | CGTTCCGAAGGGACGGGCGATG | 62.0 | -9.0 | 1.7 | -5.3 | 1.3E-07 | 9.1E-07 | BT20/MB468 | MP |
| >hg38_21_+_8433172_8446622_RNA45SN1_45S_50ntFlanks@12874.12892.19\|GCTGGAAAGGCGGCCGCCC | >hg38_21_+_8433172_8446622_RNA45SN1_45S_50ntFlanks | 12874 | 12892 | 19 | GCTGGAAAGGCGGCCGCCC | 74.8 | -9.3 | 1.8 | -5.3 | 1.3E-07 | 9.1E-07 | BT20/MB468 | MP |
| >hg38_21_+_8433172_8446622_RNA45SN1_45S_50ntFlanks@12644.12661.18\|TCTGTGGGATTATGACTG | >hg38_21_+_8433172_8446622_RNA45SN1_45S_50ntFlanks | 12644 | 12661 | 18 | TCTGTGGGATTATGACTG | 561.8 | -6.3 | 1.3 | -5.0 | 5.9E-07 | 9.1E-07 | BT20/MB468 | NUC |
| >MI0003589\|hsa-mir-582&WithFlank&5\|-\|59703600\|59703709@59.81.23&[MIMAT0004797&hsa-miR-582-3p&offsets\|0\|+1;m-147&5\|-\|59703630\|59703651&offsets\|0\|+1]\|TAACTGGTTGAACAACTGAACCC | >MI0003589\|hsa-mir-582&WithFlank&5\|-\|59703600\|59703709 | 59 | 81 | 23 | TAACTGGTTGAACAACTGAACCC | 83.1 | 10.1 | 1.9 | 5.3 | 1.3E-07 | 9.1E-07 | BT20/MB468 | MP |
| >MI0006444\|hsa-mir-1307&WithFlank&10\|-\|103394247\|103394407@87.107.21&[MIMAT0005951&hsa-miR-1307-3p&offsets\|+1\|0;m-66&10\|-\|103394300\|103394322&offsets\|+1\|-1]\|CTCGGCGTGGCGTCGGTCGTG | >MI0006444\|hsa-mir-1307&WithFlank&10\|-\|103394247\|103394407 | 87 | 107 | 21 | CTCGGCGTGGCGTCGGTCGTG | 282.0 | 4.4 | 0.9 | 5.0 | 5.4E-07 | 9.1E-07 | MB231/MB468 | NUC |

| | | | | | | | | | | | | | |
|---|---|---|---|---|---|---|---|---|---|---|---|---|---|
| >hg38_21_+_8433172_8446622_RNA45SN1_45S_50ntFlanks@9483.9505.23\|CGTGTTAGGACCCGAAAGATGGT | >hg38_21_+_8433172_8446622_RNA45SN1_45S_50ntFlanks | 9483 | 9505 | 23 | CGTGTTAGGACCCGAAAGATGGT | 287.7 | 11.5 | 2.1 | 5.6 | 2.8E-08 | 9.1E-07 | BT20/MB231 | MP |
| >hg38_21_+_8433172_8446622_RNA45SN1_45S_50ntFlanks@7977.8011.35\|CGACCTCAGATCAGACGTGGCGACCCGCTGAATTT | >hg38_21_+_8433172_8446622_RNA45SN1_45S_50ntFlanks | 7977 | 8011 | 35 | CGACCTCAGATCAGACGTGGCGACCCGCTGAATTT | 1277.4 | 3.7 | 0.7 | 5.0 | 5.4E-07 | 9.1E-07 | MB231/MB468 | NUC |
| >hg38_21_+_8433172_8446622_RNA45SN1_45S_50ntFlanks@12476.12493.18\|CCACTAATAGGGAACGTG | >hg38_21_+_8433172_8446622_RNA45SN1_45S_50ntFlanks | 12476 | 12493 | 18 | CCACTAATAGGGAACGTG | 263.5 | -5.2 | 1.0 | -5.0 | 5.4E-07 | 9.1E-07 | MB231/MB468 | NUC |
| >hg38_21_+_8433172_8446622_RNA45SN1_45S_50ntFlanks@8184.8206.23\|CGAGGCCCAGCCCGTGGACGGTG | >hg38_21_+_8433172_8446622_RNA45SN1_45S_50ntFlanks | 8184 | 8206 | 23 | CGAGGCCCAGCCCGTGGACGGTG | 381.2 | -7.0 | 1.4 | -5.0 | 5.7E-07 | 9.1E-07 | MB231/MB468 | MITO |
| >hg38_21_+_8433172_8446622_RNA45SN1_45S_50ntFlanks@3777.3800.24\|CGGTACAGTGAAACTGCGAATGGC | >hg38_21_+_8433172_8446622_RNA45SN1_45S_50ntFlanks | 3777 | 3800 | 24 | CGGTACAGTGAAACTGCGAATGGC | 246.6 | -7.6 | 1.5 | -5.0 | 4.7E-07 | 9.1E-07 | MB231/MB468 | MP |
| >hg38_21_+_8433172_8446622_RNA45SN1_45S_50ntFlanks@10927.10946.20\|CGAGGCGCCGCCGCCCCCCC | >hg38_21_+_8433172_8446622_RNA45SN1_45S_50ntFlanks | 10927 | 10946 | 20 | CGAGGCGCCGCCGCCCCCCC | 209.0 | -6.1 | 1.2 | -5.0 | 4.6E-07 | 9.1E-07 | BT20/MB468 | CYTO |
| >hg38_21_+_8433172_8446622_RNA45SN1_45S_50ntFlanks@6647.6676.30\|CGTACGACTCTTAGCGGTGGATCACTCGGC | >hg38_21_+_8433172_8446622_RNA45SN1_45S_50ntFlanks | 6647 | 6676 | 30 | CGTACGACTCTTAGCGGTGGATCACTCGGC | 765.8 | 4.2 | 0.8 | 5.3 | 1.3E-07 | 9.1E-07 | BT20/MB468 | MP |
| >hg38_21_+_8433172_8446622_RNA45SN1_45S_50ntFlanks@9913.9946.34\|AGAAAAGGTGTTGGTTGATATAGACAGCAGGACG | >hg38_21_+_8433172_8446622_RNA45SN1_45S_50ntFlanks | 9913 | 9946 | 34 | AGAAAAGGTGTTGGTTGATATAGACAGCAGGACG | 56.6 | 9.6 | 1.8 | 5.3 | 1.3E-07 | 9.1E-07 | BT20/MB468 | MP |
| >hg38_21_+_8433172_8446622_RNA45SN1_45S_50ntFlanks@11535.11553.19\|TTCCGCGCGGCGCCTCGCC | >hg38_21_+_8433172_8446622_RNA45SN1_45S_50ntFlanks | 11535 | 11553 | 19 | TTCCGCGCGGCGCCTCGCC | 196.1 | -6.0 | 1.2 | -5.0 | 4.6E-07 | 9.2E-07 | BT20/MB468 | CYTO |
| >hg38_21_+_8433172_8446622_RNA45SN1_45S_50ntFlanks@10821.10839.19\|GGAAGTCGGCAAGCCGGAT | >hg38_21_+_8433172_8446622_RNA45SN1_45S_50ntFlanks | 10821 | 10839 | 19 | GGAAGTCGGCAAGCCGGAT | 729.0 | -6.3 | 1.2 | -5.0 | 4.7E-07 | 9.2E-07 | MB231/MB468 | MP |
| >hg38_21_+_8433172_8446622_RNA45SN1_45S_50ntFlanks@7974.7997.24\|ACGCGACCTCAGATCAGACGTGGC | >hg38_21_+_8433172_8446622_RNA45SN1_45S_50ntFlanks | 7974 | 7997 | 24 | ACGCGACCTCAGATCAGACGTGGC | 3329.8 | 3.7 | 0.7 | 5.0 | 6.0E-07 | 9.2E-07 | BT20/MB468 | NUC |
| >hg38_21_+_8433172_8446622_RNA45SN1_45S_50ntFlanks@5077.5097.21\|CCGATAACGAACGAGACTCTG | >hg38_21_+_8433172_8446622_RNA45SN1_45S_50ntFlanks | 5077 | 5097 | 21 | CCGATAACGAACGAGACTCTG | 429.2 | -8.1 | 1.6 | -5.0 | 4.7E-07 | 9.2E-07 | MB231/MB468 | MP |
| >MI0000289\|hsa-mir-181a-1&WithFlank&1\|-\|198859038\|198859159@30.53.24;MI0000269\|hsa-mir-181a-2&WithFlank&9\|+\|124692436\|124692557@45.68.24&[MIMAT0000256&hsa-miR-181a-5p&offsets\|0\|+1;m-32&1\|-\|198859108\|198859130&offsets\|0\|+1];[MIMAT0000256_1&hsa-miR-181a-5p&offsets\|0\|+1;m-32&9\|+\|124692480\|124692502&offsets\|0\|+1]\|AACATTCAACGCTGTCGGTGAGTT | >MI0000289\|hsa-mir-181a-1&WithFlank&1\|-\|198859038\|198859159 | 30 | 53 | 24 | AACATTCAACGCTGTCGGTGAGTT | 434.6 | 5.0 | 1.0 | 5.0 | 5.3E-07 | 9.2E-07 | MB231/MB468 | CYTO |
| >am_trna10_ValCAC_5_-_180649395_180649467@1.34.34,trna18_ValCAC_5_-_180529253_180529325@1.34.34,trna2_ValCAC_5_+_180524070_180524142@1.34.34,trna6_ValCAC_5_+_180600650_180600722@1.34.34,trna85_ValCAC_1_-_161369490_161369562@1.34.34,trna90_ValCAC_1_-_149684088_149684161@1.34.34,trna98_ValCAC_1_-_149298555_149298627@1.34.34,trna9_ValCAC_6_+_26538282_26538354@1.34.34\|GTTTCCGTAGTGTAGTGGTTATCACGTTCGCCTC | >am_trna10_ValCAC_5_-_180649395_180649467 | 1 | 34 | 34 | GTTTCCGTAGTGTAGTGGTTATCACGTTCGCCTC | 626.6 | 7.6 | 1.5 | 5.0 | 5.7E-07 | 9.2E-07 | MB231/MB468 | MITO |
| >hg38_21_+_8433172_8446622_RNA45SN1_45S_50ntFlanks@476.497.22\|TTCGTGATCGATGTGGTGACGT | >hg38_21_+_8433172_8446622_RNA45SN1_45S_50ntFlanks | 476 | 497 | 22 | TTCGTGATCGATGTGGTGACGT | 1420.3 | 8.8 | 1.8 | 5.0 | 5.7E-07 | 9.2E-07 | MB231/MB468 | MITO |
| >hg38_21_+_8433172_8446622_RNA45SN1_45S_50ntFlanks@4007.4042.36\|CCTCGGGCCGATCGCACGCCCCCCGTGGCGGCGACG | >hg38_21_+_8433172_8446622_RNA45SN1_45S_50ntFlanks | 4007 | 4042 | 36 | CCTCGGGCCGATCGCACGCCCCCCGTGGCGGCGACG | 363.7 | -3.0 | 0.6 | -5.0 | 4.7E-07 | 9.2E-07 | BT20/MB468 | CYTO |
| >hg38_21_+_8433172_8446622_RNA45SN1_45S_50ntFlanks@11318.11342.25\|CCCCACGTCTCGTCGCGCGCGCGTC | >hg38_21_+_8433172_8446622_RNA45SN1_45S_50ntFlanks | 11318 | 11342 | 25 | CCCCACGTCTCGTCGCGCGCGCGTC | 1235.0 | -5.3 | 1.1 | -5.0 | 4.7E-07 | 9.2E-07 | BT20/MB468 | CYTO |
| >hg38_21_+_8433172_8446622_RNA45SN1_45S_50ntFlanks@12405.12425.21\|CTTTTTGATCCTTCGATGTCG | >hg38_21_+_8433172_8446622_RNA45SN1_45S_50ntFlanks | 12405 | 12425 | 21 | CTTTTTGATCCTTCGATGTCG | 1405.0 | -7.3 | 1.4 | -5.0 | 4.7E-07 | 9.2E-07 | BT20/MB468 | CYTO |
| >hg38_21_+_8433172_8446622_RNA45SN1_45S_50ntFlanks@6644.6672.29\|CCTCGTACGACTCTTAGCGGTGGATCACT | >hg38_21_+_8433172_8446622_RNA45SN1_45S_50ntFlanks | 6644 | 6672 | 29 | CCTCGTACGACTCTTAGCGGTGGATCACT | 227.0 | 4.4 | 0.9 | 5.0 | 5.7E-07 | 9.2E-07 | MB231/MB468 | MITO |
| >hg38_21_+_8433172_8446622_RNA45SN1_45S_50ntFlanks@3711.3739.29\|GTTGATCCTGCCAGTAGCATATGCTTGTC | >hg38_21_+_8433172_8446622_RNA45SN1_45S_50ntFlanks | 3711 | 3739 | 29 | GTTGATCCTGCCAGTAGCATATGCTTGTC | 317.5 | -3.4 | 0.7 | -5.0 | 5.5E-07 | 9.2E-07 | MB231/MB468 | NUC |
| >MI0000272\|hsa-mir-182&WithFlank&7\|-\|129770377\|129770498@29.53.25&[MIMAT0000259&hsa-miR-182-5p&offsets\|0\|+1;m-22&7\|-\|129770449\|129770470&offsets\|0\|+3]\|TTTGGCAATGGTAGAACTCACACTG | >MI0000272\|hsa-mir-182&WithFlank&7\|-\|129770377\|129770498 | 29 | 53 | 25 | TTTGGCAATGGTAGAACTCACACTG | 1931.1 | 3.0 | 0.6 | 5.0 | 5.5E-07 | 9.2E-07 | MB231/MB468 | NUC |
| >hg38_21_+_8433172_8446622_RNA45SN1_45S_50ntFlanks@12942.12969.28\|TTCGTAGACGACCTGCTTCTGGGTCGGG | >hg38_21_+_8433172_8446622_RNA45SN1_45S_50ntFlanks | 12942 | 12969 | 28 | TTCGTAGACGACCTGCTTCTGGGTCGGG | 61.8 | -9.0 | 1.7 | -5.3 | 1.3E-07 | 9.2E-07 | BT20/MB468 | MP |
| >hg38_21_+_8433172_8446622_RNA45SN1_45S_50ntFlanks@6758.6806.49\|ACTTGCGGCCCCGGGTTCCTCCCGGGGCTACGCCTGTCTGAGCGTCGCT | >hg38_21_+_8433172_8446622_RNA45SN1_45S_50ntFlanks | 6758 | 6806 | 49 | ACTTGCGGCCCCGGGTTCCTCCCGGGGCTACGCCTGTCTGAGCGTCGCT | 388.5 | 7.0 | 1.4 | 5.0 | 5.4E-07 | 9.2E-07 | MB231/MB468 | CYTO |
| >am_trna162_MetCAT_6_-_26758550_26758622@37.54.18,trna164_MetCAT_6_-_26735574_26735646@37.54.18,trna20_MetCAT_16_+_71460396_71460468@37.54.18,trna21_MetCAT_6_+_26701712_26701784@37.54.18,trna27_MetCAT_6_+_26766444_26766516@37.54.18,trna75_MetCAT_6_+_28912352_28912424@37.54.18,trna97_MetCAT_6_-_28921042_28921114@37.54.18\|AATCTGAAGGTCCTGAGT | >am_trna162_MetCAT_6_-_26758550_26758622 | 37 | 54 | 18 | AATCTGAAGGTCCTGAGT | 75.7 | -9.3 | 1.8 | -5.3 | 1.3E-07 | 9.3E-07 | BT20/MB468 | MP |
| >hg38_21_+_8433172_8446622_RNA45SN1_45S_50ntFlanks@12396.12418.23\|GCGACGTCGCTTTTTGATCCTTC | >hg38_21_+_8433172_8446622_RNA45SN1_45S_50ntFlanks | 12396 | 12418 | 23 | GCGACGTCGCTTTTTGATCCTTC | 708.2 | 4.4 | 0.9 | 5.0 | 5.4E-07 | 9.3E-07 | MB231/MB468 | CYTO |
| >hg38_21_+_8433172_8446622_RNA45SN1_45S_50ntFlanks@10737.10767.31\|TAAATCTCGCGCCGGGCCGTACCCATATCCG | >hg38_21_+_8433172_8446622_RNA45SN1_45S_50ntFlanks | 10737 | 10767 | 31 | TAAATCTCGCGCCGGGCCGTACCCATATCCG | 572.8 | -7.1 | 1.4 | -5.0 | 5.5E-07 | 9.3E-07 | MB231/MB468 | NUC |
| >hg38_21_+_8433172_8446622_RNA45SN1_45S_50ntFlanks@475.517.43\|CTTCGTGATCGATGTGGTGACGTCGTGCTCTCCCGGGCCGGGT | >hg38_21_+_8433172_8446622_RNA45SN1_45S_50ntFlanks | 475 | 517 | 43 | CTTCGTGATCGATGTGGTGACGTCGTGCTCTCCCGGGCCGGGT | 69.3 | 9.3 | 1.9 | 5.0 | 4.8E-07 | 9.3E-07 | MB231/MB468 | MP |
| >trnaMT_GluTTC_MT_-_14674_14742@44.72.29,trnalookalike8_GluTTC_5_-_93905172_93905240@44.72.29\|TCGTGGTTGTAGTCCGTGCGAGAATACCA | >trnaMT_GluTTC_MT_-_14674_14742 | 44 | 72 | 29 | TCGTGGTTGTAGTCCGTGCGAGAATACCA | 1145.8 | 4.3 | 0.9 | 5.0 | 5.5E-07 | 9.3E-07 | MB231/MB468 | NUC |
| >trnaMT_ProTGG_MT_-_15956_16023@41.67.27\|ATGGTGGAGTTAAAGACTTTTTCTCTG | >trnaMT_ProTGG_MT_-_15956_16023 | 41 | 67 | 27 | ATGGTGGAGTTAAAGACTTTTTCTCTG | 139.8 | -6.4 | 1.3 | -5.0 | 5.4E-07 | 9.3E-07 | MB231/MB468 | CYTO |
| >hg38_1_-_228634819_228635039_RNA5S12_5S_50ntFlanks@141.169.29\|CCGCCTGGGAATACCGGGTGCTGTAGGCT | >hg38_1_-_228634819_228635039_RNA5S12_5S_50ntFlanks | 141 | 169 | 29 | CCGCCTGGGAATACCGGGTGCTGTAGGCT | 1451.7 | 3.8 | 0.8 | 5.0 | 5.4E-07 | 9.3E-07 | MB231/MB468 | CYTO |
| >hg38_21_+_8433172_8446622_RNA45SN1_45S_50ntFlanks@5489.5515.27\|CTGAGAAGACGGTCGAACTTGACTATC | >hg38_21_+_8433172_8446622_RNA45SN1_45S_50ntFlanks | 5489 | 5515 | 27 | CTGAGAAGACGGTCGAACTTGACTATC | 327.5 | -4.0 | 0.8 | -5.0 | 5.8E-07 | 9.3E-07 | MB231/MB468 | MITO |
| >hg38_21_+_8433172_8446622_RNA45SN1_45S_50ntFlanks@12567.12587.21\|CTCAGTACGAGAGGAACCGCA | >hg38_21_+_8433172_8446622_RNA45SN1_45S_50ntFlanks | 12567 | 12587 | 21 | CTCAGTACGAGAGGAACCGCA | 351.4 | -7.3 | 1.4 | -5.3 | 1.3E-07 | 9.3E-07 | BT20/MB468 | MP |
| >trnaMT_PheGAA_MT_+_577_647@28.56.29\|ATACACTGAAAATGTTTAGACGGGCTCAC | >trnaMT_PheGAA_MT_+_577_647 | 28 | 56 | 29 | ATACACTGAAAATGTTTAGACGGGCTCAC | 83.9 | 10.1 | 1.9 | 5.3 | 1.3E-07 | 9.3E-07 | BT20/MB468 | MP |
| >hg38_21_+_8433172_8446622_RNA45SN1_45S_50ntFlanks@6732.6773.42\|ACATTGATCATCGACACTTCGAACGCACTTGCGGCCCCGGGT | >hg38_21_+_8433172_8446622_RNA45SN1_45S_50ntFlanks | 6732 | 6773 | 42 | ACATTGATCATCGACACTTCGAACGCACTTGCGGCCCCGGGT | 825.9 | 4.8 | 1.0 | 5.0 | 5.6E-07 | 9.3E-07 | MB231/MB468 | NUC |
| >hg38_21_+_8433172_8446622_RNA45SN1_45S_50ntFlanks@12628.12661.34\|TGGGGCGAAGCTACCATCTGTGGGATTATGACTG | >hg38_21_+_8433172_8446622_RNA45SN1_45S_50ntFlanks | 12628 | 12661 | 34 | TGGGGCGAAGCTACCATCTGTGGGATTATGACTG | 3113.0 | -5.1 | 1.0 | -5.1 | 4.1E-07 | 9.3E-07 | BT20/MB468 | MITO |
| >hg38_21_+_8433172_8446622_RNA45SN1_45S_50ntFlanks@8157.8178.22\|CTCGTGGGGGGCCCAAGTCCTT | >hg38_21_+_8433172_8446622_RNA45SN1_45S_50ntFlanks | 8157 | 8178 | 22 | CTCGTGGGGGGCCCAAGTCCTT | 629.0 | -7.6 | 1.4 | -5.3 | 1.3E-07 | 9.3E-07 | BT20/MB468 | MP |
| >hg38_21_+_8433172_8446622_RNA45SN1_45S_50ntFlanks@11583.11600.18\|CTGGTGCGGACCAGGGGA | >hg38_21_+_8433172_8446622_RNA45SN1_45S_50ntFlanks | 11583 | 11600 | 18 | CTGGTGCGGACCAGGGGA | 960.7 | -3.5 | 0.7 | -5.0 | 5.4E-07 | 9.3E-07 | MB231/MB468 | CYTO |
| >am_trnaMT_ArgTCG_MT_+_10405_10469@31.48.18\|TCGACTCATTAAATTATG | >am_trnaMT_ArgTCG_MT_+_10405_10469 | 31 | 48 | 18 | TCGACTCATTAAATTATG | 179.3 | -6.8 | 1.4 | -5.0 | 4.8E-07 | 9.3E-07 | MB231/MB468 | MP |
| >hg38_MT_+_598_1651_RNR1_12S_50ntFlanks@851.876.26\|TGCTTAGTTGAACAGGGCCCTGAAGC | >hg38_MT_+_598_1651_RNR1_12S_50ntFlanks | 851 | 876 | 26 | TGCTTAGTTGAACAGGGCCCTGAAGC | 79.1 | 10.1 | 1.9 | 5.3 | 1.4E-07 | 9.3E-07 | BT20/MB468 | MP |
| >hg38_21_+_8433172_8446622_RNA45SN1_45S_50ntFlanks@11641.11660.20\|CGGCGGGTGTTGACGCGATG | >hg38_21_+_8433172_8446622_RNA45SN1_45S_50ntFlanks | 11641 | 11660 | 20 | CGGCGGGTGTTGACGCGATG | 2722.0 | -7.8 | 1.6 | -5.0 | 5.6E-07 | 9.3E-07 | MB231/MB468 | NUC |
| >hg38_21_+_8433172_8446622_RNA45SN1_45S_50ntFlanks@12539.12568.30\|ATGATGTGTTGTTGCCATGGTAATCCTGCT | >hg38_21_+_8433172_8446622_RNA45SN1_45S_50ntFlanks | 12539 | 12568 | 30 | ATGATGTGTTGTTGCCATGGTAATCCTGCT | 337.9 | -7.5 | 1.5 | -5.0 | 6.1E-07 | 9.3E-07 | BT20/MB468 | NUC |
| >MI0016005\|hsa-mir-3615&WithFlank&17\|+\|74748607\|74748705@57.78.22&[MIMAT0017994&hsa-miR-3615&offsets\|0\|+1;m-271&17\|+\|74748663\|74748684&offsets\|0\|0]\|TCTCTCGGCTCCTCGCGGCTCG | >MI0016005\|hsa-mir-3615&WithFlank&17\|+\|74748607\|74748705 | 57 | 78 | 22 | TCTCTCGGCTCCTCGCGGCTCG | 922.2 | 3.6 | 0.7 | 5.0 | 5.6E-07 | 9.3E-07 | MB231/MB468 | NUC |
| >hg38_21_+_8433172_8446622_RNA45SN1_45S_50ntFlanks@3874.3899.26\|AATACATGCCGACGGGCGCTGACCCC | >hg38_21_+_8433172_8446622_RNA45SN1_45S_50ntFlanks | 3874 | 3899 | 26 | AATACATGCCGACGGGCGCTGACCCC | 772.8 | -3.8 | 0.8 | -5.0 | 5.4E-07 | 9.3E-07 | MB231/MB468 | CYTO |
| >MI0000063\|hsa-let-7b&WithFlank&22\|+\|46113680\|46113774@12.33.22&[MIMAT0000063&hsa-let-7b-5p&offsets\|0\|0;m-14&22\|+\|46113691\|46113712&offsets\|0\|0]\|TGAGGTAGTAGGTTGTGTGGTT | >MI0000063\|hsa-let-7b&WithFlank&22\|+\|46113680\|46113774 | 12 | 33 | 22 | TGAGGTAGTAGGTTGTGTGGTT | 54926.5 | 5.0 | 0.9 | 5.3 | 1.4E-07 | 9.4E-07 | BT20/MB468 | MP |
| >hg38_21_+_8433172_8446622_RNA45SN1_45S_50ntFlanks@10384.10405.22\|CCATGTGAACAGCAGTTGAACA | >hg38_21_+_8433172_8446622_RNA45SN1_45S_50ntFlanks | 10384 | 10405 | 22 | CCATGTGAACAGCAGTTGAACA | 657.7 | -8.5 | 1.6 | -5.3 | 1.4E-07 | 9.4E-07 | BT20/MB468 | MP |
| >hg38_21_+_8433172_8446622_RNA45SN1_45S_50ntFlanks@10684.10708.25\|CGGCGGCGTCCGGTGAGCTCTCGCT | >hg38_21_+_8433172_8446622_RNA45SN1_45S_50ntFlanks | 10684 | 10708 | 25 | CGGCGGCGTCCGGTGAGCTCTCGCT | 1877.4 | -3.8 | 0.8 | -5.0 | 5.8E-07 | 9.4E-07 | MB231/MB468 | MITO |
| >hg38_21_+_8433172_8446622_RNA45SN1_45S_50ntFlanks@12617.12661.45\|TGAGGAGCCAATGGGGCGAAGCTACCATCTGTGGGATTATGACTG | >hg38_21_+_8433172_8446622_RNA45SN1_45S_50ntFlanks | 12617 | 12661 | 45 | TGAGGAGCCAATGGGGCGAAGCTACCATCTGTGGGATTATGACTG | 80.1 | -10.2 | 2.0 | -5.0 | 4.9E-07 | 9.4E-07 | MB231/MB468 | MP |
| >hg38_21_+_8433172_8446622_RNA45SN1_45S_50ntFlanks@3837.3868.32\|CCTCTCCTACTTGGATAACTGTGGTAATTCTA | >hg38_21_+_8433172_8446622_RNA45SN1_45S_50ntFlanks | 3837 | 3868 | 32 | CCTCTCCTACTTGGATAACTGTGGTAATTCTA | 391.2 | -5.7 | 1.1 | -5.0 | 5.8E-07 | 9.4E-07 | MB231/MB468 | MITO |
| >hg38_21_+_8433172_8446622_RNA45SN1_45S_50ntFlanks@8113.8135.23\|GGCGCGGGACATGTGGCGTACGG | >hg38_21_+_8433172_8446622_RNA45SN1_45S_50ntFlanks | 8113 | 8135 | 23 | GGCGCGGGACATGTGGCGTACGG | 282.2 | 11.5 | 2.1 | 5.5 | 2.9E-08 | 9.4E-07 | BT20/MB231 | MP |

| | | | | | | | | | | | | | |
|---|---|---|---|---|---|---|---|---|---|---|---|---|---|
| >hg38_21_+_8433172_8446622_RNA45SN1_45S_50ntFlanks@9789.9819.31\|GCTGGCGTGGAGCCGGGCGTGGAATGCGAGT | >hg38_21_+_8433172_8446622_RNA45SN1_45S_50ntFlanks | 9789 | 9819 | 31 | GCTGGCGTGGAGCCGGGCGTGGAATGCGAGT | 451.8 | 12.2 | 2.2 | 5.5 | 2.9E-08 | 9.4E-07 | BT20/MB231 | MP |
| >hg38_21_+_8433172_8446622_RNA45SN1_45S_50ntFlanks@9805.9830.26\|GCGTGGAATGCGAGTGCCTAGTGGGC | >hg38_21_+_8433172_8446622_RNA45SN1_45S_50ntFlanks | 9805 | 9830 | 26 | GCGTGGAATGCGAGTGCCTAGTGGGC | 397.9 | 12.0 | 2.2 | 5.5 | 2.9E-08 | 9.4E-07 | BT20/MB231 | MP |
| >hg38_21_+_8433172_8446622_RNA45SN1_45S_50ntFlanks@4072.4089.18\|TCGATGGTAGTCGCCGTG | >hg38_21_+_8433172_8446622_RNA45SN1_45S_50ntFlanks | 4072 | 4089 | 18 | TCGATGGTAGTCGCCGTG | 60.4 | -9.0 | 1.7 | -5.3 | 1.4E-07 | 9.4E-07 | BT20/MB468 | MP |
| >hg38_21_+_8433172_8446622_RNA45SN1_45S_50ntFlanks@10408.10456.49\|GGTCAGTCGGTCCTGAGAGATGGGCGAGCGCCGTTCCGAAGGGACGGGC | >hg38_21_+_8433172_8446622_RNA45SN1_45S_50ntFlanks | 10408 | 10456 | 49 | GGTCAGTCGGTCCTGAGAGATGGGCGAGCGCCGTTCCGAAGGGACGGGC | 268.6 | -5.9 | 1.2 | -5.0 | 5.9E-07 | 9.4E-07 | MB231/MB468 | MITO |
| >hg38_21_+_8433172_8446622_RNA45SN1_45S_50ntFlanks@6729.6771.43\|GACACATTGATCATCGACACTTCGAACGCACTTGCGGCCCCGG | >hg38_21_+_8433172_8446622_RNA45SN1_45S_50ntFlanks | 6729 | 6771 | 43 | GACACATTGATCATCGACACTTCGAACGCACTTGCGGCCCCGG | 1202.7 | -5.9 | 1.2 | -5.0 | 5.9E-07 | 9.4E-07 | MB231/MB468 | MITO |
| >MI0000815\|hsa-mir-339&WithFlank&7\|-\|1022927\|1023032@56.76.21&[MIMAT0004702&hsa-miR-339-3p&offsets\|0\|-2;m-235&7\|-\|1022957\|1022977&offsets\|0\|0]\|TGAGCGCCTCGACGACAGAGC | >MI0000815\|hsa-mir-339&WithFlank&7\|-\|1022927\|1023032 | 56 | 76 | 21 | TGAGCGCCTCGACGACAGAGC | 1308.7 | 5.5 | 1.0 | 5.3 | 1.4E-07 | 9.4E-07 | BT20/MB468 | MP |
| >MI0000440\|hsa-mir-27b&WithFlank&9\|+\|95085439\|95085547@67.86.20&[MIMAT0000419&hsa-miR-27b-3p&offsets\|0\|-1;m-39&9\|+\|95085505\|95085525&offsets\|0\|-1]\|TTCACAGTGGCTAAGTTCTG | >MI0000440\|hsa-mir-27b&WithFlank&9\|+\|95085439\|95085547 | 67 | 86 | 20 | TTCACAGTGGCTAAGTTCTG | 4630.5 | 3.3 | 0.7 | 5.0 | 4.9E-07 | 9.4E-07 | MB231/MB468 | MP |
| >hg38_21_+_8433172_8446622_RNA45SN1_45S_50ntFlanks@12477.12497.21\|CACTAATAGGGAACGTGAGCT | >hg38_21_+_8433172_8446622_RNA45SN1_45S_50ntFlanks | 12477 | 12497 | 21 | CACTAATAGGGAACGTGAGCT | 3062.6 | 2.9 | 0.6 | 5.0 | 5.5E-07 | 9.4E-07 | MB231/MB468 | CYTO |
| >hg38_21_+_8433172_8446622_RNA45SN1_45S_50ntFlanks@3782.3814.33\|CAGTGAAACTGCGAATGGCTCATTAAATCAGTT | >hg38_21_+_8433172_8446622_RNA45SN1_45S_50ntFlanks | 3782 | 3814 | 33 | CAGTGAAACTGCGAATGGCTCATTAAATCAGTT | 888.8 | -8.3 | 1.7 | -5.0 | 6.1E-07 | 9.4E-07 | BT20/MB468 | NUC |
| >hg38_21_+_8433172_8446622_RNA45SN1_45S_50ntFlanks@11571.11602.32\|GCCGACTTAGAACTGGTGCGGACCAGGGGAAT | >hg38_21_+_8433172_8446622_RNA45SN1_45S_50ntFlanks | 11571 | 11602 | 32 | GCCGACTTAGAACTGGTGCGGACCAGGGGAAT | 1217.0 | -4.6 | 0.9 | -5.0 | 6.1E-07 | 9.4E-07 | BT20/MB468 | NUC |
| >hg38_21_+_8433172_8446622_RNA45SN1_45S_50ntFlanks@9737.9765.29\|CGAAACGATCTCAACCTATTCTCAAACTT | >hg38_21_+_8433172_8446622_RNA45SN1_45S_50ntFlanks | 9737 | 9765 | 29 | CGAAACGATCTCAACCTATTCTCAAACTT | 554.0 | 4.4 | 0.9 | 5.0 | 5.5E-07 | 9.4E-07 | MB231/MB468 | CYTO |
| >hg38_21_+_8433172_8446622_RNA45SN1_45S_50ntFlanks@11783.11802.20\|CATGAATGGATGAACGAGAT | >hg38_21_+_8433172_8446622_RNA45SN1_45S_50ntFlanks | 11783 | 11802 | 20 | CATGAATGGATGAACGAGAT | 872.2 | -8.2 | 1.6 | -5.0 | 6.1E-07 | 9.4E-07 | BT20/MB468 | NUC |
| >hg38_21_+_8433172_8446622_RNA45SN1_45S_50ntFlanks@8104.8141.38\|CCGCGGCGGGGCGCGGGACATGTGGCGTACGGAAGACC | >hg38_21_+_8433172_8446622_RNA45SN1_45S_50ntFlanks | 8104 | 8141 | 38 | CCGCGGCGGGGCGCGGGACATGTGGCGTACGGAAGACC | 3495.9 | -7.6 | 1.5 | -5.0 | 4.9E-07 | 9.4E-07 | MB231/MB468 | MP |
| >hg38_21_+_8433172_8446622_RNA45SN1_45S_50ntFlanks@12455.12476.22\|ACCAAGCGTTGGATTGTTCACC | >hg38_21_+_8433172_8446622_RNA45SN1_45S_50ntFlanks | 12455 | 12476 | 22 | ACCAAGCGTTGGATTGTTCACC | 857.4 | -4.1 | 0.8 | -5.0 | 4.9E-07 | 9.4E-07 | MB231/MB468 | MP |
| >hg38_21_+_8433172_8446622_RNA45SN1_45S_50ntFlanks@4557.4577.21\|CAGCTAGGAATAATGGAATAG | >hg38_21_+_8433172_8446622_RNA45SN1_45S_50ntFlanks | 4557 | 4577 | 21 | CAGCTAGGAATAATGGAATAG | 139.7 | -2.8 | 0.6 | -5.0 | 5.6E-07 | 9.4E-07 | MB231/MB468 | NUC |
| >hg38_21_+_8433172_8446622_RNA45SN1_45S_50ntFlanks@11581.11612.32\|AACTGGTGCGGACCAGGGGAATCCGACTGTTT | >hg38_21_+_8433172_8446622_RNA45SN1_45S_50ntFlanks | 11581 | 11612 | 32 | AACTGGTGCGGACCAGGGGAATCCGACTGTTT | 1023.4 | -2.7 | 0.5 | -5.0 | 5.6E-07 | 9.5E-07 | MB231/MB468 | NUC |
| >hg38_21_+_8433172_8446622_RNA45SN1_45S_50ntFlanks@12522.12546.25\|TTAGTTTTACCCTACTGATGATGTG | >hg38_21_+_8433172_8446622_RNA45SN1_45S_50ntFlanks | 12522 | 12546 | 25 | TTAGTTTTACCCTACTGATGATGTG | 70.7 | -9.2 | 1.8 | -5.3 | 1.4E-07 | 9.5E-07 | BT20/MB468 | MP |
| >hg38_21_+_8433172_8446622_RNA45SN1_45S_50ntFlanks@3854.3872.19\|ACTGTGGTAATTCTAGAGC | >hg38_21_+_8433172_8446622_RNA45SN1_45S_50ntFlanks | 3854 | 3872 | 19 | ACTGTGGTAATTCTAGAGC | 858.0 | -8.0 | 1.5 | -5.3 | 1.4E-07 | 9.5E-07 | BT20/MB468 | MP |
| >hg38_21_+_8433172_8446622_RNA45SN1_45S_50ntFlanks@5045.5072.28\|CTTAGTTGGTGGAGCGATTTGTCTGGTT | >hg38_21_+_8433172_8446622_RNA45SN1_45S_50ntFlanks | 5045 | 5072 | 28 | CTTAGTTGGTGGAGCGATTTGTCTGGTT | 186.3 | -9.6 | 1.9 | -5.1 | 4.2E-07 | 9.5E-07 | BT20/MB468 | MITO |
| >hg38_21_+_8433172_8446622_RNA45SN1_45S_50ntFlanks@7989.8017.29\|AGACGTGGCGACCCGCTGAATTTAAGCAT | >hg38_21_+_8433172_8446622_RNA45SN1_45S_50ntFlanks | 7989 | 8017 | 29 | AGACGTGGCGACCCGCTGAATTTAAGCAT | 462.9 | -3.9 | 0.8 | -5.0 | 4.9E-07 | 9.5E-07 | MB231/MB468 | MP |
| >hg38_21_+_8433172_8446622_RNA45SN1_45S_50ntFlanks@12083.12107.25\|CTCGCTTCTGGCGCCAAGCGCCCGG | >hg38_21_+_8433172_8446622_RNA45SN1_45S_50ntFlanks | 12083 | 12107 | 25 | CTCGCTTCTGGCGCCAAGCGCCCGG | 485.1 | -7.1 | 1.4 | -5.0 | 4.9E-07 | 9.5E-07 | MB231/MB468 | MP |
| >hg38_21_+_8433172_8446622_RNA45SN1_45S_50ntFlanks@6721.6754.34\|AATTGCAGGACACATTGATCATCGACACTTCGAA | >hg38_21_+_8433172_8446622_RNA45SN1_45S_50ntFlanks | 6721 | 6754 | 34 | AATTGCAGGACACATTGATCATCGACACTTCGAA | 273.1 | -6.8 | 1.4 | -5.0 | 5.9E-07 | 9.5E-07 | MB231/MB468 | MITO |
| >hg38_21_+_8433172_8446622_RNA45SN1_45S_50ntFlanks@10797.10817.21\|GCATGTTGGAACAATGTAGGT | >hg38_21_+_8433172_8446622_RNA45SN1_45S_50ntFlanks | 10797 | 10817 | 21 | GCATGTTGGAACAATGTAGGT | 459.5 | -3.5 | 0.7 | -5.0 | 5.5E-07 | 9.5E-07 | MB231/MB468 | CYTO |
| >hg38_21_+_8433172_8446622_RNA45SN1_45S_50ntFlanks@10403.10430.28\|ACATGGGTCAGTCGGTCCTGAGAGATGG | >hg38_21_+_8433172_8446622_RNA45SN1_45S_50ntFlanks | 10403 | 10430 | 28 | ACATGGGTCAGTCGGTCCTGAGAGATGG | 436.3 | -5.3 | 1.1 | -5.0 | 5.7E-07 | 9.5E-07 | MB231/MB468 | NUC |
| >hg38_21_+_8433172_8446622_RNA45SN1_45S_50ntFlanks@9932.9950.19\|ATAGACAGCAGGACGGTGG | >hg38_21_+_8433172_8446622_RNA45SN1_45S_50ntFlanks | 9932 | 9950 | 19 | ATAGACAGCAGGACGGTGG | 242.3 | 11.3 | 2.0 | 5.5 | 2.9E-08 | 9.5E-07 | BT20/MB231 | MP |
| >hg38_21_+_8433172_8446622_RNA45SN1_45S_50ntFlanks@10434.10453.20\|AGCGCCGTTCCGAAGGGACG | >hg38_21_+_8433172_8446622_RNA45SN1_45S_50ntFlanks | 10434 | 10453 | 20 | AGCGCCGTTCCGAAGGGACG | 79.6 | -9.4 | 1.8 | -5.3 | 1.4E-07 | 9.5E-07 | BT20/MB468 | MP |
| >hg38_21_+_8433172_8446622_RNA45SN1_45S_50ntFlanks@5472.5511.40\|ACGGCCCTGGCGGAGCGCTGAGAAGACGGTCGAACTTGAC | >hg38_21_+_8433172_8446622_RNA45SN1_45S_50ntFlanks | 5472 | 5511 | 40 | ACGGCCCTGGCGGAGCGCTGAGAAGACGGTCGAACTTGAC | 81.2 | 10.1 | 1.9 | 5.3 | 1.4E-07 | 9.5E-07 | BT20/MB468 | MP |
| >hg38_21_+_8433172_8446622_RNA45SN1_45S_50ntFlanks@9027.9057.31\|CGTCTCCTCGTGGGGGGGCCGGGCCACCCCT | >hg38_21_+_8433172_8446622_RNA45SN1_45S_50ntFlanks | 9027 | 9057 | 31 | CGTCTCCTCGTGGGGGGGCCGGGCCACCCCT | 74.7 | -9.3 | 1.8 | -5.3 | 1.4E-07 | 9.5E-07 | BT20/MB468 | MP |
| >hg38_21_+_8433172_8446622_RNA45SN1_45S_50ntFlanks@11657.11687.31\|GATGTGATTTCTGCCCAGTGCTCTGAATGTC | >hg38_21_+_8433172_8446622_RNA45SN1_45S_50ntFlanks | 11657 | 11687 | 31 | GATGTGATTTCTGCCCAGTGCTCTGAATGTC | 981.2 | 10.6 | 2.0 | 5.3 | 1.4E-07 | 9.5E-07 | BT20/MB468 | MP |
| >hg38_21_+_8433172_8446622_RNA45SN1_45S_50ntFlanks@8472.8490.19\|CGGCCCGGCGGATCTTTCC | >hg38_21_+_8433172_8446622_RNA45SN1_45S_50ntFlanks | 8472 | 8490 | 19 | CGGCCCGGCGGATCTTTCC | 165.2 | -5.5 | 1.1 | -5.0 | 5.5E-07 | 9.5E-07 | MB231/MB468 | CYTO |
| >hg38_21_+_8433172_8446622_RNA45SN1_45S_50ntFlanks@8118.8141.24\|GGGACATGTGGCGTACGGAAGACC | >hg38_21_+_8433172_8446622_RNA45SN1_45S_50ntFlanks | 8118 | 8141 | 24 | GGGACATGTGGCGTACGGAAGACC | 991.7 | 5.8 | 1.1 | 5.3 | 1.4E-07 | 9.5E-07 | BT20/MB468 | MP |
| >hg38_MT_+_1621_3279_RNR2_16S_50ntFlanks@1508.1529.22\|AAAGGACAAGAGAAATAAGGCC | >hg38_MT_+_1621_3279_RNR2_16S_50ntFlanks | 1508 | 1529 | 22 | AAAGGACAAGAGAAATAAGGCC | 75.5 | 10.0 | 1.9 | 5.3 | 1.4E-07 | 9.5E-07 | BT20/MB468 | MP |
| >MI0000734\|hsa-mir-106b&WithFlank&7\|-\|100093987\|100094080@18.39.22&[MIMAT0000680&hsa-miR-106b-5p&offsets\|0\|+1;m-103&7\|-\|100094043\|100094063&offsets\|0\|+1]\|TAAAGTGCTGACAGTGCAGATA | >MI0000734\|hsa-mir-106b&WithFlank&7\|-\|100093987\|100094080 | 18 | 39 | 22 | TAAAGTGCTGACAGTGCAGATA | 1841.9 | 5.5 | 1.1 | 5.0 | 5.9E-07 | 9.5E-07 | MB231/MB468 | MITO |
| >hg38_21_+_8433172_8446622_RNA45SN1_45S_50ntFlanks@9548.9576.29\|GAGGTCCGTAGCGGTCCTGACGTGCAAAT | >hg38_21_+_8433172_8446622_RNA45SN1_45S_50ntFlanks | 9548 | 9576 | 29 | GAGGTCCGTAGCGGTCCTGACGTGCAAAT | 159.9 | 4.0 | 0.8 | 5.0 | 5.7E-07 | 9.5E-07 | MB231/MB468 | NUC |
| >trnaMT_ValTAC_MT_+_1602_1670@42.71.30\|AGATTTCAACTTAACTTGACCGCTCTGACC | >trnaMT_ValTAC_MT_+_1602_1670 | 42 | 71 | 30 | AGATTTCAACTTAACTTGACCGCTCTGACC | 8732.9 | 5.2 | 1.0 | 5.1 | 4.2E-07 | 9.5E-07 | BT20/MB468 | MITO |
| >hg38_21_+_8433172_8446622_RNA45SN1_45S_50ntFlanks@12251.12284.34\|AAAGCTCGCTTGATCTTGATTTTCAGTACGAATA | >hg38_21_+_8433172_8446622_RNA45SN1_45S_50ntFlanks | 12251 | 12284 | 34 | AAAGCTCGCTTGATCTTGATTTTCAGTACGAATA | 862.9 | -6.3 | 1.3 | -5.0 | 5.9E-07 | 9.5E-07 | MB231/MB468 | MITO |
| >hg38_21_+_8433172_8446622_RNA45SN1_45S_50ntFlanks@6742.6771.30\|TCGACACTTCGAACGCACTTGCGGCCCCGG | >hg38_21_+_8433172_8446622_RNA45SN1_45S_50ntFlanks | 6742 | 6771 | 30 | TCGACACTTCGAACGCACTTGCGGCCCCGG | 383.2 | -7.6 | 1.5 | -5.0 | 6.2E-07 | 9.5E-07 | BT20/MB468 | NUC |
| >hg38_21_+_8433172_8446622_RNA45SN1_45S_50ntFlanks@11675.11706.32\|TGCTCTGAATGTCAAAGTGAAGAAATTCAATG | >hg38_21_+_8433172_8446622_RNA45SN1_45S_50ntFlanks | 11675 | 11706 | 32 | TGCTCTGAATGTCAAAGTGAAGAAATTCAATG | 690.6 | 3.7 | 0.7 | 5.0 | 4.8E-07 | 9.5E-07 | BT20/MB468 | CYTO |
| >hg38_21_+_8433172_8446622_RNA45SN1_45S_50ntFlanks@10342.10382.41\|ATATTCAAACGAGAACTTTGAAGGCCGAAGTGGAGAAGGGT | >hg38_21_+_8433172_8446622_RNA45SN1_45S_50ntFlanks | 10342 | 10382 | 41 | ATATTCAAACGAGAACTTTGAAGGCCGAAGTGGAGAAGGGT | 254.7 | 6.4 | 1.3 | 5.0 | 5.5E-07 | 9.5E-07 | MB231/MB468 | CYTO |
| >hg38_1_-_228634819_228635039_RNA5S12_5S_50ntFlanks@51.78.28\|GTCTACGGCCATACCACCCTGAACGCGC | >hg38_1_-_228634819_228635039_RNA5S12_5S_50ntFlanks | 51 | 78 | 28 | GTCTACGGCCATACCACCCTGAACGCGC | 266.9 | 6.3 | 1.3 | 5.0 | 5.6E-07 | 9.6E-07 | MB231/MB468 | CYTO |
| >hg38_21_+_8433172_8446622_RNA45SN1_45S_50ntFlanks@11208.11237.30\|GCGGCGACTCTGGACGCGAGCCGGGCCCTT | >hg38_21_+_8433172_8446622_RNA45SN1_45S_50ntFlanks | 11208 | 11237 | 30 | GCGGCGACTCTGGACGCGAGCCGGGCCCTT | 140.7 | -4.1 | 0.8 | -5.0 | 5.6E-07 | 9.6E-07 | MB231/MB468 | CYTO |
| >hg38_21_+_8433172_8446622_RNA45SN1_45S_50ntFlanks@6657.6681.25\|TTAGCGGTGGATCACTCGGCTCGTG | >hg38_21_+_8433172_8446622_RNA45SN1_45S_50ntFlanks | 6657 | 6681 | 25 | TTAGCGGTGGATCACTCGGCTCGTG | 713.9 | -2.0 | 0.4 | -5.1 | 4.3E-07 | 9.6E-07 | BT20/MB468 | MITO |
| >hg38_21_+_8433172_8446622_RNA45SN1_45S_50ntFlanks@10821.10838.18\|GGAAGTCGGCAAGCCGGA | >hg38_21_+_8433172_8446622_RNA45SN1_45S_50ntFlanks | 10821 | 10838 | 18 | GGAAGTCGGCAAGCCGGA | 511.7 | -5.5 | 1.1 | -5.0 | 5.7E-07 | 9.6E-07 | MB231/MB468 | NUC |
| >am_trna162_MetCAT_6_-_26758550_26758622@31.53.23,trna164_MetCAT_6_-_26735574_26735646@31.53.23,trna20_MetCAT_16_+_71460396_71460468@31.53.23,trna21_MetCAT_6_+_26701712_26701784@31.53.23,trna27_MetCAT_6_+_26766444_26766516@31.53.23,trna75_MetCAT_6_+_28912352_28912424@31.53.23,trna97_MetCAT_6_-_28921042_28921114@31.53.23\|TCTCATAATCTGAAGGTCCTGAG | >am_trna162_MetCAT_6_-_26758550_26758622 | 31 | 53 | 23 | TCTCATAATCTGAAGGTCCTGAG | 231.4 | -6.6 | 1.3 | -5.0 | 4.9E-07 | 9.6E-07 | BT20/MB468 | CYTO |
| >MI0001733\|hsa-mir-452&WithFlank&X\|-\|151959622\|151959718@59.79.21&[MIMAT0001636&hsa-miR-452-3p&offsets\|-5\|-6;m-457&X\|-\|151959639\|151959660&offsets\|0\|-1]\|TCAGTCTCATCTGCAAAGAAG | >MI0001733\|hsa-mir-452&WithFlank&X\|-\|151959622\|151959718 | 59 | 79 | 21 | TCAGTCTCATCTGCAAAGAAG | 106.0 | 10.5 | 2.0 | 5.3 | 1.4E-07 | 9.6E-07 | BT20/MB468 | MP |
| >hg38_MT_+_598_1651_RNR1_12S_50ntFlanks@447.469.23\|TGAACACACAATAGCTAAGACCC | >hg38_MT_+_598_1651_RNR1_12S_50ntFlanks | 447 | 469 | 23 | TGAACACACAATAGCTAAGACCC | 131.2 | 10.8 | 2.0 | 5.3 | 1.4E-07 | 9.6E-07 | BT20/MB468 | MP |
| >MI0000809\|hsa-mir-151a&WithFlank&8\|-\|140732558\|140732659@51.72.22(+1A)&[MIMAT0000757&hsa-miR-151a-3p&offsets\|-2\|-1(+1A);m-30&8\|-\|140732587\|140732607&offsets\|-2\|-1(+1A)]\|TACTAGACTGAAGCTCCTTGAGA | >MI0000809\|hsa-mir-151a&WithFlank&8\|-\|140732558\|140732659 | 51 | 72 | 22(+1A) | TACTAGACTGAAGCTCCTTGAGA | 355.8 | 4.6 | 0.9 | 5.3 | 1.4E-07 | 9.6E-07 | BT20/MB468 | MP |
| >hg38_21_+_8433172_8446622_RNA45SN1_45S_50ntFlanks@10795.10818.24\|TGGCATGTTGGAACAATGTAGGTA | >hg38_21_+_8433172_8446622_RNA45SN1_45S_50ntFlanks | 10795 | 10818 | 24 | TGGCATGTTGGAACAATGTAGGTA | 419.9 | -3.0 | 0.6 | -5.1 | 4.3E-07 | 9.6E-07 | BT20/MB468 | MITO |
| >hg38_21_+_8433172_8446622_RNA45SN1_45S_50ntFlanks@11212.11252.41\|CGACTCTGGACGCGAGCCGGGCCCTTCCCGTGGATCGCCCC | >hg38_21_+_8433172_8446622_RNA45SN1_45S_50ntFlanks | 11212 | 11252 | 41 | CGACTCTGGACGCGAGCCGGGCCCTTCCCGTGGATCGCCCC | 2398.2 | -7.2 | 1.5 | -5.0 | 6.3E-07 | 9.6E-07 | BT20/MB468 | NUC |
| >hg38_21_+_8433172_8446622_RNA45SN1_45S_50ntFlanks@10683.10712.30\|CCGGCGGCGTCCGGTGAGCTCTCGCTGGCC | >hg38_21_+_8433172_8446622_RNA45SN1_45S_50ntFlanks | 10683 | 10712 | 30 | CCGGCGGCGTCCGGTGAGCTCTCGCTGGCC | 2462.4 | -3.8 | 0.8 | -5.0 | 5.6E-07 | 9.6E-07 | MB231/MB468 | CYTO |
| >hg38_21_+_8433172_8446622_RNA45SN1_45S_50ntFlanks@5491.5514.24\|GAGAAGACGGTCGAACTTGACTAT | >hg38_21_+_8433172_8446622_RNA45SN1_45S_50ntFlanks | 5491 | 5514 | 24 | GAGAAGACGGTCGAACTTGACTAT | 263.3 | -3.6 | 0.7 | -5.0 | 5.0E-07 | 9.6E-07 | MB231/MB468 | MP |
| >hg38_21_+_8433172_8446622_RNA45SN1_45S_50ntFlanks@11569.11603.35\|CAGCCGACTTAGAACTGGTGCGGACCAGGGGAATC | >hg38_21_+_8433172_8446622_RNA45SN1_45S_50ntFlanks | 11569 | 11603 | 35 | CAGCCGACTTAGAACTGGTGCGGACCAGGGGAATC | 4379.6 | -4.7 | 0.9 | -5.3 | 1.4E-07 | 9.6E-07 | BT20/MB468 | MP |

<table>
<tr><td>>hg38_21_+_8433172_8446622_RNA45SN1_45S_50ntFlanks@11892.11922.31|TGACTCTAGTCTGGCACGGTGAAGAGACATG</td><td>>hg38_21_+_8433172_8446622_RNA45SN1_45S_50ntFlanks</td><td>11892</td><td>11922</td><td>31</td><td>TGACTCTAGTCTGGCACGGTGAAGAGACATG</td><td>293.4</td><td>11.5</td><td>2.1</td><td>5.5</td><td>3.0E-08</td><td>9.7E-07</td><td>BT20/MB231</td><td>MP</td></tr>
<tr><td>>hg38_21_+_8433172_8446622_RNA45SN1_45S_50ntFlanks@11552.11585.34|CCTCGGCCGGCGCCTAGCAGCCGACTTAGAACTG</td><td>>hg38_21_+_8433172_8446622_RNA45SN1_45S_50ntFlanks</td><td>11552</td><td>11585</td><td>34</td><td>CCTCGGCCGGCGCCTAGCAGCCGACTTAGAACTG</td><td>343.1</td><td>-3.7</td><td>0.7</td><td>-5.0</td><td>6.0E-07</td><td>9.7E-07</td><td>MB231/MB468</td><td>MITO</td></tr>
<tr><td>>hg38_21_+_8433172_8446622_RNA45SN1_45S_50ntFlanks@5469.5514.46|CCCACGGCCCTGGCGGAGCGCTGAGAAGACGGTCGAACTTGACTAT</td><td>>hg38_21_+_8433172_8446622_RNA45SN1_45S_50ntFlanks</td><td>5469</td><td>5514</td><td>46</td><td>CCCACGGCCCTGGCGGAGCGCTGAGAAGACGGTCGAACTTGACTAT</td><td>182.8</td><td>-6.9</td><td>1.4</td><td>-5.0</td><td>4.9E-07</td><td>9.7E-07</td><td>BT20/MB468</td><td>CYTO</td></tr>
<tr><td>>hg38_21_+_8433172_8446622_RNA45SN1_45S_50ntFlanks@9859.9885.27|GGGATGAACCGAACGCCGGGTTAAGGC</td><td>>hg38_21_+_8433172_8446622_RNA45SN1_45S_50ntFlanks</td><td>9859</td><td>9885</td><td>27</td><td>GGGATGAACCGAACGCCGGGTTAAGGC</td><td>62.8</td><td>9.7</td><td>1.8</td><td>5.3</td><td>1.4E-07</td><td>9.7E-07</td><td>BT20/MB468</td><td>MP</td></tr>
<tr><td>>hg38_21_+_8433172_8446622_RNA45SN1_45S_50ntFlanks@6722.6739.18|ATTGCAGGACACATTGAT</td><td>>hg38_21_+_8433172_8446622_RNA45SN1_45S_50ntFlanks</td><td>6722</td><td>6739</td><td>18</td><td>ATTGCAGGACACATTGAT</td><td>68.3</td><td>-9.2</td><td>1.7</td><td>-5.3</td><td>1.4E-07</td><td>9.7E-07</td><td>BT20/MB468</td><td>MP</td></tr>
<tr><td>>hg38_21_+_8433172_8446622_RNA45SN1_45S_50ntFlanks@9533.9566.34|GAGGAAACTCTGGTGGAGGTCCGTAGCGGTCCTG</td><td>>hg38_21_+_8433172_8446622_RNA45SN1_45S_50ntFlanks</td><td>9533</td><td>9566</td><td>34</td><td>GAGGAAACTCTGGTGGAGGTCCGTAGCGGTCCTG</td><td>187.8</td><td>-7.1</td><td>1.4</td><td>-5.0</td><td>5.0E-07</td><td>9.7E-07</td><td>MB231/MB468</td><td>MP</td></tr>
<tr><td>>am_trna116_GluCTC_1_-_145399233_145399304@1.28.28,trna59_GluCTC_1_+_249168447_249168518@1.28.28,trna71_GluCTC_1_-_161439189_161439260@1.28.28,trna74_GluCTC_1_-_161431809_161431880@1.28.28,trna77_GluCTC_1_-_161424398_161424469@1.28.28,trna77_GluCTC_6_+_28949976_28950047@1.28.28,trna80_GluCTC_1_-_161417018_161417089@1.28.28,trna87_GluCTC_6_-_126101393_126101464@1.28.28|TCCCTGGTGGTCTAGTGGTTAGGATTCG</td><td>>am_trna116_GluCTC_1_-_145399233_145399304</td><td>1</td><td>28</td><td>28</td><td>TCCCTGGTGGTCTAGTGGTTAGGATTCG</td><td>23349.0</td><td>-3.5</td><td>0.7</td><td>-5.0</td><td>5.6E-07</td><td>9.7E-07</td><td>MB231/MB468</td><td>CYTO</td></tr>
<tr><td>>hg38_21_+_8433172_8446622_RNA45SN1_45S_50ntFlanks@10834.10862.29|CCGGATCCGTAACTTCGGGATAAGGATTG</td><td>>hg38_21_+_8433172_8446622_RNA45SN1_45S_50ntFlanks</td><td>10834</td><td>10862</td><td>29</td><td>CCGGATCCGTAACTTCGGGATAAGGATTG</td><td>84.6</td><td>-9.5</td><td>1.8</td><td>-5.3</td><td>1.4E-07</td><td>9.7E-07</td><td>BT20/MB468</td><td>MP</td></tr>
<tr><td>>hg38_21_+_8433172_8446622_RNA45SN1_45S_50ntFlanks@10058.10078.21|CGGCAGTCGAGAGTGGACGGG</td><td>>hg38_21_+_8433172_8446622_RNA45SN1_45S_50ntFlanks</td><td>10058</td><td>10078</td><td>21</td><td>CGGCAGTCGAGAGTGGACGGG</td><td>283.8</td><td>11.5</td><td>2.1</td><td>5.5</td><td>3.0E-08</td><td>9.7E-07</td><td>BT20/MB231</td><td>MP</td></tr>
<tr><td>>hg38_21_+_8433172_8446622_RNA45SN1_45S_50ntFlanks@6698.6717.20|AGCTAGCTGCGAGAATTAAT</td><td>>hg38_21_+_8433172_8446622_RNA45SN1_45S_50ntFlanks</td><td>6698</td><td>6717</td><td>20</td><td>AGCTAGCTGCGAGAATTAAT</td><td>616.6</td><td>-7.6</td><td>1.5</td><td>-5.0</td><td>5.0E-07</td><td>9.7E-07</td><td>MB231/MB468</td><td>MP</td></tr>
<tr><td>>am_trna128_GlyGCC_6_-_27870686_27870756@1.31.31,trna133_GlyCCC_1_-_16872434_16872504@1.31.31,trna18_GlyGCC_16_+_70822597_70822667@1.31.31,trna19_GlyGCC_16_+_70823410_70823480@1.31.31,trna19_GlyGCC_2_-_157257659_157257729@1.31.31,trna24_GlyGCC_16_-_70812942_70813012@1.31.31,trna25_GlyGCC_16_-_70812114_70812184@1.31.31,trna4_GlyCCC_1_+_17188416_17188486@1.31.31,trna5_GlyGCC_17_+_8029064_8029134@1.31.31,trna68_GlyGCC_1_-_161493637_161493707@1.31.31|GCATTGGTGGTTCAGTGGTAGAATTCTCGCC</td><td>>am_trna128_GlyGCC_6_-_27870686_27870756</td><td>1</td><td>31</td><td>31</td><td>GCATTGGTGGTTCAGTGGTAGAATTCTCGCC</td><td>25327.6</td><td>4.9</td><td>1.0</td><td>5.0</td><td>5.1E-07</td><td>9.7E-07</td><td>MB231/MB468</td><td>MP</td></tr>
<tr><td>>hg38_21_+_8433172_8446622_RNA45SN1_45S_50ntFlanks@9539.9563.25|ACTCTGGTGGAGGTCCGTAGCGGTC</td><td>>hg38_21_+_8433172_8446622_RNA45SN1_45S_50ntFlanks</td><td>9539</td><td>9563</td><td>25</td><td>ACTCTGGTGGAGGTCCGTAGCGGTC</td><td>310.2</td><td>-4.9</td><td>1.0</td><td>-5.0</td><td>5.7E-07</td><td>9.8E-07</td><td>MB231/MB468</td><td>CYTO</td></tr>
<tr><td>>hg38_21_+_8433172_8446622_RNA45SN1_45S_50ntFlanks@3846.3881.36|CTTGGATAACTGTGGTAATTCTAGAGCTAATACATG</td><td>>hg38_21_+_8433172_8446622_RNA45SN1_45S_50ntFlanks</td><td>3846</td><td>3881</td><td>36</td><td>CTTGGATAACTGTGGTAATTCTAGAGCTAATACATG</td><td>341.0</td><td>-7.5</td><td>1.5</td><td>-5.0</td><td>5.1E-07</td><td>9.8E-07</td><td>MB231/MB468</td><td>MP</td></tr>
<tr><td>>hg38_21_+_8433172_8446622_RNA45SN1_45S_50ntFlanks@10535.10562.28|CGCGAGGCGTCCAGTGCGGTAACGCGAC</td><td>>hg38_21_+_8433172_8446622_RNA45SN1_45S_50ntFlanks</td><td>10535</td><td>10562</td><td>28</td><td>CGCGAGGCGTCCAGTGCGGTAACGCGAC</td><td>87.0</td><td>-9.5</td><td>1.8</td><td>-5.3</td><td>1.4E-07</td><td>9.8E-07</td><td>BT20/MB468</td><td>MP</td></tr>
<tr><td>>hg38_21_+_8433172_8446622_RNA45SN1_45S_50ntFlanks@10683.10711.29|CCGGCGGCGTCCGGTGAGCTCTCGCTGGC</td><td>>hg38_21_+_8433172_8446622_RNA45SN1_45S_50ntFlanks</td><td>10683</td><td>10711</td><td>29</td><td>CCGGCGGCGTCCGGTGAGCTCTCGCTGGC</td><td>2193.8</td><td>-8.9</td><td>1.8</td><td>-5.0</td><td>6.4E-07</td><td>9.8E-07</td><td>BT20/MB468</td><td>NUC</td></tr>
<tr><td>>hg38_21_+_8433172_8446622_RNA45SN1_45S_50ntFlanks@12633.12659.27|CGAAGCTACCATCTGTGGGATTATGAC</td><td>>hg38_21_+_8433172_8446622_RNA45SN1_45S_50ntFlanks</td><td>12633</td><td>12659</td><td>27</td><td>CGAAGCTACCATCTGTGGGATTATGAC</td><td>596.3</td><td>12.6</td><td>2.3</td><td>5.5</td><td>3.1E-08</td><td>9.8E-07</td><td>BT20/MB231</td><td>MP</td></tr>
<tr><td>>hg38_21_+_8433172_8446622_RNA45SN1_45S_50ntFlanks@12993.13015.23|CCCTCGCTGCGATCTATTGAAAG</td><td>>hg38_21_+_8433172_8446622_RNA45SN1_45S_50ntFlanks</td><td>12993</td><td>13015</td><td>23</td><td>CCCTCGCTGCGATCTATTGAAAG</td><td>1352.2</td><td>-8.5</td><td>1.7</td><td>-5.0</td><td>6.4E-07</td><td>9.8E-07</td><td>BT20/MB468</td><td>NUC</td></tr>
<tr><td>>hg38_21_+_8433172_8446622_RNA45SN1_45S_50ntFlanks@5446.5466.21|CGGATCGGCCCCGCCGGGGTC</td><td>>hg38_21_+_8433172_8446622_RNA45SN1_45S_50ntFlanks</td><td>5446</td><td>5466</td><td>21</td><td>CGGATCGGCCCCGCCGGGGTC</td><td>10545.7</td><td>-7.0</td><td>1.4</td><td>-5.1</td><td>4.4E-07</td><td>9.8E-07</td><td>BT20/MB468</td><td>MITO</td></tr>
<tr><td>>MI0000088|hsa-mir-30a&WithFlank&6|-|71403545|71403627@12.35.24(+1U)&[MIMAT0000087&hsa-miR-30a-5p&offsets|0|+2(+1U);m-15&6|-|71403595|71403616&offsets|0|+2(+1U)]|TGTAAACATCCTCGACTGGAAGCTT</td><td>>MI0000088|hsa-mir-30a&WithFlank&6|-|71403545|71403627</td><td>12</td><td>35</td><td>24(+1U)</td><td>TGTAAACATCCTCGACTGGAAGCTT</td><td>8949.5</td><td>-4.9</td><td>0.9</td><td>-5.5</td><td>4.8E-08</td><td>9.8E-07</td><td>BT20/MB231</td><td>MITO</td></tr>
<tr><td>>MI0000263|hsa-mir-7-1&WithFlank&9|-|83969742|83969863@30.54.25&[MIMAT0000252_2&hsa-miR-7-5p&offsets|0|+1;m-401&9|-|83969811|83969834&offsets|0|+1]|TGGAAGACTAGTGATTTTGTTGTTT</td><td>>MI0000263|hsa-mir-7-1&WithFlank&9|-|83969742|83969863</td><td>30</td><td>54</td><td>25</td><td>TGGAAGACTAGTGATTTTGTTGTTT</td><td>357.5</td><td>4.0</td><td>0.8</td><td>5.0</td><td>5.7E-07</td><td>9.8E-07</td><td>MB231/MB468</td><td>CYTO</td></tr>
<tr><td>>hg38_21_+_8433172_8446622_RNA45SN1_45S_50ntFlanks@10806.10827.22|AACAATGTAGGTAAGGGAAGTC</td><td>>hg38_21_+_8433172_8446622_RNA45SN1_45S_50ntFlanks</td><td>10806</td><td>10827</td><td>22</td><td>AACAATGTAGGTAAGGGAAGTC</td><td>362.0</td><td>-6.8</td><td>1.4</td><td>-5.0</td><td>6.1E-07</td><td>9.8E-07</td><td>MB231/MB468</td><td>MITO</td></tr>
<tr><td>>hg38_21_+_8433172_8446622_RNA45SN1_45S_50ntFlanks@12242.12273.32|AGAAGGGCAAAAGCTCGCTTGATCTTGATTTT</td><td>>hg38_21_+_8433172_8446622_RNA45SN1_45S_50ntFlanks</td><td>12242</td><td>12273</td><td>32</td><td>AGAAGGGCAAAAGCTCGCTTGATCTTGATTTT</td><td>838.4</td><td>-7.3</td><td>1.5</td><td>-5.0</td><td>6.1E-07</td><td>9.8E-07</td><td>MB231/MB468</td><td>MITO</td></tr>
<tr><td>>MI0000076|hsa-mir-20a&WithFlank&13|+|91351059|91351141@14.33.20(+1A)&[MIMAT0000075&hsa-miR-20a-5p&offsets|0|-3(+1A);m-96&13|+|91351072|91351094&offsets|0|-3(+1A)]|TAAAGTGCTTATAGTGCAGGA</td><td>>MI0000076|hsa-mir-20a&WithFlank&13|+|91351059|91351141</td><td>14</td><td>33</td><td>20(+1A)</td><td>TAAAGTGCTTATAGTGCAGGA</td><td>144.7</td><td>-10.6</td><td>1.9</td><td>-5.5</td><td>3.1E-08</td><td>9.8E-07</td><td>BT20/MB231</td><td>MP</td></tr>
<tr><td>>hg38_21_+_8433172_8446622_RNA45SN1_45S_50ntFlanks@4163.4181.19|CATCCAAGGAAGGCAGCAG</td><td>>hg38_21_+_8433172_8446622_RNA45SN1_45S_50ntFlanks</td><td>4163</td><td>4181</td><td>19</td><td>CATCCAAGGAAGGCAGCAG</td><td>98.4</td><td>-9.7</td><td>1.8</td><td>-5.3</td><td>1.4E-07</td><td>9.8E-07</td><td>BT20/MB468</td><td>MP</td></tr>
<tr><td>>hg38_21_+_8433172_8446622_RNA45SN1_45S_50ntFlanks@11233.11253.21|CCCTTCCCGTGGATCGCCCCA</td><td>>hg38_21_+_8433172_8446622_RNA45SN1_45S_50ntFlanks</td><td>11233</td><td>11253</td><td>21</td><td>CCCTTCCCGTGGATCGCCCCA</td><td>357.4</td><td>-6.8</td><td>1.3</td><td>-5.0</td><td>5.0E-07</td><td>9.8E-07</td><td>BT20/MB468</td><td>CYTO</td></tr>
<tr><td>>MI0000063|hsa-let-7b&WithFlank&22|+|46113680|46113774@12.32.21(+1G)&[MIMAT0000063&hsa-let-7b-5p&offsets|0|-1(+1G);m-14&22|+|46113691|46113712&offsets|0|-1(+1G)]|TGAGGTAGTAGGTTGTGTGGTG</td><td>>MI0000063|hsa-let-7b&WithFlank&22|+|46113680|46113774</td><td>12</td><td>32</td><td>21(+1G)</td><td>TGAGGTAGTAGGTTGTGTGGTG</td><td>320.6</td><td>3.4</td><td>0.7</td><td>5.0</td><td>6.4E-07</td><td>9.8E-07</td><td>BT20/MB468</td><td>NUC</td></tr>
<tr><td>>hg38_1_-_228634819_228635039_RNA5S12_5S_50ntFlanks@84.103.20|CTCGTCTGATCTCGGAAGCT</td><td>>hg38_1_-_228634819_228635039_RNA5S12_5S_50ntFlanks</td><td>84</td><td>103</td><td>20</td><td>CTCGTCTGATCTCGGAAGCT</td><td>1170.6</td><td>-7.5</td><td>1.5</td><td>-5.0</td><td>5.9E-07</td><td>9.8E-07</td><td>MB231/MB468</td><td>NUC</td></tr>
<tr><td>>trnaMT_LeuTAG_MT_+_12266_12336@37.74.38|CCCCAAAAAATTTTGGTGCAACTCCAAATAAAAGTACCA</td><td>>trnaMT_LeuTAG_MT_+_12266_12336</td><td>37</td><td>74</td><td>38</td><td>CCCCAAAAAATTTTGGTGCAACTCCAAATAAAAGTACCA</td><td>237.3</td><td>-11.3</td><td>2.0</td><td>-5.5</td><td>3.1E-08</td><td>9.8E-07</td><td>BT20/MB231</td><td>MP</td></tr>
<tr><td>>hg38_21_+_8433172_8446622_RNA45SN1_45S_50ntFlanks@4731.4764.34|AAGAACGAAAGTCGGAGGTTCGAAGACGATCAGA</td><td>>hg38_21_+_8433172_8446622_RNA45SN1_45S_50ntFlanks</td><td>4731</td><td>4764</td><td>34</td><td>AAGAACGAAAGTCGGAGGTTCGAAGACGATCAGA</td><td>978.0</td><td>7.9</td><td>1.6</td><td>5.0</td><td>5.0E-07</td><td>9.8E-07</td><td>BT20/MB468</td><td>CYTO</td></tr>
<tr><td>>hg38_21_+_8433172_8446622_RNA45SN1_45S_50ntFlanks@12003.12043.41|GCGGGCCGCCGGTGAAATACCACTACTCTGATCGTTTTTTC</td><td>>hg38_21_+_8433172_8446622_RNA45SN1_45S_50ntFlanks</td><td>12003</td><td>12043</td><td>41</td><td>GCGGGCCGCCGGTGAAATACCACTACTCTGATCGTTTTTTC</td><td>246.1</td><td>-5.4</td><td>1.1</td><td>-5.0</td><td>5.7E-07</td><td>9.8E-07</td><td>MB231/MB468</td><td>CYTO</td></tr>
<tr><td>>hg38_21_+_8433172_8446622_RNA45SN1_45S_50ntFlanks@12437.12474.38|CATTGTGAAGCAGAATTCACCAAGCGTTGGATTGTTCA</td><td>>hg38_21_+_8433172_8446622_RNA45SN1_45S_50ntFlanks</td><td>12437</td><td>12474</td><td>38</td><td>CATTGTGAAGCAGAATTCACCAAGCGTTGGATTGTTCA</td><td>536.4</td><td>-4.2</td><td>0.8</td><td>-5.0</td><td>5.0E-07</td><td>9.8E-07</td><td>BT20/MB468</td><td>CYTO</td></tr>
<tr><td>>hg38_21_+_8433172_8446622_RNA45SN1_45S_50ntFlanks@11569.11601.33|CAGCCGACTTAGAACTGGTGCGGACCAGGGGAA</td><td>>hg38_21_+_8433172_8446622_RNA45SN1_45S_50ntFlanks</td><td>11569</td><td>11601</td><td>33</td><td>CAGCCGACTTAGAACTGGTGCGGACCAGGGGAA</td><td>3688.6</td><td>-4.4</td><td>0.9</td><td>-5.0</td><td>5.7E-07</td><td>9.8E-07</td><td>MB231/MB468</td><td>CYTO</td></tr>
<tr><td>>hg38_21_+_8433172_8446622_RNA45SN1_45S_50ntFlanks@12633.12657.25|CGAAGCTACCATCTGTGGGATTATG</td><td>>hg38_21_+_8433172_8446622_RNA45SN1_45S_50ntFlanks</td><td>12633</td><td>12657</td><td>25</td><td>CGAAGCTACCATCTGTGGGATTATG</td><td>177.3</td><td>-4.8</td><td>1.0</td><td>-5.0</td><td>5.7E-07</td><td>9.8E-07</td><td>MB231/MB468</td><td>CYTO</td></tr>
<tr><td>>hg38_21_+_8433172_8446622_RNA45SN1_45S_50ntFlanks@3903.3935.33|CGCGGGGGGGATGCGTGCATTTATCAGATCAAA</td><td>>hg38_21_+_8433172_8446622_RNA45SN1_45S_50ntFlanks</td><td>3903</td><td>3935</td><td>33</td><td>CGCGGGGGGGATGCGTGCATTTATCAGATCAAA</td><td>57.3</td><td>-9.7</td><td>1.9</td><td>-5.0</td><td>5.1E-07</td><td>9.8E-07</td><td>MB231/MB468</td><td>MP</td></tr>
<tr><td>>am_MI0003639|hsa-mir-625&WithFlank&14|+|65471096|65471192@58.79.22(+1C)&[MIMAT0004808&hsa-miR-625-3p&offsets|0|0(+1C);m-137&14|+|65471153|65471174&offsets|0|0(+1C)]|GACTATAGAACTTTCCCCCTCAC</td><td>>am_MI0003639|hsa-mir-625&WithFlank&14|+|65471096|65471192</td><td>58</td><td>79</td><td>22(+1C)</td><td>GACTATAGAACTTTCCCCCTCAC</td><td>47.7</td><td>8.8</td><td>1.8</td><td>5.0</td><td>5.1E-07</td><td>9.8E-07</td><td>MB231/MB468</td><td>MP</td></tr>
<tr><td>>trnaMT_PheGAA_MT_+_577_647@36.54.19|AAAATGTTTAGACGGGCTC</td><td>>trnaMT_PheGAA_MT_+_577_647</td><td>36</td><td>54</td><td>19</td><td>AAAATGTTTAGACGGGCTC</td><td>537.1</td><td>5.4</td><td>1.0</td><td>5.3</td><td>1.4E-07</td><td>9.9E-07</td><td>BT20/MB468</td><td>MP</td></tr>
<tr><td>>hg38_21_+_8433172_8446622_RNA45SN1_45S_50ntFlanks@11675.11702.28|TGCTCTGAATGTCAAAGTGAAGAAATTC</td><td>>hg38_21_+_8433172_8446622_RNA45SN1_45S_50ntFlanks</td><td>11675</td><td>11702</td><td>28</td><td>TGCTCTGAATGTCAAAGTGAAGAAATTC</td><td>1180.5</td><td>2.9</td><td>0.6</td><td>5.0</td><td>5.8E-07</td><td>9.9E-07</td><td>MB231/MB468</td><td>CYTO</td></tr>
<tr><td>>hg38_21_+_8433172_8446622_RNA45SN1_45S_50ntFlanks@8182.8204.23|ATCGAGGCCCAGCCCGTGGACGG</td><td>>hg38_21_+_8433172_8446622_RNA45SN1_45S_50ntFlanks</td><td>8182</td><td>8204</td><td>23</td><td>ATCGAGGCCCAGCCCGTGGACGG</td><td>553.7</td><td>-7.3</td><td>1.5</td><td>-5.0</td><td>5.1E-07</td><td>9.9E-07</td><td>MB231/MB468</td><td>MP</td></tr>
<tr><td>>MI0000298|hsa-mir-221&WithFlank&X|-|45746151|45746272@71.93.23(+1U)&[MIMAT0000278&hsa-miR-221-3p&offsets|0|0(+1U);m-91&X|-|45746181|45746202&offsets|0|+1(+1U)]|AGCTACATTGTCTGCTGGGTTTCT</td><td>>MI0000298|hsa-mir-221&WithFlank&X|-|45746151|45746272</td><td>71</td><td>93</td><td>23(+1U)</td><td>AGCTACATTGTCTGCTGGGTTTCT</td><td>6567.8</td><td>3.7</td><td>0.7</td><td>5.0</td><td>5.1E-07</td><td>9.9E-07</td><td>MB231/MB468</td><td>MP</td></tr>
<tr><td>>hg38_21_+_8433172_8446622_RNA45SN1_45S_50ntFlanks@12412.12434.23|ATCCTTCGATGTCGGCTCTTCCT</td><td>>hg38_21_+_8433172_8446622_RNA45SN1_45S_50ntFlanks</td><td>12412</td><td>12434</td><td>23</td><td>ATCCTTCGATGTCGGCTCTTCCT</td><td>59.8</td><td>-9.0</td><td>1.7</td><td>-5.3</td><td>1.5E-07</td><td>9.9E-07</td><td>BT20/MB468</td><td>MP</td></tr>
<tr><td>>hg38_21_+_8433172_8446622_RNA45SN1_45S_50ntFlanks@12712.12733.22|CGGAGCCTCGGTTGGCCTCGGA</td><td>>hg38_21_+_8433172_8446622_RNA45SN1_45S_50ntFlanks</td><td>12712</td><td>12733</td><td>22</td><td>CGGAGCCTCGGTTGGCCTCGGA</td><td>428.9</td><td>-7.3</td><td>1.5</td><td>-5.0</td><td>6.2E-07</td><td>9.9E-07</td><td>MB231/MB468</td><td>MITO</td></tr>
<tr><td>>hg38_21_+_8433172_8446622_RNA45SN1_45S_50ntFlanks@12998.13039.42|GCTGCGATCTATTGAAAGTCAGCCCTCGACACAAGGGTTTGT</td><td>>hg38_21_+_8433172_8446622_RNA45SN1_45S_50ntFlanks</td><td>12998</td><td>13039</td><td>42</td><td>GCTGCGATCTATTGAAAGTCAGCCCTCGACACAAGGGTTTGT</td><td>1843.2</td><td>6.9</td><td>1.3</td><td>5.3</td><td>1.5E-07</td><td>9.9E-07</td><td>BT20/MB468</td><td>MP</td></tr>
<tr><td>>hg38_21_+_8433172_8446622_RNA45SN1_45S_50ntFlanks@10866.10894.29|CTAAGGGCTGGGTCGGTCGGGCTGGGGCG</td><td>>hg38_21_+_8433172_8446622_RNA45SN1_45S_50ntFlanks</td><td>10866</td><td>10894</td><td>29</td><td>CTAAGGGCTGGGTCGGTCGGGCTGGGGCG</td><td>828.1</td><td>-7.2</td><td>1.5</td><td>-5.0</td><td>5.9E-07</td><td>9.9E-07</td><td>MB231/MB468</td><td>NUC</td></tr>
<tr><td>>hg38_21_+_8433172_8446622_RNA45SN1_45S_50ntFlanks@12785.12805.21|CCCCGCGCGCGCGGGAGGGCG</td><td>>hg38_21_+_8433172_8446622_RNA45SN1_45S_50ntFlanks</td><td>12785</td><td>12805</td><td>21</td><td>CCCCGCGCGCGCGGGAGGGCG</td><td>3923.4</td><td>-9.2</td><td>1.8</td><td>-5.0</td><td>6.5E-07</td><td>9.9E-07</td><td>BT20/MB468</td><td>NUC</td></tr>
<tr><td>>hg38_21_+_8433172_8446622_RNA45SN1_45S_50ntFlanks@9347.9370.24|AGCCGCCGTGGCGCAATGAAGGTG</td><td>>hg38_21_+_8433172_8446622_RNA45SN1_45S_50ntFlanks</td><td>9347</td><td>9370</td><td>24</td><td>AGCCGCCGTGGCGCAATGAAGGTG</td><td>67.6</td><td>-9.2</td><td>1.7</td><td>-5.3</td><td>1.5E-07</td><td>9.9E-07</td><td>BT20/MB468</td><td>MP</td></tr>
</table>

<table>
<tr><th></th><th></th><th></th><th></th><th></th><th></th><th></th><th></th><th></th><th></th><th></th><th></th><th></th><th></th></tr>
<tr><td>>hg38_21_+_8433172_8446622_RNA45SN1_45S_50ntFlanks@12404.12436.33|GCTTTTTGATCCTTCGATGTCGGCTCTTCCTAT</td><td>>hg38_21_+_8433172_8446622_RNA45SN1_45S_50ntFlanks</td><td>12404</td><td>12436</td><td>33</td><td>GCTTTTTGATCCTTCGATGTCGGCTCTTCCTAT</td><td>268.2</td><td>-5.5</td><td>1.1</td><td>-5.0</td><td>6.2E-07</td><td>9.9E-07</td><td>MB231/MB468</td><td>MITO</td></tr>
<tr><td>>hg38_21_+_8433172_8446622_RNA45SN1_45S_50ntFlanks@8111.8138.28|GGGGCGCGGGACATGTGGCGTACGGAAG</td><td>>hg38_21_+_8433172_8446622_RNA45SN1_45S_50ntFlanks</td><td>8111</td><td>8138</td><td>28</td><td>GGGGCGCGGGACATGTGGCGTACGGAAG</td><td>430.8</td><td>12.1</td><td>2.2</td><td>5.5</td><td>3.1E-08</td><td>9.9E-07</td><td>BT20/MB231</td><td>MP</td></tr>
<tr><td>>hg38_21_+_8433172_8446622_RNA45SN1_45S_50ntFlanks@10786.10823.38|AACAGCCTCTGGCATGTTGGAACAATGTAGGTAAGGGA</td><td>>hg38_21_+_8433172_8446622_RNA45SN1_45S_50ntFlanks</td><td>10786</td><td>10823</td><td>38</td><td>AACAGCCTCTGGCATGTTGGAACAATGTAGGTAAGGGA</td><td>1168.0</td><td>-3.8</td><td>0.7</td><td>-5.0</td><td>4.4E-07</td><td>9.9E-07</td><td>BT20/MB468</td><td>MITO</td></tr>
<tr><td>>hg38_21_+_8433172_8446622_RNA45SN1_45S_50ntFlanks@6775.6803.29|CCTCCCGGGGCTACGCCTGTCTGAGCGTC</td><td>>hg38_21_+_8433172_8446622_RNA45SN1_45S_50ntFlanks</td><td>6775</td><td>6803</td><td>29</td><td>CCTCCCGGGGCTACGCCTGTCTGAGCGTC</td><td>1689.7</td><td>-5.8</td><td>1.2</td><td>-5.0</td><td>6.2E-07</td><td>1.0E-06</td><td>MB231/MB468</td><td>MITO</td></tr>
<tr><td>>hg38_21_+_8433172_8446622_RNA45SN1_45S_50ntFlanks@8381.8405.25|AAGAGGGCGTGAAACCGTTAAGAGG</td><td>>hg38_21_+_8433172_8446622_RNA45SN1_45S_50ntFlanks</td><td>8381</td><td>8405</td><td>25</td><td>AAGAGGGCGTGAAACCGTTAAGAGG</td><td>2157.5</td><td>-4.4</td><td>0.8</td><td>-5.3</td><td>1.5E-07</td><td>1.0E-06</td><td>BT20/MB468</td><td>MP</td></tr>
<tr><td>>hg38_21_+_8433172_8446622_RNA45SN1_45S_50ntFlanks@10877.10901.25|GTCGGTCGGGCTGGGGCGCGAAGCG</td><td>>hg38_21_+_8433172_8446622_RNA45SN1_45S_50ntFlanks</td><td>10877</td><td>10901</td><td>25</td><td>GTCGGTCGGGCTGGGGCGCGAAGCG</td><td>182.0</td><td>-6.3</td><td>1.3</td><td>-5.0</td><td>6.0E-07</td><td>1.0E-06</td><td>MB231/MB468</td><td>NUC</td></tr>
<tr><td>>hg38_21_+_8433172_8446622_RNA45SN1_45S_50ntFlanks@11573.11599.27|CGACTTAGAACTGGTGCGGACCAGGGG</td><td>>hg38_21_+_8433172_8446622_RNA45SN1_45S_50ntFlanks</td><td>11573</td><td>11599</td><td>27</td><td>CGACTTAGAACTGGTGCGGACCAGGGG</td><td>224.9</td><td>11.1</td><td>2.0</td><td>5.5</td><td>3.2E-08</td><td>1.0E-06</td><td>BT20/MB231</td><td>MP</td></tr>
<tr><td>>hg38_21_+_8433172_8446622_RNA45SN1_45S_50ntFlanks@11662.11685.24|GATTTCTGCCCAGTGCTCTGAATG</td><td>>hg38_21_+_8433172_8446622_RNA45SN1_45S_50ntFlanks</td><td>11662</td><td>11685</td><td>24</td><td>GATTTCTGCCCAGTGCTCTGAATG</td><td>257.7</td><td>-5.7</td><td>1.1</td><td>-5.0</td><td>5.2E-07</td><td>1.0E-06</td><td>MB231/MB468</td><td>MP</td></tr>
<tr><td>>trnaMT_HisGTG_MT_+_12138_12206@27.63.37|GATTGTGAATCTGACAACAGAGGCTTACGACCCCTTA</td><td>>trnaMT_HisGTG_MT_+_12138_12206</td><td>27</td><td>63</td><td>37</td><td>GATTGTGAATCTGACAACAGAGGCTTACGACCCCTTA</td><td>53.5</td><td>9.0</td><td>1.8</td><td>5.0</td><td>5.2E-07</td><td>1.0E-06</td><td>MB231/MB468</td><td>MP</td></tr>
<tr><td>>hg38_21_+_8433172_8446622_RNA45SN1_45S_50ntFlanks@9578.9607.30|GGTCGTCCGACCTGGGTATAGGGGCGAAAG</td><td>>hg38_21_+_8433172_8446622_RNA45SN1_45S_50ntFlanks</td><td>9578</td><td>9607</td><td>30</td><td>GGTCGTCCGACCTGGGTATAGGGGCGAAAG</td><td>84.4</td><td>10.1</td><td>1.9</td><td>5.3</td><td>1.5E-07</td><td>1.0E-06</td><td>BT20/MB468</td><td>MP</td></tr>
<tr><td>>hg38_21_+_8433172_8446622_RNA45SN1_45S_50ntFlanks@4557.4576.20|CAGCTAGGAATAATGGAATA</td><td>>hg38_21_+_8433172_8446622_RNA45SN1_45S_50ntFlanks</td><td>4557</td><td>4576</td><td>20</td><td>CAGCTAGGAATAATGGAATA</td><td>300.5</td><td>-7.4</td><td>1.5</td><td>-5.0</td><td>6.5E-07</td><td>1.0E-06</td><td>BT20/MB468</td><td>NUC</td></tr>
<tr><td>>MI0000542|hsa-mir-320a&WithFlank&8|-|22244960|22245043@48.70.23(+3U)&[MIMAT0000510&hsa-miR-320a-3p&offsets|0|+1(+3U);m-60&8|-|22244975|22244996&offsets|0|+1(+3U)]|AAAAGCTGGGTTGAGAGGGCGAATTT</td><td>>MI0000542|hsa-mir-320a&WithFlank&8|-|22244960|22245043</td><td>48</td><td>70</td><td>23(+3U)</td><td>AAAAGCTGGGTTGAGAGGGCGAATTT</td><td>76.1</td><td>10.0</td><td>1.9</td><td>5.3</td><td>1.5E-07</td><td>1.0E-06</td><td>BT20/MB468</td><td>MP</td></tr>
<tr><td>>hg38_21_+_8433172_8446622_RNA45SN1_45S_50ntFlanks@11846.11867.22|GAACGGGCTTGGCGGAATCAGC</td><td>>hg38_21_+_8433172_8446622_RNA45SN1_45S_50ntFlanks</td><td>11846</td><td>11867</td><td>22</td><td>GAACGGGCTTGGCGGAATCAGC</td><td>154.4</td><td>5.7</td><td>1.1</td><td>5.0</td><td>5.8E-07</td><td>1.0E-06</td><td>MB231/MB468</td><td>CYTO</td></tr>
<tr><td>>hg38_21_+_8433172_8446622_RNA45SN1_45S_50ntFlanks@10781.10805.25|AGGTGAACAGCCTCTGGCATGTTGG</td><td>>hg38_21_+_8433172_8446622_RNA45SN1_45S_50ntFlanks</td><td>10781</td><td>10805</td><td>25</td><td>AGGTGAACAGCCTCTGGCATGTTGG</td><td>259.9</td><td>-7.2</td><td>1.4</td><td>-5.3</td><td>1.5E-07</td><td>1.0E-06</td><td>BT20/MB468</td><td>MP</td></tr>
<tr><td>>hg38_21_+_8433172_8446622_RNA45SN1_45S_50ntFlanks@10548.10577.30|GTGCGGTAACGCGACCGATCCCGGAGAAGC</td><td>>hg38_21_+_8433172_8446622_RNA45SN1_45S_50ntFlanks</td><td>10548</td><td>10577</td><td>30</td><td>GTGCGGTAACGCGACCGATCCCGGAGAAGC</td><td>405.7</td><td>9.1</td><td>1.8</td><td>5.0</td><td>4.5E-07</td><td>1.0E-06</td><td>BT20/MB468</td><td>MITO</td></tr>
<tr><td>>hg38_21_+_8433172_8446622_RNA45SN1_45S_50ntFlanks@11566.11586.21|TAGCAGCCGACTTAGAACTGG</td><td>>hg38_21_+_8433172_8446622_RNA45SN1_45S_50ntFlanks</td><td>11566</td><td>11586</td><td>21</td><td>TAGCAGCCGACTTAGAACTGG</td><td>315.4</td><td>3.9</td><td>0.8</td><td>5.0</td><td>6.0E-07</td><td>1.0E-06</td><td>MB231/MB468</td><td>NUC</td></tr>
<tr><td>>MI0000079|hsa-mir-23a&WithFlank&19|-|13836581|13836665@52.74.23&[MIMAT0000078&hsa-miR-23a-3p&offsets|+1|+3;m-31&19|-|13836595|13836615&offsets|+1|+3]|TCACATTGCCAGGGATTTCCAAC</td><td>>MI0000079|hsa-mir-23a&WithFlank&19|-|13836581|13836665</td><td>52</td><td>74</td><td>23</td><td>TCACATTGCCAGGGATTTCCAAC</td><td>263.9</td><td>5.1</td><td>1.0</td><td>5.0</td><td>6.3E-07</td><td>1.0E-06</td><td>MB231/MB468</td><td>MITO</td></tr>
<tr><td>>hg38_21_+_8433172_8446622_RNA45SN1_45S_50ntFlanks@12946.12964.19|TAGACGACCTGCTTCTGGG</td><td>>hg38_21_+_8433172_8446622_RNA45SN1_45S_50ntFlanks</td><td>12946</td><td>12964</td><td>19</td><td>TAGACGACCTGCTTCTGGG</td><td>320.5</td><td>-7.4</td><td>1.5</td><td>-5.0</td><td>6.6E-07</td><td>1.0E-06</td><td>BT20/MB468</td><td>NUC</td></tr>
<tr><td>>hg38_21_+_8433172_8446622_RNA45SN1_45S_50ntFlanks@12274.12318.45|CAGTACGAATACAGACCGTGAAAGCGGGGCCTCACGATCCTTCTG</td><td>>hg38_21_+_8433172_8446622_RNA45SN1_45S_50ntFlanks</td><td>12274</td><td>12318</td><td>45</td><td>CAGTACGAATACAGACCGTGAAAGCGGGGCCTCACGATCCTTCTG</td><td>109.2</td><td>-8.9</td><td>1.8</td><td>-5.0</td><td>4.5E-07</td><td>1.0E-06</td><td>BT20/MB468</td><td>MITO</td></tr>
<tr><td>>hg38_21_+_8433172_8446622_RNA45SN1_45S_50ntFlanks@8191.8212.22|CAGCCCGTGGACGGTGTGAGGC</td><td>>hg38_21_+_8433172_8446622_RNA45SN1_45S_50ntFlanks</td><td>8191</td><td>8212</td><td>22</td><td>CAGCCCGTGGACGGTGTGAGGC</td><td>671.1</td><td>-6.0</td><td>1.2</td><td>-5.0</td><td>6.0E-07</td><td>1.0E-06</td><td>MB231/MB468</td><td>NUC</td></tr>
<tr><td>>hg38_21_+_8433172_8446622_RNA45SN1_45S_50ntFlanks@10313.10340.28|GCAGGTGCAGATCTTGGTGGTAGTAGCA</td><td>>hg38_21_+_8433172_8446622_RNA45SN1_45S_50ntFlanks</td><td>10313</td><td>10340</td><td>28</td><td>GCAGGTGCAGATCTTGGTGGTAGTAGCA</td><td>469.6</td><td>12.2</td><td>2.2</td><td>5.5</td><td>3.2E-08</td><td>1.0E-06</td><td>BT20/MB231</td><td>MP</td></tr>
<tr><td>>hg38_21_+_8433172_8446622_RNA45SN1_45S_50ntFlanks@11491.11516.26|CCGCGGGCCGGTCCCCCCCGCCGGGT</td><td>>hg38_21_+_8433172_8446622_RNA45SN1_45S_50ntFlanks</td><td>11491</td><td>11516</td><td>26</td><td>CCGCGGGCCGGTCCCCCCCGCCGGGT</td><td>102.7</td><td>-9.8</td><td>1.9</td><td>-5.3</td><td>1.5E-07</td><td>1.0E-06</td><td>BT20/MB468</td><td>MP</td></tr>
<tr><td>>trna146_GlnCTG_6_-_27515531_27515602@15.43.29,trna1_GlnCTG_6_+_18836402_18836473@15.43.29,trna3_GlnCTG_17_+_8023070_8023141@15.43.29,trna49_GlnCTG_6_+_27487308_27487379@15.43.29,trna7_GlnCTG_15_-_66161400_66161471@15.43.29,trna99_GlnCTG_6_-_28909378_28909449@15.43.29|ATGGTTAGCACTCTGGACTCTGAATCCAG</td><td>>trna146_GlnCTG_6_-_27515531_27515602</td><td>15</td><td>43</td><td>29</td><td>ATGGTTAGCACTCTGGACTCTGAATCCAG</td><td>68.3</td><td>-9.2</td><td>1.7</td><td>-5.3</td><td>1.5E-07</td><td>1.0E-06</td><td>BT20/MB468</td><td>MP</td></tr>
<tr><td>>hg38_21_+_8433172_8446622_RNA45SN1_45S_50ntFlanks@10687.10705.19|CGGCGTCCGGTGAGCTCTC</td><td>>hg38_21_+_8433172_8446622_RNA45SN1_45S_50ntFlanks</td><td>10687</td><td>10705</td><td>19</td><td>CGGCGTCCGGTGAGCTCTC</td><td>873.7</td><td>-8.2</td><td>1.6</td><td>-5.0</td><td>6.6E-07</td><td>1.0E-06</td><td>BT20/MB468</td><td>NUC</td></tr>
<tr><td>>MI0000070|hsa-mir-16-1&WithFlank&13|-|50048967|50049067@20.42.23;MI0000115|hsa-mir-16-2&WithFlank&3|+|160404739|160404831@16.38.23&[MIMAT0000069&hsa-miR-16-5p&offsets|0|+1;m-62&13|-|50049027|50049048&offsets|0|+1];[MIMAT0000069_1&hsa-miR-16-5p&offsets|0|+1;m-61&3|+|160404754|160404775&offsets|0|+1]|TAGCAGCACGTAAATATTGGCGT</td><td>>MI0000070|hsa-mir-16-1&WithFlank&13|-|50048967|50049067</td><td>20</td><td>42</td><td>23</td><td>TAGCAGCACGTAAATATTGGCGT</td><td>1004.8</td><td>4.4</td><td>0.9</td><td>5.0</td><td>6.6E-07</td><td>1.0E-06</td><td>BT20/MB468</td><td>NUC</td></tr>
<tr><td>>MI0000434|hsa-let-7i&WithFlank&12|+|62603680|62603775@12.30.19(+1A)&[MIMAT0000415&hsa-let-7i-5p&offsets|0|-3(+1A);m-49&12|+|62603691|62603712&offsets|0|-3(+1A)]|TGAGGTAGTAGTTTGTGCTA</td><td>>MI0000434|hsa-let-7i&WithFlank&12|+|62603680|62603775</td><td>12</td><td>30</td><td>19(+1A)</td><td>TGAGGTAGTAGTTTGTGCTA</td><td>66.9</td><td>9.3</td><td>1.9</td><td>5.0</td><td>5.2E-07</td><td>1.0E-06</td><td>MB231/MB468</td><td>MP</td></tr>
<tr><td>>hg38_21_+_8433172_8446622_RNA45SN1_45S_50ntFlanks@11584.11623.40|TGGTGCGGACCAGGGGAATCCGACTGTTTAATTAAAACAA</td><td>>hg38_21_+_8433172_8446622_RNA45SN1_45S_50ntFlanks</td><td>11584</td><td>11623</td><td>40</td><td>TGGTGCGGACCAGGGGAATCCGACTGTTTAATTAAAACAA</td><td>885.7</td><td>9.4</td><td>1.8</td><td>5.3</td><td>1.5E-07</td><td>1.0E-06</td><td>BT20/MB468</td><td>MP</td></tr>
<tr><td>>hg38_21_+_8433172_8446622_RNA45SN1_45S_50ntFlanks@11999.12043.45|CCCTGCGGGCCGCCGGTGAAATACCACTACTCTGATCGTTTTTTC</td><td>>hg38_21_+_8433172_8446622_RNA45SN1_45S_50ntFlanks</td><td>11999</td><td>12043</td><td>45</td><td>CCCTGCGGGCCGCCGGTGAAATACCACTACTCTGATCGTTTTTTC</td><td>1763.2</td><td>-7.4</td><td>1.5</td><td>-5.0</td><td>4.5E-07</td><td>1.0E-06</td><td>BT20/MB468</td><td>MITO</td></tr>
<tr><td>>hg38_21_+_8433172_8446622_RNA45SN1_45S_50ntFlanks@9925.9951.27|GGTTGATATAGACAGCAGGACGGTGGC</td><td>>hg38_21_+_8433172_8446622_RNA45SN1_45S_50ntFlanks</td><td>9925</td><td>9951</td><td>27</td><td>GGTTGATATAGACAGCAGGACGGTGGC</td><td>136.0</td><td>10.4</td><td>1.9</td><td>5.5</td><td>3.2E-08</td><td>1.0E-06</td><td>BT20/MB231</td><td>MP</td></tr>
<tr><td>>hg38_21_+_8433172_8446622_RNA45SN1_45S_50ntFlanks@11199.11220.22|CCGGCGGCGGCGGCGACTCTGG</td><td>>hg38_21_+_8433172_8446622_RNA45SN1_45S_50ntFlanks</td><td>11199</td><td>11220</td><td>22</td><td>CCGGCGGCGGCGGCGACTCTGG</td><td>1491.9</td><td>-8.7</td><td>1.7</td><td>-5.0</td><td>6.6E-07</td><td>1.0E-06</td><td>BT20/MB468</td><td>NUC</td></tr>
<tr><td>>hg38_1_-_228634819_228635039_RNA5S12_5S_50ntFlanks@85.123.39|TCGTCTGATCTCGGAAGCTAAGCAGGGTCGGGCCTGGTT</td><td>>hg38_1_-_228634819_228635039_RNA5S12_5S_50ntFlanks</td><td>85</td><td>123</td><td>39</td><td>TCGTCTGATCTCGGAAGCTAAGCAGGGTCGGGCCTGGTT</td><td>120.6</td><td>10.2</td><td>1.9</td><td>5.5</td><td>3.3E-08</td><td>1.0E-06</td><td>BT20/MB231</td><td>MP</td></tr>
<tr><td>>MI0000446|hsa-mir-125b-1&WithFlank&11|-|122099751|122099850@61.80.20(+3U)&[MIMAT0004592&hsa-miR-125b-1-3p&offsets|0|-2(+3U);m-485&11|-|122099769|122099790&offsets|0|-2(+3U)]|ACGGGTTAGGCTCTTGGGAGTTT</td><td>>MI0000446|hsa-mir-125b-1&WithFlank&11|-|122099751|122099850</td><td>61</td><td>80</td><td>20(+3U)</td><td>ACGGGTTAGGCTCTTGGGAGTTT</td><td>141.7</td><td>-10.5</td><td>1.9</td><td>-5.5</td><td>3.3E-08</td><td>1.0E-06</td><td>BT20/MB231</td><td>MP</td></tr>
<tr><td>>hg38_21_+_8433172_8446622_RNA45SN1_45S_50ntFlanks@9653.9682.30|AGGATAGCTGGCGCTCTCGCAGACCCGACG</td><td>>hg38_21_+_8433172_8446622_RNA45SN1_45S_50ntFlanks</td><td>9653</td><td>9682</td><td>30</td><td>AGGATAGCTGGCGCTCTCGCAGACCCGACG</td><td>131.3</td><td>-6.6</td><td>1.3</td><td>-5.0</td><td>5.3E-07</td><td>1.0E-06</td><td>MB231/MB468</td><td>MP</td></tr>
<tr><td>>hg38_21_+_8433172_8446622_RNA45SN1_45S_50ntFlanks@11572.11602.31|CCGACTTAGAACTGGTGCGGACCAGGGGAAT</td><td>>hg38_21_+_8433172_8446622_RNA45SN1_45S_50ntFlanks</td><td>11572</td><td>11602</td><td>31</td><td>CCGACTTAGAACTGGTGCGGACCAGGGGAAT</td><td>7184.6</td><td>-4.2</td><td>0.8</td><td>-5.3</td><td>1.5E-07</td><td>1.0E-06</td><td>BT20/MB468</td><td>MP</td></tr>
<tr><td>>MI0000081|hsa-mir-24-2&WithFlank&19|-|13836281|13836365@56.74.19(+1U);MI0000080|hsa-mir-24-1&WithFlank&9|+|95086015|95086094@50.68.19(+1U)&[MIMAT0000080&hsa-miR-24-3p&offsets|0|-3(+1U);m-36&19|-|13836288|13836310&offsets|0|-4(+1U)];[MIMAT0000080_1&hsa-miR-24-3p&offsets|0|-3(+1U);m-35&9|+|95086064|95086086&offsets|0|-4(+1U)]|TGGCTCAGTTCAGCAGGAAT</td><td>>MI0000081|hsa-mir-24-2&WithFlank&19|-|13836281|13836365</td><td>56</td><td>74</td><td>19(+1U)</td><td>TGGCTCAGTTCAGCAGGAAT</td><td>1087.3</td><td>5.1</td><td>1.0</td><td>5.0</td><td>6.4E-07</td><td>1.0E-06</td><td>MB231/MB468</td><td>MITO</td></tr>
<tr><td>>hg38_21_+_8433172_8446622_RNA45SN1_45S_50ntFlanks@11718.11740.23|AACGGCGGGAGTAACTATGACTC</td><td>>hg38_21_+_8433172_8446622_RNA45SN1_45S_50ntFlanks</td><td>11718</td><td>11740</td><td>23</td><td>AACGGCGGGAGTAACTATGACTC</td><td>521.5</td><td>-3.7</td><td>0.7</td><td>-5.0</td><td>4.6E-07</td><td>1.0E-06</td><td>BT20/MB468</td><td>MITO</td></tr>
<tr><td>>hg38_21_+_8433172_8446622_RNA45SN1_45S_50ntFlanks@12415.12438.24|CTTCGATGTCGGCTCTTCCTATCA</td><td>>hg38_21_+_8433172_8446622_RNA45SN1_45S_50ntFlanks</td><td>12415</td><td>12438</td><td>24</td><td>CTTCGATGTCGGCTCTTCCTATCA</td><td>621.1</td><td>2.9</td><td>0.6</td><td>5.0</td><td>5.2E-07</td><td>1.0E-06</td><td>BT20/MB468</td><td>CYTO</td></tr>
<tr><td>>hg38_21_+_8433172_8446622_RNA45SN1_45S_50ntFlanks@9924.9949.26|TGGTTGATATAGACAGCAGGACGGTG</td><td>>hg38_21_+_8433172_8446622_RNA45SN1_45S_50ntFlanks</td><td>9924</td><td>9949</td><td>26</td><td>TGGTTGATATAGACAGCAGGACGGTG</td><td>56.8</td><td>9.6</td><td>1.8</td><td>5.3</td><td>1.5E-07</td><td>1.0E-06</td><td>BT20/MB468</td><td>MP</td></tr>
<tr><td>>hg38_21_+_8433172_8446622_RNA45SN1_45S_50ntFlanks@9352.9378.27|CCGTGGCGCAATGAAGGTGAAGGCCGG</td><td>>hg38_21_+_8433172_8446622_RNA45SN1_45S_50ntFlanks</td><td>9352</td><td>9378</td><td>27</td><td>CCGTGGCGCAATGAAGGTGAAGGCCGG</td><td>118.0</td><td>-10.0</td><td>1.9</td><td>-5.3</td><td>1.5E-07</td><td>1.0E-06</td><td>BT20/MB468</td><td>MP</td></tr>
<tr><td>>hg38_21_+_8433172_8446622_RNA45SN1_45S_50ntFlanks@6652.6687.36|GACTCTTAGCGGTGGATCACTCGGCTCGTGCGTCGA</td><td>>hg38_21_+_8433172_8446622_RNA45SN1_45S_50ntFlanks</td><td>6652</td><td>6687</td><td>36</td><td>GACTCTTAGCGGTGGATCACTCGGCTCGTGCGTCGA</td><td>286.2</td><td>4.0</td><td>0.8</td><td>5.0</td><td>6.0E-07</td><td>1.0E-06</td><td>MB231/MB468</td><td>CYTO</td></tr>
<tr><td>>MI0000434|hsa-let-7i&WithFlank&12|+|62603680|62603775@12.32.21(+1A)&[MIMAT0000415&hsa-let-7i-5p&offsets|0|-1(+1A);m-49&12|+|62603691|62603712&offsets|0|-1(+1A)]|TGAGGTAGTAGTTTGTGCTGTA</td><td>>MI0000434|hsa-let-7i&WithFlank&12|+|62603680|62603775</td><td>12</td><td>32</td><td>21(+1A)</td><td>TGAGGTAGTAGTTTGTGCTGTA</td><td>6619.0</td><td>4.7</td><td>0.9</td><td>5.0</td><td>5.3E-07</td><td>1.0E-06</td><td>MB231/MB468</td><td>MP</td></tr>
<tr><td>>hg38_1_-_228634819_228635039_RNA5S12_5S_50ntFlanks@149.171.23|GAATACCGGGTGCTGTAGGCTTT</td><td>>hg38_1_-_228634819_228635039_RNA5S12_5S_50ntFlanks</td><td>149</td><td>171</td><td>23</td><td>GAATACCGGGTGCTGTAGGCTTT</td><td>5212.3</td><td>-4.1</td><td>0.8</td><td>-5.0</td><td>5.3E-07</td><td>1.0E-06</td><td>MB231/MB468</td><td>MP</td></tr>
<tr><td>>hg38_21_+_8433172_8446622_RNA45SN1_45S_50ntFlanks@10347.10368.22|CAAACGAGAACTTTGAAGGCCG</td><td>>hg38_21_+_8433172_8446622_RNA45SN1_45S_50ntFlanks</td><td>10347</td><td>10368</td><td>22</td><td>CAAACGAGAACTTTGAAGGCCG</td><td>628.7</td><td>-2.9</td><td>0.6</td><td>-5.0</td><td>6.0E-07</td><td>1.0E-06</td><td>MB231/MB468</td><td>CYTO</td></tr>
<tr><td>>MI0000746|hsa-mir-99b&WithFlank&19|+|51692606|51692687@13.32.20&[MIMAT0000689&hsa-miR-99b-5p&offsets|0|-2;m-10&19|+|51692618|51692639&offsets|0|-2]|CACCCGTAGAACCGACCTTG</td><td>>MI0000746|hsa-mir-99b&WithFlank&19|+|51692606|51692687</td><td>13</td><td>32</td><td>20</td><td>CACCCGTAGAACCGACCTTG</td><td>296.7</td><td>4.5</td><td>0.9</td><td>5.0</td><td>6.4E-07</td><td>1.0E-06</td><td>MB231/MB468</td><td>MITO</td></tr>
<tr><td>>hg38_21_+_8433172_8446622_RNA45SN1_45S_50ntFlanks@8196.8213.18|CGTGGACGGTGTGAGGCC</td><td>>hg38_21_+_8433172_8446622_RNA45SN1_45S_50ntFlanks</td><td>8196</td><td>8213</td><td>18</td><td>CGTGGACGGTGTGAGGCC</td><td>3221.7</td><td>-3.6</td><td>0.7</td><td>-5.0</td><td>5.3E-07</td><td>1.0E-06</td><td>MB231/MB468</td><td>MP</td></tr>
<tr><td>>hg38_21_+_8433172_8446622_RNA45SN1_45S_50ntFlanks@10681.10712.32|TTCCGGCGGCGTCCGGTGAGCTCTCGCTGGCC</td><td>>hg38_21_+_8433172_8446622_RNA45SN1_45S_50ntFlanks</td><td>10681</td><td>10712</td><td>32</td><td>TTCCGGCGGCGTCCGGTGAGCTCTCGCTGGCC</td><td>2106.0</td><td>-4.4</td><td>0.9</td><td>-5.0</td><td>5.2E-07</td><td>1.0E-06</td><td>BT20/MB468</td><td>CYTO</td></tr>
<tr><td>>hg38_21_+_8433172_8446622_RNA45SN1_45S_50ntFlanks@9794.9816.23|CGTGGAGCCGGGCGTGGAATGCG</td><td>>hg38_21_+_8433172_8446622_RNA45SN1_45S_50ntFlanks</td><td>9794</td><td>9816</td><td>23</td><td>CGTGGAGCCGGGCGTGGAATGCG</td><td>509.1</td><td>12.3</td><td>2.2</td><td>5.5</td><td>3.3E-08</td><td>1.0E-06</td><td>BT20/MB231</td><td>MP</td></tr>
</table>

| | | | | | | | | | | | | | |
|---|---|---|---|---|---|---|---|---|---|---|---|---|---|
| >hg38_21_+_8433172_8446622_RNA45SN1_45S_50ntFlanks@12419.12452.34\|GATGTCGGCTCTTCCTATCATTGTGAAGCAGAAT | >hg38_21_+_8433172_8446622_RNA45SN1_45S_50ntFlanks | 12419 | 12452 | 34 | GATGTCGGCTCTTCCTATCATTGTGAAGCAGAAT | 338.6 | 5.3 | 1.1 | 5.0 | 5.2E-07 | 1.0E-06 | BT20/MB468 | CYTO |
| >hg38_21_+_8433172_8446622_RNA45SN1_45S_50ntFlanks@12932.12956.25\|CGCTAAACCATTCGTAGACGACCTG | >hg38_21_+_8433172_8446622_RNA45SN1_45S_50ntFlanks | 12932 | 12956 | 25 | CGCTAAACCATTCGTAGACGACCTG | 594.5 | -2.6 | 0.5 | -5.0 | 5.2E-07 | 1.0E-06 | BT20/MB468 | CYTO |
| >hg38_21_+_8433172_8446622_RNA45SN1_45S_50ntFlanks@12588.12606.19\|GGTTCAGACATTTGGTGTA | >hg38_21_+_8433172_8446622_RNA45SN1_45S_50ntFlanks | 12588 | 12606 | 19 | GGTTCAGACATTTGGTGTA | 61.3 | -9.0 | 1.7 | -5.3 | 1.5E-07 | 1.0E-06 | BT20/MB468 | MP |
| >hg38_21_+_8433172_8446622_RNA45SN1_45S_50ntFlanks@9500.9525.26\|GATGGTGAACTATGCCTGGGCAGGGC | >hg38_21_+_8433172_8446622_RNA45SN1_45S_50ntFlanks | 9500 | 9525 | 26 | GATGGTGAACTATGCCTGGGCAGGGC | 382.3 | 11.9 | 2.2 | 5.5 | 3.3E-08 | 1.0E-06 | BT20/MB231 | MP |
| >hg38_21_+_8433172_8446622_RNA45SN1_45S_50ntFlanks@10873.10891.19\|CTGGGTCGGTCGGGCTGGG | >hg38_21_+_8433172_8446622_RNA45SN1_45S_50ntFlanks | 10873 | 10891 | 19 | CTGGGTCGGTCGGGCTGGG | 704.9 | -7.3 | 1.5 | -5.0 | 6.2E-07 | 1.0E-06 | MB231/MB468 | NUC |
| >hg38_21_+_8433172_8446622_RNA45SN1_45S_50ntFlanks@8340.8373.34\|ACCGTAAGGGAAAGTTGAAAAGAACTTTGAAGAG | >hg38_21_+_8433172_8446622_RNA45SN1_45S_50ntFlanks | 8340 | 8373 | 34 | ACCGTAAGGGAAAGTTGAAAAGAACTTTGAAGAG | 243.3 | 8.7 | 1.7 | 5.3 | 1.5E-07 | 1.0E-06 | BT20/MB468 | MP |
| >hg38_21_+_8433172_8446622_RNA45SN1_45S_50ntFlanks@10786.10820.35\|AACAGCCTCTGGCATGTTGGAACAATGTAGGTAAG | >hg38_21_+_8433172_8446622_RNA45SN1_45S_50ntFlanks | 10786 | 10820 | 35 | AACAGCCTCTGGCATGTTGGAACAATGTAGGTAAG | 350.4 | -3.0 | 0.6 | -5.0 | 6.0E-07 | 1.0E-06 | MB231/MB468 | CYTO |
| >am_trna119_LysCTT_1_-_145395522_145395594@7.36.30,trna11_LysCTT_5_-_180648979_180649051@7.36.30,trna128_LysCTT_1_-_55423542_55423614@7.36.30,trna13_LysCTT_6_+_26556774_26556846@7.36.30,trna6_LysCTT_19_-_52425396_52425468@7.36.30,trna7_LysCTT_16_+_3225692_3225764@7.36.30,trna9_LysCTT_5_+_180634755_180634827@7.36.30\|CTAGCTCAGTCGGTAGAGCATGAGACTCTT | >am_trna119_LysCTT_1_-_145395522_145395594 | 7 | 36 | 30 | CTAGCTCAGTCGGTAGAGCATGAGACTCTT | 591.4 | -5.7 | 1.0 | -5.4 | 5.1E-08 | 1.0E-06 | BT20/MB231 | MITO |
| >hg38_21_+_8433172_8446622_RNA45SN1_45S_50ntFlanks@12457.12484.28\|CAAGCGTTGGATTGTTCACCCACTAATA | >hg38_21_+_8433172_8446622_RNA45SN1_45S_50ntFlanks | 12457 | 12484 | 28 | CAAGCGTTGGATTGTTCACCCACTAATA | 597.4 | -4.2 | 0.8 | -5.0 | 4.6E-07 | 1.0E-06 | BT20/MB468 | MITO |
| >hg38_21_+_8433172_8446622_RNA45SN1_45S_50ntFlanks@11718.11751.34\|AACGGCGGGAGTAACTATGACTCTCTTAAGGTAG | >hg38_21_+_8433172_8446622_RNA45SN1_45S_50ntFlanks | 11718 | 11751 | 34 | AACGGCGGGAGTAACTATGACTCTCTTAAGGTAG | 823.0 | -6.7 | 1.3 | -5.0 | 6.0E-07 | 1.0E-06 | MB231/MB468 | CYTO |
| >MI0003628\|hsa-mir-615&WithFlank&12\|+\|54033944\|54034051@67.85.19&[MIMAT0003283&hsa-miR-615-3p&offsets\|0\|-3;m-495&12\|+\|54034010\|54034030&offsets\|0\|-2]\|TCCGAGCCTGGGTCTCCCT | >MI0003628\|hsa-mir-615&WithFlank&12\|+\|54033944\|54034051 | 67 | 85 | 19 | TCCGAGCCTGGGTCTCCCT | 62.4 | 9.7 | 1.8 | 5.2 | 1.5E-07 | 1.0E-06 | BT20/MB468 | MP |
| >hg38_21_+_8433172_8446622_RNA45SN1_45S_50ntFlanks@11874.11906.33\|AGAAGACCCTGTTGAGCTTGACTCTAGTCTGGC | >hg38_21_+_8433172_8446622_RNA45SN1_45S_50ntFlanks | 11874 | 11906 | 33 | AGAAGACCCTGTTGAGCTTGACTCTAGTCTGGC | 1118.3 | -5.7 | 1.1 | -5.0 | 5.3E-07 | 1.0E-06 | BT20/MB468 | CYTO |
| >MI0000088\|hsa-mir-30a&WithFlank&6\|-\|71403545\|71403627@53.73.21(+2U)&[MIMAT0000088&hsa-miR-30a-3p&offsets\|0\|-1(+2U);m-38&6\|-\|71403554\|71403575&offsets\|0\|-1(+2U)]\|CTTTCAGTCGGATGTTTGCAGTT | >MI0000088\|hsa-mir-30a&WithFlank&6\|-\|71403545\|71403627 | 53 | 73 | 21(+2U) | CTTTCAGTCGGATGTTTGCAGTT | 344.6 | 7.9 | 1.6 | 5.0 | 6.5E-07 | 1.0E-06 | MB231/MB468 | MITO |
| >am_trna133_GlyCCC_1_-_16872434_16872504@1.33.33,trna4_GlyCCC_1_+_17188416_17188486@1.33.33\|GCATTGGTGGTTCAGTGGTAGAATTCTCGCCTC | >am_trna133_GlyCCC_1_-_16872434_16872504 | 1 | 33 | 33 | GCATTGGTGGTTCAGTGGTAGAATTCTCGCCTC | 167.7 | 11.1 | 2.1 | 5.2 | 1.5E-07 | 1.0E-06 | BT20/MB468 | MP |
| >MI0000459\|hsa-mir-143&WithFlank&5\|+\|149428912\|149429029@67.86.20&[MIMAT0000435&hsa-miR-143-3p&offsets\|0\|-1;m-3&5\|+\|149428978\|149428997&offsets\|0\|0]\|TGAGATGAAGCACTGTAGCT | >MI0000459\|hsa-mir-143&WithFlank&5\|+\|149428912\|149429029 | 67 | 86 | 20 | TGAGATGAAGCACTGTAGCT | 66.6 | -9.1 | 1.7 | -5.2 | 1.5E-07 | 1.0E-06 | BT20/MB468 | MP |
| >hg38_21_+_8433172_8446622_RNA45SN1_45S_50ntFlanks@9343.9368.26\|CGAAAGCCGCCGTGGCGCAATGAAGG | >hg38_21_+_8433172_8446622_RNA45SN1_45S_50ntFlanks | 9343 | 9368 | 26 | CGAAAGCCGCCGTGGCGCAATGAAGG | 72.2 | -9.3 | 1.8 | -5.2 | 1.5E-07 | 1.0E-06 | BT20/MB468 | MP |
| >hg38_21_+_8433172_8446622_RNA45SN1_45S_50ntFlanks@10351.10371.21\|CGAGAACTTTGAAGGCCGAAG | >hg38_21_+_8433172_8446622_RNA45SN1_45S_50ntFlanks | 10351 | 10371 | 21 | CGAGAACTTTGAAGGCCGAAG | 1017.2 | -3.2 | 0.6 | -5.0 | 6.2E-07 | 1.0E-06 | MB231/MB468 | NUC |
| >hg38_21_+_8433172_8446622_RNA45SN1_45S_50ntFlanks@12203.12221.19\|TAAGGCGAGCTCAGGGAGG | >hg38_21_+_8433172_8446622_RNA45SN1_45S_50ntFlanks | 12203 | 12221 | 19 | TAAGGCGAGCTCAGGGAGG | 142.0 | -6.9 | 1.4 | -5.0 | 5.4E-07 | 1.0E-06 | MB231/MB468 | MP |
| >trnaMT_PheGAA_MT_+_577_647@23.57.35\|AAGCAATACACTGAAAATGTTTAGACGGGCTCACA | >trnaMT_PheGAA_MT_+_577_647 | 23 | 57 | 35 | AAGCAATACACTGAAAATGTTTAGACGGGCTCACA | 125.0 | 10.7 | 2.0 | 5.2 | 1.5E-07 | 1.0E-06 | BT20/MB468 | MP |
| >hg38_21_+_8433172_8446622_RNA45SN1_45S_50ntFlanks@4472.4493.22\|CTTAGCTGAGTGTCCCGCGGGG | >hg38_21_+_8433172_8446622_RNA45SN1_45S_50ntFlanks | 4472 | 4493 | 22 | CTTAGCTGAGTGTCCCGCGGGG | 449.5 | -6.1 | 1.2 | -5.0 | 6.1E-07 | 1.0E-06 | MB231/MB468 | CYTO |
| >MI0000086\|hsa-mir-28&WithFlank&3\|+\|188688775\|188688872@60.82.23(+1C)&[MIMAT0004502&hsa-miR-28-3p&offsets\|0\|+1(+1C);m-28&3\|+\|188688834\|188688855&offsets\|0\|+1(+1C)]\|CACTAGATTGTGAGCTCCTGGAGC | >MI0000086\|hsa-mir-28&WithFlank&3\|+\|188688775\|188688872 | 60 | 82 | 23(+1C) | CACTAGATTGTGAGCTCCTGGAGC | 49.8 | 8.9 | 1.8 | 5.0 | 5.4E-07 | 1.0E-06 | MB231/MB468 | MP |
| >hg38_21_+_8433172_8446622_RNA45SN1_45S_50ntFlanks@12415.12443.29\|CTTCGATGTCGGCTCTTCCTATCATTGTG | >hg38_21_+_8433172_8446622_RNA45SN1_45S_50ntFlanks | 12415 | 12443 | 29 | CTTCGATGTCGGCTCTTCCTATCATTGTG | 3925.2 | -2.2 | 0.4 | -5.0 | 5.3E-07 | 1.0E-06 | BT20/MB468 | CYTO |
| >hg38_21_+_8433172_8446622_RNA45SN1_45S_50ntFlanks@10885.10902.18\|GGCTGGGGCGCGAAGCGG | >hg38_21_+_8433172_8446622_RNA45SN1_45S_50ntFlanks | 10885 | 10902 | 18 | GGCTGGGGCGCGAAGCGG | 403.8 | -7.6 | 1.5 | -5.0 | 6.8E-07 | 1.0E-06 | BT20/MB468 | NUC |
| >hg38_21_+_8433172_8446622_RNA45SN1_45S_50ntFlanks@12627.12653.27\|ATGGGGCGAAGCTACCATCTGTGGGAT | >hg38_21_+_8433172_8446622_RNA45SN1_45S_50ntFlanks | 12627 | 12653 | 27 | ATGGGGCGAAGCTACCATCTGTGGGAT | 2168.3 | -5.7 | 1.1 | -5.0 | 5.3E-07 | 1.0E-06 | BT20/MB468 | CYTO |
| >hg38_21_+_8433172_8446622_RNA45SN1_45S_50ntFlanks@10234.10268.35\|CGGACGCTACGCCGCGACGAGTAGGAGGGCCGCTG | >hg38_21_+_8433172_8446622_RNA45SN1_45S_50ntFlanks | 10234 | 10268 | 35 | CGGACGCTACGCCGCGACGAGTAGGAGGGCCGCTG | 61.3 | -9.0 | 1.7 | -5.2 | 1.5E-07 | 1.0E-06 | BT20/MB468 | MP |
| >hg38_21_+_8433172_8446622_RNA45SN1_45S_50ntFlanks@8196.8222.27\|CGTGGACGGTGTGAGGCCGGTAGCGGC | >hg38_21_+_8433172_8446622_RNA45SN1_45S_50ntFlanks | 8196 | 8222 | 27 | CGTGGACGGTGTGAGGCCGGTAGCGGC | 378.6 | -4.1 | 0.8 | -5.2 | 1.5E-07 | 1.0E-06 | BT20/MB468 | MP |
| >hg38_21_+_8433172_8446622_RNA45SN1_45S_50ntFlanks@3831.3860.30\|CTCGCTCCTCTCCTACTTGGATAACTGTGG | >hg38_21_+_8433172_8446622_RNA45SN1_45S_50ntFlanks | 3831 | 3860 | 30 | CTCGCTCCTCTCCTACTTGGATAACTGTGG | 430.9 | -2.0 | 0.4 | -5.0 | 5.3E-07 | 1.0E-06 | BT20/MB468 | CYTO |
| >hg38_21_+_8433172_8446622_RNA45SN1_45S_50ntFlanks@11918.11954.37\|ACATGAGAGGTGTAGAATAAGTGGGAGGCCCCCGGCG | >hg38_21_+_8433172_8446622_RNA45SN1_45S_50ntFlanks | 11918 | 11954 | 37 | ACATGAGAGGTGTAGAATAAGTGGGAGGCCCCCGGCG | 340.2 | -6.2 | 1.2 | -5.0 | 6.1E-07 | 1.0E-06 | MB231/MB468 | CYTO |
| >hg38_21_+_8433172_8446622_RNA45SN1_45S_50ntFlanks@7977.8003.27\|CGACCTCAGATCAGACGTGGCGACCCG | >hg38_21_+_8433172_8446622_RNA45SN1_45S_50ntFlanks | 7977 | 8003 | 27 | CGACCTCAGATCAGACGTGGCGACCCG | 429.4 | -5.8 | 1.2 | -5.0 | 6.9E-07 | 1.1E-06 | BT20/MB468 | NUC |
| >hg38_21_+_8433172_8446622_RNA45SN1_45S_50ntFlanks@5084.5108.25\|CGAACGAGACTCTGGCATGCTAACT | >hg38_21_+_8433172_8446622_RNA45SN1_45S_50ntFlanks | 5084 | 5108 | 25 | CGAACGAGACTCTGGCATGCTAACT | 1396.3 | -7.5 | 1.5 | -5.0 | 6.6E-07 | 1.1E-06 | MB231/MB468 | MITO |
| >hg38_21_+_8433172_8446622_RNA45SN1_45S_50ntFlanks@12407.12443.37\|TTTTGATCCTTCGATGTCGGCTCTTCCTATCATTGTG | >hg38_21_+_8433172_8446622_RNA45SN1_45S_50ntFlanks | 12407 | 12443 | 37 | TTTTGATCCTTCGATGTCGGCTCTTCCTATCATTGTG | 317.4 | -6.1 | 1.2 | -5.0 | 5.5E-07 | 1.1E-06 | MB231/MB468 | MP |
| >hg38_21_+_8433172_8446622_RNA45SN1_45S_50ntFlanks@8836.8855.20\|CGGCGGTGGCGGCGGCGGCG | >hg38_21_+_8433172_8446622_RNA45SN1_45S_50ntFlanks | 8836 | 8855 | 20 | CGGCGGTGGCGGCGGCGGCG | 69.4 | -9.2 | 1.8 | -5.2 | 1.6E-07 | 1.1E-06 | BT20/MB468 | MP |
| >hg38_21_+_8433172_8446622_RNA45SN1_45S_50ntFlanks@4730.4751.22\|CAAGAACGAAAGTCGGAGGTTC | >hg38_21_+_8433172_8446622_RNA45SN1_45S_50ntFlanks | 4730 | 4751 | 22 | CAAGAACGAAAGTCGGAGGTTC | 398.8 | -5.5 | 1.1 | -5.0 | 6.6E-07 | 1.1E-06 | MB231/MB468 | MITO |
| >hg38_21_+_8433172_8446622_RNA45SN1_45S_50ntFlanks@9166.9197.32\|CCGGGCGGGTCGCGCCGTCGGGCCCGGGGGAG | >hg38_21_+_8433172_8446622_RNA45SN1_45S_50ntFlanks | 9166 | 9197 | 32 | CCGGGCGGGTCGCGCCGTCGGGCCCGGGGGAG | 2117.7 | -7.9 | 1.6 | -5.0 | 4.7E-07 | 1.1E-06 | BT20/MB468 | MITO |
| >MI0000088\|hsa-mir-30a&WithFlank&6\|-\|71403545\|71403627@12.32.21&[MIMAT0000087&hsa-miR-30a-5p&offsets\|0\|-1;m-15&6\|-\|71403595\|71403616&offsets\|0\|-1]\|TGTAAACATCCTCGACTGGAA | >MI0000088\|hsa-mir-30a&WithFlank&6\|-\|71403545\|71403627 | 12 | 32 | 21 | TGTAAACATCCTCGACTGGAA | 433.9 | 7.3 | 1.5 | 5.0 | 5.4E-07 | 1.1E-06 | BT20/MB468 | CYTO |
| >hg38_21_+_8433172_8446622_RNA45SN1_45S_50ntFlanks@12662.12703.42\|AACGCCTCTAAGTCAGAATCCCGCCCAGGCGGAACGATACGG | >hg38_21_+_8433172_8446622_RNA45SN1_45S_50ntFlanks | 12662 | 12703 | 42 | AACGCCTCTAAGTCAGAATCCCGCCCAGGCGGAACGATACGG | 113.0 | -8.9 | 1.8 | -5.0 | 4.7E-07 | 1.1E-06 | BT20/MB468 | MITO |
| >hg38_21_+_8433172_8446622_RNA45SN1_45S_50ntFlanks@11873.11901.29\|AAGAAGACCCTGTTGAGCTTGACTCTAGT | >hg38_21_+_8433172_8446622_RNA45SN1_45S_50ntFlanks | 11873 | 11901 | 29 | AAGAAGACCCTGTTGAGCTTGACTCTAGT | 203.2 | -2.9 | 0.6 | -5.0 | 6.3E-07 | 1.1E-06 | MB231/MB468 | NUC |
| >hg38_21_+_8433172_8446622_RNA45SN1_45S_50ntFlanks@8382.8405.24\|AGAGGGCGTGAAACCGTTAAGAGG | >hg38_21_+_8433172_8446622_RNA45SN1_45S_50ntFlanks | 8382 | 8405 | 24 | AGAGGGCGTGAAACCGTTAAGAGG | 306.2 | -4.8 | 1.0 | -5.0 | 5.5E-07 | 1.1E-06 | MB231/MB468 | MP |
| >hg38_21_+_8433172_8446622_RNA45SN1_45S_50ntFlanks@10392.10435.44\|ACAGCAGTTGAACATGGGTCAGTCGGTCCTGAGAGATGGGCGAG | >hg38_21_+_8433172_8446622_RNA45SN1_45S_50ntFlanks | 10392 | 10435 | 44 | ACAGCAGTTGAACATGGGTCAGTCGGTCCTGAGAGATGGGCGAG | 753.6 | -7.5 | 1.5 | -5.0 | 5.4E-07 | 1.1E-06 | BT20/MB468 | CYTO |
| >trnaMT_GluTTC_MT_-_14674_14742@44.70.27,trnalookalike8_GluTTC_5_-_93905172_93905240@44.70.27\|TCGTGGTTGTAGTCCGTGCGAGAATAC | >trnaMT_GluTTC_MT_-_14674_14742 | 44 | 70 | 27 | TCGTGGTTGTAGTCCGTGCGAGAATAC | 110.0 | 10.5 | 2.0 | 5.2 | 1.6E-07 | 1.1E-06 | BT20/MB468 | MP |
| >hg38_21_+_8433172_8446622_RNA45SN1_45S_50ntFlanks@12001.12031.31\|CTGCGGGCCGCCGGTGAAATACCACTACTCT | >hg38_21_+_8433172_8446622_RNA45SN1_45S_50ntFlanks | 12001 | 12031 | 31 | CTGCGGGCCGCCGGTGAAATACCACTACTCT | 354.7 | -2.6 | 0.5 | -5.0 | 4.7E-07 | 1.1E-06 | BT20/MB468 | MITO |
| >hg38_21_+_8433172_8446622_RNA45SN1_45S_50ntFlanks@8079.8103.25\|AAGAGCCCAGCGCCGAATCCCCGCC | >hg38_21_+_8433172_8446622_RNA45SN1_45S_50ntFlanks | 8079 | 8103 | 25 | AAGAGCCCAGCGCCGAATCCCCGCC | 681.1 | -3.6 | 0.7 | -5.0 | 4.7E-07 | 1.1E-06 | BT20/MB468 | MITO |
| >hg38_21_+_8433172_8446622_RNA45SN1_45S_50ntFlanks@5084.5104.21\|CGAACGAGACTCTGGCATGCT | >hg38_21_+_8433172_8446622_RNA45SN1_45S_50ntFlanks | 5084 | 5104 | 21 | CGAACGAGACTCTGGCATGCT | 950.8 | -7.1 | 1.4 | -5.0 | 6.7E-07 | 1.1E-06 | MB231/MB468 | MITO |
| >hg38_21_+_8433172_8446622_RNA45SN1_45S_50ntFlanks@4291.4317.27\|AGGATCCATTGGAGGGCAAGTCTGGTG | >hg38_21_+_8433172_8446622_RNA45SN1_45S_50ntFlanks | 4291 | 4317 | 27 | AGGATCCATTGGAGGGCAAGTCTGGTG | 197.6 | -6.7 | 1.3 | -5.0 | 6.4E-07 | 1.1E-06 | MB231/MB468 | NUC |
| >hg38_21_+_8433172_8446622_RNA45SN1_45S_50ntFlanks@8170.8213.44\|CAAGTCCTTCTGATCGAGGCCCAGCCCGTGGACGGTGTGAGGCC | >hg38_21_+_8433172_8446622_RNA45SN1_45S_50ntFlanks | 8170 | 8213 | 44 | CAAGTCCTTCTGATCGAGGCCCAGCCCGTGGACGGTGTGAGGCC | 817.8 | -3.9 | 0.8 | -5.0 | 6.4E-07 | 1.1E-06 | MB231/MB468 | NUC |
| >hg38_21_+_8433172_8446622_RNA45SN1_45S_50ntFlanks@3993.4024.32\|GTGACTCTAGATAACCTCGGGCCGATCGCACG | >hg38_21_+_8433172_8446622_RNA45SN1_45S_50ntFlanks | 3993 | 4024 | 32 | GTGACTCTAGATAACCTCGGGCCGATCGCACG | 3003.2 | -4.6 | 0.9 | -5.0 | 6.7E-07 | 1.1E-06 | MB231/MB468 | MITO |
| >hg38_21_+_8433172_8446622_RNA45SN1_45S_50ntFlanks@9739.9762.24\|AAACGATCTCAACCTATTCTCAAA | >hg38_21_+_8433172_8446622_RNA45SN1_45S_50ntFlanks | 9739 | 9762 | 24 | AAACGATCTCAACCTATTCTCAAA | 387.7 | -7.6 | 1.5 | -5.0 | 6.9E-07 | 1.1E-06 | BT20/MB468 | NUC |
| >hg38_21_+_8433172_8446622_RNA45SN1_45S_50ntFlanks@8340.8375.36\|ACCGTAAGGGAAAGTTGAAAAGAACTTTGAAGAGAG | >hg38_21_+_8433172_8446622_RNA45SN1_45S_50ntFlanks | 8340 | 8375 | 36 | ACCGTAAGGGAAAGTTGAAAAGAACTTTGAAGAGAG | 781.6 | 9.2 | 1.8 | 5.2 | 1.6E-07 | 1.1E-06 | BT20/MB468 | MP |

<table>
<tr><td>>hg38_21_+_8433172_8446622_RNA45SN1_45S_50ntFlanks@9792.9813.22|GGCGTGGAGCCGGGCGTGGAAT</td><td>>hg38_21_+_8433172_8446622_RNA45SN1_45S_50ntFlanks</td><td>9792</td><td>9813</td><td>22</td><td>GGCGTGGAGCCGGGCGTGGAAT</td><td>64.4</td><td>9.8</td><td>1.9</td><td>5.2</td><td>1.6E-07</td><td>1.1E-06</td><td>BT20/MB468</td><td>MP</td></tr>
<tr><td>>MI0000438|hsa-mir-15b&WithFlank&3|+|160404582|160404691@26.47.22&[MIMAT0000417&hsa-miR-15b-5p&offsets|0|0;m-110&3|+|160404607|160404628&offsets|0|0]|TAGCAGCACATCATGGTTTACA</td><td>>MI0000438|hsa-mir-15b&WithFlank&3|+|160404582|160404691</td><td>26</td><td>47</td><td>22</td><td>TAGCAGCACATCATGGTTTACA</td><td>1234.8</td><td>4.9</td><td>1.0</td><td>5.0</td><td>6.7E-07</td><td>1.1E-06</td><td>MB231/MB468</td><td>MITO</td></tr>
<tr><td>>MI0000477|hsa-mir-146a&WithFlank&5|+|160485346|160485456@27.48.22&[MIMAT0000449&hsa-miR-146a-5p&offsets|0|0;m-129&5|+|160485372|160485393&offsets|0|0]|TGAGAACTGAATTCCATGGGTT</td><td>>MI0000477|hsa-mir-146a&WithFlank&5|+|160485346|160485456</td><td>27</td><td>48</td><td>22</td><td>TGAGAACTGAATTCCATGGGTT</td><td>23365.4</td><td>-10.2</td><td>1.9</td><td>-5.4</td><td>5.3E-08</td><td>1.1E-06</td><td>BT20/MB231</td><td>MITO</td></tr>
<tr><td>>hg38_21_+_8433172_8446622_RNA45SN1_45S_50ntFlanks@12893.12933.41|CCTCGCCCGTCACGCACCGCACGTTCGTGGGGAACCTGGCG</td><td>>hg38_21_+_8433172_8446622_RNA45SN1_45S_50ntFlanks</td><td>12893</td><td>12933</td><td>41</td><td>CCTCGCCCGTCACGCACCGCACGTTCGTGGGGAACCTGGCG</td><td>1584.5</td><td>-8.7</td><td>1.7</td><td>-5.2</td><td>1.6E-07</td><td>1.1E-06</td><td>BT20/MB468</td><td>MP</td></tr>
<tr><td>>hg38_21_+_8433172_8446622_RNA45SN1_45S_50ntFlanks@10375.10424.50|AGAAGGGTTCCATGTGAACAGCAGTTGAACATGGGTCAGTCGGTCCTGAG</td><td>>hg38_21_+_8433172_8446622_RNA45SN1_45S_50ntFlanks</td><td>10375</td><td>10424</td><td>50</td><td>AGAAGGGTTCCATGTGAACAGCAGTTGAACATGGGTCAGTCGGTCCTGAG</td><td>3048.8</td><td>6.2</td><td>1.2</td><td>5.0</td><td>4.8E-07</td><td>1.1E-06</td><td>BT20/MB468</td><td>MITO</td></tr>
<tr><td>>hg38_21_+_8433172_8446622_RNA45SN1_45S_50ntFlanks@4069.4093.25|CTTTCGATGGTAGTCGCCGTGCCTA</td><td>>hg38_21_+_8433172_8446622_RNA45SN1_45S_50ntFlanks</td><td>4069</td><td>4093</td><td>25</td><td>CTTTCGATGGTAGTCGCCGTGCCTA</td><td>116.6</td><td>-5.5</td><td>1.1</td><td>-5.0</td><td>5.4E-07</td><td>1.1E-06</td><td>BT20/MB468</td><td>CYTO</td></tr>
<tr><td>>trnaMT_TyrGTA_MT_-_5826_5891@23.57.35|TTGGACTGTAAATCTAAAGACAGGGGTTAGGCCTC</td><td>>trnaMT_TyrGTA_MT_-_5826_5891</td><td>23</td><td>57</td><td>35</td><td>TTGGACTGTAAATCTAAAGACAGGGGTTAGGCCTC</td><td>99.6</td><td>10.4</td><td>2.0</td><td>5.2</td><td>1.6E-07</td><td>1.1E-06</td><td>BT20/MB468</td><td>MP</td></tr>
<tr><td>>hg38_21_+_8433172_8446622_RNA45SN1_45S_50ntFlanks@4122.4142.21|AGGGTTCGATTCCGGAGAGGG</td><td>>hg38_21_+_8433172_8446622_RNA45SN1_45S_50ntFlanks</td><td>4122</td><td>4142</td><td>21</td><td>AGGGTTCGATTCCGGAGAGGG</td><td>215.5</td><td>-6.8</td><td>1.4</td><td>-5.0</td><td>5.5E-07</td><td>1.1E-06</td><td>MB231/MB468</td><td>MP</td></tr>
<tr><td>>MI0000289|hsa-mir-181a-1&WithFlank&1|-|198859038|198859159@30.53.24(+1C);MI0000269|hsa-mir-181a-2&WithFlank&9|+|124692436|124692557@45.68.24(+1C)&[MIMAT0000256&hsa-miR-181a-5p&offsets|0|+1(+1C);m-32&1|-|198859108|198859130&offsets|0|+1(+1C)];[MIMAT0000256_1&hsa-miR-181a-5p&offsets|0|+1(+1C);m-32&9|+|124692480|124692502&offsets|0|+1(+1C)]|AACATTCAACGCTGTCGGTGAGTTC</td><td>>MI0000289|hsa-mir-181a-1&WithFlank&1|-|198859038|198859159</td><td>30</td><td>53</td><td>24(+1C)</td><td>AACATTCAACGCTGTCGGTGAGTTC</td><td>531.3</td><td>8.8</td><td>1.8</td><td>5.0</td><td>6.7E-07</td><td>1.1E-06</td><td>MB231/MB468</td><td>MITO</td></tr>
<tr><td>>hg38_21_+_8433172_8446622_RNA45SN1_45S_50ntFlanks@10402.10429.28|AACATGGGTCAGTCGGTCCTGAGAGATG</td><td>>hg38_21_+_8433172_8446622_RNA45SN1_45S_50ntFlanks</td><td>10402</td><td>10429</td><td>28</td><td>AACATGGGTCAGTCGGTCCTGAGAGATG</td><td>2068.7</td><td>-2.4</td><td>0.5</td><td>-5.0</td><td>4.8E-07</td><td>1.1E-06</td><td>BT20/MB468</td><td>MITO</td></tr>
<tr><td>>hg38_21_+_8433172_8446622_RNA45SN1_45S_50ntFlanks@11822.11844.23|TATCCAGCGAAACCACAGCCAAG</td><td>>hg38_21_+_8433172_8446622_RNA45SN1_45S_50ntFlanks</td><td>11822</td><td>11844</td><td>23</td><td>TATCCAGCGAAACCACAGCCAAG</td><td>77.9</td><td>-9.4</td><td>1.8</td><td>-5.2</td><td>1.6E-07</td><td>1.1E-06</td><td>BT20/MB468</td><td>MP</td></tr>
<tr><td>>hg38_21_+_8433172_8446622_RNA45SN1_45S_50ntFlanks@475.501.27|CTTCGTGATCGATGTGGTGACGTCGTG</td><td>>hg38_21_+_8433172_8446622_RNA45SN1_45S_50ntFlanks</td><td>475</td><td>501</td><td>27</td><td>CTTCGTGATCGATGTGGTGACGTCGTG</td><td>1262.3</td><td>7.0</td><td>1.4</td><td>5.0</td><td>4.8E-07</td><td>1.1E-06</td><td>BT20/MB468</td><td>MITO</td></tr>
<tr><td>>MI0001445|hsa-mir-423&WithFlank&17|+|30117073|30117178@59.80.22(+1C)&[MIMAT0001340&hsa-miR-423-3p&offsets|0|-1(+1C);m-100&17|+|30117131|30117153&offsets|0|-1(+1C)]|AGCTCGGTCTGAGGCCCCTCAGC</td><td>>MI0001445|hsa-mir-423&WithFlank&17|+|30117073|30117178</td><td>59</td><td>80</td><td>22(+1C)</td><td>AGCTCGGTCTGAGGCCCCTCAGC</td><td>966.4</td><td>4.3</td><td>0.9</td><td>5.0</td><td>5.6E-07</td><td>1.1E-06</td><td>MB231/MB468</td><td>MP</td></tr>
<tr><td>>MI0005544|hsa-mir-147b&WithFlank&15|+|45433044|45433135@55.73.19(+1U)&[MIMAT0004928&hsa-miR-147b-3p&offsets|0|-2(+1U);m-575&15|+|45433098|45433118&offsets|0|-2(+1U)]|GTGTGCGGAAATGCTTCTGT</td><td>>MI0005544|hsa-mir-147b&WithFlank&15|+|45433044|45433135</td><td>55</td><td>73</td><td>19(+1U)</td><td>GTGTGCGGAAATGCTTCTGT</td><td>126.6</td><td>-10.4</td><td>1.9</td><td>-5.5</td><td>3.5E-08</td><td>1.1E-06</td><td>BT20/MB231</td><td>MP</td></tr>
<tr><td>>hg38_21_+_8433172_8446622_RNA45SN1_45S_50ntFlanks@11572.11607.36|CCGACTTAGAACTGGTGCGGACCAGGGGAATCCGAC</td><td>>hg38_21_+_8433172_8446622_RNA45SN1_45S_50ntFlanks</td><td>11572</td><td>11607</td><td>36</td><td>CCGACTTAGAACTGGTGCGGACCAGGGGAATCCGAC</td><td>231.9</td><td>-7.2</td><td>1.4</td><td>-5.0</td><td>5.6E-07</td><td>1.1E-06</td><td>MB231/MB468</td><td>MP</td></tr>
<tr><td>>hg38_21_+_8433172_8446622_RNA45SN1_45S_50ntFlanks@8168.8203.36|CCCAAGTCCTTCTGATCGAGGCCCAGCCCGTGGACG</td><td>>hg38_21_+_8433172_8446622_RNA45SN1_45S_50ntFlanks</td><td>8168</td><td>8203</td><td>36</td><td>CCCAAGTCCTTCTGATCGAGGCCCAGCCCGTGGACG</td><td>225.1</td><td>-6.6</td><td>1.3</td><td>-5.0</td><td>6.4E-07</td><td>1.1E-06</td><td>MB231/MB468</td><td>NUC</td></tr>
<tr><td>>hg38_21_+_8433172_8446622_RNA45SN1_45S_50ntFlanks@6649.6675.27|TACGACTCTTAGCGGTGGATCACTCGG</td><td>>hg38_21_+_8433172_8446622_RNA45SN1_45S_50ntFlanks</td><td>6649</td><td>6675</td><td>27</td><td>TACGACTCTTAGCGGTGGATCACTCGG</td><td>364.5</td><td>-2.3</td><td>0.5</td><td>-5.0</td><td>7.0E-07</td><td>1.1E-06</td><td>BT20/MB468</td><td>NUC</td></tr>
<tr><td>>hg38_21_+_8433172_8446622_RNA45SN1_45S_50ntFlanks@6670.6697.28|ACTCGGCTCGTGCGTCGATGAAGAACGC</td><td>>hg38_21_+_8433172_8446622_RNA45SN1_45S_50ntFlanks</td><td>6670</td><td>6697</td><td>28</td><td>ACTCGGCTCGTGCGTCGATGAAGAACGC</td><td>146.3</td><td>-6.7</td><td>1.3</td><td>-5.0</td><td>5.6E-07</td><td>1.1E-06</td><td>MB231/MB468</td><td>MP</td></tr>
<tr><td>>hg38_21_+_8433172_8446622_RNA45SN1_45S_50ntFlanks@10116.10139.24|TGCGTCGGAGGGCGGCGGCGGCGG</td><td>>hg38_21_+_8433172_8446622_RNA45SN1_45S_50ntFlanks</td><td>10116</td><td>10139</td><td>24</td><td>TGCGTCGGAGGGCGGCGGCGGCGG</td><td>247.5</td><td>-6.3</td><td>1.3</td><td>-5.0</td><td>5.5E-07</td><td>1.1E-06</td><td>BT20/MB468</td><td>CYTO</td></tr>
<tr><td>>hg38_21_+_8433172_8446622_RNA45SN1_45S_50ntFlanks@10819.10836.18|AGGGAAGTCGGCAAGCCG</td><td>>hg38_21_+_8433172_8446622_RNA45SN1_45S_50ntFlanks</td><td>10819</td><td>10836</td><td>18</td><td>AGGGAAGTCGGCAAGCCG</td><td>183.4</td><td>-4.2</td><td>0.8</td><td>-5.0</td><td>6.3E-07</td><td>1.1E-06</td><td>MB231/MB468</td><td>CYTO</td></tr>
<tr><td>>MI0003776|hsa-mir-320b-1&WithFlank&1|+|116671740|116671823@48.66.19;MI0003839|hsa-mir-320b-2&WithFlank&1|-|224257034|224257116@47.65.19;MI0003778|hsa-mir-320c-1&WithFlank&18|+|21683512|21683595@48.66.19;MI0008191|hsa-mir-320c-2&WithFlank&18|+|24321669|24321752@48.66.19;MI0000542|hsa-mir-320a&WithFlank&8|-|22244960|22245043@48.66.19&[MIMAT0005792&hsa-miR-320b&offsets|0|-3;m-183&1|+|116671787|116671806&offsets|0|-1];[MIMAT0005792_1&hsa-miR-320b&offsets|0|-3;m-183&1|-|224257051|224257070&offsets|0|-1];[MIMAT0005793&hsa-miR-320c&offsets|0|-1;m-222&18|+|21683559|21683578&offsets|0|-1];[MIMAT0005793_1&hsa-miR-320c&offsets|0|-1;m-219&18|+|24321716|24321735&offsets|0|-1];[MIMAT0000510&hsa-miR-320a-3p&offsets|0|-3;m-60&8|-|22244975|22244996&offsets|0|-3]|AAAAGCTGGGTTGAGAGGG</td><td>>MI0003776|hsa-mir-320b-1&WithFlank&1|+|116671740|116671823</td><td>48</td><td>66</td><td>19</td><td>AAAAGCTGGGTTGAGAGGG</td><td>2635.0</td><td>7.8</td><td>1.5</td><td>5.2</td><td>1.6E-07</td><td>1.1E-06</td><td>BT20/MB468</td><td>MP</td></tr>
<tr><td>>am_trnaMT_TyrGTA_MT_-_5826_5891@1.31.31|GGTAAAATGGCTGAGTGAAGCATTGGACTGT</td><td>>am_trnaMT_TyrGTA_MT_-_5826_5891</td><td>1</td><td>31</td><td>31</td><td>GGTAAAATGGCTGAGTGAAGCATTGGACTGT</td><td>1351.3</td><td>6.3</td><td>1.3</td><td>5.0</td><td>6.7E-07</td><td>1.1E-06</td><td>MB231/MB468</td><td>MITO</td></tr>
<tr><td>>hg38_21_+_8433172_8446622_RNA45SN1_45S_50ntFlanks@10420.10442.23|CTGAGAGATGGGCGAGCGCCGTT</td><td>>hg38_21_+_8433172_8446622_RNA45SN1_45S_50ntFlanks</td><td>10420</td><td>10442</td><td>23</td><td>CTGAGAGATGGGCGAGCGCCGTT</td><td>181.3</td><td>-3.1</td><td>0.6</td><td>-5.0</td><td>6.3E-07</td><td>1.1E-06</td><td>MB231/MB468</td><td>CYTO</td></tr>
<tr><td>>MI0001445|hsa-mir-423&WithFlank&17|+|30117073|30117178@59.81.23&[MIMAT0001340&hsa-miR-423-3p&offsets|0|0;m-100&17|+|30117131|30117153&offsets|0|0]|AGCTCGGTCTGAGGCCCCTCAGT</td><td>>MI0001445|hsa-mir-423&WithFlank&17|+|30117073|30117178</td><td>59</td><td>81</td><td>23</td><td>AGCTCGGTCTGAGGCCCCTCAGT</td><td>5438.5</td><td>3.0</td><td>0.6</td><td>5.0</td><td>7.0E-07</td><td>1.1E-06</td><td>BT20/MB468</td><td>NUC</td></tr>
<tr><td>>hg38_21_+_8433172_8446622_RNA45SN1_45S_50ntFlanks@4574.4611.38|ATAGGACCGCGGTTCTATTTTGTTGGTTTTCGGAACTG</td><td>>hg38_21_+_8433172_8446622_RNA45SN1_45S_50ntFlanks</td><td>4574</td><td>4611</td><td>38</td><td>ATAGGACCGCGGTTCTATTTTGTTGGTTTTCGGAACTG</td><td>131.0</td><td>-9.1</td><td>1.8</td><td>-5.0</td><td>4.8E-07</td><td>1.1E-06</td><td>BT20/MB468</td><td>MITO</td></tr>
<tr><td>>am_trna128_GlyGCC_6_-_27870686_27870756@10.29.20,trna133_GlyCCC_1_-_16872434_16872504@10.29.20,trna17_GluTTC_2_-_204229131_204229201@10.29.20,trna18_GlyGCC_16_+_70822597_70822667@10.29.20,trna19_GlyGCC_16_+_70823410_70823480@10.29.20,trna19_GlyGCC_2_-_157257659_157257729@10.29.20,trna24_GlyGCC_16_-_70812942_70813012@10.29.20,trna25_GlyGCC_16_-_70812114_70812184@10.29.20,trna2_GlyGCC_21_-_18827107_18827177@10.29.20,trna35_GlyGCC_1_+_161413094_161413164@10.29.20,trna37_GlyGCC_1_+_161420467_161420537@10.29.20,trna39_GlyGCC_1_+_161427898_161427968@10.29.20,trna41_GlyGCC_1_+_161435258_161435328@10.29.20,trna4_GlyCCC_1_+_17188416_17188486@10.29.20,trna5_GlyGCC_17_+_8029064_8029134@10.29.20,trna68_GlyGCC_1_-_161493637_161493707@10.29.20,trna91_GlyCCC_1_-_149680210_149680280@10.29.20,trna99_ValCAC_1_-_149294666_149294736@10.29.20|GTTCAGTGGTAGAATTCTCG</td><td>>am_trna128_GlyGCC_6_-_27870686_27870756</td><td>10</td><td>29</td><td>20</td><td>GTTCAGTGGTAGAATTCTCG</td><td>113.8</td><td>-9.9</td><td>1.9</td><td>-5.2</td><td>1.6E-07</td><td>1.1E-06</td><td>BT20/MB468</td><td>MP</td></tr>
<tr><td>>hg38_21_+_8433172_8446622_RNA45SN1_45S_50ntFlanks@11592.11612.21|ACCAGGGGAATCCGACTGTTT</td><td>>hg38_21_+_8433172_8446622_RNA45SN1_45S_50ntFlanks</td><td>11592</td><td>11612</td><td>21</td><td>ACCAGGGGAATCCGACTGTTT</td><td>223.7</td><td>-6.4</td><td>1.3</td><td>-5.0</td><td>6.4E-07</td><td>1.1E-06</td><td>MB231/MB468</td><td>NUC</td></tr>
<tr><td>>MI0000650|hsa-mir-200c&WithFlank&12|+|6963693|6963772@50.72.23(+1U)&[MIMAT0000617&hsa-miR-200c-3p&offsets|0|0(+1U);m-54&12|+|6963742|6963764&offsets|0|0(+1U)]|TAATACTGCCGGGTAATGATGGAT</td><td>>MI0000650|hsa-mir-200c&WithFlank&12|+|6963693|6963772</td><td>50</td><td>72</td><td>23(+1U)</td><td>TAATACTGCCGGGTAATGATGGAT</td><td>1985.3</td><td>4.3</td><td>0.8</td><td>5.2</td><td>1.6E-07</td><td>1.1E-06</td><td>BT20/MB468</td><td>MP</td></tr>
<tr><td>>MI0000067|hsa-let-7f-1&WithFlank&9|+|94176341|94176439@13.33.21(+1A);MI0000068|hsa-let-7f-2&WithFlank&X|-|53557186|53557280@14.34.21(+1A)&[MIMAT0000067&hsa-let-7f-5p&offsets|0|-1(+1A);m-20&9|+|94176353|94176374&offsets|0|-1(+1A)];[MIMAT0000067_1&hsa-let-7f-5p&offsets|0|-1(+1A);m-19&X|-|53557246|53557267&offsets|0|-1(+1A)]|TGAGGTAGTAGATTGTATAGTA</td><td>>MI0000067|hsa-let-7f-1&WithFlank&9|+|94176341|94176439</td><td>13</td><td>33</td><td>21(+1A)</td><td>TGAGGTAGTAGATTGTATAGTA</td><td>1321.3</td><td>4.4</td><td>0.9</td><td>5.0</td><td>6.3E-07</td><td>1.1E-06</td><td>MB231/MB468</td><td>CYTO</td></tr>
<tr><td>>hg38_1_-_228634819_228635039_RNA5S12_5S_50ntFlanks@135.171.37|GGGAGACCGCCTGGGAATACCGGGTGCTGTAGGCTTT</td><td>>hg38_1_-_228634819_228635039_RNA5S12_5S_50ntFlanks</td><td>135</td><td>171</td><td>37</td><td>GGGAGACCGCCTGGGAATACCGGGTGCTGTAGGCTTT</td><td>9099.8</td><td>4.9</td><td>1.0</td><td>5.0</td><td>4.8E-07</td><td>1.1E-06</td><td>BT20/MB468</td><td>MITO</td></tr>
<tr><td>>hg38_21_+_8433172_8446622_RNA45SN1_45S_50ntFlanks@7985.8015.31|GATCAGACGTGGCGACCCGCTGAATTTAAGC</td><td>>hg38_21_+_8433172_8446622_RNA45SN1_45S_50ntFlanks</td><td>7985</td><td>8015</td><td>31</td><td>GATCAGACGTGGCGACCCGCTGAATTTAAGC</td><td>62.8</td><td>-9.1</td><td>1.7</td><td>-5.2</td><td>1.6E-07</td><td>1.1E-06</td><td>BT20/MB468</td><td>MP</td></tr>
<tr><td>>hg38_21_+_8433172_8446622_RNA45SN1_45S_50ntFlanks@4064.4089.26|ATCAACTTTCGATGGTAGTCGCCGTG</td><td>>hg38_21_+_8433172_8446622_RNA45SN1_45S_50ntFlanks</td><td>4064</td><td>4089</td><td>26</td><td>ATCAACTTTCGATGGTAGTCGCCGTG</td><td>57.9</td><td>-8.9</td><td>1.7</td><td>-5.2</td><td>1.6E-07</td><td>1.1E-06</td><td>BT20/MB468</td><td>MP</td></tr>
<tr><td>>trna2_GlyGCC_21_-_18827107_18827177@1.29.29,trna35_GlyGCC_1_+_161413094_161413164@1.29.29,trna37_GlyGCC_1_+_161420467_161420537@1.29.29,trna39_GlyGCC_1_+_161427898_161427968@1.29.29,trna41_GlyGCC_1_+_161435258_161435328@1.29.29|GCATGGGTGGTTCAGTGGTAGAATTCTCG</td><td>>trna2_GlyGCC_21_-_18827107_18827177</td><td>1</td><td>29</td><td>29</td><td>GCATGGGTGGTTCAGTGGTAGAATTCTCG</td><td>40137.3</td><td>-3.4</td><td>0.7</td><td>-5.0</td><td>5.5E-07</td><td>1.1E-06</td><td>BT20/MB468</td><td>CYTO</td></tr>
<tr><td>>hg38_21_+_8433172_8446622_RNA45SN1_45S_50ntFlanks@12588.12615.28|GGTTCAGACATTTGGTGTATGTGCTTGG</td><td>>hg38_21_+_8433172_8446622_RNA45SN1_45S_50ntFlanks</td><td>12588</td><td>12615</td><td>28</td><td>GGTTCAGACATTTGGTGTATGTGCTTGG</td><td>69.4</td><td>-9.2</td><td>1.8</td><td>-5.2</td><td>1.6E-07</td><td>1.1E-06</td><td>BT20/MB468</td><td>MP</td></tr>
<tr><td>>hg38_21_+_8433172_8446622_RNA45SN1_45S_50ntFlanks@12709.12739.31|CCGCGGAGCCTCGGTTGGCCTCGGATAGCCG</td><td>>hg38_21_+_8433172_8446622_RNA45SN1_45S_50ntFlanks</td><td>12709</td><td>12739</td><td>31</td><td>CCGCGGAGCCTCGGTTGGCCTCGGATAGCCG</td><td>63.0</td><td>-9.1</td><td>1.7</td><td>-5.2</td><td>1.6E-07</td><td>1.1E-06</td><td>BT20/MB468</td><td>MP</td></tr>
</table>

| | | | | | | | | | | | | | |
|---|---|---|---|---|---|---|---|---|---|---|---|---|---|
| >MI0000088\|hsa-mir-30a&WithFlank&6\|-\|71403545\|71403627@12.35.24(+1C)&[MIMAT0000087&hsa-miR-30a-5p&offsets\|0\|+2(+1C);m-15&6\|-\|71403595\|71403616&offsets\|0\|+2(+1C)]\|TGTAAACATCCTCGACTGGAAGCTC | >MI0000088\|hsa-mir-30a&WithFlank&6\|-\|71403545\|71403627 | 12 | 35 | 24(+1C) | TGTAAACATCCTCGACTGGAAGCTC | 1280.9 | -4.9 | 0.9 | -5.4 | 5.4E-08 | 1.1E-06 | BT20/MB231 | MITO |
| >am_trna128_GlyGCC_6_-_27870686_27870756@3.34.32,trna18_GlyGCC_16_+_70822597_70822667@3.34.32,trna19_GlyGCC_16_+_70823410_70823480@3.34.32,trna19_GlyGCC_2_-_157257659_157257729@3.34.32,trna24_GlyGCC_16_-_70812942_70813012@3.34.32,trna25_GlyGCC_16_-_70812114_70812184@3.34.32,trna5_GlyGCC_17_+_8029064_8029134@3.34.32,trna68_GlyGCC_1_-_161493637_161493707@3.34.32\|ATTGGTGGTTCAGTGGTAGAATTCTCGCCTGC | >am_trna128_GlyGCC_6_-_27870686_27870756 | 3 | 34 | 32 | ATTGGTGGTTCAGTGGTAGAATTCTCGCCTGC | 556.4 | -5.4 | 1.0 | -5.4 | 5.5E-08 | 1.1E-06 | BT20/MB231 | MITO |
| >MI0001446\|hsa-mir-424&WithFlank&X\|-\|134546608\|134546717@54.73.20&[MIMAT0004749&hsa-miR-424-3p&offsets\|0\|-1;m-305&X\|-\|134546644\|134546664&offsets\|0\|-1]\|CAAAACGTGAGGCGCTGCTA | >MI0001446\|hsa-mir-424&WithFlank&X\|-\|134546608\|134546717 | 54 | 73 | 20 | CAAAACGTGAGGCGCTGCTA | 96.3 | 10.3 | 2.0 | 5.2 | 1.6E-07 | 1.1E-06 | BT20/MB468 | MP |
| >hg38_21_+_8433172_8446622_RNA45SN1_45S_50ntFlanks@10866.10886.21\|CTAAGGGCTGGGTCGGTCGGG | >hg38_21_+_8433172_8446622_RNA45SN1_45S_50ntFlanks | 10866 | 10886 | 21 | CTAAGGGCTGGGTCGGTCGGG | 1063.7 | -8.4 | 1.7 | -5.0 | 7.1E-07 | 1.1E-06 | BT20/MB468 | NUC |
| >hg38_21_+_8433172_8446622_RNA45SN1_45S_50ntFlanks@8472.8492.21\|CGGCCCGGCGGATCTTTCCCG | >hg38_21_+_8433172_8446622_RNA45SN1_45S_50ntFlanks | 8472 | 8492 | 21 | CGGCCCGGCGGATCTTTCCCG | 335.9 | -6.2 | 1.2 | -5.0 | 5.5E-07 | 1.1E-06 | BT20/MB468 | CYTO |
| >MI0000809\|hsa-mir-151a&WithFlank&8\|-\|140732558\|140732659@51.72.22&[MIMAT0000757&hsa-miR-151a-3p&offsets\|-2\|-1;m-30&8\|-\|140732587\|140732607&offsets\|-2\|-1]\|TACTAGACTGAAGCTCCTTGAG | >MI0000809\|hsa-mir-151a&WithFlank&8\|-\|140732558\|140732659 | 51 | 72 | 22 | TACTAGACTGAAGCTCCTTGAG | 521.7 | 3.7 | 0.7 | 5.0 | 5.6E-07 | 1.1E-06 | MB231/MB468 | MP |
| >hg38_21_+_8433172_8446622_RNA45SN1_45S_50ntFlanks@12073.12097.25\|CCGAGGGGCTCTCGCTTCTGGCGCC | >hg38_21_+_8433172_8446622_RNA45SN1_45S_50ntFlanks | 12073 | 12097 | 25 | CCGAGGGGCTCTCGCTTCTGGCGCC | 2655.2 | -3.1 | 0.6 | -5.0 | 6.3E-07 | 1.1E-06 | MB231/MB468 | CYTO |
| >hg38_21_+_8433172_8446622_RNA45SN1_45S_50ntFlanks@9789.9830.42\|GCTGGCGTGGAGCCGGGCGTGGAATGCGAGTGCCTAGTGGGC | >hg38_21_+_8433172_8446622_RNA45SN1_45S_50ntFlanks | 9789 | 9830 | 42 | GCTGGCGTGGAGCCGGGCGTGGAATGCGAGTGCCTAGTGGGC | 430.6 | 12.1 | 2.2 | 5.5 | 3.5E-08 | 1.1E-06 | BT20/MB231 | MP |
| >hg38_21_+_8433172_8446622_RNA45SN1_45S_50ntFlanks@12306.12327.22\|CACGATCCTTCTGACCTTTTGG | >hg38_21_+_8433172_8446622_RNA45SN1_45S_50ntFlanks | 12306 | 12327 | 22 | CACGATCCTTCTGACCTTTTGG | 530.5 | -6.5 | 1.3 | -5.0 | 6.8E-07 | 1.1E-06 | MB231/MB468 | MITO |
| >hg38_21_+_8433172_8446622_RNA45SN1_45S_50ntFlanks@5414.5445.32\|CTACTACCGATTGGATGGTTTAGTGAGGCCCT | >hg38_21_+_8433172_8446622_RNA45SN1_45S_50ntFlanks | 5414 | 5445 | 32 | CTACTACCGATTGGATGGTTTAGTGAGGCCCT | 140.0 | -5.6 | 1.1 | -5.0 | 5.5E-07 | 1.1E-06 | BT20/MB468 | CYTO |
| >hg38_21_+_8433172_8446622_RNA45SN1_45S_50ntFlanks@4734.4767.34\|AACGAAAGTCGGAGGTTCGAAGACGATCAGATAC | >hg38_21_+_8433172_8446622_RNA45SN1_45S_50ntFlanks | 4734 | 4767 | 34 | AACGAAAGTCGGAGGTTCGAAGACGATCAGATAC | 49.5 | 8.9 | 1.8 | 5.0 | 5.7E-07 | 1.1E-06 | MB231/MB468 | MP |
| >MI0000077\|hsa-mir-21&WithFlank&17\|+\|59841260\|59841343@14.35.22(+1A)&[MIMAT0000076&hsa-miR-21-5p&offsets\|0\|0(+1A);m-1&17\|+\|59841273\|59841294&offsets\|0\|0(+1A)]\|TAGCTTATCAGACTGATGTTGAA | >MI0000077\|hsa-mir-21&WithFlank&17\|+\|59841260\|59841343 | 14 | 35 | 22(+1A) | TAGCTTATCAGACTGATGTTGAA | 8772.4 | 5.6 | 1.1 | 5.0 | 6.3E-07 | 1.1E-06 | MB231/MB468 | CYTO |
| >hg38_21_+_8433172_8446622_RNA45SN1_45S_50ntFlanks@10382.10424.43\|TTCCATGTGAACAGCAGTTGAACATGGGTCAGTCGGTCCTGAG | >hg38_21_+_8433172_8446622_RNA45SN1_45S_50ntFlanks | 10382 | 10424 | 43 | TTCCATGTGAACAGCAGTTGAACATGGGTCAGTCGGTCCTGAG | 315.5 | 4.3 | 0.9 | 5.0 | 6.8E-07 | 1.1E-06 | MB231/MB468 | MITO |
| >hg38_21_+_8433172_8446622_RNA45SN1_45S_50ntFlanks@10786.10825.40\|AACAGCCTCTGGCATGTTGGAACAATGTAGGTAAGGGAAG | >hg38_21_+_8433172_8446622_RNA45SN1_45S_50ntFlanks | 10786 | 10825 | 40 | AACAGCCTCTGGCATGTTGGAACAATGTAGGTAAGGGAAG | 533.6 | -7.2 | 1.4 | -5.0 | 6.5E-07 | 1.1E-06 | MB231/MB468 | NUC |
| >hg38_21_+_8433172_8446622_RNA45SN1_45S_50ntFlanks@8194.8216.23\|CCCGTGGACGGTGTGAGGCCGGT | >hg38_21_+_8433172_8446622_RNA45SN1_45S_50ntFlanks | 8194 | 8216 | 23 | CCCGTGGACGGTGTGAGGCCGGT | 369.7 | -7.1 | 1.4 | -5.0 | 5.7E-07 | 1.1E-06 | MB231/MB468 | MP |
| >hg38_21_+_8433172_8446622_RNA45SN1_45S_50ntFlanks@12580.12604.25\|GAACCGCAGGTTCAGACATTTGGTG | >hg38_21_+_8433172_8446622_RNA45SN1_45S_50ntFlanks | 12580 | 12604 | 25 | GAACCGCAGGTTCAGACATTTGGTG | 64.1 | -9.1 | 1.7 | -5.2 | 1.6E-07 | 1.1E-06 | BT20/MB468 | MP |
| >hg38_MT_+_1621_3279_RNR2_16S_50ntFlanks@721.740.20\|CTGCGTCAGATTAAAACACT | >hg38_MT_+_1621_3279_RNR2_16S_50ntFlanks | 721 | 740 | 20 | CTGCGTCAGATTAAAACACT | 59.0 | 9.6 | 1.8 | 5.2 | 1.6E-07 | 1.1E-06 | BT20/MB468 | MP |
| >MI0003513\|hsa-mir-455&WithFlank&9\|+\|114209428\|114209535@22.45.24(+1U)&[MIMAT0003150&hsa-miR-455-5p&offsets\|0\|+2(+1U);m-232&9\|+\|114209449\|114209470&offsets\|0\|+2(+1U)]\|TATGTGCCTTTGGACTACATCGTGT | >MI0003513\|hsa-mir-455&WithFlank&9\|+\|114209428\|114209535 | 22 | 45 | 24(+1U) | TATGTGCCTTTGGACTACATCGTGT | 48.0 | 8.8 | 1.8 | 5.0 | 5.7E-07 | 1.1E-06 | MB231/MB468 | MP |
| >am_MI0000809\|hsa-mir-151a&WithFlank&8\|-\|140732558\|140732659@52.73.22&[MIMAT0000757&hsa-miR-151a-3p&offsets\|-1\|0;m-30&8\|-\|140732587\|140732607&offsets\|-1\|0]\|ACTAGACTGAAGCTCCTTGAGG | >am_MI0000809\|hsa-mir-151a&WithFlank&8\|-\|140732558\|140732659 | 52 | 73 | 22 | ACTAGACTGAAGCTCCTTGAGG | 420.3 | 2.2 | 0.4 | 5.0 | 7.1E-07 | 1.1E-06 | BT20/MB468 | NUC |
| >hg38_21_+_8433172_8446622_RNA45SN1_45S_50ntFlanks@10348.10382.35\|AAACGAGAACTTTGAAGGCCGAAGTGGAGAAGGGT | >hg38_21_+_8433172_8446622_RNA45SN1_45S_50ntFlanks | 10348 | 10382 | 35 | AAACGAGAACTTTGAAGGCCGAAGTGGAGAAGGGT | 502.9 | -3.4 | 0.7 | -5.0 | 7.1E-07 | 1.1E-06 | BT20/MB468 | NUC |
| >hg38_21_+_8433172_8446622_RNA45SN1_45S_50ntFlanks@4724.4748.25\|ATTAATCAAGAACGAAAGTCGGAGG | >hg38_21_+_8433172_8446622_RNA45SN1_45S_50ntFlanks | 4724 | 4748 | 25 | ATTAATCAAGAACGAAAGTCGGAGG | 463.2 | -4.1 | 0.8 | -5.0 | 4.9E-07 | 1.1E-06 | BT20/MB468 | MITO |
| >hg38_21_+_8433172_8446622_RNA45SN1_45S_50ntFlanks@12997.13039.43\|CGCTGCGATCTATTGAAAGTCAGCCCTCGACACAAGGGTTTGT | >hg38_21_+_8433172_8446622_RNA45SN1_45S_50ntFlanks | 12997 | 13039 | 43 | CGCTGCGATCTATTGAAAGTCAGCCCTCGACACAAGGGTTTGT | 6561.3 | 5.8 | 1.1 | 5.2 | 1.6E-07 | 1.1E-06 | BT20/MB468 | MP |
| >hg38_21_+_8433172_8446622_RNA45SN1_45S_50ntFlanks@12942.12970.29\|TTCGTAGACGACCTGCTTCTGGGTCGGGG | >hg38_21_+_8433172_8446622_RNA45SN1_45S_50ntFlanks | 12942 | 12970 | 29 | TTCGTAGACGACCTGCTTCTGGGTCGGGG | 96.5 | -9.7 | 1.8 | -5.2 | 1.6E-07 | 1.1E-06 | BT20/MB468 | MP |
| >hg38_21_+_8433172_8446622_RNA45SN1_45S_50ntFlanks@6646.6680.35\|TCGTACGACTCTTAGCGGTGGATCACTCGGCTCGT | >hg38_21_+_8433172_8446622_RNA45SN1_45S_50ntFlanks | 6646 | 6680 | 35 | TCGTACGACTCTTAGCGGTGGATCACTCGGCTCGT | 22415.1 | 2.7 | 0.5 | 5.0 | 4.9E-07 | 1.1E-06 | BT20/MB468 | MITO |
| >hg38_21_+_8433172_8446622_RNA45SN1_45S_50ntFlanks@5237.5282.46\|ACTGACTGGCTCAGCGTGTGCCTACCCTACGCCGGCAGGCGCGGGT | >hg38_21_+_8433172_8446622_RNA45SN1_45S_50ntFlanks | 5237 | 5282 | 46 | ACTGACTGGCTCAGCGTGTGCCTACCCTACGCCGGCAGGCGCGGGT | 1068.2 | 10.3 | 2.0 | 5.2 | 1.6E-07 | 1.1E-06 | BT20/MB468 | MP |
| >hg38_21_+_8433172_8446622_RNA45SN1_45S_50ntFlanks@3776.3814.39\|CCGGTACAGTGAAACTGCGAATGGCTCATTAAATCAGTT | >hg38_21_+_8433172_8446622_RNA45SN1_45S_50ntFlanks | 3776 | 3814 | 39 | CCGGTACAGTGAAACTGCGAATGGCTCATTAAATCAGTT | 226.6 | -3.5 | 0.7 | -5.0 | 6.6E-07 | 1.1E-06 | MB231/MB468 | NUC |
| >hg38_21_+_8433172_8446622_RNA45SN1_45S_50ntFlanks@10846.10866.21\|CTTCGGGATAAGGATTGGCTC | >hg38_21_+_8433172_8446622_RNA45SN1_45S_50ntFlanks | 10846 | 10866 | 21 | CTTCGGGATAAGGATTGGCTC | 535.2 | -7.9 | 1.6 | -5.0 | 7.2E-07 | 1.1E-06 | BT20/MB468 | NUC |
| >hg38_21_+_8433172_8446622_RNA45SN1_45S_50ntFlanks@12936.12966.31\|AAACCATTCGTAGACGACCTGCTTCTGGGTC | >hg38_21_+_8433172_8446622_RNA45SN1_45S_50ntFlanks | 12936 | 12966 | 31 | AAACCATTCGTAGACGACCTGCTTCTGGGTC | 536.0 | -6.8 | 1.4 | -5.0 | 5.6E-07 | 1.1E-06 | BT20/MB468 | CYTO |
| >hg38_21_+_8433172_8446622_RNA45SN1_45S_50ntFlanks@485.511.27\|GATGTGGTGACGTCGTGCTCTCCCGGG | >hg38_21_+_8433172_8446622_RNA45SN1_45S_50ntFlanks | 485 | 511 | 27 | GATGTGGTGACGTCGTGCTCTCCCGGG | 78.7 | 10.0 | 1.9 | 5.2 | 1.6E-07 | 1.1E-06 | BT20/MB468 | MP |
| >MI0000342\|hsa-mir-200b&WithFlank&1\|+\|1167098\|1167204@63.85.23(+1C)&[MIMAT0000318&hsa-miR-200b-3p&offsets\|0\|+1(+1C);m-93&1\|+\|1167160\|1167182&offsets\|0\|0(+1C)]\|TAATACTGCCTGGTAATGATGACC | >MI0000342\|hsa-mir-200b&WithFlank&1\|+\|1167098\|1167204 | 63 | 85 | 23(+1C) | TAATACTGCCTGGTAATGATGACC | 617.1 | -7.6 | 1.5 | -5.0 | 7.2E-07 | 1.1E-06 | BT20/MB468 | NUC |
| >hg38_21_+_8433172_8446622_RNA45SN1_45S_50ntFlanks@10402.10419.18\|AACATGGGTCAGTCGGTC | >hg38_21_+_8433172_8446622_RNA45SN1_45S_50ntFlanks | 10402 | 10419 | 18 | AACATGGGTCAGTCGGTC | 1184.6 | -4.9 | 1.0 | -5.0 | 6.4E-07 | 1.1E-06 | MB231/MB468 | CYTO |
| >hg38_1_-_228634819_228635039_RNA5S12_5S_50ntFlanks@130.171.42\|TGGATGGGAGACCGCCTGGGAATACCGGGTGCTGTAGGCTTT | >hg38_1_-_228634819_228635039_RNA5S12_5S_50ntFlanks | 130 | 171 | 42 | TGGATGGGAGACCGCCTGGGAATACCGGGTGCTGTAGGCTTT | 841.8 | 7.2 | 1.4 | 5.2 | 1.6E-07 | 1.1E-06 | BT20/MB468 | MP |
| >MI0000273\|hsa-mir-183&WithFlank&7\|-\|129774899\|129775020@34.53.20(+1C)&[MIMAT0000261&hsa-miR-183-5p&offsets\|+1\|-1(+1C);m-41&7\|-\|129774966\|129774987&offsets\|0\|-2(+1C)]\|ATGGCACTGGTAGAATTCACC | >MI0000273\|hsa-mir-183&WithFlank&7\|-\|129774899\|129775020 | 34 | 53 | 20(+1C) | ATGGCACTGGTAGAATTCACC | 634.3 | 4.0 | 0.7 | 5.5 | 4.5E-08 | 1.1E-06 | BT20/MB231 | CYTO |
| >MI0033423\|hsa-mir-10399&WithFlank&7\|-\|139044094\|139044163@43.64.22&[MIMAT0041630&hsa-miR-10399-3p&offsets\|-1\|0;TJU_CMC_MD2.ID03061.3p-miR&offsets\|+3\|+3;m-2657&7\|-\|139044103\|139044124&offsets\|+3\|+3]\|TCTCTCGGACAAGCTGTAGGTC | >MI0033423\|hsa-mir-10399&WithFlank&7\|-\|139044094\|139044163 | 43 | 64 | 22 | TCTCTCGGACAAGCTGTAGGTC | 47.2 | 8.8 | 1.8 | 5.0 | 5.7E-07 | 1.1E-06 | MB231/MB468 | MP |
| >hg38_21_+_8433172_8446622_RNA45SN1_45S_50ntFlanks@6651.6674.24\|CGACTCTTAGCGGTGGATCACTCG | >hg38_21_+_8433172_8446622_RNA45SN1_45S_50ntFlanks | 6651 | 6674 | 24 | CGACTCTTAGCGGTGGATCACTCG | 2231.1 | 3.7 | 0.7 | 5.0 | 6.4E-07 | 1.1E-06 | MB231/MB468 | CYTO |
| >MI0000084\|hsa-mir-26b&WithFlank&2\|+\|218402640\|218402728@18.39.22&[MIMAT0000083&hsa-miR-26b-5p&offsets\|0\|+1;m-53&2\|+\|218402657\|218402678&offsets\|0\|0]\|TTCAAGTAATTCAGGATAGGTT | >MI0000084\|hsa-mir-26b&WithFlank&2\|+\|218402640\|218402728 | 18 | 39 | 22 | TTCAAGTAATTCAGGATAGGTT | 6410.8 | 4.8 | 1.0 | 5.0 | 6.9E-07 | 1.1E-06 | MB231/MB468 | MITO |
| >hg38_21_+_8433172_8446622_RNA45SN1_45S_50ntFlanks@4892.4911.20\|AAACTTAAAGGAATTGACGG | >hg38_21_+_8433172_8446622_RNA45SN1_45S_50ntFlanks | 4892 | 4911 | 20 | AAACTTAAAGGAATTGACGG | 367.0 | -7.7 | 1.6 | -5.0 | 6.9E-07 | 1.1E-06 | MB231/MB468 | MITO |
| >hg38_21_+_8433172_8446622_RNA45SN1_45S_50ntFlanks@9565.9588.24\|TGACGTGCAAATCGGTCGTCCGAC | >hg38_21_+_8433172_8446622_RNA45SN1_45S_50ntFlanks | 9565 | 9588 | 24 | TGACGTGCAAATCGGTCGTCCGAC | 164.9 | -5.5 | 1.1 | -5.0 | 5.7E-07 | 1.1E-06 | MB231/MB468 | MP |
| >hg38_21_+_8433172_8446622_RNA45SN1_45S_50ntFlanks@5492.5525.34\|AGAAGACGGTCGAACTTGACTATCTAGAGGAAGT | >hg38_21_+_8433172_8446622_RNA45SN1_45S_50ntFlanks | 5492 | 5525 | 34 | AGAAGACGGTCGAACTTGACTATCTAGAGGAAGT | 492.9 | -3.8 | 0.8 | -5.0 | 5.7E-07 | 1.1E-06 | MB231/MB468 | MP |
| >hg38_21_+_8433172_8446622_RNA45SN1_45S_50ntFlanks@9934.9960.27\|AGACAGCAGGACGGTGGCCATGGAAGT | >hg38_21_+_8433172_8446622_RNA45SN1_45S_50ntFlanks | 9934 | 9960 | 27 | AGACAGCAGGACGGTGGCCATGGAAGT | 59.0 | 9.6 | 1.8 | 5.2 | 1.7E-07 | 1.1E-06 | BT20/MB468 | MP |
| >hg38_21_+_8433172_8446622_RNA45SN1_45S_50ntFlanks@5188.5218.31\|ATAACAGGTCTGTGATGCCCTTAGATGTCCG | >hg38_21_+_8433172_8446622_RNA45SN1_45S_50ntFlanks | 5188 | 5218 | 31 | ATAACAGGTCTGTGATGCCCTTAGATGTCCG | 219.2 | -9.9 | 2.0 | -5.0 | 5.0E-07 | 1.1E-06 | BT20/MB468 | MITO |
| >hg38_21_+_8433172_8446622_RNA45SN1_45S_50ntFlanks@9738.9760.23\|GAAACGATCTCAACCTATTCTCA | >hg38_21_+_8433172_8446622_RNA45SN1_45S_50ntFlanks | 9738 | 9760 | 23 | GAAACGATCTCAACCTATTCTCA | 382.5 | -7.0 | 1.4 | -5.0 | 5.0E-07 | 1.1E-06 | BT20/MB468 | MITO |
| >hg38_21_+_8433172_8446622_RNA45SN1_45S_50ntFlanks@11729.11751.23\|TAACTATGACTCTCTTAAGGTAG | >hg38_21_+_8433172_8446622_RNA45SN1_45S_50ntFlanks | 11729 | 11751 | 23 | TAACTATGACTCTCTTAAGGTAG | 38105.0 | -7.5 | 1.5 | -5.0 | 5.6E-07 | 1.1E-06 | BT20/MB468 | CYTO |
| >hg38_21_+_8433172_8446622_RNA45SN1_45S_50ntFlanks@12892.12927.36\|CCCTCGCCCGTCACGCACCGCACGTTCGTGGGGAAC | >hg38_21_+_8433172_8446622_RNA45SN1_45S_50ntFlanks | 12892 | 12927 | 36 | CCCTCGCCCGTCACGCACCGCACGTTCGTGGGGAAC | 66.2 | -9.1 | 1.7 | -5.2 | 1.7E-07 | 1.1E-06 | BT20/MB468 | MP |
| >hg38_21_+_8433172_8446622_RNA45SN1_45S_50ntFlanks@8369.8387.19\|AAGAGAGAGTTCAAGAGGG | >hg38_21_+_8433172_8446622_RNA45SN1_45S_50ntFlanks | 8369 | 8387 | 19 | AAGAGAGAGTTCAAGAGGG | 61.7 | -9.0 | 1.7 | -5.2 | 1.7E-07 | 1.1E-06 | BT20/MB468 | MP |
| >hg38_21_+_8433172_8446622_RNA45SN1_45S_50ntFlanks@9143.9162.20\|GGGCGGACTGTCCCCAGTGC | >hg38_21_+_8433172_8446622_RNA45SN1_45S_50ntFlanks | 9143 | 9162 | 20 | GGGCGGACTGTCCCCAGTGC | 74.3 | -9.3 | 1.8 | -5.2 | 1.7E-07 | 1.1E-06 | BT20/MB468 | MP |

| | | | | | | | | | | | | | |
|---|---|---|---|---|---|---|---|---|---|---|---|---|---|
| >hg38_21_+_8433172_8446622_RNA45SN1_45S_50ntFlanks@10729.10755.27\|AGAGGGTGTAAATCTCGCGCCGGGCCG | >hg38_21_+_8433172_8446622_RNA45SN1_45S_50ntFlanks | 10729 | 10755 | 27 | AGAGGGTGTAAATCTCGCGCCGGGCCG | 323.0 | -5.8 | 1.2 | -5.0 | 6.5E-07 | 1.1E-06 | MB231/MB468 | CYTO |
| >hg38_21_+_8433172_8446622_RNA45SN1_45S_50ntFlanks@10473.10495.23\|CTCGGCCGATCGAAAGGGAGTCG | >hg38_21_+_8433172_8446622_RNA45SN1_45S_50ntFlanks | 10473 | 10495 | 23 | CTCGGCCGATCGAAAGGGAGTCG | 194.3 | -7.1 | 1.4 | -5.0 | 5.8E-07 | 1.1E-06 | MB231/MB468 | MP |
| >hg38_21_+_8433172_8446622_RNA45SN1_45S_50ntFlanks@6697.6720.24\|CAGCTAGCTGCGAGAATTAATGTG | >hg38_21_+_8433172_8446622_RNA45SN1_45S_50ntFlanks | 6697 | 6720 | 24 | CAGCTAGCTGCGAGAATTAATGTG | 1719.1 | -4.0 | 0.8 | -5.0 | 5.8E-07 | 1.1E-06 | MB231/MB468 | MP |
| >hg38_21_+_8433172_8446622_RNA45SN1_45S_50ntFlanks@11562.11601.40\|CGCCTAGCAGCCGACTTAGAACTGGTGCGGACCAGGGGAA | >hg38_21_+_8433172_8446622_RNA45SN1_45S_50ntFlanks | 11562 | 11601 | 40 | CGCCTAGCAGCCGACTTAGAACTGGTGCGGACCAGGGGAA | 855.3 | -8.3 | 1.7 | -5.0 | 7.3E-07 | 1.1E-06 | BT20/MB468 | NUC |
| >hg38_21_+_8433172_8446622_RNA45SN1_45S_50ntFlanks@10869.10889.21\|AGGGCTGGGTCGGTCGGGCTG | >hg38_21_+_8433172_8446622_RNA45SN1_45S_50ntFlanks | 10869 | 10889 | 21 | AGGGCTGGGTCGGTCGGGCTG | 3098.6 | -9.3 | 1.9 | -5.0 | 7.3E-07 | 1.1E-06 | BT20/MB468 | NUC |
| >hg38_21_+_8433172_8446622_RNA45SN1_45S_50ntFlanks@3802.3829.28\|CATTAAATCAGTTATGGTTCCTTTGGTC | >hg38_21_+_8433172_8446622_RNA45SN1_45S_50ntFlanks | 3802 | 3829 | 28 | CATTAAATCAGTTATGGTTCCTTTGGTC | 274.3 | -3.7 | 0.7 | -5.0 | 5.7E-07 | 1.1E-06 | BT20/MB468 | CYTO |
| >hg38_21_+_8433172_8446622_RNA45SN1_45S_50ntFlanks@12006.12032.27\|GGCCGCCGGTGAAATACCACTACTCTG | >hg38_21_+_8433172_8446622_RNA45SN1_45S_50ntFlanks | 12006 | 12032 | 27 | GGCCGCCGGTGAAATACCACTACTCTG | 1518.1 | -7.0 | 1.4 | -5.0 | 7.3E-07 | 1.1E-06 | BT20/MB468 | NUC |
| >hg38_21_+_8433172_8446622_RNA45SN1_45S_50ntFlanks@5078.5110.33\|CGATAACGAACGAGACTCTGGCATGCTAACTAG | >hg38_21_+_8433172_8446622_RNA45SN1_45S_50ntFlanks | 5078 | 5110 | 33 | CGATAACGAACGAGACTCTGGCATGCTAACTAG | 399.2 | -8.2 | 1.6 | -5.0 | 5.8E-07 | 1.1E-06 | MB231/MB468 | MP |
| >hg38_21_+_8433172_8446622_RNA45SN1_45S_50ntFlanks@11552.11572.21\|CCTCGGCCGGCGCCTAGCAGC | >hg38_21_+_8433172_8446622_RNA45SN1_45S_50ntFlanks | 11552 | 11572 | 21 | CCTCGGCCGGCGCCTAGCAGC | 423.2 | -6.1 | 1.2 | -5.0 | 7.4E-07 | 1.1E-06 | BT20/MB468 | NUC |
| >hg38_21_+_8433172_8446622_RNA45SN1_45S_50ntFlanks@10466.10497.32\|CGTTGCCCTCGGCCGATCGAAAGGGAGTCGGG | >hg38_21_+_8433172_8446622_RNA45SN1_45S_50ntFlanks | 10466 | 10497 | 32 | CGTTGCCCTCGGCCGATCGAAAGGGAGTCGGG | 925.7 | -5.5 | 1.1 | -5.0 | 6.6E-07 | 1.1E-06 | MB231/MB468 | CYTO |
| >MI0000811\|hsa-mir-148b&WithFlank&12\|+\|54337210\|54337320@69.89.21(+1G)&[MIMAT0000759&hsa-miR-148b-3p&offsets\|0\|-1(+1G);m-77&12\|+\|54337278\|54337299&offsets\|0\|-1(+1G)]\|TCAGTGCATCACAGAACTTTGG | >MI0000811\|hsa-mir-148b&WithFlank&12\|+\|54337210\|54337320 | 69 | 89 | 21(+1G) | TCAGTGCATCACAGAACTTTGG | 288.5 | 6.1 | 1.2 | 5.0 | 5.7E-07 | 1.1E-06 | BT20/MB468 | CYTO |
| >hg38_21_+_8433172_8446622_RNA45SN1_45S_50ntFlanks@6661.6698.38\|CGGTGGATCACTCGGCTCGTGCGTCGATGAAGAACGCA | >hg38_21_+_8433172_8446622_RNA45SN1_45S_50ntFlanks | 6661 | 6698 | 38 | CGGTGGATCACTCGGCTCGTGCGTCGATGAAGAACGCA | 1324.0 | -3.5 | 0.7 | -5.0 | 7.1E-07 | 1.1E-06 | MB231/MB468 | MITO |
| >MI0000266\|hsa-mir-10a&WithFlank&17\|-\|48579832\|48579953@28.50.23(+1C)&[MIMAT0000253&hsa-miR-10a-5p&offsets\|0\|0(+1C);m-18&17\|-\|48579905\|48579926&offsets\|0\|+1(+1C)]\|TACCCTGTAGATCCGAATTTGTGC | >MI0000266\|hsa-mir-10a&WithFlank&17\|-\|48579832\|48579953 | 28 | 50 | 23(+1C) | TACCCTGTAGATCCGAATTTGTGC | 46.3 | 8.8 | 1.8 | 5.0 | 5.9E-07 | 1.1E-06 | MB231/MB468 | MP |
| >hg38_21_+_8433172_8446622_RNA45SN1_45S_50ntFlanks@5444.5467.24\|CTCGGATCGGCCCCGCCGGGGTCG | >hg38_21_+_8433172_8446622_RNA45SN1_45S_50ntFlanks | 5444 | 5467 | 24 | CTCGGATCGGCCCCGCCGGGGTCG | 1502.6 | -8.5 | 1.7 | -5.0 | 7.4E-07 | 1.1E-06 | BT20/MB468 | NUC |
| >hg38_21_+_8433172_8446622_RNA45SN1_45S_50ntFlanks@12161.12179.19\|TGACTGGGGCGGTACACCT | >hg38_21_+_8433172_8446622_RNA45SN1_45S_50ntFlanks | 12161 | 12179 | 19 | TGACTGGGGCGGTACACCT | 463.6 | -4.2 | 0.8 | -5.0 | 5.1E-07 | 1.1E-06 | BT20/MB468 | MITO |
| >trna10_LysCTT_16_+_3241501_3241573@14.31.18,trna13_LysCTT_14_-_58706613_58706685@14.31.18,trna2_LysCTT_15_+_79152904_79152976@14.31.18,trna4_LysCTT_18_-_43669271_43669343@14.31.18\|AGTCGGTAGAGCATGGGA | >trna10_LysCTT_16_+_3241501_3241573 | 14 | 31 | 18 | AGTCGGTAGAGCATGGGA | 616.2 | -3.8 | 0.8 | -5.0 | 5.9E-07 | 1.1E-06 | MB231/MB468 | MP |
| >MI0000542\|hsa-mir-320a&WithFlank&8\|-\|22244960\|22245043@48.68.21(+2U)&[MIMAT0000510&hsa-miR-320a-3p&offsets\|0\|-1(+2U);m-60&8\|-\|22244975\|22244996&offsets\|0\|-1(+2U)]\|AAAAGCTGGGTTGAGAGGGCGTT | >MI0000542\|hsa-mir-320a&WithFlank&8\|-\|22244960\|22245043 | 48 | 68 | 21(+2U) | AAAAGCTGGGTTGAGAGGGCGTT | 10989.9 | 4.3 | 0.8 | 5.5 | 4.7E-08 | 1.1E-06 | BT20/MB231 | CYTO |
| >hg38_21_+_8433172_8446622_RNA45SN1_45S_50ntFlanks@10401.10431.31\|GAACATGGGTCAGTCGGTCCTGAGAGATGGG | >hg38_21_+_8433172_8446622_RNA45SN1_45S_50ntFlanks | 10401 | 10431 | 31 | GAACATGGGTCAGTCGGTCCTGAGAGATGGG | 232.8 | -6.2 | 1.2 | -5.0 | 5.9E-07 | 1.1E-06 | MB231/MB468 | MP |
| >trna10_AspGTC_12_-_125424193_125424264@52.70.19,trna12_AspGTC_12_-_125411891_125411962@52.70.19,trna144_AspGTC_6_-_27551236_27551307@52.70.19,trna38_AspGTC_17_-_8125556_8125627@52.70.19,trna45_AspGTC_6_+_27447453_27447524@52.70.19,trna48_AspGTC_6_+_27471523_27471594@52.70.19,trna4_AspGTC_12_+_96429799_96429870@52.70.19,trna69_AspGTC_1_-_161440205_161440276@52.70.19,trna72_AspGTC_1_-_161432824_161432895@52.70.19,trna75_AspGTC_1_-_161425414_161425485@52.70.19,trna78_AspGTC_1_-_161418033_161418104@52.70.19,trna81_AspGTC_1_-_161410615_161410686@52.70.19\|GTTCGATTCCCCGACGGGG | >trna10_AspGTC_12_-_125424193_125424264 | 52 | 70 | 19 | GTTCGATTCCCCGACGGGG | 85.9 | -9.5 | 1.8 | -5.2 | 1.7E-07 | 1.1E-06 | BT20/MB468 | MP |
| >hg38_21_+_8433172_8446622_RNA45SN1_45S_50ntFlanks@10572.10589.18\|AGAAGCCGGCGGGAGCCC | >hg38_21_+_8433172_8446622_RNA45SN1_45S_50ntFlanks | 10572 | 10589 | 18 | AGAAGCCGGCGGGAGCCC | 256.3 | -6.0 | 1.2 | -5.0 | 5.8E-07 | 1.1E-06 | BT20/MB468 | CYTO |
| >hg38_21_+_8433172_8446622_RNA45SN1_45S_50ntFlanks@10432.10456.25\|CGAGCGCCGTTCCGAAGGGACGGGC | >hg38_21_+_8433172_8446622_RNA45SN1_45S_50ntFlanks | 10432 | 10456 | 25 | CGAGCGCCGTTCCGAAGGGACGGGC | 61.0 | -9.0 | 1.7 | -5.2 | 1.7E-07 | 1.1E-06 | BT20/MB468 | MP |
| >hg38_21_+_8433172_8446622_RNA45SN1_45S_50ntFlanks@11321.11340.20\|CACGTCTCGTCGCGCGCGCG | >hg38_21_+_8433172_8446622_RNA45SN1_45S_50ntFlanks | 11321 | 11340 | 20 | CACGTCTCGTCGCGCGCGCG | 399.1 | -6.8 | 1.4 | -5.0 | 6.8E-07 | 1.1E-06 | MB231/MB468 | NUC |
| >hg38_21_+_8433172_8446622_RNA45SN1_45S_50ntFlanks@10427.10453.27\|ATGGGCGAGCGCCGTTCCGAAGGGACG | >hg38_21_+_8433172_8446622_RNA45SN1_45S_50ntFlanks | 10427 | 10453 | 27 | ATGGGCGAGCGCCGTTCCGAAGGGACG | 396.2 | -7.2 | 1.4 | -5.0 | 5.9E-07 | 1.1E-06 | MB231/MB468 | MP |
| >hg38_21_+_8433172_8446622_RNA45SN1_45S_50ntFlanks@12934.12957.24\|CTAAACCATTCGTAGACGACCTGC | >hg38_21_+_8433172_8446622_RNA45SN1_45S_50ntFlanks | 12934 | 12957 | 24 | CTAAACCATTCGTAGACGACCTGC | 234.6 | -5.2 | 1.0 | -5.0 | 7.1E-07 | 1.1E-06 | MB231/MB468 | MITO |
| >trna91_GlyCCC_1_-_149680210_149680280@1.31.31,trna99_ValCAC_1_-_149294666_149294736@1.31.31\|GCACTGGTGGTTCAGTGGTAGAATTCTCGCC | >trna91_GlyCCC_1_-_149680210_149680280 | 1 | 31 | 31 | GCACTGGTGGTTCAGTGGTAGAATTCTCGCC | 53.3 | 9.0 | 1.8 | 5.0 | 5.9E-07 | 1.1E-06 | MB231/MB468 | MP |
| >hg38_21_+_8433172_8446622_RNA45SN1_45S_50ntFlanks@10236.10259.24\|GACGCTACGCCGCGACGAGTAGGA | >hg38_21_+_8433172_8446622_RNA45SN1_45S_50ntFlanks | 10236 | 10259 | 24 | GACGCTACGCCGCGACGAGTAGGA | 288.4 | 11.5 | 2.1 | 5.5 | 3.7E-08 | 1.1E-06 | BT20/MB231 | MP |
| >hg38_MT_+_1621_3279_RNR2_16S_50ntFlanks@51.92.42\|GCTAAACCTAGCCCCAAACCCACTCCACCTTACTACCAGACA | >hg38_MT_+_1621_3279_RNR2_16S_50ntFlanks | 51 | 92 | 42 | GCTAAACCTAGCCCCAAACCCACTCCACCTTACTACCAGACA | 855.9 | 5.9 | 1.2 | 5.0 | 6.8E-07 | 1.1E-06 | MB231/MB468 | NUC |
| >hg38_21_+_8433172_8446622_RNA45SN1_45S_50ntFlanks@12255.12272.18\|CTCGCTTGATCTTGATTT | >hg38_21_+_8433172_8446622_RNA45SN1_45S_50ntFlanks | 12255 | 12272 | 18 | CTCGCTTGATCTTGATTT | 399.2 | -5.0 | 1.0 | -5.0 | 7.1E-07 | 1.1E-06 | MB231/MB468 | MITO |
| >hg38_21_+_8433172_8446622_RNA45SN1_45S_50ntFlanks@8349.8375.27\|GAAAGTTGAAAAGAACTTTGAAGAGAG | >hg38_21_+_8433172_8446622_RNA45SN1_45S_50ntFlanks | 8349 | 8375 | 27 | GAAAGTTGAAAAGAACTTTGAAGAGAG | 3059.3 | 10.0 | 1.9 | 5.2 | 1.7E-07 | 1.1E-06 | BT20/MB468 | MP |
| >trna129_MetCAT_6_-_27870271_27870342@31.50.20,trna142_MetCAT_6_-_27560600_27560671@31.50.20,trna150_MetCAT_6_-_27300764_27300835@31.50.20,trna169_MetCAT_6_-_26330529_26330600@31.50.20,trna171_MetCAT_6_-_26313352_26313423@31.50.20,trna20_MetCAT_17_-_80452597_80452668@31.50.20,trna2_MetCAT_6_+_26286754_26286825@31.50.20,trna32_MetCAT_1_+_153643726_153643797@31.50.20,trna61_MetCAT_6_+_27745664_27745735@31.50.20\|CCCATAACCCAGAGGTCGAT | >trna129_MetCAT_6_-_27870271_27870342 | 31 | 50 | 20 | CCCATAACCCAGAGGTCGAT | 62.1 | -9.0 | 1.7 | -5.2 | 1.7E-07 | 1.1E-06 | BT20/MB468 | MP |
| >hg38_21_+_8433172_8446622_RNA45SN1_45S_50ntFlanks@12124.12150.27\|CGCGACCCGCTCCGGGGACAGTGCCAG | >hg38_21_+_8433172_8446622_RNA45SN1_45S_50ntFlanks | 12124 | 12150 | 27 | CGCGACCCGCTCCGGGGACAGTGCCAG | 70.8 | -9.2 | 1.8 | -5.2 | 1.7E-07 | 1.1E-06 | BT20/MB468 | MP |
| >hg38_21_+_8433172_8446622_RNA45SN1_45S_50ntFlanks@4760.4782.23\|TCAGATACCGTCGTAGTTCCGAC | >hg38_21_+_8433172_8446622_RNA45SN1_45S_50ntFlanks | 4760 | 4782 | 23 | TCAGATACCGTCGTAGTTCCGAC | 1140.6 | -7.8 | 1.5 | -5.2 | 1.7E-07 | 1.1E-06 | BT20/MB468 | MP |
| >hg38_21_+_8433172_8446622_RNA45SN1_45S_50ntFlanks@8196.8224.29\|CGTGGACGGTGTGAGGCCGGTAGCGGCCC | >hg38_21_+_8433172_8446622_RNA45SN1_45S_50ntFlanks | 8196 | 8224 | 29 | CGTGGACGGTGTGAGGCCGGTAGCGGCCC | 551.6 | -5.7 | 1.1 | -5.0 | 7.2E-07 | 1.1E-06 | MB231/MB468 | MITO |
| >hg38_21_+_8433172_8446622_RNA45SN1_45S_50ntFlanks@10462.10497.36\|CCTCCGTTGCCCTCGGCCGATCGAAAGGGAGTCGGG | >hg38_21_+_8433172_8446622_RNA45SN1_45S_50ntFlanks | 10462 | 10497 | 36 | CCTCCGTTGCCCTCGGCCGATCGAAAGGGAGTCGGG | 71.5 | -9.2 | 1.8 | -5.2 | 1.7E-07 | 1.1E-06 | BT20/MB468 | MP |
| >hg38_21_+_8433172_8446622_RNA45SN1_45S_50ntFlanks@10350.10378.29\|ACGAGAACTTTGAAGGCCGAAGTGGAGAA | >hg38_21_+_8433172_8446622_RNA45SN1_45S_50ntFlanks | 10350 | 10378 | 29 | ACGAGAACTTTGAAGGCCGAAGTGGAGAA | 321.7 | -5.3 | 1.1 | -5.0 | 7.2E-07 | 1.1E-06 | MB231/MB468 | MITO |
| >hg38_21_+_8433172_8446622_RNA45SN1_45S_50ntFlanks@10351.10389.39\|CGAGAACTTTGAAGGCCGAAGTGGAGAAGGGTTCCATGT | >hg38_21_+_8433172_8446622_RNA45SN1_45S_50ntFlanks | 10351 | 10389 | 39 | CGAGAACTTTGAAGGCCGAAGTGGAGAAGGGTTCCATGT | 351.3 | -5.9 | 1.2 | -5.0 | 6.9E-07 | 1.1E-06 | MB231/MB468 | NUC |
| >hg38_21_+_8433172_8446622_RNA45SN1_45S_50ntFlanks@3851.3871.21\|ATAACTGTGGTAATTCTAGAG | >hg38_21_+_8433172_8446622_RNA45SN1_45S_50ntFlanks | 3851 | 3871 | 21 | ATAACTGTGGTAATTCTAGAG | 73.7 | -9.3 | 1.8 | -5.2 | 1.7E-07 | 1.1E-06 | BT20/MB468 | MP |
| >hg38_21_+_8433172_8446622_RNA45SN1_45S_50ntFlanks@10789.10817.29\|AGCCTCTGGCATGTTGGAACAATGTAGGT | >hg38_21_+_8433172_8446622_RNA45SN1_45S_50ntFlanks | 10789 | 10817 | 29 | AGCCTCTGGCATGTTGGAACAATGTAGGT | 104.6 | -5.1 | 1.0 | -5.0 | 6.0E-07 | 1.1E-06 | MB231/MB468 | MP |
| >hg38_21_+_8433172_8446622_RNA45SN1_45S_50ntFlanks@11209.11232.24\|CGGCGACTCTGGACGCGAGCCGGG | >hg38_21_+_8433172_8446622_RNA45SN1_45S_50ntFlanks | 11209 | 11232 | 24 | CGGCGACTCTGGACGCGAGCCGGG | 35529.3 | -8.6 | 1.7 | -4.9 | 7.5E-07 | 1.1E-06 | BT20/MB468 | NUC |
| >hg38_21_+_8433172_8446622_RNA45SN1_45S_50ntFlanks@11783.11803.21\|CATGAATGGATGAACGAGATT | >hg38_21_+_8433172_8446622_RNA45SN1_45S_50ntFlanks | 11783 | 11803 | 21 | CATGAATGGATGAACGAGATT | 3476.6 | -4.1 | 0.8 | -5.0 | 5.8E-07 | 1.1E-06 | BT20/MB468 | CYTO |
| >hg38_21_+_8433172_8446622_RNA45SN1_45S_50ntFlanks@11582.11603.22\|ACTGGTGCGGACCAGGGGAATC | >hg38_21_+_8433172_8446622_RNA45SN1_45S_50ntFlanks | 11582 | 11603 | 22 | ACTGGTGCGGACCAGGGGAATC | 3278.1 | -4.4 | 0.9 | -5.0 | 6.0E-07 | 1.1E-06 | MB231/MB468 | MP |
| >hg38_21_+_8433172_8446622_RNA45SN1_45S_50ntFlanks@12456.12499.44\|CCAAGCGTTGGATTGTTCACCCACTAATAGGGAACGTGAGCTGG | >hg38_21_+_8433172_8446622_RNA45SN1_45S_50ntFlanks | 12456 | 12499 | 44 | CCAAGCGTTGGATTGTTCACCCACTAATAGGGAACGTGAGCTGG | 747.6 | -4.4 | 0.9 | -5.0 | 6.7E-07 | 1.1E-06 | MB231/MB468 | CYTO |

<table>
<tr><th></th><th></th><th></th><th></th><th></th><th></th><th></th><th></th><th></th><th></th><th></th><th></th><th></th><th></th></tr>
<tr><td>>hg38_21_+_8433172_8446622_RNA45SN1_45S_50ntFlanks@9556.9582.27|TAGCGGTCCTGACGTGCAAATCGGTCG</td><td>>hg38_21_+_8433172_8446622_RNA45SN1_45S_50ntFlanks</td><td>9556</td><td>9582</td><td>27</td><td>TAGCGGTCCTGACGTGCAAATCGGTCG</td><td>587.9</td><td>-5.0</td><td>1.0</td><td>-5.0</td><td>6.7E-07</td><td>1.1E-06</td><td>MB231/MB468</td><td>CYTO</td></tr>
<tr><td>>hg38_21_+_8433172_8446622_RNA45SN1_45S_50ntFlanks@8356.8386.31|GAAAAGAACTTTGAAGAGAGAGTTCAAGAGG</td><td>>hg38_21_+_8433172_8446622_RNA45SN1_45S_50ntFlanks</td><td>8356</td><td>8386</td><td>31</td><td>GAAAAGAACTTTGAAGAGAGAGTTCAAGAGG</td><td>238.0</td><td>-4.3</td><td>0.9</td><td>-5.0</td><td>6.0E-07</td><td>1.1E-06</td><td>MB231/MB468</td><td>MP</td></tr>
<tr><td>>hg38_21_+_8433172_8446622_RNA45SN1_45S_50ntFlanks@12991.13008.18|CTCCCTCGCTGCGATCTA</td><td>>hg38_21_+_8433172_8446622_RNA45SN1_45S_50ntFlanks</td><td>12991</td><td>13008</td><td>18</td><td>CTCCCTCGCTGCGATCTA</td><td>404.8</td><td>-5.3</td><td>1.1</td><td>-5.0</td><td>6.7E-07</td><td>1.1E-06</td><td>MB231/MB468</td><td>CYTO</td></tr>
<tr><td>>hg38_21_+_8433172_8446622_RNA45SN1_45S_50ntFlanks@8184.8211.28|CGAGGCCCAGCCCGTGGACGGTGTGAGG</td><td>>hg38_21_+_8433172_8446622_RNA45SN1_45S_50ntFlanks</td><td>8184</td><td>8211</td><td>28</td><td>CGAGGCCCAGCCCGTGGACGGTGTGAGG</td><td>405.5</td><td>-7.2</td><td>1.5</td><td>-5.0</td><td>6.0E-07</td><td>1.1E-06</td><td>MB231/MB468</td><td>MP</td></tr>
<tr><td>>trna111_HisGTG_1_-_147774845_147774916@-1A.34.35,trna118_HisGTG_1_-_145396881_145396952@-1A.34.35,trna16_HisGTG_1_+_146544773_146544844@-1A.34.35,trna1_HisGTG_15_+_45493349_45493420@-1A.34.35,trna21_HisGTG_1_+_147753471_147753542@-1A.34.35,trna33_HisGTG_6_+_27125906_27125977@-1A.34.35,trna7_HisGTG_9_-_14433938_14434009@-1A.34.35,trna8_HisGTG_15_-_45492611_45492682@-1A.34.35,trna9_HisGTG_15_-_45490804_45490875@-1A.34.35|AGCCGTGATCGTATAGTGGTTAGTACTCTGCGTTG</td><td>>trna111_HisGTG_1_-_147774845_147774916</td><td>-1A</td><td>34</td><td>35</td><td>AGCCGTGATCGTATAGTGGTTAGTACTCTGCGTTG</td><td>50.9</td><td>8.9</td><td>1.8</td><td>5.0</td><td>6.0E-07</td><td>1.1E-06</td><td>MB231/MB468</td><td>MP</td></tr>
<tr><td>>MI0000102|hsa-mir-100&WithFlank&11|-|122152223|122152314@20.40.21&[MIMAT0000098&hsa-miR-100-5p&offsets|+1|0;m-27&11|-|122152276|122152296&offsets|+1|+1]|ACCCGTAGATCCGAACTTGTG</td><td>>MI0000102|hsa-mir-100&WithFlank&11|-|122152223|122152314</td><td>20</td><td>40</td><td>21</td><td>ACCCGTAGATCCGAACTTGTG</td><td>1869.6</td><td>-7.7</td><td>1.4</td><td>-5.5</td><td>3.8E-08</td><td>1.1E-06</td><td>BT20/MB231</td><td>MP</td></tr>
<tr><td>>MI0000298|hsa-mir-221&WithFlank&X|-|45746151|45746272@71.92.22(+1A)&[MIMAT0000278&hsa-miR-221-3p&offsets|0|-1(+1A);m-91&X|-|45746181|45746202&offsets|0|0(+1A)]|AGCTACATTGTCTGCTGGGTTTA</td><td>>MI0000298|hsa-mir-221&WithFlank&X|-|45746151|45746272</td><td>71</td><td>92</td><td>22(+1A)</td><td>AGCTACATTGTCTGCTGGGTTTA</td><td>1990.2</td><td>4.2</td><td>0.8</td><td>5.0</td><td>6.0E-07</td><td>1.1E-06</td><td>MB231/MB468</td><td>MP</td></tr>
<tr><td>>hg38_21_+_8433172_8446622_RNA45SN1_45S_50ntFlanks@8117.8140.24|CGGGACATGTGGCGTACGGAAGAC</td><td>>hg38_21_+_8433172_8446622_RNA45SN1_45S_50ntFlanks</td><td>8117</td><td>8140</td><td>24</td><td>CGGGACATGTGGCGTACGGAAGAC</td><td>288.6</td><td>11.5</td><td>2.1</td><td>5.5</td><td>3.8E-08</td><td>1.1E-06</td><td>BT20/MB231</td><td>MP</td></tr>
<tr><td>>hg38_21_+_8433172_8446622_RNA45SN1_45S_50ntFlanks@6647.6668.22|CGTACGACTCTTAGCGGTGGAT</td><td>>hg38_21_+_8433172_8446622_RNA45SN1_45S_50ntFlanks</td><td>6647</td><td>6668</td><td>22</td><td>CGTACGACTCTTAGCGGTGGAT</td><td>787.3</td><td>3.3</td><td>0.7</td><td>5.0</td><td>6.9E-07</td><td>1.1E-06</td><td>MB231/MB468</td><td>NUC</td></tr>
<tr><td>>MI0000108|hsa-mir-103a-2&WithFlank&20|+|3917488|3917577@54.76.23(+1C);MI0000109|hsa-mir-103a-1&WithFlank&5|-|168560890|168560979@54.76.23(+1C)&[MIMAT0000101&hsa-miR-103a-3p&offsets|0|0(+1C);m-12&20|+|3917541|3917563&offsets|0|0(+1C)];[MIMAT0000101_1&hsa-miR-103a-3p&offsets|0|0(+1C);m-13&5|-|168560904|168560926&offsets|0|0(+1C);m-44220&5|-|168560908|168560926&offsets|0|+4(+1C)]|AGCAGCATTGTACAGGGCTATGAC</td><td>>MI0000108|hsa-mir-103a-2&WithFlank&20|+|3917488|3917577</td><td>54</td><td>76</td><td>23(+1C)</td><td>AGCAGCATTGTACAGGGCTATGAC</td><td>666.5</td><td>7.6</td><td>1.5</td><td>5.0</td><td>7.2E-07</td><td>1.2E-06</td><td>MB231/MB468</td><td>MITO</td></tr>
<tr><td>>am_trna111_HisGTG_1_-_147774845_147774916@1.29.29,trna118_HisGTG_1_-_145396881_145396952@1.29.29,trna16_HisGTG_1_+_146544773_146544844@1.29.29,trna1_HisGTG_15_+_45493349_45493420@1.29.29,trna21_HisGTG_1_+_147753471_147753542@1.29.29,trna33_HisGTG_6_+_27125906_27125977@1.29.29,trna7_HisGTG_9_-_14433938_14434009@1.29.29,trna8_HisGTG_15_-_45492611_45492682@1.29.29,trna9_HisGTG_15_-_45490804_45490875@1.29.29|GCCGTGATCGTATAGTGGTTAGTACTCTG</td><td>>am_trna111_HisGTG_1_-_147774845_147774916</td><td>1</td><td>29</td><td>29</td><td>GCCGTGATCGTATAGTGGTTAGTACTCTG</td><td>4974.5</td><td>-4.5</td><td>0.9</td><td>-5.0</td><td>6.0E-07</td><td>1.2E-06</td><td>MB231/MB468</td><td>MP</td></tr>
<tr><td>>MI0000299|hsa-mir-222&WithFlank&X|-|45747009|45747130@75.96.22(+1C)&[MIMAT0000279&hsa-miR-222-3p&offsets|0|+1(+1C);m-140&X|-|45747033|45747056&offsets|0|-2(+1C)]|AGCTACATCTGGCTACTGGGTCC</td><td>>MI0000299|hsa-mir-222&WithFlank&X|-|45747009|45747130</td><td>75</td><td>96</td><td>22(+1C)</td><td>AGCTACATCTGGCTACTGGGTCC</td><td>1246.3</td><td>4.8</td><td>1.0</td><td>5.0</td><td>6.0E-07</td><td>1.2E-06</td><td>MB231/MB468</td><td>MP</td></tr>
<tr><td>>hg38_21_+_8433172_8446622_RNA45SN1_45S_50ntFlanks@10248.10270.23|CGACGAGTAGGAGGGCCGCTGCG</td><td>>hg38_21_+_8433172_8446622_RNA45SN1_45S_50ntFlanks</td><td>10248</td><td>10270</td><td>23</td><td>CGACGAGTAGGAGGGCCGCTGCG</td><td>224.6</td><td>11.1</td><td>2.0</td><td>5.5</td><td>3.8E-08</td><td>1.2E-06</td><td>BT20/MB231</td><td>MP</td></tr>
<tr><td>>hg38_21_+_8433172_8446622_RNA45SN1_45S_50ntFlanks@11919.11958.40|CATGAGAGGTGTAGAATAAGTGGGAGGCCCCCGGCGCCCC</td><td>>hg38_21_+_8433172_8446622_RNA45SN1_45S_50ntFlanks</td><td>11919</td><td>11958</td><td>40</td><td>CATGAGAGGTGTAGAATAAGTGGGAGGCCCCCGGCGCCCC</td><td>49.9</td><td>-9.5</td><td>1.9</td><td>-5.0</td><td>6.0E-07</td><td>1.2E-06</td><td>MB231/MB468</td><td>MP</td></tr>
<tr><td>>hg38_21_+_8433172_8446622_RNA45SN1_45S_50ntFlanks@4069.4094.26|CTTTCGATGGTAGTCGCCGTGCCTAC</td><td>>hg38_21_+_8433172_8446622_RNA45SN1_45S_50ntFlanks</td><td>4069</td><td>4094</td><td>26</td><td>CTTTCGATGGTAGTCGCCGTGCCTAC</td><td>489.8</td><td>-7.2</td><td>1.4</td><td>-5.0</td><td>6.0E-07</td><td>1.2E-06</td><td>MB231/MB468</td><td>MP</td></tr>
<tr><td>>hg38_21_+_8433172_8446622_RNA45SN1_45S_50ntFlanks@10410.10431.22|TCAGTCGGTCCTGAGAGATGGG</td><td>>hg38_21_+_8433172_8446622_RNA45SN1_45S_50ntFlanks</td><td>10410</td><td>10431</td><td>22</td><td>TCAGTCGGTCCTGAGAGATGGG</td><td>521.3</td><td>-3.6</td><td>0.7</td><td>-5.0</td><td>5.2E-07</td><td>1.2E-06</td><td>BT20/MB468</td><td>MITO</td></tr>
<tr><td>>hg38_21_+_8433172_8446622_RNA45SN1_45S_50ntFlanks@11235.11269.35|CTTCCCGTGGATCGCCCCAGCTGCGGCGGGCGTCG</td><td>>hg38_21_+_8433172_8446622_RNA45SN1_45S_50ntFlanks</td><td>11235</td><td>11269</td><td>35</td><td>CTTCCCGTGGATCGCCCCAGCTGCGGCGGGCGTCG</td><td>500.5</td><td>-6.6</td><td>1.3</td><td>-5.0</td><td>5.9E-07</td><td>1.2E-06</td><td>BT20/MB468</td><td>CYTO</td></tr>
<tr><td>>hg38_21_+_8433172_8446622_RNA45SN1_45S_50ntFlanks@4736.4758.23|CGAAAGTCGGAGGTTCGAAGACG</td><td>>hg38_21_+_8433172_8446622_RNA45SN1_45S_50ntFlanks</td><td>4736</td><td>4758</td><td>23</td><td>CGAAAGTCGGAGGTTCGAAGACG</td><td>1005.7</td><td>-6.1</td><td>1.2</td><td>-5.0</td><td>6.0E-07</td><td>1.2E-06</td><td>MB231/MB468</td><td>MP</td></tr>
<tr><td>>hg38_21_+_8433172_8446622_RNA45SN1_45S_50ntFlanks@5414.5443.30|CTACTACCGATTGGATGGTTTAGTGAGGCC</td><td>>hg38_21_+_8433172_8446622_RNA45SN1_45S_50ntFlanks</td><td>5414</td><td>5443</td><td>30</td><td>CTACTACCGATTGGATGGTTTAGTGAGGCC</td><td>67.4</td><td>-9.2</td><td>1.8</td><td>-5.2</td><td>1.7E-07</td><td>1.2E-06</td><td>BT20/MB468</td><td>MP</td></tr>
<tr><td>>hg38_21_+_8433172_8446622_RNA45SN1_45S_50ntFlanks@11668.11702.35|TGCCCAGTGCTCTGAATGTCAAAGTGAAGAAATTC</td><td>>hg38_21_+_8433172_8446622_RNA45SN1_45S_50ntFlanks</td><td>11668</td><td>11702</td><td>35</td><td>TGCCCAGTGCTCTGAATGTCAAAGTGAAGAAATTC</td><td>7162.4</td><td>4.1</td><td>0.8</td><td>5.0</td><td>6.8E-07</td><td>1.2E-06</td><td>MB231/MB468</td><td>CYTO</td></tr>
<tr><td>>hg38_21_+_8433172_8446622_RNA45SN1_45S_50ntFlanks@10808.10837.30|CAATGTAGGTAAGGGAAGTCGGCAAGCCGG</td><td>>hg38_21_+_8433172_8446622_RNA45SN1_45S_50ntFlanks</td><td>10808</td><td>10837</td><td>30</td><td>CAATGTAGGTAAGGGAAGTCGGCAAGCCGG</td><td>209.9</td><td>-4.3</td><td>0.9</td><td>-5.0</td><td>5.9E-07</td><td>1.2E-06</td><td>BT20/MB468</td><td>CYTO</td></tr>
<tr><td>>hg38_21_+_8433172_8446622_RNA45SN1_45S_50ntFlanks@11729.11746.18|TAACTATGACTCTCTTAA</td><td>>hg38_21_+_8433172_8446622_RNA45SN1_45S_50ntFlanks</td><td>11729</td><td>11746</td><td>18</td><td>TAACTATGACTCTCTTAA</td><td>246.8</td><td>-2.6</td><td>0.5</td><td>-5.0</td><td>5.9E-07</td><td>1.2E-06</td><td>BT20/MB468</td><td>CYTO</td></tr>
<tr><td>>hg38_21_+_8433172_8446622_RNA45SN1_45S_50ntFlanks@12935.12959.25|TAAACCATTCGTAGACGACCTGCTT</td><td>>hg38_21_+_8433172_8446622_RNA45SN1_45S_50ntFlanks</td><td>12935</td><td>12959</td><td>25</td><td>TAAACCATTCGTAGACGACCTGCTT</td><td>727.6</td><td>-8.0</td><td>1.6</td><td>-4.9</td><td>7.6E-07</td><td>1.2E-06</td><td>BT20/MB468</td><td>NUC</td></tr>
<tr><td>>hg38_21_+_8433172_8446622_RNA45SN1_45S_50ntFlanks@12625.12659.35|CAATGGGGCGAAGCTACCATCTGTGGGATTATGAC</td><td>>hg38_21_+_8433172_8446622_RNA45SN1_45S_50ntFlanks</td><td>12625</td><td>12659</td><td>35</td><td>CAATGGGGCGAAGCTACCATCTGTGGGATTATGAC</td><td>289.6</td><td>-5.1</td><td>1.0</td><td>-5.0</td><td>7.0E-07</td><td>1.2E-06</td><td>MB231/MB468</td><td>NUC</td></tr>
<tr><td>>hg38_21_+_8433172_8446622_RNA45SN1_45S_50ntFlanks@12417.12455.39|TCGATGTCGGCTCTTCCTATCATTGTGAAGCAGAATTCA</td><td>>hg38_21_+_8433172_8446622_RNA45SN1_45S_50ntFlanks</td><td>12417</td><td>12455</td><td>39</td><td>TCGATGTCGGCTCTTCCTATCATTGTGAAGCAGAATTCA</td><td>1814.1</td><td>-4.2</td><td>0.8</td><td>-5.0</td><td>5.2E-07</td><td>1.2E-06</td><td>BT20/MB468</td><td>MITO</td></tr>
<tr><td>>hg38_21_+_8433172_8446622_RNA45SN1_45S_50ntFlanks@4733.4768.36|GAACGAAAGTCGGAGGTTCGAAGACGATCAGATACC</td><td>>hg38_21_+_8433172_8446622_RNA45SN1_45S_50ntFlanks</td><td>4733</td><td>4768</td><td>36</td><td>GAACGAAAGTCGGAGGTTCGAAGACGATCAGATACC</td><td>185.3</td><td>6.9</td><td>1.4</td><td>5.0</td><td>7.0E-07</td><td>1.2E-06</td><td>MB231/MB468</td><td>NUC</td></tr>
<tr><td>>MI0000266|hsa-mir-10a&WithFlank&17|-|48579832|48579953@28.49.22(+1U)&[MIMAT0000253&hsa-miR-10a-5p&offsets|0|-1(+1U);m-18&17|-|48579905|48579926&offsets|0|0(+1U)]|TACCCTGTAGATCCGAATTTGTT</td><td>>MI0000266|hsa-mir-10a&WithFlank&17|-|48579832|48579953</td><td>28</td><td>49</td><td>22(+1U)</td><td>TACCCTGTAGATCCGAATTTGTT</td><td>45.0</td><td>8.7</td><td>1.7</td><td>5.0</td><td>6.1E-07</td><td>1.2E-06</td><td>MB231/MB468</td><td>MP</td></tr>
<tr><td>>hg38_21_+_8433172_8446622_RNA45SN1_45S_50ntFlanks@11876.11900.25|AAGACCCTGTTGAGCTTGACTCTAG</td><td>>hg38_21_+_8433172_8446622_RNA45SN1_45S_50ntFlanks</td><td>11876</td><td>11900</td><td>25</td><td>AAGACCCTGTTGAGCTTGACTCTAG</td><td>564.8</td><td>-4.3</td><td>0.9</td><td>-5.0</td><td>6.8E-07</td><td>1.2E-06</td><td>MB231/MB468</td><td>CYTO</td></tr>
<tr><td>>trna20_GluTTC_2_-_131094701_131094772@1.36.36,trna5_GluTTC_13_-_41634874_41634945@1.36.36|TCCCATATGGTCTAGCGGTTAGGATTCCTGGTTTTC</td><td>>trna20_GluTTC_2_-_131094701_131094772</td><td>1</td><td>36</td><td>36</td><td>TCCCATATGGTCTAGCGGTTAGGATTCCTGGTTTTC</td><td>66.3</td><td>9.3</td><td>1.9</td><td>5.0</td><td>6.1E-07</td><td>1.2E-06</td><td>MB231/MB468</td><td>MP</td></tr>
<tr><td>>hg38_21_+_8433172_8446622_RNA45SN1_45S_50ntFlanks@5073.5115.43|AATTCCGATAACGAACGAGACTCTGGCATGCTAACTAGTTACG</td><td>>hg38_21_+_8433172_8446622_RNA45SN1_45S_50ntFlanks</td><td>5073</td><td>5115</td><td>43</td><td>AATTCCGATAACGAACGAGACTCTGGCATGCTAACTAGTTACG</td><td>324.9</td><td>-6.0</td><td>1.2</td><td>-4.9</td><td>7.6E-07</td><td>1.2E-06</td><td>BT20/MB468</td><td>NUC</td></tr>
<tr><td>>hg38_21_+_8433172_8446622_RNA45SN1_45S_50ntFlanks@3993.4026.34|GTGACTCTAGATAACCTCGGGCCGATCGCACGCC</td><td>>hg38_21_+_8433172_8446622_RNA45SN1_45S_50ntFlanks</td><td>3993</td><td>4026</td><td>34</td><td>GTGACTCTAGATAACCTCGGGCCGATCGCACGCC</td><td>706.5</td><td>-4.1</td><td>0.8</td><td>-5.0</td><td>7.0E-07</td><td>1.2E-06</td><td>MB231/MB468</td><td>NUC</td></tr>
<tr><td>>hg38_21_+_8433172_8446622_RNA45SN1_45S_50ntFlanks@6670.6697.28|ACTCGGCTCGTGCGTCGATGAAGAACGC</td><td>>hg38_21_+_8433172_8446622_RNA45SN1_45S_50ntFlanks</td><td>6670</td><td>6697</td><td>28</td><td>ACTCGGCTCGTGCGTCGATGAAGAACGC</td><td>636.1</td><td>-3.9</td><td>0.8</td><td>-5.0</td><td>7.3E-07</td><td>1.2E-06</td><td>MB231/MB468</td><td>MITO</td></tr>
<tr><td>>hg38_21_+_8433172_8446622_RNA45SN1_45S_50ntFlanks@11665.11697.33|TTCTGCCCAGTGCTCTGAATGTCAAAGTGAAGA</td><td>>hg38_21_+_8433172_8446622_RNA45SN1_45S_50ntFlanks</td><td>11665</td><td>11697</td><td>33</td><td>TTCTGCCCAGTGCTCTGAATGTCAAAGTGAAGA</td><td>56.3</td><td>9.6</td><td>1.8</td><td>5.2</td><td>1.8E-07</td><td>1.2E-06</td><td>BT20/MB468</td><td>MP</td></tr>
<tr><td>>hg38_21_+_8433172_8446622_RNA45SN1_45S_50ntFlanks@4102.4129.28|ACCACGGGTGACGGGGAATCAGGGTTCG</td><td>>hg38_21_+_8433172_8446622_RNA45SN1_45S_50ntFlanks</td><td>4102</td><td>4129</td><td>28</td><td>ACCACGGGTGACGGGGAATCAGGGTTCG</td><td>144.9</td><td>-6.8</td><td>1.4</td><td>-5.0</td><td>6.1E-07</td><td>1.2E-06</td><td>MB231/MB468</td><td>MP</td></tr>
<tr><td>>hg38_21_+_8433172_8446622_RNA45SN1_45S_50ntFlanks@10317.10341.25|GTGCAGATCTTGGTGGTAGTAGCAA</td><td>>hg38_21_+_8433172_8446622_RNA45SN1_45S_50ntFlanks</td><td>10317</td><td>10341</td><td>25</td><td>GTGCAGATCTTGGTGGTAGTAGCAA</td><td>289.4</td><td>11.5</td><td>2.1</td><td>5.5</td><td>3.9E-08</td><td>1.2E-06</td><td>BT20/MB231</td><td>MP</td></tr>
<tr><td>>hg38_21_+_8433172_8446622_RNA45SN1_45S_50ntFlanks@10248.10280.33|CGACGAGTAGGAGGGCCGCTGCGGTGAGCCTTG</td><td>>hg38_21_+_8433172_8446622_RNA45SN1_45S_50ntFlanks</td><td>10248</td><td>10280</td><td>33</td><td>CGACGAGTAGGAGGGCCGCTGCGGTGAGCCTTG</td><td>229.7</td><td>11.2</td><td>2.0</td><td>5.5</td><td>3.9E-08</td><td>1.2E-06</td><td>BT20/MB231</td><td>MP</td></tr>
<tr><td>>hg38_21_+_8433172_8446622_RNA45SN1_45S_50ntFlanks@3765.3792.28|AGTACGCACGGCCGGTACAGTGAAACTG</td><td>>hg38_21_+_8433172_8446622_RNA45SN1_45S_50ntFlanks</td><td>3765</td><td>3792</td><td>28</td><td>AGTACGCACGGCCGGTACAGTGAAACTG</td><td>317.2</td><td>-5.3</td><td>1.1</td><td>-5.0</td><td>7.0E-07</td><td>1.2E-06</td><td>MB231/MB468</td><td>NUC</td></tr>
<tr><td>>hg38_21_+_8433172_8446622_RNA45SN1_45S_50ntFlanks@10794.10822.29|CTGGCATGTTGGAACAATGTAGGTAAGGG</td><td>>hg38_21_+_8433172_8446622_RNA45SN1_45S_50ntFlanks</td><td>10794</td><td>10822</td><td>29</td><td>CTGGCATGTTGGAACAATGTAGGTAAGGG</td><td>527.5</td><td>-4.1</td><td>0.8</td><td>-5.2</td><td>1.8E-07</td><td>1.2E-06</td><td>BT20/MB468</td><td>MP</td></tr>
<tr><td>>hg38_21_+_8433172_8446622_RNA45SN1_45S_50ntFlanks@12458.12486.29|AAGCGTTGGATTGTTCACCCACTAATAGG</td><td>>hg38_21_+_8433172_8446622_RNA45SN1_45S_50ntFlanks</td><td>12458</td><td>12486</td><td>29</td><td>AAGCGTTGGATTGTTCACCCACTAATAGG</td><td>393.7</td><td>-6.0</td><td>1.2</td><td>-4.9</td><td>7.7E-07</td><td>1.2E-06</td><td>BT20/MB468</td><td>NUC</td></tr>
<tr><td>>MI0003188|hsa-mir-503&WithFlank&X|-|134546322|134546404@12.30.19(+1A)&[MIMAT0002874&hsa-miR-503-5p&offsets|0|-4(+1A);m-297&X|-|134546371|134546393&offsets|0|-4(+1A)]|TAGCAGCGGGAACAGTTCTA</td><td>>MI0003188|hsa-mir-503&WithFlank&X|-|134546322|134546404</td><td>12</td><td>30</td><td>19(+1A)</td><td>TAGCAGCGGGAACAGTTCTA</td><td>52.4</td><td>8.9</td><td>1.8</td><td>5.0</td><td>6.1E-07</td><td>1.2E-06</td><td>MB231/MB468</td><td>MP</td></tr>
</table>

| | | | | | | | | | | | | | |
|---|---|---|---|---|---|---|---|---|---|---|---|---|---|
| >trna10_AspGTC_12_-_125424193_125424264@31.72.42,trna12_AspGTC_12_-_125411891_125411962@31.72.42,trna38_AspGTC_17_-_8125556_8125627@31.72.42,trna45_AspGTC_6_+_27447453_27447524@31.72.42,trna48_AspGTC_6_+_27471523_27471594@31.72.42,trna4_AspGTC_12_+_96429799_96429870@31.72.42,trna69_AspGTC_1_-_161440205_161440276@31.72.42,trna72_AspGTC_1_-_161432824_161432895@31.72.42,trna75_AspGTC_1_-_161425414_161425485@31.72.42,trna78_AspGTC_1_-_161418033_161418104@31.72.42,trna81_AspGTC_1_-_161410615_161410686@31.72.42\|CCTGTCACGCGGGAGACCGGGGTTCGATTCCCCGACGGGGAG | >trna10_AspGTC_12_-_125424193_125424264 | 31 | 72 | 42 | CCTGTCACGCGGGAGACCGGGGTTCGATTCCCCGACGGGGAG | 92.2 | -9.6 | 1.8 | -5.2 | 1.8E-07 | 1.2E-06 | BT20/MB468 | MP |
| >hg38_21_+_8433172_8446622_RNA45SN1_45S_50ntFlanks@12896.12931.36\|CGCCCGTCACGCACCGCACGTTCGTGGGGAACCTGG | >hg38_21_+_8433172_8446622_RNA45SN1_45S_50ntFlanks | 12896 | 12931 | 36 | CGCCCGTCACGCACCGCACGTTCGTGGGGAACCTGG | 213.1 | -7.0 | 1.4 | -5.0 | 7.4E-07 | 1.2E-06 | MB231/MB468 | MITO |
| >hg38_21_+_8433172_8446622_RNA45SN1_45S_50ntFlanks@9927.9952.26\|TTGATATAGACAGCAGGACGGTGGCC | >hg38_21_+_8433172_8446622_RNA45SN1_45S_50ntFlanks | 9927 | 9952 | 26 | TTGATATAGACAGCAGGACGGTGGCC | 63.2 | 9.7 | 1.9 | 5.2 | 1.8E-07 | 1.2E-06 | BT20/MB468 | MP |
| >hg38_21_+_8433172_8446622_RNA45SN1_45S_50ntFlanks@4005.4042.38\|AACCTCGGGCCGATCGCACGCCCCCCGTGGCGGCGACG | >hg38_21_+_8433172_8446622_RNA45SN1_45S_50ntFlanks | 4005 | 4042 | 38 | AACCTCGGGCCGATCGCACGCCCCCCGTGGCGGCGACG | 686.9 | -3.8 | 0.8 | -5.0 | 6.9E-07 | 1.2E-06 | MB231/MB468 | CYTO |
| >hg38_21_+_8433172_8446622_RNA45SN1_45S_50ntFlanks@12936.12961.26\|AAACCATTCGTAGACGACCTGCTTCT | >hg38_21_+_8433172_8446622_RNA45SN1_45S_50ntFlanks | 12936 | 12961 | 26 | AAACCATTCGTAGACGACCTGCTTCT | 260.7 | -5.2 | 1.0 | -5.0 | 7.4E-07 | 1.2E-06 | MB231/MB468 | MITO |
| >hg38_21_+_8433172_8446622_RNA45SN1_45S_50ntFlanks@12941.12965.25\|ATTCGTAGACGACCTGCTTCTGGGT | >hg38_21_+_8433172_8446622_RNA45SN1_45S_50ntFlanks | 12941 | 12965 | 25 | ATTCGTAGACGACCTGCTTCTGGGT | 274.8 | -7.0 | 1.4 | -5.0 | 6.1E-07 | 1.2E-06 | MB231/MB468 | MP |
| >hg38_21_+_8433172_8446622_RNA45SN1_45S_50ntFlanks@6776.6803.28\|CTCCCGGGGCTACGCCTGTCTGAGCGTC | >hg38_21_+_8433172_8446622_RNA45SN1_45S_50ntFlanks | 6776 | 6803 | 28 | CTCCCGGGGCTACGCCTGTCTGAGCGTC | 605.4 | -2.6 | 0.5 | -5.0 | 6.9E-07 | 1.2E-06 | MB231/MB468 | CYTO |
| >am_trnaMT_TyrGTA_MT_-_5826_5891@13.38.26\|GAGTGAAGCATTGGACTGTAAATCTA | >am_trnaMT_TyrGTA_MT_-_5826_5891 | 13 | 38 | 26 | GAGTGAAGCATTGGACTGTAAATCTA | 71.3 | 9.9 | 1.9 | 5.2 | 1.8E-07 | 1.2E-06 | BT20/MB468 | MP |
| >hg38_21_+_8433172_8446622_RNA45SN1_45S_50ntFlanks@10393.10433.41\|CAGCAGTTGAACATGGGTCAGTCGGTCCTGAGAGATGGGCG | >hg38_21_+_8433172_8446622_RNA45SN1_45S_50ntFlanks | 10393 | 10433 | 41 | CAGCAGTTGAACATGGGTCAGTCGGTCCTGAGAGATGGGCG | 1025.5 | -8.6 | 1.7 | -4.9 | 7.7E-07 | 1.2E-06 | BT20/MB468 | NUC |
| >trnaMT_ProTGG_MT_-_15956_16023@44.70.27\|GTGGAGTTAAAGACTTTTTCTCTGACC | >trnaMT_ProTGG_MT_-_15956_16023 | 44 | 70 | 27 | GTGGAGTTAAAGACTTTTTCTCTGACC | 485.2 | 3.9 | 0.8 | 5.0 | 6.2E-07 | 1.2E-06 | MB231/MB468 | MP |
| >MI0000449\|hsa-mir-132&WithFlank&17\|-\|2049902\|2050014@65.86.22(+1U)&[MIMAT0000426&hsa-miR-132-3p&offsets\|0\|0(+1U);m-138&17\|-\|2049929\|2049950&offsets\|0\|0(+1U)]\|TAACAGTCTACAGCCATGGTCGT | >MI0000449\|hsa-mir-132&WithFlank&17\|-\|2049902\|2050014 | 65 | 86 | 22(+1U) | TAACAGTCTACAGCCATGGTCGT | 206.1 | 6.7 | 1.3 | 5.0 | 7.1E-07 | 1.2E-06 | MB231/MB468 | NUC |
| >hg38_21_+_8433172_8446622_RNA45SN1_45S_50ntFlanks@8198.8224.27\|TGGACGGTGTGAGGCCGGTAGCGGCCC | >hg38_21_+_8433172_8446622_RNA45SN1_45S_50ntFlanks | 8198 | 8224 | 27 | TGGACGGTGTGAGGCCGGTAGCGGCCC | 544.3 | -4.6 | 0.9 | -5.0 | 6.9E-07 | 1.2E-06 | MB231/MB468 | CYTO |
| >am_MI0005763\|hsa-mir-941-1&WithFlank&20\|+\|63919443\|63919526@53.75.23;MI0005764\|hsa-mir-941-2&WithFlank&20\|+\|63919499\|63919582@53.75.23;MI0005765\|hsa-mir-941-3&WithFlank&20\|+\|63919555\|63919638@53.75.23;MI0005766\|hsa-mir-941-4&WithFlank&20\|+\|63919750\|63919833@53.75.23;MI0031520\|hsa-mir-941-5&WithFlank&20\|+\|63919862\|63919945@53.75.23&[MIMAT0004984&hsa-miR-941&offsets\|0\|0;m-263&20\|+\|63919495\|63919517&offsets\|0\|0];[MIMAT0004984_1&hsa-miR-941&offsets\|0\|0;m-263&20\|+\|63919551\|63919573&offsets\|0\|0];[MIMAT0004984_2&hsa-miR-941&offsets\|0\|0;m-263&20\|+\|63919607\|63919629&offsets\|0\|0];[MIMAT0004984_3&hsa-miR-941&offsets\|0\|0;m-263&20\|+\|63919802\|63919824&offsets\|0\|0];[MIMAT0004984_4&hsa-miR-941&offsets\|0\|0;m-263&20\|+\|63919914\|63919936&offsets\|0\|0]\|CACCCGGCTGTGTGCACATGTGC | >am_MI0005763\|hsa-mir-941-1&WithFlank&20\|+\|63919443\|63919526 | 53 | 75 | 23 | CACCCGGCTGTGTGCACATGTGC | 3173.4 | 3.4 | 0.7 | 5.0 | 6.2E-07 | 1.2E-06 | MB231/MB468 | MP |
| >hg38_21_+_8433172_8446622_RNA45SN1_45S_50ntFlanks@12746.12765.20\|CGCCTGTCCCCGCCGGCGGG | >hg38_21_+_8433172_8446622_RNA45SN1_45S_50ntFlanks | 12746 | 12765 | 20 | CGCCTGTCCCCGCCGGCGGG | 145.6 | -10.3 | 2.0 | -5.2 | 1.8E-07 | 1.2E-06 | BT20/MB468 | MP |
| >am_trna116_GluCTC_1_-_145399233_145399304@1.29.29,trna59_GluCTC_1_+_249168447_249168518@1.29.29,trna71_GluCTC_1_-_161439189_161439260@1.29.29,trna74_GluCTC_1_-_161431809_161431880@1.29.29,trna77_GluCTC_1_-_161424398_161424469@1.29.29,trna77_GluCTC_6_+_28949976_28950047@1.29.29,trna80_GluCTC_1_-_161417018_161417089@1.29.29,trna87_GluCTC_6_-_126101393_126101464@1.29.29\|TCCCTGGTGGTCTAGTGGTTAGGATTCGG | >am_trna116_GluCTC_1_-_145399233_145399304 | 1 | 29 | 29 | TCCCTGGTGGTCTAGTGGTTAGGATTCGG | 10377.3 | -4.2 | 0.8 | -4.9 | 7.5E-07 | 1.2E-06 | MB231/MB468 | MITO |
| >hg38_21_+_8433172_8446622_RNA45SN1_45S_50ntFlanks@11906.11931.26\|CACGGTGAAGAGACATGAGAGGTGTA | >hg38_21_+_8433172_8446622_RNA45SN1_45S_50ntFlanks | 11906 | 11931 | 26 | CACGGTGAAGAGACATGAGAGGTGTA | 224.1 | 11.1 | 2.0 | 5.5 | 4.0E-08 | 1.2E-06 | BT20/MB231 | MP |
| >hg38_21_+_8433172_8446622_RNA45SN1_45S_50ntFlanks@8176.8201.26\|CTTCTGATCGAGGCCCAGCCCGTGGA | >hg38_21_+_8433172_8446622_RNA45SN1_45S_50ntFlanks | 8176 | 8201 | 26 | CTTCTGATCGAGGCCCAGCCCGTGGA | 1214.6 | -6.3 | 1.3 | -5.0 | 6.2E-07 | 1.2E-06 | MB231/MB468 | MP |
| >hg38_21_+_8433172_8446622_RNA45SN1_45S_50ntFlanks@6652.6687.36\|GACTCTTAGCGGTGGATCACTCGGCTCGTGCGTCGA | >hg38_21_+_8433172_8446622_RNA45SN1_45S_50ntFlanks | 6652 | 6687 | 36 | GACTCTTAGCGGTGGATCACTCGGCTCGTGCGTCGA | 3688.4 | 4.1 | 0.8 | 5.0 | 7.2E-07 | 1.2E-06 | MB231/MB468 | NUC |
| >hg38_21_+_8433172_8446622_RNA45SN1_45S_50ntFlanks@8555.8581.27\|CCCGGAGGGGGCGGGCTCCGGCGGGTG | >hg38_21_+_8433172_8446622_RNA45SN1_45S_50ntFlanks | 8555 | 8581 | 27 | CCCGGAGGGGGCGGGCTCCGGCGGGTG | 71.9 | -9.2 | 1.8 | -5.2 | 1.8E-07 | 1.2E-06 | BT20/MB468 | MP |
| >trnaMT_GluTTC_MT_-_14674_14742@40.63.24,trnalookalike8_GluTTC_5_-_93905172_93905240@40.63.24\|TTGGTCGTGGTTGTAGTCCGTGCG | >trnaMT_GluTTC_MT_-_14674_14742 | 40 | 63 | 24 | TTGGTCGTGGTTGTAGTCCGTGCG | 785.9 | 5.5 | 1.1 | 5.2 | 1.8E-07 | 1.2E-06 | BT20/MB468 | MP |
| >hg38_21_+_8433172_8446622_RNA45SN1_45S_50ntFlanks@9120.9140.21\|TGCCGCGCGCGGGTCGGGGGG | >hg38_21_+_8433172_8446622_RNA45SN1_45S_50ntFlanks | 9120 | 9140 | 21 | TGCCGCGCGCGGGTCGGGGGG | 74.8 | -9.3 | 1.8 | -5.2 | 1.8E-07 | 1.2E-06 | BT20/MB468 | MP |
| >hg38_21_+_8433172_8446622_RNA45SN1_45S_50ntFlanks@8153.8181.29\|GCCGCTCGTGGGGGGCCCAAGTCCTTCTG | >hg38_21_+_8433172_8446622_RNA45SN1_45S_50ntFlanks | 8153 | 8181 | 29 | GCCGCTCGTGGGGGGCCCAAGTCCTTCTG | 387.9 | -4.0 | 0.8 | -5.0 | 7.2E-07 | 1.2E-06 | MB231/MB468 | NUC |
| >hg38_21_+_8433172_8446622_RNA45SN1_45S_50ntFlanks@11579.11602.24\|AGAACTGGTGCGGACCAGGGGAAT | >hg38_21_+_8433172_8446622_RNA45SN1_45S_50ntFlanks | 11579 | 11602 | 24 | AGAACTGGTGCGGACCAGGGGAAT | 539.5 | -6.1 | 1.2 | -4.9 | 7.5E-07 | 1.2E-06 | MB231/MB468 | MITO |
| >trnaMT_HisGTG_MT_+_12138_12206@25.63.39\|CAGATTGTGAATCTGACAACAGAGGCTTACGACCCCTTA | >trnaMT_HisGTG_MT_+_12138_12206 | 25 | 63 | 39 | CAGATTGTGAATCTGACAACAGAGGCTTACGACCCCTTA | 115.7 | 10.6 | 2.0 | 5.2 | 1.8E-07 | 1.2E-06 | BT20/MB468 | MP |
| >hg38_21_+_8433172_8446622_RNA45SN1_45S_50ntFlanks@11315.11332.18\|CCACCCCACGTCTCGTCG | >hg38_21_+_8433172_8446622_RNA45SN1_45S_50ntFlanks | 11315 | 11332 | 18 | CCACCCCACGTCTCGTCG | 2445.2 | -9.5 | 1.9 | -4.9 | 7.8E-07 | 1.2E-06 | BT20/MB468 | NUC |
| >hg38_21_+_8433172_8446622_RNA45SN1_45S_50ntFlanks@12003.12043.41\|GCGGGCCGCCGGTGAAATACCACTACTCTGATCGTTTTTTC | >hg38_21_+_8433172_8446622_RNA45SN1_45S_50ntFlanks | 12003 | 12043 | 41 | GCGGGCCGCCGGTGAAATACCACTACTCTGATCGTTTTTTC | 208.8 | -10.8 | 2.1 | -5.2 | 1.8E-07 | 1.2E-06 | BT20/MB468 | MP |
| >hg38_21_+_8433172_8446622_RNA45SN1_45S_50ntFlanks@12605.12626.22\|TATGTGCTTGGCTGAGGAGCCA | >hg38_21_+_8433172_8446622_RNA45SN1_45S_50ntFlanks | 12605 | 12626 | 22 | TATGTGCTTGGCTGAGGAGCCA | 1910.9 | -6.7 | 1.3 | -5.2 | 1.8E-07 | 1.2E-06 | BT20/MB468 | MP |
| >MI0000434\|hsa-let-7i&WithFlank&12\|+\|62603680\|62603775@12.33.22&[MIMAT0000415&hsa-let-7i-5p&offsets\|0\|0;m-49&12\|+\|62603691\|62603712&offsets\|0\|0]\|TGAGGTAGTAGTTTGTGCTGTT | >MI0000434\|hsa-let-7i&WithFlank&12\|+\|62603680\|62603775 | 12 | 33 | 22 | TGAGGTAGTAGTTTGTGCTGTT | 142405.3 | 4.2 | 0.8 | 5.0 | 6.2E-07 | 1.2E-06 | MB231/MB468 | MP |
| >hg38_21_+_8433172_8446622_RNA45SN1_45S_50ntFlanks@10349.10380.32\|AACGAGAACTTTGAAGGCCGAAGTGGAGAAGG | >hg38_21_+_8433172_8446622_RNA45SN1_45S_50ntFlanks | 10349 | 10380 | 32 | AACGAGAACTTTGAAGGCCGAAGTGGAGAAGG | 380.5 | -3.6 | 0.7 | -4.9 | 7.8E-07 | 1.2E-06 | BT20/MB468 | NUC |
| >hg38_21_+_8433172_8446622_RNA45SN1_45S_50ntFlanks@12675.12703.29\|CAGAATCCCGCCCAGGCGGAACGATACGG | >hg38_21_+_8433172_8446622_RNA45SN1_45S_50ntFlanks | 12675 | 12703 | 29 | CAGAATCCCGCCCAGGCGGAACGATACGG | 89.0 | -9.6 | 1.8 | -5.2 | 1.8E-07 | 1.2E-06 | BT20/MB468 | MP |
| >hg38_21_+_8433172_8446622_RNA45SN1_45S_50ntFlanks@11453.11477.25\|TCCTCGGCGCGCGGCGGCGGCGGCG | >hg38_21_+_8433172_8446622_RNA45SN1_45S_50ntFlanks | 11453 | 11477 | 25 | TCCTCGGCGCGCGGCGGCGGCGGCG | 60.8 | -9.0 | 1.7 | -5.2 | 1.8E-07 | 1.2E-06 | BT20/MB468 | MP |
| >trna129_MetCAT_6_-_27870271_27870342@33.51.19,trna142_MetCAT_6_-_27560600_27560671@33.51.19,trna150_MetCAT_6_-_27300764_27300835@33.51.19,trna169_MetCAT_6_-_26330529_26330600@33.51.19,trna171_MetCAT_6_-_26313352_26313423@33.51.19,trna20_MetCAT_17_-_80452597_80452668@33.51.19,trna2_MetCAT_6_+_26286754_26286825@33.51.19,trna32_MetCAT_1_+_153643726_153643797@33.51.19,trna61_MetCAT_6_+_27745664_27745735@33.51.19\|CATAACCCAGAGGTCGATG | >trna129_MetCAT_6_-_27870271_27870342 | 33 | 51 | 19 | CATAACCCAGAGGTCGATG | 187.3 | -11.4 | 2.3 | -5.0 | 6.3E-07 | 1.2E-06 | MB231/MB468 | MP |
| >hg38_21_+_8433172_8446622_RNA45SN1_45S_50ntFlanks@12243.12265.23\|GAAGGGCAAAAGCTCGCTTGATC | >hg38_21_+_8433172_8446622_RNA45SN1_45S_50ntFlanks | 12243 | 12265 | 23 | GAAGGGCAAAAGCTCGCTTGATC | 280.4 | -3.4 | 0.7 | -5.0 | 6.3E-07 | 1.2E-06 | MB231/MB468 | MP |
| >MI0000087\|hsa-mir-29a&WithFlank&7\|-\|130876741\|130876816@48.70.23&[MIMAT0000086&hsa-miR-29a-3p&offsets\|0\|+1;m-26&7\|-\|130876749\|130876769&offsets\|0\|+2]\|TAGCACCATCTGAAATCGGTTAT | >MI0000087\|hsa-mir-29a&WithFlank&7\|-\|130876741\|130876816 | 48 | 70 | 23 | TAGCACCATCTGAAATCGGTTAT | 685.3 | 6.1 | 1.2 | 5.0 | 6.3E-07 | 1.2E-06 | MB231/MB468 | MP |
| >hg38_21_+_8433172_8446622_RNA45SN1_45S_50ntFlanks@4727.4765.39\|AATCAAGAACGAAAGTCGGAGGTTCGAAGACGATCAGAT | >hg38_21_+_8433172_8446622_RNA45SN1_45S_50ntFlanks | 4727 | 4765 | 39 | AATCAAGAACGAAAGTCGGAGGTTCGAAGACGATCAGAT | 2831.9 | 5.9 | 1.2 | 5.0 | 6.1E-07 | 1.2E-06 | BT20/MB468 | CYTO |
| >MI0000078\|hsa-mir-22&WithFlank&17\|-\|1713897\|1713993@21.41.21&[MIMAT0004495&hsa-miR-22-5p&offsets\|0\|-1;m-217&17\|-\|1713953\|1713973&offsets\|0\|0]\|AGTTCTTCAGTGGCAAGCTTT | >MI0000078\|hsa-mir-22&WithFlank&17\|-\|1713897\|1713993 | 21 | 41 | 21 | AGTTCTTCAGTGGCAAGCTTT | 139.5 | 3.4 | 0.7 | 4.9 | 7.9E-07 | 1.2E-06 | BT20/MB468 | NUC |
| >MI0000299\|hsa-mir-222&WithFlank&X\|-\|45747009\|45747130@75.98.24&[MIMAT0000279&hsa-miR-222-3p&offsets\|0\|+3;m-140&X\|-\|45747033\|45747056&offsets\|0\|0]\|AGCTACATCTGGCTACTGGGTCTC | >MI0000299\|hsa-mir-222&WithFlank&X\|-\|45747009\|45747130 | 75 | 98 | 24 | AGCTACATCTGGCTACTGGGTCTC | 6511.3 | 4.1 | 0.8 | 5.0 | 6.3E-07 | 1.2E-06 | MB231/MB468 | MP |
| >hg38_21_+_8433172_8446622_RNA45SN1_45S_50ntFlanks@10350.10382.33\|ACGAGAACTTTGAAGGCCGAAGTGGAGAAGGGT | >hg38_21_+_8433172_8446622_RNA45SN1_45S_50ntFlanks | 10350 | 10382 | 33 | ACGAGAACTTTGAAGGCCGAAGTGGAGAAGGGT | 1387.9 | -3.4 | 0.7 | -5.0 | 7.2E-07 | 1.2E-06 | MB231/MB468 | NUC |
| >hg38_21_+_8433172_8446622_RNA45SN1_45S_50ntFlanks@10797.10826.30\|GCATGTTGGAACAATGTAGGTAAGGGAAGT | >hg38_21_+_8433172_8446622_RNA45SN1_45S_50ntFlanks | 10797 | 10826 | 30 | GCATGTTGGAACAATGTAGGTAAGGGAAGT | 692.6 | -4.0 | 0.8 | -5.2 | 1.8E-07 | 1.2E-06 | BT20/MB468 | MP |

<table>
<tr><td>>hg38_MT_+_1621_3279_RNR2_16S_50ntFlanks@51.90.40|GCTAAACCTAGCCCCAAACCCACTCCACCTTACTACCAGA</td><td>>hg38_MT_+_1621_3279_RNR2_16S_50ntFlanks</td><td>51</td><td>90</td><td>40</td><td>GCTAAACCTAGCCCCAAACCCACTCCACCTTACTACCAGA</td><td>3855.6</td><td>5.3</td><td>1.1</td><td>4.9</td><td>7.6E-07</td><td>1.2E-06</td><td>MB231/MB468</td><td>MITO</td></tr>
<tr><td>>hg38_21_+_8433172_8446622_RNA45SN1_45S_50ntFlanks@12993.13012.20|CCCTCGCTGCGATCTATTGA</td><td>>hg38_21_+_8433172_8446622_RNA45SN1_45S_50ntFlanks</td><td>12993</td><td>13012</td><td>20</td><td>CCCTCGCTGCGATCTATTGA</td><td>1044.5</td><td>-8.3</td><td>1.7</td><td>-4.9</td><td>7.9E-07</td><td>1.2E-06</td><td>BT20/MB468</td><td>NUC</td></tr>
<tr><td>>hg38_21_+_8433172_8446622_RNA45SN1_45S_50ntFlanks@11874.11892.19|AGAAGACCCTGTTGAGCTT</td><td>>hg38_21_+_8433172_8446622_RNA45SN1_45S_50ntFlanks</td><td>11874</td><td>11892</td><td>19</td><td>AGAAGACCCTGTTGAGCTT</td><td>571.9</td><td>-6.1</td><td>1.2</td><td>-4.9</td><td>7.6E-07</td><td>1.2E-06</td><td>MB231/MB468</td><td>MITO</td></tr>
<tr><td>>hg38_21_+_8433172_8446622_RNA45SN1_45S_50ntFlanks@4224.4242.19|AAAATAACAATACAGGACT</td><td>>hg38_21_+_8433172_8446622_RNA45SN1_45S_50ntFlanks</td><td>4224</td><td>4242</td><td>19</td><td>AAAATAACAATACAGGACT</td><td>794.9</td><td>-6.7</td><td>1.4</td><td>-4.9</td><td>7.9E-07</td><td>1.2E-06</td><td>BT20/MB468</td><td>NUC</td></tr>
<tr><td>>hg38_21_+_8433172_8446622_RNA45SN1_45S_50ntFlanks@8176.8218.43|CTTCTGATCGAGGCCCAGCCCGTGGACGGTGTGAGGCCGGTAG</td><td>>hg38_21_+_8433172_8446622_RNA45SN1_45S_50ntFlanks</td><td>8176</td><td>8218</td><td>43</td><td>CTTCTGATCGAGGCCCAGCCCGTGGACGGTGTGAGGCCGGTAG</td><td>500.0</td><td>-6.0</td><td>1.2</td><td>-4.9</td><td>7.6E-07</td><td>1.2E-06</td><td>MB231/MB468</td><td>MITO</td></tr>
<tr><td>>hg38_21_+_8433172_8446622_RNA45SN1_45S_50ntFlanks@7987.8006.20|TCAGACGTGGCGACCCGCTG</td><td>>hg38_21_+_8433172_8446622_RNA45SN1_45S_50ntFlanks</td><td>7987</td><td>8006</td><td>20</td><td>TCAGACGTGGCGACCCGCTG</td><td>819.2</td><td>-8.0</td><td>1.5</td><td>-5.2</td><td>1.8E-07</td><td>1.2E-06</td><td>BT20/MB468</td><td>MP</td></tr>
<tr><td>>hg38_21_+_8433172_8446622_RNA45SN1_45S_50ntFlanks@11784.11807.24|ATGAATGGATGAACGAGATTCCCA</td><td>>hg38_21_+_8433172_8446622_RNA45SN1_45S_50ntFlanks</td><td>11784</td><td>11807</td><td>24</td><td>ATGAATGGATGAACGAGATTCCCA</td><td>100.1</td><td>-9.7</td><td>1.9</td><td>-5.2</td><td>1.8E-07</td><td>1.2E-06</td><td>BT20/MB468</td><td>MP</td></tr>
<tr><td>>hg38_21_+_8433172_8446622_RNA45SN1_45S_50ntFlanks@8423.8467.45|CGCAGTCCGCCCGGAGGATTCAACCCGGCGGCGGGTCCGGCCGTG</td><td>>hg38_21_+_8433172_8446622_RNA45SN1_45S_50ntFlanks</td><td>8423</td><td>8467</td><td>45</td><td>CGCAGTCCGCCCGGAGGATTCAACCCGGCGGCGGGTCCGGCCGTG</td><td>156.3</td><td>-10.4</td><td>2.0</td><td>-5.2</td><td>1.8E-07</td><td>1.2E-06</td><td>BT20/MB468</td><td>MP</td></tr>
<tr><td>>hg38_21_+_8433172_8446622_RNA45SN1_45S_50ntFlanks@10404.10429.26|CATGGGTCAGTCGGTCCTGAGAGATG</td><td>>hg38_21_+_8433172_8446622_RNA45SN1_45S_50ntFlanks</td><td>10404</td><td>10429</td><td>26</td><td>CATGGGTCAGTCGGTCCTGAGAGATG</td><td>1447.3</td><td>-3.0</td><td>0.6</td><td>-5.0</td><td>7.1E-07</td><td>1.2E-06</td><td>MB231/MB468</td><td>CYTO</td></tr>
<tr><td>>MI0000085|hsa-mir-27a&WithFlank&19|-|13836434|13836523@57.77.21&[MIMAT0000084&hsa-miR-27a-3p&offsets|0|0;m-43&19|-|13836447|13836467&offsets|0|0]|TTCACAGTGGCTAAGTTCCGC</td><td>>MI0000085|hsa-mir-27a&WithFlank&19|-|13836434|13836523</td><td>57</td><td>77</td><td>21</td><td>TTCACAGTGGCTAAGTTCCGC</td><td>3516.2</td><td>5.1</td><td>1.0</td><td>4.9</td><td>7.6E-07</td><td>1.2E-06</td><td>MB231/MB468</td><td>MITO</td></tr>
<tr><td>>trna10_GluCTC_3_-_103879524_103879595@16.35.20,trna116_GluCTC_1_-_145399233_145399304@16.35.20,trna25_GluCTC_2_-_71273488_71273560@16.35.20,trna59_GluCTC_1_+_249168447_249168518@16.35.20,trna71_GluCTC_1_-_161439189_161439260@16.35.20,trna74_GluCTC_1_-_161431809_161431880@16.35.20,trna77_GluCTC_1_-_161424398_161424469@16.35.20,trna77_GluCTC_6_+_28949976_28950047@16.35.20,trna80_GluCTC_1_-_161417018_161417089@16.35.20,trna87_GluCTC_6_-_126101393_126101464@16.35.20|TGGTTAGGATTCGGCGCTCT</td><td>>trna10_GluCTC_3_-_103879524_103879595</td><td>16</td><td>35</td><td>20</td><td>TGGTTAGGATTCGGCGCTCT</td><td>300.9</td><td>7.1</td><td>1.4</td><td>5.0</td><td>6.2E-07</td><td>1.2E-06</td><td>BT20/MB468</td><td>CYTO</td></tr>
<tr><td>>am_trna117_GlyTCC_1_-_145397864_145397935@1.28.28,trna45_GlyTCC_1_+_161500903_161500974@1.28.28,trna70_GlyTCC_1_-_161439547_161439618@1.28.28,trna73_GlyTCC_1_-_161432166_161432237@1.28.28,trna76_GlyTCC_1_-_161424756_161424827@1.28.28,trna79_GlyTCC_1_-_161417375_161417446@1.28.28|GCGTTGGTGGTATAGTGGTGAGCATAGC</td><td>>am_trna117_GlyTCC_1_-_145397864_145397935</td><td>1</td><td>28</td><td>28</td><td>GCGTTGGTGGTATAGTGGTGAGCATAGC</td><td>620.8</td><td>-8.3</td><td>1.7</td><td>-4.9</td><td>8.0E-07</td><td>1.2E-06</td><td>BT20/MB468</td><td>NUC</td></tr>
<tr><td>>trnaMT_SerGCT_MT_+_12207_12265@1.25.25|GAGAAAGCTCACAAGAACTGCTAAC</td><td>>trnaMT_SerGCT_MT_+_12207_12265</td><td>1</td><td>25</td><td>25</td><td>GAGAAAGCTCACAAGAACTGCTAAC</td><td>59796.7</td><td>5.0</td><td>1.0</td><td>4.9</td><td>7.7E-07</td><td>1.2E-06</td><td>MB231/MB468</td><td>MITO</td></tr>
<tr><td>>MI0000746|hsa-mir-99b&WithFlank&19|+|51692606|51692687@13.34.22(+1A)&[MIMAT0000689&hsa-miR-99b-5p&offsets|0|0(+1A);m-10&19|+|51692618|51692639&offsets|0|0(+1A)]|CACCCGTAGAACCGACCTTGCGA</td><td>>MI0000746|hsa-mir-99b&WithFlank&19|+|51692606|51692687</td><td>13</td><td>34</td><td>22(+1A)</td><td>CACCCGTAGAACCGACCTTGCGA</td><td>444.5</td><td>1.9</td><td>0.4</td><td>4.9</td><td>8.0E-07</td><td>1.2E-06</td><td>BT20/MB468</td><td>NUC</td></tr>
<tr><td>>hg38_21_+_8433172_8446622_RNA45SN1_45S_50ntFlanks@9339.9368.30|CGCACGAAAGCCGCCGTGGCGCAATGAAGG</td><td>>hg38_21_+_8433172_8446622_RNA45SN1_45S_50ntFlanks</td><td>9339</td><td>9368</td><td>30</td><td>CGCACGAAAGCCGCCGTGGCGCAATGAAGG</td><td>527.2</td><td>-8.0</td><td>1.5</td><td>-5.2</td><td>1.8E-07</td><td>1.2E-06</td><td>BT20/MB468</td><td>MP</td></tr>
<tr><td>>hg38_21_+_8433172_8446622_RNA45SN1_45S_50ntFlanks@11570.11605.36|AGCCGACTTAGAACTGGTGCGGACCAGGGGAATCCG</td><td>>hg38_21_+_8433172_8446622_RNA45SN1_45S_50ntFlanks</td><td>11570</td><td>11605</td><td>36</td><td>AGCCGACTTAGAACTGGTGCGGACCAGGGGAATCCG</td><td>330.0</td><td>-6.6</td><td>1.3</td><td>-5.0</td><td>7.3E-07</td><td>1.2E-06</td><td>MB231/MB468</td><td>NUC</td></tr>
<tr><td>>MI0025242|hsa-mir-7706&WithFlank&15|+|85380590|85380668@50.72.23&[MIMAT0030021&hsa-miR-7706&offsets|0|-1;m-390&15|+|85380639|85380662&offsets|0|-1]|TGAAGCGCCTGTGCTCTGCCGAG</td><td>>MI0025242|hsa-mir-7706&WithFlank&15|+|85380590|85380668</td><td>50</td><td>72</td><td>23</td><td>TGAAGCGCCTGTGCTCTGCCGAG</td><td>557.4</td><td>7.4</td><td>1.5</td><td>5.0</td><td>6.2E-07</td><td>1.2E-06</td><td>BT20/MB468</td><td>CYTO</td></tr>
<tr><td>>hg38_21_+_8433172_8446622_RNA45SN1_45S_50ntFlanks@10355.10374.20|AACTTTGAAGGCCGAAGTGG</td><td>>hg38_21_+_8433172_8446622_RNA45SN1_45S_50ntFlanks</td><td>10355</td><td>10374</td><td>20</td><td>AACTTTGAAGGCCGAAGTGG</td><td>116.9</td><td>-9.9</td><td>1.9</td><td>-5.2</td><td>1.9E-07</td><td>1.2E-06</td><td>BT20/MB468</td><td>MP</td></tr>
<tr><td>>hg38_21_+_8433172_8446622_RNA45SN1_45S_50ntFlanks@9760.9788.29|AAACTTTAAATGGGTAAGAAGCCCGGCTC</td><td>>hg38_21_+_8433172_8446622_RNA45SN1_45S_50ntFlanks</td><td>9760</td><td>9788</td><td>29</td><td>AAACTTTAAATGGGTAAGAAGCCCGGCTC</td><td>92.5</td><td>-9.6</td><td>1.8</td><td>-5.2</td><td>1.9E-07</td><td>1.2E-06</td><td>BT20/MB468</td><td>MP</td></tr>
<tr><td>>hg38_21_+_8433172_8446622_RNA45SN1_45S_50ntFlanks@11316.11342.27|CACCCCACGTCTCGTCGCGCGCGCGTC</td><td>>hg38_21_+_8433172_8446622_RNA45SN1_45S_50ntFlanks</td><td>11316</td><td>11342</td><td>27</td><td>CACCCCACGTCTCGTCGCGCGCGCGTC</td><td>4322.9</td><td>-7.4</td><td>1.5</td><td>-5.0</td><td>7.2E-07</td><td>1.2E-06</td><td>MB231/MB468</td><td>CYTO</td></tr>
<tr><td>>hg38_21_+_8433172_8446622_RNA45SN1_45S_50ntFlanks@12458.12487.30|AAGCGTTGGATTGTTCACCCACTAATAGGG</td><td>>hg38_21_+_8433172_8446622_RNA45SN1_45S_50ntFlanks</td><td>12458</td><td>12487</td><td>30</td><td>AAGCGTTGGATTGTTCACCCACTAATAGGG</td><td>725.6</td><td>-4.9</td><td>1.0</td><td>-5.0</td><td>7.2E-07</td><td>1.2E-06</td><td>MB231/MB468</td><td>CYTO</td></tr>
<tr><td>>hg38_21_+_8433172_8446622_RNA45SN1_45S_50ntFlanks@11211.11250.40|GCGACTCTGGACGCGAGCCGGGCCCTTCCCGTGGATCGCC</td><td>>hg38_21_+_8433172_8446622_RNA45SN1_45S_50ntFlanks</td><td>11211</td><td>11250</td><td>40</td><td>GCGACTCTGGACGCGAGCCGGGCCCTTCCCGTGGATCGCC</td><td>319.2</td><td>4.0</td><td>0.8</td><td>5.0</td><td>5.5E-07</td><td>1.2E-06</td><td>BT20/MB468</td><td>MITO</td></tr>
<tr><td>>hg38_21_+_8433172_8446622_RNA45SN1_45S_50ntFlanks@12479.12497.19|CTAATAGGGAACGTGAGCT</td><td>>hg38_21_+_8433172_8446622_RNA45SN1_45S_50ntFlanks</td><td>12479</td><td>12497</td><td>19</td><td>CTAATAGGGAACGTGAGCT</td><td>1473.6</td><td>-4.1</td><td>0.8</td><td>-4.9</td><td>8.1E-07</td><td>1.2E-06</td><td>BT20/MB468</td><td>NUC</td></tr>
<tr><td>>hg38_21_+_8433172_8446622_RNA45SN1_45S_50ntFlanks@10690.10710.21|CGTCCGGTGAGCTCTCGCTGG</td><td>>hg38_21_+_8433172_8446622_RNA45SN1_45S_50ntFlanks</td><td>10690</td><td>10710</td><td>21</td><td>CGTCCGGTGAGCTCTCGCTGG</td><td>1213.9</td><td>-8.2</td><td>1.6</td><td>-5.2</td><td>1.9E-07</td><td>1.2E-06</td><td>BT20/MB468</td><td>MP</td></tr>
<tr><td>>hg38_21_+_8433172_8446622_RNA45SN1_45S_50ntFlanks@9505.9527.23|TGAACTATGCCTGGGCAGGGCGA</td><td>>hg38_21_+_8433172_8446622_RNA45SN1_45S_50ntFlanks</td><td>9505</td><td>9527</td><td>23</td><td>TGAACTATGCCTGGGCAGGGCGA</td><td>95.8</td><td>-9.7</td><td>1.9</td><td>-5.2</td><td>1.9E-07</td><td>1.2E-06</td><td>BT20/MB468</td><td>MP</td></tr>
<tr><td>>hg38_21_+_8433172_8446622_RNA45SN1_45S_50ntFlanks@11663.11697.35|ATTTCTGCCCAGTGCTCTGAATGTCAAAGTGAAGA</td><td>>hg38_21_+_8433172_8446622_RNA45SN1_45S_50ntFlanks</td><td>11663</td><td>11697</td><td>35</td><td>ATTTCTGCCCAGTGCTCTGAATGTCAAAGTGAAGA</td><td>73.1</td><td>9.9</td><td>1.9</td><td>5.2</td><td>1.9E-07</td><td>1.2E-06</td><td>BT20/MB468</td><td>MP</td></tr>
<tr><td>>hg38_21_+_8433172_8446622_RNA45SN1_45S_50ntFlanks@9133.9151.19|TCGGGGGGCGGGGCGGACT</td><td>>hg38_21_+_8433172_8446622_RNA45SN1_45S_50ntFlanks</td><td>9133</td><td>9151</td><td>19</td><td>TCGGGGGGCGGGGCGGACT</td><td>79.2</td><td>-9.4</td><td>1.8</td><td>-5.2</td><td>1.9E-07</td><td>1.2E-06</td><td>BT20/MB468</td><td>MP</td></tr>
<tr><td>>hg38_21_+_8433172_8446622_RNA45SN1_45S_50ntFlanks@6653.6685.33|ACTCTTAGCGGTGGATCACTCGGCTCGTGCGTC</td><td>>hg38_21_+_8433172_8446622_RNA45SN1_45S_50ntFlanks</td><td>6653</td><td>6685</td><td>33</td><td>ACTCTTAGCGGTGGATCACTCGGCTCGTGCGTC</td><td>304.6</td><td>4.3</td><td>0.8</td><td>5.2</td><td>1.9E-07</td><td>1.2E-06</td><td>BT20/MB468</td><td>MP</td></tr>
<tr><td>>hg38_21_+_8433172_8446622_RNA45SN1_45S_50ntFlanks@12002.12043.42|TGCGGGCCGCCGGTGAAATACCACTACTCTGATCGTTTTTTC</td><td>>hg38_21_+_8433172_8446622_RNA45SN1_45S_50ntFlanks</td><td>12002</td><td>12043</td><td>42</td><td>TGCGGGCCGCCGGTGAAATACCACTACTCTGATCGTTTTTTC</td><td>617.4</td><td>-7.0</td><td>1.4</td><td>-5.0</td><td>7.2E-07</td><td>1.2E-06</td><td>MB231/MB468</td><td>CYTO</td></tr>
<tr><td>>hg38_21_+_8433172_8446622_RNA45SN1_45S_50ntFlanks@9536.9571.36|GAAACTCTGGTGGAGGTCCGTAGCGGTCCTGACGTG</td><td>>hg38_21_+_8433172_8446622_RNA45SN1_45S_50ntFlanks</td><td>9536</td><td>9571</td><td>36</td><td>GAAACTCTGGTGGAGGTCCGTAGCGGTCCTGACGTG</td><td>155.0</td><td>-5.7</td><td>1.1</td><td>-5.0</td><td>6.4E-07</td><td>1.2E-06</td><td>MB231/MB468</td><td>MP</td></tr>
<tr><td>>MI0000446|hsa-mir-125b-1&WithFlank&11|-|122099751|122099850@60.80.21(+1U)&[MIMAT0004592&hsa-miR-125b-1-3p&offsets|-1|-2(+1U);m-485&11|-|122099769|122099790&offsets|-1|-2(+1U)]|CACGGGTTAGGCTCTTGGGAGT</td><td>>MI0000446|hsa-mir-125b-1&WithFlank&11|-|122099751|122099850</td><td>60</td><td>80</td><td>21(+1U)</td><td>CACGGGTTAGGCTCTTGGGAGT</td><td>124.9</td><td>-10.4</td><td>1.9</td><td>-5.5</td><td>4.2E-08</td><td>1.2E-06</td><td>BT20/MB231</td><td>MP</td></tr>
<tr><td>>MI0003560|hsa-mir-92b&WithFlank&1|+|155195171|155195278@67.88.22(+3U)&[MIMAT0003218&hsa-miR-92b-3p&offsets|0|0(+3U);m-115&1|+|155195237|155195258&offsets|0|0(+3U)]|TATTGCACTCGTCCCGGCCTCCTTT</td><td>>MI0003560|hsa-mir-92b&WithFlank&1|+|155195171|155195278</td><td>67</td><td>88</td><td>22(+3U)</td><td>TATTGCACTCGTCCCGGCCTCCTTT</td><td>63.1</td><td>9.2</td><td>1.8</td><td>5.0</td><td>6.5E-07</td><td>1.2E-06</td><td>MB231/MB468</td><td>MP</td></tr>
<tr><td>>hg38_21_+_8433172_8446622_RNA45SN1_45S_50ntFlanks@12658.12675.18|ACTGAACGCCTCTAAGTC</td><td>>hg38_21_+_8433172_8446622_RNA45SN1_45S_50ntFlanks</td><td>12658</td><td>12675</td><td>18</td><td>ACTGAACGCCTCTAAGTC</td><td>787.0</td><td>-4.9</td><td>1.0</td><td>-5.0</td><td>7.2E-07</td><td>1.2E-06</td><td>MB231/MB468</td><td>CYTO</td></tr>
<tr><td>>hg38_21_+_8433172_8446622_RNA45SN1_45S_50ntFlanks@11670.11704.35|CCCAGTGCTCTGAATGTCAAAGTGAAGAAATTCAA</td><td>>hg38_21_+_8433172_8446622_RNA45SN1_45S_50ntFlanks</td><td>11670</td><td>11704</td><td>35</td><td>CCCAGTGCTCTGAATGTCAAAGTGAAGAAATTCAA</td><td>1521.8</td><td>-8.5</td><td>1.7</td><td>-4.9</td><td>8.1E-07</td><td>1.2E-06</td><td>BT20/MB468</td><td>NUC</td></tr>
<tr><td>>MI0000439|hsa-mir-23b&WithFlank&9|+|95085202|95085310@65.86.22(+1U)&[MIMAT0000418&hsa-miR-23b-3p&offsets|+1|0(+1U);m-42&9|+|95085265|95085285&offsets|+1|+2(+1U)]|TCACATTGCCAGGGATTACCACT</td><td>>MI0000439|hsa-mir-23b&WithFlank&9|+|95085202|95085310</td><td>65</td><td>86</td><td>22(+1U)</td><td>TCACATTGCCAGGGATTACCACT</td><td>76.5</td><td>10.0</td><td>1.9</td><td>5.2</td><td>1.9E-07</td><td>1.2E-06</td><td>BT20/MB468</td><td>MP</td></tr>
<tr><td>>hg38_21_+_8433172_8446622_RNA45SN1_45S_50ntFlanks@9570.9596.27|TGCAAATCGGTCGTCCGACCTGGGTAT</td><td>>hg38_21_+_8433172_8446622_RNA45SN1_45S_50ntFlanks</td><td>9570</td><td>9596</td><td>27</td><td>TGCAAATCGGTCGTCCGACCTGGGTAT</td><td>3439.7</td><td>-8.7</td><td>1.7</td><td>-5.2</td><td>1.9E-07</td><td>1.2E-06</td><td>BT20/MB468</td><td>MP</td></tr>
<tr><td>>hg38_21_+_8433172_8446622_RNA45SN1_45S_50ntFlanks@11573.11592.20|CGACTTAGAACTGGTGCGGA</td><td>>hg38_21_+_8433172_8446622_RNA45SN1_45S_50ntFlanks</td><td>11573</td><td>11592</td><td>20</td><td>CGACTTAGAACTGGTGCGGA</td><td>403.6</td><td>-3.3</td><td>0.7</td><td>-5.0</td><td>6.3E-07</td><td>1.2E-06</td><td>BT20/MB468</td><td>CYTO</td></tr>
<tr><td>>trnaMT_PheGAA_MT_+_577_647@33.56.24|CTGAAAATGTTTAGACGGGCTCAC</td><td>>trnaMT_PheGAA_MT_+_577_647</td><td>33</td><td>56</td><td>24</td><td>CTGAAAATGTTTAGACGGGCTCAC</td><td>87.2</td><td>10.2</td><td>2.0</td><td>5.2</td><td>1.9E-07</td><td>1.2E-06</td><td>BT20/MB468</td><td>MP</td></tr>
<tr><td>>MI0000298|hsa-mir-221&WithFlank&X|-|45746151|45746272@31.55.25(+1C)&[MIMAT0004568&hsa-miR-221-5p&offsets|0|+3(+1C);m-405&X|-|45746220|45746242&offsets|0|+2(+1C)]|ACCTGGCATACAATGTAGATTTCTGC</td><td>>MI0000298|hsa-mir-221&WithFlank&X|-|45746151|45746272</td><td>31</td><td>55</td><td>25(+1C)</td><td>ACCTGGCATACAATGTAGATTTCTGC</td><td>198.2</td><td>7.9</td><td>1.6</td><td>4.9</td><td>7.8E-07</td><td>1.2E-06</td><td>MB231/MB468</td><td>MITO</td></tr>
<tr><td>>hg38_21_+_8433172_8446622_RNA45SN1_45S_50ntFlanks@5494.5514.21|AAGACGGTCGAACTTGACTAT</td><td>>hg38_21_+_8433172_8446622_RNA45SN1_45S_50ntFlanks</td><td>5494</td><td>5514</td><td>21</td><td>AAGACGGTCGAACTTGACTAT</td><td>2147.6</td><td>-8.3</td><td>1.6</td><td>-5.2</td><td>1.9E-07</td><td>1.2E-06</td><td>BT20/MB468</td><td>MP</td></tr>
<tr><td>>MI0000082|hsa-mir-25&WithFlank&7|-|100093554|100093649@58.79.22(+2A)&[MIMAT0000081&hsa-miR-25-3p&offsets|0|0(+2A);m-23&7|-|100093571|100093592&offsets|0|0(+2A)]|CATTGCACTTGTCTCGGTCTGAAA</td><td>>MI0000082|hsa-mir-25&WithFlank&7|-|100093554|100093649</td><td>58</td><td>79</td><td>22(+2A)</td><td>CATTGCACTTGTCTCGGTCTGAAA</td><td>299.6</td><td>5.1</td><td>1.0</td><td>5.0</td><td>7.3E-07</td><td>1.2E-06</td><td>MB231/MB468</td><td>CYTO</td></tr>
<tr><td>>hg38_21_+_8433172_8446622_RNA45SN1_45S_50ntFlanks@10375.10424.50|AGAAGGGTTCCATGTGAACAGCAGTTGAACATGGGTCAGTCGGTCCTGAG</td><td>>hg38_21_+_8433172_8446622_RNA45SN1_45S_50ntFlanks</td><td>10375</td><td>10424</td><td>50</td><td>AGAAGGGTTCCATGTGAACAGCAGTTGAACATGGGTCAGTCGGTCCTGAG</td><td>139.5</td><td>10.5</td><td>1.9</td><td>5.5</td><td>4.2E-08</td><td>1.2E-06</td><td>BT20/MB231</td><td>MP</td></tr>
<tr><td>>MI0000063|hsa-let-7b&WithFlank&22|+|46113680|46113774@12.32.21(+1C)&[MIMAT0000063&hsa-let-7b-5p&offsets|0|-1(+1C);m-14&22|+|46113691|46113712&offsets|0|-1(+1C)]|TGAGGTAGTAGGTTGTGTGGTC</td><td>>MI0000063|hsa-let-7b&WithFlank&22|+|46113680|46113774</td><td>12</td><td>32</td><td>21(+1C)</td><td>TGAGGTAGTAGGTTGTGTGGTC</td><td>7807.4</td><td>5.1</td><td>1.0</td><td>5.2</td><td>1.9E-07</td><td>1.2E-06</td><td>BT20/MB468</td><td>MP</td></tr>
<tr><td>>trnaMT_GluTTC_MT_-_14674_14742@41.72.32,trnalookalike8_GluTTC_5_-_93905172_93905240@41.72.32|TGGTCGTGGTTGTAGTCCGTGCGAGAATACCA</td><td>>trnaMT_GluTTC_MT_-_14674_14742</td><td>41</td><td>72</td><td>32</td><td>TGGTCGTGGTTGTAGTCCGTGCGAGAATACCA</td><td>12315.6</td><td>4.4</td><td>0.9</td><td>5.0</td><td>5.6E-07</td><td>1.2E-06</td><td>BT20/MB468</td><td>MITO</td></tr>
<tr><td>>hg38_21_+_8433172_8446622_RNA45SN1_45S_50ntFlanks@12007.12032.26|GCCGCCGGTGAAATACCACTACTCTG</td><td>>hg38_21_+_8433172_8446622_RNA45SN1_45S_50ntFlanks</td><td>12007</td><td>12032</td><td>26</td><td>GCCGCCGGTGAAATACCACTACTCTG</td><td>9943.5</td><td>-4.6</td><td>0.9</td><td>-5.0</td><td>5.6E-07</td><td>1.2E-06</td><td>BT20/MB468</td><td>MITO</td></tr>
</table>

<table>
<tr><td>>hg38_21_+_8433172_8446622_RNA45SN1_45S_50ntFlanks@6721.6758.38|AATTGCAGGACACATTGATCATCGACACTTCGAACGCA</td><td>>hg38_21_+_8433172_8446622_RNA45SN1_45S_50ntFlanks</td><td>6721</td><td>6758</td><td>38</td><td>AATTGCAGGACACATTGATCATCGACACTTCGAACGCA</td><td>1620.3</td><td>-3.7</td><td>0.7</td><td>-5.0</td><td>5.6E-07</td><td>1.2E-06</td><td>BT20/MB468</td><td>MITO</td></tr>
<tr><td>>MI0000072|hsa-mir-18a&WithFlank&13|+|91350745|91350827@12.34.23&[MIMAT0000072&hsa-miR-18a-5p&offsets|0|0;m-262&13|+|91350756|91350778&offsets|0|0]|TAAGGTGCATCTAGTGCAGATAG</td><td>>MI0000072|hsa-mir-18a&WithFlank&13|+|91350745|91350827</td><td>12</td><td>34</td><td>23</td><td>TAAGGTGCATCTAGTGCAGATAG</td><td>531.2</td><td>5.2</td><td>1.1</td><td>4.9</td><td>7.8E-07</td><td>1.2E-06</td><td>MB231/MB468</td><td>MITO</td></tr>
<tr><td>>MI0000446|hsa-mir-125b-1&WithFlank&11|-|122099751|122099850@21.40.20(+1U);MI0000470|hsa-mir-125b-2&WithFlank&21|+|16590231|16590331@23.42.20(+1U)&[MIMAT0000423&hsa-miR-125b-5p&offsets|0|-2(+1U);m-67&11|-|122099809|122099830&offsets|0|-2(+1U)];[MIMAT0000423_1&hsa-miR-125b-5p&offsets|0|-2(+1U);m-68&21|+|16590253|16590274&offsets|0|-2(+1U)]|TCCCTGAGACCCTAACTTGTT</td><td>>MI0000446|hsa-mir-125b-1&WithFlank&11|-|122099751|122099850</td><td>21</td><td>40</td><td>20(+1U)</td><td>TCCCTGAGACCCTAACTTGTT</td><td>148.3</td><td>-10.6</td><td>1.9</td><td>-5.5</td><td>4.2E-08</td><td>1.2E-06</td><td>BT20/MB231</td><td>MP</td></tr>
<tr><td>>hg38_21_+_8433172_8446622_RNA45SN1_45S_50ntFlanks@3913.3937.25|ATGCGTGCATTTATCAGATCAAAAC</td><td>>hg38_21_+_8433172_8446622_RNA45SN1_45S_50ntFlanks</td><td>3913</td><td>3937</td><td>25</td><td>ATGCGTGCATTTATCAGATCAAAAC</td><td>107.9</td><td>-9.8</td><td>1.9</td><td>-5.2</td><td>1.9E-07</td><td>1.2E-06</td><td>BT20/MB468</td><td>MP</td></tr>
<tr><td>>MI0000082|hsa-mir-25&WithFlank&7|-|100093554|100093649@58.79.22(+1A)&[MIMAT0000081&hsa-miR-25-3p&offsets|0|0(+1A);m-23&7|-|100093571|100093592&offsets|0|0(+1A)]|CATTGCACTTGTCTCGGTCTGAA</td><td>>MI0000082|hsa-mir-25&WithFlank&7|-|100093554|100093649</td><td>58</td><td>79</td><td>22(+1A)</td><td>CATTGCACTTGTCTCGGTCTGAA</td><td>368.7</td><td>4.7</td><td>1.0</td><td>4.9</td><td>7.8E-07</td><td>1.2E-06</td><td>MB231/MB468</td><td>MITO</td></tr>
<tr><td>>hg38_21_+_8433172_8446622_RNA45SN1_45S_50ntFlanks@9563.9596.34|CCTGACGTGCAAATCGGTCGTCCGACCTGGGTAT</td><td>>hg38_21_+_8433172_8446622_RNA45SN1_45S_50ntFlanks</td><td>9563</td><td>9596</td><td>34</td><td>CCTGACGTGCAAATCGGTCGTCCGACCTGGGTAT</td><td>999.9</td><td>-7.0</td><td>1.4</td><td>-5.0</td><td>6.4E-07</td><td>1.2E-06</td><td>BT20/MB468</td><td>CYTO</td></tr>
<tr><td>>trna2_GlyGCC_21_-_18827107_18827177@1.37.37,trna35_GlyGCC_1_+_161413094_161413164@1.37.37,trna37_GlyGCC_1_+_161420467_161420537@1.37.37,trna39_GlyGCC_1_+_161427898_161427968@1.37.37,trna41_GlyGCC_1_+_161435258_161435328@1.37.37|GCATGGGTGGTTCAGTGGTAGAATTCTCGCCTGCCAC</td><td>>trna2_GlyGCC_21_-_18827107_18827177</td><td>1</td><td>37</td><td>37</td><td>GCATGGGTGGTTCAGTGGTAGAATTCTCGCCTGCCAC</td><td>344.5</td><td>8.7</td><td>1.7</td><td>5.0</td><td>6.4E-07</td><td>1.2E-06</td><td>BT20/MB468</td><td>CYTO</td></tr>
<tr><td>>hg38_21_+_8433172_8446622_RNA45SN1_45S_50ntFlanks@10411.10430.20|CAGTCGGTCCTGAGAGATGG</td><td>>hg38_21_+_8433172_8446622_RNA45SN1_45S_50ntFlanks</td><td>10411</td><td>10430</td><td>20</td><td>CAGTCGGTCCTGAGAGATGG</td><td>10593.7</td><td>-8.4</td><td>1.7</td><td>-5.0</td><td>6.4E-07</td><td>1.2E-06</td><td>BT20/MB468</td><td>CYTO</td></tr>
<tr><td>>hg38_21_+_8433172_8446622_RNA45SN1_45S_50ntFlanks@12626.12667.42|AATGGGGCGAAGCTACCATCTGTGGGATTATGACTGAACGCC</td><td>>hg38_21_+_8433172_8446622_RNA45SN1_45S_50ntFlanks</td><td>12626</td><td>12667</td><td>42</td><td>AATGGGGCGAAGCTACCATCTGTGGGATTATGACTGAACGCC</td><td>291.0</td><td>6.0</td><td>1.2</td><td>5.0</td><td>6.4E-07</td><td>1.2E-06</td><td>BT20/MB468</td><td>CYTO</td></tr>
<tr><td>>hg38_21_+_8433172_8446622_RNA45SN1_45S_50ntFlanks@10350.10388.39|ACGAGAACTTTGAAGGCCGAAGTGGAGAAGGGTTCCATG</td><td>>hg38_21_+_8433172_8446622_RNA45SN1_45S_50ntFlanks</td><td>10350</td><td>10388</td><td>39</td><td>ACGAGAACTTTGAAGGCCGAAGTGGAGAAGGGTTCCATG</td><td>2667.0</td><td>-4.4</td><td>0.9</td><td>-5.0</td><td>6.4E-07</td><td>1.2E-06</td><td>BT20/MB468</td><td>CYTO</td></tr>
<tr><td>>hg38_21_+_8433172_8446622_RNA45SN1_45S_50ntFlanks@6661.6679.19|CGGTGGATCACTCGGCTCG</td><td>>hg38_21_+_8433172_8446622_RNA45SN1_45S_50ntFlanks</td><td>6661</td><td>6679</td><td>19</td><td>CGGTGGATCACTCGGCTCG</td><td>788.4</td><td>3.2</td><td>0.6</td><td>5.0</td><td>5.6E-07</td><td>1.2E-06</td><td>BT20/MB468</td><td>MITO</td></tr>
<tr><td>>hg38_21_+_8433172_8446622_RNA45SN1_45S_50ntFlanks@4220.4242.23|ACGAAAAATAACAATACAGGACT</td><td>>hg38_21_+_8433172_8446622_RNA45SN1_45S_50ntFlanks</td><td>4220</td><td>4242</td><td>23</td><td>ACGAAAAATAACAATACAGGACT</td><td>895.7</td><td>-6.7</td><td>1.4</td><td>-4.9</td><td>8.2E-07</td><td>1.2E-06</td><td>BT20/MB468</td><td>NUC</td></tr>
<tr><td>>hg38_21_+_8433172_8446622_RNA45SN1_45S_50ntFlanks@10766.10797.32|CGCAGCAGGTCTCCAAGGTGAACAGCCTCTGG</td><td>>hg38_21_+_8433172_8446622_RNA45SN1_45S_50ntFlanks</td><td>10766</td><td>10797</td><td>32</td><td>CGCAGCAGGTCTCCAAGGTGAACAGCCTCTGG</td><td>79.0</td><td>-8.4</td><td>1.7</td><td>-5.0</td><td>5.6E-07</td><td>1.3E-06</td><td>BT20/MB468</td><td>MITO</td></tr>
<tr><td>>hg38_21_+_8433172_8446622_RNA45SN1_45S_50ntFlanks@10348.10377.30|AAACGAGAACTTTGAAGGCCGAAGTGGAGA</td><td>>hg38_21_+_8433172_8446622_RNA45SN1_45S_50ntFlanks</td><td>10348</td><td>10377</td><td>30</td><td>AAACGAGAACTTTGAAGGCCGAAGTGGAGA</td><td>372.7</td><td>-2.4</td><td>0.5</td><td>-4.9</td><td>7.6E-07</td><td>1.3E-06</td><td>MB231/MB468</td><td>NUC</td></tr>
<tr><td>>hg38_21_+_8433172_8446622_RNA45SN1_45S_50ntFlanks@9343.9367.25|CGAAAGCCGCCGTGGCGCAATGAAG</td><td>>hg38_21_+_8433172_8446622_RNA45SN1_45S_50ntFlanks</td><td>9343</td><td>9367</td><td>25</td><td>CGAAAGCCGCCGTGGCGCAATGAAG</td><td>72.8</td><td>-9.3</td><td>1.8</td><td>-5.2</td><td>1.9E-07</td><td>1.3E-06</td><td>BT20/MB468</td><td>MP</td></tr>
<tr><td>>MI0000081|hsa-mir-24-2&WithFlank&19|-|13836281|13836365@56.76.21(+1A);MI0000080|hsa-mir-24-1&WithFlank&9|+|95086015|95086094@50.70.21(+1A)&[MIMAT0000080&hsa-miR-24-3p&offsets|0|-1(+1A);m-36&19|-|13836288|13836310&offsets|0|-2(+1A)];[MIMAT0000080_1&hsa-miR-24-3p&offsets|0|-1(+1A);m-35&9|+|95086064|95086086&offsets|0|-2(+1A)]|TGGCTCAGTTCAGCAGGAACAA</td><td>>MI0000081|hsa-mir-24-2&WithFlank&19|-|13836281|13836365</td><td>56</td><td>76</td><td>21(+1A)</td><td>TGGCTCAGTTCAGCAGGAACAA</td><td>1267.2</td><td>4.3</td><td>0.9</td><td>4.9</td><td>7.9E-07</td><td>1.3E-06</td><td>MB231/MB468</td><td>MITO</td></tr>
<tr><td>>hg38_21_+_8433172_8446622_RNA45SN1_45S_50ntFlanks@10351.10379.29|CGAGAACTTTGAAGGCCGAAGTGGAGAAG</td><td>>hg38_21_+_8433172_8446622_RNA45SN1_45S_50ntFlanks</td><td>10351</td><td>10379</td><td>29</td><td>CGAGAACTTTGAAGGCCGAAGTGGAGAAG</td><td>674.2</td><td>-3.1</td><td>0.6</td><td>-5.0</td><td>7.4E-07</td><td>1.3E-06</td><td>MB231/MB468</td><td>CYTO</td></tr>
<tr><td>>hg38_21_+_8433172_8446622_RNA45SN1_45S_50ntFlanks@12997.13038.42|CGCTGCGATCTATTGAAAGTCAGCCCTCGACACAAGGGTTTG</td><td>>hg38_21_+_8433172_8446622_RNA45SN1_45S_50ntFlanks</td><td>12997</td><td>13038</td><td>42</td><td>CGCTGCGATCTATTGAAAGTCAGCCCTCGACACAAGGGTTTG</td><td>115.0</td><td>10.6</td><td>2.0</td><td>5.2</td><td>1.9E-07</td><td>1.3E-06</td><td>BT20/MB468</td><td>MP</td></tr>
<tr><td>>hg38_21_+_8433172_8446622_RNA45SN1_45S_50ntFlanks@6646.6676.31|TCGTACGACTCTTAGCGGTGGATCACTCGGC</td><td>>hg38_21_+_8433172_8446622_RNA45SN1_45S_50ntFlanks</td><td>6646</td><td>6676</td><td>31</td><td>TCGTACGACTCTTAGCGGTGGATCACTCGGC</td><td>3189.4</td><td>4.0</td><td>0.8</td><td>5.2</td><td>1.9E-07</td><td>1.3E-06</td><td>BT20/MB468</td><td>MP</td></tr>
<tr><td>>MI0000736|hsa-mir-30c-1&WithFlank&1|+|40757278|40757378@23.45.23(+1A);MI0000254|hsa-mir-30c-2&WithFlank&6|-|71376954|71377037@13.35.23(+1A)&[MIMAT0000244&hsa-miR-30c-5p&offsets|0|0(+1A);m-57&1|+|40757300|40757323&offsets|0|-1(+1A)];[MIMAT0000244_1&hsa-miR-30c-5p&offsets|0|0(+1A);m-56&6|-|71377002|71377025&offsets|0|-1(+1A)]|TGTAAACATCCTACACTCTCAGCA</td><td>>MI0000736|hsa-mir-30c-1&WithFlank&1|+|40757278|40757378</td><td>23</td><td>45</td><td>23(+1A)</td><td>TGTAAACATCCTACACTCTCAGCA</td><td>1300.3</td><td>4.8</td><td>1.0</td><td>5.0</td><td>5.7E-07</td><td>1.3E-06</td><td>BT20/MB468</td><td>MITO</td></tr>
<tr><td>>trna111_HisGTG_1_-_147774845_147774916@1.33.33,trna118_HisGTG_1_-_145396881_145396952@1.33.33,trna16_HisGTG_1_+_146544773_146544844@1.33.33,trna1_HisGTG_15_+_45493349_45493420@1.33.33,trna21_HisGTG_1_+_147753471_147753542@1.33.33,trna33_HisGTG_6_+_27125906_27125977@1.33.33,trna7_HisGTG_9_-_14433938_14434009@1.33.33,trna8_HisGTG_15_-_45492611_45492682@1.33.33,trna9_HisGTG_15_-_45490804_45490875@1.33.33|GCCGTGATCGTATAGTGGTTAGTACTCTGCGTT</td><td>>trna111_HisGTG_1_-_147774845_147774916</td><td>1</td><td>33</td><td>33</td><td>GCCGTGATCGTATAGTGGTTAGTACTCTGCGTT</td><td>724.6</td><td>4.6</td><td>0.9</td><td>4.9</td><td>7.9E-07</td><td>1.3E-06</td><td>MB231/MB468</td><td>MITO</td></tr>
<tr><td>>MI0000650|hsa-mir-200c&WithFlank&12|+|6963693|6963772@50.72.23(+2A)&[MIMAT0000617&hsa-miR-200c-3p&offsets|0|0(+2A);m-54&12|+|6963742|6963764&offsets|0|0(+2A)]|TAATACTGCCGGGTAATGATGGAAA</td><td>>MI0000650|hsa-mir-200c&WithFlank&12|+|6963693|6963772</td><td>50</td><td>72</td><td>23(+2A)</td><td>TAATACTGCCGGGTAATGATGGAAA</td><td>159.9</td><td>10.7</td><td>1.9</td><td>5.5</td><td>4.3E-08</td><td>1.3E-06</td><td>BT20/MB231</td><td>MP</td></tr>
<tr><td>>hg38_21_+_8433172_8446622_RNA45SN1_45S_50ntFlanks@8116.8142.27|GCGGGACATGTGGCGTACGGAAGACCC</td><td>>hg38_21_+_8433172_8446622_RNA45SN1_45S_50ntFlanks</td><td>8116</td><td>8142</td><td>27</td><td>GCGGGACATGTGGCGTACGGAAGACCC</td><td>260.9</td><td>11.4</td><td>2.1</td><td>5.5</td><td>4.3E-08</td><td>1.3E-06</td><td>BT20/MB231</td><td>MP</td></tr>
<tr><td>>hg38_21_+_8433172_8446622_RNA45SN1_45S_50ntFlanks@4772.4793.22|GTAGTTCCGACCATAAACGATG</td><td>>hg38_21_+_8433172_8446622_RNA45SN1_45S_50ntFlanks</td><td>4772</td><td>4793</td><td>22</td><td>GTAGTTCCGACCATAAACGATG</td><td>374.3</td><td>-3.1</td><td>0.6</td><td>-4.9</td><td>7.6E-07</td><td>1.3E-06</td><td>MB231/MB468</td><td>NUC</td></tr>
<tr><td>>hg38_MT_+_1621_3279_RNR2_16S_50ntFlanks@341.373.33|AAATAGTGGGAAGATTTATAGGTAGAGGCGACA</td><td>>hg38_MT_+_1621_3279_RNR2_16S_50ntFlanks</td><td>341</td><td>373</td><td>33</td><td>AAATAGTGGGAAGATTTATAGGTAGAGGCGACA</td><td>49.2</td><td>8.8</td><td>1.8</td><td>5.0</td><td>6.6E-07</td><td>1.3E-06</td><td>MB231/MB468</td><td>MP</td></tr>
<tr><td>>hg38_21_+_8433172_8446622_RNA45SN1_45S_50ntFlanks@12638.12660.23|CTACCATCTGTGGGATTATGACT</td><td>>hg38_21_+_8433172_8446622_RNA45SN1_45S_50ntFlanks</td><td>12638</td><td>12660</td><td>23</td><td>CTACCATCTGTGGGATTATGACT</td><td>65.4</td><td>-9.1</td><td>1.8</td><td>-5.2</td><td>1.9E-07</td><td>1.3E-06</td><td>BT20/MB468</td><td>MP</td></tr>
<tr><td>>hg38_21_+_8433172_8446622_RNA45SN1_45S_50ntFlanks@12386.12414.29|AGCGTTCATAGCGACGTCGCTTTTTGATC</td><td>>hg38_21_+_8433172_8446622_RNA45SN1_45S_50ntFlanks</td><td>12386</td><td>12414</td><td>29</td><td>AGCGTTCATAGCGACGTCGCTTTTTGATC</td><td>374.6</td><td>-6.6</td><td>1.3</td><td>-5.0</td><td>6.5E-07</td><td>1.3E-06</td><td>BT20/MB468</td><td>CYTO</td></tr>
<tr><td>>MI0000081|hsa-mir-24-2&WithFlank&19|-|13836281|13836365@56.77.22;MI0000080|hsa-mir-24-1&WithFlank&9|+|95086015|95086094@50.71.22&[MIMAT0000080&hsa-miR-24-3p&offsets|0|0;m-36&19|-|13836288|13836310&offsets|0|-1];[MIMAT0000080_1&hsa-miR-24-3p&offsets|0|0;m-35&9|+|95086064|95086086&offsets|0|-1]|TGGCTCAGTTCAGCAGGAACAG</td><td>>MI0000081|hsa-mir-24-2&WithFlank&19|-|13836281|13836365</td><td>56</td><td>77</td><td>22</td><td>TGGCTCAGTTCAGCAGGAACAG</td><td>3920.1</td><td>3.5</td><td>0.7</td><td>5.0</td><td>7.4E-07</td><td>1.3E-06</td><td>MB231/MB468</td><td>CYTO</td></tr>
<tr><td>>hg38_21_+_8433172_8446622_RNA45SN1_45S_50ntFlanks@3899.3920.22|CCTTCGCGGGGGGGATGCGTGC</td><td>>hg38_21_+_8433172_8446622_RNA45SN1_45S_50ntFlanks</td><td>3899</td><td>3920</td><td>22</td><td>CCTTCGCGGGGGGGATGCGTGC</td><td>588.9</td><td>-6.8</td><td>1.4</td><td>-5.0</td><td>5.7E-07</td><td>1.3E-06</td><td>BT20/MB468</td><td>MITO</td></tr>
<tr><td>>hg38_21_+_8433172_8446622_RNA45SN1_45S_50ntFlanks@3823.3859.37|TTTGGTCGCTCGCTCCTCTCCTACTTGGATAACTGTG</td><td>>hg38_21_+_8433172_8446622_RNA45SN1_45S_50ntFlanks</td><td>3823</td><td>3859</td><td>37</td><td>TTTGGTCGCTCGCTCCTCTCCTACTTGGATAACTGTG</td><td>262.5</td><td>3.7</td><td>0.8</td><td>5.0</td><td>7.4E-07</td><td>1.3E-06</td><td>MB231/MB468</td><td>CYTO</td></tr>
<tr><td>>hg38_21_+_8433172_8446622_RNA45SN1_45S_50ntFlanks@9533.9563.31|GAGGAAACTCTGGTGGAGGTCCGTAGCGGTC</td><td>>hg38_21_+_8433172_8446622_RNA45SN1_45S_50ntFlanks</td><td>9533</td><td>9563</td><td>31</td><td>GAGGAAACTCTGGTGGAGGTCCGTAGCGGTC</td><td>9741.5</td><td>-9.7</td><td>2.0</td><td>-4.9</td><td>8.3E-07</td><td>1.3E-06</td><td>BT20/MB468</td><td>NUC</td></tr>
<tr><td>>hg38_21_+_8433172_8446622_RNA45SN1_45S_50ntFlanks@9268.9285.18|ACGTCGGCTACCCACCCG</td><td>>hg38_21_+_8433172_8446622_RNA45SN1_45S_50ntFlanks</td><td>9268</td><td>9285</td><td>18</td><td>ACGTCGGCTACCCACCCG</td><td>102.4</td><td>-6.4</td><td>1.3</td><td>-4.9</td><td>7.4E-07</td><td>1.3E-06</td><td>MB231/MB468</td><td>CYTO</td></tr>
<tr><td>>hg38_21_+_8433172_8446622_RNA45SN1_45S_50ntFlanks@10786.10819.34|AACAGCCTCTGGCATGTTGGAACAATGTAGGTAA</td><td>>hg38_21_+_8433172_8446622_RNA45SN1_45S_50ntFlanks</td><td>10786</td><td>10819</td><td>34</td><td>AACAGCCTCTGGCATGTTGGAACAATGTAGGTAA</td><td>284.3</td><td>-6.7</td><td>1.4</td><td>-4.9</td><td>8.0E-07</td><td>1.3E-06</td><td>MB231/MB468</td><td>MITO</td></tr>
<tr><td>>hg38_21_+_8433172_8446622_RNA45SN1_45S_50ntFlanks@4975.5001.27|CCCGGACACGGACAGGATTGACAGATT</td><td>>hg38_21_+_8433172_8446622_RNA45SN1_45S_50ntFlanks</td><td>4975</td><td>5001</td><td>27</td><td>CCCGGACACGGACAGGATTGACAGATT</td><td>134.2</td><td>-5.2</td><td>1.0</td><td>-5.0</td><td>6.6E-07</td><td>1.3E-06</td><td>MB231/MB468</td><td>MP</td></tr>
<tr><td>>trna116_GluCTC_1_-_145399233_145399304@1.33.33,trna59_GluCTC_1_+_249168447_249168518@1.33.33,trna71_GluCTC_1_-_161439189_161439260@1.33.33,trna74_GluCTC_1_-_161431809_161431880@1.33.33,trna77_GluCTC_1_-_161424398_161424469@1.33.33,trna77_GluCTC_6_+_28949976_28950047@1.33.33,trna80_GluCTC_1_-_161417018_161417089@1.33.33,trna87_GluCTC_6_-_126101393_126101464@1.33.33|TCCCTGGTGGTCTAGTGGTTAGGATTCGGCGCT</td><td>>trna116_GluCTC_1_-_145399233_145399304</td><td>1</td><td>33</td><td>33</td><td>TCCCTGGTGGTCTAGTGGTTAGGATTCGGCGCT</td><td>8168.9</td><td>4.2</td><td>0.9</td><td>4.9</td><td>8.0E-07</td><td>1.3E-06</td><td>MB231/MB468</td><td>MITO</td></tr>
<tr><td>>hg38_21_+_8433172_8446622_RNA45SN1_45S_50ntFlanks@10425.10456.32|AGATGGGCGAGCGCCGTTCCGAAGGGACGGGC</td><td>>hg38_21_+_8433172_8446622_RNA45SN1_45S_50ntFlanks</td><td>10425</td><td>10456</td><td>32</td><td>AGATGGGCGAGCGCCGTTCCGAAGGGACGGGC</td><td>197.6</td><td>-7.0</td><td>1.4</td><td>-5.0</td><td>6.6E-07</td><td>1.3E-06</td><td>MB231/MB468</td><td>MP</td></tr>
<tr><td>>hg38_21_+_8433172_8446622_RNA45SN1_45S_50ntFlanks@12408.12431.24|TTTGATCCTTCGATGTCGGCTCTT</td><td>>hg38_21_+_8433172_8446622_RNA45SN1_45S_50ntFlanks</td><td>12408</td><td>12431</td><td>24</td><td>TTTGATCCTTCGATGTCGGCTCTT</td><td>240.2</td><td>-7.1</td><td>1.4</td><td>-5.0</td><td>6.6E-07</td><td>1.3E-06</td><td>MB231/MB468</td><td>MP</td></tr>
<tr><td>>MI0000272|hsa-mir-182&WithFlank&7|-|129770377|129770498@29.51.23(+1C)&[MIMAT0000259&hsa-miR-182-5p&offsets|0|-1(+1C);m-22&7|-|129770449|129770470&offsets|0|+1(+1C)]|TTTGGCAATGGTAGAACTCACACC</td><td>>MI0000272|hsa-mir-182&WithFlank&7|-|129770377|129770498</td><td>29</td><td>51</td><td>23(+1C)</td><td>TTTGGCAATGGTAGAACTCACACC</td><td>785.5</td><td>2.8</td><td>0.6</td><td>4.9</td><td>7.6E-07</td><td>1.3E-06</td><td>MB231/MB468</td><td>NUC</td></tr>
<tr><td>>hg38_21_+_8433172_8446622_RNA45SN1_45S_50ntFlanks@11233.11250.18|CCCTTCCCGTGGATCGCC</td><td>>hg38_21_+_8433172_8446622_RNA45SN1_45S_50ntFlanks</td><td>11233</td><td>11250</td><td>18</td><td>CCCTTCCCGTGGATCGCC</td><td>1244.4</td><td>-7.6</td><td>1.5</td><td>-5.0</td><td>6.5E-07</td><td>1.3E-06</td><td>BT20/MB468</td><td>CYTO</td></tr>
</table>

<table>
<tr><td></td><td></td><td></td><td></td><td></td><td></td><td></td><td></td><td></td><td></td><td></td><td></td><td></td><td></td></tr>
<tr><td>>trna116_GluCTC_1_-_145399233_145399304@32.75.44,trna71_GluCTC_1_-_161439189_161439260@32.75.44,trna74_GluCTC_1_-_161431809_161431880@32.75.44,trna77_GluCTC_1_-_161424398_161424469@32.75.44,trna77_GluCTC_6_+_28949976_28950047@32.75.44,trna80_GluCTC_1_-_161417018_161417089@32.75.44,trna87_GluCTC_6_-_126101393_126101464@32.75.44|CTCTCACCGCCGCGGCCCGGGTTCGATTCCCGGTCAGGGAACCA</td><td>>trna116_GluCTC_1_-_145399233_145399304</td><td>32</td><td>75</td><td>44</td><td>CTCTCACCGCCGCGGCCCGGGTTCGATTCCCGGTCAGGGAACCA</td><td>1079.2</td><td>-5.6</td><td>1.1</td><td>-4.9</td><td>7.5E-07</td><td>1.3E-06</td><td>MB231/MB468</td><td>CYTO</td></tr>
<tr><td>>hg38_21_+_8433172_8446622_RNA45SN1_45S_50ntFlanks@8352.8379.28|AGTTGAAAAGAACTTTGAAGAGAGAGTT</td><td>>hg38_21_+_8433172_8446622_RNA45SN1_45S_50ntFlanks</td><td>8352</td><td>8379</td><td>28</td><td>AGTTGAAAAGAACTTTGAAGAGAGAGTT</td><td>436.7</td><td>-8.0</td><td>1.6</td><td>-4.9</td><td>8.4E-07</td><td>1.3E-06</td><td>BT20/MB468</td><td>NUC</td></tr>
<tr><td>>hg38_21_+_8433172_8446622_RNA45SN1_45S_50ntFlanks@10458.10484.27|ATGGCCTCCGTTGCCCTCGGCCGATCG</td><td>>hg38_21_+_8433172_8446622_RNA45SN1_45S_50ntFlanks</td><td>10458</td><td>10484</td><td>27</td><td>ATGGCCTCCGTTGCCCTCGGCCGATCG</td><td>692.5</td><td>-8.0</td><td>1.6</td><td>-4.9</td><td>8.4E-07</td><td>1.3E-06</td><td>BT20/MB468</td><td>NUC</td></tr>
<tr><td>>hg38_21_+_8433172_8446622_RNA45SN1_45S_50ntFlanks@7974.8018.45|ACGCGACCTCAGATCAGACGTGGCGACCCGCTGAATTTAAGCATA</td><td>>hg38_21_+_8433172_8446622_RNA45SN1_45S_50ntFlanks</td><td>7974</td><td>8018</td><td>45</td><td>ACGCGACCTCAGATCAGACGTGGCGACCCGCTGAATTTAAGCATA</td><td>3540.0</td><td>5.2</td><td>1.0</td><td>5.0</td><td>5.7E-07</td><td>1.3E-06</td><td>BT20/MB468</td><td>MITO</td></tr>
<tr><td>>MI0000650|hsa-mir-200c&WithFlank&12|+|6963693|6963772@50.70.21(+1C)&[MIMAT0000617&hsa-miR-200c-3p&offsets|0|-2(+1C);m-54&12|+|6963742|6963764&offsets|0|-2(+1C)]|TAATACTGCCGGGTAATGATGC</td><td>>MI0000650|hsa-mir-200c&WithFlank&12|+|6963693|6963772</td><td>50</td><td>70</td><td>21(+1C)</td><td>TAATACTGCCGGGTAATGATGC</td><td>132.7</td><td>10.4</td><td>1.9</td><td>5.5</td><td>4.4E-08</td><td>1.3E-06</td><td>BT20/MB231</td><td>MP</td></tr>
<tr><td>>hg38_21_+_8433172_8446622_RNA45SN1_45S_50ntFlanks@10016.10033.18|AATGGATGGCGCTGGAGC</td><td>>hg38_21_+_8433172_8446622_RNA45SN1_45S_50ntFlanks</td><td>10016</td><td>10033</td><td>18</td><td>AATGGATGGCGCTGGAGC</td><td>508.5</td><td>12.3</td><td>2.3</td><td>5.5</td><td>4.4E-08</td><td>1.3E-06</td><td>BT20/MB231</td><td>MP</td></tr>
<tr><td>>hg38_21_+_8433172_8446622_RNA45SN1_45S_50ntFlanks@8113.8141.29|GGCGCGGGACATGTGGCGTACGGAAGACC</td><td>>hg38_21_+_8433172_8446622_RNA45SN1_45S_50ntFlanks</td><td>8113</td><td>8141</td><td>29</td><td>GGCGCGGGACATGTGGCGTACGGAAGACC</td><td>292.6</td><td>11.5</td><td>2.1</td><td>5.5</td><td>4.4E-08</td><td>1.3E-06</td><td>BT20/MB231</td><td>MP</td></tr>
<tr><td>>hg38_21_+_8433172_8446622_RNA45SN1_45S_50ntFlanks@7977.7995.19|CGACCTCAGATCAGACGTG</td><td>>hg38_21_+_8433172_8446622_RNA45SN1_45S_50ntFlanks</td><td>7977</td><td>7995</td><td>19</td><td>CGACCTCAGATCAGACGTG</td><td>474.2</td><td>-7.7</td><td>1.6</td><td>-4.9</td><td>8.4E-07</td><td>1.3E-06</td><td>BT20/MB468</td><td>NUC</td></tr>
<tr><td>>hg38_21_+_8433172_8446622_RNA45SN1_45S_50ntFlanks@11578.11605.28|TAGAACTGGTGCGGACCAGGGGAATCCG</td><td>>hg38_21_+_8433172_8446622_RNA45SN1_45S_50ntFlanks</td><td>11578</td><td>11605</td><td>28</td><td>TAGAACTGGTGCGGACCAGGGGAATCCG</td><td>1159.1</td><td>-8.3</td><td>1.7</td><td>-4.9</td><td>8.4E-07</td><td>1.3E-06</td><td>BT20/MB468</td><td>NUC</td></tr>
<tr><td>>hg38_21_+_8433172_8446622_RNA45SN1_45S_50ntFlanks@10391.10431.41|AACAGCAGTTGAACATGGGTCAGTCGGTCCTGAGAGATGGG</td><td>>hg38_21_+_8433172_8446622_RNA45SN1_45S_50ntFlanks</td><td>10391</td><td>10431</td><td>41</td><td>AACAGCAGTTGAACATGGGTCAGTCGGTCCTGAGAGATGGG</td><td>216.5</td><td>-4.4</td><td>0.9</td><td>-4.9</td><td>7.5E-07</td><td>1.3E-06</td><td>MB231/MB468</td><td>CYTO</td></tr>
<tr><td>>hg38_21_+_8433172_8446622_RNA45SN1_45S_50ntFlanks@7976.8009.34|GCGACCTCAGATCAGACGTGGCGACCCGCTGAAT</td><td>>hg38_21_+_8433172_8446622_RNA45SN1_45S_50ntFlanks</td><td>7976</td><td>8009</td><td>34</td><td>GCGACCTCAGATCAGACGTGGCGACCCGCTGAAT</td><td>1769.4</td><td>6.1</td><td>1.2</td><td>5.2</td><td>2.0E-07</td><td>1.3E-06</td><td>BT20/MB468</td><td>MP</td></tr>
<tr><td>>MI0000477|hsa-mir-146a&WithFlank&5|+|160485346|160485456@27.45.19&[MIMAT0000449&hsa-miR-146a-5p&offsets|0|-3;m-129&5|+|160485372|160485393&offsets|0|-3]|TGAGAACTGAATTCCATGG</td><td>>MI0000477|hsa-mir-146a&WithFlank&5|+|160485346|160485456</td><td>27</td><td>45</td><td>19</td><td>TGAGAACTGAATTCCATGG</td><td>60.8</td><td>9.1</td><td>1.8</td><td>5.0</td><td>6.7E-07</td><td>1.3E-06</td><td>MB231/MB468</td><td>MP</td></tr>
<tr><td>>trna116_GluCTC_1_-_145399233_145399304@4.35.32,trna59_GluCTC_1_+_249168447_249168518@4.35.32,trna71_GluCTC_1_-_161439189_161439260@4.35.32,trna74_GluCTC_1_-_161431809_161431880@4.35.32,trna77_GluCTC_1_-_161424398_161424469@4.35.32,trna77_GluCTC_6_+_28949976_28950047@4.35.32,trna80_GluCTC_1_-_161417018_161417089@4.35.32,trna87_GluCTC_6_-_126101393_126101464@4.35.32|CTGGTGGTCTAGTGGTTAGGATTCGGCGCTCT</td><td>>trna116_GluCTC_1_-_145399233_145399304</td><td>4</td><td>35</td><td>32</td><td>CTGGTGGTCTAGTGGTTAGGATTCGGCGCTCT</td><td>198.8</td><td>4.8</td><td>1.0</td><td>4.9</td><td>8.1E-07</td><td>1.3E-06</td><td>MB231/MB468</td><td>MITO</td></tr>
<tr><td>>hg38_21_+_8433172_8446622_RNA45SN1_45S_50ntFlanks@9545.9569.25|GTGGAGGTCCGTAGCGGTCCTGACG</td><td>>hg38_21_+_8433172_8446622_RNA45SN1_45S_50ntFlanks</td><td>9545</td><td>9569</td><td>25</td><td>GTGGAGGTCCGTAGCGGTCCTGACG</td><td>228.3</td><td>-3.3</td><td>0.7</td><td>-4.9</td><td>7.7E-07</td><td>1.3E-06</td><td>MB231/MB468</td><td>NUC</td></tr>
<tr><td>>hg38_21_+_8433172_8446622_RNA45SN1_45S_50ntFlanks@12907.12933.27|CACCGCACGTTCGTGGGGAACCTGGCG</td><td>>hg38_21_+_8433172_8446622_RNA45SN1_45S_50ntFlanks</td><td>12907</td><td>12933</td><td>27</td><td>CACCGCACGTTCGTGGGGAACCTGGCG</td><td>1115.7</td><td>-6.5</td><td>1.3</td><td>-4.9</td><td>7.7E-07</td><td>1.3E-06</td><td>MB231/MB468</td><td>NUC</td></tr>
<tr><td>>MI0000103|hsa-mir-101-1&WithFlank&1|-|65058428|65058514@52.73.22(+1A);MI0000739|hsa-mir-101-2&WithFlank&9|+|4850291|4850381@54.75.22(+1A)&[MIMAT0000099&hsa-miR-101-3p&offsets|-1|0(+1A);m-17&1|-|65058443|65058462&offsets|-1|+1(+1A)];[MIMAT0000099_1&hsa-miR-101-3p&offsets|-1|0(+1A);m-16&9|+|4850345|4850364&offsets|-1|+1(+1A)]|GTACAGTACTGTGATAACTGAAA</td><td>>MI0000103|hsa-mir-101-1&WithFlank&1|-|65058428|65058514</td><td>52</td><td>73</td><td>22(+1A)</td><td>GTACAGTACTGTGATAACTGAAA</td><td>226.5</td><td>5.2</td><td>1.1</td><td>4.9</td><td>7.7E-07</td><td>1.3E-06</td><td>MB231/MB468</td><td>NUC</td></tr>
<tr><td>>hg38_21_+_8433172_8446622_RNA45SN1_45S_50ntFlanks@9554.9577.24|CGTAGCGGTCCTGACGTGCAAATC</td><td>>hg38_21_+_8433172_8446622_RNA45SN1_45S_50ntFlanks</td><td>9554</td><td>9577</td><td>24</td><td>CGTAGCGGTCCTGACGTGCAAATC</td><td>2939.5</td><td>2.8</td><td>0.6</td><td>4.9</td><td>7.5E-07</td><td>1.3E-06</td><td>MB231/MB468</td><td>CYTO</td></tr>
<tr><td>>MI0003137|hsa-mir-193b&WithFlank&16|+|14303961|14304055@57.77.21(+1C)&[MIMAT0002819&hsa-miR-193b-3p&offsets|0|-1(+1C);m-166&16|+|14304017|14304038&offsets|0|-1(+1C)]|AACTGGCCCTCAAAGTCCCGCC</td><td>>MI0003137|hsa-mir-193b&WithFlank&16|+|14303961|14304055</td><td>57</td><td>77</td><td>21(+1C)</td><td>AACTGGCCCTCAAAGTCCCGCC</td><td>398.9</td><td>4.5</td><td>0.8</td><td>5.4</td><td>5.4E-08</td><td>1.3E-06</td><td>BT20/MB231</td><td>CYTO</td></tr>
<tr><td>>trna134_GluTTC_1_-_16861774_16861845@1.34.34,trna5_GluTTC_1_+_17199078_17199149@1.34.34,trna84_GluTTC_1_-_161391883_161391954@1.34.34,trna94_GluTTC_1_-_149664355_149664427@1.34.34|TCCCTGGTGGTCTAGTGGCTAGGATTCGGCGCTT</td><td>>trna134_GluTTC_1_-_16861774_16861845</td><td>1</td><td>34</td><td>34</td><td>TCCCTGGTGGTCTAGTGGCTAGGATTCGGCGCTT</td><td>531.4</td><td>4.5</td><td>0.8</td><td>5.4</td><td>5.4E-08</td><td>1.3E-06</td><td>BT20/MB231</td><td>CYTO</td></tr>
<tr><td>>hg38_21_+_8433172_8446622_RNA45SN1_45S_50ntFlanks@6722.6746.25|ATTGCAGGACACATTGATCATCGAC</td><td>>hg38_21_+_8433172_8446622_RNA45SN1_45S_50ntFlanks</td><td>6722</td><td>6746</td><td>25</td><td>ATTGCAGGACACATTGATCATCGAC</td><td>212.3</td><td>-3.9</td><td>0.8</td><td>-4.9</td><td>7.5E-07</td><td>1.3E-06</td><td>MB231/MB468</td><td>CYTO</td></tr>
<tr><td>>hg38_21_+_8433172_8446622_RNA45SN1_45S_50ntFlanks@10461.10497.37|GCCTCCGTTGCCCTCGGCCGATCGAAAGGGAGTCGGG</td><td>>hg38_21_+_8433172_8446622_RNA45SN1_45S_50ntFlanks</td><td>10461</td><td>10497</td><td>37</td><td>GCCTCCGTTGCCCTCGGCCGATCGAAAGGGAGTCGGG</td><td>152.8</td><td>-5.9</td><td>1.2</td><td>-5.0</td><td>6.6E-07</td><td>1.3E-06</td><td>BT20/MB468</td><td>CYTO</td></tr>
<tr><td>>MI0000086|hsa-mir-28&WithFlank&3|+|188688775|188688872@60.80.21&[MIMAT0004502&hsa-miR-28-3p&offsets|0|-1;m-28&3|+|188688834|188688855&offsets|0|-1]|CACTAGATTGTGAGCTCCTGG</td><td>>MI0000086|hsa-mir-28&WithFlank&3|+|188688775|188688872</td><td>60</td><td>80</td><td>21</td><td>CACTAGATTGTGAGCTCCTGG</td><td>529.0</td><td>8.0</td><td>1.6</td><td>4.9</td><td>8.1E-07</td><td>1.3E-06</td><td>MB231/MB468</td><td>MITO</td></tr>
<tr><td>>hg38_21_+_8433172_8446622_RNA45SN1_45S_50ntFlanks@6654.6689.36|CTCTTAGCGGTGGATCACTCGGCTCGTGCGTCGATG</td><td>>hg38_21_+_8433172_8446622_RNA45SN1_45S_50ntFlanks</td><td>6654</td><td>6689</td><td>36</td><td>CTCTTAGCGGTGGATCACTCGGCTCGTGCGTCGATG</td><td>456.7</td><td>-6.1</td><td>1.2</td><td>-5.0</td><td>6.7E-07</td><td>1.3E-06</td><td>MB231/MB468</td><td>MP</td></tr>
<tr><td>>MI0000063|hsa-let-7b&WithFlank&22|+|46113680|46113774@12.33.22(+1A)&[MIMAT0000063&hsa-let-7b-5p&offsets|0|0(+1A);m-14&22|+|46113691|46113712&offsets|0|0(+1A)]|TGAGGTAGTAGGTTGTGTGGTTA</td><td>>MI0000063|hsa-let-7b&WithFlank&22|+|46113680|46113774</td><td>12</td><td>33</td><td>22(+1A)</td><td>TGAGGTAGTAGGTTGTGTGGTTA</td><td>6545.5</td><td>5.5</td><td>1.1</td><td>5.2</td><td>2.0E-07</td><td>1.3E-06</td><td>BT20/MB468</td><td>MP</td></tr>
<tr><td>>hg38_21_+_8433172_8446622_RNA45SN1_45S_50ntFlanks@8114.8140.27|GCGCGGGACATGTGGCGTACGGAAGAC</td><td>>hg38_21_+_8433172_8446622_RNA45SN1_45S_50ntFlanks</td><td>8114</td><td>8140</td><td>27</td><td>GCGCGGGACATGTGGCGTACGGAAGAC</td><td>10089.8</td><td>-5.1</td><td>1.0</td><td>-5.2</td><td>2.0E-07</td><td>1.3E-06</td><td>BT20/MB468</td><td>MP</td></tr>
<tr><td>>hg38_21_+_8433172_8446622_RNA45SN1_45S_50ntFlanks@9032.9060.29|CCTCGTGGGGGGGCCGGGCCACCCCTCCC</td><td>>hg38_21_+_8433172_8446622_RNA45SN1_45S_50ntFlanks</td><td>9032</td><td>9060</td><td>29</td><td>CCTCGTGGGGGGGCCGGGCCACCCCTCCC</td><td>65.2</td><td>-9.1</td><td>1.8</td><td>-5.2</td><td>2.0E-07</td><td>1.3E-06</td><td>BT20/MB468</td><td>MP</td></tr>
<tr><td>>hg38_21_+_8433172_8446622_RNA45SN1_45S_50ntFlanks@10389.10433.45|TGAACAGCAGTTGAACATGGGTCAGTCGGTCCTGAGAGATGGGCG</td><td>>hg38_21_+_8433172_8446622_RNA45SN1_45S_50ntFlanks</td><td>10389</td><td>10433</td><td>45</td><td>TGAACAGCAGTTGAACATGGGTCAGTCGGTCCTGAGAGATGGGCG</td><td>274.4</td><td>-7.8</td><td>1.6</td><td>-4.9</td><td>7.7E-07</td><td>1.3E-06</td><td>MB231/MB468</td><td>NUC</td></tr>
<tr><td>>hg38_21_+_8433172_8446622_RNA45SN1_45S_50ntFlanks@4724.4765.42|ATTAATCAAGAACGAAAGTCGGAGGTTCGAAGACGATCAGAT</td><td>>hg38_21_+_8433172_8446622_RNA45SN1_45S_50ntFlanks</td><td>4724</td><td>4765</td><td>42</td><td>ATTAATCAAGAACGAAAGTCGGAGGTTCGAAGACGATCAGAT</td><td>497.6</td><td>5.3</td><td>1.1</td><td>4.9</td><td>7.5E-07</td><td>1.3E-06</td><td>MB231/MB468</td><td>CYTO</td></tr>
<tr><td>>hg38_21_+_8433172_8446622_RNA45SN1_45S_50ntFlanks@5074.5094.21|ATTCCGATAACGAACGAGACT</td><td>>hg38_21_+_8433172_8446622_RNA45SN1_45S_50ntFlanks</td><td>5074</td><td>5094</td><td>21</td><td>ATTCCGATAACGAACGAGACT</td><td>368.1</td><td>-8.0</td><td>1.6</td><td>-5.0</td><td>6.7E-07</td><td>1.3E-06</td><td>MB231/MB468</td><td>MP</td></tr>
<tr><td>>hg38_21_+_8433172_8446622_RNA45SN1_45S_50ntFlanks@10392.10428.37|ACAGCAGTTGAACATGGGTCAGTCGGTCCTGAGAGAT</td><td>>hg38_21_+_8433172_8446622_RNA45SN1_45S_50ntFlanks</td><td>10392</td><td>10428</td><td>37</td><td>ACAGCAGTTGAACATGGGTCAGTCGGTCCTGAGAGAT</td><td>757.9</td><td>-8.1</td><td>1.7</td><td>-4.9</td><td>8.5E-07</td><td>1.3E-06</td><td>BT20/MB468</td><td>NUC</td></tr>
<tr><td>>hg38_21_+_8433172_8446622_RNA45SN1_45S_50ntFlanks@6758.6806.49|ACTTGCGGCCCCGGGTTCCTCCCGGGGCTACGCCTGTCTGAGCGTCGCT</td><td>>hg38_21_+_8433172_8446622_RNA45SN1_45S_50ntFlanks</td><td>6758</td><td>6806</td><td>49</td><td>ACTTGCGGCCCCGGGTTCCTCCCGGGGCTACGCCTGTCTGAGCGTCGCT</td><td>1075.7</td><td>6.5</td><td>1.3</td><td>5.0</td><td>5.8E-07</td><td>1.3E-06</td><td>BT20/MB468</td><td>MITO</td></tr>
<tr><td>>hg38_21_+_8433172_8446622_RNA45SN1_45S_50ntFlanks@11200.11227.28|CGGCGGCGGCGGCGACTCTGGACGCGAG</td><td>>hg38_21_+_8433172_8446622_RNA45SN1_45S_50ntFlanks</td><td>11200</td><td>11227</td><td>28</td><td>CGGCGGCGGCGGCGACTCTGGACGCGAG</td><td>323.5</td><td>-4.6</td><td>0.9</td><td>-5.0</td><td>6.6E-07</td><td>1.3E-06</td><td>BT20/MB468</td><td>CYTO</td></tr>
<tr><td>>hg38_21_+_8433172_8446622_RNA45SN1_45S_50ntFlanks@10873.10892.20|CTGGGTCGGTCGGGCTGGGG</td><td>>hg38_21_+_8433172_8446622_RNA45SN1_45S_50ntFlanks</td><td>10873</td><td>10892</td><td>20</td><td>CTGGGTCGGTCGGGCTGGGG</td><td>1800.7</td><td>-8.6</td><td>1.8</td><td>-4.9</td><td>8.5E-07</td><td>1.3E-06</td><td>BT20/MB468</td><td>NUC</td></tr>
<tr><td>>hg38_21_+_8433172_8446622_RNA45SN1_45S_50ntFlanks@4738.4761.24|AAAGTCGGAGGTTCGAAGACGATC</td><td>>hg38_21_+_8433172_8446622_RNA45SN1_45S_50ntFlanks</td><td>4738</td><td>4761</td><td>24</td><td>AAAGTCGGAGGTTCGAAGACGATC</td><td>451.4</td><td>-2.6</td><td>0.5</td><td>-4.9</td><td>8.5E-07</td><td>1.3E-06</td><td>BT20/MB468</td><td>NUC</td></tr>
<tr><td>>hg38_MT_+_598_1651_RNR1_12S_50ntFlanks@416.455.40|AAATAGACTACGAAAGTGGCTTTAACATATCTGAACACAC</td><td>>hg38_MT_+_598_1651_RNR1_12S_50ntFlanks</td><td>416</td><td>455</td><td>40</td><td>AAATAGACTACGAAAGTGGCTTTAACATATCTGAACACAC</td><td>163.2</td><td>11.1</td><td>2.1</td><td>5.2</td><td>2.0E-07</td><td>1.3E-06</td><td>BT20/MB468</td><td>MP</td></tr>
<tr><td>>hg38_21_+_8433172_8446622_RNA45SN1_45S_50ntFlanks@4003.4023.21|ATAACCTCGGGCCGATCGCAC</td><td>>hg38_21_+_8433172_8446622_RNA45SN1_45S_50ntFlanks</td><td>4003</td><td>4023</td><td>21</td><td>ATAACCTCGGGCCGATCGCAC</td><td>1073.1</td><td>-5.7</td><td>1.2</td><td>-4.9</td><td>8.1E-07</td><td>1.3E-06</td><td>MB231/MB468</td><td>MITO</td></tr>
<tr><td>>hg38_21_+_8433172_8446622_RNA45SN1_45S_50ntFlanks@12941.12966.26|ATTCGTAGACGACCTGCTTCTGGGTC</td><td>>hg38_21_+_8433172_8446622_RNA45SN1_45S_50ntFlanks</td><td>12941</td><td>12966</td><td>26</td><td>ATTCGTAGACGACCTGCTTCTGGGTC</td><td>534.6</td><td>-7.7</td><td>1.5</td><td>-5.2</td><td>2.0E-07</td><td>1.3E-06</td><td>BT20/MB468</td><td>MP</td></tr>
<tr><td>>MI0000542|hsa-mir-320a&WithFlank&8|-|22244960|22245043@50.70.21(+1G)&[MIMAT0000510&hsa-miR-320a-3p&offsets|+2|+1(+1G);m-60&8|-|22244975|22244996&offsets|+2|+1(+1G)]|AAGCTGGGTTGAGAGGGCGAAG</td><td>>MI0000542|hsa-mir-320a&WithFlank&8|-|22244960|22245043</td><td>50</td><td>70</td><td>21(+1G)</td><td>AAGCTGGGTTGAGAGGGCGAAG</td><td>79.7</td><td>10.1</td><td>1.9</td><td>5.2</td><td>2.0E-07</td><td>1.3E-06</td><td>BT20/MB468</td><td>MP</td></tr>
<tr><td>>hg38_21_+_8433172_8446622_RNA45SN1_45S_50ntFlanks@3850.3871.22|GATAACTGTGGTAATTCTAGAG</td><td>>hg38_21_+_8433172_8446622_RNA45SN1_45S_50ntFlanks</td><td>3850</td><td>3871</td><td>22</td><td>GATAACTGTGGTAATTCTAGAG</td><td>273.9</td><td>-6.9</td><td>1.4</td><td>-5.0</td><td>6.8E-07</td><td>1.3E-06</td><td>MB231/MB468</td><td>MP</td></tr>
<tr><td>>hg38_21_+_8433172_8446622_RNA45SN1_45S_50ntFlanks@12418.12457.40|CGATGTCGGCTCTTCCTATCATTGTGAAGCAGAATTCACC</td><td>>hg38_21_+_8433172_8446622_RNA45SN1_45S_50ntFlanks</td><td>12418</td><td>12457</td><td>40</td><td>CGATGTCGGCTCTTCCTATCATTGTGAAGCAGAATTCACC</td><td>987.0</td><td>3.8</td><td>0.8</td><td>4.9</td><td>7.6E-07</td><td>1.3E-06</td><td>MB231/MB468</td><td>CYTO</td></tr>
<tr><td>>hg38_21_+_8433172_8446622_RNA45SN1_45S_50ntFlanks@10389.10424.36|TGAACAGCAGTTGAACATGGGTCAGTCGGTCCTGAG</td><td>>hg38_21_+_8433172_8446622_RNA45SN1_45S_50ntFlanks</td><td>10389</td><td>10424</td><td>36</td><td>TGAACAGCAGTTGAACATGGGTCAGTCGGTCCTGAG</td><td>4565.2</td><td>-3.2</td><td>0.7</td><td>-4.9</td><td>8.5E-07</td><td>1.3E-06</td><td>BT20/MB468</td><td>NUC</td></tr>
<tr><td>>hg38_21_+_8433172_8446622_RNA45SN1_45S_50ntFlanks@12639.12659.21|TACCATCTGTGGGATTATGAC</td><td>>hg38_21_+_8433172_8446622_RNA45SN1_45S_50ntFlanks</td><td>12639</td><td>12659</td><td>21</td><td>TACCATCTGTGGGATTATGAC</td><td>282.1</td><td>-7.3</td><td>1.5</td><td>-5.0</td><td>6.8E-07</td><td>1.3E-06</td><td>MB231/MB468</td><td>MP</td></tr>
</table>

| | | | | | | | | | | | | | |
|---|---|---|---|---|---|---|---|---|---|---|---|---|---|
| >hg38_21_+_8433172_8446622_RNA45SN1_45S_50ntFlanks@12416.12436.21\|TTCGATGTCGGCTCTTCCTAT | >hg38_21_+_8433172_8446622_RNA45SN1_45S_50ntFlanks | 12416 | 12436 | 21 | TTCGATGTCGGCTCTTCCTAT | 1418.8 | -3.6 | 0.7 | -4.9 | 8.2E-07 | 1.3E-06 | MB231/MB468 | MITO |
| >hg38_21_+_8433172_8446622_RNA45SN1_45S_50ntFlanks@3853.3873.21\|AACTGTGGTAATTCTAGAGCT | >hg38_21_+_8433172_8446622_RNA45SN1_45S_50ntFlanks | 3853 | 3873 | 21 | AACTGTGGTAATTCTAGAGCT | 773.9 | -5.6 | 1.1 | -5.0 | 6.8E-07 | 1.3E-06 | MB231/MB468 | MP |
| >hg38_21_+_8433172_8446622_RNA45SN1_45S_50ntFlanks@8176.8206.31\|CTTCTGATCGAGGCCCAGCCCGTGGACGGTG | >hg38_21_+_8433172_8446622_RNA45SN1_45S_50ntFlanks | 8176 | 8206 | 31 | CTTCTGATCGAGGCCCAGCCCGTGGACGGTG | 949.9 | -3.8 | 0.7 | -5.2 | 2.0E-07 | 1.3E-06 | BT20/MB468 | MP |
| >hg38_21_+_8433172_8446622_RNA45SN1_45S_50ntFlanks@8358.8394.37\|AAAGAACTTTGAAGAGAGAGTTCAAGAGGGCGTGAAA | >hg38_21_+_8433172_8446622_RNA45SN1_45S_50ntFlanks | 8358 | 8394 | 37 | AAAGAACTTTGAAGAGAGAGTTCAAGAGGGCGTGAAA | 55.3 | 9.5 | 1.8 | 5.2 | 2.0E-07 | 1.3E-06 | BT20/MB468 | MP |
| >hg38_21_+_8433172_8446622_RNA45SN1_45S_50ntFlanks@9808.9830.23\|TGGAATGCGAGTGCCTAGTGGGC | >hg38_21_+_8433172_8446622_RNA45SN1_45S_50ntFlanks | 9808 | 9830 | 23 | TGGAATGCGAGTGCCTAGTGGGC | 256.7 | 11.3 | 2.1 | 5.5 | 4.5E-08 | 1.3E-06 | BT20/MB231 | MP |
| >trna32_LysCTT_16_-_3207406_3207478@4.33.30\|CGGCTAGCTCAGTCGGTAGAGCATGAGACC | >trna32_LysCTT_16_-_3207406_3207478 | 4 | 33 | 30 | CGGCTAGCTCAGTCGGTAGAGCATGAGACC | 150.6 | -10.6 | 1.9 | -5.5 | 4.5E-08 | 1.3E-06 | BT20/MB231 | MP |
| >am_trna10_ValCAC_5_-_180649395_180649467@2.32.31,trna12_ValAAC_5_-_180645270_180645342@2.32.31,trna132_ValAAC_6_-_27721179_27721251@2.32.31,trna136_ValAAC_6_-_27648885_27648957@2.32.31,trna139_ValAAC_6_-_27618707_27618779@2.32.31,trna18_ValCAC_5_-_180529253_180529325@2.32.31,trna2_ValAAC_3_+_169490018_169490090@2.32.31,trna2_ValCAC_5_+_180524070_180524142@2.32.31,trna4_ValAAC_5_+_180591154_180591226@2.32.31,trna5_ValAAC_5_+_180596610_180596682@2.32.31,trna6_ValCAC_5_+_180600650_180600722@2.32.31,trna85_ValCAC_1_-_161369490_161369562@2.32.31,trna90_ValCAC_1_-_149684088_149684161@2.32.31,trna98_ValCAC_1_-_149298555_149298627@2.32.31,trna9_ValCAC_6_+_26538282_26538354@2.32.31\|TTTCCGTAGTGTAGTGGTTATCACGTTCGCC | >am_trna10_ValCAC_5_-_180649395_180649467 | 2 | 32 | 31 | TTTCCGTAGTGTAGTGGTTATCACGTTCGCC | 71.1 | 9.4 | 1.9 | 5.0 | 6.8E-07 | 1.3E-06 | MB231/MB468 | MP |
| >MI0000272\|hsa-mir-182&WithFlank&7\|-\|129770377\|129770498@29.54.26&[MIMAT0000259&hsa-miR-182-5p&offsets\|0\|+2;m-22&7\|-\|129770449\|129770470&offsets\|0\|+4]\|TTTGGCAATGGTAGAACTCACACTGG | >MI0000272\|hsa-mir-182&WithFlank&7\|-\|129770377\|129770498 | 29 | 54 | 26 | TTTGGCAATGGTAGAACTCACACTGG | 943.7 | 2.9 | 0.6 | 4.9 | 7.9E-07 | 1.3E-06 | MB231/MB468 | NUC |
| >hg38_21_+_8433172_8446622_RNA45SN1_45S_50ntFlanks@10392.10411.20\|ACAGCAGTTGAACATGGGTC | >hg38_21_+_8433172_8446622_RNA45SN1_45S_50ntFlanks | 10392 | 10411 | 20 | ACAGCAGTTGAACATGGGTC | 284.4 | -5.6 | 1.1 | -5.0 | 6.8E-07 | 1.3E-06 | MB231/MB468 | MP |
| >hg38_21_+_8433172_8446622_RNA45SN1_45S_50ntFlanks@12408.12440.33\|TTTGATCCTTCGATGTCGGCTCTTCCTATCATT | >hg38_21_+_8433172_8446622_RNA45SN1_45S_50ntFlanks | 12408 | 12440 | 33 | TTTGATCCTTCGATGTCGGCTCTTCCTATCATT | 206.1 | 5.4 | 1.1 | 4.9 | 7.7E-07 | 1.3E-06 | MB231/MB468 | CYTO |
| >hg38_MT_+_1621_3279_RNR2_16S_50ntFlanks@483.518.36\|AAGAGGAACAGCTCTTTGGACACTAGGAAAAAACCT | >hg38_MT_+_1621_3279_RNR2_16S_50ntFlanks | 483 | 518 | 36 | AAGAGGAACAGCTCTTTGGACACTAGGAAAAAACCT | 61.8 | 9.2 | 1.8 | 5.0 | 6.8E-07 | 1.3E-06 | MB231/MB468 | MP |
| >hg38_21_+_8433172_8446622_RNA45SN1_45S_50ntFlanks@3991.4013.23\|TGGTGACTCTAGATAACCTCGGG | >hg38_21_+_8433172_8446622_RNA45SN1_45S_50ntFlanks | 3991 | 4013 | 23 | TGGTGACTCTAGATAACCTCGGG | 311.6 | -6.6 | 1.3 | -4.9 | 7.9E-07 | 1.3E-06 | MB231/MB468 | NUC |
| >hg38_21_+_8433172_8446622_RNA45SN1_45S_50ntFlanks@10794.10822.29\|CTGGCATGTTGGAACAATGTAGGTAAGGG | >hg38_21_+_8433172_8446622_RNA45SN1_45S_50ntFlanks | 10794 | 10822 | 29 | CTGGCATGTTGGAACAATGTAGGTAAGGG | 308.2 | -2.7 | 0.5 | -5.0 | 5.9E-07 | 1.3E-06 | BT20/MB468 | MITO |
| >MI0000063\|hsa-let-7b&WithFlank&22\|+\|46113680\|46113774@12.32.21(+1G)&[MIMAT0000063&hsa-let-7b-5p&offsets\|0\|-1(+1G);m-14&22\|+\|46113691\|46113712&offsets\|0\|-1(+1G)]\|TGAGGTAGTAGGTTGTGTGGTG | >MI0000063\|hsa-let-7b&WithFlank&22\|+\|46113680\|46113774 | 12 | 32 | 21(+1G) | TGAGGTAGTAGGTTGTGTGGTG | 1031.7 | 3.4 | 0.6 | 5.4 | 8.6E-08 | 1.3E-06 | BT20/MB231 | NUC |
| >MI0000542\|hsa-mir-320a&WithFlank&8\|-\|22244960\|22245043@48.72.25&[MIMAT0000510&hsa-miR-320a-3p&offsets\|0\|+3;m-60&8\|-\|22244975\|22244996&offsets\|0\|+3]\|AAAAGCTGGGTTGAGAGGGCGAAAA | >MI0000542\|hsa-mir-320a&WithFlank&8\|-\|22244960\|22245043 | 48 | 72 | 25 | AAAAGCTGGGTTGAGAGGGCGAAAA | 275.1 | 8.5 | 1.6 | 5.2 | 2.0E-07 | 1.3E-06 | BT20/MB468 | MP |
| >hg38_21_+_8433172_8446622_RNA45SN1_45S_50ntFlanks@3738.3756.19\|TCTCAAAGATTAAGCCATG | >hg38_21_+_8433172_8446622_RNA45SN1_45S_50ntFlanks | 3738 | 3756 | 19 | TCTCAAAGATTAAGCCATG | 1071.4 | -7.8 | 1.5 | -5.2 | 2.0E-07 | 1.3E-06 | BT20/MB468 | MP |
| >MI0000095\|hsa-mir-93&WithFlank&7\|-\|100093762\|100093853@17.38.22&[MIMAT0000093&hsa-miR-93-5p&offsets\|0\|-1;m-29&7\|-\|100093815\|100093837&offsets\|0\|-1]\|CAAAGTGCTGTTCGTGCAGGTA | >MI0000095\|hsa-mir-93&WithFlank&7\|-\|100093762\|100093853 | 17 | 38 | 22 | CAAAGTGCTGTTCGTGCAGGTA | 160.9 | 2.5 | 0.5 | 4.9 | 8.6E-07 | 1.3E-06 | BT20/MB468 | NUC |
| >hg38_21_+_8433172_8446622_RNA45SN1_45S_50ntFlanks@12360.12384.25\|CAGGGATAACTGGCTTGTGGCGGCC | >hg38_21_+_8433172_8446622_RNA45SN1_45S_50ntFlanks | 12360 | 12384 | 25 | CAGGGATAACTGGCTTGTGGCGGCC | 191.8 | -5.0 | 1.0 | -4.9 | 7.9E-07 | 1.3E-06 | MB231/MB468 | NUC |
| >hg38_1_-_228634819_228635039_RNA5S12_5S_50ntFlanks@149.170.22\|GAATACCGGGTGCTGTAGGCTT | >hg38_1_-_228634819_228635039_RNA5S12_5S_50ntFlanks | 149 | 170 | 22 | GAATACCGGGTGCTGTAGGCTT | 2124.5 | 2.6 | 0.5 | 5.0 | 5.9E-07 | 1.3E-06 | BT20/MB468 | MITO |
| >hg38_21_+_8433172_8446622_RNA45SN1_45S_50ntFlanks@9713.9737.25\|GCGAATGATTAGAGGTCTTGGGGCC | >hg38_21_+_8433172_8446622_RNA45SN1_45S_50ntFlanks | 9713 | 9737 | 25 | GCGAATGATTAGAGGTCTTGGGGCC | 687.7 | -5.9 | 1.2 | -5.0 | 5.9E-07 | 1.3E-06 | BT20/MB468 | MITO |
| >hg38_21_+_8433172_8446622_RNA45SN1_45S_50ntFlanks@11670.11704.35\|CCCAGTGCTCTGAATGTCAAAGTGAAGAAATTCAA | >hg38_21_+_8433172_8446622_RNA45SN1_45S_50ntFlanks | 11670 | 11704 | 35 | CCCAGTGCTCTGAATGTCAAAGTGAAGAAATTCAA | 235.3 | -2.2 | 0.4 | -5.0 | 6.7E-07 | 1.3E-06 | BT20/MB468 | CYTO |
| >am_trnaMT_LeuTAG_MT_+_12266_12336@43.61.19\|AAATTTTGGTGCAACTCCA | >am_trnaMT_LeuTAG_MT_+_12266_12336 | 43 | 61 | 19 | AAATTTTGGTGCAACTCCA | 67.3 | -9.2 | 1.8 | -5.2 | 2.0E-07 | 1.3E-06 | BT20/MB468 | MP |
| >hg38_21_+_8433172_8446622_RNA45SN1_45S_50ntFlanks@5486.5515.30\|GCGCTGAGAAGACGGTCGAACTTGACTATC | >hg38_21_+_8433172_8446622_RNA45SN1_45S_50ntFlanks | 5486 | 5515 | 30 | GCGCTGAGAAGACGGTCGAACTTGACTATC | 689.4 | 4.0 | 0.8 | 5.0 | 6.7E-07 | 1.3E-06 | BT20/MB468 | CYTO |
| >MI0000289\|hsa-mir-181a-1&WithFlank&1\|-\|198859038\|198859159@30.50.21;MI0000269\|hsa-mir-181a-2&WithFlank&9\|+\|124692436\|124692557@45.65.21&[MIMAT0000256&hsa-miR-181a-5p&offsets\|0\|-2;m-32&1\|-\|198859108\|198859130&offsets\|0\|-2];[MIMAT0000256_1&hsa-miR-181a-5p&offsets\|0\|-2;m-32&9\|+\|124692480\|124692502&offsets\|0\|-2]\|AACATTCAACGCTGTCGGTGA | >MI0000289\|hsa-mir-181a-1&WithFlank&1\|-\|198859038\|198859159 | 30 | 50 | 21 | AACATTCAACGCTGTCGGTGA | 578.6 | 4.6 | 0.9 | 4.9 | 7.9E-07 | 1.3E-06 | MB231/MB468 | NUC |
| >MI0000087\|hsa-mir-29a&WithFlank&7\|-\|130876741\|130876816@10.34.25&[MIMAT0004503&hsa-miR-29a-5p&offsets\|0\|+3;m-315&7\|-\|130876785\|130876807&offsets\|0\|+2]\|ACTGATTTCTTTTGGTGTTCAGAGT | >MI0000087\|hsa-mir-29a&WithFlank&7\|-\|130876741\|130876816 | 10 | 34 | 25 | ACTGATTTCTTTTGGTGTTCAGAGT | 129.3 | -10.4 | 1.9 | -5.5 | 4.6E-08 | 1.3E-06 | BT20/MB231 | MP |
| >hg38_21_+_8433172_8446622_RNA45SN1_45S_50ntFlanks@11660.11693.34\|GTGATTTCTGCCCAGTGCTCTGAATGTCAAAGTG | >hg38_21_+_8433172_8446622_RNA45SN1_45S_50ntFlanks | 11660 | 11693 | 34 | GTGATTTCTGCCCAGTGCTCTGAATGTCAAAGTG | 481.3 | 12.2 | 2.2 | 5.5 | 4.6E-08 | 1.3E-06 | BT20/MB231 | MP |
| >MI0003129\|hsa-mir-146b&WithFlank&10\|+\|102436506\|102436590@15.34.20(+2U)&[MIMAT0002809&hsa-miR-146b-5p&offsets\|0\|-3(+2U);m-70&10\|+\|102436520\|102436543&offsets\|0\|-4(+2U)]\|TGAGAACTGAATTCCATAGGTT | >MI0003129\|hsa-mir-146b&WithFlank&10\|+\|102436506\|102436590 | 15 | 34 | 20(+2U) | TGAGAACTGAATTCCATAGGTT | 152.3 | -10.6 | 1.9 | -5.5 | 4.6E-08 | 1.3E-06 | BT20/MB231 | MP |
| >hg38_21_+_8433172_8446622_RNA45SN1_45S_50ntFlanks@11517.11539.23\|CCGCCCCCGGGGCCGCGGTTCCG | >hg38_21_+_8433172_8446622_RNA45SN1_45S_50ntFlanks | 11517 | 11539 | 23 | CCGCCCCCGGGGCCGCGGTTCCG | 203.7 | -6.1 | 1.2 | -5.0 | 5.9E-07 | 1.3E-06 | BT20/MB468 | MITO |
| >hg38_21_+_8433172_8446622_RNA45SN1_45S_50ntFlanks@8191.8220.30\|CAGCCCGTGGACGGTGTGAGGCCGGTAGCG | >hg38_21_+_8433172_8446622_RNA45SN1_45S_50ntFlanks | 8191 | 8220 | 30 | CAGCCCGTGGACGGTGTGAGGCCGGTAGCG | 89.9 | -9.6 | 1.8 | -5.2 | 2.0E-07 | 1.3E-06 | BT20/MB468 | MP |
| >hg38_21_+_8433172_8446622_RNA45SN1_45S_50ntFlanks@6733.6773.41\|CATTGATCATCGACACTTCGAACGCACTTGCGGCCCCGGGT | >hg38_21_+_8433172_8446622_RNA45SN1_45S_50ntFlanks | 6733 | 6773 | 41 | CATTGATCATCGACACTTCGAACGCACTTGCGGCCCCGGGT | 68.6 | 9.8 | 1.9 | 5.2 | 2.0E-07 | 1.3E-06 | BT20/MB468 | MP |
| >hg38_21_+_8433172_8446622_RNA45SN1_45S_50ntFlanks@4102.4127.26\|ACCACGGGTGACGGGGAATCAGGGTT | >hg38_21_+_8433172_8446622_RNA45SN1_45S_50ntFlanks | 4102 | 4127 | 26 | ACCACGGGTGACGGGGAATCAGGGTT | 690.6 | -5.6 | 1.1 | -4.9 | 7.9E-07 | 1.3E-06 | MB231/MB468 | NUC |
| >hg38_21_+_8433172_8446622_RNA45SN1_45S_50ntFlanks@10458.10484.27\|ATGGCCTCCGTTGCCCTCGGCCGATCG | >hg38_21_+_8433172_8446622_RNA45SN1_45S_50ntFlanks | 10458 | 10484 | 27 | ATGGCCTCCGTTGCCCTCGGCCGATCG | 613.9 | -4.8 | 1.0 | -4.9 | 7.7E-07 | 1.3E-06 | MB231/MB468 | CYTO |
| >hg38_21_+_8433172_8446622_RNA45SN1_45S_50ntFlanks@10872.10902.31\|GCTGGGTCGGTCGGGCTGGGGCGCGAAGCGG | >hg38_21_+_8433172_8446622_RNA45SN1_45S_50ntFlanks | 10872 | 10902 | 31 | GCTGGGTCGGTCGGGCTGGGGCGCGAAGCGG | 332.0 | -7.5 | 1.5 | -4.9 | 8.7E-07 | 1.3E-06 | BT20/MB468 | NUC |
| >MI0000440\|hsa-mir-27b&WithFlank&9\|+\|95085439\|95085547@67.86.20(+1A)&[MIMAT0000419&hsa-miR-27b-3p&offsets\|0\|-1(+1A);m-39&9\|+\|95085505\|95085525&offsets\|0\|-1(+1A)]\|TTCACAGTGGCTAAGTTCTGA | >MI0000440\|hsa-mir-27b&WithFlank&9\|+\|95085439\|95085547 | 67 | 86 | 20(+1A) | TTCACAGTGGCTAAGTTCTGA | 1997.5 | 3.9 | 0.8 | 5.0 | 6.9E-07 | 1.3E-06 | MB231/MB468 | MP |
| >hg38_21_+_8433172_8446622_RNA45SN1_45S_50ntFlanks@6648.6696.49\|GTACGACTCTTAGCGGTGGATCACTCGGCTCGTGCGTCGATGAAGAACG | >hg38_21_+_8433172_8446622_RNA45SN1_45S_50ntFlanks | 6648 | 6696 | 49 | GTACGACTCTTAGCGGTGGATCACTCGGCTCGTGCGTCGATGAAGAACG | 573.8 | 5.5 | 1.1 | 4.9 | 7.8E-07 | 1.3E-06 | MB231/MB468 | CYTO |
| >hg38_21_+_8433172_8446622_RNA45SN1_45S_50ntFlanks@11322.11342.21\|ACGTCTCGTCGCGCGCGCGTC | >hg38_21_+_8433172_8446622_RNA45SN1_45S_50ntFlanks | 11322 | 11342 | 21 | ACGTCTCGTCGCGCGCGCGTC | 2620.8 | -4.9 | 1.0 | -4.9 | 8.0E-07 | 1.3E-06 | MB231/MB468 | NUC |
| >hg38_21_+_8433172_8446622_RNA45SN1_45S_50ntFlanks@10395.10424.30\|GCAGTTGAACATGGGTCAGTCGGTCCTGAG | >hg38_21_+_8433172_8446622_RNA45SN1_45S_50ntFlanks | 10395 | 10424 | 30 | GCAGTTGAACATGGGTCAGTCGGTCCTGAG | 5364.7 | 5.8 | 1.1 | 5.2 | 2.0E-07 | 1.3E-06 | BT20/MB468 | MP |
| >MI0000746\|hsa-mir-99b&WithFlank&19\|+\|51692606\|51692687@51.72.22&[MIMAT0004678&hsa-miR-99b-3p&offsets\|0\|0;m-151&19\|+\|51692656\|51692678&offsets\|0\|-1]\|CAAGCTCGTGTCTGTGGGTCCG | >MI0000746\|hsa-mir-99b&WithFlank&19\|+\|51692606\|51692687 | 51 | 72 | 22 | CAAGCTCGTGTCTGTGGGTCCG | 1366.3 | 4.6 | 0.9 | 5.2 | 2.0E-07 | 1.3E-06 | BT20/MB468 | MP |
| >hg38_21_+_8433172_8446622_RNA45SN1_45S_50ntFlanks@13000.13039.40\|TGCGATCTATTGAAAGTCAGCCCTCGACACAAGGGTTTGT | >hg38_21_+_8433172_8446622_RNA45SN1_45S_50ntFlanks | 13000 | 13039 | 40 | TGCGATCTATTGAAAGTCAGCCCTCGACACAAGGGTTTGT | 556.8 | 4.6 | 0.9 | 4.9 | 7.8E-07 | 1.3E-06 | MB231/MB468 | CYTO |
| >hg38_21_+_8433172_8446622_RNA45SN1_45S_50ntFlanks@4724.4748.25\|ATTAATCAAGAACGAAAGTCGGAGG | >hg38_21_+_8433172_8446622_RNA45SN1_45S_50ntFlanks | 4724 | 4748 | 25 | ATTAATCAAGAACGAAAGTCGGAGG | 377.4 | -6.7 | 1.4 | -4.9 | 8.0E-07 | 1.3E-06 | MB231/MB468 | NUC |
| >MI0000809\|hsa-mir-151a&WithFlank&8\|-\|140732558\|140732659@51.72.22&[MIMAT0000757&hsa-miR-151a-3p&offsets\|-2\|-1;m-30&8\|-\|140732587\|140732607&offsets\|-2\|-1]\|TACTAGACTGAAGCTCCTTGAG | >MI0000809\|hsa-mir-151a&WithFlank&8\|-\|140732558\|140732659 | 51 | 72 | 22 | TACTAGACTGAAGCTCCTTGAG | 966.6 | 5.3 | 1.0 | 5.2 | 2.0E-07 | 1.3E-06 | BT20/MB468 | MP |
| >trnaMT_PheGAA_MT_+_577_647@37.56.20\|AAATGTTTAGACGGGCTCAC | >trnaMT_PheGAA_MT_+_577_647 | 37 | 56 | 20 | AAATGTTTAGACGGGCTCAC | 240.1 | 6.7 | 1.4 | 4.9 | 8.3E-07 | 1.3E-06 | MB231/MB468 | MITO |
| >MI0023622\|hsa-mir-486-2&WithFlank&8\|+\|41660438\|41660513@7.29.23&[MIMAT0002177_1&hsa-miR-486-5p&offsets\|0\|+1;TJU_CMC_MD2.ID03138.5p-miR&offsets\|0\|-1;m-106&8\|+\|41660444\|41660465&offsets\|0\|+1]\|TCCTGTACTGAGCTGCCCCGAGC | >MI0023622\|hsa-mir-486-2&WithFlank&8\|+\|41660438\|41660513 | 7 | 29 | 23 | TCCTGTACTGAGCTGCCCCGAGC | 59.6 | 9.1 | 1.8 | 5.0 | 6.9E-07 | 1.3E-06 | MB231/MB468 | MP |

| | | | | | | | | | | | | | |
|---|---|---|---|---|---|---|---|---|---|---|---|---|---|
| >hg38_21_+_8433172_8446622_RNA45SN1_45S_50ntFlanks@11769.11794.26\|TAATTAGTGACGCGCATGAATGGATG | >hg38_21_+_8433172_8446622_RNA45SN1_45S_50ntFlanks | 11769 | 11794 | 26 | TAATTAGTGACGCGCATGAATGGATG | 674.3 | -4.5 | 0.9 | -5.0 | 6.9E-07 | 1.3E-06 | MB231/MB468 | MP |
| >hg38_21_+_8433172_8446622_RNA45SN1_45S_50ntFlanks@11778.11806.29\|ACGCGCATGAATGGATGAACGAGATTCCC | >hg38_21_+_8433172_8446622_RNA45SN1_45S_50ntFlanks | 11778 | 11806 | 29 | ACGCGCATGAATGGATGAACGAGATTCCC | 253.2 | -6.9 | 1.4 | -5.0 | 6.9E-07 | 1.3E-06 | MB231/MB468 | MP |
| >hg38_21_+_8433172_8446622_RNA45SN1_45S_50ntFlanks@8836.8862.27\|CGGCGGTGGCGGCGGCGGCGGCGGCGG | >hg38_21_+_8433172_8446622_RNA45SN1_45S_50ntFlanks | 8836 | 8862 | 27 | CGGCGGTGGCGGCGGCGGCGGCGGCGG | 72.3 | -9.3 | 1.8 | -5.2 | 2.0E-07 | 1.3E-06 | BT20/MB468 | MP |
| >hg38_21_+_8433172_8446622_RNA45SN1_45S_50ntFlanks@11270.11294.25\|CGGCCGCCCCCGGGGAGCCCGGCGG | >hg38_21_+_8433172_8446622_RNA45SN1_45S_50ntFlanks | 11270 | 11294 | 25 | CGGCCGCCCCCGGGGAGCCCGGCGG | 88.1 | -9.5 | 1.8 | -5.2 | 2.0E-07 | 1.3E-06 | BT20/MB468 | MP |
| >hg38_21_+_8433172_8446622_RNA45SN1_45S_50ntFlanks@4328.4352.25\|CGGTAATTCCAGCTCCAATAGCGTA | >hg38_21_+_8433172_8446622_RNA45SN1_45S_50ntFlanks | 4328 | 4352 | 25 | CGGTAATTCCAGCTCCAATAGCGTA | 77.3 | -9.4 | 1.8 | -5.2 | 2.0E-07 | 1.3E-06 | BT20/MB468 | MP |
| >hg38_21_+_8433172_8446622_RNA45SN1_45S_50ntFlanks@6659.6681.23\|AGCGGTGGATCACTCGGCTCGTG | >hg38_21_+_8433172_8446622_RNA45SN1_45S_50ntFlanks | 6659 | 6681 | 23 | AGCGGTGGATCACTCGGCTCGTG | 1303.5 | -2.6 | 0.5 | -5.0 | 6.0E-07 | 1.3E-06 | BT20/MB468 | MITO |
| >hg38_21_+_8433172_8446622_RNA45SN1_45S_50ntFlanks@10574.10612.39\|AAGCCGGCGGGAGCCCCGGGGAGAGTTCTCTTTTCTTTG | >hg38_21_+_8433172_8446622_RNA45SN1_45S_50ntFlanks | 10574 | 10612 | 39 | AAGCCGGCGGGAGCCCCGGGGAGAGTTCTCTTTTCTTTG | 220.5 | -6.5 | 1.3 | -5.0 | 6.8E-07 | 1.3E-06 | BT20/MB468 | CYTO |
| >hg38_21_+_8433172_8446622_RNA45SN1_45S_50ntFlanks@6649.6668.20\|TACGACTCTTAGCGGTGGAT | >hg38_21_+_8433172_8446622_RNA45SN1_45S_50ntFlanks | 6649 | 6668 | 20 | TACGACTCTTAGCGGTGGAT | 236.8 | 3.1 | 0.6 | 4.9 | 8.0E-07 | 1.3E-06 | MB231/MB468 | NUC |
| >hg38_21_+_8433172_8446622_RNA45SN1_45S_50ntFlanks@4753.4782.30\|AAGACGATCAGATACCGTCGTAGTTCCGAC | >hg38_21_+_8433172_8446622_RNA45SN1_45S_50ntFlanks | 4753 | 4782 | 30 | AAGACGATCAGATACCGTCGTAGTTCCGAC | 452.2 | -5.6 | 1.1 | -4.9 | 8.0E-07 | 1.3E-06 | MB231/MB468 | NUC |
| >hg38_21_+_8433172_8446622_RNA45SN1_45S_50ntFlanks@9569.9597.29\|GTGCAAATCGGTCGTCCGACCTGGGTATA | >hg38_21_+_8433172_8446622_RNA45SN1_45S_50ntFlanks | 9569 | 9597 | 29 | GTGCAAATCGGTCGTCCGACCTGGGTATA | 492.8 | -3.2 | 0.6 | -4.9 | 8.0E-07 | 1.3E-06 | MB231/MB468 | NUC |
| >hg38_21_+_8433172_8446622_RNA45SN1_45S_50ntFlanks@11582.11603.22\|ACTGGTGCGGACCAGGGGAATC | >hg38_21_+_8433172_8446622_RNA45SN1_45S_50ntFlanks | 11582 | 11603 | 22 | ACTGGTGCGGACCAGGGGAATC | 727.0 | -4.0 | 0.8 | -4.9 | 8.4E-07 | 1.3E-06 | MB231/MB468 | MITO |
| >hg38_21_+_8433172_8446622_RNA45SN1_45S_50ntFlanks@8118.8142.25\|GGGACATGTGGCGTACGGAAGACCC | >hg38_21_+_8433172_8446622_RNA45SN1_45S_50ntFlanks | 8118 | 8142 | 25 | GGGACATGTGGCGTACGGAAGACCC | 233.4 | 11.2 | 2.0 | 5.5 | 4.7E-08 | 1.3E-06 | BT20/MB231 | MP |
| >hg38_21_+_8433172_8446622_RNA45SN1_45S_50ntFlanks@3853.3872.20\|AACTGTGGTAATTCTAGAGC | >hg38_21_+_8433172_8446622_RNA45SN1_45S_50ntFlanks | 3853 | 3872 | 20 | AACTGTGGTAATTCTAGAGC | 491.4 | -7.0 | 1.4 | -5.0 | 7.0E-07 | 1.3E-06 | MB231/MB468 | MP |
| >hg38_21_+_8433172_8446622_RNA45SN1_45S_50ntFlanks@6661.6709.49\|CGGTGGATCACTCGGCTCGTGCGTCGATGAAGAACGCAGCTAGCTGCGA | >hg38_21_+_8433172_8446622_RNA45SN1_45S_50ntFlanks | 6661 | 6709 | 49 | CGGTGGATCACTCGGCTCGTGCGTCGATGAAGAACGCAGCTAGCTGCGA | 2389.6 | 6.0 | 1.1 | 5.4 | 6.8E-08 | 1.3E-06 | BT20/MB231 | MITO |
| >hg38_21_+_8433172_8446622_RNA45SN1_45S_50ntFlanks@10423.10443.21\|AGAGATGGGCGAGCGCCGTTC | >hg38_21_+_8433172_8446622_RNA45SN1_45S_50ntFlanks | 10423 | 10443 | 21 | AGAGATGGGCGAGCGCCGTTC | 209.4 | -11.6 | 2.3 | -5.0 | 7.0E-07 | 1.3E-06 | MB231/MB468 | MP |
| >am_trnaMT_GluTTC_MT_-_14674_14742@24.54.31,trnalookalike8_GluTTC_5_-_93905172_93905240@24.54.31\|ATGGTTTTTCATATCATTGGTCGTGGTTGTA | >am_trnaMT_GluTTC_MT_-_14674_14742 | 24 | 54 | 31 | ATGGTTTTTCATATCATTGGTCGTGGTTGTA | 443.0 | 9.0 | 1.7 | 5.2 | 2.1E-07 | 1.3E-06 | BT20/MB468 | MP |
| >trna10_LysCTT_16_+_3241501_3241573@13.33.21,trna13_LysCTT_14_-_58706613_58706685@13.33.21,trna2_LysCTT_15_+_79152904_79152976@13.33.21,trna4_LysCTT_18_-_43669271_43669343@13.33.21\|CAGTCGGTAGAGCATGGGACT | >trna10_LysCTT_16_+_3241501_3241573 | 13 | 33 | 21 | CAGTCGGTAGAGCATGGGACT | 83.0 | 10.1 | 1.9 | 5.2 | 2.1E-07 | 1.3E-06 | BT20/MB468 | MP |
| >MI0005756\|hsa-mir-934&WithFlank&X\|+\|136550872\|136550966@21.41.21(+1A)&[MIMAT0004977&hsa-miR-934&offsets\|0\|-1(+1A);m-943&X\|+\|136550892\|136550913&offsets\|0\|-1(+1A)]\|TGTCTACTACTGGAGACACTGA | >MI0005756\|hsa-mir-934&WithFlank&X\|+\|136550872\|136550966 | 21 | 41 | 21(+1A) | TGTCTACTACTGGAGACACTGA | 206.4 | 11.0 | 2.0 | 5.5 | 4.7E-08 | 1.3E-06 | BT20/MB231 | MP |
| >hg38_21_+_8433172_8446622_RNA45SN1_45S_50ntFlanks@10791.10816.26\|CCTCTGGCATGTTGGAACAATGTAGG | >hg38_21_+_8433172_8446622_RNA45SN1_45S_50ntFlanks | 10791 | 10816 | 26 | CCTCTGGCATGTTGGAACAATGTAGG | 176.4 | 10.8 | 2.0 | 5.5 | 4.7E-08 | 1.3E-06 | BT20/MB231 | MP |
| >hg38_21_+_8433172_8446622_RNA45SN1_45S_50ntFlanks@475.507.33\|CTTCGTGATCGATGTGGTGACGTCGTGCTCTCC | >hg38_21_+_8433172_8446622_RNA45SN1_45S_50ntFlanks | 475 | 507 | 33 | CTTCGTGATCGATGTGGTGACGTCGTGCTCTCC | 1126.6 | 4.8 | 1.0 | 4.9 | 8.4E-07 | 1.3E-06 | MB231/MB468 | MITO |
| >hg38_1_-_228634819_228635039_RNA5S12_5S_50ntFlanks@84.106.23\|CTCGTCTGATCTCGGAAGCTAAG | >hg38_1_-_228634819_228635039_RNA5S12_5S_50ntFlanks | 84 | 106 | 23 | CTCGTCTGATCTCGGAAGCTAAG | 282.4 | -7.3 | 1.5 | -4.9 | 8.8E-07 | 1.3E-06 | BT20/MB468 | NUC |
| >hg38_21_+_8433172_8446622_RNA45SN1_45S_50ntFlanks@9307.9330.24\|AAGGAGTCTAACACGTGCGCGAGT | >hg38_21_+_8433172_8446622_RNA45SN1_45S_50ntFlanks | 9307 | 9330 | 24 | AAGGAGTCTAACACGTGCGCGAGT | 445.7 | 6.6 | 1.3 | 4.9 | 8.1E-07 | 1.3E-06 | MB231/MB468 | NUC |
| >MI0000088\|hsa-mir-30a&WithFlank&6\|-\|71403545\|71403627@12.35.24(+1A)&[MIMAT0000087&hsa-miR-30a-5p&offsets\|0\|+2(+1A);m-15&6\|-\|71403595\|71403616&offsets\|0\|+2(+1A)]\|TGTAAACATCCTCGACTGGAAGCTA | >MI0000088\|hsa-mir-30a&WithFlank&6\|-\|71403545\|71403627 | 12 | 35 | 24(+1A) | TGTAAACATCCTCGACTGGAAGCTA | 1426.8 | 4.4 | 0.9 | 5.0 | 7.0E-07 | 1.3E-06 | MB231/MB468 | MP |
| >hg38_21_+_8433172_8446622_RNA45SN1_45S_50ntFlanks@11540.11584.45\|CGCGGCGCCTCGCCTCGGCCGGCGCCTAGCAGCCGACTTAGAACT | >hg38_21_+_8433172_8446622_RNA45SN1_45S_50ntFlanks | 11540 | 11584 | 45 | CGCGGCGCCTCGCCTCGGCCGGCGCCTAGCAGCCGACTTAGAACT | 4433.6 | -4.6 | 0.9 | -5.4 | 5.7E-08 | 1.3E-06 | BT20/MB231 | CYTO |
| >hg38_21_+_8433172_8446622_RNA45SN1_45S_50ntFlanks@11709.11728.20\|GCGCGGGTAAACGGCGGGAG | >hg38_21_+_8433172_8446622_RNA45SN1_45S_50ntFlanks | 11709 | 11728 | 20 | GCGCGGGTAAACGGCGGGAG | 60.6 | -9.0 | 1.7 | -5.2 | 2.1E-07 | 1.3E-06 | BT20/MB468 | MP |
| >hg38_21_+_8433172_8446622_RNA45SN1_45S_50ntFlanks@11583.11605.23\|CTGGTGCGGACCAGGGGAATCCG | >hg38_21_+_8433172_8446622_RNA45SN1_45S_50ntFlanks | 11583 | 11605 | 23 | CTGGTGCGGACCAGGGGAATCCG | 1466.6 | -8.4 | 1.7 | -4.9 | 8.8E-07 | 1.3E-06 | BT20/MB468 | NUC |
| >MI0000081\|hsa-mir-24-2&WithFlank&19\|-\|13836281\|13836365@56.77.22;MI0000080\|hsa-mir-24-1&WithFlank&9\|+\|95086015\|95086094@50.71.22&[MIMAT0000080&hsa-miR-24-3p&offsets\|0\|0;m-36&19\|-\|13836288\|13836310&offsets\|0\|-1];[MIMAT0000080_1&hsa-miR-24-3p&offsets\|0\|0;m-35&9\|+\|95086064\|95086086&offsets\|0\|-1]\|TGGCTCAGTTCAGCAGGAACAG | >MI0000081\|hsa-mir-24-2&WithFlank&19\|-\|13836281\|13836365 | 56 | 77 | 22 | TGGCTCAGTTCAGCAGGAACAG | 9917.2 | 3.9 | 0.8 | 4.9 | 8.5E-07 | 1.3E-06 | MB231/MB468 | MITO |
| >hg38_21_+_8433172_8446622_RNA45SN1_45S_50ntFlanks@10351.10389.39\|CGAGAACTTTGAAGGCCGAAGTGGAGAAGGGTTCCATGT | >hg38_21_+_8433172_8446622_RNA45SN1_45S_50ntFlanks | 10351 | 10389 | 39 | CGAGAACTTTGAAGGCCGAAGTGGAGAAGGGTTCCATGT | 466.6 | -7.2 | 1.5 | -4.9 | 8.5E-07 | 1.3E-06 | MB231/MB468 | MITO |
| >hg38_21_+_8433172_8446622_RNA45SN1_45S_50ntFlanks@6646.6667.22\|TCGTACGACTCTTAGCGGTGGA | >hg38_21_+_8433172_8446622_RNA45SN1_45S_50ntFlanks | 6646 | 6667 | 22 | TCGTACGACTCTTAGCGGTGGA | 8759.6 | 4.5 | 0.9 | 5.2 | 2.1E-07 | 1.3E-06 | BT20/MB468 | MP |
| >hg38_21_+_8433172_8446622_RNA45SN1_45S_50ntFlanks@6729.6772.44\|GACACATTGATCATCGACACTTCGAACGCACTTGCGGCCCCGGG | >hg38_21_+_8433172_8446622_RNA45SN1_45S_50ntFlanks | 6729 | 6772 | 44 | GACACATTGATCATCGACACTTCGAACGCACTTGCGGCCCCGGG | 303.2 | -5.6 | 1.1 | -4.9 | 8.5E-07 | 1.3E-06 | MB231/MB468 | MITO |
| >trnaMT_LeuTAG_MT_+_12266_12336@42.74.33\|AAAATTTTGGTGCAACTCCAAATAAAAGTACCA | >trnaMT_LeuTAG_MT_+_12266_12336 | 42 | 74 | 33 | AAAATTTTGGTGCAACTCCAAATAAAAGTACCA | 465.8 | -4.4 | 0.9 | -4.9 | 8.5E-07 | 1.3E-06 | MB231/MB468 | MITO |
| >MI0000446\|hsa-mir-125b-1&WithFlank&11\|-\|122099751\|122099850@21.43.23(+1U)&[MIMAT0000423&hsa-miR-125b-5p&offsets\|0\|+1(+1U);m-67&11\|-\|122099809\|122099830&offsets\|0\|+1(+1U)]\|TCCCTGAGACCCTAACTTGTGATT | >MI0000446\|hsa-mir-125b-1&WithFlank&11\|-\|122099751\|122099850 | 21 | 43 | 23(+1U) | TCCCTGAGACCCTAACTTGTGATT | 50.8 | 8.9 | 1.8 | 5.0 | 7.1E-07 | 1.3E-06 | MB231/MB468 | MP |
| >trnaMT_GluTTC_MT_-_14674_14742@35.71.37,trnalookalike8_GluTTC_5_-_93905172_93905240@35.71.37\|TATCATTGGTCGTGGTTGTAGTCCGTGCGAGAATACC | >trnaMT_GluTTC_MT_-_14674_14742 | 35 | 71 | 37 | TATCATTGGTCGTGGTTGTAGTCCGTGCGAGAATACC | 1484.8 | -4.9 | 0.9 | -5.4 | 5.8E-08 | 1.3E-06 | BT20/MB231 | CYTO |
| >hg38_MT_+_1621_3279_RNR2_16S_50ntFlanks@493.517.25\|GCTCTTTGGACACTAGGAAAAAACC | >hg38_MT_+_1621_3279_RNR2_16S_50ntFlanks | 493 | 517 | 25 | GCTCTTTGGACACTAGGAAAAAACC | 366.5 | -4.1 | 0.8 | -5.4 | 5.8E-08 | 1.3E-06 | BT20/MB231 | CYTO |
| >hg38_21_+_8433172_8446622_RNA45SN1_45S_50ntFlanks@12635.12657.23\|AAGCTACCATCTGTGGGATTATG | >hg38_21_+_8433172_8446622_RNA45SN1_45S_50ntFlanks | 12635 | 12657 | 23 | AAGCTACCATCTGTGGGATTATG | 171.2 | -6.8 | 1.4 | -5.0 | 6.9E-07 | 1.3E-06 | BT20/MB468 | CYTO |
| >hg38_21_+_8433172_8446622_RNA45SN1_45S_50ntFlanks@11603.11620.18\|CCGACTGTTTAATTAAAA | >hg38_21_+_8433172_8446622_RNA45SN1_45S_50ntFlanks | 11603 | 11620 | 18 | CCGACTGTTTAATTAAAA | 435.9 | -5.3 | 1.1 | -4.9 | 8.5E-07 | 1.3E-06 | MB231/MB468 | MITO |
| >hg38_21_+_8433172_8446622_RNA45SN1_45S_50ntFlanks@10821.10838.18\|GGAAGTCGGCAAGCCGGA | >hg38_21_+_8433172_8446622_RNA45SN1_45S_50ntFlanks | 10821 | 10838 | 18 | GGAAGTCGGCAAGCCGGA | 1031.9 | -8.3 | 1.7 | -4.9 | 8.9E-07 | 1.3E-06 | BT20/MB468 | NUC |
| >hg38_21_+_8433172_8446622_RNA45SN1_45S_50ntFlanks@10841.10869.29\|CGTAACTTCGGGATAAGGATTGGCTCTAA | >hg38_21_+_8433172_8446622_RNA45SN1_45S_50ntFlanks | 10841 | 10869 | 29 | CGTAACTTCGGGATAAGGATTGGCTCTAA | 621.9 | -6.2 | 1.3 | -4.9 | 8.9E-07 | 1.3E-06 | BT20/MB468 | NUC |
| >hg38_21_+_8433172_8446622_RNA45SN1_45S_50ntFlanks@6735.6770.36\|TTGATCATCGACACTTCGAACGCACTTGCGGCCCCG | >hg38_21_+_8433172_8446622_RNA45SN1_45S_50ntFlanks | 6735 | 6770 | 36 | TTGATCATCGACACTTCGAACGCACTTGCGGCCCCG | 232.3 | -3.9 | 0.8 | -4.9 | 7.9E-07 | 1.3E-06 | MB231/MB468 | CYTO |
| >hg38_21_+_8433172_8446622_RNA45SN1_45S_50ntFlanks@10840.10868.29\|CCGTAACTTCGGGATAAGGATTGGCTCTA | >hg38_21_+_8433172_8446622_RNA45SN1_45S_50ntFlanks | 10840 | 10868 | 29 | CCGTAACTTCGGGATAAGGATTGGCTCTA | 574.3 | -3.8 | 0.8 | -5.0 | 6.9E-07 | 1.3E-06 | BT20/MB468 | CYTO |
| >hg38_21_+_8433172_8446622_RNA45SN1_45S_50ntFlanks@10786.10827.42\|AACAGCCTCTGGCATGTTGGAACAATGTAGGTAAGGGAAGTC | >hg38_21_+_8433172_8446622_RNA45SN1_45S_50ntFlanks | 10786 | 10827 | 42 | AACAGCCTCTGGCATGTTGGAACAATGTAGGTAAGGGAAGTC | 237.6 | -7.3 | 1.5 | -5.0 | 7.1E-07 | 1.3E-06 | MB231/MB468 | MP |
| >hg38_21_+_8433172_8446622_RNA45SN1_45S_50ntFlanks@11729.11759.31\|TAACTATGACTCTCTTAAGGTAGCCAAATGC | >hg38_21_+_8433172_8446622_RNA45SN1_45S_50ntFlanks | 11729 | 11759 | 31 | TAACTATGACTCTCTTAAGGTAGCCAAATGC | 103.3 | -9.8 | 1.9 | -5.2 | 2.1E-07 | 1.3E-06 | BT20/MB468 | MP |
| >MI0000289\|hsa-mir-181a-1&WithFlank&1\|-\|198859038\|198859159@30.53.24;MI0000269\|hsa-mir-181a-2&WithFlank&9\|+\|124692436\|124692557@45.68.24&[MIMAT0000256&hsa-miR-181a-5p&offsets\|0\|+1;m-32&1\|-\|198859108\|198859130&offsets\|0\|+1];[MIMAT0000256_1&hsa-miR-181a-5p&offsets\|0\|+1;m-32&9\|+\|124692480\|124692502&offsets\|0\|+1]\|AACATTCAACGCTGTCGGTGAGTT | >MI0000289\|hsa-mir-181a-1&WithFlank&1\|-\|198859038\|198859159 | 30 | 53 | 24 | AACATTCAACGCTGTCGGTGAGTT | 3192.8 | -5.1 | 0.9 | -5.4 | 6.9E-08 | 1.3E-06 | BT20/MB231 | MITO |
| >hg38_21_+_8433172_8446622_RNA45SN1_45S_50ntFlanks@10237.10258.22\|ACGCTACGCCGCGACGAGTAGG | >hg38_21_+_8433172_8446622_RNA45SN1_45S_50ntFlanks | 10237 | 10258 | 22 | ACGCTACGCCGCGACGAGTAGG | 155.2 | 10.6 | 1.9 | 5.5 | 4.8E-08 | 1.3E-06 | BT20/MB231 | MP |
| >MI0000294\|hsa-mir-218-1&WithFlank&4\|+\|20528269\|20528390@31.52.22(+1A);MI0000295\|hsa-mir-218-2&WithFlank&5\|-\|168768140\|168768261@31.52.22(+1A)&[MIMAT0000275&hsa-miR-218-5p&offsets\|0\|+1(+1A);m-185&4\|+\|20528299\|20528319&offsets\|0\|+1(+1A)];[MIMAT0000275_1&hsa-miR-218-5p&offsets\|0\|+1(+1A);m-187&5\|-\|168768211\|168768231&offsets\|0\|+1(+1A)]\|TTGTGCTTGATCTAACCATGTGA | >MI0000294\|hsa-mir-218-1&WithFlank&4\|+\|20528269\|20528390 | 31 | 52 | 22(+1A) | TTGTGCTTGATCTAACCATGTGA | 144.0 | -10.6 | 1.9 | -5.5 | 4.8E-08 | 1.3E-06 | BT20/MB231 | MP |
| >hg38_21_+_8433172_8446622_RNA45SN1_45S_50ntFlanks@4103.4120.18\|CCACGGGTGACGGGGAAT | >hg38_21_+_8433172_8446622_RNA45SN1_45S_50ntFlanks | 4103 | 4120 | 18 | CCACGGGTGACGGGGAAT | 209.9 | -3.1 | 0.6 | -4.9 | 8.0E-07 | 1.4E-06 | MB231/MB468 | CYTO |
| >hg38_21_+_8433172_8446622_RNA45SN1_45S_50ntFlanks@5084.5115.32\|CGAACGAGACTCTGGCATGCTAACTAGTTACG | >hg38_21_+_8433172_8446622_RNA45SN1_45S_50ntFlanks | 5084 | 5115 | 32 | CGAACGAGACTCTGGCATGCTAACTAGTTACG | 1062.8 | -6.7 | 1.4 | -4.9 | 8.6E-07 | 1.4E-06 | MB231/MB468 | MITO |
| >hg38_21_+_8433172_8446622_RNA45SN1_45S_50ntFlanks@10771.10806.36\|CAGGTCTCCAAGGTGAACAGCCTCTGGCATGTTGGA | >hg38_21_+_8433172_8446622_RNA45SN1_45S_50ntFlanks | 10771 | 10806 | 36 | CAGGTCTCCAAGGTGAACAGCCTCTGGCATGTTGGA | 584.1 | -6.9 | 1.4 | -5.0 | 7.0E-07 | 1.4E-06 | BT20/MB468 | CYTO |
| >hg38_21_+_8433172_8446622_RNA45SN1_45S_50ntFlanks@9556.9597.42\|TAGCGGTCCTGACGTGCAAATCGGTCGTCCGACCTGGGTATA | >hg38_21_+_8433172_8446622_RNA45SN1_45S_50ntFlanks | 9556 | 9597 | 42 | TAGCGGTCCTGACGTGCAAATCGGTCGTCCGACCTGGGTATA | 127.6 | -9.1 | 1.8 | -5.0 | 6.1E-07 | 1.4E-06 | BT20/MB468 | MITO |

| | | | | | | | | | | | | | |
|---|---|---|---|---|---|---|---|---|---|---|---|---|---|
| >am_trna134_GluTTC_1_-_16861774_16861845@1.30.30,trna5_GluTTC_1_+_17199078_17199149@1.30.30,trna84_GluTTC_1_-_161391883_161391954@1.30.30,trna94_GluTTC_1_-_149664355_149664427@1.30.30\|TCCCTGGTGGTCTAGTGGCTAGGATTCGGC | >am_trna134_GluTTC_1_-_16861774_16861845 | 1 | 30 | 30 | TCCCTGGTGGTCTAGTGGCTAGGATTCGGC | 1247.4 | -3.5 | 0.7 | -4.9 | 8.0E-07 | 1.4E-06 | MB231/MB468 | CYTO |
| >hg38_21_+_8433172_8446622_RNA45SN1_45S_50ntFlanks@12417.12438.22\|TCGATGTCGGCTCTTCCTATCA | >hg38_21_+_8433172_8446622_RNA45SN1_45S_50ntFlanks | 12417 | 12438 | 22 | TCGATGTCGGCTCTTCCTATCA | 1605.8 | 2.9 | 0.6 | 4.9 | 8.0E-07 | 1.4E-06 | MB231/MB468 | CYTO |
| >hg38_21_+_8433172_8446622_RNA45SN1_45S_50ntFlanks@9822.9850.29\|CTAGTGGGCCACTTTTGGTAAGCAGAACT | >hg38_21_+_8433172_8446622_RNA45SN1_45S_50ntFlanks | 9822 | 9850 | 29 | CTAGTGGGCCACTTTTGGTAAGCAGAACT | 265.1 | 11.4 | 2.1 | 5.5 | 4.9E-08 | 1.4E-06 | BT20/MB231 | MP |
| >hg38_21_+_8433172_8446622_RNA45SN1_45S_50ntFlanks@8382.8400.19\|AGAGGGCGTGAAACCGTTA | >hg38_21_+_8433172_8446622_RNA45SN1_45S_50ntFlanks | 8382 | 8400 | 19 | AGAGGGCGTGAAACCGTTA | 280.1 | -7.2 | 1.5 | -5.0 | 7.1E-07 | 1.4E-06 | MB231/MB468 | MP |
| >hg38_MT_+_1621_3279_RNR2_16S_50ntFlanks@719.745.27\|GCCTGCGTCAGATTAAAACACTGAACT | >hg38_MT_+_1621_3279_RNR2_16S_50ntFlanks | 719 | 745 | 27 | GCCTGCGTCAGATTAAAACACTGAACT | 51.6 | 8.9 | 1.8 | 5.0 | 7.1E-07 | 1.4E-06 | MB231/MB468 | MP |
| >hg38_21_+_8433172_8446622_RNA45SN1_45S_50ntFlanks@11669.11691.23\|GCCCAGTGCTCTGAATGTCAAAG | >hg38_21_+_8433172_8446622_RNA45SN1_45S_50ntFlanks | 11669 | 11691 | 23 | GCCCAGTGCTCTGAATGTCAAAG | 324.7 | 3.0 | 0.6 | 4.9 | 8.2E-07 | 1.4E-06 | MB231/MB468 | NUC |
| >hg38_21_+_8433172_8446622_RNA45SN1_45S_50ntFlanks@6664.6688.25\|TGGATCACTCGGCTCGTGCGTCGAT | >hg38_21_+_8433172_8446622_RNA45SN1_45S_50ntFlanks | 6664 | 6688 | 25 | TGGATCACTCGGCTCGTGCGTCGAT | 624.5 | -5.6 | 1.1 | -4.9 | 8.6E-07 | 1.4E-06 | MB231/MB468 | MITO |
| >MI0000433\|hsa-let-7g&WithFlank&3\|-\|52268272\|52268367@11.32.22(+1A)&[MIMAT0000414&hsa-let-7g-5p&offsets\|0\|0(+1A);m-58&3\|-\|52268336\|52268357&offsets\|0\|0(+1A)]\|TGAGGTAGTAGTTTGTACAGTTA | >MI0000433\|hsa-let-7g&WithFlank&3\|-\|52268272\|52268367 | 11 | 32 | 22(+1A) | TGAGGTAGTAGTTTGTACAGTTA | 472.0 | 4.0 | 0.8 | 4.9 | 8.0E-07 | 1.4E-06 | MB231/MB468 | CYTO |
| >MI0000749\|hsa-mir-30e&WithFlank&1\|+\|40754349\|40754452@65.83.19&[MIMAT0000693&hsa-miR-30e-3p&offsets\|0\|-3;m-25&1\|+\|40754413\|40754434&offsets\|0\|-3]\|CTTTCAGTCGGATGTTTAC | >MI0000749\|hsa-mir-30e&WithFlank&1\|+\|40754349\|40754452 | 65 | 83 | 19 | CTTTCAGTCGGATGTTTAC | 800.7 | 4.0 | 0.8 | 4.9 | 8.0E-07 | 1.4E-06 | MB231/MB468 | CYTO |
| >hg38_21_+_8433172_8446622_RNA45SN1_45S_50ntFlanks@9551.9569.19\|GTCCGTAGCGGTCCTGACG | >hg38_21_+_8433172_8446622_RNA45SN1_45S_50ntFlanks | 9551 | 9569 | 19 | GTCCGTAGCGGTCCTGACG | 789.4 | 3.5 | 0.7 | 4.9 | 8.0E-07 | 1.4E-06 | MB231/MB468 | CYTO |
| >hg38_21_+_8433172_8446622_RNA45SN1_45S_50ntFlanks@9954.9983.30\|TGGAAGTCGGAATCCGCTAAGGAGTGTGTA | >hg38_21_+_8433172_8446622_RNA45SN1_45S_50ntFlanks | 9954 | 9983 | 30 | TGGAAGTCGGAATCCGCTAAGGAGTGTGTA | 65.2 | 9.8 | 1.9 | 5.2 | 2.1E-07 | 1.4E-06 | BT20/MB468 | MP |
| >hg38_21_+_8433172_8446622_RNA45SN1_45S_50ntFlanks@4759.4785.27\|ATCAGATACCGTCGTAGTTCCGACCAT | >hg38_21_+_8433172_8446622_RNA45SN1_45S_50ntFlanks | 4759 | 4785 | 27 | ATCAGATACCGTCGTAGTTCCGACCAT | 60.6 | -9.0 | 1.7 | -5.2 | 2.1E-07 | 1.4E-06 | BT20/MB468 | MP |
| >hg38_21_+_8433172_8446622_RNA45SN1_45S_50ntFlanks@11576.11607.32\|CTTAGAACTGGTGCGGACCAGGGGAATCCGAC | >hg38_21_+_8433172_8446622_RNA45SN1_45S_50ntFlanks | 11576 | 11607 | 32 | CTTAGAACTGGTGCGGACCAGGGGAATCCGAC | 366.2 | -5.3 | 1.1 | -4.9 | 8.2E-07 | 1.4E-06 | MB231/MB468 | NUC |
| >hg38_21_+_8433172_8446622_RNA45SN1_45S_50ntFlanks@10781.10801.21\|AGGTGAACAGCCTCTGGCATG | >hg38_21_+_8433172_8446622_RNA45SN1_45S_50ntFlanks | 10781 | 10801 | 21 | AGGTGAACAGCCTCTGGCATG | 715.0 | -5.3 | 1.1 | -5.0 | 7.0E-07 | 1.4E-06 | BT20/MB468 | CYTO |
| >hg38_21_+_8433172_8446622_RNA45SN1_45S_50ntFlanks@5421.5441.21\|CGATTGGATGGTTTAGTGAGG | >hg38_21_+_8433172_8446622_RNA45SN1_45S_50ntFlanks | 5421 | 5441 | 21 | CGATTGGATGGTTTAGTGAGG | 82.4 | -9.4 | 1.8 | -5.2 | 2.1E-07 | 1.4E-06 | BT20/MB468 | MP |
| >trnaMT_GluTTC_MT_-_14674_14742@46.71.26,trnalookalike8_GluTTC_5_-_93905172_93905240@46.71.26\|GTGGTTGTAGTCCGTGCGAGAATACC | >trnaMT_GluTTC_MT_-_14674_14742 | 46 | 71 | 26 | GTGGTTGTAGTCCGTGCGAGAATACC | 156.2 | 3.9 | 0.8 | 4.9 | 8.6E-07 | 1.4E-06 | MB231/MB468 | MITO |
| >hg38_21_+_8433172_8446622_RNA45SN1_45S_50ntFlanks@9459.9503.45\|CCGGGGAGGTGGAGCACGAGCGCACGTGTTAGGACCCGAAAGATG | >hg38_21_+_8433172_8446622_RNA45SN1_45S_50ntFlanks | 9459 | 9503 | 45 | CCGGGGAGGTGGAGCACGAGCGCACGTGTTAGGACCCGAAAGATG | 1246.7 | -7.2 | 1.5 | -5.0 | 6.2E-07 | 1.4E-06 | BT20/MB468 | MITO |
| >MI0000266\|hsa-mir-10a&WithFlank&17\|-\|48579832\|48579953@28.49.22(+1U)&[MIMAT0000253&hsa-miR-10a-5p&offsets\|0\|-1(+1U);m-18&17\|-\|48579905\|48579926&offsets\|0\|0(+1U)]\|TACCCTGTAGATCCGAATTTGTT | >MI0000266\|hsa-mir-10a&WithFlank&17\|-\|48579832\|48579953 | 28 | 49 | 22(+1U) | TACCCTGTAGATCCGAATTTGTT | 88.4 | 10.2 | 2.0 | 5.2 | 2.1E-07 | 1.4E-06 | BT20/MB468 | MP |
| >hg38_21_+_8433172_8446622_RNA45SN1_45S_50ntFlanks@12995.13038.44\|CTCGCTGCGATCTATTGAAAGTCAGCCCTCGACACAAGGGTTTG | >hg38_21_+_8433172_8446622_RNA45SN1_45S_50ntFlanks | 12995 | 13038 | 44 | CTCGCTGCGATCTATTGAAAGTCAGCCCTCGACACAAGGGTTTG | 93.9 | 10.3 | 2.0 | 5.2 | 2.1E-07 | 1.4E-06 | BT20/MB468 | MP |
| >hg38_21_+_8433172_8446622_RNA45SN1_45S_50ntFlanks@5020.5050.31\|CCGTGGGTGGTGGTGCATGGCCGTTCTTAGT | >hg38_21_+_8433172_8446622_RNA45SN1_45S_50ntFlanks | 5020 | 5050 | 31 | CCGTGGGTGGTGGTGCATGGCCGTTCTTAGT | 104.2 | 10.4 | 2.0 | 5.2 | 2.1E-07 | 1.4E-06 | BT20/MB468 | MP |
| >trna111_HisGTG_1_-_147774845_147774916@1.32.32,trna118_HisGTG_1_-_145396881_145396952@1.32.32,trna16_HisGTG_1_+_146544773_146544844@1.32.32,trna1_HisGTG_15_+_45493349_45493420@1.32.32,trna21_HisGTG_1_+_147753471_147753542@1.32.32,trna33_HisGTG_6_+_27125906_27125977@1.32.32,trna7_HisGTG_9_-_14433938_14434009@1.32.32,trna8_HisGTG_15_-_45492611_45492682@1.32.32,trna9_HisGTG_15_-_45490804_45490875@1.32.32\|GCCGTGATCGTATAGTGGTTAGTACTCTGCGT | >trna111_HisGTG_1_-_147774845_147774916 | 1 | 32 | 32 | GCCGTGATCGTATAGTGGTTAGTACTCTGCGT | 297.2 | 8.0 | 1.6 | 4.9 | 8.6E-07 | 1.4E-06 | MB231/MB468 | MITO |
| >hg38_21_+_8433172_8446622_RNA45SN1_45S_50ntFlanks@7974.8011.38\|ACGCGACCTCAGATCAGACGTGGCGACCCGCTGAATTT | >hg38_21_+_8433172_8446622_RNA45SN1_45S_50ntFlanks | 7974 | 8011 | 38 | ACGCGACCTCAGATCAGACGTGGCGACCCGCTGAATTT | 15670.8 | 3.0 | 0.6 | 4.9 | 9.0E-07 | 1.4E-06 | BT20/MB468 | NUC |
| >am_trna10_ValCAC_5_-_180649395_180649467@1.36.36,trna18_ValCAC_5_-_180529253_180529325@1.36.36,trna2_ValCAC_5_+_180524070_180524142@1.36.36,trna6_ValCAC_5_+_180600650_180600722@1.36.36,trna85_ValCAC_1_-_161369490_161369562@1.36.36,trna90_ValCAC_1_-_149684088_149684161@1.36.36,trna98_ValCAC_1_-_149298555_149298627@1.36.36,trna9_ValCAC_6_+_26538282_26538354@1.36.36\|GTTTCCGTAGTGTAGTGGTTATCACGTTCGCCTCAC | >am_trna10_ValCAC_5_-_180649395_180649467 | 1 | 36 | 36 | GTTTCCGTAGTGTAGTGGTTATCACGTTCGCCTCAC | 361.8 | 7.4 | 1.5 | 4.9 | 8.3E-07 | 1.4E-06 | MB231/MB468 | NUC |
| >hg38_21_+_8433172_8446622_RNA45SN1_45S_50ntFlanks@13008.13027.20\|ATTGAAAGTCAGCCCTCGAC | >hg38_21_+_8433172_8446622_RNA45SN1_45S_50ntFlanks | 13008 | 13027 | 20 | ATTGAAAGTCAGCCCTCGAC | 992.7 | -5.0 | 1.0 | -4.9 | 8.1E-07 | 1.4E-06 | MB231/MB468 | CYTO |
| >hg38_21_+_8433172_8446622_RNA45SN1_45S_50ntFlanks@3984.4026.43\|GCGGCTTTGGTGACTCTAGATAACCTCGGGCCGATCGCACGCC | >hg38_21_+_8433172_8446622_RNA45SN1_45S_50ntFlanks | 3984 | 4026 | 43 | GCGGCTTTGGTGACTCTAGATAACCTCGGGCCGATCGCACGCC | 933.8 | -8.2 | 1.6 | -5.2 | 2.1E-07 | 1.4E-06 | BT20/MB468 | MP |
| >hg38_21_+_8433172_8446622_RNA45SN1_45S_50ntFlanks@11568.11597.30\|GCAGCCGACTTAGAACTGGTGCGGACCAGG | >hg38_21_+_8433172_8446622_RNA45SN1_45S_50ntFlanks | 11568 | 11597 | 30 | GCAGCCGACTTAGAACTGGTGCGGACCAGG | 186.4 | 10.9 | 2.0 | 5.5 | 5.0E-08 | 1.4E-06 | BT20/MB231 | MP |
| >hg38_21_+_8433172_8446622_RNA45SN1_45S_50ntFlanks@11902.11931.30\|CTGGCACGGTGAAGAGACATGAGAGGTGTA | >hg38_21_+_8433172_8446622_RNA45SN1_45S_50ntFlanks | 11902 | 11931 | 30 | CTGGCACGGTGAAGAGACATGAGAGGTGTA | 288.3 | 11.5 | 2.1 | 5.5 | 5.0E-08 | 1.4E-06 | BT20/MB231 | MP |
| >hg38_21_+_8433172_8446622_RNA45SN1_45S_50ntFlanks@9969.9989.21\|GCTAAGGAGTGTGTAACAACT | >hg38_21_+_8433172_8446622_RNA45SN1_45S_50ntFlanks | 9969 | 9989 | 21 | GCTAAGGAGTGTGTAACAACT | 258.0 | 11.3 | 2.1 | 5.5 | 5.0E-08 | 1.4E-06 | BT20/MB231 | MP |
| >MI0000102\|hsa-mir-100&WithFlank&11\|-\|122152223\|122152314@19.39.21(+1U)&[MIMAT0000098&hsa-miR-100-5p&offsets\|0\|-1(+1U);m-27&11\|-\|122152276\|122152296&offsets\|0\|0(+1U)]\|AACCCGTAGATCCGAACTTGTT | >MI0000102\|hsa-mir-100&WithFlank&11\|-\|122152223\|122152314 | 19 | 39 | 21(+1U) | AACCCGTAGATCCGAACTTGTT | 11547.9 | -8.4 | 1.5 | -5.5 | 5.0E-08 | 1.4E-06 | BT20/MB231 | MP |
| >hg38_21_+_8433172_8446622_RNA45SN1_45S_50ntFlanks@9505.9526.22\|TGAACTATGCCTGGGCAGGGCG | >hg38_21_+_8433172_8446622_RNA45SN1_45S_50ntFlanks | 9505 | 9526 | 22 | TGAACTATGCCTGGGCAGGGCG | 1170.7 | -8.4 | 1.6 | -5.2 | 2.1E-07 | 1.4E-06 | BT20/MB468 | MP |
| >hg38_21_+_8433172_8446622_RNA45SN1_45S_50ntFlanks@12995.13039.45\|CTCGCTGCGATCTATTGAAAGTCAGCCCTCGACACAAGGGTTTGT | >hg38_21_+_8433172_8446622_RNA45SN1_45S_50ntFlanks | 12995 | 13039 | 45 | CTCGCTGCGATCTATTGAAAGTCAGCCCTCGACACAAGGGTTTGT | 5282.0 | 5.3 | 1.0 | 5.2 | 2.1E-07 | 1.4E-06 | BT20/MB468 | MP |
| >hg38_MT_+_1621_3279_RNR2_16S_50ntFlanks@1349.1389.41\|ACAATAGGGTTTACGACCTCGATGTTGGATCAGGACATCCC | >hg38_MT_+_1621_3279_RNR2_16S_50ntFlanks | 1349 | 1389 | 41 | ACAATAGGGTTTACGACCTCGATGTTGGATCAGGACATCCC | 112.1 | 10.6 | 2.0 | 5.2 | 2.1E-07 | 1.4E-06 | BT20/MB468 | MP |
| >hg38_21_+_8433172_8446622_RNA45SN1_45S_50ntFlanks@6649.6696.48\|TACGACTCTTAGCGGTGGATCACTCGGCTCGTGCGTCGATGAAGAACG | >hg38_21_+_8433172_8446622_RNA45SN1_45S_50ntFlanks | 6649 | 6696 | 48 | TACGACTCTTAGCGGTGGATCACTCGGCTCGTGCGTCGATGAAGAACG | 178.6 | 4.7 | 1.0 | 4.9 | 8.1E-07 | 1.4E-06 | MB231/MB468 | CYTO |
| >hg38_21_+_8433172_8446622_RNA45SN1_45S_50ntFlanks@10878.10901.24\|TCGGTCGGGCTGGGGCGCGAAGCG | >hg38_21_+_8433172_8446622_RNA45SN1_45S_50ntFlanks | 10878 | 10901 | 24 | TCGGTCGGGCTGGGGCGCGAAGCG | 1240.0 | -8.4 | 1.7 | -4.9 | 9.1E-07 | 1.4E-06 | BT20/MB468 | NUC |
| >hg38_21_+_8433172_8446622_RNA45SN1_45S_50ntFlanks@10681.10708.28\|TTCCGGCGGCGTCCGGTGAGCTCTCGCT | >hg38_21_+_8433172_8446622_RNA45SN1_45S_50ntFlanks | 10681 | 10708 | 28 | TTCCGGCGGCGTCCGGTGAGCTCTCGCT | 315.0 | -6.7 | 1.4 | -4.9 | 8.3E-07 | 1.4E-06 | MB231/MB468 | NUC |
| >hg38_21_+_8433172_8446622_RNA45SN1_45S_50ntFlanks@4220.4242.23\|ACGAAAAATAACAATACAGGACT | >hg38_21_+_8433172_8446622_RNA45SN1_45S_50ntFlanks | 4220 | 4242 | 23 | ACGAAAAATAACAATACAGGACT | 459.1 | -4.3 | 0.9 | -4.9 | 8.3E-07 | 1.4E-06 | MB231/MB468 | NUC |
| >hg38_21_+_8433172_8446622_RNA45SN1_45S_50ntFlanks@12407.12440.34\|TTTTGATCCTTCGATGTCGGCTCTTCCTATCATT | >hg38_21_+_8433172_8446622_RNA45SN1_45S_50ntFlanks | 12407 | 12440 | 34 | TTTTGATCCTTCGATGTCGGCTCTTCCTATCATT | 671.9 | -4.3 | 0.9 | -4.9 | 9.1E-07 | 1.4E-06 | BT20/MB468 | NUC |
| >hg38_21_+_8433172_8446622_RNA45SN1_45S_50ntFlanks@7989.8006.18\|AGACGTGGCGACCCGCTG | >hg38_21_+_8433172_8446622_RNA45SN1_45S_50ntFlanks | 7989 | 8006 | 18 | AGACGTGGCGACCCGCTG | 607.3 | -3.2 | 0.7 | -4.9 | 8.3E-07 | 1.4E-06 | MB231/MB468 | NUC |
| >hg38_21_+_8433172_8446622_RNA45SN1_45S_50ntFlanks@12991.13013.23\|CTCCCTCGCTGCGATCTATTGAA | >hg38_21_+_8433172_8446622_RNA45SN1_45S_50ntFlanks | 12991 | 13013 | 23 | CTCCCTCGCTGCGATCTATTGAA | 9022.8 | -4.2 | 0.9 | -4.9 | 8.1E-07 | 1.4E-06 | MB231/MB468 | CYTO |
| >hg38_21_+_8433172_8446622_RNA45SN1_45S_50ntFlanks@9554.9573.20\|CGTAGCGGTCCTGACGTGCA | >hg38_21_+_8433172_8446622_RNA45SN1_45S_50ntFlanks | 9554 | 9573 | 20 | CGTAGCGGTCCTGACGTGCA | 4028.3 | -2.7 | 0.5 | -5.0 | 6.2E-07 | 1.4E-06 | BT20/MB468 | MITO |
| >hg38_21_+_8433172_8446622_RNA45SN1_45S_50ntFlanks@10835.10867.33\|CGGATCCGTAACTTCGGGATAAGGATTGGCTCT | >hg38_21_+_8433172_8446622_RNA45SN1_45S_50ntFlanks | 10835 | 10867 | 33 | CGGATCCGTAACTTCGGGATAAGGATTGGCTCT | 507.8 | 3.5 | 0.7 | 5.0 | 7.1E-07 | 1.4E-06 | BT20/MB468 | CYTO |
| >hg38_21_+_8433172_8446622_RNA45SN1_45S_50ntFlanks@12457.12476.20\|CAAGCGTTGGATTGTTCACC | >hg38_21_+_8433172_8446622_RNA45SN1_45S_50ntFlanks | 12457 | 12476 | 20 | CAAGCGTTGGATTGTTCACC | 9431.8 | -4.2 | 0.9 | -5.0 | 7.1E-07 | 1.4E-06 | BT20/MB468 | CYTO |
| >hg38_21_+_8433172_8446622_RNA45SN1_45S_50ntFlanks@6730.6756.27\|ACACATTGATCATCGACACTTCGAACG | >hg38_21_+_8433172_8446622_RNA45SN1_45S_50ntFlanks | 6730 | 6756 | 27 | ACACATTGATCATCGACACTTCGAACG | 297.1 | -3.2 | 0.7 | -4.9 | 8.7E-07 | 1.4E-06 | MB231/MB468 | MITO |
| >hg38_21_+_8433172_8446622_RNA45SN1_45S_50ntFlanks@4002.4024.23\|GATAACCTCGGGCCGATCGCACG | >hg38_21_+_8433172_8446622_RNA45SN1_45S_50ntFlanks | 4002 | 4024 | 23 | GATAACCTCGGGCCGATCGCACG | 1012.5 | -5.4 | 1.1 | -5.0 | 6.2E-07 | 1.4E-06 | BT20/MB468 | MITO |
| >hg38_21_+_8433172_8446622_RNA45SN1_45S_50ntFlanks@8170.8195.26\|CAAGTCCTTCTGATCGAGGCCCAGCC | >hg38_21_+_8433172_8446622_RNA45SN1_45S_50ntFlanks | 8170 | 8195 | 26 | CAAGTCCTTCTGATCGAGGCCCAGCC | 1842.5 | -6.9 | 1.3 | -5.2 | 2.1E-07 | 1.4E-06 | BT20/MB468 | MP |
| >hg38_MT_+_1621_3279_RNR2_16S_50ntFlanks@51.92.42\|GCTAAACCTAGCCCCAAACCCACTCCACCTTACTACCAGACA | >hg38_MT_+_1621_3279_RNR2_16S_50ntFlanks | 51 | 92 | 42 | GCTAAACCTAGCCCCAAACCCACTCCACCTTACTACCAGACA | 5361.4 | 5.7 | 1.2 | 4.9 | 8.7E-07 | 1.4E-06 | MB231/MB468 | MITO |

| | | | | | | | | | | | | | |
|---|---|---|---|---|---|---|---|---|---|---|---|---|---|
| >hg38_21_+_8433172_8446622_RNA45SN1_45S_50ntFlanks@12420.12460.41\|ATGTCGGCTCTTCCTATCATTGTGAAGCAGAATTCACCAAG | >hg38_21_+_8433172_8446622_RNA45SN1_45S_50ntFlanks | 12420 | 12460 | 41 | ATGTCGGCTCTTCCTATCATTGTGAAGCAGAATTCACCAAG | 879.2 | -8.5 | 1.7 | -4.9 | 9.1E-07 | 1.4E-06 | BT20/MB468 | NUC |
| >hg38_21_+_8433172_8446622_RNA45SN1_45S_50ntFlanks@12640.12669.30\|ACCATCTGTGGGATTATGACTGAACGCCTC | >hg38_21_+_8433172_8446622_RNA45SN1_45S_50ntFlanks | 12640 | 12669 | 30 | ACCATCTGTGGGATTATGACTGAACGCCTC | 557.5 | 7.7 | 1.6 | 5.0 | 7.1E-07 | 1.4E-06 | BT20/MB468 | CYTO |
| >hg38_21_+_8433172_8446622_RNA45SN1_45S_50ntFlanks@4008.4042.35\|CTCGGGCCGATCGCACGCCCCCCGTGGCGGCGACG | >hg38_21_+_8433172_8446622_RNA45SN1_45S_50ntFlanks | 4008 | 4042 | 35 | CTCGGGCCGATCGCACGCCCCCCGTGGCGGCGACG | 185.7 | -5.0 | 1.0 | -4.9 | 8.4E-07 | 1.4E-06 | MB231/MB468 | NUC |
| >hg38_21_+_8433172_8446622_RNA45SN1_45S_50ntFlanks@6650.6673.24\|ACGACTCTTAGCGGTGGATCACTC | >hg38_21_+_8433172_8446622_RNA45SN1_45S_50ntFlanks | 6650 | 6673 | 24 | ACGACTCTTAGCGGTGGATCACTC | 246.2 | 4.5 | 0.9 | 5.2 | 2.2E-07 | 1.4E-06 | BT20/MB468 | MP |
| >hg38_21_+_8433172_8446622_RNA45SN1_45S_50ntFlanks@12539.12568.30\|ATGATGTGTTGTTGCCATGGTAATCCTGCT | >hg38_21_+_8433172_8446622_RNA45SN1_45S_50ntFlanks | 12539 | 12568 | 30 | ATGATGTGTTGTTGCCATGGTAATCCTGCT | 583.7 | -4.2 | 0.8 | -5.2 | 2.2E-07 | 1.4E-06 | BT20/MB468 | MP |
| >hg38_21_+_8433172_8446622_RNA45SN1_45S_50ntFlanks@6660.6681.22\|GCGGTGGATCACTCGGCTCGTG | >hg38_21_+_8433172_8446622_RNA45SN1_45S_50ntFlanks | 6660 | 6681 | 22 | GCGGTGGATCACTCGGCTCGTG | 325.4 | -3.2 | 0.6 | -4.9 | 8.2E-07 | 1.4E-06 | MB231/MB468 | CYTO |
| >hg38_21_+_8433172_8446622_RNA45SN1_45S_50ntFlanks@3837.3872.36\|CCTCTCCTACTTGGATAACTGTGGTAATTCTAGAGC | >hg38_21_+_8433172_8446622_RNA45SN1_45S_50ntFlanks | 3837 | 3872 | 36 | CCTCTCCTACTTGGATAACTGTGGTAATTCTAGAGC | 245.7 | -3.2 | 0.6 | -4.9 | 8.4E-07 | 1.4E-06 | MB231/MB468 | NUC |
| >hg38_21_+_8433172_8446622_RNA45SN1_45S_50ntFlanks@12624.12657.34\|CCAATGGGGCGAAGCTACCATCTGTGGGATTATG | >hg38_21_+_8433172_8446622_RNA45SN1_45S_50ntFlanks | 12624 | 12657 | 34 | CCAATGGGGCGAAGCTACCATCTGTGGGATTATG | 578.6 | -5.5 | 1.1 | -4.9 | 8.4E-07 | 1.4E-06 | MB231/MB468 | NUC |
| >MI0003643\|hsa-mir-629&WithFlank&15\|-\|70079366\|70079474@28.47.20&[MIMAT0004810&hsa-miR-629-5p&offsets\|0\|-1;m-150&15\|-\|70079427\|70079447&offsets\|0\|-1]\|TGGGTTTACGTTGGGAGAAC | >MI0003643\|hsa-mir-629&WithFlank&15\|-\|70079366\|70079474 | 28 | 47 | 20 | TGGGTTTACGTTGGGAGAAC | 48.0 | 8.8 | 1.8 | 5.0 | 7.3E-07 | 1.4E-06 | MB231/MB468 | MP |
| >trnaMT_LeuTAA_MT_+_3230_3304@1.37.37\|GTTAAGATGGCAGAGCCCGGTAATCGCATAAAACTTA | >trnaMT_LeuTAA_MT_+_3230_3304 | 1 | 37 | 37 | GTTAAGATGGCAGAGCCCGGTAATCGCATAAAACTTA | 410.0 | 7.2 | 1.5 | 4.9 | 8.8E-07 | 1.4E-06 | MB231/MB468 | MITO |
| >hg38_21_+_8433172_8446622_RNA45SN1_45S_50ntFlanks@11571.11600.30\|GCCGACTTAGAACTGGTGCGGACCAGGGGA | >hg38_21_+_8433172_8446622_RNA45SN1_45S_50ntFlanks | 11571 | 11600 | 30 | GCCGACTTAGAACTGGTGCGGACCAGGGGA | 479.1 | 2.7 | 0.5 | 5.0 | 6.3E-07 | 1.4E-06 | BT20/MB468 | MITO |
| >hg38_21_+_8433172_8446622_RNA45SN1_45S_50ntFlanks@6647.6695.49\|CGTACGACTCTTAGCGGTGGATCACTCGGCTCGTGCGTCGATGAAGAAC | >hg38_21_+_8433172_8446622_RNA45SN1_45S_50ntFlanks | 6647 | 6695 | 49 | CGTACGACTCTTAGCGGTGGATCACTCGGCTCGTGCGTCGATGAAGAAC | 2804.5 | 7.0 | 1.4 | 5.0 | 6.3E-07 | 1.4E-06 | BT20/MB468 | MITO |
| >hg38_21_+_8433172_8446622_RNA45SN1_45S_50ntFlanks@9180.9197.18\|CCGTCGGGCCCGGGGGAG | >hg38_21_+_8433172_8446622_RNA45SN1_45S_50ntFlanks | 9180 | 9197 | 18 | CCGTCGGGCCCGGGGGAG | 1319.5 | -8.2 | 1.6 | -5.2 | 2.2E-07 | 1.4E-06 | BT20/MB468 | MP |
| >hg38_21_+_8433172_8446622_RNA45SN1_45S_50ntFlanks@11890.11933.44\|CTTGACTCTAGTCTGGCACGGTGAAGAGACATGAGAGGTGTAGA | >hg38_21_+_8433172_8446622_RNA45SN1_45S_50ntFlanks | 11890 | 11933 | 44 | CTTGACTCTAGTCTGGCACGGTGAAGAGACATGAGAGGTGTAGA | 62.4 | 9.7 | 1.9 | 5.2 | 2.2E-07 | 1.4E-06 | BT20/MB468 | MP |
| >hg38_21_+_8433172_8446622_RNA45SN1_45S_50ntFlanks@6677.6714.38\|TCGTGCGTCGATGAAGAACGCAGCTAGCTGCGAGAATT | >hg38_21_+_8433172_8446622_RNA45SN1_45S_50ntFlanks | 6677 | 6714 | 38 | TCGTGCGTCGATGAAGAACGCAGCTAGCTGCGAGAATT | 62.8 | 9.7 | 1.9 | 5.2 | 2.2E-07 | 1.4E-06 | BT20/MB468 | MP |
| >hg38_21_+_8433172_8446622_RNA45SN1_45S_50ntFlanks@4733.4759.27\|GAACGAAAGTCGGAGGTTCGAAGACGA | >hg38_21_+_8433172_8446622_RNA45SN1_45S_50ntFlanks | 4733 | 4759 | 27 | GAACGAAAGTCGGAGGTTCGAAGACGA | 602.4 | -5.9 | 1.2 | -4.9 | 9.2E-07 | 1.4E-06 | BT20/MB468 | NUC |
| >hg38_21_+_8433172_8446622_RNA45SN1_45S_50ntFlanks@11325.11344.20\|TCTCGTCGCGCGCGCGTCCG | >hg38_21_+_8433172_8446622_RNA45SN1_45S_50ntFlanks | 11325 | 11344 | 20 | TCTCGTCGCGCGCGCGTCCG | 845.4 | -4.4 | 0.9 | -4.9 | 8.2E-07 | 1.4E-06 | MB231/MB468 | CYTO |
| >hg38_21_+_8433172_8446622_RNA45SN1_45S_50ntFlanks@4733.4761.29\|GAACGAAAGTCGGAGGTTCGAAGACGATC | >hg38_21_+_8433172_8446622_RNA45SN1_45S_50ntFlanks | 4733 | 4761 | 29 | GAACGAAAGTCGGAGGTTCGAAGACGATC | 392.9 | 3.3 | 0.7 | 4.9 | 8.5E-07 | 1.4E-06 | MB231/MB468 | NUC |
| >hg38_21_+_8433172_8446622_RNA45SN1_45S_50ntFlanks@9567.9603.37\|ACGTGCAAATCGGTCGTCCGACCTGGGTATAGGGGCG | >hg38_21_+_8433172_8446622_RNA45SN1_45S_50ntFlanks | 9567 | 9603 | 37 | ACGTGCAAATCGGTCGTCCGACCTGGGTATAGGGGCG | 2272.8 | -4.9 | 1.0 | -4.9 | 8.8E-07 | 1.4E-06 | MB231/MB468 | MITO |
| >hg38_21_+_8433172_8446622_RNA45SN1_45S_50ntFlanks@6663.6689.27\|GTGGATCACTCGGCTCGTGCGTCGATG | >hg38_21_+_8433172_8446622_RNA45SN1_45S_50ntFlanks | 6663 | 6689 | 27 | GTGGATCACTCGGCTCGTGCGTCGATG | 3586.4 | -2.7 | 0.5 | -5.0 | 6.3E-07 | 1.4E-06 | BT20/MB468 | MITO |
| >MI0000071\|hsa-mir-17&WithFlank&13\|+\|91350599\|91350694@20.42.23(+1C)&[MIMAT0000070&hsa-miR-17-5p&offsets\|0\|0(+1C);m-88&13\|+\|91350618\|91350640&offsets\|0\|0(+1C)]\|CAAAGTGCTTACAGTGCAGGTAGC | >MI0000071\|hsa-mir-17&WithFlank&13\|+\|91350599\|91350694 | 20 | 42 | 23(+1C) | CAAAGTGCTTACAGTGCAGGTAGC | 564.1 | 6.8 | 1.4 | 4.9 | 8.9E-07 | 1.4E-06 | MB231/MB468 | MITO |
| >hg38_21_+_8433172_8446622_RNA45SN1_45S_50ntFlanks@10778.10818.41\|CCAAGGTGAACAGCCTCTGGCATGTTGGAACAATGTAGGTA | >hg38_21_+_8433172_8446622_RNA45SN1_45S_50ntFlanks | 10778 | 10818 | 41 | CCAAGGTGAACAGCCTCTGGCATGTTGGAACAATGTAGGTA | 425.3 | -4.9 | 1.0 | -4.9 | 8.3E-07 | 1.4E-06 | MB231/MB468 | CYTO |
| >hg38_21_+_8433172_8446622_RNA45SN1_45S_50ntFlanks@4560.4579.20\|CTAGGAATAATGGAATAGGA | >hg38_21_+_8433172_8446622_RNA45SN1_45S_50ntFlanks | 4560 | 4579 | 20 | CTAGGAATAATGGAATAGGA | 223.7 | -6.6 | 1.3 | -4.9 | 8.5E-07 | 1.4E-06 | MB231/MB468 | NUC |
| >hg38_21_+_8433172_8446622_RNA45SN1_45S_50ntFlanks@9923.9946.24\|TTGGTTGATATAGACAGCAGGACG | >hg38_21_+_8433172_8446622_RNA45SN1_45S_50ntFlanks | 9923 | 9946 | 24 | TTGGTTGATATAGACAGCAGGACG | 55.1 | 9.5 | 1.8 | 5.2 | 2.2E-07 | 1.4E-06 | BT20/MB468 | MP |
| >hg38_1_-_228634819_228635039_RNA5S12_5S_50ntFlanks@84.102.19\|CTCGTCTGATCTCGGAAGC | >hg38_1_-_228634819_228635039_RNA5S12_5S_50ntFlanks | 84 | 102 | 19 | CTCGTCTGATCTCGGAAGC | 963.0 | -8.1 | 1.6 | -4.9 | 9.3E-07 | 1.4E-06 | BT20/MB468 | NUC |
| >hg38_21_+_8433172_8446622_RNA45SN1_45S_50ntFlanks@6778.6802.25\|CCCGGGGCTACGCCTGTCTGAGCGT | >hg38_21_+_8433172_8446622_RNA45SN1_45S_50ntFlanks | 6778 | 6802 | 25 | CCCGGGGCTACGCCTGTCTGAGCGT | 1656.0 | 4.2 | 0.9 | 4.9 | 8.5E-07 | 1.4E-06 | MB231/MB468 | NUC |
| >hg38_21_+_8433172_8446622_RNA45SN1_45S_50ntFlanks@9936.9968.33\|ACAGCAGGACGGTGGCCATGGAAGTCGGAATCC | >hg38_21_+_8433172_8446622_RNA45SN1_45S_50ntFlanks | 9936 | 9968 | 33 | ACAGCAGGACGGTGGCCATGGAAGTCGGAATCC | 320.5 | 11.7 | 2.1 | 5.4 | 5.1E-08 | 1.4E-06 | BT20/MB231 | MP |
| >hg38_21_+_8433172_8446622_RNA45SN1_45S_50ntFlanks@12007.12031.25\|GCCGCCGGTGAAATACCACTACTCT | >hg38_21_+_8433172_8446622_RNA45SN1_45S_50ntFlanks | 12007 | 12031 | 25 | GCCGCCGGTGAAATACCACTACTCT | 1465.0 | -6.4 | 1.3 | -5.0 | 7.4E-07 | 1.4E-06 | MB231/MB468 | MP |
| >am_trna119_LysCTT_1_-_145395522_145395594@15.32.18,trna11_LysCTT_5_-_180648979_180649051@15.32.18,trna128_LysCTT_1_-_55423542_55423614@15.32.18,trna13_LysCTT_6_+_26556774_26556846@15.32.18,trna32_LysCTT_16_-_3207406_3207478@15.32.18,trna6_LysCTT_19_-_52425396_52425468@15.32.18,trna7_LysCTT_16_+_3225692_3225764@15.32.18,trna9_LysCTT_5_+_180634755_180634827@15.32.18\|GTCGGTAGAGCATGAGAC | >am_trna119_LysCTT_1_-_145395522_145395594 | 15 | 32 | 18 | GTCGGTAGAGCATGAGAC | 639.9 | 6.7 | 1.3 | 5.0 | 7.2E-07 | 1.4E-06 | BT20/MB468 | CYTO |
| >trna116_GluCTC_1_-_145399233_145399304@31.70.40,trna71_GluCTC_1_-_161439189_161439260@31.70.40,trna74_GluCTC_1_-_161431809_161431880@31.70.40,trna77_GluCTC_1_-_161424398_161424469@31.70.40,trna77_GluCTC_6_+_28949976_28950047@31.70.40,trna80_GluCTC_1_-_161417018_161417089@31.70.40,trna87_GluCTC_6_-_126101393_126101464@31.70.40\|GCTCTCACCGCCGCGGCCCGGGTTCGATTCCCGGTCAGGG | >trna116_GluCTC_1_-_145399233_145399304 | 31 | 70 | 40 | GCTCTCACCGCCGCGGCCCGGGTTCGATTCCCGGTCAGGG | 901.6 | -8.1 | 1.7 | -4.9 | 9.3E-07 | 1.4E-06 | BT20/MB468 | NUC |
| >trna14_LysTTT_11_-_59327808_59327880@1.30.30,trna2_LysTTT_17_+_8022473_8022545@1.30.30,trna54_LysTTT_1_+_204475655_204475727@1.30.30,trna5_LysTTT_11_+_59323902_59323974@1.30.30,trna62_LysTTT_1_-_204476158_204476230@1.30.30,trna76_LysTTT_6_+_28918806_28918878@1.30.30\|GCCCGGATAGCTCAGTCGGTAGAGCATCAG | >trna14_LysTTT_11_-_59327808_59327880 | 1 | 30 | 30 | GCCCGGATAGCTCAGTCGGTAGAGCATCAG | 241.4 | -6.8 | 1.4 | -4.9 | 8.9E-07 | 1.4E-06 | MB231/MB468 | MITO |
| >MI0000542\|hsa-mir-320a&WithFlank&8\|-\|22244960\|22245043@48.69.22(+1C)&[MIMAT0000510&hsa-miR-320a-3p&offsets\|0\|0(+1C);m-60&8\|-\|22244975\|22244996&offsets\|0\|0(+1C)]\|AAAAGCTGGGTTGAGAGGGCGAC | >MI0000542\|hsa-mir-320a&WithFlank&8\|-\|22244960\|22245043 | 48 | 69 | 22(+1C) | AAAAGCTGGGTTGAGAGGGCGAC | 1404.6 | 7.0 | 1.4 | 5.2 | 2.2E-07 | 1.4E-06 | BT20/MB468 | MP |
| >hg38_21_+_8433172_8446622_RNA45SN1_45S_50ntFlanks@12360.12384.25\|CAGGGATAACTGGCTTGTGGCGGCC | >hg38_21_+_8433172_8446622_RNA45SN1_45S_50ntFlanks | 12360 | 12384 | 25 | CAGGGATAACTGGCTTGTGGCGGCC | 256.7 | -4.2 | 0.9 | -5.0 | 7.2E-07 | 1.4E-06 | BT20/MB468 | CYTO |
| >hg38_21_+_8433172_8446622_RNA45SN1_45S_50ntFlanks@9554.9592.39\|CGTAGCGGTCCTGACGTGCAAATCGGTCGTCCGACCTGG | >hg38_21_+_8433172_8446622_RNA45SN1_45S_50ntFlanks | 9554 | 9592 | 39 | CGTAGCGGTCCTGACGTGCAAATCGGTCGTCCGACCTGG | 242.0 | -7.4 | 1.5 | -4.9 | 8.9E-07 | 1.4E-06 | MB231/MB468 | MITO |
| >MI0000542\|hsa-mir-320a&WithFlank&8\|-\|22244960\|22245043@48.69.22&[MIMAT0000510&hsa-miR-320a-3p&offsets\|0\|0;m-60&8\|-\|22244975\|22244996&offsets\|0\|0]\|AAAAGCTGGGTTGAGAGGGCGA | >MI0000542\|hsa-mir-320a&WithFlank&8\|-\|22244960\|22245043 | 48 | 69 | 22 | AAAAGCTGGGTTGAGAGGGCGA | 49908.5 | 4.5 | 0.8 | 5.4 | 6.1E-08 | 1.4E-06 | BT20/MB231 | CYTO |
| >hg38_21_+_8433172_8446622_RNA45SN1_45S_50ntFlanks@12938.12959.22\|ACCATTCGTAGACGACCTGCTT | >hg38_21_+_8433172_8446622_RNA45SN1_45S_50ntFlanks | 12938 | 12959 | 22 | ACCATTCGTAGACGACCTGCTT | 278.6 | -2.6 | 0.5 | -4.9 | 8.6E-07 | 1.4E-06 | MB231/MB468 | NUC |
| >hg38_21_+_8433172_8446622_RNA45SN1_45S_50ntFlanks@10362.10388.27\|AAGGCCGAAGTGGAGAAGGGTTCCATG | >hg38_21_+_8433172_8446622_RNA45SN1_45S_50ntFlanks | 10362 | 10388 | 27 | AAGGCCGAAGTGGAGAAGGGTTCCATG | 809.5 | -6.8 | 1.4 | -5.0 | 6.4E-07 | 1.4E-06 | BT20/MB468 | MITO |
| >hg38_21_+_8433172_8446622_RNA45SN1_45S_50ntFlanks@11561.11594.34\|GCGCCTAGCAGCCGACTTAGAACTGGTGCGGACC | >hg38_21_+_8433172_8446622_RNA45SN1_45S_50ntFlanks | 11561 | 11594 | 34 | GCGCCTAGCAGCCGACTTAGAACTGGTGCGGACC | 477.3 | -4.7 | 0.9 | -4.9 | 8.4E-07 | 1.4E-06 | MB231/MB468 | CYTO |
| >hg38_21_+_8433172_8446622_RNA45SN1_45S_50ntFlanks@11672.11710.39\|CAGTGCTCTGAATGTCAAAGTGAAGAAATTCAATGAAGC | >hg38_21_+_8433172_8446622_RNA45SN1_45S_50ntFlanks | 11672 | 11710 | 39 | CAGTGCTCTGAATGTCAAAGTGAAGAAATTCAATGAAGC | 60.4 | 9.7 | 1.9 | 5.2 | 2.2E-07 | 1.4E-06 | BT20/MB468 | MP |
| >hg38_21_+_8433172_8446622_RNA45SN1_45S_50ntFlanks@12627.12657.31\|ATGGGGCGAAGCTACCATCTGTGGGATTATG | >hg38_21_+_8433172_8446622_RNA45SN1_45S_50ntFlanks | 12627 | 12657 | 31 | ATGGGGCGAAGCTACCATCTGTGGGATTATG | 920.6 | -7.7 | 1.5 | -5.2 | 2.2E-07 | 1.4E-06 | BT20/MB468 | MP |
| >hg38_21_+_8433172_8446622_RNA45SN1_45S_50ntFlanks@8154.8188.35\|CCGCTCGTGGGGGGCCCAAGTCCTTCTGATCGAGG | >hg38_21_+_8433172_8446622_RNA45SN1_45S_50ntFlanks | 8154 | 8188 | 35 | CCGCTCGTGGGGGGCCCAAGTCCTTCTGATCGAGG | 63.8 | -9.1 | 1.8 | -5.2 | 2.2E-07 | 1.4E-06 | BT20/MB468 | MP |
| >hg38_21_+_8433172_8446622_RNA45SN1_45S_50ntFlanks@10798.10825.28\|CATGTTGGAACAATGTAGGTAAGGGAAG | >hg38_21_+_8433172_8446622_RNA45SN1_45S_50ntFlanks | 10798 | 10825 | 28 | CATGTTGGAACAATGTAGGTAAGGGAAG | 267.0 | -5.3 | 1.1 | -4.9 | 8.6E-07 | 1.4E-06 | MB231/MB468 | NUC |
| >trnaMT_GluTTC_MT_-_14674_14742@24.63.40,trnalookalike8_GluTTC_5_-_93905172_93905240@24.63.40\|ATGGTTTTTCATATCATTGGTCGTGGTTGTAGTCCGTGCG | >trnaMT_GluTTC_MT_-_14674_14742 | 24 | 63 | 40 | ATGGTTTTTCATATCATTGGTCGTGGTTGTAGTCCGTGCG | 122.2 | 10.7 | 2.1 | 5.2 | 2.2E-07 | 1.4E-06 | BT20/MB468 | MP |
| >hg38_21_+_8433172_8446622_RNA45SN1_45S_50ntFlanks@5175.5207.33\|CCGAGATTGAGCAATAACAGGTCTGTGATGCCC | >hg38_21_+_8433172_8446622_RNA45SN1_45S_50ntFlanks | 5175 | 5207 | 33 | CCGAGATTGAGCAATAACAGGTCTGTGATGCCC | 56.5 | -9.7 | 2.0 | -4.9 | 7.4E-07 | 1.4E-06 | MB231/MB468 | MP |
| >hg38_21_+_8433172_8446622_RNA45SN1_45S_50ntFlanks@10678.10706.29\|CGGTTCCGGCGGCGTCCGGTGAGCTCTCG | >hg38_21_+_8433172_8446622_RNA45SN1_45S_50ntFlanks | 10678 | 10706 | 29 | CGGTTCCGGCGGCGTCCGGTGAGCTCTCG | 17079.6 | -4.9 | 1.0 | -5.0 | 6.4E-07 | 1.4E-06 | BT20/MB468 | MITO |
| >hg38_21_+_8433172_8446622_RNA45SN1_45S_50ntFlanks@5445.5466.22\|TCGGATCGGCCCCGCCGGGGTC | >hg38_21_+_8433172_8446622_RNA45SN1_45S_50ntFlanks | 5445 | 5466 | 22 | TCGGATCGGCCCCGCCGGGGTC | 5189.8 | -3.2 | 0.7 | -5.0 | 7.3E-07 | 1.4E-06 | BT20/MB468 | CYTO |

| | | | | | | | | | | | | | |
|---|---|---|---|---|---|---|---|---|---|---|---|---|---|
| >hg38_21_+_8433172_8446622_RNA45SN1_45S_50ntFlanks@9604.9622.19\|AAAGACTAATCGAACCATC | >hg38_21_+_8433172_8446622_RNA45SN1_45S_50ntFlanks | 9604 | 9622 | 19 | AAAGACTAATCGAACCATC | 520.6 | -2.9 | 0.6 | -5.0 | 6.4E-07 | 1.4E-06 | BT20/MB468 | MITO |
| >hg38_21_+_8433172_8446622_RNA45SN1_45S_50ntFlanks@10458.10480.23\|ATGGCCTCCGTTGCCCTCGGCCG | >hg38_21_+_8433172_8446622_RNA45SN1_45S_50ntFlanks | 10458 | 10480 | 23 | ATGGCCTCCGTTGCCCTCGGCCG | 99.6 | -9.7 | 1.9 | -5.2 | 2.2E-07 | 1.4E-06 | BT20/MB468 | MP |
| >hg38_21_+_8433172_8446622_RNA45SN1_45S_50ntFlanks@10537.10571.35\|CGAGGCGTCCAGTGCGGTAACGCGACCGATCCCGG | >hg38_21_+_8433172_8446622_RNA45SN1_45S_50ntFlanks | 10537 | 10571 | 35 | CGAGGCGTCCAGTGCGGTAACGCGACCGATCCCGG | 57.2 | -8.9 | 1.7 | -5.2 | 2.2E-07 | 1.4E-06 | BT20/MB468 | MP |
| >MI0000449\|hsa-mir-132&WithFlank&17\|-\|2049902\|2050014@65.85.21(+1A)&[MIMAT0000426&hsa-miR-132-3p&offsets\|0\|-1(+1A);m-138&17\|-\|2049929\|2049950&offsets\|0\|-1(+1A)]\|TAACAGTCTACAGCCATGGTCA | >MI0000449\|hsa-mir-132&WithFlank&17\|-\|2049902\|2050014 | 65 | 85 | 21(+1A) | TAACAGTCTACAGCCATGGTCA | 57.3 | 9.1 | 1.8 | 4.9 | 7.5E-07 | 1.4E-06 | MB231/MB468 | MP |
| >MI0005566\|hsa-mir-374b&WithFlank&X\|-\|74218541\|74218624@17.37.21&[MIMAT0004955&hsa-miR-374b-5p&offsets\|0\|-1;m-153&X\|-\|74218587\|74218608&offsets\|0\|-1]\|ATATAATACAACCTGCTAAGT | >MI0005566\|hsa-mir-374b&WithFlank&X\|-\|74218541\|74218624 | 17 | 37 | 21 | ATATAATACAACCTGCTAAGT | 1466.1 | 5.4 | 1.1 | 4.9 | 9.0E-07 | 1.4E-06 | MB231/MB468 | MITO |
| >hg38_21_+_8433172_8446622_RNA45SN1_45S_50ntFlanks@12942.12962.21\|TTCGTAGACGACCTGCTTCTG | >hg38_21_+_8433172_8446622_RNA45SN1_45S_50ntFlanks | 12942 | 12962 | 21 | TTCGTAGACGACCTGCTTCTG | 568.2 | -7.8 | 1.6 | -4.9 | 9.4E-07 | 1.4E-06 | BT20/MB468 | NUC |
| >hg38_21_+_8433172_8446622_RNA45SN1_45S_50ntFlanks@9736.9754.19\|CCGAAACGATCTCAACCTA | >hg38_21_+_8433172_8446622_RNA45SN1_45S_50ntFlanks | 9736 | 9754 | 19 | CCGAAACGATCTCAACCTA | 497.4 | -5.5 | 1.1 | -5.0 | 6.5E-07 | 1.4E-06 | BT20/MB468 | MITO |
| >hg38_21_+_8433172_8446622_RNA45SN1_45S_50ntFlanks@10401.10420.20\|GAACATGGGTCAGTCGGTCC | >hg38_21_+_8433172_8446622_RNA45SN1_45S_50ntFlanks | 10401 | 10420 | 20 | GAACATGGGTCAGTCGGTCC | 364.6 | -4.6 | 0.9 | -4.9 | 7.5E-07 | 1.4E-06 | MB231/MB468 | MP |
| >hg38_21_+_8433172_8446622_RNA45SN1_45S_50ntFlanks@12419.12453.35\|GATGTCGGCTCTTCCTATCATTGTGAAGCAGAATT | >hg38_21_+_8433172_8446622_RNA45SN1_45S_50ntFlanks | 12419 | 12453 | 35 | GATGTCGGCTCTTCCTATCATTGTGAAGCAGAATT | 830.3 | 5.8 | 1.2 | 4.9 | 8.4E-07 | 1.4E-06 | MB231/MB468 | CYTO |
| >hg38_21_+_8433172_8446622_RNA45SN1_45S_50ntFlanks@4657.4680.24\|CTAGAGGTGAAATTCTTGGACCGG | >hg38_21_+_8433172_8446622_RNA45SN1_45S_50ntFlanks | 4657 | 4680 | 24 | CTAGAGGTGAAATTCTTGGACCGG | 156.0 | -6.1 | 1.2 | -4.9 | 8.7E-07 | 1.4E-06 | MB231/MB468 | NUC |
| >MI0000085\|hsa-mir-27a&WithFlank&19\|-\|13836434\|13836523@57.76.20&[MIMAT0000084&hsa-miR-27a-3p&offsets\|0\|-1;m-43&19\|-\|13836447\|13836467&offsets\|0\|-1]\|TTCACAGTGGCTAAGTTCCG | >MI0000085\|hsa-mir-27a&WithFlank&19\|-\|13836434\|13836523 | 57 | 76 | 20 | TTCACAGTGGCTAAGTTCCG | 5270.7 | 4.7 | 1.0 | 4.9 | 8.4E-07 | 1.4E-06 | MB231/MB468 | CYTO |
| >hg38_21_+_8433172_8446622_RNA45SN1_45S_50ntFlanks@8381.8405.25\|AAGAGGGCGTGAAACCGTTAAGAGG | >hg38_21_+_8433172_8446622_RNA45SN1_45S_50ntFlanks | 8381 | 8405 | 25 | AAGAGGGCGTGAAACCGTTAAGAGG | 444.2 | -3.8 | 0.8 | -5.0 | 6.5E-07 | 1.4E-06 | BT20/MB468 | MITO |
| >MI0000255\|hsa-mir-30d&WithFlank&8\|-\|134804870\|134804951@12.35.24&[MIMAT0000245&hsa-miR-30d-5p&offsets\|0\|+2;m-21&8\|-\|134804919\|134804940&offsets\|0\|+2]\|TGTAAACATCCCCGACTGGAAGCT | >MI0000255\|hsa-mir-30d&WithFlank&8\|-\|134804870\|134804951 | 12 | 35 | 24 | TGTAAACATCCCCGACTGGAAGCT | 21167.8 | 4.5 | 0.9 | 5.2 | 2.2E-07 | 1.4E-06 | BT20/MB468 | MP |
| >hg38_21_+_8433172_8446622_RNA45SN1_45S_50ntFlanks@5492.5512.21\|AGAAGACGGTCGAACTTGACT | >hg38_21_+_8433172_8446622_RNA45SN1_45S_50ntFlanks | 5492 | 5512 | 21 | AGAAGACGGTCGAACTTGACT | 542.8 | -3.7 | 0.8 | -4.9 | 7.5E-07 | 1.4E-06 | MB231/MB468 | MP |
| >MI0005756\|hsa-mir-934&WithFlank&X\|+\|136550872\|136550966@21.40.20(+1A)&[MIMAT0004977&hsa-miR-934&offsets\|0\|-2(+1A);m-943&X\|+\|136550892\|136550913&offsets\|0\|-2(+1A)]\|TGTCTACTACTGGAGACACTA | >MI0005756\|hsa-mir-934&WithFlank&X\|+\|136550872\|136550966 | 21 | 40 | 20(+1A) | TGTCTACTACTGGAGACACTA | 67.8 | 5.8 | 1.2 | 4.9 | 9.4E-07 | 1.4E-06 | BT20/MB468 | NUC |
| >hg38_21_+_8433172_8446622_RNA45SN1_45S_50ntFlanks@12405.12446.42\|CTTTTTGATCCTTCGATGTCGGCTCTTCCTATCATTGTGAAG | >hg38_21_+_8433172_8446622_RNA45SN1_45S_50ntFlanks | 12405 | 12446 | 42 | CTTTTTGATCCTTCGATGTCGGCTCTTCCTATCATTGTGAAG | 1267.5 | -4.1 | 0.8 | -5.0 | 6.5E-07 | 1.4E-06 | BT20/MB468 | MITO |
| >hg38_MT_+_1621_3279_RNR2_16S_50ntFlanks@51.84.34\|GCTAAACCTAGCCCCAAACCCACTCCACCTTACT | >hg38_MT_+_1621_3279_RNR2_16S_50ntFlanks | 51 | 84 | 34 | GCTAAACCTAGCCCCAAACCCACTCCACCTTACT | 469.1 | 4.2 | 0.9 | 4.9 | 8.7E-07 | 1.4E-06 | MB231/MB468 | NUC |
| >MI0000273\|hsa-mir-183&WithFlank&7\|-\|129774899\|129775020@33.53.21(+1A)&[MIMAT0000261&hsa-miR-183-5p&offsets\|0\|-1(+1A);m-41&7\|-\|129774966\|129774987&offsets\|-1\|-2(+1A)]\|TATGGCACTGGTAGAATTCACA | >MI0000273\|hsa-mir-183&WithFlank&7\|-\|129774899\|129775020 | 33 | 53 | 21(+1A) | TATGGCACTGGTAGAATTCACA | 199.2 | 2.3 | 0.5 | 4.9 | 9.5E-07 | 1.4E-06 | BT20/MB468 | NUC |
| >hg38_21_+_8433172_8446622_RNA45SN1_45S_50ntFlanks@11668.11701.34\|TGCCCAGTGCTCTGAATGTCAAAGTGAAGAAATT | >hg38_21_+_8433172_8446622_RNA45SN1_45S_50ntFlanks | 11668 | 11701 | 34 | TGCCCAGTGCTCTGAATGTCAAAGTGAAGAAATT | 1752.4 | -6.8 | 1.4 | -4.9 | 9.5E-07 | 1.4E-06 | BT20/MB468 | NUC |
| >MI0000081\|hsa-mir-24-2&WithFlank&19\|-\|13836281\|13836365@56.77.22(+1U);MI0000080\|hsa-mir-24-1&WithFlank&9\|+\|95086015\|95086094@50.71.22(+1U)&[MIMAT0000080&hsa-miR-24-3p&offsets\|0\|0(+1U);m-36&19\|-\|13836288\|13836310&offsets\|0\|-1(+1U)];[MIMAT0000080_1&hsa-miR-24-3p&offsets\|0\|0(+1U);m-35&9\|+\|95086064\|95086086&offsets\|0\|-1(+1U)]\|TGGCTCAGTTCAGCAGGAACAGT | >MI0000081\|hsa-mir-24-2&WithFlank&19\|-\|13836281\|13836365 | 56 | 77 | 22(+1U) | TGGCTCAGTTCAGCAGGAACAGT | 2135.3 | 4.7 | 1.0 | 4.9 | 8.5E-07 | 1.4E-06 | MB231/MB468 | CYTO |
| >hg38_21_+_8433172_8446622_RNA45SN1_45S_50ntFlanks@11848.11872.25\|ACGGGCTTGGCGGAATCAGCGGGGA | >hg38_21_+_8433172_8446622_RNA45SN1_45S_50ntFlanks | 11848 | 11872 | 25 | ACGGGCTTGGCGGAATCAGCGGGGA | 302.3 | -6.5 | 1.3 | -4.9 | 8.7E-07 | 1.4E-06 | MB231/MB468 | NUC |
| >hg38_21_+_8433172_8446622_RNA45SN1_45S_50ntFlanks@12415.12438.24\|CTTCGATGTCGGCTCTTCCTATCA | >hg38_21_+_8433172_8446622_RNA45SN1_45S_50ntFlanks | 12415 | 12438 | 24 | CTTCGATGTCGGCTCTTCCTATCA | 896.5 | 3.3 | 0.7 | 4.9 | 8.5E-07 | 1.4E-06 | MB231/MB468 | CYTO |
| >hg38_21_+_8433172_8446622_RNA45SN1_45S_50ntFlanks@6661.6701.41\|CGGTGGATCACTCGGCTCGTGCGTCGATGAAGAACGCAGCT | >hg38_21_+_8433172_8446622_RNA45SN1_45S_50ntFlanks | 6661 | 6701 | 41 | CGGTGGATCACTCGGCTCGTGCGTCGATGAAGAACGCAGCT | 1455.4 | 5.2 | 1.0 | 5.2 | 2.2E-07 | 1.4E-06 | BT20/MB468 | MP |
| >hg38_21_+_8433172_8446622_RNA45SN1_45S_50ntFlanks@12636.12659.24\|AGCTACCATCTGTGGGATTATGAC | >hg38_21_+_8433172_8446622_RNA45SN1_45S_50ntFlanks | 12636 | 12659 | 24 | AGCTACCATCTGTGGGATTATGAC | 2940.2 | -4.8 | 1.0 | -5.0 | 6.5E-07 | 1.4E-06 | BT20/MB468 | MITO |
| >hg38_21_+_8433172_8446622_RNA45SN1_45S_50ntFlanks@7981.8008.28\|CTCAGATCAGACGTGGCGACCCGCTGAA | >hg38_21_+_8433172_8446622_RNA45SN1_45S_50ntFlanks | 7981 | 8008 | 28 | CTCAGATCAGACGTGGCGACCCGCTGAA | 1487.5 | -2.5 | 0.5 | -5.0 | 6.5E-07 | 1.4E-06 | BT20/MB468 | MITO |
| >hg38_21_+_8433172_8446622_RNA45SN1_45S_50ntFlanks@11572.11609.38\|CCGACTTAGAACTGGTGCGGACCAGGGGAATCCGACTG | >hg38_21_+_8433172_8446622_RNA45SN1_45S_50ntFlanks | 11572 | 11609 | 38 | CCGACTTAGAACTGGTGCGGACCAGGGGAATCCGACTG | 553.5 | -3.1 | 0.6 | -4.9 | 8.7E-07 | 1.4E-06 | MB231/MB468 | NUC |
| >hg38_21_+_8433172_8446622_RNA45SN1_45S_50ntFlanks@11704.11728.25\|ATGAAGCGCGGGTAAACGGCGGGAG | >hg38_21_+_8433172_8446622_RNA45SN1_45S_50ntFlanks | 11704 | 11728 | 25 | ATGAAGCGCGGGTAAACGGCGGGAG | 851.9 | -6.7 | 1.3 | -5.0 | 6.5E-07 | 1.4E-06 | BT20/MB468 | MITO |
| >MI0000273\|hsa-mir-183&WithFlank&7\|-\|129774899\|129775020@34.55.22&[MIMAT0000261&hsa-miR-183-5p&offsets\|+1\|+1;m-41&7\|-\|129774966\|129774987&offsets\|0\|0]\|ATGGCACTGGTAGAATTCACTG | >MI0000273\|hsa-mir-183&WithFlank&7\|-\|129774899\|129775020 | 34 | 55 | 22 | ATGGCACTGGTAGAATTCACTG | 1125.2 | 2.4 | 0.5 | 4.9 | 9.5E-07 | 1.4E-06 | BT20/MB468 | NUC |
| >hg38_21_+_8433172_8446622_RNA45SN1_45S_50ntFlanks@10411.10439.29\|CAGTCGGTCCTGAGAGATGGGCGAGCGCC | >hg38_21_+_8433172_8446622_RNA45SN1_45S_50ntFlanks | 10411 | 10439 | 29 | CAGTCGGTCCTGAGAGATGGGCGAGCGCC | 193.6 | -5.6 | 1.1 | -4.9 | 7.6E-07 | 1.4E-06 | MB231/MB468 | MP |
| >am_trna2_GlyGCC_21_-_18827107_18827177@4.26.23,trna35_GlyGCC_1_+_161413094_161413164@4.26.23,trna37_GlyGCC_1_+_161420467_161420537@4.26.23,trna39_GlyGCC_1_+_161427898_161427968@4.26.23,trna41_GlyGCC_1_+_161435258_161435328@4.26.23\|TGGGTGGTTCAGTGGTAGAATTC | >am_trna2_GlyGCC_21_-_18827107_18827177 | 4 | 26 | 23 | TGGGTGGTTCAGTGGTAGAATTC | 73.2 | 9.9 | 1.9 | 5.2 | 2.3E-07 | 1.4E-06 | BT20/MB468 | MP |
| >hg38_21_+_8433172_8446622_RNA45SN1_45S_50ntFlanks@11161.11181.21\|CACGGTCCCCCGCGAGGGGGG | >hg38_21_+_8433172_8446622_RNA45SN1_45S_50ntFlanks | 11161 | 11181 | 21 | CACGGTCCCCCGCGAGGGGGG | 125.5 | -10.1 | 1.9 | -5.2 | 2.3E-07 | 1.4E-06 | BT20/MB468 | MP |
| >hg38_21_+_8433172_8446622_RNA45SN1_45S_50ntFlanks@10726.10751.26\|GGGAGAGGGTGTAAATCTCGCGCCGG | >hg38_21_+_8433172_8446622_RNA45SN1_45S_50ntFlanks | 10726 | 10751 | 26 | GGGAGAGGGTGTAAATCTCGCGCCGG | 79.7 | -9.4 | 1.8 | -5.2 | 2.3E-07 | 1.4E-06 | BT20/MB468 | MP |
| >hg38_21_+_8433172_8446622_RNA45SN1_45S_50ntFlanks@11641.11665.25\|CGGCGGGTGTTGACGCGATGTGATT | >hg38_21_+_8433172_8446622_RNA45SN1_45S_50ntFlanks | 11641 | 11665 | 25 | CGGCGGGTGTTGACGCGATGTGATT | 9112.1 | -8.9 | 1.7 | -5.2 | 2.3E-07 | 1.4E-06 | BT20/MB468 | MP |
| >hg38_21_+_8433172_8446622_RNA45SN1_45S_50ntFlanks@7992.8018.27\|CGTGGCGACCCGCTGAATTTAAGCATA | >hg38_21_+_8433172_8446622_RNA45SN1_45S_50ntFlanks | 7992 | 8018 | 27 | CGTGGCGACCCGCTGAATTTAAGCATA | 153.0 | 5.9 | 1.2 | 4.9 | 8.5E-07 | 1.4E-06 | MB231/MB468 | CYTO |
| >trnaMT_PheGAA_MT_+_577_647@43.74.32\|TTAGACGGGCTCACATCACCCCATAAACACCA | >trnaMT_PheGAA_MT_+_577_647 | 43 | 74 | 32 | TTAGACGGGCTCACATCACCCCATAAACACCA | 396.4 | 8.1 | 1.6 | 4.9 | 8.8E-07 | 1.4E-06 | MB231/MB468 | NUC |
| >hg38_21_+_8433172_8446622_RNA45SN1_45S_50ntFlanks@8359.8378.20\|AAGAACTTTGAAGAGAGAGT | >hg38_21_+_8433172_8446622_RNA45SN1_45S_50ntFlanks | 8359 | 8378 | 20 | AAGAACTTTGAAGAGAGAGT | 852.6 | -6.3 | 1.3 | -4.9 | 7.6E-07 | 1.4E-06 | MB231/MB468 | MP |
| >hg38_MT_+_1621_3279_RNR2_16S_50ntFlanks@622.652.31\|TATAACTGAACTCCTCACACCCAATTGGACC | >hg38_MT_+_1621_3279_RNR2_16S_50ntFlanks | 622 | 652 | 31 | TATAACTGAACTCCTCACACCCAATTGGACC | 226.2 | 5.9 | 1.2 | 4.9 | 9.2E-07 | 1.4E-06 | MB231/MB468 | MITO |
| >hg38_21_+_8433172_8446622_RNA45SN1_45S_50ntFlanks@10808.10837.30\|CAATGTAGGTAAGGGAAGTCGGCAAGCCGG | >hg38_21_+_8433172_8446622_RNA45SN1_45S_50ntFlanks | 10808 | 10837 | 30 | CAATGTAGGTAAGGGAAGTCGGCAAGCCGG | 286.2 | -5.4 | 1.1 | -4.9 | 8.8E-07 | 1.4E-06 | MB231/MB468 | NUC |
| >hg38_21_+_8433172_8446622_RNA45SN1_45S_50ntFlanks@12321.12344.24\|CTTTTGGGTTTTAAGCAGGAGGTG | >hg38_21_+_8433172_8446622_RNA45SN1_45S_50ntFlanks | 12321 | 12344 | 24 | CTTTTGGGTTTTAAGCAGGAGGTG | 56.7 | -8.9 | 1.7 | -5.2 | 2.3E-07 | 1.4E-06 | BT20/MB468 | MP |
| >am_trnaMT_PheGAA_MT_+_577_647@41.62.22\|GTTTAGACGGGCTCACATCACC | >am_trnaMT_PheGAA_MT_+_577_647 | 41 | 62 | 22 | GTTTAGACGGGCTCACATCACC | 533.9 | 6.6 | 1.3 | 4.9 | 8.6E-07 | 1.4E-06 | MB231/MB468 | CYTO |
| >hg38_21_+_8433172_8446622_RNA45SN1_45S_50ntFlanks@10348.10372.25\|AAACGAGAACTTTGAAGGCCGAAGT | >hg38_21_+_8433172_8446622_RNA45SN1_45S_50ntFlanks | 10348 | 10372 | 25 | AAACGAGAACTTTGAAGGCCGAAGT | 544.7 | -2.3 | 0.5 | -4.9 | 9.6E-07 | 1.4E-06 | BT20/MB468 | NUC |
| >hg38_21_+_8433172_8446622_RNA45SN1_45S_50ntFlanks@11562.11591.30\|CGCCTAGCAGCCGACTTAGAACTGGTGCGG | >hg38_21_+_8433172_8446622_RNA45SN1_45S_50ntFlanks | 11562 | 11591 | 30 | CGCCTAGCAGCCGACTTAGAACTGGTGCGG | 112.2 | -5.5 | 1.1 | -4.9 | 7.5E-07 | 1.4E-06 | BT20/MB468 | CYTO |
| >hg38_21_+_8433172_8446622_RNA45SN1_45S_50ntFlanks@12971.12989.19\|TTTCGTACGTAGCAGAGCA | >hg38_21_+_8433172_8446622_RNA45SN1_45S_50ntFlanks | 12971 | 12989 | 19 | TTTCGTACGTAGCAGAGCA | 310.1 | -4.5 | 0.9 | -4.9 | 7.5E-07 | 1.4E-06 | BT20/MB468 | CYTO |
| >hg38_21_+_8433172_8446622_RNA45SN1_45S_50ntFlanks@4733.4761.29\|GAACGAAAGTCGGAGGTTCGAAGACGATC | >hg38_21_+_8433172_8446622_RNA45SN1_45S_50ntFlanks | 4733 | 4761 | 29 | GAACGAAAGTCGGAGGTTCGAAGACGATC | 1046.6 | 6.0 | 1.2 | 4.9 | 7.5E-07 | 1.4E-06 | BT20/MB468 | CYTO |
| >hg38_21_+_8433172_8446622_RNA45SN1_45S_50ntFlanks@11595.11626.32\|AGGGGAATCCGACTGTTTAATTAAAACAAAGC | >hg38_21_+_8433172_8446622_RNA45SN1_45S_50ntFlanks | 11595 | 11626 | 32 | AGGGGAATCCGACTGTTTAATTAAAACAAAGC | 65.8 | 9.8 | 1.9 | 5.2 | 2.3E-07 | 1.5E-06 | BT20/MB468 | MP |
| >MI0000077\|hsa-mir-21&WithFlank&17\|+\|59841260\|59841343@13.37.25&[MIMAT0000076&hsa-miR-21-5p&offsets\|-1\|+2;m-1&17\|+\|59841273\|59841294&offsets\|-1\|+2]\|GTAGCTTATCAGACTGATGTTGACT | >MI0000077\|hsa-mir-21&WithFlank&17\|+\|59841260\|59841343 | 13 | 37 | 25 | GTAGCTTATCAGACTGATGTTGACT | 262.0 | 7.9 | 1.6 | 4.9 | 9.2E-07 | 1.5E-06 | MB231/MB468 | MITO |
| >hg38_21_+_8433172_8446622_RNA45SN1_45S_50ntFlanks@10410.10453.44\|TCAGTCGGTCCTGAGAGATGGGCGAGCGCCGTTCCGAAGGGACG | >hg38_21_+_8433172_8446622_RNA45SN1_45S_50ntFlanks | 10410 | 10453 | 44 | TCAGTCGGTCCTGAGAGATGGGCGAGCGCCGTTCCGAAGGGACG | 1113.7 | -6.1 | 1.2 | -4.9 | 9.2E-07 | 1.5E-06 | MB231/MB468 | MITO |
| >hg38_21_+_8433172_8446622_RNA45SN1_45S_50ntFlanks@7992.8012.21\|CGTGGCGACCCGCTGAATTTA | >hg38_21_+_8433172_8446622_RNA45SN1_45S_50ntFlanks | 7992 | 8012 | 21 | CGTGGCGACCCGCTGAATTTA | 618.9 | -7.2 | 1.4 | -4.9 | 7.6E-07 | 1.5E-06 | MB231/MB468 | MP |
| >hg38_21_+_8433172_8446622_RNA45SN1_45S_50ntFlanks@12420.12455.36\|ATGTCGGCTCTTCCTATCATTGTGAAGCAGAATTCA | >hg38_21_+_8433172_8446622_RNA45SN1_45S_50ntFlanks | 12420 | 12455 | 36 | ATGTCGGCTCTTCCTATCATTGTGAAGCAGAATTCA | 8616.8 | -4.7 | 1.0 | -4.9 | 9.2E-07 | 1.5E-06 | MB231/MB468 | MITO |
| >hg38_21_+_8433172_8446622_RNA45SN1_45S_50ntFlanks@7975.7992.18\|CGCGACCTCAGATCAGAC | >hg38_21_+_8433172_8446622_RNA45SN1_45S_50ntFlanks | 7975 | 7992 | 18 | CGCGACCTCAGATCAGAC | 15754.5 | 4.3 | 0.9 | 4.9 | 8.6E-07 | 1.5E-06 | MB231/MB468 | CYTO |
| >hg38_21_+_8433172_8446622_RNA45SN1_45S_50ntFlanks@6670.6699.30\|ACTCGGCTCGTGCGTCGATGAAGAACGCAG | >hg38_21_+_8433172_8446622_RNA45SN1_45S_50ntFlanks | 6670 | 6699 | 30 | ACTCGGCTCGTGCGTCGATGAAGAACGCAG | 1535.2 | -2.9 | 0.6 | -5.0 | 6.6E-07 | 1.5E-06 | BT20/MB468 | MITO |

<table>
<tr><td>>hg38_21_+_8433172_8446622_RNA45SN1_45S_50ntFlanks@4570.4587.18|TGGAATAGGACCGCGGTT</td><td>>hg38_21_+_8433172_8446622_RNA45SN1_45S_50ntFlanks</td><td>4570</td><td>4587</td><td>18</td><td>TGGAATAGGACCGCGGTT</td><td>698.7</td><td>-7.1</td><td>1.4</td><td>-4.9</td><td>8.9E-07</td><td>1.5E-06</td><td>MB231/MB468</td><td>NUC</td></tr>
<tr><td>>hg38_21_+_8433172_8446622_RNA45SN1_45S_50ntFlanks@11672.11691.20|CAGTGCTCTGAATGTCAAAG</td><td>>hg38_21_+_8433172_8446622_RNA45SN1_45S_50ntFlanks</td><td>11672</td><td>11691</td><td>20</td><td>CAGTGCTCTGAATGTCAAAG</td><td>641.9</td><td>-4.2</td><td>0.8</td><td>-4.9</td><td>9.6E-07</td><td>1.5E-06</td><td>BT20/MB468</td><td>NUC</td></tr>
<tr><td>>MI0000067|hsa-let-7f-1&WithFlank&9|+|94176341|94176439@69.89.21(+1U)&[MIMAT0004486&hsa-let-7f-1-3p&offsets|0|-1(+1U);m-275&9|+|94176409|94176430&offsets|0|-1(+1U)]|CTATACAATCTATTGCCTTCCT</td><td>>MI0000067|hsa-let-7f-1&WithFlank&9|+|94176341|94176439</td><td>69</td><td>89</td><td>21(+1U)</td><td>CTATACAATCTATTGCCTTCCT</td><td>426.8</td><td>3.8</td><td>0.8</td><td>4.9</td><td>7.6E-07</td><td>1.5E-06</td><td>MB231/MB468</td><td>MP</td></tr>
<tr><td>>hg38_21_+_8433172_8446622_RNA45SN1_45S_50ntFlanks@12007.12038.32|GCCGCCGGTGAAATACCACTACTCTGATCGTT</td><td>>hg38_21_+_8433172_8446622_RNA45SN1_45S_50ntFlanks</td><td>12007</td><td>12038</td><td>32</td><td>GCCGCCGGTGAAATACCACTACTCTGATCGTT</td><td>167.6</td><td>-5.7</td><td>1.1</td><td>-4.9</td><td>7.7E-07</td><td>1.5E-06</td><td>MB231/MB468</td><td>MP</td></tr>
<tr><td>>hg38_21_+_8433172_8446622_RNA45SN1_45S_50ntFlanks@4567.4589.23|TAATGGAATAGGACCGCGGTTCT</td><td>>hg38_21_+_8433172_8446622_RNA45SN1_45S_50ntFlanks</td><td>4567</td><td>4589</td><td>23</td><td>TAATGGAATAGGACCGCGGTTCT</td><td>648.0</td><td>-5.2</td><td>1.1</td><td>-4.9</td><td>9.2E-07</td><td>1.5E-06</td><td>MB231/MB468</td><td>MITO</td></tr>
<tr><td>>hg38_21_+_8433172_8446622_RNA45SN1_45S_50ntFlanks@5472.5491.20|ACGGCCCTGGCGGAGCGCTG</td><td>>hg38_21_+_8433172_8446622_RNA45SN1_45S_50ntFlanks</td><td>5472</td><td>5491</td><td>20</td><td>ACGGCCCTGGCGGAGCGCTG</td><td>1072.1</td><td>3.2</td><td>0.6</td><td>4.9</td><td>8.6E-07</td><td>1.5E-06</td><td>MB231/MB468</td><td>CYTO</td></tr>
<tr><td>>hg38_21_+_8433172_8446622_RNA45SN1_45S_50ntFlanks@11210.11233.24|GGCGACTCTGGACGCGAGCCGGGC</td><td>>hg38_21_+_8433172_8446622_RNA45SN1_45S_50ntFlanks</td><td>11210</td><td>11233</td><td>24</td><td>GGCGACTCTGGACGCGAGCCGGGC</td><td>354.5</td><td>-5.4</td><td>1.1</td><td>-4.9</td><td>8.9E-07</td><td>1.5E-06</td><td>MB231/MB468</td><td>NUC</td></tr>
<tr><td>>hg38_21_+_8433172_8446622_RNA45SN1_45S_50ntFlanks@12637.12661.25|GCTACCATCTGTGGGATTATGACTG</td><td>>hg38_21_+_8433172_8446622_RNA45SN1_45S_50ntFlanks</td><td>12637</td><td>12661</td><td>25</td><td>GCTACCATCTGTGGGATTATGACTG</td><td>675.4</td><td>-6.3</td><td>1.3</td><td>-4.9</td><td>9.7E-07</td><td>1.5E-06</td><td>BT20/MB468</td><td>NUC</td></tr>
<tr><td>>hg38_21_+_8433172_8446622_RNA45SN1_45S_50ntFlanks@7991.8013.23|ACGTGGCGACCCGCTGAATTTAA</td><td>>hg38_21_+_8433172_8446622_RNA45SN1_45S_50ntFlanks</td><td>7991</td><td>8013</td><td>23</td><td>ACGTGGCGACCCGCTGAATTTAA</td><td>646.7</td><td>-7.3</td><td>1.5</td><td>-4.9</td><td>7.7E-07</td><td>1.5E-06</td><td>MB231/MB468</td><td>MP</td></tr>
<tr><td>>hg38_21_+_8433172_8446622_RNA45SN1_45S_50ntFlanks@12624.12651.28|CCAATGGGGCGAAGCTACCATCTGTGGG</td><td>>hg38_21_+_8433172_8446622_RNA45SN1_45S_50ntFlanks</td><td>12624</td><td>12651</td><td>28</td><td>CCAATGGGGCGAAGCTACCATCTGTGGG</td><td>97.3</td><td>-9.7</td><td>1.9</td><td>-5.2</td><td>2.3E-07</td><td>1.5E-06</td><td>BT20/MB468</td><td>MP</td></tr>
<tr><td>>MI0001446|hsa-mir-424&WithFlank&X|-|134546608|134546717@54.74.21&[MIMAT0004749&hsa-miR-424-3p&offsets|0|0;m-305&X|-|134546644|134546664&offsets|0|0]|CAAAACGTGAGGCGCTGCTAT</td><td>>MI0001446|hsa-mir-424&WithFlank&X|-|134546608|134546717</td><td>54</td><td>74</td><td>21</td><td>CAAAACGTGAGGCGCTGCTAT</td><td>132.7</td><td>4.5</td><td>0.9</td><td>4.9</td><td>9.7E-07</td><td>1.5E-06</td><td>BT20/MB468</td><td>NUC</td></tr>
<tr><td>>MI0000298|hsa-mir-221&WithFlank&X|-|45746151|45746272@71.90.20(+1C)&[MIMAT0000278&hsa-miR-221-3p&offsets|0|-3(+1C);m-91&X|-|45746181|45746202&offsets|0|-2(+1C)]|AGCTACATTGTCTGCTGGGTC</td><td>>MI0000298|hsa-mir-221&WithFlank&X|-|45746151|45746272</td><td>71</td><td>90</td><td>20(+1C)</td><td>AGCTACATTGTCTGCTGGGTC</td><td>348.6</td><td>8.1</td><td>1.7</td><td>4.9</td><td>9.3E-07</td><td>1.5E-06</td><td>MB231/MB468</td><td>MITO</td></tr>
<tr><td>>am_MI0000809|hsa-mir-151a&WithFlank&8|-|140732558|140732659@53.73.21&[MIMAT0000757&hsa-miR-151a-3p&offsets|0|0;m-30&8|-|140732587|140732607&offsets|0|0]|CTAGACTGAAGCTCCTTGAGG</td><td>>am_MI0000809|hsa-mir-151a&WithFlank&8|-|140732558|140732659</td><td>53</td><td>73</td><td>21</td><td>CTAGACTGAAGCTCCTTGAGG</td><td>33732.6</td><td>3.6</td><td>0.7</td><td>4.9</td><td>7.7E-07</td><td>1.5E-06</td><td>MB231/MB468</td><td>MP</td></tr>
<tr><td>>hg38_21_+_8433172_8446622_RNA45SN1_45S_50ntFlanks@4512.4532.21|AAAAAATTAGAGTGTTCAAAG</td><td>>hg38_21_+_8433172_8446622_RNA45SN1_45S_50ntFlanks</td><td>4512</td><td>4532</td><td>21</td><td>AAAAAATTAGAGTGTTCAAAG</td><td>63.8</td><td>-9.1</td><td>1.8</td><td>-5.2</td><td>2.3E-07</td><td>1.5E-06</td><td>BT20/MB468</td><td>MP</td></tr>
<tr><td>>hg38_21_+_8433172_8446622_RNA45SN1_45S_50ntFlanks@10850.10868.19|GGGATAAGGATTGGCTCTA</td><td>>hg38_21_+_8433172_8446622_RNA45SN1_45S_50ntFlanks</td><td>10850</td><td>10868</td><td>19</td><td>GGGATAAGGATTGGCTCTA</td><td>189.7</td><td>-4.9</td><td>1.0</td><td>-4.9</td><td>8.9E-07</td><td>1.5E-06</td><td>MB231/MB468</td><td>NUC</td></tr>
<tr><td>>hg38_21_+_8433172_8446622_RNA45SN1_45S_50ntFlanks@11567.11603.37|AGCAGCCGACTTAGAACTGGTGCGGACCAGGGGAATC</td><td>>hg38_21_+_8433172_8446622_RNA45SN1_45S_50ntFlanks</td><td>11567</td><td>11603</td><td>37</td><td>AGCAGCCGACTTAGAACTGGTGCGGACCAGGGGAATC</td><td>629.0</td><td>-5.9</td><td>1.2</td><td>-4.9</td><td>7.7E-07</td><td>1.5E-06</td><td>MB231/MB468</td><td>MP</td></tr>
<tr><td>>hg38_1_-_228634819_228635039_RNA5S12_5S_50ntFlanks@142.169.28|CGCCTGGGAATACCGGGTGCTGTAGGCT</td><td>>hg38_1_-_228634819_228635039_RNA5S12_5S_50ntFlanks</td><td>142</td><td>169</td><td>28</td><td>CGCCTGGGAATACCGGGTGCTGTAGGCT</td><td>212.6</td><td>-3.1</td><td>0.6</td><td>-4.9</td><td>7.7E-07</td><td>1.5E-06</td><td>MB231/MB468</td><td>MP</td></tr>
<tr><td>>hg38_21_+_8433172_8446622_RNA45SN1_45S_50ntFlanks@12405.12428.24|CTTTTTGATCCTTCGATGTCGGCT</td><td>>hg38_21_+_8433172_8446622_RNA45SN1_45S_50ntFlanks</td><td>12405</td><td>12428</td><td>24</td><td>CTTTTTGATCCTTCGATGTCGGCT</td><td>198.6</td><td>-6.3</td><td>1.3</td><td>-4.9</td><td>8.9E-07</td><td>1.5E-06</td><td>MB231/MB468</td><td>NUC</td></tr>
<tr><td>>hg38_21_+_8433172_8446622_RNA45SN1_45S_50ntFlanks@5485.5510.26|AGCGCTGAGAAGACGGTCGAACTTGA</td><td>>hg38_21_+_8433172_8446622_RNA45SN1_45S_50ntFlanks</td><td>5485</td><td>5510</td><td>26</td><td>AGCGCTGAGAAGACGGTCGAACTTGA</td><td>156.0</td><td>-4.0</td><td>0.8</td><td>-4.9</td><td>8.7E-07</td><td>1.5E-06</td><td>MB231/MB468</td><td>CYTO</td></tr>
<tr><td>>MI0003188|hsa-mir-503&WithFlank&X|-|134546322|134546404@12.30.19(+1A)&[MIMAT0002874&hsa-miR-503-5p&offsets|0|-4(+1A);m-297&X|-|134546371|134546393&offsets|0|-4(+1A)]|TAGCAGCGGGAACAGTTCTA</td><td>>MI0003188|hsa-mir-503&WithFlank&X|-|134546322|134546404</td><td>12</td><td>30</td><td>19(+1A)</td><td>TAGCAGCGGGAACAGTTCTA</td><td>86.7</td><td>10.2</td><td>2.0</td><td>5.2</td><td>2.3E-07</td><td>1.5E-06</td><td>BT20/MB468</td><td>MP</td></tr>
<tr><td>>hg38_21_+_8433172_8446622_RNA45SN1_45S_50ntFlanks@11572.11612.41|CCGACTTAGAACTGGTGCGGACCAGGGGAATCCGACTGTTT</td><td>>hg38_21_+_8433172_8446622_RNA45SN1_45S_50ntFlanks</td><td>11572</td><td>11612</td><td>41</td><td>CCGACTTAGAACTGGTGCGGACCAGGGGAATCCGACTGTTT</td><td>1840.7</td><td>-5.9</td><td>1.2</td><td>-4.9</td><td>7.7E-07</td><td>1.5E-06</td><td>MB231/MB468</td><td>MP</td></tr>
<tr><td>>hg38_MT_+_1621_3279_RNR2_16S_50ntFlanks@749.777.29|AATTAACAGCCCAATATCTACAATCAACC</td><td>>hg38_MT_+_1621_3279_RNR2_16S_50ntFlanks</td><td>749</td><td>777</td><td>29</td><td>AATTAACAGCCCAATATCTACAATCAACC</td><td>43.0</td><td>8.6</td><td>1.7</td><td>4.9</td><td>7.7E-07</td><td>1.5E-06</td><td>MB231/MB468</td><td>MP</td></tr>
<tr><td>>hg38_21_+_8433172_8446622_RNA45SN1_45S_50ntFlanks@10410.10453.44|TCAGTCGGTCCTGAGAGATGGGCGAGCGCCGTTCCGAAGGGACG</td><td>>hg38_21_+_8433172_8446622_RNA45SN1_45S_50ntFlanks</td><td>10410</td><td>10453</td><td>44</td><td>TCAGTCGGTCCTGAGAGATGGGCGAGCGCCGTTCCGAAGGGACG</td><td>705.4</td><td>-3.7</td><td>0.8</td><td>-4.9</td><td>8.7E-07</td><td>1.5E-06</td><td>MB231/MB468</td><td>CYTO</td></tr>
<tr><td>>am_trnaMT_ArgTCG_MT_+_10405_10469@28.48.21|ATTTCGACTCATTAAATTATG</td><td>>am_trnaMT_ArgTCG_MT_+_10405_10469</td><td>28</td><td>48</td><td>21</td><td>ATTTCGACTCATTAAATTATG</td><td>704.5</td><td>-6.2</td><td>1.3</td><td>-4.9</td><td>8.7E-07</td><td>1.5E-06</td><td>MB231/MB468</td><td>CYTO</td></tr>
<tr><td>>MI0001445|hsa-mir-423&WithFlank&17|+|30117073|30117178@59.80.22(+1A)&[MIMAT0001340&hsa-miR-423-3p&offsets|0|-1(+1A);m-100&17|+|30117131|30117153&offsets|0|-1(+1A)]|AGCTCGGTCTGAGGCCCCTCAGA</td><td>>MI0001445|hsa-mir-423&WithFlank&17|+|30117073|30117178</td><td>59</td><td>80</td><td>22(+1A)</td><td>AGCTCGGTCTGAGGCCCCTCAGA</td><td>683.3</td><td>4.7</td><td>0.9</td><td>4.9</td><td>9.4E-07</td><td>1.5E-06</td><td>MB231/MB468</td><td>MITO</td></tr>
<tr><td>>hg38_21_+_8433172_8446622_RNA45SN1_45S_50ntFlanks@11569.11591.23|CAGCCGACTTAGAACTGGTGCGG</td><td>>hg38_21_+_8433172_8446622_RNA45SN1_45S_50ntFlanks</td><td>11569</td><td>11591</td><td>23</td><td>CAGCCGACTTAGAACTGGTGCGG</td><td>256.2</td><td>-6.7</td><td>1.4</td><td>-4.9</td><td>7.6E-07</td><td>1.5E-06</td><td>BT20/MB468</td><td>CYTO</td></tr>
<tr><td>>hg38_21_+_8433172_8446622_RNA45SN1_45S_50ntFlanks@12242.12268.27|AGAAGGGCAAAAGCTCGCTTGATCTTG</td><td>>hg38_21_+_8433172_8446622_RNA45SN1_45S_50ntFlanks</td><td>12242</td><td>12268</td><td>27</td><td>AGAAGGGCAAAAGCTCGCTTGATCTTG</td><td>390.9</td><td>-7.7</td><td>1.6</td><td>-4.9</td><td>9.8E-07</td><td>1.5E-06</td><td>BT20/MB468</td><td>NUC</td></tr>
<tr><td>>hg38_21_+_8433172_8446622_RNA45SN1_45S_50ntFlanks@4444.4472.29|CCTTGCCTCTCGGCGCCCCCTCGATGCTC</td><td>>hg38_21_+_8433172_8446622_RNA45SN1_45S_50ntFlanks</td><td>4444</td><td>4472</td><td>29</td><td>CCTTGCCTCTCGGCGCCCCCTCGATGCTC</td><td>201.0</td><td>-5.7</td><td>1.2</td><td>-4.9</td><td>8.8E-07</td><td>1.5E-06</td><td>MB231/MB468</td><td>CYTO</td></tr>
<tr><td>>hg38_21_+_8433172_8446622_RNA45SN1_45S_50ntFlanks@12252.12288.37|AAGCTCGCTTGATCTTGATTTTCAGTACGAATACAGA</td><td>>hg38_21_+_8433172_8446622_RNA45SN1_45S_50ntFlanks</td><td>12252</td><td>12288</td><td>37</td><td>AAGCTCGCTTGATCTTGATTTTCAGTACGAATACAGA</td><td>273.2</td><td>-6.8</td><td>1.4</td><td>-4.9</td><td>9.4E-07</td><td>1.5E-06</td><td>MB231/MB468</td><td>MITO</td></tr>
<tr><td>>MI0000065|hsa-let-7d&WithFlank&9|+|94178828|94178926@68.88.21&[MIMAT0004484&hsa-let-7d-3p&offsets|0|-1;m-98&9|+|94178895|94178916&offsets|0|-1]|CTATACGACCTGCTGCCTTTC</td><td>>MI0000065|hsa-let-7d&WithFlank&9|+|94178828|94178926</td><td>68</td><td>88</td><td>21</td><td>CTATACGACCTGCTGCCTTTC</td><td>734.1</td><td>7.5</td><td>1.5</td><td>4.9</td><td>7.7E-07</td><td>1.5E-06</td><td>BT20/MB468</td><td>CYTO</td></tr>
<tr><td>>hg38_21_+_8433172_8446622_RNA45SN1_45S_50ntFlanks@9533.9553.21|GAGGAAACTCTGGTGGAGGTC</td><td>>hg38_21_+_8433172_8446622_RNA45SN1_45S_50ntFlanks</td><td>9533</td><td>9553</td><td>21</td><td>GAGGAAACTCTGGTGGAGGTC</td><td>20099.9</td><td>-3.9</td><td>0.8</td><td>-4.9</td><td>8.8E-07</td><td>1.5E-06</td><td>MB231/MB468</td><td>CYTO</td></tr>
<tr><td>>hg38_21_+_8433172_8446622_RNA45SN1_45S_50ntFlanks@12973.12992.20|TCGTACGTAGCAGAGCAGCT</td><td>>hg38_21_+_8433172_8446622_RNA45SN1_45S_50ntFlanks</td><td>12973</td><td>12992</td><td>20</td><td>TCGTACGTAGCAGAGCAGCT</td><td>1105.1</td><td>-5.1</td><td>1.0</td><td>-4.9</td><td>7.7E-07</td><td>1.5E-06</td><td>BT20/MB468</td><td>CYTO</td></tr>
<tr><td>>hg38_21_+_8433172_8446622_RNA45SN1_45S_50ntFlanks@12273.12299.27|TCAGTACGAATACAGACCGTGAAAGCG</td><td>>hg38_21_+_8433172_8446622_RNA45SN1_45S_50ntFlanks</td><td>12273</td><td>12299</td><td>27</td><td>TCAGTACGAATACAGACCGTGAAAGCG</td><td>61.0</td><td>-9.0</td><td>1.7</td><td>-5.2</td><td>2.3E-07</td><td>1.5E-06</td><td>BT20/MB468</td><td>MP</td></tr>
<tr><td>>hg38_21_+_8433172_8446622_RNA45SN1_45S_50ntFlanks@11573.11593.21|CGACTTAGAACTGGTGCGGAC</td><td>>hg38_21_+_8433172_8446622_RNA45SN1_45S_50ntFlanks</td><td>11573</td><td>11593</td><td>21</td><td>CGACTTAGAACTGGTGCGGAC</td><td>255.6</td><td>-3.7</td><td>0.7</td><td>-4.9</td><td>9.4E-07</td><td>1.5E-06</td><td>MB231/MB468</td><td>MITO</td></tr>
<tr><td>>hg38_MT_+_1621_3279_RNR2_16S_50ntFlanks@1348.1389.42|AACAATAGGGTTTACGACCTCGATGTTGGATCAGGACATCCC</td><td>>hg38_MT_+_1621_3279_RNR2_16S_50ntFlanks</td><td>1348</td><td>1389</td><td>42</td><td>AACAATAGGGTTTACGACCTCGATGTTGGATCAGGACATCCC</td><td>46.5</td><td>8.8</td><td>1.8</td><td>4.9</td><td>7.8E-07</td><td>1.5E-06</td><td>MB231/MB468</td><td>MP</td></tr>
<tr><td>>hg38_21_+_8433172_8446622_RNA45SN1_45S_50ntFlanks@9567.9584.18|ACGTGCAAATCGGTCGTC</td><td>>hg38_21_+_8433172_8446622_RNA45SN1_45S_50ntFlanks</td><td>9567</td><td>9584</td><td>18</td><td>ACGTGCAAATCGGTCGTC</td><td>581.4</td><td>-7.0</td><td>1.4</td><td>-4.9</td><td>7.7E-07</td><td>1.5E-06</td><td>BT20/MB468</td><td>CYTO</td></tr>
<tr><td>>hg38_21_+_8433172_8446622_RNA45SN1_45S_50ntFlanks@12957.12991.35|CTTCTGGGTCGGGGTTTCGTACGTAGCAGAGCAGC</td><td>>hg38_21_+_8433172_8446622_RNA45SN1_45S_50ntFlanks</td><td>12957</td><td>12991</td><td>35</td><td>CTTCTGGGTCGGGGTTTCGTACGTAGCAGAGCAGC</td><td>123.6</td><td>-4.2</td><td>0.8</td><td>-4.9</td><td>8.8E-07</td><td>1.5E-06</td><td>MB231/MB468</td><td>CYTO</td></tr>
<tr><td>>hg38_21_+_8433172_8446622_RNA45SN1_45S_50ntFlanks@6661.6699.39|CGGTGGATCACTCGGCTCGTGCGTCGATGAAGAACGCAG</td><td>>hg38_21_+_8433172_8446622_RNA45SN1_45S_50ntFlanks</td><td>6661</td><td>6699</td><td>39</td><td>CGGTGGATCACTCGGCTCGTGCGTCGATGAAGAACGCAG</td><td>3585.9</td><td>-2.8</td><td>0.6</td><td>-4.9</td><td>9.9E-07</td><td>1.5E-06</td><td>BT20/MB468</td><td>NUC</td></tr>
<tr><td>>hg38_21_+_8433172_8446622_RNA45SN1_45S_50ntFlanks@9554.9572.19|CGTAGCGGTCCTGACGTGC</td><td>>hg38_21_+_8433172_8446622_RNA45SN1_45S_50ntFlanks</td><td>9554</td><td>9572</td><td>19</td><td>CGTAGCGGTCCTGACGTGC</td><td>20343.7</td><td>-2.8</td><td>0.6</td><td>-4.9</td><td>8.9E-07</td><td>1.5E-06</td><td>MB231/MB468</td><td>CYTO</td></tr>
<tr><td>>hg38_MT_+_1621_3279_RNR2_16S_50ntFlanks@716.740.25|TAAGCCTGCGTCAGATTAAAACACT</td><td>>hg38_MT_+_1621_3279_RNR2_16S_50ntFlanks</td><td>716</td><td>740</td><td>25</td><td>TAAGCCTGCGTCAGATTAAAACACT</td><td>50.3</td><td>8.9</td><td>1.8</td><td>4.9</td><td>7.9E-07</td><td>1.5E-06</td><td>MB231/MB468</td><td>MP</td></tr>
<tr><td>>hg38_21_+_8433172_8446622_RNA45SN1_45S_50ntFlanks@8921.8940.20|CCGGGGCCGAGGGAGCGAGA</td><td>>hg38_21_+_8433172_8446622_RNA45SN1_45S_50ntFlanks</td><td>8921</td><td>8940</td><td>20</td><td>CCGGGGCCGAGGGAGCGAGA</td><td>66.7</td><td>9.8</td><td>1.9</td><td>5.2</td><td>2.4E-07</td><td>1.5E-06</td><td>BT20/MB468</td><td>MP</td></tr>
<tr><td>>MI0000076|hsa-mir-20a&WithFlank&13|+|91351059|91351141@14.35.22(+1A)&[MIMAT0000075&hsa-miR-20a-5p&offsets|0|-1(+1A);m-96&13|+|91351072|91351094&offsets|0|-1(+1A)]|TAAAGTGCTTATAGTGCAGGTAA</td><td>>MI0000076|hsa-mir-20a&WithFlank&13|+|91351059|91351141</td><td>14</td><td>35</td><td>22(+1A)</td><td>TAAAGTGCTTATAGTGCAGGTAA</td><td>637.0</td><td>5.1</td><td>1.0</td><td>4.9</td><td>7.7E-07</td><td>1.5E-06</td><td>BT20/MB468</td><td>CYTO</td></tr>
<tr><td>>hg38_21_+_8433172_8446622_RNA45SN1_45S_50ntFlanks@11205.11232.28|GCGGCGGCGACTCTGGACGCGAGCCGGG</td><td>>hg38_21_+_8433172_8446622_RNA45SN1_45S_50ntFlanks</td><td>11205</td><td>11232</td><td>28</td><td>GCGGCGGCGACTCTGGACGCGAGCCGGG</td><td>1032.3</td><td>-3.7</td><td>0.8</td><td>-4.9</td><td>7.7E-07</td><td>1.5E-06</td><td>BT20/MB468</td><td>CYTO</td></tr>
<tr><td>>hg38_21_+_8433172_8446622_RNA45SN1_45S_50ntFlanks@3989.4024.36|TTTGGTGACTCTAGATAACCTCGGGCCGATCGCACG</td><td>>hg38_21_+_8433172_8446622_RNA45SN1_45S_50ntFlanks</td><td>3989</td><td>4024</td><td>36</td><td>TTTGGTGACTCTAGATAACCTCGGGCCGATCGCACG</td><td>477.7</td><td>-6.0</td><td>1.2</td><td>-4.9</td><td>9.9E-07</td><td>1.5E-06</td><td>BT20/MB468</td><td>NUC</td></tr>
<tr><td>>hg38_21_+_8433172_8446622_RNA45SN1_45S_50ntFlanks@9574.9597.24|AATCGGTCGTCCGACCTGGGTATA</td><td>>hg38_21_+_8433172_8446622_RNA45SN1_45S_50ntFlanks</td><td>9574</td><td>9597</td><td>24</td><td>AATCGGTCGTCCGACCTGGGTATA</td><td>885.1</td><td>-3.6</td><td>0.7</td><td>-4.9</td><td>9.1E-07</td><td>1.5E-06</td><td>MB231/MB468</td><td>NUC</td></tr>
<tr><td>>hg38_21_+_8433172_8446622_RNA45SN1_45S_50ntFlanks@6743.6771.29|CGACACTTCGAACGCACTTGCGGCCCCGG</td><td>>hg38_21_+_8433172_8446622_RNA45SN1_45S_50ntFlanks</td><td>6743</td><td>6771</td><td>29</td><td>CGACACTTCGAACGCACTTGCGGCCCCGG</td><td>949.4</td><td>-5.5</td><td>1.1</td><td>-4.9</td><td>7.7E-07</td><td>1.5E-06</td><td>BT20/MB468</td><td>CYTO</td></tr>
<tr><td>>hg38_MT_+_1621_3279_RNR2_16S_50ntFlanks@999.1034.36|AGGGACCTGTATGAATGGCTCCACGAGGGTTCAGCT</td><td>>hg38_MT_+_1621_3279_RNR2_16S_50ntFlanks</td><td>999</td><td>1034</td><td>36</td><td>AGGGACCTGTATGAATGGCTCCACGAGGGTTCAGCT</td><td>40.3</td><td>8.6</td><td>1.7</td><td>4.9</td><td>7.9E-07</td><td>1.5E-06</td><td>MB231/MB468</td><td>MP</td></tr>
<tr><td>>hg38_21_+_8433172_8446622_RNA45SN1_45S_50ntFlanks@10346.10374.29|TCAAACGAGAACTTTGAAGGCCGAAGTGG</td><td>>hg38_21_+_8433172_8446622_RNA45SN1_45S_50ntFlanks</td><td>10346</td><td>10374</td><td>29</td><td>TCAAACGAGAACTTTGAAGGCCGAAGTGG</td><td>2341.5</td><td>-4.6</td><td>0.9</td><td>-5.2</td><td>2.4E-07</td><td>1.5E-06</td><td>BT20/MB468</td><td>MP</td></tr>
<tr><td>>hg38_21_+_8433172_8446622_RNA45SN1_45S_50ntFlanks@9545.9574.30|GTGGAGGTCCGTAGCGGTCCTGACGTGCAA</td><td>>hg38_21_+_8433172_8446622_RNA45SN1_45S_50ntFlanks</td><td>9545</td><td>9574</td><td>30</td><td>GTGGAGGTCCGTAGCGGTCCTGACGTGCAA</td><td>438.0</td><td>-2.5</td><td>0.5</td><td>-5.0</td><td>6.8E-07</td><td>1.5E-06</td><td>BT20/MB468</td><td>MITO</td></tr>
<tr><td>>hg38_21_+_8433172_8446622_RNA45SN1_45S_50ntFlanks@4919.4940.22|CCACCAGGAGTGGAGCCTGCGG</td><td>>hg38_21_+_8433172_8446622_RNA45SN1_45S_50ntFlanks</td><td>4919</td><td>4940</td><td>22</td><td>CCACCAGGAGTGGAGCCTGCGG</td><td>252.2</td><td>-7.1</td><td>1.4</td><td>-5.2</td><td>2.4E-07</td><td>1.5E-06</td><td>BT20/MB468</td><td>MP</td></tr>
<tr><td>>trna10_LysCTT_16_+_3241501_3241573@5.35.31,trna13_LysCTT_14_-_58706613_58706685@5.35.31,trna2_LysCTT_15_+_79152904_79152976@5.35.31|GGCTAGCTCAGTCGGTAGAGCATGGGACTCT</td><td>>trna10_LysCTT_16_+_3241501_3241573</td><td>5</td><td>35</td><td>31</td><td>GGCTAGCTCAGTCGGTAGAGCATGGGACTCT</td><td>713.0</td><td>7.1</td><td>1.4</td><td>4.9</td><td>9.5E-07</td><td>1.5E-06</td><td>MB231/MB468</td><td>MITO</td></tr>
</table>

<table>
<tr><td>>MI0000446|hsa-mir-125b-1&WithFlank&11|-|122099751|122099850@21.39.19(+1C);MI0000470|hsa-mir-125b-2&WithFlank&21|+|16590231|16590331@23.41.19(+1C)&[MIMAT0000423&hsa-miR-125b-5p&offsets|0|-3(+1C);m-67&11|-|122099809|122099830&offsets|0|-3(+1C)];[MIMAT0000423_1&hsa-miR-125b-5p&offsets|0|-3(+1C);m-68&21|+|16590253|16590274&offsets|0|-3(+1C)]|TCCCTGAGACCCTAACTTGC</td><td>>MI0000446|hsa-mir-125b-1&WithFlank&11|-|122099751|122099850</td><td>21</td><td>39</td><td>19(+1C)</td><td>TCCCTGAGACCCTAACTTGC</td><td>176.6</td><td>-10.9</td><td>2.0</td><td>-5.4</td><td>5.5E-08</td><td>1.5E-06</td><td>BT20/MB231</td><td>MP</td></tr>
<tr><td>>hg38_21_+_8433172_8446622_RNA45SN1_45S_50ntFlanks@7988.8017.30|CAGACGTGGCGACCCGCTGAATTTAAGCAT</td><td>>hg38_21_+_8433172_8446622_RNA45SN1_45S_50ntFlanks</td><td>7988</td><td>8017</td><td>30</td><td>CAGACGTGGCGACCCGCTGAATTTAAGCAT</td><td>324.8</td><td>3.2</td><td>0.6</td><td>4.9</td><td>8.9E-07</td><td>1.5E-06</td><td>MB231/MB468</td><td>CYTO</td></tr>
<tr><td>>MI0000273|hsa-mir-183&WithFlank&7|-|129774899|129775020@34.56.23&[MIMAT0000261&hsa-miR-183-5p&offsets|+1|+2;m-41&7|-|129774966|129774987&offsets|0|+1]|ATGGCACTGGTAGAATTCACTGT</td><td>>MI0000273|hsa-mir-183&WithFlank&7|-|129774899|129775020</td><td>34</td><td>56</td><td>23</td><td>ATGGCACTGGTAGAATTCACTGT</td><td>322.3</td><td>3.9</td><td>0.8</td><td>4.9</td><td>8.9E-07</td><td>1.5E-06</td><td>MB231/MB468</td><td>CYTO</td></tr>
<tr><td>>hg38_21_+_8433172_8446622_RNA45SN1_45S_50ntFlanks@5470.5516.47|CCACGGCCCTGGCGGAGCGCTGAGAAGACGGTCGAACTTGACTATCT</td><td>>hg38_21_+_8433172_8446622_RNA45SN1_45S_50ntFlanks</td><td>5470</td><td>5516</td><td>47</td><td>CCACGGCCCTGGCGGAGCGCTGAGAAGACGGTCGAACTTGACTATCT</td><td>382.4</td><td>6.7</td><td>1.3</td><td>5.0</td><td>6.9E-07</td><td>1.5E-06</td><td>BT20/MB468</td><td>MITO</td></tr>
<tr><td>>am_trna10_ValCAC_5_-_180649395_180649467@6.29.24,trna12_ValAAC_5_-_180645270_180645342@6.29.24,trna132_ValAAC_6_-_27721179_27721251@6.29.24,trna136_ValAAC_6_-_27648885_27648957@6.29.24,trna139_ValAAC_6_-_27618707_27618779@6.29.24,trna18_ValCAC_5_-_180529253_180529325@6.29.24,trna2_ValAAC_3_+_169490018_169490090@6.29.24,trna2_ValCAC_5_+_180524070_180524142@6.29.24,trna4_ValAAC_5_+_180591154_180591226@6.29.24,trna5_ValAAC_5_+_180596610_180596682@6.29.24,trna6_ValCAC_5_+_180600650_180600722@6.29.24,trna85_ValCAC_1_-_161369490_161369562@6.29.24,trna90_ValCAC_1_-_149684088_149684161@6.29.24,trna98_ValCAC_1_-_149298555_149298627@6.29.24,trna9_ValCAC_6_+_26538282_26538354@6.29.24|CGTAGTGTAGTGGTTATCACGTTC</td><td>>am_trna10_ValCAC_5_-_180649395_180649467</td><td>6</td><td>29</td><td>24</td><td>CGTAGTGTAGTGGTTATCACGTTC</td><td>187.2</td><td>-7.3</td><td>1.5</td><td>-4.9</td><td>7.9E-07</td><td>1.5E-06</td><td>MB231/MB468</td><td>MP</td></tr>
<tr><td>>hg38_21_+_8433172_8446622_RNA45SN1_45S_50ntFlanks@9537.9563.27|AAACTCTGGTGGAGGTCCGTAGCGGTC</td><td>>hg38_21_+_8433172_8446622_RNA45SN1_45S_50ntFlanks</td><td>9537</td><td>9563</td><td>27</td><td>AAACTCTGGTGGAGGTCCGTAGCGGTC</td><td>178.5</td><td>-6.0</td><td>1.2</td><td>-4.9</td><td>7.8E-07</td><td>1.5E-06</td><td>BT20/MB468</td><td>CYTO</td></tr>
<tr><td>>hg38_21_+_8433172_8446622_RNA45SN1_45S_50ntFlanks@10856.10873.18|AGGATTGGCTCTAAGGGC</td><td>>hg38_21_+_8433172_8446622_RNA45SN1_45S_50ntFlanks</td><td>10856</td><td>10873</td><td>18</td><td>AGGATTGGCTCTAAGGGC</td><td>192.5</td><td>-5.6</td><td>1.1</td><td>-4.9</td><td>8.0E-07</td><td>1.5E-06</td><td>MB231/MB468</td><td>MP</td></tr>
<tr><td>>hg38_21_+_8433172_8446622_RNA45SN1_45S_50ntFlanks@6651.6691.41|CGACTCTTAGCGGTGGATCACTCGGCTCGTGCGTCGATGAA</td><td>>hg38_21_+_8433172_8446622_RNA45SN1_45S_50ntFlanks</td><td>6651</td><td>6691</td><td>41</td><td>CGACTCTTAGCGGTGGATCACTCGGCTCGTGCGTCGATGAA</td><td>805.8</td><td>4.5</td><td>0.9</td><td>4.9</td><td>9.2E-07</td><td>1.5E-06</td><td>MB231/MB468</td><td>NUC</td></tr>
<tr><td>>hg38_21_+_8433172_8446622_RNA45SN1_45S_50ntFlanks@5467.5499.33|GGCCCACGGCCCTGGCGGAGCGCTGAGAAGACG</td><td>>hg38_21_+_8433172_8446622_RNA45SN1_45S_50ntFlanks</td><td>5467</td><td>5499</td><td>33</td><td>GGCCCACGGCCCTGGCGGAGCGCTGAGAAGACG</td><td>2015.8</td><td>-4.4</td><td>0.9</td><td>-4.9</td><td>9.0E-07</td><td>1.5E-06</td><td>MB231/MB468</td><td>CYTO</td></tr>
<tr><td>>hg38_21_+_8433172_8446622_RNA45SN1_45S_50ntFlanks@6649.6689.41|TACGACTCTTAGCGGTGGATCACTCGGCTCGTGCGTCGATG</td><td>>hg38_21_+_8433172_8446622_RNA45SN1_45S_50ntFlanks</td><td>6649</td><td>6689</td><td>41</td><td>TACGACTCTTAGCGGTGGATCACTCGGCTCGTGCGTCGATG</td><td>1351.7</td><td>-4.6</td><td>0.9</td><td>-4.9</td><td>8.0E-07</td><td>1.5E-06</td><td>MB231/MB468</td><td>MP</td></tr>
<tr><td>>MI0000076|hsa-mir-20a&WithFlank&13|+|91351059|91351141@14.36.23(+1C)&[MIMAT0000075&hsa-miR-20a-5p&offsets|0|0(+1C);m-96&13|+|91351072|91351094&offsets|0|0(+1C)]|TAAAGTGCTTATAGTGCAGGTAGC</td><td>>MI0000076|hsa-mir-20a&WithFlank&13|+|91351059|91351141</td><td>14</td><td>36</td><td>23(+1C)</td><td>TAAAGTGCTTATAGTGCAGGTAGC</td><td>121.6</td><td>-10.3</td><td>1.9</td><td>-5.4</td><td>5.6E-08</td><td>1.5E-06</td><td>BT20/MB231</td><td>MP</td></tr>
<tr><td>>MI0000102|hsa-mir-100&WithFlank&11|-|122152223|122152314@20.40.21(+1U)&[MIMAT0000098&hsa-miR-100-5p&offsets|+1|0(+1U);m-27&11|-|122152276|122152296&offsets|+1|+1(+1U)]|ACCCGTAGATCCGAACTTGTGT</td><td>>MI0000102|hsa-mir-100&WithFlank&11|-|122152223|122152314</td><td>20</td><td>40</td><td>21(+1U)</td><td>ACCCGTAGATCCGAACTTGTGT</td><td>117.1</td><td>-10.3</td><td>1.9</td><td>-5.4</td><td>5.6E-08</td><td>1.5E-06</td><td>BT20/MB231</td><td>MP</td></tr>
<tr><td>>hg38_21_+_8433172_8446622_RNA45SN1_45S_50ntFlanks@8105.8139.35|CGCGGCGGGGCGCGGGACATGTGGCGTACGGAAGA</td><td>>hg38_21_+_8433172_8446622_RNA45SN1_45S_50ntFlanks</td><td>8105</td><td>8139</td><td>35</td><td>CGCGGCGGGGCGCGGGACATGTGGCGTACGGAAGA</td><td>3347.7</td><td>-7.1</td><td>1.4</td><td>-4.9</td><td>8.0E-07</td><td>1.5E-06</td><td>MB231/MB468</td><td>MP</td></tr>
<tr><td>>hg38_21_+_8433172_8446622_RNA45SN1_45S_50ntFlanks@6786.6807.22|TACGCCTGTCTGAGCGTCGCTT</td><td>>hg38_21_+_8433172_8446622_RNA45SN1_45S_50ntFlanks</td><td>6786</td><td>6807</td><td>22</td><td>TACGCCTGTCTGAGCGTCGCTT</td><td>189.9</td><td>4.1</td><td>0.8</td><td>5.2</td><td>2.4E-07</td><td>1.5E-06</td><td>BT20/MB468</td><td>MP</td></tr>
<tr><td>>hg38_21_+_8433172_8446622_RNA45SN1_45S_50ntFlanks@13001.13027.27|GCGATCTATTGAAAGTCAGCCCTCGAC</td><td>>hg38_21_+_8433172_8446622_RNA45SN1_45S_50ntFlanks</td><td>13001</td><td>13027</td><td>27</td><td>GCGATCTATTGAAAGTCAGCCCTCGAC</td><td>55.5</td><td>-8.9</td><td>1.7</td><td>-5.2</td><td>2.4E-07</td><td>1.5E-06</td><td>BT20/MB468</td><td>MP</td></tr>
<tr><td>>hg38_21_+_8433172_8446622_RNA45SN1_45S_50ntFlanks@13002.13032.31|CGATCTATTGAAAGTCAGCCCTCGACACAAG</td><td>>hg38_21_+_8433172_8446622_RNA45SN1_45S_50ntFlanks</td><td>13002</td><td>13032</td><td>31</td><td>CGATCTATTGAAAGTCAGCCCTCGACACAAG</td><td>67.4</td><td>-9.2</td><td>1.8</td><td>-5.2</td><td>2.4E-07</td><td>1.5E-06</td><td>BT20/MB468</td><td>MP</td></tr>
<tr><td>>MI0003513|hsa-mir-455&WithFlank&9|+|114209428|114209535@22.44.23&[MIMAT0003150&hsa-miR-455-5p&offsets|0|+1;m-232&9|+|114209449|114209470&offsets|0|+1]|TATGTGCCTTTGGACTACATCGT</td><td>>MI0003513|hsa-mir-455&WithFlank&9|+|114209428|114209535</td><td>22</td><td>44</td><td>23</td><td>TATGTGCCTTTGGACTACATCGT</td><td>741.3</td><td>-3.4</td><td>0.6</td><td>-5.3</td><td>1.0E-07</td><td>1.5E-06</td><td>BT20/MB231</td><td>NUC</td></tr>
<tr><td>>hg38_21_+_8433172_8446622_RNA45SN1_45S_50ntFlanks@10393.10426.34|CAGCAGTTGAACATGGGTCAGTCGGTCCTGAGAG</td><td>>hg38_21_+_8433172_8446622_RNA45SN1_45S_50ntFlanks</td><td>10393</td><td>10426</td><td>34</td><td>CAGCAGTTGAACATGGGTCAGTCGGTCCTGAGAG</td><td>436.5</td><td>-5.4</td><td>1.1</td><td>-4.9</td><td>9.7E-07</td><td>1.5E-06</td><td>MB231/MB468</td><td>MITO</td></tr>
<tr><td>>hg38_21_+_8433172_8446622_RNA45SN1_45S_50ntFlanks@9541.9571.31|TCTGGTGGAGGTCCGTAGCGGTCCTGACGTG</td><td>>hg38_21_+_8433172_8446622_RNA45SN1_45S_50ntFlanks</td><td>9541</td><td>9571</td><td>31</td><td>TCTGGTGGAGGTCCGTAGCGGTCCTGACGTG</td><td>279.2</td><td>-2.8</td><td>0.6</td><td>-4.9</td><td>9.3E-07</td><td>1.5E-06</td><td>MB231/MB468</td><td>NUC</td></tr>
<tr><td>>hg38_21_+_8433172_8446622_RNA45SN1_45S_50ntFlanks@9937.9955.19|CAGCAGGACGGTGGCCATG</td><td>>hg38_21_+_8433172_8446622_RNA45SN1_45S_50ntFlanks</td><td>9937</td><td>9955</td><td>19</td><td>CAGCAGGACGGTGGCCATG</td><td>473.8</td><td>5.9</td><td>1.1</td><td>5.4</td><td>6.6E-08</td><td>1.5E-06</td><td>BT20/MB231</td><td>CYTO</td></tr>
<tr><td>>hg38_21_+_8433172_8446622_RNA45SN1_45S_50ntFlanks@11594.11611.18|CAGGGGAATCCGACTGTT</td><td>>hg38_21_+_8433172_8446622_RNA45SN1_45S_50ntFlanks</td><td>11594</td><td>11611</td><td>18</td><td>CAGGGGAATCCGACTGTT</td><td>1810.1</td><td>-3.1</td><td>0.6</td><td>-5.0</td><td>7.0E-07</td><td>1.5E-06</td><td>BT20/MB468</td><td>MITO</td></tr>
<tr><td>>hg38_21_+_8433172_8446622_RNA45SN1_45S_50ntFlanks@6721.6752.32|AATTGCAGGACACATTGATCATCGACACTTCG</td><td>>hg38_21_+_8433172_8446622_RNA45SN1_45S_50ntFlanks</td><td>6721</td><td>6752</td><td>32</td><td>AATTGCAGGACACATTGATCATCGACACTTCG</td><td>948.5</td><td>-5.5</td><td>1.1</td><td>-4.9</td><td>9.7E-07</td><td>1.5E-06</td><td>MB231/MB468</td><td>MITO</td></tr>
<tr><td>>MI0000487|hsa-mir-193a&WithFlank&17|+|31559990|31560089@27.47.21&[MIMAT0004614&hsa-miR-193a-5p&offsets|0|-1;m-123&17|+|31560016|31560037&offsets|0|-1]|TGGGTCTTTGCGGGCGAGATG</td><td>>MI0000487|hsa-mir-193a&WithFlank&17|+|31559990|31560089</td><td>27</td><td>47</td><td>21</td><td>TGGGTCTTTGCGGGCGAGATG</td><td>203.2</td><td>3.7</td><td>0.7</td><td>5.4</td><td>7.9E-08</td><td>1.5E-06</td><td>BT20/MB231</td><td>MITO</td></tr>
<tr><td>>hg38_21_+_8433172_8446622_RNA45SN1_45S_50ntFlanks@10391.10435.45|AACAGCAGTTGAACATGGGTCAGTCGGTCCTGAGAGATGGGCGAG</td><td>>hg38_21_+_8433172_8446622_RNA45SN1_45S_50ntFlanks</td><td>10391</td><td>10435</td><td>45</td><td>AACAGCAGTTGAACATGGGTCAGTCGGTCCTGAGAGATGGGCGAG</td><td>104.0</td><td>-9.8</td><td>1.9</td><td>-5.2</td><td>2.4E-07</td><td>1.5E-06</td><td>BT20/MB468</td><td>MP</td></tr>
<tr><td>>MI0000070|hsa-mir-16-1&WithFlank&13|-|50048967|50049067@20.40.21(+1A);MI0000115|hsa-mir-16-2&WithFlank&3|+|160404739|160404831@16.36.21(+1A)&[MIMAT0000069&hsa-miR-16-5p&offsets|0|-1(+1A);m-62&13|-|50049027|50049048&offsets|0|-1(+1A)];[MIMAT0000069_1&hsa-miR-16-5p&offsets|0|-1(+1A);m-61&3|+|160404754|160404775&offsets|0|-1(+1A)]|TAGCAGCACGTAAATATTGGCA</td><td>>MI0000070|hsa-mir-16-1&WithFlank&13|-|50048967|50049067</td><td>20</td><td>40</td><td>21(+1A)</td><td>TAGCAGCACGTAAATATTGGCA</td><td>653.9</td><td>5.9</td><td>1.2</td><td>4.9</td><td>9.7E-07</td><td>1.5E-06</td><td>MB231/MB468</td><td>MITO</td></tr>
<tr><td>>hg38_21_+_8433172_8446622_RNA45SN1_45S_50ntFlanks@12654.12671.18|TATGACTGAACGCCTCTA</td><td>>hg38_21_+_8433172_8446622_RNA45SN1_45S_50ntFlanks</td><td>12654</td><td>12671</td><td>18</td><td>TATGACTGAACGCCTCTA</td><td>142.3</td><td>-5.1</td><td>1.0</td><td>-4.9</td><td>9.1E-07</td><td>1.5E-06</td><td>MB231/MB468</td><td>CYTO</td></tr>
<tr><td>>hg38_21_+_8433172_8446622_RNA45SN1_45S_50ntFlanks@11323.11344.22|CGTCTCGTCGCGCGCGCGTCCG</td><td>>hg38_21_+_8433172_8446622_RNA45SN1_45S_50ntFlanks</td><td>11323</td><td>11344</td><td>22</td><td>CGTCTCGTCGCGCGCGCGTCCG</td><td>15196.5</td><td>-6.4</td><td>1.3</td><td>-4.9</td><td>7.9E-07</td><td>1.5E-06</td><td>BT20/MB468</td><td>CYTO</td></tr>
<tr><td>>hg38_21_+_8433172_8446622_RNA45SN1_45S_50ntFlanks@11581.11604.24|AACTGGTGCGGACCAGGGGAATCC</td><td>>hg38_21_+_8433172_8446622_RNA45SN1_45S_50ntFlanks</td><td>11581</td><td>11604</td><td>24</td><td>AACTGGTGCGGACCAGGGGAATCC</td><td>253.9</td><td>-5.9</td><td>1.2</td><td>-4.9</td><td>8.1E-07</td><td>1.5E-06</td><td>MB231/MB468</td><td>MP</td></tr>
<tr><td>>hg38_21_+_8433172_8446622_RNA45SN1_45S_50ntFlanks@4772.4793.22|GTAGTTCCGACCATAAACGATG</td><td>>hg38_21_+_8433172_8446622_RNA45SN1_45S_50ntFlanks</td><td>4772</td><td>4793</td><td>22</td><td>GTAGTTCCGACCATAAACGATG</td><td>375.0</td><td>-2.2</td><td>0.4</td><td>-4.9</td><td>8.0E-07</td><td>1.5E-06</td><td>BT20/MB468</td><td>CYTO</td></tr>
<tr><td>>hg38_21_+_8433172_8446622_RNA45SN1_45S_50ntFlanks@11592.11611.20|ACCAGGGGAATCCGACTGTT</td><td>>hg38_21_+_8433172_8446622_RNA45SN1_45S_50ntFlanks</td><td>11592</td><td>11611</td><td>20</td><td>ACCAGGGGAATCCGACTGTT</td><td>423.2</td><td>-3.6</td><td>0.7</td><td>-5.0</td><td>7.0E-07</td><td>1.5E-06</td><td>BT20/MB468</td><td>MITO</td></tr>
<tr><td>>hg38_21_+_8433172_8446622_RNA45SN1_45S_50ntFlanks@8196.8226.31|CGTGGACGGTGTGAGGCCGGTAGCGGCCCCC</td><td>>hg38_21_+_8433172_8446622_RNA45SN1_45S_50ntFlanks</td><td>8196</td><td>8226</td><td>31</td><td>CGTGGACGGTGTGAGGCCGGTAGCGGCCCCC</td><td>674.8</td><td>-7.9</td><td>1.6</td><td>-4.9</td><td>1.0E-06</td><td>1.5E-06</td><td>BT20/MB468</td><td>NUC</td></tr>
<tr><td>>hg38_21_+_8433172_8446622_RNA45SN1_45S_50ntFlanks@9533.9562.30|GAGGAAACTCTGGTGGAGGTCCGTAGCGGT</td><td>>hg38_21_+_8433172_8446622_RNA45SN1_45S_50ntFlanks</td><td>9533</td><td>9562</td><td>30</td><td>GAGGAAACTCTGGTGGAGGTCCGTAGCGGT</td><td>435.4</td><td>-5.7</td><td>1.2</td><td>-4.9</td><td>8.1E-07</td><td>1.5E-06</td><td>MB231/MB468</td><td>MP</td></tr>
<tr><td>>hg38_21_+_8433172_8446622_RNA45SN1_45S_50ntFlanks@11847.11872.26|AACGGGCTTGGCGGAATCAGCGGGGA</td><td>>hg38_21_+_8433172_8446622_RNA45SN1_45S_50ntFlanks</td><td>11847</td><td>11872</td><td>26</td><td>AACGGGCTTGGCGGAATCAGCGGGGA</td><td>503.6</td><td>-2.7</td><td>0.5</td><td>-5.0</td><td>7.0E-07</td><td>1.5E-06</td><td>BT20/MB468</td><td>MITO</td></tr>
<tr><td>>hg38_21_+_8433172_8446622_RNA45SN1_45S_50ntFlanks@5482.5514.33|CGGAGCGCTGAGAAGACGGTCGAACTTGACTAT</td><td>>hg38_21_+_8433172_8446622_RNA45SN1_45S_50ntFlanks</td><td>5482</td><td>5514</td><td>33</td><td>CGGAGCGCTGAGAAGACGGTCGAACTTGACTAT</td><td>266.0</td><td>-3.7</td><td>0.8</td><td>-5.0</td><td>7.0E-07</td><td>1.5E-06</td><td>BT20/MB468</td><td>MITO</td></tr>
<tr><td>>hg38_21_+_8433172_8446622_RNA45SN1_45S_50ntFlanks@9788.9830.43|CGCTGGCGTGGAGCCGGGCGTGGAATGCGAGTGCCTAGTGGGC</td><td>>hg38_21_+_8433172_8446622_RNA45SN1_45S_50ntFlanks</td><td>9788</td><td>9830</td><td>43</td><td>CGCTGGCGTGGAGCCGGGCGTGGAATGCGAGTGCCTAGTGGGC</td><td>404.8</td><td>12.0</td><td>2.2</td><td>5.4</td><td>5.7E-08</td><td>1.5E-06</td><td>BT20/MB231</td><td>MP</td></tr>
<tr><td>>MI0000455|hsa-mir-138-2&WithFlank&16|+|56858512|56858607@16.37.22;MI0000476|hsa-mir-138-1&WithFlank&3|+|44114206|44114316@29.50.22&[MIMAT0000430&hsa-miR-138-5p&offsets|0|-1;m-509&16|+|56858527|56858549&offsets|0|-1];[MIMAT0000430_1&hsa-miR-138-5p&offsets|0|-1;m-497&3|+|44114234|44114256&offsets|0|-1]|AGCTGGTGTTGTGAATCAGGCC</td><td>>MI0000455|hsa-mir-138-2&WithFlank&16|+|56858512|56858607</td><td>16</td><td>37</td><td>22</td><td>AGCTGGTGTTGTGAATCAGGCC</td><td>115.4</td><td>-10.2</td><td>1.9</td><td>-5.4</td><td>5.7E-08</td><td>1.5E-06</td><td>BT20/MB231</td><td>MP</td></tr>
<tr><td>>hg38_21_+_8433172_8446622_RNA45SN1_45S_50ntFlanks@10736.10755.20|GTAAATCTCGCGCCGGGCCG</td><td>>hg38_21_+_8433172_8446622_RNA45SN1_45S_50ntFlanks</td><td>10736</td><td>10755</td><td>20</td><td>GTAAATCTCGCGCCGGGCCG</td><td>135.8</td><td>-5.8</td><td>1.2</td><td>-4.9</td><td>8.0E-07</td><td>1.5E-06</td><td>BT20/MB468</td><td>CYTO</td></tr>
<tr><td>>hg38_21_+_8433172_8446622_RNA45SN1_45S_50ntFlanks@4498.4519.22|AAGCGTTTACTTTGAAAAAATT</td><td>>hg38_21_+_8433172_8446622_RNA45SN1_45S_50ntFlanks</td><td>4498</td><td>4519</td><td>22</td><td>AAGCGTTTACTTTGAAAAAATT</td><td>573.9</td><td>4.0</td><td>0.8</td><td>4.9</td><td>8.0E-07</td><td>1.5E-06</td><td>BT20/MB468</td><td>CYTO</td></tr>
<tr><td>>trna10_AlaCGC_6_+_26553731_26553802@58.75.18,trna119_AlaCGC_6_-_28641613_28641684@58.75.18,trna13_AlaCGC_2_+_157257281_157257352@58.75.18,trna13_AlaTGC_12_-_125406301_125406372@58.75.18,trna66_AlaTGC_6_+_28611222_28611293@58.75.18,trna8_AlaTGC_12_+_125424512_125424583@58.75.18,trna8_AlaTGC_5_+_180633868_180633939@58.75.18|TCCCCGGCATCTCCACCA</td><td>>trna10_AlaCGC_6_+_26553731_26553802</td><td>58</td><td>75</td><td>18</td><td>TCCCCGGCATCTCCACCA</td><td>559.1</td><td>4.0</td><td>0.8</td><td>4.9</td><td>9.8E-07</td><td>1.5E-06</td><td>MB231/MB468</td><td>MITO</td></tr>
<tr><td>>hg38_21_+_8433172_8446622_RNA45SN1_45S_50ntFlanks@4732.4768.37|AGAACGAAAGTCGGAGGTTCGAAGACGATCAGATACC</td><td>>hg38_21_+_8433172_8446622_RNA45SN1_45S_50ntFlanks</td><td>4732</td><td>4768</td><td>37</td><td>AGAACGAAAGTCGGAGGTTCGAAGACGATCAGATACC</td><td>217.8</td><td>6.0</td><td>1.2</td><td>4.9</td><td>9.4E-07</td><td>1.5E-06</td><td>MB231/MB468</td><td>NUC</td></tr>
<tr><td>>hg38_21_+_8433172_8446622_RNA45SN1_45S_50ntFlanks@5237.5255.19|ACTGACTGGCTCAGCGTGT</td><td>>hg38_21_+_8433172_8446622_RNA45SN1_45S_50ntFlanks</td><td>5237</td><td>5255</td><td>19</td><td>ACTGACTGGCTCAGCGTGT</td><td>944.0</td><td>5.9</td><td>1.1</td><td>5.2</td><td>2.4E-07</td><td>1.5E-06</td><td>BT20/MB468</td><td>MP</td></tr>
<tr><td>>hg38_21_+_8433172_8446622_RNA45SN1_45S_50ntFlanks@12957.12975.19|CTTCTGGGTCGGGGTTTCG</td><td>>hg38_21_+_8433172_8446622_RNA45SN1_45S_50ntFlanks</td><td>12957</td><td>12975</td><td>19</td><td>CTTCTGGGTCGGGGTTTCG</td><td>344.2</td><td>-6.7</td><td>1.4</td><td>-4.9</td><td>9.4E-07</td><td>1.5E-06</td><td>MB231/MB468</td><td>NUC</td></tr>
</table>

<table>
<tr><td>>hg38_MT_+_1621_3279_RNR2_16S_50ntFlanks@1376.1396.21|GATCAGGACATCCCGATGGTG</td><td>>hg38_MT_+_1621_3279_RNR2_16S_50ntFlanks</td><td>1376</td><td>1396</td><td>21</td><td>GATCAGGACATCCCGATGGTG</td><td>67.7</td><td>-9.2</td><td>1.8</td><td>-5.2</td><td>2.4E-07</td><td>1.5E-06</td><td>BT20/MB468</td><td>MP</td></tr>
<tr><td>>hg38_21_+_8433172_8446622_RNA45SN1_45S_50ntFlanks@11783.11805.23|CATGAATGGATGAACGAGATTCC</td><td>>hg38_21_+_8433172_8446622_RNA45SN1_45S_50ntFlanks</td><td>11783</td><td>11805</td><td>23</td><td>CATGAATGGATGAACGAGATTCC</td><td>2330.8</td><td>-4.2</td><td>0.9</td><td>-4.9</td><td>9.2E-07</td><td>1.5E-06</td><td>MB231/MB468</td><td>CYTO</td></tr>
<tr><td>>MI0000815|hsa-mir-339&WithFlank&7|-|1022927|1023032@56.77.22(+1U)&[MIMAT0004702&hsa-miR-339-3p&offsets|0|-1(+1U);m-235&7|-|1022957|1022977&offsets|0|+1(+1U)]|TGAGCGCCTCGACGACAGAGCCT</td><td>>MI0000815|hsa-mir-339&WithFlank&7|-|1022927|1023032</td><td>56</td><td>77</td><td>22(+1U)</td><td>TGAGCGCCTCGACGACAGAGCCT</td><td>105.5</td><td>10.5</td><td>2.0</td><td>5.2</td><td>2.4E-07</td><td>1.5E-06</td><td>BT20/MB468</td><td>MP</td></tr>
<tr><td>>hg38_21_+_8433172_8446622_RNA45SN1_45S_50ntFlanks@10781.10801.21|AGGTGAACAGCCTCTGGCATG</td><td>>hg38_21_+_8433172_8446622_RNA45SN1_45S_50ntFlanks</td><td>10781</td><td>10801</td><td>21</td><td>AGGTGAACAGCCTCTGGCATG</td><td>939.7</td><td>-6.0</td><td>1.2</td><td>-5.2</td><td>2.4E-07</td><td>1.5E-06</td><td>BT20/MB468</td><td>MP</td></tr>
<tr><td>>hg38_21_+_8433172_8446622_RNA45SN1_45S_50ntFlanks@11190.11209.20|CCCGGGGGGCCGGCGGCGGC</td><td>>hg38_21_+_8433172_8446622_RNA45SN1_45S_50ntFlanks</td><td>11190</td><td>11209</td><td>20</td><td>CCCGGGGGGCCGGCGGCGGC</td><td>101.3</td><td>-9.7</td><td>1.9</td><td>-5.2</td><td>2.5E-07</td><td>1.5E-06</td><td>BT20/MB468</td><td>MP</td></tr>
<tr><td>>am_trnaMT_ArgTCG_MT_+_10405_10469@31.48.18|TCGACTCATTAAATTATG</td><td>>am_trnaMT_ArgTCG_MT_+_10405_10469</td><td>31</td><td>48</td><td>18</td><td>TCGACTCATTAAATTATG</td><td>155.4</td><td>-7.0</td><td>1.4</td><td>-4.9</td><td>9.5E-07</td><td>1.5E-06</td><td>MB231/MB468</td><td>NUC</td></tr>
<tr><td>>hg38_21_+_8433172_8446622_RNA45SN1_45S_50ntFlanks@10382.10424.43|TTCCATGTGAACAGCAGTTGAACATGGGTCAGTCGGTCCTGAG</td><td>>hg38_21_+_8433172_8446622_RNA45SN1_45S_50ntFlanks</td><td>10382</td><td>10424</td><td>43</td><td>TTCCATGTGAACAGCAGTTGAACATGGGTCAGTCGGTCCTGAG</td><td>1067.3</td><td>8.0</td><td>1.6</td><td>5.2</td><td>2.5E-07</td><td>1.5E-06</td><td>BT20/MB468</td><td>MP</td></tr>
<tr><td>>hg38_21_+_8433172_8446622_RNA45SN1_45S_50ntFlanks@8112.8141.30|GGGCGCGGGACATGTGGCGTACGGAAGACC</td><td>>hg38_21_+_8433172_8446622_RNA45SN1_45S_50ntFlanks</td><td>8112</td><td>8141</td><td>30</td><td>GGGCGCGGGACATGTGGCGTACGGAAGACC</td><td>227.2</td><td>11.2</td><td>2.1</td><td>5.4</td><td>5.8E-08</td><td>1.6E-06</td><td>BT20/MB231</td><td>MP</td></tr>
<tr><td>>MI0000063|hsa-let-7b&WithFlank&22|+|46113680|46113774@12.33.22(+1A)&[MIMAT0000063&hsa-let-7b-5p&offsets|0|0(+1A);m-14&22|+|46113691|46113712&offsets|0|0(+1A)]|TGAGGTAGTAGGTTGTGTGGTTA</td><td>>MI0000063|hsa-let-7b&WithFlank&22|+|46113680|46113774</td><td>12</td><td>33</td><td>22(+1A)</td><td>TGAGGTAGTAGGTTGTGTGGTTA</td><td>590.7</td><td>3.1</td><td>0.6</td><td>4.9</td><td>9.5E-07</td><td>1.6E-06</td><td>MB231/MB468</td><td>NUC</td></tr>
<tr><td>>hg38_21_+_8433172_8446622_RNA45SN1_45S_50ntFlanks@12461.12478.18|CGTTGGATTGTTCACCCA</td><td>>hg38_21_+_8433172_8446622_RNA45SN1_45S_50ntFlanks</td><td>12461</td><td>12478</td><td>18</td><td>CGTTGGATTGTTCACCCA</td><td>64.8</td><td>-9.1</td><td>1.8</td><td>-5.2</td><td>2.5E-07</td><td>1.6E-06</td><td>BT20/MB468</td><td>MP</td></tr>
<tr><td>>hg38_21_+_8433172_8446622_RNA45SN1_45S_50ntFlanks@12644.12675.32|TCTGTGGGATTATGACTGAACGCCTCTAAGTC</td><td>>hg38_21_+_8433172_8446622_RNA45SN1_45S_50ntFlanks</td><td>12644</td><td>12675</td><td>32</td><td>TCTGTGGGATTATGACTGAACGCCTCTAAGTC</td><td>384.4</td><td>-7.6</td><td>1.6</td><td>-4.9</td><td>1.0E-06</td><td>1.6E-06</td><td>BT20/MB468</td><td>NUC</td></tr>
<tr><td>>hg38_21_+_8433172_8446622_RNA45SN1_45S_50ntFlanks@6652.6672.21|GACTCTTAGCGGTGGATCACT</td><td>>hg38_21_+_8433172_8446622_RNA45SN1_45S_50ntFlanks</td><td>6652</td><td>6672</td><td>21</td><td>GACTCTTAGCGGTGGATCACT</td><td>305.2</td><td>6.9</td><td>1.4</td><td>4.9</td><td>8.1E-07</td><td>1.6E-06</td><td>BT20/MB468</td><td>CYTO</td></tr>
<tr><td>>trnaMT_SerGCT_MT_+_12207_12265@1.48.48|GAGAAAGCTCACAAGAACTGCTAACTCATGCCCCCATGTCTAACAACA</td><td>>trnaMT_SerGCT_MT_+_12207_12265</td><td>1</td><td>48</td><td>48</td><td>GAGAAAGCTCACAAGAACTGCTAACTCATGCCCCCATGTCTAACAACA</td><td>307.5</td><td>12.3</td><td>2.5</td><td>4.9</td><td>9.3E-07</td><td>1.6E-06</td><td>MB231/MB468</td><td>CYTO</td></tr>
<tr><td>>hg38_21_+_8433172_8446622_RNA45SN1_45S_50ntFlanks@3843.3874.32|CTACTTGGATAACTGTGGTAATTCTAGAGCTA</td><td>>hg38_21_+_8433172_8446622_RNA45SN1_45S_50ntFlanks</td><td>3843</td><td>3874</td><td>32</td><td>CTACTTGGATAACTGTGGTAATTCTAGAGCTA</td><td>280.3</td><td>-7.3</td><td>1.5</td><td>-4.9</td><td>8.2E-07</td><td>1.6E-06</td><td>MB231/MB468</td><td>MP</td></tr>
<tr><td>>hg38_21_+_8433172_8446622_RNA45SN1_45S_50ntFlanks@11792.11814.23|ATGAACGAGATTCCCACTGTCCC</td><td>>hg38_21_+_8433172_8446622_RNA45SN1_45S_50ntFlanks</td><td>11792</td><td>11814</td><td>23</td><td>ATGAACGAGATTCCCACTGTCCC</td><td>63.4</td><td>-9.1</td><td>1.8</td><td>-5.2</td><td>2.5E-07</td><td>1.6E-06</td><td>BT20/MB468</td><td>MP</td></tr>
<tr><td>>hg38_21_+_8433172_8446622_RNA45SN1_45S_50ntFlanks@12937.12958.22|AACCATTCGTAGACGACCTGCT</td><td>>hg38_21_+_8433172_8446622_RNA45SN1_45S_50ntFlanks</td><td>12937</td><td>12958</td><td>22</td><td>AACCATTCGTAGACGACCTGCT</td><td>394.5</td><td>-2.5</td><td>0.5</td><td>-4.9</td><td>9.5E-07</td><td>1.6E-06</td><td>MB231/MB468</td><td>NUC</td></tr>
<tr><td>>hg38_MT_+_1621_3279_RNR2_16S_50ntFlanks@1176.1215.40|GCATTAAAAATTTCGGTTGGGGCGACCTCGGAGCAGAACC</td><td>>hg38_MT_+_1621_3279_RNR2_16S_50ntFlanks</td><td>1176</td><td>1215</td><td>40</td><td>GCATTAAAAATTTCGGTTGGGGCGACCTCGGAGCAGAACC</td><td>129.0</td><td>10.8</td><td>2.1</td><td>5.2</td><td>2.5E-07</td><td>1.6E-06</td><td>BT20/MB468</td><td>MP</td></tr>
<tr><td>>hg38_1_-_228634819_228635039_RNA5S12_5S_50ntFlanks@134.170.37|TGGGAGACCGCCTGGGAATACCGGGTGCTGTAGGCTT</td><td>>hg38_1_-_228634819_228635039_RNA5S12_5S_50ntFlanks</td><td>134</td><td>170</td><td>37</td><td>TGGGAGACCGCCTGGGAATACCGGGTGCTGTAGGCTT</td><td>19106.3</td><td>4.0</td><td>0.8</td><td>4.9</td><td>9.9E-07</td><td>1.6E-06</td><td>MB231/MB468</td><td>MITO</td></tr>
<tr><td>>hg38_21_+_8433172_8446622_RNA45SN1_45S_50ntFlanks@8177.8206.30|TTCTGATCGAGGCCCAGCCCGTGGACGGTG</td><td>>hg38_21_+_8433172_8446622_RNA45SN1_45S_50ntFlanks</td><td>8177</td><td>8206</td><td>30</td><td>TTCTGATCGAGGCCCAGCCCGTGGACGGTG</td><td>128.2</td><td>-5.5</td><td>1.1</td><td>-4.9</td><td>8.2E-07</td><td>1.6E-06</td><td>MB231/MB468</td><td>MP</td></tr>
<tr><td>>am_trna10_ValCAC_5_-_180649395_180649467@1.36.36,trna18_ValCAC_5_-_180529253_180529325@1.36.36,trna2_ValCAC_5_+_180524070_180524142@1.36.36,trna6_ValCAC_5_+_180600650_180600722@1.36.36,trna85_ValCAC_1_-_161369490_161369562@1.36.36,trna90_ValCAC_1_-_149684088_149684161@1.36.36,trna98_ValCAC_1_-_149298555_149298627@1.36.36,trna9_ValCAC_6_+_26538282_26538354@1.36.36|GTTTCCGTAGTGTAGTGGTTATCACGTTCGCCTCAC</td><td>>am_trna10_ValCAC_5_-_180649395_180649467</td><td>1</td><td>36</td><td>36</td><td>GTTTCCGTAGTGTAGTGGTTATCACGTTCGCCTCAC</td><td>445.9</td><td>6.9</td><td>1.4</td><td>4.9</td><td>1.0E-06</td><td>1.6E-06</td><td>MB231/MB468</td><td>MITO</td></tr>
<tr><td>>hg38_MT_+_598_1651_RNR1_12S_50ntFlanks@52.87.36|ATAGGTTTGGTCCTAGCCTTTCTATTAGCTCTTAGT</td><td>>hg38_MT_+_598_1651_RNR1_12S_50ntFlanks</td><td>52</td><td>87</td><td>36</td><td>ATAGGTTTGGTCCTAGCCTTTCTATTAGCTCTTAGT</td><td>7040.4</td><td>5.3</td><td>1.1</td><td>5.0</td><td>7.1E-07</td><td>1.6E-06</td><td>BT20/MB468</td><td>MITO</td></tr>
<tr><td>>trnaMT_PheGAA_MT_+_577_647@31.55.25|CACTGAAAATGTTTAGACGGGCTCA</td><td>>trnaMT_PheGAA_MT_+_577_647</td><td>31</td><td>55</td><td>25</td><td>CACTGAAAATGTTTAGACGGGCTCA</td><td>77.0</td><td>10.0</td><td>1.9</td><td>5.2</td><td>2.5E-07</td><td>1.6E-06</td><td>BT20/MB468</td><td>MP</td></tr>
<tr><td>>hg38_21_+_8433172_8446622_RNA45SN1_45S_50ntFlanks@12247.12272.26|GGCAAAAGCTCGCTTGATCTTGATTT</td><td>>hg38_21_+_8433172_8446622_RNA45SN1_45S_50ntFlanks</td><td>12247</td><td>12272</td><td>26</td><td>GGCAAAAGCTCGCTTGATCTTGATTT</td><td>299.6</td><td>-7.2</td><td>1.5</td><td>-4.9</td><td>1.0E-06</td><td>1.6E-06</td><td>MB231/MB468</td><td>MITO</td></tr>
<tr><td>>MI0000542|hsa-mir-320a&WithFlank&8|-|22244960|22245043@49.70.22&[MIMAT0000510&hsa-miR-320a-3p&offsets|+1|+1;m-60&8|-|22244975|22244996&offsets|+1|+1]|AAAGCTGGGTTGAGAGGGCGAA</td><td>>MI0000542|hsa-mir-320a&WithFlank&8|-|22244960|22245043</td><td>49</td><td>70</td><td>22</td><td>AAAGCTGGGTTGAGAGGGCGAA</td><td>1078.9</td><td>6.8</td><td>1.3</td><td>5.2</td><td>2.5E-07</td><td>1.6E-06</td><td>BT20/MB468</td><td>MP</td></tr>
<tr><td>>hg38_21_+_8433172_8446622_RNA45SN1_45S_50ntFlanks@4734.4751.18|AACGAAAGTCGGAGGTTC</td><td>>hg38_21_+_8433172_8446622_RNA45SN1_45S_50ntFlanks</td><td>4734</td><td>4751</td><td>18</td><td>AACGAAAGTCGGAGGTTC</td><td>888.6</td><td>-3.3</td><td>0.7</td><td>-4.9</td><td>9.6E-07</td><td>1.6E-06</td><td>MB231/MB468</td><td>NUC</td></tr>
<tr><td>>trnaMT_SerGCT_MT_+_12207_12265@1.26.26|GAGAAAGCTCACAAGAACTGCTAACT</td><td>>trnaMT_SerGCT_MT_+_12207_12265</td><td>1</td><td>26</td><td>26</td><td>GAGAAAGCTCACAAGAACTGCTAACT</td><td>39958.6</td><td>6.1</td><td>1.2</td><td>5.0</td><td>7.2E-07</td><td>1.6E-06</td><td>BT20/MB468</td><td>MITO</td></tr>
<tr><td>>MI0000743|hsa-mir-34c&WithFlank&11|+|111513433|111513521@52.73.22&[MIMAT0004677&hsa-miR-34c-3p&offsets|0|0;m-474&11|+|111513484|111513505&offsets|0|0]|AATCACTAACCACACGGCCAGG</td><td>>MI0000743|hsa-mir-34c&WithFlank&11|+|111513433|111513521</td><td>52</td><td>73</td><td>22</td><td>AATCACTAACCACACGGCCAGG</td><td>46.5</td><td>8.8</td><td>1.8</td><td>4.9</td><td>8.3E-07</td><td>1.6E-06</td><td>MB231/MB468</td><td>MP</td></tr>
<tr><td>>MI0000815|hsa-mir-339&WithFlank&7|-|1022927|1023032@56.77.22(+1A)&[MIMAT0004702&hsa-miR-339-3p&offsets|0|-1(+1A);m-235&7|-|1022957|1022977&offsets|0|+1(+1A)]|TGAGCGCCTCGACGACAGAGCCA</td><td>>MI0000815|hsa-mir-339&WithFlank&7|-|1022927|1023032</td><td>56</td><td>77</td><td>22(+1A)</td><td>TGAGCGCCTCGACGACAGAGCCA</td><td>46.7</td><td>8.8</td><td>1.8</td><td>4.9</td><td>8.3E-07</td><td>1.6E-06</td><td>MB231/MB468</td><td>MP</td></tr>
<tr><td>>trnaMT_MetCAT_MT_+_4402_4469@33.71.39|TACCCCGAAAATGTTGGTTATACCCTTCCCGTACTACCA</td><td>>trnaMT_MetCAT_MT_+_4402_4469</td><td>33</td><td>71</td><td>39</td><td>TACCCCGAAAATGTTGGTTATACCCTTCCCGTACTACCA</td><td>559.3</td><td>6.4</td><td>1.3</td><td>4.9</td><td>1.0E-06</td><td>1.6E-06</td><td>MB231/MB468</td><td>MITO</td></tr>
<tr><td>>hg38_21_+_8433172_8446622_RNA45SN1_45S_50ntFlanks@7983.8009.27|CAGATCAGACGTGGCGACCCGCTGAAT</td><td>>hg38_21_+_8433172_8446622_RNA45SN1_45S_50ntFlanks</td><td>7983</td><td>8009</td><td>27</td><td>CAGATCAGACGTGGCGACCCGCTGAAT</td><td>1068.7</td><td>-4.2</td><td>0.8</td><td>-4.9</td><td>8.3E-07</td><td>1.6E-06</td><td>MB231/MB468</td><td>MP</td></tr>
<tr><td>>hg38_21_+_8433172_8446622_RNA45SN1_45S_50ntFlanks@12416.12434.19|TTCGATGTCGGCTCTTCCT</td><td>>hg38_21_+_8433172_8446622_RNA45SN1_45S_50ntFlanks</td><td>12416</td><td>12434</td><td>19</td><td>TTCGATGTCGGCTCTTCCT</td><td>1538.4</td><td>-3.1</td><td>0.6</td><td>-4.9</td><td>1.0E-06</td><td>1.6E-06</td><td>BT20/MB468</td><td>NUC</td></tr>
<tr><td>>hg38_21_+_8433172_8446622_RNA45SN1_45S_50ntFlanks@9920.9951.32|GTGTTGGTTGATATAGACAGCAGGACGGTGGC</td><td>>hg38_21_+_8433172_8446622_RNA45SN1_45S_50ntFlanks</td><td>9920</td><td>9951</td><td>32</td><td>GTGTTGGTTGATATAGACAGCAGGACGGTGGC</td><td>57.8</td><td>9.6</td><td>1.9</td><td>5.2</td><td>2.5E-07</td><td>1.6E-06</td><td>BT20/MB468</td><td>MP</td></tr>
<tr><td>>hg38_21_+_8433172_8446622_RNA45SN1_45S_50ntFlanks@10726.10757.32|GGGAGAGGGTGTAAATCTCGCGCCGGGCCGTA</td><td>>hg38_21_+_8433172_8446622_RNA45SN1_45S_50ntFlanks</td><td>10726</td><td>10757</td><td>32</td><td>GGGAGAGGGTGTAAATCTCGCGCCGGGCCGTA</td><td>55.0</td><td>-8.9</td><td>1.7</td><td>-5.2</td><td>2.5E-07</td><td>1.6E-06</td><td>BT20/MB468</td><td>MP</td></tr>
<tr><td>>hg38_21_+_8433172_8446622_RNA45SN1_45S_50ntFlanks@11785.11803.19|TGAATGGATGAACGAGATT</td><td>>hg38_21_+_8433172_8446622_RNA45SN1_45S_50ntFlanks</td><td>11785</td><td>11803</td><td>19</td><td>TGAATGGATGAACGAGATT</td><td>1922.9</td><td>-4.7</td><td>0.9</td><td>-4.9</td><td>9.4E-07</td><td>1.6E-06</td><td>MB231/MB468</td><td>CYTO</td></tr>
<tr><td>>hg38_21_+_8433172_8446622_RNA45SN1_45S_50ntFlanks@6732.6772.41|ACATTGATCATCGACACTTCGAACGCACTTGCGGCCCCGGG</td><td>>hg38_21_+_8433172_8446622_RNA45SN1_45S_50ntFlanks</td><td>6732</td><td>6772</td><td>41</td><td>ACATTGATCATCGACACTTCGAACGCACTTGCGGCCCCGGG</td><td>177.3</td><td>-11.3</td><td>2.3</td><td>-4.9</td><td>8.3E-07</td><td>1.6E-06</td><td>MB231/MB468</td><td>MP</td></tr>
<tr><td>>hg38_MT_+_598_1651_RNR1_12S_50ntFlanks@568.611.44|CAAAGGACCTGGCGGTGCTTCATATCCCTCTAGAGGAGCCTGTT</td><td>>hg38_MT_+_598_1651_RNR1_12S_50ntFlanks</td><td>568</td><td>611</td><td>44</td><td>CAAAGGACCTGGCGGTGCTTCATATCCCTCTAGAGGAGCCTGTT</td><td>41.8</td><td>8.6</td><td>1.7</td><td>4.9</td><td>8.4E-07</td><td>1.6E-06</td><td>MB231/MB468</td><td>MP</td></tr>
<tr><td>>hg38_21_+_8433172_8446622_RNA45SN1_45S_50ntFlanks@12416.12435.20|TTCGATGTCGGCTCTTCCTA</td><td>>hg38_21_+_8433172_8446622_RNA45SN1_45S_50ntFlanks</td><td>12416</td><td>12435</td><td>20</td><td>TTCGATGTCGGCTCTTCCTA</td><td>595.8</td><td>-3.4</td><td>0.7</td><td>-4.9</td><td>1.0E-06</td><td>1.6E-06</td><td>MB231/MB468</td><td>MITO</td></tr>
<tr><td>>hg38_21_+_8433172_8446622_RNA45SN1_45S_50ntFlanks@6650.6679.30|ACGACTCTTAGCGGTGGATCACTCGGCTCG</td><td>>hg38_21_+_8433172_8446622_RNA45SN1_45S_50ntFlanks</td><td>6650</td><td>6679</td><td>30</td><td>ACGACTCTTAGCGGTGGATCACTCGGCTCG</td><td>2059.2</td><td>-2.5</td><td>0.5</td><td>-4.9</td><td>1.1E-06</td><td>1.6E-06</td><td>BT20/MB468</td><td>NUC</td></tr>
<tr><td>>hg38_21_+_8433172_8446622_RNA45SN1_45S_50ntFlanks@9930.9950.21|ATATAGACAGCAGGACGGTGG</td><td>>hg38_21_+_8433172_8446622_RNA45SN1_45S_50ntFlanks</td><td>9930</td><td>9950</td><td>21</td><td>ATATAGACAGCAGGACGGTGG</td><td>137.4</td><td>10.4</td><td>1.9</td><td>5.4</td><td>5.9E-08</td><td>1.6E-06</td><td>BT20/MB231</td><td>MP</td></tr>
<tr><td>>hg38_21_+_8433172_8446622_RNA45SN1_45S_50ntFlanks@12635.12660.26|AAGCTACCATCTGTGGGATTATGACT</td><td>>hg38_21_+_8433172_8446622_RNA45SN1_45S_50ntFlanks</td><td>12635</td><td>12660</td><td>26</td><td>AAGCTACCATCTGTGGGATTATGACT</td><td>412.2</td><td>-2.9</td><td>0.6</td><td>-5.0</td><td>7.3E-07</td><td>1.6E-06</td><td>BT20/MB468</td><td>MITO</td></tr>
<tr><td>>hg38_21_+_8433172_8446622_RNA45SN1_45S_50ntFlanks@12632.12661.30|GCGAAGCTACCATCTGTGGGATTATGACTG</td><td>>hg38_21_+_8433172_8446622_RNA45SN1_45S_50ntFlanks</td><td>12632</td><td>12661</td><td>30</td><td>GCGAAGCTACCATCTGTGGGATTATGACTG</td><td>866.3</td><td>-6.5</td><td>1.3</td><td>-4.9</td><td>1.1E-06</td><td>1.6E-06</td><td>BT20/MB468</td><td>NUC</td></tr>
<tr><td>>am_trnaMT_HisGTG_MT_+_12138_12206@24.51.28|TCAGATTGTGAATCTGACAACAGAGGCT</td><td>>am_trnaMT_HisGTG_MT_+_12138_12206</td><td>24</td><td>51</td><td>28</td><td>TCAGATTGTGAATCTGACAACAGAGGCT</td><td>75.2</td><td>10.0</td><td>1.9</td><td>5.2</td><td>2.5E-07</td><td>1.6E-06</td><td>BT20/MB468</td><td>MP</td></tr>
<tr><td>>hg38_21_+_8433172_8446622_RNA45SN1_45S_50ntFlanks@12634.12652.19|GAAGCTACCATCTGTGGGA</td><td>>hg38_21_+_8433172_8446622_RNA45SN1_45S_50ntFlanks</td><td>12634</td><td>12652</td><td>19</td><td>GAAGCTACCATCTGTGGGA</td><td>69.3</td><td>-9.2</td><td>1.8</td><td>-5.2</td><td>2.5E-07</td><td>1.6E-06</td><td>BT20/MB468</td><td>MP</td></tr>
<tr><td>>MI0000102|hsa-mir-100&WithFlank&11|-|122152223|122152314@19.38.20&[MIMAT0000098&hsa-miR-100-5p&offsets|0|-2;m-27&11|-|122152276|122152296&offsets|0|-1]|AACCCGTAGATCCGAACTTG</td><td>>MI0000102|hsa-mir-100&WithFlank&11|-|122152223|122152314</td><td>19</td><td>38</td><td>20</td><td>AACCCGTAGATCCGAACTTG</td><td>22851.2</td><td>-5.9</td><td>1.1</td><td>-5.3</td><td>1.1E-07</td><td>1.6E-06</td><td>BT20/MB231</td><td>NUC</td></tr>
<tr><td>>MI0000102|hsa-mir-100&WithFlank&11|-|122152223|122152314@19.38.20(+1G)&[MIMAT0000098&hsa-miR-100-5p&offsets|0|-2(+1G);m-27&11|-|122152276|122152296&offsets|0|-1(+1G)]|AACCCGTAGATCCGAACTTGG</td><td>>MI0000102|hsa-mir-100&WithFlank&11|-|122152223|122152314</td><td>19</td><td>38</td><td>20(+1G)</td><td>AACCCGTAGATCCGAACTTGG</td><td>4117.3</td><td>-8.0</td><td>1.5</td><td>-5.4</td><td>5.9E-08</td><td>1.6E-06</td><td>BT20/MB231</td><td>MP</td></tr>
<tr><td>>hg38_21_+_8433172_8446622_RNA45SN1_45S_50ntFlanks@9554.9573.20|CGTAGCGGTCCTGACGTGCA</td><td>>hg38_21_+_8433172_8446622_RNA45SN1_45S_50ntFlanks</td><td>9554</td><td>9573</td><td>20</td><td>CGTAGCGGTCCTGACGTGCA</td><td>11885.7</td><td>-2.1</td><td>0.4</td><td>-4.9</td><td>8.2E-07</td><td>1.6E-06</td><td>BT20/MB468</td><td>CYTO</td></tr>
<tr><td>>trna116_GluCTC_1_-_145399233_145399304@36.75.40,trna134_GluTTC_1_-_16861774_16861845@36.75.40,trna71_GluCTC_1_-_161439189_161439260@36.75.40,trna74_GluCTC_1_-_161431809_161431880@36.75.40,trna77_GluCTC_1_-_161424398_161424469@36.75.40,trna77_GluCTC_6_+_28949976_28950047@36.75.40,trna80_GluCTC_1_-_161417018_161417089@36.75.40,trna84_GluTTC_1_-_161391883_161391954@36.75.40,trna87_GluCTC_6_-_126101393_126101464@36.75.40|CACCGCCGCGGCCCGGGTTCGATTCCCGGTCAGGGAACCA</td><td>>trna116_GluCTC_1_-_145399233_145399304</td><td>36</td><td>75</td><td>40</td><td>CACCGCCGCGGCCCGGGTTCGATTCCCGGTCAGGGAACCA</td><td>2930.3</td><td>5.7</td><td>1.2</td><td>4.9</td><td>8.2E-07</td><td>1.6E-06</td><td>BT20/MB468</td><td>CYTO</td></tr>
</table>

<table>
<tr><td></td><td></td><td></td><td></td><td></td><td></td><td></td><td></td><td></td><td></td><td></td><td></td><td></td><td></td></tr>
<tr><td>>hg38_21_+_8433172_8446622_RNA45SN1_45S_50ntFlanks@6699.6722.24|GCTAGCTGCGAGAATTAATGTGAA</td><td>>hg38_21_+_8433172_8446622_RNA45SN1_45S_50ntFlanks</td><td>6699</td><td>6722</td><td>24</td><td>GCTAGCTGCGAGAATTAATGTGAA</td><td>140.3</td><td>-5.0</td><td>1.0</td><td>-4.9</td><td>8.4E-07</td><td>1.6E-06</td><td>MB231/MB468</td><td>MP</td></tr>
<tr><td>>hg38_21_+_8433172_8446622_RNA45SN1_45S_50ntFlanks@11999.12043.45|CCCTGCGGGCCGCCGGTGAAATACCACTACTCTGATCGTTTTTTC</td><td>>hg38_21_+_8433172_8446622_RNA45SN1_45S_50ntFlanks</td><td>11999</td><td>12043</td><td>45</td><td>CCCTGCGGGCCGCCGGTGAAATACCACTACTCTGATCGTTTTTTC</td><td>249.1</td><td>-11.0</td><td>2.1</td><td>-5.2</td><td>2.5E-07</td><td>1.6E-06</td><td>BT20/MB468</td><td>MP</td></tr>
<tr><td>>hg38_21_+_8433172_8446622_RNA45SN1_45S_50ntFlanks@10410.10428.19|TCAGTCGGTCCTGAGAGAT</td><td>>hg38_21_+_8433172_8446622_RNA45SN1_45S_50ntFlanks</td><td>10410</td><td>10428</td><td>19</td><td>TCAGTCGGTCCTGAGAGAT</td><td>875.3</td><td>-5.3</td><td>1.1</td><td>-4.9</td><td>9.5E-07</td><td>1.6E-06</td><td>MB231/MB468</td><td>CYTO</td></tr>
<tr><td>>hg38_21_+_8433172_8446622_RNA45SN1_45S_50ntFlanks@9141.9164.24|CGGGGCGGACTGTCCCCAGTGCGC</td><td>>hg38_21_+_8433172_8446622_RNA45SN1_45S_50ntFlanks</td><td>9141</td><td>9164</td><td>24</td><td>CGGGGCGGACTGTCCCCAGTGCGC</td><td>220.6</td><td>-7.0</td><td>1.4</td><td>-5.2</td><td>2.5E-07</td><td>1.6E-06</td><td>BT20/MB468</td><td>MP</td></tr>
<tr><td>>hg38_1_-_228634819_228635039_RNA5S12_5S_50ntFlanks@51.89.39|GTCTACGGCCATACCACCCTGAACGCGCCCGATCTCGTC</td><td>>hg38_1_-_228634819_228635039_RNA5S12_5S_50ntFlanks</td><td>51</td><td>89</td><td>39</td><td>GTCTACGGCCATACCACCCTGAACGCGCCCGATCTCGTC</td><td>15520.2</td><td>4.3</td><td>0.9</td><td>4.9</td><td>1.0E-06</td><td>1.6E-06</td><td>MB231/MB468</td><td>MITO</td></tr>
<tr><td>>hg38_21_+_8433172_8446622_RNA45SN1_45S_50ntFlanks@12641.12661.21|CCATCTGTGGGATTATGACTG</td><td>>hg38_21_+_8433172_8446622_RNA45SN1_45S_50ntFlanks</td><td>12641</td><td>12661</td><td>21</td><td>CCATCTGTGGGATTATGACTG</td><td>802.8</td><td>-6.3</td><td>1.3</td><td>-4.9</td><td>1.1E-06</td><td>1.6E-06</td><td>BT20/MB468</td><td>NUC</td></tr>
<tr><td>>hg38_21_+_8433172_8446622_RNA45SN1_45S_50ntFlanks@10906.10936.31|TGGGCGCGCGCCGCGGCTGGACGAGGCGCCG</td><td>>hg38_21_+_8433172_8446622_RNA45SN1_45S_50ntFlanks</td><td>10906</td><td>10936</td><td>31</td><td>TGGGCGCGCGCCGCGGCTGGACGAGGCGCCG</td><td>61.1</td><td>-9.0</td><td>1.7</td><td>-5.2</td><td>2.6E-07</td><td>1.6E-06</td><td>BT20/MB468</td><td>MP</td></tr>
<tr><td>>hg38_21_+_8433172_8446622_RNA45SN1_45S_50ntFlanks@12222.12241.20|ACAGAAACCTCCCGTGGAGC</td><td>>hg38_21_+_8433172_8446622_RNA45SN1_45S_50ntFlanks</td><td>12222</td><td>12241</td><td>20</td><td>ACAGAAACCTCCCGTGGAGC</td><td>88.1</td><td>-9.5</td><td>1.9</td><td>-5.2</td><td>2.6E-07</td><td>1.6E-06</td><td>BT20/MB468</td><td>MP</td></tr>
<tr><td>>hg38_21_+_8433172_8446622_RNA45SN1_45S_50ntFlanks@11570.11592.23|AGCCGACTTAGAACTGGTGCGGA</td><td>>hg38_21_+_8433172_8446622_RNA45SN1_45S_50ntFlanks</td><td>11570</td><td>11592</td><td>23</td><td>AGCCGACTTAGAACTGGTGCGGA</td><td>271.3</td><td>-5.5</td><td>1.1</td><td>-4.9</td><td>1.1E-06</td><td>1.6E-06</td><td>BT20/MB468</td><td>NUC</td></tr>
<tr><td>>MI0000749|hsa-mir-30e&WithFlank&1|+|40754349|40754452@65.86.22&[MIMAT0000693&hsa-miR-30e-3p&offsets|0|0;m-25&1|+|40754413|40754434&offsets|0|0]|CTTTCAGTCGGATGTTTACAGC</td><td>>MI0000749|hsa-mir-30e&WithFlank&1|+|40754349|40754452</td><td>65</td><td>86</td><td>22</td><td>CTTTCAGTCGGATGTTTACAGC</td><td>3088.9</td><td>2.9</td><td>0.6</td><td>4.9</td><td>9.8E-07</td><td>1.6E-06</td><td>MB231/MB468</td><td>NUC</td></tr>
<tr><td>>trna10_GlyTCC_17_+_8124866_8124937@31.51.21,trna117_GlyTCC_1_-_145397864_145397935@31.51.21,trna2_GlyTCC_19_+_4724082_4724153@31.51.21,trna45_GlyTCC_1_+_161500903_161500974@31.51.21,trna70_GlyTCC_1_-_161439547_161439618@31.51.21,trna73_GlyTCC_1_-_161432166_161432237@31.51.21,trna76_GlyTCC_1_-_161424756_161424827@31.51.21,trna79_GlyTCC_1_-_161417375_161417446@31.51.21,trna82_GlyTCC_1_-_161409961_161410032@31.51.21|CCTTCCAAGCAGTTGACCCGG</td><td>>trna10_GlyTCC_17_+_8124866_8124937</td><td>31</td><td>51</td><td>21</td><td>CCTTCCAAGCAGTTGACCCGG</td><td>90.7</td><td>-9.6</td><td>1.9</td><td>-5.2</td><td>2.6E-07</td><td>1.6E-06</td><td>BT20/MB468</td><td>MP</td></tr>
<tr><td>>am_trna111_HisGTG_1_-_147774845_147774916@-1G.23.24,trna118_HisGTG_1_-_145396881_145396952@-1G.23.24,trna16_HisGTG_1_+_146544773_146544844@-1G.23.24,trna1_HisGTG_15_+_45493349_45493420@-1G.23.24,trna21_HisGTG_1_+_147753471_147753542@-1G.23.24,trna33_HisGTG_6_+_27125906_27125977@-1G.23.24,trna7_HisGTG_9_-_14433938_14434009@-1G.23.24,trna8_HisGTG_15_-_45492611_45492682@-1G.23.24,trna9_HisGTG_15_-_45490804_45490875@-1G.23.24|GGCCGTGATCGTATAGTGGTTAGT</td><td>>am_trna111_HisGTG_1_-_147774845_147774916</td><td>-1G</td><td>23</td><td>24</td><td>GGCCGTGATCGTATAGTGGTTAGT</td><td>1289.1</td><td>-5.6</td><td>1.0</td><td>-5.4</td><td>8.3E-08</td><td>1.6E-06</td><td>BT20/MB231</td><td>MITO</td></tr>
<tr><td>>hg38_1_-_228634819_228635039_RNA5S12_5S_50ntFlanks@129.171.43|TTGGATGGGAGACCGCCTGGGAATACCGGGTGCTGTAGGCTTT</td><td>>hg38_1_-_228634819_228635039_RNA5S12_5S_50ntFlanks</td><td>129</td><td>171</td><td>43</td><td>TTGGATGGGAGACCGCCTGGGAATACCGGGTGCTGTAGGCTTT</td><td>1050.6</td><td>7.8</td><td>1.6</td><td>5.0</td><td>7.3E-07</td><td>1.6E-06</td><td>BT20/MB468</td><td>MITO</td></tr>
<tr><td>>MI0000086|hsa-mir-28&WithFlank&3|+|188688775|188688872@60.79.20(+1U)&[MIMAT0004502&hsa-miR-28-3p&offsets|0|-2(+1U);m-28&3|+|188688834|188688855&offsets|0|-2(+1U)]|CACTAGATTGTGAGCTCCTGT</td><td>>MI0000086|hsa-mir-28&WithFlank&3|+|188688775|188688872</td><td>60</td><td>79</td><td>20(+1U)</td><td>CACTAGATTGTGAGCTCCTGT</td><td>44.9</td><td>8.7</td><td>1.8</td><td>4.9</td><td>8.5E-07</td><td>1.6E-06</td><td>MB231/MB468</td><td>MP</td></tr>
<tr><td>>hg38_21_+_8433172_8446622_RNA45SN1_45S_50ntFlanks@6732.6771.40|ACATTGATCATCGACACTTCGAACGCACTTGCGGCCCCGG</td><td>>hg38_21_+_8433172_8446622_RNA45SN1_45S_50ntFlanks</td><td>6732</td><td>6771</td><td>40</td><td>ACATTGATCATCGACACTTCGAACGCACTTGCGGCCCCGG</td><td>1648.0</td><td>-4.2</td><td>0.9</td><td>-4.9</td><td>9.9E-07</td><td>1.6E-06</td><td>MB231/MB468</td><td>NUC</td></tr>
<tr><td>>hg38_21_+_8433172_8446622_RNA45SN1_45S_50ntFlanks@12008.12033.26|CCGCCGGTGAAATACCACTACTCTGA</td><td>>hg38_21_+_8433172_8446622_RNA45SN1_45S_50ntFlanks</td><td>12008</td><td>12033</td><td>26</td><td>CCGCCGGTGAAATACCACTACTCTGA</td><td>706.7</td><td>-5.6</td><td>1.1</td><td>-4.9</td><td>9.6E-07</td><td>1.6E-06</td><td>MB231/MB468</td><td>CYTO</td></tr>
<tr><td>>hg38_21_+_8433172_8446622_RNA45SN1_45S_50ntFlanks@10382.10424.43|TTCCATGTGAACAGCAGTTGAACATGGGTCAGTCGGTCCTGAG</td><td>>hg38_21_+_8433172_8446622_RNA45SN1_45S_50ntFlanks</td><td>10382</td><td>10424</td><td>43</td><td>TTCCATGTGAACAGCAGTTGAACATGGGTCAGTCGGTCCTGAG</td><td>1404.9</td><td>5.2</td><td>1.1</td><td>4.9</td><td>9.9E-07</td><td>1.6E-06</td><td>MB231/MB468</td><td>NUC</td></tr>
<tr><td>>hg38_21_+_8433172_8446622_RNA45SN1_45S_50ntFlanks@3853.3881.29|AACTGTGGTAATTCTAGAGCTAATACATG</td><td>>hg38_21_+_8433172_8446622_RNA45SN1_45S_50ntFlanks</td><td>3853</td><td>3881</td><td>29</td><td>AACTGTGGTAATTCTAGAGCTAATACATG</td><td>510.2</td><td>-4.1</td><td>0.8</td><td>-4.9</td><td>8.4E-07</td><td>1.6E-06</td><td>BT20/MB468</td><td>CYTO</td></tr>
<tr><td>>MI0000476|hsa-mir-138-1&WithFlank&3|+|44114206|44114316@29.53.25(+1U)&[MIMAT0000430_1&hsa-miR-138-5p&offsets|0|+2(+1U);m-497&3|+|44114234|44114256&offsets|0|+2(+1U)]|AGCTGGTGTTGTGAATCAGGCCGTTT</td><td>>MI0000476|hsa-mir-138-1&WithFlank&3|+|44114206|44114316</td><td>29</td><td>53</td><td>25(+1U)</td><td>AGCTGGTGTTGTGAATCAGGCCGTTT</td><td>41.6</td><td>8.6</td><td>1.7</td><td>4.9</td><td>8.5E-07</td><td>1.6E-06</td><td>MB231/MB468</td><td>MP</td></tr>
<tr><td>>hg38_21_+_8433172_8446622_RNA45SN1_45S_50ntFlanks@10411.10456.46|CAGTCGGTCCTGAGAGATGGGCGAGCGCCGTTCCGAAGGGACGGGC</td><td>>hg38_21_+_8433172_8446622_RNA45SN1_45S_50ntFlanks</td><td>10411</td><td>10456</td><td>46</td><td>CAGTCGGTCCTGAGAGATGGGCGAGCGCCGTTCCGAAGGGACGGGC</td><td>291.6</td><td>-7.5</td><td>1.5</td><td>-4.9</td><td>9.9E-07</td><td>1.6E-06</td><td>MB231/MB468</td><td>NUC</td></tr>
<tr><td>>hg38_21_+_8433172_8446622_RNA45SN1_45S_50ntFlanks@6730.6756.27|ACACATTGATCATCGACACTTCGAACG</td><td>>hg38_21_+_8433172_8446622_RNA45SN1_45S_50ntFlanks</td><td>6730</td><td>6756</td><td>27</td><td>ACACATTGATCATCGACACTTCGAACG</td><td>288.4</td><td>-2.6</td><td>0.5</td><td>-4.9</td><td>9.9E-07</td><td>1.6E-06</td><td>MB231/MB468</td><td>NUC</td></tr>
<tr><td>>hg38_21_+_8433172_8446622_RNA45SN1_45S_50ntFlanks@10351.10377.27|CGAGAACTTTGAAGGCCGAAGTGGAGA</td><td>>hg38_21_+_8433172_8446622_RNA45SN1_45S_50ntFlanks</td><td>10351</td><td>10377</td><td>27</td><td>CGAGAACTTTGAAGGCCGAAGTGGAGA</td><td>2576.0</td><td>-3.3</td><td>0.7</td><td>-5.0</td><td>7.4E-07</td><td>1.6E-06</td><td>BT20/MB468</td><td>MITO</td></tr>
<tr><td>>MI0000542|hsa-mir-320a&WithFlank&8|-|22244960|22245043@48.68.21(+3U)&[MIMAT0000510&hsa-miR-320a-3p&offsets|0|-1(+3U);m-60&8|-|22244975|22244996&offsets|0|-1(+3U)]|AAAAGCTGGGTTGAGAGGGCGTTT</td><td>>MI0000542|hsa-mir-320a&WithFlank&8|-|22244960|22245043</td><td>48</td><td>68</td><td>21(+3U)</td><td>AAAAGCTGGGTTGAGAGGGCGTTT</td><td>1656.8</td><td>8.8</td><td>1.7</td><td>5.2</td><td>2.6E-07</td><td>1.6E-06</td><td>BT20/MB468</td><td>MP</td></tr>
<tr><td>>hg38_21_+_8433172_8446622_RNA45SN1_45S_50ntFlanks@9579.9596.18|GTCGTCCGACCTGGGTAT</td><td>>hg38_21_+_8433172_8446622_RNA45SN1_45S_50ntFlanks</td><td>9579</td><td>9596</td><td>18</td><td>GTCGTCCGACCTGGGTAT</td><td>64.7</td><td>-9.1</td><td>1.8</td><td>-5.2</td><td>2.6E-07</td><td>1.6E-06</td><td>BT20/MB468</td><td>MP</td></tr>
<tr><td>>MI0000736|hsa-mir-30c-1&WithFlank&1|+|40757278|40757378@23.46.24(+3U);MI0000254|hsa-mir-30c-2&WithFlank&6|-|71376954|71377037@13.36.24(+3U)&[MIMAT0000244&hsa-miR-30c-5p&offsets|0|+1(+3U);m-57&1|+|40757300|40757323&offsets|0|0(+3U)];[MIMAT0000244_1&hsa-miR-30c-5p&offsets|0|+1(+3U);m-56&6|-|71377002|71377025&offsets|0|0(+3U)]|TGTAAACATCCTACACTCTCAGCTTTT</td><td>>MI0000736|hsa-mir-30c-1&WithFlank&1|+|40757278|40757378</td><td>23</td><td>46</td><td>24(+3U)</td><td>TGTAAACATCCTACACTCTCAGCTTTT</td><td>309.3</td><td>8.3</td><td>1.7</td><td>4.9</td><td>1.0E-06</td><td>1.6E-06</td><td>MB231/MB468</td><td>MITO</td></tr>
<tr><td>>hg38_21_+_8433172_8446622_RNA45SN1_45S_50ntFlanks@11217.11239.23|CTGGACGCGAGCCGGGCCCTTCC</td><td>>hg38_21_+_8433172_8446622_RNA45SN1_45S_50ntFlanks</td><td>11217</td><td>11239</td><td>23</td><td>CTGGACGCGAGCCGGGCCCTTCC</td><td>563.5</td><td>-3.8</td><td>0.8</td><td>-4.9</td><td>1.0E-06</td><td>1.6E-06</td><td>MB231/MB468</td><td>MITO</td></tr>
<tr><td>>MI0000088|hsa-mir-30a&WithFlank&6|-|71403545|71403627@12.37.26&[MIMAT0000087&hsa-miR-30a-5p&offsets|0|+4;m-15&6|-|71403595|71403616&offsets|0|+4]|TGTAAACATCCTCGACTGGAAGCTGT</td><td>>MI0000088|hsa-mir-30a&WithFlank&6|-|71403545|71403627</td><td>12</td><td>37</td><td>26</td><td>TGTAAACATCCTCGACTGGAAGCTGT</td><td>117.7</td><td>-10.3</td><td>1.9</td><td>-5.4</td><td>6.1E-08</td><td>1.6E-06</td><td>BT20/MB231</td><td>MP</td></tr>
<tr><td>>hg38_21_+_8433172_8446622_RNA45SN1_45S_50ntFlanks@483.515.33|TCGATGTGGTGACGTCGTGCTCTCCCGGGCCGG</td><td>>hg38_21_+_8433172_8446622_RNA45SN1_45S_50ntFlanks</td><td>483</td><td>515</td><td>33</td><td>TCGATGTGGTGACGTCGTGCTCTCCCGGGCCGG</td><td>445.4</td><td>4.6</td><td>0.9</td><td>4.9</td><td>1.0E-06</td><td>1.6E-06</td><td>MB231/MB468</td><td>NUC</td></tr>
<tr><td>>hg38_21_+_8433172_8446622_RNA45SN1_45S_50ntFlanks@10236.10272.37|GACGCTACGCCGCGACGAGTAGGAGGGCCGCTGCGGT</td><td>>hg38_21_+_8433172_8446622_RNA45SN1_45S_50ntFlanks</td><td>10236</td><td>10272</td><td>37</td><td>GACGCTACGCCGCGACGAGTAGGAGGGCCGCTGCGGT</td><td>58.4</td><td>9.6</td><td>1.9</td><td>5.2</td><td>2.6E-07</td><td>1.6E-06</td><td>BT20/MB468</td><td>MP</td></tr>
<tr><td>>hg38_21_+_8433172_8446622_RNA45SN1_45S_50ntFlanks@11874.11893.20|AGAAGACCCTGTTGAGCTTG</td><td>>hg38_21_+_8433172_8446622_RNA45SN1_45S_50ntFlanks</td><td>11874</td><td>11893</td><td>20</td><td>AGAAGACCCTGTTGAGCTTG</td><td>5637.6</td><td>-5.2</td><td>1.0</td><td>-5.1</td><td>2.6E-07</td><td>1.6E-06</td><td>BT20/MB468</td><td>MP</td></tr>
<tr><td>>hg38_21_+_8433172_8446622_RNA45SN1_45S_50ntFlanks@11575.11593.19|ACTTAGAACTGGTGCGGAC</td><td>>hg38_21_+_8433172_8446622_RNA45SN1_45S_50ntFlanks</td><td>11575</td><td>11593</td><td>19</td><td>ACTTAGAACTGGTGCGGAC</td><td>416.4</td><td>-2.8</td><td>0.6</td><td>-4.9</td><td>7.5E-07</td><td>1.6E-06</td><td>BT20/MB468</td><td>MITO</td></tr>
<tr><td>>hg38_21_+_8433172_8446622_RNA45SN1_45S_50ntFlanks@11729.11747.19|TAACTATGACTCTCTTAAG</td><td>>hg38_21_+_8433172_8446622_RNA45SN1_45S_50ntFlanks</td><td>11729</td><td>11747</td><td>19</td><td>TAACTATGACTCTCTTAAG</td><td>1326.5</td><td>-3.3</td><td>0.7</td><td>-4.9</td><td>7.5E-07</td><td>1.6E-06</td><td>BT20/MB468</td><td>MITO</td></tr>
<tr><td>>hg38_21_+_8433172_8446622_RNA45SN1_45S_50ntFlanks@12000.12032.33|CCTGCGGGCCGCCGGTGAAATACCACTACTCTG</td><td>>hg38_21_+_8433172_8446622_RNA45SN1_45S_50ntFlanks</td><td>12000</td><td>12032</td><td>33</td><td>CCTGCGGGCCGCCGGTGAAATACCACTACTCTG</td><td>603.3</td><td>-4.1</td><td>0.8</td><td>-4.9</td><td>9.7E-07</td><td>1.6E-06</td><td>MB231/MB468</td><td>CYTO</td></tr>
<tr><td>>hg38_21_+_8433172_8446622_RNA45SN1_45S_50ntFlanks@12510.12527.18|TCGTGAGACAGGTTAGTT</td><td>>hg38_21_+_8433172_8446622_RNA45SN1_45S_50ntFlanks</td><td>12510</td><td>12527</td><td>18</td><td>TCGTGAGACAGGTTAGTT</td><td>353.2</td><td>-7.2</td><td>1.4</td><td>-5.1</td><td>2.6E-07</td><td>1.6E-06</td><td>BT20/MB468</td><td>MP</td></tr>
<tr><td>>hg38_21_+_8433172_8446622_RNA45SN1_45S_50ntFlanks@8189.8211.23|CCCAGCCCGTGGACGGTGTGAGG</td><td>>hg38_21_+_8433172_8446622_RNA45SN1_45S_50ntFlanks</td><td>8189</td><td>8211</td><td>23</td><td>CCCAGCCCGTGGACGGTGTGAGG</td><td>120.5</td><td>-10.0</td><td>1.9</td><td>-5.1</td><td>2.6E-07</td><td>1.6E-06</td><td>BT20/MB468</td><td>MP</td></tr>
<tr><td>>hg38_21_+_8433172_8446622_RNA45SN1_45S_50ntFlanks@8741.8758.18|TCCGTCCGTCCGTCCGTC</td><td>>hg38_21_+_8433172_8446622_RNA45SN1_45S_50ntFlanks</td><td>8741</td><td>8758</td><td>18</td><td>TCCGTCCGTCCGTCCGTC</td><td>92.9</td><td>-9.6</td><td>1.9</td><td>-5.1</td><td>2.6E-07</td><td>1.6E-06</td><td>BT20/MB468</td><td>MP</td></tr>
<tr><td>>MI0001445|hsa-mir-423&WithFlank&17|+|30117073|30117178@23.43.21&[MIMAT0004748&hsa-miR-423-5p&offsets|0|-2;m-141&17|+|30117095|30117117&offsets|0|-2]|TGAGGGGCAGAGAGCGAGACT</td><td>>MI0001445|hsa-mir-423&WithFlank&17|+|30117073|30117178</td><td>23</td><td>43</td><td>21</td><td>TGAGGGGCAGAGAGCGAGACT</td><td>2424.1</td><td>4.2</td><td>0.9</td><td>4.9</td><td>8.6E-07</td><td>1.6E-06</td><td>MB231/MB468</td><td>MP</td></tr>
<tr><td>>MI0000102|hsa-mir-100&WithFlank&11|-|122152223|122152314@19.39.21(+1A)&[MIMAT0000098&hsa-miR-100-5p&offsets|0|-1(+1A);m-27&11|-|122152276|122152296&offsets|0|0(+1A)]|AACCCGTAGATCCGAACTTGTA</td><td>>MI0000102|hsa-mir-100&WithFlank&11|-|122152223|122152314</td><td>19</td><td>39</td><td>21(+1A)</td><td>AACCCGTAGATCCGAACTTGTA</td><td>88755.6</td><td>13.7</td><td>2.8</td><td>4.9</td><td>1.0E-06</td><td>1.6E-06</td><td>MB231/MB468</td><td>MITO</td></tr>
<tr><td>>hg38_21_+_8433172_8446622_RNA45SN1_45S_50ntFlanks@12415.12450.36|CTTCGATGTCGGCTCTTCCTATCATTGTGAAGCAGA</td><td>>hg38_21_+_8433172_8446622_RNA45SN1_45S_50ntFlanks</td><td>12415</td><td>12450</td><td>36</td><td>CTTCGATGTCGGCTCTTCCTATCATTGTGAAGCAGA</td><td>210.9</td><td>2.6</td><td>0.5</td><td>4.9</td><td>9.7E-07</td><td>1.6E-06</td><td>MB231/MB468</td><td>CYTO</td></tr>
<tr><td>>MI0000065|hsa-let-7d&WithFlank&9|+|94178828|94178926@68.88.21(+1C)&[MIMAT0004484&hsa-let-7d-3p&offsets|0|-1(+1C);m-98&9|+|94178895|94178916&offsets|0|-1(+1C)]|CTATACGACCTGCTGCCTTTCC</td><td>>MI0000065|hsa-let-7d&WithFlank&9|+|94178828|94178926</td><td>68</td><td>88</td><td>21(+1C)</td><td>CTATACGACCTGCTGCCTTTCC</td><td>180.4</td><td>2.6</td><td>0.5</td><td>4.9</td><td>1.1E-06</td><td>1.6E-06</td><td>BT20/MB468</td><td>NUC</td></tr>
<tr><td>>hg38_21_+_8433172_8446622_RNA45SN1_45S_50ntFlanks@9554.9577.24|CGTAGCGGTCCTGACGTGCAAATC</td><td>>hg38_21_+_8433172_8446622_RNA45SN1_45S_50ntFlanks</td><td>9554</td><td>9577</td><td>24</td><td>CGTAGCGGTCCTGACGTGCAAATC</td><td>1424.8</td><td>1.8</td><td>0.4</td><td>4.9</td><td>8.5E-07</td><td>1.6E-06</td><td>BT20/MB468</td><td>CYTO</td></tr>
<tr><td>>hg38_21_+_8433172_8446622_RNA45SN1_45S_50ntFlanks@12589.12606.18|GTTCAGACATTTGGTGTA</td><td>>hg38_21_+_8433172_8446622_RNA45SN1_45S_50ntFlanks</td><td>12589</td><td>12606</td><td>18</td><td>GTTCAGACATTTGGTGTA</td><td>56.5</td><td>-8.9</td><td>1.7</td><td>-5.1</td><td>2.6E-07</td><td>1.6E-06</td><td>BT20/MB468</td><td>MP</td></tr>
</table>

| | | | | | | | | | | | | | |
|---|---|---|---|---|---|---|---|---|---|---|---|---|---|
| >hg38_21_+_8433172_8446622_RNA45SN1_45S_50ntFlanks@4934.4951.18\|CCTGCGGCTTAATTTGAC | >hg38_21_+_8433172_8446622_RNA45SN1_45S_50ntFlanks | 4934 | 4951 | 18 | CCTGCGGCTTAATTTGAC | 170.3 | -9.5 | 1.9 | -4.9 | 7.5E-07 | 1.6E-06 | BT20/MB468 | MITO |
| >MI0005756\|hsa-mir-934&WithFlank&X\|+\|136550872\|136550966@21.43.23&[MIMAT0004977&hsa-miR-934&offsets\|0\|+1;m-943&X\|+\|136550892\|136550913&offsets\|0\|+1]\|TGTCTACTACTGGAGACACTGGT | >MI0005756\|hsa-mir-934&WithFlank&X\|+\|136550872\|136550966 | 21 | 43 | 23 | TGTCTACTACTGGAGACACTGGT | 463.3 | 3.2 | 0.7 | 4.9 | 1.1E-06 | 1.6E-06 | BT20/MB468 | NUC |
| >hg38_21_+_8433172_8446622_RNA45SN1_45S_50ntFlanks@10392.10419.28\|ACAGCAGTTGAACATGGGTCAGTCGGTC | >hg38_21_+_8433172_8446622_RNA45SN1_45S_50ntFlanks | 10392 | 10419 | 28 | ACAGCAGTTGAACATGGGTCAGTCGGTC | 309.0 | -6.8 | 1.4 | -4.9 | 1.0E-06 | 1.6E-06 | MB231/MB468 | MITO |
| >hg38_21_+_8433172_8446622_RNA45SN1_45S_50ntFlanks@12149.12180.32\|AGGTGGGGAGTTTGACTGGGGCGGTACACCTG | >hg38_21_+_8433172_8446622_RNA45SN1_45S_50ntFlanks | 12149 | 12180 | 32 | AGGTGGGGAGTTTGACTGGGGCGGTACACCTG | 58.4 | -8.9 | 1.7 | -5.1 | 2.6E-07 | 1.6E-06 | BT20/MB468 | MP |
| >hg38_21_+_8433172_8446622_RNA45SN1_45S_50ntFlanks@10513.10534.22\|CCGGAGTGGCGGAGATGGGCGC | >hg38_21_+_8433172_8446622_RNA45SN1_45S_50ntFlanks | 10513 | 10534 | 22 | CCGGAGTGGCGGAGATGGGCGC | 349.3 | -6.5 | 1.3 | -4.9 | 7.5E-07 | 1.6E-06 | BT20/MB468 | MITO |
| >hg38_21_+_8433172_8446622_RNA45SN1_45S_50ntFlanks@475.498.24\|CTTCGTGATCGATGTGGTGACGTC | >hg38_21_+_8433172_8446622_RNA45SN1_45S_50ntFlanks | 475 | 498 | 24 | CTTCGTGATCGATGTGGTGACGTC | 4507.6 | 4.5 | 0.9 | 4.9 | 8.7E-07 | 1.6E-06 | MB231/MB468 | MP |
| >hg38_21_+_8433172_8446622_RNA45SN1_45S_50ntFlanks@11552.11592.41\|CCTCGGCCGGCGCCTAGCAGCCGACTTAGAACTGGTGCGGA | >hg38_21_+_8433172_8446622_RNA45SN1_45S_50ntFlanks | 11552 | 11592 | 41 | CCTCGGCCGGCGCCTAGCAGCCGACTTAGAACTGGTGCGGA | 610.1 | -4.6 | 0.9 | -4.9 | 1.0E-06 | 1.6E-06 | MB231/MB468 | NUC |
| >MI0003205\|hsa-mir-532&WithFlank&X\|+\|50003142\|50003244@26.47.22&[MIMAT0002888&hsa-miR-532-5p&offsets\|0\|0;m-65&X\|+\|50003167\|50003188&offsets\|0\|0]\|CATGCCTTGAGTGTAGGACCGT | >MI0003205\|hsa-mir-532&WithFlank&X\|+\|50003142\|50003244 | 26 | 47 | 22 | CATGCCTTGAGTGTAGGACCGT | 1477.0 | 3.4 | 0.7 | 4.9 | 9.8E-07 | 1.6E-06 | MB231/MB468 | CYTO |
| >hg38_21_+_8433172_8446622_RNA45SN1_45S_50ntFlanks@3842.3883.42\|CCTACTTGGATAACTGTGGTAATTCTAGAGCTAATACATGCC | >hg38_21_+_8433172_8446622_RNA45SN1_45S_50ntFlanks | 3842 | 3883 | 42 | CCTACTTGGATAACTGTGGTAATTCTAGAGCTAATACATGCC | 391.0 | 6.7 | 1.4 | 4.9 | 9.8E-07 | 1.6E-06 | MB231/MB468 | CYTO |
| >hg38_21_+_8433172_8446622_RNA45SN1_45S_50ntFlanks@12488.12519.32\|AACGTGAGCTGGGTTTAGACCGTCGTGAGACA | >hg38_21_+_8433172_8446622_RNA45SN1_45S_50ntFlanks | 12488 | 12519 | 32 | AACGTGAGCTGGGTTTAGACCGTCGTGAGACA | 272.6 | -6.7 | 1.4 | -4.9 | 1.0E-06 | 1.6E-06 | MB231/MB468 | MITO |
| >hg38_21_+_8433172_8446622_RNA45SN1_45S_50ntFlanks@10677.10710.34\|GCGGTTCCGGCGGCGTCCGGTGAGCTCTCGCTGG | >hg38_21_+_8433172_8446622_RNA45SN1_45S_50ntFlanks | 10677 | 10710 | 34 | GCGGTTCCGGCGGCGTCCGGTGAGCTCTCGCTGG | 653.1 | -5.1 | 1.0 | -4.9 | 8.7E-07 | 1.6E-06 | MB231/MB468 | MP |
| >hg38_21_+_8433172_8446622_RNA45SN1_45S_50ntFlanks@6651.6677.27\|CGACTCTTAGCGGTGGATCACTCGGCT | >hg38_21_+_8433172_8446622_RNA45SN1_45S_50ntFlanks | 6651 | 6677 | 27 | CGACTCTTAGCGGTGGATCACTCGGCT | 55706.1 | 4.0 | 0.8 | 4.9 | 7.5E-07 | 1.6E-06 | BT20/MB468 | MITO |
| >hg38_21_+_8433172_8446622_RNA45SN1_45S_50ntFlanks@10680.10709.30\|GTTCCGGCGGCGTCCGGTGAGCTCTCGCTG | >hg38_21_+_8433172_8446622_RNA45SN1_45S_50ntFlanks | 10680 | 10709 | 30 | GTTCCGGCGGCGTCCGGTGAGCTCTCGCTG | 4380.8 | -4.1 | 0.8 | -4.9 | 7.5E-07 | 1.6E-06 | BT20/MB468 | MITO |
| >hg38_21_+_8433172_8446622_RNA45SN1_45S_50ntFlanks@8196.8223.28\|CGTGGACGGTGTGAGGCCGGTAGCGGCC | >hg38_21_+_8433172_8446622_RNA45SN1_45S_50ntFlanks | 8196 | 8223 | 28 | CGTGGACGGTGTGAGGCCGGTAGCGGCC | 1188.1 | -6.2 | 1.3 | -4.9 | 8.7E-07 | 1.6E-06 | MB231/MB468 | MP |
| >hg38_21_+_8433172_8446622_RNA45SN1_45S_50ntFlanks@10731.10752.22\|AGGGTGTAAATCTCGCGCCGGG | >hg38_21_+_8433172_8446622_RNA45SN1_45S_50ntFlanks | 10731 | 10752 | 22 | AGGGTGTAAATCTCGCGCCGGG | 301.7 | -6.1 | 1.2 | -4.9 | 8.6E-07 | 1.6E-06 | BT20/MB468 | CYTO |
| >hg38_21_+_8433172_8446622_RNA45SN1_45S_50ntFlanks@9914.9947.34\|GAAAAGGTGTTGGTTGATATAGACAGCAGGACGG | >hg38_21_+_8433172_8446622_RNA45SN1_45S_50ntFlanks | 9914 | 9947 | 34 | GAAAAGGTGTTGGTTGATATAGACAGCAGGACGG | 56.7 | 9.6 | 1.9 | 5.1 | 2.6E-07 | 1.6E-06 | BT20/MB468 | MP |
| >hg38_21_+_8433172_8446622_RNA45SN1_45S_50ntFlanks@12087.12110.24\|CTTCTGGCGCCAAGCGCCCGGCCG | >hg38_21_+_8433172_8446622_RNA45SN1_45S_50ntFlanks | 12087 | 12110 | 24 | CTTCTGGCGCCAAGCGCCCGGCCG | 88.4 | -9.5 | 1.9 | -5.1 | 2.6E-07 | 1.6E-06 | BT20/MB468 | MP |
| >MI0001448\|hsa-mir-425&WithFlank&3\|-\|49020142\|49020240@20.39.20&[MIMAT0003393&hsa-miR-425-5p&offsets\|0\|-3;m-95&3\|-\|49020200\|49020221&offsets\|0\|-2]\|AATGACACGATCACTCCCGT | >MI0001448\|hsa-mir-425&WithFlank&3\|-\|49020142\|49020240 | 20 | 39 | 20 | AATGACACGATCACTCCCGT | 206.4 | 5.8 | 1.2 | 4.9 | 1.0E-06 | 1.6E-06 | MB231/MB468 | NUC |
| >hg38_21_+_8433172_8446622_RNA45SN1_45S_50ntFlanks@6651.6687.37\|CGACTCTTAGCGGTGGATCACTCGGCTCGTGCGTCGA | >hg38_21_+_8433172_8446622_RNA45SN1_45S_50ntFlanks | 6651 | 6687 | 37 | CGACTCTTAGCGGTGGATCACTCGGCTCGTGCGTCGA | 718.8 | 4.7 | 1.0 | 4.9 | 8.6E-07 | 1.6E-06 | BT20/MB468 | CYTO |
| >hg38_21_+_8433172_8446622_RNA45SN1_45S_50ntFlanks@12935.12958.24\|TAAACCATTCGTAGACGACCTGCT | >hg38_21_+_8433172_8446622_RNA45SN1_45S_50ntFlanks | 12935 | 12958 | 24 | TAAACCATTCGTAGACGACCTGCT | 354.2 | -7.4 | 1.5 | -4.9 | 1.1E-06 | 1.7E-06 | BT20/MB468 | NUC |
| >hg38_21_+_8433172_8446622_RNA45SN1_45S_50ntFlanks@476.518.43\|TTCGTGATCGATGTGGTGACGTCGTGCTCTCCCGGGCCGGGTC | >hg38_21_+_8433172_8446622_RNA45SN1_45S_50ntFlanks | 476 | 518 | 43 | TTCGTGATCGATGTGGTGACGTCGTGCTCTCCCGGGCCGGGTC | 72.1 | 9.9 | 1.9 | 5.1 | 2.7E-07 | 1.7E-06 | BT20/MB468 | MP |
| >hg38_21_+_8433172_8446622_RNA45SN1_45S_50ntFlanks@4364.4389.26\|CTGCAGTTAAAAAGCTCGTAGTTGGA | >hg38_21_+_8433172_8446622_RNA45SN1_45S_50ntFlanks | 4364 | 4389 | 26 | CTGCAGTTAAAAAGCTCGTAGTTGGA | 54.5 | -8.8 | 1.7 | -5.1 | 2.7E-07 | 1.7E-06 | BT20/MB468 | MP |
| >hg38_21_+_8433172_8446622_RNA45SN1_45S_50ntFlanks@10926.10946.21\|ACGAGGCGCCGCCGCCCCCCC | >hg38_21_+_8433172_8446622_RNA45SN1_45S_50ntFlanks | 10926 | 10946 | 21 | ACGAGGCGCCGCCGCCCCCCC | 222.0 | -6.1 | 1.2 | -4.9 | 8.6E-07 | 1.7E-06 | BT20/MB468 | CYTO |
| >am_trna116_GluCTC_1_-_145399233_145399304@1.26.26,trna59_GluCTC_1_+_249168447_249168518@1.26.26,trna71_GluCTC_1_-_161439189_161439260@1.26.26,trna74_GluCTC_1_-_161431809_161431880@1.26.26,trna77_GluCTC_1_-_161424398_161424469@1.26.26,trna77_GluCTC_6_+_28949976_28950047@1.26.26,trna80_GluCTC_1_-_161417018_161417089@1.26.26,trna87_GluCTC_6_-_126101393_126101464@1.26.26\|TCCCTGGTGGTCTAGTGGTTAGGATT | >am_trna116_GluCTC_1_-_145399233_145399304 | 1 | 26 | 26 | TCCCTGGTGGTCTAGTGGTTAGGATT | 942.9 | 4.0 | 0.8 | 4.9 | 7.6E-07 | 1.7E-06 | BT20/MB468 | MITO |
| >MI0000433\|hsa-let-7g&WithFlank&3\|-\|52268272\|52268367@11.31.21(+1A)&[MIMAT0000414&hsa-let-7g-5p&offsets\|0\|-1(+1A);m-58&3\|-\|52268336\|52268357&offsets\|0\|-1(+1A)]\|TGAGGTAGTAGTTTGTACAGTA | >MI0000433\|hsa-let-7g&WithFlank&3\|-\|52268272\|52268367 | 11 | 31 | 21(+1A) | TGAGGTAGTAGTTTGTACAGTA | 1768.6 | 4.8 | 1.0 | 4.9 | 8.7E-07 | 1.7E-06 | MB231/MB468 | MP |
| >hg38_21_+_8433172_8446622_RNA45SN1_45S_50ntFlanks@10401.10430.30\|GAACATGGGTCAGTCGGTCCTGAGAGATGG | >hg38_21_+_8433172_8446622_RNA45SN1_45S_50ntFlanks | 10401 | 10430 | 30 | GAACATGGGTCAGTCGGTCCTGAGAGATGG | 414.8 | -3.7 | 0.7 | -4.9 | 1.0E-06 | 1.7E-06 | MB231/MB468 | NUC |
| >trna12_TrpCCA_17_+_19411494_19411565@30.52.23,trna168_TrpCCA_6_-_26331672_26331743@30.52.23,trna170_TrpCCA_6_-_26319330_26319401@30.52.23,trna1_TrpCCA_7_+_99067307_99067378@30.52.23,trna39_TrpCCA_17_-_8124187_8124258@30.52.23,trna6_TrpCCA_17_+_8089676_8089747@30.52.23\|ACTCCAGATCAGAAGGTTGCGTG | >trna12_TrpCCA_17_+_19411494_19411565 | 30 | 52 | 23 | ACTCCAGATCAGAAGGTTGCGTG | 121.6 | -5.4 | 1.1 | -4.9 | 8.6E-07 | 1.7E-06 | BT20/MB468 | CYTO |
| >am_trnaMT_ValTAC_MT_+_1602_1670@30.68.39\|CTTACACTTAGGAGATTTCAACTTAACTTGACCGCTCTG | >am_trnaMT_ValTAC_MT_+_1602_1670 | 30 | 68 | 39 | CTTACACTTAGGAGATTTCAACTTAACTTGACCGCTCTG | 816.5 | -8.1 | 1.7 | -4.9 | 1.0E-06 | 1.7E-06 | MB231/MB468 | NUC |
| >hg38_21_+_8433172_8446622_RNA45SN1_45S_50ntFlanks@7991.8012.22\|ACGTGGCGACCCGCTGAATTTA | >hg38_21_+_8433172_8446622_RNA45SN1_45S_50ntFlanks | 7991 | 8012 | 22 | ACGTGGCGACCCGCTGAATTTA | 4436.9 | -4.4 | 0.9 | -5.1 | 2.7E-07 | 1.7E-06 | BT20/MB468 | MP |
| >hg38_21_+_8433172_8446622_RNA45SN1_45S_50ntFlanks@10410.10433.24\|TCAGTCGGTCCTGAGAGATGGGCG | >hg38_21_+_8433172_8446622_RNA45SN1_45S_50ntFlanks | 10410 | 10433 | 24 | TCAGTCGGTCCTGAGAGATGGGCG | 1488.9 | -6.3 | 1.2 | -5.1 | 2.7E-07 | 1.7E-06 | BT20/MB468 | MP |
| >MI0000301\|hsa-mir-224&WithFlank&X\|-\|151958572\|151958664@14.37.24&[MIMAT0000281&hsa-miR-224-5p&offsets\|+1\|0;m-238&X\|-\|151958630\|151958651&offsets\|0\|+2]\|CAAGTCACTAGTGGTTCCGTTTAG | >MI0000301\|hsa-mir-224&WithFlank&X\|-\|151958572\|151958664 | 14 | 37 | 24 | CAAGTCACTAGTGGTTCCGTTTAG | 286.5 | 7.9 | 1.6 | 4.9 | 8.8E-07 | 1.7E-06 | MB231/MB468 | MP |
| >hg38_21_+_8433172_8446622_RNA45SN1_45S_50ntFlanks@5071.5110.40\|TTAATTCCGATAACGAACGAGACTCTGGCATGCTAACTAG | >hg38_21_+_8433172_8446622_RNA45SN1_45S_50ntFlanks | 5071 | 5110 | 40 | TTAATTCCGATAACGAACGAGACTCTGGCATGCTAACTAG | 237.4 | -3.7 | 0.8 | -4.9 | 1.0E-06 | 1.7E-06 | MB231/MB468 | NUC |
| >trna111_HisGTG_1_-_147774845_147774916@-1A.20.21,trna118_HisGTG_1_-_145396881_145396952@-1A.20.21,trna16_HisGTG_1_+_146544773_146544844@-1A.20.21,trna1_HisGTG_15_+_45493349_45493420@-1A.20.21,trna21_HisGTG_1_+_147753471_147753542@-1A.20.21,trna33_HisGTG_6_+_27125906_27125977@-1A.20.21,trna7_HisGTG_9_-_14433938_14434009@-1A.20.21,trna8_HisGTG_15_-_45492611_45492682@-1A.20.21,trna9_HisGTG_15_-_45490804_45490875@-1A.20.21\|AGCCGTGATCGTATAGTGGTT | >trna111_HisGTG_1_-_147774845_147774916 | -1A | 20 | 21 | AGCCGTGATCGTATAGTGGTT | 139.7 | -10.5 | 1.9 | -5.4 | 6.3E-08 | 1.7E-06 | BT20/MB231 | MP |
| >hg38_21_+_8433172_8446622_RNA45SN1_45S_50ntFlanks@11209.11229.21\|CGGCGACTCTGGACGCGAGCC | >hg38_21_+_8433172_8446622_RNA45SN1_45S_50ntFlanks | 11209 | 11229 | 21 | CGGCGACTCTGGACGCGAGCC | 768.8 | -3.9 | 0.8 | -4.9 | 9.9E-07 | 1.7E-06 | MB231/MB468 | CYTO |
| >hg38_21_+_8433172_8446622_RNA45SN1_45S_50ntFlanks@6733.6770.38\|CATTGATCATCGACACTTCGAACGCACTTGCGGCCCCG | >hg38_21_+_8433172_8446622_RNA45SN1_45S_50ntFlanks | 6733 | 6770 | 38 | CATTGATCATCGACACTTCGAACGCACTTGCGGCCCCG | 5894.5 | -9.4 | 1.9 | -4.9 | 1.1E-06 | 1.7E-06 | BT20/MB468 | NUC |
| >trnaMT_PheGAA_MT_+_577_647@28.57.30\|ATACACTGAAAATGTTTAGACGGGCTCACA | >trnaMT_PheGAA_MT_+_577_647 | 28 | 57 | 30 | ATACACTGAAAATGTTTAGACGGGCTCACA | 458.3 | 7.6 | 1.5 | 4.9 | 7.6E-07 | 1.7E-06 | BT20/MB468 | MITO |
| >hg38_21_+_8433172_8446622_RNA45SN1_45S_50ntFlanks@3843.3873.31\|CTACTTGGATAACTGTGGTAATTCTAGAGCT | >hg38_21_+_8433172_8446622_RNA45SN1_45S_50ntFlanks | 3843 | 3873 | 31 | CTACTTGGATAACTGTGGTAATTCTAGAGCT | 237.8 | -5.6 | 1.1 | -4.9 | 8.8E-07 | 1.7E-06 | MB231/MB468 | MP |
| >hg38_21_+_8433172_8446622_RNA45SN1_45S_50ntFlanks@10351.10380.30\|CGAGAACTTTGAAGGCCGAAGTGGAGAAGG | >hg38_21_+_8433172_8446622_RNA45SN1_45S_50ntFlanks | 10351 | 10380 | 30 | CGAGAACTTTGAAGGCCGAAGTGGAGAAGG | 536.6 | -2.9 | 0.6 | -4.9 | 9.9E-07 | 1.7E-06 | MB231/MB468 | CYTO |
| >hg38_21_+_8433172_8446622_RNA45SN1_45S_50ntFlanks@3850.3873.24\|GATAACTGTGGTAATTCTAGAGCT | >hg38_21_+_8433172_8446622_RNA45SN1_45S_50ntFlanks | 3850 | 3873 | 24 | GATAACTGTGGTAATTCTAGAGCT | 416.5 | -4.3 | 0.9 | -4.9 | 1.1E-06 | 1.7E-06 | MB231/MB468 | MITO |
| >MI0000095\|hsa-mir-93&WithFlank&7\|-\|100093762\|100093853@17.38.22&[MIMAT0000093&hsa-miR-93-5p&offsets\|0\|-1;m-29&7\|-\|100093815\|100093837&offsets\|0\|-1]\|CAAAGTGCTGTTCGTGCAGGTA | >MI0000095\|hsa-mir-93&WithFlank&7\|-\|100093762\|100093853 | 17 | 38 | 22 | CAAAGTGCTGTTCGTGCAGGTA | 481.4 | 5.3 | 1.1 | 4.9 | 9.9E-07 | 1.7E-06 | MB231/MB468 | CYTO |
| >hg38_21_+_8433172_8446622_RNA45SN1_45S_50ntFlanks@9534.9569.36\|AGGAAACTCTGGTGGAGGTCCGTAGCGGTCCTGACG | >hg38_21_+_8433172_8446622_RNA45SN1_45S_50ntFlanks | 9534 | 9569 | 36 | AGGAAACTCTGGTGGAGGTCCGTAGCGGTCCTGACG | 4478.5 | -3.9 | 0.8 | -4.9 | 8.7E-07 | 1.7E-06 | BT20/MB468 | CYTO |
| >hg38_21_+_8433172_8446622_RNA45SN1_45S_50ntFlanks@11569.11607.39\|CAGCCGACTTAGAACTGGTGCGGACCAGGGGAATCCGAC | >hg38_21_+_8433172_8446622_RNA45SN1_45S_50ntFlanks | 11569 | 11607 | 39 | CAGCCGACTTAGAACTGGTGCGGACCAGGGGAATCCGAC | 378.0 | -6.0 | 1.2 | -4.9 | 1.1E-06 | 1.7E-06 | BT20/MB468 | NUC |
| >hg38_21_+_8433172_8446622_RNA45SN1_45S_50ntFlanks@10798.10817.20\|CATGTTGGAACAATGTAGGT | >hg38_21_+_8433172_8446622_RNA45SN1_45S_50ntFlanks | 10798 | 10817 | 20 | CATGTTGGAACAATGTAGGT | 627.5 | -3.3 | 0.7 | -4.9 | 8.7E-07 | 1.7E-06 | BT20/MB468 | CYTO |
| >am_trnaMT_PheGAA_MT_+_577_647@40.62.23\|TGTTTAGACGGGCTCACATCACC | >am_trnaMT_PheGAA_MT_+_577_647 | 40 | 62 | 23 | TGTTTAGACGGGCTCACATCACC | 435.2 | 8.6 | 1.7 | 4.9 | 8.7E-07 | 1.7E-06 | BT20/MB468 | CYTO |
| >hg38_MT_+_1621_3279_RNR2_16S_50ntFlanks@51.88.38\|GCTAAACCTAGCCCCAAACCCACTCCACCTTACTACCA | >hg38_MT_+_1621_3279_RNR2_16S_50ntFlanks | 51 | 88 | 38 | GCTAAACCTAGCCCCAAACCCACTCCACCTTACTACCA | 4971.5 | 5.1 | 1.0 | 4.9 | 7.7E-07 | 1.7E-06 | BT20/MB468 | MITO |

<table>
<tr><th></th><th></th><th></th><th></th><th></th><th></th><th></th><th></th><th></th><th></th><th></th><th></th><th></th><th></th></tr>
<tr><td>>MI0000264|hsa-mir-7-2&WithFlank&15|+|88611819|88611940@38.61.24;MI0000265|hsa-mir-7-3&WithFlank&19|+|4770664|4770785@37.60.24;MI0000263|hsa-mir-7-1&WithFlank&9|-|83969742|83969863@30.53.24&[MIMAT0000252&hsa-miR-7-5p&offsets|0|0;m-400&15|+|88611856|88611879&offsets|0|0];[MIMAT0000252_1&hsa-miR-7-5p&offsets|0|0;m-402&19|+|4770700|4770723&offsets|0|0];[MIMAT0000252_2&hsa-miR-7-5p&offsets|0|0;m-401&9|-|83969811|83969834&offsets|0|0]|TGGAAGACTAGTGATTTTGTTGTT</td><td>>MI0000264|hsa-mir-7-2&WithFlank&15|+|88611819|88611940</td><td>38</td><td>61</td><td>24</td><td>TGGAAGACTAGTGATTTTGTTGTT</td><td>22825.8</td><td>3.0</td><td>0.6</td><td>4.9</td><td>1.1E-06</td><td>1.7E-06</td><td>BT20/MB468</td><td>NUC</td></tr>
<tr><td>>MI0000449|hsa-mir-132&WithFlank&17|-|2049902|2050014@65.84.20&[MIMAT0000426&hsa-miR-132-3p&offsets|0|-2;m-138&17|-|2049929|2049950&offsets|0|-2]|TAACAGTCTACAGCCATGGT</td><td>>MI0000449|hsa-mir-132&WithFlank&17|-|2049902|2050014</td><td>65</td><td>84</td><td>20</td><td>TAACAGTCTACAGCCATGGT</td><td>100.4</td><td>10.4</td><td>2.0</td><td>5.1</td><td>2.7E-07</td><td>1.7E-06</td><td>BT20/MB468</td><td>MP</td></tr>
<tr><td>>hg38_21_+_8433172_8446622_RNA45SN1_45S_50ntFlanks@8168.8187.20|CCCAAGTCCTTCTGATCGAG</td><td>>hg38_21_+_8433172_8446622_RNA45SN1_45S_50ntFlanks</td><td>8168</td><td>8187</td><td>20</td><td>CCCAAGTCCTTCTGATCGAG</td><td>398.8</td><td>-4.9</td><td>1.0</td><td>-4.9</td><td>1.0E-06</td><td>1.7E-06</td><td>MB231/MB468</td><td>CYTO</td></tr>
<tr><td>>hg38_21_+_8433172_8446622_RNA45SN1_45S_50ntFlanks@11324.11344.21|GTCTCGTCGCGCGCGCGTCCG</td><td>>hg38_21_+_8433172_8446622_RNA45SN1_45S_50ntFlanks</td><td>11324</td><td>11344</td><td>21</td><td>GTCTCGTCGCGCGCGCGTCCG</td><td>352.1</td><td>-4.2</td><td>0.9</td><td>-4.9</td><td>1.0E-06</td><td>1.7E-06</td><td>MB231/MB468</td><td>CYTO</td></tr>
<tr><td>>trnaMT_ProTGG_MT_-_15956_16023@50.71.22|TTAAAGACTTTTTCTCTGACCA</td><td>>trnaMT_ProTGG_MT_-_15956_16023</td><td>50</td><td>71</td><td>22</td><td>TTAAAGACTTTTTCTCTGACCA</td><td>710.9</td><td>-3.4</td><td>0.6</td><td>-5.3</td><td>1.1E-07</td><td>1.7E-06</td><td>BT20/MB231</td><td>NUC</td></tr>
<tr><td>>MI0000272|hsa-mir-182&WithFlank&7|-|129770377|129770498@29.50.22&[MIMAT0000259&hsa-miR-182-5p&offsets|0|-2;m-22&7|-|129770449|129770470&offsets|0|0]|TTTGGCAATGGTAGAACTCACA</td><td>>MI0000272|hsa-mir-182&WithFlank&7|-|129770377|129770498</td><td>29</td><td>50</td><td>22</td><td>TTTGGCAATGGTAGAACTCACA</td><td>5596.8</td><td>4.6</td><td>0.8</td><td>5.4</td><td>7.3E-08</td><td>1.7E-06</td><td>BT20/MB231</td><td>CYTO</td></tr>
<tr><td>>hg38_21_+_8433172_8446622_RNA45SN1_45S_50ntFlanks@7981.7998.18|CTCAGATCAGACGTGGCG</td><td>>hg38_21_+_8433172_8446622_RNA45SN1_45S_50ntFlanks</td><td>7981</td><td>7998</td><td>18</td><td>CTCAGATCAGACGTGGCG</td><td>113.0</td><td>-5.1</td><td>1.0</td><td>-4.9</td><td>1.0E-06</td><td>1.7E-06</td><td>MB231/MB468</td><td>CYTO</td></tr>
<tr><td>>trna10_AspGTC_12_-_125424193_125424264@38.71.34,trna12_AspGTC_12_-_125411891_125411962@38.71.34,trna144_AspGTC_6_-_27551236_27551307@38.71.34,trna38_AspGTC_17_-_8125556_8125627@38.71.34,trna45_AspGTC_6_+_27447453_27447524@38.71.34,trna48_AspGTC_6_+_27471523_27471594@38.71.34,trna4_AspGTC_12_+_96429799_96429870@38.71.34,trna69_AspGTC_1_-_161440205_161440276@38.71.34,trna72_AspGTC_1_-_161432824_161432895@38.71.34,trna75_AspGTC_1_-_161425414_161425485@38.71.34,trna78_AspGTC_1_-_161418033_161418104@38.71.34,trna81_AspGTC_1_-_161410615_161410686@38.71.34|CGCGGGAGACCGGGGTTCGATTCCCCGACGGGGA</td><td>>trna10_AspGTC_12_-_125424193_125424264</td><td>38</td><td>71</td><td>34</td><td>CGCGGGAGACCGGGGTTCGATTCCCCGACGGGGA</td><td>393.2</td><td>-3.9</td><td>0.8</td><td>-4.9</td><td>8.9E-07</td><td>1.7E-06</td><td>MB231/MB468</td><td>MP</td></tr>
<tr><td>>hg38_21_+_8433172_8446622_RNA45SN1_45S_50ntFlanks@3841.3872.32|TCCTACTTGGATAACTGTGGTAATTCTAGAGC</td><td>>hg38_21_+_8433172_8446622_RNA45SN1_45S_50ntFlanks</td><td>3841</td><td>3872</td><td>32</td><td>TCCTACTTGGATAACTGTGGTAATTCTAGAGC</td><td>336.2</td><td>3.5</td><td>0.7</td><td>4.9</td><td>8.7E-07</td><td>1.7E-06</td><td>BT20/MB468</td><td>CYTO</td></tr>
<tr><td>>hg38_21_+_8433172_8446622_RNA45SN1_45S_50ntFlanks@9569.9592.24|GTGCAAATCGGTCGTCCGACCTGG</td><td>>hg38_21_+_8433172_8446622_RNA45SN1_45S_50ntFlanks</td><td>9569</td><td>9592</td><td>24</td><td>GTGCAAATCGGTCGTCCGACCTGG</td><td>326.4</td><td>-4.5</td><td>0.9</td><td>-4.9</td><td>1.0E-06</td><td>1.7E-06</td><td>MB231/MB468</td><td>CYTO</td></tr>
<tr><td>>hg38_21_+_8433172_8446622_RNA45SN1_45S_50ntFlanks@6719.6750.32|TGAATTGCAGGACACATTGATCATCGACACTT</td><td>>hg38_21_+_8433172_8446622_RNA45SN1_45S_50ntFlanks</td><td>6719</td><td>6750</td><td>32</td><td>TGAATTGCAGGACACATTGATCATCGACACTT</td><td>208.6</td><td>-5.0</td><td>1.0</td><td>-4.9</td><td>1.0E-06</td><td>1.7E-06</td><td>MB231/MB468</td><td>NUC</td></tr>
<tr><td>>hg38_21_+_8433172_8446622_RNA45SN1_45S_50ntFlanks@11874.11905.32|AGAAGACCCTGTTGAGCTTGACTCTAGTCTGG</td><td>>hg38_21_+_8433172_8446622_RNA45SN1_45S_50ntFlanks</td><td>11874</td><td>11905</td><td>32</td><td>AGAAGACCCTGTTGAGCTTGACTCTAGTCTGG</td><td>170.5</td><td>-6.2</td><td>1.3</td><td>-4.9</td><td>1.0E-06</td><td>1.7E-06</td><td>MB231/MB468</td><td>NUC</td></tr>
<tr><td>>MI0000078|hsa-mir-22&WithFlank&17|-|1713897|1713993@21.40.20(+1C)&[MIMAT0004495&hsa-miR-22-5p&offsets|0|-2(+1C);m-217&17|-|1713953|1713973&offsets|0|-1(+1C)]|AGTTCTTCAGTGGCAAGCTTC</td><td>>MI0000078|hsa-mir-22&WithFlank&17|-|1713897|1713993</td><td>21</td><td>40</td><td>20(+1C)</td><td>AGTTCTTCAGTGGCAAGCTTC</td><td>376.3</td><td>6.0</td><td>1.2</td><td>4.9</td><td>8.9E-07</td><td>1.7E-06</td><td>MB231/MB468</td><td>MP</td></tr>
<tr><td>>hg38_21_+_8433172_8446622_RNA45SN1_45S_50ntFlanks@11727.11745.19|AGTAACTATGACTCTCTTA</td><td>>hg38_21_+_8433172_8446622_RNA45SN1_45S_50ntFlanks</td><td>11727</td><td>11745</td><td>19</td><td>AGTAACTATGACTCTCTTA</td><td>464.7</td><td>3.2</td><td>0.7</td><td>4.9</td><td>1.0E-06</td><td>1.7E-06</td><td>MB231/MB468</td><td>CYTO</td></tr>
<tr><td>>hg38_21_+_8433172_8446622_RNA45SN1_45S_50ntFlanks@3843.3861.19|CTACTTGGATAACTGTGGT</td><td>>hg38_21_+_8433172_8446622_RNA45SN1_45S_50ntFlanks</td><td>3843</td><td>3861</td><td>19</td><td>CTACTTGGATAACTGTGGT</td><td>615.7</td><td>-2.7</td><td>0.6</td><td>-4.9</td><td>1.0E-06</td><td>1.7E-06</td><td>MB231/MB468</td><td>NUC</td></tr>
<tr><td>>hg38_21_+_8433172_8446622_RNA45SN1_45S_50ntFlanks@12712.12757.46|CGGAGCCTCGGTTGGCCTCGGATAGCCGGTCCCCCGCCTGTCCCCG</td><td>>hg38_21_+_8433172_8446622_RNA45SN1_45S_50ntFlanks</td><td>12712</td><td>12757</td><td>46</td><td>CGGAGCCTCGGTTGGCCTCGGATAGCCGGTCCCCCGCCTGTCCCCG</td><td>179.2</td><td>-10.6</td><td>2.1</td><td>-5.1</td><td>2.7E-07</td><td>1.7E-06</td><td>BT20/MB468</td><td>MP</td></tr>
<tr><td>>MI0000285|hsa-mir-205&WithFlank&1|+|209432127|209432248@40.61.22(+1G)&[MIMAT0000266&hsa-miR-205-5p&offsets|0|0(+1G);m-351&1|+|209432166|209432187&offsets|0|0(+1G)]|TCCTTCATTCCACCGGAGTCTGG</td><td>>MI0000285|hsa-mir-205&WithFlank&1|+|209432127|209432248</td><td>40</td><td>61</td><td>22(+1G)</td><td>TCCTTCATTCCACCGGAGTCTGG</td><td>114.6</td><td>10.2</td><td>1.9</td><td>5.4</td><td>6.4E-08</td><td>1.7E-06</td><td>BT20/MB231</td><td>MP</td></tr>
<tr><td>>hg38_21_+_8433172_8446622_RNA45SN1_45S_50ntFlanks@13006.13027.22|CTATTGAAAGTCAGCCCTCGAC</td><td>>hg38_21_+_8433172_8446622_RNA45SN1_45S_50ntFlanks</td><td>13006</td><td>13027</td><td>22</td><td>CTATTGAAAGTCAGCCCTCGAC</td><td>1366.3</td><td>-7.6</td><td>1.5</td><td>-4.9</td><td>8.8E-07</td><td>1.7E-06</td><td>BT20/MB468</td><td>CYTO</td></tr>
<tr><td>>MI0000070|hsa-mir-16-1&WithFlank&13|-|50048967|50049067@20.40.21(+1A);MI0000115|hsa-mir-16-2&WithFlank&3|+|160404739|160404831@16.36.21(+1A)&[MIMAT0000069&hsa-miR-16-5p&offsets|0|-1(+1A);m-62&13|-|50049027|50049048&offsets|0|-1(+1A)];[MIMAT0000069_1&hsa-miR-16-5p&offsets|0|-1(+1A);m-61&3|+|160404754|160404775&offsets|0|-1(+1A)]|TAGCAGCACGTAAATATTGGCA</td><td>>MI0000070|hsa-mir-16-1&WithFlank&13|-|50048967|50049067</td><td>20</td><td>40</td><td>21(+1A)</td><td>TAGCAGCACGTAAATATTGGCA</td><td>307.5</td><td>5.0</td><td>1.0</td><td>4.9</td><td>1.0E-06</td><td>1.7E-06</td><td>MB231/MB468</td><td>CYTO</td></tr>
<tr><td>>hg38_21_+_8433172_8446622_RNA45SN1_45S_50ntFlanks@12935.12955.21|TAAACCATTCGTAGACGACCT</td><td>>hg38_21_+_8433172_8446622_RNA45SN1_45S_50ntFlanks</td><td>12935</td><td>12955</td><td>21</td><td>TAAACCATTCGTAGACGACCT</td><td>213.0</td><td>-3.8</td><td>0.8</td><td>-4.9</td><td>1.0E-06</td><td>1.7E-06</td><td>MB231/MB468</td><td>CYTO</td></tr>
<tr><td>>hg38_21_+_8433172_8446622_RNA45SN1_45S_50ntFlanks@12488.12509.22|AACGTGAGCTGGGTTTAGACCG</td><td>>hg38_21_+_8433172_8446622_RNA45SN1_45S_50ntFlanks</td><td>12488</td><td>12509</td><td>22</td><td>AACGTGAGCTGGGTTTAGACCG</td><td>11565.1</td><td>-9.6</td><td>2.0</td><td>-4.9</td><td>1.1E-06</td><td>1.7E-06</td><td>BT20/MB468</td><td>NUC</td></tr>
<tr><td>>hg38_21_+_8433172_8446622_RNA45SN1_45S_50ntFlanks@11784.11806.23|ATGAATGGATGAACGAGATTCCC</td><td>>hg38_21_+_8433172_8446622_RNA45SN1_45S_50ntFlanks</td><td>11784</td><td>11806</td><td>23</td><td>ATGAATGGATGAACGAGATTCCC</td><td>646.9</td><td>-6.7</td><td>1.4</td><td>-4.9</td><td>1.0E-06</td><td>1.7E-06</td><td>MB231/MB468</td><td>CYTO</td></tr>
<tr><td>>hg38_21_+_8433172_8446622_RNA45SN1_45S_50ntFlanks@10369.10401.33|AAGTGGAGAAGGGTTCCATGTGAACAGCAGTTG</td><td>>hg38_21_+_8433172_8446622_RNA45SN1_45S_50ntFlanks</td><td>10369</td><td>10401</td><td>33</td><td>AAGTGGAGAAGGGTTCCATGTGAACAGCAGTTG</td><td>200.9</td><td>-6.1</td><td>1.2</td><td>-4.9</td><td>1.0E-06</td><td>1.7E-06</td><td>MB231/MB468</td><td>NUC</td></tr>
<tr><td>>hg38_21_+_8433172_8446622_RNA45SN1_45S_50ntFlanks@12995.13016.22|CTCGCTGCGATCTATTGAAAGT</td><td>>hg38_21_+_8433172_8446622_RNA45SN1_45S_50ntFlanks</td><td>12995</td><td>13016</td><td>22</td><td>CTCGCTGCGATCTATTGAAAGT</td><td>248.2</td><td>4.2</td><td>0.9</td><td>4.9</td><td>1.1E-06</td><td>1.7E-06</td><td>MB231/MB468</td><td>MITO</td></tr>
<tr><td>>hg38_21_+_8433172_8446622_RNA45SN1_45S_50ntFlanks@11703.11727.25|AATGAAGCGCGGGTAAACGGCGGGA</td><td>>hg38_21_+_8433172_8446622_RNA45SN1_45S_50ntFlanks</td><td>11703</td><td>11727</td><td>25</td><td>AATGAAGCGCGGGTAAACGGCGGGA</td><td>274.3</td><td>-7.2</td><td>1.5</td><td>-4.9</td><td>9.0E-07</td><td>1.7E-06</td><td>MB231/MB468</td><td>MP</td></tr>
<tr><td>>trna6_TrpCCA_12_+_98898030_98898101@30.52.23|ACTCCAGATCAGAAGGCTGCGTG</td><td>>trna6_TrpCCA_12_+_98898030_98898101</td><td>30</td><td>52</td><td>23</td><td>ACTCCAGATCAGAAGGCTGCGTG</td><td>109.1</td><td>-5.4</td><td>1.1</td><td>-4.9</td><td>8.8E-07</td><td>1.7E-06</td><td>BT20/MB468</td><td>CYTO</td></tr>
<tr><td>>hg38_21_+_8433172_8446622_RNA45SN1_45S_50ntFlanks@7991.8014.24|ACGTGGCGACCCGCTGAATTTAAG</td><td>>hg38_21_+_8433172_8446622_RNA45SN1_45S_50ntFlanks</td><td>7991</td><td>8014</td><td>24</td><td>ACGTGGCGACCCGCTGAATTTAAG</td><td>1753.2</td><td>-4.4</td><td>0.8</td><td>-5.1</td><td>2.7E-07</td><td>1.7E-06</td><td>BT20/MB468</td><td>MP</td></tr>
<tr><td>>hg38_MT_+_598_1651_RNR1_12S_50ntFlanks@427.469.43|GAAAGTGGCTTTAACATATCTGAACACACAATAGCTAAGACCC</td><td>>hg38_MT_+_598_1651_RNR1_12S_50ntFlanks</td><td>427</td><td>469</td><td>43</td><td>GAAAGTGGCTTTAACATATCTGAACACACAATAGCTAAGACCC</td><td>122.2</td><td>10.7</td><td>2.1</td><td>5.1</td><td>2.7E-07</td><td>1.7E-06</td><td>BT20/MB468</td><td>MP</td></tr>
<tr><td>>hg38_21_+_8433172_8446622_RNA45SN1_45S_50ntFlanks@12072.12097.26|CCCGAGGGGCTCTCGCTTCTGGCGCC</td><td>>hg38_21_+_8433172_8446622_RNA45SN1_45S_50ntFlanks</td><td>12072</td><td>12097</td><td>26</td><td>CCCGAGGGGCTCTCGCTTCTGGCGCC</td><td>1253.2</td><td>-8.4</td><td>1.7</td><td>-4.9</td><td>1.1E-06</td><td>1.7E-06</td><td>BT20/MB468</td><td>NUC</td></tr>
<tr><td>>hg38_21_+_8433172_8446622_RNA45SN1_45S_50ntFlanks@11808.11827.20|CTGTCCCTACCTACTATCCA</td><td>>hg38_21_+_8433172_8446622_RNA45SN1_45S_50ntFlanks</td><td>11808</td><td>11827</td><td>20</td><td>CTGTCCCTACCTACTATCCA</td><td>103.8</td><td>-9.8</td><td>1.9</td><td>-5.1</td><td>2.7E-07</td><td>1.7E-06</td><td>BT20/MB468</td><td>MP</td></tr>
<tr><td>>MI0000650|hsa-mir-200c&WithFlank&12|+|6963693|6963772@50.71.22(+1U)&[MIMAT0000617&hsa-miR-200c-3p&offsets|0|-1(+1U);m-54&12|+|6963742|6963764&offsets|0|-1(+1U)]|TAATACTGCCGGGTAATGATGGT</td><td>>MI0000650|hsa-mir-200c&WithFlank&12|+|6963693|6963772</td><td>50</td><td>71</td><td>22(+1U)</td><td>TAATACTGCCGGGTAATGATGGT</td><td>149.0</td><td>-5.6</td><td>1.1</td><td>-4.9</td><td>9.0E-07</td><td>1.7E-06</td><td>MB231/MB468</td><td>MP</td></tr>
<tr><td>>hg38_21_+_8433172_8446622_RNA45SN1_45S_50ntFlanks@4358.4391.34|AAGTTGCTGCAGTTAAAAAGCTCGTAGTTGGATC</td><td>>hg38_21_+_8433172_8446622_RNA45SN1_45S_50ntFlanks</td><td>4358</td><td>4391</td><td>34</td><td>AAGTTGCTGCAGTTAAAAAGCTCGTAGTTGGATC</td><td>155.8</td><td>-5.5</td><td>1.1</td><td>-4.9</td><td>9.0E-07</td><td>1.7E-06</td><td>MB231/MB468</td><td>MP</td></tr>
<tr><td>>hg38_21_+_8433172_8446622_RNA45SN1_45S_50ntFlanks@6785.6806.22|CTACGCCTGTCTGAGCGTCGCT</td><td>>hg38_21_+_8433172_8446622_RNA45SN1_45S_50ntFlanks</td><td>6785</td><td>6806</td><td>22</td><td>CTACGCCTGTCTGAGCGTCGCT</td><td>5692.8</td><td>3.6</td><td>0.7</td><td>4.9</td><td>7.8E-07</td><td>1.7E-06</td><td>BT20/MB468</td><td>MITO</td></tr>
<tr><td>>hg38_21_+_8433172_8446622_RNA45SN1_45S_50ntFlanks@10678.10698.21|CGGTTCCGGCGGCGTCCGGTG</td><td>>hg38_21_+_8433172_8446622_RNA45SN1_45S_50ntFlanks</td><td>10678</td><td>10698</td><td>21</td><td>CGGTTCCGGCGGCGTCCGGTG</td><td>81.0</td><td>-9.4</td><td>1.8</td><td>-5.1</td><td>2.7E-07</td><td>1.7E-06</td><td>BT20/MB468</td><td>MP</td></tr>
<tr><td>>MI0000063|hsa-let-7b&WithFlank&22|+|46113680|46113774@12.31.20(+1C)&[MIMAT0000063&hsa-let-7b-5p&offsets|0|-2(+1C);m-14&22|+|46113691|46113712&offsets|0|-2(+1C)]|TGAGGTAGTAGGTTGTGTGGC</td><td>>MI0000063|hsa-let-7b&WithFlank&22|+|46113680|46113774</td><td>12</td><td>31</td><td>20(+1C)</td><td>TGAGGTAGTAGGTTGTGTGGC</td><td>861.8</td><td>2.8</td><td>0.5</td><td>5.3</td><td>1.2E-07</td><td>1.7E-06</td><td>BT20/MB231</td><td>NUC</td></tr>
<tr><td>>hg38_21_+_8433172_8446622_RNA45SN1_45S_50ntFlanks@10842.10865.24|GTAACTTCGGGATAAGGATTGGCT</td><td>>hg38_21_+_8433172_8446622_RNA45SN1_45S_50ntFlanks</td><td>10842</td><td>10865</td><td>24</td><td>GTAACTTCGGGATAAGGATTGGCT</td><td>131.3</td><td>-6.7</td><td>1.4</td><td>-4.9</td><td>9.0E-07</td><td>1.7E-06</td><td>MB231/MB468</td><td>MP</td></tr>
<tr><td>>trna119_LysCTT_1_-_145395522_145395594@1.35.35,trna11_LysCTT_5_-_180648979_180649051@1.35.35,trna13_LysCTT_6_+_26556774_26556846@1.35.35,trna7_LysCTT_16_+_3225692_3225764@1.35.35,trna9_LysCTT_5_+_180634755_180634827@1.35.35|GCCCGGCTAGCTCAGTCGGTAGAGCATGAGACTCT</td><td>>trna119_LysCTT_1_-_145395522_145395594</td><td>1</td><td>35</td><td>35</td><td>GCCCGGCTAGCTCAGTCGGTAGAGCATGAGACTCT</td><td>338.8</td><td>3.3</td><td>0.7</td><td>4.9</td><td>1.1E-06</td><td>1.7E-06</td><td>BT20/MB468</td><td>NUC</td></tr>
<tr><td>>hg38_21_+_8433172_8446622_RNA45SN1_45S_50ntFlanks@11670.11692.23|CCCAGTGCTCTGAATGTCAAAGT</td><td>>hg38_21_+_8433172_8446622_RNA45SN1_45S_50ntFlanks</td><td>11670</td><td>11692</td><td>23</td><td>CCCAGTGCTCTGAATGTCAAAGT</td><td>165.5</td><td>-3.6</td><td>0.7</td><td>-4.9</td><td>9.0E-07</td><td>1.7E-06</td><td>MB231/MB468</td><td>MP</td></tr>
<tr><td>>hg38_21_+_8433172_8446622_RNA45SN1_45S_50ntFlanks@3875.3894.20|ATACATGCCGACGGGCGCTG</td><td>>hg38_21_+_8433172_8446622_RNA45SN1_45S_50ntFlanks</td><td>3875</td><td>3894</td><td>20</td><td>ATACATGCCGACGGGCGCTG</td><td>57.5</td><td>-8.9</td><td>1.7</td><td>-5.1</td><td>2.8E-07</td><td>1.7E-06</td><td>BT20/MB468</td><td>MP</td></tr>
<tr><td>>hg38_21_+_8433172_8446622_RNA45SN1_45S_50ntFlanks@5484.5526.43|GAGCGCTGAGAAGACGGTCGAACTTGACTATCTAGAGGAAGTA</td><td>>hg38_21_+_8433172_8446622_RNA45SN1_45S_50ntFlanks</td><td>5484</td><td>5526</td><td>43</td><td>GAGCGCTGAGAAGACGGTCGAACTTGACTATCTAGAGGAAGTA</td><td>399.7</td><td>-7.8</td><td>1.6</td><td>-4.9</td><td>1.1E-06</td><td>1.7E-06</td><td>BT20/MB468</td><td>NUC</td></tr>
<tr><td>>MI0000811|hsa-mir-148b&WithFlank&12|+|54337210|54337320@69.90.22(+1A)&[MIMAT0000759&hsa-miR-148b-3p&offsets|0|0(+1A);m-77&12|+|54337278|54337299&offsets|0|0(+1A)]|TCAGTGCATCACAGAACTTTGTA</td><td>>MI0000811|hsa-mir-148b&WithFlank&12|+|54337210|54337320</td><td>69</td><td>90</td><td>22(+1A)</td><td>TCAGTGCATCACAGAACTTTGTA</td><td>75.7</td><td>1.9</td><td>0.4</td><td>4.9</td><td>1.1E-06</td><td>1.7E-06</td><td>BT20/MB468</td><td>NUC</td></tr>
<tr><td>>hg38_21_+_8433172_8446622_RNA45SN1_45S_50ntFlanks@4472.4497.26|CTTAGCTGAGTGTCCCGCGGGGCCCG</td><td>>hg38_21_+_8433172_8446622_RNA45SN1_45S_50ntFlanks</td><td>4472</td><td>4497</td><td>26</td><td>CTTAGCTGAGTGTCCCGCGGGGCCCG</td><td>373.8</td><td>-6.4</td><td>1.3</td><td>-4.9</td><td>8.9E-07</td><td>1.7E-06</td><td>BT20/MB468</td><td>CYTO</td></tr>
<tr><td>>hg38_21_+_8433172_8446622_RNA45SN1_45S_50ntFlanks@10725.10755.31|GGGGAGAGGGTGTAAATCTCGCGCCGGGCCG</td><td>>hg38_21_+_8433172_8446622_RNA45SN1_45S_50ntFlanks</td><td>10725</td><td>10755</td><td>31</td><td>GGGGAGAGGGTGTAAATCTCGCGCCGGGCCG</td><td>85.5</td><td>-9.5</td><td>1.8</td><td>-5.1</td><td>2.8E-07</td><td>1.7E-06</td><td>BT20/MB468</td><td>MP</td></tr>
</table>

<table>
<tr><th></th><th></th><th></th><th></th><th></th><th></th><th></th><th></th><th></th><th></th><th></th><th></th><th></th><th></th></tr>
<tr><td>>hg38_21_+_8433172_8446622_RNA45SN1_45S_50ntFlanks@12415.12436.22|CTTCGATGTCGGCTCTTCCTAT</td><td>>hg38_21_+_8433172_8446622_RNA45SN1_45S_50ntFlanks</td><td>12415</td><td>12436</td><td>22</td><td>CTTCGATGTCGGCTCTTCCTAT</td><td>3944.5</td><td>-3.6</td><td>0.7</td><td>-4.9</td><td>8.9E-07</td><td>1.7E-06</td><td>BT20/MB468</td><td>CYTO</td></tr>
<tr><td>>hg38_21_+_8433172_8446622_RNA45SN1_45S_50ntFlanks@12539.12579.41|ATGATGTGTTGTTGCCATGGTAATCCTGCTCAGTACGAGAG</td><td>>hg38_21_+_8433172_8446622_RNA45SN1_45S_50ntFlanks</td><td>12539</td><td>12579</td><td>41</td><td>ATGATGTGTTGTTGCCATGGTAATCCTGCTCAGTACGAGAG</td><td>86.9</td><td>-8.5</td><td>1.7</td><td>-4.9</td><td>7.9E-07</td><td>1.7E-06</td><td>BT20/MB468</td><td>MITO</td></tr>
<tr><td>>hg38_21_+_8433172_8446622_RNA45SN1_45S_50ntFlanks@4449.4472.24|CCTCTCGGCGCCCCCTCGATGCTC</td><td>>hg38_21_+_8433172_8446622_RNA45SN1_45S_50ntFlanks</td><td>4449</td><td>4472</td><td>24</td><td>CCTCTCGGCGCCCCCTCGATGCTC</td><td>2267.9</td><td>-8.5</td><td>1.7</td><td>-4.9</td><td>1.1E-06</td><td>1.7E-06</td><td>MB231/MB468</td><td>NUC</td></tr>
<tr><td>>hg38_21_+_8433172_8446622_RNA45SN1_45S_50ntFlanks@7969.7988.20|CCGAGACGCGACCTCAGATC</td><td>>hg38_21_+_8433172_8446622_RNA45SN1_45S_50ntFlanks</td><td>7969</td><td>7988</td><td>20</td><td>CCGAGACGCGACCTCAGATC</td><td>54.9</td><td>9.5</td><td>1.9</td><td>5.1</td><td>2.8E-07</td><td>1.7E-06</td><td>BT20/MB468</td><td>MP</td></tr>
<tr><td>>MI0000433|hsa-let-7g&WithFlank&3|-|52268272|52268367@11.31.21(+1C)&[MIMAT0000414&hsa-let-7g-5p&offsets|0|-1(+1C);m-58&3|-|52268336|52268357&offsets|0|-1(+1C)]|TGAGGTAGTAGTTTGTACAGTC</td><td>>MI0000433|hsa-let-7g&WithFlank&3|-|52268272|52268367</td><td>11</td><td>31</td><td>21(+1C)</td><td>TGAGGTAGTAGTTTGTACAGTC</td><td>7124.9</td><td>4.8</td><td>1.0</td><td>4.9</td><td>9.1E-07</td><td>1.7E-06</td><td>MB231/MB468</td><td>MP</td></tr>
<tr><td>>hg38_21_+_8433172_8446622_RNA45SN1_45S_50ntFlanks@12420.12454.35|ATGTCGGCTCTTCCTATCATTGTGAAGCAGAATTC</td><td>>hg38_21_+_8433172_8446622_RNA45SN1_45S_50ntFlanks</td><td>12420</td><td>12454</td><td>35</td><td>ATGTCGGCTCTTCCTATCATTGTGAAGCAGAATTC</td><td>661.6</td><td>-6.3</td><td>1.3</td><td>-4.9</td><td>1.1E-06</td><td>1.7E-06</td><td>BT20/MB468</td><td>NUC</td></tr>
<tr><td>>hg38_21_+_8433172_8446622_RNA45SN1_45S_50ntFlanks@9545.9571.27|GTGGAGGTCCGTAGCGGTCCTGACGTG</td><td>>hg38_21_+_8433172_8446622_RNA45SN1_45S_50ntFlanks</td><td>9545</td><td>9571</td><td>27</td><td>GTGGAGGTCCGTAGCGGTCCTGACGTG</td><td>6505.9</td><td>-4.7</td><td>0.9</td><td>-5.1</td><td>2.8E-07</td><td>1.7E-06</td><td>BT20/MB468</td><td>MP</td></tr>
<tr><td>>hg38_21_+_8433172_8446622_RNA45SN1_45S_50ntFlanks@6776.6803.28|CTCCCGGGGCTACGCCTGTCTGAGCGTC</td><td>>hg38_21_+_8433172_8446622_RNA45SN1_45S_50ntFlanks</td><td>6776</td><td>6803</td><td>28</td><td>CTCCCGGGGCTACGCCTGTCTGAGCGTC</td><td>715.5</td><td>-2.8</td><td>0.6</td><td>-4.9</td><td>1.1E-06</td><td>1.7E-06</td><td>MB231/MB468</td><td>NUC</td></tr>
<tr><td>>am_trna119_LysCTT_1_-_145395522_145395594@7.36.30,trna11_LysCTT_5_-_180648979_180649051@7.36.30,trna128_LysCTT_1_-_55423542_55423614@7.36.30,trna13_LysCTT_6_+_26556774_26556846@7.36.30,trna6_LysCTT_19_-_52425396_52425468@7.36.30,trna7_LysCTT_16_+_3225692_3225764@7.36.30,trna9_LysCTT_5_+_180634755_180634827@7.36.30|CTAGCTCAGTCGGTAGAGCATGAGACTCTT</td><td>>am_trna119_LysCTT_1_-_145395522_145395594</td><td>7</td><td>36</td><td>30</td><td>CTAGCTCAGTCGGTAGAGCATGAGACTCTT</td><td>545.2</td><td>8.1</td><td>1.7</td><td>4.9</td><td>1.1E-06</td><td>1.7E-06</td><td>MB231/MB468</td><td>MITO</td></tr>
<tr><td>>hg38_21_+_8433172_8446622_RNA45SN1_45S_50ntFlanks@12444.12484.41|AAGCAGAATTCACCAAGCGTTGGATTGTTCACCCACTAATA</td><td>>hg38_21_+_8433172_8446622_RNA45SN1_45S_50ntFlanks</td><td>12444</td><td>12484</td><td>41</td><td>AAGCAGAATTCACCAAGCGTTGGATTGTTCACCCACTAATA</td><td>571.7</td><td>-6.0</td><td>1.2</td><td>-4.9</td><td>1.1E-06</td><td>1.7E-06</td><td>MB231/MB468</td><td>MITO</td></tr>
<tr><td>>hg38_21_+_8433172_8446622_RNA45SN1_45S_50ntFlanks@12991.13010.20|CTCCCTCGCTGCGATCTATT</td><td>>hg38_21_+_8433172_8446622_RNA45SN1_45S_50ntFlanks</td><td>12991</td><td>13010</td><td>20</td><td>CTCCCTCGCTGCGATCTATT</td><td>534.6</td><td>-3.9</td><td>0.8</td><td>-4.9</td><td>9.0E-07</td><td>1.7E-06</td><td>BT20/MB468</td><td>CYTO</td></tr>
<tr><td>>MI0003137|hsa-mir-193b&WithFlank&16|+|14303961|14304055@57.79.23&[MIMAT0002819&hsa-miR-193b-3p&offsets|0|+1;m-166&16|+|14304017|14304038&offsets|0|+1]|AACTGGCCCTCAAAGTCCCGCTT</td><td>>MI0003137|hsa-mir-193b&WithFlank&16|+|14303961|14304055</td><td>57</td><td>79</td><td>23</td><td>AACTGGCCCTCAAAGTCCCGCTT</td><td>189.0</td><td>5.0</td><td>0.9</td><td>5.4</td><td>7.6E-08</td><td>1.7E-06</td><td>BT20/MB231</td><td>CYTO</td></tr>
<tr><td>>hg38_21_+_8433172_8446622_RNA45SN1_45S_50ntFlanks@483.514.32|TCGATGTGGTGACGTCGTGCTCTCCCGGGCCG</td><td>>hg38_21_+_8433172_8446622_RNA45SN1_45S_50ntFlanks</td><td>483</td><td>514</td><td>32</td><td>TCGATGTGGTGACGTCGTGCTCTCCCGGGCCG</td><td>255.3</td><td>4.1</td><td>0.8</td><td>4.9</td><td>1.1E-06</td><td>1.7E-06</td><td>MB231/MB468</td><td>NUC</td></tr>
<tr><td>>trnaMT_PheGAA_MT_+_577_647@21.53.33|CAAAGCAATACACTGAAAATGTTTAGACGGGCT</td><td>>trnaMT_PheGAA_MT_+_577_647</td><td>21</td><td>53</td><td>33</td><td>CAAAGCAATACACTGAAAATGTTTAGACGGGCT</td><td>86.9</td><td>10.2</td><td>2.0</td><td>5.1</td><td>2.8E-07</td><td>1.7E-06</td><td>BT20/MB468</td><td>MP</td></tr>
<tr><td>>hg38_21_+_8433172_8446622_RNA45SN1_45S_50ntFlanks@4882.4911.30|TGCAAAGCTGAAACTTAAAGGAATTGACGG</td><td>>hg38_21_+_8433172_8446622_RNA45SN1_45S_50ntFlanks</td><td>4882</td><td>4911</td><td>30</td><td>TGCAAAGCTGAAACTTAAAGGAATTGACGG</td><td>55.1</td><td>9.5</td><td>1.9</td><td>5.1</td><td>2.8E-07</td><td>1.7E-06</td><td>BT20/MB468</td><td>MP</td></tr>
<tr><td>>hg38_21_+_8433172_8446622_RNA45SN1_45S_50ntFlanks@12671.12703.33|AAGTCAGAATCCCGCCCAGGCGGAACGATACGG</td><td>>hg38_21_+_8433172_8446622_RNA45SN1_45S_50ntFlanks</td><td>12671</td><td>12703</td><td>33</td><td>AAGTCAGAATCCCGCCCAGGCGGAACGATACGG</td><td>91.9</td><td>-9.6</td><td>1.9</td><td>-5.1</td><td>2.8E-07</td><td>1.7E-06</td><td>BT20/MB468</td><td>MP</td></tr>
<tr><td>>hg38_21_+_8433172_8446622_RNA45SN1_45S_50ntFlanks@12477.12497.21|CACTAATAGGGAACGTGAGCT</td><td>>hg38_21_+_8433172_8446622_RNA45SN1_45S_50ntFlanks</td><td>12477</td><td>12497</td><td>21</td><td>CACTAATAGGGAACGTGAGCT</td><td>447.3</td><td>2.8</td><td>0.6</td><td>4.9</td><td>8.0E-07</td><td>1.7E-06</td><td>BT20/MB468</td><td>MITO</td></tr>
<tr><td>>hg38_21_+_8433172_8446622_RNA45SN1_45S_50ntFlanks@11874.11901.28|AGAAGACCCTGTTGAGCTTGACTCTAGT</td><td>>hg38_21_+_8433172_8446622_RNA45SN1_45S_50ntFlanks</td><td>11874</td><td>11901</td><td>28</td><td>AGAAGACCCTGTTGAGCTTGACTCTAGT</td><td>322.6</td><td>-7.3</td><td>1.5</td><td>-4.9</td><td>1.2E-06</td><td>1.7E-06</td><td>BT20/MB468</td><td>NUC</td></tr>
<tr><td>>hg38_21_+_8433172_8446622_RNA45SN1_45S_50ntFlanks@12414.12448.35|CCTTCGATGTCGGCTCTTCCTATCATTGTGAAGCA</td><td>>hg38_21_+_8433172_8446622_RNA45SN1_45S_50ntFlanks</td><td>12414</td><td>12448</td><td>35</td><td>CCTTCGATGTCGGCTCTTCCTATCATTGTGAAGCA</td><td>222.7</td><td>5.3</td><td>1.1</td><td>4.9</td><td>1.1E-06</td><td>1.7E-06</td><td>MB231/MB468</td><td>NUC</td></tr>
<tr><td>>hg38_21_+_8433172_8446622_RNA45SN1_45S_50ntFlanks@6779.6806.28|CCGGGGCTACGCCTGTCTGAGCGTCGCT</td><td>>hg38_21_+_8433172_8446622_RNA45SN1_45S_50ntFlanks</td><td>6779</td><td>6806</td><td>28</td><td>CCGGGGCTACGCCTGTCTGAGCGTCGCT</td><td>3227.8</td><td>3.9</td><td>0.8</td><td>4.9</td><td>8.0E-07</td><td>1.7E-06</td><td>BT20/MB468</td><td>MITO</td></tr>
<tr><td>>hg38_21_+_8433172_8446622_RNA45SN1_45S_50ntFlanks@12046.12068.23|TGACCCGGTGAGGCGGGGGGGCG</td><td>>hg38_21_+_8433172_8446622_RNA45SN1_45S_50ntFlanks</td><td>12046</td><td>12068</td><td>23</td><td>TGACCCGGTGAGGCGGGGGGGCG</td><td>60.4</td><td>-9.0</td><td>1.8</td><td>-5.1</td><td>2.8E-07</td><td>1.7E-06</td><td>BT20/MB468</td><td>MP</td></tr>
<tr><td>>hg38_21_+_8433172_8446622_RNA45SN1_45S_50ntFlanks@9534.9570.37|AGGAAACTCTGGTGGAGGTCCGTAGCGGTCCTGACGT</td><td>>hg38_21_+_8433172_8446622_RNA45SN1_45S_50ntFlanks</td><td>9534</td><td>9570</td><td>37</td><td>AGGAAACTCTGGTGGAGGTCCGTAGCGGTCCTGACGT</td><td>2154.9</td><td>-2.8</td><td>0.6</td><td>-4.9</td><td>9.1E-07</td><td>1.7E-06</td><td>BT20/MB468</td><td>CYTO</td></tr>
<tr><td>>hg38_21_+_8433172_8446622_RNA45SN1_45S_50ntFlanks@10389.10408.20|TGAACAGCAGTTGAACATGG</td><td>>hg38_21_+_8433172_8446622_RNA45SN1_45S_50ntFlanks</td><td>10389</td><td>10408</td><td>20</td><td>TGAACAGCAGTTGAACATGG</td><td>286.0</td><td>-5.0</td><td>1.0</td><td>-4.9</td><td>1.1E-06</td><td>1.7E-06</td><td>MB231/MB468</td><td>NUC</td></tr>
<tr><td>>hg38_21_+_8433172_8446622_RNA45SN1_45S_50ntFlanks@10343.10369.27|TATTCAAACGAGAACTTTGAAGGCCGA</td><td>>hg38_21_+_8433172_8446622_RNA45SN1_45S_50ntFlanks</td><td>10343</td><td>10369</td><td>27</td><td>TATTCAAACGAGAACTTTGAAGGCCGA</td><td>192.6</td><td>10.9</td><td>2.0</td><td>5.4</td><td>6.6E-08</td><td>1.7E-06</td><td>BT20/MB231</td><td>MP</td></tr>
<tr><td>>hg38_21_+_8433172_8446622_RNA45SN1_45S_50ntFlanks@11906.11929.24|CACGGTGAAGAGACATGAGAGGTG</td><td>>hg38_21_+_8433172_8446622_RNA45SN1_45S_50ntFlanks</td><td>11906</td><td>11929</td><td>24</td><td>CACGGTGAAGAGACATGAGAGGTG</td><td>255.4</td><td>11.3</td><td>2.1</td><td>5.4</td><td>6.6E-08</td><td>1.7E-06</td><td>BT20/MB231</td><td>MP</td></tr>
<tr><td>>trnaMT_PheGAA_MT_+_577_647@33.54.22|CTGAAAATGTTTAGACGGGCTC</td><td>>trnaMT_PheGAA_MT_+_577_647</td><td>33</td><td>54</td><td>22</td><td>CTGAAAATGTTTAGACGGGCTC</td><td>89.3</td><td>10.2</td><td>2.0</td><td>5.1</td><td>2.8E-07</td><td>1.7E-06</td><td>BT20/MB468</td><td>MP</td></tr>
<tr><td>>hg38_21_+_8433172_8446622_RNA45SN1_45S_50ntFlanks@9559.9596.38|CGGTCCTGACGTGCAAATCGGTCGTCCGACCTGGGTAT</td><td>>hg38_21_+_8433172_8446622_RNA45SN1_45S_50ntFlanks</td><td>9559</td><td>9596</td><td>38</td><td>CGGTCCTGACGTGCAAATCGGTCGTCCGACCTGGGTAT</td><td>116.1</td><td>-8.9</td><td>1.8</td><td>-4.9</td><td>8.0E-07</td><td>1.7E-06</td><td>BT20/MB468</td><td>MITO</td></tr>
<tr><td>>hg38_21_+_8433172_8446622_RNA45SN1_45S_50ntFlanks@9569.9596.28|GTGCAAATCGGTCGTCCGACCTGGGTAT</td><td>>hg38_21_+_8433172_8446622_RNA45SN1_45S_50ntFlanks</td><td>9569</td><td>9596</td><td>28</td><td>GTGCAAATCGGTCGTCCGACCTGGGTAT</td><td>501.7</td><td>-3.2</td><td>0.6</td><td>-4.9</td><td>8.0E-07</td><td>1.7E-06</td><td>BT20/MB468</td><td>MITO</td></tr>
<tr><td>>hg38_21_+_8433172_8446622_RNA45SN1_45S_50ntFlanks@10685.10706.22|GGCGGCGTCCGGTGAGCTCTCG</td><td>>hg38_21_+_8433172_8446622_RNA45SN1_45S_50ntFlanks</td><td>10685</td><td>10706</td><td>22</td><td>GGCGGCGTCCGGTGAGCTCTCG</td><td>259.5</td><td>-7.0</td><td>1.4</td><td>-4.9</td><td>9.3E-07</td><td>1.7E-06</td><td>MB231/MB468</td><td>MP</td></tr>
<tr><td>>MI0000255|hsa-mir-30d&WithFlank&8|-|134804870|134804951@12.33.22(+1A)&[MIMAT0000245&hsa-miR-30d-5p&offsets|0|0(+1A);m-21&8|-|134804919|134804940&offsets|0|0(+1A)]|TGTAAACATCCCCGACTGGAAGA</td><td>>MI0000255|hsa-mir-30d&WithFlank&8|-|134804870|134804951</td><td>12</td><td>33</td><td>22(+1A)</td><td>TGTAAACATCCCCGACTGGAAGA</td><td>792.9</td><td>4.9</td><td>1.0</td><td>5.1</td><td>2.8E-07</td><td>1.7E-06</td><td>BT20/MB468</td><td>MP</td></tr>
<tr><td>>am_trnaMT_LeuTAG_MT_+_12266_12336@38.70.33|CCCAAAAATTTTGGTGCAACTCCAAATAAAAGT</td><td>>am_trnaMT_LeuTAG_MT_+_12266_12336</td><td>38</td><td>70</td><td>33</td><td>CCCAAAAATTTTGGTGCAACTCCAAATAAAAGT</td><td>293.8</td><td>-11.6</td><td>2.1</td><td>-5.4</td><td>6.7E-08</td><td>1.8E-06</td><td>BT20/MB231</td><td>MP</td></tr>
<tr><td>>hg38_21_+_8433172_8446622_RNA45SN1_45S_50ntFlanks@6774.6806.33|TCCTCCCGGGGCTACGCCTGTCTGAGCGTCGCT</td><td>>hg38_21_+_8433172_8446622_RNA45SN1_45S_50ntFlanks</td><td>6774</td><td>6806</td><td>33</td><td>TCCTCCCGGGGCTACGCCTGTCTGAGCGTCGCT</td><td>418.7</td><td>4.8</td><td>0.9</td><td>5.1</td><td>2.8E-07</td><td>1.8E-06</td><td>BT20/MB468</td><td>MP</td></tr>
<tr><td>>hg38_21_+_8433172_8446622_RNA45SN1_45S_50ntFlanks@8168.8212.45|CCCAAGTCCTTCTGATCGAGGCCCAGCCCGTGGACGGTGTGAGGC</td><td>>hg38_21_+_8433172_8446622_RNA45SN1_45S_50ntFlanks</td><td>8168</td><td>8212</td><td>45</td><td>CCCAAGTCCTTCTGATCGAGGCCCAGCCCGTGGACGGTGTGAGGC</td><td>1245.1</td><td>-5.1</td><td>1.0</td><td>-4.9</td><td>1.1E-06</td><td>1.8E-06</td><td>MB231/MB468</td><td>NUC</td></tr>
<tr><td>>hg38_21_+_8433172_8446622_RNA45SN1_45S_50ntFlanks@11593.11612.20|CCAGGGGAATCCGACTGTTT</td><td>>hg38_21_+_8433172_8446622_RNA45SN1_45S_50ntFlanks</td><td>11593</td><td>11612</td><td>20</td><td>CCAGGGGAATCCGACTGTTT</td><td>1191.3</td><td>-4.3</td><td>0.9</td><td>-4.9</td><td>9.2E-07</td><td>1.8E-06</td><td>BT20/MB468</td><td>CYTO</td></tr>
<tr><td>>hg38_21_+_8433172_8446622_RNA45SN1_45S_50ntFlanks@3878.3898.21|CATGCCGACGGGCGCTGACCC</td><td>>hg38_21_+_8433172_8446622_RNA45SN1_45S_50ntFlanks</td><td>3878</td><td>3898</td><td>21</td><td>CATGCCGACGGGCGCTGACCC</td><td>368.5</td><td>-3.4</td><td>0.7</td><td>-4.9</td><td>1.1E-06</td><td>1.8E-06</td><td>MB231/MB468</td><td>NUC</td></tr>
<tr><td>>hg38_21_+_8433172_8446622_RNA45SN1_45S_50ntFlanks@10877.10901.25|GTCGGTCGGGCTGGGGCGCGAAGCG</td><td>>hg38_21_+_8433172_8446622_RNA45SN1_45S_50ntFlanks</td><td>10877</td><td>10901</td><td>25</td><td>GTCGGTCGGGCTGGGGCGCGAAGCG</td><td>373.9</td><td>-7.4</td><td>1.5</td><td>-4.9</td><td>1.2E-06</td><td>1.8E-06</td><td>BT20/MB468</td><td>NUC</td></tr>
<tr><td>>hg38_21_+_8433172_8446622_RNA45SN1_45S_50ntFlanks@3783.3801.19|AGTGAAACTGCGAATGGCT</td><td>>hg38_21_+_8433172_8446622_RNA45SN1_45S_50ntFlanks</td><td>3783</td><td>3801</td><td>19</td><td>AGTGAAACTGCGAATGGCT</td><td>1386.9</td><td>-6.6</td><td>1.4</td><td>-4.9</td><td>1.1E-06</td><td>1.8E-06</td><td>MB231/MB468</td><td>MITO</td></tr>
<tr><td>>am_trna111_HisGTG_1_-_147774845_147774916@-1G.29.30,trna118_HisGTG_1_-_145396881_145396952@-1G.29.30,trna16_HisGTG_1_+_146544773_146544844@-1G.29.30,trna1_HisGTG_15_+_45493349_45493420@-1G.29.30,trna21_HisGTG_1_+_147753471_147753542@-1G.29.30,trna33_HisGTG_6_+_27125906_27125977@-1G.29.30,trna7_HisGTG_9_-_14433938_14434009@-1G.29.30,trna8_HisGTG_15_-_45492611_45492682@-1G.29.30,trna9_HisGTG_15_-_45490804_45490875@-1G.29.30|GGCCGTGATCGTATAGTGGTTAGTACTCTG</td><td>>am_trna111_HisGTG_1_-_147774845_147774916</td><td>-1G</td><td>29</td><td>30</td><td>GGCCGTGATCGTATAGTGGTTAGTACTCTG</td><td>1718.5</td><td>-5.6</td><td>1.1</td><td>-4.9</td><td>8.1E-07</td><td>1.8E-06</td><td>BT20/MB468</td><td>MITO</td></tr>
<tr><td>>hg38_21_+_8433172_8446622_RNA45SN1_45S_50ntFlanks@12458.12478.21|AAGCGTTGGATTGTTCACCCA</td><td>>hg38_21_+_8433172_8446622_RNA45SN1_45S_50ntFlanks</td><td>12458</td><td>12478</td><td>21</td><td>AAGCGTTGGATTGTTCACCCA</td><td>785.7</td><td>-4.2</td><td>0.9</td><td>-4.9</td><td>1.1E-06</td><td>1.8E-06</td><td>MB231/MB468</td><td>MITO</td></tr>
<tr><td>>trnaMT_GluTTC_MT_-_14674_14742@38.72.35,trnalookalike8_GluTTC_5_-_93905172_93905240@38.72.35|CATTGGTCGTGGTTGTAGTCCGTGCGAGAATACCA</td><td>>trnaMT_GluTTC_MT_-_14674_14742</td><td>38</td><td>72</td><td>35</td><td>CATTGGTCGTGGTTGTAGTCCGTGCGAGAATACCA</td><td>3834.6</td><td>3.3</td><td>0.7</td><td>4.9</td><td>1.1E-06</td><td>1.8E-06</td><td>MB231/MB468</td><td>MITO</td></tr>
<tr><td>>hg38_21_+_8433172_8446622_RNA45SN1_45S_50ntFlanks@8357.8395.39|AAAAGAACTTTGAAGAGAGAGTTCAAGAGGGCGTGAAAC</td><td>>hg38_21_+_8433172_8446622_RNA45SN1_45S_50ntFlanks</td><td>8357</td><td>8395</td><td>39</td><td>AAAAGAACTTTGAAGAGAGAGTTCAAGAGGGCGTGAAAC</td><td>212.3</td><td>11.1</td><td>2.0</td><td>5.4</td><td>6.7E-08</td><td>1.8E-06</td><td>BT20/MB231</td><td>MP</td></tr>
<tr><td>>hg38_21_+_8433172_8446622_RNA45SN1_45S_50ntFlanks@11727.11751.25|AGTAACTATGACTCTCTTAAGGTAG</td><td>>hg38_21_+_8433172_8446622_RNA45SN1_45S_50ntFlanks</td><td>11727</td><td>11751</td><td>25</td><td>AGTAACTATGACTCTCTTAAGGTAG</td><td>1356.0</td><td>-3.1</td><td>0.6</td><td>-4.9</td><td>1.1E-06</td><td>1.8E-06</td><td>MB231/MB468</td><td>NUC</td></tr>
<tr><td>>hg38_21_+_8433172_8446622_RNA45SN1_45S_50ntFlanks@5055.5081.27|GGAGCGATTTGTCTGGTTAATTCCGAT</td><td>>hg38_21_+_8433172_8446622_RNA45SN1_45S_50ntFlanks</td><td>5055</td><td>5081</td><td>27</td><td>GGAGCGATTTGTCTGGTTAATTCCGAT</td><td>339.1</td><td>7.3</td><td>1.5</td><td>4.9</td><td>8.1E-07</td><td>1.8E-06</td><td>BT20/MB468</td><td>MITO</td></tr>
<tr><td>>MI0000071|hsa-mir-17&WithFlank&13|+|91350599|91350694@20.41.22(+1A)&[MIMAT0000070&hsa-miR-17-5p&offsets|0|-1(+1A);m-88&13|+|91350618|91350640&offsets|0|-1(+1A)]|CAAAGTGCTTACAGTGCAGGTAA</td><td>>MI0000071|hsa-mir-17&WithFlank&13|+|91350599|91350694</td><td>20</td><td>41</td><td>22(+1A)</td><td>CAAAGTGCTTACAGTGCAGGTAA</td><td>1414.7</td><td>5.5</td><td>1.1</td><td>4.9</td><td>1.1E-06</td><td>1.8E-06</td><td>MB231/MB468</td><td>MITO</td></tr>
<tr><td>>hg38_21_+_8433172_8446622_RNA45SN1_45S_50ntFlanks@8912.8933.22|CGCCGAATCCCGGGGCCGAGGG</td><td>>hg38_21_+_8433172_8446622_RNA45SN1_45S_50ntFlanks</td><td>8912</td><td>8933</td><td>22</td><td>CGCCGAATCCCGGGGCCGAGGG</td><td>73.0</td><td>9.9</td><td>1.9</td><td>5.1</td><td>2.9E-07</td><td>1.8E-06</td><td>BT20/MB468</td><td>MP</td></tr>
<tr><td>>hg38_21_+_8433172_8446622_RNA45SN1_45S_50ntFlanks@7976.8010.35|GCGACCTCAGATCAGACGTGGCGACCCGCTGAATT</td><td>>hg38_21_+_8433172_8446622_RNA45SN1_45S_50ntFlanks</td><td>7976</td><td>8010</td><td>35</td><td>GCGACCTCAGATCAGACGTGGCGACCCGCTGAATT</td><td>432.3</td><td>3.3</td><td>0.7</td><td>4.9</td><td>1.2E-06</td><td>1.8E-06</td><td>BT20/MB468</td><td>NUC</td></tr>
<tr><td>>hg38_21_+_8433172_8446622_RNA45SN1_45S_50ntFlanks@10787.10825.39|ACAGCCTCTGGCATGTTGGAACAATGTAGGTAAGGGAAG</td><td>>hg38_21_+_8433172_8446622_RNA45SN1_45S_50ntFlanks</td><td>10787</td><td>10825</td><td>39</td><td>ACAGCCTCTGGCATGTTGGAACAATGTAGGTAAGGGAAG</td><td>87.9</td><td>-9.5</td><td>1.9</td><td>-5.1</td><td>2.9E-07</td><td>1.8E-06</td><td>BT20/MB468</td><td>MP</td></tr>
<tr><td>>hg38_21_+_8433172_8446622_RNA45SN1_45S_50ntFlanks@8159.8178.20|CGTGGGGGGCCCAAGTCCTT</td><td>>hg38_21_+_8433172_8446622_RNA45SN1_45S_50ntFlanks</td><td>8159</td><td>8178</td><td>20</td><td>CGTGGGGGGCCCAAGTCCTT</td><td>157.8</td><td>-7.0</td><td>1.4</td><td>-4.9</td><td>9.4E-07</td><td>1.8E-06</td><td>MB231/MB468</td><td>MP</td></tr>
</table>

<table>
<tr><th></th><th></th><th></th><th></th><th></th><th></th><th></th><th></th><th></th><th></th><th></th><th></th><th></th><th></th><th></th></tr>
<tr><td>>hg38_21_+_8433172_8446622_RNA45SN1_45S_50ntFlanks@8356.8375.20|GAAAAGAACTTTGAAGAGAG</td><td>>hg38_21_+_8433172_8446622_RNA45SN1_45S_50ntFlanks</td><td>8356</td><td>8375</td><td>20</td><td>GAAAAGAACTTTGAAGAGAG</td><td>357.5</td><td>-3.3</td><td>0.7</td><td>-4.9</td><td>1.1E-06</td><td>1.8E-06</td><td>MB231/MB468</td><td>NUC</td><td></td></tr>
<tr><td>>hg38_21_+_8433172_8446622_RNA45SN1_45S_50ntFlanks@10396.10424.29|CAGTTGAACATGGGTCAGTCGGTCCTGAG</td><td>>hg38_21_+_8433172_8446622_RNA45SN1_45S_50ntFlanks</td><td>10396</td><td>10424</td><td>29</td><td>CAGTTGAACATGGGTCAGTCGGTCCTGAG</td><td>8274.0</td><td>5.3</td><td>1.0</td><td>5.1</td><td>2.9E-07</td><td>1.8E-06</td><td>BT20/MB468</td><td>MP</td><td></td></tr>
<tr><td>>hg38_21_+_8433172_8446622_RNA45SN1_45S_50ntFlanks@12717.12741.25|CCTCGGTTGGCCTCGGATAGCCGGT</td><td>>hg38_21_+_8433172_8446622_RNA45SN1_45S_50ntFlanks</td><td>12717</td><td>12741</td><td>25</td><td>CCTCGGTTGGCCTCGGATAGCCGGT</td><td>59.2</td><td>-9.0</td><td>1.7</td><td>-5.1</td><td>2.9E-07</td><td>1.8E-06</td><td>BT20/MB468</td><td>MP</td><td></td></tr>
<tr><td>>hg38_21_+_8433172_8446622_RNA45SN1_45S_50ntFlanks@12499.12518.20|GGTTTAGACCGTCGTGAGAC</td><td>>hg38_21_+_8433172_8446622_RNA45SN1_45S_50ntFlanks</td><td>12499</td><td>12518</td><td>20</td><td>GGTTTAGACCGTCGTGAGAC</td><td>185.2</td><td>-6.4</td><td>1.3</td><td>-4.9</td><td>9.3E-07</td><td>1.8E-06</td><td>BT20/MB468</td><td>CYTO</td><td></td></tr>
<tr><td>>hg38_21_+_8433172_8446622_RNA45SN1_45S_50ntFlanks@6661.6696.36|CGGTGGATCACTCGGCTCGTGCGTCGATGAAGAACG</td><td>>hg38_21_+_8433172_8446622_RNA45SN1_45S_50ntFlanks</td><td>6661</td><td>6696</td><td>36</td><td>CGGTGGATCACTCGGCTCGTGCGTCGATGAAGAACG</td><td>507.0</td><td>3.6</td><td>0.7</td><td>4.9</td><td>9.3E-07</td><td>1.8E-06</td><td>BT20/MB468</td><td>CYTO</td><td></td></tr>
<tr><td>>MI0000077|hsa-mir-21&WithFlank&17|+|59841260|59841343@13.35.23&[MIMAT0000076&hsa-miR-21-5p&offsets|-1|0;m-1&17|+|59841273|59841294&offsets|-1|0]|GTAGCTTATCAGACTGATGTTGA</td><td>>MI0000077|hsa-mir-21&WithFlank&17|+|59841260|59841343</td><td>13</td><td>35</td><td>23</td><td>GTAGCTTATCAGACTGATGTTGA</td><td>2221.3</td><td>3.1</td><td>0.6</td><td>4.9</td><td>9.4E-07</td><td>1.8E-06</td><td>MB231/MB468</td><td>MP</td><td></td></tr>
<tr><td>>MI0000455|hsa-mir-138-2&WithFlank&16|+|56858512|56858607@16.39.24&[MIMAT0000430&hsa-miR-138-5p&offsets|0|+1;m-509&16|+|56858527|56858549&offsets|0|+1]|AGCTGGTGTTGTGAATCAGGCCGA</td><td>>MI0000455|hsa-mir-138-2&WithFlank&16|+|56858512|56858607</td><td>16</td><td>39</td><td>24</td><td>AGCTGGTGTTGTGAATCAGGCCGA</td><td>111.0</td><td>-10.2</td><td>1.9</td><td>-5.4</td><td>6.8E-08</td><td>1.8E-06</td><td>BT20/MB231</td><td>MP</td><td></td></tr>
<tr><td>>hg38_21_+_8433172_8446622_RNA45SN1_45S_50ntFlanks@4588.4621.34|CTATTTTGTTGGTTTTCGGAACTGAGGCCATGAT</td><td>>hg38_21_+_8433172_8446622_RNA45SN1_45S_50ntFlanks</td><td>4588</td><td>4621</td><td>34</td><td>CTATTTTGTTGGTTTTCGGAACTGAGGCCATGAT</td><td>294.6</td><td>-5.9</td><td>1.2</td><td>-4.9</td><td>9.4E-07</td><td>1.8E-06</td><td>MB231/MB468</td><td>MP</td><td></td></tr>
<tr><td>>trnaMT_GluTTC_MT_-_14674_14742@31.72.42,trnalookalike8_GluTTC_5_-_93905172_93905240@31.72.42|TTCATATCATTGGTCGTGGTTGTAGTCCGTGCGAGAATACCA</td><td>>trnaMT_GluTTC_MT_-_14674_14742</td><td>31</td><td>72</td><td>42</td><td>TTCATATCATTGGTCGTGGTTGTAGTCCGTGCGAGAATACCA</td><td>1139.9</td><td>5.1</td><td>1.0</td><td>4.9</td><td>1.1E-06</td><td>1.8E-06</td><td>MB231/MB468</td><td>NUC</td><td></td></tr>
<tr><td>>hg38_21_+_8433172_8446622_RNA45SN1_45S_50ntFlanks@4003.4039.37|ATAACCTCGGGCCGATCGCACGCCCCCCGTGGCGGCG</td><td>>hg38_21_+_8433172_8446622_RNA45SN1_45S_50ntFlanks</td><td>4003</td><td>4039</td><td>37</td><td>ATAACCTCGGGCCGATCGCACGCCCCCCGTGGCGGCG</td><td>398.1</td><td>-5.3</td><td>1.1</td><td>-4.9</td><td>1.1E-06</td><td>1.8E-06</td><td>MB231/MB468</td><td>NUC</td><td></td></tr>
<tr><td>>MI0000446|hsa-mir-125b-1&WithFlank&11|-|122099751|122099850@21.42.22(+1C);MI0000470|hsa-mir-125b-2&WithFlank&21|+|16590231|16590331@23.44.22(+1C)&[MIMAT0000423&hsa-miR-125b-5p&offsets|0|0(+1C);m-67&11|-|122099809|122099830&offsets|0|0(+1C)];[MIMAT0000423_1&hsa-miR-125b-5p&offsets|0|0(+1C);m-68&21|+|16590253|16590274&offsets|0|0(+1C)]|TCCCTGAGACCCTAACTTGTGAC</td><td>>MI0000446|hsa-mir-125b-1&WithFlank&11|-|122099751|122099850</td><td>21</td><td>42</td><td>22(+1C)</td><td>TCCCTGAGACCCTAACTTGTGAC</td><td>122.1</td><td>-10.3</td><td>1.9</td><td>-5.4</td><td>6.9E-08</td><td>1.8E-06</td><td>BT20/MB231</td><td>MP</td><td></td></tr>
<tr><td>>am_trnaMT_PheGAA_MT_+_577_647@41.62.22|GTTTAGACGGGCTCACATCACC</td><td>>am_trnaMT_PheGAA_MT_+_577_647</td><td>41</td><td>62</td><td>22</td><td>GTTTAGACGGGCTCACATCACC</td><td>1050.8</td><td>5.7</td><td>1.1</td><td>5.1</td><td>2.9E-07</td><td>1.8E-06</td><td>BT20/MB468</td><td>MP</td><td></td></tr>
<tr><td>>hg38_21_+_8433172_8446622_RNA45SN1_45S_50ntFlanks@12678.12711.34|AATCCCGCCCAGGCGGAACGATACGGCAGCGCCG</td><td>>hg38_21_+_8433172_8446622_RNA45SN1_45S_50ntFlanks</td><td>12678</td><td>12711</td><td>34</td><td>AATCCCGCCCAGGCGGAACGATACGGCAGCGCCG</td><td>66.1</td><td>-9.1</td><td>1.8</td><td>-5.1</td><td>2.9E-07</td><td>1.8E-06</td><td>BT20/MB468</td><td>MP</td><td></td></tr>
<tr><td>>hg38_21_+_8433172_8446622_RNA45SN1_45S_50ntFlanks@12942.12979.38|TTCGTAGACGACCTGCTTCTGGGTCGGGGTTTCGTACG</td><td>>hg38_21_+_8433172_8446622_RNA45SN1_45S_50ntFlanks</td><td>12942</td><td>12979</td><td>38</td><td>TTCGTAGACGACCTGCTTCTGGGTCGGGGTTTCGTACG</td><td>296.6</td><td>-5.0</td><td>1.0</td><td>-4.9</td><td>1.1E-06</td><td>1.8E-06</td><td>MB231/MB468</td><td>CYTO</td><td></td></tr>
<tr><td>>MI0000114|hsa-mir-107&WithFlank&10|-|89592741|89592833@56.77.22&[MIMAT0000104&hsa-miR-107&offsets|0|-1;m-78&10|-|89592758|89592778&offsets|0|+1]|AGCAGCATTGTACAGGGCTATC</td><td>>MI0000114|hsa-mir-107&WithFlank&10|-|89592741|89592833</td><td>56</td><td>77</td><td>22</td><td>AGCAGCATTGTACAGGGCTATC</td><td>548.8</td><td>5.0</td><td>1.0</td><td>4.9</td><td>1.1E-06</td><td>1.8E-06</td><td>MB231/MB468</td><td>CYTO</td><td></td></tr>
<tr><td>>hg38_21_+_8433172_8446622_RNA45SN1_45S_50ntFlanks@10727.10755.29|GGAGAGGGTGTAAATCTCGCGCCGGGCCG</td><td>>hg38_21_+_8433172_8446622_RNA45SN1_45S_50ntFlanks</td><td>10727</td><td>10755</td><td>29</td><td>GGAGAGGGTGTAAATCTCGCGCCGGGCCG</td><td>709.2</td><td>-6.3</td><td>1.3</td><td>-4.9</td><td>1.1E-06</td><td>1.8E-06</td><td>MB231/MB468</td><td>CYTO</td><td></td></tr>
<tr><td>>hg38_21_+_8433172_8446622_RNA45SN1_45S_50ntFlanks@11682.11703.22|AATGTCAAAGTGAAGAAATTCA</td><td>>hg38_21_+_8433172_8446622_RNA45SN1_45S_50ntFlanks</td><td>11682</td><td>11703</td><td>22</td><td>AATGTCAAAGTGAAGAAATTCA</td><td>63.9</td><td>-9.1</td><td>1.8</td><td>-5.1</td><td>2.9E-07</td><td>1.8E-06</td><td>BT20/MB468</td><td>MP</td><td></td></tr>
<tr><td>>hg38_21_+_8433172_8446622_RNA45SN1_45S_50ntFlanks@10101.10126.26|CGCGCGCGTGTGGTGTGCGTCGGAGG</td><td>>hg38_21_+_8433172_8446622_RNA45SN1_45S_50ntFlanks</td><td>10101</td><td>10126</td><td>26</td><td>CGCGCGCGTGTGGTGTGCGTCGGAGG</td><td>303.0</td><td>-6.4</td><td>1.3</td><td>-4.9</td><td>9.4E-07</td><td>1.8E-06</td><td>BT20/MB468</td><td>CYTO</td><td></td></tr>
<tr><td>>hg38_21_+_8433172_8446622_RNA45SN1_45S_50ntFlanks@475.499.25|CTTCGTGATCGATGTGGTGACGTCG</td><td>>hg38_21_+_8433172_8446622_RNA45SN1_45S_50ntFlanks</td><td>475</td><td>499</td><td>25</td><td>CTTCGTGATCGATGTGGTGACGTCG</td><td>498.6</td><td>6.3</td><td>1.3</td><td>4.9</td><td>1.1E-06</td><td>1.8E-06</td><td>MB231/MB468</td><td>MITO</td><td></td></tr>
<tr><td>>hg38_21_+_8433172_8446622_RNA45SN1_45S_50ntFlanks@3776.3802.27|CCGGTACAGTGAAACTGCGAATGGCTC</td><td>>hg38_21_+_8433172_8446622_RNA45SN1_45S_50ntFlanks</td><td>3776</td><td>3802</td><td>27</td><td>CCGGTACAGTGAAACTGCGAATGGCTC</td><td>62.2</td><td>-9.0</td><td>1.8</td><td>-5.1</td><td>2.9E-07</td><td>1.8E-06</td><td>BT20/MB468</td><td>MP</td><td></td></tr>
<tr><td>>hg38_21_+_8433172_8446622_RNA45SN1_45S_50ntFlanks@4283.4301.19|CTTTAACGAGGATCCATTG</td><td>>hg38_21_+_8433172_8446622_RNA45SN1_45S_50ntFlanks</td><td>4283</td><td>4301</td><td>19</td><td>CTTTAACGAGGATCCATTG</td><td>53.2</td><td>-8.8</td><td>1.7</td><td>-5.1</td><td>2.9E-07</td><td>1.8E-06</td><td>BT20/MB468</td><td>MP</td><td></td></tr>
<tr><td>>MI0000076|hsa-mir-20a&WithFlank&13|+|91351059|91351141@14.35.22(+1A)&[MIMAT0000075&hsa-miR-20a-5p&offsets|0|-1(+1A);m-96&13|+|91351072|91351094&offsets|0|-1(+1A)]|TAAAGTGCTTATAGTGCAGGTAA</td><td>>MI0000076|hsa-mir-20a&WithFlank&13|+|91351059|91351141</td><td>14</td><td>35</td><td>22(+1A)</td><td>TAAAGTGCTTATAGTGCAGGTAA</td><td>3052.1</td><td>-5.0</td><td>0.9</td><td>-5.3</td><td>9.4E-08</td><td>1.8E-06</td><td>BT20/MB231</td><td>MITO</td><td></td></tr>
<tr><td>>MI0000542|hsa-mir-320a&WithFlank&8|-|22244960|22245043@50.70.21(+1C)&[MIMAT0000510&hsa-miR-320a-3p&offsets|+2|+1(+1C);m-60&8|-|22244975|22244996&offsets|+2|+1(+1C)]|AAGCTGGGTTGAGAGGGCGAAC</td><td>>MI0000542|hsa-mir-320a&WithFlank&8|-|22244960|22245043</td><td>50</td><td>70</td><td>21(+1C)</td><td>AAGCTGGGTTGAGAGGGCGAAC</td><td>84.9</td><td>10.2</td><td>2.0</td><td>5.1</td><td>2.9E-07</td><td>1.8E-06</td><td>BT20/MB468</td><td>MP</td><td></td></tr>
<tr><td>>MI0001733|hsa-mir-452&WithFlank&X|-|151959622|151959718@23.43.21&[MIMAT0001635&hsa-miR-452-5p&offsets|+3|+2;m-188&X|-|151959677|151959699&offsets|+3|+1]|TGTTTGCAGAGGAAACTGAGA</td><td>>MI0001733|hsa-mir-452&WithFlank&X|-|151959622|151959718</td><td>23</td><td>43</td><td>21</td><td>TGTTTGCAGAGGAAACTGAGA</td><td>750.4</td><td>4.7</td><td>0.9</td><td>5.4</td><td>8.0E-08</td><td>1.8E-06</td><td>BT20/MB231</td><td>CYTO</td><td></td></tr>
<tr><td>>hg38_21_+_8433172_8446622_RNA45SN1_45S_50ntFlanks@3823.3853.31|TTTGGTCGCTCGCTCCTCTCCTACTTGGATA</td><td>>hg38_21_+_8433172_8446622_RNA45SN1_45S_50ntFlanks</td><td>3823</td><td>3853</td><td>31</td><td>TTTGGTCGCTCGCTCCTCTCCTACTTGGATA</td><td>269.3</td><td>-3.9</td><td>0.8</td><td>-4.9</td><td>1.1E-06</td><td>1.8E-06</td><td>MB231/MB468</td><td>NUC</td><td></td></tr>
<tr><td>>hg38_21_+_8433172_8446622_RNA45SN1_45S_50ntFlanks@10393.10422.30|CAGCAGTTGAACATGGGTCAGTCGGTCCTG</td><td>>hg38_21_+_8433172_8446622_RNA45SN1_45S_50ntFlanks</td><td>10393</td><td>10422</td><td>30</td><td>CAGCAGTTGAACATGGGTCAGTCGGTCCTG</td><td>1346.8</td><td>-8.4</td><td>1.7</td><td>-4.9</td><td>1.2E-06</td><td>1.8E-06</td><td>BT20/MB468</td><td>NUC</td><td></td></tr>
<tr><td>>MI0000469|hsa-mir-125a&WithFlank&19|+|51693248|51693345@21.43.23&[MIMAT0000443&hsa-miR-125a-5p&offsets|0|-1;m-63&19|+|51693268|51693289&offsets|0|+1]|TCCCTGAGACCCTTTAACCTGTG</td><td>>MI0000469|hsa-mir-125a&WithFlank&19|+|51693248|51693345</td><td>21</td><td>43</td><td>23</td><td>TCCCTGAGACCCTTTAACCTGTG</td><td>9537.3</td><td>4.1</td><td>0.8</td><td>4.9</td><td>1.2E-06</td><td>1.8E-06</td><td>MB231/MB468</td><td>MITO</td><td></td></tr>
<tr><td>>hg38_21_+_8433172_8446622_RNA45SN1_45S_50ntFlanks@12656.12674.19|TGACTGAACGCCTCTAAGT</td><td>>hg38_21_+_8433172_8446622_RNA45SN1_45S_50ntFlanks</td><td>12656</td><td>12674</td><td>19</td><td>TGACTGAACGCCTCTAAGT</td><td>191.6</td><td>-2.5</td><td>0.5</td><td>-4.9</td><td>9.4E-07</td><td>1.8E-06</td><td>BT20/MB468</td><td>CYTO</td><td></td></tr>
<tr><td>>hg38_21_+_8433172_8446622_RNA45SN1_45S_50ntFlanks@11643.11660.18|GCGGGTGTTGACGCGATG</td><td>>hg38_21_+_8433172_8446622_RNA45SN1_45S_50ntFlanks</td><td>11643</td><td>11660</td><td>18</td><td>GCGGGTGTTGACGCGATG</td><td>4475.8</td><td>-5.6</td><td>1.1</td><td>-5.1</td><td>2.9E-07</td><td>1.8E-06</td><td>BT20/MB468</td><td>MP</td><td></td></tr>
<tr><td>>hg38_21_+_8433172_8446622_RNA45SN1_45S_50ntFlanks@12244.12272.29|AAGGGCAAAAGCTCGCTTGATCTTGATTT</td><td>>hg38_21_+_8433172_8446622_RNA45SN1_45S_50ntFlanks</td><td>12244</td><td>12272</td><td>29</td><td>AAGGGCAAAAGCTCGCTTGATCTTGATTT</td><td>993.5</td><td>-6.1</td><td>1.3</td><td>-4.9</td><td>1.2E-06</td><td>1.8E-06</td><td>MB231/MB468</td><td>MITO</td><td></td></tr>
<tr><td>>hg38_21_+_8433172_8446622_RNA45SN1_45S_50ntFlanks@5445.5467.23|TCGGATCGGCCCCGCCGGGGTCG</td><td>>hg38_21_+_8433172_8446622_RNA45SN1_45S_50ntFlanks</td><td>5445</td><td>5467</td><td>23</td><td>TCGGATCGGCCCCGCCGGGGTCG</td><td>234.8</td><td>-4.5</td><td>0.9</td><td>-4.9</td><td>9.4E-07</td><td>1.8E-06</td><td>BT20/MB468</td><td>CYTO</td><td></td></tr>
<tr><td>>hg38_21_+_8433172_8446622_RNA45SN1_45S_50ntFlanks@11786.11804.19|GAATGGATGAACGAGATTC</td><td>>hg38_21_+_8433172_8446622_RNA45SN1_45S_50ntFlanks</td><td>11786</td><td>11804</td><td>19</td><td>GAATGGATGAACGAGATTC</td><td>1530.1</td><td>-6.4</td><td>1.2</td><td>-5.1</td><td>2.9E-07</td><td>1.8E-06</td><td>BT20/MB468</td><td>MP</td><td></td></tr>
<tr><td>>hg38_21_+_8433172_8446622_RNA45SN1_45S_50ntFlanks@11190.11220.31|CCCGGGGGGCCGGCGGCGGCGGCGACTCTGG</td><td>>hg38_21_+_8433172_8446622_RNA45SN1_45S_50ntFlanks</td><td>11190</td><td>11220</td><td>31</td><td>CCCGGGGGGCCGGCGGCGGCGGCGACTCTGG</td><td>88.5</td><td>-9.5</td><td>1.9</td><td>-5.1</td><td>2.9E-07</td><td>1.8E-06</td><td>BT20/MB468</td><td>MP</td><td></td></tr>
<tr><td>>hg38_21_+_8433172_8446622_RNA45SN1_45S_50ntFlanks@4449.4470.22|CCTCTCGGCGCCCCCTCGATGC</td><td>>hg38_21_+_8433172_8446622_RNA45SN1_45S_50ntFlanks</td><td>4449</td><td>4470</td><td>22</td><td>CCTCTCGGCGCCCCCTCGATGC</td><td>232.7</td><td>-6.7</td><td>1.4</td><td>-4.9</td><td>1.1E-06</td><td>1.8E-06</td><td>MB231/MB468</td><td>NUC</td><td></td></tr>
<tr><td>>MI0000261|hsa-mir-139&WithFlank&11|-|72615057|72615136@13.35.23&[MIMAT0000250&hsa-miR-139-5p&offsets|0|0;m-186&11|-|72615102|72615124&offsets|0|0]|TCTACAGTGCACGTGTCTCCAGT</td><td>>MI0000261|hsa-mir-139&WithFlank&11|-|72615057|72615136</td><td>13</td><td>35</td><td>23</td><td>TCTACAGTGCACGTGTCTCCAGT</td><td>1206.1</td><td>8.5</td><td>1.7</td><td>4.9</td><td>9.6E-07</td><td>1.8E-06</td><td>MB231/MB468</td><td>MP</td><td></td></tr>
<tr><td>>hg38_21_+_8433172_8446622_RNA45SN1_45S_50ntFlanks@11896.11929.34|TCTAGTCTGGCACGGTGAAGAGACATGAGAGGTG</td><td>>hg38_21_+_8433172_8446622_RNA45SN1_45S_50ntFlanks</td><td>11896</td><td>11929</td><td>34</td><td>TCTAGTCTGGCACGGTGAAGAGACATGAGAGGTG</td><td>268.8</td><td>11.4</td><td>2.1</td><td>5.4</td><td>7.0E-08</td><td>1.8E-06</td><td>BT20/MB231</td><td>MP</td><td></td></tr>
<tr><td>>MI0000070|hsa-mir-16-1&WithFlank&13|-|50048967|50049067@20.42.23;MI0000115|hsa-mir-16-2&WithFlank&3|+|160404739|160404831@16.38.23&[MIMAT0000069&hsa-miR-16-5p&offsets|0|+1;m-62&13|-|50049027|50049048&offsets|0|+1];[MIMAT0000069_1&hsa-miR-16-5p&offsets|0|+1;m-61&3|+|160404754|160404775&offsets|0|+1]|TAGCAGCACGTAAATATTGGCGT</td><td>>MI0000070|hsa-mir-16-1&WithFlank&13|-|50048967|50049067</td><td>20</td><td>42</td><td>23</td><td>TAGCAGCACGTAAATATTGGCGT</td><td>703.8</td><td>4.8</td><td>1.0</td><td>4.9</td><td>1.1E-06</td><td>1.8E-06</td><td>MB231/MB468</td><td>CYTO</td><td></td></tr>
<tr><td>>hg38_21_+_8433172_8446622_RNA45SN1_45S_50ntFlanks@11600.11625.26|AATCCGACTGTTTAATTAAAACAAAG</td><td>>hg38_21_+_8433172_8446622_RNA45SN1_45S_50ntFlanks</td><td>11600</td><td>11625</td><td>26</td><td>AATCCGACTGTTTAATTAAAACAAAG</td><td>65.4</td><td>9.8</td><td>1.9</td><td>5.1</td><td>3.0E-07</td><td>1.8E-06</td><td>BT20/MB468</td><td>MP</td><td></td></tr>
<tr><td>>hg38_21_+_8433172_8446622_RNA45SN1_45S_50ntFlanks@9914.9946.33|GAAAAGGTGTTGGTTGATATAGACAGCAGGACG</td><td>>hg38_21_+_8433172_8446622_RNA45SN1_45S_50ntFlanks</td><td>9914</td><td>9946</td><td>33</td><td>GAAAAGGTGTTGGTTGATATAGACAGCAGGACG</td><td>57.9</td><td>9.6</td><td>1.9</td><td>5.1</td><td>3.0E-07</td><td>1.8E-06</td><td>BT20/MB468</td><td>MP</td><td></td></tr>
<tr><td>>hg38_21_+_8433172_8446622_RNA45SN1_45S_50ntFlanks@9202.9221.20|TCTCGGGGCCACGCGCGCGT</td><td>>hg38_21_+_8433172_8446622_RNA45SN1_45S_50ntFlanks</td><td>9202</td><td>9221</td><td>20</td><td>TCTCGGGGCCACGCGCGCGT</td><td>60.8</td><td>-9.0</td><td>1.8</td><td>-5.1</td><td>3.0E-07</td><td>1.8E-06</td><td>BT20/MB468</td><td>MP</td><td></td></tr>
<tr><td>>hg38_MT_+_1621_3279_RNR2_16S_50ntFlanks@1176.1215.40|GCATTAAAAATTTCGGTTGGGGCGACCTCGGAGCAGAACC</td><td>>hg38_MT_+_1621_3279_RNR2_16S_50ntFlanks</td><td>1176</td><td>1215</td><td>40</td><td>GCATTAAAAATTTCGGTTGGGGCGACCTCGGAGCAGAACC</td><td>43.7</td><td>8.7</td><td>1.8</td><td>4.9</td><td>9.6E-07</td><td>1.8E-06</td><td>MB231/MB468</td><td>MP</td><td></td></tr>
<tr><td>>MI0000094|hsa-mir-92a-2&WithFlank&X|-|134169532|134169618@54.76.23&[MIMAT0000092_1&hsa-miR-92a-3p&offsets|0|+1;m-11&X|-|134169544|134169565&offsets|0|+1]|TATTGCACTTGTCCCGGCCTGTG</td><td>>MI0000094|hsa-mir-92a-2&WithFlank&X|-|134169532|134169618</td><td>54</td><td>76</td><td>23</td><td>TATTGCACTTGTCCCGGCCTGTG</td><td>458.4</td><td>5.7</td><td>1.2</td><td>4.9</td><td>9.5E-07</td><td>1.8E-06</td><td>BT20/MB468</td><td>CYTO</td><td></td></tr>
<tr><td>>MI0000074|hsa-mir-19b-1&WithFlank&13|+|91351186|91351284@60.81.22;MI0000075|hsa-mir-19b-2&WithFlank&X|-|134169665|134169772@68.89.22&[MIMAT0000074&hsa-miR-19b-3p&offsets|0|-1;m-122&13|+|91351245|91351267&offsets|0|-1];[MIMAT0000074_1&hsa-miR-19b-3p&offsets|0|-1;m-121&X|-|134169683|134169705&offsets|0|-1]|TGTGCAAATCCATGCAAAACTG</td><td>>MI0000074|hsa-mir-19b-1&WithFlank&13|+|91351186|91351284</td><td>60</td><td>81</td><td>22</td><td>TGTGCAAATCCATGCAAAACTG</td><td>563.1</td><td>5.5</td><td>1.1</td><td>4.9</td><td>1.2E-06</td><td>1.8E-06</td><td>MB231/MB468</td><td>MITO</td><td></td></tr>
</table>

| | | | | | | | | | | | | |
|---|---|---|---|---|---|---|---|---|---|---|---|---|
| >am_MI0005763\|hsa-mir-941-1&WithFlank&20\|+\|63919443\|63919526@53.74.22;MI0005764\|hsa-mir-941-2&WithFlank&20\|+\|63919499\|63919582@53.74.22;MI0005765\|hsa-mir-941-3&WithFlank&20\|+\|63919555\|63919638@53.74.22;MI0005766\|hsa-mir-941-4&WithFlank&20\|+\|63919750\|63919833@53.74.22;MI0031520\|hsa-mir-941-5&WithFlank&20\|+\|63919862\|63919945@53.74.22&[MIMAT0004984&hsa-miR-941&offsets\|0\|-1;m-263&20\|+\|63919495\|63919517&offsets\|0\|-1];[MIMAT0004984_1&hsa-miR-941&offsets\|0\|-1;m-263&20\|+\|63919551\|63919573&offsets\|0\|-1];[MIMAT0004984_2&hsa-miR-941&offsets\|0\|-1;m-263&20\|+\|63919607\|63919629&offsets\|0\|-1];[MIMAT0004984_3&hsa-miR-941&offsets\|0\|-1;m-263&20\|+\|63919802\|63919824&offsets\|0\|-1];[MIMAT0004984_4&hsa-miR-941&offsets\|0\|-1;m-263&20\|+\|63919914\|63919936&offsets\|0\|-1]\|CACCCGGCTGTGTGCACATGTG | >am_MI0005763\|hsa-mir-941-1&WithFlank&20\|+\|63919443\|63919526 | 53 | 74 | 22 | CACCCGGCTGTGTGCACATGTG | 418.4 | 7.0 | 1.4 | 4.9 | 1.2E-06 | 1.8E-06 | MB231/MB468 | MITO |
| >hg38_MT_+_1621_3279_RNR2_16S_50ntFlanks@713.749.37\|GCATAAGCCTGCGTCAGATTAAAACACTGAACTGACA | >hg38_MT_+_1621_3279_RNR2_16S_50ntFlanks | 713 | 749 | 37 | GCATAAGCCTGCGTCAGATTAAAACACTGAACTGACA | 85.0 | 10.2 | 2.0 | 5.1 | 3.0E-07 | 1.8E-06 | BT20/MB468 | MP |
| >hg38_21_+_8433172_8446622_RNA45SN1_45S_50ntFlanks@9534.9554.21\|AGGAAACTCTGGTGGAGGTCC | >hg38_21_+_8433172_8446622_RNA45SN1_45S_50ntFlanks | 9534 | 9554 | 21 | AGGAAACTCTGGTGGAGGTCC | 501.8 | -7.3 | 1.5 | -4.9 | 9.5E-07 | 1.8E-06 | BT20/MB468 | CYTO |
| >hg38_21_+_8433172_8446622_RNA45SN1_45S_50ntFlanks@10411.10434.24\|CAGTCGGTCCTGAGAGATGGGCGA | >hg38_21_+_8433172_8446622_RNA45SN1_45S_50ntFlanks | 10411 | 10434 | 24 | CAGTCGGTCCTGAGAGATGGGCGA | 85.0 | -9.5 | 1.9 | -5.1 | 3.0E-07 | 1.8E-06 | BT20/MB468 | MP |
| >MI0000746\|hsa-mir-99b&WithFlank&19\|+\|51692606\|51692687@13.34.22(+2U)&[MIMAT0000689&hsa-miR-99b-5p&offsets\|0\|0(+2U);m-10&19\|+\|51692618\|51692639&offsets\|0\|0(+2U)]\|CACCCGTAGAACCGACCTTGCGTT | >MI0000746\|hsa-mir-99b&WithFlank&19\|+\|51692606\|51692687 | 13 | 34 | 22(+2U) | CACCCGTAGAACCGACCTTGCGTT | 224.2 | 5.5 | 1.1 | 4.9 | 1.1E-06 | 1.8E-06 | MB231/MB468 | NUC |
| >hg38_21_+_8433172_8446622_RNA45SN1_45S_50ntFlanks@7977.8017.41\|CGACCTCAGATCAGACGTGGCGACCCGCTGAATTTAAGCAT | >hg38_21_+_8433172_8446622_RNA45SN1_45S_50ntFlanks | 7977 | 8017 | 41 | CGACCTCAGATCAGACGTGGCGACCCGCTGAATTTAAGCAT | 782.3 | -5.2 | 1.0 | -5.4 | 8.2E-08 | 1.8E-06 | BT20/MB231 | CYTO |
| >hg38_21_+_8433172_8446622_RNA45SN1_45S_50ntFlanks@5084.5121.38\|CGAACGAGACTCTGGCATGCTAACTAGTTACGCGACCC | >hg38_21_+_8433172_8446622_RNA45SN1_45S_50ntFlanks | 5084 | 5121 | 38 | CGAACGAGACTCTGGCATGCTAACTAGTTACGCGACCC | 551.6 | -6.3 | 1.3 | -4.9 | 1.2E-06 | 1.8E-06 | MB231/MB468 | MITO |
| >MI0000456\|hsa-mir-140&WithFlank&16\|+\|69933075\|69933186@69.90.22(+1U)&[MIMAT0004597&hsa-miR-140-3p&offsets\|+1\|+2(+1U);m-45&16\|+\|69933143\|69933164&offsets\|0\|0(+1U)]\|ACCACAGGGTAGAACCACGGACT | >MI0000456\|hsa-mir-140&WithFlank&16\|+\|69933075\|69933186 | 69 | 90 | 22(+1U) | ACCACAGGGTAGAACCACGGACT | 1172.1 | 5.9 | 1.2 | 4.9 | 9.7E-07 | 1.8E-06 | MB231/MB468 | MP |
| >hg38_21_+_8433172_8446622_RNA45SN1_45S_50ntFlanks@12159.12180.22\|TTTGACTGGGGCGGTACACCTG | >hg38_21_+_8433172_8446622_RNA45SN1_45S_50ntFlanks | 12159 | 12180 | 22 | TTTGACTGGGGCGGTACACCTG | 726.4 | -6.3 | 1.3 | -4.9 | 1.1E-06 | 1.8E-06 | MB231/MB468 | CYTO |
| >trna10_LysCTT_16_+_3241501_3241573@4.36.33,trna13_LysCTT_14_-_58706613_58706685@4.36.33,trna2_LysCTT_15_+_79152904_79152976@4.36.33\|CGGCTAGCTCAGTCGGTAGAGCATGGGACTCTT | >trna10_LysCTT_16_+_3241501_3241573 | 4 | 36 | 33 | CGGCTAGCTCAGTCGGTAGAGCATGGGACTCTT | 904.7 | 7.3 | 1.5 | 4.9 | 1.2E-06 | 1.8E-06 | MB231/MB468 | MITO |
| >hg38_21_+_8433172_8446622_RNA45SN1_45S_50ntFlanks@10248.10273.26\|CGACGAGTAGGAGGGCCGCTGCGGTG | >hg38_21_+_8433172_8446622_RNA45SN1_45S_50ntFlanks | 10248 | 10273 | 26 | CGACGAGTAGGAGGGCCGCTGCGGTG | 202.0 | 11.0 | 2.0 | 5.4 | 7.2E-08 | 1.8E-06 | BT20/MB231 | MP |
| >hg38_21_+_8433172_8446622_RNA45SN1_45S_50ntFlanks@11569.11597.29\|CAGCCGACTTAGAACTGGTGCGGACCAGG | >hg38_21_+_8433172_8446622_RNA45SN1_45S_50ntFlanks | 11569 | 11597 | 29 | CAGCCGACTTAGAACTGGTGCGGACCAGG | 200.4 | 11.0 | 2.0 | 5.4 | 7.2E-08 | 1.8E-06 | BT20/MB231 | MP |
| >hg38_21_+_8433172_8446622_RNA45SN1_45S_50ntFlanks@8118.8141.24\|GGGACATGTGGCGTACGGAAGACC | >hg38_21_+_8433172_8446622_RNA45SN1_45S_50ntFlanks | 8118 | 8141 | 24 | GGGACATGTGGCGTACGGAAGACC | 2450.8 | 7.8 | 1.4 | 5.4 | 7.2E-08 | 1.8E-06 | BT20/MB231 | MP |
| >hg38_MT_+_598_1651_RNR1_12S_50ntFlanks@569.611.43\|AAAGGACCTGGCGGTGCTTCATATCCCTCTAGAGGAGCCTGTT | >hg38_MT_+_598_1651_RNR1_12S_50ntFlanks | 569 | 611 | 43 | AAAGGACCTGGCGGTGCTTCATATCCCTCTAGAGGAGCCTGTT | 108.1 | -10.2 | 1.9 | -5.4 | 7.2E-08 | 1.8E-06 | BT20/MB231 | MP |
| >hg38_21_+_8433172_8446622_RNA45SN1_45S_50ntFlanks@5496.5516.21\|GACGGTCGAACTTGACTATCT | >hg38_21_+_8433172_8446622_RNA45SN1_45S_50ntFlanks | 5496 | 5516 | 21 | GACGGTCGAACTTGACTATCT | 203.4 | 8.3 | 1.6 | 5.1 | 3.0E-07 | 1.8E-06 | BT20/MB468 | MP |
| >hg38_21_+_8433172_8446622_RNA45SN1_45S_50ntFlanks@4261.4282.22\|TGGAATGAGTCCACTTTAAATC | >hg38_21_+_8433172_8446622_RNA45SN1_45S_50ntFlanks | 4261 | 4282 | 22 | TGGAATGAGTCCACTTTAAATC | 57.0 | -8.9 | 1.7 | -5.1 | 3.0E-07 | 1.8E-06 | BT20/MB468 | MP |
| >hg38_21_+_8433172_8446622_RNA45SN1_45S_50ntFlanks@4728.4761.34\|ATCAAGAACGAAAGTCGGAGGTTCGAAGACGATC | >hg38_21_+_8433172_8446622_RNA45SN1_45S_50ntFlanks | 4728 | 4761 | 34 | ATCAAGAACGAAAGTCGGAGGTTCGAAGACGATC | 62.3 | -9.0 | 1.8 | -5.1 | 3.0E-07 | 1.8E-06 | BT20/MB468 | MP |
| >hg38_21_+_8433172_8446622_RNA45SN1_45S_50ntFlanks@9240.9275.36\|GCGGAGCGAGCGCACGGGGTCGGCGGCGACGTCGGC | >hg38_21_+_8433172_8446622_RNA45SN1_45S_50ntFlanks | 9240 | 9275 | 36 | GCGGAGCGAGCGCACGGGGTCGGCGGCGACGTCGGC | 67.9 | -9.2 | 1.8 | -5.1 | 3.0E-07 | 1.8E-06 | BT20/MB468 | MP |
| >MI0000446\|hsa-mir-125b-1&WithFlank&11\|-\|122099751\|122099850@21.43.23&[MIMAT0000423&hsa-miR-125b-5p&offsets\|0\|+1;m-67&11\|-\|122099809\|122099830&offsets\|0\|+1]\|TCCCTGAGACCCTAACTTGTGAT | >MI0000446\|hsa-mir-125b-1&WithFlank&11\|-\|122099751\|122099850 | 21 | 43 | 23 | TCCCTGAGACCCTAACTTGTGAT | 304.0 | 7.9 | 1.6 | 4.9 | 9.8E-07 | 1.8E-06 | MB231/MB468 | MP |
| >hg38_21_+_8433172_8446622_RNA45SN1_45S_50ntFlanks@10392.10429.38\|ACAGCAGTTGAACATGGGTCAGTCGGTCCTGAGAGATG | >hg38_21_+_8433172_8446622_RNA45SN1_45S_50ntFlanks | 10392 | 10429 | 38 | ACAGCAGTTGAACATGGGTCAGTCGGTCCTGAGAGATG | 2825.8 | -6.1 | 1.2 | -4.9 | 1.2E-06 | 1.8E-06 | MB231/MB468 | MITO |
| >hg38_21_+_8433172_8446622_RNA45SN1_45S_50ntFlanks@12971.12990.20\|TTTCGTACGTAGCAGAGCAG | >hg38_21_+_8433172_8446622_RNA45SN1_45S_50ntFlanks | 12971 | 12990 | 20 | TTTCGTACGTAGCAGAGCAG | 500.3 | -6.4 | 1.3 | -4.9 | 9.7E-07 | 1.8E-06 | BT20/MB468 | CYTO |
| >hg38_21_+_8433172_8446622_RNA45SN1_45S_50ntFlanks@8406.8442.37\|TAAACGGGTGGGGTCCGCGCAGTCCGCCCGGAGGATT | >hg38_21_+_8433172_8446622_RNA45SN1_45S_50ntFlanks | 8406 | 8442 | 37 | TAAACGGGTGGGGTCCGCGCAGTCCGCCCGGAGGATT | 66.6 | -9.1 | 1.8 | -5.1 | 3.0E-07 | 1.9E-06 | BT20/MB468 | MP |
| >MI0003776\|hsa-mir-320b-1&WithFlank&1\|+\|116671740\|116671823@48.66.19(+1A);MI0003839\|hsa-mir-320b-2&WithFlank&1\|-\|224257034\|224257116@47.65.19(+1A);MI0003778\|hsa-mir-320c-1&WithFlank&18\|+\|21683512\|21683595@48.66.19(+1A);MI0008191\|hsa-mir-320c-2&WithFlank&18\|+\|24321669\|24321752@48.66.19(+1A);MI0000542\|hsa-mir-320a&WithFlank&8\|-\|22244960\|22245043@48.66.19(+1A)&[MIMAT0005792&hsa-miR-320b&offsets\|0\|-3(+1A);m-183&1\|+\|116671787\|116671806&offsets\|0\|-1(+1A)];[MIMAT0005792_1&hsa-miR-320b&offsets\|0\|-3(+1A);m-183&1\|-\|224257051\|224257070&offsets\|0\|-1(+1A)];[MIMAT0005793&hsa-miR-320c&offsets\|0\|-1(+1A);m-222&18\|+\|21683559\|21683578&offsets\|0\|-1(+1A)];[MIMAT0005793_1&hsa-miR-320c&offsets\|0\|-1(+1A);m-219&18\|+\|24321716\|24321735&offsets\|0\|-1(+1A)];[MIMAT0000510&hsa-miR-320a-3p&offsets\|0\|-3(+1A);m-60&8\|-\|22244975\|22244996&offsets\|0\|-3(+1A)]\|AAAAGCTGGGTTGAGAGGGA | >MI0003776\|hsa-mir-320b-1&WithFlank&1\|+\|116671740\|116671823 | 48 | 66 | 19(+1A) | AAAAGCTGGGTTGAGAGGGA | 869.5 | 4.7 | 0.9 | 5.4 | 8.3E-08 | 1.9E-06 | BT20/MB231 | CYTO |
| >hg38_21_+_8433172_8446622_RNA45SN1_45S_50ntFlanks@4367.4391.25\|CAGTTAAAAAGCTCGTAGTTGGATC | >hg38_21_+_8433172_8446622_RNA45SN1_45S_50ntFlanks | 4367 | 4391 | 25 | CAGTTAAAAAGCTCGTAGTTGGATC | 355.7 | -5.0 | 1.0 | -4.9 | 8.5E-07 | 1.9E-06 | BT20/MB468 | MITO |
| >hg38_21_+_8433172_8446622_RNA45SN1_45S_50ntFlanks@10681.10708.28\|TTCCGGCGGCGTCCGGTGAGCTCTCGCT | >hg38_21_+_8433172_8446622_RNA45SN1_45S_50ntFlanks | 10681 | 10708 | 28 | TTCCGGCGGCGTCCGGTGAGCTCTCGCT | 662.1 | -8.0 | 1.7 | -4.8 | 1.2E-06 | 1.9E-06 | BT20/MB468 | NUC |
| >MI0000734\|hsa-mir-106b&WithFlank&7\|-\|100093987\|100094080@58.78.21&[MIMAT0004672&hsa-miR-106b-3p&offsets\|0\|-1;m-87&7\|-\|100094003\|100094023&offsets\|0\|0]\|CCGCACTGTGGGTACTTGCTG | >MI0000734\|hsa-mir-106b&WithFlank&7\|-\|100093987\|100094080 | 58 | 78 | 21 | CCGCACTGTGGGTACTTGCTG | 2480.0 | 5.6 | 1.1 | 5.1 | 3.0E-07 | 1.9E-06 | BT20/MB468 | MP |
| >hg38_21_+_8433172_8446622_RNA45SN1_45S_50ntFlanks@12474.12498.25\|ACCCACTAATAGGGAACGTGAGCTG | >hg38_21_+_8433172_8446622_RNA45SN1_45S_50ntFlanks | 12474 | 12498 | 25 | ACCCACTAATAGGGAACGTGAGCTG | 168.4 | -6.6 | 1.4 | -4.9 | 9.8E-07 | 1.9E-06 | MB231/MB468 | MP |
| >hg38_1_-_228634819_228635039_RNA5S12_5S_50ntFlanks@150.169.20\|AATACCGGGTGCTGTAGGCT | >hg38_1_-_228634819_228635039_RNA5S12_5S_50ntFlanks | 150 | 169 | 20 | AATACCGGGTGCTGTAGGCT | 953.6 | 3.7 | 0.8 | 4.9 | 1.1E-06 | 1.9E-06 | MB231/MB468 | CYTO |
| >hg38_21_+_8433172_8446622_RNA45SN1_45S_50ntFlanks@9523.9571.49\|GGCGAAGCCAGAGGAAACTCTGGTGGAGGTCCGTAGCGGTCCTGACGTG | >hg38_21_+_8433172_8446622_RNA45SN1_45S_50ntFlanks | 9523 | 9571 | 49 | GGCGAAGCCAGAGGAAACTCTGGTGGAGGTCCGTAGCGGTCCTGACGTG | 257.9 | 6.3 | 1.3 | 4.9 | 1.1E-06 | 1.9E-06 | MB231/MB468 | CYTO |
| >hg38_21_+_8433172_8446622_RNA45SN1_45S_50ntFlanks@12445.12472.28\|AGCAGAATTCACCAAGCGTTGGATTGTT | >hg38_21_+_8433172_8446622_RNA45SN1_45S_50ntFlanks | 12445 | 12472 | 28 | AGCAGAATTCACCAAGCGTTGGATTGTT | 57.6 | -8.9 | 1.7 | -5.1 | 3.0E-07 | 1.9E-06 | BT20/MB468 | MP |
| >trna13_ValCAC_19_-_4724647_4724719@1.32.32\|GTTTCCGTAGTGTAGCGGTTATCACATTCGCC | >trna13_ValCAC_19_-_4724647_4724719 | 1 | 32 | 32 | GTTTCCGTAGTGTAGCGGTTATCACATTCGCC | 235.3 | 7.9 | 1.6 | 4.9 | 9.9E-07 | 1.9E-06 | MB231/MB468 | MP |
| >hg38_21_+_8433172_8446622_RNA45SN1_45S_50ntFlanks@10398.10424.27\|GTTGAACATGGGTCAGTCGGTCCTGAG | >hg38_21_+_8433172_8446622_RNA45SN1_45S_50ntFlanks | 10398 | 10424 | 27 | GTTGAACATGGGTCAGTCGGTCCTGAG | 5569.7 | 7.5 | 1.5 | 5.1 | 3.1E-07 | 1.9E-06 | BT20/MB468 | MP |
| >hg38_21_+_8433172_8446622_RNA45SN1_45S_50ntFlanks@10685.10710.26\|GGCGGCGTCCGGTGAGCTCTCGCTGG | >hg38_21_+_8433172_8446622_RNA45SN1_45S_50ntFlanks | 10685 | 10710 | 26 | GGCGGCGTCCGGTGAGCTCTCGCTGG | 455.0 | -2.7 | 0.6 | -4.9 | 1.1E-06 | 1.9E-06 | MB231/MB468 | CYTO |
| >hg38_21_+_8433172_8446622_RNA45SN1_45S_50ntFlanks@7983.8008.26\|CAGATCAGACGTGGCGACCCGCTGAA | >hg38_21_+_8433172_8446622_RNA45SN1_45S_50ntFlanks | 7983 | 8008 | 26 | CAGATCAGACGTGGCGACCCGCTGAA | 4057.5 | -5.5 | 1.1 | -5.1 | 3.1E-07 | 1.9E-06 | BT20/MB468 | MP |
| >hg38_MT_+_598_1651_RNR1_12S_50ntFlanks@972.1002.31\|GGTAAGTGTACTGGAAAGTGCACTTGGACGA | >hg38_MT_+_598_1651_RNR1_12S_50ntFlanks | 972 | 1002 | 31 | GGTAAGTGTACTGGAAAGTGCACTTGGACGA | 58.8 | 9.1 | 1.9 | 4.9 | 9.9E-07 | 1.9E-06 | MB231/MB468 | MP |
| >hg38_21_+_8433172_8446622_RNA45SN1_45S_50ntFlanks@10387.10407.21\|TGTGAACAGCAGTTGAACATG | >hg38_21_+_8433172_8446622_RNA45SN1_45S_50ntFlanks | 10387 | 10407 | 21 | TGTGAACAGCAGTTGAACATG | 511.7 | -6.4 | 1.3 | -4.9 | 9.9E-07 | 1.9E-06 | MB231/MB468 | MP |
| >hg38_21_+_8433172_8446622_RNA45SN1_45S_50ntFlanks@12949.12990.42\|ACGACCTGCTTCTGGGTCGGGGTTTCGTACGTAGCAGAGCAG | >hg38_21_+_8433172_8446622_RNA45SN1_45S_50ntFlanks | 12949 | 12990 | 42 | ACGACCTGCTTCTGGGTCGGGGTTTCGTACGTAGCAGAGCAG | 200.9 | -4.7 | 1.0 | -4.9 | 1.1E-06 | 1.9E-06 | MB231/MB468 | CYTO |
| >hg38_21_+_8433172_8446622_RNA45SN1_45S_50ntFlanks@4006.4042.37\|ACCTCGGGCCGATCGCACGCCCCCCGTGGCGGCGACG | >hg38_21_+_8433172_8446622_RNA45SN1_45S_50ntFlanks | 4006 | 4042 | 37 | ACCTCGGGCCGATCGCACGCCCCCCGTGGCGGCGACG | 835.8 | -5.2 | 1.1 | -4.9 | 1.1E-06 | 1.9E-06 | MB231/MB468 | CYTO |
| >hg38_21_+_8433172_8446622_RNA45SN1_45S_50ntFlanks@12418.12436.19\|CGATGTCGGCTCTTCCTAT | >hg38_21_+_8433172_8446622_RNA45SN1_45S_50ntFlanks | 12418 | 12436 | 19 | CGATGTCGGCTCTTCCTAT | 69064.6 | -7.2 | 1.5 | -4.8 | 1.3E-06 | 1.9E-06 | BT20/MB468 | NUC |
| >hg38_21_+_8433172_8446622_RNA45SN1_45S_50ntFlanks@11718.11739.22\|AACGGCGGGAGTAACTATGACT | >hg38_21_+_8433172_8446622_RNA45SN1_45S_50ntFlanks | 11718 | 11739 | 22 | AACGGCGGGAGTAACTATGACT | 569.4 | -5.7 | 1.2 | -4.9 | 1.2E-06 | 1.9E-06 | MB231/MB468 | NUC |
| >hg38_21_+_8433172_8446622_RNA45SN1_45S_50ntFlanks@10107.10131.25\|CGTGTGGTGTGCGTCGGAGGGCGGC | >hg38_21_+_8433172_8446622_RNA45SN1_45S_50ntFlanks | 10107 | 10131 | 25 | CGTGTGGTGTGCGTCGGAGGGCGGC | 207.6 | 11.0 | 2.0 | 5.4 | 7.3E-08 | 1.9E-06 | BT20/MB231 | MP |
| >am_trnaMT_ValTAC_MT_+_1602_1670@46.67.22\|TTCAACTTAACTTGACCGCTCT | >am_trnaMT_ValTAC_MT_+_1602_1670 | 46 | 67 | 22 | TTCAACTTAACTTGACCGCTCT | 814.4 | 9.1 | 1.9 | 4.9 | 9.8E-07 | 1.9E-06 | BT20/MB468 | CYTO |

| | | | | | | | | | | | | | |
|---|---|---|---|---|---|---|---|---|---|---|---|---|---|
| >hg38_21_+_8433172_8446622_RNA45SN1_45S_50ntFlanks@6646.6668.23\|TCGTACGACTCTTAGCGGTGGAT | >hg38_21_+_8433172_8446622_RNA45SN1_45S_50ntFlanks | 6646 | 6668 | 23 | TCGTACGACTCTTAGCGGTGGAT | 3471.9 | 3.1 | 0.6 | 4.9 | 1.2E-06 | 1.9E-06 | MB231/MB468 | NUC |
| >hg38_21_+_8433172_8446622_RNA45SN1_45S_50ntFlanks@11572.11604.33\|CCGACTTAGAACTGGTGCGGACCAGGGGAATCC | >hg38_21_+_8433172_8446622_RNA45SN1_45S_50ntFlanks | 11572 | 11604 | 33 | CCGACTTAGAACTGGTGCGGACCAGGGGAATCC | 563.7 | -3.7 | 0.8 | -4.9 | 1.2E-06 | 1.9E-06 | MB231/MB468 | MITO |
| >hg38_21_+_8433172_8446622_RNA45SN1_45S_50ntFlanks@9621.9650.30\|TCTAGTAGCTGGTTCCCTCCGAAGTTTCCC | >hg38_21_+_8433172_8446622_RNA45SN1_45S_50ntFlanks | 9621 | 9650 | 30 | TCTAGTAGCTGGTTCCCTCCGAAGTTTCCC | 160.9 | -6.2 | 1.3 | -4.9 | 1.2E-06 | 1.9E-06 | MB231/MB468 | NUC |
| >hg38_21_+_8433172_8446622_RNA45SN1_45S_50ntFlanks@9533.9569.37\|GAGGAAACTCTGGTGGAGGTCCGTAGCGGTCCTGACG | >hg38_21_+_8433172_8446622_RNA45SN1_45S_50ntFlanks | 9533 | 9569 | 37 | GAGGAAACTCTGGTGGAGGTCCGTAGCGGTCCTGACG | 905.7 | -3.7 | 0.8 | -4.9 | 1.2E-06 | 1.9E-06 | MB231/MB468 | NUC |
| >hg38_21_+_8433172_8446622_RNA45SN1_45S_50ntFlanks@11918.11959.42\|ACATGAGAGGTGTAGAATAAGTGGGAGGCCCCCGGCGCCCCC | >hg38_21_+_8433172_8446622_RNA45SN1_45S_50ntFlanks | 11918 | 11959 | 42 | ACATGAGAGGTGTAGAATAAGTGGGAGGCCCCCGGCGCCCCC | 51.8 | -9.6 | 2.0 | -4.9 | 9.9E-07 | 1.9E-06 | MB231/MB468 | MP |
| >hg38_21_+_8433172_8446622_RNA45SN1_45S_50ntFlanks@12997.13014.18\|CGCTGCGATCTATTGAAA | >hg38_21_+_8433172_8446622_RNA45SN1_45S_50ntFlanks | 12997 | 13014 | 18 | CGCTGCGATCTATTGAAA | 1592.7 | -3.4 | 0.7 | -4.8 | 1.3E-06 | 1.9E-06 | BT20/MB468 | NUC |
| >hg38_21_+_8433172_8446622_RNA45SN1_45S_50ntFlanks@12366.12383.18\|TAACTGGCTTGTGGCGGC | >hg38_21_+_8433172_8446622_RNA45SN1_45S_50ntFlanks | 12366 | 12383 | 18 | TAACTGGCTTGTGGCGGC | 455.8 | -6.7 | 1.4 | -4.9 | 1.2E-06 | 1.9E-06 | MB231/MB468 | NUC |
| >hg38_21_+_8433172_8446622_RNA45SN1_45S_50ntFlanks@11612.11660.49\|TAATTAAAACAAAGCATCGCGAAGGCCCGCGGCGGGTGTTGACGCGATG | >hg38_21_+_8433172_8446622_RNA45SN1_45S_50ntFlanks | 11612 | 11660 | 49 | TAATTAAAACAAAGCATCGCGAAGGCCCGCGGCGGGTGTTGACGCGATG | 108.9 | -8.9 | 1.8 | -4.9 | 8.7E-07 | 1.9E-06 | BT20/MB468 | MITO |
| >hg38_21_+_8433172_8446622_RNA45SN1_45S_50ntFlanks@12568.12587.20\|TCAGTACGAGAGGAACCGCA | >hg38_21_+_8433172_8446622_RNA45SN1_45S_50ntFlanks | 12568 | 12587 | 20 | TCAGTACGAGAGGAACCGCA | 90.2 | -5.6 | 1.1 | -4.9 | 9.9E-07 | 1.9E-06 | BT20/MB468 | CYTO |
| >hg38_21_+_8433172_8446622_RNA45SN1_45S_50ntFlanks@10788.10806.19\|CAGCCTCTGGCATGTTGGA | >hg38_21_+_8433172_8446622_RNA45SN1_45S_50ntFlanks | 10788 | 10806 | 19 | CAGCCTCTGGCATGTTGGA | 275.8 | -5.3 | 1.1 | -4.9 | 9.9E-07 | 1.9E-06 | BT20/MB468 | CYTO |
| >hg38_21_+_8433172_8446622_RNA45SN1_45S_50ntFlanks@6789.6806.18\|GCCTGTCTGAGCGTCGCT | >hg38_21_+_8433172_8446622_RNA45SN1_45S_50ntFlanks | 6789 | 6806 | 18 | GCCTGTCTGAGCGTCGCT | 3468.1 | 3.4 | 0.7 | 4.9 | 1.2E-06 | 1.9E-06 | MB231/MB468 | MITO |
| >hg38_21_+_8433172_8446622_RNA45SN1_45S_50ntFlanks@10402.10435.34\|AACATGGGTCAGTCGGTCCTGAGAGATGGGCGAG | >hg38_21_+_8433172_8446622_RNA45SN1_45S_50ntFlanks | 10402 | 10435 | 34 | AACATGGGTCAGTCGGTCCTGAGAGATGGGCGAG | 746.4 | -6.6 | 1.4 | -4.9 | 9.9E-07 | 1.9E-06 | BT20/MB468 | CYTO |
| >hg38_21_+_8433172_8446622_RNA45SN1_45S_50ntFlanks@6760.6806.47\|TTGCGGCCCCGGGTTCCTCCCGGGGCTACGCCTGTCTGAGCGTCGCT | >hg38_21_+_8433172_8446622_RNA45SN1_45S_50ntFlanks | 6760 | 6806 | 47 | TTGCGGCCCCGGGTTCCTCCCGGGGCTACGCCTGTCTGAGCGTCGCT | 60.4 | 9.7 | 1.9 | 5.1 | 3.1E-07 | 1.9E-06 | BT20/MB468 | MP |
| >MI0000825\|hsa-mir-345&WithFlank&14\|+\|100307853\|100307962@24.45.22&[MIMAT0000772&hsa-miR-345-5p&offsets\|0\|0;m-193&14\|+\|100307876\|100307896&offsets\|0\|+1]\|GCTGACTCCTAGTCCAGGGCTC | >MI0000825\|hsa-mir-345&WithFlank&14\|+\|100307853\|100307962 | 24 | 45 | 22 | GCTGACTCCTAGTCCAGGGCTC | 324.2 | 7.0 | 1.4 | 4.9 | 9.9E-07 | 1.9E-06 | BT20/MB468 | CYTO |
| >trnaMT_LeuTAA_MT_+_3230_3304@16.45.30\|CCCGGTAATCGCATAAAACTTAAAACTTTA | >trnaMT_LeuTAA_MT_+_3230_3304 | 16 | 45 | 30 | CCCGGTAATCGCATAAAACTTAAAACTTTA | 114.7 | -9.9 | 1.9 | -5.1 | 3.1E-07 | 1.9E-06 | BT20/MB468 | MP |
| >hg38_21_+_8433172_8446622_RNA45SN1_45S_50ntFlanks@12419.12462.44\|GATGTCGGCTCTTCCTATCATTGTGAAGCAGAATTCACCAAGCG | >hg38_21_+_8433172_8446622_RNA45SN1_45S_50ntFlanks | 12419 | 12462 | 44 | GATGTCGGCTCTTCCTATCATTGTGAAGCAGAATTCACCAAGCG | 1541.0 | -5.1 | 1.0 | -5.4 | 8.5E-08 | 1.9E-06 | BT20/MB231 | CYTO |
| >hg38_21_+_8433172_8446622_RNA45SN1_45S_50ntFlanks@8175.8216.42\|CCTTCTGATCGAGGCCCAGCCCGTGGACGGTGTGAGGCCGGT | >hg38_21_+_8433172_8446622_RNA45SN1_45S_50ntFlanks | 8175 | 8216 | 42 | CCTTCTGATCGAGGCCCAGCCCGTGGACGGTGTGAGGCCGGT | 1008.0 | 8.3 | 1.6 | 5.1 | 3.1E-07 | 1.9E-06 | BT20/MB468 | MP |
| >trnaMT_HisGTG_MT_+_12138_12206@18.55.38\|AAAACATCAGATTGTGAATCTGACAACAGAGGCTTACG | >trnaMT_HisGTG_MT_+_12138_12206 | 18 | 55 | 38 | AAAACATCAGATTGTGAATCTGACAACAGAGGCTTACG | 50.4 | 8.9 | 1.8 | 4.9 | 1.0E-06 | 1.9E-06 | MB231/MB468 | MP |
| >hg38_21_+_8433172_8446622_RNA45SN1_45S_50ntFlanks@6738.6757.20\|ATCATCGACACTTCGAACGC | >hg38_21_+_8433172_8446622_RNA45SN1_45S_50ntFlanks | 6738 | 6757 | 20 | ATCATCGACACTTCGAACGC | 61.6 | -9.0 | 1.8 | -5.1 | 3.1E-07 | 1.9E-06 | BT20/MB468 | MP |
| >hg38_21_+_8433172_8446622_RNA45SN1_45S_50ntFlanks@4202.4223.22\|CGACCCGGGGAGGTAGTGACGA | >hg38_21_+_8433172_8446622_RNA45SN1_45S_50ntFlanks | 4202 | 4223 | 22 | CGACCCGGGGAGGTAGTGACGA | 419.4 | -7.2 | 1.5 | -4.9 | 1.0E-06 | 1.9E-06 | MB231/MB468 | MP |
| >hg38_21_+_8433172_8446622_RNA45SN1_45S_50ntFlanks@11778.11803.26\|ACGCGCATGAATGGATGAACGAGATT | >hg38_21_+_8433172_8446622_RNA45SN1_45S_50ntFlanks | 11778 | 11803 | 26 | ACGCGCATGAATGGATGAACGAGATT | 624.6 | -7.8 | 1.6 | -4.8 | 1.3E-06 | 1.9E-06 | BT20/MB468 | NUC |
| >MI0000065\|hsa-let-7d&WithFlank&9\|+\|94178828\|94178926@68.89.22&[MIMAT0004484&hsa-let-7d-3p&offsets\|0\|0;m-98&9\|+\|94178895\|94178916&offsets\|0\|0]\|CTATACGACCTGCTGCCTTTCT | >MI0000065\|hsa-let-7d&WithFlank&9\|+\|94178828\|94178926 | 68 | 89 | 22 | CTATACGACCTGCTGCCTTTCT | 1169.1 | 2.4 | 0.5 | 4.8 | 1.3E-06 | 1.9E-06 | BT20/MB468 | NUC |
| >hg38_21_+_8433172_8446622_RNA45SN1_45S_50ntFlanks@12073.12090.18\|CCGAGGGGCTCTCGCTTC | >hg38_21_+_8433172_8446622_RNA45SN1_45S_50ntFlanks | 12073 | 12090 | 18 | CCGAGGGGCTCTCGCTTC | 164.8 | -6.2 | 1.3 | -4.9 | 1.2E-06 | 1.9E-06 | MB231/MB468 | NUC |
| >hg38_21_+_8433172_8446622_RNA45SN1_45S_50ntFlanks@476.515.40\|TTCGTGATCGATGTGGTGACGTCGTGCTCTCCCGGGCCGG | >hg38_21_+_8433172_8446622_RNA45SN1_45S_50ntFlanks | 476 | 515 | 40 | TTCGTGATCGATGTGGTGACGTCGTGCTCTCCCGGGCCGG | 1108.3 | 4.6 | 0.9 | 4.9 | 1.2E-06 | 1.9E-06 | MB231/MB468 | NUC |
| >hg38_21_+_8433172_8446622_RNA45SN1_45S_50ntFlanks@4082.4101.20\|TCGCCGTGCCTACCATGGTG | >hg38_21_+_8433172_8446622_RNA45SN1_45S_50ntFlanks | 4082 | 4101 | 20 | TCGCCGTGCCTACCATGGTG | 78.5 | -9.4 | 1.8 | -5.1 | 3.1E-07 | 1.9E-06 | BT20/MB468 | MP |
| >trna116_GluCTC_1_-_145399233_145399304@32.68.37,trna59_GluCTC_1_+_249168447_249168518@32.68.37,trna71_GluCTC_1_-_161439189_161439260@32.68.37,trna74_GluCTC_1_-_161431809_161431880@32.68.37,trna77_GluCTC_1_-_161424398_161424469@32.68.37,trna77_GluCTC_6_+_28949976_28950047@32.68.37,trna80_GluCTC_1_-_161417018_161417089@32.68.37,trna87_GluCTC_6_-_126101393_126101464@32.68.37\|CTCTCACCGCCGCGGCCCGGGTTCGATTCCCGGTCAG | >trna116_GluCTC_1_-_145399233_145399304 | 32 | 68 | 37 | CTCTCACCGCCGCGGCCCGGGTTCGATTCCCGGTCAG | 108.2 | -6.1 | 1.2 | -4.9 | 9.9E-07 | 1.9E-06 | BT20/MB468 | CYTO |
| >hg38_21_+_8433172_8446622_RNA45SN1_45S_50ntFlanks@10394.10424.31\|AGCAGTTGAACATGGGTCAGTCGGTCCTGAG | >hg38_21_+_8433172_8446622_RNA45SN1_45S_50ntFlanks | 10394 | 10424 | 31 | AGCAGTTGAACATGGGTCAGTCGGTCCTGAG | 986.8 | 3.6 | 0.7 | 4.9 | 9.9E-07 | 1.9E-06 | BT20/MB468 | CYTO |
| >hg38_21_+_8433172_8446622_RNA45SN1_45S_50ntFlanks@11844.11861.18\|GGGAACGGGCTTGGCGGA | >hg38_21_+_8433172_8446622_RNA45SN1_45S_50ntFlanks | 11844 | 11861 | 18 | GGGAACGGGCTTGGCGGA | 163.3 | -6.9 | 1.4 | -4.9 | 1.0E-06 | 1.9E-06 | MB231/MB468 | MP |
| >hg38_21_+_8433172_8446622_RNA45SN1_45S_50ntFlanks@10369.10393.25\|AAGTGGAGAAGGGTTCCATGTGAAC | >hg38_21_+_8433172_8446622_RNA45SN1_45S_50ntFlanks | 10369 | 10393 | 25 | AAGTGGAGAAGGGTTCCATGTGAAC | 59.3 | -9.0 | 1.8 | -5.1 | 3.1E-07 | 1.9E-06 | BT20/MB468 | MP |
| >hg38_21_+_8433172_8446622_RNA45SN1_45S_50ntFlanks@6646.6671.26\|TCGTACGACTCTTAGCGGTGGATCAC | >hg38_21_+_8433172_8446622_RNA45SN1_45S_50ntFlanks | 6646 | 6671 | 26 | TCGTACGACTCTTAGCGGTGGATCAC | 348.3 | 3.9 | 0.8 | 4.9 | 1.1E-06 | 1.9E-06 | MB231/MB468 | CYTO |
| >hg38_21_+_8433172_8446622_RNA45SN1_45S_50ntFlanks@7990.8011.22\|GACGTGGCGACCCGCTGAATTT | >hg38_21_+_8433172_8446622_RNA45SN1_45S_50ntFlanks | 7990 | 8011 | 22 | GACGTGGCGACCCGCTGAATTT | 544.7 | 5.8 | 1.2 | 4.9 | 1.0E-06 | 1.9E-06 | BT20/MB468 | CYTO |
| >hg38_21_+_8433172_8446622_RNA45SN1_45S_50ntFlanks@10807.10840.34\|ACAATGTAGGTAAGGGAAGTCGGCAAGCCGGATC | >hg38_21_+_8433172_8446622_RNA45SN1_45S_50ntFlanks | 10807 | 10840 | 34 | ACAATGTAGGTAAGGGAAGTCGGCAAGCCGGATC | 223.6 | -4.1 | 0.8 | -4.9 | 1.0E-06 | 1.9E-06 | BT20/MB468 | CYTO |
| >hg38_21_+_8433172_8446622_RNA45SN1_45S_50ntFlanks@9951.9979.29\|CCATGGAAGTCGGAATCCGCTAAGGAGTG | >hg38_21_+_8433172_8446622_RNA45SN1_45S_50ntFlanks | 9951 | 9979 | 29 | CCATGGAAGTCGGAATCCGCTAAGGAGTG | 182.1 | -6.8 | 1.4 | -4.9 | 1.0E-06 | 1.9E-06 | MB231/MB468 | MP |
| >hg38_21_+_8433172_8446622_RNA45SN1_45S_50ntFlanks@10245.10262.18\|CCGCGACGAGTAGGAGGG | >hg38_21_+_8433172_8446622_RNA45SN1_45S_50ntFlanks | 10245 | 10262 | 18 | CCGCGACGAGTAGGAGGG | 151.9 | 10.6 | 2.0 | 5.4 | 7.6E-08 | 1.9E-06 | BT20/MB231 | MP |
| >hg38_21_+_8433172_8446622_RNA45SN1_45S_50ntFlanks@10341.10369.29\|AATATTCAAACGAGAACTTTGAAGGCCGA | >hg38_21_+_8433172_8446622_RNA45SN1_45S_50ntFlanks | 10341 | 10369 | 29 | AATATTCAAACGAGAACTTTGAAGGCCGA | 197.1 | 11.0 | 2.0 | 5.4 | 7.6E-08 | 1.9E-06 | BT20/MB231 | MP |
| >hg38_21_+_8433172_8446622_RNA45SN1_45S_50ntFlanks@11913.11940.28\|AAGAGACATGAGAGGTGTAGAATAAGTG | >hg38_21_+_8433172_8446622_RNA45SN1_45S_50ntFlanks | 11913 | 11940 | 28 | AAGAGACATGAGAGGTGTAGAATAAGTG | 184.4 | 10.9 | 2.0 | 5.4 | 7.6E-08 | 1.9E-06 | BT20/MB231 | MP |
| >hg38_21_+_8433172_8446622_RNA45SN1_45S_50ntFlanks@6744.6772.29\|GACACTTCGAACGCACTTGCGGCCCCGGG | >hg38_21_+_8433172_8446622_RNA45SN1_45S_50ntFlanks | 6744 | 6772 | 29 | GACACTTCGAACGCACTTGCGGCCCCGGG | 582.3 | -7.0 | 1.5 | -4.9 | 1.2E-06 | 1.9E-06 | MB231/MB468 | MITO |
| >hg38_21_+_8433172_8446622_RNA45SN1_45S_50ntFlanks@12420.12441.22\|ATGTCGGCTCTTCCTATCATTG | >hg38_21_+_8433172_8446622_RNA45SN1_45S_50ntFlanks | 12420 | 12441 | 22 | ATGTCGGCTCTTCCTATCATTG | 2894.7 | -3.4 | 0.7 | -4.9 | 1.0E-06 | 1.9E-06 | BT20/MB468 | CYTO |
| >hg38_21_+_8433172_8446622_RNA45SN1_45S_50ntFlanks@4724.4751.28\|ATTAATCAAGAACGAAAGTCGGAGGTTC | >hg38_21_+_8433172_8446622_RNA45SN1_45S_50ntFlanks | 4724 | 4751 | 28 | ATTAATCAAGAACGAAAGTCGGAGGTTC | 565.9 | -3.7 | 0.8 | -4.9 | 1.0E-06 | 1.9E-06 | BT20/MB468 | CYTO |
| >hg38_21_+_8433172_8446622_RNA45SN1_45S_50ntFlanks@11199.11223.25\|CCGGCGGCGGCGGCGACTCTGGACG | >hg38_21_+_8433172_8446622_RNA45SN1_45S_50ntFlanks | 11199 | 11223 | 25 | CCGGCGGCGGCGGCGACTCTGGACG | 521.2 | -6.4 | 1.3 | -4.9 | 1.0E-06 | 1.9E-06 | BT20/MB468 | CYTO |
| >hg38_21_+_8433172_8446622_RNA45SN1_45S_50ntFlanks@7977.8012.36\|CGACCTCAGATCAGACGTGGCGACCCGCTGAATTTA | >hg38_21_+_8433172_8446622_RNA45SN1_45S_50ntFlanks | 7977 | 8012 | 36 | CGACCTCAGATCAGACGTGGCGACCCGCTGAATTTA | 572.3 | -3.8 | 0.8 | -4.8 | 1.3E-06 | 1.9E-06 | BT20/MB468 | NUC |
| >hg38_21_+_8433172_8446622_RNA45SN1_45S_50ntFlanks@10392.10424.33\|ACAGCAGTTGAACATGGGTCAGTCGGTCCTGAG | >hg38_21_+_8433172_8446622_RNA45SN1_45S_50ntFlanks | 10392 | 10424 | 33 | ACAGCAGTTGAACATGGGTCAGTCGGTCCTGAG | 15640.8 | 3.5 | 0.7 | 4.9 | 8.9E-07 | 1.9E-06 | BT20/MB468 | MITO |
| >hg38_21_+_8433172_8446622_RNA45SN1_45S_50ntFlanks@4730.4758.29\|CAAGAACGAAAGTCGGAGGTTCGAAGACG | >hg38_21_+_8433172_8446622_RNA45SN1_45S_50ntFlanks | 4730 | 4758 | 29 | CAAGAACGAAAGTCGGAGGTTCGAAGACG | 2392.0 | -3.7 | 0.7 | -4.9 | 8.9E-07 | 1.9E-06 | BT20/MB468 | MITO |
| >hg38_21_+_8433172_8446622_RNA45SN1_45S_50ntFlanks@4735.4759.25\|ACGAAAGTCGGAGGTTCGAAGACGA | >hg38_21_+_8433172_8446622_RNA45SN1_45S_50ntFlanks | 4735 | 4759 | 25 | ACGAAAGTCGGAGGTTCGAAGACGA | 685.8 | -3.4 | 0.7 | -4.9 | 1.0E-06 | 1.9E-06 | MB231/MB468 | MP |
| >hg38_21_+_8433172_8446622_RNA45SN1_45S_50ntFlanks@11217.11238.22\|CTGGACGCGAGCCGGGCCCTTC | >hg38_21_+_8433172_8446622_RNA45SN1_45S_50ntFlanks | 11217 | 11238 | 22 | CTGGACGCGAGCCGGGCCCTTC | 666.8 | -4.1 | 0.8 | -4.9 | 1.0E-06 | 1.9E-06 | MB231/MB468 | MP |
| >hg38_21_+_8433172_8446622_RNA45SN1_45S_50ntFlanks@6645.6662.18\|CTCGTACGACTCTTAGCG | >hg38_21_+_8433172_8446622_RNA45SN1_45S_50ntFlanks | 6645 | 6662 | 18 | CTCGTACGACTCTTAGCG | 6216.2 | 3.2 | 0.6 | 4.9 | 1.0E-06 | 1.9E-06 | BT20/MB468 | CYTO |
| >MI0000750\|hsa-mir-26a-2&WithFlank&12\|-\|57824603\|57824698@20.41.22(+2U);MI0000083\|hsa-mir-26a-1&WithFlank&3\|+\|37969398\|37969486@16.37.22(+2U)&[MIMAT0000082&hsa-miR-26a-5p&offsets\|0\|0(+2U);m-33&12\|-\|57824658\|57824679&offsets\|0\|0(+2U)];[MIMAT0000082_1&hsa-miR-26a-5p&offsets\|0\|0(+2U);m-34&3\|+\|37969413\|37969434&offsets\|0\|0(+2U)]\|TTCAAGTAATCCAGGATAGGCTTT | >MI0000750\|hsa-mir-26a-2&WithFlank&12\|-\|57824603\|57824698 | 20 | 41 | 22(+2U) | TTCAAGTAATCCAGGATAGGCTTT | 341.9 | 2.4 | 0.5 | 4.8 | 1.3E-06 | 1.9E-06 | BT20/MB468 | NUC |
| >hg38_21_+_8433172_8446622_RNA45SN1_45S_50ntFlanks@8403.8422.20\|AGGTAAACGGGTGGGGTCCG | >hg38_21_+_8433172_8446622_RNA45SN1_45S_50ntFlanks | 8403 | 8422 | 20 | AGGTAAACGGGTGGGGTCCG | 308.0 | -7.2 | 1.4 | -5.1 | 3.2E-07 | 1.9E-06 | BT20/MB468 | MP |
| >hg38_21_+_8433172_8446622_RNA45SN1_45S_50ntFlanks@11775.11794.20\|GTGACGCGCATGAATGGATG | >hg38_21_+_8433172_8446622_RNA45SN1_45S_50ntFlanks | 11775 | 11794 | 20 | GTGACGCGCATGAATGGATG | 428.2 | 3.3 | 0.7 | 4.9 | 1.2E-06 | 1.9E-06 | MB231/MB468 | CYTO |
| >hg38_MT_+_598_1651_RNR1_12S_50ntFlanks@966.1004.39\|TAACATGGTAAGTGTACTGGAAAGTGCACTTGGACGAAC | >hg38_MT_+_598_1651_RNR1_12S_50ntFlanks | 966 | 1004 | 39 | TAACATGGTAAGTGTACTGGAAAGTGCACTTGGACGAAC | 200.6 | 6.5 | 1.3 | 4.9 | 1.2E-06 | 1.9E-06 | MB231/MB468 | NUC |
| >hg38_21_+_8433172_8446622_RNA45SN1_45S_50ntFlanks@10420.10443.24\|CTGAGAGATGGGCGAGCGCCGTTC | >hg38_21_+_8433172_8446622_RNA45SN1_45S_50ntFlanks | 10420 | 10443 | 24 | CTGAGAGATGGGCGAGCGCCGTTC | 808.6 | -4.9 | 1.0 | -4.9 | 1.2E-06 | 1.9E-06 | MB231/MB468 | CYTO |

| | | | | | | | | | | | | | |
|---|---|---|---|---|---|---|---|---|---|---|---|---|---|
| >hg38_21_+_8433172_8446622_RNA45SN1_45S_50ntFlanks@11208.11228.21\|GCGGCGACTCTGGACGCGAGC | >hg38_21_+_8433172_8446622_RNA45SN1_45S_50ntFlanks | 11208 | 11228 | 21 | GCGGCGACTCTGGACGCGAGC | 1209.6 | -6.3 | 1.3 | -4.9 | 1.0E-06 | 1.9E-06 | MB231/MB468 | MP |
| >hg38_21_+_8433172_8446622_RNA45SN1_45S_50ntFlanks@12394.12419.26\|TAGCGACGTCGCTTTTTGATCCTTCG | >hg38_21_+_8433172_8446622_RNA45SN1_45S_50ntFlanks | 12394 | 12419 | 26 | TAGCGACGTCGCTTTTTGATCCTTCG | 218.0 | -7.1 | 1.5 | -4.9 | 1.0E-06 | 1.9E-06 | MB231/MB468 | MP |
| >MI0000440\|hsa-mir-27b&WithFlank&9\|+\|95085439\|95085547@67.87.21&[MIMAT0000419&hsa-miR-27b-3p&offsets\|0\|0;m-39&9\|+\|95085505\|95085525&offsets\|0\|0]\|TTCACAGTGGCTAAGTTCTGC | >MI0000440\|hsa-mir-27b&WithFlank&9\|+\|95085439\|95085547 | 67 | 87 | 21 | TTCACAGTGGCTAAGTTCTGC | 24429.7 | 3.4 | 0.7 | 4.9 | 1.0E-06 | 1.9E-06 | MB231/MB468 | MP |
| >hg38_21_+_8433172_8446622_RNA45SN1_45S_50ntFlanks@4919.4938.20\|CCACCAGGAGTGGAGCCTGC | >hg38_21_+_8433172_8446622_RNA45SN1_45S_50ntFlanks | 4919 | 4938 | 20 | CCACCAGGAGTGGAGCCTGC | 190.7 | -7.0 | 1.4 | -4.9 | 1.0E-06 | 1.9E-06 | MB231/MB468 | MP |
| >hg38_21_+_8433172_8446622_RNA45SN1_45S_50ntFlanks@8077.8104.28\|GGAAGAGCCCAGCGCCGAATCCCCGCCC | >hg38_21_+_8433172_8446622_RNA45SN1_45S_50ntFlanks | 8077 | 8104 | 28 | GGAAGAGCCCAGCGCCGAATCCCCGCCC | 364.4 | -7.8 | 1.6 | -4.9 | 1.0E-06 | 1.9E-06 | MB231/MB468 | MP |
| >hg38_21_+_8433172_8446622_RNA45SN1_45S_50ntFlanks@11540.11582.43\|CGCGGCGCCTCGCCTCGGCCGGCGCCTAGCAGCCGACTTAGAA | >hg38_21_+_8433172_8446622_RNA45SN1_45S_50ntFlanks | 11540 | 11582 | 43 | CGCGGCGCCTCGCCTCGGCCGGCGCCTAGCAGCCGACTTAGAA | 933.4 | 3.7 | 0.8 | 4.9 | 1.2E-06 | 1.9E-06 | MB231/MB468 | CYTO |
| >hg38_21_+_8433172_8446622_RNA45SN1_45S_50ntFlanks@11318.11336.19\|CCCCACGTCTCGTCGCGCG | >hg38_21_+_8433172_8446622_RNA45SN1_45S_50ntFlanks | 11318 | 11336 | 19 | CCCCACGTCTCGTCGCGCG | 691.4 | -5.4 | 1.1 | -4.9 | 1.0E-06 | 1.9E-06 | BT20/MB468 | CYTO |
| >hg38_21_+_8433172_8446622_RNA45SN1_45S_50ntFlanks@11571.11597.27\|GCCGACTTAGAACTGGTGCGGACCAGG | >hg38_21_+_8433172_8446622_RNA45SN1_45S_50ntFlanks | 11571 | 11597 | 27 | GCCGACTTAGAACTGGTGCGGACCAGG | 184.3 | 10.9 | 2.0 | 5.4 | 7.7E-08 | 1.9E-06 | BT20/MB231 | MP |
| >trna4_ArgTCG_15_+_89878304_89878376@31.53.23\|ACTTCGGATCAGAAGATTGCAGG | >trna4_ArgTCG_15_+_89878304_89878376 | 31 | 53 | 23 | ACTTCGGATCAGAAGATTGCAGG | 121.9 | -10.0 | 2.0 | -5.1 | 3.2E-07 | 1.9E-06 | BT20/MB468 | MP |
| >am_trna119_LysCTT_1_-_145395522_145395594@9.36.28,trna11_LysCTT_5_-_180648979_180649051@9.36.28,trna128_LysCTT_1_-_55423542_55423614@9.36.28,trna13_LysCTT_6_+_26556774_26556846@9.36.28,trna6_LysCTT_19_-_52425396_52425468@9.36.28,trna7_LysCTT_16_+_3225692_3225764@9.36.28,trna9_LysCTT_5_+_180634755_180634827@9.36.28\|AGCTCAGTCGGTAGAGCATGAGACTCTT | >am_trna119_LysCTT_1_-_145395522_145395594 | 9 | 36 | 28 | AGCTCAGTCGGTAGAGCATGAGACTCTT | 43.6 | 8.7 | 1.8 | 4.9 | 1.0E-06 | 1.9E-06 | MB231/MB468 | MP |
| >hg38_21_+_8433172_8446622_RNA45SN1_45S_50ntFlanks@10544.10563.20\|TCCAGTGCGGTAACGCGACC | >hg38_21_+_8433172_8446622_RNA45SN1_45S_50ntFlanks | 10544 | 10563 | 20 | TCCAGTGCGGTAACGCGACC | 56.2 | -8.9 | 1.7 | -5.1 | 3.2E-07 | 2.0E-06 | BT20/MB468 | MP |
| >MI0000542\|hsa-mir-320a&WithFlank&8\|-\|22244960\|22245043@47.68.22(+1U)&[MIMAT0000510&hsa-miR-320a-3p&offsets\|-1\|-1(+1U);m-60&8\|-\|22244975\|22244996&offsets\|-1\|-1(+1U)]\|GAAAAGCTGGGTTGAGAGGGCGT | >MI0000542\|hsa-mir-320a&WithFlank&8\|-\|22244960\|22245043 | 47 | 68 | 22(+1U) | GAAAAGCTGGGTTGAGAGGGCGT | 298.7 | 4.3 | 0.8 | 5.3 | 8.9E-08 | 2.0E-06 | BT20/MB231 | CYTO |
| >hg38_21_+_8433172_8446622_RNA45SN1_45S_50ntFlanks@9553.9573.21\|CCGTAGCGGTCCTGACGTGCA | >hg38_21_+_8433172_8446622_RNA45SN1_45S_50ntFlanks | 9553 | 9573 | 21 | CCGTAGCGGTCCTGACGTGCA | 410.9 | 2.5 | 0.5 | 4.9 | 1.2E-06 | 2.0E-06 | MB231/MB468 | CYTO |
| >hg38_21_+_8433172_8446622_RNA45SN1_45S_50ntFlanks@4005.4022.18\|AACCTCGGGCCGATCGCA | >hg38_21_+_8433172_8446622_RNA45SN1_45S_50ntFlanks | 4005 | 4022 | 18 | AACCTCGGGCCGATCGCA | 548.1 | -5.5 | 1.1 | -4.8 | 1.3E-06 | 2.0E-06 | MB231/MB468 | MITO |
| >hg38_21_+_8433172_8446622_RNA45SN1_45S_50ntFlanks@12006.12035.30\|GGCCGCCGGTGAAATACCACTACTCTGATC | >hg38_21_+_8433172_8446622_RNA45SN1_45S_50ntFlanks | 12006 | 12035 | 30 | GGCCGCCGGTGAAATACCACTACTCTGATC | 752.5 | 3.5 | 0.7 | 5.3 | 1.0E-07 | 2.0E-06 | BT20/MB231 | MITO |
| >MI0000071\|hsa-mir-17&WithFlank&13\|+\|91350599\|91350694@20.42.23&[MIMAT0000070&hsa-miR-17-5p&offsets\|0\|0;m-88&13\|+\|91350618\|91350640&offsets\|0\|0]\|CAAAGTGCTTACAGTGCAGGTAG | >MI0000071\|hsa-mir-17&WithFlank&13\|+\|91350599\|91350694 | 20 | 42 | 23 | CAAAGTGCTTACAGTGCAGGTAG | 2496.0 | 4.0 | 0.8 | 4.9 | 9.0E-07 | 2.0E-06 | BT20/MB468 | MITO |
| >MI0000764\|hsa-mir-363&WithFlank&X\|-\|134169372\|134169458@56.76.21&[MIMAT0000707&hsa-miR-363-3p&offsets\|0\|-1;m-178&X\|-\|134169381\|134169403&offsets\|0\|-2]\|AATTGCACGGTATCCATCTGT | >MI0000764\|hsa-mir-363&WithFlank&X\|-\|134169372\|134169458 | 56 | 76 | 21 | AATTGCACGGTATCCATCTGT | 145.8 | 10.5 | 2.0 | 5.4 | 7.8E-08 | 2.0E-06 | BT20/MB231 | MP |
| >hg38_21_+_8433172_8446622_RNA45SN1_45S_50ntFlanks@4003.4020.18\|ATAACCTCGGGCCGATCG | >hg38_21_+_8433172_8446622_RNA45SN1_45S_50ntFlanks | 4003 | 4020 | 18 | ATAACCTCGGGCCGATCG | 569.2 | -4.7 | 1.0 | -4.9 | 1.0E-06 | 2.0E-06 | BT20/MB468 | CYTO |
| >hg38_21_+_8433172_8446622_RNA45SN1_45S_50ntFlanks@12633.12651.19\|CGAAGCTACCATCTGTGGG | >hg38_21_+_8433172_8446622_RNA45SN1_45S_50ntFlanks | 12633 | 12651 | 19 | CGAAGCTACCATCTGTGGG | 510.4 | -5.0 | 1.0 | -4.9 | 1.0E-06 | 2.0E-06 | BT20/MB468 | CYTO |
| >hg38_MT_+_598_1651_RNR1_12S_50ntFlanks@911.946.36\|CTTCAAAGGACATTTAACTAAAACCCCTACGCATTT | >hg38_MT_+_598_1651_RNR1_12S_50ntFlanks | 911 | 946 | 36 | CTTCAAAGGACATTTAACTAAAACCCCTACGCATTT | 322.4 | 3.7 | 0.8 | 4.9 | 9.0E-07 | 2.0E-06 | BT20/MB468 | MITO |
| >hg38_21_+_8433172_8446622_RNA45SN1_45S_50ntFlanks@11718.11742.25\|AACGGCGGGAGTAACTATGACTCTC | >hg38_21_+_8433172_8446622_RNA45SN1_45S_50ntFlanks | 11718 | 11742 | 25 | AACGGCGGGAGTAACTATGACTCTC | 1466.4 | -8.3 | 1.7 | -4.8 | 1.3E-06 | 2.0E-06 | BT20/MB468 | NUC |
| >hg38_21_+_8433172_8446622_RNA45SN1_45S_50ntFlanks@9554.9596.43\|CGTAGCGGTCCTGACGTGCAAATCGGTCGTCCGACCTGGGTAT | >hg38_21_+_8433172_8446622_RNA45SN1_45S_50ntFlanks | 9554 | 9596 | 43 | CGTAGCGGTCCTGACGTGCAAATCGGTCGTCCGACCTGGGTAT | 112.8 | -9.9 | 1.9 | -5.1 | 3.2E-07 | 2.0E-06 | BT20/MB468 | MP |
| >trna116_GluCTC_1_-_145399233_145399304@35.74.40,trna134_GluTTC_1_-_16861774_16861845@35.74.40,trna71_GluCTC_1_-_161439189_161439260@35.74.40,trna74_GluCTC_1_-_161431809_161431880@35.74.40,trna77_GluCTC_1_-_161424398_161424469@35.74.40,trna77_GluCTC_6_+_28949976_28950047@35.74.40,trna80_GluCTC_1_-_161417018_161417089@35.74.40,trna84_GluTTC_1_-_161391883_161391954@35.74.40,trna87_GluCTC_6_-_126101393_126101464@35.74.40\|TCACCGCCGCGGCCCGGGTTCGATTCCCGGTCAGGGAACC | >trna116_GluCTC_1_-_145399233_145399304 | 35 | 74 | 40 | TCACCGCCGCGGCCCGGGTTCGATTCCCGGTCAGGGAACC | 1933.5 | -5.0 | 0.9 | -5.3 | 9.0E-08 | 2.0E-06 | BT20/MB231 | CYTO |
| >hg38_21_+_8433172_8446622_RNA45SN1_45S_50ntFlanks@10389.10407.19\|TGAACAGCAGTTGAACATG | >hg38_21_+_8433172_8446622_RNA45SN1_45S_50ntFlanks | 10389 | 10407 | 19 | TGAACAGCAGTTGAACATG | 2593.7 | -8.1 | 1.7 | -4.9 | 1.0E-06 | 2.0E-06 | BT20/MB468 | CYTO |
| >hg38_21_+_8433172_8446622_RNA45SN1_45S_50ntFlanks@10411.10453.43\|CAGTCGGTCCTGAGAGATGGGCGAGCGCCGTTCCGAAGGGACG | >hg38_21_+_8433172_8446622_RNA45SN1_45S_50ntFlanks | 10411 | 10453 | 43 | CAGTCGGTCCTGAGAGATGGGCGAGCGCCGTTCCGAAGGGACG | 1129.4 | -7.2 | 1.5 | -4.9 | 1.0E-06 | 2.0E-06 | BT20/MB468 | CYTO |
| >MI0000683\|hsa-mir-181b-2&WithFlank&9\|+\|124693704\|124693804@22.46.25&[MIMAT0000257_1&hsa-miR-181b-5p&offsets\|0\|+2;m-72&9\|+\|124693725\|124693747&offsets\|0\|+2]\|AACATTCATTGCTGTCGGTGGGTTT | >MI0000683\|hsa-mir-181b-2&WithFlank&9\|+\|124693704\|124693804 | 22 | 46 | 25 | AACATTCATTGCTGTCGGTGGGTTT | 966.8 | 4.1 | 0.8 | 4.9 | 1.0E-06 | 2.0E-06 | MB231/MB468 | MP |
| >hg38_21_+_8433172_8446622_RNA45SN1_45S_50ntFlanks@12181.12200.20\|TCAAACGGTAACGCAGGTGT | >hg38_21_+_8433172_8446622_RNA45SN1_45S_50ntFlanks | 12181 | 12200 | 20 | TCAAACGGTAACGCAGGTGT | 93.7 | -9.6 | 1.9 | -5.1 | 3.2E-07 | 2.0E-06 | BT20/MB468 | MP |
| >hg38_21_+_8433172_8446622_RNA45SN1_45S_50ntFlanks@9471.9503.33\|AGCACGAGCGCACGTGTTAGGACCCGAAAGATG | >hg38_21_+_8433172_8446622_RNA45SN1_45S_50ntFlanks | 9471 | 9503 | 33 | AGCACGAGCGCACGTGTTAGGACCCGAAAGATG | 893.5 | -2.6 | 0.5 | -4.9 | 9.1E-07 | 2.0E-06 | BT20/MB468 | MITO |
| >hg38_21_+_8433172_8446622_RNA45SN1_45S_50ntFlanks@12617.12654.38\|TGAGGAGCCAATGGGGCGAAGCTACCATCTGTGGGATT | >hg38_21_+_8433172_8446622_RNA45SN1_45S_50ntFlanks | 12617 | 12654 | 38 | TGAGGAGCCAATGGGGCGAAGCTACCATCTGTGGGATT | 804.0 | -3.3 | 0.7 | -4.9 | 1.0E-06 | 2.0E-06 | BT20/MB468 | CYTO |
| >MI0000734\|hsa-mir-106b&WithFlank&7\|-\|100093987\|100094080@56.76.21(+1A)&[MIMAT0004672&hsa-miR-106b-3p&offsets\|-2\|-3(+1A);m-87&7\|-\|100094003\|100094023&offsets\|-2\|-2(+1A)]\|TACCGCACTGTGGGTACTTGCA | >MI0000734\|hsa-mir-106b&WithFlank&7\|-\|100093987\|100094080 | 56 | 76 | 21(+1A) | TACCGCACTGTGGGTACTTGCA | 59.0 | 9.1 | 1.9 | 4.9 | 1.0E-06 | 2.0E-06 | MB231/MB468 | MP |
| >hg38_21_+_8433172_8446622_RNA45SN1_45S_50ntFlanks@10784.10818.35\|TGAACAGCCTCTGGCATGTTGGAACAATGTAGGTA | >hg38_21_+_8433172_8446622_RNA45SN1_45S_50ntFlanks | 10784 | 10818 | 35 | TGAACAGCCTCTGGCATGTTGGAACAATGTAGGTA | 338.9 | -7.3 | 1.5 | -4.8 | 1.3E-06 | 2.0E-06 | BT20/MB468 | NUC |
| >hg38_21_+_8433172_8446622_RNA45SN1_45S_50ntFlanks@8170.8202.33\|CAAGTCCTTCTGATCGAGGCCCAGCCCGTGGAC | >hg38_21_+_8433172_8446622_RNA45SN1_45S_50ntFlanks | 8170 | 8202 | 33 | CAAGTCCTTCTGATCGAGGCCCAGCCCGTGGAC | 70.0 | -9.2 | 1.8 | -5.1 | 3.3E-07 | 2.0E-06 | BT20/MB468 | MP |
| >MI0000081\|hsa-mir-24-2&WithFlank&19\|-\|13836281\|13836365@56.74.19(+1U);MI0000080\|hsa-mir-24-1&WithFlank&9\|+\|95086015\|95086094@50.68.19(+1U)&[MIMAT0000080&hsa-miR-24-3p&offsets\|0\|-3(+1U);m-36&19\|-\|13836288\|13836310&offsets\|0\|-4(+1U)];[MIMAT0000080_1&hsa-miR-24-3p&offsets\|0\|-3(+1U);m-35&9\|+\|95086064\|95086086&offsets\|0\|-4(+1U)]\|TGGCTCAGTTCAGCAGGAAT | >MI0000081\|hsa-mir-24-2&WithFlank&19\|-\|13836281\|13836365 | 56 | 74 | 19(+1U) | TGGCTCAGTTCAGCAGGAAT | 354.4 | 4.6 | 0.9 | 4.9 | 1.2E-06 | 2.0E-06 | MB231/MB468 | CYTO |
| >hg38_21_+_8433172_8446622_RNA45SN1_45S_50ntFlanks@9552.9571.20\|TCCGTAGCGGTCCTGACGTG | >hg38_21_+_8433172_8446622_RNA45SN1_45S_50ntFlanks | 9552 | 9571 | 20 | TCCGTAGCGGTCCTGACGTG | 287.1 | -7.0 | 1.4 | -4.9 | 1.1E-06 | 2.0E-06 | MB231/MB468 | MP |
| >hg38_21_+_8433172_8446622_RNA45SN1_45S_50ntFlanks@7975.8004.30\|CGCGACCTCAGATCAGACGTGGCGACCCGC | >hg38_21_+_8433172_8446622_RNA45SN1_45S_50ntFlanks | 7975 | 8004 | 30 | CGCGACCTCAGATCAGACGTGGCGACCCGC | 9516.0 | 3.0 | 0.6 | 4.8 | 1.3E-06 | 2.0E-06 | BT20/MB468 | NUC |
| >MI0000811\|hsa-mir-148b&WithFlank&12\|+\|54337210\|54337320@69.90.22(+1U)&[MIMAT0000759&hsa-miR-148b-3p&offsets\|0\|0(+1U);m-77&12\|+\|54337278\|54337299&offsets\|0\|0(+1U)]\|TCAGTGCATCACAGAACTTTGTT | >MI0000811\|hsa-mir-148b&WithFlank&12\|+\|54337210\|54337320 | 69 | 90 | 22(+1U) | TCAGTGCATCACAGAACTTTGTT | 1011.2 | 2.6 | 0.5 | 4.8 | 1.3E-06 | 2.0E-06 | BT20/MB468 | NUC |
| >hg38_21_+_8433172_8446622_RNA45SN1_45S_50ntFlanks@10873.10904.32\|CTGGGTCGGTCGGGCTGGGGCGCGAAGCGGGG | >hg38_21_+_8433172_8446622_RNA45SN1_45S_50ntFlanks | 10873 | 10904 | 32 | CTGGGTCGGTCGGGCTGGGGCGCGAAGCGGGG | 484.2 | -6.9 | 1.4 | -4.9 | 1.2E-06 | 2.0E-06 | MB231/MB468 | NUC |
| >hg38_21_+_8433172_8446622_RNA45SN1_45S_50ntFlanks@7974.8006.33\|ACGCGACCTCAGATCAGACGTGGCGACCCGCTG | >hg38_21_+_8433172_8446622_RNA45SN1_45S_50ntFlanks | 7974 | 8006 | 33 | ACGCGACCTCAGATCAGACGTGGCGACCCGCTG | 27048.3 | -3.8 | 0.8 | -4.8 | 1.3E-06 | 2.0E-06 | MB231/MB468 | MITO |
| >hg38_21_+_8433172_8446622_RNA45SN1_45S_50ntFlanks@12456.12474.19\|CCAAGCGTTGGATTGTTCA | >hg38_21_+_8433172_8446622_RNA45SN1_45S_50ntFlanks | 12456 | 12474 | 19 | CCAAGCGTTGGATTGTTCA | 824.4 | -6.7 | 1.4 | -4.9 | 9.2E-07 | 2.0E-06 | BT20/MB468 | MITO |
| >MI0000285\|hsa-mir-205&WithFlank&1\|+\|209432127\|209432248@40.62.23(+1U)&[MIMAT0000266&hsa-miR-205-5p&offsets\|0\|+1(+1U);m-351&1\|+\|209432166\|209432187&offsets\|0\|+1(+1U)]\|TCCTTCATTCCACCGGAGTCTGTT | >MI0000285\|hsa-mir-205&WithFlank&1\|+\|209432127\|209432248 | 40 | 62 | 23(+1U) | TCCTTCATTCCACCGGAGTCTGTT | 108.3 | 10.1 | 1.9 | 5.4 | 8.0E-08 | 2.0E-06 | BT20/MB231 | MP |
| >trna10_AspGTC_12_-_125424193_125424264@33.69.37,trna12_AspGTC_12_-_125411891_125411962@33.69.37,trna144_AspGTC_6_-_27551236_27551307@33.69.37,trna38_AspGTC_17_-_8125556_8125627@33.69.37,trna45_AspGTC_6_+_27447453_27447524@33.69.37,trna48_AspGTC_6_+_27471523_27471594@33.69.37,trna4_AspGTC_12_+_96429799_96429870@33.69.37,trna69_AspGTC_1_-_161440205_161440276@33.69.37,trna72_AspGTC_1_-_161432824_161432895@33.69.37,trna75_AspGTC_1_-_161425414_161425485@33.69.37,trna78_AspGTC_1_-_161418033_161418104@33.69.37,trna81_AspGTC_1_-_161410615_161410686@33.69.37\|TGTCACGCGGGAGACCGGGGTTCGATTCCCCGACGGG | >trna10_AspGTC_12_-_125424193_125424264 | 33 | 69 | 37 | TGTCACGCGGGAGACCGGGGTTCGATTCCCCGACGGG | 172.5 | 10.8 | 2.0 | 5.4 | 8.0E-08 | 2.0E-06 | BT20/MB231 | MP |

| | | | | | | | | | | | | | |
|---|---|---|---|---|---|---|---|---|---|---|---|---|---|
| >hg38_21_+_8433172_8446622_RNA45SN1_45S_50ntFlanks@10234.10259.26\|CGGACGCTACGCCGCGACGAGTAGGA | >hg38_21_+_8433172_8446622_RNA45SN1_45S_50ntFlanks | 10234 | 10259 | 26 | CGGACGCTACGCCGCGACGAGTAGGA | 226.6 | 11.2 | 2.1 | 5.4 | 8.0E-08 | 2.0E-06 | BT20/MB231 | MP |
| >hg38_21_+_8433172_8446622_RNA45SN1_45S_50ntFlanks@4681.4699.19\|CGCAAGACGGACCAGAGCG | >hg38_21_+_8433172_8446622_RNA45SN1_45S_50ntFlanks | 4681 | 4699 | 19 | CGCAAGACGGACCAGAGCG | 151.5 | -6.7 | 1.4 | -4.9 | 1.1E-06 | 2.0E-06 | MB231/MB468 | MP |
| >hg38_MT_+_598_1651_RNR1_12S_50ntFlanks@628.665.38\|CCGATCAACCTCACCACCTCTTGCTCAGCCTATATACC | >hg38_MT_+_598_1651_RNR1_12S_50ntFlanks | 628 | 665 | 38 | CCGATCAACCTCACCACCTCTTGCTCAGCCTATATACC | 85.7 | 10.2 | 2.0 | 5.1 | 3.3E-07 | 2.0E-06 | BT20/MB468 | MP |
| >hg38_21_+_8433172_8446622_RNA45SN1_45S_50ntFlanks@12629.12654.26\|GGGGCGAAGCTACCATCTGTGGGATT | >hg38_21_+_8433172_8446622_RNA45SN1_45S_50ntFlanks | 12629 | 12654 | 26 | GGGGCGAAGCTACCATCTGTGGGATT | 63.8 | -9.1 | 1.8 | -5.1 | 3.3E-07 | 2.0E-06 | BT20/MB468 | MP |
| >trnaMT_PheGAA_MT_+_577_647@39.59.21\|ATGTTTAGACGGGCTCACATC | >trnaMT_PheGAA_MT_+_577_647 | 39 | 59 | 21 | ATGTTTAGACGGGCTCACATC | 240.9 | 8.2 | 1.7 | 4.9 | 1.0E-06 | 2.0E-06 | BT20/MB468 | CYTO |
| >hg38_21_+_8433172_8446622_RNA45SN1_45S_50ntFlanks@11670.11703.34\|CCCAGTGCTCTGAATGTCAAAGTGAAGAAATTCA | >hg38_21_+_8433172_8446622_RNA45SN1_45S_50ntFlanks | 11670 | 11703 | 34 | CCCAGTGCTCTGAATGTCAAAGTGAAGAAATTCA | 289.0 | -7.1 | 1.4 | -4.9 | 1.1E-06 | 2.0E-06 | MB231/MB468 | MP |
| >hg38_21_+_8433172_8446622_RNA45SN1_45S_50ntFlanks@10806.10828.23\|AACAATGTAGGTAAGGGAAGTCG | >hg38_21_+_8433172_8446622_RNA45SN1_45S_50ntFlanks | 10806 | 10828 | 23 | AACAATGTAGGTAAGGGAAGTCG | 308.4 | -4.6 | 0.9 | -4.9 | 1.1E-06 | 2.0E-06 | BT20/MB468 | CYTO |
| >hg38_21_+_8433172_8446622_RNA45SN1_45S_50ntFlanks@483.521.39\|TCGATGTGGTGACGTCGTGCTCTCCCGGGCCGGGTCCGA | >hg38_21_+_8433172_8446622_RNA45SN1_45S_50ntFlanks | 483 | 521 | 39 | TCGATGTGGTGACGTCGTGCTCTCCCGGGCCGGGTCCGA | 221.6 | 4.2 | 0.9 | 4.8 | 1.2E-06 | 2.0E-06 | MB231/MB468 | NUC |
| >hg38_21_+_8433172_8446622_RNA45SN1_45S_50ntFlanks@9556.9573.18\|TAGCGGTCCTGACGTGCA | >hg38_21_+_8433172_8446622_RNA45SN1_45S_50ntFlanks | 9556 | 9573 | 18 | TAGCGGTCCTGACGTGCA | 468.7 | -5.8 | 1.2 | -4.9 | 1.1E-06 | 2.0E-06 | MB231/MB468 | MP |
| >hg38_21_+_8433172_8446622_RNA45SN1_45S_50ntFlanks@5485.5514.30\|AGCGCTGAGAAGACGGTCGAACTTGACTAT | >hg38_21_+_8433172_8446622_RNA45SN1_45S_50ntFlanks | 5485 | 5514 | 30 | AGCGCTGAGAAGACGGTCGAACTTGACTAT | 5456.1 | -7.1 | 1.5 | -4.8 | 1.3E-06 | 2.0E-06 | BT20/MB468 | NUC |
| >hg38_21_+_8433172_8446622_RNA45SN1_45S_50ntFlanks@11540.11563.24\|CGCGGCGCCTCGCCTCGGCCGGCG | >hg38_21_+_8433172_8446622_RNA45SN1_45S_50ntFlanks | 11540 | 11563 | 24 | CGCGGCGCCTCGCCTCGGCCGGCG | 158.5 | -5.7 | 1.2 | -4.9 | 1.1E-06 | 2.0E-06 | BT20/MB468 | CYTO |
| >hg38_21_+_8433172_8446622_RNA45SN1_45S_50ntFlanks@4995.5034.40\|ACAGATTGATAGCTCTTTCTCGATTCCGTGGGTGGTGGTG | >hg38_21_+_8433172_8446622_RNA45SN1_45S_50ntFlanks | 4995 | 5034 | 40 | ACAGATTGATAGCTCTTTCTCGATTCCGTGGGTGGTGGTG | 171.0 | -11.3 | 2.3 | -4.9 | 1.1E-06 | 2.0E-06 | MB231/MB468 | MP |
| >MI0000462\|hsa-mir-152&WithFlank&17\|-\|48037155\|48037253@60.81.22(+1U)&[MIMAT0000438&hsa-miR-152-3p&offsets\|0\|+1(+1U);m-97&17\|-\|48037174\|48037194&offsets\|0\|+1(+1U)]\|TCAGTGCATGACAGAACTTGGGT | >MI0000462\|hsa-mir-152&WithFlank&17\|-\|48037155\|48037253 | 60 | 81 | 22(+1U) | TCAGTGCATGACAGAACTTGGGT | 301.7 | 8.9 | 1.7 | 5.1 | 3.3E-07 | 2.0E-06 | BT20/MB468 | MP |
| >hg38_21_+_8433172_8446622_RNA45SN1_45S_50ntFlanks@10392.10418.27\|ACAGCAGTTGAACATGGGTCAGTCGGT | >hg38_21_+_8433172_8446622_RNA45SN1_45S_50ntFlanks | 10392 | 10418 | 27 | ACAGCAGTTGAACATGGGTCAGTCGGT | 381.1 | -5.2 | 1.1 | -4.8 | 1.2E-06 | 2.0E-06 | MB231/MB468 | NUC |
| >hg38_21_+_8433172_8446622_RNA45SN1_45S_50ntFlanks@8812.8835.24\|CGCGGGTCGGGGCGGCGGCGGCGG | >hg38_21_+_8433172_8446622_RNA45SN1_45S_50ntFlanks | 8812 | 8835 | 24 | CGCGGGTCGGGGCGGCGGCGGCGG | 84.0 | -9.5 | 1.9 | -5.1 | 3.3E-07 | 2.0E-06 | BT20/MB468 | MP |
| >hg38_21_+_8433172_8446622_RNA45SN1_45S_50ntFlanks@12244.12273.30\|AAGGGCAAAAGCTCGCTTGATCTTGATTTT | >hg38_21_+_8433172_8446622_RNA45SN1_45S_50ntFlanks | 12244 | 12273 | 30 | AAGGGCAAAAGCTCGCTTGATCTTGATTTT | 190.9 | -6.9 | 1.4 | -4.9 | 1.1E-06 | 2.0E-06 | MB231/MB468 | MP |
| >hg38_21_+_8433172_8446622_RNA45SN1_45S_50ntFlanks@5035.5061.27\|CATGGCCGTTCTTAGTTGGTGGAGCGA | >hg38_21_+_8433172_8446622_RNA45SN1_45S_50ntFlanks | 5035 | 5061 | 27 | CATGGCCGTTCTTAGTTGGTGGAGCGA | 179.0 | -9.6 | 2.0 | -4.9 | 9.3E-07 | 2.0E-06 | BT20/MB468 | MITO |
| >hg38_21_+_8433172_8446622_RNA45SN1_45S_50ntFlanks@3993.4013.21\|GTGACTCTAGATAACCTCGGG | >hg38_21_+_8433172_8446622_RNA45SN1_45S_50ntFlanks | 3993 | 4013 | 21 | GTGACTCTAGATAACCTCGGG | 881.1 | -7.2 | 1.5 | -4.8 | 1.3E-06 | 2.0E-06 | MB231/MB468 | MITO |
| >hg38_21_+_8433172_8446622_RNA45SN1_45S_50ntFlanks@10669.10706.38\|AAAGCGTCGCGGTTCCGGCGGCGTCCGGTGAGCTCTCG | >hg38_21_+_8433172_8446622_RNA45SN1_45S_50ntFlanks | 10669 | 10706 | 38 | AAAGCGTCGCGGTTCCGGCGGCGTCCGGTGAGCTCTCG | 2702.6 | -9.3 | 1.9 | -4.8 | 1.4E-06 | 2.0E-06 | BT20/MB468 | NUC |
| >hg38_21_+_8433172_8446622_RNA45SN1_45S_50ntFlanks@12418.12456.39\|CGATGTCGGCTCTTCCTATCATTGTGAAGCAGAATTCAC | >hg38_21_+_8433172_8446622_RNA45SN1_45S_50ntFlanks | 12418 | 12456 | 39 | CGATGTCGGCTCTTCCTATCATTGTGAAGCAGAATTCAC | 2327.1 | -4.2 | 0.9 | -4.8 | 1.2E-06 | 2.0E-06 | MB231/MB468 | NUC |
| >hg38_21_+_8433172_8446622_RNA45SN1_45S_50ntFlanks@12993.13014.22\|CCCTCGCTGCGATCTATTGAAA | >hg38_21_+_8433172_8446622_RNA45SN1_45S_50ntFlanks | 12993 | 13014 | 22 | CCCTCGCTGCGATCTATTGAAA | 592.1 | -3.3 | 0.7 | -4.8 | 1.2E-06 | 2.0E-06 | MB231/MB468 | NUC |
| >hg38_21_+_8433172_8446622_RNA45SN1_45S_50ntFlanks@10396.10430.35\|CAGTTGAACATGGGTCAGTCGGTCCTGAGAGATGG | >hg38_21_+_8433172_8446622_RNA45SN1_45S_50ntFlanks | 10396 | 10430 | 35 | CAGTTGAACATGGGTCAGTCGGTCCTGAGAGATGG | 162.6 | -6.9 | 1.4 | -4.9 | 1.1E-06 | 2.0E-06 | MB231/MB468 | MP |
| >hg38_21_+_8433172_8446622_RNA45SN1_45S_50ntFlanks@11575.11592.18\|ACTTAGAACTGGTGCGGA | >hg38_21_+_8433172_8446622_RNA45SN1_45S_50ntFlanks | 11575 | 11592 | 18 | ACTTAGAACTGGTGCGGA | 3580.5 | -6.3 | 1.3 | -4.9 | 1.1E-06 | 2.0E-06 | BT20/MB468 | CYTO |
| >hg38_21_+_8433172_8446622_RNA45SN1_45S_50ntFlanks@10389.10429.41\|TGAACAGCAGTTGAACATGGGTCAGTCGGTCCTGAGAGATG | >hg38_21_+_8433172_8446622_RNA45SN1_45S_50ntFlanks | 10389 | 10429 | 41 | TGAACAGCAGTTGAACATGGGTCAGTCGGTCCTGAGAGATG | 443.5 | -8.0 | 1.6 | -4.9 | 1.1E-06 | 2.0E-06 | MB231/MB468 | MP |
| >MI0000082\|hsa-mir-25&WithFlank&7\|-\|100093554\|100093649@58.77.20&[MIMAT0000081&hsa-miR-25-3p&offsets\|0\|-2;m-23&7\|-\|100093571\|100093592&offsets\|0\|-2]\|CATTGCACTTGTCTCGGTCT | >MI0000082\|hsa-mir-25&WithFlank&7\|-\|100093554\|100093649 | 58 | 77 | 20 | CATTGCACTTGTCTCGGTCT | 277.5 | 6.1 | 1.3 | 4.8 | 1.3E-06 | 2.0E-06 | MB231/MB468 | MITO |
| >hg38_21_+_8433172_8446622_RNA45SN1_45S_50ntFlanks@8687.8705.19\|CGGCTCCGGGACGGCTGGG | >hg38_21_+_8433172_8446622_RNA45SN1_45S_50ntFlanks | 8687 | 8705 | 19 | CGGCTCCGGGACGGCTGGG | 203.6 | 6.2 | 1.3 | 4.8 | 1.2E-06 | 2.0E-06 | MB231/MB468 | NUC |
| >hg38_21_+_8433172_8446622_RNA45SN1_45S_50ntFlanks@12948.12982.35\|GACGACCTGCTTCTGGGTCGGGGTTTCGTACGTAG | >hg38_21_+_8433172_8446622_RNA45SN1_45S_50ntFlanks | 12948 | 12982 | 35 | GACGACCTGCTTCTGGGTCGGGGTTTCGTACGTAG | 244.8 | -6.4 | 1.3 | -4.8 | 1.2E-06 | 2.0E-06 | MB231/MB468 | NUC |
| >hg38_21_+_8433172_8446622_RNA45SN1_45S_50ntFlanks@11919.11950.32\|CATGAGAGGTGTAGAATAAGTGGGAGGCCCCC | >hg38_21_+_8433172_8446622_RNA45SN1_45S_50ntFlanks | 11919 | 11950 | 32 | CATGAGAGGTGTAGAATAAGTGGGAGGCCCCC | 269.9 | -4.7 | 1.0 | -4.9 | 1.1E-06 | 2.0E-06 | BT20/MB468 | CYTO |
| >hg38_21_+_8433172_8446622_RNA45SN1_45S_50ntFlanks@9789.9820.32\|GCTGGCGTGGAGCCGGGCGTGGAATGCGAGTG | >hg38_21_+_8433172_8446622_RNA45SN1_45S_50ntFlanks | 9789 | 9820 | 32 | GCTGGCGTGGAGCCGGGCGTGGAATGCGAGTG | 361.4 | 11.8 | 2.2 | 5.4 | 8.2E-08 | 2.0E-06 | BT20/MB231 | MP |
| >MI0000082\|hsa-mir-25&WithFlank&7\|-\|100093554\|100093649@58.79.22&[MIMAT0000081&hsa-miR-25-3p&offsets\|0\|0;m-23&7\|-\|100093571\|100093592&offsets\|0\|0]\|CATTGCACTTGTCTCGGTCTGA | >MI0000082\|hsa-mir-25&WithFlank&7\|-\|100093554\|100093649 | 58 | 79 | 22 | CATTGCACTTGTCTCGGTCTGA | 37377.0 | 4.8 | 0.9 | 5.1 | 3.3E-07 | 2.0E-06 | BT20/MB468 | MP |
| >MI0003685\|hsa-mir-421&WithFlank&X\|-\|74218371\|74218467@57.75.19(+1U)&[MIMAT0003339&hsa-miR-421&offsets\|+3\|-1(+1U);m-309&X\|-\|74218392\|74218414&offsets\|+3\|-1(+1U)]\|AACAGACATTAATTGGGCGT | >MI0003685\|hsa-mir-421&WithFlank&X\|-\|74218371\|74218467 | 57 | 75 | 19(+1U) | AACAGACATTAATTGGGCGT | 87.7 | 10.2 | 2.0 | 5.1 | 3.3E-07 | 2.0E-06 | BT20/MB468 | MP |
| >hg38_21_+_8433172_8446622_RNA45SN1_45S_50ntFlanks@10131.10149.19\|CGGCGGCGGCGGCGGCGGG | >hg38_21_+_8433172_8446622_RNA45SN1_45S_50ntFlanks | 10131 | 10149 | 19 | CGGCGGCGGCGGCGGCGGG | 392.6 | -7.5 | 1.5 | -5.1 | 3.4E-07 | 2.0E-06 | BT20/MB468 | MP |
| >hg38_21_+_8433172_8446622_RNA45SN1_45S_50ntFlanks@8848.8866.19\|CGGCGGCGGCGGCGGCGGG | >hg38_21_+_8433172_8446622_RNA45SN1_45S_50ntFlanks | 8848 | 8866 | 19 | CGGCGGCGGCGGCGGCGGG | 392.6 | -7.5 | 1.5 | -5.1 | 3.4E-07 | 2.0E-06 | BT20/MB468 | MP |
| >hg38_21_+_8433172_8446622_RNA45SN1_45S_50ntFlanks@9574.9592.19\|AATCGGTCGTCCGACCTGG | >hg38_21_+_8433172_8446622_RNA45SN1_45S_50ntFlanks | 9574 | 9592 | 19 | AATCGGTCGTCCGACCTGG | 1062.9 | -5.6 | 1.1 | -5.1 | 3.4E-07 | 2.0E-06 | BT20/MB468 | MP |
| >hg38_21_+_8433172_8446622_RNA45SN1_45S_50ntFlanks@10317.10353.37\|GTGCAGATCTTGGTGGTAGTAGCAAATATTCAAACGA | >hg38_21_+_8433172_8446622_RNA45SN1_45S_50ntFlanks | 10317 | 10353 | 37 | GTGCAGATCTTGGTGGTAGTAGCAAATATTCAAACGA | 52.0 | 9.4 | 1.9 | 5.1 | 3.4E-07 | 2.0E-06 | BT20/MB468 | MP |
| >hg38_21_+_8433172_8446622_RNA45SN1_45S_50ntFlanks@10716.10736.21\|GAAAATCCGGGGGAGAGGGTG | >hg38_21_+_8433172_8446622_RNA45SN1_45S_50ntFlanks | 10716 | 10736 | 21 | GAAAATCCGGGGGAGAGGGTG | 380.0 | -4.1 | 0.8 | -4.9 | 1.1E-06 | 2.0E-06 | MB231/MB468 | MP |
| >hg38_21_+_8433172_8446622_RNA45SN1_45S_50ntFlanks@9739.9766.28\|AAACGATCTCAACCTATTCTCAAACTTT | >hg38_21_+_8433172_8446622_RNA45SN1_45S_50ntFlanks | 9739 | 9766 | 28 | AAACGATCTCAACCTATTCTCAAACTTT | 970.2 | -3.4 | 0.7 | -4.9 | 9.4E-07 | 2.0E-06 | BT20/MB468 | MITO |
| >trna10_LysCTT_16_+_3241501_3241573@1.29.29,trna13_LysCTT_14_-_58706613_58706685@1.29.29,trna2_LysCTT_15_+_79152904_79152976@1.29.29\|GCCCGGCTAGCTCAGTCGGTAGAGCATGG | >trna10_LysCTT_16_+_3241501_3241573 | 1 | 29 | 29 | GCCCGGCTAGCTCAGTCGGTAGAGCATGG | 368.0 | -4.1 | 0.8 | -4.8 | 1.3E-06 | 2.0E-06 | MB231/MB468 | MITO |
| >MI0001445\|hsa-mir-423&WithFlank&17\|+\|30117073\|30117178@58.81.24&[MIMAT0001340&hsa-miR-423-3p&offsets\|-1\|0;m-100&17\|+\|30117131\|30117153&offsets\|-1\|0]\|AAGCTCGGTCTGAGGCCCCTCAGT | >MI0001445\|hsa-mir-423&WithFlank&17\|+\|30117073\|30117178 | 58 | 81 | 24 | AAGCTCGGTCTGAGGCCCCTCAGT | 741.4 | 3.8 | 0.8 | 4.9 | 1.1E-06 | 2.0E-06 | MB231/MB468 | MP |
| >hg38_21_+_8433172_8446622_RNA45SN1_45S_50ntFlanks@3835.3871.37\|CTCCTCTCCTACTTGGATAACTGTGGTAATTCTAGAG | >hg38_21_+_8433172_8446622_RNA45SN1_45S_50ntFlanks | 3835 | 3871 | 37 | CTCCTCTCCTACTTGGATAACTGTGGTAATTCTAGAG | 389.3 | -2.6 | 0.5 | -4.9 | 9.4E-07 | 2.0E-06 | BT20/MB468 | MITO |
| >am_MI0000786\|hsa-mir-378a&WithFlank&5\|+\|149732819\|149732896@49.70.22&[MIMAT0000732&hsa-miR-378a-3p&offsets\|0\|0;m-44&5\|+\|149732867\|149732888&offsets\|0\|0]\|ACTGGACTTGGAGTCAGAAGGC | >am_MI0000786\|hsa-mir-378a&WithFlank&5\|+\|149732819\|149732896 | 49 | 70 | 22 | ACTGGACTTGGAGTCAGAAGGC | 11705.4 | 2.9 | 0.6 | 4.8 | 1.3E-06 | 2.0E-06 | MB231/MB468 | NUC |
| >hg38_21_+_8433172_8446622_RNA45SN1_45S_50ntFlanks@12408.12435.28\|TTTGATCCTTCGATGTCGGCTCTTCCTA | >hg38_21_+_8433172_8446622_RNA45SN1_45S_50ntFlanks | 12408 | 12435 | 28 | TTTGATCCTTCGATGTCGGCTCTTCCTA | 258.9 | -3.7 | 0.8 | -4.8 | 1.3E-06 | 2.0E-06 | MB231/MB468 | NUC |
| >trna130_GlnTTG_6_-_27763640_27763711@30.52.23,trna16_GlnTTG_17_+_47269890_47269961@30.52.23,trna173_GlnTTG_6_-_26311975_26312046@30.52.23,trna174_GlnTTG_6_-_26311424_26311495@30.52.23\|GACTTTGAATCCAGCGATCCGAG | >trna130_GlnTTG_6_-_27763640_27763711 | 30 | 52 | 23 | GACTTTGAATCCAGCGATCCGAG | 83.5 | -9.5 | 1.9 | -5.1 | 3.4E-07 | 2.0E-06 | BT20/MB468 | MP |
| >hg38_21_+_8433172_8446622_RNA45SN1_45S_50ntFlanks@12638.12661.24\|CTACCATCTGTGGGATTATGACTG | >hg38_21_+_8433172_8446622_RNA45SN1_45S_50ntFlanks | 12638 | 12661 | 24 | CTACCATCTGTGGGATTATGACTG | 4307.9 | -5.2 | 1.0 | -5.1 | 3.4E-07 | 2.0E-06 | BT20/MB468 | MP |
| >MI0000456\|hsa-mir-140&WithFlank&16\|+\|69933075\|69933186@68.90.23(+1C)&[MIMAT0004597&hsa-miR-140-3p&offsets\|0\|+2(+1C);m-45&16\|+\|69933143\|69933164&offsets\|-1\|0(+1C)]\|TACCACAGGGTAGAACCACGGACC | >MI0000456\|hsa-mir-140&WithFlank&16\|+\|69933075\|69933186 | 68 | 90 | 23(+1C) | TACCACAGGGTAGAACCACGGACC | 51.0 | 8.9 | 1.8 | 4.9 | 1.1E-06 | 2.0E-06 | MB231/MB468 | MP |
| >hg38_21_+_8433172_8446622_RNA45SN1_45S_50ntFlanks@8178.8212.35\|TCTGATCGAGGCCCAGCCCGTGGACGGTGTGAGGC | >hg38_21_+_8433172_8446622_RNA45SN1_45S_50ntFlanks | 8178 | 8212 | 35 | TCTGATCGAGGCCCAGCCCGTGGACGGTGTGAGGC | 152.4 | -7.0 | 1.4 | -4.9 | 1.1E-06 | 2.0E-06 | MB231/MB468 | MP |
| >hg38_21_+_8433172_8446622_RNA45SN1_45S_50ntFlanks@9536.9563.28\|GAAACTCTGGTGGAGGTCCGTAGCGGTC | >hg38_21_+_8433172_8446622_RNA45SN1_45S_50ntFlanks | 9536 | 9563 | 28 | GAAACTCTGGTGGAGGTCCGTAGCGGTC | 178.9 | -6.9 | 1.4 | -4.9 | 1.1E-06 | 2.0E-06 | MB231/MB468 | MP |
| >hg38_21_+_8433172_8446622_RNA45SN1_45S_50ntFlanks@11323.11342.20\|CGTCTCGTCGCGCGCGCGTC | >hg38_21_+_8433172_8446622_RNA45SN1_45S_50ntFlanks | 11323 | 11342 | 20 | CGTCTCGTCGCGCGCGCGTC | 9434.7 | -4.7 | 1.0 | -4.9 | 1.1E-06 | 2.0E-06 | BT20/MB468 | CYTO |
| >hg38_21_+_8433172_8446622_RNA45SN1_45S_50ntFlanks@11124.11146.23\|CCGGCGGCGGCCCGCCGCGGGGC | >hg38_21_+_8433172_8446622_RNA45SN1_45S_50ntFlanks | 11124 | 11146 | 23 | CCGGCGGCGGCCCGCCGCGGGGC | 125.3 | -10.0 | 2.0 | -5.1 | 3.4E-07 | 2.0E-06 | BT20/MB468 | MP |
| >hg38_21_+_8433172_8446622_RNA45SN1_45S_50ntFlanks@5485.5519.35\|AGCGCTGAGAAGACGGTCGAACTTGACTATCTAGA | >hg38_21_+_8433172_8446622_RNA45SN1_45S_50ntFlanks | 5485 | 5519 | 35 | AGCGCTGAGAAGACGGTCGAACTTGACTATCTAGA | 474.1 | 3.9 | 0.8 | 4.9 | 1.1E-06 | 2.0E-06 | BT20/MB468 | CYTO |

<table>
<tr><td>>trna10_GluCTC_3_-_103879524_103879595@16.34.19,trna116_GluCTC_1_-_145399233_145399304@16.34.19,trna25_GluCTC_2_-_71273488_71273560@16.34.19,trna59_GluCTC_1_+_249168447_249168518@16.34.19,trna71_GluCTC_1_-_161439189_161439260@16.34.19,trna74_GluCTC_1_-_161431809_161431880@16.34.19,trna77_GluCTC_1_-_161424398_161424469@16.34.19,trna77_GluCTC_6_+_28949976_28950047@16.34.19,trna80_GluCTC_1_-_161417018_161417089@16.34.19,trna87_GluCTC_6_-_126101393_126101464@16.34.19|TGGTTAGGATTCGGCGCTC</td><td>>trna10_GluCTC_3_-_103879524_103879595</td><td>16</td><td>34</td><td>19</td><td>TGGTTAGGATTCGGCGCTC</td><td>825.0</td><td>6.2</td><td>1.3</td><td>4.9</td><td>1.1E-06</td><td>2.0E-06</td><td>BT20/MB468</td><td>CYTO</td></tr>
<tr><td>>hg38_21_+_8433172_8446622_RNA45SN1_45S_50ntFlanks@12615.12653.39|GCTGAGGAGCCAATGGGGCGAAGCTACCATCTGTGGGAT</td><td>>hg38_21_+_8433172_8446622_RNA45SN1_45S_50ntFlanks</td><td>12615</td><td>12653</td><td>39</td><td>GCTGAGGAGCCAATGGGGCGAAGCTACCATCTGTGGGAT</td><td>1100.9</td><td>-5.0</td><td>1.0</td><td>-4.9</td><td>1.2E-06</td><td>2.0E-06</td><td>MB231/MB468</td><td>CYTO</td></tr>
<tr><td>>hg38_21_+_8433172_8446622_RNA45SN1_45S_50ntFlanks@10690.10709.20|CGTCCGGTGAGCTCTCGCTG</td><td>>hg38_21_+_8433172_8446622_RNA45SN1_45S_50ntFlanks</td><td>10690</td><td>10709</td><td>20</td><td>CGTCCGGTGAGCTCTCGCTG</td><td>3740.4</td><td>-9.0</td><td>1.9</td><td>-4.8</td><td>1.4E-06</td><td>2.0E-06</td><td>BT20/MB468</td><td>NUC</td></tr>
<tr><td>>trna116_GluCTC_1_-_145399233_145399304@32.69.38,trna59_GluCTC_1_+_249168447_249168518@32.69.38,trna71_GluCTC_1_-_161439189_161439260@32.69.38,trna74_GluCTC_1_-_161431809_161431880@32.69.38,trna77_GluCTC_1_-_161424398_161424469@32.69.38,trna77_GluCTC_6_+_28949976_28950047@32.69.38,trna80_GluCTC_1_-_161417018_161417089@32.69.38,trna87_GluCTC_6_-_126101393_126101464@32.69.38|CTCTCACCGCCGCGGCCCGGGTTCGATTCCCGGTCAGG</td><td>>trna116_GluCTC_1_-_145399233_145399304</td><td>32</td><td>69</td><td>38</td><td>CTCTCACCGCCGCGGCCCGGGTTCGATTCCCGGTCAGG</td><td>517.9</td><td>-3.3</td><td>0.7</td><td>-4.9</td><td>1.1E-06</td><td>2.0E-06</td><td>BT20/MB468</td><td>CYTO</td></tr>
<tr><td>>hg38_21_+_8433172_8446622_RNA45SN1_45S_50ntFlanks@10798.10818.21|CATGTTGGAACAATGTAGGTA</td><td>>hg38_21_+_8433172_8446622_RNA45SN1_45S_50ntFlanks</td><td>10798</td><td>10818</td><td>21</td><td>CATGTTGGAACAATGTAGGTA</td><td>1519.3</td><td>-6.2</td><td>1.2</td><td>-5.1</td><td>3.4E-07</td><td>2.1E-06</td><td>BT20/MB468</td><td>MP</td></tr>
<tr><td>>MI0000103|hsa-mir-101-1&WithFlank&1|-|65058428|65058514@53.73.21(+1A);MI0000739|hsa-mir-101-2&WithFlank&9|+|4850291|4850381@55.75.21(+1A)&[MIMAT0000099&hsa-miR-101-3p&offsets|0|0(+1A);m-17&1|-|65058443|65058462&offsets|0|+1(+1A)];[MIMAT0000099_1&hsa-miR-101-3p&offsets|0|0(+1A);m-16&9|+|4850345|4850364&offsets|0|+1(+1A)]|TACAGTACTGTGATAACTGAAA</td><td>>MI0000103|hsa-mir-101-1&WithFlank&1|-|65058428|65058514</td><td>53</td><td>73</td><td>21(+1A)</td><td>TACAGTACTGTGATAACTGAAA</td><td>228.8</td><td>5.5</td><td>1.1</td><td>4.8</td><td>1.3E-06</td><td>2.1E-06</td><td>MB231/MB468</td><td>NUC</td></tr>
<tr><td>>hg38_21_+_8433172_8446622_RNA45SN1_45S_50ntFlanks@10725.10752.28|GGGGAGAGGGTGTAAATCTCGCGCCGGG</td><td>>hg38_21_+_8433172_8446622_RNA45SN1_45S_50ntFlanks</td><td>10725</td><td>10752</td><td>28</td><td>GGGGAGAGGGTGTAAATCTCGCGCCGGG</td><td>107.9</td><td>-5.6</td><td>1.2</td><td>-4.9</td><td>1.1E-06</td><td>2.1E-06</td><td>BT20/MB468</td><td>CYTO</td></tr>
<tr><td>>hg38_21_+_8433172_8446622_RNA45SN1_45S_50ntFlanks@8343.8365.23|GTAAGGGAAAGTTGAAAAGAACT</td><td>>hg38_21_+_8433172_8446622_RNA45SN1_45S_50ntFlanks</td><td>8343</td><td>8365</td><td>23</td><td>GTAAGGGAAAGTTGAAAAGAACT</td><td>57.8</td><td>9.6</td><td>1.9</td><td>5.1</td><td>3.4E-07</td><td>2.1E-06</td><td>BT20/MB468</td><td>MP</td></tr>
<tr><td>>hg38_21_+_8433172_8446622_RNA45SN1_45S_50ntFlanks@12456.12481.26|CCAAGCGTTGGATTGTTCACCCACTA</td><td>>hg38_21_+_8433172_8446622_RNA45SN1_45S_50ntFlanks</td><td>12456</td><td>12481</td><td>26</td><td>CCAAGCGTTGGATTGTTCACCCACTA</td><td>395.9</td><td>-4.9</td><td>1.0</td><td>-4.9</td><td>9.5E-07</td><td>2.1E-06</td><td>BT20/MB468</td><td>MITO</td></tr>
<tr><td>>hg38_21_+_8433172_8446622_RNA45SN1_45S_50ntFlanks@7986.8020.35|ATCAGACGTGGCGACCCGCTGAATTTAAGCATATT</td><td>>hg38_21_+_8433172_8446622_RNA45SN1_45S_50ntFlanks</td><td>7986</td><td>8020</td><td>35</td><td>ATCAGACGTGGCGACCCGCTGAATTTAAGCATATT</td><td>90.6</td><td>-8.6</td><td>1.8</td><td>-4.9</td><td>9.5E-07</td><td>2.1E-06</td><td>BT20/MB468</td><td>MITO</td></tr>
<tr><td>>hg38_21_+_8433172_8446622_RNA45SN1_45S_50ntFlanks@7981.8005.25|CTCAGATCAGACGTGGCGACCCGCT</td><td>>hg38_21_+_8433172_8446622_RNA45SN1_45S_50ntFlanks</td><td>7981</td><td>8005</td><td>25</td><td>CTCAGATCAGACGTGGCGACCCGCT</td><td>141.0</td><td>10.5</td><td>2.0</td><td>5.4</td><td>8.3E-08</td><td>2.1E-06</td><td>BT20/MB231</td><td>MP</td></tr>
<tr><td>>hg38_21_+_8433172_8446622_RNA45SN1_45S_50ntFlanks@8113.8140.28|GGCGCGGGACATGTGGCGTACGGAAGAC</td><td>>hg38_21_+_8433172_8446622_RNA45SN1_45S_50ntFlanks</td><td>8113</td><td>8140</td><td>28</td><td>GGCGCGGGACATGTGGCGTACGGAAGAC</td><td>331.2</td><td>11.7</td><td>2.2</td><td>5.4</td><td>8.3E-08</td><td>2.1E-06</td><td>BT20/MB231</td><td>MP</td></tr>
<tr><td>>hg38_21_+_8433172_8446622_RNA45SN1_45S_50ntFlanks@5078.5110.33|CGATAACGAACGAGACTCTGGCATGCTAACTAG</td><td>>hg38_21_+_8433172_8446622_RNA45SN1_45S_50ntFlanks</td><td>5078</td><td>5110</td><td>33</td><td>CGATAACGAACGAGACTCTGGCATGCTAACTAG</td><td>606.4</td><td>-8.1</td><td>1.7</td><td>-4.8</td><td>1.4E-06</td><td>2.1E-06</td><td>BT20/MB468</td><td>NUC</td></tr>
<tr><td>>hg38_21_+_8433172_8446622_RNA45SN1_45S_50ntFlanks@10402.10429.28|AACATGGGTCAGTCGGTCCTGAGAGATG</td><td>>hg38_21_+_8433172_8446622_RNA45SN1_45S_50ntFlanks</td><td>10402</td><td>10429</td><td>28</td><td>AACATGGGTCAGTCGGTCCTGAGAGATG</td><td>3875.8</td><td>-4.2</td><td>0.8</td><td>-5.1</td><td>3.4E-07</td><td>2.1E-06</td><td>BT20/MB468</td><td>MP</td></tr>
<tr><td>>hg38_21_+_8433172_8446622_RNA45SN1_45S_50ntFlanks@4590.4615.26|ATTTTGTTGGTTTTCGGAACTGAGGC</td><td>>hg38_21_+_8433172_8446622_RNA45SN1_45S_50ntFlanks</td><td>4590</td><td>4615</td><td>26</td><td>ATTTTGTTGGTTTTCGGAACTGAGGC</td><td>263.9</td><td>-7.0</td><td>1.5</td><td>-4.8</td><td>1.4E-06</td><td>2.1E-06</td><td>BT20/MB468</td><td>NUC</td></tr>
<tr><td>>hg38_MT_+_1621_3279_RNR2_16S_50ntFlanks@51.79.29|GCTAAACCTAGCCCCAAACCCACTCCACC</td><td>>hg38_MT_+_1621_3279_RNR2_16S_50ntFlanks</td><td>51</td><td>79</td><td>29</td><td>GCTAAACCTAGCCCCAAACCCACTCCACC</td><td>475.3</td><td>3.5</td><td>0.7</td><td>4.8</td><td>1.3E-06</td><td>2.1E-06</td><td>MB231/MB468</td><td>NUC</td></tr>
<tr><td>>hg38_MT_+_1621_3279_RNR2_16S_50ntFlanks@1095.1132.38|AGACGAGAAGACCCTATGGAGCTTTAATTTATTAATGC</td><td>>hg38_MT_+_1621_3279_RNR2_16S_50ntFlanks</td><td>1095</td><td>1132</td><td>38</td><td>AGACGAGAAGACCCTATGGAGCTTTAATTTATTAATGC</td><td>97.2</td><td>10.3</td><td>2.0</td><td>5.1</td><td>3.4E-07</td><td>2.1E-06</td><td>BT20/MB468</td><td>MP</td></tr>
<tr><td>>hg38_21_+_8433172_8446622_RNA45SN1_45S_50ntFlanks@9568.9601.34|CGTGCAAATCGGTCGTCCGACCTGGGTATAGGGG</td><td>>hg38_21_+_8433172_8446622_RNA45SN1_45S_50ntFlanks</td><td>9568</td><td>9601</td><td>34</td><td>CGTGCAAATCGGTCGTCCGACCTGGGTATAGGGG</td><td>199.2</td><td>-7.3</td><td>1.5</td><td>-4.9</td><td>1.1E-06</td><td>2.1E-06</td><td>MB231/MB468</td><td>MP</td></tr>
<tr><td>>hg38_21_+_8433172_8446622_RNA45SN1_45S_50ntFlanks@6721.6756.36|AATTGCAGGACACATTGATCATCGACACTTCGAACG</td><td>>hg38_21_+_8433172_8446622_RNA45SN1_45S_50ntFlanks</td><td>6721</td><td>6756</td><td>36</td><td>AATTGCAGGACACATTGATCATCGACACTTCGAACG</td><td>1557.8</td><td>-8.5</td><td>1.8</td><td>-4.8</td><td>1.4E-06</td><td>2.1E-06</td><td>BT20/MB468</td><td>NUC</td></tr>
<tr><td>>trnaMT_GluTTC_MT_-_14674_14742@26.65.40,trnalookalike8_GluTTC_5_-_93905172_93905240@26.65.40|GGTTTTTCATATCATTGGTCGTGGTTGTAGTCCGTGCGAG</td><td>>trnaMT_GluTTC_MT_-_14674_14742</td><td>26</td><td>65</td><td>40</td><td>GGTTTTTCATATCATTGGTCGTGGTTGTAGTCCGTGCGAG</td><td>88.0</td><td>10.2</td><td>2.0</td><td>5.1</td><td>3.4E-07</td><td>2.1E-06</td><td>BT20/MB468</td><td>MP</td></tr>
<tr><td>>hg38_21_+_8433172_8446622_RNA45SN1_45S_50ntFlanks@9570.9597.28|TGCAAATCGGTCGTCCGACCTGGGTATA</td><td>>hg38_21_+_8433172_8446622_RNA45SN1_45S_50ntFlanks</td><td>9570</td><td>9597</td><td>28</td><td>TGCAAATCGGTCGTCCGACCTGGGTATA</td><td>2477.2</td><td>-7.1</td><td>1.4</td><td>-5.1</td><td>3.4E-07</td><td>2.1E-06</td><td>BT20/MB468</td><td>MP</td></tr>
<tr><td>>trna10_IleTAT_19_-_39902808_39902900@33.52.20,trna29_IleTAT_6_+_26988125_26988218@33.52.20,trna55_IleTAT_6_+_27599200_27599293@33.52.20,trna5_IleTAT_2_+_43037676_43037768@33.52.20,trna63_IleTAT_6_+_28505367_28505460@33.52.20|CTTATAATGCCGAGGTTGTG</td><td>>trna10_IleTAT_19_-_39902808_39902900</td><td>33</td><td>52</td><td>20</td><td>CTTATAATGCCGAGGTTGTG</td><td>69.1</td><td>-9.2</td><td>1.8</td><td>-5.1</td><td>3.4E-07</td><td>2.1E-06</td><td>BT20/MB468</td><td>MP</td></tr>
<tr><td>>trnaMT_GluTTC_MT_-_14674_14742@39.72.34,trnalookalike8_GluTTC_5_-_93905172_93905240@39.72.34|ATTGGTCGTGGTTGTAGTCCGTGCGAGAATACCA</td><td>>trnaMT_GluTTC_MT_-_14674_14742</td><td>39</td><td>72</td><td>34</td><td>ATTGGTCGTGGTTGTAGTCCGTGCGAGAATACCA</td><td>5017.1</td><td>7.1</td><td>1.5</td><td>4.9</td><td>1.1E-06</td><td>2.1E-06</td><td>BT20/MB468</td><td>CYTO</td></tr>
<tr><td>>hg38_21_+_8433172_8446622_RNA45SN1_45S_50ntFlanks@3842.3862.21|CCTACTTGGATAACTGTGGTA</td><td>>hg38_21_+_8433172_8446622_RNA45SN1_45S_50ntFlanks</td><td>3842</td><td>3862</td><td>21</td><td>CCTACTTGGATAACTGTGGTA</td><td>698.3</td><td>3.8</td><td>0.8</td><td>4.9</td><td>1.1E-06</td><td>2.1E-06</td><td>BT20/MB468</td><td>CYTO</td></tr>
<tr><td>>hg38_21_+_8433172_8446622_RNA45SN1_45S_50ntFlanks@4722.4752.31|TCATTAATCAAGAACGAAAGTCGGAGGTTCG</td><td>>hg38_21_+_8433172_8446622_RNA45SN1_45S_50ntFlanks</td><td>4722</td><td>4752</td><td>31</td><td>TCATTAATCAAGAACGAAAGTCGGAGGTTCG</td><td>953.7</td><td>-7.3</td><td>1.5</td><td>-4.9</td><td>1.1E-06</td><td>2.1E-06</td><td>BT20/MB468</td><td>CYTO</td></tr>
<tr><td>>hg38_MT_+_1621_3279_RNR2_16S_50ntFlanks@253.282.30|AACTTTGCAAGGAGAGCCAAAGCTAAGACC</td><td>>hg38_MT_+_1621_3279_RNR2_16S_50ntFlanks</td><td>253</td><td>282</td><td>30</td><td>AACTTTGCAAGGAGAGCCAAAGCTAAGACC</td><td>50.2</td><td>8.9</td><td>1.8</td><td>4.9</td><td>1.1E-06</td><td>2.1E-06</td><td>MB231/MB468</td><td>MP</td></tr>
<tr><td>>hg38_21_+_8433172_8446622_RNA45SN1_45S_50ntFlanks@12413.12431.19|TCCTTCGATGTCGGCTCTT</td><td>>hg38_21_+_8433172_8446622_RNA45SN1_45S_50ntFlanks</td><td>12413</td><td>12431</td><td>19</td><td>TCCTTCGATGTCGGCTCTT</td><td>343.4</td><td>-2.9</td><td>0.6</td><td>-4.8</td><td>1.2E-06</td><td>2.1E-06</td><td>MB231/MB468</td><td>CYTO</td></tr>
<tr><td>>hg38_21_+_8433172_8446622_RNA45SN1_45S_50ntFlanks@8113.8138.26|GGCGCGGGACATGTGGCGTACGGAAG</td><td>>hg38_21_+_8433172_8446622_RNA45SN1_45S_50ntFlanks</td><td>8113</td><td>8138</td><td>26</td><td>GGCGCGGGACATGTGGCGTACGGAAG</td><td>351.8</td><td>-7.7</td><td>1.6</td><td>-4.8</td><td>1.4E-06</td><td>2.1E-06</td><td>BT20/MB468</td><td>NUC</td></tr>
<tr><td>>hg38_21_+_8433172_8446622_RNA45SN1_45S_50ntFlanks@11573.11613.41|CGACTTAGAACTGGTGCGGACCAGGGGAATCCGACTGTTTA</td><td>>hg38_21_+_8433172_8446622_RNA45SN1_45S_50ntFlanks</td><td>11573</td><td>11613</td><td>41</td><td>CGACTTAGAACTGGTGCGGACCAGGGGAATCCGACTGTTTA</td><td>1898.8</td><td>4.8</td><td>1.0</td><td>4.9</td><td>9.6E-07</td><td>2.1E-06</td><td>BT20/MB468</td><td>MITO</td></tr>
<tr><td>>MI0000103|hsa-mir-101-1&WithFlank&1|-|65058428|65058514@53.72.20;MI0000739|hsa-mir-101-2&WithFlank&9|+|4850291|4850381@55.74.20&[MIMAT0000099&hsa-miR-101-3p&offsets|0|-1;m-17&1|-|65058443|65058462&offsets|0|0];[MIMAT0000099_1&hsa-miR-101-3p&offsets|0|-1;m-16&9|+|4850345|4850364&offsets|0|0]|TACAGTACTGTGATAACTGA</td><td>>MI0000103|hsa-mir-101-1&WithFlank&1|-|65058428|65058514</td><td>53</td><td>72</td><td>20</td><td>TACAGTACTGTGATAACTGA</td><td>819.9</td><td>5.0</td><td>1.0</td><td>4.8</td><td>1.3E-06</td><td>2.1E-06</td><td>MB231/MB468</td><td>MITO</td></tr>
<tr><td>>hg38_21_+_8433172_8446622_RNA45SN1_45S_50ntFlanks@12822.12859.38|CCGGGACCGGGGTCCGGTGCGGAGTGCCCTTCGTCCTG</td><td>>hg38_21_+_8433172_8446622_RNA45SN1_45S_50ntFlanks</td><td>12822</td><td>12859</td><td>38</td><td>CCGGGACCGGGGTCCGGTGCGGAGTGCCCTTCGTCCTG</td><td>56.9</td><td>-8.9</td><td>1.7</td><td>-5.1</td><td>3.5E-07</td><td>2.1E-06</td><td>BT20/MB468</td><td>MP</td></tr>
<tr><td>>hg38_21_+_8433172_8446622_RNA45SN1_45S_50ntFlanks@5491.5516.26|GAGAAGACGGTCGAACTTGACTATCT</td><td>>hg38_21_+_8433172_8446622_RNA45SN1_45S_50ntFlanks</td><td>5491</td><td>5516</td><td>26</td><td>GAGAAGACGGTCGAACTTGACTATCT</td><td>368.3</td><td>3.1</td><td>0.6</td><td>4.8</td><td>1.3E-06</td><td>2.1E-06</td><td>MB231/MB468</td><td>NUC</td></tr>
<tr><td>>MI0000446|hsa-mir-125b-1&WithFlank&11|-|122099751|122099850@21.41.21;MI0000470|hsa-mir-125b-2&WithFlank&21|+|16590231|16590331@23.43.21&[MIMAT0000423&hsa-miR-125b-5p&offsets|0|-1;m-67&11|-|122099809|122099830&offsets|0|-1];[MIMAT0000423_1&hsa-miR-125b-5p&offsets|0|-1;m-68&21|+|16590253|16590274&offsets|0|-1]|TCCCTGAGACCCTAACTTGTG</td><td>>MI0000446|hsa-mir-125b-1&WithFlank&11|-|122099751|122099850</td><td>21</td><td>41</td><td>21</td><td>TCCCTGAGACCCTAACTTGTG</td><td>1426.6</td><td>5.1</td><td>1.0</td><td>4.9</td><td>1.1E-06</td><td>2.1E-06</td><td>BT20/MB468</td><td>CYTO</td></tr>
<tr><td>>hg38_21_+_8433172_8446622_RNA45SN1_45S_50ntFlanks@10843.10867.25|TAACTTCGGGATAAGGATTGGCTCT</td><td>>hg38_21_+_8433172_8446622_RNA45SN1_45S_50ntFlanks</td><td>10843</td><td>10867</td><td>25</td><td>TAACTTCGGGATAAGGATTGGCTCT</td><td>709.7</td><td>-7.9</td><td>1.6</td><td>-4.8</td><td>1.4E-06</td><td>2.1E-06</td><td>BT20/MB468</td><td>NUC</td></tr>
<tr><td>>hg38_21_+_8433172_8446622_RNA45SN1_45S_50ntFlanks@9533.9574.42|GAGGAAACTCTGGTGGAGGTCCGTAGCGGTCCTGACGTGCAA</td><td>>hg38_21_+_8433172_8446622_RNA45SN1_45S_50ntFlanks</td><td>9533</td><td>9574</td><td>42</td><td>GAGGAAACTCTGGTGGAGGTCCGTAGCGGTCCTGACGTGCAA</td><td>698.3</td><td>-8.4</td><td>1.7</td><td>-4.8</td><td>1.4E-06</td><td>2.1E-06</td><td>BT20/MB468</td><td>NUC</td></tr>
<tr><td>>MI0000650|hsa-mir-200c&WithFlank&12|+|6963693|6963772@49.72.24&[MIMAT0000617&hsa-miR-200c-3p&offsets|-1|0;m-54&12|+|6963742|6963764&offsets|-1|0]|CTAATACTGCCGGGTAATGATGGA</td><td>>MI0000650|hsa-mir-200c&WithFlank&12|+|6963693|6963772</td><td>49</td><td>72</td><td>24</td><td>CTAATACTGCCGGGTAATGATGGA</td><td>126.1</td><td>10.3</td><td>1.9</td><td>5.4</td><td>8.5E-08</td><td>2.1E-06</td><td>BT20/MB231</td><td>MP</td></tr>
<tr><td>>MI0000088|hsa-mir-30a&WithFlank&6|-|71403545|71403627@53.73.21(+1A)&[MIMAT0000088&hsa-miR-30a-3p&offsets|0|-1(+1A);m-38&6|-|71403554|71403575&offsets|0|-1(+1A)]|CTTTCAGTCGGATGTTTGCAGA</td><td>>MI0000088|hsa-mir-30a&WithFlank&6|-|71403545|71403627</td><td>53</td><td>73</td><td>21(+1A)</td><td>CTTTCAGTCGGATGTTTGCAGA</td><td>492.9</td><td>7.0</td><td>1.5</td><td>4.8</td><td>1.3E-06</td><td>2.1E-06</td><td>MB231/MB468</td><td>MITO</td></tr>
<tr><td>>hg38_21_+_8433172_8446622_RNA45SN1_45S_50ntFlanks@12001.12031.31|CTGCGGGCCGCCGGTGAAATACCACTACTCT</td><td>>hg38_21_+_8433172_8446622_RNA45SN1_45S_50ntFlanks</td><td>12001</td><td>12031</td><td>31</td><td>CTGCGGGCCGCCGGTGAAATACCACTACTCT</td><td>767.1</td><td>-5.8</td><td>1.2</td><td>-4.8</td><td>1.3E-06</td><td>2.1E-06</td><td>MB231/MB468</td><td>MITO</td></tr>
<tr><td>>hg38_21_+_8433172_8446622_RNA45SN1_45S_50ntFlanks@9504.9525.22|GTGAACTATGCCTGGGCAGGGC</td><td>>hg38_21_+_8433172_8446622_RNA45SN1_45S_50ntFlanks</td><td>9504</td><td>9525</td><td>22</td><td>GTGAACTATGCCTGGGCAGGGC</td><td>2746.2</td><td>-5.0</td><td>1.0</td><td>-5.1</td><td>3.5E-07</td><td>2.1E-06</td><td>BT20/MB468</td><td>MP</td></tr>
<tr><td>>hg38_21_+_8433172_8446622_RNA45SN1_45S_50ntFlanks@9576.9594.19|TCGGTCGTCCGACCTGGGT</td><td>>hg38_21_+_8433172_8446622_RNA45SN1_45S_50ntFlanks</td><td>9576</td><td>9594</td><td>19</td><td>TCGGTCGTCCGACCTGGGT</td><td>248.8</td><td>-6.9</td><td>1.4</td><td>-4.9</td><td>1.1E-06</td><td>2.1E-06</td><td>MB231/MB468</td><td>MP</td></tr>
<tr><td>>hg38_21_+_8433172_8446622_RNA45SN1_45S_50ntFlanks@5078.5117.40|CGATAACGAACGAGACTCTGGCATGCTAACTAGTTACGCG</td><td>>hg38_21_+_8433172_8446622_RNA45SN1_45S_50ntFlanks</td><td>5078</td><td>5117</td><td>40</td><td>CGATAACGAACGAGACTCTGGCATGCTAACTAGTTACGCG</td><td>397.2</td><td>-7.7</td><td>1.6</td><td>-4.9</td><td>1.1E-06</td><td>2.1E-06</td><td>MB231/MB468</td><td>MP</td></tr>
<tr><td>>hg38_21_+_8433172_8446622_RNA45SN1_45S_50ntFlanks@12996.13036.41|TCGCTGCGATCTATTGAAAGTCAGCCCTCGACACAAGGGTT</td><td>>hg38_21_+_8433172_8446622_RNA45SN1_45S_50ntFlanks</td><td>12996</td><td>13036</td><td>41</td><td>TCGCTGCGATCTATTGAAAGTCAGCCCTCGACACAAGGGTT</td><td>2721.5</td><td>6.8</td><td>1.4</td><td>4.9</td><td>9.7E-07</td><td>2.1E-06</td><td>BT20/MB468</td><td>MITO</td></tr>
<tr><td>>hg38_21_+_8433172_8446622_RNA45SN1_45S_50ntFlanks@12457.12486.30|CAAGCGTTGGATTGTTCACCCACTAATAGG</td><td>>hg38_21_+_8433172_8446622_RNA45SN1_45S_50ntFlanks</td><td>12457</td><td>12486</td><td>30</td><td>CAAGCGTTGGATTGTTCACCCACTAATAGG</td><td>169.9</td><td>-6.7</td><td>1.4</td><td>-4.9</td><td>1.1E-06</td><td>2.1E-06</td><td>MB231/MB468</td><td>MP</td></tr>
</table>

| | | | | | | | | | | | | | |
|---|---|---|---|---|---|---|---|---|---|---|---|---|---|
| >trna2_GlyGCC_21_-_18827107_18827177@3.31.29,trna35_GlyGCC_1_+_161413094_161413164@3.31.29,trna37_GlyGCC_1_+_161420467_161420537@3.31.29,trna39_GlyGCC_1_+_161427898_161427968@3.31.29,trna41_GlyGCC_1_+_161435258_161435328@3.31.29\|ATGGGTGGTTCAGTGGTAGAATTCTCGCC | >trna2_GlyGCC_21_-_18827107_18827177 | 3 | 31 | 29 | ATGGGTGGTTCAGTGGTAGAATTCTCGCC | 106.6 | 10.1 | 1.9 | 5.4 | 8.6E-08 | 2.1E-06 | BT20/MB231 | MP |
| >MI0000101\|hsa-mir-99a&WithFlank&21\|+\|16539083\|16539175@18.39.22&[MIMAT0000097&hsa-miR-99a-5p&offsets\|-1\|-1;m-64&21\|+\|16539101\|16539121&offsets\|-1\|0]\|AAACCCGTAGATCCGATCTTGT | >MI0000101\|hsa-mir-99a&WithFlank&21\|+\|16539083\|16539175 | 18 | 39 | 22 | AAACCCGTAGATCCGATCTTGT | 114.2 | -10.2 | 1.9 | -5.4 | 8.6E-08 | 2.1E-06 | BT20/MB231 | MP |
| >MI0003599\|hsa-mir-589&WithFlank&7\|-\|5495813\|5495923@30.49.20&[MIMAT0004799&hsa-miR-589-5p&offsets\|0\|-2;m-162&7\|-\|5495874\|5495894&offsets\|0\|-1]\|TGAGAACCACGTCTGCTCTG | >MI0003599\|hsa-mir-589&WithFlank&7\|-\|5495813\|5495923 | 30 | 49 | 20 | TGAGAACCACGTCTGCTCTG | 77.9 | 10.0 | 2.0 | 5.1 | 3.5E-07 | 2.1E-06 | BT20/MB468 | MP |
| >hg38_21_+_8433172_8446622_RNA45SN1_45S_50ntFlanks@11233.11269.37\|CCCTTCCCGTGGATCGCCCCAGCTGCGGCGGGCGTCG | >hg38_21_+_8433172_8446622_RNA45SN1_45S_50ntFlanks | 11233 | 11269 | 37 | CCCTTCCCGTGGATCGCCCCAGCTGCGGCGGGCGTCG | 69.3 | -9.2 | 1.8 | -5.1 | 3.5E-07 | 2.1E-06 | BT20/MB468 | MP |
| >trna10_AspGTC_12_-_125424193_125424264@40.69.30,trna12_AspGTC_12_-_125411891_125411962@40.69.30,trna144_AspGTC_6_-_27551236_27551307@40.69.30,trna38_AspGTC_17_-_8125556_8125627@40.69.30,trna45_AspGTC_6_+_27447453_27447524@40.69.30,trna48_AspGTC_6_+_27471523_27471594@40.69.30,trna4_AspGTC_12_+_96429799_96429870@40.69.30,trna69_AspGTC_1_-_161440205_161440276@40.69.30,trna72_AspGTC_1_-_161432824_161432895@40.69.30,trna75_AspGTC_1_-_161425414_161425485@40.69.30,trna78_AspGTC_1_-_161418033_161418104@40.69.30,trna81_AspGTC_1_-_161410615_161410686@40.69.30\|CGGGAGACCGGGGTTCGATTCCCCGACGGG | >trna10_AspGTC_12_-_125424193_125424264 | 40 | 69 | 30 | CGGGAGACCGGGGTTCGATTCCCCGACGGG | 372.2 | -3.4 | 0.7 | -4.9 | 1.1E-06 | 2.1E-06 | MB231/MB468 | MP |
| >hg38_21_+_8433172_8446622_RNA45SN1_45S_50ntFlanks@10792.10822.31\|CTCTGGCATGTTGGAACAATGTAGGTAAGGG | >hg38_21_+_8433172_8446622_RNA45SN1_45S_50ntFlanks | 10792 | 10822 | 31 | CTCTGGCATGTTGGAACAATGTAGGTAAGGG | 365.9 | -5.3 | 1.1 | -4.8 | 1.4E-06 | 2.1E-06 | MB231/MB468 | MITO |
| >MI0000815\|hsa-mir-339&WithFlank&7\|-\|1022927\|1023032@56.76.21&[MIMAT0004702&hsa-miR-339-3p&offsets\|0\|-2;m-235&7\|-\|1022957\|1022977&offsets\|0\|0]\|TGAGCGCCTCGACGACAGAGC | >MI0000815\|hsa-mir-339&WithFlank&7\|-\|1022927\|1023032 | 56 | 76 | 21 | TGAGCGCCTCGACGACAGAGC | 86.3 | 2.6 | 0.5 | 4.8 | 1.4E-06 | 2.1E-06 | BT20/MB468 | NUC |
| >hg38_21_+_8433172_8446622_RNA45SN1_45S_50ntFlanks@9544.9566.23\|GGTGGAGGTCCGTAGCGGTCCTG | >hg38_21_+_8433172_8446622_RNA45SN1_45S_50ntFlanks | 9544 | 9566 | 23 | GGTGGAGGTCCGTAGCGGTCCTG | 210.5 | -4.0 | 0.8 | -4.8 | 1.3E-06 | 2.1E-06 | MB231/MB468 | CYTO |
| >hg38_21_+_8433172_8446622_RNA45SN1_45S_50ntFlanks@12029.12049.21\|TCTGATCGTTTTTTCACTGAC | >hg38_21_+_8433172_8446622_RNA45SN1_45S_50ntFlanks | 12029 | 12049 | 21 | TCTGATCGTTTTTTCACTGAC | 74.9 | 10.0 | 2.0 | 5.1 | 3.5E-07 | 2.1E-06 | BT20/MB468 | MP |
| >hg38_21_+_8433172_8446622_RNA45SN1_45S_50ntFlanks@7979.8006.28\|ACCTCAGATCAGACGTGGCGACCCGCTG | >hg38_21_+_8433172_8446622_RNA45SN1_45S_50ntFlanks | 7979 | 8006 | 28 | ACCTCAGATCAGACGTGGCGACCCGCTG | 2643.3 | -3.9 | 0.8 | -4.8 | 1.3E-06 | 2.1E-06 | MB231/MB468 | CYTO |
| >hg38_21_+_8433172_8446622_RNA45SN1_45S_50ntFlanks@10797.10825.29\|GCATGTTGGAACAATGTAGGTAAGGGAAG | >hg38_21_+_8433172_8446622_RNA45SN1_45S_50ntFlanks | 10797 | 10825 | 29 | GCATGTTGGAACAATGTAGGTAAGGGAAG | 480.8 | -6.7 | 1.4 | -4.8 | 1.4E-06 | 2.1E-06 | MB231/MB468 | MITO |
| >MI0000085\|hsa-mir-27a&WithFlank&19\|-\|13836434\|13836523@57.76.20(+3U)&[MIMAT0000084&hsa-miR-27a-3p&offsets\|0\|-1(+3U);m-43&19\|-\|13836447\|13836467&offsets\|0\|-1(+3U)]\|TTCACAGTGGCTAAGTTCCGTTT | >MI0000085\|hsa-mir-27a&WithFlank&19\|-\|13836434\|13836523 | 57 | 76 | 20(+3U) | TTCACAGTGGCTAAGTTCCGTTT | 343.7 | 4.9 | 1.0 | 4.8 | 1.3E-06 | 2.1E-06 | MB231/MB468 | CYTO |
| >hg38_21_+_8433172_8446622_RNA45SN1_45S_50ntFlanks@12907.12932.26\|CACCGCACGTTCGTGGGGAACCTGGC | >hg38_21_+_8433172_8446622_RNA45SN1_45S_50ntFlanks | 12907 | 12932 | 26 | CACCGCACGTTCGTGGGGAACCTGGC | 504.7 | -8.3 | 1.7 | -4.8 | 1.4E-06 | 2.1E-06 | BT20/MB468 | NUC |
| >MI0000089\|hsa-mir-31&WithFlank&9\|-\|21512109\|21512191@14.35.22(+1C)&[MIMAT0000089&hsa-miR-31-5p&offsets\|0\|+1(+1C);m-360&9\|-\|21512157\|21512178&offsets\|0\|0(+1C)]\|AGGCAAGATGCTGGCATAGCTGC | >MI0000089\|hsa-mir-31&WithFlank&9\|-\|21512109\|21512191 | 14 | 35 | 22(+1C) | AGGCAAGATGCTGGCATAGCTGC | 139.3 | 10.5 | 2.0 | 5.4 | 8.7E-08 | 2.1E-06 | BT20/MB231 | MP |
| >MI0001445\|hsa-mir-423&WithFlank&17\|+\|30117073\|30117178@23.44.22&[MIMAT0004748&hsa-miR-423-5p&offsets\|0\|-1;m-141&17\|+\|30117095\|30117117&offsets\|0\|-1]\|TGAGGGGCAGAGAGCGAGACTT | >MI0001445\|hsa-mir-423&WithFlank&17\|+\|30117073\|30117178 | 23 | 44 | 22 | TGAGGGGCAGAGAGCGAGACTT | 5989.4 | 7.5 | 1.5 | 5.1 | 3.5E-07 | 2.1E-06 | BT20/MB468 | MP |
| >hg38_21_+_8433172_8446622_RNA45SN1_45S_50ntFlanks@10666.10706.41\|TGGAAAGCGTCGCGGTTCCGGCGGCGTCCGGTGAGCTCTCG | >hg38_21_+_8433172_8446622_RNA45SN1_45S_50ntFlanks | 10666 | 10706 | 41 | TGGAAAGCGTCGCGGTTCCGGCGGCGTCCGGTGAGCTCTCG | 562.3 | -8.1 | 1.7 | -4.8 | 1.4E-06 | 2.1E-06 | BT20/MB468 | NUC |
| >MI0003137\|hsa-mir-193b&WithFlank&16\|+\|14303961\|14304055@57.77.21(+1A)&[MIMAT0002819&hsa-miR-193b-3p&offsets\|0\|-1(+1A);m-166&16\|+\|14304017\|14304038&offsets\|0\|-1(+1A)]\|AACTGGCCCTCAAAGTCCCGCA | >MI0003137\|hsa-mir-193b&WithFlank&16\|+\|14303961\|14304055 | 57 | 77 | 21(+1A) | AACTGGCCCTCAAAGTCCCGCA | 507.3 | 6.3 | 1.2 | 5.3 | 1.4E-07 | 2.1E-06 | BT20/MB231 | NUC |
| >hg38_21_+_8433172_8446622_RNA45SN1_45S_50ntFlanks@10412.10429.18\|AGTCGGTCCTGAGAGATG | >hg38_21_+_8433172_8446622_RNA45SN1_45S_50ntFlanks | 10412 | 10429 | 18 | AGTCGGTCCTGAGAGATG | 262.8 | -5.2 | 1.1 | -4.8 | 1.3E-06 | 2.1E-06 | MB231/MB468 | NUC |
| >hg38_21_+_8433172_8446622_RNA45SN1_45S_50ntFlanks@6652.6679.28\|GACTCTTAGCGGTGGATCACTCGGCTCG | >hg38_21_+_8433172_8446622_RNA45SN1_45S_50ntFlanks | 6652 | 6679 | 28 | GACTCTTAGCGGTGGATCACTCGGCTCG | 2916.6 | -3.4 | 0.7 | -4.8 | 1.4E-06 | 2.1E-06 | BT20/MB468 | NUC |
| >hg38_21_+_8433172_8446622_RNA45SN1_45S_50ntFlanks@12404.12441.38\|GCTTTTTGATCCTTCGATGTCGGCTCTTCCTATCATTG | >hg38_21_+_8433172_8446622_RNA45SN1_45S_50ntFlanks | 12404 | 12441 | 38 | GCTTTTTGATCCTTCGATGTCGGCTCTTCCTATCATTG | 836.1 | -8.5 | 1.8 | -4.8 | 1.4E-06 | 2.1E-06 | BT20/MB468 | NUC |
| >hg38_21_+_8433172_8446622_RNA45SN1_45S_50ntFlanks@6647.6667.21\|CGTACGACTCTTAGCGGTGGA | >hg38_21_+_8433172_8446622_RNA45SN1_45S_50ntFlanks | 6647 | 6667 | 21 | CGTACGACTCTTAGCGGTGGA | 2477.8 | 4.4 | 0.9 | 5.1 | 3.5E-07 | 2.1E-06 | BT20/MB468 | MP |
| >hg38_21_+_8433172_8446622_RNA45SN1_45S_50ntFlanks@9556.9573.18\|TAGCGGTCCTGACGTGCA | >hg38_21_+_8433172_8446622_RNA45SN1_45S_50ntFlanks | 9556 | 9573 | 18 | TAGCGGTCCTGACGTGCA | 302.1 | -5.5 | 1.1 | -4.8 | 1.4E-06 | 2.1E-06 | MB231/MB468 | MITO |
| >am_trnaMT_HisGTG_MT_+_12138_12206@22.51.30\|CATCAGATTGTGAATCTGACAACAGAGGCT | >am_trnaMT_HisGTG_MT_+_12138_12206 | 22 | 51 | 30 | CATCAGATTGTGAATCTGACAACAGAGGCT | 85.6 | 10.2 | 2.0 | 5.1 | 3.5E-07 | 2.1E-06 | BT20/MB468 | MP |
| >hg38_21_+_8433172_8446622_RNA45SN1_45S_50ntFlanks@4367.4393.27\|CAGTTAAAAAGCTCGTAGTTGGATCTT | >hg38_21_+_8433172_8446622_RNA45SN1_45S_50ntFlanks | 4367 | 4393 | 27 | CAGTTAAAAAGCTCGTAGTTGGATCTT | 114.0 | -6.6 | 1.4 | -4.8 | 1.3E-06 | 2.1E-06 | MB231/MB468 | NUC |
| >hg38_21_+_8433172_8446622_RNA45SN1_45S_50ntFlanks@12004.12032.29\|CGGGCCGCCGGTGAAATACCACTACTCTG | >hg38_21_+_8433172_8446622_RNA45SN1_45S_50ntFlanks | 12004 | 12032 | 29 | CGGGCCGCCGGTGAAATACCACTACTCTG | 941.6 | -6.1 | 1.3 | -4.8 | 1.4E-06 | 2.1E-06 | MB231/MB468 | MITO |
| >hg38_21_+_8433172_8446622_RNA45SN1_45S_50ntFlanks@10357.10388.32\|CTTTGAAGGCCGAAGTGGAGAAGGGTTCCATG | >hg38_21_+_8433172_8446622_RNA45SN1_45S_50ntFlanks | 10357 | 10388 | 32 | CTTTGAAGGCCGAAGTGGAGAAGGGTTCCATG | 683.9 | -5.5 | 1.1 | -4.8 | 1.3E-06 | 2.1E-06 | MB231/MB468 | CYTO |
| >hg38_21_+_8433172_8446622_RNA45SN1_45S_50ntFlanks@9561.9596.36\|GTCCTGACGTGCAAATCGGTCGTCCGACCTGGGTAT | >hg38_21_+_8433172_8446622_RNA45SN1_45S_50ntFlanks | 9561 | 9596 | 36 | GTCCTGACGTGCAAATCGGTCGTCCGACCTGGGTAT | 365.3 | -3.5 | 0.7 | -4.9 | 9.9E-07 | 2.1E-06 | BT20/MB468 | MITO |
| >hg38_21_+_8433172_8446622_RNA45SN1_45S_50ntFlanks@10921.10945.25\|GCTGGACGAGGCGCCGCCGCCCCCC | >hg38_21_+_8433172_8446622_RNA45SN1_45S_50ntFlanks | 10921 | 10945 | 25 | GCTGGACGAGGCGCCGCCGCCCCCC | 203.2 | -7.2 | 1.5 | -4.9 | 1.1E-06 | 2.1E-06 | MB231/MB468 | MP |
| >MI0001445\|hsa-mir-423&WithFlank&17\|+\|30117073\|30117178@23.43.21(+1C)&[MIMAT0004748&hsa-miR-423-5p&offsets\|0\|-2(+1C);m-141&17\|+\|30117095\|30117117&offsets\|0\|-2(+1C)]\|TGAGGGGCAGAGAGCGAGACTC | >MI0001445\|hsa-mir-423&WithFlank&17\|+\|30117073\|30117178 | 23 | 43 | 21(+1C) | TGAGGGGCAGAGAGCGAGACTC | 824.3 | 8.1 | 1.6 | 5.1 | 3.6E-07 | 2.1E-06 | BT20/MB468 | MP |
| >hg38_21_+_8433172_8446622_RNA45SN1_45S_50ntFlanks@12420.12441.22\|ATGTCGGCTCTTCCTATCATTG | >hg38_21_+_8433172_8446622_RNA45SN1_45S_50ntFlanks | 12420 | 12441 | 22 | ATGTCGGCTCTTCCTATCATTG | 483.1 | -7.1 | 1.5 | -4.9 | 1.1E-06 | 2.1E-06 | MB231/MB468 | MP |
| >hg38_21_+_8433172_8446622_RNA45SN1_45S_50ntFlanks@12539.12569.31\|ATGATGTGTTGTTGCCATGGTAATCCTGCTC | >hg38_21_+_8433172_8446622_RNA45SN1_45S_50ntFlanks | 12539 | 12569 | 31 | ATGATGTGTTGTTGCCATGGTAATCCTGCTC | 1671.2 | -5.4 | 1.1 | -4.9 | 1.1E-06 | 2.1E-06 | BT20/MB468 | CYTO |
| >hg38_21_+_8433172_8446622_RNA45SN1_45S_50ntFlanks@10351.10385.35\|CGAGAACTTTGAAGGCCGAAGTGGAGAAGGGTTCC | >hg38_21_+_8433172_8446622_RNA45SN1_45S_50ntFlanks | 10351 | 10385 | 35 | CGAGAACTTTGAAGGCCGAAGTGGAGAAGGGTTCC | 356.8 | -5.3 | 1.1 | -4.8 | 1.3E-06 | 2.1E-06 | MB231/MB468 | NUC |
| >hg38_21_+_8433172_8446622_RNA45SN1_45S_50ntFlanks@10805.10840.36\|GAACAATGTAGGTAAGGGAAGTCGGCAAGCCGGATC | >hg38_21_+_8433172_8446622_RNA45SN1_45S_50ntFlanks | 10805 | 10840 | 36 | GAACAATGTAGGTAAGGGAAGTCGGCAAGCCGGATC | 497.2 | -7.7 | 1.6 | -4.8 | 1.4E-06 | 2.1E-06 | BT20/MB468 | NUC |
| >hg38_21_+_8433172_8446622_RNA45SN1_45S_50ntFlanks@8159.8181.23\|CGTGGGGGGCCCAAGTCCTTCTG | >hg38_21_+_8433172_8446622_RNA45SN1_45S_50ntFlanks | 8159 | 8181 | 23 | CGTGGGGGGCCCAAGTCCTTCTG | 155.3 | -9.4 | 1.9 | -4.9 | 9.9E-07 | 2.1E-06 | BT20/MB468 | MITO |
| >hg38_21_+_8433172_8446622_RNA45SN1_45S_50ntFlanks@3835.3861.27\|CTCCTCTCCTACTTGGATAACTGTGGT | >hg38_21_+_8433172_8446622_RNA45SN1_45S_50ntFlanks | 3835 | 3861 | 27 | CTCCTCTCCTACTTGGATAACTGTGGT | 1162.1 | -8.2 | 1.7 | -4.8 | 1.4E-06 | 2.1E-06 | BT20/MB468 | NUC |
| >hg38_21_+_8433172_8446622_RNA45SN1_45S_50ntFlanks@10463.10484.22\|CTCCGTTGCCCTCGGCCGATCG | >hg38_21_+_8433172_8446622_RNA45SN1_45S_50ntFlanks | 10463 | 10484 | 22 | CTCCGTTGCCCTCGGCCGATCG | 59.2 | -9.0 | 1.8 | -5.1 | 3.6E-07 | 2.1E-06 | BT20/MB468 | MP |
| >hg38_21_+_8433172_8446622_RNA45SN1_45S_50ntFlanks@4130.4150.21\|ATTCCGGAGAGGGAGCCTGAG | >hg38_21_+_8433172_8446622_RNA45SN1_45S_50ntFlanks | 4130 | 4150 | 21 | ATTCCGGAGAGGGAGCCTGAG | 435.3 | -5.1 | 1.1 | -4.8 | 1.4E-06 | 2.1E-06 | MB231/MB468 | MITO |
| >hg38_21_+_8433172_8446622_RNA45SN1_45S_50ntFlanks@13008.13028.21\|ATTGAAAGTCAGCCCTCGACA | >hg38_21_+_8433172_8446622_RNA45SN1_45S_50ntFlanks | 13008 | 13028 | 21 | ATTGAAAGTCAGCCCTCGACA | 232.3 | -6.8 | 1.4 | -4.8 | 1.4E-06 | 2.1E-06 | MB231/MB468 | MITO |
| >MI0003560\|hsa-mir-92b&WithFlank&1\|+\|155195171\|155195278@67.86.20(+1U)&[MIMAT0003218&hsa-miR-92b-3p&offsets\|0\|-2(+1U);m-115&1\|+\|155195237\|155195258&offsets\|0\|-2(+1U)]\|TATTGCACTCGTCCCGGCCTT | >MI0003560\|hsa-mir-92b&WithFlank&1\|+\|155195171\|155195278 | 67 | 86 | 20(+1U) | TATTGCACTCGTCCCGGCCTT | 464.5 | 9.5 | 1.9 | 5.1 | 3.6E-07 | 2.2E-06 | BT20/MB468 | MP |
| >hg38_21_+_8433172_8446622_RNA45SN1_45S_50ntFlanks@4727.4764.38\|AATCAAGAACGAAAGTCGGAGGTTCGAAGACGATCAGA | >hg38_21_+_8433172_8446622_RNA45SN1_45S_50ntFlanks | 4727 | 4764 | 38 | AATCAAGAACGAAAGTCGGAGGTTCGAAGACGATCAGA | 632.3 | 7.9 | 1.6 | 4.9 | 1.1E-06 | 2.2E-06 | BT20/MB468 | CYTO |
| >hg38_21_+_8433172_8446622_RNA45SN1_45S_50ntFlanks@12635.12661.27\|AAGCTACCATCTGTGGGATTATGACTG | >hg38_21_+_8433172_8446622_RNA45SN1_45S_50ntFlanks | 12635 | 12661 | 27 | AAGCTACCATCTGTGGGATTATGACTG | 767.5 | -3.0 | 0.6 | -4.9 | 1.1E-06 | 2.2E-06 | BT20/MB468 | CYTO |
| >MI0003560\|hsa-mir-92b&WithFlank&1\|+\|155195171\|155195278@67.85.19&[MIMAT0003218&hsa-miR-92b-3p&offsets\|0\|-3;m-115&1\|+\|155195237\|155195258&offsets\|0\|-3]\|TATTGCACTCGTCCCGGCC | >MI0003560\|hsa-mir-92b&WithFlank&1\|+\|155195171\|155195278 | 67 | 85 | 19 | TATTGCACTCGTCCCGGCC | 55.2 | 9.0 | 1.9 | 4.9 | 1.1E-06 | 2.2E-06 | MB231/MB468 | MP |

<table>
<tr><td></td><td></td><td></td><td></td><td></td><td></td><td></td><td></td><td></td><td></td><td></td><td></td><td></td><td></td></tr>
<tr><td>>MI0000264|hsa-mir-7-2&WithFlank&15|+|88611819|88611940@38.60.23(+1A);MI0000265|hsa-mir-7-3&WithFlank&19|+|4770664|4770785@37.59.23(+1A);MI0000263|hsa-mir-7-1&WithFlank&9|-|83969742|83969863@30.52.23(+1A)&[MIMAT0000252&hsa-miR-7-5p&offsets|0|-1(+1A);m-400&15|+|88611856|88611879&offsets|0|-1(+1A)];[MIMAT0000252_1&hsa-miR-7-5p&offsets|0|-1(+1A);m-402&19|+|4770700|4770723&offsets|0|-1(+1A)];[MIMAT0000252_2&hsa-miR-7-5p&offsets|0|-1(+1A);m-401&9|-|83969811|83969834&offsets|0|-1(+1A)]|TGGAAGACTAGTGATTTTGTTGTA</td><td>>MI0000264|hsa-mir-7-2&WithFlank&15|+|88611819|88611940</td><td>38</td><td>60</td><td>23(+1A)</td><td>TGGAAGACTAGTGATTTTGTTGTA</td><td>495.3</td><td>4.0</td><td>0.8</td><td>4.8</td><td>1.3E-06</td><td>2.2E-06</td><td>MB231/MB468</td><td>CYTO</td></tr>
<tr><td>>hg38_21_+_8433172_8446622_RNA45SN1_45S_50ntFlanks@12163.12181.19|ACTGGGGCGGTACACCTGT</td><td>>hg38_21_+_8433172_8446622_RNA45SN1_45S_50ntFlanks</td><td>12163</td><td>12181</td><td>19</td><td>ACTGGGGCGGTACACCTGT</td><td>294.1</td><td>-5.8</td><td>1.2</td><td>-4.8</td><td>1.3E-06</td><td>2.2E-06</td><td>MB231/MB468</td><td>CYTO</td></tr>
<tr><td>>hg38_21_+_8433172_8446622_RNA45SN1_45S_50ntFlanks@6650.6680.31|ACGACTCTTAGCGGTGGATCACTCGGCTCGT</td><td>>hg38_21_+_8433172_8446622_RNA45SN1_45S_50ntFlanks</td><td>6650</td><td>6680</td><td>31</td><td>ACGACTCTTAGCGGTGGATCACTCGGCTCGT</td><td>359.0</td><td>-3.1</td><td>0.6</td><td>-4.9</td><td>1.2E-06</td><td>2.2E-06</td><td>MB231/MB468</td><td>MP</td></tr>
<tr><td>>hg38_21_+_8433172_8446622_RNA45SN1_45S_50ntFlanks@9573.9595.23|AAATCGGTCGTCCGACCTGGGTA</td><td>>hg38_21_+_8433172_8446622_RNA45SN1_45S_50ntFlanks</td><td>9573</td><td>9595</td><td>23</td><td>AAATCGGTCGTCCGACCTGGGTA</td><td>1259.8</td><td>-4.6</td><td>0.9</td><td>-4.9</td><td>1.0E-06</td><td>2.2E-06</td><td>BT20/MB468</td><td>MITO</td></tr>
<tr><td>>hg38_21_+_8433172_8446622_RNA45SN1_45S_50ntFlanks@3777.3805.29|CGGTACAGTGAAACTGCGAATGGCTCATT</td><td>>hg38_21_+_8433172_8446622_RNA45SN1_45S_50ntFlanks</td><td>3777</td><td>3805</td><td>29</td><td>CGGTACAGTGAAACTGCGAATGGCTCATT</td><td>726.0</td><td>-6.1</td><td>1.3</td><td>-4.8</td><td>1.5E-06</td><td>2.2E-06</td><td>BT20/MB468</td><td>NUC</td></tr>
<tr><td>>hg38_21_+_8433172_8446622_RNA45SN1_45S_50ntFlanks@12324.12365.42|TTGGGTTTTAAGCAGGAGGTGTCAGAAAAGTTACCACAGGGA</td><td>>hg38_21_+_8433172_8446622_RNA45SN1_45S_50ntFlanks</td><td>12324</td><td>12365</td><td>42</td><td>TTGGGTTTTAAGCAGGAGGTGTCAGAAAAGTTACCACAGGGA</td><td>124.4</td><td>-10.0</td><td>2.0</td><td>-5.1</td><td>3.6E-07</td><td>2.2E-06</td><td>BT20/MB468</td><td>MP</td></tr>
<tr><td>>hg38_21_+_8433172_8446622_RNA45SN1_45S_50ntFlanks@12160.12181.22|TTGACTGGGGCGGTACACCTGT</td><td>>hg38_21_+_8433172_8446622_RNA45SN1_45S_50ntFlanks</td><td>12160</td><td>12181</td><td>22</td><td>TTGACTGGGGCGGTACACCTGT</td><td>242.0</td><td>-7.0</td><td>1.4</td><td>-5.1</td><td>3.6E-07</td><td>2.2E-06</td><td>BT20/MB468</td><td>MP</td></tr>
<tr><td>>hg38_21_+_8433172_8446622_RNA45SN1_45S_50ntFlanks@10841.10863.23|CGTAACTTCGGGATAAGGATTGG</td><td>>hg38_21_+_8433172_8446622_RNA45SN1_45S_50ntFlanks</td><td>10841</td><td>10863</td><td>23</td><td>CGTAACTTCGGGATAAGGATTGG</td><td>394.8</td><td>-2.7</td><td>0.5</td><td>-4.9</td><td>1.0E-06</td><td>2.2E-06</td><td>BT20/MB468</td><td>MITO</td></tr>
<tr><td>>hg38_21_+_8433172_8446622_RNA45SN1_45S_50ntFlanks@12456.12487.32|CCAAGCGTTGGATTGTTCACCCACTAATAGGG</td><td>>hg38_21_+_8433172_8446622_RNA45SN1_45S_50ntFlanks</td><td>12456</td><td>12487</td><td>32</td><td>CCAAGCGTTGGATTGTTCACCCACTAATAGGG</td><td>34370.0</td><td>-7.5</td><td>1.5</td><td>-4.9</td><td>1.1E-06</td><td>2.2E-06</td><td>BT20/MB468</td><td>CYTO</td></tr>
<tr><td>>hg38_21_+_8433172_8446622_RNA45SN1_45S_50ntFlanks@8342.8365.24|CGTAAGGGAAAGTTGAAAAGAACT</td><td>>hg38_21_+_8433172_8446622_RNA45SN1_45S_50ntFlanks</td><td>8342</td><td>8365</td><td>24</td><td>CGTAAGGGAAAGTTGAAAAGAACT</td><td>3384.8</td><td>6.4</td><td>1.3</td><td>4.8</td><td>1.3E-06</td><td>2.2E-06</td><td>MB231/MB468</td><td>CYTO</td></tr>
<tr><td>>trnaMT_GluTTC_MT_-_14674_14742@24.58.35,trnalookalike8_GluTTC_5_-_93905172_93905240@24.58.35|ATGGTTTTTCATATCATTGGTCGTGGTTGTAGTCC</td><td>>trnaMT_GluTTC_MT_-_14674_14742</td><td>24</td><td>58</td><td>35</td><td>ATGGTTTTTCATATCATTGGTCGTGGTTGTAGTCC</td><td>609.3</td><td>9.2</td><td>1.8</td><td>5.1</td><td>3.6E-07</td><td>2.2E-06</td><td>BT20/MB468</td><td>MP</td></tr>
<tr><td>>hg38_21_+_8433172_8446622_RNA45SN1_45S_50ntFlanks@10798.10822.25|CATGTTGGAACAATGTAGGTAAGGG</td><td>>hg38_21_+_8433172_8446622_RNA45SN1_45S_50ntFlanks</td><td>10798</td><td>10822</td><td>25</td><td>CATGTTGGAACAATGTAGGTAAGGG</td><td>400.6</td><td>-7.4</td><td>1.5</td><td>-4.8</td><td>1.5E-06</td><td>2.2E-06</td><td>BT20/MB468</td><td>NUC</td></tr>
<tr><td>>hg38_21_+_8433172_8446622_RNA45SN1_45S_50ntFlanks@3903.3938.36|CGCGGGGGGGATGCGTGCATTTATCAGATCAAAACC</td><td>>hg38_21_+_8433172_8446622_RNA45SN1_45S_50ntFlanks</td><td>3903</td><td>3938</td><td>36</td><td>CGCGGGGGGGATGCGTGCATTTATCAGATCAAAACC</td><td>116.8</td><td>-9.9</td><td>2.0</td><td>-5.1</td><td>3.6E-07</td><td>2.2E-06</td><td>BT20/MB468</td><td>MP</td></tr>
<tr><td>>hg38_1_-_228634819_228635039_RNA5S12_5S_50ntFlanks@130.171.42|TGGATGGGAGACCGCCTGGGAATACCGGGTGCTGTAGGCTTT</td><td>>hg38_1_-_228634819_228635039_RNA5S12_5S_50ntFlanks</td><td>130</td><td>171</td><td>42</td><td>TGGATGGGAGACCGCCTGGGAATACCGGGTGCTGTAGGCTTT</td><td>1416.9</td><td>7.9</td><td>1.6</td><td>4.9</td><td>1.0E-06</td><td>2.2E-06</td><td>BT20/MB468</td><td>MITO</td></tr>
<tr><td>>hg38_21_+_8433172_8446622_RNA45SN1_45S_50ntFlanks@10789.10816.28|AGCCTCTGGCATGTTGGAACAATGTAGG</td><td>>hg38_21_+_8433172_8446622_RNA45SN1_45S_50ntFlanks</td><td>10789</td><td>10816</td><td>28</td><td>AGCCTCTGGCATGTTGGAACAATGTAGG</td><td>294.7</td><td>-5.5</td><td>1.1</td><td>-4.8</td><td>1.5E-06</td><td>2.2E-06</td><td>BT20/MB468</td><td>NUC</td></tr>
<tr><td>>hg38_21_+_8433172_8446622_RNA45SN1_45S_50ntFlanks@7992.8017.26|CGTGGCGACCCGCTGAATTTAAGCAT</td><td>>hg38_21_+_8433172_8446622_RNA45SN1_45S_50ntFlanks</td><td>7992</td><td>8017</td><td>26</td><td>CGTGGCGACCCGCTGAATTTAAGCAT</td><td>181.9</td><td>5.9</td><td>1.2</td><td>4.8</td><td>1.3E-06</td><td>2.2E-06</td><td>MB231/MB468</td><td>CYTO</td></tr>
<tr><td>>hg38_21_+_8433172_8446622_RNA45SN1_45S_50ntFlanks@12488.12524.37|AACGTGAGCTGGGTTTAGACCGTCGTGAGACAGGTTA</td><td>>hg38_21_+_8433172_8446622_RNA45SN1_45S_50ntFlanks</td><td>12488</td><td>12524</td><td>37</td><td>AACGTGAGCTGGGTTTAGACCGTCGTGAGACAGGTTA</td><td>444.8</td><td>-7.1</td><td>1.5</td><td>-4.8</td><td>1.4E-06</td><td>2.2E-06</td><td>MB231/MB468</td><td>NUC</td></tr>
<tr><td>>am_trna128_GlyGCC_6_-_27870686_27870756@1.34.34,trna18_GlyGCC_16_+_70822597_70822667@1.34.34,trna19_GlyGCC_16_+_70823410_70823480@1.34.34,trna19_GlyGCC_2_-_157257659_157257729@1.34.34,trna24_GlyGCC_16_-_70812942_70813012@1.34.34,trna25_GlyGCC_16_-_70812114_70812184@1.34.34,trna5_GlyGCC_17_+_8029064_8029134@1.34.34,trna68_GlyGCC_1_-_161493637_161493707@1.34.34|GCATTGGTGGTTCAGTGGTAGAATTCTCGCCTGC</td><td>>am_trna128_GlyGCC_6_-_27870686_27870756</td><td>1</td><td>34</td><td>34</td><td>GCATTGGTGGTTCAGTGGTAGAATTCTCGCCTGC</td><td>3005.6</td><td>-4.4</td><td>0.8</td><td>-5.3</td><td>1.2E-07</td><td>2.2E-06</td><td>BT20/MB231</td><td>MITO</td></tr>
<tr><td>>trna10_AspGTC_12_-_125424193_125424264@39.73.35,trna12_AspGTC_12_-_125411891_125411962@39.73.35,trna144_AspGTC_6_-_27551236_27551307@39.73.35,trna38_AspGTC_17_-_8125556_8125627@39.73.35,trna45_AspGTC_6_+_27447453_27447524@39.73.35,trna48_AspGTC_6_+_27471523_27471594@39.73.35,trna4_AspGTC_12_+_96429799_96429870@39.73.35,trna69_AspGTC_1_-_161440205_161440276@39.73.35,trna72_AspGTC_1_-_161432824_161432895@39.73.35,trna75_AspGTC_1_-_161425414_161425485@39.73.35,trna78_AspGTC_1_-_161418033_161418104@39.73.35,trna81_AspGTC_1_-_161410615_161410686@39.73.35|GCGGGAGACCGGGGTTCGATTCCCCGACGGGGAGC</td><td>>trna10_AspGTC_12_-_125424193_125424264</td><td>39</td><td>73</td><td>35</td><td>GCGGGAGACCGGGGTTCGATTCCCCGACGGGGAGC</td><td>4053.7</td><td>4.9</td><td>1.0</td><td>4.8</td><td>1.3E-06</td><td>2.2E-06</td><td>MB231/MB468</td><td>CYTO</td></tr>
<tr><td>>hg38_21_+_8433172_8446622_RNA45SN1_45S_50ntFlanks@6650.6679.30|ACGACTCTTAGCGGTGGATCACTCGGCTCG</td><td>>hg38_21_+_8433172_8446622_RNA45SN1_45S_50ntFlanks</td><td>6650</td><td>6679</td><td>30</td><td>ACGACTCTTAGCGGTGGATCACTCGGCTCG</td><td>822.5</td><td>-4.1</td><td>0.8</td><td>-4.9</td><td>1.2E-06</td><td>2.2E-06</td><td>MB231/MB468</td><td>MP</td></tr>
<tr><td>>hg38_21_+_8433172_8446622_RNA45SN1_45S_50ntFlanks@12420.12437.18|ATGTCGGCTCTTCCTATC</td><td>>hg38_21_+_8433172_8446622_RNA45SN1_45S_50ntFlanks</td><td>12420</td><td>12437</td><td>18</td><td>ATGTCGGCTCTTCCTATC</td><td>1413.7</td><td>-6.2</td><td>1.3</td><td>-4.9</td><td>1.2E-06</td><td>2.2E-06</td><td>MB231/MB468</td><td>MP</td></tr>
<tr><td>>hg38_21_+_8433172_8446622_RNA45SN1_45S_50ntFlanks@8437.8467.31|AGGATTCAACCCGGCGGCGGGTCCGGCCGTG</td><td>>hg38_21_+_8433172_8446622_RNA45SN1_45S_50ntFlanks</td><td>8437</td><td>8467</td><td>31</td><td>AGGATTCAACCCGGCGGCGGGTCCGGCCGTG</td><td>366.6</td><td>-7.4</td><td>1.5</td><td>-5.1</td><td>3.7E-07</td><td>2.2E-06</td><td>BT20/MB468</td><td>MP</td></tr>
<tr><td>>MI0002470|hsa-mir-486-1&WithFlank&8|-|41660435|41660514@10.31.22(+1A);MI0023622|hsa-mir-486-2&WithFlank&8|+|41660438|41660513@7.28.22(+1A)&[MIMAT0002177&hsa-miR-486-5p&offsets|0|0(+1A);m-107&8|-|41660484|41660505&offsets|0|0(+1A)];[MIMAT0002177_1&hsa-miR-486-5p&offsets|0|0(+1A);TJU_CMC_MD2.ID03138.5p-miR&offsets|0|-2(+1A);m-106&8|+|41660444|41660465&offsets|0|0(+1A)]|TCCTGTACTGAGCTGCCCCGAGA</td><td>>MI0002470|hsa-mir-486-1&WithFlank&8|-|41660435|41660514</td><td>10</td><td>31</td><td>22(+1A)</td><td>TCCTGTACTGAGCTGCCCCGAGA</td><td>41.7</td><td>8.6</td><td>1.8</td><td>4.9</td><td>1.2E-06</td><td>2.2E-06</td><td>MB231/MB468</td><td>MP</td></tr>
<tr><td>>hg38_21_+_8433172_8446622_RNA45SN1_45S_50ntFlanks@8337.8378.42|AGTACCGTAAGGGAAAGTTGAAAAGAACTTTGAAGAGAGAGT</td><td>>hg38_21_+_8433172_8446622_RNA45SN1_45S_50ntFlanks</td><td>8337</td><td>8378</td><td>42</td><td>AGTACCGTAAGGGAAAGTTGAAAAGAACTTTGAAGAGAGAGT</td><td>59.0</td><td>9.6</td><td>1.9</td><td>5.1</td><td>3.7E-07</td><td>2.2E-06</td><td>BT20/MB468</td><td>MP</td></tr>
<tr><td>>hg38_21_+_8433172_8446622_RNA45SN1_45S_50ntFlanks@10807.10840.34|ACAATGTAGGTAAGGGAAGTCGGCAAGCCGGATC</td><td>>hg38_21_+_8433172_8446622_RNA45SN1_45S_50ntFlanks</td><td>10807</td><td>10840</td><td>34</td><td>ACAATGTAGGTAAGGGAAGTCGGCAAGCCGGATC</td><td>481.5</td><td>-6.3</td><td>1.3</td><td>-4.9</td><td>1.0E-06</td><td>2.2E-06</td><td>BT20/MB468</td><td>MITO</td></tr>
<tr><td>>hg38_MT_+_1621_3279_RNR2_16S_50ntFlanks@1431.1451.21|AAAGTCCTACGTGATCTGAGT</td><td>>hg38_MT_+_1621_3279_RNR2_16S_50ntFlanks</td><td>1431</td><td>1451</td><td>21</td><td>AAAGTCCTACGTGATCTGAGT</td><td>212.3</td><td>4.6</td><td>0.9</td><td>4.8</td><td>1.4E-06</td><td>2.2E-06</td><td>MB231/MB468</td><td>MITO</td></tr>
<tr><td>>trna12_ValAAC_5_-_180645270_180645342@1.34.34,trna132_ValAAC_6_-_27721179_27721251@1.34.34,trna136_ValAAC_6_-_27648885_27648957@1.34.34,trna139_ValAAC_6_-_27618707_27618779@1.34.34,trna2_ValAAC_3_+_169490018_169490090@1.34.34,trna4_ValAAC_5_+_180591154_180591226@1.34.34,trna5_ValAAC_5_+_180596610_180596682@1.34.34|GTTTCCGTAGTGTAGTGGTTATCACGTTCGCCTA</td><td>>trna12_ValAAC_5_-_180645270_180645342</td><td>1</td><td>34</td><td>34</td><td>GTTTCCGTAGTGTAGTGGTTATCACGTTCGCCTA</td><td>72.6</td><td>9.4</td><td>1.9</td><td>4.9</td><td>1.2E-06</td><td>2.2E-06</td><td>MB231/MB468</td><td>MP</td></tr>
<tr><td>>hg38_21_+_8433172_8446622_RNA45SN1_45S_50ntFlanks@12933.12955.23|GCTAAACCATTCGTAGACGACCT</td><td>>hg38_21_+_8433172_8446622_RNA45SN1_45S_50ntFlanks</td><td>12933</td><td>12955</td><td>23</td><td>GCTAAACCATTCGTAGACGACCT</td><td>202.2</td><td>-3.5</td><td>0.7</td><td>-4.8</td><td>1.4E-06</td><td>2.2E-06</td><td>MB231/MB468</td><td>NUC</td></tr>
<tr><td>>hg38_21_+_8433172_8446622_RNA45SN1_45S_50ntFlanks@12417.12443.27|TCGATGTCGGCTCTTCCTATCATTGTG</td><td>>hg38_21_+_8433172_8446622_RNA45SN1_45S_50ntFlanks</td><td>12417</td><td>12443</td><td>27</td><td>TCGATGTCGGCTCTTCCTATCATTGTG</td><td>1068.5</td><td>-7.6</td><td>1.6</td><td>-4.9</td><td>1.2E-06</td><td>2.2E-06</td><td>MB231/MB468</td><td>MP</td></tr>
<tr><td>>hg38_21_+_8433172_8446622_RNA45SN1_45S_50ntFlanks@10482.10509.28|TCGAAAGGGAGTCGGGTTCAGATCCCCG</td><td>>hg38_21_+_8433172_8446622_RNA45SN1_45S_50ntFlanks</td><td>10482</td><td>10509</td><td>28</td><td>TCGAAAGGGAGTCGGGTTCAGATCCCCG</td><td>175.8</td><td>-6.3</td><td>1.3</td><td>-4.9</td><td>1.2E-06</td><td>2.2E-06</td><td>BT20/MB468</td><td>CYTO</td></tr>
<tr><td>>hg38_21_+_8433172_8446622_RNA45SN1_45S_50ntFlanks@3705.3740.36|TACCTGGTTGATCCTGCCAGTAGCATATGCTTGTCT</td><td>>hg38_21_+_8433172_8446622_RNA45SN1_45S_50ntFlanks</td><td>3705</td><td>3740</td><td>36</td><td>TACCTGGTTGATCCTGCCAGTAGCATATGCTTGTCT</td><td>77.3</td><td>10.0</td><td>2.0</td><td>5.1</td><td>3.7E-07</td><td>2.2E-06</td><td>BT20/MB468</td><td>MP</td></tr>
<tr><td>>trnaMT_PheGAA_MT_+_577_647@38.60.23|AATGTTTAGACGGGCTCACATCA</td><td>>trnaMT_PheGAA_MT_+_577_647</td><td>38</td><td>60</td><td>23</td><td>AATGTTTAGACGGGCTCACATCA</td><td>80.0</td><td>10.1</td><td>2.0</td><td>5.1</td><td>3.7E-07</td><td>2.2E-06</td><td>BT20/MB468</td><td>MP</td></tr>
<tr><td>>MI0000299|hsa-mir-222&WithFlank&X|-|45747009|45747130@75.99.25(+3U)&[MIMAT0000279&hsa-miR-222-3p&offsets|0|+4(+3U);m-140&X|-|45747033|45747056&offsets|0|+1(+3U)]|AGCTACATCTGGCTACTGGGTCTCTTTT</td><td>>MI0000299|hsa-mir-222&WithFlank&X|-|45747009|45747130</td><td>75</td><td>99</td><td>25(+3U)</td><td>AGCTACATCTGGCTACTGGGTCTCTTTT</td><td>1170.7</td><td>-3.9</td><td>0.7</td><td>-5.3</td><td>1.5E-07</td><td>2.2E-06</td><td>BT20/MB231</td><td>NUC</td></tr>
<tr><td>>hg38_21_+_8433172_8446622_RNA45SN1_45S_50ntFlanks@5472.5510.39|ACGGCCCTGGCGGAGCGCTGAGAAGACGGTCGAACTTGA</td><td>>hg38_21_+_8433172_8446622_RNA45SN1_45S_50ntFlanks</td><td>5472</td><td>5510</td><td>39</td><td>ACGGCCCTGGCGGAGCGCTGAGAAGACGGTCGAACTTGA</td><td>1058.9</td><td>4.7</td><td>1.0</td><td>4.9</td><td>1.2E-06</td><td>2.2E-06</td><td>BT20/MB468</td><td>CYTO</td></tr>
<tr><td>>hg38_21_+_8433172_8446622_RNA45SN1_45S_50ntFlanks@12638.12661.24|CTACCATCTGTGGGATTATGACTG</td><td>>hg38_21_+_8433172_8446622_RNA45SN1_45S_50ntFlanks</td><td>12638</td><td>12661</td><td>24</td><td>CTACCATCTGTGGGATTATGACTG</td><td>1028.8</td><td>-8.2</td><td>1.7</td><td>-4.8</td><td>1.5E-06</td><td>2.2E-06</td><td>BT20/MB468</td><td>NUC</td></tr>
<tr><td>>hg38_21_+_8433172_8446622_RNA45SN1_45S_50ntFlanks@5482.5514.33|CGGAGCGCTGAGAAGACGGTCGAACTTGACTAT</td><td>>hg38_21_+_8433172_8446622_RNA45SN1_45S_50ntFlanks</td><td>5482</td><td>5514</td><td>33</td><td>CGGAGCGCTGAGAAGACGGTCGAACTTGACTAT</td><td>286.6</td><td>-2.2</td><td>0.4</td><td>-4.9</td><td>1.2E-06</td><td>2.2E-06</td><td>BT20/MB468</td><td>CYTO</td></tr>
<tr><td>>hg38_21_+_8433172_8446622_RNA45SN1_45S_50ntFlanks@11873.11909.37|AAGAAGACCCTGTTGAGCTTGACTCTAGTCTGGCACG</td><td>>hg38_21_+_8433172_8446622_RNA45SN1_45S_50ntFlanks</td><td>11873</td><td>11909</td><td>37</td><td>AAGAAGACCCTGTTGAGCTTGACTCTAGTCTGGCACG</td><td>357.2</td><td>-3.7</td><td>0.8</td><td>-4.9</td><td>1.0E-06</td><td>2.2E-06</td><td>BT20/MB468</td><td>MITO</td></tr>
<tr><td>>hg38_21_+_8433172_8446622_RNA45SN1_45S_50ntFlanks@10916.10939.24|CCGCGGCTGGACGAGGCGCCGCCG</td><td>>hg38_21_+_8433172_8446622_RNA45SN1_45S_50ntFlanks</td><td>10916</td><td>10939</td><td>24</td><td>CCGCGGCTGGACGAGGCGCCGCCG</td><td>80.3</td><td>-9.4</td><td>1.9</td><td>-5.1</td><td>3.7E-07</td><td>2.2E-06</td><td>BT20/MB468</td><td>MP</td></tr>
<tr><td>>hg38_21_+_8433172_8446622_RNA45SN1_45S_50ntFlanks@9240.9271.32|GCGGAGCGAGCGCACGGGGTCGGCGGCGACGT</td><td>>hg38_21_+_8433172_8446622_RNA45SN1_45S_50ntFlanks</td><td>9240</td><td>9271</td><td>32</td><td>GCGGAGCGAGCGCACGGGGTCGGCGGCGACGT</td><td>57.6</td><td>-8.9</td><td>1.8</td><td>-5.1</td><td>3.7E-07</td><td>2.2E-06</td><td>BT20/MB468</td><td>MP</td></tr>
</table>

<table>
<tr><td>>hg38_21_+_8433172_8446622_RNA45SN1_45S_50ntFlanks@12996.13017.22|TCGCTGCGATCTATTGAAAGTC</td><td>>hg38_21_+_8433172_8446622_RNA45SN1_45S_50ntFlanks</td><td>12996</td><td>13017</td><td>22</td><td>TCGCTGCGATCTATTGAAAGTC</td><td>195.9</td><td>3.8</td><td>0.8</td><td>4.8</td><td>1.4E-06</td><td>2.2E-06</td><td>MB231/MB468</td><td>MITO</td></tr>
<tr><td>>hg38_21_+_8433172_8446622_RNA45SN1_45S_50ntFlanks@10733.10752.20|GGTGTAAATCTCGCGCCGGG</td><td>>hg38_21_+_8433172_8446622_RNA45SN1_45S_50ntFlanks</td><td>10733</td><td>10752</td><td>20</td><td>GGTGTAAATCTCGCGCCGGG</td><td>54.3</td><td>-8.8</td><td>1.7</td><td>-5.1</td><td>3.7E-07</td><td>2.2E-06</td><td>BT20/MB468</td><td>MP</td></tr>
<tr><td>>hg38_21_+_8433172_8446622_RNA45SN1_45S_50ntFlanks@12419.12460.42|GATGTCGGCTCTTCCTATCATTGTGAAGCAGAATTCACCAAG</td><td>>hg38_21_+_8433172_8446622_RNA45SN1_45S_50ntFlanks</td><td>12419</td><td>12460</td><td>42</td><td>GATGTCGGCTCTTCCTATCATTGTGAAGCAGAATTCACCAAG</td><td>390.1</td><td>-4.1</td><td>0.8</td><td>-5.3</td><td>1.0E-07</td><td>2.2E-06</td><td>BT20/MB231</td><td>CYTO</td></tr>
<tr><td>>hg38_21_+_8433172_8446622_RNA45SN1_45S_50ntFlanks@3840.3879.40|CTCCTACTTGGATAACTGTGGTAATTCTAGAGCTAATACA</td><td>>hg38_21_+_8433172_8446622_RNA45SN1_45S_50ntFlanks</td><td>3840</td><td>3879</td><td>40</td><td>CTCCTACTTGGATAACTGTGGTAATTCTAGAGCTAATACA</td><td>616.7</td><td>4.4</td><td>0.9</td><td>4.9</td><td>1.0E-06</td><td>2.2E-06</td><td>BT20/MB468</td><td>MITO</td></tr>
<tr><td>>MI0000266|hsa-mir-10a&WithFlank&17|-|48579832|48579953@27.49.23&[MIMAT0000253&hsa-miR-10a-5p&offsets|-1|-1;m-18&17|-|48579905|48579926&offsets|-1|0]|ATACCCTGTAGATCCGAATTTGT</td><td>>MI0000266|hsa-mir-10a&WithFlank&17|-|48579832|48579953</td><td>27</td><td>49</td><td>23</td><td>ATACCCTGTAGATCCGAATTTGT</td><td>81.4</td><td>10.1</td><td>2.0</td><td>5.1</td><td>3.7E-07</td><td>2.2E-06</td><td>BT20/MB468</td><td>MP</td></tr>
<tr><td>>hg38_MT_+_598_1651_RNR1_12S_50ntFlanks@52.84.33|ATAGGTTTGGTCCTAGCCTTTCTATTAGCTCTT</td><td>>hg38_MT_+_598_1651_RNR1_12S_50ntFlanks</td><td>52</td><td>84</td><td>33</td><td>ATAGGTTTGGTCCTAGCCTTTCTATTAGCTCTT</td><td>3358.0</td><td>5.1</td><td>1.0</td><td>4.9</td><td>1.0E-06</td><td>2.2E-06</td><td>BT20/MB468</td><td>MITO</td></tr>
<tr><td>>MI0003776|hsa-mir-320b-1&WithFlank&1|+|116671740|116671823@48.67.20(+1U);MI0003839|hsa-mir-320b-2&WithFlank&1|-|224257034|224257116@47.66.20(+1U);MI0000542|hsa-mir-320a&WithFlank&8|-|22244960|22245043@48.67.20(+1U)&[MIMAT0005792&hsa-miR-320b&offsets|0|-2(+1U);m-183&1|+|116671787|116671806&offsets|0|0(+1U)];[MIMAT0005792_1&hsa-miR-320b&offsets|0|-2(+1U);m-183&1|-|224257051|224257070&offsets|0|0(+1U)];[MIMAT0000510&hsa-miR-320a-3p&offsets|0|-2(+1U);m-60&8|-|22244975|22244996&offsets|0|-2(+1U)]|AAAAGCTGGGTTGAGAGGGCT</td><td>>MI0003776|hsa-mir-320b-1&WithFlank&1|+|116671740|116671823</td><td>48</td><td>67</td><td>20(+1U)</td><td>AAAAGCTGGGTTGAGAGGGCT</td><td>1841.1</td><td>7.3</td><td>1.4</td><td>5.1</td><td>3.7E-07</td><td>2.2E-06</td><td>BT20/MB468</td><td>MP</td></tr>
<tr><td>>hg38_21_+_8433172_8446622_RNA45SN1_45S_50ntFlanks@11821.11853.33|CTATCCAGCGAAACCACAGCCAAGGGAACGGGC</td><td>>hg38_21_+_8433172_8446622_RNA45SN1_45S_50ntFlanks</td><td>11821</td><td>11853</td><td>33</td><td>CTATCCAGCGAAACCACAGCCAAGGGAACGGGC</td><td>54.0</td><td>-8.8</td><td>1.7</td><td>-5.1</td><td>3.7E-07</td><td>2.2E-06</td><td>BT20/MB468</td><td>MP</td></tr>
<tr><td>>hg38_21_+_8433172_8446622_RNA45SN1_45S_50ntFlanks@12539.12568.30|ATGATGTGTTGTTGCCATGGTAATCCTGCT</td><td>>hg38_21_+_8433172_8446622_RNA45SN1_45S_50ntFlanks</td><td>12539</td><td>12568</td><td>30</td><td>ATGATGTGTTGTTGCCATGGTAATCCTGCT</td><td>162.2</td><td>-6.2</td><td>1.3</td><td>-4.8</td><td>1.4E-06</td><td>2.2E-06</td><td>MB231/MB468</td><td>NUC</td></tr>
<tr><td>>hg38_21_+_8433172_8446622_RNA45SN1_45S_50ntFlanks@4243.4263.21|CTTTCGAGGCCCTGTAATTGG</td><td>>hg38_21_+_8433172_8446622_RNA45SN1_45S_50ntFlanks</td><td>4243</td><td>4263</td><td>21</td><td>CTTTCGAGGCCCTGTAATTGG</td><td>523.1</td><td>-5.5</td><td>1.1</td><td>-4.8</td><td>1.4E-06</td><td>2.2E-06</td><td>MB231/MB468</td><td>NUC</td></tr>
<tr><td>>hg38_21_+_8433172_8446622_RNA45SN1_45S_50ntFlanks@8180.8211.32|TGATCGAGGCCCAGCCCGTGGACGGTGTGAGG</td><td>>hg38_21_+_8433172_8446622_RNA45SN1_45S_50ntFlanks</td><td>8180</td><td>8211</td><td>32</td><td>TGATCGAGGCCCAGCCCGTGGACGGTGTGAGG</td><td>346.0</td><td>-6.6</td><td>1.4</td><td>-4.8</td><td>1.4E-06</td><td>2.2E-06</td><td>MB231/MB468</td><td>NUC</td></tr>
<tr><td>>hg38_MT_+_1621_3279_RNR2_16S_50ntFlanks@1428.1458.31|ATTAAAGTCCTACGTGATCTGAGTTCAGACC</td><td>>hg38_MT_+_1621_3279_RNR2_16S_50ntFlanks</td><td>1428</td><td>1458</td><td>31</td><td>ATTAAAGTCCTACGTGATCTGAGTTCAGACC</td><td>377.3</td><td>3.4</td><td>0.7</td><td>4.8</td><td>1.4E-06</td><td>2.2E-06</td><td>MB231/MB468</td><td>MITO</td></tr>
<tr><td>>hg38_21_+_8433172_8446622_RNA45SN1_45S_50ntFlanks@11316.11338.23|CACCCCACGTCTCGTCGCGCGCG</td><td>>hg38_21_+_8433172_8446622_RNA45SN1_45S_50ntFlanks</td><td>11316</td><td>11338</td><td>23</td><td>CACCCCACGTCTCGTCGCGCGCG</td><td>351.1</td><td>-7.3</td><td>1.4</td><td>-5.1</td><td>3.7E-07</td><td>2.2E-06</td><td>BT20/MB468</td><td>MP</td></tr>
<tr><td>>hg38_21_+_8433172_8446622_RNA45SN1_45S_50ntFlanks@6669.6699.31|CACTCGGCTCGTGCGTCGATGAAGAACGCAG</td><td>>hg38_21_+_8433172_8446622_RNA45SN1_45S_50ntFlanks</td><td>6669</td><td>6699</td><td>31</td><td>CACTCGGCTCGTGCGTCGATGAAGAACGCAG</td><td>957.9</td><td>-3.7</td><td>0.8</td><td>-4.8</td><td>1.4E-06</td><td>2.2E-06</td><td>MB231/MB468</td><td>MITO</td></tr>
<tr><td>>hg38_21_+_8433172_8446622_RNA45SN1_45S_50ntFlanks@12641.12661.21|CCATCTGTGGGATTATGACTG</td><td>>hg38_21_+_8433172_8446622_RNA45SN1_45S_50ntFlanks</td><td>12641</td><td>12661</td><td>21</td><td>CCATCTGTGGGATTATGACTG</td><td>379.4</td><td>11.9</td><td>2.2</td><td>5.3</td><td>9.2E-08</td><td>2.2E-06</td><td>BT20/MB231</td><td>MP</td></tr>
<tr><td>>MI0000456|hsa-mir-140&WithFlank&16|+|69933075|69933186@70.90.21(+1U)&[MIMAT0004597&hsa-miR-140-3p&offsets|+2|+2(+1U);m-45&16|+|69933143|69933164&offsets|+1|0(+1U)]|CCACAGGGTAGAACCACGGACT</td><td>>MI0000456|hsa-mir-140&WithFlank&16|+|69933075|69933186</td><td>70</td><td>90</td><td>21(+1U)</td><td>CCACAGGGTAGAACCACGGACT</td><td>41.2</td><td>8.6</td><td>1.8</td><td>4.9</td><td>1.2E-06</td><td>2.2E-06</td><td>MB231/MB468</td><td>MP</td></tr>
<tr><td>>MI0000650|hsa-mir-200c&WithFlank&12|+|6963693|6963772@50.69.20&[MIMAT0000617&hsa-miR-200c-3p&offsets|0|-3;m-54&12|+|6963742|6963764&offsets|0|-3]|TAATACTGCCGGGTAATGAT</td><td>>MI0000650|hsa-mir-200c&WithFlank&12|+|6963693|6963772</td><td>50</td><td>69</td><td>20</td><td>TAATACTGCCGGGTAATGAT</td><td>1489.7</td><td>6.9</td><td>1.4</td><td>5.1</td><td>3.7E-07</td><td>2.2E-06</td><td>BT20/MB468</td><td>MP</td></tr>
<tr><td>>hg38_21_+_8433172_8446622_RNA45SN1_45S_50ntFlanks@10866.10901.36|CTAAGGGCTGGGTCGGTCGGGCTGGGGCGCGAAGCG</td><td>>hg38_21_+_8433172_8446622_RNA45SN1_45S_50ntFlanks</td><td>10866</td><td>10901</td><td>36</td><td>CTAAGGGCTGGGTCGGTCGGGCTGGGGCGCGAAGCG</td><td>743.6</td><td>-3.8</td><td>0.8</td><td>-4.9</td><td>1.0E-06</td><td>2.2E-06</td><td>BT20/MB468</td><td>MITO</td></tr>
<tr><td>>hg38_21_+_8433172_8446622_RNA45SN1_45S_50ntFlanks@10687.10710.24|CGGCGTCCGGTGAGCTCTCGCTGG</td><td>>hg38_21_+_8433172_8446622_RNA45SN1_45S_50ntFlanks</td><td>10687</td><td>10710</td><td>24</td><td>CGGCGTCCGGTGAGCTCTCGCTGG</td><td>6292.7</td><td>-6.3</td><td>1.3</td><td>-4.9</td><td>1.2E-06</td><td>2.2E-06</td><td>BT20/MB468</td><td>CYTO</td></tr>
<tr><td>>hg38_21_+_8433172_8446622_RNA45SN1_45S_50ntFlanks@10409.10432.24|GTCAGTCGGTCCTGAGAGATGGGC</td><td>>hg38_21_+_8433172_8446622_RNA45SN1_45S_50ntFlanks</td><td>10409</td><td>10432</td><td>24</td><td>GTCAGTCGGTCCTGAGAGATGGGC</td><td>267.9</td><td>-2.8</td><td>0.6</td><td>-4.8</td><td>1.4E-06</td><td>2.2E-06</td><td>MB231/MB468</td><td>NUC</td></tr>
<tr><td>>hg38_21_+_8433172_8446622_RNA45SN1_45S_50ntFlanks@10841.10859.19|CGTAACTTCGGGATAAGGA</td><td>>hg38_21_+_8433172_8446622_RNA45SN1_45S_50ntFlanks</td><td>10841</td><td>10859</td><td>19</td><td>CGTAACTTCGGGATAAGGA</td><td>215.9</td><td>-5.3</td><td>1.1</td><td>-4.8</td><td>1.4E-06</td><td>2.2E-06</td><td>MB231/MB468</td><td>NUC</td></tr>
<tr><td>>MI0000767|hsa-mir-365a&WithFlank&16|+|14309279|14309377@62.82.21;MI0000769|hsa-mir-365b&WithFlank&17|+|31575405|31575527@74.94.21&[MIMAT0000710&hsa-miR-365a-3p&offsets|0|-1;m-176&16|+|14309340|14309361&offsets|0|-1];[MIMAT0022834&hsa-miR-365b-3p&offsets|0|-1;m-175&17|+|31575478|31575499&offsets|0|-1]|TAATGCCCCTAAAAATCCTTA</td><td>>MI0000767|hsa-mir-365a&WithFlank&16|+|14309279|14309377</td><td>62</td><td>82</td><td>21</td><td>TAATGCCCCTAAAAATCCTTA</td><td>537.6</td><td>4.6</td><td>0.9</td><td>5.3</td><td>1.0E-07</td><td>2.2E-06</td><td>BT20/MB231</td><td>CYTO</td></tr>
<tr><td>>hg38_21_+_8433172_8446622_RNA45SN1_45S_50ntFlanks@12995.13038.44|CTCGCTGCGATCTATTGAAAGTCAGCCCTCGACACAAGGGTTTG</td><td>>hg38_21_+_8433172_8446622_RNA45SN1_45S_50ntFlanks</td><td>12995</td><td>13038</td><td>44</td><td>CTCGCTGCGATCTATTGAAAGTCAGCCCTCGACACAAGGGTTTG</td><td>1029.7</td><td>5.8</td><td>1.2</td><td>4.8</td><td>1.4E-06</td><td>2.2E-06</td><td>MB231/MB468</td><td>NUC</td></tr>
<tr><td>>hg38_21_+_8433172_8446622_RNA45SN1_45S_50ntFlanks@10922.10941.20|CTGGACGAGGCGCCGCCGCC</td><td>>hg38_21_+_8433172_8446622_RNA45SN1_45S_50ntFlanks</td><td>10922</td><td>10941</td><td>20</td><td>CTGGACGAGGCGCCGCCGCC</td><td>705.0</td><td>-5.8</td><td>1.2</td><td>-4.8</td><td>1.3E-06</td><td>2.2E-06</td><td>MB231/MB468</td><td>CYTO</td></tr>
<tr><td>>hg38_21_+_8433172_8446622_RNA45SN1_45S_50ntFlanks@10681.10712.32|TTCCGGCGGCGTCCGGTGAGCTCTCGCTGGCC</td><td>>hg38_21_+_8433172_8446622_RNA45SN1_45S_50ntFlanks</td><td>10681</td><td>10712</td><td>32</td><td>TTCCGGCGGCGTCCGGTGAGCTCTCGCTGGCC</td><td>6670.7</td><td>-9.3</td><td>1.9</td><td>-4.8</td><td>1.5E-06</td><td>2.2E-06</td><td>BT20/MB468</td><td>NUC</td></tr>
<tr><td>>hg38_21_+_8433172_8446622_RNA45SN1_45S_50ntFlanks@9956.9994.39|GAAGTCGGAATCCGCTAAGGAGTGTGTAACAACTCACCT</td><td>>hg38_21_+_8433172_8446622_RNA45SN1_45S_50ntFlanks</td><td>9956</td><td>9994</td><td>39</td><td>GAAGTCGGAATCCGCTAAGGAGTGTGTAACAACTCACCT</td><td>60.4</td><td>9.7</td><td>1.9</td><td>5.1</td><td>3.8E-07</td><td>2.2E-06</td><td>BT20/MB468</td><td>MP</td></tr>
<tr><td>>hg38_MT_+_598_1651_RNR1_12S_50ntFlanks@912.950.39|TTCAAAGGACATTTAACTAAAACCCCTACGCATTTATAT</td><td>>hg38_MT_+_598_1651_RNR1_12S_50ntFlanks</td><td>912</td><td>950</td><td>39</td><td>TTCAAAGGACATTTAACTAAAACCCCTACGCATTTATAT</td><td>90.6</td><td>10.2</td><td>2.0</td><td>5.1</td><td>3.8E-07</td><td>2.2E-06</td><td>BT20/MB468</td><td>MP</td></tr>
<tr><td>>hg38_21_+_8433172_8446622_RNA45SN1_45S_50ntFlanks@13009.13028.20|TTGAAAGTCAGCCCTCGACA</td><td>>hg38_21_+_8433172_8446622_RNA45SN1_45S_50ntFlanks</td><td>13009</td><td>13028</td><td>20</td><td>TTGAAAGTCAGCCCTCGACA</td><td>63.5</td><td>-9.1</td><td>1.8</td><td>-5.1</td><td>3.8E-07</td><td>2.2E-06</td><td>BT20/MB468</td><td>MP</td></tr>
<tr><td>>hg38_21_+_8433172_8446622_RNA45SN1_45S_50ntFlanks@5489.5521.33|CTGAGAAGACGGTCGAACTTGACTATCTAGAGG</td><td>>hg38_21_+_8433172_8446622_RNA45SN1_45S_50ntFlanks</td><td>5489</td><td>5521</td><td>33</td><td>CTGAGAAGACGGTCGAACTTGACTATCTAGAGG</td><td>117.1</td><td>-5.1</td><td>1.1</td><td>-4.9</td><td>1.2E-06</td><td>2.2E-06</td><td>MB231/MB468</td><td>MP</td></tr>
<tr><td>>hg38_21_+_8433172_8446622_RNA45SN1_45S_50ntFlanks@12407.12437.31|TTTTGATCCTTCGATGTCGGCTCTTCCTATC</td><td>>hg38_21_+_8433172_8446622_RNA45SN1_45S_50ntFlanks</td><td>12407</td><td>12437</td><td>31</td><td>TTTTGATCCTTCGATGTCGGCTCTTCCTATC</td><td>175.4</td><td>-5.1</td><td>1.0</td><td>-4.9</td><td>1.2E-06</td><td>2.2E-06</td><td>MB231/MB468</td><td>MP</td></tr>
<tr><td>>trna11_GluTTC_15_-_26327381_26327452@41.75.35,trna3_GluTTC_13_-_45492062_45492133@41.75.35|AGGCGGCCCGGGTTCGACTCCCGGTGTGGGAACCA</td><td>>trna11_GluTTC_15_-_26327381_26327452</td><td>41</td><td>75</td><td>35</td><td>AGGCGGCCCGGGTTCGACTCCCGGTGTGGGAACCA</td><td>65.3</td><td>9.2</td><td>1.9</td><td>4.9</td><td>1.2E-06</td><td>2.2E-06</td><td>MB231/MB468</td><td>MP</td></tr>
<tr><td>>hg38_21_+_8433172_8446622_RNA45SN1_45S_50ntFlanks@9930.9952.23|ATATAGACAGCAGGACGGTGGCC</td><td>>hg38_21_+_8433172_8446622_RNA45SN1_45S_50ntFlanks</td><td>9930</td><td>9952</td><td>23</td><td>ATATAGACAGCAGGACGGTGGCC</td><td>262.8</td><td>-7.0</td><td>1.4</td><td>-4.9</td><td>1.2E-06</td><td>2.3E-06</td><td>MB231/MB468</td><td>MP</td></tr>
<tr><td>>hg38_21_+_8433172_8446622_RNA45SN1_45S_50ntFlanks@12417.12458.42|TCGATGTCGGCTCTTCCTATCATTGTGAAGCAGAATTCACCA</td><td>>hg38_21_+_8433172_8446622_RNA45SN1_45S_50ntFlanks</td><td>12417</td><td>12458</td><td>42</td><td>TCGATGTCGGCTCTTCCTATCATTGTGAAGCAGAATTCACCA</td><td>219.2</td><td>-4.3</td><td>0.8</td><td>-5.3</td><td>1.0E-07</td><td>2.3E-06</td><td>BT20/MB231</td><td>CYTO</td></tr>
<tr><td>>hg38_21_+_8433172_8446622_RNA45SN1_45S_50ntFlanks@4729.4761.33|TCAAGAACGAAAGTCGGAGGTTCGAAGACGATC</td><td>>hg38_21_+_8433172_8446622_RNA45SN1_45S_50ntFlanks</td><td>4729</td><td>4761</td><td>33</td><td>TCAAGAACGAAAGTCGGAGGTTCGAAGACGATC</td><td>240.8</td><td>4.6</td><td>1.0</td><td>4.8</td><td>1.4E-06</td><td>2.3E-06</td><td>MB231/MB468</td><td>CYTO</td></tr>
<tr><td>>hg38_21_+_8433172_8446622_RNA45SN1_45S_50ntFlanks@11582.11601.20|ACTGGTGCGGACCAGGGGAA</td><td>>hg38_21_+_8433172_8446622_RNA45SN1_45S_50ntFlanks</td><td>11582</td><td>11601</td><td>20</td><td>ACTGGTGCGGACCAGGGGAA</td><td>462.4</td><td>-7.3</td><td>1.5</td><td>-4.9</td><td>1.2E-06</td><td>2.3E-06</td><td>MB231/MB468</td><td>MP</td></tr>
<tr><td>>trna14_LysTTT_11_-_59327808_59327880@1.35.35,trna2_LysTTT_17_+_8022473_8022545@1.35.35,trna54_LysTTT_1_+_204475655_204475727@1.35.35,trna5_LysTTT_11_+_59323902_59323974@1.35.35,trna62_LysTTT_1_-_204476158_204476230@1.35.35,trna76_LysTTT_6_+_28918806_28918878@1.35.35|GCCCGGATAGCTCAGTCGGTAGAGCATCAGACTTT</td><td>>trna14_LysTTT_11_-_59327808_59327880</td><td>1</td><td>35</td><td>35</td><td>GCCCGGATAGCTCAGTCGGTAGAGCATCAGACTTT</td><td>56.8</td><td>9.0</td><td>1.9</td><td>4.9</td><td>1.2E-06</td><td>2.3E-06</td><td>MB231/MB468</td><td>MP</td></tr>
<tr><td>>hg38_21_+_8433172_8446622_RNA45SN1_45S_50ntFlanks@11322.11342.21|ACGTCTCGTCGCGCGCGCGTC</td><td>>hg38_21_+_8433172_8446622_RNA45SN1_45S_50ntFlanks</td><td>11322</td><td>11342</td><td>21</td><td>ACGTCTCGTCGCGCGCGCGTC</td><td>2383.7</td><td>-4.6</td><td>0.9</td><td>-4.8</td><td>1.4E-06</td><td>2.3E-06</td><td>MB231/MB468</td><td>CYTO</td></tr>
<tr><td>>hg38_21_+_8433172_8446622_RNA45SN1_45S_50ntFlanks@11894.11916.23|ACTCTAGTCTGGCACGGTGAAGA</td><td>>hg38_21_+_8433172_8446622_RNA45SN1_45S_50ntFlanks</td><td>11894</td><td>11916</td><td>23</td><td>ACTCTAGTCTGGCACGGTGAAGA</td><td>660.2</td><td>-7.0</td><td>1.5</td><td>-4.8</td><td>1.4E-06</td><td>2.3E-06</td><td>MB231/MB468</td><td>NUC</td></tr>
<tr><td>>hg38_21_+_8433172_8446622_RNA45SN1_45S_50ntFlanks@11731.11751.21|ACTATGACTCTCTTAAGGTAG</td><td>>hg38_21_+_8433172_8446622_RNA45SN1_45S_50ntFlanks</td><td>11731</td><td>11751</td><td>21</td><td>ACTATGACTCTCTTAAGGTAG</td><td>157.0</td><td>-6.6</td><td>1.4</td><td>-4.9</td><td>1.2E-06</td><td>2.3E-06</td><td>MB231/MB468</td><td>MP</td></tr>
<tr><td>>MI0000078|hsa-mir-22&WithFlank&17|-|1713897|1713993@59.80.22&[MIMAT0000077&hsa-miR-22-3p&offsets|0|0;m-2&17|-|1713914|1713935&offsets|0|0]|AAGCTGCCAGTTGAAGAACTGT</td><td>>MI0000078|hsa-mir-22&WithFlank&17|-|1713897|1713993</td><td>59</td><td>80</td><td>22</td><td>AAGCTGCCAGTTGAAGAACTGT</td><td>24956.5</td><td>5.3</td><td>1.0</td><td>5.1</td><td>3.8E-07</td><td>2.3E-06</td><td>BT20/MB468</td><td>MP</td></tr>
<tr><td>>MI0000478|hsa-mir-149&WithFlank&2|+|240455995|240456095@21.42.22&[MIMAT0000450&hsa-miR-149-5p&offsets|0|-1;m-195&2|+|240456015|240456037&offsets|0|-1]|TCTGGCTCCGTGTCTTCACTCC</td><td>>MI0000478|hsa-mir-149&WithFlank&2|+|240455995|240456095</td><td>21</td><td>42</td><td>22</td><td>TCTGGCTCCGTGTCTTCACTCC</td><td>688.8</td><td>7.8</td><td>1.5</td><td>5.1</td><td>3.8E-07</td><td>2.3E-06</td><td>BT20/MB468</td><td>MP</td></tr>
<tr><td>>hg38_21_+_8433172_8446622_RNA45SN1_45S_50ntFlanks@12300.12318.19|GGGCCTCACGATCCTTCTG</td><td>>hg38_21_+_8433172_8446622_RNA45SN1_45S_50ntFlanks</td><td>12300</td><td>12318</td><td>19</td><td>GGGCCTCACGATCCTTCTG</td><td>1384.4</td><td>-7.4</td><td>1.5</td><td>-4.9</td><td>1.0E-06</td><td>2.3E-06</td><td>BT20/MB468</td><td>MITO</td></tr>
<tr><td>>hg38_21_+_8433172_8446622_RNA45SN1_45S_50ntFlanks@6726.6743.18|CAGGACACATTGATCATC</td><td>>hg38_21_+_8433172_8446622_RNA45SN1_45S_50ntFlanks</td><td>6726</td><td>6743</td><td>18</td><td>CAGGACACATTGATCATC</td><td>177.9</td><td>-6.8</td><td>1.4</td><td>-4.9</td><td>1.2E-06</td><td>2.3E-06</td><td>MB231/MB468</td><td>MP</td></tr>
<tr><td>>hg38_1_-_228634819_228635039_RNA5S12_5S_50ntFlanks@133.171.39|ATGGGAGACCGCCTGGGAATACCGGGTGCTGTAGGCTTT</td><td>>hg38_1_-_228634819_228635039_RNA5S12_5S_50ntFlanks</td><td>133</td><td>171</td><td>39</td><td>ATGGGAGACCGCCTGGGAATACCGGGTGCTGTAGGCTTT</td><td>3894.1</td><td>4.1</td><td>0.8</td><td>4.8</td><td>1.4E-06</td><td>2.3E-06</td><td>MB231/MB468</td><td>NUC</td></tr>
<tr><td>>hg38_21_+_8433172_8446622_RNA45SN1_45S_50ntFlanks@7971.7988.18|GAGACGCGACCTCAGATC</td><td>>hg38_21_+_8433172_8446622_RNA45SN1_45S_50ntFlanks</td><td>7971</td><td>7988</td><td>18</td><td>GAGACGCGACCTCAGATC</td><td>379.3</td><td>3.7</td><td>0.8</td><td>4.8</td><td>1.4E-06</td><td>2.3E-06</td><td>MB231/MB468</td><td>NUC</td></tr>
</table>

<table>
<tr><td>>MI0002470|hsa-mir-486-1&WithFlank&8|-|41660435|41660514@10.29.20;MI0023622|hsa-mir-486-2&WithFlank&8|+|41660438|41660513@7.26.20&[MIMAT0002177&hsa-miR-486-5p&offsets|0|-2;m-107&8|-|41660484|41660505&offsets|0|-2];[MIMAT0002177_1&hsa-miR-486-5p&offsets|0|-2;TJU_CMC_MD2.ID03138.5p-miR&offsets|0|-4;m-106&8|+|41660444|41660465&offsets|0|-2]|TCCTGTACTGAGCTGCCCCG</td><td>>MI0002470|hsa-mir-486-1&WithFlank&8|-|41660435|41660514</td><td>10</td><td>29</td><td>20</td><td>TCCTGTACTGAGCTGCCCCG</td><td>43.5</td><td>8.7</td><td>1.8</td><td>4.9</td><td>1.2E-06</td><td>2.3E-06</td><td>MB231/MB468</td><td>MP</td></tr>
<tr><td>>hg38_21_+_8433172_8446622_RNA45SN1_45S_50ntFlanks@10391.10422.32|AACAGCAGTTGAACATGGGTCAGTCGGTCCTG</td><td>>hg38_21_+_8433172_8446622_RNA45SN1_45S_50ntFlanks</td><td>10391</td><td>10422</td><td>32</td><td>AACAGCAGTTGAACATGGGTCAGTCGGTCCTG</td><td>311.0</td><td>-6.9</td><td>1.4</td><td>-4.9</td><td>1.2E-06</td><td>2.3E-06</td><td>MB231/MB468</td><td>MP</td></tr>
<tr><td>>hg38_21_+_8433172_8446622_RNA45SN1_45S_50ntFlanks@10466.10486.21|CGTTGCCCTCGGCCGATCGAA</td><td>>hg38_21_+_8433172_8446622_RNA45SN1_45S_50ntFlanks</td><td>10466</td><td>10486</td><td>21</td><td>CGTTGCCCTCGGCCGATCGAA</td><td>314.1</td><td>-6.8</td><td>1.4</td><td>-4.8</td><td>1.5E-06</td><td>2.3E-06</td><td>MB231/MB468</td><td>MITO</td></tr>
<tr><td>>hg38_21_+_8433172_8446622_RNA45SN1_45S_50ntFlanks@6732.6752.21|ACATTGATCATCGACACTTCG</td><td>>hg38_21_+_8433172_8446622_RNA45SN1_45S_50ntFlanks</td><td>6732</td><td>6752</td><td>21</td><td>ACATTGATCATCGACACTTCG</td><td>470.0</td><td>-5.3</td><td>1.1</td><td>-4.9</td><td>1.2E-06</td><td>2.3E-06</td><td>MB231/MB468</td><td>MP</td></tr>
<tr><td>>hg38_21_+_8433172_8446622_RNA45SN1_45S_50ntFlanks@9311.9336.26|AGTCTAACACGTGCGCGAGTCGGGGG</td><td>>hg38_21_+_8433172_8446622_RNA45SN1_45S_50ntFlanks</td><td>9311</td><td>9336</td><td>26</td><td>AGTCTAACACGTGCGCGAGTCGGGGG</td><td>172.4</td><td>-6.8</td><td>1.4</td><td>-4.9</td><td>1.2E-06</td><td>2.3E-06</td><td>MB231/MB468</td><td>MP</td></tr>
<tr><td>>trnaMT_ProTGG_MT_-_15956_16023@50.71.22|TTAAAGACTTTTTCTCTGACCA</td><td>>trnaMT_ProTGG_MT_-_15956_16023</td><td>50</td><td>71</td><td>22</td><td>TTAAAGACTTTTTCTCTGACCA</td><td>750.8</td><td>5.1</td><td>1.0</td><td>4.9</td><td>1.1E-06</td><td>2.3E-06</td><td>BT20/MB468</td><td>MITO</td></tr>
<tr><td>>hg38_21_+_8433172_8446622_RNA45SN1_45S_50ntFlanks@10412.10429.18|AGTCGGTCCTGAGAGATG</td><td>>hg38_21_+_8433172_8446622_RNA45SN1_45S_50ntFlanks</td><td>10412</td><td>10429</td><td>18</td><td>AGTCGGTCCTGAGAGATG</td><td>521.4</td><td>-7.7</td><td>1.6</td><td>-4.8</td><td>1.5E-06</td><td>2.3E-06</td><td>BT20/MB468</td><td>NUC</td></tr>
<tr><td>>hg38_21_+_8433172_8446622_RNA45SN1_45S_50ntFlanks@7992.8012.21|CGTGGCGACCCGCTGAATTTA</td><td>>hg38_21_+_8433172_8446622_RNA45SN1_45S_50ntFlanks</td><td>7992</td><td>8012</td><td>21</td><td>CGTGGCGACCCGCTGAATTTA</td><td>1470.0</td><td>-3.8</td><td>0.8</td><td>-4.9</td><td>1.1E-06</td><td>2.3E-06</td><td>BT20/MB468</td><td>MITO</td></tr>
<tr><td>>hg38_21_+_8433172_8446622_RNA45SN1_45S_50ntFlanks@4731.4751.21|AAGAACGAAAGTCGGAGGTTC</td><td>>hg38_21_+_8433172_8446622_RNA45SN1_45S_50ntFlanks</td><td>4731</td><td>4751</td><td>21</td><td>AAGAACGAAAGTCGGAGGTTC</td><td>729.2</td><td>-3.5</td><td>0.7</td><td>-4.8</td><td>1.5E-06</td><td>2.3E-06</td><td>MB231/MB468</td><td>MITO</td></tr>
<tr><td>>MI0000441|hsa-mir-30b&WithFlank&8|-|134800514|134800613@23.43.21(+1A)&[MIMAT0000420&hsa-miR-30b-5p&offsets|0|-1(+1A);m-76&8|-|134800570|134800591&offsets|0|-1(+1A)]|TGTAAACATCCTACACTCAGCA</td><td>>MI0000441|hsa-mir-30b&WithFlank&8|-|134800514|134800613</td><td>23</td><td>43</td><td>21(+1A)</td><td>TGTAAACATCCTACACTCAGCA</td><td>1118.4</td><td>9.0</td><td>1.9</td><td>4.8</td><td>1.5E-06</td><td>2.3E-06</td><td>MB231/MB468</td><td>MITO</td></tr>
<tr><td>>hg38_21_+_8433172_8446622_RNA45SN1_45S_50ntFlanks@8112.8138.27|GGGCGCGGGACATGTGGCGTACGGAAG</td><td>>hg38_21_+_8433172_8446622_RNA45SN1_45S_50ntFlanks</td><td>8112</td><td>8138</td><td>27</td><td>GGGCGCGGGACATGTGGCGTACGGAAG</td><td>206.4</td><td>11.0</td><td>2.1</td><td>5.3</td><td>9.4E-08</td><td>2.3E-06</td><td>BT20/MB231</td><td>MP</td></tr>
<tr><td>>hg38_21_+_8433172_8446622_RNA45SN1_45S_50ntFlanks@3780.3801.22|TACAGTGAAACTGCGAATGGCT</td><td>>hg38_21_+_8433172_8446622_RNA45SN1_45S_50ntFlanks</td><td>3780</td><td>3801</td><td>22</td><td>TACAGTGAAACTGCGAATGGCT</td><td>389.6</td><td>-6.0</td><td>1.2</td><td>-4.8</td><td>1.4E-06</td><td>2.3E-06</td><td>MB231/MB468</td><td>NUC</td></tr>
<tr><td>>hg38_21_+_8433172_8446622_RNA45SN1_45S_50ntFlanks@10827.10844.18|CGGCAAGCCGGATCCGTA</td><td>>hg38_21_+_8433172_8446622_RNA45SN1_45S_50ntFlanks</td><td>10827</td><td>10844</td><td>18</td><td>CGGCAAGCCGGATCCGTA</td><td>60.5</td><td>-9.0</td><td>1.8</td><td>-5.1</td><td>3.8E-07</td><td>2.3E-06</td><td>BT20/MB468</td><td>MP</td></tr>
<tr><td>>hg38_21_+_8433172_8446622_RNA45SN1_45S_50ntFlanks@12713.12742.30|GGAGCCTCGGTTGGCCTCGGATAGCCGGTC</td><td>>hg38_21_+_8433172_8446622_RNA45SN1_45S_50ntFlanks</td><td>12713</td><td>12742</td><td>30</td><td>GGAGCCTCGGTTGGCCTCGGATAGCCGGTC</td><td>63.9</td><td>-9.1</td><td>1.8</td><td>-5.1</td><td>3.8E-07</td><td>2.3E-06</td><td>BT20/MB468</td><td>MP</td></tr>
<tr><td>>hg38_21_+_8433172_8446622_RNA45SN1_45S_50ntFlanks@10780.10801.22|AAGGTGAACAGCCTCTGGCATG</td><td>>hg38_21_+_8433172_8446622_RNA45SN1_45S_50ntFlanks</td><td>10780</td><td>10801</td><td>22</td><td>AAGGTGAACAGCCTCTGGCATG</td><td>304.7</td><td>-5.4</td><td>1.1</td><td>-4.8</td><td>1.5E-06</td><td>2.3E-06</td><td>BT20/MB468</td><td>NUC</td></tr>
<tr><td>>hg38_21_+_8433172_8446622_RNA45SN1_45S_50ntFlanks@10926.10947.22|ACGAGGCGCCGCCGCCCCCCCC</td><td>>hg38_21_+_8433172_8446622_RNA45SN1_45S_50ntFlanks</td><td>10926</td><td>10947</td><td>22</td><td>ACGAGGCGCCGCCGCCCCCCCC</td><td>218.6</td><td>-11.6</td><td>2.4</td><td>-4.9</td><td>1.2E-06</td><td>2.3E-06</td><td>MB231/MB468</td><td>MP</td></tr>
<tr><td>>MI0003599|hsa-mir-589&WithFlank&7|-|5495813|5495923@30.51.22(+1U)&[MIMAT0004799&hsa-miR-589-5p&offsets|0|0(+1U);m-162&7|-|5495874|5495894&offsets|0|+1(+1U)]|TGAGAACCACGTCTGCTCTGAGT</td><td>>MI0003599|hsa-mir-589&WithFlank&7|-|5495813|5495923</td><td>30</td><td>51</td><td>22(+1U)</td><td>TGAGAACCACGTCTGCTCTGAGT</td><td>53.1</td><td>9.0</td><td>1.8</td><td>4.9</td><td>1.2E-06</td><td>2.3E-06</td><td>MB231/MB468</td><td>MP</td></tr>
<tr><td>>hg38_21_+_8433172_8446622_RNA45SN1_45S_50ntFlanks@11848.11871.24|ACGGGCTTGGCGGAATCAGCGGGG</td><td>>hg38_21_+_8433172_8446622_RNA45SN1_45S_50ntFlanks</td><td>11848</td><td>11871</td><td>24</td><td>ACGGGCTTGGCGGAATCAGCGGGG</td><td>344.5</td><td>-5.7</td><td>1.2</td><td>-4.9</td><td>1.2E-06</td><td>2.3E-06</td><td>MB231/MB468</td><td>MP</td></tr>
<tr><td>>trna10_AspGTC_12_-_125424193_125424264@37.70.34,trna12_AspGTC_12_-_125411891_125411962@37.70.34,trna144_AspGTC_6_-_27551236_27551307@37.70.34,trna38_AspGTC_17_-_8125556_8125627@37.70.34,trna45_AspGTC_6_+_27447453_27447524@37.70.34,trna48_AspGTC_6_+_27471523_27471594@37.70.34,trna4_AspGTC_12_+_96429799_96429870@37.70.34,trna69_AspGTC_1_-_161440205_161440276@37.70.34,trna72_AspGTC_1_-_161432824_161432895@37.70.34,trna75_AspGTC_1_-_161425414_161425485@37.70.34,trna78_AspGTC_1_-_161418033_161418104@37.70.34,trna81_AspGTC_1_-_161410615_161410686@37.70.34|ACGCGGGAGACCGGGGTTCGATTCCCCGACGGGG</td><td>>trna10_AspGTC_12_-_125424193_125424264</td><td>37</td><td>70</td><td>34</td><td>ACGCGGGAGACCGGGGTTCGATTCCCCGACGGGG</td><td>371.1</td><td>-5.7</td><td>1.2</td><td>-4.8</td><td>1.5E-06</td><td>2.3E-06</td><td>BT20/MB468</td><td>NUC</td></tr>
<tr><td>>hg38_21_+_8433172_8446622_RNA45SN1_45S_50ntFlanks@7975.8001.27|CGCGACCTCAGATCAGACGTGGCGACC</td><td>>hg38_21_+_8433172_8446622_RNA45SN1_45S_50ntFlanks</td><td>7975</td><td>8001</td><td>27</td><td>CGCGACCTCAGATCAGACGTGGCGACC</td><td>2726.0</td><td>3.3</td><td>0.6</td><td>5.2</td><td>1.6E-07</td><td>2.3E-06</td><td>BT20/MB231</td><td>NUC</td></tr>
<tr><td>>trna10_LysCTT_16_+_3241501_3241573@1.29.29,trna13_LysCTT_14_-_58706613_58706685@1.29.29,trna2_LysCTT_15_+_79152904_79152976@1.29.29|GCCCGGCTAGCTCAGTCGGTAGAGCATGG</td><td>>trna10_LysCTT_16_+_3241501_3241573</td><td>1</td><td>29</td><td>29</td><td>GCCCGGCTAGCTCAGTCGGTAGAGCATGG</td><td>383.7</td><td>-5.8</td><td>1.2</td><td>-4.9</td><td>1.2E-06</td><td>2.3E-06</td><td>MB231/MB468</td><td>MP</td></tr>
<tr><td>>am_trna116_GluCTC_1_-_145399233_145399304@11.28.18,trna25_GluCTC_2_-_71273488_71273560@11.28.18,trna59_GluCTC_1_+_249168447_249168518@11.28.18,trna71_GluCTC_1_-_161439189_161439260@11.28.18,trna74_GluCTC_1_-_161431809_161431880@11.28.18,trna77_GluCTC_1_-_161424398_161424469@11.28.18,trna77_GluCTC_6_+_28949976_28950047@11.28.18,trna80_GluCTC_1_-_161417018_161417089@11.28.18,trna87_GluCTC_6_-_126101393_126101464@11.28.18|TCTAGTGGTTAGGATTCG</td><td>>am_trna116_GluCTC_1_-_145399233_145399304</td><td>11</td><td>28</td><td>18</td><td>TCTAGTGGTTAGGATTCG</td><td>107.8</td><td>-9.8</td><td>1.9</td><td>-5.1</td><td>3.9E-07</td><td>2.3E-06</td><td>BT20/MB468</td><td>MP</td></tr>
<tr><td>>hg38_21_+_8433172_8446622_RNA45SN1_45S_50ntFlanks@12412.12436.25|ATCCTTCGATGTCGGCTCTTCCTAT</td><td>>hg38_21_+_8433172_8446622_RNA45SN1_45S_50ntFlanks</td><td>12412</td><td>12436</td><td>25</td><td>ATCCTTCGATGTCGGCTCTTCCTAT</td><td>2220.4</td><td>-7.7</td><td>1.6</td><td>-4.8</td><td>1.4E-06</td><td>2.3E-06</td><td>MB231/MB468</td><td>NUC</td></tr>
<tr><td>>hg38_21_+_8433172_8446622_RNA45SN1_45S_50ntFlanks@11580.11601.22|GAACTGGTGCGGACCAGGGGAA</td><td>>hg38_21_+_8433172_8446622_RNA45SN1_45S_50ntFlanks</td><td>11580</td><td>11601</td><td>22</td><td>GAACTGGTGCGGACCAGGGGAA</td><td>674.5</td><td>-5.6</td><td>1.2</td><td>-4.8</td><td>1.4E-06</td><td>2.3E-06</td><td>MB231/MB468</td><td>NUC</td></tr>
<tr><td>>hg38_21_+_8433172_8446622_RNA45SN1_45S_50ntFlanks@12634.12661.28|GAAGCTACCATCTGTGGGATTATGACTG</td><td>>hg38_21_+_8433172_8446622_RNA45SN1_45S_50ntFlanks</td><td>12634</td><td>12661</td><td>28</td><td>GAAGCTACCATCTGTGGGATTATGACTG</td><td>1360.1</td><td>-5.9</td><td>1.2</td><td>-4.8</td><td>1.5E-06</td><td>2.3E-06</td><td>MB231/MB468</td><td>MITO</td></tr>
<tr><td>>hg38_MT_+_1621_3279_RNR2_16S_50ntFlanks@717.740.24|AAGCCTGCGTCAGATTAAAACACT</td><td>>hg38_MT_+_1621_3279_RNR2_16S_50ntFlanks</td><td>717</td><td>740</td><td>24</td><td>AAGCCTGCGTCAGATTAAAACACT</td><td>44.9</td><td>8.7</td><td>1.8</td><td>4.9</td><td>1.2E-06</td><td>2.3E-06</td><td>MB231/MB468</td><td>MP</td></tr>
<tr><td>>hg38_21_+_8433172_8446622_RNA45SN1_45S_50ntFlanks@11452.11474.23|CTCCTCGGCGCGCGGCGGCGGCG</td><td>>hg38_21_+_8433172_8446622_RNA45SN1_45S_50ntFlanks</td><td>11452</td><td>11474</td><td>23</td><td>CTCCTCGGCGCGCGGCGGCGGCG</td><td>54.9</td><td>-8.9</td><td>1.7</td><td>-5.1</td><td>3.9E-07</td><td>2.3E-06</td><td>BT20/MB468</td><td>MP</td></tr>
<tr><td>>hg38_21_+_8433172_8446622_RNA45SN1_45S_50ntFlanks@12936.12960.25|AAACCATTCGTAGACGACCTGCTTC</td><td>>hg38_21_+_8433172_8446622_RNA45SN1_45S_50ntFlanks</td><td>12936</td><td>12960</td><td>25</td><td>AAACCATTCGTAGACGACCTGCTTC</td><td>919.9</td><td>-3.1</td><td>0.6</td><td>-4.8</td><td>1.4E-06</td><td>2.3E-06</td><td>MB231/MB468</td><td>NUC</td></tr>
<tr><td>>MI0000095|hsa-mir-93&WithFlank&7|-|100093762|100093853@17.39.23(+1A)&[MIMAT0000093&hsa-miR-93-5p&offsets|0|0(+1A);m-29&7|-|100093815|100093837&offsets|0|0(+1A)]|CAAAGTGCTGTTCGTGCAGGTAGA</td><td>>MI0000095|hsa-mir-93&WithFlank&7|-|100093762|100093853</td><td>17</td><td>39</td><td>23(+1A)</td><td>CAAAGTGCTGTTCGTGCAGGTAGA</td><td>1340.4</td><td>4.4</td><td>0.9</td><td>4.8</td><td>1.5E-06</td><td>2.3E-06</td><td>MB231/MB468</td><td>MITO</td></tr>
<tr><td>>hg38_21_+_8433172_8446622_RNA45SN1_45S_50ntFlanks@8191.8208.18|CAGCCCGTGGACGGTGTG</td><td>>hg38_21_+_8433172_8446622_RNA45SN1_45S_50ntFlanks</td><td>8191</td><td>8208</td><td>18</td><td>CAGCCCGTGGACGGTGTG</td><td>83.3</td><td>-9.5</td><td>1.9</td><td>-5.1</td><td>3.9E-07</td><td>2.3E-06</td><td>BT20/MB468</td><td>MP</td></tr>
<tr><td>>hg38_21_+_8433172_8446622_RNA45SN1_45S_50ntFlanks@12631.12659.29|GGCGAAGCTACCATCTGTGGGATTATGAC</td><td>>hg38_21_+_8433172_8446622_RNA45SN1_45S_50ntFlanks</td><td>12631</td><td>12659</td><td>29</td><td>GGCGAAGCTACCATCTGTGGGATTATGAC</td><td>336.5</td><td>-7.5</td><td>1.6</td><td>-4.8</td><td>1.6E-06</td><td>2.3E-06</td><td>BT20/MB468</td><td>NUC</td></tr>
<tr><td>>MI0025242|hsa-mir-7706&WithFlank&15|+|85380590|85380668@50.72.23(+1U)&[MIMAT0030021&hsa-miR-7706&offsets|0|-1(+1U);m-390&15|+|85380639|85380662&offsets|0|-1(+1U)]|TGAAGCGCCTGTGCTCTGCCGAGT</td><td>>MI0025242|hsa-mir-7706&WithFlank&15|+|85380590|85380668</td><td>50</td><td>72</td><td>23(+1U)</td><td>TGAAGCGCCTGTGCTCTGCCGAGT</td><td>228.8</td><td>3.6</td><td>0.8</td><td>4.8</td><td>1.4E-06</td><td>2.3E-06</td><td>MB231/MB468</td><td>NUC</td></tr>
<tr><td>>hg38_21_+_8433172_8446622_RNA45SN1_45S_50ntFlanks@10348.10388.41|AAACGAGAACTTTGAAGGCCGAAGTGGAGAAGGGTTCCATG</td><td>>hg38_21_+_8433172_8446622_RNA45SN1_45S_50ntFlanks</td><td>10348</td><td>10388</td><td>41</td><td>AAACGAGAACTTTGAAGGCCGAAGTGGAGAAGGGTTCCATG</td><td>829.6</td><td>-6.6</td><td>1.4</td><td>-4.8</td><td>1.6E-06</td><td>2.3E-06</td><td>BT20/MB468</td><td>NUC</td></tr>
<tr><td>>am_trnaMT_PheGAA_MT_+_577_647@40.57.18|TGTTTAGACGGGCTCACA</td><td>>am_trnaMT_PheGAA_MT_+_577_647</td><td>40</td><td>57</td><td>18</td><td>TGTTTAGACGGGCTCACA</td><td>573.5</td><td>4.4</td><td>0.9</td><td>5.1</td><td>3.9E-07</td><td>2.3E-06</td><td>BT20/MB468</td><td>MP</td></tr>
<tr><td>>hg38_21_+_8433172_8446622_RNA45SN1_45S_50ntFlanks@9552.9572.21|TCCGTAGCGGTCCTGACGTGC</td><td>>hg38_21_+_8433172_8446622_RNA45SN1_45S_50ntFlanks</td><td>9552</td><td>9572</td><td>21</td><td>TCCGTAGCGGTCCTGACGTGC</td><td>480.2</td><td>-7.5</td><td>1.6</td><td>-4.8</td><td>1.6E-06</td><td>2.3E-06</td><td>BT20/MB468</td><td>NUC</td></tr>
<tr><td>>hg38_21_+_8433172_8446622_RNA45SN1_45S_50ntFlanks@9653.9679.27|AGGATAGCTGGCGCTCTCGCAGACCCG</td><td>>hg38_21_+_8433172_8446622_RNA45SN1_45S_50ntFlanks</td><td>9653</td><td>9679</td><td>27</td><td>AGGATAGCTGGCGCTCTCGCAGACCCG</td><td>150.7</td><td>-6.5</td><td>1.3</td><td>-4.9</td><td>1.2E-06</td><td>2.3E-06</td><td>MB231/MB468</td><td>MP</td></tr>
<tr><td>>hg38_21_+_8433172_8446622_RNA45SN1_45S_50ntFlanks@10821.10838.18|GGAAGTCGGCAAGCCGGA</td><td>>hg38_21_+_8433172_8446622_RNA45SN1_45S_50ntFlanks</td><td>10821</td><td>10838</td><td>18</td><td>GGAAGTCGGCAAGCCGGA</td><td>371.3</td><td>-6.7</td><td>1.4</td><td>-4.8</td><td>1.5E-06</td><td>2.3E-06</td><td>MB231/MB468</td><td>MITO</td></tr>
<tr><td>>hg38_21_+_8433172_8446622_RNA45SN1_45S_50ntFlanks@11573.11612.40|CGACTTAGAACTGGTGCGGACCAGGGGAATCCGACTGTTT</td><td>>hg38_21_+_8433172_8446622_RNA45SN1_45S_50ntFlanks</td><td>11573</td><td>11612</td><td>40</td><td>CGACTTAGAACTGGTGCGGACCAGGGGAATCCGACTGTTT</td><td>478.5</td><td>-4.1</td><td>0.9</td><td>-4.8</td><td>1.6E-06</td><td>2.3E-06</td><td>BT20/MB468</td><td>NUC</td></tr>
<tr><td>>am_trnaMT_TyrGTA_MT_-_5826_5891@13.37.25|GAGTGAAGCATTGGACTGTAAATCT</td><td>>am_trnaMT_TyrGTA_MT_-_5826_5891</td><td>13</td><td>37</td><td>25</td><td>GAGTGAAGCATTGGACTGTAAATCT</td><td>83.6</td><td>10.1</td><td>2.0</td><td>5.1</td><td>3.9E-07</td><td>2.3E-06</td><td>BT20/MB468</td><td>MP</td></tr>
<tr><td>>MI0001445|hsa-mir-423&WithFlank&17|+|30117073|30117178@58.78.21&[MIMAT0001340&hsa-miR-423-3p&offsets|-1|-3;m-100&17|+|30117131|30117153&offsets|-1|-3]|AAGCTCGGTCTGAGGCCCCTC</td><td>>MI0001445|hsa-mir-423&WithFlank&17|+|30117073|30117178</td><td>58</td><td>78</td><td>21</td><td>AAGCTCGGTCTGAGGCCCCTC</td><td>542.0</td><td>6.7</td><td>1.4</td><td>4.8</td><td>1.4E-06</td><td>2.3E-06</td><td>MB231/MB468</td><td>CYTO</td></tr>
<tr><td>>hg38_21_+_8433172_8446622_RNA45SN1_45S_50ntFlanks@12626.12657.32|AATGGGGCGAAGCTACCATCTGTGGGATTATG</td><td>>hg38_21_+_8433172_8446622_RNA45SN1_45S_50ntFlanks</td><td>12626</td><td>12657</td><td>32</td><td>AATGGGGCGAAGCTACCATCTGTGGGATTATG</td><td>1179.4</td><td>-8.3</td><td>1.7</td><td>-4.8</td><td>1.6E-06</td><td>2.3E-06</td><td>BT20/MB468</td><td>NUC</td></tr>
</table>

<table>
<tr><td></td><td></td><td></td><td></td><td></td><td></td><td></td><td></td><td></td><td></td><td></td><td></td><td></td><td></td></tr>
<tr><td>>trna116_GluCTC_1_-_145399233_145399304@37.74.38,trna134_GluTTC_1_-_16861774_16861845@37.74.38,trna71_GluCTC_1_-_161439189_161439260@37.74.38,trna74_GluCTC_1_-_161431809_161431880@37.74.38,trna77_GluCTC_1_-_161424398_161424469@37.74.38,trna77_GluCTC_6_+_28949976_28950047@37.74.38,trna80_GluCTC_1_-_161417018_161417089@37.74.38,trna84_GluTTC_1_-_161391883_161391954@37.74.38,trna87_GluCTC_6_-_126101393_126101464@37.74.38|ACCGCCGCGGCCCGGGTTCGATTCCCGGTCAGGGAACC</td><td>>trna116_GluCTC_1_-_145399233_145399304</td><td>37</td><td>74</td><td>38</td><td>ACCGCCGCGGCCCGGGTTCGATTCCCGGTCAGGGAACC</td><td>612.6</td><td>5.6</td><td>1.2</td><td>4.8</td><td>1.5E-06</td><td>2.3E-06</td><td>MB231/MB468</td><td>MITO</td></tr>
<tr><td>>hg38_21_+_8433172_8446622_RNA45SN1_45S_50ntFlanks@9794.9819.26|CGTGGAGCCGGGCGTGGAATGCGAGT</td><td>>hg38_21_+_8433172_8446622_RNA45SN1_45S_50ntFlanks</td><td>9794</td><td>9819</td><td>26</td><td>CGTGGAGCCGGGCGTGGAATGCGAGT</td><td>185.0</td><td>10.9</td><td>2.0</td><td>5.3</td><td>9.6E-08</td><td>2.3E-06</td><td>BT20/MB231</td><td>MP</td></tr>
<tr><td>>hg38_21_+_8433172_8446622_RNA45SN1_45S_50ntFlanks@8171.8215.45|AAGTCCTTCTGATCGAGGCCCAGCCCGTGGACGGTGTGAGGCCGG</td><td>>hg38_21_+_8433172_8446622_RNA45SN1_45S_50ntFlanks</td><td>8171</td><td>8215</td><td>45</td><td>AAGTCCTTCTGATCGAGGCCCAGCCCGTGGACGGTGTGAGGCCGG</td><td>130.6</td><td>-10.9</td><td>2.2</td><td>-4.8</td><td>1.2E-06</td><td>2.3E-06</td><td>MB231/MB468</td><td>MP</td></tr>
<tr><td>>MI0001445|hsa-mir-423&WithFlank&17|+|30117073|30117178@59.81.23(+1A)&[MIMAT0001340&hsa-miR-423-3p&offsets|0|0(+1A);m-100&17|+|30117131|30117153&offsets|0|0(+1A)]|AGCTCGGTCTGAGGCCCCTCAGTA</td><td>>MI0001445|hsa-mir-423&WithFlank&17|+|30117073|30117178</td><td>59</td><td>81</td><td>23(+1A)</td><td>AGCTCGGTCTGAGGCCCCTCAGTA</td><td>719.4</td><td>4.2</td><td>0.9</td><td>4.8</td><td>1.4E-06</td><td>2.3E-06</td><td>MB231/MB468</td><td>NUC</td></tr>
<tr><td>>hg38_21_+_8433172_8446622_RNA45SN1_45S_50ntFlanks@9621.9652.32|TCTAGTAGCTGGTTCCCTCCGAAGTTTCCCTC</td><td>>hg38_21_+_8433172_8446622_RNA45SN1_45S_50ntFlanks</td><td>9621</td><td>9652</td><td>32</td><td>TCTAGTAGCTGGTTCCCTCCGAAGTTTCCCTC</td><td>309.4</td><td>-3.3</td><td>0.7</td><td>-4.9</td><td>1.1E-06</td><td>2.3E-06</td><td>BT20/MB468</td><td>MITO</td></tr>
<tr><td>>hg38_21_+_8433172_8446622_RNA45SN1_45S_50ntFlanks@10922.10943.22|CTGGACGAGGCGCCGCCGCCCC</td><td>>hg38_21_+_8433172_8446622_RNA45SN1_45S_50ntFlanks</td><td>10922</td><td>10943</td><td>22</td><td>CTGGACGAGGCGCCGCCGCCCC</td><td>161.2</td><td>-6.4</td><td>1.3</td><td>-4.8</td><td>1.4E-06</td><td>2.3E-06</td><td>MB231/MB468</td><td>NUC</td></tr>
<tr><td>>hg38_21_+_8433172_8446622_RNA45SN1_45S_50ntFlanks@4283.4316.34|CTTTAACGAGGATCCATTGGAGGGCAAGTCTGGT</td><td>>hg38_21_+_8433172_8446622_RNA45SN1_45S_50ntFlanks</td><td>4283</td><td>4316</td><td>34</td><td>CTTTAACGAGGATCCATTGGAGGGCAAGTCTGGT</td><td>243.1</td><td>-7.2</td><td>1.5</td><td>-4.8</td><td>1.2E-06</td><td>2.3E-06</td><td>MB231/MB468</td><td>MP</td></tr>
<tr><td>>hg38_21_+_8433172_8446622_RNA45SN1_45S_50ntFlanks@4588.4619.32|CTATTTTGTTGGTTTTCGGAACTGAGGCCATG</td><td>>hg38_21_+_8433172_8446622_RNA45SN1_45S_50ntFlanks</td><td>4588</td><td>4619</td><td>32</td><td>CTATTTTGTTGGTTTTCGGAACTGAGGCCATG</td><td>595.7</td><td>-6.1</td><td>1.3</td><td>-4.8</td><td>1.2E-06</td><td>2.3E-06</td><td>MB231/MB468</td><td>MP</td></tr>
<tr><td>>MI0000746|hsa-mir-99b&WithFlank&19|+|51692606|51692687@51.70.20&[MIMAT0004678&hsa-miR-99b-3p&offsets|0|-2;m-151&19|+|51692656|51692678&offsets|0|-3]|CAAGCTCGTGTCTGTGGGTC</td><td>>MI0000746|hsa-mir-99b&WithFlank&19|+|51692606|51692687</td><td>51</td><td>70</td><td>20</td><td>CAAGCTCGTGTCTGTGGGTC</td><td>455.5</td><td>5.1</td><td>1.0</td><td>5.1</td><td>3.9E-07</td><td>2.3E-06</td><td>BT20/MB468</td><td>MP</td></tr>
<tr><td>>hg38_21_+_8433172_8446622_RNA45SN1_45S_50ntFlanks@10238.10259.22|CGCTACGCCGCGACGAGTAGGA</td><td>>hg38_21_+_8433172_8446622_RNA45SN1_45S_50ntFlanks</td><td>10238</td><td>10259</td><td>22</td><td>CGCTACGCCGCGACGAGTAGGA</td><td>720.3</td><td>4.6</td><td>0.9</td><td>5.1</td><td>3.9E-07</td><td>2.3E-06</td><td>BT20/MB468</td><td>MP</td></tr>
<tr><td>>trnaMT_GluTTC_MT_-_14674_14742@41.72.32,trnalookalike8_GluTTC_5_-_93905172_93905240@41.72.32|TGGTCGTGGTTGTAGTCCGTGCGAGAATACCA</td><td>>trnaMT_GluTTC_MT_-_14674_14742</td><td>41</td><td>72</td><td>32</td><td>TGGTCGTGGTTGTAGTCCGTGCGAGAATACCA</td><td>9683.8</td><td>3.9</td><td>0.8</td><td>4.8</td><td>1.5E-06</td><td>2.3E-06</td><td>MB231/MB468</td><td>NUC</td></tr>
<tr><td>>hg38_21_+_8433172_8446622_RNA45SN1_45S_50ntFlanks@7989.8019.31|AGACGTGGCGACCCGCTGAATTTAAGCATAT</td><td>>hg38_21_+_8433172_8446622_RNA45SN1_45S_50ntFlanks</td><td>7989</td><td>8019</td><td>31</td><td>AGACGTGGCGACCCGCTGAATTTAAGCATAT</td><td>257.1</td><td>8.3</td><td>1.7</td><td>4.9</td><td>1.2E-06</td><td>2.3E-06</td><td>BT20/MB468</td><td>CYTO</td></tr>
<tr><td>>am_trna117_GlyTCC_1_-_145397864_145397935@1.28.28,trna45_GlyTCC_1_+_161500903_161500974@1.28.28,trna70_GlyTCC_1_-_161439547_161439618@1.28.28,trna73_GlyTCC_1_-_161432166_161432237@1.28.28,trna76_GlyTCC_1_-_161424756_161424827@1.28.28,trna79_GlyTCC_1_-_161417375_161417446@1.28.28|GCGTTGGTGGTATAGTGGTGAGCATAGC</td><td>>am_trna117_GlyTCC_1_-_145397864_145397935</td><td>1</td><td>28</td><td>28</td><td>GCGTTGGTGGTATAGTGGTGAGCATAGC</td><td>184.2</td><td>-3.5</td><td>0.7</td><td>-4.8</td><td>1.4E-06</td><td>2.3E-06</td><td>MB231/MB468</td><td>CYTO</td></tr>
<tr><td>>hg38_21_+_8433172_8446622_RNA45SN1_45S_50ntFlanks@12628.12659.32|TGGGGCGAAGCTACCATCTGTGGGATTATGAC</td><td>>hg38_21_+_8433172_8446622_RNA45SN1_45S_50ntFlanks</td><td>12628</td><td>12659</td><td>32</td><td>TGGGGCGAAGCTACCATCTGTGGGATTATGAC</td><td>1344.5</td><td>-6.5</td><td>1.3</td><td>-4.9</td><td>1.1E-06</td><td>2.3E-06</td><td>BT20/MB468</td><td>MITO</td></tr>
<tr><td>>hg38_21_+_8433172_8446622_RNA45SN1_45S_50ntFlanks@4050.4081.32|CGAACGTCTGCCCTATCAACTTTCGATGGTAG</td><td>>hg38_21_+_8433172_8446622_RNA45SN1_45S_50ntFlanks</td><td>4050</td><td>4081</td><td>32</td><td>CGAACGTCTGCCCTATCAACTTTCGATGGTAG</td><td>221.5</td><td>-6.0</td><td>1.2</td><td>-4.9</td><td>1.1E-06</td><td>2.3E-06</td><td>BT20/MB468</td><td>MITO</td></tr>
<tr><td>>hg38_21_+_8433172_8446622_RNA45SN1_45S_50ntFlanks@12407.12446.40|TTTTGATCCTTCGATGTCGGCTCTTCCTATCATTGTGAAG</td><td>>hg38_21_+_8433172_8446622_RNA45SN1_45S_50ntFlanks</td><td>12407</td><td>12446</td><td>40</td><td>TTTTGATCCTTCGATGTCGGCTCTTCCTATCATTGTGAAG</td><td>2567.3</td><td>-8.9</td><td>1.9</td><td>-4.8</td><td>1.6E-06</td><td>2.3E-06</td><td>BT20/MB468</td><td>NUC</td></tr>
<tr><td>>hg38_21_+_8433172_8446622_RNA45SN1_45S_50ntFlanks@11365.11387.23|GGCGGCGGCGGTCGGCGGGCGGC</td><td>>hg38_21_+_8433172_8446622_RNA45SN1_45S_50ntFlanks</td><td>11365</td><td>11387</td><td>23</td><td>GGCGGCGGCGGTCGGCGGGCGGC</td><td>211.4</td><td>11.1</td><td>2.1</td><td>5.3</td><td>9.7E-08</td><td>2.3E-06</td><td>BT20/MB231</td><td>MP</td></tr>
<tr><td>>MI0000683|hsa-mir-181b-2&WithFlank&9|+|124693704|124693804@22.46.25(+1C)&[MIMAT0000257_1&hsa-miR-181b-5p&offsets|0|+2(+1C);m-72&9|+|124693725|124693747&offsets|0|+2(+1C)]|AACATTCATTGCTGTCGGTGGGTTTC</td><td>>MI0000683|hsa-mir-181b-2&WithFlank&9|+|124693704|124693804</td><td>22</td><td>46</td><td>25(+1C)</td><td>AACATTCATTGCTGTCGGTGGGTTTC</td><td>118.9</td><td>-10.3</td><td>1.9</td><td>-5.3</td><td>9.8E-08</td><td>2.3E-06</td><td>BT20/MB231</td><td>MP</td></tr>
<tr><td>>hg38_21_+_8433172_8446622_RNA45SN1_45S_50ntFlanks@12073.12091.19|CCGAGGGGCTCTCGCTTCT</td><td>>hg38_21_+_8433172_8446622_RNA45SN1_45S_50ntFlanks</td><td>12073</td><td>12091</td><td>19</td><td>CCGAGGGGCTCTCGCTTCT</td><td>200.0</td><td>-6.2</td><td>1.3</td><td>-4.8</td><td>1.2E-06</td><td>2.3E-06</td><td>BT20/MB468</td><td>CYTO</td></tr>
<tr><td>>hg38_21_+_8433172_8446622_RNA45SN1_45S_50ntFlanks@8527.8551.25|CCTTCCCCCGCCGCCCCTCCTCCTC</td><td>>hg38_21_+_8433172_8446622_RNA45SN1_45S_50ntFlanks</td><td>8527</td><td>8551</td><td>25</td><td>CCTTCCCCCGCCGCCCCTCCTCCTC</td><td>427.1</td><td>-7.7</td><td>1.6</td><td>-4.8</td><td>1.6E-06</td><td>2.3E-06</td><td>BT20/MB468</td><td>NUC</td></tr>
<tr><td>>hg38_21_+_8433172_8446622_RNA45SN1_45S_50ntFlanks@7975.8006.32|CGCGACCTCAGATCAGACGTGGCGACCCGCTG</td><td>>hg38_21_+_8433172_8446622_RNA45SN1_45S_50ntFlanks</td><td>7975</td><td>8006</td><td>32</td><td>CGCGACCTCAGATCAGACGTGGCGACCCGCTG</td><td>269557.6</td><td>-4.3</td><td>0.9</td><td>-4.8</td><td>1.5E-06</td><td>2.3E-06</td><td>MB231/MB468</td><td>MITO</td></tr>
<tr><td>>hg38_MT_+_1621_3279_RNR2_16S_50ntFlanks@113.146.34|CCAAATAAAGTATAGGCGATAGAAATTGAAACCT</td><td>>hg38_MT_+_1621_3279_RNR2_16S_50ntFlanks</td><td>113</td><td>146</td><td>34</td><td>CCAAATAAAGTATAGGCGATAGAAATTGAAACCT</td><td>49.9</td><td>8.9</td><td>1.8</td><td>4.8</td><td>1.3E-06</td><td>2.3E-06</td><td>MB231/MB468</td><td>MP</td></tr>
<tr><td>>MI0000477|hsa-mir-146a&WithFlank&5|+|160485346|160485456@27.45.19(+3U)&[MIMAT0000449&hsa-miR-146a-5p&offsets|0|-3(+3U);m-129&5|+|160485372|160485393&offsets|0|-3(+3U)]|TGAGAACTGAATTCCATGGTTT</td><td>>MI0000477|hsa-mir-146a&WithFlank&5|+|160485346|160485456</td><td>27</td><td>45</td><td>19(+3U)</td><td>TGAGAACTGAATTCCATGGTTT</td><td>166.3</td><td>-10.8</td><td>2.0</td><td>-5.3</td><td>9.8E-08</td><td>2.4E-06</td><td>BT20/MB231</td><td>MP</td></tr>
<tr><td>>hg38_21_+_8433172_8446622_RNA45SN1_45S_50ntFlanks@7972.8010.39|AGACGCGACCTCAGATCAGACGTGGCGACCCGCTGAATT</td><td>>hg38_21_+_8433172_8446622_RNA45SN1_45S_50ntFlanks</td><td>7972</td><td>8010</td><td>39</td><td>AGACGCGACCTCAGATCAGACGTGGCGACCCGCTGAATT</td><td>429.4</td><td>4.5</td><td>0.9</td><td>4.8</td><td>1.6E-06</td><td>2.4E-06</td><td>BT20/MB468</td><td>NUC</td></tr>
<tr><td>>hg38_21_+_8433172_8446622_RNA45SN1_45S_50ntFlanks@12488.12518.31|AACGTGAGCTGGGTTTAGACCGTCGTGAGAC</td><td>>hg38_21_+_8433172_8446622_RNA45SN1_45S_50ntFlanks</td><td>12488</td><td>12518</td><td>31</td><td>AACGTGAGCTGGGTTTAGACCGTCGTGAGAC</td><td>215.3</td><td>-4.8</td><td>1.0</td><td>-4.8</td><td>1.5E-06</td><td>2.4E-06</td><td>MB231/MB468</td><td>NUC</td></tr>
<tr><td>>hg38_21_+_8433172_8446622_RNA45SN1_45S_50ntFlanks@8169.8186.18|CCAAGTCCTTCTGATCGA</td><td>>hg38_21_+_8433172_8446622_RNA45SN1_45S_50ntFlanks</td><td>8169</td><td>8186</td><td>18</td><td>CCAAGTCCTTCTGATCGA</td><td>437.8</td><td>-7.0</td><td>1.5</td><td>-4.8</td><td>1.5E-06</td><td>2.4E-06</td><td>MB231/MB468</td><td>MITO</td></tr>
<tr><td>>hg38_MT_+_598_1651_RNR1_12S_50ntFlanks@52.77.26|ATAGGTTTGGTCCTAGCCTTTCTATT</td><td>>hg38_MT_+_598_1651_RNR1_12S_50ntFlanks</td><td>52</td><td>77</td><td>26</td><td>ATAGGTTTGGTCCTAGCCTTTCTATT</td><td>203.3</td><td>4.1</td><td>0.8</td><td>4.8</td><td>1.5E-06</td><td>2.4E-06</td><td>MB231/MB468</td><td>NUC</td></tr>
<tr><td>>MI0001445|hsa-mir-423&WithFlank&17|+|30117073|30117178@23.45.23&[MIMAT0004748&hsa-miR-423-5p&offsets|0|0;m-141&17|+|30117095|30117117&offsets|0|0]|TGAGGGGCAGAGAGCGAGACTTT</td><td>>MI0001445|hsa-mir-423&WithFlank&17|+|30117073|30117178</td><td>23</td><td>45</td><td>23</td><td>TGAGGGGCAGAGAGCGAGACTTT</td><td>13386.5</td><td>4.3</td><td>0.9</td><td>4.8</td><td>1.3E-06</td><td>2.4E-06</td><td>MB231/MB468</td><td>MP</td></tr>
<tr><td>>hg38_21_+_8433172_8446622_RNA45SN1_45S_50ntFlanks@9496.9526.31|GAAAGATGGTGAACTATGCCTGGGCAGGGCG</td><td>>hg38_21_+_8433172_8446622_RNA45SN1_45S_50ntFlanks</td><td>9496</td><td>9526</td><td>31</td><td>GAAAGATGGTGAACTATGCCTGGGCAGGGCG</td><td>260.3</td><td>11.4</td><td>2.1</td><td>5.3</td><td>9.9E-08</td><td>2.4E-06</td><td>BT20/MB231</td><td>MP</td></tr>
<tr><td>>hg38_21_+_8433172_8446622_RNA45SN1_45S_50ntFlanks@7974.8005.32|ACGCGACCTCAGATCAGACGTGGCGACCCGCT</td><td>>hg38_21_+_8433172_8446622_RNA45SN1_45S_50ntFlanks</td><td>7974</td><td>8005</td><td>32</td><td>ACGCGACCTCAGATCAGACGTGGCGACCCGCT</td><td>6700.8</td><td>4.5</td><td>0.9</td><td>5.1</td><td>4.0E-07</td><td>2.4E-06</td><td>BT20/MB468</td><td>MP</td></tr>
<tr><td>>hg38_21_+_8433172_8446622_RNA45SN1_45S_50ntFlanks@11576.11607.32|CTTAGAACTGGTGCGGACCAGGGGAATCCGAC</td><td>>hg38_21_+_8433172_8446622_RNA45SN1_45S_50ntFlanks</td><td>11576</td><td>11607</td><td>32</td><td>CTTAGAACTGGTGCGGACCAGGGGAATCCGAC</td><td>545.3</td><td>-3.1</td><td>0.6</td><td>-4.8</td><td>1.4E-06</td><td>2.4E-06</td><td>MB231/MB468</td><td>CYTO</td></tr>
<tr><td>>hg38_21_+_8433172_8446622_RNA45SN1_45S_50ntFlanks@11578.11602.25|TAGAACTGGTGCGGACCAGGGGAAT</td><td>>hg38_21_+_8433172_8446622_RNA45SN1_45S_50ntFlanks</td><td>11578</td><td>11602</td><td>25</td><td>TAGAACTGGTGCGGACCAGGGGAAT</td><td>242.6</td><td>-3.0</td><td>0.6</td><td>-4.8</td><td>1.4E-06</td><td>2.4E-06</td><td>MB231/MB468</td><td>CYTO</td></tr>
<tr><td>>hg38_21_+_8433172_8446622_RNA45SN1_45S_50ntFlanks@12416.12449.34|TTCGATGTCGGCTCTTCCTATCATTGTGAAGCAG</td><td>>hg38_21_+_8433172_8446622_RNA45SN1_45S_50ntFlanks</td><td>12416</td><td>12449</td><td>34</td><td>TTCGATGTCGGCTCTTCCTATCATTGTGAAGCAG</td><td>371.1</td><td>3.5</td><td>0.7</td><td>4.8</td><td>1.5E-06</td><td>2.4E-06</td><td>MB231/MB468</td><td>MITO</td></tr>
<tr><td>>hg38_21_+_8433172_8446622_RNA45SN1_45S_50ntFlanks@9504.9524.21|GTGAACTATGCCTGGGCAGGG</td><td>>hg38_21_+_8433172_8446622_RNA45SN1_45S_50ntFlanks</td><td>9504</td><td>9524</td><td>21</td><td>GTGAACTATGCCTGGGCAGGG</td><td>576.4</td><td>-6.6</td><td>1.4</td><td>-4.8</td><td>1.3E-06</td><td>2.4E-06</td><td>BT20/MB468</td><td>CYTO</td></tr>
<tr><td>>MI0000085|hsa-mir-27a&WithFlank&19|-|13836434|13836523@16.36.21&[MIMAT0004501&hsa-miR-27a-5p&offsets|0|-1;m-270&19|-|13836487|13836508&offsets|0|-1]|AGGGCTTAGCTGCTTGTGAGC</td><td>>MI0000085|hsa-mir-27a&WithFlank&19|-|13836434|13836523</td><td>16</td><td>36</td><td>21</td><td>AGGGCTTAGCTGCTTGTGAGC</td><td>483.9</td><td>6.1</td><td>1.3</td><td>4.8</td><td>1.5E-06</td><td>2.4E-06</td><td>MB231/MB468</td><td>NUC</td></tr>
<tr><td>>hg38_21_+_8433172_8446622_RNA45SN1_45S_50ntFlanks@3823.3852.30|TTTGGTCGCTCGCTCCTCTCCTACTTGGAT</td><td>>hg38_21_+_8433172_8446622_RNA45SN1_45S_50ntFlanks</td><td>3823</td><td>3852</td><td>30</td><td>TTTGGTCGCTCGCTCCTCTCCTACTTGGAT</td><td>622.2</td><td>-4.0</td><td>0.8</td><td>-4.8</td><td>1.5E-06</td><td>2.4E-06</td><td>MB231/MB468</td><td>NUC</td></tr>
<tr><td>>am_trnaMT_ValTAC_MT_+_1602_1670@49.68.20|AACTTAACTTGACCGCTCTG</td><td>>am_trnaMT_ValTAC_MT_+_1602_1670</td><td>49</td><td>68</td><td>20</td><td>AACTTAACTTGACCGCTCTG</td><td>1049.2</td><td>-4.4</td><td>0.9</td><td>-4.8</td><td>1.5E-06</td><td>2.4E-06</td><td>MB231/MB468</td><td>MITO</td></tr>
<tr><td>>hg38_21_+_8433172_8446622_RNA45SN1_45S_50ntFlanks@3885.3915.31|ACGGGCGCTGACCCCCTTCGCGGGGGGGATG</td><td>>hg38_21_+_8433172_8446622_RNA45SN1_45S_50ntFlanks</td><td>3885</td><td>3915</td><td>31</td><td>ACGGGCGCTGACCCCCTTCGCGGGGGGGATG</td><td>547.8</td><td>-3.4</td><td>0.7</td><td>-4.8</td><td>1.3E-06</td><td>2.4E-06</td><td>BT20/MB468</td><td>CYTO</td></tr>
<tr><td>>hg38_21_+_8433172_8446622_RNA45SN1_45S_50ntFlanks@10346.10370.25|TCAAACGAGAACTTTGAAGGCCGAA</td><td>>hg38_21_+_8433172_8446622_RNA45SN1_45S_50ntFlanks</td><td>10346</td><td>10370</td><td>25</td><td>TCAAACGAGAACTTTGAAGGCCGAA</td><td>283.3</td><td>-3.7</td><td>0.8</td><td>-4.8</td><td>1.5E-06</td><td>2.4E-06</td><td>MB231/MB468</td><td>MITO</td></tr>
<tr><td>>hg38_21_+_8433172_8446622_RNA45SN1_45S_50ntFlanks@10778.10818.41|CCAAGGTGAACAGCCTCTGGCATGTTGGAACAATGTAGGTA</td><td>>hg38_21_+_8433172_8446622_RNA45SN1_45S_50ntFlanks</td><td>10778</td><td>10818</td><td>41</td><td>CCAAGGTGAACAGCCTCTGGCATGTTGGAACAATGTAGGTA</td><td>100.4</td><td>-9.7</td><td>1.9</td><td>-5.1</td><td>4.0E-07</td><td>2.4E-06</td><td>BT20/MB468</td><td>MP</td></tr>
<tr><td>>hg38_21_+_8433172_8446622_RNA45SN1_45S_50ntFlanks@12456.12493.38|CCAAGCGTTGGATTGTTCACCCACTAATAGGGAACGTG</td><td>>hg38_21_+_8433172_8446622_RNA45SN1_45S_50ntFlanks</td><td>12456</td><td>12493</td><td>38</td><td>CCAAGCGTTGGATTGTTCACCCACTAATAGGGAACGTG</td><td>451.4</td><td>-3.1</td><td>0.6</td><td>-4.8</td><td>1.3E-06</td><td>2.4E-06</td><td>BT20/MB468</td><td>CYTO</td></tr>
<tr><td>>hg38_21_+_8433172_8446622_RNA45SN1_45S_50ntFlanks@11553.11591.39|CTCGGCCGGCGCCTAGCAGCCGACTTAGAACTGGTGCGG</td><td>>hg38_21_+_8433172_8446622_RNA45SN1_45S_50ntFlanks</td><td>11553</td><td>11591</td><td>39</td><td>CTCGGCCGGCGCCTAGCAGCCGACTTAGAACTGGTGCGG</td><td>118.8</td><td>-3.6</td><td>0.8</td><td>-4.8</td><td>1.4E-06</td><td>2.4E-06</td><td>MB231/MB468</td><td>CYTO</td></tr>
<tr><td>>hg38_21_+_8433172_8446622_RNA45SN1_45S_50ntFlanks@10510.10533.24|AATCCGGAGTGGCGGAGATGGGCG</td><td>>hg38_21_+_8433172_8446622_RNA45SN1_45S_50ntFlanks</td><td>10510</td><td>10533</td><td>24</td><td>AATCCGGAGTGGCGGAGATGGGCG</td><td>1409.8</td><td>-5.6</td><td>1.1</td><td>-4.9</td><td>1.1E-06</td><td>2.4E-06</td><td>BT20/MB468</td><td>MITO</td></tr>
<tr><td>>hg38_21_+_8433172_8446622_RNA45SN1_45S_50ntFlanks@12415.12442.28|CTTCGATGTCGGCTCTTCCTATCATTGT</td><td>>hg38_21_+_8433172_8446622_RNA45SN1_45S_50ntFlanks</td><td>12415</td><td>12442</td><td>28</td><td>CTTCGATGTCGGCTCTTCCTATCATTGT</td><td>643.1</td><td>2.4</td><td>0.5</td><td>4.8</td><td>1.3E-06</td><td>2.4E-06</td><td>BT20/MB468</td><td>CYTO</td></tr>
<tr><td>>hg38_21_+_8433172_8446622_RNA45SN1_45S_50ntFlanks@12499.12517.19|GGTTTAGACCGTCGTGAGA</td><td>>hg38_21_+_8433172_8446622_RNA45SN1_45S_50ntFlanks</td><td>12499</td><td>12517</td><td>19</td><td>GGTTTAGACCGTCGTGAGA</td><td>81.5</td><td>-9.4</td><td>1.9</td><td>-5.1</td><td>4.0E-07</td><td>2.4E-06</td><td>BT20/MB468</td><td>MP</td></tr>
</table>

| | | | | | | | | | | | | | |
|---|---|---|---|---|---|---|---|---|---|---|---|---|---|
| >hg38_21_+_8433172_8446622_RNA45SN1_45S_50ntFlanks@9740.9758.19\|AACGATCTCAACCTATTCT | >hg38_21_+_8433172_8446622_RNA45SN1_45S_50ntFlanks | 9740 | 9758 | 19 | AACGATCTCAACCTATTCT | 267.2 | -2.5 | 0.5 | -4.8 | 1.5E-06 | 2.4E-06 | MB231/MB468 | NUC |
| >MI0000255\|hsa-mir-30d&WithFlank&8\|-\|134804870\|134804951@12.35.24&[MIMAT0000245&hsa-miR-30d-5p&offsets\|0\|+2;m-21&8\|-\|134804919\|134804940&offsets\|0\|+2]\|TGTAAACATCCCCGACTGGAAGCT | >MI0000255\|hsa-mir-30d&WithFlank&8\|-\|134804870\|134804951 | 12 | 35 | 24 | TGTAAACATCCCCGACTGGAAGCT | 15272.4 | 3.4 | 0.7 | 4.8 | 1.3E-06 | 2.4E-06 | MB231/MB468 | MP |
| >hg38_21_+_8433172_8446622_RNA45SN1_45S_50ntFlanks@6649.6679.31\|TACGACTCTTAGCGGTGGATCACTCGGCTCG | >hg38_21_+_8433172_8446622_RNA45SN1_45S_50ntFlanks | 6649 | 6679 | 31 | TACGACTCTTAGCGGTGGATCACTCGGCTCG | 511.2 | -4.7 | 1.0 | -4.8 | 1.3E-06 | 2.4E-06 | MB231/MB468 | MP |
| >hg38_21_+_8433172_8446622_RNA45SN1_45S_50ntFlanks@9533.9571.39\|GAGGAAACTCTGGTGGAGGTCCGTAGCGGTCCTGACGTG | >hg38_21_+_8433172_8446622_RNA45SN1_45S_50ntFlanks | 9533 | 9571 | 39 | GAGGAAACTCTGGTGGAGGTCCGTAGCGGTCCTGACGTG | 1960.8 | -5.1 | 1.1 | -4.8 | 1.3E-06 | 2.4E-06 | MB231/MB468 | MP |
| >hg38_21_+_8433172_8446622_RNA45SN1_45S_50ntFlanks@3793.3810.18\|CGAATGGCTCATTAAATC | >hg38_21_+_8433172_8446622_RNA45SN1_45S_50ntFlanks | 3793 | 3810 | 18 | CGAATGGCTCATTAAATC | 147.9 | -4.9 | 1.0 | -4.8 | 1.5E-06 | 2.4E-06 | MB231/MB468 | NUC |
| >MI0003560\|hsa-mir-92b&WithFlank&1\|+\|155195171\|155195278@67.86.20(+1U)&[MIMAT0003218&hsa-miR-92b-3p&offsets\|0\|-2(+1U);m-115&1\|+\|155195237\|155195258&offsets\|0\|-2(+1U)]\|TATTGCACTCGTCCCGGCCTT | >MI0003560\|hsa-mir-92b&WithFlank&1\|+\|155195171\|155195278 | 67 | 86 | 20(+1U) | TATTGCACTCGTCCCGGCCTT | 300.0 | 8.0 | 1.7 | 4.8 | 1.3E-06 | 2.4E-06 | MB231/MB468 | MP |
| >hg38_21_+_8433172_8446622_RNA45SN1_45S_50ntFlanks@12418.12443.26\|CGATGTCGGCTCTTCCTATCATTGTG | >hg38_21_+_8433172_8446622_RNA45SN1_45S_50ntFlanks | 12418 | 12443 | 26 | CGATGTCGGCTCTTCCTATCATTGTG | 11242.1 | -3.2 | 0.7 | -4.8 | 1.5E-06 | 2.4E-06 | MB231/MB468 | CYTO |
| >hg38_21_+_8433172_8446622_RNA45SN1_45S_50ntFlanks@12654.12674.21\|TATGACTGAACGCCTCTAAGT | >hg38_21_+_8433172_8446622_RNA45SN1_45S_50ntFlanks | 12654 | 12674 | 21 | TATGACTGAACGCCTCTAAGT | 722.0 | -6.6 | 1.4 | -4.8 | 1.3E-06 | 2.4E-06 | BT20/MB468 | CYTO |
| >MI0000273\|hsa-mir-183&WithFlank&7\|-\|129774899\|129775020@34.56.23&[MIMAT0000261&hsa-miR-183-5p&offsets\|+1\|+2;m-41&7\|-\|129774966\|129774987&offsets\|0\|+1]\|ATGGCACTGGTAGAATTCACTGT | >MI0000273\|hsa-mir-183&WithFlank&7\|-\|129774899\|129775020 | 34 | 56 | 23 | ATGGCACTGGTAGAATTCACTGT | 288.2 | 1.9 | 0.4 | 4.8 | 1.6E-06 | 2.4E-06 | BT20/MB468 | NUC |
| >hg38_21_+_8433172_8446622_RNA45SN1_45S_50ntFlanks@12416.12444.29\|TTCGATGTCGGCTCTTCCTATCATTGTGA | >hg38_21_+_8433172_8446622_RNA45SN1_45S_50ntFlanks | 12416 | 12444 | 29 | TTCGATGTCGGCTCTTCCTATCATTGTGA | 1405.6 | 2.6 | 0.5 | 4.8 | 1.3E-06 | 2.4E-06 | BT20/MB468 | CYTO |
| >hg38_21_+_8433172_8446622_RNA45SN1_45S_50ntFlanks@10797.10822.26\|GCATGTTGGAACAATGTAGGTAAGGG | >hg38_21_+_8433172_8446622_RNA45SN1_45S_50ntFlanks | 10797 | 10822 | 26 | GCATGTTGGAACAATGTAGGTAAGGG | 235.6 | -2.8 | 0.6 | -4.8 | 1.3E-06 | 2.4E-06 | BT20/MB468 | CYTO |
| >hg38_21_+_8433172_8446622_RNA45SN1_45S_50ntFlanks@12944.12982.39\|CGTAGACGACCTGCTTCTGGGTCGGGGTTTCGTACGTAG | >hg38_21_+_8433172_8446622_RNA45SN1_45S_50ntFlanks | 12944 | 12982 | 39 | CGTAGACGACCTGCTTCTGGGTCGGGGTTTCGTACGTAG | 104.3 | -6.0 | 1.2 | -4.8 | 1.3E-06 | 2.4E-06 | BT20/MB468 | CYTO |
| >hg38_21_+_8433172_8446622_RNA45SN1_45S_50ntFlanks@11318.11343.26\|CCCCACGTCTCGTCGCGCGCGCGTCC | >hg38_21_+_8433172_8446622_RNA45SN1_45S_50ntFlanks | 11318 | 11343 | 26 | CCCCACGTCTCGTCGCGCGCGCGTCC | 445.1 | -4.7 | 1.0 | -4.8 | 1.3E-06 | 2.4E-06 | BT20/MB468 | CYTO |
| >hg38_21_+_8433172_8446622_RNA45SN1_45S_50ntFlanks@12944.12964.21\|CGTAGACGACCTGCTTCTGGG | >hg38_21_+_8433172_8446622_RNA45SN1_45S_50ntFlanks | 12944 | 12964 | 21 | CGTAGACGACCTGCTTCTGGG | 191.0 | -5.7 | 1.2 | -4.8 | 1.3E-06 | 2.4E-06 | BT20/MB468 | CYTO |
| >hg38_21_+_8433172_8446622_RNA45SN1_45S_50ntFlanks@11580.11602.23\|GAACTGGTGCGGACCAGGGGAAT | >hg38_21_+_8433172_8446622_RNA45SN1_45S_50ntFlanks | 11580 | 11602 | 23 | GAACTGGTGCGGACCAGGGGAAT | 824.6 | -6.3 | 1.3 | -4.8 | 1.6E-06 | 2.4E-06 | BT20/MB468 | NUC |
| >hg38_21_+_8433172_8446622_RNA45SN1_45S_50ntFlanks@12416.12457.42\|TTCGATGTCGGCTCTTCCTATCATTGTGAAGCAGAATTCACC | >hg38_21_+_8433172_8446622_RNA45SN1_45S_50ntFlanks | 12416 | 12457 | 42 | TTCGATGTCGGCTCTTCCTATCATTGTGAAGCAGAATTCACC | 2388.8 | -4.5 | 0.8 | -5.3 | 1.1E-07 | 2.4E-06 | BT20/MB231 | CYTO |
| >hg38_21_+_8433172_8446622_RNA45SN1_45S_50ntFlanks@10829.10858.30\|GCAAGCCGGATCCGTAACTTCGGGATAAGG | >hg38_21_+_8433172_8446622_RNA45SN1_45S_50ntFlanks | 10829 | 10858 | 30 | GCAAGCCGGATCCGTAACTTCGGGATAAGG | 275.4 | -6.9 | 1.4 | -4.8 | 1.6E-06 | 2.4E-06 | MB231/MB468 | MITO |
| >MI0000746\|hsa-mir-99b&WithFlank&19\|+\|51692606\|51692687@14.32.19(+1U)&[MIMAT0000689&hsa-miR-99b-5p&offsets\|+1\|-2(+1U);m-10&19\|+\|51692618\|51692639&offsets\|+1\|-2(+1U)]\|ACCCGTAGAACCGACCTTGT | >MI0000746\|hsa-mir-99b&WithFlank&19\|+\|51692606\|51692687 | 14 | 32 | 19(+1U) | ACCCGTAGAACCGACCTTGT | 57.7 | 9.6 | 1.9 | 5.1 | 4.1E-07 | 2.4E-06 | BT20/MB468 | MP |
| >hg38_21_+_8433172_8446622_RNA45SN1_45S_50ntFlanks@10787.10818.32\|ACAGCCTCTGGCATGTTGGAACAATGTAGGTA | >hg38_21_+_8433172_8446622_RNA45SN1_45S_50ntFlanks | 10787 | 10818 | 32 | ACAGCCTCTGGCATGTTGGAACAATGTAGGTA | 1013.8 | -3.1 | 0.6 | -4.8 | 1.3E-06 | 2.4E-06 | BT20/MB468 | CYTO |
| >hg38_1_-_228634819_228635039_RNA5S12_5S_50ntFlanks@148.169.22\|GGAATACCGGGTGCTGTAGGCT | >hg38_1_-_228634819_228635039_RNA5S12_5S_50ntFlanks | 148 | 169 | 22 | GGAATACCGGGTGCTGTAGGCT | 71661.6 | 9.1 | 1.7 | 5.3 | 1.0E-07 | 2.4E-06 | BT20/MB231 | MP |
| >hg38_21_+_8433172_8446622_RNA45SN1_45S_50ntFlanks@4367.4394.28\|CAGTTAAAAAGCTCGTAGTTGGATCTTG | >hg38_21_+_8433172_8446622_RNA45SN1_45S_50ntFlanks | 4367 | 4394 | 28 | CAGTTAAAAAGCTCGTAGTTGGATCTTG | 222.5 | -5.9 | 1.2 | -4.8 | 1.3E-06 | 2.4E-06 | BT20/MB468 | CYTO |
| >hg38_21_+_8433172_8446622_RNA45SN1_45S_50ntFlanks@5417.5438.22\|CTACCGATTGGATGGTTTAGTG | >hg38_21_+_8433172_8446622_RNA45SN1_45S_50ntFlanks | 5417 | 5438 | 22 | CTACCGATTGGATGGTTTAGTG | 55.9 | -8.9 | 1.8 | -5.1 | 4.1E-07 | 2.4E-06 | BT20/MB468 | MP |
| >hg38_21_+_8433172_8446622_RNA45SN1_45S_50ntFlanks@7991.8009.19\|ACGTGGCGACCCGCTGAAT | >hg38_21_+_8433172_8446622_RNA45SN1_45S_50ntFlanks | 7991 | 8009 | 19 | ACGTGGCGACCCGCTGAAT | 9334.4 | -3.9 | 0.8 | -5.1 | 4.1E-07 | 2.4E-06 | BT20/MB468 | MP |
| >hg38_21_+_8433172_8446622_RNA45SN1_45S_50ntFlanks@9534.9563.30\|AGGAAACTCTGGTGGAGGTCCGTAGCGGTC | >hg38_21_+_8433172_8446622_RNA45SN1_45S_50ntFlanks | 9534 | 9563 | 30 | AGGAAACTCTGGTGGAGGTCCGTAGCGGTC | 13233.8 | -5.9 | 1.2 | -5.1 | 4.1E-07 | 2.4E-06 | BT20/MB468 | MP |
| >hg38_21_+_8433172_8446622_RNA45SN1_45S_50ntFlanks@9485.9510.26\|TGTTAGGACCCGAAAGATGGTGAACT | >hg38_21_+_8433172_8446622_RNA45SN1_45S_50ntFlanks | 9485 | 9510 | 26 | TGTTAGGACCCGAAAGATGGTGAACT | 50.5 | 9.4 | 1.9 | 5.1 | 4.1E-07 | 2.4E-06 | BT20/MB468 | MP |
| >am_trna116_GluCTC_1_-_145399233_145399304@11.29.19,trna25_GluCTC_2_-_71273488_71273560@11.29.19,trna59_GluCTC_1_+_249168447_249168518@11.29.19,trna71_GluCTC_1_-_161439189_161439260@11.29.19,trna74_GluCTC_1_-_161431809_161431880@11.29.19,trna77_GluCTC_1_-_161424398_161424469@11.29.19,trna77_GluCTC_6_+_28949976_28950047@11.29.19,trna80_GluCTC_1_-_161417018_161417089@11.29.19,trna87_GluCTC_6_-_126101393_126101464@11.29.19\|TCTAGTGGTTAGGATTCGG | >am_trna116_GluCTC_1_-_145399233_145399304 | 11 | 29 | 19 | TCTAGTGGTTAGGATTCGG | 64.7 | -9.1 | 1.8 | -5.1 | 4.1E-07 | 2.4E-06 | BT20/MB468 | MP |
| >hg38_21_+_8433172_8446622_RNA45SN1_45S_50ntFlanks@10791.10816.26\|CCTCTGGCATGTTGGAACAATGTAGG | >hg38_21_+_8433172_8446622_RNA45SN1_45S_50ntFlanks | 10791 | 10816 | 26 | CCTCTGGCATGTTGGAACAATGTAGG | 284.4 | -5.6 | 1.2 | -4.8 | 1.7E-06 | 2.4E-06 | BT20/MB468 | NUC |
| >hg38_21_+_8433172_8446622_RNA45SN1_45S_50ntFlanks@3811.3837.27\|AGTTATGGTTCCTTTGGTCGCTCGCTC | >hg38_21_+_8433172_8446622_RNA45SN1_45S_50ntFlanks | 3811 | 3837 | 27 | AGTTATGGTTCCTTTGGTCGCTCGCTC | 416.2 | -7.5 | 1.6 | -4.8 | 1.7E-06 | 2.4E-06 | BT20/MB468 | NUC |
| >hg38_21_+_8433172_8446622_RNA45SN1_45S_50ntFlanks@8183.8223.41\|TCGAGGCCCAGCCCGTGGACGGTGTGAGGCCGGTAGCGGCC | >hg38_21_+_8433172_8446622_RNA45SN1_45S_50ntFlanks | 8183 | 8223 | 41 | TCGAGGCCCAGCCCGTGGACGGTGTGAGGCCGGTAGCGGCC | 239.2 | -6.9 | 1.4 | -4.8 | 1.6E-06 | 2.4E-06 | MB231/MB468 | MITO |
| >hg38_21_+_8433172_8446622_RNA45SN1_45S_50ntFlanks@10388.10424.37\|GTGAACAGCAGTTGAACATGGGTCAGTCGGTCCTGAG | >hg38_21_+_8433172_8446622_RNA45SN1_45S_50ntFlanks | 10388 | 10424 | 37 | GTGAACAGCAGTTGAACATGGGTCAGTCGGTCCTGAG | 20514.3 | 4.8 | 0.9 | 5.3 | 1.3E-07 | 2.4E-06 | BT20/MB231 | MITO |
| >hg38_21_+_8433172_8446622_RNA45SN1_45S_50ntFlanks@8339.8375.37\|TACCGTAAGGGAAAGTTGAAAAGAACTTTGAAGAGAG | >hg38_21_+_8433172_8446622_RNA45SN1_45S_50ntFlanks | 8339 | 8375 | 37 | TACCGTAAGGGAAAGTTGAAAAGAACTTTGAAGAGAG | 430.3 | 5.8 | 1.1 | 5.3 | 1.3E-07 | 2.4E-06 | BT20/MB231 | MITO |
| >MI0003820\|hsa-mir-454&WithFlank&17\|-\|59137752\|59137878@70.95.26&[MIMAT0003885&hsa-miR-454-3p&offsets\|0\|+3;m-272&17\|-\|59137787\|59137809&offsets\|0\|+3]\|TAGTGCAATATTGCTTATAGGGTTTT | >MI0003820\|hsa-mir-454&WithFlank&17\|-\|59137752\|59137878 | 70 | 95 | 26 | TAGTGCAATATTGCTTATAGGGTTTT | 278.1 | -5.3 | 1.0 | -5.3 | 1.3E-07 | 2.4E-06 | BT20/MB231 | MITO |
| >hg38_21_+_8433172_8446622_RNA45SN1_45S_50ntFlanks@4722.4759.38\|TCATTAATCAAGAACGAAAGTCGGAGGTTCGAAGACGA | >hg38_21_+_8433172_8446622_RNA45SN1_45S_50ntFlanks | 4722 | 4759 | 38 | TCATTAATCAAGAACGAAAGTCGGAGGTTCGAAGACGA | 3007.1 | -4.5 | 0.9 | -4.8 | 1.3E-06 | 2.4E-06 | BT20/MB468 | CYTO |
| >hg38_21_+_8433172_8446622_RNA45SN1_45S_50ntFlanks@11781.11805.25\|CGCATGAATGGATGAACGAGATTCC | >hg38_21_+_8433172_8446622_RNA45SN1_45S_50ntFlanks | 11781 | 11805 | 25 | CGCATGAATGGATGAACGAGATTCC | 480.1 | -4.8 | 1.0 | -4.8 | 1.5E-06 | 2.4E-06 | MB231/MB468 | CYTO |
| >hg38_21_+_8433172_8446622_RNA45SN1_45S_50ntFlanks@12694.12711.18\|AACGATACGGCAGCGCCG | >hg38_21_+_8433172_8446622_RNA45SN1_45S_50ntFlanks | 12694 | 12711 | 18 | AACGATACGGCAGCGCCG | 96.6 | -9.7 | 1.9 | -5.1 | 4.2E-07 | 2.4E-06 | BT20/MB468 | MP |
| >hg38_21_+_8433172_8446622_RNA45SN1_45S_50ntFlanks@12073.12092.20\|CCGAGGGGCTCTCGCTTCTG | >hg38_21_+_8433172_8446622_RNA45SN1_45S_50ntFlanks | 12073 | 12092 | 20 | CCGAGGGGCTCTCGCTTCTG | 787.3 | -5.1 | 1.0 | -4.8 | 1.3E-06 | 2.4E-06 | BT20/MB468 | CYTO |
| >hg38_MT_+_1621_3279_RNR2_16S_50ntFlanks@721.740.20\|CTGCGTCAGATTAAAACACT | >hg38_MT_+_1621_3279_RNR2_16S_50ntFlanks | 721 | 740 | 20 | CTGCGTCAGATTAAAACACT | 40.9 | 8.6 | 1.8 | 4.8 | 1.3E-06 | 2.4E-06 | MB231/MB468 | MP |
| >hg38_MT_+_1621_3279_RNR2_16S_50ntFlanks@459.492.34\|CTTGTAAATTTAACTGTTAGTCCAAAGAGGAACA | >hg38_MT_+_1621_3279_RNR2_16S_50ntFlanks | 459 | 492 | 34 | CTTGTAAATTTAACTGTTAGTCCAAAGAGGAACA | 262.9 | 7.7 | 1.6 | 4.8 | 1.5E-06 | 2.4E-06 | MB231/MB468 | NUC |
| >MI0000067\|hsa-let-7f-1&WithFlank&9\|+\|94176341\|94176439@69.89.21(+1U)&[MIMAT0004486&hsa-let-7f-1-3p&offsets\|0\|-1(+1U);m-275&9\|+\|94176409\|94176430&offsets\|0\|-1(+1U)]\|CTATACAATCTATTGCCTTCCT | >MI0000067\|hsa-let-7f-1&WithFlank&9\|+\|94176341\|94176439 | 69 | 89 | 21(+1U) | CTATACAATCTATTGCCTTCCT | 311.6 | 7.9 | 1.6 | 4.8 | 1.6E-06 | 2.4E-06 | MB231/MB468 | MITO |
| >trna111_HisGTG_1_-_147774845_147774916@-1T.31.32,trna118_HisGTG_1_-_145396881_145396952@-1T.31.32,trna16_HisGTG_1_+_146544773_146544844@-1T.31.32,trna1_HisGTG_15_+_45493349_45493420@-1T.31.32,trna21_HisGTG_1_+_147753471_147753542@-1T.31.32,trna33_HisGTG_6_+_27125906_27125977@-1T.31.32,trna7_HisGTG_9_-_14433938_14434009@-1T.31.32,trna8_HisGTG_15_-_45492611_45492682@-1T.31.32,trna9_HisGTG_15_-_45490804_45490875@-1T.31.32\|TGCCGTGATCGTATAGTGGTTAGTACTCTGCG | >trna111_HisGTG_1_-_147774845_147774916 | -1T | 31 | 32 | TGCCGTGATCGTATAGTGGTTAGTACTCTGCG | 9689.2 | -4.0 | 0.8 | -4.8 | 1.5E-06 | 2.5E-06 | MB231/MB468 | NUC |
| >hg38_21_+_8433172_8446622_RNA45SN1_45S_50ntFlanks@10869.10891.23\|AGGGCTGGGTCGGTCGGGCTGGG | >hg38_21_+_8433172_8446622_RNA45SN1_45S_50ntFlanks | 10869 | 10891 | 23 | AGGGCTGGGTCGGTCGGGCTGGG | 1081.4 | -5.1 | 1.1 | -4.8 | 1.5E-06 | 2.5E-06 | MB231/MB468 | CYTO |
| >hg38_21_+_8433172_8446622_RNA45SN1_45S_50ntFlanks@10349.10381.33\|AACGAGAACTTTGAAGGCCGAAGTGGAGAAGGG | >hg38_21_+_8433172_8446622_RNA45SN1_45S_50ntFlanks | 10349 | 10381 | 33 | AACGAGAACTTTGAAGGCCGAAGTGGAGAAGGG | 308.8 | -3.1 | 0.6 | -4.8 | 1.5E-06 | 2.5E-06 | MB231/MB468 | NUC |
| >hg38_21_+_8433172_8446622_RNA45SN1_45S_50ntFlanks@9482.9509.28\|ACGTGTTAGGACCCGAAAGATGGTGAAC | >hg38_21_+_8433172_8446622_RNA45SN1_45S_50ntFlanks | 9482 | 9509 | 28 | ACGTGTTAGGACCCGAAAGATGGTGAAC | 256.9 | 11.3 | 2.1 | 5.3 | 1.0E-07 | 2.5E-06 | BT20/MB231 | MP |
| >MI0000743\|hsa-mir-34c&WithFlank&11\|+\|111513433\|111513521@19.39.21&[MIMAT0000686&hsa-miR-34c-5p&offsets\|0\|-2;m-336&11\|+\|111513451\|111513473&offsets\|0\|-2]\|AGGCAGTGTAGTTAGCTGATT | >MI0000743\|hsa-mir-34c&WithFlank&11\|+\|111513433\|111513521 | 19 | 39 | 21 | AGGCAGTGTAGTTAGCTGATT | 118.0 | -10.3 | 1.9 | -5.3 | 1.0E-07 | 2.5E-06 | BT20/MB231 | MP |

<table>
<tr><td>>MI0000093|hsa-mir-92a-1&WithFlank&13|+|91351308|91351397@54.75.22(+1A);MI0000094|hsa-mir-92a-2&WithFlank&X|-|134169532|134169618@54.75.22(+1A)&[MIMAT0000092&hsa-miR-92a-3p&offsets|0|0(+1A);m-9&13|+|91351361|91351382&offsets|0|0(+1A)];[MIMAT0000092_1&hsa-miR-92a-3p&offsets|0|0(+1A);m-11&X|-|134169544|134169565&offsets|0|0(+1A)]|TATTGCACTTGTCCCGGCCTGTA</td><td>>MI0000093|hsa-mir-92a-1&WithFlank&13|+|91351308|91351397</td><td>54</td><td>75</td><td>22(+1A)</td><td>TATTGCACTTGTCCCGGCCTGTA</td><td>7350.1</td><td>4.1</td><td>0.8</td><td>4.8</td><td>1.6E-06</td><td>2.5E-06</td><td>MB231/MB468</td><td>MITO</td></tr>
<tr><td>>MI0000093|hsa-mir-92a-1&WithFlank&13|+|91351308|91351397@55.75.21;MI0000094|hsa-mir-92a-2&WithFlank&X|-|134169532|134169618@55.75.21&[MIMAT0000092&hsa-miR-92a-3p&offsets|+1|0;m-9&13|+|91351361|91351382&offsets|+1|0];[MIMAT0000092_1&hsa-miR-92a-3p&offsets|+1|0;m-11&X|-|134169544|134169565&offsets|+1|0]|ATTGCACTTGTCCCGGCCTGT</td><td>>MI0000093|hsa-mir-92a-1&WithFlank&13|+|91351308|91351397</td><td>55</td><td>75</td><td>21</td><td>ATTGCACTTGTCCCGGCCTGT</td><td>1148.3</td><td>3.4</td><td>0.7</td><td>4.8</td><td>1.5E-06</td><td>2.5E-06</td><td>MB231/MB468</td><td>NUC</td></tr>
<tr><td>>hg38_21_+_8433172_8446622_RNA45SN1_45S_50ntFlanks@12934.12955.22|CTAAACCATTCGTAGACGACCT</td><td>>hg38_21_+_8433172_8446622_RNA45SN1_45S_50ntFlanks</td><td>12934</td><td>12955</td><td>22</td><td>CTAAACCATTCGTAGACGACCT</td><td>673.1</td><td>-4.6</td><td>0.9</td><td>-4.8</td><td>1.5E-06</td><td>2.5E-06</td><td>MB231/MB468</td><td>CYTO</td></tr>
<tr><td>>hg38_21_+_8433172_8446622_RNA45SN1_45S_50ntFlanks@8158.8181.24|TCGTGGGGGGCCCAAGTCCTTCTG</td><td>>hg38_21_+_8433172_8446622_RNA45SN1_45S_50ntFlanks</td><td>8158</td><td>8181</td><td>24</td><td>TCGTGGGGGGCCCAAGTCCTTCTG</td><td>52.6</td><td>-8.8</td><td>1.7</td><td>-5.1</td><td>4.2E-07</td><td>2.5E-06</td><td>BT20/MB468</td><td>MP</td></tr>
<tr><td>>trna14_LysTTT_11_-_59327808_59327880@3.33.31,trna2_LysTTT_17_+_8022473_8022545@3.33.31,trna54_LysTTT_1_+_204475655_204475727@3.33.31,trna5_LysTTT_11_+_59323902_59323974@3.33.31,trna62_LysTTT_1_-_204476158_204476230@3.33.31,trna76_LysTTT_6_+_28918806_28918878@3.33.31|CCGGATAGCTCAGTCGGTAGAGCATCAGACT</td><td>>trna14_LysTTT_11_-_59327808_59327880</td><td>3</td><td>33</td><td>31</td><td>CCGGATAGCTCAGTCGGTAGAGCATCAGACT</td><td>438.3</td><td>-5.3</td><td>1.0</td><td>-5.3</td><td>1.3E-07</td><td>2.5E-06</td><td>BT20/MB231</td><td>MITO</td></tr>
<tr><td>>hg38_21_+_8433172_8446622_RNA45SN1_45S_50ntFlanks@9622.9650.29|CTAGTAGCTGGTTCCCTCCGAAGTTTCCC</td><td>>hg38_21_+_8433172_8446622_RNA45SN1_45S_50ntFlanks</td><td>9622</td><td>9650</td><td>29</td><td>CTAGTAGCTGGTTCCCTCCGAAGTTTCCC</td><td>237.1</td><td>-7.1</td><td>1.4</td><td>-5.1</td><td>4.2E-07</td><td>2.5E-06</td><td>BT20/MB468</td><td>MP</td></tr>
<tr><td>>trna14_LysTTT_11_-_59327808_59327880@3.33.31,trna2_LysTTT_17_+_8022473_8022545@3.33.31,trna54_LysTTT_1_+_204475655_204475727@3.33.31,trna5_LysTTT_11_+_59323902_59323974@3.33.31,trna62_LysTTT_1_-_204476158_204476230@3.33.31,trna76_LysTTT_6_+_28918806_28918878@3.33.31|CCGGATAGCTCAGTCGGTAGAGCATCAGACT</td><td>>trna14_LysTTT_11_-_59327808_59327880</td><td>3</td><td>33</td><td>31</td><td>CCGGATAGCTCAGTCGGTAGAGCATCAGACT</td><td>385.5</td><td>6.9</td><td>1.4</td><td>4.8</td><td>1.6E-06</td><td>2.5E-06</td><td>MB231/MB468</td><td>MITO</td></tr>
<tr><td>>hg38_21_+_8433172_8446622_RNA45SN1_45S_50ntFlanks@9736.9753.18|CCGAAACGATCTCAACCT</td><td>>hg38_21_+_8433172_8446622_RNA45SN1_45S_50ntFlanks</td><td>9736</td><td>9753</td><td>18</td><td>CCGAAACGATCTCAACCT</td><td>334.1</td><td>-7.6</td><td>1.6</td><td>-4.8</td><td>1.7E-06</td><td>2.5E-06</td><td>BT20/MB468</td><td>NUC</td></tr>
<tr><td>>MI0000066|hsa-let-7e&WithFlank&19|+|51692780|51692870@14.32.19&[MIMAT0000066&hsa-let-7e-5p&offsets|0|-3;m-50&19|+|51692793|51692814&offsets|0|-3]|TGAGGTAGGAGGTTGTATA</td><td>>MI0000066|hsa-let-7e&WithFlank&19|+|51692780|51692870</td><td>14</td><td>32</td><td>19</td><td>TGAGGTAGGAGGTTGTATA</td><td>668.4</td><td>4.3</td><td>0.8</td><td>5.3</td><td>1.2E-07</td><td>2.5E-06</td><td>BT20/MB231</td><td>CYTO</td></tr>
<tr><td>>hg38_21_+_8433172_8446622_RNA45SN1_45S_50ntFlanks@4733.4751.19|GAACGAAAGTCGGAGGTTC</td><td>>hg38_21_+_8433172_8446622_RNA45SN1_45S_50ntFlanks</td><td>4733</td><td>4751</td><td>19</td><td>GAACGAAAGTCGGAGGTTC</td><td>678.8</td><td>-3.6</td><td>0.7</td><td>-4.8</td><td>1.3E-06</td><td>2.5E-06</td><td>MB231/MB468</td><td>MP</td></tr>
<tr><td>>hg38_21_+_8433172_8446622_RNA45SN1_45S_50ntFlanks@6654.6688.35|CTCTTAGCGGTGGATCACTCGGCTCGTGCGTCGAT</td><td>>hg38_21_+_8433172_8446622_RNA45SN1_45S_50ntFlanks</td><td>6654</td><td>6688</td><td>35</td><td>CTCTTAGCGGTGGATCACTCGGCTCGTGCGTCGAT</td><td>382.6</td><td>-3.6</td><td>0.8</td><td>-4.8</td><td>1.6E-06</td><td>2.5E-06</td><td>MB231/MB468</td><td>MITO</td></tr>
<tr><td>>trnaMT_GluTTC_MT_-_14674_14742@26.58.33,trnalookalike8_GluTTC_5_-_93905172_93905240@26.58.33|GGTTTTTCATATCATTGGTCGTGGTTGTAGTCC</td><td>>trnaMT_GluTTC_MT_-_14674_14742</td><td>26</td><td>58</td><td>33</td><td>GGTTTTTCATATCATTGGTCGTGGTTGTAGTCC</td><td>567.9</td><td>9.4</td><td>1.9</td><td>5.1</td><td>4.2E-07</td><td>2.5E-06</td><td>BT20/MB468</td><td>MP</td></tr>
<tr><td>>hg38_21_+_8433172_8446622_RNA45SN1_45S_50ntFlanks@10389.10429.41|TGAACAGCAGTTGAACATGGGTCAGTCGGTCCTGAGAGATG</td><td>>hg38_21_+_8433172_8446622_RNA45SN1_45S_50ntFlanks</td><td>10389</td><td>10429</td><td>41</td><td>TGAACAGCAGTTGAACATGGGTCAGTCGGTCCTGAGAGATG</td><td>628.0</td><td>-5.7</td><td>1.2</td><td>-4.8</td><td>1.6E-06</td><td>2.5E-06</td><td>MB231/MB468</td><td>MITO</td></tr>
<tr><td>>hg38_21_+_8433172_8446622_RNA45SN1_45S_50ntFlanks@10840.10864.25|CCGTAACTTCGGGATAAGGATTGGC</td><td>>hg38_21_+_8433172_8446622_RNA45SN1_45S_50ntFlanks</td><td>10840</td><td>10864</td><td>25</td><td>CCGTAACTTCGGGATAAGGATTGGC</td><td>239.9</td><td>-4.1</td><td>0.9</td><td>-4.8</td><td>1.5E-06</td><td>2.5E-06</td><td>MB231/MB468</td><td>CYTO</td></tr>
<tr><td>>hg38_21_+_8433172_8446622_RNA45SN1_45S_50ntFlanks@3899.3936.38|CCTTCGCGGGGGGGATGCGTGCATTTATCAGATCAAAA</td><td>>hg38_21_+_8433172_8446622_RNA45SN1_45S_50ntFlanks</td><td>3899</td><td>3936</td><td>38</td><td>CCTTCGCGGGGGGGATGCGTGCATTTATCAGATCAAAA</td><td>121.9</td><td>-10.0</td><td>2.0</td><td>-5.1</td><td>4.2E-07</td><td>2.5E-06</td><td>BT20/MB468</td><td>MP</td></tr>
<tr><td>>MI0000078|hsa-mir-22&WithFlank&17|-|1713897|1713993@59.79.21(+1C)&[MIMAT0000077&hsa-miR-22-3p&offsets|0|-1(+1C);m-2&17|-|1713914|1713935&offsets|0|-1(+1C)]|AAGCTGCCAGTTGAAGAACTGC</td><td>>MI0000078|hsa-mir-22&WithFlank&17|-|1713897|1713993</td><td>59</td><td>79</td><td>21(+1C)</td><td>AAGCTGCCAGTTGAAGAACTGC</td><td>3397.8</td><td>5.3</td><td>1.0</td><td>5.1</td><td>4.2E-07</td><td>2.5E-06</td><td>BT20/MB468</td><td>MP</td></tr>
<tr><td>>hg38_21_+_8433172_8446622_RNA45SN1_45S_50ntFlanks@4369.4391.23|GTTAAAAAGCTCGTAGTTGGATC</td><td>>hg38_21_+_8433172_8446622_RNA45SN1_45S_50ntFlanks</td><td>4369</td><td>4391</td><td>23</td><td>GTTAAAAAGCTCGTAGTTGGATC</td><td>395.9</td><td>-5.5</td><td>1.1</td><td>-4.8</td><td>1.3E-06</td><td>2.5E-06</td><td>MB231/MB468</td><td>MP</td></tr>
<tr><td>>hg38_21_+_8433172_8446622_RNA45SN1_45S_50ntFlanks@12712.12730.19|CGGAGCCTCGGTTGGCCTC</td><td>>hg38_21_+_8433172_8446622_RNA45SN1_45S_50ntFlanks</td><td>12712</td><td>12730</td><td>19</td><td>CGGAGCCTCGGTTGGCCTC</td><td>100.3</td><td>-9.7</td><td>1.9</td><td>-5.1</td><td>4.2E-07</td><td>2.5E-06</td><td>BT20/MB468</td><td>MP</td></tr>
<tr><td>>hg38_21_+_8433172_8446622_RNA45SN1_45S_50ntFlanks@3773.3799.27|CGGCCGGTACAGTGAAACTGCGAATGG</td><td>>hg38_21_+_8433172_8446622_RNA45SN1_45S_50ntFlanks</td><td>3773</td><td>3799</td><td>27</td><td>CGGCCGGTACAGTGAAACTGCGAATGG</td><td>173.7</td><td>-6.7</td><td>1.4</td><td>-4.8</td><td>1.3E-06</td><td>2.5E-06</td><td>MB231/MB468</td><td>MP</td></tr>
<tr><td>>hg38_21_+_8433172_8446622_RNA45SN1_45S_50ntFlanks@6721.6757.37|AATTGCAGGACACATTGATCATCGACACTTCGAACGC</td><td>>hg38_21_+_8433172_8446622_RNA45SN1_45S_50ntFlanks</td><td>6721</td><td>6757</td><td>37</td><td>AATTGCAGGACACATTGATCATCGACACTTCGAACGC</td><td>435.1</td><td>-5.6</td><td>1.2</td><td>-4.8</td><td>1.6E-06</td><td>2.5E-06</td><td>MB231/MB468</td><td>NUC</td></tr>
<tr><td>>MI0003188|hsa-mir-503&WithFlank&X|-|134546322|134546404@12.30.19(+2A)&[MIMAT0002874&hsa-miR-503-5p&offsets|0|-4(+2A);m-297&X|-|134546371|134546393&offsets|0|-4(+2A)]|TAGCAGCGGGAACAGTTCTAA</td><td>>MI0003188|hsa-mir-503&WithFlank&X|-|134546322|134546404</td><td>12</td><td>30</td><td>19(+2A)</td><td>TAGCAGCGGGAACAGTTCTAA</td><td>45.5</td><td>8.7</td><td>1.8</td><td>4.8</td><td>1.3E-06</td><td>2.5E-06</td><td>MB231/MB468</td><td>MP</td></tr>
<tr><td>>hg38_21_+_8433172_8446622_RNA45SN1_45S_50ntFlanks@13002.13030.29|CGATCTATTGAAAGTCAGCCCTCGACACA</td><td>>hg38_21_+_8433172_8446622_RNA45SN1_45S_50ntFlanks</td><td>13002</td><td>13030</td><td>29</td><td>CGATCTATTGAAAGTCAGCCCTCGACACA</td><td>58.9</td><td>-9.0</td><td>1.8</td><td>-5.1</td><td>4.3E-07</td><td>2.5E-06</td><td>BT20/MB468</td><td>MP</td></tr>
<tr><td>>hg38_21_+_8433172_8446622_RNA45SN1_45S_50ntFlanks@9574.9603.30|AATCGGTCGTCCGACCTGGGTATAGGGGCG</td><td>>hg38_21_+_8433172_8446622_RNA45SN1_45S_50ntFlanks</td><td>9574</td><td>9603</td><td>30</td><td>AATCGGTCGTCCGACCTGGGTATAGGGGCG</td><td>6910.1</td><td>-4.2</td><td>0.9</td><td>-4.8</td><td>1.3E-06</td><td>2.5E-06</td><td>MB231/MB468</td><td>MP</td></tr>
<tr><td>>hg38_21_+_8433172_8446622_RNA45SN1_45S_50ntFlanks@9203.9220.18|CTCGGGGCCACGCGCGCG</td><td>>hg38_21_+_8433172_8446622_RNA45SN1_45S_50ntFlanks</td><td>9203</td><td>9220</td><td>18</td><td>CTCGGGGCCACGCGCGCG</td><td>1046.0</td><td>-6.4</td><td>1.3</td><td>-5.1</td><td>4.3E-07</td><td>2.5E-06</td><td>BT20/MB468</td><td>MP</td></tr>
<tr><td>>hg38_21_+_8433172_8446622_RNA45SN1_45S_50ntFlanks@9760.9789.30|AAACTTTAAATGGGTAAGAAGCCCGGCTCG</td><td>>hg38_21_+_8433172_8446622_RNA45SN1_45S_50ntFlanks</td><td>9760</td><td>9789</td><td>30</td><td>AAACTTTAAATGGGTAAGAAGCCCGGCTCG</td><td>73.9</td><td>-9.3</td><td>1.8</td><td>-5.1</td><td>4.3E-07</td><td>2.5E-06</td><td>BT20/MB468</td><td>MP</td></tr>
<tr><td>>am_trna2_GlyTCC_19_+_4724082_4724153@1.29.29|GCGTTGGTGGTATAGTGGTTAGCATAGCT</td><td>>am_trna2_GlyTCC_19_+_4724082_4724153</td><td>1</td><td>29</td><td>29</td><td>GCGTTGGTGGTATAGTGGTTAGCATAGCT</td><td>441.5</td><td>-3.4</td><td>0.7</td><td>-4.8</td><td>1.3E-06</td><td>2.5E-06</td><td>BT20/MB468</td><td>CYTO</td></tr>
<tr><td>>hg38_21_+_8433172_8446622_RNA45SN1_45S_50ntFlanks@3777.3805.29|CGGTACAGTGAAACTGCGAATGGCTCATT</td><td>>hg38_21_+_8433172_8446622_RNA45SN1_45S_50ntFlanks</td><td>3777</td><td>3805</td><td>29</td><td>CGGTACAGTGAAACTGCGAATGGCTCATT</td><td>382.0</td><td>-3.2</td><td>0.7</td><td>-4.8</td><td>1.6E-06</td><td>2.5E-06</td><td>MB231/MB468</td><td>NUC</td></tr>
<tr><td>>hg38_21_+_8433172_8446622_RNA45SN1_45S_50ntFlanks@10458.10475.18|ATGGCCTCCGTTGCCCTC</td><td>>hg38_21_+_8433172_8446622_RNA45SN1_45S_50ntFlanks</td><td>10458</td><td>10475</td><td>18</td><td>ATGGCCTCCGTTGCCCTC</td><td>79.4</td><td>-9.4</td><td>1.9</td><td>-5.1</td><td>4.3E-07</td><td>2.5E-06</td><td>BT20/MB468</td><td>MP</td></tr>
<tr><td>>hg38_21_+_8433172_8446622_RNA45SN1_45S_50ntFlanks@10680.10705.26|GTTCCGGCGGCGTCCGGTGAGCTCTC</td><td>>hg38_21_+_8433172_8446622_RNA45SN1_45S_50ntFlanks</td><td>10680</td><td>10705</td><td>26</td><td>GTTCCGGCGGCGTCCGGTGAGCTCTC</td><td>440.1</td><td>-5.7</td><td>1.2</td><td>-4.8</td><td>1.6E-06</td><td>2.5E-06</td><td>MB231/MB468</td><td>MITO</td></tr>
<tr><td>>hg38_21_+_8433172_8446622_RNA45SN1_45S_50ntFlanks@9503.9523.21|GGTGAACTATGCCTGGGCAGG</td><td>>hg38_21_+_8433172_8446622_RNA45SN1_45S_50ntFlanks</td><td>9503</td><td>9523</td><td>21</td><td>GGTGAACTATGCCTGGGCAGG</td><td>174.7</td><td>-7.1</td><td>1.5</td><td>-4.8</td><td>1.3E-06</td><td>2.5E-06</td><td>MB231/MB468</td><td>MP</td></tr>
<tr><td>>hg38_21_+_8433172_8446622_RNA45SN1_45S_50ntFlanks@11566.11584.19|TAGCAGCCGACTTAGAACT</td><td>>hg38_21_+_8433172_8446622_RNA45SN1_45S_50ntFlanks</td><td>11566</td><td>11584</td><td>19</td><td>TAGCAGCCGACTTAGAACT</td><td>195.6</td><td>4.3</td><td>0.9</td><td>4.8</td><td>1.6E-06</td><td>2.5E-06</td><td>MB231/MB468</td><td>MITO</td></tr>
<tr><td>>hg38_21_+_8433172_8446622_RNA45SN1_45S_50ntFlanks@10846.10872.27|CTTCGGGATAAGGATTGGCTCTAAGGG</td><td>>hg38_21_+_8433172_8446622_RNA45SN1_45S_50ntFlanks</td><td>10846</td><td>10872</td><td>27</td><td>CTTCGGGATAAGGATTGGCTCTAAGGG</td><td>424.8</td><td>-7.5</td><td>1.6</td><td>-4.8</td><td>1.7E-06</td><td>2.5E-06</td><td>BT20/MB468</td><td>NUC</td></tr>
<tr><td>>hg38_21_+_8433172_8446622_RNA45SN1_45S_50ntFlanks@4069.4098.30|CTTTCGATGGTAGTCGCCGTGCCTACCATG</td><td>>hg38_21_+_8433172_8446622_RNA45SN1_45S_50ntFlanks</td><td>4069</td><td>4098</td><td>30</td><td>CTTTCGATGGTAGTCGCCGTGCCTACCATG</td><td>403.9</td><td>-5.1</td><td>1.0</td><td>-4.8</td><td>1.3E-06</td><td>2.5E-06</td><td>BT20/MB468</td><td>CYTO</td></tr>
<tr><td>>MI0000301|hsa-mir-224&WithFlank&X|-|151958572|151958664@14.35.22(+1U)&[MIMAT0000281&hsa-miR-224-5p&offsets|+1|-2(+1U);m-238&X|-|151958630|151958651&offsets|0|0(+1U)]|CAAGTCACTAGTGGTTCCGTTTT</td><td>>MI0000301|hsa-mir-224&WithFlank&X|-|151958572|151958664</td><td>14</td><td>35</td><td>22(+1U)</td><td>CAAGTCACTAGTGGTTCCGTTTT</td><td>64.0</td><td>9.7</td><td>1.9</td><td>5.1</td><td>4.3E-07</td><td>2.5E-06</td><td>BT20/MB468</td><td>MP</td></tr>
<tr><td>>hg38_21_+_8433172_8446622_RNA45SN1_45S_50ntFlanks@7989.8011.23|AGACGTGGCGACCCGCTGAATTT</td><td>>hg38_21_+_8433172_8446622_RNA45SN1_45S_50ntFlanks</td><td>7989</td><td>8011</td><td>23</td><td>AGACGTGGCGACCCGCTGAATTT</td><td>3722.7</td><td>4.3</td><td>0.9</td><td>4.8</td><td>1.6E-06</td><td>2.5E-06</td><td>MB231/MB468</td><td>NUC</td></tr>
<tr><td>>hg38_21_+_8433172_8446622_RNA45SN1_45S_50ntFlanks@4439.4471.33|CCGCCCCTTGCCTCTCGGCGCCCCCTCGATGCT</td><td>>hg38_21_+_8433172_8446622_RNA45SN1_45S_50ntFlanks</td><td>4439</td><td>4471</td><td>33</td><td>CCGCCCCTTGCCTCTCGGCGCCCCCTCGATGCT</td><td>132.5</td><td>-4.1</td><td>0.8</td><td>-4.8</td><td>1.3E-06</td><td>2.5E-06</td><td>BT20/MB468</td><td>CYTO</td></tr>
<tr><td>>hg38_21_+_8433172_8446622_RNA45SN1_45S_50ntFlanks@12996.13013.18|TCGCTGCGATCTATTGAA</td><td>>hg38_21_+_8433172_8446622_RNA45SN1_45S_50ntFlanks</td><td>12996</td><td>13013</td><td>18</td><td>TCGCTGCGATCTATTGAA</td><td>5088.6</td><td>3.5</td><td>0.7</td><td>4.8</td><td>1.5E-06</td><td>2.5E-06</td><td>MB231/MB468</td><td>CYTO</td></tr>
<tr><td>>hg38_21_+_8433172_8446622_RNA45SN1_45S_50ntFlanks@6648.6678.31|GTACGACTCTTAGCGGTGGATCACTCGGCTC</td><td>>hg38_21_+_8433172_8446622_RNA45SN1_45S_50ntFlanks</td><td>6648</td><td>6678</td><td>31</td><td>GTACGACTCTTAGCGGTGGATCACTCGGCTC</td><td>198.1</td><td>2.5</td><td>0.5</td><td>4.8</td><td>1.5E-06</td><td>2.5E-06</td><td>MB231/MB468</td><td>CYTO</td></tr>
<tr><td>>hg38_21_+_8433172_8446622_RNA45SN1_45S_50ntFlanks@10867.10902.36|TAAGGGCTGGGTCGGTCGGGCTGGGGCGCGAAGCGG</td><td>>hg38_21_+_8433172_8446622_RNA45SN1_45S_50ntFlanks</td><td>10867</td><td>10902</td><td>36</td><td>TAAGGGCTGGGTCGGTCGGGCTGGGGCGCGAAGCGG</td><td>399.0</td><td>-7.5</td><td>1.6</td><td>-4.8</td><td>1.7E-06</td><td>2.5E-06</td><td>BT20/MB468</td><td>NUC</td></tr>
<tr><td>>trna111_HisGTG_1_-_147774845_147774916@-1T.34.35,trna118_HisGTG_1_-_145396881_145396952@-1T.34.35,trna16_HisGTG_1_+_146544773_146544844@-1T.34.35,trna1_HisGTG_15_+_45493349_45493420@-1T.34.35,trna21_HisGTG_1_+_147753471_147753542@-1T.34.35,trna33_HisGTG_6_+_27125906_27125977@-1T.34.35,trna7_HisGTG_9_-_14433938_14434009@-1T.34.35,trna8_HisGTG_15_-_45492611_45492682@-1T.34.35,trna9_HisGTG_15_-_45490804_45490875@-1T.34.35|TGCCGTGATCGTATAGTGGTTAGTACTCTGCGTTG</td><td>>trna111_HisGTG_1_-_147774845_147774916</td><td>-1T</td><td>34</td><td>35</td><td>TGCCGTGATCGTATAGTGGTTAGTACTCTGCGTTG</td><td>708.7</td><td>4.5</td><td>0.9</td><td>4.8</td><td>1.6E-06</td><td>2.5E-06</td><td>MB231/MB468</td><td>MITO</td></tr>
<tr><td>>hg38_21_+_8433172_8446622_RNA45SN1_45S_50ntFlanks@11534.11552.19|GTTCCGCGCGGCGCCTCGC</td><td>>hg38_21_+_8433172_8446622_RNA45SN1_45S_50ntFlanks</td><td>11534</td><td>11552</td><td>19</td><td>GTTCCGCGCGGCGCCTCGC</td><td>631.4</td><td>-6.8</td><td>1.4</td><td>-4.8</td><td>1.3E-06</td><td>2.5E-06</td><td>BT20/MB468</td><td>CYTO</td></tr>
<tr><td>>hg38_21_+_8433172_8446622_RNA45SN1_45S_50ntFlanks@6722.6746.25|ATTGCAGGACACATTGATCATCGAC</td><td>>hg38_21_+_8433172_8446622_RNA45SN1_45S_50ntFlanks</td><td>6722</td><td>6746</td><td>25</td><td>ATTGCAGGACACATTGATCATCGAC</td><td>194.8</td><td>-4.7</td><td>1.0</td><td>-4.8</td><td>1.6E-06</td><td>2.5E-06</td><td>MB231/MB468</td><td>NUC</td></tr>
<tr><td>>hg38_21_+_8433172_8446622_RNA45SN1_45S_50ntFlanks@12990.13015.26|GCTCCCTCGCTGCGATCTATTGAAAG</td><td>>hg38_21_+_8433172_8446622_RNA45SN1_45S_50ntFlanks</td><td>12990</td><td>13015</td><td>26</td><td>GCTCCCTCGCTGCGATCTATTGAAAG</td><td>1052.1</td><td>-4.1</td><td>0.9</td><td>-4.8</td><td>1.6E-06</td><td>2.5E-06</td><td>MB231/MB468</td><td>NUC</td></tr>
<tr><td>>hg38_21_+_8433172_8446622_RNA45SN1_45S_50ntFlanks@9738.9769.32|GAAACGATCTCAACCTATTCTCAAACTTTAAA</td><td>>hg38_21_+_8433172_8446622_RNA45SN1_45S_50ntFlanks</td><td>9738</td><td>9769</td><td>32</td><td>GAAACGATCTCAACCTATTCTCAAACTTTAAA</td><td>982.3</td><td>-2.7</td><td>0.6</td><td>-4.9</td><td>1.2E-06</td><td>2.5E-06</td><td>BT20/MB468</td><td>MITO</td></tr>
<tr><td>>MI0005756|hsa-mir-934&WithFlank&X|+|136550872|136550966@21.40.20&[MIMAT0004977&hsa-miR-934&offsets|0|-2;m-943&X|+|136550892|136550913&offsets|0|-2]|TGTCTACTACTGGAGACACT</td><td>>MI0005756|hsa-mir-934&WithFlank&X|+|136550872|136550966</td><td>21</td><td>40</td><td>20</td><td>TGTCTACTACTGGAGACACT</td><td>143.5</td><td>10.5</td><td>2.0</td><td>5.3</td><td>1.1E-07</td><td>2.5E-06</td><td>BT20/MB231</td><td>MP</td></tr>
<tr><td>>hg38_21_+_8433172_8446622_RNA45SN1_45S_50ntFlanks@4045.4066.22|CCATTCGAACGTCTGCCCTATC</td><td>>hg38_21_+_8433172_8446622_RNA45SN1_45S_50ntFlanks</td><td>4045</td><td>4066</td><td>22</td><td>CCATTCGAACGTCTGCCCTATC</td><td>1023.4</td><td>-6.2</td><td>1.3</td><td>-4.9</td><td>1.2E-06</td><td>2.5E-06</td><td>BT20/MB468</td><td>MITO</td></tr>
</table>

<table>
<tr><td></td><td></td><td></td><td></td><td></td><td></td><td></td><td></td><td></td><td></td><td></td><td></td><td></td><td></td></tr>
<tr><td>>trnaMT_HisGTG_MT_+_12138_12206@26.55.30|AGATTGTGAATCTGACAACAGAGGCTTACG</td><td>>trnaMT_HisGTG_MT_+_12138_12206</td><td>26</td><td>55</td><td>30</td><td>AGATTGTGAATCTGACAACAGAGGCTTACG</td><td>84.8</td><td>10.2</td><td>2.0</td><td>5.1</td><td>4.3E-07</td><td>2.5E-06</td><td>BT20/MB468</td><td>MP</td></tr>
<tr><td>>hg38_21_+_8433172_8446622_RNA45SN1_45S_50ntFlanks@11564.11590.27|CCTAGCAGCCGACTTAGAACTGGTGCG</td><td>>hg38_21_+_8433172_8446622_RNA45SN1_45S_50ntFlanks</td><td>11564</td><td>11590</td><td>27</td><td>CCTAGCAGCCGACTTAGAACTGGTGCG</td><td>214.5</td><td>-4.2</td><td>0.9</td><td>-4.8</td><td>1.5E-06</td><td>2.5E-06</td><td>MB231/MB468</td><td>CYTO</td></tr>
<tr><td>>hg38_21_+_8433172_8446622_RNA45SN1_45S_50ntFlanks@10458.10481.24|ATGGCCTCCGTTGCCCTCGGCCGA</td><td>>hg38_21_+_8433172_8446622_RNA45SN1_45S_50ntFlanks</td><td>10458</td><td>10481</td><td>24</td><td>ATGGCCTCCGTTGCCCTCGGCCGA</td><td>109.4</td><td>-9.9</td><td>1.9</td><td>-5.1</td><td>4.3E-07</td><td>2.5E-06</td><td>BT20/MB468</td><td>MP</td></tr>
<tr><td>>hg38_21_+_8433172_8446622_RNA45SN1_45S_50ntFlanks@12631.12659.29|GGCGAAGCTACCATCTGTGGGATTATGAC</td><td>>hg38_21_+_8433172_8446622_RNA45SN1_45S_50ntFlanks</td><td>12631</td><td>12659</td><td>29</td><td>GGCGAAGCTACCATCTGTGGGATTATGAC</td><td>1158.5</td><td>-6.1</td><td>1.3</td><td>-4.8</td><td>1.6E-06</td><td>2.5E-06</td><td>MB231/MB468</td><td>MITO</td></tr>
<tr><td>>hg38_1_-_228634819_228635039_RNA5S12_5S_50ntFlanks@51.87.37|GTCTACGGCCATACCACCCTGAACGCGCCCGATCTCG</td><td>>hg38_1_-_228634819_228635039_RNA5S12_5S_50ntFlanks</td><td>51</td><td>87</td><td>37</td><td>GTCTACGGCCATACCACCCTGAACGCGCCCGATCTCG</td><td>5237.6</td><td>8.2</td><td>1.7</td><td>4.8</td><td>1.3E-06</td><td>2.5E-06</td><td>BT20/MB468</td><td>CYTO</td></tr>
<tr><td>>hg38_21_+_8433172_8446622_RNA45SN1_45S_50ntFlanks@10664.10710.47|CTTGGAAAGCGTCGCGGTTCCGGCGGCGTCCGGTGAGCTCTCGCTGG</td><td>>hg38_21_+_8433172_8446622_RNA45SN1_45S_50ntFlanks</td><td>10664</td><td>10710</td><td>47</td><td>CTTGGAAAGCGTCGCGGTTCCGGCGGCGTCCGGTGAGCTCTCGCTGG</td><td>358.9</td><td>-7.4</td><td>1.5</td><td>-4.8</td><td>1.3E-06</td><td>2.5E-06</td><td>BT20/MB468</td><td>CYTO</td></tr>
<tr><td>>hg38_21_+_8433172_8446622_RNA45SN1_45S_50ntFlanks@3994.4013.20|TGACTCTAGATAACCTCGGG</td><td>>hg38_21_+_8433172_8446622_RNA45SN1_45S_50ntFlanks</td><td>3994</td><td>4013</td><td>20</td><td>TGACTCTAGATAACCTCGGG</td><td>252.9</td><td>-3.3</td><td>0.7</td><td>-4.8</td><td>1.6E-06</td><td>2.5E-06</td><td>MB231/MB468</td><td>NUC</td></tr>
<tr><td>>hg38_21_+_8433172_8446622_RNA45SN1_45S_50ntFlanks@11316.11340.25|CACCCCACGTCTCGTCGCGCGCGCG</td><td>>hg38_21_+_8433172_8446622_RNA45SN1_45S_50ntFlanks</td><td>11316</td><td>11340</td><td>25</td><td>CACCCCACGTCTCGTCGCGCGCGCG</td><td>414.4</td><td>-4.6</td><td>1.0</td><td>-4.8</td><td>1.3E-06</td><td>2.5E-06</td><td>BT20/MB468</td><td>CYTO</td></tr>
<tr><td>>hg38_21_+_8433172_8446622_RNA45SN1_45S_50ntFlanks@10802.10819.18|TTGGAACAATGTAGGTAA</td><td>>hg38_21_+_8433172_8446622_RNA45SN1_45S_50ntFlanks</td><td>10802</td><td>10819</td><td>18</td><td>TTGGAACAATGTAGGTAA</td><td>53.7</td><td>-8.8</td><td>1.7</td><td>-5.1</td><td>4.3E-07</td><td>2.5E-06</td><td>BT20/MB468</td><td>MP</td></tr>
<tr><td>>MI0000103|hsa-mir-101-1&WithFlank&1|-|65058428|65058514@52.73.22;MI0000739|hsa-mir-101-2&WithFlank&9|+|4850291|4850381@54.75.22&[MIMAT0000099&hsa-miR-101-3p&offsets|-1|0;m-17&1|-|65058443|65058462&offsets|-1|+1];[MIMAT0000099_1&hsa-miR-101-3p&offsets|-1|0;m-16&9|+|4850345|4850364&offsets|-1|+1]|GTACAGTACTGTGATAACTGAA</td><td>>MI0000103|hsa-mir-101-1&WithFlank&1|-|65058428|65058514</td><td>52</td><td>73</td><td>22</td><td>GTACAGTACTGTGATAACTGAA</td><td>1448.2</td><td>5.1</td><td>1.1</td><td>4.8</td><td>1.6E-06</td><td>2.5E-06</td><td>MB231/MB468</td><td>MITO</td></tr>
<tr><td>>hg38_21_+_8433172_8446622_RNA45SN1_45S_50ntFlanks@11576.11602.27|CTTAGAACTGGTGCGGACCAGGGGAAT</td><td>>hg38_21_+_8433172_8446622_RNA45SN1_45S_50ntFlanks</td><td>11576</td><td>11602</td><td>27</td><td>CTTAGAACTGGTGCGGACCAGGGGAAT</td><td>1456.2</td><td>-4.3</td><td>0.9</td><td>-4.8</td><td>1.4E-06</td><td>2.5E-06</td><td>BT20/MB468</td><td>CYTO</td></tr>
<tr><td>>hg38_21_+_8433172_8446622_RNA45SN1_45S_50ntFlanks@12242.12268.27|AGAAGGGCAAAAGCTCGCTTGATCTTG</td><td>>hg38_21_+_8433172_8446622_RNA45SN1_45S_50ntFlanks</td><td>12242</td><td>12268</td><td>27</td><td>AGAAGGGCAAAAGCTCGCTTGATCTTG</td><td>502.8</td><td>-6.9</td><td>1.4</td><td>-4.8</td><td>1.6E-06</td><td>2.5E-06</td><td>MB231/MB468</td><td>MITO</td></tr>
<tr><td>>hg38_21_+_8433172_8446622_RNA45SN1_45S_50ntFlanks@12510.12530.21|TCGTGAGACAGGTTAGTTTTA</td><td>>hg38_21_+_8433172_8446622_RNA45SN1_45S_50ntFlanks</td><td>12510</td><td>12530</td><td>21</td><td>TCGTGAGACAGGTTAGTTTTA</td><td>56.5</td><td>-8.9</td><td>1.8</td><td>-5.1</td><td>4.3E-07</td><td>2.5E-06</td><td>BT20/MB468</td><td>MP</td></tr>
<tr><td>>hg38_21_+_8433172_8446622_RNA45SN1_45S_50ntFlanks@12438.12455.18|ATTGTGAAGCAGAATTCA</td><td>>hg38_21_+_8433172_8446622_RNA45SN1_45S_50ntFlanks</td><td>12438</td><td>12455</td><td>18</td><td>ATTGTGAAGCAGAATTCA</td><td>2794.2</td><td>-8.0</td><td>1.7</td><td>-4.8</td><td>1.4E-06</td><td>2.5E-06</td><td>BT20/MB468</td><td>CYTO</td></tr>
<tr><td>>trnaMT_GluTTC_MT_-_14674_14742@39.72.34,trnalookalike8_GluTTC_5_-_93905172_93905240@39.72.34|ATTGGTCGTGGTTGTAGTCCGTGCGAGAATACCA</td><td>>trnaMT_GluTTC_MT_-_14674_14742</td><td>39</td><td>72</td><td>34</td><td>ATTGGTCGTGGTTGTAGTCCGTGCGAGAATACCA</td><td>11051.2</td><td>3.6</td><td>0.8</td><td>4.8</td><td>1.6E-06</td><td>2.5E-06</td><td>MB231/MB468</td><td>MITO</td></tr>
<tr><td>>hg38_21_+_8433172_8446622_RNA45SN1_45S_50ntFlanks@4160.4177.18|CCACATCCAAGGAAGGCA</td><td>>hg38_21_+_8433172_8446622_RNA45SN1_45S_50ntFlanks</td><td>4160</td><td>4177</td><td>18</td><td>CCACATCCAAGGAAGGCA</td><td>135.6</td><td>-10.9</td><td>2.3</td><td>-4.8</td><td>1.4E-06</td><td>2.5E-06</td><td>MB231/MB468</td><td>MP</td></tr>
<tr><td>>hg38_21_+_8433172_8446622_RNA45SN1_45S_50ntFlanks@11580.11601.22|GAACTGGTGCGGACCAGGGGAA</td><td>>hg38_21_+_8433172_8446622_RNA45SN1_45S_50ntFlanks</td><td>11580</td><td>11601</td><td>22</td><td>GAACTGGTGCGGACCAGGGGAA</td><td>302.2</td><td>-3.1</td><td>0.6</td><td>-4.9</td><td>1.2E-06</td><td>2.5E-06</td><td>BT20/MB468</td><td>MITO</td></tr>
<tr><td>>hg38_21_+_8433172_8446622_RNA45SN1_45S_50ntFlanks@12993.13014.22|CCCTCGCTGCGATCTATTGAAA</td><td>>hg38_21_+_8433172_8446622_RNA45SN1_45S_50ntFlanks</td><td>12993</td><td>13014</td><td>22</td><td>CCCTCGCTGCGATCTATTGAAA</td><td>1121.8</td><td>-8.1</td><td>1.7</td><td>-4.8</td><td>1.7E-06</td><td>2.5E-06</td><td>BT20/MB468</td><td>NUC</td></tr>
<tr><td>>hg38_21_+_8433172_8446622_RNA45SN1_45S_50ntFlanks@6664.6688.25|TGGATCACTCGGCTCGTGCGTCGAT</td><td>>hg38_21_+_8433172_8446622_RNA45SN1_45S_50ntFlanks</td><td>6664</td><td>6688</td><td>25</td><td>TGGATCACTCGGCTCGTGCGTCGAT</td><td>188.1</td><td>10.9</td><td>2.0</td><td>5.3</td><td>1.1E-07</td><td>2.5E-06</td><td>BT20/MB231</td><td>MP</td></tr>
<tr><td>>MI0000270|hsa-mir-181b-1&WithFlank&1|-|198858867|198858988@42.63.22(+1C);MI0000683|hsa-mir-181b-2&WithFlank&9|+|124693704|124693804@22.43.22(+1C)&[MIMAT0000257&hsa-miR-181b-5p&offsets|0|-1(+1C);m-73&1|-|198858925|198858947&offsets|0|-1(+1C)];[MIMAT0000257_1&hsa-miR-181b-5p&offsets|0|-1(+1C);m-72&9|+|124693725|124693747&offsets|0|-1(+1C)]|AACATTCATTGCTGTCGGTGGGC</td><td>>MI0000270|hsa-mir-181b-1&WithFlank&1|-|198858867|198858988</td><td>42</td><td>63</td><td>22(+1C)</td><td>AACATTCATTGCTGTCGGTGGGC</td><td>357.5</td><td>6.2</td><td>1.3</td><td>4.8</td><td>1.7E-06</td><td>2.5E-06</td><td>MB231/MB468</td><td>MITO</td></tr>
<tr><td>>hg38_21_+_8433172_8446622_RNA45SN1_45S_50ntFlanks@11455.11472.18|CTCGGCGCGCGGCGGCGG</td><td>>hg38_21_+_8433172_8446622_RNA45SN1_45S_50ntFlanks</td><td>11455</td><td>11472</td><td>18</td><td>CTCGGCGCGCGGCGGCGG</td><td>63.3</td><td>-9.1</td><td>1.8</td><td>-5.1</td><td>4.4E-07</td><td>2.6E-06</td><td>BT20/MB468</td><td>MP</td></tr>
<tr><td>>hg38_21_+_8433172_8446622_RNA45SN1_45S_50ntFlanks@5487.5516.30|CGCTGAGAAGACGGTCGAACTTGACTATCT</td><td>>hg38_21_+_8433172_8446622_RNA45SN1_45S_50ntFlanks</td><td>5487</td><td>5516</td><td>30</td><td>CGCTGAGAAGACGGTCGAACTTGACTATCT</td><td>1528.0</td><td>3.7</td><td>0.7</td><td>5.3</td><td>1.4E-07</td><td>2.6E-06</td><td>BT20/MB231</td><td>MITO</td></tr>
<tr><td>>hg38_21_+_8433172_8446622_RNA45SN1_45S_50ntFlanks@10458.10483.26|ATGGCCTCCGTTGCCCTCGGCCGATC</td><td>>hg38_21_+_8433172_8446622_RNA45SN1_45S_50ntFlanks</td><td>10458</td><td>10483</td><td>26</td><td>ATGGCCTCCGTTGCCCTCGGCCGATC</td><td>219.5</td><td>-4.7</td><td>1.0</td><td>-4.8</td><td>1.5E-06</td><td>2.6E-06</td><td>MB231/MB468</td><td>CYTO</td></tr>
<tr><td>>hg38_21_+_8433172_8446622_RNA45SN1_45S_50ntFlanks@11571.11600.30|GCCGACTTAGAACTGGTGCGGACCAGGGGA</td><td>>hg38_21_+_8433172_8446622_RNA45SN1_45S_50ntFlanks</td><td>11571</td><td>11600</td><td>30</td><td>GCCGACTTAGAACTGGTGCGGACCAGGGGA</td><td>476.6</td><td>-3.1</td><td>0.7</td><td>-4.8</td><td>1.4E-06</td><td>2.6E-06</td><td>MB231/MB468</td><td>MP</td></tr>
<tr><td>>MI0001445|hsa-mir-423&WithFlank&17|+|30117073|30117178@59.81.23&[MIMAT0001340&hsa-miR-423-3p&offsets|0|0;m-100&17|+|30117131|30117153&offsets|0|0]|AGCTCGGTCTGAGGCCCCTCAGT</td><td>>MI0001445|hsa-mir-423&WithFlank&17|+|30117073|30117178</td><td>59</td><td>81</td><td>23</td><td>AGCTCGGTCTGAGGCCCCTCAGT</td><td>6264.2</td><td>3.8</td><td>0.8</td><td>4.8</td><td>1.4E-06</td><td>2.6E-06</td><td>MB231/MB468</td><td>MP</td></tr>
<tr><td>>hg38_21_+_8433172_8446622_RNA45SN1_45S_50ntFlanks@9790.9830.41|CTGGCGTGGAGCCGGGCGTGGAATGCGAGTGCCTAGTGGGC</td><td>>hg38_21_+_8433172_8446622_RNA45SN1_45S_50ntFlanks</td><td>9790</td><td>9830</td><td>41</td><td>CTGGCGTGGAGCCGGGCGTGGAATGCGAGTGCCTAGTGGGC</td><td>278.2</td><td>11.5</td><td>2.2</td><td>5.3</td><td>1.1E-07</td><td>2.6E-06</td><td>BT20/MB231</td><td>MP</td></tr>
<tr><td>>hg38_MT_+_598_1651_RNR1_12S_50ntFlanks@850.877.28|GTGCTTAGTTGAACAGGGCCCTGAAGCG</td><td>>hg38_MT_+_598_1651_RNR1_12S_50ntFlanks</td><td>850</td><td>877</td><td>28</td><td>GTGCTTAGTTGAACAGGGCCCTGAAGCG</td><td>65.7</td><td>9.8</td><td>1.9</td><td>5.1</td><td>4.4E-07</td><td>2.6E-06</td><td>BT20/MB468</td><td>MP</td></tr>
<tr><td>>hg38_21_+_8433172_8446622_RNA45SN1_45S_50ntFlanks@4047.4066.20|ATTCGAACGTCTGCCCTATC</td><td>>hg38_21_+_8433172_8446622_RNA45SN1_45S_50ntFlanks</td><td>4047</td><td>4066</td><td>20</td><td>ATTCGAACGTCTGCCCTATC</td><td>1711.3</td><td>-2.1</td><td>0.4</td><td>-4.8</td><td>1.4E-06</td><td>2.6E-06</td><td>BT20/MB468</td><td>CYTO</td></tr>
<tr><td>>MI0003137|hsa-mir-193b&WithFlank&16|+|14303961|14304055@19.40.22(+1U)&[MIMAT0004767&hsa-miR-193b-5p&offsets|-1|-1(+1U);m-343&16|+|14303980|14304001&offsets|-1|-1(+1U)]|TCGGGGTTTTGAGGGCGAGATGT</td><td>>MI0003137|hsa-mir-193b&WithFlank&16|+|14303961|14304055</td><td>19</td><td>40</td><td>22(+1U)</td><td>TCGGGGTTTTGAGGGCGAGATGT</td><td>72.6</td><td>9.9</td><td>2.0</td><td>5.1</td><td>4.4E-07</td><td>2.6E-06</td><td>BT20/MB468</td><td>MP</td></tr>
<tr><td>>hg38_21_+_8433172_8446622_RNA45SN1_45S_50ntFlanks@11452.11469.18|CTCCTCGGCGCGCGGCGG</td><td>>hg38_21_+_8433172_8446622_RNA45SN1_45S_50ntFlanks</td><td>11452</td><td>11469</td><td>18</td><td>CTCCTCGGCGCGCGGCGG</td><td>53.3</td><td>-8.8</td><td>1.7</td><td>-5.1</td><td>4.4E-07</td><td>2.6E-06</td><td>BT20/MB468</td><td>MP</td></tr>
<tr><td>>hg38_21_+_8433172_8446622_RNA45SN1_45S_50ntFlanks@10392.10411.20|ACAGCAGTTGAACATGGGTC</td><td>>hg38_21_+_8433172_8446622_RNA45SN1_45S_50ntFlanks</td><td>10392</td><td>10411</td><td>20</td><td>ACAGCAGTTGAACATGGGTC</td><td>322.8</td><td>-2.7</td><td>0.6</td><td>-4.8</td><td>1.6E-06</td><td>2.6E-06</td><td>MB231/MB468</td><td>CYTO</td></tr>
<tr><td>>MI0000750|hsa-mir-26a-2&WithFlank&12|-|57824603|57824698@20.39.20(+1U);MI0000083|hsa-mir-26a-1&WithFlank&3|+|37969398|37969486@16.35.20(+1U)&[MIMAT0000082&hsa-miR-26a-5p&offsets|0|-2(+1U);m-33&12|-|57824658|57824679&offsets|0|-2(+1U)];[MIMAT0000082_1&hsa-miR-26a-5p&offsets|0|-2(+1U);m-34&3|+|37969413|37969434&offsets|0|-2(+1U)]|TTCAAGTAATCCAGGATAGGT</td><td>>MI0000750|hsa-mir-26a-2&WithFlank&12|-|57824603|57824698</td><td>20</td><td>39</td><td>20(+1U)</td><td>TTCAAGTAATCCAGGATAGGT</td><td>433.4</td><td>7.3</td><td>1.5</td><td>4.8</td><td>1.7E-06</td><td>2.6E-06</td><td>MB231/MB468</td><td>MITO</td></tr>
<tr><td>>MI0006444|hsa-mir-1307&WithFlank&10|-|103394247|103394407@87.108.22&[MIMAT0005951&hsa-miR-1307-3p&offsets|+1|+1;m-66&10|-|103394300|103394322&offsets|+1|0]|CTCGGCGTGGCGTCGGTCGTGG</td><td>>MI0006444|hsa-mir-1307&WithFlank&10|-|103394247|103394407</td><td>87</td><td>108</td><td>22</td><td>CTCGGCGTGGCGTCGGTCGTGG</td><td>587.4</td><td>3.8</td><td>0.8</td><td>4.8</td><td>1.7E-06</td><td>2.6E-06</td><td>MB231/MB468</td><td>MITO</td></tr>
<tr><td>>hg38_21_+_8433172_8446622_RNA45SN1_45S_50ntFlanks@5078.5115.38|CGATAACGAACGAGACTCTGGCATGCTAACTAGTTACG</td><td>>hg38_21_+_8433172_8446622_RNA45SN1_45S_50ntFlanks</td><td>5078</td><td>5115</td><td>38</td><td>CGATAACGAACGAGACTCTGGCATGCTAACTAGTTACG</td><td>324.3</td><td>3.7</td><td>0.8</td><td>4.8</td><td>1.4E-06</td><td>2.6E-06</td><td>BT20/MB468</td><td>CYTO</td></tr>
<tr><td>>hg38_21_+_8433172_8446622_RNA45SN1_45S_50ntFlanks@10786.10829.44|AACAGCCTCTGGCATGTTGGAACAATGTAGGTAAGGGAAGTCGG</td><td>>hg38_21_+_8433172_8446622_RNA45SN1_45S_50ntFlanks</td><td>10786</td><td>10829</td><td>44</td><td>AACAGCCTCTGGCATGTTGGAACAATGTAGGTAAGGGAAGTCGG</td><td>141.1</td><td>-4.7</td><td>1.0</td><td>-4.8</td><td>1.6E-06</td><td>2.6E-06</td><td>MB231/MB468</td><td>CYTO</td></tr>
<tr><td>>hg38_21_+_8433172_8446622_RNA45SN1_45S_50ntFlanks@6650.6669.20|ACGACTCTTAGCGGTGGATC</td><td>>hg38_21_+_8433172_8446622_RNA45SN1_45S_50ntFlanks</td><td>6650</td><td>6669</td><td>20</td><td>ACGACTCTTAGCGGTGGATC</td><td>820.5</td><td>1.8</td><td>0.4</td><td>4.8</td><td>1.7E-06</td><td>2.6E-06</td><td>BT20/MB468</td><td>NUC</td></tr>
<tr><td>>hg38_21_+_8433172_8446622_RNA45SN1_45S_50ntFlanks@10787.10818.32|ACAGCCTCTGGCATGTTGGAACAATGTAGGTA</td><td>>hg38_21_+_8433172_8446622_RNA45SN1_45S_50ntFlanks</td><td>10787</td><td>10818</td><td>32</td><td>ACAGCCTCTGGCATGTTGGAACAATGTAGGTA</td><td>742.6</td><td>-7.9</td><td>1.7</td><td>-4.8</td><td>1.7E-06</td><td>2.6E-06</td><td>BT20/MB468</td><td>NUC</td></tr>
<tr><td>>hg38_21_+_8433172_8446622_RNA45SN1_45S_50ntFlanks@6662.6688.27|GGTGGATCACTCGGCTCGTGCGTCGAT</td><td>>hg38_21_+_8433172_8446622_RNA45SN1_45S_50ntFlanks</td><td>6662</td><td>6688</td><td>27</td><td>GGTGGATCACTCGGCTCGTGCGTCGAT</td><td>230.6</td><td>5.0</td><td>0.9</td><td>5.3</td><td>1.4E-07</td><td>2.6E-06</td><td>BT20/MB231</td><td>MITO</td></tr>
<tr><td>>hg38_21_+_8433172_8446622_RNA45SN1_45S_50ntFlanks@10786.10826.41|AACAGCCTCTGGCATGTTGGAACAATGTAGGTAAGGGAAGT</td><td>>hg38_21_+_8433172_8446622_RNA45SN1_45S_50ntFlanks</td><td>10786</td><td>10826</td><td>41</td><td>AACAGCCTCTGGCATGTTGGAACAATGTAGGTAAGGGAAGT</td><td>635.4</td><td>-4.1</td><td>0.9</td><td>-4.8</td><td>1.6E-06</td><td>2.6E-06</td><td>MB231/MB468</td><td>CYTO</td></tr>
<tr><td>>hg38_21_+_8433172_8446622_RNA45SN1_45S_50ntFlanks@12636.12654.19|AGCTACCATCTGTGGGATT</td><td>>hg38_21_+_8433172_8446622_RNA45SN1_45S_50ntFlanks</td><td>12636</td><td>12654</td><td>19</td><td>AGCTACCATCTGTGGGATT</td><td>561.6</td><td>-6.2</td><td>1.3</td><td>-4.8</td><td>1.6E-06</td><td>2.6E-06</td><td>MB231/MB468</td><td>CYTO</td></tr>
<tr><td>>hg38_21_+_8433172_8446622_RNA45SN1_45S_50ntFlanks@13002.13031.30|CGATCTATTGAAAGTCAGCCCTCGACACAA</td><td>>hg38_21_+_8433172_8446622_RNA45SN1_45S_50ntFlanks</td><td>13002</td><td>13031</td><td>30</td><td>CGATCTATTGAAAGTCAGCCCTCGACACAA</td><td>146.2</td><td>-5.0</td><td>1.0</td><td>-4.8</td><td>1.6E-06</td><td>2.6E-06</td><td>MB231/MB468</td><td>CYTO</td></tr>
<tr><td>>hg38_21_+_8433172_8446622_RNA45SN1_45S_50ntFlanks@6729.6770.42|GACACATTGATCATCGACACTTCGAACGCACTTGCGGCCCCG</td><td>>hg38_21_+_8433172_8446622_RNA45SN1_45S_50ntFlanks</td><td>6729</td><td>6770</td><td>42</td><td>GACACATTGATCATCGACACTTCGAACGCACTTGCGGCCCCG</td><td>1039.2</td><td>-5.5</td><td>1.2</td><td>-4.8</td><td>1.7E-06</td><td>2.6E-06</td><td>MB231/MB468</td><td>MITO</td></tr>
<tr><td>>hg38_21_+_8433172_8446622_RNA45SN1_45S_50ntFlanks@10806.10840.35|AACAATGTAGGTAAGGGAAGTCGGCAAGCCGGATC</td><td>>hg38_21_+_8433172_8446622_RNA45SN1_45S_50ntFlanks</td><td>10806</td><td>10840</td><td>35</td><td>AACAATGTAGGTAAGGGAAGTCGGCAAGCCGGATC</td><td>1130.1</td><td>-8.2</td><td>1.7</td><td>-4.8</td><td>1.8E-06</td><td>2.6E-06</td><td>BT20/MB468</td><td>NUC</td></tr>
<tr><td>>hg38_21_+_8433172_8446622_RNA45SN1_45S_50ntFlanks@12084.12108.25|TCGCTTCTGGCGCCAAGCGCCCGGC</td><td>>hg38_21_+_8433172_8446622_RNA45SN1_45S_50ntFlanks</td><td>12084</td><td>12108</td><td>25</td><td>TCGCTTCTGGCGCCAAGCGCCCGGC</td><td>113.7</td><td>-9.9</td><td>2.0</td><td>-5.0</td><td>4.4E-07</td><td>2.6E-06</td><td>BT20/MB468</td><td>MP</td></tr>
<tr><td>>hg38_21_+_8433172_8446622_RNA45SN1_45S_50ntFlanks@6739.6770.32|TCATCGACACTTCGAACGCACTTGCGGCCCCG</td><td>>hg38_21_+_8433172_8446622_RNA45SN1_45S_50ntFlanks</td><td>6739</td><td>6770</td><td>32</td><td>TCATCGACACTTCGAACGCACTTGCGGCCCCG</td><td>338.1</td><td>-6.7</td><td>1.4</td><td>-4.8</td><td>1.6E-06</td><td>2.6E-06</td><td>MB231/MB468</td><td>NUC</td></tr>
<tr><td>>hg38_21_+_8433172_8446622_RNA45SN1_45S_50ntFlanks@10929.10946.18|AGGCGCCGCCGCCCCCCC</td><td>>hg38_21_+_8433172_8446622_RNA45SN1_45S_50ntFlanks</td><td>10929</td><td>10946</td><td>18</td><td>AGGCGCCGCCGCCCCCCC</td><td>137.3</td><td>-10.2</td><td>2.0</td><td>-5.0</td><td>4.4E-07</td><td>2.6E-06</td><td>BT20/MB468</td><td>MP</td></tr>
<tr><td>>hg38_21_+_8433172_8446622_RNA45SN1_45S_50ntFlanks@10391.10408.18|AACAGCAGTTGAACATGG</td><td>>hg38_21_+_8433172_8446622_RNA45SN1_45S_50ntFlanks</td><td>10391</td><td>10408</td><td>18</td><td>AACAGCAGTTGAACATGG</td><td>396.6</td><td>-7.3</td><td>1.4</td><td>-5.0</td><td>4.5E-07</td><td>2.6E-06</td><td>BT20/MB468</td><td>MP</td></tr>
<tr><td>>hg38_21_+_8433172_8446622_RNA45SN1_45S_50ntFlanks@3993.4024.32|GTGACTCTAGATAACCTCGGGCCGATCGCACG</td><td>>hg38_21_+_8433172_8446622_RNA45SN1_45S_50ntFlanks</td><td>3993</td><td>4024</td><td>32</td><td>GTGACTCTAGATAACCTCGGGCCGATCGCACG</td><td>832.3</td><td>-5.7</td><td>1.2</td><td>-4.8</td><td>1.4E-06</td><td>2.6E-06</td><td>MB231/MB468</td><td>MP</td></tr>
</table>

| | | | | | | | | | | | | | |
|---|---|---|---|---|---|---|---|---|---|---|---|---|---|
| >trna16_LeuAAG_16_+_22308461_22308542@14.32.19,trna16_LeuAAG_5_-_180601044_180601125@14.32.19,trna19_LeuAAG_5_-_180524474_180524555@14.32.19,trna1_LeuAAG_14_+_21078291_21078372@14.32.19,trna2_LeuTAG_14_+_21093529_21093610@14.32.19,trna3_LeuAAG_5_+_180528840_180528921@14.32.19,trna42_LeuTAG_17_-_8023632_8023713@14.32.19,trna78_LeuAAG_6_+_28956779_28956860@14.32.19,trna7_LeuAAG_5_+_180614701_180614782@14.32.19,trna98_LeuAAG_6_-_28911399_28911480@14.32.19\|AGCGGTCTAAGGCGCTGGA | >trna16_LeuAAG_16_+_22308461_22308542 | 14 | 32 | 19 | AGCGGTCTAAGGCGCTGGA | 65.4 | -9.1 | 1.8 | -5.0 | 4.5E-07 | 2.6E-06 | BT20/MB468 | MP |
| >hg38_21_+_8433172_8446622_RNA45SN1_45S_50ntFlanks@12941.12965.25\|ATTCGTAGACGACCTGCTTCTGGGT | >hg38_21_+_8433172_8446622_RNA45SN1_45S_50ntFlanks | 12941 | 12965 | 25 | ATTCGTAGACGACCTGCTTCTGGGT | 402.9 | -7.5 | 1.6 | -4.8 | 1.8E-06 | 2.6E-06 | BT20/MB468 | NUC |
| >hg38_1_-_228634819_228635039_RNA5S12_5S_50ntFlanks@150.170.21\|AATACCGGGTGCTGTAGGCTT | >hg38_1_-_228634819_228635039_RNA5S12_5S_50ntFlanks | 150 | 170 | 21 | AATACCGGGTGCTGTAGGCTT | 4095.5 | -3.6 | 0.8 | -4.8 | 1.7E-06 | 2.6E-06 | MB231/MB468 | MITO |
| >hg38_21_+_8433172_8446622_RNA45SN1_45S_50ntFlanks@10396.10426.31\|CAGTTGAACATGGGTCAGTCGGTCCTGAGAG | >hg38_21_+_8433172_8446622_RNA45SN1_45S_50ntFlanks | 10396 | 10426 | 31 | CAGTTGAACATGGGTCAGTCGGTCCTGAGAG | 166.0 | -6.9 | 1.4 | -4.8 | 1.4E-06 | 2.6E-06 | MB231/MB468 | MP |
| >MI0001733\|hsa-mir-452&WithFlank&X\|-\|151959622\|151959718@23.43.21(+1U)&[MIMAT0001635&hsa-miR-452-5p&offsets\|+3\|+2(+1U);m-188&X\|-\|151959677\|151959699&offsets\|+3\|+1(+1U)]\|TGTTTGCAGAGGAAACTGAGAT | >MI0001733\|hsa-mir-452&WithFlank&X\|-\|151959622\|151959718 | 23 | 43 | 21(+1U) | TGTTTGCAGAGGAAACTGAGAT | 52.4 | 8.9 | 1.9 | 4.8 | 1.4E-06 | 2.6E-06 | MB231/MB468 | MP |
| >hg38_21_+_8433172_8446622_RNA45SN1_45S_50ntFlanks@10693.10711.19\|CCGGTGAGCTCTCGCTGGC | >hg38_21_+_8433172_8446622_RNA45SN1_45S_50ntFlanks | 10693 | 10711 | 19 | CCGGTGAGCTCTCGCTGGC | 255.9 | -7.3 | 1.5 | -4.8 | 1.4E-06 | 2.6E-06 | MB231/MB468 | MP |
| >trna10_AspGTC_12_-_125424193_125424264@32.70.39,trna12_AspGTC_12_-_125411891_125411962@32.70.39,trna144_AspGTC_6_-_27551236_27551307@32.70.39,trna38_AspGTC_17_-_8125556_8125627@32.70.39,trna45_AspGTC_6_+_27447453_27447524@32.70.39,trna48_AspGTC_6_+_27471523_27471594@32.70.39,trna4_AspGTC_12_+_96429799_96429870@32.70.39,trna69_AspGTC_1_-_161440205_161440276@32.70.39,trna72_AspGTC_1_-_161432824_161432895@32.70.39,trna75_AspGTC_1_-_161425414_161425485@32.70.39,trna78_AspGTC_1_-_161418033_161418104@32.70.39,trna81_AspGTC_1_-_161410615_161410686@32.70.39\|CTGTCACGCGGGAGACCGGGGTTCGATTCCCCGACGGGG | >trna10_AspGTC_12_-_125424193_125424264 | 32 | 70 | 39 | CTGTCACGCGGGAGACCGGGGTTCGATTCCCCGACGGGG | 477.7 | -6.3 | 1.3 | -4.8 | 1.4E-06 | 2.6E-06 | BT20/MB468 | CYTO |
| >hg38_21_+_8433172_8446622_RNA45SN1_45S_50ntFlanks@12941.12964.24\|ATTCGTAGACGACCTGCTTCTGGG | >hg38_21_+_8433172_8446622_RNA45SN1_45S_50ntFlanks | 12941 | 12964 | 24 | ATTCGTAGACGACCTGCTTCTGGG | 114.6 | -5.3 | 1.1 | -4.8 | 1.4E-06 | 2.6E-06 | BT20/MB468 | CYTO |
| >hg38_21_+_8433172_8446622_RNA45SN1_45S_50ntFlanks@4555.4574.20\|CGCAGCTAGGAATAATGGAA | >hg38_21_+_8433172_8446622_RNA45SN1_45S_50ntFlanks | 4555 | 4574 | 20 | CGCAGCTAGGAATAATGGAA | 289.7 | -6.9 | 1.4 | -4.8 | 1.4E-06 | 2.6E-06 | MB231/MB468 | MP |
| >hg38_21_+_8433172_8446622_RNA45SN1_45S_50ntFlanks@4759.4793.35\|ATCAGATACCGTCGTAGTTCCGACCATAAACGATG | >hg38_21_+_8433172_8446622_RNA45SN1_45S_50ntFlanks | 4759 | 4793 | 35 | ATCAGATACCGTCGTAGTTCCGACCATAAACGATG | 660.7 | -7.6 | 1.6 | -4.8 | 1.4E-06 | 2.6E-06 | MB231/MB468 | MP |
| >hg38_21_+_8433172_8446622_RNA45SN1_45S_50ntFlanks@11896.11931.36\|TCTAGTCTGGCACGGTGAAGAGACATGAGAGGTGTA | >hg38_21_+_8433172_8446622_RNA45SN1_45S_50ntFlanks | 11896 | 11931 | 36 | TCTAGTCTGGCACGGTGAAGAGACATGAGAGGTGTA | 205.6 | 11.0 | 2.1 | 5.3 | 1.1E-07 | 2.6E-06 | BT20/MB231 | MP |
| >hg38_21_+_8433172_8446622_RNA45SN1_45S_50ntFlanks@9482.9503.22\|ACGTGTTAGGACCCGAAAGATG | >hg38_21_+_8433172_8446622_RNA45SN1_45S_50ntFlanks | 9482 | 9503 | 22 | ACGTGTTAGGACCCGAAAGATG | 221.9 | 11.1 | 2.1 | 5.3 | 1.1E-07 | 2.6E-06 | BT20/MB231 | MP |
| >hg38_21_+_8433172_8446622_RNA45SN1_45S_50ntFlanks@9954.9982.29\|TGGAAGTCGGAATCCGCTAAGGAGTGTGT | >hg38_21_+_8433172_8446622_RNA45SN1_45S_50ntFlanks | 9954 | 9982 | 29 | TGGAAGTCGGAATCCGCTAAGGAGTGTGT | 60.7 | 9.7 | 1.9 | 5.0 | 4.5E-07 | 2.6E-06 | BT20/MB468 | MP |
| >hg38_MT_+_1621_3279_RNR2_16S_50ntFlanks@112.145.34\|CCCAAATAAAGTATAGGCGATAGAAATTGAAACC | >hg38_MT_+_1621_3279_RNR2_16S_50ntFlanks | 112 | 145 | 34 | CCCAAATAAAGTATAGGCGATAGAAATTGAAACC | 89.7 | 10.2 | 2.0 | 5.0 | 4.5E-07 | 2.6E-06 | BT20/MB468 | MP |
| >hg38_21_+_8433172_8446622_RNA45SN1_45S_50ntFlanks@5471.5499.29\|CACGGCCCTGGCGGAGCGCTGAGAAGACG | >hg38_21_+_8433172_8446622_RNA45SN1_45S_50ntFlanks | 5471 | 5499 | 29 | CACGGCCCTGGCGGAGCGCTGAGAAGACG | 2049.4 | -6.2 | 1.3 | -4.8 | 1.7E-06 | 2.6E-06 | MB231/MB468 | MITO |
| >hg38_21_+_8433172_8446622_RNA45SN1_45S_50ntFlanks@6748.6770.23\|CTTCGAACGCACTTGCGGCCCCG | >hg38_21_+_8433172_8446622_RNA45SN1_45S_50ntFlanks | 6748 | 6770 | 23 | CTTCGAACGCACTTGCGGCCCCG | 325.9 | -6.1 | 1.3 | -4.8 | 1.4E-06 | 2.6E-06 | BT20/MB468 | CYTO |
| >hg38_MT_+_598_1651_RNR1_12S_50ntFlanks@844.870.27\|AGTAGAGTGCTTAGTTGAACAGGGCCC | >hg38_MT_+_598_1651_RNR1_12S_50ntFlanks | 844 | 870 | 27 | AGTAGAGTGCTTAGTTGAACAGGGCCC | 92.5 | 10.3 | 2.0 | 5.0 | 4.5E-07 | 2.6E-06 | BT20/MB468 | MP |
| >am_trna111_HisGTG_1_-_147774845_147774916@1.29.29,trna118_HisGTG_1_-_145396881_145396952@1.29.29,trna16_HisGTG_1_+_146544773_146544844@1.29.29,trna1_HisGTG_15_+_45493349_45493420@1.29.29,trna21_HisGTG_1_+_147753471_147753542@1.29.29,trna33_HisGTG_6_+_27125906_27125977@1.29.29,trna7_HisGTG_9_-_14433938_14434009@1.29.29,trna8_HisGTG_15_-_45492611_45492682@1.29.29,trna9_HisGTG_15_-_45490804_45490875@1.29.29\|GCCGTGATCGTATAGTGGTTAGTACTCTG | >am_trna111_HisGTG_1_-_147774845_147774916 | 1 | 29 | 29 | GCCGTGATCGTATAGTGGTTAGTACTCTG | 19176.5 | -9.7 | 2.0 | -4.8 | 1.8E-06 | 2.6E-06 | BT20/MB468 | NUC |
| >hg38_21_+_8433172_8446622_RNA45SN1_45S_50ntFlanks@12999.13028.30\|CTGCGATCTATTGAAAGTCAGCCCTCGACA | >hg38_21_+_8433172_8446622_RNA45SN1_45S_50ntFlanks | 12999 | 13028 | 30 | CTGCGATCTATTGAAAGTCAGCCCTCGACA | 1185.2 | -8.3 | 1.7 | -4.8 | 1.8E-06 | 2.6E-06 | BT20/MB468 | NUC |
| >hg38_21_+_8433172_8446622_RNA45SN1_45S_50ntFlanks@11580.11601.22\|GAACTGGTGCGGACCAGGGGAA | >hg38_21_+_8433172_8446622_RNA45SN1_45S_50ntFlanks | 11580 | 11601 | 22 | GAACTGGTGCGGACCAGGGGAA | 1408.9 | -8.3 | 1.7 | -4.8 | 1.8E-06 | 2.6E-06 | BT20/MB468 | NUC |
| >hg38_21_+_8433172_8446622_RNA45SN1_45S_50ntFlanks@10414.10443.30\|TCGGTCCTGAGAGATGGGCGAGCGCCGTTC | >hg38_21_+_8433172_8446622_RNA45SN1_45S_50ntFlanks | 10414 | 10443 | 30 | TCGGTCCTGAGAGATGGGCGAGCGCCGTTC | 150.2 | -4.2 | 0.9 | -4.8 | 1.4E-06 | 2.6E-06 | BT20/MB468 | CYTO |
| >hg38_21_+_8433172_8446622_RNA45SN1_45S_50ntFlanks@11317.11338.22\|ACCCCACGTCTCGTCGCGCGCG | >hg38_21_+_8433172_8446622_RNA45SN1_45S_50ntFlanks | 11317 | 11338 | 22 | ACCCCACGTCTCGTCGCGCGCG | 56.3 | -8.9 | 1.8 | -5.0 | 4.5E-07 | 2.6E-06 | BT20/MB468 | MP |
| >MI0000085\|hsa-mir-27a&WithFlank&19\|-\|13836434\|13836523@57.76.20&[MIMAT0000084&hsa-miR-27a-3p&offsets\|0\|-1;m-43&19\|-\|13836447\|13836467&offsets\|0\|-1]\|TTCACAGTGGCTAAGTTCCG | >MI0000085\|hsa-mir-27a&WithFlank&19\|-\|13836434\|13836523 | 57 | 76 | 20 | TTCACAGTGGCTAAGTTCCG | 5022.0 | 4.0 | 0.8 | 4.9 | 1.2E-06 | 2.6E-06 | BT20/MB468 | MITO |
| >hg38_21_+_8433172_8446622_RNA45SN1_45S_50ntFlanks@11876.11906.31\|AAGACCCTGTTGAGCTTGACTCTAGTCTGGC | >hg38_21_+_8433172_8446622_RNA45SN1_45S_50ntFlanks | 11876 | 11906 | 31 | AAGACCCTGTTGAGCTTGACTCTAGTCTGGC | 333.6 | -6.5 | 1.4 | -4.8 | 1.4E-06 | 2.6E-06 | BT20/MB468 | CYTO |
| >hg38_21_+_8433172_8446622_RNA45SN1_45S_50ntFlanks@12619.12654.36\|AGGAGCCAATGGGGCGAAGCTACCATCTGTGGGATT | >hg38_21_+_8433172_8446622_RNA45SN1_45S_50ntFlanks | 12619 | 12654 | 36 | AGGAGCCAATGGGGCGAAGCTACCATCTGTGGGATT | 1188.8 | -3.4 | 0.7 | -4.8 | 1.4E-06 | 2.6E-06 | BT20/MB468 | CYTO |
| >hg38_21_+_8433172_8446622_RNA45SN1_45S_50ntFlanks@3771.3791.21\|CACGGCCGGTACAGTGAAACT | >hg38_21_+_8433172_8446622_RNA45SN1_45S_50ntFlanks | 3771 | 3791 | 21 | CACGGCCGGTACAGTGAAACT | 700.1 | -7.8 | 1.6 | -4.8 | 1.8E-06 | 2.6E-06 | BT20/MB468 | NUC |
| >hg38_21_+_8433172_8446622_RNA45SN1_45S_50ntFlanks@6779.6799.21\|CCGGGGCTACGCCTGTCTGAG | >hg38_21_+_8433172_8446622_RNA45SN1_45S_50ntFlanks | 6779 | 6799 | 21 | CCGGGGCTACGCCTGTCTGAG | 78.6 | -9.4 | 1.9 | -5.0 | 4.5E-07 | 2.6E-06 | BT20/MB468 | MP |
| >hg38_21_+_8433172_8446622_RNA45SN1_45S_50ntFlanks@8198.8222.25\|TGGACGGTGTGAGGCCGGTAGCGGC | >hg38_21_+_8433172_8446622_RNA45SN1_45S_50ntFlanks | 8198 | 8222 | 25 | TGGACGGTGTGAGGCCGGTAGCGGC | 444.7 | -7.5 | 1.6 | -4.8 | 1.8E-06 | 2.6E-06 | BT20/MB468 | NUC |
| >hg38_21_+_8433172_8446622_RNA45SN1_45S_50ntFlanks@4102.4127.26\|ACCACGGGTGACGGGGAATCAGGGTT | >hg38_21_+_8433172_8446622_RNA45SN1_45S_50ntFlanks | 4102 | 4127 | 26 | ACCACGGGTGACGGGGAATCAGGGTT | 268.5 | -2.3 | 0.5 | -4.8 | 1.6E-06 | 2.6E-06 | MB231/MB468 | CYTO |
| >hg38_21_+_8433172_8446622_RNA45SN1_45S_50ntFlanks@7990.8014.25\|GACGTGGCGACCCGCTGAATTTAAG | >hg38_21_+_8433172_8446622_RNA45SN1_45S_50ntFlanks | 7990 | 8014 | 25 | GACGTGGCGACCCGCTGAATTTAAG | 255.1 | -5.5 | 1.1 | -4.8 | 1.4E-06 | 2.6E-06 | MB231/MB468 | MP |
| >hg38_21_+_8433172_8446622_RNA45SN1_45S_50ntFlanks@8198.8224.27\|TGGACGGTGTGAGGCCGGTAGCGGCCC | >hg38_21_+_8433172_8446622_RNA45SN1_45S_50ntFlanks | 8198 | 8224 | 27 | TGGACGGTGTGAGGCCGGTAGCGGCCC | 457.1 | -4.6 | 1.0 | -4.8 | 1.4E-06 | 2.6E-06 | BT20/MB468 | CYTO |
| >hg38_21_+_8433172_8446622_RNA45SN1_45S_50ntFlanks@5216.5233.18\|CCGGGGCTGCACGCGCGC | >hg38_21_+_8433172_8446622_RNA45SN1_45S_50ntFlanks | 5216 | 5233 | 18 | CCGGGGCTGCACGCGCGC | 512.6 | 7.4 | 1.5 | 4.8 | 1.7E-06 | 2.6E-06 | MB231/MB468 | NUC |
| >hg38_21_+_8433172_8446622_RNA45SN1_45S_50ntFlanks@8363.8391.29\|ACTTTGAAGAGAGAGTTCAAGAGGGCGTG | >hg38_21_+_8433172_8446622_RNA45SN1_45S_50ntFlanks | 8363 | 8391 | 29 | ACTTTGAAGAGAGAGTTCAAGAGGGCGTG | 78.9 | 10.0 | 2.0 | 5.0 | 4.5E-07 | 2.6E-06 | BT20/MB468 | MP |
| >hg38_21_+_8433172_8446622_RNA45SN1_45S_50ntFlanks@12202.12222.21\|CTAAGGCGAGCTCAGGGAGGA | >hg38_21_+_8433172_8446622_RNA45SN1_45S_50ntFlanks | 12202 | 12222 | 21 | CTAAGGCGAGCTCAGGGAGGA | 282.2 | -6.6 | 1.4 | -4.8 | 1.7E-06 | 2.6E-06 | MB231/MB468 | NUC |
| >hg38_21_+_8433172_8446622_RNA45SN1_45S_50ntFlanks@6645.6665.21\|CTCGTACGACTCTTAGCGGTG | >hg38_21_+_8433172_8446622_RNA45SN1_45S_50ntFlanks | 6645 | 6665 | 21 | CTCGTACGACTCTTAGCGGTG | 4061.1 | 3.4 | 0.7 | 4.8 | 1.6E-06 | 2.6E-06 | MB231/MB468 | CYTO |
| >MI0000487\|hsa-mir-193a&WithFlank&17\|+\|31559990\|31560089@27.48.22&[MIMAT0004614&hsa-miR-193a-5p&offsets\|0\|0;m-123&17\|+\|31560016\|31560037&offsets\|0\|0]\|TGGGTCTTTGCGGGCGAGATGA | >MI0000487\|hsa-mir-193a&WithFlank&17\|+\|31559990\|31560089 | 27 | 48 | 22 | TGGGTCTTTGCGGGCGAGATGA | 1339.9 | 5.8 | 1.1 | 5.0 | 4.6E-07 | 2.6E-06 | BT20/MB468 | MP |
| >hg38_21_+_8433172_8446622_RNA45SN1_45S_50ntFlanks@9930.9956.27\|ATATAGACAGCAGGACGGTGGCCATGG | >hg38_21_+_8433172_8446622_RNA45SN1_45S_50ntFlanks | 9930 | 9956 | 27 | ATATAGACAGCAGGACGGTGGCCATGG | 319.7 | -7.7 | 1.6 | -4.8 | 1.4E-06 | 2.6E-06 | MB231/MB468 | MP |
| >hg38_21_+_8433172_8446622_RNA45SN1_45S_50ntFlanks@6733.6774.42\|CATTGATCATCGACACTTCGAACGCACTTGCGGCCCCGGGTT | >hg38_21_+_8433172_8446622_RNA45SN1_45S_50ntFlanks | 6733 | 6774 | 42 | CATTGATCATCGACACTTCGAACGCACTTGCGGCCCCGGGTT | 57.7 | 9.6 | 1.9 | 5.0 | 4.6E-07 | 2.6E-06 | BT20/MB468 | MP |
| >hg38_21_+_8433172_8446622_RNA45SN1_45S_50ntFlanks@12934.12954.21\|CTAAACCATTCGTAGACGACC | >hg38_21_+_8433172_8446622_RNA45SN1_45S_50ntFlanks | 12934 | 12954 | 21 | CTAAACCATTCGTAGACGACC | 124.6 | -6.7 | 1.4 | -4.8 | 1.4E-06 | 2.6E-06 | MB231/MB468 | MP |
| >hg38_21_+_8433172_8446622_RNA45SN1_45S_50ntFlanks@12633.12661.29\|CGAAGCTACCATCTGTGGGATTATGACTG | >hg38_21_+_8433172_8446622_RNA45SN1_45S_50ntFlanks | 12633 | 12661 | 29 | CGAAGCTACCATCTGTGGGATTATGACTG | 586.6 | -3.1 | 0.6 | -4.8 | 1.6E-06 | 2.6E-06 | MB231/MB468 | CYTO |
| >hg38_21_+_8433172_8446622_RNA45SN1_45S_50ntFlanks@6740.6771.32\|CATCGACACTTCGAACGCACTTGCGGCCCCGG | >hg38_21_+_8433172_8446622_RNA45SN1_45S_50ntFlanks | 6740 | 6771 | 32 | CATCGACACTTCGAACGCACTTGCGGCCCCGG | 485.4 | -4.5 | 0.9 | -4.8 | 1.4E-06 | 2.6E-06 | BT20/MB468 | CYTO |
| >hg38_21_+_8433172_8446622_RNA45SN1_45S_50ntFlanks@5019.5047.29\|TCCGTGGGTGGTGGTGCATGGCCGTTCTT | >hg38_21_+_8433172_8446622_RNA45SN1_45S_50ntFlanks | 5019 | 5047 | 29 | TCCGTGGGTGGTGGTGCATGGCCGTTCTT | 77.1 | 10.0 | 2.0 | 5.0 | 4.6E-07 | 2.6E-06 | BT20/MB468 | MP |
| >hg38_MT_+_1621_3279_RNR2_16S_50ntFlanks@729.753.25\|GATTAAAACACTGAACTGACAATTA | >hg38_MT_+_1621_3279_RNR2_16S_50ntFlanks | 729 | 753 | 25 | GATTAAAACACTGAACTGACAATTA | 61.8 | 9.2 | 1.9 | 4.8 | 1.4E-06 | 2.6E-06 | MB231/MB468 | MP |
| >hg38_MT_+_598_1651_RNR1_12S_50ntFlanks@843.881.39\|GAGTAGAGTGCTTAGTTGAACAGGGCCCTGAAGCGCGTA | >hg38_MT_+_598_1651_RNR1_12S_50ntFlanks | 843 | 881 | 39 | GAGTAGAGTGCTTAGTTGAACAGGGCCCTGAAGCGCGTA | 92.7 | 10.3 | 2.0 | 5.0 | 4.6E-07 | 2.7E-06 | BT20/MB468 | MP |

<table>
<tr><td></td><td></td><td></td><td></td><td></td><td></td><td></td><td></td><td></td><td></td><td></td><td></td><td></td><td></td></tr>
<tr><td>>hg38_21_+_8433172_8446622_RNA45SN1_45S_50ntFlanks@11755.11780.26|AATGCCTCGTCATCTAATTAGTGACG</td><td>>hg38_21_+_8433172_8446622_RNA45SN1_45S_50ntFlanks</td><td>11755</td><td>11780</td><td>26</td><td>AATGCCTCGTCATCTAATTAGTGACG</td><td>58.5</td><td>-9.0</td><td>1.8</td><td>-5.0</td><td>4.6E-07</td><td>2.7E-06</td><td>BT20/MB468</td><td>MP</td></tr>
<tr><td>>hg38_21_+_8433172_8446622_RNA45SN1_45S_50ntFlanks@8159.8181.23|CGTGGGGGGCCCAAGTCCTTCTG</td><td>>hg38_21_+_8433172_8446622_RNA45SN1_45S_50ntFlanks</td><td>8159</td><td>8181</td><td>23</td><td>CGTGGGGGGCCCAAGTCCTTCTG</td><td>373.4</td><td>-7.8</td><td>1.6</td><td>-4.8</td><td>1.7E-06</td><td>2.7E-06</td><td>MB231/MB468</td><td>MITO</td></tr>
<tr><td>>hg38_21_+_8433172_8446622_RNA45SN1_45S_50ntFlanks@11209.11230.22|CGGCGACTCTGGACGCGAGCCG</td><td>>hg38_21_+_8433172_8446622_RNA45SN1_45S_50ntFlanks</td><td>11209</td><td>11230</td><td>22</td><td>CGGCGACTCTGGACGCGAGCCG</td><td>1177.7</td><td>-3.8</td><td>0.8</td><td>-4.8</td><td>1.4E-06</td><td>2.7E-06</td><td>BT20/MB468</td><td>CYTO</td></tr>
<tr><td>>hg38_21_+_8433172_8446622_RNA45SN1_45S_50ntFlanks@5486.5515.30|GCGCTGAGAAGACGGTCGAACTTGACTATC</td><td>>hg38_21_+_8433172_8446622_RNA45SN1_45S_50ntFlanks</td><td>5486</td><td>5515</td><td>30</td><td>GCGCTGAGAAGACGGTCGAACTTGACTATC</td><td>426.5</td><td>-7.0</td><td>1.5</td><td>-4.8</td><td>1.4E-06</td><td>2.7E-06</td><td>MB231/MB468</td><td>MP</td></tr>
<tr><td>>hg38_21_+_8433172_8446622_RNA45SN1_45S_50ntFlanks@12997.13017.21|CGCTGCGATCTATTGAAAGTC</td><td>>hg38_21_+_8433172_8446622_RNA45SN1_45S_50ntFlanks</td><td>12997</td><td>13017</td><td>21</td><td>CGCTGCGATCTATTGAAAGTC</td><td>286.3</td><td>3.5</td><td>0.7</td><td>4.8</td><td>1.7E-06</td><td>2.7E-06</td><td>MB231/MB468</td><td>MITO</td></tr>
<tr><td>>hg38_21_+_8433172_8446622_RNA45SN1_45S_50ntFlanks@10869.10892.24|AGGGCTGGGTCGGTCGGGCTGGGG</td><td>>hg38_21_+_8433172_8446622_RNA45SN1_45S_50ntFlanks</td><td>10869</td><td>10892</td><td>24</td><td>AGGGCTGGGTCGGTCGGGCTGGGG</td><td>1177.1</td><td>-8.3</td><td>1.7</td><td>-4.8</td><td>1.8E-06</td><td>2.7E-06</td><td>BT20/MB468</td><td>NUC</td></tr>
<tr><td>>MI0000088|hsa-mir-30a&WithFlank&6|-|71403545|71403627@54.75.22(+1U)&[MIMAT0000088&hsa-miR-30a-3p&offsets|+1|+1(+1U);m-38&6|-|71403554|71403575&offsets|+1|+1(+1U)]|TTTCAGTCGGATGTTTGCAGCTT</td><td>>MI0000088|hsa-mir-30a&WithFlank&6|-|71403545|71403627</td><td>54</td><td>75</td><td>22(+1U)</td><td>TTTCAGTCGGATGTTTGCAGCTT</td><td>45.9</td><td>8.7</td><td>1.8</td><td>4.8</td><td>1.4E-06</td><td>2.7E-06</td><td>MB231/MB468</td><td>MP</td></tr>
<tr><td>>hg38_1_-_228634819_228635039_RNA5S12_5S_50ntFlanks@150.168.19|AATACCGGGTGCTGTAGGC</td><td>>hg38_1_-_228634819_228635039_RNA5S12_5S_50ntFlanks</td><td>150</td><td>168</td><td>19</td><td>AATACCGGGTGCTGTAGGC</td><td>161.6</td><td>-6.6</td><td>1.4</td><td>-4.8</td><td>1.4E-06</td><td>2.7E-06</td><td>MB231/MB468</td><td>MP</td></tr>
<tr><td>>am_trnaMT_PheGAA_MT_+_577_647@40.63.24|TGTTTAGACGGGCTCACATCACCC</td><td>>am_trnaMT_PheGAA_MT_+_577_647</td><td>40</td><td>63</td><td>24</td><td>TGTTTAGACGGGCTCACATCACCC</td><td>111.3</td><td>10.5</td><td>2.1</td><td>5.0</td><td>4.6E-07</td><td>2.7E-06</td><td>BT20/MB468</td><td>MP</td></tr>
<tr><td>>hg38_21_+_8433172_8446622_RNA45SN1_45S_50ntFlanks@4764.4783.20|ATACCGTCGTAGTTCCGACC</td><td>>hg38_21_+_8433172_8446622_RNA45SN1_45S_50ntFlanks</td><td>4764</td><td>4783</td><td>20</td><td>ATACCGTCGTAGTTCCGACC</td><td>56.0</td><td>-8.9</td><td>1.8</td><td>-5.0</td><td>4.6E-07</td><td>2.7E-06</td><td>BT20/MB468</td><td>MP</td></tr>
<tr><td>>hg38_21_+_8433172_8446622_RNA45SN1_45S_50ntFlanks@7980.8009.30|CCTCAGATCAGACGTGGCGACCCGCTGAAT</td><td>>hg38_21_+_8433172_8446622_RNA45SN1_45S_50ntFlanks</td><td>7980</td><td>8009</td><td>30</td><td>CCTCAGATCAGACGTGGCGACCCGCTGAAT</td><td>231.4</td><td>-2.6</td><td>0.5</td><td>-4.8</td><td>1.7E-06</td><td>2.7E-06</td><td>MB231/MB468</td><td>NUC</td></tr>
<tr><td>>hg38_21_+_8433172_8446622_RNA45SN1_45S_50ntFlanks@10348.10376.29|AAACGAGAACTTTGAAGGCCGAAGTGGAG</td><td>>hg38_21_+_8433172_8446622_RNA45SN1_45S_50ntFlanks</td><td>10348</td><td>10376</td><td>29</td><td>AAACGAGAACTTTGAAGGCCGAAGTGGAG</td><td>310.4</td><td>-2.6</td><td>0.5</td><td>-4.8</td><td>1.7E-06</td><td>2.7E-06</td><td>MB231/MB468</td><td>NUC</td></tr>
<tr><td>>hg38_21_+_8433172_8446622_RNA45SN1_45S_50ntFlanks@12988.13013.26|CAGCTCCCTCGCTGCGATCTATTGAA</td><td>>hg38_21_+_8433172_8446622_RNA45SN1_45S_50ntFlanks</td><td>12988</td><td>13013</td><td>26</td><td>CAGCTCCCTCGCTGCGATCTATTGAA</td><td>881.9</td><td>-6.1</td><td>1.3</td><td>-4.8</td><td>1.8E-06</td><td>2.7E-06</td><td>BT20/MB468</td><td>NUC</td></tr>
<tr><td>>hg38_21_+_8433172_8446622_RNA45SN1_45S_50ntFlanks@3790.3814.25|CTGCGAATGGCTCATTAAATCAGTT</td><td>>hg38_21_+_8433172_8446622_RNA45SN1_45S_50ntFlanks</td><td>3790</td><td>3814</td><td>25</td><td>CTGCGAATGGCTCATTAAATCAGTT</td><td>65.4</td><td>-9.1</td><td>1.8</td><td>-5.0</td><td>4.6E-07</td><td>2.7E-06</td><td>BT20/MB468</td><td>MP</td></tr>
<tr><td>>hg38_21_+_8433172_8446622_RNA45SN1_45S_50ntFlanks@11873.11893.21|AAGAAGACCCTGTTGAGCTTG</td><td>>hg38_21_+_8433172_8446622_RNA45SN1_45S_50ntFlanks</td><td>11873</td><td>11893</td><td>21</td><td>AAGAAGACCCTGTTGAGCTTG</td><td>247.6</td><td>11.3</td><td>2.1</td><td>5.3</td><td>1.2E-07</td><td>2.7E-06</td><td>BT20/MB231</td><td>MP</td></tr>
<tr><td>>am_trna134_GluTTC_1_-_16861774_16861845@1.30.30,trna5_GluTTC_1_+_17199078_17199149@1.30.30,trna84_GluTTC_1_-_161391883_161391954@1.30.30,trna94_GluTTC_1_-_149664355_149664427@1.30.30|TCCCTGGTGGTCTAGTGGCTAGGATTCGGC</td><td>>am_trna134_GluTTC_1_-_16861774_16861845</td><td>1</td><td>30</td><td>30</td><td>TCCCTGGTGGTCTAGTGGCTAGGATTCGGC</td><td>1464.1</td><td>-7.7</td><td>1.6</td><td>-4.8</td><td>1.7E-06</td><td>2.7E-06</td><td>MB231/MB468</td><td>NUC</td></tr>
<tr><td>>hg38_21_+_8433172_8446622_RNA45SN1_45S_50ntFlanks@11920.11940.21|ATGAGAGGTGTAGAATAAGTG</td><td>>hg38_21_+_8433172_8446622_RNA45SN1_45S_50ntFlanks</td><td>11920</td><td>11940</td><td>21</td><td>ATGAGAGGTGTAGAATAAGTG</td><td>222.1</td><td>11.1</td><td>2.1</td><td>5.3</td><td>1.2E-07</td><td>2.7E-06</td><td>BT20/MB231</td><td>MP</td></tr>
<tr><td>>hg38_21_+_8433172_8446622_RNA45SN1_45S_50ntFlanks@11117.11134.18|GCAGGGGCCGGCGGCGGC</td><td>>hg38_21_+_8433172_8446622_RNA45SN1_45S_50ntFlanks</td><td>11117</td><td>11134</td><td>18</td><td>GCAGGGGCCGGCGGCGGC</td><td>67.9</td><td>-9.2</td><td>1.8</td><td>-5.0</td><td>4.6E-07</td><td>2.7E-06</td><td>BT20/MB468</td><td>MP</td></tr>
<tr><td>>hg38_21_+_8433172_8446622_RNA45SN1_45S_50ntFlanks@8196.8223.28|CGTGGACGGTGTGAGGCCGGTAGCGGCC</td><td>>hg38_21_+_8433172_8446622_RNA45SN1_45S_50ntFlanks</td><td>8196</td><td>8223</td><td>28</td><td>CGTGGACGGTGTGAGGCCGGTAGCGGCC</td><td>1267.1</td><td>-8.3</td><td>1.7</td><td>-4.8</td><td>1.8E-06</td><td>2.7E-06</td><td>BT20/MB468</td><td>NUC</td></tr>
<tr><td>>hg38_21_+_8433172_8446622_RNA45SN1_45S_50ntFlanks@12202.12221.20|CTAAGGCGAGCTCAGGGAGG</td><td>>hg38_21_+_8433172_8446622_RNA45SN1_45S_50ntFlanks</td><td>12202</td><td>12221</td><td>20</td><td>CTAAGGCGAGCTCAGGGAGG</td><td>489.4</td><td>-4.5</td><td>0.9</td><td>-4.8</td><td>1.2E-06</td><td>2.7E-06</td><td>BT20/MB468</td><td>MITO</td></tr>
<tr><td>>hg38_21_+_8433172_8446622_RNA45SN1_45S_50ntFlanks@9951.9982.32|CCATGGAAGTCGGAATCCGCTAAGGAGTGTGT</td><td>>hg38_21_+_8433172_8446622_RNA45SN1_45S_50ntFlanks</td><td>9951</td><td>9982</td><td>32</td><td>CCATGGAAGTCGGAATCCGCTAAGGAGTGTGT</td><td>55.0</td><td>9.5</td><td>1.9</td><td>5.0</td><td>4.6E-07</td><td>2.7E-06</td><td>BT20/MB468</td><td>MP</td></tr>
<tr><td>>hg38_21_+_8433172_8446622_RNA45SN1_45S_50ntFlanks@8176.8201.26|CTTCTGATCGAGGCCCAGCCCGTGGA</td><td>>hg38_21_+_8433172_8446622_RNA45SN1_45S_50ntFlanks</td><td>8176</td><td>8201</td><td>26</td><td>CTTCTGATCGAGGCCCAGCCCGTGGA</td><td>860.0</td><td>-4.5</td><td>0.9</td><td>-5.0</td><td>4.6E-07</td><td>2.7E-06</td><td>BT20/MB468</td><td>MP</td></tr>
<tr><td>>hg38_21_+_8433172_8446622_RNA45SN1_45S_50ntFlanks@6647.6667.21|CGTACGACTCTTAGCGGTGGA</td><td>>hg38_21_+_8433172_8446622_RNA45SN1_45S_50ntFlanks</td><td>6647</td><td>6667</td><td>21</td><td>CGTACGACTCTTAGCGGTGGA</td><td>390.8</td><td>3.2</td><td>0.7</td><td>4.8</td><td>1.6E-06</td><td>2.7E-06</td><td>MB231/MB468</td><td>CYTO</td></tr>
<tr><td>>hg38_21_+_8433172_8446622_RNA45SN1_45S_50ntFlanks@5084.5110.27|CGAACGAGACTCTGGCATGCTAACTAG</td><td>>hg38_21_+_8433172_8446622_RNA45SN1_45S_50ntFlanks</td><td>5084</td><td>5110</td><td>27</td><td>CGAACGAGACTCTGGCATGCTAACTAG</td><td>2384.7</td><td>-5.5</td><td>1.1</td><td>-4.8</td><td>1.2E-06</td><td>2.7E-06</td><td>BT20/MB468</td><td>MITO</td></tr>
<tr><td>>hg38_21_+_8433172_8446622_RNA45SN1_45S_50ntFlanks@5472.5509.38|ACGGCCCTGGCGGAGCGCTGAGAAGACGGTCGAACTTG</td><td>>hg38_21_+_8433172_8446622_RNA45SN1_45S_50ntFlanks</td><td>5472</td><td>5509</td><td>38</td><td>ACGGCCCTGGCGGAGCGCTGAGAAGACGGTCGAACTTG</td><td>872.6</td><td>4.3</td><td>0.9</td><td>4.8</td><td>1.7E-06</td><td>2.7E-06</td><td>MB231/MB468</td><td>NUC</td></tr>
<tr><td>>am_MI0000786|hsa-mir-378a&WithFlank&5|+|149732819|149732896@49.70.22(+1U)&[MIMAT0000732&hsa-miR-378a-3p&offsets|0|0(+1U);m-44&5|+|149732867|149732888&offsets|0|0(+1U)]|ACTGGACTTGGAGTCAGAAGGCT</td><td>>am_MI0000786|hsa-mir-378a&WithFlank&5|+|149732819|149732896</td><td>49</td><td>70</td><td>22(+1U)</td><td>ACTGGACTTGGAGTCAGAAGGCT</td><td>1143.2</td><td>3.5</td><td>0.7</td><td>4.8</td><td>1.6E-06</td><td>2.7E-06</td><td>MB231/MB468</td><td>CYTO</td></tr>
<tr><td>>hg38_21_+_8433172_8446622_RNA45SN1_45S_50ntFlanks@11771.11790.20|ATTAGTGACGCGCATGAATG</td><td>>hg38_21_+_8433172_8446622_RNA45SN1_45S_50ntFlanks</td><td>11771</td><td>11790</td><td>20</td><td>ATTAGTGACGCGCATGAATG</td><td>399.8</td><td>-4.1</td><td>0.9</td><td>-4.8</td><td>1.8E-06</td><td>2.7E-06</td><td>BT20/MB468</td><td>NUC</td></tr>
<tr><td>>hg38_21_+_8433172_8446622_RNA45SN1_45S_50ntFlanks@10690.10712.23|CGTCCGGTGAGCTCTCGCTGGCC</td><td>>hg38_21_+_8433172_8446622_RNA45SN1_45S_50ntFlanks</td><td>10690</td><td>10712</td><td>23</td><td>CGTCCGGTGAGCTCTCGCTGGCC</td><td>91.5</td><td>-9.6</td><td>1.9</td><td>-5.0</td><td>4.6E-07</td><td>2.7E-06</td><td>BT20/MB468</td><td>MP</td></tr>
<tr><td>>MI0000078|hsa-mir-22&WithFlank&17|-|1713897|1713993@59.79.21(+1A)&[MIMAT0000077&hsa-miR-22-3p&offsets|0|-1(+1A);m-2&17|-|1713914|1713935&offsets|0|-1(+1A)]|AAGCTGCCAGTTGAAGAACTGA</td><td>>MI0000078|hsa-mir-22&WithFlank&17|-|1713897|1713993</td><td>59</td><td>79</td><td>21(+1A)</td><td>AAGCTGCCAGTTGAAGAACTGA</td><td>759.5</td><td>5.5</td><td>1.1</td><td>5.0</td><td>4.7E-07</td><td>2.7E-06</td><td>BT20/MB468</td><td>MP</td></tr>
<tr><td>>hg38_21_+_8433172_8446622_RNA45SN1_45S_50ntFlanks@10445.10463.19|GAAGGGACGGGCGATGGCC</td><td>>hg38_21_+_8433172_8446622_RNA45SN1_45S_50ntFlanks</td><td>10445</td><td>10463</td><td>19</td><td>GAAGGGACGGGCGATGGCC</td><td>52.3</td><td>9.5</td><td>1.9</td><td>5.0</td><td>4.7E-07</td><td>2.7E-06</td><td>BT20/MB468</td><td>MP</td></tr>
<tr><td>>hg38_21_+_8433172_8446622_RNA45SN1_45S_50ntFlanks@10348.10383.36|AAACGAGAACTTTGAAGGCCGAAGTGGAGAAGGGTT</td><td>>hg38_21_+_8433172_8446622_RNA45SN1_45S_50ntFlanks</td><td>10348</td><td>10383</td><td>36</td><td>AAACGAGAACTTTGAAGGCCGAAGTGGAGAAGGGTT</td><td>1274.0</td><td>-4.0</td><td>0.8</td><td>-4.8</td><td>1.4E-06</td><td>2.7E-06</td><td>MB231/MB468</td><td>MP</td></tr>
<tr><td>>hg38_21_+_8433172_8446622_RNA45SN1_45S_50ntFlanks@9534.9574.41|AGGAAACTCTGGTGGAGGTCCGTAGCGGTCCTGACGTGCAA</td><td>>hg38_21_+_8433172_8446622_RNA45SN1_45S_50ntFlanks</td><td>9534</td><td>9574</td><td>41</td><td>AGGAAACTCTGGTGGAGGTCCGTAGCGGTCCTGACGTGCAA</td><td>263.0</td><td>-7.3</td><td>1.5</td><td>-4.8</td><td>1.4E-06</td><td>2.7E-06</td><td>MB231/MB468</td><td>MP</td></tr>
<tr><td>>hg38_21_+_8433172_8446622_RNA45SN1_45S_50ntFlanks@4732.4752.21|AGAACGAAAGTCGGAGGTTCG</td><td>>hg38_21_+_8433172_8446622_RNA45SN1_45S_50ntFlanks</td><td>4732</td><td>4752</td><td>21</td><td>AGAACGAAAGTCGGAGGTTCG</td><td>512.9</td><td>-5.4</td><td>1.1</td><td>-4.8</td><td>1.4E-06</td><td>2.7E-06</td><td>MB231/MB468</td><td>MP</td></tr>
<tr><td>>MI0001445|hsa-mir-423&WithFlank&17|+|30117073|30117178@59.79.21&[MIMAT0001340&hsa-miR-423-3p&offsets|0|-2;m-100&17|+|30117131|30117153&offsets|0|-2]|AGCTCGGTCTGAGGCCCCTCA</td><td>>MI0001445|hsa-mir-423&WithFlank&17|+|30117073|30117178</td><td>59</td><td>79</td><td>21</td><td>AGCTCGGTCTGAGGCCCCTCA</td><td>607.6</td><td>4.5</td><td>0.9</td><td>4.8</td><td>1.7E-06</td><td>2.7E-06</td><td>MB231/MB468</td><td>NUC</td></tr>
<tr><td>>hg38_21_+_8433172_8446622_RNA45SN1_45S_50ntFlanks@10808.10844.37|CAATGTAGGTAAGGGAAGTCGGCAAGCCGGATCCGTA</td><td>>hg38_21_+_8433172_8446622_RNA45SN1_45S_50ntFlanks</td><td>10808</td><td>10844</td><td>37</td><td>CAATGTAGGTAAGGGAAGTCGGCAAGCCGGATCCGTA</td><td>143.5</td><td>-5.5</td><td>1.1</td><td>-4.8</td><td>1.4E-06</td><td>2.7E-06</td><td>BT20/MB468</td><td>CYTO</td></tr>
<tr><td>>hg38_21_+_8433172_8446622_RNA45SN1_45S_50ntFlanks@11707.11728.22|AAGCGCGGGTAAACGGCGGGAG</td><td>>hg38_21_+_8433172_8446622_RNA45SN1_45S_50ntFlanks</td><td>11707</td><td>11728</td><td>22</td><td>AAGCGCGGGTAAACGGCGGGAG</td><td>586.3</td><td>-6.8</td><td>1.4</td><td>-4.8</td><td>1.7E-06</td><td>2.7E-06</td><td>MB231/MB468</td><td>NUC</td></tr>
<tr><td>>hg38_21_+_8433172_8446622_RNA45SN1_45S_50ntFlanks@8182.8199.18|ATCGAGGCCCAGCCCGTG</td><td>>hg38_21_+_8433172_8446622_RNA45SN1_45S_50ntFlanks</td><td>8182</td><td>8199</td><td>18</td><td>ATCGAGGCCCAGCCCGTG</td><td>167.4</td><td>-11.3</td><td>2.3</td><td>-4.8</td><td>1.5E-06</td><td>2.7E-06</td><td>MB231/MB468</td><td>MP</td></tr>
<tr><td>>hg38_21_+_8433172_8446622_RNA45SN1_45S_50ntFlanks@10684.10708.25|CGGCGGCGTCCGGTGAGCTCTCGCT</td><td>>hg38_21_+_8433172_8446622_RNA45SN1_45S_50ntFlanks</td><td>10684</td><td>10708</td><td>25</td><td>CGGCGGCGTCCGGTGAGCTCTCGCT</td><td>1016.4</td><td>-3.2</td><td>0.7</td><td>-4.8</td><td>1.7E-06</td><td>2.7E-06</td><td>MB231/MB468</td><td>NUC</td></tr>
<tr><td>>hg38_21_+_8433172_8446622_RNA45SN1_45S_50ntFlanks@12415.12442.28|CTTCGATGTCGGCTCTTCCTATCATTGT</td><td>>hg38_21_+_8433172_8446622_RNA45SN1_45S_50ntFlanks</td><td>12415</td><td>12442</td><td>28</td><td>CTTCGATGTCGGCTCTTCCTATCATTGT</td><td>1279.1</td><td>3.3</td><td>0.7</td><td>4.8</td><td>1.6E-06</td><td>2.7E-06</td><td>MB231/MB468</td><td>CYTO</td></tr>
<tr><td>>hg38_21_+_8433172_8446622_RNA45SN1_45S_50ntFlanks@12624.12659.36|CCAATGGGGCGAAGCTACCATCTGTGGGATTATGAC</td><td>>hg38_21_+_8433172_8446622_RNA45SN1_45S_50ntFlanks</td><td>12624</td><td>12659</td><td>36</td><td>CCAATGGGGCGAAGCTACCATCTGTGGGATTATGAC</td><td>316.6</td><td>-5.2</td><td>1.1</td><td>-4.8</td><td>1.7E-06</td><td>2.7E-06</td><td>MB231/MB468</td><td>NUC</td></tr>
<tr><td>>hg38_21_+_8433172_8446622_RNA45SN1_45S_50ntFlanks@11783.11806.24|CATGAATGGATGAACGAGATTCCC</td><td>>hg38_21_+_8433172_8446622_RNA45SN1_45S_50ntFlanks</td><td>11783</td><td>11806</td><td>24</td><td>CATGAATGGATGAACGAGATTCCC</td><td>1800.7</td><td>-7.7</td><td>1.6</td><td>-4.8</td><td>1.5E-06</td><td>2.7E-06</td><td>BT20/MB468</td><td>CYTO</td></tr>
<tr><td>>hg38_21_+_8433172_8446622_RNA45SN1_45S_50ntFlanks@8342.8364.23|CGTAAGGGAAAGTTGAAAAGAAC</td><td>>hg38_21_+_8433172_8446622_RNA45SN1_45S_50ntFlanks</td><td>8342</td><td>8364</td><td>23</td><td>CGTAAGGGAAAGTTGAAAAGAAC</td><td>804.6</td><td>6.8</td><td>1.4</td><td>4.8</td><td>1.6E-06</td><td>2.7E-06</td><td>MB231/MB468</td><td>CYTO</td></tr>
<tr><td>>hg38_21_+_8433172_8446622_RNA45SN1_45S_50ntFlanks@10407.10426.20|GGGTCAGTCGGTCCTGAGAG</td><td>>hg38_21_+_8433172_8446622_RNA45SN1_45S_50ntFlanks</td><td>10407</td><td>10426</td><td>20</td><td>GGGTCAGTCGGTCCTGAGAG</td><td>57.9</td><td>-8.9</td><td>1.8</td><td>-5.0</td><td>4.7E-07</td><td>2.7E-06</td><td>BT20/MB468</td><td>MP</td></tr>
<tr><td>>hg38_21_+_8433172_8446622_RNA45SN1_45S_50ntFlanks@10391.10411.21|AACAGCAGTTGAACATGGGTC</td><td>>hg38_21_+_8433172_8446622_RNA45SN1_45S_50ntFlanks</td><td>10391</td><td>10411</td><td>21</td><td>AACAGCAGTTGAACATGGGTC</td><td>264.0</td><td>-2.3</td><td>0.5</td><td>-4.8</td><td>1.7E-06</td><td>2.7E-06</td><td>MB231/MB468</td><td>CYTO</td></tr>
<tr><td>>hg38_21_+_8433172_8446622_RNA45SN1_45S_50ntFlanks@6658.6689.32|TAGCGGTGGATCACTCGGCTCGTGCGTCGATG</td><td>>hg38_21_+_8433172_8446622_RNA45SN1_45S_50ntFlanks</td><td>6658</td><td>6689</td><td>32</td><td>TAGCGGTGGATCACTCGGCTCGTGCGTCGATG</td><td>520.0</td><td>-5.2</td><td>1.1</td><td>-4.8</td><td>1.8E-06</td><td>2.7E-06</td><td>MB231/MB468</td><td>MITO</td></tr>
<tr><td>>hg38_21_+_8433172_8446622_RNA45SN1_45S_50ntFlanks@4760.4779.20|TCAGATACCGTCGTAGTTCC</td><td>>hg38_21_+_8433172_8446622_RNA45SN1_45S_50ntFlanks</td><td>4760</td><td>4779</td><td>20</td><td>TCAGATACCGTCGTAGTTCC</td><td>156.9</td><td>-6.9</td><td>1.4</td><td>-4.8</td><td>1.5E-06</td><td>2.7E-06</td><td>MB231/MB468</td><td>MP</td></tr>
<tr><td>>hg38_21_+_8433172_8446622_RNA45SN1_45S_50ntFlanks@10725.10752.28|GGGGAGAGGGTGTAAATCTCGCGCCGGG</td><td>>hg38_21_+_8433172_8446622_RNA45SN1_45S_50ntFlanks</td><td>10725</td><td>10752</td><td>28</td><td>GGGGAGAGGGTGTAAATCTCGCGCCGGG</td><td>82.6</td><td>-9.4</td><td>1.9</td><td>-5.0</td><td>4.7E-07</td><td>2.7E-06</td><td>BT20/MB468</td><td>MP</td></tr>
<tr><td>>hg38_21_+_8433172_8446622_RNA45SN1_45S_50ntFlanks@8212.8235.24|CCGGTAGCGGCCCCCGGCGCGCCG</td><td>>hg38_21_+_8433172_8446622_RNA45SN1_45S_50ntFlanks</td><td>8212</td><td>8235</td><td>24</td><td>CCGGTAGCGGCCCCCGGCGCGCCG</td><td>57.1</td><td>-8.9</td><td>1.8</td><td>-5.0</td><td>4.7E-07</td><td>2.7E-06</td><td>BT20/MB468</td><td>MP</td></tr>
<tr><td>>hg38_21_+_8433172_8446622_RNA45SN1_45S_50ntFlanks@11641.11664.24|CGGCGGGTGTTGACGCGATGTGAT</td><td>>hg38_21_+_8433172_8446622_RNA45SN1_45S_50ntFlanks</td><td>11641</td><td>11664</td><td>24</td><td>CGGCGGGTGTTGACGCGATGTGAT</td><td>2739.7</td><td>-6.8</td><td>1.4</td><td>-5.0</td><td>4.7E-07</td><td>2.7E-06</td><td>BT20/MB468</td><td>MP</td></tr>
<tr><td>>hg38_21_+_8433172_8446622_RNA45SN1_45S_50ntFlanks@12393.12414.22|ATAGCGACGTCGCTTTTTGATC</td><td>>hg38_21_+_8433172_8446622_RNA45SN1_45S_50ntFlanks</td><td>12393</td><td>12414</td><td>22</td><td>ATAGCGACGTCGCTTTTTGATC</td><td>475.2</td><td>-3.8</td><td>0.8</td><td>-4.8</td><td>1.9E-06</td><td>2.7E-06</td><td>BT20/MB468</td><td>NUC</td></tr>
</table>

<table>
<tr><td>>am_trnaMT_MetCAT_MT_+_4402_4469@1.47.47|AGTAAGGTCAGCTAAATAAGCTATCGGGCCCATACCCCGAAAATGTT</td><td>>am_trnaMT_MetCAT_MT_+_4402_4469</td><td>1</td><td>47</td><td>47</td><td>AGTAAGGTCAGCTAAATAAGCTATCGGGCCCATACCCCGAAAATGTT</td><td>700.9</td><td>7.5</td><td>1.5</td><td>4.8</td><td>1.3E-06</td><td>2.7E-06</td><td>BT20/MB468</td><td>MITO</td></tr>
<tr><td>>hg38_21_+_8433172_8446622_RNA45SN1_45S_50ntFlanks@10840.10867.28|CCGTAACTTCGGGATAAGGATTGGCTCT</td><td>>hg38_21_+_8433172_8446622_RNA45SN1_45S_50ntFlanks</td><td>10840</td><td>10867</td><td>28</td><td>CCGTAACTTCGGGATAAGGATTGGCTCT</td><td>444.1</td><td>-2.5</td><td>0.5</td><td>-4.8</td><td>1.7E-06</td><td>2.7E-06</td><td>MB231/MB468</td><td>NUC</td></tr>
<tr><td>>hg38_21_+_8433172_8446622_RNA45SN1_45S_50ntFlanks@10922.10946.25|CTGGACGAGGCGCCGCCGCCCCCCC</td><td>>hg38_21_+_8433172_8446622_RNA45SN1_45S_50ntFlanks</td><td>10922</td><td>10946</td><td>25</td><td>CTGGACGAGGCGCCGCCGCCCCCCC</td><td>2071.7</td><td>-8.6</td><td>1.8</td><td>-4.8</td><td>1.9E-06</td><td>2.7E-06</td><td>BT20/MB468</td><td>NUC</td></tr>
<tr><td>>MI0000465|hsa-mir-191&WithFlank&3|-|49020612|49020715@22.43.22(+1A)&[MIMAT0000440&hsa-miR-191-5p&offsets|0|-1(+1A);m-48&3|-|49020672|49020694&offsets|0|-1(+1A)]|CAACGGAATCCCAAAAGCAGCTA</td><td>>MI0000465|hsa-mir-191&WithFlank&3|-|49020612|49020715</td><td>22</td><td>43</td><td>22(+1A)</td><td>CAACGGAATCCCAAAAGCAGCTA</td><td>532.1</td><td>2.4</td><td>0.5</td><td>4.8</td><td>1.9E-06</td><td>2.7E-06</td><td>BT20/MB468</td><td>NUC</td></tr>
<tr><td>>hg38_21_+_8433172_8446622_RNA45SN1_45S_50ntFlanks@11668.11691.24|TGCCCAGTGCTCTGAATGTCAAAG</td><td>>hg38_21_+_8433172_8446622_RNA45SN1_45S_50ntFlanks</td><td>11668</td><td>11691</td><td>24</td><td>TGCCCAGTGCTCTGAATGTCAAAG</td><td>381.9</td><td>3.8</td><td>0.8</td><td>4.8</td><td>1.3E-06</td><td>2.7E-06</td><td>BT20/MB468</td><td>MITO</td></tr>
<tr><td>>MI0000456|hsa-mir-140&WithFlank&16|+|69933075|69933186@69.90.22(+1U)&[MIMAT0004597&hsa-miR-140-3p&offsets|+1|+2(+1U);m-45&16|+|69933143|69933164&offsets|0|0(+1U)]|ACCACAGGGTAGAACCACGGACT</td><td>>MI0000456|hsa-mir-140&WithFlank&16|+|69933075|69933186</td><td>69</td><td>90</td><td>22(+1U)</td><td>ACCACAGGGTAGAACCACGGACT</td><td>111.6</td><td>4.1</td><td>0.9</td><td>4.8</td><td>1.9E-06</td><td>2.7E-06</td><td>BT20/MB468</td><td>NUC</td></tr>
<tr><td>>hg38_21_+_8433172_8446622_RNA45SN1_45S_50ntFlanks@10680.10711.32|GTTCCGGCGGCGTCCGGTGAGCTCTCGCTGGC</td><td>>hg38_21_+_8433172_8446622_RNA45SN1_45S_50ntFlanks</td><td>10680</td><td>10711</td><td>32</td><td>GTTCCGGCGGCGTCCGGTGAGCTCTCGCTGGC</td><td>21360.0</td><td>-10.0</td><td>2.1</td><td>-4.8</td><td>1.9E-06</td><td>2.7E-06</td><td>BT20/MB468</td><td>NUC</td></tr>
<tr><td>>hg38_MT_+_1621_3279_RNR2_16S_50ntFlanks@1348.1384.37|AACAATAGGGTTTACGACCTCGATGTTGGATCAGGAC</td><td>>hg38_MT_+_1621_3279_RNR2_16S_50ntFlanks</td><td>1348</td><td>1384</td><td>37</td><td>AACAATAGGGTTTACGACCTCGATGTTGGATCAGGAC</td><td>42.0</td><td>8.6</td><td>1.8</td><td>4.8</td><td>1.5E-06</td><td>2.7E-06</td><td>MB231/MB468</td><td>MP</td></tr>
<tr><td>>hg38_21_+_8433172_8446622_RNA45SN1_45S_50ntFlanks@4367.4398.32|CAGTTAAAAAGCTCGTAGTTGGATCTTGGGAG</td><td>>hg38_21_+_8433172_8446622_RNA45SN1_45S_50ntFlanks</td><td>4367</td><td>4398</td><td>32</td><td>CAGTTAAAAAGCTCGTAGTTGGATCTTGGGAG</td><td>319.2</td><td>-7.4</td><td>1.5</td><td>-5.0</td><td>4.8E-07</td><td>2.7E-06</td><td>BT20/MB468</td><td>MP</td></tr>
<tr><td>>hg38_MT_+_1621_3279_RNR2_16S_50ntFlanks@488.526.39|GAACAGCTCTTTGGACACTAGGAAAAAACCTTGTAGAGA</td><td>>hg38_MT_+_1621_3279_RNR2_16S_50ntFlanks</td><td>488</td><td>526</td><td>39</td><td>GAACAGCTCTTTGGACACTAGGAAAAAACCTTGTAGAGA</td><td>52.4</td><td>9.5</td><td>1.9</td><td>5.0</td><td>4.8E-07</td><td>2.7E-06</td><td>BT20/MB468</td><td>MP</td></tr>
<tr><td>>hg38_21_+_8433172_8446622_RNA45SN1_45S_50ntFlanks@8348.8375.28|GGAAAGTTGAAAAGAACTTTGAAGAGAG</td><td>>hg38_21_+_8433172_8446622_RNA45SN1_45S_50ntFlanks</td><td>8348</td><td>8375</td><td>28</td><td>GGAAAGTTGAAAAGAACTTTGAAGAGAG</td><td>3127.4</td><td>9.7</td><td>1.9</td><td>5.0</td><td>4.8E-07</td><td>2.7E-06</td><td>BT20/MB468</td><td>MP</td></tr>
<tr><td>>hg38_21_+_8433172_8446622_RNA45SN1_45S_50ntFlanks@11562.11611.50|CGCCTAGCAGCCGACTTAGAACTGGTGCGGACCAGGGGAATCCGACTGTT</td><td>>hg38_21_+_8433172_8446622_RNA45SN1_45S_50ntFlanks</td><td>11562</td><td>11611</td><td>50</td><td>CGCCTAGCAGCCGACTTAGAACTGGTGCGGACCAGGGGAATCCGACTGTT</td><td>267.9</td><td>-7.1</td><td>1.5</td><td>-4.8</td><td>1.8E-06</td><td>2.7E-06</td><td>MB231/MB468</td><td>MITO</td></tr>
<tr><td>>hg38_MT_+_1621_3279_RNR2_16S_50ntFlanks@712.740.29|CGCATAAGCCTGCGTCAGATTAAAACACT</td><td>>hg38_MT_+_1621_3279_RNR2_16S_50ntFlanks</td><td>712</td><td>740</td><td>29</td><td>CGCATAAGCCTGCGTCAGATTAAAACACT</td><td>38.3</td><td>8.5</td><td>1.8</td><td>4.8</td><td>1.5E-06</td><td>2.7E-06</td><td>MB231/MB468</td><td>MP</td></tr>
<tr><td>>hg38_21_+_8433172_8446622_RNA45SN1_45S_50ntFlanks@4727.4751.25|AATCAAGAACGAAAGTCGGAGGTTC</td><td>>hg38_21_+_8433172_8446622_RNA45SN1_45S_50ntFlanks</td><td>4727</td><td>4751</td><td>25</td><td>AATCAAGAACGAAAGTCGGAGGTTC</td><td>640.0</td><td>-4.1</td><td>0.8</td><td>-4.8</td><td>1.5E-06</td><td>2.7E-06</td><td>MB231/MB468</td><td>MP</td></tr>
<tr><td>>hg38_21_+_8433172_8446622_RNA45SN1_45S_50ntFlanks@12991.13017.27|CTCCCTCGCTGCGATCTATTGAAAGTC</td><td>>hg38_21_+_8433172_8446622_RNA45SN1_45S_50ntFlanks</td><td>12991</td><td>13017</td><td>27</td><td>CTCCCTCGCTGCGATCTATTGAAAGTC</td><td>459.8</td><td>-5.2</td><td>1.1</td><td>-4.8</td><td>1.8E-06</td><td>2.7E-06</td><td>MB231/MB468</td><td>MITO</td></tr>
<tr><td>>hg38_21_+_8433172_8446622_RNA45SN1_45S_50ntFlanks@12456.12499.44|CCAAGCGTTGGATTGTTCACCCACTAATAGGGAACGTGAGCTGG</td><td>>hg38_21_+_8433172_8446622_RNA45SN1_45S_50ntFlanks</td><td>12456</td><td>12499</td><td>44</td><td>CCAAGCGTTGGATTGTTCACCCACTAATAGGGAACGTGAGCTGG</td><td>414.4</td><td>-7.3</td><td>1.5</td><td>-4.8</td><td>1.7E-06</td><td>2.8E-06</td><td>MB231/MB468</td><td>NUC</td></tr>
<tr><td>>hg38_21_+_8433172_8446622_RNA45SN1_45S_50ntFlanks@6721.6741.21|AATTGCAGGACACATTGATCA</td><td>>hg38_21_+_8433172_8446622_RNA45SN1_45S_50ntFlanks</td><td>6721</td><td>6741</td><td>21</td><td>AATTGCAGGACACATTGATCA</td><td>457.6</td><td>-6.7</td><td>1.4</td><td>-4.8</td><td>1.5E-06</td><td>2.8E-06</td><td>BT20/MB468</td><td>CYTO</td></tr>
<tr><td>>hg38_21_+_8433172_8446622_RNA45SN1_45S_50ntFlanks@12416.12446.31|TTCGATGTCGGCTCTTCCTATCATTGTGAAG</td><td>>hg38_21_+_8433172_8446622_RNA45SN1_45S_50ntFlanks</td><td>12416</td><td>12446</td><td>31</td><td>TTCGATGTCGGCTCTTCCTATCATTGTGAAG</td><td>3133.7</td><td>3.3</td><td>0.7</td><td>4.8</td><td>1.7E-06</td><td>2.8E-06</td><td>MB231/MB468</td><td>CYTO</td></tr>
<tr><td>>hg38_21_+_8433172_8446622_RNA45SN1_45S_50ntFlanks@10677.10711.35|GCGGTTCCGGCGGCGTCCGGTGAGCTCTCGCTGGC</td><td>>hg38_21_+_8433172_8446622_RNA45SN1_45S_50ntFlanks</td><td>10677</td><td>10711</td><td>35</td><td>GCGGTTCCGGCGGCGTCCGGTGAGCTCTCGCTGGC</td><td>1462.9</td><td>-3.8</td><td>0.8</td><td>-4.8</td><td>1.7E-06</td><td>2.8E-06</td><td>MB231/MB468</td><td>NUC</td></tr>
<tr><td>>hg38_21_+_8433172_8446622_RNA45SN1_45S_50ntFlanks@12420.12448.29|ATGTCGGCTCTTCCTATCATTGTGAAGCA</td><td>>hg38_21_+_8433172_8446622_RNA45SN1_45S_50ntFlanks</td><td>12420</td><td>12448</td><td>29</td><td>ATGTCGGCTCTTCCTATCATTGTGAAGCA</td><td>2673.5</td><td>3.3</td><td>0.7</td><td>4.8</td><td>1.7E-06</td><td>2.8E-06</td><td>MB231/MB468</td><td>CYTO</td></tr>
<tr><td>>hg38_21_+_8433172_8446622_RNA45SN1_45S_50ntFlanks@11824.11844.21|TCCAGCGAAACCACAGCCAAG</td><td>>hg38_21_+_8433172_8446622_RNA45SN1_45S_50ntFlanks</td><td>11824</td><td>11844</td><td>21</td><td>TCCAGCGAAACCACAGCCAAG</td><td>62.8</td><td>-9.1</td><td>1.8</td><td>-5.0</td><td>4.8E-07</td><td>2.8E-06</td><td>BT20/MB468</td><td>MP</td></tr>
<tr><td>>hg38_21_+_8433172_8446622_RNA45SN1_45S_50ntFlanks@475.508.34|CTTCGTGATCGATGTGGTGACGTCGTGCTCTCCC</td><td>>hg38_21_+_8433172_8446622_RNA45SN1_45S_50ntFlanks</td><td>475</td><td>508</td><td>34</td><td>CTTCGTGATCGATGTGGTGACGTCGTGCTCTCCC</td><td>412.2</td><td>4.5</td><td>0.9</td><td>4.8</td><td>1.8E-06</td><td>2.8E-06</td><td>MB231/MB468</td><td>MITO</td></tr>
<tr><td>>trna152_ValCAC_6_-_27248049_27248121@1.24.24|GCTTCTGTAGTGTAGTGGTTATCA</td><td>>trna152_ValCAC_6_-_27248049_27248121</td><td>1</td><td>24</td><td>24</td><td>GCTTCTGTAGTGTAGTGGTTATCA</td><td>221.0</td><td>-11.7</td><td>2.4</td><td>-4.8</td><td>1.5E-06</td><td>2.8E-06</td><td>MB231/MB468</td><td>MP</td></tr>
<tr><td>>hg38_21_+_8433172_8446622_RNA45SN1_45S_50ntFlanks@7974.7992.19|ACGCGACCTCAGATCAGAC</td><td>>hg38_21_+_8433172_8446622_RNA45SN1_45S_50ntFlanks</td><td>7974</td><td>7992</td><td>19</td><td>ACGCGACCTCAGATCAGAC</td><td>936.4</td><td>3.3</td><td>0.7</td><td>4.8</td><td>1.5E-06</td><td>2.8E-06</td><td>BT20/MB468</td><td>CYTO</td></tr>
<tr><td>>MI0000483|hsa-mir-186&WithFlank&1|-|71067625|71067722@21.42.22&[MIMAT0000456&hsa-miR-186-5p&offsets|0|0;m-84&1|-|71067681|71067702&offsets|0|0]|CAAAGAATTCTCCTTTTGGGCT</td><td>>MI0000483|hsa-mir-186&WithFlank&1|-|71067625|71067722</td><td>21</td><td>42</td><td>22</td><td>CAAAGAATTCTCCTTTTGGGCT</td><td>1389.8</td><td>4.5</td><td>0.9</td><td>4.8</td><td>1.8E-06</td><td>2.8E-06</td><td>MB231/MB468</td><td>MITO</td></tr>
<tr><td>>am_trna111_HisGTG_1_-_147774845_147774916@-1G.29.30,trna118_HisGTG_1_-_145396881_145396952@-1G.29.30,trna16_HisGTG_1_+_146544773_146544844@-1G.29.30,trna1_HisGTG_15_+_45493349_45493420@-1G.29.30,trna21_HisGTG_1_+_147753471_147753542@-1G.29.30,trna33_HisGTG_6_+_27125906_27125977@-1G.29.30,trna7_HisGTG_9_-_14433938_14434009@-1G.29.30,trna8_HisGTG_15_-_45492611_45492682@-1G.29.30,trna9_HisGTG_15_-_45490804_45490875@-1G.29.30|GGCCGTGATCGTATAGTGGTTAGTACTCTG</td><td>>am_trna111_HisGTG_1_-_147774845_147774916</td><td>-1G</td><td>29</td><td>30</td><td>GGCCGTGATCGTATAGTGGTTAGTACTCTG</td><td>12901.9</td><td>-3.5</td><td>0.7</td><td>-4.8</td><td>1.7E-06</td><td>2.8E-06</td><td>MB231/MB468</td><td>CYTO</td></tr>
<tr><td>>hg38_21_+_8433172_8446622_RNA45SN1_45S_50ntFlanks@11565.11600.36|CTAGCAGCCGACTTAGAACTGGTGCGGACCAGGGGA</td><td>>hg38_21_+_8433172_8446622_RNA45SN1_45S_50ntFlanks</td><td>11565</td><td>11600</td><td>36</td><td>CTAGCAGCCGACTTAGAACTGGTGCGGACCAGGGGA</td><td>267.7</td><td>-3.2</td><td>0.7</td><td>-4.8</td><td>1.7E-06</td><td>2.8E-06</td><td>MB231/MB468</td><td>NUC</td></tr>
<tr><td>>trna10_LysCTT_16_+_3241501_3241573@4.33.30,trna13_LysCTT_14_-_58706613_58706685@4.33.30,trna2_LysCTT_15_+_79152904_79152976@4.33.30|CGGCTAGCTCAGTCGGTAGAGCATGGGACT</td><td>>trna10_LysCTT_16_+_3241501_3241573</td><td>4</td><td>33</td><td>30</td><td>CGGCTAGCTCAGTCGGTAGAGCATGGGACT</td><td>510.8</td><td>4.1</td><td>0.9</td><td>4.8</td><td>1.5E-06</td><td>2.8E-06</td><td>MB231/MB468</td><td>MP</td></tr>
<tr><td>>hg38_21_+_8433172_8446622_RNA45SN1_45S_50ntFlanks@6662.6689.28|GGTGGATCACTCGGCTCGTGCGTCGATG</td><td>>hg38_21_+_8433172_8446622_RNA45SN1_45S_50ntFlanks</td><td>6662</td><td>6689</td><td>28</td><td>GGTGGATCACTCGGCTCGTGCGTCGATG</td><td>232.1</td><td>-6.8</td><td>1.4</td><td>-4.8</td><td>1.5E-06</td><td>2.8E-06</td><td>MB231/MB468</td><td>MP</td></tr>
<tr><td>>hg38_MT_+_598_1651_RNR1_12S_50ntFlanks@570.612.43|AAGGACCTGGCGGTGCTTCATATCCCTCTAGAGGAGCCTGTTC</td><td>>hg38_MT_+_598_1651_RNR1_12S_50ntFlanks</td><td>570</td><td>612</td><td>43</td><td>AAGGACCTGGCGGTGCTTCATATCCCTCTAGAGGAGCCTGTTC</td><td>105.4</td><td>-10.1</td><td>1.9</td><td>-5.3</td><td>1.2E-07</td><td>2.8E-06</td><td>BT20/MB231</td><td>MP</td></tr>
<tr><td>>hg38_21_+_8433172_8446622_RNA45SN1_45S_50ntFlanks@10680.10705.26|GTTCCGGCGGCGTCCGGTGAGCTCTC</td><td>>hg38_21_+_8433172_8446622_RNA45SN1_45S_50ntFlanks</td><td>10680</td><td>10705</td><td>26</td><td>GTTCCGGCGGCGTCCGGTGAGCTCTC</td><td>711.3</td><td>-5.3</td><td>1.1</td><td>-4.8</td><td>1.7E-06</td><td>2.8E-06</td><td>MB231/MB468</td><td>NUC</td></tr>
<tr><td>>hg38_21_+_8433172_8446622_RNA45SN1_45S_50ntFlanks@3854.3898.45|ACTGTGGTAATTCTAGAGCTAATACATGCCGACGGGCGCTGACCC</td><td>>hg38_21_+_8433172_8446622_RNA45SN1_45S_50ntFlanks</td><td>3854</td><td>3898</td><td>45</td><td>ACTGTGGTAATTCTAGAGCTAATACATGCCGACGGGCGCTGACCC</td><td>1633.0</td><td>-7.9</td><td>1.7</td><td>-4.8</td><td>1.8E-06</td><td>2.8E-06</td><td>MB231/MB468</td><td>NUC</td></tr>
<tr><td>>hg38_MT_+_1621_3279_RNR2_16S_50ntFlanks@484.518.35|AGAGGAACAGCTCTTTGGACACTAGGAAAAAACCT</td><td>>hg38_MT_+_1621_3279_RNR2_16S_50ntFlanks</td><td>484</td><td>518</td><td>35</td><td>AGAGGAACAGCTCTTTGGACACTAGGAAAAAACCT</td><td>54.4</td><td>9.5</td><td>1.9</td><td>5.0</td><td>4.8E-07</td><td>2.8E-06</td><td>BT20/MB468</td><td>MP</td></tr>
<tr><td>>hg38_21_+_8433172_8446622_RNA45SN1_45S_50ntFlanks@11571.11605.35|GCCGACTTAGAACTGGTGCGGACCAGGGGAATCCG</td><td>>hg38_21_+_8433172_8446622_RNA45SN1_45S_50ntFlanks</td><td>11571</td><td>11605</td><td>35</td><td>GCCGACTTAGAACTGGTGCGGACCAGGGGAATCCG</td><td>270.4</td><td>-3.0</td><td>0.6</td><td>-4.8</td><td>1.8E-06</td><td>2.8E-06</td><td>MB231/MB468</td><td>NUC</td></tr>
<tr><td>>hg38_21_+_8433172_8446622_RNA45SN1_45S_50ntFlanks@8476.8497.22|CCGGCGGATCTTTCCCGCCCCC</td><td>>hg38_21_+_8433172_8446622_RNA45SN1_45S_50ntFlanks</td><td>8476</td><td>8497</td><td>22</td><td>CCGGCGGATCTTTCCCGCCCCC</td><td>90.4</td><td>-9.6</td><td>1.9</td><td>-5.0</td><td>4.9E-07</td><td>2.8E-06</td><td>BT20/MB468</td><td>MP</td></tr>
<tr><td>>hg38_21_+_8433172_8446622_RNA45SN1_45S_50ntFlanks@11671.11698.28|CCAGTGCTCTGAATGTCAAAGTGAAGAA</td><td>>hg38_21_+_8433172_8446622_RNA45SN1_45S_50ntFlanks</td><td>11671</td><td>11698</td><td>28</td><td>CCAGTGCTCTGAATGTCAAAGTGAAGAA</td><td>114.2</td><td>-6.5</td><td>1.4</td><td>-4.8</td><td>1.5E-06</td><td>2.8E-06</td><td>MB231/MB468</td><td>MP</td></tr>
<tr><td>>hg38_21_+_8433172_8446622_RNA45SN1_45S_50ntFlanks@12616.12661.46|CTGAGGAGCCAATGGGGCGAAGCTACCATCTGTGGGATTATGACTG</td><td>>hg38_21_+_8433172_8446622_RNA45SN1_45S_50ntFlanks</td><td>12616</td><td>12661</td><td>46</td><td>CTGAGGAGCCAATGGGGCGAAGCTACCATCTGTGGGATTATGACTG</td><td>155.1</td><td>-11.1</td><td>2.3</td><td>-4.8</td><td>1.5E-06</td><td>2.8E-06</td><td>MB231/MB468</td><td>MP</td></tr>
<tr><td>>hg38_21_+_8433172_8446622_RNA45SN1_45S_50ntFlanks@8382.8402.21|AGAGGGCGTGAAACCGTTAAG</td><td>>hg38_21_+_8433172_8446622_RNA45SN1_45S_50ntFlanks</td><td>8382</td><td>8402</td><td>21</td><td>AGAGGGCGTGAAACCGTTAAG</td><td>719.2</td><td>-5.7</td><td>1.2</td><td>-4.8</td><td>1.5E-06</td><td>2.8E-06</td><td>MB231/MB468</td><td>MP</td></tr>
<tr><td>>hg38_21_+_8433172_8446622_RNA45SN1_45S_50ntFlanks@11739.11765.27|TCTCTTAAGGTAGCCAAATGCCTCGTC</td><td>>hg38_21_+_8433172_8446622_RNA45SN1_45S_50ntFlanks</td><td>11739</td><td>11765</td><td>27</td><td>TCTCTTAAGGTAGCCAAATGCCTCGTC</td><td>53.7</td><td>-8.8</td><td>1.8</td><td>-5.0</td><td>4.9E-07</td><td>2.8E-06</td><td>BT20/MB468</td><td>MP</td></tr>
<tr><td>>hg38_21_+_8433172_8446622_RNA45SN1_45S_50ntFlanks@11027.11075.49|CTCTCTCTCTCTCTCCCCCGCTCCCCGTCCTCCCCCCTCCCCGGGGGAG</td><td>>hg38_21_+_8433172_8446622_RNA45SN1_45S_50ntFlanks</td><td>11027</td><td>11075</td><td>49</td><td>CTCTCTCTCTCTCTCCCCCGCTCCCCGTCCTCCCCCCTCCCCGGGGGAG</td><td>258.9</td><td>-6.2</td><td>1.3</td><td>-4.8</td><td>1.5E-06</td><td>2.8E-06</td><td>BT20/MB468</td><td>CYTO</td></tr>
<tr><td>>hg38_21_+_8433172_8446622_RNA45SN1_45S_50ntFlanks@13004.13023.20|ATCTATTGAAAGTCAGCCCT</td><td>>hg38_21_+_8433172_8446622_RNA45SN1_45S_50ntFlanks</td><td>13004</td><td>13023</td><td>20</td><td>ATCTATTGAAAGTCAGCCCT</td><td>59.7</td><td>-9.0</td><td>1.8</td><td>-5.0</td><td>4.9E-07</td><td>2.8E-06</td><td>BT20/MB468</td><td>MP</td></tr>
<tr><td>>MI0000081|hsa-mir-24-2&WithFlank&19|-|13836281|13836365@56.76.21(+1A);MI0000080|hsa-mir-24-1&WithFlank&9|+|95086015|95086094@50.70.21(+1A)&[MIMAT0000080&hsa-miR-24-3p&offsets|0|-1(+1A);m-36&19|-|13836288|13836310&offsets|0|-2(+1A)];[MIMAT0000080_1&hsa-miR-24-3p&offsets|0|-1(+1A);m-35&9|+|95086064|95086086&offsets|0|-2(+1A)]|TGGCTCAGTTCAGCAGGAACAA</td><td>>MI0000081|hsa-mir-24-2&WithFlank&19|-|13836281|13836365</td><td>56</td><td>76</td><td>21(+1A)</td><td>TGGCTCAGTTCAGCAGGAACAA</td><td>443.3</td><td>3.9</td><td>0.8</td><td>4.8</td><td>1.7E-06</td><td>2.8E-06</td><td>MB231/MB468</td><td>CYTO</td></tr>
<tr><td>>hg38_21_+_8433172_8446622_RNA45SN1_45S_50ntFlanks@10357.10377.21|CTTTGAAGGCCGAAGTGGAGA</td><td>>hg38_21_+_8433172_8446622_RNA45SN1_45S_50ntFlanks</td><td>10357</td><td>10377</td><td>21</td><td>CTTTGAAGGCCGAAGTGGAGA</td><td>308.5</td><td>-5.0</td><td>1.0</td><td>-4.8</td><td>1.8E-06</td><td>2.8E-06</td><td>MB231/MB468</td><td>NUC</td></tr>
<tr><td>>MI0000087|hsa-mir-29a&WithFlank&7|-|130876741|130876816@48.66.19(+1C)&[MIMAT0000086&hsa-miR-29a-3p&offsets|0|-3(+1C);m-26&7|-|130876749|130876769&offsets|0|-2(+1C)]|TAGCACCATCTGAAATCGGC</td><td>>MI0000087|hsa-mir-29a&WithFlank&7|-|130876741|130876816</td><td>48</td><td>66</td><td>19(+1C)</td><td>TAGCACCATCTGAAATCGGC</td><td>198.7</td><td>7.8</td><td>1.6</td><td>4.8</td><td>1.5E-06</td><td>2.8E-06</td><td>MB231/MB468</td><td>MP</td></tr>
<tr><td>>hg38_21_+_8433172_8446622_RNA45SN1_45S_50ntFlanks@9554.9582.29|CGTAGCGGTCCTGACGTGCAAATCGGTCG</td><td>>hg38_21_+_8433172_8446622_RNA45SN1_45S_50ntFlanks</td><td>9554</td><td>9582</td><td>29</td><td>CGTAGCGGTCCTGACGTGCAAATCGGTCG</td><td>164.3</td><td>-6.9</td><td>1.4</td><td>-4.8</td><td>1.5E-06</td><td>2.8E-06</td><td>MB231/MB468</td><td>MP</td></tr>
<tr><td>>hg38_21_+_8433172_8446622_RNA45SN1_45S_50ntFlanks@10343.10380.38|TATTCAAACGAGAACTTTGAAGGCCGAAGTGGAGAAGG</td><td>>hg38_21_+_8433172_8446622_RNA45SN1_45S_50ntFlanks</td><td>10343</td><td>10380</td><td>38</td><td>TATTCAAACGAGAACTTTGAAGGCCGAAGTGGAGAAGG</td><td>60.0</td><td>9.7</td><td>1.9</td><td>5.0</td><td>4.9E-07</td><td>2.8E-06</td><td>BT20/MB468</td><td>MP</td></tr>
</table>

| | | | | | | | | | | | | | |
|---|---|---|---|---|---|---|---|---|---|---|---|---|---|
| >hg38_MT_+_1621_3279_RNR2_16S_50ntFlanks@618.656.39\|AACATATAACTGAACTCCTCACACCCAATTGGACCAATC | >hg38_MT_+_1621_3279_RNR2_16S_50ntFlanks | 618 | 656 | 39 | AACATATAACTGAACTCCTCACACCCAATTGGACCAATC | 91.8 | 10.3 | 2.0 | 5.0 | 4.9E-07 | 2.8E-06 | BT20/MB468 | MP |
| >hg38_21_+_8433172_8446622_RNA45SN1_45S_50ntFlanks@10845.10867.23\|ACTTCGGGATAAGGATTGGCTCT | >hg38_21_+_8433172_8446622_RNA45SN1_45S_50ntFlanks | 10845 | 10867 | 23 | ACTTCGGGATAAGGATTGGCTCT | 226.4 | -6.9 | 1.4 | -4.8 | 1.5E-06 | 2.8E-06 | MB231/MB468 | MP |
| >hg38_21_+_8433172_8446622_RNA45SN1_45S_50ntFlanks@12456.12489.34\|CCAAGCGTTGGATTGTTCACCCACTAATAGGGAA | >hg38_21_+_8433172_8446622_RNA45SN1_45S_50ntFlanks | 12456 | 12489 | 34 | CCAAGCGTTGGATTGTTCACCCACTAATAGGGAA | 686.8 | -6.8 | 1.4 | -4.8 | 1.3E-06 | 2.8E-06 | BT20/MB468 | MITO |
| >hg38_21_+_8433172_8446622_RNA45SN1_45S_50ntFlanks@10686.10712.27\|GCGGCGTCCGGTGAGCTCTCGCTGGCC | >hg38_21_+_8433172_8446622_RNA45SN1_45S_50ntFlanks | 10686 | 10712 | 27 | GCGGCGTCCGGTGAGCTCTCGCTGGCC | 435.5 | -2.6 | 0.5 | -4.8 | 1.7E-06 | 2.8E-06 | MB231/MB468 | CYTO |
| >trna116_GluCTC_1_-_145399233_145399304@7.31.25,trna59_GluCTC_1_+_249168447_249168518@7.31.25,trna71_GluCTC_1_-_161439189_161439260@7.31.25,trna74_GluCTC_1_-_161431809_161431880@7.31.25,trna77_GluCTC_1_-_161424398_161424469@7.31.25,trna77_GluCTC_6_+_28949976_28950047@7.31.25,trna80_GluCTC_1_-_161417018_161417089@7.31.25,trna87_GluCTC_6_-_126101393_126101464@7.31.25\|GTGGTCTAGTGGTTAGGATTCGGCG | >trna116_GluCTC_1_-_145399233_145399304 | 7 | 31 | 25 | GTGGTCTAGTGGTTAGGATTCGGCG | 150.4 | -6.1 | 1.3 | -4.8 | 1.8E-06 | 2.8E-06 | MB231/MB468 | NUC |
| >hg38_21_+_8433172_8446622_RNA45SN1_45S_50ntFlanks@7974.8010.37\|ACGCGACCTCAGATCAGACGTGGCGACCCGCTGAATT | >hg38_21_+_8433172_8446622_RNA45SN1_45S_50ntFlanks | 7974 | 8010 | 37 | ACGCGACCTCAGATCAGACGTGGCGACCCGCTGAATT | 17308.5 | 3.1 | 0.7 | 4.8 | 1.8E-06 | 2.8E-06 | MB231/MB468 | MITO |
| >hg38_21_+_8433172_8446622_RNA45SN1_45S_50ntFlanks@10681.10710.30\|TTCCGGCGGCGTCCGGTGAGCTCTCGCTGG | >hg38_21_+_8433172_8446622_RNA45SN1_45S_50ntFlanks | 10681 | 10710 | 30 | TTCCGGCGGCGTCCGGTGAGCTCTCGCTGG | 1349.7 | -5.3 | 1.1 | -4.8 | 1.3E-06 | 2.8E-06 | BT20/MB468 | MITO |
| >MI0000077\|hsa-mir-21&WithFlank&17\|+\|59841260\|59841343@15.37.23&[MIMAT0000076&hsa-miR-21-5p&offsets\|+1\|+2;m-1&17\|+\|59841273\|59841294&offsets\|+1\|+2]\|AGCTTATCAGACTGATGTTGACT | >MI0000077\|hsa-mir-21&WithFlank&17\|+\|59841260\|59841343 | 15 | 37 | 23 | AGCTTATCAGACTGATGTTGACT | 310.9 | 8.0 | 1.7 | 4.8 | 1.8E-06 | 2.8E-06 | MB231/MB468 | MITO |
| >MI0000743\|hsa-mir-34c&WithFlank&11\|+\|111513433\|111513521@19.40.22&[MIMAT0000686&hsa-miR-34c-5p&offsets\|0\|-1;m-336&11\|+\|111513451\|111513473&offsets\|0\|-1]\|AGGCAGTGTAGTTAGCTGATTG | >MI0000743\|hsa-mir-34c&WithFlank&11\|+\|111513433\|111513521 | 19 | 40 | 22 | AGGCAGTGTAGTTAGCTGATTG | 110.3 | -10.2 | 1.9 | -5.3 | 1.2E-07 | 2.8E-06 | BT20/MB231 | MP |
| >hg38_21_+_8433172_8446622_RNA45SN1_45S_50ntFlanks@12404.12432.29\|GCTTTTTGATCCTTCGATGTCGGCTCTTC | >hg38_21_+_8433172_8446622_RNA45SN1_45S_50ntFlanks | 12404 | 12432 | 29 | GCTTTTTGATCCTTCGATGTCGGCTCTTC | 138.4 | -6.6 | 1.4 | -4.8 | 1.5E-06 | 2.8E-06 | MB231/MB468 | MP |
| >am_trna10_ValCAC_5_-_180649395_180649467@3.23.21,trna12_ValAAC_5_-_180645270_180645342@3.23.21,trna132_ValAAC_6_-_27721179_27721251@3.23.21,trna136_ValAAC_6_-_27648885_27648957@3.23.21,trna139_ValAAC_6_-_27618707_27618779@3.23.21,trna18_ValCAC_5_-_180529253_180529325@3.23.21,trna2_ValAAC_3_+_169490018_169490090@3.23.21,trna2_ValCAC_5_+_180524070_180524142@3.23.21,trna37_ValAAC_6_+_27203288_27203360@3.23.21,trna4_ValAAC_5_+_180591154_180591226@3.23.21,trna5_ValAAC_5_+_180596610_180596682@3.23.21,trna6_ValCAC_5_+_180600650_180600722@3.23.21,trna85_ValCAC_1_-_161369490_161369562@3.23.21,trna90_ValCAC_1_-_149684088_149684161@3.23.21,trna98_ValCAC_1_-_149298555_149298627@3.23.21,trna9_ValCAC_6_+_26538282_26538354@3.23.21\|TTCCGTAGTGTAGTGGTTATC | >am_trna10_ValCAC_5_-_180649395_180649467 | 3 | 23 | 21 | TTCCGTAGTGTAGTGGTTATC | 255.9 | 7.4 | 1.5 | 4.8 | 1.8E-06 | 2.8E-06 | MB231/MB468 | MITO |
| >hg38_21_+_8433172_8446622_RNA45SN1_45S_50ntFlanks@5470.5514.45\|CCACGGCCCTGGCGGAGCGCTGAGAAGACGGTCGAACTTGACTAT | >hg38_21_+_8433172_8446622_RNA45SN1_45S_50ntFlanks | 5470 | 5514 | 45 | CCACGGCCCTGGCGGAGCGCTGAGAAGACGGTCGAACTTGACTAT | 576.1 | -6.2 | 1.3 | -4.8 | 1.8E-06 | 2.8E-06 | MB231/MB468 | NUC |
| >hg38_21_+_8433172_8446622_RNA45SN1_45S_50ntFlanks@11573.11601.29\|CGACTTAGAACTGGTGCGGACCAGGGGAA | >hg38_21_+_8433172_8446622_RNA45SN1_45S_50ntFlanks | 11573 | 11601 | 29 | CGACTTAGAACTGGTGCGGACCAGGGGAA | 1066.6 | -3.7 | 0.8 | -4.8 | 1.7E-06 | 2.8E-06 | MB231/MB468 | CYTO |
| >hg38_21_+_8433172_8446622_RNA45SN1_45S_50ntFlanks@6721.6751.31\|AATTGCAGGACACATTGATCATCGACACTTC | >hg38_21_+_8433172_8446622_RNA45SN1_45S_50ntFlanks | 6721 | 6751 | 31 | AATTGCAGGACACATTGATCATCGACACTTC | 173.5 | -3.3 | 0.7 | -4.8 | 1.8E-06 | 2.8E-06 | MB231/MB468 | NUC |
| >hg38_21_+_8433172_8446622_RNA45SN1_45S_50ntFlanks@11315.11332.18\|CCACCCCACGTCTCGTCG | >hg38_21_+_8433172_8446622_RNA45SN1_45S_50ntFlanks | 11315 | 11332 | 18 | CCACCCCACGTCTCGTCG | 641.8 | -6.7 | 1.4 | -4.8 | 1.7E-06 | 2.8E-06 | MB231/MB468 | CYTO |
| >hg38_21_+_8433172_8446622_RNA45SN1_45S_50ntFlanks@8182.8211.30\|ATCGAGGCCCAGCCCGTGGACGGTGTGAGG | >hg38_21_+_8433172_8446622_RNA45SN1_45S_50ntFlanks | 8182 | 8211 | 30 | ATCGAGGCCCAGCCCGTGGACGGTGTGAGG | 216.6 | -5.4 | 1.1 | -4.8 | 1.7E-06 | 2.8E-06 | MB231/MB468 | CYTO |
| >hg38_21_+_8433172_8446622_RNA45SN1_45S_50ntFlanks@6759.6807.49\|CTTGCGGCCCCGGGTTCCTCCCGGGGCTACGCCTGTCTGAGCGTCGCTT | >hg38_21_+_8433172_8446622_RNA45SN1_45S_50ntFlanks | 6759 | 6807 | 49 | CTTGCGGCCCCGGGTTCCTCCCGGGGCTACGCCTGTCTGAGCGTCGCTT | 688.7 | 5.3 | 1.1 | 4.8 | 1.3E-06 | 2.8E-06 | BT20/MB468 | MITO |
| >hg38_21_+_8433172_8446622_RNA45SN1_45S_50ntFlanks@12944.12962.19\|CGTAGACGACCTGCTTCTG | >hg38_21_+_8433172_8446622_RNA45SN1_45S_50ntFlanks | 12944 | 12962 | 19 | CGTAGACGACCTGCTTCTG | 236.0 | -2.5 | 0.5 | -4.8 | 1.8E-06 | 2.8E-06 | MB231/MB468 | NUC |
| >hg38_21_+_8433172_8446622_RNA45SN1_45S_50ntFlanks@6671.6696.26\|CTCGGCTCGTGCGTCGATGAAGAACG | >hg38_21_+_8433172_8446622_RNA45SN1_45S_50ntFlanks | 6671 | 6696 | 26 | CTCGGCTCGTGCGTCGATGAAGAACG | 3682.8 | -4.0 | 0.8 | -4.8 | 1.3E-06 | 2.8E-06 | BT20/MB468 | MITO |
| >hg38_21_+_8433172_8446622_RNA45SN1_45S_50ntFlanks@5446.5466.21\|CGGATCGGCCCCGCCGGGGTC | >hg38_21_+_8433172_8446622_RNA45SN1_45S_50ntFlanks | 5446 | 5466 | 21 | CGGATCGGCCCCGCCGGGGTC | 235.0 | 11.2 | 2.1 | 5.3 | 1.2E-07 | 2.8E-06 | BT20/MB231 | MP |
| >hg38_21_+_8433172_8446622_RNA45SN1_45S_50ntFlanks@8179.8223.45\|CTGATCGAGGCCCAGCCCGTGGACGGTGTGAGGCCGGTAGCGGCC | >hg38_21_+_8433172_8446622_RNA45SN1_45S_50ntFlanks | 8179 | 8223 | 45 | CTGATCGAGGCCCAGCCCGTGGACGGTGTGAGGCCGGTAGCGGCC | 210.3 | -6.8 | 1.4 | -4.8 | 1.8E-06 | 2.8E-06 | MB231/MB468 | NUC |
| >hg38_21_+_8433172_8446622_RNA45SN1_45S_50ntFlanks@8338.8378.41\|GTACCGTAAGGGAAAGTTGAAAAGAACTTTGAAGAGAGAGT | >hg38_21_+_8433172_8446622_RNA45SN1_45S_50ntFlanks | 8338 | 8378 | 41 | GTACCGTAAGGGAAAGTTGAAAAGAACTTTGAAGAGAGAGT | 62.7 | 9.7 | 1.9 | 5.0 | 5.0E-07 | 2.8E-06 | BT20/MB468 | MP |
| >hg38_21_+_8433172_8446622_RNA45SN1_45S_50ntFlanks@8183.8223.41\|TCGAGGCCCAGCCCGTGGACGGTGTGAGGCCGGTAGCGGCC | >hg38_21_+_8433172_8446622_RNA45SN1_45S_50ntFlanks | 8183 | 8223 | 41 | TCGAGGCCCAGCCCGTGGACGGTGTGAGGCCGGTAGCGGCC | 275.0 | -6.1 | 1.3 | -4.8 | 1.5E-06 | 2.8E-06 | BT20/MB468 | CYTO |
| >hg38_21_+_8433172_8446622_RNA45SN1_45S_50ntFlanks@12894.12928.35\|CTCGCCCGTCACGCACCGCACGTTCGTGGGGAACC | >hg38_21_+_8433172_8446622_RNA45SN1_45S_50ntFlanks | 12894 | 12928 | 35 | CTCGCCCGTCACGCACCGCACGTTCGTGGGGAACC | 52.4 | -8.8 | 1.7 | -5.0 | 5.0E-07 | 2.8E-06 | BT20/MB468 | MP |
| >hg38_21_+_8433172_8446622_RNA45SN1_45S_50ntFlanks@10393.10432.40\|CAGCAGTTGAACATGGGTCAGTCGGTCCTGAGAGATGGGC | >hg38_21_+_8433172_8446622_RNA45SN1_45S_50ntFlanks | 10393 | 10432 | 40 | CAGCAGTTGAACATGGGTCAGTCGGTCCTGAGAGATGGGC | 396.6 | -7.7 | 1.6 | -4.8 | 1.9E-06 | 2.9E-06 | BT20/MB468 | NUC |
| >hg38_21_+_8433172_8446622_RNA45SN1_45S_50ntFlanks@5035.5058.24\|CATGGCCGTTCTTAGTTGGTGGAG | >hg38_21_+_8433172_8446622_RNA45SN1_45S_50ntFlanks | 5035 | 5058 | 24 | CATGGCCGTTCTTAGTTGGTGGAG | 203.3 | -9.8 | 2.0 | -4.8 | 1.3E-06 | 2.9E-06 | BT20/MB468 | MITO |
| >hg38_21_+_8433172_8446622_RNA45SN1_45S_50ntFlanks@11584.11612.29\|TGGTGCGGACCAGGGGAATCCGACTGTTT | >hg38_21_+_8433172_8446622_RNA45SN1_45S_50ntFlanks | 11584 | 11612 | 29 | TGGTGCGGACCAGGGGAATCCGACTGTTT | 766.5 | 4.9 | 0.9 | 5.2 | 2.0E-07 | 2.9E-06 | BT20/MB231 | NUC |
| >hg38_21_+_8433172_8446622_RNA45SN1_45S_50ntFlanks@10786.10831.46\|AACAGCCTCTGGCATGTTGGAACAATGTAGGTAAGGGAAGTCGGCA | >hg38_21_+_8433172_8446622_RNA45SN1_45S_50ntFlanks | 10786 | 10831 | 46 | AACAGCCTCTGGCATGTTGGAACAATGTAGGTAAGGGAAGTCGGCA | 208.2 | -6.1 | 1.3 | -4.8 | 1.9E-06 | 2.9E-06 | MB231/MB468 | MITO |
| >hg38_21_+_8433172_8446622_RNA45SN1_45S_50ntFlanks@4730.4752.23\|CAAGAACGAAAGTCGGAGGTTCG | >hg38_21_+_8433172_8446622_RNA45SN1_45S_50ntFlanks | 4730 | 4752 | 23 | CAAGAACGAAAGTCGGAGGTTCG | 155.3 | -5.7 | 1.2 | -4.8 | 1.3E-06 | 2.9E-06 | BT20/MB468 | MITO |
| >hg38_21_+_8433172_8446622_RNA45SN1_45S_50ntFlanks@12409.12438.30\|TTGATCCTTCGATGTCGGCTCTTCCTATCA | >hg38_21_+_8433172_8446622_RNA45SN1_45S_50ntFlanks | 12409 | 12438 | 30 | TTGATCCTTCGATGTCGGCTCTTCCTATCA | 226.8 | 3.5 | 0.7 | 4.8 | 1.8E-06 | 2.9E-06 | MB231/MB468 | NUC |
| >hg38_21_+_8433172_8446622_RNA45SN1_45S_50ntFlanks@3990.4026.37\|TTGGTGACTCTAGATAACCTCGGGCCGATCGCACGCC | >hg38_21_+_8433172_8446622_RNA45SN1_45S_50ntFlanks | 3990 | 4026 | 37 | TTGGTGACTCTAGATAACCTCGGGCCGATCGCACGCC | 470.5 | -3.6 | 0.8 | -4.8 | 1.8E-06 | 2.9E-06 | MB231/MB468 | NUC |
| >MI0000261\|hsa-mir-139&WithFlank&11\|-\|72615057\|72615136@49.71.23&[MIMAT0004552&hsa-miR-139-3p&offsets\|0\|0;m-291&11\|-\|72615066\|72615088&offsets\|0\|0]\|TGGAGACGCGGCCCTGTTGGAGT | >MI0000261\|hsa-mir-139&WithFlank&11\|-\|72615057\|72615136 | 49 | 71 | 23 | TGGAGACGCGGCCCTGTTGGAGT | 42.7 | 8.6 | 1.8 | 4.8 | 1.5E-06 | 2.9E-06 | MB231/MB468 | MP |
| >hg38_21_+_8433172_8446622_RNA45SN1_45S_50ntFlanks@3815.3842.28\|ATGGTTCCTTTGGTCGCTCGCTCCTCTC | >hg38_21_+_8433172_8446622_RNA45SN1_45S_50ntFlanks | 3815 | 3842 | 28 | ATGGTTCCTTTGGTCGCTCGCTCCTCTC | 154.3 | -6.7 | 1.4 | -4.8 | 1.5E-06 | 2.9E-06 | MB231/MB468 | MP |
| >hg38_21_+_8433172_8446622_RNA45SN1_45S_50ntFlanks@10349.10368.20\|AACGAGAACTTTGAAGGCCG | >hg38_21_+_8433172_8446622_RNA45SN1_45S_50ntFlanks | 10349 | 10368 | 20 | AACGAGAACTTTGAAGGCCG | 2838.6 | -3.9 | 0.8 | -4.8 | 1.8E-06 | 2.9E-06 | MB231/MB468 | NUC |
| >hg38_21_+_8433172_8446622_RNA45SN1_45S_50ntFlanks@3705.3736.32\|TACCTGGTTGATCCTGCCAGTAGCATATGCTT | >hg38_21_+_8433172_8446622_RNA45SN1_45S_50ntFlanks | 3705 | 3736 | 32 | TACCTGGTTGATCCTGCCAGTAGCATATGCTT | 2430.6 | 5.7 | 1.2 | 4.8 | 1.3E-06 | 2.9E-06 | BT20/MB468 | MITO |
| >hg38_21_+_8433172_8446622_RNA45SN1_45S_50ntFlanks@12045.12063.19\|CTGACCCGGTGAGGCGGGG | >hg38_21_+_8433172_8446622_RNA45SN1_45S_50ntFlanks | 12045 | 12063 | 19 | CTGACCCGGTGAGGCGGGG | 68.6 | -9.2 | 1.8 | -5.0 | 5.0E-07 | 2.9E-06 | BT20/MB468 | MP |
| >hg38_21_+_8433172_8446622_RNA45SN1_45S_50ntFlanks@4764.4782.19\|ATACCGTCGTAGTTCCGAC | >hg38_21_+_8433172_8446622_RNA45SN1_45S_50ntFlanks | 4764 | 4782 | 19 | ATACCGTCGTAGTTCCGAC | 68.6 | -9.2 | 1.8 | -5.0 | 5.0E-07 | 2.9E-06 | BT20/MB468 | MP |
| >MI0003776\|hsa-mir-320b-1&WithFlank&1\|+\|116671740\|116671823@49.67.19;MI0003839\|hsa-mir-320b-2&WithFlank&1\|-\|224257034\|224257116@48.66.19;MI0000542\|hsa-mir-320a&WithFlank&8\|-\|22244960\|22245043@49.67.19&[MIMAT0005792&hsa-miR-320b&offsets\|+1\|-2;m-183&1\|+\|116671787\|116671806&offsets\|+1\|0];[MIMAT0005792_1&hsa-miR-320b&offsets\|+1\|-2;m-183&1\|-\|224257051\|224257070&offsets\|+1\|0];[MIMAT0000510&hsa-miR-320a-3p&offsets\|+1\|-2;m-60&8\|-\|22244975\|22244996&offsets\|+1\|-2]\|AAAGCTGGGTTGAGAGGGC | >MI0003776\|hsa-mir-320b-1&WithFlank&1\|+\|116671740\|116671823 | 49 | 67 | 19 | AAAGCTGGGTTGAGAGGGC | 494.5 | 7.9 | 1.6 | 5.0 | 5.0E-07 | 2.9E-06 | BT20/MB468 | MP |
| >hg38_21_+_8433172_8446622_RNA45SN1_45S_50ntFlanks@10574.10607.34\|AAGCCGGCGGGAGCCCCGGGGAGAGTTCTCTTTT | >hg38_21_+_8433172_8446622_RNA45SN1_45S_50ntFlanks | 10574 | 10607 | 34 | AAGCCGGCGGGAGCCCCGGGGAGAGTTCTCTTTT | 385.0 | -6.5 | 1.4 | -4.8 | 1.5E-06 | 2.9E-06 | BT20/MB468 | CYTO |
| >hg38_21_+_8433172_8446622_RNA45SN1_45S_50ntFlanks@12242.12265.24\|AGAAGGGCAAAAGCTCGCTTGATC | >hg38_21_+_8433172_8446622_RNA45SN1_45S_50ntFlanks | 12242 | 12265 | 24 | AGAAGGGCAAAAGCTCGCTTGATC | 409.8 | -5.4 | 1.1 | -4.8 | 1.5E-06 | 2.9E-06 | MB231/MB468 | MP |
| >hg38_21_+_8433172_8446622_RNA45SN1_45S_50ntFlanks@10342.10373.32\|ATATTCAAACGAGAACTTTGAAGGCCGAAGTG | >hg38_21_+_8433172_8446622_RNA45SN1_45S_50ntFlanks | 10342 | 10373 | 32 | ATATTCAAACGAGAACTTTGAAGGCCGAAGTG | 323.5 | 3.3 | 0.7 | 4.8 | 1.7E-06 | 2.9E-06 | MB231/MB468 | CYTO |
| >hg38_21_+_8433172_8446622_RNA45SN1_45S_50ntFlanks@11875.11898.24\|GAAGACCCTGTTGAGCTTGACTCT | >hg38_21_+_8433172_8446622_RNA45SN1_45S_50ntFlanks | 11875 | 11898 | 24 | GAAGACCCTGTTGAGCTTGACTCT | 710.3 | -4.3 | 0.9 | -4.8 | 2.0E-06 | 2.9E-06 | BT20/MB468 | NUC |

<table>
<tr><th></th><th></th><th></th><th></th><th></th><th></th><th></th><th></th><th></th><th></th><th></th><th></th><th></th><th></th></tr>
<tr><td>>hg38_21_+_8433172_8446622_RNA45SN1_45S_50ntFlanks@11665.11685.21|TTCTGCCCAGTGCTCTGAATG</td><td>>hg38_21_+_8433172_8446622_RNA45SN1_45S_50ntFlanks</td><td>11665</td><td>11685</td><td>21</td><td>TTCTGCCCAGTGCTCTGAATG</td><td>427.3</td><td>-3.5</td><td>0.7</td><td>-4.8</td><td>1.9E-06</td><td>2.9E-06</td><td>MB231/MB468</td><td>MITO</td></tr>
<tr><td>>hg38_21_+_8433172_8446622_RNA45SN1_45S_50ntFlanks@12008.12042.35|CCGCCGGTGAAATACCACTACTCTGATCGTTTTTT</td><td>>hg38_21_+_8433172_8446622_RNA45SN1_45S_50ntFlanks</td><td>12008</td><td>12042</td><td>35</td><td>CCGCCGGTGAAATACCACTACTCTGATCGTTTTTT</td><td>1081.0</td><td>-7.1</td><td>1.5</td><td>-4.8</td><td>1.3E-06</td><td>2.9E-06</td><td>BT20/MB468</td><td>MITO</td></tr>
<tr><td>>MI0000457|hsa-mir-141&WithFlank&12|+|6964091|6964197@65.84.20&[MIMAT0000432&hsa-miR-141-3p&offsets|0|-2;m-134&12|+|6964155|6964175&offsets|0|-1]|TAACACTGTCTGGTAAAGAT</td><td>>MI0000457|hsa-mir-141&WithFlank&12|+|6964091|6964197</td><td>65</td><td>84</td><td>20</td><td>TAACACTGTCTGGTAAAGAT</td><td>107.5</td><td>10.1</td><td>1.9</td><td>5.3</td><td>1.3E-07</td><td>2.9E-06</td><td>BT20/MB231</td><td>MP</td></tr>
<tr><td>>hg38_21_+_8433172_8446622_RNA45SN1_45S_50ntFlanks@11235.11255.21|CTTCCCGTGGATCGCCCCAGC</td><td>>hg38_21_+_8433172_8446622_RNA45SN1_45S_50ntFlanks</td><td>11235</td><td>11255</td><td>21</td><td>CTTCCCGTGGATCGCCCCAGC</td><td>851.1</td><td>-6.3</td><td>1.3</td><td>-4.8</td><td>2.0E-06</td><td>2.9E-06</td><td>BT20/MB468</td><td>NUC</td></tr>
<tr><td>>hg38_21_+_8433172_8446622_RNA45SN1_45S_50ntFlanks@10361.10388.28|GAAGGCCGAAGTGGAGAAGGGTTCCATG</td><td>>hg38_21_+_8433172_8446622_RNA45SN1_45S_50ntFlanks</td><td>10361</td><td>10388</td><td>28</td><td>GAAGGCCGAAGTGGAGAAGGGTTCCATG</td><td>766.1</td><td>-6.1</td><td>1.3</td><td>-4.8</td><td>1.6E-06</td><td>2.9E-06</td><td>BT20/MB468</td><td>CYTO</td></tr>
<tr><td>>hg38_21_+_8433172_8446622_RNA45SN1_45S_50ntFlanks@6644.6667.24|CCTCGTACGACTCTTAGCGGTGGA</td><td>>hg38_21_+_8433172_8446622_RNA45SN1_45S_50ntFlanks</td><td>6644</td><td>6667</td><td>24</td><td>CCTCGTACGACTCTTAGCGGTGGA</td><td>305.8</td><td>3.7</td><td>0.8</td><td>4.8</td><td>1.9E-06</td><td>2.9E-06</td><td>MB231/MB468</td><td>MITO</td></tr>
<tr><td>>hg38_21_+_8433172_8446622_RNA45SN1_45S_50ntFlanks@12625.12661.37|CAATGGGGCGAAGCTACCATCTGTGGGATTATGACTG</td><td>>hg38_21_+_8433172_8446622_RNA45SN1_45S_50ntFlanks</td><td>12625</td><td>12661</td><td>37</td><td>CAATGGGGCGAAGCTACCATCTGTGGGATTATGACTG</td><td>1061.8</td><td>-3.5</td><td>0.7</td><td>-4.8</td><td>1.4E-06</td><td>2.9E-06</td><td>BT20/MB468</td><td>MITO</td></tr>
<tr><td>>MI0000255|hsa-mir-30d&WithFlank&8|-|134804870|134804951@12.35.24(+1A)&[MIMAT0000245&hsa-miR-30d-5p&offsets|0|+2(+1A);m-21&8|-|134804919|134804940&offsets|0|+2(+1A)]|TGTAAACATCCCCGACTGGAAGCTA</td><td>>MI0000255|hsa-mir-30d&WithFlank&8|-|134804870|134804951</td><td>12</td><td>35</td><td>24(+1A)</td><td>TGTAAACATCCCCGACTGGAAGCTA</td><td>309.5</td><td>8.0</td><td>1.7</td><td>4.8</td><td>1.9E-06</td><td>2.9E-06</td><td>MB231/MB468</td><td>MITO</td></tr>
<tr><td>>hg38_21_+_8433172_8446622_RNA45SN1_45S_50ntFlanks@12456.12496.41|CCAAGCGTTGGATTGTTCACCCACTAATAGGGAACGTGAGC</td><td>>hg38_21_+_8433172_8446622_RNA45SN1_45S_50ntFlanks</td><td>12456</td><td>12496</td><td>41</td><td>CCAAGCGTTGGATTGTTCACCCACTAATAGGGAACGTGAGC</td><td>964.3</td><td>-3.8</td><td>0.8</td><td>-4.8</td><td>1.8E-06</td><td>2.9E-06</td><td>MB231/MB468</td><td>NUC</td></tr>
<tr><td>>hg38_21_+_8433172_8446622_RNA45SN1_45S_50ntFlanks@4975.4999.25|CCCGGACACGGACAGGATTGACAGA</td><td>>hg38_21_+_8433172_8446622_RNA45SN1_45S_50ntFlanks</td><td>4975</td><td>4999</td><td>25</td><td>CCCGGACACGGACAGGATTGACAGA</td><td>1042.0</td><td>-5.4</td><td>1.1</td><td>-4.8</td><td>1.9E-06</td><td>2.9E-06</td><td>MB231/MB468</td><td>MITO</td></tr>
<tr><td>>hg38_21_+_8433172_8446622_RNA45SN1_45S_50ntFlanks@11210.11233.24|GGCGACTCTGGACGCGAGCCGGGC</td><td>>hg38_21_+_8433172_8446622_RNA45SN1_45S_50ntFlanks</td><td>11210</td><td>11233</td><td>24</td><td>GGCGACTCTGGACGCGAGCCGGGC</td><td>459.5</td><td>-3.5</td><td>0.7</td><td>-4.8</td><td>1.6E-06</td><td>2.9E-06</td><td>BT20/MB468</td><td>CYTO</td></tr>
<tr><td>>hg38_21_+_8433172_8446622_RNA45SN1_45S_50ntFlanks@10385.10424.40|CATGTGAACAGCAGTTGAACATGGGTCAGTCGGTCCTGAG</td><td>>hg38_21_+_8433172_8446622_RNA45SN1_45S_50ntFlanks</td><td>10385</td><td>10424</td><td>40</td><td>CATGTGAACAGCAGTTGAACATGGGTCAGTCGGTCCTGAG</td><td>696.1</td><td>4.2</td><td>0.9</td><td>4.8</td><td>1.8E-06</td><td>2.9E-06</td><td>MB231/MB468</td><td>CYTO</td></tr>
<tr><td>>hg38_21_+_8433172_8446622_RNA45SN1_45S_50ntFlanks@12420.12443.24|ATGTCGGCTCTTCCTATCATTGTG</td><td>>hg38_21_+_8433172_8446622_RNA45SN1_45S_50ntFlanks</td><td>12420</td><td>12443</td><td>24</td><td>ATGTCGGCTCTTCCTATCATTGTG</td><td>1764.1</td><td>-3.1</td><td>0.6</td><td>-4.8</td><td>1.4E-06</td><td>2.9E-06</td><td>BT20/MB468</td><td>MITO</td></tr>
<tr><td>>MI0000434|hsa-let-7i&WithFlank&12|+|62603680|62603775@12.32.21(+1C)&[MIMAT0000415&hsa-let-7i-5p&offsets|0|-1(+1C);m-49&12|+|62603691|62603712&offsets|0|-1(+1C)]|TGAGGTAGTAGTTTGTGCTGTC</td><td>>MI0000434|hsa-let-7i&WithFlank&12|+|62603680|62603775</td><td>12</td><td>32</td><td>21(+1C)</td><td>TGAGGTAGTAGTTTGTGCTGTC</td><td>5896.2</td><td>5.0</td><td>1.1</td><td>4.8</td><td>1.9E-06</td><td>2.9E-06</td><td>MB231/MB468</td><td>MITO</td></tr>
<tr><td>>hg38_21_+_8433172_8446622_RNA45SN1_45S_50ntFlanks@8184.8206.23|CGAGGCCCAGCCCGTGGACGGTG</td><td>>hg38_21_+_8433172_8446622_RNA45SN1_45S_50ntFlanks</td><td>8184</td><td>8206</td><td>23</td><td>CGAGGCCCAGCCCGTGGACGGTG</td><td>404.3</td><td>-7.0</td><td>1.5</td><td>-4.8</td><td>1.6E-06</td><td>2.9E-06</td><td>MB231/MB468</td><td>MP</td></tr>
<tr><td>>hg38_21_+_8433172_8446622_RNA45SN1_45S_50ntFlanks@6699.6717.19|GCTAGCTGCGAGAATTAAT</td><td>>hg38_21_+_8433172_8446622_RNA45SN1_45S_50ntFlanks</td><td>6699</td><td>6717</td><td>19</td><td>GCTAGCTGCGAGAATTAAT</td><td>499.9</td><td>-4.7</td><td>1.0</td><td>-4.8</td><td>1.8E-06</td><td>2.9E-06</td><td>MB231/MB468</td><td>CYTO</td></tr>
<tr><td>>hg38_21_+_8433172_8446622_RNA45SN1_45S_50ntFlanks@9454.9500.47|CCGCGCCGGGGAGGTGGAGCACGAGCGCACGTGTTAGGACCCGAAAG</td><td>>hg38_21_+_8433172_8446622_RNA45SN1_45S_50ntFlanks</td><td>9454</td><td>9500</td><td>47</td><td>CCGCGCCGGGGAGGTGGAGCACGAGCGCACGTGTTAGGACCCGAAAG</td><td>124.3</td><td>-10.0</td><td>2.0</td><td>-5.0</td><td>5.1E-07</td><td>2.9E-06</td><td>BT20/MB468</td><td>MP</td></tr>
<tr><td>>hg38_21_+_8433172_8446622_RNA45SN1_45S_50ntFlanks@11577.11594.18|TTAGAACTGGTGCGGACC</td><td>>hg38_21_+_8433172_8446622_RNA45SN1_45S_50ntFlanks</td><td>11577</td><td>11594</td><td>18</td><td>TTAGAACTGGTGCGGACC</td><td>707.5</td><td>-6.0</td><td>1.3</td><td>-4.8</td><td>2.0E-06</td><td>2.9E-06</td><td>BT20/MB468</td><td>NUC</td></tr>
<tr><td>>hg38_21_+_8433172_8446622_RNA45SN1_45S_50ntFlanks@10693.10712.20|CCGGTGAGCTCTCGCTGGCC</td><td>>hg38_21_+_8433172_8446622_RNA45SN1_45S_50ntFlanks</td><td>10693</td><td>10712</td><td>20</td><td>CCGGTGAGCTCTCGCTGGCC</td><td>79.4</td><td>-9.4</td><td>1.9</td><td>-5.0</td><td>5.1E-07</td><td>2.9E-06</td><td>BT20/MB468</td><td>MP</td></tr>
<tr><td>>hg38_21_+_8433172_8446622_RNA45SN1_45S_50ntFlanks@12635.12652.18|AAGCTACCATCTGTGGGA</td><td>>hg38_21_+_8433172_8446622_RNA45SN1_45S_50ntFlanks</td><td>12635</td><td>12652</td><td>18</td><td>AAGCTACCATCTGTGGGA</td><td>123.8</td><td>-5.5</td><td>1.1</td><td>-4.8</td><td>1.6E-06</td><td>2.9E-06</td><td>BT20/MB468</td><td>CYTO</td></tr>
<tr><td>>hg38_21_+_8433172_8446622_RNA45SN1_45S_50ntFlanks@11769.11791.23|TAATTAGTGACGCGCATGAATGG</td><td>>hg38_21_+_8433172_8446622_RNA45SN1_45S_50ntFlanks</td><td>11769</td><td>11791</td><td>23</td><td>TAATTAGTGACGCGCATGAATGG</td><td>279.4</td><td>-6.5</td><td>1.4</td><td>-4.8</td><td>1.9E-06</td><td>2.9E-06</td><td>MB231/MB468</td><td>MITO</td></tr>
<tr><td>>hg38_21_+_8433172_8446622_RNA45SN1_45S_50ntFlanks@3861.3898.38|TAATTCTAGAGCTAATACATGCCGACGGGCGCTGACCC</td><td>>hg38_21_+_8433172_8446622_RNA45SN1_45S_50ntFlanks</td><td>3861</td><td>3898</td><td>38</td><td>TAATTCTAGAGCTAATACATGCCGACGGGCGCTGACCC</td><td>128.7</td><td>-10.9</td><td>2.3</td><td>-4.8</td><td>1.6E-06</td><td>2.9E-06</td><td>MB231/MB468</td><td>MP</td></tr>
<tr><td>>MI0000273|hsa-mir-183&WithFlank&7|-|129774899|129775020@34.53.20(+1A)&[MIMAT0000261&hsa-miR-183-5p&offsets|+1|-1(+1A);m-41&7|-|129774966|129774987&offsets|0|-2(+1A)]|ATGGCACTGGTAGAATTCACA</td><td>>MI0000273|hsa-mir-183&WithFlank&7|-|129774899|129775020</td><td>34</td><td>53</td><td>20(+1A)</td><td>ATGGCACTGGTAGAATTCACA</td><td>356.9</td><td>4.4</td><td>0.8</td><td>5.3</td><td>1.4E-07</td><td>2.9E-06</td><td>BT20/MB231</td><td>CYTO</td></tr>
<tr><td>>trnaMT_PheGAA_MT_+_577_647@37.54.18|AAATGTTTAGACGGGCTC</td><td>>trnaMT_PheGAA_MT_+_577_647</td><td>37</td><td>54</td><td>18</td><td>AAATGTTTAGACGGGCTC</td><td>38.5</td><td>8.5</td><td>1.8</td><td>4.8</td><td>1.6E-06</td><td>2.9E-06</td><td>MB231/MB468</td><td>MP</td></tr>
<tr><td>>hg38_21_+_8433172_8446622_RNA45SN1_45S_50ntFlanks@6652.6677.26|GACTCTTAGCGGTGGATCACTCGGCT</td><td>>hg38_21_+_8433172_8446622_RNA45SN1_45S_50ntFlanks</td><td>6652</td><td>6677</td><td>26</td><td>GACTCTTAGCGGTGGATCACTCGGCT</td><td>929.2</td><td>3.2</td><td>0.7</td><td>4.8</td><td>1.6E-06</td><td>2.9E-06</td><td>BT20/MB468</td><td>CYTO</td></tr>
<tr><td>>MI0000542|hsa-mir-320a&WithFlank&8|-|22244960|22245043@47.68.22(+1C)&[MIMAT0000510&hsa-miR-320a-3p&offsets|-1|-1(+1C);m-60&8|-|22244975|22244996&offsets|-1|-1(+1C)]|GAAAAGCTGGGTTGAGAGGGCGC</td><td>>MI0000542|hsa-mir-320a&WithFlank&8|-|22244960|22245043</td><td>47</td><td>68</td><td>22(+1C)</td><td>GAAAAGCTGGGTTGAGAGGGCGC</td><td>73.6</td><td>9.9</td><td>2.0</td><td>5.0</td><td>5.1E-07</td><td>2.9E-06</td><td>BT20/MB468</td><td>MP</td></tr>
<tr><td>>hg38_1_-_228634819_228635039_RNA5S12_5S_50ntFlanks@134.170.37|TGGGAGACCGCCTGGGAATACCGGGTGCTGTAGGCTT</td><td>>hg38_1_-_228634819_228635039_RNA5S12_5S_50ntFlanks</td><td>134</td><td>170</td><td>37</td><td>TGGGAGACCGCCTGGGAATACCGGGTGCTGTAGGCTT</td><td>33715.1</td><td>4.9</td><td>1.0</td><td>5.0</td><td>5.1E-07</td><td>2.9E-06</td><td>BT20/MB468</td><td>MP</td></tr>
<tr><td>>hg38_21_+_8433172_8446622_RNA45SN1_45S_50ntFlanks@10687.10712.26|CGGCGTCCGGTGAGCTCTCGCTGGCC</td><td>>hg38_21_+_8433172_8446622_RNA45SN1_45S_50ntFlanks</td><td>10687</td><td>10712</td><td>26</td><td>CGGCGTCCGGTGAGCTCTCGCTGGCC</td><td>2055.1</td><td>-8.6</td><td>1.8</td><td>-4.8</td><td>2.0E-06</td><td>2.9E-06</td><td>BT20/MB468</td><td>NUC</td></tr>
<tr><td>>hg38_21_+_8433172_8446622_RNA45SN1_45S_50ntFlanks@11664.11692.29|TTTCTGCCCAGTGCTCTGAATGTCAAAGT</td><td>>hg38_21_+_8433172_8446622_RNA45SN1_45S_50ntFlanks</td><td>11664</td><td>11692</td><td>29</td><td>TTTCTGCCCAGTGCTCTGAATGTCAAAGT</td><td>54.2</td><td>9.5</td><td>1.9</td><td>5.0</td><td>5.1E-07</td><td>2.9E-06</td><td>BT20/MB468</td><td>MP</td></tr>
<tr><td>>hg38_21_+_8433172_8446622_RNA45SN1_45S_50ntFlanks@11729.11764.36|TAACTATGACTCTCTTAAGGTAGCCAAATGCCTCGT</td><td>>hg38_21_+_8433172_8446622_RNA45SN1_45S_50ntFlanks</td><td>11729</td><td>11764</td><td>36</td><td>TAACTATGACTCTCTTAAGGTAGCCAAATGCCTCGT</td><td>1617.3</td><td>-7.5</td><td>1.6</td><td>-4.8</td><td>1.6E-06</td><td>2.9E-06</td><td>BT20/MB468</td><td>CYTO</td></tr>
<tr><td>>hg38_21_+_8433172_8446622_RNA45SN1_45S_50ntFlanks@9572.9594.23|CAAATCGGTCGTCCGACCTGGGT</td><td>>hg38_21_+_8433172_8446622_RNA45SN1_45S_50ntFlanks</td><td>9572</td><td>9594</td><td>23</td><td>CAAATCGGTCGTCCGACCTGGGT</td><td>925.2</td><td>-4.2</td><td>0.9</td><td>-4.8</td><td>1.4E-06</td><td>2.9E-06</td><td>BT20/MB468</td><td>MITO</td></tr>
<tr><td>>hg38_21_+_8433172_8446622_RNA45SN1_45S_50ntFlanks@8183.8204.22|TCGAGGCCCAGCCCGTGGACGG</td><td>>hg38_21_+_8433172_8446622_RNA45SN1_45S_50ntFlanks</td><td>8183</td><td>8204</td><td>22</td><td>TCGAGGCCCAGCCCGTGGACGG</td><td>119.3</td><td>-5.2</td><td>1.1</td><td>-4.8</td><td>1.6E-06</td><td>2.9E-06</td><td>MB231/MB468</td><td>MP</td></tr>
<tr><td>>hg38_21_+_8433172_8446622_RNA45SN1_45S_50ntFlanks@12419.12437.19|GATGTCGGCTCTTCCTATC</td><td>>hg38_21_+_8433172_8446622_RNA45SN1_45S_50ntFlanks</td><td>12419</td><td>12437</td><td>19</td><td>GATGTCGGCTCTTCCTATC</td><td>3532.2</td><td>2.0</td><td>0.4</td><td>4.8</td><td>1.6E-06</td><td>2.9E-06</td><td>BT20/MB468</td><td>CYTO</td></tr>
<tr><td>>hg38_21_+_8433172_8446622_RNA45SN1_45S_50ntFlanks@10830.10851.22|CAAGCCGGATCCGTAACTTCGG</td><td>>hg38_21_+_8433172_8446622_RNA45SN1_45S_50ntFlanks</td><td>10830</td><td>10851</td><td>22</td><td>CAAGCCGGATCCGTAACTTCGG</td><td>242.6</td><td>-3.8</td><td>0.8</td><td>-4.8</td><td>1.9E-06</td><td>2.9E-06</td><td>MB231/MB468</td><td>MITO</td></tr>
<tr><td>>hg38_21_+_8433172_8446622_RNA45SN1_45S_50ntFlanks@12936.12953.18|AAACCATTCGTAGACGAC</td><td>>hg38_21_+_8433172_8446622_RNA45SN1_45S_50ntFlanks</td><td>12936</td><td>12953</td><td>18</td><td>AAACCATTCGTAGACGAC</td><td>232.3</td><td>-2.7</td><td>0.6</td><td>-4.8</td><td>1.4E-06</td><td>2.9E-06</td><td>BT20/MB468</td><td>MITO</td></tr>
<tr><td>>hg38_21_+_8433172_8446622_RNA45SN1_45S_50ntFlanks@12655.12673.19|ATGACTGAACGCCTCTAAG</td><td>>hg38_21_+_8433172_8446622_RNA45SN1_45S_50ntFlanks</td><td>12655</td><td>12673</td><td>19</td><td>ATGACTGAACGCCTCTAAG</td><td>1207.0</td><td>-6.0</td><td>1.3</td><td>-4.8</td><td>1.8E-06</td><td>2.9E-06</td><td>MB231/MB468</td><td>NUC</td></tr>
<tr><td>>hg38_21_+_8433172_8446622_RNA45SN1_45S_50ntFlanks@11881.11898.18|CCTGTTGAGCTTGACTCT</td><td>>hg38_21_+_8433172_8446622_RNA45SN1_45S_50ntFlanks</td><td>11881</td><td>11898</td><td>18</td><td>CCTGTTGAGCTTGACTCT</td><td>171.3</td><td>-3.1</td><td>0.6</td><td>-4.8</td><td>1.8E-06</td><td>2.9E-06</td><td>MB231/MB468</td><td>CYTO</td></tr>
<tr><td>>hg38_21_+_8433172_8446622_RNA45SN1_45S_50ntFlanks@6652.6673.22|GACTCTTAGCGGTGGATCACTC</td><td>>hg38_21_+_8433172_8446622_RNA45SN1_45S_50ntFlanks</td><td>6652</td><td>6673</td><td>22</td><td>GACTCTTAGCGGTGGATCACTC</td><td>289.3</td><td>4.4</td><td>0.9</td><td>4.8</td><td>1.8E-06</td><td>2.9E-06</td><td>MB231/MB468</td><td>NUC</td></tr>
<tr><td>>hg38_21_+_8433172_8446622_RNA45SN1_45S_50ntFlanks@4069.4095.27|CTTTCGATGGTAGTCGCCGTGCCTACC</td><td>>hg38_21_+_8433172_8446622_RNA45SN1_45S_50ntFlanks</td><td>4069</td><td>4095</td><td>27</td><td>CTTTCGATGGTAGTCGCCGTGCCTACC</td><td>276.5</td><td>-4.0</td><td>0.8</td><td>-4.8</td><td>1.8E-06</td><td>2.9E-06</td><td>MB231/MB468</td><td>CYTO</td></tr>
<tr><td>>hg38_21_+_8433172_8446622_RNA45SN1_45S_50ntFlanks@9571.9595.25|GCAAATCGGTCGTCCGACCTGGGTA</td><td>>hg38_21_+_8433172_8446622_RNA45SN1_45S_50ntFlanks</td><td>9571</td><td>9595</td><td>25</td><td>GCAAATCGGTCGTCCGACCTGGGTA</td><td>236.5</td><td>-4.0</td><td>0.8</td><td>-4.8</td><td>1.8E-06</td><td>2.9E-06</td><td>MB231/MB468</td><td>CYTO</td></tr>
<tr><td>>hg38_21_+_8433172_8446622_RNA45SN1_45S_50ntFlanks@8177.8204.28|TTCTGATCGAGGCCCAGCCCGTGGACGG</td><td>>hg38_21_+_8433172_8446622_RNA45SN1_45S_50ntFlanks</td><td>8177</td><td>8204</td><td>28</td><td>TTCTGATCGAGGCCCAGCCCGTGGACGG</td><td>964.9</td><td>-6.3</td><td>1.3</td><td>-4.8</td><td>2.0E-06</td><td>2.9E-06</td><td>BT20/MB468</td><td>NUC</td></tr>
<tr><td>>am_trnaMT_ValTAC_MT_+_1602_1670@48.67.20|CAACTTAACTTGACCGCTCT</td><td>>am_trnaMT_ValTAC_MT_+_1602_1670</td><td>48</td><td>67</td><td>20</td><td>CAACTTAACTTGACCGCTCT</td><td>3743.6</td><td>6.6</td><td>1.4</td><td>4.8</td><td>1.8E-06</td><td>2.9E-06</td><td>MB231/MB468</td><td>CYTO</td></tr>
<tr><td>>hg38_21_+_8433172_8446622_RNA45SN1_45S_50ntFlanks@4785.4803.19|TAAACGATGCCGACCGGCG</td><td>>hg38_21_+_8433172_8446622_RNA45SN1_45S_50ntFlanks</td><td>4785</td><td>4803</td><td>19</td><td>TAAACGATGCCGACCGGCG</td><td>65.8</td><td>-9.1</td><td>1.8</td><td>-5.0</td><td>5.1E-07</td><td>2.9E-06</td><td>BT20/MB468</td><td>MP</td></tr>
<tr><td>>MI0000095|hsa-mir-93&WithFlank&7|-|100093762|100093853@17.37.21&[MIMAT0000093&hsa-miR-93-5p&offsets|0|-2;m-29&7|-|100093815|100093837&offsets|0|-2]|CAAAGTGCTGTTCGTGCAGGT</td><td>>MI0000095|hsa-mir-93&WithFlank&7|-|100093762|100093853</td><td>17</td><td>37</td><td>21</td><td>CAAAGTGCTGTTCGTGCAGGT</td><td>595.9</td><td>5.0</td><td>1.0</td><td>4.8</td><td>1.8E-06</td><td>2.9E-06</td><td>MB231/MB468</td><td>CYTO</td></tr>
<tr><td>>MI0000266|hsa-mir-10a&WithFlank&17|-|48579832|48579953@29.50.22(+1C)&[MIMAT0000253&hsa-miR-10a-5p&offsets|+1|0(+1C);m-18&17|-|48579905|48579926&offsets|+1|+1(+1C)]|ACCCTGTAGATCCGAATTTGTGC</td><td>>MI0000266|hsa-mir-10a&WithFlank&17|-|48579832|48579953</td><td>29</td><td>50</td><td>22(+1C)</td><td>ACCCTGTAGATCCGAATTTGTGC</td><td>42.2</td><td>8.6</td><td>1.8</td><td>4.8</td><td>1.6E-06</td><td>2.9E-06</td><td>MB231/MB468</td><td>MP</td></tr>
<tr><td>>MI0000439|hsa-mir-23b&WithFlank&9|+|95085202|95085310@65.86.22(+1U)&[MIMAT0000418&hsa-miR-23b-3p&offsets|+1|0(+1U);m-42&9|+|95085265|95085285&offsets|+1|+2(+1U)]|TCACATTGCCAGGGATTACCACT</td><td>>MI0000439|hsa-mir-23b&WithFlank&9|+|95085202|95085310</td><td>65</td><td>86</td><td>22(+1U)</td><td>TCACATTGCCAGGGATTACCACT</td><td>344.1</td><td>6.9</td><td>1.5</td><td>4.8</td><td>1.9E-06</td><td>2.9E-06</td><td>MB231/MB468</td><td>MITO</td></tr>
<tr><td>>MI0003776|hsa-mir-320b-1&WithFlank&1|+|116671740|116671823@48.67.20;MI0003839|hsa-mir-320b-2&WithFlank&1|-|224257034|224257116@47.66.20;MI0000542|hsa-mir-320a&WithFlank&8|-|22244960|22245043@48.67.20&[MIMAT0005792&hsa-miR-320b&offsets|0|-2;m-183&1|+|116671787|116671806&offsets|0|0];[MIMAT0005792_1&hsa-miR-320b&offsets|0|-2;m-183&1|-|224257051|224257070&offsets|0|0];[MIMAT0000510&hsa-miR-320a-3p&offsets|0|-2;m-60&8|-|22244975|22244996&offsets|0|-2]|AAAAGCTGGGTTGAGAGGGC</td><td>>MI0003776|hsa-mir-320b-1&WithFlank&1|+|116671740|116671823</td><td>48</td><td>67</td><td>20</td><td>AAAAGCTGGGTTGAGAGGGC</td><td>4391.6</td><td>3.8</td><td>0.7</td><td>5.3</td><td>1.4E-07</td><td>2.9E-06</td><td>BT20/MB231</td><td>CYTO</td></tr>
<tr><td>>hg38_21_+_8433172_8446622_RNA45SN1_45S_50ntFlanks@4003.4025.23|ATAACCTCGGGCCGATCGCACGC</td><td>>hg38_21_+_8433172_8446622_RNA45SN1_45S_50ntFlanks</td><td>4003</td><td>4025</td><td>23</td><td>ATAACCTCGGGCCGATCGCACGC</td><td>776.9</td><td>-4.9</td><td>1.0</td><td>-4.8</td><td>1.6E-06</td><td>2.9E-06</td><td>BT20/MB468</td><td>CYTO</td></tr>
<tr><td>>hg38_21_+_8433172_8446622_RNA45SN1_45S_50ntFlanks@10365.10388.24|GCCGAAGTGGAGAAGGGTTCCATG</td><td>>hg38_21_+_8433172_8446622_RNA45SN1_45S_50ntFlanks</td><td>10365</td><td>10388</td><td>24</td><td>GCCGAAGTGGAGAAGGGTTCCATG</td><td>57.7</td><td>-8.9</td><td>1.8</td><td>-5.0</td><td>5.2E-07</td><td>2.9E-06</td><td>BT20/MB468</td><td>MP</td></tr>
<tr><td>>hg38_21_+_8433172_8446622_RNA45SN1_45S_50ntFlanks@4102.4127.26|ACCACGGGTGACGGGGAATCAGGGTT</td><td>>hg38_21_+_8433172_8446622_RNA45SN1_45S_50ntFlanks</td><td>4102</td><td>4127</td><td>26</td><td>ACCACGGGTGACGGGGAATCAGGGTT</td><td>680.3</td><td>-2.9</td><td>0.6</td><td>-4.8</td><td>1.6E-06</td><td>2.9E-06</td><td>MB231/MB468</td><td>MP</td></tr>
</table>

| | | | | | | | | | | | | | |
|---|---|---|---|---|---|---|---|---|---|---|---|---|---|
| >hg38_21_+_8433172_8446622_RNA45SN1_45S_50ntFlanks@8170.8217.48\|CAAGTCCTTCTGATCGAGGCCCAGCCCGTGGACGGTGTGAGGCCGGTA | >hg38_21_+_8433172_8446622_RNA45SN1_45S_50ntFlanks | 8170 | 8217 | 48 | CAAGTCCTTCTGATCGAGGCCCAGCCCGTGGACGGTGTGAGGCCGGTA | 285.2 | -6.0 | 1.3 | -4.8 | 1.9E-06 | 2.9E-06 | MB231/MB468 | MITO |
| >MI0000542\|hsa-mir-320a&WithFlank&8\|-\|22244960\|22245043@49.68.20(+2U)&[MIMAT0000510&hsa-miR-320a-3p&offsets\|+1\|-1(+2U);m-60&8\|-\|22244975\|22244996&offsets\|+1\|-1(+2U)]\|AAAGCTGGGTTGAGAGGGCGTT | >MI0000542\|hsa-mir-320a&WithFlank&8\|-\|22244960\|22245043 | 49 | 68 | 20(+2U) | AAAGCTGGGTTGAGAGGGCGTT | 464.4 | 4.1 | 0.8 | 5.3 | 1.4E-07 | 2.9E-06 | BT20/MB231 | CYTO |
| >hg38_21_+_8433172_8446622_RNA45SN1_45S_50ntFlanks@7983.8012.30\|CAGATCAGACGTGGCGACCCGCTGAATTTA | >hg38_21_+_8433172_8446622_RNA45SN1_45S_50ntFlanks | 7983 | 8012 | 30 | CAGATCAGACGTGGCGACCCGCTGAATTTA | 761.1 | -4.3 | 0.9 | -4.8 | 1.4E-06 | 2.9E-06 | BT20/MB468 | MITO |
| >hg38_21_+_8433172_8446622_RNA45SN1_45S_50ntFlanks@3876.3898.23\|TACATGCCGACGGGCGCTGACCC | >hg38_21_+_8433172_8446622_RNA45SN1_45S_50ntFlanks | 3876 | 3898 | 23 | TACATGCCGACGGGCGCTGACCC | 1439.4 | -3.5 | 0.7 | -4.8 | 1.6E-06 | 2.9E-06 | BT20/MB468 | CYTO |
| >hg38_21_+_8433172_8446622_RNA45SN1_45S_50ntFlanks@10248.10282.35\|CGACGAGTAGGAGGGCCGCTGCGGTGAGCCTTGAA | >hg38_21_+_8433172_8446622_RNA45SN1_45S_50ntFlanks | 10248 | 10282 | 35 | CGACGAGTAGGAGGGCCGCTGCGGTGAGCCTTGAA | 56.8 | 9.6 | 1.9 | 5.0 | 5.2E-07 | 3.0E-06 | BT20/MB468 | MP |
| >hg38_21_+_8433172_8446622_RNA45SN1_45S_50ntFlanks@3985.4024.40\|CGGCTTTGGTGACTCTAGATAACCTCGGGCCGATCGCACG | >hg38_21_+_8433172_8446622_RNA45SN1_45S_50ntFlanks | 3985 | 4024 | 40 | CGGCTTTGGTGACTCTAGATAACCTCGGGCCGATCGCACG | 1071.2 | -6.7 | 1.4 | -4.8 | 1.6E-06 | 3.0E-06 | BT20/MB468 | CYTO |
| >hg38_21_+_8433172_8446622_RNA45SN1_45S_50ntFlanks@11563.11611.49\|GCCTAGCAGCCGACTTAGAACTGGTGCGGACCAGGGGAATCCGACTGTT | >hg38_21_+_8433172_8446622_RNA45SN1_45S_50ntFlanks | 11563 | 11611 | 49 | GCCTAGCAGCCGACTTAGAACTGGTGCGGACCAGGGGAATCCGACTGTT | 980.4 | 6.9 | 1.4 | 4.8 | 1.4E-06 | 3.0E-06 | BT20/MB468 | MITO |
| >hg38_21_+_8433172_8446622_RNA45SN1_45S_50ntFlanks@11771.11794.24\|ATTAGTGACGCGCATGAATGGATG | >hg38_21_+_8433172_8446622_RNA45SN1_45S_50ntFlanks | 11771 | 11794 | 24 | ATTAGTGACGCGCATGAATGGATG | 1020.9 | -3.2 | 0.7 | -4.8 | 1.9E-06 | 3.0E-06 | MB231/MB468 | NUC |
| >hg38_MT_+_598_1651_RNR1_12S_50ntFlanks@849.872.24\|AGTGCTTAGTTGAACAGGGCCCTG | >hg38_MT_+_598_1651_RNR1_12S_50ntFlanks | 849 | 872 | 24 | AGTGCTTAGTTGAACAGGGCCCTG | 79.8 | 10.1 | 2.0 | 5.0 | 5.2E-07 | 3.0E-06 | BT20/MB468 | MP |
| >hg38_21_+_8433172_8446622_RNA45SN1_45S_50ntFlanks@12359.12376.18\|ACAGGGATAACTGGCTTG | >hg38_21_+_8433172_8446622_RNA45SN1_45S_50ntFlanks | 12359 | 12376 | 18 | ACAGGGATAACTGGCTTG | 52.4 | -8.8 | 1.8 | -5.0 | 5.2E-07 | 3.0E-06 | BT20/MB468 | MP |
| >hg38_21_+_8433172_8446622_RNA45SN1_45S_50ntFlanks@11821.11846.26\|CTATCCAGCGAAACCACAGCCAAGGG | >hg38_21_+_8433172_8446622_RNA45SN1_45S_50ntFlanks | 11821 | 11846 | 26 | CTATCCAGCGAAACCACAGCCAAGGG | 69.9 | -9.2 | 1.8 | -5.0 | 5.2E-07 | 3.0E-06 | BT20/MB468 | MP |
| >hg38_21_+_8433172_8446622_RNA45SN1_45S_50ntFlanks@9929.9951.23\|GATATAGACAGCAGGACGGTGGC | >hg38_21_+_8433172_8446622_RNA45SN1_45S_50ntFlanks | 9929 | 9951 | 23 | GATATAGACAGCAGGACGGTGGC | 708.1 | -7.4 | 1.5 | -4.8 | 1.6E-06 | 3.0E-06 | MB231/MB468 | MP |
| >hg38_21_+_8433172_8446622_RNA45SN1_45S_50ntFlanks@4000.4025.26\|TAGATAACCTCGGGCCGATCGCACGC | >hg38_21_+_8433172_8446622_RNA45SN1_45S_50ntFlanks | 4000 | 4025 | 26 | TAGATAACCTCGGGCCGATCGCACGC | 295.6 | -6.6 | 1.4 | -4.8 | 1.9E-06 | 3.0E-06 | MB231/MB468 | NUC |
| >MI0000441\|hsa-mir-30b&WithFlank&8\|-\|134800514\|134800613@23.44.22(+1U)&[MIMAT0000420&hsa-miR-30b-5p&offsets\|0\|0(+1U);m-76&8\|-\|134800570\|134800591&offsets\|0\|0(+1U)]\|TGTAAACATCCTACACTCAGCTT | >MI0000441\|hsa-mir-30b&WithFlank&8\|-\|134800514\|134800613 | 23 | 44 | 22(+1U) | TGTAAACATCCTACACTCAGCTT | 41.1 | 8.6 | 1.8 | 4.8 | 1.6E-06 | 3.0E-06 | MB231/MB468 | MP |
| >hg38_21_+_8433172_8446622_RNA45SN1_45S_50ntFlanks@3984.4024.41\|GCGGCTTTGGTGACTCTAGATAACCTCGGGCCGATCGCACG | >hg38_21_+_8433172_8446622_RNA45SN1_45S_50ntFlanks | 3984 | 4024 | 41 | GCGGCTTTGGTGACTCTAGATAACCTCGGGCCGATCGCACG | 841.5 | -7.9 | 1.6 | -4.8 | 1.6E-06 | 3.0E-06 | BT20/MB468 | CYTO |
| >MI0000239\|hsa-mir-197&WithFlank&1\|+\|109598887\|109598973@54.72.19&[MIMAT0000227&hsa-miR-197-3p&offsets\|0\|-3;m-99&1\|+\|109598940\|109598961&offsets\|0\|-3]\|TTCACCACCTTCTCCACCC | >MI0000239\|hsa-mir-197&WithFlank&1\|+\|109598887\|109598973 | 54 | 72 | 19 | TTCACCACCTTCTCCACCC | 349.3 | 3.2 | 0.6 | 5.2 | 2.1E-07 | 3.0E-06 | BT20/MB231 | NUC |
| >hg38_21_+_8433172_8446622_RNA45SN1_45S_50ntFlanks@12152.12180.29\|TGGGGAGTTTGACTGGGGCGGTACACCTG | >hg38_21_+_8433172_8446622_RNA45SN1_45S_50ntFlanks | 12152 | 12180 | 29 | TGGGGAGTTTGACTGGGGCGGTACACCTG | 304.7 | -7.0 | 1.4 | -5.0 | 5.2E-07 | 3.0E-06 | BT20/MB468 | MP |
| >MI0000734\|hsa-mir-106b&WithFlank&7\|-\|100093987\|100094080@18.38.21&[MIMAT0000680&hsa-miR-106b-5p&offsets\|0\|0;m-103&7\|-\|100094043\|100094063&offsets\|0\|0]\|TAAAGTGCTGACAGTGCAGAT | >MI0000734\|hsa-mir-106b&WithFlank&7\|-\|100093987\|100094080 | 18 | 38 | 21 | TAAAGTGCTGACAGTGCAGAT | 529.9 | 4.1 | 0.9 | 4.8 | 2.0E-06 | 3.0E-06 | BT20/MB468 | NUC |
| >hg38_21_+_8433172_8446622_RNA45SN1_45S_50ntFlanks@9926.9950.25\|GTTGATATAGACAGCAGGACGGTGG | >hg38_21_+_8433172_8446622_RNA45SN1_45S_50ntFlanks | 9926 | 9950 | 25 | GTTGATATAGACAGCAGGACGGTGG | 54.0 | 9.5 | 1.9 | 5.0 | 5.2E-07 | 3.0E-06 | BT20/MB468 | MP |
| >hg38_MT_+_598_1651_RNR1_12S_50ntFlanks@843.875.33\|GAGTAGAGTGCTTAGTTGAACAGGGCCCTGAAG | >hg38_MT_+_598_1651_RNR1_12S_50ntFlanks | 843 | 875 | 33 | GAGTAGAGTGCTTAGTTGAACAGGGCCCTGAAG | 101.8 | 10.4 | 2.1 | 5.0 | 5.2E-07 | 3.0E-06 | BT20/MB468 | MP |
| >hg38_21_+_8433172_8446622_RNA45SN1_45S_50ntFlanks@13008.13028.21\|ATTGAAAGTCAGCCCTCGACA | >hg38_21_+_8433172_8446622_RNA45SN1_45S_50ntFlanks | 13008 | 13028 | 21 | ATTGAAAGTCAGCCCTCGACA | 905.8 | -4.4 | 0.9 | -4.8 | 1.8E-06 | 3.0E-06 | MB231/MB468 | CYTO |
| >MI0000803\|hsa-mir-330&WithFlank&19\|-\|45638988\|45639093@63.86.24(+1U)&[MIMAT0000751&hsa-miR-330-3p&offsets\|0\|+1(+1U);m-347&19\|-\|45639009\|45639030&offsets\|-1\|+1(+1U)]\|GCAAAGCACACGGCCTGCAGAGAGT | >MI0000803\|hsa-mir-330&WithFlank&19\|-\|45638988\|45639093 | 63 | 86 | 24(+1U) | GCAAAGCACACGGCCTGCAGAGAGT | 287.8 | 3.7 | 0.8 | 4.8 | 1.6E-06 | 3.0E-06 | MB231/MB468 | MP |
| >hg38_21_+_8433172_8446622_RNA45SN1_45S_50ntFlanks@10571.10594.24\|GAGAAGCCGGCGGGAGCCCCGGGG | >hg38_21_+_8433172_8446622_RNA45SN1_45S_50ntFlanks | 10571 | 10594 | 24 | GAGAAGCCGGCGGGAGCCCCGGGG | 56.7 | -8.9 | 1.8 | -5.0 | 5.2E-07 | 3.0E-06 | BT20/MB468 | MP |
| >hg38_21_+_8433172_8446622_RNA45SN1_45S_50ntFlanks@5470.5499.30\|CCACGGCCCTGGCGGAGCGCTGAGAAGACG | >hg38_21_+_8433172_8446622_RNA45SN1_45S_50ntFlanks | 5470 | 5499 | 30 | CCACGGCCCTGGCGGAGCGCTGAGAAGACG | 2027.1 | -8.5 | 1.8 | -4.7 | 2.0E-06 | 3.0E-06 | BT20/MB468 | NUC |
| >hg38_21_+_8433172_8446622_RNA45SN1_45S_50ntFlanks@6647.6676.30\|CGTACGACTCTTAGCGGTGGATCACTCGGC | >hg38_21_+_8433172_8446622_RNA45SN1_45S_50ntFlanks | 6647 | 6676 | 30 | CGTACGACTCTTAGCGGTGGATCACTCGGC | 609.8 | 4.3 | 0.9 | 4.8 | 1.9E-06 | 3.0E-06 | MB231/MB468 | NUC |
| >hg38_21_+_8433172_8446622_RNA45SN1_45S_50ntFlanks@9536.9563.28\|GAAACTCTGGTGGAGGTCCGTAGCGGTC | >hg38_21_+_8433172_8446622_RNA45SN1_45S_50ntFlanks | 9536 | 9563 | 28 | GAAACTCTGGTGGAGGTCCGTAGCGGTC | 303.6 | -5.4 | 1.1 | -4.8 | 1.9E-06 | 3.0E-06 | MB231/MB468 | NUC |
| >hg38_1_-_228634819_228635039_RNA5S12_5S_50ntFlanks@51.97.47\|GTCTACGGCCATACCACCCTGAACGCGCCCGATCTCGTCTGATCTCG | >hg38_1_-_228634819_228635039_RNA5S12_5S_50ntFlanks | 51 | 97 | 47 | GTCTACGGCCATACCACCCTGAACGCGCCCGATCTCGTCTGATCTCG | 229.2 | 6.1 | 1.3 | 4.8 | 1.8E-06 | 3.0E-06 | MB231/MB468 | CYTO |
| >hg38_21_+_8433172_8446622_RNA45SN1_45S_50ntFlanks@8196.8215.20\|CGTGGACGGTGTGAGGCCGG | >hg38_21_+_8433172_8446622_RNA45SN1_45S_50ntFlanks | 8196 | 8215 | 20 | CGTGGACGGTGTGAGGCCGG | 946.4 | -7.0 | 1.5 | -4.8 | 2.0E-06 | 3.0E-06 | MB231/MB468 | MITO |
| >hg38_21_+_8433172_8446622_RNA45SN1_45S_50ntFlanks@4414.4438.25\|CCGCGAGGCGAGCCACCGCCCGTCC | >hg38_21_+_8433172_8446622_RNA45SN1_45S_50ntFlanks | 4414 | 4438 | 25 | CCGCGAGGCGAGCCACCGCCCGTCC | 158.7 | -11.2 | 2.3 | -4.8 | 1.6E-06 | 3.0E-06 | MB231/MB468 | MP |
| >trnaMT_PheGAA_MT_+_577_647@27.53.27\|AATACACTGAAAATGTTTAGACGGGCT | >trnaMT_PheGAA_MT_+_577_647 | 27 | 53 | 27 | AATACACTGAAAATGTTTAGACGGGCT | 89.3 | 10.2 | 2.0 | 5.0 | 5.3E-07 | 3.0E-06 | BT20/MB468 | MP |
| >hg38_21_+_8433172_8446622_RNA45SN1_45S_50ntFlanks@10420.10442.23\|CTGAGAGATGGGCGAGCGCCGTT | >hg38_21_+_8433172_8446622_RNA45SN1_45S_50ntFlanks | 10420 | 10442 | 23 | CTGAGAGATGGGCGAGCGCCGTT | 91.0 | -9.6 | 1.9 | -5.0 | 5.3E-07 | 3.0E-06 | BT20/MB468 | MP |
| >hg38_21_+_8433172_8446622_RNA45SN1_45S_50ntFlanks@9564.9588.25\|CTGACGTGCAAATCGGTCGTCCGAC | >hg38_21_+_8433172_8446622_RNA45SN1_45S_50ntFlanks | 9564 | 9588 | 25 | CTGACGTGCAAATCGGTCGTCCGAC | 55.2 | -8.9 | 1.8 | -5.0 | 5.3E-07 | 3.0E-06 | BT20/MB468 | MP |
| >MI0000285\|hsa-mir-205&WithFlank&1\|+\|209432127\|209432248@40.60.21(+1C)&[MIMAT0000266&hsa-miR-205-5p&offsets\|0\|-1(+1C);m-351&1\|+\|209432166\|209432187&offsets\|0\|-1(+1C)]\|TCCTTCATTCCACCGGAGTCTC | >MI0000285\|hsa-mir-205&WithFlank&1\|+\|209432127\|209432248 | 40 | 60 | 21(+1C) | TCCTTCATTCCACCGGAGTCTC | 127.4 | 10.3 | 2.0 | 5.3 | 1.3E-07 | 3.0E-06 | BT20/MB231 | MP |
| >MI0005566\|hsa-mir-374b&WithFlank&X\|-\|74218541\|74218624@17.38.22(+1C)&[MIMAT0004955&hsa-miR-374b-5p&offsets\|0\|0(+1C);m-153&X\|-\|74218587\|74218608&offsets\|0\|0(+1C)]\|ATATAATACAACCTGCTAAGTGC | >MI0005566\|hsa-mir-374b&WithFlank&X\|-\|74218541\|74218624 | 17 | 38 | 22(+1C) | ATATAATACAACCTGCTAAGTGC | 124.4 | -10.4 | 2.0 | -5.3 | 1.3E-07 | 3.0E-06 | BT20/MB231 | MP |
| >hg38_21_+_8433172_8446622_RNA45SN1_45S_50ntFlanks@4073.4095.23\|CGATGGTAGTCGCCGTGCCTACC | >hg38_21_+_8433172_8446622_RNA45SN1_45S_50ntFlanks | 4073 | 4095 | 23 | CGATGGTAGTCGCCGTGCCTACC | 60.1 | -9.0 | 1.8 | -5.0 | 5.3E-07 | 3.0E-06 | BT20/MB468 | MP |
| >hg38_21_+_8433172_8446622_RNA45SN1_45S_50ntFlanks@7973.7993.21\|GACGCGACCTCAGATCAGACG | >hg38_21_+_8433172_8446622_RNA45SN1_45S_50ntFlanks | 7973 | 7993 | 21 | GACGCGACCTCAGATCAGACG | 510.5 | 3.5 | 0.7 | 4.8 | 1.8E-06 | 3.0E-06 | MB231/MB468 | CYTO |
| >hg38_21_+_8433172_8446622_RNA45SN1_45S_50ntFlanks@8198.8223.26\|TGGACGGTGTGAGGCCGGTAGCGGCC | >hg38_21_+_8433172_8446622_RNA45SN1_45S_50ntFlanks | 8198 | 8223 | 26 | TGGACGGTGTGAGGCCGGTAGCGGCC | 687.7 | -4.7 | 1.0 | -4.8 | 1.8E-06 | 3.0E-06 | MB231/MB468 | CYTO |
| >hg38_21_+_8433172_8446622_RNA45SN1_45S_50ntFlanks@12936.12960.25\|AAACCATTCGTAGACGACCTGCTTC | >hg38_21_+_8433172_8446622_RNA45SN1_45S_50ntFlanks | 12936 | 12960 | 25 | AAACCATTCGTAGACGACCTGCTTC | 362.3 | -3.4 | 0.7 | -4.8 | 2.0E-06 | 3.0E-06 | MB231/MB468 | MITO |
| >hg38_21_+_8433172_8446622_RNA45SN1_45S_50ntFlanks@3900.3919.20\|CTTCGCGGGGGGGATGCGTG | >hg38_21_+_8433172_8446622_RNA45SN1_45S_50ntFlanks | 3900 | 3919 | 20 | CTTCGCGGGGGGGATGCGTG | 302.0 | -2.6 | 0.5 | -4.8 | 1.6E-06 | 3.0E-06 | BT20/MB468 | CYTO |
| >hg38_21_+_8433172_8446622_RNA45SN1_45S_50ntFlanks@3837.3860.24\|CCTCTCCTACTTGGATAACTGTGG | >hg38_21_+_8433172_8446622_RNA45SN1_45S_50ntFlanks | 3837 | 3860 | 24 | CCTCTCCTACTTGGATAACTGTGG | 236.8 | -7.3 | 1.5 | -4.8 | 1.6E-06 | 3.0E-06 | MB231/MB468 | MP |
| >hg38_21_+_8433172_8446622_RNA45SN1_45S_50ntFlanks@4159.4178.20\|ACCACATCCAAGGAAGGCAG | >hg38_21_+_8433172_8446622_RNA45SN1_45S_50ntFlanks | 4159 | 4178 | 20 | ACCACATCCAAGGAAGGCAG | 63.0 | -9.1 | 1.8 | -5.0 | 5.3E-07 | 3.0E-06 | BT20/MB468 | MP |
| >hg38_21_+_8433172_8446622_RNA45SN1_45S_50ntFlanks@3782.3814.33\|CAGTGAAACTGCGAATGGCTCATTAAATCAGTT | >hg38_21_+_8433172_8446622_RNA45SN1_45S_50ntFlanks | 3782 | 3814 | 33 | CAGTGAAACTGCGAATGGCTCATTAAATCAGTT | 213.2 | -7.5 | 1.6 | -4.8 | 1.6E-06 | 3.0E-06 | MB231/MB468 | MP |
| >hg38_21_+_8433172_8446622_RNA45SN1_45S_50ntFlanks@11930.11958.29\|TAGAATAAGTGGGAGGCCCCCGGCGCCCC | >hg38_21_+_8433172_8446622_RNA45SN1_45S_50ntFlanks | 11930 | 11958 | 29 | TAGAATAAGTGGGAGGCCCCCGGCGCCCC | 52.1 | -8.8 | 1.8 | -5.0 | 5.3E-07 | 3.0E-06 | BT20/MB468 | MP |
| >hg38_MT_+_598_1651_RNR1_12S_50ntFlanks@429.469.41\|AAGTGGCTTTAACATATCTGAACACACAATAGCTAAGACCC | >hg38_MT_+_598_1651_RNR1_12S_50ntFlanks | 429 | 469 | 41 | AAGTGGCTTTAACATATCTGAACACACAATAGCTAAGACCC | 40.0 | 8.5 | 1.8 | 4.8 | 1.6E-06 | 3.0E-06 | MB231/MB468 | MP |
| >hg38_21_+_8433172_8446622_RNA45SN1_45S_50ntFlanks@8196.8219.24\|CGTGGACGGTGTGAGGCCGGTAGC | >hg38_21_+_8433172_8446622_RNA45SN1_45S_50ntFlanks | 8196 | 8219 | 24 | CGTGGACGGTGTGAGGCCGGTAGC | 400.7 | 2.5 | 0.5 | 4.8 | 1.6E-06 | 3.0E-06 | BT20/MB468 | CYTO |
| >hg38_21_+_8433172_8446622_RNA45SN1_45S_50ntFlanks@12161.12181.21\|TGACTGGGGCGGTACACCTGT | >hg38_21_+_8433172_8446622_RNA45SN1_45S_50ntFlanks | 12161 | 12181 | 21 | TGACTGGGGCGGTACACCTGT | 213.9 | -6.1 | 1.3 | -4.8 | 1.4E-06 | 3.0E-06 | BT20/MB468 | MITO |
| >hg38_21_+_8433172_8446622_RNA45SN1_45S_50ntFlanks@6660.6688.29\|GCGGTGGATCACTCGGCTCGTGCGTCGAT | >hg38_21_+_8433172_8446622_RNA45SN1_45S_50ntFlanks | 6660 | 6688 | 29 | GCGGTGGATCACTCGGCTCGTGCGTCGAT | 389.5 | -3.9 | 0.8 | -4.8 | 2.0E-06 | 3.0E-06 | MB231/MB468 | MITO |
| >hg38_21_+_8433172_8446622_RNA45SN1_45S_50ntFlanks@8170.8190.21\|CAAGTCCTTCTGATCGAGGCC | >hg38_21_+_8433172_8446622_RNA45SN1_45S_50ntFlanks | 8170 | 8190 | 21 | CAAGTCCTTCTGATCGAGGCC | 59.6 | -9.0 | 1.8 | -5.0 | 5.3E-07 | 3.0E-06 | BT20/MB468 | MP |
| >MI0000086\|hsa-mir-28&WithFlank&3\|+\|188688775\|188688872@60.80.21(+1G)&[MIMAT0004502&hsa-miR-28-3p&offsets\|0\|-1(+1G);m-28&3\|+\|188688834\|188688855&offsets\|0\|-1(+1G)]\|CACTAGATTGTGAGCTCCTGGG | >MI0000086\|hsa-mir-28&WithFlank&3\|+\|188688775\|188688872 | 60 | 80 | 21(+1G) | CACTAGATTGTGAGCTCCTGGG | 47.6 | 8.8 | 1.8 | 4.8 | 1.6E-06 | 3.0E-06 | MB231/MB468 | MP |

<table>
<tr><td>>hg38_21_+_8433172_8446622_RNA45SN1_45S_50ntFlanks@4367.4394.28|CAGTTAAAAAGCTCGTAGTTGGATCTTG</td><td>>hg38_21_+_8433172_8446622_RNA45SN1_45S_50ntFlanks</td><td>4367</td><td>4394</td><td>28</td><td>CAGTTAAAAAGCTCGTAGTTGGATCTTG</td><td>199.6</td><td>-6.9</td><td>1.4</td><td>-5.0</td><td>5.3E-07</td><td>3.0E-06</td><td>BT20/MB468</td><td>MP</td></tr>
<tr><td>>hg38_21_+_8433172_8446622_RNA45SN1_45S_50ntFlanks@3815.3837.23|ATGGTTCCTTTGGTCGCTCGCTC</td><td>>hg38_21_+_8433172_8446622_RNA45SN1_45S_50ntFlanks</td><td>3815</td><td>3837</td><td>23</td><td>ATGGTTCCTTTGGTCGCTCGCTC</td><td>143.2</td><td>-6.6</td><td>1.4</td><td>-4.8</td><td>1.6E-06</td><td>3.0E-06</td><td>MB231/MB468</td><td>MP</td></tr>
<tr><td>>hg38_21_+_8433172_8446622_RNA45SN1_45S_50ntFlanks@12274.12307.34|CAGTACGAATACAGACCGTGAAAGCGGGGCCTCA</td><td>>hg38_21_+_8433172_8446622_RNA45SN1_45S_50ntFlanks</td><td>12274</td><td>12307</td><td>34</td><td>CAGTACGAATACAGACCGTGAAAGCGGGGCCTCA</td><td>88.8</td><td>-8.6</td><td>1.8</td><td>-4.8</td><td>1.4E-06</td><td>3.0E-06</td><td>BT20/MB468</td><td>MITO</td></tr>
<tr><td>>hg38_21_+_8433172_8446622_RNA45SN1_45S_50ntFlanks@4588.4615.28|CTATTTTGTTGGTTTTCGGAACTGAGGC</td><td>>hg38_21_+_8433172_8446622_RNA45SN1_45S_50ntFlanks</td><td>4588</td><td>4615</td><td>28</td><td>CTATTTTGTTGGTTTTCGGAACTGAGGC</td><td>727.9</td><td>-7.7</td><td>1.6</td><td>-4.7</td><td>2.1E-06</td><td>3.0E-06</td><td>BT20/MB468</td><td>NUC</td></tr>
<tr><td>>hg38_21_+_8433172_8446622_RNA45SN1_45S_50ntFlanks@3831.3872.42|CTCGCTCCTCTCCTACTTGGATAACTGTGGTAATTCTAGAGC</td><td>>hg38_21_+_8433172_8446622_RNA45SN1_45S_50ntFlanks</td><td>3831</td><td>3872</td><td>42</td><td>CTCGCTCCTCTCCTACTTGGATAACTGTGGTAATTCTAGAGC</td><td>258.5</td><td>-4.2</td><td>0.9</td><td>-4.8</td><td>1.9E-06</td><td>3.0E-06</td><td>MB231/MB468</td><td>NUC</td></tr>
<tr><td>>hg38_21_+_8433172_8446622_RNA45SN1_45S_50ntFlanks@9561.9596.36|GTCCTGACGTGCAAATCGGTCGTCCGACCTGGGTAT</td><td>>hg38_21_+_8433172_8446622_RNA45SN1_45S_50ntFlanks</td><td>9561</td><td>9596</td><td>36</td><td>GTCCTGACGTGCAAATCGGTCGTCCGACCTGGGTAT</td><td>301.5</td><td>-3.8</td><td>0.8</td><td>-4.8</td><td>1.9E-06</td><td>3.0E-06</td><td>MB231/MB468</td><td>NUC</td></tr>
<tr><td>>trna12_ValAAC_5_-_180645270_180645342@1.34.34,trna132_ValAAC_6_-_27721179_27721251@1.34.34,trna136_ValAAC_6_-_27648885_27648957@1.34.34,trna139_ValAAC_6_-_27618707_27618779@1.34.34,trna2_ValAAC_3_+_169490018_169490090@1.34.34,trna4_ValAAC_5_+_180591154_180591226@1.34.34,trna5_ValAAC_5_+_180596610_180596682@1.34.34|GTTTCCGTAGTGTAGTGGTTATCACGTTCGCCTA</td><td>>trna12_ValAAC_5_-_180645270_180645342</td><td>1</td><td>34</td><td>34</td><td>GTTTCCGTAGTGTAGTGGTTATCACGTTCGCCTA</td><td>857.2</td><td>13.7</td><td>2.9</td><td>4.8</td><td>1.9E-06</td><td>3.0E-06</td><td>MB231/MB468</td><td>CYTO</td></tr>
<tr><td>>hg38_21_+_8433172_8446622_RNA45SN1_45S_50ntFlanks@10348.10368.21|AAACGAGAACTTTGAAGGCCG</td><td>>hg38_21_+_8433172_8446622_RNA45SN1_45S_50ntFlanks</td><td>10348</td><td>10368</td><td>21</td><td>AAACGAGAACTTTGAAGGCCG</td><td>1357.2</td><td>-3.6</td><td>0.7</td><td>-4.8</td><td>1.9E-06</td><td>3.0E-06</td><td>MB231/MB468</td><td>CYTO</td></tr>
<tr><td>>hg38_21_+_8433172_8446622_RNA45SN1_45S_50ntFlanks@11586.11611.26|GTGCGGACCAGGGGAATCCGACTGTT</td><td>>hg38_21_+_8433172_8446622_RNA45SN1_45S_50ntFlanks</td><td>11586</td><td>11611</td><td>26</td><td>GTGCGGACCAGGGGAATCCGACTGTT</td><td>2398.8</td><td>-3.3</td><td>0.7</td><td>-4.8</td><td>1.6E-06</td><td>3.0E-06</td><td>BT20/MB468</td><td>CYTO</td></tr>
<tr><td>>MI0001448|hsa-mir-425&WithFlank&3|-|49020142|49020240@20.43.24&[MIMAT0003393&hsa-miR-425-5p&offsets|0|+1;m-95&3|-|49020200|49020221&offsets|0|+2]|AATGACACGATCACTCCCGTTGAG</td><td>>MI0001448|hsa-mir-425&WithFlank&3|-|49020142|49020240</td><td>20</td><td>43</td><td>24</td><td>AATGACACGATCACTCCCGTTGAG</td><td>365.1</td><td>1.8</td><td>0.4</td><td>4.7</td><td>2.1E-06</td><td>3.0E-06</td><td>BT20/MB468</td><td>NUC</td></tr>
<tr><td>>MI0000542|hsa-mir-320a&WithFlank&8|-|22244960|22245043@48.71.24&[MIMAT0000510&hsa-miR-320a-3p&offsets|0|+2;m-60&8|-|22244975|22244996&offsets|0|+2]|AAAAGCTGGGTTGAGAGGGCGAAA</td><td>>MI0000542|hsa-mir-320a&WithFlank&8|-|22244960|22245043</td><td>48</td><td>71</td><td>24</td><td>AAAAGCTGGGTTGAGAGGGCGAAA</td><td>10259.6</td><td>4.3</td><td>0.8</td><td>5.3</td><td>1.5E-07</td><td>3.0E-06</td><td>BT20/MB231</td><td>CYTO</td></tr>
<tr><td>>hg38_21_+_8433172_8446622_RNA45SN1_45S_50ntFlanks@6646.6685.40|TCGTACGACTCTTAGCGGTGGATCACTCGGCTCGTGCGTC</td><td>>hg38_21_+_8433172_8446622_RNA45SN1_45S_50ntFlanks</td><td>6646</td><td>6685</td><td>40</td><td>TCGTACGACTCTTAGCGGTGGATCACTCGGCTCGTGCGTC</td><td>6709.1</td><td>4.5</td><td>0.9</td><td>5.3</td><td>1.5E-07</td><td>3.0E-06</td><td>BT20/MB231</td><td>CYTO</td></tr>
<tr><td>>hg38_21_+_8433172_8446622_RNA45SN1_45S_50ntFlanks@5075.5097.23|TTCCGATAACGAACGAGACTCTG</td><td>>hg38_21_+_8433172_8446622_RNA45SN1_45S_50ntFlanks</td><td>5075</td><td>5097</td><td>23</td><td>TTCCGATAACGAACGAGACTCTG</td><td>206.5</td><td>-11.6</td><td>2.4</td><td>-4.8</td><td>1.6E-06</td><td>3.0E-06</td><td>MB231/MB468</td><td>MP</td></tr>
<tr><td>>hg38_21_+_8433172_8446622_RNA45SN1_45S_50ntFlanks@7989.8016.28|AGACGTGGCGACCCGCTGAATTTAAGCA</td><td>>hg38_21_+_8433172_8446622_RNA45SN1_45S_50ntFlanks</td><td>7989</td><td>8016</td><td>28</td><td>AGACGTGGCGACCCGCTGAATTTAAGCA</td><td>522.4</td><td>6.0</td><td>1.3</td><td>4.8</td><td>1.6E-06</td><td>3.0E-06</td><td>BT20/MB468</td><td>CYTO</td></tr>
<tr><td>>hg38_21_+_8433172_8446622_RNA45SN1_45S_50ntFlanks@9620.9648.29|ATCTAGTAGCTGGTTCCCTCCGAAGTTTC</td><td>>hg38_21_+_8433172_8446622_RNA45SN1_45S_50ntFlanks</td><td>9620</td><td>9648</td><td>29</td><td>ATCTAGTAGCTGGTTCCCTCCGAAGTTTC</td><td>54.5</td><td>-8.8</td><td>1.8</td><td>-5.0</td><td>5.4E-07</td><td>3.0E-06</td><td>BT20/MB468</td><td>MP</td></tr>
<tr><td>>hg38_21_+_8433172_8446622_RNA45SN1_45S_50ntFlanks@10853.10873.21|ATAAGGATTGGCTCTAAGGGC</td><td>>hg38_21_+_8433172_8446622_RNA45SN1_45S_50ntFlanks</td><td>10853</td><td>10873</td><td>21</td><td>ATAAGGATTGGCTCTAAGGGC</td><td>178.7</td><td>10.8</td><td>2.1</td><td>5.3</td><td>1.3E-07</td><td>3.0E-06</td><td>BT20/MB231</td><td>MP</td></tr>
<tr><td>>hg38_21_+_8433172_8446622_RNA45SN1_45S_50ntFlanks@10771.10818.48|CAGGTCTCCAAGGTGAACAGCCTCTGGCATGTTGGAACAATGTAGGTA</td><td>>hg38_21_+_8433172_8446622_RNA45SN1_45S_50ntFlanks</td><td>10771</td><td>10818</td><td>48</td><td>CAGGTCTCCAAGGTGAACAGCCTCTGGCATGTTGGAACAATGTAGGTA</td><td>165.0</td><td>-11.2</td><td>2.3</td><td>-4.8</td><td>1.6E-06</td><td>3.0E-06</td><td>MB231/MB468</td><td>MP</td></tr>
<tr><td>>MI0000061|hsa-let-7a-2&WithFlank&11|-|122146516|122146599@11.29.19;MI0000062|hsa-let-7a-3&WithFlank&22|+|46112743|46112828@10.28.19;MI0000060|hsa-let-7a-1&WithFlank&9|+|94175951|94176042@12.30.19&[MIMAT0000062&hsa-let-7a-5p&offsets|0|-3;m-6&11|-|122146568|122146589&offsets|0|-3];[MIMAT0000062_1&hsa-let-7a-5p&offsets|0|-3;m-5&22|+|46112752|46112773&offsets|0|-3];[MIMAT0000062_2&hsa-let-7a-5p&offsets|0|-3;m-4&9|+|94175962|94175983&offsets|0|-3]|TGAGGTAGTAGGTTGTATA</td><td>>MI0000061|hsa-let-7a-2&WithFlank&11|-|122146516|122146599</td><td>11</td><td>29</td><td>19</td><td>TGAGGTAGTAGGTTGTATA</td><td>264.1</td><td>2.2</td><td>0.5</td><td>4.7</td><td>2.1E-06</td><td>3.0E-06</td><td>BT20/MB468</td><td>NUC</td></tr>
<tr><td>>hg38_21_+_8433172_8446622_RNA45SN1_45S_50ntFlanks@9623.9649.27|TAGTAGCTGGTTCCCTCCGAAGTTTCC</td><td>>hg38_21_+_8433172_8446622_RNA45SN1_45S_50ntFlanks</td><td>9623</td><td>9649</td><td>27</td><td>TAGTAGCTGGTTCCCTCCGAAGTTTCC</td><td>262.3</td><td>-7.0</td><td>1.5</td><td>-4.7</td><td>2.1E-06</td><td>3.0E-06</td><td>BT20/MB468</td><td>NUC</td></tr>
<tr><td>>hg38_21_+_8433172_8446622_RNA45SN1_45S_50ntFlanks@6651.6691.41|CGACTCTTAGCGGTGGATCACTCGGCTCGTGCGTCGATGAA</td><td>>hg38_21_+_8433172_8446622_RNA45SN1_45S_50ntFlanks</td><td>6651</td><td>6691</td><td>41</td><td>CGACTCTTAGCGGTGGATCACTCGGCTCGTGCGTCGATGAA</td><td>1261.4</td><td>3.1</td><td>0.6</td><td>4.8</td><td>1.4E-06</td><td>3.1E-06</td><td>BT20/MB468</td><td>MITO</td></tr>
<tr><td>>hg38_21_+_8433172_8446622_RNA45SN1_45S_50ntFlanks@5497.5514.18|ACGGTCGAACTTGACTAT</td><td>>hg38_21_+_8433172_8446622_RNA45SN1_45S_50ntFlanks</td><td>5497</td><td>5514</td><td>18</td><td>ACGGTCGAACTTGACTAT</td><td>138.2</td><td>-11.0</td><td>2.3</td><td>-4.8</td><td>1.6E-06</td><td>3.1E-06</td><td>MB231/MB468</td><td>MP</td></tr>
<tr><td>>hg38_21_+_8433172_8446622_RNA45SN1_45S_50ntFlanks@8179.8206.28|CTGATCGAGGCCCAGCCCGTGGACGGTG</td><td>>hg38_21_+_8433172_8446622_RNA45SN1_45S_50ntFlanks</td><td>8179</td><td>8206</td><td>28</td><td>CTGATCGAGGCCCAGCCCGTGGACGGTG</td><td>320.9</td><td>-6.7</td><td>1.4</td><td>-4.8</td><td>1.9E-06</td><td>3.1E-06</td><td>MB231/MB468</td><td>NUC</td></tr>
<tr><td>>hg38_21_+_8433172_8446622_RNA45SN1_45S_50ntFlanks@11588.11611.24|GCGGACCAGGGGAATCCGACTGTT</td><td>>hg38_21_+_8433172_8446622_RNA45SN1_45S_50ntFlanks</td><td>11588</td><td>11611</td><td>24</td><td>GCGGACCAGGGGAATCCGACTGTT</td><td>280.9</td><td>2.4</td><td>0.5</td><td>4.8</td><td>1.7E-06</td><td>3.1E-06</td><td>BT20/MB468</td><td>CYTO</td></tr>
<tr><td>>hg38_21_+_8433172_8446622_RNA45SN1_45S_50ntFlanks@12420.12443.24|ATGTCGGCTCTTCCTATCATTGTG</td><td>>hg38_21_+_8433172_8446622_RNA45SN1_45S_50ntFlanks</td><td>12420</td><td>12443</td><td>24</td><td>ATGTCGGCTCTTCCTATCATTGTG</td><td>5082.4</td><td>-3.6</td><td>0.8</td><td>-4.8</td><td>1.7E-06</td><td>3.1E-06</td><td>BT20/MB468</td><td>CYTO</td></tr>
<tr><td>>hg38_21_+_8433172_8446622_RNA45SN1_45S_50ntFlanks@11316.11334.19|CACCCCACGTCTCGTCGCG</td><td>>hg38_21_+_8433172_8446622_RNA45SN1_45S_50ntFlanks</td><td>11316</td><td>11334</td><td>19</td><td>CACCCCACGTCTCGTCGCG</td><td>1517.8</td><td>-5.8</td><td>1.2</td><td>-4.8</td><td>1.7E-06</td><td>3.1E-06</td><td>BT20/MB468</td><td>CYTO</td></tr>
<tr><td>>hg38_21_+_8433172_8446622_RNA45SN1_45S_50ntFlanks@4588.4613.26|CTATTTTGTTGGTTTTCGGAACTGAG</td><td>>hg38_21_+_8433172_8446622_RNA45SN1_45S_50ntFlanks</td><td>4588</td><td>4613</td><td>26</td><td>CTATTTTGTTGGTTTTCGGAACTGAG</td><td>184.6</td><td>-5.8</td><td>1.2</td><td>-4.8</td><td>1.7E-06</td><td>3.1E-06</td><td>BT20/MB468</td><td>CYTO</td></tr>
<tr><td>>hg38_21_+_8433172_8446622_RNA45SN1_45S_50ntFlanks@9493.9525.33|CCCGAAAGATGGTGAACTATGCCTGGGCAGGGC</td><td>>hg38_21_+_8433172_8446622_RNA45SN1_45S_50ntFlanks</td><td>9493</td><td>9525</td><td>33</td><td>CCCGAAAGATGGTGAACTATGCCTGGGCAGGGC</td><td>291.9</td><td>11.5</td><td>2.2</td><td>5.3</td><td>1.4E-07</td><td>3.1E-06</td><td>BT20/MB231</td><td>MP</td></tr>
<tr><td>>hg38_21_+_8433172_8446622_RNA45SN1_45S_50ntFlanks@6744.6771.28|GACACTTCGAACGCACTTGCGGCCCCGG</td><td>>hg38_21_+_8433172_8446622_RNA45SN1_45S_50ntFlanks</td><td>6744</td><td>6771</td><td>28</td><td>GACACTTCGAACGCACTTGCGGCCCCGG</td><td>519.6</td><td>-5.3</td><td>1.1</td><td>-4.8</td><td>1.9E-06</td><td>3.1E-06</td><td>MB231/MB468</td><td>NUC</td></tr>
<tr><td>>hg38_21_+_8433172_8446622_RNA45SN1_45S_50ntFlanks@6655.6673.19|TCTTAGCGGTGGATCACTC</td><td>>hg38_21_+_8433172_8446622_RNA45SN1_45S_50ntFlanks</td><td>6655</td><td>6673</td><td>19</td><td>TCTTAGCGGTGGATCACTC</td><td>52.8</td><td>9.5</td><td>1.9</td><td>5.0</td><td>5.4E-07</td><td>3.1E-06</td><td>BT20/MB468</td><td>MP</td></tr>
<tr><td>>hg38_21_+_8433172_8446622_RNA45SN1_45S_50ntFlanks@11795.11819.25|AACGAGATTCCCACTGTCCCTACCT</td><td>>hg38_21_+_8433172_8446622_RNA45SN1_45S_50ntFlanks</td><td>11795</td><td>11819</td><td>25</td><td>AACGAGATTCCCACTGTCCCTACCT</td><td>248.8</td><td>-5.2</td><td>1.1</td><td>-4.8</td><td>2.0E-06</td><td>3.1E-06</td><td>MB231/MB468</td><td>MITO</td></tr>
<tr><td>>hg38_21_+_8433172_8446622_RNA45SN1_45S_50ntFlanks@6723.6743.21|TTGCAGGACACATTGATCATC</td><td>>hg38_21_+_8433172_8446622_RNA45SN1_45S_50ntFlanks</td><td>6723</td><td>6743</td><td>21</td><td>TTGCAGGACACATTGATCATC</td><td>1310.8</td><td>-8.4</td><td>1.8</td><td>-4.7</td><td>2.1E-06</td><td>3.1E-06</td><td>BT20/MB468</td><td>NUC</td></tr>
<tr><td>>MI0000342|hsa-mir-200b&WithFlank&1|+|1167098|1167204@27.48.22&[MIMAT0004571&hsa-miR-200b-5p&offsets|0|0;m-312&1|+|1167124|1167145&offsets|0|0]|CATCTTACTGGGCAGCATTGGA</td><td>>MI0000342|hsa-mir-200b&WithFlank&1|+|1167098|1167204</td><td>27</td><td>48</td><td>22</td><td>CATCTTACTGGGCAGCATTGGA</td><td>51.9</td><td>-8.8</td><td>1.8</td><td>-5.0</td><td>5.4E-07</td><td>3.1E-06</td><td>BT20/MB468</td><td>MP</td></tr>
<tr><td>>hg38_21_+_8433172_8446622_RNA45SN1_45S_50ntFlanks@12626.12647.22|AATGGGGCGAAGCTACCATCTG</td><td>>hg38_21_+_8433172_8446622_RNA45SN1_45S_50ntFlanks</td><td>12626</td><td>12647</td><td>22</td><td>AATGGGGCGAAGCTACCATCTG</td><td>62.7</td><td>-9.1</td><td>1.8</td><td>-5.0</td><td>5.4E-07</td><td>3.1E-06</td><td>BT20/MB468</td><td>MP</td></tr>
<tr><td>>hg38_21_+_8433172_8446622_RNA45SN1_45S_50ntFlanks@12430.12455.26|TTCCTATCATTGTGAAGCAGAATTCA</td><td>>hg38_21_+_8433172_8446622_RNA45SN1_45S_50ntFlanks</td><td>12430</td><td>12455</td><td>26</td><td>TTCCTATCATTGTGAAGCAGAATTCA</td><td>52.3</td><td>-8.8</td><td>1.8</td><td>-5.0</td><td>5.4E-07</td><td>3.1E-06</td><td>BT20/MB468</td><td>MP</td></tr>
<tr><td>>hg38_21_+_8433172_8446622_RNA45SN1_45S_50ntFlanks@5080.5120.41|ATAACGAACGAGACTCTGGCATGCTAACTAGTTACGCGACC</td><td>>hg38_21_+_8433172_8446622_RNA45SN1_45S_50ntFlanks</td><td>5080</td><td>5120</td><td>41</td><td>ATAACGAACGAGACTCTGGCATGCTAACTAGTTACGCGACC</td><td>141.3</td><td>-11.0</td><td>2.3</td><td>-4.8</td><td>1.7E-06</td><td>3.1E-06</td><td>MB231/MB468</td><td>MP</td></tr>
<tr><td>>hg38_21_+_8433172_8446622_RNA45SN1_45S_50ntFlanks@4557.4577.21|CAGCTAGGAATAATGGAATAG</td><td>>hg38_21_+_8433172_8446622_RNA45SN1_45S_50ntFlanks</td><td>4557</td><td>4577</td><td>21</td><td>CAGCTAGGAATAATGGAATAG</td><td>50.7</td><td>-8.7</td><td>1.7</td><td>-5.0</td><td>5.5E-07</td><td>3.1E-06</td><td>BT20/MB468</td><td>MP</td></tr>
<tr><td>>hg38_21_+_8433172_8446622_RNA45SN1_45S_50ntFlanks@5527.5548.22|AAAGTCGTAACAAGGTTTCCGT</td><td>>hg38_21_+_8433172_8446622_RNA45SN1_45S_50ntFlanks</td><td>5527</td><td>5548</td><td>22</td><td>AAAGTCGTAACAAGGTTTCCGT</td><td>385.5</td><td>-7.5</td><td>1.6</td><td>-4.7</td><td>2.1E-06</td><td>3.1E-06</td><td>BT20/MB468</td><td>NUC</td></tr>
<tr><td>>hg38_21_+_8433172_8446622_RNA45SN1_45S_50ntFlanks@12001.12032.32|CTGCGGGCCGCCGGTGAAATACCACTACTCTG</td><td>>hg38_21_+_8433172_8446622_RNA45SN1_45S_50ntFlanks</td><td>12001</td><td>12032</td><td>32</td><td>CTGCGGGCCGCCGGTGAAATACCACTACTCTG</td><td>4339.2</td><td>-6.6</td><td>1.4</td><td>-4.8</td><td>2.0E-06</td><td>3.1E-06</td><td>MB231/MB468</td><td>NUC</td></tr>
<tr><td>>MI0000471|hsa-mir-126&WithFlank&9|+|136670596|136670692@58.78.21(+1C)&[MIMAT0000445&hsa-miR-126-3p&offsets|0|-1(+1C);m-40&9|+|136670653|136670674&offsets|0|-1(+1C)]|TCGTACCGTGAGTAATAATGCC</td><td>>MI0000471|hsa-mir-126&WithFlank&9|+|136670596|136670692</td><td>58</td><td>78</td><td>21(+1C)</td><td>TCGTACCGTGAGTAATAATGCC</td><td>57.4</td><td>9.1</td><td>1.9</td><td>4.8</td><td>1.7E-06</td><td>3.1E-06</td><td>MB231/MB468</td><td>MP</td></tr>
<tr><td>>hg38_21_+_8433172_8446622_RNA45SN1_45S_50ntFlanks@12071.12096.26|CCCCGAGGGGCTCTCGCTTCTGGCGC</td><td>>hg38_21_+_8433172_8446622_RNA45SN1_45S_50ntFlanks</td><td>12071</td><td>12096</td><td>26</td><td>CCCCGAGGGGCTCTCGCTTCTGGCGC</td><td>267.7</td><td>-5.5</td><td>1.2</td><td>-4.8</td><td>2.0E-06</td><td>3.1E-06</td><td>MB231/MB468</td><td>NUC</td></tr>
<tr><td>>hg38_21_+_8433172_8446622_RNA45SN1_45S_50ntFlanks@3921.3959.39|ATTTATCAGATCAAAACCAACCCGGTCAGCCCCTCTCCG</td><td>>hg38_21_+_8433172_8446622_RNA45SN1_45S_50ntFlanks</td><td>3921</td><td>3959</td><td>39</td><td>ATTTATCAGATCAAAACCAACCCGGTCAGCCCCTCTCCG</td><td>309.7</td><td>-6.0</td><td>1.3</td><td>-4.8</td><td>1.9E-06</td><td>3.1E-06</td><td>MB231/MB468</td><td>CYTO</td></tr>
<tr><td>>hg38_21_+_8433172_8446622_RNA45SN1_45S_50ntFlanks@11483.11502.20|CGGAGGGGCCGCGGGCCGGT</td><td>>hg38_21_+_8433172_8446622_RNA45SN1_45S_50ntFlanks</td><td>11483</td><td>11502</td><td>20</td><td>CGGAGGGGCCGCGGGCCGGT</td><td>69.2</td><td>-9.2</td><td>1.8</td><td>-5.0</td><td>5.5E-07</td><td>3.1E-06</td><td>BT20/MB468</td><td>MP</td></tr>
<tr><td>>MI0000816|hsa-mir-335&WithFlank&7|+|130496105|130496210@58.78.21&[MIMAT0004703&hsa-miR-335-3p&offsets|0|-1;m-159&7|+|130496162|130496183&offsets|0|-1]|TTTTTCATTATTGCTCCTGAC</td><td>>MI0000816|hsa-mir-335&WithFlank&7|+|130496105|130496210</td><td>58</td><td>78</td><td>21</td><td>TTTTTCATTATTGCTCCTGAC</td><td>424.8</td><td>8.1</td><td>1.7</td><td>4.8</td><td>2.0E-06</td><td>3.1E-06</td><td>MB231/MB468</td><td>MITO</td></tr>
<tr><td>>hg38_21_+_8433172_8446622_RNA45SN1_45S_50ntFlanks@12997.13037.41|CGCTGCGATCTATTGAAAGTCAGCCCTCGACACAAGGGTTT</td><td>>hg38_21_+_8433172_8446622_RNA45SN1_45S_50ntFlanks</td><td>12997</td><td>13037</td><td>41</td><td>CGCTGCGATCTATTGAAAGTCAGCCCTCGACACAAGGGTTT</td><td>258.6</td><td>6.2</td><td>1.3</td><td>4.8</td><td>2.0E-06</td><td>3.1E-06</td><td>MB231/MB468</td><td>MITO</td></tr>
<tr><td>>hg38_21_+_8433172_8446622_RNA45SN1_45S_50ntFlanks@7984.8004.21|AGATCAGACGTGGCGACCCGC</td><td>>hg38_21_+_8433172_8446622_RNA45SN1_45S_50ntFlanks</td><td>7984</td><td>8004</td><td>21</td><td>AGATCAGACGTGGCGACCCGC</td><td>812.8</td><td>-3.4</td><td>0.7</td><td>-4.8</td><td>1.5E-06</td><td>3.1E-06</td><td>BT20/MB468</td><td>MITO</td></tr>
<tr><td>>hg38_21_+_8433172_8446622_RNA45SN1_45S_50ntFlanks@9307.9338.32|AAGGAGTCTAACACGTGCGCGAGTCGGGGGCT</td><td>>hg38_21_+_8433172_8446622_RNA45SN1_45S_50ntFlanks</td><td>9307</td><td>9338</td><td>32</td><td>AAGGAGTCTAACACGTGCGCGAGTCGGGGGCT</td><td>2590.0</td><td>-6.2</td><td>1.3</td><td>-4.8</td><td>1.7E-06</td><td>3.1E-06</td><td>MB231/MB468</td><td>MP</td></tr>
<tr><td>>hg38_21_+_8433172_8446622_RNA45SN1_45S_50ntFlanks@10797.10840.44|GCATGTTGGAACAATGTAGGTAAGGGAAGTCGGCAAGCCGGATC</td><td>>hg38_21_+_8433172_8446622_RNA45SN1_45S_50ntFlanks</td><td>10797</td><td>10840</td><td>44</td><td>GCATGTTGGAACAATGTAGGTAAGGGAAGTCGGCAAGCCGGATC</td><td>280.0</td><td>3.3</td><td>0.7</td><td>4.8</td><td>1.9E-06</td><td>3.1E-06</td><td>MB231/MB468</td><td>CYTO</td></tr>
</table>

| | | | | | | | | | | | | | |
|---|---|---|---|---|---|---|---|---|---|---|---|---|---|
| >hg38_21_+_8433172_8446622_RNA45SN1_45S_50ntFlanks@11565.11601.37\|CTAGCAGCCGACTTAGAACTGGTGCGGACCAGGGGAA | >hg38_21_+_8433172_8446622_RNA45SN1_45S_50ntFlanks | 11565 | 11601 | 37 | CTAGCAGCCGACTTAGAACTGGTGCGGACCAGGGGAA | 965.3 | -8.1 | 1.7 | -4.7 | 2.1E-06 | 3.1E-06 | BT20/MB468 | NUC |
| >hg38_21_+_8433172_8446622_RNA45SN1_45S_50ntFlanks@4681.4707.27\|CGCAAGACGGACCAGAGCGAAAGCATT | >hg38_21_+_8433172_8446622_RNA45SN1_45S_50ntFlanks | 4681 | 4707 | 27 | CGCAAGACGGACCAGAGCGAAAGCATT | 102.7 | -5.0 | 1.0 | -4.8 | 1.7E-06 | 3.1E-06 | MB231/MB468 | MP |
| >hg38_21_+_8433172_8446622_RNA45SN1_45S_50ntFlanks@11705.11740.36\|TGAAGCGCGGGTAAACGGCGGGAGTAACTATGACTC | >hg38_21_+_8433172_8446622_RNA45SN1_45S_50ntFlanks | 11705 | 11740 | 36 | TGAAGCGCGGGTAAACGGCGGGAGTAACTATGACTC | 337.9 | -6.4 | 1.3 | -4.8 | 2.0E-06 | 3.1E-06 | MB231/MB468 | NUC |
| >hg38_21_+_8433172_8446622_RNA45SN1_45S_50ntFlanks@5084.5112.29\|CGAACGAGACTCTGGCATGCTAACTAGTT | >hg38_21_+_8433172_8446622_RNA45SN1_45S_50ntFlanks | 5084 | 5112 | 29 | CGAACGAGACTCTGGCATGCTAACTAGTT | 1052.3 | -5.5 | 1.2 | -4.8 | 2.0E-06 | 3.1E-06 | MB231/MB468 | NUC |
| >hg38_21_+_8433172_8446622_RNA45SN1_45S_50ntFlanks@10830.10853.24\|CAAGCCGGATCCGTAACTTCGGGA | >hg38_21_+_8433172_8446622_RNA45SN1_45S_50ntFlanks | 10830 | 10853 | 24 | CAAGCCGGATCCGTAACTTCGGGA | 220.1 | -4.2 | 0.9 | -4.8 | 1.9E-06 | 3.1E-06 | MB231/MB468 | CYTO |
| >hg38_21_+_8433172_8446622_RNA45SN1_45S_50ntFlanks@3815.3837.23\|ATGGTTCCTTTGGTCGCTCGCTC | >hg38_21_+_8433172_8446622_RNA45SN1_45S_50ntFlanks | 3815 | 3837 | 23 | ATGGTTCCTTTGGTCGCTCGCTC | 834.7 | -6.2 | 1.3 | -4.7 | 2.1E-06 | 3.1E-06 | BT20/MB468 | NUC |
| >hg38_MT_+_1621_3279_RNR2_16S_50ntFlanks@728.759.32\|AGATTAAAACACTGAACTGACAATTAACAGCC | >hg38_MT_+_1621_3279_RNR2_16S_50ntFlanks | 728 | 759 | 32 | AGATTAAAACACTGAACTGACAATTAACAGCC | 44.7 | 8.7 | 1.8 | 4.8 | 1.7E-06 | 3.1E-06 | MB231/MB468 | MP |
| >hg38_21_+_8433172_8446622_RNA45SN1_45S_50ntFlanks@12363.12382.20\|GGATAACTGGCTTGTGGCGG | >hg38_21_+_8433172_8446622_RNA45SN1_45S_50ntFlanks | 12363 | 12382 | 20 | GGATAACTGGCTTGTGGCGG | 399.4 | -6.6 | 1.4 | -4.8 | 1.7E-06 | 3.1E-06 | BT20/MB468 | CYTO |
| >hg38_21_+_8433172_8446622_RNA45SN1_45S_50ntFlanks@9554.9596.43\|CGTAGCGGTCCTGACGTGCAAATCGGTCGTCCGACCTGGGTAT | >hg38_21_+_8433172_8446622_RNA45SN1_45S_50ntFlanks | 9554 | 9596 | 43 | CGTAGCGGTCCTGACGTGCAAATCGGTCGTCCGACCTGGGTAT | 1283.0 | -7.6 | 1.6 | -4.8 | 1.5E-06 | 3.1E-06 | BT20/MB468 | MITO |
| >hg38_21_+_8433172_8446622_RNA45SN1_45S_50ntFlanks@9341.9368.28\|CACGAAAGCCGCCGTGGCGCAATGAAGG | >hg38_21_+_8433172_8446622_RNA45SN1_45S_50ntFlanks | 9341 | 9368 | 28 | CACGAAAGCCGCCGTGGCGCAATGAAGG | 63.3 | -9.1 | 1.8 | -5.0 | 5.5E-07 | 3.1E-06 | BT20/MB468 | MP |
| >hg38_21_+_8433172_8446622_RNA45SN1_45S_50ntFlanks@6731.6770.40\|CACATTGATCATCGACACTTCGAACGCACTTGCGGCCCCG | >hg38_21_+_8433172_8446622_RNA45SN1_45S_50ntFlanks | 6731 | 6770 | 40 | CACATTGATCATCGACACTTCGAACGCACTTGCGGCCCCG | 972.4 | -5.0 | 1.1 | -4.8 | 1.9E-06 | 3.1E-06 | MB231/MB468 | CYTO |
| >hg38_21_+_8433172_8446622_RNA45SN1_45S_50ntFlanks@11784.11805.22\|ATGAATGGATGAACGAGATTCC | >hg38_21_+_8433172_8446622_RNA45SN1_45S_50ntFlanks | 11784 | 11805 | 22 | ATGAATGGATGAACGAGATTCC | 283.7 | -5.2 | 1.1 | -4.7 | 2.0E-06 | 3.1E-06 | MB231/MB468 | MITO |
| >hg38_21_+_8433172_8446622_RNA45SN1_45S_50ntFlanks@474.498.25\|TCTTCGTGATCGATGTGGTGACGTC | >hg38_21_+_8433172_8446622_RNA45SN1_45S_50ntFlanks | 474 | 498 | 25 | TCTTCGTGATCGATGTGGTGACGTC | 546.0 | 7.8 | 1.6 | 4.7 | 2.0E-06 | 3.1E-06 | MB231/MB468 | MITO |
| >hg38_21_+_8433172_8446622_RNA45SN1_45S_50ntFlanks@12936.12962.27\|AAACCATTCGTAGACGACCTGCTTCTG | >hg38_21_+_8433172_8446622_RNA45SN1_45S_50ntFlanks | 12936 | 12962 | 27 | AAACCATTCGTAGACGACCTGCTTCTG | 570.5 | -5.4 | 1.1 | -4.8 | 2.0E-06 | 3.1E-06 | MB231/MB468 | NUC |
| >am_trna119_LysCTT_1_-_145395522_145395594@7.34.28,trna11_LysCTT_5_-_180648979_180649051@7.34.28,trna128_LysCTT_1_-_55423542_55423614@7.34.28,trna13_LysCTT_6_+_26556774_26556846@7.34.28,trna6_LysCTT_19_-_52425396_52425468@7.34.28,trna7_LysCTT_16_+_3225692_3225764@7.34.28,trna9_LysCTT_5_+_180634755_180634827@7.34.28\|CTAGCTCAGTCGGTAGAGCATGAGACTC | >am_trna119_LysCTT_1_-_145395522_145395594 | 7 | 34 | 28 | CTAGCTCAGTCGGTAGAGCATGAGACTC | 44.0 | 8.7 | 1.8 | 4.8 | 1.7E-06 | 3.1E-06 | MB231/MB468 | MP |
| >trnaMT_GluTTC_MT_-_14674_14742@21.58.38,trnalookalike8_GluTTC_5_-_93905172_93905240@21.58.38\|ACGATGGTTTTTCATATCATTGGTCGTGGTTGTAGTCC | >trnaMT_GluTTC_MT_-_14674_14742 | 21 | 58 | 38 | ACGATGGTTTTTCATATCATTGGTCGTGGTTGTAGTCC | 877.5 | 9.6 | 1.9 | 5.0 | 5.5E-07 | 3.1E-06 | BT20/MB468 | MP |
| >hg38_21_+_8433172_8446622_RNA45SN1_45S_50ntFlanks@11767.11803.37\|TCTAATTAGTGACGCGCATGAATGGATGAACGAGATT | >hg38_21_+_8433172_8446622_RNA45SN1_45S_50ntFlanks | 11767 | 11803 | 37 | TCTAATTAGTGACGCGCATGAATGGATGAACGAGATT | 399.2 | -3.3 | 0.7 | -4.8 | 1.7E-06 | 3.1E-06 | BT20/MB468 | CYTO |
| >hg38_21_+_8433172_8446622_RNA45SN1_45S_50ntFlanks@6658.6689.32\|TAGCGGTGGATCACTCGGCTCGTGCGTCGATG | >hg38_21_+_8433172_8446622_RNA45SN1_45S_50ntFlanks | 6658 | 6689 | 32 | TAGCGGTGGATCACTCGGCTCGTGCGTCGATG | 263.0 | -6.9 | 1.4 | -4.8 | 1.7E-06 | 3.1E-06 | MB231/MB468 | MP |
| >MI0000085\|hsa-mir-27a&WithFlank&19\|-\|13836434\|13836523@57.75.19(+1A)&[MIMAT0000084&hsa-miR-27a-3p&offsets\|0\|-2(+1A);m-43&19\|-\|13836447\|13836467&offsets\|0\|-2(+1A)]\|TTCACAGTGGCTAAGTTCCA | >MI0000085\|hsa-mir-27a&WithFlank&19\|-\|13836434\|13836523 | 57 | 75 | 19(+1A) | TTCACAGTGGCTAAGTTCCA | 911.1 | 4.9 | 1.0 | 4.8 | 1.9E-06 | 3.1E-06 | MB231/MB468 | CYTO |
| >hg38_21_+_8433172_8446622_RNA45SN1_45S_50ntFlanks@8189.8212.24\|CCCAGCCCGTGGACGGTGTGAGGC | >hg38_21_+_8433172_8446622_RNA45SN1_45S_50ntFlanks | 8189 | 8212 | 24 | CCCAGCCCGTGGACGGTGTGAGGC | 514.0 | -5.8 | 1.2 | -4.8 | 1.9E-06 | 3.1E-06 | MB231/MB468 | CYTO |
| >hg38_21_+_8433172_8446622_RNA45SN1_45S_50ntFlanks@8372.8406.35\|AGAGAGTTCAAGAGGGCGTGAAACCGTTAAGAGGT | >hg38_21_+_8433172_8446622_RNA45SN1_45S_50ntFlanks | 8372 | 8406 | 35 | AGAGAGTTCAAGAGGGCGTGAAACCGTTAAGAGGT | 49.9 | 9.4 | 1.9 | 5.0 | 5.6E-07 | 3.1E-06 | BT20/MB468 | MP |
| >hg38_21_+_8433172_8446622_RNA45SN1_45S_50ntFlanks@11682.11702.21\|AATGTCAAAGTGAAGAAATTC | >hg38_21_+_8433172_8446622_RNA45SN1_45S_50ntFlanks | 11682 | 11702 | 21 | AATGTCAAAGTGAAGAAATTC | 315.1 | -6.9 | 1.4 | -4.7 | 2.2E-06 | 3.1E-06 | BT20/MB468 | NUC |
| >MI0000471\|hsa-mir-126&WithFlank&9\|+\|136670596\|136670692@58.78.21(+1U)&[MIMAT0000445&hsa-miR-126-3p&offsets\|0\|-1(+1U);m-40&9\|+\|136670653\|136670674&offsets\|0\|-1(+1U)]\|TCGTACCGTGAGTAATAATGCT | >MI0000471\|hsa-mir-126&WithFlank&9\|+\|136670596\|136670692 | 58 | 78 | 21(+1U) | TCGTACCGTGAGTAATAATGCT | 76.8 | 10.0 | 2.0 | 5.0 | 5.6E-07 | 3.1E-06 | BT20/MB468 | MP |
| >hg38_21_+_8433172_8446622_RNA45SN1_45S_50ntFlanks@10841.10872.32\|CGTAACTTCGGGATAAGGATTGGCTCTAAGGG | >hg38_21_+_8433172_8446622_RNA45SN1_45S_50ntFlanks | 10841 | 10872 | 32 | CGTAACTTCGGGATAAGGATTGGCTCTAAGGG | 156.1 | -6.9 | 1.4 | -4.8 | 1.7E-06 | 3.1E-06 | MB231/MB468 | MP |
| >hg38_21_+_8433172_8446622_RNA45SN1_45S_50ntFlanks@11778.11795.18\|ACGCGCATGAATGGATGA | >hg38_21_+_8433172_8446622_RNA45SN1_45S_50ntFlanks | 11778 | 11795 | 18 | ACGCGCATGAATGGATGA | 156.1 | -6.1 | 1.3 | -4.8 | 2.0E-06 | 3.1E-06 | MB231/MB468 | NUC |
| >hg38_21_+_8433172_8446622_RNA45SN1_45S_50ntFlanks@6662.6699.38\|GGTGGATCACTCGGCTCGTGCGTCGATGAAGAACGCAG | >hg38_21_+_8433172_8446622_RNA45SN1_45S_50ntFlanks | 6662 | 6699 | 38 | GGTGGATCACTCGGCTCGTGCGTCGATGAAGAACGCAG | 132.5 | 10.4 | 2.0 | 5.3 | 1.4E-07 | 3.2E-06 | BT20/MB231 | MP |
| >hg38_21_+_8433172_8446622_RNA45SN1_45S_50ntFlanks@12914.12932.19\|CGTTCGTGGGGAACCTGGC | >hg38_21_+_8433172_8446622_RNA45SN1_45S_50ntFlanks | 12914 | 12932 | 19 | CGTTCGTGGGGAACCTGGC | 157.4 | -6.7 | 1.4 | -4.8 | 1.7E-06 | 3.2E-06 | MB231/MB468 | MP |
| >hg38_21_+_8433172_8446622_RNA45SN1_45S_50ntFlanks@12457.12488.32\|CAAGCGTTGGATTGTTCACCCACTAATAGGGA | >hg38_21_+_8433172_8446622_RNA45SN1_45S_50ntFlanks | 12457 | 12488 | 32 | CAAGCGTTGGATTGTTCACCCACTAATAGGGA | 419.5 | -5.8 | 1.2 | -4.7 | 2.1E-06 | 3.2E-06 | MB231/MB468 | MITO |
| >hg38_21_+_8433172_8446622_RNA45SN1_45S_50ntFlanks@475.493.19\|CTTCGTGATCGATGTGGTG | >hg38_21_+_8433172_8446622_RNA45SN1_45S_50ntFlanks | 475 | 493 | 19 | CTTCGTGATCGATGTGGTG | 450.9 | 3.6 | 0.8 | 4.8 | 2.0E-06 | 3.2E-06 | MB231/MB468 | NUC |
| >hg38_21_+_8433172_8446622_RNA45SN1_45S_50ntFlanks@6772.6801.30\|GTTCCTCCCGGGGCTACGCCTGTCTGAGCG | >hg38_21_+_8433172_8446622_RNA45SN1_45S_50ntFlanks | 6772 | 6801 | 30 | GTTCCTCCCGGGGCTACGCCTGTCTGAGCG | 330.0 | -5.1 | 1.1 | -4.7 | 2.1E-06 | 3.2E-06 | MB231/MB468 | MITO |
| >hg38_21_+_8433172_8446622_RNA45SN1_45S_50ntFlanks@6721.6770.50\|AATTGCAGGACACATTGATCATCGACACTTCGAACGCACTTGCGGCCCCG | >hg38_21_+_8433172_8446622_RNA45SN1_45S_50ntFlanks | 6721 | 6770 | 50 | AATTGCAGGACACATTGATCATCGACACTTCGAACGCACTTGCGGCCCCG | 351.9 | -4.9 | 1.0 | -4.8 | 1.9E-06 | 3.2E-06 | MB231/MB468 | CYTO |
| >hg38_21_+_8433172_8446622_RNA45SN1_45S_50ntFlanks@7974.7993.20\|ACGCGACCTCAGATCAGACG | >hg38_21_+_8433172_8446622_RNA45SN1_45S_50ntFlanks | 7974 | 7993 | 20 | ACGCGACCTCAGATCAGACG | 5156.2 | 4.5 | 0.9 | 4.8 | 1.7E-06 | 3.2E-06 | BT20/MB468 | CYTO |
| >hg38_21_+_8433172_8446622_RNA45SN1_45S_50ntFlanks@6760.6806.47\|TTGCGGCCCCGGGTTCCTCCCGGGGCTACGCCTGTCTGAGCGTCGCT | >hg38_21_+_8433172_8446622_RNA45SN1_45S_50ntFlanks | 6760 | 6806 | 47 | TTGCGGCCCCGGGTTCCTCCCGGGGCTACGCCTGTCTGAGCGTCGCT | 1244.8 | 7.7 | 1.6 | 4.8 | 1.5E-06 | 3.2E-06 | BT20/MB468 | MITO |
| >hg38_21_+_8433172_8446622_RNA45SN1_45S_50ntFlanks@9564.9603.40\|CTGACGTGCAAATCGGTCGTCCGACCTGGGTATAGGGGCG | >hg38_21_+_8433172_8446622_RNA45SN1_45S_50ntFlanks | 9564 | 9603 | 40 | CTGACGTGCAAATCGGTCGTCCGACCTGGGTATAGGGGCG | 410.0 | -6.2 | 1.3 | -4.7 | 2.2E-06 | 3.2E-06 | BT20/MB468 | NUC |
| >hg38_21_+_8433172_8446622_RNA45SN1_45S_50ntFlanks@4657.4686.30\|CTAGAGGTGAAATTCTTGGACCGGCGCAAG | >hg38_21_+_8433172_8446622_RNA45SN1_45S_50ntFlanks | 4657 | 4686 | 30 | CTAGAGGTGAAATTCTTGGACCGGCGCAAG | 236.3 | -4.7 | 1.0 | -4.8 | 1.7E-06 | 3.2E-06 | BT20/MB468 | CYTO |
| >hg38_21_+_8433172_8446622_RNA45SN1_45S_50ntFlanks@6646.6695.50\|TCGTACGACTCTTAGCGGTGGATCACTCGGCTCGTGCGTCGATGAAGAAC | >hg38_21_+_8433172_8446622_RNA45SN1_45S_50ntFlanks | 6646 | 6695 | 50 | TCGTACGACTCTTAGCGGTGGATCACTCGGCTCGTGCGTCGATGAAGAAC | 398.7 | 6.2 | 1.3 | 4.8 | 1.9E-06 | 3.2E-06 | MB231/MB468 | CYTO |
| >hg38_21_+_8433172_8446622_RNA45SN1_45S_50ntFlanks@8191.8212.22\|CAGCCCGTGGACGGTGTGAGGC | >hg38_21_+_8433172_8446622_RNA45SN1_45S_50ntFlanks | 8191 | 8212 | 22 | CAGCCCGTGGACGGTGTGAGGC | 558.2 | -4.5 | 0.9 | -4.8 | 1.9E-06 | 3.2E-06 | MB231/MB468 | CYTO |
| >hg38_21_+_8433172_8446622_RNA45SN1_45S_50ntFlanks@7981.8009.29\|CTCAGATCAGACGTGGCGACCCGCTGAAT | >hg38_21_+_8433172_8446622_RNA45SN1_45S_50ntFlanks | 7981 | 8009 | 29 | CTCAGATCAGACGTGGCGACCCGCTGAAT | 597.0 | -3.1 | 0.6 | -4.7 | 2.1E-06 | 3.2E-06 | MB231/MB468 | MITO |
| >MI0000082\|hsa-mir-25&WithFlank&7\|-\|100093554\|100093649@58.78.21&[MIMAT0000081&hsa-miR-25-3p&offsets\|0\|-1;m-23&7\|-\|100093571\|100093592&offsets\|0\|-1]\|CATTGCACTTGTCTCGGTCTG | >MI0000082\|hsa-mir-25&WithFlank&7\|-\|100093554\|100093649 | 58 | 78 | 21 | CATTGCACTTGTCTCGGTCTG | 6276.2 | 4.6 | 0.9 | 5.0 | 5.6E-07 | 3.2E-06 | BT20/MB468 | MP |
| >MI0000063\|hsa-let-7b&WithFlank&22\|+\|46113680\|46113774@12.34.23(+1U)&[MIMAT0000063&hsa-let-7b-5p&offsets\|0\|+1(+1U);m-14&22\|+\|46113691\|46113712&offsets\|0\|+1(+1U)]\|TGAGGTAGTAGGTTGTGTGGTTTT | >MI0000063\|hsa-let-7b&WithFlank&22\|+\|46113680\|46113774 | 12 | 34 | 23(+1U) | TGAGGTAGTAGGTTGTGTGGTTTT | 733.3 | 4.8 | 1.0 | 5.0 | 5.6E-07 | 3.2E-06 | BT20/MB468 | MP |
| >hg38_21_+_8433172_8446622_RNA45SN1_45S_50ntFlanks@12417.12446.30\|TCGATGTCGGCTCTTCCTATCATTGTGAAG | >hg38_21_+_8433172_8446622_RNA45SN1_45S_50ntFlanks | 12417 | 12446 | 30 | TCGATGTCGGCTCTTCCTATCATTGTGAAG | 571.7 | -2.1 | 0.4 | -4.8 | 1.5E-06 | 3.2E-06 | BT20/MB468 | MITO |
| >hg38_21_+_8433172_8446622_RNA45SN1_45S_50ntFlanks@4853.4883.31\|AAGTCTTTGGGTTCCGGGGGGAGTATGGTTG | >hg38_21_+_8433172_8446622_RNA45SN1_45S_50ntFlanks | 4853 | 4883 | 31 | AAGTCTTTGGGTTCCGGGGGGAGTATGGTTG | 200.9 | -2.9 | 0.6 | -4.8 | 1.7E-06 | 3.2E-06 | BT20/MB468 | CYTO |
| >MI0000457\|hsa-mir-141&WithFlank&12\|+\|6964091\|6964197@65.85.21&[MIMAT0000432&hsa-miR-141-3p&offsets\|0\|-1;m-134&12\|+\|6964155\|6964175&offsets\|0\|0]\|TAACACTGTCTGGTAAAGATG | >MI0000457\|hsa-mir-141&WithFlank&12\|+\|6964091\|6964197 | 65 | 85 | 21 | TAACACTGTCTGGTAAAGATG | 172.8 | 4.8 | 1.0 | 4.7 | 2.2E-06 | 3.2E-06 | BT20/MB468 | NUC |
| >trna111_HisGTG_1_-_147774845_147774916@-1T.22.23,trna118_HisGTG_1_-_145396881_145396952@-1T.22.23,trna16_HisGTG_1_+_146544773_146544844@-1T.22.23,trna1_HisGTG_15_+_45493349_45493420@-1T.22.23,trna21_HisGTG_1_+_147753471_147753542@-1T.22.23,trna33_HisGTG_6_+_27125906_27125977@-1T.22.23,trna7_HisGTG_9_-_14433938_14434009@-1T.22.23,trna8_HisGTG_15_-_45492611_45492682@-1T.22.23,trna9_HisGTG_15_-_45490804_45490875@-1T.22.23\|TGCCGTGATCGTATAGTGGTTAG | >trna111_HisGTG_1_-_147774845_147774916 | -1T | 22 | 23 | TGCCGTGATCGTATAGTGGTTAG | 256.5 | 7.8 | 1.6 | 4.8 | 1.7E-06 | 3.2E-06 | MB231/MB468 | MP |
| >hg38_21_+_8433172_8446622_RNA45SN1_45S_50ntFlanks@4723.4757.35\|CATTAATCAAGAACGAAAGTCGGAGGTTCGAAGAC | >hg38_21_+_8433172_8446622_RNA45SN1_45S_50ntFlanks | 4723 | 4757 | 35 | CATTAATCAAGAACGAAAGTCGGAGGTTCGAAGAC | 183.2 | -4.7 | 1.0 | -4.8 | 2.0E-06 | 3.2E-06 | MB231/MB468 | NUC |
| >hg38_21_+_8433172_8446622_RNA45SN1_45S_50ntFlanks@9778.9819.42\|AAGCCCGGCTCGCTGGCGTGGAGCCGGGCGTGGAATGCGAGT | >hg38_21_+_8433172_8446622_RNA45SN1_45S_50ntFlanks | 9778 | 9819 | 42 | AAGCCCGGCTCGCTGGCGTGGAGCCGGGCGTGGAATGCGAGT | 55.4 | 9.5 | 1.9 | 5.0 | 5.7E-07 | 3.2E-06 | BT20/MB468 | MP |

<table>
<tr><td>>hg38_21_+_8433172_8446622_RNA45SN1_45S_50ntFlanks@10375.10424.50|AGAAGGGTTCCATGTGAACAGCAGTTGAACATGGGTCAGTCGGTCCTGAG</td><td>>hg38_21_+_8433172_8446622_RNA45SN1_45S_50ntFlanks</td><td>10375</td><td>10424</td><td>50</td><td>AGAAGGGTTCCATGTGAACAGCAGTTGAACATGGGTCAGTCGGTCCTGAG</td><td>2617.1</td><td>6.0</td><td>1.2</td><td>5.2</td><td>1.8E-07</td><td>3.2E-06</td><td>BT20/MB231</td><td>MITO</td></tr>
<tr><td>>hg38_21_+_8433172_8446622_RNA45SN1_45S_50ntFlanks@11581.11600.20|AACTGGTGCGGACCAGGGGA</td><td>>hg38_21_+_8433172_8446622_RNA45SN1_45S_50ntFlanks</td><td>11581</td><td>11600</td><td>20</td><td>AACTGGTGCGGACCAGGGGA</td><td>307.0</td><td>-7.2</td><td>1.5</td><td>-4.7</td><td>2.2E-06</td><td>3.2E-06</td><td>BT20/MB468</td><td>NUC</td></tr>
<tr><td>>hg38_21_+_8433172_8446622_RNA45SN1_45S_50ntFlanks@12618.12654.37|GAGGAGCCAATGGGGCGAAGCTACCATCTGTGGGATT</td><td>>hg38_21_+_8433172_8446622_RNA45SN1_45S_50ntFlanks</td><td>12618</td><td>12654</td><td>37</td><td>GAGGAGCCAATGGGGCGAAGCTACCATCTGTGGGATT</td><td>173.9</td><td>-5.3</td><td>1.1</td><td>-4.8</td><td>1.7E-06</td><td>3.2E-06</td><td>MB231/MB468</td><td>MP</td></tr>
<tr><td>>hg38_21_+_8433172_8446622_RNA45SN1_45S_50ntFlanks@3776.3801.26|CCGGTACAGTGAAACTGCGAATGGCT</td><td>>hg38_21_+_8433172_8446622_RNA45SN1_45S_50ntFlanks</td><td>3776</td><td>3801</td><td>26</td><td>CCGGTACAGTGAAACTGCGAATGGCT</td><td>621.6</td><td>-3.3</td><td>0.7</td><td>-4.8</td><td>2.0E-06</td><td>3.2E-06</td><td>MB231/MB468</td><td>CYTO</td></tr>
<tr><td>>hg38_21_+_8433172_8446622_RNA45SN1_45S_50ntFlanks@5217.5240.24|CGGGGCTGCACGCGCGCTACACTG</td><td>>hg38_21_+_8433172_8446622_RNA45SN1_45S_50ntFlanks</td><td>5217</td><td>5240</td><td>24</td><td>CGGGGCTGCACGCGCGCTACACTG</td><td>194.1</td><td>-9.7</td><td>2.0</td><td>-4.8</td><td>1.5E-06</td><td>3.2E-06</td><td>BT20/MB468</td><td>MITO</td></tr>
<tr><td>>hg38_21_+_8433172_8446622_RNA45SN1_45S_50ntFlanks@5083.5104.22|ACGAACGAGACTCTGGCATGCT</td><td>>hg38_21_+_8433172_8446622_RNA45SN1_45S_50ntFlanks</td><td>5083</td><td>5104</td><td>22</td><td>ACGAACGAGACTCTGGCATGCT</td><td>264.1</td><td>3.9</td><td>0.8</td><td>4.8</td><td>2.0E-06</td><td>3.2E-06</td><td>MB231/MB468</td><td>CYTO</td></tr>
<tr><td>>hg38_21_+_8433172_8446622_RNA45SN1_45S_50ntFlanks@7971.8005.35|GAGACGCGACCTCAGATCAGACGTGGCGACCCGCT</td><td>>hg38_21_+_8433172_8446622_RNA45SN1_45S_50ntFlanks</td><td>7971</td><td>8005</td><td>35</td><td>GAGACGCGACCTCAGATCAGACGTGGCGACCCGCT</td><td>895.0</td><td>5.4</td><td>1.0</td><td>5.2</td><td>2.2E-07</td><td>3.2E-06</td><td>BT20/MB231</td><td>NUC</td></tr>
<tr><td>>hg38_21_+_8433172_8446622_RNA45SN1_45S_50ntFlanks@12043.12063.21|CACTGACCCGGTGAGGCGGGG</td><td>>hg38_21_+_8433172_8446622_RNA45SN1_45S_50ntFlanks</td><td>12043</td><td>12063</td><td>21</td><td>CACTGACCCGGTGAGGCGGGG</td><td>61.6</td><td>-9.0</td><td>1.8</td><td>-5.0</td><td>5.7E-07</td><td>3.2E-06</td><td>BT20/MB468</td><td>MP</td></tr>
<tr><td>>hg38_21_+_8433172_8446622_RNA45SN1_45S_50ntFlanks@6784.6804.21|GCTACGCCTGTCTGAGCGTCG</td><td>>hg38_21_+_8433172_8446622_RNA45SN1_45S_50ntFlanks</td><td>6784</td><td>6804</td><td>21</td><td>GCTACGCCTGTCTGAGCGTCG</td><td>65.9</td><td>-9.1</td><td>1.8</td><td>-5.0</td><td>5.7E-07</td><td>3.2E-06</td><td>BT20/MB468</td><td>MP</td></tr>
<tr><td>>hg38_21_+_8433172_8446622_RNA45SN1_45S_50ntFlanks@3802.3834.33|CATTAAATCAGTTATGGTTCCTTTGGTCGCTCG</td><td>>hg38_21_+_8433172_8446622_RNA45SN1_45S_50ntFlanks</td><td>3802</td><td>3834</td><td>33</td><td>CATTAAATCAGTTATGGTTCCTTTGGTCGCTCG</td><td>63.0</td><td>-9.1</td><td>1.8</td><td>-5.0</td><td>5.7E-07</td><td>3.2E-06</td><td>BT20/MB468</td><td>MP</td></tr>
<tr><td>>MI0000063|hsa-let-7b&WithFlank&22|+|46113680|46113774@12.32.21&[MIMAT0000063&hsa-let-7b-5p&offsets|0|-1;m-14&22|+|46113691|46113712&offsets|0|-1]|TGAGGTAGTAGGTTGTGTGGT</td><td>>MI0000063|hsa-let-7b&WithFlank&22|+|46113680|46113774</td><td>12</td><td>32</td><td>21</td><td>TGAGGTAGTAGGTTGTGTGGT</td><td>2075.3</td><td>2.2</td><td>0.5</td><td>4.7</td><td>2.2E-06</td><td>3.2E-06</td><td>BT20/MB468</td><td>NUC</td></tr>
<tr><td>>am_MI0039740|hsa-mir-12136&WithFlank&1|-|632609|632691@7.32.26(+1G)&[MIMAT0049032&hsa-miR-12136&offsets|0|+8(+1G)]|GAAAAAGTCATGGAGGCCATGGGGTTG</td><td>>am_MI0039740|hsa-mir-12136&WithFlank&1|-|632609|632691</td><td>7</td><td>32</td><td>26(+1G)</td><td>GAAAAAGTCATGGAGGCCATGGGGTTG</td><td>412.8</td><td>-3.0</td><td>0.6</td><td>-4.7</td><td>2.2E-06</td><td>3.2E-06</td><td>BT20/MB468</td><td>NUC</td></tr>
<tr><td>>trnaMT_SerTGA_MT_-_7446_7514@1.27.27,trnalookalike6_SerTGA_1_-_567997_568065@1.27.27|GAAAAAGTCATGGAGGCCATGGGGTTG</td><td>>trnaMT_SerTGA_MT_-_7446_7514</td><td>1</td><td>27</td><td>27</td><td>GAAAAAGTCATGGAGGCCATGGGGTTG</td><td>412.8</td><td>-3.0</td><td>0.6</td><td>-4.7</td><td>2.2E-06</td><td>3.2E-06</td><td>BT20/MB468</td><td>NUC</td></tr>
<tr><td>>hg38_21_+_8433172_8446622_RNA45SN1_45S_50ntFlanks@10783.10817.35|GTGAACAGCCTCTGGCATGTTGGAACAATGTAGGT</td><td>>hg38_21_+_8433172_8446622_RNA45SN1_45S_50ntFlanks</td><td>10783</td><td>10817</td><td>35</td><td>GTGAACAGCCTCTGGCATGTTGGAACAATGTAGGT</td><td>321.9</td><td>-3.2</td><td>0.7</td><td>-4.7</td><td>2.0E-06</td><td>3.2E-06</td><td>MB231/MB468</td><td>NUC</td></tr>
<tr><td>>hg38_21_+_8433172_8446622_RNA45SN1_45S_50ntFlanks@9266.9285.20|CGACGTCGGCTACCCACCCG</td><td>>hg38_21_+_8433172_8446622_RNA45SN1_45S_50ntFlanks</td><td>9266</td><td>9285</td><td>20</td><td>CGACGTCGGCTACCCACCCG</td><td>698.3</td><td>-6.9</td><td>1.5</td><td>-4.7</td><td>2.2E-06</td><td>3.2E-06</td><td>BT20/MB468</td><td>NUC</td></tr>
<tr><td>>hg38_21_+_8433172_8446622_RNA45SN1_45S_50ntFlanks@11573.11592.20|CGACTTAGAACTGGTGCGGA</td><td>>hg38_21_+_8433172_8446622_RNA45SN1_45S_50ntFlanks</td><td>11573</td><td>11592</td><td>20</td><td>CGACTTAGAACTGGTGCGGA</td><td>351.1</td><td>-5.2</td><td>1.1</td><td>-4.7</td><td>2.1E-06</td><td>3.2E-06</td><td>MB231/MB468</td><td>MITO</td></tr>
<tr><td>>hg38_21_+_8433172_8446622_RNA45SN1_45S_50ntFlanks@7988.8010.23|CAGACGTGGCGACCCGCTGAATT</td><td>>hg38_21_+_8433172_8446622_RNA45SN1_45S_50ntFlanks</td><td>7988</td><td>8010</td><td>23</td><td>CAGACGTGGCGACCCGCTGAATT</td><td>1353.9</td><td>-3.7</td><td>0.8</td><td>-4.7</td><td>2.1E-06</td><td>3.2E-06</td><td>MB231/MB468</td><td>MITO</td></tr>
<tr><td>>hg38_21_+_8433172_8446622_RNA45SN1_45S_50ntFlanks@476.512.37|TTCGTGATCGATGTGGTGACGTCGTGCTCTCCCGGGC</td><td>>hg38_21_+_8433172_8446622_RNA45SN1_45S_50ntFlanks</td><td>476</td><td>512</td><td>37</td><td>TTCGTGATCGATGTGGTGACGTCGTGCTCTCCCGGGC</td><td>360.4</td><td>7.0</td><td>1.5</td><td>4.8</td><td>1.5E-06</td><td>3.2E-06</td><td>BT20/MB468</td><td>MITO</td></tr>
<tr><td>>hg38_21_+_8433172_8446622_RNA45SN1_45S_50ntFlanks@5227.5246.20|CGCGCGCTACACTGACTGGC</td><td>>hg38_21_+_8433172_8446622_RNA45SN1_45S_50ntFlanks</td><td>5227</td><td>5246</td><td>20</td><td>CGCGCGCTACACTGACTGGC</td><td>207.3</td><td>-9.8</td><td>2.0</td><td>-4.8</td><td>1.5E-06</td><td>3.2E-06</td><td>BT20/MB468</td><td>MITO</td></tr>
<tr><td>>hg38_21_+_8433172_8446622_RNA45SN1_45S_50ntFlanks@10239.10258.20|GCTACGCCGCGACGAGTAGG</td><td>>hg38_21_+_8433172_8446622_RNA45SN1_45S_50ntFlanks</td><td>10239</td><td>10258</td><td>20</td><td>GCTACGCCGCGACGAGTAGG</td><td>126.7</td><td>10.3</td><td>2.0</td><td>5.3</td><td>1.4E-07</td><td>3.2E-06</td><td>BT20/MB231</td><td>MP</td></tr>
<tr><td>>hg38_21_+_8433172_8446622_RNA45SN1_45S_50ntFlanks@9004.9026.23|TCTCCCCCGCGGGGGCGCGCCGG</td><td>>hg38_21_+_8433172_8446622_RNA45SN1_45S_50ntFlanks</td><td>9004</td><td>9026</td><td>23</td><td>TCTCCCCCGCGGGGGCGCGCCGG</td><td>235.4</td><td>-5.5</td><td>1.2</td><td>-4.8</td><td>2.0E-06</td><td>3.2E-06</td><td>MB231/MB468</td><td>CYTO</td></tr>
<tr><td>>hg38_21_+_8433172_8446622_RNA45SN1_45S_50ntFlanks@6731.6772.42|CACATTGATCATCGACACTTCGAACGCACTTGCGGCCCCGGG</td><td>>hg38_21_+_8433172_8446622_RNA45SN1_45S_50ntFlanks</td><td>6731</td><td>6772</td><td>42</td><td>CACATTGATCATCGACACTTCGAACGCACTTGCGGCCCCGGG</td><td>571.8</td><td>-3.2</td><td>0.7</td><td>-4.8</td><td>1.5E-06</td><td>3.2E-06</td><td>BT20/MB468</td><td>MITO</td></tr>
<tr><td>>hg38_21_+_8433172_8446622_RNA45SN1_45S_50ntFlanks@6672.6699.28|TCGGCTCGTGCGTCGATGAAGAACGCAG</td><td>>hg38_21_+_8433172_8446622_RNA45SN1_45S_50ntFlanks</td><td>6672</td><td>6699</td><td>28</td><td>TCGGCTCGTGCGTCGATGAAGAACGCAG</td><td>115.3</td><td>-5.0</td><td>1.0</td><td>-4.8</td><td>1.8E-06</td><td>3.2E-06</td><td>MB231/MB468</td><td>MP</td></tr>
<tr><td>>hg38_21_+_8433172_8446622_RNA45SN1_45S_50ntFlanks@3835.3869.35|CTCCTCTCCTACTTGGATAACTGTGGTAATTCTAG</td><td>>hg38_21_+_8433172_8446622_RNA45SN1_45S_50ntFlanks</td><td>3835</td><td>3869</td><td>35</td><td>CTCCTCTCCTACTTGGATAACTGTGGTAATTCTAG</td><td>280.5</td><td>-6.4</td><td>1.3</td><td>-4.7</td><td>2.1E-06</td><td>3.2E-06</td><td>MB231/MB468</td><td>NUC</td></tr>
<tr><td>>hg38_21_+_8433172_8446622_RNA45SN1_45S_50ntFlanks@6669.6686.18|CACTCGGCTCGTGCGTCG</td><td>>hg38_21_+_8433172_8446622_RNA45SN1_45S_50ntFlanks</td><td>6669</td><td>6686</td><td>18</td><td>CACTCGGCTCGTGCGTCG</td><td>726.8</td><td>-5.3</td><td>1.1</td><td>-4.7</td><td>2.1E-06</td><td>3.2E-06</td><td>MB231/MB468</td><td>MITO</td></tr>
<tr><td>>hg38_21_+_8433172_8446622_RNA45SN1_45S_50ntFlanks@12449.12480.32|GAATTCACCAAGCGTTGGATTGTTCACCCACT</td><td>>hg38_21_+_8433172_8446622_RNA45SN1_45S_50ntFlanks</td><td>12449</td><td>12480</td><td>32</td><td>GAATTCACCAAGCGTTGGATTGTTCACCCACT</td><td>174.7</td><td>6.2</td><td>1.3</td><td>4.7</td><td>2.1E-06</td><td>3.2E-06</td><td>MB231/MB468</td><td>NUC</td></tr>
<tr><td>>hg38_21_+_8433172_8446622_RNA45SN1_45S_50ntFlanks@6653.6681.29|ACTCTTAGCGGTGGATCACTCGGCTCGTG</td><td>>hg38_21_+_8433172_8446622_RNA45SN1_45S_50ntFlanks</td><td>6653</td><td>6681</td><td>29</td><td>ACTCTTAGCGGTGGATCACTCGGCTCGTG</td><td>1860.8</td><td>-3.2</td><td>0.7</td><td>-4.8</td><td>2.0E-06</td><td>3.2E-06</td><td>MB231/MB468</td><td>CYTO</td></tr>
<tr><td>>hg38_21_+_8433172_8446622_RNA45SN1_45S_50ntFlanks@8355.8389.35|TGAAAAGAACTTTGAAGAGAGAGTTCAAGAGGGCG</td><td>>hg38_21_+_8433172_8446622_RNA45SN1_45S_50ntFlanks</td><td>8355</td><td>8389</td><td>35</td><td>TGAAAAGAACTTTGAAGAGAGAGTTCAAGAGGGCG</td><td>49.8</td><td>9.4</td><td>1.9</td><td>5.0</td><td>5.8E-07</td><td>3.2E-06</td><td>BT20/MB468</td><td>MP</td></tr>
<tr><td>>hg38_21_+_8433172_8446622_RNA45SN1_45S_50ntFlanks@11885.11905.21|TTGAGCTTGACTCTAGTCTGG</td><td>>hg38_21_+_8433172_8446622_RNA45SN1_45S_50ntFlanks</td><td>11885</td><td>11905</td><td>21</td><td>TTGAGCTTGACTCTAGTCTGG</td><td>980.9</td><td>-7.0</td><td>1.5</td><td>-4.8</td><td>1.8E-06</td><td>3.3E-06</td><td>BT20/MB468</td><td>CYTO</td></tr>
<tr><td>>am_MI0003581|hsa-mir-574&WithFlank&4|+|38868026|38868133@31.51.21(+1A)&[MIMAT0004795&hsa-miR-574-5p&offsets|0|-2(+1A);m-213&4|+|38868056|38868078&offsets|0|-2(+1A)]|TGAGTGTGTGTGTGTGAGTGTA</td><td>>am_MI0003581|hsa-mir-574&WithFlank&4|+|38868026|38868133</td><td>31</td><td>51</td><td>21(+1A)</td><td>TGAGTGTGTGTGTGTGAGTGTA</td><td>44.8</td><td>8.7</td><td>1.8</td><td>4.8</td><td>1.8E-06</td><td>3.3E-06</td><td>MB231/MB468</td><td>MP</td></tr>
<tr><td>>hg38_21_+_8433172_8446622_RNA45SN1_45S_50ntFlanks@10822.10840.19|GAAGTCGGCAAGCCGGATC</td><td>>hg38_21_+_8433172_8446622_RNA45SN1_45S_50ntFlanks</td><td>10822</td><td>10840</td><td>19</td><td>GAAGTCGGCAAGCCGGATC</td><td>529.9</td><td>-5.3</td><td>1.1</td><td>-4.8</td><td>2.0E-06</td><td>3.3E-06</td><td>MB231/MB468</td><td>CYTO</td></tr>
<tr><td>>hg38_21_+_8433172_8446622_RNA45SN1_45S_50ntFlanks@11580.11600.21|GAACTGGTGCGGACCAGGGGA</td><td>>hg38_21_+_8433172_8446622_RNA45SN1_45S_50ntFlanks</td><td>11580</td><td>11600</td><td>21</td><td>GAACTGGTGCGGACCAGGGGA</td><td>469.0</td><td>-7.9</td><td>1.6</td><td>-5.0</td><td>5.8E-07</td><td>3.3E-06</td><td>BT20/MB468</td><td>MP</td></tr>
<tr><td>>trna116_GluCTC_1_-_145399233_145399304@1.42.42,trna59_GluCTC_1_+_249168447_249168518@1.42.42,trna71_GluCTC_1_-_161439189_161439260@1.42.42,trna74_GluCTC_1_-_161431809_161431880@1.42.42,trna77_GluCTC_1_-_161424398_161424469@1.42.42,trna77_GluCTC_6_+_28949976_28950047@1.42.42,trna80_GluCTC_1_-_161417018_161417089@1.42.42,trna87_GluCTC_6_-_126101393_126101464@1.42.42|TCCCTGGTGGTCTAGTGGTTAGGATTCGGCGCTCTCACCGCC</td><td>>trna116_GluCTC_1_-_145399233_145399304</td><td>1</td><td>42</td><td>42</td><td>TCCCTGGTGGTCTAGTGGTTAGGATTCGGCGCTCTCACCGCC</td><td>1446.3</td><td>4.9</td><td>1.0</td><td>4.8</td><td>2.0E-06</td><td>3.3E-06</td><td>MB231/MB468</td><td>CYTO</td></tr>
<tr><td>>hg38_21_+_8433172_8446622_RNA45SN1_45S_50ntFlanks@4726.4758.33|TAATCAAGAACGAAAGTCGGAGGTTCGAAGACG</td><td>>hg38_21_+_8433172_8446622_RNA45SN1_45S_50ntFlanks</td><td>4726</td><td>4758</td><td>33</td><td>TAATCAAGAACGAAAGTCGGAGGTTCGAAGACG</td><td>2337.8</td><td>-2.9</td><td>0.6</td><td>-4.8</td><td>2.0E-06</td><td>3.3E-06</td><td>MB231/MB468</td><td>CYTO</td></tr>
<tr><td>>trna10_LysCTT_16_+_3241501_3241573@4.35.32,trna13_LysCTT_14_-_58706613_58706685@4.35.32,trna2_LysCTT_15_+_79152904_79152976@4.35.32|CGGCTAGCTCAGTCGGTAGAGCATGGGACTCT</td><td>>trna10_LysCTT_16_+_3241501_3241573</td><td>4</td><td>35</td><td>32</td><td>CGGCTAGCTCAGTCGGTAGAGCATGGGACTCT</td><td>1981.0</td><td>-4.3</td><td>0.8</td><td>-5.2</td><td>1.8E-07</td><td>3.3E-06</td><td>BT20/MB231</td><td>MITO</td></tr>
<tr><td>>hg38_21_+_8433172_8446622_RNA45SN1_45S_50ntFlanks@11785.11805.21|TGAATGGATGAACGAGATTCC</td><td>>hg38_21_+_8433172_8446622_RNA45SN1_45S_50ntFlanks</td><td>11785</td><td>11805</td><td>21</td><td>TGAATGGATGAACGAGATTCC</td><td>535.6</td><td>-3.8</td><td>0.8</td><td>-4.7</td><td>2.1E-06</td><td>3.3E-06</td><td>MB231/MB468</td><td>NUC</td></tr>
<tr><td>>hg38_21_+_8433172_8446622_RNA45SN1_45S_50ntFlanks@4752.4782.31|GAAGACGATCAGATACCGTCGTAGTTCCGAC</td><td>>hg38_21_+_8433172_8446622_RNA45SN1_45S_50ntFlanks</td><td>4752</td><td>4782</td><td>31</td><td>GAAGACGATCAGATACCGTCGTAGTTCCGAC</td><td>298.1</td><td>-7.1</td><td>1.5</td><td>-4.8</td><td>1.8E-06</td><td>3.3E-06</td><td>MB231/MB468</td><td>MP</td></tr>
<tr><td>>hg38_21_+_8433172_8446622_RNA45SN1_45S_50ntFlanks@11537.11584.48|CCGCGCGGCGCCTCGCCTCGGCCGGCGCCTAGCAGCCGACTTAGAACT</td><td>>hg38_21_+_8433172_8446622_RNA45SN1_45S_50ntFlanks</td><td>11537</td><td>11584</td><td>48</td><td>CCGCGCGGCGCCTCGCCTCGGCCGGCGCCTAGCAGCCGACTTAGAACT</td><td>151.9</td><td>5.5</td><td>1.2</td><td>4.8</td><td>2.0E-06</td><td>3.3E-06</td><td>MB231/MB468</td><td>CYTO</td></tr>
<tr><td>>hg38_21_+_8433172_8446622_RNA45SN1_45S_50ntFlanks@9489.9525.37|AGGACCCGAAAGATGGTGAACTATGCCTGGGCAGGGC</td><td>>hg38_21_+_8433172_8446622_RNA45SN1_45S_50ntFlanks</td><td>9489</td><td>9525</td><td>37</td><td>AGGACCCGAAAGATGGTGAACTATGCCTGGGCAGGGC</td><td>217.5</td><td>11.1</td><td>2.1</td><td>5.3</td><td>1.5E-07</td><td>3.3E-06</td><td>BT20/MB231</td><td>MP</td></tr>
<tr><td>>MI0000298|hsa-mir-221&WithFlank&X|-|45746151|45746272@71.88.18(+4U)&[MIMAT0000278&hsa-miR-221-3p&offsets|0|-5(+4U);m-91&X|-|45746181|45746202&offsets|0|-4(+4U)]|AGCTACATTGTCTGCTGGTTTT</td><td>>MI0000298|hsa-mir-221&WithFlank&X|-|45746151|45746272</td><td>71</td><td>88</td><td>18(+4U)</td><td>AGCTACATTGTCTGCTGGTTTT</td><td>52.1</td><td>8.9</td><td>1.9</td><td>4.8</td><td>1.8E-06</td><td>3.3E-06</td><td>MB231/MB468</td><td>MP</td></tr>
<tr><td>>hg38_21_+_8433172_8446622_RNA45SN1_45S_50ntFlanks@11789.11806.18|TGGATGAACGAGATTCCC</td><td>>hg38_21_+_8433172_8446622_RNA45SN1_45S_50ntFlanks</td><td>11789</td><td>11806</td><td>18</td><td>TGGATGAACGAGATTCCC</td><td>152.3</td><td>-5.7</td><td>1.2</td><td>-4.8</td><td>1.8E-06</td><td>3.3E-06</td><td>MB231/MB468</td><td>MP</td></tr>
<tr><td>>hg38_21_+_8433172_8446622_RNA45SN1_45S_50ntFlanks@3782.3805.24|CAGTGAAACTGCGAATGGCTCATT</td><td>>hg38_21_+_8433172_8446622_RNA45SN1_45S_50ntFlanks</td><td>3782</td><td>3805</td><td>24</td><td>CAGTGAAACTGCGAATGGCTCATT</td><td>438.0</td><td>-3.7</td><td>0.8</td><td>-4.7</td><td>2.1E-06</td><td>3.3E-06</td><td>MB231/MB468</td><td>NUC</td></tr>
<tr><td>>trna15_ValAAC_5_-_180615416_180615488@1.29.29|GTTTCCGTAGTGTAGTGGTCATCACGTTC</td><td>>trna15_ValAAC_5_-_180615416_180615488</td><td>1</td><td>29</td><td>29</td><td>GTTTCCGTAGTGTAGTGGTCATCACGTTC</td><td>578.8</td><td>-5.5</td><td>1.2</td><td>-4.7</td><td>2.1E-06</td><td>3.3E-06</td><td>MB231/MB468</td><td>NUC</td></tr>
<tr><td>>hg38_21_+_8433172_8446622_RNA45SN1_45S_50ntFlanks@12420.12460.41|ATGTCGGCTCTTCCTATCATTGTGAAGCAGAATTCACCAAG</td><td>>hg38_21_+_8433172_8446622_RNA45SN1_45S_50ntFlanks</td><td>12420</td><td>12460</td><td>41</td><td>ATGTCGGCTCTTCCTATCATTGTGAAGCAGAATTCACCAAG</td><td>2220.3</td><td>-4.4</td><td>0.8</td><td>-5.2</td><td>1.8E-07</td><td>3.3E-06</td><td>BT20/MB231</td><td>MITO</td></tr>
<tr><td>>MI0003560|hsa-mir-92b&WithFlank&1|+|155195171|155195278@67.88.22&[MIMAT0003218&hsa-miR-92b-3p&offsets|0|0;m-115&1|+|155195237|155195258&offsets|0|0]|TATTGCACTCGTCCCGGCCTCC</td><td>>MI0003560|hsa-mir-92b&WithFlank&1|+|155195171|155195278</td><td>67</td><td>88</td><td>22</td><td>TATTGCACTCGTCCCGGCCTCC</td><td>1234.1</td><td>2.7</td><td>0.6</td><td>4.7</td><td>2.2E-06</td><td>3.3E-06</td><td>BT20/MB468</td><td>NUC</td></tr>
<tr><td>>hg38_21_+_8433172_8446622_RNA45SN1_45S_50ntFlanks@12482.12499.18|ATAGGGAACGTGAGCTGG</td><td>>hg38_21_+_8433172_8446622_RNA45SN1_45S_50ntFlanks</td><td>12482</td><td>12499</td><td>18</td><td>ATAGGGAACGTGAGCTGG</td><td>145.1</td><td>-5.4</td><td>1.1</td><td>-4.8</td><td>1.8E-06</td><td>3.3E-06</td><td>MB231/MB468</td><td>MP</td></tr>
<tr><td>>MI0000477|hsa-mir-146a&WithFlank&5|+|160485346|160485456@27.45.19(+1A)&[MIMAT0000449&hsa-miR-146a-5p&offsets|0|-3(+1A);m-129&5|+|160485372|160485393&offsets|0|-3(+1A)]|TGAGAACTGAATTCCATGGA</td><td>>MI0000477|hsa-mir-146a&WithFlank&5|+|160485346|160485456</td><td>27</td><td>45</td><td>19(+1A)</td><td>TGAGAACTGAATTCCATGGA</td><td>130.2</td><td>-10.4</td><td>2.0</td><td>-5.3</td><td>1.5E-07</td><td>3.3E-06</td><td>BT20/MB231</td><td>MP</td></tr>
</table>

<table>
<tr><td>>am_trna2_GlyGCC_21_-_18827107_18827177@1.24.24,trna35_GlyGCC_1_+_161413094_161413164@1.24.24,trna37_GlyGCC_1_+_161420467_161420537@1.24.24,trna39_GlyGCC_1_+_161427898_161427968@1.24.24,trna41_GlyGCC_1_+_161435258_161435328@1.24.24|GCATGGGTGGTTCAGTGGTAGAAT</td><td>>am_trna2_GlyGCC_21_-_18827107_18827177</td><td>1</td><td>24</td><td>24</td><td>GCATGGGTGGTTCAGTGGTAGAAT</td><td>107.8</td><td>10.1</td><td>1.9</td><td>5.3</td><td>1.5E-07</td><td>3.3E-06</td><td>BT20/MB231</td><td>MP</td></tr>
<tr><td>>hg38_21_+_8433172_8446622_RNA45SN1_45S_50ntFlanks@9795.9819.25|GTGGAGCCGGGCGTGGAATGCGAGT</td><td>>hg38_21_+_8433172_8446622_RNA45SN1_45S_50ntFlanks</td><td>9795</td><td>9819</td><td>25</td><td>GTGGAGCCGGGCGTGGAATGCGAGT</td><td>203.5</td><td>11.0</td><td>2.1</td><td>5.3</td><td>1.5E-07</td><td>3.3E-06</td><td>BT20/MB231</td><td>MP</td></tr>
<tr><td>>hg38_21_+_8433172_8446622_RNA45SN1_45S_50ntFlanks@10807.10826.20|ACAATGTAGGTAAGGGAAGT</td><td>>hg38_21_+_8433172_8446622_RNA45SN1_45S_50ntFlanks</td><td>10807</td><td>10826</td><td>20</td><td>ACAATGTAGGTAAGGGAAGT</td><td>342.6</td><td>-4.7</td><td>1.0</td><td>-4.8</td><td>1.5E-06</td><td>3.3E-06</td><td>BT20/MB468</td><td>MITO</td></tr>
<tr><td>>hg38_21_+_8433172_8446622_RNA45SN1_45S_50ntFlanks@12008.12036.29|CCGCCGGTGAAATACCACTACTCTGATCG</td><td>>hg38_21_+_8433172_8446622_RNA45SN1_45S_50ntFlanks</td><td>12008</td><td>12036</td><td>29</td><td>CCGCCGGTGAAATACCACTACTCTGATCG</td><td>652.6</td><td>-7.8</td><td>1.7</td><td>-4.7</td><td>2.3E-06</td><td>3.3E-06</td><td>BT20/MB468</td><td>NUC</td></tr>
<tr><td>>hg38_21_+_8433172_8446622_RNA45SN1_45S_50ntFlanks@12222.12248.27|ACAGAAACCTCCCGTGGAGCAGAAGGG</td><td>>hg38_21_+_8433172_8446622_RNA45SN1_45S_50ntFlanks</td><td>12222</td><td>12248</td><td>27</td><td>ACAGAAACCTCCCGTGGAGCAGAAGGG</td><td>64.3</td><td>-9.1</td><td>1.8</td><td>-5.0</td><td>5.8E-07</td><td>3.3E-06</td><td>BT20/MB468</td><td>MP</td></tr>
<tr><td>>hg38_21_+_8433172_8446622_RNA45SN1_45S_50ntFlanks@6647.6665.19|CGTACGACTCTTAGCGGTG</td><td>>hg38_21_+_8433172_8446622_RNA45SN1_45S_50ntFlanks</td><td>6647</td><td>6665</td><td>19</td><td>CGTACGACTCTTAGCGGTG</td><td>2551.9</td><td>3.4</td><td>0.7</td><td>4.7</td><td>2.1E-06</td><td>3.3E-06</td><td>MB231/MB468</td><td>NUC</td></tr>
<tr><td>>MI0001445|hsa-mir-423&WithFlank&17|+|30117073|30117178@23.43.21&[MIMAT0004748&hsa-miR-423-5p&offsets|0|-2;m-141&17|+|30117095|30117117&offsets|0|-2]|TGAGGGGCAGAGAGCGAGACT</td><td>>MI0001445|hsa-mir-423&WithFlank&17|+|30117073|30117178</td><td>23</td><td>43</td><td>21</td><td>TGAGGGGCAGAGAGCGAGACT</td><td>201.6</td><td>1.8</td><td>0.4</td><td>4.7</td><td>2.3E-06</td><td>3.3E-06</td><td>BT20/MB468</td><td>NUC</td></tr>
<tr><td>>hg38_21_+_8433172_8446622_RNA45SN1_45S_50ntFlanks@11890.11907.18|CTTGACTCTAGTCTGGCA</td><td>>hg38_21_+_8433172_8446622_RNA45SN1_45S_50ntFlanks</td><td>11890</td><td>11907</td><td>18</td><td>CTTGACTCTAGTCTGGCA</td><td>1046.3</td><td>-5.5</td><td>1.2</td><td>-4.8</td><td>1.8E-06</td><td>3.3E-06</td><td>BT20/MB468</td><td>CYTO</td></tr>
<tr><td>>MI0000103|hsa-mir-101-1&WithFlank&1|-|65058428|65058514@52.73.22(+2U);MI0000739|hsa-mir-101-2&WithFlank&9|+|4850291|4850381@54.75.22(+2U)&[MIMAT0000099&hsa-miR-101-3p&offsets|-1|0(+2U);m-17&1|-|65058443|65058462&offsets|-1|+1(+2U)];[MIMAT0000099_1&hsa-miR-101-3p&offsets|-1|0(+2U);m-16&9|+|4850345|4850364&offsets|-1|+1(+2U)]|GTACAGTACTGTGATAACTGAATT</td><td>>MI0000103|hsa-mir-101-1&WithFlank&1|-|65058428|65058514</td><td>52</td><td>73</td><td>22(+2U)</td><td>GTACAGTACTGTGATAACTGAATT</td><td>98.9</td><td>-10.0</td><td>1.9</td><td>-5.3</td><td>1.5E-07</td><td>3.3E-06</td><td>BT20/MB231</td><td>MP</td></tr>
<tr><td>>hg38_21_+_8433172_8446622_RNA45SN1_45S_50ntFlanks@8169.8195.27|CCAAGTCCTTCTGATCGAGGCCCAGCC</td><td>>hg38_21_+_8433172_8446622_RNA45SN1_45S_50ntFlanks</td><td>8169</td><td>8195</td><td>27</td><td>CCAAGTCCTTCTGATCGAGGCCCAGCC</td><td>317.3</td><td>-6.9</td><td>1.4</td><td>-4.8</td><td>1.8E-06</td><td>3.3E-06</td><td>MB231/MB468</td><td>MP</td></tr>
<tr><td>>hg38_21_+_8433172_8446622_RNA45SN1_45S_50ntFlanks@12934.12952.19|CTAAACCATTCGTAGACGA</td><td>>hg38_21_+_8433172_8446622_RNA45SN1_45S_50ntFlanks</td><td>12934</td><td>12952</td><td>19</td><td>CTAAACCATTCGTAGACGA</td><td>315.1</td><td>-5.2</td><td>1.1</td><td>-4.7</td><td>2.1E-06</td><td>3.3E-06</td><td>MB231/MB468</td><td>NUC</td></tr>
<tr><td>>MI0000066|hsa-let-7e&WithFlank&19|+|51692780|51692870@14.35.22(+1U)&[MIMAT0000066&hsa-let-7e-5p&offsets|0|0(+1U);m-50&19|+|51692793|51692814&offsets|0|0(+1U)]|TGAGGTAGGAGGTTGTATAGTTT</td><td>>MI0000066|hsa-let-7e&WithFlank&19|+|51692780|51692870</td><td>14</td><td>35</td><td>22(+1U)</td><td>TGAGGTAGGAGGTTGTATAGTTT</td><td>425.5</td><td>4.5</td><td>0.9</td><td>4.7</td><td>2.2E-06</td><td>3.3E-06</td><td>MB231/MB468</td><td>MITO</td></tr>
<tr><td>>hg38_21_+_8433172_8446622_RNA45SN1_45S_50ntFlanks@9567.9592.26|ACGTGCAAATCGGTCGTCCGACCTGG</td><td>>hg38_21_+_8433172_8446622_RNA45SN1_45S_50ntFlanks</td><td>9567</td><td>9592</td><td>26</td><td>ACGTGCAAATCGGTCGTCCGACCTGG</td><td>325.7</td><td>-5.1</td><td>1.1</td><td>-4.7</td><td>2.1E-06</td><td>3.3E-06</td><td>MB231/MB468</td><td>NUC</td></tr>
<tr><td>>hg38_21_+_8433172_8446622_RNA45SN1_45S_50ntFlanks@9481.9503.23|CACGTGTTAGGACCCGAAAGATG</td><td>>hg38_21_+_8433172_8446622_RNA45SN1_45S_50ntFlanks</td><td>9481</td><td>9503</td><td>23</td><td>CACGTGTTAGGACCCGAAAGATG</td><td>228.7</td><td>11.2</td><td>2.1</td><td>5.3</td><td>1.5E-07</td><td>3.3E-06</td><td>BT20/MB231</td><td>MP</td></tr>
<tr><td>>hg38_21_+_8433172_8446622_RNA45SN1_45S_50ntFlanks@9928.9946.19|TGATATAGACAGCAGGACG</td><td>>hg38_21_+_8433172_8446622_RNA45SN1_45S_50ntFlanks</td><td>9928</td><td>9946</td><td>19</td><td>TGATATAGACAGCAGGACG</td><td>106.9</td><td>10.1</td><td>1.9</td><td>5.3</td><td>1.5E-07</td><td>3.3E-06</td><td>BT20/MB231</td><td>MP</td></tr>
<tr><td>>hg38_21_+_8433172_8446622_RNA45SN1_45S_50ntFlanks@3927.3959.33|CAGATCAAAACCAACCCGGTCAGCCCCTCTCCG</td><td>>hg38_21_+_8433172_8446622_RNA45SN1_45S_50ntFlanks</td><td>3927</td><td>3959</td><td>33</td><td>CAGATCAAAACCAACCCGGTCAGCCCCTCTCCG</td><td>145.6</td><td>-3.8</td><td>0.8</td><td>-4.8</td><td>1.8E-06</td><td>3.3E-06</td><td>BT20/MB468</td><td>CYTO</td></tr>
<tr><td>>hg38_21_+_8433172_8446622_RNA45SN1_45S_50ntFlanks@12973.13001.29|TCGTACGTAGCAGAGCAGCTCCCTCGCTG</td><td>>hg38_21_+_8433172_8446622_RNA45SN1_45S_50ntFlanks</td><td>12973</td><td>13001</td><td>29</td><td>TCGTACGTAGCAGAGCAGCTCCCTCGCTG</td><td>349.4</td><td>-6.1</td><td>1.3</td><td>-4.8</td><td>1.8E-06</td><td>3.3E-06</td><td>BT20/MB468</td><td>CYTO</td></tr>
<tr><td>>hg38_MT_+_1621_3279_RNR2_16S_50ntFlanks@826.856.31|AAGGAAAGGTTAAAAAAAGTAAAAGGAACTC</td><td>>hg38_MT_+_1621_3279_RNR2_16S_50ntFlanks</td><td>826</td><td>856</td><td>31</td><td>AAGGAAAGGTTAAAAAAAGTAAAAGGAACTC</td><td>40.3</td><td>8.6</td><td>1.8</td><td>4.8</td><td>1.8E-06</td><td>3.3E-06</td><td>MB231/MB468</td><td>MP</td></tr>
<tr><td>>hg38_21_+_8433172_8446622_RNA45SN1_45S_50ntFlanks@11539.11583.45|GCGCGGCGCCTCGCCTCGGCCGGCGCCTAGCAGCCGACTTAGAAC</td><td>>hg38_21_+_8433172_8446622_RNA45SN1_45S_50ntFlanks</td><td>11539</td><td>11583</td><td>45</td><td>GCGCGGCGCCTCGCCTCGGCCGGCGCCTAGCAGCCGACTTAGAAC</td><td>466.9</td><td>-4.5</td><td>0.8</td><td>-5.2</td><td>1.6E-07</td><td>3.3E-06</td><td>BT20/MB231</td><td>CYTO</td></tr>
<tr><td>>hg38_21_+_8433172_8446622_RNA45SN1_45S_50ntFlanks@11561.11587.27|GCGCCTAGCAGCCGACTTAGAACTGGT</td><td>>hg38_21_+_8433172_8446622_RNA45SN1_45S_50ntFlanks</td><td>11561</td><td>11587</td><td>27</td><td>GCGCCTAGCAGCCGACTTAGAACTGGT</td><td>166.6</td><td>-3.7</td><td>0.8</td><td>-4.7</td><td>2.0E-06</td><td>3.3E-06</td><td>MB231/MB468</td><td>CYTO</td></tr>
<tr><td>>hg38_21_+_8433172_8446622_RNA45SN1_45S_50ntFlanks@11208.11232.25|GCGGCGACTCTGGACGCGAGCCGGG</td><td>>hg38_21_+_8433172_8446622_RNA45SN1_45S_50ntFlanks</td><td>11208</td><td>11232</td><td>25</td><td>GCGGCGACTCTGGACGCGAGCCGGG</td><td>15546.2</td><td>-5.2</td><td>1.1</td><td>-4.8</td><td>1.8E-06</td><td>3.3E-06</td><td>BT20/MB468</td><td>CYTO</td></tr>
<tr><td>>hg38_21_+_8433172_8446622_RNA45SN1_45S_50ntFlanks@4546.4563.18|CCTGGATACCGCAGCTAG</td><td>>hg38_21_+_8433172_8446622_RNA45SN1_45S_50ntFlanks</td><td>4546</td><td>4563</td><td>18</td><td>CCTGGATACCGCAGCTAG</td><td>63.5</td><td>-9.1</td><td>1.8</td><td>-5.0</td><td>5.9E-07</td><td>3.3E-06</td><td>BT20/MB468</td><td>MP</td></tr>
<tr><td>>hg38_21_+_8433172_8446622_RNA45SN1_45S_50ntFlanks@6645.6675.31|CTCGTACGACTCTTAGCGGTGGATCACTCGG</td><td>>hg38_21_+_8433172_8446622_RNA45SN1_45S_50ntFlanks</td><td>6645</td><td>6675</td><td>31</td><td>CTCGTACGACTCTTAGCGGTGGATCACTCGG</td><td>530.0</td><td>-2.3</td><td>0.5</td><td>-4.7</td><td>2.3E-06</td><td>3.3E-06</td><td>BT20/MB468</td><td>NUC</td></tr>
<tr><td>>hg38_21_+_8433172_8446622_RNA45SN1_45S_50ntFlanks@11319.11344.26|CCCACGTCTCGTCGCGCGCGCGTCCG</td><td>>hg38_21_+_8433172_8446622_RNA45SN1_45S_50ntFlanks</td><td>11319</td><td>11344</td><td>26</td><td>CCCACGTCTCGTCGCGCGCGCGTCCG</td><td>10772.8</td><td>-9.4</td><td>2.0</td><td>-4.7</td><td>2.3E-06</td><td>3.3E-06</td><td>BT20/MB468</td><td>NUC</td></tr>
<tr><td>>hg38_21_+_8433172_8446622_RNA45SN1_45S_50ntFlanks@12874.12891.18|GCTGGAAAGGCGGCCGCC</td><td>>hg38_21_+_8433172_8446622_RNA45SN1_45S_50ntFlanks</td><td>12874</td><td>12891</td><td>18</td><td>GCTGGAAAGGCGGCCGCC</td><td>334.2</td><td>-6.2</td><td>1.3</td><td>-4.8</td><td>1.8E-06</td><td>3.3E-06</td><td>BT20/MB468</td><td>CYTO</td></tr>
<tr><td>>hg38_MT_+_1621_3279_RNR2_16S_50ntFlanks@51.87.37|GCTAAACCTAGCCCCAAACCCACTCCACCTTACTACC</td><td>>hg38_MT_+_1621_3279_RNR2_16S_50ntFlanks</td><td>51</td><td>87</td><td>37</td><td>GCTAAACCTAGCCCCAAACCCACTCCACCTTACTACC</td><td>4539.2</td><td>4.5</td><td>0.9</td><td>4.7</td><td>2.2E-06</td><td>3.3E-06</td><td>MB231/MB468</td><td>MITO</td></tr>
<tr><td>>hg38_21_+_8433172_8446622_RNA45SN1_45S_50ntFlanks@8403.8422.20|AGGTAAACGGGTGGGGTCCG</td><td>>hg38_21_+_8433172_8446622_RNA45SN1_45S_50ntFlanks</td><td>8403</td><td>8422</td><td>20</td><td>AGGTAAACGGGTGGGGTCCG</td><td>231.2</td><td>-5.9</td><td>1.2</td><td>-4.8</td><td>1.6E-06</td><td>3.3E-06</td><td>BT20/MB468</td><td>MITO</td></tr>
<tr><td>>MI0000077|hsa-mir-21&WithFlank&17|+|59841260|59841343@14.34.21(+1G)&[MIMAT0000076&hsa-miR-21-5p&offsets|0|-1(+1G);m-1&17|+|59841273|59841294&offsets|0|-1(+1G)]|TAGCTTATCAGACTGATGTTGG</td><td>>MI0000077|hsa-mir-21&WithFlank&17|+|59841260|59841343</td><td>14</td><td>34</td><td>21(+1G)</td><td>TAGCTTATCAGACTGATGTTGG</td><td>1401.4</td><td>5.3</td><td>1.1</td><td>4.7</td><td>2.2E-06</td><td>3.3E-06</td><td>MB231/MB468</td><td>MITO</td></tr>
<tr><td>>am_trnaMT_TyrGTA_MT_-_5826_5891@38.55.18|AAAGACAGGGGTTAGGCC</td><td>>am_trnaMT_TyrGTA_MT_-_5826_5891</td><td>38</td><td>55</td><td>18</td><td>AAAGACAGGGGTTAGGCC</td><td>66.0</td><td>9.8</td><td>2.0</td><td>5.0</td><td>5.9E-07</td><td>3.3E-06</td><td>BT20/MB468</td><td>MP</td></tr>
<tr><td>>hg38_21_+_8433172_8446622_RNA45SN1_45S_50ntFlanks@10410.10443.34|TCAGTCGGTCCTGAGAGATGGGCGAGCGCCGTTC</td><td>>hg38_21_+_8433172_8446622_RNA45SN1_45S_50ntFlanks</td><td>10410</td><td>10443</td><td>34</td><td>TCAGTCGGTCCTGAGAGATGGGCGAGCGCCGTTC</td><td>315.1</td><td>-6.7</td><td>1.4</td><td>-4.8</td><td>1.6E-06</td><td>3.3E-06</td><td>BT20/MB468</td><td>MITO</td></tr>
<tr><td>>hg38_21_+_8433172_8446622_RNA45SN1_45S_50ntFlanks@11890.11912.23|CTTGACTCTAGTCTGGCACGGTG</td><td>>hg38_21_+_8433172_8446622_RNA45SN1_45S_50ntFlanks</td><td>11890</td><td>11912</td><td>23</td><td>CTTGACTCTAGTCTGGCACGGTG</td><td>477.5</td><td>-3.4</td><td>0.7</td><td>-4.8</td><td>1.6E-06</td><td>3.3E-06</td><td>BT20/MB468</td><td>MITO</td></tr>
<tr><td>>hg38_21_+_8433172_8446622_RNA45SN1_45S_50ntFlanks@11217.11247.31|CTGGACGCGAGCCGGGCCCTTCCCGTGGATC</td><td>>hg38_21_+_8433172_8446622_RNA45SN1_45S_50ntFlanks</td><td>11217</td><td>11247</td><td>31</td><td>CTGGACGCGAGCCGGGCCCTTCCCGTGGATC</td><td>449.0</td><td>3.8</td><td>0.8</td><td>4.8</td><td>1.6E-06</td><td>3.3E-06</td><td>BT20/MB468</td><td>MITO</td></tr>
<tr><td>>hg38_21_+_8433172_8446622_RNA45SN1_45S_50ntFlanks@10402.10426.25|AACATGGGTCAGTCGGTCCTGAGAG</td><td>>hg38_21_+_8433172_8446622_RNA45SN1_45S_50ntFlanks</td><td>10402</td><td>10426</td><td>25</td><td>AACATGGGTCAGTCGGTCCTGAGAG</td><td>621.8</td><td>-7.3</td><td>1.5</td><td>-5.0</td><td>5.9E-07</td><td>3.3E-06</td><td>BT20/MB468</td><td>MP</td></tr>
<tr><td>>trna2_GlyGCC_21_-_18827107_18827177@4.31.28,trna35_GlyGCC_1_+_161413094_161413164@4.31.28,trna37_GlyGCC_1_+_161420467_161420537@4.31.28,trna39_GlyGCC_1_+_161427898_161427968@4.31.28,trna41_GlyGCC_1_+_161435258_161435328@4.31.28|TGGGTGGTTCAGTGGTAGAATTCTCGCC</td><td>>trna2_GlyGCC_21_-_18827107_18827177</td><td>4</td><td>31</td><td>28</td><td>TGGGTGGTTCAGTGGTAGAATTCTCGCC</td><td>83.3</td><td>10.1</td><td>2.0</td><td>5.0</td><td>5.9E-07</td><td>3.3E-06</td><td>BT20/MB468</td><td>MP</td></tr>
<tr><td>>hg38_21_+_8433172_8446622_RNA45SN1_45S_50ntFlanks@485.515.31|GATGTGGTGACGTCGTGCTCTCCCGGGCCGG</td><td>>hg38_21_+_8433172_8446622_RNA45SN1_45S_50ntFlanks</td><td>485</td><td>515</td><td>31</td><td>GATGTGGTGACGTCGTGCTCTCCCGGGCCGG</td><td>60.8</td><td>9.7</td><td>1.9</td><td>5.0</td><td>6.0E-07</td><td>3.3E-06</td><td>BT20/MB468</td><td>MP</td></tr>
<tr><td>>hg38_21_+_8433172_8446622_RNA45SN1_45S_50ntFlanks@7990.8010.21|GACGTGGCGACCCGCTGAATT</td><td>>hg38_21_+_8433172_8446622_RNA45SN1_45S_50ntFlanks</td><td>7990</td><td>8010</td><td>21</td><td>GACGTGGCGACCCGCTGAATT</td><td>355.4</td><td>4.2</td><td>0.9</td><td>4.7</td><td>2.1E-06</td><td>3.3E-06</td><td>MB231/MB468</td><td>NUC</td></tr>
<tr><td>>hg38_21_+_8433172_8446622_RNA45SN1_45S_50ntFlanks@8196.8225.30|CGTGGACGGTGTGAGGCCGGTAGCGGCCCC</td><td>>hg38_21_+_8433172_8446622_RNA45SN1_45S_50ntFlanks</td><td>8196</td><td>8225</td><td>30</td><td>CGTGGACGGTGTGAGGCCGGTAGCGGCCCC</td><td>398.8</td><td>-5.4</td><td>1.1</td><td>-4.7</td><td>2.2E-06</td><td>3.3E-06</td><td>MB231/MB468</td><td>MITO</td></tr>
<tr><td>>hg38_21_+_8433172_8446622_RNA45SN1_45S_50ntFlanks@10513.10532.20|CCGGAGTGGCGGAGATGGGC</td><td>>hg38_21_+_8433172_8446622_RNA45SN1_45S_50ntFlanks</td><td>10513</td><td>10532</td><td>20</td><td>CCGGAGTGGCGGAGATGGGC</td><td>795.9</td><td>-5.8</td><td>1.2</td><td>-4.8</td><td>1.8E-06</td><td>3.4E-06</td><td>MB231/MB468</td><td>MP</td></tr>
<tr><td>>hg38_21_+_8433172_8446622_RNA45SN1_45S_50ntFlanks@4444.4472.29|CCTTGCCTCTCGGCGCCCCCTCGATGCTC</td><td>>hg38_21_+_8433172_8446622_RNA45SN1_45S_50ntFlanks</td><td>4444</td><td>4472</td><td>29</td><td>CCTTGCCTCTCGGCGCCCCCTCGATGCTC</td><td>170.9</td><td>-6.1</td><td>1.3</td><td>-4.8</td><td>1.8E-06</td><td>3.4E-06</td><td>BT20/MB468</td><td>CYTO</td></tr>
<tr><td>>hg38_21_+_8433172_8446622_RNA45SN1_45S_50ntFlanks@4370.4394.25|TTAAAAAGCTCGTAGTTGGATCTTG</td><td>>hg38_21_+_8433172_8446622_RNA45SN1_45S_50ntFlanks</td><td>4370</td><td>4394</td><td>25</td><td>TTAAAAAGCTCGTAGTTGGATCTTG</td><td>51.9</td><td>-8.8</td><td>1.8</td><td>-5.0</td><td>6.0E-07</td><td>3.4E-06</td><td>BT20/MB468</td><td>MP</td></tr>
<tr><td>>am_trna10_ValCAC_5_-_180649395_180649467@1.34.34,trna18_ValCAC_5_-_180529253_180529325@1.34.34,trna2_ValCAC_5_+_180524070_180524142@1.34.34,trna6_ValCAC_5_+_180600650_180600722@1.34.34,trna85_ValCAC_1_-_161369490_161369562@1.34.34,trna90_ValCAC_1_-_149684088_149684161@1.34.34,trna98_ValCAC_1_-_149298555_149298627@1.34.34,trna9_ValCAC_6_+_26538282_26538354@1.34.34|GTTTCCGTAGTGTAGTGGTTATCACGTTCGCCTC</td><td>>am_trna10_ValCAC_5_-_180649395_180649467</td><td>1</td><td>34</td><td>34</td><td>GTTTCCGTAGTGTAGTGGTTATCACGTTCGCCTC</td><td>491.5</td><td>8.5</td><td>1.8</td><td>4.8</td><td>1.8E-06</td><td>3.4E-06</td><td>MB231/MB468</td><td>MP</td></tr>
<tr><td>>hg38_21_+_8433172_8446622_RNA45SN1_45S_50ntFlanks@10355.10384.30|AACTTTGAAGGCCGAAGTGGAGAAGGGTTC</td><td>>hg38_21_+_8433172_8446622_RNA45SN1_45S_50ntFlanks</td><td>10355</td><td>10384</td><td>30</td><td>AACTTTGAAGGCCGAAGTGGAGAAGGGTTC</td><td>146.2</td><td>-6.9</td><td>1.4</td><td>-4.8</td><td>1.8E-06</td><td>3.4E-06</td><td>MB231/MB468</td><td>MP</td></tr>
<tr><td>>hg38_21_+_8433172_8446622_RNA45SN1_45S_50ntFlanks@10394.10424.31|AGCAGTTGAACATGGGTCAGTCGGTCCTGAG</td><td>>hg38_21_+_8433172_8446622_RNA45SN1_45S_50ntFlanks</td><td>10394</td><td>10424</td><td>31</td><td>AGCAGTTGAACATGGGTCAGTCGGTCCTGAG</td><td>6424.6</td><td>5.7</td><td>1.1</td><td>5.0</td><td>6.0E-07</td><td>3.4E-06</td><td>BT20/MB468</td><td>MP</td></tr>
<tr><td>>hg38_21_+_8433172_8446622_RNA45SN1_45S_50ntFlanks@11881.11905.25|CCTGTTGAGCTTGACTCTAGTCTGG</td><td>>hg38_21_+_8433172_8446622_RNA45SN1_45S_50ntFlanks</td><td>11881</td><td>11905</td><td>25</td><td>CCTGTTGAGCTTGACTCTAGTCTGG</td><td>211.0</td><td>-6.9</td><td>1.4</td><td>-5.0</td><td>6.0E-07</td><td>3.4E-06</td><td>BT20/MB468</td><td>MP</td></tr>
<tr><td>>MI0000270|hsa-mir-181b-1&WithFlank&1|-|198858867|198858988@42.60.19;MI0000683|hsa-mir-181b-2&WithFlank&9|+|124693704|124693804@22.40.19&[MIMAT0000257&hsa-miR-181b-5p&offsets|0|-4;m-73&1|-|198858925|198858947&offsets|0|-4];[MIMAT0000257_1&hsa-miR-181b-5p&offsets|0|-4;m-72&9|+|124693725|124693747&offsets|0|-4]|AACATTCATTGCTGTCGGT</td><td>>MI0000270|hsa-mir-181b-1&WithFlank&1|-|198858867|198858988</td><td>42</td><td>60</td><td>19</td><td>AACATTCATTGCTGTCGGT</td><td>181.3</td><td>5.6</td><td>1.2</td><td>4.7</td><td>2.1E-06</td><td>3.4E-06</td><td>MB231/MB468</td><td>NUC</td></tr>
<tr><td>>hg38_21_+_8433172_8446622_RNA45SN1_45S_50ntFlanks@11668.11703.36|TGCCCAGTGCTCTGAATGTCAAAGTGAAGAAATTCA</td><td>>hg38_21_+_8433172_8446622_RNA45SN1_45S_50ntFlanks</td><td>11668</td><td>11703</td><td>36</td><td>TGCCCAGTGCTCTGAATGTCAAAGTGAAGAAATTCA</td><td>482.6</td><td>-4.0</td><td>0.8</td><td>-4.7</td><td>2.3E-06</td><td>3.4E-06</td><td>BT20/MB468</td><td>NUC</td></tr>
</table>

<table>
<tr><td>>hg38_21_+_8433172_8446622_RNA45SN1_45S_50ntFlanks@9533.9564.32|GAGGAAACTCTGGTGGAGGTCCGTAGCGGTCC</td><td>>hg38_21_+_8433172_8446622_RNA45SN1_45S_50ntFlanks</td><td>9533</td><td>9564</td><td>32</td><td>GAGGAAACTCTGGTGGAGGTCCGTAGCGGTCC</td><td>1707.9</td><td>-8.5</td><td>1.8</td><td>-4.7</td><td>2.3E-06</td><td>3.4E-06</td><td>BT20/MB468</td><td>NUC</td></tr>
<tr><td>>MI0000082|hsa-mir-25&WithFlank&7|-|100093554|100093649@58.79.22(+2A)&[MIMAT0000081&hsa-miR-25-3p&offsets|0|0(+2A);m-23&7|-|100093571|100093592&offsets|0|0(+2A)]|CATTGCACTTGTCTCGGTCTGAAA</td><td>>MI0000082|hsa-mir-25&WithFlank&7|-|100093554|100093649</td><td>58</td><td>79</td><td>22(+2A)</td><td>CATTGCACTTGTCTCGGTCTGAAA</td><td>388.6</td><td>3.3</td><td>0.7</td><td>4.7</td><td>2.1E-06</td><td>3.4E-06</td><td>MB231/MB468</td><td>NUC</td></tr>
<tr><td>>hg38_21_+_8433172_8446622_RNA45SN1_45S_50ntFlanks@11896.11922.27|TCTAGTCTGGCACGGTGAAGAGACATG</td><td>>hg38_21_+_8433172_8446622_RNA45SN1_45S_50ntFlanks</td><td>11896</td><td>11922</td><td>27</td><td>TCTAGTCTGGCACGGTGAAGAGACATG</td><td>245.3</td><td>11.3</td><td>2.1</td><td>5.2</td><td>1.5E-07</td><td>3.4E-06</td><td>BT20/MB231</td><td>MP</td></tr>
<tr><td>>hg38_21_+_8433172_8446622_RNA45SN1_45S_50ntFlanks@3840.3865.26|CTCCTACTTGGATAACTGTGGTAATT</td><td>>hg38_21_+_8433172_8446622_RNA45SN1_45S_50ntFlanks</td><td>3840</td><td>3865</td><td>26</td><td>CTCCTACTTGGATAACTGTGGTAATT</td><td>304.4</td><td>4.0</td><td>0.8</td><td>5.2</td><td>2.4E-07</td><td>3.4E-06</td><td>BT20/MB231</td><td>NUC</td></tr>
<tr><td>>hg38_21_+_8433172_8446622_RNA45SN1_45S_50ntFlanks@8171.8211.41|AAGTCCTTCTGATCGAGGCCCAGCCCGTGGACGGTGTGAGG</td><td>>hg38_21_+_8433172_8446622_RNA45SN1_45S_50ntFlanks</td><td>8171</td><td>8211</td><td>41</td><td>AAGTCCTTCTGATCGAGGCCCAGCCCGTGGACGGTGTGAGG</td><td>99.4</td><td>-10.5</td><td>2.2</td><td>-4.8</td><td>1.8E-06</td><td>3.4E-06</td><td>MB231/MB468</td><td>MP</td></tr>
<tr><td>>hg38_21_+_8433172_8446622_RNA45SN1_45S_50ntFlanks@11894.11920.27|ACTCTAGTCTGGCACGGTGAAGAGACA</td><td>>hg38_21_+_8433172_8446622_RNA45SN1_45S_50ntFlanks</td><td>11894</td><td>11920</td><td>27</td><td>ACTCTAGTCTGGCACGGTGAAGAGACA</td><td>53.8</td><td>-8.8</td><td>1.8</td><td>-5.0</td><td>6.0E-07</td><td>3.4E-06</td><td>BT20/MB468</td><td>MP</td></tr>
<tr><td>>hg38_21_+_8433172_8446622_RNA45SN1_45S_50ntFlanks@7976.8006.31|GCGACCTCAGATCAGACGTGGCGACCCGCTG</td><td>>hg38_21_+_8433172_8446622_RNA45SN1_45S_50ntFlanks</td><td>7976</td><td>8006</td><td>31</td><td>GCGACCTCAGATCAGACGTGGCGACCCGCTG</td><td>527.0</td><td>-2.8</td><td>0.6</td><td>-4.7</td><td>2.3E-06</td><td>3.4E-06</td><td>BT20/MB468</td><td>NUC</td></tr>
<tr><td>>hg38_21_+_8433172_8446622_RNA45SN1_45S_50ntFlanks@9416.9453.38|CAGTCCGCCGAGGGCGCACCACCGGCCCGTCTCGCCCG</td><td>>hg38_21_+_8433172_8446622_RNA45SN1_45S_50ntFlanks</td><td>9416</td><td>9453</td><td>38</td><td>CAGTCCGCCGAGGGCGCACCACCGGCCCGTCTCGCCCG</td><td>52.5</td><td>-8.8</td><td>1.8</td><td>-5.0</td><td>6.0E-07</td><td>3.4E-06</td><td>BT20/MB468</td><td>MP</td></tr>
<tr><td>>hg38_21_+_8433172_8446622_RNA45SN1_45S_50ntFlanks@10387.10410.24|TGTGAACAGCAGTTGAACATGGGT</td><td>>hg38_21_+_8433172_8446622_RNA45SN1_45S_50ntFlanks</td><td>10387</td><td>10410</td><td>24</td><td>TGTGAACAGCAGTTGAACATGGGT</td><td>253.6</td><td>-3.1</td><td>0.7</td><td>-4.7</td><td>2.1E-06</td><td>3.4E-06</td><td>MB231/MB468</td><td>CYTO</td></tr>
<tr><td>>hg38_MT_+_598_1651_RNR1_12S_50ntFlanks@52.90.39|ATAGGTTTGGTCCTAGCCTTTCTATTAGCTCTTAGTAAG</td><td>>hg38_MT_+_598_1651_RNR1_12S_50ntFlanks</td><td>52</td><td>90</td><td>39</td><td>ATAGGTTTGGTCCTAGCCTTTCTATTAGCTCTTAGTAAG</td><td>1109.6</td><td>4.7</td><td>1.0</td><td>4.8</td><td>1.6E-06</td><td>3.4E-06</td><td>BT20/MB468</td><td>MITO</td></tr>
<tr><td>>hg38_21_+_8433172_8446622_RNA45SN1_45S_50ntFlanks@11667.11688.22|CTGCCCAGTGCTCTGAATGTCA</td><td>>hg38_21_+_8433172_8446622_RNA45SN1_45S_50ntFlanks</td><td>11667</td><td>11688</td><td>22</td><td>CTGCCCAGTGCTCTGAATGTCA</td><td>352.2</td><td>5.3</td><td>1.1</td><td>4.7</td><td>2.1E-06</td><td>3.4E-06</td><td>MB231/MB468</td><td>CYTO</td></tr>
<tr><td>>hg38_21_+_8433172_8446622_RNA45SN1_45S_50ntFlanks@10905.10933.29|CTGGGCGCGCGCCGCGGCTGGACGAGGCG</td><td>>hg38_21_+_8433172_8446622_RNA45SN1_45S_50ntFlanks</td><td>10905</td><td>10933</td><td>29</td><td>CTGGGCGCGCGCCGCGGCTGGACGAGGCG</td><td>1456.2</td><td>-8.6</td><td>1.8</td><td>-4.7</td><td>2.3E-06</td><td>3.4E-06</td><td>BT20/MB468</td><td>NUC</td></tr>
<tr><td>>trnaMT_MetCAT_MT_+_4402_4469@32.71.40|ATACCCCGAAAATGTTGGTTATACCCTTCCCGTACTACCA</td><td>>trnaMT_MetCAT_MT_+_4402_4469</td><td>32</td><td>71</td><td>40</td><td>ATACCCCGAAAATGTTGGTTATACCCTTCCCGTACTACCA</td><td>320.7</td><td>8.2</td><td>1.7</td><td>4.7</td><td>2.1E-06</td><td>3.4E-06</td><td>MB231/MB468</td><td>NUC</td></tr>
<tr><td>>hg38_21_+_8433172_8446622_RNA45SN1_45S_50ntFlanks@11027.11060.34|CTCTCTCTCTCTCTCCCCCGCTCCCCGTCCTCCC</td><td>>hg38_21_+_8433172_8446622_RNA45SN1_45S_50ntFlanks</td><td>11027</td><td>11060</td><td>34</td><td>CTCTCTCTCTCTCTCCCCCGCTCCCCGTCCTCCC</td><td>6480.9</td><td>-9.5</td><td>2.0</td><td>-4.7</td><td>2.3E-06</td><td>3.4E-06</td><td>BT20/MB468</td><td>NUC</td></tr>
<tr><td>>hg38_21_+_8433172_8446622_RNA45SN1_45S_50ntFlanks@8169.8215.47|CCAAGTCCTTCTGATCGAGGCCCAGCCCGTGGACGGTGTGAGGCCGG</td><td>>hg38_21_+_8433172_8446622_RNA45SN1_45S_50ntFlanks</td><td>8169</td><td>8215</td><td>47</td><td>CCAAGTCCTTCTGATCGAGGCCCAGCCCGTGGACGGTGTGAGGCCGG</td><td>183.6</td><td>-4.7</td><td>1.0</td><td>-4.7</td><td>2.1E-06</td><td>3.4E-06</td><td>MB231/MB468</td><td>CYTO</td></tr>
<tr><td>>trna10_LysCTT_16_+_3241501_3241573@7.35.29,trna13_LysCTT_14_-_58706613_58706685@7.35.29,trna2_LysCTT_15_+_79152904_79152976@7.35.29,trna4_LysCTT_18_-_43669271_43669343@7.35.29|CTAGCTCAGTCGGTAGAGCATGGGACTCT</td><td>>trna10_LysCTT_16_+_3241501_3241573</td><td>7</td><td>35</td><td>29</td><td>CTAGCTCAGTCGGTAGAGCATGGGACTCT</td><td>282.7</td><td>8.0</td><td>1.7</td><td>4.7</td><td>2.2E-06</td><td>3.4E-06</td><td>MB231/MB468</td><td>MITO</td></tr>
<tr><td>>hg38_21_+_8433172_8446622_RNA45SN1_45S_50ntFlanks@12635.12658.24|AAGCTACCATCTGTGGGATTATGA</td><td>>hg38_21_+_8433172_8446622_RNA45SN1_45S_50ntFlanks</td><td>12635</td><td>12658</td><td>24</td><td>AAGCTACCATCTGTGGGATTATGA</td><td>54.8</td><td>-8.9</td><td>1.8</td><td>-5.0</td><td>6.0E-07</td><td>3.4E-06</td><td>BT20/MB468</td><td>MP</td></tr>
<tr><td>>hg38_21_+_8433172_8446622_RNA45SN1_45S_50ntFlanks@12418.12451.34|CGATGTCGGCTCTTCCTATCATTGTGAAGCAGAA</td><td>>hg38_21_+_8433172_8446622_RNA45SN1_45S_50ntFlanks</td><td>12418</td><td>12451</td><td>34</td><td>CGATGTCGGCTCTTCCTATCATTGTGAAGCAGAA</td><td>420.1</td><td>-7.5</td><td>1.6</td><td>-4.7</td><td>2.3E-06</td><td>3.4E-06</td><td>BT20/MB468</td><td>NUC</td></tr>
<tr><td>>hg38_21_+_8433172_8446622_RNA45SN1_45S_50ntFlanks@6646.6676.31|TCGTACGACTCTTAGCGGTGGATCACTCGGC</td><td>>hg38_21_+_8433172_8446622_RNA45SN1_45S_50ntFlanks</td><td>6646</td><td>6676</td><td>31</td><td>TCGTACGACTCTTAGCGGTGGATCACTCGGC</td><td>875.0</td><td>4.0</td><td>0.8</td><td>4.7</td><td>2.1E-06</td><td>3.4E-06</td><td>MB231/MB468</td><td>CYTO</td></tr>
<tr><td>>am_trna10_ValCAC_5_-_180649395_180649467@1.29.29,trna12_ValAAC_5_-_180645270_180645342@1.29.29,trna132_ValAAC_6_-_27721179_27721251@1.29.29,trna136_ValAAC_6_-_27648885_27648957@1.29.29,trna139_ValAAC_6_-_27618707_27618779@1.29.29,trna18_ValCAC_5_-_180529253_180529325@1.29.29,trna2_ValAAC_3_+_169490018_169490090@1.29.29,trna2_ValCAC_5_+_180524070_180524142@1.29.29,trna4_ValAAC_5_+_180591154_180591226@1.29.29,trna5_ValAAC_5_+_180596610_180596682@1.29.29,trna6_ValCAC_5_+_180600650_180600722@1.29.29,trna85_ValCAC_1_-_161369490_161369562@1.29.29,trna90_ValCAC_1_-_149684088_149684161@1.29.29,trna98_ValCAC_1_-_149298555_149298627@1.29.29,trna9_ValCAC_6_+_26538282_26538354@1.29.29|GTTTCCGTAGTGTAGTGGTTATCACGTTC</td><td>>am_trna10_ValCAC_5_-_180649395_180649467</td><td>1</td><td>29</td><td>29</td><td>GTTTCCGTAGTGTAGTGGTTATCACGTTC</td><td>9054.5</td><td>-4.5</td><td>1.0</td><td>-4.7</td><td>2.2E-06</td><td>3.4E-06</td><td>MB231/MB468</td><td>MITO</td></tr>
<tr><td>>am_MI0000086|hsa-mir-28&WithFlank&3|+|188688775|188688872@60.78.19&[MIMAT0004502&hsa-miR-28-3p&offsets|0|-3;m-28&3|+|188688834|188688855&offsets|0|-3]|CACTAGATTGTGAGCTCCT</td><td>>am_MI0000086|hsa-mir-28&WithFlank&3|+|188688775|188688872</td><td>60</td><td>78</td><td>19</td><td>CACTAGATTGTGAGCTCCT</td><td>336.9</td><td>7.7</td><td>1.6</td><td>5.0</td><td>6.1E-07</td><td>3.4E-06</td><td>BT20/MB468</td><td>MP</td></tr>
<tr><td>>hg38_21_+_8433172_8446622_RNA45SN1_45S_50ntFlanks@11771.11809.39|ATTAGTGACGCGCATGAATGGATGAACGAGATTCCCACT</td><td>>hg38_21_+_8433172_8446622_RNA45SN1_45S_50ntFlanks</td><td>11771</td><td>11809</td><td>39</td><td>ATTAGTGACGCGCATGAATGGATGAACGAGATTCCCACT</td><td>48.4</td><td>9.3</td><td>1.9</td><td>5.0</td><td>6.1E-07</td><td>3.4E-06</td><td>BT20/MB468</td><td>MP</td></tr>
<tr><td>>hg38_21_+_8433172_8446622_RNA45SN1_45S_50ntFlanks@8359.8377.19|AAGAACTTTGAAGAGAGAG</td><td>>hg38_21_+_8433172_8446622_RNA45SN1_45S_50ntFlanks</td><td>8359</td><td>8377</td><td>19</td><td>AAGAACTTTGAAGAGAGAG</td><td>461.2</td><td>-7.3</td><td>1.5</td><td>-4.8</td><td>1.8E-06</td><td>3.4E-06</td><td>MB231/MB468</td><td>MP</td></tr>
<tr><td>>MI0000077|hsa-mir-21&WithFlank&17|+|59841260|59841343@14.31.18(+1C)&[MIMAT0000076&hsa-miR-21-5p&offsets|0|-4(+1C);m-1&17|+|59841273|59841294&offsets|0|-4(+1C)]|TAGCTTATCAGACTGATGC</td><td>>MI0000077|hsa-mir-21&WithFlank&17|+|59841260|59841343</td><td>14</td><td>31</td><td>18(+1C)</td><td>TAGCTTATCAGACTGATGC</td><td>481.9</td><td>4.5</td><td>0.9</td><td>4.8</td><td>1.8E-06</td><td>3.4E-06</td><td>MB231/MB468</td><td>MP</td></tr>
<tr><td>>MI0000102|hsa-mir-100&WithFlank&11|-|122152223|122152314@19.40.22(+1U)&[MIMAT0000098&hsa-miR-100-5p&offsets|0|0(+1U);m-27&11|-|122152276|122152296&offsets|0|+1(+1U)]|AACCCGTAGATCCGAACTTGTGT</td><td>>MI0000102|hsa-mir-100&WithFlank&11|-|122152223|122152314</td><td>19</td><td>40</td><td>22(+1U)</td><td>AACCCGTAGATCCGAACTTGTGT</td><td>53466.7</td><td>13.1</td><td>2.8</td><td>4.7</td><td>2.2E-06</td><td>3.4E-06</td><td>MB231/MB468</td><td>MITO</td></tr>
<tr><td>>hg38_21_+_8433172_8446622_RNA45SN1_45S_50ntFlanks@10049.10081.33|GGCCGTCGCCGGCAGTCGAGAGTGGACGGGAGC</td><td>>hg38_21_+_8433172_8446622_RNA45SN1_45S_50ntFlanks</td><td>10049</td><td>10081</td><td>33</td><td>GGCCGTCGCCGGCAGTCGAGAGTGGACGGGAGC</td><td>48.2</td><td>9.3</td><td>1.9</td><td>5.0</td><td>6.1E-07</td><td>3.4E-06</td><td>BT20/MB468</td><td>MP</td></tr>
<tr><td>>hg38_21_+_8433172_8446622_RNA45SN1_45S_50ntFlanks@10421.10453.33|TGAGAGATGGGCGAGCGCCGTTCCGAAGGGACG</td><td>>hg38_21_+_8433172_8446622_RNA45SN1_45S_50ntFlanks</td><td>10421</td><td>10453</td><td>33</td><td>TGAGAGATGGGCGAGCGCCGTTCCGAAGGGACG</td><td>634.0</td><td>-4.9</td><td>1.0</td><td>-4.8</td><td>1.8E-06</td><td>3.4E-06</td><td>BT20/MB468</td><td>CYTO</td></tr>
<tr><td>>hg38_21_+_8433172_8446622_RNA45SN1_45S_50ntFlanks@10350.10377.28|ACGAGAACTTTGAAGGCCGAAGTGGAGA</td><td>>hg38_21_+_8433172_8446622_RNA45SN1_45S_50ntFlanks</td><td>10350</td><td>10377</td><td>28</td><td>ACGAGAACTTTGAAGGCCGAAGTGGAGA</td><td>6358.1</td><td>-5.5</td><td>1.1</td><td>-5.0</td><td>6.1E-07</td><td>3.4E-06</td><td>BT20/MB468</td><td>MP</td></tr>
<tr><td>>hg38_21_+_8433172_8446622_RNA45SN1_45S_50ntFlanks@4243.4264.22|CTTTCGAGGCCCTGTAATTGGA</td><td>>hg38_21_+_8433172_8446622_RNA45SN1_45S_50ntFlanks</td><td>4243</td><td>4264</td><td>22</td><td>CTTTCGAGGCCCTGTAATTGGA</td><td>183.3</td><td>-6.9</td><td>1.5</td><td>-4.8</td><td>1.8E-06</td><td>3.4E-06</td><td>MB231/MB468</td><td>MP</td></tr>
<tr><td>>trnaMT_SerGCT_MT_+_12207_12265@12.61.50|CAAGAACTGCTAACTCATGCCCCCATGTCTAACAACATGGCTTTCTCACC</td><td>>trnaMT_SerGCT_MT_+_12207_12265</td><td>12</td><td>61</td><td>50</td><td>CAAGAACTGCTAACTCATGCCCCCATGTCTAACAACATGGCTTTCTCACC</td><td>480.1</td><td>12.7</td><td>2.5</td><td>5.0</td><td>6.1E-07</td><td>3.4E-06</td><td>BT20/MB468</td><td>MP</td></tr>
<tr><td>>hg38_21_+_8433172_8446622_RNA45SN1_45S_50ntFlanks@9572.9590.19|CAAATCGGTCGTCCGACCT</td><td>>hg38_21_+_8433172_8446622_RNA45SN1_45S_50ntFlanks</td><td>9572</td><td>9590</td><td>19</td><td>CAAATCGGTCGTCCGACCT</td><td>69.3</td><td>-9.2</td><td>1.8</td><td>-5.0</td><td>6.1E-07</td><td>3.4E-06</td><td>BT20/MB468</td><td>MP</td></tr>
<tr><td>>hg38_21_+_8433172_8446622_RNA45SN1_45S_50ntFlanks@4978.4999.22|GGACACGGACAGGATTGACAGA</td><td>>hg38_21_+_8433172_8446622_RNA45SN1_45S_50ntFlanks</td><td>4978</td><td>4999</td><td>22</td><td>GGACACGGACAGGATTGACAGA</td><td>216.8</td><td>-6.9</td><td>1.4</td><td>-4.8</td><td>1.8E-06</td><td>3.4E-06</td><td>MB231/MB468</td><td>MP</td></tr>
<tr><td>>hg38_21_+_8433172_8446622_RNA45SN1_45S_50ntFlanks@12413.12436.24|TCCTTCGATGTCGGCTCTTCCTAT</td><td>>hg38_21_+_8433172_8446622_RNA45SN1_45S_50ntFlanks</td><td>12413</td><td>12436</td><td>24</td><td>TCCTTCGATGTCGGCTCTTCCTAT</td><td>1042.6</td><td>-5.8</td><td>1.2</td><td>-4.7</td><td>2.2E-06</td><td>3.4E-06</td><td>MB231/MB468</td><td>NUC</td></tr>
<tr><td>>hg38_21_+_8433172_8446622_RNA45SN1_45S_50ntFlanks@4472.4494.23|CTTAGCTGAGTGTCCCGCGGGGC</td><td>>hg38_21_+_8433172_8446622_RNA45SN1_45S_50ntFlanks</td><td>4472</td><td>4494</td><td>23</td><td>CTTAGCTGAGTGTCCCGCGGGGC</td><td>1014.9</td><td>-6.9</td><td>1.4</td><td>-4.8</td><td>1.9E-06</td><td>3.4E-06</td><td>BT20/MB468</td><td>CYTO</td></tr>
<tr><td>>hg38_21_+_8433172_8446622_RNA45SN1_45S_50ntFlanks@11579.11605.27|AGAACTGGTGCGGACCAGGGGAATCCG</td><td>>hg38_21_+_8433172_8446622_RNA45SN1_45S_50ntFlanks</td><td>11579</td><td>11605</td><td>27</td><td>AGAACTGGTGCGGACCAGGGGAATCCG</td><td>1616.9</td><td>-6.4</td><td>1.3</td><td>-5.0</td><td>6.2E-07</td><td>3.4E-06</td><td>BT20/MB468</td><td>MP</td></tr>
<tr><td>>hg38_21_+_8433172_8446622_RNA45SN1_45S_50ntFlanks@8359.8377.19|AAGAACTTTGAAGAGAGAG</td><td>>hg38_21_+_8433172_8446622_RNA45SN1_45S_50ntFlanks</td><td>8359</td><td>8377</td><td>19</td><td>AAGAACTTTGAAGAGAGAG</td><td>329.8</td><td>-7.6</td><td>1.5</td><td>-5.0</td><td>6.2E-07</td><td>3.4E-06</td><td>BT20/MB468</td><td>MP</td></tr>
<tr><td>>hg38_21_+_8433172_8446622_RNA45SN1_45S_50ntFlanks@9131.9152.22|GGTCGGGGGGCGGGGCGGACTG</td><td>>hg38_21_+_8433172_8446622_RNA45SN1_45S_50ntFlanks</td><td>9131</td><td>9152</td><td>22</td><td>GGTCGGGGGGCGGGGCGGACTG</td><td>727.7</td><td>-8.0</td><td>1.6</td><td>-5.0</td><td>6.2E-07</td><td>3.4E-06</td><td>BT20/MB468</td><td>MP</td></tr>
<tr><td>>hg38_21_+_8433172_8446622_RNA45SN1_45S_50ntFlanks@10399.10424.26|TTGAACATGGGTCAGTCGGTCCTGAG</td><td>>hg38_21_+_8433172_8446622_RNA45SN1_45S_50ntFlanks</td><td>10399</td><td>10424</td><td>26</td><td>TTGAACATGGGTCAGTCGGTCCTGAG</td><td>423.8</td><td>3.7</td><td>0.8</td><td>4.7</td><td>2.2E-06</td><td>3.5E-06</td><td>MB231/MB468</td><td>NUC</td></tr>
<tr><td>>hg38_21_+_8433172_8446622_RNA45SN1_45S_50ntFlanks@8168.8203.36|CCCAAGTCCTTCTGATCGAGGCCCAGCCCGTGGACG</td><td>>hg38_21_+_8433172_8446622_RNA45SN1_45S_50ntFlanks</td><td>8168</td><td>8203</td><td>36</td><td>CCCAAGTCCTTCTGATCGAGGCCCAGCCCGTGGACG</td><td>284.6</td><td>-6.7</td><td>1.4</td><td>-4.8</td><td>1.9E-06</td><td>3.5E-06</td><td>BT20/MB468</td><td>CYTO</td></tr>
<tr><td>>hg38_21_+_8433172_8446622_RNA45SN1_45S_50ntFlanks@12627.12660.34|ATGGGGCGAAGCTACCATCTGTGGGATTATGACT</td><td>>hg38_21_+_8433172_8446622_RNA45SN1_45S_50ntFlanks</td><td>12627</td><td>12660</td><td>34</td><td>ATGGGGCGAAGCTACCATCTGTGGGATTATGACT</td><td>159.9</td><td>-6.6</td><td>1.4</td><td>-4.8</td><td>1.9E-06</td><td>3.5E-06</td><td>MB231/MB468</td><td>MP</td></tr>
<tr><td>>hg38_21_+_8433172_8446622_RNA45SN1_45S_50ntFlanks@6733.6774.42|CATTGATCATCGACACTTCGAACGCACTTGCGGCCCCGGGTT</td><td>>hg38_21_+_8433172_8446622_RNA45SN1_45S_50ntFlanks</td><td>6733</td><td>6774</td><td>42</td><td>CATTGATCATCGACACTTCGAACGCACTTGCGGCCCCGGGTT</td><td>260.2</td><td>4.8</td><td>1.0</td><td>4.7</td><td>2.2E-06</td><td>3.5E-06</td><td>MB231/MB468</td><td>NUC</td></tr>
<tr><td>>MI0000063|hsa-let-7b&WithFlank&22|+|46113680|46113774@12.33.22&[MIMAT0000063&hsa-let-7b-5p&offsets|0|0;m-14&22|+|46113691|46113712&offsets|0|0]|TGAGGTAGTAGGTTGTGTGGTT</td><td>>MI0000063|hsa-let-7b&WithFlank&22|+|46113680|46113774</td><td>12</td><td>33</td><td>22</td><td>TGAGGTAGTAGGTTGTGTGGTT</td><td>53382.4</td><td>2.7</td><td>0.5</td><td>5.2</td><td>2.5E-07</td><td>3.5E-06</td><td>BT20/MB231</td><td>NUC</td></tr>
<tr><td>>hg38_21_+_8433172_8446622_RNA45SN1_45S_50ntFlanks@9736.9767.32|CCGAAACGATCTCAACCTATTCTCAAACTTTA</td><td>>hg38_21_+_8433172_8446622_RNA45SN1_45S_50ntFlanks</td><td>9736</td><td>9767</td><td>32</td><td>CCGAAACGATCTCAACCTATTCTCAAACTTTA</td><td>351.4</td><td>-5.2</td><td>1.1</td><td>-4.7</td><td>2.2E-06</td><td>3.5E-06</td><td>MB231/MB468</td><td>NUC</td></tr>
<tr><td>>hg38_21_+_8433172_8446622_RNA45SN1_45S_50ntFlanks@10425.10453.29|AGATGGGCGAGCGCCGTTCCGAAGGGACG</td><td>>hg38_21_+_8433172_8446622_RNA45SN1_45S_50ntFlanks</td><td>10425</td><td>10453</td><td>29</td><td>AGATGGGCGAGCGCCGTTCCGAAGGGACG</td><td>539.8</td><td>-4.8</td><td>1.0</td><td>-4.8</td><td>1.9E-06</td><td>3.5E-06</td><td>BT20/MB468</td><td>CYTO</td></tr>
<tr><td>>MI0000273|hsa-mir-183&WithFlank&7|-|129774899|129775020@34.55.22&[MIMAT0000261&hsa-miR-183-5p&offsets|+1|+1;m-41&7|-|129774966|129774987&offsets|0|0]|ATGGCACTGGTAGAATTCACTG</td><td>>MI0000273|hsa-mir-183&WithFlank&7|-|129774899|129775020</td><td>34</td><td>55</td><td>22</td><td>ATGGCACTGGTAGAATTCACTG</td><td>3844.7</td><td>4.0</td><td>0.8</td><td>5.2</td><td>1.7E-07</td><td>3.5E-06</td><td>BT20/MB231</td><td>CYTO</td></tr>
<tr><td>>trna5_AspGTC_12_+_98897281_98897352@36.74.39|CACGCGGGAGACCGGGGTTCAATTCCCCGACGGGGAGCC</td><td>>trna5_AspGTC_12_+_98897281_98897352</td><td>36</td><td>74</td><td>39</td><td>CACGCGGGAGACCGGGGTTCAATTCCCCGACGGGGAGCC</td><td>1759.2</td><td>-5.4</td><td>1.0</td><td>-5.2</td><td>1.7E-07</td><td>3.5E-06</td><td>BT20/MB231</td><td>CYTO</td></tr>
</table>

| | | | | | | | | | | | | | |
|---|---|---|---|---|---|---|---|---|---|---|---|---|---|
| >hg38_MT_+_1621_3279_RNR2_16S_50ntFlanks@1425.1449.25\|ACGATTAAAGTCCTACGTGATCTGA | >hg38_MT_+_1621_3279_RNR2_16S_50ntFlanks | 1425 | 1449 | 25 | ACGATTAAAGTCCTACGTGATCTGA | 54.1 | 9.0 | 1.9 | 4.8 | 1.9E-06 | 3.5E-06 | MB231/MB468 | MP |
| >hg38_21_+_8433172_8446622_RNA45SN1_45S_50ntFlanks@4723.4758.36\|CATTAATCAAGAACGAAAGTCGGAGGTTCGAAGACG | >hg38_21_+_8433172_8446622_RNA45SN1_45S_50ntFlanks | 4723 | 4758 | 36 | CATTAATCAAGAACGAAAGTCGGAGGTTCGAAGACG | 3498.7 | -3.5 | 0.7 | -4.8 | 1.9E-06 | 3.5E-06 | BT20/MB468 | CYTO |
| >hg38_21_+_8433172_8446622_RNA45SN1_45S_50ntFlanks@11663.11681.19\|ATTTCTGCCCAGTGCTCTG | >hg38_21_+_8433172_8446622_RNA45SN1_45S_50ntFlanks | 11663 | 11681 | 19 | ATTTCTGCCCAGTGCTCTG | 138.0 | -6.8 | 1.4 | -4.8 | 1.9E-06 | 3.5E-06 | MB231/MB468 | MP |
| >am_MI0000809\|hsa-mir-151a&WithFlank&8\|-\|140732558\|140732659@53.71.19&[MIMAT0000757&hsa-miR-151a-3p&offsets\|0\|-2;m-30&8\|-\|140732587\|140732607&offsets\|0\|-2]\|CTAGACTGAAGCTCCTTGA | >am_MI0000809\|hsa-mir-151a&WithFlank&8\|-\|140732558\|140732659 | 53 | 71 | 19 | CTAGACTGAAGCTCCTTGA | 337.5 | 6.8 | 1.4 | 4.7 | 2.3E-06 | 3.5E-06 | MB231/MB468 | MITO |
| >trna111_HisGTG_1_-_147774845_147774916@-1G.32.33,trna118_HisGTG_1_-_145396881_145396952@-1G.32.33,trna16_HisGTG_1_+_146544773_146544844@-1G.32.33,trna1_HisGTG_15_+_45493349_45493420@-1G.32.33,trna21_HisGTG_1_+_147753471_147753542@-1G.32.33,trna33_HisGTG_6_+_27125906_27125977@-1G.32.33,trna7_HisGTG_9_-_14433938_14434009@-1G.32.33,trna8_HisGTG_15_-_45492611_45492682@-1G.32.33,trna9_HisGTG_15_-_45490804_45490875@-1G.32.33\|GGCCGTGATCGTATAGTGGTTAGTACTCTGCGT | >trna111_HisGTG_1_-_147774845_147774916 | -1G | 32 | 33 | GGCCGTGATCGTATAGTGGTTAGTACTCTGCGT | 1853.7 | -5.3 | 1.0 | -5.2 | 2.5E-07 | 3.5E-06 | BT20/MB231 | NUC |
| >hg38_21_+_8433172_8446622_RNA45SN1_45S_50ntFlanks@12001.12032.32\|CTGCGGGCCGCCGGTGAAATACCACTACTCTG | >hg38_21_+_8433172_8446622_RNA45SN1_45S_50ntFlanks | 12001 | 12032 | 32 | CTGCGGGCCGCCGGTGAAATACCACTACTCTG | 3870.5 | -4.9 | 1.0 | -4.7 | 2.1E-06 | 3.5E-06 | MB231/MB468 | CYTO |
| >hg38_21_+_8433172_8446622_RNA45SN1_45S_50ntFlanks@4995.5024.30\|ACAGATTGATAGCTCTTTCTCGATTCCGTG | >hg38_21_+_8433172_8446622_RNA45SN1_45S_50ntFlanks | 4995 | 5024 | 30 | ACAGATTGATAGCTCTTTCTCGATTCCGTG | 900.5 | -4.5 | 0.9 | -4.8 | 1.6E-06 | 3.5E-06 | BT20/MB468 | MITO |
| >hg38_1_-_228634819_228635039_RNA5S12_5S_50ntFlanks@139.169.31\|GACCGCCTGGGAATACCGGGTGCTGTAGGCT | >hg38_1_-_228634819_228635039_RNA5S12_5S_50ntFlanks | 139 | 169 | 31 | GACCGCCTGGGAATACCGGGTGCTGTAGGCT | 660.9 | 4.3 | 0.9 | 4.7 | 2.2E-06 | 3.5E-06 | MB231/MB468 | NUC |
| >hg38_21_+_8433172_8446622_RNA45SN1_45S_50ntFlanks@11573.11590.18\|CGACTTAGAACTGGTGCG | >hg38_21_+_8433172_8446622_RNA45SN1_45S_50ntFlanks | 11573 | 11590 | 18 | CGACTTAGAACTGGTGCG | 702.8 | -3.8 | 0.8 | -4.8 | 1.9E-06 | 3.5E-06 | MB231/MB468 | MP |
| >hg38_21_+_8433172_8446622_RNA45SN1_45S_50ntFlanks@12539.12568.30\|ATGATGTGTTGTTGCCATGGTAATCCTGCT | >hg38_21_+_8433172_8446622_RNA45SN1_45S_50ntFlanks | 12539 | 12568 | 30 | ATGATGTGTTGTTGCCATGGTAATCCTGCT | 210.7 | -2.5 | 0.5 | -4.8 | 1.6E-06 | 3.5E-06 | BT20/MB468 | MITO |
| >hg38_21_+_8433172_8446622_RNA45SN1_45S_50ntFlanks@10385.10406.22\|CATGTGAACAGCAGTTGAACAT | >hg38_21_+_8433172_8446622_RNA45SN1_45S_50ntFlanks | 10385 | 10406 | 22 | CATGTGAACAGCAGTTGAACAT | 76.6 | -9.3 | 1.9 | -5.0 | 6.2E-07 | 3.5E-06 | BT20/MB468 | MP |
| >hg38_21_+_8433172_8446622_RNA45SN1_45S_50ntFlanks@5444.5469.26\|CTCGGATCGGCCCCGCCGGGGTCGGC | >hg38_21_+_8433172_8446622_RNA45SN1_45S_50ntFlanks | 5444 | 5469 | 26 | CTCGGATCGGCCCCGCCGGGGTCGGC | 54.2 | -8.8 | 1.8 | -5.0 | 6.3E-07 | 3.5E-06 | BT20/MB468 | MP |
| >hg38_21_+_8433172_8446622_RNA45SN1_45S_50ntFlanks@10771.10818.48\|CAGGTCTCCAAGGTGAACAGCCTCTGGCATGTTGGAACAATGTAGGTA | >hg38_21_+_8433172_8446622_RNA45SN1_45S_50ntFlanks | 10771 | 10818 | 48 | CAGGTCTCCAAGGTGAACAGCCTCTGGCATGTTGGAACAATGTAGGTA | 945.3 | -5.6 | 1.2 | -4.7 | 2.1E-06 | 3.5E-06 | MB231/MB468 | CYTO |
| >hg38_21_+_8433172_8446622_RNA45SN1_45S_50ntFlanks@6719.6751.33\|TGAATTGCAGGACACATTGATCATCGACACTTC | >hg38_21_+_8433172_8446622_RNA45SN1_45S_50ntFlanks | 6719 | 6751 | 33 | TGAATTGCAGGACACATTGATCATCGACACTTC | 588.0 | -7.2 | 1.5 | -4.8 | 1.9E-06 | 3.5E-06 | MB231/MB468 | MP |
| >hg38_21_+_8433172_8446622_RNA45SN1_45S_50ntFlanks@9546.9563.18\|TGGAGGTCCGTAGCGGTC | >hg38_21_+_8433172_8446622_RNA45SN1_45S_50ntFlanks | 9546 | 9563 | 18 | TGGAGGTCCGTAGCGGTC | 606.4 | -7.7 | 1.6 | -4.7 | 2.4E-06 | 3.5E-06 | BT20/MB468 | NUC |
| >hg38_21_+_8433172_8446622_RNA45SN1_45S_50ntFlanks@8183.8223.41\|TCGAGGCCCAGCCCGTGGACGGTGTGAGGCCGGTAGCGGCC | >hg38_21_+_8433172_8446622_RNA45SN1_45S_50ntFlanks | 8183 | 8223 | 41 | TCGAGGCCCAGCCCGTGGACGGTGTGAGGCCGGTAGCGGCC | 333.3 | -4.3 | 0.9 | -4.7 | 2.2E-06 | 3.5E-06 | MB231/MB468 | CYTO |
| >hg38_21_+_8433172_8446622_RNA45SN1_45S_50ntFlanks@9553.9570.18\|CCGTAGCGGTCCTGACGT | >hg38_21_+_8433172_8446622_RNA45SN1_45S_50ntFlanks | 9553 | 9570 | 18 | CCGTAGCGGTCCTGACGT | 530.9 | 2.3 | 0.5 | 4.7 | 2.2E-06 | 3.5E-06 | MB231/MB468 | CYTO |
| >hg38_1_-_228634819_228635039_RNA5S12_5S_50ntFlanks@85.108.24\|TCGTCTGATCTCGGAAGCTAAGCA | >hg38_1_-_228634819_228635039_RNA5S12_5S_50ntFlanks | 85 | 108 | 24 | TCGTCTGATCTCGGAAGCTAAGCA | 263.3 | 11.4 | 2.2 | 5.2 | 1.6E-07 | 3.5E-06 | BT20/MB231 | MP |
| >MI0000102\|hsa-mir-100&WithFlank&11\|-\|122152223\|122152314@20.40.21(+1A)&[MIMAT0000098&hsa-miR-100-5p&offsets\|+1\|0(+1A);m-27&11\|-\|122152276\|122152296&offsets\|+1\|+1(+1A)]\|ACCCGTAGATCCGAACTTGTGA | >MI0000102\|hsa-mir-100&WithFlank&11\|-\|122152223\|122152314 | 20 | 40 | 21(+1A) | ACCCGTAGATCCGAACTTGTGA | 102.8 | -10.1 | 1.9 | -5.2 | 1.6E-07 | 3.5E-06 | BT20/MB231 | MP |
| >hg38_21_+_8433172_8446622_RNA45SN1_45S_50ntFlanks@4069.4101.33\|CTTTCGATGGTAGTCGCCGTGCCTACCATGGTG | >hg38_21_+_8433172_8446622_RNA45SN1_45S_50ntFlanks | 4069 | 4101 | 33 | CTTTCGATGGTAGTCGCCGTGCCTACCATGGTG | 670.3 | -3.0 | 0.6 | -4.7 | 2.2E-06 | 3.5E-06 | MB231/MB468 | CYTO |
| >MI0003820\|hsa-mir-454&WithFlank&17\|-\|59137752\|59137878@70.92.23&[MIMAT0003885&hsa-miR-454-3p&offsets\|0\|0;m-272&17\|-\|59137787\|59137809&offsets\|0\|0]\|TAGTGCAATATTGCTTATAGGGT | >MI0003820\|hsa-mir-454&WithFlank&17\|-\|59137752\|59137878 | 70 | 92 | 23 | TAGTGCAATATTGCTTATAGGGT | 318.7 | 7.0 | 1.5 | 4.7 | 2.3E-06 | 3.5E-06 | MB231/MB468 | MITO |
| >trna10_AspGTC_12_-_125424193_125424264@37.73.37,trna12_AspGTC_12_-_125411891_125411962@37.73.37,trna144_AspGTC_6_-_27551236_27551307@37.73.37,trna38_AspGTC_17_-_8125556_8125627@37.73.37,trna45_AspGTC_6_+_27447453_27447524@37.73.37,trna48_AspGTC_6_+_27471523_27471594@37.73.37,trna4_AspGTC_12_+_96429799_96429870@37.73.37,trna69_AspGTC_1_-_161440205_161440276@37.73.37,trna72_AspGTC_1_-_161432824_161432895@37.73.37,trna75_AspGTC_1_-_161425414_161425485@37.73.37,trna78_AspGTC_1_-_161418033_161418104@37.73.37,trna81_AspGTC_1_-_161410615_161410686@37.73.37\|ACGCGGGAGACCGGGGTTCGATTCCCCGACGGGGAGC | >trna10_AspGTC_12_-_125424193_125424264 | 37 | 73 | 37 | ACGCGGGAGACCGGGGTTCGATTCCCCGACGGGGAGC | 23659.1 | -5.5 | 1.0 | -5.2 | 1.7E-07 | 3.5E-06 | BT20/MB231 | CYTO |
| >hg38_21_+_8433172_8446622_RNA45SN1_45S_50ntFlanks@10365.10390.26\|GCCGAAGTGGAGAAGGGTTCCATGTG | >hg38_21_+_8433172_8446622_RNA45SN1_45S_50ntFlanks | 10365 | 10390 | 26 | GCCGAAGTGGAGAAGGGTTCCATGTG | 48.4 | -8.7 | 1.7 | -5.0 | 6.3E-07 | 3.5E-06 | BT20/MB468 | MP |
| >hg38_21_+_8433172_8446622_RNA45SN1_45S_50ntFlanks@12938.12972.35\|ACCATTCGTAGACGACCTGCTTCTGGGTCGGGGTT | >hg38_21_+_8433172_8446622_RNA45SN1_45S_50ntFlanks | 12938 | 12972 | 35 | ACCATTCGTAGACGACCTGCTTCTGGGTCGGGGTT | 59.1 | -9.0 | 1.8 | -5.0 | 6.3E-07 | 3.5E-06 | BT20/MB468 | MP |
| >hg38_21_+_8433172_8446622_RNA45SN1_45S_50ntFlanks@12604.12624.21\|GTATGTGCTTGGCTGAGGAGC | >hg38_21_+_8433172_8446622_RNA45SN1_45S_50ntFlanks | 12604 | 12624 | 21 | GTATGTGCTTGGCTGAGGAGC | 308.1 | 5.4 | 1.1 | 4.7 | 2.2E-06 | 3.5E-06 | MB231/MB468 | CYTO |
| >hg38_21_+_8433172_8446622_RNA45SN1_45S_50ntFlanks@12997.13038.42\|CGCTGCGATCTATTGAAAGTCAGCCCTCGACACAAGGGTTTG | >hg38_21_+_8433172_8446622_RNA45SN1_45S_50ntFlanks | 12997 | 13038 | 42 | CGCTGCGATCTATTGAAAGTCAGCCCTCGACACAAGGGTTTG | 1100.7 | 6.6 | 1.4 | 4.7 | 2.2E-06 | 3.5E-06 | MB231/MB468 | NUC |
| >hg38_21_+_8433172_8446622_RNA45SN1_45S_50ntFlanks@4567.4589.23\|TAATGGAATAGGACCGCGGTTCT | >hg38_21_+_8433172_8446622_RNA45SN1_45S_50ntFlanks | 4567 | 4589 | 23 | TAATGGAATAGGACCGCGGTTCT | 312.8 | -3.6 | 0.8 | -4.8 | 1.9E-06 | 3.5E-06 | MB231/MB468 | MP |
| >hg38_21_+_8433172_8446622_RNA45SN1_45S_50ntFlanks@12156.12180.25\|GAGTTTGACTGGGGCGGTACACCTG | >hg38_21_+_8433172_8446622_RNA45SN1_45S_50ntFlanks | 12156 | 12180 | 25 | GAGTTTGACTGGGGCGGTACACCTG | 313.7 | -7.1 | 1.4 | -5.0 | 6.3E-07 | 3.5E-06 | BT20/MB468 | MP |
| >hg38_21_+_8433172_8446622_RNA45SN1_45S_50ntFlanks@11729.11746.18\|TAACTATGACTCTCTTAA | >hg38_21_+_8433172_8446622_RNA45SN1_45S_50ntFlanks | 11729 | 11746 | 18 | TAACTATGACTCTCTTAA | 202.4 | -3.3 | 0.7 | -4.7 | 2.2E-06 | 3.5E-06 | MB231/MB468 | NUC |
| >hg38_21_+_8433172_8446622_RNA45SN1_45S_50ntFlanks@5489.5506.18\|CTGAGAAGACGGTCGAAC | >hg38_21_+_8433172_8446622_RNA45SN1_45S_50ntFlanks | 5489 | 5506 | 18 | CTGAGAAGACGGTCGAAC | 258.4 | -6.8 | 1.4 | -4.8 | 1.9E-06 | 3.5E-06 | MB231/MB468 | MP |
| >hg38_21_+_8433172_8446622_RNA45SN1_45S_50ntFlanks@6652.6683.32\|GACTCTTAGCGGTGGATCACTCGGCTCGTGCG | >hg38_21_+_8433172_8446622_RNA45SN1_45S_50ntFlanks | 6652 | 6683 | 32 | GACTCTTAGCGGTGGATCACTCGGCTCGTGCG | 360.2 | 2.9 | 0.6 | 4.7 | 2.2E-06 | 3.5E-06 | MB231/MB468 | CYTO |
| >hg38_21_+_8433172_8446622_RNA45SN1_45S_50ntFlanks@11577.11612.36\|TTAGAACTGGTGCGGACCAGGGGAATCCGACTGTTT | >hg38_21_+_8433172_8446622_RNA45SN1_45S_50ntFlanks | 11577 | 11612 | 36 | TTAGAACTGGTGCGGACCAGGGGAATCCGACTGTTT | 187.5 | -5.6 | 1.2 | -4.7 | 2.3E-06 | 3.5E-06 | MB231/MB468 | MITO |
| >hg38_21_+_8433172_8446622_RNA45SN1_45S_50ntFlanks@9566.9592.27\|GACGTGCAAATCGGTCGTCCGACCTGG | >hg38_21_+_8433172_8446622_RNA45SN1_45S_50ntFlanks | 9566 | 9592 | 27 | GACGTGCAAATCGGTCGTCCGACCTGG | 486.1 | -2.8 | 0.6 | -4.7 | 2.2E-06 | 3.5E-06 | MB231/MB468 | CYTO |
| >hg38_21_+_8433172_8446622_RNA45SN1_45S_50ntFlanks@9739.9767.29\|AAACGATCTCAACCTATTCTCAAACTTTA | >hg38_21_+_8433172_8446622_RNA45SN1_45S_50ntFlanks | 9739 | 9767 | 29 | AAACGATCTCAACCTATTCTCAAACTTTA | 860.3 | -3.3 | 0.7 | -4.7 | 2.3E-06 | 3.5E-06 | MB231/MB468 | NUC |
| >hg38_21_+_8433172_8446622_RNA45SN1_45S_50ntFlanks@11575.11594.20\|ACTTAGAACTGGTGCGGACC | >hg38_21_+_8433172_8446622_RNA45SN1_45S_50ntFlanks | 11575 | 11594 | 20 | ACTTAGAACTGGTGCGGACC | 1118.0 | -5.8 | 1.2 | -4.8 | 1.9E-06 | 3.5E-06 | MB231/MB468 | MP |
| >hg38_21_+_8433172_8446622_RNA45SN1_45S_50ntFlanks@9554.9597.44\|CGTAGCGGTCCTGACGTGCAAATCGGTCGTCCGACCTGGGTATA | >hg38_21_+_8433172_8446622_RNA45SN1_45S_50ntFlanks | 9554 | 9597 | 44 | CGTAGCGGTCCTGACGTGCAAATCGGTCGTCCGACCTGGGTATA | 2806.8 | -6.2 | 1.3 | -4.7 | 2.3E-06 | 3.5E-06 | MB231/MB468 | NUC |
| >MI0000456\|hsa-mir-140&WithFlank&16\|+\|69933075\|69933186@68.87.20(+1A)&[MIMAT0004597&hsa-miR-140-3p&offsets\|0\|-1(+1A);m-45&16\|+\|69933143\|69933164&offsets\|-1\|-3(+1A)]\|TACCACAGGGTAGAACCACGA | >MI0000456\|hsa-mir-140&WithFlank&16\|+\|69933075\|69933186 | 68 | 87 | 20(+1A) | TACCACAGGGTAGAACCACGA | 42.5 | 8.6 | 1.8 | 4.8 | 1.9E-06 | 3.5E-06 | MB231/MB468 | MP |
| >hg38_21_+_8433172_8446622_RNA45SN1_45S_50ntFlanks@6734.6752.19\|ATTGATCATCGACACTTCG | >hg38_21_+_8433172_8446622_RNA45SN1_45S_50ntFlanks | 6734 | 6752 | 19 | ATTGATCATCGACACTTCG | 509.4 | 2.7 | 0.6 | 4.8 | 1.7E-06 | 3.5E-06 | BT20/MB468 | MITO |
| >MI0000087\|hsa-mir-29a&WithFlank&7\|-\|130876741\|130876816@48.66.19(+1A)&[MIMAT0000086&hsa-miR-29a-3p&offsets\|0\|-3(+1A);m-26&7\|-\|130876749\|130876769&offsets\|0\|-2(+1A)]\|TAGCACCATCTGAAATCGGA | >MI0000087\|hsa-mir-29a&WithFlank&7\|-\|130876741\|130876816 | 48 | 66 | 19(+1A) | TAGCACCATCTGAAATCGGA | 52.9 | 8.9 | 1.9 | 4.8 | 1.9E-06 | 3.5E-06 | MB231/MB468 | MP |
| >hg38_21_+_8433172_8446622_RNA45SN1_45S_50ntFlanks@10410.10434.25\|TCAGTCGGTCCTGAGAGATGGGCGA | >hg38_21_+_8433172_8446622_RNA45SN1_45S_50ntFlanks | 10410 | 10434 | 25 | TCAGTCGGTCCTGAGAGATGGGCGA | 61.2 | -9.0 | 1.8 | -5.0 | 6.4E-07 | 3.5E-06 | BT20/MB468 | MP |
| >hg38_21_+_8433172_8446622_RNA45SN1_45S_50ntFlanks@9545.9573.29\|GTGGAGGTCCGTAGCGGTCCTGACGTGCA | >hg38_21_+_8433172_8446622_RNA45SN1_45S_50ntFlanks | 9545 | 9573 | 29 | GTGGAGGTCCGTAGCGGTCCTGACGTGCA | 2225.3 | -3.6 | 0.8 | -4.7 | 2.3E-06 | 3.5E-06 | MB231/MB468 | NUC |
| >hg38_21_+_8433172_8446622_RNA45SN1_45S_50ntFlanks@12991.13015.25\|CTCCCTCGCTGCGATCTATTGAAAG | >hg38_21_+_8433172_8446622_RNA45SN1_45S_50ntFlanks | 12991 | 13015 | 25 | CTCCCTCGCTGCGATCTATTGAAAG | 2834.1 | -3.7 | 0.8 | -4.7 | 2.2E-06 | 3.5E-06 | MB231/MB468 | CYTO |
| >hg38_21_+_8433172_8446622_RNA45SN1_45S_50ntFlanks@11752.11780.29\|CCAAATGCCTCGTCATCTAATTAGTGACG | >hg38_21_+_8433172_8446622_RNA45SN1_45S_50ntFlanks | 11752 | 11780 | 29 | CCAAATGCCTCGTCATCTAATTAGTGACG | 52.2 | -8.8 | 1.8 | -5.0 | 6.4E-07 | 3.6E-06 | BT20/MB468 | MP |
| >hg38_21_+_8433172_8446622_RNA45SN1_45S_50ntFlanks@8365.8395.31\|TTTGAAGAGAGAGTTCAAGAGGGCGTGAAAC | >hg38_21_+_8433172_8446622_RNA45SN1_45S_50ntFlanks | 8365 | 8395 | 31 | TTTGAAGAGAGAGTTCAAGAGGGCGTGAAAC | 53.8 | 9.5 | 1.9 | 5.0 | 6.4E-07 | 3.6E-06 | BT20/MB468 | MP |
| >hg38_21_+_8433172_8446622_RNA45SN1_45S_50ntFlanks@11573.11591.19\|CGACTTAGAACTGGTGCGG | >hg38_21_+_8433172_8446622_RNA45SN1_45S_50ntFlanks | 11573 | 11591 | 19 | CGACTTAGAACTGGTGCGG | 721.8 | -4.9 | 1.0 | -5.0 | 6.4E-07 | 3.6E-06 | BT20/MB468 | MP |

| | | | | | | | | | | | | | |
|---|---|---|---|---|---|---|---|---|---|---|---|---|---|
| >hg38_21_+_8433172_8446622_RNA45SN1_45S_50ntFlanks@11886.11906.21\|TGAGCTTGACTCTAGTCTGGC | >hg38_21_+_8433172_8446622_RNA45SN1_45S_50ntFlanks | 11886 | 11906 | 21 | TGAGCTTGACTCTAGTCTGGC | 63.2 | -9.1 | 1.8 | -5.0 | 6.4E-07 | 3.6E-06 | BT20/MB468 | MP |
| >hg38_21_+_8433172_8446622_RNA45SN1_45S_50ntFlanks@3870.3898.29\|AGCTAATACATGCCGACGGGCGCTGACCC | >hg38_21_+_8433172_8446622_RNA45SN1_45S_50ntFlanks | 3870 | 3898 | 29 | AGCTAATACATGCCGACGGGCGCTGACCC | 349.5 | -5.0 | 1.0 | -4.8 | 1.9E-06 | 3.6E-06 | BT20/MB468 | CYTO |
| >trna10_LysCTT_16_+_3241501_3241573@1.36.36,trna13_LysCTT_14_-_58706613_58706685@1.36.36,trna2_LysCTT_15_+_79152904_79152976@1.36.36\|GCCCGGCTAGCTCAGTCGGTAGAGCATGGGACTCTT | >trna10_LysCTT_16_+_3241501_3241573 | 1 | 36 | 36 | GCCCGGCTAGCTCAGTCGGTAGAGCATGGGACTCTT | 5094.1 | 5.9 | 1.2 | 4.7 | 2.3E-06 | 3.6E-06 | MB231/MB468 | MITO |
| >hg38_21_+_8433172_8446622_RNA45SN1_45S_50ntFlanks@11729.11768.40\|TAACTATGACTCTCTTAAGGTAGCCAAATGCCTCGTCATC | >hg38_21_+_8433172_8446622_RNA45SN1_45S_50ntFlanks | 11729 | 11768 | 40 | TAACTATGACTCTCTTAAGGTAGCCAAATGCCTCGTCATC | 57.7 | -8.9 | 1.8 | -5.0 | 6.4E-07 | 3.6E-06 | BT20/MB468 | MP |
| >hg38_21_+_8433172_8446622_RNA45SN1_45S_50ntFlanks@8104.8139.36\|CCGCGGCGGGGCGCGGGACATGTGGCGTACGGAAGA | >hg38_21_+_8433172_8446622_RNA45SN1_45S_50ntFlanks | 8104 | 8139 | 36 | CCGCGGCGGGGCGCGGGACATGTGGCGTACGGAAGA | 1411.3 | -6.6 | 1.4 | -4.8 | 1.9E-06 | 3.6E-06 | MB231/MB468 | MP |
| >MI0000469\|hsa-mir-125a&WithFlank&19\|+\|51693248\|51693345@21.43.23&[MIMAT0000443&hsa-miR-125a-5p&offsets\|0\|-1;m-63&19\|+\|51693268\|51693289&offsets\|0\|+1]\|TCCCTGAGACCCTTTAACCTGTG | >MI0000469\|hsa-mir-125a&WithFlank&19\|+\|51693248\|51693345 | 21 | 43 | 23 | TCCCTGAGACCCTTTAACCTGTG | 11327.1 | 3.5 | 0.7 | 4.7 | 2.3E-06 | 3.6E-06 | MB231/MB468 | NUC |
| >hg38_21_+_8433172_8446622_RNA45SN1_45S_50ntFlanks@8184.8204.21\|CGAGGCCCAGCCCGTGGACGG | >hg38_21_+_8433172_8446622_RNA45SN1_45S_50ntFlanks | 8184 | 8204 | 21 | CGAGGCCCAGCCCGTGGACGG | 343.1 | -6.7 | 1.4 | -4.7 | 2.3E-06 | 3.6E-06 | MB231/MB468 | MITO |
| >hg38_21_+_8433172_8446622_RNA45SN1_45S_50ntFlanks@4555.4577.23\|CGCAGCTAGGAATAATGGAATAG | >hg38_21_+_8433172_8446622_RNA45SN1_45S_50ntFlanks | 4555 | 4577 | 23 | CGCAGCTAGGAATAATGGAATAG | 207.8 | -5.9 | 1.2 | -4.7 | 2.3E-06 | 3.6E-06 | MB231/MB468 | NUC |
| >trna119_LysCTT_1_-_145395522_145395594@1.29.29,trna11_LysCTT_5_-_180648979_180649051@1.29.29,trna13_LysCTT_6_+_26556774_26556846@1.29.29,trna32_LysCTT_16_-_3207406_3207478@1.29.29,trna7_LysCTT_16_+_3225692_3225764@1.29.29,trna9_LysCTT_5_+_180634755_180634827@1.29.29\|GCCCGGCTAGCTCAGTCGGTAGAGCATGA | >trna119_LysCTT_1_-_145395522_145395594 | 1 | 29 | 29 | GCCCGGCTAGCTCAGTCGGTAGAGCATGA | 2819.0 | -3.8 | 0.8 | -4.7 | 2.2E-06 | 3.6E-06 | MB231/MB468 | CYTO |
| >hg38_21_+_8433172_8446622_RNA45SN1_45S_50ntFlanks@12934.12965.32\|CTAAACCATTCGTAGACGACCTGCTTCTGGGT | >hg38_21_+_8433172_8446622_RNA45SN1_45S_50ntFlanks | 12934 | 12965 | 32 | CTAAACCATTCGTAGACGACCTGCTTCTGGGT | 979.5 | -5.8 | 1.2 | -4.8 | 1.9E-06 | 3.6E-06 | BT20/MB468 | CYTO |
| >hg38_21_+_8433172_8446622_RNA45SN1_45S_50ntFlanks@12626.12652.27\|AATGGGGCGAAGCTACCATCTGTGGGA | >hg38_21_+_8433172_8446622_RNA45SN1_45S_50ntFlanks | 12626 | 12652 | 27 | AATGGGGCGAAGCTACCATCTGTGGGA | 286.2 | -3.1 | 0.7 | -4.7 | 2.2E-06 | 3.6E-06 | MB231/MB468 | CYTO |
| >hg38_21_+_8433172_8446622_RNA45SN1_45S_50ntFlanks@4380.4410.31\|CGTAGTTGGATCTTGGGAGCGGGCGGGCGGT | >hg38_21_+_8433172_8446622_RNA45SN1_45S_50ntFlanks | 4380 | 4410 | 31 | CGTAGTTGGATCTTGGGAGCGGGCGGGCGGT | 398.7 | -5.6 | 1.2 | -4.7 | 2.5E-06 | 3.6E-06 | BT20/MB468 | NUC |
| >MI0000542\|hsa-mir-320a&WithFlank&8\|-\|22244960\|22245043@49.69.21&[MIMAT0000510&hsa-miR-320a-3p&offsets\|+1\|0;m-60&8\|-\|22244975\|22244996&offsets\|+1\|0]\|AAAGCTGGGTTGAGAGGGCGA | >MI0000542\|hsa-mir-320a&WithFlank&8\|-\|22244960\|22245043 | 49 | 69 | 21 | AAAGCTGGGTTGAGAGGGCGA | 3800.1 | 7.2 | 1.4 | 5.0 | 6.4E-07 | 3.6E-06 | BT20/MB468 | MP |
| >hg38_21_+_8433172_8446622_RNA45SN1_45S_50ntFlanks@11206.11232.27\|CGGCGGCGACTCTGGACGCGAGCCGGG | >hg38_21_+_8433172_8446622_RNA45SN1_45S_50ntFlanks | 11206 | 11232 | 27 | CGGCGGCGACTCTGGACGCGAGCCGGG | 440.6 | -5.7 | 1.2 | -4.7 | 2.4E-06 | 3.6E-06 | MB231/MB468 | MITO |
| >hg38_21_+_8433172_8446622_RNA45SN1_45S_50ntFlanks@12420.12446.27\|ATGTCGGCTCTTCCTATCATTGTGAAG | >hg38_21_+_8433172_8446622_RNA45SN1_45S_50ntFlanks | 12420 | 12446 | 27 | ATGTCGGCTCTTCCTATCATTGTGAAG | 3536.1 | -3.4 | 0.7 | -4.7 | 2.4E-06 | 3.6E-06 | MB231/MB468 | MITO |
| >hg38_21_+_8433172_8446622_RNA45SN1_45S_50ntFlanks@7988.8011.24\|CAGACGTGGCGACCCGCTGAATTT | >hg38_21_+_8433172_8446622_RNA45SN1_45S_50ntFlanks | 7988 | 8011 | 24 | CAGACGTGGCGACCCGCTGAATTT | 2835.4 | -3.7 | 0.8 | -4.7 | 2.4E-06 | 3.6E-06 | MB231/MB468 | MITO |
| >hg38_21_+_8433172_8446622_RNA45SN1_45S_50ntFlanks@11668.11706.39\|TGCCCAGTGCTCTGAATGTCAAAGTGAAGAAATTCAATG | >hg38_21_+_8433172_8446622_RNA45SN1_45S_50ntFlanks | 11668 | 11706 | 39 | TGCCCAGTGCTCTGAATGTCAAAGTGAAGAAATTCAATG | 690.2 | 4.2 | 0.9 | 4.7 | 2.3E-06 | 3.6E-06 | MB231/MB468 | NUC |
| >hg38_21_+_8433172_8446622_RNA45SN1_45S_50ntFlanks@8112.8135.24\|GGGCGCGGGACATGTGGCGTACGG | >hg38_21_+_8433172_8446622_RNA45SN1_45S_50ntFlanks | 8112 | 8135 | 24 | GGGCGCGGGACATGTGGCGTACGG | 272.0 | 11.4 | 2.2 | 5.2 | 1.6E-07 | 3.6E-06 | BT20/MB231 | MP |
| >hg38_21_+_8433172_8446622_RNA45SN1_45S_50ntFlanks@10798.10825.28\|CATGTTGGAACAATGTAGGTAAGGGAAG | >hg38_21_+_8433172_8446622_RNA45SN1_45S_50ntFlanks | 10798 | 10825 | 28 | CATGTTGGAACAATGTAGGTAAGGGAAG | 229.3 | -4.3 | 0.9 | -4.7 | 2.2E-06 | 3.6E-06 | MB231/MB468 | CYTO |
| >hg38_21_+_8433172_8446622_RNA45SN1_45S_50ntFlanks@13008.13025.18\|ATTGAAAGTCAGCCCTCG | >hg38_21_+_8433172_8446622_RNA45SN1_45S_50ntFlanks | 13008 | 13025 | 18 | ATTGAAAGTCAGCCCTCG | 1449.6 | -5.9 | 1.2 | -4.8 | 2.0E-06 | 3.6E-06 | BT20/MB468 | CYTO |
| >MI0000459\|hsa-mir-143&WithFlank&5\|+\|149428912\|149429029@67.86.20(+1U)&[MIMAT0000435&hsa-miR-143-3p&offsets\|0\|-1(+1U);m-3&5\|+\|149428978\|149428997&offsets\|0\|0(+1U)]\|TGAGATGAAGCACTGTAGCTT | >MI0000459\|hsa-mir-143&WithFlank&5\|+\|149428912\|149429029 | 67 | 86 | 20(+1U) | TGAGATGAAGCACTGTAGCTT | 123.3 | -6.5 | 1.4 | -4.8 | 2.0E-06 | 3.6E-06 | MB231/MB468 | MP |
| >hg38_21_+_8433172_8446622_RNA45SN1_45S_50ntFlanks@12073.12098.26\|CCGAGGGGCTCTCGCTTCTGGCGCCA | >hg38_21_+_8433172_8446622_RNA45SN1_45S_50ntFlanks | 12073 | 12098 | 26 | CCGAGGGGCTCTCGCTTCTGGCGCCA | 366.3 | -4.3 | 0.9 | -4.7 | 2.2E-06 | 3.6E-06 | MB231/MB468 | CYTO |
| >hg38_21_+_8433172_8446622_RNA45SN1_45S_50ntFlanks@3705.3741.37\|TACCTGGTTGATCCTGCCAGTAGCATATGCTTGTCTC | >hg38_21_+_8433172_8446622_RNA45SN1_45S_50ntFlanks | 3705 | 3741 | 37 | TACCTGGTTGATCCTGCCAGTAGCATATGCTTGTCTC | 1906.6 | 3.1 | 0.6 | 5.2 | 2.6E-07 | 3.6E-06 | BT20/MB231 | NUC |
| >hg38_MT_+_1621_3279_RNR2_16S_50ntFlanks@1347.1388.42\|CAACAATAGGGTTTACGACCTCGATGTTGGATCAGGACATCC | >hg38_MT_+_1621_3279_RNR2_16S_50ntFlanks | 1347 | 1388 | 42 | CAACAATAGGGTTTACGACCTCGATGTTGGATCAGGACATCC | 37.7 | 8.5 | 1.8 | 4.8 | 2.0E-06 | 3.6E-06 | MB231/MB468 | MP |
| >hg38_1_-_228634819_228635039_RNA5S12_5S_50ntFlanks@148.168.21\|GGAATACCGGGTGCTGTAGGC | >hg38_1_-_228634819_228635039_RNA5S12_5S_50ntFlanks | 148 | 168 | 21 | GGAATACCGGGTGCTGTAGGC | 185.5 | -6.8 | 1.4 | -4.8 | 2.0E-06 | 3.6E-06 | MB231/MB468 | MP |
| >hg38_21_+_8433172_8446622_RNA45SN1_45S_50ntFlanks@10385.10424.40\|CATGTGAACAGCAGTTGAACATGGGTCAGTCGGTCCTGAG | >hg38_21_+_8433172_8446622_RNA45SN1_45S_50ntFlanks | 10385 | 10424 | 40 | CATGTGAACAGCAGTTGAACATGGGTCAGTCGGTCCTGAG | 995.2 | -3.6 | 0.8 | -4.7 | 2.5E-06 | 3.6E-06 | BT20/MB468 | NUC |
| >hg38_21_+_8433172_8446622_RNA45SN1_45S_50ntFlanks@12935.12952.18\|TAAACCATTCGTAGACGA | >hg38_21_+_8433172_8446622_RNA45SN1_45S_50ntFlanks | 12935 | 12952 | 18 | TAAACCATTCGTAGACGA | 227.1 | -6.1 | 1.3 | -4.8 | 2.0E-06 | 3.6E-06 | BT20/MB468 | CYTO |
| >hg38_21_+_8433172_8446622_RNA45SN1_45S_50ntFlanks@4726.4751.26\|TAATCAAGAACGAAAGTCGGAGGTTC | >hg38_21_+_8433172_8446622_RNA45SN1_45S_50ntFlanks | 4726 | 4751 | 26 | TAATCAAGAACGAAAGTCGGAGGTTC | 203.1 | -5.9 | 1.2 | -4.7 | 2.3E-06 | 3.6E-06 | MB231/MB468 | NUC |
| >hg38_21_+_8433172_8446622_RNA45SN1_45S_50ntFlanks@4736.4759.24\|CGAAAGTCGGAGGTTCGAAGACGA | >hg38_21_+_8433172_8446622_RNA45SN1_45S_50ntFlanks | 4736 | 4759 | 24 | CGAAAGTCGGAGGTTCGAAGACGA | 184.5 | -4.7 | 1.0 | -4.7 | 2.3E-06 | 3.6E-06 | MB231/MB468 | NUC |
| >hg38_21_+_8433172_8446622_RNA45SN1_45S_50ntFlanks@11785.11806.22\|TGAATGGATGAACGAGATTCCC | >hg38_21_+_8433172_8446622_RNA45SN1_45S_50ntFlanks | 11785 | 11806 | 22 | TGAATGGATGAACGAGATTCCC | 749.5 | -6.9 | 1.5 | -4.7 | 2.2E-06 | 3.6E-06 | MB231/MB468 | CYTO |
| >hg38_21_+_8433172_8446622_RNA45SN1_45S_50ntFlanks@3985.4009.25\|CGGCTTTGGTGACTCTAGATAACCT | >hg38_21_+_8433172_8446622_RNA45SN1_45S_50ntFlanks | 3985 | 4009 | 25 | CGGCTTTGGTGACTCTAGATAACCT | 232.5 | -4.8 | 1.0 | -4.7 | 2.3E-06 | 3.6E-06 | MB231/MB468 | NUC |
| >MI0000299\|hsa-mir-222&WithFlank&X\|-\|45747009\|45747130@75.99.25(+4U)&[MIMAT0000279&hsa-miR-222-3p&offsets\|0\|+4(+4U);m-140&X\|-\|45747033\|45747056&offsets\|0\|+1(+4U)]\|AGCTACATCTGGCTACTGGGTCTCTTTTT | >MI0000299\|hsa-mir-222&WithFlank&X\|-\|45747009\|45747130 | 75 | 99 | 25(+4U) | AGCTACATCTGGCTACTGGGTCTCTTTTT | 569.5 | -4.4 | 0.9 | -5.2 | 2.0E-07 | 3.6E-06 | BT20/MB231 | MITO |
| >am_trna119_LysCTT_1_-_145395522_145395594@7.36.30,trna11_LysCTT_5_-_180648979_180649051@7.36.30,trna128_LysCTT_1_-_55423542_55423614@7.36.30,trna13_LysCTT_6_+_26556774_26556846@7.36.30,trna6_LysCTT_19_-_52425396_52425468@7.36.30,trna7_LysCTT_16_+_3225692_3225764@7.36.30,trna9_LysCTT_5_+_180634755_180634827@7.36.30\|CTAGCTCAGTCGGTAGAGCATGAGACTCTT | >am_trna119_LysCTT_1_-_145395522_145395594 | 7 | 36 | 30 | CTAGCTCAGTCGGTAGAGCATGAGACTCTT | 43.5 | 8.7 | 1.8 | 4.8 | 2.0E-06 | 3.6E-06 | MB231/MB468 | MP |
| >hg38_21_+_8433172_8446622_RNA45SN1_45S_50ntFlanks@11669.11692.24\|GCCCAGTGCTCTGAATGTCAAAGT | >hg38_21_+_8433172_8446622_RNA45SN1_45S_50ntFlanks | 11669 | 11692 | 24 | GCCCAGTGCTCTGAATGTCAAAGT | 269.1 | 6.8 | 1.4 | 4.8 | 2.0E-06 | 3.6E-06 | BT20/MB468 | CYTO |
| >hg38_21_+_8433172_8446622_RNA45SN1_45S_50ntFlanks@12420.12446.27\|ATGTCGGCTCTTCCTATCATTGTGAAG | >hg38_21_+_8433172_8446622_RNA45SN1_45S_50ntFlanks | 12420 | 12446 | 27 | ATGTCGGCTCTTCCTATCATTGTGAAG | 10755.0 | -2.8 | 0.6 | -4.7 | 2.3E-06 | 3.6E-06 | MB231/MB468 | CYTO |
| >hg38_21_+_8433172_8446622_RNA45SN1_45S_50ntFlanks@10786.10815.30\|AACAGCCTCTGGCATGTTGGAACAATGTAG | >hg38_21_+_8433172_8446622_RNA45SN1_45S_50ntFlanks | 10786 | 10815 | 30 | AACAGCCTCTGGCATGTTGGAACAATGTAG | 487.6 | -5.2 | 1.1 | -4.7 | 2.3E-06 | 3.6E-06 | MB231/MB468 | CYTO |
| >hg38_21_+_8433172_8446622_RNA45SN1_45S_50ntFlanks@5186.5204.19\|CAATAACAGGTCTGTGATG | >hg38_21_+_8433172_8446622_RNA45SN1_45S_50ntFlanks | 5186 | 5204 | 19 | CAATAACAGGTCTGTGATG | 367.5 | -4.8 | 1.0 | -5.0 | 6.6E-07 | 3.6E-06 | BT20/MB468 | MP |
| >hg38_21_+_8433172_8446622_RNA45SN1_45S_50ntFlanks@3769.3794.26\|CGCACGGCCGGTACAGTGAAACTGCG | >hg38_21_+_8433172_8446622_RNA45SN1_45S_50ntFlanks | 3769 | 3794 | 26 | CGCACGGCCGGTACAGTGAAACTGCG | 694.9 | -5.8 | 1.2 | -4.7 | 2.4E-06 | 3.6E-06 | MB231/MB468 | MITO |
| >MI0000077\|hsa-mir-21&WithFlank&17\|+\|59841260\|59841343@14.31.18(+1C)&[MIMAT0000076&hsa-miR-21-5p&offsets\|0\|-4(+1C);m-1&17\|+\|59841273\|59841294&offsets\|0\|-4(+1C)]\|TAGCTTATCAGACTGATGC | >MI0000077\|hsa-mir-21&WithFlank&17\|+\|59841260\|59841343 | 14 | 31 | 18(+1C) | TAGCTTATCAGACTGATGC | 348.2 | 8.6 | 1.8 | 4.8 | 2.0E-06 | 3.6E-06 | BT20/MB468 | CYTO |
| >hg38_21_+_8433172_8446622_RNA45SN1_45S_50ntFlanks@10386.10410.25\|ATGTGAACAGCAGTTGAACATGGGT | >hg38_21_+_8433172_8446622_RNA45SN1_45S_50ntFlanks | 10386 | 10410 | 25 | ATGTGAACAGCAGTTGAACATGGGT | 286.7 | -4.9 | 1.0 | -4.7 | 2.3E-06 | 3.6E-06 | MB231/MB468 | NUC |
| >hg38_21_+_8433172_8446622_RNA45SN1_45S_50ntFlanks@12928.12956.29\|CTGGCGCTAAACCATTCGTAGACGACCTG | >hg38_21_+_8433172_8446622_RNA45SN1_45S_50ntFlanks | 12928 | 12956 | 29 | CTGGCGCTAAACCATTCGTAGACGACCTG | 292.4 | -5.9 | 1.2 | -4.8 | 1.7E-06 | 3.7E-06 | BT20/MB468 | MITO |
| >trna14_LysTTT_11_-_59327808_59327880@4.36.33,trna2_LysTTT_17_+_8022473_8022545@4.36.33,trna54_LysTTT_1_+_204475655_204475727@4.36.33,trna5_LysTTT_11_+_59323902_59323974@4.36.33,trna62_LysTTT_1_-_204476158_204476230@4.36.33,trna76_LysTTT_6_+_28918806_28918878@4.36.33\|CGGATAGCTCAGTCGGTAGAGCATCAGACTTTT | >trna14_LysTTT_11_-_59327808_59327880 | 4 | 36 | 33 | CGGATAGCTCAGTCGGTAGAGCATCAGACTTTT | 48.5 | 8.8 | 1.9 | 4.8 | 2.0E-06 | 3.7E-06 | MB231/MB468 | MP |
| >hg38_21_+_8433172_8446622_RNA45SN1_45S_50ntFlanks@8197.8223.27\|GTGGACGGTGTGAGGCCGGTAGCGGCC | >hg38_21_+_8433172_8446622_RNA45SN1_45S_50ntFlanks | 8197 | 8223 | 27 | GTGGACGGTGTGAGGCCGGTAGCGGCC | 180.1 | -7.0 | 1.5 | -4.8 | 2.0E-06 | 3.7E-06 | MB231/MB468 | MP |
| >hg38_21_+_8433172_8446622_RNA45SN1_45S_50ntFlanks@3994.4023.30\|TGACTCTAGATAACCTCGGGCCGATCGCAC | >hg38_21_+_8433172_8446622_RNA45SN1_45S_50ntFlanks | 3994 | 4023 | 30 | TGACTCTAGATAACCTCGGGCCGATCGCAC | 188.3 | 3.3 | 0.7 | 4.7 | 2.3E-06 | 3.7E-06 | MB231/MB468 | NUC |
| >hg38_21_+_8433172_8446622_RNA45SN1_45S_50ntFlanks@5236.5257.22\|CACTGACTGGCTCAGCGTGTGC | >hg38_21_+_8433172_8446622_RNA45SN1_45S_50ntFlanks | 5236 | 5257 | 22 | CACTGACTGGCTCAGCGTGTGC | 1423.9 | -7.5 | 1.6 | -4.7 | 2.4E-06 | 3.7E-06 | MB231/MB468 | MITO |
| >hg38_21_+_8433172_8446622_RNA45SN1_45S_50ntFlanks@10726.10750.25\|GGGAGAGGGTGTAAATCTCGCGCCG | >hg38_21_+_8433172_8446622_RNA45SN1_45S_50ntFlanks | 10726 | 10750 | 25 | GGGAGAGGGTGTAAATCTCGCGCCG | 96.6 | -9.7 | 1.9 | -5.0 | 6.6E-07 | 3.7E-06 | BT20/MB468 | MP |

<table>
<tr><td>>hg38_21_+_8433172_8446622_RNA45SN1_45S_50ntFlanks@11874.11900.27|AGAAGACCCTGTTGAGCTTGACTCTAG</td><td>>hg38_21_+_8433172_8446622_RNA45SN1_45S_50ntFlanks</td><td>11874</td><td>11900</td><td>27</td><td>AGAAGACCCTGTTGAGCTTGACTCTAG</td><td>931.2</td><td>-7.0</td><td>1.5</td><td>-4.7</td><td>2.3E-06</td><td>3.7E-06</td><td>MB231/MB468</td><td>NUC</td></tr>
<tr><td>>hg38_21_+_8433172_8446622_RNA45SN1_45S_50ntFlanks@7982.8009.28|TCAGATCAGACGTGGCGACCCGCTGAAT</td><td>>hg38_21_+_8433172_8446622_RNA45SN1_45S_50ntFlanks</td><td>7982</td><td>8009</td><td>28</td><td>TCAGATCAGACGTGGCGACCCGCTGAAT</td><td>325.9</td><td>-3.2</td><td>0.7</td><td>-4.8</td><td>2.0E-06</td><td>3.7E-06</td><td>MB231/MB468</td><td>MP</td></tr>
<tr><td>>hg38_21_+_8433172_8446622_RNA45SN1_45S_50ntFlanks@13002.13019.18|CGATCTATTGAAAGTCAG</td><td>>hg38_21_+_8433172_8446622_RNA45SN1_45S_50ntFlanks</td><td>13002</td><td>13019</td><td>18</td><td>CGATCTATTGAAAGTCAG</td><td>71.3</td><td>-9.2</td><td>1.9</td><td>-5.0</td><td>6.6E-07</td><td>3.7E-06</td><td>BT20/MB468</td><td>MP</td></tr>
<tr><td>>hg38_21_+_8433172_8446622_RNA45SN1_45S_50ntFlanks@11567.11601.35|AGCAGCCGACTTAGAACTGGTGCGGACCAGGGGAA</td><td>>hg38_21_+_8433172_8446622_RNA45SN1_45S_50ntFlanks</td><td>11567</td><td>11601</td><td>35</td><td>AGCAGCCGACTTAGAACTGGTGCGGACCAGGGGAA</td><td>684.6</td><td>-5.6</td><td>1.2</td><td>-4.8</td><td>2.0E-06</td><td>3.7E-06</td><td>MB231/MB468</td><td>MP</td></tr>
<tr><td>>hg38_21_+_8433172_8446622_RNA45SN1_45S_50ntFlanks@11561.11587.27|GCGCCTAGCAGCCGACTTAGAACTGGT</td><td>>hg38_21_+_8433172_8446622_RNA45SN1_45S_50ntFlanks</td><td>11561</td><td>11587</td><td>27</td><td>GCGCCTAGCAGCCGACTTAGAACTGGT</td><td>388.3</td><td>-5.3</td><td>1.1</td><td>-4.7</td><td>2.4E-06</td><td>3.7E-06</td><td>MB231/MB468</td><td>MITO</td></tr>
<tr><td>>hg38_21_+_8433172_8446622_RNA45SN1_45S_50ntFlanks@12712.12733.22|CGGAGCCTCGGTTGGCCTCGGA</td><td>>hg38_21_+_8433172_8446622_RNA45SN1_45S_50ntFlanks</td><td>12712</td><td>12733</td><td>22</td><td>CGGAGCCTCGGTTGGCCTCGGA</td><td>579.1</td><td>-4.7</td><td>0.9</td><td>-5.0</td><td>6.6E-07</td><td>3.7E-06</td><td>BT20/MB468</td><td>MP</td></tr>
<tr><td>>hg38_21_+_8433172_8446622_RNA45SN1_45S_50ntFlanks@4208.4242.35|GGGGAGGTAGTGACGAAAAATAACAATACAGGACT</td><td>>hg38_21_+_8433172_8446622_RNA45SN1_45S_50ntFlanks</td><td>4208</td><td>4242</td><td>35</td><td>GGGGAGGTAGTGACGAAAAATAACAATACAGGACT</td><td>219.1</td><td>8.6</td><td>1.7</td><td>5.0</td><td>6.6E-07</td><td>3.7E-06</td><td>BT20/MB468</td><td>MP</td></tr>
<tr><td>>hg38_21_+_8433172_8446622_RNA45SN1_45S_50ntFlanks@9539.9569.31|ACTCTGGTGGAGGTCCGTAGCGGTCCTGACG</td><td>>hg38_21_+_8433172_8446622_RNA45SN1_45S_50ntFlanks</td><td>9539</td><td>9569</td><td>31</td><td>ACTCTGGTGGAGGTCCGTAGCGGTCCTGACG</td><td>634.6</td><td>-4.0</td><td>0.8</td><td>-4.7</td><td>2.4E-06</td><td>3.7E-06</td><td>MB231/MB468</td><td>NUC</td></tr>
<tr><td>>MI0000749|hsa-mir-30e&WithFlank&1|+|40754349|40754452@65.83.19&[MIMAT0000693&hsa-miR-30e-3p&offsets|0|-3;m-25&1|+|40754413|40754434&offsets|0|-3]|CTTTCAGTCGGATGTTTAC</td><td>>MI0000749|hsa-mir-30e&WithFlank&1|+|40754349|40754452</td><td>65</td><td>83</td><td>19</td><td>CTTTCAGTCGGATGTTTAC</td><td>623.8</td><td>3.8</td><td>0.8</td><td>4.8</td><td>1.7E-06</td><td>3.7E-06</td><td>BT20/MB468</td><td>MITO</td></tr>
<tr><td>>hg38_21_+_8433172_8446622_RNA45SN1_45S_50ntFlanks@8355.8388.34|TGAAAAGAACTTTGAAGAGAGAGTTCAAGAGGGC</td><td>>hg38_21_+_8433172_8446622_RNA45SN1_45S_50ntFlanks</td><td>8355</td><td>8388</td><td>34</td><td>TGAAAAGAACTTTGAAGAGAGAGTTCAAGAGGGC</td><td>50.3</td><td>9.4</td><td>1.9</td><td>5.0</td><td>6.6E-07</td><td>3.7E-06</td><td>BT20/MB468</td><td>MP</td></tr>
<tr><td>>hg38_21_+_8433172_8446622_RNA45SN1_45S_50ntFlanks@4130.4158.29|ATTCCGGAGAGGGAGCCTGAGAAACGGCT</td><td>>hg38_21_+_8433172_8446622_RNA45SN1_45S_50ntFlanks</td><td>4130</td><td>4158</td><td>29</td><td>ATTCCGGAGAGGGAGCCTGAGAAACGGCT</td><td>116.5</td><td>-6.6</td><td>1.4</td><td>-4.8</td><td>2.0E-06</td><td>3.7E-06</td><td>MB231/MB468</td><td>MP</td></tr>
<tr><td>>trna119_LysCTT_1_-_145395522_145395594@5.36.32,trna11_LysCTT_5_-_180648979_180649051@5.36.32,trna13_LysCTT_6_+_26556774_26556846@5.36.32,trna7_LysCTT_16_+_3225692_3225764@5.36.32,trna9_LysCTT_5_+_180634755_180634827@5.36.32|GGCTAGCTCAGTCGGTAGAGCATGAGACTCTT</td><td>>trna119_LysCTT_1_-_145395522_145395594</td><td>5</td><td>36</td><td>32</td><td>GGCTAGCTCAGTCGGTAGAGCATGAGACTCTT</td><td>1368.2</td><td>6.4</td><td>1.4</td><td>4.7</td><td>2.4E-06</td><td>3.7E-06</td><td>MB231/MB468</td><td>MITO</td></tr>
<tr><td>>hg38_21_+_8433172_8446622_RNA45SN1_45S_50ntFlanks@7974.8004.31|ACGCGACCTCAGATCAGACGTGGCGACCCGC</td><td>>hg38_21_+_8433172_8446622_RNA45SN1_45S_50ntFlanks</td><td>7974</td><td>8004</td><td>31</td><td>ACGCGACCTCAGATCAGACGTGGCGACCCGC</td><td>640.0</td><td>2.9</td><td>0.6</td><td>4.7</td><td>2.4E-06</td><td>3.7E-06</td><td>MB231/MB468</td><td>NUC</td></tr>
<tr><td>>hg38_21_+_8433172_8446622_RNA45SN1_45S_50ntFlanks@12423.12445.23|TCGGCTCTTCCTATCATTGTGAA</td><td>>hg38_21_+_8433172_8446622_RNA45SN1_45S_50ntFlanks</td><td>12423</td><td>12445</td><td>23</td><td>TCGGCTCTTCCTATCATTGTGAA</td><td>268.9</td><td>-2.4</td><td>0.5</td><td>-4.7</td><td>2.4E-06</td><td>3.7E-06</td><td>MB231/MB468</td><td>NUC</td></tr>
<tr><td>>hg38_21_+_8433172_8446622_RNA45SN1_45S_50ntFlanks@11561.11583.23|GCGCCTAGCAGCCGACTTAGAAC</td><td>>hg38_21_+_8433172_8446622_RNA45SN1_45S_50ntFlanks</td><td>11561</td><td>11583</td><td>23</td><td>GCGCCTAGCAGCCGACTTAGAAC</td><td>771.1</td><td>-5.6</td><td>1.2</td><td>-4.7</td><td>2.4E-06</td><td>3.7E-06</td><td>MB231/MB468</td><td>MITO</td></tr>
<tr><td>>hg38_21_+_8433172_8446622_RNA45SN1_45S_50ntFlanks@9540.9570.31|CTCTGGTGGAGGTCCGTAGCGGTCCTGACGT</td><td>>hg38_21_+_8433172_8446622_RNA45SN1_45S_50ntFlanks</td><td>9540</td><td>9570</td><td>31</td><td>CTCTGGTGGAGGTCCGTAGCGGTCCTGACGT</td><td>221.9</td><td>11.1</td><td>2.1</td><td>5.2</td><td>1.7E-07</td><td>3.7E-06</td><td>BT20/MB231</td><td>MP</td></tr>
<tr><td>>MI0000446|hsa-mir-125b-1&WithFlank&11|-|122099751|122099850@21.41.21(+1C);MI0000470|hsa-mir-125b-2&WithFlank&21|+|16590231|16590331@23.43.21(+1C)&[MIMAT0000423&hsa-miR-125b-5p&offsets|0|-1(+1C);m-67&11|-|122099809|122099830&offsets|0|-1(+1C)];[MIMAT0000423_1&hsa-miR-125b-5p&offsets|0|-1(+1C);m-68&21|+|16590253|16590274&offsets|0|-1(+1C)]|TCCCTGAGACCCTAACTTGTGC</td><td>>MI0000446|hsa-mir-125b-1&WithFlank&11|-|122099751|122099850</td><td>21</td><td>41</td><td>21(+1C)</td><td>TCCCTGAGACCCTAACTTGTGC</td><td>123.0</td><td>-10.3</td><td>2.0</td><td>-5.2</td><td>1.7E-07</td><td>3.7E-06</td><td>BT20/MB231</td><td>MP</td></tr>
<tr><td>>hg38_21_+_8433172_8446622_RNA45SN1_45S_50ntFlanks@12157.12181.25|AGTTTGACTGGGGCGGTACACCTGT</td><td>>hg38_21_+_8433172_8446622_RNA45SN1_45S_50ntFlanks</td><td>12157</td><td>12181</td><td>25</td><td>AGTTTGACTGGGGCGGTACACCTGT</td><td>2103.8</td><td>-7.9</td><td>1.6</td><td>-5.0</td><td>6.7E-07</td><td>3.7E-06</td><td>BT20/MB468</td><td>MP</td></tr>
<tr><td>>hg38_21_+_8433172_8446622_RNA45SN1_45S_50ntFlanks@3830.3852.23|GCTCGCTCCTCTCCTACTTGGAT</td><td>>hg38_21_+_8433172_8446622_RNA45SN1_45S_50ntFlanks</td><td>3830</td><td>3852</td><td>23</td><td>GCTCGCTCCTCTCCTACTTGGAT</td><td>247.9</td><td>-5.4</td><td>1.1</td><td>-4.7</td><td>2.4E-06</td><td>3.7E-06</td><td>MB231/MB468</td><td>NUC</td></tr>
<tr><td>>hg38_21_+_8433172_8446622_RNA45SN1_45S_50ntFlanks@10481.10499.19|ATCGAAAGGGAGTCGGGTT</td><td>>hg38_21_+_8433172_8446622_RNA45SN1_45S_50ntFlanks</td><td>10481</td><td>10499</td><td>19</td><td>ATCGAAAGGGAGTCGGGTT</td><td>58.4</td><td>-8.9</td><td>1.8</td><td>-5.0</td><td>6.7E-07</td><td>3.7E-06</td><td>BT20/MB468</td><td>MP</td></tr>
<tr><td>>hg38_21_+_8433172_8446622_RNA45SN1_45S_50ntFlanks@10798.10816.19|CATGTTGGAACAATGTAGG</td><td>>hg38_21_+_8433172_8446622_RNA45SN1_45S_50ntFlanks</td><td>10798</td><td>10816</td><td>19</td><td>CATGTTGGAACAATGTAGG</td><td>522.2</td><td>-4.6</td><td>1.0</td><td>-4.7</td><td>2.3E-06</td><td>3.7E-06</td><td>MB231/MB468</td><td>CYTO</td></tr>
<tr><td>>MI0000471|hsa-mir-126&WithFlank&9|+|136670596|136670692@58.79.22&[MIMAT0000445&hsa-miR-126-3p&offsets|0|0;m-40&9|+|136670653|136670674&offsets|0|0]|TCGTACCGTGAGTAATAATGCG</td><td>>MI0000471|hsa-mir-126&WithFlank&9|+|136670596|136670692</td><td>58</td><td>79</td><td>22</td><td>TCGTACCGTGAGTAATAATGCG</td><td>3016.0</td><td>7.2</td><td>1.5</td><td>5.0</td><td>6.7E-07</td><td>3.7E-06</td><td>BT20/MB468</td><td>MP</td></tr>
<tr><td>>hg38_21_+_8433172_8446622_RNA45SN1_45S_50ntFlanks@6652.6686.35|GACTCTTAGCGGTGGATCACTCGGCTCGTGCGTCG</td><td>>hg38_21_+_8433172_8446622_RNA45SN1_45S_50ntFlanks</td><td>6652</td><td>6686</td><td>35</td><td>GACTCTTAGCGGTGGATCACTCGGCTCGTGCGTCG</td><td>1642.7</td><td>-1.8</td><td>0.4</td><td>-4.7</td><td>2.5E-06</td><td>3.7E-06</td><td>BT20/MB468</td><td>NUC</td></tr>
<tr><td>>MI0000071|hsa-mir-17&WithFlank&13|+|91350599|91350694@20.43.24&[MIMAT0000070&hsa-miR-17-5p&offsets|0|+1;m-88&13|+|91350618|91350640&offsets|0|+1]|CAAAGTGCTTACAGTGCAGGTAGT</td><td>>MI0000071|hsa-mir-17&WithFlank&13|+|91350599|91350694</td><td>20</td><td>43</td><td>24</td><td>CAAAGTGCTTACAGTGCAGGTAGT</td><td>3836.1</td><td>-5.0</td><td>1.0</td><td>-5.2</td><td>2.1E-07</td><td>3.7E-06</td><td>BT20/MB231</td><td>MITO</td></tr>
<tr><td>>MI0003776|hsa-mir-320b-1&WithFlank&1|+|116671740|116671823@48.66.19(+3A);MI0003839|hsa-mir-320b-2&WithFlank&1|-|224257034|224257116@47.65.19(+3A);MI0003778|hsa-mir-320c-1&WithFlank&18|+|21683512|21683595@48.66.19(+3A);MI0008191|hsa-mir-320c-2&WithFlank&18|+|24321669|24321752@48.66.19(+3A);MI0000542|hsa-mir-320a&WithFlank&8|-|22244960|22245043@48.66.19(+3A)&[MIMAT0005792&hsa-miR-320b&offsets|0|-3(+3A);m-183&1|+|116671787|116671806&offsets|0|-1(+3A)];[MIMAT0005792_1&hsa-miR-320b&offsets|0|-3(+3A);m-183&1|-|224257051|224257070&offsets|0|-1(+3A)];[MIMAT0005793&hsa-miR-320c&offsets|0|-1(+3A);m-222&18|+|21683559|21683578&offsets|0|-1(+3A)];[MIMAT0005793_1&hsa-miR-320c&offsets|0|-1(+3A);m-219&18|+|24321716|24321735&offsets|0|-1(+3A)];[MIMAT0000510&hsa-miR-320a-3p&offsets|0|-3(+3A);m-60&8|-|22244975|22244996&offsets|0|-3(+3A)]|AAAAGCTGGGTTGAGAGGGAAA</td><td>>MI0003776|hsa-mir-320b-1&WithFlank&1|+|116671740|116671823</td><td>48</td><td>66</td><td>19(+3A)</td><td>AAAAGCTGGGTTGAGAGGGAAA</td><td>85.2</td><td>10.2</td><td>2.0</td><td>5.0</td><td>6.7E-07</td><td>3.7E-06</td><td>BT20/MB468</td><td>MP</td></tr>
<tr><td>>hg38_21_+_8433172_8446622_RNA45SN1_45S_50ntFlanks@10392.10422.31|ACAGCAGTTGAACATGGGTCAGTCGGTCCTG</td><td>>hg38_21_+_8433172_8446622_RNA45SN1_45S_50ntFlanks</td><td>10392</td><td>10422</td><td>31</td><td>ACAGCAGTTGAACATGGGTCAGTCGGTCCTG</td><td>404.2</td><td>-5.3</td><td>1.1</td><td>-4.7</td><td>2.4E-06</td><td>3.7E-06</td><td>MB231/MB468</td><td>MITO</td></tr>
<tr><td>>trna27_GlyCCC_2_-_70476123_70476193@1.29.29,trna34_GlyCCC_16_-_686736_686806@1.29.29|GCGCCGCTGGTGTAGTGGTATCATGCAAG</td><td>>trna27_GlyCCC_2_-_70476123_70476193</td><td>1</td><td>29</td><td>29</td><td>GCGCCGCTGGTGTAGTGGTATCATGCAAG</td><td>611.4</td><td>-8.4</td><td>1.8</td><td>-4.8</td><td>2.0E-06</td><td>3.7E-06</td><td>MB231/MB468</td><td>MP</td></tr>
<tr><td>>hg38_MT_+_1621_3279_RNR2_16S_50ntFlanks@1349.1387.39|ACAATAGGGTTTACGACCTCGATGTTGGATCAGGACATC</td><td>>hg38_MT_+_1621_3279_RNR2_16S_50ntFlanks</td><td>1349</td><td>1387</td><td>39</td><td>ACAATAGGGTTTACGACCTCGATGTTGGATCAGGACATC</td><td>39.7</td><td>8.5</td><td>1.8</td><td>4.8</td><td>2.0E-06</td><td>3.7E-06</td><td>MB231/MB468</td><td>MP</td></tr>
<tr><td>>am_MI0000809|hsa-mir-151a&WithFlank&8|-|140732558|140732659@53.72.20(+1C)&[MIMAT0000757&hsa-miR-151a-3p&offsets|0|-1(+1C);m-30&8|-|140732587|140732607&offsets|0|-1(+1C)]|CTAGACTGAAGCTCCTTGAGC</td><td>>am_MI0000809|hsa-mir-151a&WithFlank&8|-|140732558|140732659</td><td>53</td><td>72</td><td>20(+1C)</td><td>CTAGACTGAAGCTCCTTGAGC</td><td>460.3</td><td>6.9</td><td>1.5</td><td>4.7</td><td>2.4E-06</td><td>3.7E-06</td><td>MB231/MB468</td><td>MITO</td></tr>
<tr><td>>hg38_21_+_8433172_8446622_RNA45SN1_45S_50ntFlanks@9203.9225.23|CTCGGGGCCACGCGCGCGTCCCC</td><td>>hg38_21_+_8433172_8446622_RNA45SN1_45S_50ntFlanks</td><td>9203</td><td>9225</td><td>23</td><td>CTCGGGGCCACGCGCGCGTCCCC</td><td>602.2</td><td>-5.9</td><td>1.2</td><td>-5.0</td><td>6.7E-07</td><td>3.7E-06</td><td>BT20/MB468</td><td>MP</td></tr>
<tr><td>>hg38_21_+_8433172_8446622_RNA45SN1_45S_50ntFlanks@3813.3830.18|TTATGGTTCCTTTGGTCG</td><td>>hg38_21_+_8433172_8446622_RNA45SN1_45S_50ntFlanks</td><td>3813</td><td>3830</td><td>18</td><td>TTATGGTTCCTTTGGTCG</td><td>48.0</td><td>-8.7</td><td>1.7</td><td>-5.0</td><td>6.7E-07</td><td>3.7E-06</td><td>BT20/MB468</td><td>MP</td></tr>
<tr><td>>hg38_21_+_8433172_8446622_RNA45SN1_45S_50ntFlanks@8105.8138.34|CGCGGCGGGGCGCGGGACATGTGGCGTACGGAAG</td><td>>hg38_21_+_8433172_8446622_RNA45SN1_45S_50ntFlanks</td><td>8105</td><td>8138</td><td>34</td><td>CGCGGCGGGGCGCGGGACATGTGGCGTACGGAAG</td><td>8000.7</td><td>-4.3</td><td>0.9</td><td>-5.0</td><td>6.7E-07</td><td>3.7E-06</td><td>BT20/MB468</td><td>MP</td></tr>
<tr><td>>hg38_21_+_8433172_8446622_RNA45SN1_45S_50ntFlanks@11567.11603.37|AGCAGCCGACTTAGAACTGGTGCGGACCAGGGGAATC</td><td>>hg38_21_+_8433172_8446622_RNA45SN1_45S_50ntFlanks</td><td>11567</td><td>11603</td><td>37</td><td>AGCAGCCGACTTAGAACTGGTGCGGACCAGGGGAATC</td><td>190.5</td><td>-2.0</td><td>0.4</td><td>-4.8</td><td>2.0E-06</td><td>3.7E-06</td><td>BT20/MB468</td><td>CYTO</td></tr>
<tr><td>>hg38_21_+_8433172_8446622_RNA45SN1_45S_50ntFlanks@6722.6743.22|ATTGCAGGACACATTGATCATC</td><td>>hg38_21_+_8433172_8446622_RNA45SN1_45S_50ntFlanks</td><td>6722</td><td>6743</td><td>22</td><td>ATTGCAGGACACATTGATCATC</td><td>557.0</td><td>-5.5</td><td>1.2</td><td>-4.8</td><td>2.0E-06</td><td>3.7E-06</td><td>MB231/MB468</td><td>MP</td></tr>
<tr><td>>MI0000683|hsa-mir-181b-2&WithFlank&9|+|124693704|124693804@22.46.25&[MIMAT0000257_1&hsa-miR-181b-5p&offsets|0|+2;m-72&9|+|124693725|124693747&offsets|0|+2]|AACATTCATTGCTGTCGGTGGGTTT</td><td>>MI0000683|hsa-mir-181b-2&WithFlank&9|+|124693704|124693804</td><td>22</td><td>46</td><td>25</td><td>AACATTCATTGCTGTCGGTGGGTTT</td><td>68.5</td><td>3.0</td><td>0.6</td><td>4.7</td><td>2.6E-06</td><td>3.7E-06</td><td>BT20/MB468</td><td>NUC</td></tr>
<tr><td>>hg38_21_+_8433172_8446622_RNA45SN1_45S_50ntFlanks@12043.12068.26|CACTGACCCGGTGAGGCGGGGGGGCG</td><td>>hg38_21_+_8433172_8446622_RNA45SN1_45S_50ntFlanks</td><td>12043</td><td>12068</td><td>26</td><td>CACTGACCCGGTGAGGCGGGGGGGCG</td><td>66.1</td><td>-9.1</td><td>1.8</td><td>-5.0</td><td>6.7E-07</td><td>3.7E-06</td><td>BT20/MB468</td><td>MP</td></tr>
<tr><td>>MI0000093|hsa-mir-92a-1&WithFlank&13|+|91351308|91351397@55.75.21;MI0000094|hsa-mir-92a-2&WithFlank&X|-|134169532|134169618@55.75.21&[MIMAT0000092&hsa-miR-92a-3p&offsets|+1|0;m-9&13|+|91351361|91351382&offsets|+1|0];[MIMAT0000092_1&hsa-miR-92a-3p&offsets|+1|0;m-11&X|-|134169544|134169565&offsets|+1|0]|ATTGCACTTGTCCCGGCCTGT</td><td>>MI0000093|hsa-mir-92a-1&WithFlank&13|+|91351308|91351397</td><td>55</td><td>75</td><td>21</td><td>ATTGCACTTGTCCCGGCCTGT</td><td>914.8</td><td>3.9</td><td>0.8</td><td>4.7</td><td>2.5E-06</td><td>3.7E-06</td><td>MB231/MB468</td><td>MITO</td></tr>
<tr><td>>hg38_21_+_8433172_8446622_RNA45SN1_45S_50ntFlanks@3861.3881.21|TAATTCTAGAGCTAATACATG</td><td>>hg38_21_+_8433172_8446622_RNA45SN1_45S_50ntFlanks</td><td>3861</td><td>3881</td><td>21</td><td>TAATTCTAGAGCTAATACATG</td><td>460.5</td><td>-7.1</td><td>1.5</td><td>-4.7</td><td>2.3E-06</td><td>3.7E-06</td><td>MB231/MB468</td><td>CYTO</td></tr>
<tr><td>>hg38_21_+_8433172_8446622_RNA45SN1_45S_50ntFlanks@10841.10868.28|CGTAACTTCGGGATAAGGATTGGCTCTA</td><td>>hg38_21_+_8433172_8446622_RNA45SN1_45S_50ntFlanks</td><td>10841</td><td>10868</td><td>28</td><td>CGTAACTTCGGGATAAGGATTGGCTCTA</td><td>3392.7</td><td>-4.5</td><td>0.9</td><td>-5.0</td><td>6.8E-07</td><td>3.7E-06</td><td>BT20/MB468</td><td>MP</td></tr>
<tr><td>>hg38_21_+_8433172_8446622_RNA45SN1_45S_50ntFlanks@11314.11342.29|CCCACCCCACGTCTCGTCGCGCGCGCGTC</td><td>>hg38_21_+_8433172_8446622_RNA45SN1_45S_50ntFlanks</td><td>11314</td><td>11342</td><td>29</td><td>CCCACCCCACGTCTCGTCGCGCGCGCGTC</td><td>171.9</td><td>-4.6</td><td>1.0</td><td>-4.8</td><td>2.0E-06</td><td>3.7E-06</td><td>BT20/MB468</td><td>CYTO</td></tr>
<tr><td>>MI0000081|hsa-mir-24-2&WithFlank&19|-|13836281|13836365@56.76.21(+1U);MI0000080|hsa-mir-24-1&WithFlank&9|+|95086015|95086094@50.70.21(+1U)&[MIMAT0000080&hsa-miR-24-3p&offsets|0|-1(+1U);m-36&19|-|13836288|13836310&offsets|0|-2(+1U)];[MIMAT0000080_1&hsa-miR-24-3p&offsets|0|-1(+1U);m-35&9|+|95086064|95086086&offsets|0|-2(+1U)]|TGGCTCAGTTCAGCAGGAACAT</td><td>>MI0000081|hsa-mir-24-2&WithFlank&19|-|13836281|13836365</td><td>56</td><td>76</td><td>21(+1U)</td><td>TGGCTCAGTTCAGCAGGAACAT</td><td>711.5</td><td>4.2</td><td>0.9</td><td>4.8</td><td>2.0E-06</td><td>3.7E-06</td><td>MB231/MB468</td><td>MP</td></tr>
<tr><td>>trnaMT_PheGAA_MT_+_577_647@39.57.19|ATGTTTAGACGGGCTCACA</td><td>>trnaMT_PheGAA_MT_+_577_647</td><td>39</td><td>57</td><td>19</td><td>ATGTTTAGACGGGCTCACA</td><td>121.2</td><td>3.2</td><td>0.7</td><td>4.7</td><td>2.6E-06</td><td>3.7E-06</td><td>BT20/MB468</td><td>NUC</td></tr>
</table>

| | | | | | | | | | | | | | |
|---|---|---|---|---|---|---|---|---|---|---|---|---|---|
| >hg38_21_+_8433172_8446622_RNA45SN1_45S_50ntFlanks@12431.12455.25\|TCCTATCATTGTGAAGCAGAATTCA | >hg38_21_+_8433172_8446622_RNA45SN1_45S_50ntFlanks | 12431 | 12455 | 25 | TCCTATCATTGTGAAGCAGAATTCA | 229.1 | -3.3 | 0.7 | -4.8 | 1.8E-06 | 3.7E-06 | BT20/MB468 | MITO |
| >MI0000063\|hsa-let-7b&WithFlank&22\|+\|46113680\|46113774@12.31.20&[MIMAT0000063&hsa-let-7b-5p&offsets\|0\|-2;m-14&22\|+\|46113691\|46113712&offsets\|0\|-2]\|TGAGGTAGTAGGTTGTGTGG | >MI0000063\|hsa-let-7b&WithFlank&22\|+\|46113680\|46113774 | 12 | 31 | 20 | TGAGGTAGTAGGTTGTGTGG | 830.4 | 2.8 | 0.5 | 5.1 | 2.7E-07 | 3.7E-06 | BT20/MB231 | NUC |
| >hg38_21_+_8433172_8446622_RNA45SN1_45S_50ntFlanks@6651.6679.29\|CGACTCTTAGCGGTGGATCACTCGGCTCG | >hg38_21_+_8433172_8446622_RNA45SN1_45S_50ntFlanks | 6651 | 6679 | 29 | CGACTCTTAGCGGTGGATCACTCGGCTCG | 7716.1 | -2.6 | 0.5 | -4.7 | 2.6E-06 | 3.7E-06 | BT20/MB468 | NUC |
| >hg38_21_+_8433172_8446622_RNA45SN1_45S_50ntFlanks@12436.12473.38\|TCATTGTGAAGCAGAATTCACCAAGCGTTGGATTGTTC | >hg38_21_+_8433172_8446622_RNA45SN1_45S_50ntFlanks | 12436 | 12473 | 38 | TCATTGTGAAGCAGAATTCACCAAGCGTTGGATTGTTC | 482.4 | -3.7 | 0.8 | -4.7 | 2.4E-06 | 3.7E-06 | MB231/MB468 | NUC |
| >hg38_21_+_8433172_8446622_RNA45SN1_45S_50ntFlanks@10353.10383.31\|AGAACTTTGAAGGCCGAAGTGGAGAAGGGTT | >hg38_21_+_8433172_8446622_RNA45SN1_45S_50ntFlanks | 10353 | 10383 | 31 | AGAACTTTGAAGGCCGAAGTGGAGAAGGGTT | 603.4 | -3.1 | 0.7 | -4.7 | 2.4E-06 | 3.7E-06 | MB231/MB468 | NUC |
| >hg38_21_+_8433172_8446622_RNA45SN1_45S_50ntFlanks@10771.10806.36\|CAGGTCTCCAAGGTGAACAGCCTCTGGCATGTTGGA | >hg38_21_+_8433172_8446622_RNA45SN1_45S_50ntFlanks | 10771 | 10806 | 36 | CAGGTCTCCAAGGTGAACAGCCTCTGGCATGTTGGA | 664.7 | -4.9 | 1.0 | -4.8 | 1.8E-06 | 3.7E-06 | BT20/MB468 | MITO |
| >am_trnaMT_ValTAC_MT_+_1602_1670@28.66.39\|AACTTACACTTAGGAGATTTCAACTTAACTTGACCGCTC | >am_trnaMT_ValTAC_MT_+_1602_1670 | 28 | 66 | 39 | AACTTACACTTAGGAGATTTCAACTTAACTTGACCGCTC | 567.8 | 12.9 | 2.6 | 5.0 | 6.8E-07 | 3.7E-06 | BT20/MB468 | MP |
| >hg38_21_+_8433172_8446622_RNA45SN1_45S_50ntFlanks@4685.4722.38\|AGACGGACCAGAGCGAAAGCATTTGCCAAGAATGTTTT | >hg38_21_+_8433172_8446622_RNA45SN1_45S_50ntFlanks | 4685 | 4722 | 38 | AGACGGACCAGAGCGAAAGCATTTGCCAAGAATGTTTT | 282.7 | -5.5 | 1.2 | -4.7 | 2.6E-06 | 3.7E-06 | BT20/MB468 | NUC |
| >hg38_21_+_8433172_8446622_RNA45SN1_45S_50ntFlanks@5473.5516.44\|CGGCCCTGGCGGAGCGCTGAGAAGACGGTCGAACTTGACTATCT | >hg38_21_+_8433172_8446622_RNA45SN1_45S_50ntFlanks | 5473 | 5516 | 44 | CGGCCCTGGCGGAGCGCTGAGAAGACGGTCGAACTTGACTATCT | 432.8 | 4.3 | 0.9 | 4.7 | 2.4E-06 | 3.7E-06 | MB231/MB468 | NUC |
| >hg38_21_+_8433172_8446622_RNA45SN1_45S_50ntFlanks@3705.3735.31\|TACCTGGTTGATCCTGCCAGTAGCATATGCT | >hg38_21_+_8433172_8446622_RNA45SN1_45S_50ntFlanks | 3705 | 3735 | 31 | TACCTGGTTGATCCTGCCAGTAGCATATGCT | 382.0 | 3.3 | 0.6 | 5.1 | 2.7E-07 | 3.7E-06 | BT20/MB231 | NUC |
| >trna116_GluCTC_1_-_145399233_145399304@1.42.42,trna59_GluCTC_1_+_249168447_249168518@1.42.42,trna71_GluCTC_1_-_161439189_161439260@1.42.42,trna74_GluCTC_1_-_161431809_161431880@1.42.42,trna77_GluCTC_1_-_161424398_161424469@1.42.42,trna77_GluCTC_6_+_28949976_28950047@1.42.42,trna80_GluCTC_1_-_161417018_161417089@1.42.42,trna87_GluCTC_6_-_126101393_126101464@1.42.42\|TCCCTGGTGGTCTAGTGGTTAGGATTCGGCGCTCTCACCGCC | >trna116_GluCTC_1_-_145399233_145399304 | 1 | 42 | 42 | TCCCTGGTGGTCTAGTGGTTAGGATTCGGCGCTCTCACCGCC | 2317.2 | 5.8 | 1.2 | 4.7 | 2.0E-06 | 3.7E-06 | BT20/MB468 | CYTO |
| >hg38_1_-_228634819_228635039_RNA5S12_5S_50ntFlanks@123.169.47\|TAGTACTTGGATGGGAGACCGCCTGGGAATACCGGGTGCTGTAGGCT | >hg38_1_-_228634819_228635039_RNA5S12_5S_50ntFlanks | 123 | 169 | 47 | TAGTACTTGGATGGGAGACCGCCTGGGAATACCGGGTGCTGTAGGCT | 528.6 | 6.6 | 1.4 | 4.8 | 1.8E-06 | 3.7E-06 | BT20/MB468 | MITO |
| >hg38_21_+_8433172_8446622_RNA45SN1_45S_50ntFlanks@9539.9572.34\|ACTCTGGTGGAGGTCCGTAGCGGTCCTGACGTGC | >hg38_21_+_8433172_8446622_RNA45SN1_45S_50ntFlanks | 9539 | 9572 | 34 | ACTCTGGTGGAGGTCCGTAGCGGTCCTGACGTGC | 318.4 | 5.5 | 1.1 | 5.2 | 2.1E-07 | 3.8E-06 | BT20/MB231 | MITO |
| >hg38_21_+_8433172_8446622_RNA45SN1_45S_50ntFlanks@476.516.41\|TTCGTGATCGATGTGGTGACGTCGTGCTCTCCCGGGCCGGG | >hg38_21_+_8433172_8446622_RNA45SN1_45S_50ntFlanks | 476 | 516 | 41 | TTCGTGATCGATGTGGTGACGTCGTGCTCTCCCGGGCCGGG | 714.6 | 5.5 | 1.2 | 4.7 | 2.5E-06 | 3.8E-06 | MB231/MB468 | MITO |
| >hg38_21_+_8433172_8446622_RNA45SN1_45S_50ntFlanks@12932.12973.42\|CGCTAAACCATTCGTAGACGACCTGCTTCTGGGTCGGGGTTT | >hg38_21_+_8433172_8446622_RNA45SN1_45S_50ntFlanks | 12932 | 12973 | 42 | CGCTAAACCATTCGTAGACGACCTGCTTCTGGGTCGGGGTTT | 243.7 | -5.7 | 1.2 | -4.7 | 2.5E-06 | 3.8E-06 | MB231/MB468 | MITO |
| >hg38_21_+_8433172_8446622_RNA45SN1_45S_50ntFlanks@12407.12448.42\|TTTTGATCCTTCGATGTCGGCTCTTCCTATCATTGTGAAGCA | >hg38_21_+_8433172_8446622_RNA45SN1_45S_50ntFlanks | 12407 | 12448 | 42 | TTTTGATCCTTCGATGTCGGCTCTTCCTATCATTGTGAAGCA | 389.8 | 2.9 | 0.6 | 4.8 | 1.8E-06 | 3.8E-06 | BT20/MB468 | MITO |
| >hg38_21_+_8433172_8446622_RNA45SN1_45S_50ntFlanks@12569.12587.19\|CAGTACGAGAGGAACCGCA | >hg38_21_+_8433172_8446622_RNA45SN1_45S_50ntFlanks | 12569 | 12587 | 19 | CAGTACGAGAGGAACCGCA | 333.5 | -7.0 | 1.5 | -4.7 | 2.1E-06 | 3.8E-06 | MB231/MB468 | MP |
| >hg38_21_+_8433172_8446622_RNA45SN1_45S_50ntFlanks@4160.4180.21\|CCACATCCAAGGAAGGCAGCA | >hg38_21_+_8433172_8446622_RNA45SN1_45S_50ntFlanks | 4160 | 4180 | 21 | CCACATCCAAGGAAGGCAGCA | 72.2 | -9.3 | 1.9 | -5.0 | 6.8E-07 | 3.8E-06 | BT20/MB468 | MP |
| >hg38_21_+_8433172_8446622_RNA45SN1_45S_50ntFlanks@4514.4537.24\|AAAATTAGAGTGTTCAAAGCAGGC | >hg38_21_+_8433172_8446622_RNA45SN1_45S_50ntFlanks | 4514 | 4537 | 24 | AAAATTAGAGTGTTCAAAGCAGGC | 52.0 | -8.8 | 1.8 | -5.0 | 6.8E-07 | 3.8E-06 | BT20/MB468 | MP |
| >hg38_MT_+_1621_3279_RNR2_16S_50ntFlanks@1348.1385.38\|AACAATAGGGTTTACGACCTCGATGTTGGATCAGGACA | >hg38_MT_+_1621_3279_RNR2_16S_50ntFlanks | 1348 | 1385 | 38 | AACAATAGGGTTTACGACCTCGATGTTGGATCAGGACA | 208.5 | 5.1 | 1.1 | 4.7 | 2.5E-06 | 3.8E-06 | MB231/MB468 | MITO |
| >hg38_21_+_8433172_8446622_RNA45SN1_45S_50ntFlanks@10389.10408.20\|TGAACAGCAGTTGAACATGG | >hg38_21_+_8433172_8446622_RNA45SN1_45S_50ntFlanks | 10389 | 10408 | 20 | TGAACAGCAGTTGAACATGG | 169.1 | -5.9 | 1.3 | -4.7 | 2.1E-06 | 3.8E-06 | BT20/MB468 | CYTO |
| >hg38_21_+_8433172_8446622_RNA45SN1_45S_50ntFlanks@11598.11627.30\|GGAATCCGACTGTTTAATTAAAACAAAGCA | >hg38_21_+_8433172_8446622_RNA45SN1_45S_50ntFlanks | 11598 | 11627 | 30 | GGAATCCGACTGTTTAATTAAAACAAAGCA | 51.8 | 9.4 | 1.9 | 5.0 | 6.9E-07 | 3.8E-06 | BT20/MB468 | MP |
| >hg38_21_+_8433172_8446622_RNA45SN1_45S_50ntFlanks@11255.11282.28\|CTGCGGCGGGCGTCGCGGCCGCCCCCGG | >hg38_21_+_8433172_8446622_RNA45SN1_45S_50ntFlanks | 11255 | 11282 | 28 | CTGCGGCGGGCGTCGCGGCCGCCCCCGG | 70.7 | -9.2 | 1.9 | -5.0 | 6.9E-07 | 3.8E-06 | BT20/MB468 | MP |
| >hg38_21_+_8433172_8446622_RNA45SN1_45S_50ntFlanks@8737.8756.20\|CCCGTCCGTCCGTCCGTCCG | >hg38_21_+_8433172_8446622_RNA45SN1_45S_50ntFlanks | 8737 | 8756 | 20 | CCCGTCCGTCCGTCCGTCCG | 54.2 | -8.8 | 1.8 | -5.0 | 6.9E-07 | 3.8E-06 | BT20/MB468 | MP |
| >hg38_21_+_8433172_8446622_RNA45SN1_45S_50ntFlanks@12938.12966.29\|ACCATTCGTAGACGACCTGCTTCTGGGTC | >hg38_21_+_8433172_8446622_RNA45SN1_45S_50ntFlanks | 12938 | 12966 | 29 | ACCATTCGTAGACGACCTGCTTCTGGGTC | 242.1 | -6.1 | 1.3 | -4.7 | 2.1E-06 | 3.8E-06 | BT20/MB468 | CYTO |
| >hg38_21_+_8433172_8446622_RNA45SN1_45S_50ntFlanks@3789.3810.22\|ACTGCGAATGGCTCATTAAATC | >hg38_21_+_8433172_8446622_RNA45SN1_45S_50ntFlanks | 3789 | 3810 | 22 | ACTGCGAATGGCTCATTAAATC | 668.4 | -4.0 | 0.8 | -4.7 | 2.4E-06 | 3.8E-06 | MB231/MB468 | NUC |
| >am_trna111_HisGTG_1_-_147774845_147774916@-1G.29.30,trna118_HisGTG_1_-_145396881_145396952@-1G.29.30,trna16_HisGTG_1_+_146544773_146544844@-1G.29.30,trna1_HisGTG_15_+_45493349_45493420@-1G.29.30,trna21_HisGTG_1_+_147753471_147753542@-1G.29.30,trna33_HisGTG_6_+_27125906_27125977@-1G.29.30,trna7_HisGTG_9_-_14433938_14434009@-1G.29.30,trna8_HisGTG_15_-_45492611_45492682@-1G.29.30,trna9_HisGTG_15_-_45490804_45490875@-1G.29.30\|GGCCGTGATCGTATAGTGGTTAGTACTCTG | >am_trna111_HisGTG_1_-_147774845_147774916 | -1G | 29 | 30 | GGCCGTGATCGTATAGTGGTTAGTACTCTG | 5771.1 | -4.2 | 0.9 | -4.7 | 2.1E-06 | 3.8E-06 | MB231/MB468 | MP |
| >hg38_1_-_228634819_228635039_RNA5S12_5S_50ntFlanks@85.126.42\|TCGTCTGATCTCGGAAGCTAAGCAGGGTCGGGCCTGGTTAGT | >hg38_1_-_228634819_228635039_RNA5S12_5S_50ntFlanks | 85 | 126 | 42 | TCGTCTGATCTCGGAAGCTAAGCAGGGTCGGGCCTGGTTAGT | 53.5 | 9.5 | 1.9 | 5.0 | 6.9E-07 | 3.8E-06 | BT20/MB468 | MP |
| >hg38_21_+_8433172_8446622_RNA45SN1_45S_50ntFlanks@8814.8843.30\|CGGGTCGGGGCGGCGGCGGCGGCGGCGGTG | >hg38_21_+_8433172_8446622_RNA45SN1_45S_50ntFlanks | 8814 | 8843 | 30 | CGGGTCGGGGCGGCGGCGGCGGCGGCGGTG | 55.0 | -8.9 | 1.8 | -5.0 | 6.9E-07 | 3.8E-06 | BT20/MB468 | MP |
| >hg38_21_+_8433172_8446622_RNA45SN1_45S_50ntFlanks@8176.8204.29\|CTTCTGATCGAGGCCCAGCCCGTGGACGG | >hg38_21_+_8433172_8446622_RNA45SN1_45S_50ntFlanks | 8176 | 8204 | 29 | CTTCTGATCGAGGCCCAGCCCGTGGACGG | 786.3 | -2.9 | 0.6 | -4.8 | 1.8E-06 | 3.8E-06 | BT20/MB468 | MITO |
| >hg38_21_+_8433172_8446622_RNA45SN1_45S_50ntFlanks@9307.9335.29\|AAGGAGTCTAACACGTGCGCGAGTCGGGG | >hg38_21_+_8433172_8446622_RNA45SN1_45S_50ntFlanks | 9307 | 9335 | 29 | AAGGAGTCTAACACGTGCGCGAGTCGGGG | 751.2 | -7.3 | 1.5 | -4.7 | 2.1E-06 | 3.8E-06 | MB231/MB468 | MP |
| >hg38_21_+_8433172_8446622_RNA45SN1_45S_50ntFlanks@10471.10497.27\|CCCTCGGCCGATCGAAAGGGAGTCGGG | >hg38_21_+_8433172_8446622_RNA45SN1_45S_50ntFlanks | 10471 | 10497 | 27 | CCCTCGGCCGATCGAAAGGGAGTCGGG | 71.6 | -9.2 | 1.9 | -5.0 | 6.9E-07 | 3.8E-06 | BT20/MB468 | MP |
| >hg38_21_+_8433172_8446622_RNA45SN1_45S_50ntFlanks@10351.10369.19\|CGAGAACTTTGAAGGCCGA | >hg38_21_+_8433172_8446622_RNA45SN1_45S_50ntFlanks | 10351 | 10369 | 19 | CGAGAACTTTGAAGGCCGA | 569.7 | -2.3 | 0.5 | -4.7 | 2.1E-06 | 3.8E-06 | BT20/MB468 | CYTO |
| >hg38_21_+_8433172_8446622_RNA45SN1_45S_50ntFlanks@10872.10891.20\|GCTGGGTCGGTCGGGCTGGG | >hg38_21_+_8433172_8446622_RNA45SN1_45S_50ntFlanks | 10872 | 10891 | 20 | GCTGGGTCGGTCGGGCTGGG | 198.4 | -6.8 | 1.4 | -4.7 | 2.1E-06 | 3.8E-06 | MB231/MB468 | MP |
| >hg38_21_+_8433172_8446622_RNA45SN1_45S_50ntFlanks@10350.10370.21\|ACGAGAACTTTGAAGGCCGAA | >hg38_21_+_8433172_8446622_RNA45SN1_45S_50ntFlanks | 10350 | 10370 | 21 | ACGAGAACTTTGAAGGCCGAA | 957.2 | -4.3 | 0.9 | -4.7 | 2.1E-06 | 3.8E-06 | MB231/MB468 | MP |
| >hg38_21_+_8433172_8446622_RNA45SN1_45S_50ntFlanks@3854.3872.19\|ACTGTGGTAATTCTAGAGC | >hg38_21_+_8433172_8446622_RNA45SN1_45S_50ntFlanks | 3854 | 3872 | 19 | ACTGTGGTAATTCTAGAGC | 579.2 | -3.4 | 0.7 | -4.7 | 2.4E-06 | 3.8E-06 | MB231/MB468 | CYTO |
| >MI0000063\|hsa-let-7b&WithFlank&22\|+\|46113680\|46113774@66.86.21(+1U)&[MIMAT0004482&hsa-let-7b-3p&offsets\|0\|-1(+1U);m-171&22\|+\|46113745\|46113766&offsets\|0\|-1(+1U)]\|CTATACAACCTACTGCCTTCCT | >MI0000063\|hsa-let-7b&WithFlank&22\|+\|46113680\|46113774 | 66 | 86 | 21(+1U) | CTATACAACCTACTGCCTTCCT | 546.0 | 5.5 | 1.1 | 5.0 | 7.0E-07 | 3.8E-06 | BT20/MB468 | MP |
| >hg38_21_+_8433172_8446622_RNA45SN1_45S_50ntFlanks@12641.12661.21\|CCATCTGTGGGATTATGACTG | >hg38_21_+_8433172_8446622_RNA45SN1_45S_50ntFlanks | 12641 | 12661 | 21 | CCATCTGTGGGATTATGACTG | 397.2 | -3.3 | 0.7 | -4.7 | 2.5E-06 | 3.8E-06 | MB231/MB468 | NUC |
| >MI0000255\|hsa-mir-30d&WithFlank&8\|-\|134804870\|134804951@12.34.23(+1G)&[MIMAT0000245&hsa-miR-30d-5p&offsets\|0\|+1(+1G);m-21&8\|-\|134804919\|134804940&offsets\|0\|+1(+1G)]\|TGTAAACATCCCCGACTGGAAGCG | >MI0000255\|hsa-mir-30d&WithFlank&8\|-\|134804870\|134804951 | 12 | 34 | 23(+1G) | TGTAAACATCCCCGACTGGAAGCG | 682.8 | 5.1 | 1.1 | 4.7 | 2.4E-06 | 3.8E-06 | MB231/MB468 | CYTO |
| >hg38_21_+_8433172_8446622_RNA45SN1_45S_50ntFlanks@11675.11706.32\|TGCTCTGAATGTCAAAGTGAAGAAATTCAATG | >hg38_21_+_8433172_8446622_RNA45SN1_45S_50ntFlanks | 11675 | 11706 | 32 | TGCTCTGAATGTCAAAGTGAAGAAATTCAATG | 752.0 | 3.6 | 0.8 | 4.7 | 2.4E-06 | 3.8E-06 | MB231/MB468 | CYTO |
| >hg38_21_+_8433172_8446622_RNA45SN1_45S_50ntFlanks@12404.12432.29\|GCTTTTTGATCCTTCGATGTCGGCTCTTC | >hg38_21_+_8433172_8446622_RNA45SN1_45S_50ntFlanks | 12404 | 12432 | 29 | GCTTTTTGATCCTTCGATGTCGGCTCTTC | 606.6 | -5.9 | 1.3 | -4.7 | 2.6E-06 | 3.8E-06 | BT20/MB468 | NUC |
| >hg38_21_+_8433172_8446622_RNA45SN1_45S_50ntFlanks@11710.11727.18\|CGCGGGTAAACGGCGGGA | >hg38_21_+_8433172_8446622_RNA45SN1_45S_50ntFlanks | 11710 | 11727 | 18 | CGCGGGTAAACGGCGGGA | 52.2 | -8.8 | 1.8 | -5.0 | 7.0E-07 | 3.8E-06 | BT20/MB468 | MP |
| >MI0000542\|hsa-mir-320a&WithFlank&8\|-\|22244960\|22245043@50.71.22&[MIMAT0000510&hsa-miR-320a-3p&offsets\|+2\|+2;m-60&8\|-\|22244975\|22244996&offsets\|+2\|+2]\|AAGCTGGGTTGAGAGGGCGAAA | >MI0000542\|hsa-mir-320a&WithFlank&8\|-\|22244960\|22245043 | 50 | 71 | 22 | AAGCTGGGTTGAGAGGGCGAAA | 275.1 | 4.0 | 0.8 | 5.2 | 1.9E-07 | 3.8E-06 | BT20/MB231 | CYTO |
| >MI0000087\|hsa-mir-29a&WithFlank&7\|-\|130876741\|130876816@47.69.23(+1A)&[MIMAT0000086&hsa-miR-29a-3p&offsets\|-1\|0(+1A);m-26&7\|-\|130876749\|130876769&offsets\|-1\|+1(+1A)]\|CTAGCACCATCTGAAATCGGTTAA | >MI0000087\|hsa-mir-29a&WithFlank&7\|-\|130876741\|130876816 | 47 | 69 | 23(+1A) | CTAGCACCATCTGAAATCGGTTAA | 44.4 | 8.7 | 1.8 | 4.7 | 2.1E-06 | 3.8E-06 | MB231/MB468 | MP |

<table>
<tr><td>>MI0000087|hsa-mir-29a&WithFlank&7|-|130876741|130876816@47.68.22(+2U)&[MIMAT0000086&hsa-miR-29a-3p&offsets|-1|-1(+2U);m-26&7|-|130876749|130876769&offsets|-1|0(+2U)]|CTAGCACCATCTGAAATCGGTTTT</td><td>>MI0000087|hsa-mir-29a&WithFlank&7|-|130876741|130876816</td><td>47</td><td>68</td><td>22(+2U)</td><td>CTAGCACCATCTGAAATCGGTTTT</td><td>72.9</td><td>9.9</td><td>2.0</td><td>5.0</td><td>7.0E-07</td><td>3.9E-06</td><td>BT20/MB468</td><td>MP</td></tr>
<tr><td>>hg38_21_+_8433172_8446622_RNA45SN1_45S_50ntFlanks@11873.11905.33|AAGAAGACCCTGTTGAGCTTGACTCTAGTCTGG</td><td>>hg38_21_+_8433172_8446622_RNA45SN1_45S_50ntFlanks</td><td>11873</td><td>11905</td><td>33</td><td>AAGAAGACCCTGTTGAGCTTGACTCTAGTCTGG</td><td>709.4</td><td>-5.0</td><td>1.0</td><td>-4.8</td><td>1.8E-06</td><td>3.9E-06</td><td>BT20/MB468</td><td>MITO</td></tr>
<tr><td>>hg38_21_+_8433172_8446622_RNA45SN1_45S_50ntFlanks@10349.10374.26|AACGAGAACTTTGAAGGCCGAAGTGG</td><td>>hg38_21_+_8433172_8446622_RNA45SN1_45S_50ntFlanks</td><td>10349</td><td>10374</td><td>26</td><td>AACGAGAACTTTGAAGGCCGAAGTGG</td><td>741.6</td><td>-2.5</td><td>0.5</td><td>-4.7</td><td>2.1E-06</td><td>3.9E-06</td><td>BT20/MB468</td><td>CYTO</td></tr>
<tr><td>>hg38_21_+_8433172_8446622_RNA45SN1_45S_50ntFlanks@10682.10709.28|TCCGGCGGCGTCCGGTGAGCTCTCGCTG</td><td>>hg38_21_+_8433172_8446622_RNA45SN1_45S_50ntFlanks</td><td>10682</td><td>10709</td><td>28</td><td>TCCGGCGGCGTCCGGTGAGCTCTCGCTG</td><td>48.0</td><td>-8.7</td><td>1.7</td><td>-5.0</td><td>7.0E-07</td><td>3.9E-06</td><td>BT20/MB468</td><td>MP</td></tr>
<tr><td>>MI0003560|hsa-mir-92b&WithFlank&1|+|155195171|155195278@67.87.21(+1A)&[MIMAT0003218&hsa-miR-92b-3p&offsets|0|-1(+1A);m-115&1|+|155195237|155195258&offsets|0|-1(+1A)]|TATTGCACTCGTCCCGGCCTCA</td><td>>MI0003560|hsa-mir-92b&WithFlank&1|+|155195171|155195278</td><td>67</td><td>87</td><td>21(+1A)</td><td>TATTGCACTCGTCCCGGCCTCA</td><td>1074.4</td><td>7.5</td><td>1.5</td><td>5.0</td><td>7.0E-07</td><td>3.9E-06</td><td>BT20/MB468</td><td>MP</td></tr>
<tr><td>>hg38_21_+_8433172_8446622_RNA45SN1_45S_50ntFlanks@12308.12328.21|CGATCCTTCTGACCTTTTGGG</td><td>>hg38_21_+_8433172_8446622_RNA45SN1_45S_50ntFlanks</td><td>12308</td><td>12328</td><td>21</td><td>CGATCCTTCTGACCTTTTGGG</td><td>49.2</td><td>-8.7</td><td>1.8</td><td>-5.0</td><td>7.0E-07</td><td>3.9E-06</td><td>BT20/MB468</td><td>MP</td></tr>
<tr><td>>MI0000456|hsa-mir-140&WithFlank&16|+|69933075|69933186@69.90.22(+1C)&[MIMAT0004597&hsa-miR-140-3p&offsets|+1|+2(+1C);m-45&16|+|69933143|69933164&offsets|0|0(+1C)]|ACCACAGGGTAGAACCACGGACC</td><td>>MI0000456|hsa-mir-140&WithFlank&16|+|69933075|69933186</td><td>69</td><td>90</td><td>22(+1C)</td><td>ACCACAGGGTAGAACCACGGACC</td><td>190.1</td><td>7.5</td><td>1.6</td><td>4.7</td><td>2.1E-06</td><td>3.9E-06</td><td>MB231/MB468</td><td>MP</td></tr>
<tr><td>>hg38_21_+_8433172_8446622_RNA45SN1_45S_50ntFlanks@10477.10494.18|GCCGATCGAAAGGGAGTC</td><td>>hg38_21_+_8433172_8446622_RNA45SN1_45S_50ntFlanks</td><td>10477</td><td>10494</td><td>18</td><td>GCCGATCGAAAGGGAGTC</td><td>142.6</td><td>-6.8</td><td>1.4</td><td>-4.7</td><td>2.1E-06</td><td>3.9E-06</td><td>MB231/MB468</td><td>MP</td></tr>
<tr><td>>MI0001445|hsa-mir-423&WithFlank&17|+|30117073|30117178@23.42.20(+1A)&[MIMAT0004748&hsa-miR-423-5p&offsets|0|-3(+1A);m-141&17|+|30117095|30117117&offsets|0|-3(+1A)]|TGAGGGGCAGAGAGCGAGACA</td><td>>MI0001445|hsa-mir-423&WithFlank&17|+|30117073|30117178</td><td>23</td><td>42</td><td>20(+1A)</td><td>TGAGGGGCAGAGAGCGAGACA</td><td>1217.5</td><td>6.9</td><td>1.4</td><td>5.0</td><td>7.1E-07</td><td>3.9E-06</td><td>BT20/MB468</td><td>MP</td></tr>
<tr><td>>MI0000650|hsa-mir-200c&WithFlank&12|+|6963693|6963772@50.69.20(+1A)&[MIMAT0000617&hsa-miR-200c-3p&offsets|0|-3(+1A);m-54&12|+|6963742|6963764&offsets|0|-3(+1A)]|TAATACTGCCGGGTAATGATA</td><td>>MI0000650|hsa-mir-200c&WithFlank&12|+|6963693|6963772</td><td>50</td><td>69</td><td>20(+1A)</td><td>TAATACTGCCGGGTAATGATA</td><td>504.1</td><td>6.6</td><td>1.3</td><td>5.0</td><td>7.1E-07</td><td>3.9E-06</td><td>BT20/MB468</td><td>MP</td></tr>
<tr><td>>hg38_21_+_8433172_8446622_RNA45SN1_45S_50ntFlanks@10414.10431.18|TCGGTCCTGAGAGATGGG</td><td>>hg38_21_+_8433172_8446622_RNA45SN1_45S_50ntFlanks</td><td>10414</td><td>10431</td><td>18</td><td>TCGGTCCTGAGAGATGGG</td><td>841.1</td><td>-6.9</td><td>1.4</td><td>-4.7</td><td>2.1E-06</td><td>3.9E-06</td><td>BT20/MB468</td><td>CYTO</td></tr>
<tr><td>>hg38_21_+_8433172_8446622_RNA45SN1_45S_50ntFlanks@9459.9500.42|CCGGGGAGGTGGAGCACGAGCGCACGTGTTAGGACCCGAAAG</td><td>>hg38_21_+_8433172_8446622_RNA45SN1_45S_50ntFlanks</td><td>9459</td><td>9500</td><td>42</td><td>CCGGGGAGGTGGAGCACGAGCGCACGTGTTAGGACCCGAAAG</td><td>79.8</td><td>-8.4</td><td>1.8</td><td>-4.8</td><td>1.8E-06</td><td>3.9E-06</td><td>BT20/MB468</td><td>MITO</td></tr>
<tr><td>>hg38_21_+_8433172_8446622_RNA45SN1_45S_50ntFlanks@7989.8010.22|AGACGTGGCGACCCGCTGAATT</td><td>>hg38_21_+_8433172_8446622_RNA45SN1_45S_50ntFlanks</td><td>7989</td><td>8010</td><td>22</td><td>AGACGTGGCGACCCGCTGAATT</td><td>2035.6</td><td>4.5</td><td>1.0</td><td>4.7</td><td>2.5E-06</td><td>3.9E-06</td><td>MB231/MB468</td><td>NUC</td></tr>
<tr><td>>hg38_21_+_8433172_8446622_RNA45SN1_45S_50ntFlanks@3921.3959.39|ATTTATCAGATCAAAACCAACCCGGTCAGCCCCTCTCCG</td><td>>hg38_21_+_8433172_8446622_RNA45SN1_45S_50ntFlanks</td><td>3921</td><td>3959</td><td>39</td><td>ATTTATCAGATCAAAACCAACCCGGTCAGCCCCTCTCCG</td><td>259.5</td><td>-6.4</td><td>1.4</td><td>-4.7</td><td>2.1E-06</td><td>3.9E-06</td><td>BT20/MB468</td><td>CYTO</td></tr>
<tr><td>>hg38_21_+_8433172_8446622_RNA45SN1_45S_50ntFlanks@10681.10706.26|TTCCGGCGGCGTCCGGTGAGCTCTCG</td><td>>hg38_21_+_8433172_8446622_RNA45SN1_45S_50ntFlanks</td><td>10681</td><td>10706</td><td>26</td><td>TTCCGGCGGCGTCCGGTGAGCTCTCG</td><td>8156.2</td><td>-9.2</td><td>2.0</td><td>-4.7</td><td>2.7E-06</td><td>3.9E-06</td><td>BT20/MB468</td><td>NUC</td></tr>
<tr><td>>hg38_21_+_8433172_8446622_RNA45SN1_45S_50ntFlanks@11671.11693.23|CCAGTGCTCTGAATGTCAAAGTG</td><td>>hg38_21_+_8433172_8446622_RNA45SN1_45S_50ntFlanks</td><td>11671</td><td>11693</td><td>23</td><td>CCAGTGCTCTGAATGTCAAAGTG</td><td>1897.3</td><td>-8.6</td><td>1.8</td><td>-4.7</td><td>2.7E-06</td><td>3.9E-06</td><td>BT20/MB468</td><td>NUC</td></tr>
<tr><td>>hg38_21_+_8433172_8446622_RNA45SN1_45S_50ntFlanks@9923.9947.25|TTGGTTGATATAGACAGCAGGACGG</td><td>>hg38_21_+_8433172_8446622_RNA45SN1_45S_50ntFlanks</td><td>9923</td><td>9947</td><td>25</td><td>TTGGTTGATATAGACAGCAGGACGG</td><td>46.9</td><td>9.3</td><td>1.9</td><td>5.0</td><td>7.1E-07</td><td>3.9E-06</td><td>BT20/MB468</td><td>MP</td></tr>
<tr><td>>MI0000462|hsa-mir-152&WithFlank&17|-|48037155|48037253@60.80.21(+2U)&[MIMAT0000438&hsa-miR-152-3p&offsets|0|0(+2U);m-97&17|-|48037174|48037194&offsets|0|0(+2U)]|TCAGTGCATGACAGAACTTGGTT</td><td>>MI0000462|hsa-mir-152&WithFlank&17|-|48037155|48037253</td><td>60</td><td>80</td><td>21(+2U)</td><td>TCAGTGCATGACAGAACTTGGTT</td><td>38.0</td><td>8.5</td><td>1.8</td><td>4.7</td><td>2.1E-06</td><td>3.9E-06</td><td>MB231/MB468</td><td>MP</td></tr>
<tr><td>>hg38_21_+_8433172_8446622_RNA45SN1_45S_50ntFlanks@9180.9197.18|CCGTCGGGCCCGGGGGAG</td><td>>hg38_21_+_8433172_8446622_RNA45SN1_45S_50ntFlanks</td><td>9180</td><td>9197</td><td>18</td><td>CCGTCGGGCCCGGGGGAG</td><td>2030.1</td><td>-7.3</td><td>1.5</td><td>-4.8</td><td>1.8E-06</td><td>3.9E-06</td><td>BT20/MB468</td><td>MITO</td></tr>
<tr><td>>hg38_21_+_8433172_8446622_RNA45SN1_45S_50ntFlanks@10922.10941.20|CTGGACGAGGCGCCGCCGCC</td><td>>hg38_21_+_8433172_8446622_RNA45SN1_45S_50ntFlanks</td><td>10922</td><td>10941</td><td>20</td><td>CTGGACGAGGCGCCGCCGCC</td><td>632.6</td><td>-3.4</td><td>0.7</td><td>-4.7</td><td>2.1E-06</td><td>3.9E-06</td><td>BT20/MB468</td><td>CYTO</td></tr>
<tr><td>>trna106_HisGTG_1_-_149155828_149155899@13.31.19,trna111_HisGTG_1_-_147774845_147774916@13.31.19,trna118_HisGTG_1_-_145396881_145396952@13.31.19,trna16_HisGTG_1_+_146544773_146544844@13.31.19,trna1_HisGTG_15_+_45493349_45493420@13.31.19,trna21_HisGTG_1_+_147753471_147753542@13.31.19,trna33_HisGTG_6_+_27125906_27125977@13.31.19,trna7_HisGTG_9_-_14433938_14434009@13.31.19,trna8_HisGTG_15_-_45492611_45492682@13.31.19,trna9_HisGTG_15_-_45490804_45490875@13.31.19|TAGTGGTTAGTACTCTGCG</td><td>>trna106_HisGTG_1_-_149155828_149155899</td><td>13</td><td>31</td><td>19</td><td>TAGTGGTTAGTACTCTGCG</td><td>146.8</td><td>-5.0</td><td>1.0</td><td>-4.7</td><td>2.1E-06</td><td>3.9E-06</td><td>MB231/MB468</td><td>MP</td></tr>
<tr><td>>hg38_21_+_8433172_8446622_RNA45SN1_45S_50ntFlanks@13009.13028.20|TTGAAAGTCAGCCCTCGACA</td><td>>hg38_21_+_8433172_8446622_RNA45SN1_45S_50ntFlanks</td><td>13009</td><td>13028</td><td>20</td><td>TTGAAAGTCAGCCCTCGACA</td><td>297.1</td><td>-6.9</td><td>1.5</td><td>-4.7</td><td>2.1E-06</td><td>3.9E-06</td><td>BT20/MB468</td><td>CYTO</td></tr>
<tr><td>>MI0003778|hsa-mir-320c-1&WithFlank&18|+|21683512|21683595@48.67.20;MI0008191|hsa-mir-320c-2&WithFlank&18|+|24321669|24321752@48.67.20&[MIMAT0005793&hsa-miR-320c&offsets|0|0;m-222&18|+|21683559|21683578&offsets|0|0];[MIMAT0005793_1&hsa-miR-320c&offsets|0|0;m-219&18|+|24321716|24321735&offsets|0|0]|AAAAGCTGGGTTGAGAGGGT</td><td>>MI0003778|hsa-mir-320c-1&WithFlank&18|+|21683512|21683595</td><td>48</td><td>67</td><td>20</td><td>AAAAGCTGGGTTGAGAGGGT</td><td>1730.9</td><td>4.1</td><td>0.8</td><td>5.2</td><td>2.0E-07</td><td>3.9E-06</td><td>BT20/MB231</td><td>CYTO</td></tr>
<tr><td>>hg38_21_+_8433172_8446622_RNA45SN1_45S_50ntFlanks@7978.8008.31|GACCTCAGATCAGACGTGGCGACCCGCTGAA</td><td>>hg38_21_+_8433172_8446622_RNA45SN1_45S_50ntFlanks</td><td>7978</td><td>8008</td><td>31</td><td>GACCTCAGATCAGACGTGGCGACCCGCTGAA</td><td>350.4</td><td>-3.4</td><td>0.7</td><td>-4.7</td><td>2.7E-06</td><td>3.9E-06</td><td>BT20/MB468</td><td>NUC</td></tr>
<tr><td>>hg38_21_+_8433172_8446622_RNA45SN1_45S_50ntFlanks@11235.11255.21|CTTCCCGTGGATCGCCCCAGC</td><td>>hg38_21_+_8433172_8446622_RNA45SN1_45S_50ntFlanks</td><td>11235</td><td>11255</td><td>21</td><td>CTTCCCGTGGATCGCCCCAGC</td><td>365.1</td><td>-4.2</td><td>0.9</td><td>-4.7</td><td>2.4E-06</td><td>3.9E-06</td><td>MB231/MB468</td><td>CYTO</td></tr>
<tr><td>>hg38_21_+_8433172_8446622_RNA45SN1_45S_50ntFlanks@12003.12026.24|GCGGGCCGCCGGTGAAATACCACT</td><td>>hg38_21_+_8433172_8446622_RNA45SN1_45S_50ntFlanks</td><td>12003</td><td>12026</td><td>24</td><td>GCGGGCCGCCGGTGAAATACCACT</td><td>102.3</td><td>10.0</td><td>1.9</td><td>5.2</td><td>1.8E-07</td><td>3.9E-06</td><td>BT20/MB231</td><td>MP</td></tr>
<tr><td>>hg38_21_+_8433172_8446622_RNA45SN1_45S_50ntFlanks@11807.11830.24|ACTGTCCCTACCTACTATCCAGCG</td><td>>hg38_21_+_8433172_8446622_RNA45SN1_45S_50ntFlanks</td><td>11807</td><td>11830</td><td>24</td><td>ACTGTCCCTACCTACTATCCAGCG</td><td>176.8</td><td>-6.9</td><td>1.5</td><td>-4.7</td><td>2.1E-06</td><td>3.9E-06</td><td>MB231/MB468</td><td>MP</td></tr>
<tr><td>>hg38_21_+_8433172_8446622_RNA45SN1_45S_50ntFlanks@11727.11753.27|AGTAACTATGACTCTCTTAAGGTAGCC</td><td>>hg38_21_+_8433172_8446622_RNA45SN1_45S_50ntFlanks</td><td>11727</td><td>11753</td><td>27</td><td>AGTAACTATGACTCTCTTAAGGTAGCC</td><td>166.8</td><td>-5.3</td><td>1.1</td><td>-4.7</td><td>2.1E-06</td><td>3.9E-06</td><td>MB231/MB468</td><td>MP</td></tr>
<tr><td>>hg38_21_+_8433172_8446622_RNA45SN1_45S_50ntFlanks@9926.9948.23|GTTGATATAGACAGCAGGACGGT</td><td>>hg38_21_+_8433172_8446622_RNA45SN1_45S_50ntFlanks</td><td>9926</td><td>9948</td><td>23</td><td>GTTGATATAGACAGCAGGACGGT</td><td>125.4</td><td>10.3</td><td>2.0</td><td>5.2</td><td>1.8E-07</td><td>3.9E-06</td><td>BT20/MB231</td><td>MP</td></tr>
<tr><td>>hg38_21_+_8433172_8446622_RNA45SN1_45S_50ntFlanks@5212.5255.44|ATGTCCGGGGCTGCACGCGCGCTACACTGACTGGCTCAGCGTGT</td><td>>hg38_21_+_8433172_8446622_RNA45SN1_45S_50ntFlanks</td><td>5212</td><td>5255</td><td>44</td><td>ATGTCCGGGGCTGCACGCGCGCTACACTGACTGGCTCAGCGTGT</td><td>72.4</td><td>9.9</td><td>2.0</td><td>5.0</td><td>7.2E-07</td><td>3.9E-06</td><td>BT20/MB468</td><td>MP</td></tr>
<tr><td>>hg38_21_+_8433172_8446622_RNA45SN1_45S_50ntFlanks@12419.12444.26|GATGTCGGCTCTTCCTATCATTGTGA</td><td>>hg38_21_+_8433172_8446622_RNA45SN1_45S_50ntFlanks</td><td>12419</td><td>12444</td><td>26</td><td>GATGTCGGCTCTTCCTATCATTGTGA</td><td>927.4</td><td>2.7</td><td>0.6</td><td>4.7</td><td>2.1E-06</td><td>3.9E-06</td><td>BT20/MB468</td><td>CYTO</td></tr>
<tr><td>>hg38_21_+_8433172_8446622_RNA45SN1_45S_50ntFlanks@11918.11949.32|ACATGAGAGGTGTAGAATAAGTGGGAGGCCCC</td><td>>hg38_21_+_8433172_8446622_RNA45SN1_45S_50ntFlanks</td><td>11918</td><td>11949</td><td>32</td><td>ACATGAGAGGTGTAGAATAAGTGGGAGGCCCC</td><td>499.8</td><td>-6.6</td><td>1.4</td><td>-4.7</td><td>2.1E-06</td><td>3.9E-06</td><td>BT20/MB468</td><td>CYTO</td></tr>
<tr><td>>hg38_21_+_8433172_8446622_RNA45SN1_45S_50ntFlanks@11641.11663.23|CGGCGGGTGTTGACGCGATGTGA</td><td>>hg38_21_+_8433172_8446622_RNA45SN1_45S_50ntFlanks</td><td>11641</td><td>11663</td><td>23</td><td>CGGCGGGTGTTGACGCGATGTGA</td><td>1405.7</td><td>-7.7</td><td>1.6</td><td>-5.0</td><td>7.2E-07</td><td>3.9E-06</td><td>BT20/MB468</td><td>MP</td></tr>
<tr><td>>hg38_21_+_8433172_8446622_RNA45SN1_45S_50ntFlanks@9914.9948.35|GAAAAGGTGTTGGTTGATATAGACAGCAGGACGGT</td><td>>hg38_21_+_8433172_8446622_RNA45SN1_45S_50ntFlanks</td><td>9914</td><td>9948</td><td>35</td><td>GAAAAGGTGTTGGTTGATATAGACAGCAGGACGGT</td><td>146.8</td><td>10.5</td><td>2.0</td><td>5.2</td><td>1.8E-07</td><td>3.9E-06</td><td>BT20/MB231</td><td>MP</td></tr>
<tr><td>>am_trna119_LysCTT_1_-_145395522_145395594@13.33.21,trna11_LysCTT_5_-_180648979_180649051@13.33.21,trna128_LysCTT_1_-_55423542_55423614@13.33.21,trna13_LysCTT_6_+_26556774_26556846@13.33.21,trna6_LysCTT_19_-_52425396_52425468@13.33.21,trna7_LysCTT_16_+_3225692_3225764@13.33.21,trna9_LysCTT_5_+_180634755_180634827@13.33.21|CAGTCGGTAGAGCATGAGACT</td><td>>am_trna119_LysCTT_1_-_145395522_145395594</td><td>13</td><td>33</td><td>21</td><td>CAGTCGGTAGAGCATGAGACT</td><td>993.1</td><td>-4.5</td><td>0.9</td><td>-5.2</td><td>2.2E-07</td><td>3.9E-06</td><td>BT20/MB231</td><td>MITO</td></tr>
<tr><td>>hg38_21_+_8433172_8446622_RNA45SN1_45S_50ntFlanks@7983.8014.32|CAGATCAGACGTGGCGACCCGCTGAATTTAAG</td><td>>hg38_21_+_8433172_8446622_RNA45SN1_45S_50ntFlanks</td><td>7983</td><td>8014</td><td>32</td><td>CAGATCAGACGTGGCGACCCGCTGAATTTAAG</td><td>55.9</td><td>-8.9</td><td>1.8</td><td>-5.0</td><td>7.2E-07</td><td>3.9E-06</td><td>BT20/MB468</td><td>MP</td></tr>
<tr><td>>hg38_21_+_8433172_8446622_RNA45SN1_45S_50ntFlanks@9792.9830.39|GGCGTGGAGCCGGGCGTGGAATGCGAGTGCCTAGTGGGC</td><td>>hg38_21_+_8433172_8446622_RNA45SN1_45S_50ntFlanks</td><td>9792</td><td>9830</td><td>39</td><td>GGCGTGGAGCCGGGCGTGGAATGCGAGTGCCTAGTGGGC</td><td>207.1</td><td>11.0</td><td>2.1</td><td>5.2</td><td>1.8E-07</td><td>3.9E-06</td><td>BT20/MB231</td><td>MP</td></tr>
<tr><td>>hg38_21_+_8433172_8446622_RNA45SN1_45S_50ntFlanks@10544.10567.24|TCCAGTGCGGTAACGCGACCGATC</td><td>>hg38_21_+_8433172_8446622_RNA45SN1_45S_50ntFlanks</td><td>10544</td><td>10567</td><td>24</td><td>TCCAGTGCGGTAACGCGACCGATC</td><td>607.6</td><td>-4.0</td><td>0.8</td><td>-5.0</td><td>7.2E-07</td><td>3.9E-06</td><td>BT20/MB468</td><td>MP</td></tr>
<tr><td>>hg38_21_+_8433172_8446622_RNA45SN1_45S_50ntFlanks@10781.10818.38|AGGTGAACAGCCTCTGGCATGTTGGAACAATGTAGGTA</td><td>>hg38_21_+_8433172_8446622_RNA45SN1_45S_50ntFlanks</td><td>10781</td><td>10818</td><td>38</td><td>AGGTGAACAGCCTCTGGCATGTTGGAACAATGTAGGTA</td><td>974.0</td><td>-7.0</td><td>1.4</td><td>-5.0</td><td>7.2E-07</td><td>3.9E-06</td><td>BT20/MB468</td><td>MP</td></tr>
<tr><td>>hg38_21_+_8433172_8446622_RNA45SN1_45S_50ntFlanks@8169.8186.18|CCAAGTCCTTCTGATCGA</td><td>>hg38_21_+_8433172_8446622_RNA45SN1_45S_50ntFlanks</td><td>8169</td><td>8186</td><td>18</td><td>CCAAGTCCTTCTGATCGA</td><td>356.6</td><td>-4.7</td><td>1.0</td><td>-4.7</td><td>2.4E-06</td><td>3.9E-06</td><td>MB231/MB468</td><td>CYTO</td></tr>
<tr><td>>hg38_21_+_8433172_8446622_RNA45SN1_45S_50ntFlanks@10927.10947.21|CGAGGCGCCGCCGCCCCCCCC</td><td>>hg38_21_+_8433172_8446622_RNA45SN1_45S_50ntFlanks</td><td>10927</td><td>10947</td><td>21</td><td>CGAGGCGCCGCCGCCCCCCCC</td><td>176.2</td><td>-11.3</td><td>2.4</td><td>-4.7</td><td>2.1E-06</td><td>3.9E-06</td><td>MB231/MB468</td><td>MP</td></tr>
<tr><td>>hg38_21_+_8433172_8446622_RNA45SN1_45S_50ntFlanks@10912.10931.20|CGCGCCGCGGCTGGACGAGG</td><td>>hg38_21_+_8433172_8446622_RNA45SN1_45S_50ntFlanks</td><td>10912</td><td>10931</td><td>20</td><td>CGCGCCGCGGCTGGACGAGG</td><td>220.6</td><td>-5.6</td><td>1.2</td><td>-4.7</td><td>2.4E-06</td><td>3.9E-06</td><td>MB231/MB468</td><td>CYTO</td></tr>
<tr><td>>MI0003656|hsa-mir-641&WithFlank&19|-|40282537|40282647@22.46.25&[MIMAT0003311&hsa-miR-641&offsets|0|+1;m-786&19|-|40282603|40282626&offsets|0|+1]|AAAGACATAGGATAGAGTCACCTCT</td><td>>MI0003656|hsa-mir-641&WithFlank&19|-|40282537|40282647</td><td>22</td><td>46</td><td>25</td><td>AAAGACATAGGATAGAGTCACCTCT</td><td>43.6</td><td>8.7</td><td>1.8</td><td>4.7</td><td>2.1E-06</td><td>3.9E-06</td><td>MB231/MB468</td><td>MP</td></tr>
<tr><td>>hg38_21_+_8433172_8446622_RNA45SN1_45S_50ntFlanks@5186.5204.19|CAATAACAGGTCTGTGATG</td><td>>hg38_21_+_8433172_8446622_RNA45SN1_45S_50ntFlanks</td><td>5186</td><td>5204</td><td>19</td><td>CAATAACAGGTCTGTGATG</td><td>1094.8</td><td>-6.9</td><td>1.5</td><td>-4.7</td><td>2.6E-06</td><td>3.9E-06</td><td>MB231/MB468</td><td>MITO</td></tr>
<tr><td>>hg38_21_+_8433172_8446622_RNA45SN1_45S_50ntFlanks@11571.11602.32|GCCGACTTAGAACTGGTGCGGACCAGGGGAAT</td><td>>hg38_21_+_8433172_8446622_RNA45SN1_45S_50ntFlanks</td><td>11571</td><td>11602</td><td>32</td><td>GCCGACTTAGAACTGGTGCGGACCAGGGGAAT</td><td>509.1</td><td>-3.5</td><td>0.7</td><td>-4.7</td><td>2.2E-06</td><td>3.9E-06</td><td>MB231/MB468</td><td>MP</td></tr>
</table>

| | | | | | | | | | | | | | |
|---|---|---|---|---|---|---|---|---|---|---|---|---|---|
| >hg38_21_+_8433172_8446622_RNA45SN1_45S_50ntFlanks@9538.9571.34\|AACTCTGGTGGAGGTCCGTAGCGGTCCTGACGTG | >hg38_21_+_8433172_8446622_RNA45SN1_45S_50ntFlanks | 9538 | 9571 | 34 | AACTCTGGTGGAGGTCCGTAGCGGTCCTGACGTG | 167.3 | -6.8 | 1.4 | -4.7 | 2.2E-06 | 3.9E-06 | MB231/MB468 | MP |
| >hg38_21_+_8433172_8446622_RNA45SN1_45S_50ntFlanks@11210.11232.23\|GGCGACTCTGGACGCGAGCCGGG | >hg38_21_+_8433172_8446622_RNA45SN1_45S_50ntFlanks | 11210 | 11232 | 23 | GGCGACTCTGGACGCGAGCCGGG | 3028.6 | -8.8 | 1.9 | -4.7 | 2.7E-06 | 3.9E-06 | BT20/MB468 | NUC |
| >trna10_LysCTT_16_+_3241501_3241573@4.36.33,trna13_LysCTT_14_-_58706613_58706685@4.36.33,trna2_LysCTT_15_+_79152904_79152976@4.36.33\|CGGCTAGCTCAGTCGGTAGAGCATGGGACTCTT | >trna10_LysCTT_16_+_3241501_3241573 | 4 | 36 | 33 | CGGCTAGCTCAGTCGGTAGAGCATGGGACTCTT | 1002.2 | -4.5 | 0.9 | -5.2 | 2.3E-07 | 4.0E-06 | BT20/MB231 | MITO |
| >hg38_21_+_8433172_8446622_RNA45SN1_45S_50ntFlanks@11885.11904.20\|TTGAGCTTGACTCTAGTCTG | >hg38_21_+_8433172_8446622_RNA45SN1_45S_50ntFlanks | 11885 | 11904 | 20 | TTGAGCTTGACTCTAGTCTG | 1816.8 | -6.2 | 1.3 | -5.0 | 7.2E-07 | 4.0E-06 | BT20/MB468 | MP |
| >hg38_21_+_8433172_8446622_RNA45SN1_45S_50ntFlanks@10547.10564.18\|AGTGCGGTAACGCGACCG | >hg38_21_+_8433172_8446622_RNA45SN1_45S_50ntFlanks | 10547 | 10564 | 18 | AGTGCGGTAACGCGACCG | 73.7 | -9.3 | 1.9 | -5.0 | 7.2E-07 | 4.0E-06 | BT20/MB468 | MP |
| >hg38_21_+_8433172_8446622_RNA45SN1_45S_50ntFlanks@6719.6750.32\|TGAATTGCAGGACACATTGATCATCGACACTT | >hg38_21_+_8433172_8446622_RNA45SN1_45S_50ntFlanks | 6719 | 6750 | 32 | TGAATTGCAGGACACATTGATCATCGACACTT | 114.1 | -3.7 | 0.8 | -4.7 | 2.5E-06 | 4.0E-06 | MB231/MB468 | CYTO |
| >hg38_21_+_8433172_8446622_RNA45SN1_45S_50ntFlanks@10394.10424.31\|AGCAGTTGAACATGGGTCAGTCGGTCCTGAG | >hg38_21_+_8433172_8446622_RNA45SN1_45S_50ntFlanks | 10394 | 10424 | 31 | AGCAGTTGAACATGGGTCAGTCGGTCCTGAG | 1152.0 | 3.6 | 0.8 | 4.7 | 2.5E-06 | 4.0E-06 | MB231/MB468 | CYTO |
| >hg38_21_+_8433172_8446622_RNA45SN1_45S_50ntFlanks@9720.9746.27\|ATTAGAGGTCTTGGGGCCGAAACGATC | >hg38_21_+_8433172_8446622_RNA45SN1_45S_50ntFlanks | 9720 | 9746 | 27 | ATTAGAGGTCTTGGGGCCGAAACGATC | 95.1 | -9.7 | 1.9 | -5.0 | 7.3E-07 | 4.0E-06 | BT20/MB468 | MP |
| >hg38_21_+_8433172_8446622_RNA45SN1_45S_50ntFlanks@11874.11902.29\|AGAAGACCCTGTTGAGCTTGACTCTAGTC | >hg38_21_+_8433172_8446622_RNA45SN1_45S_50ntFlanks | 11874 | 11902 | 29 | AGAAGACCCTGTTGAGCTTGACTCTAGTC | 403.8 | -3.3 | 0.7 | -4.7 | 2.5E-06 | 4.0E-06 | MB231/MB468 | CYTO |
| >hg38_21_+_8433172_8446622_RNA45SN1_45S_50ntFlanks@4221.4242.22\|CGAAAAATAACAATACAGGACT | >hg38_21_+_8433172_8446622_RNA45SN1_45S_50ntFlanks | 4221 | 4242 | 22 | CGAAAAATAACAATACAGGACT | 266.6 | -5.3 | 1.1 | -4.7 | 2.2E-06 | 4.0E-06 | MB231/MB468 | MP |
| >hg38_21_+_8433172_8446622_RNA45SN1_45S_50ntFlanks@9352.9374.23\|CCGTGGCGCAATGAAGGTGAAGG | >hg38_21_+_8433172_8446622_RNA45SN1_45S_50ntFlanks | 9352 | 9374 | 23 | CCGTGGCGCAATGAAGGTGAAGG | 62.8 | -9.1 | 1.8 | -5.0 | 7.3E-07 | 4.0E-06 | BT20/MB468 | MP |
| >hg38_21_+_8433172_8446622_RNA45SN1_45S_50ntFlanks@12417.12437.21\|TCGATGTCGGCTCTTCCTATC | >hg38_21_+_8433172_8446622_RNA45SN1_45S_50ntFlanks | 12417 | 12437 | 21 | TCGATGTCGGCTCTTCCTATC | 471.1 | -5.5 | 1.2 | -4.7 | 2.2E-06 | 4.0E-06 | MB231/MB468 | MP |
| >hg38_21_+_8433172_8446622_RNA45SN1_45S_50ntFlanks@12324.12355.32\|TTGGGTTTTAAGCAGGAGGTGTCAGAAAAGTT | >hg38_21_+_8433172_8446622_RNA45SN1_45S_50ntFlanks | 12324 | 12355 | 32 | TTGGGTTTTAAGCAGGAGGTGTCAGAAAAGTT | 176.9 | -5.7 | 1.2 | -4.7 | 2.2E-06 | 4.0E-06 | BT20/MB468 | CYTO |
| >hg38_21_+_8433172_8446622_RNA45SN1_45S_50ntFlanks@3855.3874.20\|CTGTGGTAATTCTAGAGCTA | >hg38_21_+_8433172_8446622_RNA45SN1_45S_50ntFlanks | 3855 | 3874 | 20 | CTGTGGTAATTCTAGAGCTA | 133.4 | -6.4 | 1.3 | -4.7 | 2.2E-06 | 4.0E-06 | MB231/MB468 | MP |
| >MI0000086\|hsa-mir-28&WithFlank&3\|+\|188688775\|188688872@60.81.22(+1C)&[MIMAT0004502&hsa-miR-28-3p&offsets\|0\|0(+1C);m-28&3\|+\|188688834\|188688855&offsets\|0\|0(+1C)]\|CACTAGATTGTGAGCTCCTGGAC | >MI0000086\|hsa-mir-28&WithFlank&3\|+\|188688775\|188688872 | 60 | 81 | 22(+1C) | CACTAGATTGTGAGCTCCTGGAC | 51.3 | 8.9 | 1.9 | 4.7 | 2.2E-06 | 4.0E-06 | MB231/MB468 | MP |
| >hg38_21_+_8433172_8446622_RNA45SN1_45S_50ntFlanks@9791.9811.21\|TGGCGTGGAGCCGGGCGTGGA | >hg38_21_+_8433172_8446622_RNA45SN1_45S_50ntFlanks | 9791 | 9811 | 21 | TGGCGTGGAGCCGGGCGTGGA | 62.2 | 9.7 | 2.0 | 5.0 | 7.3E-07 | 4.0E-06 | BT20/MB468 | MP |
| >hg38_21_+_8433172_8446622_RNA45SN1_45S_50ntFlanks@6721.6760.40\|AATTGCAGGACACATTGATCATCGACACTTCGAACGCACT | >hg38_21_+_8433172_8446622_RNA45SN1_45S_50ntFlanks | 6721 | 6760 | 40 | AATTGCAGGACACATTGATCATCGACACTTCGAACGCACT | 1313.9 | -4.1 | 0.9 | -4.7 | 2.6E-06 | 4.0E-06 | MB231/MB468 | NUC |
| >trnaMT_HisGTG_MT_+_12138_12206@27.55.29\|GATTGTGAATCTGACAACAGAGGCTTACG | >trnaMT_HisGTG_MT_+_12138_12206 | 27 | 55 | 29 | GATTGTGAATCTGACAACAGAGGCTTACG | 90.8 | 10.3 | 2.1 | 5.0 | 7.3E-07 | 4.0E-06 | BT20/MB468 | MP |
| >MI0000087\|hsa-mir-29a&WithFlank&7\|-\|130876741\|130876816@47.64.18&[MIMAT0000086&hsa-miR-29a-3p&offsets\|-1\|-5;m-26&7\|-\|130876749\|130876769&offsets\|-1\|-4]\|CTAGCACCATCTGAAATC | >MI0000087\|hsa-mir-29a&WithFlank&7\|-\|130876741\|130876816 | 47 | 64 | 18 | CTAGCACCATCTGAAATC | 50.4 | 8.9 | 1.9 | 4.7 | 2.2E-06 | 4.0E-06 | MB231/MB468 | MP |
| >hg38_21_+_8433172_8446622_RNA45SN1_45S_50ntFlanks@4740.4759.20\|AGTCGGAGGTTCGAAGACGA | >hg38_21_+_8433172_8446622_RNA45SN1_45S_50ntFlanks | 4740 | 4759 | 20 | AGTCGGAGGTTCGAAGACGA | 129.0 | -6.5 | 1.4 | -4.7 | 2.2E-06 | 4.0E-06 | MB231/MB468 | MP |
| >hg38_21_+_8433172_8446622_RNA45SN1_45S_50ntFlanks@11575.11607.33\|ACTTAGAACTGGTGCGGACCAGGGGAATCCGAC | >hg38_21_+_8433172_8446622_RNA45SN1_45S_50ntFlanks | 11575 | 11607 | 33 | ACTTAGAACTGGTGCGGACCAGGGGAATCCGAC | 562.3 | -5.5 | 1.2 | -4.7 | 2.6E-06 | 4.0E-06 | MB231/MB468 | MITO |
| >hg38_21_+_8433172_8446622_RNA45SN1_45S_50ntFlanks@6671.6697.27\|CTCGGCTCGTGCGTCGATGAAGAACGC | >hg38_21_+_8433172_8446622_RNA45SN1_45S_50ntFlanks | 6671 | 6697 | 27 | CTCGGCTCGTGCGTCGATGAAGAACGC | 934.5 | -8.0 | 1.7 | -4.7 | 2.8E-06 | 4.0E-06 | BT20/MB468 | NUC |
| >hg38_21_+_8433172_8446622_RNA45SN1_45S_50ntFlanks@6653.6687.35\|ACTCTTAGCGGTGGATCACTCGGCTCGTGCGTCGA | >hg38_21_+_8433172_8446622_RNA45SN1_45S_50ntFlanks | 6653 | 6687 | 35 | ACTCTTAGCGGTGGATCACTCGGCTCGTGCGTCGA | 915.4 | 3.2 | 0.7 | 4.7 | 2.6E-06 | 4.0E-06 | MB231/MB468 | NUC |
| >hg38_21_+_8433172_8446622_RNA45SN1_45S_50ntFlanks@11542.11584.43\|CGGCGCCTCGCCTCGGCCGGCGCCTAGCAGCCGACTTAGAACT | >hg38_21_+_8433172_8446622_RNA45SN1_45S_50ntFlanks | 11542 | 11584 | 43 | CGGCGCCTCGCCTCGGCCGGCGCCTAGCAGCCGACTTAGAACT | 514.3 | 7.0 | 1.5 | 4.8 | 1.9E-06 | 4.0E-06 | BT20/MB468 | MITO |
| >hg38_21_+_8433172_8446622_RNA45SN1_45S_50ntFlanks@7973.8006.34\|GACGCGACCTCAGATCAGACGTGGCGACCCGCTG | >hg38_21_+_8433172_8446622_RNA45SN1_45S_50ntFlanks | 7973 | 8006 | 34 | GACGCGACCTCAGATCAGACGTGGCGACCCGCTG | 2413.6 | -3.1 | 0.7 | -4.7 | 2.6E-06 | 4.0E-06 | MB231/MB468 | MITO |
| >hg38_21_+_8433172_8446622_RNA45SN1_45S_50ntFlanks@9835.9868.34\|TTTGGTAAGCAGAACTGGCGCTGCGGGATGAACC | >hg38_21_+_8433172_8446622_RNA45SN1_45S_50ntFlanks | 9835 | 9868 | 34 | TTTGGTAAGCAGAACTGGCGCTGCGGGATGAACC | 160.9 | 10.7 | 2.0 | 5.2 | 1.9E-07 | 4.0E-06 | BT20/MB231 | MP |
| >hg38_21_+_8433172_8446622_RNA45SN1_45S_50ntFlanks@11561.11586.26\|GCGCCTAGCAGCCGACTTAGAACTGG | >hg38_21_+_8433172_8446622_RNA45SN1_45S_50ntFlanks | 11561 | 11586 | 26 | GCGCCTAGCAGCCGACTTAGAACTGG | 1148.8 | -3.0 | 0.6 | -4.8 | 1.9E-06 | 4.0E-06 | BT20/MB468 | MITO |
| >hg38_21_+_8433172_8446622_RNA45SN1_45S_50ntFlanks@5485.5526.42\|AGCGCTGAGAAGACGGTCGAACTTGACTATCTAGAGGAAGTA | >hg38_21_+_8433172_8446622_RNA45SN1_45S_50ntFlanks | 5485 | 5526 | 42 | AGCGCTGAGAAGACGGTCGAACTTGACTATCTAGAGGAAGTA | 485.0 | -4.5 | 0.9 | -4.7 | 2.5E-06 | 4.0E-06 | MB231/MB468 | CYTO |
| >hg38_21_+_8433172_8446622_RNA45SN1_45S_50ntFlanks@11602.11623.22\|TCCGACTGTTTAATTAAAACAA | >hg38_21_+_8433172_8446622_RNA45SN1_45S_50ntFlanks | 11602 | 11623 | 22 | TCCGACTGTTTAATTAAAACAA | 1258.9 | 4.0 | 0.8 | 5.2 | 2.3E-07 | 4.0E-06 | BT20/MB231 | MITO |
| >hg38_21_+_8433172_8446622_RNA45SN1_45S_50ntFlanks@10679.10709.31\|GGTTCCGGCGGCGTCCGGTGAGCTCTCGCTG | >hg38_21_+_8433172_8446622_RNA45SN1_45S_50ntFlanks | 10679 | 10709 | 31 | GGTTCCGGCGGCGTCCGGTGAGCTCTCGCTG | 4545.6 | -4.4 | 0.9 | -4.7 | 2.2E-06 | 4.0E-06 | BT20/MB468 | CYTO |
| >hg38_21_+_8433172_8446622_RNA45SN1_45S_50ntFlanks@11882.11904.23\|CTGTTGAGCTTGACTCTAGTCTG | >hg38_21_+_8433172_8446622_RNA45SN1_45S_50ntFlanks | 11882 | 11904 | 23 | CTGTTGAGCTTGACTCTAGTCTG | 919.1 | -5.6 | 1.2 | -4.7 | 2.2E-06 | 4.0E-06 | BT20/MB468 | CYTO |
| >hg38_21_+_8433172_8446622_RNA45SN1_45S_50ntFlanks@6661.6683.23\|CGGTGGATCACTCGGCTCGTGCG | >hg38_21_+_8433172_8446622_RNA45SN1_45S_50ntFlanks | 6661 | 6683 | 23 | CGGTGGATCACTCGGCTCGTGCG | 203.4 | 2.7 | 0.6 | 4.7 | 2.5E-06 | 4.0E-06 | MB231/MB468 | CYTO |
| >hg38_21_+_8433172_8446622_RNA45SN1_45S_50ntFlanks@11576.11611.36\|CTTAGAACTGGTGCGGACCAGGGGAATCCGACTGTT | >hg38_21_+_8433172_8446622_RNA45SN1_45S_50ntFlanks | 11576 | 11611 | 36 | CTTAGAACTGGTGCGGACCAGGGGAATCCGACTGTT | 422.3 | -3.3 | 0.7 | -4.7 | 2.5E-06 | 4.0E-06 | MB231/MB468 | CYTO |
| >hg38_21_+_8433172_8446622_RNA45SN1_45S_50ntFlanks@3776.3805.30\|CCGGTACAGTGAAACTGCGAATGGCTCATT | >hg38_21_+_8433172_8446622_RNA45SN1_45S_50ntFlanks | 3776 | 3805 | 30 | CCGGTACAGTGAAACTGCGAATGGCTCATT | 595.5 | -5.3 | 1.1 | -4.7 | 2.6E-06 | 4.0E-06 | MB231/MB468 | MITO |
| >hg38_21_+_8433172_8446622_RNA45SN1_45S_50ntFlanks@9716.9737.22\|AATGATTAGAGGTCTTGGGGCC | >hg38_21_+_8433172_8446622_RNA45SN1_45S_50ntFlanks | 9716 | 9737 | 22 | AATGATTAGAGGTCTTGGGGCC | 653.8 | -7.7 | 1.6 | -4.7 | 2.8E-06 | 4.0E-06 | BT20/MB468 | NUC |
| >hg38_21_+_8433172_8446622_RNA45SN1_45S_50ntFlanks@8157.8181.25\|CTCGTGGGGGGCCCAAGTCCTTCTG | >hg38_21_+_8433172_8446622_RNA45SN1_45S_50ntFlanks | 8157 | 8181 | 25 | CTCGTGGGGGGCCCAAGTCCTTCTG | 2027.9 | -4.3 | 0.9 | -4.8 | 1.9E-06 | 4.0E-06 | BT20/MB468 | MITO |
| >hg38_21_+_8433172_8446622_RNA45SN1_45S_50ntFlanks@12606.12626.21\|ATGTGCTTGGCTGAGGAGCCA | >hg38_21_+_8433172_8446622_RNA45SN1_45S_50ntFlanks | 12606 | 12626 | 21 | ATGTGCTTGGCTGAGGAGCCA | 334.6 | -5.3 | 1.1 | -4.7 | 2.6E-06 | 4.0E-06 | MB231/MB468 | NUC |
| >hg38_21_+_8433172_8446622_RNA45SN1_45S_50ntFlanks@10351.10370.20\|CGAGAACTTTGAAGGCCGAA | >hg38_21_+_8433172_8446622_RNA45SN1_45S_50ntFlanks | 10351 | 10370 | 20 | CGAGAACTTTGAAGGCCGAA | 430.4 | -6.0 | 1.3 | -4.7 | 2.8E-06 | 4.0E-06 | BT20/MB468 | NUC |
| >hg38_21_+_8433172_8446622_RNA45SN1_45S_50ntFlanks@12934.12971.38\|CTAAACCATTCGTAGACGACCTGCTTCTGGGTCGGGGT | >hg38_21_+_8433172_8446622_RNA45SN1_45S_50ntFlanks | 12934 | 12971 | 38 | CTAAACCATTCGTAGACGACCTGCTTCTGGGTCGGGGT | 119.9 | -6.3 | 1.3 | -4.7 | 2.2E-06 | 4.0E-06 | BT20/MB468 | CYTO |
| >hg38_21_+_8433172_8446622_RNA45SN1_45S_50ntFlanks@12083.12107.25\|CTCGCTTCTGGCGCCAAGCGCCCGG | >hg38_21_+_8433172_8446622_RNA45SN1_45S_50ntFlanks | 12083 | 12107 | 25 | CTCGCTTCTGGCGCCAAGCGCCCGG | 218.9 | -6.0 | 1.3 | -4.7 | 2.2E-06 | 4.0E-06 | BT20/MB468 | CYTO |
| >hg38_21_+_8433172_8446622_RNA45SN1_45S_50ntFlanks@10513.10533.21\|CCGGAGTGGCGGAGATGGGCG | >hg38_21_+_8433172_8446622_RNA45SN1_45S_50ntFlanks | 10513 | 10533 | 21 | CCGGAGTGGCGGAGATGGGCG | 2619.9 | -7.5 | 1.6 | -4.8 | 1.9E-06 | 4.0E-06 | BT20/MB468 | MITO |
| >hg38_21_+_8433172_8446622_RNA45SN1_45S_50ntFlanks@12941.12965.25\|ATTCGTAGACGACCTGCTTCTGGGT | >hg38_21_+_8433172_8446622_RNA45SN1_45S_50ntFlanks | 12941 | 12965 | 25 | ATTCGTAGACGACCTGCTTCTGGGT | 301.7 | -4.3 | 0.9 | -4.7 | 2.2E-06 | 4.0E-06 | BT20/MB468 | CYTO |
| >hg38_21_+_8433172_8446622_RNA45SN1_45S_50ntFlanks@10362.10390.29\|AAGGCCGAAGTGGAGAAGGGTTCCATGTG | >hg38_21_+_8433172_8446622_RNA45SN1_45S_50ntFlanks | 10362 | 10390 | 29 | AAGGCCGAAGTGGAGAAGGGTTCCATGTG | 1939.7 | -6.7 | 1.4 | -4.9 | 7.4E-07 | 4.0E-06 | BT20/MB468 | MP |
| >hg38_21_+_8433172_8446622_RNA45SN1_45S_50ntFlanks@12475.12498.24\|CCCACTAATAGGGAACGTGAGCTG | >hg38_21_+_8433172_8446622_RNA45SN1_45S_50ntFlanks | 12475 | 12498 | 24 | CCCACTAATAGGGAACGTGAGCTG | 553.1 | -4.4 | 0.9 | -4.7 | 2.5E-06 | 4.0E-06 | MB231/MB468 | CYTO |
| >MI0000811\|hsa-mir-148b&WithFlank&12\|+\|54337210\|54337320@69.90.22(+1A)&[MIMAT0000759&hsa-miR-148b-3p&offsets\|0\|0(+1A);m-77&12\|+\|54337278\|54337299&offsets\|0\|0(+1A)]\|TCAGTGCATCACAGAACTTTGTA | >MI0000811\|hsa-mir-148b&WithFlank&12\|+\|54337210\|54337320 | 69 | 90 | 22(+1A) | TCAGTGCATCACAGAACTTTGTA | 270.2 | 5.1 | 1.1 | 4.7 | 2.7E-06 | 4.0E-06 | MB231/MB468 | MITO |
| >hg38_21_+_8433172_8446622_RNA45SN1_45S_50ntFlanks@5067.5102.36\|CTGGTTAATTCCGATAACGAACGAGACTCTGGCATG | >hg38_21_+_8433172_8446622_RNA45SN1_45S_50ntFlanks | 5067 | 5102 | 36 | CTGGTTAATTCCGATAACGAACGAGACTCTGGCATG | 129.4 | -10.9 | 2.3 | -4.7 | 2.2E-06 | 4.0E-06 | MB231/MB468 | MP |
| >MI0000767\|hsa-mir-365a&WithFlank&16\|+\|14309279\|14309377@62.81.20;MI0000769\|hsa-mir-365b&WithFlank&17\|+\|31575405\|31575527@74.93.20&[MIMAT0000710&hsa-miR-365a-3p&offsets\|0\|-2;m-176&16\|+\|14309340\|14309361&offsets\|0\|-2];[MIMAT0022834&hsa-miR-365b-3p&offsets\|0\|-2;m-175&17\|+\|31575478\|31575499&offsets\|0\|-2]\|TAATGCCCCTAAAAATCCTT | >MI0000767\|hsa-mir-365a&WithFlank&16\|+\|14309279\|14309377 | 62 | 81 | 20 | TAATGCCCCTAAAAATCCTT | 276.2 | 4.5 | 0.9 | 5.2 | 2.0E-07 | 4.1E-06 | BT20/MB231 | CYTO |
| >hg38_21_+_8433172_8446622_RNA45SN1_45S_50ntFlanks@12974.12991.18\|CGTACGTAGCAGAGCAGC | >hg38_21_+_8433172_8446622_RNA45SN1_45S_50ntFlanks | 12974 | 12991 | 18 | CGTACGTAGCAGAGCAGC | 1727.3 | -7.5 | 1.6 | -4.7 | 2.2E-06 | 4.1E-06 | MB231/MB468 | MP |
| >hg38_21_+_8433172_8446622_RNA45SN1_45S_50ntFlanks@3790.3810.21\|CTGCGAATGGCTCATTAAATC | >hg38_21_+_8433172_8446622_RNA45SN1_45S_50ntFlanks | 3790 | 3810 | 21 | CTGCGAATGGCTCATTAAATC | 184.8 | -6.9 | 1.5 | -4.7 | 2.2E-06 | 4.1E-06 | MB231/MB468 | MP |
| >hg38_21_+_8433172_8446622_RNA45SN1_45S_50ntFlanks@11212.11251.40\|CGACTCTGGACGCGAGCCGGGCCCTTCCCGTGGATCGCCC | >hg38_21_+_8433172_8446622_RNA45SN1_45S_50ntFlanks | 11212 | 11251 | 40 | CGACTCTGGACGCGAGCCGGGCCCTTCCCGTGGATCGCCC | 289.9 | -4.6 | 1.0 | -4.7 | 2.7E-06 | 4.1E-06 | MB231/MB468 | MITO |

<table>
<tr><td></td><td></td><td></td><td></td><td></td><td></td><td></td><td></td><td></td><td></td><td></td><td></td><td></td><td></td></tr>
<tr><td>>hg38_21_+_8433172_8446622_RNA45SN1_45S_50ntFlanks@10400.10429.30|TGAACATGGGTCAGTCGGTCCTGAGAGATG</td><td>>hg38_21_+_8433172_8446622_RNA45SN1_45S_50ntFlanks</td><td>10400</td><td>10429</td><td>30</td><td>TGAACATGGGTCAGTCGGTCCTGAGAGATG</td><td>179.4</td><td>10.8</td><td>2.1</td><td>5.2</td><td>1.9E-07</td><td>4.1E-06</td><td>BT20/MB231</td><td>MP</td></tr>
<tr><td>>MI0000746|hsa-mir-99b&WithFlank&19|+|51692606|51692687@51.72.22&[MIMAT0004678&hsa-miR-99b-3p&offsets|0|0;m-151&19|+|51692656|51692678&offsets|0|-1]|CAAGCTCGTGTCTGTGGGTCCG</td><td>>MI0000746|hsa-mir-99b&WithFlank&19|+|51692606|51692687</td><td>51</td><td>72</td><td>22</td><td>CAAGCTCGTGTCTGTGGGTCCG</td><td>702.0</td><td>3.3</td><td>0.7</td><td>4.7</td><td>2.6E-06</td><td>4.1E-06</td><td>MB231/MB468</td><td>NUC</td></tr>
<tr><td>>hg38_21_+_8433172_8446622_RNA45SN1_45S_50ntFlanks@3863.3898.36|ATTCTAGAGCTAATACATGCCGACGGGCGCTGACCC</td><td>>hg38_21_+_8433172_8446622_RNA45SN1_45S_50ntFlanks</td><td>3863</td><td>3898</td><td>36</td><td>ATTCTAGAGCTAATACATGCCGACGGGCGCTGACCC</td><td>239.4</td><td>-4.3</td><td>0.9</td><td>-4.7</td><td>2.5E-06</td><td>4.1E-06</td><td>MB231/MB468</td><td>CYTO</td></tr>
<tr><td>>hg38_21_+_8433172_8446622_RNA45SN1_45S_50ntFlanks@5071.5094.24|TTAATTCCGATAACGAACGAGACT</td><td>>hg38_21_+_8433172_8446622_RNA45SN1_45S_50ntFlanks</td><td>5071</td><td>5094</td><td>24</td><td>TTAATTCCGATAACGAACGAGACT</td><td>401.1</td><td>-8.2</td><td>1.7</td><td>-4.7</td><td>2.2E-06</td><td>4.1E-06</td><td>MB231/MB468</td><td>MP</td></tr>
<tr><td>>hg38_21_+_8433172_8446622_RNA45SN1_45S_50ntFlanks@9481.9510.30|CACGTGTTAGGACCCGAAAGATGGTGAACT</td><td>>hg38_21_+_8433172_8446622_RNA45SN1_45S_50ntFlanks</td><td>9481</td><td>9510</td><td>30</td><td>CACGTGTTAGGACCCGAAAGATGGTGAACT</td><td>242.9</td><td>11.3</td><td>2.2</td><td>5.2</td><td>1.9E-07</td><td>4.1E-06</td><td>BT20/MB231</td><td>MP</td></tr>
<tr><td>>MI0002468|hsa-mir-484&WithFlank&16|+|15643288|15643378@14.35.22(+1U)&[MIMAT0002174&hsa-miR-484&offsets|0|0(+1U);m-127&16|+|15643301|15643321&offsets|0|+1(+1U)]|TCAGGCTCAGTCCCCTCCCGATT</td><td>>MI0002468|hsa-mir-484&WithFlank&16|+|15643288|15643378</td><td>14</td><td>35</td><td>22(+1U)</td><td>TCAGGCTCAGTCCCCTCCCGATT</td><td>489.5</td><td>4.1</td><td>0.9</td><td>4.7</td><td>2.6E-06</td><td>4.1E-06</td><td>MB231/MB468</td><td>NUC</td></tr>
<tr><td>>hg38_21_+_8433172_8446622_RNA45SN1_45S_50ntFlanks@10401.10424.24|GAACATGGGTCAGTCGGTCCTGAG</td><td>>hg38_21_+_8433172_8446622_RNA45SN1_45S_50ntFlanks</td><td>10401</td><td>10424</td><td>24</td><td>GAACATGGGTCAGTCGGTCCTGAG</td><td>7566.5</td><td>4.6</td><td>0.9</td><td>4.9</td><td>7.5E-07</td><td>4.1E-06</td><td>BT20/MB468</td><td>MP</td></tr>
<tr><td>>hg38_21_+_8433172_8446622_RNA45SN1_45S_50ntFlanks@4724.4758.35|ATTAATCAAGAACGAAAGTCGGAGGTTCGAAGACG</td><td>>hg38_21_+_8433172_8446622_RNA45SN1_45S_50ntFlanks</td><td>4724</td><td>4758</td><td>35</td><td>ATTAATCAAGAACGAAAGTCGGAGGTTCGAAGACG</td><td>5404.8</td><td>-3.4</td><td>0.7</td><td>-4.7</td><td>2.2E-06</td><td>4.1E-06</td><td>BT20/MB468</td><td>CYTO</td></tr>
<tr><td>>hg38_21_+_8433172_8446622_RNA45SN1_45S_50ntFlanks@12641.12659.19|CCATCTGTGGGATTATGAC</td><td>>hg38_21_+_8433172_8446622_RNA45SN1_45S_50ntFlanks</td><td>12641</td><td>12659</td><td>19</td><td>CCATCTGTGGGATTATGAC</td><td>501.8</td><td>-4.0</td><td>0.9</td><td>-4.7</td><td>2.6E-06</td><td>4.1E-06</td><td>MB231/MB468</td><td>NUC</td></tr>
<tr><td>>hg38_21_+_8433172_8446622_RNA45SN1_45S_50ntFlanks@12008.12035.28|CCGCCGGTGAAATACCACTACTCTGATC</td><td>>hg38_21_+_8433172_8446622_RNA45SN1_45S_50ntFlanks</td><td>12008</td><td>12035</td><td>28</td><td>CCGCCGGTGAAATACCACTACTCTGATC</td><td>486.3</td><td>-7.5</td><td>1.6</td><td>-4.7</td><td>2.8E-06</td><td>4.1E-06</td><td>BT20/MB468</td><td>NUC</td></tr>
<tr><td>>hg38_21_+_8433172_8446622_RNA45SN1_45S_50ntFlanks@6651.6670.20|CGACTCTTAGCGGTGGATCA</td><td>>hg38_21_+_8433172_8446622_RNA45SN1_45S_50ntFlanks</td><td>6651</td><td>6670</td><td>20</td><td>CGACTCTTAGCGGTGGATCA</td><td>10875.1</td><td>-3.8</td><td>0.8</td><td>-4.7</td><td>2.7E-06</td><td>4.1E-06</td><td>MB231/MB468</td><td>MITO</td></tr>
<tr><td>>hg38_21_+_8433172_8446622_RNA45SN1_45S_50ntFlanks@5059.5080.22|CGATTTGTCTGGTTAATTCCGA</td><td>>hg38_21_+_8433172_8446622_RNA45SN1_45S_50ntFlanks</td><td>5059</td><td>5080</td><td>22</td><td>CGATTTGTCTGGTTAATTCCGA</td><td>105.2</td><td>-9.8</td><td>2.0</td><td>-4.9</td><td>7.5E-07</td><td>4.1E-06</td><td>BT20/MB468</td><td>MP</td></tr>
<tr><td>>hg38_21_+_8433172_8446622_RNA45SN1_45S_50ntFlanks@6731.6772.42|CACATTGATCATCGACACTTCGAACGCACTTGCGGCCCCGGG</td><td>>hg38_21_+_8433172_8446622_RNA45SN1_45S_50ntFlanks</td><td>6731</td><td>6772</td><td>42</td><td>CACATTGATCATCGACACTTCGAACGCACTTGCGGCCCCGGG</td><td>457.7</td><td>-5.7</td><td>1.2</td><td>-4.7</td><td>2.6E-06</td><td>4.1E-06</td><td>MB231/MB468</td><td>NUC</td></tr>
<tr><td>>MI0000456|hsa-mir-140&WithFlank&16|+|69933075|69933186@70.90.21&[MIMAT0004597&hsa-miR-140-3p&offsets|+2|+2;m-45&16|+|69933143|69933164&offsets|+1|0]|CCACAGGGTAGAACCACGGAC</td><td>>MI0000456|hsa-mir-140&WithFlank&16|+|69933075|69933186</td><td>70</td><td>90</td><td>21</td><td>CCACAGGGTAGAACCACGGAC</td><td>80.5</td><td>10.1</td><td>2.0</td><td>4.9</td><td>7.5E-07</td><td>4.1E-06</td><td>BT20/MB468</td><td>MP</td></tr>
<tr><td>>hg38_21_+_8433172_8446622_RNA45SN1_45S_50ntFlanks@12418.12448.31|CGATGTCGGCTCTTCCTATCATTGTGAAGCA</td><td>>hg38_21_+_8433172_8446622_RNA45SN1_45S_50ntFlanks</td><td>12418</td><td>12448</td><td>31</td><td>CGATGTCGGCTCTTCCTATCATTGTGAAGCA</td><td>1791.7</td><td>3.4</td><td>0.7</td><td>4.7</td><td>2.5E-06</td><td>4.1E-06</td><td>MB231/MB468</td><td>CYTO</td></tr>
<tr><td>>hg38_21_+_8433172_8446622_RNA45SN1_45S_50ntFlanks@12999.13028.30|CTGCGATCTATTGAAAGTCAGCCCTCGACA</td><td>>hg38_21_+_8433172_8446622_RNA45SN1_45S_50ntFlanks</td><td>12999</td><td>13028</td><td>30</td><td>CTGCGATCTATTGAAAGTCAGCCCTCGACA</td><td>504.6</td><td>-3.5</td><td>0.7</td><td>-4.7</td><td>2.7E-06</td><td>4.1E-06</td><td>MB231/MB468</td><td>MITO</td></tr>
<tr><td>>hg38_21_+_8433172_8446622_RNA45SN1_45S_50ntFlanks@11035.11060.26|CTCTCTCCCCCGCTCCCCGTCCTCCC</td><td>>hg38_21_+_8433172_8446622_RNA45SN1_45S_50ntFlanks</td><td>11035</td><td>11060</td><td>26</td><td>CTCTCTCCCCCGCTCCCCGTCCTCCC</td><td>460.3</td><td>-7.1</td><td>1.5</td><td>-4.7</td><td>2.6E-06</td><td>4.1E-06</td><td>MB231/MB468</td><td>NUC</td></tr>
<tr><td>>hg38_21_+_8433172_8446622_RNA45SN1_45S_50ntFlanks@12419.12440.22|GATGTCGGCTCTTCCTATCATT</td><td>>hg38_21_+_8433172_8446622_RNA45SN1_45S_50ntFlanks</td><td>12419</td><td>12440</td><td>22</td><td>GATGTCGGCTCTTCCTATCATT</td><td>596.9</td><td>3.6</td><td>0.8</td><td>4.7</td><td>2.6E-06</td><td>4.1E-06</td><td>MB231/MB468</td><td>NUC</td></tr>
<tr><td>>am_trna10_ValCAC_5_-_180649395_180649467@1.29.29,trna12_ValAAC_5_-_180645270_180645342@1.29.29,trna132_ValAAC_6_-_27721179_27721251@1.29.29,trna136_ValAAC_6_-_27648885_27648957@1.29.29,trna139_ValAAC_6_-_27618707_27618779@1.29.29,trna18_ValCAC_5_-_180529253_180529325@1.29.29,trna2_ValAAC_3_+_169490018_169490090@1.29.29,trna2_ValCAC_5_+_180524070_180524142@1.29.29,trna4_ValAAC_5_+_180591154_180591226@1.29.29,trna5_ValAAC_5_+_180596610_180596682@1.29.29,trna6_ValCAC_5_+_180600650_180600722@1.29.29,trna85_ValCAC_1_-_161369490_161369562@1.29.29,trna90_ValCAC_1_-_149684088_149684161@1.29.29,trna98_ValCAC_1_-_149298555_149298627@1.29.29,trna9_ValCAC_6_+_26538282_26538354@1.29.29|GTTTCCGTAGTGTAGTGGTTATCACGTTC</td><td>>am_trna10_ValCAC_5_-_180649395_180649467</td><td>1</td><td>29</td><td>29</td><td>GTTTCCGTAGTGTAGTGGTTATCACGTTC</td><td>6037.1</td><td>-4.5</td><td>1.0</td><td>-4.7</td><td>2.6E-06</td><td>4.1E-06</td><td>MB231/MB468</td><td>NUC</td></tr>
<tr><td>>hg38_21_+_8433172_8446622_RNA45SN1_45S_50ntFlanks@9544.9573.30|GGTGGAGGTCCGTAGCGGTCCTGACGTGCA</td><td>>hg38_21_+_8433172_8446622_RNA45SN1_45S_50ntFlanks</td><td>9544</td><td>9573</td><td>30</td><td>GGTGGAGGTCCGTAGCGGTCCTGACGTGCA</td><td>1061.7</td><td>-6.2</td><td>1.3</td><td>-4.7</td><td>2.8E-06</td><td>4.1E-06</td><td>BT20/MB468</td><td>NUC</td></tr>
<tr><td>>hg38_21_+_8433172_8446622_RNA45SN1_45S_50ntFlanks@9567.9593.27|ACGTGCAAATCGGTCGTCCGACCTGGG</td><td>>hg38_21_+_8433172_8446622_RNA45SN1_45S_50ntFlanks</td><td>9567</td><td>9593</td><td>27</td><td>ACGTGCAAATCGGTCGTCCGACCTGGG</td><td>196.8</td><td>-4.0</td><td>0.9</td><td>-4.7</td><td>2.6E-06</td><td>4.1E-06</td><td>MB231/MB468</td><td>CYTO</td></tr>
<tr><td>>hg38_21_+_8433172_8446622_RNA45SN1_45S_50ntFlanks@10819.10837.19|AGGGAAGTCGGCAAGCCGG</td><td>>hg38_21_+_8433172_8446622_RNA45SN1_45S_50ntFlanks</td><td>10819</td><td>10837</td><td>19</td><td>AGGGAAGTCGGCAAGCCGG</td><td>489.1</td><td>-3.6</td><td>0.8</td><td>-4.7</td><td>2.3E-06</td><td>4.1E-06</td><td>BT20/MB468</td><td>CYTO</td></tr>
<tr><td>>MI0000073|hsa-mir-19a&WithFlank&13|+|91350885|91350978@55.77.23(+1C)&[MIMAT0000073&hsa-miR-19a-3p&offsets|0|0(+1C);m-201&13|+|91350939|91350961&offsets|0|0(+1C)]|TGTGCAAATCTATGCAAAACTGAC</td><td>>MI0000073|hsa-mir-19a&WithFlank&13|+|91350885|91350978</td><td>55</td><td>77</td><td>23(+1C)</td><td>TGTGCAAATCTATGCAAAACTGAC</td><td>110.7</td><td>-10.2</td><td>2.0</td><td>-5.2</td><td>1.9E-07</td><td>4.1E-06</td><td>BT20/MB231</td><td>MP</td></tr>
<tr><td>>hg38_21_+_8433172_8446622_RNA45SN1_45S_50ntFlanks@8357.8376.20|AAAAGAACTTTGAAGAGAGA</td><td>>hg38_21_+_8433172_8446622_RNA45SN1_45S_50ntFlanks</td><td>8357</td><td>8376</td><td>20</td><td>AAAAGAACTTTGAAGAGAGA</td><td>1913.0</td><td>-5.4</td><td>1.1</td><td>-4.9</td><td>7.6E-07</td><td>4.1E-06</td><td>BT20/MB468</td><td>MP</td></tr>
<tr><td>>hg38_21_+_8433172_8446622_RNA45SN1_45S_50ntFlanks@4006.4042.37|ACCTCGGGCCGATCGCACGCCCCCCGTGGCGGCGACG</td><td>>hg38_21_+_8433172_8446622_RNA45SN1_45S_50ntFlanks</td><td>4006</td><td>4042</td><td>37</td><td>ACCTCGGGCCGATCGCACGCCCCCCGTGGCGGCGACG</td><td>693.0</td><td>-6.5</td><td>1.4</td><td>-4.7</td><td>2.3E-06</td><td>4.1E-06</td><td>BT20/MB468</td><td>CYTO</td></tr>
<tr><td>>MI0000433|hsa-let-7g&WithFlank&3|-|52268272|52268367@11.31.21(+1A)&[MIMAT0000414&hsa-let-7g-5p&offsets|0|-1(+1A);m-58&3|-|52268336|52268357&offsets|0|-1(+1A)]|TGAGGTAGTAGTTTGTACAGTA</td><td>>MI0000433|hsa-let-7g&WithFlank&3|-|52268272|52268367</td><td>11</td><td>31</td><td>21(+1A)</td><td>TGAGGTAGTAGTTTGTACAGTA</td><td>924.4</td><td>2.6</td><td>0.6</td><td>4.7</td><td>2.8E-06</td><td>4.1E-06</td><td>BT20/MB468</td><td>NUC</td></tr>
<tr><td>>hg38_21_+_8433172_8446622_RNA45SN1_45S_50ntFlanks@3855.3881.27|CTGTGGTAATTCTAGAGCTAATACATG</td><td>>hg38_21_+_8433172_8446622_RNA45SN1_45S_50ntFlanks</td><td>3855</td><td>3881</td><td>27</td><td>CTGTGGTAATTCTAGAGCTAATACATG</td><td>426.9</td><td>-7.5</td><td>1.6</td><td>-4.7</td><td>2.9E-06</td><td>4.1E-06</td><td>BT20/MB468</td><td>NUC</td></tr>
<tr><td>>hg38_21_+_8433172_8446622_RNA45SN1_45S_50ntFlanks@12912.12943.32|CACGTTCGTGGGGAACCTGGCGCTAAACCATT</td><td>>hg38_21_+_8433172_8446622_RNA45SN1_45S_50ntFlanks</td><td>12912</td><td>12943</td><td>32</td><td>CACGTTCGTGGGGAACCTGGCGCTAAACCATT</td><td>373.7</td><td>-5.2</td><td>1.1</td><td>-4.7</td><td>2.9E-06</td><td>4.1E-06</td><td>BT20/MB468</td><td>NUC</td></tr>
<tr><td>>trnaMT_TrpTCA_MT_+_5512_5579@18.35.18,trnalookalike3_TrpTCA_1_+_566062_566129@18.35.18|ACAGACCAAGAGCCTTCA</td><td>>trnaMT_TrpTCA_MT_+_5512_5579</td><td>18</td><td>35</td><td>18</td><td>ACAGACCAAGAGCCTTCA</td><td>365.8</td><td>6.9</td><td>1.5</td><td>4.7</td><td>2.7E-06</td><td>4.1E-06</td><td>MB231/MB468</td><td>MITO</td></tr>
<tr><td>>hg38_21_+_8433172_8446622_RNA45SN1_45S_50ntFlanks@10402.10428.27|AACATGGGTCAGTCGGTCCTGAGAGAT</td><td>>hg38_21_+_8433172_8446622_RNA45SN1_45S_50ntFlanks</td><td>10402</td><td>10428</td><td>27</td><td>AACATGGGTCAGTCGGTCCTGAGAGAT</td><td>167.6</td><td>-2.7</td><td>0.6</td><td>-4.7</td><td>2.3E-06</td><td>4.1E-06</td><td>BT20/MB468</td><td>CYTO</td></tr>
<tr><td>>hg38_21_+_8433172_8446622_RNA45SN1_45S_50ntFlanks@9027.9064.38|CGTCTCCTCGTGGGGGGGCCGGGCCACCCCTCCCACGG</td><td>>hg38_21_+_8433172_8446622_RNA45SN1_45S_50ntFlanks</td><td>9027</td><td>9064</td><td>38</td><td>CGTCTCCTCGTGGGGGGGCCGGGCCACCCCTCCCACGG</td><td>1155.1</td><td>-6.9</td><td>1.5</td><td>-4.8</td><td>2.0E-06</td><td>4.1E-06</td><td>BT20/MB468</td><td>MITO</td></tr>
<tr><td>>hg38_21_+_8433172_8446622_RNA45SN1_45S_50ntFlanks@9862.9898.37|ATGAACCGAACGCCGGGTTAAGGCGCCCGATGCCGAC</td><td>>hg38_21_+_8433172_8446622_RNA45SN1_45S_50ntFlanks</td><td>9862</td><td>9898</td><td>37</td><td>ATGAACCGAACGCCGGGTTAAGGCGCCCGATGCCGAC</td><td>209.2</td><td>-7.7</td><td>1.6</td><td>-4.7</td><td>2.7E-06</td><td>4.1E-06</td><td>MB231/MB468</td><td>MITO</td></tr>
<tr><td>>hg38_21_+_8433172_8446622_RNA45SN1_45S_50ntFlanks@3874.3894.21|AATACATGCCGACGGGCGCTG</td><td>>hg38_21_+_8433172_8446622_RNA45SN1_45S_50ntFlanks</td><td>3874</td><td>3894</td><td>21</td><td>AATACATGCCGACGGGCGCTG</td><td>48.0</td><td>-8.7</td><td>1.8</td><td>-4.9</td><td>7.6E-07</td><td>4.1E-06</td><td>BT20/MB468</td><td>MP</td></tr>
<tr><td>>am_trnaMT_HisGTG_MT_+_12138_12206@18.51.34|AAAACATCAGATTGTGAATCTGACAACAGAGGCT</td><td>>am_trnaMT_HisGTG_MT_+_12138_12206</td><td>18</td><td>51</td><td>34</td><td>AAAACATCAGATTGTGAATCTGACAACAGAGGCT</td><td>1023.0</td><td>8.2</td><td>1.7</td><td>4.9</td><td>7.6E-07</td><td>4.2E-06</td><td>BT20/MB468</td><td>MP</td></tr>
<tr><td>>hg38_21_+_8433172_8446622_RNA45SN1_45S_50ntFlanks@10869.10893.25|AGGGCTGGGTCGGTCGGGCTGGGGC</td><td>>hg38_21_+_8433172_8446622_RNA45SN1_45S_50ntFlanks</td><td>10869</td><td>10893</td><td>25</td><td>AGGGCTGGGTCGGTCGGGCTGGGGC</td><td>227.2</td><td>-4.3</td><td>0.9</td><td>-4.7</td><td>2.3E-06</td><td>4.2E-06</td><td>MB231/MB468</td><td>MP</td></tr>
<tr><td>>hg38_21_+_8433172_8446622_RNA45SN1_45S_50ntFlanks@10360.10383.24|TGAAGGCCGAAGTGGAGAAGGGTT</td><td>>hg38_21_+_8433172_8446622_RNA45SN1_45S_50ntFlanks</td><td>10360</td><td>10383</td><td>24</td><td>TGAAGGCCGAAGTGGAGAAGGGTT</td><td>249.2</td><td>-4.8</td><td>1.0</td><td>-4.7</td><td>2.7E-06</td><td>4.2E-06</td><td>MB231/MB468</td><td>NUC</td></tr>
<tr><td>>hg38_21_+_8433172_8446622_RNA45SN1_45S_50ntFlanks@6667.6698.32|ATCACTCGGCTCGTGCGTCGATGAAGAACGCA</td><td>>hg38_21_+_8433172_8446622_RNA45SN1_45S_50ntFlanks</td><td>6667</td><td>6698</td><td>32</td><td>ATCACTCGGCTCGTGCGTCGATGAAGAACGCA</td><td>47.7</td><td>-8.7</td><td>1.8</td><td>-4.9</td><td>7.6E-07</td><td>4.2E-06</td><td>BT20/MB468</td><td>MP</td></tr>
<tr><td>>trnaMT_IleGAT_MT_+_4263_4331@42.71.30|AATAGGAGCTTAAACCCCCTTATTTCTACC</td><td>>trnaMT_IleGAT_MT_+_4263_4331</td><td>42</td><td>71</td><td>30</td><td>AATAGGAGCTTAAACCCCCTTATTTCTACC</td><td>37.8</td><td>8.5</td><td>1.8</td><td>4.7</td><td>2.3E-06</td><td>4.2E-06</td><td>MB231/MB468</td><td>MP</td></tr>
<tr><td>>MI0000737|hsa-mir-200a&WithFlank&1|+|1167857|1167958@60.81.22(+1C)&[MIMAT0000682&hsa-miR-200a-3p&offsets|0|0(+1C);m-142&1|+|1167916|1167938&offsets|0|-1(+1C)]|TAACACTGTCTGGTAACGATGTC</td><td>>MI0000737|hsa-mir-200a&WithFlank&1|+|1167857|1167958</td><td>60</td><td>81</td><td>22(+1C)</td><td>TAACACTGTCTGGTAACGATGTC</td><td>1116.2</td><td>-6.0</td><td>1.3</td><td>-4.7</td><td>2.3E-06</td><td>4.2E-06</td><td>MB231/MB468</td><td>MP</td></tr>
<tr><td>>hg38_21_+_8433172_8446622_RNA45SN1_45S_50ntFlanks@5083.5120.38|ACGAACGAGACTCTGGCATGCTAACTAGTTACGCGACC</td><td>>hg38_21_+_8433172_8446622_RNA45SN1_45S_50ntFlanks</td><td>5083</td><td>5120</td><td>38</td><td>ACGAACGAGACTCTGGCATGCTAACTAGTTACGCGACC</td><td>461.4</td><td>-5.5</td><td>1.2</td><td>-4.7</td><td>2.3E-06</td><td>4.2E-06</td><td>MB231/MB468</td><td>MP</td></tr>
<tr><td>>hg38_21_+_8433172_8446622_RNA45SN1_45S_50ntFlanks@10481.10498.18|ATCGAAAGGGAGTCGGGT</td><td>>hg38_21_+_8433172_8446622_RNA45SN1_45S_50ntFlanks</td><td>10481</td><td>10498</td><td>18</td><td>ATCGAAAGGGAGTCGGGT</td><td>335.2</td><td>-5.7</td><td>1.2</td><td>-4.7</td><td>2.6E-06</td><td>4.2E-06</td><td>MB231/MB468</td><td>CYTO</td></tr>
<tr><td>>hg38_21_+_8433172_8446622_RNA45SN1_45S_50ntFlanks@9545.9566.22|GTGGAGGTCCGTAGCGGTCCTG</td><td>>hg38_21_+_8433172_8446622_RNA45SN1_45S_50ntFlanks</td><td>9545</td><td>9566</td><td>22</td><td>GTGGAGGTCCGTAGCGGTCCTG</td><td>2281.8</td><td>-7.2</td><td>1.5</td><td>-4.7</td><td>2.3E-06</td><td>4.2E-06</td><td>BT20/MB468</td><td>CYTO</td></tr>
<tr><td>>hg38_21_+_8433172_8446622_RNA45SN1_45S_50ntFlanks@7968.7988.21|TCCGAGACGCGACCTCAGATC</td><td>>hg38_21_+_8433172_8446622_RNA45SN1_45S_50ntFlanks</td><td>7968</td><td>7988</td><td>21</td><td>TCCGAGACGCGACCTCAGATC</td><td>647.6</td><td>4.1</td><td>0.9</td><td>4.7</td><td>2.7E-06</td><td>4.2E-06</td><td>MB231/MB468</td><td>NUC</td></tr>
<tr><td>>MI0000737|hsa-mir-200a&WithFlank&1|+|1167857|1167958@60.80.21(+1C)&[MIMAT0000682&hsa-miR-200a-3p&offsets|0|-1(+1C);m-142&1|+|1167916|1167938&offsets|0|-2(+1C)]|TAACACTGTCTGGTAACGATGC</td><td>>MI0000737|hsa-mir-200a&WithFlank&1|+|1167857|1167958</td><td>60</td><td>80</td><td>21(+1C)</td><td>TAACACTGTCTGGTAACGATGC</td><td>238.6</td><td>-6.6</td><td>1.4</td><td>-4.7</td><td>2.3E-06</td><td>4.2E-06</td><td>MB231/MB468</td><td>MP</td></tr>
<tr><td>>hg38_21_+_8433172_8446622_RNA45SN1_45S_50ntFlanks@8196.8224.29|CGTGGACGGTGTGAGGCCGGTAGCGGCCC</td><td>>hg38_21_+_8433172_8446622_RNA45SN1_45S_50ntFlanks</td><td>8196</td><td>8224</td><td>29</td><td>CGTGGACGGTGTGAGGCCGGTAGCGGCCC</td><td>2686.5</td><td>-8.6</td><td>1.8</td><td>-4.7</td><td>2.9E-06</td><td>4.2E-06</td><td>BT20/MB468</td><td>NUC</td></tr>
<tr><td>>hg38_21_+_8433172_8446622_RNA45SN1_45S_50ntFlanks@8172.8212.41|AGTCCTTCTGATCGAGGCCCAGCCCGTGGACGGTGTGAGGC</td><td>>hg38_21_+_8433172_8446622_RNA45SN1_45S_50ntFlanks</td><td>8172</td><td>8212</td><td>41</td><td>AGTCCTTCTGATCGAGGCCCAGCCCGTGGACGGTGTGAGGC</td><td>2196.4</td><td>-8.6</td><td>1.8</td><td>-4.7</td><td>2.9E-06</td><td>4.2E-06</td><td>BT20/MB468</td><td>NUC</td></tr>
</table>

<table>
<tr><th></th><th></th><th></th><th></th><th></th><th></th><th></th><th></th><th></th><th></th><th></th><th></th><th></th><th></th></tr>
<tr><td>>MI0000108|hsa-mir-103a-2&WithFlank&20|+|3917488|3917577@54.77.24;MI0000109|hsa-mir-103a-1&WithFlank&5|-|168560890|168560979@54.77.24&[MIMAT0000101&hsa-miR-103a-3p&offsets|0|+1;m-12&20|+|3917541|3917563&offsets|0|+1];[MIMAT0000101_1&hsa-miR-103a-3p&offsets|0|+1;m-13&5|-|168560904|168560926&offsets|0|+1;m-44220&5|-|168560908|168560926&offsets|0|+5]|AGCAGCATTGTACAGGGCTATGAA</td><td>>MI0000108|hsa-mir-103a-2&WithFlank&20|+|3917488|3917577</td><td>54</td><td>77</td><td>24</td><td>AGCAGCATTGTACAGGGCTATGAA</td><td>337.3</td><td>2.2</td><td>0.5</td><td>4.7</td><td>2.9E-06</td><td>4.2E-06</td><td>BT20/MB468</td><td>NUC</td></tr>
<tr><td>>hg38_21_+_8433172_8446622_RNA45SN1_45S_50ntFlanks@3853.3898.46|AACTGTGGTAATTCTAGAGCTAATACATGCCGACGGGCGCTGACCC</td><td>>hg38_21_+_8433172_8446622_RNA45SN1_45S_50ntFlanks</td><td>3853</td><td>3898</td><td>46</td><td>AACTGTGGTAATTCTAGAGCTAATACATGCCGACGGGCGCTGACCC</td><td>161.7</td><td>-11.2</td><td>2.4</td><td>-4.7</td><td>2.3E-06</td><td>4.2E-06</td><td>MB231/MB468</td><td>MP</td></tr>
<tr><td>>hg38_21_+_8433172_8446622_RNA45SN1_45S_50ntFlanks@11874.11909.36|AGAAGACCCTGTTGAGCTTGACTCTAGTCTGGCACG</td><td>>hg38_21_+_8433172_8446622_RNA45SN1_45S_50ntFlanks</td><td>11874</td><td>11909</td><td>36</td><td>AGAAGACCCTGTTGAGCTTGACTCTAGTCTGGCACG</td><td>577.3</td><td>-3.3</td><td>0.7</td><td>-4.8</td><td>2.0E-06</td><td>4.2E-06</td><td>BT20/MB468</td><td>MITO</td></tr>
<tr><td>>hg38_21_+_8433172_8446622_RNA45SN1_45S_50ntFlanks@6731.6772.42|CACATTGATCATCGACACTTCGAACGCACTTGCGGCCCCGGG</td><td>>hg38_21_+_8433172_8446622_RNA45SN1_45S_50ntFlanks</td><td>6731</td><td>6772</td><td>42</td><td>CACATTGATCATCGACACTTCGAACGCACTTGCGGCCCCGGG</td><td>145.8</td><td>-11.1</td><td>2.3</td><td>-4.7</td><td>2.3E-06</td><td>4.2E-06</td><td>MB231/MB468</td><td>MP</td></tr>
<tr><td>>MI0000087|hsa-mir-29a&WithFlank&7|-|130876741|130876816@48.67.20(+1C)&[MIMAT0000086&hsa-miR-29a-3p&offsets|0|-2(+1C);m-26&7|-|130876749|130876769&offsets|0|-1(+1C)]|TAGCACCATCTGAAATCGGTC</td><td>>MI0000087|hsa-mir-29a&WithFlank&7|-|130876741|130876816</td><td>48</td><td>67</td><td>20(+1C)</td><td>TAGCACCATCTGAAATCGGTC</td><td>348.5</td><td>8.0</td><td>1.7</td><td>4.7</td><td>2.9E-06</td><td>4.2E-06</td><td>BT20/MB468</td><td>NUC</td></tr>
<tr><td>>hg38_21_+_8433172_8446622_RNA45SN1_45S_50ntFlanks@10871.10891.21|GGCTGGGTCGGTCGGGCTGGG</td><td>>hg38_21_+_8433172_8446622_RNA45SN1_45S_50ntFlanks</td><td>10871</td><td>10891</td><td>21</td><td>GGCTGGGTCGGTCGGGCTGGG</td><td>296.9</td><td>11.5</td><td>2.2</td><td>5.2</td><td>2.0E-07</td><td>4.2E-06</td><td>BT20/MB231</td><td>MP</td></tr>
<tr><td>>hg38_21_+_8433172_8446622_RNA45SN1_45S_50ntFlanks@8112.8140.29|GGGCGCGGGACATGTGGCGTACGGAAGAC</td><td>>hg38_21_+_8433172_8446622_RNA45SN1_45S_50ntFlanks</td><td>8112</td><td>8140</td><td>29</td><td>GGGCGCGGGACATGTGGCGTACGGAAGAC</td><td>289.8</td><td>11.5</td><td>2.2</td><td>5.2</td><td>2.0E-07</td><td>4.2E-06</td><td>BT20/MB231</td><td>MP</td></tr>
<tr><td>>hg38_21_+_8433172_8446622_RNA45SN1_45S_50ntFlanks@10402.10420.19|AACATGGGTCAGTCGGTCC</td><td>>hg38_21_+_8433172_8446622_RNA45SN1_45S_50ntFlanks</td><td>10402</td><td>10420</td><td>19</td><td>AACATGGGTCAGTCGGTCC</td><td>1564.1</td><td>-7.0</td><td>1.5</td><td>-4.7</td><td>2.3E-06</td><td>4.2E-06</td><td>BT20/MB468</td><td>CYTO</td></tr>
<tr><td>>hg38_21_+_8433172_8446622_RNA45SN1_45S_50ntFlanks@8196.8214.19|CGTGGACGGTGTGAGGCCG</td><td>>hg38_21_+_8433172_8446622_RNA45SN1_45S_50ntFlanks</td><td>8196</td><td>8214</td><td>19</td><td>CGTGGACGGTGTGAGGCCG</td><td>2073.3</td><td>-8.4</td><td>1.8</td><td>-4.7</td><td>2.9E-06</td><td>4.2E-06</td><td>BT20/MB468</td><td>NUC</td></tr>
<tr><td>>hg38_21_+_8433172_8446622_RNA45SN1_45S_50ntFlanks@9937.9980.44|CAGCAGGACGGTGGCCATGGAAGTCGGAATCCGCTAAGGAGTGT</td><td>>hg38_21_+_8433172_8446622_RNA45SN1_45S_50ntFlanks</td><td>9937</td><td>9980</td><td>44</td><td>CAGCAGGACGGTGGCCATGGAAGTCGGAATCCGCTAAGGAGTGT</td><td>49.5</td><td>9.4</td><td>1.9</td><td>4.9</td><td>7.7E-07</td><td>4.2E-06</td><td>BT20/MB468</td><td>MP</td></tr>
<tr><td>>hg38_21_+_8433172_8446622_RNA45SN1_45S_50ntFlanks@10466.10484.19|CGTTGCCCTCGGCCGATCG</td><td>>hg38_21_+_8433172_8446622_RNA45SN1_45S_50ntFlanks</td><td>10466</td><td>10484</td><td>19</td><td>CGTTGCCCTCGGCCGATCG</td><td>58.2</td><td>-8.9</td><td>1.8</td><td>-4.9</td><td>7.7E-07</td><td>4.2E-06</td><td>BT20/MB468</td><td>MP</td></tr>
<tr><td>>hg38_MT_+_1621_3279_RNR2_16S_50ntFlanks@826.859.34|AAGGAAAGGTTAAAAAAAGTAAAAGGAACTCGGC</td><td>>hg38_MT_+_1621_3279_RNR2_16S_50ntFlanks</td><td>826</td><td>859</td><td>34</td><td>AAGGAAAGGTTAAAAAAAGTAAAAGGAACTCGGC</td><td>61.0</td><td>9.7</td><td>2.0</td><td>4.9</td><td>7.7E-07</td><td>4.2E-06</td><td>BT20/MB468</td><td>MP</td></tr>
<tr><td>>MI0000081|hsa-mir-24-2&WithFlank&19|-|13836281|13836365@56.74.19(+1U);MI0000080|hsa-mir-24-1&WithFlank&9|+|95086015|95086094@50.68.19(+1U)&[MIMAT0000080&hsa-miR-24-3p&offsets|0|-3(+1U);m-36&19|-|13836288|13836310&offsets|0|-4(+1U)];[MIMAT0000080_1&hsa-miR-24-3p&offsets|0|-3(+1U);m-35&9|+|95086064|95086086&offsets|0|-4(+1U)]|TGGCTCAGTTCAGCAGGAAT</td><td>>MI0000081|hsa-mir-24-2&WithFlank&19|-|13836281|13836365</td><td>56</td><td>74</td><td>19(+1U)</td><td>TGGCTCAGTTCAGCAGGAAT</td><td>313.3</td><td>3.1</td><td>0.7</td><td>4.7</td><td>2.9E-06</td><td>4.2E-06</td><td>BT20/MB468</td><td>NUC</td></tr>
<tr><td>>hg38_21_+_8433172_8446622_RNA45SN1_45S_50ntFlanks@12405.12426.22|CTTTTTGATCCTTCGATGTCGG</td><td>>hg38_21_+_8433172_8446622_RNA45SN1_45S_50ntFlanks</td><td>12405</td><td>12426</td><td>22</td><td>CTTTTTGATCCTTCGATGTCGG</td><td>5593.4</td><td>-7.7</td><td>1.6</td><td>-4.7</td><td>2.3E-06</td><td>4.2E-06</td><td>BT20/MB468</td><td>CYTO</td></tr>
<tr><td>>hg38_21_+_8433172_8446622_RNA45SN1_45S_50ntFlanks@6772.6799.28|GTTCCTCCCGGGGCTACGCCTGTCTGAG</td><td>>hg38_21_+_8433172_8446622_RNA45SN1_45S_50ntFlanks</td><td>6772</td><td>6799</td><td>28</td><td>GTTCCTCCCGGGGCTACGCCTGTCTGAG</td><td>1849.8</td><td>-5.5</td><td>1.2</td><td>-4.8</td><td>2.0E-06</td><td>4.2E-06</td><td>BT20/MB468</td><td>MITO</td></tr>
<tr><td>>hg38_21_+_8433172_8446622_RNA45SN1_45S_50ntFlanks@5472.5494.23|ACGGCCCTGGCGGAGCGCTGAGA</td><td>>hg38_21_+_8433172_8446622_RNA45SN1_45S_50ntFlanks</td><td>5472</td><td>5494</td><td>23</td><td>ACGGCCCTGGCGGAGCGCTGAGA</td><td>282.3</td><td>3.9</td><td>0.8</td><td>4.7</td><td>2.7E-06</td><td>4.2E-06</td><td>MB231/MB468</td><td>NUC</td></tr>
<tr><td>>hg38_21_+_8433172_8446622_RNA45SN1_45S_50ntFlanks@10410.10443.34|TCAGTCGGTCCTGAGAGATGGGCGAGCGCCGTTC</td><td>>hg38_21_+_8433172_8446622_RNA45SN1_45S_50ntFlanks</td><td>10410</td><td>10443</td><td>34</td><td>TCAGTCGGTCCTGAGAGATGGGCGAGCGCCGTTC</td><td>397.7</td><td>-5.1</td><td>1.1</td><td>-4.7</td><td>2.7E-06</td><td>4.2E-06</td><td>MB231/MB468</td><td>NUC</td></tr>
<tr><td>>hg38_21_+_8433172_8446622_RNA45SN1_45S_50ntFlanks@9622.9649.28|CTAGTAGCTGGTTCCCTCCGAAGTTTCC</td><td>>hg38_21_+_8433172_8446622_RNA45SN1_45S_50ntFlanks</td><td>9622</td><td>9649</td><td>28</td><td>CTAGTAGCTGGTTCCCTCCGAAGTTTCC</td><td>255.2</td><td>-2.9</td><td>0.6</td><td>-4.8</td><td>2.0E-06</td><td>4.2E-06</td><td>BT20/MB468</td><td>MITO</td></tr>
<tr><td>>trna10_AspGTC_12_-_125424193_125424264@35.74.40,trna12_AspGTC_12_-_125411891_125411962@35.74.40,trna144_AspGTC_6_-_27551236_27551307@35.74.40,trna38_AspGTC_17_-_8125556_8125627@35.74.40,trna45_AspGTC_6_+_27447453_27447524@35.74.40,trna48_AspGTC_6_+_27471523_27471594@35.74.40,trna4_AspGTC_12_+_96429799_96429870@35.74.40,trna69_AspGTC_1_-_161440205_161440276@35.74.40,trna72_AspGTC_1_-_161432824_161432895@35.74.40,trna75_AspGTC_1_-_161425414_161425485@35.74.40,trna78_AspGTC_1_-_161418033_161418104@35.74.40,trna81_AspGTC_1_-_161410615_161410686@35.74.40|TCACGCGGGAGACCGGGGTTCGATTCCCCGACGGGGAGCC</td><td>>trna10_AspGTC_12_-_125424193_125424264</td><td>35</td><td>74</td><td>40</td><td>TCACGCGGGAGACCGGGGTTCGATTCCCCGACGGGGAGCC</td><td>178.6</td><td>7.7</td><td>1.7</td><td>4.7</td><td>2.8E-06</td><td>4.2E-06</td><td>MB231/MB468</td><td>MITO</td></tr>
<tr><td>>hg38_MT_+_1621_3279_RNR2_16S_50ntFlanks@1508.1529.22|AAAGGACAAGAGAAATAAGGCC</td><td>>hg38_MT_+_1621_3279_RNR2_16S_50ntFlanks</td><td>1508</td><td>1529</td><td>22</td><td>AAAGGACAAGAGAAATAAGGCC</td><td>45.7</td><td>8.7</td><td>1.8</td><td>4.7</td><td>2.3E-06</td><td>4.2E-06</td><td>MB231/MB468</td><td>MP</td></tr>
<tr><td>>hg38_21_+_8433172_8446622_RNA45SN1_45S_50ntFlanks@6647.6672.26|CGTACGACTCTTAGCGGTGGATCACT</td><td>>hg38_21_+_8433172_8446622_RNA45SN1_45S_50ntFlanks</td><td>6647</td><td>6672</td><td>26</td><td>CGTACGACTCTTAGCGGTGGATCACT</td><td>448.9</td><td>2.5</td><td>0.5</td><td>4.7</td><td>2.9E-06</td><td>4.2E-06</td><td>BT20/MB468</td><td>NUC</td></tr>
<tr><td>>am_MI0000809|hsa-mir-151a&WithFlank&8|-|140732558|140732659@53.73.21(+3U)&[MIMAT0000757&hsa-miR-151a-3p&offsets|0|0(+3U);m-30&8|-|140732587|140732607&offsets|0|0(+3U)]|CTAGACTGAAGCTCCTTGAGGTTT</td><td>>am_MI0000809|hsa-mir-151a&WithFlank&8|-|140732558|140732659</td><td>53</td><td>73</td><td>21(+3U)</td><td>CTAGACTGAAGCTCCTTGAGGTTT</td><td>762.4</td><td>5.5</td><td>1.2</td><td>4.7</td><td>2.3E-06</td><td>4.2E-06</td><td>MB231/MB468</td><td>MP</td></tr>
<tr><td>>hg38_21_+_8433172_8446622_RNA45SN1_45S_50ntFlanks@4002.4042.41|GATAACCTCGGGCCGATCGCACGCCCCCCGTGGCGGCGACG</td><td>>hg38_21_+_8433172_8446622_RNA45SN1_45S_50ntFlanks</td><td>4002</td><td>4042</td><td>41</td><td>GATAACCTCGGGCCGATCGCACGCCCCCCGTGGCGGCGACG</td><td>1057.9</td><td>-6.1</td><td>1.3</td><td>-4.7</td><td>2.3E-06</td><td>4.2E-06</td><td>BT20/MB468</td><td>CYTO</td></tr>
<tr><td>>hg38_21_+_8433172_8446622_RNA45SN1_45S_50ntFlanks@4962.4985.24|AAACCTCACCCGGCCCGGACACGG</td><td>>hg38_21_+_8433172_8446622_RNA45SN1_45S_50ntFlanks</td><td>4962</td><td>4985</td><td>24</td><td>AAACCTCACCCGGCCCGGACACGG</td><td>365.0</td><td>-6.1</td><td>1.3</td><td>-4.7</td><td>2.8E-06</td><td>4.2E-06</td><td>MB231/MB468</td><td>MITO</td></tr>
<tr><td>>MI0000085|hsa-mir-27a&WithFlank&19|-|13836434|13836523@57.75.19&[MIMAT0000084&hsa-miR-27a-3p&offsets|0|-2;m-43&19|-|13836447|13836467&offsets|0|-2]|TTCACAGTGGCTAAGTTCC</td><td>>MI0000085|hsa-mir-27a&WithFlank&19|-|13836434|13836523</td><td>57</td><td>75</td><td>19</td><td>TTCACAGTGGCTAAGTTCC</td><td>658.8</td><td>4.7</td><td>1.0</td><td>4.7</td><td>2.8E-06</td><td>4.2E-06</td><td>MB231/MB468</td><td>MITO</td></tr>
<tr><td>>hg38_21_+_8433172_8446622_RNA45SN1_45S_50ntFlanks@4453.4472.20|TCGGCGCCCCCTCGATGCTC</td><td>>hg38_21_+_8433172_8446622_RNA45SN1_45S_50ntFlanks</td><td>4453</td><td>4472</td><td>20</td><td>TCGGCGCCCCCTCGATGCTC</td><td>197.2</td><td>-6.6</td><td>1.4</td><td>-4.7</td><td>2.3E-06</td><td>4.2E-06</td><td>MB231/MB468</td><td>MP</td></tr>
<tr><td>>hg38_21_+_8433172_8446622_RNA45SN1_45S_50ntFlanks@8105.8143.39|CGCGGCGGGGCGCGGGACATGTGGCGTACGGAAGACCCG</td><td>>hg38_21_+_8433172_8446622_RNA45SN1_45S_50ntFlanks</td><td>8105</td><td>8143</td><td>39</td><td>CGCGGCGGGGCGCGGGACATGTGGCGTACGGAAGACCCG</td><td>1671.5</td><td>-5.8</td><td>1.2</td><td>-4.7</td><td>2.6E-06</td><td>4.2E-06</td><td>MB231/MB468</td><td>CYTO</td></tr>
<tr><td>>hg38_21_+_8433172_8446622_RNA45SN1_45S_50ntFlanks@11581.11604.24|AACTGGTGCGGACCAGGGGAATCC</td><td>>hg38_21_+_8433172_8446622_RNA45SN1_45S_50ntFlanks</td><td>11581</td><td>11604</td><td>24</td><td>AACTGGTGCGGACCAGGGGAATCC</td><td>203.1</td><td>-5.1</td><td>1.1</td><td>-4.7</td><td>2.8E-06</td><td>4.2E-06</td><td>MB231/MB468</td><td>MITO</td></tr>
<tr><td>>trna116_GluCTC_1_-_145399233_145399304@39.75.37,trna134_GluTTC_1_-_16861774_16861845@39.75.37,trna71_GluCTC_1_-_161439189_161439260@39.75.37,trna74_GluCTC_1_-_161431809_161431880@39.75.37,trna77_GluCTC_1_-_161424398_161424469@39.75.37,trna77_GluCTC_6_+_28949976_28950047@39.75.37,trna80_GluCTC_1_-_161417018_161417089@39.75.37,trna84_GluTTC_1_-_161391883_161391954@39.75.37,trna87_GluCTC_6_-_126101393_126101464@39.75.37|CGCCGCGGCCCGGGTTCGATTCCCGGTCAGGGAACCA</td><td>>trna116_GluCTC_1_-_145399233_145399304</td><td>39</td><td>75</td><td>37</td><td>CGCCGCGGCCCGGGTTCGATTCCCGGTCAGGGAACCA</td><td>570.3</td><td>4.3</td><td>0.9</td><td>4.7</td><td>2.3E-06</td><td>4.2E-06</td><td>BT20/MB468</td><td>CYTO</td></tr>
<tr><td>>hg38_21_+_8433172_8446622_RNA45SN1_45S_50ntFlanks@11326.11345.20|CTCGTCGCGCGCGCGTCCGC</td><td>>hg38_21_+_8433172_8446622_RNA45SN1_45S_50ntFlanks</td><td>11326</td><td>11345</td><td>20</td><td>CTCGTCGCGCGCGCGTCCGC</td><td>708.2</td><td>8.3</td><td>1.8</td><td>4.7</td><td>2.8E-06</td><td>4.3E-06</td><td>MB231/MB468</td><td>MITO</td></tr>
<tr><td>>hg38_21_+_8433172_8446622_RNA45SN1_45S_50ntFlanks@12479.12498.20|CTAATAGGGAACGTGAGCTG</td><td>>hg38_21_+_8433172_8446622_RNA45SN1_45S_50ntFlanks</td><td>12479</td><td>12498</td><td>20</td><td>CTAATAGGGAACGTGAGCTG</td><td>1380.6</td><td>-2.9</td><td>0.6</td><td>-4.7</td><td>2.6E-06</td><td>4.3E-06</td><td>MB231/MB468</td><td>CYTO</td></tr>
<tr><td>>hg38_21_+_8433172_8446622_RNA45SN1_45S_50ntFlanks@6649.6693.45|TACGACTCTTAGCGGTGGATCACTCGGCTCGTGCGTCGATGAAGA</td><td>>hg38_21_+_8433172_8446622_RNA45SN1_45S_50ntFlanks</td><td>6649</td><td>6693</td><td>45</td><td>TACGACTCTTAGCGGTGGATCACTCGGCTCGTGCGTCGATGAAGA</td><td>512.9</td><td>4.3</td><td>0.9</td><td>4.8</td><td>2.0E-06</td><td>4.3E-06</td><td>BT20/MB468</td><td>MITO</td></tr>
<tr><td>>hg38_21_+_8433172_8446622_RNA45SN1_45S_50ntFlanks@8196.8225.30|CGTGGACGGTGTGAGGCCGGTAGCGGCCCC</td><td>>hg38_21_+_8433172_8446622_RNA45SN1_45S_50ntFlanks</td><td>8196</td><td>8225</td><td>30</td><td>CGTGGACGGTGTGAGGCCGGTAGCGGCCCC</td><td>984.8</td><td>-3.2</td><td>0.7</td><td>-4.7</td><td>2.3E-06</td><td>4.3E-06</td><td>BT20/MB468</td><td>CYTO</td></tr>
<tr><td>>hg38_21_+_8433172_8446622_RNA45SN1_45S_50ntFlanks@12539.12587.49|ATGATGTGTTGTTGCCATGGTAATCCTGCTCAGTACGAGAGGAACCGCA</td><td>>hg38_21_+_8433172_8446622_RNA45SN1_45S_50ntFlanks</td><td>12539</td><td>12587</td><td>49</td><td>ATGATGTGTTGTTGCCATGGTAATCCTGCTCAGTACGAGAGGAACCGCA</td><td>210.4</td><td>-5.8</td><td>1.2</td><td>-4.7</td><td>2.6E-06</td><td>4.3E-06</td><td>MB231/MB468</td><td>CYTO</td></tr>
<tr><td>>hg38_21_+_8433172_8446622_RNA45SN1_45S_50ntFlanks@11575.11604.30|ACTTAGAACTGGTGCGGACCAGGGGAATCC</td><td>>hg38_21_+_8433172_8446622_RNA45SN1_45S_50ntFlanks</td><td>11575</td><td>11604</td><td>30</td><td>ACTTAGAACTGGTGCGGACCAGGGGAATCC</td><td>548.4</td><td>-7.7</td><td>1.7</td><td>-4.7</td><td>3.0E-06</td><td>4.3E-06</td><td>BT20/MB468</td><td>NUC</td></tr>
<tr><td>>hg38_21_+_8433172_8446622_RNA45SN1_45S_50ntFlanks@12332.12350.19|TAAGCAGGAGGTGTCAGAA</td><td>>hg38_21_+_8433172_8446622_RNA45SN1_45S_50ntFlanks</td><td>12332</td><td>12350</td><td>19</td><td>TAAGCAGGAGGTGTCAGAA</td><td>150.1</td><td>-6.5</td><td>1.4</td><td>-4.7</td><td>2.3E-06</td><td>4.3E-06</td><td>MB231/MB468</td><td>MP</td></tr>
<tr><td>>hg38_21_+_8433172_8446622_RNA45SN1_45S_50ntFlanks@12071.12094.24|CCCCGAGGGGCTCTCGCTTCTGGC</td><td>>hg38_21_+_8433172_8446622_RNA45SN1_45S_50ntFlanks</td><td>12071</td><td>12094</td><td>24</td><td>CCCCGAGGGGCTCTCGCTTCTGGC</td><td>543.5</td><td>-3.2</td><td>0.7</td><td>-4.7</td><td>2.3E-06</td><td>4.3E-06</td><td>BT20/MB468</td><td>CYTO</td></tr>
<tr><td>>hg38_21_+_8433172_8446622_RNA45SN1_45S_50ntFlanks@6676.6696.21|CTCGTGCGTCGATGAAGAACG</td><td>>hg38_21_+_8433172_8446622_RNA45SN1_45S_50ntFlanks</td><td>6676</td><td>6696</td><td>21</td><td>CTCGTGCGTCGATGAAGAACG</td><td>366.8</td><td>-5.4</td><td>1.1</td><td>-4.7</td><td>2.3E-06</td><td>4.3E-06</td><td>MB231/MB468</td><td>MP</td></tr>
<tr><td>>hg38_21_+_8433172_8446622_RNA45SN1_45S_50ntFlanks@4588.4615.28|CTATTTTGTTGGTTTTCGGAACTGAGGC</td><td>>hg38_21_+_8433172_8446622_RNA45SN1_45S_50ntFlanks</td><td>4588</td><td>4615</td><td>28</td><td>CTATTTTGTTGGTTTTCGGAACTGAGGC</td><td>295.7</td><td>-4.3</td><td>0.9</td><td>-4.7</td><td>2.7E-06</td><td>4.3E-06</td><td>MB231/MB468</td><td>CYTO</td></tr>
<tr><td>>MI0000087|hsa-mir-29a&WithFlank&7|-|130876741|130876816@47.66.20(+1C)&[MIMAT0000086&hsa-miR-29a-3p&offsets|-1|-3(+1C);m-26&7|-|130876749|130876769&offsets|-1|-2(+1C)]|CTAGCACCATCTGAAATCGGC</td><td>>MI0000087|hsa-mir-29a&WithFlank&7|-|130876741|130876816</td><td>47</td><td>66</td><td>20(+1C)</td><td>CTAGCACCATCTGAAATCGGC</td><td>50.7</td><td>8.9</td><td>1.9</td><td>4.7</td><td>2.3E-06</td><td>4.3E-06</td><td>MB231/MB468</td><td>MP</td></tr>
<tr><td>>MI0000433|hsa-let-7g&WithFlank&3|-|52268272|52268367@11.32.22(+1A)&[MIMAT0000414&hsa-let-7g-5p&offsets|0|0(+1A);m-58&3|-|52268336|52268357&offsets|0|0(+1A)]|TGAGGTAGTAGTTTGTACAGTTA</td><td>>MI0000433|hsa-let-7g&WithFlank&3|-|52268272|52268367</td><td>11</td><td>32</td><td>22(+1A)</td><td>TGAGGTAGTAGTTTGTACAGTTA</td><td>1543.9</td><td>4.6</td><td>1.0</td><td>4.7</td><td>2.3E-06</td><td>4.3E-06</td><td>MB231/MB468</td><td>MP</td></tr>
</table>

| | | | | | | | | | | | | | |
|---|---|---|---|---|---|---|---|---|---|---|---|---|---|
| >hg38_21_+_8433172_8446622_RNA45SN1_45S_50ntFlanks@12625.12654.30\|CAATGGGGCGAAGCTACCATCTGTGGGATT | >hg38_21_+_8433172_8446622_RNA45SN1_45S_50ntFlanks | 12625 | 12654 | 30 | CAATGGGGCGAAGCTACCATCTGTGGGATT | 3793.9 | -4.8 | 1.0 | -4.9 | 7.9E-07 | 4.3E-06 | BT20/MB468 | MP |
| >hg38_21_+_8433172_8446622_RNA45SN1_45S_50ntFlanks@12714.12745.32\|GAGCCTCGGTTGGCCTCGGATAGCCGGTCCCC | >hg38_21_+_8433172_8446622_RNA45SN1_45S_50ntFlanks | 12714 | 12745 | 32 | GAGCCTCGGTTGGCCTCGGATAGCCGGTCCCC | 396.0 | -4.3 | 0.9 | -4.9 | 7.9E-07 | 4.3E-06 | BT20/MB468 | MP |
| >MI0015866\|hsa-mir-4326&WithFlank&20\|+\|63286802\|63286872@17.40.24&[MIMAT0016888&hsa-miR-4326&offsets\|0\|+4;m-339&20\|+\|63286818\|63286841&offsets\|0\|0]\|TGTTCCTCTGTCTCCCAGACTCTG | >MI0015866\|hsa-mir-4326&WithFlank&20\|+\|63286802\|63286872 | 17 | 40 | 24 | TGTTCCTCTGTCTCCCAGACTCTG | 220.4 | 7.6 | 1.6 | 4.7 | 2.8E-06 | 4.3E-06 | MB231/MB468 | MITO |
| >hg38_21_+_8433172_8446622_RNA45SN1_45S_50ntFlanks@11320.11344.25\|CCACGTCTCGTCGCGCGCGCGTCCG | >hg38_21_+_8433172_8446622_RNA45SN1_45S_50ntFlanks | 11320 | 11344 | 25 | CCACGTCTCGTCGCGCGCGCGTCCG | 4074.7 | -5.7 | 1.2 | -4.7 | 2.4E-06 | 4.3E-06 | BT20/MB468 | CYTO |
| >hg38_21_+_8433172_8446622_RNA45SN1_45S_50ntFlanks@6719.6741.23\|TGAATTGCAGGACACATTGATCA | >hg38_21_+_8433172_8446622_RNA45SN1_45S_50ntFlanks | 6719 | 6741 | 23 | TGAATTGCAGGACACATTGATCA | 260.4 | -4.4 | 0.9 | -4.7 | 2.4E-06 | 4.3E-06 | BT20/MB468 | CYTO |
| >hg38_21_+_8433172_8446622_RNA45SN1_45S_50ntFlanks@9573.9595.23\|AAATCGGTCGTCCGACCTGGGTA | >hg38_21_+_8433172_8446622_RNA45SN1_45S_50ntFlanks | 9573 | 9595 | 23 | AAATCGGTCGTCCGACCTGGGTA | 1935.7 | -4.2 | 0.9 | -4.7 | 2.7E-06 | 4.3E-06 | MB231/MB468 | CYTO |
| >hg38_21_+_8433172_8446622_RNA45SN1_45S_50ntFlanks@11582.11616.35\|ACTGGTGCGGACCAGGGGAATCCGACTGTTTAATT | >hg38_21_+_8433172_8446622_RNA45SN1_45S_50ntFlanks | 11582 | 11616 | 35 | ACTGGTGCGGACCAGGGGAATCCGACTGTTTAATT | 466.9 | 5.8 | 1.2 | 4.9 | 7.9E-07 | 4.3E-06 | BT20/MB468 | MP |
| >hg38_21_+_8433172_8446622_RNA45SN1_45S_50ntFlanks@5494.5522.29\|AAGACGGTCGAACTTGACTATCTAGAGGA | >hg38_21_+_8433172_8446622_RNA45SN1_45S_50ntFlanks | 5494 | 5522 | 29 | AAGACGGTCGAACTTGACTATCTAGAGGA | 129.6 | -6.5 | 1.4 | -4.7 | 2.4E-06 | 4.3E-06 | MB231/MB468 | MP |
| >MI0000273\|hsa-mir-183&WithFlank&7\|-\|129774899\|129775020@33.55.23&[MIMAT0000261&hsa-miR-183-5p&offsets\|0\|+1;m-41&7\|-\|129774966\|129774987&offsets\|-1\|0]\|TATGGCACTGGTAGAATTCACTG | >MI0000273\|hsa-mir-183&WithFlank&7\|-\|129774899\|129775020 | 33 | 55 | 23 | TATGGCACTGGTAGAATTCACTG | 4284.0 | 3.8 | 0.7 | 5.2 | 2.2E-07 | 4.3E-06 | BT20/MB231 | CYTO |
| >hg38_1_-_228634819_228635039_RNA5S12_5S_50ntFlanks@51.85.35\|GTCTACGGCCATACCACCCTGAACGCGCCCGATCT | >hg38_1_-_228634819_228635039_RNA5S12_5S_50ntFlanks | 51 | 85 | 35 | GTCTACGGCCATACCACCCTGAACGCGCCCGATCT | 710.7 | 4.3 | 0.9 | 4.7 | 2.1E-06 | 4.3E-06 | BT20/MB468 | MITO |
| >hg38_21_+_8433172_8446622_RNA45SN1_45S_50ntFlanks@10118.10138.21\|CGTCGGAGGGCGGCGGCGGCG | >hg38_21_+_8433172_8446622_RNA45SN1_45S_50ntFlanks | 10118 | 10138 | 21 | CGTCGGAGGGCGGCGGCGGCG | 85.5 | -9.5 | 1.9 | -4.9 | 7.9E-07 | 4.3E-06 | BT20/MB468 | MP |
| >am_trnaMT_AlaTGC_MT_-_5587_5655@1.36.36,trnalookalike1_AlaTGC_14_+_32954252_32954320@1.36.36,trnalookalike4_AlaTGC_1_-_566137_566205@1.36.36\|AAGGGCTTAGCTTAATTAAAGTGGCTGATTTGCGTT | >am_trnaMT_AlaTGC_MT_-_5587_5655 | 1 | 36 | 36 | AAGGGCTTAGCTTAATTAAAGTGGCTGATTTGCGTT | 44.6 | 8.7 | 1.8 | 4.7 | 2.4E-06 | 4.3E-06 | MB231/MB468 | MP |
| >hg38_21_+_8433172_8446622_RNA45SN1_45S_50ntFlanks@11771.11792.22\|ATTAGTGACGCGCATGAATGGA | >hg38_21_+_8433172_8446622_RNA45SN1_45S_50ntFlanks | 11771 | 11792 | 22 | ATTAGTGACGCGCATGAATGGA | 537.7 | -7.6 | 1.6 | -4.7 | 3.0E-06 | 4.3E-06 | BT20/MB468 | NUC |
| >MI0000298\|hsa-mir-221&WithFlank&X\|-\|45746151\|45746272@31.53.23(+1C)&[MIMAT0004568&hsa-miR-221-5p&offsets\|0\|+1(+1C);m-405&X\|-\|45746220\|45746242&offsets\|0\|0(+1C)]\|ACCTGGCATACAATGTAGATTTCC | >MI0000298\|hsa-mir-221&WithFlank&X\|-\|45746151\|45746272 | 31 | 53 | 23(+1C) | ACCTGGCATACAATGTAGATTTCC | 279.0 | 7.8 | 1.7 | 4.7 | 2.9E-06 | 4.3E-06 | MB231/MB468 | MITO |
| >MI0000065\|hsa-let-7d&WithFlank&9\|+\|94178828\|94178926@68.89.22&[MIMAT0004484&hsa-let-7d-3p&offsets\|0\|0;m-98&9\|+\|94178895\|94178916&offsets\|0\|0]\|CTATACGACCTGCTGCCTTTCT | >MI0000065\|hsa-let-7d&WithFlank&9\|+\|94178828\|94178926 | 68 | 89 | 22 | CTATACGACCTGCTGCCTTTCT | 2314.2 | 3.7 | 0.8 | 4.7 | 2.4E-06 | 4.3E-06 | MB231/MB468 | MP |
| >MI0003776\|hsa-mir-320b-1&WithFlank&1\|+\|116671740\|116671823@48.66.19(+1A);MI0003839\|hsa-mir-320b-2&WithFlank&1\|-\|224257034\|224257116@47.65.19(+1A);MI0003778\|hsa-mir-320c-1&WithFlank&18\|+\|21683512\|21683595@48.66.19(+1A);MI0008191\|hsa-mir-320c-2&WithFlank&18\|+\|24321669\|24321752@48.66.19(+1A);MI0000542\|hsa-mir-320a&WithFlank&8\|-\|22244960\|22245043@48.66.19(+1A)&[MIMAT0005792&hsa-miR-320b&offsets\|0\|-3(+1A);m-183&1\|+\|116671787\|116671806&offsets\|0\|-1(+1A)];[MIMAT0005792_1&hsa-miR-320b&offsets\|0\|-3(+1A);m-183&1\|-\|224257051\|224257070&offsets\|0\|-1(+1A)];[MIMAT0005793&hsa-miR-320c&offsets\|0\|-1(+1A);m-222&18\|+\|21683559\|21683578&offsets\|0\|-1(+1A)];[MIMAT0005793_1&hsa-miR-320c&offsets\|0\|-1(+1A);m-219&18\|+\|24321716\|24321735&offsets\|0\|-1(+1A)];[MIMAT0000510&hsa-miR-320a-3p&offsets\|0\|-3(+1A);m-60&8\|-\|22244975\|22244996&offsets\|0\|-3(+1A)]\|AAAAGCTGGGTTGAGAGGGA | >MI0003776\|hsa-mir-320b-1&WithFlank&1\|+\|116671740\|116671823 | 48 | 66 | 19(+1A) | AAAAGCTGGGTTGAGAGGGA | 263.9 | 3.3 | 0.7 | 5.1 | 3.1E-07 | 4.3E-06 | BT20/MB231 | NUC |
| >hg38_21_+_8433172_8446622_RNA45SN1_45S_50ntFlanks@8380.8406.27\|CAAGAGGGCGTGAAACCGTTAAGAGGT | >hg38_21_+_8433172_8446622_RNA45SN1_45S_50ntFlanks | 8380 | 8406 | 27 | CAAGAGGGCGTGAAACCGTTAAGAGGT | 1163.0 | -3.3 | 0.7 | -4.7 | 2.1E-06 | 4.3E-06 | BT20/MB468 | MITO |
| >hg38_21_+_8433172_8446622_RNA45SN1_45S_50ntFlanks@10689.10708.20\|GCGTCCGGTGAGCTCTCGCT | >hg38_21_+_8433172_8446622_RNA45SN1_45S_50ntFlanks | 10689 | 10708 | 20 | GCGTCCGGTGAGCTCTCGCT | 1332.6 | -5.7 | 1.2 | -4.7 | 2.8E-06 | 4.3E-06 | MB231/MB468 | NUC |
| >hg38_21_+_8433172_8446622_RNA45SN1_45S_50ntFlanks@10905.10936.32\|CTGGGCGCGCGCCGCGGCTGGACGAGGCGCCG | >hg38_21_+_8433172_8446622_RNA45SN1_45S_50ntFlanks | 10905 | 10936 | 32 | CTGGGCGCGCGCCGCGGCTGGACGAGGCGCCG | 506.6 | -7.8 | 1.7 | -4.7 | 3.0E-06 | 4.3E-06 | BT20/MB468 | NUC |
| >hg38_21_+_8433172_8446622_RNA45SN1_45S_50ntFlanks@5078.5097.20\|CGATAACGAACGAGACTCTG | >hg38_21_+_8433172_8446622_RNA45SN1_45S_50ntFlanks | 5078 | 5097 | 20 | CGATAACGAACGAGACTCTG | 697.9 | -6.0 | 1.3 | -4.7 | 2.9E-06 | 4.3E-06 | MB231/MB468 | MITO |
| >hg38_21_+_8433172_8446622_RNA45SN1_45S_50ntFlanks@7974.7992.19\|ACGCGACCTCAGATCAGAC | >hg38_21_+_8433172_8446622_RNA45SN1_45S_50ntFlanks | 7974 | 7992 | 19 | ACGCGACCTCAGATCAGAC | 2993.0 | 4.5 | 0.9 | 4.9 | 8.0E-07 | 4.3E-06 | BT20/MB468 | MP |
| >hg38_21_+_8433172_8446622_RNA45SN1_45S_50ntFlanks@10784.10826.43\|TGAACAGCCTCTGGCATGTTGGAACAATGTAGGTAAGGGAAGT | >hg38_21_+_8433172_8446622_RNA45SN1_45S_50ntFlanks | 10784 | 10826 | 43 | TGAACAGCCTCTGGCATGTTGGAACAATGTAGGTAAGGGAAGT | 264.0 | -6.0 | 1.3 | -4.7 | 2.9E-06 | 4.3E-06 | MB231/MB468 | MITO |
| >trnaMT_SerGCT_MT_+_12207_12265@10.38.29\|CACAAGAACTGCTAACTCATGCCCCCATG | >trnaMT_SerGCT_MT_+_12207_12265 | 10 | 38 | 29 | CACAAGAACTGCTAACTCATGCCCCCATG | 213.2 | -6.7 | 1.4 | -4.7 | 2.4E-06 | 4.3E-06 | MB231/MB468 | MP |
| >trna111_HisGTG_1_-_147774845_147774916@-1T.26.27,trna118_HisGTG_1_-_145396881_145396952@-1T.26.27,trna16_HisGTG_1_+_146544773_146544844@-1T.26.27,trna1_HisGTG_15_+_45493349_45493420@-1T.26.27,trna21_HisGTG_1_+_147753471_147753542@-1T.26.27,trna33_HisGTG_6_+_27125906_27125977@-1T.26.27,trna7_HisGTG_9_-_14433938_14434009@-1T.26.27,trna8_HisGTG_15_-_45492611_45492682@-1T.26.27,trna9_HisGTG_15_-_45490804_45490875@-1T.26.27\|TGCCGTGATCGTATAGTGGTTAGTACT | >trna111_HisGTG_1_-_147774845_147774916 | -1T | 26 | 27 | TGCCGTGATCGTATAGTGGTTAGTACT | 1354.8 | 3.6 | 0.8 | 4.7 | 2.4E-06 | 4.3E-06 | BT20/MB468 | CYTO |
| >hg38_21_+_8433172_8446622_RNA45SN1_45S_50ntFlanks@7991.8017.27\|ACGTGGCGACCCGCTGAATTTAAGCAT | >hg38_21_+_8433172_8446622_RNA45SN1_45S_50ntFlanks | 7991 | 8017 | 27 | ACGTGGCGACCCGCTGAATTTAAGCAT | 416.4 | -6.9 | 1.5 | -4.7 | 2.4E-06 | 4.3E-06 | MB231/MB468 | MP |
| >hg38_21_+_8433172_8446622_RNA45SN1_45S_50ntFlanks@6729.6771.43\|GACACATTGATCATCGACACTTCGAACGCACTTGCGGCCCCGG | >hg38_21_+_8433172_8446622_RNA45SN1_45S_50ntFlanks | 6729 | 6771 | 43 | GACACATTGATCATCGACACTTCGAACGCACTTGCGGCCCCGG | 124.5 | -10.8 | 2.3 | -4.7 | 2.4E-06 | 4.3E-06 | MB231/MB468 | MP |
| >am_trna13_ThrTGT_5_-_180618687_180618758@1.29.29,trna20_ThrTGT_14_-_21099319_21099391@1.29.29\|GGCTCCATAGCTCAGGGGTTAGAGCACTG | >am_trna13_ThrTGT_5_-_180618687_180618758 | 1 | 29 | 29 | GGCTCCATAGCTCAGGGGTTAGAGCACTG | 106.8 | -5.6 | 1.2 | -4.7 | 2.4E-06 | 4.3E-06 | BT20/MB468 | CYTO |
| >MI0000650\|hsa-mir-200c&WithFlank&12\|+\|6963693\|6963772@50.71.22(+1G)&[MIMAT0000617&hsa-miR-200c-3p&offsets\|0\|-1(+1G);m-54&12\|+\|6963742\|6963764&offsets\|0\|-1(+1G)]\|TAATACTGCCGGGTAATGATGGG | >MI0000650\|hsa-mir-200c&WithFlank&12\|+\|6963693\|6963772 | 50 | 71 | 22(+1G) | TAATACTGCCGGGTAATGATGGG | 1599.5 | 4.9 | 1.0 | 4.9 | 8.0E-07 | 4.3E-06 | BT20/MB468 | MP |
| >hg38_21_+_8433172_8446622_RNA45SN1_45S_50ntFlanks@5525.5548.24\|TAAAAGTCGTAACAAGGTTTCCGT | >hg38_21_+_8433172_8446622_RNA45SN1_45S_50ntFlanks | 5525 | 5548 | 24 | TAAAAGTCGTAACAAGGTTTCCGT | 50.2 | -8.7 | 1.8 | -4.9 | 8.0E-07 | 4.3E-06 | BT20/MB468 | MP |
| >hg38_21_+_8433172_8446622_RNA45SN1_45S_50ntFlanks@12994.13039.46\|CCTCGCTGCGATCTATTGAAAGTCAGCCCTCGACACAAGGGTTTGT | >hg38_21_+_8433172_8446622_RNA45SN1_45S_50ntFlanks | 12994 | 13039 | 46 | CCTCGCTGCGATCTATTGAAAGTCAGCCCTCGACACAAGGGTTTGT | 18966.8 | 5.0 | 1.0 | 4.7 | 2.1E-06 | 4.3E-06 | BT20/MB468 | MITO |
| >am_trna11_ArgACG_3_-_45730491_45730563@39.56.18,trna138_ArgACG_6_-_27638344_27638416@39.56.18,trna156_ArgACG_6_-_27181623_27181695@39.56.18,trna36_ArgACG_6_+_27182952_27183024@39.56.18\|TCAGAAGATTCTAGGTTC | >am_trna11_ArgACG_3_-_45730491_45730563 | 39 | 56 | 18 | TCAGAAGATTCTAGGTTC | 137.9 | -6.2 | 1.3 | -4.7 | 2.8E-06 | 4.3E-06 | MB231/MB468 | NUC |
| >MI0000650\|hsa-mir-200c&WithFlank&12\|+\|6963693\|6963772@50.70.21(+1A)&[MIMAT0000617&hsa-miR-200c-3p&offsets\|0\|-2(+1A);m-54&12\|+\|6963742\|6963764&offsets\|0\|-2(+1A)]\|TAATACTGCCGGGTAATGATGA | >MI0000650\|hsa-mir-200c&WithFlank&12\|+\|6963693\|6963772 | 50 | 70 | 21(+1A) | TAATACTGCCGGGTAATGATGA | 3847.6 | 10.2 | 2.0 | 5.2 | 2.2E-07 | 4.3E-06 | BT20/MB231 | CYTO |
| >hg38_21_+_8433172_8446622_RNA45SN1_45S_50ntFlanks@10402.10419.18\|AACATGGGTCAGTCGGTC | >hg38_21_+_8433172_8446622_RNA45SN1_45S_50ntFlanks | 10402 | 10419 | 18 | AACATGGGTCAGTCGGTC | 981.2 | -5.3 | 1.1 | -4.7 | 2.4E-06 | 4.3E-06 | BT20/MB468 | CYTO |
| >hg38_21_+_8433172_8446622_RNA45SN1_45S_50ntFlanks@13005.13028.24\|TCTATTGAAAGTCAGCCCTCGACA | >hg38_21_+_8433172_8446622_RNA45SN1_45S_50ntFlanks | 13005 | 13028 | 24 | TCTATTGAAAGTCAGCCCTCGACA | 332.2 | -6.4 | 1.4 | -4.7 | 2.4E-06 | 4.4E-06 | BT20/MB468 | CYTO |
| >hg38_21_+_8433172_8446622_RNA45SN1_45S_50ntFlanks@11737.11768.32\|ACTCTCTTAAGGTAGCCAAATGCCTCGTCATC | >hg38_21_+_8433172_8446622_RNA45SN1_45S_50ntFlanks | 11737 | 11768 | 32 | ACTCTCTTAAGGTAGCCAAATGCCTCGTCATC | 140.7 | -5.6 | 1.2 | -4.7 | 2.8E-06 | 4.4E-06 | MB231/MB468 | NUC |
| >MI0000065\|hsa-let-7d&WithFlank&9\|+\|94178828\|94178926@68.89.22&[MIMAT0004484&hsa-let-7d-3p&offsets\|0\|0;m-98&9\|+\|94178895\|94178916&offsets\|0\|0]\|CTATACGACCTGCTGCCTTTCT | >MI0000065\|hsa-let-7d&WithFlank&9\|+\|94178828\|94178926 | 68 | 89 | 22 | CTATACGACCTGCTGCCTTTCT | 4255.0 | 5.2 | 1.1 | 4.9 | 8.1E-07 | 4.4E-06 | BT20/MB468 | MP |
| >hg38_21_+_8433172_8446622_RNA45SN1_45S_50ntFlanks@10784.10801.18\|TGAACAGCCTCTGGCATG | >hg38_21_+_8433172_8446622_RNA45SN1_45S_50ntFlanks | 10784 | 10801 | 18 | TGAACAGCCTCTGGCATG | 112.1 | -10.7 | 2.3 | -4.7 | 2.4E-06 | 4.4E-06 | MB231/MB468 | MP |
| >hg38_21_+_8433172_8446622_RNA45SN1_45S_50ntFlanks@12938.12960.23\|ACCATTCGTAGACGACCTGCTTC | >hg38_21_+_8433172_8446622_RNA45SN1_45S_50ntFlanks | 12938 | 12960 | 23 | ACCATTCGTAGACGACCTGCTTC | 305.5 | 3.3 | 0.7 | 4.7 | 2.7E-06 | 4.4E-06 | MB231/MB468 | CYTO |
| >MI0000095\|hsa-mir-93&WithFlank&7\|-\|100093762\|100093853@17.39.23&[MIMAT0000093&hsa-miR-93-5p&offsets\|0\|0;m-29&7\|-\|100093815\|100093837&offsets\|0\|0]\|CAAAGTGCTGTTCGTGCAGGTAG | >MI0000095\|hsa-mir-93&WithFlank&7\|-\|100093762\|100093853 | 17 | 39 | 23 | CAAAGTGCTGTTCGTGCAGGTAG | 5704.2 | 4.6 | 1.0 | 4.7 | 2.7E-06 | 4.4E-06 | MB231/MB468 | CYTO |
| >hg38_21_+_8433172_8446622_RNA45SN1_45S_50ntFlanks@10916.10936.21\|CCGCGGCTGGACGAGGCGCCG | >hg38_21_+_8433172_8446622_RNA45SN1_45S_50ntFlanks | 10916 | 10936 | 21 | CCGCGGCTGGACGAGGCGCCG | 289.5 | -6.9 | 1.4 | -4.9 | 8.1E-07 | 4.4E-06 | BT20/MB468 | MP |
| >hg38_21_+_8433172_8446622_RNA45SN1_45S_50ntFlanks@6719.6739.21\|TGAATTGCAGGACACATTGAT | >hg38_21_+_8433172_8446622_RNA45SN1_45S_50ntFlanks | 6719 | 6739 | 21 | TGAATTGCAGGACACATTGAT | 325.4 | -7.1 | 1.4 | -4.9 | 8.1E-07 | 4.4E-06 | BT20/MB468 | MP |
| >hg38_21_+_8433172_8446622_RNA45SN1_45S_50ntFlanks@9544.9570.27\|GGTGGAGGTCCGTAGCGGTCCTGACGT | >hg38_21_+_8433172_8446622_RNA45SN1_45S_50ntFlanks | 9544 | 9570 | 27 | GGTGGAGGTCCGTAGCGGTCCTGACGT | 517.6 | -2.8 | 0.6 | -4.7 | 2.8E-06 | 4.4E-06 | MB231/MB468 | NUC |
| >hg38_21_+_8433172_8446622_RNA45SN1_45S_50ntFlanks@12973.12992.20\|TCGTACGTAGCAGAGCAGCT | >hg38_21_+_8433172_8446622_RNA45SN1_45S_50ntFlanks | 12973 | 12992 | 20 | TCGTACGTAGCAGAGCAGCT | 391.3 | -5.5 | 1.2 | -4.7 | 2.4E-06 | 4.4E-06 | MB231/MB468 | MP |

<table>
<tr><th></th><th></th><th></th><th></th><th></th><th></th><th></th><th></th><th></th><th></th><th></th><th></th><th></th><th></th></tr>
<tr><td>>am_trna162_MetCAT_6_-_26758550_26758622@31.54.24,trna164_MetCAT_6_-_26735574_26735646@31.54.24,trna20_MetCAT_16_+_71460396_71460468@31.54.24,trna21_MetCAT_6_+_26701712_26701784@31.54.24,trna27_MetCAT_6_+_26766444_26766516@31.54.24,trna75_MetCAT_6_+_28912352_28912424@31.54.24,trna97_MetCAT_6_-_28921042_28921114@31.54.24|TCTCATAATCTGAAGGTCCTGAGT</td><td>>am_trna162_MetCAT_6_-_26758550_26758622</td><td>31</td><td>54</td><td>24</td><td>TCTCATAATCTGAAGGTCCTGAGT</td><td>480.8</td><td>-7.2</td><td>1.5</td><td>-4.7</td><td>3.0E-06</td><td>4.4E-06</td><td>BT20/MB468</td><td>NUC</td></tr>
<tr><td>>am_trnaMT_PheGAA_MT_+_577_647@40.62.23|TGTTTAGACGGGCTCACATCACC</td><td>>am_trnaMT_PheGAA_MT_+_577_647</td><td>40</td><td>62</td><td>23</td><td>TGTTTAGACGGGCTCACATCACC</td><td>182.0</td><td>5.4</td><td>1.2</td><td>4.7</td><td>2.8E-06</td><td>4.4E-06</td><td>MB231/MB468</td><td>NUC</td></tr>
<tr><td>>hg38_21_+_8433172_8446622_RNA45SN1_45S_50ntFlanks@12625.12657.33|CAATGGGGCGAAGCTACCATCTGTGGGATTATG</td><td>>hg38_21_+_8433172_8446622_RNA45SN1_45S_50ntFlanks</td><td>12625</td><td>12657</td><td>33</td><td>CAATGGGGCGAAGCTACCATCTGTGGGATTATG</td><td>555.1</td><td>-3.7</td><td>0.8</td><td>-4.7</td><td>2.1E-06</td><td>4.4E-06</td><td>BT20/MB468</td><td>MITO</td></tr>
<tr><td>>hg38_21_+_8433172_8446622_RNA45SN1_45S_50ntFlanks@11584.11613.30|TGGTGCGGACCAGGGGAATCCGACTGTTTA</td><td>>hg38_21_+_8433172_8446622_RNA45SN1_45S_50ntFlanks</td><td>11584</td><td>11613</td><td>30</td><td>TGGTGCGGACCAGGGGAATCCGACTGTTTA</td><td>197.0</td><td>3.7</td><td>0.7</td><td>5.2</td><td>2.6E-07</td><td>4.4E-06</td><td>BT20/MB231</td><td>MITO</td></tr>
<tr><td>>MI0000815|hsa-mir-339&WithFlank&7|-|1022927|1023032@21.44.24&[MIMAT0000764&hsa-miR-339-5p&offsets|0|+1;m-163&7|-|1022990|1023012&offsets|0|+1]|TCCCTGTCCTCCAGGAGCTCACGT</td><td>>MI0000815|hsa-mir-339&WithFlank&7|-|1022927|1023032</td><td>21</td><td>44</td><td>24</td><td>TCCCTGTCCTCCAGGAGCTCACGT</td><td>291.7</td><td>6.1</td><td>1.3</td><td>4.7</td><td>2.9E-06</td><td>4.4E-06</td><td>MB231/MB468</td><td>MITO</td></tr>
<tr><td>>hg38_21_+_8433172_8446622_RNA45SN1_45S_50ntFlanks@12934.12956.23|CTAAACCATTCGTAGACGACCTG</td><td>>hg38_21_+_8433172_8446622_RNA45SN1_45S_50ntFlanks</td><td>12934</td><td>12956</td><td>23</td><td>CTAAACCATTCGTAGACGACCTG</td><td>10445.8</td><td>-5.9</td><td>1.3</td><td>-4.7</td><td>2.7E-06</td><td>4.4E-06</td><td>MB231/MB468</td><td>CYTO</td></tr>
<tr><td>>hg38_21_+_8433172_8446622_RNA45SN1_45S_50ntFlanks@10362.10379.18|AAGGCCGAAGTGGAGAAG</td><td>>hg38_21_+_8433172_8446622_RNA45SN1_45S_50ntFlanks</td><td>10362</td><td>10379</td><td>18</td><td>AAGGCCGAAGTGGAGAAG</td><td>242.2</td><td>-5.4</td><td>1.2</td><td>-4.7</td><td>2.8E-06</td><td>4.4E-06</td><td>MB231/MB468</td><td>NUC</td></tr>
<tr><td>>hg38_21_+_8433172_8446622_RNA45SN1_45S_50ntFlanks@12458.12487.30|AAGCGTTGGATTGTTCACCCACTAATAGGG</td><td>>hg38_21_+_8433172_8446622_RNA45SN1_45S_50ntFlanks</td><td>12458</td><td>12487</td><td>30</td><td>AAGCGTTGGATTGTTCACCCACTAATAGGG</td><td>231.4</td><td>-6.9</td><td>1.5</td><td>-4.7</td><td>2.4E-06</td><td>4.4E-06</td><td>MB231/MB468</td><td>MP</td></tr>
<tr><td>>MI0001445|hsa-mir-423&WithFlank&17|+|30117073|30117178@59.82.24&[MIMAT0001340&hsa-miR-423-3p&offsets|0|+1;m-100&17|+|30117131|30117153&offsets|0|+1]|AGCTCGGTCTGAGGCCCCTCAGTC</td><td>>MI0001445|hsa-mir-423&WithFlank&17|+|30117073|30117178</td><td>59</td><td>82</td><td>24</td><td>AGCTCGGTCTGAGGCCCCTCAGTC</td><td>291.5</td><td>6.1</td><td>1.3</td><td>4.7</td><td>2.9E-06</td><td>4.4E-06</td><td>MB231/MB468</td><td>MITO</td></tr>
<tr><td>>hg38_21_+_8433172_8446622_RNA45SN1_45S_50ntFlanks@5080.5112.33|ATAACGAACGAGACTCTGGCATGCTAACTAGTT</td><td>>hg38_21_+_8433172_8446622_RNA45SN1_45S_50ntFlanks</td><td>5080</td><td>5112</td><td>33</td><td>ATAACGAACGAGACTCTGGCATGCTAACTAGTT</td><td>243.8</td><td>-6.1</td><td>1.3</td><td>-4.7</td><td>2.4E-06</td><td>4.4E-06</td><td>MB231/MB468</td><td>MP</td></tr>
<tr><td>>hg38_21_+_8433172_8446622_RNA45SN1_45S_50ntFlanks@12681.12708.28|CCCGCCCAGGCGGAACGATACGGCAGCG</td><td>>hg38_21_+_8433172_8446622_RNA45SN1_45S_50ntFlanks</td><td>12681</td><td>12708</td><td>28</td><td>CCCGCCCAGGCGGAACGATACGGCAGCG</td><td>58.7</td><td>-9.0</td><td>1.8</td><td>-4.9</td><td>8.2E-07</td><td>4.4E-06</td><td>BT20/MB468</td><td>MP</td></tr>
<tr><td>>hg38_21_+_8433172_8446622_RNA45SN1_45S_50ntFlanks@12995.13037.43|CTCGCTGCGATCTATTGAAAGTCAGCCCTCGACACAAGGGTTT</td><td>>hg38_21_+_8433172_8446622_RNA45SN1_45S_50ntFlanks</td><td>12995</td><td>13037</td><td>43</td><td>CTCGCTGCGATCTATTGAAAGTCAGCCCTCGACACAAGGGTTT</td><td>3787.4</td><td>11.1</td><td>2.4</td><td>4.7</td><td>2.8E-06</td><td>4.4E-06</td><td>MB231/MB468</td><td>CYTO</td></tr>
<tr><td>>trna10_AspGTC_12_-_125424193_125424264@38.75.38,trna12_AspGTC_12_-_125411891_125411962@38.75.38,trna144_AspGTC_6_-_27551236_27551307@38.75.38,trna38_AspGTC_17_-_8125556_8125627@38.75.38,trna45_AspGTC_6_+_27447453_27447524@38.75.38,trna48_AspGTC_6_+_27471523_27471594@38.75.38,trna4_AspGTC_12_+_96429799_96429870@38.75.38,trna69_AspGTC_1_-_161440205_161440276@38.75.38,trna72_AspGTC_1_-_161432824_161432895@38.75.38,trna75_AspGTC_1_-_161425414_161425485@38.75.38,trna78_AspGTC_1_-_161418033_161418104@38.75.38,trna81_AspGTC_1_-_161410615_161410686@38.75.38|CGCGGGAGACCGGGGTTCGATTCCCCGACGGGGAGCCA</td><td>>trna10_AspGTC_12_-_125424193_125424264</td><td>38</td><td>75</td><td>38</td><td>CGCGGGAGACCGGGGTTCGATTCCCCGACGGGGAGCCA</td><td>775.0</td><td>3.2</td><td>0.7</td><td>4.7</td><td>2.4E-06</td><td>4.4E-06</td><td>BT20/MB468</td><td>CYTO</td></tr>
<tr><td>>MI0000065|hsa-let-7d&WithFlank&9|+|94178828|94178926@68.88.21&[MIMAT0004484&hsa-let-7d-3p&offsets|0|-1;m-98&9|+|94178895|94178916&offsets|0|-1]|CTATACGACCTGCTGCCTTTC</td><td>>MI0000065|hsa-let-7d&WithFlank&9|+|94178828|94178926</td><td>68</td><td>88</td><td>21</td><td>CTATACGACCTGCTGCCTTTC</td><td>452.2</td><td>6.5</td><td>1.4</td><td>4.7</td><td>2.8E-06</td><td>4.5E-06</td><td>MB231/MB468</td><td>CYTO</td></tr>
<tr><td>>hg38_21_+_8433172_8446622_RNA45SN1_45S_50ntFlanks@12447.12485.39|CAGAATTCACCAAGCGTTGGATTGTTCACCCACTAATAG</td><td>>hg38_21_+_8433172_8446622_RNA45SN1_45S_50ntFlanks</td><td>12447</td><td>12485</td><td>39</td><td>CAGAATTCACCAAGCGTTGGATTGTTCACCCACTAATAG</td><td>227.0</td><td>-3.6</td><td>0.8</td><td>-4.7</td><td>2.9E-06</td><td>4.5E-06</td><td>MB231/MB468</td><td>NUC</td></tr>
<tr><td>>hg38_21_+_8433172_8446622_RNA45SN1_45S_50ntFlanks@8358.8396.39|AAAGAACTTTGAAGAGAGAGTTCAAGAGGGCGTGAAACC</td><td>>hg38_21_+_8433172_8446622_RNA45SN1_45S_50ntFlanks</td><td>8358</td><td>8396</td><td>39</td><td>AAAGAACTTTGAAGAGAGAGTTCAAGAGGGCGTGAAACC</td><td>51.4</td><td>9.4</td><td>1.9</td><td>4.9</td><td>8.3E-07</td><td>4.5E-06</td><td>BT20/MB468</td><td>MP</td></tr>
<tr><td>>hg38_21_+_8433172_8446622_RNA45SN1_45S_50ntFlanks@12935.12966.32|TAAACCATTCGTAGACGACCTGCTTCTGGGTC</td><td>>hg38_21_+_8433172_8446622_RNA45SN1_45S_50ntFlanks</td><td>12935</td><td>12966</td><td>32</td><td>TAAACCATTCGTAGACGACCTGCTTCTGGGTC</td><td>348.5</td><td>-6.3</td><td>1.3</td><td>-4.7</td><td>2.5E-06</td><td>4.5E-06</td><td>BT20/MB468</td><td>CYTO</td></tr>
<tr><td>>trna14_LysTTT_11_-_59327808_59327880@1.30.30,trna2_LysTTT_17_+_8022473_8022545@1.30.30,trna54_LysTTT_1_+_204475655_204475727@1.30.30,trna5_LysTTT_11_+_59323902_59323974@1.30.30,trna62_LysTTT_1_-_204476158_204476230@1.30.30,trna76_LysTTT_6_+_28918806_28918878@1.30.30|GCCCGGATAGCTCAGTCGGTAGAGCATCAG</td><td>>trna14_LysTTT_11_-_59327808_59327880</td><td>1</td><td>30</td><td>30</td><td>GCCCGGATAGCTCAGTCGGTAGAGCATCAG</td><td>569.9</td><td>-5.8</td><td>1.2</td><td>-4.7</td><td>2.4E-06</td><td>4.5E-06</td><td>MB231/MB468</td><td>MP</td></tr>
<tr><td>>trnaMT_HisGTG_MT_+_12138_12206@32.70.39|TGAATCTGACAACAGAGGCTTACGACCCCTTATTTACCC</td><td>>trnaMT_HisGTG_MT_+_12138_12206</td><td>32</td><td>70</td><td>39</td><td>TGAATCTGACAACAGAGGCTTACGACCCCTTATTTACCC</td><td>1658.7</td><td>4.6</td><td>1.0</td><td>4.7</td><td>2.9E-06</td><td>4.5E-06</td><td>MB231/MB468</td><td>NUC</td></tr>
<tr><td>>hg38_1_-_228634819_228635039_RNA5S12_5S_50ntFlanks@129.171.43|TTGGATGGGAGACCGCCTGGGAATACCGGGTGCTGTAGGCTTT</td><td>>hg38_1_-_228634819_228635039_RNA5S12_5S_50ntFlanks</td><td>129</td><td>171</td><td>43</td><td>TTGGATGGGAGACCGCCTGGGAATACCGGGTGCTGTAGGCTTT</td><td>1056.8</td><td>5.9</td><td>1.3</td><td>4.7</td><td>2.9E-06</td><td>4.5E-06</td><td>MB231/MB468</td><td>NUC</td></tr>
<tr><td>>am_trnaMT_ValTAC_MT_+_1602_1670@44.68.25|ATTTCAACTTAACTTGACCGCTCTG</td><td>>am_trnaMT_ValTAC_MT_+_1602_1670</td><td>44</td><td>68</td><td>25</td><td>ATTTCAACTTAACTTGACCGCTCTG</td><td>875.1</td><td>-6.5</td><td>1.4</td><td>-4.7</td><td>2.9E-06</td><td>4.5E-06</td><td>MB231/MB468</td><td>NUC</td></tr>
<tr><td>>hg38_21_+_8433172_8446622_RNA45SN1_45S_50ntFlanks@10781.10801.21|AGGTGAACAGCCTCTGGCATG</td><td>>hg38_21_+_8433172_8446622_RNA45SN1_45S_50ntFlanks</td><td>10781</td><td>10801</td><td>21</td><td>AGGTGAACAGCCTCTGGCATG</td><td>282.0</td><td>-4.8</td><td>1.0</td><td>-4.7</td><td>2.9E-06</td><td>4.5E-06</td><td>MB231/MB468</td><td>NUC</td></tr>
<tr><td>>MI0001445|hsa-mir-423&WithFlank&17|+|30117073|30117178@59.80.22(+1G)&[MIMAT0001340&hsa-miR-423-3p&offsets|0|-1(+1G);m-100&17|+|30117131|30117153&offsets|0|-1(+1G)]|AGCTCGGTCTGAGGCCCCTCAGG</td><td>>MI0001445|hsa-mir-423&WithFlank&17|+|30117073|30117178</td><td>59</td><td>80</td><td>22(+1G)</td><td>AGCTCGGTCTGAGGCCCCTCAGG</td><td>295.6</td><td>5.1</td><td>1.1</td><td>4.7</td><td>3.0E-06</td><td>4.5E-06</td><td>MB231/MB468</td><td>MITO</td></tr>
<tr><td>>hg38_21_+_8433172_8446622_RNA45SN1_45S_50ntFlanks@10684.10709.26|CGGCGGCGTCCGGTGAGCTCTCGCTG</td><td>>hg38_21_+_8433172_8446622_RNA45SN1_45S_50ntFlanks</td><td>10684</td><td>10709</td><td>26</td><td>CGGCGGCGTCCGGTGAGCTCTCGCTG</td><td>2404.7</td><td>-6.9</td><td>1.5</td><td>-4.7</td><td>3.1E-06</td><td>4.5E-06</td><td>BT20/MB468</td><td>NUC</td></tr>
<tr><td>>hg38_21_+_8433172_8446622_RNA45SN1_45S_50ntFlanks@9534.9571.38|AGGAAACTCTGGTGGAGGTCCGTAGCGGTCCTGACGTG</td><td>>hg38_21_+_8433172_8446622_RNA45SN1_45S_50ntFlanks</td><td>9534</td><td>9571</td><td>38</td><td>AGGAAACTCTGGTGGAGGTCCGTAGCGGTCCTGACGTG</td><td>7719.2</td><td>-3.8</td><td>0.8</td><td>-4.7</td><td>2.1E-06</td><td>4.5E-06</td><td>BT20/MB468</td><td>MITO</td></tr>
<tr><td>>MI0000767|hsa-mir-365a&WithFlank&16|+|14309279|14309377@62.83.22;MI0000769|hsa-mir-365b&WithFlank&17|+|31575405|31575527@74.95.22&[MIMAT0000710&hsa-miR-365a-3p&offsets|0|0;m-176&16|+|14309340|14309361&offsets|0|0];[MIMAT0022834&hsa-miR-365b-3p&offsets|0|0;m-175&17|+|31575478|31575499&offsets|0|0]|TAATGCCCCTAAAAATCCTTAT</td><td>>MI0000767|hsa-mir-365a&WithFlank&16|+|14309279|14309377</td><td>62</td><td>83</td><td>22</td><td>TAATGCCCCTAAAAATCCTTAT</td><td>551.1</td><td>5.5</td><td>1.2</td><td>4.7</td><td>3.0E-06</td><td>4.5E-06</td><td>MB231/MB468</td><td>MITO</td></tr>
<tr><td>>hg38_21_+_8433172_8446622_RNA45SN1_45S_50ntFlanks@11211.11254.44|GCGACTCTGGACGCGAGCCGGGCCCTTCCCGTGGATCGCCCCAG</td><td>>hg38_21_+_8433172_8446622_RNA45SN1_45S_50ntFlanks</td><td>11211</td><td>11254</td><td>44</td><td>GCGACTCTGGACGCGAGCCGGGCCCTTCCCGTGGATCGCCCCAG</td><td>1691.1</td><td>-8.9</td><td>1.9</td><td>-4.7</td><td>3.1E-06</td><td>4.5E-06</td><td>BT20/MB468</td><td>NUC</td></tr>
<tr><td>>hg38_21_+_8433172_8446622_RNA45SN1_45S_50ntFlanks@10684.10712.29|CGGCGGCGTCCGGTGAGCTCTCGCTGGCC</td><td>>hg38_21_+_8433172_8446622_RNA45SN1_45S_50ntFlanks</td><td>10684</td><td>10712</td><td>29</td><td>CGGCGGCGTCCGGTGAGCTCTCGCTGGCC</td><td>1240.4</td><td>-8.2</td><td>1.8</td><td>-4.7</td><td>3.1E-06</td><td>4.5E-06</td><td>BT20/MB468</td><td>NUC</td></tr>
<tr><td>>hg38_21_+_8433172_8446622_RNA45SN1_45S_50ntFlanks@8194.8216.23|CCCGTGGACGGTGTGAGGCCGGT</td><td>>hg38_21_+_8433172_8446622_RNA45SN1_45S_50ntFlanks</td><td>8194</td><td>8216</td><td>23</td><td>CCCGTGGACGGTGTGAGGCCGGT</td><td>232.2</td><td>-2.8</td><td>0.6</td><td>-4.7</td><td>2.8E-06</td><td>4.5E-06</td><td>MB231/MB468</td><td>CYTO</td></tr>
<tr><td>>MI0001445|hsa-mir-423&WithFlank&17|+|30117073|30117178@59.79.21&[MIMAT0001340&hsa-miR-423-3p&offsets|0|-2;m-100&17|+|30117131|30117153&offsets|0|-2]|AGCTCGGTCTGAGGCCCCTCA</td><td>>MI0001445|hsa-mir-423&WithFlank&17|+|30117073|30117178</td><td>59</td><td>79</td><td>21</td><td>AGCTCGGTCTGAGGCCCCTCA</td><td>398.8</td><td>4.2</td><td>0.9</td><td>4.7</td><td>3.0E-06</td><td>4.5E-06</td><td>MB231/MB468</td><td>MITO</td></tr>
<tr><td>>hg38_21_+_8433172_8446622_RNA45SN1_45S_50ntFlanks@5445.5466.22|TCGGATCGGCCCCGCCGGGGTC</td><td>>hg38_21_+_8433172_8446622_RNA45SN1_45S_50ntFlanks</td><td>5445</td><td>5466</td><td>22</td><td>TCGGATCGGCCCCGCCGGGGTC</td><td>12854.5</td><td>-7.0</td><td>1.4</td><td>-4.9</td><td>8.3E-07</td><td>4.5E-06</td><td>BT20/MB468</td><td>MP</td></tr>
<tr><td>>hg38_21_+_8433172_8446622_RNA45SN1_45S_50ntFlanks@10802.10820.19|TTGGAACAATGTAGGTAAG</td><td>>hg38_21_+_8433172_8446622_RNA45SN1_45S_50ntFlanks</td><td>10802</td><td>10820</td><td>19</td><td>TTGGAACAATGTAGGTAAG</td><td>172.3</td><td>-6.6</td><td>1.4</td><td>-4.7</td><td>2.5E-06</td><td>4.5E-06</td><td>MB231/MB468</td><td>MP</td></tr>
<tr><td>>hg38_21_+_8433172_8446622_RNA45SN1_45S_50ntFlanks@12028.12058.31|CTCTGATCGTTTTTTCACTGACCCGGTGAGG</td><td>>hg38_21_+_8433172_8446622_RNA45SN1_45S_50ntFlanks</td><td>12028</td><td>12058</td><td>31</td><td>CTCTGATCGTTTTTTCACTGACCCGGTGAGG</td><td>222.9</td><td>11.1</td><td>2.1</td><td>5.2</td><td>2.1E-07</td><td>4.5E-06</td><td>BT20/MB231</td><td>MP</td></tr>
<tr><td>>hg38_21_+_8433172_8446622_RNA45SN1_45S_50ntFlanks@11234.11254.21|CCTTCCCGTGGATCGCCCCAG</td><td>>hg38_21_+_8433172_8446622_RNA45SN1_45S_50ntFlanks</td><td>11234</td><td>11254</td><td>21</td><td>CCTTCCCGTGGATCGCCCCAG</td><td>300.5</td><td>-6.0</td><td>1.3</td><td>-4.7</td><td>2.1E-06</td><td>4.5E-06</td><td>BT20/MB468</td><td>MITO</td></tr>
<tr><td>>hg38_21_+_8433172_8446622_RNA45SN1_45S_50ntFlanks@10391.10422.32|AACAGCAGTTGAACATGGGTCAGTCGGTCCTG</td><td>>hg38_21_+_8433172_8446622_RNA45SN1_45S_50ntFlanks</td><td>10391</td><td>10422</td><td>32</td><td>AACAGCAGTTGAACATGGGTCAGTCGGTCCTG</td><td>256.9</td><td>-5.1</td><td>1.1</td><td>-4.7</td><td>3.0E-06</td><td>4.5E-06</td><td>MB231/MB468</td><td>MITO</td></tr>
<tr><td>>hg38_21_+_8433172_8446622_RNA45SN1_45S_50ntFlanks@7980.8011.32|CCTCAGATCAGACGTGGCGACCCGCTGAATTT</td><td>>hg38_21_+_8433172_8446622_RNA45SN1_45S_50ntFlanks</td><td>7980</td><td>8011</td><td>32</td><td>CCTCAGATCAGACGTGGCGACCCGCTGAATTT</td><td>190.9</td><td>-5.2</td><td>1.1</td><td>-4.7</td><td>2.5E-06</td><td>4.5E-06</td><td>MB231/MB468</td><td>MP</td></tr>
<tr><td>>hg38_21_+_8433172_8446622_RNA45SN1_45S_50ntFlanks@11649.11670.22|GTTGACGCGATGTGATTTCTGC</td><td>>hg38_21_+_8433172_8446622_RNA45SN1_45S_50ntFlanks</td><td>11649</td><td>11670</td><td>22</td><td>GTTGACGCGATGTGATTTCTGC</td><td>149.6</td><td>10.6</td><td>2.0</td><td>5.2</td><td>2.1E-07</td><td>4.5E-06</td><td>BT20/MB231</td><td>MP</td></tr>
<tr><td>>hg38_21_+_8433172_8446622_RNA45SN1_45S_50ntFlanks@11603.11623.21|CCGACTGTTTAATTAAAACAA</td><td>>hg38_21_+_8433172_8446622_RNA45SN1_45S_50ntFlanks</td><td>11603</td><td>11623</td><td>21</td><td>CCGACTGTTTAATTAAAACAA</td><td>2192.5</td><td>4.3</td><td>0.8</td><td>5.1</td><td>2.6E-07</td><td>4.5E-06</td><td>BT20/MB231</td><td>MITO</td></tr>
<tr><td>>MI0000070|hsa-mir-16-1&WithFlank&13|-|50048967|50049067@20.39.20(+1U);MI0000115|hsa-mir-16-2&WithFlank&3|+|160404739|160404831@16.35.20(+1U)&[MIMAT0000069&hsa-miR-16-5p&offsets|0|-2(+1U);m-62&13|-|50049027|50049048&offsets|0|-2(+1U)];[MIMAT0000069_1&hsa-miR-16-5p&offsets|0|-2(+1U);m-61&3|+|160404754|160404775&offsets|0|-2(+1U)]|TAGCAGCACGTAAATATTGGT</td><td>>MI0000070|hsa-mir-16-1&WithFlank&13|-|50048967|50049067</td><td>20</td><td>39</td><td>20(+1U)</td><td>TAGCAGCACGTAAATATTGGT</td><td>160.1</td><td>3.2</td><td>0.7</td><td>4.7</td><td>3.1E-06</td><td>4.5E-06</td><td>BT20/MB468</td><td>NUC</td></tr>
<tr><td>>MI0003560|hsa-mir-92b&WithFlank&1|+|155195171|155195278@67.85.19(+1C)&[MIMAT0003218&hsa-miR-92b-3p&offsets|0|-3(+1C);m-115&1|+|155195237|155195258&offsets|0|-3(+1C)]|TATTGCACTCGTCCCGGCCC</td><td>>MI0003560|hsa-mir-92b&WithFlank&1|+|155195171|155195278</td><td>67</td><td>85</td><td>19(+1C)</td><td>TATTGCACTCGTCCCGGCCC</td><td>56.4</td><td>9.0</td><td>1.9</td><td>4.7</td><td>2.5E-06</td><td>4.5E-06</td><td>MB231/MB468</td><td>MP</td></tr>
</table>

<table>
<tr><td>>hg38_21_+_8433172_8446622_RNA45SN1_45S_50ntFlanks@6651.6679.29|CGACTCTTAGCGGTGGATCACTCGGCTCG</td><td>>hg38_21_+_8433172_8446622_RNA45SN1_45S_50ntFlanks</td><td>6651</td><td>6679</td><td>29</td><td>CGACTCTTAGCGGTGGATCACTCGGCTCG</td><td>8835.8</td><td>-4.2</td><td>0.9</td><td>-4.7</td><td>2.5E-06</td><td>4.5E-06</td><td>MB231/MB468</td><td>MP</td></tr>
<tr><td>>hg38_21_+_8433172_8446622_RNA45SN1_45S_50ntFlanks@12957.12988.32|CTTCTGGGTCGGGGTTTCGTACGTAGCAGAGC</td><td>>hg38_21_+_8433172_8446622_RNA45SN1_45S_50ntFlanks</td><td>12957</td><td>12988</td><td>32</td><td>CTTCTGGGTCGGGGTTTCGTACGTAGCAGAGC</td><td>911.2</td><td>-7.9</td><td>1.7</td><td>-4.7</td><td>3.1E-06</td><td>4.5E-06</td><td>BT20/MB468</td><td>NUC</td></tr>
<tr><td>>MI0003776|hsa-mir-320b-1&WithFlank&1|+|116671740|116671823@48.66.19(+1G);MI0003839|hsa-mir-320b-2&WithFlank&1|-|224257034|224257116@47.65.19(+1G);MI0003778|hsa-mir-320c-1&WithFlank&18|+|21683512|21683595@48.66.19(+1G);MI0008191|hsa-mir-320c-2&WithFlank&18|+|24321669|24321752@48.66.19(+1G);MI0000542|hsa-mir-320a&WithFlank&8|-|22244960|22245043@48.66.19(+1G)&[MIMAT0005792&hsa-miR-320b&offsets|0|-3(+1G);m-183&1|+|116671787|116671806&offsets|0|-1(+1G)];[MIMAT0005792_1&hsa-miR-320b&offsets|0|-3(+1G);m-183&1|-|224257051|224257070&offsets|0|-1(+1G)];[MIMAT0005793&hsa-miR-320c&offsets|0|-1(+1G);m-222&18|+|21683559|21683578&offsets|0|-1(+1G)];[MIMAT0005793_1&hsa-miR-320c&offsets|0|-1(+1G);m-219&18|+|24321716|24321735&offsets|0|-1(+1G)];[MIMAT0000510&hsa-miR-320a-3p&offsets|0|-3(+1G);m-60&8|-|22244975|22244996&offsets|0|-3(+1G)]|AAAAGCTGGGTTGAGAGGGG</td><td>>MI0003776|hsa-mir-320b-1&WithFlank&1|+|116671740|116671823</td><td>48</td><td>66</td><td>19(+1G)</td><td>AAAAGCTGGGTTGAGAGGGG</td><td>70.2</td><td>9.9</td><td>2.0</td><td>4.9</td><td>8.4E-07</td><td>4.5E-06</td><td>BT20/MB468</td><td>MP</td></tr>
<tr><td>>hg38_21_+_8433172_8446622_RNA45SN1_45S_50ntFlanks@12405.12436.32|CTTTTTGATCCTTCGATGTCGGCTCTTCCTAT</td><td>>hg38_21_+_8433172_8446622_RNA45SN1_45S_50ntFlanks</td><td>12405</td><td>12436</td><td>32</td><td>CTTTTTGATCCTTCGATGTCGGCTCTTCCTAT</td><td>7047.2</td><td>-6.8</td><td>1.4</td><td>-4.7</td><td>2.5E-06</td><td>4.5E-06</td><td>BT20/MB468</td><td>CYTO</td></tr>
<tr><td>>hg38_21_+_8433172_8446622_RNA45SN1_45S_50ntFlanks@11226.11254.29|AGCCGGGCCCTTCCCGTGGATCGCCCCAG</td><td>>hg38_21_+_8433172_8446622_RNA45SN1_45S_50ntFlanks</td><td>11226</td><td>11254</td><td>29</td><td>AGCCGGGCCCTTCCCGTGGATCGCCCCAG</td><td>57.2</td><td>-8.9</td><td>1.8</td><td>-4.9</td><td>8.4E-07</td><td>4.5E-06</td><td>BT20/MB468</td><td>MP</td></tr>
<tr><td>>hg38_MT_+_598_1651_RNR1_12S_50ntFlanks@853.879.27|CTTAGTTGAACAGGGCCCTGAAGCGCG</td><td>>hg38_MT_+_598_1651_RNR1_12S_50ntFlanks</td><td>853</td><td>879</td><td>27</td><td>CTTAGTTGAACAGGGCCCTGAAGCGCG</td><td>65.2</td><td>9.8</td><td>2.0</td><td>4.9</td><td>8.4E-07</td><td>4.5E-06</td><td>BT20/MB468</td><td>MP</td></tr>
<tr><td>>hg38_21_+_8433172_8446622_RNA45SN1_45S_50ntFlanks@5473.5514.42|CGGCCCTGGCGGAGCGCTGAGAAGACGGTCGAACTTGACTAT</td><td>>hg38_21_+_8433172_8446622_RNA45SN1_45S_50ntFlanks</td><td>5473</td><td>5514</td><td>42</td><td>CGGCCCTGGCGGAGCGCTGAGAAGACGGTCGAACTTGACTAT</td><td>852.0</td><td>-5.8</td><td>1.2</td><td>-4.7</td><td>2.5E-06</td><td>4.5E-06</td><td>MB231/MB468</td><td>MP</td></tr>
<tr><td>>hg38_21_+_8433172_8446622_RNA45SN1_45S_50ntFlanks@9913.9949.37|AGAAAAGGTGTTGGTTGATATAGACAGCAGGACGGTG</td><td>>hg38_21_+_8433172_8446622_RNA45SN1_45S_50ntFlanks</td><td>9913</td><td>9949</td><td>37</td><td>AGAAAAGGTGTTGGTTGATATAGACAGCAGGACGGTG</td><td>47.2</td><td>9.3</td><td>1.9</td><td>4.9</td><td>8.4E-07</td><td>4.5E-06</td><td>BT20/MB468</td><td>MP</td></tr>
<tr><td>>hg38_21_+_8433172_8446622_RNA45SN1_45S_50ntFlanks@12405.12428.24|CTTTTTGATCCTTCGATGTCGGCT</td><td>>hg38_21_+_8433172_8446622_RNA45SN1_45S_50ntFlanks</td><td>12405</td><td>12428</td><td>24</td><td>CTTTTTGATCCTTCGATGTCGGCT</td><td>260.3</td><td>-4.1</td><td>0.9</td><td>-4.7</td><td>2.5E-06</td><td>4.5E-06</td><td>BT20/MB468</td><td>CYTO</td></tr>
<tr><td>>hg38_21_+_8433172_8446622_RNA45SN1_45S_50ntFlanks@10342.10374.33|ATATTCAAACGAGAACTTTGAAGGCCGAAGTGG</td><td>>hg38_21_+_8433172_8446622_RNA45SN1_45S_50ntFlanks</td><td>10342</td><td>10374</td><td>33</td><td>ATATTCAAACGAGAACTTTGAAGGCCGAAGTGG</td><td>252.7</td><td>-4.4</td><td>0.9</td><td>-4.7</td><td>2.5E-06</td><td>4.5E-06</td><td>MB231/MB468</td><td>MP</td></tr>
<tr><td>>trna111_HisGTG_1_-_147774845_147774916@-1T.29.30,trna118_HisGTG_1_-_145396881_145396952@-1T.29.30,trna16_HisGTG_1_+_146544773_146544844@-1T.29.30,trna1_HisGTG_15_+_45493349_45493420@-1T.29.30,trna21_HisGTG_1_+_147753471_147753542@-1T.29.30,trna33_HisGTG_6_+_27125906_27125977@-1T.29.30,trna7_HisGTG_9_-_14433938_14434009@-1T.29.30,trna8_HisGTG_15_-_45492611_45492682@-1T.29.30,trna9_HisGTG_15_-_45490804_45490875@-1T.29.30|TGCCGTGATCGTATAGTGGTTAGTACTCTG</td><td>>trna111_HisGTG_1_-_147774845_147774916</td><td>-1T</td><td>29</td><td>30</td><td>TGCCGTGATCGTATAGTGGTTAGTACTCTG</td><td>1086.1</td><td>-5.4</td><td>1.1</td><td>-4.7</td><td>2.2E-06</td><td>4.5E-06</td><td>BT20/MB468</td><td>MITO</td></tr>
<tr><td>>hg38_21_+_8433172_8446622_RNA45SN1_45S_50ntFlanks@10664.10706.43|CTTGGAAAGCGTCGCGGTTCCGGCGGCGTCCGGTGAGCTCTCG</td><td>>hg38_21_+_8433172_8446622_RNA45SN1_45S_50ntFlanks</td><td>10664</td><td>10706</td><td>43</td><td>CTTGGAAAGCGTCGCGGTTCCGGCGGCGTCCGGTGAGCTCTCG</td><td>662.3</td><td>-5.1</td><td>1.1</td><td>-4.7</td><td>2.8E-06</td><td>4.6E-06</td><td>MB231/MB468</td><td>CYTO</td></tr>
<tr><td>>hg38_21_+_8433172_8446622_RNA45SN1_45S_50ntFlanks@10787.10826.40|ACAGCCTCTGGCATGTTGGAACAATGTAGGTAAGGGAAGT</td><td>>hg38_21_+_8433172_8446622_RNA45SN1_45S_50ntFlanks</td><td>10787</td><td>10826</td><td>40</td><td>ACAGCCTCTGGCATGTTGGAACAATGTAGGTAAGGGAAGT</td><td>214.5</td><td>-7.1</td><td>1.5</td><td>-4.7</td><td>2.5E-06</td><td>4.6E-06</td><td>MB231/MB468</td><td>MP</td></tr>
<tr><td>>trnaMT_CysGCA_MT_-_5761_5826@47.69.23|CTTCAAACCTGCCGGGGCTTCCA</td><td>>trnaMT_CysGCA_MT_-_5761_5826</td><td>47</td><td>69</td><td>23</td><td>CTTCAAACCTGCCGGGGCTTCCA</td><td>631.5</td><td>-4.7</td><td>1.0</td><td>-4.7</td><td>3.0E-06</td><td>4.6E-06</td><td>MB231/MB468</td><td>MITO</td></tr>
<tr><td>>hg38_21_+_8433172_8446622_RNA45SN1_45S_50ntFlanks@12717.12747.31|CCTCGGTTGGCCTCGGATAGCCGGTCCCCCG</td><td>>hg38_21_+_8433172_8446622_RNA45SN1_45S_50ntFlanks</td><td>12717</td><td>12747</td><td>31</td><td>CCTCGGTTGGCCTCGGATAGCCGGTCCCCCG</td><td>103.5</td><td>-6.4</td><td>1.4</td><td>-4.7</td><td>2.8E-06</td><td>4.6E-06</td><td>MB231/MB468</td><td>CYTO</td></tr>
<tr><td>>hg38_21_+_8433172_8446622_RNA45SN1_45S_50ntFlanks@10414.10443.30|TCGGTCCTGAGAGATGGGCGAGCGCCGTTC</td><td>>hg38_21_+_8433172_8446622_RNA45SN1_45S_50ntFlanks</td><td>10414</td><td>10443</td><td>30</td><td>TCGGTCCTGAGAGATGGGCGAGCGCCGTTC</td><td>182.2</td><td>-3.6</td><td>0.8</td><td>-4.7</td><td>2.8E-06</td><td>4.6E-06</td><td>MB231/MB468</td><td>CYTO</td></tr>
<tr><td>>hg38_21_+_8433172_8446622_RNA45SN1_45S_50ntFlanks@4145.4168.24|CCTGAGAAACGGCTACCACATCCA</td><td>>hg38_21_+_8433172_8446622_RNA45SN1_45S_50ntFlanks</td><td>4145</td><td>4168</td><td>24</td><td>CCTGAGAAACGGCTACCACATCCA</td><td>63.5</td><td>-9.1</td><td>1.8</td><td>-4.9</td><td>8.5E-07</td><td>4.6E-06</td><td>BT20/MB468</td><td>MP</td></tr>
<tr><td>>trna20_GluTTC_2_-_131094701_131094772@1.35.35,trna5_GluTTC_13_-_41634874_41634945@1.35.35|TCCCATATGGTCTAGCGGTTAGGATTCCTGGTTTT</td><td>>trna20_GluTTC_2_-_131094701_131094772</td><td>1</td><td>35</td><td>35</td><td>TCCCATATGGTCTAGCGGTTAGGATTCCTGGTTTT</td><td>52.1</td><td>8.9</td><td>1.9</td><td>4.7</td><td>2.5E-06</td><td>4.6E-06</td><td>MB231/MB468</td><td>MP</td></tr>
<tr><td>>hg38_21_+_8433172_8446622_RNA45SN1_45S_50ntFlanks@4005.4023.19|AACCTCGGGCCGATCGCAC</td><td>>hg38_21_+_8433172_8446622_RNA45SN1_45S_50ntFlanks</td><td>4005</td><td>4023</td><td>19</td><td>AACCTCGGGCCGATCGCAC</td><td>1413.2</td><td>3.3</td><td>0.7</td><td>4.7</td><td>2.9E-06</td><td>4.6E-06</td><td>MB231/MB468</td><td>CYTO</td></tr>
<tr><td>>hg38_21_+_8433172_8446622_RNA45SN1_45S_50ntFlanks@9923.9955.33|TTGGTTGATATAGACAGCAGGACGGTGGCCATG</td><td>>hg38_21_+_8433172_8446622_RNA45SN1_45S_50ntFlanks</td><td>9923</td><td>9955</td><td>33</td><td>TTGGTTGATATAGACAGCAGGACGGTGGCCATG</td><td>48.2</td><td>9.3</td><td>1.9</td><td>4.9</td><td>8.5E-07</td><td>4.6E-06</td><td>BT20/MB468</td><td>MP</td></tr>
<tr><td>>hg38_21_+_8433172_8446622_RNA45SN1_45S_50ntFlanks@11092.11116.25|CGGCGGGGGGAGAAGGGTCGGGGCG</td><td>>hg38_21_+_8433172_8446622_RNA45SN1_45S_50ntFlanks</td><td>11092</td><td>11116</td><td>25</td><td>CGGCGGGGGGAGAAGGGTCGGGGCG</td><td>66.2</td><td>-9.1</td><td>1.9</td><td>-4.9</td><td>8.5E-07</td><td>4.6E-06</td><td>BT20/MB468</td><td>MP</td></tr>
<tr><td>>hg38_21_+_8433172_8446622_RNA45SN1_45S_50ntFlanks@4974.4999.26|GCCCGGACACGGACAGGATTGACAGA</td><td>>hg38_21_+_8433172_8446622_RNA45SN1_45S_50ntFlanks</td><td>4974</td><td>4999</td><td>26</td><td>GCCCGGACACGGACAGGATTGACAGA</td><td>220.0</td><td>-6.8</td><td>1.5</td><td>-4.7</td><td>2.5E-06</td><td>4.6E-06</td><td>MB231/MB468</td><td>MP</td></tr>
<tr><td>>trnaMT_ValTAC_MT_+_1602_1670@44.72.29|ATTTCAACTTAACTTGACCGCTCTGACCA</td><td>>trnaMT_ValTAC_MT_+_1602_1670</td><td>44</td><td>72</td><td>29</td><td>ATTTCAACTTAACTTGACCGCTCTGACCA</td><td>9948.9</td><td>5.4</td><td>1.1</td><td>4.7</td><td>2.2E-06</td><td>4.6E-06</td><td>BT20/MB468</td><td>MITO</td></tr>
<tr><td>>hg38_21_+_8433172_8446622_RNA45SN1_45S_50ntFlanks@9571.9595.25|GCAAATCGGTCGTCCGACCTGGGTA</td><td>>hg38_21_+_8433172_8446622_RNA45SN1_45S_50ntFlanks</td><td>9571</td><td>9595</td><td>25</td><td>GCAAATCGGTCGTCCGACCTGGGTA</td><td>278.3</td><td>-7.0</td><td>1.5</td><td>-4.7</td><td>2.5E-06</td><td>4.6E-06</td><td>MB231/MB468</td><td>MP</td></tr>
<tr><td>>hg38_21_+_8433172_8446622_RNA45SN1_45S_50ntFlanks@12347.12375.29|AGAAAAGTTACCACAGGGATAACTGGCTT</td><td>>hg38_21_+_8433172_8446622_RNA45SN1_45S_50ntFlanks</td><td>12347</td><td>12375</td><td>29</td><td>AGAAAAGTTACCACAGGGATAACTGGCTT</td><td>176.9</td><td>-11.3</td><td>2.4</td><td>-4.7</td><td>2.5E-06</td><td>4.6E-06</td><td>MB231/MB468</td><td>MP</td></tr>
<tr><td>>hg38_21_+_8433172_8446622_RNA45SN1_45S_50ntFlanks@3985.4008.24|CGGCTTTGGTGACTCTAGATAACC</td><td>>hg38_21_+_8433172_8446622_RNA45SN1_45S_50ntFlanks</td><td>3985</td><td>4008</td><td>24</td><td>CGGCTTTGGTGACTCTAGATAACC</td><td>384.7</td><td>-7.3</td><td>1.6</td><td>-4.7</td><td>3.2E-06</td><td>4.6E-06</td><td>BT20/MB468</td><td>NUC</td></tr>
<tr><td>>hg38_21_+_8433172_8446622_RNA45SN1_45S_50ntFlanks@9569.9594.26|GTGCAAATCGGTCGTCCGACCTGGGT</td><td>>hg38_21_+_8433172_8446622_RNA45SN1_45S_50ntFlanks</td><td>9569</td><td>9594</td><td>26</td><td>GTGCAAATCGGTCGTCCGACCTGGGT</td><td>146.8</td><td>-5.1</td><td>1.1</td><td>-4.7</td><td>2.5E-06</td><td>4.6E-06</td><td>MB231/MB468</td><td>MP</td></tr>
<tr><td>>hg38_21_+_8433172_8446622_RNA45SN1_45S_50ntFlanks@8111.8132.22|GGGGCGCGGGACATGTGGCGTA</td><td>>hg38_21_+_8433172_8446622_RNA45SN1_45S_50ntFlanks</td><td>8111</td><td>8132</td><td>22</td><td>GGGGCGCGGGACATGTGGCGTA</td><td>144.5</td><td>10.5</td><td>2.0</td><td>5.2</td><td>2.2E-07</td><td>4.6E-06</td><td>BT20/MB231</td><td>MP</td></tr>
<tr><td>>hg38_21_+_8433172_8446622_RNA45SN1_45S_50ntFlanks@11768.11792.25|CTAATTAGTGACGCGCATGAATGGA</td><td>>hg38_21_+_8433172_8446622_RNA45SN1_45S_50ntFlanks</td><td>11768</td><td>11792</td><td>25</td><td>CTAATTAGTGACGCGCATGAATGGA</td><td>2069.9</td><td>-5.7</td><td>1.2</td><td>-4.7</td><td>2.5E-06</td><td>4.6E-06</td><td>BT20/MB468</td><td>CYTO</td></tr>
<tr><td>>hg38_21_+_8433172_8446622_RNA45SN1_45S_50ntFlanks@10458.10497.40|ATGGCCTCCGTTGCCCTCGGCCGATCGAAAGGGAGTCGGG</td><td>>hg38_21_+_8433172_8446622_RNA45SN1_45S_50ntFlanks</td><td>10458</td><td>10497</td><td>40</td><td>ATGGCCTCCGTTGCCCTCGGCCGATCGAAAGGGAGTCGGG</td><td>762.8</td><td>-7.6</td><td>1.5</td><td>-4.9</td><td>8.6E-07</td><td>4.6E-06</td><td>BT20/MB468</td><td>MP</td></tr>
<tr><td>>hg38_21_+_8433172_8446622_RNA45SN1_45S_50ntFlanks@9930.9959.30|ATATAGACAGCAGGACGGTGGCCATGGAAG</td><td>>hg38_21_+_8433172_8446622_RNA45SN1_45S_50ntFlanks</td><td>9930</td><td>9959</td><td>30</td><td>ATATAGACAGCAGGACGGTGGCCATGGAAG</td><td>1856.1</td><td>-7.4</td><td>1.6</td><td>-4.7</td><td>2.2E-06</td><td>4.6E-06</td><td>BT20/MB468</td><td>MITO</td></tr>
<tr><td>>hg38_21_+_8433172_8446622_RNA45SN1_45S_50ntFlanks@12420.12451.32|ATGTCGGCTCTTCCTATCATTGTGAAGCAGAA</td><td>>hg38_21_+_8433172_8446622_RNA45SN1_45S_50ntFlanks</td><td>12420</td><td>12451</td><td>32</td><td>ATGTCGGCTCTTCCTATCATTGTGAAGCAGAA</td><td>373.9</td><td>-7.3</td><td>1.6</td><td>-4.7</td><td>3.2E-06</td><td>4.6E-06</td><td>BT20/MB468</td><td>NUC</td></tr>
<tr><td>>MI0000457|hsa-mir-141&WithFlank&12|+|6964091|6964197@23.43.21&[MIMAT0004598&hsa-miR-141-5p&offsets|0|-1;m-221&12|+|6964113|6964133&offsets|0|0]|CATCTTCCAGTACAGTGTTGG</td><td>>MI0000457|hsa-mir-141&WithFlank&12|+|6964091|6964197</td><td>23</td><td>43</td><td>21</td><td>CATCTTCCAGTACAGTGTTGG</td><td>123.0</td><td>10.3</td><td>2.0</td><td>5.2</td><td>2.2E-07</td><td>4.6E-06</td><td>BT20/MB231</td><td>MP</td></tr>
<tr><td>>hg38_21_+_8433172_8446622_RNA45SN1_45S_50ntFlanks@4684.4720.37|AAGACGGACCAGAGCGAAAGCATTTGCCAAGAATGTT</td><td>>hg38_21_+_8433172_8446622_RNA45SN1_45S_50ntFlanks</td><td>4684</td><td>4720</td><td>37</td><td>AAGACGGACCAGAGCGAAAGCATTTGCCAAGAATGTT</td><td>498.5</td><td>5.4</td><td>1.1</td><td>4.7</td><td>2.9E-06</td><td>4.6E-06</td><td>MB231/MB468</td><td>CYTO</td></tr>
<tr><td>>hg38_21_+_8433172_8446622_RNA45SN1_45S_50ntFlanks@3911.3937.27|GGATGCGTGCATTTATCAGATCAAAAC</td><td>>hg38_21_+_8433172_8446622_RNA45SN1_45S_50ntFlanks</td><td>3911</td><td>3937</td><td>27</td><td>GGATGCGTGCATTTATCAGATCAAAAC</td><td>62.4</td><td>-9.0</td><td>1.8</td><td>-4.9</td><td>8.6E-07</td><td>4.6E-06</td><td>BT20/MB468</td><td>MP</td></tr>
<tr><td>>hg38_21_+_8433172_8446622_RNA45SN1_45S_50ntFlanks@4723.4748.26|CATTAATCAAGAACGAAAGTCGGAGG</td><td>>hg38_21_+_8433172_8446622_RNA45SN1_45S_50ntFlanks</td><td>4723</td><td>4748</td><td>26</td><td>CATTAATCAAGAACGAAAGTCGGAGG</td><td>291.9</td><td>-6.3</td><td>1.4</td><td>-4.7</td><td>3.0E-06</td><td>4.6E-06</td><td>MB231/MB468</td><td>NUC</td></tr>
<tr><td>>hg38_21_+_8433172_8446622_RNA45SN1_45S_50ntFlanks@11874.11912.39|AGAAGACCCTGTTGAGCTTGACTCTAGTCTGGCACGGTG</td><td>>hg38_21_+_8433172_8446622_RNA45SN1_45S_50ntFlanks</td><td>11874</td><td>11912</td><td>39</td><td>AGAAGACCCTGTTGAGCTTGACTCTAGTCTGGCACGGTG</td><td>184.5</td><td>-6.1</td><td>1.3</td><td>-4.7</td><td>2.6E-06</td><td>4.6E-06</td><td>BT20/MB468</td><td>CYTO</td></tr>
<tr><td>>hg38_21_+_8433172_8446622_RNA45SN1_45S_50ntFlanks@7984.8003.20|AGATCAGACGTGGCGACCCG</td><td>>hg38_21_+_8433172_8446622_RNA45SN1_45S_50ntFlanks</td><td>7984</td><td>8003</td><td>20</td><td>AGATCAGACGTGGCGACCCG</td><td>510.5</td><td>-6.7</td><td>1.4</td><td>-4.7</td><td>3.0E-06</td><td>4.6E-06</td><td>MB231/MB468</td><td>NUC</td></tr>
<tr><td>>hg38_21_+_8433172_8446622_RNA45SN1_45S_50ntFlanks@10683.10712.30|CCGGCGGCGTCCGGTGAGCTCTCGCTGGCC</td><td>>hg38_21_+_8433172_8446622_RNA45SN1_45S_50ntFlanks</td><td>10683</td><td>10712</td><td>30</td><td>CCGGCGGCGTCCGGTGAGCTCTCGCTGGCC</td><td>1187.3</td><td>-7.3</td><td>1.5</td><td>-4.7</td><td>2.5E-06</td><td>4.6E-06</td><td>MB231/MB468</td><td>MP</td></tr>
<tr><td>>MI0000254|hsa-mir-30c-2&WithFlank&6|-|71376954|71377037@53.74.22&[MIMAT0004550&hsa-miR-30c-2-3p&offsets|0|0;m-206&6|-|71376964|71376985&offsets|0|0]|CTGGGAGAAGGCTGTTTACTCT</td><td>>MI0000254|hsa-mir-30c-2&WithFlank&6|-|71376954|71377037</td><td>53</td><td>74</td><td>22</td><td>CTGGGAGAAGGCTGTTTACTCT</td><td>360.0</td><td>3.2</td><td>0.7</td><td>4.7</td><td>2.5E-06</td><td>4.6E-06</td><td>MB231/MB468</td><td>MP</td></tr>
<tr><td>>hg38_21_+_8433172_8446622_RNA45SN1_45S_50ntFlanks@9925.9949.25|GGTTGATATAGACAGCAGGACGGTG</td><td>>hg38_21_+_8433172_8446622_RNA45SN1_45S_50ntFlanks</td><td>9925</td><td>9949</td><td>25</td><td>GGTTGATATAGACAGCAGGACGGTG</td><td>120.5</td><td>10.2</td><td>2.0</td><td>5.2</td><td>2.2E-07</td><td>4.6E-06</td><td>BT20/MB231</td><td>MP</td></tr>
<tr><td>>hg38_21_+_8433172_8446622_RNA45SN1_45S_50ntFlanks@11568.11593.26|GCAGCCGACTTAGAACTGGTGCGGAC</td><td>>hg38_21_+_8433172_8446622_RNA45SN1_45S_50ntFlanks</td><td>11568</td><td>11593</td><td>26</td><td>GCAGCCGACTTAGAACTGGTGCGGAC</td><td>337.5</td><td>-6.2</td><td>1.3</td><td>-4.7</td><td>2.6E-06</td><td>4.7E-06</td><td>MB231/MB468</td><td>MP</td></tr>
<tr><td>>hg38_21_+_8433172_8446622_RNA45SN1_45S_50ntFlanks@11317.11343.27|ACCCCACGTCTCGTCGCGCGCGCGTCC</td><td>>hg38_21_+_8433172_8446622_RNA45SN1_45S_50ntFlanks</td><td>11317</td><td>11343</td><td>27</td><td>ACCCCACGTCTCGTCGCGCGCGCGTCC</td><td>451.0</td><td>-4.6</td><td>1.0</td><td>-4.7</td><td>2.6E-06</td><td>4.7E-06</td><td>BT20/MB468</td><td>CYTO</td></tr>
<tr><td>>hg38_21_+_8433172_8446622_RNA45SN1_45S_50ntFlanks@6734.6771.38|ATTGATCATCGACACTTCGAACGCACTTGCGGCCCCGG</td><td>>hg38_21_+_8433172_8446622_RNA45SN1_45S_50ntFlanks</td><td>6734</td><td>6771</td><td>38</td><td>ATTGATCATCGACACTTCGAACGCACTTGCGGCCCCGG</td><td>898.8</td><td>-3.0</td><td>0.6</td><td>-4.7</td><td>3.0E-06</td><td>4.7E-06</td><td>MB231/MB468</td><td>NUC</td></tr>
<tr><td>>hg38_21_+_8433172_8446622_RNA45SN1_45S_50ntFlanks@6661.6697.37|CGGTGGATCACTCGGCTCGTGCGTCGATGAAGAACGC</td><td>>hg38_21_+_8433172_8446622_RNA45SN1_45S_50ntFlanks</td><td>6661</td><td>6697</td><td>37</td><td>CGGTGGATCACTCGGCTCGTGCGTCGATGAAGAACGC</td><td>781.2</td><td>2.7</td><td>0.6</td><td>4.7</td><td>2.6E-06</td><td>4.7E-06</td><td>BT20/MB468</td><td>CYTO</td></tr>
<tr><td>>hg38_21_+_8433172_8446622_RNA45SN1_45S_50ntFlanks@5477.5516.40|CCTGGCGGAGCGCTGAGAAGACGGTCGAACTTGACTATCT</td><td>>hg38_21_+_8433172_8446622_RNA45SN1_45S_50ntFlanks</td><td>5477</td><td>5516</td><td>40</td><td>CCTGGCGGAGCGCTGAGAAGACGGTCGAACTTGACTATCT</td><td>1237.7</td><td>4.6</td><td>0.9</td><td>5.1</td><td>2.7E-07</td><td>4.7E-06</td><td>BT20/MB231</td><td>MITO</td></tr>
</table>

<table>
<tr><td>>MI0000279|hsa-mir-196a-2&WithFlank&12|+|53991732|53991853@31.52.22;MI0000238|hsa-mir-196a-1&WithFlank&17|-|48632484|48632565@13.34.22&[MIMAT0000226&hsa-miR-196a-5p&offsets|0|0;m-261&12|+|53991762|53991783&offsets|0|0];[MIMAT0000226_1&hsa-miR-196a-5p&offsets|0|0;m-268&17|-|48632532|48632553&offsets|0|0]|TAGGTAGTTTCATGTTGTTGGG</td><td>>MI0000279|hsa-mir-196a-2&WithFlank&12|+|53991732|53991853</td><td>31</td><td>52</td><td>22</td><td>TAGGTAGTTTCATGTTGTTGGG</td><td>5166.3</td><td>10.6</td><td>2.0</td><td>5.2</td><td>2.4E-07</td><td>4.7E-06</td><td>BT20/MB231</td><td>CYTO</td></tr>
<tr><td>>hg38_21_+_8433172_8446622_RNA45SN1_45S_50ntFlanks@12388.12434.47|CGTTCATAGCGACGTCGCTTTTTGATCCTTCGATGTCGGCTCTTCCT</td><td>>hg38_21_+_8433172_8446622_RNA45SN1_45S_50ntFlanks</td><td>12388</td><td>12434</td><td>47</td><td>CGTTCATAGCGACGTCGCTTTTTGATCCTTCGATGTCGGCTCTTCCT</td><td>741.1</td><td>-5.0</td><td>1.0</td><td>-5.2</td><td>2.4E-07</td><td>4.7E-06</td><td>BT20/MB231</td><td>CYTO</td></tr>
<tr><td>>hg38_21_+_8433172_8446622_RNA45SN1_45S_50ntFlanks@11212.11231.20|CGACTCTGGACGCGAGCCGG</td><td>>hg38_21_+_8433172_8446622_RNA45SN1_45S_50ntFlanks</td><td>11212</td><td>11231</td><td>20</td><td>CGACTCTGGACGCGAGCCGG</td><td>366.4</td><td>-5.0</td><td>1.1</td><td>-4.7</td><td>2.6E-06</td><td>4.7E-06</td><td>BT20/MB468</td><td>CYTO</td></tr>
<tr><td>>hg38_21_+_8433172_8446622_RNA45SN1_45S_50ntFlanks@6653.6681.29|ACTCTTAGCGGTGGATCACTCGGCTCGTG</td><td>>hg38_21_+_8433172_8446622_RNA45SN1_45S_50ntFlanks</td><td>6653</td><td>6681</td><td>29</td><td>ACTCTTAGCGGTGGATCACTCGGCTCGTG</td><td>1588.6</td><td>-3.1</td><td>0.7</td><td>-4.7</td><td>2.6E-06</td><td>4.7E-06</td><td>BT20/MB468</td><td>CYTO</td></tr>
<tr><td>>MI0000782|hsa-mir-374a&WithFlank&X|-|74287280|74287363@18.38.21(+1A)&[MIMAT0000727&hsa-miR-374a-5p&offsets|0|-1(+1A);m-184&X|-|74287325|74287346&offsets|0|-1(+1A)]|TTATAATACAACCTGATAAGTA</td><td>>MI0000782|hsa-mir-374a&WithFlank&X|-|74287280|74287363</td><td>18</td><td>38</td><td>21(+1A)</td><td>TTATAATACAACCTGATAAGTA</td><td>116.1</td><td>-10.3</td><td>2.0</td><td>-5.2</td><td>2.2E-07</td><td>4.7E-06</td><td>BT20/MB231</td><td>MP</td></tr>
<tr><td>>MI0000542|hsa-mir-320a&WithFlank&8|-|22244960|22245043@48.68.21(+1U)&[MIMAT0000510&hsa-miR-320a-3p&offsets|0|-1(+1U);m-60&8|-|22244975|22244996&offsets|0|-1(+1U)]|AAAAGCTGGGTTGAGAGGGCGT</td><td>>MI0000542|hsa-mir-320a&WithFlank&8|-|22244960|22245043</td><td>48</td><td>68</td><td>21(+1U)</td><td>AAAAGCTGGGTTGAGAGGGCGT</td><td>26397.1</td><td>4.0</td><td>0.8</td><td>5.2</td><td>2.4E-07</td><td>4.7E-06</td><td>BT20/MB231</td><td>CYTO</td></tr>
<tr><td>>am_trna134_GluTTC_1_-_16861774_16861845@1.30.30,trna5_GluTTC_1_+_17199078_17199149@1.30.30,trna84_GluTTC_1_-_161391883_161391954@1.30.30,trna94_GluTTC_1_-_149664355_149664427@1.30.30|TCCCTGGTGGTCTAGTGGCTAGGATTCGGC</td><td>>am_trna134_GluTTC_1_-_16861774_16861845</td><td>1</td><td>30</td><td>30</td><td>TCCCTGGTGGTCTAGTGGCTAGGATTCGGC</td><td>902.2</td><td>3.9</td><td>0.8</td><td>5.2</td><td>2.4E-07</td><td>4.7E-06</td><td>BT20/MB231</td><td>CYTO</td></tr>
<tr><td>>MI0003776|hsa-mir-320b-1&WithFlank&1|+|116671740|116671823@49.68.20;MI0003839|hsa-mir-320b-2&WithFlank&1|-|224257034|224257116@48.67.20&[MIMAT0005792&hsa-miR-320b&offsets|+1|-1;m-183&1|+|116671787|116671806&offsets|+1|+1];[MIMAT0005792_1&hsa-miR-320b&offsets|+1|-1;m-183&1|-|224257051|224257070&offsets|+1|+1]|AAAGCTGGGTTGAGAGGGCA</td><td>>MI0003776|hsa-mir-320b-1&WithFlank&1|+|116671740|116671823</td><td>49</td><td>68</td><td>20</td><td>AAAGCTGGGTTGAGAGGGCA</td><td>242.1</td><td>8.5</td><td>1.7</td><td>4.9</td><td>8.7E-07</td><td>4.7E-06</td><td>BT20/MB468</td><td>MP</td></tr>
<tr><td>>hg38_21_+_8433172_8446622_RNA45SN1_45S_50ntFlanks@9005.9026.22|CTCCCCCGCGGGGGCGCGCCGG</td><td>>hg38_21_+_8433172_8446622_RNA45SN1_45S_50ntFlanks</td><td>9005</td><td>9026</td><td>22</td><td>CTCCCCCGCGGGGGCGCGCCGG</td><td>57.4</td><td>-8.9</td><td>1.8</td><td>-4.9</td><td>8.7E-07</td><td>4.7E-06</td><td>BT20/MB468</td><td>MP</td></tr>
<tr><td>>hg38_21_+_8433172_8446622_RNA45SN1_45S_50ntFlanks@8105.8141.37|CGCGGCGGGGCGCGGGACATGTGGCGTACGGAAGACC</td><td>>hg38_21_+_8433172_8446622_RNA45SN1_45S_50ntFlanks</td><td>8105</td><td>8141</td><td>37</td><td>CGCGGCGGGGCGCGGGACATGTGGCGTACGGAAGACC</td><td>243.3</td><td>-4.8</td><td>1.0</td><td>-4.7</td><td>3.0E-06</td><td>4.7E-06</td><td>MB231/MB468</td><td>NUC</td></tr>
<tr><td>>hg38_21_+_8433172_8446622_RNA45SN1_45S_50ntFlanks@11208.11232.25|GCGGCGACTCTGGACGCGAGCCGGG</td><td>>hg38_21_+_8433172_8446622_RNA45SN1_45S_50ntFlanks</td><td>11208</td><td>11232</td><td>25</td><td>GCGGCGACTCTGGACGCGAGCCGGG</td><td>260.2</td><td>11.4</td><td>2.2</td><td>5.2</td><td>2.3E-07</td><td>4.7E-06</td><td>BT20/MB231</td><td>MP</td></tr>
<tr><td>>hg38_21_+_8433172_8446622_RNA45SN1_45S_50ntFlanks@10351.10382.32|CGAGAACTTTGAAGGCCGAAGTGGAGAAGGGT</td><td>>hg38_21_+_8433172_8446622_RNA45SN1_45S_50ntFlanks</td><td>10351</td><td>10382</td><td>32</td><td>CGAGAACTTTGAAGGCCGAAGTGGAGAAGGGT</td><td>1176.8</td><td>-2.1</td><td>0.4</td><td>-4.7</td><td>2.6E-06</td><td>4.7E-06</td><td>BT20/MB468</td><td>CYTO</td></tr>
<tr><td>>hg38_21_+_8433172_8446622_RNA45SN1_45S_50ntFlanks@11785.11805.21|TGAATGGATGAACGAGATTCC</td><td>>hg38_21_+_8433172_8446622_RNA45SN1_45S_50ntFlanks</td><td>11785</td><td>11805</td><td>21</td><td>TGAATGGATGAACGAGATTCC</td><td>307.6</td><td>-5.2</td><td>1.1</td><td>-4.7</td><td>3.1E-06</td><td>4.7E-06</td><td>MB231/MB468</td><td>MITO</td></tr>
<tr><td>>hg38_21_+_8433172_8446622_RNA45SN1_45S_50ntFlanks@10783.10817.35|GTGAACAGCCTCTGGCATGTTGGAACAATGTAGGT</td><td>>hg38_21_+_8433172_8446622_RNA45SN1_45S_50ntFlanks</td><td>10783</td><td>10817</td><td>35</td><td>GTGAACAGCCTCTGGCATGTTGGAACAATGTAGGT</td><td>428.4</td><td>-3.4</td><td>0.7</td><td>-4.7</td><td>2.9E-06</td><td>4.7E-06</td><td>MB231/MB468</td><td>CYTO</td></tr>
<tr><td>>hg38_21_+_8433172_8446622_RNA45SN1_45S_50ntFlanks@10394.10430.37|AGCAGTTGAACATGGGTCAGTCGGTCCTGAGAGATGG</td><td>>hg38_21_+_8433172_8446622_RNA45SN1_45S_50ntFlanks</td><td>10394</td><td>10430</td><td>37</td><td>AGCAGTTGAACATGGGTCAGTCGGTCCTGAGAGATGG</td><td>389.0</td><td>-7.3</td><td>1.6</td><td>-4.7</td><td>3.3E-06</td><td>4.7E-06</td><td>BT20/MB468</td><td>NUC</td></tr>
<tr><td>>hg38_21_+_8433172_8446622_RNA45SN1_45S_50ntFlanks@12108.12125.18|CCGCGCGCCGGCCGGGCG</td><td>>hg38_21_+_8433172_8446622_RNA45SN1_45S_50ntFlanks</td><td>12108</td><td>12125</td><td>18</td><td>CCGCGCGCCGGCCGGGCG</td><td>304.7</td><td>-6.0</td><td>1.3</td><td>-4.7</td><td>2.6E-06</td><td>4.7E-06</td><td>BT20/MB468</td><td>CYTO</td></tr>
<tr><td>>am_MI0015825|hsa-mir-378c&WithFlank&10|-|130962582|130962674@17.35.19;MI0016749|hsa-mir-378d-1&WithFlank&4|-|5923269|5923334@37.55.19;MI0000786|hsa-mir-378a&WithFlank&5|+|149732819|149732896@49.67.19;MI0003840|hsa-mir-378d-2&WithFlank&8|-|93916016|93916125@28.46.19&[MIMAT0016847&hsa-miR-378c&offsets|0|-6;m-130&10|-|130962639|130962658&offsets|0|-1];[MIMAT0018926&hsa-miR-378d&offsets|0|-1;m-228&4|-|5923281|5923298&offsets|0|+1];[MIMAT0000732&hsa-miR-378a-3p&offsets|0|-3;m-44&5|+|149732867|149732888&offsets|0|-3];[MIMAT0018926_1&hsa-miR-378d&offsets|0|-1;m-227&8|-|93916081|93916098&offsets|0|+1]|ACTGGACTTGGAGTCAGAA</td><td>>am_MI0015825|hsa-mir-378c&WithFlank&10|-|130962582|130962674</td><td>17</td><td>35</td><td>19</td><td>ACTGGACTTGGAGTCAGAA</td><td>163.4</td><td>4.4</td><td>0.9</td><td>4.7</td><td>3.1E-06</td><td>4.7E-06</td><td>MB231/MB468</td><td>MITO</td></tr>
<tr><td>>hg38_21_+_8433172_8446622_RNA45SN1_45S_50ntFlanks@3984.4002.19|GCGGCTTTGGTGACTCTAG</td><td>>hg38_21_+_8433172_8446622_RNA45SN1_45S_50ntFlanks</td><td>3984</td><td>4002</td><td>19</td><td>GCGGCTTTGGTGACTCTAG</td><td>556.4</td><td>-6.0</td><td>1.3</td><td>-4.7</td><td>2.9E-06</td><td>4.7E-06</td><td>MB231/MB468</td><td>CYTO</td></tr>
<tr><td>>MI0000285|hsa-mir-205&WithFlank&1|+|209432127|209432248@40.60.21&[MIMAT0000266&hsa-miR-205-5p&offsets|0|-1;m-351&1|+|209432166|209432187&offsets|0|-1]|TCCTTCATTCCACCGGAGTCT</td><td>>MI0000285|hsa-mir-205&WithFlank&1|+|209432127|209432248</td><td>40</td><td>60</td><td>21</td><td>TCCTTCATTCCACCGGAGTCT</td><td>2110.3</td><td>3.1</td><td>0.7</td><td>4.7</td><td>3.3E-06</td><td>4.7E-06</td><td>BT20/MB468</td><td>NUC</td></tr>
<tr><td>>hg38_21_+_8433172_8446622_RNA45SN1_45S_50ntFlanks@6772.6807.36|GTTCCTCCCGGGGCTACGCCTGTCTGAGCGTCGCTT</td><td>>hg38_21_+_8433172_8446622_RNA45SN1_45S_50ntFlanks</td><td>6772</td><td>6807</td><td>36</td><td>GTTCCTCCCGGGGCTACGCCTGTCTGAGCGTCGCTT</td><td>461.6</td><td>-6.8</td><td>1.5</td><td>-4.7</td><td>2.6E-06</td><td>4.7E-06</td><td>MB231/MB468</td><td>MP</td></tr>
<tr><td>>hg38_21_+_8433172_8446622_RNA45SN1_45S_50ntFlanks@11567.11603.37|AGCAGCCGACTTAGAACTGGTGCGGACCAGGGGAATC</td><td>>hg38_21_+_8433172_8446622_RNA45SN1_45S_50ntFlanks</td><td>11567</td><td>11603</td><td>37</td><td>AGCAGCCGACTTAGAACTGGTGCGGACCAGGGGAATC</td><td>1734.7</td><td>-5.7</td><td>1.2</td><td>-4.7</td><td>3.1E-06</td><td>4.7E-06</td><td>MB231/MB468</td><td>MITO</td></tr>
<tr><td>>hg38_21_+_8433172_8446622_RNA45SN1_45S_50ntFlanks@3769.3794.26|CGCACGGCCGGTACAGTGAAACTGCG</td><td>>hg38_21_+_8433172_8446622_RNA45SN1_45S_50ntFlanks</td><td>3769</td><td>3794</td><td>26</td><td>CGCACGGCCGGTACAGTGAAACTGCG</td><td>400.5</td><td>-3.9</td><td>0.8</td><td>-4.7</td><td>3.1E-06</td><td>4.7E-06</td><td>MB231/MB468</td><td>NUC</td></tr>
<tr><td>>MI0000063|hsa-let-7b&WithFlank&22|+|46113680|46113774@12.34.23(+1U)&[MIMAT0000063&hsa-let-7b-5p&offsets|0|+1(+1U);m-14&22|+|46113691|46113712&offsets|0|+1(+1U)]|TGAGGTAGTAGGTTGTGTGGTTTT</td><td>>MI0000063|hsa-let-7b&WithFlank&22|+|46113680|46113774</td><td>12</td><td>34</td><td>23(+1U)</td><td>TGAGGTAGTAGGTTGTGTGGTTTT</td><td>516.0</td><td>3.6</td><td>0.8</td><td>4.7</td><td>2.6E-06</td><td>4.7E-06</td><td>MB231/MB468</td><td>MP</td></tr>
<tr><td>>MI0000298|hsa-mir-221&WithFlank&X|-|45746151|45746272@71.90.20(+1A)&[MIMAT0000278&hsa-miR-221-3p&offsets|0|-3(+1A);m-91&X|-|45746181|45746202&offsets|0|-2(+1A)]|AGCTACATTGTCTGCTGGGTA</td><td>>MI0000298|hsa-mir-221&WithFlank&X|-|45746151|45746272</td><td>71</td><td>90</td><td>20(+1A)</td><td>AGCTACATTGTCTGCTGGGTA</td><td>259.5</td><td>5.5</td><td>1.2</td><td>4.7</td><td>2.6E-06</td><td>4.7E-06</td><td>MB231/MB468</td><td>MP</td></tr>
<tr><td>>trnaMT_LeuTAA_MT_+_3230_3304@19.42.24|GGTAATCGCATAAAACTTAAAACT</td><td>>trnaMT_LeuTAA_MT_+_3230_3304</td><td>19</td><td>42</td><td>24</td><td>GGTAATCGCATAAAACTTAAAACT</td><td>305.0</td><td>4.3</td><td>0.8</td><td>5.2</td><td>2.5E-07</td><td>4.7E-06</td><td>BT20/MB231</td><td>CYTO</td></tr>
<tr><td>>hg38_21_+_8433172_8446622_RNA45SN1_45S_50ntFlanks@6666.6698.33|GATCACTCGGCTCGTGCGTCGATGAAGAACGCA</td><td>>hg38_21_+_8433172_8446622_RNA45SN1_45S_50ntFlanks</td><td>6666</td><td>6698</td><td>33</td><td>GATCACTCGGCTCGTGCGTCGATGAAGAACGCA</td><td>49.2</td><td>-8.7</td><td>1.8</td><td>-4.9</td><td>8.8E-07</td><td>4.7E-06</td><td>BT20/MB468</td><td>MP</td></tr>
<tr><td>>hg38_21_+_8433172_8446622_RNA45SN1_45S_50ntFlanks@5492.5521.30|AGAAGACGGTCGAACTTGACTATCTAGAGG</td><td>>hg38_21_+_8433172_8446622_RNA45SN1_45S_50ntFlanks</td><td>5492</td><td>5521</td><td>30</td><td>AGAAGACGGTCGAACTTGACTATCTAGAGG</td><td>281.7</td><td>-5.4</td><td>1.1</td><td>-4.7</td><td>2.6E-06</td><td>4.7E-06</td><td>MB231/MB468</td><td>MP</td></tr>
<tr><td>>hg38_21_+_8433172_8446622_RNA45SN1_45S_50ntFlanks@11594.11612.19|CAGGGGAATCCGACTGTTT</td><td>>hg38_21_+_8433172_8446622_RNA45SN1_45S_50ntFlanks</td><td>11594</td><td>11612</td><td>19</td><td>CAGGGGAATCCGACTGTTT</td><td>844.3</td><td>-7.1</td><td>1.5</td><td>-4.7</td><td>2.6E-06</td><td>4.7E-06</td><td>MB231/MB468</td><td>MP</td></tr>
<tr><td>>hg38_21_+_8433172_8446622_RNA45SN1_45S_50ntFlanks@9913.9947.35|AGAAAAGGTGTTGGTTGATATAGACAGCAGGACGG</td><td>>hg38_21_+_8433172_8446622_RNA45SN1_45S_50ntFlanks</td><td>9913</td><td>9947</td><td>35</td><td>AGAAAAGGTGTTGGTTGATATAGACAGCAGGACGG</td><td>59.5</td><td>9.6</td><td>2.0</td><td>4.9</td><td>8.8E-07</td><td>4.7E-06</td><td>BT20/MB468</td><td>MP</td></tr>
<tr><td>>hg38_21_+_8433172_8446622_RNA45SN1_45S_50ntFlanks@12307.12326.20|ACGATCCTTCTGACCTTTTG</td><td>>hg38_21_+_8433172_8446622_RNA45SN1_45S_50ntFlanks</td><td>12307</td><td>12326</td><td>20</td><td>ACGATCCTTCTGACCTTTTG</td><td>48.4</td><td>-8.7</td><td>1.8</td><td>-4.9</td><td>8.8E-07</td><td>4.7E-06</td><td>BT20/MB468</td><td>MP</td></tr>
<tr><td>>hg38_21_+_8433172_8446622_RNA45SN1_45S_50ntFlanks@3996.4022.27|ACTCTAGATAACCTCGGGCCGATCGCA</td><td>>hg38_21_+_8433172_8446622_RNA45SN1_45S_50ntFlanks</td><td>3996</td><td>4022</td><td>27</td><td>ACTCTAGATAACCTCGGGCCGATCGCA</td><td>871.2</td><td>-8.1</td><td>1.7</td><td>-4.7</td><td>3.3E-06</td><td>4.7E-06</td><td>BT20/MB468</td><td>NUC</td></tr>
<tr><td>>hg38_21_+_8433172_8446622_RNA45SN1_45S_50ntFlanks@6645.6673.29|CTCGTACGACTCTTAGCGGTGGATCACTC</td><td>>hg38_21_+_8433172_8446622_RNA45SN1_45S_50ntFlanks</td><td>6645</td><td>6673</td><td>29</td><td>CTCGTACGACTCTTAGCGGTGGATCACTC</td><td>265.1</td><td>4.0</td><td>0.9</td><td>4.7</td><td>3.1E-06</td><td>4.7E-06</td><td>MB231/MB468</td><td>NUC</td></tr>
<tr><td>>hg38_21_+_8433172_8446622_RNA45SN1_45S_50ntFlanks@9714.9737.24|CGAATGATTAGAGGTCTTGGGGCC</td><td>>hg38_21_+_8433172_8446622_RNA45SN1_45S_50ntFlanks</td><td>9714</td><td>9737</td><td>24</td><td>CGAATGATTAGAGGTCTTGGGGCC</td><td>830.6</td><td>-4.9</td><td>1.0</td><td>-4.7</td><td>3.0E-06</td><td>4.7E-06</td><td>MB231/MB468</td><td>CYTO</td></tr>
<tr><td>>hg38_21_+_8433172_8446622_RNA45SN1_45S_50ntFlanks@10355.10377.23|AACTTTGAAGGCCGAAGTGGAGA</td><td>>hg38_21_+_8433172_8446622_RNA45SN1_45S_50ntFlanks</td><td>10355</td><td>10377</td><td>23</td><td>AACTTTGAAGGCCGAAGTGGAGA</td><td>179.9</td><td>-7.2</td><td>1.5</td><td>-4.7</td><td>2.6E-06</td><td>4.7E-06</td><td>MB231/MB468</td><td>MP</td></tr>
<tr><td>>hg38_21_+_8433172_8446622_RNA45SN1_45S_50ntFlanks@4730.4765.36|CAAGAACGAAAGTCGGAGGTTCGAAGACGATCAGAT</td><td>>hg38_21_+_8433172_8446622_RNA45SN1_45S_50ntFlanks</td><td>4730</td><td>4765</td><td>36</td><td>CAAGAACGAAAGTCGGAGGTTCGAAGACGATCAGAT</td><td>268.1</td><td>4.4</td><td>0.9</td><td>4.9</td><td>8.8E-07</td><td>4.8E-06</td><td>BT20/MB468</td><td>MP</td></tr>
<tr><td>>MI0001445|hsa-mir-423&WithFlank&17|+|30117073|30117178@23.45.23(+1C)&[MIMAT0004748&hsa-miR-423-5p&offsets|0|0(+1C);m-141&17|+|30117095|30117117&offsets|0|0(+1C)]|TGAGGGGCAGAGAGCGAGACTTTC</td><td>>MI0001445|hsa-mir-423&WithFlank&17|+|30117073|30117178</td><td>23</td><td>45</td><td>23(+1C)</td><td>TGAGGGGCAGAGAGCGAGACTTTC</td><td>565.5</td><td>6.7</td><td>1.4</td><td>4.9</td><td>8.9E-07</td><td>4.8E-06</td><td>BT20/MB468</td><td>MP</td></tr>
<tr><td>>hg38_21_+_8433172_8446622_RNA45SN1_45S_50ntFlanks@7975.8009.35|CGCGACCTCAGATCAGACGTGGCGACCCGCTGAAT</td><td>>hg38_21_+_8433172_8446622_RNA45SN1_45S_50ntFlanks</td><td>7975</td><td>8009</td><td>35</td><td>CGCGACCTCAGATCAGACGTGGCGACCCGCTGAAT</td><td>51592.7</td><td>3.1</td><td>0.7</td><td>4.7</td><td>3.1E-06</td><td>4.8E-06</td><td>MB231/MB468</td><td>NUC</td></tr>
<tr><td>>hg38_21_+_8433172_8446622_RNA45SN1_45S_50ntFlanks@9942.9978.37|GGACGGTGGCCATGGAAGTCGGAATCCGCTAAGGAGT</td><td>>hg38_21_+_8433172_8446622_RNA45SN1_45S_50ntFlanks</td><td>9942</td><td>9978</td><td>37</td><td>GGACGGTGGCCATGGAAGTCGGAATCCGCTAAGGAGT</td><td>45.9</td><td>9.3</td><td>1.9</td><td>4.9</td><td>8.9E-07</td><td>4.8E-06</td><td>BT20/MB468</td><td>MP</td></tr>
<tr><td>>hg38_21_+_8433172_8446622_RNA45SN1_45S_50ntFlanks@4722.4748.27|TCATTAATCAAGAACGAAAGTCGGAGG</td><td>>hg38_21_+_8433172_8446622_RNA45SN1_45S_50ntFlanks</td><td>4722</td><td>4748</td><td>27</td><td>TCATTAATCAAGAACGAAAGTCGGAGG</td><td>385.9</td><td>-7.1</td><td>1.4</td><td>-4.9</td><td>8.9E-07</td><td>4.8E-06</td><td>BT20/MB468</td><td>MP</td></tr>
<tr><td>>hg38_21_+_8433172_8446622_RNA45SN1_45S_50ntFlanks@12993.13039.47|CCCTCGCTGCGATCTATTGAAAGTCAGCCCTCGACACAAGGGTTTGT</td><td>>hg38_21_+_8433172_8446622_RNA45SN1_45S_50ntFlanks</td><td>12993</td><td>13039</td><td>47</td><td>CCCTCGCTGCGATCTATTGAAAGTCAGCCCTCGACACAAGGGTTTGT</td><td>4838.0</td><td>-5.9</td><td>1.3</td><td>-4.7</td><td>3.3E-06</td><td>4.8E-06</td><td>BT20/MB468</td><td>NUC</td></tr>
<tr><td>>hg38_21_+_8433172_8446622_RNA45SN1_45S_50ntFlanks@5424.5466.43|TTGGATGGTTTAGTGAGGCCCTCGGATCGGCCCCGCCGGGGTC</td><td>>hg38_21_+_8433172_8446622_RNA45SN1_45S_50ntFlanks</td><td>5424</td><td>5466</td><td>43</td><td>TTGGATGGTTTAGTGAGGCCCTCGGATCGGCCCCGCCGGGGTC</td><td>388.9</td><td>-6.9</td><td>1.5</td><td>-4.7</td><td>3.1E-06</td><td>4.8E-06</td><td>MB231/MB468</td><td>NUC</td></tr>
<tr><td>>hg38_21_+_8433172_8446622_RNA45SN1_45S_50ntFlanks@11236.11254.19|TTCCCGTGGATCGCCCCAG</td><td>>hg38_21_+_8433172_8446622_RNA45SN1_45S_50ntFlanks</td><td>11236</td><td>11254</td><td>19</td><td>TTCCCGTGGATCGCCCCAG</td><td>56.6</td><td>-8.9</td><td>1.8</td><td>-4.9</td><td>8.9E-07</td><td>4.8E-06</td><td>BT20/MB468</td><td>MP</td></tr>
<tr><td>>hg38_21_+_8433172_8446622_RNA45SN1_45S_50ntFlanks@12155.12182.28|GGAGTTTGACTGGGGCGGTACACCTGTC</td><td>>hg38_21_+_8433172_8446622_RNA45SN1_45S_50ntFlanks</td><td>12155</td><td>12182</td><td>28</td><td>GGAGTTTGACTGGGGCGGTACACCTGTC</td><td>516.9</td><td>-7.2</td><td>1.5</td><td>-4.7</td><td>2.6E-06</td><td>4.8E-06</td><td>MB231/MB468</td><td>MP</td></tr>
<tr><td>>hg38_21_+_8433172_8446622_RNA45SN1_45S_50ntFlanks@4752.4786.35|GAAGACGATCAGATACCGTCGTAGTTCCGACCATA</td><td>>hg38_21_+_8433172_8446622_RNA45SN1_45S_50ntFlanks</td><td>4752</td><td>4786</td><td>35</td><td>GAAGACGATCAGATACCGTCGTAGTTCCGACCATA</td><td>412.6</td><td>-5.4</td><td>1.2</td><td>-4.7</td><td>3.2E-06</td><td>4.8E-06</td><td>MB231/MB468</td><td>MITO</td></tr>
<tr><td>>hg38_21_+_8433172_8446622_RNA45SN1_45S_50ntFlanks@8241.8261.21|GGGTCTTCCCGGAGTCGGGTT</td><td>>hg38_21_+_8433172_8446622_RNA45SN1_45S_50ntFlanks</td><td>8241</td><td>8261</td><td>21</td><td>GGGTCTTCCCGGAGTCGGGTT</td><td>155.8</td><td>10.6</td><td>2.1</td><td>5.2</td><td>2.3E-07</td><td>4.8E-06</td><td>BT20/MB231</td><td>MP</td></tr>
</table>

<table>
<tr><td>>hg38_21_+_8433172_8446622_RNA45SN1_45S_50ntFlanks@10878.10904.27|TCGGTCGGGCTGGGGCGCGAAGCGGGG</td><td>>hg38_21_+_8433172_8446622_RNA45SN1_45S_50ntFlanks</td><td>10878</td><td>10904</td><td>27</td><td>TCGGTCGGGCTGGGGCGCGAAGCGGGG</td><td>422.9</td><td>-7.4</td><td>1.6</td><td>-4.6</td><td>3.3E-06</td><td>4.8E-06</td><td>BT20/MB468</td><td>NUC</td></tr>
<tr><td>>MI0000477|hsa-mir-146a&WithFlank&5|+|160485346|160485456@27.48.22(+2A)&[MIMAT0000449&hsa-miR-146a-5p&offsets|0|0(+2A);m-129&5|+|160485372|160485393&offsets|0|0(+2A)]|TGAGAACTGAATTCCATGGGTTAA</td><td>>MI0000477|hsa-mir-146a&WithFlank&5|+|160485346|160485456</td><td>27</td><td>48</td><td>22(+2A)</td><td>TGAGAACTGAATTCCATGGGTTAA</td><td>45.5</td><td>8.7</td><td>1.9</td><td>4.7</td><td>2.6E-06</td><td>4.8E-06</td><td>MB231/MB468</td><td>MP</td></tr>
<tr><td>>MI0000477|hsa-mir-146a&WithFlank&5|+|160485346|160485456@26.47.22(+1C)&[MIMAT0000449&hsa-miR-146a-5p&offsets|-1|-1(+1C);m-129&5|+|160485372|160485393&offsets|-1|-1(+1C)]|TTGAGAACTGAATTCCATGGGTC</td><td>>MI0000477|hsa-mir-146a&WithFlank&5|+|160485346|160485456</td><td>26</td><td>47</td><td>22(+1C)</td><td>TTGAGAACTGAATTCCATGGGTC</td><td>55.5</td><td>9.0</td><td>1.9</td><td>4.7</td><td>2.6E-06</td><td>4.8E-06</td><td>MB231/MB468</td><td>MP</td></tr>
<tr><td>>MI0005561|hsa-mir-877&WithFlank&6|+|30584326|30584423@7.26.20(+1U)&[MIMAT0004949&hsa-miR-877-5p&offsets|0|0(+1U);m-382&6|+|30584332|30584354&offsets|0|-3(+1U)]|GTAGAGGAGATGGCGCAGGGT</td><td>>MI0005561|hsa-mir-877&WithFlank&6|+|30584326|30584423</td><td>7</td><td>26</td><td>20(+1U)</td><td>GTAGAGGAGATGGCGCAGGGT</td><td>42.5</td><td>8.6</td><td>1.8</td><td>4.7</td><td>2.6E-06</td><td>4.8E-06</td><td>MB231/MB468</td><td>MP</td></tr>
<tr><td>>hg38_21_+_8433172_8446622_RNA45SN1_45S_50ntFlanks@12184.12201.18|AACGGTAACGCAGGTGTC</td><td>>hg38_21_+_8433172_8446622_RNA45SN1_45S_50ntFlanks</td><td>12184</td><td>12201</td><td>18</td><td>AACGGTAACGCAGGTGTC</td><td>59.3</td><td>-9.0</td><td>1.8</td><td>-4.9</td><td>8.9E-07</td><td>4.8E-06</td><td>BT20/MB468</td><td>MP</td></tr>
<tr><td>>hg38_21_+_8433172_8446622_RNA45SN1_45S_50ntFlanks@8176.8195.20|CTTCTGATCGAGGCCCAGCC</td><td>>hg38_21_+_8433172_8446622_RNA45SN1_45S_50ntFlanks</td><td>8176</td><td>8195</td><td>20</td><td>CTTCTGATCGAGGCCCAGCC</td><td>1453.2</td><td>-3.8</td><td>0.8</td><td>-4.7</td><td>2.6E-06</td><td>4.8E-06</td><td>MB231/MB468</td><td>MP</td></tr>
<tr><td>>hg38_21_+_8433172_8446622_RNA45SN1_45S_50ntFlanks@3862.3898.37|AATTCTAGAGCTAATACATGCCGACGGGCGCTGACCC</td><td>>hg38_21_+_8433172_8446622_RNA45SN1_45S_50ntFlanks</td><td>3862</td><td>3898</td><td>37</td><td>AATTCTAGAGCTAATACATGCCGACGGGCGCTGACCC</td><td>105.3</td><td>-10.6</td><td>2.3</td><td>-4.7</td><td>2.6E-06</td><td>4.8E-06</td><td>MB231/MB468</td><td>MP</td></tr>
<tr><td>>hg38_21_+_8433172_8446622_RNA45SN1_45S_50ntFlanks@12163.12181.19|ACTGGGGCGGTACACCTGT</td><td>>hg38_21_+_8433172_8446622_RNA45SN1_45S_50ntFlanks</td><td>12163</td><td>12181</td><td>19</td><td>ACTGGGGCGGTACACCTGT</td><td>249.1</td><td>-4.5</td><td>1.0</td><td>-4.7</td><td>2.6E-06</td><td>4.8E-06</td><td>BT20/MB468</td><td>CYTO</td></tr>
<tr><td>>hg38_21_+_8433172_8446622_RNA45SN1_45S_50ntFlanks@11207.11254.48|GGCGGCGACTCTGGACGCGAGCCGGGCCCTTCCCGTGGATCGCCCCAG</td><td>>hg38_21_+_8433172_8446622_RNA45SN1_45S_50ntFlanks</td><td>11207</td><td>11254</td><td>48</td><td>GGCGGCGACTCTGGACGCGAGCCGGGCCCTTCCCGTGGATCGCCCCAG</td><td>1480.7</td><td>-7.8</td><td>1.7</td><td>-4.7</td><td>3.1E-06</td><td>4.8E-06</td><td>MB231/MB468</td><td>NUC</td></tr>
<tr><td>>hg38_21_+_8433172_8446622_RNA45SN1_45S_50ntFlanks@9534.9558.25|AGGAAACTCTGGTGGAGGTCCGTAG</td><td>>hg38_21_+_8433172_8446622_RNA45SN1_45S_50ntFlanks</td><td>9534</td><td>9558</td><td>25</td><td>AGGAAACTCTGGTGGAGGTCCGTAG</td><td>211.3</td><td>-5.0</td><td>1.1</td><td>-4.7</td><td>3.1E-06</td><td>4.8E-06</td><td>MB231/MB468</td><td>NUC</td></tr>
<tr><td>>hg38_21_+_8433172_8446622_RNA45SN1_45S_50ntFlanks@9835.9864.30|TTTGGTAAGCAGAACTGGCGCTGCGGGATG</td><td>>hg38_21_+_8433172_8446622_RNA45SN1_45S_50ntFlanks</td><td>9835</td><td>9864</td><td>30</td><td>TTTGGTAAGCAGAACTGGCGCTGCGGGATG</td><td>182.1</td><td>10.8</td><td>2.1</td><td>5.2</td><td>2.3E-07</td><td>4.8E-06</td><td>BT20/MB231</td><td>MP</td></tr>
<tr><td>>trna17_ValTAC_11_-_59318102_59318174@3.36.34,trna4_ValTAC_X_-_18693029_18693101@3.36.34|TTCCATAGTGTAGTGGTTATCACGTCTGCTTTAC</td><td>>trna17_ValTAC_11_-_59318102_59318174</td><td>3</td><td>36</td><td>34</td><td>TTCCATAGTGTAGTGGTTATCACGTCTGCTTTAC</td><td>196.2</td><td>-4.8</td><td>0.9</td><td>-5.1</td><td>2.8E-07</td><td>4.8E-06</td><td>BT20/MB231</td><td>MITO</td></tr>
<tr><td>>hg38_21_+_8433172_8446622_RNA45SN1_45S_50ntFlanks@5495.5516.22|AGACGGTCGAACTTGACTATCT</td><td>>hg38_21_+_8433172_8446622_RNA45SN1_45S_50ntFlanks</td><td>5495</td><td>5516</td><td>22</td><td>AGACGGTCGAACTTGACTATCT</td><td>1075.1</td><td>4.3</td><td>0.8</td><td>5.1</td><td>2.9E-07</td><td>4.8E-06</td><td>BT20/MB231</td><td>MITO</td></tr>
<tr><td>>trnaMT_PheGAA_MT_+_577_647@35.60.26|GAAAATGTTTAGACGGGCTCACATCA</td><td>>trnaMT_PheGAA_MT_+_577_647</td><td>35</td><td>60</td><td>26</td><td>GAAAATGTTTAGACGGGCTCACATCA</td><td>543.3</td><td>6.5</td><td>1.3</td><td>4.9</td><td>9.0E-07</td><td>4.8E-06</td><td>BT20/MB468</td><td>MP</td></tr>
<tr><td>>MI0025242|hsa-mir-7706&WithFlank&15|+|85380590|85380668@50.71.22&[MIMAT0030021&hsa-miR-7706&offsets|0|-2;m-390&15|+|85380639|85380662&offsets|0|-2]|TGAAGCGCCTGTGCTCTGCCGA</td><td>>MI0025242|hsa-mir-7706&WithFlank&15|+|85380590|85380668</td><td>50</td><td>71</td><td>22</td><td>TGAAGCGCCTGTGCTCTGCCGA</td><td>578.0</td><td>7.2</td><td>1.5</td><td>4.7</td><td>2.7E-06</td><td>4.8E-06</td><td>BT20/MB468</td><td>CYTO</td></tr>
<tr><td>>MI0000063|hsa-let-7b&WithFlank&22|+|46113680|46113774@12.32.21(+1A)&[MIMAT0000063&hsa-let-7b-5p&offsets|0|-1(+1A);m-14&22|+|46113691|46113712&offsets|0|-1(+1A)]|TGAGGTAGTAGGTTGTGTGGTA</td><td>>MI0000063|hsa-let-7b&WithFlank&22|+|46113680|46113774</td><td>12</td><td>32</td><td>21(+1A)</td><td>TGAGGTAGTAGGTTGTGTGGTA</td><td>325.1</td><td>3.1</td><td>0.7</td><td>4.7</td><td>3.1E-06</td><td>4.8E-06</td><td>MB231/MB468</td><td>NUC</td></tr>
<tr><td>>hg38_21_+_8433172_8446622_RNA45SN1_45S_50ntFlanks@475.509.35|CTTCGTGATCGATGTGGTGACGTCGTGCTCTCCCG</td><td>>hg38_21_+_8433172_8446622_RNA45SN1_45S_50ntFlanks</td><td>475</td><td>509</td><td>35</td><td>CTTCGTGATCGATGTGGTGACGTCGTGCTCTCCCG</td><td>9696.6</td><td>3.4</td><td>0.7</td><td>4.7</td><td>3.1E-06</td><td>4.8E-06</td><td>MB231/MB468</td><td>NUC</td></tr>
<tr><td>>hg38_21_+_8433172_8446622_RNA45SN1_45S_50ntFlanks@8552.8579.28|CTCCCCGGAGGGGGCGGGCTCCGGCGGG</td><td>>hg38_21_+_8433172_8446622_RNA45SN1_45S_50ntFlanks</td><td>8552</td><td>8579</td><td>28</td><td>CTCCCCGGAGGGGGCGGGCTCCGGCGGG</td><td>114.8</td><td>-9.9</td><td>2.0</td><td>-4.9</td><td>9.0E-07</td><td>4.8E-06</td><td>BT20/MB468</td><td>MP</td></tr>
<tr><td>>hg38_21_+_8433172_8446622_RNA45SN1_45S_50ntFlanks@12410.12433.24|TGATCCTTCGATGTCGGCTCTTCC</td><td>>hg38_21_+_8433172_8446622_RNA45SN1_45S_50ntFlanks</td><td>12410</td><td>12433</td><td>24</td><td>TGATCCTTCGATGTCGGCTCTTCC</td><td>516.1</td><td>4.2</td><td>0.9</td><td>4.7</td><td>3.0E-06</td><td>4.8E-06</td><td>MB231/MB468</td><td>CYTO</td></tr>
<tr><td>>hg38_21_+_8433172_8446622_RNA45SN1_45S_50ntFlanks@5083.5115.33|ACGAACGAGACTCTGGCATGCTAACTAGTTACG</td><td>>hg38_21_+_8433172_8446622_RNA45SN1_45S_50ntFlanks</td><td>5083</td><td>5115</td><td>33</td><td>ACGAACGAGACTCTGGCATGCTAACTAGTTACG</td><td>159.5</td><td>-6.8</td><td>1.5</td><td>-4.7</td><td>2.7E-06</td><td>4.8E-06</td><td>MB231/MB468</td><td>MP</td></tr>
<tr><td>>hg38_21_+_8433172_8446622_RNA45SN1_45S_50ntFlanks@9562.9596.35|TCCTGACGTGCAAATCGGTCGTCCGACCTGGGTAT</td><td>>hg38_21_+_8433172_8446622_RNA45SN1_45S_50ntFlanks</td><td>9562</td><td>9596</td><td>35</td><td>TCCTGACGTGCAAATCGGTCGTCCGACCTGGGTAT</td><td>942.5</td><td>-6.9</td><td>1.5</td><td>-4.7</td><td>2.7E-06</td><td>4.8E-06</td><td>BT20/MB468</td><td>CYTO</td></tr>
<tr><td>>hg38_21_+_8433172_8446622_RNA45SN1_45S_50ntFlanks@12510.12546.37|TCGTGAGACAGGTTAGTTTTACCCTACTGATGATGTG</td><td>>hg38_21_+_8433172_8446622_RNA45SN1_45S_50ntFlanks</td><td>12510</td><td>12546</td><td>37</td><td>TCGTGAGACAGGTTAGTTTTACCCTACTGATGATGTG</td><td>61.4</td><td>-9.0</td><td>1.8</td><td>-4.9</td><td>9.0E-07</td><td>4.8E-06</td><td>BT20/MB468</td><td>MP</td></tr>
<tr><td>>hg38_MT_+_1621_3279_RNR2_16S_50ntFlanks@1095.1120.26|AGACGAGAAGACCCTATGGAGCTTTA</td><td>>hg38_MT_+_1621_3279_RNR2_16S_50ntFlanks</td><td>1095</td><td>1120</td><td>26</td><td>AGACGAGAAGACCCTATGGAGCTTTA</td><td>584.0</td><td>6.1</td><td>1.3</td><td>4.7</td><td>2.3E-06</td><td>4.8E-06</td><td>BT20/MB468</td><td>MITO</td></tr>
<tr><td>>MI0000482|hsa-mir-185&WithFlank&22|+|20033133|20033226@21.42.22(+1A)&[MIMAT0000455&hsa-miR-185-5p&offsets|0|0(+1A);m-145&22|+|20033153|20033174&offsets|0|0(+1A)]|TGGAGAGAAAGGCAGTTCCTGAA</td><td>>MI0000482|hsa-mir-185&WithFlank&22|+|20033133|20033226</td><td>21</td><td>42</td><td>22(+1A)</td><td>TGGAGAGAAAGGCAGTTCCTGAA</td><td>400.2</td><td>4.1</td><td>0.8</td><td>5.2</td><td>2.5E-07</td><td>4.8E-06</td><td>BT20/MB231</td><td>CYTO</td></tr>
<tr><td>>hg38_21_+_8433172_8446622_RNA45SN1_45S_50ntFlanks@10572.10598.27|AGAAGCCGGCGGGAGCCCCGGGGAGAG</td><td>>hg38_21_+_8433172_8446622_RNA45SN1_45S_50ntFlanks</td><td>10572</td><td>10598</td><td>27</td><td>AGAAGCCGGCGGGAGCCCCGGGGAGAG</td><td>51.8</td><td>-8.8</td><td>1.8</td><td>-4.9</td><td>9.0E-07</td><td>4.8E-06</td><td>BT20/MB468</td><td>MP</td></tr>
<tr><td>>hg38_21_+_8433172_8446622_RNA45SN1_45S_50ntFlanks@11565.11600.36|CTAGCAGCCGACTTAGAACTGGTGCGGACCAGGGGA</td><td>>hg38_21_+_8433172_8446622_RNA45SN1_45S_50ntFlanks</td><td>11565</td><td>11600</td><td>36</td><td>CTAGCAGCCGACTTAGAACTGGTGCGGACCAGGGGA</td><td>127.8</td><td>-5.3</td><td>1.1</td><td>-4.7</td><td>2.7E-06</td><td>4.8E-06</td><td>MB231/MB468</td><td>MP</td></tr>
<tr><td>>trnaMT_PheGAA_MT_+_577_647@35.54.20|GAAAATGTTTAGACGGGCTC</td><td>>trnaMT_PheGAA_MT_+_577_647</td><td>35</td><td>54</td><td>20</td><td>GAAAATGTTTAGACGGGCTC</td><td>873.4</td><td>5.1</td><td>1.0</td><td>4.9</td><td>9.1E-07</td><td>4.8E-06</td><td>BT20/MB468</td><td>MP</td></tr>
<tr><td>>hg38_21_+_8433172_8446622_RNA45SN1_45S_50ntFlanks@7975.8016.42|CGCGACCTCAGATCAGACGTGGCGACCCGCTGAATTTAAGCA</td><td>>hg38_21_+_8433172_8446622_RNA45SN1_45S_50ntFlanks</td><td>7975</td><td>8016</td><td>42</td><td>CGCGACCTCAGATCAGACGTGGCGACCCGCTGAATTTAAGCA</td><td>13615.6</td><td>5.0</td><td>1.1</td><td>4.7</td><td>3.1E-06</td><td>4.9E-06</td><td>MB231/MB468</td><td>NUC</td></tr>
<tr><td>>hg38_21_+_8433172_8446622_RNA45SN1_45S_50ntFlanks@6661.6689.29|CGGTGGATCACTCGGCTCGTGCGTCGATG</td><td>>hg38_21_+_8433172_8446622_RNA45SN1_45S_50ntFlanks</td><td>6661</td><td>6689</td><td>29</td><td>CGGTGGATCACTCGGCTCGTGCGTCGATG</td><td>13471.3</td><td>4.2</td><td>0.8</td><td>5.1</td><td>2.9E-07</td><td>4.9E-06</td><td>BT20/MB231</td><td>MITO</td></tr>
<tr><td>>hg38_21_+_8433172_8446622_RNA45SN1_45S_50ntFlanks@11670.11700.31|CCCAGTGCTCTGAATGTCAAAGTGAAGAAAT</td><td>>hg38_21_+_8433172_8446622_RNA45SN1_45S_50ntFlanks</td><td>11670</td><td>11700</td><td>31</td><td>CCCAGTGCTCTGAATGTCAAAGTGAAGAAAT</td><td>150.0</td><td>-2.2</td><td>0.5</td><td>-4.7</td><td>2.7E-06</td><td>4.9E-06</td><td>BT20/MB468</td><td>CYTO</td></tr>
<tr><td>>hg38_21_+_8433172_8446622_RNA45SN1_45S_50ntFlanks@8323.8354.32|ACCGATAGTCAACAAGTACCGTAAGGGAAAGT</td><td>>hg38_21_+_8433172_8446622_RNA45SN1_45S_50ntFlanks</td><td>8323</td><td>8354</td><td>32</td><td>ACCGATAGTCAACAAGTACCGTAAGGGAAAGT</td><td>146.7</td><td>-5.4</td><td>1.2</td><td>-4.7</td><td>2.7E-06</td><td>4.9E-06</td><td>BT20/MB468</td><td>CYTO</td></tr>
<tr><td>>hg38_MT_+_1621_3279_RNR2_16S_50ntFlanks@498.530.33|TTGGACACTAGGAAAAAACCTTGTAGAGAGAGT</td><td>>hg38_MT_+_1621_3279_RNR2_16S_50ntFlanks</td><td>498</td><td>530</td><td>33</td><td>TTGGACACTAGGAAAAAACCTTGTAGAGAGAGT</td><td>53.7</td><td>9.5</td><td>1.9</td><td>4.9</td><td>9.1E-07</td><td>4.9E-06</td><td>BT20/MB468</td><td>MP</td></tr>
<tr><td>>hg38_21_+_8433172_8446622_RNA45SN1_45S_50ntFlanks@8354.8388.35|TTGAAAAGAACTTTGAAGAGAGAGTTCAAGAGGGC</td><td>>hg38_21_+_8433172_8446622_RNA45SN1_45S_50ntFlanks</td><td>8354</td><td>8388</td><td>35</td><td>TTGAAAAGAACTTTGAAGAGAGAGTTCAAGAGGGC</td><td>45.9</td><td>9.3</td><td>1.9</td><td>4.9</td><td>9.1E-07</td><td>4.9E-06</td><td>BT20/MB468</td><td>MP</td></tr>
<tr><td>>trna10_LysCTT_16_+_3241501_3241573@1.30.30,trna13_LysCTT_14_-_58706613_58706685@1.30.30,trna2_LysCTT_15_+_79152904_79152976@1.30.30|GCCCGGCTAGCTCAGTCGGTAGAGCATGGG</td><td>>trna10_LysCTT_16_+_3241501_3241573</td><td>1</td><td>30</td><td>30</td><td>GCCCGGCTAGCTCAGTCGGTAGAGCATGGG</td><td>70.8</td><td>-9.2</td><td>1.9</td><td>-4.9</td><td>9.1E-07</td><td>4.9E-06</td><td>BT20/MB468</td><td>MP</td></tr>
<tr><td>>hg38_1_-_228634819_228635039_RNA5S12_5S_50ntFlanks@136.170.35|GGAGACCGCCTGGGAATACCGGGTGCTGTAGGCTT</td><td>>hg38_1_-_228634819_228635039_RNA5S12_5S_50ntFlanks</td><td>136</td><td>170</td><td>35</td><td>GGAGACCGCCTGGGAATACCGGGTGCTGTAGGCTT</td><td>2246.7</td><td>4.8</td><td>1.0</td><td>4.7</td><td>2.7E-06</td><td>4.9E-06</td><td>BT20/MB468</td><td>CYTO</td></tr>
<tr><td>>hg38_21_+_8433172_8446622_RNA45SN1_45S_50ntFlanks@10408.10434.27|GGTCAGTCGGTCCTGAGAGATGGGCGA</td><td>>hg38_21_+_8433172_8446622_RNA45SN1_45S_50ntFlanks</td><td>10408</td><td>10434</td><td>27</td><td>GGTCAGTCGGTCCTGAGAGATGGGCGA</td><td>375.1</td><td>-6.8</td><td>1.5</td><td>-4.7</td><td>3.2E-06</td><td>4.9E-06</td><td>MB231/MB468</td><td>NUC</td></tr>
<tr><td>>trna11_GluTTC_15_-_26327381_26327452@1.30.30,trna3_GluTTC_13_-_45492062_45492133@1.30.30|TCCCACATGGTCTAGCGGTTAGGATTCCTG</td><td>>trna11_GluTTC_15_-_26327381_26327452</td><td>1</td><td>30</td><td>30</td><td>TCCCACATGGTCTAGCGGTTAGGATTCCTG</td><td>2059.8</td><td>-4.3</td><td>0.9</td><td>-4.7</td><td>2.7E-06</td><td>4.9E-06</td><td>MB231/MB468</td><td>MP</td></tr>
<tr><td>>hg38_21_+_8433172_8446622_RNA45SN1_45S_50ntFlanks@4047.4064.18|ATTCGAACGTCTGCCCTA</td><td>>hg38_21_+_8433172_8446622_RNA45SN1_45S_50ntFlanks</td><td>4047</td><td>4064</td><td>18</td><td>ATTCGAACGTCTGCCCTA</td><td>428.8</td><td>-6.8</td><td>1.4</td><td>-4.7</td><td>2.3E-06</td><td>4.9E-06</td><td>BT20/MB468</td><td>MITO</td></tr>
<tr><td>>hg38_21_+_8433172_8446622_RNA45SN1_45S_50ntFlanks@9563.9597.35|CCTGACGTGCAAATCGGTCGTCCGACCTGGGTATA</td><td>>hg38_21_+_8433172_8446622_RNA45SN1_45S_50ntFlanks</td><td>9563</td><td>9597</td><td>35</td><td>CCTGACGTGCAAATCGGTCGTCCGACCTGGGTATA</td><td>530.3</td><td>-4.6</td><td>1.0</td><td>-4.7</td><td>3.0E-06</td><td>4.9E-06</td><td>MB231/MB468</td><td>CYTO</td></tr>
<tr><td>>hg38_21_+_8433172_8446622_RNA45SN1_45S_50ntFlanks@11209.11252.44|CGGCGACTCTGGACGCGAGCCGGGCCCTTCCCGTGGATCGCCCC</td><td>>hg38_21_+_8433172_8446622_RNA45SN1_45S_50ntFlanks</td><td>11209</td><td>11252</td><td>44</td><td>CGGCGACTCTGGACGCGAGCCGGGCCCTTCCCGTGGATCGCCCC</td><td>3699.3</td><td>-7.2</td><td>1.5</td><td>-4.6</td><td>3.4E-06</td><td>4.9E-06</td><td>BT20/MB468</td><td>NUC</td></tr>
<tr><td>>hg38_21_+_8433172_8446622_RNA45SN1_45S_50ntFlanks@8841.8859.19|GTGGCGGCGGCGGCGGCGG</td><td>>hg38_21_+_8433172_8446622_RNA45SN1_45S_50ntFlanks</td><td>8841</td><td>8859</td><td>19</td><td>GTGGCGGCGGCGGCGGCGG</td><td>57.8</td><td>-8.9</td><td>1.8</td><td>-4.9</td><td>9.1E-07</td><td>4.9E-06</td><td>BT20/MB468</td><td>MP</td></tr>
<tr><td>>hg38_21_+_8433172_8446622_RNA45SN1_45S_50ntFlanks@10238.10272.35|CGCTACGCCGCGACGAGTAGGAGGGCCGCTGCGGT</td><td>>hg38_21_+_8433172_8446622_RNA45SN1_45S_50ntFlanks</td><td>10238</td><td>10272</td><td>35</td><td>CGCTACGCCGCGACGAGTAGGAGGGCCGCTGCGGT</td><td>267.6</td><td>7.8</td><td>1.6</td><td>4.9</td><td>9.1E-07</td><td>4.9E-06</td><td>BT20/MB468</td><td>MP</td></tr>
<tr><td>>hg38_21_+_8433172_8446622_RNA45SN1_45S_50ntFlanks@11783.11802.20|CATGAATGGATGAACGAGAT</td><td>>hg38_21_+_8433172_8446622_RNA45SN1_45S_50ntFlanks</td><td>11783</td><td>11802</td><td>20</td><td>CATGAATGGATGAACGAGAT</td><td>699.6</td><td>-7.5</td><td>1.5</td><td>-4.9</td><td>9.1E-07</td><td>4.9E-06</td><td>BT20/MB468</td><td>MP</td></tr>
<tr><td>>hg38_21_+_8433172_8446622_RNA45SN1_45S_50ntFlanks@12162.12182.21|GACTGGGGCGGTACACCTGTC</td><td>>hg38_21_+_8433172_8446622_RNA45SN1_45S_50ntFlanks</td><td>12162</td><td>12182</td><td>21</td><td>GACTGGGGCGGTACACCTGTC</td><td>56.2</td><td>-8.9</td><td>1.8</td><td>-4.9</td><td>9.1E-07</td><td>4.9E-06</td><td>BT20/MB468</td><td>MP</td></tr>
<tr><td>>hg38_21_+_8433172_8446622_RNA45SN1_45S_50ntFlanks@5487.5515.29|CGCTGAGAAGACGGTCGAACTTGACTATC</td><td>>hg38_21_+_8433172_8446622_RNA45SN1_45S_50ntFlanks</td><td>5487</td><td>5515</td><td>29</td><td>CGCTGAGAAGACGGTCGAACTTGACTATC</td><td>250.3</td><td>11.3</td><td>2.2</td><td>5.2</td><td>2.4E-07</td><td>4.9E-06</td><td>BT20/MB231</td><td>MP</td></tr>
<tr><td>>hg38_21_+_8433172_8446622_RNA45SN1_45S_50ntFlanks@11317.11343.27|ACCCCACGTCTCGTCGCGCGCGTCC</td><td>>hg38_21_+_8433172_8446622_RNA45SN1_45S_50ntFlanks</td><td>11317</td><td>11343</td><td>27</td><td>ACCCCACGTCTCGTCGCGCGCGCGTCC</td><td>536.2</td><td>-4.4</td><td>1.0</td><td>-4.7</td><td>3.1E-06</td><td>4.9E-06</td><td>MB231/MB468</td><td>CYTO</td></tr>
<tr><td>>MI0001445|hsa-mir-423&WithFlank&17|+|30117073|30117178@59.81.23(+1A)&[MIMAT0001340&hsa-miR-423-3p&offsets|0|0(+1A);m-100&17|+|30117131|30117153&offsets|0|0(+1A)]|AGCTCGGTCTGAGGCCCCTCAGTA</td><td>>MI0001445|hsa-mir-423&WithFlank&17|+|30117073|30117178</td><td>59</td><td>81</td><td>23(+1A)</td><td>AGCTCGGTCTGAGGCCCCTCAGTA</td><td>644.9</td><td>4.2</td><td>0.9</td><td>4.7</td><td>3.2E-06</td><td>4.9E-06</td><td>MB231/MB468</td><td>MITO</td></tr>
<tr><td>>MI0000076|hsa-mir-20a&WithFlank&13|+|91351059|91351141@14.36.23(+1A)&[MIMAT0000075&hsa-miR-20a-5p&offsets|0|0(+1A);m-96&13|+|91351072|91351094&offsets|0|0(+1A)]|TAAAGTGCTTATAGTGCAGGTAGA</td><td>>MI0000076|hsa-mir-20a&WithFlank&13|+|91351059|91351141</td><td>14</td><td>36</td><td>23(+1A)</td><td>TAAAGTGCTTATAGTGCAGGTAGA</td><td>1624.2</td><td>-5.2</td><td>1.0</td><td>-5.2</td><td>2.4E-07</td><td>4.9E-06</td><td>BT20/MB231</td><td>MP</td></tr>
<tr><td>>MI0000289|hsa-mir-181a-1&WithFlank&1|-|198859038|198859159@70.88.19&[MIMAT0000270&hsa-miR-181a-3p&offsets|0|-3;m-125&1|-|198859069|198859090&offsets|0|-3]|ACCATCGACCGTTGATTGT</td><td>>MI0000289|hsa-mir-181a-1&WithFlank&1|-|198859038|198859159</td><td>70</td><td>88</td><td>19</td><td>ACCATCGACCGTTGATTGT</td><td>92.6</td><td>-9.9</td><td>1.9</td><td>-5.2</td><td>2.4E-07</td><td>4.9E-06</td><td>BT20/MB231</td><td>MP</td></tr>
</table>

| | | | | | | | | | | | | | |
|---|---|---|---|---|---|---|---|---|---|---|---|---|---|
| >hg38_21_+_8433172_8446622_RNA45SN1_45S_50ntFlanks@4047.4071.25\|ATTCGAACGTCTGCCCTATCAACTT | >hg38_21_+_8433172_8446622_RNA45SN1_45S_50ntFlanks | 4047 | 4071 | 25 | ATTCGAACGTCTGCCCTATCAACTT | 271.1 | -4.9 | 1.1 | -4.7 | 3.2E-06 | 4.9E-06 | MB231/MB468 | NUC |
| >hg38_21_+_8433172_8446622_RNA45SN1_45S_50ntFlanks@3860.3898.39\|GTAATTCTAGAGCTAATACATGCCGACGGGCGCTGACCC | >hg38_21_+_8433172_8446622_RNA45SN1_45S_50ntFlanks | 3860 | 3898 | 39 | GTAATTCTAGAGCTAATACATGCCGACGGGCGCTGACCC | 197.0 | -6.3 | 1.3 | -4.7 | 2.7E-06 | 4.9E-06 | BT20/MB468 | CYTO |
| >hg38_21_+_8433172_8446622_RNA45SN1_45S_50ntFlanks@12045.12072.28\|CTGACCCGGTGAGGCGGGGGGGCGAGCC | >hg38_21_+_8433172_8446622_RNA45SN1_45S_50ntFlanks | 12045 | 12072 | 28 | CTGACCCGGTGAGGCGGGGGGGCGAGCC | 328.1 | -6.4 | 1.4 | -4.7 | 2.3E-06 | 4.9E-06 | BT20/MB468 | MITO |
| >hg38_21_+_8433172_8446622_RNA45SN1_45S_50ntFlanks@4526.4564.39\|TTCAAAGCAGGCCCGAGCCGCCTGGATACCGCAGCTAGG | >hg38_21_+_8433172_8446622_RNA45SN1_45S_50ntFlanks | 4526 | 4564 | 39 | TTCAAAGCAGGCCCGAGCCGCCTGGATACCGCAGCTAGG | 36.9 | 8.4 | 1.8 | 4.7 | 2.7E-06 | 4.9E-06 | MB231/MB468 | MP |
| >hg38_21_+_8433172_8446622_RNA45SN1_45S_50ntFlanks@9862.9885.24\|ATGAACCGAACGCCGGGTTAAGGC | >hg38_21_+_8433172_8446622_RNA45SN1_45S_50ntFlanks | 9862 | 9885 | 24 | ATGAACCGAACGCCGGGTTAAGGC | 142.5 | 10.5 | 2.0 | 5.2 | 2.4E-07 | 4.9E-06 | BT20/MB231 | MP |
| >hg38_21_+_8433172_8446622_RNA45SN1_45S_50ntFlanks@10186.10209.24\|TCCTCCCCTCCTCCCGCCCACGCC | >hg38_21_+_8433172_8446622_RNA45SN1_45S_50ntFlanks | 10186 | 10209 | 24 | TCCTCCCCTCCTCCCGCCCACGCC | 173.0 | -11.3 | 2.4 | -4.7 | 2.7E-06 | 4.9E-06 | MB231/MB468 | MP |
| >hg38_21_+_8433172_8446622_RNA45SN1_45S_50ntFlanks@12539.12567.29\|ATGATGTGTTGTTGCCATGGTAATCCTGC | >hg38_21_+_8433172_8446622_RNA45SN1_45S_50ntFlanks | 12539 | 12567 | 29 | ATGATGTGTTGTTGCCATGGTAATCCTGC | 514.1 | -2.9 | 0.6 | -4.7 | 2.3E-06 | 4.9E-06 | BT20/MB468 | MITO |
| >hg38_21_+_8433172_8446622_RNA45SN1_45S_50ntFlanks@7986.8015.30\|ATCAGACGTGGCGACCCGCTGAATTTAAGC | >hg38_21_+_8433172_8446622_RNA45SN1_45S_50ntFlanks | 7986 | 8015 | 30 | ATCAGACGTGGCGACCCGCTGAATTTAAGC | 682.7 | -6.9 | 1.5 | -4.7 | 3.3E-06 | 4.9E-06 | MB231/MB468 | MITO |
| >hg38_21_+_8433172_8446622_RNA45SN1_45S_50ntFlanks@8198.8225.28\|TGGACGGTGTGAGGCCGGTAGCGGCCCC | >hg38_21_+_8433172_8446622_RNA45SN1_45S_50ntFlanks | 8198 | 8225 | 28 | TGGACGGTGTGAGGCCGGTAGCGGCCCC | 790.9 | -4.6 | 1.0 | -4.7 | 3.1E-06 | 4.9E-06 | MB231/MB468 | CYTO |
| >MI0000286\|hsa-mir-210&WithFlank&11\|-\|568083\|568204@72.93.22(+1A)&[MIMAT0000267&hsa-miR-210-3p&offsets\|0\|0(+1A);m-102&11\|-\|568112\|568133&offsets\|0\|0(+1A)]\|CTGTGCGTGTGACAGCGGCTGAA | >MI0000286\|hsa-mir-210&WithFlank&11\|-\|568083\|568204 | 72 | 93 | 22(+1A) | CTGTGCGTGTGACAGCGGCTGAA | 300.9 | 8.0 | 1.7 | 4.7 | 3.3E-06 | 4.9E-06 | MB231/MB468 | MITO |
| >trnaMT_HisGTG_MT_+_12138_12206@43.72.30\|ACAGAGGCTTACGACCCCTTATTTACCCCA | >trnaMT_HisGTG_MT_+_12138_12206 | 43 | 72 | 30 | ACAGAGGCTTACGACCCCTTATTTACCCCA | 463.5 | 7.4 | 1.5 | 4.9 | 9.2E-07 | 4.9E-06 | BT20/MB468 | MP |
| >MI0000088\|hsa-mir-30a&WithFlank&6\|-\|71403545\|71403627@53.70.18(+1U)&[MIMAT0000088&hsa-miR-30a-3p&offsets\|0\|-4(+1U);m-38&6\|-\|71403554\|71403575&offsets\|0\|-4(+1U)]\|CTTTCAGTCGGATGTTTGT | >MI0000088\|hsa-mir-30a&WithFlank&6\|-\|71403545\|71403627 | 53 | 70 | 18(+1U) | CTTTCAGTCGGATGTTTGT | 298.9 | 6.7 | 1.4 | 4.7 | 3.3E-06 | 4.9E-06 | MB231/MB468 | MITO |
| >hg38_21_+_8433172_8446622_RNA45SN1_45S_50ntFlanks@12632.12661.30\|GCGAAGCTACCATCTGTGGGATTATGACTG | >hg38_21_+_8433172_8446622_RNA45SN1_45S_50ntFlanks | 12632 | 12661 | 30 | GCGAAGCTACCATCTGTGGGATTATGACTG | 463.4 | -3.0 | 0.6 | -4.7 | 3.2E-06 | 4.9E-06 | MB231/MB468 | NUC |
| >hg38_21_+_8433172_8446622_RNA45SN1_45S_50ntFlanks@4128.4150.23\|CGATTCCGGAGAGGGAGCCTGAG | >hg38_21_+_8433172_8446622_RNA45SN1_45S_50ntFlanks | 4128 | 4150 | 23 | CGATTCCGGAGAGGGAGCCTGAG | 142.2 | -3.0 | 0.6 | -4.7 | 2.7E-06 | 4.9E-06 | MB231/MB468 | MP |
| >hg38_21_+_8433172_8446622_RNA45SN1_45S_50ntFlanks@9482.9516.35\|ACGTGTTAGGACCCGAAAGATGGTGAACTATGCCT | >hg38_21_+_8433172_8446622_RNA45SN1_45S_50ntFlanks | 9482 | 9516 | 35 | ACGTGTTAGGACCCGAAAGATGGTGAACTATGCCT | 46.2 | 9.3 | 1.9 | 4.9 | 9.2E-07 | 4.9E-06 | BT20/MB468 | MP |
| >hg38_21_+_8433172_8446622_RNA45SN1_45S_50ntFlanks@8195.8212.18\|CCGTGGACGGTGTGAGGC | >hg38_21_+_8433172_8446622_RNA45SN1_45S_50ntFlanks | 8195 | 8212 | 18 | CCGTGGACGGTGTGAGGC | 479.8 | 4.2 | 0.9 | 4.7 | 2.7E-06 | 4.9E-06 | BT20/MB468 | CYTO |
| >hg38_1_-_228634819_228635039_RNA5S12_5S_50ntFlanks@140.170.31\|ACCGCCTGGGAATACCGGGTGCTGTAGGCTT | >hg38_1_-_228634819_228635039_RNA5S12_5S_50ntFlanks | 140 | 170 | 31 | ACCGCCTGGGAATACCGGGTGCTGTAGGCTT | 24460.9 | 4.4 | 0.9 | 4.7 | 2.7E-06 | 4.9E-06 | BT20/MB468 | CYTO |
| >hg38_21_+_8433172_8446622_RNA45SN1_45S_50ntFlanks@6719.6739.21\|TGAATTGCAGGACACATTGAT | >hg38_21_+_8433172_8446622_RNA45SN1_45S_50ntFlanks | 6719 | 6739 | 21 | TGAATTGCAGGACACATTGAT | 646.3 | -6.6 | 1.4 | -4.7 | 2.7E-06 | 4.9E-06 | BT20/MB468 | CYTO |
| >MI0002470\|hsa-mir-486-1&WithFlank&8\|-\|41660435\|41660514@10.31.22;MI0023622\|hsa-mir-486-2&WithFlank&8\|+\|41660438\|41660513@7.28.22&[MIMAT0002177&hsa-miR-486-5p&offsets\|0\|0;m-107&8\|-\|41660484\|41660505&offsets\|0\|0];[MIMAT0002177_1&hsa-miR-486-5p&offsets\|0\|0;TJU_CMC_MD2.ID03138.5p-miR&offsets\|0\|-2;m-106&8\|+\|41660444\|41660465&offsets\|0\|0]\|TCCTGTACTGAGCTGCCCCGAG | >MI0002470\|hsa-mir-486-1&WithFlank&8\|-\|41660435\|41660514 | 10 | 31 | 22 | TCCTGTACTGAGCTGCCCCGAG | 154.4 | 7.3 | 1.5 | 4.7 | 2.7E-06 | 4.9E-06 | MB231/MB468 | MP |
| >hg38_21_+_8433172_8446622_RNA45SN1_45S_50ntFlanks@9913.9951.39\|AGAAAAGGTGTTGGTTGATATAGACAGCAGGACGGTGGC | >hg38_21_+_8433172_8446622_RNA45SN1_45S_50ntFlanks | 9913 | 9951 | 39 | AGAAAAGGTGTTGGTTGATATAGACAGCAGGACGGTGGC | 45.4 | 9.3 | 1.9 | 4.9 | 9.3E-07 | 4.9E-06 | BT20/MB468 | MP |
| >MI0001445\|hsa-mir-423&WithFlank&17\|+\|30117073\|30117178@23.44.22(+1G)&[MIMAT0004748&hsa-miR-423-5p&offsets\|0\|-1(+1G);m-141&17\|+\|30117095\|30117117&offsets\|0\|-1(+1G)]\|TGAGGGGCAGAGAGCGAGACTTG | >MI0001445\|hsa-mir-423&WithFlank&17\|+\|30117073\|30117178 | 23 | 44 | 22(+1G) | TGAGGGGCAGAGAGCGAGACTTG | 428.3 | 6.9 | 1.4 | 4.9 | 9.3E-07 | 4.9E-06 | BT20/MB468 | MP |
| >MI0000809\|hsa-mir-151a&WithFlank&8\|-\|140732558\|140732659@51.72.22(+1A)&[MIMAT0000757&hsa-miR-151a-3p&offsets\|-2\|-1(+1A);m-30&8\|-\|140732587\|140732607&offsets\|-2\|-1(+1A)]\|TACTAGACTGAAGCTCCTTGAGA | >MI0000809\|hsa-mir-151a&WithFlank&8\|-\|140732558\|140732659 | 51 | 72 | 22(+1A) | TACTAGACTGAAGCTCCTTGAGA | 463.8 | 6.8 | 1.5 | 4.7 | 3.3E-06 | 4.9E-06 | MB231/MB468 | MITO |
| >hg38_21_+_8433172_8446622_RNA45SN1_45S_50ntFlanks@3823.3850.28\|TTTGGTCGCTCGCTCCTCTCCTACTTGG | >hg38_21_+_8433172_8446622_RNA45SN1_45S_50ntFlanks | 3823 | 3850 | 28 | TTTGGTCGCTCGCTCCTCTCCTACTTGG | 602.7 | -7.7 | 1.6 | -4.6 | 3.4E-06 | 4.9E-06 | BT20/MB468 | NUC |
| >hg38_21_+_8433172_8446622_RNA45SN1_45S_50ntFlanks@4972.4999.28\|CGGCCCGGACACGGACAGGATTGACAGA | >hg38_21_+_8433172_8446622_RNA45SN1_45S_50ntFlanks | 4972 | 4999 | 28 | CGGCCCGGACACGGACAGGATTGACAGA | 565.6 | -7.2 | 1.5 | -4.7 | 2.7E-06 | 4.9E-06 | MB231/MB468 | MP |
| >hg38_21_+_8433172_8446622_RNA45SN1_45S_50ntFlanks@12420.12457.38\|ATGTCGGCTCTTCCTATCATTGTGAAGCAGAATTCACC | >hg38_21_+_8433172_8446622_RNA45SN1_45S_50ntFlanks | 12420 | 12457 | 38 | ATGTCGGCTCTTCCTATCATTGTGAAGCAGAATTCACC | 456.1 | -3.9 | 0.8 | -4.6 | 3.4E-06 | 5.0E-06 | BT20/MB468 | NUC |
| >hg38_21_+_8433172_8446622_RNA45SN1_45S_50ntFlanks@6719.6737.19\|TGAATTGCAGGACACATTG | >hg38_21_+_8433172_8446622_RNA45SN1_45S_50ntFlanks | 6719 | 6737 | 19 | TGAATTGCAGGACACATTG | 1403.7 | -7.6 | 1.6 | -4.9 | 9.3E-07 | 5.0E-06 | BT20/MB468 | MP |
| >hg38_MT_+_1621_3279_RNR2_16S_50ntFlanks@838.864.27\|AAAAAAGTAAAAGGAACTCGGCAAATC | >hg38_MT_+_1621_3279_RNR2_16S_50ntFlanks | 838 | 864 | 27 | AAAAAAGTAAAAGGAACTCGGCAAATC | 45.1 | 8.7 | 1.9 | 4.7 | 2.7E-06 | 5.0E-06 | MB231/MB468 | MP |
| >trna111_HisGTG_1_-_147774845_147774916@-1G.36.37,trna118_HisGTG_1_-_145396881_145396952@-1G.36.37,trna16_HisGTG_1_+_146544773_146544844@-1G.36.37,trna1_HisGTG_15_+_45493349_45493420@-1G.36.37,trna21_HisGTG_1_+_147753471_147753542@-1G.36.37,trna33_HisGTG_6_+_27125906_27125977@-1G.36.37,trna7_HisGTG_9_-_14433938_14434009@-1G.36.37,trna8_HisGTG_15_-_45492611_45492682@-1G.36.37,trna9_HisGTG_15_-_45490804_45490875@-1G.36.37\|GGCCGTGATCGTATAGTGGTTAGTACTCTGCGTTGTG | >trna111_HisGTG_1_-_147774845_147774916 | -1G | 36 | 37 | GGCCGTGATCGTATAGTGGTTAGTACTCTGCGTTGTG | 355.1 | 8.2 | 1.7 | 4.7 | 2.7E-06 | 5.0E-06 | BT20/MB468 | CYTO |
| >hg38_21_+_8433172_8446622_RNA45SN1_45S_50ntFlanks@3783.3800.18\|AGTGAAACTGCGAATGGC | >hg38_21_+_8433172_8446622_RNA45SN1_45S_50ntFlanks | 3783 | 3800 | 18 | AGTGAAACTGCGAATGGC | 387.0 | -4.7 | 1.0 | -4.7 | 3.3E-06 | 5.0E-06 | MB231/MB468 | MITO |
| >MI0000077\|hsa-mir-21&WithFlank&17\|+\|59841260\|59841343@14.34.21&[MIMAT0000076&hsa-miR-21-5p&offsets\|0\|-1;m-1&17\|+\|59841273\|59841294&offsets\|0\|-1]\|TAGCTTATCAGACTGATGTTG | >MI0000077\|hsa-mir-21&WithFlank&17\|+\|59841260\|59841343 | 14 | 34 | 21 | TAGCTTATCAGACTGATGTTG | 4245.7 | 4.9 | 1.0 | 4.7 | 3.1E-06 | 5.0E-06 | MB231/MB468 | CYTO |
| >hg38_21_+_8433172_8446622_RNA45SN1_45S_50ntFlanks@6645.6663.19\|CTCGTACGACTCTTAGCGG | >hg38_21_+_8433172_8446622_RNA45SN1_45S_50ntFlanks | 6645 | 6663 | 19 | CTCGTACGACTCTTAGCGG | 3884.2 | 3.2 | 0.7 | 4.7 | 3.1E-06 | 5.0E-06 | MB231/MB468 | CYTO |
| >hg38_21_+_8433172_8446622_RNA45SN1_45S_50ntFlanks@3991.4023.33\|TGGTGACTCTAGATAACCTCGGGCCGATCGCAC | >hg38_21_+_8433172_8446622_RNA45SN1_45S_50ntFlanks | 3991 | 4023 | 33 | TGGTGACTCTAGATAACCTCGGGCCGATCGCAC | 429.7 | 2.6 | 0.6 | 4.7 | 3.1E-06 | 5.0E-06 | MB231/MB468 | CYTO |
| >hg38_21_+_8433172_8446622_RNA45SN1_45S_50ntFlanks@12257.12284.28\|CGCTTGATCTTGATTTTCAGTACGAATA | >hg38_21_+_8433172_8446622_RNA45SN1_45S_50ntFlanks | 12257 | 12284 | 28 | CGCTTGATCTTGATTTTCAGTACGAATA | 502.1 | -6.1 | 1.3 | -4.7 | 3.3E-06 | 5.0E-06 | MB231/MB468 | MITO |
| >hg38_21_+_8433172_8446622_RNA45SN1_45S_50ntFlanks@3854.3872.19\|ACTGTGGTAATTCTAGAGC | >hg38_21_+_8433172_8446622_RNA45SN1_45S_50ntFlanks | 3854 | 3872 | 19 | ACTGTGGTAATTCTAGAGC | 505.0 | -2.9 | 0.6 | -4.7 | 2.8E-06 | 5.0E-06 | BT20/MB468 | CYTO |
| >trna116_GluCTC_1_-_145399233_145399304@39.73.35,trna134_GluTTC_1_-_16861774_16861845@39.73.35,trna71_GluCTC_1_-_161439189_161439260@39.73.35,trna74_GluCTC_1_-_161431809_161431880@39.73.35,trna77_GluCTC_1_-_161424398_161424469@39.73.35,trna77_GluCTC_6_+_28949976_28950047@39.73.35,trna80_GluCTC_1_-_161417018_161417089@39.73.35,trna84_GluTTC_1_-_161391883_161391954@39.73.35,trna87_GluCTC_6_-_126101393_126101464@39.73.35\|CGCCGCGGCCCGGGTTCGATTCCCGGTCAGGGAAC | >trna116_GluCTC_1_-_145399233_145399304 | 39 | 73 | 35 | CGCCGCGGCCCGGGTTCGATTCCCGGTCAGGGAAC | 637.4 | 6.1 | 1.3 | 4.7 | 3.1E-06 | 5.0E-06 | MB231/MB468 | CYTO |
| >hg38_21_+_8433172_8446622_RNA45SN1_45S_50ntFlanks@5421.5466.46\|CGATTGGATGGTTTAGTGAGGCCCTCGGATCGGCCCCGCCGGGGTC | >hg38_21_+_8433172_8446622_RNA45SN1_45S_50ntFlanks | 5421 | 5466 | 46 | CGATTGGATGGTTTAGTGAGGCCCTCGGATCGGCCCCGCCGGGGTC | 75.9 | -9.3 | 1.9 | -4.9 | 9.4E-07 | 5.0E-06 | BT20/MB468 | MP |
| >hg38_21_+_8433172_8446622_RNA45SN1_45S_50ntFlanks@12717.12757.41\|CCTCGGTTGGCCTCGGATAGCCGGTCCCCCGCCTGTCCCCG | >hg38_21_+_8433172_8446622_RNA45SN1_45S_50ntFlanks | 12717 | 12757 | 41 | CCTCGGTTGGCCTCGGATAGCCGGTCCCCCGCCTGTCCCCG | 142.1 | -11.0 | 2.3 | -4.7 | 2.8E-06 | 5.0E-06 | MB231/MB468 | MP |
| >hg38_21_+_8433172_8446622_RNA45SN1_45S_50ntFlanks@6677.6700.24\|TCGTGCGTCGATGAAGAACGCAGC | >hg38_21_+_8433172_8446622_RNA45SN1_45S_50ntFlanks | 6677 | 6700 | 24 | TCGTGCGTCGATGAAGAACGCAGC | 83.8 | 10.1 | 2.1 | 4.9 | 9.4E-07 | 5.0E-06 | BT20/MB468 | MP |
| >hg38_21_+_8433172_8446622_RNA45SN1_45S_50ntFlanks@10546.10564.19\|CAGTGCGGTAACGCGACCG | >hg38_21_+_8433172_8446622_RNA45SN1_45S_50ntFlanks | 10546 | 10564 | 19 | CAGTGCGGTAACGCGACCG | 81.4 | -9.4 | 1.9 | -4.9 | 9.4E-07 | 5.0E-06 | BT20/MB468 | MP |
| >hg38_21_+_8433172_8446622_RNA45SN1_45S_50ntFlanks@8380.8402.23\|CAAGAGGGCGTGAAACCGTTAAG | >hg38_21_+_8433172_8446622_RNA45SN1_45S_50ntFlanks | 8380 | 8402 | 23 | CAAGAGGGCGTGAAACCGTTAAG | 6969.8 | -7.2 | 1.5 | -4.9 | 9.4E-07 | 5.0E-06 | BT20/MB468 | MP |
| >hg38_21_+_8433172_8446622_RNA45SN1_45S_50ntFlanks@7983.8005.23\|CAGATCAGACGTGGCGACCCGCT | >hg38_21_+_8433172_8446622_RNA45SN1_45S_50ntFlanks | 7983 | 8005 | 23 | CAGATCAGACGTGGCGACCCGCT | 1250.4 | -2.5 | 0.5 | -4.7 | 2.4E-06 | 5.0E-06 | BT20/MB468 | MITO |
| >MI0000478\|hsa-mir-149&WithFlank&2\|+\|240455995\|240456095@21.43.23&[MIMAT0000450&hsa-miR-149-5p&offsets\|0\|0;m-195&2\|+\|240456015\|240456037&offsets\|0\|0]\|TCTGGCTCCGTGTCTTCACTCCC | >MI0000478\|hsa-mir-149&WithFlank&2\|+\|240455995\|240456095 | 21 | 43 | 23 | TCTGGCTCCGTGTCTTCACTCCC | 663.4 | 8.8 | 1.8 | 4.9 | 9.5E-07 | 5.0E-06 | BT20/MB468 | MP |
| >hg38_21_+_8433172_8446622_RNA45SN1_45S_50ntFlanks@11212.11229.18\|CGACTCTGGACGCGAGCC | >hg38_21_+_8433172_8446622_RNA45SN1_45S_50ntFlanks | 11212 | 11229 | 18 | CGACTCTGGACGCGAGCC | 478.7 | -5.1 | 1.1 | -4.7 | 3.1E-06 | 5.0E-06 | MB231/MB468 | CYTO |
| >hg38_21_+_8433172_8446622_RNA45SN1_45S_50ntFlanks@7974.8005.32\|ACGCGACCTCAGATCAGACGTGGCGACCCGCT | >hg38_21_+_8433172_8446622_RNA45SN1_45S_50ntFlanks | 7974 | 8005 | 32 | ACGCGACCTCAGATCAGACGTGGCGACCCGCT | 37192.6 | 4.8 | 0.9 | 5.1 | 3.7E-07 | 5.0E-06 | BT20/MB231 | NUC |
| >hg38_21_+_8433172_8446622_RNA45SN1_45S_50ntFlanks@12397.12419.23\|CGACGTCGCTTTTTGATCCTTCG | >hg38_21_+_8433172_8446622_RNA45SN1_45S_50ntFlanks | 12397 | 12419 | 23 | CGACGTCGCTTTTTGATCCTTCG | 395.4 | -6.7 | 1.4 | -4.6 | 3.3E-06 | 5.0E-06 | MB231/MB468 | MITO |

<table>
<tr><td></td><td></td><td></td><td></td><td></td><td></td><td></td><td></td><td></td><td></td><td></td><td></td><td></td><td></td></tr>
<tr><td>>MI0000063|hsa-let-7b&WithFlank&22|+|46113680|46113774@12.31.20&[MIMAT0000063&hsa-let-7b-5p&offsets|0|-2;m-14&22|+|46113691|46113712&offsets|0|-2]|TGAGGTAGTAGGTTGTGTGG</td><td>>MI0000063|hsa-let-7b&WithFlank&22|+|46113680|46113774</td><td>12</td><td>31</td><td>20</td><td>TGAGGTAGTAGGTTGTGTGG</td><td>289.7</td><td>2.0</td><td>0.4</td><td>4.6</td><td>3.5E-06</td><td>5.0E-06</td><td>BT20/MB468</td><td>NUC</td></tr>
<tr><td>>hg38_21_+_8433172_8446622_RNA45SN1_45S_50ntFlanks@10381.10424.44|GTTCCATGTGAACAGCAGTTGAACATGGGTCAGTCGGTCCTGAG</td><td>>hg38_21_+_8433172_8446622_RNA45SN1_45S_50ntFlanks</td><td>10381</td><td>10424</td><td>44</td><td>GTTCCATGTGAACAGCAGTTGAACATGGGTCAGTCGGTCCTGAG</td><td>983.5</td><td>5.0</td><td>1.1</td><td>4.7</td><td>3.3E-06</td><td>5.0E-06</td><td>MB231/MB468</td><td>NUC</td></tr>
<tr><td>>hg38_21_+_8433172_8446622_RNA45SN1_45S_50ntFlanks@11905.11929.25|GCACGGTGAAGAGACATGAGAGGTG</td><td>>hg38_21_+_8433172_8446622_RNA45SN1_45S_50ntFlanks</td><td>11905</td><td>11929</td><td>25</td><td>GCACGGTGAAGAGACATGAGAGGTG</td><td>230.9</td><td>11.2</td><td>2.2</td><td>5.2</td><td>2.5E-07</td><td>5.0E-06</td><td>BT20/MB231</td><td>MP</td></tr>
<tr><td>>am_trna128_GlyGCC_6_-_27870686_27870756@1.33.33,trna18_GlyGCC_16_+_70822597_70822667@1.33.33,trna19_GlyGCC_16_+_70823410_70823480@1.33.33,trna19_GlyGCC_2_-_157257659_157257729@1.33.33,trna24_GlyGCC_16_-_70812942_70813012@1.33.33,trna25_GlyGCC_16_-_70812114_70812184@1.33.33,trna5_GlyGCC_17_+_8029064_8029134@1.33.33,trna68_GlyGCC_1_-_161493637_161493707@1.33.33|GCATTGGTGGTTCAGTGGTAGAATTCTCGCCTG</td><td>>am_trna128_GlyGCC_6_-_27870686_27870756</td><td>1</td><td>33</td><td>33</td><td>GCATTGGTGGTTCAGTGGTAGAATTCTCGCCTG</td><td>3688.3</td><td>5.2</td><td>1.1</td><td>4.7</td><td>2.8E-06</td><td>5.0E-06</td><td>MB231/MB468</td><td>MP</td></tr>
<tr><td>>trnaMT_SerGCT_MT_+_12207_12265@1.41.41|GAGAAAGCTCACAAGAACTGCTAACTCATGCCCCCATGTCT</td><td>>trnaMT_SerGCT_MT_+_12207_12265</td><td>1</td><td>41</td><td>41</td><td>GAGAAAGCTCACAAGAACTGCTAACTCATGCCCCCATGTCT</td><td>37.3</td><td>8.4</td><td>1.8</td><td>4.7</td><td>2.8E-06</td><td>5.0E-06</td><td>MB231/MB468</td><td>MP</td></tr>
<tr><td>>hg38_21_+_8433172_8446622_RNA45SN1_45S_50ntFlanks@11999.12037.39|CCCTGCGGGCCGCCGGTGAAATACCACTACTCTGATCGT</td><td>>hg38_21_+_8433172_8446622_RNA45SN1_45S_50ntFlanks</td><td>11999</td><td>12037</td><td>39</td><td>CCCTGCGGGCCGCCGGTGAAATACCACTACTCTGATCGT</td><td>178.0</td><td>-6.3</td><td>1.4</td><td>-4.7</td><td>3.3E-06</td><td>5.0E-06</td><td>MB231/MB468</td><td>NUC</td></tr>
<tr><td>>hg38_21_+_8433172_8446622_RNA45SN1_45S_50ntFlanks@11671.11701.31|CCAGTGCTCTGAATGTCAAAGTGAAGAAATT</td><td>>hg38_21_+_8433172_8446622_RNA45SN1_45S_50ntFlanks</td><td>11671</td><td>11701</td><td>31</td><td>CCAGTGCTCTGAATGTCAAAGTGAAGAAATT</td><td>255.2</td><td>-6.9</td><td>1.5</td><td>-4.7</td><td>2.8E-06</td><td>5.0E-06</td><td>MB231/MB468</td><td>MP</td></tr>
<tr><td>>hg38_21_+_8433172_8446622_RNA45SN1_45S_50ntFlanks@11728.11748.21|GTAACTATGACTCTCTTAAGG</td><td>>hg38_21_+_8433172_8446622_RNA45SN1_45S_50ntFlanks</td><td>11728</td><td>11748</td><td>21</td><td>GTAACTATGACTCTCTTAAGG</td><td>662.2</td><td>-6.2</td><td>1.3</td><td>-4.6</td><td>3.5E-06</td><td>5.0E-06</td><td>BT20/MB468</td><td>NUC</td></tr>
<tr><td>>hg38_21_+_8433172_8446622_RNA45SN1_45S_50ntFlanks@3779.3801.23|GTACAGTGAAACTGCGAATGGCT</td><td>>hg38_21_+_8433172_8446622_RNA45SN1_45S_50ntFlanks</td><td>3779</td><td>3801</td><td>23</td><td>GTACAGTGAAACTGCGAATGGCT</td><td>158.8</td><td>-6.8</td><td>1.5</td><td>-4.7</td><td>2.8E-06</td><td>5.0E-06</td><td>MB231/MB468</td><td>MP</td></tr>
<tr><td>>hg38_21_+_8433172_8446622_RNA45SN1_45S_50ntFlanks@10404.10431.28|CATGGGTCAGTCGGTCCTGAGAGATGGG</td><td>>hg38_21_+_8433172_8446622_RNA45SN1_45S_50ntFlanks</td><td>10404</td><td>10431</td><td>28</td><td>CATGGGTCAGTCGGTCCTGAGAGATGGG</td><td>172.9</td><td>-5.7</td><td>1.2</td><td>-4.7</td><td>2.8E-06</td><td>5.0E-06</td><td>BT20/MB468</td><td>CYTO</td></tr>
<tr><td>>hg38_21_+_8433172_8446622_RNA45SN1_45S_50ntFlanks@11641.11685.45|CGGCGGGTGTTGACGCGATGTGATTTCTGCCCAGTGCTCTGAATG</td><td>>hg38_21_+_8433172_8446622_RNA45SN1_45S_50ntFlanks</td><td>11641</td><td>11685</td><td>45</td><td>CGGCGGGTGTTGACGCGATGTGATTTCTGCCCAGTGCTCTGAATG</td><td>4797.2</td><td>-4.8</td><td>1.0</td><td>-4.7</td><td>2.4E-06</td><td>5.1E-06</td><td>BT20/MB468</td><td>MITO</td></tr>
<tr><td>>trna116_GluCTC_1_-_145399233_145399304@1.39.39,trna59_GluCTC_1_+_249168447_249168518@1.39.39,trna71_GluCTC_1_-_161439189_161439260@1.39.39,trna74_GluCTC_1_-_161431809_161431880@1.39.39,trna77_GluCTC_1_-_161424398_161424469@1.39.39,trna77_GluCTC_6_+_28949976_28950047@1.39.39,trna80_GluCTC_1_-_161417018_161417089@1.39.39,trna87_GluCTC_6_-_126101393_126101464@1.39.39|TCCCTGGTGGTCTAGTGGTTAGGATTCGGCGCTCTCACC</td><td>>trna116_GluCTC_1_-_145399233_145399304</td><td>1</td><td>39</td><td>39</td><td>TCCCTGGTGGTCTAGTGGTTAGGATTCGGCGCTCTCACC</td><td>2695.7</td><td>4.1</td><td>0.9</td><td>4.7</td><td>3.3E-06</td><td>5.1E-06</td><td>MB231/MB468</td><td>NUC</td></tr>
<tr><td>>hg38_21_+_8433172_8446622_RNA45SN1_45S_50ntFlanks@6730.6771.42|ACACATTGATCATCGACACTTCGAACGCACTTGCGGCCCCGG</td><td>>hg38_21_+_8433172_8446622_RNA45SN1_45S_50ntFlanks</td><td>6730</td><td>6771</td><td>42</td><td>ACACATTGATCATCGACACTTCGAACGCACTTGCGGCCCCGG</td><td>601.3</td><td>-6.0</td><td>1.3</td><td>-4.7</td><td>3.3E-06</td><td>5.1E-06</td><td>MB231/MB468</td><td>NUC</td></tr>
<tr><td>>trnaMT_GluTTC_MT_-_14674_14742@20.58.39,trnalookalike8_GluTTC_5_-_93905172_93905240@20.58.39|AACGATGGTTTTTCATATCATTGGTCGTGGTTGTAGTCC</td><td>>trnaMT_GluTTC_MT_-_14674_14742</td><td>20</td><td>58</td><td>39</td><td>AACGATGGTTTTTCATATCATTGGTCGTGGTTGTAGTCC</td><td>36.3</td><td>8.4</td><td>1.8</td><td>4.7</td><td>2.8E-06</td><td>5.1E-06</td><td>MB231/MB468</td><td>MP</td></tr>
<tr><td>>hg38_21_+_8433172_8446622_RNA45SN1_45S_50ntFlanks@11539.11584.46|GCGCGGCGCCTCGCCTCGGCCGGCGCCTAGCAGCCGACTTAGAACT</td><td>>hg38_21_+_8433172_8446622_RNA45SN1_45S_50ntFlanks</td><td>11539</td><td>11584</td><td>46</td><td>GCGCGGCGCCTCGCCTCGGCCGGCGCCTAGCAGCCGACTTAGAACT</td><td>827.9</td><td>-4.5</td><td>0.9</td><td>-5.1</td><td>2.7E-07</td><td>5.1E-06</td><td>BT20/MB231</td><td>CYTO</td></tr>
<tr><td>>MI0003776|hsa-mir-320b-1&WithFlank&1|+|116671740|116671823@48.68.21;MI0003839|hsa-mir-320b-2&WithFlank&1|-|224257034|224257116@47.67.21&[MIMAT0005792&hsa-miR-320b&offsets|0|-1;m-183&1|+|116671787|116671806&offsets|0|+1];[MIMAT0005792_1&hsa-miR-320b&offsets|0|-1;m-183&1|-|224257051|224257070&offsets|0|+1]|AAAAGCTGGGTTGAGAGGGCA</td><td>>MI0003776|hsa-mir-320b-1&WithFlank&1|+|116671740|116671823</td><td>48</td><td>68</td><td>21</td><td>AAAAGCTGGGTTGAGAGGGCA</td><td>4732.3</td><td>4.5</td><td>0.9</td><td>5.1</td><td>2.7E-07</td><td>5.1E-06</td><td>BT20/MB231</td><td>CYTO</td></tr>
<tr><td>>hg38_21_+_8433172_8446622_RNA45SN1_45S_50ntFlanks@10869.10903.35|AGGGCTGGGTCGGTCGGGCTGGGGCGCGAAGCGGG</td><td>>hg38_21_+_8433172_8446622_RNA45SN1_45S_50ntFlanks</td><td>10869</td><td>10903</td><td>35</td><td>AGGGCTGGGTCGGTCGGGCTGGGGCGCGAAGCGGG</td><td>301.4</td><td>-6.5</td><td>1.4</td><td>-4.7</td><td>3.3E-06</td><td>5.1E-06</td><td>MB231/MB468</td><td>NUC</td></tr>
<tr><td>>hg38_1_-_228634819_228635039_RNA5S12_5S_50ntFlanks@51.81.31|GTCTACGGCCATACCACCCTGAACGCGCCCG</td><td>>hg38_1_-_228634819_228635039_RNA5S12_5S_50ntFlanks</td><td>51</td><td>81</td><td>31</td><td>GTCTACGGCCATACCACCCTGAACGCGCCCG</td><td>2502.9</td><td>4.4</td><td>0.9</td><td>4.7</td><td>3.2E-06</td><td>5.1E-06</td><td>MB231/MB468</td><td>CYTO</td></tr>
<tr><td>>hg38_21_+_8433172_8446622_RNA45SN1_45S_50ntFlanks@3842.3859.18|CCTACTTGGATAACTGTG</td><td>>hg38_21_+_8433172_8446622_RNA45SN1_45S_50ntFlanks</td><td>3842</td><td>3859</td><td>18</td><td>CCTACTTGGATAACTGTG</td><td>413.4</td><td>-6.1</td><td>1.3</td><td>-4.7</td><td>2.8E-06</td><td>5.1E-06</td><td>MB231/MB468</td><td>MP</td></tr>
<tr><td>>hg38_21_+_8433172_8446622_RNA45SN1_45S_50ntFlanks@476.517.42|TTCGTGATCGATGTGGTGACGTCGTGCTCTCCCGGGCCGGGT</td><td>>hg38_21_+_8433172_8446622_RNA45SN1_45S_50ntFlanks</td><td>476</td><td>517</td><td>42</td><td>TTCGTGATCGATGTGGTGACGTCGTGCTCTCCCGGGCCGGGT</td><td>11131.8</td><td>4.3</td><td>0.9</td><td>4.7</td><td>3.3E-06</td><td>5.1E-06</td><td>MB231/MB468</td><td>NUC</td></tr>
<tr><td>>hg38_21_+_8433172_8446622_RNA45SN1_45S_50ntFlanks@5074.5110.37|ATTCCGATAACGAACGAGACTCTGGCATGCTAACTAG</td><td>>hg38_21_+_8433172_8446622_RNA45SN1_45S_50ntFlanks</td><td>5074</td><td>5110</td><td>37</td><td>ATTCCGATAACGAACGAGACTCTGGCATGCTAACTAG</td><td>319.1</td><td>-4.0</td><td>0.9</td><td>-4.7</td><td>3.3E-06</td><td>5.1E-06</td><td>MB231/MB468</td><td>NUC</td></tr>
<tr><td>>hg38_21_+_8433172_8446622_RNA45SN1_45S_50ntFlanks@10835.10867.33|CGGATCCGTAACTTCGGGATAAGGATTGGCTCT</td><td>>hg38_21_+_8433172_8446622_RNA45SN1_45S_50ntFlanks</td><td>10835</td><td>10867</td><td>33</td><td>CGGATCCGTAACTTCGGGATAAGGATTGGCTCT</td><td>673.8</td><td>3.6</td><td>0.8</td><td>4.7</td><td>3.2E-06</td><td>5.1E-06</td><td>MB231/MB468</td><td>CYTO</td></tr>
<tr><td>>MI0000477|hsa-mir-146a&WithFlank&5|+|160485346|160485456@27.46.20(+2C)&[MIMAT0000449&hsa-miR-146a-5p&offsets|0|-2(+2C);m-129&5|+|160485372|160485393&offsets|0|-2(+2C)]|TGAGAACTGAATTCCATGGGCC</td><td>>MI0000477|hsa-mir-146a&WithFlank&5|+|160485346|160485456</td><td>27</td><td>46</td><td>20(+2C)</td><td>TGAGAACTGAATTCCATGGGCC</td><td>54.2</td><td>9.0</td><td>1.9</td><td>4.7</td><td>2.8E-06</td><td>5.1E-06</td><td>MB231/MB468</td><td>MP</td></tr>
<tr><td>>hg38_21_+_8433172_8446622_RNA45SN1_45S_50ntFlanks@11703.11726.24|AATGAAGCGCGGGTAAACGGCGGG</td><td>>hg38_21_+_8433172_8446622_RNA45SN1_45S_50ntFlanks</td><td>11703</td><td>11726</td><td>24</td><td>AATGAAGCGCGGGTAAACGGCGGG</td><td>830.7</td><td>-7.8</td><td>1.7</td><td>-4.6</td><td>3.6E-06</td><td>5.1E-06</td><td>BT20/MB468</td><td>NUC</td></tr>
<tr><td>>MI0000289|hsa-mir-181a-1&WithFlank&1|-|198859038|198859159@30.54.25;MI0000269|hsa-mir-181a-2&WithFlank&9|+|124692436|124692557@45.69.25&[MIMAT0000256&hsa-miR-181a-5p&offsets|0|+2;m-32&1|-|198859108|198859130&offsets|0|+2];[MIMAT0000256_1&hsa-miR-181a-5p&offsets|0|+2;m-32&9|+|124692480|124692502&offsets|0|+2]|AACATTCAACGCTGTCGGTGAGTTT</td><td>>MI0000289|hsa-mir-181a-1&WithFlank&1|-|198859038|198859159</td><td>30</td><td>54</td><td>25</td><td>AACATTCAACGCTGTCGGTGAGTTT</td><td>7671.1</td><td>-5.4</td><td>1.1</td><td>-5.2</td><td>2.5E-07</td><td>5.1E-06</td><td>BT20/MB231</td><td>MP</td></tr>
<tr><td>>hg38_21_+_8433172_8446622_RNA45SN1_45S_50ntFlanks@11597.11621.25|GGGAATCCGACTGTTTAATTAAAAC</td><td>>hg38_21_+_8433172_8446622_RNA45SN1_45S_50ntFlanks</td><td>11597</td><td>11621</td><td>25</td><td>GGGAATCCGACTGTTTAATTAAAAC</td><td>167.7</td><td>10.7</td><td>2.1</td><td>5.2</td><td>2.5E-07</td><td>5.1E-06</td><td>BT20/MB231</td><td>MP</td></tr>
<tr><td>>hg38_21_+_8433172_8446622_RNA45SN1_45S_50ntFlanks@5485.5513.29|AGCGCTGAGAAGACGGTCGAACTTGACTA</td><td>>hg38_21_+_8433172_8446622_RNA45SN1_45S_50ntFlanks</td><td>5485</td><td>5513</td><td>29</td><td>AGCGCTGAGAAGACGGTCGAACTTGACTA</td><td>187.8</td><td>10.9</td><td>2.1</td><td>5.2</td><td>2.5E-07</td><td>5.1E-06</td><td>BT20/MB231</td><td>MP</td></tr>
<tr><td>>hg38_21_+_8433172_8446622_RNA45SN1_45S_50ntFlanks@9802.9819.18|CGGGCGTGGAATGCGAGT</td><td>>hg38_21_+_8433172_8446622_RNA45SN1_45S_50ntFlanks</td><td>9802</td><td>9819</td><td>18</td><td>CGGGCGTGGAATGCGAGT</td><td>129.5</td><td>10.3</td><td>2.0</td><td>5.2</td><td>2.5E-07</td><td>5.1E-06</td><td>BT20/MB231</td><td>MP</td></tr>
<tr><td>>MI0000085|hsa-mir-27a&WithFlank&19|-|13836434|13836523@57.77.21(+2U)&[MIMAT0000084&hsa-miR-27a-3p&offsets|0|0(+2U);m-43&19|-|13836447|13836467&offsets|0|0(+2U)]|TTCACAGTGGCTAAGTTCCGCTT</td><td>>MI0000085|hsa-mir-27a&WithFlank&19|-|13836434|13836523</td><td>57</td><td>77</td><td>21(+2U)</td><td>TTCACAGTGGCTAAGTTCCGCTT</td><td>98.3</td><td>-10.0</td><td>1.9</td><td>-5.2</td><td>2.5E-07</td><td>5.1E-06</td><td>BT20/MB231</td><td>MP</td></tr>
<tr><td>>MI0000471|hsa-mir-126&WithFlank&9|+|136670596|136670692@59.78.20(+1A)&[MIMAT0000445&hsa-miR-126-3p&offsets|+1|-1(+1A);m-40&9|+|136670653|136670674&offsets|+1|-1(+1A)]|CGTACCGTGAGTAATAATGCA</td><td>>MI0000471|hsa-mir-126&WithFlank&9|+|136670596|136670692</td><td>59</td><td>78</td><td>20(+1A)</td><td>CGTACCGTGAGTAATAATGCA</td><td>80.0</td><td>10.1</td><td>2.1</td><td>4.9</td><td>9.6E-07</td><td>5.1E-06</td><td>BT20/MB468</td><td>MP</td></tr>
<tr><td>>hg38_21_+_8433172_8446622_RNA45SN1_45S_50ntFlanks@9624.9648.25|AGTAGCTGGTTCCCTCCGAAGTTTC</td><td>>hg38_21_+_8433172_8446622_RNA45SN1_45S_50ntFlanks</td><td>9624</td><td>9648</td><td>25</td><td>AGTAGCTGGTTCCCTCCGAAGTTTC</td><td>93.3</td><td>-9.6</td><td>2.0</td><td>-4.9</td><td>9.6E-07</td><td>5.1E-06</td><td>BT20/MB468</td><td>MP</td></tr>
<tr><td>>hg38_21_+_8433172_8446622_RNA45SN1_45S_50ntFlanks@6775.6806.32|CCTCCCGGGGCTACGCCTGTCTGAGCGTCGCT</td><td>>hg38_21_+_8433172_8446622_RNA45SN1_45S_50ntFlanks</td><td>6775</td><td>6806</td><td>32</td><td>CCTCCCGGGGCTACGCCTGTCTGAGCGTCGCT</td><td>179.6</td><td>2.3</td><td>0.5</td><td>4.6</td><td>3.6E-06</td><td>5.1E-06</td><td>BT20/MB468</td><td>NUC</td></tr>
<tr><td>>trnaMT_PheGAA_MT_+_577_647@15.35.21|CCTCCTCAAAGCAATACACTG</td><td>>trnaMT_PheGAA_MT_+_577_647</td><td>15</td><td>35</td><td>21</td><td>CCTCCTCAAAGCAATACACTG</td><td>215.6</td><td>-6.6</td><td>1.4</td><td>-4.6</td><td>3.3E-06</td><td>5.1E-06</td><td>MB231/MB468</td><td>NUC</td></tr>
<tr><td>>hg38_21_+_8433172_8446622_RNA45SN1_45S_50ntFlanks@3985.4014.30|CGGCTTTGGTGACTCTAGATAACCTCGGGC</td><td>>hg38_21_+_8433172_8446622_RNA45SN1_45S_50ntFlanks</td><td>3985</td><td>4014</td><td>30</td><td>CGGCTTTGGTGACTCTAGATAACCTCGGGC</td><td>382.8</td><td>-6.4</td><td>1.4</td><td>-4.6</td><td>3.3E-06</td><td>5.1E-06</td><td>MB231/MB468</td><td>NUC</td></tr>
<tr><td>>MI0000750|hsa-mir-26a-2&WithFlank&12|-|57824603|57824698@20.42.23;MI0000083|hsa-mir-26a-1&WithFlank&3|+|37969398|37969486@16.38.23&[MIMAT0000082&hsa-miR-26a-5p&offsets|0|+1;m-33&12|-|57824658|57824679&offsets|0|+1];[MIMAT0000082_1&hsa-miR-26a-5p&offsets|0|+1;m-34&3|+|37969413|37969434&offsets|0|+1]|TTCAAGTAATCCAGGATAGGCTG</td><td>>MI0000750|hsa-mir-26a-2&WithFlank&12|-|57824603|57824698</td><td>20</td><td>42</td><td>23</td><td>TTCAAGTAATCCAGGATAGGCTG</td><td>197.2</td><td>4.6</td><td>1.0</td><td>4.6</td><td>3.4E-06</td><td>5.1E-06</td><td>MB231/MB468</td><td>MITO</td></tr>
<tr><td>>hg38_21_+_8433172_8446622_RNA45SN1_45S_50ntFlanks@10385.10404.20|CATGTGAACAGCAGTTGAAC</td><td>>hg38_21_+_8433172_8446622_RNA45SN1_45S_50ntFlanks</td><td>10385</td><td>10404</td><td>20</td><td>CATGTGAACAGCAGTTGAAC</td><td>76.4</td><td>-9.3</td><td>1.9</td><td>-4.9</td><td>9.6E-07</td><td>5.1E-06</td><td>BT20/MB468</td><td>MP</td></tr>
<tr><td>>hg38_21_+_8433172_8446622_RNA45SN1_45S_50ntFlanks@6651.6690.40|CGACTCTTAGCGGTGGATCACTCGGCTCGTGCGTCGATGA</td><td>>hg38_21_+_8433172_8446622_RNA45SN1_45S_50ntFlanks</td><td>6651</td><td>6690</td><td>40</td><td>CGACTCTTAGCGGTGGATCACTCGGCTCGTGCGTCGATGA</td><td>9395.7</td><td>2.8</td><td>0.6</td><td>4.7</td><td>2.5E-06</td><td>5.1E-06</td><td>BT20/MB468</td><td>MITO</td></tr>
<tr><td>>hg38_21_+_8433172_8446622_RNA45SN1_45S_50ntFlanks@4727.4767.41|AATCAAGAACGAAAGTCGGAGGTTCGAAGACGATCAGATAC</td><td>>hg38_21_+_8433172_8446622_RNA45SN1_45S_50ntFlanks</td><td>4727</td><td>4767</td><td>41</td><td>AATCAAGAACGAAAGTCGGAGGTTCGAAGACGATCAGATAC</td><td>4406.6</td><td>11.2</td><td>2.4</td><td>4.7</td><td>3.2E-06</td><td>5.1E-06</td><td>MB231/MB468</td><td>CYTO</td></tr>
<tr><td>>hg38_21_+_8433172_8446622_RNA45SN1_45S_50ntFlanks@4002.4023.22|GATAACCTCGGGCCGATCGCAC</td><td>>hg38_21_+_8433172_8446622_RNA45SN1_45S_50ntFlanks</td><td>4002</td><td>4023</td><td>22</td><td>GATAACCTCGGGCCGATCGCAC</td><td>133.9</td><td>-6.6</td><td>1.4</td><td>-4.7</td><td>2.8E-06</td><td>5.1E-06</td><td>MB231/MB468</td><td>MP</td></tr>
<tr><td>>hg38_21_+_8433172_8446622_RNA45SN1_45S_50ntFlanks@4129.4162.34|GATTCCGGAGAGGGAGCCTGAGAAACGGCTACCA</td><td>>hg38_21_+_8433172_8446622_RNA45SN1_45S_50ntFlanks</td><td>4129</td><td>4162</td><td>34</td><td>GATTCCGGAGAGGGAGCCTGAGAAACGGCTACCA</td><td>476.9</td><td>-6.7</td><td>1.4</td><td>-4.6</td><td>3.3E-06</td><td>5.1E-06</td><td>MB231/MB468</td><td>NUC</td></tr>
<tr><td>>hg38_21_+_8433172_8446622_RNA45SN1_45S_50ntFlanks@12283.12306.24|TACAGACCGTGAAAGCGGGGCCTC</td><td>>hg38_21_+_8433172_8446622_RNA45SN1_45S_50ntFlanks</td><td>12283</td><td>12306</td><td>24</td><td>TACAGACCGTGAAAGCGGGGCCTC</td><td>66.2</td><td>-9.1</td><td>1.9</td><td>-4.9</td><td>9.7E-07</td><td>5.1E-06</td><td>BT20/MB468</td><td>MP</td></tr>
<tr><td>>trna152_ValCAC_6_-_27248049_27248121@1.29.29|GCTTCTGTAGTGTAGTGGTTATCACGTTC</td><td>>trna152_ValCAC_6_-_27248049_27248121</td><td>1</td><td>29</td><td>29</td><td>GCTTCTGTAGTGTAGTGGTTATCACGTTC</td><td>3726.9</td><td>-4.2</td><td>0.9</td><td>-4.6</td><td>3.4E-06</td><td>5.1E-06</td><td>MB231/MB468</td><td>MITO</td></tr>
<tr><td>>hg38_21_+_8433172_8446622_RNA45SN1_45S_50ntFlanks@6721.6742.22|AATTGCAGGACACATTGATCAT</td><td>>hg38_21_+_8433172_8446622_RNA45SN1_45S_50ntFlanks</td><td>6721</td><td>6742</td><td>22</td><td>AATTGCAGGACACATTGATCAT</td><td>612.1</td><td>-5.0</td><td>1.1</td><td>-4.7</td><td>3.2E-06</td><td>5.1E-06</td><td>MB231/MB468</td><td>CYTO</td></tr>
</table>

<table>
<tr><td>>trnaMT_PheGAA_MT_+_577_647@32.62.31|ACTGAAAATGTTTAGACGGGCTCACATCACC</td><td>>trnaMT_PheGAA_MT_+_577_647</td><td>32</td><td>62</td><td>31</td><td>ACTGAAAATGTTTAGACGGGCTCACATCACC</td><td>77.4</td><td>10.0</td><td>2.0</td><td>4.9</td><td>9.7E-07</td><td>5.2E-06</td><td>BT20/MB468</td><td>MP</td></tr>
<tr><td>>hg38_21_+_8433172_8446622_RNA45SN1_45S_50ntFlanks@10816.10840.25|GTAAGGGAAGTCGGCAAGCCGGATC</td><td>>hg38_21_+_8433172_8446622_RNA45SN1_45S_50ntFlanks</td><td>10816</td><td>10840</td><td>25</td><td>GTAAGGGAAGTCGGCAAGCCGGATC</td><td>330.0</td><td>-5.0</td><td>1.1</td><td>-4.6</td><td>3.4E-06</td><td>5.2E-06</td><td>MB231/MB468</td><td>NUC</td></tr>
<tr><td>>MI0000084|hsa-mir-26b&WithFlank&2|+|218402640|218402728@18.38.21(+1C)&[MIMAT0000083&hsa-miR-26b-5p&offsets|0|0(+1C);m-53&2|+|218402657|218402678&offsets|0|-1(+1C)]|TTCAAGTAATTCAGGATAGGTC</td><td>>MI0000084|hsa-mir-26b&WithFlank&2|+|218402640|218402728</td><td>18</td><td>38</td><td>21(+1C)</td><td>TTCAAGTAATTCAGGATAGGTC</td><td>904.5</td><td>4.8</td><td>1.0</td><td>4.6</td><td>3.4E-06</td><td>5.2E-06</td><td>MB231/MB468</td><td>MITO</td></tr>
<tr><td>>MI0001445|hsa-mir-423&WithFlank&17|+|30117073|30117178@59.81.23(+1U)&[MIMAT0001340&hsa-miR-423-3p&offsets|0|0(+1U);m-100&17|+|30117131|30117153&offsets|0|0(+1U)]|AGCTCGGTCTGAGGCCCCTCAGTT</td><td>>MI0001445|hsa-mir-423&WithFlank&17|+|30117073|30117178</td><td>59</td><td>81</td><td>23(+1U)</td><td>AGCTCGGTCTGAGGCCCCTCAGTT</td><td>375.2</td><td>2.2</td><td>0.5</td><td>4.6</td><td>3.6E-06</td><td>5.2E-06</td><td>BT20/MB468</td><td>NUC</td></tr>
<tr><td>>hg38_21_+_8433172_8446622_RNA45SN1_45S_50ntFlanks@10682.10706.25|TCCGGCGGCGTCCGGTGAGCTCTCG</td><td>>hg38_21_+_8433172_8446622_RNA45SN1_45S_50ntFlanks</td><td>10682</td><td>10706</td><td>25</td><td>TCCGGCGGCGTCCGGTGAGCTCTCG</td><td>5838.3</td><td>-9.0</td><td>1.9</td><td>-4.6</td><td>3.6E-06</td><td>5.2E-06</td><td>BT20/MB468</td><td>NUC</td></tr>
<tr><td>>hg38_21_+_8433172_8446622_RNA45SN1_45S_50ntFlanks@12456.12475.20|CCAAGCGTTGGATTGTTCAC</td><td>>hg38_21_+_8433172_8446622_RNA45SN1_45S_50ntFlanks</td><td>12456</td><td>12475</td><td>20</td><td>CCAAGCGTTGGATTGTTCAC</td><td>9574.1</td><td>-8.8</td><td>1.8</td><td>-4.9</td><td>9.8E-07</td><td>5.2E-06</td><td>BT20/MB468</td><td>MP</td></tr>
<tr><td>>trna143_LysTTT_6_-_27559593_27559665@13.33.21,trna149_LysTTT_6_-_27302769_27302841@13.33.21,trna14_LysTTT_11_-_59327808_59327880@13.33.21,trna2_LysTTT_17_+_8022473_8022545@13.33.21,trna54_LysTTT_1_+_204475655_204475727@13.33.21,trna5_LysTTT_11_+_59323902_59323974@13.33.21,trna62_LysTTT_1_-_204476158_204476230@13.33.21,trna76_LysTTT_6_+_28918806_28918878@13.33.21|CAGTCGGTAGAGCATCAGACT</td><td>>trna143_LysTTT_6_-_27559593_27559665</td><td>13</td><td>33</td><td>21</td><td>CAGTCGGTAGAGCATCAGACT</td><td>35.4</td><td>8.4</td><td>1.8</td><td>4.7</td><td>2.9E-06</td><td>5.2E-06</td><td>MB231/MB468</td><td>MP</td></tr>
<tr><td>>hg38_21_+_8433172_8446622_RNA45SN1_45S_50ntFlanks@10369.10407.39|AAGTGGAGAAGGGTTCCATGTGAACAGCAGTTGAACATG</td><td>>hg38_21_+_8433172_8446622_RNA45SN1_45S_50ntFlanks</td><td>10369</td><td>10407</td><td>39</td><td>AAGTGGAGAAGGGTTCCATGTGAACAGCAGTTGAACATG</td><td>943.0</td><td>-6.8</td><td>1.4</td><td>-4.7</td><td>2.9E-06</td><td>5.2E-06</td><td>BT20/MB468</td><td>CYTO</td></tr>
<tr><td>>hg38_21_+_8433172_8446622_RNA45SN1_45S_50ntFlanks@8327.8375.49|ATAGTCAACAAGTACCGTAAGGGAAAGTTGAAAAGAACTTTGAAGAGAG</td><td>>hg38_21_+_8433172_8446622_RNA45SN1_45S_50ntFlanks</td><td>8327</td><td>8375</td><td>49</td><td>ATAGTCAACAAGTACCGTAAGGGAAAGTTGAAAAGAACTTTGAAGAGAG</td><td>114.7</td><td>10.2</td><td>2.0</td><td>5.2</td><td>2.6E-07</td><td>5.2E-06</td><td>BT20/MB231</td><td>MP</td></tr>
<tr><td>>hg38_21_+_8433172_8446622_RNA45SN1_45S_50ntFlanks@8403.8420.18|AGGTAAACGGGTGGGGTC</td><td>>hg38_21_+_8433172_8446622_RNA45SN1_45S_50ntFlanks</td><td>8403</td><td>8420</td><td>18</td><td>AGGTAAACGGGTGGGGTC</td><td>54.6</td><td>-8.9</td><td>1.8</td><td>-4.9</td><td>9.8E-07</td><td>5.2E-06</td><td>BT20/MB468</td><td>MP</td></tr>
<tr><td>>MI0000273|hsa-mir-183&WithFlank&7|-|129774899|129775020@33.53.21(+1A)&[MIMAT0000261&hsa-miR-183-5p&offsets|0|-1(+1A);m-41&7|-|129774966|129774987&offsets|-1|-2(+1A)]|TATGGCACTGGTAGAATTCACA</td><td>>MI0000273|hsa-mir-183&WithFlank&7|-|129774899|129775020</td><td>33</td><td>53</td><td>21(+1A)</td><td>TATGGCACTGGTAGAATTCACA</td><td>1176.8</td><td>4.0</td><td>0.8</td><td>5.1</td><td>2.8E-07</td><td>5.2E-06</td><td>BT20/MB231</td><td>CYTO</td></tr>
<tr><td>>hg38_21_+_8433172_8446622_RNA45SN1_45S_50ntFlanks@6654.6700.47|CTCTTAGCGGTGGATCACTCGGCTCGTGCGTCGATGAAGAACGCAGC</td><td>>hg38_21_+_8433172_8446622_RNA45SN1_45S_50ntFlanks</td><td>6654</td><td>6700</td><td>47</td><td>CTCTTAGCGGTGGATCACTCGGCTCGTGCGTCGATGAAGAACGCAGC</td><td>78.8</td><td>10.0</td><td>2.1</td><td>4.9</td><td>9.8E-07</td><td>5.2E-06</td><td>BT20/MB468</td><td>MP</td></tr>
<tr><td>>hg38_21_+_8433172_8446622_RNA45SN1_45S_50ntFlanks@12418.12435.18|CGATGTCGGCTCTTCCTA</td><td>>hg38_21_+_8433172_8446622_RNA45SN1_45S_50ntFlanks</td><td>12418</td><td>12435</td><td>18</td><td>CGATGTCGGCTCTTCCTA</td><td>11962.4</td><td>-4.5</td><td>1.0</td><td>-4.7</td><td>2.9E-06</td><td>5.2E-06</td><td>BT20/MB468</td><td>CYTO</td></tr>
<tr><td>>MI0001445|hsa-mir-423&WithFlank&17|+|30117073|30117178@59.78.20&[MIMAT0001340&hsa-miR-423-3p&offsets|0|-3;m-100&17|+|30117131|30117153&offsets|0|-3]|AGCTCGGTCTGAGGCCCCTC</td><td>>MI0001445|hsa-mir-423&WithFlank&17|+|30117073|30117178</td><td>59</td><td>78</td><td>20</td><td>AGCTCGGTCTGAGGCCCCTC</td><td>2196.4</td><td>4.3</td><td>0.9</td><td>4.6</td><td>3.5E-06</td><td>5.2E-06</td><td>MB231/MB468</td><td>MITO</td></tr>
<tr><td>>hg38_21_+_8433172_8446622_RNA45SN1_45S_50ntFlanks@12420.12459.40|ATGTCGGCTCTTCCTATCATTGTGAAGCAGAATTCACCAA</td><td>>hg38_21_+_8433172_8446622_RNA45SN1_45S_50ntFlanks</td><td>12420</td><td>12459</td><td>40</td><td>ATGTCGGCTCTTCCTATCATTGTGAAGCAGAATTCACCAA</td><td>2518.4</td><td>-4.3</td><td>0.8</td><td>-5.1</td><td>3.1E-07</td><td>5.2E-06</td><td>BT20/MB231</td><td>MITO</td></tr>
<tr><td>>hg38_21_+_8433172_8446622_RNA45SN1_45S_50ntFlanks@10680.10708.29|GTTCCGGCGGCGTCCGGTGAGCTCTCGCT</td><td>>hg38_21_+_8433172_8446622_RNA45SN1_45S_50ntFlanks</td><td>10680</td><td>10708</td><td>29</td><td>GTTCCGGCGGCGTCCGGTGAGCTCTCGCT</td><td>14065.5</td><td>-4.7</td><td>1.0</td><td>-4.7</td><td>2.9E-06</td><td>5.2E-06</td><td>BT20/MB468</td><td>CYTO</td></tr>
<tr><td>>hg38_21_+_8433172_8446622_RNA45SN1_45S_50ntFlanks@12002.12031.30|TGCGGGCCGCCGGTGAAATACCACTACTCT</td><td>>hg38_21_+_8433172_8446622_RNA45SN1_45S_50ntFlanks</td><td>12002</td><td>12031</td><td>30</td><td>TGCGGGCCGCCGGTGAAATACCACTACTCT</td><td>964.6</td><td>-5.4</td><td>1.2</td><td>-4.6</td><td>3.4E-06</td><td>5.2E-06</td><td>MB231/MB468</td><td>NUC</td></tr>
<tr><td>>hg38_21_+_8433172_8446622_RNA45SN1_45S_50ntFlanks@10350.10380.31|ACGAGAACTTTGAAGGCCGAAGTGGAGAAGG</td><td>>hg38_21_+_8433172_8446622_RNA45SN1_45S_50ntFlanks</td><td>10350</td><td>10380</td><td>31</td><td>ACGAGAACTTTGAAGGCCGAAGTGGAGAAGG</td><td>651.9</td><td>-5.8</td><td>1.2</td><td>-4.7</td><td>2.9E-06</td><td>5.2E-06</td><td>MB231/MB468</td><td>MP</td></tr>
<tr><td>>hg38_21_+_8433172_8446622_RNA45SN1_45S_50ntFlanks@9568.9592.25|CGTGCAAATCGGTCGTCCGACCTGG</td><td>>hg38_21_+_8433172_8446622_RNA45SN1_45S_50ntFlanks</td><td>9568</td><td>9592</td><td>25</td><td>CGTGCAAATCGGTCGTCCGACCTGG</td><td>310.3</td><td>-2.8</td><td>0.6</td><td>-4.7</td><td>2.5E-06</td><td>5.2E-06</td><td>BT20/MB468</td><td>MITO</td></tr>
<tr><td>>hg38_21_+_8433172_8446622_RNA45SN1_45S_50ntFlanks@7988.8016.29|CAGACGTGGCGACCCGCTGAATTTAAGCA</td><td>>hg38_21_+_8433172_8446622_RNA45SN1_45S_50ntFlanks</td><td>7988</td><td>8016</td><td>29</td><td>CAGACGTGGCGACCCGCTGAATTTAAGCA</td><td>519.0</td><td>3.7</td><td>0.8</td><td>4.7</td><td>3.3E-06</td><td>5.2E-06</td><td>MB231/MB468</td><td>CYTO</td></tr>
<tr><td>>MI0000255|hsa-mir-30d&WithFlank&8|-|134804870|134804951@12.34.23(+1C)&[MIMAT0000245&hsa-miR-30d-5p&offsets|0|+1(+1C);m-21&8|-|134804919|134804940&offsets|0|+1(+1C)]|TGTAAACATCCCCGACTGGAAGCC</td><td>>MI0000255|hsa-mir-30d&WithFlank&8|-|134804870|134804951</td><td>12</td><td>34</td><td>23(+1C)</td><td>TGTAAACATCCCCGACTGGAAGCC</td><td>3266.5</td><td>4.4</td><td>0.9</td><td>4.9</td><td>9.9E-07</td><td>5.2E-06</td><td>BT20/MB468</td><td>MP</td></tr>
<tr><td>>hg38_21_+_8433172_8446622_RNA45SN1_45S_50ntFlanks@6661.6685.25|CGGTGGATCACTCGGCTCGTGCGTC</td><td>>hg38_21_+_8433172_8446622_RNA45SN1_45S_50ntFlanks</td><td>6661</td><td>6685</td><td>25</td><td>CGGTGGATCACTCGGCTCGTGCGTC</td><td>385.0</td><td>2.2</td><td>0.5</td><td>4.6</td><td>3.4E-06</td><td>5.2E-06</td><td>MB231/MB468</td><td>NUC</td></tr>
<tr><td>>hg38_21_+_8433172_8446622_RNA45SN1_45S_50ntFlanks@8177.8206.30|TTCTGATCGAGGCCCAGCCCGTGGACGGTG</td><td>>hg38_21_+_8433172_8446622_RNA45SN1_45S_50ntFlanks</td><td>8177</td><td>8206</td><td>30</td><td>TTCTGATCGAGGCCCAGCCCGTGGACGGTG</td><td>455.5</td><td>-3.1</td><td>0.7</td><td>-4.6</td><td>3.4E-06</td><td>5.2E-06</td><td>MB231/MB468</td><td>NUC</td></tr>
<tr><td>>hg38_21_+_8433172_8446622_RNA45SN1_45S_50ntFlanks@12892.12931.40|CCCTCGCCCGTCACGCACCGCACGTTCGTGGGGAACCTGG</td><td>>hg38_21_+_8433172_8446622_RNA45SN1_45S_50ntFlanks</td><td>12892</td><td>12931</td><td>40</td><td>CCCTCGCCCGTCACGCACCGCACGTTCGTGGGGAACCTGG</td><td>92.3</td><td>-9.6</td><td>2.0</td><td>-4.9</td><td>9.9E-07</td><td>5.2E-06</td><td>BT20/MB468</td><td>MP</td></tr>
<tr><td>>hg38_21_+_8433172_8446622_RNA45SN1_45S_50ntFlanks@12004.12038.35|CGGGCCGCCGGTGAAATACCACTACTCTGATCGTT</td><td>>hg38_21_+_8433172_8446622_RNA45SN1_45S_50ntFlanks</td><td>12004</td><td>12038</td><td>35</td><td>CGGGCCGCCGGTGAAATACCACTACTCTGATCGTT</td><td>172.0</td><td>10.8</td><td>2.1</td><td>5.1</td><td>2.6E-07</td><td>5.2E-06</td><td>BT20/MB231</td><td>MP</td></tr>
<tr><td>>MI0000077|hsa-mir-21&WithFlank&17|+|59841260|59841343@14.35.22(+2U)&[MIMAT0000076&hsa-miR-21-5p&offsets|0|0(+2U);m-1&17|+|59841273|59841294&offsets|0|0(+2U)]|TAGCTTATCAGACTGATGTTGATT</td><td>>MI0000077|hsa-mir-21&WithFlank&17|+|59841260|59841343</td><td>14</td><td>35</td><td>22(+2U)</td><td>TAGCTTATCAGACTGATGTTGATT</td><td>325.7</td><td>7.9</td><td>1.7</td><td>4.6</td><td>3.5E-06</td><td>5.2E-06</td><td>MB231/MB468</td><td>MITO</td></tr>
<tr><td>>hg38_21_+_8433172_8446622_RNA45SN1_45S_50ntFlanks@11592.11616.25|ACCAGGGGAATCCGACTGTTTAATT</td><td>>hg38_21_+_8433172_8446622_RNA45SN1_45S_50ntFlanks</td><td>11592</td><td>11616</td><td>25</td><td>ACCAGGGGAATCCGACTGTTTAATT</td><td>334.0</td><td>-3.4</td><td>0.7</td><td>-4.6</td><td>3.5E-06</td><td>5.2E-06</td><td>MB231/MB468</td><td>MITO</td></tr>
<tr><td>>hg38_21_+_8433172_8446622_RNA45SN1_45S_50ntFlanks@7969.8005.37|CCGAGACGCGACCTCAGATCAGACGTGGCGACCCGCT</td><td>>hg38_21_+_8433172_8446622_RNA45SN1_45S_50ntFlanks</td><td>7969</td><td>8005</td><td>37</td><td>CCGAGACGCGACCTCAGATCAGACGTGGCGACCCGCT</td><td>242.1</td><td>5.4</td><td>1.2</td><td>4.6</td><td>3.7E-06</td><td>5.2E-06</td><td>BT20/MB468</td><td>NUC</td></tr>
<tr><td>>hg38_21_+_8433172_8446622_RNA45SN1_45S_50ntFlanks@6650.6690.41|ACGACTCTTAGCGGTGGATCACTCGGCTCGTGCGTCGATGA</td><td>>hg38_21_+_8433172_8446622_RNA45SN1_45S_50ntFlanks</td><td>6650</td><td>6690</td><td>41</td><td>ACGACTCTTAGCGGTGGATCACTCGGCTCGTGCGTCGATGA</td><td>761.7</td><td>3.0</td><td>0.6</td><td>4.7</td><td>2.5E-06</td><td>5.3E-06</td><td>BT20/MB468</td><td>MITO</td></tr>
<tr><td>>hg38_21_+_8433172_8446622_RNA45SN1_45S_50ntFlanks@12841.12869.29|CGGAGTGCCCTTCGTCCTGGGAAACGGGG</td><td>>hg38_21_+_8433172_8446622_RNA45SN1_45S_50ntFlanks</td><td>12841</td><td>12869</td><td>29</td><td>CGGAGTGCCCTTCGTCCTGGGAAACGGGG</td><td>143.5</td><td>-11.0</td><td>2.4</td><td>-4.7</td><td>2.9E-06</td><td>5.3E-06</td><td>MB231/MB468</td><td>MP</td></tr>
<tr><td>>hg38_21_+_8433172_8446622_RNA45SN1_45S_50ntFlanks@12503.12521.19|TAGACCGTCGTGAGACAGG</td><td>>hg38_21_+_8433172_8446622_RNA45SN1_45S_50ntFlanks</td><td>12503</td><td>12521</td><td>19</td><td>TAGACCGTCGTGAGACAGG</td><td>82.5</td><td>-9.4</td><td>1.9</td><td>-4.9</td><td>9.9E-07</td><td>5.3E-06</td><td>BT20/MB468</td><td>MP</td></tr>
<tr><td>>MI0000449|hsa-mir-132&WithFlank&17|-|2049902|2050014@29.49.21&[MIMAT0004594&hsa-miR-132-5p&offsets|0|-1;m-414&17|-|2049965|2049986&offsets|0|-1]|ACCGTGGCTTTCGATTGTTAC</td><td>>MI0000449|hsa-mir-132&WithFlank&17|-|2049902|2050014</td><td>29</td><td>49</td><td>21</td><td>ACCGTGGCTTTCGATTGTTAC</td><td>62.5</td><td>9.7</td><td>2.0</td><td>4.9</td><td>1.0E-06</td><td>5.3E-06</td><td>BT20/MB468</td><td>MP</td></tr>
<tr><td>>MI0000078|hsa-mir-22&WithFlank&17|-|1713897|1713993@59.77.19(+1C)&[MIMAT0000077&hsa-miR-22-3p&offsets|0|-3(+1C);m-2&17|-|1713914|1713935&offsets|0|-3(+1C)]|AAGCTGCCAGTTGAAGAACC</td><td>>MI0000078|hsa-mir-22&WithFlank&17|-|1713897|1713993</td><td>59</td><td>77</td><td>19(+1C)</td><td>AAGCTGCCAGTTGAAGAACC</td><td>51.6</td><td>9.4</td><td>1.9</td><td>4.9</td><td>1.0E-06</td><td>5.3E-06</td><td>BT20/MB468</td><td>MP</td></tr>
<tr><td>>MI0000255|hsa-mir-30d&WithFlank&8|-|134804870|134804951@12.34.23(+1C)&[MIMAT0000245&hsa-miR-30d-5p&offsets|0|+1(+1C);m-21&8|-|134804919|134804940&offsets|0|+1(+1C)]|TGTAAACATCCCCGACTGGAAGCC</td><td>>MI0000255|hsa-mir-30d&WithFlank&8|-|134804870|134804951</td><td>12</td><td>34</td><td>23(+1C)</td><td>TGTAAACATCCCCGACTGGAAGCC</td><td>2760.9</td><td>3.6</td><td>0.8</td><td>4.7</td><td>2.9E-06</td><td>5.3E-06</td><td>MB231/MB468</td><td>MP</td></tr>
<tr><td>>hg38_21_+_8433172_8446622_RNA45SN1_45S_50ntFlanks@4722.4755.34|TCATTAATCAAGAACGAAAGTCGGAGGTTCGAAG</td><td>>hg38_21_+_8433172_8446622_RNA45SN1_45S_50ntFlanks</td><td>4722</td><td>4755</td><td>34</td><td>TCATTAATCAAGAACGAAAGTCGGAGGTTCGAAG</td><td>133.0</td><td>-4.4</td><td>0.9</td><td>-4.7</td><td>3.3E-06</td><td>5.3E-06</td><td>MB231/MB468</td><td>CYTO</td></tr>
<tr><td>>MI0000542|hsa-mir-320a&WithFlank&8|-|22244960|22245043@49.68.20(+1C)&[MIMAT0000510&hsa-miR-320a-3p&offsets|+1|-1(+1C);m-60&8|-|22244975|22244996&offsets|+1|-1(+1C)]|AAAGCTGGGTTGAGAGGGCGC</td><td>>MI0000542|hsa-mir-320a&WithFlank&8|-|22244960|22245043</td><td>49</td><td>68</td><td>20(+1C)</td><td>AAAGCTGGGTTGAGAGGGCGC</td><td>267.6</td><td>8.7</td><td>1.8</td><td>4.9</td><td>1.0E-06</td><td>5.3E-06</td><td>BT20/MB468</td><td>MP</td></tr>
<tr><td>>hg38_21_+_8433172_8446622_RNA45SN1_45S_50ntFlanks@5475.5506.32|GCCCTGGCGGAGCGCTGAGAAGACGGTCGAAC</td><td>>hg38_21_+_8433172_8446622_RNA45SN1_45S_50ntFlanks</td><td>5475</td><td>5506</td><td>32</td><td>GCCCTGGCGGAGCGCTGAGAAGACGGTCGAAC</td><td>55.9</td><td>9.6</td><td>2.0</td><td>4.9</td><td>1.0E-06</td><td>5.3E-06</td><td>BT20/MB468</td><td>MP</td></tr>
<tr><td>>hg38_21_+_8433172_8446622_RNA45SN1_45S_50ntFlanks@3802.3841.40|CATTAAATCAGTTATGGTTCCTTTGGTCGCTCGCTCCTCT</td><td>>hg38_21_+_8433172_8446622_RNA45SN1_45S_50ntFlanks</td><td>3802</td><td>3841</td><td>40</td><td>CATTAAATCAGTTATGGTTCCTTTGGTCGCTCGCTCCTCT</td><td>197.1</td><td>-4.0</td><td>0.9</td><td>-4.6</td><td>3.4E-06</td><td>5.3E-06</td><td>MB231/MB468</td><td>NUC</td></tr>
<tr><td>>hg38_21_+_8433172_8446622_RNA45SN1_45S_50ntFlanks@11208.11251.44|GCGGCGACTCTGGACGCGAGCCGGGCCCTTCCCGTGGATCGCCC</td><td>>hg38_21_+_8433172_8446622_RNA45SN1_45S_50ntFlanks</td><td>11208</td><td>11251</td><td>44</td><td>GCGGCGACTCTGGACGCGAGCCGGGCCCTTCCCGTGGATCGCCC</td><td>307.0</td><td>-4.7</td><td>1.0</td><td>-4.7</td><td>3.3E-06</td><td>5.3E-06</td><td>MB231/MB468</td><td>CYTO</td></tr>
<tr><td>>hg38_21_+_8433172_8446622_RNA45SN1_45S_50ntFlanks@11211.11243.33|GCGACTCTGGACGCGAGCCGGGCCCTTCCCGTG</td><td>>hg38_21_+_8433172_8446622_RNA45SN1_45S_50ntFlanks</td><td>11211</td><td>11243</td><td>33</td><td>GCGACTCTGGACGCGAGCCGGGCCCTTCCCGTG</td><td>193.9</td><td>10.9</td><td>2.1</td><td>5.1</td><td>2.6E-07</td><td>5.3E-06</td><td>BT20/MB231</td><td>MP</td></tr>
<tr><td>>MI0000102|hsa-mir-100&WithFlank&11|-|122152223|122152314@19.41.23&[MIMAT0000098&hsa-miR-100-5p&offsets|0|+1;m-27&11|-|122152276|122152296&offsets|0|+2]|AACCCGTAGATCCGAACTTGTGG</td><td>>MI0000102|hsa-mir-100&WithFlank&11|-|122152223|122152314</td><td>19</td><td>41</td><td>23</td><td>AACCCGTAGATCCGAACTTGTGG</td><td>3314.4</td><td>-7.5</td><td>1.5</td><td>-5.1</td><td>2.6E-07</td><td>5.3E-06</td><td>BT20/MB231</td><td>MP</td></tr>
<tr><td>>MI0003137|hsa-mir-193b&WithFlank&16|+|14303961|14304055@57.78.22&[MIMAT0002819&hsa-miR-193b-3p&offsets|0|0;m-166&16|+|14304017|14304038&offsets|0|0]|AACTGGCCCTCAAAGTCCCGCT</td><td>>MI0003137|hsa-mir-193b&WithFlank&16|+|14303961|14304055</td><td>57</td><td>78</td><td>22</td><td>AACTGGCCCTCAAAGTCCCGCT</td><td>6320.0</td><td>5.5</td><td>1.1</td><td>4.9</td><td>1.0E-06</td><td>5.3E-06</td><td>BT20/MB468</td><td>MP</td></tr>
<tr><td>>MI0000077|hsa-mir-21&WithFlank&17|+|59841260|59841343@14.34.21(+2C)&[MIMAT0000076&hsa-miR-21-5p&offsets|0|-1(+2C);m-1&17|+|59841273|59841294&offsets|0|-1(+2C)]|TAGCTTATCAGACTGATGTTGCC</td><td>>MI0000077|hsa-mir-21&WithFlank&17|+|59841260|59841343</td><td>14</td><td>34</td><td>21(+2C)</td><td>TAGCTTATCAGACTGATGTTGCC</td><td>955.5</td><td>4.2</td><td>0.9</td><td>4.7</td><td>2.9E-06</td><td>5.3E-06</td><td>MB231/MB468</td><td>MP</td></tr>
<tr><td>>hg38_21_+_8433172_8446622_RNA45SN1_45S_50ntFlanks@12437.12474.38|CATTGTGAAGCAGAATTCACCAAGCGTTGGATTGTTCA</td><td>>hg38_21_+_8433172_8446622_RNA45SN1_45S_50ntFlanks</td><td>12437</td><td>12474</td><td>38</td><td>CATTGTGAAGCAGAATTCACCAAGCGTTGGATTGTTCA</td><td>311.6</td><td>-7.4</td><td>1.6</td><td>-4.7</td><td>2.9E-06</td><td>5.3E-06</td><td>MB231/MB468</td><td>MP</td></tr>
<tr><td>>hg38_21_+_8433172_8446622_RNA45SN1_45S_50ntFlanks@10692.10709.18|TCCGGTGAGCTCTCGCTG</td><td>>hg38_21_+_8433172_8446622_RNA45SN1_45S_50ntFlanks</td><td>10692</td><td>10709</td><td>18</td><td>TCCGGTGAGCTCTCGCTG</td><td>144.0</td><td>-5.7</td><td>1.2</td><td>-4.7</td><td>2.9E-06</td><td>5.3E-06</td><td>BT20/MB468</td><td>CYTO</td></tr>
<tr><td>>trnaMT_ValTAC_MT_+_1602_1670@42.71.30|AGATTTCAACTTAACTTGACCGCTCTGACC</td><td>>trnaMT_ValTAC_MT_+_1602_1670</td><td>42</td><td>71</td><td>30</td><td>AGATTTCAACTTAACTTGACCGCTCTGACC</td><td>11810.8</td><td>4.3</td><td>0.9</td><td>4.6</td><td>3.5E-06</td><td>5.3E-06</td><td>MB231/MB468</td><td>MITO</td></tr>
</table>

<table>
<tr><td></td><td></td><td></td><td></td><td></td><td></td><td></td><td></td><td></td><td></td><td></td><td></td><td></td><td></td></tr>
<tr><td>>hg38_21_+_8433172_8446622_RNA45SN1_45S_50ntFlanks@10346.10374.29|TCAAACGAGAACTTTGAAGGCCGAAGTGG</td><td>>hg38_21_+_8433172_8446622_RNA45SN1_45S_50ntFlanks</td><td>10346</td><td>10374</td><td>29</td><td>TCAAACGAGAACTTTGAAGGCCGAAGTGG</td><td>678.2</td><td>-2.7</td><td>0.6</td><td>-4.7</td><td>2.9E-06</td><td>5.3E-06</td><td>BT20/MB468</td><td>CYTO</td></tr>
<tr><td>>hg38_21_+_8433172_8446622_RNA45SN1_45S_50ntFlanks@9526.9569.44|GAAGCCAGAGGAAACTCTGGTGGAGGTCCGTAGCGGTCCTGACG</td><td>>hg38_21_+_8433172_8446622_RNA45SN1_45S_50ntFlanks</td><td>9526</td><td>9569</td><td>44</td><td>GAAGCCAGAGGAAACTCTGGTGGAGGTCCGTAGCGGTCCTGACG</td><td>320.2</td><td>6.1</td><td>1.3</td><td>4.6</td><td>3.3E-06</td><td>5.3E-06</td><td>MB231/MB468</td><td>CYTO</td></tr>
<tr><td>>hg38_21_+_8433172_8446622_RNA45SN1_45S_50ntFlanks@11817.11844.28|CCTACTATCCAGCGAAACCACAGCCAAG</td><td>>hg38_21_+_8433172_8446622_RNA45SN1_45S_50ntFlanks</td><td>11817</td><td>11844</td><td>28</td><td>CCTACTATCCAGCGAAACCACAGCCAAG</td><td>51.1</td><td>-8.8</td><td>1.8</td><td>-4.9</td><td>1.0E-06</td><td>5.3E-06</td><td>BT20/MB468</td><td>MP</td></tr>
<tr><td>>hg38_21_+_8433172_8446622_RNA45SN1_45S_50ntFlanks@6709.6731.23|AGAATTAATGTGAATTGCAGGAC</td><td>>hg38_21_+_8433172_8446622_RNA45SN1_45S_50ntFlanks</td><td>6709</td><td>6731</td><td>23</td><td>AGAATTAATGTGAATTGCAGGAC</td><td>54.9</td><td>-8.9</td><td>1.8</td><td>-4.9</td><td>1.0E-06</td><td>5.3E-06</td><td>BT20/MB468</td><td>MP</td></tr>
<tr><td>>hg38_21_+_8433172_8446622_RNA45SN1_45S_50ntFlanks@11572.11592.21|CCGACTTAGAACTGGTGCGGA</td><td>>hg38_21_+_8433172_8446622_RNA45SN1_45S_50ntFlanks</td><td>11572</td><td>11592</td><td>21</td><td>CCGACTTAGAACTGGTGCGGA</td><td>142.8</td><td>-4.7</td><td>1.0</td><td>-4.6</td><td>3.5E-06</td><td>5.3E-06</td><td>MB231/MB468</td><td>NUC</td></tr>
<tr><td>>MI0000542|hsa-mir-320a&WithFlank&8|-|22244960|22245043@48.71.24(+1C)&[MIMAT0000510&hsa-miR-320a-3p&offsets|0|+2(+1C);m-60&8|-|22244975|22244996&offsets|0|+2(+1C)]|AAAAGCTGGGTTGAGAGGGCGAAAC</td><td>>MI0000542|hsa-mir-320a&WithFlank&8|-|22244960|22245043</td><td>48</td><td>71</td><td>24(+1C)</td><td>AAAAGCTGGGTTGAGAGGGCGAAAC</td><td>246.2</td><td>3.9</td><td>0.8</td><td>5.1</td><td>2.9E-07</td><td>5.3E-06</td><td>BT20/MB231</td><td>CYTO</td></tr>
<tr><td>>MI0001448|hsa-mir-425&WithFlank&3|-|49020142|49020240@64.81.18(+1U)&[MIMAT0001343&hsa-miR-425-3p&offsets|+3|-1(+1U);m-258&3|-|49020160|49020181&offsets|+4|0(+1U)]|GGGAATGTCGTGTCCGCCT</td><td>>MI0001448|hsa-mir-425&WithFlank&3|-|49020142|49020240</td><td>64</td><td>81</td><td>18(+1U)</td><td>GGGAATGTCGTGTCCGCCT</td><td>75.4</td><td>10.0</td><td>2.0</td><td>4.9</td><td>1.0E-06</td><td>5.3E-06</td><td>BT20/MB468</td><td>MP</td></tr>
<tr><td>>MI0000088|hsa-mir-30a&WithFlank&6|-|71403545|71403627@53.71.19&[MIMAT0000088&hsa-miR-30a-3p&offsets|0|-3;m-38&6|-|71403554|71403575&offsets|0|-3]|CTTTCAGTCGGATGTTTGC</td><td>>MI0000088|hsa-mir-30a&WithFlank&6|-|71403545|71403627</td><td>53</td><td>71</td><td>19</td><td>CTTTCAGTCGGATGTTTGC</td><td>2140.3</td><td>4.0</td><td>0.9</td><td>4.7</td><td>2.9E-06</td><td>5.3E-06</td><td>MB231/MB468</td><td>MP</td></tr>
<tr><td>>hg38_MT_+_1621_3279_RNR2_16S_50ntFlanks@382.411.30|GAGCCTGGTGATAGCTGGTTGTCCAAGATA</td><td>>hg38_MT_+_1621_3279_RNR2_16S_50ntFlanks</td><td>382</td><td>411</td><td>30</td><td>GAGCCTGGTGATAGCTGGTTGTCCAAGATA</td><td>39.9</td><td>8.5</td><td>1.8</td><td>4.7</td><td>2.9E-06</td><td>5.3E-06</td><td>MB231/MB468</td><td>MP</td></tr>
<tr><td>>MI0000085|hsa-mir-27a&WithFlank&19|-|13836434|13836523@57.76.20&[MIMAT0000084&hsa-miR-27a-3p&offsets|0|-1;m-43&19|-|13836447|13836467&offsets|0|-1]|TTCACAGTGGCTAAGTTCCG</td><td>>MI0000085|hsa-mir-27a&WithFlank&19|-|13836434|13836523</td><td>57</td><td>76</td><td>20</td><td>TTCACAGTGGCTAAGTTCCG</td><td>13717.1</td><td>4.2</td><td>0.9</td><td>4.6</td><td>3.6E-06</td><td>5.3E-06</td><td>MB231/MB468</td><td>MITO</td></tr>
<tr><td>>MI0000299|hsa-mir-222&WithFlank&X|-|45747009|45747130@75.95.21(+1U)&[MIMAT0000279&hsa-miR-222-3p&offsets|0|0(+1U);m-140&X|-|45747033|45747056&offsets|0|-3(+1U)]|AGCTACATCTGGCTACTGGGTT</td><td>>MI0000299|hsa-mir-222&WithFlank&X|-|45747009|45747130</td><td>75</td><td>95</td><td>21(+1U)</td><td>AGCTACATCTGGCTACTGGGTT</td><td>348.8</td><td>6.8</td><td>1.5</td><td>4.6</td><td>3.6E-06</td><td>5.3E-06</td><td>MB231/MB468</td><td>MITO</td></tr>
<tr><td>>hg38_21_+_8433172_8446622_RNA45SN1_45S_50ntFlanks@6660.6688.29|GCGGTGGATCACTCGGCTCGTGCGTCGAT</td><td>>hg38_21_+_8433172_8446622_RNA45SN1_45S_50ntFlanks</td><td>6660</td><td>6688</td><td>29</td><td>GCGGTGGATCACTCGGCTCGTGCGTCGAT</td><td>493.7</td><td>3.8</td><td>0.8</td><td>5.1</td><td>3.2E-07</td><td>5.3E-06</td><td>BT20/MB231</td><td>MITO</td></tr>
<tr><td>>am_trna111_HisGTG_1_-_147774845_147774916@-1G.26.27,trna118_HisGTG_1_-_145396881_145396952@-1G.26.27,trna16_HisGTG_1_+_146544773_146544844@-1G.26.27,trna1_HisGTG_15_+_45493349_45493420@-1G.26.27,trna21_HisGTG_1_+_147753471_147753542@-1G.26.27,trna33_HisGTG_6_+_27125906_27125977@-1G.26.27,trna7_HisGTG_9_-_14433938_14434009@-1G.26.27,trna8_HisGTG_15_-_45492611_45492682@-1G.26.27,trna9_HisGTG_15_-_45490804_45490875@-1G.26.27|GGCCGTGATCGTATAGTGGTTAGTACT</td><td>>am_trna111_HisGTG_1_-_147774845_147774916</td><td>-1G</td><td>26</td><td>27</td><td>GGCCGTGATCGTATAGTGGTTAGTACT</td><td>970.7</td><td>4.3</td><td>0.9</td><td>4.6</td><td>3.6E-06</td><td>5.3E-06</td><td>MB231/MB468</td><td>MITO</td></tr>
<tr><td>>hg38_21_+_8433172_8446622_RNA45SN1_45S_50ntFlanks@4224.4241.18|AAAATAACAATACAGGAC</td><td>>hg38_21_+_8433172_8446622_RNA45SN1_45S_50ntFlanks</td><td>4224</td><td>4241</td><td>18</td><td>AAAATAACAATACAGGAC</td><td>51.5</td><td>-8.8</td><td>1.8</td><td>-4.9</td><td>1.0E-06</td><td>5.3E-06</td><td>BT20/MB468</td><td>MP</td></tr>
<tr><td>>hg38_1_-_228634819_228635039_RNA5S12_5S_50ntFlanks@151.169.19|ATACCGGGTGCTGTAGGCT</td><td>>hg38_1_-_228634819_228635039_RNA5S12_5S_50ntFlanks</td><td>151</td><td>169</td><td>19</td><td>ATACCGGGTGCTGTAGGCT</td><td>549.9</td><td>-3.4</td><td>0.7</td><td>-4.6</td><td>3.6E-06</td><td>5.3E-06</td><td>MB231/MB468</td><td>MITO</td></tr>
<tr><td>>hg38_21_+_8433172_8446622_RNA45SN1_45S_50ntFlanks@5444.5461.18|CTCGGATCGGCCCCGCCG</td><td>>hg38_21_+_8433172_8446622_RNA45SN1_45S_50ntFlanks</td><td>5444</td><td>5461</td><td>18</td><td>CTCGGATCGGCCCCGCCG</td><td>105.9</td><td>-6.4</td><td>1.4</td><td>-4.7</td><td>3.0E-06</td><td>5.4E-06</td><td>MB231/MB468</td><td>MP</td></tr>
<tr><td>>trna10_MetCAT_8_-_124169470_124169542@31.51.21,trna22_MetCAT_16_-_87417628_87417700@31.51.21|TCTCATAATCTGAAGGTCGTG</td><td>>trna10_MetCAT_8_-_124169470_124169542</td><td>31</td><td>51</td><td>21</td><td>TCTCATAATCTGAAGGTCGTG</td><td>732.4</td><td>-7.4</td><td>1.5</td><td>-4.9</td><td>1.0E-06</td><td>5.4E-06</td><td>BT20/MB468</td><td>MP</td></tr>
<tr><td>>hg38_21_+_8433172_8446622_RNA45SN1_45S_50ntFlanks@10808.10828.21|CAATGTAGGTAAGGGAAGTCG</td><td>>hg38_21_+_8433172_8446622_RNA45SN1_45S_50ntFlanks</td><td>10808</td><td>10828</td><td>21</td><td>CAATGTAGGTAAGGGAAGTCG</td><td>962.7</td><td>-6.7</td><td>1.4</td><td>-4.7</td><td>3.0E-06</td><td>5.4E-06</td><td>BT20/MB468</td><td>CYTO</td></tr>
<tr><td>>hg38_21_+_8433172_8446622_RNA45SN1_45S_50ntFlanks@3875.3899.25|ATACATGCCGACGGGCGCTGACCCC</td><td>>hg38_21_+_8433172_8446622_RNA45SN1_45S_50ntFlanks</td><td>3875</td><td>3899</td><td>25</td><td>ATACATGCCGACGGGCGCTGACCCC</td><td>1540.9</td><td>-6.8</td><td>1.5</td><td>-4.6</td><td>3.7E-06</td><td>5.4E-06</td><td>BT20/MB468</td><td>NUC</td></tr>
<tr><td>>hg38_21_+_8433172_8446622_RNA45SN1_45S_50ntFlanks@12540.12569.30|TGATGTGTTGTTGCCATGGTAATCCTGCTC</td><td>>hg38_21_+_8433172_8446622_RNA45SN1_45S_50ntFlanks</td><td>12540</td><td>12569</td><td>30</td><td>TGATGTGTTGTTGCCATGGTAATCCTGCTC</td><td>280.3</td><td>-6.4</td><td>1.4</td><td>-4.6</td><td>3.5E-06</td><td>5.4E-06</td><td>MB231/MB468</td><td>NUC</td></tr>
<tr><td>>MI0000285|hsa-mir-205&WithFlank&1|+|209432127|209432248@40.61.22(+1C)&[MIMAT0000266&hsa-miR-205-5p&offsets|0|0(+1C);m-351&1|+|209432166|209432187&offsets|0|0(+1C)]|TCCTTCATTCCACCGGAGTCTGC</td><td>>MI0000285|hsa-mir-205&WithFlank&1|+|209432127|209432248</td><td>40</td><td>61</td><td>22(+1C)</td><td>TCCTTCATTCCACCGGAGTCTGC</td><td>9774.5</td><td>11.1</td><td>2.2</td><td>5.1</td><td>2.9E-07</td><td>5.4E-06</td><td>BT20/MB231</td><td>CYTO</td></tr>
<tr><td>>MI0000478|hsa-mir-149&WithFlank&2|+|240455995|240456095@21.41.21(+1A)&[MIMAT0000450&hsa-miR-149-5p&offsets|0|-2(+1A);m-195&2|+|240456015|240456037&offsets|0|-2(+1A)]|TCTGGCTCCGTGTCTTCACTCA</td><td>>MI0000478|hsa-mir-149&WithFlank&2|+|240455995|240456095</td><td>21</td><td>41</td><td>21(+1A)</td><td>TCTGGCTCCGTGTCTTCACTCA</td><td>490.1</td><td>4.4</td><td>0.9</td><td>5.1</td><td>2.9E-07</td><td>5.4E-06</td><td>BT20/MB231</td><td>CYTO</td></tr>
<tr><td>>trna13_ValCAC_19_-_4724647_4724719@1.34.34|GTTTCCGTAGTGTAGCGGTTATCACATTCGCCTC</td><td>>trna13_ValCAC_19_-_4724647_4724719</td><td>1</td><td>34</td><td>34</td><td>GTTTCCGTAGTGTAGCGGTTATCACATTCGCCTC</td><td>1043.2</td><td>-5.1</td><td>1.0</td><td>-5.1</td><td>2.9E-07</td><td>5.4E-06</td><td>BT20/MB231</td><td>CYTO</td></tr>
<tr><td>>hg38_21_+_8433172_8446622_RNA45SN1_45S_50ntFlanks@7977.8016.40|CGACCTCAGATCAGACGTGGCGACCCGCTGAATTTAAGCA</td><td>>hg38_21_+_8433172_8446622_RNA45SN1_45S_50ntFlanks</td><td>7977</td><td>8016</td><td>40</td><td>CGACCTCAGATCAGACGTGGCGACCCGCTGAATTTAAGCA</td><td>2507.8</td><td>-5.2</td><td>1.0</td><td>-5.1</td><td>2.9E-07</td><td>5.4E-06</td><td>BT20/MB231</td><td>CYTO</td></tr>
<tr><td>>MI0000063|hsa-let-7b&WithFlank&22|+|46113680|46113774@12.34.23&[MIMAT0000063&hsa-let-7b-5p&offsets|0|+1;m-14&22|+|46113691|46113712&offsets|0|+1]|TGAGGTAGTAGGTTGTGTGGTTT</td><td>>MI0000063|hsa-let-7b&WithFlank&22|+|46113680|46113774</td><td>12</td><td>34</td><td>23</td><td>TGAGGTAGTAGGTTGTGTGGTTT</td><td>5146.1</td><td>3.5</td><td>0.8</td><td>4.7</td><td>3.0E-06</td><td>5.4E-06</td><td>MB231/MB468</td><td>MP</td></tr>
<tr><td>>hg38_21_+_8433172_8446622_RNA45SN1_45S_50ntFlanks@12997.13015.19|CGCTGCGATCTATTGAAAG</td><td>>hg38_21_+_8433172_8446622_RNA45SN1_45S_50ntFlanks</td><td>12997</td><td>13015</td><td>19</td><td>CGCTGCGATCTATTGAAAG</td><td>932.7</td><td>2.4</td><td>0.5</td><td>4.6</td><td>3.5E-06</td><td>5.4E-06</td><td>MB231/MB468</td><td>NUC</td></tr>
<tr><td>>hg38_21_+_8433172_8446622_RNA45SN1_45S_50ntFlanks@485.530.46|GATGTGGTGACGTCGTGCTCTCCCGGGCCGGGTCCGAGCCGCGACG</td><td>>hg38_21_+_8433172_8446622_RNA45SN1_45S_50ntFlanks</td><td>485</td><td>530</td><td>46</td><td>GATGTGGTGACGTCGTGCTCTCCCGGGCCGGGTCCGAGCCGCGACG</td><td>78.2</td><td>10.0</td><td>2.1</td><td>4.9</td><td>1.0E-06</td><td>5.4E-06</td><td>BT20/MB468</td><td>MP</td></tr>
<tr><td>>hg38_21_+_8433172_8446622_RNA45SN1_45S_50ntFlanks@8192.8212.21|AGCCCGTGGACGGTGTGAGGC</td><td>>hg38_21_+_8433172_8446622_RNA45SN1_45S_50ntFlanks</td><td>8192</td><td>8212</td><td>21</td><td>AGCCCGTGGACGGTGTGAGGC</td><td>384.0</td><td>-3.5</td><td>0.8</td><td>-4.7</td><td>3.0E-06</td><td>5.4E-06</td><td>MB231/MB468</td><td>MP</td></tr>
<tr><td>>hg38_21_+_8433172_8446622_RNA45SN1_45S_50ntFlanks@11570.11612.43|AGCCGACTTAGAACTGGTGCGGACCAGGGGAATCCGACTGTTT</td><td>>hg38_21_+_8433172_8446622_RNA45SN1_45S_50ntFlanks</td><td>11570</td><td>11612</td><td>43</td><td>AGCCGACTTAGAACTGGTGCGGACCAGGGGAATCCGACTGTTT</td><td>1566.4</td><td>-4.4</td><td>0.9</td><td>-4.6</td><td>3.6E-06</td><td>5.4E-06</td><td>MB231/MB468</td><td>MITO</td></tr>
<tr><td>>hg38_21_+_8433172_8446622_RNA45SN1_45S_50ntFlanks@11829.11852.24|CGAAACCACAGCCAAGGGAACGGG</td><td>>hg38_21_+_8433172_8446622_RNA45SN1_45S_50ntFlanks</td><td>11829</td><td>11852</td><td>24</td><td>CGAAACCACAGCCAAGGGAACGGG</td><td>49.8</td><td>-8.7</td><td>1.8</td><td>-4.9</td><td>1.0E-06</td><td>5.4E-06</td><td>BT20/MB468</td><td>MP</td></tr>
<tr><td>>hg38_21_+_8433172_8446622_RNA45SN1_45S_50ntFlanks@12606.12625.20|ATGTGCTTGGCTGAGGAGCC</td><td>>hg38_21_+_8433172_8446622_RNA45SN1_45S_50ntFlanks</td><td>12606</td><td>12625</td><td>20</td><td>ATGTGCTTGGCTGAGGAGCC</td><td>863.2</td><td>-3.1</td><td>0.7</td><td>-4.6</td><td>3.5E-06</td><td>5.4E-06</td><td>MB231/MB468</td><td>NUC</td></tr>
<tr><td>>hg38_21_+_8433172_8446622_RNA45SN1_45S_50ntFlanks@7980.8005.26|CCTCAGATCAGACGTGGCGACCCGCT</td><td>>hg38_21_+_8433172_8446622_RNA45SN1_45S_50ntFlanks</td><td>7980</td><td>8005</td><td>26</td><td>CCTCAGATCAGACGTGGCGACCCGCT</td><td>147.2</td><td>10.5</td><td>2.0</td><td>5.1</td><td>2.7E-07</td><td>5.4E-06</td><td>BT20/MB231</td><td>MP</td></tr>
<tr><td>>hg38_21_+_8433172_8446622_RNA45SN1_45S_50ntFlanks@475.504.30|CTTCGTGATCGATGTGGTGACGTCGTGCTC</td><td>>hg38_21_+_8433172_8446622_RNA45SN1_45S_50ntFlanks</td><td>475</td><td>504</td><td>30</td><td>CTTCGTGATCGATGTGGTGACGTCGTGCTC</td><td>467.5</td><td>3.6</td><td>0.8</td><td>4.7</td><td>3.0E-06</td><td>5.4E-06</td><td>MB231/MB468</td><td>MP</td></tr>
<tr><td>>hg38_21_+_8433172_8446622_RNA45SN1_45S_50ntFlanks@9621.9646.26|TCTAGTAGCTGGTTCCCTCCGAAGTT</td><td>>hg38_21_+_8433172_8446622_RNA45SN1_45S_50ntFlanks</td><td>9621</td><td>9646</td><td>26</td><td>TCTAGTAGCTGGTTCCCTCCGAAGTT</td><td>254.8</td><td>-6.4</td><td>1.4</td><td>-4.6</td><td>3.6E-06</td><td>5.4E-06</td><td>MB231/MB468</td><td>MITO</td></tr>
<tr><td>>hg38_21_+_8433172_8446622_RNA45SN1_45S_50ntFlanks@5236.5256.21|CACTGACTGGCTCAGCGTGTG</td><td>>hg38_21_+_8433172_8446622_RNA45SN1_45S_50ntFlanks</td><td>5236</td><td>5256</td><td>21</td><td>CACTGACTGGCTCAGCGTGTG</td><td>171.1</td><td>-3.8</td><td>0.8</td><td>-4.7</td><td>3.0E-06</td><td>5.4E-06</td><td>MB231/MB468</td><td>MP</td></tr>
<tr><td>>hg38_21_+_8433172_8446622_RNA45SN1_45S_50ntFlanks@10246.10279.34|CGCGACGAGTAGGAGGGCCGCTGCGGTGAGCCTT</td><td>>hg38_21_+_8433172_8446622_RNA45SN1_45S_50ntFlanks</td><td>10246</td><td>10279</td><td>34</td><td>CGCGACGAGTAGGAGGGCCGCTGCGGTGAGCCTT</td><td>199.0</td><td>11.0</td><td>2.1</td><td>5.1</td><td>2.7E-07</td><td>5.4E-06</td><td>BT20/MB231</td><td>MP</td></tr>
<tr><td>>hg38_21_+_8433172_8446622_RNA45SN1_45S_50ntFlanks@6774.6807.34|TCCTCCCGGGGCTACGCCTGTCTGAGCGTCGCTT</td><td>>hg38_21_+_8433172_8446622_RNA45SN1_45S_50ntFlanks</td><td>6774</td><td>6807</td><td>34</td><td>TCCTCCCGGGGCTACGCCTGTCTGAGCGTCGCTT</td><td>87.7</td><td>2.4</td><td>0.5</td><td>4.6</td><td>3.8E-06</td><td>5.4E-06</td><td>BT20/MB468</td><td>NUC</td></tr>
<tr><td>>hg38_21_+_8433172_8446622_RNA45SN1_45S_50ntFlanks@12615.12661.47|GCTGAGGAGCCAATGGGGCGAAGCTACCATCTGTGGGATTATGACTG</td><td>>hg38_21_+_8433172_8446622_RNA45SN1_45S_50ntFlanks</td><td>12615</td><td>12661</td><td>47</td><td>GCTGAGGAGCCAATGGGGCGAAGCTACCATCTGTGGGATTATGACTG</td><td>107.1</td><td>-5.8</td><td>1.2</td><td>-4.7</td><td>3.0E-06</td><td>5.4E-06</td><td>BT20/MB468</td><td>CYTO</td></tr>
<tr><td>>MI0000273|hsa-mir-183&WithFlank&7|-|129774899|129775020@33.54.22&[MIMAT0000261&hsa-miR-183-5p&offsets|0|0;m-41&7|-|129774966|129774987&offsets|-1|-1]|TATGGCACTGGTAGAATTCACT</td><td>>MI0000273|hsa-mir-183&WithFlank&7|-|129774899|129775020</td><td>33</td><td>54</td><td>22</td><td>TATGGCACTGGTAGAATTCACT</td><td>2307.1</td><td>3.5</td><td>0.7</td><td>4.6</td><td>3.4E-06</td><td>5.4E-06</td><td>MB231/MB468</td><td>CYTO</td></tr>
<tr><td>>MI0000811|hsa-mir-148b&WithFlank&12|+|54337210|54337320@69.90.22(+2U)&[MIMAT0000759&hsa-miR-148b-3p&offsets|0|0(+2U);m-77&12|+|54337278|54337299&offsets|0|0(+2U)]|TCAGTGCATCACAGAACTTTGTTT</td><td>>MI0000811|hsa-mir-148b&WithFlank&12|+|54337210|54337320</td><td>69</td><td>90</td><td>22(+2U)</td><td>TCAGTGCATCACAGAACTTTGTTT</td><td>619.8</td><td>7.2</td><td>1.6</td><td>4.6</td><td>3.6E-06</td><td>5.4E-06</td><td>MB231/MB468</td><td>MITO</td></tr>
<tr><td>>hg38_21_+_8433172_8446622_RNA45SN1_45S_50ntFlanks@8169.8214.46|CCAAGTCCTTCTGATCGAGGCCCAGCCCGTGGACGGTGTGAGGCCG</td><td>>hg38_21_+_8433172_8446622_RNA45SN1_45S_50ntFlanks</td><td>8169</td><td>8214</td><td>46</td><td>CCAAGTCCTTCTGATCGAGGCCCAGCCCGTGGACGGTGTGAGGCCG</td><td>689.4</td><td>-7.4</td><td>1.6</td><td>-4.6</td><td>3.6E-06</td><td>5.4E-06</td><td>MB231/MB468</td><td>MITO</td></tr>
<tr><td>>hg38_21_+_8433172_8446622_RNA45SN1_45S_50ntFlanks@12642.12659.18|CATCTGTGGGATTATGAC</td><td>>hg38_21_+_8433172_8446622_RNA45SN1_45S_50ntFlanks</td><td>12642</td><td>12659</td><td>18</td><td>CATCTGTGGGATTATGAC</td><td>364.7</td><td>-3.3</td><td>0.7</td><td>-4.6</td><td>3.4E-06</td><td>5.4E-06</td><td>MB231/MB468</td><td>CYTO</td></tr>
<tr><td>>trnaMT_MetCAT_MT_+_4402_4469@49.71.23|GTTATACCCTTCCCGTACTACCA</td><td>>trnaMT_MetCAT_MT_+_4402_4469</td><td>49</td><td>71</td><td>23</td><td>GTTATACCCTTCCCGTACTACCA</td><td>243.0</td><td>6.0</td><td>1.3</td><td>4.6</td><td>3.4E-06</td><td>5.4E-06</td><td>MB231/MB468</td><td>CYTO</td></tr>
<tr><td>>hg38_21_+_8433172_8446622_RNA45SN1_45S_50ntFlanks@11104.11133.30|AAGGGTCGGGGCGGCAGGGGCCGGCGGCGG</td><td>>hg38_21_+_8433172_8446622_RNA45SN1_45S_50ntFlanks</td><td>11104</td><td>11133</td><td>30</td><td>AAGGGTCGGGGCGGCAGGGGCCGGCGGCGG</td><td>397.3</td><td>-7.4</td><td>1.5</td><td>-4.9</td><td>1.0E-06</td><td>5.4E-06</td><td>BT20/MB468</td><td>MP</td></tr>
<tr><td>>hg38_21_+_8433172_8446622_RNA45SN1_45S_50ntFlanks@8198.8226.29|TGGACGGTGTGAGGCCGGTAGCGGCCCCC</td><td>>hg38_21_+_8433172_8446622_RNA45SN1_45S_50ntFlanks</td><td>8198</td><td>8226</td><td>29</td><td>TGGACGGTGTGAGGCCGGTAGCGGCCCCC</td><td>136.7</td><td>-5.5</td><td>1.2</td><td>-4.7</td><td>3.0E-06</td><td>5.4E-06</td><td>BT20/MB468</td><td>CYTO</td></tr>
<tr><td>>MI0001448|hsa-mir-425&WithFlank&3|-|49020142|49020240@20.43.24(+1C)&[MIMAT0003393&hsa-miR-425-5p&offsets|0|+1(+1C);m-95&3|-|49020200|49020221&offsets|0|+2(+1C)]|AATGACACGATCACTCCCGTTGAGC</td><td>>MI0001448|hsa-mir-425&WithFlank&3|-|49020142|49020240</td><td>20</td><td>43</td><td>24(+1C)</td><td>AATGACACGATCACTCCCGTTGAGC</td><td>142.2</td><td>2.6</td><td>0.6</td><td>4.6</td><td>3.8E-06</td><td>5.4E-06</td><td>BT20/MB468</td><td>NUC</td></tr>
<tr><td>>hg38_21_+_8433172_8446622_RNA45SN1_45S_50ntFlanks@10362.10380.19|AAGGCCGAAGTGGAGAAGG</td><td>>hg38_21_+_8433172_8446622_RNA45SN1_45S_50ntFlanks</td><td>10362</td><td>10380</td><td>19</td><td>AAGGCCGAAGTGGAGAAGG</td><td>182.1</td><td>-6.5</td><td>1.4</td><td>-4.7</td><td>3.0E-06</td><td>5.4E-06</td><td>MB231/MB468</td><td>MP</td></tr>
</table>

<table>
<tr><td></td><td></td><td></td><td></td><td></td><td></td><td></td><td></td><td></td><td></td><td></td><td></td><td></td><td></td></tr>
<tr><td>>hg38_21_+_8433172_8446622_RNA45SN1_45S_50ntFlanks@10312.10340.29|CGCAGGTGCAGATCTTGGTGGTAGTAGCA</td><td>>hg38_21_+_8433172_8446622_RNA45SN1_45S_50ntFlanks</td><td>10312</td><td>10340</td><td>29</td><td>CGCAGGTGCAGATCTTGGTGGTAGTAGCA</td><td>223.1</td><td>11.1</td><td>2.2</td><td>5.1</td><td>2.7E-07</td><td>5.5E-06</td><td>BT20/MB231</td><td>MP</td></tr>
<tr><td>>hg38_21_+_8433172_8446622_RNA45SN1_45S_50ntFlanks@12249.12271.23|CAAAAGCTCGCTTGATCTTGATT</td><td>>hg38_21_+_8433172_8446622_RNA45SN1_45S_50ntFlanks</td><td>12249</td><td>12271</td><td>23</td><td>CAAAAGCTCGCTTGATCTTGATT</td><td>481.9</td><td>-5.6</td><td>1.2</td><td>-4.6</td><td>3.6E-06</td><td>5.5E-06</td><td>MB231/MB468</td><td>MITO</td></tr>
<tr><td>>trnaMT_LysTTT_MT_+_8295_8364@1.47.47,trnalookalike7_LysTTT_1_+_568844_568913@1.47.47|CACTGTAAAGCTAACTTAGCATTAACCTTTTAAGTTAAAGATTAAGA</td><td>>trnaMT_LysTTT_MT_+_8295_8364</td><td>1</td><td>47</td><td>47</td><td>CACTGTAAAGCTAACTTAGCATTAACCTTTTAAGTTAAAGATTAAGA</td><td>452.5</td><td>-12.1</td><td>2.4</td><td>-5.1</td><td>3.3E-07</td><td>5.5E-06</td><td>BT20/MB231</td><td>MITO</td></tr>
<tr><td>>hg38_21_+_8433172_8446622_RNA45SN1_45S_50ntFlanks@9573.9595.23|AAATCGGTCGTCCGACCTGGGTA</td><td>>hg38_21_+_8433172_8446622_RNA45SN1_45S_50ntFlanks</td><td>9573</td><td>9595</td><td>23</td><td>AAATCGGTCGTCCGACCTGGGTA</td><td>543.1</td><td>-6.0</td><td>1.3</td><td>-4.6</td><td>3.8E-06</td><td>5.5E-06</td><td>BT20/MB468</td><td>NUC</td></tr>
<tr><td>>hg38_21_+_8433172_8446622_RNA45SN1_45S_50ntFlanks@9567.9603.37|ACGTGCAAATCGGTCGTCCGACCTGGGTATAGGGGCG</td><td>>hg38_21_+_8433172_8446622_RNA45SN1_45S_50ntFlanks</td><td>9567</td><td>9603</td><td>37</td><td>ACGTGCAAATCGGTCGTCCGACCTGGGTATAGGGGCG</td><td>491.3</td><td>-6.0</td><td>1.3</td><td>-4.7</td><td>3.0E-06</td><td>5.5E-06</td><td>MB231/MB468</td><td>MP</td></tr>
<tr><td>>am_trna128_GlyGCC_6_-_27870686_27870756@7.29.23,trna133_GlyCCC_1_-_16872434_16872504@7.29.23,trna17_GluTTC_2_-_204229131_204229201@7.29.23,trna18_GlyGCC_16_+_70822597_70822667@7.29.23,trna19_GlyGCC_16_+_70823410_70823480@7.29.23,trna19_GlyGCC_2_-_157257659_157257729@7.29.23,trna24_GlyGCC_16_-_70812942_70813012@7.29.23,trna25_GlyGCC_16_-_70812114_70812184@7.29.23,trna2_GlyGCC_21_-_18827107_18827177@7.29.23,trna35_GlyGCC_1_+_161413094_161413164@7.29.23,trna37_GlyGCC_1_+_161420467_161420537@7.29.23,trna39_GlyGCC_1_+_161427898_161427968@7.29.23,trna41_GlyGCC_1_+_161435258_161435328@7.29.23,trna4_GlyCCC_1_+_17188416_17188486@7.29.23,trna5_GlyGCC_17_+_8029064_8029134@7.29.23,trna68_GlyGCC_1_-_161493637_161493707@7.29.23,trna91_GlyCCC_1_-_149680210_149680280@7.29.23,trna99_ValCAC_1_-_149294666_149294736@7.29.23|GTGGTTCAGTGGTAGAATTCTCG</td><td>>am_trna128_GlyGCC_6_-_27870686_27870756</td><td>7</td><td>29</td><td>23</td><td>GTGGTTCAGTGGTAGAATTCTCG</td><td>62.8</td><td>-9.1</td><td>1.9</td><td>-4.9</td><td>1.0E-06</td><td>5.5E-06</td><td>BT20/MB468</td><td>MP</td></tr>
<tr><td>>hg38_21_+_8433172_8446622_RNA45SN1_45S_50ntFlanks@11933.11969.37|AATAAGTGGGAGGCCCCCGGCGCCCCCCCGGTGTCCC</td><td>>hg38_21_+_8433172_8446622_RNA45SN1_45S_50ntFlanks</td><td>11933</td><td>11969</td><td>37</td><td>AATAAGTGGGAGGCCCCCGGCGCCCCCCCGGTGTCCC</td><td>74.4</td><td>-8.3</td><td>1.8</td><td>-4.7</td><td>2.6E-06</td><td>5.5E-06</td><td>BT20/MB468</td><td>MITO</td></tr>
<tr><td>>hg38_21_+_8433172_8446622_RNA45SN1_45S_50ntFlanks@6733.6773.41|CATTGATCATCGACACTTCGAACGCACTTGCGGCCCCGGGT</td><td>>hg38_21_+_8433172_8446622_RNA45SN1_45S_50ntFlanks</td><td>6733</td><td>6773</td><td>41</td><td>CATTGATCATCGACACTTCGAACGCACTTGCGGCCCCGGGT</td><td>450.8</td><td>4.4</td><td>0.9</td><td>4.6</td><td>3.6E-06</td><td>5.5E-06</td><td>MB231/MB468</td><td>NUC</td></tr>
<tr><td>>hg38_21_+_8433172_8446622_RNA45SN1_45S_50ntFlanks@8176.8215.40|CTTCTGATCGAGGCCCAGCCCGTGGACGGTGTGAGGCCGG</td><td>>hg38_21_+_8433172_8446622_RNA45SN1_45S_50ntFlanks</td><td>8176</td><td>8215</td><td>40</td><td>CTTCTGATCGAGGCCCAGCCCGTGGACGGTGTGAGGCCGG</td><td>340.6</td><td>-7.3</td><td>1.6</td><td>-4.6</td><td>3.8E-06</td><td>5.5E-06</td><td>BT20/MB468</td><td>NUC</td></tr>
<tr><td>>hg38_21_+_8433172_8446622_RNA45SN1_45S_50ntFlanks@12997.13035.39|CGCTGCGATCTATTGAAAGTCAGCCCTCGACACAAGGGT</td><td>>hg38_21_+_8433172_8446622_RNA45SN1_45S_50ntFlanks</td><td>12997</td><td>13035</td><td>39</td><td>CGCTGCGATCTATTGAAAGTCAGCCCTCGACACAAGGGT</td><td>404.0</td><td>7.6</td><td>1.6</td><td>4.7</td><td>3.0E-06</td><td>5.5E-06</td><td>BT20/MB468</td><td>CYTO</td></tr>
<tr><td>>hg38_21_+_8433172_8446622_RNA45SN1_45S_50ntFlanks@10247.10278.32|GCGACGAGTAGGAGGGCCGCTGCGGTGAGCCT</td><td>>hg38_21_+_8433172_8446622_RNA45SN1_45S_50ntFlanks</td><td>10247</td><td>10278</td><td>32</td><td>GCGACGAGTAGGAGGGCCGCTGCGGTGAGCCT</td><td>155.9</td><td>10.6</td><td>2.1</td><td>5.1</td><td>2.8E-07</td><td>5.5E-06</td><td>BT20/MB231</td><td>MP</td></tr>
<tr><td>>hg38_21_+_8433172_8446622_RNA45SN1_45S_50ntFlanks@12004.12035.32|CGGGCCGCCGGTGAAATACCACTACTCTGATC</td><td>>hg38_21_+_8433172_8446622_RNA45SN1_45S_50ntFlanks</td><td>12004</td><td>12035</td><td>32</td><td>CGGGCCGCCGGTGAAATACCACTACTCTGATC</td><td>2087.1</td><td>7.2</td><td>1.4</td><td>5.1</td><td>2.8E-07</td><td>5.5E-06</td><td>BT20/MB231</td><td>MP</td></tr>
<tr><td>>hg38_21_+_8433172_8446622_RNA45SN1_45S_50ntFlanks@9483.9509.27|CGTGTTAGGACCCGAAAGATGGTGAAC</td><td>>hg38_21_+_8433172_8446622_RNA45SN1_45S_50ntFlanks</td><td>9483</td><td>9509</td><td>27</td><td>CGTGTTAGGACCCGAAAGATGGTGAAC</td><td>234.9</td><td>11.2</td><td>2.2</td><td>5.1</td><td>2.8E-07</td><td>5.5E-06</td><td>BT20/MB231</td><td>MP</td></tr>
<tr><td>>MI0000446|hsa-mir-125b-1&WithFlank&11|-|122099751|122099850@61.80.20&[MIMAT0004592&hsa-miR-125b-1-3p&offsets|0|-2;m-485&11|-|122099769|122099790&offsets|0|-2]|ACGGGTTAGGCTCTTGGGAG</td><td>>MI0000446|hsa-mir-125b-1&WithFlank&11|-|122099751|122099850</td><td>61</td><td>80</td><td>20</td><td>ACGGGTTAGGCTCTTGGGAG</td><td>103.9</td><td>-10.1</td><td>2.0</td><td>-5.1</td><td>2.8E-07</td><td>5.5E-06</td><td>BT20/MB231</td><td>MP</td></tr>
<tr><td>>MI0000469|hsa-mir-125a&WithFlank&19|+|51693248|51693345@21.44.24&[MIMAT0000443&hsa-miR-125a-5p&offsets|0|0;m-63&19|+|51693268|51693289&offsets|0|+2]|TCCCTGAGACCCTTTAACCTGTGA</td><td>>MI0000469|hsa-mir-125a&WithFlank&19|+|51693248|51693345</td><td>21</td><td>44</td><td>24</td><td>TCCCTGAGACCCTTTAACCTGTGA</td><td>3166.4</td><td>4.4</td><td>1.0</td><td>4.6</td><td>3.6E-06</td><td>5.5E-06</td><td>MB231/MB468</td><td>MITO</td></tr>
<tr><td>>hg38_21_+_8433172_8446622_RNA45SN1_45S_50ntFlanks@10402.10426.25|AACATGGGTCAGTCGGTCCTGAGAG</td><td>>hg38_21_+_8433172_8446622_RNA45SN1_45S_50ntFlanks</td><td>10402</td><td>10426</td><td>25</td><td>AACATGGGTCAGTCGGTCCTGAGAG</td><td>294.4</td><td>-2.4</td><td>0.5</td><td>-4.7</td><td>2.6E-06</td><td>5.5E-06</td><td>BT20/MB468</td><td>MITO</td></tr>
<tr><td>>MI0000088|hsa-mir-30a&WithFlank&6|-|71403545|71403627@12.35.24(+1U)&[MIMAT0000087&hsa-miR-30a-5p&offsets|0|+2(+1U);m-15&6|-|71403595|71403616&offsets|0|+2(+1U)]|TGTAAACATCCTCGACTGGAAGCTT</td><td>>MI0000088|hsa-mir-30a&WithFlank&6|-|71403545|71403627</td><td>12</td><td>35</td><td>24(+1U)</td><td>TGTAAACATCCTCGACTGGAAGCTT</td><td>702.2</td><td>5.9</td><td>1.3</td><td>4.7</td><td>3.0E-06</td><td>5.5E-06</td><td>BT20/MB468</td><td>CYTO</td></tr>
<tr><td>>trna152_ValCAC_6_-_27248049_27248121@1.35.35|GCTTCTGTAGTGTAGTGGTTATCACGTTCGCCTCA</td><td>>trna152_ValCAC_6_-_27248049_27248121</td><td>1</td><td>35</td><td>35</td><td>GCTTCTGTAGTGTAGTGGTTATCACGTTCGCCTCA</td><td>74.9</td><td>10.0</td><td>2.0</td><td>4.9</td><td>1.0E-06</td><td>5.5E-06</td><td>BT20/MB468</td><td>MP</td></tr>
<tr><td>>hg38_MT_+_1621_3279_RNR2_16S_50ntFlanks@1415.1452.38|CGTTTGTTCAACGATTAAAGTCCTACGTGATCTGAGTT</td><td>>hg38_MT_+_1621_3279_RNR2_16S_50ntFlanks</td><td>1415</td><td>1452</td><td>38</td><td>CGTTTGTTCAACGATTAAAGTCCTACGTGATCTGAGTT</td><td>39.6</td><td>8.5</td><td>1.8</td><td>4.7</td><td>3.0E-06</td><td>5.5E-06</td><td>MB231/MB468</td><td>MP</td></tr>
<tr><td>>hg38_21_+_8433172_8446622_RNA45SN1_45S_50ntFlanks@9938.9968.31|AGCAGGACGGTGGCCATGGAAGTCGGAATCC</td><td>>hg38_21_+_8433172_8446622_RNA45SN1_45S_50ntFlanks</td><td>9938</td><td>9968</td><td>31</td><td>AGCAGGACGGTGGCCATGGAAGTCGGAATCC</td><td>1033.8</td><td>4.7</td><td>1.0</td><td>4.9</td><td>1.0E-06</td><td>5.5E-06</td><td>BT20/MB468</td><td>MP</td></tr>
<tr><td>>MI0000650|hsa-mir-200c&WithFlank&12|+|6963693|6963772@50.72.23(+1C)&[MIMAT0000617&hsa-miR-200c-3p&offsets|0|0(+1C);m-54&12|+|6963742|6963764&offsets|0|0(+1C)]|TAATACTGCCGGGTAATGATGGAC</td><td>>MI0000650|hsa-mir-200c&WithFlank&12|+|6963693|6963772</td><td>50</td><td>72</td><td>23(+1C)</td><td>TAATACTGCCGGGTAATGATGGAC</td><td>303.5</td><td>4.2</td><td>0.9</td><td>4.9</td><td>1.0E-06</td><td>5.5E-06</td><td>BT20/MB468</td><td>MP</td></tr>
<tr><td>>hg38_21_+_8433172_8446622_RNA45SN1_45S_50ntFlanks@3790.3810.21|CTGCGAATGGCTCATTAAATC</td><td>>hg38_21_+_8433172_8446622_RNA45SN1_45S_50ntFlanks</td><td>3790</td><td>3810</td><td>21</td><td>CTGCGAATGGCTCATTAAATC</td><td>408.2</td><td>-6.8</td><td>1.5</td><td>-4.6</td><td>3.7E-06</td><td>5.5E-06</td><td>MB231/MB468</td><td>MITO</td></tr>
<tr><td>>hg38_21_+_8433172_8446622_RNA45SN1_45S_50ntFlanks@9923.9951.29|TTGGTTGATATAGACAGCAGGACGGTGGC</td><td>>hg38_21_+_8433172_8446622_RNA45SN1_45S_50ntFlanks</td><td>9923</td><td>9951</td><td>29</td><td>TTGGTTGATATAGACAGCAGGACGGTGGC</td><td>45.1</td><td>9.2</td><td>1.9</td><td>4.9</td><td>1.0E-06</td><td>5.5E-06</td><td>BT20/MB468</td><td>MP</td></tr>
<tr><td>>hg38_21_+_8433172_8446622_RNA45SN1_45S_50ntFlanks@12007.12035.29|GCCGCCGGTGAAATACCACTACTCTGATC</td><td>>hg38_21_+_8433172_8446622_RNA45SN1_45S_50ntFlanks</td><td>12007</td><td>12035</td><td>29</td><td>GCCGCCGGTGAAATACCACTACTCTGATC</td><td>1451.4</td><td>3.6</td><td>0.8</td><td>4.7</td><td>2.6E-06</td><td>5.5E-06</td><td>BT20/MB468</td><td>MITO</td></tr>
<tr><td>>hg38_21_+_8433172_8446622_RNA45SN1_45S_50ntFlanks@12337.12370.34|AGGAGGTGTCAGAAAAGTTACCACAGGGATAACT</td><td>>hg38_21_+_8433172_8446622_RNA45SN1_45S_50ntFlanks</td><td>12337</td><td>12370</td><td>34</td><td>AGGAGGTGTCAGAAAAGTTACCACAGGGATAACT</td><td>50.6</td><td>9.4</td><td>1.9</td><td>4.9</td><td>1.0E-06</td><td>5.5E-06</td><td>BT20/MB468</td><td>MP</td></tr>
<tr><td>>hg38_21_+_8433172_8446622_RNA45SN1_45S_50ntFlanks@11480.11502.23|CGGCGGAGGGGCCGCGGGCCGGT</td><td>>hg38_21_+_8433172_8446622_RNA45SN1_45S_50ntFlanks</td><td>11480</td><td>11502</td><td>23</td><td>CGGCGGAGGGGCCGCGGGCCGGT</td><td>129.6</td><td>-5.4</td><td>1.2</td><td>-4.7</td><td>3.1E-06</td><td>5.5E-06</td><td>BT20/MB468</td><td>CYTO</td></tr>
<tr><td>>trna119_LysCTT_1_-_145395522_145395594@1.31.31,trna11_LysCTT_5_-_180648979_180649051@1.31.31,trna13_LysCTT_6_+_26556774_26556846@1.31.31,trna32_LysCTT_16_-_3207406_3207478@1.31.31,trna7_LysCTT_16_+_3225692_3225764@1.31.31,trna9_LysCTT_5_+_180634755_180634827@1.31.31|GCCCGGCTAGCTCAGTCGGTAGAGCATGAGA</td><td>>trna119_LysCTT_1_-_145395522_145395594</td><td>1</td><td>31</td><td>31</td><td>GCCCGGCTAGCTCAGTCGGTAGAGCATGAGA</td><td>1793.1</td><td>-3.9</td><td>0.8</td><td>-4.7</td><td>3.0E-06</td><td>5.5E-06</td><td>MB231/MB468</td><td>MP</td></tr>
<tr><td>>hg38_21_+_8433172_8446622_RNA45SN1_45S_50ntFlanks@9554.9571.18|CGTAGCGGTCCTGACGTG</td><td>>hg38_21_+_8433172_8446622_RNA45SN1_45S_50ntFlanks</td><td>9554</td><td>9571</td><td>18</td><td>CGTAGCGGTCCTGACGTG</td><td>55084.7</td><td>-5.8</td><td>1.2</td><td>-4.9</td><td>1.1E-06</td><td>5.5E-06</td><td>BT20/MB468</td><td>MP</td></tr>
<tr><td>>hg38_21_+_8433172_8446622_RNA45SN1_45S_50ntFlanks@5491.5516.26|GAGAAGACGGTCGAACTTGACTATCT</td><td>>hg38_21_+_8433172_8446622_RNA45SN1_45S_50ntFlanks</td><td>5491</td><td>5516</td><td>26</td><td>GAGAAGACGGTCGAACTTGACTATCT</td><td>19255.9</td><td>5.4</td><td>1.1</td><td>5.1</td><td>3.4E-07</td><td>5.5E-06</td><td>BT20/MB231</td><td>MITO</td></tr>
<tr><td>>hg38_21_+_8433172_8446622_RNA45SN1_45S_50ntFlanks@3787.3810.24|AAACTGCGAATGGCTCATTAAATC</td><td>>hg38_21_+_8433172_8446622_RNA45SN1_45S_50ntFlanks</td><td>3787</td><td>3810</td><td>24</td><td>AAACTGCGAATGGCTCATTAAATC</td><td>468.3</td><td>-5.7</td><td>1.2</td><td>-4.7</td><td>3.1E-06</td><td>5.5E-06</td><td>MB231/MB468</td><td>MP</td></tr>
<tr><td>>hg38_21_+_8433172_8446622_RNA45SN1_45S_50ntFlanks@11641.11684.44|CGGCGGGTGTTGACGCGATGTGATTTCTGCCCAGTGCTCTGAAT</td><td>>hg38_21_+_8433172_8446622_RNA45SN1_45S_50ntFlanks</td><td>11641</td><td>11684</td><td>44</td><td>CGGCGGGTGTTGACGCGATGTGATTTCTGCCCAGTGCTCTGAAT</td><td>538.2</td><td>-5.9</td><td>1.3</td><td>-4.6</td><td>3.7E-06</td><td>5.5E-06</td><td>MB231/MB468</td><td>MITO</td></tr>
<tr><td>>trnaMT_SerGCT_MT_+_12207_12265@1.22.22|GAGAAAGCTCACAAGAACTGCT</td><td>>trnaMT_SerGCT_MT_+_12207_12265</td><td>1</td><td>22</td><td>22</td><td>GAGAAAGCTCACAAGAACTGCT</td><td>332.8</td><td>4.3</td><td>0.9</td><td>4.6</td><td>3.5E-06</td><td>5.5E-06</td><td>MB231/MB468</td><td>CYTO</td></tr>
<tr><td>>hg38_21_+_8433172_8446622_RNA45SN1_45S_50ntFlanks@5472.5516.45|ACGGCCCTGGCGGAGCGCTGAGAAGACGGTCGAACTTGACTATCT</td><td>>hg38_21_+_8433172_8446622_RNA45SN1_45S_50ntFlanks</td><td>5472</td><td>5516</td><td>45</td><td>ACGGCCCTGGCGGAGCGCTGAGAAGACGGTCGAACTTGACTATCT</td><td>1378.3</td><td>5.7</td><td>1.2</td><td>4.6</td><td>3.7E-06</td><td>5.5E-06</td><td>MB231/MB468</td><td>MITO</td></tr>
<tr><td>>hg38_21_+_8433172_8446622_RNA45SN1_45S_50ntFlanks@6769.6806.38|CGGGTTCCTCCCGGGGCTACGCCTGTCTGAGCGTCGCT</td><td>>hg38_21_+_8433172_8446622_RNA45SN1_45S_50ntFlanks</td><td>6769</td><td>6806</td><td>38</td><td>CGGGTTCCTCCCGGGGCTACGCCTGTCTGAGCGTCGCT</td><td>41.8</td><td>8.6</td><td>1.8</td><td>4.7</td><td>3.1E-06</td><td>5.5E-06</td><td>MB231/MB468</td><td>MP</td></tr>
<tr><td>>hg38_21_+_8433172_8446622_RNA45SN1_45S_50ntFlanks@11561.11594.34|GCGCCTAGCAGCCGACTTAGAACTGGTGCGGACC</td><td>>hg38_21_+_8433172_8446622_RNA45SN1_45S_50ntFlanks</td><td>11561</td><td>11594</td><td>34</td><td>GCGCCTAGCAGCCGACTTAGAACTGGTGCGGACC</td><td>280.5</td><td>-2.9</td><td>0.6</td><td>-4.6</td><td>3.6E-06</td><td>5.5E-06</td><td>MB231/MB468</td><td>NUC</td></tr>
<tr><td>>hg38_21_+_8433172_8446622_RNA45SN1_45S_50ntFlanks@11999.12026.28|CCCTGCGGGCCGCCGGTGAAATACCACT</td><td>>hg38_21_+_8433172_8446622_RNA45SN1_45S_50ntFlanks</td><td>11999</td><td>12026</td><td>28</td><td>CCCTGCGGGCCGCCGGTGAAATACCACT</td><td>117.0</td><td>-10.0</td><td>2.0</td><td>-4.9</td><td>1.1E-06</td><td>5.5E-06</td><td>BT20/MB468</td><td>MP</td></tr>
<tr><td>>hg38_21_+_8433172_8446622_RNA45SN1_45S_50ntFlanks@10678.10706.29|CGGTTCCGGCGGCGTCCGGTGAGCTCTCG</td><td>>hg38_21_+_8433172_8446622_RNA45SN1_45S_50ntFlanks</td><td>10678</td><td>10706</td><td>29</td><td>CGGTTCCGGCGGCGTCCGGTGAGCTCTCG</td><td>35448.2</td><td>-8.3</td><td>1.8</td><td>-4.6</td><td>3.9E-06</td><td>5.6E-06</td><td>BT20/MB468</td><td>NUC</td></tr>
<tr><td>>hg38_21_+_8433172_8446622_RNA45SN1_45S_50ntFlanks@12328.12346.19|GTTTTAAGCAGGAGGTGTC</td><td>>hg38_21_+_8433172_8446622_RNA45SN1_45S_50ntFlanks</td><td>12328</td><td>12346</td><td>19</td><td>GTTTTAAGCAGGAGGTGTC</td><td>181.6</td><td>-3.6</td><td>0.8</td><td>-4.6</td><td>3.5E-06</td><td>5.6E-06</td><td>MB231/MB468</td><td>CYTO</td></tr>
<tr><td>>hg38_21_+_8433172_8446622_RNA45SN1_45S_50ntFlanks@8153.8188.36|GCCGCTCGTGGGGGGCCCAAGTCCTTCTGATCGAGG</td><td>>hg38_21_+_8433172_8446622_RNA45SN1_45S_50ntFlanks</td><td>8153</td><td>8188</td><td>36</td><td>GCCGCTCGTGGGGGGCCCAAGTCCTTCTGATCGAGG</td><td>55.5</td><td>-8.9</td><td>1.8</td><td>-4.9</td><td>1.1E-06</td><td>5.6E-06</td><td>BT20/MB468</td><td>MP</td></tr>
<tr><td>>MI0000750|hsa-mir-26a-2&WithFlank&12|-|57824603|57824698@20.41.22(+1A);MI0000083|hsa-mir-26a-1&WithFlank&3|+|37969398|37969486@16.37.22(+1A)&[MIMAT0000082&hsa-miR-26a-5p&offsets|0|0(+1A);m-33&12|-|57824658|57824679&offsets|0|0(+1A)];[MIMAT0000082_1&hsa-miR-26a-5p&offsets|0|0(+1A);m-34&3|+|37969413|37969434&offsets|0|0(+1A)]|TTCAAGTAATCCAGGATAGGCTA</td><td>>MI0000750|hsa-mir-26a-2&WithFlank&12|-|57824603|57824698</td><td>20</td><td>41</td><td>22(+1A)</td><td>TTCAAGTAATCCAGGATAGGCTA</td><td>302.0</td><td>1.9</td><td>0.4</td><td>4.6</td><td>3.9E-06</td><td>5.6E-06</td><td>BT20/MB468</td><td>NUC</td></tr>
<tr><td>>hg38_MT_+_598_1651_RNR1_12S_50ntFlanks@634.653.20|AACCTCACCACCTCTTGCTC</td><td>>hg38_MT_+_598_1651_RNR1_12S_50ntFlanks</td><td>634</td><td>653</td><td>20</td><td>AACCTCACCACCTCTTGCTC</td><td>57.4</td><td>9.6</td><td>2.0</td><td>4.9</td><td>1.1E-06</td><td>5.6E-06</td><td>BT20/MB468</td><td>MP</td></tr>
<tr><td>>hg38_21_+_8433172_8446622_RNA45SN1_45S_50ntFlanks@9533.9550.18|GAGGAAACTCTGGTGGAG</td><td>>hg38_21_+_8433172_8446622_RNA45SN1_45S_50ntFlanks</td><td>9533</td><td>9550</td><td>18</td><td>GAGGAAACTCTGGTGGAG</td><td>281.7</td><td>-3.9</td><td>0.8</td><td>-4.7</td><td>3.1E-06</td><td>5.6E-06</td><td>MB231/MB468</td><td>MP</td></tr>
<tr><td>>hg38_21_+_8433172_8446622_RNA45SN1_45S_50ntFlanks@8155.8181.27|CGCTCGTGGGGGGCCCAAGTCCTTCTG</td><td>>hg38_21_+_8433172_8446622_RNA45SN1_45S_50ntFlanks</td><td>8155</td><td>8181</td><td>27</td><td>CGCTCGTGGGGGGCCCAAGTCCTTCTG</td><td>1089.5</td><td>-8.1</td><td>1.7</td><td>-4.6</td><td>3.9E-06</td><td>5.6E-06</td><td>BT20/MB468</td><td>NUC</td></tr>
<tr><td>>hg38_21_+_8433172_8446622_RNA45SN1_45S_50ntFlanks@12539.12571.33|ATGATGTGTTGTTGCCATGGTAATCCTGCTCAG</td><td>>hg38_21_+_8433172_8446622_RNA45SN1_45S_50ntFlanks</td><td>12539</td><td>12571</td><td>33</td><td>ATGATGTGTTGTTGCCATGGTAATCCTGCTCAG</td><td>59.9</td><td>-9.0</td><td>1.8</td><td>-4.9</td><td>1.1E-06</td><td>5.6E-06</td><td>BT20/MB468</td><td>MP</td></tr>
</table>

<table>
<tr><td>>hg38_MT_+_1621_3279_RNR2_16S_50ntFlanks@51.93.43|GCTAAACCTAGCCCCAAACCCACTCCACCTTACTACCAGACAA</td><td>>hg38_MT_+_1621_3279_RNR2_16S_50ntFlanks</td><td>51</td><td>93</td><td>43</td><td>GCTAAACCTAGCCCCAAACCCACTCCACCTTACTACCAGACAA</td><td>503.4</td><td>5.8</td><td>1.2</td><td>4.7</td><td>2.7E-06</td><td>5.6E-06</td><td>BT20/MB468</td><td>MITO</td></tr>
<tr><td>>hg38_21_+_8433172_8446622_RNA45SN1_45S_50ntFlanks@3984.4004.21|GCGGCTTTGGTGACTCTAGAT</td><td>>hg38_21_+_8433172_8446622_RNA45SN1_45S_50ntFlanks</td><td>3984</td><td>4004</td><td>21</td><td>GCGGCTTTGGTGACTCTAGAT</td><td>357.9</td><td>-6.6</td><td>1.4</td><td>-4.6</td><td>3.6E-06</td><td>5.6E-06</td><td>MB231/MB468</td><td>NUC</td></tr>
<tr><td>>hg38_21_+_8433172_8446622_RNA45SN1_45S_50ntFlanks@5078.5098.21|CGATAACGAACGAGACTCTGG</td><td>>hg38_21_+_8433172_8446622_RNA45SN1_45S_50ntFlanks</td><td>5078</td><td>5098</td><td>21</td><td>CGATAACGAACGAGACTCTGG</td><td>926.6</td><td>-6.9</td><td>1.5</td><td>-4.6</td><td>3.9E-06</td><td>5.6E-06</td><td>BT20/MB468</td><td>NUC</td></tr>
<tr><td>>hg38_1_-_228634819_228635039_RNA5S12_5S_50ntFlanks@131.169.39|GGATGGGAGACCGCCTGGGAATACCGGGTGCTGTAGGCT</td><td>>hg38_1_-_228634819_228635039_RNA5S12_5S_50ntFlanks</td><td>131</td><td>169</td><td>39</td><td>GGATGGGAGACCGCCTGGGAATACCGGGTGCTGTAGGCT</td><td>420.5</td><td>4.5</td><td>0.9</td><td>5.1</td><td>3.4E-07</td><td>5.6E-06</td><td>BT20/MB231</td><td>MITO</td></tr>
<tr><td>>hg38_21_+_8433172_8446622_RNA45SN1_45S_50ntFlanks@11589.11607.19|CGGACCAGGGGAATCCGAC</td><td>>hg38_21_+_8433172_8446622_RNA45SN1_45S_50ntFlanks</td><td>11589</td><td>11607</td><td>19</td><td>CGGACCAGGGGAATCCGAC</td><td>443.8</td><td>4.0</td><td>0.8</td><td>5.1</td><td>3.4E-07</td><td>5.6E-06</td><td>BT20/MB231</td><td>MITO</td></tr>
<tr><td>>hg38_MT_+_1621_3279_RNR2_16S_50ntFlanks@626.663.38|ACTGAACTCCTCACACCCAATTGGACCAATCTATCACC</td><td>>hg38_MT_+_1621_3279_RNR2_16S_50ntFlanks</td><td>626</td><td>663</td><td>38</td><td>ACTGAACTCCTCACACCCAATTGGACCAATCTATCACC</td><td>82.4</td><td>10.1</td><td>2.1</td><td>4.9</td><td>1.1E-06</td><td>5.6E-06</td><td>BT20/MB468</td><td>MP</td></tr>
<tr><td>>hg38_21_+_8433172_8446622_RNA45SN1_45S_50ntFlanks@12618.12654.37|GAGGAGCCAATGGGGCGAAGCTACCATCTGTGGGATT</td><td>>hg38_21_+_8433172_8446622_RNA45SN1_45S_50ntFlanks</td><td>12618</td><td>12654</td><td>37</td><td>GAGGAGCCAATGGGGCGAAGCTACCATCTGTGGGATT</td><td>372.7</td><td>-6.8</td><td>1.5</td><td>-4.6</td><td>3.7E-06</td><td>5.6E-06</td><td>MB231/MB468</td><td>MITO</td></tr>
<tr><td>>hg38_21_+_8433172_8446622_RNA45SN1_45S_50ntFlanks@11930.11967.38|TAGAATAAGTGGGAGGCCCCCGGCGCCCCCCCGGTGTC</td><td>>hg38_21_+_8433172_8446622_RNA45SN1_45S_50ntFlanks</td><td>11930</td><td>11967</td><td>38</td><td>TAGAATAAGTGGGAGGCCCCCGGCGCCCCCCCGGTGTC</td><td>88.9</td><td>-8.6</td><td>1.8</td><td>-4.7</td><td>2.7E-06</td><td>5.6E-06</td><td>BT20/MB468</td><td>MITO</td></tr>
<tr><td>>hg38_21_+_8433172_8446622_RNA45SN1_45S_50ntFlanks@9952.9982.31|CATGGAAGTCGGAATCCGCTAAGGAGTGTGT</td><td>>hg38_21_+_8433172_8446622_RNA45SN1_45S_50ntFlanks</td><td>9952</td><td>9982</td><td>31</td><td>CATGGAAGTCGGAATCCGCTAAGGAGTGTGT</td><td>55.1</td><td>9.5</td><td>2.0</td><td>4.9</td><td>1.1E-06</td><td>5.6E-06</td><td>BT20/MB468</td><td>MP</td></tr>
<tr><td>>hg38_21_+_8433172_8446622_RNA45SN1_45S_50ntFlanks@6723.6744.22|TTGCAGGACACATTGATCATCG</td><td>>hg38_21_+_8433172_8446622_RNA45SN1_45S_50ntFlanks</td><td>6723</td><td>6744</td><td>22</td><td>TTGCAGGACACATTGATCATCG</td><td>1696.3</td><td>-5.2</td><td>1.1</td><td>-4.6</td><td>3.5E-06</td><td>5.6E-06</td><td>MB231/MB468</td><td>CYTO</td></tr>
<tr><td>>hg38_21_+_8433172_8446622_RNA45SN1_45S_50ntFlanks@5486.5514.29|GCGCTGAGAAGACGGTCGAACTTGACTAT</td><td>>hg38_21_+_8433172_8446622_RNA45SN1_45S_50ntFlanks</td><td>5486</td><td>5514</td><td>29</td><td>GCGCTGAGAAGACGGTCGAACTTGACTAT</td><td>1149.1</td><td>-3.5</td><td>0.8</td><td>-4.6</td><td>3.5E-06</td><td>5.6E-06</td><td>MB231/MB468</td><td>CYTO</td></tr>
<tr><td>>hg38_21_+_8433172_8446622_RNA45SN1_45S_50ntFlanks@11593.11612.20|CCAGGGGAATCCGACTGTTT</td><td>>hg38_21_+_8433172_8446622_RNA45SN1_45S_50ntFlanks</td><td>11593</td><td>11612</td><td>20</td><td>CCAGGGGAATCCGACTGTTT</td><td>868.3</td><td>-3.1</td><td>0.7</td><td>-4.7</td><td>2.7E-06</td><td>5.6E-06</td><td>BT20/MB468</td><td>MITO</td></tr>
<tr><td>>hg38_21_+_8433172_8446622_RNA45SN1_45S_50ntFlanks@9806.9838.33|CGTGGAATGCGAGTGCCTAGTGGGCCACTTTTG</td><td>>hg38_21_+_8433172_8446622_RNA45SN1_45S_50ntFlanks</td><td>9806</td><td>9838</td><td>33</td><td>CGTGGAATGCGAGTGCCTAGTGGGCCACTTTTG</td><td>490.7</td><td>-4.7</td><td>1.0</td><td>-4.7</td><td>2.7E-06</td><td>5.6E-06</td><td>BT20/MB468</td><td>MITO</td></tr>
<tr><td>>MI0000542|hsa-mir-320a&WithFlank&8|-|22244960|22245043@49.69.21(+1U)&[MIMAT0000510&hsa-miR-320a-3p&offsets|+1|0(+1U);m-60&8|-|22244975|22244996&offsets|+1|0(+1U)]|AAAGCTGGGTTGAGAGGGCGAT</td><td>>MI0000542|hsa-mir-320a&WithFlank&8|-|22244960|22245043</td><td>49</td><td>69</td><td>21(+1U)</td><td>AAAGCTGGGTTGAGAGGGCGAT</td><td>910.5</td><td>6.8</td><td>1.4</td><td>4.9</td><td>1.1E-06</td><td>5.6E-06</td><td>BT20/MB468</td><td>MP</td></tr>
<tr><td>>hg38_21_+_8433172_8446622_RNA45SN1_45S_50ntFlanks@475.512.38|CTTCGTGATCGATGTGGTGACGTCGTGCTCTCCCGGGC</td><td>>hg38_21_+_8433172_8446622_RNA45SN1_45S_50ntFlanks</td><td>475</td><td>512</td><td>38</td><td>CTTCGTGATCGATGTGGTGACGTCGTGCTCTCCCGGGC</td><td>448.0</td><td>8.4</td><td>1.8</td><td>4.7</td><td>2.7E-06</td><td>5.6E-06</td><td>BT20/MB468</td><td>MITO</td></tr>
<tr><td>>hg38_21_+_8433172_8446622_RNA45SN1_45S_50ntFlanks@11572.11594.23|CCGACTTAGAACTGGTGCGGACC</td><td>>hg38_21_+_8433172_8446622_RNA45SN1_45S_50ntFlanks</td><td>11572</td><td>11594</td><td>23</td><td>CCGACTTAGAACTGGTGCGGACC</td><td>326.4</td><td>-3.2</td><td>0.7</td><td>-4.6</td><td>3.7E-06</td><td>5.6E-06</td><td>MB231/MB468</td><td>MITO</td></tr>
<tr><td>>hg38_21_+_8433172_8446622_RNA45SN1_45S_50ntFlanks@5472.5489.18|ACGGCCCTGGCGGAGCGC</td><td>>hg38_21_+_8433172_8446622_RNA45SN1_45S_50ntFlanks</td><td>5472</td><td>5489</td><td>18</td><td>ACGGCCCTGGCGGAGCGC</td><td>147.1</td><td>4.8</td><td>1.0</td><td>4.6</td><td>3.9E-06</td><td>5.6E-06</td><td>BT20/MB468</td><td>NUC</td></tr>
<tr><td>>hg38_21_+_8433172_8446622_RNA45SN1_45S_50ntFlanks@6740.6770.31|CATCGACACTTCGAACGCACTTGCGGCCCCG</td><td>>hg38_21_+_8433172_8446622_RNA45SN1_45S_50ntFlanks</td><td>6740</td><td>6770</td><td>31</td><td>CATCGACACTTCGAACGCACTTGCGGCCCCG</td><td>850.1</td><td>-6.6</td><td>1.4</td><td>-4.7</td><td>3.1E-06</td><td>5.6E-06</td><td>BT20/MB468</td><td>CYTO</td></tr>
<tr><td>>MI0003188|hsa-mir-503&WithFlank&X|-|134546322|134546404@12.33.22&[MIMAT0002874&hsa-miR-503-5p&offsets|0|-1;m-297&X|-|134546371|134546393&offsets|0|-1]|TAGCAGCGGGAACAGTTCTGCA</td><td>>MI0003188|hsa-mir-503&WithFlank&X|-|134546322|134546404</td><td>12</td><td>33</td><td>22</td><td>TAGCAGCGGGAACAGTTCTGCA</td><td>39.6</td><td>8.5</td><td>1.8</td><td>4.7</td><td>3.1E-06</td><td>5.6E-06</td><td>MB231/MB468</td><td>MP</td></tr>
<tr><td>>hg38_21_+_8433172_8446622_RNA45SN1_45S_50ntFlanks@12408.12443.36|TTTGATCCTTCGATGTCGGCTCTTCCTATCATTGTG</td><td>>hg38_21_+_8433172_8446622_RNA45SN1_45S_50ntFlanks</td><td>12408</td><td>12443</td><td>36</td><td>TTTGATCCTTCGATGTCGGCTCTTCCTATCATTGTG</td><td>1600.9</td><td>-6.6</td><td>1.4</td><td>-4.6</td><td>3.9E-06</td><td>5.6E-06</td><td>BT20/MB468</td><td>NUC</td></tr>
<tr><td>>hg38_21_+_8433172_8446622_RNA45SN1_45S_50ntFlanks@12567.12604.38|CTCAGTACGAGAGGAACCGCAGGTTCAGACATTTGGTG</td><td>>hg38_21_+_8433172_8446622_RNA45SN1_45S_50ntFlanks</td><td>12567</td><td>12604</td><td>38</td><td>CTCAGTACGAGAGGAACCGCAGGTTCAGACATTTGGTG</td><td>64.4</td><td>-9.1</td><td>1.9</td><td>-4.9</td><td>1.1E-06</td><td>5.6E-06</td><td>BT20/MB468</td><td>MP</td></tr>
<tr><td>>hg38_21_+_8433172_8446622_RNA45SN1_45S_50ntFlanks@4445.4470.26|CTTGCCTCTCGGCGCCCCCTCGATGC</td><td>>hg38_21_+_8433172_8446622_RNA45SN1_45S_50ntFlanks</td><td>4445</td><td>4470</td><td>26</td><td>CTTGCCTCTCGGCGCCCCCTCGATGC</td><td>131.2</td><td>-3.8</td><td>0.8</td><td>-4.7</td><td>3.1E-06</td><td>5.6E-06</td><td>BT20/MB468</td><td>CYTO</td></tr>
<tr><td>>MI0000102|hsa-mir-100&WithFlank&11|-|122152223|122152314@20.38.19&[MIMAT0000098&hsa-miR-100-5p&offsets|+1|-2;m-27&11|-|122152276|122152296&offsets|+1|-1]|ACCCGTAGATCCGAACTTG</td><td>>MI0000102|hsa-mir-100&WithFlank&11|-|122152223|122152314</td><td>20</td><td>38</td><td>19</td><td>ACCCGTAGATCCGAACTTG</td><td>59.2</td><td>9.1</td><td>2.0</td><td>4.7</td><td>3.1E-06</td><td>5.6E-06</td><td>MB231/MB468</td><td>MP</td></tr>
<tr><td>>hg38_21_+_8433172_8446622_RNA45SN1_45S_50ntFlanks@12457.12484.28|CAAGCGTTGGATTGTTCACCCACTAATA</td><td>>hg38_21_+_8433172_8446622_RNA45SN1_45S_50ntFlanks</td><td>12457</td><td>12484</td><td>28</td><td>CAAGCGTTGGATTGTTCACCCACTAATA</td><td>526.3</td><td>-5.4</td><td>1.2</td><td>-4.6</td><td>3.7E-06</td><td>5.6E-06</td><td>MB231/MB468</td><td>NUC</td></tr>
<tr><td>>trnaMT_GluTTC_MT_-_14674_14742@40.72.33,trnalookalike8_GluTTC_5_-_93905172_93905240@40.72.33|TTGGTCGTGGTTGTAGTCCGTGCGAGAATACCA</td><td>>trnaMT_GluTTC_MT_-_14674_14742</td><td>40</td><td>72</td><td>33</td><td>TTGGTCGTGGTTGTAGTCCGTGCGAGAATACCA</td><td>22184.0</td><td>3.8</td><td>0.8</td><td>4.6</td><td>3.7E-06</td><td>5.6E-06</td><td>MB231/MB468</td><td>NUC</td></tr>
<tr><td>>trnaMT_SerGCT_MT_+_12207_12265@1.21.21|GAGAAAGCTCACAAGAACTGC</td><td>>trnaMT_SerGCT_MT_+_12207_12265</td><td>1</td><td>21</td><td>21</td><td>GAGAAAGCTCACAAGAACTGC</td><td>405.2</td><td>5.5</td><td>1.2</td><td>4.6</td><td>3.7E-06</td><td>5.6E-06</td><td>MB231/MB468</td><td>NUC</td></tr>
<tr><td>>hg38_21_+_8433172_8446622_RNA45SN1_45S_50ntFlanks@11577.11601.25|TTAGAACTGGTGCGGACCAGGGGAA</td><td>>hg38_21_+_8433172_8446622_RNA45SN1_45S_50ntFlanks</td><td>11577</td><td>11601</td><td>25</td><td>TTAGAACTGGTGCGGACCAGGGGAA</td><td>368.9</td><td>-6.4</td><td>1.4</td><td>-4.6</td><td>3.7E-06</td><td>5.6E-06</td><td>MB231/MB468</td><td>NUC</td></tr>
<tr><td>>hg38_21_+_8433172_8446622_RNA45SN1_45S_50ntFlanks@10830.10874.45|CAAGCCGGATCCGTAACTTCGGGATAAGGATTGGCTCTAAGGGCT</td><td>>hg38_21_+_8433172_8446622_RNA45SN1_45S_50ntFlanks</td><td>10830</td><td>10874</td><td>45</td><td>CAAGCCGGATCCGTAACTTCGGGATAAGGATTGGCTCTAAGGGCT</td><td>1235.6</td><td>-6.9</td><td>1.5</td><td>-4.6</td><td>3.7E-06</td><td>5.6E-06</td><td>MB231/MB468</td><td>NUC</td></tr>
<tr><td>>MI0000078|hsa-mir-22&WithFlank&17|-|1713897|1713993@59.80.22(+1A)&[MIMAT0000077&hsa-miR-22-3p&offsets|0|0(+1A);m-2&17|-|1713914|1713935&offsets|0|0(+1A)]|AAGCTGCCAGTTGAAGAACTGTA</td><td>>MI0000078|hsa-mir-22&WithFlank&17|-|1713897|1713993</td><td>59</td><td>80</td><td>22(+1A)</td><td>AAGCTGCCAGTTGAAGAACTGTA</td><td>281.8</td><td>6.9</td><td>1.5</td><td>4.6</td><td>3.8E-06</td><td>5.6E-06</td><td>MB231/MB468</td><td>MITO</td></tr>
<tr><td>>hg38_21_+_8433172_8446622_RNA45SN1_45S_50ntFlanks@5519.5545.27|AGGAAGTAAAAGTCGTAACAAGGTTTC</td><td>>hg38_21_+_8433172_8446622_RNA45SN1_45S_50ntFlanks</td><td>5519</td><td>5545</td><td>27</td><td>AGGAAGTAAAAGTCGTAACAAGGTTTC</td><td>73.6</td><td>-9.3</td><td>1.9</td><td>-4.9</td><td>1.1E-06</td><td>5.6E-06</td><td>BT20/MB468</td><td>MP</td></tr>
<tr><td>>hg38_21_+_8433172_8446622_RNA45SN1_45S_50ntFlanks@10866.10883.18|CTAAGGGCTGGGTCGGTC</td><td>>hg38_21_+_8433172_8446622_RNA45SN1_45S_50ntFlanks</td><td>10866</td><td>10883</td><td>18</td><td>CTAAGGGCTGGGTCGGTC</td><td>513.1</td><td>-7.5</td><td>1.6</td><td>-4.6</td><td>4.0E-06</td><td>5.7E-06</td><td>BT20/MB468</td><td>NUC</td></tr>
<tr><td>>hg38_21_+_8433172_8446622_RNA45SN1_45S_50ntFlanks@11793.11810.18|TGAACGAGATTCCCACTG</td><td>>hg38_21_+_8433172_8446622_RNA45SN1_45S_50ntFlanks</td><td>11793</td><td>11810</td><td>18</td><td>TGAACGAGATTCCCACTG</td><td>66.6</td><td>-9.1</td><td>1.9</td><td>-4.9</td><td>1.1E-06</td><td>5.7E-06</td><td>BT20/MB468</td><td>MP</td></tr>
<tr><td>>hg38_21_+_8433172_8446622_RNA45SN1_45S_50ntFlanks@11891.11909.19|TTGACTCTAGTCTGGCACG</td><td>>hg38_21_+_8433172_8446622_RNA45SN1_45S_50ntFlanks</td><td>11891</td><td>11909</td><td>19</td><td>TTGACTCTAGTCTGGCACG</td><td>361.5</td><td>-3.6</td><td>0.8</td><td>-4.6</td><td>3.6E-06</td><td>5.7E-06</td><td>MB231/MB468</td><td>CYTO</td></tr>
<tr><td>>hg38_21_+_8433172_8446622_RNA45SN1_45S_50ntFlanks@6645.6663.19|CTCGTACGACTCTTAGCGG</td><td>>hg38_21_+_8433172_8446622_RNA45SN1_45S_50ntFlanks</td><td>6645</td><td>6663</td><td>19</td><td>CTCGTACGACTCTTAGCGG</td><td>3171.0</td><td>3.2</td><td>0.7</td><td>4.7</td><td>3.1E-06</td><td>5.7E-06</td><td>BT20/MB468</td><td>CYTO</td></tr>
<tr><td>>hg38_21_+_8433172_8446622_RNA45SN1_45S_50ntFlanks@10689.10708.20|GCGTCCGGTGAGCTCTCGCT</td><td>>hg38_21_+_8433172_8446622_RNA45SN1_45S_50ntFlanks</td><td>10689</td><td>10708</td><td>20</td><td>GCGTCCGGTGAGCTCTCGCT</td><td>1224.3</td><td>-5.1</td><td>1.1</td><td>-4.7</td><td>3.1E-06</td><td>5.7E-06</td><td>BT20/MB468</td><td>CYTO</td></tr>
<tr><td>>hg38_21_+_8433172_8446622_RNA45SN1_45S_50ntFlanks@12475.12498.24|CCCACTAATAGGGAACGTGAGCTG</td><td>>hg38_21_+_8433172_8446622_RNA45SN1_45S_50ntFlanks</td><td>12475</td><td>12498</td><td>24</td><td>CCCACTAATAGGGAACGTGAGCTG</td><td>299.3</td><td>-2.7</td><td>0.6</td><td>-4.7</td><td>2.7E-06</td><td>5.7E-06</td><td>BT20/MB468</td><td>MITO</td></tr>
<tr><td>>hg38_21_+_8433172_8446622_RNA45SN1_45S_50ntFlanks@4002.4042.41|GATAACCTCGGGCCGATCGCACGCCCCCCGTGGCGGCGACG</td><td>>hg38_21_+_8433172_8446622_RNA45SN1_45S_50ntFlanks</td><td>4002</td><td>4042</td><td>41</td><td>GATAACCTCGGGCCGATCGCACGCCCCCCGTGGCGGCGACG</td><td>1562.7</td><td>-8.1</td><td>1.8</td><td>-4.6</td><td>4.0E-06</td><td>5.7E-06</td><td>BT20/MB468</td><td>NUC</td></tr>
<tr><td>>MI0003776|hsa-mir-320b-1&WithFlank&1|+|116671740|116671823@48.67.20;MI0003839|hsa-mir-320b-2&WithFlank&1|-|224257034|224257116@47.66.20;MI0000542|hsa-mir-320a&WithFlank&8|-|22244960|22245043@48.67.20&[MIMAT0005792&hsa-miR-320b&offsets|0|-2;m-183&1|+|116671787|116671806&offsets|0|0];[MIMAT0005792_1&hsa-miR-320b&offsets|0|-2;m-183&1|-|224257051|224257070&offsets|0|0];[MIMAT0000510&hsa-miR-320a-3p&offsets|0|-2;m-60&8|-|22244975|22244996&offsets|0|-2]|AAAAGCTGGGTTGAGAGGGC</td><td>>MI0003776|hsa-mir-320b-1&WithFlank&1|+|116671740|116671823</td><td>48</td><td>67</td><td>20</td><td>AAAAGCTGGGTTGAGAGGGC</td><td>1297.0</td><td>2.8</td><td>0.6</td><td>5.1</td><td>4.2E-07</td><td>5.7E-06</td><td>BT20/MB231</td><td>NUC</td></tr>
<tr><td>>hg38_21_+_8433172_8446622_RNA45SN1_45S_50ntFlanks@11207.11225.19|GGCGGCGACTCTGGACGCG</td><td>>hg38_21_+_8433172_8446622_RNA45SN1_45S_50ntFlanks</td><td>11207</td><td>11225</td><td>19</td><td>GGCGGCGACTCTGGACGCG</td><td>206.0</td><td>-6.1</td><td>1.3</td><td>-4.6</td><td>3.7E-06</td><td>5.7E-06</td><td>MB231/MB468</td><td>NUC</td></tr>
<tr><td>>hg38_21_+_8433172_8446622_RNA45SN1_45S_50ntFlanks@6650.6678.29|ACGACTCTTAGCGGTGGATCACTCGGCTC</td><td>>hg38_21_+_8433172_8446622_RNA45SN1_45S_50ntFlanks</td><td>6650</td><td>6678</td><td>29</td><td>ACGACTCTTAGCGGTGGATCACTCGGCTC</td><td>294.9</td><td>2.5</td><td>0.5</td><td>4.6</td><td>3.6E-06</td><td>5.7E-06</td><td>MB231/MB468</td><td>CYTO</td></tr>
<tr><td>>hg38_21_+_8433172_8446622_RNA45SN1_45S_50ntFlanks@12319.12346.28|ACCTTTTGGGTTTTAAGCAGGAGGTGTC</td><td>>hg38_21_+_8433172_8446622_RNA45SN1_45S_50ntFlanks</td><td>12319</td><td>12346</td><td>28</td><td>ACCTTTTGGGTTTTAAGCAGGAGGTGTC</td><td>109.3</td><td>-10.6</td><td>2.3</td><td>-4.7</td><td>3.1E-06</td><td>5.7E-06</td><td>MB231/MB468</td><td>MP</td></tr>
<tr><td>>hg38_21_+_8433172_8446622_RNA45SN1_45S_50ntFlanks@10350.10381.32|ACGAGAACTTTGAAGGCCGAAGTGGAGAAGGG</td><td>>hg38_21_+_8433172_8446622_RNA45SN1_45S_50ntFlanks</td><td>10350</td><td>10381</td><td>32</td><td>ACGAGAACTTTGAAGGCCGAAGTGGAGAAGGG</td><td>411.1</td><td>-5.1</td><td>1.1</td><td>-4.6</td><td>3.8E-06</td><td>5.7E-06</td><td>MB231/MB468</td><td>MITO</td></tr>
<tr><td>>hg38_21_+_8433172_8446622_RNA45SN1_45S_50ntFlanks@3878.3898.21|CATGCCGACGGGCGCTGACCC</td><td>>hg38_21_+_8433172_8446622_RNA45SN1_45S_50ntFlanks</td><td>3878</td><td>3898</td><td>21</td><td>CATGCCGACGGGCGCTGACCC</td><td>419.2</td><td>-6.6</td><td>1.4</td><td>-4.6</td><td>3.8E-06</td><td>5.7E-06</td><td>MB231/MB468</td><td>MITO</td></tr>
<tr><td>>trnaMT_ValTAC_MT_+_1602_1670@50.71.22|ACTTAACTTGACCGCTCTGACC</td><td>>trnaMT_ValTAC_MT_+_1602_1670</td><td>50</td><td>71</td><td>22</td><td>ACTTAACTTGACCGCTCTGACC</td><td>343.8</td><td>4.9</td><td>1.1</td><td>4.6</td><td>3.6E-06</td><td>5.7E-06</td><td>MB231/MB468</td><td>CYTO</td></tr>
<tr><td>>trnaMT_PheGAA_MT_+_577_647@21.55.35|CAAAGCAATACACTGAAAATGTTTAGACGGGCTCA</td><td>>trnaMT_PheGAA_MT_+_577_647</td><td>21</td><td>55</td><td>35</td><td>CAAAGCAATACACTGAAAATGTTTAGACGGGCTCA</td><td>65.1</td><td>9.8</td><td>2.0</td><td>4.9</td><td>1.1E-06</td><td>5.7E-06</td><td>BT20/MB468</td><td>MP</td></tr>
<tr><td>>hg38_21_+_8433172_8446622_RNA45SN1_45S_50ntFlanks@11640.11660.21|GCGGCGGGTGTTGACGCGATG</td><td>>hg38_21_+_8433172_8446622_RNA45SN1_45S_50ntFlanks</td><td>11640</td><td>11660</td><td>21</td><td>GCGGCGGGTGTTGACGCGATG</td><td>499.1</td><td>-7.0</td><td>1.5</td><td>-4.7</td><td>3.1E-06</td><td>5.7E-06</td><td>MB231/MB468</td><td>MP</td></tr>
<tr><td>>hg38_21_+_8433172_8446622_RNA45SN1_45S_50ntFlanks@11561.11588.28|GCGCCTAGCAGCCGACTTAGAACTGGTG</td><td>>hg38_21_+_8433172_8446622_RNA45SN1_45S_50ntFlanks</td><td>11561</td><td>11588</td><td>28</td><td>GCGCCTAGCAGCCGACTTAGAACTGGTG</td><td>1001.4</td><td>-4.0</td><td>0.9</td><td>-4.7</td><td>3.2E-06</td><td>5.7E-06</td><td>BT20/MB468</td><td>CYTO</td></tr>
<tr><td>>hg38_21_+_8433172_8446622_RNA45SN1_45S_50ntFlanks@11569.11586.18|CAGCCGACTTAGAACTGG</td><td>>hg38_21_+_8433172_8446622_RNA45SN1_45S_50ntFlanks</td><td>11569</td><td>11586</td><td>18</td><td>CAGCCGACTTAGAACTGG</td><td>1040.6</td><td>-3.2</td><td>0.7</td><td>-4.7</td><td>3.2E-06</td><td>5.7E-06</td><td>BT20/MB468</td><td>CYTO</td></tr>
<tr><td>>hg38_21_+_8433172_8446622_RNA45SN1_45S_50ntFlanks@6689.6720.32|GAAGAACGCAGCTAGCTGCGAGAATTAATGTG</td><td>>hg38_21_+_8433172_8446622_RNA45SN1_45S_50ntFlanks</td><td>6689</td><td>6720</td><td>32</td><td>GAAGAACGCAGCTAGCTGCGAGAATTAATGTG</td><td>329.4</td><td>-6.1</td><td>1.3</td><td>-4.7</td><td>3.2E-06</td><td>5.7E-06</td><td>BT20/MB468</td><td>CYTO</td></tr>
<tr><td>>hg38_21_+_8433172_8446622_RNA45SN1_45S_50ntFlanks@8198.8237.40|TGGACGGTGTGAGGCCGGTAGCGGCCCCCGGCGCGCCGGG</td><td>>hg38_21_+_8433172_8446622_RNA45SN1_45S_50ntFlanks</td><td>8198</td><td>8237</td><td>40</td><td>TGGACGGTGTGAGGCCGGTAGCGGCCCCCGGCGCGCCGGG</td><td>112.0</td><td>-5.7</td><td>1.2</td><td>-4.7</td><td>3.2E-06</td><td>5.7E-06</td><td>BT20/MB468</td><td>CYTO</td></tr>
<tr><td>>MI0003599|hsa-mir-589&WithFlank&7|-|5495813|5495923@30.52.23&[MIMAT0004799&hsa-miR-589-5p&offsets|0|+1;m-162&7|-|5495874|5495894&offsets|0|+2]|TGAGAACCACGTCTGCTCTGAGC</td><td>>MI0003599|hsa-mir-589&WithFlank&7|-|5495813|5495923</td><td>30</td><td>52</td><td>23</td><td>TGAGAACCACGTCTGCTCTGAGC</td><td>285.6</td><td>7.5</td><td>1.6</td><td>4.6</td><td>3.8E-06</td><td>5.7E-06</td><td>MB231/MB468</td><td>MITO</td></tr>
</table>

<table>
<tr><td></td><td></td><td></td><td></td><td></td><td></td><td></td><td></td><td></td><td></td><td></td><td></td><td></td><td></td></tr>
<tr><td>>hg38_21_+_8433172_8446622_RNA45SN1_45S_50ntFlanks@3789.3810.22|ACTGCGAATGGCTCATTAAATC</td><td>>hg38_21_+_8433172_8446622_RNA45SN1_45S_50ntFlanks</td><td>3789</td><td>3810</td><td>22</td><td>ACTGCGAATGGCTCATTAAATC</td><td>461.1</td><td>-7.1</td><td>1.5</td><td>-4.7</td><td>3.2E-06</td><td>5.7E-06</td><td>MB231/MB468</td><td>MP</td></tr>
<tr><td>>hg38_21_+_8433172_8446622_RNA45SN1_45S_50ntFlanks@11565.11599.35|CTAGCAGCCGACTTAGAACTGGTGCGGACCAGGGG</td><td>>hg38_21_+_8433172_8446622_RNA45SN1_45S_50ntFlanks</td><td>11565</td><td>11599</td><td>35</td><td>CTAGCAGCCGACTTAGAACTGGTGCGGACCAGGGG</td><td>214.1</td><td>11.1</td><td>2.2</td><td>5.1</td><td>2.9E-07</td><td>5.7E-06</td><td>BT20/MB231</td><td>MP</td></tr>
<tr><td>>hg38_21_+_8433172_8446622_RNA45SN1_45S_50ntFlanks@11912.11940.29|GAAGAGACATGAGAGGTGTAGAATAAGTG</td><td>>hg38_21_+_8433172_8446622_RNA45SN1_45S_50ntFlanks</td><td>11912</td><td>11940</td><td>29</td><td>GAAGAGACATGAGAGGTGTAGAATAAGTG</td><td>164.9</td><td>10.7</td><td>2.1</td><td>5.1</td><td>2.9E-07</td><td>5.7E-06</td><td>BT20/MB231</td><td>MP</td></tr>
<tr><td>>hg38_21_+_8433172_8446622_RNA45SN1_45S_50ntFlanks@11552.11592.41|CCTCGGCCGGCGCCTAGCAGCCGACTTAGAACTGGTGCGGA</td><td>>hg38_21_+_8433172_8446622_RNA45SN1_45S_50ntFlanks</td><td>11552</td><td>11592</td><td>41</td><td>CCTCGGCCGGCGCCTAGCAGCCGACTTAGAACTGGTGCGGA</td><td>1257.1</td><td>-8.6</td><td>1.9</td><td>-4.6</td><td>4.0E-06</td><td>5.7E-06</td><td>BT20/MB468</td><td>NUC</td></tr>
<tr><td>>hg38_21_+_8433172_8446622_RNA45SN1_45S_50ntFlanks@11675.11709.35|TGCTCTGAATGTCAAAGTGAAGAAATTCAATGAAG</td><td>>hg38_21_+_8433172_8446622_RNA45SN1_45S_50ntFlanks</td><td>11675</td><td>11709</td><td>35</td><td>TGCTCTGAATGTCAAAGTGAAGAAATTCAATGAAG</td><td>482.9</td><td>3.9</td><td>0.8</td><td>4.6</td><td>3.6E-06</td><td>5.7E-06</td><td>MB231/MB468</td><td>CYTO</td></tr>
<tr><td>>hg38_21_+_8433172_8446622_RNA45SN1_45S_50ntFlanks@9483.9520.38|CGTGTTAGGACCCGAAAGATGGTGAACTATGCCTGGGC</td><td>>hg38_21_+_8433172_8446622_RNA45SN1_45S_50ntFlanks</td><td>9483</td><td>9520</td><td>38</td><td>CGTGTTAGGACCCGAAAGATGGTGAACTATGCCTGGGC</td><td>46.3</td><td>9.3</td><td>1.9</td><td>4.9</td><td>1.1E-06</td><td>5.7E-06</td><td>BT20/MB468</td><td>MP</td></tr>
<tr><td>>hg38_21_+_8433172_8446622_RNA45SN1_45S_50ntFlanks@6656.6686.31|CTTAGCGGTGGATCACTCGGCTCGTGCGTCG</td><td>>hg38_21_+_8433172_8446622_RNA45SN1_45S_50ntFlanks</td><td>6656</td><td>6686</td><td>31</td><td>CTTAGCGGTGGATCACTCGGCTCGTGCGTCG</td><td>524.3</td><td>-6.6</td><td>1.4</td><td>-4.6</td><td>3.8E-06</td><td>5.7E-06</td><td>MB231/MB468</td><td>MITO</td></tr>
<tr><td>>MI0001445|hsa-mir-423&WithFlank&17|+|30117073|30117178@23.46.24&[MIMAT0004748&hsa-miR-423-5p&offsets|0|+1;m-141&17|+|30117095|30117117&offsets|0|+1]|TGAGGGGCAGAGAGCGAGACTTTT</td><td>>MI0001445|hsa-mir-423&WithFlank&17|+|30117073|30117178</td><td>23</td><td>46</td><td>24</td><td>TGAGGGGCAGAGAGCGAGACTTTT</td><td>2102.9</td><td>4.2</td><td>0.9</td><td>4.7</td><td>3.2E-06</td><td>5.7E-06</td><td>MB231/MB468</td><td>MP</td></tr>
<tr><td>>MI0000456|hsa-mir-140&WithFlank&16|+|69933075|69933186@69.91.23&[MIMAT0004597&hsa-miR-140-3p&offsets|+1|+3;m-45&16|+|69933143|69933164&offsets|0|+1]|ACCACAGGGTAGAACCACGGACA</td><td>>MI0000456|hsa-mir-140&WithFlank&16|+|69933075|69933186</td><td>69</td><td>91</td><td>23</td><td>ACCACAGGGTAGAACCACGGACA</td><td>327.1</td><td>7.9</td><td>1.7</td><td>4.6</td><td>3.8E-06</td><td>5.7E-06</td><td>MB231/MB468</td><td>MITO</td></tr>
<tr><td>>hg38_21_+_8433172_8446622_RNA45SN1_45S_50ntFlanks@9929.9948.20|GATATAGACAGCAGGACGGT</td><td>>hg38_21_+_8433172_8446622_RNA45SN1_45S_50ntFlanks</td><td>9929</td><td>9948</td><td>20</td><td>GATATAGACAGCAGGACGGT</td><td>318.4</td><td>-5.5</td><td>1.2</td><td>-4.6</td><td>3.8E-06</td><td>5.7E-06</td><td>MB231/MB468</td><td>MITO</td></tr>
<tr><td>>trnaMT_LeuTAA_MT_+_3230_3304@1.34.34|GTTAAGATGGCAGAGCCCGGTAATCGCATAAAAC</td><td>>trnaMT_LeuTAA_MT_+_3230_3304</td><td>1</td><td>34</td><td>34</td><td>GTTAAGATGGCAGAGCCCGGTAATCGCATAAAAC</td><td>43.3</td><td>8.7</td><td>1.9</td><td>4.7</td><td>3.2E-06</td><td>5.7E-06</td><td>MB231/MB468</td><td>MP</td></tr>
<tr><td>>hg38_21_+_8433172_8446622_RNA45SN1_45S_50ntFlanks@9178.9197.20|CGCCGTCGGGCCCGGGGGAG</td><td>>hg38_21_+_8433172_8446622_RNA45SN1_45S_50ntFlanks</td><td>9178</td><td>9197</td><td>20</td><td>CGCCGTCGGGCCCGGGGGAG</td><td>887.7</td><td>-4.1</td><td>0.9</td><td>-4.7</td><td>2.7E-06</td><td>5.7E-06</td><td>BT20/MB468</td><td>MITO</td></tr>
<tr><td>>hg38_21_+_8433172_8446622_RNA45SN1_45S_50ntFlanks@10357.10388.32|CTTTGAAGGCCGAAGTGGAGAAGGGTTCCATG</td><td>>hg38_21_+_8433172_8446622_RNA45SN1_45S_50ntFlanks</td><td>10357</td><td>10388</td><td>32</td><td>CTTTGAAGGCCGAAGTGGAGAAGGGTTCCATG</td><td>586.3</td><td>-5.7</td><td>1.2</td><td>-4.7</td><td>3.2E-06</td><td>5.7E-06</td><td>BT20/MB468</td><td>CYTO</td></tr>
<tr><td>>hg38_21_+_8433172_8446622_RNA45SN1_45S_50ntFlanks@12836.12859.24|CGGTGCGGAGTGCCCTTCGTCCTG</td><td>>hg38_21_+_8433172_8446622_RNA45SN1_45S_50ntFlanks</td><td>12836</td><td>12859</td><td>24</td><td>CGGTGCGGAGTGCCCTTCGTCCTG</td><td>75.9</td><td>-9.3</td><td>1.9</td><td>-4.9</td><td>1.1E-06</td><td>5.7E-06</td><td>BT20/MB468</td><td>MP</td></tr>
<tr><td>>hg38_21_+_8433172_8446622_RNA45SN1_45S_50ntFlanks@9533.9567.35|GAGGAAACTCTGGTGGAGGTCCGTAGCGGTCCTGA</td><td>>hg38_21_+_8433172_8446622_RNA45SN1_45S_50ntFlanks</td><td>9533</td><td>9567</td><td>35</td><td>GAGGAAACTCTGGTGGAGGTCCGTAGCGGTCCTGA</td><td>1149.3</td><td>-8.2</td><td>1.8</td><td>-4.6</td><td>4.0E-06</td><td>5.7E-06</td><td>BT20/MB468</td><td>NUC</td></tr>
<tr><td>>hg38_21_+_8433172_8446622_RNA45SN1_45S_50ntFlanks@11885.11904.20|TTGAGCTTGACTCTAGTCTG</td><td>>hg38_21_+_8433172_8446622_RNA45SN1_45S_50ntFlanks</td><td>11885</td><td>11904</td><td>20</td><td>TTGAGCTTGACTCTAGTCTG</td><td>804.6</td><td>-3.5</td><td>0.8</td><td>-4.7</td><td>2.8E-06</td><td>5.7E-06</td><td>BT20/MB468</td><td>MITO</td></tr>
<tr><td>>hg38_21_+_8433172_8446622_RNA45SN1_45S_50ntFlanks@8183.8212.30|TCGAGGCCCAGCCCGTGGACGGTGTGAGGC</td><td>>hg38_21_+_8433172_8446622_RNA45SN1_45S_50ntFlanks</td><td>8183</td><td>8212</td><td>30</td><td>TCGAGGCCCAGCCCGTGGACGGTGTGAGGC</td><td>182.6</td><td>-7.0</td><td>1.5</td><td>-4.7</td><td>3.2E-06</td><td>5.7E-06</td><td>MB231/MB468</td><td>MP</td></tr>
<tr><td>>hg38_21_+_8433172_8446622_RNA45SN1_45S_50ntFlanks@12407.12434.28|TTTTGATCCTTCGATGTCGGCTCTTCCT</td><td>>hg38_21_+_8433172_8446622_RNA45SN1_45S_50ntFlanks</td><td>12407</td><td>12434</td><td>28</td><td>TTTTGATCCTTCGATGTCGGCTCTTCCT</td><td>292.3</td><td>-5.3</td><td>1.1</td><td>-4.7</td><td>3.2E-06</td><td>5.7E-06</td><td>MB231/MB468</td><td>MP</td></tr>
<tr><td>>MI0000270|hsa-mir-181b-1&WithFlank&1|-|198858867|198858988@42.65.24;MI0000683|hsa-mir-181b-2&WithFlank&9|+|124693704|124693804@22.45.24&[MIMAT0000257&hsa-miR-181b-5p&offsets|0|+1;m-73&1|-|198858925|198858947&offsets|0|+1];[MIMAT0000257_1&hsa-miR-181b-5p&offsets|0|+1;m-72&9|+|124693725|124693747&offsets|0|+1]|AACATTCATTGCTGTCGGTGGGTT</td><td>>MI0000270|hsa-mir-181b-1&WithFlank&1|-|198858867|198858988</td><td>42</td><td>65</td><td>24</td><td>AACATTCATTGCTGTCGGTGGGTT</td><td>3439.9</td><td>3.6</td><td>0.8</td><td>4.7</td><td>3.2E-06</td><td>5.7E-06</td><td>MB231/MB468</td><td>MP</td></tr>
<tr><td>>MI0000805|hsa-mir-342&WithFlank&14|+|100109649|100109759@67.90.24&[MIMAT0000753&hsa-miR-342-3p&offsets|0|+1;m-128&14|+|100109715|100109737&offsets|0|+1]|TCTCACACAGAAATCGCACCCGTC</td><td>>MI0000805|hsa-mir-342&WithFlank&14|+|100109649|100109759</td><td>67</td><td>90</td><td>24</td><td>TCTCACACAGAAATCGCACCCGTC</td><td>314.0</td><td>4.6</td><td>1.0</td><td>4.6</td><td>3.8E-06</td><td>5.7E-06</td><td>MB231/MB468</td><td>MITO</td></tr>
<tr><td>>hg38_21_+_8433172_8446622_RNA45SN1_45S_50ntFlanks@10411.10431.21|CAGTCGGTCCTGAGAGATGGG</td><td>>hg38_21_+_8433172_8446622_RNA45SN1_45S_50ntFlanks</td><td>10411</td><td>10431</td><td>21</td><td>CAGTCGGTCCTGAGAGATGGG</td><td>218.3</td><td>11.1</td><td>2.2</td><td>5.1</td><td>2.9E-07</td><td>5.7E-06</td><td>BT20/MB231</td><td>MP</td></tr>
<tr><td>>hg38_21_+_8433172_8446622_RNA45SN1_45S_50ntFlanks@9925.9952.28|GGTTGATATAGACAGCAGGACGGTGGCC</td><td>>hg38_21_+_8433172_8446622_RNA45SN1_45S_50ntFlanks</td><td>9925</td><td>9952</td><td>28</td><td>GGTTGATATAGACAGCAGGACGGTGGCC</td><td>105.9</td><td>10.1</td><td>2.0</td><td>5.1</td><td>2.9E-07</td><td>5.7E-06</td><td>BT20/MB231</td><td>MP</td></tr>
<tr><td>>hg38_1_-_228634819_228635039_RNA5S12_5S_50ntFlanks@148.171.24|GGAATACCGGGTGCTGTAGGCTTT</td><td>>hg38_1_-_228634819_228635039_RNA5S12_5S_50ntFlanks</td><td>148</td><td>171</td><td>24</td><td>GGAATACCGGGTGCTGTAGGCTTT</td><td>6922.5</td><td>3.4</td><td>0.7</td><td>4.6</td><td>3.6E-06</td><td>5.7E-06</td><td>MB231/MB468</td><td>CYTO</td></tr>
<tr><td>>hg38_21_+_8433172_8446622_RNA45SN1_45S_50ntFlanks@9567.9586.20|ACGTGCAAATCGGTCGTCCG</td><td>>hg38_21_+_8433172_8446622_RNA45SN1_45S_50ntFlanks</td><td>9567</td><td>9586</td><td>20</td><td>ACGTGCAAATCGGTCGTCCG</td><td>559.0</td><td>-6.5</td><td>1.4</td><td>-4.7</td><td>3.2E-06</td><td>5.8E-06</td><td>BT20/MB468</td><td>CYTO</td></tr>
<tr><td>>hg38_21_+_8433172_8446622_RNA45SN1_45S_50ntFlanks@12478.12497.20|ACTAATAGGGAACGTGAGCT</td><td>>hg38_21_+_8433172_8446622_RNA45SN1_45S_50ntFlanks</td><td>12478</td><td>12497</td><td>20</td><td>ACTAATAGGGAACGTGAGCT</td><td>844.1</td><td>-3.1</td><td>0.7</td><td>-4.6</td><td>4.0E-06</td><td>5.8E-06</td><td>BT20/MB468</td><td>NUC</td></tr>
<tr><td>>hg38_21_+_8433172_8446622_RNA45SN1_45S_50ntFlanks@3903.3937.35|CGCGGGGGGGATGCGTGCATTTATCAGATCAAAAC</td><td>>hg38_21_+_8433172_8446622_RNA45SN1_45S_50ntFlanks</td><td>3903</td><td>3937</td><td>35</td><td>CGCGGGGGGGATGCGTGCATTTATCAGATCAAAAC</td><td>114.4</td><td>-10.7</td><td>2.3</td><td>-4.7</td><td>3.2E-06</td><td>5.8E-06</td><td>MB231/MB468</td><td>MP</td></tr>
<tr><td>>hg38_21_+_8433172_8446622_RNA45SN1_45S_50ntFlanks@12539.12573.35|ATGATGTGTTGTTGCCATGGTAATCCTGCTCAGTA</td><td>>hg38_21_+_8433172_8446622_RNA45SN1_45S_50ntFlanks</td><td>12539</td><td>12573</td><td>35</td><td>ATGATGTGTTGTTGCCATGGTAATCCTGCTCAGTA</td><td>349.6</td><td>-6.6</td><td>1.4</td><td>-4.6</td><td>3.8E-06</td><td>5.8E-06</td><td>MB231/MB468</td><td>NUC</td></tr>
<tr><td>>hg38_21_+_8433172_8446622_RNA45SN1_45S_50ntFlanks@6700.6720.21|CTAGCTGCGAGAATTAATGTG</td><td>>hg38_21_+_8433172_8446622_RNA45SN1_45S_50ntFlanks</td><td>6700</td><td>6720</td><td>21</td><td>CTAGCTGCGAGAATTAATGTG</td><td>8844.8</td><td>-3.8</td><td>0.8</td><td>-4.6</td><td>3.8E-06</td><td>5.8E-06</td><td>MB231/MB468</td><td>NUC</td></tr>
<tr><td>>hg38_21_+_8433172_8446622_RNA45SN1_45S_50ntFlanks@10350.10375.26|ACGAGAACTTTGAAGGCCGAAGTGGA</td><td>>hg38_21_+_8433172_8446622_RNA45SN1_45S_50ntFlanks</td><td>10350</td><td>10375</td><td>26</td><td>ACGAGAACTTTGAAGGCCGAAGTGGA</td><td>347.0</td><td>-7.3</td><td>1.6</td><td>-4.7</td><td>3.2E-06</td><td>5.8E-06</td><td>MB231/MB468</td><td>MP</td></tr>
<tr><td>>am_MI0000809|hsa-mir-151a&WithFlank&8|-|140732558|140732659@53.71.19&[MIMAT0000757&hsa-miR-151a-3p&offsets|0|-2;m-30&8|-|140732587|140732607&offsets|0|-2]|CTAGACTGAAGCTCCTTGA</td><td>>am_MI0000809|hsa-mir-151a&WithFlank&8|-|140732558|140732659</td><td>53</td><td>71</td><td>19</td><td>CTAGACTGAAGCTCCTTGA</td><td>1622.2</td><td>6.7</td><td>1.4</td><td>4.9</td><td>1.1E-06</td><td>5.8E-06</td><td>BT20/MB468</td><td>MP</td></tr>
<tr><td>>am_trnaMT_PheGAA_MT_+_577_647@44.62.19|TAGACGGGCTCACATCACC</td><td>>am_trnaMT_PheGAA_MT_+_577_647</td><td>44</td><td>62</td><td>19</td><td>TAGACGGGCTCACATCACC</td><td>560.0</td><td>2.8</td><td>0.6</td><td>4.6</td><td>3.9E-06</td><td>5.8E-06</td><td>MB231/MB468</td><td>MITO</td></tr>
<tr><td>>MI0000285|hsa-mir-205&WithFlank&1|+|209432127|209432248@40.61.22&[MIMAT0000266&hsa-miR-205-5p&offsets|0|0;m-351&1|+|209432166|209432187&offsets|0|0]|TCCTTCATTCCACCGGAGTCTG</td><td>>MI0000285|hsa-mir-205&WithFlank&1|+|209432127|209432248</td><td>40</td><td>61</td><td>22</td><td>TCCTTCATTCCACCGGAGTCTG</td><td>817.5</td><td>-3.6</td><td>0.8</td><td>-4.6</td><td>3.6E-06</td><td>5.8E-06</td><td>MB231/MB468</td><td>CYTO</td></tr>
<tr><td>>hg38_21_+_8433172_8446622_RNA45SN1_45S_50ntFlanks@10735.10757.23|TGTAAATCTCGCGCCGGGCCGTA</td><td>>hg38_21_+_8433172_8446622_RNA45SN1_45S_50ntFlanks</td><td>10735</td><td>10757</td><td>23</td><td>TGTAAATCTCGCGCCGGGCCGTA</td><td>244.2</td><td>-4.2</td><td>0.9</td><td>-4.6</td><td>3.6E-06</td><td>5.8E-06</td><td>MB231/MB468</td><td>CYTO</td></tr>
<tr><td>>hg38_21_+_8433172_8446622_RNA45SN1_45S_50ntFlanks@9740.9767.28|AACGATCTCAACCTATTCTCAAACTTTA</td><td>>hg38_21_+_8433172_8446622_RNA45SN1_45S_50ntFlanks</td><td>9740</td><td>9767</td><td>28</td><td>AACGATCTCAACCTATTCTCAAACTTTA</td><td>726.0</td><td>-3.6</td><td>0.8</td><td>-4.6</td><td>3.9E-06</td><td>5.8E-06</td><td>MB231/MB468</td><td>MITO</td></tr>
<tr><td>>hg38_21_+_8433172_8446622_RNA45SN1_45S_50ntFlanks@10315.10336.22|AGGTGCAGATCTTGGTGGTAGT</td><td>>hg38_21_+_8433172_8446622_RNA45SN1_45S_50ntFlanks</td><td>10315</td><td>10336</td><td>22</td><td>AGGTGCAGATCTTGGTGGTAGT</td><td>54.2</td><td>9.5</td><td>2.0</td><td>4.9</td><td>1.1E-06</td><td>5.8E-06</td><td>BT20/MB468</td><td>MP</td></tr>
<tr><td>>trna58_LeuCAA_1_+_249168054_249168159@48.65.18|CCGGATGGAGGCGTGGGT</td><td>>trna58_LeuCAA_1_+_249168054_249168159</td><td>48</td><td>65</td><td>18</td><td>CCGGATGGAGGCGTGGGT</td><td>605.5</td><td>7.4</td><td>1.6</td><td>4.7</td><td>3.2E-06</td><td>5.8E-06</td><td>BT20/MB468</td><td>CYTO</td></tr>
<tr><td>>MI0001445|hsa-mir-423&WithFlank&17|+|30117073|30117178@58.77.20(+1U)&[MIMAT0001340&hsa-miR-423-3p&offsets|-1|-4(+1U);m-100&17|+|30117131|30117153&offsets|-1|-4(+1U)]|AAGCTCGGTCTGAGGCCCCTT</td><td>>MI0001445|hsa-mir-423&WithFlank&17|+|30117073|30117178</td><td>58</td><td>77</td><td>20(+1U)</td><td>AAGCTCGGTCTGAGGCCCCTT</td><td>51.3</td><td>8.9</td><td>1.9</td><td>4.7</td><td>3.2E-06</td><td>5.8E-06</td><td>MB231/MB468</td><td>MP</td></tr>
<tr><td>>hg38_21_+_8433172_8446622_RNA45SN1_45S_50ntFlanks@12418.12454.37|CGATGTCGGCTCTTCCTATCATTGTGAAGCAGAATTC</td><td>>hg38_21_+_8433172_8446622_RNA45SN1_45S_50ntFlanks</td><td>12418</td><td>12454</td><td>37</td><td>CGATGTCGGCTCTTCCTATCATTGTGAAGCAGAATTC</td><td>883.1</td><td>-8.1</td><td>1.8</td><td>-4.6</td><td>4.1E-06</td><td>5.8E-06</td><td>BT20/MB468</td><td>NUC</td></tr>
<tr><td>>hg38_21_+_8433172_8446622_RNA45SN1_45S_50ntFlanks@5469.5516.48|CCCACGGCCCTGGCGGAGCGCTGAGAAGACGGTCGAACTTGACTATCT</td><td>>hg38_21_+_8433172_8446622_RNA45SN1_45S_50ntFlanks</td><td>5469</td><td>5516</td><td>48</td><td>CCCACGGCCCTGGCGGAGCGCTGAGAAGACGGTCGAACTTGACTATCT</td><td>1542.7</td><td>8.8</td><td>1.9</td><td>4.7</td><td>2.8E-06</td><td>5.8E-06</td><td>BT20/MB468</td><td>MITO</td></tr>
<tr><td>>hg38_MT_+_598_1651_RNR1_12S_50ntFlanks@850.873.24|GTGCTTAGTTGAACAGGGCCCTGA</td><td>>hg38_MT_+_598_1651_RNR1_12S_50ntFlanks</td><td>850</td><td>873</td><td>24</td><td>GTGCTTAGTTGAACAGGGCCCTGA</td><td>65.4</td><td>9.8</td><td>2.0</td><td>4.9</td><td>1.1E-06</td><td>5.8E-06</td><td>BT20/MB468</td><td>MP</td></tr>
<tr><td>>hg38_21_+_8433172_8446622_RNA45SN1_45S_50ntFlanks@6652.6694.43|GACTCTTAGCGGTGGATCACTCGGCTCGTGCGTCGATGAAGAA</td><td>>hg38_21_+_8433172_8446622_RNA45SN1_45S_50ntFlanks</td><td>6652</td><td>6694</td><td>43</td><td>GACTCTTAGCGGTGGATCACTCGGCTCGTGCGTCGATGAAGAA</td><td>867.4</td><td>4.1</td><td>0.9</td><td>4.6</td><td>3.8E-06</td><td>5.8E-06</td><td>MB231/MB468</td><td>NUC</td></tr>
<tr><td>>hg38_21_+_8433172_8446622_RNA45SN1_45S_50ntFlanks@9932.9960.29|ATAGACAGCAGGACGGTGGCCATGGAAGT</td><td>>hg38_21_+_8433172_8446622_RNA45SN1_45S_50ntFlanks</td><td>9932</td><td>9960</td><td>29</td><td>ATAGACAGCAGGACGGTGGCCATGGAAGT</td><td>189.0</td><td>10.9</td><td>2.1</td><td>5.1</td><td>3.0E-07</td><td>5.8E-06</td><td>BT20/MB231</td><td>MP</td></tr>
<tr><td>>hg38_21_+_8433172_8446622_RNA45SN1_45S_50ntFlanks@6742.6771.30|TCGACACTTCGAACGCACTTGCGGCCCCGG</td><td>>hg38_21_+_8433172_8446622_RNA45SN1_45S_50ntFlanks</td><td>6742</td><td>6771</td><td>30</td><td>TCGACACTTCGAACGCACTTGCGGCCCCGG</td><td>193.2</td><td>-4.8</td><td>1.0</td><td>-4.6</td><td>3.8E-06</td><td>5.8E-06</td><td>MB231/MB468</td><td>NUC</td></tr>
<tr><td>>hg38_21_+_8433172_8446622_RNA45SN1_45S_50ntFlanks@12991.13011.21|CTCCCTCGCTGCGATCTATTG</td><td>>hg38_21_+_8433172_8446622_RNA45SN1_45S_50ntFlanks</td><td>12991</td><td>13011</td><td>21</td><td>CTCCCTCGCTGCGATCTATTG</td><td>1390.1</td><td>-5.7</td><td>1.2</td><td>-4.7</td><td>3.2E-06</td><td>5.8E-06</td><td>BT20/MB468</td><td>CYTO</td></tr>
<tr><td>>hg38_21_+_8433172_8446622_RNA45SN1_45S_50ntFlanks@11581.11611.31|AACTGGTGCGGACCAGGGGAATCCGACTGTT</td><td>>hg38_21_+_8433172_8446622_RNA45SN1_45S_50ntFlanks</td><td>11581</td><td>11611</td><td>31</td><td>AACTGGTGCGGACCAGGGGAATCCGACTGTT</td><td>1998.0</td><td>-4.4</td><td>0.9</td><td>-4.9</td><td>1.1E-06</td><td>5.8E-06</td><td>BT20/MB468</td><td>MP</td></tr>
<tr><td>>MI0000086|hsa-mir-28&WithFlank&3|+|188688775|188688872@61.82.22(+1U)&[MIMAT0004502&hsa-miR-28-3p&offsets|+1|+1(+1U);m-28&3|+|188688834|188688855&offsets|+1|+1(+1U)]|ACTAGATTGTGAGCTCCTGGAGT</td><td>>MI0000086|hsa-mir-28&WithFlank&3|+|188688775|188688872</td><td>61</td><td>82</td><td>22(+1U)</td><td>ACTAGATTGTGAGCTCCTGGAGT</td><td>546.6</td><td>6.4</td><td>1.3</td><td>4.9</td><td>1.1E-06</td><td>5.8E-06</td><td>BT20/MB468</td><td>MP</td></tr>
<tr><td>>hg38_21_+_8433172_8446622_RNA45SN1_45S_50ntFlanks@11600.11629.30|AATCCGACTGTTTAATTAAAACAAAGCATC</td><td>>hg38_21_+_8433172_8446622_RNA45SN1_45S_50ntFlanks</td><td>11600</td><td>11629</td><td>30</td><td>AATCCGACTGTTTAATTAAAACAAAGCATC</td><td>152.8</td><td>10.6</td><td>2.1</td><td>5.1</td><td>3.0E-07</td><td>5.8E-06</td><td>BT20/MB231</td><td>MP</td></tr>
<tr><td>>hg38_21_+_8433172_8446622_RNA45SN1_45S_50ntFlanks@10868.10892.25|AAGGGCTGGGTCGGTCGGGCTGGGG</td><td>>hg38_21_+_8433172_8446622_RNA45SN1_45S_50ntFlanks</td><td>10868</td><td>10892</td><td>25</td><td>AAGGGCTGGGTCGGTCGGGCTGGGG</td><td>227.7</td><td>-6.8</td><td>1.5</td><td>-4.7</td><td>3.2E-06</td><td>5.8E-06</td><td>MB231/MB468</td><td>MP</td></tr>
<tr><td>>hg38_1_-_228634819_228635039_RNA5S12_5S_50ntFlanks@136.170.35|GGAGACCGCCTGGGAATACCGGGTGCTGTAGGCTT</td><td>>hg38_1_-_228634819_228635039_RNA5S12_5S_50ntFlanks</td><td>136</td><td>170</td><td>35</td><td>GGAGACCGCCTGGGAATACCGGGTGCTGTAGGCTT</td><td>4425.7</td><td>4.0</td><td>0.9</td><td>4.7</td><td>2.8E-06</td><td>5.8E-06</td><td>BT20/MB468</td><td>MITO</td></tr>
<tr><td>>hg38_21_+_8433172_8446622_RNA45SN1_45S_50ntFlanks@6661.6707.47|CGGTGGATCACTCGGCTCGTGCGTCGATGAAGAACGCAGCTAGCTGC</td><td>>hg38_21_+_8433172_8446622_RNA45SN1_45S_50ntFlanks</td><td>6661</td><td>6707</td><td>47</td><td>CGGTGGATCACTCGGCTCGTGCGTCGATGAAGAACGCAGCTAGCTGC</td><td>692.3</td><td>7.6</td><td>1.6</td><td>4.7</td><td>3.3E-06</td><td>5.8E-06</td><td>BT20/MB468</td><td>CYTO</td></tr>
</table>

<table>
<tr><td>>trnaMT_SerGCT_MT_+_12207_12265@14.61.48|AGAACTGCTAACTCATGCCCCCATGTCTAACAACATGGCTTTCTCACC</td><td>>trnaMT_SerGCT_MT_+_12207_12265</td><td>14</td><td>61</td><td>48</td><td>AGAACTGCTAACTCATGCCCCCATGTCTAACAACATGGCTTTCTCACC</td><td>384.1</td><td>12.3</td><td>2.5</td><td>4.9</td><td>1.1E-06</td><td>5.8E-06</td><td>BT20/MB468</td><td>MP</td></tr>
<tr><td>>hg38_21_+_8433172_8446622_RNA45SN1_45S_50ntFlanks@4328.4346.19|CGGTAATTCCAGCTCCAAT</td><td>>hg38_21_+_8433172_8446622_RNA45SN1_45S_50ntFlanks</td><td>4328</td><td>4346</td><td>19</td><td>CGGTAATTCCAGCTCCAAT</td><td>44.9</td><td>-8.6</td><td>1.8</td><td>-4.9</td><td>1.1E-06</td><td>5.8E-06</td><td>BT20/MB468</td><td>MP</td></tr>
<tr><td>>MI0000298|hsa-mir-221&WithFlank&X|-|45746151|45746272@71.92.22&[MIMAT0000278&hsa-miR-221-3p&offsets|0|-1;m-91&X|-|45746181|45746202&offsets|0|0]|AGCTACATTGTCTGCTGGGTTT</td><td>>MI0000298|hsa-mir-221&WithFlank&X|-|45746151|45746272</td><td>71</td><td>92</td><td>22</td><td>AGCTACATTGTCTGCTGGGTTT</td><td>1527.9</td><td>2.3</td><td>0.5</td><td>4.6</td><td>4.1E-06</td><td>5.8E-06</td><td>BT20/MB468</td><td>NUC</td></tr>
<tr><td>>MI0000077|hsa-mir-21&WithFlank&17|+|59841260|59841343@14.34.21(+3C)&[MIMAT0000076&hsa-miR-21-5p&offsets|0|-1(+3C);m-1&17|+|59841273|59841294&offsets|0|-1(+3C)]|TAGCTTATCAGACTGATGTTGCCC</td><td>>MI0000077|hsa-mir-21&WithFlank&17|+|59841260|59841343</td><td>14</td><td>34</td><td>21(+3C)</td><td>TAGCTTATCAGACTGATGTTGCCC</td><td>42.2</td><td>8.6</td><td>1.9</td><td>4.7</td><td>3.2E-06</td><td>5.8E-06</td><td>MB231/MB468</td><td>MP</td></tr>
<tr><td>>hg38_21_+_8433172_8446622_RNA45SN1_45S_50ntFlanks@5107.5136.30|CTAGTTACGCGACCCCCGAGCGGTCGGCGT</td><td>>hg38_21_+_8433172_8446622_RNA45SN1_45S_50ntFlanks</td><td>5107</td><td>5136</td><td>30</td><td>CTAGTTACGCGACCCCCGAGCGGTCGGCGT</td><td>82.3</td><td>-9.4</td><td>1.9</td><td>-4.9</td><td>1.1E-06</td><td>5.9E-06</td><td>BT20/MB468</td><td>MP</td></tr>
<tr><td>>hg38_21_+_8433172_8446622_RNA45SN1_45S_50ntFlanks@6645.6685.41|CTCGTACGACTCTTAGCGGTGGATCACTCGGCTCGTGCGTC</td><td>>hg38_21_+_8433172_8446622_RNA45SN1_45S_50ntFlanks</td><td>6645</td><td>6685</td><td>41</td><td>CTCGTACGACTCTTAGCGGTGGATCACTCGGCTCGTGCGTC</td><td>1358.4</td><td>5.5</td><td>1.1</td><td>4.9</td><td>1.1E-06</td><td>5.9E-06</td><td>BT20/MB468</td><td>MP</td></tr>
<tr><td>>hg38_21_+_8433172_8446622_RNA45SN1_45S_50ntFlanks@6653.6695.43|ACTCTTAGCGGTGGATCACTCGGCTCGTGCGTCGATGAAGAAC</td><td>>hg38_21_+_8433172_8446622_RNA45SN1_45S_50ntFlanks</td><td>6653</td><td>6695</td><td>43</td><td>ACTCTTAGCGGTGGATCACTCGGCTCGTGCGTCGATGAAGAAC</td><td>457.8</td><td>6.9</td><td>1.5</td><td>4.7</td><td>2.8E-06</td><td>5.9E-06</td><td>BT20/MB468</td><td>MITO</td></tr>
<tr><td>>hg38_21_+_8433172_8446622_RNA45SN1_45S_50ntFlanks@3939.3959.21|AACCCGGTCAGCCCCTCTCCG</td><td>>hg38_21_+_8433172_8446622_RNA45SN1_45S_50ntFlanks</td><td>3939</td><td>3959</td><td>21</td><td>AACCCGGTCAGCCCCTCTCCG</td><td>1762.5</td><td>-6.9</td><td>1.5</td><td>-4.6</td><td>4.1E-06</td><td>5.9E-06</td><td>BT20/MB468</td><td>NUC</td></tr>
<tr><td>>hg38_21_+_8433172_8446622_RNA45SN1_45S_50ntFlanks@6653.6689.37|ACTCTTAGCGGTGGATCACTCGGCTCGTGCGTCGATG</td><td>>hg38_21_+_8433172_8446622_RNA45SN1_45S_50ntFlanks</td><td>6653</td><td>6689</td><td>37</td><td>ACTCTTAGCGGTGGATCACTCGGCTCGTGCGTCGATG</td><td>1040.4</td><td>-4.9</td><td>1.1</td><td>-4.7</td><td>3.3E-06</td><td>5.9E-06</td><td>MB231/MB468</td><td>MP</td></tr>
<tr><td>>hg38_21_+_8433172_8446622_RNA45SN1_45S_50ntFlanks@6682.6718.37|CGTCGATGAAGAACGCAGCTAGCTGCGAGAATTAATG</td><td>>hg38_21_+_8433172_8446622_RNA45SN1_45S_50ntFlanks</td><td>6682</td><td>6718</td><td>37</td><td>CGTCGATGAAGAACGCAGCTAGCTGCGAGAATTAATG</td><td>98.4</td><td>-9.7</td><td>2.0</td><td>-4.9</td><td>1.1E-06</td><td>5.9E-06</td><td>BT20/MB468</td><td>MP</td></tr>
<tr><td>>hg38_21_+_8433172_8446622_RNA45SN1_45S_50ntFlanks@7978.8006.29|GACCTCAGATCAGACGTGGCGACCCGCTG</td><td>>hg38_21_+_8433172_8446622_RNA45SN1_45S_50ntFlanks</td><td>7978</td><td>8006</td><td>29</td><td>GACCTCAGATCAGACGTGGCGACCCGCTG</td><td>732.4</td><td>-5.2</td><td>1.1</td><td>-4.6</td><td>3.9E-06</td><td>5.9E-06</td><td>MB231/MB468</td><td>MITO</td></tr>
<tr><td>>MI0000103|hsa-mir-101-1&WithFlank&1|-|65058428|65058514@53.73.21;MI0000739|hsa-mir-101-2&WithFlank&9|+|4850291|4850381@55.75.21&[MIMAT0000099&hsa-miR-101-3p&offsets|0|0;m-17&1|-|65058443|65058462&offsets|0|+1];[MIMAT0000099_1&hsa-miR-101-3p&offsets|0|0;m-16&9|+|4850345|4850364&offsets|0|+1]|TACAGTACTGTGATAACTGAA</td><td>>MI0000103|hsa-mir-101-1&WithFlank&1|-|65058428|65058514</td><td>53</td><td>73</td><td>21</td><td>TACAGTACTGTGATAACTGAA</td><td>456.2</td><td>5.6</td><td>1.2</td><td>4.6</td><td>3.8E-06</td><td>5.9E-06</td><td>MB231/MB468</td><td>NUC</td></tr>
<tr><td>>MI0000272|hsa-mir-182&WithFlank&7|-|129770377|129770498@29.52.24&[MIMAT0000259&hsa-miR-182-5p&offsets|0|0;m-22&7|-|129770449|129770470&offsets|0|+2]|TTTGGCAATGGTAGAACTCACACT</td><td>>MI0000272|hsa-mir-182&WithFlank&7|-|129770377|129770498</td><td>29</td><td>52</td><td>24</td><td>TTTGGCAATGGTAGAACTCACACT</td><td>7223.5</td><td>2.7</td><td>0.6</td><td>4.6</td><td>3.8E-06</td><td>5.9E-06</td><td>MB231/MB468</td><td>NUC</td></tr>
<tr><td>>hg38_21_+_8433172_8446622_RNA45SN1_45S_50ntFlanks@5487.5516.30|CGCTGAGAAGACGGTCGAACTTGACTATCT</td><td>>hg38_21_+_8433172_8446622_RNA45SN1_45S_50ntFlanks</td><td>5487</td><td>5516</td><td>30</td><td>CGCTGAGAAGACGGTCGAACTTGACTATCT</td><td>8814.6</td><td>3.6</td><td>0.8</td><td>4.7</td><td>3.3E-06</td><td>5.9E-06</td><td>BT20/MB468</td><td>CYTO</td></tr>
<tr><td>>hg38_21_+_8433172_8446622_RNA45SN1_45S_50ntFlanks@8154.8182.29|CCGCTCGTGGGGGGCCCAAGTCCTTCTGA</td><td>>hg38_21_+_8433172_8446622_RNA45SN1_45S_50ntFlanks</td><td>8154</td><td>8182</td><td>29</td><td>CCGCTCGTGGGGGGCCCAAGTCCTTCTGA</td><td>668.0</td><td>-5.2</td><td>1.1</td><td>-4.7</td><td>2.8E-06</td><td>5.9E-06</td><td>BT20/MB468</td><td>MITO</td></tr>
<tr><td>>hg38_21_+_8433172_8446622_RNA45SN1_45S_50ntFlanks@4000.4024.25|TAGATAACCTCGGGCCGATCGCACG</td><td>>hg38_21_+_8433172_8446622_RNA45SN1_45S_50ntFlanks</td><td>4000</td><td>4024</td><td>25</td><td>TAGATAACCTCGGGCCGATCGCACG</td><td>516.3</td><td>-4.5</td><td>1.0</td><td>-4.7</td><td>3.3E-06</td><td>5.9E-06</td><td>BT20/MB468</td><td>CYTO</td></tr>
<tr><td>>MI0000266|hsa-mir-10a&WithFlank&17|-|48579832|48579953@29.48.20&[MIMAT0000253&hsa-miR-10a-5p&offsets|+1|-2;m-18&17|-|48579905|48579926&offsets|+1|-1]|ACCCTGTAGATCCGAATTTG</td><td>>MI0000266|hsa-mir-10a&WithFlank&17|-|48579832|48579953</td><td>29</td><td>48</td><td>20</td><td>ACCCTGTAGATCCGAATTTG</td><td>68.9</td><td>9.9</td><td>2.0</td><td>4.9</td><td>1.1E-06</td><td>5.9E-06</td><td>BT20/MB468</td><td>MP</td></tr>
<tr><td>>hg38_21_+_8433172_8446622_RNA45SN1_45S_50ntFlanks@10381.10424.44|GTTCCATGTGAACAGCAGTTGAACATGGGTCAGTCGGTCCTGAG</td><td>>hg38_21_+_8433172_8446622_RNA45SN1_45S_50ntFlanks</td><td>10381</td><td>10424</td><td>44</td><td>GTTCCATGTGAACAGCAGTTGAACATGGGTCAGTCGGTCCTGAG</td><td>301.0</td><td>3.9</td><td>0.9</td><td>4.6</td><td>4.0E-06</td><td>5.9E-06</td><td>MB231/MB468</td><td>MITO</td></tr>
<tr><td>>trnaMT_PheGAA_MT_+_577_647@28.57.30|ATACACTGAAAATGTTTAGACGGGCTCACA</td><td>>trnaMT_PheGAA_MT_+_577_647</td><td>28</td><td>57</td><td>30</td><td>ATACACTGAAAATGTTTAGACGGGCTCACA</td><td>247.3</td><td>8.6</td><td>1.8</td><td>4.9</td><td>1.1E-06</td><td>5.9E-06</td><td>BT20/MB468</td><td>MP</td></tr>
<tr><td>>hg38_21_+_8433172_8446622_RNA45SN1_45S_50ntFlanks@475.506.32|CTTCGTGATCGATGTGGTGACGTCGTGCTCTC</td><td>>hg38_21_+_8433172_8446622_RNA45SN1_45S_50ntFlanks</td><td>475</td><td>506</td><td>32</td><td>CTTCGTGATCGATGTGGTGACGTCGTGCTCTC</td><td>36160.9</td><td>3.5</td><td>0.8</td><td>4.6</td><td>3.9E-06</td><td>5.9E-06</td><td>MB231/MB468</td><td>NUC</td></tr>
<tr><td>>MI0000736|hsa-mir-30c-1&WithFlank&1|+|40757278|40757378@23.41.19(+1A);MI0000254|hsa-mir-30c-2&WithFlank&6|-|71376954|71377037@13.31.19(+1A)&[MIMAT0000244&hsa-miR-30c-5p&offsets|0|-4(+1A);m-57&1|+|40757300|40757323&offsets|0|-5(+1A)];[MIMAT0000244_1&hsa-miR-30c-5p&offsets|0|-4(+1A);m-56&6|-|71377002|71377025&offsets|0|-5(+1A)]|TGTAAACATCCTACACTCTA</td><td>>MI0000736|hsa-mir-30c-1&WithFlank&1|+|40757278|40757378</td><td>23</td><td>41</td><td>19(+1A)</td><td>TGTAAACATCCTACACTCTA</td><td>58.2</td><td>9.1</td><td>2.0</td><td>4.7</td><td>3.3E-06</td><td>5.9E-06</td><td>MB231/MB468</td><td>MP</td></tr>
<tr><td>>hg38_21_+_8433172_8446622_RNA45SN1_45S_50ntFlanks@12971.12991.21|TTTCGTACGTAGCAGAGCAGC</td><td>>hg38_21_+_8433172_8446622_RNA45SN1_45S_50ntFlanks</td><td>12971</td><td>12991</td><td>21</td><td>TTTCGTACGTAGCAGAGCAGC</td><td>494.0</td><td>-5.0</td><td>1.1</td><td>-4.7</td><td>3.3E-06</td><td>5.9E-06</td><td>BT20/MB468</td><td>CYTO</td></tr>
<tr><td>>hg38_21_+_8433172_8446622_RNA45SN1_45S_50ntFlanks@11570.11593.24|AGCCGACTTAGAACTGGTGCGGAC</td><td>>hg38_21_+_8433172_8446622_RNA45SN1_45S_50ntFlanks</td><td>11570</td><td>11593</td><td>24</td><td>AGCCGACTTAGAACTGGTGCGGAC</td><td>165.4</td><td>-5.5</td><td>1.2</td><td>-4.7</td><td>3.3E-06</td><td>5.9E-06</td><td>MB231/MB468</td><td>MP</td></tr>
<tr><td>>hg38_21_+_8433172_8446622_RNA45SN1_45S_50ntFlanks@9940.9978.39|CAGGACGGTGGCCATGGAAGTCGGAATCCGCTAAGGAGT</td><td>>hg38_21_+_8433172_8446622_RNA45SN1_45S_50ntFlanks</td><td>9940</td><td>9978</td><td>39</td><td>CAGGACGGTGGCCATGGAAGTCGGAATCCGCTAAGGAGT</td><td>326.7</td><td>8.4</td><td>1.7</td><td>4.9</td><td>1.1E-06</td><td>5.9E-06</td><td>BT20/MB468</td><td>MP</td></tr>
<tr><td>>hg38_21_+_8433172_8446622_RNA45SN1_45S_50ntFlanks@11561.11601.41|GCGCCTAGCAGCCGACTTAGAACTGGTGCGGACCAGGGGAA</td><td>>hg38_21_+_8433172_8446622_RNA45SN1_45S_50ntFlanks</td><td>11561</td><td>11601</td><td>41</td><td>GCGCCTAGCAGCCGACTTAGAACTGGTGCGGACCAGGGGAA</td><td>2682.1</td><td>-3.9</td><td>0.8</td><td>-4.7</td><td>2.9E-06</td><td>5.9E-06</td><td>BT20/MB468</td><td>MITO</td></tr>
<tr><td>>hg38_21_+_8433172_8446622_RNA45SN1_45S_50ntFlanks@10821.10865.45|GGAAGTCGGCAAGCCGGATCCGTAACTTCGGGATAAGGATTGGCT</td><td>>hg38_21_+_8433172_8446622_RNA45SN1_45S_50ntFlanks</td><td>10821</td><td>10865</td><td>45</td><td>GGAAGTCGGCAAGCCGGATCCGTAACTTCGGGATAAGGATTGGCT</td><td>1127.3</td><td>-6.8</td><td>1.5</td><td>-4.6</td><td>3.9E-06</td><td>5.9E-06</td><td>MB231/MB468</td><td>NUC</td></tr>
<tr><td>>trnaMT_GluTTC_MT_-_14674_14742@41.72.32,trnalookalike8_GluTTC_5_-_93905172_93905240@41.72.32|TGGTCGTGGTTGTAGTCCGTGCGAGAATACCA</td><td>>trnaMT_GluTTC_MT_-_14674_14742</td><td>41</td><td>72</td><td>32</td><td>TGGTCGTGGTTGTAGTCCGTGCGAGAATACCA</td><td>1029.2</td><td>4.6</td><td>1.0</td><td>4.7</td><td>3.3E-06</td><td>5.9E-06</td><td>BT20/MB468</td><td>CYTO</td></tr>
<tr><td>>hg38_21_+_8433172_8446622_RNA45SN1_45S_50ntFlanks@4738.4766.29|AAAGTCGGAGGTTCGAAGACGATCAGATA</td><td>>hg38_21_+_8433172_8446622_RNA45SN1_45S_50ntFlanks</td><td>4738</td><td>4766</td><td>29</td><td>AAAGTCGGAGGTTCGAAGACGATCAGATA</td><td>503.5</td><td>5.0</td><td>1.1</td><td>4.6</td><td>3.7E-06</td><td>5.9E-06</td><td>MB231/MB468</td><td>CYTO</td></tr>
<tr><td>>hg38_21_+_8433172_8446622_RNA45SN1_45S_50ntFlanks@12946.12988.43|TAGACGACCTGCTTCTGGGTCGGGGTTTCGTACGTAGCAGAGC</td><td>>hg38_21_+_8433172_8446622_RNA45SN1_45S_50ntFlanks</td><td>12946</td><td>12988</td><td>43</td><td>TAGACGACCTGCTTCTGGGTCGGGGTTTCGTACGTAGCAGAGC</td><td>594.3</td><td>-7.9</td><td>1.7</td><td>-4.6</td><td>4.2E-06</td><td>5.9E-06</td><td>BT20/MB468</td><td>NUC</td></tr>
<tr><td>>hg38_21_+_8433172_8446622_RNA45SN1_45S_50ntFlanks@12163.12182.20|ACTGGGGCGGTACACCTGTC</td><td>>hg38_21_+_8433172_8446622_RNA45SN1_45S_50ntFlanks</td><td>12163</td><td>12182</td><td>20</td><td>ACTGGGGCGGTACACCTGTC</td><td>605.4</td><td>-4.0</td><td>0.9</td><td>-4.6</td><td>3.7E-06</td><td>6.0E-06</td><td>MB231/MB468</td><td>CYTO</td></tr>
<tr><td>>hg38_21_+_8433172_8446622_RNA45SN1_45S_50ntFlanks@8169.8214.46|CCAAGTCCTTCTGATCGAGGCCCAGCCCGTGGACGGTGTGAGGCCG</td><td>>hg38_21_+_8433172_8446622_RNA45SN1_45S_50ntFlanks</td><td>8169</td><td>8214</td><td>46</td><td>CCAAGTCCTTCTGATCGAGGCCCAGCCCGTGGACGGTGTGAGGCCG</td><td>88.2</td><td>-10.3</td><td>2.2</td><td>-4.7</td><td>3.3E-06</td><td>6.0E-06</td><td>MB231/MB468</td><td>MP</td></tr>
<tr><td>>hg38_21_+_8433172_8446622_RNA45SN1_45S_50ntFlanks@12944.12982.39|CGTAGACGACCTGCTTCTGGGTCGGGGTTTCGTACGTAG</td><td>>hg38_21_+_8433172_8446622_RNA45SN1_45S_50ntFlanks</td><td>12944</td><td>12982</td><td>39</td><td>CGTAGACGACCTGCTTCTGGGTCGGGGTTTCGTACGTAG</td><td>127.0</td><td>-4.1</td><td>0.9</td><td>-4.6</td><td>3.8E-06</td><td>6.0E-06</td><td>MB231/MB468</td><td>CYTO</td></tr>
<tr><td>>hg38_21_+_8433172_8446622_RNA45SN1_45S_50ntFlanks@9792.9821.30|GGCGTGGAGCCGGGCGTGGAATGCGAGTGC</td><td>>hg38_21_+_8433172_8446622_RNA45SN1_45S_50ntFlanks</td><td>9792</td><td>9821</td><td>30</td><td>GGCGTGGAGCCGGGCGTGGAATGCGAGTGC</td><td>209.8</td><td>11.0</td><td>2.2</td><td>5.1</td><td>3.1E-07</td><td>6.0E-06</td><td>BT20/MB231</td><td>MP</td></tr>
<tr><td>>am_trna162_MetCAT_6_-_26758550_26758622@31.54.24,trna164_MetCAT_6_-_26735574_26735646@31.54.24,trna20_MetCAT_16_+_71460396_71460468@31.54.24,trna21_MetCAT_6_+_26701712_26701784@31.54.24,trna27_MetCAT_6_+_26766444_26766516@31.54.24,trna75_MetCAT_6_+_28912352_28912424@31.54.24,trna97_MetCAT_6_-_28921042_28921114@31.54.24|TCTCATAATCTGAAGGTCCTGAGT</td><td>>am_trna162_MetCAT_6_-_26758550_26758622</td><td>31</td><td>54</td><td>24</td><td>TCTCATAATCTGAAGGTCCTGAGT</td><td>528.4</td><td>-4.3</td><td>0.9</td><td>-4.6</td><td>3.8E-06</td><td>6.0E-06</td><td>MB231/MB468</td><td>CYTO</td></tr>
<tr><td>>hg38_21_+_8433172_8446622_RNA45SN1_45S_50ntFlanks@5425.5450.26|TGGATGGTTTAGTGAGGCCCTCGGAT</td><td>>hg38_21_+_8433172_8446622_RNA45SN1_45S_50ntFlanks</td><td>5425</td><td>5450</td><td>26</td><td>TGGATGGTTTAGTGAGGCCCTCGGAT</td><td>176.8</td><td>6.4</td><td>1.4</td><td>4.6</td><td>3.9E-06</td><td>6.0E-06</td><td>MB231/MB468</td><td>NUC</td></tr>
<tr><td>>hg38_21_+_8433172_8446622_RNA45SN1_45S_50ntFlanks@12655.12675.21|ATGACTGAACGCCTCTAAGTC</td><td>>hg38_21_+_8433172_8446622_RNA45SN1_45S_50ntFlanks</td><td>12655</td><td>12675</td><td>21</td><td>ATGACTGAACGCCTCTAAGTC</td><td>1071.8</td><td>-7.4</td><td>1.5</td><td>-4.9</td><td>1.2E-06</td><td>6.0E-06</td><td>BT20/MB468</td><td>MP</td></tr>
<tr><td>>hg38_21_+_8433172_8446622_RNA45SN1_45S_50ntFlanks@11124.11145.22|CCGGCGGCGGCCCGCCGCGGGG</td><td>>hg38_21_+_8433172_8446622_RNA45SN1_45S_50ntFlanks</td><td>11124</td><td>11145</td><td>22</td><td>CCGGCGGCGGCCCGCCGCGGGG</td><td>133.4</td><td>-10.1</td><td>2.1</td><td>-4.9</td><td>1.2E-06</td><td>6.0E-06</td><td>BT20/MB468</td><td>MP</td></tr>
<tr><td>>hg38_21_+_8433172_8446622_RNA45SN1_45S_50ntFlanks@10798.10826.29|CATGTTGGAACAATGTAGGTAAGGGAAGT</td><td>>hg38_21_+_8433172_8446622_RNA45SN1_45S_50ntFlanks</td><td>10798</td><td>10826</td><td>29</td><td>CATGTTGGAACAATGTAGGTAAGGGAAGT</td><td>284.1</td><td>-2.2</td><td>0.5</td><td>-4.7</td><td>2.9E-06</td><td>6.0E-06</td><td>BT20/MB468</td><td>MITO</td></tr>
<tr><td>>trna10_LysCTT_16_+_3241501_3241573@1.29.29,trna13_LysCTT_14_-_58706613_58706685@1.29.29,trna2_LysCTT_15_+_79152904_79152976@1.29.29|GCCCGGCTAGCTCAGTCGGTAGAGCATGG</td><td>>trna10_LysCTT_16_+_3241501_3241573</td><td>1</td><td>29</td><td>29</td><td>GCCCGGCTAGCTCAGTCGGTAGAGCATGG</td><td>965.0</td><td>-5.4</td><td>1.2</td><td>-4.6</td><td>3.3E-06</td><td>6.0E-06</td><td>BT20/MB468</td><td>CYTO</td></tr>
<tr><td>>hg38_21_+_8433172_8446622_RNA45SN1_45S_50ntFlanks@12420.12461.42|ATGTCGGCTCTTCCTATCATTGTGAAGCAGAATTCACCAAGC</td><td>>hg38_21_+_8433172_8446622_RNA45SN1_45S_50ntFlanks</td><td>12420</td><td>12461</td><td>42</td><td>ATGTCGGCTCTTCCTATCATTGTGAAGCAGAATTCACCAAGC</td><td>249.7</td><td>-4.5</td><td>0.9</td><td>-5.1</td><td>3.3E-07</td><td>6.0E-06</td><td>BT20/MB231</td><td>CYTO</td></tr>
<tr><td>>MI0000469|hsa-mir-125a&WithFlank&19|+|51693248|51693345@21.42.22(+1A)&[MIMAT0000443&hsa-miR-125a-5p&offsets|0|-2(+1A);m-63&19|+|51693268|51693289&offsets|0|0(+1A)]|TCCCTGAGACCCTTTAACCTGTA</td><td>>MI0000469|hsa-mir-125a&WithFlank&19|+|51693248|51693345</td><td>21</td><td>42</td><td>22(+1A)</td><td>TCCCTGAGACCCTTTAACCTGTA</td><td>1228.1</td><td>4.4</td><td>1.0</td><td>4.6</td><td>4.0E-06</td><td>6.0E-06</td><td>MB231/MB468</td><td>MITO</td></tr>
<tr><td>>MI0000279|hsa-mir-196a-2&WithFlank&12|+|53991732|53991853@31.52.22;MI0000238|hsa-mir-196a-1&WithFlank&17|-|48632484|48632565@13.34.22&[MIMAT0000226&hsa-miR-196a-5p&offsets|0|0;m-261&12|+|53991762|53991783&offsets|0|0];[MIMAT0000226_1&hsa-miR-196a-5p&offsets|0|0;m-268&17|-|48632532|48632553&offsets|0|0]|TAGGTAGTTTCATGTTGTTGGG</td><td>>MI0000279|hsa-mir-196a-2&WithFlank&12|+|53991732|53991853</td><td>31</td><td>52</td><td>22</td><td>TAGGTAGTTTCATGTTGTTGGG</td><td>3756.6</td><td>4.9</td><td>1.0</td><td>4.9</td><td>1.2E-06</td><td>6.0E-06</td><td>BT20/MB468</td><td>MP</td></tr>
<tr><td>>hg38_21_+_8433172_8446622_RNA45SN1_45S_50ntFlanks@6722.6740.19|ATTGCAGGACACATTGATC</td><td>>hg38_21_+_8433172_8446622_RNA45SN1_45S_50ntFlanks</td><td>6722</td><td>6740</td><td>19</td><td>ATTGCAGGACACATTGATC</td><td>531.2</td><td>-3.8</td><td>0.8</td><td>-4.9</td><td>1.2E-06</td><td>6.0E-06</td><td>BT20/MB468</td><td>MP</td></tr>
<tr><td>>hg38_21_+_8433172_8446622_RNA45SN1_45S_50ntFlanks@12008.12029.22|CCGCCGGTGAAATACCACTACT</td><td>>hg38_21_+_8433172_8446622_RNA45SN1_45S_50ntFlanks</td><td>12008</td><td>12029</td><td>22</td><td>CCGCCGGTGAAATACCACTACT</td><td>1566.5</td><td>-7.7</td><td>1.6</td><td>-4.9</td><td>1.2E-06</td><td>6.0E-06</td><td>BT20/MB468</td><td>MP</td></tr>
<tr><td>>hg38_21_+_8433172_8446622_RNA45SN1_45S_50ntFlanks@4160.4182.23|CCACATCCAAGGAAGGCAGCAGG</td><td>>hg38_21_+_8433172_8446622_RNA45SN1_45S_50ntFlanks</td><td>4160</td><td>4182</td><td>23</td><td>CCACATCCAAGGAAGGCAGCAGG</td><td>349.6</td><td>-7.7</td><td>1.6</td><td>-4.9</td><td>1.2E-06</td><td>6.0E-06</td><td>BT20/MB468</td><td>MP</td></tr>
</table>

<table>
<tr><td>>MI0000077|hsa-mir-21&WithFlank&17|+|59841260|59841343@14.34.21(+3C)&[MIMAT0000076&hsa-miR-21-5p&offsets|0|-1(+3C);m-1&17|+|59841273|59841294&offsets|0|-1(+3C)]|TAGCTTATCAGACTGATGTTGCCC</td><td>>MI0000077|hsa-mir-21&WithFlank&17|+|59841260|59841343</td><td>14</td><td>34</td><td>21(+3C)</td><td>TAGCTTATCAGACTGATGTTGCCC</td><td>263.4</td><td>-4.3</td><td>0.9</td><td>-5.1</td><td>3.7E-07</td><td>6.0E-06</td><td>BT20/MB231</td><td>MITO</td></tr>
<tr><td>>MI0000085|hsa-mir-27a&WithFlank&19|-|13836434|13836523@57.76.20(+1A)&[MIMAT0000084&hsa-miR-27a-3p&offsets|0|-1(+1A);m-43&19|-|13836447|13836467&offsets|0|-1(+1A)]|TTCACAGTGGCTAAGTTCCGA</td><td>>MI0000085|hsa-mir-27a&WithFlank&19|-|13836434|13836523</td><td>57</td><td>76</td><td>20(+1A)</td><td>TTCACAGTGGCTAAGTTCCGA</td><td>1163.5</td><td>4.5</td><td>1.0</td><td>4.6</td><td>3.8E-06</td><td>6.0E-06</td><td>MB231/MB468</td><td>CYTO</td></tr>
<tr><td>>hg38_21_+_8433172_8446622_RNA45SN1_45S_50ntFlanks@11569.11606.38|CAGCCGACTTAGAACTGGTGCGGACCAGGGGAATCCGA</td><td>>hg38_21_+_8433172_8446622_RNA45SN1_45S_50ntFlanks</td><td>11569</td><td>11606</td><td>38</td><td>CAGCCGACTTAGAACTGGTGCGGACCAGGGGAATCCGA</td><td>473.5</td><td>-6.8</td><td>1.5</td><td>-4.6</td><td>4.0E-06</td><td>6.0E-06</td><td>MB231/MB468</td><td>MITO</td></tr>
<tr><td>>hg38_21_+_8433172_8446622_RNA45SN1_45S_50ntFlanks@11783.11800.18|CATGAATGGATGAACGAG</td><td>>hg38_21_+_8433172_8446622_RNA45SN1_45S_50ntFlanks</td><td>11783</td><td>11800</td><td>18</td><td>CATGAATGGATGAACGAG</td><td>349.2</td><td>-3.6</td><td>0.8</td><td>-4.6</td><td>3.4E-06</td><td>6.0E-06</td><td>BT20/MB468</td><td>CYTO</td></tr>
<tr><td>>MI0033423|hsa-mir-10399&WithFlank&7|-|139044094|139044163@45.64.20(+1U)&[MIMAT0041630&hsa-miR-10399-3p&offsets|+1|0(+1U);TJU_CMC_MD2.ID03061.3p-miR&offsets|+5|+3(+1U);m-2657&7|-|139044103|139044124&offsets|+5|+3(+1U)]|TCTCGGACAAGCTGTAGGTCT</td><td>>MI0033423|hsa-mir-10399&WithFlank&7|-|139044094|139044163</td><td>45</td><td>64</td><td>20(+1U)</td><td>TCTCGGACAAGCTGTAGGTCT</td><td>36.6</td><td>8.4</td><td>1.8</td><td>4.6</td><td>3.3E-06</td><td>6.0E-06</td><td>MB231/MB468</td><td>MP</td></tr>
<tr><td>>hg38_21_+_8433172_8446622_RNA45SN1_45S_50ntFlanks@6650.6675.26|ACGACTCTTAGCGGTGGATCACTCGG</td><td>>hg38_21_+_8433172_8446622_RNA45SN1_45S_50ntFlanks</td><td>6650</td><td>6675</td><td>26</td><td>ACGACTCTTAGCGGTGGATCACTCGG</td><td>2673.8</td><td>3.2</td><td>0.7</td><td>4.7</td><td>2.9E-06</td><td>6.0E-06</td><td>BT20/MB468</td><td>MITO</td></tr>
<tr><td>>hg38_21_+_8433172_8446622_RNA45SN1_45S_50ntFlanks@3776.3805.30|CCGGTACAGTGAAACTGCGAATGGCTCATT</td><td>>hg38_21_+_8433172_8446622_RNA45SN1_45S_50ntFlanks</td><td>3776</td><td>3805</td><td>30</td><td>CCGGTACAGTGAAACTGCGAATGGCTCATT</td><td>618.7</td><td>-6.4</td><td>1.4</td><td>-4.6</td><td>3.3E-06</td><td>6.0E-06</td><td>MB231/MB468</td><td>MP</td></tr>
<tr><td>>hg38_21_+_8433172_8446622_RNA45SN1_45S_50ntFlanks@6644.6664.21|CCTCGTACGACTCTTAGCGGT</td><td>>hg38_21_+_8433172_8446622_RNA45SN1_45S_50ntFlanks</td><td>6644</td><td>6664</td><td>21</td><td>CCTCGTACGACTCTTAGCGGT</td><td>154.5</td><td>4.2</td><td>0.9</td><td>4.9</td><td>1.2E-06</td><td>6.0E-06</td><td>BT20/MB468</td><td>MP</td></tr>
<tr><td>>hg38_21_+_8433172_8446622_RNA45SN1_45S_50ntFlanks@9590.9610.21|TGGGTATAGGGGCGAAAGACT</td><td>>hg38_21_+_8433172_8446622_RNA45SN1_45S_50ntFlanks</td><td>9590</td><td>9610</td><td>21</td><td>TGGGTATAGGGGCGAAAGACT</td><td>49.1</td><td>9.4</td><td>1.9</td><td>4.9</td><td>1.2E-06</td><td>6.0E-06</td><td>BT20/MB468</td><td>MP</td></tr>
<tr><td>>hg38_21_+_8433172_8446622_RNA45SN1_45S_50ntFlanks@6660.6708.49|GCGGTGGATCACTCGGCTCGTGCGTCGATGAAGAACGCAGCTAGCTGCG</td><td>>hg38_21_+_8433172_8446622_RNA45SN1_45S_50ntFlanks</td><td>6660</td><td>6708</td><td>49</td><td>GCGGTGGATCACTCGGCTCGTGCGTCGATGAAGAACGCAGCTAGCTGCG</td><td>775.4</td><td>4.5</td><td>1.0</td><td>4.7</td><td>2.9E-06</td><td>6.0E-06</td><td>BT20/MB468</td><td>MITO</td></tr>
<tr><td>>hg38_21_+_8433172_8446622_RNA45SN1_45S_50ntFlanks@9579.9597.19|GTCGTCCGACCTGGGTATA</td><td>>hg38_21_+_8433172_8446622_RNA45SN1_45S_50ntFlanks</td><td>9579</td><td>9597</td><td>19</td><td>GTCGTCCGACCTGGGTATA</td><td>53.4</td><td>-8.8</td><td>1.8</td><td>-4.9</td><td>1.2E-06</td><td>6.0E-06</td><td>BT20/MB468</td><td>MP</td></tr>
<tr><td>>MI0000433|hsa-let-7g&WithFlank&3|-|52268272|52268367@11.33.23&[MIMAT0000414&hsa-let-7g-5p&offsets|0|+1;m-58&3|-|52268336|52268357&offsets|0|+1]|TGAGGTAGTAGTTTGTACAGTTT</td><td>>MI0000433|hsa-let-7g&WithFlank&3|-|52268272|52268367</td><td>11</td><td>33</td><td>23</td><td>TGAGGTAGTAGTTTGTACAGTTT</td><td>1094.5</td><td>5.0</td><td>1.0</td><td>4.9</td><td>1.2E-06</td><td>6.0E-06</td><td>BT20/MB468</td><td>MP</td></tr>
<tr><td>>hg38_21_+_8433172_8446622_RNA45SN1_45S_50ntFlanks@9534.9552.19|AGGAAACTCTGGTGGAGGT</td><td>>hg38_21_+_8433172_8446622_RNA45SN1_45S_50ntFlanks</td><td>9534</td><td>9552</td><td>19</td><td>AGGAAACTCTGGTGGAGGT</td><td>2091.1</td><td>-4.2</td><td>0.9</td><td>-4.9</td><td>1.2E-06</td><td>6.0E-06</td><td>BT20/MB468</td><td>MP</td></tr>
<tr><td>>MI0000081|hsa-mir-24-2&WithFlank&19|-|13836281|13836365@56.76.21(+1A);MI0000080|hsa-mir-24-1&WithFlank&9|+|95086015|95086094@50.70.21(+1A)&[MIMAT0000080&hsa-miR-24-3p&offsets|0|-1(+1A);m-36&19|-|13836288|13836310&offsets|0|-2(+1A)];[MIMAT0000080_1&hsa-miR-24-3p&offsets|0|-1(+1A);m-35&9|+|95086064|95086086&offsets|0|-2(+1A)]|TGGCTCAGTTCAGCAGGAACAA</td><td>>MI0000081|hsa-mir-24-2&WithFlank&19|-|13836281|13836365</td><td>56</td><td>76</td><td>21(+1A)</td><td>TGGCTCAGTTCAGCAGGAACAA</td><td>5304.9</td><td>3.8</td><td>0.8</td><td>4.6</td><td>3.3E-06</td><td>6.0E-06</td><td>MB231/MB468</td><td>MP</td></tr>
<tr><td>>hg38_21_+_8433172_8446622_RNA45SN1_45S_50ntFlanks@6651.6670.20|CGACTCTTAGCGGTGGATCA</td><td>>hg38_21_+_8433172_8446622_RNA45SN1_45S_50ntFlanks</td><td>6651</td><td>6670</td><td>20</td><td>CGACTCTTAGCGGTGGATCA</td><td>1500.9</td><td>-4.0</td><td>0.8</td><td>-4.9</td><td>1.2E-06</td><td>6.0E-06</td><td>BT20/MB468</td><td>MP</td></tr>
<tr><td>>hg38_21_+_8433172_8446622_RNA45SN1_45S_50ntFlanks@10402.10422.21|AACATGGGTCAGTCGGTCCTG</td><td>>hg38_21_+_8433172_8446622_RNA45SN1_45S_50ntFlanks</td><td>10402</td><td>10422</td><td>21</td><td>AACATGGGTCAGTCGGTCCTG</td><td>946.5</td><td>-4.2</td><td>0.9</td><td>-4.9</td><td>1.2E-06</td><td>6.0E-06</td><td>BT20/MB468</td><td>MP</td></tr>
<tr><td>>hg38_21_+_8433172_8446622_RNA45SN1_45S_50ntFlanks@6719.6752.34|TGAATTGCAGGACACATTGATCATCGACACTTCG</td><td>>hg38_21_+_8433172_8446622_RNA45SN1_45S_50ntFlanks</td><td>6719</td><td>6752</td><td>34</td><td>TGAATTGCAGGACACATTGATCATCGACACTTCG</td><td>390.0</td><td>-7.5</td><td>1.5</td><td>-4.9</td><td>1.2E-06</td><td>6.0E-06</td><td>BT20/MB468</td><td>MP</td></tr>
<tr><td>>hg38_21_+_8433172_8446622_RNA45SN1_45S_50ntFlanks@4102.4121.20|ACCACGGGTGACGGGGAATC</td><td>>hg38_21_+_8433172_8446622_RNA45SN1_45S_50ntFlanks</td><td>4102</td><td>4121</td><td>20</td><td>ACCACGGGTGACGGGGAATC</td><td>1334.1</td><td>-4.2</td><td>0.9</td><td>-4.6</td><td>3.4E-06</td><td>6.0E-06</td><td>BT20/MB468</td><td>CYTO</td></tr>
<tr><td>>am_trnaMT_ProTGG_MT_-_15956_16023@52.70.19|AAAGACTTTTTCTCTGACC</td><td>>am_trnaMT_ProTGG_MT_-_15956_16023</td><td>52</td><td>70</td><td>19</td><td>AAAGACTTTTTCTCTGACC</td><td>856.6</td><td>4.4</td><td>1.0</td><td>4.6</td><td>4.0E-06</td><td>6.0E-06</td><td>MB231/MB468</td><td>MITO</td></tr>
<tr><td>>hg38_21_+_8433172_8446622_RNA45SN1_45S_50ntFlanks@11579.11606.28|AGAACTGGTGCGGACCAGGGGAATCCGA</td><td>>hg38_21_+_8433172_8446622_RNA45SN1_45S_50ntFlanks</td><td>11579</td><td>11606</td><td>28</td><td>AGAACTGGTGCGGACCAGGGGAATCCGA</td><td>275.9</td><td>-6.4</td><td>1.4</td><td>-4.6</td><td>4.0E-06</td><td>6.0E-06</td><td>MB231/MB468</td><td>MITO</td></tr>
<tr><td>>hg38_21_+_8433172_8446622_RNA45SN1_45S_50ntFlanks@12285.12306.22|CAGACCGTGAAAGCGGGGCCTC</td><td>>hg38_21_+_8433172_8446622_RNA45SN1_45S_50ntFlanks</td><td>12285</td><td>12306</td><td>22</td><td>CAGACCGTGAAAGCGGGGCCTC</td><td>69.0</td><td>-9.2</td><td>1.9</td><td>-4.9</td><td>1.2E-06</td><td>6.0E-06</td><td>BT20/MB468</td><td>MP</td></tr>
<tr><td>>hg38_21_+_8433172_8446622_RNA45SN1_45S_50ntFlanks@7984.8011.28|AGATCAGACGTGGCGACCCGCTGAATTT</td><td>>hg38_21_+_8433172_8446622_RNA45SN1_45S_50ntFlanks</td><td>7984</td><td>8011</td><td>28</td><td>AGATCAGACGTGGCGACCCGCTGAATTT</td><td>203.5</td><td>-6.9</td><td>1.5</td><td>-4.6</td><td>3.4E-06</td><td>6.0E-06</td><td>MB231/MB468</td><td>MP</td></tr>
<tr><td>>MI0000471|hsa-mir-126&WithFlank&9|+|136670596|136670692@58.78.21&[MIMAT0000445&hsa-miR-126-3p&offsets|0|-1;m-40&9|+|136670653|136670674&offsets|0|-1]|TCGTACCGTGAGTAATAATGC</td><td>>MI0000471|hsa-mir-126&WithFlank&9|+|136670596|136670692</td><td>58</td><td>78</td><td>21</td><td>TCGTACCGTGAGTAATAATGC</td><td>1583.6</td><td>7.0</td><td>1.4</td><td>4.9</td><td>1.2E-06</td><td>6.0E-06</td><td>BT20/MB468</td><td>MP</td></tr>
<tr><td>>hg38_21_+_8433172_8446622_RNA45SN1_45S_50ntFlanks@10351.10380.30|CGAGAACTTTGAAGGCCGAAGTGGAGAAGG</td><td>>hg38_21_+_8433172_8446622_RNA45SN1_45S_50ntFlanks</td><td>10351</td><td>10380</td><td>30</td><td>CGAGAACTTTGAAGGCCGAAGTGGAGAAGG</td><td>644.9</td><td>-5.3</td><td>1.1</td><td>-4.6</td><td>4.1E-06</td><td>6.1E-06</td><td>MB231/MB468</td><td>MITO</td></tr>
<tr><td>>hg38_21_+_8433172_8446622_RNA45SN1_45S_50ntFlanks@12007.12042.36|GCCGCCGGTGAAATACCACTACTCTGATCGTTTTTT</td><td>>hg38_21_+_8433172_8446622_RNA45SN1_45S_50ntFlanks</td><td>12007</td><td>12042</td><td>36</td><td>GCCGCCGGTGAAATACCACTACTCTGATCGTTTTTT</td><td>176.4</td><td>-4.8</td><td>1.0</td><td>-4.6</td><td>3.8E-06</td><td>6.1E-06</td><td>MB231/MB468</td><td>CYTO</td></tr>
<tr><td>>hg38_21_+_8433172_8446622_RNA45SN1_45S_50ntFlanks@9569.9592.24|GTGCAAATCGGTCGTCCGACCTGG</td><td>>hg38_21_+_8433172_8446622_RNA45SN1_45S_50ntFlanks</td><td>9569</td><td>9592</td><td>24</td><td>GTGCAAATCGGTCGTCCGACCTGG</td><td>271.5</td><td>-5.9</td><td>1.3</td><td>-4.6</td><td>3.4E-06</td><td>6.1E-06</td><td>BT20/MB468</td><td>CYTO</td></tr>
<tr><td>>hg38_21_+_8433172_8446622_RNA45SN1_45S_50ntFlanks@3910.3937.28|GGGATGCGTGCATTTATCAGATCAAAAC</td><td>>hg38_21_+_8433172_8446622_RNA45SN1_45S_50ntFlanks</td><td>3910</td><td>3937</td><td>28</td><td>GGGATGCGTGCATTTATCAGATCAAAAC</td><td>89.4</td><td>-9.6</td><td>2.0</td><td>-4.9</td><td>1.2E-06</td><td>6.1E-06</td><td>BT20/MB468</td><td>MP</td></tr>
<tr><td>>hg38_21_+_8433172_8446622_RNA45SN1_45S_50ntFlanks@8357.8389.33|AAAAGAACTTTGAAGAGAGAGTTCAAGAGGGCG</td><td>>hg38_21_+_8433172_8446622_RNA45SN1_45S_50ntFlanks</td><td>8357</td><td>8389</td><td>33</td><td>AAAAGAACTTTGAAGAGAGAGTTCAAGAGGGCG</td><td>461.6</td><td>-7.5</td><td>1.6</td><td>-4.6</td><td>4.3E-06</td><td>6.1E-06</td><td>BT20/MB468</td><td>NUC</td></tr>
<tr><td>>hg38_21_+_8433172_8446622_RNA45SN1_45S_50ntFlanks@10351.10370.20|CGAGAACTTTGAAGGCCGAA</td><td>>hg38_21_+_8433172_8446622_RNA45SN1_45S_50ntFlanks</td><td>10351</td><td>10370</td><td>20</td><td>CGAGAACTTTGAAGGCCGAA</td><td>279.2</td><td>-1.9</td><td>0.4</td><td>-4.6</td><td>3.4E-06</td><td>6.1E-06</td><td>BT20/MB468</td><td>CYTO</td></tr>
<tr><td>>hg38_21_+_8433172_8446622_RNA45SN1_45S_50ntFlanks@3705.3744.40|TACCTGGTTGATCCTGCCAGTAGCATATGCTTGTCTCAAA</td><td>>hg38_21_+_8433172_8446622_RNA45SN1_45S_50ntFlanks</td><td>3705</td><td>3744</td><td>40</td><td>TACCTGGTTGATCCTGCCAGTAGCATATGCTTGTCTCAAA</td><td>444.4</td><td>2.4</td><td>0.5</td><td>4.6</td><td>4.3E-06</td><td>6.1E-06</td><td>BT20/MB468</td><td>NUC</td></tr>
<tr><td>>hg38_21_+_8433172_8446622_RNA45SN1_45S_50ntFlanks@9739.9764.26|AAACGATCTCAACCTATTCTCAAACT</td><td>>hg38_21_+_8433172_8446622_RNA45SN1_45S_50ntFlanks</td><td>9739</td><td>9764</td><td>26</td><td>AAACGATCTCAACCTATTCTCAAACT</td><td>533.4</td><td>-6.0</td><td>1.3</td><td>-4.6</td><td>4.3E-06</td><td>6.1E-06</td><td>BT20/MB468</td><td>NUC</td></tr>
<tr><td>>hg38_21_+_8433172_8446622_RNA45SN1_45S_50ntFlanks@8357.8377.21|AAAAGAACTTTGAAGAGAGAG</td><td>>hg38_21_+_8433172_8446622_RNA45SN1_45S_50ntFlanks</td><td>8357</td><td>8377</td><td>21</td><td>AAAAGAACTTTGAAGAGAGAG</td><td>1702.3</td><td>-5.8</td><td>1.2</td><td>-4.6</td><td>3.4E-06</td><td>6.1E-06</td><td>BT20/MB468</td><td>CYTO</td></tr>
<tr><td>>hg38_21_+_8433172_8446622_RNA45SN1_45S_50ntFlanks@11208.11228.21|GCGGCGACTCTGGACGCGAGC</td><td>>hg38_21_+_8433172_8446622_RNA45SN1_45S_50ntFlanks</td><td>11208</td><td>11228</td><td>21</td><td>GCGGCGACTCTGGACGCGAGC</td><td>762.8</td><td>-4.0</td><td>0.9</td><td>-4.6</td><td>3.8E-06</td><td>6.1E-06</td><td>MB231/MB468</td><td>CYTO</td></tr>
<tr><td>>hg38_21_+_8433172_8446622_RNA45SN1_45S_50ntFlanks@3816.3852.37|TGGTTCCTTTGGTCGCTCGCTCCTCTCCTACTTGGAT</td><td>>hg38_21_+_8433172_8446622_RNA45SN1_45S_50ntFlanks</td><td>3816</td><td>3852</td><td>37</td><td>TGGTTCCTTTGGTCGCTCGCTCCTCTCCTACTTGGAT</td><td>220.3</td><td>-5.2</td><td>1.1</td><td>-4.6</td><td>4.0E-06</td><td>6.1E-06</td><td>MB231/MB468</td><td>NUC</td></tr>
<tr><td>>MI0000273|hsa-mir-183&WithFlank&7|-|129774899|129775020@33.55.23&[MIMAT0000261&hsa-miR-183-5p&offsets|0|+1;m-41&7|-|129774966|129774987&offsets|-1|0]|TATGGCACTGGTAGAATTCACTG</td><td>>MI0000273|hsa-mir-183&WithFlank&7|-|129774899|129775020</td><td>33</td><td>55</td><td>23</td><td>TATGGCACTGGTAGAATTCACTG</td><td>3777.0</td><td>2.6</td><td>0.5</td><td>5.0</td><td>4.5E-07</td><td>6.1E-06</td><td>BT20/MB231</td><td>NUC</td></tr>
<tr><td>>hg38_21_+_8433172_8446622_RNA45SN1_45S_50ntFlanks@11208.11226.19|GCGGCGACTCTGGACGCGA</td><td>>hg38_21_+_8433172_8446622_RNA45SN1_45S_50ntFlanks</td><td>11208</td><td>11226</td><td>19</td><td>GCGGCGACTCTGGACGCGA</td><td>182.2</td><td>-5.6</td><td>1.2</td><td>-4.6</td><td>3.4E-06</td><td>6.1E-06</td><td>BT20/MB468</td><td>CYTO</td></tr>
<tr><td>>hg38_21_+_8433172_8446622_RNA45SN1_45S_50ntFlanks@8353.8392.40|GTTGAAAAGAACTTTGAAGAGAGAGTTCAAGAGGGCGTGA</td><td>>hg38_21_+_8433172_8446622_RNA45SN1_45S_50ntFlanks</td><td>8353</td><td>8392</td><td>40</td><td>GTTGAAAAGAACTTTGAAGAGAGAGTTCAAGAGGGCGTGA</td><td>44.9</td><td>9.2</td><td>1.9</td><td>4.9</td><td>1.2E-06</td><td>6.1E-06</td><td>BT20/MB468</td><td>MP</td></tr>
<tr><td>>hg38_21_+_8433172_8446622_RNA45SN1_45S_50ntFlanks@12932.12972.41|CGCTAAACCATTCGTAGACGACCTGCTTCTGGGTCGGGGTT</td><td>>hg38_21_+_8433172_8446622_RNA45SN1_45S_50ntFlanks</td><td>12932</td><td>12972</td><td>41</td><td>CGCTAAACCATTCGTAGACGACCTGCTTCTGGGTCGGGGTT</td><td>460.0</td><td>-3.5</td><td>0.7</td><td>-4.7</td><td>3.0E-06</td><td>6.1E-06</td><td>BT20/MB468</td><td>MITO</td></tr>
<tr><td>>hg38_MT_+_598_1651_RNR1_12S_50ntFlanks@569.612.44|AAAGGACCTGGCGGTGCTTCATATCCCTCTAGAGGAGCCTGTTC</td><td>>hg38_MT_+_598_1651_RNR1_12S_50ntFlanks</td><td>569</td><td>612</td><td>44</td><td>AAAGGACCTGGCGGTGCTTCATATCCCTCTAGAGGAGCCTGTTC</td><td>93.3</td><td>-9.9</td><td>1.9</td><td>-5.1</td><td>3.2E-07</td><td>6.1E-06</td><td>BT20/MB231</td><td>MP</td></tr>
<tr><td>>MI0000077|hsa-mir-21&WithFlank&17|+|59841260|59841343@14.32.19&[MIMAT0000076&hsa-miR-21-5p&offsets|0|-3;m-1&17|+|59841273|59841294&offsets|0|-3]|TAGCTTATCAGACTGATGT</td><td>>MI0000077|hsa-mir-21&WithFlank&17|+|59841260|59841343</td><td>14</td><td>32</td><td>19</td><td>TAGCTTATCAGACTGATGT</td><td>760.6</td><td>6.1</td><td>1.3</td><td>4.6</td><td>4.1E-06</td><td>6.1E-06</td><td>MB231/MB468</td><td>MITO</td></tr>
<tr><td>>hg38_21_+_8433172_8446622_RNA45SN1_45S_50ntFlanks@7973.8005.33|GACGCGACCTCAGATCAGACGTGGCGACCCGCT</td><td>>hg38_21_+_8433172_8446622_RNA45SN1_45S_50ntFlanks</td><td>7973</td><td>8005</td><td>33</td><td>GACGCGACCTCAGATCAGACGTGGCGACCCGCT</td><td>1044.8</td><td>4.4</td><td>0.9</td><td>4.9</td><td>1.2E-06</td><td>6.1E-06</td><td>BT20/MB468</td><td>MP</td></tr>
<tr><td>>hg38_MT_+_1621_3279_RNR2_16S_50ntFlanks@496.519.24|CTTTGGACACTAGGAAAAAACCTT</td><td>>hg38_MT_+_1621_3279_RNR2_16S_50ntFlanks</td><td>496</td><td>519</td><td>24</td><td>CTTTGGACACTAGGAAAAAACCTT</td><td>53.2</td><td>9.5</td><td>2.0</td><td>4.9</td><td>1.2E-06</td><td>6.1E-06</td><td>BT20/MB468</td><td>MP</td></tr>
<tr><td>>hg38_21_+_8433172_8446622_RNA45SN1_45S_50ntFlanks@9339.9379.41|CGCACGAAAGCCGCCGTGGCGCAATGAAGGTGAAGGCCGGC</td><td>>hg38_21_+_8433172_8446622_RNA45SN1_45S_50ntFlanks</td><td>9339</td><td>9379</td><td>41</td><td>CGCACGAAAGCCGCCGTGGCGCAATGAAGGTGAAGGCCGGC</td><td>58.9</td><td>-9.0</td><td>1.8</td><td>-4.9</td><td>1.2E-06</td><td>6.1E-06</td><td>BT20/MB468</td><td>MP</td></tr>
<tr><td>>hg38_21_+_8433172_8446622_RNA45SN1_45S_50ntFlanks@7975.8004.30|CGCGACCTCAGATCAGACGTGGCGACCCGC</td><td>>hg38_21_+_8433172_8446622_RNA45SN1_45S_50ntFlanks</td><td>7975</td><td>8004</td><td>30</td><td>CGCGACCTCAGATCAGACGTGGCGACCCGC</td><td>29789.9</td><td>3.4</td><td>0.7</td><td>5.0</td><td>4.6E-07</td><td>6.1E-06</td><td>BT20/MB231</td><td>NUC</td></tr>
<tr><td>>trna10_AspGTC_12_-_125424193_125424264@38.69.32,trna12_AspGTC_12_-_125411891_125411962@38.69.32,trna144_AspGTC_6_-_27551236_27551307@38.69.32,trna38_AspGTC_17_-_8125556_8125627@38.69.32,trna45_AspGTC_6_+_27447453_27447524@38.69.32,trna48_AspGTC_6_+_27471523_27471594@38.69.32,trna4_AspGTC_12_+_96429799_96429870@38.69.32,trna69_AspGTC_1_-_161440205_161440276@38.69.32,trna72_AspGTC_1_-_161432824_161432895@38.69.32,trna75_AspGTC_1_-_161425414_161425485@38.69.32,trna78_AspGTC_1_-_161418033_161418104@38.69.32,trna81_AspGTC_1_-_161410615_161410686@38.69.32|CGCGGGAGACCGGGGTTCGATTCCCCGACGGG</td><td>>trna10_AspGTC_12_-_125424193_125424264</td><td>38</td><td>69</td><td>32</td><td>CGCGGGAGACCGGGGTTCGATTCCCCGACGGG</td><td>1171.5</td><td>-3.2</td><td>0.7</td><td>-4.6</td><td>3.4E-06</td><td>6.2E-06</td><td>MB231/MB468</td><td>MP</td></tr>
<tr><td>>hg38_21_+_8433172_8446622_RNA45SN1_45S_50ntFlanks@11199.11227.29|CCGGCGGCGGCGGCGACTCTGGACGCGAG</td><td>>hg38_21_+_8433172_8446622_RNA45SN1_45S_50ntFlanks</td><td>11199</td><td>11227</td><td>29</td><td>CCGGCGGCGGCGGCGACTCTGGACGCGAG</td><td>905.6</td><td>-7.0</td><td>1.5</td><td>-4.6</td><td>4.0E-06</td><td>6.2E-06</td><td>MB231/MB468</td><td>NUC</td></tr>
<tr><td>>hg38_21_+_8433172_8446622_RNA45SN1_45S_50ntFlanks@9604.9621.18|AAAGACTAATCGAACCAT</td><td>>hg38_21_+_8433172_8446622_RNA45SN1_45S_50ntFlanks</td><td>9604</td><td>9621</td><td>18</td><td>AAAGACTAATCGAACCAT</td><td>609.9</td><td>-7.7</td><td>1.7</td><td>-4.6</td><td>4.3E-06</td><td>6.2E-06</td><td>BT20/MB468</td><td>NUC</td></tr>
<tr><td>>hg38_21_+_8433172_8446622_RNA45SN1_45S_50ntFlanks@10319.10336.18|GCAGATCTTGGTGGTAGT</td><td>>hg38_21_+_8433172_8446622_RNA45SN1_45S_50ntFlanks</td><td>10319</td><td>10336</td><td>18</td><td>GCAGATCTTGGTGGTAGT</td><td>176.3</td><td>10.8</td><td>2.1</td><td>5.1</td><td>3.2E-07</td><td>6.2E-06</td><td>BT20/MB231</td><td>MP</td></tr>
</table>

<table>
<tr><td>>hg38_21_+_8433172_8446622_RNA45SN1_45S_50ntFlanks@11483.11502.20|CGGAGGGGCCGCGGGCCGGT</td><td>>hg38_21_+_8433172_8446622_RNA45SN1_45S_50ntFlanks</td><td>11483</td><td>11502</td><td>20</td><td>CGGAGGGGCCGCGGGCCGGT</td><td>113.3</td><td>-3.9</td><td>0.8</td><td>-4.6</td><td>3.4E-06</td><td>6.2E-06</td><td>BT20/MB468</td><td>CYTO</td></tr>
<tr><td>>hg38_21_+_8433172_8446622_RNA45SN1_45S_50ntFlanks@7975.7992.18|CGCGACCTCAGATCAGAC</td><td>>hg38_21_+_8433172_8446622_RNA45SN1_45S_50ntFlanks</td><td>7975</td><td>7992</td><td>18</td><td>CGCGACCTCAGATCAGAC</td><td>20077.3</td><td>4.6</td><td>0.9</td><td>4.9</td><td>1.2E-06</td><td>6.2E-06</td><td>BT20/MB468</td><td>MP</td></tr>
<tr><td>>hg38_21_+_8433172_8446622_RNA45SN1_45S_50ntFlanks@4027.4046.20|CCCCGTGGCGGCGACGACCC</td><td>>hg38_21_+_8433172_8446622_RNA45SN1_45S_50ntFlanks</td><td>4027</td><td>4046</td><td>20</td><td>CCCCGTGGCGGCGACGACCC</td><td>51.2</td><td>-8.8</td><td>1.8</td><td>-4.9</td><td>1.2E-06</td><td>6.2E-06</td><td>BT20/MB468</td><td>MP</td></tr>
<tr><td>>hg38_21_+_8433172_8446622_RNA45SN1_45S_50ntFlanks@6734.6770.37|ATTGATCATCGACACTTCGAACGCACTTGCGGCCCCG</td><td>>hg38_21_+_8433172_8446622_RNA45SN1_45S_50ntFlanks</td><td>6734</td><td>6770</td><td>37</td><td>ATTGATCATCGACACTTCGAACGCACTTGCGGCCCCG</td><td>388.0</td><td>-2.4</td><td>0.5</td><td>-4.6</td><td>3.5E-06</td><td>6.2E-06</td><td>BT20/MB468</td><td>CYTO</td></tr>
<tr><td>>hg38_21_+_8433172_8446622_RNA45SN1_45S_50ntFlanks@10768.10793.26|CAGCAGGTCTCCAAGGTGAACAGCCT</td><td>>hg38_21_+_8433172_8446622_RNA45SN1_45S_50ntFlanks</td><td>10768</td><td>10793</td><td>26</td><td>CAGCAGGTCTCCAAGGTGAACAGCCT</td><td>51.8</td><td>-8.8</td><td>1.8</td><td>-4.9</td><td>1.2E-06</td><td>6.2E-06</td><td>BT20/MB468</td><td>MP</td></tr>
<tr><td>>MI0000433|hsa-let-7g&WithFlank&3|-|52268272|52268367@11.32.22&[MIMAT0000414&hsa-let-7g-5p&offsets|0|0;m-58&3|-|52268336|52268357&offsets|0|0]|TGAGGTAGTAGTTTGTACAGTT</td><td>>MI0000433|hsa-let-7g&WithFlank&3|-|52268272|52268367</td><td>11</td><td>32</td><td>22</td><td>TGAGGTAGTAGTTTGTACAGTT</td><td>41019.5</td><td>4.3</td><td>0.9</td><td>4.6</td><td>3.4E-06</td><td>6.2E-06</td><td>MB231/MB468</td><td>MP</td></tr>
<tr><td>>hg38_21_+_8433172_8446622_RNA45SN1_45S_50ntFlanks@10186.10207.22|TCCTCCCCTCCTCCCGCCCACG</td><td>>hg38_21_+_8433172_8446622_RNA45SN1_45S_50ntFlanks</td><td>10186</td><td>10207</td><td>22</td><td>TCCTCCCCTCCTCCCGCCCACG</td><td>162.6</td><td>-11.2</td><td>2.4</td><td>-4.6</td><td>3.4E-06</td><td>6.2E-06</td><td>MB231/MB468</td><td>MP</td></tr>
<tr><td>>hg38_21_+_8433172_8446622_RNA45SN1_45S_50ntFlanks@11882.11904.23|CTGTTGAGCTTGACTCTAGTCTG</td><td>>hg38_21_+_8433172_8446622_RNA45SN1_45S_50ntFlanks</td><td>11882</td><td>11904</td><td>23</td><td>CTGTTGAGCTTGACTCTAGTCTG</td><td>500.8</td><td>-5.0</td><td>1.1</td><td>-4.6</td><td>4.1E-06</td><td>6.2E-06</td><td>MB231/MB468</td><td>NUC</td></tr>
<tr><td>>hg38_21_+_8433172_8446622_RNA45SN1_45S_50ntFlanks@3822.3853.32|CTTTGGTCGCTCGCTCCTCTCCTACTTGGATA</td><td>>hg38_21_+_8433172_8446622_RNA45SN1_45S_50ntFlanks</td><td>3822</td><td>3853</td><td>32</td><td>CTTTGGTCGCTCGCTCCTCTCCTACTTGGATA</td><td>218.4</td><td>-6.4</td><td>1.4</td><td>-4.6</td><td>4.1E-06</td><td>6.2E-06</td><td>MB231/MB468</td><td>NUC</td></tr>
<tr><td>>trna152_ValCAC_6_-_27248049_27248121@1.32.32|GCTTCTGTAGTGTAGTGGTTATCACGTTCGCC</td><td>>trna152_ValCAC_6_-_27248049_27248121</td><td>1</td><td>32</td><td>32</td><td>GCTTCTGTAGTGTAGTGGTTATCACGTTCGCC</td><td>769.7</td><td>5.9</td><td>1.2</td><td>4.9</td><td>1.2E-06</td><td>6.2E-06</td><td>BT20/MB468</td><td>MP</td></tr>
<tr><td>>hg38_21_+_8433172_8446622_RNA45SN1_45S_50ntFlanks@11906.11928.23|CACGGTGAAGAGACATGAGAGGT</td><td>>hg38_21_+_8433172_8446622_RNA45SN1_45S_50ntFlanks</td><td>11906</td><td>11928</td><td>23</td><td>CACGGTGAAGAGACATGAGAGGT</td><td>185.6</td><td>10.9</td><td>2.1</td><td>5.1</td><td>3.2E-07</td><td>6.2E-06</td><td>BT20/MB231</td><td>MP</td></tr>
<tr><td>>hg38_21_+_8433172_8446622_RNA45SN1_45S_50ntFlanks@11771.11794.24|ATTAGTGACGCGCATGAATGGATG</td><td>>hg38_21_+_8433172_8446622_RNA45SN1_45S_50ntFlanks</td><td>11771</td><td>11794</td><td>24</td><td>ATTAGTGACGCGCATGAATGGATG</td><td>495.8</td><td>-3.7</td><td>0.8</td><td>-4.6</td><td>3.5E-06</td><td>6.2E-06</td><td>MB231/MB468</td><td>MP</td></tr>
<tr><td>>hg38_21_+_8433172_8446622_RNA45SN1_45S_50ntFlanks@5084.5120.37|CGAACGAGACTCTGGCATGCTAACTAGTTACGCGACC</td><td>>hg38_21_+_8433172_8446622_RNA45SN1_45S_50ntFlanks</td><td>5084</td><td>5120</td><td>37</td><td>CGAACGAGACTCTGGCATGCTAACTAGTTACGCGACC</td><td>1109.5</td><td>-4.6</td><td>1.0</td><td>-4.6</td><td>4.1E-06</td><td>6.2E-06</td><td>MB231/MB468</td><td>NUC</td></tr>
<tr><td>>hg38_21_+_8433172_8446622_RNA45SN1_45S_50ntFlanks@11847.11886.40|AACGGGCTTGGCGGAATCAGCGGGGAAAGAAGACCCTGTT</td><td>>hg38_21_+_8433172_8446622_RNA45SN1_45S_50ntFlanks</td><td>11847</td><td>11886</td><td>40</td><td>AACGGGCTTGGCGGAATCAGCGGGGAAAGAAGACCCTGTT</td><td>148.3</td><td>10.5</td><td>2.1</td><td>5.1</td><td>3.2E-07</td><td>6.2E-06</td><td>BT20/MB231</td><td>MP</td></tr>
<tr><td>>hg38_21_+_8433172_8446622_RNA45SN1_45S_50ntFlanks@11571.11605.35|GCCGACTTAGAACTGGTGCGGACCAGGGGAATCCG</td><td>>hg38_21_+_8433172_8446622_RNA45SN1_45S_50ntFlanks</td><td>11571</td><td>11605</td><td>35</td><td>GCCGACTTAGAACTGGTGCGGACCAGGGGAATCCG</td><td>172.1</td><td>-6.9</td><td>1.5</td><td>-4.6</td><td>3.5E-06</td><td>6.2E-06</td><td>MB231/MB468</td><td>MP</td></tr>
<tr><td>>MI0000446|hsa-mir-125b-1&WithFlank&11|-|122099751|122099850@61.78.18&[MIMAT0004592&hsa-miR-125b-1-3p&offsets|0|-4;m-485&11|-|122099769|122099790&offsets|0|-4]|ACGGGTTAGGCTCTTGGG</td><td>>MI0000446|hsa-mir-125b-1&WithFlank&11|-|122099751|122099850</td><td>61</td><td>78</td><td>18</td><td>ACGGGTTAGGCTCTTGGG</td><td>43.4</td><td>8.7</td><td>1.9</td><td>4.6</td><td>3.5E-06</td><td>6.2E-06</td><td>MB231/MB468</td><td>MP</td></tr>
<tr><td>>hg38_21_+_8433172_8446622_RNA45SN1_45S_50ntFlanks@6680.6718.39|TGCGTCGATGAAGAACGCAGCTAGCTGCGAGAATTAATG</td><td>>hg38_21_+_8433172_8446622_RNA45SN1_45S_50ntFlanks</td><td>6680</td><td>6718</td><td>39</td><td>TGCGTCGATGAAGAACGCAGCTAGCTGCGAGAATTAATG</td><td>132.8</td><td>-5.6</td><td>1.2</td><td>-4.6</td><td>3.5E-06</td><td>6.2E-06</td><td>BT20/MB468</td><td>CYTO</td></tr>
<tr><td>>hg38_21_+_8433172_8446622_RNA45SN1_45S_50ntFlanks@3776.3801.26|CCGGTACAGTGAAACTGCGAATGGCT</td><td>>hg38_21_+_8433172_8446622_RNA45SN1_45S_50ntFlanks</td><td>3776</td><td>3801</td><td>26</td><td>CCGGTACAGTGAAACTGCGAATGGCT</td><td>537.3</td><td>-2.9</td><td>0.6</td><td>-4.6</td><td>3.5E-06</td><td>6.2E-06</td><td>BT20/MB468</td><td>CYTO</td></tr>
<tr><td>>hg38_21_+_8433172_8446622_RNA45SN1_45S_50ntFlanks@9716.9737.22|AATGATTAGAGGTCTTGGGGCC</td><td>>hg38_21_+_8433172_8446622_RNA45SN1_45S_50ntFlanks</td><td>9716</td><td>9737</td><td>22</td><td>AATGATTAGAGGTCTTGGGGCC</td><td>655.7</td><td>-4.0</td><td>0.9</td><td>-4.7</td><td>3.0E-06</td><td>6.2E-06</td><td>BT20/MB468</td><td>MITO</td></tr>
<tr><td>>MI0003188|hsa-mir-503&WithFlank&X|-|134546322|134546404@12.30.19(+2A)&[MIMAT0002874&hsa-miR-503-5p&offsets|0|-4(+2A);m-297&X|-|134546371|134546393&offsets|0|-4(+2A)]|TAGCAGCGGGAACAGTTCTAA</td><td>>MI0003188|hsa-mir-503&WithFlank&X|-|134546322|134546404</td><td>12</td><td>30</td><td>19(+2A)</td><td>TAGCAGCGGGAACAGTTCTAA</td><td>60.7</td><td>9.7</td><td>2.0</td><td>4.9</td><td>1.2E-06</td><td>6.2E-06</td><td>BT20/MB468</td><td>MP</td></tr>
<tr><td>>hg38_21_+_8433172_8446622_RNA45SN1_45S_50ntFlanks@11317.11342.26|ACCCCACGTCTCGTCGCGCGCGCGTC</td><td>>hg38_21_+_8433172_8446622_RNA45SN1_45S_50ntFlanks</td><td>11317</td><td>11342</td><td>26</td><td>ACCCCACGTCTCGTCGCGCGCGCGTC</td><td>1495.0</td><td>-5.2</td><td>1.1</td><td>-4.6</td><td>3.5E-06</td><td>6.2E-06</td><td>BT20/MB468</td><td>CYTO</td></tr>
<tr><td>>hg38_21_+_8433172_8446622_RNA45SN1_45S_50ntFlanks@12248.12268.21|GCAAAAGCTCGCTTGATCTTG</td><td>>hg38_21_+_8433172_8446622_RNA45SN1_45S_50ntFlanks</td><td>12248</td><td>12268</td><td>21</td><td>GCAAAAGCTCGCTTGATCTTG</td><td>52.5</td><td>-8.8</td><td>1.8</td><td>-4.9</td><td>1.2E-06</td><td>6.2E-06</td><td>BT20/MB468</td><td>MP</td></tr>
<tr><td>>hg38_21_+_8433172_8446622_RNA45SN1_45S_50ntFlanks@12033.12070.38|ATCGTTTTTTCACTGACCCGGTGAGGCGGGGGGGCGAG</td><td>>hg38_21_+_8433172_8446622_RNA45SN1_45S_50ntFlanks</td><td>12033</td><td>12070</td><td>38</td><td>ATCGTTTTTTCACTGACCCGGTGAGGCGGGGGGGCGAG</td><td>61.7</td><td>-9.0</td><td>1.9</td><td>-4.9</td><td>1.2E-06</td><td>6.2E-06</td><td>BT20/MB468</td><td>MP</td></tr>
<tr><td>>trna10_LysCTT_16_+_3241501_3241573@1.35.35,trna13_LysCTT_14_-_58706613_58706685@1.35.35,trna2_LysCTT_15_+_79152904_79152976@1.35.35|GCCCGGCTAGCTCAGTCGGTAGAGCATGGGACTCT</td><td>>trna10_LysCTT_16_+_3241501_3241573</td><td>1</td><td>35</td><td>35</td><td>GCCCGGCTAGCTCAGTCGGTAGAGCATGGGACTCT</td><td>6938.7</td><td>6.5</td><td>1.3</td><td>4.9</td><td>1.2E-06</td><td>6.2E-06</td><td>BT20/MB468</td><td>MP</td></tr>
<tr><td>>hg38_21_+_8433172_8446622_RNA45SN1_45S_50ntFlanks@4568.4589.22|AATGGAATAGGACCGCGGTTCT</td><td>>hg38_21_+_8433172_8446622_RNA45SN1_45S_50ntFlanks</td><td>4568</td><td>4589</td><td>22</td><td>AATGGAATAGGACCGCGGTTCT</td><td>389.3</td><td>3.3</td><td>0.7</td><td>4.6</td><td>3.9E-06</td><td>6.2E-06</td><td>MB231/MB468</td><td>CYTO</td></tr>
<tr><td>>hg38_21_+_8433172_8446622_RNA45SN1_45S_50ntFlanks@12945.12972.28|GTAGACGACCTGCTTCTGGGTCGGGGTT</td><td>>hg38_21_+_8433172_8446622_RNA45SN1_45S_50ntFlanks</td><td>12945</td><td>12972</td><td>28</td><td>GTAGACGACCTGCTTCTGGGTCGGGGTT</td><td>50.2</td><td>-8.7</td><td>1.8</td><td>-4.9</td><td>1.2E-06</td><td>6.2E-06</td><td>BT20/MB468</td><td>MP</td></tr>
<tr><td>>hg38_21_+_8433172_8446622_RNA45SN1_45S_50ntFlanks@8179.8203.25|CTGATCGAGGCCCAGCCCGTGGACG</td><td>>hg38_21_+_8433172_8446622_RNA45SN1_45S_50ntFlanks</td><td>8179</td><td>8203</td><td>25</td><td>CTGATCGAGGCCCAGCCCGTGGACG</td><td>437.9</td><td>-7.3</td><td>1.6</td><td>-4.6</td><td>4.4E-06</td><td>6.3E-06</td><td>BT20/MB468</td><td>NUC</td></tr>
<tr><td>>hg38_21_+_8433172_8446622_RNA45SN1_45S_50ntFlanks@11518.11537.20|CGCCCCCGGGGCCGCGGTTC</td><td>>hg38_21_+_8433172_8446622_RNA45SN1_45S_50ntFlanks</td><td>11518</td><td>11537</td><td>20</td><td>CGCCCCCGGGGCCGCGGTTC</td><td>193.1</td><td>10.8</td><td>2.4</td><td>4.6</td><td>4.2E-06</td><td>6.3E-06</td><td>MB231/MB468</td><td>MITO</td></tr>
<tr><td>>hg38_21_+_8433172_8446622_RNA45SN1_45S_50ntFlanks@4044.4063.20|CCCATTCGAACGTCTGCCCT</td><td>>hg38_21_+_8433172_8446622_RNA45SN1_45S_50ntFlanks</td><td>4044</td><td>4063</td><td>20</td><td>CCCATTCGAACGTCTGCCCT</td><td>236.8</td><td>-4.0</td><td>0.9</td><td>-4.6</td><td>3.9E-06</td><td>6.3E-06</td><td>MB231/MB468</td><td>CYTO</td></tr>
<tr><td>>hg38_21_+_8433172_8446622_RNA45SN1_45S_50ntFlanks@6733.6756.24|CATTGATCATCGACACTTCGAACG</td><td>>hg38_21_+_8433172_8446622_RNA45SN1_45S_50ntFlanks</td><td>6733</td><td>6756</td><td>24</td><td>CATTGATCATCGACACTTCGAACG</td><td>1133.4</td><td>-3.5</td><td>0.8</td><td>-4.6</td><td>4.2E-06</td><td>6.3E-06</td><td>MB231/MB468</td><td>MITO</td></tr>
<tr><td>>MI0003628|hsa-mir-615&WithFlank&12|+|54033944|54034051@67.86.20&[MIMAT0003283&hsa-miR-615-3p&offsets|0|-2;m-495&12|+|54034010|54034030&offsets|0|-1]|TCCGAGCCTGGGTCTCCCTC</td><td>>MI0003628|hsa-mir-615&WithFlank&12|+|54033944|54034051</td><td>67</td><td>86</td><td>20</td><td>TCCGAGCCTGGGTCTCCCTC</td><td>103.2</td><td>10.0</td><td>2.0</td><td>5.1</td><td>3.3E-07</td><td>6.3E-06</td><td>BT20/MB231</td><td>MP</td></tr>
<tr><td>>hg38_21_+_8433172_8446622_RNA45SN1_45S_50ntFlanks@10248.10289.42|CGACGAGTAGGAGGGCCGCTGCGGTGAGCCTTGAAGCCTAGG</td><td>>hg38_21_+_8433172_8446622_RNA45SN1_45S_50ntFlanks</td><td>10248</td><td>10289</td><td>42</td><td>CGACGAGTAGGAGGGCCGCTGCGGTGAGCCTTGAAGCCTAGG</td><td>141.5</td><td>10.5</td><td>2.1</td><td>5.1</td><td>3.3E-07</td><td>6.3E-06</td><td>BT20/MB231</td><td>MP</td></tr>
<tr><td>>hg38_21_+_8433172_8446622_RNA45SN1_45S_50ntFlanks@8105.8134.30|CGCGGCGGGGCGCGGGACATGTGGCGTACG</td><td>>hg38_21_+_8433172_8446622_RNA45SN1_45S_50ntFlanks</td><td>8105</td><td>8134</td><td>30</td><td>CGCGGCGGGGCGCGGGACATGTGGCGTACG</td><td>119.5</td><td>10.2</td><td>2.0</td><td>5.1</td><td>3.3E-07</td><td>6.3E-06</td><td>BT20/MB231</td><td>MP</td></tr>
<tr><td>>hg38_21_+_8433172_8446622_RNA45SN1_45S_50ntFlanks@10912.10931.20|CGCGCCGCGGCTGGACGAGG</td><td>>hg38_21_+_8433172_8446622_RNA45SN1_45S_50ntFlanks</td><td>10912</td><td>10931</td><td>20</td><td>CGCGCCGCGGCTGGACGAGG</td><td>601.5</td><td>-5.8</td><td>1.3</td><td>-4.6</td><td>4.1E-06</td><td>6.3E-06</td><td>MB231/MB468</td><td>NUC</td></tr>
<tr><td>>hg38_21_+_8433172_8446622_RNA45SN1_45S_50ntFlanks@11729.11753.25|TAACTATGACTCTCTTAAGGTAGCC</td><td>>hg38_21_+_8433172_8446622_RNA45SN1_45S_50ntFlanks</td><td>11729</td><td>11753</td><td>25</td><td>TAACTATGACTCTCTTAAGGTAGCC</td><td>1436.6</td><td>-7.7</td><td>1.6</td><td>-4.9</td><td>1.2E-06</td><td>6.3E-06</td><td>BT20/MB468</td><td>MP</td></tr>
<tr><td>>hg38_21_+_8433172_8446622_RNA45SN1_45S_50ntFlanks@11564.11587.24|CCTAGCAGCCGACTTAGAACTGGT</td><td>>hg38_21_+_8433172_8446622_RNA45SN1_45S_50ntFlanks</td><td>11564</td><td>11587</td><td>24</td><td>CCTAGCAGCCGACTTAGAACTGGT</td><td>379.0</td><td>-7.2</td><td>1.6</td><td>-4.6</td><td>4.4E-06</td><td>6.3E-06</td><td>BT20/MB468</td><td>NUC</td></tr>
<tr><td>>hg38_21_+_8433172_8446622_RNA45SN1_45S_50ntFlanks@5468.5499.32|GCCCACGGCCCTGGCGGAGCGCTGAGAAGACG</td><td>>hg38_21_+_8433172_8446622_RNA45SN1_45S_50ntFlanks</td><td>5468</td><td>5499</td><td>32</td><td>GCCCACGGCCCTGGCGGAGCGCTGAGAAGACG</td><td>735.0</td><td>-7.1</td><td>1.5</td><td>-4.6</td><td>4.1E-06</td><td>6.3E-06</td><td>MB231/MB468</td><td>NUC</td></tr>
<tr><td>>hg38_21_+_8433172_8446622_RNA45SN1_45S_50ntFlanks@12991.13019.29|CTCCCTCGCTGCGATCTATTGAAAGTCAG</td><td>>hg38_21_+_8433172_8446622_RNA45SN1_45S_50ntFlanks</td><td>12991</td><td>13019</td><td>29</td><td>CTCCCTCGCTGCGATCTATTGAAAGTCAG</td><td>310.6</td><td>-5.9</td><td>1.3</td><td>-4.6</td><td>3.5E-06</td><td>6.3E-06</td><td>BT20/MB468</td><td>CYTO</td></tr>
<tr><td>>hg38_21_+_8433172_8446622_RNA45SN1_45S_50ntFlanks@10248.10281.34|CGACGAGTAGGAGGGCCGCTGCGGTGAGCCTTGA</td><td>>hg38_21_+_8433172_8446622_RNA45SN1_45S_50ntFlanks</td><td>10248</td><td>10281</td><td>34</td><td>CGACGAGTAGGAGGGCCGCTGCGGTGAGCCTTGA</td><td>204.2</td><td>8.1</td><td>1.7</td><td>4.9</td><td>1.2E-06</td><td>6.3E-06</td><td>BT20/MB468</td><td>MP</td></tr>
<tr><td>>hg38_21_+_8433172_8446622_RNA45SN1_45S_50ntFlanks@12456.12493.38|CCAAGCGTTGGATTGTTCACCCACTAATAGGGAACGTG</td><td>>hg38_21_+_8433172_8446622_RNA45SN1_45S_50ntFlanks</td><td>12456</td><td>12493</td><td>38</td><td>CCAAGCGTTGGATTGTTCACCCACTAATAGGGAACGTG</td><td>843.2</td><td>-3.9</td><td>0.8</td><td>-4.6</td><td>4.1E-06</td><td>6.3E-06</td><td>MB231/MB468</td><td>NUC</td></tr>
<tr><td>>MI0000542|hsa-mir-320a&WithFlank&8|-|22244960|22245043@48.68.21&[MIMAT0000510&hsa-miR-320a-3p&offsets|0|-1;m-60&8|-|22244975|22244996&offsets|0|-1]|AAAAGCTGGGTTGAGAGGGCG</td><td>>MI0000542|hsa-mir-320a&WithFlank&8|-|22244960|22245043</td><td>48</td><td>68</td><td>21</td><td>AAAAGCTGGGTTGAGAGGGCG</td><td>1075.9</td><td>3.6</td><td>0.8</td><td>4.6</td><td>4.1E-06</td><td>6.3E-06</td><td>MB231/MB468</td><td>NUC</td></tr>
<tr><td>>hg38_21_+_8433172_8446622_RNA45SN1_45S_50ntFlanks@12990.13013.24|GCTCCCTCGCTGCGATCTATTGAA</td><td>>hg38_21_+_8433172_8446622_RNA45SN1_45S_50ntFlanks</td><td>12990</td><td>13013</td><td>24</td><td>GCTCCCTCGCTGCGATCTATTGAA</td><td>7789.3</td><td>-4.3</td><td>0.9</td><td>-4.6</td><td>4.1E-06</td><td>6.3E-06</td><td>MB231/MB468</td><td>NUC</td></tr>
<tr><td>>hg38_21_+_8433172_8446622_RNA45SN1_45S_50ntFlanks@4538.4559.22|CCGAGCCGCCTGGATACCGCAG</td><td>>hg38_21_+_8433172_8446622_RNA45SN1_45S_50ntFlanks</td><td>4538</td><td>4559</td><td>22</td><td>CCGAGCCGCCTGGATACCGCAG</td><td>58.0</td><td>-8.9</td><td>1.8</td><td>-4.9</td><td>1.2E-06</td><td>6.3E-06</td><td>BT20/MB468</td><td>MP</td></tr>
<tr><td>>hg38_21_+_8433172_8446622_RNA45SN1_45S_50ntFlanks@12605.12622.18|TATGTGCTTGGCTGAGGA</td><td>>hg38_21_+_8433172_8446622_RNA45SN1_45S_50ntFlanks</td><td>12605</td><td>12622</td><td>18</td><td>TATGTGCTTGGCTGAGGA</td><td>170.2</td><td>-3.5</td><td>0.8</td><td>-4.6</td><td>4.0E-06</td><td>6.3E-06</td><td>MB231/MB468</td><td>CYTO</td></tr>
<tr><td>>hg38_21_+_8433172_8446622_RNA45SN1_45S_50ntFlanks@12081.12108.28|CTCTCGCTTCTGGCGCCAAGCGCCCGGC</td><td>>hg38_21_+_8433172_8446622_RNA45SN1_45S_50ntFlanks</td><td>12081</td><td>12108</td><td>28</td><td>CTCTCGCTTCTGGCGCCAAGCGCCCGGC</td><td>550.1</td><td>-6.6</td><td>1.4</td><td>-4.6</td><td>3.5E-06</td><td>6.3E-06</td><td>BT20/MB468</td><td>CYTO</td></tr>
<tr><td>>hg38_21_+_8433172_8446622_RNA45SN1_45S_50ntFlanks@12894.12926.33|CTCGCCCGTCACGCACCGCACGTTCGTGGGGAA</td><td>>hg38_21_+_8433172_8446622_RNA45SN1_45S_50ntFlanks</td><td>12894</td><td>12926</td><td>33</td><td>CTCGCCCGTCACGCACCGCACGTTCGTGGGGAA</td><td>61.8</td><td>-9.0</td><td>1.9</td><td>-4.9</td><td>1.2E-06</td><td>6.3E-06</td><td>BT20/MB468</td><td>MP</td></tr>
<tr><td>>hg38_21_+_8433172_8446622_RNA45SN1_45S_50ntFlanks@12656.12675.20|TGACTGAACGCCTCTAAGTC</td><td>>hg38_21_+_8433172_8446622_RNA45SN1_45S_50ntFlanks</td><td>12656</td><td>12675</td><td>20</td><td>TGACTGAACGCCTCTAAGTC</td><td>279.7</td><td>-5.1</td><td>1.1</td><td>-4.6</td><td>4.1E-06</td><td>6.3E-06</td><td>MB231/MB468</td><td>NUC</td></tr>
<tr><td>>hg38_21_+_8433172_8446622_RNA45SN1_45S_50ntFlanks@3771.3799.29|CACGGCCGGTACAGTGAAACTGCGAATGG</td><td>>hg38_21_+_8433172_8446622_RNA45SN1_45S_50ntFlanks</td><td>3771</td><td>3799</td><td>29</td><td>CACGGCCGGTACAGTGAAACTGCGAATGG</td><td>264.0</td><td>-6.7</td><td>1.4</td><td>-4.6</td><td>3.5E-06</td><td>6.3E-06</td><td>MB231/MB468</td><td>MP</td></tr>
<tr><td>>hg38_21_+_8433172_8446622_RNA45SN1_45S_50ntFlanks@5059.5093.35|CGATTTGTCTGGTTAATTCCGATAACGAACGAGAC</td><td>>hg38_21_+_8433172_8446622_RNA45SN1_45S_50ntFlanks</td><td>5059</td><td>5093</td><td>35</td><td>CGATTTGTCTGGTTAATTCCGATAACGAACGAGAC</td><td>108.8</td><td>-10.6</td><td>2.3</td><td>-4.6</td><td>3.5E-06</td><td>6.3E-06</td><td>MB231/MB468</td><td>MP</td></tr>
<tr><td>>trna119_LysCTT_1_-_145395522_145395594@1.33.33,trna11_LysCTT_5_-_180648979_180649051@1.33.33,trna13_LysCTT_6_+_26556774_26556846@1.33.33,trna7_LysCTT_16_+_3225692_3225764@1.33.33,trna9_LysCTT_5_+_180634755_180634827@1.33.33|GCCCGGCTAGCTCAGTCGGTAGAGCATGAGACT</td><td>>trna119_LysCTT_1_-_145395522_145395594</td><td>1</td><td>33</td><td>33</td><td>GCCCGGCTAGCTCAGTCGGTAGAGCATGAGACT</td><td>2695.9</td><td>4.3</td><td>0.9</td><td>4.6</td><td>3.5E-06</td><td>6.3E-06</td><td>MB231/MB468</td><td>MP</td></tr>
</table>

| | | | | | | | | | | | | | |
|---|---|---|---|---|---|---|---|---|---|---|---|---|---|
| >hg38_21_+_8433172_8446622_RNA45SN1_45S_50ntFlanks@10906.10925.20\|TGGGCGCGCGCCGCGGCTGG | >hg38_21_+_8433172_8446622_RNA45SN1_45S_50ntFlanks | 10906 | 10925 | 20 | TGGGCGCGCGCCGCGGCTGG | 241.8 | -4.3 | 0.9 | -4.6 | 4.0E-06 | 6.3E-06 | MB231/MB468 | CYTO |
| >hg38_21_+_8433172_8446622_RNA45SN1_45S_50ntFlanks@9777.9819.43\|GAAGCCCGGCTCGCTGGCGTGGAGCCGGGCGTGGAATGCGAGT | >hg38_21_+_8433172_8446622_RNA45SN1_45S_50ntFlanks | 9777 | 9819 | 43 | GAAGCCCGGCTCGCTGGCGTGGAGCCGGGCGTGGAATGCGAGT | 59.4 | 9.6 | 2.0 | 4.9 | 1.2E-06 | 6.3E-06 | BT20/MB468 | MP |
| >hg38_21_+_8433172_8446622_RNA45SN1_45S_50ntFlanks@5496.5526.31\|GACGGTCGAACTTGACTATCTAGAGGAAGTA | >hg38_21_+_8433172_8446622_RNA45SN1_45S_50ntFlanks | 5496 | 5526 | 31 | GACGGTCGAACTTGACTATCTAGAGGAAGTA | 143.3 | -5.6 | 1.2 | -4.6 | 3.5E-06 | 6.3E-06 | MB231/MB468 | MP |
| >am_trna2_GlyGCC_21_-_18827107_18827177@1.18.18,trna35_GlyGCC_1_+_161413094_161413164@1.18.18,trna37_GlyGCC_1_+_161420467_161420537@1.18.18,trna39_GlyGCC_1_+_161427898_161427968@1.18.18,trna41_GlyGCC_1_+_161435258_161435328@1.18.18\|GCATGGGTGGTTCAGTGG | >am_trna2_GlyGCC_21_-_18827107_18827177 | 1 | 18 | 18 | GCATGGGTGGTTCAGTGG | 112.7 | 10.2 | 2.0 | 5.1 | 3.3E-07 | 6.3E-06 | BT20/MB231 | MP |
| >hg38_21_+_8433172_8446622_RNA45SN1_45S_50ntFlanks@9915.9947.33\|AAAAGGTGTTGGTTGATATAGACAGCAGGACGG | >hg38_21_+_8433172_8446622_RNA45SN1_45S_50ntFlanks | 9915 | 9947 | 33 | AAAAGGTGTTGGTTGATATAGACAGCAGGACGG | 145.6 | 10.5 | 2.1 | 5.1 | 3.3E-07 | 6.3E-06 | BT20/MB231 | MP |
| >MI0000061\|hsa-let-7a-2&WithFlank&11\|-\|122146516\|122146599@11.31.21(+1C);MI0000062\|hsa-let-7a-3&WithFlank&22\|+\|46112743\|46112828@10.30.21(+1C);MI0000060\|hsa-let-7a-1&WithFlank&9\|+\|94175951\|94176042@12.32.21(+1C)&[MIMAT0000062&hsa-let-7a-5p&offsets\|0\|-1(+1C);m-6&11\|-\|122146568\|122146589&offsets\|0\|-1(+1C)];[MIMAT0000062_1&hsa-let-7a-5p&offsets\|0\|-1(+1C);m-5&22\|+\|46112752\|46112773&offsets\|0\|-1(+1C)];[MIMAT0000062_2&hsa-let-7a-5p&offsets\|0\|-1(+1C);m-4&9\|+\|94175962\|94175983&offsets\|0\|-1(+1C)]\|TGAGGTAGTAGGTTGTATAGTC | >MI0000061\|hsa-let-7a-2&WithFlank&11\|-\|122146516\|122146599 | 11 | 31 | 21(+1C) | TGAGGTAGTAGGTTGTATAGTC | 8269.4 | 4.4 | 0.9 | 4.6 | 3.5E-06 | 6.3E-06 | MB231/MB468 | MP |
| >hg38_21_+_8433172_8446622_RNA45SN1_45S_50ntFlanks@12445.12475.31\|AGCAGAATTCACCAAGCGTTGGATTGTTCAC | >hg38_21_+_8433172_8446622_RNA45SN1_45S_50ntFlanks | 12445 | 12475 | 31 | AGCAGAATTCACCAAGCGTTGGATTGTTCAC | 71.7 | -9.2 | 1.9 | -4.9 | 1.2E-06 | 6.3E-06 | BT20/MB468 | MP |
| >hg38_21_+_8433172_8446622_RNA45SN1_45S_50ntFlanks@9534.9575.42\|AGGAAACTCTGGTGGAGGTCCGTAGCGGTCCTGACGTGCAAA | >hg38_21_+_8433172_8446622_RNA45SN1_45S_50ntFlanks | 9534 | 9575 | 42 | AGGAAACTCTGGTGGAGGTCCGTAGCGGTCCTGACGTGCAAA | 446.2 | -4.4 | 0.9 | -4.6 | 4.2E-06 | 6.3E-06 | MB231/MB468 | NUC |
| >MI0003513\|hsa-mir-455&WithFlank&9\|+\|114209428\|114209535@24.45.22(+1U)&[MIMAT0003150&hsa-miR-455-5p&offsets\|+2\|+2(+1U);m-232&9\|+\|114209449\|114209470&offsets\|+2\|+2(+1U)]\|TGTGCCTTTGGACTACATCGTGT | >MI0003513\|hsa-mir-455&WithFlank&9\|+\|114209428\|114209535 | 24 | 45 | 22(+1U) | TGTGCCTTTGGACTACATCGTGT | 92.3 | -9.9 | 1.9 | -5.1 | 3.3E-07 | 6.3E-06 | BT20/MB231 | MP |
| >hg38_21_+_8433172_8446622_RNA45SN1_45S_50ntFlanks@12164.12181.18\|CTGGGGCGGTACACCTGT | >hg38_21_+_8433172_8446622_RNA45SN1_45S_50ntFlanks | 12164 | 12181 | 18 | CTGGGGCGGTACACCTGT | 270.5 | -6.9 | 1.4 | -4.9 | 1.2E-06 | 6.3E-06 | BT20/MB468 | MP |
| >trna10_LysCTT_16_+_3241501_3241573@13.33.21,trna13_LysCTT_14_-_58706613_58706685@13.33.21,trna2_LysCTT_15_+_79152904_79152976@13.33.21,trna4_LysCTT_18_-_43669271_43669343@13.33.21\|CAGTCGGTAGAGCATGGGACT | >trna10_LysCTT_16_+_3241501_3241573 | 13 | 33 | 21 | CAGTCGGTAGAGCATGGGACT | 43.0 | 8.6 | 1.9 | 4.6 | 3.5E-06 | 6.3E-06 | MB231/MB468 | MP |
| >hg38_21_+_8433172_8446622_RNA45SN1_45S_50ntFlanks@11578.11601.24\|TAGAACTGGTGCGGACCAGGGGAA | >hg38_21_+_8433172_8446622_RNA45SN1_45S_50ntFlanks | 11578 | 11601 | 24 | TAGAACTGGTGCGGACCAGGGGAA | 166.7 | -6.6 | 1.4 | -4.6 | 3.5E-06 | 6.3E-06 | MB231/MB468 | MP |
| >hg38_21_+_8433172_8446622_RNA45SN1_45S_50ntFlanks@9572.9596.25\|CAAATCGGTCGTCCGACCTGGGTAT | >hg38_21_+_8433172_8446622_RNA45SN1_45S_50ntFlanks | 9572 | 9596 | 25 | CAAATCGGTCGTCCGACCTGGGTAT | 4945.6 | -7.7 | 1.7 | -4.6 | 3.6E-06 | 6.3E-06 | BT20/MB468 | CYTO |
| >hg38_21_+_8433172_8446622_RNA45SN1_45S_50ntFlanks@10786.10823.38\|AACAGCCTCTGGCATGTTGGAACAATGTAGGTAAGGGA | >hg38_21_+_8433172_8446622_RNA45SN1_45S_50ntFlanks | 10786 | 10823 | 38 | AACAGCCTCTGGCATGTTGGAACAATGTAGGTAAGGGA | 406.1 | -6.5 | 1.4 | -4.6 | 4.2E-06 | 6.3E-06 | MB231/MB468 | NUC |
| >hg38_21_+_8433172_8446622_RNA45SN1_45S_50ntFlanks@11537.11580.44\|CCGCGCGGCGCCTCGCCTCGGCCGGCGCCTAGCAGCCGACTTAG | >hg38_21_+_8433172_8446622_RNA45SN1_45S_50ntFlanks | 11537 | 11580 | 44 | CCGCGCGGCGCCTCGCCTCGGCCGGCGCCTAGCAGCCGACTTAG | 2660.3 | -9.6 | 2.1 | -4.6 | 4.5E-06 | 6.3E-06 | BT20/MB468 | NUC |
| >hg38_21_+_8433172_8446622_RNA45SN1_45S_50ntFlanks@9569.9595.27\|GTGCAAATCGGTCGTCCGACCTGGGTA | >hg38_21_+_8433172_8446622_RNA45SN1_45S_50ntFlanks | 9569 | 9595 | 27 | GTGCAAATCGGTCGTCCGACCTGGGTA | 396.1 | -7.3 | 1.6 | -4.6 | 4.5E-06 | 6.3E-06 | BT20/MB468 | NUC |
| >hg38_21_+_8433172_8446622_RNA45SN1_45S_50ntFlanks@12624.12659.36\|CCAATGGGGCGAAGCTACCATCTGTGGGATTATGAC | >hg38_21_+_8433172_8446622_RNA45SN1_45S_50ntFlanks | 12624 | 12659 | 36 | CCAATGGGGCGAAGCTACCATCTGTGGGATTATGAC | 360.7 | -7.2 | 1.5 | -4.8 | 1.2E-06 | 6.3E-06 | BT20/MB468 | MP |
| >hg38_21_+_8433172_8446622_RNA45SN1_45S_50ntFlanks@12481.12499.19\|AATAGGGAACGTGAGCTGG | >hg38_21_+_8433172_8446622_RNA45SN1_45S_50ntFlanks | 12481 | 12499 | 19 | AATAGGGAACGTGAGCTGG | 250.5 | -2.6 | 0.6 | -4.6 | 4.2E-06 | 6.3E-06 | MB231/MB468 | NUC |
| >hg38_21_+_8433172_8446622_RNA45SN1_45S_50ntFlanks@9533.9572.40\|GAGGAAACTCTGGTGGAGGTCCGTAGCGGTCCTGACGTGC | >hg38_21_+_8433172_8446622_RNA45SN1_45S_50ntFlanks | 9533 | 9572 | 40 | GAGGAAACTCTGGTGGAGGTCCGTAGCGGTCCTGACGTGC | 1538.5 | -3.8 | 0.8 | -4.6 | 4.2E-06 | 6.3E-06 | MB231/MB468 | NUC |
| >hg38_21_+_8433172_8446622_RNA45SN1_45S_50ntFlanks@6738.6756.19\|ATCATCGACACTTCGAACG | >hg38_21_+_8433172_8446622_RNA45SN1_45S_50ntFlanks | 6738 | 6756 | 19 | ATCATCGACACTTCGAACG | 415.8 | -6.6 | 1.4 | -4.6 | 4.3E-06 | 6.3E-06 | MB231/MB468 | MITO |
| >trnaMT_GluTTC_MT_-_14674_14742@32.72.41,trnalookalike8_GluTTC_5_-_93905172_93905240@32.72.41\|TCATATCATTGGTCGTGGTTGTAGTCCGTGCGAGAATACCA | >trnaMT_GluTTC_MT_-_14674_14742 | 32 | 72 | 41 | TCATATCATTGGTCGTGGTTGTAGTCCGTGCGAGAATACCA | 549.5 | 4.0 | 0.9 | 4.6 | 4.2E-06 | 6.3E-06 | MB231/MB468 | NUC |
| >hg38_21_+_8433172_8446622_RNA45SN1_45S_50ntFlanks@8192.8214.23\|AGCCCGTGGACGGTGTGAGGCCG | >hg38_21_+_8433172_8446622_RNA45SN1_45S_50ntFlanks | 8192 | 8214 | 23 | AGCCCGTGGACGGTGTGAGGCCG | 336.8 | -3.7 | 0.8 | -4.6 | 4.2E-06 | 6.3E-06 | MB231/MB468 | NUC |
| >hg38_21_+_8433172_8446622_RNA45SN1_45S_50ntFlanks@9156.9188.33\|CCAGTGCGCCCCGGGCGGGTCGCGCCGTCGGGC | >hg38_21_+_8433172_8446622_RNA45SN1_45S_50ntFlanks | 9156 | 9188 | 33 | CCAGTGCGCCCCGGGCGGGTCGCGCCGTCGGGC | 896.5 | -5.9 | 1.1 | -5.1 | 3.5E-07 | 6.3E-06 | BT20/MB231 | CYTO |
| >hg38_21_+_8433172_8446622_RNA45SN1_45S_50ntFlanks@6726.6744.19\|CAGGACACATTGATCATCG | >hg38_21_+_8433172_8446622_RNA45SN1_45S_50ntFlanks | 6726 | 6744 | 19 | CAGGACACATTGATCATCG | 456.8 | -7.1 | 1.5 | -4.6 | 3.6E-06 | 6.4E-06 | BT20/MB468 | CYTO |
| >hg38_21_+_8433172_8446622_RNA45SN1_45S_50ntFlanks@10240.10260.21\|CTACGCCGCGACGAGTAGGAG | >hg38_21_+_8433172_8446622_RNA45SN1_45S_50ntFlanks | 10240 | 10260 | 21 | CTACGCCGCGACGAGTAGGAG | 167.4 | -6.5 | 1.4 | -4.6 | 3.5E-06 | 6.4E-06 | MB231/MB468 | MP |
| >trnaMT_PheGAA_MT_+_577_647@28.62.35\|ATACACTGAAAATGTTTAGACGGGCTCACATCACC | >trnaMT_PheGAA_MT_+_577_647 | 28 | 62 | 35 | ATACACTGAAAATGTTTAGACGGGCTCACATCACC | 71.0 | 9.9 | 2.0 | 4.8 | 1.2E-06 | 6.4E-06 | BT20/MB468 | MP |
| >hg38_MT_+_598_1651_RNR1_12S_50ntFlanks@811.828.18\|AAGGGTCGAAGGTGGATT | >hg38_MT_+_598_1651_RNR1_12S_50ntFlanks | 811 | 828 | 18 | AAGGGTCGAAGGTGGATT | 48.9 | 9.4 | 1.9 | 4.8 | 1.2E-06 | 6.4E-06 | BT20/MB468 | MP |
| >hg38_21_+_8433172_8446622_RNA45SN1_45S_50ntFlanks@9941.9981.41\|AGGACGGTGGCCATGGAAGTCGGAATCCGCTAAGGAGTGTG | >hg38_21_+_8433172_8446622_RNA45SN1_45S_50ntFlanks | 9941 | 9981 | 41 | AGGACGGTGGCCATGGAAGTCGGAATCCGCTAAGGAGTGTG | 44.7 | 9.2 | 1.9 | 4.8 | 1.2E-06 | 6.4E-06 | BT20/MB468 | MP |
| >hg38_21_+_8433172_8446622_RNA45SN1_45S_50ntFlanks@11568.11588.21\|GCAGCCGACTTAGAACTGGTG | >hg38_21_+_8433172_8446622_RNA45SN1_45S_50ntFlanks | 11568 | 11588 | 21 | GCAGCCGACTTAGAACTGGTG | 562.0 | -2.3 | 0.5 | -4.6 | 4.0E-06 | 6.4E-06 | MB231/MB468 | CYTO |
| >hg38_21_+_8433172_8446622_RNA45SN1_45S_50ntFlanks@3866.3898.33\|CTAGAGCTAATACATGCCGACGGGCGCTGACCC | >hg38_21_+_8433172_8446622_RNA45SN1_45S_50ntFlanks | 3866 | 3898 | 33 | CTAGAGCTAATACATGCCGACGGGCGCTGACCC | 291.7 | -7.4 | 1.5 | -4.8 | 1.2E-06 | 6.4E-06 | BT20/MB468 | MP |
| >hg38_21_+_8433172_8446622_RNA45SN1_45S_50ntFlanks@10355.10390.36\|AACTTTGAAGGCCGAAGTGGAGAAGGGTTCCATGTG | >hg38_21_+_8433172_8446622_RNA45SN1_45S_50ntFlanks | 10355 | 10390 | 36 | AACTTTGAAGGCCGAAGTGGAGAAGGGTTCCATGTG | 312.1 | -6.6 | 1.4 | -4.6 | 4.3E-06 | 6.4E-06 | MB231/MB468 | MITO |
| >hg38_21_+_8433172_8446622_RNA45SN1_45S_50ntFlanks@9166.9194.29\|CCGGGCGGGTCGCGCCGTCGGGCCCGGGG | >hg38_21_+_8433172_8446622_RNA45SN1_45S_50ntFlanks | 9166 | 9194 | 29 | CCGGGCGGGTCGCGCCGTCGGGCCCGGGG | 54.0 | -8.8 | 1.8 | -4.8 | 1.2E-06 | 6.4E-06 | BT20/MB468 | MP |
| >hg38_21_+_8433172_8446622_RNA45SN1_45S_50ntFlanks@10787.10815.29\|ACAGCCTCTGGCATGTTGGAACAATGTAG | >hg38_21_+_8433172_8446622_RNA45SN1_45S_50ntFlanks | 10787 | 10815 | 29 | ACAGCCTCTGGCATGTTGGAACAATGTAG | 240.6 | -3.8 | 0.8 | -4.6 | 4.0E-06 | 6.4E-06 | MB231/MB468 | CYTO |
| >hg38_21_+_8433172_8446622_RNA45SN1_45S_50ntFlanks@4661.4690.30\|AGGTGAAATTCTTGGACCGGCGCAAGACGG | >hg38_21_+_8433172_8446622_RNA45SN1_45S_50ntFlanks | 4661 | 4690 | 30 | AGGTGAAATTCTTGGACCGGCGCAAGACGG | 396.3 | -3.5 | 0.8 | -4.6 | 3.6E-06 | 6.4E-06 | MB231/MB468 | MP |
| >hg38_21_+_8433172_8446622_RNA45SN1_45S_50ntFlanks@10771.10816.46\|CAGGTCTCCAAGGTGAACAGCCTCTGGCATGTTGGAACAATGTAGG | >hg38_21_+_8433172_8446622_RNA45SN1_45S_50ntFlanks | 10771 | 10816 | 46 | CAGGTCTCCAAGGTGAACAGCCTCTGGCATGTTGGAACAATGTAGG | 663.3 | -5.8 | 1.3 | -4.6 | 4.2E-06 | 6.4E-06 | MB231/MB468 | NUC |
| >hg38_21_+_8433172_8446622_RNA45SN1_45S_50ntFlanks@11540.11583.44\|CGCGGCGCCTCGCCTCGGCCGGCGCCTAGCAGCCGACTTAGAAC | >hg38_21_+_8433172_8446622_RNA45SN1_45S_50ntFlanks | 11540 | 11583 | 44 | CGCGGCGCCTCGCCTCGGCCGGCGCCTAGCAGCCGACTTAGAAC | 1884.0 | -4.3 | 0.8 | -5.1 | 3.5E-07 | 6.4E-06 | BT20/MB231 | CYTO |
| >hg38_1_-_228634819_228635039_RNA5S12_5S_50ntFlanks@51.74.24\|GTCTACGGCCATACCACCCTGAAC | >hg38_1_-_228634819_228635039_RNA5S12_5S_50ntFlanks | 51 | 74 | 24 | GTCTACGGCCATACCACCCTGAAC | 395.9 | 6.7 | 1.5 | 4.6 | 4.0E-06 | 6.4E-06 | MB231/MB468 | CYTO |
| >hg38_21_+_8433172_8446622_RNA45SN1_45S_50ntFlanks@8176.8215.40\|CTTCTGATCGAGGCCCAGCCCGTGGACGGTGTGAGGCCGG | >hg38_21_+_8433172_8446622_RNA45SN1_45S_50ntFlanks | 8176 | 8215 | 40 | CTTCTGATCGAGGCCCAGCCCGTGGACGGTGTGAGGCCGG | 442.4 | -5.2 | 1.1 | -4.6 | 4.0E-06 | 6.4E-06 | MB231/MB468 | CYTO |
| >MI0000802\|hsa-mir-340&WithFlank&5\|-\|180015297\|180015403@22.42.21(+1C)&[MIMAT0004692&hsa-miR-340-5p&offsets\|0\|-1(+1C);m-220&5\|-\|180015361\|180015382&offsets\|0\|-1(+1C)]\|TTATAAAGCAATGAGACTGATC | >MI0000802\|hsa-mir-340&WithFlank&5\|-\|180015297\|180015403 | 22 | 42 | 21(+1C) | TTATAAAGCAATGAGACTGATC | 100.3 | 3.5 | 0.8 | 4.6 | 4.5E-06 | 6.4E-06 | BT20/MB468 | NUC |
| >hg38_21_+_8433172_8446622_RNA45SN1_45S_50ntFlanks@12477.12498.22\|CACTAATAGGGAACGTGAGCTG | >hg38_21_+_8433172_8446622_RNA45SN1_45S_50ntFlanks | 12477 | 12498 | 22 | CACTAATAGGGAACGTGAGCTG | 754.1 | -5.7 | 1.2 | -4.6 | 3.6E-06 | 6.4E-06 | MB231/MB468 | MP |
| >MI0000095\|hsa-mir-93&WithFlank&7\|-\|100093762\|100093853@17.38.22(+1A)&[MIMAT0000093&hsa-miR-93-5p&offsets\|0\|-1(+1A);m-29&7\|-\|100093815\|100093837&offsets\|0\|-1(+1A)]\|CAAAGTGCTGTTCGTGCAGGTAA | >MI0000095\|hsa-mir-93&WithFlank&7\|-\|100093762\|100093853 | 17 | 38 | 22(+1A) | CAAAGTGCTGTTCGTGCAGGTAA | 350.0 | 3.0 | 0.7 | 4.6 | 4.5E-06 | 6.4E-06 | BT20/MB468 | NUC |
| >hg38_21_+_8433172_8446622_RNA45SN1_45S_50ntFlanks@11564.11583.20\|CCTAGCAGCCGACTTAGAAC | >hg38_21_+_8433172_8446622_RNA45SN1_45S_50ntFlanks | 11564 | 11583 | 20 | CCTAGCAGCCGACTTAGAAC | 507.9 | -5.7 | 1.2 | -4.6 | 4.5E-06 | 6.4E-06 | BT20/MB468 | NUC |
| >hg38_21_+_8433172_8446622_RNA45SN1_45S_50ntFlanks@12408.12426.19\|TTTGATCCTTCGATGTCGG | >hg38_21_+_8433172_8446622_RNA45SN1_45S_50ntFlanks | 12408 | 12426 | 19 | TTTGATCCTTCGATGTCGG | 128.9 | -3.9 | 0.9 | -4.6 | 3.6E-06 | 6.4E-06 | BT20/MB468 | CYTO |
| >hg38_21_+_8433172_8446622_RNA45SN1_45S_50ntFlanks@12417.12455.39\|TCGATGTCGGCTCTTCCTATCATTGTGAAGCAGAATTCA | >hg38_21_+_8433172_8446622_RNA45SN1_45S_50ntFlanks | 12417 | 12455 | 39 | TCGATGTCGGCTCTTCCTATCATTGTGAAGCAGAATTCA | 1795.7 | -3.9 | 0.8 | -4.6 | 4.2E-06 | 6.4E-06 | MB231/MB468 | NUC |
| >hg38_21_+_8433172_8446622_RNA45SN1_45S_50ntFlanks@8102.8143.42\|CCCCGCGGCGGGGCGCGGGACATGTGGCGTACGGAAGACCCG | >hg38_21_+_8433172_8446622_RNA45SN1_45S_50ntFlanks | 8102 | 8143 | 42 | CCCCGCGGCGGGGCGCGGGACATGTGGCGTACGGAAGACCCG | 91.8 | -10.4 | 2.2 | -4.6 | 3.6E-06 | 6.4E-06 | MB231/MB468 | MP |
| >MI0000440\|hsa-mir-27b&WithFlank&9\|+\|95085439\|95085547@67.85.19&[MIMAT0000419&hsa-miR-27b-3p&offsets\|0\|-2;m-39&9\|+\|95085505\|95085525&offsets\|0\|-2]\|TTCACAGTGGCTAAGTTCT | >MI0000440\|hsa-mir-27b&WithFlank&9\|+\|95085439\|95085547 | 67 | 85 | 19 | TTCACAGTGGCTAAGTTCT | 1399.1 | 4.0 | 0.8 | 4.8 | 1.3E-06 | 6.4E-06 | BT20/MB468 | MP |
| >am_trnaMT_PheGAA_MT_+_577_647@40.58.19\|TGTTTAGACGGGCTCACAT | >am_trnaMT_PheGAA_MT_+_577_647 | 40 | 58 | 19 | TGTTTAGACGGGCTCACAT | 221.3 | 4.9 | 1.0 | 4.8 | 1.3E-06 | 6.4E-06 | BT20/MB468 | MP |
| >hg38_21_+_8433172_8446622_RNA45SN1_45S_50ntFlanks@6651.6690.40\|CGACTCTTAGCGGTGGATCACTCGGCTCGTGCGTCGATGA | >hg38_21_+_8433172_8446622_RNA45SN1_45S_50ntFlanks | 6651 | 6690 | 40 | CGACTCTTAGCGGTGGATCACTCGGCTCGTGCGTCGATGA | 2745.9 | 3.0 | 0.7 | 4.6 | 4.2E-06 | 6.4E-06 | MB231/MB468 | NUC |

| | | | | | | | | | | | | | |
|---|---|---|---|---|---|---|---|---|---|---|---|---|---|
| >hg38_21_+_8433172_8446622_RNA45SN1_45S_50ntFlanks@4297.4320.24\|CATTGGAGGGCAAGTCTGGTGCCA | >hg38_21_+_8433172_8446622_RNA45SN1_45S_50ntFlanks | 4297 | 4320 | 24 | CATTGGAGGGCAAGTCTGGTGCCA | 161.2 | -5.6 | 1.2 | -4.6 | 3.6E-06 | 6.4E-06 | BT20/MB468 | CYTO |
| >hg38_21_+_8433172_8446622_RNA45SN1_45S_50ntFlanks@10727.10763.37\|GGAGAGGGTGTAAATCTCGCGCCGGGCCGTACCCATA | >hg38_21_+_8433172_8446622_RNA45SN1_45S_50ntFlanks | 10727 | 10763 | 37 | GGAGAGGGTGTAAATCTCGCGCCGGGCCGTACCCATA | 57.2 | -8.9 | 1.8 | -4.8 | 1.3E-06 | 6.4E-06 | BT20/MB468 | MP |
| >hg38_21_+_8433172_8446622_RNA45SN1_45S_50ntFlanks@12841.12860.20\|CGGAGTGCCCTTCGTCCTGG | >hg38_21_+_8433172_8446622_RNA45SN1_45S_50ntFlanks | 12841 | 12860 | 20 | CGGAGTGCCCTTCGTCCTGG | 127.9 | -10.9 | 2.3 | -4.6 | 3.6E-06 | 6.4E-06 | MB231/MB468 | MP |
| >hg38_21_+_8433172_8446622_RNA45SN1_45S_50ntFlanks@3738.3757.20\|TCTCAAAGATTAAGCCATGC | >hg38_21_+_8433172_8446622_RNA45SN1_45S_50ntFlanks | 3738 | 3757 | 20 | TCTCAAAGATTAAGCCATGC | 46.7 | -8.6 | 1.8 | -4.8 | 1.3E-06 | 6.4E-06 | BT20/MB468 | MP |
| >hg38_1_-_228634819_228635039_RNA5S12_5S_50ntFlanks@148.170.23\|GGAATACCGGGTGCTGTAGGCTT | >hg38_1_-_228634819_228635039_RNA5S12_5S_50ntFlanks | 148 | 170 | 23 | GGAATACCGGGTGCTGTAGGCTT | 4893.0 | 3.1 | 0.7 | 4.6 | 4.1E-06 | 6.5E-06 | MB231/MB468 | CYTO |
| >MI0000650\|hsa-mir-200c&WithFlank&12\|+\|6963693\|6963772@50.72.23(+2U)&[MIMAT0000617&hsa-miR-200c-3p&offsets\|0\|0(+2U);m-54&12\|+\|6963742\|6963764&offsets\|0\|0(+2U)]\|TAATACTGCCGGGTAATGATGGATT | >MI0000650\|hsa-mir-200c&WithFlank&12\|+\|6963693\|6963772 | 50 | 72 | 23(+2U) | TAATACTGCCGGGTAATGATGGATT | 113.6 | 10.2 | 2.0 | 5.1 | 3.4E-07 | 6.5E-06 | BT20/MB231 | MP |
| >hg38_21_+_8433172_8446622_RNA45SN1_45S_50ntFlanks@11781.11803.23\|CGCATGAATGGATGAACGAGATT | >hg38_21_+_8433172_8446622_RNA45SN1_45S_50ntFlanks | 11781 | 11803 | 23 | CGCATGAATGGATGAACGAGATT | 228.3 | 11.2 | 2.2 | 5.1 | 3.4E-07 | 6.5E-06 | BT20/MB231 | MP |
| >hg38_21_+_8433172_8446622_RNA45SN1_45S_50ntFlanks@8246.8268.23\|TTCCCGGAGTCGGGTTGCTTGGG | >hg38_21_+_8433172_8446622_RNA45SN1_45S_50ntFlanks | 8246 | 8268 | 23 | TTCCCGGAGTCGGGTTGCTTGGG | 160.2 | 10.7 | 2.1 | 5.1 | 3.4E-07 | 6.5E-06 | BT20/MB231 | MP |
| >hg38_21_+_8433172_8446622_RNA45SN1_45S_50ntFlanks@11670.11691.22\|CCCAGTGCTCTGAATGTCAAAG | >hg38_21_+_8433172_8446622_RNA45SN1_45S_50ntFlanks | 11670 | 11691 | 22 | CCCAGTGCTCTGAATGTCAAAG | 1592.3 | -4.9 | 1.1 | -4.6 | 4.3E-06 | 6.5E-06 | MB231/MB468 | MITO |
| >hg38_MT_+_1621_3279_RNR2_16S_50ntFlanks@201.230.30\|ATATAGCAAGGACTAACCCCTATACCTTCT | >hg38_MT_+_1621_3279_RNR2_16S_50ntFlanks | 201 | 230 | 30 | ATATAGCAAGGACTAACCCCTATACCTTCT | 40.4 | 8.6 | 1.8 | 4.6 | 3.6E-06 | 6.5E-06 | MB231/MB468 | MP |
| >trna10_AspGTC_12_-_125424193_125424264@1.36.36,trna12_AspGTC_12_-_125411891_125411962@1.36.36,trna38_AspGTC_17_-_8125556_8125627@1.36.36,trna45_AspGTC_6_+_27447453_27447524@1.36.36,trna48_AspGTC_6_+_27471523_27471594@1.36.36,trna4_AspGTC_12_+_96429799_96429870@1.36.36,trna69_AspGTC_1_-_161440205_161440276@1.36.36,trna72_AspGTC_1_-_161432824_161432895@1.36.36,trna75_AspGTC_1_-_161425414_161425485@1.36.36,trna78_AspGTC_1_-_161418033_161418104@1.36.36,trna81_AspGTC_1_-_161410615_161410686@1.36.36\|TCCTCGTTAGTATAGTGGTGAGTATCCCCGCCTGTC | >trna10_AspGTC_12_-_125424193_125424264 | 1 | 36 | 36 | TCCTCGTTAGTATAGTGGTGAGTATCCCCGCCTGTC | 67.5 | 9.3 | 2.0 | 4.6 | 3.6E-06 | 6.5E-06 | MB231/MB468 | MP |
| >hg38_21_+_8433172_8446622_RNA45SN1_45S_50ntFlanks@9598.9615.18\|GGGGCGAAAGACTAATCG | >hg38_21_+_8433172_8446622_RNA45SN1_45S_50ntFlanks | 9598 | 9615 | 18 | GGGGCGAAAGACTAATCG | 176.9 | -6.5 | 1.4 | -4.6 | 3.6E-06 | 6.5E-06 | MB231/MB468 | MP |
| >MI0000086\|hsa-mir-28&WithFlank&3\|+\|188688775\|188688872@62.82.21(+1C)&[MIMAT0004502&hsa-miR-28-3p&offsets\|+2\|+1(+1C);m-28&3\|+\|188688834\|188688855&offsets\|+2\|+1(+1C)]\|CTAGATTGTGAGCTCCTGGAGC | >MI0000086\|hsa-mir-28&WithFlank&3\|+\|188688775\|188688872 | 62 | 82 | 21(+1C) | CTAGATTGTGAGCTCCTGGAGC | 43.7 | 8.7 | 1.9 | 4.6 | 3.6E-06 | 6.5E-06 | MB231/MB468 | MP |
| >trnaMT_ValTAC_MT_+_1602_1670@54.71.18\|AACTTGACCGCTCTGACC | >trnaMT_ValTAC_MT_+_1602_1670 | 54 | 71 | 18 | AACTTGACCGCTCTGACC | 1371.5 | 4.1 | 0.9 | 4.7 | 3.1E-06 | 6.5E-06 | BT20/MB468 | MITO |
| >hg38_21_+_8433172_8446622_RNA45SN1_45S_50ntFlanks@4982.5003.22\|ACGGACAGGATTGACAGATTGA | >hg38_21_+_8433172_8446622_RNA45SN1_45S_50ntFlanks | 4982 | 5003 | 22 | ACGGACAGGATTGACAGATTGA | 184.0 | -9.6 | 2.1 | -4.7 | 3.1E-06 | 6.5E-06 | BT20/MB468 | MITO |
| >hg38_21_+_8433172_8446622_RNA45SN1_45S_50ntFlanks@10788.10806.19\|CAGCCTCTGGCATGTTGGA | >hg38_21_+_8433172_8446622_RNA45SN1_45S_50ntFlanks | 10788 | 10806 | 19 | CAGCCTCTGGCATGTTGGA | 156.9 | -6.4 | 1.4 | -4.6 | 3.6E-06 | 6.5E-06 | MB231/MB468 | MP |
| >MI0000440\|hsa-mir-27b&WithFlank&9\|+\|95085439\|95085547@67.87.21(+1U)&[MIMAT0000419&hsa-miR-27b-3p&offsets\|0\|0(+1U);m-39&9\|+\|95085505\|95085525&offsets\|0\|0(+1U)]\|TTCACAGTGGCTAAGTTCTGCT | >MI0000440\|hsa-mir-27b&WithFlank&9\|+\|95085439\|95085547 | 67 | 87 | 21(+1U) | TTCACAGTGGCTAAGTTCTGCT | 1425.3 | 4.0 | 0.9 | 4.6 | 3.6E-06 | 6.5E-06 | MB231/MB468 | MP |
| >MI0000803\|hsa-mir-330&WithFlank&19\|-\|45638988\|45639093@63.86.24(+1C)&[MIMAT0000751&hsa-miR-330-3p&offsets\|0\|+1(+1C);m-347&19\|-\|45639009\|45639030&offsets\|-1\|+1(+1C)]\|GCAAAGCACACGGCCTGCAGAGAGC | >MI0000803\|hsa-mir-330&WithFlank&19\|-\|45638988\|45639093 | 63 | 86 | 24(+1C) | GCAAAGCACACGGCCTGCAGAGAGC | 40.5 | 8.6 | 1.8 | 4.6 | 3.6E-06 | 6.5E-06 | MB231/MB468 | MP |
| >MI0000061\|hsa-let-7a-2&WithFlank&11\|-\|122146516\|122146599@11.31.21(+1G);MI0000062\|hsa-let-7a-3&WithFlank&22\|+\|46112743\|46112828@10.30.21(+1G);MI0000060\|hsa-let-7a-1&WithFlank&9\|+\|94175951\|94176042@12.32.21(+1G)&[MIMAT0000062&hsa-let-7a-5p&offsets\|0\|-1(+1G);m-6&11\|-\|122146568\|122146589&offsets\|0\|-1(+1G)];[MIMAT0000062_1&hsa-let-7a-5p&offsets\|0\|-1(+1G);m-5&22\|+\|46112752\|46112773&offsets\|0\|-1(+1G)];[MIMAT0000062_2&hsa-let-7a-5p&offsets\|0\|-1(+1G);m-4&9\|+\|94175962\|94175983&offsets\|0\|-1(+1G)]\|TGAGGTAGTAGGTTGTATAGTG | >MI0000061\|hsa-let-7a-2&WithFlank&11\|-\|122146516\|122146599 | 11 | 31 | 21(+1G) | TGAGGTAGTAGGTTGTATAGTG | 762.2 | 4.4 | 0.9 | 4.6 | 4.1E-06 | 6.5E-06 | MB231/MB468 | CYTO |
| >hg38_21_+_8433172_8446622_RNA45SN1_45S_50ntFlanks@9546.9572.27\|TGGAGGTCCGTAGCGGTCCTGACGTGC | >hg38_21_+_8433172_8446622_RNA45SN1_45S_50ntFlanks | 9546 | 9572 | 27 | TGGAGGTCCGTAGCGGTCCTGACGTGC | 624.6 | 2.6 | 0.6 | 4.6 | 4.1E-06 | 6.5E-06 | MB231/MB468 | CYTO |
| >hg38_21_+_8433172_8446622_RNA45SN1_45S_50ntFlanks@5471.5494.24\|CACGGCCCTGGCGGAGCGCTGAGA | >hg38_21_+_8433172_8446622_RNA45SN1_45S_50ntFlanks | 5471 | 5494 | 24 | CACGGCCCTGGCGGAGCGCTGAGA | 1118.4 | 5.2 | 1.1 | 4.6 | 3.7E-06 | 6.5E-06 | BT20/MB468 | CYTO |
| >hg38_21_+_8433172_8446622_RNA45SN1_45S_50ntFlanks@11666.11685.20\|TCTGCCCAGTGCTCTGAATG | >hg38_21_+_8433172_8446622_RNA45SN1_45S_50ntFlanks | 11666 | 11685 | 20 | TCTGCCCAGTGCTCTGAATG | 1615.7 | -2.6 | 0.6 | -4.6 | 4.3E-06 | 6.5E-06 | MB231/MB468 | NUC |
| >hg38_21_+_8433172_8446622_RNA45SN1_45S_50ntFlanks@4560.4578.19\|CTAGGAATAATGGAATAGG | >hg38_21_+_8433172_8446622_RNA45SN1_45S_50ntFlanks | 4560 | 4578 | 19 | CTAGGAATAATGGAATAGG | 218.7 | -4.3 | 0.9 | -4.6 | 4.1E-06 | 6.5E-06 | MB231/MB468 | CYTO |
| >MI0000273\|hsa-mir-183&WithFlank&7\|-\|129774899\|129775020@34.54.21&[MIMAT0000261&hsa-miR-183-5p&offsets\|+1\|0;m-41&7\|-\|129774966\|129774987&offsets\|0\|-1]\|ATGGCACTGGTAGAATTCACT | >MI0000273\|hsa-mir-183&WithFlank&7\|-\|129774899\|129775020 | 34 | 54 | 21 | ATGGCACTGGTAGAATTCACT | 1184.6 | 2.1 | 0.5 | 4.6 | 4.6E-06 | 6.6E-06 | BT20/MB468 | NUC |
| >hg38_21_+_8433172_8446622_RNA45SN1_45S_50ntFlanks@12416.12447.32\|TTCGATGTCGGCTCTTCCTATCATTGTGAAGC | >hg38_21_+_8433172_8446622_RNA45SN1_45S_50ntFlanks | 12416 | 12447 | 32 | TTCGATGTCGGCTCTTCCTATCATTGTGAAGC | 674.9 | 5.7 | 1.2 | 4.6 | 3.7E-06 | 6.6E-06 | BT20/MB468 | CYTO |
| >hg38_21_+_8433172_8446622_RNA45SN1_45S_50ntFlanks@11583.11600.18\|CTGGTGCGGACCAGGGGA | >hg38_21_+_8433172_8446622_RNA45SN1_45S_50ntFlanks | 11583 | 11600 | 18 | CTGGTGCGGACCAGGGGA | 823.1 | -3.4 | 0.7 | -4.6 | 3.7E-06 | 6.6E-06 | BT20/MB468 | CYTO |
| >hg38_21_+_8433172_8446622_RNA45SN1_45S_50ntFlanks@10348.10374.27\|AAACGAGAACTTTGAAGGCCGAAGTGG | >hg38_21_+_8433172_8446622_RNA45SN1_45S_50ntFlanks | 10348 | 10374 | 27 | AAACGAGAACTTTGAAGGCCGAAGTGG | 469.8 | -2.5 | 0.5 | -4.7 | 3.2E-06 | 6.6E-06 | BT20/MB468 | MITO |
| >hg38_21_+_8433172_8446622_RNA45SN1_45S_50ntFlanks@10914.10932.19\|CGCCGCGGCTGGACGAGGC | >hg38_21_+_8433172_8446622_RNA45SN1_45S_50ntFlanks | 10914 | 10932 | 19 | CGCCGCGGCTGGACGAGGC | 269.4 | -6.1 | 1.3 | -4.7 | 3.2E-06 | 6.6E-06 | BT20/MB468 | MITO |
| >hg38_21_+_8433172_8446622_RNA45SN1_45S_50ntFlanks@11694.11717.24\|AAGAAATTCAATGAAGCGCGGGTA | >hg38_21_+_8433172_8446622_RNA45SN1_45S_50ntFlanks | 11694 | 11717 | 24 | AAGAAATTCAATGAAGCGCGGGTA | 385.8 | -4.9 | 1.1 | -4.6 | 4.2E-06 | 6.6E-06 | MB231/MB468 | CYTO |
| >hg38_21_+_8433172_8446622_RNA45SN1_45S_50ntFlanks@8198.8216.19\|TGGACGGTGTGAGGCCGGT | >hg38_21_+_8433172_8446622_RNA45SN1_45S_50ntFlanks | 8198 | 8216 | 19 | TGGACGGTGTGAGGCCGGT | 252.4 | -6.6 | 1.4 | -4.6 | 3.7E-06 | 6.6E-06 | MB231/MB468 | MP |
| >MI0000085\|hsa-mir-27a&WithFlank&19\|-\|13836434\|13836523@57.76.20(+3U)&[MIMAT0000084&hsa-miR-27a-3p&offsets\|0\|-1(+3U);m-43&19\|-\|13836447\|13836467&offsets\|0\|-1(+3U)]\|TTCACAGTGGCTAAGTTCCGTTT | >MI0000085\|hsa-mir-27a&WithFlank&19\|-\|13836434\|13836523 | 57 | 76 | 20(+3U) | TTCACAGTGGCTAAGTTCCGTTT | 2020.9 | 3.9 | 0.8 | 4.6 | 3.7E-06 | 6.6E-06 | MB231/MB468 | MP |
| >hg38_21_+_8433172_8446622_RNA45SN1_45S_50ntFlanks@10392.10429.38\|ACAGCAGTTGAACATGGGTCAGTCGGTCCTGAGAGATG | >hg38_21_+_8433172_8446622_RNA45SN1_45S_50ntFlanks | 10392 | 10429 | 38 | ACAGCAGTTGAACATGGGTCAGTCGGTCCTGAGAGATG | 2849.1 | -3.8 | 0.8 | -4.6 | 4.2E-06 | 6.6E-06 | MB231/MB468 | CYTO |
| >hg38_21_+_8433172_8446622_RNA45SN1_45S_50ntFlanks@12996.13038.43\|TCGCTGCGATCTATTGAAAGTCAGCCCTCGACACAAGGGTTTG | >hg38_21_+_8433172_8446622_RNA45SN1_45S_50ntFlanks | 12996 | 13038 | 43 | TCGCTGCGATCTATTGAAAGTCAGCCCTCGACACAAGGGTTTG | 778.5 | 5.4 | 1.2 | 4.6 | 4.3E-06 | 6.6E-06 | MB231/MB468 | NUC |
| >MI0000071\|hsa-mir-17&WithFlank&13\|+\|91350599\|91350694@20.43.24&[MIMAT0000070&hsa-miR-17-5p&offsets\|0\|+1;m-88&13\|+\|91350618\|91350640&offsets\|0\|+1]\|CAAAGTGCTTACAGTGCAGGTAGT | >MI0000071\|hsa-mir-17&WithFlank&13\|+\|91350599\|91350694 | 20 | 43 | 24 | CAAAGTGCTTACAGTGCAGGTAGT | 997.5 | 4.8 | 1.0 | 4.6 | 4.2E-06 | 6.6E-06 | MB231/MB468 | CYTO |
| >hg38_21_+_8433172_8446622_RNA45SN1_45S_50ntFlanks@11345.11367.23\|CTGGGGGCGGGGAGCGGTCGGGC | >hg38_21_+_8433172_8446622_RNA45SN1_45S_50ntFlanks | 11345 | 11367 | 23 | CTGGGGGCGGGGAGCGGTCGGGC | 70.3 | -9.2 | 1.9 | -4.8 | 1.3E-06 | 6.6E-06 | BT20/MB468 | MP |
| >hg38_21_+_8433172_8446622_RNA45SN1_45S_50ntFlanks@12345.12365.21\|TCAGAAAAGTTACCACAGGGA | >hg38_21_+_8433172_8446622_RNA45SN1_45S_50ntFlanks | 12345 | 12365 | 21 | TCAGAAAAGTTACCACAGGGA | 53.6 | -8.8 | 1.8 | -4.8 | 1.3E-06 | 6.6E-06 | BT20/MB468 | MP |
| >hg38_21_+_8433172_8446622_RNA45SN1_45S_50ntFlanks@12006.12033.28\|GGCCGCCGGTGAAATACCACTACTCTGA | >hg38_21_+_8433172_8446622_RNA45SN1_45S_50ntFlanks | 12006 | 12033 | 28 | GGCCGCCGGTGAAATACCACTACTCTGA | 454.0 | 3.4 | 0.7 | 5.1 | 4.1E-07 | 6.6E-06 | BT20/MB231 | MITO |
| >MI0000095\|hsa-mir-93&WithFlank&7\|-\|100093762\|100093853@17.38.22(+1A)&[MIMAT0000093&hsa-miR-93-5p&offsets\|0\|-1(+1A);m-29&7\|-\|100093815\|100093837&offsets\|0\|-1(+1A)]\|CAAAGTGCTGTTCGTGCAGGTAA | >MI0000095\|hsa-mir-93&WithFlank&7\|-\|100093762\|100093853 | 17 | 38 | 22(+1A) | CAAAGTGCTGTTCGTGCAGGTAA | 3499.6 | 3.8 | 0.8 | 4.6 | 3.7E-06 | 6.6E-06 | MB231/MB468 | MP |
| >hg38_MT_+_1621_3279_RNR2_16S_50ntFlanks@51.88.38\|GCTAAACCTAGCCCCAAACCCACTCCACCTTACTACCA | >hg38_MT_+_1621_3279_RNR2_16S_50ntFlanks | 51 | 88 | 38 | GCTAAACCTAGCCCCAAACCCACTCCACCTTACTACCA | 5245.2 | 4.3 | 0.9 | 4.6 | 3.7E-06 | 6.6E-06 | MB231/MB468 | MP |
| >hg38_21_+_8433172_8446622_RNA45SN1_45S_50ntFlanks@10771.10801.31\|CAGGTCTCCAAGGTGAACAGCCTCTGGCATG | >hg38_21_+_8433172_8446622_RNA45SN1_45S_50ntFlanks | 10771 | 10801 | 31 | CAGGTCTCCAAGGTGAACAGCCTCTGGCATG | 61.9 | -9.0 | 1.9 | -4.8 | 1.3E-06 | 6.6E-06 | BT20/MB468 | MP |
| >hg38_21_+_8433172_8446622_RNA45SN1_45S_50ntFlanks@12407.12446.40\|TTTTGATCCTTCGATGTCGGCTCTTCCTATCATTGTGAAG | >hg38_21_+_8433172_8446622_RNA45SN1_45S_50ntFlanks | 12407 | 12446 | 40 | TTTTGATCCTTCGATGTCGGCTCTTCCTATCATTGTGAAG | 629.8 | -3.7 | 0.8 | -4.6 | 4.4E-06 | 6.6E-06 | MB231/MB468 | MITO |
| >MI0000102\|hsa-mir-100&WithFlank&11\|-\|122152223\|122152314@22.39.18(+1A)&[MIMAT0000098&hsa-miR-100-5p&offsets\|+3\|-1(+1A);m-27&11\|-\|122152276\|122152296&offsets\|+3\|0(+1A)]\|CCGTAGATCCGAACTTGTA | >MI0000102\|hsa-mir-100&WithFlank&11\|-\|122152223\|122152314 | 22 | 39 | 18(+1A) | CCGTAGATCCGAACTTGTA | 62.6 | 9.2 | 2.0 | 4.6 | 3.7E-06 | 6.6E-06 | MB231/MB468 | MP |
| >hg38_21_+_8433172_8446622_RNA45SN1_45S_50ntFlanks@5485.5516.32\|AGCGCTGAGAAGACGGTCGAACTTGACTATCT | >hg38_21_+_8433172_8446622_RNA45SN1_45S_50ntFlanks | 5485 | 5516 | 32 | AGCGCTGAGAAGACGGTCGAACTTGACTATCT | 6999.2 | 6.9 | 1.4 | 5.0 | 4.9E-07 | 6.6E-06 | BT20/MB231 | NUC |
| >hg38_21_+_8433172_8446622_RNA45SN1_45S_50ntFlanks@12044.12067.24\|ACTGACCCGGTGAGGCGGGGGGGC | >hg38_21_+_8433172_8446622_RNA45SN1_45S_50ntFlanks | 12044 | 12067 | 24 | ACTGACCCGGTGAGGCGGGGGGGC | 64.4 | -9.1 | 1.9 | -4.8 | 1.3E-06 | 6.6E-06 | BT20/MB468 | MP |
| >hg38_21_+_8433172_8446622_RNA45SN1_45S_50ntFlanks@10186.10208.23\|TCCTCCCCTCCTCCCGCCCACGC | >hg38_21_+_8433172_8446622_RNA45SN1_45S_50ntFlanks | 10186 | 10208 | 23 | TCCTCCCCTCCTCCCGCCCACGC | 166.7 | -11.2 | 2.4 | -4.6 | 3.7E-06 | 6.6E-06 | MB231/MB468 | MP |
| >hg38_21_+_8433172_8446622_RNA45SN1_45S_50ntFlanks@11667.11685.19\|CTGCCCAGTGCTCTGAATG | >hg38_21_+_8433172_8446622_RNA45SN1_45S_50ntFlanks | 11667 | 11685 | 19 | CTGCCCAGTGCTCTGAATG | 688.6 | -5.7 | 1.2 | -4.6 | 3.7E-06 | 6.6E-06 | MB231/MB468 | MP |

| | | | | | | | | | | | | | |
|---|---|---|---|---|---|---|---|---|---|---|---|---|---|
| >hg38_21_+_8433172_8446622_RNA45SN1_45S_50ntFlanks@12417.12437.21\|TCGATGTCGGCTCTTCCTATC | >hg38_21_+_8433172_8446622_RNA45SN1_45S_50ntFlanks | 12417 | 12437 | 21 | TCGATGTCGGCTCTTCCTATC | 3915.8 | -2.4 | 0.5 | -4.6 | 4.4E-06 | 6.6E-06 | MB231/MB468 | NUC |
| >hg38_21_+_8433172_8446622_RNA45SN1_45S_50ntFlanks@6646.6673.28\|TCGTACGACTCTTAGCGGTGGATCACTC | >hg38_21_+_8433172_8446622_RNA45SN1_45S_50ntFlanks | 6646 | 6673 | 28 | TCGTACGACTCTTAGCGGTGGATCACTC | 1338.6 | 3.7 | 0.8 | 4.8 | 1.3E-06 | 6.6E-06 | BT20/MB468 | MP |
| >hg38_21_+_8433172_8446622_RNA45SN1_45S_50ntFlanks@4732.4752.21\|AGAACGAAAGTCGGAGGTTCG | >hg38_21_+_8433172_8446622_RNA45SN1_45S_50ntFlanks | 4732 | 4752 | 21 | AGAACGAAAGTCGGAGGTTCG | 298.4 | -7.0 | 1.5 | -4.6 | 4.7E-06 | 6.6E-06 | BT20/MB468 | NUC |
| >MI0003560\|hsa-mir-92b&WithFlank&1\|+\|155195171\|155195278@67.88.22(+1U)&[MIMAT0003218&hsa-miR-92b-3p&offsets\|0\|0(+1U);m-115&1\|+\|155195237\|155195258&offsets\|0\|0(+1U)]\|TATTGCACTCGTCCCGGCCTCCT | >MI0003560\|hsa-mir-92b&WithFlank&1\|+\|155195171\|155195278 | 67 | 88 | 22(+1U) | TATTGCACTCGTCCCGGCCTCCT | 407.0 | 6.9 | 1.5 | 4.6 | 4.5E-06 | 6.6E-06 | MB231/MB468 | MITO |
| >hg38_21_+_8433172_8446622_RNA45SN1_45S_50ntFlanks@12405.12424.20\|CTTTTTGATCCTTCGATGTC | >hg38_21_+_8433172_8446622_RNA45SN1_45S_50ntFlanks | 12405 | 12424 | 20 | CTTTTTGATCCTTCGATGTC | 994.1 | -7.4 | 1.5 | -4.8 | 1.3E-06 | 6.6E-06 | BT20/MB468 | MP |
| >hg38_21_+_8433172_8446622_RNA45SN1_45S_50ntFlanks@11710.11728.19\|CGCGGGTAAACGGCGGGAG | >hg38_21_+_8433172_8446622_RNA45SN1_45S_50ntFlanks | 11710 | 11728 | 19 | CGCGGGTAAACGGCGGGAG | 205.9 | -5.9 | 1.3 | -4.7 | 3.2E-06 | 6.7E-06 | BT20/MB468 | MITO |
| >hg38_MT_+_1621_3279_RNR2_16S_50ntFlanks@459.492.34\|CTTGTAAATTTAACTGTTAGTCCAAAGAGGAACA | >hg38_MT_+_1621_3279_RNR2_16S_50ntFlanks | 459 | 492 | 34 | CTTGTAAATTTAACTGTTAGTCCAAAGAGGAACA | 516.2 | 3.9 | 0.9 | 4.6 | 4.5E-06 | 6.7E-06 | MB231/MB468 | MITO |
| >hg38_21_+_8433172_8446622_RNA45SN1_45S_50ntFlanks@6648.6665.18\|GTACGACTCTTAGCGGTG | >hg38_21_+_8433172_8446622_RNA45SN1_45S_50ntFlanks | 6648 | 6665 | 18 | GTACGACTCTTAGCGGTG | 1554.8 | 4.0 | 0.8 | 4.8 | 1.3E-06 | 6.7E-06 | BT20/MB468 | MP |
| >hg38_21_+_8433172_8446622_RNA45SN1_45S_50ntFlanks@475.506.32\|CTTCGTGATCGATGTGGTGACGTCGTGCTCTC | >hg38_21_+_8433172_8446622_RNA45SN1_45S_50ntFlanks | 475 | 506 | 32 | CTTCGTGATCGATGTGGTGACGTCGTGCTCTC | 777.3 | 5.2 | 1.1 | 4.8 | 1.3E-06 | 6.7E-06 | BT20/MB468 | MP |
| >MI0000088\|hsa-mir-30a&WithFlank&6\|-\|71403545\|71403627@13.35.23&[MIMAT0000087&hsa-miR-30a-5p&offsets\|+1\|+2;m-15&6\|-\|71403595\|71403616&offsets\|+1\|+2]\|GTAAACATCCTCGACTGGAAGCT | >MI0000088\|hsa-mir-30a&WithFlank&6\|-\|71403545\|71403627 | 13 | 35 | 23 | GTAAACATCCTCGACTGGAAGCT | 216.4 | 4.0 | 0.9 | 4.6 | 4.7E-06 | 6.7E-06 | BT20/MB468 | NUC |
| >hg38_21_+_8433172_8446622_RNA45SN1_45S_50ntFlanks@474.497.24\|TCTTCGTGATCGATGTGGTGACGT | >hg38_21_+_8433172_8446622_RNA45SN1_45S_50ntFlanks | 474 | 497 | 24 | TCTTCGTGATCGATGTGGTGACGT | 598.1 | 7.0 | 1.5 | 4.6 | 4.5E-06 | 6.7E-06 | MB231/MB468 | MITO |
| >hg38_21_+_8433172_8446622_RNA45SN1_45S_50ntFlanks@11643.11662.20\|GCGGGTGTTGACGCGATGTG | >hg38_21_+_8433172_8446622_RNA45SN1_45S_50ntFlanks | 11643 | 11662 | 20 | GCGGGTGTTGACGCGATGTG | 226.9 | -3.5 | 0.8 | -4.6 | 3.7E-06 | 6.7E-06 | MB231/MB468 | MP |
| >MI0033423\|hsa-mir-10399&WithFlank&7\|-\|139044094\|139044163@45.63.19(+1U)&[MIMAT0041630&hsa-miR-10399-3p&offsets\|+1\|-1(+1U);TJU_CMC_MD2.ID03061.3p-miR&offsets\|+5\|+2(+1U);m-2657&7\|-\|139044103\|139044124&offsets\|+5\|+2(+1U)]\|TCTCGGACAAGCTGTAGGTT | >MI0033423\|hsa-mir-10399&WithFlank&7\|-\|139044094\|139044163 | 45 | 63 | 19(+1U) | TCTCGGACAAGCTGTAGGTT | 37.3 | 8.4 | 1.8 | 4.6 | 3.7E-06 | 6.7E-06 | MB231/MB468 | MP |
| >hg38_21_+_8433172_8446622_RNA45SN1_45S_50ntFlanks@10344.10361.18\|ATTCAAACGAGAACTTTG | >hg38_21_+_8433172_8446622_RNA45SN1_45S_50ntFlanks | 10344 | 10361 | 18 | ATTCAAACGAGAACTTTG | 594.9 | -5.3 | 1.1 | -4.6 | 3.7E-06 | 6.7E-06 | MB231/MB468 | MP |
| >hg38_21_+_8433172_8446622_RNA45SN1_45S_50ntFlanks@9562.9592.31\|TCCTGACGTGCAAATCGGTCGTCCGACCTGG | >hg38_21_+_8433172_8446622_RNA45SN1_45S_50ntFlanks | 9562 | 9592 | 31 | TCCTGACGTGCAAATCGGTCGTCCGACCTGG | 217.7 | -5.9 | 1.3 | -4.6 | 3.8E-06 | 6.7E-06 | BT20/MB468 | CYTO |
| >hg38_21_+_8433172_8446622_RNA45SN1_45S_50ntFlanks@51.72.22\|GCTGACACGCTGTCCTCTGGCG | >hg38_21_+_8433172_8446622_RNA45SN1_45S_50ntFlanks | 51 | 72 | 22 | GCTGACACGCTGTCCTCTGGCG | 153.4 | -6.4 | 1.4 | -4.6 | 3.7E-06 | 6.7E-06 | MB231/MB468 | MP |
| >trna13_ValCAC_19_-_4724647_4724719@1.29.29\|GTTTCCGTAGTGTAGCGGTTATCACATTC | >trna13_ValCAC_19_-_4724647_4724719 | 1 | 29 | 29 | GTTTCCGTAGTGTAGCGGTTATCACATTC | 648.0 | -6.1 | 1.3 | -4.6 | 3.7E-06 | 6.7E-06 | MB231/MB468 | MP |
| >hg38_21_+_8433172_8446622_RNA45SN1_45S_50ntFlanks@8195.8213.19\|CCGTGGACGGTGTGAGGCC | >hg38_21_+_8433172_8446622_RNA45SN1_45S_50ntFlanks | 8195 | 8213 | 19 | CCGTGGACGGTGTGAGGCC | 366.8 | -3.2 | 0.7 | -4.6 | 4.7E-06 | 6.7E-06 | BT20/MB468 | NUC |
| >MI0000683\|hsa-mir-181b-2&WithFlank&9\|+\|124693704\|124693804@23.46.24&[MIMAT0000257_1&hsa-miR-181b-5p&offsets\|+1\|+2;m-72&9\|+\|124693725\|124693747&offsets\|+1\|+2]\|ACATTCATTGCTGTCGGTGGGTTT | >MI0000683\|hsa-mir-181b-2&WithFlank&9\|+\|124693704\|124693804 | 23 | 46 | 24 | ACATTCATTGCTGTCGGTGGGTTT | 41.1 | 8.6 | 1.9 | 4.6 | 3.7E-06 | 6.7E-06 | MB231/MB468 | MP |
| >hg38_21_+_8433172_8446622_RNA45SN1_45S_50ntFlanks@6651.6689.39\|CGACTCTTAGCGGTGGATCACTCGGCTCGTGCGTCGATG | >hg38_21_+_8433172_8446622_RNA45SN1_45S_50ntFlanks | 6651 | 6689 | 39 | CGACTCTTAGCGGTGGATCACTCGGCTCGTGCGTCGATG | 29288.9 | -2.5 | 0.5 | -4.6 | 4.7E-06 | 6.7E-06 | BT20/MB468 | NUC |
| >trna111_HisGTG_1_-_147774845_147774916@1.34.34,trna118_HisGTG_1_-_145396881_145396952@1.34.34,trna16_HisGTG_1_+_146544773_146544844@1.34.34,trna1_HisGTG_15_+_45493349_45493420@1.34.34,trna21_HisGTG_1_+_147753471_147753542@1.34.34,trna33_HisGTG_6_+_27125906_27125977@1.34.34,trna7_HisGTG_9_-_14433938_14434009@1.34.34,trna8_HisGTG_15_-_45492611_45492682@1.34.34,trna9_HisGTG_15_-_45490804_45490875@1.34.34\|GCCGTGATCGTATAGTGGTTAGTACTCTGCGTTG | >trna111_HisGTG_1_-_147774845_147774916 | 1 | 34 | 34 | GCCGTGATCGTATAGTGGTTAGTACTCTGCGTTG | 960.3 | 4.3 | 0.9 | 4.6 | 4.5E-06 | 6.7E-06 | MB231/MB468 | MITO |
| >hg38_21_+_8433172_8446622_RNA45SN1_45S_50ntFlanks@10907.10931.25\|GGGCGCGCGCCGCGGCTGGACGAGG | >hg38_21_+_8433172_8446622_RNA45SN1_45S_50ntFlanks | 10907 | 10931 | 25 | GGGCGCGCGCCGCGGCTGGACGAGG | 126.0 | 10.3 | 2.0 | 5.1 | 3.6E-07 | 6.7E-06 | BT20/MB231 | MP |
| >hg38_21_+_8433172_8446622_RNA45SN1_45S_50ntFlanks@11206.11232.27\|CGGCGGCGACTCTGGACGCGAGCCGGG | >hg38_21_+_8433172_8446622_RNA45SN1_45S_50ntFlanks | 11206 | 11232 | 27 | CGGCGGCGACTCTGGACGCGAGCCGGG | 143.0 | 10.5 | 2.1 | 5.1 | 3.6E-07 | 6.7E-06 | BT20/MB231 | MP |
| >hg38_21_+_8433172_8446622_RNA45SN1_45S_50ntFlanks@9790.9820.31\|CTGGCGTGGAGCCGGGCGTGGAATGCGAGTG | >hg38_21_+_8433172_8446622_RNA45SN1_45S_50ntFlanks | 9790 | 9820 | 31 | CTGGCGTGGAGCCGGGCGTGGAATGCGAGTG | 215.2 | 11.1 | 2.2 | 5.1 | 3.6E-07 | 6.7E-06 | BT20/MB231 | MP |
| >MI0002470\|hsa-mir-486-1&WithFlank&8\|-\|41660435\|41660514@10.30.21;MI0023622\|hsa-mir-486-2&WithFlank&8\|+\|41660438\|41660513@7.27.21&[MIMAT0002177&hsa-miR-486-5p&offsets\|0\|-1;m-107&8\|-\|41660484\|41660505&offsets\|0\|-1];[MIMAT0002177_1&hsa-miR-486-5p&offsets\|0\|-1;TJU_CMC_MD2.ID03138.5p-miR&offsets\|0\|-3;m-106&8\|+\|41660444\|41660465&offsets\|0\|-1]\|TCCTGTACTGAGCTGCCCCGA | >MI0002470\|hsa-mir-486-1&WithFlank&8\|-\|41660435\|41660514 | 10 | 30 | 21 | TCCTGTACTGAGCTGCCCCGA | 94.4 | -10.0 | 2.0 | -5.1 | 3.6E-07 | 6.7E-06 | BT20/MB231 | MP |
| >MI0006349\|hsa-mir-1287&WithFlank&10\|-\|98395212\|98395313@22.42.21&[MIMAT0005878&hsa-miR-1287-5p&offsets\|0\|-1;m-244&10\|-\|98395272\|98395292&offsets\|0\|0]\|TGCTGGATCAGTGGTTCGAGT | >MI0006349\|hsa-mir-1287&WithFlank&10\|-\|98395212\|98395313 | 22 | 42 | 21 | TGCTGGATCAGTGGTTCGAGT | 95.3 | -10.0 | 2.0 | -5.1 | 3.6E-07 | 6.7E-06 | BT20/MB231 | MP |
| >hg38_21_+_8433172_8446622_RNA45SN1_45S_50ntFlanks@7970.8005.36\|CGAGACGCGACCTCAGATCAGACGTGGCGACCCGCT | >hg38_21_+_8433172_8446622_RNA45SN1_45S_50ntFlanks | 7970 | 8005 | 36 | CGAGACGCGACCTCAGATCAGACGTGGCGACCCGCT | 728.7 | 4.5 | 0.9 | 5.0 | 5.0E-07 | 6.7E-06 | BT20/MB231 | NUC |
| >hg38_21_+_8433172_8446622_RNA45SN1_45S_50ntFlanks@10788.10820.33\|CAGCCTCTGGCATGTTGGAACAATGTAGGTAAG | >hg38_21_+_8433172_8446622_RNA45SN1_45S_50ntFlanks | 10788 | 10820 | 33 | CAGCCTCTGGCATGTTGGAACAATGTAGGTAAG | 104.5 | -4.8 | 1.0 | -4.6 | 3.8E-06 | 6.7E-06 | MB231/MB468 | MP |
| >hg38_21_+_8433172_8446622_RNA45SN1_45S_50ntFlanks@12420.12443.24\|ATGTCGGCTCTTCCTATCATTGTG | >hg38_21_+_8433172_8446622_RNA45SN1_45S_50ntFlanks | 12420 | 12443 | 24 | ATGTCGGCTCTTCCTATCATTGTG | 3986.7 | -4.5 | 1.0 | -4.6 | 4.5E-06 | 6.7E-06 | MB231/MB468 | MITO |
| >MI0000102\|hsa-mir-100&WithFlank&11\|-\|122152223\|122152314@20.38.19(+1C)&[MIMAT0000098&hsa-miR-100-5p&offsets\|+1\|-2(+1C);m-27&11\|-\|122152276\|122152296&offsets\|+1\|-1(+1C)]\|ACCCGTAGATCCGAACTTGC | >MI0000102\|hsa-mir-100&WithFlank&11\|-\|122152223\|122152314 | 20 | 38 | 19(+1C) | ACCCGTAGATCCGAACTTGC | 44.5 | 8.7 | 1.9 | 4.6 | 3.8E-06 | 6.7E-06 | MB231/MB468 | MP |
| >hg38_21_+_8433172_8446622_RNA45SN1_45S_50ntFlanks@10351.10370.20\|CGAGAACTTTGAAGGCCGAA | >hg38_21_+_8433172_8446622_RNA45SN1_45S_50ntFlanks | 10351 | 10370 | 20 | CGAGAACTTTGAAGGCCGAA | 316.2 | -2.5 | 0.5 | -4.6 | 4.3E-06 | 6.8E-06 | MB231/MB468 | CYTO |
| >hg38_21_+_8433172_8446622_RNA45SN1_45S_50ntFlanks@8443.8467.25\|CAACCCGGCGGCGGGTCCGGCCGTG | >hg38_21_+_8433172_8446622_RNA45SN1_45S_50ntFlanks | 8443 | 8467 | 25 | CAACCCGGCGGCGGGTCCGGCCGTG | 48.2 | -8.7 | 1.8 | -4.8 | 1.3E-06 | 6.8E-06 | BT20/MB468 | MP |
| >trnaMT_PheGAA_MT_+_577_647@38.56.19\|AATGTTTAGACGGGCTCAC | >trnaMT_PheGAA_MT_+_577_647 | 38 | 56 | 19 | AATGTTTAGACGGGCTCAC | 459.6 | 6.5 | 1.3 | 4.8 | 1.3E-06 | 6.8E-06 | BT20/MB468 | MP |
| >hg38_MT_+_1621_3279_RNR2_16S_50ntFlanks@1370.1414.45\|ATGTTGGATCAGGACATCCCGATGGTGCAGCCGCTATTAAAGGTT | >hg38_MT_+_1621_3279_RNR2_16S_50ntFlanks | 1370 | 1414 | 45 | ATGTTGGATCAGGACATCCCGATGGTGCAGCCGCTATTAAAGGTT | 339.9 | 6.9 | 1.5 | 4.7 | 3.3E-06 | 6.8E-06 | BT20/MB468 | MITO |
| >trna10_AlaCGC_6_+_26553731_26553802@58.75.18,trna119_AlaCGC_6_-_28641613_28641684@58.75.18,trna13_AlaCGC_2_+_157257281_157257352@58.75.18,trna13_AlaTGC_12_-_125406301_125406372@58.75.18,trna66_AlaTGC_6_+_28611222_28611293@58.75.18,trna8_AlaTGC_12_+_125424512_125424583@58.75.18,trna8_AlaTGC_5_+_180633868_180633939@58.75.18\|TCCCCGGCATCTCCACCA | >trna10_AlaCGC_6_+_26553731_26553802 | 58 | 75 | 18 | TCCCCGGCATCTCCACCA | 888.1 | 4.3 | 0.9 | 4.6 | 4.5E-06 | 6.8E-06 | MB231/MB468 | NUC |
| >MI0000737\|hsa-mir-200a&WithFlank&1\|+\|1167857\|1167958@60.83.24&[MIMAT0000682&hsa-miR-200a-3p&offsets\|0\|+2;m-142&1\|+\|1167916\|1167938&offsets\|0\|+1]\|TAACACTGTCTGGTAACGATGTTC | >MI0000737\|hsa-mir-200a&WithFlank&1\|+\|1167857\|1167958 | 60 | 83 | 24 | TAACACTGTCTGGTAACGATGTTC | 252.3 | -6.9 | 1.5 | -4.6 | 3.8E-06 | 6.8E-06 | MB231/MB468 | MP |
| >hg38_21_+_8433172_8446622_RNA45SN1_45S_50ntFlanks@12994.13015.22\|CCTCGCTGCGATCTATTGAAAG | >hg38_21_+_8433172_8446622_RNA45SN1_45S_50ntFlanks | 12994 | 13015 | 22 | CCTCGCTGCGATCTATTGAAAG | 1145.8 | -3.3 | 0.7 | -4.6 | 4.5E-06 | 6.8E-06 | MB231/MB468 | NUC |
| >hg38_21_+_8433172_8446622_RNA45SN1_45S_50ntFlanks@12948.12988.41\|GACGACCTGCTTCTGGGTCGGGGTTTCGTACGTAGCAGAGC | >hg38_21_+_8433172_8446622_RNA45SN1_45S_50ntFlanks | 12948 | 12988 | 41 | GACGACCTGCTTCTGGGTCGGGGTTTCGTACGTAGCAGAGC | 1193.9 | -8.1 | 1.8 | -4.6 | 4.8E-06 | 6.8E-06 | BT20/MB468 | NUC |
| >hg38_21_+_8433172_8446622_RNA45SN1_45S_50ntFlanks@12992.13016.25\|TCCCTCGCTGCGATCTATTGAAAGT | >hg38_21_+_8433172_8446622_RNA45SN1_45S_50ntFlanks | 12992 | 13016 | 25 | TCCCTCGCTGCGATCTATTGAAAGT | 642.7 | -7.6 | 1.7 | -4.6 | 4.8E-06 | 6.8E-06 | BT20/MB468 | NUC |
| >hg38_21_+_8433172_8446622_RNA45SN1_45S_50ntFlanks@3854.3875.22\|ACTGTGGTAATTCTAGAGCTAA | >hg38_21_+_8433172_8446622_RNA45SN1_45S_50ntFlanks | 3854 | 3875 | 22 | ACTGTGGTAATTCTAGAGCTAA | 45.8 | -8.6 | 1.8 | -4.8 | 1.3E-06 | 6.8E-06 | BT20/MB468 | MP |
| >hg38_21_+_8433172_8446622_RNA45SN1_45S_50ntFlanks@10391.10432.42\|AACAGCAGTTGAACATGGGTCAGTCGGTCCTGAGAGATGGGC | >hg38_21_+_8433172_8446622_RNA45SN1_45S_50ntFlanks | 10391 | 10432 | 42 | AACAGCAGTTGAACATGGGTCAGTCGGTCCTGAGAGATGGGC | 227.0 | -6.9 | 1.5 | -4.6 | 4.5E-06 | 6.8E-06 | MB231/MB468 | NUC |
| >hg38_21_+_8433172_8446622_RNA45SN1_45S_50ntFlanks@12007.12042.36\|GCCGCCGGTGAAATACCACTACTCTGATCGTTTTTT | >hg38_21_+_8433172_8446622_RNA45SN1_45S_50ntFlanks | 12007 | 12042 | 36 | GCCGCCGGTGAAATACCACTACTCTGATCGTTTTTT | 147.1 | -6.6 | 1.4 | -4.6 | 3.8E-06 | 6.8E-06 | BT20/MB468 | CYTO |
| >MI0000298\|hsa-mir-221&WithFlank&X\|-\|45746151\|45746272@71.93.23&[MIMAT0000278&hsa-miR-221-3p&offsets\|0\|0;m-91&X\|-\|45746181\|45746202&offsets\|0\|+1]\|AGCTACATTGTCTGCTGGGTTTC | >MI0000298\|hsa-mir-221&WithFlank&X\|-\|45746151\|45746272 | 71 | 93 | 23 | AGCTACATTGTCTGCTGGGTTTC | 50489.6 | 3.7 | 0.8 | 4.6 | 3.8E-06 | 6.8E-06 | MB231/MB468 | MP |
| >MI0000076\|hsa-mir-20a&WithFlank&13\|+\|91351059\|91351141@14.36.23&[MIMAT0000075&hsa-miR-20a-5p&offsets\|0\|0;m-96&13\|+\|91351072\|91351094&offsets\|0\|0]\|TAAAGTGCTTATAGTGCAGGTAG | >MI0000076\|hsa-mir-20a&WithFlank&13\|+\|91351059\|91351141 | 14 | 36 | 23 | TAAAGTGCTTATAGTGCAGGTAG | 25230.2 | -4.6 | 0.9 | -5.1 | 4.3E-07 | 6.8E-06 | BT20/MB231 | MITO |
| >MI0000805\|hsa-mir-342&WithFlank&14\|+\|100109649\|100109759@67.90.24&[MIMAT0000753&hsa-miR-342-3p&offsets\|0\|+1;m-128&14\|+\|100109715\|100109737&offsets\|0\|+1]\|TCTCACACAGAAATCGCACCCGTC | >MI0000805\|hsa-mir-342&WithFlank&14\|+\|100109649\|100109759 | 67 | 90 | 24 | TCTCACACAGAAATCGCACCCGTC | 347.8 | 2.9 | 0.6 | 4.6 | 4.5E-06 | 6.8E-06 | MB231/MB468 | NUC |

| | | | | | | | | | | | | | |
|---|---|---|---|---|---|---|---|---|---|---|---|---|---|
| >hg38_21_+_8433172_8446622_RNA45SN1_45S_50ntFlanks@8359.8394.36\|AAGAACTTTGAAGAGAGAGTTCAAGAGGGCGTGAAA | >hg38_21_+_8433172_8446622_RNA45SN1_45S_50ntFlanks | 8359 | 8394 | 36 | AAGAACTTTGAAGAGAGAGTTCAAGAGGGCGTGAAA | 46.4 | 9.3 | 1.9 | 4.8 | 1.3E-06 | 6.8E-06 | BT20/MB468 | MP |
| >hg38_MT_+_598_1651_RNR1_12S_50ntFlanks@416.453.38\|AAATAGACTACGAAAGTGGCTTTAACATATCTGAACAC | >hg38_MT_+_598_1651_RNR1_12S_50ntFlanks | 416 | 453 | 38 | AAATAGACTACGAAAGTGGCTTTAACATATCTGAACAC | 37.9 | 8.5 | 1.8 | 4.6 | 3.8E-06 | 6.8E-06 | MB231/MB468 | MP |
| >am_trna10_ValCAC_5_-_180649395_180649467@1.33.33,trna12_ValAAC_5_-_180645270_180645342@1.33.33,trna132_ValAAC_6_-_27721179_27721251@1.33.33,trna136_ValAAC_6_-_27648885_27648957@1.33.33,trna139_ValAAC_6_-_27618707_27618779@1.33.33,trna18_ValCAC_5_-_180529253_180529325@1.33.33,trna2_ValAAC_3_+_169490018_169490090@1.33.33,trna2_ValCAC_5_+_180524070_180524142@1.33.33,trna4_ValAAC_5_+_180591154_180591226@1.33.33,trna5_ValAAC_5_+_180596610_180596682@1.33.33,trna6_ValCAC_5_+_180600650_180600722@1.33.33,trna85_ValCAC_1_-_161369490_161369562@1.33.33,trna90_ValCAC_1_-_149684088_149684161@1.33.33,trna98_ValCAC_1_-_149298555_149298627@1.33.33,trna9_ValCAC_6_+_26538282_26538354@1.33.33\|GTTTCCGTAGTGTAGTGGTTATCACGTTCGCCT | >am_trna10_ValCAC_5_-_180649395_180649467 | 1 | 33 | 33 | GTTTCCGTAGTGTAGTGGTTATCACGTTCGCCT | 988.7 | 8.7 | 1.9 | 4.6 | 3.8E-06 | 6.8E-06 | MB231/MB468 | MP |
| >hg38_21_+_8433172_8446622_RNA45SN1_45S_50ntFlanks@9545.9569.25\|GTGGAGGTCCGTAGCGGTCCTGACG | >hg38_21_+_8433172_8446622_RNA45SN1_45S_50ntFlanks | 9545 | 9569 | 25 | GTGGAGGTCCGTAGCGGTCCTGACG | 1231.2 | -3.3 | 0.7 | -4.6 | 4.3E-06 | 6.8E-06 | MB231/MB468 | CYTO |
| >hg38_21_+_8433172_8446622_RNA45SN1_45S_50ntFlanks@10866.10883.18\|CTAAGGGCTGGGTCGGTC | >hg38_21_+_8433172_8446622_RNA45SN1_45S_50ntFlanks | 10866 | 10883 | 18 | CTAAGGGCTGGGTCGGTC | 252.9 | -6.2 | 1.4 | -4.6 | 4.5E-06 | 6.8E-06 | MB231/MB468 | NUC |
| >hg38_21_+_8433172_8446622_RNA45SN1_45S_50ntFlanks@11849.11886.38\|CGGGCTTGGCGGAATCAGCGGGGAAAGAAGACCCTGTT | >hg38_21_+_8433172_8446622_RNA45SN1_45S_50ntFlanks | 11849 | 11886 | 38 | CGGGCTTGGCGGAATCAGCGGGGAAAGAAGACCCTGTT | 51.4 | 9.4 | 2.0 | 4.8 | 1.3E-06 | 6.8E-06 | BT20/MB468 | MP |
| >hg38_21_+_8433172_8446622_RNA45SN1_45S_50ntFlanks@6706.6732.27\|GCGAGAATTAATGTGAATTGCAGGACA | >hg38_21_+_8433172_8446622_RNA45SN1_45S_50ntFlanks | 6706 | 6732 | 27 | GCGAGAATTAATGTGAATTGCAGGACA | 212.0 | 7.4 | 1.5 | 4.8 | 1.3E-06 | 6.8E-06 | BT20/MB468 | MP |
| >hg38_21_+_8433172_8446622_RNA45SN1_45S_50ntFlanks@12444.12474.31\|AAGCAGAATTCACCAAGCGTTGGATTGTTCA | >hg38_21_+_8433172_8446622_RNA45SN1_45S_50ntFlanks | 12444 | 12474 | 31 | AAGCAGAATTCACCAAGCGTTGGATTGTTCA | 64.0 | -9.1 | 1.9 | -4.8 | 1.3E-06 | 6.8E-06 | BT20/MB468 | MP |
| >hg38_21_+_8433172_8446622_RNA45SN1_45S_50ntFlanks@5084.5108.25\|CGAACGAGACTCTGGCATGCTAACT | >hg38_21_+_8433172_8446622_RNA45SN1_45S_50ntFlanks | 5084 | 5108 | 25 | CGAACGAGACTCTGGCATGCTAACT | 404.8 | -7.5 | 1.6 | -4.6 | 3.8E-06 | 6.8E-06 | MB231/MB468 | MP |
| >hg38_21_+_8433172_8446622_RNA45SN1_45S_50ntFlanks@9792.9819.28\|GGCGTGGAGCCGGGCGTGGAATGCGAGT | >hg38_21_+_8433172_8446622_RNA45SN1_45S_50ntFlanks | 9792 | 9819 | 28 | GGCGTGGAGCCGGGCGTGGAATGCGAGT | 174.2 | 8.1 | 1.7 | 4.8 | 1.3E-06 | 6.8E-06 | BT20/MB468 | MP |
| >hg38_1_-_228634819_228635039_RNA5S12_5S_50ntFlanks@148.169.22\|GGAATACCGGGTGCTGTAGGCT | >hg38_1_-_228634819_228635039_RNA5S12_5S_50ntFlanks | 148 | 169 | 22 | GGAATACCGGGTGCTGTAGGCT | 766.8 | 3.7 | 0.7 | 5.1 | 4.3E-07 | 6.8E-06 | BT20/MB231 | MITO |
| >hg38_21_+_8433172_8446622_RNA45SN1_45S_50ntFlanks@12637.12661.25\|GCTACCATCTGTGGGATTATGACTG | >hg38_21_+_8433172_8446622_RNA45SN1_45S_50ntFlanks | 12637 | 12661 | 25 | GCTACCATCTGTGGGATTATGACTG | 358.2 | -2.8 | 0.6 | -4.6 | 4.5E-06 | 6.8E-06 | MB231/MB468 | NUC |
| >hg38_21_+_8433172_8446622_RNA45SN1_45S_50ntFlanks@12204.12221.18\|AAGGCGAGCTCAGGGAGG | >hg38_21_+_8433172_8446622_RNA45SN1_45S_50ntFlanks | 12204 | 12221 | 18 | AAGGCGAGCTCAGGGAGG | 302.5 | -4.2 | 0.9 | -4.6 | 3.8E-06 | 6.9E-06 | MB231/MB468 | MP |
| >hg38_21_+_8433172_8446622_RNA45SN1_45S_50ntFlanks@11208.11234.27\|GCGGCGACTCTGGACGCGAGCCGGGCC | >hg38_21_+_8433172_8446622_RNA45SN1_45S_50ntFlanks | 11208 | 11234 | 27 | GCGGCGACTCTGGACGCGAGCCGGGCC | 5826.0 | -7.4 | 1.6 | -4.6 | 4.8E-06 | 6.9E-06 | BT20/MB468 | NUC |
| >hg38_21_+_8433172_8446622_RNA45SN1_45S_50ntFlanks@7990.8012.23\|GACGTGGCGACCCGCTGAATTTA | >hg38_21_+_8433172_8446622_RNA45SN1_45S_50ntFlanks | 7990 | 8012 | 23 | GACGTGGCGACCCGCTGAATTTA | 335.0 | 3.3 | 0.7 | 4.6 | 4.3E-06 | 6.9E-06 | MB231/MB468 | CYTO |
| >hg38_21_+_8433172_8446622_RNA45SN1_45S_50ntFlanks@12974.12992.19\|CGTACGTAGCAGAGCAGCT | >hg38_21_+_8433172_8446622_RNA45SN1_45S_50ntFlanks | 12974 | 12992 | 19 | CGTACGTAGCAGAGCAGCT | 370.6 | -2.5 | 0.6 | -4.6 | 4.3E-06 | 6.9E-06 | MB231/MB468 | CYTO |
| >hg38_21_+_8433172_8446622_RNA45SN1_45S_50ntFlanks@5188.5206.19\|ATAACAGGTCTGTGATGCC | >hg38_21_+_8433172_8446622_RNA45SN1_45S_50ntFlanks | 5188 | 5206 | 19 | ATAACAGGTCTGTGATGCC | 242.8 | -7.3 | 1.6 | -4.6 | 3.8E-06 | 6.9E-06 | MB231/MB468 | MP |
| >hg38_21_+_8433172_8446622_RNA45SN1_45S_50ntFlanks@9029.9066.38\|TCTCCTCGTGGGGGGGCCGGGCCACCCCTCCCACGGCG | >hg38_21_+_8433172_8446622_RNA45SN1_45S_50ntFlanks | 9029 | 9066 | 38 | TCTCCTCGTGGGGGGGCCGGGCCACCCCTCCCACGGCG | 47.4 | -8.6 | 1.8 | -4.8 | 1.4E-06 | 6.9E-06 | BT20/MB468 | MP |
| >MI0003674\|hsa-mir-653&WithFlank&7\|-\|93482754\|93482861@22.42.21(+1U)&[MIMAT0003328&hsa-miR-653-5p&offsets\|+3\|+3(+1U);m-368&7\|-\|93482819\|93482840&offsets\|0\|-1(+1U)]\|TTGAAACAATCTCTACTGAACT | >MI0003674\|hsa-mir-653&WithFlank&7\|-\|93482754\|93482861 | 22 | 42 | 21(+1U) | TTGAAACAATCTCTACTGAACT | 35.1 | 8.4 | 1.8 | 4.6 | 3.8E-06 | 6.9E-06 | MB231/MB468 | MP |
| >MI0000269\|hsa-mir-181a-2&WithFlank&9\|+\|124692436\|124692557@83.103.21(+1U)&[MIMAT0004558&hsa-miR-181a-2-3p&offsets\|0\|-1(+1U);m-71&9\|+\|124692518\|124692539&offsets\|0\|-1(+1U)]\|ACCACTGACCGTTGACTGTACT | >MI0000269\|hsa-mir-181a-2&WithFlank&9\|+\|124692436\|124692557 | 83 | 103 | 21(+1U) | ACCACTGACCGTTGACTGTACT | 427.3 | 7.7 | 1.7 | 4.6 | 4.6E-06 | 6.9E-06 | MB231/MB468 | MITO |
| >hg38_21_+_8433172_8446622_RNA45SN1_45S_50ntFlanks@5427.5466.40\|GATGGTTTAGTGAGGCCCTCGGATCGGCCCCGCCGGGGTC | >hg38_21_+_8433172_8446622_RNA45SN1_45S_50ntFlanks | 5427 | 5466 | 40 | GATGGTTTAGTGAGGCCCTCGGATCGGCCCCGCCGGGGTC | 116.4 | -10.7 | 2.3 | -4.6 | 3.8E-06 | 6.9E-06 | MB231/MB468 | MP |
| >hg38_21_+_8433172_8446622_RNA45SN1_45S_50ntFlanks@10798.10818.21\|CATGTTGGAACAATGTAGGTA | >hg38_21_+_8433172_8446622_RNA45SN1_45S_50ntFlanks | 10798 | 10818 | 21 | CATGTTGGAACAATGTAGGTA | 1105.1 | -8.0 | 1.8 | -4.6 | 4.8E-06 | 6.9E-06 | BT20/MB468 | NUC |
| >hg38_21_+_8433172_8446622_RNA45SN1_45S_50ntFlanks@6650.6695.46\|ACGACTCTTAGCGGTGGATCACTCGGCTCGTGCGTCGATGAAGAAC | >hg38_21_+_8433172_8446622_RNA45SN1_45S_50ntFlanks | 6650 | 6695 | 46 | ACGACTCTTAGCGGTGGATCACTCGGCTCGTGCGTCGATGAAGAAC | 651.4 | 5.1 | 1.1 | 4.6 | 4.5E-06 | 6.9E-06 | MB231/MB468 | NUC |
| >hg38_1_-_228634819_228635039_RNA5S12_5S_50ntFlanks@151.170.20\|ATACCGGGTGCTGTAGGCTT | >hg38_1_-_228634819_228635039_RNA5S12_5S_50ntFlanks | 151 | 170 | 20 | ATACCGGGTGCTGTAGGCTT | 9885.4 | -3.4 | 0.7 | -4.6 | 3.8E-06 | 6.9E-06 | MB231/MB468 | MP |
| >hg38_21_+_8433172_8446622_RNA45SN1_45S_50ntFlanks@10845.10872.28\|ACTTCGGGATAAGGATTGGCTCTAAGGG | >hg38_21_+_8433172_8446622_RNA45SN1_45S_50ntFlanks | 10845 | 10872 | 28 | ACTTCGGGATAAGGATTGGCTCTAAGGG | 254.7 | -6.3 | 1.4 | -4.6 | 4.5E-06 | 6.9E-06 | MB231/MB468 | NUC |
| >hg38_21_+_8433172_8446622_RNA45SN1_45S_50ntFlanks@7991.8011.21\|ACGTGGCGACCCGCTGAATTT | >hg38_21_+_8433172_8446622_RNA45SN1_45S_50ntFlanks | 7991 | 8011 | 21 | ACGTGGCGACCCGCTGAATTT | 2745.2 | -3.8 | 0.8 | -4.6 | 3.8E-06 | 6.9E-06 | MB231/MB468 | MP |
| >hg38_21_+_8433172_8446622_RNA45SN1_45S_50ntFlanks@10798.10820.23\|CATGTTGGAACAATGTAGGTAAG | >hg38_21_+_8433172_8446622_RNA45SN1_45S_50ntFlanks | 10798 | 10820 | 23 | CATGTTGGAACAATGTAGGTAAG | 113.9 | -6.4 | 1.4 | -4.6 | 3.8E-06 | 6.9E-06 | MB231/MB468 | MP |
| >MI0000272\|hsa-mir-182&WithFlank&7\|-\|129770377\|129770498@29.53.25(+1A)&[MIMAT0000259&hsa-miR-182-5p&offsets\|0\|+1(+1A);m-22&7\|-\|129770449\|129770470&offsets\|0\|+3(+1A)]\|TTTGGCAATGGTAGAACTCACACTGA | >MI0000272\|hsa-mir-182&WithFlank&7\|-\|129770377\|129770498 | 29 | 53 | 25(+1A) | TTTGGCAATGGTAGAACTCACACTGA | 1351.5 | 4.0 | 0.8 | 5.1 | 3.8E-07 | 6.9E-06 | BT20/MB231 | CYTO |
| >hg38_21_+_8433172_8446622_RNA45SN1_45S_50ntFlanks@3840.3865.26\|CTCCTACTTGGATAACTGTGGTAATT | >hg38_21_+_8433172_8446622_RNA45SN1_45S_50ntFlanks | 3840 | 3865 | 26 | CTCCTACTTGGATAACTGTGGTAATT | 276.9 | 3.0 | 0.7 | 4.6 | 4.4E-06 | 6.9E-06 | MB231/MB468 | CYTO |
| >hg38_21_+_8433172_8446622_RNA45SN1_45S_50ntFlanks@6652.6692.41\|GACTCTTAGCGGTGGATCACTCGGCTCGTGCGTCGATGAAG | >hg38_21_+_8433172_8446622_RNA45SN1_45S_50ntFlanks | 6652 | 6692 | 41 | GACTCTTAGCGGTGGATCACTCGGCTCGTGCGTCGATGAAG | 1505.9 | 4.8 | 1.0 | 4.6 | 3.9E-06 | 6.9E-06 | BT20/MB468 | CYTO |
| >hg38_21_+_8433172_8446622_RNA45SN1_45S_50ntFlanks@10843.10868.26\|TAACTTCGGGATAAGGATTGGCTCTA | >hg38_21_+_8433172_8446622_RNA45SN1_45S_50ntFlanks | 10843 | 10868 | 26 | TAACTTCGGGATAAGGATTGGCTCTA | 950.8 | -5.8 | 1.3 | -4.6 | 3.9E-06 | 6.9E-06 | BT20/MB468 | CYTO |
| >hg38_21_+_8433172_8446622_RNA45SN1_45S_50ntFlanks@4102.4121.20\|ACCACGGGTGACGGGGAATC | >hg38_21_+_8433172_8446622_RNA45SN1_45S_50ntFlanks | 4102 | 4121 | 20 | ACCACGGGTGACGGGGAATC | 1859.6 | -4.3 | 0.9 | -4.8 | 1.4E-06 | 6.9E-06 | BT20/MB468 | MP |
| >hg38_21_+_8433172_8446622_RNA45SN1_45S_50ntFlanks@11918.11956.39\|ACATGAGAGGTGTAGAATAAGTGGGAGGCCCCCGGCGCC | >hg38_21_+_8433172_8446622_RNA45SN1_45S_50ntFlanks | 11918 | 11956 | 39 | ACATGAGAGGTGTAGAATAAGTGGGAGGCCCCCGGCGCC | 145.7 | -5.9 | 1.3 | -4.6 | 3.9E-06 | 6.9E-06 | BT20/MB468 | CYTO |
| >hg38_21_+_8433172_8446622_RNA45SN1_45S_50ntFlanks@9533.9571.39\|GAGGAAACTCTGGTGGAGGTCCGTAGCGGTCCTGACGTG | >hg38_21_+_8433172_8446622_RNA45SN1_45S_50ntFlanks | 9533 | 9571 | 39 | GAGGAAACTCTGGTGGAGGTCCGTAGCGGTCCTGACGTG | 1988.4 | -4.7 | 1.0 | -4.6 | 4.6E-06 | 6.9E-06 | MB231/MB468 | MITO |
| >hg38_21_+_8433172_8446622_RNA45SN1_45S_50ntFlanks@7976.8010.35\|GCGACCTCAGATCAGACGTGGCGACCCGCTGAATT | >hg38_21_+_8433172_8446622_RNA45SN1_45S_50ntFlanks | 7976 | 8010 | 35 | GCGACCTCAGATCAGACGTGGCGACCCGCTGAATT | 1074.4 | 9.8 | 2.0 | 4.8 | 1.4E-06 | 6.9E-06 | BT20/MB468 | MP |
| >hg38_21_+_8433172_8446622_RNA45SN1_45S_50ntFlanks@6771.6799.29\|GGTTCCTCCCGGGGCTACGCCTGTCTGAG | >hg38_21_+_8433172_8446622_RNA45SN1_45S_50ntFlanks | 6771 | 6799 | 29 | GGTTCCTCCCGGGGCTACGCCTGTCTGAG | 435.7 | -4.9 | 1.1 | -4.6 | 3.3E-06 | 6.9E-06 | BT20/MB468 | MITO |
| >hg38_21_+_8433172_8446622_RNA45SN1_45S_50ntFlanks@475.503.29\|CTTCGTGATCGATGTGGTGACGTCGTGCT | >hg38_21_+_8433172_8446622_RNA45SN1_45S_50ntFlanks | 475 | 503 | 29 | CTTCGTGATCGATGTGGTGACGTCGTGCT | 7540.2 | 3.6 | 0.8 | 4.6 | 4.6E-06 | 6.9E-06 | MB231/MB468 | NUC |
| >am_MI0000748\|hsa-mir-130b&WithFlank&22\|+\|21653298\|21653391@19.39.21&[MIMAT0004680&hsa-miR-130b-5p&offsets\|0\|0;m-234&22\|+\|21653316\|21653337&offsets\|0\|-1]\|ACTCTTTCCCTGTTGCACTAC | >am_MI0000748\|hsa-mir-130b&WithFlank&22\|+\|21653298\|21653391 | 19 | 39 | 21 | ACTCTTTCCCTGTTGCACTAC | 281.0 | 5.5 | 1.2 | 4.6 | 4.6E-06 | 6.9E-06 | MB231/MB468 | MITO |
| >hg38_21_+_8433172_8446622_RNA45SN1_45S_50ntFlanks@4680.4720.41\|GCGCAAGACGGACCAGAGCGAAAGCATTTGCCAAGAATGTT | >hg38_21_+_8433172_8446622_RNA45SN1_45S_50ntFlanks | 4680 | 4720 | 41 | GCGCAAGACGGACCAGAGCGAAAGCATTTGCCAAGAATGTT | 43.6 | 9.2 | 1.9 | 4.8 | 1.4E-06 | 6.9E-06 | BT20/MB468 | MP |
| >trna124_ArgTCG_6_-_28510891_28510963@31.53.23,trna19_ArgTCG_17_+_73031208_73031280@31.53.23,trna3_ArgTCG_6_+_26299905_26299977@31.53.23,trna4_ArgTCG_6_+_26323046_26323118@31.53.23\|ACTTCGGATCAGAAGATTGAGGG | >trna124_ArgTCG_6_-_28510891_28510963 | 31 | 53 | 23 | ACTTCGGATCAGAAGATTGAGGG | 81.6 | -9.4 | 2.0 | -4.8 | 1.4E-06 | 6.9E-06 | BT20/MB468 | MP |
| >hg38_21_+_8433172_8446622_RNA45SN1_45S_50ntFlanks@10402.10428.27\|AACATGGGTCAGTCGGTCCTGAGAGAT | >hg38_21_+_8433172_8446622_RNA45SN1_45S_50ntFlanks | 10402 | 10428 | 27 | AACATGGGTCAGTCGGTCCTGAGAGAT | 673.1 | -7.6 | 1.7 | -4.6 | 4.9E-06 | 6.9E-06 | BT20/MB468 | NUC |
| >hg38_21_+_8433172_8446622_RNA45SN1_45S_50ntFlanks@10350.10384.35\|ACGAGAACTTTGAAGGCCGAAGTGGAGAAGGGTTC | >hg38_21_+_8433172_8446622_RNA45SN1_45S_50ntFlanks | 10350 | 10384 | 35 | ACGAGAACTTTGAAGGCCGAAGTGGAGAAGGGTTC | 2939.4 | -5.4 | 1.1 | -4.8 | 1.4E-06 | 6.9E-06 | BT20/MB468 | MP |
| >hg38_21_+_8433172_8446622_RNA45SN1_45S_50ntFlanks@8423.8459.37\|CGCAGTCCGCCCGGAGGATTCAACCCGGCGGCGGGTC | >hg38_21_+_8433172_8446622_RNA45SN1_45S_50ntFlanks | 8423 | 8459 | 37 | CGCAGTCCGCCCGGAGGATTCAACCCGGCGGCGGGTC | 68.8 | -9.2 | 1.9 | -4.8 | 1.4E-06 | 6.9E-06 | BT20/MB468 | MP |
| >hg38_21_+_8433172_8446622_RNA45SN1_45S_50ntFlanks@3782.3801.20\|CAGTGAAACTGCGAATGGCT | >hg38_21_+_8433172_8446622_RNA45SN1_45S_50ntFlanks | 3782 | 3801 | 20 | CAGTGAAACTGCGAATGGCT | 2234.0 | -6.1 | 1.3 | -4.6 | 3.3E-06 | 6.9E-06 | BT20/MB468 | MITO |
| >hg38_21_+_8433172_8446622_RNA45SN1_45S_50ntFlanks@9339.9377.39\|CGCACGAAAGCCGCCGTGGCGCAATGAAGGTGAAGGCCG | >hg38_21_+_8433172_8446622_RNA45SN1_45S_50ntFlanks | 9339 | 9377 | 39 | CGCACGAAAGCCGCCGTGGCGCAATGAAGGTGAAGGCCG | 340.8 | -6.8 | 1.5 | -4.6 | 3.4E-06 | 6.9E-06 | BT20/MB468 | MITO |

| | | | | | | | | | | | | | |
|---|---|---|---|---|---|---|---|---|---|---|---|---|---|
| >hg38_21_+_8433172_8446622_RNA45SN1_45S_50ntFlanks@4128.4159.32\|CGATTCCGGAGAGGGAGCCTGAGAAACGGCTA | >hg38_21_+_8433172_8446622_RNA45SN1_45S_50ntFlanks | 4128 | 4159 | 32 | CGATTCCGGAGAGGGAGCCTGAGAAACGGCTA | 60.7 | -9.0 | 1.9 | -4.8 | 1.4E-06 | 6.9E-06 | BT20/MB468 | MP |
| >hg38_21_+_8433172_8446622_RNA45SN1_45S_50ntFlanks@6702.6726.25\|AGCTGCGAGAATTAATGTGAATTGC | >hg38_21_+_8433172_8446622_RNA45SN1_45S_50ntFlanks | 6702 | 6726 | 25 | AGCTGCGAGAATTAATGTGAATTGC | 219.4 | 5.3 | 1.2 | 4.6 | 4.6E-06 | 6.9E-06 | MB231/MB468 | NUC |
| >hg38_21_+_8433172_8446622_RNA45SN1_45S_50ntFlanks@11579.11602.24\|AGAACTGGTGCGGACCAGGGGAAT | >hg38_21_+_8433172_8446622_RNA45SN1_45S_50ntFlanks | 11579 | 11602 | 24 | AGAACTGGTGCGGACCAGGGGAAT | 253.5 | -2.9 | 0.6 | -4.6 | 3.4E-06 | 6.9E-06 | BT20/MB468 | MITO |
| >hg38_21_+_8433172_8446622_RNA45SN1_45S_50ntFlanks@13002.13028.27\|CGATCTATTGAAAGTCAGCCCTCGACA | >hg38_21_+_8433172_8446622_RNA45SN1_45S_50ntFlanks | 13002 | 13028 | 27 | CGATCTATTGAAAGTCAGCCCTCGACA | 6004.5 | -7.1 | 1.5 | -4.6 | 4.4E-06 | 7.0E-06 | MB231/MB468 | CYTO |
| >hg38_21_+_8433172_8446622_RNA45SN1_45S_50ntFlanks@10343.10361.19\|TATTCAAACGAGAACTTTG | >hg38_21_+_8433172_8446622_RNA45SN1_45S_50ntFlanks | 10343 | 10361 | 19 | TATTCAAACGAGAACTTTG | 286.9 | -5.3 | 1.2 | -4.6 | 4.7E-06 | 7.0E-06 | MB231/MB468 | MITO |
| >hg38_MT_+_598_1651_RNR1_12S_50ntFlanks@972.1004.33\|GGTAAGTGTACTGGAAAGTGCACTTGGACGAAC | >hg38_MT_+_598_1651_RNR1_12S_50ntFlanks | 972 | 1004 | 33 | GGTAAGTGTACTGGAAAGTGCACTTGGACGAAC | 600.8 | 6.9 | 1.5 | 4.6 | 4.7E-06 | 7.0E-06 | MB231/MB468 | MITO |
| >hg38_21_+_8433172_8446622_RNA45SN1_45S_50ntFlanks@10351.10372.22\|CGAGAACTTTGAAGGCCGAAGT | >hg38_21_+_8433172_8446622_RNA45SN1_45S_50ntFlanks | 10351 | 10372 | 22 | CGAGAACTTTGAAGGCCGAAGT | 354.5 | -2.0 | 0.4 | -4.6 | 3.9E-06 | 7.0E-06 | BT20/MB468 | CYTO |
| >hg38_21_+_8433172_8446622_RNA45SN1_45S_50ntFlanks@10873.10891.19\|CTGGGTCGGTCGGGCTGGG | >hg38_21_+_8433172_8446622_RNA45SN1_45S_50ntFlanks | 10873 | 10891 | 19 | CTGGGTCGGTCGGGCTGGG | 1392.8 | -8.2 | 1.8 | -4.6 | 4.9E-06 | 7.0E-06 | BT20/MB468 | NUC |
| >hg38_21_+_8433172_8446622_RNA45SN1_45S_50ntFlanks@7980.8009.30\|CCTCAGATCAGACGTGGCGACCCGCTGAAT | >hg38_21_+_8433172_8446622_RNA45SN1_45S_50ntFlanks | 7980 | 8009 | 30 | CCTCAGATCAGACGTGGCGACCCGCTGAAT | 498.9 | -2.5 | 0.5 | -4.6 | 4.9E-06 | 7.0E-06 | BT20/MB468 | NUC |
| >hg38_21_+_8433172_8446622_RNA45SN1_45S_50ntFlanks@10841.10868.28\|CGTAACTTCGGGATAAGGATTGGCTCTA | >hg38_21_+_8433172_8446622_RNA45SN1_45S_50ntFlanks | 10841 | 10868 | 28 | CGTAACTTCGGGATAAGGATTGGCTCTA | 1059.2 | -2.8 | 0.6 | -4.6 | 3.4E-06 | 7.0E-06 | BT20/MB468 | MITO |
| >hg38_21_+_8433172_8446622_RNA45SN1_45S_50ntFlanks@12423.12457.35\|TCGGCTCTTCCTATCATTGTGAAGCAGAATTCACC | >hg38_21_+_8433172_8446622_RNA45SN1_45S_50ntFlanks | 12423 | 12457 | 35 | TCGGCTCTTCCTATCATTGTGAAGCAGAATTCACC | 261.6 | 3.6 | 0.8 | 4.6 | 4.6E-06 | 7.0E-06 | MB231/MB468 | NUC |
| >hg38_21_+_8433172_8446622_RNA45SN1_45S_50ntFlanks@8356.8376.21\|GAAAAGAACTTTGAAGAGAGA | >hg38_21_+_8433172_8446622_RNA45SN1_45S_50ntFlanks | 8356 | 8376 | 21 | GAAAAGAACTTTGAAGAGAGA | 487.2 | -5.6 | 1.2 | -4.6 | 3.9E-06 | 7.0E-06 | MB231/MB468 | MP |
| >hg38_21_+_8433172_8446622_RNA45SN1_45S_50ntFlanks@10344.10362.19\|ATTCAAACGAGAACTTTGA | >hg38_21_+_8433172_8446622_RNA45SN1_45S_50ntFlanks | 10344 | 10362 | 19 | ATTCAAACGAGAACTTTGA | 160.0 | 4.1 | 0.8 | 4.8 | 1.4E-06 | 7.0E-06 | BT20/MB468 | MP |
| >am_trna119_LysCTT_1_-_145395522_145395594@8.36.29,trna11_LysCTT_5_-_180648979_180649051@8.36.29,trna128_LysCTT_1_-_55423542_55423614@8.36.29,trna13_LysCTT_6_+_26556774_26556846@8.36.29,trna6_LysCTT_19_-_52425396_52425468@8.36.29,trna7_LysCTT_16_+_3225692_3225764@8.36.29,trna9_LysCTT_5_+_180634755_180634827@8.36.29\|TAGCTCAGTCGGTAGAGCATGAGACTCTT | >am_trna119_LysCTT_1_-_145395522_145395594 | 8 | 36 | 29 | TAGCTCAGTCGGTAGAGCATGAGACTCTT | 39.0 | 8.5 | 1.8 | 4.6 | 3.9E-06 | 7.0E-06 | MB231/MB468 | MP |
| >hg38_21_+_8433172_8446622_RNA45SN1_45S_50ntFlanks@6652.6685.34\|GACTCTTAGCGGTGGATCACTCGGCTCGTGCGTC | >hg38_21_+_8433172_8446622_RNA45SN1_45S_50ntFlanks | 6652 | 6685 | 34 | GACTCTTAGCGGTGGATCACTCGGCTCGTGCGTC | 468.0 | 2.5 | 0.5 | 4.6 | 3.4E-06 | 7.0E-06 | BT20/MB468 | MITO |
| >hg38_21_+_8433172_8446622_RNA45SN1_45S_50ntFlanks@12149.12181.33\|AGGTGGGGAGTTTGACTGGGGCGGTACACCTGT | >hg38_21_+_8433172_8446622_RNA45SN1_45S_50ntFlanks | 12149 | 12181 | 33 | AGGTGGGGAGTTTGACTGGGGCGGTACACCTGT | 53.4 | -8.8 | 1.8 | -4.8 | 1.4E-06 | 7.0E-06 | BT20/MB468 | MP |
| >hg38_MT_+_598_1651_RNR1_12S_50ntFlanks@431.469.39\|GTGGCTTTAACATATCTGAACACACAATAGCTAAGACCC | >hg38_MT_+_598_1651_RNR1_12S_50ntFlanks | 431 | 469 | 39 | GTGGCTTTAACATATCTGAACACACAATAGCTAAGACCC | 38.3 | 8.5 | 1.8 | 4.6 | 3.9E-06 | 7.0E-06 | MB231/MB468 | MP |
| >hg38_21_+_8433172_8446622_RNA45SN1_45S_50ntFlanks@11572.11590.19\|CCGACTTAGAACTGGTGCG | >hg38_21_+_8433172_8446622_RNA45SN1_45S_50ntFlanks | 11572 | 11590 | 19 | CCGACTTAGAACTGGTGCG | 3515.0 | -4.8 | 1.0 | -4.8 | 1.4E-06 | 7.0E-06 | BT20/MB468 | MP |
| >hg38_21_+_8433172_8446622_RNA45SN1_45S_50ntFlanks@8021.8046.26\|AGTCAGCGGAGGAGAAGAAACTAACC | >hg38_21_+_8433172_8446622_RNA45SN1_45S_50ntFlanks | 8021 | 8046 | 26 | AGTCAGCGGAGGAGAAGAAACTAACC | 172.0 | -4.1 | 0.8 | -5.1 | 3.9E-07 | 7.0E-06 | BT20/MB231 | CYTO |
| >hg38_21_+_8433172_8446622_RNA45SN1_45S_50ntFlanks@11604.11629.26\|CGACTGTTTAATTAAAACAAAGCATC | >hg38_21_+_8433172_8446622_RNA45SN1_45S_50ntFlanks | 11604 | 11629 | 26 | CGACTGTTTAATTAAAACAAAGCATC | 141.5 | 10.5 | 2.1 | 5.1 | 3.8E-07 | 7.0E-06 | BT20/MB231 | MP |
| >hg38_21_+_8433172_8446622_RNA45SN1_45S_50ntFlanks@12456.12479.24\|CCAAGCGTTGGATTGTTCACCCAC | >hg38_21_+_8433172_8446622_RNA45SN1_45S_50ntFlanks | 12456 | 12479 | 24 | CCAAGCGTTGGATTGTTCACCCAC | 381.8 | -4.7 | 1.0 | -4.6 | 3.4E-06 | 7.0E-06 | BT20/MB468 | MITO |
| >MI0003776\|hsa-mir-320b-1&WithFlank&1\|+\|116671740\|116671823@49.68.20;MI0003839\|hsa-mir-320b-2&WithFlank&1\|-\|224257034\|224257116@48.67.20&[MIMAT0005792&hsa-miR-320b&offsets\|+1\|-1;m-183&1\|+\|116671787\|116671806&offsets\|+1\|+1];[MIMAT0005792_1&hsa-miR-320b&offsets\|+1\|-1;m-183&1\|-\|224257051\|224257070&offsets\|+1\|+1]\|AAAGCTGGGTTGAGAGGGCA | >MI0003776\|hsa-mir-320b-1&WithFlank&1\|+\|116671740\|116671823 | 49 | 68 | 20 | AAAGCTGGGTTGAGAGGGCA | 235.4 | 4.3 | 0.8 | 5.1 | 4.0E-07 | 7.0E-06 | BT20/MB231 | CYTO |
| >hg38_21_+_8433172_8446622_RNA45SN1_45S_50ntFlanks@7985.8011.27\|GATCAGACGTGGCGACCCGCTGAATTT | >hg38_21_+_8433172_8446622_RNA45SN1_45S_50ntFlanks | 7985 | 8011 | 27 | GATCAGACGTGGCGACCCGCTGAATTT | 209.7 | -6.6 | 1.4 | -4.6 | 3.9E-06 | 7.0E-06 | MB231/MB468 | MP |
| >MI0000471\|hsa-mir-126&WithFlank&9\|+\|136670596\|136670692@58.78.21(+1A)&[MIMAT0000445&hsa-miR-126-3p&offsets\|0\|-1(+1A);m-40&9\|+\|136670653\|136670674&offsets\|0\|-1(+1A)]\|TCGTACCGTGAGTAATAATGCA | >MI0000471\|hsa-mir-126&WithFlank&9\|+\|136670596\|136670692 | 58 | 78 | 21(+1A) | TCGTACCGTGAGTAATAATGCA | 557.5 | 7.8 | 1.6 | 4.8 | 1.4E-06 | 7.0E-06 | BT20/MB468 | MP |
| >hg38_21_+_8433172_8446622_RNA45SN1_45S_50ntFlanks@6661.6692.32\|CGGTGGATCACTCGGCTCGTGCGTCGATGAAG | >hg38_21_+_8433172_8446622_RNA45SN1_45S_50ntFlanks | 6661 | 6692 | 32 | CGGTGGATCACTCGGCTCGTGCGTCGATGAAG | 480.2 | 4.5 | 1.0 | 4.6 | 4.0E-06 | 7.1E-06 | BT20/MB468 | CYTO |
| >hg38_21_+_8433172_8446622_RNA45SN1_45S_50ntFlanks@9926.9946.21\|GTTGATATAGACAGCAGGACG | >hg38_21_+_8433172_8446622_RNA45SN1_45S_50ntFlanks | 9926 | 9946 | 21 | GTTGATATAGACAGCAGGACG | 104.0 | 10.0 | 2.0 | 5.1 | 3.8E-07 | 7.1E-06 | BT20/MB231 | MP |
| >hg38_21_+_8433172_8446622_RNA45SN1_45S_50ntFlanks@6742.6770.29\|TCGACACTTCGAACGCACTTGCGGCCCCG | >hg38_21_+_8433172_8446622_RNA45SN1_45S_50ntFlanks | 6742 | 6770 | 29 | TCGACACTTCGAACGCACTTGCGGCCCCG | 484.9 | -6.4 | 1.4 | -4.6 | 4.0E-06 | 7.1E-06 | BT20/MB468 | CYTO |
| >hg38_21_+_8433172_8446622_RNA45SN1_45S_50ntFlanks@4922.4941.20\|CCAGGAGTGGAGCCTGCGGC | >hg38_21_+_8433172_8446622_RNA45SN1_45S_50ntFlanks | 4922 | 4941 | 20 | CCAGGAGTGGAGCCTGCGGC | 643.9 | -5.4 | 1.2 | -4.6 | 4.8E-06 | 7.1E-06 | MB231/MB468 | MITO |
| >hg38_21_+_8433172_8446622_RNA45SN1_45S_50ntFlanks@10396.10435.40\|CAGTTGAACATGGGTCAGTCGGTCCTGAGAGATGGGCGAG | >hg38_21_+_8433172_8446622_RNA45SN1_45S_50ntFlanks | 10396 | 10435 | 40 | CAGTTGAACATGGGTCAGTCGGTCCTGAGAGATGGGCGAG | 175.3 | -4.1 | 0.9 | -4.6 | 4.5E-06 | 7.1E-06 | MB231/MB468 | CYTO |
| >MI0000815\|hsa-mir-339&WithFlank&7\|-\|1022927\|1023032@56.77.22(+1U)&[MIMAT0004702&hsa-miR-339-3p&offsets\|0\|-1(+1U);m-235&7\|-\|1022957\|1022977&offsets\|0\|+1(+1U)]\|TGAGCGCCTCGACGACAGAGCCT | >MI0000815\|hsa-mir-339&WithFlank&7\|-\|1022927\|1023032 | 56 | 77 | 22(+1U) | TGAGCGCCTCGACGACAGAGCCT | 41.7 | 8.6 | 1.9 | 4.6 | 4.0E-06 | 7.1E-06 | MB231/MB468 | MP |
| >MI0000100\|hsa-mir-98&WithFlank&X\|-\|53556217\|53556347@28.49.22&[MIMAT0000096&hsa-miR-98-5p&offsets\|0\|0;m-143&X\|-\|53556299\|53556320&offsets\|0\|0]\|TGAGGTAGTAAGTTGTATTGTT | >MI0000100\|hsa-mir-98&WithFlank&X\|-\|53556217\|53556347 | 28 | 49 | 22 | TGAGGTAGTAAGTTGTATTGTT | 2654.8 | 3.9 | 0.9 | 4.6 | 4.8E-06 | 7.1E-06 | MB231/MB468 | MITO |
| >am_trnaMT_IleGAT_MT_+_4263_4331@1.32.32\|AGAAATATGTCTGATAAAAGAGTTACTTTGAT | >am_trnaMT_IleGAT_MT_+_4263_4331 | 1 | 32 | 32 | AGAAATATGTCTGATAAAAGAGTTACTTTGAT | 86.7 | -9.8 | 1.9 | -5.1 | 3.8E-07 | 7.1E-06 | BT20/MB231 | MP |
| >MI0000084\|hsa-mir-26b&WithFlank&2\|+\|218402640\|218402728@18.38.21&[MIMAT0000083&hsa-miR-26b-5p&offsets\|0\|0;m-53&2\|+\|218402657\|218402678&offsets\|0\|-1]\|TTCAAGTAATTCAGGATAGGT | >MI0000084\|hsa-mir-26b&WithFlank&2\|+\|218402640\|218402728 | 18 | 38 | 21 | TTCAAGTAATTCAGGATAGGT | 713.1 | 4.5 | 1.0 | 4.6 | 4.8E-06 | 7.1E-06 | MB231/MB468 | MITO |
| >hg38_21_+_8433172_8446622_RNA45SN1_45S_50ntFlanks@10877.10902.26\|GTCGGTCGGGCTGGGGCGCGAAGCGG | >hg38_21_+_8433172_8446622_RNA45SN1_45S_50ntFlanks | 10877 | 10902 | 26 | GTCGGTCGGGCTGGGGCGCGAAGCGG | 504.3 | -7.5 | 1.6 | -4.6 | 5.0E-06 | 7.1E-06 | BT20/MB468 | NUC |
| >hg38_21_+_8433172_8446622_RNA45SN1_45S_50ntFlanks@11769.11790.22\|TAATTAGTGACGCGCATGAATG | >hg38_21_+_8433172_8446622_RNA45SN1_45S_50ntFlanks | 11769 | 11790 | 22 | TAATTAGTGACGCGCATGAATG | 999.7 | -3.4 | 0.7 | -4.6 | 4.5E-06 | 7.1E-06 | MB231/MB468 | CYTO |
| >trna11_GluTTC_15_-_26327381_26327452@1.36.36,trna3_GluTTC_13_-_45492062_45492133@1.36.36\|TCCCACATGGTCTAGCGGTTAGGATTCCTGGTTTTC | >trna11_GluTTC_15_-_26327381_26327452 | 1 | 36 | 36 | TCCCACATGGTCTAGCGGTTAGGATTCCTGGTTTTC | 528.1 | 6.7 | 1.5 | 4.6 | 4.8E-06 | 7.1E-06 | MB231/MB468 | MITO |
| >hg38_21_+_8433172_8446622_RNA45SN1_45S_50ntFlanks@12418.12438.21\|CGATGTCGGCTCTTCCTATCA | >hg38_21_+_8433172_8446622_RNA45SN1_45S_50ntFlanks | 12418 | 12438 | 21 | CGATGTCGGCTCTTCCTATCA | 4560.9 | -4.7 | 1.0 | -4.6 | 4.8E-06 | 7.1E-06 | MB231/MB468 | MITO |
| >hg38_21_+_8433172_8446622_RNA45SN1_45S_50ntFlanks@13002.13029.28\|CGATCTATTGAAAGTCAGCCCTCGACAC | >hg38_21_+_8433172_8446622_RNA45SN1_45S_50ntFlanks | 13002 | 13029 | 28 | CGATCTATTGAAAGTCAGCCCTCGACAC | 3337.9 | -6.7 | 1.5 | -4.6 | 4.5E-06 | 7.1E-06 | MB231/MB468 | CYTO |
| >MI0000433\|hsa-let-7g&WithFlank&3\|-\|52268272\|52268367@11.28.18&[MIMAT0000414&hsa-let-7g-5p&offsets\|0\|-4;m-58&3\|-\|52268336\|52268357&offsets\|0\|-4]\|TGAGGTAGTAGTTTGTAC | >MI0000433\|hsa-let-7g&WithFlank&3\|-\|52268272\|52268367 | 11 | 28 | 18 | TGAGGTAGTAGTTTGTAC | 1055.5 | 2.0 | 0.4 | 4.6 | 5.0E-06 | 7.1E-06 | BT20/MB468 | NUC |
| >hg38_21_+_8433172_8446622_RNA45SN1_45S_50ntFlanks@5487.5512.26\|CGCTGAGAAGACGGTCGAACTTGACT | >hg38_21_+_8433172_8446622_RNA45SN1_45S_50ntFlanks | 5487 | 5512 | 26 | CGCTGAGAAGACGGTCGAACTTGACT | 550.4 | -6.9 | 1.5 | -4.6 | 4.0E-06 | 7.1E-06 | MB231/MB468 | MP |
| >hg38_21_+_8433172_8446622_RNA45SN1_45S_50ntFlanks@11593.11616.24\|CCAGGGGAATCCGACTGTTTAATT | >hg38_21_+_8433172_8446622_RNA45SN1_45S_50ntFlanks | 11593 | 11616 | 24 | CCAGGGGAATCCGACTGTTTAATT | 200.5 | -2.4 | 0.5 | -4.6 | 4.7E-06 | 7.1E-06 | MB231/MB468 | NUC |
| >MI0000734\|hsa-mir-106b&WithFlank&7\|-\|100093987\|100094080@58.77.20(+1A)&[MIMAT0004672&hsa-miR-106b-3p&offsets\|0\|-2(+1A);m-87&7\|-\|100094003\|100094023&offsets\|0\|-1(+1A)]\|CCGCACTGTGGGTACTTGCTA | >MI0000734\|hsa-mir-106b&WithFlank&7\|-\|100093987\|100094080 | 58 | 77 | 20(+1A) | CCGCACTGTGGGTACTTGCTA | 67.2 | 3.2 | 0.7 | 4.6 | 5.0E-06 | 7.1E-06 | BT20/MB468 | NUC |
| >hg38_21_+_8433172_8446622_RNA45SN1_45S_50ntFlanks@12872.12891.20\|CGGCTGGAAAGGCGGCCGCC | >hg38_21_+_8433172_8446622_RNA45SN1_45S_50ntFlanks | 12872 | 12891 | 20 | CGGCTGGAAAGGCGGCCGCC | 74.5 | -9.3 | 1.9 | -4.8 | 1.4E-06 | 7.2E-06 | BT20/MB468 | MP |
| >MI0000440\|hsa-mir-27b&WithFlank&9\|+\|95085439\|95085547@67.87.21(+2U)&[MIMAT0000419&hsa-miR-27b-3p&offsets\|0\|0(+2U);m-39&9\|+\|95085505\|95085525&offsets\|0\|0(+2U)]\|TTCACAGTGGCTAAGTTCTGCTT | >MI0000440\|hsa-mir-27b&WithFlank&9\|+\|95085439\|95085547 | 67 | 87 | 21(+2U) | TTCACAGTGGCTAAGTTCTGCTT | 667.8 | 6.0 | 1.3 | 4.6 | 4.8E-06 | 7.2E-06 | MB231/MB468 | MITO |
| >hg38_21_+_8433172_8446622_RNA45SN1_45S_50ntFlanks@12636.12654.19\|AGCTACCATCTGTGGGATT | >hg38_21_+_8433172_8446622_RNA45SN1_45S_50ntFlanks | 12636 | 12654 | 19 | AGCTACCATCTGTGGGATT | 467.7 | -6.2 | 1.4 | -4.6 | 4.0E-06 | 7.2E-06 | BT20/MB468 | CYTO |
| >MI0000809\|hsa-mir-151a&WithFlank&8\|-\|140732558\|140732659@51.73.23&[MIMAT0000757&hsa-miR-151a-3p&offsets\|-2\|0;m-30&8\|-\|140732587\|140732607&offsets\|-2\|0]\|TACTAGACTGAAGCTCCTTGAGG | >MI0000809\|hsa-mir-151a&WithFlank&8\|-\|140732558\|140732659 | 51 | 73 | 23 | TACTAGACTGAAGCTCCTTGAGG | 861.4 | 1.7 | 0.4 | 4.6 | 5.1E-06 | 7.2E-06 | BT20/MB468 | NUC |
| >hg38_21_+_8433172_8446622_RNA45SN1_45S_50ntFlanks@12417.12457.41\|TCGATGTCGGCTCTTCCTATCATTGTGAAGCAGAATTCACC | >hg38_21_+_8433172_8446622_RNA45SN1_45S_50ntFlanks | 12417 | 12457 | 41 | TCGATGTCGGCTCTTCCTATCATTGTGAAGCAGAATTCACC | 634.5 | -4.0 | 0.8 | -5.1 | 4.1E-07 | 7.2E-06 | BT20/MB231 | CYTO |
| >hg38_21_+_8433172_8446622_RNA45SN1_45S_50ntFlanks@9504.9527.24\|GTGAACTATGCCTGGGCAGGGCGA | >hg38_21_+_8433172_8446622_RNA45SN1_45S_50ntFlanks | 9504 | 9527 | 24 | GTGAACTATGCCTGGGCAGGGCGA | 1611.5 | -6.5 | 1.4 | -4.8 | 1.4E-06 | 7.2E-06 | BT20/MB468 | MP |

| | | | | | | | | | | | | | |
|---|---|---|---|---|---|---|---|---|---|---|---|---|---|
| >hg38_21_+_8433172_8446622_RNA45SN1_45S_50ntFlanks@11572.11606.35\|CCGACTTAGAACTGGTGCGGACCAGGGGAATCCGA | >hg38_21_+_8433172_8446622_RNA45SN1_45S_50ntFlanks | 11572 | 11606 | 35 | CCGACTTAGAACTGGTGCGGACCAGGGGAATCCGA | 936.0 | -7.9 | 1.7 | -4.6 | 5.1E-06 | 7.2E-06 | BT20/MB468 | NUC |
| >hg38_21_+_8433172_8446622_RNA45SN1_45S_50ntFlanks@11211.11254.44\|GCGACTCTGGACGCGAGCCGGGCCCTTCCCGTGGATCGCCCCAG | >hg38_21_+_8433172_8446622_RNA45SN1_45S_50ntFlanks | 11211 | 11254 | 44 | GCGACTCTGGACGCGAGCCGGGCCCTTCCCGTGGATCGCCCCAG | 794.6 | -7.5 | 1.6 | -4.6 | 4.7E-06 | 7.2E-06 | MB231/MB468 | NUC |
| >hg38_21_+_8433172_8446622_RNA45SN1_45S_50ntFlanks@10782.10817.36\|GGTGAACAGCCTCTGGCATGTTGGAACAATGTAGGT | >hg38_21_+_8433172_8446622_RNA45SN1_45S_50ntFlanks | 10782 | 10817 | 36 | GGTGAACAGCCTCTGGCATGTTGGAACAATGTAGGT | 48.7 | -8.7 | 1.8 | -4.8 | 1.4E-06 | 7.2E-06 | BT20/MB468 | MP |
| >hg38_21_+_8433172_8446622_RNA45SN1_45S_50ntFlanks@10846.10868.23\|CTTCGGGATAAGGATTGGCTCTA | >hg38_21_+_8433172_8446622_RNA45SN1_45S_50ntFlanks | 10846 | 10868 | 23 | CTTCGGGATAAGGATTGGCTCTA | 403.2 | -3.9 | 0.8 | -4.8 | 1.4E-06 | 7.2E-06 | BT20/MB468 | MP |
| >hg38_21_+_8433172_8446622_RNA45SN1_45S_50ntFlanks@12627.12652.26\|ATGGGGCGAAGCTACCATCTGTGGGA | >hg38_21_+_8433172_8446622_RNA45SN1_45S_50ntFlanks | 12627 | 12652 | 26 | ATGGGGCGAAGCTACCATCTGTGGGA | 1063.7 | -7.8 | 1.6 | -4.8 | 1.4E-06 | 7.2E-06 | BT20/MB468 | MP |
| >hg38_MT_+_598_1651_RNR1_12S_50ntFlanks@606.633.28\|CCTGTTCTGTAATCGATAAACCCCGATC | >hg38_MT_+_598_1651_RNR1_12S_50ntFlanks | 606 | 633 | 28 | CCTGTTCTGTAATCGATAAACCCCGATC | 278.4 | -3.6 | 0.7 | -5.1 | 4.1E-07 | 7.2E-06 | BT20/MB231 | CYTO |
| >hg38_21_+_8433172_8446622_RNA45SN1_45S_50ntFlanks@10411.10428.18\|CAGTCGGTCCTGAGAGAT | >hg38_21_+_8433172_8446622_RNA45SN1_45S_50ntFlanks | 10411 | 10428 | 18 | CAGTCGGTCCTGAGAGAT | 3844.7 | -5.6 | 1.2 | -4.6 | 4.6E-06 | 7.2E-06 | MB231/MB468 | CYTO |
| >MI0000077\|hsa-mir-21&WithFlank&17\|+\|59841260\|59841343@14.36.23(+1A)&[MIMAT0000076&hsa-miR-21-5p&offsets\|0\|+1(+1A);m-1&17\|+\|59841273\|59841294&offsets\|0\|+1(+1A)]\|TAGCTTATCAGACTGATGTTGACA | >MI0000077\|hsa-mir-21&WithFlank&17\|+\|59841260\|59841343 | 14 | 36 | 23(+1A) | TAGCTTATCAGACTGATGTTGACA | 28269.9 | 2.6 | 0.6 | 4.6 | 5.1E-06 | 7.2E-06 | BT20/MB468 | NUC |
| >MI0000273\|hsa-mir-183&WithFlank&7\|-\|129774899\|129775020@33.53.21(+1C)&[MIMAT0000261&hsa-miR-183-5p&offsets\|0\|-1(+1C);m-41&7\|-\|129774966\|129774987&offsets\|-1\|-2(+1C)]\|TATGGCACTGGTAGAATTCACC | >MI0000273\|hsa-mir-183&WithFlank&7\|-\|129774899\|129775020 | 33 | 53 | 21(+1C) | TATGGCACTGGTAGAATTCACC | 607.5 | 1.8 | 0.4 | 4.6 | 5.1E-06 | 7.3E-06 | BT20/MB468 | NUC |
| >MI0001733\|hsa-mir-452&WithFlank&X\|-\|151959622\|151959718@20.42.23(+1U)&[MIMAT0001635&hsa-miR-452-5p&offsets\|0\|+1(+1U);m-188&X\|-\|151959677\|151959699&offsets\|0\|0(+1U)]\|AACTGTTTGCAGAGGAAACTGAGT | >MI0001733\|hsa-mir-452&WithFlank&X\|-\|151959622\|151959718 | 20 | 42 | 23(+1U) | AACTGTTTGCAGAGGAAACTGAGT | 39.9 | 8.5 | 1.9 | 4.6 | 4.1E-06 | 7.3E-06 | MB231/MB468 | MP |
| >hg38_21_+_8433172_8446622_RNA45SN1_45S_50ntFlanks@6733.6752.20\|CATTGATCATCGACACTTCG | >hg38_21_+_8433172_8446622_RNA45SN1_45S_50ntFlanks | 6733 | 6752 | 20 | CATTGATCATCGACACTTCG | 935.2 | -3.4 | 0.8 | -4.6 | 4.9E-06 | 7.3E-06 | MB231/MB468 | MITO |
| >hg38_21_+_8433172_8446622_RNA45SN1_45S_50ntFlanks@5074.5112.39\|ATTCCGATAACGAACGAGACTCTGGCATGCTAACTAGTT | >hg38_21_+_8433172_8446622_RNA45SN1_45S_50ntFlanks | 5074 | 5112 | 39 | ATTCCGATAACGAACGAGACTCTGGCATGCTAACTAGTT | 128.1 | -10.9 | 2.4 | -4.6 | 4.1E-06 | 7.3E-06 | MB231/MB468 | MP |
| >hg38_21_+_8433172_8446622_RNA45SN1_45S_50ntFlanks@6647.6680.34\|CGTACGACTCTTAGCGGTGGATCACTCGGCTCGT | >hg38_21_+_8433172_8446622_RNA45SN1_45S_50ntFlanks | 6647 | 6680 | 34 | CGTACGACTCTTAGCGGTGGATCACTCGGCTCGT | 3044.6 | 2.3 | 0.5 | 4.6 | 3.5E-06 | 7.3E-06 | BT20/MB468 | MITO |
| >hg38_21_+_8433172_8446622_RNA45SN1_45S_50ntFlanks@8171.8216.46\|AAGTCCTTCTGATCGAGGCCCAGCCCGTGGACGGTGTGAGGCCGGT | >hg38_21_+_8433172_8446622_RNA45SN1_45S_50ntFlanks | 8171 | 8216 | 46 | AAGTCCTTCTGATCGAGGCCCAGCCCGTGGACGGTGTGAGGCCGGT | 508.9 | -6.0 | 1.3 | -4.6 | 5.1E-06 | 7.3E-06 | BT20/MB468 | NUC |
| >trna2_GlyGCC_21_-_18827107_18827177@1.32.32,trna35_GlyGCC_1_+_161413094_161413164@1.32.32,trna37_GlyGCC_1_+_161420467_161420537@1.32.32,trna39_GlyGCC_1_+_161427898_161427968@1.32.32,trna41_GlyGCC_1_+_161435258_161435328@1.32.32\|GCATGGGTGGTTCAGTGGTAGAATTCTCGCCT | >trna2_GlyGCC_21_-_18827107_18827177 | 1 | 32 | 32 | GCATGGGTGGTTCAGTGGTAGAATTCTCGCCT | 1583.7 | 4.7 | 0.9 | 5.0 | 5.5E-07 | 7.3E-06 | BT20/MB231 | NUC |
| >hg38_21_+_8433172_8446622_RNA45SN1_45S_50ntFlanks@12081.12106.26\|CTCTCGCTTCTGGCGCCAAGCGCCCG | >hg38_21_+_8433172_8446622_RNA45SN1_45S_50ntFlanks | 12081 | 12106 | 26 | CTCTCGCTTCTGGCGCCAAGCGCCCG | 559.4 | -6.6 | 1.4 | -4.6 | 4.1E-06 | 7.3E-06 | BT20/MB468 | CYTO |
| >trnaMT_GluTTC_MT_-_14674_14742@29.69.41,trnalookalike8_GluTTC_5_-_93905172_93905240@29.69.41\|TTTTCATATCATTGGTCGTGGTTGTAGTCCGTGCGAGAATA | >trnaMT_GluTTC_MT_-_14674_14742 | 29 | 69 | 41 | TTTTCATATCATTGGTCGTGGTTGTAGTCCGTGCGAGAATA | 73.6 | 9.9 | 2.1 | 4.8 | 1.4E-06 | 7.3E-06 | BT20/MB468 | MP |
| >am_MI0003583\|hsa-mir-576&WithFlank&4\|+\|109488692\|109488801@61.83.23&[MIMAT0004796&hsa-miR-576-3p&offsets\|0\|+1;m-524&4\|+\|109488752\|109488772&offsets\|0\|+2]\|AAGATGTGGAAAAATTGGAATCC | >am_MI0003583\|hsa-mir-576&WithFlank&4\|+\|109488692\|109488801 | 61 | 83 | 23 | AAGATGTGGAAAAATTGGAATCC | 78.0 | 3.3 | 0.7 | 4.6 | 5.2E-06 | 7.3E-06 | BT20/MB468 | NUC |
| >hg38_21_+_8433172_8446622_RNA45SN1_45S_50ntFlanks@475.515.41\|CTTCGTGATCGATGTGGTGACGTCGTGCTCTCCCGGGCCGG | >hg38_21_+_8433172_8446622_RNA45SN1_45S_50ntFlanks | 475 | 515 | 41 | CTTCGTGATCGATGTGGTGACGTCGTGCTCTCCCGGGCCGG | 579.9 | 7.2 | 1.6 | 4.6 | 3.5E-06 | 7.3E-06 | BT20/MB468 | MITO |
| >hg38_21_+_8433172_8446622_RNA45SN1_45S_50ntFlanks@9540.9571.32\|CTCTGGTGGAGGTCCGTAGCGGTCCTGACGTG | >hg38_21_+_8433172_8446622_RNA45SN1_45S_50ntFlanks | 9540 | 9571 | 32 | CTCTGGTGGAGGTCCGTAGCGGTCCTGACGTG | 802.5 | -3.1 | 0.7 | -4.6 | 4.6E-06 | 7.3E-06 | MB231/MB468 | CYTO |
| >MI0000088\|hsa-mir-30a&WithFlank&6\|-\|71403545\|71403627@12.33.22(+1U)&[MIMAT0000087&hsa-miR-30a-5p&offsets\|0\|0(+1U);m-15&6\|-\|71403595\|71403616&offsets\|0\|0(+1U)]\|TGTAAACATCCTCGACTGGAAGT | >MI0000088\|hsa-mir-30a&WithFlank&6\|-\|71403545\|71403627 | 12 | 33 | 22(+1U) | TGTAAACATCCTCGACTGGAAGT | 1525.0 | 4.3 | 0.9 | 4.6 | 5.2E-06 | 7.3E-06 | BT20/MB468 | NUC |
| >hg38_21_+_8433172_8446622_RNA45SN1_45S_50ntFlanks@10770.10805.36\|GCAGGTCTCCAAGGTGAACAGCCTCTGGCATGTTGG | >hg38_21_+_8433172_8446622_RNA45SN1_45S_50ntFlanks | 10770 | 10805 | 36 | GCAGGTCTCCAAGGTGAACAGCCTCTGGCATGTTGG | 183.8 | -4.8 | 1.1 | -4.6 | 4.8E-06 | 7.3E-06 | MB231/MB468 | NUC |
| >hg38_MT_+_598_1651_RNR1_12S_50ntFlanks@634.665.32\|AACCTCACCACCTCTTGCTCAGCCTATATACC | >hg38_MT_+_598_1651_RNR1_12S_50ntFlanks | 634 | 665 | 32 | AACCTCACCACCTCTTGCTCAGCCTATATACC | 245.5 | 7.3 | 1.6 | 4.6 | 4.9E-06 | 7.3E-06 | MB231/MB468 | MITO |
| >hg38_21_+_8433172_8446622_RNA45SN1_45S_50ntFlanks@12437.12475.39\|CATTGTGAAGCAGAATTCACCAAGCGTTGGATTGTTCAC | >hg38_21_+_8433172_8446622_RNA45SN1_45S_50ntFlanks | 12437 | 12475 | 39 | CATTGTGAAGCAGAATTCACCAAGCGTTGGATTGTTCAC | 87.1 | -9.5 | 2.0 | -4.8 | 1.5E-06 | 7.3E-06 | BT20/MB468 | MP |
| >hg38_21_+_8433172_8446622_RNA45SN1_45S_50ntFlanks@10317.10352.36\|GTGCAGATCTTGGTGGTAGTAGCAAATATTCAAACG | >hg38_21_+_8433172_8446622_RNA45SN1_45S_50ntFlanks | 10317 | 10352 | 36 | GTGCAGATCTTGGTGGTAGTAGCAAATATTCAAACG | 131.2 | 10.4 | 2.0 | 5.1 | 4.0E-07 | 7.3E-06 | BT20/MB231 | MP |
| >hg38_21_+_8433172_8446622_RNA45SN1_45S_50ntFlanks@6782.6801.20\|GGGCTACGCCTGTCTGAGCG | >hg38_21_+_8433172_8446622_RNA45SN1_45S_50ntFlanks | 6782 | 6801 | 20 | GGGCTACGCCTGTCTGAGCG | 45.7 | -8.6 | 1.8 | -4.8 | 1.5E-06 | 7.3E-06 | BT20/MB468 | MP |
| >hg38_21_+_8433172_8446622_RNA45SN1_45S_50ntFlanks@11564.11591.28\|CCTAGCAGCCGACTTAGAACTGGTGCGG | >hg38_21_+_8433172_8446622_RNA45SN1_45S_50ntFlanks | 11564 | 11591 | 28 | CCTAGCAGCCGACTTAGAACTGGTGCGG | 991.2 | -6.9 | 1.5 | -4.6 | 4.1E-06 | 7.4E-06 | BT20/MB468 | CYTO |
| >hg38_21_+_8433172_8446622_RNA45SN1_45S_50ntFlanks@3861.3881.21\|TAATTCTAGAGCTAATACATG | >hg38_21_+_8433172_8446622_RNA45SN1_45S_50ntFlanks | 3861 | 3881 | 21 | TAATTCTAGAGCTAATACATG | 395.2 | -7.8 | 1.7 | -4.6 | 4.1E-06 | 7.4E-06 | BT20/MB468 | CYTO |
| >hg38_21_+_8433172_8446622_RNA45SN1_45S_50ntFlanks@12285.12320.36\|CAGACCGTGAAAGCGGGGCCTCACGATCCTTCTGAC | >hg38_21_+_8433172_8446622_RNA45SN1_45S_50ntFlanks | 12285 | 12320 | 36 | CAGACCGTGAAAGCGGGGCCTCACGATCCTTCTGAC | 391.4 | -5.2 | 1.1 | -4.6 | 5.0E-06 | 7.4E-06 | MB231/MB468 | MITO |
| >hg38_21_+_8433172_8446622_RNA45SN1_45S_50ntFlanks@11572.11600.29\|CCGACTTAGAACTGGTGCGGACCAGGGGA | >hg38_21_+_8433172_8446622_RNA45SN1_45S_50ntFlanks | 11572 | 11600 | 29 | CCGACTTAGAACTGGTGCGGACCAGGGGA | 6896.4 | -3.8 | 0.8 | -4.8 | 1.5E-06 | 7.4E-06 | BT20/MB468 | MP |
| >hg38_21_+_8433172_8446622_RNA45SN1_45S_50ntFlanks@9342.9370.29\|ACGAAAGCCGCCGTGGCGCAATGAAGGTG | >hg38_21_+_8433172_8446622_RNA45SN1_45S_50ntFlanks | 9342 | 9370 | 29 | ACGAAAGCCGCCGTGGCGCAATGAAGGTG | 372.2 | -7.1 | 1.5 | -4.8 | 1.5E-06 | 7.4E-06 | BT20/MB468 | MP |
| >hg38_21_+_8433172_8446622_RNA45SN1_45S_50ntFlanks@6652.6700.49\|GACTCTTAGCGGTGGATCACTCGGCTCGTGCGTCGATGAAGAACGCAGC | >hg38_21_+_8433172_8446622_RNA45SN1_45S_50ntFlanks | 6652 | 6700 | 49 | GACTCTTAGCGGTGGATCACTCGGCTCGTGCGTCGATGAAGAACGCAGC | 635.2 | 6.6 | 1.4 | 4.6 | 4.7E-06 | 7.4E-06 | MB231/MB468 | CYTO |
| >hg38_21_+_8433172_8446622_RNA45SN1_45S_50ntFlanks@10677.10710.34\|GCGGTTCCGGCGGCGTCCGGTGAGCTCTCGCTGG | >hg38_21_+_8433172_8446622_RNA45SN1_45S_50ntFlanks | 10677 | 10710 | 34 | GCGGTTCCGGCGGCGTCCGGTGAGCTCTCGCTGG | 1894.1 | -2.8 | 0.6 | -4.6 | 4.7E-06 | 7.4E-06 | MB231/MB468 | CYTO |
| >hg38_21_+_8433172_8446622_RNA45SN1_45S_50ntFlanks@12638.12661.24\|CTACCATCTGTGGGATTATGACTG | >hg38_21_+_8433172_8446622_RNA45SN1_45S_50ntFlanks | 12638 | 12661 | 24 | CTACCATCTGTGGGATTATGACTG | 306.3 | -2.8 | 0.6 | -4.6 | 4.7E-06 | 7.4E-06 | MB231/MB468 | CYTO |
| >am_trnaMT_ValTAC_MT_+_1602_1670@28.65.38\|AACTTACACTTAGGAGATTTCAACTTAACTTGACCGCT | >am_trnaMT_ValTAC_MT_+_1602_1670 | 28 | 65 | 38 | AACTTACACTTAGGAGATTTCAACTTAACTTGACCGCT | 34.8 | 8.3 | 1.8 | 4.6 | 4.1E-06 | 7.4E-06 | MB231/MB468 | MP |
| >hg38_21_+_8433172_8446622_RNA45SN1_45S_50ntFlanks@7975.8015.41\|CGCGACCTCAGATCAGACGTGGCGACCCGCTGAATTTAAGC | >hg38_21_+_8433172_8446622_RNA45SN1_45S_50ntFlanks | 7975 | 8015 | 41 | CGCGACCTCAGATCAGACGTGGCGACCCGCTGAATTTAAGC | 6739.5 | 3.9 | 0.9 | 4.6 | 5.2E-06 | 7.4E-06 | BT20/MB468 | NUC |
| >hg38_21_+_8433172_8446622_RNA45SN1_45S_50ntFlanks@3961.3983.23\|CCCCGGCCGGGGGGCGGGCGCCG | >hg38_21_+_8433172_8446622_RNA45SN1_45S_50ntFlanks | 3961 | 3983 | 23 | CCCCGGCCGGGGGGCGGGCGCCG | 54.1 | -8.8 | 1.8 | -4.8 | 1.5E-06 | 7.4E-06 | BT20/MB468 | MP |
| >hg38_21_+_8433172_8446622_RNA45SN1_45S_50ntFlanks@12415.12442.28\|CTTCGATGTCGGCTCTTCCTATCATTGT | >hg38_21_+_8433172_8446622_RNA45SN1_45S_50ntFlanks | 12415 | 12442 | 28 | CTTCGATGTCGGCTCTTCCTATCATTGT | 337.8 | -2.7 | 0.6 | -4.6 | 5.2E-06 | 7.4E-06 | BT20/MB468 | NUC |
| >hg38_21_+_8433172_8446622_RNA45SN1_45S_50ntFlanks@11565.11585.21\|CTAGCAGCCGACTTAGAACTG | >hg38_21_+_8433172_8446622_RNA45SN1_45S_50ntFlanks | 11565 | 11585 | 21 | CTAGCAGCCGACTTAGAACTG | 231.5 | -2.2 | 0.5 | -4.6 | 4.9E-06 | 7.4E-06 | MB231/MB468 | NUC |
| >MI0000750\|hsa-mir-26a-2&WithFlank&12\|-\|57824603\|57824698@20.40.21(+1G);MI0000083\|hsa-mir-26a-1&WithFlank&3\|+\|37969398\|37969486@16.36.21(+1G)&[MIMAT0000082&hsa-miR-26a-5p&offsets\|0\|-1(+1G);m-33&12\|-\|57824658\|57824679&offsets\|0\|-1(+1G)];[MIMAT0000082_1&hsa-miR-26a-5p&offsets\|0\|-1(+1G);m-34&3\|+\|37969413\|37969434&offsets\|0\|-1(+1G)]\|TTCAAGTAATCCAGGATAGGCG | >MI0000750\|hsa-mir-26a-2&WithFlank&12\|-\|57824603\|57824698 | 20 | 40 | 21(+1G) | TTCAAGTAATCCAGGATAGGCG | 2204.8 | 4.4 | 1.0 | 4.6 | 5.0E-06 | 7.4E-06 | MB231/MB468 | MITO |
| >MI0000071\|hsa-mir-17&WithFlank&13\|+\|91350599\|91350694@20.40.21&[MIMAT0000070&hsa-miR-17-5p&offsets\|0\|-2;m-88&13\|+\|91350618\|91350640&offsets\|0\|-2]\|CAAAGTGCTTACAGTGCAGGT | >MI0000071\|hsa-mir-17&WithFlank&13\|+\|91350599\|91350694 | 20 | 40 | 21 | CAAAGTGCTTACAGTGCAGGT | 1325.3 | -4.4 | 0.9 | -5.0 | 4.7E-07 | 7.4E-06 | BT20/MB231 | MITO |
| >hg38_21_+_8433172_8446622_RNA45SN1_45S_50ntFlanks@9716.9737.22\|AATGATTAGAGGTCTTGGGGCC | >hg38_21_+_8433172_8446622_RNA45SN1_45S_50ntFlanks | 9716 | 9737 | 22 | AATGATTAGAGGTCTTGGGGCC | 320.9 | -2.7 | 0.6 | -4.6 | 4.2E-06 | 7.4E-06 | BT20/MB468 | CYTO |
| >trna116_GluCTC_1_-_145399233_145399304@3.34.32,trna59_GluCTC_1_+_249168447_249168518@3.34.32,trna71_GluCTC_1_-_161439189_161439260@3.34.32,trna74_GluCTC_1_-_161431809_161431880@3.34.32,trna77_GluCTC_1_-_161424398_161424469@3.34.32,trna77_GluCTC_6_+_28949976_28950047@3.34.32,trna80_GluCTC_1_-_161417018_161417089@3.34.32,trna87_GluCTC_6_-_126101393_126101464@3.34.32\|CCTGGTGGTCTAGTGGTTAGGATTCGGCGCTC | >trna116_GluCTC_1_-_145399233_145399304 | 3 | 34 | 32 | CCTGGTGGTCTAGTGGTTAGGATTCGGCGCTC | 278.9 | 4.4 | 1.0 | 4.6 | 5.0E-06 | 7.4E-06 | MB231/MB468 | MITO |
| >MI0000272\|hsa-mir-182&WithFlank&7\|-\|129770377\|129770498@29.54.26(+1U)&[MIMAT0000259&hsa-miR-182-5p&offsets\|0\|+2(+1U);m-22&7\|-\|129770449\|129770470&offsets\|0\|+4(+1U)]\|TTTGGCAATGGTAGAACTCACACTGGT | >MI0000272\|hsa-mir-182&WithFlank&7\|-\|129770377\|129770498 | 29 | 54 | 26(+1U) | TTTGGCAATGGTAGAACTCACACTGGT | 335.4 | 4.9 | 1.1 | 4.6 | 5.0E-06 | 7.4E-06 | MB231/MB468 | MITO |
| >hg38_MT_+_598_1651_RNR1_12S_50ntFlanks@854.872.19\|TTAGTTGAACAGGGCCCTG | >hg38_MT_+_598_1651_RNR1_12S_50ntFlanks | 854 | 872 | 19 | TTAGTTGAACAGGGCCCTG | 56.9 | 9.6 | 2.0 | 4.8 | 1.5E-06 | 7.4E-06 | BT20/MB468 | MP |
| >hg38_21_+_8433172_8446622_RNA45SN1_45S_50ntFlanks@3861.3898.38\|TAATTCTAGAGCTAATACATGCCGACGGGCGCTGACCC | >hg38_21_+_8433172_8446622_RNA45SN1_45S_50ntFlanks | 3861 | 3898 | 38 | TAATTCTAGAGCTAATACATGCCGACGGGCGCTGACCC | 343.3 | -6.8 | 1.5 | -4.6 | 4.9E-06 | 7.5E-06 | MB231/MB468 | NUC |
| >hg38_21_+_8433172_8446622_RNA45SN1_45S_50ntFlanks@4000.4022.23\|TAGATAACCTCGGGCCGATCGCA | >hg38_21_+_8433172_8446622_RNA45SN1_45S_50ntFlanks | 4000 | 4022 | 23 | TAGATAACCTCGGGCCGATCGCA | 547.3 | -7.6 | 1.7 | -4.6 | 5.3E-06 | 7.5E-06 | BT20/MB468 | NUC |

<table>
<tr><td>>trnaMT_GluTTC_MT_-_14674_14742@34.71.38,trnalookalike8_GluTTC_5_-_93905172_93905240@34.71.38|ATATCATTGGTCGTGGTTGTAGTCCGTGCGAGAATACC</td><td>>trnaMT_GluTTC_MT_-_14674_14742</td><td>34</td><td>71</td><td>38</td><td>ATATCATTGGTCGTGGTTGTAGTCCGTGCGAGAATACC</td><td>550.4</td><td>3.7</td><td>0.8</td><td>4.6</td><td>5.0E-06</td><td>7.5E-06</td><td>MB231/MB468</td><td>MITO</td></tr>
<tr><td>>hg38_21_+_8433172_8446622_RNA45SN1_45S_50ntFlanks@10346.10368.23|TCAAACGAGAACTTTGAAGGCCG</td><td>>hg38_21_+_8433172_8446622_RNA45SN1_45S_50ntFlanks</td><td>10346</td><td>10368</td><td>23</td><td>TCAAACGAGAACTTTGAAGGCCG</td><td>2699.2</td><td>-2.9</td><td>0.6</td><td>-4.6</td><td>3.6E-06</td><td>7.5E-06</td><td>BT20/MB468</td><td>MITO</td></tr>
<tr><td>>hg38_21_+_8433172_8446622_RNA45SN1_45S_50ntFlanks@6650.6673.24|ACGACTCTTAGCGGTGGATCACTC</td><td>>hg38_21_+_8433172_8446622_RNA45SN1_45S_50ntFlanks</td><td>6650</td><td>6673</td><td>24</td><td>ACGACTCTTAGCGGTGGATCACTC</td><td>278.6</td><td>4.6</td><td>1.0</td><td>4.6</td><td>5.0E-06</td><td>7.5E-06</td><td>MB231/MB468</td><td>NUC</td></tr>
<tr><td>>hg38_21_+_8433172_8446622_RNA45SN1_45S_50ntFlanks@9299.9331.33|CACGGACCAAGGAGTCTAACACGTGCGCGAGTC</td><td>>hg38_21_+_8433172_8446622_RNA45SN1_45S_50ntFlanks</td><td>9299</td><td>9331</td><td>33</td><td>CACGGACCAAGGAGTCTAACACGTGCGCGAGTC</td><td>44.2</td><td>9.2</td><td>1.9</td><td>4.8</td><td>1.5E-06</td><td>7.5E-06</td><td>BT20/MB468</td><td>MP</td></tr>
<tr><td>>hg38_21_+_8433172_8446622_RNA45SN1_45S_50ntFlanks@8169.8203.35|CCAAGTCCTTCTGATCGAGGCCCAGCCCGTGGACG</td><td>>hg38_21_+_8433172_8446622_RNA45SN1_45S_50ntFlanks</td><td>8169</td><td>8203</td><td>35</td><td>CCAAGTCCTTCTGATCGAGGCCCAGCCCGTGGACG</td><td>251.5</td><td>-7.0</td><td>1.5</td><td>-4.6</td><td>4.2E-06</td><td>7.5E-06</td><td>MB231/MB468</td><td>MP</td></tr>
<tr><td>>hg38_21_+_8433172_8446622_RNA45SN1_45S_50ntFlanks@10291.10320.30|CGCGGGCCCGGGTGGAGCCGCCGCAGGTGC</td><td>>hg38_21_+_8433172_8446622_RNA45SN1_45S_50ntFlanks</td><td>10291</td><td>10320</td><td>30</td><td>CGCGGGCCCGGGTGGAGCCGCCGCAGGTGC</td><td>120.1</td><td>-6.5</td><td>1.4</td><td>-4.6</td><td>4.2E-06</td><td>7.5E-06</td><td>MB231/MB468</td><td>MP</td></tr>
<tr><td>>hg38_MT_+_1621_3279_RNR2_16S_50ntFlanks@1508.1547.40|AAAGGACAAGAGAAATAAGGCCTACTTCACAAAGCGCCTT</td><td>>hg38_MT_+_1621_3279_RNR2_16S_50ntFlanks</td><td>1508</td><td>1547</td><td>40</td><td>AAAGGACAAGAGAAATAAGGCCTACTTCACAAAGCGCCTT</td><td>55.7</td><td>9.5</td><td>2.0</td><td>4.8</td><td>1.5E-06</td><td>7.5E-06</td><td>BT20/MB468</td><td>MP</td></tr>
<tr><td>>hg38_21_+_8433172_8446622_RNA45SN1_45S_50ntFlanks@11998.12032.35|GCCCTGCGGGCCGCCGGTGAAATACCACTACTCTG</td><td>>hg38_21_+_8433172_8446622_RNA45SN1_45S_50ntFlanks</td><td>11998</td><td>12032</td><td>35</td><td>GCCCTGCGGGCCGCCGGTGAAATACCACTACTCTG</td><td>57.3</td><td>-8.9</td><td>1.9</td><td>-4.8</td><td>1.5E-06</td><td>7.5E-06</td><td>BT20/MB468</td><td>MP</td></tr>
<tr><td>>hg38_21_+_8433172_8446622_RNA45SN1_45S_50ntFlanks@10786.10817.32|AACAGCCTCTGGCATGTTGGAACAATGTAGGT</td><td>>hg38_21_+_8433172_8446622_RNA45SN1_45S_50ntFlanks</td><td>10786</td><td>10817</td><td>32</td><td>AACAGCCTCTGGCATGTTGGAACAATGTAGGT</td><td>1347.6</td><td>-4.2</td><td>0.9</td><td>-4.8</td><td>1.5E-06</td><td>7.5E-06</td><td>BT20/MB468</td><td>MP</td></tr>
<tr><td>>MI0000067|hsa-let-7f-1&WithFlank&9|+|94176341|94176439@13.30.18;MI0000068|hsa-let-7f-2&WithFlank&X|-|53557186|53557280@14.31.18&[MIMAT0000067&hsa-let-7f-5p&offsets|0|-4;m-20&9|+|94176353|94176374&offsets|0|-4];[MIMAT0000067_1&hsa-let-7f-5p&offsets|0|-4;m-19&X|-|53557246|53557267&offsets|0|-4]|TGAGGTAGTAGATTGTAT</td><td>>MI0000067|hsa-let-7f-1&WithFlank&9|+|94176341|94176439</td><td>13</td><td>30</td><td>18</td><td>TGAGGTAGTAGATTGTAT</td><td>1779.9</td><td>2.0</td><td>0.4</td><td>4.6</td><td>5.3E-06</td><td>7.5E-06</td><td>BT20/MB468</td><td>NUC</td></tr>
<tr><td>>trna11_SerAGA_8_-_96281885_96281966@1.18.18,trna145_SerAGA_6_-_27521192_27521273@1.18.18,trna147_SerAGA_6_-_27509554_27509635@1.18.18,trna148_SerTGA_6_-_27473607_27473688@1.18.18,trna172_SerTGA_6_-_26312824_26312905@1.18.18,trna35_SerAGA_17_-_8129928_8130009@1.18.18,trna44_SerAGA_6_+_27446591_27446672@1.18.18,trna46_SerAGA_6_+_27463593_27463674@1.18.18,trna47_SerAGA_6_+_27470818_27470899@1.18.18,trna50_SerAGA_6_+_27499987_27500068@1.18.18,trna51_SerTGA_6_+_27513468_27513549@1.18.18,trna5_SerAGA_6_+_26327817_26327898@1.18.18|GTAGTCGTGGCCGAGTGG</td><td>>trna11_SerAGA_8_-_96281885_96281966</td><td>1</td><td>18</td><td>18</td><td>GTAGTCGTGGCCGAGTGG</td><td>324.2</td><td>4.6</td><td>1.0</td><td>4.8</td><td>1.5E-06</td><td>7.5E-06</td><td>BT20/MB468</td><td>MP</td></tr>
<tr><td>>hg38_21_+_8433172_8446622_RNA45SN1_45S_50ntFlanks@6663.6689.27|GTGGATCACTCGGCTCGTGCGTCGATG</td><td>>hg38_21_+_8433172_8446622_RNA45SN1_45S_50ntFlanks</td><td>6663</td><td>6689</td><td>27</td><td>GTGGATCACTCGGCTCGTGCGTCGATG</td><td>5426.2</td><td>-3.9</td><td>0.8</td><td>-4.8</td><td>1.5E-06</td><td>7.5E-06</td><td>BT20/MB468</td><td>MP</td></tr>
<tr><td>>hg38_MT_+_1621_3279_RNR2_16S_50ntFlanks@113.145.33|CCAAATAAAGTATAGGCGATAGAAATTGAAACC</td><td>>hg38_MT_+_1621_3279_RNR2_16S_50ntFlanks</td><td>113</td><td>145</td><td>33</td><td>CCAAATAAAGTATAGGCGATAGAAATTGAAACC</td><td>39.1</td><td>8.5</td><td>1.8</td><td>4.6</td><td>4.2E-06</td><td>7.5E-06</td><td>MB231/MB468</td><td>MP</td></tr>
<tr><td>>hg38_21_+_8433172_8446622_RNA45SN1_45S_50ntFlanks@12912.12933.22|CACGTTCGTGGGGAACCTGGCG</td><td>>hg38_21_+_8433172_8446622_RNA45SN1_45S_50ntFlanks</td><td>12912</td><td>12933</td><td>22</td><td>CACGTTCGTGGGGAACCTGGCG</td><td>613.1</td><td>-3.6</td><td>0.8</td><td>-4.6</td><td>3.6E-06</td><td>7.5E-06</td><td>BT20/MB468</td><td>MITO</td></tr>
<tr><td>>hg38_MT_+_1621_3279_RNR2_16S_50ntFlanks@341.377.37|AAATAGTGGGAAGATTTATAGGTAGAGGCGACAAACC</td><td>>hg38_MT_+_1621_3279_RNR2_16S_50ntFlanks</td><td>341</td><td>377</td><td>37</td><td>AAATAGTGGGAAGATTTATAGGTAGAGGCGACAAACC</td><td>68.1</td><td>9.8</td><td>2.0</td><td>4.8</td><td>1.5E-06</td><td>7.5E-06</td><td>BT20/MB468</td><td>MP</td></tr>
<tr><td>>MI0001733|hsa-mir-452&WithFlank&X|-|151959622|151959718@23.43.21&[MIMAT0001635&hsa-miR-452-5p&offsets|+3|+2;m-188&X|-|151959677|151959699&offsets|+3|+1]|TGTTTGCAGAGGAAACTGAGA</td><td>>MI0001733|hsa-mir-452&WithFlank&X|-|151959622|151959718</td><td>23</td><td>43</td><td>21</td><td>TGTTTGCAGAGGAAACTGAGA</td><td>603.5</td><td>9.2</td><td>1.9</td><td>4.8</td><td>1.5E-06</td><td>7.5E-06</td><td>BT20/MB468</td><td>MP</td></tr>
<tr><td>>MI0003140|hsa-mir-512-1&WithFlank&19|+|53666673|53666768@57.78.22;MI0003141|hsa-mir-512-2&WithFlank&19|+|53669151|53669260@63.84.22&[MIMAT0002823&hsa-miR-512-3p&offsets|0|0;m-849&19|+|53666729|53666751&offsets|0|-1];[MIMAT0002823_1&hsa-miR-512-3p&offsets|0|0;m-849&19|+|53669213|53669235&offsets|0|-1]|AAGTGCTGTCATAGCTGAGGTC</td><td>>MI0003140|hsa-mir-512-1&WithFlank&19|+|53666673|53666768</td><td>57</td><td>78</td><td>22</td><td>AAGTGCTGTCATAGCTGAGGTC</td><td>61.7</td><td>9.7</td><td>2.0</td><td>4.8</td><td>1.5E-06</td><td>7.5E-06</td><td>BT20/MB468</td><td>MP</td></tr>
<tr><td>>MI0003762|hsa-mir-550a-3&WithFlank&7|-|29680728|29680834@27.47.21&[MIMAT0020925&hsa-miR-550a-3-5p&offsets|0|+1;m-494&7|-|29680788|29680808&offsets|0|0]|AGTGCCTGAGGGAGTAAGAGT</td><td>>MI0003762|hsa-mir-550a-3&WithFlank&7|-|29680728|29680834</td><td>27</td><td>47</td><td>21</td><td>AGTGCCTGAGGGAGTAAGAGT</td><td>57.5</td><td>9.6</td><td>2.0</td><td>4.8</td><td>1.5E-06</td><td>7.5E-06</td><td>BT20/MB468</td><td>MP</td></tr>
<tr><td>>am_trnaMT_HisGTG_MT_+_12138_12206@25.51.27|CAGATTGTGAATCTGACAACAGAGGCT</td><td>>am_trnaMT_HisGTG_MT_+_12138_12206</td><td>25</td><td>51</td><td>27</td><td>CAGATTGTGAATCTGACAACAGAGGCT</td><td>178.3</td><td>8.3</td><td>1.7</td><td>4.8</td><td>1.5E-06</td><td>7.5E-06</td><td>BT20/MB468</td><td>MP</td></tr>
<tr><td>>trna143_LysTTT_6_-_27559593_27559665@13.30.18,trna149_LysTTT_6_-_27302769_27302841@13.30.18,trna14_LysTTT_11_-_59327808_59327880@13.30.18,trna2_LysTTT_17_+_8022473_8022545@13.30.18,trna54_LysTTT_1_+_204475655_204475727@13.30.18,trna5_LysTTT_11_+_59323902_59323974@13.30.18,trna62_LysTTT_1_-_204476158_204476230@13.30.18,trna76_LysTTT_6_+_28918806_28918878@13.30.18|CAGTCGGTAGAGCATCAG</td><td>>trna143_LysTTT_6_-_27559593_27559665</td><td>13</td><td>30</td><td>18</td><td>CAGTCGGTAGAGCATCAG</td><td>327.6</td><td>-6.5</td><td>1.4</td><td>-4.6</td><td>5.0E-06</td><td>7.5E-06</td><td>MB231/MB468</td><td>NUC</td></tr>
<tr><td>>am_trna128_GlyGCC_6_-_27870686_27870756@1.32.32,trna133_GlyCCC_1_-_16872434_16872504@1.32.32,trna18_GlyGCC_16_+_70822597_70822667@1.32.32,trna19_GlyGCC_16_+_70823410_70823480@1.32.32,trna19_GlyGCC_2_-_157257659_157257729@1.32.32,trna24_GlyGCC_16_-_70812942_70813012@1.32.32,trna25_GlyGCC_16_-_70812114_70812184@1.32.32,trna4_GlyCCC_1_+_17188416_17188486@1.32.32,trna5_GlyGCC_17_+_8029064_8029134@1.32.32,trna68_GlyGCC_1_-_161493637_161493707@1.32.32|GCATTGGTGGTTCAGTGGTAGAATTCTCGCCT</td><td>>am_trna128_GlyGCC_6_-_27870686_27870756</td><td>1</td><td>32</td><td>32</td><td>GCATTGGTGGTTCAGTGGTAGAATTCTCGCCT</td><td>18418.3</td><td>7.2</td><td>1.5</td><td>4.8</td><td>1.5E-06</td><td>7.5E-06</td><td>BT20/MB468</td><td>MP</td></tr>
<tr><td>>hg38_21_+_8433172_8446622_RNA45SN1_45S_50ntFlanks@4682.4720.39|GCAAGACGGACCAGAGCGAAAGCATTTGCCAAGAATGTT</td><td>>hg38_21_+_8433172_8446622_RNA45SN1_45S_50ntFlanks</td><td>4682</td><td>4720</td><td>39</td><td>GCAAGACGGACCAGAGCGAAAGCATTTGCCAAGAATGTT</td><td>43.5</td><td>9.2</td><td>1.9</td><td>4.8</td><td>1.5E-06</td><td>7.5E-06</td><td>BT20/MB468</td><td>MP</td></tr>
<tr><td>>hg38_21_+_8433172_8446622_RNA45SN1_45S_50ntFlanks@3991.4024.34|TGGTGACTCTAGATAACCTCGGGCCGATCGCACG</td><td>>hg38_21_+_8433172_8446622_RNA45SN1_45S_50ntFlanks</td><td>3991</td><td>4024</td><td>34</td><td>TGGTGACTCTAGATAACCTCGGGCCGATCGCACG</td><td>766.6</td><td>-5.1</td><td>1.1</td><td>-4.6</td><td>5.1E-06</td><td>7.5E-06</td><td>MB231/MB468</td><td>MITO</td></tr>
<tr><td>>MI0000481|hsa-mir-184&WithFlank&15|+|79209782|79209877@59.80.22&[MIMAT0000454&hsa-miR-184&offsets|0|0;m-460&15|+|79209840|79209861&offsets|0|0]|TGGACGGAGAACTGATAAGGGT</td><td>>MI0000481|hsa-mir-184&WithFlank&15|+|79209782|79209877</td><td>59</td><td>80</td><td>22</td><td>TGGACGGAGAACTGATAAGGGT</td><td>556.7</td><td>5.2</td><td>1.0</td><td>5.1</td><td>4.1E-07</td><td>7.5E-06</td><td>BT20/MB231</td><td>MP</td></tr>
<tr><td>>hg38_21_+_8433172_8446622_RNA45SN1_45S_50ntFlanks@4725.4751.27|TTAATCAAGAACGAAAGTCGGAGGTTC</td><td>>hg38_21_+_8433172_8446622_RNA45SN1_45S_50ntFlanks</td><td>4725</td><td>4751</td><td>27</td><td>TTAATCAAGAACGAAAGTCGGAGGTTC</td><td>229.9</td><td>-6.5</td><td>1.4</td><td>-4.6</td><td>4.2E-06</td><td>7.5E-06</td><td>MB231/MB468</td><td>MP</td></tr>
<tr><td>>hg38_21_+_8433172_8446622_RNA45SN1_45S_50ntFlanks@10679.10706.28|GGTTCCGGCGGCGTCCGGTGAGCTCTCG</td><td>>hg38_21_+_8433172_8446622_RNA45SN1_45S_50ntFlanks</td><td>10679</td><td>10706</td><td>28</td><td>GGTTCCGGCGGCGTCCGGTGAGCTCTCG</td><td>8188.1</td><td>-5.1</td><td>1.1</td><td>-4.8</td><td>1.5E-06</td><td>7.5E-06</td><td>BT20/MB468</td><td>MP</td></tr>
<tr><td>>hg38_21_+_8433172_8446622_RNA45SN1_45S_50ntFlanks@12619.12654.36|AGGAGCCAATGGGGCGAAGCTACCATCTGTGGGATT</td><td>>hg38_21_+_8433172_8446622_RNA45SN1_45S_50ntFlanks</td><td>12619</td><td>12654</td><td>36</td><td>AGGAGCCAATGGGGCGAAGCTACCATCTGTGGGATT</td><td>1078.5</td><td>-4.2</td><td>0.9</td><td>-4.8</td><td>1.5E-06</td><td>7.5E-06</td><td>BT20/MB468</td><td>MP</td></tr>
<tr><td>>hg38_21_+_8433172_8446622_RNA45SN1_45S_50ntFlanks@476.519.44|TTCGTGATCGATGTGGTGACGTCGTGCTCTCCCGGGCCGGGTCC</td><td>>hg38_21_+_8433172_8446622_RNA45SN1_45S_50ntFlanks</td><td>476</td><td>519</td><td>44</td><td>TTCGTGATCGATGTGGTGACGTCGTGCTCTCCCGGGCCGGGTCC</td><td>1945.9</td><td>4.5</td><td>1.0</td><td>4.6</td><td>5.0E-06</td><td>7.6E-06</td><td>MB231/MB468</td><td>NUC</td></tr>
<tr><td>>MI0000102|hsa-mir-100&WithFlank&11|-|122152223|122152314@19.38.20&[MIMAT0000098&hsa-miR-100-5p&offsets|0|-2;m-27&11|-|122152276|122152296&offsets|0|-1]|AACCCGTAGATCCGAACTTG</td><td>>MI0000102|hsa-mir-100&WithFlank&11|-|122152223|122152314</td><td>19</td><td>38</td><td>20</td><td>AACCCGTAGATCCGAACTTG</td><td>62213.4</td><td>-6.8</td><td>1.3</td><td>-5.1</td><td>4.1E-07</td><td>7.6E-06</td><td>BT20/MB231</td><td>MP</td></tr>
<tr><td>>trna116_GluCTC_1_-_145399233_145399304@32.70.39,trna71_GluCTC_1_-_161439189_161439260@32.70.39,trna74_GluCTC_1_-_161431809_161431880@32.70.39,trna77_GluCTC_1_-_161424398_161424469@32.70.39,trna77_GluCTC_6_+_28949976_28950047@32.70.39,trna80_GluCTC_1_-_161417018_161417089@32.70.39,trna87_GluCTC_6_-_126101393_126101464@32.70.39|CTCTCACCGCCGCGGCCCGGGTTCGATTCCCGGTCAGGG</td><td>>trna116_GluCTC_1_-_145399233_145399304</td><td>32</td><td>70</td><td>39</td><td>CTCTCACCGCCGCGGCCCGGGTTCGATTCCCGGTCAGGG</td><td>5590.9</td><td>-7.6</td><td>1.7</td><td>-4.6</td><td>4.3E-06</td><td>7.6E-06</td><td>BT20/MB468</td><td>CYTO</td></tr>
<tr><td>>MI0003178|hsa-mir-519a-1&WithFlank&19|+|53752391|53752487@59.80.22;MI0003182|hsa-mir-519a-2&WithFlank&19|+|53762338|53762436@60.81.22&[MIMAT0002869&hsa-miR-519a-3p&offsets|0|0;m-880&19|+|53752449|53752470&offsets|0|0];[MIMAT0002869_1&hsa-miR-519a-3p&offsets|0|0;m-880&19|+|53762397|53762418&offsets|0|0]|AAAGTGCATCCTTTTAGAGTGT</td><td>>MI0003178|hsa-mir-519a-1&WithFlank&19|+|53752391|53752487</td><td>59</td><td>80</td><td>22</td><td>AAAGTGCATCCTTTTAGAGTGT</td><td>34.6</td><td>8.3</td><td>1.8</td><td>4.6</td><td>4.2E-06</td><td>7.6E-06</td><td>MB231/MB468</td><td>MP</td></tr>
<tr><td>>hg38_21_+_8433172_8446622_RNA45SN1_45S_50ntFlanks@12717.12747.31|CCTCGGTTGGCCTCGGATAGCCGGTCCCCCG</td><td>>hg38_21_+_8433172_8446622_RNA45SN1_45S_50ntFlanks</td><td>12717</td><td>12747</td><td>31</td><td>CCTCGGTTGGCCTCGGATAGCCGGTCCCCCG</td><td>83.3</td><td>-8.7</td><td>1.9</td><td>-4.6</td><td>4.3E-06</td><td>7.6E-06</td><td>BT20/MB468</td><td>CYTO</td></tr>
<tr><td>>hg38_21_+_8433172_8446622_RNA45SN1_45S_50ntFlanks@10402.10433.32|AACATGGGTCAGTCGGTCCTGAGAGATGGGCG</td><td>>hg38_21_+_8433172_8446622_RNA45SN1_45S_50ntFlanks</td><td>10402</td><td>10433</td><td>32</td><td>AACATGGGTCAGTCGGTCCTGAGAGATGGGCG</td><td>601.2</td><td>-4.4</td><td>1.0</td><td>-4.6</td><td>4.8E-06</td><td>7.6E-06</td><td>MB231/MB468</td><td>CYTO</td></tr>
<tr><td>>hg38_21_+_8433172_8446622_RNA45SN1_45S_50ntFlanks@11570.11591.22|AGCCGACTTAGAACTGGTGCGG</td><td>>hg38_21_+_8433172_8446622_RNA45SN1_45S_50ntFlanks</td><td>11570</td><td>11591</td><td>22</td><td>AGCCGACTTAGAACTGGTGCGG</td><td>342.3</td><td>-4.6</td><td>1.0</td><td>-4.6</td><td>4.3E-06</td><td>7.6E-06</td><td>BT20/MB468</td><td>CYTO</td></tr>
<tr><td>>trna10_GluCTC_3_-_103879524_103879595@16.33.18,trna116_GluCTC_1_-_145399233_145399304@16.33.18,trna25_GluCTC_2_-_71273488_71273560@16.33.18,trna59_GluCTC_1_+_249168447_249168518@16.33.18,trna71_GluCTC_1_-_161439189_161439260@16.33.18,trna74_GluCTC_1_-_161431809_161431880@16.33.18,trna77_GluCTC_1_-_161424398_161424469@16.33.18,trna77_GluCTC_6_+_28949976_28950047@16.33.18,trna80_GluCTC_1_-_161417018_161417089@16.33.18,trna87_GluCTC_6_-_126101393_126101464@16.33.18|TGGTTAGGATTCGGCGCT</td><td>>trna10_GluCTC_3_-_103879524_103879595</td><td>16</td><td>33</td><td>18</td><td>TGGTTAGGATTCGGCGCT</td><td>718.2</td><td>4.5</td><td>1.0</td><td>4.6</td><td>4.8E-06</td><td>7.6E-06</td><td>MB231/MB468</td><td>CYTO</td></tr>
</table>

<table>
<tr><td>>MI0000542|hsa-mir-320a&WithFlank&8|-|22244960|22245043@49.68.20(+2U)&[MIMAT0000510&hsa-miR-320a-3p&offsets|+1|-1(+2U);m-60&8|-|22244975|22244996&offsets|+1|-1(+2U)]|AAAGCTGGGTTGAGAGGGCGTT</td><td>>MI0000542|hsa-mir-320a&WithFlank&8|-|22244960|22245043</td><td>49</td><td>68</td><td>20(+2U)</td><td>AAAGCTGGGTTGAGAGGGCGTT</td><td>620.6</td><td>7.9</td><td>1.6</td><td>4.8</td><td>1.5E-06</td><td>7.6E-06</td><td>BT20/MB468</td><td>MP</td></tr>
<tr><td>>hg38_21_+_8433172_8446622_RNA45SN1_45S_50ntFlanks@12444.12484.41|AAGCAGAATTCACCAAGCGTTGGATTGTTCACCCACTAATA</td><td>>hg38_21_+_8433172_8446622_RNA45SN1_45S_50ntFlanks</td><td>12444</td><td>12484</td><td>41</td><td>AAGCAGAATTCACCAAGCGTTGGATTGTTCACCCACTAATA</td><td>75.5</td><td>-9.3</td><td>1.9</td><td>-4.8</td><td>1.5E-06</td><td>7.6E-06</td><td>BT20/MB468</td><td>MP</td></tr>
<tr><td>>hg38_21_+_8433172_8446622_RNA45SN1_45S_50ntFlanks@11828.11856.29|GCGAAACCACAGCCAAGGGAACGGGCTTG</td><td>>hg38_21_+_8433172_8446622_RNA45SN1_45S_50ntFlanks</td><td>11828</td><td>11856</td><td>29</td><td>GCGAAACCACAGCCAAGGGAACGGGCTTG</td><td>62.2</td><td>-9.0</td><td>1.9</td><td>-4.8</td><td>1.5E-06</td><td>7.6E-06</td><td>BT20/MB468</td><td>MP</td></tr>
<tr><td>>MI0000102|hsa-mir-100&WithFlank&11|-|122152223|122152314@19.40.22(+2A)&[MIMAT0000098&hsa-miR-100-5p&offsets|0|0(+2A);m-27&11|-|122152276|122152296&offsets|0|+1(+2A)]|AACCCGTAGATCCGAACTTGTGAA</td><td>>MI0000102|hsa-mir-100&WithFlank&11|-|122152223|122152314</td><td>19</td><td>40</td><td>22(+2A)</td><td>AACCCGTAGATCCGAACTTGTGAA</td><td>1688.8</td><td>-5.8</td><td>1.2</td><td>-5.0</td><td>5.8E-07</td><td>7.6E-06</td><td>BT20/MB231</td><td>NUC</td></tr>
<tr><td>>hg38_MT_+_1621_3279_RNR2_16S_50ntFlanks@1427.1451.25|GATTAAAGTCCTACGTGATCTGAGT</td><td>>hg38_MT_+_1621_3279_RNR2_16S_50ntFlanks</td><td>1427</td><td>1451</td><td>25</td><td>GATTAAAGTCCTACGTGATCTGAGT</td><td>362.5</td><td>4.2</td><td>0.9</td><td>4.6</td><td>4.2E-06</td><td>7.6E-06</td><td>MB231/MB468</td><td>MP</td></tr>
<tr><td>>hg38_21_+_8433172_8446622_RNA45SN1_45S_50ntFlanks@4724.4759.36|ATTAATCAAGAACGAAAGTCGGAGGTTCGAAGACGA</td><td>>hg38_21_+_8433172_8446622_RNA45SN1_45S_50ntFlanks</td><td>4724</td><td>4759</td><td>36</td><td>ATTAATCAAGAACGAAAGTCGGAGGTTCGAAGACGA</td><td>2184.6</td><td>-3.8</td><td>0.8</td><td>-4.6</td><td>4.3E-06</td><td>7.6E-06</td><td>BT20/MB468</td><td>CYTO</td></tr>
<tr><td>>hg38_21_+_8433172_8446622_RNA45SN1_45S_50ntFlanks@12934.12957.24|CTAAACCATTCGTAGACGACCTGC</td><td>>hg38_21_+_8433172_8446622_RNA45SN1_45S_50ntFlanks</td><td>12934</td><td>12957</td><td>24</td><td>CTAAACCATTCGTAGACGACCTGC</td><td>411.5</td><td>-2.6</td><td>0.6</td><td>-4.6</td><td>4.8E-06</td><td>7.6E-06</td><td>MB231/MB468</td><td>CYTO</td></tr>
<tr><td>>hg38_21_+_8433172_8446622_RNA45SN1_45S_50ntFlanks@10678.10709.32|CGGTTCCGGCGGCGTCCGGTGAGCTCTCGCTG</td><td>>hg38_21_+_8433172_8446622_RNA45SN1_45S_50ntFlanks</td><td>10678</td><td>10709</td><td>32</td><td>CGGTTCCGGCGGCGTCCGGTGAGCTCTCGCTG</td><td>16457.1</td><td>-4.5</td><td>1.0</td><td>-4.6</td><td>4.3E-06</td><td>7.6E-06</td><td>BT20/MB468</td><td>CYTO</td></tr>
<tr><td>>trna10_GluCTC_3_-_103879524_103879595@16.39.24,trna116_GluCTC_1_-_145399233_145399304@16.39.24,trna59_GluCTC_1_+_249168447_249168518@16.39.24,trna71_GluCTC_1_-_161439189_161439260@16.39.24,trna74_GluCTC_1_-_161431809_161431880@16.39.24,trna77_GluCTC_1_-_161424398_161424469@16.39.24,trna77_GluCTC_6_+_28949976_28950047@16.39.24,trna80_GluCTC_1_-_161417018_161417089@16.39.24,trna87_GluCTC_6_-_126101393_126101464@16.39.24|TGGTTAGGATTCGGCGCTCTCACC</td><td>>trna10_GluCTC_3_-_103879524_103879595</td><td>16</td><td>39</td><td>24</td><td>TGGTTAGGATTCGGCGCTCTCACC</td><td>869.8</td><td>5.9</td><td>1.3</td><td>4.6</td><td>4.3E-06</td><td>7.6E-06</td><td>BT20/MB468</td><td>CYTO</td></tr>
<tr><td>>MI0000469|hsa-mir-125a&WithFlank&19|+|51693248|51693345@21.42.22(+1A)&[MIMAT0000443&hsa-miR-125a-5p&offsets|0|-2(+1A);m-63&19|+|51693268|51693289&offsets|0|0(+1A)]|TCCCTGAGACCCTTTAACCTGTA</td><td>>MI0000469|hsa-mir-125a&WithFlank&19|+|51693248|51693345</td><td>21</td><td>42</td><td>22(+1A)</td><td>TCCCTGAGACCCTTTAACCTGTA</td><td>828.7</td><td>3.5</td><td>0.8</td><td>4.6</td><td>5.0E-06</td><td>7.6E-06</td><td>MB231/MB468</td><td>NUC</td></tr>
<tr><td>>hg38_21_+_8433172_8446622_RNA45SN1_45S_50ntFlanks@8380.8402.23|CAAGAGGGCGTGAAACCGTTAAG</td><td>>hg38_21_+_8433172_8446622_RNA45SN1_45S_50ntFlanks</td><td>8380</td><td>8402</td><td>23</td><td>CAAGAGGGCGTGAAACCGTTAAG</td><td>1307.5</td><td>-6.9</td><td>1.5</td><td>-4.6</td><td>4.3E-06</td><td>7.6E-06</td><td>BT20/MB468</td><td>CYTO</td></tr>
<tr><td>>hg38_21_+_8433172_8446622_RNA45SN1_45S_50ntFlanks@5479.5506.28|TGGCGGAGCGCTGAGAAGACGGTCGAAC</td><td>>hg38_21_+_8433172_8446622_RNA45SN1_45S_50ntFlanks</td><td>5479</td><td>5506</td><td>28</td><td>TGGCGGAGCGCTGAGAAGACGGTCGAAC</td><td>53.7</td><td>9.0</td><td>2.0</td><td>4.6</td><td>4.3E-06</td><td>7.6E-06</td><td>MB231/MB468</td><td>MP</td></tr>
<tr><td>>hg38_21_+_8433172_8446622_RNA45SN1_45S_50ntFlanks@10393.10431.39|CAGCAGTTGAACATGGGTCAGTCGGTCCTGAGAGATGGG</td><td>>hg38_21_+_8433172_8446622_RNA45SN1_45S_50ntFlanks</td><td>10393</td><td>10431</td><td>39</td><td>CAGCAGTTGAACATGGGTCAGTCGGTCCTGAGAGATGGG</td><td>295.4</td><td>-6.7</td><td>1.5</td><td>-4.6</td><td>5.2E-06</td><td>7.6E-06</td><td>MB231/MB468</td><td>MITO</td></tr>
<tr><td>>hg38_21_+_8433172_8446622_RNA45SN1_45S_50ntFlanks@9567.9601.35|ACGTGCAAATCGGTCGTCCGACCTGGGTATAGGGG</td><td>>hg38_21_+_8433172_8446622_RNA45SN1_45S_50ntFlanks</td><td>9567</td><td>9601</td><td>35</td><td>ACGTGCAAATCGGTCGTCCGACCTGGGTATAGGGG</td><td>536.8</td><td>-5.2</td><td>1.1</td><td>-4.6</td><td>5.2E-06</td><td>7.6E-06</td><td>MB231/MB468</td><td>MITO</td></tr>
<tr><td>>hg38_21_+_8433172_8446622_RNA45SN1_45S_50ntFlanks@10830.10853.24|CAAGCCGGATCCGTAACTTCGGGA</td><td>>hg38_21_+_8433172_8446622_RNA45SN1_45S_50ntFlanks</td><td>10830</td><td>10853</td><td>24</td><td>CAAGCCGGATCCGTAACTTCGGGA</td><td>181.4</td><td>-5.9</td><td>1.3</td><td>-4.6</td><td>4.3E-06</td><td>7.6E-06</td><td>BT20/MB468</td><td>CYTO</td></tr>
<tr><td>>hg38_21_+_8433172_8446622_RNA45SN1_45S_50ntFlanks@11319.11342.24|CCCACGTCTCGTCGCGCGCGCGTC</td><td>>hg38_21_+_8433172_8446622_RNA45SN1_45S_50ntFlanks</td><td>11319</td><td>11342</td><td>24</td><td>CCCACGTCTCGTCGCGCGCGCGTC</td><td>775.4</td><td>-5.0</td><td>1.1</td><td>-4.6</td><td>4.3E-06</td><td>7.6E-06</td><td>BT20/MB468</td><td>CYTO</td></tr>
<tr><td>>hg38_1_-_228634819_228635039_RNA5S12_5S_50ntFlanks@90.123.34|TGATCTCGGAAGCTAAGCAGGGTCGGGCCTGGTT</td><td>>hg38_1_-_228634819_228635039_RNA5S12_5S_50ntFlanks</td><td>90</td><td>123</td><td>34</td><td>TGATCTCGGAAGCTAAGCAGGGTCGGGCCTGGTT</td><td>208.0</td><td>4.3</td><td>0.9</td><td>4.6</td><td>5.1E-06</td><td>7.6E-06</td><td>MB231/MB468</td><td>NUC</td></tr>
<tr><td>>hg38_21_+_8433172_8446622_RNA45SN1_45S_50ntFlanks@11785.11805.21|TGAATGGATGAACGAGATTCC</td><td>>hg38_21_+_8433172_8446622_RNA45SN1_45S_50ntFlanks</td><td>11785</td><td>11805</td><td>21</td><td>TGAATGGATGAACGAGATTCC</td><td>1100.5</td><td>-4.7</td><td>1.0</td><td>-4.6</td><td>4.9E-06</td><td>7.6E-06</td><td>MB231/MB468</td><td>CYTO</td></tr>
<tr><td>>MI0000071|hsa-mir-17&WithFlank&13|+|91350599|91350694@20.42.23(+1A)&[MIMAT0000070&hsa-miR-17-5p&offsets|0|0(+1A);m-88&13|+|91350618|91350640&offsets|0|0(+1A)]|CAAAGTGCTTACAGTGCAGGTAGA</td><td>>MI0000071|hsa-mir-17&WithFlank&13|+|91350599|91350694</td><td>20</td><td>42</td><td>23(+1A)</td><td>CAAAGTGCTTACAGTGCAGGTAGA</td><td>2556.0</td><td>4.5</td><td>1.0</td><td>4.6</td><td>4.9E-06</td><td>7.6E-06</td><td>MB231/MB468</td><td>CYTO</td></tr>
<tr><td>>hg38_21_+_8433172_8446622_RNA45SN1_45S_50ntFlanks@12646.12670.25|TGTGGGATTATGACTGAACGCCTCT</td><td>>hg38_21_+_8433172_8446622_RNA45SN1_45S_50ntFlanks</td><td>12646</td><td>12670</td><td>25</td><td>TGTGGGATTATGACTGAACGCCTCT</td><td>49.2</td><td>9.4</td><td>1.9</td><td>4.8</td><td>1.5E-06</td><td>7.7E-06</td><td>BT20/MB468</td><td>MP</td></tr>
<tr><td>>hg38_21_+_8433172_8446622_RNA45SN1_45S_50ntFlanks@11571.11601.31|GCCGACTTAGAACTGGTGCGGACCAGGGGAA</td><td>>hg38_21_+_8433172_8446622_RNA45SN1_45S_50ntFlanks</td><td>11571</td><td>11601</td><td>31</td><td>GCCGACTTAGAACTGGTGCGGACCAGGGGAA</td><td>714.8</td><td>-4.3</td><td>0.9</td><td>-4.6</td><td>5.2E-06</td><td>7.7E-06</td><td>MB231/MB468</td><td>MITO</td></tr>
<tr><td>>MI0000542|hsa-mir-320a&WithFlank&8|-|22244960|22245043@48.69.22(+1C)&[MIMAT0000510&hsa-miR-320a-3p&offsets|0|0(+1C);m-60&8|-|22244975|22244996&offsets|0|0(+1C)]|AAAAGCTGGGTTGAGAGGGCGAC</td><td>>MI0000542|hsa-mir-320a&WithFlank&8|-|22244960|22245043</td><td>48</td><td>69</td><td>22(+1C)</td><td>AAAAGCTGGGTTGAGAGGGCGAC</td><td>3766.5</td><td>4.3</td><td>0.9</td><td>5.1</td><td>4.4E-07</td><td>7.7E-06</td><td>BT20/MB231</td><td>CYTO</td></tr>
<tr><td>>hg38_21_+_8433172_8446622_RNA45SN1_45S_50ntFlanks@9554.9584.31|CGTAGCGGTCCTGACGTGCAAATCGGTCGTC</td><td>>hg38_21_+_8433172_8446622_RNA45SN1_45S_50ntFlanks</td><td>9554</td><td>9584</td><td>31</td><td>CGTAGCGGTCCTGACGTGCAAATCGGTCGTC</td><td>789.1</td><td>-7.2</td><td>1.6</td><td>-4.6</td><td>4.3E-06</td><td>7.7E-06</td><td>MB231/MB468</td><td>MP</td></tr>
<tr><td>>hg38_21_+_8433172_8446622_RNA45SN1_45S_50ntFlanks@4995.5022.28|ACAGATTGATAGCTCTTTCTCGATTCCG</td><td>>hg38_21_+_8433172_8446622_RNA45SN1_45S_50ntFlanks</td><td>4995</td><td>5022</td><td>28</td><td>ACAGATTGATAGCTCTTTCTCGATTCCG</td><td>996.6</td><td>-6.5</td><td>1.4</td><td>-4.6</td><td>5.1E-06</td><td>7.7E-06</td><td>MB231/MB468</td><td>NUC</td></tr>
<tr><td>>hg38_21_+_8433172_8446622_RNA45SN1_45S_50ntFlanks@8117.8142.26|CGGGACATGTGGCGTACGGAAGACCC</td><td>>hg38_21_+_8433172_8446622_RNA45SN1_45S_50ntFlanks</td><td>8117</td><td>8142</td><td>26</td><td>CGGGACATGTGGCGTACGGAAGACCC</td><td>101.0</td><td>10.0</td><td>2.0</td><td>5.1</td><td>4.2E-07</td><td>7.7E-06</td><td>BT20/MB231</td><td>MP</td></tr>
<tr><td>>hg38_21_+_8433172_8446622_RNA45SN1_45S_50ntFlanks@3775.3798.24|GCCGGTACAGTGAAACTGCGAATG</td><td>>hg38_21_+_8433172_8446622_RNA45SN1_45S_50ntFlanks</td><td>3775</td><td>3798</td><td>24</td><td>GCCGGTACAGTGAAACTGCGAATG</td><td>536.3</td><td>-7.4</td><td>1.6</td><td>-4.5</td><td>5.4E-06</td><td>7.7E-06</td><td>BT20/MB468</td><td>NUC</td></tr>
<tr><td>>am_MI0000809|hsa-mir-151a&WithFlank&8|-|140732558|140732659@53.73.21(+2U)&[MIMAT0000757&hsa-miR-151a-3p&offsets|0|0(+2U);m-30&8|-|140732587|140732607&offsets|0|0(+2U)]|CTAGACTGAAGCTCCTTGAGGTT</td><td>>am_MI0000809|hsa-mir-151a&WithFlank&8|-|140732558|140732659</td><td>53</td><td>73</td><td>21(+2U)</td><td>CTAGACTGAAGCTCCTTGAGGTT</td><td>668.7</td><td>5.5</td><td>1.2</td><td>4.6</td><td>4.3E-06</td><td>7.7E-06</td><td>MB231/MB468</td><td>MP</td></tr>
<tr><td>>hg38_21_+_8433172_8446622_RNA45SN1_45S_50ntFlanks@11769.11786.18|TAATTAGTGACGCGCATG</td><td>>hg38_21_+_8433172_8446622_RNA45SN1_45S_50ntFlanks</td><td>11769</td><td>11786</td><td>18</td><td>TAATTAGTGACGCGCATG</td><td>557.5</td><td>-6.7</td><td>1.5</td><td>-4.6</td><td>4.3E-06</td><td>7.7E-06</td><td>BT20/MB468</td><td>CYTO</td></tr>
<tr><td>>hg38_21_+_8433172_8446622_RNA45SN1_45S_50ntFlanks@10350.10376.27|ACGAGAACTTTGAAGGCCGAAGTGGAG</td><td>>hg38_21_+_8433172_8446622_RNA45SN1_45S_50ntFlanks</td><td>10350</td><td>10376</td><td>27</td><td>ACGAGAACTTTGAAGGCCGAAGTGGAG</td><td>3344.4</td><td>-5.7</td><td>1.2</td><td>-4.8</td><td>1.5E-06</td><td>7.7E-06</td><td>BT20/MB468</td><td>MP</td></tr>
<tr><td>>hg38_21_+_8433172_8446622_RNA45SN1_45S_50ntFlanks@476.502.27|TTCGTGATCGATGTGGTGACGTCGTGC</td><td>>hg38_21_+_8433172_8446622_RNA45SN1_45S_50ntFlanks</td><td>476</td><td>502</td><td>27</td><td>TTCGTGATCGATGTGGTGACGTCGTGC</td><td>280.0</td><td>5.7</td><td>1.3</td><td>4.6</td><td>5.2E-06</td><td>7.7E-06</td><td>MB231/MB468</td><td>MITO</td></tr>
<tr><td>>hg38_21_+_8433172_8446622_RNA45SN1_45S_50ntFlanks@8121.8142.22|ACATGTGGCGTACGGAAGACCC</td><td>>hg38_21_+_8433172_8446622_RNA45SN1_45S_50ntFlanks</td><td>8121</td><td>8142</td><td>22</td><td>ACATGTGGCGTACGGAAGACCC</td><td>105.6</td><td>10.1</td><td>2.0</td><td>5.1</td><td>4.2E-07</td><td>7.7E-06</td><td>BT20/MB231</td><td>MP</td></tr>
<tr><td>>hg38_21_+_8433172_8446622_RNA45SN1_45S_50ntFlanks@10073.10092.20|GACGGGAGCGGCGGGGGCGG</td><td>>hg38_21_+_8433172_8446622_RNA45SN1_45S_50ntFlanks</td><td>10073</td><td>10092</td><td>20</td><td>GACGGGAGCGGCGGGGGCGG</td><td>377.8</td><td>-7.1</td><td>1.5</td><td>-4.8</td><td>1.5E-06</td><td>7.7E-06</td><td>BT20/MB468</td><td>MP</td></tr>
<tr><td>>hg38_21_+_8433172_8446622_RNA45SN1_45S_50ntFlanks@6647.6670.24|CGTACGACTCTTAGCGGTGGATCA</td><td>>hg38_21_+_8433172_8446622_RNA45SN1_45S_50ntFlanks</td><td>6647</td><td>6670</td><td>24</td><td>CGTACGACTCTTAGCGGTGGATCA</td><td>3450.1</td><td>-3.4</td><td>0.7</td><td>-4.6</td><td>5.2E-06</td><td>7.7E-06</td><td>MB231/MB468</td><td>MITO</td></tr>
<tr><td>>hg38_21_+_8433172_8446622_RNA45SN1_45S_50ntFlanks@7991.8010.20|ACGTGGCGACCCGCTGAATT</td><td>>hg38_21_+_8433172_8446622_RNA45SN1_45S_50ntFlanks</td><td>7991</td><td>8010</td><td>20</td><td>ACGTGGCGACCCGCTGAATT</td><td>328.4</td><td>2.4</td><td>0.5</td><td>4.6</td><td>4.4E-06</td><td>7.7E-06</td><td>BT20/MB468</td><td>CYTO</td></tr>
<tr><td>>hg38_21_+_8433172_8446622_RNA45SN1_45S_50ntFlanks@11586.11611.26|GTGCGGACCAGGGGAATCCGACTGTT</td><td>>hg38_21_+_8433172_8446622_RNA45SN1_45S_50ntFlanks</td><td>11586</td><td>11611</td><td>26</td><td>GTGCGGACCAGGGGAATCCGACTGTT</td><td>2877.5</td><td>-3.0</td><td>0.7</td><td>-4.6</td><td>4.9E-06</td><td>7.7E-06</td><td>MB231/MB468</td><td>CYTO</td></tr>
<tr><td>>hg38_21_+_8433172_8446622_RNA45SN1_45S_50ntFlanks@11875.11893.19|GAAGACCCTGTTGAGCTTG</td><td>>hg38_21_+_8433172_8446622_RNA45SN1_45S_50ntFlanks</td><td>11875</td><td>11893</td><td>19</td><td>GAAGACCCTGTTGAGCTTG</td><td>708.5</td><td>-5.4</td><td>1.2</td><td>-4.6</td><td>5.1E-06</td><td>7.7E-06</td><td>MB231/MB468</td><td>NUC</td></tr>
<tr><td>>hg38_21_+_8433172_8446622_RNA45SN1_45S_50ntFlanks@12407.12438.32|TTTTGATCCTTCGATGTCGGCTCTTCCTATCA</td><td>>hg38_21_+_8433172_8446622_RNA45SN1_45S_50ntFlanks</td><td>12407</td><td>12438</td><td>32</td><td>TTTTGATCCTTCGATGTCGGCTCTTCCTATCA</td><td>300.6</td><td>3.6</td><td>0.8</td><td>4.6</td><td>4.9E-06</td><td>7.7E-06</td><td>MB231/MB468</td><td>CYTO</td></tr>
<tr><td>>hg38_21_+_8433172_8446622_RNA45SN1_45S_50ntFlanks@11874.11910.37|AGAAGACCCTGTTGAGCTTGACTCTAGTCTGGCACGG</td><td>>hg38_21_+_8433172_8446622_RNA45SN1_45S_50ntFlanks</td><td>11874</td><td>11910</td><td>37</td><td>AGAAGACCCTGTTGAGCTTGACTCTAGTCTGGCACGG</td><td>236.2</td><td>-6.3</td><td>1.4</td><td>-4.6</td><td>5.1E-06</td><td>7.7E-06</td><td>MB231/MB468</td><td>NUC</td></tr>
<tr><td>>MI0000478|hsa-mir-149&WithFlank&2|+|240455995|240456095@21.43.23&[MIMAT0000450&hsa-miR-149-5p&offsets|0|0;m-195&2|+|240456015|240456037&offsets|0|0]|TCTGGCTCCGTGTCTTCACTCCC</td><td>>MI0000478|hsa-mir-149&WithFlank&2|+|240455995|240456095</td><td>21</td><td>43</td><td>23</td><td>TCTGGCTCCGTGTCTTCACTCCC</td><td>2561.7</td><td>4.0</td><td>0.8</td><td>5.0</td><td>4.4E-07</td><td>7.7E-06</td><td>BT20/MB231</td><td>CYTO</td></tr>
<tr><td>>MI0000650|hsa-mir-200c&WithFlank&12|+|6963693|6963772@50.71.22(+1U)&[MIMAT0000617&hsa-miR-200c-3p&offsets|0|-1(+1U);m-54&12|+|6963742|6963764&offsets|0|-1(+1U)]|TAATACTGCCGGGTAATGATGGT</td><td>>MI0000650|hsa-mir-200c&WithFlank&12|+|6963693|6963772</td><td>50</td><td>71</td><td>22(+1U)</td><td>TAATACTGCCGGGTAATGATGGT</td><td>2898.3</td><td>9.9</td><td>2.0</td><td>5.0</td><td>4.5E-07</td><td>7.7E-06</td><td>BT20/MB231</td><td>CYTO</td></tr>
<tr><td>>trnaMT_MetCAT_MT_+_4402_4469@35.71.37|CCCCGAAAATGTTGGTTATACCCTTCCCGTACTACCA</td><td>>trnaMT_MetCAT_MT_+_4402_4469</td><td>35</td><td>71</td><td>37</td><td>CCCCGAAAATGTTGGTTATACCCTTCCCGTACTACCA</td><td>336.6</td><td>-4.1</td><td>0.8</td><td>-5.0</td><td>4.5E-07</td><td>7.7E-06</td><td>BT20/MB231</td><td>CYTO</td></tr>
<tr><td>>hg38_21_+_8433172_8446622_RNA45SN1_45S_50ntFlanks@12027.12049.23|ACTCTGATCGTTTTTTCACTGAC</td><td>>hg38_21_+_8433172_8446622_RNA45SN1_45S_50ntFlanks</td><td>12027</td><td>12049</td><td>23</td><td>ACTCTGATCGTTTTTTCACTGAC</td><td>200.5</td><td>11.0</td><td>2.2</td><td>5.1</td><td>4.2E-07</td><td>7.7E-06</td><td>BT20/MB231</td><td>MP</td></tr>
<tr><td>>hg38_21_+_8433172_8446622_RNA45SN1_45S_50ntFlanks@9454.9503.50|CCGCGCCGGGGAGGTGGAGCACGAGCGCACGTGTTAGGACCCGAAAGATG</td><td>>hg38_21_+_8433172_8446622_RNA45SN1_45S_50ntFlanks</td><td>9454</td><td>9503</td><td>50</td><td>CCGCGCCGGGGAGGTGGAGCACGAGCGCACGTGTTAGGACCCGAAAGATG</td><td>1819.8</td><td>-9.1</td><td>1.9</td><td>-4.8</td><td>1.5E-06</td><td>7.7E-06</td><td>BT20/MB468</td><td>MP</td></tr>
<tr><td>>MI0000102|hsa-mir-100&WithFlank&11|-|122152223|122152314@19.39.21(+1A)&[MIMAT0000098&hsa-miR-100-5p&offsets|0|-1(+1A);m-27&11|-|122152276|122152296&offsets|0|0(+1A)]|AACCCGTAGATCCGAACTTGTA</td><td>>MI0000102|hsa-mir-100&WithFlank&11|-|122152223|122152314</td><td>19</td><td>39</td><td>21(+1A)</td><td>AACCCGTAGATCCGAACTTGTA</td><td>316.9</td><td>7.8</td><td>1.7</td><td>4.5</td><td>5.5E-06</td><td>7.7E-06</td><td>BT20/MB468</td><td>NUC</td></tr>
<tr><td>>hg38_MT_+_598_1651_RNR1_12S_50ntFlanks@972.1002.31|GGTAAGTGTACTGGAAAGTGCACTTGGACGA</td><td>>hg38_MT_+_598_1651_RNR1_12S_50ntFlanks</td><td>972</td><td>1002</td><td>31</td><td>GGTAAGTGTACTGGAAAGTGCACTTGGACGA</td><td>128.6</td><td>-10.4</td><td>2.1</td><td>-5.1</td><td>4.3E-07</td><td>7.7E-06</td><td>BT20/MB231</td><td>MP</td></tr>
</table>

<table>
<tr><td>>am_MI0000786|hsa-mir-378a&WithFlank&5|+|149732819|149732896@49.69.21(+1U)&[MIMAT0000732&hsa-miR-378a-3p&offsets|0|-1(+1U);m-44&5|+|149732867|149732888&offsets|0|-1(+1U)]|ACTGGACTTGGAGTCAGAAGGT</td><td>>am_MI0000786|hsa-mir-378a&WithFlank&5|+|149732819|149732896</td><td>49</td><td>69</td><td>21(+1U)</td><td>ACTGGACTTGGAGTCAGAAGGT</td><td>1967.2</td><td>3.8</td><td>0.8</td><td>4.6</td><td>4.9E-06</td><td>7.7E-06</td><td>MB231/MB468</td><td>CYTO</td></tr>
<tr><td>>hg38_21_+_8433172_8446622_RNA45SN1_45S_50ntFlanks@4240.4258.19|ACTCTTTCGAGGCCCTGTA</td><td>>hg38_21_+_8433172_8446622_RNA45SN1_45S_50ntFlanks</td><td>4240</td><td>4258</td><td>19</td><td>ACTCTTTCGAGGCCCTGTA</td><td>540.5</td><td>-4.8</td><td>1.0</td><td>-4.6</td><td>4.4E-06</td><td>7.7E-06</td><td>BT20/MB468</td><td>CYTO</td></tr>
<tr><td>>hg38_21_+_8433172_8446622_RNA45SN1_45S_50ntFlanks@10689.10708.20|GCGTCCGGTGAGCTCTCGCT</td><td>>hg38_21_+_8433172_8446622_RNA45SN1_45S_50ntFlanks</td><td>10689</td><td>10708</td><td>20</td><td>GCGTCCGGTGAGCTCTCGCT</td><td>2759.6</td><td>-8.4</td><td>1.8</td><td>-4.5</td><td>5.5E-06</td><td>7.7E-06</td><td>BT20/MB468</td><td>NUC</td></tr>
<tr><td>>hg38_21_+_8433172_8446622_RNA45SN1_45S_50ntFlanks@4726.4751.26|TAATCAAGAACGAAAGTCGGAGGTTC</td><td>>hg38_21_+_8433172_8446622_RNA45SN1_45S_50ntFlanks</td><td>4726</td><td>4751</td><td>26</td><td>TAATCAAGAACGAAAGTCGGAGGTTC</td><td>201.2</td><td>-3.0</td><td>0.7</td><td>-4.6</td><td>4.4E-06</td><td>7.7E-06</td><td>BT20/MB468</td><td>CYTO</td></tr>
<tr><td>>MI0000239|hsa-mir-197&WithFlank&1|+|109598887|109598973@54.75.22&[MIMAT0000227&hsa-miR-197-3p&offsets|0|0;m-99&1|+|109598940|109598961&offsets|0|0]|TTCACCACCTTCTCCACCCAGC</td><td>>MI0000239|hsa-mir-197&WithFlank&1|+|109598887|109598973</td><td>54</td><td>75</td><td>22</td><td>TTCACCACCTTCTCCACCCAGC</td><td>1450.8</td><td>4.0</td><td>0.9</td><td>4.6</td><td>5.2E-06</td><td>7.7E-06</td><td>MB231/MB468</td><td>MITO</td></tr>
<tr><td>>trna10_AspGTC_12_-_125424193_125424264@1.29.29,trna12_AspGTC_12_-_125411891_125411962@1.29.29,trna38_AspGTC_17_-_8125556_8125627@1.29.29,trna45_AspGTC_6_+_27447453_27447524@1.29.29,trna48_AspGTC_6_+_27471523_27471594@1.29.29,trna4_AspGTC_12_+_96429799_96429870@1.29.29,trna69_AspGTC_1_-_161440205_161440276@1.29.29,trna72_AspGTC_1_-_161432824_161432895@1.29.29,trna75_AspGTC_1_-_161425414_161425485@1.29.29,trna78_AspGTC_1_-_161418033_161418104@1.29.29,trna81_AspGTC_1_-_161410615_161410686@1.29.29|TCCTCGTTAGTATAGTGGTGAGTATCCCC</td><td>>trna10_AspGTC_12_-_125424193_125424264</td><td>1</td><td>29</td><td>29</td><td>TCCTCGTTAGTATAGTGGTGAGTATCCCC</td><td>207.7</td><td>-6.6</td><td>1.5</td><td>-4.6</td><td>5.2E-06</td><td>7.7E-06</td><td>MB231/MB468</td><td>MITO</td></tr>
<tr><td>>MI0005559|hsa-mir-744&WithFlank&17|+|12081893|12082002@17.38.22&[MIMAT0004945&hsa-miR-744-5p&offsets|0|0;m-172&17|+|12081909|12081929&offsets|0|+1]|TGCGGGGCTAGGGCTAACAGCA</td><td>>MI0005559|hsa-mir-744&WithFlank&17|+|12081893|12082002</td><td>17</td><td>38</td><td>22</td><td>TGCGGGGCTAGGGCTAACAGCA</td><td>610.9</td><td>4.3</td><td>0.9</td><td>4.8</td><td>1.6E-06</td><td>7.7E-06</td><td>BT20/MB468</td><td>MP</td></tr>
<tr><td>>hg38_21_+_8433172_8446622_RNA45SN1_45S_50ntFlanks@12399.12433.35|ACGTCGCTTTTTGATCCTTCGATGTCGGCTCTTCC</td><td>>hg38_21_+_8433172_8446622_RNA45SN1_45S_50ntFlanks</td><td>12399</td><td>12433</td><td>35</td><td>ACGTCGCTTTTTGATCCTTCGATGTCGGCTCTTCC</td><td>366.0</td><td>3.6</td><td>0.8</td><td>4.6</td><td>4.9E-06</td><td>7.8E-06</td><td>MB231/MB468</td><td>CYTO</td></tr>
<tr><td>>hg38_21_+_8433172_8446622_RNA45SN1_45S_50ntFlanks@7986.8005.20|ATCAGACGTGGCGACCCGCT</td><td>>hg38_21_+_8433172_8446622_RNA45SN1_45S_50ntFlanks</td><td>7986</td><td>8005</td><td>20</td><td>ATCAGACGTGGCGACCCGCT</td><td>393.0</td><td>-5.7</td><td>1.2</td><td>-4.6</td><td>4.9E-06</td><td>7.8E-06</td><td>MB231/MB468</td><td>CYTO</td></tr>
<tr><td>>hg38_21_+_8433172_8446622_RNA45SN1_45S_50ntFlanks@10780.10801.22|AAGGTGAACAGCCTCTGGCATG</td><td>>hg38_21_+_8433172_8446622_RNA45SN1_45S_50ntFlanks</td><td>10780</td><td>10801</td><td>22</td><td>AAGGTGAACAGCCTCTGGCATG</td><td>240.0</td><td>-6.6</td><td>1.4</td><td>-4.6</td><td>4.4E-06</td><td>7.8E-06</td><td>MB231/MB468</td><td>MP</td></tr>
<tr><td>>MI0000272|hsa-mir-182&WithFlank&7|-|129770377|129770498@29.52.24(+1A)&[MIMAT0000259&hsa-miR-182-5p&offsets|0|0(+1A);m-22&7|-|129770449|129770470&offsets|0|+2(+1A)]|TTTGGCAATGGTAGAACTCACACTA</td><td>>MI0000272|hsa-mir-182&WithFlank&7|-|129770377|129770498</td><td>29</td><td>52</td><td>24(+1A)</td><td>TTTGGCAATGGTAGAACTCACACTA</td><td>2445.2</td><td>4.0</td><td>0.8</td><td>5.0</td><td>4.5E-07</td><td>7.8E-06</td><td>BT20/MB231</td><td>CYTO</td></tr>
<tr><td>>hg38_MT_+_1621_3279_RNR2_16S_50ntFlanks@194.230.37|AAGCATAATATAGCAAGGACTAACCCCTATACCTTCT</td><td>>hg38_MT_+_1621_3279_RNR2_16S_50ntFlanks</td><td>194</td><td>230</td><td>37</td><td>AAGCATAATATAGCAAGGACTAACCCCTATACCTTCT</td><td>34.7</td><td>8.3</td><td>1.8</td><td>4.6</td><td>4.4E-06</td><td>7.8E-06</td><td>MB231/MB468</td><td>MP</td></tr>
<tr><td>>hg38_21_+_8433172_8446622_RNA45SN1_45S_50ntFlanks@12418.12446.29|CGATGTCGGCTCTTCCTATCATTGTGAAG</td><td>>hg38_21_+_8433172_8446622_RNA45SN1_45S_50ntFlanks</td><td>12418</td><td>12446</td><td>29</td><td>CGATGTCGGCTCTTCCTATCATTGTGAAG</td><td>8333.5</td><td>-9.1</td><td>2.0</td><td>-4.5</td><td>5.5E-06</td><td>7.8E-06</td><td>BT20/MB468</td><td>NUC</td></tr>
<tr><td>>hg38_21_+_8433172_8446622_RNA45SN1_45S_50ntFlanks@8848.8867.20|CGGCGGCGGCGGCGGCGGGA</td><td>>hg38_21_+_8433172_8446622_RNA45SN1_45S_50ntFlanks</td><td>8848</td><td>8867</td><td>20</td><td>CGGCGGCGGCGGCGGCGGGA</td><td>407.5</td><td>-6.3</td><td>1.4</td><td>-4.6</td><td>3.8E-06</td><td>7.8E-06</td><td>BT20/MB468</td><td>MITO</td></tr>
<tr><td>>hg38_21_+_8433172_8446622_RNA45SN1_45S_50ntFlanks@3854.3871.18|ACTGTGGTAATTCTAGAG</td><td>>hg38_21_+_8433172_8446622_RNA45SN1_45S_50ntFlanks</td><td>3854</td><td>3871</td><td>18</td><td>ACTGTGGTAATTCTAGAG</td><td>275.4</td><td>-4.2</td><td>0.9</td><td>-4.6</td><td>5.0E-06</td><td>7.8E-06</td><td>MB231/MB468</td><td>CYTO</td></tr>
<tr><td>>MI0003185|hsa-mir-501&WithFlank&X|+|50009716|50009811@57.78.22&[MIMAT0004774&hsa-miR-501-3p&offsets|0|0;m-168&X|+|50009772|50009793&offsets|0|0]|AATGCACCCGGGCAAGGATTCT</td><td>>MI0003185|hsa-mir-501&WithFlank&X|+|50009716|50009811</td><td>57</td><td>78</td><td>22</td><td>AATGCACCCGGGCAAGGATTCT</td><td>193.3</td><td>4.5</td><td>1.0</td><td>4.6</td><td>5.3E-06</td><td>7.8E-06</td><td>MB231/MB468</td><td>MITO</td></tr>
<tr><td>>hg38_21_+_8433172_8446622_RNA45SN1_45S_50ntFlanks@11670.11707.38|CCCAGTGCTCTGAATGTCAAAGTGAAGAAATTCAATGA</td><td>>hg38_21_+_8433172_8446622_RNA45SN1_45S_50ntFlanks</td><td>11670</td><td>11707</td><td>38</td><td>CCCAGTGCTCTGAATGTCAAAGTGAAGAAATTCAATGA</td><td>393.5</td><td>-7.3</td><td>1.6</td><td>-4.5</td><td>5.5E-06</td><td>7.8E-06</td><td>BT20/MB468</td><td>NUC</td></tr>
<tr><td>>hg38_21_+_8433172_8446622_RNA45SN1_45S_50ntFlanks@4569.4589.21|ATGGAATAGGACCGCGGTTCT</td><td>>hg38_21_+_8433172_8446622_RNA45SN1_45S_50ntFlanks</td><td>4569</td><td>4589</td><td>21</td><td>ATGGAATAGGACCGCGGTTCT</td><td>239.2</td><td>-3.5</td><td>0.8</td><td>-4.6</td><td>4.4E-06</td><td>7.8E-06</td><td>MB231/MB468</td><td>MP</td></tr>
<tr><td>>hg38_21_+_8433172_8446622_RNA45SN1_45S_50ntFlanks@4588.4619.32|CTATTTTGTTGGTTTTCGGAACTGAGGCCATG</td><td>>hg38_21_+_8433172_8446622_RNA45SN1_45S_50ntFlanks</td><td>4588</td><td>4619</td><td>32</td><td>CTATTTTGTTGGTTTTCGGAACTGAGGCCATG</td><td>242.5</td><td>-2.9</td><td>0.6</td><td>-4.6</td><td>3.8E-06</td><td>7.8E-06</td><td>BT20/MB468</td><td>MITO</td></tr>
<tr><td>>hg38_21_+_8433172_8446622_RNA45SN1_45S_50ntFlanks@8171.8213.43|AAGTCCTTCTGATCGAGGCCCAGCCCGTGGACGGTGTGAGGCC</td><td>>hg38_21_+_8433172_8446622_RNA45SN1_45S_50ntFlanks</td><td>8171</td><td>8213</td><td>43</td><td>AAGTCCTTCTGATCGAGGCCCAGCCCGTGGACGGTGTGAGGCC</td><td>353.1</td><td>-3.5</td><td>0.8</td><td>-4.6</td><td>5.3E-06</td><td>7.8E-06</td><td>MB231/MB468</td><td>MITO</td></tr>
<tr><td>>hg38_21_+_8433172_8446622_RNA45SN1_45S_50ntFlanks@12937.12966.30|AACCATTCGTAGACGACCTGCTTCTGGGTC</td><td>>hg38_21_+_8433172_8446622_RNA45SN1_45S_50ntFlanks</td><td>12937</td><td>12966</td><td>30</td><td>AACCATTCGTAGACGACCTGCTTCTGGGTC</td><td>488.8</td><td>-6.3</td><td>1.4</td><td>-4.6</td><td>4.4E-06</td><td>7.8E-06</td><td>BT20/MB468</td><td>CYTO</td></tr>
<tr><td>>hg38_21_+_8433172_8446622_RNA45SN1_45S_50ntFlanks@6655.6681.27|TCTTAGCGGTGGATCACTCGGCTCGTG</td><td>>hg38_21_+_8433172_8446622_RNA45SN1_45S_50ntFlanks</td><td>6655</td><td>6681</td><td>27</td><td>TCTTAGCGGTGGATCACTCGGCTCGTG</td><td>1502.7</td><td>-2.9</td><td>0.6</td><td>-4.6</td><td>4.4E-06</td><td>7.8E-06</td><td>BT20/MB468</td><td>CYTO</td></tr>
<tr><td>>hg38_21_+_8433172_8446622_RNA45SN1_45S_50ntFlanks@6740.6758.19|CATCGACACTTCGAACGCA</td><td>>hg38_21_+_8433172_8446622_RNA45SN1_45S_50ntFlanks</td><td>6740</td><td>6758</td><td>19</td><td>CATCGACACTTCGAACGCA</td><td>57.5</td><td>-8.9</td><td>1.9</td><td>-4.8</td><td>1.6E-06</td><td>7.8E-06</td><td>BT20/MB468</td><td>MP</td></tr>
<tr><td>>hg38_1_-_228634819_228635039_RNA5S12_5S_50ntFlanks@84.111.28|CTCGTCTGATCTCGGAAGCTAAGCAGGG</td><td>>hg38_1_-_228634819_228635039_RNA5S12_5S_50ntFlanks</td><td>84</td><td>111</td><td>28</td><td>CTCGTCTGATCTCGGAAGCTAAGCAGGG</td><td>287.2</td><td>-6.3</td><td>1.4</td><td>-4.6</td><td>5.2E-06</td><td>7.9E-06</td><td>MB231/MB468</td><td>NUC</td></tr>
<tr><td>>MI0001445|hsa-mir-423&WithFlank&17|+|30117073|30117178@23.45.23(+1A)&[MIMAT0004748&hsa-miR-423-5p&offsets|0|0(+1A);m-141&17|+|30117095|30117117&offsets|0|0(+1A)]|TGAGGGGCAGAGAGCGAGACTTTA</td><td>>MI0001445|hsa-mir-423&WithFlank&17|+|30117073|30117178</td><td>23</td><td>45</td><td>23(+1A)</td><td>TGAGGGGCAGAGAGCGAGACTTTA</td><td>491.6</td><td>6.5</td><td>1.4</td><td>4.8</td><td>1.6E-06</td><td>7.9E-06</td><td>BT20/MB468</td><td>MP</td></tr>
<tr><td>>hg38_21_+_8433172_8446622_RNA45SN1_45S_50ntFlanks@9564.9593.30|CTGACGTGCAAATCGGTCGTCCGACCTGGG</td><td>>hg38_21_+_8433172_8446622_RNA45SN1_45S_50ntFlanks</td><td>9564</td><td>9593</td><td>30</td><td>CTGACGTGCAAATCGGTCGTCCGACCTGGG</td><td>104.4</td><td>-5.4</td><td>1.2</td><td>-4.6</td><td>4.5E-06</td><td>7.9E-06</td><td>BT20/MB468</td><td>CYTO</td></tr>
<tr><td>>hg38_21_+_8433172_8446622_RNA45SN1_45S_50ntFlanks@5485.5516.32|AGCGCTGAGAAGACGGTCGAACTTGACTATCT</td><td>>hg38_21_+_8433172_8446622_RNA45SN1_45S_50ntFlanks</td><td>5485</td><td>5516</td><td>32</td><td>AGCGCTGAGAAGACGGTCGAACTTGACTATCT</td><td>17337.0</td><td>5.2</td><td>1.1</td><td>4.6</td><td>3.8E-06</td><td>7.9E-06</td><td>BT20/MB468</td><td>MITO</td></tr>
<tr><td>>hg38_21_+_8433172_8446622_RNA45SN1_45S_50ntFlanks@8192.8215.24|AGCCCGTGGACGGTGTGAGGCCGG</td><td>>hg38_21_+_8433172_8446622_RNA45SN1_45S_50ntFlanks</td><td>8192</td><td>8215</td><td>24</td><td>AGCCCGTGGACGGTGTGAGGCCGG</td><td>244.9</td><td>-3.8</td><td>0.8</td><td>-4.6</td><td>5.0E-06</td><td>7.9E-06</td><td>MB231/MB468</td><td>CYTO</td></tr>
<tr><td>>hg38_21_+_8433172_8446622_RNA45SN1_45S_50ntFlanks@9736.9753.18|CCGAAACGATCTCAACCT</td><td>>hg38_21_+_8433172_8446622_RNA45SN1_45S_50ntFlanks</td><td>9736</td><td>9753</td><td>18</td><td>CCGAAACGATCTCAACCT</td><td>361.9</td><td>-4.0</td><td>0.9</td><td>-4.6</td><td>5.0E-06</td><td>7.9E-06</td><td>MB231/MB468</td><td>CYTO</td></tr>
<tr><td>>hg38_21_+_8433172_8446622_RNA45SN1_45S_50ntFlanks@9621.9649.29|TCTAGTAGCTGGTTCCCTCCGAAGTTTCC</td><td>>hg38_21_+_8433172_8446622_RNA45SN1_45S_50ntFlanks</td><td>9621</td><td>9649</td><td>29</td><td>TCTAGTAGCTGGTTCCCTCCGAAGTTTCC</td><td>258.2</td><td>-2.7</td><td>0.6</td><td>-4.6</td><td>3.8E-06</td><td>7.9E-06</td><td>BT20/MB468</td><td>MITO</td></tr>
<tr><td>>trna116_GluCTC_1_-_145399233_145399304@1.36.36,trna59_GluCTC_1_+_249168447_249168518@1.36.36,trna71_GluCTC_1_-_161439189_161439260@1.36.36,trna74_GluCTC_1_-_161431809_161431880@1.36.36,trna77_GluCTC_1_-_161424398_161424469@1.36.36,trna77_GluCTC_6_+_28949976_28950047@1.36.36,trna80_GluCTC_1_-_161417018_161417089@1.36.36,trna87_GluCTC_6_-_126101393_126101464@1.36.36|TCCCTGGTGGTCTAGTGGTTAGGATTCGGCGCTCTC</td><td>>trna116_GluCTC_1_-_145399233_145399304</td><td>1</td><td>36</td><td>36</td><td>TCCCTGGTGGTCTAGTGGTTAGGATTCGGCGCTCTC</td><td>353.6</td><td>2.6</td><td>0.6</td><td>4.5</td><td>5.6E-06</td><td>7.9E-06</td><td>BT20/MB468</td><td>NUC</td></tr>
<tr><td>>MI0000239|hsa-mir-197&WithFlank&1|+|109598887|109598973@54.74.21(+1U)&[MIMAT0000227&hsa-miR-197-3p&offsets|0|-1(+1U);m-99&1|+|109598940|109598961&offsets|0|-1(+1U)]|TTCACCACCTTCTCCACCCAGT</td><td>>MI0000239|hsa-mir-197&WithFlank&1|+|109598887|109598973</td><td>54</td><td>74</td><td>21(+1U)</td><td>TTCACCACCTTCTCCACCCAGT</td><td>268.7</td><td>3.0</td><td>0.7</td><td>4.6</td><td>5.2E-06</td><td>7.9E-06</td><td>MB231/MB468</td><td>NUC</td></tr>
<tr><td>>hg38_21_+_8433172_8446622_RNA45SN1_45S_50ntFlanks@11670.11697.28|CCCAGTGCTCTGAATGTCAAAGTGAAGA</td><td>>hg38_21_+_8433172_8446622_RNA45SN1_45S_50ntFlanks</td><td>11670</td><td>11697</td><td>28</td><td>CCCAGTGCTCTGAATGTCAAAGTGAAGA</td><td>152.1</td><td>-5.0</td><td>1.1</td><td>-4.6</td><td>4.4E-06</td><td>7.9E-06</td><td>MB231/MB468</td><td>MP</td></tr>
<tr><td>>hg38_21_+_8433172_8446622_RNA45SN1_45S_50ntFlanks@6719.6747.29|TGAATTGCAGGACACATTGATCATCGACA</td><td>>hg38_21_+_8433172_8446622_RNA45SN1_45S_50ntFlanks</td><td>6719</td><td>6747</td><td>29</td><td>TGAATTGCAGGACACATTGATCATCGACA</td><td>1076.5</td><td>-3.2</td><td>0.7</td><td>-4.6</td><td>4.5E-06</td><td>7.9E-06</td><td>BT20/MB468</td><td>CYTO</td></tr>
<tr><td>>hg38_21_+_8433172_8446622_RNA45SN1_45S_50ntFlanks@11540.11581.42|CGCGGCGCCTCGCCTCGGCCGGCGCCTAGCAGCCGACTTAGA</td><td>>hg38_21_+_8433172_8446622_RNA45SN1_45S_50ntFlanks</td><td>11540</td><td>11581</td><td>42</td><td>CGCGGCGCCTCGCCTCGGCCGGCGCCTAGCAGCCGACTTAGA</td><td>398.6</td><td>3.3</td><td>0.7</td><td>4.6</td><td>5.0E-06</td><td>7.9E-06</td><td>MB231/MB468</td><td>CYTO</td></tr>
<tr><td>>hg38_21_+_8433172_8446622_RNA45SN1_45S_50ntFlanks@11552.11592.41|CCTCGGCCGGCGCCTAGCAGCCGACTTAGAACTGGTGCGGA</td><td>>hg38_21_+_8433172_8446622_RNA45SN1_45S_50ntFlanks</td><td>11552</td><td>11592</td><td>41</td><td>CCTCGGCCGGCGCCTAGCAGCCGACTTAGAACTGGTGCGGA</td><td>331.7</td><td>-4.3</td><td>1.0</td><td>-4.6</td><td>5.0E-06</td><td>7.9E-06</td><td>MB231/MB468</td><td>CYTO</td></tr>
<tr><td>>MI0000063|hsa-let-7b&WithFlank&22|+|46113680|46113774@12.33.22(+1A)&[MIMAT0000063&hsa-let-7b-5p&offsets|0|0(+1A);m-14&22|+|46113691|46113712&offsets|0|0(+1A)]|TGAGGTAGTAGGTTGTGTGGTTA</td><td>>MI0000063|hsa-let-7b&WithFlank&22|+|46113680|46113774</td><td>12</td><td>33</td><td>22(+1A)</td><td>TGAGGTAGTAGGTTGTGTGGTTA</td><td>478.0</td><td>4.2</td><td>0.9</td><td>4.6</td><td>5.3E-06</td><td>7.9E-06</td><td>MB231/MB468</td><td>MITO</td></tr>
<tr><td>>hg38_21_+_8433172_8446622_RNA45SN1_45S_50ntFlanks@10411.10442.32|CAGTCGGTCCTGAGAGATGGGCGAGCGCCGTT</td><td>>hg38_21_+_8433172_8446622_RNA45SN1_45S_50ntFlanks</td><td>10411</td><td>10442</td><td>32</td><td>CAGTCGGTCCTGAGAGATGGGCGAGCGCCGTT</td><td>437.1</td><td>-5.2</td><td>1.1</td><td>-4.6</td><td>5.2E-06</td><td>7.9E-06</td><td>MB231/MB468</td><td>NUC</td></tr>
<tr><td>>hg38_21_+_8433172_8446622_RNA45SN1_45S_50ntFlanks@10239.10262.24|GCTACGCCGCGACGAGTAGGAGGG</td><td>>hg38_21_+_8433172_8446622_RNA45SN1_45S_50ntFlanks</td><td>10239</td><td>10262</td><td>24</td><td>GCTACGCCGCGACGAGTAGGAGGG</td><td>142.2</td><td>10.5</td><td>2.1</td><td>5.1</td><td>4.4E-07</td><td>7.9E-06</td><td>BT20/MB231</td><td>MP</td></tr>
<tr><td>>hg38_21_+_8433172_8446622_RNA45SN1_45S_50ntFlanks@7981.7998.18|CTCAGATCAGACGTGGCG</td><td>>hg38_21_+_8433172_8446622_RNA45SN1_45S_50ntFlanks</td><td>7981</td><td>7998</td><td>18</td><td>CTCAGATCAGACGTGGCG</td><td>152.7</td><td>-5.9</td><td>1.3</td><td>-4.6</td><td>5.2E-06</td><td>7.9E-06</td><td>MB231/MB468</td><td>NUC</td></tr>
<tr><td>>hg38_21_+_8433172_8446622_RNA45SN1_45S_50ntFlanks@11622.11640.19|AAAGCATCGCGAAGGCCCG</td><td>>hg38_21_+_8433172_8446622_RNA45SN1_45S_50ntFlanks</td><td>11622</td><td>11640</td><td>19</td><td>AAAGCATCGCGAAGGCCCG</td><td>604.5</td><td>-5.9</td><td>1.3</td><td>-4.6</td><td>5.4E-06</td><td>7.9E-06</td><td>MB231/MB468</td><td>MITO</td></tr>
<tr><td>>hg38_21_+_8433172_8446622_RNA45SN1_45S_50ntFlanks@7986.8018.33|ATCAGACGTGGCGACCCGCTGAATTTAAGCATA</td><td>>hg38_21_+_8433172_8446622_RNA45SN1_45S_50ntFlanks</td><td>7986</td><td>8018</td><td>33</td><td>ATCAGACGTGGCGACCCGCTGAATTTAAGCATA</td><td>1474.9</td><td>-7.1</td><td>1.6</td><td>-4.6</td><td>5.4E-06</td><td>7.9E-06</td><td>MB231/MB468</td><td>MITO</td></tr>
<tr><td>>hg38_21_+_8433172_8446622_RNA45SN1_45S_50ntFlanks@12420.12437.18|ATGTCGGCTCTTCCTATC</td><td>>hg38_21_+_8433172_8446622_RNA45SN1_45S_50ntFlanks</td><td>12420</td><td>12437</td><td>18</td><td>ATGTCGGCTCTTCCTATC</td><td>4898.0</td><td>-3.3</td><td>0.7</td><td>-4.5</td><td>5.4E-06</td><td>7.9E-06</td><td>MB231/MB468</td><td>MITO</td></tr>
<tr><td>>hg38_21_+_8433172_8446622_RNA45SN1_45S_50ntFlanks@4666.4695.30|AAATTCTTGGACCGGCGCAAGACGGACCAG</td><td>>hg38_21_+_8433172_8446622_RNA45SN1_45S_50ntFlanks</td><td>4666</td><td>4695</td><td>30</td><td>AAATTCTTGGACCGGCGCAAGACGGACCAG</td><td>48.2</td><td>-8.7</td><td>1.8</td><td>-4.8</td><td>1.6E-06</td><td>7.9E-06</td><td>BT20/MB468</td><td>MP</td></tr>
</table>

<table>
<tr><td>>MI0000446|hsa-mir-125b-1&WithFlank&11|-|122099751|122099850@21.43.23&[MIMAT0000423&hsa-miR-125b-5p&offsets|0|+1;m-67&11|-|122099809|122099830&offsets|0|+1]|TCCCTGAGACCCTAACTTGTGAT</td><td>>MI0000446|hsa-mir-125b-1&WithFlank&11|-|122099751|122099850</td><td>21</td><td>43</td><td>23</td><td>TCCCTGAGACCCTAACTTGTGAT</td><td>544.9</td><td>-8.1</td><td>1.6</td><td>-5.1</td><td>4.4E-07</td><td>7.9E-06</td><td>BT20/MB231</td><td>MP</td></tr>
<tr><td>>trnaMT_GluTTC_MT_-_14674_14742@39.64.26,trnalookalike8_GluTTC_5_-_93905172_93905240@39.64.26|ATTGGTCGTGGTTGTAGTCCGTGCGA</td><td>>trnaMT_GluTTC_MT_-_14674_14742</td><td>39</td><td>64</td><td>26</td><td>ATTGGTCGTGGTTGTAGTCCGTGCGA</td><td>165.2</td><td>8.2</td><td>1.7</td><td>4.8</td><td>1.6E-06</td><td>7.9E-06</td><td>BT20/MB468</td><td>MP</td></tr>
<tr><td>>MI0000263|hsa-mir-7-1&WithFlank&9|-|83969742|83969863@30.54.25&[MIMAT0000252_2&hsa-miR-7-5p&offsets|0|+1;m-401&9|-|83969811|83969834&offsets|0|+1]|TGGAAGACTAGTGATTTTGTTGTTT</td><td>>MI0000263|hsa-mir-7-1&WithFlank&9|-|83969742|83969863</td><td>30</td><td>54</td><td>25</td><td>TGGAAGACTAGTGATTTTGTTGTTT</td><td>267.7</td><td>2.4</td><td>0.5</td><td>4.5</td><td>5.6E-06</td><td>8.0E-06</td><td>BT20/MB468</td><td>NUC</td></tr>
<tr><td>>hg38_21_+_8433172_8446622_RNA45SN1_45S_50ntFlanks@11571.11592.22|GCCGACTTAGAACTGGTGCGGA</td><td>>hg38_21_+_8433172_8446622_RNA45SN1_45S_50ntFlanks</td><td>11571</td><td>11592</td><td>22</td><td>GCCGACTTAGAACTGGTGCGGA</td><td>986.9</td><td>-5.7</td><td>1.2</td><td>-4.6</td><td>4.5E-06</td><td>8.0E-06</td><td>MB231/MB468</td><td>MP</td></tr>
<tr><td>>hg38_21_+_8433172_8446622_RNA45SN1_45S_50ntFlanks@4225.4243.19|AAATAACAATACAGGACTC</td><td>>hg38_21_+_8433172_8446622_RNA45SN1_45S_50ntFlanks</td><td>4225</td><td>4243</td><td>19</td><td>AAATAACAATACAGGACTC</td><td>365.9</td><td>-6.8</td><td>1.5</td><td>-4.6</td><td>4.5E-06</td><td>8.0E-06</td><td>MB231/MB468</td><td>MP</td></tr>
<tr><td>>hg38_21_+_8433172_8446622_RNA45SN1_45S_50ntFlanks@9736.9765.30|CCGAAACGATCTCAACCTATTCTCAAACTT</td><td>>hg38_21_+_8433172_8446622_RNA45SN1_45S_50ntFlanks</td><td>9736</td><td>9765</td><td>30</td><td>CCGAAACGATCTCAACCTATTCTCAAACTT</td><td>132.5</td><td>-5.3</td><td>1.2</td><td>-4.6</td><td>5.1E-06</td><td>8.0E-06</td><td>MB231/MB468</td><td>CYTO</td></tr>
<tr><td>>MI0003513|hsa-mir-455&WithFlank&9|+|114209428|114209535@22.45.24&[MIMAT0003150&hsa-miR-455-5p&offsets|0|+2;m-232&9|+|114209449|114209470&offsets|0|+2]|TATGTGCCTTTGGACTACATCGTG</td><td>>MI0003513|hsa-mir-455&WithFlank&9|+|114209428|114209535</td><td>22</td><td>45</td><td>24</td><td>TATGTGCCTTTGGACTACATCGTG</td><td>253.4</td><td>-3.7</td><td>0.7</td><td>-5.0</td><td>5.1E-07</td><td>8.0E-06</td><td>BT20/MB231</td><td>MITO</td></tr>
<tr><td>>MI0000456|hsa-mir-140&WithFlank&16|+|69933075|69933186@68.89.22&[MIMAT0004597&hsa-miR-140-3p&offsets|0|+1;m-45&16|+|69933143|69933164&offsets|-1|-1]|TACCACAGGGTAGAACCACGGA</td><td>>MI0000456|hsa-mir-140&WithFlank&16|+|69933075|69933186</td><td>68</td><td>89</td><td>22</td><td>TACCACAGGGTAGAACCACGGA</td><td>684.7</td><td>5.4</td><td>1.2</td><td>4.6</td><td>4.5E-06</td><td>8.0E-06</td><td>MB231/MB468</td><td>MP</td></tr>
<tr><td>>hg38_21_+_8433172_8446622_RNA45SN1_45S_50ntFlanks@10349.10377.29|AACGAGAACTTTGAAGGCCGAAGTGGAGA</td><td>>hg38_21_+_8433172_8446622_RNA45SN1_45S_50ntFlanks</td><td>10349</td><td>10377</td><td>29</td><td>AACGAGAACTTTGAAGGCCGAAGTGGAGA</td><td>3334.0</td><td>-6.3</td><td>1.4</td><td>-4.6</td><td>4.5E-06</td><td>8.0E-06</td><td>MB231/MB468</td><td>MP</td></tr>
<tr><td>>am_trna111_HisGTG_1_-_147774845_147774916@-1G.29.30,trna118_HisGTG_1_-_145396881_145396952@-1G.29.30,trna16_HisGTG_1_+_146544773_146544844@-1G.29.30,trna1_HisGTG_15_+_45493349_45493420@-1G.29.30,trna21_HisGTG_1_+_147753471_147753542@-1G.29.30,trna33_HisGTG_6_+_27125906_27125977@-1G.29.30,trna7_HisGTG_9_-_14433938_14434009@-1G.29.30,trna8_HisGTG_15_-_45492611_45492682@-1G.29.30,trna9_HisGTG_15_-_45490804_45490875@-1G.29.30|GGCCGTGATCGTATAGTGGTTAGTACTCTG</td><td>>am_trna111_HisGTG_1_-_147774845_147774916</td><td>-1G</td><td>29</td><td>30</td><td>GGCCGTGATCGTATAGTGGTTAGTACTCTG</td><td>3722.4</td><td>-4.9</td><td>1.0</td><td>-4.8</td><td>1.6E-06</td><td>8.0E-06</td><td>BT20/MB468</td><td>MP</td></tr>
<tr><td>>hg38_21_+_8433172_8446622_RNA45SN1_45S_50ntFlanks@11569.11593.25|CAGCCGACTTAGAACTGGTGCGGAC</td><td>>hg38_21_+_8433172_8446622_RNA45SN1_45S_50ntFlanks</td><td>11569</td><td>11593</td><td>25</td><td>CAGCCGACTTAGAACTGGTGCGGAC</td><td>153.9</td><td>-2.8</td><td>0.6</td><td>-4.6</td><td>3.9E-06</td><td>8.0E-06</td><td>BT20/MB468</td><td>MITO</td></tr>
<tr><td>>hg38_21_+_8433172_8446622_RNA45SN1_45S_50ntFlanks@10391.10424.34|AACAGCAGTTGAACATGGGTCAGTCGGTCCTGAG</td><td>>hg38_21_+_8433172_8446622_RNA45SN1_45S_50ntFlanks</td><td>10391</td><td>10424</td><td>34</td><td>AACAGCAGTTGAACATGGGTCAGTCGGTCCTGAG</td><td>6473.3</td><td>-2.9</td><td>0.6</td><td>-4.5</td><td>5.7E-06</td><td>8.0E-06</td><td>BT20/MB468</td><td>NUC</td></tr>
<tr><td>>hg38_21_+_8433172_8446622_RNA45SN1_45S_50ntFlanks@5472.5515.44|ACGGCCCTGGCGGAGCGCTGAGAAGACGGTCGAACTTGACTATC</td><td>>hg38_21_+_8433172_8446622_RNA45SN1_45S_50ntFlanks</td><td>5472</td><td>5515</td><td>44</td><td>ACGGCCCTGGCGGAGCGCTGAGAAGACGGTCGAACTTGACTATC</td><td>326.0</td><td>4.5</td><td>1.0</td><td>4.6</td><td>5.3E-06</td><td>8.0E-06</td><td>MB231/MB468</td><td>NUC</td></tr>
<tr><td>>hg38_21_+_8433172_8446622_RNA45SN1_45S_50ntFlanks@10481.10500.20|ATCGAAAGGGAGTCGGGTTC</td><td>>hg38_21_+_8433172_8446622_RNA45SN1_45S_50ntFlanks</td><td>10481</td><td>10500</td><td>20</td><td>ATCGAAAGGGAGTCGGGTTC</td><td>101.2</td><td>-10.5</td><td>2.3</td><td>-4.6</td><td>4.5E-06</td><td>8.0E-06</td><td>MB231/MB468</td><td>MP</td></tr>
<tr><td>>hg38_21_+_8433172_8446622_RNA45SN1_45S_50ntFlanks@3882.3915.34|CCGACGGGCGCTGACCCCCTTCGCGGGGGGGATG</td><td>>hg38_21_+_8433172_8446622_RNA45SN1_45S_50ntFlanks</td><td>3882</td><td>3915</td><td>34</td><td>CCGACGGGCGCTGACCCCCTTCGCGGGGGGGATG</td><td>316.0</td><td>-2.9</td><td>0.6</td><td>-4.6</td><td>4.5E-06</td><td>8.0E-06</td><td>BT20/MB468</td><td>CYTO</td></tr>
<tr><td>>hg38_21_+_8433172_8446622_RNA45SN1_45S_50ntFlanks@11317.11338.22|ACCCCACGTCTCGTCGCGCGCG</td><td>>hg38_21_+_8433172_8446622_RNA45SN1_45S_50ntFlanks</td><td>11317</td><td>11338</td><td>22</td><td>ACCCCACGTCTCGTCGCGCGCG</td><td>1079.1</td><td>-5.0</td><td>1.1</td><td>-4.6</td><td>4.5E-06</td><td>8.0E-06</td><td>BT20/MB468</td><td>CYTO</td></tr>
<tr><td>>hg38_21_+_8433172_8446622_RNA45SN1_45S_50ntFlanks@9548.9571.24|GAGGTCCGTAGCGGTCCTGACGTG</td><td>>hg38_21_+_8433172_8446622_RNA45SN1_45S_50ntFlanks</td><td>9548</td><td>9571</td><td>24</td><td>GAGGTCCGTAGCGGTCCTGACGTG</td><td>627.5</td><td>-5.4</td><td>1.2</td><td>-4.6</td><td>4.5E-06</td><td>8.0E-06</td><td>MB231/MB468</td><td>MP</td></tr>
<tr><td>>hg38_21_+_8433172_8446622_RNA45SN1_45S_50ntFlanks@6733.6769.37|CATTGATCATCGACACTTCGAACGCACTTGCGGCCCC</td><td>>hg38_21_+_8433172_8446622_RNA45SN1_45S_50ntFlanks</td><td>6733</td><td>6769</td><td>37</td><td>CATTGATCATCGACACTTCGAACGCACTTGCGGCCCC</td><td>266.3</td><td>-4.9</td><td>1.1</td><td>-4.5</td><td>5.4E-06</td><td>8.0E-06</td><td>MB231/MB468</td><td>MITO</td></tr>
<tr><td>>MI0000298|hsa-mir-221&WithFlank&X|-|45746151|45746272@31.56.26&[MIMAT0004568&hsa-miR-221-5p&offsets|0|+4;m-405&X|-|45746220|45746242&offsets|0|+3]|ACCTGGCATACAATGTAGATTTCTGT</td><td>>MI0000298|hsa-mir-221&WithFlank&X|-|45746151|45746272</td><td>31</td><td>56</td><td>26</td><td>ACCTGGCATACAATGTAGATTTCTGT</td><td>1761.9</td><td>-3.4</td><td>0.7</td><td>-5.0</td><td>5.1E-07</td><td>8.0E-06</td><td>BT20/MB231</td><td>MITO</td></tr>
<tr><td>>hg38_21_+_8433172_8446622_RNA45SN1_45S_50ntFlanks@7974.8006.33|ACGCGACCTCAGATCAGACGTGGCGACCCGCTG</td><td>>hg38_21_+_8433172_8446622_RNA45SN1_45S_50ntFlanks</td><td>7974</td><td>8006</td><td>33</td><td>ACGCGACCTCAGATCAGACGTGGCGACCCGCTG</td><td>11000.5</td><td>4.2</td><td>0.8</td><td>5.0</td><td>6.1E-07</td><td>8.0E-06</td><td>BT20/MB231</td><td>NUC</td></tr>
<tr><td>>MI0002468|hsa-mir-484&WithFlank&16|+|15643288|15643378@14.35.22(+1U)&[MIMAT0002174&hsa-miR-484&offsets|0|0(+1U);m-127&16|+|15643301|15643321&offsets|0|+1(+1U)]|TCAGGCTCAGTCCCCTCCCGATT</td><td>>MI0002468|hsa-mir-484&WithFlank&16|+|15643288|15643378</td><td>14</td><td>35</td><td>22(+1U)</td><td>TCAGGCTCAGTCCCCTCCCGATT</td><td>772.0</td><td>4.3</td><td>1.0</td><td>4.5</td><td>5.4E-06</td><td>8.0E-06</td><td>MB231/MB468</td><td>MITO</td></tr>
<tr><td>>hg38_21_+_8433172_8446622_RNA45SN1_45S_50ntFlanks@7971.8010.40|GAGACGCGACCTCAGATCAGACGTGGCGACCCGCTGAATT</td><td>>hg38_21_+_8433172_8446622_RNA45SN1_45S_50ntFlanks</td><td>7971</td><td>8010</td><td>40</td><td>GAGACGCGACCTCAGATCAGACGTGGCGACCCGCTGAATT</td><td>383.1</td><td>4.5</td><td>1.0</td><td>4.5</td><td>5.7E-06</td><td>8.0E-06</td><td>BT20/MB468</td><td>NUC</td></tr>
<tr><td>>hg38_21_+_8433172_8446622_RNA45SN1_45S_50ntFlanks@8115.8141.27|CGCGGGACATGTGGCGTACGGAAGACC</td><td>>hg38_21_+_8433172_8446622_RNA45SN1_45S_50ntFlanks</td><td>8115</td><td>8141</td><td>27</td><td>CGCGGGACATGTGGCGTACGGAAGACC</td><td>160.7</td><td>-5.9</td><td>1.3</td><td>-4.6</td><td>5.3E-06</td><td>8.0E-06</td><td>MB231/MB468</td><td>NUC</td></tr>
<tr><td>>hg38_21_+_8433172_8446622_RNA45SN1_45S_50ntFlanks@5492.5512.21|AGAAGACGGTCGAACTTGACT</td><td>>hg38_21_+_8433172_8446622_RNA45SN1_45S_50ntFlanks</td><td>5492</td><td>5512</td><td>21</td><td>AGAAGACGGTCGAACTTGACT</td><td>3075.6</td><td>6.0</td><td>1.3</td><td>4.6</td><td>4.6E-06</td><td>8.0E-06</td><td>BT20/MB468</td><td>CYTO</td></tr>
<tr><td>>hg38_21_+_8433172_8446622_RNA45SN1_45S_50ntFlanks@6649.6675.27|TACGACTCTTAGCGGTGGATCACTCGG</td><td>>hg38_21_+_8433172_8446622_RNA45SN1_45S_50ntFlanks</td><td>6649</td><td>6675</td><td>27</td><td>TACGACTCTTAGCGGTGGATCACTCGG</td><td>1131.6</td><td>3.0</td><td>0.6</td><td>4.6</td><td>3.9E-06</td><td>8.0E-06</td><td>BT20/MB468</td><td>MITO</td></tr>
<tr><td>>hg38_21_+_8433172_8446622_RNA45SN1_45S_50ntFlanks@10385.10424.40|CATGTGAACAGCAGTTGAACATGGGTCAGTCGGTCCTGAG</td><td>>hg38_21_+_8433172_8446622_RNA45SN1_45S_50ntFlanks</td><td>10385</td><td>10424</td><td>40</td><td>CATGTGAACAGCAGTTGAACATGGGTCAGTCGGTCCTGAG</td><td>2501.5</td><td>3.8</td><td>0.8</td><td>4.6</td><td>3.9E-06</td><td>8.0E-06</td><td>BT20/MB468</td><td>MITO</td></tr>
<tr><td>>hg38_21_+_8433172_8446622_RNA45SN1_45S_50ntFlanks@3846.3869.24|CTTGGATAACTGTGGTAATTCTAG</td><td>>hg38_21_+_8433172_8446622_RNA45SN1_45S_50ntFlanks</td><td>3846</td><td>3869</td><td>24</td><td>CTTGGATAACTGTGGTAATTCTAG</td><td>641.0</td><td>-2.7</td><td>0.6</td><td>-4.6</td><td>4.6E-06</td><td>8.0E-06</td><td>BT20/MB468</td><td>CYTO</td></tr>
<tr><td>>hg38_21_+_8433172_8446622_RNA45SN1_45S_50ntFlanks@11362.11379.18|TCGGGCGGCGGCGGTCGG</td><td>>hg38_21_+_8433172_8446622_RNA45SN1_45S_50ntFlanks</td><td>11362</td><td>11379</td><td>18</td><td>TCGGGCGGCGGCGGTCGG</td><td>48.8</td><td>-8.7</td><td>1.8</td><td>-4.8</td><td>1.6E-06</td><td>8.0E-06</td><td>BT20/MB468</td><td>MP</td></tr>
<tr><td>>hg38_1_-_228634819_228635039_RNA5S12_5S_50ntFlanks@51.87.37|GTCTACGGCCATACCACCCTGAACGCGCCCGATCTCG</td><td>>hg38_1_-_228634819_228635039_RNA5S12_5S_50ntFlanks</td><td>51</td><td>87</td><td>37</td><td>GTCTACGGCCATACCACCCTGAACGCGCCCGATCTCG</td><td>502.6</td><td>3.6</td><td>0.8</td><td>4.5</td><td>5.5E-06</td><td>8.1E-06</td><td>MB231/MB468</td><td>MITO</td></tr>
<tr><td>>hg38_MT_+_1621_3279_RNR2_16S_50ntFlanks@1431.1451.21|AAAGTCCTACGTGATCTGAGT</td><td>>hg38_MT_+_1621_3279_RNR2_16S_50ntFlanks</td><td>1431</td><td>1451</td><td>21</td><td>AAAGTCCTACGTGATCTGAGT</td><td>241.9</td><td>3.2</td><td>0.7</td><td>4.6</td><td>4.5E-06</td><td>8.1E-06</td><td>MB231/MB468</td><td>MP</td></tr>
<tr><td>>hg38_21_+_8433172_8446622_RNA45SN1_45S_50ntFlanks@10771.10808.38|CAGGTCTCCAAGGTGAACAGCCTCTGGCATGTTGGAAC</td><td>>hg38_21_+_8433172_8446622_RNA45SN1_45S_50ntFlanks</td><td>10771</td><td>10808</td><td>38</td><td>CAGGTCTCCAAGGTGAACAGCCTCTGGCATGTTGGAAC</td><td>71.8</td><td>-9.2</td><td>1.9</td><td>-4.8</td><td>1.6E-06</td><td>8.1E-06</td><td>BT20/MB468</td><td>MP</td></tr>
<tr><td>>hg38_21_+_8433172_8446622_RNA45SN1_45S_50ntFlanks@13002.13021.20|CGATCTATTGAAAGTCAGCC</td><td>>hg38_21_+_8433172_8446622_RNA45SN1_45S_50ntFlanks</td><td>13002</td><td>13021</td><td>20</td><td>CGATCTATTGAAAGTCAGCC</td><td>48.7</td><td>-8.7</td><td>1.8</td><td>-4.8</td><td>1.6E-06</td><td>8.1E-06</td><td>BT20/MB468</td><td>MP</td></tr>
<tr><td>>MI0000542|hsa-mir-320a&WithFlank&8|-|22244960|22245043@48.70.23&[MIMAT0000510&hsa-miR-320a-3p&offsets|0|+1;m-60&8|-|22244975|22244996&offsets|0|+1]|AAAAGCTGGGTTGAGAGGGCGAA</td><td>>MI0000542|hsa-mir-320a&WithFlank&8|-|22244960|22245043</td><td>48</td><td>70</td><td>23</td><td>AAAAGCTGGGTTGAGAGGGCGAA</td><td>25020.2</td><td>4.0</td><td>0.8</td><td>5.0</td><td>4.7E-07</td><td>8.1E-06</td><td>BT20/MB231</td><td>CYTO</td></tr>
<tr><td>>MI0016003|hsa-mir-3613&WithFlank&13|-|49996409|49996507@22.43.22&[MIMAT0017990&hsa-miR-3613-5p&offsets|0|0;m-259&13|-|49996465|49996486&offsets|0|0]|TGTTGTACTTTTTTTTTTGTTC</td><td>>MI0016003|hsa-mir-3613&WithFlank&13|-|49996409|49996507</td><td>22</td><td>43</td><td>22</td><td>TGTTGTACTTTTTTTTTTGTTC</td><td>236.6</td><td>4.8</td><td>0.9</td><td>5.0</td><td>4.8E-07</td><td>8.1E-06</td><td>BT20/MB231</td><td>CYTO</td></tr>
<tr><td>>MI0000273|hsa-mir-183&WithFlank&7|-|129774899|129775020@34.54.21&[MIMAT0000261&hsa-miR-183-5p&offsets|+1|0;m-41&7|-|129774966|129774987&offsets|0|-1]|ATGGCACTGGTAGAATTCACT</td><td>>MI0000273|hsa-mir-183&WithFlank&7|-|129774899|129775020</td><td>34</td><td>54</td><td>21</td><td>ATGGCACTGGTAGAATTCACT</td><td>533.8</td><td>3.7</td><td>0.8</td><td>4.6</td><td>5.2E-06</td><td>8.1E-06</td><td>MB231/MB468</td><td>CYTO</td></tr>
<tr><td>>hg38_21_+_8433172_8446622_RNA45SN1_45S_50ntFlanks@12642.12669.28|CATCTGTGGGATTATGACTGAACGCCTC</td><td>>hg38_21_+_8433172_8446622_RNA45SN1_45S_50ntFlanks</td><td>12642</td><td>12669</td><td>28</td><td>CATCTGTGGGATTATGACTGAACGCCTC</td><td>276.5</td><td>5.4</td><td>1.2</td><td>4.6</td><td>5.2E-06</td><td>8.1E-06</td><td>MB231/MB468</td><td>CYTO</td></tr>
<tr><td>>hg38_21_+_8433172_8446622_RNA45SN1_45S_50ntFlanks@12644.12661.18|TCTGTGGGATTATGACTG</td><td>>hg38_21_+_8433172_8446622_RNA45SN1_45S_50ntFlanks</td><td>12644</td><td>12661</td><td>18</td><td>TCTGTGGGATTATGACTG</td><td>385.3</td><td>-3.0</td><td>0.7</td><td>-4.6</td><td>5.2E-06</td><td>8.1E-06</td><td>MB231/MB468</td><td>CYTO</td></tr>
<tr><td>>hg38_21_+_8433172_8446622_RNA45SN1_45S_50ntFlanks@4983.5003.21|CGGACAGGATTGACAGATTGA</td><td>>hg38_21_+_8433172_8446622_RNA45SN1_45S_50ntFlanks</td><td>4983</td><td>5003</td><td>21</td><td>CGGACAGGATTGACAGATTGA</td><td>131.4</td><td>-10.9</td><td>2.4</td><td>-4.6</td><td>4.6E-06</td><td>8.1E-06</td><td>MB231/MB468</td><td>MP</td></tr>
<tr><td>>trnaMT_GluTTC_MT_-_14674_14742@40.58.19,trnalookalike8_GluTTC_5_-_93905172_93905240@40.58.19|TTGGTCGTGGTTGTAGTCC</td><td>>trnaMT_GluTTC_MT_-_14674_14742</td><td>40</td><td>58</td><td>19</td><td>TTGGTCGTGGTTGTAGTCC</td><td>250.9</td><td>4.3</td><td>0.9</td><td>4.5</td><td>5.5E-06</td><td>8.1E-06</td><td>MB231/MB468</td><td>MITO</td></tr>
<tr><td>>trna10_LysCTT_16_+_3241501_3241573@11.35.25,trna13_LysCTT_14_-_58706613_58706685@11.35.25,trna2_LysCTT_15_+_79152904_79152976@11.35.25,trna4_LysCTT_18_-_43669271_43669343@11.35.25|CTCAGTCGGTAGAGCATGGGACTCT</td><td>>trna10_LysCTT_16_+_3241501_3241573</td><td>11</td><td>35</td><td>25</td><td>CTCAGTCGGTAGAGCATGGGACTCT</td><td>38.5</td><td>8.5</td><td>1.9</td><td>4.6</td><td>4.6E-06</td><td>8.1E-06</td><td>MB231/MB468</td><td>MP</td></tr>
<tr><td>>trna116_GluCTC_1_-_145399233_145399304@39.75.37,trna134_GluTTC_1_-_16861774_16861845@39.75.37,trna71_GluCTC_1_-_161439189_161439260@39.75.37,trna74_GluCTC_1_-_161431809_161431880@39.75.37,trna77_GluCTC_1_-_161424398_161424469@39.75.37,trna77_GluCTC_6_+_28949976_28950047@39.75.37,trna80_GluCTC_1_-_161417018_161417089@39.75.37,trna84_GluTTC_1_-_161391883_161391954@39.75.37,trna87_GluCTC_6_-_126101393_126101464@39.75.37|CGCCGCGGCCCGGGTTCGATTCCCGGTCAGGGAACCA</td><td>>trna116_GluCTC_1_-_145399233_145399304</td><td>39</td><td>75</td><td>37</td><td>CGCCGCGGCCCGGGTTCGATTCCCGGTCAGGGAACCA</td><td>1665.1</td><td>5.7</td><td>1.2</td><td>4.6</td><td>5.2E-06</td><td>8.1E-06</td><td>MB231/MB468</td><td>CYTO</td></tr>
<tr><td>>hg38_21_+_8433172_8446622_RNA45SN1_45S_50ntFlanks@11592.11611.20|ACCAGGGGAATCCGACTGTT</td><td>>hg38_21_+_8433172_8446622_RNA45SN1_45S_50ntFlanks</td><td>11592</td><td>11611</td><td>20</td><td>ACCAGGGGAATCCGACTGTT</td><td>280.5</td><td>-6.2</td><td>1.4</td><td>-4.5</td><td>5.4E-06</td><td>8.1E-06</td><td>MB231/MB468</td><td>NUC</td></tr>
</table>

<table>
<tr><td>>trna143_LysTTT_6_-_27559593_27559665@14.32.19,trna149_LysTTT_6_-_27302769_27302841@14.32.19,trna14_LysTTT_11_-_59327808_59327880@14.32.19,trna2_LysTTT_17_+_8022473_8022545@14.32.19,trna54_LysTTT_1_+_204475655_204475727@14.32.19,trna5_LysTTT_11_+_59323902_59323974@14.32.19,trna62_LysTTT_1_-_204476158_204476230@14.32.19,trna76_LysTTT_6_+_28918806_28918878@14.32.19|AGTCGGTAGAGCATCAGAC</td><td>>trna143_LysTTT_6_-_27559593_27559665</td><td>14</td><td>32</td><td>19</td><td>AGTCGGTAGAGCATCAGAC</td><td>567.7</td><td>3.7</td><td>0.8</td><td>4.6</td><td>4.6E-06</td><td>8.1E-06</td><td>BT20/MB468</td><td>CYTO</td></tr>
<tr><td>>trnaMT_GluTTC_MT_-_14674_14742@40.61.22,trnalookalike8_GluTTC_5_-_93905172_93905240@40.61.22|TTGGTCGTGGTTGTAGTCCGTG</td><td>>trnaMT_GluTTC_MT_-_14674_14742</td><td>40</td><td>61</td><td>22</td><td>TTGGTCGTGGTTGTAGTCCGTG</td><td>656.0</td><td>4.7</td><td>1.0</td><td>4.8</td><td>1.6E-06</td><td>8.1E-06</td><td>BT20/MB468</td><td>MP</td></tr>
<tr><td>>hg38_21_+_8433172_8446622_RNA45SN1_45S_50ntFlanks@11210.11231.22|GGCGACTCTGGACGCGAGCCGG</td><td>>hg38_21_+_8433172_8446622_RNA45SN1_45S_50ntFlanks</td><td>11210</td><td>11231</td><td>22</td><td>GGCGACTCTGGACGCGAGCCGG</td><td>216.7</td><td>-4.9</td><td>1.1</td><td>-4.5</td><td>5.4E-06</td><td>8.1E-06</td><td>MB231/MB468</td><td>NUC</td></tr>
<tr><td>>hg38_21_+_8433172_8446622_RNA45SN1_45S_50ntFlanks@10871.10888.18|GGCTGGGTCGGTCGGGCT</td><td>>hg38_21_+_8433172_8446622_RNA45SN1_45S_50ntFlanks</td><td>10871</td><td>10888</td><td>18</td><td>GGCTGGGTCGGTCGGGCT</td><td>211.4</td><td>11.1</td><td>2.2</td><td>5.0</td><td>4.5E-07</td><td>8.1E-06</td><td>BT20/MB231</td><td>MP</td></tr>
<tr><td>>hg38_21_+_8433172_8446622_RNA45SN1_45S_50ntFlanks@9572.9598.27|CAAATCGGTCGTCCGACCTGGGTATAG</td><td>>hg38_21_+_8433172_8446622_RNA45SN1_45S_50ntFlanks</td><td>9572</td><td>9598</td><td>27</td><td>CAAATCGGTCGTCCGACCTGGGTATAG</td><td>264.9</td><td>-5.8</td><td>1.3</td><td>-4.6</td><td>4.0E-06</td><td>8.2E-06</td><td>BT20/MB468</td><td>MITO</td></tr>
<tr><td>>MI0000750|hsa-mir-26a-2&WithFlank&12|-|57824603|57824698@20.40.21;MI0000083|hsa-mir-26a-1&WithFlank&3|+|37969398|37969486@16.36.21&[MIMAT0000082&hsa-miR-26a-5p&offsets|0|-1;m-33&12|-|57824658|57824679&offsets|0|-1];[MIMAT0000082_1&hsa-miR-26a-5p&offsets|0|-1;m-34&3|+|37969413|37969434&offsets|0|-1]|TTCAAGTAATCCAGGATAGGC</td><td>>MI0000750|hsa-mir-26a-2&WithFlank&12|-|57824603|57824698</td><td>20</td><td>40</td><td>21</td><td>TTCAAGTAATCCAGGATAGGC</td><td>394.2</td><td>2.0</td><td>0.4</td><td>4.5</td><td>5.8E-06</td><td>8.2E-06</td><td>BT20/MB468</td><td>NUC</td></tr>
<tr><td>>hg38_21_+_8433172_8446622_RNA45SN1_45S_50ntFlanks@10512.10535.24|TCCGGAGTGGCGGAGATGGGCGCC</td><td>>hg38_21_+_8433172_8446622_RNA45SN1_45S_50ntFlanks</td><td>10512</td><td>10535</td><td>24</td><td>TCCGGAGTGGCGGAGATGGGCGCC</td><td>158.8</td><td>-3.8</td><td>0.8</td><td>-4.6</td><td>4.6E-06</td><td>8.2E-06</td><td>MB231/MB468</td><td>MP</td></tr>
<tr><td>>hg38_21_+_8433172_8446622_RNA45SN1_45S_50ntFlanks@11589.11623.35|CGGACCAGGGGAATCCGACTGTTTAATTAAAACAA</td><td>>hg38_21_+_8433172_8446622_RNA45SN1_45S_50ntFlanks</td><td>11589</td><td>11623</td><td>35</td><td>CGGACCAGGGGAATCCGACTGTTTAATTAAAACAA</td><td>1851.7</td><td>4.6</td><td>0.9</td><td>5.0</td><td>5.2E-07</td><td>8.2E-06</td><td>BT20/MB231</td><td>MITO</td></tr>
<tr><td>>hg38_21_+_8433172_8446622_RNA45SN1_45S_50ntFlanks@7991.8016.26|ACGTGGCGACCCGCTGAATTTAAGCA</td><td>>hg38_21_+_8433172_8446622_RNA45SN1_45S_50ntFlanks</td><td>7991</td><td>8016</td><td>26</td><td>ACGTGGCGACCCGCTGAATTTAAGCA</td><td>521.8</td><td>3.6</td><td>0.8</td><td>4.6</td><td>5.2E-06</td><td>8.2E-06</td><td>MB231/MB468</td><td>CYTO</td></tr>
<tr><td>>hg38_21_+_8433172_8446622_RNA45SN1_45S_50ntFlanks@10783.10817.35|GTGAACAGCCTCTGGCATGTTGGAACAATGTAGGT</td><td>>hg38_21_+_8433172_8446622_RNA45SN1_45S_50ntFlanks</td><td>10783</td><td>10817</td><td>35</td><td>GTGAACAGCCTCTGGCATGTTGGAACAATGTAGGT</td><td>622.8</td><td>-7.5</td><td>1.7</td><td>-4.5</td><td>5.8E-06</td><td>8.2E-06</td><td>BT20/MB468</td><td>NUC</td></tr>
<tr><td>>hg38_21_+_8433172_8446622_RNA45SN1_45S_50ntFlanks@10727.10759.33|GGAGAGGGTGTAAATCTCGCGCCGGGCCGTACC</td><td>>hg38_21_+_8433172_8446622_RNA45SN1_45S_50ntFlanks</td><td>10727</td><td>10759</td><td>33</td><td>GGAGAGGGTGTAAATCTCGCGCCGGGCCGTACC</td><td>126.6</td><td>-6.4</td><td>1.4</td><td>-4.6</td><td>4.6E-06</td><td>8.2E-06</td><td>MB231/MB468</td><td>MP</td></tr>
<tr><td>>hg38_21_+_8433172_8446622_RNA45SN1_45S_50ntFlanks@5420.5445.26|CCGATTGGATGGTTTAGTGAGGCCCT</td><td>>hg38_21_+_8433172_8446622_RNA45SN1_45S_50ntFlanks</td><td>5420</td><td>5445</td><td>26</td><td>CCGATTGGATGGTTTAGTGAGGCCCT</td><td>1855.8</td><td>-7.0</td><td>1.5</td><td>-4.6</td><td>4.7E-06</td><td>8.2E-06</td><td>BT20/MB468</td><td>CYTO</td></tr>
<tr><td>>MI0000478|hsa-mir-149&WithFlank&2|+|240455995|240456095@21.42.22&[MIMAT0000450&hsa-miR-149-5p&offsets|0|-1;m-195&2|+|240456015|240456037&offsets|0|-1]|TCTGGCTCCGTGTCTTCACTCC</td><td>>MI0000478|hsa-mir-149&WithFlank&2|+|240455995|240456095</td><td>21</td><td>42</td><td>22</td><td>TCTGGCTCCGTGTCTTCACTCC</td><td>1927.0</td><td>3.8</td><td>0.8</td><td>5.0</td><td>4.8E-07</td><td>8.2E-06</td><td>BT20/MB231</td><td>CYTO</td></tr>
<tr><td>>MI0005756|hsa-mir-934&WithFlank&X|+|136550872|136550966@21.42.22(+2A)&[MIMAT0004977&hsa-miR-934&offsets|0|0(+2A);m-943&X|+|136550892|136550913&offsets|0|0(+2A)]|TGTCTACTACTGGAGACACTGGAA</td><td>>MI0005756|hsa-mir-934&WithFlank&X|+|136550872|136550966</td><td>21</td><td>42</td><td>22(+2A)</td><td>TGTCTACTACTGGAGACACTGGAA</td><td>110.6</td><td>-10.7</td><td>2.3</td><td>-4.6</td><td>4.6E-06</td><td>8.2E-06</td><td>MB231/MB468</td><td>MP</td></tr>
<tr><td>>MI0016005|hsa-mir-3615&WithFlank&17|+|74748607|74748705@57.78.22&[MIMAT0017994&hsa-miR-3615&offsets|0|+1;m-271&17|+|74748663|74748684&offsets|0|0]|TCTCTCGGCTCCTCGCGGCTCG</td><td>>MI0016005|hsa-mir-3615&WithFlank&17|+|74748607|74748705</td><td>57</td><td>78</td><td>22</td><td>TCTCTCGGCTCCTCGCGGCTCG</td><td>402.6</td><td>3.9</td><td>0.9</td><td>4.5</td><td>5.6E-06</td><td>8.2E-06</td><td>MB231/MB468</td><td>MITO</td></tr>
<tr><td>>hg38_21_+_8433172_8446622_RNA45SN1_45S_50ntFlanks@11874.11892.19|AGAAGACCCTGTTGAGCTT</td><td>>hg38_21_+_8433172_8446622_RNA45SN1_45S_50ntFlanks</td><td>11874</td><td>11892</td><td>19</td><td>AGAAGACCCTGTTGAGCTT</td><td>467.7</td><td>-5.2</td><td>1.1</td><td>-4.5</td><td>5.5E-06</td><td>8.2E-06</td><td>MB231/MB468</td><td>NUC</td></tr>
<tr><td>>hg38_21_+_8433172_8446622_RNA45SN1_45S_50ntFlanks@10678.10710.33|CGGTTCCGGCGGCGTCCGGTGAGCTCTCGCTGG</td><td>>hg38_21_+_8433172_8446622_RNA45SN1_45S_50ntFlanks</td><td>10678</td><td>10710</td><td>33</td><td>CGGTTCCGGCGGCGTCCGGTGAGCTCTCGCTGG</td><td>119282.6</td><td>-6.0</td><td>1.3</td><td>-4.8</td><td>1.7E-06</td><td>8.2E-06</td><td>BT20/MB468</td><td>MP</td></tr>
<tr><td>>MI0000239|hsa-mir-197&WithFlank&1|+|109598887|109598973@54.74.21&[MIMAT0000227&hsa-miR-197-3p&offsets|0|-1;m-99&1|+|109598940|109598961&offsets|0|-1]|TTCACCACCTTCTCCACCCAG</td><td>>MI0000239|hsa-mir-197&WithFlank&1|+|109598887|109598973</td><td>54</td><td>74</td><td>21</td><td>TTCACCACCTTCTCCACCCAG</td><td>401.2</td><td>3.3</td><td>0.7</td><td>5.0</td><td>6.3E-07</td><td>8.2E-06</td><td>BT20/MB231</td><td>NUC</td></tr>
<tr><td>>MI0000063|hsa-let-7b&WithFlank&22|+|46113680|46113774@12.34.23(+1U)&[MIMAT0000063&hsa-let-7b-5p&offsets|0|+1(+1U);m-14&22|+|46113691|46113712&offsets|0|+1(+1U)]|TGAGGTAGTAGGTTGTGTGGTTTT</td><td>>MI0000063|hsa-let-7b&WithFlank&22|+|46113680|46113774</td><td>12</td><td>34</td><td>23(+1U)</td><td>TGAGGTAGTAGGTTGTGTGGTTTT</td><td>174.1</td><td>2.3</td><td>0.5</td><td>4.5</td><td>5.9E-06</td><td>8.3E-06</td><td>BT20/MB468</td><td>NUC</td></tr>
<tr><td>>hg38_21_+_8433172_8446622_RNA45SN1_45S_50ntFlanks@11567.11605.39|AGCAGCCGACTTAGAACTGGTGCGGACCAGGGGAATCCG</td><td>>hg38_21_+_8433172_8446622_RNA45SN1_45S_50ntFlanks</td><td>11567</td><td>11605</td><td>39</td><td>AGCAGCCGACTTAGAACTGGTGCGGACCAGGGGAATCCG</td><td>439.9</td><td>-7.3</td><td>1.6</td><td>-4.5</td><td>5.9E-06</td><td>8.3E-06</td><td>BT20/MB468</td><td>NUC</td></tr>
<tr><td>>hg38_21_+_8433172_8446622_RNA45SN1_45S_50ntFlanks@12949.12989.41|ACGACCTGCTTCTGGGTCGGGGTTTCGTACGTAGCAGAGCA</td><td>>hg38_21_+_8433172_8446622_RNA45SN1_45S_50ntFlanks</td><td>12949</td><td>12989</td><td>41</td><td>ACGACCTGCTTCTGGGTCGGGGTTTCGTACGTAGCAGAGCA</td><td>181.9</td><td>-5.0</td><td>1.1</td><td>-4.5</td><td>5.5E-06</td><td>8.3E-06</td><td>MB231/MB468</td><td>NUC</td></tr>
<tr><td>>MI0003776|hsa-mir-320b-1&WithFlank&1|+|116671740|116671823@48.66.19(+1A);MI0003839|hsa-mir-320b-2&WithFlank&1|-|224257034|224257116@47.65.19(+1A);MI0003778|hsa-mir-320c-1&WithFlank&18|+|21683512|21683595@48.66.19(+1A);MI0008191|hsa-mir-320c-2&WithFlank&18|+|24321669|24321752@48.66.19(+1A);MI0000542|hsa-mir-320a&WithFlank&8|-|22244960|22245043@48.66.19(+1A)&[MIMAT0005792&hsa-miR-320b&offsets|0|-3(+1A);m-183&1|+|116671787|116671806&offsets|0|-1(+1A)];[MIMAT0005792_1&hsa-miR-320b&offsets|0|-3(+1A);m-183&1|-|224257051|224257070&offsets|0|-1(+1A)];[MIMAT0005793&hsa-miR-320c&offsets|0|-1(+1A);m-222&18|+|21683559|21683578&offsets|0|-1(+1A)];[MIMAT0005793_1&hsa-miR-320c&offsets|0|-1(+1A);m-219&18|+|24321716|24321735&offsets|0|-1(+1A)];[MIMAT0000510&hsa-miR-320a-3p&offsets|0|-3(+1A);m-60&8|-|22244975|22244996&offsets|0|-3(+1A)]|AAAAGCTGGGTTGAGAGGGA</td><td>>MI0003776|hsa-mir-320b-1&WithFlank&1|+|116671740|116671823</td><td>48</td><td>66</td><td>19(+1A)</td><td>AAAAGCTGGGTTGAGAGGGA</td><td>1761.7</td><td>8.2</td><td>1.7</td><td>4.8</td><td>1.7E-06</td><td>8.3E-06</td><td>BT20/MB468</td><td>MP</td></tr>
<tr><td>>hg38_21_+_8433172_8446622_RNA45SN1_45S_50ntFlanks@12949.12982.34|ACGACCTGCTTCTGGGTCGGGGTTTCGTACGTAG</td><td>>hg38_21_+_8433172_8446622_RNA45SN1_45S_50ntFlanks</td><td>12949</td><td>12982</td><td>34</td><td>ACGACCTGCTTCTGGGTCGGGGTTTCGTACGTAG</td><td>55.9</td><td>-8.9</td><td>1.9</td><td>-4.8</td><td>1.7E-06</td><td>8.3E-06</td><td>BT20/MB468</td><td>MP</td></tr>
<tr><td>>hg38_21_+_8433172_8446622_RNA45SN1_45S_50ntFlanks@12912.12934.23|CACGTTCGTGGGGAACCTGGCGC</td><td>>hg38_21_+_8433172_8446622_RNA45SN1_45S_50ntFlanks</td><td>12912</td><td>12934</td><td>23</td><td>CACGTTCGTGGGGAACCTGGCGC</td><td>101.9</td><td>-10.5</td><td>2.3</td><td>-4.6</td><td>4.6E-06</td><td>8.3E-06</td><td>MB231/MB468</td><td>MP</td></tr>
<tr><td>>MI0000298|hsa-mir-221&WithFlank&X|-|45746151|45746272@31.55.25&[MIMAT0004568&hsa-miR-221-5p&offsets|0|+3;m-405&X|-|45746220|45746242&offsets|0|+2]|ACCTGGCATACAATGTAGATTTCTG</td><td>>MI0000298|hsa-mir-221&WithFlank&X|-|45746151|45746272</td><td>31</td><td>55</td><td>25</td><td>ACCTGGCATACAATGTAGATTTCTG</td><td>198.0</td><td>6.4</td><td>1.4</td><td>4.5</td><td>5.6E-06</td><td>8.3E-06</td><td>MB231/MB468</td><td>MITO</td></tr>
<tr><td>>hg38_21_+_8433172_8446622_RNA45SN1_45S_50ntFlanks@10402.10419.18|AACATGGGTCAGTCGGTC</td><td>>hg38_21_+_8433172_8446622_RNA45SN1_45S_50ntFlanks</td><td>10402</td><td>10419</td><td>18</td><td>AACATGGGTCAGTCGGTC</td><td>1017.6</td><td>-4.7</td><td>1.0</td><td>-4.8</td><td>1.7E-06</td><td>8.3E-06</td><td>BT20/MB468</td><td>MP</td></tr>
<tr><td>>hg38_21_+_8433172_8446622_RNA45SN1_45S_50ntFlanks@10234.10261.28|CGGACGCTACGCCGCGACGAGTAGGAGG</td><td>>hg38_21_+_8433172_8446622_RNA45SN1_45S_50ntFlanks</td><td>10234</td><td>10261</td><td>28</td><td>CGGACGCTACGCCGCGACGAGTAGGAGG</td><td>130.1</td><td>10.4</td><td>2.1</td><td>5.0</td><td>4.6E-07</td><td>8.3E-06</td><td>BT20/MB231</td><td>MP</td></tr>
<tr><td>>hg38_21_+_8433172_8446622_RNA45SN1_45S_50ntFlanks@8120.8140.21|GACATGTGGCGTACGGAAGAC</td><td>>hg38_21_+_8433172_8446622_RNA45SN1_45S_50ntFlanks</td><td>8120</td><td>8140</td><td>21</td><td>GACATGTGGCGTACGGAAGAC</td><td>97.4</td><td>9.9</td><td>2.0</td><td>5.0</td><td>4.6E-07</td><td>8.3E-06</td><td>BT20/MB231</td><td>MP</td></tr>
<tr><td>>trnaMT_SerGCT_MT_+_12207_12265@1.48.48|GAGAAAGCTCACAAGAACTGCTAACTCATGCCCCCATGTCTAACAACA</td><td>>trnaMT_SerGCT_MT_+_12207_12265</td><td>1</td><td>48</td><td>48</td><td>GAGAAAGCTCACAAGAACTGCTAACTCATGCCCCCATGTCTAACAACA</td><td>1413.0</td><td>6.7</td><td>1.5</td><td>4.5</td><td>5.6E-06</td><td>8.3E-06</td><td>MB231/MB468</td><td>MITO</td></tr>
<tr><td>>MI0000285|hsa-mir-205&WithFlank&1|+|209432127|209432248@40.61.22(+1C)&[MIMAT0000266&hsa-miR-205-5p&offsets|0|0(+1C);m-351&1|+|209432166|209432187&offsets|0|0(+1C)]|TCCTTCATTCCACCGGAGTCTGC</td><td>>MI0000285|hsa-mir-205&WithFlank&1|+|209432127|209432248</td><td>40</td><td>61</td><td>22(+1C)</td><td>TCCTTCATTCCACCGGAGTCTGC</td><td>2263.7</td><td>3.3</td><td>0.7</td><td>4.5</td><td>5.9E-06</td><td>8.3E-06</td><td>BT20/MB468</td><td>NUC</td></tr>
<tr><td>>hg38_21_+_8433172_8446622_RNA45SN1_45S_50ntFlanks@3838.3868.31|CTCTCCTACTTGGATAACTGTGGTAATTCTA</td><td>>hg38_21_+_8433172_8446622_RNA45SN1_45S_50ntFlanks</td><td>3838</td><td>3868</td><td>31</td><td>CTCTCCTACTTGGATAACTGTGGTAATTCTA</td><td>355.6</td><td>-5.3</td><td>1.2</td><td>-4.5</td><td>5.6E-06</td><td>8.3E-06</td><td>MB231/MB468</td><td>MITO</td></tr>
<tr><td>>hg38_21_+_8433172_8446622_RNA45SN1_45S_50ntFlanks@10726.10745.20|GGGAGAGGGTGTAAATCTCG</td><td>>hg38_21_+_8433172_8446622_RNA45SN1_45S_50ntFlanks</td><td>10726</td><td>10745</td><td>20</td><td>GGGAGAGGGTGTAAATCTCG</td><td>61.6</td><td>-9.0</td><td>1.9</td><td>-4.8</td><td>1.7E-06</td><td>8.3E-06</td><td>BT20/MB468</td><td>MP</td></tr>
<tr><td>>hg38_21_+_8433172_8446622_RNA45SN1_45S_50ntFlanks@12934.12972.39|CTAAACCATTCGTAGACGACCTGCTTCTGGGTCGGGGTT</td><td>>hg38_21_+_8433172_8446622_RNA45SN1_45S_50ntFlanks</td><td>12934</td><td>12972</td><td>39</td><td>CTAAACCATTCGTAGACGACCTGCTTCTGGGTCGGGGTT</td><td>1182.0</td><td>-8.0</td><td>1.7</td><td>-4.8</td><td>1.7E-06</td><td>8.3E-06</td><td>BT20/MB468</td><td>MP</td></tr>
<tr><td>>hg38_MT_+_598_1651_RNR1_12S_50ntFlanks@844.871.28|AGTAGAGTGCTTAGTTGAACAGGGCCCT</td><td>>hg38_MT_+_598_1651_RNR1_12S_50ntFlanks</td><td>844</td><td>871</td><td>28</td><td>AGTAGAGTGCTTAGTTGAACAGGGCCCT</td><td>251.1</td><td>8.8</td><td>1.8</td><td>4.8</td><td>1.7E-06</td><td>8.3E-06</td><td>BT20/MB468</td><td>MP</td></tr>
<tr><td>>hg38_21_+_8433172_8446622_RNA45SN1_45S_50ntFlanks@5472.5491.20|ACGGCCCTGGCGGAGCGCTG</td><td>>hg38_21_+_8433172_8446622_RNA45SN1_45S_50ntFlanks</td><td>5472</td><td>5491</td><td>20</td><td>ACGGCCCTGGCGGAGCGCTG</td><td>3036.9</td><td>5.1</td><td>1.1</td><td>4.6</td><td>4.7E-06</td><td>8.3E-06</td><td>BT20/MB468</td><td>CYTO</td></tr>
<tr><td>>hg38_21_+_8433172_8446622_RNA45SN1_45S_50ntFlanks@11718.11738.21|AACGGCGGGAGTAACTATGAC</td><td>>hg38_21_+_8433172_8446622_RNA45SN1_45S_50ntFlanks</td><td>11718</td><td>11738</td><td>21</td><td>AACGGCGGGAGTAACTATGAC</td><td>643.2</td><td>-5.7</td><td>1.2</td><td>-4.6</td><td>4.7E-06</td><td>8.3E-06</td><td>BT20/MB468</td><td>CYTO</td></tr>
<tr><td>>hg38_21_+_8433172_8446622_RNA45SN1_45S_50ntFlanks@11233.11253.21|CCCTTCCCGTGGATCGCCCCA</td><td>>hg38_21_+_8433172_8446622_RNA45SN1_45S_50ntFlanks</td><td>11233</td><td>11253</td><td>21</td><td>CCCTTCCCGTGGATCGCCCCA</td><td>371.4</td><td>-6.2</td><td>1.3</td><td>-4.6</td><td>4.0E-06</td><td>8.3E-06</td><td>BT20/MB468</td><td>MITO</td></tr>
<tr><td>>am_trna2_GlyGCC_21_-_18827107_18827177@3.26.24,trna35_GlyGCC_1_+_161413094_161413164@3.26.24,trna37_GlyGCC_1_+_161420467_161420537@3.26.24,trna39_GlyGCC_1_+_161427898_161427968@3.26.24,trna41_GlyGCC_1_+_161435258_161435328@3.26.24|ATGGGTGGTTCAGTGGTAGAATTC</td><td>>am_trna2_GlyGCC_21_-_18827107_18827177</td><td>3</td><td>26</td><td>24</td><td>ATGGGTGGTTCAGTGGTAGAATTC</td><td>44.6</td><td>9.2</td><td>1.9</td><td>4.8</td><td>1.7E-06</td><td>8.3E-06</td><td>BT20/MB468</td><td>MP</td></tr>
<tr><td>>hg38_21_+_8433172_8446622_RNA45SN1_45S_50ntFlanks@12436.12473.38|TCATTGTGAAGCAGAATTCACCAAGCGTTGGATTGTTC</td><td>>hg38_21_+_8433172_8446622_RNA45SN1_45S_50ntFlanks</td><td>12436</td><td>12473</td><td>38</td><td>TCATTGTGAAGCAGAATTCACCAAGCGTTGGATTGTTC</td><td>335.9</td><td>-5.5</td><td>1.2</td><td>-4.6</td><td>4.7E-06</td><td>8.3E-06</td><td>MB231/MB468</td><td>MP</td></tr>
<tr><td>>hg38_21_+_8433172_8446622_RNA45SN1_45S_50ntFlanks@5083.5110.28|ACGAACGAGACTCTGGCATGCTAACTAG</td><td>>hg38_21_+_8433172_8446622_RNA45SN1_45S_50ntFlanks</td><td>5083</td><td>5110</td><td>28</td><td>ACGAACGAGACTCTGGCATGCTAACTAG</td><td>895.3</td><td>-6.7</td><td>1.5</td><td>-4.6</td><td>4.7E-06</td><td>8.3E-06</td><td>MB231/MB468</td><td>MP</td></tr>
<tr><td>>hg38_21_+_8433172_8446622_RNA45SN1_45S_50ntFlanks@10853.10873.21|ATAAGGATTGGCTCTAAGGGC</td><td>>hg38_21_+_8433172_8446622_RNA45SN1_45S_50ntFlanks</td><td>10853</td><td>10873</td><td>21</td><td>ATAAGGATTGGCTCTAAGGGC</td><td>318.0</td><td>-2.9</td><td>0.6</td><td>-4.6</td><td>4.7E-06</td><td>8.3E-06</td><td>BT20/MB468</td><td>CYTO</td></tr>
<tr><td>>hg38_21_+_8433172_8446622_RNA45SN1_45S_50ntFlanks@6661.6679.19|CGGTGGATCACTCGGCTCG</td><td>>hg38_21_+_8433172_8446622_RNA45SN1_45S_50ntFlanks</td><td>6661</td><td>6679</td><td>19</td><td>CGGTGGATCACTCGGCTCG</td><td>207.4</td><td>-3.4</td><td>0.7</td><td>-4.6</td><td>4.7E-06</td><td>8.3E-06</td><td>MB231/MB468</td><td>MP</td></tr>
</table>

| | | | | | | | | | | | | | |
|---|---|---|---|---|---|---|---|---|---|---|---|---|---|
| >MI0000456\|hsa-mir-140&WithFlank&16\|+\|69933075\|69933186@29.47.19(+1C)&[MIMAT0000431&hsa-miR-140-5p&offsets\|0\|-3(+1C);m-173&16\|+\|69933103\|69933124&offsets\|0\|-3(+1C)]\|CAGTGGTTTTACCCTATGGC | >MI0000456\|hsa-mir-140&WithFlank&16\|+\|69933075\|69933186 | 29 | 47 | 19(+1C) | CAGTGGTTTTACCCTATGGC | 56.6 | 9.0 | 2.0 | 4.6 | 4.7E-06 | 8.3E-06 | MB231/MB468 | MP |
| >hg38_21_+_8433172_8446622_RNA45SN1_45S_50ntFlanks@7979.8006.28\|ACCTCAGATCAGACGTGGCGACCCGCTG | >hg38_21_+_8433172_8446622_RNA45SN1_45S_50ntFlanks | 7979 | 8006 | 28 | ACCTCAGATCAGACGTGGCGACCCGCTG | 2215.0 | -4.1 | 0.9 | -4.6 | 4.7E-06 | 8.3E-06 | BT20/MB468 | CYTO |
| >hg38_21_+_8433172_8446622_RNA45SN1_45S_50ntFlanks@10350.10381.32\|ACGAGAACTTTGAAGGCCGAAGTGGAGAAGGG | >hg38_21_+_8433172_8446622_RNA45SN1_45S_50ntFlanks | 10350 | 10381 | 32 | ACGAGAACTTTGAAGGCCGAAGTGGAGAAGGG | 321.6 | -2.5 | 0.6 | -4.6 | 5.3E-06 | 8.4E-06 | MB231/MB468 | CYTO |
| >hg38_21_+_8433172_8446622_RNA45SN1_45S_50ntFlanks@6646.6667.22\|TCGTACGACTCTTAGCGGTGGA | >hg38_21_+_8433172_8446622_RNA45SN1_45S_50ntFlanks | 6646 | 6667 | 22 | TCGTACGACTCTTAGCGGTGGA | 2169.3 | 3.1 | 0.7 | 4.6 | 5.3E-06 | 8.4E-06 | MB231/MB468 | CYTO |
| >MI0000736\|hsa-mir-30c-1&WithFlank&1\|+\|40757278\|40757378@23.48.26;MI0000254\|hsa-mir-30c-2&WithFlank&6\|-\|71376954\|71377037@13.38.26&[MIMAT0000244&hsa-miR-30c-5p&offsets\|0\|+3;m-57&1\|+\|40757300\|40757323&offsets\|0\|+2];[MIMAT0000244_1&hsa-miR-30c-5p&offsets\|0\|+3;m-56&6\|-\|71377002\|71377025&offsets\|0\|+2]\|TGTAAACATCCTACACTCTCAGCTGT | >MI0000736\|hsa-mir-30c-1&WithFlank&1\|+\|40757278\|40757378 | 23 | 48 | 26 | TGTAAACATCCTACACTCTCAGCTGT | 503.2 | -4.1 | 0.8 | -5.0 | 5.4E-07 | 8.4E-06 | BT20/MB231 | MITO |
| >hg38_21_+_8433172_8446622_RNA45SN1_45S_50ntFlanks@10404.10424.21\|CATGGGTCAGTCGGTCCTGAG | >hg38_21_+_8433172_8446622_RNA45SN1_45S_50ntFlanks | 10404 | 10424 | 21 | CATGGGTCAGTCGGTCCTGAG | 1746.1 | -2.3 | 0.5 | -4.5 | 5.9E-06 | 8.4E-06 | BT20/MB468 | NUC |
| >hg38_21_+_8433172_8446622_RNA45SN1_45S_50ntFlanks@12669.12703.35\|CTAAGTCAGAATCCCGCCCAGGCGGAACGATACGG | >hg38_21_+_8433172_8446622_RNA45SN1_45S_50ntFlanks | 12669 | 12703 | 35 | CTAAGTCAGAATCCCGCCCAGGCGGAACGATACGG | 60.4 | -9.0 | 1.9 | -4.8 | 1.7E-06 | 8.4E-06 | BT20/MB468 | MP |
| >hg38_21_+_8433172_8446622_RNA45SN1_45S_50ntFlanks@6700.6718.19\|CTAGCTGCGAGAATTAATG | >hg38_21_+_8433172_8446622_RNA45SN1_45S_50ntFlanks | 6700 | 6718 | 19 | CTAGCTGCGAGAATTAATG | 7051.9 | -8.5 | 1.8 | -4.8 | 1.7E-06 | 8.4E-06 | BT20/MB468 | MP |
| >MI0000736\|hsa-mir-30c-1&WithFlank&1\|+\|40757278\|40757378@23.48.26;MI0000254\|hsa-mir-30c-2&WithFlank&6\|-\|71376954\|71377037@13.38.26&[MIMAT0000244&hsa-miR-30c-5p&offsets\|0\|+3;m-57&1\|+\|40757300\|40757323&offsets\|0\|+2];[MIMAT0000244_1&hsa-miR-30c-5p&offsets\|0\|+3;m-56&6\|-\|71377002\|71377025&offsets\|0\|+2]\|TGTAAACATCCTACACTCTCAGCTGT | >MI0000736\|hsa-mir-30c-1&WithFlank&1\|+\|40757278\|40757378 | 23 | 48 | 26 | TGTAAACATCCTACACTCTCAGCTGT | 437.8 | 6.7 | 1.5 | 4.5 | 5.7E-06 | 8.4E-06 | MB231/MB468 | MITO |
| >hg38_21_+_8433172_8446622_RNA45SN1_45S_50ntFlanks@9574.9594.21\|AATCGGTCGTCCGACCTGGGT | >hg38_21_+_8433172_8446622_RNA45SN1_45S_50ntFlanks | 9574 | 9594 | 21 | AATCGGTCGTCCGACCTGGGT | 424.4 | -2.5 | 0.6 | -4.6 | 4.8E-06 | 8.4E-06 | BT20/MB468 | CYTO |
| >hg38_21_+_8433172_8446622_RNA45SN1_45S_50ntFlanks@10402.10426.25\|AACATGGGTCAGTCGGTCCTGAGAG | >hg38_21_+_8433172_8446622_RNA45SN1_45S_50ntFlanks | 10402 | 10426 | 25 | AACATGGGTCAGTCGGTCCTGAGAG | 414.9 | -5.1 | 1.1 | -4.5 | 5.6E-06 | 8.4E-06 | MB231/MB468 | NUC |
| >hg38_21_+_8433172_8446622_RNA45SN1_45S_50ntFlanks@11641.11679.39\|CGGCGGGTGTTGACGCGATGTGATTTCTGCCCAGTGCTC | >hg38_21_+_8433172_8446622_RNA45SN1_45S_50ntFlanks | 11641 | 11679 | 39 | CGGCGGGTGTTGACGCGATGTGATTTCTGCCCAGTGCTC | 327.4 | -7.5 | 1.6 | -4.8 | 1.7E-06 | 8.4E-06 | BT20/MB468 | MP |
| >hg38_21_+_8433172_8446622_RNA45SN1_45S_50ntFlanks@3846.3868.23\|CTTGGATAACTGTGGTAATTCTA | >hg38_21_+_8433172_8446622_RNA45SN1_45S_50ntFlanks | 3846 | 3868 | 23 | CTTGGATAACTGTGGTAATTCTA | 493.6 | -2.6 | 0.6 | -4.5 | 5.6E-06 | 8.4E-06 | MB231/MB468 | NUC |
| >hg38_21_+_8433172_8446622_RNA45SN1_45S_50ntFlanks@10878.10901.24\|TCGGTCGGGCTGGGGCGCGAAGCG | >hg38_21_+_8433172_8446622_RNA45SN1_45S_50ntFlanks | 10878 | 10901 | 24 | TCGGTCGGGCTGGGGCGCGAAGCG | 611.6 | -6.7 | 1.5 | -4.5 | 5.6E-06 | 8.4E-06 | MB231/MB468 | NUC |
| >hg38_21_+_8433172_8446622_RNA45SN1_45S_50ntFlanks@11671.11703.33\|CCAGTGCTCTGAATGTCAAAGTGAAGAAATTCA | >hg38_21_+_8433172_8446622_RNA45SN1_45S_50ntFlanks | 11671 | 11703 | 33 | CCAGTGCTCTGAATGTCAAAGTGAAGAAATTCA | 485.4 | -5.9 | 1.3 | -4.6 | 4.7E-06 | 8.4E-06 | MB231/MB468 | MP |
| >hg38_21_+_8433172_8446622_RNA45SN1_45S_50ntFlanks@10793.10817.25\|TCTGGCATGTTGGAACAATGTAGGT | >hg38_21_+_8433172_8446622_RNA45SN1_45S_50ntFlanks | 10793 | 10817 | 25 | TCTGGCATGTTGGAACAATGTAGGT | 115.4 | -6.2 | 1.4 | -4.6 | 4.7E-06 | 8.4E-06 | MB231/MB468 | MP |
| >hg38_21_+_8433172_8446622_RNA45SN1_45S_50ntFlanks@11217.11246.30\|CTGGACGCGAGCCGGGCCCTTCCCGTGGAT | >hg38_21_+_8433172_8446622_RNA45SN1_45S_50ntFlanks | 11217 | 11246 | 30 | CTGGACGCGAGCCGGGCCCTTCCCGTGGAT | 151.0 | 3.7 | 0.8 | 4.5 | 5.7E-06 | 8.4E-06 | MB231/MB468 | MITO |
| >hg38_21_+_8433172_8446622_RNA45SN1_45S_50ntFlanks@10349.10374.26\|AACGAGAACTTTGAAGGCCGAAGTGG | >hg38_21_+_8433172_8446622_RNA45SN1_45S_50ntFlanks | 10349 | 10374 | 26 | AACGAGAACTTTGAAGGCCGAAGTGG | 2169.0 | -4.9 | 1.0 | -4.8 | 1.7E-06 | 8.4E-06 | BT20/MB468 | MP |
| >MI0000483\|hsa-mir-186&WithFlank&1\|-\|71067625\|71067722@21.41.21&[MIMAT0000456&hsa-miR-186-5p&offsets\|0\|-1;m-84&1\|-\|71067681\|71067702&offsets\|0\|-1]\|CAAAGAATTCTCCTTTTGGGC | >MI0000483\|hsa-mir-186&WithFlank&1\|-\|71067625\|71067722 | 21 | 41 | 21 | CAAAGAATTCTCCTTTTGGGC | 205.6 | 4.6 | 1.0 | 4.5 | 5.7E-06 | 8.4E-06 | MB231/MB468 | MITO |
| >MI0006444\|hsa-mir-1307&WithFlank&10\|-\|103394247\|103394407@86.107.22(+1U)&[MIMAT0005951&hsa-miR-1307-3p&offsets\|0\|0(+1U);m-66&10\|-\|103394300\|103394322&offsets\|0\|-1(+1U)]\|ACTCGGCGTGGCGTCGGTCGTGT | >MI0006444\|hsa-mir-1307&WithFlank&10\|-\|103394247\|103394407 | 86 | 107 | 22(+1U) | ACTCGGCGTGGCGTCGGTCGTGT | 142.5 | 2.1 | 0.5 | 4.5 | 6.0E-06 | 8.4E-06 | BT20/MB468 | NUC |
| >hg38_21_+_8433172_8446622_RNA45SN1_45S_50ntFlanks@11319.11342.24\|CCCACGTCTCGTCGCGCGCGCGTC | >hg38_21_+_8433172_8446622_RNA45SN1_45S_50ntFlanks | 11319 | 11342 | 24 | CCCACGTCTCGTCGCGCGCGCGTC | 978.4 | -6.9 | 1.5 | -4.5 | 5.6E-06 | 8.4E-06 | MB231/MB468 | NUC |
| >hg38_21_+_8433172_8446622_RNA45SN1_45S_50ntFlanks@10317.10335.19\|GTGCAGATCTTGGTGGTAG | >hg38_21_+_8433172_8446622_RNA45SN1_45S_50ntFlanks | 10317 | 10335 | 19 | GTGCAGATCTTGGTGGTAG | 219.9 | 11.1 | 2.2 | 5.0 | 4.7E-07 | 8.4E-06 | BT20/MB231 | MP |
| >hg38_21_+_8433172_8446622_RNA45SN1_45S_50ntFlanks@3846.3872.27\|CTTGGATAACTGTGGTAATTCTAGAGC | >hg38_21_+_8433172_8446622_RNA45SN1_45S_50ntFlanks | 3846 | 3872 | 27 | CTTGGATAACTGTGGTAATTCTAGAGC | 230.5 | -6.5 | 1.4 | -4.6 | 4.8E-06 | 8.5E-06 | MB231/MB468 | MP |
| >trna119_LysCTT_1_-_145395522_145395594@4.34.31,trna11_LysCTT_5_-_180648979_180649051@4.34.31,trna13_LysCTT_6_+_26556774_26556846@4.34.31,trna7_LysCTT_16_+_3225692_3225764@4.34.31,trna9_LysCTT_5_+_180634755_180634827@4.34.31\|CGGCTAGCTCAGTCGGTAGAGCATGAGACTC | >trna119_LysCTT_1_-_145395522_145395594 | 4 | 34 | 31 | CGGCTAGCTCAGTCGGTAGAGCATGAGACTC | 784.7 | -5.2 | 1.0 | -5.0 | 5.5E-07 | 8.5E-06 | BT20/MB231 | MITO |
| >trna119_LysCTT_1_-_145395522_145395594@3.32.30,trna11_LysCTT_5_-_180648979_180649051@3.32.30,trna13_LysCTT_6_+_26556774_26556846@3.32.30,trna32_LysCTT_16_-_3207406_3207478@3.32.30,trna7_LysCTT_16_+_3225692_3225764@3.32.30,trna9_LysCTT_5_+_180634755_180634827@3.32.30\|CCGGCTAGCTCAGTCGGTAGAGCATGAGAC | >trna119_LysCTT_1_-_145395522_145395594 | 3 | 32 | 30 | CCGGCTAGCTCAGTCGGTAGAGCATGAGAC | 209.3 | 5.7 | 1.2 | 4.5 | 5.7E-06 | 8.5E-06 | MB231/MB468 | MITO |
| >hg38_21_+_8433172_8446622_RNA45SN1_45S_50ntFlanks@9605.9622.18\|AAGACTAATCGAACCATC | >hg38_21_+_8433172_8446622_RNA45SN1_45S_50ntFlanks | 9605 | 9622 | 18 | AAGACTAATCGAACCATC | 1160.8 | -4.6 | 1.0 | -4.8 | 1.7E-06 | 8.5E-06 | BT20/MB468 | MP |
| >MI0000764\|hsa-mir-363&WithFlank&X\|-\|134169372\|134169458@56.78.23&[MIMAT0000707&hsa-miR-363-3p&offsets\|0\|+1;m-178&X\|-\|134169381\|134169403&offsets\|0\|0]\|AATTGCACGGTATCCATCTGTAA | >MI0000764\|hsa-mir-363&WithFlank&X\|-\|134169372\|134169458 | 56 | 78 | 23 | AATTGCACGGTATCCATCTGTAA | 87.3 | 9.8 | 1.9 | 5.0 | 4.7E-07 | 8.5E-06 | BT20/MB231 | MP |
| >hg38_21_+_8433172_8446622_RNA45SN1_45S_50ntFlanks@12539.12563.25\|ATGATGTGTTGTTGCCATGGTAATC | >hg38_21_+_8433172_8446622_RNA45SN1_45S_50ntFlanks | 12539 | 12563 | 25 | ATGATGTGTTGTTGCCATGGTAATC | 370.4 | -6.0 | 1.3 | -4.6 | 4.8E-06 | 8.5E-06 | BT20/MB468 | CYTO |
| >hg38_21_+_8433172_8446622_RNA45SN1_45S_50ntFlanks@6671.6697.27\|CTCGGCTCGTGCGTCGATGAAGAACGC | >hg38_21_+_8433172_8446622_RNA45SN1_45S_50ntFlanks | 6671 | 6697 | 27 | CTCGGCTCGTGCGTCGATGAAGAACGC | 366.8 | -4.2 | 0.9 | -4.6 | 4.8E-06 | 8.5E-06 | BT20/MB468 | CYTO |
| >hg38_21_+_8433172_8446622_RNA45SN1_45S_50ntFlanks@10787.10816.30\|ACAGCCTCTGGCATGTTGGAACAATGTAGG | >hg38_21_+_8433172_8446622_RNA45SN1_45S_50ntFlanks | 10787 | 10816 | 30 | ACAGCCTCTGGCATGTTGGAACAATGTAGG | 577.8 | -3.2 | 0.7 | -4.5 | 5.4E-06 | 8.5E-06 | MB231/MB468 | CYTO |
| >hg38_21_+_8433172_8446622_RNA45SN1_45S_50ntFlanks@5487.5523.37\|CGCTGAGAAGACGGTCGAACTTGACTATCTAGAGGAA | >hg38_21_+_8433172_8446622_RNA45SN1_45S_50ntFlanks | 5487 | 5523 | 37 | CGCTGAGAAGACGGTCGAACTTGACTATCTAGAGGAA | 61.3 | -9.0 | 1.9 | -4.8 | 1.7E-06 | 8.5E-06 | BT20/MB468 | MP |
| >hg38_21_+_8433172_8446622_RNA45SN1_45S_50ntFlanks@12405.12445.41\|CTTTTTGATCCTTCGATGTCGGCTCTTCCTATCATTGTGAA | >hg38_21_+_8433172_8446622_RNA45SN1_45S_50ntFlanks | 12405 | 12445 | 41 | CTTTTTGATCCTTCGATGTCGGCTCTTCCTATCATTGTGAA | 686.8 | -5.3 | 1.2 | -4.5 | 5.8E-06 | 8.5E-06 | MB231/MB468 | MITO |
| >hg38_1_-_228634819_228635039_RNA5S12_5S_50ntFlanks@51.94.44\|GTCTACGGCCATACCACCCTGAACGCGCCCGATCTCGTCTGATC | >hg38_1_-_228634819_228635039_RNA5S12_5S_50ntFlanks | 51 | 94 | 44 | GTCTACGGCCATACCACCCTGAACGCGCCCGATCTCGTCTGATC | 247.7 | 3.5 | 0.8 | 4.5 | 5.6E-06 | 8.5E-06 | MB231/MB468 | NUC |
| >hg38_MT_+_1621_3279_RNR2_16S_50ntFlanks@51.95.45\|GCTAAACCTAGCCCCAAACCCACTCCACCTTACTACCAGACAACC | >hg38_MT_+_1621_3279_RNR2_16S_50ntFlanks | 51 | 95 | 45 | GCTAAACCTAGCCCCAAACCCACTCCACCTTACTACCAGACAACC | 1661.4 | 6.1 | 1.3 | 4.5 | 5.6E-06 | 8.5E-06 | MB231/MB468 | NUC |
| >hg38_21_+_8433172_8446622_RNA45SN1_45S_50ntFlanks@11672.11699.28\|CAGTGCTCTGAATGTCAAAGTGAAGAAA | >hg38_21_+_8433172_8446622_RNA45SN1_45S_50ntFlanks | 11672 | 11699 | 28 | CAGTGCTCTGAATGTCAAAGTGAAGAAA | 207.7 | -2.2 | 0.5 | -4.5 | 5.6E-06 | 8.5E-06 | MB231/MB468 | NUC |
| >hg38_21_+_8433172_8446622_RNA45SN1_45S_50ntFlanks@6648.6670.23\|GTACGACTCTTAGCGGTGGATCA | >hg38_21_+_8433172_8446622_RNA45SN1_45S_50ntFlanks | 6648 | 6670 | 23 | GTACGACTCTTAGCGGTGGATCA | 899.7 | -3.4 | 0.8 | -4.5 | 5.8E-06 | 8.5E-06 | MB231/MB468 | MITO |
| >MI0000811\|hsa-mir-148b&WithFlank&12\|+\|54337210\|54337320@69.91.23&[MIMAT0000759&hsa-miR-148b-3p&offsets\|0\|+1;m-77&12\|+\|54337278\|54337299&offsets\|0\|+1]\|TCAGTGCATCACAGAACTTTGTC | >MI0000811\|hsa-mir-148b&WithFlank&12\|+\|54337210\|54337320 | 69 | 91 | 23 | TCAGTGCATCACAGAACTTTGTC | 753.6 | 7.3 | 1.6 | 4.5 | 5.8E-06 | 8.5E-06 | MB231/MB468 | MITO |
| >hg38_21_+_8433172_8446622_RNA45SN1_45S_50ntFlanks@12907.12932.26\|CACCGCACGTTCGTGGGGAACCTGGC | >hg38_21_+_8433172_8446622_RNA45SN1_45S_50ntFlanks | 12907 | 12932 | 26 | CACCGCACGTTCGTGGGGAACCTGGC | 516.7 | -5.3 | 1.2 | -4.5 | 5.8E-06 | 8.5E-06 | MB231/MB468 | MITO |
| >hg38_21_+_8433172_8446622_RNA45SN1_45S_50ntFlanks@12416.12445.30\|TTCGATGTCGGCTCTTCCTATCATTGTGAA | >hg38_21_+_8433172_8446622_RNA45SN1_45S_50ntFlanks | 12416 | 12445 | 30 | TTCGATGTCGGCTCTTCCTATCATTGTGAA | 593.8 | 2.7 | 0.6 | 4.6 | 4.8E-06 | 8.5E-06 | BT20/MB468 | CYTO |
| >hg38_21_+_8433172_8446622_RNA45SN1_45S_50ntFlanks@12526.12559.34\|TTTTACCCTACTGATGATGTGTTGTTGCCATGGT | >hg38_21_+_8433172_8446622_RNA45SN1_45S_50ntFlanks | 12526 | 12559 | 34 | TTTTACCCTACTGATGATGTGTTGTTGCCATGGT | 49.7 | -8.7 | 1.8 | -4.8 | 1.7E-06 | 8.5E-06 | BT20/MB468 | MP |
| >hg38_21_+_8433172_8446622_RNA45SN1_45S_50ntFlanks@12437.12474.38\|CATTGTGAAGCAGAATTCACCAAGCGTTGGATTGTTCA | >hg38_21_+_8433172_8446622_RNA45SN1_45S_50ntFlanks | 12437 | 12474 | 38 | CATTGTGAAGCAGAATTCACCAAGCGTTGGATTGTTCA | 314.5 | -6.8 | 1.5 | -4.5 | 5.8E-06 | 8.5E-06 | MB231/MB468 | MITO |
| >trnaMT_LeuTAA_MT_+_3230_3304@1.42.42\|GTTAAGATGGCAGAGCCCGGTAATCGCATAAAACTTAAAACT | >trnaMT_LeuTAA_MT_+_3230_3304 | 1 | 42 | 42 | GTTAAGATGGCAGAGCCCGGTAATCGCATAAAACTTAAAACT | 556.1 | 8.2 | 1.8 | 4.5 | 5.8E-06 | 8.5E-06 | MB231/MB468 | MITO |
| >hg38_21_+_8433172_8446622_RNA45SN1_45S_50ntFlanks@5188.5207.20\|ATAACAGGTCTGTGATGCCC | >hg38_21_+_8433172_8446622_RNA45SN1_45S_50ntFlanks | 5188 | 5207 | 20 | ATAACAGGTCTGTGATGCCC | 62.1 | -9.0 | 1.9 | -4.8 | 1.7E-06 | 8.5E-06 | BT20/MB468 | MP |
| >MI0000087\|hsa-mir-29a&WithFlank&7\|-\|130876741\|130876816@10.30.21&[MIMAT0004503&hsa-miR-29a-5p&offsets\|0\|-1;m-315&7\|-\|130876785\|130876807&offsets\|0\|-2]\|ACTGATTTCTTTTGGTGTTCA | >MI0000087\|hsa-mir-29a&WithFlank&7\|-\|130876741\|130876816 | 10 | 30 | 21 | ACTGATTTCTTTTGGTGTTCA | 34.8 | 8.3 | 1.8 | 4.6 | 4.8E-06 | 8.5E-06 | MB231/MB468 | MP |
| >hg38_21_+_8433172_8446622_RNA45SN1_45S_50ntFlanks@10320.10340.21\|CAGATCTTGGTGGTAGTAGCA | >hg38_21_+_8433172_8446622_RNA45SN1_45S_50ntFlanks | 10320 | 10340 | 21 | CAGATCTTGGTGGTAGTAGCA | 151.4 | -6.6 | 1.5 | -4.6 | 4.8E-06 | 8.5E-06 | MB231/MB468 | MP |
| >hg38_1_-_228634819_228635039_RNA5S12_5S_50ntFlanks@149.170.22\|GAATACCGGGTGCTGTAGGCTT | >hg38_1_-_228634819_228635039_RNA5S12_5S_50ntFlanks | 149 | 170 | 22 | GAATACCGGGTGCTGTAGGCTT | 4287.1 | -3.4 | 0.7 | -4.6 | 4.8E-06 | 8.5E-06 | MB231/MB468 | MP |
| >hg38_21_+_8433172_8446622_RNA45SN1_45S_50ntFlanks@9571.9597.27\|GCAAATCGGTCGTCCGACCTGGGTATA | >hg38_21_+_8433172_8446622_RNA45SN1_45S_50ntFlanks | 9571 | 9597 | 27 | GCAAATCGGTCGTCCGACCTGGGTATA | 352.1 | -6.2 | 1.3 | -4.6 | 4.2E-06 | 8.5E-06 | BT20/MB468 | MITO |

| | | | | | | | | | | | | | |
|---|---|---|---|---|---|---|---|---|---|---|---|---|---|
| >hg38_21_+_8433172_8446622_RNA45SN1_45S_50ntFlanks@3835.3860.26\|CTCCTCTCCTACTTGGATAACTGTGG | >hg38_21_+_8433172_8446622_RNA45SN1_45S_50ntFlanks | 3835 | 3860 | 26 | CTCCTCTCCTACTTGGATAACTGTGG | 2052.8 | -2.8 | 0.6 | -4.6 | 4.9E-06 | 8.5E-06 | BT20/MB468 | CYTO |
| >hg38_21_+_8433172_8446622_RNA45SN1_45S_50ntFlanks@11209.11243.35\|CGGCGACTCTGGACGCGAGCCGGGCCCTTCCCGTG | >hg38_21_+_8433172_8446622_RNA45SN1_45S_50ntFlanks | 11209 | 11243 | 35 | CGGCGACTCTGGACGCGAGCCGGGCCCTTCCCGTG | 165.6 | 10.7 | 2.1 | 5.0 | 4.8E-07 | 8.5E-06 | BT20/MB231 | MP |
| >hg38_21_+_8433172_8446622_RNA45SN1_45S_50ntFlanks@11575.11607.33\|ACTTAGAACTGGTGCGGACCAGGGGAATCCGAC | >hg38_21_+_8433172_8446622_RNA45SN1_45S_50ntFlanks | 11575 | 11607 | 33 | ACTTAGAACTGGTGCGGACCAGGGGAATCCGAC | 716.5 | -2.5 | 0.6 | -4.5 | 5.5E-06 | 8.5E-06 | MB231/MB468 | CYTO |
| >hg38_21_+_8433172_8446622_RNA45SN1_45S_50ntFlanks@10808.10827.20\|CAATGTAGGTAAGGGAAGTC | >hg38_21_+_8433172_8446622_RNA45SN1_45S_50ntFlanks | 10808 | 10827 | 20 | CAATGTAGGTAAGGGAAGTC | 301.5 | -6.8 | 1.4 | -4.8 | 1.7E-06 | 8.6E-06 | BT20/MB468 | MP |
| >hg38_21_+_8433172_8446622_RNA45SN1_45S_50ntFlanks@12616.12654.39\|CTGAGGAGCCAATGGGGCGAAGCTACCATCTGTGGGATT | >hg38_21_+_8433172_8446622_RNA45SN1_45S_50ntFlanks | 12616 | 12654 | 39 | CTGAGGAGCCAATGGGGCGAAGCTACCATCTGTGGGATT | 2101.7 | -3.7 | 0.8 | -4.6 | 4.2E-06 | 8.6E-06 | BT20/MB468 | MITO |
| >hg38_21_+_8433172_8446622_RNA45SN1_45S_50ntFlanks@10362.10385.24\|AAGGCCGAAGTGGAGAAGGGTTCC | >hg38_21_+_8433172_8446622_RNA45SN1_45S_50ntFlanks | 10362 | 10385 | 24 | AAGGCCGAAGTGGAGAAGGGTTCC | 172.8 | -6.9 | 1.5 | -4.6 | 4.8E-06 | 8.6E-06 | MB231/MB468 | MP |
| >hg38_21_+_8433172_8446622_RNA45SN1_45S_50ntFlanks@8102.8140.39\|CCCCGCGGCGGGGCGCGGGACATGTGGCGTACGGAAGAC | >hg38_21_+_8433172_8446622_RNA45SN1_45S_50ntFlanks | 8102 | 8140 | 39 | CCCCGCGGCGGGGCGCGGGACATGTGGCGTACGGAAGAC | 64.4 | -9.1 | 1.9 | -4.8 | 1.7E-06 | 8.6E-06 | BT20/MB468 | MP |
| >MI0000077\|hsa-mir-21&WithFlank&17\|+\|59841260\|59841343@14.35.22&[MIMAT0000076&hsa-miR-21-5p&offsets\|0\|0;m-1&17\|+\|59841273\|59841294&offsets\|0\|0]\|TAGCTTATCAGACTGATGTTGA | >MI0000077\|hsa-mir-21&WithFlank&17\|+\|59841260\|59841343 | 14 | 35 | 22 | TAGCTTATCAGACTGATGTTGA | 24919.5 | 4.7 | 1.0 | 4.5 | 5.5E-06 | 8.6E-06 | MB231/MB468 | CYTO |
| >am_trna10_ValCAC_5_-_180649395_180649467@1.32.32,trna12_ValAAC_5_-_180645270_180645342@1.32.32,trna132_ValAAC_6_-_27721179_27721251@1.32.32,trna136_ValAAC_6_-_27648885_27648957@1.32.32,trna139_ValAAC_6_-_27618707_27618779@1.32.32,trna18_ValCAC_5_-_180529253_180529325@1.32.32,trna2_ValAAC_3_+_169490018_169490090@1.32.32,trna2_ValCAC_5_+_180524070_180524142@1.32.32,trna4_ValAAC_5_+_180591154_180591226@1.32.32,trna5_ValAAC_5_+_180596610_180596682@1.32.32,trna6_ValCAC_5_+_180600650_180600722@1.32.32,trna85_ValCAC_1_-_161369490_161369562@1.32.32,trna90_ValCAC_1_-_149684088_149684161@1.32.32,trna98_ValCAC_1_-_149298555_149298627@1.32.32,trna9_ValCAC_6_+_26538282_26538354@1.32.32\|GTTTCCGTAGTGTAGTGGTTATCACGTTCGCC | >am_trna10_ValCAC_5_-_180649395_180649467 | 1 | 32 | 32 | GTTTCCGTAGTGTAGTGGTTATCACGTTCGCC | 959.9 | 5.8 | 1.2 | 4.8 | 1.7E-06 | 8.6E-06 | BT20/MB468 | MP |
| >trnaMT_PheGAA_MT_+_577_647@42.74.33\|TTTAGACGGGCTCACATCACCCCATAAACACCA | >trnaMT_PheGAA_MT_+_577_647 | 42 | 74 | 33 | TTTAGACGGGCTCACATCACCCCATAAACACCA | 192.8 | 7.3 | 1.6 | 4.5 | 5.7E-06 | 8.6E-06 | MB231/MB468 | NUC |
| >hg38_21_+_8433172_8446622_RNA45SN1_45S_50ntFlanks@11581.11600.20\|AACTGGTGCGGACCAGGGGA | >hg38_21_+_8433172_8446622_RNA45SN1_45S_50ntFlanks | 11581 | 11600 | 20 | AACTGGTGCGGACCAGGGGA | 629.7 | -7.7 | 1.6 | -4.8 | 1.7E-06 | 8.6E-06 | BT20/MB468 | MP |
| >MI0000088\|hsa-mir-30a&WithFlank&6\|-\|71403545\|71403627@13.31.19&[MIMAT0000087&hsa-miR-30a-5p&offsets\|+1\|-2;m-15&6\|-\|71403595\|71403616&offsets\|+1\|-2]\|GTAAACATCCTCGACTGGA | >MI0000088\|hsa-mir-30a&WithFlank&6\|-\|71403545\|71403627 | 13 | 31 | 19 | GTAAACATCCTCGACTGGA | 76.0 | 10.0 | 2.1 | 4.8 | 1.7E-06 | 8.6E-06 | BT20/MB468 | MP |
| >trna20_GluTTC_2_-_131094701_131094772@1.33.33,trna5_GluTTC_13_-_41634874_41634945@1.33.33\|TCCCATATGGTCTAGCGGTTAGGATTCCTGGTT | >trna20_GluTTC_2_-_131094701_131094772 | 1 | 33 | 33 | TCCCATATGGTCTAGCGGTTAGGATTCCTGGTT | 483.9 | 9.5 | 2.0 | 4.8 | 1.7E-06 | 8.6E-06 | BT20/MB468 | MP |
| >hg38_21_+_8433172_8446622_RNA45SN1_45S_50ntFlanks@12434.12454.21\|TATCATTGTGAAGCAGAATTC | >hg38_21_+_8433172_8446622_RNA45SN1_45S_50ntFlanks | 12434 | 12454 | 21 | TATCATTGTGAAGCAGAATTC | 50.0 | -8.7 | 1.8 | -4.8 | 1.7E-06 | 8.6E-06 | BT20/MB468 | MP |
| >hg38_21_+_8433172_8446622_RNA45SN1_45S_50ntFlanks@8552.8577.26\|CTCCCCGGAGGGGGCGGGCTCCGGCG | >hg38_21_+_8433172_8446622_RNA45SN1_45S_50ntFlanks | 8552 | 8577 | 26 | CTCCCCGGAGGGGGCGGGCTCCGGCG | 359.5 | -7.5 | 1.6 | -4.8 | 1.7E-06 | 8.6E-06 | BT20/MB468 | MP |
| >hg38_21_+_8433172_8446622_RNA45SN1_45S_50ntFlanks@5061.5093.33\|ATTTGTCTGGTTAATTCCGATAACGAACGAGAC | >hg38_21_+_8433172_8446622_RNA45SN1_45S_50ntFlanks | 5061 | 5093 | 33 | ATTTGTCTGGTTAATTCCGATAACGAACGAGAC | 86.8 | -9.5 | 2.0 | -4.8 | 1.7E-06 | 8.6E-06 | BT20/MB468 | MP |
| >hg38_21_+_8433172_8446622_RNA45SN1_45S_50ntFlanks@474.506.33\|TCTTCGTGATCGATGTGGTGACGTCGTGCTCTC | >hg38_21_+_8433172_8446622_RNA45SN1_45S_50ntFlanks | 474 | 506 | 33 | TCTTCGTGATCGATGTGGTGACGTCGTGCTCTC | 1257.0 | 3.5 | 0.8 | 4.5 | 5.7E-06 | 8.6E-06 | MB231/MB468 | NUC |
| >hg38_21_+_8433172_8446622_RNA45SN1_45S_50ntFlanks@6696.6720.25\|GCAGCTAGCTGCGAGAATTAATGTG | >hg38_21_+_8433172_8446622_RNA45SN1_45S_50ntFlanks | 6696 | 6720 | 25 | GCAGCTAGCTGCGAGAATTAATGTG | 171.2 | 4.0 | 0.9 | 4.5 | 5.7E-06 | 8.6E-06 | MB231/MB468 | NUC |
| >hg38_21_+_8433172_8446622_RNA45SN1_45S_50ntFlanks@3843.3881.39\|CTACTTGGATAACTGTGGTAATTCTAGAGCTAATACATG | >hg38_21_+_8433172_8446622_RNA45SN1_45S_50ntFlanks | 3843 | 3881 | 39 | CTACTTGGATAACTGTGGTAATTCTAGAGCTAATACATG | 86.3 | -10.3 | 2.3 | -4.6 | 4.8E-06 | 8.6E-06 | MB231/MB468 | MP |
| >hg38_21_+_8433172_8446622_RNA45SN1_45S_50ntFlanks@12927.12953.27\|CCTGGCGCTAAACCATTCGTAGACGAC | >hg38_21_+_8433172_8446622_RNA45SN1_45S_50ntFlanks | 12927 | 12953 | 27 | CCTGGCGCTAAACCATTCGTAGACGAC | 363.2 | -3.3 | 0.7 | -4.6 | 4.2E-06 | 8.6E-06 | BT20/MB468 | MITO |
| >hg38_21_+_8433172_8446622_RNA45SN1_45S_50ntFlanks@12161.12180.20\|TGACTGGGGCGGTACACCTG | >hg38_21_+_8433172_8446622_RNA45SN1_45S_50ntFlanks | 12161 | 12180 | 20 | TGACTGGGGCGGTACACCTG | 1233.2 | -7.4 | 1.6 | -4.8 | 1.7E-06 | 8.6E-06 | BT20/MB468 | MP |
| >hg38_21_+_8433172_8446622_RNA45SN1_45S_50ntFlanks@12072.12091.20\|CCCGAGGGGCTCTCGCTTCT | >hg38_21_+_8433172_8446622_RNA45SN1_45S_50ntFlanks | 12072 | 12091 | 20 | CCCGAGGGGCTCTCGCTTCT | 108.0 | -5.4 | 1.2 | -4.6 | 4.9E-06 | 8.6E-06 | BT20/MB468 | CYTO |
| >hg38_21_+_8433172_8446622_RNA45SN1_45S_50ntFlanks@6651.6674.24\|CGACTCTTAGCGGTGGATCACTCG | >hg38_21_+_8433172_8446622_RNA45SN1_45S_50ntFlanks | 6651 | 6674 | 24 | CGACTCTTAGCGGTGGATCACTCG | 2573.5 | 4.2 | 0.9 | 4.6 | 4.9E-06 | 8.6E-06 | BT20/MB468 | CYTO |
| >hg38_21_+_8433172_8446622_RNA45SN1_45S_50ntFlanks@6661.6707.47\|CGGTGGATCACTCGGCTCGTGCGTCGATGAAGAACGCAGCTAGCTGC | >hg38_21_+_8433172_8446622_RNA45SN1_45S_50ntFlanks | 6661 | 6707 | 47 | CGGTGGATCACTCGGCTCGTGCGTCGATGAAGAACGCAGCTAGCTGC | 166.1 | 5.3 | 1.2 | 4.5 | 5.5E-06 | 8.6E-06 | MB231/MB468 | CYTO |
| >hg38_21_+_8433172_8446622_RNA45SN1_45S_50ntFlanks@3815.3841.27\|ATGGTTCCTTTGGTCGCTCGCTCCTCT | >hg38_21_+_8433172_8446622_RNA45SN1_45S_50ntFlanks | 3815 | 3841 | 27 | ATGGTTCCTTTGGTCGCTCGCTCCTCT | 513.2 | -5.7 | 1.3 | -4.5 | 6.1E-06 | 8.6E-06 | BT20/MB468 | NUC |
| >hg38_21_+_8433172_8446622_RNA45SN1_45S_50ntFlanks@9538.9571.34\|AACTCTGGTGGAGGTCCGTAGCGGTCCTGACGTG | >hg38_21_+_8433172_8446622_RNA45SN1_45S_50ntFlanks | 9538 | 9571 | 34 | AACTCTGGTGGAGGTCCGTAGCGGTCCTGACGTG | 3772.8 | -8.5 | 1.9 | -4.5 | 6.1E-06 | 8.6E-06 | BT20/MB468 | NUC |
| >hg38_21_+_8433172_8446622_RNA45SN1_45S_50ntFlanks@10682.10712.31\|TCCGGCGGCGTCCGGTGAGCTCTCGCTGGCC | >hg38_21_+_8433172_8446622_RNA45SN1_45S_50ntFlanks | 10682 | 10712 | 31 | TCCGGCGGCGTCCGGTGAGCTCTCGCTGGCC | 804.0 | -7.7 | 1.7 | -4.5 | 6.1E-06 | 8.6E-06 | BT20/MB468 | NUC |
| >hg38_21_+_8433172_8446622_RNA45SN1_45S_50ntFlanks@12044.12061.18\|ACTGACCCGGTGAGGCGG | >hg38_21_+_8433172_8446622_RNA45SN1_45S_50ntFlanks | 12044 | 12061 | 18 | ACTGACCCGGTGAGGCGG | 157.9 | -11.2 | 2.4 | -4.6 | 4.9E-06 | 8.6E-06 | MB231/MB468 | MP |
| >MI0003776\|hsa-mir-320b-1&WithFlank&1\|+\|116671740\|116671823@48.67.20;MI0003839\|hsa-mir-320b-2&WithFlank&1\|-\|224257034\|224257116@47.66.20;MI0000542\|hsa-mir-320a&WithFlank&8\|-\|22244960\|22245043@48.67.20&[MIMAT0005792&hsa-miR-320b&offsets\|0\|-2;m-183&1\|+\|116671787\|116671806&offsets\|0\|0];[MIMAT0005792_1&hsa-miR-320b&offsets\|0\|-2;m-183&1\|-\|224257051\|224257070&offsets\|0\|0];[MIMAT0000510&hsa-miR-320a-3p&offsets\|0\|-2;m-60&8\|-\|22244975\|22244996&offsets\|0\|-2]\|AAAAGCTGGGTTGAGAGGGC | >MI0003776\|hsa-mir-320b-1&WithFlank&1\|+\|116671740\|116671823 | 48 | 67 | 20 | AAAAGCTGGGTTGAGAGGGC | 903.1 | 3.3 | 0.7 | 5.0 | 5.6E-07 | 8.6E-06 | BT20/MB231 | MITO |
| >hg38_1_-_228634819_228635039_RNA5S12_5S_50ntFlanks@129.170.42\|TTGGATGGGAGACCGCCTGGGAATACCGGGTGCTGTAGGCTT | >hg38_1_-_228634819_228635039_RNA5S12_5S_50ntFlanks | 129 | 170 | 42 | TTGGATGGGAGACCGCCTGGGAATACCGGGTGCTGTAGGCTT | 1432.3 | 5.3 | 1.2 | 4.5 | 5.8E-06 | 8.6E-06 | MB231/MB468 | NUC |
| >hg38_21_+_8433172_8446622_RNA45SN1_45S_50ntFlanks@4555.4572.18\|CGCAGCTAGGAATAATGG | >hg38_21_+_8433172_8446622_RNA45SN1_45S_50ntFlanks | 4555 | 4572 | 18 | CGCAGCTAGGAATAATGG | 473.8 | -3.3 | 0.7 | -4.5 | 5.5E-06 | 8.6E-06 | MB231/MB468 | CYTO |
| >hg38_21_+_8433172_8446622_RNA45SN1_45S_50ntFlanks@3816.3837.22\|TGGTTCCTTTGGTCGCTCGCTC | >hg38_21_+_8433172_8446622_RNA45SN1_45S_50ntFlanks | 3816 | 3837 | 22 | TGGTTCCTTTGGTCGCTCGCTC | 454.5 | -7.3 | 1.6 | -4.5 | 6.2E-06 | 8.7E-06 | BT20/MB468 | NUC |
| >MI0000066\|hsa-let-7e&WithFlank&19\|+\|51692780\|51692870@14.33.20&[MIMAT0000066&hsa-let-7e-5p&offsets\|0\|-2;m-50&19\|+\|51692793\|51692814&offsets\|0\|-2]\|TGAGGTAGGAGGTTGTATAG | >MI0000066\|hsa-let-7e&WithFlank&19\|+\|51692780\|51692870 | 14 | 33 | 20 | TGAGGTAGGAGGTTGTATAG | 53.0 | 9.5 | 2.0 | 4.8 | 1.8E-06 | 8.7E-06 | BT20/MB468 | MP |
| >hg38_21_+_8433172_8446622_RNA45SN1_45S_50ntFlanks@8358.8375.18\|AAAGAACTTTGAAGAGAG | >hg38_21_+_8433172_8446622_RNA45SN1_45S_50ntFlanks | 8358 | 8375 | 18 | AAAGAACTTTGAAGAGAG | 4772.2 | 4.2 | 0.9 | 4.8 | 1.8E-06 | 8.7E-06 | BT20/MB468 | MP |
| >MI0000477\|hsa-mir-146a&WithFlank&5\|+\|160485346\|160485456@27.45.19&[MIMAT0000449&hsa-miR-146a-5p&offsets\|0\|-3;m-129&5\|+\|160485372\|160485393&offsets\|0\|-3]\|TGAGAACTGAATTCCATGG | >MI0000477\|hsa-mir-146a&WithFlank&5\|+\|160485346\|160485456 | 27 | 45 | 19 | TGAGAACTGAATTCCATGG | 105.8 | -10.1 | 2.0 | -5.0 | 4.9E-07 | 8.7E-06 | BT20/MB231 | MP |
| >hg38_21_+_8433172_8446622_RNA45SN1_45S_50ntFlanks@11920.11950.31\|ATGAGAGGTGTAGAATAAGTGGGAGGCCCCC | >hg38_21_+_8433172_8446622_RNA45SN1_45S_50ntFlanks | 11920 | 11950 | 31 | ATGAGAGGTGTAGAATAAGTGGGAGGCCCCC | 890.1 | -6.2 | 1.4 | -4.6 | 4.9E-06 | 8.7E-06 | MB231/MB468 | MP |
| >hg38_21_+_8433172_8446622_RNA45SN1_45S_50ntFlanks@12553.12580.28\|CCATGGTAATCCTGCTCAGTACGAGAGG | >hg38_21_+_8433172_8446622_RNA45SN1_45S_50ntFlanks | 12553 | 12580 | 28 | CCATGGTAATCCTGCTCAGTACGAGAGG | 81.9 | -8.4 | 1.8 | -4.6 | 4.2E-06 | 8.7E-06 | BT20/MB468 | MITO |
| >hg38_21_+_8433172_8446622_RNA45SN1_45S_50ntFlanks@3802.3838.37\|CATTAAATCAGTTATGGTTCCTTTGGTCGCTCGCTCC | >hg38_21_+_8433172_8446622_RNA45SN1_45S_50ntFlanks | 3802 | 3838 | 37 | CATTAAATCAGTTATGGTTCCTTTGGTCGCTCGCTCC | 206.3 | -5.3 | 1.2 | -4.5 | 5.8E-06 | 8.7E-06 | MB231/MB468 | NUC |
| >hg38_21_+_8433172_8446622_RNA45SN1_45S_50ntFlanks@5074.5091.18\|ATTCCGATAACGAACGAG | >hg38_21_+_8433172_8446622_RNA45SN1_45S_50ntFlanks | 5074 | 5091 | 18 | ATTCCGATAACGAACGAG | 77.2 | -9.4 | 2.0 | -4.8 | 1.8E-06 | 8.7E-06 | BT20/MB468 | MP |
| >MI0000067\|hsa-let-7f-1&WithFlank&9\|+\|94176341\|94176439@13.30.18;MI0000068\|hsa-let-7f-2&WithFlank&X\|-\|53557186\|53557280@14.31.18&[MIMAT0000067&hsa-let-7f-5p&offsets\|0\|-4;m-20&9\|+\|94176353\|94176374&offsets\|0\|-4];[MIMAT0000067_1&hsa-let-7f-5p&offsets\|0\|-4;m-19&X\|-\|53557246\|53557267&offsets\|0\|-4]\|TGAGGTAGTAGATTGTAT | >MI0000067\|hsa-let-7f-1&WithFlank&9\|+\|94176341\|94176439 | 13 | 30 | 18 | TGAGGTAGTAGATTGTAT | 3879.3 | 4.8 | 1.0 | 4.6 | 4.9E-06 | 8.7E-06 | MB231/MB468 | MP |
| >hg38_21_+_8433172_8446622_RNA45SN1_45S_50ntFlanks@5419.5443.25\|ACCGATTGGATGGTTTAGTGAGGCC | >hg38_21_+_8433172_8446622_RNA45SN1_45S_50ntFlanks | 5419 | 5443 | 25 | ACCGATTGGATGGTTTAGTGAGGCC | 135.9 | -3.6 | 0.8 | -4.5 | 5.6E-06 | 8.7E-06 | MB231/MB468 | CYTO |
| >hg38_21_+_8433172_8446622_RNA45SN1_45S_50ntFlanks@7966.8005.40\|CCTCCGAGACGCGACCTCAGATCAGACGTGGCGACCCGCT | >hg38_21_+_8433172_8446622_RNA45SN1_45S_50ntFlanks | 7966 | 8005 | 40 | CCTCCGAGACGCGACCTCAGATCAGACGTGGCGACCCGCT | 192.8 | 5.5 | 1.2 | 4.5 | 6.2E-06 | 8.7E-06 | BT20/MB468 | NUC |
| >hg38_21_+_8433172_8446622_RNA45SN1_45S_50ntFlanks@6721.6751.31\|AATTGCAGGACACATTGATCATCGACACTTC | >hg38_21_+_8433172_8446622_RNA45SN1_45S_50ntFlanks | 6721 | 6751 | 31 | AATTGCAGGACACATTGATCATCGACACTTC | 230.4 | -5.1 | 1.1 | -4.6 | 4.9E-06 | 8.7E-06 | MB231/MB468 | MP |
| >MI0001519\|hsa-mir-20b&WithFlank&X\|-\|134169803\|134169883@12.34.23&[MIMAT0001413&hsa-miR-20b-5p&offsets\|0\|0;m-241&X\|-\|134169850\|134169872&offsets\|0\|0]\|CAAAGTGCTCATAGTGCAGGTAG | >MI0001519\|hsa-mir-20b&WithFlank&X\|-\|134169803\|134169883 | 12 | 34 | 23 | CAAAGTGCTCATAGTGCAGGTAG | 125.3 | 10.3 | 2.0 | 5.0 | 4.9E-07 | 8.7E-06 | BT20/MB231 | MP |

<table>
<tr><th></th><th></th><th></th><th></th><th></th><th></th><th></th><th></th><th></th><th></th><th></th><th></th><th></th><th></th></tr>
<tr><td>>MI0000061|hsa-let-7a-2&WithFlank&11|-|122146516|122146599@11.30.20(+1G);MI0000062|hsa-let-7a-3&WithFlank&22|+|46112743|46112828@10.29.20(+1G);MI0000060|hsa-let-7a-1&WithFlank&9|+|94175951|94176042@12.31.20(+1G)&[MIMAT0000062&hsa-let-7a-5p&offsets|0|-2(+1G);m-6&11|-|122146568|122146589&offsets|0|-2(+1G)];[MIMAT0000062_1&hsa-let-7a-5p&offsets|0|-2(+1G);m-5&22|+|46112752|46112773&offsets|0|-2(+1G)];[MIMAT0000062_2&hsa-let-7a-5p&offsets|0|-2(+1G);m-4&9|+|94175962|94175983&offsets|0|-2(+1G)]|TGAGGTAGTAGGTTGTATAGG</td><td>>MI0000061|hsa-let-7a-2&WithFlank&11|-|122146516|122146599</td><td>11</td><td>30</td><td>20(+1G)</td><td>TGAGGTAGTAGGTTGTATAGG</td><td>320.3</td><td>6.9</td><td>1.5</td><td>4.6</td><td>5.0E-06</td><td>8.7E-06</td><td>BT20/MB468</td><td>CYTO</td></tr>
<tr><td>>hg38_21_+_8433172_8446622_RNA45SN1_45S_50ntFlanks@12932.12969.38|CGCTAAACCATTCGTAGACGACCTGCTTCTGGGTCGGG</td><td>>hg38_21_+_8433172_8446622_RNA45SN1_45S_50ntFlanks</td><td>12932</td><td>12969</td><td>38</td><td>CGCTAAACCATTCGTAGACGACCTGCTTCTGGGTCGGG</td><td>93.8</td><td>-10.4</td><td>2.3</td><td>-4.6</td><td>4.9E-06</td><td>8.7E-06</td><td>MB231/MB468</td><td>MP</td></tr>
<tr><td>>MI0000082|hsa-mir-25&WithFlank&7|-|100093554|100093649@58.78.21(+1C)&[MIMAT0000081&hsa-miR-25-3p&offsets|0|-1(+1C);m-23&7|-|100093571|100093592&offsets|0|-1(+1C)]|CATTGCACTTGTCTCGGTCTGC</td><td>>MI0000082|hsa-mir-25&WithFlank&7|-|100093554|100093649</td><td>58</td><td>78</td><td>21(+1C)</td><td>CATTGCACTTGTCTCGGTCTGC</td><td>248.0</td><td>7.6</td><td>1.7</td><td>4.5</td><td>5.9E-06</td><td>8.7E-06</td><td>MB231/MB468</td><td>MITO</td></tr>
<tr><td>>hg38_21_+_8433172_8446622_RNA45SN1_45S_50ntFlanks@4729.4751.23|TCAAGAACGAAAGTCGGAGGTTC</td><td>>hg38_21_+_8433172_8446622_RNA45SN1_45S_50ntFlanks</td><td>4729</td><td>4751</td><td>23</td><td>TCAAGAACGAAAGTCGGAGGTTC</td><td>234.8</td><td>-6.3</td><td>1.4</td><td>-4.5</td><td>5.9E-06</td><td>8.7E-06</td><td>MB231/MB468</td><td>MITO</td></tr>
<tr><td>>hg38_21_+_8433172_8446622_RNA45SN1_45S_50ntFlanks@11663.11692.30|ATTTCTGCCCAGTGCTCTGAATGTCAAAGT</td><td>>hg38_21_+_8433172_8446622_RNA45SN1_45S_50ntFlanks</td><td>11663</td><td>11692</td><td>30</td><td>ATTTCTGCCCAGTGCTCTGAATGTCAAAGT</td><td>421.3</td><td>3.5</td><td>0.7</td><td>5.0</td><td>5.7E-07</td><td>8.8E-06</td><td>BT20/MB231</td><td>MITO</td></tr>
<tr><td>>hg38_21_+_8433172_8446622_RNA45SN1_45S_50ntFlanks@3930.3959.30|ATCAAAACCAACCCGGTCAGCCCCTCTCCG</td><td>>hg38_21_+_8433172_8446622_RNA45SN1_45S_50ntFlanks</td><td>3930</td><td>3959</td><td>30</td><td>ATCAAAACCAACCCGGTCAGCCCCTCTCCG</td><td>751.5</td><td>-6.0</td><td>1.3</td><td>-4.5</td><td>5.6E-06</td><td>8.8E-06</td><td>MB231/MB468</td><td>CYTO</td></tr>
<tr><td>>trna10_LysCTT_16_+_3241501_3241573@1.29.29,trna13_LysCTT_14_-_58706613_58706685@1.29.29,trna2_LysCTT_15_+_79152904_79152976@1.29.29|GCCCGGCTAGCTCAGTCGGTAGAGCATGG</td><td>>trna10_LysCTT_16_+_3241501_3241573</td><td>1</td><td>29</td><td>29</td><td>GCCCGGCTAGCTCAGTCGGTAGAGCATGG</td><td>322.3</td><td>-5.0</td><td>1.1</td><td>-4.5</td><td>5.8E-06</td><td>8.8E-06</td><td>MB231/MB468</td><td>NUC</td></tr>
<tr><td>>hg38_21_+_8433172_8446622_RNA45SN1_45S_50ntFlanks@10866.10892.27|CTAAGGGCTGGGTCGGTCGGGCTGGGG</td><td>>hg38_21_+_8433172_8446622_RNA45SN1_45S_50ntFlanks</td><td>10866</td><td>10892</td><td>27</td><td>CTAAGGGCTGGGTCGGTCGGGCTGGGG</td><td>1075.8</td><td>-6.0</td><td>1.3</td><td>-4.8</td><td>1.8E-06</td><td>8.8E-06</td><td>BT20/MB468</td><td>MP</td></tr>
<tr><td>>MI0016005|hsa-mir-3615&WithFlank&17|+|74748607|74748705@57.79.23&[MIMAT0017994&hsa-miR-3615&offsets|0|+2;m-271&17|+|74748663|74748684&offsets|0|+1]|TCTCTCGGCTCCTCGCGGCTCGC</td><td>>MI0016005|hsa-mir-3615&WithFlank&17|+|74748607|74748705</td><td>57</td><td>79</td><td>23</td><td>TCTCTCGGCTCCTCGCGGCTCGC</td><td>213.6</td><td>4.0</td><td>0.9</td><td>4.5</td><td>6.0E-06</td><td>8.8E-06</td><td>MB231/MB468</td><td>MITO</td></tr>
<tr><td>>hg38_21_+_8433172_8446622_RNA45SN1_45S_50ntFlanks@7988.8011.24|CAGACGTGGCGACCCGCTGAATTT</td><td>>hg38_21_+_8433172_8446622_RNA45SN1_45S_50ntFlanks</td><td>7988</td><td>8011</td><td>24</td><td>CAGACGTGGCGACCCGCTGAATTT</td><td>476.3</td><td>2.9</td><td>0.6</td><td>4.6</td><td>5.0E-06</td><td>8.8E-06</td><td>BT20/MB468</td><td>CYTO</td></tr>
<tr><td>>MI0000478|hsa-mir-149&WithFlank&2|+|240455995|240456095@21.43.23&[MIMAT0000450&hsa-miR-149-5p&offsets|0|0;m-195&2|+|240456015|240456037&offsets|0|0]|TCTGGCTCCGTGTCTTCACTCCC</td><td>>MI0000478|hsa-mir-149&WithFlank&2|+|240455995|240456095</td><td>21</td><td>43</td><td>23</td><td>TCTGGCTCCGTGTCTTCACTCCC</td><td>2187.6</td><td>3.9</td><td>0.8</td><td>5.0</td><td>6.7E-07</td><td>8.8E-06</td><td>BT20/MB231</td><td>NUC</td></tr>
<tr><td>>hg38_21_+_8433172_8446622_RNA45SN1_45S_50ntFlanks@9562.9595.34|TCCTGACGTGCAAATCGGTCGTCCGACCTGGGTA</td><td>>hg38_21_+_8433172_8446622_RNA45SN1_45S_50ntFlanks</td><td>9562</td><td>9595</td><td>34</td><td>TCCTGACGTGCAAATCGGTCGTCCGACCTGGGTA</td><td>250.7</td><td>-5.8</td><td>1.3</td><td>-4.6</td><td>5.0E-06</td><td>8.8E-06</td><td>BT20/MB468</td><td>CYTO</td></tr>
<tr><td>>hg38_21_+_8433172_8446622_RNA45SN1_45S_50ntFlanks@10099.10125.27|CGCGCGCGCGTGTGGTGTGCGTCGGAG</td><td>>hg38_21_+_8433172_8446622_RNA45SN1_45S_50ntFlanks</td><td>10099</td><td>10125</td><td>27</td><td>CGCGCGCGCGTGTGGTGTGCGTCGGAG</td><td>78.1</td><td>-9.4</td><td>2.0</td><td>-4.8</td><td>1.8E-06</td><td>8.8E-06</td><td>BT20/MB468</td><td>MP</td></tr>
<tr><td>>hg38_21_+_8433172_8446622_RNA45SN1_45S_50ntFlanks@9747.9767.21|TCAACCTATTCTCAAACTTTA</td><td>>hg38_21_+_8433172_8446622_RNA45SN1_45S_50ntFlanks</td><td>9747</td><td>9767</td><td>21</td><td>TCAACCTATTCTCAAACTTTA</td><td>467.3</td><td>8.3</td><td>1.8</td><td>4.6</td><td>4.3E-06</td><td>8.8E-06</td><td>BT20/MB468</td><td>MITO</td></tr>
<tr><td>>hg38_21_+_8433172_8446622_RNA45SN1_45S_50ntFlanks@8836.8866.31|CGGCGGTGGCGGCGGCGGCGGCGGCGGCGGG</td><td>>hg38_21_+_8433172_8446622_RNA45SN1_45S_50ntFlanks</td><td>8836</td><td>8866</td><td>31</td><td>CGGCGGTGGCGGCGGCGGCGGCGGCGGCGGG</td><td>90.4</td><td>-10.4</td><td>2.3</td><td>-4.6</td><td>4.9E-06</td><td>8.8E-06</td><td>MB231/MB468</td><td>MP</td></tr>
<tr><td>>hg38_21_+_8433172_8446622_RNA45SN1_45S_50ntFlanks@11452.11472.21|CTCCTCGGCGCGCGGCGGCGG</td><td>>hg38_21_+_8433172_8446622_RNA45SN1_45S_50ntFlanks</td><td>11452</td><td>11472</td><td>21</td><td>CTCCTCGGCGCGCGGCGGCGG</td><td>2375.1</td><td>-8.7</td><td>1.9</td><td>-4.5</td><td>6.3E-06</td><td>8.8E-06</td><td>BT20/MB468</td><td>NUC</td></tr>
<tr><td>>hg38_21_+_8433172_8446622_RNA45SN1_45S_50ntFlanks@12456.12498.43|CCAAGCGTTGGATTGTTCACCCACTAATAGGGAACGTGAGCTG</td><td>>hg38_21_+_8433172_8446622_RNA45SN1_45S_50ntFlanks</td><td>12456</td><td>12498</td><td>43</td><td>CCAAGCGTTGGATTGTTCACCCACTAATAGGGAACGTGAGCTG</td><td>240.1</td><td>-6.6</td><td>1.5</td><td>-4.6</td><td>5.0E-06</td><td>8.8E-06</td><td>BT20/MB468</td><td>CYTO</td></tr>
<tr><td>>hg38_21_+_8433172_8446622_RNA45SN1_45S_50ntFlanks@9538.9569.32|AACTCTGGTGGAGGTCCGTAGCGGTCCTGACG</td><td>>hg38_21_+_8433172_8446622_RNA45SN1_45S_50ntFlanks</td><td>9538</td><td>9569</td><td>32</td><td>AACTCTGGTGGAGGTCCGTAGCGGTCCTGACG</td><td>605.6</td><td>-7.7</td><td>1.7</td><td>-4.5</td><td>6.3E-06</td><td>8.8E-06</td><td>BT20/MB468</td><td>NUC</td></tr>
<tr><td>>hg38_21_+_8433172_8446622_RNA45SN1_45S_50ntFlanks@12419.12457.39|GATGTCGGCTCTTCCTATCATTGTGAAGCAGAATTCACC</td><td>>hg38_21_+_8433172_8446622_RNA45SN1_45S_50ntFlanks</td><td>12419</td><td>12457</td><td>39</td><td>GATGTCGGCTCTTCCTATCATTGTGAAGCAGAATTCACC</td><td>535.1</td><td>5.7</td><td>1.3</td><td>4.6</td><td>5.0E-06</td><td>8.8E-06</td><td>BT20/MB468</td><td>CYTO</td></tr>
<tr><td>>hg38_21_+_8433172_8446622_RNA45SN1_45S_50ntFlanks@9035.9058.24|CGTGGGGGGGCCGGGCCACCCCTC</td><td>>hg38_21_+_8433172_8446622_RNA45SN1_45S_50ntFlanks</td><td>9035</td><td>9058</td><td>24</td><td>CGTGGGGGGGCCGGGCCACCCCTC</td><td>327.2</td><td>-7.3</td><td>1.5</td><td>-4.8</td><td>1.8E-06</td><td>8.8E-06</td><td>BT20/MB468</td><td>MP</td></tr>
<tr><td>>hg38_21_+_8433172_8446622_RNA45SN1_45S_50ntFlanks@3813.3834.22|TTATGGTTCCTTTGGTCGCTCG</td><td>>hg38_21_+_8433172_8446622_RNA45SN1_45S_50ntFlanks</td><td>3813</td><td>3834</td><td>22</td><td>TTATGGTTCCTTTGGTCGCTCG</td><td>1046.8</td><td>-7.9</td><td>1.7</td><td>-4.5</td><td>6.3E-06</td><td>8.8E-06</td><td>BT20/MB468</td><td>NUC</td></tr>
<tr><td>>hg38_21_+_8433172_8446622_RNA45SN1_45S_50ntFlanks@11580.11599.20|GAACTGGTGCGGACCAGGGG</td><td>>hg38_21_+_8433172_8446622_RNA45SN1_45S_50ntFlanks</td><td>11580</td><td>11599</td><td>20</td><td>GAACTGGTGCGGACCAGGGG</td><td>63.6</td><td>-9.1</td><td>1.9</td><td>-4.8</td><td>1.8E-06</td><td>8.8E-06</td><td>BT20/MB468</td><td>MP</td></tr>
<tr><td>>hg38_21_+_8433172_8446622_RNA45SN1_45S_50ntFlanks@9545.9571.27|GTGGAGGTCCGTAGCGGTCCTGACGTG</td><td>>hg38_21_+_8433172_8446622_RNA45SN1_45S_50ntFlanks</td><td>9545</td><td>9571</td><td>27</td><td>GTGGAGGTCCGTAGCGGTCCTGACGTG</td><td>28391.9</td><td>-3.5</td><td>0.8</td><td>-4.5</td><td>5.7E-06</td><td>8.8E-06</td><td>MB231/MB468</td><td>CYTO</td></tr>
<tr><td>>hg38_21_+_8433172_8446622_RNA45SN1_45S_50ntFlanks@8323.8354.32|ACCGATAGTCAACAAGTACCGTAAGGGAAAGT</td><td>>hg38_21_+_8433172_8446622_RNA45SN1_45S_50ntFlanks</td><td>8323</td><td>8354</td><td>32</td><td>ACCGATAGTCAACAAGTACCGTAAGGGAAAGT</td><td>54.2</td><td>-8.8</td><td>1.9</td><td>-4.8</td><td>1.8E-06</td><td>8.8E-06</td><td>BT20/MB468</td><td>MP</td></tr>
<tr><td>>trna111_HisGTG_1_-_147774845_147774916@-1G.35.36,trna118_HisGTG_1_-_145396881_145396952@-1G.35.36,trna16_HisGTG_1_+_146544773_146544844@-1G.35.36,trna1_HisGTG_15_+_45493349_45493420@-1G.35.36,trna21_HisGTG_1_+_147753471_147753542@-1G.35.36,trna33_HisGTG_6_+_27125906_27125977@-1G.35.36,trna7_HisGTG_9_-_14433938_14434009@-1G.35.36,trna8_HisGTG_15_-_45492611_45492682@-1G.35.36,trna9_HisGTG_15_-_45490804_45490875@-1G.35.36|GGCCGTGATCGTATAGTGGTTAGTACTCTGCGTTGT</td><td>>trna111_HisGTG_1_-_147774845_147774916</td><td>-1G</td><td>35</td><td>36</td><td>GGCCGTGATCGTATAGTGGTTAGTACTCTGCGTTGT</td><td>81.1</td><td>5.3</td><td>1.2</td><td>4.5</td><td>6.3E-06</td><td>8.8E-06</td><td>BT20/MB468</td><td>NUC</td></tr>
<tr><td>>hg38_21_+_8433172_8446622_RNA45SN1_45S_50ntFlanks@10787.10826.40|ACAGCCTCTGGCATGTTGGAACAATGTAGGTAAGGGAAGT</td><td>>hg38_21_+_8433172_8446622_RNA45SN1_45S_50ntFlanks</td><td>10787</td><td>10826</td><td>40</td><td>ACAGCCTCTGGCATGTTGGAACAATGTAGGTAAGGGAAGT</td><td>348.2</td><td>-7.2</td><td>1.6</td><td>-4.5</td><td>6.3E-06</td><td>8.9E-06</td><td>BT20/MB468</td><td>NUC</td></tr>
<tr><td>>hg38_21_+_8433172_8446622_RNA45SN1_45S_50ntFlanks@4224.4242.19|AAAATAACAATACAGGACT</td><td>>hg38_21_+_8433172_8446622_RNA45SN1_45S_50ntFlanks</td><td>4224</td><td>4242</td><td>19</td><td>AAAATAACAATACAGGACT</td><td>394.0</td><td>-5.4</td><td>1.2</td><td>-4.5</td><td>5.9E-06</td><td>8.9E-06</td><td>MB231/MB468</td><td>NUC</td></tr>
<tr><td>>trnaMT_PheGAA_MT_+_577_647@39.56.18|ATGTTTAGACGGGCTCAC</td><td>>trnaMT_PheGAA_MT_+_577_647</td><td>39</td><td>56</td><td>18</td><td>ATGTTTAGACGGGCTCAC</td><td>294.6</td><td>2.9</td><td>0.7</td><td>4.5</td><td>6.3E-06</td><td>8.9E-06</td><td>BT20/MB468</td><td>NUC</td></tr>
<tr><td>>hg38_21_+_8433172_8446622_RNA45SN1_45S_50ntFlanks@13007.13028.22|TATTGAAAGTCAGCCCTCGACA</td><td>>hg38_21_+_8433172_8446622_RNA45SN1_45S_50ntFlanks</td><td>13007</td><td>13028</td><td>22</td><td>TATTGAAAGTCAGCCCTCGACA</td><td>1025.0</td><td>-3.9</td><td>0.9</td><td>-4.5</td><td>5.7E-06</td><td>8.9E-06</td><td>MB231/MB468</td><td>CYTO</td></tr>
<tr><td>>hg38_21_+_8433172_8446622_RNA45SN1_45S_50ntFlanks@4567.4607.41|TAATGGAATAGGACCGCGGTTCTATTTTGTTGGTTTTCGGA</td><td>>hg38_21_+_8433172_8446622_RNA45SN1_45S_50ntFlanks</td><td>4567</td><td>4607</td><td>41</td><td>TAATGGAATAGGACCGCGGTTCTATTTTGTTGGTTTTCGGA</td><td>354.2</td><td>12.2</td><td>2.6</td><td>4.8</td><td>1.8E-06</td><td>8.9E-06</td><td>BT20/MB468</td><td>MP</td></tr>
<tr><td>>MI0016887|hsa-mir-4521&WithFlank&17|+|8186939|8187010@10.34.25&[MIMAT0019058&hsa-miR-4521&offsets|0|+3;m-765&17|+|8186948|8186969&offsets|0|+3]|GCTAAGGAAGTCCTGTGCTCAGTTT</td><td>>MI0016887|hsa-mir-4521&WithFlank&17|+|8186939|8187010</td><td>10</td><td>34</td><td>25</td><td>GCTAAGGAAGTCCTGTGCTCAGTTT</td><td>169.7</td><td>5.1</td><td>1.1</td><td>4.5</td><td>6.3E-06</td><td>8.9E-06</td><td>BT20/MB468</td><td>NUC</td></tr>
<tr><td>>hg38_21_+_8433172_8446622_RNA45SN1_45S_50ntFlanks@11568.11605.38|GCAGCCGACTTAGAACTGGTGCGGACCAGGGGAATCCG</td><td>>hg38_21_+_8433172_8446622_RNA45SN1_45S_50ntFlanks</td><td>11568</td><td>11605</td><td>38</td><td>GCAGCCGACTTAGAACTGGTGCGGACCAGGGGAATCCG</td><td>969.9</td><td>-7.9</td><td>1.7</td><td>-4.5</td><td>6.3E-06</td><td>8.9E-06</td><td>BT20/MB468</td><td>NUC</td></tr>
<tr><td>>MI0023622|hsa-mir-486-2&WithFlank&8|+|41660438|41660513@7.29.23&[MIMAT0002177_1&hsa-miR-486-5p&offsets|0|+1;TJU_CMC_MD2.ID03138.5p-miR&offsets|0|-1;m-106&8|+|41660444|41660465&offsets|0|+1]|TCCTGTACTGAGCTGCCCCGAGC</td><td>>MI0023622|hsa-mir-486-2&WithFlank&8|+|41660438|41660513</td><td>7</td><td>29</td><td>23</td><td>TCCTGTACTGAGCTGCCCCGAGC</td><td>104.8</td><td>-10.1</td><td>2.0</td><td>-5.0</td><td>5.0E-07</td><td>8.9E-06</td><td>BT20/MB231</td><td>MP</td></tr>
<tr><td>>MI0000088|hsa-mir-30a&WithFlank&6|-|71403545|71403627@12.33.22(+1A)&[MIMAT0000087&hsa-miR-30a-5p&offsets|0|0(+1A);m-15&6|-|71403595|71403616&offsets|0|0(+1A)]|TGTAAACATCCTCGACTGGAAGA</td><td>>MI0000088|hsa-mir-30a&WithFlank&6|-|71403545|71403627</td><td>12</td><td>33</td><td>22(+1A)</td><td>TGTAAACATCCTCGACTGGAAGA</td><td>339.2</td><td>3.9</td><td>0.9</td><td>4.5</td><td>6.3E-06</td><td>8.9E-06</td><td>BT20/MB468</td><td>NUC</td></tr>
<tr><td>>hg38_21_+_8433172_8446622_RNA45SN1_45S_50ntFlanks@12479.12498.20|CTAATAGGGAACGTGAGCTG</td><td>>hg38_21_+_8433172_8446622_RNA45SN1_45S_50ntFlanks</td><td>12479</td><td>12498</td><td>20</td><td>CTAATAGGGAACGTGAGCTG</td><td>2304.6</td><td>-8.3</td><td>1.8</td><td>-4.5</td><td>6.3E-06</td><td>8.9E-06</td><td>BT20/MB468</td><td>NUC</td></tr>
<tr><td>>hg38_21_+_8433172_8446622_RNA45SN1_45S_50ntFlanks@10846.10863.18|CTTCGGGATAAGGATTGG</td><td>>hg38_21_+_8433172_8446622_RNA45SN1_45S_50ntFlanks</td><td>10846</td><td>10863</td><td>18</td><td>CTTCGGGATAAGGATTGG</td><td>862.6</td><td>-6.7</td><td>1.5</td><td>-4.6</td><td>5.1E-06</td><td>8.9E-06</td><td>BT20/MB468</td><td>CYTO</td></tr>
<tr><td>>hg38_21_+_8433172_8446622_RNA45SN1_45S_50ntFlanks@6649.6681.33|TACGACTCTTAGCGGTGGATCACTCGGCTCGTG</td><td>>hg38_21_+_8433172_8446622_RNA45SN1_45S_50ntFlanks</td><td>6649</td><td>6681</td><td>33</td><td>TACGACTCTTAGCGGTGGATCACTCGGCTCGTG</td><td>1836.6</td><td>-3.9</td><td>0.8</td><td>-4.8</td><td>1.8E-06</td><td>8.9E-06</td><td>BT20/MB468</td><td>MP</td></tr>
<tr><td>>hg38_21_+_8433172_8446622_RNA45SN1_45S_50ntFlanks@10572.10607.36|AGAAGCCGGCGGGAGCCCCGGGGAGAGTTCTCTTTT</td><td>>hg38_21_+_8433172_8446622_RNA45SN1_45S_50ntFlanks</td><td>10572</td><td>10607</td><td>36</td><td>AGAAGCCGGCGGGAGCCCCGGGGAGAGTTCTCTTTT</td><td>400.1</td><td>-6.1</td><td>1.3</td><td>-4.6</td><td>5.1E-06</td><td>8.9E-06</td><td>BT20/MB468</td><td>CYTO</td></tr>
<tr><td>>hg38_21_+_8433172_8446622_RNA45SN1_45S_50ntFlanks@12523.12546.24|TAGTTTTACCCTACTGATGATGTG</td><td>>hg38_21_+_8433172_8446622_RNA45SN1_45S_50ntFlanks</td><td>12523</td><td>12546</td><td>24</td><td>TAGTTTTACCCTACTGATGATGTG</td><td>48.4</td><td>-8.7</td><td>1.8</td><td>-4.8</td><td>1.8E-06</td><td>8.9E-06</td><td>BT20/MB468</td><td>MP</td></tr>
<tr><td>>am_trnaMT_ValTAC_MT_+_1602_1670@30.68.39|CTTACACTTAGGAGATTTCAACTTAACTTGACCGCTCTG</td><td>>am_trnaMT_ValTAC_MT_+_1602_1670</td><td>30</td><td>68</td><td>39</td><td>CTTACACTTAGGAGATTTCAACTTAACTTGACCGCTCTG</td><td>92.2</td><td>-10.4</td><td>2.3</td><td>-4.6</td><td>5.0E-06</td><td>8.9E-06</td><td>MB231/MB468</td><td>MP</td></tr>
<tr><td>>hg38_21_+_8433172_8446622_RNA45SN1_45S_50ntFlanks@8198.8222.25|TGGACGGTGTGAGGCCGGTAGCGGC</td><td>>hg38_21_+_8433172_8446622_RNA45SN1_45S_50ntFlanks</td><td>8198</td><td>8222</td><td>25</td><td>TGGACGGTGTGAGGCCGGTAGCGGC</td><td>218.8</td><td>-4.7</td><td>1.0</td><td>-4.5</td><td>6.0E-06</td><td>8.9E-06</td><td>MB231/MB468</td><td>NUC</td></tr>
<tr><td>>MI0000102|hsa-mir-100&WithFlank&11|-|122152223|122152314@21.40.20&[MIMAT0000098&hsa-miR-100-5p&offsets|+2|0;m-27&11|-|122152276|122152296&offsets|+2|+1]|CCCGTAGATCCGAACTTGTG</td><td>>MI0000102|hsa-mir-100&WithFlank&11|-|122152223|122152314</td><td>21</td><td>40</td><td>20</td><td>CCCGTAGATCCGAACTTGTG</td><td>37.6</td><td>8.5</td><td>1.9</td><td>4.6</td><td>5.0E-06</td><td>8.9E-06</td><td>MB231/MB468</td><td>MP</td></tr>
<tr><td>>hg38_21_+_8433172_8446622_RNA45SN1_45S_50ntFlanks@12035.12053.19|CGTTTTTTCACTGACCCGG</td><td>>hg38_21_+_8433172_8446622_RNA45SN1_45S_50ntFlanks</td><td>12035</td><td>12053</td><td>19</td><td>CGTTTTTTCACTGACCCGG</td><td>130.7</td><td>-5.6</td><td>1.2</td><td>-4.6</td><td>5.1E-06</td><td>8.9E-06</td><td>BT20/MB468</td><td>CYTO</td></tr>
<tr><td>>hg38_21_+_8433172_8446622_RNA45SN1_45S_50ntFlanks@8194.8230.37|CCCGTGGACGGTGTGAGGCCGGTAGCGGCCCCCGGCG</td><td>>hg38_21_+_8433172_8446622_RNA45SN1_45S_50ntFlanks</td><td>8194</td><td>8230</td><td>37</td><td>CCCGTGGACGGTGTGAGGCCGGTAGCGGCCCCCGGCG</td><td>48.8</td><td>-8.7</td><td>1.8</td><td>-4.8</td><td>1.8E-06</td><td>8.9E-06</td><td>BT20/MB468</td><td>MP</td></tr>
<tr><td>>hg38_21_+_8433172_8446622_RNA45SN1_45S_50ntFlanks@11584.11611.28|TGGTGCGGACCAGGGGAATCCGACTGTT</td><td>>hg38_21_+_8433172_8446622_RNA45SN1_45S_50ntFlanks</td><td>11584</td><td>11611</td><td>28</td><td>TGGTGCGGACCAGGGGAATCCGACTGTT</td><td>899.8</td><td>3.4</td><td>0.7</td><td>5.0</td><td>5.9E-07</td><td>8.9E-06</td><td>BT20/MB231</td><td>MITO</td></tr>
<tr><td>>hg38_21_+_8433172_8446622_RNA45SN1_45S_50ntFlanks@9481.9515.35|CACGTGTTAGGACCCGAAAGATGGTGAACTATGCC</td><td>>hg38_21_+_8433172_8446622_RNA45SN1_45S_50ntFlanks</td><td>9481</td><td>9515</td><td>35</td><td>CACGTGTTAGGACCCGAAAGATGGTGAACTATGCC</td><td>49.3</td><td>9.4</td><td>2.0</td><td>4.8</td><td>1.8E-06</td><td>8.9E-06</td><td>BT20/MB468</td><td>MP</td></tr>
</table>

<table>
<thead>
<tr><th></th><th></th><th></th><th></th><th></th><th></th><th></th><th></th><th></th><th></th><th></th><th></th><th></th><th></th></tr>
</thead>
<tbody>
<tr><td>>hg38_21_+_8433172_8446622_RNA45SN1_45S_50ntFlanks@10351.10376.26|CGAGAACTTTGAAGGCCGAAGTGGAG</td><td>>hg38_21_+_8433172_8446622_RNA45SN1_45S_50ntFlanks</td><td>10351</td><td>10376</td><td>26</td><td>CGAGAACTTTGAAGGCCGAAGTGGAG</td><td>7559.1</td><td>-7.7</td><td>1.6</td><td>-4.8</td><td>1.8E-06</td><td>8.9E-06</td><td>BT20/MB468</td><td>MP</td></tr>
<tr><td>>hg38_21_+_8433172_8446622_RNA45SN1_45S_50ntFlanks@10353.10374.22|AGAACTTTGAAGGCCGAAGTGG</td><td>>hg38_21_+_8433172_8446622_RNA45SN1_45S_50ntFlanks</td><td>10353</td><td>10374</td><td>22</td><td>AGAACTTTGAAGGCCGAAGTGG</td><td>283.7</td><td>-6.8</td><td>1.5</td><td>-4.6</td><td>5.0E-06</td><td>8.9E-06</td><td>MB231/MB468</td><td>MP</td></tr>
<tr><td>>hg38_21_+_8433172_8446622_RNA45SN1_45S_50ntFlanks@6722.6756.35|ATTGCAGGACACATTGATCATCGACACTTCGAACG</td><td>>hg38_21_+_8433172_8446622_RNA45SN1_45S_50ntFlanks</td><td>6722</td><td>6756</td><td>35</td><td>ATTGCAGGACACATTGATCATCGACACTTCGAACG</td><td>118.9</td><td>-5.2</td><td>1.1</td><td>-4.6</td><td>5.0E-06</td><td>8.9E-06</td><td>MB231/MB468</td><td>MP</td></tr>
<tr><td>>hg38_21_+_8433172_8446622_RNA45SN1_45S_50ntFlanks@9133.9152.20|TCGGGGGGCGGGGCGGACTG</td><td>>hg38_21_+_8433172_8446622_RNA45SN1_45S_50ntFlanks</td><td>9133</td><td>9152</td><td>20</td><td>TCGGGGGGCGGGGCGGACTG</td><td>204.6</td><td>-5.9</td><td>1.3</td><td>-4.6</td><td>4.4E-06</td><td>9.0E-06</td><td>BT20/MB468</td><td>MITO</td></tr>
<tr><td>>hg38_21_+_8433172_8446622_RNA45SN1_45S_50ntFlanks@10234.10263.30|CGGACGCTACGCCGCGACGAGTAGGAGGGC</td><td>>hg38_21_+_8433172_8446622_RNA45SN1_45S_50ntFlanks</td><td>10234</td><td>10263</td><td>30</td><td>CGGACGCTACGCCGCGACGAGTAGGAGGGC</td><td>133.8</td><td>10.4</td><td>2.1</td><td>5.0</td><td>5.1E-07</td><td>9.0E-06</td><td>BT20/MB231</td><td>MP</td></tr>
<tr><td>>hg38_21_+_8433172_8446622_RNA45SN1_45S_50ntFlanks@12416.12453.38|TTCGATGTCGGCTCTTCCTATCATTGTGAAGCAGAATT</td><td>>hg38_21_+_8433172_8446622_RNA45SN1_45S_50ntFlanks</td><td>12416</td><td>12453</td><td>38</td><td>TTCGATGTCGGCTCTTCCTATCATTGTGAAGCAGAATT</td><td>250.2</td><td>5.4</td><td>1.2</td><td>4.5</td><td>6.0E-06</td><td>9.0E-06</td><td>MB231/MB468</td><td>NUC</td></tr>
<tr><td>>trna111_HisGTG_1_-_147774845_147774916@-1G.34.35,trna118_HisGTG_1_-_145396881_145396952@-1G.34.35,trna16_HisGTG_1_+_146544773_146544844@-1G.34.35,trna1_HisGTG_15_+_45493349_45493420@-1G.34.35,trna21_HisGTG_1_+_147753471_147753542@-1G.34.35,trna33_HisGTG_6_+_27125906_27125977@-1G.34.35,trna7_HisGTG_9_-_14433938_14434009@-1G.34.35,trna8_HisGTG_15_-_45492611_45492682@-1G.34.35,trna9_HisGTG_15_-_45490804_45490875@-1G.34.35|GGCCGTGATCGTATAGTGGTTAGTACTCTGCGTTG</td><td>>trna111_HisGTG_1_-_147774845_147774916</td><td>-1G</td><td>34</td><td>35</td><td>GGCCGTGATCGTATAGTGGTTAGTACTCTGCGTTG</td><td>7042.1</td><td>-5.0</td><td>1.0</td><td>-5.0</td><td>6.9E-07</td><td>9.0E-06</td><td>BT20/MB231</td><td>NUC</td></tr>
<tr><td>>MI0000736|hsa-mir-30c-1&WithFlank&1|+|40757278|40757378@23.44.22(+1U);MI0000254|hsa-mir-30c-2&WithFlank&6|-|71376954|71377037@13.34.22(+1U)&[MIMAT0000244&hsa-miR-30c-5p&offsets|0|-1(+1U);m-57&1|+|40757300|40757323&offsets|0|-2(+1U)];[MIMAT0000244_1&hsa-miR-30c-5p&offsets|0|-1(+1U);m-56&6|-|71377002|71377025&offsets|0|-2(+1U)]|TGTAAACATCCTACACTCTCAGT</td><td>>MI0000736|hsa-mir-30c-1&WithFlank&1|+|40757278|40757378</td><td>23</td><td>44</td><td>22(+1U)</td><td>TGTAAACATCCTACACTCTCAGT</td><td>2056.7</td><td>4.4</td><td>1.0</td><td>4.6</td><td>5.1E-06</td><td>9.0E-06</td><td>MB231/MB468</td><td>MP</td></tr>
<tr><td>>hg38_21_+_8433172_8446622_RNA45SN1_45S_50ntFlanks@10390.10424.35|GAACAGCAGTTGAACATGGGTCAGTCGGTCCTGAG</td><td>>hg38_21_+_8433172_8446622_RNA45SN1_45S_50ntFlanks</td><td>10390</td><td>10424</td><td>35</td><td>GAACAGCAGTTGAACATGGGTCAGTCGGTCCTGAG</td><td>2112.8</td><td>8.0</td><td>1.7</td><td>4.8</td><td>1.8E-06</td><td>9.0E-06</td><td>BT20/MB468</td><td>MP</td></tr>
<tr><td>>hg38_21_+_8433172_8446622_RNA45SN1_45S_50ntFlanks@9542.9571.30|CTGGTGGAGGTCCGTAGCGGTCCTGACGTG</td><td>>hg38_21_+_8433172_8446622_RNA45SN1_45S_50ntFlanks</td><td>9542</td><td>9571</td><td>30</td><td>CTGGTGGAGGTCCGTAGCGGTCCTGACGTG</td><td>164.0</td><td>-2.8</td><td>0.6</td><td>-4.5</td><td>6.0E-06</td><td>9.0E-06</td><td>MB231/MB468</td><td>NUC</td></tr>
<tr><td>>hg38_21_+_8433172_8446622_RNA45SN1_45S_50ntFlanks@12624.12657.34|CCAATGGGGCGAAGCTACCATCTGTGGGATTATG</td><td>>hg38_21_+_8433172_8446622_RNA45SN1_45S_50ntFlanks</td><td>12624</td><td>12657</td><td>34</td><td>CCAATGGGGCGAAGCTACCATCTGTGGGATTATG</td><td>349.9</td><td>-3.7</td><td>0.8</td><td>-4.5</td><td>5.8E-06</td><td>9.0E-06</td><td>MB231/MB468</td><td>CYTO</td></tr>
<tr><td>>MI0000067|hsa-let-7f-1&WithFlank&9|+|94176341|94176439@13.34.22(+1A);MI0000068|hsa-let-7f-2&WithFlank&X|-|53557186|53557280@14.35.22(+1A)&[MIMAT0000067&hsa-let-7f-5p&offsets|0|0(+1A);m-20&9|+|94176353|94176374&offsets|0|0(+1A)];[MIMAT0000067_1&hsa-let-7f-5p&offsets|0|0(+1A);m-19&X|-|53557246|53557267&offsets|0|0(+1A)]|TGAGGTAGTAGATTGTATAGTTA</td><td>>MI0000067|hsa-let-7f-1&WithFlank&9|+|94176341|94176439</td><td>13</td><td>34</td><td>22(+1A)</td><td>TGAGGTAGTAGATTGTATAGTTA</td><td>529.4</td><td>4.2</td><td>0.9</td><td>4.5</td><td>6.1E-06</td><td>9.0E-06</td><td>MB231/MB468</td><td>MITO</td></tr>
<tr><td>>trnaMT_SerGCT_MT_+_12207_12265@1.19.19|GAGAAAGCTCACAAGAACT</td><td>>trnaMT_SerGCT_MT_+_12207_12265</td><td>1</td><td>19</td><td>19</td><td>GAGAAAGCTCACAAGAACT</td><td>1506.4</td><td>4.0</td><td>0.9</td><td>4.5</td><td>6.4E-06</td><td>9.0E-06</td><td>BT20/MB468</td><td>NUC</td></tr>
<tr><td>>hg38_21_+_8433172_8446622_RNA45SN1_45S_50ntFlanks@8335.8375.41|CAAGTACCGTAAGGGAAAGTTGAAAAGAACTTTGAAGAGAG</td><td>>hg38_21_+_8433172_8446622_RNA45SN1_45S_50ntFlanks</td><td>8335</td><td>8375</td><td>41</td><td>CAAGTACCGTAAGGGAAAGTTGAAAAGAACTTTGAAGAGAG</td><td>454.2</td><td>5.8</td><td>1.2</td><td>5.0</td><td>6.0E-07</td><td>9.0E-06</td><td>BT20/MB231</td><td>MITO</td></tr>
<tr><td>>am_trna128_GlyGCC_6_-_27870686_27870756@1.29.29,trna133_GlyCCC_1_-_16872434_16872504@1.29.29,trna18_GlyGCC_16_+_70822597_70822667@1.29.29,trna19_GlyGCC_16_+_70823410_70823480@1.29.29,trna19_GlyGCC_2_-_157257659_157257729@1.29.29,trna24_GlyGCC_16_-_70812942_70813012@1.29.29,trna25_GlyGCC_16_-_70812114_70812184@1.29.29,trna4_GlyCCC_1_+_17188416_17188486@1.29.29,trna5_GlyGCC_17_+_8029064_8029134@1.29.29,trna68_GlyGCC_1_-_161493637_161493707@1.29.29|GCATTGGTGGTTCAGTGGTAGAATTCTCG</td><td>>am_trna128_GlyGCC_6_-_27870686_27870756</td><td>1</td><td>29</td><td>29</td><td>GCATTGGTGGTTCAGTGGTAGAATTCTCG</td><td>7893.9</td><td>-4.0</td><td>0.9</td><td>-4.5</td><td>6.1E-06</td><td>9.0E-06</td><td>MB231/MB468</td><td>MITO</td></tr>
<tr><td>>hg38_21_+_8433172_8446622_RNA45SN1_45S_50ntFlanks@6722.6770.49|ATTGCAGGACACATTGATCATCGACACTTCGAACGCACTTGCGGCCCCG</td><td>>hg38_21_+_8433172_8446622_RNA45SN1_45S_50ntFlanks</td><td>6722</td><td>6770</td><td>49</td><td>ATTGCAGGACACATTGATCATCGACACTTCGAACGCACTTGCGGCCCCG</td><td>434.3</td><td>-7.0</td><td>1.5</td><td>-4.5</td><td>6.1E-06</td><td>9.0E-06</td><td>MB231/MB468</td><td>MITO</td></tr>
<tr><td>>hg38_21_+_8433172_8446622_RNA45SN1_45S_50ntFlanks@10064.10082.19|TCGAGAGTGGACGGGAGCG</td><td>>hg38_21_+_8433172_8446622_RNA45SN1_45S_50ntFlanks</td><td>10064</td><td>10082</td><td>19</td><td>TCGAGAGTGGACGGGAGCG</td><td>112.6</td><td>10.1</td><td>2.0</td><td>5.0</td><td>5.1E-07</td><td>9.0E-06</td><td>BT20/MB231</td><td>MP</td></tr>
<tr><td>>trnaMT_HisGTG_MT_+_12138_12206@20.55.36|AACATCAGATTGTGAATCTGACAACAGAGGCTTACG</td><td>>trnaMT_HisGTG_MT_+_12138_12206</td><td>20</td><td>55</td><td>36</td><td>AACATCAGATTGTGAATCTGACAACAGAGGCTTACG</td><td>461.7</td><td>8.6</td><td>1.9</td><td>4.6</td><td>4.4E-06</td><td>9.0E-06</td><td>BT20/MB468</td><td>MITO</td></tr>
<tr><td>>hg38_MT_+_598_1651_RNR1_12S_50ntFlanks@698.730.33|AAGTAAGCGCAAGTACCCACGTAAAGACGTTAG</td><td>>hg38_MT_+_598_1651_RNR1_12S_50ntFlanks</td><td>698</td><td>730</td><td>33</td><td>AAGTAAGCGCAAGTACCCACGTAAAGACGTTAG</td><td>36.8</td><td>8.4</td><td>1.8</td><td>4.6</td><td>5.1E-06</td><td>9.0E-06</td><td>MB231/MB468</td><td>MP</td></tr>
<tr><td>>hg38_21_+_8433172_8446622_RNA45SN1_45S_50ntFlanks@10729.10752.24|AGAGGGTGTAAATCTCGCGCCGGG</td><td>>hg38_21_+_8433172_8446622_RNA45SN1_45S_50ntFlanks</td><td>10729</td><td>10752</td><td>24</td><td>AGAGGGTGTAAATCTCGCGCCGGG</td><td>175.3</td><td>-5.5</td><td>1.2</td><td>-4.6</td><td>5.1E-06</td><td>9.0E-06</td><td>BT20/MB468</td><td>CYTO</td></tr>
<tr><td>>trna111_HisGTG_1_-_147774845_147774916@2.30.29,trna118_HisGTG_1_-_145396881_145396952@2.30.29,trna16_HisGTG_1_+_146544773_146544844@2.30.29,trna1_HisGTG_15_+_45493349_45493420@2.30.29,trna21_HisGTG_1_+_147753471_147753542@2.30.29,trna33_HisGTG_6_+_27125906_27125977@2.30.29,trna7_HisGTG_9_-_14433938_14434009@2.30.29,trna8_HisGTG_15_-_45492611_45492682@2.30.29,trna9_HisGTG_15_-_45490804_45490875@2.30.29|CCGTGATCGTATAGTGGTTAGTACTCTGC</td><td>>trna111_HisGTG_1_-_147774845_147774916</td><td>2</td><td>30</td><td>29</td><td>CCGTGATCGTATAGTGGTTAGTACTCTGC</td><td>1041.7</td><td>3.7</td><td>0.8</td><td>4.5</td><td>5.8E-06</td><td>9.0E-06</td><td>MB231/MB468</td><td>CYTO</td></tr>
<tr><td>>hg38_21_+_8433172_8446622_RNA45SN1_45S_50ntFlanks@8190.8212.23|CCAGCCCGTGGACGGTGTGAGGC</td><td>>hg38_21_+_8433172_8446622_RNA45SN1_45S_50ntFlanks</td><td>8190</td><td>8212</td><td>23</td><td>CCAGCCCGTGGACGGTGTGAGGC</td><td>352.7</td><td>-3.0</td><td>0.7</td><td>-4.6</td><td>5.2E-06</td><td>9.0E-06</td><td>BT20/MB468</td><td>CYTO</td></tr>
<tr><td>>hg38_21_+_8433172_8446622_RNA45SN1_45S_50ntFlanks@6649.6669.21|TACGACTCTTAGCGGTGGATC</td><td>>hg38_21_+_8433172_8446622_RNA45SN1_45S_50ntFlanks</td><td>6649</td><td>6669</td><td>21</td><td>TACGACTCTTAGCGGTGGATC</td><td>782.2</td><td>2.7</td><td>0.6</td><td>4.6</td><td>5.2E-06</td><td>9.0E-06</td><td>BT20/MB468</td><td>CYTO</td></tr>
<tr><td>>MI0000114|hsa-mir-107&WithFlank&10|-|89592741|89592833@56.76.21(+1U);MI0000108|hsa-mir-103a-2&WithFlank&20|+|3917488|3917577@54.74.21(+1U);MI0000109|hsa-mir-103a-1&WithFlank&5|-|168560890|168560979@54.74.21(+1U)&[MIMAT0000104&hsa-miR-107&offsets|0|-2(+1U);m-78&10|-|89592758|89592778&offsets|0|0(+1U)];[MIMAT0000101&hsa-miR-103a-3p&offsets|0|-2(+1U);m-12&20|+|3917541|3917563&offsets|0|-2(+1U)];[MIMAT0000101_1&hsa-miR-103a-3p&offsets|0|-2(+1U);m-13&5|-|168560904|168560926&offsets|0|-2(+1U);m-44220&5|-|168560908|168560926&offsets|0|+2(+1U)]|AGCAGCATTGTACAGGGCTATT</td><td>>MI0000114|hsa-mir-107&WithFlank&10|-|89592741|89592833</td><td>56</td><td>76</td><td>21(+1U)</td><td>AGCAGCATTGTACAGGGCTATT</td><td>346.3</td><td>7.8</td><td>1.7</td><td>4.5</td><td>6.2E-06</td><td>9.0E-06</td><td>MB231/MB468</td><td>MITO</td></tr>
<tr><td>>hg38_21_+_8433172_8446622_RNA45SN1_45S_50ntFlanks@10794.10822.29|CTGGCATGTTGGAACAATGTAGGTAAGGG</td><td>>hg38_21_+_8433172_8446622_RNA45SN1_45S_50ntFlanks</td><td>10794</td><td>10822</td><td>29</td><td>CTGGCATGTTGGAACAATGTAGGTAAGGG</td><td>302.4</td><td>-6.4</td><td>1.4</td><td>-4.5</td><td>6.0E-06</td><td>9.0E-06</td><td>MB231/MB468</td><td>NUC</td></tr>
<tr><td>>MI0000078|hsa-mir-22&WithFlank&17|-|1713897|1713993@21.40.20(+1A)&[MIMAT0004495&hsa-miR-22-5p&offsets|0|-2(+1A);m-217&17|-|1713953|1713973&offsets|0|-1(+1A)]|AGTTCTTCAGTGGCAAGCTTA</td><td>>MI0000078|hsa-mir-22&WithFlank&17|-|1713897|1713993</td><td>21</td><td>40</td><td>20(+1A)</td><td>AGTTCTTCAGTGGCAAGCTTA</td><td>44.0</td><td>8.7</td><td>1.9</td><td>4.6</td><td>5.1E-06</td><td>9.0E-06</td><td>MB231/MB468</td><td>MP</td></tr>
<tr><td>>hg38_21_+_8433172_8446622_RNA45SN1_45S_50ntFlanks@6682.6699.18|CGTCGATGAAGAACGCAG</td><td>>hg38_21_+_8433172_8446622_RNA45SN1_45S_50ntFlanks</td><td>6682</td><td>6699</td><td>18</td><td>CGTCGATGAAGAACGCAG</td><td>62.0</td><td>-9.0</td><td>1.9</td><td>-4.8</td><td>1.8E-06</td><td>9.0E-06</td><td>BT20/MB468</td><td>MP</td></tr>
<tr><td>>hg38_21_+_8433172_8446622_RNA45SN1_45S_50ntFlanks@12477.12500.24|CACTAATAGGGAACGTGAGCTGGG</td><td>>hg38_21_+_8433172_8446622_RNA45SN1_45S_50ntFlanks</td><td>12477</td><td>12500</td><td>24</td><td>CACTAATAGGGAACGTGAGCTGGG</td><td>373.0</td><td>-7.1</td><td>1.6</td><td>-4.5</td><td>6.5E-06</td><td>9.1E-06</td><td>BT20/MB468</td><td>NUC</td></tr>
<tr><td>>hg38_21_+_8433172_8446622_RNA45SN1_45S_50ntFlanks@11795.11820.26|AACGAGATTCCCACTGTCCCTACCTA</td><td>>hg38_21_+_8433172_8446622_RNA45SN1_45S_50ntFlanks</td><td>11795</td><td>11820</td><td>26</td><td>AACGAGATTCCCACTGTCCCTACCTA</td><td>69.6</td><td>-9.2</td><td>1.9</td><td>-4.8</td><td>1.9E-06</td><td>9.1E-06</td><td>BT20/MB468</td><td>MP</td></tr>
<tr><td>>hg38_21_+_8433172_8446622_RNA45SN1_45S_50ntFlanks@11594.11612.19|CAGGGGAATCCGACTGTTT</td><td>>hg38_21_+_8433172_8446622_RNA45SN1_45S_50ntFlanks</td><td>11594</td><td>11612</td><td>19</td><td>CAGGGGAATCCGACTGTTT</td><td>943.0</td><td>-3.2</td><td>0.7</td><td>-4.5</td><td>5.8E-06</td><td>9.1E-06</td><td>MB231/MB468</td><td>CYTO</td></tr>
<tr><td>>hg38_21_+_8433172_8446622_RNA45SN1_45S_50ntFlanks@474.509.36|TCTTCGTGATCGATGTGGTGACGTCGTGCTCTCCCG</td><td>>hg38_21_+_8433172_8446622_RNA45SN1_45S_50ntFlanks</td><td>474</td><td>509</td><td>36</td><td>TCTTCGTGATCGATGTGGTGACGTCGTGCTCTCCCG</td><td>308.3</td><td>3.2</td><td>0.7</td><td>4.5</td><td>6.1E-06</td><td>9.1E-06</td><td>MB231/MB468</td><td>NUC</td></tr>
<tr><td>>am_trna111_HisGTG_1_-_147774845_147774916@3.29.27,trna118_HisGTG_1_-_145396881_145396952@3.29.27,trna16_HisGTG_1_+_146544773_146544844@3.29.27,trna1_HisGTG_15_+_45493349_45493420@3.29.27,trna21_HisGTG_1_+_147753471_147753542@3.29.27,trna33_HisGTG_6_+_27125906_27125977@3.29.27,trna7_HisGTG_9_-_14433938_14434009@3.29.27,trna8_HisGTG_15_-_45492611_45492682@3.29.27,trna9_HisGTG_15_-_45490804_45490875@3.29.27|CGTGATCGTATAGTGGTTAGTACTCTG</td><td>>am_trna111_HisGTG_1_-_147774845_147774916</td><td>3</td><td>29</td><td>27</td><td>CGTGATCGTATAGTGGTTAGTACTCTG</td><td>172.5</td><td>-5.6</td><td>1.2</td><td>-4.6</td><td>4.4E-06</td><td>9.1E-06</td><td>BT20/MB468</td><td>MITO</td></tr>
<tr><td>>hg38_MT_+_598_1651_RNR1_12S_50ntFlanks@843.871.29|GAGTAGAGTGCTTAGTTGAACAGGGCCCT</td><td>>hg38_MT_+_598_1651_RNR1_12S_50ntFlanks</td><td>843</td><td>871</td><td>29</td><td>GAGTAGAGTGCTTAGTTGAACAGGGCCCT</td><td>435.1</td><td>8.7</td><td>1.8</td><td>4.8</td><td>1.9E-06</td><td>9.1E-06</td><td>BT20/MB468</td><td>MP</td></tr>
<tr><td>>am_trna111_HisGTG_1_-_147774845_147774916@4.29.26,trna118_HisGTG_1_-_145396881_145396952@4.29.26,trna16_HisGTG_1_+_146544773_146544844@4.29.26,trna1_HisGTG_15_+_45493349_45493420@4.29.26,trna21_HisGTG_1_+_147753471_147753542@4.29.26,trna33_HisGTG_6_+_27125906_27125977@4.29.26,trna7_HisGTG_9_-_14433938_14434009@4.29.26,trna8_HisGTG_15_-_45492611_45492682@4.29.26,trna9_HisGTG_15_-_45490804_45490875@4.29.26|GTGATCGTATAGTGGTTAGTACTCTG</td><td>>am_trna111_HisGTG_1_-_147774845_147774916</td><td>4</td><td>29</td><td>26</td><td>GTGATCGTATAGTGGTTAGTACTCTG</td><td>46.1</td><td>-8.6</td><td>1.8</td><td>-4.8</td><td>1.9E-06</td><td>9.1E-06</td><td>BT20/MB468</td><td>MP</td></tr>
</tbody>
</table>

<table>
<tr><th></th><th></th><th></th><th></th><th></th><th></th><th></th><th></th><th></th><th></th><th></th><th></th><th></th><th></th></tr>
<tr><td>>MI0000095|hsa-mir-93&WithFlank&7|-|100093762|100093853@17.38.22(+1A)&[MIMAT0000093&hsa-miR-93-5p&offsets|0|-1(+1A);m-29&7|-|100093815|100093837&offsets|0|-1(+1A)]|CAAAGTGCTGTTCGTGCAGGTAA</td><td>>MI0000095|hsa-mir-93&WithFlank&7|-|100093762|100093853</td><td>17</td><td>38</td><td>22(+1A)</td><td>CAAAGTGCTGTTCGTGCAGGTAA</td><td>619.1</td><td>4.9</td><td>1.1</td><td>4.5</td><td>5.8E-06</td><td>9.1E-06</td><td>MB231/MB468</td><td>CYTO</td></tr>
<tr><td>>hg38_21_+_8433172_8446622_RNA45SN1_45S_50ntFlanks@474.505.32|TCTTCGTGATCGATGTGGTGACGTCGTGCTCT</td><td>>hg38_21_+_8433172_8446622_RNA45SN1_45S_50ntFlanks</td><td>474</td><td>505</td><td>32</td><td>TCTTCGTGATCGATGTGGTGACGTCGTGCTCT</td><td>169.7</td><td>7.2</td><td>1.6</td><td>4.5</td><td>6.2E-06</td><td>9.1E-06</td><td>MB231/MB468</td><td>MITO</td></tr>
<tr><td>>hg38_21_+_8433172_8446622_RNA45SN1_45S_50ntFlanks@7976.8005.30|GCGACCTCAGATCAGACGTGGCGACCCGCT</td><td>>hg38_21_+_8433172_8446622_RNA45SN1_45S_50ntFlanks</td><td>7976</td><td>8005</td><td>30</td><td>GCGACCTCAGATCAGACGTGGCGACCCGCT</td><td>324.9</td><td>6.0</td><td>1.3</td><td>4.8</td><td>1.9E-06</td><td>9.1E-06</td><td>BT20/MB468</td><td>MP</td></tr>
<tr><td>>trna111_HisGTG_1_-_147774845_147774916@3.30.28,trna118_HisGTG_1_-_145396881_145396952@3.30.28,trna16_HisGTG_1_+_146544773_146544844@3.30.28,trna1_HisGTG_15_+_45493349_45493420@3.30.28,trna21_HisGTG_1_+_147753471_147753542@3.30.28,trna33_HisGTG_6_+_27125906_27125977@3.30.28,trna7_HisGTG_9_-_14433938_14434009@3.30.28,trna8_HisGTG_15_-_45492611_45492682@3.30.28,trna9_HisGTG_15_-_45490804_45490875@3.30.28|CGTGATCGTATAGTGGTTAGTACTCTGC</td><td>>trna111_HisGTG_1_-_147774845_147774916</td><td>3</td><td>30</td><td>28</td><td>CGTGATCGTATAGTGGTTAGTACTCTGC</td><td>195.9</td><td>4.2</td><td>0.9</td><td>4.8</td><td>1.9E-06</td><td>9.1E-06</td><td>BT20/MB468</td><td>MP</td></tr>
<tr><td>>hg38_21_+_8433172_8446622_RNA45SN1_45S_50ntFlanks@11319.11343.25|CCCACGTCTCGTCGCGCGCGCGTCC</td><td>>hg38_21_+_8433172_8446622_RNA45SN1_45S_50ntFlanks</td><td>11319</td><td>11343</td><td>25</td><td>CCCACGTCTCGTCGCGCGCGCGTCC</td><td>296.8</td><td>-5.6</td><td>1.2</td><td>-4.5</td><td>6.2E-06</td><td>9.1E-06</td><td>MB231/MB468</td><td>MITO</td></tr>
<tr><td>>hg38_21_+_8433172_8446622_RNA45SN1_45S_50ntFlanks@6719.6745.27|TGAATTGCAGGACACATTGATCATCGA</td><td>>hg38_21_+_8433172_8446622_RNA45SN1_45S_50ntFlanks</td><td>6719</td><td>6745</td><td>27</td><td>TGAATTGCAGGACACATTGATCATCGA</td><td>924.5</td><td>-6.6</td><td>1.4</td><td>-4.6</td><td>5.2E-06</td><td>9.1E-06</td><td>BT20/MB468</td><td>CYTO</td></tr>
<tr><td>>hg38_21_+_8433172_8446622_RNA45SN1_45S_50ntFlanks@12615.12634.20|GCTGAGGAGCCAATGGGGCG</td><td>>hg38_21_+_8433172_8446622_RNA45SN1_45S_50ntFlanks</td><td>12615</td><td>12634</td><td>20</td><td>GCTGAGGAGCCAATGGGGCG</td><td>141.2</td><td>-3.6</td><td>0.8</td><td>-4.5</td><td>5.8E-06</td><td>9.1E-06</td><td>MB231/MB468</td><td>CYTO</td></tr>
<tr><td>>hg38_21_+_8433172_8446622_RNA45SN1_45S_50ntFlanks@12714.12745.32|GAGCCTCGGTTGGCCTCGGATAGCCGGTCCCC</td><td>>hg38_21_+_8433172_8446622_RNA45SN1_45S_50ntFlanks</td><td>12714</td><td>12745</td><td>32</td><td>GAGCCTCGGTTGGCCTCGGATAGCCGGTCCCC</td><td>524.3</td><td>-7.0</td><td>1.5</td><td>-4.6</td><td>5.1E-06</td><td>9.1E-06</td><td>MB231/MB468</td><td>MP</td></tr>
<tr><td>>hg38_21_+_8433172_8446622_RNA45SN1_45S_50ntFlanks@10387.10409.23|TGTGAACAGCAGTTGAACATGGG</td><td>>hg38_21_+_8433172_8446622_RNA45SN1_45S_50ntFlanks</td><td>10387</td><td>10409</td><td>23</td><td>TGTGAACAGCAGTTGAACATGGG</td><td>107.0</td><td>-10.6</td><td>2.3</td><td>-4.6</td><td>5.1E-06</td><td>9.1E-06</td><td>MB231/MB468</td><td>MP</td></tr>
<tr><td>>hg38_21_+_8433172_8446622_RNA45SN1_45S_50ntFlanks@9478.9497.20|GCGCACGTGTTAGGACCCGA</td><td>>hg38_21_+_8433172_8446622_RNA45SN1_45S_50ntFlanks</td><td>9478</td><td>9497</td><td>20</td><td>GCGCACGTGTTAGGACCCGA</td><td>348.1</td><td>5.7</td><td>1.3</td><td>4.5</td><td>6.1E-06</td><td>9.1E-06</td><td>MB231/MB468</td><td>NUC</td></tr>
<tr><td>>trna119_LysCTT_1_-_145395522_145395594@3.33.31,trna11_LysCTT_5_-_180648979_180649051@3.33.31,trna13_LysCTT_6_+_26556774_26556846@3.33.31,trna7_LysCTT_16_+_3225692_3225764@3.33.31,trna9_LysCTT_5_+_180634755_180634827@3.33.31|CCGGCTAGCTCAGTCGGTAGAGCATGAGACT</td><td>>trna119_LysCTT_1_-_145395522_145395594</td><td>3</td><td>33</td><td>31</td><td>CCGGCTAGCTCAGTCGGTAGAGCATGAGACT</td><td>1354.9</td><td>-5.4</td><td>1.1</td><td>-5.0</td><td>6.1E-07</td><td>9.1E-06</td><td>BT20/MB231</td><td>MITO</td></tr>
<tr><td>>hg38_21_+_8433172_8446622_RNA45SN1_45S_50ntFlanks@3837.3860.24|CCTCTCCTACTTGGATAACTGTGG</td><td>>hg38_21_+_8433172_8446622_RNA45SN1_45S_50ntFlanks</td><td>3837</td><td>3860</td><td>24</td><td>CCTCTCCTACTTGGATAACTGTGG</td><td>219.0</td><td>-2.6</td><td>0.6</td><td>-4.6</td><td>5.2E-06</td><td>9.2E-06</td><td>BT20/MB468</td><td>CYTO</td></tr>
<tr><td>>hg38_1_-_228634819_228635039_RNA5S12_5S_50ntFlanks@133.170.38|ATGGGAGACCGCCTGGGAATACCGGGTGCTGTAGGCTT</td><td>>hg38_1_-_228634819_228635039_RNA5S12_5S_50ntFlanks</td><td>133</td><td>170</td><td>38</td><td>ATGGGAGACCGCCTGGGAATACCGGGTGCTGTAGGCTT</td><td>20791.0</td><td>4.2</td><td>0.9</td><td>4.6</td><td>5.2E-06</td><td>9.2E-06</td><td>BT20/MB468</td><td>CYTO</td></tr>
<tr><td>>MI0000465|hsa-mir-191&WithFlank&3|-|49020612|49020715@22.43.22(+1U)&[MIMAT0000440&hsa-miR-191-5p&offsets|0|-1(+1U);m-48&3|-|49020672|49020694&offsets|0|-1(+1U)]|CAACGGAATCCCAAAAGCAGCTT</td><td>>MI0000465|hsa-mir-191&WithFlank&3|-|49020612|49020715</td><td>22</td><td>43</td><td>22(+1U)</td><td>CAACGGAATCCCAAAAGCAGCTT</td><td>298.7</td><td>5.8</td><td>1.3</td><td>4.5</td><td>6.2E-06</td><td>9.2E-06</td><td>MB231/MB468</td><td>MITO</td></tr>
<tr><td>>am_trna128_GlyGCC_6_-_27870686_27870756@1.30.30,trna133_GlyCCC_1_-_16872434_16872504@1.30.30,trna18_GlyGCC_16_+_70822597_70822667@1.30.30,trna19_GlyGCC_16_+_70823410_70823480@1.30.30,trna19_GlyGCC_2_-_157257659_157257729@1.30.30,trna24_GlyGCC_16_-_70812942_70813012@1.30.30,trna25_GlyGCC_16_-_70812114_70812184@1.30.30,trna4_GlyCCC_1_+_17188416_17188486@1.30.30,trna5_GlyGCC_17_+_8029064_8029134@1.30.30,trna68_GlyGCC_1_-_161493637_161493707@1.30.30|GCATTGGTGGTTCAGTGGTAGAATTCTCGC</td><td>>am_trna128_GlyGCC_6_-_27870686_27870756</td><td>1</td><td>30</td><td>30</td><td>GCATTGGTGGTTCAGTGGTAGAATTCTCGC</td><td>6218.4</td><td>-4.1</td><td>0.8</td><td>-5.0</td><td>6.2E-07</td><td>9.2E-06</td><td>BT20/MB231</td><td>MITO</td></tr>
<tr><td>>hg38_21_+_8433172_8446622_RNA45SN1_45S_50ntFlanks@10403.10424.22|ACATGGGTCAGTCGGTCCTGAG</td><td>>hg38_21_+_8433172_8446622_RNA45SN1_45S_50ntFlanks</td><td>10403</td><td>10424</td><td>22</td><td>ACATGGGTCAGTCGGTCCTGAG</td><td>10086.7</td><td>4.2</td><td>0.8</td><td>5.0</td><td>6.2E-07</td><td>9.2E-06</td><td>BT20/MB231</td><td>MITO</td></tr>
<tr><td>>hg38_21_+_8433172_8446622_RNA45SN1_45S_50ntFlanks@12420.12456.37|ATGTCGGCTCTTCCTATCATTGTGAAGCAGAATTCAC</td><td>>hg38_21_+_8433172_8446622_RNA45SN1_45S_50ntFlanks</td><td>12420</td><td>12456</td><td>37</td><td>ATGTCGGCTCTTCCTATCATTGTGAAGCAGAATTCAC</td><td>1476.5</td><td>-3.9</td><td>0.8</td><td>-5.0</td><td>6.1E-07</td><td>9.2E-06</td><td>BT20/MB231</td><td>MITO</td></tr>
<tr><td>>hg38_21_+_8433172_8446622_RNA45SN1_45S_50ntFlanks@12458.12475.18|AAGCGTTGGATTGTTCAC</td><td>>hg38_21_+_8433172_8446622_RNA45SN1_45S_50ntFlanks</td><td>12458</td><td>12475</td><td>18</td><td>AAGCGTTGGATTGTTCAC</td><td>796.6</td><td>-3.6</td><td>0.8</td><td>-4.5</td><td>5.9E-06</td><td>9.2E-06</td><td>MB231/MB468</td><td>CYTO</td></tr>
<tr><td>>MI0000093|hsa-mir-92a-1&WithFlank&13|+|91351308|91351397@54.74.21(+1G);MI0000094|hsa-mir-92a-2&WithFlank&X|-|134169532|134169618@54.74.21(+1G)&[MIMAT0000092&hsa-miR-92a-3p&offsets|0|-1(+1G);m-9&13|+|91351361|91351382&offsets|0|-1(+1G)];[MIMAT0000092_1&hsa-miR-92a-3p&offsets|0|-1(+1G);m-11&X|-|134169544|134169565&offsets|0|-1(+1G)]|TATTGCACTTGTCCCGGCCTGG</td><td>>MI0000093|hsa-mir-92a-1&WithFlank&13|+|91351308|91351397</td><td>54</td><td>74</td><td>21(+1G)</td><td>TATTGCACTTGTCCCGGCCTGG</td><td>1030.8</td><td>4.6</td><td>1.0</td><td>4.5</td><td>6.2E-06</td><td>9.2E-06</td><td>MB231/MB468</td><td>MITO</td></tr>
<tr><td>>hg38_21_+_8433172_8446622_RNA45SN1_45S_50ntFlanks@9741.9765.25|ACGATCTCAACCTATTCTCAAACTT</td><td>>hg38_21_+_8433172_8446622_RNA45SN1_45S_50ntFlanks</td><td>9741</td><td>9765</td><td>25</td><td>ACGATCTCAACCTATTCTCAAACTT</td><td>48.5</td><td>-8.7</td><td>1.8</td><td>-4.8</td><td>1.9E-06</td><td>9.2E-06</td><td>BT20/MB468</td><td>MP</td></tr>
<tr><td>>hg38_21_+_8433172_8446622_RNA45SN1_45S_50ntFlanks@10771.10816.46|CAGGTCTCCAAGGTGAACAGCCTCTGGCATGTTGGAACAATGTAGG</td><td>>hg38_21_+_8433172_8446622_RNA45SN1_45S_50ntFlanks</td><td>10771</td><td>10816</td><td>46</td><td>CAGGTCTCCAAGGTGAACAGCCTCTGGCATGTTGGAACAATGTAGG</td><td>103.0</td><td>-10.6</td><td>2.3</td><td>-4.6</td><td>5.2E-06</td><td>9.2E-06</td><td>MB231/MB468</td><td>MP</td></tr>
<tr><td>>hg38_21_+_8433172_8446622_RNA45SN1_45S_50ntFlanks@10905.10926.22|CTGGGCGCGCGCCGCGGCTGGA</td><td>>hg38_21_+_8433172_8446622_RNA45SN1_45S_50ntFlanks</td><td>10905</td><td>10926</td><td>22</td><td>CTGGGCGCGCGCCGCGGCTGGA</td><td>269.0</td><td>-6.0</td><td>1.3</td><td>-4.6</td><td>5.3E-06</td><td>9.2E-06</td><td>BT20/MB468</td><td>CYTO</td></tr>
<tr><td>>hg38_21_+_8433172_8446622_RNA45SN1_45S_50ntFlanks@8156.8181.26|GCTCGTGGGGGGCCCAAGTCCTTCTG</td><td>>hg38_21_+_8433172_8446622_RNA45SN1_45S_50ntFlanks</td><td>8156</td><td>8181</td><td>26</td><td>GCTCGTGGGGGGCCCAAGTCCTTCTG</td><td>209.9</td><td>-6.6</td><td>1.5</td><td>-4.6</td><td>5.2E-06</td><td>9.2E-06</td><td>MB231/MB468</td><td>MP</td></tr>
<tr><td>>trna32_LysCTT_16_-_3207406_3207478@13.33.21|CAGTCGGTAGAGCATGAGACC</td><td>>trna32_LysCTT_16_-_3207406_3207478</td><td>13</td><td>33</td><td>21</td><td>CAGTCGGTAGAGCATGAGACC</td><td>38.6</td><td>8.5</td><td>1.9</td><td>4.6</td><td>5.2E-06</td><td>9.2E-06</td><td>MB231/MB468</td><td>MP</td></tr>
<tr><td>>hg38_21_+_8433172_8446622_RNA45SN1_45S_50ntFlanks@12417.12441.25|TCGATGTCGGCTCTTCCTATCATTG</td><td>>hg38_21_+_8433172_8446622_RNA45SN1_45S_50ntFlanks</td><td>12417</td><td>12441</td><td>25</td><td>TCGATGTCGGCTCTTCCTATCATTG</td><td>1530.9</td><td>-3.1</td><td>0.7</td><td>-4.5</td><td>6.2E-06</td><td>9.2E-06</td><td>MB231/MB468</td><td>NUC</td></tr>
<tr><td>>hg38_21_+_8433172_8446622_RNA45SN1_45S_50ntFlanks@7974.8009.36|ACGCGACCTCAGATCAGACGTGGCGACCCGCTGAAT</td><td>>hg38_21_+_8433172_8446622_RNA45SN1_45S_50ntFlanks</td><td>7974</td><td>8009</td><td>36</td><td>ACGCGACCTCAGATCAGACGTGGCGACCCGCTGAAT</td><td>13629.8</td><td>2.9</td><td>0.6</td><td>4.5</td><td>6.6E-06</td><td>9.2E-06</td><td>BT20/MB468</td><td>NUC</td></tr>
<tr><td>>hg38_21_+_8433172_8446622_RNA45SN1_45S_50ntFlanks@9940.9961.22|CAGGACGGTGGCCATGGAAGTC</td><td>>hg38_21_+_8433172_8446622_RNA45SN1_45S_50ntFlanks</td><td>9940</td><td>9961</td><td>22</td><td>CAGGACGGTGGCCATGGAAGTC</td><td>269.3</td><td>5.7</td><td>1.2</td><td>4.8</td><td>1.9E-06</td><td>9.2E-06</td><td>BT20/MB468</td><td>MP</td></tr>
<tr><td>>hg38_21_+_8433172_8446622_RNA45SN1_45S_50ntFlanks@10808.10842.35|CAATGTAGGTAAGGGAAGTCGGCAAGCCGGATCCG</td><td>>hg38_21_+_8433172_8446622_RNA45SN1_45S_50ntFlanks</td><td>10808</td><td>10842</td><td>35</td><td>CAATGTAGGTAAGGGAAGTCGGCAAGCCGGATCCG</td><td>48.3</td><td>-8.7</td><td>1.8</td><td>-4.8</td><td>1.9E-06</td><td>9.2E-06</td><td>BT20/MB468</td><td>MP</td></tr>
<tr><td>>hg38_21_+_8433172_8446622_RNA45SN1_45S_50ntFlanks@12423.12443.21|TCGGCTCTTCCTATCATTGTG</td><td>>hg38_21_+_8433172_8446622_RNA45SN1_45S_50ntFlanks</td><td>12423</td><td>12443</td><td>21</td><td>TCGGCTCTTCCTATCATTGTG</td><td>659.7</td><td>-3.2</td><td>0.7</td><td>-4.5</td><td>6.2E-06</td><td>9.2E-06</td><td>MB231/MB468</td><td>NUC</td></tr>
<tr><td>>trnaMT_PheGAA_MT_+_577_647@35.66.32|GAAAATGTTTAGACGGGCTCACATCACCCCAT</td><td>>trnaMT_PheGAA_MT_+_577_647</td><td>35</td><td>66</td><td>32</td><td>GAAAATGTTTAGACGGGCTCACATCACCCCAT</td><td>33.4</td><td>8.3</td><td>1.8</td><td>4.6</td><td>5.2E-06</td><td>9.2E-06</td><td>MB231/MB468</td><td>MP</td></tr>
<tr><td>>hg38_21_+_8433172_8446622_RNA45SN1_45S_50ntFlanks@11564.11582.19|CCTAGCAGCCGACTTAGAA</td><td>>hg38_21_+_8433172_8446622_RNA45SN1_45S_50ntFlanks</td><td>11564</td><td>11582</td><td>19</td><td>CCTAGCAGCCGACTTAGAA</td><td>949.4</td><td>-6.7</td><td>1.5</td><td>-4.6</td><td>5.3E-06</td><td>9.2E-06</td><td>BT20/MB468</td><td>CYTO</td></tr>
<tr><td>>hg38_21_+_8433172_8446622_RNA45SN1_45S_50ntFlanks@6649.6698.50|TACGACTCTTAGCGGTGGATCACTCGGCTCGTGCGTCGATGAAGAACGCA</td><td>>hg38_21_+_8433172_8446622_RNA45SN1_45S_50ntFlanks</td><td>6649</td><td>6698</td><td>50</td><td>TACGACTCTTAGCGGTGGATCACTCGGCTCGTGCGTCGATGAAGAACGCA</td><td>1464.3</td><td>-5.4</td><td>1.2</td><td>-4.5</td><td>6.6E-06</td><td>9.2E-06</td><td>BT20/MB468</td><td>NUC</td></tr>
<tr><td>>hg38_21_+_8433172_8446622_RNA45SN1_45S_50ntFlanks@8190.8212.23|CCAGCCCGTGGACGGTGTGAGGC</td><td>>hg38_21_+_8433172_8446622_RNA45SN1_45S_50ntFlanks</td><td>8190</td><td>8212</td><td>23</td><td>CCAGCCCGTGGACGGTGTGAGGC</td><td>747.5</td><td>-6.6</td><td>1.5</td><td>-4.5</td><td>6.3E-06</td><td>9.2E-06</td><td>MB231/MB468</td><td>MITO</td></tr>
<tr><td>>hg38_21_+_8433172_8446622_RNA45SN1_45S_50ntFlanks@4443.4472.30|CCCTTGCCTCTCGGCGCCCCCTCGATGCTC</td><td>>hg38_21_+_8433172_8446622_RNA45SN1_45S_50ntFlanks</td><td>4443</td><td>4472</td><td>30</td><td>CCCTTGCCTCTCGGCGCCCCCTCGATGCTC</td><td>352.7</td><td>-7.0</td><td>1.5</td><td>-4.8</td><td>1.9E-06</td><td>9.2E-06</td><td>BT20/MB468</td><td>MP</td></tr>
<tr><td>>MI0000456|hsa-mir-140&WithFlank&16|+|69933075|69933186@29.50.22&[MIMAT0000431&hsa-miR-140-5p&offsets|0|0;m-173&16|+|69933103|69933124&offsets|0|0]|CAGTGGTTTTACCCTATGGTAG</td><td>>MI0000456|hsa-mir-140&WithFlank&16|+|69933075|69933186</td><td>29</td><td>50</td><td>22</td><td>CAGTGGTTTTACCCTATGGTAG</td><td>348.2</td><td>6.7</td><td>1.5</td><td>4.5</td><td>6.3E-06</td><td>9.2E-06</td><td>MB231/MB468</td><td>MITO</td></tr>
<tr><td>>trna10_ValCAC_5_-_180649395_180649467@37.54.18,trna12_ValAAC_5_-_180645270_180645342@37.54.18,trna132_ValAAC_6_-_27721179_27721251@37.54.18,trna13_ValCAC_19_-_4724647_4724719@37.54.18,trna152_ValCAC_6_-_27248049_27248121@37.54.18,trna157_ValCAC_6_-_27173867_27173939@37.54.18,trna15_ValAAC_5_-_180615416_180615488@37.54.18,trna18_ValCAC_5_-_180529253_180529325@37.54.18,trna2_ValAAC_3_+_169490018_169490090@37.54.18,trna2_ValCAC_5_+_180524070_180524142@37.54.18,trna37_ValAAC_6_+_27203288_27203360@37.54.18,trna4_ValAAC_5_+_180591154_180591226@37.54.18,trna5_ValAAC_5_+_180596610_180596682@37.54.18,trna6_ValCAC_5_+_180600650_180600722@37.54.18,trna85_ValCAC_1_-_161369490_161369562@37.54.18,trna98_ValCAC_1_-_149298555_149298627@37.54.18,trna9_ValCAC_6_+_26538282_26538354@37.54.18|ACGCGAAAGGTCCCCGGT</td><td>>trna10_ValCAC_5_-_180649395_180649467</td><td>37</td><td>54</td><td>18</td><td>ACGCGAAAGGTCCCCGGT</td><td>106.0</td><td>-10.6</td><td>2.3</td><td>-4.6</td><td>5.2E-06</td><td>9.2E-06</td><td>MB231/MB468</td><td>MP</td></tr>
<tr><td>>hg38_21_+_8433172_8446622_RNA45SN1_45S_50ntFlanks@10664.10711.48|CTTGGAAAGCGTCGCGGTTCCGGCGGCGTCCGGTGAGCTCTCGCTGGC</td><td>>hg38_21_+_8433172_8446622_RNA45SN1_45S_50ntFlanks</td><td>10664</td><td>10711</td><td>48</td><td>CTTGGAAAGCGTCGCGGTTCCGGCGGCGTCCGGTGAGCTCTCGCTGGC</td><td>367.6</td><td>-6.6</td><td>1.4</td><td>-4.6</td><td>4.5E-06</td><td>9.2E-06</td><td>BT20/MB468</td><td>MITO</td></tr>
<tr><td>>hg38_21_+_8433172_8446622_RNA45SN1_45S_50ntFlanks@5420.5443.24|CCGATTGGATGGTTTAGTGAGGCC</td><td>>hg38_21_+_8433172_8446622_RNA45SN1_45S_50ntFlanks</td><td>5420</td><td>5443</td><td>24</td><td>CCGATTGGATGGTTTAGTGAGGCC</td><td>1640.0</td><td>-7.7</td><td>1.6</td><td>-4.8</td><td>1.9E-06</td><td>9.2E-06</td><td>BT20/MB468</td><td>MP</td></tr>
</table>

| | | | | | | | | | | | | | |
|---|---|---|---|---|---|---|---|---|---|---|---|---|---|
| >trna111_HisGTG_1_-_147774845_147774916@-1T.31.32,trna118_HisGTG_1_-_145396881_145396952@-1T.31.32,trna16_HisGTG_1_+_146544773_146544844@-1T.31.32,trna1_HisGTG_15_+_45493349_45493420@-1T.31.32,trna21_HisGTG_1_+_147753471_147753542@-1T.31.32,trna33_HisGTG_6_+_27125906_27125977@-1T.31.32,trna7_HisGTG_9_-_14433938_14434009@-1T.31.32,trna8_HisGTG_15_-_45492611_45492682@-1T.31.32,trna9_HisGTG_15_-_45490804_45490875@-1T.31.32\|TGCCGTGATCGTATAGTGGTTAGTACTCTGCG | >trna111_HisGTG_1_-_147774845_147774916 | -1T | 31 | 32 | TGCCGTGATCGTATAGTGGTTAGTACTCTGCG | 6326.0 | -4.9 | 1.0 | -4.8 | 1.9E-06 | 9.3E-06 | BT20/MB468 | MP |
| >hg38_21_+_8433172_8446622_RNA45SN1_45S_50ntFlanks@11682.11702.21\|AATGTCAAAGTGAAGAAATTC | >hg38_21_+_8433172_8446622_RNA45SN1_45S_50ntFlanks | 11682 | 11702 | 21 | AATGTCAAAGTGAAGAAATTC | 1055.9 | -5.8 | 1.3 | -4.6 | 5.2E-06 | 9.3E-06 | MB231/MB468 | MP |
| >hg38_21_+_8433172_8446622_RNA45SN1_45S_50ntFlanks@9929.9950.22\|GATATAGACAGCAGGACGGTGG | >hg38_21_+_8433172_8446622_RNA45SN1_45S_50ntFlanks | 9929 | 9950 | 22 | GATATAGACAGCAGGACGGTGG | 151.5 | -6.5 | 1.4 | -4.6 | 5.2E-06 | 9.3E-06 | MB231/MB468 | MP |
| >hg38_21_+_8433172_8446622_RNA45SN1_45S_50ntFlanks@12399.12434.36\|ACGTCGCTTTTTGATCCTTCGATGTCGGCTCTTCCT | >hg38_21_+_8433172_8446622_RNA45SN1_45S_50ntFlanks | 12399 | 12434 | 36 | ACGTCGCTTTTTGATCCTTCGATGTCGGCTCTTCCT | 577.5 | -3.3 | 0.7 | -4.5 | 6.3E-06 | 9.3E-06 | MB231/MB468 | MITO |
| >MI0000087\|hsa-mir-29a&WithFlank&7\|-\|130876741\|130876816@47.64.18&[MIMAT0000086&hsa-miR-29a-3p&offsets\|-1\|-5;m-26&7\|-\|130876749\|130876769&offsets\|-1\|-4]\|CTAGCACCATCTGAAATC | >MI0000087\|hsa-mir-29a&WithFlank&7\|-\|130876741\|130876816 | 47 | 64 | 18 | CTAGCACCATCTGAAATC | 73.9 | 10.0 | 2.1 | 4.8 | 1.9E-06 | 9.3E-06 | BT20/MB468 | MP |
| >hg38_21_+_8433172_8446622_RNA45SN1_45S_50ntFlanks@11579.11601.23\|AGAACTGGTGCGGACCAGGGGAA | >hg38_21_+_8433172_8446622_RNA45SN1_45S_50ntFlanks | 11579 | 11601 | 23 | AGAACTGGTGCGGACCAGGGGAA | 575.0 | -5.3 | 1.2 | -4.5 | 6.2E-06 | 9.3E-06 | MB231/MB468 | NUC |
| >hg38_21_+_8433172_8446622_RNA45SN1_45S_50ntFlanks@12274.12291.18\|CAGTACGAATACAGACCG | >hg38_21_+_8433172_8446622_RNA45SN1_45S_50ntFlanks | 12274 | 12291 | 18 | CAGTACGAATACAGACCG | 44.0 | -8.5 | 1.8 | -4.8 | 1.9E-06 | 9.3E-06 | BT20/MB468 | MP |
| >hg38_21_+_8433172_8446622_RNA45SN1_45S_50ntFlanks@3775.3798.24\|GCCGGTACAGTGAAACTGCGAATG | >hg38_21_+_8433172_8446622_RNA45SN1_45S_50ntFlanks | 3775 | 3798 | 24 | GCCGGTACAGTGAAACTGCGAATG | 366.8 | -7.1 | 1.6 | -4.6 | 5.2E-06 | 9.3E-06 | MB231/MB468 | MP |
| >hg38_21_+_8433172_8446622_RNA45SN1_45S_50ntFlanks@5487.5524.38\|CGCTGAGAAGACGGTCGAACTTGACTATCTAGAGGAAG | >hg38_21_+_8433172_8446622_RNA45SN1_45S_50ntFlanks | 5487 | 5524 | 38 | CGCTGAGAAGACGGTCGAACTTGACTATCTAGAGGAAG | 63.4 | -9.1 | 1.9 | -4.8 | 1.9E-06 | 9.3E-06 | BT20/MB468 | MP |
| >hg38_21_+_8433172_8446622_RNA45SN1_45S_50ntFlanks@9532.9571.40\|AGAGGAAACTCTGGTGGAGGTCCGTAGCGGTCCTGACGTG | >hg38_21_+_8433172_8446622_RNA45SN1_45S_50ntFlanks | 9532 | 9571 | 40 | AGAGGAAACTCTGGTGGAGGTCCGTAGCGGTCCTGACGTG | 45.1 | 8.7 | 1.9 | 4.6 | 5.2E-06 | 9.3E-06 | MB231/MB468 | MP |
| >hg38_21_+_8433172_8446622_RNA45SN1_45S_50ntFlanks@12394.12417.24\|TAGCGACGTCGCTTTTTGATCCTT | >hg38_21_+_8433172_8446622_RNA45SN1_45S_50ntFlanks | 12394 | 12417 | 24 | TAGCGACGTCGCTTTTTGATCCTT | 783.0 | -4.2 | 0.9 | -4.6 | 4.5E-06 | 9.3E-06 | BT20/MB468 | MITO |
| >hg38_21_+_8433172_8446622_RNA45SN1_45S_50ntFlanks@10377.10424.48\|AAGGGTTCCATGTGAACAGCAGTTGAACATGGGTCAGTCGGTCCTGAG | >hg38_21_+_8433172_8446622_RNA45SN1_45S_50ntFlanks | 10377 | 10424 | 48 | AAGGGTTCCATGTGAACAGCAGTTGAACATGGGTCAGTCGGTCCTGAG | 3936.7 | 5.3 | 1.1 | 5.0 | 6.3E-07 | 9.3E-06 | BT20/MB231 | MITO |
| >hg38_21_+_8433172_8446622_RNA45SN1_45S_50ntFlanks@10400.10424.25\|TGAACATGGGTCAGTCGGTCCTGAG | >hg38_21_+_8433172_8446622_RNA45SN1_45S_50ntFlanks | 10400 | 10424 | 25 | TGAACATGGGTCAGTCGGTCCTGAG | 12056.1 | 4.1 | 0.8 | 5.0 | 6.3E-07 | 9.3E-06 | BT20/MB231 | MITO |
| >hg38_21_+_8433172_8446622_RNA45SN1_45S_50ntFlanks@4567.4607.41\|TAATGGAATAGGACCGCGGTTCTATTTTGTTGGTTTTCGGA | >hg38_21_+_8433172_8446622_RNA45SN1_45S_50ntFlanks | 4567 | 4607 | 41 | TAATGGAATAGGACCGCGGTTCTATTTTGTTGGTTTTCGGA | 39.5 | 8.5 | 1.9 | 4.6 | 5.3E-06 | 9.3E-06 | MB231/MB468 | MP |
| >MI0000342\|hsa-mir-200b&WithFlank&1\|+\|1167098\|1167204@63.85.23(+1A)&[MIMAT0000318&hsa-miR-200b-3p&offsets\|0\|+1(+1A);m-93&1\|+\|1167160\|1167182&offsets\|0\|0(+1A)]\|TAATACTGCCTGGTAATGATGACA | >MI0000342\|hsa-mir-200b&WithFlank&1\|+\|1167098\|1167204 | 63 | 85 | 23(+1A) | TAATACTGCCTGGTAATGATGACA | 626.7 | -5.1 | 1.1 | -4.5 | 6.2E-06 | 9.3E-06 | MB231/MB468 | NUC |
| >hg38_21_+_8433172_8446622_RNA45SN1_45S_50ntFlanks@4734.4752.19\|AACGAAAGTCGGAGGTTCG | >hg38_21_+_8433172_8446622_RNA45SN1_45S_50ntFlanks | 4734 | 4752 | 19 | AACGAAAGTCGGAGGTTCG | 333.0 | -3.7 | 0.8 | -4.5 | 6.2E-06 | 9.3E-06 | MB231/MB468 | NUC |
| >MI0000264\|hsa-mir-7-2&WithFlank&15\|+\|88611819\|88611940@38.58.21;MI0000265\|hsa-mir-7-3&WithFlank&19\|+\|4770664\|4770785@37.57.21;MI0000263\|hsa-mir-7-1&WithFlank&9\|-\|83969742\|83969863@30.50.21&[MIMAT0000252&hsa-miR-7-5p&offsets\|0\|-3;m-400&15\|+\|88611856\|88611879&offsets\|0\|-3];[MIMAT0000252_1&hsa-miR-7-5p&offsets\|0\|-3;m-402&19\|+\|4770700\|4770723&offsets\|0\|-3];[MIMAT0000252_2&hsa-miR-7-5p&offsets\|0\|-3;m-401&9\|-\|83969811\|83969834&offsets\|0\|-3]\|TGGAAGACTAGTGATTTTGTT | >MI0000264\|hsa-mir-7-2&WithFlank&15\|+\|88611819\|88611940 | 38 | 58 | 21 | TGGAAGACTAGTGATTTTGTT | 339.3 | 3.0 | 0.7 | 4.5 | 6.7E-06 | 9.3E-06 | BT20/MB468 | NUC |
| >hg38_21_+_8433172_8446622_RNA45SN1_45S_50ntFlanks@4761.4780.20\|CAGATACCGTCGTAGTTCCG | >hg38_21_+_8433172_8446622_RNA45SN1_45S_50ntFlanks | 4761 | 4780 | 20 | CAGATACCGTCGTAGTTCCG | 293.4 | -3.9 | 0.9 | -4.5 | 6.0E-06 | 9.3E-06 | MB231/MB468 | CYTO |
| >hg38_21_+_8433172_8446622_RNA45SN1_45S_50ntFlanks@5481.5499.19\|GCGGAGCGCTGAGAAGACG | >hg38_21_+_8433172_8446622_RNA45SN1_45S_50ntFlanks | 5481 | 5499 | 19 | GCGGAGCGCTGAGAAGACG | 183.3 | -5.0 | 1.1 | -4.6 | 5.3E-06 | 9.3E-06 | MB231/MB468 | MP |
| >hg38_21_+_8433172_8446622_RNA45SN1_45S_50ntFlanks@11235.11252.18\|CTTCCCGTGGATCGCCCC | >hg38_21_+_8433172_8446622_RNA45SN1_45S_50ntFlanks | 11235 | 11252 | 18 | CTTCCCGTGGATCGCCCC | 1079.6 | -3.3 | 0.7 | -4.6 | 4.6E-06 | 9.3E-06 | BT20/MB468 | MITO |
| >hg38_21_+_8433172_8446622_RNA45SN1_45S_50ntFlanks@11669.11691.23\|GCCCAGTGCTCTGAATGTCAAAG | >hg38_21_+_8433172_8446622_RNA45SN1_45S_50ntFlanks | 11669 | 11691 | 23 | GCCCAGTGCTCTGAATGTCAAAG | 456.1 | 4.1 | 0.9 | 4.6 | 4.6E-06 | 9.3E-06 | BT20/MB468 | MITO |
| >hg38_21_+_8433172_8446622_RNA45SN1_45S_50ntFlanks@10788.10818.31\|CAGCCTCTGGCATGTTGGAACAATGTAGGTA | >hg38_21_+_8433172_8446622_RNA45SN1_45S_50ntFlanks | 10788 | 10818 | 31 | CAGCCTCTGGCATGTTGGAACAATGTAGGTA | 1239.2 | -3.3 | 0.7 | -4.6 | 5.4E-06 | 9.4E-06 | BT20/MB468 | CYTO |
| >hg38_21_+_8433172_8446622_RNA45SN1_45S_50ntFlanks@11200.11234.35\|CGGCGGCGGCGGCGACTCTGGACGCGAGCCGGGCC | >hg38_21_+_8433172_8446622_RNA45SN1_45S_50ntFlanks | 11200 | 11234 | 35 | CGGCGGCGGCGGCGACTCTGGACGCGAGCCGGGCC | 427.2 | -4.9 | 1.1 | -4.5 | 6.0E-06 | 9.4E-06 | MB231/MB468 | CYTO |
| >hg38_21_+_8433172_8446622_RNA45SN1_45S_50ntFlanks@11586.11613.28\|GTGCGGACCAGGGGAATCCGACTGTTTA | >hg38_21_+_8433172_8446622_RNA45SN1_45S_50ntFlanks | 11586 | 11613 | 28 | GTGCGGACCAGGGGAATCCGACTGTTTA | 344.3 | -5.4 | 1.2 | -4.6 | 5.3E-06 | 9.4E-06 | MB231/MB468 | MP |
| >trnaMT_MetCAT_MT_+_4402_4469@38.71.34\|CGAAAATGTTGGTTATACCCTTCCCGTACTACCA | >trnaMT_MetCAT_MT_+_4402_4469 | 38 | 71 | 34 | CGAAAATGTTGGTTATACCCTTCCCGTACTACCA | 439.3 | 4.4 | 1.0 | 4.5 | 6.4E-06 | 9.4E-06 | MB231/MB468 | MITO |
| >hg38_21_+_8433172_8446622_RNA45SN1_45S_50ntFlanks@11586.11612.27\|GTGCGGACCAGGGGAATCCGACTGTTT | >hg38_21_+_8433172_8446622_RNA45SN1_45S_50ntFlanks | 11586 | 11612 | 27 | GTGCGGACCAGGGGAATCCGACTGTTT | 1299.0 | -4.1 | 0.9 | -4.8 | 1.9E-06 | 9.4E-06 | BT20/MB468 | MP |
| >hg38_21_+_8433172_8446622_RNA45SN1_45S_50ntFlanks@6646.6673.28\|TCGTACGACTCTTAGCGGTGGATCACTC | >hg38_21_+_8433172_8446622_RNA45SN1_45S_50ntFlanks | 6646 | 6673 | 28 | TCGTACGACTCTTAGCGGTGGATCACTC | 1241.9 | 3.8 | 0.8 | 4.5 | 6.3E-06 | 9.4E-06 | MB231/MB468 | NUC |
| >hg38_21_+_8433172_8446622_RNA45SN1_45S_50ntFlanks@9940.9957.18\|CAGGACGGTGGCCATGGA | >hg38_21_+_8433172_8446622_RNA45SN1_45S_50ntFlanks | 9940 | 9957 | 18 | CAGGACGGTGGCCATGGA | 444.6 | 4.4 | 0.9 | 4.8 | 1.9E-06 | 9.4E-06 | BT20/MB468 | MP |
| >MI0000063\|hsa-let-7b&WithFlank&22\|+\|46113680\|46113774@12.32.21&[MIMAT0000063&hsa-let-7b-5p&offsets\|0\|-1;m-14&22\|+\|46113691\|46113712&offsets\|0\|-1]\|TGAGGTAGTAGGTTGTGTGGT | >MI0000063\|hsa-let-7b&WithFlank&22\|+\|46113680\|46113774 | 12 | 32 | 21 | TGAGGTAGTAGGTTGTGTGGT | 6353.5 | 2.5 | 0.5 | 5.0 | 7.3E-07 | 9.4E-06 | BT20/MB231 | NUC |
| >MI0000446\|hsa-mir-125b-1&WithFlank&11\|-\|122099751\|122099850@21.41.21(+1G);MI0000470\|hsa-mir-125b-2&WithFlank&21\|+\|16590231\|16590331@23.43.21(+1G)&[MIMAT0000423&hsa-miR-125b-5p&offsets\|0\|-1(+1G);m-67&11\|-\|122099809\|122099830&offsets\|0\|-1(+1G)];[MIMAT0000423_1&hsa-miR-125b-5p&offsets\|0\|-1(+1G);m-68&21\|+\|16590253\|16590274&offsets\|0\|-1(+1G)]\|TCCCTGAGACCCTAACTTGTGG | >MI0000446\|hsa-mir-125b-1&WithFlank&11\|-\|122099751\|122099850 | 21 | 41 | 21(+1G) | TCCCTGAGACCCTAACTTGTGG | 351.6 | 7.9 | 1.7 | 4.5 | 6.4E-06 | 9.4E-06 | MB231/MB468 | MITO |
| >hg38_21_+_8433172_8446622_RNA45SN1_45S_50ntFlanks@10391.10432.42\|AACAGCAGTTGAACATGGGTCAGTCGGTCCTGAGAGATGGGC | >hg38_21_+_8433172_8446622_RNA45SN1_45S_50ntFlanks | 10391 | 10432 | 42 | AACAGCAGTTGAACATGGGTCAGTCGGTCCTGAGAGATGGGC | 196.0 | -6.8 | 1.5 | -4.5 | 6.4E-06 | 9.4E-06 | MB231/MB468 | MITO |
| >MI0000446\|hsa-mir-125b-1&WithFlank&11\|-\|122099751\|122099850@61.82.22(+1C)&[MIMAT0004592&hsa-miR-125b-1-3p&offsets\|0\|0(+1C);m-485&11\|-\|122099769\|122099790&offsets\|0\|0(+1C)]\|ACGGGTTAGGCTCTTGGGAGCTC | >MI0000446\|hsa-mir-125b-1&WithFlank&11\|-\|122099751\|122099850 | 61 | 82 | 22(+1C) | ACGGGTTAGGCTCTTGGGAGCTC | 37.8 | 8.5 | 1.9 | 4.6 | 5.3E-06 | 9.4E-06 | MB231/MB468 | MP |
| >MI0000433\|hsa-let-7g&WithFlank&3\|-\|52268272\|52268367@11.32.22&[MIMAT0000414&hsa-let-7g-5p&offsets\|0\|0;m-58&3\|-\|52268336\|52268357&offsets\|0\|0]\|TGAGGTAGTAGTTTGTACAGTT | >MI0000433\|hsa-let-7g&WithFlank&3\|-\|52268272\|52268367 | 11 | 32 | 22 | TGAGGTAGTAGTTTGTACAGTT | 33887.8 | 1.8 | 0.4 | 4.5 | 6.7E-06 | 9.4E-06 | BT20/MB468 | NUC |
| >hg38_21_+_8433172_8446622_RNA45SN1_45S_50ntFlanks@5073.5113.41\|AATTCCGATAACGAACGAGACTCTGGCATGCTAACTAGTTA | >hg38_21_+_8433172_8446622_RNA45SN1_45S_50ntFlanks | 5073 | 5113 | 41 | AATTCCGATAACGAACGAGACTCTGGCATGCTAACTAGTTA | 307.6 | -6.7 | 1.5 | -4.5 | 6.3E-06 | 9.4E-06 | MB231/MB468 | NUC |
| >hg38_21_+_8433172_8446622_RNA45SN1_45S_50ntFlanks@11741.11759.19\|TCTTAAGGTAGCCAAATGC | >hg38_21_+_8433172_8446622_RNA45SN1_45S_50ntFlanks | 11741 | 11759 | 19 | TCTTAAGGTAGCCAAATGC | 280.9 | -6.7 | 1.5 | -4.5 | 5.4E-06 | 9.4E-06 | BT20/MB468 | CYTO |
| >hg38_21_+_8433172_8446622_RNA45SN1_45S_50ntFlanks@6721.6743.23\|AATTGCAGGACACATTGATCATC | >hg38_21_+_8433172_8446622_RNA45SN1_45S_50ntFlanks | 6721 | 6743 | 23 | AATTGCAGGACACATTGATCATC | 3087.7 | -5.0 | 1.1 | -4.8 | 1.9E-06 | 9.4E-06 | BT20/MB468 | MP |
| >hg38_21_+_8433172_8446622_RNA45SN1_45S_50ntFlanks@11255.11272.18\|CTGCGGCGGGCGTCGCGG | >hg38_21_+_8433172_8446622_RNA45SN1_45S_50ntFlanks | 11255 | 11272 | 18 | CTGCGGCGGGCGTCGCGG | 62.7 | -9.1 | 1.9 | -4.8 | 1.9E-06 | 9.4E-06 | BT20/MB468 | MP |
| >hg38_21_+_8433172_8446622_RNA45SN1_45S_50ntFlanks@8913.8936.24\|GCCGAATCCCGGGGCCGAGGGAGC | >hg38_21_+_8433172_8446622_RNA45SN1_45S_50ntFlanks | 8913 | 8936 | 24 | GCCGAATCCCGGGGCCGAGGGAGC | 52.9 | 9.5 | 2.0 | 4.8 | 1.9E-06 | 9.4E-06 | BT20/MB468 | MP |
| >MI0000433\|hsa-let-7g&WithFlank&3\|-\|52268272\|52268367@11.28.18&[MIMAT0000414&hsa-let-7g-5p&offsets\|0\|-4;m-58&3\|-\|52268336\|52268357&offsets\|0\|-4]\|TGAGGTAGTAGTTTGTAC | >MI0000433\|hsa-let-7g&WithFlank&3\|-\|52268272\|52268367 | 11 | 28 | 18 | TGAGGTAGTAGTTTGTAC | 2541.2 | 4.9 | 1.1 | 4.6 | 5.3E-06 | 9.4E-06 | MB231/MB468 | MP |
| >hg38_21_+_8433172_8446622_RNA45SN1_45S_50ntFlanks@9481.9509.29\|CACGTGTTAGGACCCGAAAGATGGTGAAC | >hg38_21_+_8433172_8446622_RNA45SN1_45S_50ntFlanks | 9481 | 9509 | 29 | CACGTGTTAGGACCCGAAAGATGGTGAAC | 51.8 | 9.4 | 2.0 | 4.8 | 1.9E-06 | 9.4E-06 | BT20/MB468 | MP |
| >hg38_21_+_8433172_8446622_RNA45SN1_45S_50ntFlanks@12159.12179.21\|TTTGACTGGGGCGGTACACCT | >hg38_21_+_8433172_8446622_RNA45SN1_45S_50ntFlanks | 12159 | 12179 | 21 | TTTGACTGGGGCGGTACACCT | 886.2 | -6.5 | 1.4 | -4.5 | 5.4E-06 | 9.5E-06 | BT20/MB468 | CYTO |
| >hg38_21_+_8433172_8446622_RNA45SN1_45S_50ntFlanks@4153.4184.32\|ACGGCTACCACATCCAAGGAAGGCAGCAGGCG | >hg38_21_+_8433172_8446622_RNA45SN1_45S_50ntFlanks | 4153 | 4184 | 32 | ACGGCTACCACATCCAAGGAAGGCAGCAGGCG | 53.0 | -8.8 | 1.9 | -4.8 | 1.9E-06 | 9.5E-06 | BT20/MB468 | MP |
| >MI0003820\|hsa-mir-454&WithFlank&17\|-\|59137752\|59137878@70.92.23&[MIMAT0003885&hsa-miR-454-3p&offsets\|0\|0;m-272&17\|-\|59137787\|59137809&offsets\|0\|0]\|TAGTGCAATATTGCTTATAGGGT | >MI0003820\|hsa-mir-454&WithFlank&17\|-\|59137752\|59137878 | 70 | 92 | 23 | TAGTGCAATATTGCTTATAGGGT | 113.1 | 2.7 | 0.6 | 4.5 | 6.8E-06 | 9.5E-06 | BT20/MB468 | NUC |
| >MI0000446\|hsa-mir-125b-1&WithFlank&11\|-\|122099751\|122099850@21.41.21(+1U);MI0000470\|hsa-mir-125b-2&WithFlank&21\|+\|16590231\|16590331@23.43.21(+1U)&[MIMAT0000423&hsa-miR-125b-5p&offsets\|0\|-1(+1U);m-67&11\|-\|122099809\|122099830&offsets\|0\|-1(+1U)];[MIMAT0000423_1&hsa-miR-125b-5p&offsets\|0\|-1(+1U);m-68&21\|+\|16590253\|16590274&offsets\|0\|-1(+1U)]\|TCCCTGAGACCCTAACTTGTGT | >MI0000446\|hsa-mir-125b-1&WithFlank&11\|-\|122099751\|122099850 | 21 | 41 | 21(+1U) | TCCCTGAGACCCTAACTTGTGT | 475.8 | 5.8 | 1.3 | 4.6 | 5.4E-06 | 9.5E-06 | MB231/MB468 | MP |
| >hg38_21_+_8433172_8446622_RNA45SN1_45S_50ntFlanks@6723.6740.18\|TTGCAGGACACATTGATC | >hg38_21_+_8433172_8446622_RNA45SN1_45S_50ntFlanks | 6723 | 6740 | 18 | TTGCAGGACACATTGATC | 985.8 | -8.1 | 1.8 | -4.5 | 6.8E-06 | 9.5E-06 | BT20/MB468 | NUC |
| >hg38_21_+_8433172_8446622_RNA45SN1_45S_50ntFlanks@8154.8179.26\|CCGCTCGTGGGGGGCCCAAGTCCTTC | >hg38_21_+_8433172_8446622_RNA45SN1_45S_50ntFlanks | 8154 | 8179 | 26 | CCGCTCGTGGGGGGCCCAAGTCCTTC | 180.7 | -5.9 | 1.3 | -4.6 | 4.6E-06 | 9.5E-06 | BT20/MB468 | MITO |
| >hg38_21_+_8433172_8446622_RNA45SN1_45S_50ntFlanks@12488.12524.37\|AACGTGAGCTGGGTTTAGACCGTCGTGAGACAGGTTA | >hg38_21_+_8433172_8446622_RNA45SN1_45S_50ntFlanks | 12488 | 12524 | 37 | AACGTGAGCTGGGTTTAGACCGTCGTGAGACAGGTTA | 237.9 | -6.8 | 1.5 | -4.5 | 6.5E-06 | 9.5E-06 | MB231/MB468 | MITO |

<table>
<tr><td>>MI0000103|hsa-mir-101-1&WithFlank&1|-|65058428|65058514@52.72.21;MI0000739|hsa-mir-101-2&WithFlank&9|+|4850291|4850381@54.74.21&[MIMAT0000099&hsa-miR-101-3p&offsets|-1|-1;m-17&1|-|65058443|65058462&offsets|-1|0];[MIMAT0000099_1&hsa-miR-101-3p&offsets|-1|-1;m-16&9|+|4850345|4850364&offsets|-1|0]|GTACAGTACTGTGATAACTGA</td><td>>MI0000103|hsa-mir-101-1&WithFlank&1|-|65058428|65058514</td><td>52</td><td>72</td><td>21</td><td>GTACAGTACTGTGATAACTGA</td><td>546.2</td><td>5.1</td><td>1.1</td><td>4.5</td><td>6.5E-06</td><td>9.5E-06</td><td>MB231/MB468</td><td>MITO</td></tr>
<tr><td>>MI0003188|hsa-mir-503&WithFlank&X|-|134546322|134546404@12.31.20(+1A)&[MIMAT0002874&hsa-miR-503-5p&offsets|0|-3(+1A);m-297&X|-|134546371|134546393&offsets|0|-3(+1A)]|TAGCAGCGGGAACAGTTCTGA</td><td>>MI0003188|hsa-mir-503&WithFlank&X|-|134546322|134546404</td><td>12</td><td>31</td><td>20(+1A)</td><td>TAGCAGCGGGAACAGTTCTGA</td><td>366.1</td><td>8.9</td><td>1.9</td><td>4.8</td><td>2.0E-06</td><td>9.5E-06</td><td>BT20/MB468</td><td>MP</td></tr>
<tr><td>>MI0003763|hsa-mir-767&WithFlank&X|-|152393415|152393535@33.55.23&[MIMAT0003882&hsa-miR-767-5p&offsets|0|0;m-788&X|-|152393481|152393503&offsets|0|0]|TGCACCATGGTTGTCTGAGCATG</td><td>>MI0003763|hsa-mir-767&WithFlank&X|-|152393415|152393535</td><td>33</td><td>55</td><td>23</td><td>TGCACCATGGTTGTCTGAGCATG</td><td>86.6</td><td>9.8</td><td>1.9</td><td>5.0</td><td>5.4E-07</td><td>9.5E-06</td><td>BT20/MB231</td><td>MP</td></tr>
<tr><td>>hg38_21_+_8433172_8446622_RNA45SN1_45S_50ntFlanks@11785.11807.23|TGAATGGATGAACGAGATTCCCA</td><td>>hg38_21_+_8433172_8446622_RNA45SN1_45S_50ntFlanks</td><td>11785</td><td>11807</td><td>23</td><td>TGAATGGATGAACGAGATTCCCA</td><td>66.4</td><td>-9.1</td><td>1.9</td><td>-4.8</td><td>2.0E-06</td><td>9.5E-06</td><td>BT20/MB468</td><td>MP</td></tr>
<tr><td>>hg38_1_-_228634819_228635039_RNA5S12_5S_50ntFlanks@128.169.42|CTTGGATGGGAGACCGCCTGGGAATACCGGGTGCTGTAGGCT</td><td>>hg38_1_-_228634819_228635039_RNA5S12_5S_50ntFlanks</td><td>128</td><td>169</td><td>42</td><td>CTTGGATGGGAGACCGCCTGGGAATACCGGGTGCTGTAGGCT</td><td>291.6</td><td>4.2</td><td>0.9</td><td>5.0</td><td>6.5E-07</td><td>9.5E-06</td><td>BT20/MB231</td><td>MITO</td></tr>
<tr><td>>hg38_21_+_8433172_8446622_RNA45SN1_45S_50ntFlanks@11208.11254.47|GCGGCGACTCTGGACGCGAGCCGGGCCCTTCCCGTGGATCGCCCCAG</td><td>>hg38_21_+_8433172_8446622_RNA45SN1_45S_50ntFlanks</td><td>11208</td><td>11254</td><td>47</td><td>GCGGCGACTCTGGACGCGAGCCGGGCCCTTCCCGTGGATCGCCCCAG</td><td>293.4</td><td>5.9</td><td>1.2</td><td>5.0</td><td>6.5E-07</td><td>9.5E-06</td><td>BT20/MB231</td><td>MITO</td></tr>
<tr><td>>hg38_21_+_8433172_8446622_RNA45SN1_45S_50ntFlanks@12243.12273.31|GAAGGGCAAAAGCTCGCTTGATCTTGATTTT</td><td>>hg38_21_+_8433172_8446622_RNA45SN1_45S_50ntFlanks</td><td>12243</td><td>12273</td><td>31</td><td>GAAGGGCAAAAGCTCGCTTGATCTTGATTTT</td><td>539.7</td><td>-5.5</td><td>1.2</td><td>-4.5</td><td>6.5E-06</td><td>9.5E-06</td><td>MB231/MB468</td><td>MITO</td></tr>
<tr><td>>am_MI0003639|hsa-mir-625&WithFlank&14|+|65471096|65471192@58.79.22(+1C)&[MIMAT0004808&hsa-miR-625-3p&offsets|0|0(+1C);m-137&14|+|65471153|65471174&offsets|0|0(+1C)]|GACTATAGAACTTTCCCCCTCAC</td><td>>am_MI0003639|hsa-mir-625&WithFlank&14|+|65471096|65471192</td><td>58</td><td>79</td><td>22(+1C)</td><td>GACTATAGAACTTTCCCCCTCAC</td><td>87.0</td><td>-9.8</td><td>2.0</td><td>-5.0</td><td>5.4E-07</td><td>9.5E-06</td><td>BT20/MB231</td><td>MP</td></tr>
<tr><td>>am_trna134_GluTTC_1_-_16861774_16861845@1.28.28,trna5_GluTTC_1_+_17199078_17199149@1.28.28,trna84_GluTTC_1_-_161391883_161391954@1.28.28,trna94_GluTTC_1_-_149664355_149664427@1.28.28|TCCCTGGTGGTCTAGTGGCTAGGATTCG</td><td>>am_trna134_GluTTC_1_-_16861774_16861845</td><td>1</td><td>28</td><td>28</td><td>TCCCTGGTGGTCTAGTGGCTAGGATTCG</td><td>1093.6</td><td>-2.4</td><td>0.5</td><td>-4.5</td><td>5.5E-06</td><td>9.5E-06</td><td>BT20/MB468</td><td>CYTO</td></tr>
<tr><td>>hg38_21_+_8433172_8446622_RNA45SN1_45S_50ntFlanks@3852.3873.22|TAACTGTGGTAATTCTAGAGCT</td><td>>hg38_21_+_8433172_8446622_RNA45SN1_45S_50ntFlanks</td><td>3852</td><td>3873</td><td>22</td><td>TAACTGTGGTAATTCTAGAGCT</td><td>141.6</td><td>-4.9</td><td>1.1</td><td>-4.5</td><td>5.4E-06</td><td>9.5E-06</td><td>MB231/MB468</td><td>MP</td></tr>
<tr><td>>hg38_21_+_8433172_8446622_RNA45SN1_45S_50ntFlanks@3873.3899.27|TAATACATGCCGACGGGCGCTGACCCC</td><td>>hg38_21_+_8433172_8446622_RNA45SN1_45S_50ntFlanks</td><td>3873</td><td>3899</td><td>27</td><td>TAATACATGCCGACGGGCGCTGACCCC</td><td>384.5</td><td>-6.6</td><td>1.5</td><td>-4.5</td><td>6.4E-06</td><td>9.5E-06</td><td>MB231/MB468</td><td>NUC</td></tr>
<tr><td>>hg38_21_+_8433172_8446622_RNA45SN1_45S_50ntFlanks@10664.10712.49|CTTGGAAAGCGTCGCGGTTCCGGCGGCGTCCGGTGAGCTCTCGCTGGCC</td><td>>hg38_21_+_8433172_8446622_RNA45SN1_45S_50ntFlanks</td><td>10664</td><td>10712</td><td>49</td><td>CTTGGAAAGCGTCGCGGTTCCGGCGGCGTCCGGTGAGCTCTCGCTGGCC</td><td>158.3</td><td>-5.0</td><td>1.1</td><td>-4.5</td><td>6.1E-06</td><td>9.5E-06</td><td>MB231/MB468</td><td>CYTO</td></tr>
<tr><td>>hg38_21_+_8433172_8446622_RNA45SN1_45S_50ntFlanks@12419.12439.21|GATGTCGGCTCTTCCTATCAT</td><td>>hg38_21_+_8433172_8446622_RNA45SN1_45S_50ntFlanks</td><td>12419</td><td>12439</td><td>21</td><td>GATGTCGGCTCTTCCTATCAT</td><td>1099.2</td><td>3.1</td><td>0.7</td><td>4.5</td><td>6.4E-06</td><td>9.5E-06</td><td>MB231/MB468</td><td>NUC</td></tr>
<tr><td>>hg38_21_+_8433172_8446622_RNA45SN1_45S_50ntFlanks@6733.6774.42|CATTGATCATCGACACTTCGAACGCACTTGCGGCCCCGGGTT</td><td>>hg38_21_+_8433172_8446622_RNA45SN1_45S_50ntFlanks</td><td>6733</td><td>6774</td><td>42</td><td>CATTGATCATCGACACTTCGAACGCACTTGCGGCCCCGGGTT</td><td>428.6</td><td>4.3</td><td>0.9</td><td>4.5</td><td>6.5E-06</td><td>9.5E-06</td><td>MB231/MB468</td><td>MITO</td></tr>
<tr><td>>hg38_21_+_8433172_8446622_RNA45SN1_45S_50ntFlanks@11668.11691.24|TGCCCAGTGCTCTGAATGTCAAAG</td><td>>hg38_21_+_8433172_8446622_RNA45SN1_45S_50ntFlanks</td><td>11668</td><td>11691</td><td>24</td><td>TGCCCAGTGCTCTGAATGTCAAAG</td><td>369.0</td><td>2.9</td><td>0.6</td><td>4.5</td><td>6.4E-06</td><td>9.5E-06</td><td>MB231/MB468</td><td>NUC</td></tr>
<tr><td>>hg38_21_+_8433172_8446622_RNA45SN1_45S_50ntFlanks@6645.6674.30|CTCGTACGACTCTTAGCGGTGGATCACTCG</td><td>>hg38_21_+_8433172_8446622_RNA45SN1_45S_50ntFlanks</td><td>6645</td><td>6674</td><td>30</td><td>CTCGTACGACTCTTAGCGGTGGATCACTCG</td><td>572.3</td><td>3.7</td><td>0.8</td><td>4.5</td><td>5.5E-06</td><td>9.5E-06</td><td>BT20/MB468</td><td>CYTO</td></tr>
<tr><td>>hg38_21_+_8433172_8446622_RNA45SN1_45S_50ntFlanks@6721.6740.20|AATTGCAGGACACATTGATC</td><td>>hg38_21_+_8433172_8446622_RNA45SN1_45S_50ntFlanks</td><td>6721</td><td>6740</td><td>20</td><td>AATTGCAGGACACATTGATC</td><td>1474.2</td><td>-2.9</td><td>0.6</td><td>-4.6</td><td>4.7E-06</td><td>9.5E-06</td><td>BT20/MB468</td><td>MITO</td></tr>
<tr><td>>hg38_1_-_228634819_228635039_RNA5S12_5S_50ntFlanks@51.87.37|GTCTACGGCCATACCACCCTGAACGCGCCCGATCTCG</td><td>>hg38_1_-_228634819_228635039_RNA5S12_5S_50ntFlanks</td><td>51</td><td>87</td><td>37</td><td>GTCTACGGCCATACCACCCTGAACGCGCCCGATCTCG</td><td>1904.2</td><td>6.5</td><td>1.4</td><td>4.5</td><td>6.1E-06</td><td>9.6E-06</td><td>MB231/MB468</td><td>CYTO</td></tr>
<tr><td>>am_trna10_ValCAC_5_-_180649395_180649467@1.33.33,trna12_ValAAC_5_-_180645270_180645342@1.33.33,trna132_ValAAC_6_-_27721179_27721251@1.33.33,trna136_ValAAC_6_-_27648885_27648957@1.33.33,trna139_ValAAC_6_-_27618707_27618779@1.33.33,trna18_ValCAC_5_-_180529253_180529325@1.33.33,trna2_ValAAC_3_+_169490018_169490090@1.33.33,trna2_ValCAC_5_+_180524070_180524142@1.33.33,trna4_ValAAC_5_+_180591154_180591226@1.33.33,trna5_ValAAC_5_+_180596610_180596682@1.33.33,trna6_ValCAC_5_+_180600650_180600722@1.33.33,trna85_ValCAC_1_-_161369490_161369562@1.33.33,trna90_ValCAC_1_-_149684088_149684161@1.33.33,trna98_ValCAC_1_-_149298555_149298627@1.33.33,trna9_ValCAC_6_+_26538282_26538354@1.33.33|GTTTCCGTAGTGTAGTGGTTATCACGTTCGCCT</td><td>>am_trna10_ValCAC_5_-_180649395_180649467</td><td>1</td><td>33</td><td>33</td><td>GTTTCCGTAGTGTAGTGGTTATCACGTTCGCCT</td><td>1373.8</td><td>10.1</td><td>2.1</td><td>4.8</td><td>2.0E-06</td><td>9.6E-06</td><td>BT20/MB468</td><td>MP</td></tr>
<tr><td>>MI0000542|hsa-mir-320a&WithFlank&8|-|22244960|22245043@50.70.21(+1U)&[MIMAT0000510&hsa-miR-320a-3p&offsets|+2|+1(+1U);m-60&8|-|22244975|22244996&offsets|+2|+1(+1U)]|AAGCTGGGTTGAGAGGGCGAAT</td><td>>MI0000542|hsa-mir-320a&WithFlank&8|-|22244960|22245043</td><td>50</td><td>70</td><td>21(+1U)</td><td>AAGCTGGGTTGAGAGGGCGAAT</td><td>682.9</td><td>6.6</td><td>1.4</td><td>4.8</td><td>2.0E-06</td><td>9.6E-06</td><td>BT20/MB468</td><td>MP</td></tr>
<tr><td>>hg38_21_+_8433172_8446622_RNA45SN1_45S_50ntFlanks@9482.9512.31|ACGTGTTAGGACCCGAAAGATGGTGAACTAT</td><td>>hg38_21_+_8433172_8446622_RNA45SN1_45S_50ntFlanks</td><td>9482</td><td>9512</td><td>31</td><td>ACGTGTTAGGACCCGAAAGATGGTGAACTAT</td><td>52.0</td><td>9.5</td><td>2.0</td><td>4.8</td><td>2.0E-06</td><td>9.6E-06</td><td>BT20/MB468</td><td>MP</td></tr>
<tr><td>>hg38_21_+_8433172_8446622_RNA45SN1_45S_50ntFlanks@11211.11232.22|GCGACTCTGGACGCGAGCCGGG</td><td>>hg38_21_+_8433172_8446622_RNA45SN1_45S_50ntFlanks</td><td>11211</td><td>11232</td><td>22</td><td>GCGACTCTGGACGCGAGCCGGG</td><td>1040.2</td><td>-4.2</td><td>0.9</td><td>-4.8</td><td>2.0E-06</td><td>9.6E-06</td><td>BT20/MB468</td><td>MP</td></tr>
<tr><td>>hg38_21_+_8433172_8446622_RNA45SN1_45S_50ntFlanks@12307.12325.19|ACGATCCTTCTGACCTTTT</td><td>>hg38_21_+_8433172_8446622_RNA45SN1_45S_50ntFlanks</td><td>12307</td><td>12325</td><td>19</td><td>ACGATCCTTCTGACCTTTT</td><td>43.6</td><td>-8.5</td><td>1.8</td><td>-4.8</td><td>2.0E-06</td><td>9.6E-06</td><td>BT20/MB468</td><td>MP</td></tr>
<tr><td>>hg38_21_+_8433172_8446622_RNA45SN1_45S_50ntFlanks@4130.4151.22|ATTCCGGAGAGGGAGCCTGAGA</td><td>>hg38_21_+_8433172_8446622_RNA45SN1_45S_50ntFlanks</td><td>4130</td><td>4151</td><td>22</td><td>ATTCCGGAGAGGGAGCCTGAGA</td><td>148.1</td><td>-2.5</td><td>0.5</td><td>-4.5</td><td>6.4E-06</td><td>9.6E-06</td><td>MB231/MB468</td><td>NUC</td></tr>
<tr><td>>hg38_21_+_8433172_8446622_RNA45SN1_45S_50ntFlanks@12044.12072.29|ACTGACCCGGTGAGGCGGGGGGGCGAGCC</td><td>>hg38_21_+_8433172_8446622_RNA45SN1_45S_50ntFlanks</td><td>12044</td><td>12072</td><td>29</td><td>ACTGACCCGGTGAGGCGGGGGGGCGAGCC</td><td>603.9</td><td>-7.3</td><td>1.5</td><td>-4.8</td><td>2.0E-06</td><td>9.6E-06</td><td>BT20/MB468</td><td>MP</td></tr>
<tr><td>>hg38_21_+_8433172_8446622_RNA45SN1_45S_50ntFlanks@12397.12416.20|CGACGTCGCTTTTTGATCCT</td><td>>hg38_21_+_8433172_8446622_RNA45SN1_45S_50ntFlanks</td><td>12397</td><td>12416</td><td>20</td><td>CGACGTCGCTTTTTGATCCT</td><td>107.6</td><td>-6.5</td><td>1.4</td><td>-4.5</td><td>5.4E-06</td><td>9.6E-06</td><td>MB231/MB468</td><td>MP</td></tr>
<tr><td>>am_trna111_HisGTG_1_-_147774845_147774916@1.26.26,trna118_HisGTG_1_-_145396881_145396952@1.26.26,trna16_HisGTG_1_+_146544773_146544844@1.26.26,trna1_HisGTG_15_+_45493349_45493420@1.26.26,trna21_HisGTG_1_+_147753471_147753542@1.26.26,trna33_HisGTG_6_+_27125906_27125977@1.26.26,trna7_HisGTG_9_-_14433938_14434009@1.26.26,trna8_HisGTG_15_-_45492611_45492682@1.26.26,trna9_HisGTG_15_-_45490804_45490875@1.26.26|GCCGTGATCGTATAGTGGTTAGTACT</td><td>>am_trna111_HisGTG_1_-_147774845_147774916</td><td>1</td><td>26</td><td>26</td><td>GCCGTGATCGTATAGTGGTTAGTACT</td><td>1346.7</td><td>3.7</td><td>0.8</td><td>4.5</td><td>5.5E-06</td><td>9.6E-06</td><td>BT20/MB468</td><td>CYTO</td></tr>
<tr><td>>MI0005756|hsa-mir-934&WithFlank&X|+|136550872|136550966@21.41.21(+1U)&[MIMAT0004977&hsa-miR-934&offsets|0|-1(+1U);m-943&X|+|136550892|136550913&offsets|0|-1(+1U)]|TGTCTACTACTGGAGACACTGT</td><td>>MI0005756|hsa-mir-934&WithFlank&X|+|136550872|136550966</td><td>21</td><td>41</td><td>21(+1U)</td><td>TGTCTACTACTGGAGACACTGT</td><td>134.4</td><td>10.4</td><td>2.1</td><td>5.0</td><td>5.5E-07</td><td>9.6E-06</td><td>BT20/MB231</td><td>MP</td></tr>
<tr><td>>hg38_21_+_8433172_8446622_RNA45SN1_45S_50ntFlanks@9925.9948.24|GGTTGATATAGACAGCAGGACGGT</td><td>>hg38_21_+_8433172_8446622_RNA45SN1_45S_50ntFlanks</td><td>9925</td><td>9948</td><td>24</td><td>GGTTGATATAGACAGCAGGACGGT</td><td>87.7</td><td>9.8</td><td>2.0</td><td>5.0</td><td>5.5E-07</td><td>9.6E-06</td><td>BT20/MB231</td><td>MP</td></tr>
<tr><td>>MI0000448|hsa-mir-130a&WithFlank&11|+|57641192|57641292@61.82.22(+1A)&[MIMAT0000425&hsa-miR-130a-3p&offsets|0|0(+1A);m-139&11|+|57641252|57641273&offsets|0|0(+1A)]|CAGTGCAATGTTAAAAGGGCATA</td><td>>MI0000448|hsa-mir-130a&WithFlank&11|+|57641192|57641292</td><td>61</td><td>82</td><td>22(+1A)</td><td>CAGTGCAATGTTAAAAGGGCATA</td><td>90.6</td><td>-9.9</td><td>2.0</td><td>-5.0</td><td>5.5E-07</td><td>9.6E-06</td><td>BT20/MB231</td><td>MP</td></tr>
<tr><td>>MI0003513|hsa-mir-455&WithFlank&9|+|114209428|114209535@22.45.24(+1U)&[MIMAT0003150&hsa-miR-455-5p&offsets|0|+2(+1U);m-232&9|+|114209449|114209470&offsets|0|+2(+1U)]|TATGTGCCTTTGGACTACATCGTGT</td><td>>MI0003513|hsa-mir-455&WithFlank&9|+|114209428|114209535</td><td>22</td><td>45</td><td>24(+1U)</td><td>TATGTGCCTTTGGACTACATCGTGT</td><td>89.4</td><td>-9.9</td><td>2.0</td><td>-5.0</td><td>5.5E-07</td><td>9.6E-06</td><td>BT20/MB231</td><td>MP</td></tr>
<tr><td>>hg38_21_+_8433172_8446622_RNA45SN1_45S_50ntFlanks@9941.9959.19|AGGACGGTGGCCATGGAAG</td><td>>hg38_21_+_8433172_8446622_RNA45SN1_45S_50ntFlanks</td><td>9941</td><td>9959</td><td>19</td><td>AGGACGGTGGCCATGGAAG</td><td>122.9</td><td>10.3</td><td>2.1</td><td>5.0</td><td>5.5E-07</td><td>9.6E-06</td><td>BT20/MB231</td><td>MP</td></tr>
<tr><td>>hg38_21_+_8433172_8446622_RNA45SN1_45S_50ntFlanks@4096.4121.26|ATGGTGACCACGGGTGACGGGGAATC</td><td>>hg38_21_+_8433172_8446622_RNA45SN1_45S_50ntFlanks</td><td>4096</td><td>4121</td><td>26</td><td>ATGGTGACCACGGGTGACGGGGAATC</td><td>149.2</td><td>-6.6</td><td>1.5</td><td>-4.5</td><td>5.4E-06</td><td>9.6E-06</td><td>MB231/MB468</td><td>MP</td></tr>
<tr><td>>hg38_21_+_8433172_8446622_RNA45SN1_45S_50ntFlanks@9459.9500.42|CCGGGGAGGTGGAGCACGAGCGCACGTGTTAGGACCCGAAAG</td><td>>hg38_21_+_8433172_8446622_RNA45SN1_45S_50ntFlanks</td><td>9459</td><td>9500</td><td>42</td><td>CCGGGGAGGTGGAGCACGAGCGCACGTGTTAGGACCCGAAAG</td><td>65.6</td><td>-9.1</td><td>1.9</td><td>-4.8</td><td>2.0E-06</td><td>9.6E-06</td><td>BT20/MB468</td><td>MP</td></tr>
<tr><td>>hg38_21_+_8433172_8446622_RNA45SN1_45S_50ntFlanks@6648.6676.29|GTACGACTCTTAGCGGTGGATCACTCGGC</td><td>>hg38_21_+_8433172_8446622_RNA45SN1_45S_50ntFlanks</td><td>6648</td><td>6676</td><td>29</td><td>GTACGACTCTTAGCGGTGGATCACTCGGC</td><td>1137.0</td><td>6.4</td><td>1.4</td><td>4.5</td><td>5.5E-06</td><td>9.6E-06</td><td>BT20/MB468</td><td>CYTO</td></tr>
<tr><td>>hg38_21_+_8433172_8446622_RNA45SN1_45S_50ntFlanks@11565.11602.38|CTAGCAGCCGACTTAGAACTGGTGCGGACCAGGGGAAT</td><td>>hg38_21_+_8433172_8446622_RNA45SN1_45S_50ntFlanks</td><td>11565</td><td>11602</td><td>38</td><td>CTAGCAGCCGACTTAGAACTGGTGCGGACCAGGGGAAT</td><td>226.9</td><td>-3.3</td><td>0.7</td><td>-4.5</td><td>6.4E-06</td><td>9.6E-06</td><td>MB231/MB468</td><td>NUC</td></tr>
<tr><td>>hg38_21_+_8433172_8446622_RNA45SN1_45S_50ntFlanks@10798.10827.30|CATGTTGGAACAATGTAGGTAAGGGAAGTC</td><td>>hg38_21_+_8433172_8446622_RNA45SN1_45S_50ntFlanks</td><td>10798</td><td>10827</td><td>30</td><td>CATGTTGGAACAATGTAGGTAAGGGAAGTC</td><td>212.0</td><td>-6.4</td><td>1.4</td><td>-4.5</td><td>5.4E-06</td><td>9.6E-06</td><td>MB231/MB468</td><td>MP</td></tr>
</table>

| | | | | | | | | | | | | | |
|---|---|---|---|---|---|---|---|---|---|---|---|---|---|
| >am_trna128_GlyGCC_6_-_27870686_27870756@1.34.34,trna18_GlyGCC_16_+_70822597_70822667@1.34.34,trna19_GlyGCC_16_+_70823410_70823480@1.34.34,trna19_GlyGCC_2_-_157257659_157257729@1.34.34,trna24_GlyGCC_16_-_70812942_70813012@1.34.34,trna25_GlyGCC_16_-_70812114_70812184@1.34.34,trna5_GlyGCC_17_+_8029064_8029134@1.34.34,trna68_GlyGCC_1_-_161493637_161493707@1.34.34\|GCATTGGTGGTTCAGTGGTAGAATTCTCGCCTGC | >am_trna128_GlyGCC_6_-_27870686_27870756 | 1 | 34 | 34 | GCATTGGTGGTTCAGTGGTAGAATTCTCGCCTGC | 3841.0 | -4.9 | 1.0 | -5.0 | 5.7E-07 | 9.6E-06 | BT20/MB231 | CYTO |
| >hg38_21_+_8433172_8446622_RNA45SN1_45S_50ntFlanks@10387.10424.38\|TGTGAACAGCAGTTGAACATGGGTCAGTCGGTCCTGAG | >hg38_21_+_8433172_8446622_RNA45SN1_45S_50ntFlanks | 10387 | 10424 | 38 | TGTGAACAGCAGTTGAACATGGGTCAGTCGGTCCTGAG | 522.6 | 3.5 | 0.8 | 4.5 | 6.6E-06 | 9.6E-06 | MB231/MB468 | MITO |
| >hg38_21_+_8433172_8446622_RNA45SN1_45S_50ntFlanks@11572.11599.28\|CCGACTTAGAACTGGTGCGGACCAGGGG | >hg38_21_+_8433172_8446622_RNA45SN1_45S_50ntFlanks | 11572 | 11599 | 28 | CCGACTTAGAACTGGTGCGGACCAGGGG | 332.2 | -4.8 | 1.1 | -4.5 | 6.6E-06 | 9.7E-06 | MB231/MB468 | MITO |
| >hg38_21_+_8433172_8446622_RNA45SN1_45S_50ntFlanks@5485.5526.42\|AGCGCTGAGAAGACGGTCGAACTTGACTATCTAGAGGAAGTA | >hg38_21_+_8433172_8446622_RNA45SN1_45S_50ntFlanks | 5485 | 5526 | 42 | AGCGCTGAGAAGACGGTCGAACTTGACTATCTAGAGGAAGTA | 1391.3 | -6.7 | 1.5 | -4.5 | 6.9E-06 | 9.7E-06 | BT20/MB468 | NUC |
| >hg38_MT_+_1621_3279_RNR2_16S_50ntFlanks@715.740.26\|ATAAGCCTGCGTCAGATTAAAACACT | >hg38_MT_+_1621_3279_RNR2_16S_50ntFlanks | 715 | 740 | 26 | ATAAGCCTGCGTCAGATTAAAACACT | 38.2 | 8.5 | 1.9 | 4.5 | 5.5E-06 | 9.7E-06 | MB231/MB468 | MP |
| >hg38_21_+_8433172_8446622_RNA45SN1_45S_50ntFlanks@5482.5514.33\|CGGAGCGCTGAGAAGACGGTCGAACTTGACTAT | >hg38_21_+_8433172_8446622_RNA45SN1_45S_50ntFlanks | 5482 | 5514 | 33 | CGGAGCGCTGAGAAGACGGTCGAACTTGACTAT | 392.4 | -6.6 | 1.5 | -4.5 | 6.5E-06 | 9.7E-06 | MB231/MB468 | NUC |
| >hg38_21_+_8433172_8446622_RNA45SN1_45S_50ntFlanks@4731.4759.29\|AAGAACGAAAGTCGGAGGTTCGAAGACGA | >hg38_21_+_8433172_8446622_RNA45SN1_45S_50ntFlanks | 4731 | 4759 | 29 | AAGAACGAAAGTCGGAGGTTCGAAGACGA | 1132.8 | -2.8 | 0.6 | -4.6 | 4.7E-06 | 9.7E-06 | BT20/MB468 | MITO |
| >hg38_21_+_8433172_8446622_RNA45SN1_45S_50ntFlanks@10240.10259.20\|CTACGCCGCGACGAGTAGGA | >hg38_21_+_8433172_8446622_RNA45SN1_45S_50ntFlanks | 10240 | 10259 | 20 | CTACGCCGCGACGAGTAGGA | 439.4 | 5.4 | 1.2 | 4.5 | 5.6E-06 | 9.7E-06 | BT20/MB468 | CYTO |
| >hg38_21_+_8433172_8446622_RNA45SN1_45S_50ntFlanks@5083.5110.28\|ACGAACGAGACTCTGGCATGCTAACTAG | >hg38_21_+_8433172_8446622_RNA45SN1_45S_50ntFlanks | 5083 | 5110 | 28 | ACGAACGAGACTCTGGCATGCTAACTAG | 774.8 | -8.0 | 1.8 | -4.5 | 6.9E-06 | 9.7E-06 | BT20/MB468 | NUC |
| >hg38_MT_+_598_1651_RNR1_12S_50ntFlanks@428.455.28\|AAAGTGGCTTTAACATATCTGAACACAC | >hg38_MT_+_598_1651_RNR1_12S_50ntFlanks | 428 | 455 | 28 | AAAGTGGCTTTAACATATCTGAACACAC | 341.4 | 3.8 | 0.8 | 4.5 | 6.6E-06 | 9.7E-06 | MB231/MB468 | MITO |
| >hg38_21_+_8433172_8446622_RNA45SN1_45S_50ntFlanks@12634.12654.21\|GAAGCTACCATCTGTGGGATT | >hg38_21_+_8433172_8446622_RNA45SN1_45S_50ntFlanks | 12634 | 12654 | 21 | GAAGCTACCATCTGTGGGATT | 843.4 | -7.2 | 1.5 | -4.8 | 2.0E-06 | 9.7E-06 | BT20/MB468 | MP |
| >MI0000746\|hsa-mir-99b&WithFlank&19\|+\|51692606\|51692687@13.33.21(+1U)&[MIMAT0000689&hsa-miR-99b-5p&offsets\|0\|-1(+1U);m-10&19\|+\|51692618\|51692639&offsets\|0\|-1(+1U)]\|CACCCGTAGAACCGACCTTGCT | >MI0000746\|hsa-mir-99b&WithFlank&19\|+\|51692606\|51692687 | 13 | 33 | 21(+1U) | CACCCGTAGAACCGACCTTGCT | 193.9 | 2.0 | 0.5 | 4.5 | 7.0E-06 | 9.7E-06 | BT20/MB468 | NUC |
| >hg38_21_+_8433172_8446622_RNA45SN1_45S_50ntFlanks@4243.4263.21\|CTTTCGAGGCCCTGTAATTGG | >hg38_21_+_8433172_8446622_RNA45SN1_45S_50ntFlanks | 4243 | 4263 | 21 | CTTTCGAGGCCCTGTAATTGG | 260.7 | -6.9 | 1.5 | -4.5 | 5.5E-06 | 9.7E-06 | MB231/MB468 | MP |
| >hg38_21_+_8433172_8446622_RNA45SN1_45S_50ntFlanks@3775.3798.24\|GCCGGTACAGTGAAACTGCGAATG | >hg38_21_+_8433172_8446622_RNA45SN1_45S_50ntFlanks | 3775 | 3798 | 24 | GCCGGTACAGTGAAACTGCGAATG | 308.9 | -6.6 | 1.5 | -4.5 | 6.7E-06 | 9.7E-06 | MB231/MB468 | MITO |
| >hg38_21_+_8433172_8446622_RNA45SN1_45S_50ntFlanks@11781.11804.24\|CGCATGAATGGATGAACGAGATTC | >hg38_21_+_8433172_8446622_RNA45SN1_45S_50ntFlanks | 11781 | 11804 | 24 | CGCATGAATGGATGAACGAGATTC | 192.1 | 10.9 | 2.2 | 5.0 | 5.6E-07 | 9.7E-06 | BT20/MB231 | MP |
| >hg38_21_+_8433172_8446622_RNA45SN1_45S_50ntFlanks@3988.4024.37\|CTTTGGTGACTCTAGATAACCTCGGGCCGATCGCACG | >hg38_21_+_8433172_8446622_RNA45SN1_45S_50ntFlanks | 3988 | 4024 | 37 | CTTTGGTGACTCTAGATAACCTCGGGCCGATCGCACG | 682.9 | -3.5 | 0.8 | -4.5 | 6.5E-06 | 9.7E-06 | MB231/MB468 | NUC |
| >hg38_MT_+_1621_3279_RNR2_16S_50ntFlanks@1216.1248.33\|CAACCTCCGAGCAGTACATGCTAAGACTTCACC | >hg38_MT_+_1621_3279_RNR2_16S_50ntFlanks | 1216 | 1248 | 33 | CAACCTCCGAGCAGTACATGCTAAGACTTCACC | 555.3 | 4.3 | 1.0 | 4.5 | 6.7E-06 | 9.7E-06 | MB231/MB468 | MITO |
| >hg38_21_+_8433172_8446622_RNA45SN1_45S_50ntFlanks@4728.4758.31\|ATCAAGAACGAAAGTCGGAGGTTCGAAGACG | >hg38_21_+_8433172_8446622_RNA45SN1_45S_50ntFlanks | 4728 | 4758 | 31 | ATCAAGAACGAAAGTCGGAGGTTCGAAGACG | 3132.0 | -7.2 | 1.6 | -4.5 | 6.7E-06 | 9.7E-06 | MB231/MB468 | MITO |
| >hg38_21_+_8433172_8446622_RNA45SN1_45S_50ntFlanks@10516.10533.18\|GAGTGGCGGAGATGGGCG | >hg38_21_+_8433172_8446622_RNA45SN1_45S_50ntFlanks | 10516 | 10533 | 18 | GAGTGGCGGAGATGGGCG | 3888.4 | -5.1 | 1.1 | -4.8 | 2.0E-06 | 9.8E-06 | BT20/MB468 | MP |
| >MI0000279\|hsa-mir-196a-2&WithFlank&12\|+\|53991732\|53991853@31.51.21(+1U);MI0000238\|hsa-mir-196a-1&WithFlank&17\|-\|48632484\|48632565@13.33.21(+1U)&[MIMAT0000226&hsa-miR-196a-5p&offsets\|0\|-1(+1U);m-261&12\|+\|53991762\|53991783&offsets\|0\|-1(+1U)];[MIMAT0000226_1&hsa-miR-196a-5p&offsets\|0\|-1(+1U);m-268&17\|-\|48632532\|48632553&offsets\|0\|-1(+1U)]\|TAGGTAGTTTCATGTTGTTGGT | >MI0000279\|hsa-mir-196a-2&WithFlank&12\|+\|53991732\|53991853 | 31 | 51 | 21(+1U) | TAGGTAGTTTCATGTTGTTGGT | 112.2 | 10.1 | 2.0 | 5.0 | 5.6E-07 | 9.8E-06 | BT20/MB231 | MP |
| >hg38_21_+_8433172_8446622_RNA45SN1_45S_50ntFlanks@8114.8136.23\|GCGCGGGACATGTGGCGTACGGA | >hg38_21_+_8433172_8446622_RNA45SN1_45S_50ntFlanks | 8114 | 8136 | 23 | GCGCGGGACATGTGGCGTACGGA | 1959.0 | 10.2 | 2.0 | 5.0 | 5.7E-07 | 9.8E-06 | BT20/MB231 | MP |
| >MI0000298\|hsa-mir-221&WithFlank&X\|-\|45746151\|45746272@31.54.24&[MIMAT0004568&hsa-miR-221-5p&offsets\|0\|+2;m-405&X\|-\|45746220\|45746242&offsets\|0\|+1]\|ACCTGGCATACAATGTAGATTTCT | >MI0000298\|hsa-mir-221&WithFlank&X\|-\|45746151\|45746272 | 31 | 54 | 24 | ACCTGGCATACAATGTAGATTTCT | 513.5 | -4.6 | 0.9 | -5.0 | 5.7E-07 | 9.8E-06 | BT20/MB231 | MP |
| >hg38_21_+_8433172_8446622_RNA45SN1_45S_50ntFlanks@10361.10390.30\|GAAGGCCGAAGTGGAGAAGGGTTCCATGTG | >hg38_21_+_8433172_8446622_RNA45SN1_45S_50ntFlanks | 10361 | 10390 | 30 | GAAGGCCGAAGTGGAGAAGGGTTCCATGTG | 530.4 | -4.8 | 1.1 | -4.5 | 6.3E-06 | 9.8E-06 | MB231/MB468 | CYTO |
| >MI0000063\|hsa-let-7b&WithFlank&22\|+\|46113680\|46113774@12.32.21(+1A)&[MIMAT0000063&hsa-let-7b-5p&offsets\|0\|-1(+1A);m-14&22\|+\|46113691\|46113712&offsets\|0\|-1(+1A)]\|TGAGGTAGTAGGTTGTGTGGTA | >MI0000063\|hsa-let-7b&WithFlank&22\|+\|46113680\|46113774 | 12 | 32 | 21(+1A) | TGAGGTAGTAGGTTGTGTGGTA | 2421.2 | 5.0 | 1.1 | 4.8 | 2.0E-06 | 9.8E-06 | BT20/MB468 | MP |
| >hg38_1_-_228634819_228635039_RNA5S12_5S_50ntFlanks@127.170.44\|ACTTGGATGGGAGACCGCCTGGGAATACCGGGTGCTGTAGGCTT | >hg38_1_-_228634819_228635039_RNA5S12_5S_50ntFlanks | 127 | 170 | 44 | ACTTGGATGGGAGACCGCCTGGGAATACCGGGTGCTGTAGGCTT | 941.8 | 5.4 | 1.2 | 4.5 | 6.6E-06 | 9.8E-06 | MB231/MB468 | NUC |
| >trna134_GluTTC_1_-_16861774_16861845@1.31.31,trna5_GluTTC_1_+_17199078_17199149@1.31.31,trna84_GluTTC_1_-_161391883_161391954@1.31.31,trna94_GluTTC_1_-_149664355_149664427@1.31.31\|TCCCTGGTGGTCTAGTGGCTAGGATTCGGCG | >trna134_GluTTC_1_-_16861774_16861845 | 1 | 31 | 31 | TCCCTGGTGGTCTAGTGGCTAGGATTCGGCG | 4936.2 | -3.0 | 0.7 | -4.5 | 5.6E-06 | 9.8E-06 | BT20/MB468 | CYTO |
| >hg38_21_+_8433172_8446622_RNA45SN1_45S_50ntFlanks@12157.12181.25\|AGTTTGACTGGGGCGGTACACCTGT | >hg38_21_+_8433172_8446622_RNA45SN1_45S_50ntFlanks | 12157 | 12181 | 25 | AGTTTGACTGGGGCGGTACACCTGT | 214.3 | -5.7 | 1.2 | -4.5 | 5.6E-06 | 9.8E-06 | BT20/MB468 | CYTO |
| >MI0000342\|hsa-mir-200b&WithFlank&1\|+\|1167098\|1167204@63.84.22(+1U)&[MIMAT0000318&hsa-miR-200b-3p&offsets\|0\|0(+1U);m-93&1\|+\|1167160\|1167182&offsets\|0\|-1(+1U)]\|TAATACTGCCTGGTAATGATGAT | >MI0000342\|hsa-mir-200b&WithFlank&1\|+\|1167098\|1167204 | 63 | 84 | 22(+1U) | TAATACTGCCTGGTAATGATGAT | 348.2 | -5.0 | 1.1 | -4.5 | 6.6E-06 | 9.8E-06 | MB231/MB468 | NUC |
| >MI0001445\|hsa-mir-423&WithFlank&17\|+\|30117073\|30117178@23.46.24&[MIMAT0004748&hsa-miR-423-5p&offsets\|0\|+1;m-141&17\|+\|30117095\|30117117&offsets\|0\|+1]\|TGAGGGGCAGAGAGCGAGACTTTT | >MI0001445\|hsa-mir-423&WithFlank&17\|+\|30117073\|30117178 | 23 | 46 | 24 | TGAGGGGCAGAGAGCGAGACTTTT | 238.8 | 6.5 | 1.5 | 4.5 | 6.7E-06 | 9.8E-06 | MB231/MB468 | MITO |
| >trnaMT_HisGTG_MT_+_12138_12206@27.62.36\|GATTGTGAATCTGACAACAGAGGCTTACGACCCCTT | >trnaMT_HisGTG_MT_+_12138_12206 | 27 | 62 | 36 | GATTGTGAATCTGACAACAGAGGCTTACGACCCCTT | 64.7 | 9.8 | 2.1 | 4.8 | 2.0E-06 | 9.8E-06 | BT20/MB468 | MP |
| >trna10_AspGTC_12_-_125424193_125424264@1.34.34,trna12_AspGTC_12_-_125411891_125411962@1.34.34,trna38_AspGTC_17_-_8125556_8125627@1.34.34,trna45_AspGTC_6_+_27447453_27447524@1.34.34,trna48_AspGTC_6_+_27471523_27471594@1.34.34,trna4_AspGTC_12_+_96429799_96429870@1.34.34,trna69_AspGTC_1_-_161440205_161440276@1.34.34,trna72_AspGTC_1_-_161432824_161432895@1.34.34,trna75_AspGTC_1_-_161425414_161425485@1.34.34,trna78_AspGTC_1_-_161418033_161418104@1.34.34,trna81_AspGTC_1_-_161410615_161410686@1.34.34\|TCCTCGTTAGTATAGTGGTGAGTATCCCCGCCTG | >trna10_AspGTC_12_-_125424193_125424264 | 1 | 34 | 34 | TCCTCGTTAGTATAGTGGTGAGTATCCCCGCCTG | 409.4 | 4.6 | 1.0 | 4.5 | 6.7E-06 | 9.8E-06 | MB231/MB468 | MITO |
| >hg38_21_+_8433172_8446622_RNA45SN1_45S_50ntFlanks@12081.12106.26\|CTCTCGCTTCTGGCGCCAAGCGCCCG | >hg38_21_+_8433172_8446622_RNA45SN1_45S_50ntFlanks | 12081 | 12106 | 26 | CTCTCGCTTCTGGCGCCAAGCGCCCG | 483.1 | -7.3 | 1.6 | -4.5 | 7.0E-06 | 9.8E-06 | BT20/MB468 | NUC |
| >MI0000734\|hsa-mir-106b&WithFlank&7\|-\|100093987\|100094080@58.77.20&[MIMAT0004672&hsa-miR-106b-3p&offsets\|0\|-2;m-87&7\|-\|100094003\|100094023&offsets\|0\|-1]\|CCGCACTGTGGGTACTTGCT | >MI0000734\|hsa-mir-106b&WithFlank&7\|-\|100093987\|100094080 | 58 | 77 | 20 | CCGCACTGTGGGTACTTGCT | 1137.4 | 5.5 | 1.2 | 4.8 | 2.0E-06 | 9.8E-06 | BT20/MB468 | MP |
| >MI0000809\|hsa-mir-151a&WithFlank&8\|-\|140732558\|140732659@51.72.22(+1U)&[MIMAT0000757&hsa-miR-151a-3p&offsets\|-2\|-1(+1U);m-30&8\|-\|140732587\|140732607&offsets\|-2\|-1(+1U)]\|TACTAGACTGAAGCTCCTTGAGT | >MI0000809\|hsa-mir-151a&WithFlank&8\|-\|140732558\|140732659 | 51 | 72 | 22(+1U) | TACTAGACTGAAGCTCCTTGAGT | 221.1 | 4.2 | 0.9 | 4.8 | 2.0E-06 | 9.8E-06 | BT20/MB468 | MP |
| >hg38_21_+_8433172_8446622_RNA45SN1_45S_50ntFlanks@11682.11700.19\|AATGTCAAAGTGAAGAAAT | >hg38_21_+_8433172_8446622_RNA45SN1_45S_50ntFlanks | 11682 | 11700 | 19 | AATGTCAAAGTGAAGAAAT | 63.4 | -9.1 | 1.9 | -4.8 | 2.0E-06 | 9.8E-06 | BT20/MB468 | MP |
| >hg38_21_+_8433172_8446622_RNA45SN1_45S_50ntFlanks@5073.5094.22\|AATTCCGATAACGAACGAGACT | >hg38_21_+_8433172_8446622_RNA45SN1_45S_50ntFlanks | 5073 | 5094 | 22 | AATTCCGATAACGAACGAGACT | 1147.7 | -6.9 | 1.5 | -4.5 | 6.7E-06 | 9.9E-06 | MB231/MB468 | MITO |
| >hg38_21_+_8433172_8446622_RNA45SN1_45S_50ntFlanks@9939.9979.41\|GCAGGACGGTGGCCATGGAAGTCGGAATCCGCTAAGGAGTG | >hg38_21_+_8433172_8446622_RNA45SN1_45S_50ntFlanks | 9939 | 9979 | 41 | GCAGGACGGTGGCCATGGAAGTCGGAATCCGCTAAGGAGTG | 140.7 | 10.5 | 2.1 | 5.0 | 5.7E-07 | 9.9E-06 | BT20/MB231 | MP |
| >MI0000266\|hsa-mir-10a&WithFlank&17\|-\|48579832\|48579953@28.50.23(+1A)&[MIMAT0000253&hsa-miR-10a-5p&offsets\|0\|0(+1A);m-18&17\|-\|48579905\|48579926&offsets\|0\|+1(+1A)]\|TACCCTGTAGATCCGAATTTGTGA | >MI0000266\|hsa-mir-10a&WithFlank&17\|-\|48579832\|48579953 | 28 | 50 | 23(+1A) | TACCCTGTAGATCCGAATTTGTGA | 308.2 | 8.2 | 1.7 | 4.7 | 2.0E-06 | 9.9E-06 | BT20/MB468 | MP |
| >hg38_21_+_8433172_8446622_RNA45SN1_45S_50ntFlanks@10571.10591.21\|GAGAAGCCGGCGGGAGCCCCG | >hg38_21_+_8433172_8446622_RNA45SN1_45S_50ntFlanks | 10571 | 10591 | 21 | GAGAAGCCGGCGGGAGCCCCG | 612.1 | -6.0 | 1.3 | -4.5 | 6.3E-06 | 9.9E-06 | MB231/MB468 | CYTO |
| >MI0000234\|hsa-mir-192&WithFlank&11\|-\|64891131\|64891252@30.49.20(+1U)&[MIMAT0000222&hsa-miR-192-5p&offsets\|0\|-1(+1U);m-59&11\|-\|64891203\|64891223&offsets\|0\|-1(+1U)]\|CTGACCTATGAATTGACAGCT | >MI0000234\|hsa-mir-192&WithFlank&11\|-\|64891131\|64891252 | 30 | 49 | 20(+1U) | CTGACCTATGAATTGACAGCT | 507.9 | 5.0 | 1.1 | 4.5 | 6.8E-06 | 9.9E-06 | MB231/MB468 | MITO |

<table>
<tr><td>>hg38_21_+_8433172_8446622_RNA45SN1_45S_50ntFlanks@11641.11667.27|CGGCGGGTGTTGACGCGATGTGATTTC</td><td>>hg38_21_+_8433172_8446622_RNA45SN1_45S_50ntFlanks</td><td>11641</td><td>11667</td><td>27</td><td>CGGCGGGTGTTGACGCGATGTGATTTC</td><td>1365.4</td><td>-8.1</td><td>1.8</td><td>-4.5</td><td>7.1E-06</td><td>9.9E-06</td><td>BT20/MB468</td><td>NUC</td></tr>
<tr><td>>MI0000077|hsa-mir-21&WithFlank&17|+|59841260|59841343@14.34.21(+2U)&[MIMAT0000076&hsa-miR-21-5p&offsets|0|-1(+2U);m-1&17|+|59841273|59841294&offsets|0|-1(+2U)]|TAGCTTATCAGACTGATGTTGTT</td><td>>MI0000077|hsa-mir-21&WithFlank&17|+|59841260|59841343</td><td>14</td><td>34</td><td>21(+2U)</td><td>TAGCTTATCAGACTGATGTTGTT</td><td>467.0</td><td>8.6</td><td>1.9</td><td>4.5</td><td>5.7E-06</td><td>9.9E-06</td><td>BT20/MB468</td><td>CYTO</td></tr>
<tr><td>>hg38_21_+_8433172_8446622_RNA45SN1_45S_50ntFlanks@12273.12318.46|TCAGTACGAATACAGACCGTGAAAGCGGGGCCTCACGATCCTTCTG</td><td>>hg38_21_+_8433172_8446622_RNA45SN1_45S_50ntFlanks</td><td>12273</td><td>12318</td><td>46</td><td>TCAGTACGAATACAGACCGTGAAAGCGGGGCCTCACGATCCTTCTG</td><td>75.2</td><td>-8.3</td><td>1.8</td><td>-4.6</td><td>4.8E-06</td><td>9.9E-06</td><td>BT20/MB468</td><td>MITO</td></tr>
<tr><td>>hg38_MT_+_1621_3279_RNR2_16S_50ntFlanks@623.656.34|ATAACTGAACTCCTCACACCCAATTGGACCAATC</td><td>>hg38_MT_+_1621_3279_RNR2_16S_50ntFlanks</td><td>623</td><td>656</td><td>34</td><td>ATAACTGAACTCCTCACACCCAATTGGACCAATC</td><td>56.9</td><td>9.6</td><td>2.0</td><td>4.7</td><td>2.0E-06</td><td>9.9E-06</td><td>BT20/MB468</td><td>MP</td></tr>
<tr><td>>hg38_21_+_8433172_8446622_RNA45SN1_45S_50ntFlanks@12622.12654.33|AGCCAATGGGGCGAAGCTACCATCTGTGGGATT</td><td>>hg38_21_+_8433172_8446622_RNA45SN1_45S_50ntFlanks</td><td>12622</td><td>12654</td><td>33</td><td>AGCCAATGGGGCGAAGCTACCATCTGTGGGATT</td><td>440.5</td><td>-7.3</td><td>1.6</td><td>-4.5</td><td>7.1E-06</td><td>9.9E-06</td><td>BT20/MB468</td><td>NUC</td></tr>
<tr><td>>hg38_21_+_8433172_8446622_RNA45SN1_45S_50ntFlanks@11534.11553.20|GTTCCGCGCGGCGCCTCGCC</td><td>>hg38_21_+_8433172_8446622_RNA45SN1_45S_50ntFlanks</td><td>11534</td><td>11553</td><td>20</td><td>GTTCCGCGCGGCGCCTCGCC</td><td>1055.0</td><td>-6.7</td><td>1.5</td><td>-4.5</td><td>5.7E-06</td><td>9.9E-06</td><td>BT20/MB468</td><td>CYTO</td></tr>
<tr><td>>MI0003560|hsa-mir-92b&WithFlank&1|+|155195171|155195278@67.85.19(+1A)&[MIMAT0003218&hsa-miR-92b-3p&offsets|0|-3(+1A);m-115&1|+|155195237|155195258&offsets|0|-3(+1A)]|TATTGCACTCGTCCCGGCCA</td><td>>MI0003560|hsa-mir-92b&WithFlank&1|+|155195171|155195278</td><td>67</td><td>85</td><td>19(+1A)</td><td>TATTGCACTCGTCCCGGCCA</td><td>51.2</td><td>8.9</td><td>2.0</td><td>4.5</td><td>5.6E-06</td><td>9.9E-06</td><td>MB231/MB468</td><td>MP</td></tr>
<tr><td>>hg38_21_+_8433172_8446622_RNA45SN1_45S_50ntFlanks@9539.9569.31|ACTCTGGTGGAGGTCCGTAGCGGTCCTGACG</td><td>>hg38_21_+_8433172_8446622_RNA45SN1_45S_50ntFlanks</td><td>9539</td><td>9569</td><td>31</td><td>ACTCTGGTGGAGGTCCGTAGCGGTCCTGACG</td><td>1187.5</td><td>-8.1</td><td>1.8</td><td>-4.5</td><td>7.1E-06</td><td>9.9E-06</td><td>BT20/MB468</td><td>NUC</td></tr>
<tr><td>>hg38_21_+_8433172_8446622_RNA45SN1_45S_50ntFlanks@10678.10695.18|CGGTTCCGGCGGCGTCCG</td><td>>hg38_21_+_8433172_8446622_RNA45SN1_45S_50ntFlanks</td><td>10678</td><td>10695</td><td>18</td><td>CGGTTCCGGCGGCGTCCG</td><td>129.1</td><td>-5.3</td><td>1.2</td><td>-4.5</td><td>5.7E-06</td><td>9.9E-06</td><td>BT20/MB468</td><td>CYTO</td></tr>
<tr><td>>trnaMT_PheGAA_MT_+_577_647@36.60.25|AAAATGTTTAGACGGGCTCACATCA</td><td>>trnaMT_PheGAA_MT_+_577_647</td><td>36</td><td>60</td><td>25</td><td>AAAATGTTTAGACGGGCTCACATCA</td><td>33.0</td><td>8.3</td><td>1.8</td><td>4.5</td><td>5.6E-06</td><td>9.9E-06</td><td>MB231/MB468</td><td>MP</td></tr>
<tr><td>>hg38_21_+_8433172_8446622_RNA45SN1_45S_50ntFlanks@5492.5514.23|AGAAGACGGTCGAACTTGACTAT</td><td>>hg38_21_+_8433172_8446622_RNA45SN1_45S_50ntFlanks</td><td>5492</td><td>5514</td><td>23</td><td>AGAAGACGGTCGAACTTGACTAT</td><td>1819.8</td><td>-4.7</td><td>1.0</td><td>-4.5</td><td>6.4E-06</td><td>9.9E-06</td><td>MB231/MB468</td><td>CYTO</td></tr>
<tr><td>>hg38_21_+_8433172_8446622_RNA45SN1_45S_50ntFlanks@11580.11612.33|GAACTGGTGCGGACCAGGGGAATCCGACTGTTT</td><td>>hg38_21_+_8433172_8446622_RNA45SN1_45S_50ntFlanks</td><td>11580</td><td>11612</td><td>33</td><td>GAACTGGTGCGGACCAGGGGAATCCGACTGTTT</td><td>265.3</td><td>-4.9</td><td>1.1</td><td>-4.5</td><td>6.8E-06</td><td>9.9E-06</td><td>MB231/MB468</td><td>MITO</td></tr>
<tr><td>>hg38_21_+_8433172_8446622_RNA45SN1_45S_50ntFlanks@10866.10890.25|CTAAGGGCTGGGTCGGTCGGGCTGG</td><td>>hg38_21_+_8433172_8446622_RNA45SN1_45S_50ntFlanks</td><td>10866</td><td>10890</td><td>25</td><td>CTAAGGGCTGGGTCGGTCGGGCTGG</td><td>1773.5</td><td>-8.1</td><td>1.8</td><td>-4.5</td><td>7.1E-06</td><td>9.9E-06</td><td>BT20/MB468</td><td>NUC</td></tr>
<tr><td>>hg38_21_+_8433172_8446622_RNA45SN1_45S_50ntFlanks@12589.12616.28|GTTCAGACATTTGGTGTATGTGCTTGGC</td><td>>hg38_21_+_8433172_8446622_RNA45SN1_45S_50ntFlanks</td><td>12589</td><td>12616</td><td>28</td><td>GTTCAGACATTTGGTGTATGTGCTTGGC</td><td>55.1</td><td>-8.9</td><td>1.9</td><td>-4.7</td><td>2.1E-06</td><td>9.9E-06</td><td>BT20/MB468</td><td>MP</td></tr>
<tr><td>>hg38_21_+_8433172_8446622_RNA45SN1_45S_50ntFlanks@6651.6687.37|CGACTCTTAGCGGTGGATCACTCGGCTCGTGCGTCGA</td><td>>hg38_21_+_8433172_8446622_RNA45SN1_45S_50ntFlanks</td><td>6651</td><td>6687</td><td>37</td><td>CGACTCTTAGCGGTGGATCACTCGGCTCGTGCGTCGA</td><td>782.7</td><td>4.6</td><td>1.0</td><td>4.5</td><td>6.4E-06</td><td>9.9E-06</td><td>MB231/MB468</td><td>CYTO</td></tr>
<tr><td>>MI0000746|hsa-mir-99b&WithFlank&19|+|51692606|51692687@13.33.21&[MIMAT0000689&hsa-miR-99b-5p&offsets|0|-1;m-10&19|+|51692618|51692639&offsets|0|-1]|CACCCGTAGAACCGACCTTGC</td><td>>MI0000746|hsa-mir-99b&WithFlank&19|+|51692606|51692687</td><td>13</td><td>33</td><td>21</td><td>CACCCGTAGAACCGACCTTGC</td><td>7528.8</td><td>4.4</td><td>0.9</td><td>4.7</td><td>2.1E-06</td><td>9.9E-06</td><td>BT20/MB468</td><td>MP</td></tr>
<tr><td>>trna10_LysCTT_16_+_3241501_3241573@3.33.31,trna13_LysCTT_14_-_58706613_58706685@3.33.31,trna2_LysCTT_15_+_79152904_79152976@3.33.31|CCGGCTAGCTCAGTCGGTAGAGCATGGGACT</td><td>>trna10_LysCTT_16_+_3241501_3241573</td><td>3</td><td>33</td><td>31</td><td>CCGGCTAGCTCAGTCGGTAGAGCATGGGACT</td><td>238.8</td><td>6.6</td><td>1.5</td><td>4.5</td><td>5.7E-06</td><td>1.0E-05</td><td>BT20/MB468</td><td>CYTO</td></tr>
<tr><td>>MI0000101|hsa-mir-99a&WithFlank&21|+|16539083|16539175@19.38.20&[MIMAT0000097&hsa-miR-99a-5p&offsets|0|-2;m-64&21|+|16539101|16539121&offsets|0|-1]|AACCCGTAGATCCGATCTTG</td><td>>MI0000101|hsa-mir-99a&WithFlank&21|+|16539083|16539175</td><td>19</td><td>38</td><td>20</td><td>AACCCGTAGATCCGATCTTG</td><td>46.5</td><td>-8.6</td><td>1.8</td><td>-4.7</td><td>2.1E-06</td><td>1.0E-05</td><td>BT20/MB468</td><td>MP</td></tr>
<tr><td>>hg38_21_+_8433172_8446622_RNA45SN1_45S_50ntFlanks@10441.10460.20|TTCCGAAGGGACGGGCGATG</td><td>>hg38_21_+_8433172_8446622_RNA45SN1_45S_50ntFlanks</td><td>10441</td><td>10460</td><td>20</td><td>TTCCGAAGGGACGGGCGATG</td><td>328.5</td><td>-7.0</td><td>1.5</td><td>-4.5</td><td>5.6E-06</td><td>1.0E-05</td><td>MB231/MB468</td><td>MP</td></tr>
<tr><td>>hg38_21_+_8433172_8446622_RNA45SN1_45S_50ntFlanks@11598.11622.25|GGAATCCGACTGTTTAATTAAAACA</td><td>>hg38_21_+_8433172_8446622_RNA45SN1_45S_50ntFlanks</td><td>11598</td><td>11622</td><td>25</td><td>GGAATCCGACTGTTTAATTAAAACA</td><td>136.1</td><td>10.4</td><td>2.1</td><td>5.0</td><td>5.8E-07</td><td>1.0E-05</td><td>BT20/MB231</td><td>MP</td></tr>
<tr><td>>hg38_21_+_8433172_8446622_RNA45SN1_45S_50ntFlanks@4297.4314.18|CATTGGAGGGCAAGTCTG</td><td>>hg38_21_+_8433172_8446622_RNA45SN1_45S_50ntFlanks</td><td>4297</td><td>4314</td><td>18</td><td>CATTGGAGGGCAAGTCTG</td><td>263.6</td><td>-6.4</td><td>1.4</td><td>-4.5</td><td>6.8E-06</td><td>1.0E-05</td><td>MB231/MB468</td><td>MITO</td></tr>
<tr><td>>am_MI0000809|hsa-mir-151a&WithFlank&8|-|140732558|140732659@53.72.20&[MIMAT0000757&hsa-miR-151a-3p&offsets|0|-1;m-30&8|-|140732587|140732607&offsets|0|-1]|CTAGACTGAAGCTCCTTGAG</td><td>>am_MI0000809|hsa-mir-151a&WithFlank&8|-|140732558|140732659</td><td>53</td><td>72</td><td>20</td><td>CTAGACTGAAGCTCCTTGAG</td><td>1642.9</td><td>3.3</td><td>0.7</td><td>4.5</td><td>5.6E-06</td><td>1.0E-05</td><td>MB231/MB468</td><td>MP</td></tr>
<tr><td>>hg38_21_+_8433172_8446622_RNA45SN1_45S_50ntFlanks@5420.5441.22|CCGATTGGATGGTTTAGTGAGG</td><td>>hg38_21_+_8433172_8446622_RNA45SN1_45S_50ntFlanks</td><td>5420</td><td>5441</td><td>22</td><td>CCGATTGGATGGTTTAGTGAGG</td><td>48.1</td><td>-8.7</td><td>1.8</td><td>-4.7</td><td>2.1E-06</td><td>1.0E-05</td><td>BT20/MB468</td><td>MP</td></tr>
<tr><td>>hg38_21_+_8433172_8446622_RNA45SN1_45S_50ntFlanks@3838.3860.23|CTCTCCTACTTGGATAACTGTGG</td><td>>hg38_21_+_8433172_8446622_RNA45SN1_45S_50ntFlanks</td><td>3838</td><td>3860</td><td>23</td><td>CTCTCCTACTTGGATAACTGTGG</td><td>1162.1</td><td>-2.1</td><td>0.5</td><td>-4.5</td><td>5.7E-06</td><td>1.0E-05</td><td>BT20/MB468</td><td>CYTO</td></tr>
<tr><td>>hg38_21_+_8433172_8446622_RNA45SN1_45S_50ntFlanks@7979.8007.29|ACCTCAGATCAGACGTGGCGACCCGCTGA</td><td>>hg38_21_+_8433172_8446622_RNA45SN1_45S_50ntFlanks</td><td>7979</td><td>8007</td><td>29</td><td>ACCTCAGATCAGACGTGGCGACCCGCTGA</td><td>486.9</td><td>-5.5</td><td>1.2</td><td>-4.5</td><td>7.2E-06</td><td>1.0E-05</td><td>BT20/MB468</td><td>NUC</td></tr>
<tr><td>>hg38_21_+_8433172_8446622_RNA45SN1_45S_50ntFlanks@10357.10376.20|CTTTGAAGGCCGAAGTGGAG</td><td>>hg38_21_+_8433172_8446622_RNA45SN1_45S_50ntFlanks</td><td>10357</td><td>10376</td><td>20</td><td>CTTTGAAGGCCGAAGTGGAG</td><td>148.2</td><td>-3.3</td><td>0.7</td><td>-4.5</td><td>6.4E-06</td><td>1.0E-05</td><td>MB231/MB468</td><td>CYTO</td></tr>
<tr><td>>am_trna111_HisGTG_1_-_147774845_147774916@1.29.29,trna118_HisGTG_1_-_145396881_145396952@1.29.29,trna16_HisGTG_1_+_146544773_146544844@1.29.29,trna1_HisGTG_15_+_45493349_45493420@1.29.29,trna21_HisGTG_1_+_147753471_147753542@1.29.29,trna33_HisGTG_6_+_27125906_27125977@1.29.29,trna7_HisGTG_9_-_14433938_14434009@1.29.29,trna8_HisGTG_15_-_45492611_45492682@1.29.29,trna9_HisGTG_15_-_45490804_45490875@1.29.29|GCCGTGATCGTATAGTGGTTAGTACTCTG</td><td>>am_trna111_HisGTG_1_-_147774845_147774916</td><td>1</td><td>29</td><td>29</td><td>GCCGTGATCGTATAGTGGTTAGTACTCTG</td><td>2803.8</td><td>-3.9</td><td>0.9</td><td>-4.5</td><td>6.9E-06</td><td>1.0E-05</td><td>MB231/MB468</td><td>MITO</td></tr>
<tr><td>>hg38_21_+_8433172_8446622_RNA45SN1_45S_50ntFlanks@10783.10823.41|GTGAACAGCCTCTGGCATGTTGGAACAATGTAGGTAAGGGA</td><td>>hg38_21_+_8433172_8446622_RNA45SN1_45S_50ntFlanks</td><td>10783</td><td>10823</td><td>41</td><td>GTGAACAGCCTCTGGCATGTTGGAACAATGTAGGTAAGGGA</td><td>78.3</td><td>-10.2</td><td>2.2</td><td>-4.5</td><td>5.7E-06</td><td>1.0E-05</td><td>MB231/MB468</td><td>MP</td></tr>
<tr><td>>trnaMT_ValTAC_MT_+_1602_1670@29.72.44|ACTTACACTTAGGAGATTTCAACTTAACTTGACCGCTCTGACCA</td><td>>trnaMT_ValTAC_MT_+_1602_1670</td><td>29</td><td>72</td><td>44</td><td>ACTTACACTTAGGAGATTTCAACTTAACTTGACCGCTCTGACCA</td><td>1128.0</td><td>4.5</td><td>1.0</td><td>4.6</td><td>4.9E-06</td><td>1.0E-05</td><td>BT20/MB468</td><td>MITO</td></tr>
<tr><td>>hg38_21_+_8433172_8446622_RNA45SN1_45S_50ntFlanks@4224.4242.19|AAAATAACAATACAGGACT</td><td>>hg38_21_+_8433172_8446622_RNA45SN1_45S_50ntFlanks</td><td>4224</td><td>4242</td><td>19</td><td>AAAATAACAATACAGGACT</td><td>914.8</td><td>-3.9</td><td>0.9</td><td>-4.5</td><td>6.4E-06</td><td>1.0E-05</td><td>MB231/MB468</td><td>CYTO</td></tr>
<tr><td>>hg38_21_+_8433172_8446622_RNA45SN1_45S_50ntFlanks@8176.8201.26|CTTCTGATCGAGGCCCAGCCCGTGGA</td><td>>hg38_21_+_8433172_8446622_RNA45SN1_45S_50ntFlanks</td><td>8176</td><td>8201</td><td>26</td><td>CTTCTGATCGAGGCCCAGCCCGTGGA</td><td>950.1</td><td>-7.7</td><td>1.7</td><td>-4.5</td><td>7.2E-06</td><td>1.0E-05</td><td>BT20/MB468</td><td>NUC</td></tr>
<tr><td>>hg38_21_+_8433172_8446622_RNA45SN1_45S_50ntFlanks@10572.10591.20|AGAAGCCGGCGGGAGCCCCG</td><td>>hg38_21_+_8433172_8446622_RNA45SN1_45S_50ntFlanks</td><td>10572</td><td>10591</td><td>20</td><td>AGAAGCCGGCGGGAGCCCCG</td><td>344.4</td><td>-4.7</td><td>1.0</td><td>-4.6</td><td>4.9E-06</td><td>1.0E-05</td><td>BT20/MB468</td><td>MITO</td></tr>
<tr><td>>MI0000090|hsa-mir-32&WithFlank&9|-|109046223|109046304@12.33.22&[MIMAT0000090&hsa-miR-32-5p&offsets|0|0;m-208&9|-|109046273|109046293&offsets|0|+1]|TATTGCACATTACTAAGTTGCA</td><td>>MI0000090|hsa-mir-32&WithFlank&9|-|109046223|109046304</td><td>12</td><td>33</td><td>22</td><td>TATTGCACATTACTAAGTTGCA</td><td>487.5</td><td>5.0</td><td>1.1</td><td>4.5</td><td>6.9E-06</td><td>1.0E-05</td><td>MB231/MB468</td><td>MITO</td></tr>
<tr><td>>hg38_21_+_8433172_8446622_RNA45SN1_45S_50ntFlanks@11583.11601.19|CTGGTGCGGACCAGGGGAA</td><td>>hg38_21_+_8433172_8446622_RNA45SN1_45S_50ntFlanks</td><td>11583</td><td>11601</td><td>19</td><td>CTGGTGCGGACCAGGGGAA</td><td>2595.3</td><td>-8.3</td><td>1.9</td><td>-4.5</td><td>7.2E-06</td><td>1.0E-05</td><td>BT20/MB468</td><td>NUC</td></tr>
<tr><td>>hg38_21_+_8433172_8446622_RNA45SN1_45S_50ntFlanks@3850.3883.34|GATAACTGTGGTAATTCTAGAGCTAATACATGCC</td><td>>hg38_21_+_8433172_8446622_RNA45SN1_45S_50ntFlanks</td><td>3850</td><td>3883</td><td>34</td><td>GATAACTGTGGTAATTCTAGAGCTAATACATGCC</td><td>403.3</td><td>3.7</td><td>0.8</td><td>4.5</td><td>6.5E-06</td><td>1.0E-05</td><td>MB231/MB468</td><td>CYTO</td></tr>
<tr><td>>hg38_21_+_8433172_8446622_RNA45SN1_45S_50ntFlanks@8347.8378.32|GGGAAAGTTGAAAAGAACTTTGAAGAGAGAGT</td><td>>hg38_21_+_8433172_8446622_RNA45SN1_45S_50ntFlanks</td><td>8347</td><td>8378</td><td>32</td><td>GGGAAAGTTGAAAAGAACTTTGAAGAGAGAGT</td><td>196.7</td><td>6.9</td><td>1.5</td><td>4.7</td><td>2.1E-06</td><td>1.0E-05</td><td>BT20/MB468</td><td>MP</td></tr>
<tr><td>>hg38_21_+_8433172_8446622_RNA45SN1_45S_50ntFlanks@10412.10433.22|AGTCGGTCCTGAGAGATGGGCG</td><td>>hg38_21_+_8433172_8446622_RNA45SN1_45S_50ntFlanks</td><td>10412</td><td>10433</td><td>22</td><td>AGTCGGTCCTGAGAGATGGGCG</td><td>61.5</td><td>-9.0</td><td>1.9</td><td>-4.7</td><td>2.1E-06</td><td>1.0E-05</td><td>BT20/MB468</td><td>MP</td></tr>
<tr><td>>hg38_21_+_8433172_8446622_RNA45SN1_45S_50ntFlanks@11704.11724.21|ATGAAGCGCGGGTAAACGGCG</td><td>>hg38_21_+_8433172_8446622_RNA45SN1_45S_50ntFlanks</td><td>11704</td><td>11724</td><td>21</td><td>ATGAAGCGCGGGTAAACGGCG</td><td>65.9</td><td>-9.1</td><td>1.9</td><td>-4.7</td><td>2.1E-06</td><td>1.0E-05</td><td>BT20/MB468</td><td>MP</td></tr>
<tr><td>>hg38_21_+_8433172_8446622_RNA45SN1_45S_50ntFlanks@10401.10439.39|GAACATGGGTCAGTCGGTCCTGAGAGATGGGCGAGCGCC</td><td>>hg38_21_+_8433172_8446622_RNA45SN1_45S_50ntFlanks</td><td>10401</td><td>10439</td><td>39</td><td>GAACATGGGTCAGTCGGTCCTGAGAGATGGGCGAGCGCC</td><td>366.0</td><td>6.2</td><td>1.4</td><td>4.5</td><td>6.5E-06</td><td>1.0E-05</td><td>MB231/MB468</td><td>CYTO</td></tr>
<tr><td>>hg38_21_+_8433172_8446622_RNA45SN1_45S_50ntFlanks@12970.12991.22|GTTTCGTACGTAGCAGAGCAGC</td><td>>hg38_21_+_8433172_8446622_RNA45SN1_45S_50ntFlanks</td><td>12970</td><td>12991</td><td>22</td><td>GTTTCGTACGTAGCAGAGCAGC</td><td>169.8</td><td>-3.9</td><td>0.9</td><td>-4.5</td><td>6.5E-06</td><td>1.0E-05</td><td>MB231/MB468</td><td>CYTO</td></tr>
<tr><td>>MI0000439|hsa-mir-23b&WithFlank&9|+|95085202|95085310@67.84.18&[MIMAT0000418&hsa-miR-23b-3p&offsets|+3|-2;m-42&9|+|95085265|95085285&offsets|+3|0]|ACATTGCCAGGGATTACC</td><td>>MI0000439|hsa-mir-23b&WithFlank&9|+|95085202|95085310</td><td>67</td><td>84</td><td>18</td><td>ACATTGCCAGGGATTACC</td><td>235.8</td><td>11.1</td><td>2.4</td><td>4.5</td><td>5.7E-06</td><td>1.0E-05</td><td>MB231/MB468</td><td>MP</td></tr>
<tr><td>>hg38_21_+_8433172_8446622_RNA45SN1_45S_50ntFlanks@12829.12847.19|CGGGGTCCGGTGCGGAGTG</td><td>>hg38_21_+_8433172_8446622_RNA45SN1_45S_50ntFlanks</td><td>12829</td><td>12847</td><td>19</td><td>CGGGGTCCGGTGCGGAGTG</td><td>97.3</td><td>-10.5</td><td>2.3</td><td>-4.5</td><td>5.7E-06</td><td>1.0E-05</td><td>MB231/MB468</td><td>MP</td></tr>
<tr><td>>trnaMT_SerGCT_MT_+_12207_12265@29.62.34|TGCCCCCATGTCTAACAACATGGCTTTCTCACCA</td><td>>trnaMT_SerGCT_MT_+_12207_12265</td><td>29</td><td>62</td><td>34</td><td>TGCCCCCATGTCTAACAACATGGCTTTCTCACCA</td><td>548.1</td><td>4.7</td><td>1.0</td><td>4.5</td><td>6.9E-06</td><td>1.0E-05</td><td>MB231/MB468</td><td>MITO</td></tr>
<tr><td>>hg38_21_+_8433172_8446622_RNA45SN1_45S_50ntFlanks@12081.12110.30|CTCTCGCTTCTGGCGCCAAGCGCCCGGCCG</td><td>>hg38_21_+_8433172_8446622_RNA45SN1_45S_50ntFlanks</td><td>12081</td><td>12110</td><td>30</td><td>CTCTCGCTTCTGGCGCCAAGCGCCCGGCCG</td><td>831.6</td><td>-6.5</td><td>1.4</td><td>-4.5</td><td>5.8E-06</td><td>1.0E-05</td><td>BT20/MB468</td><td>CYTO</td></tr>
<tr><td>>hg38_21_+_8433172_8446622_RNA45SN1_45S_50ntFlanks@4733.4758.26|GAACGAAAGTCGGAGGTTCGAAGACG</td><td>>hg38_21_+_8433172_8446622_RNA45SN1_45S_50ntFlanks</td><td>4733</td><td>4758</td><td>26</td><td>GAACGAAAGTCGGAGGTTCGAAGACG</td><td>2533.9</td><td>-3.6</td><td>0.8</td><td>-4.5</td><td>5.7E-06</td><td>1.0E-05</td><td>MB231/MB468</td><td>MP</td></tr>
<tr><td>>hg38_21_+_8433172_8446622_RNA45SN1_45S_50ntFlanks@12444.12485.42|AAGCAGAATTCACCAAGCGTTGGATTGTTCACCCACTAATAG</td><td>>hg38_21_+_8433172_8446622_RNA45SN1_45S_50ntFlanks</td><td>12444</td><td>12485</td><td>42</td><td>AAGCAGAATTCACCAAGCGTTGGATTGTTCACCCACTAATAG</td><td>717.1</td><td>-4.3</td><td>0.9</td><td>-4.5</td><td>6.8E-06</td><td>1.0E-05</td><td>MB231/MB468</td><td>NUC</td></tr>
<tr><td>>hg38_21_+_8433172_8446622_RNA45SN1_45S_50ntFlanks@11707.11728.22|AAGCGCGGGTAAACGGCGGGAG</td><td>>hg38_21_+_8433172_8446622_RNA45SN1_45S_50ntFlanks</td><td>11707</td><td>11728</td><td>22</td><td>AAGCGCGGGTAAACGGCGGGAG</td><td>368.4</td><td>-6.9</td><td>1.5</td><td>-4.7</td><td>2.1E-06</td><td>1.0E-05</td><td>BT20/MB468</td><td>MP</td></tr>
<tr><td>>hg38_21_+_8433172_8446622_RNA45SN1_45S_50ntFlanks@6671.6699.29|CTCGGCTCGTGCGTCGATGAAGAACGCAG</td><td>>hg38_21_+_8433172_8446622_RNA45SN1_45S_50ntFlanks</td><td>6671</td><td>6699</td><td>29</td><td>CTCGGCTCGTGCGTCGATGAAGAACGCAG</td><td>2252.9</td><td>-5.3</td><td>1.2</td><td>-4.5</td><td>6.5E-06</td><td>1.0E-05</td><td>MB231/MB468</td><td>CYTO</td></tr>
</table>

<table>
<tr><th></th><th></th><th></th><th></th><th></th><th></th><th></th><th></th><th></th><th></th><th></th><th></th><th></th><th></th></tr>
<tr><td>>hg38_21_+_8433172_8446622_RNA45SN1_45S_50ntFlanks@11577.11603.27|TTAGAACTGGTGCGGACCAGGGGAATC</td><td>>hg38_21_+_8433172_8446622_RNA45SN1_45S_50ntFlanks</td><td>11577</td><td>11603</td><td>27</td><td>TTAGAACTGGTGCGGACCAGGGGAATC</td><td>428.2</td><td>-3.3</td><td>0.7</td><td>-4.6</td><td>5.0E-06</td><td>1.0E-05</td><td>BT20/MB468</td><td>MITO</td></tr>
<tr><td>>hg38_21_+_8433172_8446622_RNA45SN1_45S_50ntFlanks@5486.5514.29|GCGCTGAGAAGACGGTCGAACTTGACTAT</td><td>>hg38_21_+_8433172_8446622_RNA45SN1_45S_50ntFlanks</td><td>5486</td><td>5514</td><td>29</td><td>GCGCTGAGAAGACGGTCGAACTTGACTAT</td><td>906.1</td><td>-3.0</td><td>0.6</td><td>-4.6</td><td>5.0E-06</td><td>1.0E-05</td><td>BT20/MB468</td><td>MITO</td></tr>
<tr><td>>hg38_21_+_8433172_8446622_RNA45SN1_45S_50ntFlanks@8169.8212.44|CCAAGTCCTTCTGATCGAGGCCCAGCCCGTGGACGGTGTGAGGC</td><td>>hg38_21_+_8433172_8446622_RNA45SN1_45S_50ntFlanks</td><td>8169</td><td>8212</td><td>44</td><td>CCAAGTCCTTCTGATCGAGGCCCAGCCCGTGGACGGTGTGAGGC</td><td>257.1</td><td>-7.6</td><td>1.7</td><td>-4.5</td><td>5.8E-06</td><td>1.0E-05</td><td>MB231/MB468</td><td>MP</td></tr>
<tr><td>>trna10_AspGTC_12_-_125424193_125424264@37.73.37,trna12_AspGTC_12_-_125411891_125411962@37.73.37,trna144_AspGTC_6_-_27551236_27551307@37.73.37,trna38_AspGTC_17_-_8125556_8125627@37.73.37,trna45_AspGTC_6_+_27447453_27447524@37.73.37,trna48_AspGTC_6_+_27471523_27471594@37.73.37,trna4_AspGTC_12_+_96429799_96429870@37.73.37,trna69_AspGTC_1_-_161440205_161440276@37.73.37,trna72_AspGTC_1_-_161432824_161432895@37.73.37,trna75_AspGTC_1_-_161425414_161425485@37.73.37,trna78_AspGTC_1_-_161418033_161418104@37.73.37,trna81_AspGTC_1_-_161410615_161410686@37.73.37|ACGCGGGAGACCGGGGTTCGATTCCCCGACGGGGAGC</td><td>>trna10_AspGTC_12_-_125424193_125424264</td><td>37</td><td>73</td><td>37</td><td>ACGCGGGAGACCGGGGTTCGATTCCCCGACGGGGAGC</td><td>996.6</td><td>3.0</td><td>0.7</td><td>4.5</td><td>5.9E-06</td><td>1.0E-05</td><td>BT20/MB468</td><td>CYTO</td></tr>
<tr><td>>trna10_AspGTC_12_-_125424193_125424264@32.73.42,trna12_AspGTC_12_-_125411891_125411962@32.73.42,trna144_AspGTC_6_-_27551236_27551307@32.73.42,trna38_AspGTC_17_-_8125556_8125627@32.73.42,trna45_AspGTC_6_+_27447453_27447524@32.73.42,trna48_AspGTC_6_+_27471523_27471594@32.73.42,trna4_AspGTC_12_+_96429799_96429870@32.73.42,trna69_AspGTC_1_-_161440205_161440276@32.73.42,trna72_AspGTC_1_-_161432824_161432895@32.73.42,trna75_AspGTC_1_-_161425414_161425485@32.73.42,trna78_AspGTC_1_-_161418033_161418104@32.73.42,trna81_AspGTC_1_-_161410615_161410686@32.73.42|CTGTCACGCGGGAGACCGGGGTTCGATTCCCCGACGGGGAGC</td><td>>trna10_AspGTC_12_-_125424193_125424264</td><td>32</td><td>73</td><td>42</td><td>CTGTCACGCGGGAGACCGGGGTTCGATTCCCCGACGGGGAGC</td><td>339.5</td><td>-4.4</td><td>1.0</td><td>-4.5</td><td>5.9E-06</td><td>1.0E-05</td><td>BT20/MB468</td><td>CYTO</td></tr>
<tr><td>>MI0033423|hsa-mir-10399&WithFlank&7|-|139044094|139044163@43.64.22&[MIMAT0041630&hsa-miR-10399-3p&offsets|-1|0;TJU_CMC_MD2.ID03061.3p-miR&offsets|+3|+3;m-2657&7|-|139044103|139044124&offsets|+3|+3]|TCTCTCGGACAAGCTGTAGGTC</td><td>>MI0033423|hsa-mir-10399&WithFlank&7|-|139044094|139044163</td><td>43</td><td>64</td><td>22</td><td>TCTCTCGGACAAGCTGTAGGTC</td><td>86.2</td><td>-9.8</td><td>2.0</td><td>-5.0</td><td>5.9E-07</td><td>1.0E-05</td><td>BT20/MB231</td><td>MP</td></tr>
<tr><td>>hg38_21_+_8433172_8446622_RNA45SN1_45S_50ntFlanks@9569.9603.35|GTGCAAATCGGTCGTCCGACCTGGGTATAGGGGCG</td><td>>hg38_21_+_8433172_8446622_RNA45SN1_45S_50ntFlanks</td><td>9569</td><td>9603</td><td>35</td><td>GTGCAAATCGGTCGTCCGACCTGGGTATAGGGGCG</td><td>419.3</td><td>3.7</td><td>0.8</td><td>4.5</td><td>6.5E-06</td><td>1.0E-05</td><td>MB231/MB468</td><td>CYTO</td></tr>
<tr><td>>hg38_21_+_8433172_8446622_RNA45SN1_45S_50ntFlanks@12407.12435.29|TTTTGATCCTTCGATGTCGGCTCTTCCTA</td><td>>hg38_21_+_8433172_8446622_RNA45SN1_45S_50ntFlanks</td><td>12407</td><td>12435</td><td>29</td><td>TTTTGATCCTTCGATGTCGGCTCTTCCTA</td><td>490.8</td><td>-3.1</td><td>0.7</td><td>-4.6</td><td>5.0E-06</td><td>1.0E-05</td><td>BT20/MB468</td><td>MITO</td></tr>
<tr><td>>hg38_21_+_8433172_8446622_RNA45SN1_45S_50ntFlanks@11578.11602.25|TAGAACTGGTGCGGACCAGGGGAAT</td><td>>hg38_21_+_8433172_8446622_RNA45SN1_45S_50ntFlanks</td><td>11578</td><td>11602</td><td>25</td><td>TAGAACTGGTGCGGACCAGGGGAAT</td><td>100.0</td><td>10.0</td><td>2.0</td><td>5.0</td><td>6.0E-07</td><td>1.0E-05</td><td>BT20/MB231</td><td>MP</td></tr>
<tr><td>>hg38_21_+_8433172_8446622_RNA45SN1_45S_50ntFlanks@12893.12932.40|CCTCGCCCGTCACGCACCGCACGTTCGTGGGGAACCTGGC</td><td>>hg38_21_+_8433172_8446622_RNA45SN1_45S_50ntFlanks</td><td>12893</td><td>12932</td><td>40</td><td>CCTCGCCCGTCACGCACCGCACGTTCGTGGGGAACCTGGC</td><td>188.0</td><td>-3.8</td><td>0.8</td><td>-4.5</td><td>6.6E-06</td><td>1.0E-05</td><td>MB231/MB468</td><td>CYTO</td></tr>
<tr><td>>MI0000065|hsa-let-7d&WithFlank&9|+|94178828|94178926@68.88.21(+1C)&[MIMAT0004484&hsa-let-7d-3p&offsets|0|-1(+1C);m-98&9|+|94178895|94178916&offsets|0|-1(+1C)]|CTATACGACCTGCTGCCTTTCC</td><td>>MI0000065|hsa-let-7d&WithFlank&9|+|94178828|94178926</td><td>68</td><td>88</td><td>21(+1C)</td><td>CTATACGACCTGCTGCCTTTCC</td><td>449.4</td><td>3.9</td><td>0.9</td><td>4.5</td><td>5.8E-06</td><td>1.0E-05</td><td>MB231/MB468</td><td>MP</td></tr>
<tr><td>>hg38_21_+_8433172_8446622_RNA45SN1_45S_50ntFlanks@4574.4616.43|ATAGGACCGCGGTTCTATTTTGTTGGTTTTCGGAACTGAGGCC</td><td>>hg38_21_+_8433172_8446622_RNA45SN1_45S_50ntFlanks</td><td>4574</td><td>4616</td><td>43</td><td>ATAGGACCGCGGTTCTATTTTGTTGGTTTTCGGAACTGAGGCC</td><td>185.2</td><td>-7.2</td><td>1.6</td><td>-4.5</td><td>5.9E-06</td><td>1.0E-05</td><td>BT20/MB468</td><td>CYTO</td></tr>
<tr><td>>hg38_21_+_8433172_8446622_RNA45SN1_45S_50ntFlanks@12223.12240.18|CAGAAACCTCCCGTGGAG</td><td>>hg38_21_+_8433172_8446622_RNA45SN1_45S_50ntFlanks</td><td>12223</td><td>12240</td><td>18</td><td>CAGAAACCTCCCGTGGAG</td><td>49.6</td><td>-8.7</td><td>1.8</td><td>-4.7</td><td>2.1E-06</td><td>1.0E-05</td><td>BT20/MB468</td><td>MP</td></tr>
<tr><td>>MI0003776|hsa-mir-320b-1&WithFlank&1|+|116671740|116671823@48.68.21;MI0003839|hsa-mir-320b-2&WithFlank&1|-|224257034|224257116@47.67.21&[MIMAT0005792&hsa-miR-320b&offsets|0|-1;m-183&1|+|116671787|116671806&offsets|0|+1];[MIMAT0005792_1&hsa-miR-320b&offsets|0|-1;m-183&1|-|224257051|224257070&offsets|0|+1]|AAAAGCTGGGTTGAGAGGGCA</td><td>>MI0003776|hsa-mir-320b-1&WithFlank&1|+|116671740|116671823</td><td>48</td><td>68</td><td>21</td><td>AAAAGCTGGGTTGAGAGGGCA</td><td>890.8</td><td>2.9</td><td>0.6</td><td>4.9</td><td>8.0E-07</td><td>1.0E-05</td><td>BT20/MB231</td><td>NUC</td></tr>
<tr><td>>hg38_21_+_8433172_8446622_RNA45SN1_45S_50ntFlanks@4726.4752.27|TAATCAAGAACGAAAGTCGGAGGTTCG</td><td>>hg38_21_+_8433172_8446622_RNA45SN1_45S_50ntFlanks</td><td>4726</td><td>4752</td><td>27</td><td>TAATCAAGAACGAAAGTCGGAGGTTCG</td><td>324.6</td><td>-4.7</td><td>1.0</td><td>-4.5</td><td>5.9E-06</td><td>1.0E-05</td><td>BT20/MB468</td><td>CYTO</td></tr>
<tr><td>>hg38_21_+_8433172_8446622_RNA45SN1_45S_50ntFlanks@12992.13013.22|TCCCTCGCTGCGATCTATTGAA</td><td>>hg38_21_+_8433172_8446622_RNA45SN1_45S_50ntFlanks</td><td>12992</td><td>13013</td><td>22</td><td>TCCCTCGCTGCGATCTATTGAA</td><td>246.4</td><td>-6.4</td><td>1.4</td><td>-4.5</td><td>5.8E-06</td><td>1.0E-05</td><td>MB231/MB468</td><td>MP</td></tr>
<tr><td>>MI0000434|hsa-let-7i&WithFlank&12|+|62603680|62603775@12.31.20&[MIMAT0000415&hsa-let-7i-5p&offsets|0|-2;m-49&12|+|62603691|62603712&offsets|0|-2]|TGAGGTAGTAGTTTGTGCTG</td><td>>MI0000434|hsa-let-7i&WithFlank&12|+|62603680|62603775</td><td>12</td><td>31</td><td>20</td><td>TGAGGTAGTAGTTTGTGCTG</td><td>76.7</td><td>1.7</td><td>0.4</td><td>4.5</td><td>7.4E-06</td><td>1.0E-05</td><td>BT20/MB468</td><td>NUC</td></tr>
<tr><td>>hg38_21_+_8433172_8446622_RNA45SN1_45S_50ntFlanks@10840.10868.29|CCGTAACTTCGGGATAAGGATTGGCTCTA</td><td>>hg38_21_+_8433172_8446622_RNA45SN1_45S_50ntFlanks</td><td>10840</td><td>10868</td><td>29</td><td>CCGTAACTTCGGGATAAGGATTGGCTCTA</td><td>269.8</td><td>-2.6</td><td>0.6</td><td>-4.6</td><td>5.0E-06</td><td>1.0E-05</td><td>BT20/MB468</td><td>MITO</td></tr>
<tr><td>>hg38_21_+_8433172_8446622_RNA45SN1_45S_50ntFlanks@10786.10829.44|AACAGCCTCTGGCATGTTGGAACAATGTAGGTAAGGGAAGTCGG</td><td>>hg38_21_+_8433172_8446622_RNA45SN1_45S_50ntFlanks</td><td>10786</td><td>10829</td><td>44</td><td>AACAGCCTCTGGCATGTTGGAACAATGTAGGTAAGGGAAGTCGG</td><td>116.9</td><td>-6.2</td><td>1.4</td><td>-4.5</td><td>5.9E-06</td><td>1.0E-05</td><td>BT20/MB468</td><td>CYTO</td></tr>
<tr><td>>hg38_21_+_8433172_8446622_RNA45SN1_45S_50ntFlanks@5045.5066.22|CTTAGTTGGTGGAGCGATTTGT</td><td>>hg38_21_+_8433172_8446622_RNA45SN1_45S_50ntFlanks</td><td>5045</td><td>5066</td><td>22</td><td>CTTAGTTGGTGGAGCGATTTGT</td><td>130.0</td><td>-10.9</td><td>2.4</td><td>-4.5</td><td>5.8E-06</td><td>1.0E-05</td><td>MB231/MB468</td><td>MP</td></tr>
<tr><td>>hg38_21_+_8433172_8446622_RNA45SN1_45S_50ntFlanks@9572.9603.32|CAAATCGGTCGTCCGACCTGGGTATAGGGGCG</td><td>>hg38_21_+_8433172_8446622_RNA45SN1_45S_50ntFlanks</td><td>9572</td><td>9603</td><td>32</td><td>CAAATCGGTCGTCCGACCTGGGTATAGGGGCG</td><td>1464.6</td><td>-2.6</td><td>0.6</td><td>-4.5</td><td>6.6E-06</td><td>1.0E-05</td><td>MB231/MB468</td><td>CYTO</td></tr>
<tr><td>>hg38_21_+_8433172_8446622_RNA45SN1_45S_50ntFlanks@12488.12519.32|AACGTGAGCTGGGTTTAGACCGTCGTGAGACA</td><td>>hg38_21_+_8433172_8446622_RNA45SN1_45S_50ntFlanks</td><td>12488</td><td>12519</td><td>32</td><td>AACGTGAGCTGGGTTTAGACCGTCGTGAGACA</td><td>280.9</td><td>-6.9</td><td>1.5</td><td>-4.5</td><td>7.4E-06</td><td>1.0E-05</td><td>BT20/MB468</td><td>NUC</td></tr>
<tr><td>>hg38_21_+_8433172_8446622_RNA45SN1_45S_50ntFlanks@6661.6683.23|CGGTGGATCACTCGGCTCGTGCG</td><td>>hg38_21_+_8433172_8446622_RNA45SN1_45S_50ntFlanks</td><td>6661</td><td>6683</td><td>23</td><td>CGGTGGATCACTCGGCTCGTGCG</td><td>1461.8</td><td>3.5</td><td>0.8</td><td>4.6</td><td>5.1E-06</td><td>1.0E-05</td><td>BT20/MB468</td><td>MITO</td></tr>
<tr><td>>MI0000239|hsa-mir-197&WithFlank&1|+|109598887|109598973@54.74.21(+1U)&[MIMAT0000227&hsa-miR-197-3p&offsets|0|-1(+1U);m-99&1|+|109598940|109598961&offsets|0|-1(+1U)]|TTCACCACCTTCTCCACCCAGT</td><td>>MI0000239|hsa-mir-197&WithFlank&1|+|109598887|109598973</td><td>54</td><td>74</td><td>21(+1U)</td><td>TTCACCACCTTCTCCACCCAGT</td><td>448.6</td><td>4.7</td><td>1.0</td><td>4.5</td><td>7.1E-06</td><td>1.0E-05</td><td>MB231/MB468</td><td>MITO</td></tr>
<tr><td>>hg38_21_+_8433172_8446622_RNA45SN1_45S_50ntFlanks@8078.8104.27|GAAGAGCCCAGCGCCGAATCCCCGCCC</td><td>>hg38_21_+_8433172_8446622_RNA45SN1_45S_50ntFlanks</td><td>8078</td><td>8104</td><td>27</td><td>GAAGAGCCCAGCGCCGAATCCCCGCCC</td><td>155.8</td><td>-7.0</td><td>1.5</td><td>-4.5</td><td>5.8E-06</td><td>1.0E-05</td><td>MB231/MB468</td><td>MP</td></tr>
<tr><td>>MI0000071|hsa-mir-17&WithFlank&13|+|91350599|91350694@20.41.22&[MIMAT0000070&hsa-miR-17-5p&offsets|0|-1;m-88&13|+|91350618|91350640&offsets|0|-1]|CAAAGTGCTTACAGTGCAGGTA</td><td>>MI0000071|hsa-mir-17&WithFlank&13|+|91350599|91350694</td><td>20</td><td>41</td><td>22</td><td>CAAAGTGCTTACAGTGCAGGTA</td><td>719.3</td><td>4.7</td><td>1.0</td><td>4.5</td><td>7.1E-06</td><td>1.0E-05</td><td>MB231/MB468</td><td>MITO</td></tr>
<tr><td>>hg38_21_+_8433172_8446622_RNA45SN1_45S_50ntFlanks@4995.5021.27|ACAGATTGATAGCTCTTTCTCGATTCC</td><td>>hg38_21_+_8433172_8446622_RNA45SN1_45S_50ntFlanks</td><td>4995</td><td>5021</td><td>27</td><td>ACAGATTGATAGCTCTTTCTCGATTCC</td><td>61.9</td><td>-9.0</td><td>1.9</td><td>-4.7</td><td>2.1E-06</td><td>1.0E-05</td><td>BT20/MB468</td><td>MP</td></tr>
<tr><td>>hg38_21_+_8433172_8446622_RNA45SN1_45S_50ntFlanks@3882.3899.18|CCGACGGGCGCTGACCCC</td><td>>hg38_21_+_8433172_8446622_RNA45SN1_45S_50ntFlanks</td><td>3882</td><td>3899</td><td>18</td><td>CCGACGGGCGCTGACCCC</td><td>914.3</td><td>-7.6</td><td>1.7</td><td>-4.5</td><td>7.4E-06</td><td>1.0E-05</td><td>BT20/MB468</td><td>NUC</td></tr>
<tr><td>>hg38_21_+_8433172_8446622_RNA45SN1_45S_50ntFlanks@8356.8375.20|GAAAAGAACTTTGAAGAGAG</td><td>>hg38_21_+_8433172_8446622_RNA45SN1_45S_50ntFlanks</td><td>8356</td><td>8375</td><td>20</td><td>GAAAAGAACTTTGAAGAGAG</td><td>429.9</td><td>-5.4</td><td>1.2</td><td>-4.5</td><td>7.1E-06</td><td>1.0E-05</td><td>MB231/MB468</td><td>MITO</td></tr>
<tr><td>>hg38_21_+_8433172_8446622_RNA45SN1_45S_50ntFlanks@9298.9338.41|ACACGGACCAAGGAGTCTAACACGTGCGCGAGTCGGGGGCT</td><td>>hg38_21_+_8433172_8446622_RNA45SN1_45S_50ntFlanks</td><td>9298</td><td>9338</td><td>41</td><td>ACACGGACCAAGGAGTCTAACACGTGCGCGAGTCGGGGGCT</td><td>269.3</td><td>-6.6</td><td>1.5</td><td>-4.5</td><td>7.1E-06</td><td>1.0E-05</td><td>MB231/MB468</td><td>MITO</td></tr>
<tr><td>>MI0000078|hsa-mir-22&WithFlank&17|-|1713897|1713993@59.79.21(+1G)&[MIMAT0000077&hsa-miR-22-3p&offsets|0|-1(+1G);m-2&17|-|1713914|1713935&offsets|0|-1(+1G)]|AAGCTGCCAGTTGAAGAACTGG</td><td>>MI0000078|hsa-mir-22&WithFlank&17|-|1713897|1713993</td><td>59</td><td>79</td><td>21(+1G)</td><td>AAGCTGCCAGTTGAAGAACTGG</td><td>596.8</td><td>8.7</td><td>1.9</td><td>4.5</td><td>6.0E-06</td><td>1.0E-05</td><td>BT20/MB468</td><td>CYTO</td></tr>
<tr><td>>am_trnaMT_PheGAA_MT_+_577_647@43.60.18|TTAGACGGGCTCACATCA</td><td>>am_trnaMT_PheGAA_MT_+_577_647</td><td>43</td><td>60</td><td>18</td><td>TTAGACGGGCTCACATCA</td><td>66.0</td><td>9.8</td><td>2.1</td><td>4.7</td><td>2.2E-06</td><td>1.0E-05</td><td>BT20/MB468</td><td>MP</td></tr>
<tr><td>>hg38_21_+_8433172_8446622_RNA45SN1_45S_50ntFlanks@10140.10158.19|CGGCGGCGGGGGTGTGGGG</td><td>>hg38_21_+_8433172_8446622_RNA45SN1_45S_50ntFlanks</td><td>10140</td><td>10158</td><td>19</td><td>CGGCGGCGGGGGTGTGGGG</td><td>48.0</td><td>-8.7</td><td>1.8</td><td>-4.7</td><td>2.2E-06</td><td>1.0E-05</td><td>BT20/MB468</td><td>MP</td></tr>
<tr><td>>hg38_21_+_8433172_8446622_RNA45SN1_45S_50ntFlanks@4507.4528.22|CTTTGAAAAAATTAGAGTGTTC</td><td>>hg38_21_+_8433172_8446622_RNA45SN1_45S_50ntFlanks</td><td>4507</td><td>4528</td><td>22</td><td>CTTTGAAAAAATTAGAGTGTTC</td><td>60.6</td><td>-9.0</td><td>1.9</td><td>-4.7</td><td>2.2E-06</td><td>1.0E-05</td><td>BT20/MB468</td><td>MP</td></tr>
<tr><td>>MI0000273|hsa-mir-183&WithFlank&7|-|129774899|129775020@33.56.24&[MIMAT0000261&hsa-miR-183-5p&offsets|0|+2;m-41&7|-|129774966|129774987&offsets|-1|+1]|TATGGCACTGGTAGAATTCACTGT</td><td>>MI0000273|hsa-mir-183&WithFlank&7|-|129774899|129775020</td><td>33</td><td>56</td><td>24</td><td>TATGGCACTGGTAGAATTCACTGT</td><td>278.8</td><td>3.4</td><td>0.7</td><td>4.5</td><td>6.9E-06</td><td>1.0E-05</td><td>MB231/MB468</td><td>NUC</td></tr>
<tr><td>>hg38_21_+_8433172_8446622_RNA45SN1_45S_50ntFlanks@4733.4752.20|GAACGAAAGTCGGAGGTTCG</td><td>>hg38_21_+_8433172_8446622_RNA45SN1_45S_50ntFlanks</td><td>4733</td><td>4752</td><td>20</td><td>GAACGAAAGTCGGAGGTTCG</td><td>237.6</td><td>-5.0</td><td>1.1</td><td>-4.5</td><td>5.9E-06</td><td>1.0E-05</td><td>MB231/MB468</td><td>MP</td></tr>
<tr><td>>hg38_21_+_8433172_8446622_RNA45SN1_45S_50ntFlanks@10314.10333.20|CAGGTGCAGATCTTGGTGGT</td><td>>hg38_21_+_8433172_8446622_RNA45SN1_45S_50ntFlanks</td><td>10314</td><td>10333</td><td>20</td><td>CAGGTGCAGATCTTGGTGGT</td><td>167.6</td><td>10.7</td><td>2.1</td><td>5.0</td><td>6.1E-07</td><td>1.0E-05</td><td>BT20/MB231</td><td>MP</td></tr>
<tr><td>>hg38_21_+_8433172_8446622_RNA45SN1_45S_50ntFlanks@10916.10933.18|CCGCGGCTGGACGAGGCG</td><td>>hg38_21_+_8433172_8446622_RNA45SN1_45S_50ntFlanks</td><td>10916</td><td>10933</td><td>18</td><td>CCGCGGCTGGACGAGGCG</td><td>490.6</td><td>-4.8</td><td>1.1</td><td>-4.5</td><td>6.0E-06</td><td>1.0E-05</td><td>BT20/MB468</td><td>CYTO</td></tr>
<tr><td>>hg38_MT_+_598_1651_RNR1_12S_50ntFlanks@907.946.40|TATACTTCAAAGGACATTTAACTAAAACCCCTACGCATTT</td><td>>hg38_MT_+_598_1651_RNR1_12S_50ntFlanks</td><td>907</td><td>946</td><td>40</td><td>TATACTTCAAAGGACATTTAACTAAAACCCCTACGCATTT</td><td>267.1</td><td>8.7</td><td>1.8</td><td>4.7</td><td>2.2E-06</td><td>1.0E-05</td><td>BT20/MB468</td><td>MP</td></tr>
<tr><td>>hg38_21_+_8433172_8446622_RNA45SN1_45S_50ntFlanks@4201.4219.19|CCGACCCGGGGAGGTAGTG</td><td>>hg38_21_+_8433172_8446622_RNA45SN1_45S_50ntFlanks</td><td>4201</td><td>4219</td><td>19</td><td>CCGACCCGGGGAGGTAGTG</td><td>492.4</td><td>-7.0</td><td>1.5</td><td>-4.7</td><td>2.2E-06</td><td>1.0E-05</td><td>BT20/MB468</td><td>MP</td></tr>
</table>

| | | | | | | | | | | | | | |
|---|---|---|---|---|---|---|---|---|---|---|---|---|---|
| >MI0000108\|hsa-mir-103a-2&WithFlank&20\|+\|3917488\|3917577@54.75.22(+1G);MI0000109\|hsa-mir-103a-1&WithFlank&5\|-\|168560890\|168560979@54.75.22(+1G)&[MIMAT0000101&hsa-miR-103a-3p&offsets\|0\|-1(+1G);m-12&20\|+\|3917541\|3917563&offsets\|0\|-1(+1G)];[MIMAT0000101_1&hsa-miR-103a-3p&offsets\|0\|-1(+1G);m-13&5\|-\|168560904\|168560926&offsets\|0\|-1(+1G);m-44220&5\|-\|168560908\|168560926&offsets\|0\|+3(+1G)]\|AGCAGCATTGTACAGGGCTATGG | >MI0000108\|hsa-mir-103a-2&WithFlank&20\|+\|3917488\|3917577 | 54 | 75 | 22(+1G) | AGCAGCATTGTACAGGGCTATGG | 464.2 | 5.0 | 1.1 | 4.5 | 7.1E-06 | 1.0E-05 | MB231/MB468 | MITO |
| >hg38_21_+_8433172_8446622_RNA45SN1_45S_50ntFlanks@12633.12669.37\|CGAAGCTACCATCTGTGGGATTATGACTGAACGCCTC | >hg38_21_+_8433172_8446622_RNA45SN1_45S_50ntFlanks | 12633 | 12669 | 37 | CGAAGCTACCATCTGTGGGATTATGACTGAACGCCTC | 171.5 | 4.7 | 1.0 | 4.5 | 6.7E-06 | 1.0E-05 | MB231/MB468 | CYTO |
| >hg38_21_+_8433172_8446622_RNA45SN1_45S_50ntFlanks@10866.10900.35\|CTAAGGGCTGGGTCGGTCGGGCTGGGGCGCGAAGC | >hg38_21_+_8433172_8446622_RNA45SN1_45S_50ntFlanks | 10866 | 10900 | 35 | CTAAGGGCTGGGTCGGTCGGGCTGGGGCGCGAAGC | 179.7 | -6.1 | 1.4 | -4.5 | 7.0E-06 | 1.0E-05 | MB231/MB468 | NUC |
| >hg38_21_+_8433172_8446622_RNA45SN1_45S_50ntFlanks@8176.8223.48\|CTTCTGATCGAGGCCCAGCCCGTGGACGGTGTGAGGCCGGTAGCGGCC | >hg38_21_+_8433172_8446622_RNA45SN1_45S_50ntFlanks | 8176 | 8223 | 48 | CTTCTGATCGAGGCCCAGCCCGTGGACGGTGTGAGGCCGGTAGCGGCC | 102.2 | -10.5 | 2.3 | -4.5 | 5.9E-06 | 1.0E-05 | MB231/MB468 | MP |
| >MI0000061\|hsa-let-7a-2&WithFlank&11\|-\|122146516\|122146599@11.32.22(+1C);MI0000062\|hsa-let-7a-3&WithFlank&22\|+\|46112743\|46112828@10.31.22(+1C);MI0000060\|hsa-let-7a-1&WithFlank&9\|+\|94175951\|94176042@12.33.22(+1C)&[MIMAT0000062&hsa-let-7a-5p&offsets\|0\|0(+1C);m-6&11\|-\|122146568\|122146589&offsets\|0\|0(+1C)];[MIMAT0000062_1&hsa-let-7a-5p&offsets\|0\|0(+1C);m-5&22\|+\|46112752\|46112773&offsets\|0\|0(+1C)];[MIMAT0000062_2&hsa-let-7a-5p&offsets\|0\|0(+1C);m-4&9\|+\|94175962\|94175983&offsets\|0\|0(+1C)]\|TGAGGTAGTAGGTTGTATAGTTC | >MI0000061\|hsa-let-7a-2&WithFlank&11\|-\|122146516\|122146599 | 11 | 32 | 22(+1C) | TGAGGTAGTAGGTTGTATAGTTC | 323.3 | 6.6 | 1.5 | 4.5 | 7.1E-06 | 1.0E-05 | MB231/MB468 | MITO |
| >hg38_21_+_8433172_8446622_RNA45SN1_45S_50ntFlanks@5487.5509.23\|CGCTGAGAAGACGGTCGAACTTG | >hg38_21_+_8433172_8446622_RNA45SN1_45S_50ntFlanks | 5487 | 5509 | 23 | CGCTGAGAAGACGGTCGAACTTG | 1394.0 | -4.5 | 1.0 | -4.7 | 2.2E-06 | 1.0E-05 | BT20/MB468 | MP |
| >hg38_21_+_8433172_8446622_RNA45SN1_45S_50ntFlanks@12417.12441.25\|TCGATGTCGGCTCTTCCTATCATTG | >hg38_21_+_8433172_8446622_RNA45SN1_45S_50ntFlanks | 12417 | 12441 | 25 | TCGATGTCGGCTCTTCCTATCATTG | 970.3 | -2.7 | 0.6 | -4.6 | 5.1E-06 | 1.0E-05 | BT20/MB468 | MITO |
| >hg38_21_+_8433172_8446622_RNA45SN1_45S_50ntFlanks@3866.3899.34\|CTAGAGCTAATACATGCCGACGGGCGCTGACCCC | >hg38_21_+_8433172_8446622_RNA45SN1_45S_50ntFlanks | 3866 | 3899 | 34 | CTAGAGCTAATACATGCCGACGGGCGCTGACCCC | 170.3 | -2.9 | 0.6 | -4.5 | 6.0E-06 | 1.0E-05 | BT20/MB468 | CYTO |
| >hg38_21_+_8433172_8446622_RNA45SN1_45S_50ntFlanks@11480.11500.21\|CGGCGGAGGGGCCGCGGGCCG | >hg38_21_+_8433172_8446622_RNA45SN1_45S_50ntFlanks | 11480 | 11500 | 21 | CGGCGGAGGGGCCGCGGGCCG | 62.6 | -9.0 | 1.9 | -4.7 | 2.2E-06 | 1.0E-05 | BT20/MB468 | MP |
| >hg38_21_+_8433172_8446622_RNA45SN1_45S_50ntFlanks@10411.10428.18\|CAGTCGGTCCTGAGAGAT | >hg38_21_+_8433172_8446622_RNA45SN1_45S_50ntFlanks | 10411 | 10428 | 18 | CAGTCGGTCCTGAGAGAT | 1849.9 | -4.1 | 0.9 | -4.6 | 5.1E-06 | 1.0E-05 | BT20/MB468 | MITO |
| >trna5_AspGTC_12_+_98897281_98897352@39.75.37\|GCGGGAGACCGGGGTTCAATTCCCCGACGGGGAGCCA | >trna5_AspGTC_12_+_98897281_98897352 | 39 | 75 | 37 | GCGGGAGACCGGGGTTCAATTCCCCGACGGGGAGCCA | 1306.4 | 6.2 | 1.4 | 4.5 | 6.7E-06 | 1.0E-05 | MB231/MB468 | CYTO |
| >hg38_21_+_8433172_8446622_RNA45SN1_45S_50ntFlanks@3837.3860.24\|CCTCTCCTACTTGGATAACTGTGG | >hg38_21_+_8433172_8446622_RNA45SN1_45S_50ntFlanks | 3837 | 3860 | 24 | CCTCTCCTACTTGGATAACTGTGG | 263.5 | -2.5 | 0.6 | -4.5 | 6.7E-06 | 1.0E-05 | MB231/MB468 | CYTO |
| >hg38_21_+_8433172_8446622_RNA45SN1_45S_50ntFlanks@4045.4062.18\|CCATTCGAACGTCTGCCC | >hg38_21_+_8433172_8446622_RNA45SN1_45S_50ntFlanks | 4045 | 4062 | 18 | CCATTCGAACGTCTGCCC | 796.9 | -2.9 | 0.7 | -4.5 | 6.7E-06 | 1.0E-05 | MB231/MB468 | CYTO |
| >hg38_21_+_8433172_8446622_RNA45SN1_45S_50ntFlanks@11604.11623.20\|CGACTGTTTAATTAAAACAA | >hg38_21_+_8433172_8446622_RNA45SN1_45S_50ntFlanks | 11604 | 11623 | 20 | CGACTGTTTAATTAAAACAA | 634.5 | 4.4 | 1.0 | 4.5 | 6.7E-06 | 1.0E-05 | MB231/MB468 | CYTO |
| >hg38_21_+_8433172_8446622_RNA45SN1_45S_50ntFlanks@12477.12496.20\|CACTAATAGGGAACGTGAGC | >hg38_21_+_8433172_8446622_RNA45SN1_45S_50ntFlanks | 12477 | 12496 | 20 | CACTAATAGGGAACGTGAGC | 3013.7 | 2.6 | 0.6 | 4.5 | 6.7E-06 | 1.0E-05 | MB231/MB468 | CYTO |
| >hg38_21_+_8433172_8446622_RNA45SN1_45S_50ntFlanks@12076.12096.21\|AGGGGCTCTCGCTTCTGGCGC | >hg38_21_+_8433172_8446622_RNA45SN1_45S_50ntFlanks | 12076 | 12096 | 21 | AGGGGCTCTCGCTTCTGGCGC | 596.4 | -3.2 | 0.7 | -4.5 | 6.7E-06 | 1.0E-05 | MB231/MB468 | CYTO |
| >MI0000809\|hsa-mir-151a&WithFlank&8\|-\|140732558\|140732659@51.73.23(+1U)&[MIMAT0000757&hsa-miR-151a-3p&offsets\|-2\|0(+1U);m-30&8\|-\|140732587\|140732607&offsets\|-2\|0(+1U)]\|TACTAGACTGAAGCTCCTTGAGGT | >MI0000809\|hsa-mir-151a&WithFlank&8\|-\|140732558\|140732659 | 51 | 73 | 23(+1U) | TACTAGACTGAAGCTCCTTGAGGT | 433.0 | 6.6 | 1.5 | 4.5 | 7.2E-06 | 1.0E-05 | MB231/MB468 | MITO |
| >hg38_21_+_8433172_8446622_RNA45SN1_45S_50ntFlanks@12045.12070.26\|CTGACCCGGTGAGGCGGGGGGGCGAG | >hg38_21_+_8433172_8446622_RNA45SN1_45S_50ntFlanks | 12045 | 12070 | 26 | CTGACCCGGTGAGGCGGGGGGGCGAG | 288.8 | -6.9 | 1.5 | -4.7 | 2.2E-06 | 1.0E-05 | BT20/MB468 | MP |
| >hg38_21_+_8433172_8446622_RNA45SN1_45S_50ntFlanks@12996.13039.44\|TCGCTGCGATCTATTGAAAGTCAGCCCTCGACACAAGGGTTTGT | >hg38_21_+_8433172_8446622_RNA45SN1_45S_50ntFlanks | 12996 | 13039 | 44 | TCGCTGCGATCTATTGAAAGTCAGCCCTCGACACAAGGGTTTGT | 5274.9 | 5.3 | 1.1 | 4.7 | 2.2E-06 | 1.0E-05 | BT20/MB468 | MP |
| >hg38_21_+_8433172_8446622_RNA45SN1_45S_50ntFlanks@4064.4101.38\|ATCAACTTTCGATGGTAGTCGCCGTGCCTACCATGGTG | >hg38_21_+_8433172_8446622_RNA45SN1_45S_50ntFlanks | 4064 | 4101 | 38 | ATCAACTTTCGATGGTAGTCGCCGTGCCTACCATGGTG | 669.8 | -6.0 | 1.3 | -4.5 | 6.8E-06 | 1.0E-05 | MB231/MB468 | CYTO |
| >hg38_21_+_8433172_8446622_RNA45SN1_45S_50ntFlanks@5186.5226.41\|CAATAACAGGTCTGTGATGCCCTTAGATGTCCGGGGCTGCA | >hg38_21_+_8433172_8446622_RNA45SN1_45S_50ntFlanks | 5186 | 5226 | 41 | CAATAACAGGTCTGTGATGCCCTTAGATGTCCGGGGCTGCA | 118.2 | -10.7 | 2.4 | -4.5 | 5.9E-06 | 1.0E-05 | MB231/MB468 | MP |
| >hg38_21_+_8433172_8446622_RNA45SN1_45S_50ntFlanks@4752.4782.31\|GAAGACGATCAGATACCGTCGTAGTTCCGAC | >hg38_21_+_8433172_8446622_RNA45SN1_45S_50ntFlanks | 4752 | 4782 | 31 | GAAGACGATCAGATACCGTCGTAGTTCCGAC | 225.2 | -2.4 | 0.5 | -4.5 | 6.8E-06 | 1.0E-05 | MB231/MB468 | CYTO |
| >hg38_21_+_8433172_8446622_RNA45SN1_45S_50ntFlanks@10574.10596.23\|AAGCCGGCGGGAGCCCCGGGGAG | >hg38_21_+_8433172_8446622_RNA45SN1_45S_50ntFlanks | 10574 | 10596 | 23 | AAGCCGGCGGGAGCCCCGGGGAG | 250.5 | -5.9 | 1.3 | -4.5 | 6.1E-06 | 1.1E-05 | BT20/MB468 | CYTO |
| >hg38_21_+_8433172_8446622_RNA45SN1_45S_50ntFlanks@7984.8003.20\|AGATCAGACGTGGCGACCCG | >hg38_21_+_8433172_8446622_RNA45SN1_45S_50ntFlanks | 7984 | 8003 | 20 | AGATCAGACGTGGCGACCCG | 325.9 | -4.8 | 1.1 | -4.5 | 6.8E-06 | 1.1E-05 | MB231/MB468 | CYTO |
| >hg38_21_+_8433172_8446622_RNA45SN1_45S_50ntFlanks@8345.8378.34\|AAGGGAAAGTTGAAAAGAACTTTGAAGAGAGAGT | >hg38_21_+_8433172_8446622_RNA45SN1_45S_50ntFlanks | 8345 | 8378 | 34 | AAGGGAAAGTTGAAAAGAACTTTGAAGAGAGAGT | 144.4 | 8.0 | 1.7 | 4.7 | 2.2E-06 | 1.1E-05 | BT20/MB468 | MP |
| >MI0000446\|hsa-mir-125b-1&WithFlank&11\|-\|122099751\|122099850@21.43.23(+1U)&[MIMAT0000423&hsa-miR-125b-5p&offsets\|0\|+1(+1U);m-67&11\|-\|122099809\|122099830&offsets\|0\|+1(+1U)]\|TCCCTGAGACCCTAACTTGTGATT | >MI0000446\|hsa-mir-125b-1&WithFlank&11\|-\|122099751\|122099850 | 21 | 43 | 23(+1U) | TCCCTGAGACCCTAACTTGTGATT | 90.9 | -9.9 | 2.0 | -5.0 | 6.2E-07 | 1.1E-05 | BT20/MB231 | MP |
| >hg38_21_+_8433172_8446622_RNA45SN1_45S_50ntFlanks@10404.10430.27\|CATGGGTCAGTCGGTCCTGAGAGATGG | >hg38_21_+_8433172_8446622_RNA45SN1_45S_50ntFlanks | 10404 | 10430 | 27 | CATGGGTCAGTCGGTCCTGAGAGATGG | 366.4 | -6.0 | 1.3 | -4.5 | 6.1E-06 | 1.1E-05 | BT20/MB468 | CYTO |
| >MI0005757\|hsa-mir-935&WithFlank&19\|+\|53982301\|53982403@63.84.22&[MIMAT0004978&hsa-miR-935&offsets\|+1\|0;m-549&19\|+\|53982363\|53982384&offsets\|0\|0]\|CAGTTACCGCTTCCGCTACCGC | >MI0005757\|hsa-mir-935&WithFlank&19\|+\|53982301\|53982403 | 63 | 84 | 22 | CAGTTACCGCTTCCGCTACCGC | 85.9 | 9.8 | 2.0 | 5.0 | 6.2E-07 | 1.1E-05 | BT20/MB231 | MP |
| >hg38_21_+_8433172_8446622_RNA45SN1_45S_50ntFlanks@6663.6688.26\|GTGGATCACTCGGCTCGTGCGTCGAT | >hg38_21_+_8433172_8446622_RNA45SN1_45S_50ntFlanks | 6663 | 6688 | 26 | GTGGATCACTCGGCTCGTGCGTCGAT | 152.2 | -2.5 | 0.6 | -4.5 | 7.1E-06 | 1.1E-05 | MB231/MB468 | NUC |
| >hg38_21_+_8433172_8446622_RNA45SN1_45S_50ntFlanks@11316.11337.22\|CACCCCACGTCTCGTCGCGCGC | >hg38_21_+_8433172_8446622_RNA45SN1_45S_50ntFlanks | 11316 | 11337 | 22 | CACCCCACGTCTCGTCGCGCGC | 153.3 | -5.4 | 1.2 | -4.5 | 6.1E-06 | 1.1E-05 | BT20/MB468 | CYTO |
| >hg38_21_+_8433172_8446622_RNA45SN1_45S_50ntFlanks@9198.9218.21\|GTTCTCTCGGGGCCACGCGCG | >hg38_21_+_8433172_8446622_RNA45SN1_45S_50ntFlanks | 9198 | 9218 | 21 | GTTCTCTCGGGGCCACGCGCG | 616.2 | -7.2 | 1.5 | -4.7 | 2.2E-06 | 1.1E-05 | BT20/MB468 | MP |
| >trnaMT_ValTAC_MT_+_1602_1670@46.71.26\|TTCAACTTAACTTGACCGCTCTGACC | >trnaMT_ValTAC_MT_+_1602_1670 | 46 | 71 | 26 | TTCAACTTAACTTGACCGCTCTGACC | 747.3 | 3.9 | 0.9 | 4.6 | 5.2E-06 | 1.1E-05 | BT20/MB468 | MITO |
| >hg38_21_+_8433172_8446622_RNA45SN1_45S_50ntFlanks@4983.5016.34\|CGGACAGGATTGACAGATTGATAGCTCTTTCTCG | >hg38_21_+_8433172_8446622_RNA45SN1_45S_50ntFlanks | 4983 | 5016 | 34 | CGGACAGGATTGACAGATTGATAGCTCTTTCTCG | 1357.2 | -5.3 | 1.2 | -4.6 | 5.2E-06 | 1.1E-05 | BT20/MB468 | MITO |
| >trna119_LysCTT_1_-_145395522_145395594@1.36.36,trna11_LysCTT_5_-_180648979_180649051@1.36.36,trna13_LysCTT_6_+_26556774_26556846@1.36.36,trna7_LysCTT_16_+_3225692_3225764@1.36.36,trna9_LysCTT_5_+_180634755_180634827@1.36.36\|GCCCGGCTAGCTCAGTCGGTAGAGCATGAGACTCTT | >trna119_LysCTT_1_-_145395522_145395594 | 1 | 36 | 36 | GCCCGGCTAGCTCAGTCGGTAGAGCATGAGACTCTT | 448.7 | 3.6 | 0.8 | 4.5 | 6.0E-06 | 1.1E-05 | MB231/MB468 | MP |
| >MI0000102\|hsa-mir-100&WithFlank&11\|-\|122152223\|122152314@19.41.23(+1A)&[MIMAT0000098&hsa-miR-100-5p&offsets\|0\|+1(+1A);m-27&11\|-\|122152276\|122152296&offsets\|0\|+2(+1A)]\|AACCCGTAGATCCGAACTTGTGGA | >MI0000102\|hsa-mir-100&WithFlank&11\|-\|122152223\|122152314 | 19 | 41 | 23(+1A) | AACCCGTAGATCCGAACTTGTGGA | 34.5 | 8.3 | 1.8 | 4.5 | 6.0E-06 | 1.1E-05 | MB231/MB468 | MP |
| >hg38_21_+_8433172_8446622_RNA45SN1_45S_50ntFlanks@10443.10460.18\|CCGAAGGGACGGGCGATG | >hg38_21_+_8433172_8446622_RNA45SN1_45S_50ntFlanks | 10443 | 10460 | 18 | CCGAAGGGACGGGCGATG | 251.8 | -3.1 | 0.7 | -4.5 | 7.1E-06 | 1.1E-05 | MB231/MB468 | NUC |
| >hg38_21_+_8433172_8446622_RNA45SN1_45S_50ntFlanks@11999.12029.31\|CCCTGCGGGCCGCCGGTGAAATACCACTACT | >hg38_21_+_8433172_8446622_RNA45SN1_45S_50ntFlanks | 11999 | 12029 | 31 | CCCTGCGGGCCGCCGGTGAAATACCACTACT | 205.4 | -3.2 | 0.7 | -4.5 | 6.8E-06 | 1.1E-05 | MB231/MB468 | CYTO |
| >hg38_21_+_8433172_8446622_RNA45SN1_45S_50ntFlanks@5515.5534.20\|CTAGAGGAAGTAAAAGTCGT | >hg38_21_+_8433172_8446622_RNA45SN1_45S_50ntFlanks | 5515 | 5534 | 20 | CTAGAGGAAGTAAAAGTCGT | 201.2 | -4.1 | 0.9 | -4.5 | 6.8E-06 | 1.1E-05 | MB231/MB468 | CYTO |
| >hg38_21_+_8433172_8446622_RNA45SN1_45S_50ntFlanks@6772.6804.33\|GTTCCTCCCGGGGCTACGCCTGTCTGAGCGTCG | >hg38_21_+_8433172_8446622_RNA45SN1_45S_50ntFlanks | 6772 | 6804 | 33 | GTTCCTCCCGGGGCTACGCCTGTCTGAGCGTCG | 89.9 | -6.2 | 1.4 | -4.5 | 6.0E-06 | 1.1E-05 | MB231/MB468 | MP |
| >hg38_21_+_8433172_8446622_RNA45SN1_45S_50ntFlanks@8176.8223.48\|CTTCTGATCGAGGCCCAGCCCGTGGACGGTGTGAGGCCGGTAGCGGCC | >hg38_21_+_8433172_8446622_RNA45SN1_45S_50ntFlanks | 8176 | 8223 | 48 | CTTCTGATCGAGGCCCAGCCCGTGGACGGTGTGAGGCCGGTAGCGGCC | 1075.0 | -6.3 | 1.4 | -4.5 | 7.1E-06 | 1.1E-05 | MB231/MB468 | NUC |
| >hg38_21_+_8433172_8446622_RNA45SN1_45S_50ntFlanks@10678.10709.32\|CGGTTCCGGCGGCGTCCGGTGAGCTCTCGCTG | >hg38_21_+_8433172_8446622_RNA45SN1_45S_50ntFlanks | 10678 | 10709 | 32 | CGGTTCCGGCGGCGTCCGGTGAGCTCTCGCTG | 42263.6 | -4.9 | 1.1 | -4.6 | 5.2E-06 | 1.1E-05 | BT20/MB468 | MITO |
| >trna152_ValCAC_6_-_27248049_27248121@1.29.29\|GCTTCTGTAGTGTAGTGGTTATCACGTTC | >trna152_ValCAC_6_-_27248049_27248121 | 1 | 29 | 29 | GCTTCTGTAGTGTAGTGGTTATCACGTTC | 6815.1 | -4.7 | 1.0 | -4.5 | 7.1E-06 | 1.1E-05 | MB231/MB468 | NUC |
| >hg38_21_+_8433172_8446622_RNA45SN1_45S_50ntFlanks@12419.12442.24\|GATGTCGGCTCTTCCTATCATTGT | >hg38_21_+_8433172_8446622_RNA45SN1_45S_50ntFlanks | 12419 | 12442 | 24 | GATGTCGGCTCTTCCTATCATTGT | 848.5 | 3.6 | 0.8 | 4.5 | 7.1E-06 | 1.1E-05 | MB231/MB468 | NUC |
| >hg38_21_+_8433172_8446622_RNA45SN1_45S_50ntFlanks@7975.8007.33\|CGCGACCTCAGATCAGACGTGGCGACCCGCTGA | >hg38_21_+_8433172_8446622_RNA45SN1_45S_50ntFlanks | 7975 | 8007 | 33 | CGCGACCTCAGATCAGACGTGGCGACCCGCTGA | 115184.2 | 3.8 | 0.8 | 4.6 | 5.2E-06 | 1.1E-05 | BT20/MB468 | MITO |
| >hg38_21_+_8433172_8446622_RNA45SN1_45S_50ntFlanks@11668.11697.30\|TGCCCAGTGCTCTGAATGTCAAAGTGAAGA | >hg38_21_+_8433172_8446622_RNA45SN1_45S_50ntFlanks | 11668 | 11697 | 30 | TGCCCAGTGCTCTGAATGTCAAAGTGAAGA | 195.2 | 4.2 | 0.9 | 4.5 | 6.9E-06 | 1.1E-05 | MB231/MB468 | CYTO |
| >hg38_21_+_8433172_8446622_RNA45SN1_45S_50ntFlanks@10845.10872.28\|ACTTCGGGATAAGGATTGGCTCTAAGGG | >hg38_21_+_8433172_8446622_RNA45SN1_45S_50ntFlanks | 10845 | 10872 | 28 | ACTTCGGGATAAGGATTGGCTCTAAGGG | 245.4 | -3.9 | 0.9 | -4.5 | 6.9E-06 | 1.1E-05 | MB231/MB468 | CYTO |
| >hg38_21_+_8433172_8446622_RNA45SN1_45S_50ntFlanks@8189.8212.24\|CCCAGCCCGTGGACGGTGTGAGGC | >hg38_21_+_8433172_8446622_RNA45SN1_45S_50ntFlanks | 8189 | 8212 | 24 | CCCAGCCCGTGGACGGTGTGAGGC | 1040.5 | -7.0 | 1.5 | -4.5 | 6.0E-06 | 1.1E-05 | MB231/MB468 | MP |

<table>
<tr><td>>trna111_HisGTG_1_-_147774845_147774916@-1G.30.31,trna118_HisGTG_1_-_145396881_145396952@-1G.30.31,trna16_HisGTG_1_+_146544773_146544844@-1G.30.31,trna1_HisGTG_15_+_45493349_45493420@-1G.30.31,trna21_HisGTG_1_+_147753471_147753542@-1G.30.31,trna33_HisGTG_6_+_27125906_27125977@-1G.30.31,trna7_HisGTG_9_-_14433938_14434009@-1G.30.31,trna8_HisGTG_15_-_45492611_45492682@-1G.30.31,trna9_HisGTG_15_-_45490804_45490875@-1G.30.31|GGCCGTGATCGTATAGTGGTTAGTACTCTGC</td><td>>trna111_HisGTG_1_-_147774845_147774916</td><td>-1G</td><td>30</td><td>31</td><td>GGCCGTGATCGTATAGTGGTTAGTACTCTGC</td><td>1618.5</td><td>-4.0</td><td>0.8</td><td>-5.0</td><td>7.4E-07</td><td>1.1E-05</td><td>BT20/MB231</td><td>MITO</td></tr>
<tr><td>>hg38_21_+_8433172_8446622_RNA45SN1_45S_50ntFlanks@10422.10456.35|GAGAGATGGGCGAGCGCCGTTCCGAAGGGACGGGC</td><td>>hg38_21_+_8433172_8446622_RNA45SN1_45S_50ntFlanks</td><td>10422</td><td>10456</td><td>35</td><td>GAGAGATGGGCGAGCGCCGTTCCGAAGGGACGGGC</td><td>1052.1</td><td>3.1</td><td>0.7</td><td>4.5</td><td>6.9E-06</td><td>1.1E-05</td><td>MB231/MB468</td><td>CYTO</td></tr>
<tr><td>>MI0000542|hsa-mir-320a&WithFlank&8|-|22244960|22245043@49.71.23(+1U)&[MIMAT0000510&hsa-miR-320a-3p&offsets|+1|+2(+1U);m-60&8|-|22244975|22244996&offsets|+1|+2(+1U)]|AAAGCTGGGTTGAGAGGGCGAAAT</td><td>>MI0000542|hsa-mir-320a&WithFlank&8|-|22244960|22245043</td><td>49</td><td>71</td><td>23(+1U)</td><td>AAAGCTGGGTTGAGAGGGCGAAAT</td><td>214.7</td><td>8.2</td><td>1.7</td><td>4.7</td><td>2.2E-06</td><td>1.1E-05</td><td>BT20/MB468</td><td>MP</td></tr>
<tr><td>>MI0000272|hsa-mir-182&WithFlank&7|-|129770377|129770498@29.53.25&[MIMAT0000259&hsa-miR-182-5p&offsets|0|+1;m-22&7|-|129770449|129770470&offsets|0|+3]|TTTGGCAATGGTAGAACTCACACTG</td><td>>MI0000272|hsa-mir-182&WithFlank&7|-|129770377|129770498</td><td>29</td><td>53</td><td>25</td><td>TTTGGCAATGGTAGAACTCACACTG</td><td>1479.2</td><td>4.0</td><td>0.9</td><td>4.5</td><td>7.3E-06</td><td>1.1E-05</td><td>MB231/MB468</td><td>MITO</td></tr>
<tr><td>>hg38_21_+_8433172_8446622_RNA45SN1_45S_50ntFlanks@12456.12481.26|CCAAGCGTTGGATTGTTCACCCACTA</td><td>>hg38_21_+_8433172_8446622_RNA45SN1_45S_50ntFlanks</td><td>12456</td><td>12481</td><td>26</td><td>CCAAGCGTTGGATTGTTCACCCACTA</td><td>908.1</td><td>-7.4</td><td>1.6</td><td>-4.7</td><td>2.2E-06</td><td>1.1E-05</td><td>BT20/MB468</td><td>MP</td></tr>
<tr><td>>MI0000433|hsa-let-7g&WithFlank&3|-|52268272|52268367@11.33.23(+1U)&[MIMAT0000414&hsa-let-7g-5p&offsets|0|+1(+1U);m-58&3|-|52268336|52268357&offsets|0|+1(+1U)]|TGAGGTAGTAGTTTGTACAGTTTT</td><td>>MI0000433|hsa-let-7g&WithFlank&3|-|52268272|52268367</td><td>11</td><td>33</td><td>23(+1U)</td><td>TGAGGTAGTAGTTTGTACAGTTTT</td><td>48.1</td><td>8.8</td><td>1.9</td><td>4.5</td><td>6.1E-06</td><td>1.1E-05</td><td>MB231/MB468</td><td>MP</td></tr>
<tr><td>>trnaMT_SerGCT_MT_+_12207_12265@1.22.22|GAGAAAGCTCACAAGAACTGCT</td><td>>trnaMT_SerGCT_MT_+_12207_12265</td><td>1</td><td>22</td><td>22</td><td>GAGAAAGCTCACAAGAACTGCT</td><td>32597.0</td><td>5.8</td><td>1.3</td><td>4.6</td><td>5.3E-06</td><td>1.1E-05</td><td>BT20/MB468</td><td>MITO</td></tr>
<tr><td>>hg38_21_+_8433172_8446622_RNA45SN1_45S_50ntFlanks@12615.12659.45|GCTGAGGAGCCAATGGGGCGAAGCTACCATCTGTGGGATTATGAC</td><td>>hg38_21_+_8433172_8446622_RNA45SN1_45S_50ntFlanks</td><td>12615</td><td>12659</td><td>45</td><td>GCTGAGGAGCCAATGGGGCGAAGCTACCATCTGTGGGATTATGAC</td><td>123.2</td><td>-10.8</td><td>2.4</td><td>-4.5</td><td>6.1E-06</td><td>1.1E-05</td><td>MB231/MB468</td><td>MP</td></tr>
<tr><td>>hg38_21_+_8433172_8446622_RNA45SN1_45S_50ntFlanks@11571.11600.30|GCCGACTTAGAACTGGTGCGGACCAGGGGA</td><td>>hg38_21_+_8433172_8446622_RNA45SN1_45S_50ntFlanks</td><td>11571</td><td>11600</td><td>30</td><td>GCCGACTTAGAACTGGTGCGGACCAGGGGA</td><td>1347.8</td><td>-4.6</td><td>1.0</td><td>-4.5</td><td>7.7E-06</td><td>1.1E-05</td><td>BT20/MB468</td><td>NUC</td></tr>
<tr><td>>hg38_21_+_8433172_8446622_RNA45SN1_45S_50ntFlanks@12259.12285.27|CTTGATCTTGATTTTCAGTACGAATAC</td><td>>hg38_21_+_8433172_8446622_RNA45SN1_45S_50ntFlanks</td><td>12259</td><td>12285</td><td>27</td><td>CTTGATCTTGATTTTCAGTACGAATAC</td><td>390.1</td><td>-4.7</td><td>1.0</td><td>-4.5</td><td>7.4E-06</td><td>1.1E-05</td><td>MB231/MB468</td><td>MITO</td></tr>
<tr><td>>hg38_21_+_8433172_8446622_RNA45SN1_45S_50ntFlanks@8114.8138.25|GCGCGGGACATGTGGCGTACGGAAG</td><td>>hg38_21_+_8433172_8446622_RNA45SN1_45S_50ntFlanks</td><td>8114</td><td>8138</td><td>25</td><td>GCGCGGGACATGTGGCGTACGGAAG</td><td>15345.9</td><td>-5.2</td><td>1.1</td><td>-4.7</td><td>2.2E-06</td><td>1.1E-05</td><td>BT20/MB468</td><td>MP</td></tr>
<tr><td>>MI0000081|hsa-mir-24-2&WithFlank&19|-|13836281|13836365@56.74.19(+1A);MI0000080|hsa-mir-24-1&WithFlank&9|+|95086015|95086094@50.68.19(+1A)&[MIMAT0000080&hsa-miR-24-3p&offsets|0|-3(+1A);m-36&19|-|13836288|13836310&offsets|0|-4(+1A)];[MIMAT0000080_1&hsa-miR-24-3p&offsets|0|-3(+1A);m-35&9|+|95086064|95086086&offsets|0|-4(+1A)]|TGGCTCAGTTCAGCAGGAAA</td><td>>MI0000081|hsa-mir-24-2&WithFlank&19|-|13836281|13836365</td><td>56</td><td>74</td><td>19(+1A)</td><td>TGGCTCAGTTCAGCAGGAAA</td><td>300.6</td><td>8.2</td><td>1.8</td><td>4.5</td><td>6.2E-06</td><td>1.1E-05</td><td>BT20/MB468</td><td>CYTO</td></tr>
<tr><td>>trna10_AspGTC_12_-_125424193_125424264@39.70.32,trna12_AspGTC_12_-_125411891_125411962@39.70.32,trna144_AspGTC_6_-_27551236_27551307@39.70.32,trna38_AspGTC_17_-_8125556_8125627@39.70.32,trna45_AspGTC_6_+_27447453_27447524@39.70.32,trna48_AspGTC_6_+_27471523_27471594@39.70.32,trna4_AspGTC_12_+_96429799_96429870@39.70.32,trna69_AspGTC_1_-_161440205_161440276@39.70.32,trna72_AspGTC_1_-_161432824_161432895@39.70.32,trna75_AspGTC_1_-_161425414_161425485@39.70.32,trna78_AspGTC_1_-_161418033_161418104@39.70.32,trna81_AspGTC_1_-_161410615_161410686@39.70.32|GCGGGAGACCGGGGTTCGATTCCCCGACGGGG</td><td>>trna10_AspGTC_12_-_125424193_125424264</td><td>39</td><td>70</td><td>32</td><td>GCGGGAGACCGGGGTTCGATTCCCCGACGGGG</td><td>308.2</td><td>-3.2</td><td>0.7</td><td>-4.5</td><td>6.1E-06</td><td>1.1E-05</td><td>MB231/MB468</td><td>MP</td></tr>
<tr><td>>hg38_21_+_8433172_8446622_RNA45SN1_45S_50ntFlanks@7982.8012.31|TCAGATCAGACGTGGCGACCCGCTGAATTTA</td><td>>hg38_21_+_8433172_8446622_RNA45SN1_45S_50ntFlanks</td><td>7982</td><td>8012</td><td>31</td><td>TCAGATCAGACGTGGCGACCCGCTGAATTTA</td><td>231.6</td><td>-6.5</td><td>1.4</td><td>-4.5</td><td>6.1E-06</td><td>1.1E-05</td><td>MB231/MB468</td><td>MP</td></tr>
<tr><td>>hg38_21_+_8433172_8446622_RNA45SN1_45S_50ntFlanks@11583.11607.25|CTGGTGCGGACCAGGGGAATCCGAC</td><td>>hg38_21_+_8433172_8446622_RNA45SN1_45S_50ntFlanks</td><td>11583</td><td>11607</td><td>25</td><td>CTGGTGCGGACCAGGGGAATCCGAC</td><td>296.4</td><td>-6.6</td><td>1.5</td><td>-4.5</td><td>6.1E-06</td><td>1.1E-05</td><td>MB231/MB468</td><td>MP</td></tr>
<tr><td>>hg38_21_+_8433172_8446622_RNA45SN1_45S_50ntFlanks@11564.11592.29|CCTAGCAGCCGACTTAGAACTGGTGCGGA</td><td>>hg38_21_+_8433172_8446622_RNA45SN1_45S_50ntFlanks</td><td>11564</td><td>11592</td><td>29</td><td>CCTAGCAGCCGACTTAGAACTGGTGCGGA</td><td>152.3</td><td>-5.9</td><td>1.3</td><td>-4.5</td><td>7.2E-06</td><td>1.1E-05</td><td>MB231/MB468</td><td>NUC</td></tr>
<tr><td>>hg38_21_+_8433172_8446622_RNA45SN1_45S_50ntFlanks@4588.4615.28|CTATTTTGTTGGTTTTCGGAACTGAGGC</td><td>>hg38_21_+_8433172_8446622_RNA45SN1_45S_50ntFlanks</td><td>4588</td><td>4615</td><td>28</td><td>CTATTTTGTTGGTTTTCGGAACTGAGGC</td><td>248.4</td><td>-4.5</td><td>1.0</td><td>-4.5</td><td>6.2E-06</td><td>1.1E-05</td><td>BT20/MB468</td><td>CYTO</td></tr>
<tr><td>>hg38_21_+_8433172_8446622_RNA45SN1_45S_50ntFlanks@8354.8389.36|TTGAAAAGAACTTTGAAGAGAGAGTTCAAGAGGGCG</td><td>>hg38_21_+_8433172_8446622_RNA45SN1_45S_50ntFlanks</td><td>8354</td><td>8389</td><td>36</td><td>TTGAAAAGAACTTTGAAGAGAGAGTTCAAGAGGGCG</td><td>130.5</td><td>10.4</td><td>2.1</td><td>5.0</td><td>6.3E-07</td><td>1.1E-05</td><td>BT20/MB231</td><td>MP</td></tr>
<tr><td>>hg38_21_+_8433172_8446622_RNA45SN1_45S_50ntFlanks@12399.12419.21|ACGTCGCTTTTTGATCCTTCG</td><td>>hg38_21_+_8433172_8446622_RNA45SN1_45S_50ntFlanks</td><td>12399</td><td>12419</td><td>21</td><td>ACGTCGCTTTTTGATCCTTCG</td><td>279.7</td><td>-4.4</td><td>1.0</td><td>-4.5</td><td>6.2E-06</td><td>1.1E-05</td><td>BT20/MB468</td><td>CYTO</td></tr>
<tr><td>>hg38_21_+_8433172_8446622_RNA45SN1_45S_50ntFlanks@11235.11269.35|CTTCCCGTGGATCGCCCCAGCTGCGGCGGGCGTCG</td><td>>hg38_21_+_8433172_8446622_RNA45SN1_45S_50ntFlanks</td><td>11235</td><td>11269</td><td>35</td><td>CTTCCCGTGGATCGCCCCAGCTGCGGCGGGCGTCG</td><td>1003.6</td><td>-6.4</td><td>1.4</td><td>-4.6</td><td>5.3E-06</td><td>1.1E-05</td><td>BT20/MB468</td><td>MITO</td></tr>
<tr><td>>trna10_AlaCGC_6_+_26553731_26553802@1.18.18,trna119_AlaCGC_6_-_28641613_28641684@1.18.18,trna120_AlaAGC_6_-_28626014_28626085@1.18.18,trna13_AlaCGC_2_+_157257281_157257352@1.18.18,trna13_AlaTGC_12_-_125406301_125406372@1.18.18,trna66_AlaTGC_6_+_28611222_28611293@1.18.18,trna8_AlaTGC_12_+_125424512_125424583@1.18.18,trna8_AlaTGC_5_+_180633868_180633939@1.18.18|GGGGATGTAGCTCAGTGG</td><td>>trna10_AlaCGC_6_+_26553731_26553802</td><td>1</td><td>18</td><td>18</td><td>GGGGATGTAGCTCAGTGG</td><td>626.5</td><td>6.6</td><td>1.5</td><td>4.5</td><td>7.0E-06</td><td>1.1E-05</td><td>MB231/MB468</td><td>CYTO</td></tr>
<tr><td>>hg38_21_+_8433172_8446622_RNA45SN1_45S_50ntFlanks@9381.9418.38|CGCTCGCCGGCCGAGGTGGGATCCCGAGGCCTCTCCAG</td><td>>hg38_21_+_8433172_8446622_RNA45SN1_45S_50ntFlanks</td><td>9381</td><td>9418</td><td>38</td><td>CGCTCGCCGGCCGAGGTGGGATCCCGAGGCCTCTCCAG</td><td>53.9</td><td>-8.8</td><td>1.9</td><td>-4.7</td><td>2.3E-06</td><td>1.1E-05</td><td>BT20/MB468</td><td>MP</td></tr>
<tr><td>>hg38_21_+_8433172_8446622_RNA45SN1_45S_50ntFlanks@12157.12174.18|AGTTTGACTGGGGCGGTA</td><td>>hg38_21_+_8433172_8446622_RNA45SN1_45S_50ntFlanks</td><td>12157</td><td>12174</td><td>18</td><td>AGTTTGACTGGGGCGGTA</td><td>49.4</td><td>-8.7</td><td>1.8</td><td>-4.7</td><td>2.3E-06</td><td>1.1E-05</td><td>BT20/MB468</td><td>MP</td></tr>
<tr><td>>hg38_21_+_8433172_8446622_RNA45SN1_45S_50ntFlanks@7974.7993.20|ACGCGACCTCAGATCAGACG</td><td>>hg38_21_+_8433172_8446622_RNA45SN1_45S_50ntFlanks</td><td>7974</td><td>7993</td><td>20</td><td>ACGCGACCTCAGATCAGACG</td><td>2124.5</td><td>2.8</td><td>0.6</td><td>4.5</td><td>7.0E-06</td><td>1.1E-05</td><td>MB231/MB468</td><td>CYTO</td></tr>
<tr><td>>hg38_21_+_8433172_8446622_RNA45SN1_45S_50ntFlanks@12642.12659.18|CATCTGTGGGATTATGAC</td><td>>hg38_21_+_8433172_8446622_RNA45SN1_45S_50ntFlanks</td><td>12642</td><td>12659</td><td>18</td><td>CATCTGTGGGATTATGAC</td><td>431.4</td><td>-7.1</td><td>1.6</td><td>-4.5</td><td>7.8E-06</td><td>1.1E-05</td><td>BT20/MB468</td><td>NUC</td></tr>
<tr><td>>trnaMT_GluTTC_MT_-_14674_14742@44.72.29,trnalookalike8_GluTTC_5_-_93905172_93905240@44.72.29|TCGTGGTTGTAGTCCGTGCGAGAATACCA</td><td>>trnaMT_GluTTC_MT_-_14674_14742</td><td>44</td><td>72</td><td>29</td><td>TCGTGGTTGTAGTCCGTGCGAGAATACCA</td><td>1855.8</td><td>6.2</td><td>1.4</td><td>4.5</td><td>7.0E-06</td><td>1.1E-05</td><td>MB231/MB468</td><td>CYTO</td></tr>
<tr><td>>hg38_21_+_8433172_8446622_RNA45SN1_45S_50ntFlanks@6645.6676.32|CTCGTACGACTCTTAGCGGTGGATCACTCGGC</td><td>>hg38_21_+_8433172_8446622_RNA45SN1_45S_50ntFlanks</td><td>6645</td><td>6676</td><td>32</td><td>CTCGTACGACTCTTAGCGGTGGATCACTCGGC</td><td>1195.9</td><td>7.7</td><td>1.7</td><td>4.5</td><td>6.3E-06</td><td>1.1E-05</td><td>BT20/MB468</td><td>CYTO</td></tr>
<tr><td>>hg38_MT_+_1621_3279_RNR2_16S_50ntFlanks@51.94.44|GCTAAACCTAGCCCCAAACCCACTCCACCTTACTACCAGACAAC</td><td>>hg38_MT_+_1621_3279_RNR2_16S_50ntFlanks</td><td>51</td><td>94</td><td>44</td><td>GCTAAACCTAGCCCCAAACCCACTCCACCTTACTACCAGACAAC</td><td>759.1</td><td>5.8</td><td>1.3</td><td>4.5</td><td>7.3E-06</td><td>1.1E-05</td><td>MB231/MB468</td><td>NUC</td></tr>
<tr><td>>hg38_21_+_8433172_8446622_RNA45SN1_45S_50ntFlanks@10855.10873.19|AAGGATTGGCTCTAAGGGC</td><td>>hg38_21_+_8433172_8446622_RNA45SN1_45S_50ntFlanks</td><td>10855</td><td>10873</td><td>19</td><td>AAGGATTGGCTCTAAGGGC</td><td>201.1</td><td>-3.7</td><td>0.8</td><td>-4.5</td><td>6.2E-06</td><td>1.1E-05</td><td>MB231/MB468</td><td>MP</td></tr>
<tr><td>>trnaMT_GluTTC_MT_-_14674_14742@40.66.27,trnalookalike8_GluTTC_5_-_93905172_93905240@40.66.27|TTGGTCGTGGTTGTAGTCCGTGCGAGA</td><td>>trnaMT_GluTTC_MT_-_14674_14742</td><td>40</td><td>66</td><td>27</td><td>TTGGTCGTGGTTGTAGTCCGTGCGAGA</td><td>230.2</td><td>3.7</td><td>0.8</td><td>4.5</td><td>7.5E-06</td><td>1.1E-05</td><td>MB231/MB468</td><td>MITO</td></tr>
<tr><td>>hg38_21_+_8433172_8446622_RNA45SN1_45S_50ntFlanks@10408.10425.18|GGTCAGTCGGTCCTGAGA</td><td>>hg38_21_+_8433172_8446622_RNA45SN1_45S_50ntFlanks</td><td>10408</td><td>10425</td><td>18</td><td>GGTCAGTCGGTCCTGAGA</td><td>307.7</td><td>-5.8</td><td>1.3</td><td>-4.5</td><td>6.3E-06</td><td>1.1E-05</td><td>BT20/MB468</td><td>CYTO</td></tr>
<tr><td>>hg38_21_+_8433172_8446622_RNA45SN1_45S_50ntFlanks@11208.11230.23|GCGGCGACTCTGGACGCGAGCCG</td><td>>hg38_21_+_8433172_8446622_RNA45SN1_45S_50ntFlanks</td><td>11208</td><td>11230</td><td>23</td><td>GCGGCGACTCTGGACGCGAGCCG</td><td>1418.1</td><td>-3.6</td><td>0.8</td><td>-4.5</td><td>6.3E-06</td><td>1.1E-05</td><td>BT20/MB468</td><td>CYTO</td></tr>
<tr><td>>hg38_21_+_8433172_8446622_RNA45SN1_45S_50ntFlanks@9839.9857.19|GTAAGCAGAACTGGCGCTG</td><td>>hg38_21_+_8433172_8446622_RNA45SN1_45S_50ntFlanks</td><td>9839</td><td>9857</td><td>19</td><td>GTAAGCAGAACTGGCGCTG</td><td>53.8</td><td>-8.8</td><td>1.9</td><td>-4.7</td><td>2.3E-06</td><td>1.1E-05</td><td>BT20/MB468</td><td>MP</td></tr>
<tr><td>>hg38_21_+_8433172_8446622_RNA45SN1_45S_50ntFlanks@11575.11609.35|ACTTAGAACTGGTGCGGACCAGGGGAATCCGACTG</td><td>>hg38_21_+_8433172_8446622_RNA45SN1_45S_50ntFlanks</td><td>11575</td><td>11609</td><td>35</td><td>ACTTAGAACTGGTGCGGACCAGGGGAATCCGACTG</td><td>431.6</td><td>-3.2</td><td>0.7</td><td>-4.5</td><td>6.3E-06</td><td>1.1E-05</td><td>BT20/MB468</td><td>CYTO</td></tr>
<tr><td>>MI0003763|hsa-mir-767&WithFlank&X|-|152393415|152393535@33.54.22&[MIMAT0003882&hsa-miR-767-5p&offsets|0|-1;m-788&X|-|152393481|152393503&offsets|0|-1]|TGCACCATGGTTGTCTGAGCAT</td><td>>MI0003763|hsa-mir-767&WithFlank&X|-|152393415|152393535</td><td>33</td><td>54</td><td>22</td><td>TGCACCATGGTTGTCTGAGCAT</td><td>90.2</td><td>9.8</td><td>2.0</td><td>5.0</td><td>6.4E-07</td><td>1.1E-05</td><td>BT20/MB231</td><td>MP</td></tr>
<tr><td>>hg38_21_+_8433172_8446622_RNA45SN1_45S_50ntFlanks@12636.12659.24|AGCTACCATCTGTGGGATTATGAC</td><td>>hg38_21_+_8433172_8446622_RNA45SN1_45S_50ntFlanks</td><td>12636</td><td>12659</td><td>24</td><td>AGCTACCATCTGTGGGATTATGAC</td><td>1055.7</td><td>-7.8</td><td>1.8</td><td>-4.5</td><td>7.8E-06</td><td>1.1E-05</td><td>BT20/MB468</td><td>NUC</td></tr>
<tr><td>>hg38_21_+_8433172_8446622_RNA45SN1_45S_50ntFlanks@12628.12654.27|TGGGGCGAAGCTACCATCTGTGGGATT</td><td>>hg38_21_+_8433172_8446622_RNA45SN1_45S_50ntFlanks</td><td>12628</td><td>12654</td><td>27</td><td>TGGGGCGAAGCTACCATCTGTGGGATT</td><td>2599.7</td><td>-7.8</td><td>1.6</td><td>-4.7</td><td>2.3E-06</td><td>1.1E-05</td><td>BT20/MB468</td><td>MP</td></tr>
<tr><td>>hg38_21_+_8433172_8446622_RNA45SN1_45S_50ntFlanks@8472.8491.20|CGGCCCGGCGGATCTTTCCC</td><td>>hg38_21_+_8433172_8446622_RNA45SN1_45S_50ntFlanks</td><td>8472</td><td>8491</td><td>20</td><td>CGGCCCGGCGGATCTTTCCC</td><td>272.9</td><td>-5.8</td><td>1.3</td><td>-4.5</td><td>6.3E-06</td><td>1.1E-05</td><td>BT20/MB468</td><td>CYTO</td></tr>
<tr><td>>MI0000449|hsa-mir-132&WithFlank&17|-|2049902|2050014@65.87.23&[MIMAT0000426&hsa-miR-132-3p&offsets|0|+1;m-138&17|-|2049929|2049950&offsets|0|+1]|TAACAGTCTACAGCCATGGTCGC</td><td>>MI0000449|hsa-mir-132&WithFlank&17|-|2049902|2050014</td><td>65</td><td>87</td><td>23</td><td>TAACAGTCTACAGCCATGGTCGC</td><td>40.4</td><td>8.6</td><td>1.9</td><td>4.5</td><td>6.2E-06</td><td>1.1E-05</td><td>MB231/MB468</td><td>MP</td></tr>
<tr><td>>trnaMT_SerGCT_MT_+_12207_12265@1.21.21|GAGAAAGCTCACAAGAACTGC</td><td>>trnaMT_SerGCT_MT_+_12207_12265</td><td>1</td><td>21</td><td>21</td><td>GAGAAAGCTCACAAGAACTGC</td><td>13066.6</td><td>5.0</td><td>1.1</td><td>4.5</td><td>7.5E-06</td><td>1.1E-05</td><td>MB231/MB468</td><td>MITO</td></tr>
<tr><td>>hg38_21_+_8433172_8446622_RNA45SN1_45S_50ntFlanks@11581.11616.36|AACTGGTGCGGACCAGGGGAATCCGACTGTTTAATT</td><td>>hg38_21_+_8433172_8446622_RNA45SN1_45S_50ntFlanks</td><td>11581</td><td>11616</td><td>36</td><td>AACTGGTGCGGACCAGGGGAATCCGACTGTTTAATT</td><td>327.7</td><td>2.8</td><td>0.6</td><td>4.5</td><td>7.1E-06</td><td>1.1E-05</td><td>MB231/MB468</td><td>CYTO</td></tr>
</table>

<table>
<tr><td>>am_trna128_GlyGCC_6_-_27870686_27870756@7.34.28,trna18_GlyGCC_16_+_70822597_70822667@7.34.28,trna19_GlyGCC_16_+_70823410_70823480@7.34.28,trna19_GlyGCC_2_-_157257659_157257729@7.34.28,trna24_GlyGCC_16_-_70812942_70813012@7.34.28,trna25_GlyGCC_16_-_70812114_70812184@7.34.28,trna2_GlyGCC_21_-_18827107_18827177@7.34.28,trna35_GlyGCC_1_+_161413094_161413164@7.34.28,trna37_GlyGCC_1_+_161420467_161420537@7.34.28,trna39_GlyGCC_1_+_161427898_161427968@7.34.28,trna41_GlyGCC_1_+_161435258_161435328@7.34.28,trna5_GlyGCC_17_+_8029064_8029134@7.34.28,trna68_GlyGCC_1_-_161493637_161493707@7.34.28|GTGGTTCAGTGGTAGAATTCTCGCCTGC</td><td>>am_trna128_GlyGCC_6_-_27870686_27870756</td><td>7</td><td>34</td><td>28</td><td>GTGGTTCAGTGGTAGAATTCTCGCCTGC</td><td>165.3</td><td>5.5</td><td>1.2</td><td>4.5</td><td>7.5E-06</td><td>1.1E-05</td><td>MB231/MB468</td><td>MITO</td></tr>
<tr><td>>MI0000455|hsa-mir-138-2&WithFlank&16|+|56858512|56858607@16.36.21(+1U);MI0000476|hsa-mir-138-1&WithFlank&3|+|44114206|44114316@29.49.21(+1U)&[MIMAT0000430&hsa-miR-138-5p&offsets|0|-2(+1U);m-509&16|+|56858527|56858549&offsets|0|-2(+1U)];[MIMAT0000430_1&hsa-miR-138-5p&offsets|0|-2(+1U);m-497&3|+|44114234|44114256&offsets|0|-2(+1U)]|AGCTGGTGTTGTGAATCAGGCT</td><td>>MI0000455|hsa-mir-138-2&WithFlank&16|+|56858512|56858607</td><td>16</td><td>36</td><td>21(+1U)</td><td>AGCTGGTGTTGTGAATCAGGCT</td><td>36.8</td><td>8.4</td><td>1.9</td><td>4.5</td><td>6.2E-06</td><td>1.1E-05</td><td>MB231/MB468</td><td>MP</td></tr>
<tr><td>>hg38_21_+_8433172_8446622_RNA45SN1_45S_50ntFlanks@6719.6743.25|TGAATTGCAGGACACATTGATCATC</td><td>>hg38_21_+_8433172_8446622_RNA45SN1_45S_50ntFlanks</td><td>6719</td><td>6743</td><td>25</td><td>TGAATTGCAGGACACATTGATCATC</td><td>1725.6</td><td>-2.7</td><td>0.6</td><td>-4.5</td><td>5.4E-06</td><td>1.1E-05</td><td>BT20/MB468</td><td>MITO</td></tr>
<tr><td>>trna10_LysCTT_16_+_3241501_3241573@4.34.31,trna13_LysCTT_14_-_58706613_58706685@4.34.31,trna2_LysCTT_15_+_79152904_79152976@4.34.31|CGGCTAGCTCAGTCGGTAGAGCATGGGACTC</td><td>>trna10_LysCTT_16_+_3241501_3241573</td><td>4</td><td>34</td><td>31</td><td>CGGCTAGCTCAGTCGGTAGAGCATGGGACTC</td><td>373.5</td><td>7.5</td><td>1.7</td><td>4.5</td><td>7.5E-06</td><td>1.1E-05</td><td>MB231/MB468</td><td>MITO</td></tr>
<tr><td>>hg38_21_+_8433172_8446622_RNA45SN1_45S_50ntFlanks@6771.6799.29|GGTTCCTCCCGGGGCTACGCCTGTCTGAG</td><td>>hg38_21_+_8433172_8446622_RNA45SN1_45S_50ntFlanks</td><td>6771</td><td>6799</td><td>29</td><td>GGTTCCTCCCGGGGCTACGCCTGTCTGAG</td><td>492.5</td><td>-7.2</td><td>1.5</td><td>-4.7</td><td>2.3E-06</td><td>1.1E-05</td><td>BT20/MB468</td><td>MP</td></tr>
<tr><td>>trna10_LysCTT_16_+_3241501_3241573@1.37.37,trna13_LysCTT_14_-_58706613_58706685@1.37.37,trna2_LysCTT_15_+_79152904_79152976@1.37.37|GCCCGGCTAGCTCAGTCGGTAGAGCATGGGACTCTTA</td><td>>trna10_LysCTT_16_+_3241501_3241573</td><td>1</td><td>37</td><td>37</td><td>GCCCGGCTAGCTCAGTCGGTAGAGCATGGGACTCTTA</td><td>383.8</td><td>8.5</td><td>1.9</td><td>4.5</td><td>6.3E-06</td><td>1.1E-05</td><td>BT20/MB468</td><td>CYTO</td></tr>
<tr><td>>hg38_21_+_8433172_8446622_RNA45SN1_45S_50ntFlanks@12329.12346.18|TTTTAAGCAGGAGGTGTC</td><td>>hg38_21_+_8433172_8446622_RNA45SN1_45S_50ntFlanks</td><td>12329</td><td>12346</td><td>18</td><td>TTTTAAGCAGGAGGTGTC</td><td>251.1</td><td>-4.2</td><td>0.9</td><td>-4.5</td><td>7.1E-06</td><td>1.1E-05</td><td>MB231/MB468</td><td>CYTO</td></tr>
<tr><td>>hg38_21_+_8433172_8446622_RNA45SN1_45S_50ntFlanks@4686.4726.41|GACGGACCAGAGCGAAAGCATTTGCCAAGAATGTTTTCATT</td><td>>hg38_21_+_8433172_8446622_RNA45SN1_45S_50ntFlanks</td><td>4686</td><td>4726</td><td>41</td><td>GACGGACCAGAGCGAAAGCATTTGCCAAGAATGTTTTCATT</td><td>47.0</td><td>9.3</td><td>2.0</td><td>4.7</td><td>2.3E-06</td><td>1.1E-05</td><td>BT20/MB468</td><td>MP</td></tr>
<tr><td>>trna116_GluCTC_1_-_145399233_145399304@4.33.30,trna59_GluCTC_1_+_249168447_249168518@4.33.30,trna71_GluCTC_1_-_161439189_161439260@4.33.30,trna74_GluCTC_1_-_161431809_161431880@4.33.30,trna77_GluCTC_1_-_161424398_161424469@4.33.30,trna77_GluCTC_6_+_28949976_28950047@4.33.30,trna80_GluCTC_1_-_161417018_161417089@4.33.30,trna87_GluCTC_6_-_126101393_126101464@4.33.30|CTGGTGGTCTAGTGGTTAGGATTCGGCGCT</td><td>>trna116_GluCTC_1_-_145399233_145399304</td><td>4</td><td>33</td><td>30</td><td>CTGGTGGTCTAGTGGTTAGGATTCGGCGCT</td><td>286.5</td><td>4.1</td><td>0.9</td><td>4.5</td><td>7.6E-06</td><td>1.1E-05</td><td>MB231/MB468</td><td>MITO</td></tr>
<tr><td>>MI0000750|hsa-mir-26a-2&WithFlank&12|-|57824603|57824698@20.43.24;MI0000083|hsa-mir-26a-1&WithFlank&3|+|37969398|37969486@16.39.24&[MIMAT0000082&hsa-miR-26a-5p&offsets|0|+2;m-33&12|-|57824658|57824679&offsets|0|+2];[MIMAT0000082_1&hsa-miR-26a-5p&offsets|0|+2;m-34&3|+|37969413|37969434&offsets|0|+2]|TTCAAGTAATCCAGGATAGGCTGT</td><td>>MI0000750|hsa-mir-26a-2&WithFlank&12|-|57824603|57824698</td><td>20</td><td>43</td><td>24</td><td>TTCAAGTAATCCAGGATAGGCTGT</td><td>248.2</td><td>7.6</td><td>1.7</td><td>4.5</td><td>7.6E-06</td><td>1.1E-05</td><td>MB231/MB468</td><td>MITO</td></tr>
<tr><td>>hg38_21_+_8433172_8446622_RNA45SN1_45S_50ntFlanks@12710.12742.33|CGCGGAGCCTCGGTTGGCCTCGGATAGCCGGTC</td><td>>hg38_21_+_8433172_8446622_RNA45SN1_45S_50ntFlanks</td><td>12710</td><td>12742</td><td>33</td><td>CGCGGAGCCTCGGTTGGCCTCGGATAGCCGGTC</td><td>563.4</td><td>-7.1</td><td>1.5</td><td>-4.7</td><td>2.3E-06</td><td>1.1E-05</td><td>BT20/MB468</td><td>MP</td></tr>
<tr><td>>hg38_21_+_8433172_8446622_RNA45SN1_45S_50ntFlanks@11610.11640.31|TTTAATTAAAACAAAGCATCGCGAAGGCCCG</td><td>>hg38_21_+_8433172_8446622_RNA45SN1_45S_50ntFlanks</td><td>11610</td><td>11640</td><td>31</td><td>TTTAATTAAAACAAAGCATCGCGAAGGCCCG</td><td>44.1</td><td>-8.5</td><td>1.8</td><td>-4.7</td><td>2.3E-06</td><td>1.1E-05</td><td>BT20/MB468</td><td>MP</td></tr>
<tr><td>>hg38_21_+_8433172_8446622_RNA45SN1_45S_50ntFlanks@6640.6683.44|TCGACCTCGTACGACTCTTAGCGGTGGATCACTCGGCTCGTGCG</td><td>>hg38_21_+_8433172_8446622_RNA45SN1_45S_50ntFlanks</td><td>6640</td><td>6683</td><td>44</td><td>TCGACCTCGTACGACTCTTAGCGGTGGATCACTCGGCTCGTGCG</td><td>294.7</td><td>7.8</td><td>1.7</td><td>4.5</td><td>7.4E-06</td><td>1.1E-05</td><td>MB231/MB468</td><td>NUC</td></tr>
<tr><td>>hg38_1_-_228634819_228635039_RNA5S12_5S_50ntFlanks@138.171.34|AGACCGCCTGGGAATACCGGGTGCTGTAGGCTTT</td><td>>hg38_1_-_228634819_228635039_RNA5S12_5S_50ntFlanks</td><td>138</td><td>171</td><td>34</td><td>AGACCGCCTGGGAATACCGGGTGCTGTAGGCTTT</td><td>514.2</td><td>3.9</td><td>0.9</td><td>4.5</td><td>5.4E-06</td><td>1.1E-05</td><td>BT20/MB468</td><td>MITO</td></tr>
<tr><td>>hg38_21_+_8433172_8446622_RNA45SN1_45S_50ntFlanks@12241.12271.31|CAGAAGGGCAAAAGCTCGCTTGATCTTGATT</td><td>>hg38_21_+_8433172_8446622_RNA45SN1_45S_50ntFlanks</td><td>12241</td><td>12271</td><td>31</td><td>CAGAAGGGCAAAAGCTCGCTTGATCTTGATT</td><td>475.9</td><td>-3.4</td><td>0.8</td><td>-4.5</td><td>7.6E-06</td><td>1.1E-05</td><td>MB231/MB468</td><td>MITO</td></tr>
<tr><td>>hg38_21_+_8433172_8446622_RNA45SN1_45S_50ntFlanks@10830.10858.29|CAAGCCGGATCCGTAACTTCGGGATAAGG</td><td>>hg38_21_+_8433172_8446622_RNA45SN1_45S_50ntFlanks</td><td>10830</td><td>10858</td><td>29</td><td>CAAGCCGGATCCGTAACTTCGGGATAAGG</td><td>403.9</td><td>-4.7</td><td>1.0</td><td>-4.5</td><td>5.4E-06</td><td>1.1E-05</td><td>BT20/MB468</td><td>MITO</td></tr>
<tr><td>>hg38_21_+_8433172_8446622_RNA45SN1_45S_50ntFlanks@12415.12457.43|CTTCGATGTCGGCTCTTCCTATCATTGTGAAGCAGAATTCACC</td><td>>hg38_21_+_8433172_8446622_RNA45SN1_45S_50ntFlanks</td><td>12415</td><td>12457</td><td>43</td><td>CTTCGATGTCGGCTCTTCCTATCATTGTGAAGCAGAATTCACC</td><td>323.0</td><td>-4.4</td><td>0.9</td><td>-5.0</td><td>6.6E-07</td><td>1.1E-05</td><td>BT20/MB231</td><td>CYTO</td></tr>
<tr><td>>hg38_21_+_8433172_8446622_RNA45SN1_45S_50ntFlanks@5492.5509.18|AGAAGACGGTCGAACTTG</td><td>>hg38_21_+_8433172_8446622_RNA45SN1_45S_50ntFlanks</td><td>5492</td><td>5509</td><td>18</td><td>AGAAGACGGTCGAACTTG</td><td>251.1</td><td>-3.3</td><td>0.7</td><td>-4.5</td><td>7.4E-06</td><td>1.1E-05</td><td>MB231/MB468</td><td>NUC</td></tr>
<tr><td>>hg38_21_+_8433172_8446622_RNA45SN1_45S_50ntFlanks@5472.5502.31|ACGGCCCTGGCGGAGCGCTGAGAAGACGGTC</td><td>>hg38_21_+_8433172_8446622_RNA45SN1_45S_50ntFlanks</td><td>5472</td><td>5502</td><td>31</td><td>ACGGCCCTGGCGGAGCGCTGAGAAGACGGTC</td><td>421.9</td><td>4.6</td><td>1.0</td><td>4.7</td><td>2.3E-06</td><td>1.1E-05</td><td>BT20/MB468</td><td>MP</td></tr>
<tr><td>>MI0000086|hsa-mir-28&WithFlank&3|+|188688775|188688872@60.81.22(+1A)&[MIMAT0004502&hsa-miR-28-3p&offsets|0|0(+1A);m-28&3|+|188688834|188688855&offsets|0|0(+1A)]|CACTAGATTGTGAGCTCCTGGAA</td><td>>MI0000086|hsa-mir-28&WithFlank&3|+|188688775|188688872</td><td>60</td><td>81</td><td>22(+1A)</td><td>CACTAGATTGTGAGCTCCTGGAA</td><td>899.6</td><td>6.4</td><td>1.4</td><td>4.7</td><td>2.3E-06</td><td>1.1E-05</td><td>BT20/MB468</td><td>MP</td></tr>
<tr><td>>hg38_21_+_8433172_8446622_RNA45SN1_45S_50ntFlanks@9603.9629.27|GAAAGACTAATCGAACCATCTAGTAGC</td><td>>hg38_21_+_8433172_8446622_RNA45SN1_45S_50ntFlanks</td><td>9603</td><td>9629</td><td>27</td><td>GAAAGACTAATCGAACCATCTAGTAGC</td><td>45.2</td><td>9.2</td><td>2.0</td><td>4.7</td><td>2.3E-06</td><td>1.1E-05</td><td>BT20/MB468</td><td>MP</td></tr>
<tr><td>>hg38_1_-_228634819_228635039_RNA5S12_5S_50ntFlanks@135.172.38|GGGAGACCGCCTGGGAATACCGGGTGCTGTAGGCTTTT</td><td>>hg38_1_-_228634819_228635039_RNA5S12_5S_50ntFlanks</td><td>135</td><td>172</td><td>38</td><td>GGGAGACCGCCTGGGAATACCGGGTGCTGTAGGCTTTT</td><td>528.2</td><td>4.5</td><td>1.0</td><td>4.5</td><td>7.2E-06</td><td>1.1E-05</td><td>MB231/MB468</td><td>CYTO</td></tr>
<tr><td>>hg38_21_+_8433172_8446622_RNA45SN1_45S_50ntFlanks@10781.10799.19|AGGTGAACAGCCTCTGGCA</td><td>>hg38_21_+_8433172_8446622_RNA45SN1_45S_50ntFlanks</td><td>10781</td><td>10799</td><td>19</td><td>AGGTGAACAGCCTCTGGCA</td><td>267.9</td><td>-4.2</td><td>0.9</td><td>-4.7</td><td>2.3E-06</td><td>1.1E-05</td><td>BT20/MB468</td><td>MP</td></tr>
<tr><td>>hg38_21_+_8433172_8446622_RNA45SN1_45S_50ntFlanks@7973.7997.25|GACGCGACCTCAGATCAGACGTGGC</td><td>>hg38_21_+_8433172_8446622_RNA45SN1_45S_50ntFlanks</td><td>7973</td><td>7997</td><td>25</td><td>GACGCGACCTCAGATCAGACGTGGC</td><td>486.7</td><td>3.9</td><td>0.9</td><td>4.5</td><td>8.0E-06</td><td>1.1E-05</td><td>BT20/MB468</td><td>NUC</td></tr>
<tr><td>>hg38_21_+_8433172_8446622_RNA45SN1_45S_50ntFlanks@11733.11751.19|TATGACTCTCTTAAGGTAG</td><td>>hg38_21_+_8433172_8446622_RNA45SN1_45S_50ntFlanks</td><td>11733</td><td>11751</td><td>19</td><td>TATGACTCTCTTAAGGTAG</td><td>337.9</td><td>-4.9</td><td>1.1</td><td>-4.5</td><td>7.4E-06</td><td>1.1E-05</td><td>MB231/MB468</td><td>NUC</td></tr>
<tr><td>>hg38_21_+_8433172_8446622_RNA45SN1_45S_50ntFlanks@8157.8178.22|CTCGTGGGGGGCCCAAGTCCTT</td><td>>hg38_21_+_8433172_8446622_RNA45SN1_45S_50ntFlanks</td><td>8157</td><td>8178</td><td>22</td><td>CTCGTGGGGGGCCCAAGTCCTT</td><td>923.5</td><td>-6.8</td><td>1.5</td><td>-4.5</td><td>5.5E-06</td><td>1.1E-05</td><td>BT20/MB468</td><td>MITO</td></tr>
<tr><td>>hg38_21_+_8433172_8446622_RNA45SN1_45S_50ntFlanks@12044.12068.25|ACTGACCCGGTGAGGCGGGGGGGCG</td><td>>hg38_21_+_8433172_8446622_RNA45SN1_45S_50ntFlanks</td><td>12044</td><td>12068</td><td>25</td><td>ACTGACCCGGTGAGGCGGGGGGGCG</td><td>894.4</td><td>-7.4</td><td>1.6</td><td>-4.7</td><td>2.3E-06</td><td>1.1E-05</td><td>BT20/MB468</td><td>MP</td></tr>
<tr><td>>hg38_21_+_8433172_8446622_RNA45SN1_45S_50ntFlanks@9837.9864.28|TGGTAAGCAGAACTGGCGCTGCGGGATG</td><td>>hg38_21_+_8433172_8446622_RNA45SN1_45S_50ntFlanks</td><td>9837</td><td>9864</td><td>28</td><td>TGGTAAGCAGAACTGGCGCTGCGGGATG</td><td>157.7</td><td>10.6</td><td>2.1</td><td>5.0</td><td>6.6E-07</td><td>1.1E-05</td><td>BT20/MB231</td><td>MP</td></tr>
<tr><td>>hg38_21_+_8433172_8446622_RNA45SN1_45S_50ntFlanks@12488.12505.18|AACGTGAGCTGGGTTTAG</td><td>>hg38_21_+_8433172_8446622_RNA45SN1_45S_50ntFlanks</td><td>12488</td><td>12505</td><td>18</td><td>AACGTGAGCTGGGTTTAG</td><td>791.8</td><td>-7.9</td><td>1.8</td><td>-4.5</td><td>8.0E-06</td><td>1.1E-05</td><td>BT20/MB468</td><td>NUC</td></tr>
<tr><td>>hg38_21_+_8433172_8446622_RNA45SN1_45S_50ntFlanks@12001.12043.43|CTGCGGGCCGCCGGTGAAATACCACTACTCTGATCGTTTTTTC</td><td>>hg38_21_+_8433172_8446622_RNA45SN1_45S_50ntFlanks</td><td>12001</td><td>12043</td><td>43</td><td>CTGCGGGCCGCCGGTGAAATACCACTACTCTGATCGTTTTTTC</td><td>112.4</td><td>-10.7</td><td>2.4</td><td>-4.5</td><td>6.3E-06</td><td>1.1E-05</td><td>MB231/MB468</td><td>MP</td></tr>
<tr><td>>hg38_21_+_8433172_8446622_RNA45SN1_45S_50ntFlanks@11561.11588.28|GCGCCTAGCAGCCGACTTAGAACTGGTG</td><td>>hg38_21_+_8433172_8446622_RNA45SN1_45S_50ntFlanks</td><td>11561</td><td>11588</td><td>28</td><td>GCGCCTAGCAGCCGACTTAGAACTGGTG</td><td>394.9</td><td>-4.9</td><td>1.1</td><td>-4.5</td><td>7.5E-06</td><td>1.1E-05</td><td>MB231/MB468</td><td>NUC</td></tr>
<tr><td>>hg38_21_+_8433172_8446622_RNA45SN1_45S_50ntFlanks@11576.11601.26|CTTAGAACTGGTGCGGACCAGGGGAA</td><td>>hg38_21_+_8433172_8446622_RNA45SN1_45S_50ntFlanks</td><td>11576</td><td>11601</td><td>26</td><td>CTTAGAACTGGTGCGGACCAGGGGAA</td><td>257.0</td><td>-2.7</td><td>0.6</td><td>-4.5</td><td>5.5E-06</td><td>1.1E-05</td><td>BT20/MB468</td><td>MITO</td></tr>
<tr><td>>hg38_21_+_8433172_8446622_RNA45SN1_45S_50ntFlanks@9855.9885.31|CTGCGGGATGAACCGAACGCCGGGTTAAGGC</td><td>>hg38_21_+_8433172_8446622_RNA45SN1_45S_50ntFlanks</td><td>9855</td><td>9885</td><td>31</td><td>CTGCGGGATGAACCGAACGCCGGGTTAAGGC</td><td>137.4</td><td>10.4</td><td>2.1</td><td>5.0</td><td>6.6E-07</td><td>1.1E-05</td><td>BT20/MB231</td><td>MP</td></tr>
<tr><td>>am_trna128_GlyGCC_6_-_27870686_27870756@3.31.29,trna133_GlyCCC_1_-_16872434_16872504@3.31.29,trna18_GlyGCC_16_+_70822597_70822667@3.31.29,trna19_GlyGCC_16_+_70823410_70823480@3.31.29,trna19_GlyGCC_2_-_157257659_157257729@3.31.29,trna24_GlyGCC_16_-_70812942_70813012@3.31.29,trna25_GlyGCC_16_-_70812114_70812184@3.31.29,trna4_GlyCCC_1_+_17188416_17188486@3.31.29,trna5_GlyGCC_17_+_8029064_8029134@3.31.29,trna68_GlyGCC_1_-_161493637_161493707@3.31.29|ATTGGTGGTTCAGTGGTAGAATTCTCGCC</td><td>>am_trna128_GlyGCC_6_-_27870686_27870756</td><td>3</td><td>31</td><td>29</td><td>ATTGGTGGTTCAGTGGTAGAATTCTCGCC</td><td>234.3</td><td>5.5</td><td>1.2</td><td>4.5</td><td>7.6E-06</td><td>1.1E-05</td><td>MB231/MB468</td><td>MITO</td></tr>
<tr><td>>hg38_21_+_8433172_8446622_RNA45SN1_45S_50ntFlanks@6731.6753.23|CACATTGATCATCGACACTTCGA</td><td>>hg38_21_+_8433172_8446622_RNA45SN1_45S_50ntFlanks</td><td>6731</td><td>6753</td><td>23</td><td>CACATTGATCATCGACACTTCGA</td><td>528.5</td><td>-3.8</td><td>0.9</td><td>-4.5</td><td>7.7E-06</td><td>1.1E-05</td><td>MB231/MB468</td><td>MITO</td></tr>
<tr><td>>MI0000440|hsa-mir-27b&WithFlank&9|+|95085439|95085547@67.85.19&[MIMAT0000419&hsa-miR-27b-3p&offsets|0|-2;m-39&9|+|95085505|95085525&offsets|0|-2]|TTCACAGTGGCTAAGTTCT</td><td>>MI0000440|hsa-mir-27b&WithFlank&9|+|95085439|95085547</td><td>67</td><td>85</td><td>19</td><td>TTCACAGTGGCTAAGTTCT</td><td>1659.2</td><td>3.7</td><td>0.8</td><td>4.5</td><td>6.3E-06</td><td>1.1E-05</td><td>MB231/MB468</td><td>MP</td></tr>
<tr><td>>hg38_21_+_8433172_8446622_RNA45SN1_45S_50ntFlanks@11567.11586.20|AGCAGCCGACTTAGAACTGG</td><td>>hg38_21_+_8433172_8446622_RNA45SN1_45S_50ntFlanks</td><td>11567</td><td>11586</td><td>20</td><td>AGCAGCCGACTTAGAACTGG</td><td>813.6</td><td>2.3</td><td>0.5</td><td>4.5</td><td>5.5E-06</td><td>1.1E-05</td><td>BT20/MB468</td><td>MITO</td></tr>
<tr><td>>hg38_21_+_8433172_8446622_RNA45SN1_45S_50ntFlanks@10797.10818.22|GCATGTTGGAACAATGTAGGTA</td><td>>hg38_21_+_8433172_8446622_RNA45SN1_45S_50ntFlanks</td><td>10797</td><td>10818</td><td>22</td><td>GCATGTTGGAACAATGTAGGTA</td><td>677.4</td><td>-3.6</td><td>0.8</td><td>-4.5</td><td>7.5E-06</td><td>1.1E-05</td><td>MB231/MB468</td><td>NUC</td></tr>
</table>

<table>
<tr><td>>MI0000542|hsa-mir-320a&WithFlank&8|-|22244960|22245043@48.68.21(+1U)&[MIMAT0000510&hsa-miR-320a-3p&offsets|0|-1(+1U);m-60&8|-|22244975|22244996&offsets|0|-1(+1U)]|AAAAGCTGGGTTGAGAGGGCGT</td><td>>MI0000542|hsa-mir-320a&WithFlank&8|-|22244960|22245043</td><td>48</td><td>68</td><td>21(+1U)</td><td>AAAAGCTGGGTTGAGAGGGCGT</td><td>3890.0</td><td>4.3</td><td>0.9</td><td>4.5</td><td>6.3E-06</td><td>1.1E-05</td><td>MB231/MB468</td><td>MP</td></tr>
<tr><td>>hg38_21_+_8433172_8446622_RNA45SN1_45S_50ntFlanks@4220.4242.23|ACGAAAAAATAACAATACAGGACT</td><td>>hg38_21_+_8433172_8446622_RNA45SN1_45S_50ntFlanks</td><td>4220</td><td>4242</td><td>23</td><td>ACGAAAAAATAACAATACAGGACT</td><td>1056.8</td><td>-3.5</td><td>0.8</td><td>-4.5</td><td>7.2E-06</td><td>1.1E-05</td><td>MB231/MB468</td><td>CYTO</td></tr>
<tr><td>>hg38_21_+_8433172_8446622_RNA45SN1_45S_50ntFlanks@12078.12095.18|GGGCTCTCGCTTCTGGCG</td><td>>hg38_21_+_8433172_8446622_RNA45SN1_45S_50ntFlanks</td><td>12078</td><td>12095</td><td>18</td><td>GGGCTCTCGCTTCTGGCG</td><td>529.4</td><td>-5.0</td><td>1.1</td><td>-4.5</td><td>6.5E-06</td><td>1.1E-05</td><td>BT20/MB468</td><td>CYTO</td></tr>
<tr><td>>MI0003776|hsa-mir-320b-1&WithFlank&1|+|116671740|116671823@48.66.19;MI0003839|hsa-mir-320b-2&WithFlank&1|-|224257034|224257116@47.65.19;MI0003778|hsa-mir-320c-1&WithFlank&18|+|21683512|21683595@48.66.19;MI0008191|hsa-mir-320c-2&WithFlank&18|+|24321669|24321752@48.66.19;MI0000542|hsa-mir-320a&WithFlank&8|-|22244960|22245043@48.66.19&[MIMAT0005792&hsa-miR-320b&offsets|0|-3;m-183&1|+|116671787|116671806&offsets|0|-1];[MIMAT0005792_1&hsa-miR-320b&offsets|0|-3;m-183&1|-|224257051|224257070&offsets|0|-1];[MIMAT0005793&hsa-miR-320c&offsets|0|-1;m-222&18|+|21683559|21683578&offsets|0|-1];[MIMAT0005793_1&hsa-miR-320c&offsets|0|-1;m-219&18|+|24321716|24321735&offsets|0|-1];[MIMAT0000510&hsa-miR-320a-3p&offsets|0|-3;m-60&8|-|22244975|22244996&offsets|0|-3]|AAAAGCTGGGTTGAGAGGG</td><td>>MI0003776|hsa-mir-320b-1&WithFlank&1|+|116671740|116671823</td><td>48</td><td>66</td><td>19</td><td>AAAAGCTGGGTTGAGAGGG</td><td>102.4</td><td>3.7</td><td>0.8</td><td>4.5</td><td>8.0E-06</td><td>1.1E-05</td><td>BT20/MB468</td><td>NUC</td></tr>
<tr><td>>hg38_21_+_8433172_8446622_RNA45SN1_45S_50ntFlanks@10772.10805.34|AGGTCTCCAAGGTGAACAGCCTCTGGCATGTTGG</td><td>>hg38_21_+_8433172_8446622_RNA45SN1_45S_50ntFlanks</td><td>10772</td><td>10805</td><td>34</td><td>AGGTCTCCAAGGTGAACAGCCTCTGGCATGTTGG</td><td>156.7</td><td>-4.4</td><td>1.0</td><td>-4.5</td><td>7.5E-06</td><td>1.1E-05</td><td>MB231/MB468</td><td>NUC</td></tr>
<tr><td>>MI0000077|hsa-mir-21&WithFlank&17|+|59841260|59841343@13.36.24&[MIMAT0000076&hsa-miR-21-5p&offsets|-1|+1;m-1&17|+|59841273|59841294&offsets|-1|+1]|GTAGCTTATCAGACTGATGTTGAC</td><td>>MI0000077|hsa-mir-21&WithFlank&17|+|59841260|59841343</td><td>13</td><td>36</td><td>24</td><td>GTAGCTTATCAGACTGATGTTGAC</td><td>314.8</td><td>5.7</td><td>1.3</td><td>4.5</td><td>7.7E-06</td><td>1.1E-05</td><td>MB231/MB468</td><td>MITO</td></tr>
<tr><td>>hg38_21_+_8433172_8446622_RNA45SN1_45S_50ntFlanks@12450.12480.31|AATTCACCAAGCGTTGGATTGTTCACCCACT</td><td>>hg38_21_+_8433172_8446622_RNA45SN1_45S_50ntFlanks</td><td>12450</td><td>12480</td><td>31</td><td>AATTCACCAAGCGTTGGATTGTTCACCCACT</td><td>89.9</td><td>3.4</td><td>0.8</td><td>4.5</td><td>8.0E-06</td><td>1.1E-05</td><td>BT20/MB468</td><td>NUC</td></tr>
<tr><td>>hg38_21_+_8433172_8446622_RNA45SN1_45S_50ntFlanks@9484.9506.23|GTGTTAGGACCCGAAAGATGGTG</td><td>>hg38_21_+_8433172_8446622_RNA45SN1_45S_50ntFlanks</td><td>9484</td><td>9506</td><td>23</td><td>GTGTTAGGACCCGAAAGATGGTG</td><td>165.6</td><td>10.7</td><td>2.2</td><td>5.0</td><td>6.6E-07</td><td>1.1E-05</td><td>BT20/MB231</td><td>MP</td></tr>
<tr><td>>MI0000085|hsa-mir-27a&WithFlank&19|-|13836434|13836523@57.76.20(+1U)&[MIMAT0000084&hsa-miR-27a-3p&offsets|0|-1(+1U);m-43&19|-|13836447|13836467&offsets|0|-1(+1U)]|TTCACAGTGGCTAAGTTCCGT</td><td>>MI0000085|hsa-mir-27a&WithFlank&19|-|13836434|13836523</td><td>57</td><td>76</td><td>20(+1U)</td><td>TTCACAGTGGCTAAGTTCCGT</td><td>2014.2</td><td>3.6</td><td>0.8</td><td>4.5</td><td>5.5E-06</td><td>1.1E-05</td><td>BT20/MB468</td><td>MITO</td></tr>
<tr><td>>MI0000078|hsa-mir-22&WithFlank&17|-|1713897|1713993@59.77.19&[MIMAT0000077&hsa-miR-22-3p&offsets|0|-3;m-2&17|-|1713914|1713935&offsets|0|-3]|AAGCTGCCAGTTGAAGAAC</td><td>>MI0000078|hsa-mir-22&WithFlank&17|-|1713897|1713993</td><td>59</td><td>77</td><td>19</td><td>AAGCTGCCAGTTGAAGAAC</td><td>206.7</td><td>8.1</td><td>1.7</td><td>4.7</td><td>2.3E-06</td><td>1.1E-05</td><td>BT20/MB468</td><td>MP</td></tr>
<tr><td>>hg38_21_+_8433172_8446622_RNA45SN1_45S_50ntFlanks@11582.11612.31|ACTGGTGCGGACCAGGGGAATCCGACTGTTT</td><td>>hg38_21_+_8433172_8446622_RNA45SN1_45S_50ntFlanks</td><td>11582</td><td>11612</td><td>31</td><td>ACTGGTGCGGACCAGGGGAATCCGACTGTTT</td><td>825.5</td><td>-3.4</td><td>0.8</td><td>-4.5</td><td>7.7E-06</td><td>1.1E-05</td><td>MB231/MB468</td><td>MITO</td></tr>
<tr><td>>am_trna2_GlyGCC_21_-_18827107_18827177@1.27.27,trna35_GlyGCC_1_+_161413094_161413164@1.27.27,trna37_GlyGCC_1_+_161420467_161420537@1.27.27,trna39_GlyGCC_1_+_161427898_161427968@1.27.27,trna41_GlyGCC_1_+_161435258_161435328@1.27.27|GCATGGGTGGTTCAGTGGTAGAATTCT</td><td>>am_trna2_GlyGCC_21_-_18827107_18827177</td><td>1</td><td>27</td><td>27</td><td>GCATGGGTGGTTCAGTGGTAGAATTCT</td><td>472.4</td><td>-5.8</td><td>1.3</td><td>-4.5</td><td>6.4E-06</td><td>1.1E-05</td><td>MB231/MB468</td><td>MP</td></tr>
<tr><td>>hg38_21_+_8433172_8446622_RNA45SN1_45S_50ntFlanks@9482.9506.25|ACGTGTTAGGACCCGAAAGATGGTG</td><td>>hg38_21_+_8433172_8446622_RNA45SN1_45S_50ntFlanks</td><td>9482</td><td>9506</td><td>25</td><td>ACGTGTTAGGACCCGAAAGATGGTG</td><td>171.2</td><td>10.8</td><td>2.2</td><td>5.0</td><td>6.7E-07</td><td>1.1E-05</td><td>BT20/MB231</td><td>MP</td></tr>
<tr><td>>hg38_21_+_8433172_8446622_RNA45SN1_45S_50ntFlanks@11580.11604.25|GAACTGGTGCGGACCAGGGGAATCC</td><td>>hg38_21_+_8433172_8446622_RNA45SN1_45S_50ntFlanks</td><td>11580</td><td>11604</td><td>25</td><td>GAACTGGTGCGGACCAGGGGAATCC</td><td>351.4</td><td>-5.6</td><td>1.2</td><td>-4.5</td><td>8.1E-06</td><td>1.1E-05</td><td>BT20/MB468</td><td>NUC</td></tr>
<tr><td>>hg38_21_+_8433172_8446622_RNA45SN1_45S_50ntFlanks@9761.9786.26|AACTTTAAATGGGTAAGAAGCCCGGC</td><td>>hg38_21_+_8433172_8446622_RNA45SN1_45S_50ntFlanks</td><td>9761</td><td>9786</td><td>26</td><td>AACTTTAAATGGGTAAGAAGCCCGGC</td><td>139.6</td><td>-11.0</td><td>2.4</td><td>-4.5</td><td>6.4E-06</td><td>1.1E-05</td><td>MB231/MB468</td><td>MP</td></tr>
<tr><td>>hg38_21_+_8433172_8446622_RNA45SN1_45S_50ntFlanks@10781.10809.29|AGGTGAACAGCCTCTGGCATGTTGGAACA</td><td>>hg38_21_+_8433172_8446622_RNA45SN1_45S_50ntFlanks</td><td>10781</td><td>10809</td><td>29</td><td>AGGTGAACAGCCTCTGGCATGTTGGAACA</td><td>221.2</td><td>-6.4</td><td>1.4</td><td>-4.5</td><td>6.4E-06</td><td>1.1E-05</td><td>MB231/MB468</td><td>MP</td></tr>
<tr><td>>hg38_21_+_8433172_8446622_RNA45SN1_45S_50ntFlanks@12414.12443.30|CCTTCGATGTCGGCTCTTCCTATCATTGTG</td><td>>hg38_21_+_8433172_8446622_RNA45SN1_45S_50ntFlanks</td><td>12414</td><td>12443</td><td>30</td><td>CCTTCGATGTCGGCTCTTCCTATCATTGTG</td><td>579.8</td><td>3.0</td><td>0.7</td><td>4.5</td><td>7.3E-06</td><td>1.1E-05</td><td>MB231/MB468</td><td>CYTO</td></tr>
<tr><td>>hg38_21_+_8433172_8446622_RNA45SN1_45S_50ntFlanks@5484.5518.35|GAGCGCTGAGAAGACGGTCGAACTTGACTATCTAG</td><td>>hg38_21_+_8433172_8446622_RNA45SN1_45S_50ntFlanks</td><td>5484</td><td>5518</td><td>35</td><td>GAGCGCTGAGAAGACGGTCGAACTTGACTATCTAG</td><td>384.1</td><td>-3.8</td><td>0.8</td><td>-4.5</td><td>6.5E-06</td><td>1.1E-05</td><td>BT20/MB468</td><td>CYTO</td></tr>
<tr><td>>hg38_21_+_8433172_8446622_RNA45SN1_45S_50ntFlanks@10783.10826.44|GTGAACAGCCTCTGGCATGTTGGAACAATGTAGGTAAGGGAAGT</td><td>>hg38_21_+_8433172_8446622_RNA45SN1_45S_50ntFlanks</td><td>10783</td><td>10826</td><td>44</td><td>GTGAACAGCCTCTGGCATGTTGGAACAATGTAGGTAAGGGAAGT</td><td>213.1</td><td>-6.8</td><td>1.5</td><td>-4.5</td><td>7.8E-06</td><td>1.1E-05</td><td>MB231/MB468</td><td>MITO</td></tr>
<tr><td>>hg38_21_+_8433172_8446622_RNA45SN1_45S_50ntFlanks@3922.3959.38|TTTATCAGATCAAAACCAACCCGGTCAGCCCCTCTCCG</td><td>>hg38_21_+_8433172_8446622_RNA45SN1_45S_50ntFlanks</td><td>3922</td><td>3959</td><td>38</td><td>TTTATCAGATCAAAACCAACCCGGTCAGCCCCTCTCCG</td><td>286.7</td><td>-4.6</td><td>1.0</td><td>-4.5</td><td>7.3E-06</td><td>1.1E-05</td><td>MB231/MB468</td><td>CYTO</td></tr>
<tr><td>>hg38_21_+_8433172_8446622_RNA45SN1_45S_50ntFlanks@6646.6670.25|TCGTACGACTCTTAGCGGTGGATCA</td><td>>hg38_21_+_8433172_8446622_RNA45SN1_45S_50ntFlanks</td><td>6646</td><td>6670</td><td>25</td><td>TCGTACGACTCTTAGCGGTGGATCA</td><td>19771.6</td><td>-3.5</td><td>0.8</td><td>-4.5</td><td>7.8E-06</td><td>1.1E-05</td><td>MB231/MB468</td><td>MITO</td></tr>
<tr><td>>hg38_21_+_8433172_8446622_RNA45SN1_45S_50ntFlanks@8911.8940.30|TCGCCGAATCCCGGGGCCGAGGGAGCGAGA</td><td>>hg38_21_+_8433172_8446622_RNA45SN1_45S_50ntFlanks</td><td>8911</td><td>8940</td><td>30</td><td>TCGCCGAATCCCGGGGCCGAGGGAGCGAGA</td><td>64.8</td><td>9.8</td><td>2.1</td><td>4.7</td><td>2.4E-06</td><td>1.1E-05</td><td>BT20/MB468</td><td>MP</td></tr>
<tr><td>>hg38_21_+_8433172_8446622_RNA45SN1_45S_50ntFlanks@9955.9989.35|GGAAGTCGGAATCCGCTAAGGAGTGTGTAACAACT</td><td>>hg38_21_+_8433172_8446622_RNA45SN1_45S_50ntFlanks</td><td>9955</td><td>9989</td><td>35</td><td>GGAAGTCGGAATCCGCTAAGGAGTGTGTAACAACT</td><td>52.1</td><td>9.5</td><td>2.0</td><td>4.7</td><td>2.4E-06</td><td>1.1E-05</td><td>BT20/MB468</td><td>MP</td></tr>
<tr><td>>hg38_21_+_8433172_8446622_RNA45SN1_45S_50ntFlanks@10477.10495.19|GCCGATCGAAAGGGAGTCG</td><td>>hg38_21_+_8433172_8446622_RNA45SN1_45S_50ntFlanks</td><td>10477</td><td>10495</td><td>19</td><td>GCCGATCGAAAGGGAGTCG</td><td>322.9</td><td>-6.7</td><td>1.5</td><td>-4.5</td><td>6.4E-06</td><td>1.1E-05</td><td>MB231/MB468</td><td>MP</td></tr>
<tr><td>>hg38_21_+_8433172_8446622_RNA45SN1_45S_50ntFlanks@12991.13014.24|CTCCCTCGCTGCGATCTATTGAAA</td><td>>hg38_21_+_8433172_8446622_RNA45SN1_45S_50ntFlanks</td><td>12991</td><td>13014</td><td>24</td><td>CTCCCTCGCTGCGATCTATTGAAA</td><td>2223.8</td><td>-3.6</td><td>0.8</td><td>-4.5</td><td>7.3E-06</td><td>1.1E-05</td><td>MB231/MB468</td><td>CYTO</td></tr>
<tr><td>>hg38_21_+_8433172_8446622_RNA45SN1_45S_50ntFlanks@8369.8391.23|AAGAGAGAGTTCAAGAGGGCGTG</td><td>>hg38_21_+_8433172_8446622_RNA45SN1_45S_50ntFlanks</td><td>8369</td><td>8391</td><td>23</td><td>AAGAGAGAGTTCAAGAGGGCGTG</td><td>110.4</td><td>-6.3</td><td>1.4</td><td>-4.5</td><td>6.4E-06</td><td>1.1E-05</td><td>MB231/MB468</td><td>MP</td></tr>
<tr><td>>hg38_21_+_8433172_8446622_RNA45SN1_45S_50ntFlanks@6645.6672.28|CTCGTACGACTCTTAGCGGTGGATCACT</td><td>>hg38_21_+_8433172_8446622_RNA45SN1_45S_50ntFlanks</td><td>6645</td><td>6672</td><td>28</td><td>CTCGTACGACTCTTAGCGGTGGATCACT</td><td>750.2</td><td>5.1</td><td>1.1</td><td>4.5</td><td>7.3E-06</td><td>1.1E-05</td><td>MB231/MB468</td><td>CYTO</td></tr>
<tr><td>>hg38_21_+_8433172_8446622_RNA45SN1_45S_50ntFlanks@9622.9648.27|CTAGTAGCTGGTTCCCTCCGAAGTTTC</td><td>>hg38_21_+_8433172_8446622_RNA45SN1_45S_50ntFlanks</td><td>9622</td><td>9648</td><td>27</td><td>CTAGTAGCTGGTTCCCTCCGAAGTTTC</td><td>562.8</td><td>-5.0</td><td>1.1</td><td>-4.5</td><td>5.6E-06</td><td>1.1E-05</td><td>BT20/MB468</td><td>MITO</td></tr>
<tr><td>>hg38_21_+_8433172_8446622_RNA45SN1_45S_50ntFlanks@12570.12587.18|AGTACGAGAGGAACCGCA</td><td>>hg38_21_+_8433172_8446622_RNA45SN1_45S_50ntFlanks</td><td>12570</td><td>12587</td><td>18</td><td>AGTACGAGAGGAACCGCA</td><td>521.8</td><td>-6.2</td><td>1.4</td><td>-4.5</td><td>6.6E-06</td><td>1.1E-05</td><td>BT20/MB468</td><td>CYTO</td></tr>
<tr><td>>hg38_21_+_8433172_8446622_RNA45SN1_45S_50ntFlanks@12942.12979.38|TTCGTAGACGACCTGCTTCTGGGTCGGGGTTTCGTACG</td><td>>hg38_21_+_8433172_8446622_RNA45SN1_45S_50ntFlanks</td><td>12942</td><td>12979</td><td>38</td><td>TTCGTAGACGACCTGCTTCTGGGTCGGGGTTTCGTACG</td><td>249.3</td><td>-6.1</td><td>1.4</td><td>-4.5</td><td>6.6E-06</td><td>1.1E-05</td><td>BT20/MB468</td><td>CYTO</td></tr>
<tr><td>>hg38_21_+_8433172_8446622_RNA45SN1_45S_50ntFlanks@474.503.30|TCTTCGTGATCGATGTGGTGACGTCGTGCT</td><td>>hg38_21_+_8433172_8446622_RNA45SN1_45S_50ntFlanks</td><td>474</td><td>503</td><td>30</td><td>TCTTCGTGATCGATGTGGTGACGTCGTGCT</td><td>308.8</td><td>4.1</td><td>0.9</td><td>4.5</td><td>7.6E-06</td><td>1.1E-05</td><td>MB231/MB468</td><td>NUC</td></tr>
<tr><td>>hg38_21_+_8433172_8446622_RNA45SN1_45S_50ntFlanks@7976.8006.31|GCGACCTCAGATCAGACGTGGCGACCCGCTG</td><td>>hg38_21_+_8433172_8446622_RNA45SN1_45S_50ntFlanks</td><td>7976</td><td>8006</td><td>31</td><td>GCGACCTCAGATCAGACGTGGCGACCCGCTG</td><td>122.7</td><td>-6.3</td><td>1.4</td><td>-4.5</td><td>6.4E-06</td><td>1.1E-05</td><td>MB231/MB468</td><td>MP</td></tr>
<tr><td>>MI0003513|hsa-mir-455&WithFlank&9|+|114209428|114209535@22.44.23&[MIMAT0003150&hsa-miR-455-5p&offsets|0|+1;m-232&9|+|114209449|114209470&offsets|0|+1]|TATGTGCCTTTGGACTACATCGT</td><td>>MI0003513|hsa-mir-455&WithFlank&9|+|114209428|114209535</td><td>22</td><td>44</td><td>23</td><td>TATGTGCCTTTGGACTACATCGT</td><td>2000.6</td><td>-4.5</td><td>0.9</td><td>-5.0</td><td>6.8E-07</td><td>1.1E-05</td><td>BT20/MB231</td><td>MP</td></tr>
<tr><td>>hg38_21_+_8433172_8446622_RNA45SN1_45S_50ntFlanks@3900.3919.20|CTTCGCGGGGGGGATGCGTG</td><td>>hg38_21_+_8433172_8446622_RNA45SN1_45S_50ntFlanks</td><td>3900</td><td>3919</td><td>20</td><td>CTTCGCGGGGGGGATGCGTG</td><td>451.3</td><td>-4.7</td><td>1.0</td><td>-4.5</td><td>5.6E-06</td><td>1.1E-05</td><td>BT20/MB468</td><td>MITO</td></tr>
<tr><td>>am_trna10_LysCTT_16_+_3241501_3241573@11.28.18,trna119_LysCTT_1_-_145395522_145395594@11.28.18,trna11_LysCTT_5_-_180648979_180649051@11.28.18,trna128_LysCTT_1_-_55423542_55423614@11.28.18,trna13_LysCTT_14_-_58706613_58706685@11.28.18,trna13_LysCTT_6_+_26556774_26556846@11.28.18,trna2_LysCTT_15_+_79152904_79152976@11.28.18,trna32_LysCTT_16_-_3207406_3207478@11.28.18,trna4_LysCTT_18_-_43669271_43669343@11.28.18,trna6_LysCTT_19_-_52425396_52425468@11.28.18,trna7_LysCTT_16_+_3225692_3225764@11.28.18,trna9_LysCTT_5_+_180634755_180634827@11.28.18|CTCAGTCGGTAGAGCATG</td><td>>am_trna10_LysCTT_16_+_3241501_3241573</td><td>11</td><td>28</td><td>18</td><td>CTCAGTCGGTAGAGCATG</td><td>658.1</td><td>-7.7</td><td>1.7</td><td>-4.5</td><td>8.2E-06</td><td>1.1E-05</td><td>BT20/MB468</td><td>NUC</td></tr>
<tr><td>>MI0000487|hsa-mir-193a&WithFlank&17|+|31559990|31560089@27.48.22&[MIMAT0004614&hsa-miR-193a-5p&offsets|0|0;m-123&17|+|31560016|31560037&offsets|0|0]|TGGGTCTTTGCGGGCGAGATGA</td><td>>MI0000487|hsa-mir-193a&WithFlank&17|+|31559990|31560089</td><td>27</td><td>48</td><td>22</td><td>TGGGTCTTTGCGGGCGAGATGA</td><td>491.7</td><td>4.6</td><td>0.9</td><td>5.0</td><td>6.8E-07</td><td>1.1E-05</td><td>BT20/MB231</td><td>CYTO</td></tr>
<tr><td>>hg38_21_+_8433172_8446622_RNA45SN1_45S_50ntFlanks@10914.10936.23|CGCCGCGGCTGGACGAGGCGCCG</td><td>>hg38_21_+_8433172_8446622_RNA45SN1_45S_50ntFlanks</td><td>10914</td><td>10936</td><td>23</td><td>CGCCGCGGCTGGACGAGGCGCCG</td><td>747.2</td><td>-6.4</td><td>1.4</td><td>-4.5</td><td>5.6E-06</td><td>1.1E-05</td><td>BT20/MB468</td><td>MITO</td></tr>
<tr><td>>MI0000542|hsa-mir-320a&WithFlank&8|-|22244960|22245043@48.68.21&[MIMAT0000510&hsa-miR-320a-3p&offsets|0|-1;m-60&8|-|22244975|22244996&offsets|0|-1]|AAAAGCTGGGTTGAGAGGGCG</td><td>>MI0000542|hsa-mir-320a&WithFlank&8|-|22244960|22245043</td><td>48</td><td>68</td><td>21</td><td>AAAAGCTGGGTTGAGAGGGCG</td><td>15222.7</td><td>3.9</td><td>0.8</td><td>5.0</td><td>6.9E-07</td><td>1.1E-05</td><td>BT20/MB231</td><td>CYTO</td></tr>
<tr><td>>hg38_21_+_8433172_8446622_RNA45SN1_45S_50ntFlanks@485.511.27|GATGTGGTGACGTCGTGCTCTCCCGGG</td><td>>hg38_21_+_8433172_8446622_RNA45SN1_45S_50ntFlanks</td><td>485</td><td>511</td><td>27</td><td>GATGTGGTGACGTCGTGCTCTCCCGGG</td><td>163.7</td><td>10.6</td><td>2.4</td><td>4.5</td><td>7.9E-06</td><td>1.1E-05</td><td>MB231/MB468</td><td>MITO</td></tr>
<tr><td>>hg38_21_+_8433172_8446622_RNA45SN1_45S_50ntFlanks@6647.6668.22|CGTACGACTCTTAGCGGTGGAT</td><td>>hg38_21_+_8433172_8446622_RNA45SN1_45S_50ntFlanks</td><td>6647</td><td>6668</td><td>22</td><td>CGTACGACTCTTAGCGGTGGAT</td><td>436.8</td><td>2.6</td><td>0.6</td><td>4.5</td><td>7.4E-06</td><td>1.1E-05</td><td>MB231/MB468</td><td>CYTO</td></tr>
<tr><td>>hg38_21_+_8433172_8446622_RNA45SN1_45S_50ntFlanks@9574.9593.20|AATCGGTCGTCCGACCTGGG</td><td>>hg38_21_+_8433172_8446622_RNA45SN1_45S_50ntFlanks</td><td>9574</td><td>9593</td><td>20</td><td>AATCGGTCGTCCGACCTGGG</td><td>377.5</td><td>-4.1</td><td>0.9</td><td>-4.5</td><td>6.5E-06</td><td>1.1E-05</td><td>MB231/MB468</td><td>MP</td></tr>
<tr><td>>hg38_21_+_8433172_8446622_RNA45SN1_45S_50ntFlanks@4006.4042.37|ACCTCGGGCCGATCGCACGCCCCCCGTGGCGGCGACG</td><td>>hg38_21_+_8433172_8446622_RNA45SN1_45S_50ntFlanks</td><td>4006</td><td>4042</td><td>37</td><td>ACCTCGGGCCGATCGCACGCCCCCCGTGGCGGCGACG</td><td>710.9</td><td>-6.7</td><td>1.5</td><td>-4.5</td><td>7.7E-06</td><td>1.1E-05</td><td>MB231/MB468</td><td>NUC</td></tr>
</table>

| | | | | | | | | | | | | | |
|---|---|---|---|---|---|---|---|---|---|---|---|---|---|
| >hg38_21_+_8433172_8446622_RNA45SN1_45S_50ntFlanks@5473.5516.44\|CGGCCCTGGCGGAGCGCTGAGAAGACGGTCGAACTTGACTATCT | >hg38_21_+_8433172_8446622_RNA45SN1_45S_50ntFlanks | 5473 | 5516 | 44 | CGGCCCTGGCGGAGCGCTGAGAAGACGGTCGAACTTGACTATCT | 13587.3 | 5.8 | 1.2 | 4.9 | 8.0E-07 | 1.1E-05 | BT20/MB231 | MITO |
| >hg38_21_+_8433172_8446622_RNA45SN1_45S_50ntFlanks@10771.10807.37\|CAGGTCTCCAAGGTGAACAGCCTCTGGCATGTTGGAA | >hg38_21_+_8433172_8446622_RNA45SN1_45S_50ntFlanks | 10771 | 10807 | 37 | CAGGTCTCCAAGGTGAACAGCCTCTGGCATGTTGGAA | 81.7 | -9.4 | 2.0 | -4.7 | 2.4E-06 | 1.1E-05 | BT20/MB468 | MP |
| >hg38_21_+_8433172_8446622_RNA45SN1_45S_50ntFlanks@12635.12669.35\|AAGCTACCATCTGTGGGATTATGACTGAACGCCTC | >hg38_21_+_8433172_8446622_RNA45SN1_45S_50ntFlanks | 12635 | 12669 | 35 | AAGCTACCATCTGTGGGATTATGACTGAACGCCTC | 358.8 | 5.4 | 1.2 | 4.5 | 7.4E-06 | 1.1E-05 | MB231/MB468 | CYTO |
| >hg38_21_+_8433172_8446622_RNA45SN1_45S_50ntFlanks@9940.9981.42\|CAGGACGGTGGCCATGGAAGTCGGAATCCGCTAAGGAGTGTG | >hg38_21_+_8433172_8446622_RNA45SN1_45S_50ntFlanks | 9940 | 9981 | 42 | CAGGACGGTGGCCATGGAAGTCGGAATCCGCTAAGGAGTGTG | 172.9 | 8.3 | 1.8 | 4.7 | 2.4E-06 | 1.1E-05 | BT20/MB468 | MP |
| >hg38_21_+_8433172_8446622_RNA45SN1_45S_50ntFlanks@11209.11231.23\|CGGCGACTCTGGACGCGAGCCGG | >hg38_21_+_8433172_8446622_RNA45SN1_45S_50ntFlanks | 11209 | 11231 | 23 | CGGCGACTCTGGACGCGAGCCGG | 2462.1 | -6.6 | 1.5 | -4.5 | 8.3E-06 | 1.1E-05 | BT20/MB468 | NUC |
| >hg38_21_+_8433172_8446622_RNA45SN1_45S_50ntFlanks@8183.8206.24\|TCGAGGCCCAGCCCGTGGACGGTG | >hg38_21_+_8433172_8446622_RNA45SN1_45S_50ntFlanks | 8183 | 8206 | 24 | TCGAGGCCCAGCCCGTGGACGGTG | 168.8 | -6.5 | 1.4 | -4.5 | 6.5E-06 | 1.1E-05 | MB231/MB468 | MP |
| >hg38_21_+_8433172_8446622_RNA45SN1_45S_50ntFlanks@9930.9951.22\|ATATAGACAGCAGGACGGTGGC | >hg38_21_+_8433172_8446622_RNA45SN1_45S_50ntFlanks | 9930 | 9951 | 22 | ATATAGACAGCAGGACGGTGGC | 189.4 | -6.5 | 1.4 | -4.5 | 6.5E-06 | 1.1E-05 | MB231/MB468 | MP |
| >trna10_LysCTT_16_+_3241501_3241573@13.35.23,trna13_LysCTT_14_-_58706613_58706685@13.35.23,trna2_LysCTT_15_+_79152904_79152976@13.35.23,trna4_LysCTT_18_-_43669271_43669343@13.35.23\|CAGTCGGTAGAGCATGGGACTCT | >trna10_LysCTT_16_+_3241501_3241573 | 13 | 35 | 23 | CAGTCGGTAGAGCATGGGACTCT | 40.2 | 8.5 | 1.9 | 4.5 | 6.5E-06 | 1.1E-05 | MB231/MB468 | MP |
| >hg38_21_+_8433172_8446622_RNA45SN1_45S_50ntFlanks@3788.3810.23\|AACTGCGAATGGCTCATTAAATC | >hg38_21_+_8433172_8446622_RNA45SN1_45S_50ntFlanks | 3788 | 3810 | 23 | AACTGCGAATGGCTCATTAAATC | 182.8 | -6.7 | 1.5 | -4.5 | 6.5E-06 | 1.1E-05 | MB231/MB468 | MP |
| >hg38_21_+_8433172_8446622_RNA45SN1_45S_50ntFlanks@6730.6770.41\|ACACATTGATCATCGACACTTCGAACGCACTTGCGGCCCCG | >hg38_21_+_8433172_8446622_RNA45SN1_45S_50ntFlanks | 6730 | 6770 | 41 | ACACATTGATCATCGACACTTCGAACGCACTTGCGGCCCCG | 270.5 | -5.8 | 1.3 | -4.5 | 6.7E-06 | 1.1E-05 | BT20/MB468 | CYTO |
| >hg38_21_+_8433172_8446622_RNA45SN1_45S_50ntFlanks@11874.11902.29\|AGAAGACCCTGTTGAGCTTGACTCTAGTC | >hg38_21_+_8433172_8446622_RNA45SN1_45S_50ntFlanks | 11874 | 11902 | 29 | AGAAGACCCTGTTGAGCTTGACTCTAGTC | 347.8 | -2.9 | 0.6 | -4.5 | 6.7E-06 | 1.1E-05 | BT20/MB468 | CYTO |
| >hg38_21_+_8433172_8446622_RNA45SN1_45S_50ntFlanks@7987.8016.30\|TCAGACGTGGCGACCCGCTGAATTTAAGCA | >hg38_21_+_8433172_8446622_RNA45SN1_45S_50ntFlanks | 7987 | 8016 | 30 | TCAGACGTGGCGACCCGCTGAATTTAAGCA | 85.1 | -4.7 | 1.0 | -4.5 | 6.5E-06 | 1.2E-05 | MB231/MB468 | MP |
| >hg38_21_+_8433172_8446622_RNA45SN1_45S_50ntFlanks@12676.12700.25\|AGAATCCCGCCCAGGCGGAACGATA | >hg38_21_+_8433172_8446622_RNA45SN1_45S_50ntFlanks | 12676 | 12700 | 25 | AGAATCCCGCCCAGGCGGAACGATA | 115.5 | -10.7 | 2.4 | -4.5 | 6.5E-06 | 1.2E-05 | MB231/MB468 | MP |
| >hg38_21_+_8433172_8446622_RNA45SN1_45S_50ntFlanks@3771.3794.24\|CACGGCCGGTACAGTGAAACTGCG | >hg38_21_+_8433172_8446622_RNA45SN1_45S_50ntFlanks | 3771 | 3794 | 24 | CACGGCCGGTACAGTGAAACTGCG | 277.8 | -7.2 | 1.6 | -4.5 | 6.5E-06 | 1.2E-05 | MB231/MB468 | MP |
| >hg38_21_+_8433172_8446622_RNA45SN1_45S_50ntFlanks@3782.3800.19\|CAGTGAAACTGCGAATGGC | >hg38_21_+_8433172_8446622_RNA45SN1_45S_50ntFlanks | 3782 | 3800 | 19 | CAGTGAAACTGCGAATGGC | 509.4 | -2.3 | 0.5 | -4.5 | 7.5E-06 | 1.2E-05 | MB231/MB468 | CYTO |
| >hg38_21_+_8433172_8446622_RNA45SN1_45S_50ntFlanks@11874.11912.39\|AGAAGACCCTGTTGAGCTTGACTCTAGTCTGGCACGGTG | >hg38_21_+_8433172_8446622_RNA45SN1_45S_50ntFlanks | 11874 | 11912 | 39 | AGAAGACCCTGTTGAGCTTGACTCTAGTCTGGCACGGTG | 423.0 | -7.3 | 1.6 | -4.7 | 2.4E-06 | 1.2E-05 | BT20/MB468 | MP |
| >hg38_21_+_8433172_8446622_RNA45SN1_45S_50ntFlanks@7992.8014.23\|CGTGGCGACCCGCTGAATTTAAG | >hg38_21_+_8433172_8446622_RNA45SN1_45S_50ntFlanks | 7992 | 8014 | 23 | CGTGGCGACCCGCTGAATTTAAG | 416.3 | -5.6 | 1.2 | -4.5 | 6.6E-06 | 1.2E-05 | MB231/MB468 | MP |
| >hg38_21_+_8433172_8446622_RNA45SN1_45S_50ntFlanks@8168.8203.36\|CCCAAGTCCTTCTGATCGAGGCCCAGCCCGTGGACG | >hg38_21_+_8433172_8446622_RNA45SN1_45S_50ntFlanks | 8168 | 8203 | 36 | CCCAAGTCCTTCTGATCGAGGCCCAGCCCGTGGACG | 598.2 | -7.4 | 1.6 | -4.7 | 2.4E-06 | 1.2E-05 | BT20/MB468 | MP |
| >hg38_21_+_8433172_8446622_RNA45SN1_45S_50ntFlanks@3852.3881.30\|TAACTGTGGTAATTCTAGAGCTAATACATG | >hg38_21_+_8433172_8446622_RNA45SN1_45S_50ntFlanks | 3852 | 3881 | 30 | TAACTGTGGTAATTCTAGAGCTAATACATG | 186.2 | -5.7 | 1.3 | -4.5 | 5.7E-06 | 1.2E-05 | BT20/MB468 | MITO |
| >hg38_21_+_8433172_8446622_RNA45SN1_45S_50ntFlanks@10816.10840.25\|GTAAGGGAAGTCGGCAAGCCGGATC | >hg38_21_+_8433172_8446622_RNA45SN1_45S_50ntFlanks | 10816 | 10840 | 25 | GTAAGGGAAGTCGGCAAGCCGGATC | 287.9 | -2.4 | 0.5 | -4.5 | 7.5E-06 | 1.2E-05 | MB231/MB468 | CYTO |
| >hg38_21_+_8433172_8446622_RNA45SN1_45S_50ntFlanks@10402.10420.19\|AACATGGGTCAGTCGGTCC | >hg38_21_+_8433172_8446622_RNA45SN1_45S_50ntFlanks | 10402 | 10420 | 19 | AACATGGGTCAGTCGGTCC | 1716.6 | -5.9 | 1.3 | -4.7 | 2.4E-06 | 1.2E-05 | BT20/MB468 | MP |
| >MI0000063\|hsa-let-7b&WithFlank&22\|+\|46113680\|46113774@12.30.19(+1A)&[MIMAT0000063&hsa-let-7b-5p&offsets\|0\|-3(+1A);m-14&22\|+\|46113691\|46113712&offsets\|0\|-3(+1A)]\|TGAGGTAGTAGGTTGTGTGA | >MI0000063\|hsa-let-7b&WithFlank&22\|+\|46113680\|46113774 | 12 | 30 | 19(+1A) | TGAGGTAGTAGGTTGTGTGA | 364.9 | 6.3 | 1.3 | 4.7 | 2.4E-06 | 1.2E-05 | BT20/MB468 | MP |
| >hg38_21_+_8433172_8446622_RNA45SN1_45S_50ntFlanks@11320.11340.21\|CCACGTCTCGTCGCGCGCGCG | >hg38_21_+_8433172_8446622_RNA45SN1_45S_50ntFlanks | 11320 | 11340 | 21 | CCACGTCTCGTCGCGCGCGCG | 146.7 | -5.8 | 1.3 | -4.5 | 7.8E-06 | 1.2E-05 | MB231/MB468 | NUC |
| >hg38_21_+_8433172_8446622_RNA45SN1_45S_50ntFlanks@10420.10462.43\|CTGAGAGATGGGCGAGCGCCGTTCCGAAGGGACGGGCGATGGC | >hg38_21_+_8433172_8446622_RNA45SN1_45S_50ntFlanks | 10420 | 10462 | 43 | CTGAGAGATGGGCGAGCGCCGTTCCGAAGGGACGGGCGATGGC | 93.5 | -5.6 | 1.3 | -4.5 | 6.7E-06 | 1.2E-05 | BT20/MB468 | CYTO |
| >hg38_21_+_8433172_8446622_RNA45SN1_45S_50ntFlanks@11905.11931.27\|GCACGGTGAAGAGACATGAGAGGTGTA | >hg38_21_+_8433172_8446622_RNA45SN1_45S_50ntFlanks | 11905 | 11931 | 27 | GCACGGTGAAGAGACATGAGAGGTGTA | 154.8 | 10.6 | 2.1 | 5.0 | 6.9E-07 | 1.2E-05 | BT20/MB231 | MP |
| >hg38_21_+_8433172_8446622_RNA45SN1_45S_50ntFlanks@10683.10711.29\|CCGGCGGCGTCCGGTGAGCTCTCGCTGGC | >hg38_21_+_8433172_8446622_RNA45SN1_45S_50ntFlanks | 10683 | 10711 | 29 | CCGGCGGCGTCCGGTGAGCTCTCGCTGGC | 1446.1 | -5.2 | 1.1 | -4.5 | 5.7E-06 | 1.2E-05 | BT20/MB468 | MITO |
| >trna11_GluTTC_15_-_26327381_26327452@1.29.29,trna3_GluTTC_13_-_45492062_45492133@1.29.29\|TCCCACATGGTCTAGCGGTTAGGATTCCT | >trna11_GluTTC_15_-_26327381_26327452 | 1 | 29 | 29 | TCCCACATGGTCTAGCGGTTAGGATTCCT | 1798.9 | -4.3 | 1.0 | -4.5 | 7.8E-06 | 1.2E-05 | MB231/MB468 | NUC |
| >hg38_21_+_8433172_8446622_RNA45SN1_45S_50ntFlanks@8118.8140.23\|GGGACATGTGGCGTACGGAAGAC | >hg38_21_+_8433172_8446622_RNA45SN1_45S_50ntFlanks | 8118 | 8140 | 23 | GGGACATGTGGCGTACGGAAGAC | 514.8 | 5.5 | 1.2 | 4.7 | 2.4E-06 | 1.2E-05 | BT20/MB468 | MP |
| >hg38_21_+_8433172_8446622_RNA45SN1_45S_50ntFlanks@11568.11592.25\|GCAGCCGACTTAGAACTGGTGCGGA | >hg38_21_+_8433172_8446622_RNA45SN1_45S_50ntFlanks | 11568 | 11592 | 25 | GCAGCCGACTTAGAACTGGTGCGGA | 389.0 | -4.0 | 0.9 | -4.5 | 8.0E-06 | 1.2E-05 | MB231/MB468 | MITO |
| >hg38_21_+_8433172_8446622_RNA45SN1_45S_50ntFlanks@6779.6797.19\|CCGGGGCTACGCCTGTCTG | >hg38_21_+_8433172_8446622_RNA45SN1_45S_50ntFlanks | 6779 | 6797 | 19 | CCGGGGCTACGCCTGTCTG | 54.6 | -8.9 | 1.9 | -4.7 | 2.4E-06 | 1.2E-05 | BT20/MB468 | MP |
| >hg38_21_+_8433172_8446622_RNA45SN1_45S_50ntFlanks@3850.3869.20\|GATAACTGTGGTAATTCTAG | >hg38_21_+_8433172_8446622_RNA45SN1_45S_50ntFlanks | 3850 | 3869 | 20 | GATAACTGTGGTAATTCTAG | 250.3 | -2.3 | 0.5 | -4.5 | 7.5E-06 | 1.2E-05 | MB231/MB468 | CYTO |
| >hg38_21_+_8433172_8446622_RNA45SN1_45S_50ntFlanks@9565.9592.28\|TGACGTGCAAATCGGTCGTCCGACCTGG | >hg38_21_+_8433172_8446622_RNA45SN1_45S_50ntFlanks | 9565 | 9592 | 28 | TGACGTGCAAATCGGTCGTCCGACCTGG | 208.5 | 11.0 | 2.2 | 5.0 | 7.0E-07 | 1.2E-05 | BT20/MB231 | MP |
| >hg38_21_+_8433172_8446622_RNA45SN1_45S_50ntFlanks@9929.9959.31\|GATATAGACAGCAGGACGGTGGCCATGGAAG | >hg38_21_+_8433172_8446622_RNA45SN1_45S_50ntFlanks | 9929 | 9959 | 31 | GATATAGACAGCAGGACGGTGGCCATGGAAG | 601.3 | -5.4 | 1.2 | -4.5 | 5.8E-06 | 1.2E-05 | BT20/MB468 | MITO |
| >trnaMT_GluTTC_MT_-_14674_14742@27.72.46,trnalookalike8_GluTTC_5_-_93905172_93905240@27.72.46\|GTTTTTCATATCATTGGTCGTGGTTGTAGTCCGTGCGAGAATACCA | >trnaMT_GluTTC_MT_-_14674_14742 | 27 | 72 | 46 | GTTTTTCATATCATTGGTCGTGGTTGTAGTCCGTGCGAGAATACCA | 1306.8 | -8.6 | 1.9 | -4.5 | 8.4E-06 | 1.2E-05 | BT20/MB468 | NUC |
| >hg38_21_+_8433172_8446622_RNA45SN1_45S_50ntFlanks@10403.10431.29\|ACATGGGTCAGTCGGTCCTGAGAGATGGG | >hg38_21_+_8433172_8446622_RNA45SN1_45S_50ntFlanks | 10403 | 10431 | 29 | ACATGGGTCAGTCGGTCCTGAGAGATGGG | 384.6 | -7.2 | 1.6 | -4.5 | 8.4E-06 | 1.2E-05 | BT20/MB468 | NUC |
| >hg38_MT_+_598_1651_RNR1_12S_50ntFlanks@914.950.37\|CAAAGGACATTTAACTAAAACCCCTACGCATTTATAT | >hg38_MT_+_598_1651_RNR1_12S_50ntFlanks | 914 | 950 | 37 | CAAAGGACATTTAACTAAAACCCCTACGCATTTATAT | 248.5 | 8.8 | 1.9 | 4.7 | 2.5E-06 | 1.2E-05 | BT20/MB468 | MP |
| >hg38_21_+_8433172_8446622_RNA45SN1_45S_50ntFlanks@10841.10874.34\|CGTAACTTCGGGATAAGGATTGGCTCTAAGGGCT | >hg38_21_+_8433172_8446622_RNA45SN1_45S_50ntFlanks | 10841 | 10874 | 34 | CGTAACTTCGGGATAAGGATTGGCTCTAAGGGCT | 273.1 | -7.0 | 1.5 | -4.5 | 6.6E-06 | 1.2E-05 | MB231/MB468 | MP |
| >hg38_21_+_8433172_8446622_RNA45SN1_45S_50ntFlanks@9950.9982.33\|GCCATGGAAGTCGGAATCCGCTAAGGAGTGTGT | >hg38_21_+_8433172_8446622_RNA45SN1_45S_50ntFlanks | 9950 | 9982 | 33 | GCCATGGAAGTCGGAATCCGCTAAGGAGTGTGT | 45.2 | 9.2 | 2.0 | 4.7 | 2.5E-06 | 1.2E-05 | BT20/MB468 | MP |
| >hg38_21_+_8433172_8446622_RNA45SN1_45S_50ntFlanks@9923.9948.26\|TTGGTTGATATAGACAGCAGGACGGT | >hg38_21_+_8433172_8446622_RNA45SN1_45S_50ntFlanks | 9923 | 9948 | 26 | TTGGTTGATATAGACAGCAGGACGGT | 122.5 | 10.3 | 2.1 | 5.0 | 7.0E-07 | 1.2E-05 | BT20/MB231 | MP |
| >hg38_21_+_8433172_8446622_RNA45SN1_45S_50ntFlanks@10714.10736.23\|TTGAAAATCCGGGGGAGAGGGTG | >hg38_21_+_8433172_8446622_RNA45SN1_45S_50ntFlanks | 10714 | 10736 | 23 | TTGAAAATCCGGGGGAGAGGGTG | 110.3 | -6.4 | 1.4 | -4.5 | 6.6E-06 | 1.2E-05 | MB231/MB468 | MP |
| >MI0000446\|hsa-mir-125b-1&WithFlank&11\|-\|122099751\|122099850@21.40.20(+1A);MI0000470\|hsa-mir-125b-2&WithFlank&21\|+\|16590231\|16590331@23.42.20(+1A)&[MIMAT0000423&hsa-miR-125b-5p&offsets\|0\|-2(+1A);m-67&11\|-\|122099809\|122099830&offsets\|0\|-2(+1A)];[MIMAT0000423_1&hsa-miR-125b-5p&offsets\|0\|-2(+1A);m-68&21\|+\|16590253\|16590274&offsets\|0\|-2(+1A)]\|TCCCTGAGACCCTAACTTGTA | >MI0000446\|hsa-mir-125b-1&WithFlank&11\|-\|122099751\|122099850 | 21 | 40 | 20(+1A) | TCCCTGAGACCCTAACTTGTA | 1455.7 | -8.5 | 1.7 | -5.0 | 7.0E-07 | 1.2E-05 | BT20/MB231 | MP |
| >hg38_21_+_8433172_8446622_RNA45SN1_45S_50ntFlanks@9533.9554.22\|GAGGAAACTCTGGTGGAGGTCC | >hg38_21_+_8433172_8446622_RNA45SN1_45S_50ntFlanks | 9533 | 9554 | 22 | GAGGAAACTCTGGTGGAGGTCC | 313.2 | -5.4 | 1.2 | -4.5 | 6.7E-06 | 1.2E-05 | MB231/MB468 | MP |
| >hg38_21_+_8433172_8446622_RNA45SN1_45S_50ntFlanks@7977.8016.40\|CGACCTCAGATCAGACGTGGCGACCCGCTGAATTTAAGCA | >hg38_21_+_8433172_8446622_RNA45SN1_45S_50ntFlanks | 7977 | 8016 | 40 | CGACCTCAGATCAGACGTGGCGACCCGCTGAATTTAAGCA | 1018.5 | 5.5 | 1.2 | 4.5 | 7.9E-06 | 1.2E-05 | MB231/MB468 | NUC |
| >hg38_21_+_8433172_8446622_RNA45SN1_45S_50ntFlanks@12992.13016.25\|TCCCTCGCTGCGATCTATTGAAAGT | >hg38_21_+_8433172_8446622_RNA45SN1_45S_50ntFlanks | 12992 | 13016 | 25 | TCCCTCGCTGCGATCTATTGAAAGT | 345.8 | -2.3 | 0.5 | -4.5 | 7.9E-06 | 1.2E-05 | MB231/MB468 | NUC |
| >hg38_21_+_8433172_8446622_RNA45SN1_45S_50ntFlanks@12633.12657.25\|CGAAGCTACCATCTGTGGGATTATG | >hg38_21_+_8433172_8446622_RNA45SN1_45S_50ntFlanks | 12633 | 12657 | 25 | CGAAGCTACCATCTGTGGGATTATG | 144.0 | -6.1 | 1.3 | -4.5 | 6.8E-06 | 1.2E-05 | BT20/MB468 | CYTO |
| >hg38_21_+_8433172_8446622_RNA45SN1_45S_50ntFlanks@6719.6746.28\|TGAATTGCAGGACACATTGATCATCGAC | >hg38_21_+_8433172_8446622_RNA45SN1_45S_50ntFlanks | 6719 | 6746 | 28 | TGAATTGCAGGACACATTGATCATCGAC | 643.2 | -3.1 | 0.7 | -4.5 | 5.8E-06 | 1.2E-05 | BT20/MB468 | MITO |
| >hg38_21_+_8433172_8446622_RNA45SN1_45S_50ntFlanks@3997.4022.26\|CTCTAGATAACCTCGGGCCGATCGCA | >hg38_21_+_8433172_8446622_RNA45SN1_45S_50ntFlanks | 3997 | 4022 | 26 | CTCTAGATAACCTCGGGCCGATCGCA | 254.7 | -6.4 | 1.4 | -4.5 | 7.9E-06 | 1.2E-05 | MB231/MB468 | NUC |

<table>
<tr><td></td><td></td><td></td><td></td><td></td><td></td><td></td><td></td><td></td><td></td><td></td><td></td><td></td><td></td><td></td></tr>
<tr><td>>trna111_HisGTG_1_-_147774845_147774916@1.33.33,trna118_HisGTG_1_-_145396881_145396952@1.33.33,trna16_HisGTG_1_+_146544773_146544844@1.33.33,trna1_HisGTG_15_+_45493349_45493420@1.33.33,trna21_HisGTG_1_+_147753471_147753542@1.33.33,trna33_HisGTG_6_+_27125906_27125977@1.33.33,trna7_HisGTG_9_-_14433938_14434009@1.33.33,trna8_HisGTG_15_-_45492611_45492682@1.33.33,trna9_HisGTG_15_-_45490804_45490875@1.33.33|GCCGTGATCGTATAGTGGTTAGTACTCTGCGTT</td><td>>trna111_HisGTG_1_-_147774845_147774916</td><td>1</td><td>33</td><td>33</td><td>GCCGTGATCGTATAGTGGTTAGTACTCTGCGTT</td><td>1331.4</td><td>4.3</td><td>1.0</td><td>4.5</td><td>8.4E-06</td><td>1.2E-05</td><td>BT20/MB468</td><td>NUC</td></tr>
<tr><td>>hg38_21_+_8433172_8446622_RNA45SN1_45S_50ntFlanks@7993.8016.24|GTGGCGACCCGCTGAATTTAAGCA</td><td>>hg38_21_+_8433172_8446622_RNA45SN1_45S_50ntFlanks</td><td>7993</td><td>8016</td><td>24</td><td>GTGGCGACCCGCTGAATTTAAGCA</td><td>178.4</td><td>3.6</td><td>0.8</td><td>4.5</td><td>7.9E-06</td><td>1.2E-05</td><td>MB231/MB468</td><td>NUC</td></tr>
<tr><td>>hg38_21_+_8433172_8446622_RNA45SN1_45S_50ntFlanks@4723.4761.39|CATTAATCAAGAACGAAAGTCGGAGGTTCGAAGACGATC</td><td>>hg38_21_+_8433172_8446622_RNA45SN1_45S_50ntFlanks</td><td>4723</td><td>4761</td><td>39</td><td>CATTAATCAAGAACGAAAGTCGGAGGTTCGAAGACGATC</td><td>314.7</td><td>-5.0</td><td>1.1</td><td>-4.5</td><td>8.1E-06</td><td>1.2E-05</td><td>MB231/MB468</td><td>MITO</td></tr>
<tr><td>>trnaMT_SerGCT_MT_+_12207_12265@1.19.19|GAGAAAGCTCACAAGAACT</td><td>>trnaMT_SerGCT_MT_+_12207_12265</td><td>1</td><td>19</td><td>19</td><td>GAGAAAGCTCACAAGAACT</td><td>1421.5</td><td>7.4</td><td>1.6</td><td>4.5</td><td>6.8E-06</td><td>1.2E-05</td><td>BT20/MB468</td><td>CYTO</td></tr>
<tr><td>>hg38_21_+_8433172_8446622_RNA45SN1_45S_50ntFlanks@10396.10435.40|CAGTTGAACATGGGTCAGTCGGTCCTGAGAGATGGGCGAG</td><td>>hg38_21_+_8433172_8446622_RNA45SN1_45S_50ntFlanks</td><td>10396</td><td>10435</td><td>40</td><td>CAGTTGAACATGGGTCAGTCGGTCCTGAGAGATGGGCGAG</td><td>360.9</td><td>-5.3</td><td>1.2</td><td>-4.5</td><td>7.9E-06</td><td>1.2E-05</td><td>MB231/MB468</td><td>NUC</td></tr>
<tr><td>>hg38_21_+_8433172_8446622_RNA45SN1_45S_50ntFlanks@9539.9572.34|ACTCTGGTGGAGGTCCGTAGCGGTCCTGACGTGC</td><td>>hg38_21_+_8433172_8446622_RNA45SN1_45S_50ntFlanks</td><td>9539</td><td>9572</td><td>34</td><td>ACTCTGGTGGAGGTCCGTAGCGGTCCTGACGTGC</td><td>612.2</td><td>-7.5</td><td>1.7</td><td>-4.5</td><td>8.5E-06</td><td>1.2E-05</td><td>BT20/MB468</td><td>NUC</td></tr>
<tr><td>>hg38_21_+_8433172_8446622_RNA45SN1_45S_50ntFlanks@12242.12262.21|AGAAGGGCAAAAGCTCGCTTG</td><td>>hg38_21_+_8433172_8446622_RNA45SN1_45S_50ntFlanks</td><td>12242</td><td>12262</td><td>21</td><td>AGAAGGGCAAAAGCTCGCTTG</td><td>54.8</td><td>-8.9</td><td>1.9</td><td>-4.7</td><td>2.5E-06</td><td>1.2E-05</td><td>BT20/MB468</td><td>MP</td></tr>
<tr><td>>hg38_21_+_8433172_8446622_RNA45SN1_45S_50ntFlanks@5152.5185.34|AGGGACAAGTGGCGTTCAGCCACCCGAGATTGAG</td><td>>hg38_21_+_8433172_8446622_RNA45SN1_45S_50ntFlanks</td><td>5152</td><td>5185</td><td>34</td><td>AGGGACAAGTGGCGTTCAGCCACCCGAGATTGAG</td><td>501.5</td><td>-7.7</td><td>1.7</td><td>-4.5</td><td>7.9E-06</td><td>1.2E-05</td><td>MB231/MB468</td><td>NUC</td></tr>
<tr><td>>hg38_21_+_8433172_8446622_RNA45SN1_45S_50ntFlanks@6652.6677.26|GACTCTTAGCGGTGGATCACTCGGCT</td><td>>hg38_21_+_8433172_8446622_RNA45SN1_45S_50ntFlanks</td><td>6652</td><td>6677</td><td>26</td><td>GACTCTTAGCGGTGGATCACTCGGCT</td><td>5412.8</td><td>3.6</td><td>0.8</td><td>4.5</td><td>5.8E-06</td><td>1.2E-05</td><td>BT20/MB468</td><td>MITO</td></tr>
<tr><td>>hg38_21_+_8433172_8446622_RNA45SN1_45S_50ntFlanks@6663.6696.34|GTGGATCACTCGGCTCGTGCGTCGATGAAGAACG</td><td>>hg38_21_+_8433172_8446622_RNA45SN1_45S_50ntFlanks</td><td>6663</td><td>6696</td><td>34</td><td>GTGGATCACTCGGCTCGTGCGTCGATGAAGAACG</td><td>493.0</td><td>2.6</td><td>0.6</td><td>4.5</td><td>5.8E-06</td><td>1.2E-05</td><td>BT20/MB468</td><td>MITO</td></tr>
<tr><td>>hg38_21_+_8433172_8446622_RNA45SN1_45S_50ntFlanks@12539.12566.28|ATGATGTGTTGTTGCCATGGTAATCCTG</td><td>>hg38_21_+_8433172_8446622_RNA45SN1_45S_50ntFlanks</td><td>12539</td><td>12566</td><td>28</td><td>ATGATGTGTTGTTGCCATGGTAATCCTG</td><td>888.9</td><td>-6.5</td><td>1.4</td><td>-4.5</td><td>6.8E-06</td><td>1.2E-05</td><td>BT20/MB468</td><td>CYTO</td></tr>
<tr><td>>hg38_21_+_8433172_8446622_RNA45SN1_45S_50ntFlanks@5472.5511.40|ACGGCCCTGGCGGAGCGCTGAGAAGACGGTCGAACTTGAC</td><td>>hg38_21_+_8433172_8446622_RNA45SN1_45S_50ntFlanks</td><td>5472</td><td>5511</td><td>40</td><td>ACGGCCCTGGCGGAGCGCTGAGAAGACGGTCGAACTTGAC</td><td>36.7</td><td>8.4</td><td>1.9</td><td>4.5</td><td>6.7E-06</td><td>1.2E-05</td><td>MB231/MB468</td><td>MP</td></tr>
<tr><td>>MI0000082|hsa-mir-25&WithFlank&7|-|100093554|100093649@58.78.21(+1G)&[MIMAT0000081&hsa-miR-25-3p&offsets|0|-1(+1G);m-23&7|-|100093571|100093592&offsets|0|-1(+1G)]|CATTGCACTTGTCTCGGTCTGG</td><td>>MI0000082|hsa-mir-25&WithFlank&7|-|100093554|100093649</td><td>58</td><td>78</td><td>21(+1G)</td><td>CATTGCACTTGTCTCGGTCTGG</td><td>318.1</td><td>4.3</td><td>1.0</td><td>4.5</td><td>8.2E-06</td><td>1.2E-05</td><td>MB231/MB468</td><td>MITO</td></tr>
<tr><td>>hg38_21_+_8433172_8446622_RNA45SN1_45S_50ntFlanks@10425.10443.19|AGATGGGCGAGCGCCGTTC</td><td>>hg38_21_+_8433172_8446622_RNA45SN1_45S_50ntFlanks</td><td>10425</td><td>10443</td><td>19</td><td>AGATGGGCGAGCGCCGTTC</td><td>124.6</td><td>-10.8</td><td>2.4</td><td>-4.5</td><td>6.7E-06</td><td>1.2E-05</td><td>MB231/MB468</td><td>MP</td></tr>
<tr><td>>hg38_21_+_8433172_8446622_RNA45SN1_45S_50ntFlanks@11564.11590.27|CCTAGCAGCCGACTTAGAACTGGTGCG</td><td>>hg38_21_+_8433172_8446622_RNA45SN1_45S_50ntFlanks</td><td>11564</td><td>11590</td><td>27</td><td>CCTAGCAGCCGACTTAGAACTGGTGCG</td><td>699.9</td><td>-4.4</td><td>0.9</td><td>-4.7</td><td>2.5E-06</td><td>1.2E-05</td><td>BT20/MB468</td><td>MP</td></tr>
<tr><td>>hg38_21_+_8433172_8446622_RNA45SN1_45S_50ntFlanks@5487.5516.30|CGCTGAGAAGACGGTCGAACTTGACTATCT</td><td>>hg38_21_+_8433172_8446622_RNA45SN1_45S_50ntFlanks</td><td>5487</td><td>5516</td><td>30</td><td>CGCTGAGAAGACGGTCGAACTTGACTATCT</td><td>1180.0</td><td>-3.5</td><td>0.8</td><td>-4.5</td><td>8.2E-06</td><td>1.2E-05</td><td>MB231/MB468</td><td>MITO</td></tr>
<tr><td>>hg38_21_+_8433172_8446622_RNA45SN1_45S_50ntFlanks@12444.12478.35|AAGCAGAATTCACCAAGCGTTGGATTGTTCACCCA</td><td>>hg38_21_+_8433172_8446622_RNA45SN1_45S_50ntFlanks</td><td>12444</td><td>12478</td><td>35</td><td>AAGCAGAATTCACCAAGCGTTGGATTGTTCACCCA</td><td>510.0</td><td>-5.7</td><td>1.3</td><td>-4.5</td><td>8.5E-06</td><td>1.2E-05</td><td>BT20/MB468</td><td>NUC</td></tr>
<tr><td>>hg38_21_+_8433172_8446622_RNA45SN1_45S_50ntFlanks@8327.8354.28|ATAGTCAACAAGTACCGTAAGGGAAAGT</td><td>>hg38_21_+_8433172_8446622_RNA45SN1_45S_50ntFlanks</td><td>8327</td><td>8354</td><td>28</td><td>ATAGTCAACAAGTACCGTAAGGGAAAGT</td><td>52.0</td><td>-8.8</td><td>1.9</td><td>-4.7</td><td>2.5E-06</td><td>1.2E-05</td><td>BT20/MB468</td><td>MP</td></tr>
<tr><td>>hg38_21_+_8433172_8446622_RNA45SN1_45S_50ntFlanks@6648.6680.33|GTACGACTCTTAGCGGTGGATCACTCGGCTCGT</td><td>>hg38_21_+_8433172_8446622_RNA45SN1_45S_50ntFlanks</td><td>6648</td><td>6680</td><td>33</td><td>GTACGACTCTTAGCGGTGGATCACTCGGCTCGT</td><td>1142.0</td><td>4.1</td><td>0.9</td><td>4.5</td><td>6.9E-06</td><td>1.2E-05</td><td>BT20/MB468</td><td>CYTO</td></tr>
<tr><td>>am_trnaMT_ArgTCG_MT_+_10405_10469@29.48.20|TTTCGACTCATTAAATTATG</td><td>>am_trnaMT_ArgTCG_MT_+_10405_10469</td><td>29</td><td>48</td><td>20</td><td>TTTCGACTCATTAAATTATG</td><td>1534.5</td><td>-9.0</td><td>2.0</td><td>-4.5</td><td>8.5E-06</td><td>1.2E-05</td><td>BT20/MB468</td><td>NUC</td></tr>
<tr><td>>hg38_21_+_8433172_8446622_RNA45SN1_45S_50ntFlanks@11573.11591.19|CGACTTAGAACTGGTGCGG</td><td>>hg38_21_+_8433172_8446622_RNA45SN1_45S_50ntFlanks</td><td>11573</td><td>11591</td><td>19</td><td>CGACTTAGAACTGGTGCGG</td><td>309.3</td><td>-4.5</td><td>1.0</td><td>-4.5</td><td>6.9E-06</td><td>1.2E-05</td><td>BT20/MB468</td><td>CYTO</td></tr>
<tr><td>>hg38_21_+_8433172_8446622_RNA45SN1_45S_50ntFlanks@4570.4587.18|TGGAATAGGACCGCGGTT</td><td>>hg38_21_+_8433172_8446622_RNA45SN1_45S_50ntFlanks</td><td>4570</td><td>4587</td><td>18</td><td>TGGAATAGGACCGCGGTT</td><td>338.8</td><td>-2.5</td><td>0.5</td><td>-4.5</td><td>6.9E-06</td><td>1.2E-05</td><td>BT20/MB468</td><td>CYTO</td></tr>
<tr><td>>hg38_21_+_8433172_8446622_RNA45SN1_45S_50ntFlanks@9604.9621.18|AAAGACTAATCGAACCAT</td><td>>hg38_21_+_8433172_8446622_RNA45SN1_45S_50ntFlanks</td><td>9604</td><td>9621</td><td>18</td><td>AAAGACTAATCGAACCAT</td><td>263.5</td><td>-4.1</td><td>0.9</td><td>-4.5</td><td>6.9E-06</td><td>1.2E-05</td><td>BT20/MB468</td><td>CYTO</td></tr>
<tr><td>>MI0000253|hsa-mir-148a&WithFlank&7|-|25949913|25949992@50.73.24&[MIMAT0000243&hsa-miR-148a-3p&offsets|0|+2;m-7&7|-|25949922|25949943&offsets|0|+2]|TCAGTGCACTACAGAACTTTGTCT</td><td>>MI0000253|hsa-mir-148a&WithFlank&7|-|25949913|25949992</td><td>50</td><td>73</td><td>24</td><td>TCAGTGCACTACAGAACTTTGTCT</td><td>287.6</td><td>5.3</td><td>1.1</td><td>5.0</td><td>7.2E-07</td><td>1.2E-05</td><td>BT20/MB231</td><td>CYTO</td></tr>
<tr><td>>MI0000061|hsa-let-7a-2&WithFlank&11|-|122146516|122146599@11.32.22;MI0000062|hsa-let-7a-3&WithFlank&22|+|46112743|46112828@10.31.22;MI0000060|hsa-let-7a-1&WithFlank&9|+|94175951|94176042@12.33.22&[MIMAT0000062&hsa-let-7a-5p&offsets|0|0;m-6&11|-|122146568|122146589&offsets|0|0];[MIMAT0000062_1&hsa-let-7a-5p&offsets|0|0;m-5&22|+|46112752|46112773&offsets|0|0];[MIMAT0000062_2&hsa-let-7a-5p&offsets|0|0;m-4&9|+|94175962|94175983&offsets|0|0]|TGAGGTAGTAGGTTGTATAGTT</td><td>>MI0000061|hsa-let-7a-2&WithFlank&11|-|122146516|122146599</td><td>11</td><td>32</td><td>22</td><td>TGAGGTAGTAGGTTGTATAGTT</td><td>50348.4</td><td>3.9</td><td>0.9</td><td>4.5</td><td>6.8E-06</td><td>1.2E-05</td><td>MB231/MB468</td><td>MP</td></tr>
<tr><td>>hg38_21_+_8433172_8446622_RNA45SN1_45S_50ntFlanks@6646.6682.37|TCGTACGACTCTTAGCGGTGGATCACTCGGCTCGTGC</td><td>>hg38_21_+_8433172_8446622_RNA45SN1_45S_50ntFlanks</td><td>6646</td><td>6682</td><td>37</td><td>TCGTACGACTCTTAGCGGTGGATCACTCGGCTCGTGC</td><td>491.7</td><td>6.1</td><td>1.4</td><td>4.5</td><td>6.9E-06</td><td>1.2E-05</td><td>BT20/MB468</td><td>CYTO</td></tr>
<tr><td>>hg38_21_+_8433172_8446622_RNA45SN1_45S_50ntFlanks@4260.4277.18|TTGGAATGAGTCCACTTT</td><td>>hg38_21_+_8433172_8446622_RNA45SN1_45S_50ntFlanks</td><td>4260</td><td>4277</td><td>18</td><td>TTGGAATGAGTCCACTTT</td><td>164.3</td><td>-5.0</td><td>1.1</td><td>-4.5</td><td>6.8E-06</td><td>1.2E-05</td><td>MB231/MB468</td><td>MP</td></tr>
<tr><td>>hg38_21_+_8433172_8446622_RNA45SN1_45S_50ntFlanks@4739.4758.20|AAGTCGGAGGTTCGAAGACG</td><td>>hg38_21_+_8433172_8446622_RNA45SN1_45S_50ntFlanks</td><td>4739</td><td>4758</td><td>20</td><td>AAGTCGGAGGTTCGAAGACG</td><td>811.0</td><td>-3.5</td><td>0.7</td><td>-4.7</td><td>2.5E-06</td><td>1.2E-05</td><td>BT20/MB468</td><td>MP</td></tr>
<tr><td>>hg38_21_+_8433172_8446622_RNA45SN1_45S_50ntFlanks@6664.6681.18|TGGATCACTCGGCTCGTG</td><td>>hg38_21_+_8433172_8446622_RNA45SN1_45S_50ntFlanks</td><td>6664</td><td>6681</td><td>18</td><td>TGGATCACTCGGCTCGTG</td><td>4503.1</td><td>-4.9</td><td>1.0</td><td>-4.7</td><td>2.5E-06</td><td>1.2E-05</td><td>BT20/MB468</td><td>MP</td></tr>
<tr><td>>hg38_21_+_8433172_8446622_RNA45SN1_45S_50ntFlanks@9935.9952.18|GACAGCAGGACGGTGGCC</td><td>>hg38_21_+_8433172_8446622_RNA45SN1_45S_50ntFlanks</td><td>9935</td><td>9952</td><td>18</td><td>GACAGCAGGACGGTGGCC</td><td>104.5</td><td>10.0</td><td>2.0</td><td>5.0</td><td>7.2E-07</td><td>1.2E-05</td><td>BT20/MB231</td><td>MP</td></tr>
<tr><td>>hg38_21_+_8433172_8446622_RNA45SN1_45S_50ntFlanks@9535.9571.37|GGAAACTCTGGTGGAGGTCCGTAGCGGTCCTGACGTG</td><td>>hg38_21_+_8433172_8446622_RNA45SN1_45S_50ntFlanks</td><td>9535</td><td>9571</td><td>37</td><td>GGAAACTCTGGTGGAGGTCCGTAGCGGTCCTGACGTG</td><td>160.5</td><td>-5.3</td><td>1.2</td><td>-4.5</td><td>6.8E-06</td><td>1.2E-05</td><td>MB231/MB468</td><td>MP</td></tr>
<tr><td>>hg38_21_+_8433172_8446622_RNA45SN1_45S_50ntFlanks@12406.12435.30|TTTTTGATCCTTCGATGTCGGCTCTTCCTA</td><td>>hg38_21_+_8433172_8446622_RNA45SN1_45S_50ntFlanks</td><td>12406</td><td>12435</td><td>30</td><td>TTTTTGATCCTTCGATGTCGGCTCTTCCTA</td><td>795.9</td><td>-6.9</td><td>1.6</td><td>-4.5</td><td>8.0E-06</td><td>1.2E-05</td><td>MB231/MB468</td><td>NUC</td></tr>
<tr><td>>hg38_21_+_8433172_8446622_RNA45SN1_45S_50ntFlanks@6745.6765.21|ACACTTCGAACGCACTTGCGG</td><td>>hg38_21_+_8433172_8446622_RNA45SN1_45S_50ntFlanks</td><td>6745</td><td>6765</td><td>21</td><td>ACACTTCGAACGCACTTGCGG</td><td>219.5</td><td>-7.0</td><td>1.5</td><td>-4.7</td><td>2.5E-06</td><td>1.2E-05</td><td>BT20/MB468</td><td>MP</td></tr>
<tr><td>>hg38_21_+_8433172_8446622_RNA45SN1_45S_50ntFlanks@11808.11830.23|CTGTCCCTACCTACTATCCAGCG</td><td>>hg38_21_+_8433172_8446622_RNA45SN1_45S_50ntFlanks</td><td>11808</td><td>11830</td><td>23</td><td>CTGTCCCTACCTACTATCCAGCG</td><td>339.2</td><td>-7.0</td><td>1.6</td><td>-4.4</td><td>8.6E-06</td><td>1.2E-05</td><td>BT20/MB468</td><td>NUC</td></tr>
<tr><td>>hg38_21_+_8433172_8446622_RNA45SN1_45S_50ntFlanks@5419.5445.27|ACCGATTGGATGGTTTAGTGAGGCCCT</td><td>>hg38_21_+_8433172_8446622_RNA45SN1_45S_50ntFlanks</td><td>5419</td><td>5445</td><td>27</td><td>ACCGATTGGATGGTTTAGTGAGGCCCT</td><td>252.2</td><td>-4.0</td><td>0.9</td><td>-4.5</td><td>7.8E-06</td><td>1.2E-05</td><td>MB231/MB468</td><td>CYTO</td></tr>
<tr><td>>MI0003137|hsa-mir-193b&WithFlank&16|+|14303961|14304055@20.41.22&[MIMAT0004767&hsa-miR-193b-5p&offsets|0|0;m-343&16|+|14303980|14304001&offsets|0|0]|CGGGGTTTTGAGGGCGAGATGA</td><td>>MI0003137|hsa-mir-193b&WithFlank&16|+|14303961|14304055</td><td>20</td><td>41</td><td>22</td><td>CGGGGTTTTGAGGGCGAGATGA</td><td>412.2</td><td>8.7</td><td>1.8</td><td>4.7</td><td>2.5E-06</td><td>1.2E-05</td><td>BT20/MB468</td><td>MP</td></tr>
<tr><td>>hg38_21_+_8433172_8446622_RNA45SN1_45S_50ntFlanks@10853.10879.27|ATAAGGATTGGCTCTAAGGGCTGGGTC</td><td>>hg38_21_+_8433172_8446622_RNA45SN1_45S_50ntFlanks</td><td>10853</td><td>10879</td><td>27</td><td>ATAAGGATTGGCTCTAAGGGCTGGGTC</td><td>194.8</td><td>-6.9</td><td>1.5</td><td>-4.5</td><td>6.8E-06</td><td>1.2E-05</td><td>MB231/MB468</td><td>MP</td></tr>
<tr><td>>hg38_21_+_8433172_8446622_RNA45SN1_45S_50ntFlanks@10319.10352.34|GCAGATCTTGGTGGTAGTAGCAAATATTCAAACG</td><td>>hg38_21_+_8433172_8446622_RNA45SN1_45S_50ntFlanks</td><td>10319</td><td>10352</td><td>34</td><td>GCAGATCTTGGTGGTAGTAGCAAATATTCAAACG</td><td>109.9</td><td>10.1</td><td>2.0</td><td>5.0</td><td>7.3E-07</td><td>1.2E-05</td><td>BT20/MB231</td><td>MP</td></tr>
<tr><td>>MI0000743|hsa-mir-34c&WithFlank&11|+|111513433|111513521@52.73.22&[MIMAT0004677&hsa-miR-34c-3p&offsets|0|0;m-474&11|+|111513484|111513505&offsets|0|0]|AATCACTAACCACACGGCCAGG</td><td>>MI0000743|hsa-mir-34c&WithFlank&11|+|111513433|111513521</td><td>52</td><td>73</td><td>22</td><td>AATCACTAACCACACGGCCAGG</td><td>87.9</td><td>-9.9</td><td>2.0</td><td>-5.0</td><td>7.3E-07</td><td>1.2E-05</td><td>BT20/MB231</td><td>MP</td></tr>
<tr><td>>hg38_21_+_8433172_8446622_RNA45SN1_45S_50ntFlanks@11728.11749.22|GTAACTATGACTCTCTTAAGGT</td><td>>hg38_21_+_8433172_8446622_RNA45SN1_45S_50ntFlanks</td><td>11728</td><td>11749</td><td>22</td><td>GTAACTATGACTCTCTTAAGGT</td><td>274.1</td><td>-3.1</td><td>0.7</td><td>-4.5</td><td>8.1E-06</td><td>1.2E-05</td><td>MB231/MB468</td><td>NUC</td></tr>
<tr><td>>hg38_21_+_8433172_8446622_RNA45SN1_45S_50ntFlanks@9504.9527.24|GTGAACTATGCCTGGGCAGGGCGA</td><td>>hg38_21_+_8433172_8446622_RNA45SN1_45S_50ntFlanks</td><td>9504</td><td>9527</td><td>24</td><td>GTGAACTATGCCTGGGCAGGGCGA</td><td>590.3</td><td>-6.9</td><td>1.5</td><td>-4.5</td><td>8.3E-06</td><td>1.2E-05</td><td>MB231/MB468</td><td>MITO</td></tr>
<tr><td>>hg38_MT_+_1621_3279_RNR2_16S_50ntFlanks@826.859.34|AAGGAAAGGTTAAAAAAAGTAAAAGGAACTCGGC</td><td>>hg38_MT_+_1621_3279_RNR2_16S_50ntFlanks</td><td>826</td><td>859</td><td>34</td><td>AAGGAAAGGTTAAAAAAAGTAAAAGGAACTCGGC</td><td>36.6</td><td>8.4</td><td>1.9</td><td>4.5</td><td>6.8E-06</td><td>1.2E-05</td><td>MB231/MB468</td><td>MP</td></tr>
<tr><td>>hg38_21_+_8433172_8446622_RNA45SN1_45S_50ntFlanks@12973.13000.28|TCGTACGTAGCAGAGCAGCTCCCTCGCT</td><td>>hg38_21_+_8433172_8446622_RNA45SN1_45S_50ntFlanks</td><td>12973</td><td>13000</td><td>28</td><td>TCGTACGTAGCAGAGCAGCTCCCTCGCT</td><td>102.3</td><td>-6.2</td><td>1.4</td><td>-4.5</td><td>6.8E-06</td><td>1.2E-05</td><td>MB231/MB468</td><td>MP</td></tr>
<tr><td>>hg38_21_+_8433172_8446622_RNA45SN1_45S_50ntFlanks@12360.12384.25|CAGGGATAACTGGCTTGTGGCGGCC</td><td>>hg38_21_+_8433172_8446622_RNA45SN1_45S_50ntFlanks</td><td>12360</td><td>12384</td><td>25</td><td>CAGGGATAACTGGCTTGTGGCGGCC</td><td>310.9</td><td>-3.8</td><td>0.9</td><td>-4.5</td><td>7.8E-06</td><td>1.2E-05</td><td>MB231/MB468</td><td>CYTO</td></tr>
<tr><td>>hg38_21_+_8433172_8446622_RNA45SN1_45S_50ntFlanks@7983.8010.28|CAGATCAGACGTGGCGACCCGCTGAATT</td><td>>hg38_21_+_8433172_8446622_RNA45SN1_45S_50ntFlanks</td><td>7983</td><td>8010</td><td>28</td><td>CAGATCAGACGTGGCGACCCGCTGAATT</td><td>204.3</td><td>-5.1</td><td>1.1</td><td>-4.5</td><td>6.9E-06</td><td>1.2E-05</td><td>MB231/MB468</td><td>MP</td></tr>
<tr><td>>hg38_21_+_8433172_8446622_RNA45SN1_45S_50ntFlanks@12072.12094.23|CCCGAGGGGCTCTCGCTTCTGGC</td><td>>hg38_21_+_8433172_8446622_RNA45SN1_45S_50ntFlanks</td><td>12072</td><td>12094</td><td>23</td><td>CCCGAGGGGCTCTCGCTTCTGGC</td><td>438.4</td><td>-3.0</td><td>0.7</td><td>-4.5</td><td>7.8E-06</td><td>1.2E-05</td><td>MB231/MB468</td><td>CYTO</td></tr>
<tr><td>>hg38_21_+_8433172_8446622_RNA45SN1_45S_50ntFlanks@4047.4081.35|ATTCGAACGTCTGCCCTATCAACTTTCGATGGTAG</td><td>>hg38_21_+_8433172_8446622_RNA45SN1_45S_50ntFlanks</td><td>4047</td><td>4081</td><td>35</td><td>ATTCGAACGTCTGCCCTATCAACTTTCGATGGTAG</td><td>307.6</td><td>-5.6</td><td>1.2</td><td>-4.5</td><td>7.8E-06</td><td>1.2E-05</td><td>MB231/MB468</td><td>CYTO</td></tr>
<tr><td>>hg38_21_+_8433172_8446622_RNA45SN1_45S_50ntFlanks@4130.4151.22|ATTCCGGAGAGGGAGCCTGAGA</td><td>>hg38_21_+_8433172_8446622_RNA45SN1_45S_50ntFlanks</td><td>4130</td><td>4151</td><td>22</td><td>ATTCCGGAGAGGGAGCCTGAGA</td><td>160.4</td><td>-2.9</td><td>0.6</td><td>-4.5</td><td>6.9E-06</td><td>1.2E-05</td><td>MB231/MB468</td><td>MP</td></tr>
<tr><td>>hg38_21_+_8433172_8446622_RNA45SN1_45S_50ntFlanks@10829.10852.24|GCAAGCCGGATCCGTAACTTCGGG</td><td>>hg38_21_+_8433172_8446622_RNA45SN1_45S_50ntFlanks</td><td>10829</td><td>10852</td><td>24</td><td>GCAAGCCGGATCCGTAACTTCGGG</td><td>384.0</td><td>-4.7</td><td>1.1</td><td>-4.5</td><td>7.8E-06</td><td>1.2E-05</td><td>MB231/MB468</td><td>CYTO</td></tr>
</table>

<table>
<tr><td>>trna2_GlyGCC_21_-_18827107_18827177@1.35.35,trna35_GlyGCC_1_+_161413094_161413164@1.35.35,trna37_GlyGCC_1_+_161420467_161420537@1.35.35,trna39_GlyGCC_1_+_161427898_161427968@1.35.35,trna41_GlyGCC_1_+_161435258_161435328@1.35.35|GCATGGGTGGTTCAGTGGTAGAATTCTCGCCTGCC</td><td>>trna2_GlyGCC_21_-_18827107_18827177</td><td>1</td><td>35</td><td>35</td><td>GCATGGGTGGTTCAGTGGTAGAATTCTCGCCTGCC</td><td>486.1</td><td>5.9</td><td>1.3</td><td>4.5</td><td>7.0E-06</td><td>1.2E-05</td><td>BT20/MB468</td><td>CYTO</td></tr>
<tr><td>>hg38_21_+_8433172_8446622_RNA45SN1_45S_50ntFlanks@11322.11345.24|ACGTCTCGTCGCGCGCGCGTCCGC</td><td>>hg38_21_+_8433172_8446622_RNA45SN1_45S_50ntFlanks</td><td>11322</td><td>11345</td><td>24</td><td>ACGTCTCGTCGCGCGCGCGTCCGC</td><td>442.1</td><td>5.1</td><td>1.2</td><td>4.5</td><td>8.2E-06</td><td>1.2E-05</td><td>MB231/MB468</td><td>NUC</td></tr>
<tr><td>>hg38_21_+_8433172_8446622_RNA45SN1_45S_50ntFlanks@12071.12096.26|CCCCGAGGGGCTCTCGCTTCTGGCGC</td><td>>hg38_21_+_8433172_8446622_RNA45SN1_45S_50ntFlanks</td><td>12071</td><td>12096</td><td>26</td><td>CCCCGAGGGGCTCTCGCTTCTGGCGC</td><td>838.8</td><td>-5.1</td><td>1.1</td><td>-4.5</td><td>7.9E-06</td><td>1.2E-05</td><td>MB231/MB468</td><td>CYTO</td></tr>
<tr><td>>hg38_21_+_8433172_8446622_RNA45SN1_45S_50ntFlanks@11786.11804.19|GAATGGATGAACGAGATTC</td><td>>hg38_21_+_8433172_8446622_RNA45SN1_45S_50ntFlanks</td><td>11786</td><td>11804</td><td>19</td><td>GAATGGATGAACGAGATTC</td><td>233.6</td><td>-2.7</td><td>0.6</td><td>-4.5</td><td>6.0E-06</td><td>1.2E-05</td><td>BT20/MB468</td><td>MITO</td></tr>
<tr><td>>hg38_21_+_8433172_8446622_RNA45SN1_45S_50ntFlanks@12539.12566.28|ATGATGTGTTGTTGCCATGGTAATCCTG</td><td>>hg38_21_+_8433172_8446622_RNA45SN1_45S_50ntFlanks</td><td>12539</td><td>12566</td><td>28</td><td>ATGATGTGTTGTTGCCATGGTAATCCTG</td><td>691.2</td><td>-4.8</td><td>1.1</td><td>-4.5</td><td>6.0E-06</td><td>1.2E-05</td><td>BT20/MB468</td><td>MITO</td></tr>
<tr><td>>hg38_21_+_8433172_8446622_RNA45SN1_45S_50ntFlanks@6646.6678.33|TCGTACGACTCTTAGCGGTGGATCACTCGGCTC</td><td>>hg38_21_+_8433172_8446622_RNA45SN1_45S_50ntFlanks</td><td>6646</td><td>6678</td><td>33</td><td>TCGTACGACTCTTAGCGGTGGATCACTCGGCTC</td><td>615.8</td><td>2.7</td><td>0.6</td><td>4.5</td><td>7.9E-06</td><td>1.2E-05</td><td>MB231/MB468</td><td>CYTO</td></tr>
<tr><td>>hg38_21_+_8433172_8446622_RNA45SN1_45S_50ntFlanks@11567.11601.35|AGCAGCCGACTTAGAACTGGTGCGGACCAGGGGAA</td><td>>hg38_21_+_8433172_8446622_RNA45SN1_45S_50ntFlanks</td><td>11567</td><td>11601</td><td>35</td><td>AGCAGCCGACTTAGAACTGGTGCGGACCAGGGGAA</td><td>769.6</td><td>-3.9</td><td>0.9</td><td>-4.5</td><td>8.4E-06</td><td>1.2E-05</td><td>MB231/MB468</td><td>MITO</td></tr>
<tr><td>>hg38_21_+_8433172_8446622_RNA45SN1_45S_50ntFlanks@5444.5464.21|CTCGGATCGGCCCCGCCGGGG</td><td>>hg38_21_+_8433172_8446622_RNA45SN1_45S_50ntFlanks</td><td>5444</td><td>5464</td><td>21</td><td>CTCGGATCGGCCCCGCCGGGG</td><td>55.0</td><td>-8.9</td><td>1.9</td><td>-4.7</td><td>2.6E-06</td><td>1.2E-05</td><td>BT20/MB468</td><td>MP</td></tr>
<tr><td>>hg38_21_+_8433172_8446622_RNA45SN1_45S_50ntFlanks@6646.6671.26|TCGTACGACTCTTAGCGGTGGATCAC</td><td>>hg38_21_+_8433172_8446622_RNA45SN1_45S_50ntFlanks</td><td>6646</td><td>6671</td><td>26</td><td>TCGTACGACTCTTAGCGGTGGATCAC</td><td>497.6</td><td>3.9</td><td>0.9</td><td>4.5</td><td>8.2E-06</td><td>1.2E-05</td><td>MB231/MB468</td><td>NUC</td></tr>
<tr><td>>hg38_21_+_8433172_8446622_RNA45SN1_45S_50ntFlanks@10684.10705.22|CGGCGGCGTCCGGTGAGCTCTC</td><td>>hg38_21_+_8433172_8446622_RNA45SN1_45S_50ntFlanks</td><td>10684</td><td>10705</td><td>22</td><td>CGGCGGCGTCCGGTGAGCTCTC</td><td>528.4</td><td>-5.0</td><td>1.1</td><td>-4.5</td><td>8.2E-06</td><td>1.2E-05</td><td>MB231/MB468</td><td>NUC</td></tr>
<tr><td>>am_MI0003639|hsa-mir-625&WithFlank&14|+|65471096|65471192@58.80.23&[MIMAT0004808&hsa-miR-625-3p&offsets|0|+1;m-137&14|+|65471153|65471174&offsets|0|+1]|GACTATAGAACTTTCCCCCTCAT</td><td>>am_MI0003639|hsa-mir-625&WithFlank&14|+|65471096|65471192</td><td>58</td><td>80</td><td>23</td><td>GACTATAGAACTTTCCCCCTCAT</td><td>314.3</td><td>6.4</td><td>1.4</td><td>4.5</td><td>6.9E-06</td><td>1.2E-05</td><td>MB231/MB468</td><td>MP</td></tr>
<tr><td>>hg38_21_+_8433172_8446622_RNA45SN1_45S_50ntFlanks@10686.10709.24|GCGGCGTCCGGTGAGCTCTCGCTG</td><td>>hg38_21_+_8433172_8446622_RNA45SN1_45S_50ntFlanks</td><td>10686</td><td>10709</td><td>24</td><td>GCGGCGTCCGGTGAGCTCTCGCTG</td><td>425.6</td><td>-6.7</td><td>1.5</td><td>-4.5</td><td>6.9E-06</td><td>1.2E-05</td><td>MB231/MB468</td><td>MP</td></tr>
<tr><td>>hg38_21_+_8433172_8446622_RNA45SN1_45S_50ntFlanks@10466.10484.19|CGTTGCCCTCGGCCGATCG</td><td>>hg38_21_+_8433172_8446622_RNA45SN1_45S_50ntFlanks</td><td>10466</td><td>10484</td><td>19</td><td>CGTTGCCCTCGGCCGATCG</td><td>115.0</td><td>-5.2</td><td>1.2</td><td>-4.5</td><td>7.1E-06</td><td>1.2E-05</td><td>BT20/MB468</td><td>CYTO</td></tr>
<tr><td>>hg38_21_+_8433172_8446622_RNA45SN1_45S_50ntFlanks@8386.8406.21|GGCGTGAAACCGTTAAGAGGT</td><td>>hg38_21_+_8433172_8446622_RNA45SN1_45S_50ntFlanks</td><td>8386</td><td>8406</td><td>21</td><td>GGCGTGAAACCGTTAAGAGGT</td><td>48.2</td><td>-8.7</td><td>1.8</td><td>-4.7</td><td>2.6E-06</td><td>1.2E-05</td><td>BT20/MB468</td><td>MP</td></tr>
<tr><td>>hg38_21_+_8433172_8446622_RNA45SN1_45S_50ntFlanks@11197.11223.27|GGCCGGCGGCGGCGGCGACTCTGGACG</td><td>>hg38_21_+_8433172_8446622_RNA45SN1_45S_50ntFlanks</td><td>11197</td><td>11223</td><td>27</td><td>GGCCGGCGGCGGCGGCGACTCTGGACG</td><td>91.9</td><td>-10.4</td><td>2.3</td><td>-4.5</td><td>6.9E-06</td><td>1.2E-05</td><td>MB231/MB468</td><td>MP</td></tr>
<tr><td>>MI0002468|hsa-mir-484&WithFlank&16|+|15643288|15643378@14.35.22&[MIMAT0002174&hsa-miR-484&offsets|0|0;m-127&16|+|15643301|15643321&offsets|0|+1]|TCAGGCTCAGTCCCCTCCCGAT</td><td>>MI0002468|hsa-mir-484&WithFlank&16|+|15643288|15643378</td><td>14</td><td>35</td><td>22</td><td>TCAGGCTCAGTCCCCTCCCGAT</td><td>1855.7</td><td>4.3</td><td>1.0</td><td>4.5</td><td>8.4E-06</td><td>1.2E-05</td><td>MB231/MB468</td><td>MITO</td></tr>
<tr><td>>hg38_21_+_8433172_8446622_RNA45SN1_45S_50ntFlanks@9565.9592.28|TGACGTGCAAATCGGTCGTCCGACCTGG</td><td>>hg38_21_+_8433172_8446622_RNA45SN1_45S_50ntFlanks</td><td>9565</td><td>9592</td><td>28</td><td>TGACGTGCAAATCGGTCGTCCGACCTGG</td><td>393.8</td><td>-5.1</td><td>1.2</td><td>-4.5</td><td>8.4E-06</td><td>1.2E-05</td><td>MB231/MB468</td><td>MITO</td></tr>
<tr><td>>hg38_21_+_8433172_8446622_RNA45SN1_45S_50ntFlanks@8182.8203.22|ATCGAGGCCCAGCCCGTGGACG</td><td>>hg38_21_+_8433172_8446622_RNA45SN1_45S_50ntFlanks</td><td>8182</td><td>8203</td><td>22</td><td>ATCGAGGCCCAGCCCGTGGACG</td><td>761.4</td><td>-7.0</td><td>1.6</td><td>-4.5</td><td>6.9E-06</td><td>1.2E-05</td><td>MB231/MB468</td><td>MP</td></tr>
<tr><td>>hg38_21_+_8433172_8446622_RNA45SN1_45S_50ntFlanks@10829.10858.30|GCAAGCCGGATCCGTAACTTCGGGATAAGG</td><td>>hg38_21_+_8433172_8446622_RNA45SN1_45S_50ntFlanks</td><td>10829</td><td>10858</td><td>30</td><td>GCAAGCCGGATCCGTAACTTCGGGATAAGG</td><td>291.7</td><td>-6.2</td><td>1.4</td><td>-4.5</td><td>8.2E-06</td><td>1.2E-05</td><td>MB231/MB468</td><td>NUC</td></tr>
<tr><td>>hg38_MT_+_598_1651_RNR1_12S_50ntFlanks@746.767.22|ATGAGGTGGCAAGAAATGGGCT</td><td>>hg38_MT_+_598_1651_RNR1_12S_50ntFlanks</td><td>746</td><td>767</td><td>22</td><td>ATGAGGTGGCAAGAAATGGGCT</td><td>49.1</td><td>9.4</td><td>2.0</td><td>4.7</td><td>2.6E-06</td><td>1.2E-05</td><td>BT20/MB468</td><td>MP</td></tr>
<tr><td>>hg38_MT_+_1621_3279_RNR2_16S_50ntFlanks@51.93.43|GCTAAACCTAGCCCCAAACCCACTCCACCTTACTACCAGACAA</td><td>>hg38_MT_+_1621_3279_RNR2_16S_50ntFlanks</td><td>51</td><td>93</td><td>43</td><td>GCTAAACCTAGCCCCAAACCCACTCCACCTTACTACCAGACAA</td><td>1535.5</td><td>6.3</td><td>1.4</td><td>4.5</td><td>8.4E-06</td><td>1.2E-05</td><td>MB231/MB468</td><td>MITO</td></tr>
<tr><td>>hg38_21_+_8433172_8446622_RNA45SN1_45S_50ntFlanks@4130.4150.21|ATTCCGGAGAGGGAGCCTGAG</td><td>>hg38_21_+_8433172_8446622_RNA45SN1_45S_50ntFlanks</td><td>4130</td><td>4150</td><td>21</td><td>ATTCCGGAGAGGGAGCCTGAG</td><td>170.7</td><td>-4.9</td><td>1.1</td><td>-4.5</td><td>6.9E-06</td><td>1.2E-05</td><td>MB231/MB468</td><td>MP</td></tr>
<tr><td>>hg38_21_+_8433172_8446622_RNA45SN1_45S_50ntFlanks@10247.10281.35|GCGACGAGTAGGAGGGCCGCTGCGGTGAGCCTTGA</td><td>>hg38_21_+_8433172_8446622_RNA45SN1_45S_50ntFlanks</td><td>10247</td><td>10281</td><td>35</td><td>GCGACGAGTAGGAGGGCCGCTGCGGTGAGCCTTGA</td><td>131.1</td><td>10.4</td><td>2.1</td><td>5.0</td><td>7.4E-07</td><td>1.2E-05</td><td>BT20/MB231</td><td>MP</td></tr>
<tr><td>>hg38_21_+_8433172_8446622_RNA45SN1_45S_50ntFlanks@3846.3881.36|CTTGGATAACTGTGGTAATTCTAGAGCTAATACATG</td><td>>hg38_21_+_8433172_8446622_RNA45SN1_45S_50ntFlanks</td><td>3846</td><td>3881</td><td>36</td><td>CTTGGATAACTGTGGTAATTCTAGAGCTAATACATG</td><td>344.3</td><td>-6.0</td><td>1.3</td><td>-4.5</td><td>6.0E-06</td><td>1.2E-05</td><td>BT20/MB468</td><td>MITO</td></tr>
<tr><td>>hg38_21_+_8433172_8446622_RNA45SN1_45S_50ntFlanks@12416.12434.19|TTCGATGTCGGCTCTTCCT</td><td>>hg38_21_+_8433172_8446622_RNA45SN1_45S_50ntFlanks</td><td>12416</td><td>12434</td><td>19</td><td>TTCGATGTCGGCTCTTCCT</td><td>582.0</td><td>2.0</td><td>0.4</td><td>4.5</td><td>6.0E-06</td><td>1.2E-05</td><td>BT20/MB468</td><td>MITO</td></tr>
<tr><td>>hg38_21_+_8433172_8446622_RNA45SN1_45S_50ntFlanks@5459.5499.41|CCGGGGTCGGCCCACGGCCCTGGCGGAGCGCTGAGAAGACG</td><td>>hg38_21_+_8433172_8446622_RNA45SN1_45S_50ntFlanks</td><td>5459</td><td>5499</td><td>41</td><td>CCGGGGTCGGCCCACGGCCCTGGCGGAGCGCTGAGAAGACG</td><td>61.2</td><td>-9.0</td><td>1.9</td><td>-4.7</td><td>2.6E-06</td><td>1.2E-05</td><td>BT20/MB468</td><td>MP</td></tr>
<tr><td>>hg38_21_+_8433172_8446622_RNA45SN1_45S_50ntFlanks@9562.9595.34|TCCTGACGTGCAAATCGGTCGTCCGACCTGGGTA</td><td>>hg38_21_+_8433172_8446622_RNA45SN1_45S_50ntFlanks</td><td>9562</td><td>9595</td><td>34</td><td>TCCTGACGTGCAAATCGGTCGTCCGACCTGGGTA</td><td>299.8</td><td>-4.9</td><td>1.1</td><td>-4.5</td><td>8.3E-06</td><td>1.2E-05</td><td>MB231/MB468</td><td>NUC</td></tr>
<tr><td>>hg38_21_+_8433172_8446622_RNA45SN1_45S_50ntFlanks@12139.12162.24|GGACAGTGCCAGGTGGGGAGTTTG</td><td>>hg38_21_+_8433172_8446622_RNA45SN1_45S_50ntFlanks</td><td>12139</td><td>12162</td><td>24</td><td>GGACAGTGCCAGGTGGGGAGTTTG</td><td>51.4</td><td>-8.8</td><td>1.9</td><td>-4.7</td><td>2.6E-06</td><td>1.2E-05</td><td>BT20/MB468</td><td>MP</td></tr>
<tr><td>>MI0000683|hsa-mir-181b-2&WithFlank&9|+|124693704|124693804@22.46.25(+1U)&[MIMAT0000257_1&hsa-miR-181b-5p&offsets|0|+2(+1U);m-72&9|+|124693725|124693747&offsets|0|+2(+1U)]|AACATTCATTGCTGTCGGTGGGTTTT</td><td>>MI0000683|hsa-mir-181b-2&WithFlank&9|+|124693704|124693804</td><td>22</td><td>46</td><td>25(+1U)</td><td>AACATTCATTGCTGTCGGTGGGTTTT</td><td>756.4</td><td>-4.4</td><td>0.9</td><td>-4.9</td><td>8.5E-07</td><td>1.2E-05</td><td>BT20/MB231</td><td>MITO</td></tr>
<tr><td>>MI0000465|hsa-mir-191&WithFlank&3|-|49020612|49020715@22.43.22&[MIMAT0000440&hsa-miR-191-5p&offsets|0|-1;m-48&3|-|49020672|49020694&offsets|0|-1]|CAACGGAATCCCAAAAGCAGCT</td><td>>MI0000465|hsa-mir-191&WithFlank&3|-|49020612|49020715</td><td>22</td><td>43</td><td>22</td><td>CAACGGAATCCCAAAAGCAGCT</td><td>3462.8</td><td>1.9</td><td>0.4</td><td>4.4</td><td>8.8E-06</td><td>1.2E-05</td><td>BT20/MB468</td><td>NUC</td></tr>
<tr><td>>hg38_21_+_8433172_8446622_RNA45SN1_45S_50ntFlanks@7976.8011.36|GCGACCTCAGATCAGACGTGGCGACCCGCTGAATTT</td><td>>hg38_21_+_8433172_8446622_RNA45SN1_45S_50ntFlanks</td><td>7976</td><td>8011</td><td>36</td><td>GCGACCTCAGATCAGACGTGGCGACCCGCTGAATTT</td><td>2382.6</td><td>8.3</td><td>1.8</td><td>4.7</td><td>2.6E-06</td><td>1.2E-05</td><td>BT20/MB468</td><td>MP</td></tr>
<tr><td>>MI0000088|hsa-mir-30a&WithFlank&6|-|71403545|71403627@12.35.24(+3U)&[MIMAT0000087&hsa-miR-30a-5p&offsets|0|+2(+3U);m-15&6|-|71403595|71403616&offsets|0|+2(+3U)]|TGTAAACATCCTCGACTGGAAGCTTTT</td><td>>MI0000088|hsa-mir-30a&WithFlank&6|-|71403545|71403627</td><td>12</td><td>35</td><td>24(+3U)</td><td>TGTAAACATCCTCGACTGGAAGCTTTT</td><td>443.9</td><td>-7.9</td><td>1.6</td><td>-4.9</td><td>7.5E-07</td><td>1.2E-05</td><td>BT20/MB231</td><td>MP</td></tr>
<tr><td>>hg38_21_+_8433172_8446622_RNA45SN1_45S_50ntFlanks@9740.9759.20|AACGATCTCAACCTATTCTC</td><td>>hg38_21_+_8433172_8446622_RNA45SN1_45S_50ntFlanks</td><td>9740</td><td>9759</td><td>20</td><td>AACGATCTCAACCTATTCTC</td><td>808.5</td><td>-4.2</td><td>0.9</td><td>-4.5</td><td>8.5E-06</td><td>1.2E-05</td><td>MB231/MB468</td><td>MITO</td></tr>
<tr><td>>MI0000086|hsa-mir-28&WithFlank&3|+|188688775|188688872@60.79.20(+1A)&[MIMAT0004502&hsa-miR-28-3p&offsets|0|-2(+1A);m-28&3|+|188688834|188688855&offsets|0|-2(+1A)]|CACTAGATTGTGAGCTCCTGA</td><td>>MI0000086|hsa-mir-28&WithFlank&3|+|188688775|188688872</td><td>60</td><td>79</td><td>20(+1A)</td><td>CACTAGATTGTGAGCTCCTGA</td><td>168.7</td><td>7.3</td><td>1.6</td><td>4.5</td><td>7.0E-06</td><td>1.2E-05</td><td>MB231/MB468</td><td>MP</td></tr>
<tr><td>>hg38_21_+_8433172_8446622_RNA45SN1_45S_50ntFlanks@12416.12437.22|TTCGATGTCGGCTCTTCCTATC</td><td>>hg38_21_+_8433172_8446622_RNA45SN1_45S_50ntFlanks</td><td>12416</td><td>12437</td><td>22</td><td>TTCGATGTCGGCTCTTCCTATC</td><td>2220.5</td><td>2.6</td><td>0.6</td><td>4.5</td><td>7.2E-06</td><td>1.2E-05</td><td>BT20/MB468</td><td>CYTO</td></tr>
<tr><td>>MI0000447|hsa-mir-128-1&WithFlank&2|+|135665391|135665484@56.75.20;MI0000727|hsa-mir-128-2&WithFlank&3|+|35744470|35744565@58.77.20&[MIMAT0000424&hsa-miR-128-3p&offsets|0|-1;m-108&2|+|135665446|135665466&offsets|0|-1];[MIMAT0000424_1&hsa-miR-128-3p&offsets|0|-1;m-117&3|+|35744527|35744547&offsets|0|-1]|TCACAGTGAACCGGTCTCTT</td><td>>MI0000447|hsa-mir-128-1&WithFlank&2|+|135665391|135665484</td><td>56</td><td>75</td><td>20</td><td>TCACAGTGAACCGGTCTCTT</td><td>1198.7</td><td>3.6</td><td>0.8</td><td>4.5</td><td>8.3E-06</td><td>1.2E-05</td><td>MB231/MB468</td><td>NUC</td></tr>
<tr><td>>hg38_21_+_8433172_8446622_RNA45SN1_45S_50ntFlanks@8360.8379.20|AGAACTTTGAAGAGAGAGTT</td><td>>hg38_21_+_8433172_8446622_RNA45SN1_45S_50ntFlanks</td><td>8360</td><td>8379</td><td>20</td><td>AGAACTTTGAAGAGAGAGTT</td><td>66.9</td><td>-9.1</td><td>1.9</td><td>-4.7</td><td>2.6E-06</td><td>1.2E-05</td><td>BT20/MB468</td><td>MP</td></tr>
<tr><td>>hg38_21_+_8433172_8446622_RNA45SN1_45S_50ntFlanks@3861.3898.38|TAATTCTAGAGCTAATACATGCCGACGGGCGCTGACCC</td><td>>hg38_21_+_8433172_8446622_RNA45SN1_45S_50ntFlanks</td><td>3861</td><td>3898</td><td>38</td><td>TAATTCTAGAGCTAATACATGCCGACGGGCGCTGACCC</td><td>331.2</td><td>-6.2</td><td>1.4</td><td>-4.5</td><td>7.2E-06</td><td>1.2E-05</td><td>BT20/MB468</td><td>CYTO</td></tr>
<tr><td>>hg38_21_+_8433172_8446622_RNA45SN1_45S_50ntFlanks@12002.12031.30|TGCGGGCCGCCGGTGAAATACCACTACTCT</td><td>>hg38_21_+_8433172_8446622_RNA45SN1_45S_50ntFlanks</td><td>12002</td><td>12031</td><td>30</td><td>TGCGGGCCGCCGGTGAAATACCACTACTCT</td><td>656.6</td><td>-2.6</td><td>0.6</td><td>-4.5</td><td>6.1E-06</td><td>1.2E-05</td><td>BT20/MB468</td><td>MITO</td></tr>
<tr><td>>hg38_21_+_8433172_8446622_RNA45SN1_45S_50ntFlanks@10477.10497.21|GCCGATCGAAAGGGAGTCGGG</td><td>>hg38_21_+_8433172_8446622_RNA45SN1_45S_50ntFlanks</td><td>10477</td><td>10497</td><td>21</td><td>GCCGATCGAAAGGGAGTCGGG</td><td>63.9</td><td>-9.1</td><td>1.9</td><td>-4.7</td><td>2.6E-06</td><td>1.2E-05</td><td>BT20/MB468</td><td>MP</td></tr>
<tr><td>>hg38_21_+_8433172_8446622_RNA45SN1_45S_50ntFlanks@9487.9510.24|TTAGGACCCGAAAGATGGTGAACT</td><td>>hg38_21_+_8433172_8446622_RNA45SN1_45S_50ntFlanks</td><td>9487</td><td>9510</td><td>24</td><td>TTAGGACCCGAAAGATGGTGAACT</td><td>41.3</td><td>9.1</td><td>1.9</td><td>4.7</td><td>2.6E-06</td><td>1.2E-05</td><td>BT20/MB468</td><td>MP</td></tr>
<tr><td>>hg38_21_+_8433172_8446622_RNA45SN1_45S_50ntFlanks@9862.9892.31|ATGAACCGAACGCCGGGTTAAGGCGCCCGAT</td><td>>hg38_21_+_8433172_8446622_RNA45SN1_45S_50ntFlanks</td><td>9862</td><td>9892</td><td>31</td><td>ATGAACCGAACGCCGGGTTAAGGCGCCCGAT</td><td>144.9</td><td>-6.5</td><td>1.4</td><td>-4.5</td><td>7.1E-06</td><td>1.2E-05</td><td>MB231/MB468</td><td>MP</td></tr>
<tr><td>>hg38_21_+_8433172_8446622_RNA45SN1_45S_50ntFlanks@10420.10457.38|CTGAGAGATGGGCGAGCGCCGTTCCGAAGGGACGGGCG</td><td>>hg38_21_+_8433172_8446622_RNA45SN1_45S_50ntFlanks</td><td>10420</td><td>10457</td><td>38</td><td>CTGAGAGATGGGCGAGCGCCGTTCCGAAGGGACGGGCG</td><td>942.9</td><td>-6.5</td><td>1.5</td><td>-4.5</td><td>7.2E-06</td><td>1.2E-05</td><td>BT20/MB468</td><td>CYTO</td></tr>
<tr><td>>MI0000078|hsa-mir-22&WithFlank&17|-|1713897|1713993@59.78.20&[MIMAT0000077&hsa-miR-22-3p&offsets|0|-2;m-2&17|-|1713914|1713935&offsets|0|-2]|AAGCTGCCAGTTGAAGAACT</td><td>>MI0000078|hsa-mir-22&WithFlank&17|-|1713897|1713993</td><td>59</td><td>78</td><td>20</td><td>AAGCTGCCAGTTGAAGAACT</td><td>373.3</td><td>6.0</td><td>1.3</td><td>4.7</td><td>2.6E-06</td><td>1.2E-05</td><td>BT20/MB468</td><td>MP</td></tr>
<tr><td>>hg38_21_+_8433172_8446622_RNA45SN1_45S_50ntFlanks@10916.10933.18|CCGCGGCTGGACGAGGCG</td><td>>hg38_21_+_8433172_8446622_RNA45SN1_45S_50ntFlanks</td><td>10916</td><td>10933</td><td>18</td><td>CCGCGGCTGGACGAGGCG</td><td>683.4</td><td>-3.9</td><td>0.8</td><td>-4.7</td><td>2.6E-06</td><td>1.2E-05</td><td>BT20/MB468</td><td>MP</td></tr>
<tr><td>>hg38_21_+_8433172_8446622_RNA45SN1_45S_50ntFlanks@5489.5526.38|CTGAGAAGACGGTCGAACTTGACTATCTAGAGGAAGTA</td><td>>hg38_21_+_8433172_8446622_RNA45SN1_45S_50ntFlanks</td><td>5489</td><td>5526</td><td>38</td><td>CTGAGAAGACGGTCGAACTTGACTATCTAGAGGAAGTA</td><td>144.5</td><td>-6.6</td><td>1.5</td><td>-4.5</td><td>7.1E-06</td><td>1.2E-05</td><td>MB231/MB468</td><td>MP</td></tr>
<tr><td>>hg38_21_+_8433172_8446622_RNA45SN1_45S_50ntFlanks@10910.10931.22|CGCGCGCCGCGGCTGGACGAGG</td><td>>hg38_21_+_8433172_8446622_RNA45SN1_45S_50ntFlanks</td><td>10910</td><td>10931</td><td>22</td><td>CGCGCGCCGCGGCTGGACGAGG</td><td>138.2</td><td>10.4</td><td>2.1</td><td>4.9</td><td>7.6E-07</td><td>1.2E-05</td><td>BT20/MB231</td><td>MP</td></tr>
<tr><td>>hg38_21_+_8433172_8446622_RNA45SN1_45S_50ntFlanks@4753.4783.31|AAGACGATCAGATACCGTCGTAGTTCCGACC</td><td>>hg38_21_+_8433172_8446622_RNA45SN1_45S_50ntFlanks</td><td>4753</td><td>4783</td><td>31</td><td>AAGACGATCAGATACCGTCGTAGTTCCGACC</td><td>164.5</td><td>-7.0</td><td>1.6</td><td>-4.5</td><td>7.1E-06</td><td>1.2E-05</td><td>MB231/MB468</td><td>MP</td></tr>
<tr><td>>hg38_21_+_8433172_8446622_RNA45SN1_45S_50ntFlanks@9171.9197.27|CGGGTCGCGCCGTCGGGCCCGGGGGAG</td><td>>hg38_21_+_8433172_8446622_RNA45SN1_45S_50ntFlanks</td><td>9171</td><td>9197</td><td>27</td><td>CGGGTCGCGCCGTCGGGCCCGGGGGAG</td><td>647.0</td><td>-6.5</td><td>1.4</td><td>-4.5</td><td>6.2E-06</td><td>1.2E-05</td><td>BT20/MB468</td><td>MITO</td></tr>
</table>

| | | | | | | | | | | | | | |
|---|---|---|---|---|---|---|---|---|---|---|---|---|---|
| >hg38_1_-_228634819_228635039_RNA5S12_5S_50ntFlanks@134.172.39\|TGGGAGACCGCCTGGGAATACCGGGTGCTGTAGGCTTTT | >hg38_1_-_228634819_228635039_RNA5S12_5S_50ntFlanks | 134 | 172 | 39 | TGGGAGACCGCCTGGGAATACCGGGTGCTGTAGGCTTTT | 338.6 | 7.7 | 1.7 | 4.4 | 8.6E-06 | 1.2E-05 | MB231/MB468 | MITO |
| >hg38_21_+_8433172_8446622_RNA45SN1_45S_50ntFlanks@10794.10816.23\|CTGGCATGTTGGAACAATGTAGG | >hg38_21_+_8433172_8446622_RNA45SN1_45S_50ntFlanks | 10794 | 10816 | 23 | CTGGCATGTTGGAACAATGTAGG | 677.0 | -5.2 | 1.2 | -4.5 | 8.4E-06 | 1.2E-05 | MB231/MB468 | NUC |
| >hg38_21_+_8433172_8446622_RNA45SN1_45S_50ntFlanks@12446.12480.35\|GCAGAATTCACCAAGCGTTGGATTGTTCACCCACT | >hg38_21_+_8433172_8446622_RNA45SN1_45S_50ntFlanks | 12446 | 12480 | 35 | GCAGAATTCACCAAGCGTTGGATTGTTCACCCACT | 230.5 | 3.3 | 0.7 | 4.5 | 8.4E-06 | 1.2E-05 | MB231/MB468 | NUC |
| >hg38_21_+_8433172_8446622_RNA45SN1_45S_50ntFlanks@11572.11609.38\|CCGACTTAGAACTGGTGCGGACCAGGGGAATCCGACTG | >hg38_21_+_8433172_8446622_RNA45SN1_45S_50ntFlanks | 11572 | 11609 | 38 | CCGACTTAGAACTGGTGCGGACCAGGGGAATCCGACTG | 353.1 | -2.7 | 0.6 | -4.5 | 8.1E-06 | 1.2E-05 | MB231/MB468 | CYTO |
| >hg38_21_+_8433172_8446622_RNA45SN1_45S_50ntFlanks@10772.10818.47\|AGGTCTCCAAGGTGAACAGCCTCTGGCATGTTGGAACAATGTAGGTA | >hg38_21_+_8433172_8446622_RNA45SN1_45S_50ntFlanks | 10772 | 10818 | 47 | AGGTCTCCAAGGTGAACAGCCTCTGGCATGTTGGAACAATGTAGGTA | 223.0 | -6.8 | 1.5 | -4.4 | 8.6E-06 | 1.2E-05 | MB231/MB468 | MITO |
| >trnaMT_ValTAC_MT_+_1602_1670@52.71.20\|TTAACTTGACCGCTCTGACC | >trnaMT_ValTAC_MT_+_1602_1670 | 52 | 71 | 20 | TTAACTTGACCGCTCTGACC | 206.5 | 5.3 | 1.2 | 4.5 | 8.4E-06 | 1.2E-05 | MB231/MB468 | NUC |
| >hg38_21_+_8433172_8446622_RNA45SN1_45S_50ntFlanks@10830.10858.29\|CAAGCCGGATCCGTAACTTCGGGATAAGG | >hg38_21_+_8433172_8446622_RNA45SN1_45S_50ntFlanks | 10830 | 10858 | 29 | CAAGCCGGATCCGTAACTTCGGGATAAGG | 860.6 | -7.7 | 1.6 | -4.7 | 2.7E-06 | 1.3E-05 | BT20/MB468 | MP |
| >trna130_GlnTTG_6_-_27763640_27763711@31.52.22,trna16_GlnTTG_17_+_47269890_47269961@31.52.22,trna173_GlnTTG_6_-_26311975_26312046@31.52.22,trna174_GlnTTG_6_-_26311424_26311495@31.52.22\|ACTTTGAATCCAGCGATCCGAG | >trna130_GlnTTG_6_-_27763640_27763711 | 31 | 52 | 22 | ACTTTGAATCCAGCGATCCGAG | 104.6 | -10.6 | 2.4 | -4.5 | 7.1E-06 | 1.3E-05 | MB231/MB468 | MP |
| >MI0000074\|hsa-mir-19b-1&WithFlank&13\|+\|91351186\|91351284@60.82.23;MI0000075\|hsa-mir-19b-2&WithFlank&X\|-\|134169665\|134169772@68.90.23&[MIMAT0000074&hsa-miR-19b-3p&offsets\|0\|0;m-122&13\|+\|91351245\|91351267&offsets\|0\|0];[MIMAT0000074_1&hsa-miR-19b-3p&offsets\|0\|0;m-121&X\|-\|134169683\|134169705&offsets\|0\|0]\|TGTGCAAATCCATGCAAAACTGA | >MI0000074\|hsa-mir-19b-1&WithFlank&13\|+\|91351186\|91351284 | 60 | 82 | 23 | TGTGCAAATCCATGCAAAACTGA | 1214.9 | -4.7 | 1.0 | -4.9 | 8.8E-07 | 1.3E-05 | BT20/MB231 | MITO |
| >MI0000266\|hsa-mir-10a&WithFlank&17\|-\|48579832\|48579953@29.48.20&[MIMAT0000253&hsa-miR-10a-5p&offsets\|+1\|-2;m-18&17\|-\|48579905\|48579926&offsets\|+1\|-1]\|ACCCTGTAGATCCGAATTTG | >MI0000266\|hsa-mir-10a&WithFlank&17\|-\|48579832\|48579953 | 29 | 48 | 20 | ACCCTGTAGATCCGAATTTG | 44.6 | 8.7 | 1.9 | 4.5 | 7.2E-06 | 1.3E-05 | MB231/MB468 | MP |
| >hg38_21_+_8433172_8446622_RNA45SN1_45S_50ntFlanks@9940.9964.25\|CAGGACGGTGGCCATGGAAGTCGGA | >hg38_21_+_8433172_8446622_RNA45SN1_45S_50ntFlanks | 9940 | 9964 | 25 | CAGGACGGTGGCCATGGAAGTCGGA | 103.0 | 10.0 | 2.0 | 4.9 | 7.7E-07 | 1.3E-05 | BT20/MB231 | MP |
| >hg38_21_+_8433172_8446622_RNA45SN1_45S_50ntFlanks@9953.9983.31\|ATGGAAGTCGGAATCCGCTAAGGAGTGTGTA | >hg38_21_+_8433172_8446622_RNA45SN1_45S_50ntFlanks | 9953 | 9983 | 31 | ATGGAAGTCGGAATCCGCTAAGGAGTGTGTA | 131.7 | 10.4 | 2.1 | 4.9 | 7.7E-07 | 1.3E-05 | BT20/MB231 | MP |
| >hg38_MT_+_598_1651_RNR1_12S_50ntFlanks@568.611.44\|CAAAGGACCTGGCGGTGCTTCATATCCCTCTAGAGGAGCCTGTT | >hg38_MT_+_598_1651_RNR1_12S_50ntFlanks | 568 | 611 | 44 | CAAAGGACCTGGCGGTGCTTCATATCCCTCTAGAGGAGCCTGTT | 86.0 | -9.8 | 2.0 | -4.9 | 7.7E-07 | 1.3E-05 | BT20/MB231 | MP |
| >hg38_21_+_8433172_8446622_RNA45SN1_45S_50ntFlanks@10383.10424.42\|TCCATGTGAACAGCAGTTGAACATGGGTCAGTCGGTCCTGAG | >hg38_21_+_8433172_8446622_RNA45SN1_45S_50ntFlanks | 10383 | 10424 | 42 | TCCATGTGAACAGCAGTTGAACATGGGTCAGTCGGTCCTGAG | 770.7 | 7.5 | 1.6 | 4.7 | 2.7E-06 | 1.3E-05 | BT20/MB468 | MP |
| >hg38_MT_+_598_1651_RNR1_12S_50ntFlanks@850.876.27\|GTGCTTAGTTGAACAGGGCCCTGAAGC | >hg38_MT_+_598_1651_RNR1_12S_50ntFlanks | 850 | 876 | 27 | GTGCTTAGTTGAACAGGGCCCTGAAGC | 54.9 | 9.5 | 2.0 | 4.7 | 2.7E-06 | 1.3E-05 | BT20/MB468 | MP |
| >hg38_21_+_8433172_8446622_RNA45SN1_45S_50ntFlanks@12997.13040.44\|CGCTGCGATCTATTGAAAGTCAGCCCTCGACACAAGGGTTTGTC | >hg38_21_+_8433172_8446622_RNA45SN1_45S_50ntFlanks | 12997 | 13040 | 44 | CGCTGCGATCTATTGAAAGTCAGCCCTCGACACAAGGGTTTGTC | 382.8 | 7.5 | 1.7 | 4.4 | 8.7E-06 | 1.3E-05 | MB231/MB468 | MITO |
| >hg38_21_+_8433172_8446622_RNA45SN1_45S_50ntFlanks@10854.10879.26\|TAAGGATTGGCTCTAAGGGCTGGGTC | >hg38_21_+_8433172_8446622_RNA45SN1_45S_50ntFlanks | 10854 | 10879 | 26 | TAAGGATTGGCTCTAAGGGCTGGGTC | 119.0 | 10.2 | 2.1 | 4.9 | 7.7E-07 | 1.3E-05 | BT20/MB231 | MP |
| >hg38_21_+_8433172_8446622_RNA45SN1_45S_50ntFlanks@5420.5443.24\|CCGATTGGATGGTTTAGTGAGGCC | >hg38_21_+_8433172_8446622_RNA45SN1_45S_50ntFlanks | 5420 | 5443 | 24 | CCGATTGGATGGTTTAGTGAGGCC | 649.4 | -6.2 | 1.4 | -4.5 | 6.2E-06 | 1.3E-05 | BT20/MB468 | MITO |
| >hg38_21_+_8433172_8446622_RNA45SN1_45S_50ntFlanks@4045.4069.25\|CCATTCGAACGTCTGCCCTATCAAC | >hg38_21_+_8433172_8446622_RNA45SN1_45S_50ntFlanks | 4045 | 4069 | 25 | CCATTCGAACGTCTGCCCTATCAAC | 325.4 | -6.0 | 1.3 | -4.4 | 8.7E-06 | 1.3E-05 | MB231/MB468 | MITO |
| >hg38_21_+_8433172_8446622_RNA45SN1_45S_50ntFlanks@12932.12960.29\|CGCTAAACCATTCGTAGACGACCTGCTTC | >hg38_21_+_8433172_8446622_RNA45SN1_45S_50ntFlanks | 12932 | 12960 | 29 | CGCTAAACCATTCGTAGACGACCTGCTTC | 121.5 | -6.1 | 1.4 | -4.5 | 7.2E-06 | 1.3E-05 | MB231/MB468 | MP |
| >MI0000439\|hsa-mir-23b&WithFlank&9\|+\|95085202\|95085310@67.86.20(+1C)&[MIMAT0000418&hsa-miR-23b-3p&offsets\|+3\|0(+1C);m-42&9\|+\|95085265\|95085285&offsets\|+3\|+2(+1C)]\|ACATTGCCAGGGATTACCACC | >MI0000439\|hsa-mir-23b&WithFlank&9\|+\|95085202\|95085310 | 67 | 86 | 20(+1C) | ACATTGCCAGGGATTACCACC | 219.4 | 11.0 | 2.5 | 4.5 | 7.2E-06 | 1.3E-05 | MB231/MB468 | MP |
| >hg38_21_+_8433172_8446622_RNA45SN1_45S_50ntFlanks@10788.10828.41\|CAGCCTCTGGCATGTTGGAACAATGTAGGTAAGGGAAGTCG | >hg38_21_+_8433172_8446622_RNA45SN1_45S_50ntFlanks | 10788 | 10828 | 41 | CAGCCTCTGGCATGTTGGAACAATGTAGGTAAGGGAAGTCG | 289.9 | -6.5 | 1.5 | -4.4 | 8.7E-06 | 1.3E-05 | MB231/MB468 | MITO |
| >hg38_21_+_8433172_8446622_RNA45SN1_45S_50ntFlanks@10409.10433.25\|GTCAGTCGGTCCTGAGAGATGGGCG | >hg38_21_+_8433172_8446622_RNA45SN1_45S_50ntFlanks | 10409 | 10433 | 25 | GTCAGTCGGTCCTGAGAGATGGGCG | 338.2 | -7.0 | 1.6 | -4.4 | 9.1E-06 | 1.3E-05 | BT20/MB468 | NUC |
| >hg38_21_+_8433172_8446622_RNA45SN1_45S_50ntFlanks@12937.12962.26\|AACCATTCGTAGACGACCTGCTTCTG | >hg38_21_+_8433172_8446622_RNA45SN1_45S_50ntFlanks | 12937 | 12962 | 26 | AACCATTCGTAGACGACCTGCTTCTG | 408.6 | -6.3 | 1.4 | -4.5 | 8.6E-06 | 1.3E-05 | MB231/MB468 | NUC |
| >hg38_21_+_8433172_8446622_RNA45SN1_45S_50ntFlanks@12413.12431.19\|TCCTTCGATGTCGGCTCTT | >hg38_21_+_8433172_8446622_RNA45SN1_45S_50ntFlanks | 12413 | 12431 | 19 | TCCTTCGATGTCGGCTCTT | 255.8 | -4.7 | 1.0 | -4.5 | 8.6E-06 | 1.3E-05 | MB231/MB468 | NUC |
| >MI0000650\|hsa-mir-200c&WithFlank&12\|+\|6963693\|6963772@50.68.19(+1C)&[MIMAT0000617&hsa-miR-200c-3p&offsets\|0\|-4(+1C);m-54&12\|+\|6963742\|6963764&offsets\|0\|-4(+1C)]\|TAATACTGCCGGGTAATGAC | >MI0000650\|hsa-mir-200c&WithFlank&12\|+\|6963693\|6963772 | 50 | 68 | 19(+1C) | TAATACTGCCGGGTAATGAC | 180.6 | 8.2 | 1.7 | 4.7 | 2.7E-06 | 1.3E-05 | BT20/MB468 | MP |
| >hg38_21_+_8433172_8446622_RNA45SN1_45S_50ntFlanks@12416.12454.39\|TTCGATGTCGGCTCTTCCTATCATTGTGAAGCAGAATTC | >hg38_21_+_8433172_8446622_RNA45SN1_45S_50ntFlanks | 12416 | 12454 | 39 | TTCGATGTCGGCTCTTCCTATCATTGTGAAGCAGAATTC | 491.9 | 5.0 | 1.1 | 4.5 | 8.2E-06 | 1.3E-05 | MB231/MB468 | CYTO |
| >hg38_21_+_8433172_8446622_RNA45SN1_45S_50ntFlanks@4570.4587.18\|TGGAATAGGACCGCGGTT | >hg38_21_+_8433172_8446622_RNA45SN1_45S_50ntFlanks | 4570 | 4587 | 18 | TGGAATAGGACCGCGGTT | 399.2 | -2.7 | 0.6 | -4.5 | 8.3E-06 | 1.3E-05 | MB231/MB468 | CYTO |
| >hg38_21_+_8433172_8446622_RNA45SN1_45S_50ntFlanks@9926.9947.22\|GTTGATATAGACAGCAGGACGG | >hg38_21_+_8433172_8446622_RNA45SN1_45S_50ntFlanks | 9926 | 9947 | 22 | GTTGATATAGACAGCAGGACGG | 111.6 | 10.1 | 2.1 | 4.9 | 7.8E-07 | 1.3E-05 | BT20/MB231 | MP |
| >hg38_21_+_8433172_8446622_RNA45SN1_45S_50ntFlanks@10349.10374.26\|AACGAGAACTTTGAAGGCCGAAGTGG | >hg38_21_+_8433172_8446622_RNA45SN1_45S_50ntFlanks | 10349 | 10374 | 26 | AACGAGAACTTTGAAGGCCGAAGTGG | 1171.4 | -5.2 | 1.2 | -4.4 | 8.8E-06 | 1.3E-05 | MB231/MB468 | MITO |
| >hg38_21_+_8433172_8446622_RNA45SN1_45S_50ntFlanks@12071.12095.25\|CCCCGAGGGGCTCTCGCTTCTGGCG | >hg38_21_+_8433172_8446622_RNA45SN1_45S_50ntFlanks | 12071 | 12095 | 25 | CCCCGAGGGGCTCTCGCTTCTGGCG | 734.9 | -7.7 | 1.6 | -4.7 | 2.7E-06 | 1.3E-05 | BT20/MB468 | MP |
| >hg38_21_+_8433172_8446622_RNA45SN1_45S_50ntFlanks@4730.4751.22\|CAAGAACGAAAGTCGGAGGTTC | >hg38_21_+_8433172_8446622_RNA45SN1_45S_50ntFlanks | 4730 | 4751 | 22 | CAAGAACGAAAGTCGGAGGTTC | 1321.8 | -5.5 | 1.2 | -4.5 | 7.3E-06 | 1.3E-05 | MB231/MB468 | MP |
| >MI0000469\|hsa-mir-125a&WithFlank&19\|+\|51693248\|51693345@21.43.23(+1U)&[MIMAT0000443&hsa-miR-125a-5p&offsets\|0\|-1(+1U);m-63&19\|+\|51693268\|51693289&offsets\|0\|+1(+1U)]\|TCCCTGAGACCCTTTAACCTGTGT | >MI0000469\|hsa-mir-125a&WithFlank&19\|+\|51693248\|51693345 | 21 | 43 | 23(+1U) | TCCCTGAGACCCTTTAACCTGTGT | 909.7 | 3.6 | 0.8 | 4.4 | 8.6E-06 | 1.3E-05 | MB231/MB468 | NUC |
| >hg38_21_+_8433172_8446622_RNA45SN1_45S_50ntFlanks@10357.10390.34\|CTTTGAAGGCCGAAGTGGAGAAGGGTTCCATGTG | >hg38_21_+_8433172_8446622_RNA45SN1_45S_50ntFlanks | 10357 | 10390 | 34 | CTTTGAAGGCCGAAGTGGAGAAGGGTTCCATGTG | 374.4 | -6.7 | 1.5 | -4.5 | 7.3E-06 | 1.3E-05 | MB231/MB468 | MP |
| >hg38_21_+_8433172_8446622_RNA45SN1_45S_50ntFlanks@9577.9597.21\|CGGTCGTCCGACCTGGGTATA | >hg38_21_+_8433172_8446622_RNA45SN1_45S_50ntFlanks | 9577 | 9597 | 21 | CGGTCGTCCGACCTGGGTATA | 109.8 | -6.4 | 1.4 | -4.5 | 7.3E-06 | 1.3E-05 | MB231/MB468 | MP |
| >hg38_21_+_8433172_8446622_RNA45SN1_45S_50ntFlanks@11876.11897.22\|AAGACCCTGTTGAGCTTGACTC | >hg38_21_+_8433172_8446622_RNA45SN1_45S_50ntFlanks | 11876 | 11897 | 22 | AAGACCCTGTTGAGCTTGACTC | 738.5 | -3.0 | 0.7 | -4.5 | 8.3E-06 | 1.3E-05 | MB231/MB468 | CYTO |
| >hg38_21_+_8433172_8446622_RNA45SN1_45S_50ntFlanks@11718.11740.23\|AACGGCGGGAGTAACTATGACTC | >hg38_21_+_8433172_8446622_RNA45SN1_45S_50ntFlanks | 11718 | 11740 | 23 | AACGGCGGGAGTAACTATGACTC | 2247.8 | -5.5 | 1.2 | -4.5 | 8.3E-06 | 1.3E-05 | MB231/MB468 | CYTO |
| >hg38_MT_+_598_1651_RNR1_12S_50ntFlanks@430.468.39\|AGTGGCTTTAACATATCTGAACACACAATAGCTAAGACC | >hg38_MT_+_598_1651_RNR1_12S_50ntFlanks | 430 | 468 | 39 | AGTGGCTTTAACATATCTGAACACACAATAGCTAAGACC | 161.5 | 7.0 | 1.6 | 4.4 | 8.8E-06 | 1.3E-05 | MB231/MB468 | MITO |
| >am_trnaMT_HisGTG_MT_+_12138_12206@18.50.33\|AAAACATCAGATTGTGAATCTGACAACAGAGGC | >am_trnaMT_HisGTG_MT_+_12138_12206 | 18 | 50 | 33 | AAAACATCAGATTGTGAATCTGACAACAGAGGC | 66.9 | 9.8 | 2.1 | 4.7 | 2.7E-06 | 1.3E-05 | BT20/MB468 | MP |
| >hg38_21_+_8433172_8446622_RNA45SN1_45S_50ntFlanks@6646.6668.23\|TCGTACGACTCTTAGCGGTGGAT | >hg38_21_+_8433172_8446622_RNA45SN1_45S_50ntFlanks | 6646 | 6668 | 23 | TCGTACGACTCTTAGCGGTGGAT | 2082.0 | 2.4 | 0.5 | 4.5 | 8.3E-06 | 1.3E-05 | MB231/MB468 | CYTO |
| >hg38_21_+_8433172_8446622_RNA45SN1_45S_50ntFlanks@10819.10838.20\|AGGGAAGTCGGCAAGCCGGA | >hg38_21_+_8433172_8446622_RNA45SN1_45S_50ntFlanks | 10819 | 10838 | 20 | AGGGAAGTCGGCAAGCCGGA | 644.7 | -4.0 | 0.8 | -4.7 | 2.7E-06 | 1.3E-05 | BT20/MB468 | MP |
| >hg38_21_+_8433172_8446622_RNA45SN1_45S_50ntFlanks@12957.12986.30\|CTTCTGGGTCGGGGTTTCGTACGTAGCAGA | >hg38_21_+_8433172_8446622_RNA45SN1_45S_50ntFlanks | 12957 | 12986 | 30 | CTTCTGGGTCGGGGTTTCGTACGTAGCAGA | 1572.1 | -8.1 | 1.8 | -4.4 | 9.2E-06 | 1.3E-05 | BT20/MB468 | NUC |
| >hg38_21_+_8433172_8446622_RNA45SN1_45S_50ntFlanks@11217.11234.18\|CTGGACGCGAGCCGGGCC | >hg38_21_+_8433172_8446622_RNA45SN1_45S_50ntFlanks | 11217 | 11234 | 18 | CTGGACGCGAGCCGGGCC | 1368.2 | -4.3 | 0.9 | -4.7 | 2.7E-06 | 1.3E-05 | BT20/MB468 | MP |
| >hg38_21_+_8433172_8446622_RNA45SN1_45S_50ntFlanks@11581.11607.27\|AACTGGTGCGGACCAGGGGAATCCGAC | >hg38_21_+_8433172_8446622_RNA45SN1_45S_50ntFlanks | 11581 | 11607 | 27 | AACTGGTGCGGACCAGGGGAATCCGAC | 179.2 | -2.5 | 0.6 | -4.4 | 8.7E-06 | 1.3E-05 | MB231/MB468 | NUC |
| >hg38_21_+_8433172_8446622_RNA45SN1_45S_50ntFlanks@9930.9947.18\|ATATAGACAGCAGGACGG | >hg38_21_+_8433172_8446622_RNA45SN1_45S_50ntFlanks | 9930 | 9947 | 18 | ATATAGACAGCAGGACGG | 153.1 | -6.6 | 1.5 | -4.5 | 7.3E-06 | 1.3E-05 | MB231/MB468 | MP |
| >hg38_21_+_8433172_8446622_RNA45SN1_45S_50ntFlanks@4752.4782.31\|GAAGACGATCAGATACCGTCGTAGTTCCGAC | >hg38_21_+_8433172_8446622_RNA45SN1_45S_50ntFlanks | 4752 | 4782 | 31 | GAAGACGATCAGATACCGTCGTAGTTCCGAC | 187.5 | -2.5 | 0.6 | -4.5 | 7.5E-06 | 1.3E-05 | BT20/MB468 | CYTO |
| >hg38_21_+_8433172_8446622_RNA45SN1_45S_50ntFlanks@10427.10455.29\|ATGGGCGAGCGCCGTTCCGAAGGGACGGG | >hg38_21_+_8433172_8446622_RNA45SN1_45S_50ntFlanks | 10427 | 10455 | 29 | ATGGGCGAGCGCCGTTCCGAAGGGACGGG | 223.2 | -3.6 | 0.8 | -4.5 | 8.4E-06 | 1.3E-05 | MB231/MB468 | CYTO |
| >hg38_21_+_8433172_8446622_RNA45SN1_45S_50ntFlanks@11890.11912.23\|CTTGACTCTAGTCTGGCACGGTG | >hg38_21_+_8433172_8446622_RNA45SN1_45S_50ntFlanks | 11890 | 11912 | 23 | CTTGACTCTAGTCTGGCACGGTG | 355.4 | -4.9 | 1.1 | -4.4 | 8.7E-06 | 1.3E-05 | MB231/MB468 | NUC |
| >hg38_21_+_8433172_8446622_RNA45SN1_45S_50ntFlanks@12397.12414.18\|CGACGTCGCTTTTTGATC | >hg38_21_+_8433172_8446622_RNA45SN1_45S_50ntFlanks | 12397 | 12414 | 18 | CGACGTCGCTTTTTGATC | 2010.3 | -2.1 | 0.5 | -4.5 | 7.5E-06 | 1.3E-05 | BT20/MB468 | CYTO |
| >hg38_21_+_8433172_8446622_RNA45SN1_45S_50ntFlanks@4567.4587.21\|TAATGGAATAGGACCGCGGTT | >hg38_21_+_8433172_8446622_RNA45SN1_45S_50ntFlanks | 4567 | 4587 | 21 | TAATGGAATAGGACCGCGGTT | 715.6 | -6.7 | 1.5 | -4.4 | 8.7E-06 | 1.3E-05 | MB231/MB468 | NUC |

<table>
<tr><td>>hg38_21_+_8433172_8446622_RNA45SN1_45S_50ntFlanks@4723.4751.29|CATTAATCAAGAACGAAAGTCGGAGGTTC</td><td>>hg38_21_+_8433172_8446622_RNA45SN1_45S_50ntFlanks</td><td>4723</td><td>4751</td><td>29</td><td>CATTAATCAAGAACGAAAGTCGGAGGTTC</td><td>625.7</td><td>-3.3</td><td>0.7</td><td>-4.5</td><td>8.4E-06</td><td>1.3E-05</td><td>MB231/MB468</td><td>CYTO</td></tr>
<tr><td>>MI0000471|hsa-mir-126&WithFlank&9|+|136670596|136670692@57.78.22&[MIMAT0000445&hsa-miR-126-3p&offsets|-1|-1;m-40&9|+|136670653|136670674&offsets|-1|-1]|CTCGTACCGTGAGTAATAATGC</td><td>>MI0000471|hsa-mir-126&WithFlank&9|+|136670596|136670692</td><td>57</td><td>78</td><td>22</td><td>CTCGTACCGTGAGTAATAATGC</td><td>35.2</td><td>8.4</td><td>1.9</td><td>4.5</td><td>7.4E-06</td><td>1.3E-05</td><td>MB231/MB468</td><td>MP</td></tr>
<tr><td>>hg38_21_+_8433172_8446622_RNA45SN1_45S_50ntFlanks@12434.12455.22|TATCATTGTGAAGCAGAATTCA</td><td>>hg38_21_+_8433172_8446622_RNA45SN1_45S_50ntFlanks</td><td>12434</td><td>12455</td><td>22</td><td>TATCATTGTGAAGCAGAATTCA</td><td>210.4</td><td>-6.3</td><td>1.4</td><td>-4.5</td><td>7.5E-06</td><td>1.3E-05</td><td>BT20/MB468</td><td>CYTO</td></tr>
<tr><td>>hg38_21_+_8433172_8446622_RNA45SN1_45S_50ntFlanks@8115.8142.28|CGCGGGACATGTGGCGTACGGAAGACCC</td><td>>hg38_21_+_8433172_8446622_RNA45SN1_45S_50ntFlanks</td><td>8115</td><td>8142</td><td>28</td><td>CGCGGGACATGTGGCGTACGGAAGACCC</td><td>401.3</td><td>-2.5</td><td>0.6</td><td>-4.5</td><td>6.4E-06</td><td>1.3E-05</td><td>BT20/MB468</td><td>MITO</td></tr>
<tr><td>>MI0001448|hsa-mir-425&WithFlank&3|-|49020142|49020240@60.79.20(+1U)&[MIMAT0001343&hsa-miR-425-3p&offsets|-1|-3(+1U);m-258&3|-|49020160|49020181&offsets|0|-2(+1U)]|CATCGGGAATGTCGTGTCCGT</td><td>>MI0001448|hsa-mir-425&WithFlank&3|-|49020142|49020240</td><td>60</td><td>79</td><td>20(+1U)</td><td>CATCGGGAATGTCGTGTCCGT</td><td>41.1</td><td>8.6</td><td>1.9</td><td>4.5</td><td>7.4E-06</td><td>1.3E-05</td><td>MB231/MB468</td><td>MP</td></tr>
<tr><td>>hg38_21_+_8433172_8446622_RNA45SN1_45S_50ntFlanks@11881.11903.23|CCTGTTGAGCTTGACTCTAGTCT</td><td>>hg38_21_+_8433172_8446622_RNA45SN1_45S_50ntFlanks</td><td>11881</td><td>11903</td><td>23</td><td>CCTGTTGAGCTTGACTCTAGTCT</td><td>407.8</td><td>-5.0</td><td>1.1</td><td>-4.5</td><td>7.5E-06</td><td>1.3E-05</td><td>BT20/MB468</td><td>CYTO</td></tr>
<tr><td>>hg38_21_+_8433172_8446622_RNA45SN1_45S_50ntFlanks@3985.4007.23|CGGCTTTGGTGACTCTAGATAAC</td><td>>hg38_21_+_8433172_8446622_RNA45SN1_45S_50ntFlanks</td><td>3985</td><td>4007</td><td>23</td><td>CGGCTTTGGTGACTCTAGATAAC</td><td>52.3</td><td>-8.8</td><td>1.9</td><td>-4.7</td><td>2.7E-06</td><td>1.3E-05</td><td>BT20/MB468</td><td>MP</td></tr>
<tr><td>>hg38_21_+_8433172_8446622_RNA45SN1_45S_50ntFlanks@10395.10435.41|GCAGTTGAACATGGGTCAGTCGGTCCTGAGAGATGGGCGAG</td><td>>hg38_21_+_8433172_8446622_RNA45SN1_45S_50ntFlanks</td><td>10395</td><td>10435</td><td>41</td><td>GCAGTTGAACATGGGTCAGTCGGTCCTGAGAGATGGGCGAG</td><td>89.7</td><td>-10.4</td><td>2.3</td><td>-4.5</td><td>7.4E-06</td><td>1.3E-05</td><td>MB231/MB468</td><td>MP</td></tr>
<tr><td>>hg38_21_+_8433172_8446622_RNA45SN1_45S_50ntFlanks@7974.8002.29|ACGCGACCTCAGATCAGACGTGGCGACCC</td><td>>hg38_21_+_8433172_8446622_RNA45SN1_45S_50ntFlanks</td><td>7974</td><td>8002</td><td>29</td><td>ACGCGACCTCAGATCAGACGTGGCGACCC</td><td>1652.8</td><td>4.0</td><td>0.8</td><td>4.9</td><td>1.0E-06</td><td>1.3E-05</td><td>BT20/MB231</td><td>NUC</td></tr>
<tr><td>>hg38_21_+_8433172_8446622_RNA45SN1_45S_50ntFlanks@6647.6665.19|CGTACGACTCTTAGCGGTG</td><td>>hg38_21_+_8433172_8446622_RNA45SN1_45S_50ntFlanks</td><td>6647</td><td>6665</td><td>19</td><td>CGTACGACTCTTAGCGGTG</td><td>1431.1</td><td>2.7</td><td>0.6</td><td>4.5</td><td>8.4E-06</td><td>1.3E-05</td><td>MB231/MB468</td><td>CYTO</td></tr>
<tr><td>>hg38_21_+_8433172_8446622_RNA45SN1_45S_50ntFlanks@10678.10705.28|CGGTTCCGGCGGCGTCCGGTGAGCTCTC</td><td>>hg38_21_+_8433172_8446622_RNA45SN1_45S_50ntFlanks</td><td>10678</td><td>10705</td><td>28</td><td>CGGTTCCGGCGGCGTCCGGTGAGCTCTC</td><td>667.6</td><td>-6.6</td><td>1.5</td><td>-4.4</td><td>8.8E-06</td><td>1.3E-05</td><td>MB231/MB468</td><td>NUC</td></tr>
<tr><td>>hg38_21_+_8433172_8446622_RNA45SN1_45S_50ntFlanks@11873.11898.26|AAGAAGACCCTGTTGAGCTTGACTCT</td><td>>hg38_21_+_8433172_8446622_RNA45SN1_45S_50ntFlanks</td><td>11873</td><td>11898</td><td>26</td><td>AAGAAGACCCTGTTGAGCTTGACTCT</td><td>1686.2</td><td>-5.6</td><td>1.3</td><td>-4.4</td><td>8.8E-06</td><td>1.3E-05</td><td>MB231/MB468</td><td>NUC</td></tr>
<tr><td>>am_MI0000086|hsa-mir-28&WithFlank&3|+|188688775|188688872@60.78.19&[MIMAT0004502&hsa-miR-28-3p&offsets|0|-3;m-28&3|+|188688834|188688855&offsets|0|-3]|CACTAGATTGTGAGCTCCT</td><td>>am_MI0000086|hsa-mir-28&WithFlank&3|+|188688775|188688872</td><td>60</td><td>78</td><td>19</td><td>CACTAGATTGTGAGCTCCT</td><td>231.2</td><td>6.5</td><td>1.5</td><td>4.5</td><td>7.4E-06</td><td>1.3E-05</td><td>MB231/MB468</td><td>MP</td></tr>
<tr><td>>hg38_21_+_8433172_8446622_RNA45SN1_45S_50ntFlanks@4046.4069.24|CATTCGAACGTCTGCCCTATCAAC</td><td>>hg38_21_+_8433172_8446622_RNA45SN1_45S_50ntFlanks</td><td>4046</td><td>4069</td><td>24</td><td>CATTCGAACGTCTGCCCTATCAAC</td><td>309.2</td><td>3.3</td><td>0.7</td><td>4.5</td><td>8.4E-06</td><td>1.3E-05</td><td>MB231/MB468</td><td>CYTO</td></tr>
<tr><td>>hg38_21_+_8433172_8446622_RNA45SN1_45S_50ntFlanks@8191.8214.24|CAGCCCGTGGACGGTGTGAGGCCG</td><td>>hg38_21_+_8433172_8446622_RNA45SN1_45S_50ntFlanks</td><td>8191</td><td>8214</td><td>24</td><td>CAGCCCGTGGACGGTGTGAGGCCG</td><td>489.7</td><td>-4.2</td><td>0.9</td><td>-4.5</td><td>8.4E-06</td><td>1.3E-05</td><td>MB231/MB468</td><td>CYTO</td></tr>
<tr><td>>MI0000465|hsa-mir-191&WithFlank&3|-|49020612|49020715@22.42.21(+1G)&[MIMAT0000440&hsa-miR-191-5p&offsets|0|-2(+1G);m-48&3|-|49020672|49020694&offsets|0|-2(+1G)]|CAACGGAATCCCAAAAGCAGCG</td><td>>MI0000465|hsa-mir-191&WithFlank&3|-|49020612|49020715</td><td>22</td><td>42</td><td>21(+1G)</td><td>CAACGGAATCCCAAAAGCAGCG</td><td>124.6</td><td>2.4</td><td>0.5</td><td>4.4</td><td>9.3E-06</td><td>1.3E-05</td><td>BT20/MB468</td><td>NUC</td></tr>
<tr><td>>hg38_MT_+_1621_3279_RNR2_16S_50ntFlanks@51.87.37|GCTAAACCTAGCCCCAAACCCACTCCACCTTACTACC</td><td>>hg38_MT_+_1621_3279_RNR2_16S_50ntFlanks</td><td>51</td><td>87</td><td>37</td><td>GCTAAACCTAGCCCCAAACCCACTCCACCTTACTACC</td><td>19768.6</td><td>6.5</td><td>1.4</td><td>4.7</td><td>2.8E-06</td><td>1.3E-05</td><td>BT20/MB468</td><td>MP</td></tr>
<tr><td>>hg38_21_+_8433172_8446622_RNA45SN1_45S_50ntFlanks@11482.11502.21|GCGGAGGGGCCGCGGGCCGGT</td><td>>hg38_21_+_8433172_8446622_RNA45SN1_45S_50ntFlanks</td><td>11482</td><td>11502</td><td>21</td><td>GCGGAGGGGCCGCGGGCCGGT</td><td>59.8</td><td>-9.0</td><td>1.9</td><td>-4.7</td><td>2.8E-06</td><td>1.3E-05</td><td>BT20/MB468</td><td>MP</td></tr>
<tr><td>>hg38_21_+_8433172_8446622_RNA45SN1_45S_50ntFlanks@10237.10262.26|ACGCTACGCCGCGACGAGTAGGAGGG</td><td>>hg38_21_+_8433172_8446622_RNA45SN1_45S_50ntFlanks</td><td>10237</td><td>10262</td><td>26</td><td>ACGCTACGCCGCGACGAGTAGGAGGG</td><td>140.4</td><td>10.5</td><td>2.1</td><td>4.9</td><td>8.0E-07</td><td>1.3E-05</td><td>BT20/MB231</td><td>MP</td></tr>
<tr><td>>hg38_21_+_8433172_8446622_RNA45SN1_45S_50ntFlanks@12417.12446.30|TCGATGTCGGCTCTTCCTATCATTGTGAAG</td><td>>hg38_21_+_8433172_8446622_RNA45SN1_45S_50ntFlanks</td><td>12417</td><td>12446</td><td>30</td><td>TCGATGTCGGCTCTTCCTATCATTGTGAAG</td><td>1253.8</td><td>-3.0</td><td>0.7</td><td>-4.4</td><td>9.0E-06</td><td>1.3E-05</td><td>MB231/MB468</td><td>MITO</td></tr>
<tr><td>>hg38_21_+_8433172_8446622_RNA45SN1_45S_50ntFlanks@11584.11623.40|TGGTGCGGACCAGGGGAATCCGACTGTTTAATTAAAACAA</td><td>>hg38_21_+_8433172_8446622_RNA45SN1_45S_50ntFlanks</td><td>11584</td><td>11623</td><td>40</td><td>TGGTGCGGACCAGGGGAATCCGACTGTTTAATTAAAACAA</td><td>147.1</td><td>3.5</td><td>0.8</td><td>4.4</td><td>9.4E-06</td><td>1.3E-05</td><td>BT20/MB468</td><td>NUC</td></tr>
<tr><td>>hg38_21_+_8433172_8446622_RNA45SN1_45S_50ntFlanks@12391.12414.24|TCATAGCGACGTCGCTTTTTGATC</td><td>>hg38_21_+_8433172_8446622_RNA45SN1_45S_50ntFlanks</td><td>12391</td><td>12414</td><td>24</td><td>TCATAGCGACGTCGCTTTTTGATC</td><td>118.0</td><td>-4.7</td><td>1.1</td><td>-4.5</td><td>7.4E-06</td><td>1.3E-05</td><td>MB231/MB468</td><td>MP</td></tr>
<tr><td>>hg38_21_+_8433172_8446622_RNA45SN1_45S_50ntFlanks@10392.10428.37|ACAGCAGTTGAACATGGGTCAGTCGGTCCTGAGAGAT</td><td>>hg38_21_+_8433172_8446622_RNA45SN1_45S_50ntFlanks</td><td>10392</td><td>10428</td><td>37</td><td>ACAGCAGTTGAACATGGGTCAGTCGGTCCTGAGAGAT</td><td>215.6</td><td>-6.9</td><td>1.5</td><td>-4.5</td><td>7.4E-06</td><td>1.3E-05</td><td>MB231/MB468</td><td>MP</td></tr>
<tr><td>>hg38_21_+_8433172_8446622_RNA45SN1_45S_50ntFlanks@6655.6683.29|TCTTAGCGGTGGATCACTCGGCTCGTGCG</td><td>>hg38_21_+_8433172_8446622_RNA45SN1_45S_50ntFlanks</td><td>6655</td><td>6683</td><td>29</td><td>TCTTAGCGGTGGATCACTCGGCTCGTGCG</td><td>622.6</td><td>3.3</td><td>0.7</td><td>4.4</td><td>8.8E-06</td><td>1.3E-05</td><td>MB231/MB468</td><td>NUC</td></tr>
<tr><td>>hg38_21_+_8433172_8446622_RNA45SN1_45S_50ntFlanks@3984.4026.43|GCGGCTTTGGTGACTCTAGATAACCTCGGGCCGATCGCACGCC</td><td>>hg38_21_+_8433172_8446622_RNA45SN1_45S_50ntFlanks</td><td>3984</td><td>4026</td><td>43</td><td>GCGGCTTTGGTGACTCTAGATAACCTCGGGCCGATCGCACGCC</td><td>640.9</td><td>-7.2</td><td>1.6</td><td>-4.5</td><td>7.6E-06</td><td>1.3E-05</td><td>BT20/MB468</td><td>CYTO</td></tr>
<tr><td>>hg38_21_+_8433172_8446622_RNA45SN1_45S_50ntFlanks@4723.4759.37|CATTAATCAAGAACGAAAGTCGGAGGTTCGAAGACGA</td><td>>hg38_21_+_8433172_8446622_RNA45SN1_45S_50ntFlanks</td><td>4723</td><td>4759</td><td>37</td><td>CATTAATCAAGAACGAAAGTCGGAGGTTCGAAGACGA</td><td>1332.0</td><td>-3.7</td><td>0.8</td><td>-4.5</td><td>7.6E-06</td><td>1.3E-05</td><td>BT20/MB468</td><td>CYTO</td></tr>
<tr><td>>hg38_MT_+_1621_3279_RNR2_16S_50ntFlanks@825.860.36|TAAGGAAAGGTTAAAAAAAGTAAAAGGAACTCGGCA</td><td>>hg38_MT_+_1621_3279_RNR2_16S_50ntFlanks</td><td>825</td><td>860</td><td>36</td><td>TAAGGAAAGGTTAAAAAAAGTAAAAGGAACTCGGCA</td><td>56.2</td><td>9.6</td><td>2.0</td><td>4.7</td><td>2.8E-06</td><td>1.3E-05</td><td>BT20/MB468</td><td>MP</td></tr>
<tr><td>>hg38_21_+_8433172_8446622_RNA45SN1_45S_50ntFlanks@10879.10902.24|CGGTCGGGCTGGGGCGCGAAGCGG</td><td>>hg38_21_+_8433172_8446622_RNA45SN1_45S_50ntFlanks</td><td>10879</td><td>10902</td><td>24</td><td>CGGTCGGGCTGGGGCGCGAAGCGG</td><td>318.1</td><td>-5.0</td><td>1.1</td><td>-4.4</td><td>8.8E-06</td><td>1.3E-05</td><td>MB231/MB468</td><td>NUC</td></tr>
<tr><td>>trna2_GlyGCC_21_-_18827107_18827177@1.32.32,trna35_GlyGCC_1_+_161413094_161413164@1.32.32,trna37_GlyGCC_1_+_161420467_161420537@1.32.32,trna39_GlyGCC_1_+_161427898_161427968@1.32.32,trna41_GlyGCC_1_+_161435258_161435328@1.32.32|GCATGGGTGGTTCAGTGGTAGAATTCTCGCCT</td><td>>trna2_GlyGCC_21_-_18827107_18827177</td><td>1</td><td>32</td><td>32</td><td>GCATGGGTGGTTCAGTGGTAGAATTCTCGCCT</td><td>10237.4</td><td>5.5</td><td>1.1</td><td>4.9</td><td>9.2E-07</td><td>1.3E-05</td><td>BT20/MB231</td><td>MITO</td></tr>
<tr><td>>hg38_21_+_8433172_8446622_RNA45SN1_45S_50ntFlanks@8796.8825.30|CTCCCTCGGGAGGGCGCGCGGGTCGGGGCG</td><td>>hg38_21_+_8433172_8446622_RNA45SN1_45S_50ntFlanks</td><td>8796</td><td>8825</td><td>30</td><td>CTCCCTCGGGAGGGCGCGCGGGTCGGGGCG</td><td>1222.5</td><td>13.9</td><td>2.8</td><td>4.9</td><td>9.3E-07</td><td>1.3E-05</td><td>BT20/MB231</td><td>MITO</td></tr>
<tr><td>>am_trna119_LysCTT_1_-_145395522_145395594@8.35.28,trna11_LysCTT_5_-_180648979_180649051@8.35.28,trna128_LysCTT_1_-_55423542_55423614@8.35.28,trna13_LysCTT_6_+_26556774_26556846@8.35.28,trna6_LysCTT_19_-_52425396_52425468@8.35.28,trna7_LysCTT_16_+_3225692_3225764@8.35.28,trna9_LysCTT_5_+_180634755_180634827@8.35.28|TAGCTCAGTCGGTAGAGCATGAGACTCT</td><td>>am_trna119_LysCTT_1_-_145395522_145395594</td><td>8</td><td>35</td><td>28</td><td>TAGCTCAGTCGGTAGAGCATGAGACTCT</td><td>212.4</td><td>-4.8</td><td>1.0</td><td>-4.9</td><td>9.3E-07</td><td>1.3E-05</td><td>BT20/MB231</td><td>MITO</td></tr>
<tr><td>>hg38_21_+_8433172_8446622_RNA45SN1_45S_50ntFlanks@10683.10705.23|CCGGCGGCGTCCGGTGAGCTCTC</td><td>>hg38_21_+_8433172_8446622_RNA45SN1_45S_50ntFlanks</td><td>10683</td><td>10705</td><td>23</td><td>CCGGCGGCGTCCGGTGAGCTCTC</td><td>1171.8</td><td>-5.4</td><td>1.2</td><td>-4.4</td><td>9.0E-06</td><td>1.3E-05</td><td>MB231/MB468</td><td>MITO</td></tr>
<tr><td>>hg38_21_+_8433172_8446622_RNA45SN1_45S_50ntFlanks@12936.12958.23|AAACCATTCGTAGACGACCTGCT</td><td>>hg38_21_+_8433172_8446622_RNA45SN1_45S_50ntFlanks</td><td>12936</td><td>12958</td><td>23</td><td>AAACCATTCGTAGACGACCTGCT</td><td>566.0</td><td>-2.8</td><td>0.6</td><td>-4.4</td><td>8.9E-06</td><td>1.3E-05</td><td>MB231/MB468</td><td>NUC</td></tr>
<tr><td>>hg38_21_+_8433172_8446622_RNA45SN1_45S_50ntFlanks@10016.10054.39|AATGGATGGCGCTGGAGCGTCGGGCCCATACCCGGCCGT</td><td>>hg38_21_+_8433172_8446622_RNA45SN1_45S_50ntFlanks</td><td>10016</td><td>10054</td><td>39</td><td>AATGGATGGCGCTGGAGCGTCGGGCCCATACCCGGCCGT</td><td>49.8</td><td>9.4</td><td>2.0</td><td>4.7</td><td>2.8E-06</td><td>1.3E-05</td><td>BT20/MB468</td><td>MP</td></tr>
<tr><td>>hg38_21_+_8433172_8446622_RNA45SN1_45S_50ntFlanks@7974.8004.31|ACGCGACCTCAGATCAGACGTGGCGACCCGC</td><td>>hg38_21_+_8433172_8446622_RNA45SN1_45S_50ntFlanks</td><td>7974</td><td>8004</td><td>31</td><td>ACGCGACCTCAGATCAGACGTGGCGACCCGC</td><td>1689.5</td><td>3.3</td><td>0.8</td><td>4.4</td><td>9.4E-06</td><td>1.3E-05</td><td>BT20/MB468</td><td>NUC</td></tr>
<tr><td>>hg38_21_+_8433172_8446622_RNA45SN1_45S_50ntFlanks@12477.12499.23|CACTAATAGGGAACGTGAGCTGG</td><td>>hg38_21_+_8433172_8446622_RNA45SN1_45S_50ntFlanks</td><td>12477</td><td>12499</td><td>23</td><td>CACTAATAGGGAACGTGAGCTGG</td><td>627.9</td><td>-7.4</td><td>1.7</td><td>-4.4</td><td>9.4E-06</td><td>1.3E-05</td><td>BT20/MB468</td><td>NUC</td></tr>
<tr><td>>hg38_21_+_8433172_8446622_RNA45SN1_45S_50ntFlanks@8144.8165.22|CTCCCCGGCGCCGCTCGTGGGG</td><td>>hg38_21_+_8433172_8446622_RNA45SN1_45S_50ntFlanks</td><td>8144</td><td>8165</td><td>22</td><td>CTCCCCGGCGCCGCTCGTGGGG</td><td>86.3</td><td>-9.5</td><td>2.0</td><td>-4.7</td><td>2.8E-06</td><td>1.3E-05</td><td>BT20/MB468</td><td>MP</td></tr>
<tr><td>>MI0005544|hsa-mir-147b&WithFlank&15|+|45433044|45433135@55.73.19(+1A)&[MIMAT0004928&hsa-miR-147b-3p&offsets|0|-2(+1A);m-575&15|+|45433098|45433118&offsets|0|-2(+1A)]|GTGTGCGGAAATGCTTCTGA</td><td>>MI0005544|hsa-mir-147b&WithFlank&15|+|45433044|45433135</td><td>55</td><td>73</td><td>19(+1A)</td><td>GTGTGCGGAAATGCTTCTGA</td><td>36.8</td><td>8.4</td><td>1.9</td><td>4.5</td><td>7.5E-06</td><td>1.3E-05</td><td>MB231/MB468</td><td>MP</td></tr>
<tr><td>>hg38_21_+_8433172_8446622_RNA45SN1_45S_50ntFlanks@12625.12653.29|CAATGGGGCGAAGCTACCATCTGTGGGAT</td><td>>hg38_21_+_8433172_8446622_RNA45SN1_45S_50ntFlanks</td><td>12625</td><td>12653</td><td>29</td><td>CAATGGGGCGAAGCTACCATCTGTGGGAT</td><td>968.6</td><td>-5.2</td><td>1.2</td><td>-4.5</td><td>7.6E-06</td><td>1.3E-05</td><td>BT20/MB468</td><td>CYTO</td></tr>
<tr><td>>hg38_21_+_8433172_8446622_RNA45SN1_45S_50ntFlanks@6722.6758.37|ATTGCAGGACACATTGATCATCGACACTTCGAACGCA</td><td>>hg38_21_+_8433172_8446622_RNA45SN1_45S_50ntFlanks</td><td>6722</td><td>6758</td><td>37</td><td>ATTGCAGGACACATTGATCATCGACACTTCGAACGCA</td><td>170.8</td><td>-2.2</td><td>0.5</td><td>-4.5</td><td>7.6E-06</td><td>1.3E-05</td><td>BT20/MB468</td><td>CYTO</td></tr>
<tr><td>>hg38_21_+_8433172_8446622_RNA45SN1_45S_50ntFlanks@3882.3915.34|CCGACGGGCGCTGACCCCCTTCGCGGGGGGGATG</td><td>>hg38_21_+_8433172_8446622_RNA45SN1_45S_50ntFlanks</td><td>3882</td><td>3915</td><td>34</td><td>CCGACGGGCGCTGACCCCCTTCGCGGGGGGGATG</td><td>222.5</td><td>-4.7</td><td>1.1</td><td>-4.4</td><td>8.9E-06</td><td>1.3E-05</td><td>MB231/MB468</td><td>NUC</td></tr>
<tr><td>>hg38_21_+_8433172_8446622_RNA45SN1_45S_50ntFlanks@4082.4099.18|TCGCCGTGCCTACCATGG</td><td>>hg38_21_+_8433172_8446622_RNA45SN1_45S_50ntFlanks</td><td>4082</td><td>4099</td><td>18</td><td>TCGCCGTGCCTACCATGG</td><td>47.6</td><td>-8.7</td><td>1.8</td><td>-4.7</td><td>2.8E-06</td><td>1.3E-05</td><td>BT20/MB468</td><td>MP</td></tr>
<tr><td>>trna1_GlnCTG_6_+_18836402_18836473@58.75.18,trna3_GlnCTG_17_+_8023070_8023141@58.75.18,trna42_GlnCTG_6_+_27263212_27263283@58.75.18,trna49_GlnCTG_6_+_27487308_27487379@58.75.18,trna7_GlnCTG_15_-_66161400_66161471@58.75.18,trna99_GlnCTG_6_-_28909378_28909449@58.75.18|ATCTCGGTGGAACCTCCA</td><td>>trna1_GlnCTG_6_+_18836402_18836473</td><td>58</td><td>75</td><td>18</td><td>ATCTCGGTGGAACCTCCA</td><td>43.8</td><td>-8.5</td><td>1.8</td><td>-4.7</td><td>2.8E-06</td><td>1.3E-05</td><td>BT20/MB468</td><td>MP</td></tr>
<tr><td>>hg38_21_+_8433172_8446622_RNA45SN1_45S_50ntFlanks@12479.12499.21|CTAATAGGGAACGTGAGCTGG</td><td>>hg38_21_+_8433172_8446622_RNA45SN1_45S_50ntFlanks</td><td>12479</td><td>12499</td><td>21</td><td>CTAATAGGGAACGTGAGCTGG</td><td>221.1</td><td>-5.1</td><td>1.1</td><td>-4.5</td><td>7.5E-06</td><td>1.3E-05</td><td>MB231/MB468</td><td>MP</td></tr>
<tr><td>>hg38_21_+_8433172_8446622_RNA45SN1_45S_50ntFlanks@11729.11768.40|TAACTATGACTCTCTTAAGGTAGCCAAATGCCTCGTCATC</td><td>>hg38_21_+_8433172_8446622_RNA45SN1_45S_50ntFlanks</td><td>11729</td><td>11768</td><td>40</td><td>TAACTATGACTCTCTTAAGGTAGCCAAATGCCTCGTCATC</td><td>248.4</td><td>-4.7</td><td>1.1</td><td>-4.5</td><td>8.6E-06</td><td>1.3E-05</td><td>MB231/MB468</td><td>CYTO</td></tr>
<tr><td>>hg38_21_+_8433172_8446622_RNA45SN1_45S_50ntFlanks@6648.6680.33|GTACGACTCTTAGCGGTGGATCACTCGGCTCGT</td><td>>hg38_21_+_8433172_8446622_RNA45SN1_45S_50ntFlanks</td><td>6648</td><td>6680</td><td>33</td><td>GTACGACTCTTAGCGGTGGATCACTCGGCTCGT</td><td>4136.4</td><td>2.4</td><td>0.5</td><td>4.5</td><td>6.5E-06</td><td>1.3E-05</td><td>BT20/MB468</td><td>MITO</td></tr>
</table>

<table>
<tr><th></th><th></th><th></th><th></th><th></th><th></th><th></th><th></th><th></th><th></th><th></th><th></th><th></th><th></th><th></th></tr>
<tr><td>>hg38_21_+_8433172_8446622_RNA45SN1_45S_50ntFlanks@6733.6752.20|CATTGATCATCGACACTTCG</td><td>>hg38_21_+_8433172_8446622_RNA45SN1_45S_50ntFlanks</td><td>6733</td><td>6752</td><td>20</td><td>CATTGATCATCGACACTTCG</td><td>846.5</td><td>-2.7</td><td>0.6</td><td>-4.5</td><td>8.6E-06</td><td>1.3E-05</td><td>MB231/MB468</td><td>CYTO</td></tr>
<tr><td>>trna10_AspGTC_12_-_125424193_125424264@35.75.41,trna12_AspGTC_12_-_125411891_125411962@35.75.41,trna144_AspGTC_6_-_27551236_27551307@35.75.41,trna38_AspGTC_17_-_8125556_8125627@35.75.41,trna45_AspGTC_6_+_27447453_27447524@35.75.41,trna48_AspGTC_6_+_27471523_27471594@35.75.41,trna4_AspGTC_12_+_96429799_96429870@35.75.41,trna69_AspGTC_1_-_161440205_161440276@35.75.41,trna72_AspGTC_1_-_161432824_161432895@35.75.41,trna75_AspGTC_1_-_161425414_161425485@35.75.41,trna78_AspGTC_1_-_161418033_161418104@35.75.41,trna81_AspGTC_1_-_161410615_161410686@35.75.41|TCACGCGGGAGACCGGGGTTCGATTCCCCGACGGGGAGCCA</td><td>>trna10_AspGTC_12_-_125424193_125424264</td><td>35</td><td>75</td><td>41</td><td>TCACGCGGGAGACCGGGGTTCGATTCCCCGACGGGGAGCCA</td><td>37056.9</td><td>-5.8</td><td>1.2</td><td>-4.9</td><td>8.0E-07</td><td>1.3E-05</td><td>BT20/MB231</td><td>CYTO</td></tr>
<tr><td>>hg38_21_+_8433172_8446622_RNA45SN1_45S_50ntFlanks@4243.4271.29|CTTTCGAGGCCCTGTAATTGGAATGAGTC</td><td>>hg38_21_+_8433172_8446622_RNA45SN1_45S_50ntFlanks</td><td>4243</td><td>4271</td><td>29</td><td>CTTTCGAGGCCCTGTAATTGGAATGAGTC</td><td>199.0</td><td>-6.6</td><td>1.5</td><td>-4.5</td><td>7.5E-06</td><td>1.3E-05</td><td>MB231/MB468</td><td>MP</td></tr>
<tr><td>>hg38_21_+_8433172_8446622_RNA45SN1_45S_50ntFlanks@11197.11232.36|GGCCGGCGGCGGCGGCGACTCTGGACGCGAGCCGGG</td><td>>hg38_21_+_8433172_8446622_RNA45SN1_45S_50ntFlanks</td><td>11197</td><td>11232</td><td>36</td><td>GGCCGGCGGCGGCGGCGACTCTGGACGCGAGCCGGG</td><td>177.6</td><td>-5.7</td><td>1.3</td><td>-4.5</td><td>7.7E-06</td><td>1.3E-05</td><td>BT20/MB468</td><td>CYTO</td></tr>
<tr><td>>hg38_21_+_8433172_8446622_RNA45SN1_45S_50ntFlanks@475.498.24|CTTCGTGATCGATGTGGTGACGTC</td><td>>hg38_21_+_8433172_8446622_RNA45SN1_45S_50ntFlanks</td><td>475</td><td>498</td><td>24</td><td>CTTCGTGATCGATGTGGTGACGTC</td><td>7692.8</td><td>5.9</td><td>1.3</td><td>4.7</td><td>2.8E-06</td><td>1.3E-05</td><td>BT20/MB468</td><td>MP</td></tr>
<tr><td>>MI0000063|hsa-let-7b&WithFlank&22|+|46113680|46113774@12.33.22(+1C)&[MIMAT0000063&hsa-let-7b-5p&offsets|0|0(+1C);m-14&22|+|46113691|46113712&offsets|0|0(+1C)]|TGAGGTAGTAGGTTGTGTGGTTC</td><td>>MI0000063|hsa-let-7b&WithFlank&22|+|46113680|46113774</td><td>12</td><td>33</td><td>22(+1C)</td><td>TGAGGTAGTAGGTTGTGTGGTTC</td><td>817.6</td><td>3.9</td><td>0.9</td><td>4.5</td><td>7.5E-06</td><td>1.3E-05</td><td>MB231/MB468</td><td>MP</td></tr>
<tr><td>>hg38_21_+_8433172_8446622_RNA45SN1_45S_50ntFlanks@8170.8196.27|CAAGTCCTTCTGATCGAGGCCCAGCCC</td><td>>hg38_21_+_8433172_8446622_RNA45SN1_45S_50ntFlanks</td><td>8170</td><td>8196</td><td>27</td><td>CAAGTCCTTCTGATCGAGGCCCAGCCC</td><td>301.6</td><td>-6.5</td><td>1.5</td><td>-4.4</td><td>8.9E-06</td><td>1.3E-05</td><td>MB231/MB468</td><td>NUC</td></tr>
<tr><td>>MI0005559|hsa-mir-744&WithFlank&17|+|12081893|12082002@17.36.20(+1U)&[MIMAT0004945&hsa-miR-744-5p&offsets|0|-2(+1U);m-172&17|+|12081909|12081929&offsets|0|-1(+1U)]|TGCGGGGCTAGGGCTAACAGT</td><td>>MI0005559|hsa-mir-744&WithFlank&17|+|12081893|12082002</td><td>17</td><td>36</td><td>20(+1U)</td><td>TGCGGGGCTAGGGCTAACAGT</td><td>626.4</td><td>7.4</td><td>1.6</td><td>4.7</td><td>2.8E-06</td><td>1.3E-05</td><td>BT20/MB468</td><td>MP</td></tr>
<tr><td>>hg38_21_+_8433172_8446622_RNA45SN1_45S_50ntFlanks@10059.10082.24|GGCAGTCGAGAGTGGACGGGAGCG</td><td>>hg38_21_+_8433172_8446622_RNA45SN1_45S_50ntFlanks</td><td>10059</td><td>10082</td><td>24</td><td>GGCAGTCGAGAGTGGACGGGAGCG</td><td>43.1</td><td>9.2</td><td>2.0</td><td>4.7</td><td>2.8E-06</td><td>1.3E-05</td><td>BT20/MB468</td><td>MP</td></tr>
<tr><td>>hg38_21_+_8433172_8446622_RNA45SN1_45S_50ntFlanks@11571.11593.23|GCCGACTTAGAACTGGTGCGGAC</td><td>>hg38_21_+_8433172_8446622_RNA45SN1_45S_50ntFlanks</td><td>11571</td><td>11593</td><td>23</td><td>GCCGACTTAGAACTGGTGCGGAC</td><td>49.2</td><td>-8.7</td><td>1.9</td><td>-4.7</td><td>2.8E-06</td><td>1.3E-05</td><td>BT20/MB468</td><td>MP</td></tr>
<tr><td>>hg38_21_+_8433172_8446622_RNA45SN1_45S_50ntFlanks@9537.9571.35|AAACTCTGGTGGAGGTCCGTAGCGGTCCTGACGTG</td><td>>hg38_21_+_8433172_8446622_RNA45SN1_45S_50ntFlanks</td><td>9537</td><td>9571</td><td>35</td><td>AAACTCTGGTGGAGGTCCGTAGCGGTCCTGACGTG</td><td>2181.4</td><td>-8.1</td><td>1.8</td><td>-4.4</td><td>9.5E-06</td><td>1.3E-05</td><td>BT20/MB468</td><td>NUC</td></tr>
<tr><td>>hg38_21_+_8433172_8446622_RNA45SN1_45S_50ntFlanks@5418.5445.28|TACCGATTGGATGGTTTAGTGAGGCCCT</td><td>>hg38_21_+_8433172_8446622_RNA45SN1_45S_50ntFlanks</td><td>5418</td><td>5445</td><td>28</td><td>TACCGATTGGATGGTTTAGTGAGGCCCT</td><td>114.1</td><td>-2.4</td><td>0.5</td><td>-4.5</td><td>7.7E-06</td><td>1.3E-05</td><td>BT20/MB468</td><td>CYTO</td></tr>
<tr><td>>hg38_21_+_8433172_8446622_RNA45SN1_45S_50ntFlanks@10797.10822.26|GCATGTTGGAACAATGTAGGTAAGGG</td><td>>hg38_21_+_8433172_8446622_RNA45SN1_45S_50ntFlanks</td><td>10797</td><td>10822</td><td>26</td><td>GCATGTTGGAACAATGTAGGTAAGGG</td><td>263.0</td><td>-3.8</td><td>0.9</td><td>-4.4</td><td>8.6E-06</td><td>1.3E-05</td><td>MB231/MB468</td><td>CYTO</td></tr>
<tr><td>>trna14_LysTTT_11_-_59327808_59327880@4.33.30,trna2_LysTTT_17_+_8022473_8022545@4.33.30,trna54_LysTTT_1_+_204475655_204475727@4.33.30,trna5_LysTTT_11_+_59323902_59323974@4.33.30,trna62_LysTTT_1_-_204476158_204476230@4.33.30,trna76_LysTTT_6_+_28918806_28918878@4.33.30|CGGATAGCTCAGTCGGTAGAGCATCAGACT</td><td>>trna14_LysTTT_11_-_59327808_59327880</td><td>4</td><td>33</td><td>30</td><td>CGGATAGCTCAGTCGGTAGAGCATCAGACT</td><td>45.5</td><td>8.7</td><td>1.9</td><td>4.5</td><td>7.6E-06</td><td>1.3E-05</td><td>MB231/MB468</td><td>MP</td></tr>
<tr><td>>hg38_MT_+_598_1651_RNR1_12S_50ntFlanks@416.454.39|AAATAGACTACGAAAGTGGCTTTAACATATCTGAACACA</td><td>>hg38_MT_+_598_1651_RNR1_12S_50ntFlanks</td><td>416</td><td>454</td><td>39</td><td>AAATAGACTACGAAAGTGGCTTTAACATATCTGAACACA</td><td>36.0</td><td>8.4</td><td>1.9</td><td>4.5</td><td>7.6E-06</td><td>1.3E-05</td><td>MB231/MB468</td><td>MP</td></tr>
<tr><td>>hg38_21_+_8433172_8446622_RNA45SN1_45S_50ntFlanks@12948.12990.43|GACGACCTGCTTCTGGGTCGGGGTTTCGTACGTAGCAGAGCAG</td><td>>hg38_21_+_8433172_8446622_RNA45SN1_45S_50ntFlanks</td><td>12948</td><td>12990</td><td>43</td><td>GACGACCTGCTTCTGGGTCGGGGTTTCGTACGTAGCAGAGCAG</td><td>709.2</td><td>-6.0</td><td>1.3</td><td>-4.4</td><td>9.0E-06</td><td>1.3E-05</td><td>MB231/MB468</td><td>NUC</td></tr>
<tr><td>>hg38_21_+_8433172_8446622_RNA45SN1_45S_50ntFlanks@11206.11232.27|CGGCGGCGACTCTGGACGCGAGCCGGG</td><td>>hg38_21_+_8433172_8446622_RNA45SN1_45S_50ntFlanks</td><td>11206</td><td>11232</td><td>27</td><td>CGGCGGCGACTCTGGACGCGAGCCGGG</td><td>956.5</td><td>-5.4</td><td>1.2</td><td>-4.4</td><td>8.6E-06</td><td>1.3E-05</td><td>MB231/MB468</td><td>CYTO</td></tr>
<tr><td>>hg38_21_+_8433172_8446622_RNA45SN1_45S_50ntFlanks@11576.11613.38|CTTAGAACTGGTGCGGACCAGGGGAATCCGACTGTTTA</td><td>>hg38_21_+_8433172_8446622_RNA45SN1_45S_50ntFlanks</td><td>11576</td><td>11613</td><td>38</td><td>CTTAGAACTGGTGCGGACCAGGGGAATCCGACTGTTTA</td><td>599.9</td><td>-2.8</td><td>0.6</td><td>-4.4</td><td>9.6E-06</td><td>1.3E-05</td><td>BT20/MB468</td><td>NUC</td></tr>
<tr><td>>hg38_21_+_8433172_8446622_RNA45SN1_45S_50ntFlanks@4472.4492.21|CTTAGCTGAGTGTCCCGCGGG</td><td>>hg38_21_+_8433172_8446622_RNA45SN1_45S_50ntFlanks</td><td>4472</td><td>4492</td><td>21</td><td>CTTAGCTGAGTGTCCCGCGGG</td><td>259.8</td><td>-5.8</td><td>1.3</td><td>-4.5</td><td>7.7E-06</td><td>1.3E-05</td><td>BT20/MB468</td><td>CYTO</td></tr>
<tr><td>>MI0000063|hsa-let-7b&WithFlank&22|+|46113680|46113774@12.32.21(+1A)&[MIMAT0000063&hsa-let-7b-5p&offsets|0|-1(+1A);m-14&22|+|46113691|46113712&offsets|0|-1(+1A)]|TGAGGTAGTAGGTTGTGTGGTA</td><td>>MI0000063|hsa-let-7b&WithFlank&22|+|46113680|46113774</td><td>12</td><td>32</td><td>21(+1A)</td><td>TGAGGTAGTAGGTTGTGTGGTA</td><td>302.4</td><td>4.4</td><td>1.0</td><td>4.4</td><td>9.2E-06</td><td>1.3E-05</td><td>MB231/MB468</td><td>MITO</td></tr>
<tr><td>>hg38_21_+_8433172_8446622_RNA45SN1_45S_50ntFlanks@10910.10931.22|CGCGCGCCGCGGCTGGACGAGG</td><td>>hg38_21_+_8433172_8446622_RNA45SN1_45S_50ntFlanks</td><td>10910</td><td>10931</td><td>22</td><td>CGCGCGCCGCGGCTGGACGAGG</td><td>380.7</td><td>-5.5</td><td>1.2</td><td>-4.4</td><td>9.0E-06</td><td>1.3E-05</td><td>MB231/MB468</td><td>NUC</td></tr>
<tr><td>>MI0000065|hsa-let-7d&WithFlank&9|+|94178828|94178926@14.36.23&[MIMAT0000065&hsa-let-7d-5p&offsets|0|+1;m-90&9|+|94178841|94178862&offsets|0|+1]|AGAGGTAGTAGGTTGCATAGTTT</td><td>>MI0000065|hsa-let-7d&WithFlank&9|+|94178828|94178926</td><td>14</td><td>36</td><td>23</td><td>AGAGGTAGTAGGTTGCATAGTTT</td><td>276.7</td><td>6.4</td><td>1.4</td><td>4.4</td><td>9.2E-06</td><td>1.3E-05</td><td>MB231/MB468</td><td>MITO</td></tr>
<tr><td>>hg38_21_+_8433172_8446622_RNA45SN1_45S_50ntFlanks@4963.4985.23|AACCTCACCCGGCCCGGACACGG</td><td>>hg38_21_+_8433172_8446622_RNA45SN1_45S_50ntFlanks</td><td>4963</td><td>4985</td><td>23</td><td>AACCTCACCCGGCCCGGACACGG</td><td>337.1</td><td>-5.0</td><td>1.1</td><td>-4.4</td><td>9.2E-06</td><td>1.3E-05</td><td>MB231/MB468</td><td>MITO</td></tr>
<tr><td>>MI0000469|hsa-mir-125a&WithFlank&19|+|51693248|51693345@59.79.21&[MIMAT0004602&hsa-miR-125a-3p&offsets|0|-1;m-333&19|+|51693306|51693326&offsets|0|0]|ACAGGTGAGGTTCTTGGGAGC</td><td>>MI0000469|hsa-mir-125a&WithFlank&19|+|51693248|51693345</td><td>59</td><td>79</td><td>21</td><td>ACAGGTGAGGTTCTTGGGAGC</td><td>681.6</td><td>6.6</td><td>1.4</td><td>4.7</td><td>2.8E-06</td><td>1.3E-05</td><td>BT20/MB468</td><td>MP</td></tr>
<tr><td>>hg38_21_+_8433172_8446622_RNA45SN1_45S_50ntFlanks@6649.6681.33|TACGACTCTTAGCGGTGGATCACTCGGCTCGTG</td><td>>hg38_21_+_8433172_8446622_RNA45SN1_45S_50ntFlanks</td><td>6649</td><td>6681</td><td>33</td><td>TACGACTCTTAGCGGTGGATCACTCGGCTCGTG</td><td>4260.9</td><td>-3.8</td><td>0.9</td><td>-4.4</td><td>9.2E-06</td><td>1.3E-05</td><td>MB231/MB468</td><td>MITO</td></tr>
<tr><td>>hg38_21_+_8433172_8446622_RNA45SN1_45S_50ntFlanks@11911.11940.30|TGAAGAGACATGAGAGGTGTAGAATAAGTG</td><td>>hg38_21_+_8433172_8446622_RNA45SN1_45S_50ntFlanks</td><td>11911</td><td>11940</td><td>30</td><td>TGAAGAGACATGAGAGGTGTAGAATAAGTG</td><td>44.3</td><td>9.2</td><td>2.0</td><td>4.7</td><td>2.9E-06</td><td>1.3E-05</td><td>BT20/MB468</td><td>MP</td></tr>
<tr><td>>hg38_21_+_8433172_8446622_RNA45SN1_45S_50ntFlanks@4724.4748.25|ATTAATCAAGAACGAAAGTCGGAGG</td><td>>hg38_21_+_8433172_8446622_RNA45SN1_45S_50ntFlanks</td><td>4724</td><td>4748</td><td>25</td><td>ATTAATCAAGAACGAAAGTCGGAGG</td><td>596.9</td><td>-7.0</td><td>1.5</td><td>-4.7</td><td>2.9E-06</td><td>1.3E-05</td><td>BT20/MB468</td><td>MP</td></tr>
<tr><td>>hg38_21_+_8433172_8446622_RNA45SN1_45S_50ntFlanks@9526.9571.46|GAAGCCAGAGGAAACTCTGGTGGAGGTCCGTAGCGGTCCTGACGTG</td><td>>hg38_21_+_8433172_8446622_RNA45SN1_45S_50ntFlanks</td><td>9526</td><td>9571</td><td>46</td><td>GAAGCCAGAGGAAACTCTGGTGGAGGTCCGTAGCGGTCCTGACGTG</td><td>1335.0</td><td>-4.8</td><td>1.0</td><td>-4.9</td><td>8.2E-07</td><td>1.3E-05</td><td>BT20/MB231</td><td>CYTO</td></tr>
<tr><td>>MI0000816|hsa-mir-335&WithFlank&7|+|130496105|130496210@22.43.22(+1C)&[MIMAT0000765&hsa-miR-335-5p&offsets|0|-1(+1C);m-249&7|+|130496126|130496147&offsets|0|0(+1C)]|TCAAGAGCAATAACGAAAAATGC</td><td>>MI0000816|hsa-mir-335&WithFlank&7|+|130496105|130496210</td><td>22</td><td>43</td><td>22(+1C)</td><td>TCAAGAGCAATAACGAAAAATGC</td><td>39.8</td><td>8.5</td><td>1.9</td><td>4.5</td><td>7.7E-06</td><td>1.3E-05</td><td>MB231/MB468</td><td>MP</td></tr>
<tr><td>>MI0033423|hsa-mir-10399&WithFlank&7|-|139044094|139044163@45.63.19(+2U)&[MIMAT0041630&hsa-miR-10399-3p&offsets|+1|-1(+2U);TJU_CMC_MD2.ID03061.3p-miR&offsets|+5|+2(+2U);m-2657&7|-|139044103|139044124&offsets|+5|+2(+2U)]|TCTCGGACAAGCTGTAGGTTT</td><td>>MI0033423|hsa-mir-10399&WithFlank&7|-|139044094|139044163</td><td>45</td><td>63</td><td>19(+2U)</td><td>TCTCGGACAAGCTGTAGGTTT</td><td>220.8</td><td>6.1</td><td>1.4</td><td>4.5</td><td>7.7E-06</td><td>1.3E-05</td><td>MB231/MB468</td><td>MP</td></tr>
<tr><td>>trnaMT_MetCAT_MT_+_4402_4469@37.70.34|CCGAAAATGTTGGTTATACCCTTCCCGTACTACC</td><td>>trnaMT_MetCAT_MT_+_4402_4469</td><td>37</td><td>70</td><td>34</td><td>CCGAAAATGTTGGTTATACCCTTCCCGTACTACC</td><td>35.0</td><td>8.3</td><td>1.9</td><td>4.5</td><td>7.7E-06</td><td>1.3E-05</td><td>MB231/MB468</td><td>MP</td></tr>
<tr><td>>hg38_21_+_8433172_8446622_RNA45SN1_45S_50ntFlanks@12456.12500.45|CCAAGCGTTGGATTGTTCACCCACTAATAGGGAACGTGAGCTGGG</td><td>>hg38_21_+_8433172_8446622_RNA45SN1_45S_50ntFlanks</td><td>12456</td><td>12500</td><td>45</td><td>CCAAGCGTTGGATTGTTCACCCACTAATAGGGAACGTGAGCTGGG</td><td>206.4</td><td>-6.4</td><td>1.4</td><td>-4.5</td><td>7.8E-06</td><td>1.3E-05</td><td>BT20/MB468</td><td>CYTO</td></tr>
<tr><td>>hg38_21_+_8433172_8446622_RNA45SN1_45S_50ntFlanks@11199.11227.29|CCGGCGGCGGCGGCGACTCTGGACGCGAG</td><td>>hg38_21_+_8433172_8446622_RNA45SN1_45S_50ntFlanks</td><td>11199</td><td>11227</td><td>29</td><td>CCGGCGGCGGCGGCGACTCTGGACGCGAG</td><td>1830.8</td><td>-8.1</td><td>1.8</td><td>-4.4</td><td>9.7E-06</td><td>1.3E-05</td><td>BT20/MB468</td><td>NUC</td></tr>
<tr><td>>hg38_21_+_8433172_8446622_RNA45SN1_45S_50ntFlanks@12973.12992.20|TCGTACGTAGCAGAGCAGCT</td><td>>hg38_21_+_8433172_8446622_RNA45SN1_45S_50ntFlanks</td><td>12973</td><td>12992</td><td>20</td><td>TCGTACGTAGCAGAGCAGCT</td><td>355.1</td><td>-2.3</td><td>0.5</td><td>-4.5</td><td>6.7E-06</td><td>1.3E-05</td><td>BT20/MB468</td><td>MITO</td></tr>
<tr><td>>hg38_21_+_8433172_8446622_RNA45SN1_45S_50ntFlanks@12410.12437.28|TGATCCTTCGATGTCGGCTCTTCCTATC</td><td>>hg38_21_+_8433172_8446622_RNA45SN1_45S_50ntFlanks</td><td>12410</td><td>12437</td><td>28</td><td>TGATCCTTCGATGTCGGCTCTTCCTATC</td><td>251.4</td><td>2.5</td><td>0.6</td><td>4.4</td><td>9.1E-06</td><td>1.3E-05</td><td>MB231/MB468</td><td>NUC</td></tr>
<tr><td>>hg38_21_+_8433172_8446622_RNA45SN1_45S_50ntFlanks@4103.4121.19|CCACGGGTGACGGGGAATC</td><td>>hg38_21_+_8433172_8446622_RNA45SN1_45S_50ntFlanks</td><td>4103</td><td>4121</td><td>19</td><td>CCACGGGTGACGGGGAATC</td><td>337.1</td><td>-6.6</td><td>1.5</td><td>-4.5</td><td>6.7E-06</td><td>1.3E-05</td><td>BT20/MB468</td><td>MITO</td></tr>
<tr><td>>hg38_21_+_8433172_8446622_RNA45SN1_45S_50ntFlanks@9567.9596.30|ACGTGCAAATCGGTCGTCCGACCTGGGTAT</td><td>>hg38_21_+_8433172_8446622_RNA45SN1_45S_50ntFlanks</td><td>9567</td><td>9596</td><td>30</td><td>ACGTGCAAATCGGTCGTCCGACCTGGGTAT</td><td>4079.1</td><td>-8.0</td><td>1.7</td><td>-4.7</td><td>2.9E-06</td><td>1.3E-05</td><td>BT20/MB468</td><td>MP</td></tr>
<tr><td>>hg38_21_+_8433172_8446622_RNA45SN1_45S_50ntFlanks@11718.11738.21|AACGGCGGGAGTAACTATGAC</td><td>>hg38_21_+_8433172_8446622_RNA45SN1_45S_50ntFlanks</td><td>11718</td><td>11738</td><td>21</td><td>AACGGCGGGAGTAACTATGAC</td><td>755.5</td><td>-5.1</td><td>1.2</td><td>-4.4</td><td>8.8E-06</td><td>1.3E-05</td><td>MB231/MB468</td><td>CYTO</td></tr>
<tr><td>>MI0000476|hsa-mir-138-1&WithFlank&3|+|44114206|44114316@29.53.25(+1U)&[MIMAT0000430_1&hsa-miR-138-5p&offsets|0|+2(+1U);m-497&3|+|44114234|44114256&offsets|0|+2(+1U)]|AGCTGGTGTTGTGAATCAGGCCGTTT</td><td>>MI0000476|hsa-mir-138-1&WithFlank&3|+|44114206|44114316</td><td>29</td><td>53</td><td>25(+1U)</td><td>AGCTGGTGTTGTGAATCAGGCCGTTT</td><td>83.4</td><td>-9.8</td><td>2.0</td><td>-4.9</td><td>8.3E-07</td><td>1.3E-05</td><td>BT20/MB231</td><td>MP</td></tr>
<tr><td>>hg38_21_+_8433172_8446622_RNA45SN1_45S_50ntFlanks@6644.6668.25|CCTCGTACGACTCTTAGCGGTGGAT</td><td>>hg38_21_+_8433172_8446622_RNA45SN1_45S_50ntFlanks</td><td>6644</td><td>6668</td><td>25</td><td>CCTCGTACGACTCTTAGCGGTGGAT</td><td>157.1</td><td>4.5</td><td>1.0</td><td>4.4</td><td>9.2E-06</td><td>1.3E-05</td><td>MB231/MB468</td><td>NUC</td></tr>
<tr><td>>hg38_21_+_8433172_8446622_RNA45SN1_45S_50ntFlanks@11538.11560.23|CGCGCGGCGCCTCGCCTCGGCCG</td><td>>hg38_21_+_8433172_8446622_RNA45SN1_45S_50ntFlanks</td><td>11538</td><td>11560</td><td>23</td><td>CGCGCGGCGCCTCGCCTCGGCCG</td><td>120.7</td><td>-5.3</td><td>1.2</td><td>-4.5</td><td>7.9E-06</td><td>1.3E-05</td><td>BT20/MB468</td><td>CYTO</td></tr>
<tr><td>>hg38_21_+_8433172_8446622_RNA45SN1_45S_50ntFlanks@11316.11342.27|CACCCCACGTCTCGTCGCGCGCGTC</td><td>>hg38_21_+_8433172_8446622_RNA45SN1_45S_50ntFlanks</td><td>11316</td><td>11342</td><td>27</td><td>CACCCCACGTCTCGTCGCGCGCGTC</td><td>18529.3</td><td>-9.3</td><td>2.1</td><td>-4.4</td><td>9.8E-06</td><td>1.4E-05</td><td>BT20/MB468</td><td>NUC</td></tr>
</table>

<table>
<tr><td>>hg38_21_+_8433172_8446622_RNA45SN1_45S_50ntFlanks@12081.12107.27|CTCTCGCTTCTGGCGCCAAGCGCCCGG</td><td>>hg38_21_+_8433172_8446622_RNA45SN1_45S_50ntFlanks</td><td>12081</td><td>12107</td><td>27</td><td>CTCTCGCTTCTGGCGCCAAGCGCCCGG</td><td>490.5</td><td>-7.2</td><td>1.6</td><td>-4.4</td><td>9.8E-06</td><td>1.4E-05</td><td>BT20/MB468</td><td>NUC</td></tr>
<tr><td>>hg38_21_+_8433172_8446622_RNA45SN1_45S_50ntFlanks@12444.12476.33|AAGCAGAATTCACCAAGCGTTGGATTGTTCACC</td><td>>hg38_21_+_8433172_8446622_RNA45SN1_45S_50ntFlanks</td><td>12444</td><td>12476</td><td>33</td><td>AAGCAGAATTCACCAAGCGTTGGATTGTTCACC</td><td>573.9</td><td>-3.5</td><td>0.8</td><td>-4.4</td><td>9.4E-06</td><td>1.4E-05</td><td>MB231/MB468</td><td>MITO</td></tr>
<tr><td>>MI0000082|hsa-mir-25&WithFlank&7|-|100093554|100093649@58.78.21(+1U)&[MIMAT0000081&hsa-miR-25-3p&offsets|0|-1(+1U);m-23&7|-|100093571|100093592&offsets|0|-1(+1U)]|CATTGCACTTGTCTCGGTCTGT</td><td>>MI0000082|hsa-mir-25&WithFlank&7|-|100093554|100093649</td><td>58</td><td>78</td><td>21(+1U)</td><td>CATTGCACTTGTCTCGGTCTGT</td><td>2745.2</td><td>4.9</td><td>1.0</td><td>4.7</td><td>2.9E-06</td><td>1.4E-05</td><td>BT20/MB468</td><td>MP</td></tr>
<tr><td>>hg38_21_+_8433172_8446622_RNA45SN1_45S_50ntFlanks@11741.11758.18|TCTTAAGGTAGCCAAATG</td><td>>hg38_21_+_8433172_8446622_RNA45SN1_45S_50ntFlanks</td><td>11741</td><td>11758</td><td>18</td><td>TCTTAAGGTAGCCAAATG</td><td>480.2</td><td>-7.2</td><td>1.6</td><td>-4.5</td><td>7.9E-06</td><td>1.4E-05</td><td>BT20/MB468</td><td>CYTO</td></tr>
<tr><td>>hg38_21_+_8433172_8446622_RNA45SN1_45S_50ntFlanks@12841.12863.23|CGGAGTGCCCTTCGTCCTGGGAA</td><td>>hg38_21_+_8433172_8446622_RNA45SN1_45S_50ntFlanks</td><td>12841</td><td>12863</td><td>23</td><td>CGGAGTGCCCTTCGTCCTGGGAA</td><td>91.4</td><td>-10.4</td><td>2.3</td><td>-4.5</td><td>7.8E-06</td><td>1.4E-05</td><td>MB231/MB468</td><td>MP</td></tr>
<tr><td>>MI0000088|hsa-mir-30a&WithFlank&6|-|71403545|71403627@12.35.24(+4U)&[MIMAT0000087&hsa-miR-30a-5p&offsets|0|+2(+4U);m-15&6|-|71403595|71403616&offsets|0|+2(+4U)]|TGTAAACATCCTCGACTGGAAGCTTTTT</td><td>>MI0000088|hsa-mir-30a&WithFlank&6|-|71403545|71403627</td><td>12</td><td>35</td><td>24(+4U)</td><td>TGTAAACATCCTCGACTGGAAGCTTTTT</td><td>87.8</td><td>-9.9</td><td>2.0</td><td>-4.9</td><td>8.4E-07</td><td>1.4E-05</td><td>BT20/MB231</td><td>MP</td></tr>
<tr><td>>hg38_21_+_8433172_8446622_RNA45SN1_45S_50ntFlanks@7988.8009.22|CAGACGTGGCGACCCGCTGAAT</td><td>>hg38_21_+_8433172_8446622_RNA45SN1_45S_50ntFlanks</td><td>7988</td><td>8009</td><td>22</td><td>CAGACGTGGCGACCCGCTGAAT</td><td>560.7</td><td>2.7</td><td>0.6</td><td>4.5</td><td>7.9E-06</td><td>1.4E-05</td><td>BT20/MB468</td><td>CYTO</td></tr>
<tr><td>>hg38_21_+_8433172_8446622_RNA45SN1_45S_50ntFlanks@10350.10371.22|ACGAGAACTTTGAAGGCCGAAG</td><td>>hg38_21_+_8433172_8446622_RNA45SN1_45S_50ntFlanks</td><td>10350</td><td>10371</td><td>22</td><td>ACGAGAACTTTGAAGGCCGAAG</td><td>1838.2</td><td>-3.7</td><td>0.8</td><td>-4.4</td><td>9.4E-06</td><td>1.4E-05</td><td>MB231/MB468</td><td>MITO</td></tr>
<tr><td>>hg38_21_+_8433172_8446622_RNA45SN1_45S_50ntFlanks@10391.10428.38|AACAGCAGTTGAACATGGGTCAGTCGGTCCTGAGAGAT</td><td>>hg38_21_+_8433172_8446622_RNA45SN1_45S_50ntFlanks</td><td>10391</td><td>10428</td><td>38</td><td>AACAGCAGTTGAACATGGGTCAGTCGGTCCTGAGAGAT</td><td>245.8</td><td>-3.3</td><td>0.7</td><td>-4.4</td><td>9.2E-06</td><td>1.4E-05</td><td>MB231/MB468</td><td>NUC</td></tr>
<tr><td>>hg38_21_+_8433172_8446622_RNA45SN1_45S_50ntFlanks@6646.6675.30|TCGTACGACTCTTAGCGGTGGATCACTCGG</td><td>>hg38_21_+_8433172_8446622_RNA45SN1_45S_50ntFlanks</td><td>6646</td><td>6675</td><td>30</td><td>TCGTACGACTCTTAGCGGTGGATCACTCGG</td><td>12761.7</td><td>2.9</td><td>0.6</td><td>4.5</td><td>6.7E-06</td><td>1.4E-05</td><td>BT20/MB468</td><td>MITO</td></tr>
<tr><td>>MI0000253|hsa-mir-148a&WithFlank&7|-|25949913|25949992@50.70.21(+1A)&[MIMAT0000243&hsa-miR-148a-3p&offsets|0|-1(+1A);m-7&7|-|25949922|25949943&offsets|0|-1(+1A)]|TCAGTGCACTACAGAACTTTGA</td><td>>MI0000253|hsa-mir-148a&WithFlank&7|-|25949913|25949992</td><td>50</td><td>70</td><td>21(+1A)</td><td>TCAGTGCACTACAGAACTTTGA</td><td>1215.6</td><td>3.3</td><td>0.7</td><td>4.4</td><td>9.8E-06</td><td>1.4E-05</td><td>BT20/MB468</td><td>NUC</td></tr>
<tr><td>>MI0001733|hsa-mir-452&WithFlank&X|-|151959622|151959718@23.44.22&[MIMAT0001635&hsa-miR-452-5p&offsets|+3|+3;m-188&X|-|151959677|151959699&offsets|+3|+2]|TGTTTGCAGAGGAAACTGAGAC</td><td>>MI0001733|hsa-mir-452&WithFlank&X|-|151959622|151959718</td><td>23</td><td>44</td><td>22</td><td>TGTTTGCAGAGGAAACTGAGAC</td><td>336.0</td><td>4.1</td><td>0.8</td><td>4.9</td><td>8.3E-07</td><td>1.4E-05</td><td>BT20/MB231</td><td>CYTO</td></tr>
<tr><td>>hg38_21_+_8433172_8446622_RNA45SN1_45S_50ntFlanks@11570.11588.19|AGCCGACTTAGAACTGGTG</td><td>>hg38_21_+_8433172_8446622_RNA45SN1_45S_50ntFlanks</td><td>11570</td><td>11588</td><td>19</td><td>AGCCGACTTAGAACTGGTG</td><td>460.0</td><td>-5.4</td><td>1.2</td><td>-4.5</td><td>7.8E-06</td><td>1.4E-05</td><td>MB231/MB468</td><td>MP</td></tr>
<tr><td>>hg38_21_+_8433172_8446622_RNA45SN1_45S_50ntFlanks@11848.11872.25|ACGGGCTTGGCGGAATCAGCGGGGA</td><td>>hg38_21_+_8433172_8446622_RNA45SN1_45S_50ntFlanks</td><td>11848</td><td>11872</td><td>25</td><td>ACGGGCTTGGCGGAATCAGCGGGGA</td><td>290.9</td><td>-3.1</td><td>0.7</td><td>-4.5</td><td>6.8E-06</td><td>1.4E-05</td><td>BT20/MB468</td><td>MITO</td></tr>
<tr><td>>hg38_21_+_8433172_8446622_RNA45SN1_45S_50ntFlanks@4287.4317.31|AACGAGGATCCATTGGAGGGCAAGTCTGGTG</td><td>>hg38_21_+_8433172_8446622_RNA45SN1_45S_50ntFlanks</td><td>4287</td><td>4317</td><td>31</td><td>AACGAGGATCCATTGGAGGGCAAGTCTGGTG</td><td>338.6</td><td>-6.9</td><td>1.6</td><td>-4.5</td><td>7.8E-06</td><td>1.4E-05</td><td>MB231/MB468</td><td>MP</td></tr>
<tr><td>>hg38_21_+_8433172_8446622_RNA45SN1_45S_50ntFlanks@10319.10340.22|GCAGATCTTGGTGGTAGTAGCA</td><td>>hg38_21_+_8433172_8446622_RNA45SN1_45S_50ntFlanks</td><td>10319</td><td>10340</td><td>22</td><td>GCAGATCTTGGTGGTAGTAGCA</td><td>326.7</td><td>5.2</td><td>1.1</td><td>4.9</td><td>9.8E-07</td><td>1.4E-05</td><td>BT20/MB231</td><td>MITO</td></tr>
<tr><td>>hg38_21_+_8433172_8446622_RNA45SN1_45S_50ntFlanks@8336.8375.40|AAGTACCGTAAGGGAAAGTTGAAAAGAACTTTGAAGAGAG</td><td>>hg38_21_+_8433172_8446622_RNA45SN1_45S_50ntFlanks</td><td>8336</td><td>8375</td><td>40</td><td>AAGTACCGTAAGGGAAAGTTGAAAAGAACTTTGAAGAGAG</td><td>360.7</td><td>5.6</td><td>1.1</td><td>4.9</td><td>9.7E-07</td><td>1.4E-05</td><td>BT20/MB231</td><td>MITO</td></tr>
<tr><td>>hg38_21_+_8433172_8446622_RNA45SN1_45S_50ntFlanks@6661.6688.28|CGGTGGATCACTCGGCTCGTGCGTCGAT</td><td>>hg38_21_+_8433172_8446622_RNA45SN1_45S_50ntFlanks</td><td>6661</td><td>6688</td><td>28</td><td>CGGTGGATCACTCGGCTCGTGCGTCGAT</td><td>2515.4</td><td>-3.2</td><td>0.7</td><td>-4.4</td><td>9.4E-06</td><td>1.4E-05</td><td>MB231/MB468</td><td>MITO</td></tr>
<tr><td>>hg38_21_+_8433172_8446622_RNA45SN1_45S_50ntFlanks@12933.12972.40|GCTAAACCATTCGTAGACGACCTGCTTCTGGGTCGGGGTT</td><td>>hg38_21_+_8433172_8446622_RNA45SN1_45S_50ntFlanks</td><td>12933</td><td>12972</td><td>40</td><td>GCTAAACCATTCGTAGACGACCTGCTTCTGGGTCGGGGTT</td><td>280.6</td><td>-4.5</td><td>1.0</td><td>-4.4</td><td>8.9E-06</td><td>1.4E-05</td><td>MB231/MB468</td><td>CYTO</td></tr>
<tr><td>>hg38_21_+_8433172_8446622_RNA45SN1_45S_50ntFlanks@10482.10499.18|TCGAAAGGGAGTCGGGTT</td><td>>hg38_21_+_8433172_8446622_RNA45SN1_45S_50ntFlanks</td><td>10482</td><td>10499</td><td>18</td><td>TCGAAAGGGAGTCGGGTT</td><td>209.4</td><td>-2.3</td><td>0.5</td><td>-4.4</td><td>8.9E-06</td><td>1.4E-05</td><td>MB231/MB468</td><td>CYTO</td></tr>
<tr><td>>hg38_21_+_8433172_8446622_RNA45SN1_45S_50ntFlanks@12308.12325.18|CGATCCTTCTGACCTTTT</td><td>>hg38_21_+_8433172_8446622_RNA45SN1_45S_50ntFlanks</td><td>12308</td><td>12325</td><td>18</td><td>CGATCCTTCTGACCTTTT</td><td>347.1</td><td>-6.6</td><td>1.5</td><td>-4.4</td><td>9.5E-06</td><td>1.4E-05</td><td>MB231/MB468</td><td>MITO</td></tr>
<tr><td>>hg38_21_+_8433172_8446622_RNA45SN1_45S_50ntFlanks@10421.10457.37|TGAGAGATGGGCGAGCGCCGTTCCGAAGGGACGGGCG</td><td>>hg38_21_+_8433172_8446622_RNA45SN1_45S_50ntFlanks</td><td>10421</td><td>10457</td><td>37</td><td>TGAGAGATGGGCGAGCGCCGTTCCGAAGGGACGGGCG</td><td>672.1</td><td>-6.3</td><td>1.4</td><td>-4.5</td><td>8.0E-06</td><td>1.4E-05</td><td>BT20/MB468</td><td>CYTO</td></tr>
<tr><td>>hg38_21_+_8433172_8446622_RNA45SN1_45S_50ntFlanks@10922.10939.18|CTGGACGAGGCGCCGCCG</td><td>>hg38_21_+_8433172_8446622_RNA45SN1_45S_50ntFlanks</td><td>10922</td><td>10939</td><td>18</td><td>CTGGACGAGGCGCCGCCG</td><td>1027.2</td><td>-4.0</td><td>0.9</td><td>-4.5</td><td>8.0E-06</td><td>1.4E-05</td><td>BT20/MB468</td><td>CYTO</td></tr>
<tr><td>>hg38_21_+_8433172_8446622_RNA45SN1_45S_50ntFlanks@9923.9949.27|TTGGTTGATATAGACAGCAGGACGGTG</td><td>>hg38_21_+_8433172_8446622_RNA45SN1_45S_50ntFlanks</td><td>9923</td><td>9949</td><td>27</td><td>TTGGTTGATATAGACAGCAGGACGGTG</td><td>102.4</td><td>10.0</td><td>2.0</td><td>4.9</td><td>8.5E-07</td><td>1.4E-05</td><td>BT20/MB231</td><td>MP</td></tr>
<tr><td>>trna143_LysTTT_6_-_27559593_27559665@5.36.32,trna14_LysTTT_11_-_59327808_59327880@5.36.32,trna2_LysTTT_17_+_8022473_8022545@5.36.32,trna54_LysTTT_1_+_204475655_204475727@5.36.32,trna5_LysTTT_11_+_59323902_59323974@5.36.32,trna62_LysTTT_1_-_204476158_204476230@5.36.32,trna76_LysTTT_6_+_28918806_28918878@5.36.32|GGATAGCTCAGTCGGTAGAGCATCAGACTTTT</td><td>>trna143_LysTTT_6_-_27559593_27559665</td><td>5</td><td>36</td><td>32</td><td>GGATAGCTCAGTCGGTAGAGCATCAGACTTTT</td><td>293.1</td><td>5.9</td><td>1.3</td><td>4.5</td><td>8.0E-06</td><td>1.4E-05</td><td>BT20/MB468</td><td>CYTO</td></tr>
<tr><td>>hg38_21_+_8433172_8446622_RNA45SN1_45S_50ntFlanks@4068.4089.22|ACTTTCGATGGTAGTCGCCGTG</td><td>>hg38_21_+_8433172_8446622_RNA45SN1_45S_50ntFlanks</td><td>4068</td><td>4089</td><td>22</td><td>ACTTTCGATGGTAGTCGCCGTG</td><td>138.3</td><td>-5.5</td><td>1.2</td><td>-4.5</td><td>8.0E-06</td><td>1.4E-05</td><td>BT20/MB468</td><td>CYTO</td></tr>
<tr><td>>MI0000542|hsa-mir-320a&WithFlank&8|-|22244960|22245043@50.71.22&[MIMAT0000510&hsa-miR-320a-3p&offsets|+2|+2;m-60&8|-|22244975|22244996&offsets|+2|+2]|AAGCTGGGTTGAGAGGGCGAAA</td><td>>MI0000542|hsa-mir-320a&WithFlank&8|-|22244960|22245043</td><td>50</td><td>71</td><td>22</td><td>AAGCTGGGTTGAGAGGGCGAAA</td><td>297.6</td><td>5.8</td><td>1.2</td><td>4.7</td><td>2.9E-06</td><td>1.4E-05</td><td>BT20/MB468</td><td>MP</td></tr>
<tr><td>>hg38_21_+_8433172_8446622_RNA45SN1_45S_50ntFlanks@11755.11777.23|AATGCCTCGTCATCTAATTAGTG</td><td>>hg38_21_+_8433172_8446622_RNA45SN1_45S_50ntFlanks</td><td>11755</td><td>11777</td><td>23</td><td>AATGCCTCGTCATCTAATTAGTG</td><td>276.0</td><td>-3.2</td><td>0.7</td><td>-4.4</td><td>9.3E-06</td><td>1.4E-05</td><td>MB231/MB468</td><td>NUC</td></tr>
<tr><td>>MI0000082|hsa-mir-25&WithFlank&7|-|100093554|100093649@58.79.22&[MIMAT0000081&hsa-miR-25-3p&offsets|0|0;m-23&7|-|100093571|100093592&offsets|0|0]|CATTGCACTTGTCTCGGTCTGA</td><td>>MI0000082|hsa-mir-25&WithFlank&7|-|100093554|100093649</td><td>58</td><td>79</td><td>22</td><td>CATTGCACTTGTCTCGGTCTGA</td><td>21064.2</td><td>3.3</td><td>0.7</td><td>4.5</td><td>7.8E-06</td><td>1.4E-05</td><td>MB231/MB468</td><td>MP</td></tr>
<tr><td>>MI0000077|hsa-mir-21&WithFlank&17|+|59841260|59841343@14.37.24(+1U)&[MIMAT0000076&hsa-miR-21-5p&offsets|0|+2(+1U);m-1&17|+|59841273|59841294&offsets|0|+2(+1U)]|TAGCTTATCAGACTGATGTTGACTT</td><td>>MI0000077|hsa-mir-21&WithFlank&17|+|59841260|59841343</td><td>14</td><td>37</td><td>24(+1U)</td><td>TAGCTTATCAGACTGATGTTGACTT</td><td>34.5</td><td>8.3</td><td>1.9</td><td>4.5</td><td>7.8E-06</td><td>1.4E-05</td><td>MB231/MB468</td><td>MP</td></tr>
<tr><td>>hg38_21_+_8433172_8446622_RNA45SN1_45S_50ntFlanks@9027.9049.23|CGTCTCCTCGTGGGGGGGCCGGG</td><td>>hg38_21_+_8433172_8446622_RNA45SN1_45S_50ntFlanks</td><td>9027</td><td>9049</td><td>23</td><td>CGTCTCCTCGTGGGGGGGCCGGG</td><td>93.4</td><td>-10.4</td><td>2.3</td><td>-4.5</td><td>7.8E-06</td><td>1.4E-05</td><td>MB231/MB468</td><td>MP</td></tr>
<tr><td>>hg38_21_+_8433172_8446622_RNA45SN1_45S_50ntFlanks@11873.11905.33|AAGAAGACCCTGTTGAGCTTGACTCTAGTCTGG</td><td>>hg38_21_+_8433172_8446622_RNA45SN1_45S_50ntFlanks</td><td>11873</td><td>11905</td><td>33</td><td>AAGAAGACCCTGTTGAGCTTGACTCTAGTCTGG</td><td>263.6</td><td>-5.7</td><td>1.3</td><td>-4.5</td><td>8.0E-06</td><td>1.4E-05</td><td>BT20/MB468</td><td>CYTO</td></tr>
<tr><td>>hg38_21_+_8433172_8446622_RNA45SN1_45S_50ntFlanks@3711.3737.27|GTTGATCCTGCCAGTAGCATATGCTTG</td><td>>hg38_21_+_8433172_8446622_RNA45SN1_45S_50ntFlanks</td><td>3711</td><td>3737</td><td>27</td><td>GTTGATCCTGCCAGTAGCATATGCTTG</td><td>512.1</td><td>-4.8</td><td>1.1</td><td>-4.5</td><td>6.8E-06</td><td>1.4E-05</td><td>BT20/MB468</td><td>MITO</td></tr>
<tr><td>>MI0003137|hsa-mir-193b&WithFlank&16|+|14303961|14304055@57.79.23&[MIMAT0002819&hsa-miR-193b-3p&offsets|0|+1;m-166&16|+|14304017|14304038&offsets|0|+1]|AACTGGCCCTCAAAGTCCCGCTT</td><td>>MI0003137|hsa-mir-193b&WithFlank&16|+|14303961|14304055</td><td>57</td><td>79</td><td>23</td><td>AACTGGCCCTCAAAGTCCCGCTT</td><td>223.2</td><td>7.0</td><td>1.4</td><td>4.9</td><td>1.1E-06</td><td>1.4E-05</td><td>BT20/MB231</td><td>NUC</td></tr>
<tr><td>>hg38_21_+_8433172_8446622_RNA45SN1_45S_50ntFlanks@9604.9621.18|AAAGACTAATCGAACCAT</td><td>>hg38_21_+_8433172_8446622_RNA45SN1_45S_50ntFlanks</td><td>9604</td><td>9621</td><td>18</td><td>AAAGACTAATCGAACCAT</td><td>422.4</td><td>-5.4</td><td>1.2</td><td>-4.4</td><td>9.5E-06</td><td>1.4E-05</td><td>MB231/MB468</td><td>MITO</td></tr>
<tr><td>>hg38_21_+_8433172_8446622_RNA45SN1_45S_50ntFlanks@9339.9377.39|CGCACGAAAGCCGCCGTGGCGCAATGAAGGTGAAGGCCG</td><td>>hg38_21_+_8433172_8446622_RNA45SN1_45S_50ntFlanks</td><td>9339</td><td>9377</td><td>39</td><td>CGCACGAAAGCCGCCGTGGCGCAATGAAGGTGAAGGCCG</td><td>305.4</td><td>-7.2</td><td>1.6</td><td>-4.7</td><td>2.9E-06</td><td>1.4E-05</td><td>BT20/MB468</td><td>MP</td></tr>
<tr><td>>hg38_21_+_8433172_8446622_RNA45SN1_45S_50ntFlanks@12912.12932.21|CACGTTCGTGGGGAACCTGGC</td><td>>hg38_21_+_8433172_8446622_RNA45SN1_45S_50ntFlanks</td><td>12912</td><td>12932</td><td>21</td><td>CACGTTCGTGGGGAACCTGGC</td><td>423.8</td><td>-6.2</td><td>1.4</td><td>-4.4</td><td>9.9E-06</td><td>1.4E-05</td><td>BT20/MB468</td><td>NUC</td></tr>
<tr><td>>hg38_21_+_8433172_8446622_RNA45SN1_45S_50ntFlanks@12892.12932.41|CCCTCGCCCGTCACGCACCGCACGTTCGTGGGGAACCTGGC</td><td>>hg38_21_+_8433172_8446622_RNA45SN1_45S_50ntFlanks</td><td>12892</td><td>12932</td><td>41</td><td>CCCTCGCCCGTCACGCACCGCACGTTCGTGGGGAACCTGGC</td><td>1333.8</td><td>-8.0</td><td>1.8</td><td>-4.4</td><td>1.0E-05</td><td>1.4E-05</td><td>BT20/MB468</td><td>NUC</td></tr>
<tr><td>>hg38_21_+_8433172_8446622_RNA45SN1_45S_50ntFlanks@12934.12955.22|CTAAACCATTCGTAGACGACCT</td><td>>hg38_21_+_8433172_8446622_RNA45SN1_45S_50ntFlanks</td><td>12934</td><td>12955</td><td>22</td><td>CTAAACCATTCGTAGACGACCT</td><td>201.0</td><td>-2.7</td><td>0.6</td><td>-4.5</td><td>6.8E-06</td><td>1.4E-05</td><td>BT20/MB468</td><td>MITO</td></tr>
<tr><td>>hg38_21_+_8433172_8446622_RNA45SN1_45S_50ntFlanks@5442.5466.25|CCCTCGGATCGGCCCCGCCGGGGTC</td><td>>hg38_21_+_8433172_8446622_RNA45SN1_45S_50ntFlanks</td><td>5442</td><td>5466</td><td>25</td><td>CCCTCGGATCGGCCCCGCCGGGGTC</td><td>500.2</td><td>-6.4</td><td>1.5</td><td>-4.4</td><td>9.4E-06</td><td>1.4E-05</td><td>MB231/MB468</td><td>NUC</td></tr>
<tr><td>>hg38_21_+_8433172_8446622_RNA45SN1_45S_50ntFlanks@3840.3861.22|CTCCTACTTGGATAACTGTGGT</td><td>>hg38_21_+_8433172_8446622_RNA45SN1_45S_50ntFlanks</td><td>3840</td><td>3861</td><td>22</td><td>CTCCTACTTGGATAACTGTGGT</td><td>462.2</td><td>3.8</td><td>0.8</td><td>4.9</td><td>1.1E-06</td><td>1.4E-05</td><td>BT20/MB231</td><td>NUC</td></tr>
<tr><td>>hg38_21_+_8433172_8446622_RNA45SN1_45S_50ntFlanks@12540.12569.30|TGATGTGTTGTTGCCATGGTAATCCTGCTC</td><td>>hg38_21_+_8433172_8446622_RNA45SN1_45S_50ntFlanks</td><td>12540</td><td>12569</td><td>30</td><td>TGATGTGTTGTTGCCATGGTAATCCTGCTC</td><td>172.6</td><td>-3.3</td><td>0.7</td><td>-4.4</td><td>9.0E-06</td><td>1.4E-05</td><td>MB231/MB468</td><td>CYTO</td></tr>
<tr><td>>hg38_21_+_8433172_8446622_RNA45SN1_45S_50ntFlanks@8179.8210.32|CTGATCGAGGCCCAGCCCGTGGACGGTGTGAG</td><td>>hg38_21_+_8433172_8446622_RNA45SN1_45S_50ntFlanks</td><td>8179</td><td>8210</td><td>32</td><td>CTGATCGAGGCCCAGCCCGTGGACGGTGTGAG</td><td>662.6</td><td>-7.6</td><td>1.7</td><td>-4.4</td><td>1.0E-05</td><td>1.4E-05</td><td>BT20/MB468</td><td>NUC</td></tr>
<tr><td>>hg38_21_+_8433172_8446622_RNA45SN1_45S_50ntFlanks@11209.11228.20|CGGCGACTCTGGACGCGAGC</td><td>>hg38_21_+_8433172_8446622_RNA45SN1_45S_50ntFlanks</td><td>11209</td><td>11228</td><td>20</td><td>CGGCGACTCTGGACGCGAGC</td><td>421.3</td><td>-5.9</td><td>1.3</td><td>-4.5</td><td>7.9E-06</td><td>1.4E-05</td><td>MB231/MB468</td><td>MP</td></tr>
<tr><td>>hg38_21_+_8433172_8446622_RNA45SN1_45S_50ntFlanks@10317.10340.24|GTGCAGATCTTGGTGGTAGTAGCA</td><td>>hg38_21_+_8433172_8446622_RNA45SN1_45S_50ntFlanks</td><td>10317</td><td>10340</td><td>24</td><td>GTGCAGATCTTGGTGGTAGTAGCA</td><td>342.2</td><td>5.3</td><td>1.1</td><td>4.9</td><td>1.0E-06</td><td>1.4E-05</td><td>BT20/MB231</td><td>MITO</td></tr>
<tr><td>>hg38_21_+_8433172_8446622_RNA45SN1_45S_50ntFlanks@12420.12449.30|ATGTCGGCTCTTCCTATCATTGTGAAGCAG</td><td>>hg38_21_+_8433172_8446622_RNA45SN1_45S_50ntFlanks</td><td>12420</td><td>12449</td><td>30</td><td>ATGTCGGCTCTTCCTATCATTGTGAAGCAG</td><td>1018.5</td><td>-4.0</td><td>0.8</td><td>-4.9</td><td>1.0E-06</td><td>1.4E-05</td><td>BT20/MB231</td><td>MITO</td></tr>
<tr><td>>hg38_21_+_8433172_8446622_RNA45SN1_45S_50ntFlanks@11316.11338.23|CACCCCACGTCTCGTCGCGCGCG</td><td>>hg38_21_+_8433172_8446622_RNA45SN1_45S_50ntFlanks</td><td>11316</td><td>11338</td><td>23</td><td>CACCCCACGTCTCGTCGCGCGCG</td><td>9579.9</td><td>-9.0</td><td>2.0</td><td>-4.4</td><td>1.0E-05</td><td>1.4E-05</td><td>BT20/MB468</td><td>NUC</td></tr>
<tr><td>>hg38_21_+_8433172_8446622_RNA45SN1_45S_50ntFlanks@6761.6806.46|TGCGGCCCCGGGTTCCTCCCGGGGCTACGCCTGTCTGAGCGTCGCT</td><td>>hg38_21_+_8433172_8446622_RNA45SN1_45S_50ntFlanks</td><td>6761</td><td>6806</td><td>46</td><td>TGCGGCCCCGGGTTCCTCCCGGGGCTACGCCTGTCTGAGCGTCGCT</td><td>52.2</td><td>9.5</td><td>2.0</td><td>4.7</td><td>3.0E-06</td><td>1.4E-05</td><td>BT20/MB468</td><td>MP</td></tr>
<tr><td>>hg38_21_+_8433172_8446622_RNA45SN1_45S_50ntFlanks@6785.6814.30|CTACGCCTGTCTGAGCGTCGCTTGCCGATC</td><td>>hg38_21_+_8433172_8446622_RNA45SN1_45S_50ntFlanks</td><td>6785</td><td>6814</td><td>30</td><td>CTACGCCTGTCTGAGCGTCGCTTGCCGATC</td><td>47.6</td><td>9.3</td><td>2.0</td><td>4.7</td><td>3.0E-06</td><td>1.4E-05</td><td>BT20/MB468</td><td>MP</td></tr>
</table>

| | | | | | | | | | | | | | |
|---|---|---|---|---|---|---|---|---|---|---|---|---|---|
| >hg38_21_+_8433172_8446622_RNA45SN1_45S_50ntFlanks@10679.10711.33\|GGTTCCGGCGGCGTCCGGTGAGCTCTCGCTGGC | >hg38_21_+_8433172_8446622_RNA45SN1_45S_50ntFlanks | 10679 | 10711 | 33 | GGTTCCGGCGGCGTCCGGTGAGCTCTCGCTGGC | 4774.0 | -4.1 | 0.9 | -4.5 | 8.1E-06 | 1.4E-05 | BT20/MB468 | CYTO |
| >hg38_21_+_8433172_8446622_RNA45SN1_45S_50ntFlanks@12874.12892.19\|GCTGGAAAGGCGGCCGCCC | >hg38_21_+_8433172_8446622_RNA45SN1_45S_50ntFlanks | 12874 | 12892 | 19 | GCTGGAAAGGCGGCCGCCC | 151.3 | -5.5 | 1.2 | -4.5 | 8.1E-06 | 1.4E-05 | BT20/MB468 | CYTO |
| >hg38_21_+_8433172_8446622_RNA45SN1_45S_50ntFlanks@3854.3884.31\|ACTGTGGTAATTCTAGAGCTAATACATGCCG | >hg38_21_+_8433172_8446622_RNA45SN1_45S_50ntFlanks | 3854 | 3884 | 31 | ACTGTGGTAATTCTAGAGCTAATACATGCCG | 262.6 | -4.8 | 1.1 | -4.4 | 9.4E-06 | 1.4E-05 | MB231/MB468 | NUC |
| >MI0000469\|hsa-mir-125a&WithFlank&19\|+\|51693248\|51693345@21.42.22(+1U)&[MIMAT0000443&hsa-miR-125a-5p&offsets\|0\|-2(+1U);m-63&19\|+\|51693268\|51693289&offsets\|0\|0(+1U)]\|TCCCTGAGACCCTTTAACCTGTT | >MI0000469\|hsa-mir-125a&WithFlank&19\|+\|51693248\|51693345 | 21 | 42 | 22(+1U) | TCCCTGAGACCCTTTAACCTGTT | 85.8 | 2.2 | 0.5 | 4.4 | 1.0E-05 | 1.4E-05 | BT20/MB468 | NUC |
| >hg38_21_+_8433172_8446622_RNA45SN1_45S_50ntFlanks@10379.10424.46\|GGGTTCCATGTGAACAGCAGTTGAACATGGGTCAGTCGGTCCTGAG | >hg38_21_+_8433172_8446622_RNA45SN1_45S_50ntFlanks | 10379 | 10424 | 46 | GGGTTCCATGTGAACAGCAGTTGAACATGGGTCAGTCGGTCCTGAG | 1833.3 | 4.6 | 0.9 | 4.9 | 1.0E-06 | 1.4E-05 | BT20/MB231 | MITO |
| >hg38_21_+_8433172_8446622_RNA45SN1_45S_50ntFlanks@9127.9152.26\|CGCGGGTCGGGGGGCGGGGCGGACTG | >hg38_21_+_8433172_8446622_RNA45SN1_45S_50ntFlanks | 9127 | 9152 | 26 | CGCGGGTCGGGGGGCGGGGCGGACTG | 203.2 | -6.9 | 1.5 | -4.7 | 3.0E-06 | 1.4E-05 | BT20/MB468 | MP |
| >hg38_21_+_8433172_8446622_RNA45SN1_45S_50ntFlanks@9570.9592.23\|TGCAAATCGGTCGTCCGACCTGG | >hg38_21_+_8433172_8446622_RNA45SN1_45S_50ntFlanks | 9570 | 9592 | 23 | TGCAAATCGGTCGTCCGACCTGG | 320.3 | -7.2 | 1.5 | -4.7 | 3.0E-06 | 1.4E-05 | BT20/MB468 | MP |
| >hg38_21_+_8433172_8446622_RNA45SN1_45S_50ntFlanks@12029.12049.21\|TCTGATCGTTTTTTCACTGAC | >hg38_21_+_8433172_8446622_RNA45SN1_45S_50ntFlanks | 12029 | 12049 | 21 | TCTGATCGTTTTTTCACTGAC | 178.7 | 10.8 | 2.2 | 4.9 | 8.6E-07 | 1.4E-05 | BT20/MB231 | MP |
| >hg38_21_+_8433172_8446622_RNA45SN1_45S_50ntFlanks@485.519.35\|GATGTGGTGACGTCGTGCTCTCCCGGGCCGGGTCC | >hg38_21_+_8433172_8446622_RNA45SN1_45S_50ntFlanks | 485 | 519 | 35 | GATGTGGTGACGTCGTGCTCTCCCGGGCCGGGTCC | 153.9 | 10.5 | 2.4 | 4.4 | 9.6E-06 | 1.4E-05 | MB231/MB468 | MITO |
| >hg38_21_+_8433172_8446622_RNA45SN1_45S_50ntFlanks@10775.10816.42\|TCTCCAAGGTGAACAGCCTCTGGCATGTTGGAACAATGTAGG | >hg38_21_+_8433172_8446622_RNA45SN1_45S_50ntFlanks | 10775 | 10816 | 42 | TCTCCAAGGTGAACAGCCTCTGGCATGTTGGAACAATGTAGG | 96.0 | 9.9 | 2.0 | 4.9 | 8.7E-07 | 1.4E-05 | BT20/MB231 | MP |
| >hg38_21_+_8433172_8446622_RNA45SN1_45S_50ntFlanks@11027.11051.25\|CTCTCTCTCTCTCTCCCCCGCTCCC | >hg38_21_+_8433172_8446622_RNA45SN1_45S_50ntFlanks | 11027 | 11051 | 25 | CTCTCTCTCTCTCTCCCCCGCTCCC | 3327.2 | -8.5 | 1.9 | -4.4 | 1.0E-05 | 1.4E-05 | BT20/MB468 | NUC |
| >hg38_21_+_8433172_8446622_RNA45SN1_45S_50ntFlanks@6661.6701.41\|CGGTGGATCACTCGGCTCGTGCGTCGATGAAGAACGCAGCT | >hg38_21_+_8433172_8446622_RNA45SN1_45S_50ntFlanks | 6661 | 6701 | 41 | CGGTGGATCACTCGGCTCGTGCGTCGATGAAGAACGCAGCT | 475.0 | 2.8 | 0.6 | 4.4 | 1.0E-05 | 1.4E-05 | BT20/MB468 | NUC |
| >hg38_21_+_8433172_8446622_RNA45SN1_45S_50ntFlanks@4099.4119.21\|GTGACCACGGGTGACGGGGAA | >hg38_21_+_8433172_8446622_RNA45SN1_45S_50ntFlanks | 4099 | 4119 | 21 | GTGACCACGGGTGACGGGGAA | 42.4 | -8.5 | 1.8 | -4.7 | 3.0E-06 | 1.4E-05 | BT20/MB468 | MP |
| >hg38_21_+_8433172_8446622_RNA45SN1_45S_50ntFlanks@10482.10512.31\|TCGAAAGGGAGTCGGGTTCAGATCCCCGAAT | >hg38_21_+_8433172_8446622_RNA45SN1_45S_50ntFlanks | 10482 | 10512 | 31 | TCGAAAGGGAGTCGGGTTCAGATCCCCGAAT | 698.0 | -6.6 | 1.5 | -4.5 | 8.2E-06 | 1.4E-05 | BT20/MB468 | CYTO |
| >hg38_21_+_8433172_8446622_RNA45SN1_45S_50ntFlanks@6660.6689.30\|GCGGTGGATCACTCGGCTCGTGCGTCGATG | >hg38_21_+_8433172_8446622_RNA45SN1_45S_50ntFlanks | 6660 | 6689 | 30 | GCGGTGGATCACTCGGCTCGTGCGTCGATG | 1465.1 | -5.9 | 1.3 | -4.5 | 8.0E-06 | 1.4E-05 | MB231/MB468 | MP |
| >trna27_GlyCCC_2_-_70476123_70476193@1.28.28,trna34_GlyCCC_16_-_686736_686806@1.28.28\|GCGCCGCTGGTGTAGTGGTATCATGCAA | >trna27_GlyCCC_2_-_70476123_70476193 | 1 | 28 | 28 | GCGCCGCTGGTGTAGTGGTATCATGCAA | 123.9 | -10.8 | 2.4 | -4.5 | 8.0E-06 | 1.4E-05 | MB231/MB468 | MP |
| >hg38_21_+_8433172_8446622_RNA45SN1_45S_50ntFlanks@10726.10747.22\|GGGAGAGGGTGTAAATCTCGCG | >hg38_21_+_8433172_8446622_RNA45SN1_45S_50ntFlanks | 10726 | 10747 | 22 | GGGAGAGGGTGTAAATCTCGCG | 51.8 | -8.8 | 1.9 | -4.7 | 3.0E-06 | 1.4E-05 | BT20/MB468 | MP |
| >hg38_21_+_8433172_8446622_RNA45SN1_45S_50ntFlanks@12021.12055.35\|ACCACTACTCTGATCGTTTTTTCACTGACCCGGTG | >hg38_21_+_8433172_8446622_RNA45SN1_45S_50ntFlanks | 12021 | 12055 | 35 | ACCACTACTCTGATCGTTTTTTCACTGACCCGGTG | 108.2 | 10.1 | 2.1 | 4.9 | 8.7E-07 | 1.4E-05 | BT20/MB231 | MP |
| >hg38_21_+_8433172_8446622_RNA45SN1_45S_50ntFlanks@9534.9577.44\|AGGAAACTCTGGTGGAGGTCCGTAGCGGTCCTGACGTGCAAATC | >hg38_21_+_8433172_8446622_RNA45SN1_45S_50ntFlanks | 9534 | 9577 | 44 | AGGAAACTCTGGTGGAGGTCCGTAGCGGTCCTGACGTGCAAATC | 266.8 | 3.4 | 0.8 | 4.4 | 9.1E-06 | 1.4E-05 | MB231/MB468 | CYTO |
| >hg38_21_+_8433172_8446622_RNA45SN1_45S_50ntFlanks@12423.12455.33\|TCGGCTCTTCCTATCATTGTGAAGCAGAATTCA | >hg38_21_+_8433172_8446622_RNA45SN1_45S_50ntFlanks | 12423 | 12455 | 33 | TCGGCTCTTCCTATCATTGTGAAGCAGAATTCA | 313.0 | -6.4 | 1.4 | -4.4 | 9.7E-06 | 1.4E-05 | MB231/MB468 | MITO |
| >hg38_21_+_8433172_8446622_RNA45SN1_45S_50ntFlanks@11581.11606.26\|AACTGGTGCGGACCAGGGGAATCCGA | >hg38_21_+_8433172_8446622_RNA45SN1_45S_50ntFlanks | 11581 | 11606 | 26 | AACTGGTGCGGACCAGGGGAATCCGA | 578.8 | -3.8 | 0.9 | -4.5 | 8.0E-06 | 1.4E-05 | MB231/MB468 | MP |
| >MI0000459\|hsa-mir-143&WithFlank&5\|+\|149428912\|149429029@67.87.21(+1U)&[MIMAT0000435&hsa-miR-143-3p&offsets\|0\|0(+1U);m-3&5\|+\|149428978\|149428997&offsets\|0\|+1(+1U)]\|TGAGATGAAGCACTGTAGCTCT | >MI0000459\|hsa-mir-143&WithFlank&5\|+\|149428912\|149429029 | 67 | 87 | 21(+1U) | TGAGATGAAGCACTGTAGCTCT | 164.2 | -6.4 | 1.4 | -4.5 | 8.0E-06 | 1.4E-05 | MB231/MB468 | MP |
| >hg38_21_+_8433172_8446622_RNA45SN1_45S_50ntFlanks@12203.12232.30\|TAAGGCGAGCTCAGGGAGGACAGAAACCTC | >hg38_21_+_8433172_8446622_RNA45SN1_45S_50ntFlanks | 12203 | 12232 | 30 | TAAGGCGAGCTCAGGGAGGACAGAAACCTC | 111.1 | -10.7 | 2.4 | -4.5 | 8.0E-06 | 1.4E-05 | MB231/MB468 | MP |
| >MI0000093\|hsa-mir-92a-1&WithFlank&13\|+\|91351308\|91351397@54.74.21;MI0000094\|hsa-mir-92a-2&WithFlank&X\|-\|134169532\|134169618@54.74.21&[MIMAT0000092&hsa-miR-92a-3p&offsets\|0\|-1;m-9&13\|+\|91351361\|91351382&offsets\|0\|-1];[MIMAT0000092_1&hsa-miR-92a-3p&offsets\|0\|-1;m-11&X\|-\|134169544\|134169565&offsets\|0\|-1]\|TATTGCACTTGTCCCGGCCTG | >MI0000093\|hsa-mir-92a-1&WithFlank&13\|+\|91351308\|91351397 | 54 | 74 | 21 | TATTGCACTTGTCCCGGCCTG | 987.5 | 4.1 | 0.9 | 4.5 | 8.0E-06 | 1.4E-05 | MB231/MB468 | MP |
| >hg38_21_+_8433172_8446622_RNA45SN1_45S_50ntFlanks@7969.8010.42\|CCGAGACGCGACCTCAGATCAGACGTGGCGACCCGCTGAATT | >hg38_21_+_8433172_8446622_RNA45SN1_45S_50ntFlanks | 7969 | 8010 | 42 | CCGAGACGCGACCTCAGATCAGACGTGGCGACCCGCTGAATT | 343.1 | 4.8 | 1.1 | 4.4 | 9.5E-06 | 1.4E-05 | MB231/MB468 | NUC |
| >trna15_ValAAC_5_-_180615416_180615488@1.29.29\|GTTTCCGTAGTGTAGTGGTCATCACGTTC | >trna15_ValAAC_5_-_180615416_180615488 | 1 | 29 | 29 | GTTTCCGTAGTGTAGTGGTCATCACGTTC | 926.4 | -5.7 | 1.3 | -4.4 | 9.7E-06 | 1.4E-05 | MB231/MB468 | MITO |
| >trna111_HisGTG_1_-_147774845_147774916@-1A.31.32,trna118_HisGTG_1_-_145396881_145396952@-1A.31.32,trna16_HisGTG_1_+_146544773_146544844@-1A.31.32,trna1_HisGTG_15_+_45493349_45493420@-1A.31.32,trna21_HisGTG_1_+_147753471_147753542@-1A.31.32,trna33_HisGTG_6_+_27125906_27125977@-1A.31.32,trna7_HisGTG_9_-_14433938_14434009@-1A.31.32,trna8_HisGTG_15_-_45492611_45492682@-1A.31.32,trna9_HisGTG_15_-_45490804_45490875@-1A.31.32\|AGCCGTGATCGTATAGTGGTTAGTACTCTGCG | >trna111_HisGTG_1_-_147774845_147774916 | -1A | 31 | 32 | AGCCGTGATCGTATAGTGGTTAGTACTCTGCG | 1275.1 | -3.8 | 0.8 | -4.5 | 8.0E-06 | 1.4E-05 | MB231/MB468 | MP |
| >hg38_21_+_8433172_8446622_RNA45SN1_45S_50ntFlanks@4007.4026.20\|CCTCGGGCCGATCGCACGCC | >hg38_21_+_8433172_8446622_RNA45SN1_45S_50ntFlanks | 4007 | 4026 | 20 | CCTCGGGCCGATCGCACGCC | 188.5 | -4.3 | 1.0 | -4.5 | 8.2E-06 | 1.4E-05 | BT20/MB468 | CYTO |
| >MI0006444\|hsa-mir-1307&WithFlank&10\|-\|103394247\|103394407@87.107.21(+1U)&[MIMAT0005951&hsa-miR-1307-3p&offsets\|+1\|0(+1U);m-66&10\|-\|103394300\|103394322&offsets\|+1\|-1(+1U)]\|CTCGGCGTGGCGTCGGTCGTGT | >MI0006444\|hsa-mir-1307&WithFlank&10\|-\|103394247\|103394407 | 87 | 107 | 21(+1U) | CTCGGCGTGGCGTCGGTCGTGT | 182.6 | 5.5 | 1.2 | 4.4 | 9.8E-06 | 1.4E-05 | MB231/MB468 | MITO |
| >MI0000737\|hsa-mir-200a&WithFlank&1\|+\|1167857\|1167958@60.82.23&[MIMAT0000682&hsa-miR-200a-3p&offsets\|0\|+1;m-142&1\|+\|1167916\|1167938&offsets\|0\|0]\|TAACACTGTCTGGTAACGATGTT | >MI0000737\|hsa-mir-200a&WithFlank&1\|+\|1167857\|1167958 | 60 | 82 | 23 | TAACACTGTCTGGTAACGATGTT | 5840.8 | -6.8 | 1.4 | -4.7 | 3.0E-06 | 1.4E-05 | BT20/MB468 | MP |
| >hg38_21_+_8433172_8446622_RNA45SN1_45S_50ntFlanks@11847.11887.41\|AACGGGCTTGGCGGAATCAGCGGGGAAAGAAGACCCTGTTG | >hg38_21_+_8433172_8446622_RNA45SN1_45S_50ntFlanks | 11847 | 11887 | 41 | AACGGGCTTGGCGGAATCAGCGGGGAAAGAAGACCCTGTTG | 95.1 | 9.9 | 2.0 | 4.9 | 8.8E-07 | 1.4E-05 | BT20/MB231 | MP |
| >hg38_21_+_8433172_8446622_RNA45SN1_45S_50ntFlanks@10775.10796.22\|TCTCCAAGGTGAACAGCCTCTG | >hg38_21_+_8433172_8446622_RNA45SN1_45S_50ntFlanks | 10775 | 10796 | 22 | TCTCCAAGGTGAACAGCCTCTG | 94.1 | -6.2 | 1.4 | -4.5 | 8.1E-06 | 1.4E-05 | MB231/MB468 | MP |
| >trna119_LysCTT_1_-_145395522_145395594@3.34.32,trna11_LysCTT_5_-_180648979_180649051@3.34.32,trna13_LysCTT_6_+_26556774_26556846@3.34.32,trna7_LysCTT_16_+_3225692_3225764@3.34.32,trna9_LysCTT_5_+_180634755_180634827@3.34.32\|CCGGCTAGCTCAGTCGGTAGAGCATGAGACTC | >trna119_LysCTT_1_-_145395522_145395594 | 3 | 34 | 32 | CCGGCTAGCTCAGTCGGTAGAGCATGAGACTC | 654.8 | -5.1 | 1.0 | -4.9 | 1.0E-06 | 1.4E-05 | BT20/MB231 | MITO |
| >trnaMT_GluTTC_MT_-_14674_14742@34.71.38,trnalookalike8_GluTTC_5_-_93905172_93905240@34.71.38\|ATATCATTGGTCGTGGTTGTAGTCCGTGCGAGAATACC | >trnaMT_GluTTC_MT_-_14674_14742 | 34 | 71 | 38 | ATATCATTGGTCGTGGTTGTAGTCCGTGCGAGAATACC | 648.6 | -4.5 | 0.9 | -4.9 | 8.7E-07 | 1.4E-05 | BT20/MB231 | CYTO |
| >MI0000478\|hsa-mir-149&WithFlank&2\|+\|240455995\|240456095@21.41.21(+1U)&[MIMAT0000450&hsa-miR-149-5p&offsets\|0\|-2(+1U);m-195&2\|+\|240456015\|240456037&offsets\|0\|-2(+1U)]\|TCTGGCTCCGTGTCTTCACTCT | >MI0000478\|hsa-mir-149&WithFlank&2\|+\|240455995\|240456095 | 21 | 41 | 21(+1U) | TCTGGCTCCGTGTCTTCACTCT | 1048.0 | 4.2 | 0.9 | 4.9 | 8.7E-07 | 1.4E-05 | BT20/MB231 | CYTO |
| >hg38_21_+_8433172_8446622_RNA45SN1_45S_50ntFlanks@12931.12956.26\|GCGCTAAACCATTCGTAGACGACCTG | >hg38_21_+_8433172_8446622_RNA45SN1_45S_50ntFlanks | 12931 | 12956 | 26 | GCGCTAAACCATTCGTAGACGACCTG | 835.8 | -2.3 | 0.5 | -4.5 | 7.0E-06 | 1.4E-05 | BT20/MB468 | MITO |
| >hg38_MT_+_1621_3279_RNR2_16S_50ntFlanks@618.656.39\|AACATATAACTGAACTCCTCACACCCAATTGGACCAATC | >hg38_MT_+_1621_3279_RNR2_16S_50ntFlanks | 618 | 656 | 39 | AACATATAACTGAACTCCTCACACCCAATTGGACCAATC | 31.5 | 8.2 | 1.8 | 4.5 | 8.1E-06 | 1.4E-05 | MB231/MB468 | MP |
| >hg38_21_+_8433172_8446622_RNA45SN1_45S_50ntFlanks@6647.6678.32\|CGTACGACTCTTAGCGGTGGATCACTCGGCTC | >hg38_21_+_8433172_8446622_RNA45SN1_45S_50ntFlanks | 6647 | 6678 | 32 | CGTACGACTCTTAGCGGTGGATCACTCGGCTC | 610.1 | 3.7 | 0.8 | 4.7 | 3.0E-06 | 1.4E-05 | BT20/MB468 | MP |
| >MI0000086\|hsa-mir-28&WithFlank&3\|+\|188688775\|188688872@20.41.22&[MIMAT0000085&hsa-miR-28-5p&offsets\|0\|0;m-133&3\|+\|188688794\|188688815&offsets\|0\|0]\|AAGGAGCTCACAGTCTATTGAG | >MI0000086\|hsa-mir-28&WithFlank&3\|+\|188688775\|188688872 | 20 | 41 | 22 | AAGGAGCTCACAGTCTATTGAG | 250.6 | 6.5 | 1.5 | 4.4 | 9.8E-06 | 1.4E-05 | MB231/MB468 | MITO |
| >MI0000102\|hsa-mir-100&WithFlank&11\|-\|122152223\|122152314@19.40.22(+1C)&[MIMAT0000098&hsa-miR-100-5p&offsets\|0\|0(+1C);m-27&11\|-\|122152276\|122152296&offsets\|0\|+1(+1C)]\|AACCCGTAGATCCGAACTTGTGC | >MI0000102\|hsa-mir-100&WithFlank&11\|-\|122152223\|122152314 | 19 | 40 | 22(+1C) | AACCCGTAGATCCGAACTTGTGC | 5239.9 | -6.3 | 1.3 | -4.9 | 1.1E-06 | 1.4E-05 | BT20/MB231 | NUC |

<table>
<tr><td>>trna10_AspGTC_12_-_125424193_125424264@38.75.38,trna12_AspGTC_12_-_125411891_125411962@38.75.38,trna144_AspGTC_6_-_27551236_27551307@38.75.38,trna38_AspGTC_17_-_8125556_8125627@38.75.38,trna45_AspGTC_6_+_27447453_27447524@38.75.38,trna48_AspGTC_6_+_27471523_27471594@38.75.38,trna4_AspGTC_12_+_96429799_96429870@38.75.38,trna69_AspGTC_1_-_161440205_161440276@38.75.38,trna72_AspGTC_1_-_161432824_161432895@38.75.38,trna75_AspGTC_1_-_161425414_161425485@38.75.38,trna78_AspGTC_1_-_161418033_161418104@38.75.38,trna81_AspGTC_1_-_161410615_161410686@38.75.38|CGCGGGAGACCGGGGTTCGATTCCCCGACGGGGAGCCA</td><td>>trna10_AspGTC_12_-_125424193_125424264</td><td>38</td><td>75</td><td>38</td><td>CGCGGGAGACCGGGGTTCGATTCCCCGACGGGGAGCCA</td><td>1945.8</td><td>4.4</td><td>1.0</td><td>4.4</td><td>9.3E-06</td><td>1.4E-05</td><td>MB231/MB468</td><td>CYTO</td></tr>
<tr><td>>hg38_21_+_8433172_8446622_RNA45SN1_45S_50ntFlanks@12658.12676.19|ACTGAACGCCTCTAAGTCA</td><td>>hg38_21_+_8433172_8446622_RNA45SN1_45S_50ntFlanks</td><td>12658</td><td>12676</td><td>19</td><td>ACTGAACGCCTCTAAGTCA</td><td>52.7</td><td>-8.8</td><td>1.9</td><td>-4.7</td><td>3.0E-06</td><td>1.4E-05</td><td>BT20/MB468</td><td>MP</td></tr>
<tr><td>>hg38_21_+_8433172_8446622_RNA45SN1_45S_50ntFlanks@12707.12742.36|CGCCGCGGAGCCTCGGTTGGCCTCGGATAGCCGGTC</td><td>>hg38_21_+_8433172_8446622_RNA45SN1_45S_50ntFlanks</td><td>12707</td><td>12742</td><td>36</td><td>CGCCGCGGAGCCTCGGTTGGCCTCGGATAGCCGGTC</td><td>48.3</td><td>-8.7</td><td>1.9</td><td>-4.7</td><td>3.0E-06</td><td>1.4E-05</td><td>BT20/MB468</td><td>MP</td></tr>
<tr><td>>MI0000650|hsa-mir-200c&WithFlank&12|+|6963693|6963772@50.71.22&[MIMAT0000617&hsa-miR-200c-3p&offsets|0|-1;m-54&12|+|6963742|6963764&offsets|0|-1]|TAATACTGCCGGGTAATGATGG</td><td>>MI0000650|hsa-mir-200c&WithFlank&12|+|6963693|6963772</td><td>50</td><td>71</td><td>22</td><td>TAATACTGCCGGGTAATGATGG</td><td>307.9</td><td>-4.7</td><td>1.1</td><td>-4.4</td><td>9.6E-06</td><td>1.4E-05</td><td>MB231/MB468</td><td>NUC</td></tr>
<tr><td>>hg38_21_+_8433172_8446622_RNA45SN1_45S_50ntFlanks@5188.5206.19|ATAACAGGTCTGTGATGCC</td><td>>hg38_21_+_8433172_8446622_RNA45SN1_45S_50ntFlanks</td><td>5188</td><td>5206</td><td>19</td><td>ATAACAGGTCTGTGATGCC</td><td>959.7</td><td>-7.2</td><td>1.6</td><td>-4.4</td><td>9.8E-06</td><td>1.4E-05</td><td>MB231/MB468</td><td>MITO</td></tr>
<tr><td>>hg38_21_+_8433172_8446622_RNA45SN1_45S_50ntFlanks@10513.10546.34|CCGGAGTGGCGGAGATGGGCGCCGCGAGGCGTCC</td><td>>hg38_21_+_8433172_8446622_RNA45SN1_45S_50ntFlanks</td><td>10513</td><td>10546</td><td>34</td><td>CCGGAGTGGCGGAGATGGGCGCCGCGAGGCGTCC</td><td>136.7</td><td>-4.2</td><td>0.9</td><td>-4.4</td><td>9.3E-06</td><td>1.4E-05</td><td>MB231/MB468</td><td>CYTO</td></tr>
<tr><td>>am_trna128_GlyGCC_6_-_27870686_27870756@1.29.29,trna133_GlyCCC_1_-_16872434_16872504@1.29.29,trna18_GlyGCC_16_+_70822597_70822667@1.29.29,trna19_GlyGCC_16_+_70823410_70823480@1.29.29,trna19_GlyGCC_2_-_157257659_157257729@1.29.29,trna24_GlyGCC_16_-_70812942_70813012@1.29.29,trna25_GlyGCC_16_-_70812114_70812184@1.29.29,trna4_GlyCCC_1_+_17188416_17188486@1.29.29,trna5_GlyGCC_17_+_8029064_8029134@1.29.29,trna68_GlyGCC_1_-_161493637_161493707@1.29.29|GCATTGGTGGTTCAGTGGTAGAATTCTCG</td><td>>am_trna128_GlyGCC_6_-_27870686_27870756</td><td>1</td><td>29</td><td>29</td><td>GCATTGGTGGTTCAGTGGTAGAATTCTCG</td><td>3455.7</td><td>-4.0</td><td>0.9</td><td>-4.5</td><td>7.1E-06</td><td>1.4E-05</td><td>BT20/MB468</td><td>MITO</td></tr>
<tr><td>>hg38_21_+_8433172_8446622_RNA45SN1_45S_50ntFlanks@11316.11344.29|CACCCCACGTCTCGTCGCGCGCGCGTCCG</td><td>>hg38_21_+_8433172_8446622_RNA45SN1_45S_50ntFlanks</td><td>11316</td><td>11344</td><td>29</td><td>CACCCCACGTCTCGTCGCGCGCGCGTCCG</td><td>12287.4</td><td>-7.3</td><td>1.6</td><td>-4.4</td><td>9.3E-06</td><td>1.4E-05</td><td>MB231/MB468</td><td>CYTO</td></tr>
<tr><td>>hg38_21_+_8433172_8446622_RNA45SN1_45S_50ntFlanks@476.496.21|TTCGTGATCGATGTGGTGACG</td><td>>hg38_21_+_8433172_8446622_RNA45SN1_45S_50ntFlanks</td><td>476</td><td>496</td><td>21</td><td>TTCGTGATCGATGTGGTGACG</td><td>806.7</td><td>6.7</td><td>1.5</td><td>4.4</td><td>9.9E-06</td><td>1.4E-05</td><td>MB231/MB468</td><td>MITO</td></tr>
<tr><td>>MI0003667|hsa-mir-652&WithFlank&X|+|110055323|110055432@67.87.21&[MIMAT0003322&hsa-miR-652-3p&offsets|0|0;m-148&X|+|110055389|110055410&offsets|0|-1]|AATGGCGCCACTAGGGTTGTG</td><td>>MI0003667|hsa-mir-652&WithFlank&X|+|110055323|110055432</td><td>67</td><td>87</td><td>21</td><td>AATGGCGCCACTAGGGTTGTG</td><td>224.8</td><td>3.3</td><td>0.7</td><td>4.9</td><td>1.1E-06</td><td>1.4E-05</td><td>BT20/MB231</td><td>NUC</td></tr>
<tr><td>>hg38_21_+_8433172_8446622_RNA45SN1_45S_50ntFlanks@8465.8497.33|GTGTCGGCGGCCCGGCGGATCTTTCCCGCCCCC</td><td>>hg38_21_+_8433172_8446622_RNA45SN1_45S_50ntFlanks</td><td>8465</td><td>8497</td><td>33</td><td>GTGTCGGCGGCCCGGCGGATCTTTCCCGCCCCC</td><td>305.3</td><td>-6.4</td><td>1.4</td><td>-4.4</td><td>9.7E-06</td><td>1.4E-05</td><td>MB231/MB468</td><td>NUC</td></tr>
<tr><td>>hg38_21_+_8433172_8446622_RNA45SN1_45S_50ntFlanks@7973.8006.34|GACGCGACCTCAGATCAGACGTGGCGACCCGCTG</td><td>>hg38_21_+_8433172_8446622_RNA45SN1_45S_50ntFlanks</td><td>7973</td><td>8006</td><td>34</td><td>GACGCGACCTCAGATCAGACGTGGCGACCCGCTG</td><td>2352.4</td><td>4.4</td><td>0.9</td><td>4.9</td><td>1.1E-06</td><td>1.4E-05</td><td>BT20/MB231</td><td>NUC</td></tr>
<tr><td>>hg38_21_+_8433172_8446622_RNA45SN1_45S_50ntFlanks@10759.10785.27|CCATATCCGCAGCAGGTCTCCAAGGTG</td><td>>hg38_21_+_8433172_8446622_RNA45SN1_45S_50ntFlanks</td><td>10759</td><td>10785</td><td>27</td><td>CCATATCCGCAGCAGGTCTCCAAGGTG</td><td>46.9</td><td>-8.6</td><td>1.8</td><td>-4.7</td><td>3.1E-06</td><td>1.4E-05</td><td>BT20/MB468</td><td>MP</td></tr>
<tr><td>>MI0000440|hsa-mir-27b&WithFlank&9|+|95085439|95085547@67.88.22(+1A)&[MIMAT0000419&hsa-miR-27b-3p&offsets|0|+1(+1A);m-39&9|+|95085505|95085525&offsets|0|+1(+1A)]|TTCACAGTGGCTAAGTTCTGCAA</td><td>>MI0000440|hsa-mir-27b&WithFlank&9|+|95085439|95085547</td><td>67</td><td>88</td><td>22(+1A)</td><td>TTCACAGTGGCTAAGTTCTGCAA</td><td>308.7</td><td>4.2</td><td>0.9</td><td>4.4</td><td>9.9E-06</td><td>1.4E-05</td><td>MB231/MB468</td><td>MITO</td></tr>
<tr><td>>hg38_21_+_8433172_8446622_RNA45SN1_45S_50ntFlanks@9852.9880.29|GCGCTGCGGGATGAACCGAACGCCGGGTT</td><td>>hg38_21_+_8433172_8446622_RNA45SN1_45S_50ntFlanks</td><td>9852</td><td>9880</td><td>29</td><td>GCGCTGCGGGATGAACCGAACGCCGGGTT</td><td>215.4</td><td>-7.3</td><td>1.6</td><td>-4.5</td><td>8.2E-06</td><td>1.4E-05</td><td>MB231/MB468</td><td>MP</td></tr>
<tr><td>>MI0000266|hsa-mir-10a&WithFlank&17|-|48579832|48579953@28.50.23(+1A)&[MIMAT0000253&hsa-miR-10a-5p&offsets|0|0(+1A);m-18&17|-|48579905|48579926&offsets|0|+1(+1A)]|TACCCTGTAGATCCGAATTTGTGA</td><td>>MI0000266|hsa-mir-10a&WithFlank&17|-|48579832|48579953</td><td>28</td><td>50</td><td>23(+1A)</td><td>TACCCTGTAGATCCGAATTTGTGA</td><td>300.6</td><td>7.3</td><td>1.6</td><td>4.5</td><td>8.2E-06</td><td>1.4E-05</td><td>MB231/MB468</td><td>MP</td></tr>
<tr><td>>trna14_LysTTT_11_-_59327808_59327880@2.33.32,trna2_LysTTT_17_+_8022473_8022545@2.33.32,trna54_LysTTT_1_+_204475655_204475727@2.33.32,trna5_LysTTT_11_+_59323902_59323974@2.33.32,trna62_LysTTT_1_-_204476158_204476230@2.33.32,trna76_LysTTT_6_+_28918806_28918878@2.33.32|CCCGGATAGCTCAGTCGGTAGAGCATCAGACT</td><td>>trna14_LysTTT_11_-_59327808_59327880</td><td>2</td><td>33</td><td>32</td><td>CCCGGATAGCTCAGTCGGTAGAGCATCAGACT</td><td>35.2</td><td>8.4</td><td>1.9</td><td>4.5</td><td>8.2E-06</td><td>1.4E-05</td><td>MB231/MB468</td><td>MP</td></tr>
<tr><td>>hg38_21_+_8433172_8446622_RNA45SN1_45S_50ntFlanks@5073.5097.25|AATTCCGATAACGAACGAGACTCTG</td><td>>hg38_21_+_8433172_8446622_RNA45SN1_45S_50ntFlanks</td><td>5073</td><td>5097</td><td>25</td><td>AATTCCGATAACGAACGAGACTCTG</td><td>1630.1</td><td>-6.6</td><td>1.5</td><td>-4.4</td><td>1.0E-05</td><td>1.4E-05</td><td>BT20/MB468</td><td>NUC</td></tr>
<tr><td>>MI0000440|hsa-mir-27b&WithFlank&9|+|95085439|95085547@70.87.18&[MIMAT0000419&hsa-miR-27b-3p&offsets|+3|0;m-39&9|+|95085505|95085525&offsets|+3|0]|ACAGTGGCTAAGTTCTGC</td><td>>MI0000440|hsa-mir-27b&WithFlank&9|+|95085439|95085547</td><td>70</td><td>87</td><td>18</td><td>ACAGTGGCTAAGTTCTGC</td><td>202.8</td><td>10.9</td><td>2.4</td><td>4.5</td><td>8.2E-06</td><td>1.4E-05</td><td>MB231/MB468</td><td>MP</td></tr>
<tr><td>>hg38_21_+_8433172_8446622_RNA45SN1_45S_50ntFlanks@10845.10868.24|ACTTCGGGATAAGGATTGGCTCTA</td><td>>hg38_21_+_8433172_8446622_RNA45SN1_45S_50ntFlanks</td><td>10845</td><td>10868</td><td>24</td><td>ACTTCGGGATAAGGATTGGCTCTA</td><td>222.6</td><td>-2.5</td><td>0.6</td><td>-4.5</td><td>7.1E-06</td><td>1.4E-05</td><td>BT20/MB468</td><td>MITO</td></tr>
<tr><td>>trnaMT_PheGAA_MT_+_577_647@37.56.20|AAATGTTTAGACGGGCTCAC</td><td>>trnaMT_PheGAA_MT_+_577_647</td><td>37</td><td>56</td><td>20</td><td>AAATGTTTAGACGGGCTCAC</td><td>288.5</td><td>6.4</td><td>1.4</td><td>4.7</td><td>3.1E-06</td><td>1.4E-05</td><td>BT20/MB468</td><td>MP</td></tr>
<tr><td>>hg38_21_+_8433172_8446622_RNA45SN1_45S_50ntFlanks@474.510.37|TCTTCGTGATCGATGTGGTGACGTCGTGCTCTCCCGG</td><td>>hg38_21_+_8433172_8446622_RNA45SN1_45S_50ntFlanks</td><td>474</td><td>510</td><td>37</td><td>TCTTCGTGATCGATGTGGTGACGTCGTGCTCTCCCGG</td><td>472.4</td><td>3.3</td><td>0.7</td><td>4.4</td><td>9.7E-06</td><td>1.4E-05</td><td>MB231/MB468</td><td>NUC</td></tr>
<tr><td>>hg38_21_+_8433172_8446622_RNA45SN1_45S_50ntFlanks@10678.10707.30|CGGTTCCGGCGGCGTCCGGTGAGCTCTCGC</td><td>>hg38_21_+_8433172_8446622_RNA45SN1_45S_50ntFlanks</td><td>10678</td><td>10707</td><td>30</td><td>CGGTTCCGGCGGCGTCCGGTGAGCTCTCGC</td><td>712.9</td><td>-5.1</td><td>1.2</td><td>-4.4</td><td>9.8E-06</td><td>1.4E-05</td><td>MB231/MB468</td><td>NUC</td></tr>
<tr><td>>MI0000270|hsa-mir-181b-1&WithFlank&1|-|198858867|198858988@42.61.20;MI0000683|hsa-mir-181b-2&WithFlank&9|+|124693704|124693804@22.41.20&[MIMAT0000257&hsa-miR-181b-5p&offsets|0|-3;m-73&1|-|198858925|198858947&offsets|0|-3];[MIMAT0000257_1&hsa-miR-181b-5p&offsets|0|-3;m-72&9|+|124693725|124693747&offsets|0|-3]|AACATTCATTGCTGTCGGTG</td><td>>MI0000270|hsa-mir-181b-1&WithFlank&1|-|198858867|198858988</td><td>42</td><td>61</td><td>20</td><td>AACATTCATTGCTGTCGGTG</td><td>277.8</td><td>5.4</td><td>1.2</td><td>4.4</td><td>9.8E-06</td><td>1.4E-05</td><td>MB231/MB468</td><td>NUC</td></tr>
<tr><td>>hg38_21_+_8433172_8446622_RNA45SN1_45S_50ntFlanks@12503.12520.18|TAGACCGTCGTGAGACAG</td><td>>hg38_21_+_8433172_8446622_RNA45SN1_45S_50ntFlanks</td><td>12503</td><td>12520</td><td>18</td><td>TAGACCGTCGTGAGACAG</td><td>82.1</td><td>-9.4</td><td>2.0</td><td>-4.7</td><td>3.1E-06</td><td>1.4E-05</td><td>BT20/MB468</td><td>MP</td></tr>
<tr><td>>hg38_21_+_8433172_8446622_RNA45SN1_45S_50ntFlanks@12416.12435.20|TTCGATGTCGGCTCTTCCTA</td><td>>hg38_21_+_8433172_8446622_RNA45SN1_45S_50ntFlanks</td><td>12416</td><td>12435</td><td>20</td><td>TTCGATGTCGGCTCTTCCTA</td><td>1153.9</td><td>-2.7</td><td>0.6</td><td>-4.4</td><td>9.8E-06</td><td>1.4E-05</td><td>MB231/MB468</td><td>NUC</td></tr>
<tr><td>>hg38_21_+_8433172_8446622_RNA45SN1_45S_50ntFlanks@12615.12657.43|GCTGAGGAGCCAATGGGGCGAAGCTACCATCTGTGGGATTATG</td><td>>hg38_21_+_8433172_8446622_RNA45SN1_45S_50ntFlanks</td><td>12615</td><td>12657</td><td>43</td><td>GCTGAGGAGCCAATGGGGCGAAGCTACCATCTGTGGGATTATG</td><td>428.5</td><td>-5.9</td><td>1.3</td><td>-4.4</td><td>9.8E-06</td><td>1.4E-05</td><td>MB231/MB468</td><td>NUC</td></tr>
<tr><td>>hg38_21_+_8433172_8446622_RNA45SN1_45S_50ntFlanks@10238.10264.27|CGCTACGCCGCGACGAGTAGGAGGGCC</td><td>>hg38_21_+_8433172_8446622_RNA45SN1_45S_50ntFlanks</td><td>10238</td><td>10264</td><td>27</td><td>CGCTACGCCGCGACGAGTAGGAGGGCC</td><td>271.0</td><td>4.8</td><td>1.0</td><td>4.7</td><td>3.1E-06</td><td>1.4E-05</td><td>BT20/MB468</td><td>MP</td></tr>
<tr><td>>trnaMT_GluTTC_MT_-_14674_14742@39.69.31,trnalookalike8_GluTTC_5_-_93905172_93905240@39.69.31|ATTGGTCGTGGTTGTAGTCCGTGCGAGAATA</td><td>>trnaMT_GluTTC_MT_-_14674_14742</td><td>39</td><td>69</td><td>31</td><td>ATTGGTCGTGGTTGTAGTCCGTGCGAGAATA</td><td>466.2</td><td>5.6</td><td>1.2</td><td>4.7</td><td>3.1E-06</td><td>1.4E-05</td><td>BT20/MB468</td><td>MP</td></tr>
<tr><td>>hg38_21_+_8433172_8446622_RNA45SN1_45S_50ntFlanks@11316.11344.29|CACCCCACGTCTCGTCGCGCGCGCGTCCG</td><td>>hg38_21_+_8433172_8446622_RNA45SN1_45S_50ntFlanks</td><td>11316</td><td>11344</td><td>29</td><td>CACCCCACGTCTCGTCGCGCGCGCGTCCG</td><td>19613.4</td><td>-6.7</td><td>1.5</td><td>-4.5</td><td>7.1E-06</td><td>1.4E-05</td><td>BT20/MB468</td><td>MITO</td></tr>
<tr><td>>hg38_21_+_8433172_8446622_RNA45SN1_45S_50ntFlanks@8176.8205.30|CTTCTGATCGAGGCCCAGCCCGTGGACGGT</td><td>>hg38_21_+_8433172_8446622_RNA45SN1_45S_50ntFlanks</td><td>8176</td><td>8205</td><td>30</td><td>CTTCTGATCGAGGCCCAGCCCGTGGACGGT</td><td>368.7</td><td>-5.1</td><td>1.1</td><td>-4.5</td><td>8.2E-06</td><td>1.4E-05</td><td>MB231/MB468</td><td>MP</td></tr>
<tr><td>>hg38_21_+_8433172_8446622_RNA45SN1_45S_50ntFlanks@10788.10815.28|CAGCCTCTGGCATGTTGGAACAATGTAG</td><td>>hg38_21_+_8433172_8446622_RNA45SN1_45S_50ntFlanks</td><td>10788</td><td>10815</td><td>28</td><td>CAGCCTCTGGCATGTTGGAACAATGTAG</td><td>450.7</td><td>-3.2</td><td>0.7</td><td>-4.5</td><td>8.4E-06</td><td>1.4E-05</td><td>BT20/MB468</td><td>CYTO</td></tr>
<tr><td>>MI0000342|hsa-mir-200b&WithFlank&1|+|1167098|1167204@63.83.21&[MIMAT0000318&hsa-miR-200b-3p&offsets|0|-1;m-93&1|+|1167160|1167182&offsets|0|-2]|TAATACTGCCTGGTAATGATG</td><td>>MI0000342|hsa-mir-200b&WithFlank&1|+|1167098|1167204</td><td>63</td><td>83</td><td>21</td><td>TAATACTGCCTGGTAATGATG</td><td>307.2</td><td>-4.8</td><td>1.1</td><td>-4.4</td><td>9.8E-06</td><td>1.4E-05</td><td>MB231/MB468</td><td>NUC</td></tr>
<tr><td>>hg38_21_+_8433172_8446622_RNA45SN1_45S_50ntFlanks@6648.6695.48|GTACGACTCTTAGCGGTGGATCACTCGGCTCGTGCGTCGATGAAGAAC</td><td>>hg38_21_+_8433172_8446622_RNA45SN1_45S_50ntFlanks</td><td>6648</td><td>6695</td><td>48</td><td>GTACGACTCTTAGCGGTGGATCACTCGGCTCGTGCGTCGATGAAGAAC</td><td>977.4</td><td>5.6</td><td>1.3</td><td>4.4</td><td>9.8E-06</td><td>1.4E-05</td><td>MB231/MB468</td><td>NUC</td></tr>
<tr><td>>MI0000088|hsa-mir-30a&WithFlank&6|-|71403545|71403627@54.73.20(+1U)&[MIMAT0000088&hsa-miR-30a-3p&offsets|+1|-1(+1U);m-38&6|-|71403554|71403575&offsets|+1|-1(+1U)]|TTTCAGTCGGATGTTTGCAGT</td><td>>MI0000088|hsa-mir-30a&WithFlank&6|-|71403545|71403627</td><td>54</td><td>73</td><td>20(+1U)</td><td>TTTCAGTCGGATGTTTGCAGT</td><td>491.7</td><td>2.6</td><td>0.6</td><td>4.4</td><td>1.0E-05</td><td>1.4E-05</td><td>BT20/MB468</td><td>NUC</td></tr>
<tr><td>>hg38_21_+_8433172_8446622_RNA45SN1_45S_50ntFlanks@10841.10867.27|CGTAACTTCGGGATAAGGATTGGCTCT</td><td>>hg38_21_+_8433172_8446622_RNA45SN1_45S_50ntFlanks</td><td>10841</td><td>10867</td><td>27</td><td>CGTAACTTCGGGATAAGGATTGGCTCT</td><td>1231.0</td><td>-4.3</td><td>1.0</td><td>-4.5</td><td>8.3E-06</td><td>1.4E-05</td><td>MB231/MB468</td><td>MP</td></tr>
<tr><td>>MI0000081|hsa-mir-24-2&WithFlank&19|-|13836281|13836365@56.74.19(+1U);MI0000080|hsa-mir-24-1&WithFlank&9|+|95086015|95086094@50.68.19(+1U)&[MIMAT0000080&hsa-miR-24-3p&offsets|0|-3(+1U);m-36&19|-|13836288|13836310&offsets|0|-4(+1U)];[MIMAT0000080_1&hsa-miR-24-3p&offsets|0|-3(+1U);m-35&9|+|95086064|95086086&offsets|0|-4(+1U)]|TGGCTCAGTTCAGCAGGAAT</td><td>>MI0000081|hsa-mir-24-2&WithFlank&19|-|13836281|13836365</td><td>56</td><td>74</td><td>19(+1U)</td><td>TGGCTCAGTTCAGCAGGAAT</td><td>3334.6</td><td>4.1</td><td>0.9</td><td>4.5</td><td>8.3E-06</td><td>1.4E-05</td><td>MB231/MB468</td><td>MP</td></tr>
<tr><td>>hg38_21_+_8433172_8446622_RNA45SN1_45S_50ntFlanks@10392.10425.34|ACAGCAGTTGAACATGGGTCAGTCGGTCCTGAGA</td><td>>hg38_21_+_8433172_8446622_RNA45SN1_45S_50ntFlanks</td><td>10392</td><td>10425</td><td>34</td><td>ACAGCAGTTGAACATGGGTCAGTCGGTCCTGAGA</td><td>226.5</td><td>-4.7</td><td>1.1</td><td>-4.4</td><td>9.8E-06</td><td>1.4E-05</td><td>MB231/MB468</td><td>NUC</td></tr>
<tr><td>>hg38_21_+_8433172_8446622_RNA45SN1_45S_50ntFlanks@9554.9597.44|CGTAGCGGTCCTGACGTGCAAATCGGTCGTCCGACCTGGGTATA</td><td>>hg38_21_+_8433172_8446622_RNA45SN1_45S_50ntFlanks</td><td>9554</td><td>9597</td><td>44</td><td>CGTAGCGGTCCTGACGTGCAAATCGGTCGTCCGACCTGGGTATA</td><td>554.2</td><td>-4.9</td><td>1.1</td><td>-4.4</td><td>9.4E-06</td><td>1.4E-05</td><td>MB231/MB468</td><td>CYTO</td></tr>
<tr><td>>hg38_21_+_8433172_8446622_RNA45SN1_45S_50ntFlanks@3854.3874.21|ACTGTGGTAATTCTAGAGCTA</td><td>>hg38_21_+_8433172_8446622_RNA45SN1_45S_50ntFlanks</td><td>3854</td><td>3874</td><td>21</td><td>ACTGTGGTAATTCTAGAGCTA</td><td>159.8</td><td>-4.6</td><td>1.0</td><td>-4.4</td><td>9.8E-06</td><td>1.4E-05</td><td>MB231/MB468</td><td>NUC</td></tr>
<tr><td>>hg38_21_+_8433172_8446622_RNA45SN1_45S_50ntFlanks@11027.11050.24|CTCTCTCTCTCTCTCCCCCGCTCC</td><td>>hg38_21_+_8433172_8446622_RNA45SN1_45S_50ntFlanks</td><td>11027</td><td>11050</td><td>24</td><td>CTCTCTCTCTCTCTCCCCCGCTCC</td><td>3769.3</td><td>-8.6</td><td>2.0</td><td>-4.4</td><td>1.0E-05</td><td>1.4E-05</td><td>BT20/MB468</td><td>NUC</td></tr>
</table>

| | | | | | | | | | | | | | |
|---|---|---|---|---|---|---|---|---|---|---|---|---|---|
| >hg38_21_+_8433172_8446622_RNA45SN1_45S_50ntFlanks@11670.11700.31\|CCCAGTGCTCTGAATGTCAAAGTGAAGAAAT | >hg38_21_+_8433172_8446622_RNA45SN1_45S_50ntFlanks | 11670 | 11700 | 31 | CCCAGTGCTCTGAATGTCAAAGTGAAGAAAT | 299.5 | -6.3 | 1.4 | -4.4 | 1.0E-05 | 1.4E-05 | MB231/MB468 | MITO |
| >trnaMT_ValTAC_MT_+_1602_1670@42.72.31\|AGATTTCAACTTAACTTGACCGCTCTGACCA | >trnaMT_ValTAC_MT_+_1602_1670 | 42 | 72 | 31 | AGATTTCAACTTAACTTGACCGCTCTGACCA | 26564.2 | 4.4 | 1.0 | 4.4 | 1.0E-05 | 1.4E-05 | MB231/MB468 | MITO |
| >hg38_21_+_8433172_8446622_RNA45SN1_45S_50ntFlanks@9162.9187.26\|CGCCCCGGGCGGGTCGCGCCGTCGGG | >hg38_21_+_8433172_8446622_RNA45SN1_45S_50ntFlanks | 9162 | 9187 | 26 | CGCCCCGGGCGGGTCGCGCCGTCGGG | 153.5 | -5.7 | 1.3 | -4.5 | 7.2E-06 | 1.5E-05 | BT20/MB468 | MITO |
| >MI0000750\|hsa-mir-26a-2&WithFlank&12\|-\|57824603\|57824698@20.40.21(+1A);MI0000083\|hsa-mir-26a-1&WithFlank&3\|+\|37969398\|37969486@16.36.21(+1A)&[MIMAT0000082&hsa-miR-26a-5p&offsets\|0\|-1(+1A);m-33&12\|-\|57824658\|57824679&offsets\|0\|-1(+1A)];[MIMAT0000082_1&hsa-miR-26a-5p&offsets\|0\|-1(+1A);m-34&3\|+\|37969413\|37969434&offsets\|0\|-1(+1A)]\|TTCAAGTAATCCAGGATAGGCA | >MI0000750\|hsa-mir-26a-2&WithFlank&12\|-\|57824603\|57824698 | 20 | 40 | 21(+1A) | TTCAAGTAATCCAGGATAGGCA | 8467.9 | 4.2 | 1.0 | 4.5 | 8.3E-06 | 1.5E-05 | MB231/MB468 | MP |
| >MI0000098\|hsa-mir-96&WithFlank&7\|-\|129774686\|129774775@15.36.22(+1A)&[MIMAT0000095&hsa-miR-96-5p&offsets\|0\|-1(+1A);m-248&7\|-\|129774739\|129774761&offsets\|0\|-1(+1A)]\|TTTGGCACTAGCACATTTTTGCA | >MI0000098\|hsa-mir-96&WithFlank&7\|-\|129774686\|129774775 | 15 | 36 | 22(+1A) | TTTGGCACTAGCACATTTTTGCA | 678.1 | 3.8 | 0.8 | 4.9 | 1.2E-06 | 1.5E-05 | BT20/MB231 | NUC |
| >hg38_21_+_8433172_8446622_RNA45SN1_45S_50ntFlanks@12416.12438.23\|TTCGATGTCGGCTCTTCCTATCA | >hg38_21_+_8433172_8446622_RNA45SN1_45S_50ntFlanks | 12416 | 12438 | 23 | TTCGATGTCGGCTCTTCCTATCA | 743.1 | 3.4 | 0.8 | 4.5 | 7.2E-06 | 1.5E-05 | BT20/MB468 | MITO |
| >hg38_21_+_8433172_8446622_RNA45SN1_45S_50ntFlanks@10458.10476.19\|ATGGCCTCCGTTGCCCTCG | >hg38_21_+_8433172_8446622_RNA45SN1_45S_50ntFlanks | 10458 | 10476 | 19 | ATGGCCTCCGTTGCCCTCG | 82.7 | -10.2 | 2.3 | -4.5 | 8.4E-06 | 1.5E-05 | MB231/MB468 | MP |
| >MI0000809\|hsa-mir-151a&WithFlank&8\|-\|140732558\|140732659@53.74.22(+2A)&[MIMAT0000757&hsa-miR-151a-3p&offsets\|0\|+1(+2A);m-30&8\|-\|140732587\|140732607&offsets\|0\|+1(+2A)]\|CTAGACTGAAGCTCCTTGAGGAAA | >MI0000809\|hsa-mir-151a&WithFlank&8\|-\|140732558\|140732659 | 53 | 74 | 22(+2A) | CTAGACTGAAGCTCCTTGAGGAAA | 267.1 | 5.1 | 1.2 | 4.4 | 1.0E-05 | 1.5E-05 | MB231/MB468 | MITO |
| >MI0000449\|hsa-mir-132&WithFlank&17\|-\|2049902\|2050014@28.48.21&[MIMAT0004594&hsa-miR-132-5p&offsets\|-1\|-2;m-414&17\|-\|2049965\|2049986&offsets\|-1\|-2]\|AACCGTGGCTTTCGATTGTTA | >MI0000449\|hsa-mir-132&WithFlank&17\|-\|2049902\|2050014 | 28 | 48 | 21 | AACCGTGGCTTTCGATTGTTA | 48.9 | 9.4 | 2.0 | 4.7 | 3.1E-06 | 1.5E-05 | BT20/MB468 | MP |
| >hg38_21_+_8433172_8446622_RNA45SN1_45S_50ntFlanks@8358.8378.21\|AAAGAACTTTGAAGAGAGAGT | >hg38_21_+_8433172_8446622_RNA45SN1_45S_50ntFlanks | 8358 | 8378 | 21 | AAAGAACTTTGAAGAGAGAGT | 1142.4 | -6.2 | 1.4 | -4.5 | 8.4E-06 | 1.5E-05 | MB231/MB468 | MP |
| >hg38_21_+_8433172_8446622_RNA45SN1_45S_50ntFlanks@12654.12671.18\|TATGACTGAACGCCTCTA | >hg38_21_+_8433172_8446622_RNA45SN1_45S_50ntFlanks | 12654 | 12671 | 18 | TATGACTGAACGCCTCTA | 271.7 | -6.1 | 1.4 | -4.4 | 1.0E-05 | 1.5E-05 | MB231/MB468 | NUC |
| >hg38_21_+_8433172_8446622_RNA45SN1_45S_50ntFlanks@3875.3899.25\|ATACATGCCGACGGGCGCTGACCCC | >hg38_21_+_8433172_8446622_RNA45SN1_45S_50ntFlanks | 3875 | 3899 | 25 | ATACATGCCGACGGGCGCTGACCCC | 783.8 | -5.6 | 1.3 | -4.4 | 1.0E-05 | 1.5E-05 | MB231/MB468 | NUC |
| >MI0000734\|hsa-mir-106b&WithFlank&7\|-\|100093987\|100094080@58.79.22&[MIMAT0004672&hsa-miR-106b-3p&offsets\|0\|0;m-87&7\|-\|100094003\|100094023&offsets\|0\|+1]\|CCGCACTGTGGGTACTTGCTGC | >MI0000734\|hsa-mir-106b&WithFlank&7\|-\|100093987\|100094080 | 58 | 79 | 22 | CCGCACTGTGGGTACTTGCTGC | 6378.1 | 5.1 | 1.1 | 4.7 | 3.1E-06 | 1.5E-05 | BT20/MB468 | MP |
| >hg38_21_+_8433172_8446622_RNA45SN1_45S_50ntFlanks@4045.4067.23\|CCATTCGAACGTCTGCCCTATCA | >hg38_21_+_8433172_8446622_RNA45SN1_45S_50ntFlanks | 4045 | 4067 | 23 | CCATTCGAACGTCTGCCCTATCA | 280.2 | -4.5 | 1.0 | -4.4 | 1.0E-05 | 1.5E-05 | MB231/MB468 | NUC |
| >hg38_21_+_8433172_8446622_RNA45SN1_45S_50ntFlanks@12446.12476.31\|GCAGAATTCACCAAGCGTTGGATTGTTCACC | >hg38_21_+_8433172_8446622_RNA45SN1_45S_50ntFlanks | 12446 | 12476 | 31 | GCAGAATTCACCAAGCGTTGGATTGTTCACC | 170.2 | 3.9 | 0.9 | 4.4 | 1.0E-05 | 1.5E-05 | MB231/MB468 | NUC |
| >am_trna128_GlyGCC_6_-_27870686_27870756@3.32.30,trna133_GlyCCC_1_-_16872434_16872504@3.32.30,trna18_GlyGCC_16_+_70822597_70822667@3.32.30,trna19_GlyGCC_16_+_70823410_70823480@3.32.30,trna19_GlyGCC_2_-_157257659_157257729@3.32.30,trna24_GlyGCC_16_-_70812942_70813012@3.32.30,trna25_GlyGCC_16_-_70812114_70812184@3.32.30,trna4_GlyCCC_1_+_17188416_17188486@3.32.30,trna5_GlyGCC_17_+_8029064_8029134@3.32.30,trna68_GlyGCC_1_-_161493637_161493707@3.32.30\|ATTGGTGGTTCAGTGGTAGAATTCTCGCCT | >am_trna128_GlyGCC_6_-_27870686_27870756 | 3 | 32 | 30 | ATTGGTGGTTCAGTGGTAGAATTCTCGCCT | 35.0 | 8.4 | 1.9 | 4.5 | 8.4E-06 | 1.5E-05 | MB231/MB468 | MP |
| >hg38_21_+_8433172_8446622_RNA45SN1_45S_50ntFlanks@10387.10424.38\|TGTGAACAGCAGTTGAACATGGGTCAGTCGGTCCTGAG | >hg38_21_+_8433172_8446622_RNA45SN1_45S_50ntFlanks | 10387 | 10424 | 38 | TGTGAACAGCAGTTGAACATGGGTCAGTCGGTCCTGAG | 3550.9 | 8.1 | 1.7 | 4.7 | 3.2E-06 | 1.5E-05 | BT20/MB468 | MP |
| >hg38_21_+_8433172_8446622_RNA45SN1_45S_50ntFlanks@11578.11601.24\|TAGAACTGGTGCGGACCAGGGGAA | >hg38_21_+_8433172_8446622_RNA45SN1_45S_50ntFlanks | 11578 | 11601 | 24 | TAGAACTGGTGCGGACCAGGGGAA | 321.2 | -6.2 | 1.4 | -4.4 | 1.0E-05 | 1.5E-05 | MB231/MB468 | NUC |
| >hg38_21_+_8433172_8446622_RNA45SN1_45S_50ntFlanks@11643.11660.18\|GCGGGTGTTGACGCGATG | >hg38_21_+_8433172_8446622_RNA45SN1_45S_50ntFlanks | 11643 | 11660 | 18 | GCGGGTGTTGACGCGATG | 620.6 | -3.8 | 0.8 | -4.5 | 7.3E-06 | 1.5E-05 | BT20/MB468 | MITO |
| >hg38_21_+_8433172_8446622_RNA45SN1_45S_50ntFlanks@9240.9283.44\|GCGGAGCGAGCGCACGGGGTCGGCGGCGACGTCGGCTACCCACC | >hg38_21_+_8433172_8446622_RNA45SN1_45S_50ntFlanks | 9240 | 9283 | 44 | GCGGAGCGAGCGCACGGGGTCGGCGGCGACGTCGGCTACCCACC | 75.5 | -8.3 | 1.9 | -4.5 | 7.3E-06 | 1.5E-05 | BT20/MB468 | MITO |
| >MI0000542\|hsa-mir-320a&WithFlank&8\|-\|22244960\|22245043@48.70.23(+1U)&[MIMAT0000510&hsa-miR-320a-3p&offsets\|0\|+1(+1U);m-60&8\|-\|22244975\|22244996&offsets\|0\|+1(+1U)]\|AAAAGCTGGGTTGAGAGGGCGAAT | >MI0000542\|hsa-mir-320a&WithFlank&8\|-\|22244960\|22245043 | 48 | 70 | 23(+1U) | AAAAGCTGGGTTGAGAGGGCGAAT | 2043.0 | 3.9 | 0.9 | 4.5 | 8.4E-06 | 1.5E-05 | MB231/MB468 | MP |
| >MI0000074\|hsa-mir-19b-1&WithFlank&13\|+\|91351186\|91351284@60.82.23;MI0000075\|hsa-mir-19b-2&WithFlank&X\|-\|134169665\|134169772@68.90.23&[MIMAT0000074&hsa-miR-19b-3p&offsets\|0\|0;m-122&13\|+\|91351245\|91351267&offsets\|0\|0];[MIMAT0000074_1&hsa-miR-19b-3p&offsets\|0\|0;m-121&X\|-\|134169683\|134169705&offsets\|0\|0]\|TGTGCAAATCCATGCAAAACTGA | >MI0000074\|hsa-mir-19b-1&WithFlank&13\|+\|91351186\|91351284 | 60 | 82 | 23 | TGTGCAAATCCATGCAAAACTGA | 1272.8 | 6.5 | 1.5 | 4.4 | 1.0E-05 | 1.5E-05 | MB231/MB468 | MITO |
| >hg38_21_+_8433172_8446622_RNA45SN1_45S_50ntFlanks@10885.10902.18\|GGCTGGGGCGCGAAGCGG | >hg38_21_+_8433172_8446622_RNA45SN1_45S_50ntFlanks | 10885 | 10902 | 18 | GGCTGGGGCGCGAAGCGG | 197.4 | -5.9 | 1.3 | -4.4 | 1.0E-05 | 1.5E-05 | MB231/MB468 | NUC |
| >hg38_21_+_8433172_8446622_RNA45SN1_45S_50ntFlanks@12996.13040.45\|TCGCTGCGATCTATTGAAAGTCAGCCCTCGACACAAGGGTTTGTC | >hg38_21_+_8433172_8446622_RNA45SN1_45S_50ntFlanks | 12996 | 13040 | 45 | TCGCTGCGATCTATTGAAAGTCAGCCCTCGACACAAGGGTTTGTC | 331.5 | 6.0 | 1.4 | 4.4 | 1.0E-05 | 1.5E-05 | MB231/MB468 | NUC |
| >hg38_21_+_8433172_8446622_RNA45SN1_45S_50ntFlanks@11572.11607.36\|CCGACTTAGAACTGGTGCGGACCAGGGGAATCCGAC | >hg38_21_+_8433172_8446622_RNA45SN1_45S_50ntFlanks | 11572 | 11607 | 36 | CCGACTTAGAACTGGTGCGGACCAGGGGAATCCGAC | 288.1 | -3.0 | 0.7 | -4.4 | 1.0E-05 | 1.5E-05 | MB231/MB468 | NUC |
| >hg38_MT_+_598_1651_RNR1_12S_50ntFlanks@428.470.43\|AAAGTGGCTTTAACATATCTGAACACACAATAGCTAAGACCCA | >hg38_MT_+_598_1651_RNR1_12S_50ntFlanks | 428 | 470 | 43 | AAAGTGGCTTTAACATATCTGAACACACAATAGCTAAGACCCA | 130.5 | 7.1 | 1.6 | 4.4 | 1.0E-05 | 1.5E-05 | MB231/MB468 | MITO |
| >trnaMT_SerGCT_MT_+_12207_12265@30.62.33\|GCCCCCATGTCTAACAACATGGCTTTCTCACCA | >trnaMT_SerGCT_MT_+_12207_12265 | 30 | 62 | 33 | GCCCCCATGTCTAACAACATGGCTTTCTCACCA | 676.1 | 5.1 | 1.1 | 4.4 | 1.0E-05 | 1.5E-05 | MB231/MB468 | MITO |
| >hg38_21_+_8433172_8446622_RNA45SN1_45S_50ntFlanks@8105.8146.42\|CGCGGCGGGGCGCGGGACATGTGGCGTACGGAAGACCCGCTC | >hg38_21_+_8433172_8446622_RNA45SN1_45S_50ntFlanks | 8105 | 8146 | 42 | CGCGGCGGGGCGCGGGACATGTGGCGTACGGAAGACCCGCTC | 85.6 | -10.3 | 2.3 | -4.5 | 8.5E-06 | 1.5E-05 | MB231/MB468 | MP |
| >hg38_21_+_8433172_8446622_RNA45SN1_45S_50ntFlanks@9481.9506.26\|CACGTGTTAGGACCCGAAAGATGGTG | >hg38_21_+_8433172_8446622_RNA45SN1_45S_50ntFlanks | 9481 | 9506 | 26 | CACGTGTTAGGACCCGAAAGATGGTG | 165.3 | 10.7 | 2.2 | 4.9 | 9.3E-07 | 1.5E-05 | BT20/MB231 | MP |
| >hg38_21_+_8433172_8446622_RNA45SN1_45S_50ntFlanks@9919.9947.29\|GGTGTTGGTTGATATAGACAGCAGGACGG | >hg38_21_+_8433172_8446622_RNA45SN1_45S_50ntFlanks | 9919 | 9947 | 29 | GGTGTTGGTTGATATAGACAGCAGGACGG | 111.5 | 10.1 | 2.1 | 4.9 | 9.3E-07 | 1.5E-05 | BT20/MB231 | MP |
| >hg38_21_+_8433172_8446622_RNA45SN1_45S_50ntFlanks@9955.9983.29\|GGAAGTCGGAATCCGCTAAGGAGTGTGTA | >hg38_21_+_8433172_8446622_RNA45SN1_45S_50ntFlanks | 9955 | 9983 | 29 | GGAAGTCGGAATCCGCTAAGGAGTGTGTA | 114.8 | 10.2 | 2.1 | 4.9 | 9.2E-07 | 1.5E-05 | BT20/MB231 | MP |
| >hg38_21_+_8433172_8446622_RNA45SN1_45S_50ntFlanks@8196.8225.30\|CGTGGACGGTGTGAGGCCGGTAGCGGCCCC | >hg38_21_+_8433172_8446622_RNA45SN1_45S_50ntFlanks | 8196 | 8225 | 30 | CGTGGACGGTGTGAGGCCGGTAGCGGCCCC | 2070.1 | -8.1 | 1.9 | -4.4 | 1.1E-05 | 1.5E-05 | BT20/MB468 | NUC |
| >hg38_21_+_8433172_8446622_RNA45SN1_45S_50ntFlanks@10355.10390.36\|AACTTTGAAGGCCGAAGTGGAGAAGGGTTCCATGTG | >hg38_21_+_8433172_8446622_RNA45SN1_45S_50ntFlanks | 10355 | 10390 | 36 | AACTTTGAAGGCCGAAGTGGAGAAGGGTTCCATGTG | 371.9 | -4.4 | 1.0 | -4.4 | 9.6E-06 | 1.5E-05 | MB231/MB468 | CYTO |
| >MI0000087\|hsa-mir-29a&WithFlank&7\|-\|130876741\|130876816@48.67.20(+1C)&[MIMAT0000086&hsa-miR-29a-3p&offsets\|0\|-2(+1C);m-26&7\|-\|130876749\|130876769&offsets\|0\|-1(+1C)]\|TAGCACCATCTGAAATCGGTC | >MI0000087\|hsa-mir-29a&WithFlank&7\|-\|130876741\|130876816 | 48 | 67 | 20(+1C) | TAGCACCATCTGAAATCGGTC | 2402.0 | 6.3 | 1.3 | 4.7 | 3.2E-06 | 1.5E-05 | BT20/MB468 | MP |
| >hg38_21_+_8433172_8446622_RNA45SN1_45S_50ntFlanks@11208.11233.26\|GCGGCGACTCTGGACGCGAGCCGGGC | >hg38_21_+_8433172_8446622_RNA45SN1_45S_50ntFlanks | 11208 | 11233 | 26 | GCGGCGACTCTGGACGCGAGCCGGGC | 9386.8 | -7.2 | 1.6 | -4.4 | 1.1E-05 | 1.5E-05 | BT20/MB468 | NUC |
| >MI0000434\|hsa-let-7i&WithFlank&12\|+\|62603680\|62603775@12.33.22&[MIMAT0000415&hsa-let-7i-5p&offsets\|0\|0;m-49&12\|+\|62603691\|62603712&offsets\|0\|0]\|TGAGGTAGTAGTTTGTGCTGTT | >MI0000434\|hsa-let-7i&WithFlank&12\|+\|62603680\|62603775 | 12 | 33 | 22 | TGAGGTAGTAGTTTGTGCTGTT | 43342.9 | 4.6 | 1.0 | 4.4 | 1.0E-05 | 1.5E-05 | MB231/MB468 | MITO |
| >hg38_21_+_8433172_8446622_RNA45SN1_45S_50ntFlanks@12412.12443.32\|ATCCTTCGATGTCGGCTCTTCCTATCATTGTG | >hg38_21_+_8433172_8446622_RNA45SN1_45S_50ntFlanks | 12412 | 12443 | 32 | ATCCTTCGATGTCGGCTCTTCCTATCATTGTG | 47.5 | -8.7 | 1.9 | -4.7 | 3.2E-06 | 1.5E-05 | BT20/MB468 | MP |
| >hg38_21_+_8433172_8446622_RNA45SN1_45S_50ntFlanks@11561.11584.24\|GCGCCTAGCAGCCGACTTAGAACT | >hg38_21_+_8433172_8446622_RNA45SN1_45S_50ntFlanks | 11561 | 11584 | 24 | GCGCCTAGCAGCCGACTTAGAACT | 277.4 | -6.5 | 1.4 | -4.5 | 8.5E-06 | 1.5E-05 | MB231/MB468 | MP |
| >MI0000065\|hsa-let-7d&WithFlank&9\|+\|94178828\|94178926@14.34.21&[MIMAT0000065&hsa-let-7d-5p&offsets\|0\|-1;m-90&9\|+\|94178841\|94178862&offsets\|0\|-1]\|AGAGGTAGTAGGTTGCATAGT | >MI0000065\|hsa-let-7d&WithFlank&9\|+\|94178828\|94178926 | 14 | 34 | 21 | AGAGGTAGTAGGTTGCATAGT | 865.0 | 4.1 | 0.9 | 4.5 | 8.5E-06 | 1.5E-05 | MB231/MB468 | MP |
| >hg38_21_+_8433172_8446622_RNA45SN1_45S_50ntFlanks@12605.12634.30\|TATGTGCTTGGCTGAGGAGCCAATGGGGCG | >hg38_21_+_8433172_8446622_RNA45SN1_45S_50ntFlanks | 12605 | 12634 | 30 | TATGTGCTTGGCTGAGGAGCCAATGGGGCG | 212.0 | -4.2 | 0.9 | -4.4 | 9.7E-06 | 1.5E-05 | MB231/MB468 | CYTO |
| >hg38_21_+_8433172_8446622_RNA45SN1_45S_50ntFlanks@9544.9569.26\|GGTGGAGGTCCGTAGCGGTCCTGACG | >hg38_21_+_8433172_8446622_RNA45SN1_45S_50ntFlanks | 9544 | 9569 | 26 | GGTGGAGGTCCGTAGCGGTCCTGACG | 324.2 | -2.3 | 0.5 | -4.4 | 1.0E-05 | 1.5E-05 | MB231/MB468 | NUC |
| >hg38_21_+_8433172_8446622_RNA45SN1_45S_50ntFlanks@8836.8867.32\|CGGCGGTGGCGGCGGCGGCGGCGGCGGCGGGA | >hg38_21_+_8433172_8446622_RNA45SN1_45S_50ntFlanks | 8836 | 8867 | 32 | CGGCGGTGGCGGCGGCGGCGGCGGCGGCGGGA | 78.5 | -10.2 | 2.3 | -4.5 | 8.5E-06 | 1.5E-05 | MB231/MB468 | MP |
| >hg38_21_+_8433172_8446622_RNA45SN1_45S_50ntFlanks@6651.6672.22\|CGACTCTTAGCGGTGGATCACT | >hg38_21_+_8433172_8446622_RNA45SN1_45S_50ntFlanks | 6651 | 6672 | 22 | CGACTCTTAGCGGTGGATCACT | 2717.4 | 2.5 | 0.6 | 4.4 | 1.1E-05 | 1.5E-05 | BT20/MB468 | NUC |
| >hg38_MT_+_1621_3279_RNR2_16S_50ntFlanks@1338.1374.37\|AGTCCATATCAACAATAGGGTTTACGACCTCGATGTT | >hg38_MT_+_1621_3279_RNR2_16S_50ntFlanks | 1338 | 1374 | 37 | AGTCCATATCAACAATAGGGTTTACGACCTCGATGTT | 34.8 | 8.3 | 1.9 | 4.5 | 8.5E-06 | 1.5E-05 | MB231/MB468 | MP |
| >hg38_21_+_8433172_8446622_RNA45SN1_45S_50ntFlanks@11873.11906.34\|AAGAAGACCCTGTTGAGCTTGACTCTAGTCTGGC | >hg38_21_+_8433172_8446622_RNA45SN1_45S_50ntFlanks | 11873 | 11906 | 34 | AAGAAGACCCTGTTGAGCTTGACTCTAGTCTGGC | 673.9 | -6.6 | 1.5 | -4.4 | 1.0E-05 | 1.5E-05 | MB231/MB468 | NUC |
| >hg38_21_+_8433172_8446622_RNA45SN1_45S_50ntFlanks@8170.8187.18\|CAAGTCCTTCTGATCGAG | >hg38_21_+_8433172_8446622_RNA45SN1_45S_50ntFlanks | 8170 | 8187 | 18 | CAAGTCCTTCTGATCGAG | 341.3 | -3.2 | 0.7 | -4.5 | 7.4E-06 | 1.5E-05 | BT20/MB468 | MITO |

| | | | | | | | | | | | | | |
|---|---|---|---|---|---|---|---|---|---|---|---|---|---|
| >hg38_21_+_8433172_8446622_RNA45SN1_45S_50ntFlanks@10685.10711.27\|GGCGGCGTCCGGTGAGCTCTCGCTGGC | >hg38_21_+_8433172_8446622_RNA45SN1_45S_50ntFlanks | 10685 | 10711 | 27 | GGCGGCGTCCGGTGAGCTCTCGCTGGC | 373.6 | -3.0 | 0.7 | -4.4 | 1.0E-05 | 1.5E-05 | MB231/MB468 | MITO |
| >MI0000082\|hsa-mir-25&WithFlank&7\|-\|100093554\|100093649@58.78.21(+1G)&[MIMAT0000081&hsa-miR-25-3p&offsets\|0\|-1(+1G);m-23&7\|-\|100093571\|100093592&offsets\|0\|-1(+1G)]\|CATTGCACTTGTCTCGGTCTGG | >MI0000082\|hsa-mir-25&WithFlank&7\|-\|100093554\|100093649 | 58 | 78 | 21(+1G) | CATTGCACTTGTCTCGGTCTGG | 540.5 | 4.2 | 0.9 | 4.4 | 9.7E-06 | 1.5E-05 | MB231/MB468 | CYTO |
| >hg38_21_+_8433172_8446622_RNA45SN1_45S_50ntFlanks@11874.11897.24\|AGAAGACCCTGTTGAGCTTGACTC | >hg38_21_+_8433172_8446622_RNA45SN1_45S_50ntFlanks | 11874 | 11897 | 24 | AGAAGACCCTGTTGAGCTTGACTC | 957.2 | -3.8 | 0.9 | -4.4 | 1.0E-05 | 1.5E-05 | MB231/MB468 | NUC |
| >hg38_21_+_8433172_8446622_RNA45SN1_45S_50ntFlanks@5485.5522.38\|AGCGCTGAGAAGACGGTCGAACTTGACTATCTAGAGGA | >hg38_21_+_8433172_8446622_RNA45SN1_45S_50ntFlanks | 5485 | 5522 | 38 | AGCGCTGAGAAGACGGTCGAACTTGACTATCTAGAGGA | 163.8 | 10.7 | 2.2 | 4.9 | 9.4E-07 | 1.5E-05 | BT20/MB231 | MP |
| >trna4_ArgTCG_15_+_89878304_89878376@31.52.22\|ACTTCGGATCAGAAGATTGCAG | >trna4_ArgTCG_15_+_89878304_89878376 | 31 | 52 | 22 | ACTTCGGATCAGAAGATTGCAG | 126.0 | -10.8 | 2.4 | -4.5 | 8.6E-06 | 1.5E-05 | MB231/MB468 | MP |
| >hg38_21_+_8433172_8446622_RNA45SN1_45S_50ntFlanks@12419.12442.24\|GATGTCGGCTCTTCCTATCATTGT | >hg38_21_+_8433172_8446622_RNA45SN1_45S_50ntFlanks | 12419 | 12442 | 24 | GATGTCGGCTCTTCCTATCATTGT | 186.4 | 3.9 | 0.9 | 4.4 | 1.0E-05 | 1.5E-05 | MB231/MB468 | MITO |
| >hg38_21_+_8433172_8446622_RNA45SN1_45S_50ntFlanks@3841.3862.22\|TCCTACTTGGATAACTGTGGTA | >hg38_21_+_8433172_8446622_RNA45SN1_45S_50ntFlanks | 3841 | 3862 | 22 | TCCTACTTGGATAACTGTGGTA | 327.8 | 3.8 | 0.8 | 4.9 | 9.3E-07 | 1.5E-05 | BT20/MB231 | CYTO |
| >hg38_21_+_8433172_8446622_RNA45SN1_45S_50ntFlanks@11874.11905.32\|AGAAGACCCTGTTGAGCTTGACTCTAGTCTGG | >hg38_21_+_8433172_8446622_RNA45SN1_45S_50ntFlanks | 11874 | 11905 | 32 | AGAAGACCCTGTTGAGCTTGACTCTAGTCTGG | 743.7 | -4.9 | 1.1 | -4.5 | 7.4E-06 | 1.5E-05 | BT20/MB468 | MITO |
| >hg38_21_+_8433172_8446622_RNA45SN1_45S_50ntFlanks@9546.9572.27\|TGGAGGTCCGTAGCGGTCCTGACGTGC | >hg38_21_+_8433172_8446622_RNA45SN1_45S_50ntFlanks | 9546 | 9572 | 27 | TGGAGGTCCGTAGCGGTCCTGACGTGC | 389.3 | 2.1 | 0.5 | 4.4 | 8.8E-06 | 1.5E-05 | BT20/MB468 | CYTO |
| >hg38_21_+_8433172_8446622_RNA45SN1_45S_50ntFlanks@3711.3737.27\|GTTGATCCTGCCAGTAGCATATGCTTG | >hg38_21_+_8433172_8446622_RNA45SN1_45S_50ntFlanks | 3711 | 3737 | 27 | GTTGATCCTGCCAGTAGCATATGCTTG | 597.6 | -4.3 | 1.0 | -4.4 | 9.8E-06 | 1.5E-05 | MB231/MB468 | CYTO |
| >MI0000085\|hsa-mir-27a&WithFlank&19\|-\|13836434\|13836523@57.76.20(+1U)&[MIMAT0000084&hsa-miR-27a-3p&offsets\|0\|-1(+1U);m-43&19\|-\|13836447\|13836467&offsets\|0\|-1(+1U)]\|TTCACAGTGGCTAAGTTCCGT | >MI0000085\|hsa-mir-27a&WithFlank&19\|-\|13836434\|13836523 | 57 | 76 | 20(+1U) | TTCACAGTGGCTAAGTTCCGT | 1937.7 | 4.3 | 1.0 | 4.4 | 9.8E-06 | 1.5E-05 | MB231/MB468 | CYTO |
| >hg38_21_+_8433172_8446622_RNA45SN1_45S_50ntFlanks@8475.8496.22\|CCCGGCGGATCTTTCCCGCCCC | >hg38_21_+_8433172_8446622_RNA45SN1_45S_50ntFlanks | 8475 | 8496 | 22 | CCCGGCGGATCTTTCCCGCCCC | 71.6 | -9.2 | 2.0 | -4.7 | 3.2E-06 | 1.5E-05 | BT20/MB468 | MP |
| >hg38_21_+_8433172_8446622_RNA45SN1_45S_50ntFlanks@10680.10705.26\|GTTCCGGCGGCGTCCGGTGAGCTCTC | >hg38_21_+_8433172_8446622_RNA45SN1_45S_50ntFlanks | 10680 | 10705 | 26 | GTTCCGGCGGCGTCCGGTGAGCTCTC | 1456.8 | -6.3 | 1.4 | -4.4 | 1.1E-05 | 1.5E-05 | BT20/MB468 | NUC |
| >hg38_21_+_8433172_8446622_RNA45SN1_45S_50ntFlanks@12071.12097.27\|CCCCGAGGGGCTCTCGCTTCTGGCGCC | >hg38_21_+_8433172_8446622_RNA45SN1_45S_50ntFlanks | 12071 | 12097 | 27 | CCCCGAGGGGCTCTCGCTTCTGGCGCC | 738.6 | -7.6 | 1.7 | -4.4 | 1.1E-05 | 1.5E-05 | BT20/MB468 | NUC |
| >hg38_21_+_8433172_8446622_RNA45SN1_45S_50ntFlanks@7984.8018.35\|AGATCAGACGTGGCGACCCGCTGAATTTAAGCATA | >hg38_21_+_8433172_8446622_RNA45SN1_45S_50ntFlanks | 7984 | 8018 | 35 | AGATCAGACGTGGCGACCCGCTGAATTTAAGCATA | 59.2 | -9.0 | 1.9 | -4.7 | 3.2E-06 | 1.5E-05 | BT20/MB468 | MP |
| >hg38_21_+_8433172_8446622_RNA45SN1_45S_50ntFlanks@3866.3898.33\|CTAGAGCTAATACATGCCGACGGGCGCTGACCC | >hg38_21_+_8433172_8446622_RNA45SN1_45S_50ntFlanks | 3866 | 3898 | 33 | CTAGAGCTAATACATGCCGACGGGCGCTGACCC | 326.6 | -6.1 | 1.4 | -4.4 | 8.8E-06 | 1.5E-05 | BT20/MB468 | CYTO |
| >am_MI0000086\|hsa-mir-28&WithFlank&3\|+\|188688775\|188688872@60.77.18(+1C)&[MIMAT0004502&hsa-miR-28-3p&offsets\|0\|-4(+1C);m-28&3\|+\|188688834\|188688855&offsets\|0\|-4(+1C)]\|CACTAGATTGTGAGCTCCC | >am_MI0000086\|hsa-mir-28&WithFlank&3\|+\|188688775\|188688872 | 60 | 77 | 18(+1C) | CACTAGATTGTGAGCTCCC | 43.4 | 8.7 | 1.9 | 4.4 | 8.6E-06 | 1.5E-05 | MB231/MB468 | MP |
| >MI0000061\|hsa-let-7a-2&WithFlank&11\|-\|122146516\|122146599@11.31.21(+1A);MI0000062\|hsa-let-7a-3&WithFlank&22\|+\|46112743\|46112828@10.30.21(+1A);MI0000060\|hsa-let-7a-1&WithFlank&9\|+\|94175951\|94176042@12.32.21(+1A)&[MIMAT0000062&hsa-let-7a-5p&offsets\|0\|-1(+1A);m-6&11\|-\|122146568\|122146589&offsets\|0\|-1(+1A)];[MIMAT0000062_1&hsa-let-7a-5p&offsets\|0\|-1(+1A);m-5&22\|+\|46112752\|46112773&offsets\|0\|-1(+1A)];[MIMAT0000062_2&hsa-let-7a-5p&offsets\|0\|-1(+1A);m-4&9\|+\|94175962\|94175983&offsets\|0\|-1(+1A)]\|TGAGGTAGTAGGTTGTATAGTA | >MI0000061\|hsa-let-7a-2&WithFlank&11\|-\|122146516\|122146599 | 11 | 31 | 21(+1A) | TGAGGTAGTAGGTTGTATAGTA | 678.7 | 2.5 | 0.6 | 4.4 | 1.1E-05 | 1.5E-05 | BT20/MB468 | NUC |
| >hg38_21_+_8433172_8446622_RNA45SN1_45S_50ntFlanks@11784.11803.20\|ATGAATGGATGAACGAGATT | >hg38_21_+_8433172_8446622_RNA45SN1_45S_50ntFlanks | 11784 | 11803 | 20 | ATGAATGGATGAACGAGATT | 1416.8 | -3.7 | 0.8 | -4.4 | 9.8E-06 | 1.5E-05 | MB231/MB468 | CYTO |
| >hg38_21_+_8433172_8446622_RNA45SN1_45S_50ntFlanks@10781.10801.21\|AGGTGAACAGCCTCTGGCATG | >hg38_21_+_8433172_8446622_RNA45SN1_45S_50ntFlanks | 10781 | 10801 | 21 | AGGTGAACAGCCTCTGGCATG | 287.5 | -3.0 | 0.7 | -4.5 | 7.5E-06 | 1.5E-05 | BT20/MB468 | MITO |
| >hg38_21_+_8433172_8446622_RNA45SN1_45S_50ntFlanks@11211.11232.22\|GCGACTCTGGACGCGAGCCGGG | >hg38_21_+_8433172_8446622_RNA45SN1_45S_50ntFlanks | 11211 | 11232 | 22 | GCGACTCTGGACGCGAGCCGGG | 1042.8 | -7.7 | 1.8 | -4.4 | 1.1E-05 | 1.5E-05 | BT20/MB468 | NUC |
| >hg38_21_+_8433172_8446622_RNA45SN1_45S_50ntFlanks@10443.10460.18\|CCGAAGGGACGGGCGATG | >hg38_21_+_8433172_8446622_RNA45SN1_45S_50ntFlanks | 10443 | 10460 | 18 | CCGAAGGGACGGGCGATG | 468.3 | -7.3 | 1.7 | -4.4 | 1.1E-05 | 1.5E-05 | BT20/MB468 | NUC |
| >hg38_21_+_8433172_8446622_RNA45SN1_45S_50ntFlanks@10768.10785.18\|CAGCAGGTCTCCAAGGTG | >hg38_21_+_8433172_8446622_RNA45SN1_45S_50ntFlanks | 10768 | 10785 | 18 | CAGCAGGTCTCCAAGGTG | 504.3 | -6.3 | 1.4 | -4.4 | 1.0E-05 | 1.5E-05 | MB231/MB468 | NUC |
| >MI0003513\|hsa-mir-455&WithFlank&9\|+\|114209428\|114209535@60.80.21&[MIMAT0004784&hsa-miR-455-3p&offsets\|0\|0;m-135&9\|+\|114209487\|114209508&offsets\|0\|-1]\|GCAGTCCATGGGCATATACAC | >MI0003513\|hsa-mir-455&WithFlank&9\|+\|114209428\|114209535 | 60 | 80 | 21 | GCAGTCCATGGGCATATACAC | 40.2 | 8.5 | 1.9 | 4.4 | 8.7E-06 | 1.5E-05 | MB231/MB468 | MP |
| >hg38_21_+_8433172_8446622_RNA45SN1_45S_50ntFlanks@7975.8004.30\|CGCGACCTCAGATCAGACGTGGCGACCCGC | >hg38_21_+_8433172_8446622_RNA45SN1_45S_50ntFlanks | 7975 | 8004 | 30 | CGCGACCTCAGATCAGACGTGGCGACCCGC | 9341.7 | 4.2 | 0.9 | 4.7 | 3.3E-06 | 1.5E-05 | BT20/MB468 | MP |
| >MI0000477\|hsa-mir-146a&WithFlank&5\|+\|160485346\|160485456@26.46.21(+1C)&[MIMAT0000449&hsa-miR-146a-5p&offsets\|-1\|-2(+1C);m-129&5\|+\|160485372\|160485393&offsets\|-1\|-2(+1C)]\|TTGAGAACTGAATTCCATGGGC | >MI0000477\|hsa-mir-146a&WithFlank&5\|+\|160485346\|160485456 | 26 | 46 | 21(+1C) | TTGAGAACTGAATTCCATGGGC | 41.1 | 8.6 | 1.9 | 4.4 | 8.7E-06 | 1.5E-05 | MB231/MB468 | MP |
| >MI0001733\|hsa-mir-452&WithFlank&X\|-\|151959622\|151959718@20.42.23&[MIMAT0001635&hsa-miR-452-5p&offsets\|0\|+1;m-188&X\|-\|151959677\|151959699&offsets\|0\|0]\|AACTGTTTGCAGAGGAAACTGAG | >MI0001733\|hsa-mir-452&WithFlank&X\|-\|151959622\|151959718 | 20 | 42 | 23 | AACTGTTTGCAGAGGAAACTGAG | 1084.9 | 4.6 | 0.9 | 4.9 | 9.4E-07 | 1.5E-05 | BT20/MB231 | CYTO |
| >hg38_21_+_8433172_8446622_RNA45SN1_45S_50ntFlanks@10905.10936.32\|CTGGGCGCGCGCCGCGGCTGGACGAGGCGCCG | >hg38_21_+_8433172_8446622_RNA45SN1_45S_50ntFlanks | 10905 | 10936 | 32 | CTGGGCGCGCGCCGCGGCTGGACGAGGCGCCG | 381.3 | -6.4 | 1.4 | -4.4 | 8.9E-06 | 1.5E-05 | BT20/MB468 | CYTO |
| >hg38_21_+_8433172_8446622_RNA45SN1_45S_50ntFlanks@3835.3859.25\|CTCCTCTCCTACTTGGATAACTGTG | >hg38_21_+_8433172_8446622_RNA45SN1_45S_50ntFlanks | 3835 | 3859 | 25 | CTCCTCTCCTACTTGGATAACTGTG | 196.9 | -2.9 | 0.7 | -4.4 | 8.9E-06 | 1.5E-05 | BT20/MB468 | CYTO |
| >hg38_21_+_8433172_8446622_RNA45SN1_45S_50ntFlanks@485.514.30\|GATGTGGTGACGTCGTGCTCTCCCGGGCCG | >hg38_21_+_8433172_8446622_RNA45SN1_45S_50ntFlanks | 485 | 514 | 30 | GATGTGGTGACGTCGTGCTCTCCCGGGCCG | 150.5 | 10.5 | 2.4 | 4.4 | 1.1E-05 | 1.5E-05 | MB231/MB468 | MITO |
| >hg38_21_+_8433172_8446622_RNA45SN1_45S_50ntFlanks@11729.11768.40\|TAACTATGACTCTCTTAAGGTAGCCAAATGCCTCGTCATC | >hg38_21_+_8433172_8446622_RNA45SN1_45S_50ntFlanks | 11729 | 11768 | 40 | TAACTATGACTCTCTTAAGGTAGCCAAATGCCTCGTCATC | 204.9 | -6.2 | 1.4 | -4.4 | 8.9E-06 | 1.5E-05 | BT20/MB468 | CYTO |
| >am_trnaMT_PheGAA_MT_+_577_647@41.59.19\|GTTTAGACGGGCTCACATC | >am_trnaMT_PheGAA_MT_+_577_647 | 41 | 59 | 19 | GTTTAGACGGGCTCACATC | 863.2 | 5.1 | 1.1 | 4.7 | 3.3E-06 | 1.5E-05 | BT20/MB468 | MP |
| >hg38_21_+_8433172_8446622_RNA45SN1_45S_50ntFlanks@10823.10865.43\|AAGTCGGCAAGCCGGATCCGTAACTTCGGGATAAGGATTGGCT | >hg38_21_+_8433172_8446622_RNA45SN1_45S_50ntFlanks | 10823 | 10865 | 43 | AAGTCGGCAAGCCGGATCCGTAACTTCGGGATAAGGATTGGCT | 243.2 | -5.7 | 1.3 | -4.5 | 7.6E-06 | 1.5E-05 | BT20/MB468 | MITO |
| >hg38_21_+_8433172_8446622_RNA45SN1_45S_50ntFlanks@10386.10424.39\|ATGTGAACAGCAGTTGAACATGGGTCAGTCGGTCCTGAG | >hg38_21_+_8433172_8446622_RNA45SN1_45S_50ntFlanks | 10386 | 10424 | 39 | ATGTGAACAGCAGTTGAACATGGGTCAGTCGGTCCTGAG | 3851.9 | 8.1 | 1.7 | 4.7 | 3.3E-06 | 1.5E-05 | BT20/MB468 | MP |
| >trna5_AspGTC_12_+_98897281_98897352@38.73.36\|CGCGGGAGACCGGGGTTCAATTCCCCGACGGGGAGC | >trna5_AspGTC_12_+_98897281_98897352 | 38 | 73 | 36 | CGCGGGAGACCGGGGTTCAATTCCCCGACGGGGAGC | 430.7 | 6.4 | 1.4 | 4.4 | 1.0E-05 | 1.5E-05 | MB231/MB468 | CYTO |
| >hg38_21_+_8433172_8446622_RNA45SN1_45S_50ntFlanks@10916.10936.21\|CCGCGGCTGGACGAGGCGCCG | >hg38_21_+_8433172_8446622_RNA45SN1_45S_50ntFlanks | 10916 | 10936 | 21 | CCGCGGCTGGACGAGGCGCCG | 375.4 | -6.1 | 1.4 | -4.4 | 8.9E-06 | 1.5E-05 | BT20/MB468 | CYTO |
| >hg38_MT_+_1621_3279_RNR2_16S_50ntFlanks@498.518.21\|TTGGACACTAGGAAAAAACCT | >hg38_MT_+_1621_3279_RNR2_16S_50ntFlanks | 498 | 518 | 21 | TTGGACACTAGGAAAAAACCT | 152.9 | 6.9 | 1.5 | 4.7 | 3.3E-06 | 1.5E-05 | BT20/MB468 | MP |
| >hg38_21_+_8433172_8446622_RNA45SN1_45S_50ntFlanks@12624.12656.33\|CCAATGGGGCGAAGCTACCATCTGTGGGATTAT | >hg38_21_+_8433172_8446622_RNA45SN1_45S_50ntFlanks | 12624 | 12656 | 33 | CCAATGGGGCGAAGCTACCATCTGTGGGATTAT | 152.3 | -6.2 | 1.4 | -4.4 | 8.8E-06 | 1.5E-05 | MB231/MB468 | MP |
| >hg38_21_+_8433172_8446622_RNA45SN1_45S_50ntFlanks@6646.6678.33\|TCGTACGACTCTTAGCGGTGGATCACTCGGCTC | >hg38_21_+_8433172_8446622_RNA45SN1_45S_50ntFlanks | 6646 | 6678 | 33 | TCGTACGACTCTTAGCGGTGGATCACTCGGCTC | 3006.0 | 3.9 | 0.8 | 4.7 | 3.3E-06 | 1.5E-05 | BT20/MB468 | MP |
| >hg38_21_+_8433172_8446622_RNA45SN1_45S_50ntFlanks@12893.12935.43\|CCTCGCCCGTCACGCACCGCACGTTCGTGGGGAACCTGGCGCT | >hg38_21_+_8433172_8446622_RNA45SN1_45S_50ntFlanks | 12893 | 12935 | 43 | CCTCGCCCGTCACGCACCGCACGTTCGTGGGGAACCTGGCGCT | 680.8 | -5.1 | 1.1 | -4.5 | 7.6E-06 | 1.5E-05 | BT20/MB468 | MITO |
| >hg38_21_+_8433172_8446622_RNA45SN1_45S_50ntFlanks@5421.5445.25\|CGATTGGATGGTTTAGTGAGGCCCT | >hg38_21_+_8433172_8446622_RNA45SN1_45S_50ntFlanks | 5421 | 5445 | 25 | CGATTGGATGGTTTAGTGAGGCCCT | 504.8 | -6.5 | 1.4 | -4.5 | 7.6E-06 | 1.5E-05 | BT20/MB468 | MITO |
| >hg38_21_+_8433172_8446622_RNA45SN1_45S_50ntFlanks@3843.3869.27\|CTACTTGGATAACTGTGGTAATTCTAG | >hg38_21_+_8433172_8446622_RNA45SN1_45S_50ntFlanks | 3843 | 3869 | 27 | CTACTTGGATAACTGTGGTAATTCTAG | 184.7 | -6.8 | 1.5 | -4.7 | 3.3E-06 | 1.5E-05 | BT20/MB468 | MP |
| >hg38_21_+_8433172_8446622_RNA45SN1_45S_50ntFlanks@11577.11594.18\|TTAGAACTGGTGCGGACC | >hg38_21_+_8433172_8446622_RNA45SN1_45S_50ntFlanks | 11577 | 11594 | 18 | TTAGAACTGGTGCGGACC | 171.9 | -4.9 | 1.1 | -4.4 | 8.8E-06 | 1.5E-05 | MB231/MB468 | MP |
| >hg38_21_+_8433172_8446622_RNA45SN1_45S_50ntFlanks@11641.11680.40\|CGGCGGGTGTTGACGCGATGTGATTTCTGCCCAGTGCTCT | >hg38_21_+_8433172_8446622_RNA45SN1_45S_50ntFlanks | 11641 | 11680 | 40 | CGGCGGGTGTTGACGCGATGTGATTTCTGCCCAGTGCTCT | 1302.1 | -4.0 | 0.9 | -4.5 | 7.6E-06 | 1.5E-05 | BT20/MB468 | MITO |
| >hg38_21_+_8433172_8446622_RNA45SN1_45S_50ntFlanks@11776.11794.19\|TGACGCGCATGAATGGATG | >hg38_21_+_8433172_8446622_RNA45SN1_45S_50ntFlanks | 11776 | 11794 | 19 | TGACGCGCATGAATGGATG | 472.4 | -5.6 | 1.3 | -4.4 | 1.1E-05 | 1.5E-05 | BT20/MB468 | NUC |
| >MI0000272\|hsa-mir-182&WithFlank&7\|-\|129770377\|129770498@29.54.26&[MIMAT0000259&hsa-miR-182-5p&offsets\|0\|+2;m-22&7\|-\|129770449\|129770470&offsets\|0\|+4]\|TTTGGCAATGGTAGAACTCACACTGG | >MI0000272\|hsa-mir-182&WithFlank&7\|-\|129770377\|129770498 | 29 | 54 | 26 | TTTGGCAATGGTAGAACTCACACTGG | 861.1 | 4.1 | 0.9 | 4.4 | 1.1E-05 | 1.5E-05 | MB231/MB468 | MITO |
| >hg38_21_+_8433172_8446622_RNA45SN1_45S_50ntFlanks@7991.8017.27\|ACGTGGCGACCCGCTGAATTTAAGCAT | >hg38_21_+_8433172_8446622_RNA45SN1_45S_50ntFlanks | 7991 | 8017 | 27 | ACGTGGCGACCCGCTGAATTTAAGCAT | 201.0 | 2.8 | 0.6 | 4.4 | 1.0E-05 | 1.5E-05 | MB231/MB468 | CYTO |
| >trnaMT_PheGAA_MT_+_577_647@1.34.34\|GTTTATGTAGCTTACCTCCTCAAAGCAATACACT | >trnaMT_PheGAA_MT_+_577_647 | 1 | 34 | 34 | GTTTATGTAGCTTACCTCCTCAAAGCAATACACT | 2059.5 | 7.1 | 1.6 | 4.5 | 7.7E-06 | 1.5E-05 | BT20/MB468 | MITO |

| | | | | | | | | | | | | | |
|---|---|---|---|---|---|---|---|---|---|---|---|---|---|
| >MI0000087\|hsa-mir-29a&WithFlank&7\|-\|130876741\|130876816@48.69.22(+1G)&[MIMAT0000086&hsa-miR-29a-3p&offsets\|0\|0(+1G);m-26&7\|-\|130876749\|130876769&offsets\|0\|+1(+1G)]\|TAGCACCATCTGAAATCGGTTAG | >MI0000087\|hsa-mir-29a&WithFlank&7\|-\|130876741\|130876816 | 48 | 69 | 22(+1G) | TAGCACCATCTGAAATCGGTTAG | 35.0 | 8.3 | 1.9 | 4.4 | 8.8E-06 | 1.5E-05 | MB231/MB468 | MP |
| >hg38_21_+_8433172_8446622_RNA45SN1_45S_50ntFlanks@11873.11904.32\|AAGAAGACCCTGTTGAGCTTGACTCTAGTCTG | >hg38_21_+_8433172_8446622_RNA45SN1_45S_50ntFlanks | 11873 | 11904 | 32 | AAGAAGACCCTGTTGAGCTTGACTCTAGTCTG | 544.9 | -6.0 | 1.4 | -4.5 | 7.7E-06 | 1.5E-05 | BT20/MB468 | MITO |
| >hg38_21_+_8433172_8446622_RNA45SN1_45S_50ntFlanks@11211.11250.40\|GCGACTCTGGACGCGAGCCGGGCCCTTCCCGTGGATCGCC | >hg38_21_+_8433172_8446622_RNA45SN1_45S_50ntFlanks | 11211 | 11250 | 40 | GCGACTCTGGACGCGAGCCGGGCCCTTCCCGTGGATCGCC | 231.0 | 4.2 | 0.9 | 4.9 | 1.1E-06 | 1.5E-05 | BT20/MB231 | MITO |
| >hg38_21_+_8433172_8446622_RNA45SN1_45S_50ntFlanks@5471.5499.29\|CACGGCCCTGGCGGAGCGCTGAGAAGACG | >hg38_21_+_8433172_8446622_RNA45SN1_45S_50ntFlanks | 5471 | 5499 | 29 | CACGGCCCTGGCGGAGCGCTGAGAAGACG | 2739.3 | -5.0 | 1.1 | -4.6 | 3.3E-06 | 1.5E-05 | BT20/MB468 | MP |
| >MI0001448\|hsa-mir-425&WithFlank&3\|-\|49020142\|49020240@20.44.25&[MIMAT0003393&hsa-miR-425-5p&offsets\|0\|+2;m-95&3\|-\|49020200\|49020221&offsets\|0\|+3]\|AATGACACGATCACTCCCGTTGAGT | >MI0001448\|hsa-mir-425&WithFlank&3\|-\|49020142\|49020240 | 20 | 44 | 25 | AATGACACGATCACTCCCGTTGAGT | 1041.4 | 2.3 | 0.5 | 4.4 | 1.1E-05 | 1.5E-05 | BT20/MB468 | NUC |
| >hg38_21_+_8433172_8446622_RNA45SN1_45S_50ntFlanks@482.504.23\|ATCGATGTGGTGACGTCGTGCTC | >hg38_21_+_8433172_8446622_RNA45SN1_45S_50ntFlanks | 482 | 504 | 23 | ATCGATGTGGTGACGTCGTGCTC | 66.0 | -9.1 | 2.0 | -4.6 | 3.3E-06 | 1.5E-05 | BT20/MB468 | MP |
| >hg38_21_+_8433172_8446622_RNA45SN1_45S_50ntFlanks@10348.10380.33\|AAACGAGAACTTTGAAGGCCGAAGTGGAGAAGG | >hg38_21_+_8433172_8446622_RNA45SN1_45S_50ntFlanks | 10348 | 10380 | 33 | AAACGAGAACTTTGAAGGCCGAAGTGGAGAAGG | 184.7 | 2.8 | 0.6 | 4.4 | 1.0E-05 | 1.5E-05 | MB231/MB468 | CYTO |
| >MI0000298\|hsa-mir-221&WithFlank&X\|-\|45746151\|45746272@71.94.24&[MIMAT0000278&hsa-miR-221-3p&offsets\|0\|+1;m-91&X\|-\|45746181\|45746202&offsets\|0\|+2]\|AGCTACATTGTCTGCTGGGTTTCA | >MI0000298\|hsa-mir-221&WithFlank&X\|-\|45746151\|45746272 | 71 | 94 | 24 | AGCTACATTGTCTGCTGGGTTTCA | 3133.1 | -4.3 | 0.9 | -4.9 | 9.8E-07 | 1.5E-05 | BT20/MB231 | MP |
| >hg38_21_+_8433172_8446622_RNA45SN1_45S_50ntFlanks@10787.10816.30\|ACAGCCTCTGGCATGTTGGAACAATGTAGG | >hg38_21_+_8433172_8446622_RNA45SN1_45S_50ntFlanks | 10787 | 10816 | 30 | ACAGCCTCTGGCATGTTGGAACAATGTAGG | 106.2 | 10.1 | 2.1 | 4.9 | 9.7E-07 | 1.5E-05 | BT20/MB231 | MP |
| >MI0000085\|hsa-mir-27a&WithFlank&19\|-\|13836434\|13836523@57.76.20(+1U)&[MIMAT0000084&hsa-miR-27a-3p&offsets\|0\|-1(+1U);m-43&19\|-\|13836447\|13836467&offsets\|0\|-1(+1U)]\|TTCACAGTGGCTAAGTTCCGT | >MI0000085\|hsa-mir-27a&WithFlank&19\|-\|13836434\|13836523 | 57 | 76 | 20(+1U) | TTCACAGTGGCTAAGTTCCGT | 7742.4 | 4.3 | 1.0 | 4.4 | 1.1E-05 | 1.5E-05 | MB231/MB468 | MITO |
| >hg38_21_+_8433172_8446622_RNA45SN1_45S_50ntFlanks@8380.8407.28\|CAAGAGGGCGTGAAACCGTTAAGAGGTA | >hg38_21_+_8433172_8446622_RNA45SN1_45S_50ntFlanks | 8380 | 8407 | 28 | CAAGAGGGCGTGAAACCGTTAAGAGGTA | 337.3 | -2.9 | 0.6 | -4.5 | 7.7E-06 | 1.5E-05 | BT20/MB468 | MITO |
| >hg38_21_+_8433172_8446622_RNA45SN1_45S_50ntFlanks@12822.12848.27\|CCGGGACCGGGGTCCGGTGCGGAGTGC | >hg38_21_+_8433172_8446622_RNA45SN1_45S_50ntFlanks | 12822 | 12848 | 27 | CCGGGACCGGGGTCCGGTGCGGAGTGC | 104.9 | -10.6 | 2.4 | -4.4 | 8.9E-06 | 1.5E-05 | MB231/MB468 | MP |
| >hg38_21_+_8433172_8446622_RNA45SN1_45S_50ntFlanks@9605.9623.19\|AAGACTAATCGAACCATCT | >hg38_21_+_8433172_8446622_RNA45SN1_45S_50ntFlanks | 9605 | 9623 | 19 | AAGACTAATCGAACCATCT | 172.7 | -6.5 | 1.5 | -4.4 | 8.9E-06 | 1.5E-05 | MB231/MB468 | MP |
| >hg38_21_+_8433172_8446622_RNA45SN1_45S_50ntFlanks@7975.8012.38\|CGCGACCTCAGATCAGACGTGGCGACCCGCTGAATTTA | >hg38_21_+_8433172_8446622_RNA45SN1_45S_50ntFlanks | 7975 | 8012 | 38 | CGCGACCTCAGATCAGACGTGGCGACCCGCTGAATTTA | 28451.9 | -2.1 | 0.5 | -4.4 | 1.1E-05 | 1.6E-05 | BT20/MB468 | NUC |
| >hg38_MT_+_598_1651_RNR1_12S_50ntFlanks@442.480.39\|ATATCTGAACACACAATAGCTAAGACCCAAACTGGGATT | >hg38_MT_+_598_1651_RNR1_12S_50ntFlanks | 442 | 480 | 39 | ATATCTGAACACACAATAGCTAAGACCCAAACTGGGATT | 478.6 | 5.5 | 1.2 | 4.4 | 1.1E-05 | 1.6E-05 | MB231/MB468 | MITO |
| >hg38_21_+_8433172_8446622_RNA45SN1_45S_50ntFlanks@4000.4042.43\|TAGATAACCTCGGGCCGATCGCACGCCCCCCGTGGCGGCGACG | >hg38_21_+_8433172_8446622_RNA45SN1_45S_50ntFlanks | 4000 | 4042 | 43 | TAGATAACCTCGGGCCGATCGCACGCCCCCCGTGGCGGCGACG | 431.7 | -5.2 | 1.2 | -4.4 | 1.1E-05 | 1.6E-05 | MB231/MB468 | NUC |
| >MI0000542\|hsa-mir-320a&WithFlank&8\|-\|22244960\|22245043@48.70.23(+2U)&[MIMAT0000510&hsa-miR-320a-3p&offsets\|0\|+1(+2U);m-60&8\|-\|22244975\|22244996&offsets\|0\|+1(+2U)]\|AAAAGCTGGGTTGAGAGGGCGAATT | >MI0000542\|hsa-mir-320a&WithFlank&8\|-\|22244960\|22245043 | 48 | 70 | 23(+2U) | AAAAGCTGGGTTGAGAGGGCGAATT | 582.7 | 6.3 | 1.4 | 4.6 | 3.4E-06 | 1.6E-05 | BT20/MB468 | MP |
| >hg38_21_+_8433172_8446622_RNA45SN1_45S_50ntFlanks@10805.10822.18\|GAACAATGTAGGTAAGGG | >hg38_21_+_8433172_8446622_RNA45SN1_45S_50ntFlanks | 10805 | 10822 | 18 | GAACAATGTAGGTAAGGG | 287.3 | -5.7 | 1.3 | -4.4 | 9.1E-06 | 1.6E-05 | BT20/MB468 | CYTO |
| >hg38_21_+_8433172_8446622_RNA45SN1_45S_50ntFlanks@11666.11685.20\|TCTGCCCAGTGCTCTGAATG | >hg38_21_+_8433172_8446622_RNA45SN1_45S_50ntFlanks | 11666 | 11685 | 20 | TCTGCCCAGTGCTCTGAATG | 3697.6 | -3.3 | 0.7 | -4.4 | 1.0E-05 | 1.6E-05 | MB231/MB468 | CYTO |
| >hg38_21_+_8433172_8446622_RNA45SN1_45S_50ntFlanks@8189.8209.21\|CCCAGCCCGTGGACGGTGTGA | >hg38_21_+_8433172_8446622_RNA45SN1_45S_50ntFlanks | 8189 | 8209 | 21 | CCCAGCCCGTGGACGGTGTGA | 97.4 | -6.1 | 1.4 | -4.4 | 9.0E-06 | 1.6E-05 | MB231/MB468 | MP |
| >hg38_21_+_8433172_8446622_RNA45SN1_45S_50ntFlanks@4241.4264.24\|CTCTTTCGAGGCCCTGTAATTGGA | >hg38_21_+_8433172_8446622_RNA45SN1_45S_50ntFlanks | 4241 | 4264 | 24 | CTCTTTCGAGGCCCTGTAATTGGA | 53.1 | -8.8 | 1.9 | -4.6 | 3.4E-06 | 1.6E-05 | BT20/MB468 | MP |
| >hg38_21_+_8433172_8446622_RNA45SN1_45S_50ntFlanks@12244.12272.29\|AAGGGCAAAAGCTCGCTTGATCTTGATTT | >hg38_21_+_8433172_8446622_RNA45SN1_45S_50ntFlanks | 12244 | 12272 | 29 | AAGGGCAAAAGCTCGCTTGATCTTGATTT | 521.4 | -5.6 | 1.3 | -4.4 | 1.1E-05 | 1.6E-05 | BT20/MB468 | NUC |
| >hg38_21_+_8433172_8446622_RNA45SN1_45S_50ntFlanks@6696.6720.25\|GCAGCTAGCTGCGAGAATTAATGTG | >hg38_21_+_8433172_8446622_RNA45SN1_45S_50ntFlanks | 6696 | 6720 | 25 | GCAGCTAGCTGCGAGAATTAATGTG | 406.2 | 3.5 | 0.8 | 4.4 | 1.0E-05 | 1.6E-05 | MB231/MB468 | CYTO |
| >hg38_21_+_8433172_8446622_RNA45SN1_45S_50ntFlanks@12072.12094.23\|CCCGAGGGGCTCTCGCTTCTGGC | >hg38_21_+_8433172_8446622_RNA45SN1_45S_50ntFlanks | 12072 | 12094 | 23 | CCCGAGGGGCTCTCGCTTCTGGC | 375.1 | -2.7 | 0.6 | -4.4 | 9.1E-06 | 1.6E-05 | BT20/MB468 | CYTO |
| >MI0000440\|hsa-mir-27b&WithFlank&9\|+\|95085439\|95085547@67.88.22(+1U)&[MIMAT0000419&hsa-miR-27b-3p&offsets\|0\|+1(+1U);m-39&9\|+\|95085505\|95085525&offsets\|0\|+1(+1U)]\|TTCACAGTGGCTAAGTTCTGCAT | >MI0000440\|hsa-mir-27b&WithFlank&9\|+\|95085439\|95085547 | 67 | 88 | 22(+1U) | TTCACAGTGGCTAAGTTCTGCAT | 348.6 | 6.5 | 1.5 | 4.4 | 1.1E-05 | 1.6E-05 | MB231/MB468 | MITO |
| >hg38_21_+_8433172_8446622_RNA45SN1_45S_50ntFlanks@8144.8175.32\|CTCCCCGGCGCCGCTCGTGGGGGGCCCAAGTC | >hg38_21_+_8433172_8446622_RNA45SN1_45S_50ntFlanks | 8144 | 8175 | 32 | CTCCCCGGCGCCGCTCGTGGGGGGCCCAAGTC | 67.4 | -9.2 | 2.0 | -4.6 | 3.4E-06 | 1.6E-05 | BT20/MB468 | MP |
| >hg38_21_+_8433172_8446622_RNA45SN1_45S_50ntFlanks@11027.11053.27\|CTCTCTCTCTCTCTCCCCCGCTCCCCG | >hg38_21_+_8433172_8446622_RNA45SN1_45S_50ntFlanks | 11027 | 11053 | 27 | CTCTCTCTCTCTCTCCCCCGCTCCCCG | 1578.8 | -6.1 | 1.4 | -4.4 | 1.1E-05 | 1.6E-05 | MB231/MB468 | MITO |
| >trna111_HisGTG_1_-_147774845_147774916@-1A.22.23,trna118_HisGTG_1_-_145396881_145396952@-1A.22.23,trna16_HisGTG_1_+_146544773_146544844@-1A.22.23,trna1_HisGTG_15_+_45493349_45493420@-1A.22.23,trna21_HisGTG_1_+_147753471_147753542@-1A.22.23,trna33_HisGTG_6_+_27125906_27125977@-1A.22.23,trna7_HisGTG_9_-_14433938_14434009@-1A.22.23,trna8_HisGTG_15_-_45492611_45492682@-1A.22.23,trna9_HisGTG_15_-_45490804_45490875@-1A.22.23\|AGCCGTGATCGTATAGTGGTTAG | >trna111_HisGTG_1_-_147774845_147774916 | -1A | 22 | 23 | AGCCGTGATCGTATAGTGGTTAG | 38.9 | 8.5 | 1.9 | 4.4 | 9.0E-06 | 1.6E-05 | MB231/MB468 | MP |
| >hg38_21_+_8433172_8446622_RNA45SN1_45S_50ntFlanks@8196.8217.22\|CGTGGACGGTGTGAGGCCGGTA | >hg38_21_+_8433172_8446622_RNA45SN1_45S_50ntFlanks | 8196 | 8217 | 22 | CGTGGACGGTGTGAGGCCGGTA | 487.6 | -3.2 | 0.7 | -4.4 | 9.0E-06 | 1.6E-05 | MB231/MB468 | MP |
| >hg38_21_+_8433172_8446622_RNA45SN1_45S_50ntFlanks@11728.11747.20\|GTAACTATGACTCTCTTAAG | >hg38_21_+_8433172_8446622_RNA45SN1_45S_50ntFlanks | 11728 | 11747 | 20 | GTAACTATGACTCTCTTAAG | 915.9 | -7.8 | 1.8 | -4.4 | 1.1E-05 | 1.6E-05 | BT20/MB468 | NUC |
| >hg38_21_+_8433172_8446622_RNA45SN1_45S_50ntFlanks@7989.8008.20\|AGACGTGGCGACCCGCTGAA | >hg38_21_+_8433172_8446622_RNA45SN1_45S_50ntFlanks | 7989 | 8008 | 20 | AGACGTGGCGACCCGCTGAA | 24780.8 | -4.3 | 0.9 | -4.6 | 3.4E-06 | 1.6E-05 | BT20/MB468 | MP |
| >MI0000434\|hsa-let-7i&WithFlank&12\|+\|62603680\|62603775@12.33.22(+2U)&[MIMAT0000415&hsa-let-7i-5p&offsets\|0\|0(+2U);m-49&12\|+\|62603691\|62603712&offsets\|0\|0(+2U)]\|TGAGGTAGTAGTTTGTGCTGTTTT | >MI0000434\|hsa-let-7i&WithFlank&12\|+\|62603680\|62603775 | 12 | 33 | 22(+2U) | TGAGGTAGTAGTTTGTGCTGTTTT | 283.0 | 5.3 | 1.2 | 4.4 | 9.0E-06 | 1.6E-05 | MB231/MB468 | MP |
| >MI0000108\|hsa-mir-103a-2&WithFlank&20\|+\|3917488\|3917577@54.76.23(+1U);MI0000109\|hsa-mir-103a-1&WithFlank&5\|-\|168560890\|168560979@54.76.23(+1U)&[MIMAT0000101&hsa-miR-103a-3p&offsets\|0\|0(+1U);m-12&20\|+\|3917541\|3917563&offsets\|0\|0(+1U)];[MIMAT0000101_1&hsa-miR-103a-3p&offsets\|0\|0(+1U);m-13&5\|-\|168560904\|168560926&offsets\|0\|0(+1U);m-44220&5\|-\|168560908\|168560926&offsets\|0\|+4(+1U)]\|AGCAGCATTGTACAGGGCTATGAT | >MI0000108\|hsa-mir-103a-2&WithFlank&20\|+\|3917488\|3917577 | 54 | 76 | 23(+1U) | AGCAGCATTGTACAGGGCTATGAT | 644.6 | 2.1 | 0.5 | 4.4 | 1.1E-05 | 1.6E-05 | BT20/MB468 | NUC |
| >trna119_LysCTT_1_-_145395522_145395594@2.36.35,trna11_LysCTT_5_-_180648979_180649051@2.36.35,trna13_LysCTT_6_+_26556774_26556846@2.36.35,trna7_LysCTT_16_+_3225692_3225764@2.36.35,trna9_LysCTT_5_+_180634755_180634827@2.36.35\|CCCGGCTAGCTCAGTCGGTAGAGCATGAGACTCTT | >trna119_LysCTT_1_-_145395522_145395594 | 2 | 36 | 35 | CCCGGCTAGCTCAGTCGGTAGAGCATGAGACTCTT | 35.5 | 8.4 | 1.9 | 4.4 | 9.0E-06 | 1.6E-05 | MB231/MB468 | MP |
| >MI0000063\|hsa-let-7b&WithFlank&22\|+\|46113680\|46113774@12.32.21&[MIMAT0000063&hsa-let-7b-5p&offsets\|0\|-1;m-14&22\|+\|46113691\|46113712&offsets\|0\|-1]\|TGAGGTAGTAGGTTGTGTGGT | >MI0000063\|hsa-let-7b&WithFlank&22\|+\|46113680\|46113774 | 12 | 32 | 21 | TGAGGTAGTAGGTTGTGTGGT | 1609.3 | 3.0 | 0.7 | 4.4 | 1.1E-05 | 1.6E-05 | MB231/MB468 | NUC |
| >hg38_21_+_8433172_8446622_RNA45SN1_45S_50ntFlanks@8359.8392.34\|AAGAACTTTGAAGAGAGAGTTCAAGAGGGCGTGA | >hg38_21_+_8433172_8446622_RNA45SN1_45S_50ntFlanks | 8359 | 8392 | 34 | AAGAACTTTGAAGAGAGAGTTCAAGAGGGCGTGA | 41.5 | 9.1 | 2.0 | 4.6 | 3.4E-06 | 1.6E-05 | BT20/MB468 | MP |
| >hg38_21_+_8433172_8446622_RNA45SN1_45S_50ntFlanks@11577.11603.27\|TTAGAACTGGTGCGGACCAGGGGAATC | >hg38_21_+_8433172_8446622_RNA45SN1_45S_50ntFlanks | 11577 | 11603 | 27 | TTAGAACTGGTGCGGACCAGGGGAATC | 991.7 | -6.0 | 1.3 | -4.6 | 3.4E-06 | 1.6E-05 | BT20/MB468 | MP |
| >hg38_21_+_8433172_8446622_RNA45SN1_45S_50ntFlanks@10771.10805.35\|CAGGTCTCCAAGGTGAACAGCCTCTGGCATGTTGG | >hg38_21_+_8433172_8446622_RNA45SN1_45S_50ntFlanks | 10771 | 10805 | 35 | CAGGTCTCCAAGGTGAACAGCCTCTGGCATGTTGG | 693.0 | -6.3 | 1.4 | -4.4 | 9.2E-06 | 1.6E-05 | BT20/MB468 | CYTO |
| >trna116_GluCTC_1_-_145399233_145399304@32.69.38,trna59_GluCTC_1_+_249168447_249168518@32.69.38,trna71_GluCTC_1_-_161439189_161439260@32.69.38,trna74_GluCTC_1_-_161431809_161431880@32.69.38,trna77_GluCTC_1_-_161424398_161424469@32.69.38,trna77_GluCTC_6_+_28949976_28950047@32.69.38,trna80_GluCTC_1_-_161417018_161417089@32.69.38,trna87_GluCTC_6_-_126101393_126101464@32.69.38\|CTCTCACCGCCGCGGCCCGGGTTCGATTCCCGGTCAGG | >trna116_GluCTC_1_-_145399233_145399304 | 32 | 69 | 38 | CTCTCACCGCCGCGGCCCGGGTTCGATTCCCGGTCAGG | 1619.8 | -8.0 | 1.8 | -4.4 | 1.1E-05 | 1.6E-05 | BT20/MB468 | NUC |
| >am_trna119_LysCTT_1_-_145395522_145395594@6.35.30,trna11_LysCTT_5_-_180648979_180649051@6.35.30,trna128_LysCTT_1_-_55423542_55423614@6.35.30,trna13_LysCTT_6_+_26556774_26556846@6.35.30,trna6_LysCTT_19_-_52425396_52425468@6.35.30,trna7_LysCTT_16_+_3225692_3225764@6.35.30,trna9_LysCTT_5_+_180634755_180634827@6.35.30\|GCTAGCTCAGTCGGTAGAGCATGAGACTCT | >am_trna119_LysCTT_1_-_145395522_145395594 | 6 | 35 | 30 | GCTAGCTCAGTCGGTAGAGCATGAGACTCT | 249.8 | -4.8 | 1.0 | -4.9 | 1.2E-06 | 1.6E-05 | BT20/MB231 | MITO |
| >hg38_21_+_8433172_8446622_RNA45SN1_45S_50ntFlanks@11598.11623.26\|GGAATCCGACTGTTTAATTAAAACAA | >hg38_21_+_8433172_8446622_RNA45SN1_45S_50ntFlanks | 11598 | 11623 | 26 | GGAATCCGACTGTTTAATTAAAACAA | 209.7 | 3.8 | 0.8 | 4.9 | 1.2E-06 | 1.6E-05 | BT20/MB231 | MITO |

| | | | | | | | | | | | | | |
|---|---|---|---|---|---|---|---|---|---|---|---|---|---|
| >hg38_21_+_8433172_8446622_RNA45SN1_45S_50ntFlanks@6661.6708.48\|CGGTGGATCACTCGGCTCGTGCGTCGATGAAGAACGCAGCTAGCTGCG | >hg38_21_+_8433172_8446622_RNA45SN1_45S_50ntFlanks | 6661 | 6708 | 48 | CGGTGGATCACTCGGCTCGTGCGTCGATGAAGAACGCAGCTAGCTGCG | 9479.1 | 4.9 | 1.0 | 4.9 | 1.2E-06 | 1.6E-05 | BT20/MB231 | MITO |
| >hg38_21_+_8433172_8446622_RNA45SN1_45S_50ntFlanks@6648.6665.18\|GTACGACTCTTAGCGGTG | >hg38_21_+_8433172_8446622_RNA45SN1_45S_50ntFlanks | 6648 | 6665 | 18 | GTACGACTCTTAGCGGTG | 333.2 | 3.5 | 0.8 | 4.4 | 1.1E-05 | 1.6E-05 | MB231/MB468 | NUC |
| >hg38_21_+_8433172_8446622_RNA45SN1_45S_50ntFlanks@10841.10866.26\|CGTAACTTCGGGATAAGGATTGGCTC | >hg38_21_+_8433172_8446622_RNA45SN1_45S_50ntFlanks | 10841 | 10866 | 26 | CGTAACTTCGGGATAAGGATTGGCTC | 1499.9 | -2.8 | 0.6 | -4.4 | 9.2E-06 | 1.6E-05 | BT20/MB468 | CYTO |
| >hg38_21_+_8433172_8446622_RNA45SN1_45S_50ntFlanks@4040.4063.24\|ACGACCCATTCGAACGTCTGCCCT | >hg38_21_+_8433172_8446622_RNA45SN1_45S_50ntFlanks | 4040 | 4063 | 24 | ACGACCCATTCGAACGTCTGCCCT | 142.4 | -2.3 | 0.5 | -4.4 | 9.2E-06 | 1.6E-05 | BT20/MB468 | CYTO |
| >MI0003776\|hsa-mir-320b-1&WithFlank&1\|+\|116671740\|116671823@48.67.20(+1U);MI0003839\|hsa-mir-320b-2&WithFlank&1\|-\|224257034\|224257116@47.66.20(+1U);MI0000542\|hsa-mir-320a&WithFlank&8\|-\|22244960\|22245043@48.67.20(+1U)&[MIMAT0005792&hsa-miR-320b&offsets\|0\|-2(+1U);m-183&1\|+\|116671787\|116671806&offsets\|0\|0(+1U)];[MIMAT0005792_1&hsa-miR-320b&offsets\|0\|-2(+1U);m-183&1\|-\|224257051\|224257070&offsets\|0\|0(+1U)];[MIMAT0000510&hsa-miR-320a-3p&offsets\|0\|-2(+1U);m-60&8\|-\|22244975\|22244996&offsets\|0\|-2(+1U)]\|AAAAGCTGGGTTGAGAGGGCT | >MI0003776\|hsa-mir-320b-1&WithFlank&1\|+\|116671740\|116671823 | 48 | 67 | 20(+1U) | AAAAGCTGGGTTGAGAGGGCT | 1197.0 | 4.0 | 0.8 | 4.9 | 9.9E-07 | 1.6E-05 | BT20/MB231 | CYTO |
| >hg38_21_+_8433172_8446622_RNA45SN1_45S_50ntFlanks@8105.8139.35\|CGCGGCGGGGCGCGGGACATGTGGCGTACGGAAGA | >hg38_21_+_8433172_8446622_RNA45SN1_45S_50ntFlanks | 8105 | 8139 | 35 | CGCGGCGGGGCGCGGGACATGTGGCGTACGGAAGA | 171.5 | -5.8 | 1.3 | -4.4 | 1.1E-05 | 1.6E-05 | MB231/MB468 | NUC |
| >MI0005559\|hsa-mir-744&WithFlank&17\|+\|12081893\|12082002@17.38.22&[MIMAT0004945&hsa-miR-744-5p&offsets\|0\|0;m-172&17\|+\|12081909\|12081929&offsets\|0\|+1]\|TGCGGGGCTAGGGCTAACAGCA | >MI0005559\|hsa-mir-744&WithFlank&17\|+\|12081893\|12082002 | 17 | 38 | 22 | TGCGGGGCTAGGGCTAACAGCA | 242.6 | 4.0 | 0.9 | 4.4 | 1.1E-05 | 1.6E-05 | MB231/MB468 | MITO |
| >hg38_21_+_8433172_8446622_RNA45SN1_45S_50ntFlanks@12300.12325.26\|GGGCCTCACGATCCTTCTGACCTTTT | >hg38_21_+_8433172_8446622_RNA45SN1_45S_50ntFlanks | 12300 | 12325 | 26 | GGGCCTCACGATCCTTCTGACCTTTT | 357.3 | -5.3 | 1.2 | -4.4 | 1.1E-05 | 1.6E-05 | MB231/MB468 | MITO |
| >hg38_21_+_8433172_8446622_RNA45SN1_45S_50ntFlanks@6730.6756.27\|ACACATTGATCATCGACACTTCGAACG | >hg38_21_+_8433172_8446622_RNA45SN1_45S_50ntFlanks | 6730 | 6756 | 27 | ACACATTGATCATCGACACTTCGAACG | 185.9 | -6.3 | 1.4 | -4.4 | 9.1E-06 | 1.6E-05 | MB231/MB468 | MP |
| >trnaMT_PheGAA_MT_+_577_647@38.57.20\|AATGTTTAGACGGGCTCACA | >trnaMT_PheGAA_MT_+_577_647 | 38 | 57 | 20 | AATGTTTAGACGGGCTCACA | 213.4 | 3.9 | 0.9 | 4.4 | 1.1E-05 | 1.6E-05 | MB231/MB468 | MITO |
| >hg38_21_+_8433172_8446622_RNA45SN1_45S_50ntFlanks@11564.11584.21\|CCTAGCAGCCGACTTAGAACT | >hg38_21_+_8433172_8446622_RNA45SN1_45S_50ntFlanks | 11564 | 11584 | 21 | CCTAGCAGCCGACTTAGAACT | 440.5 | -3.1 | 0.7 | -4.4 | 1.1E-05 | 1.6E-05 | MB231/MB468 | MITO |
| >hg38_21_+_8433172_8446622_RNA45SN1_45S_50ntFlanks@10411.10443.33\|CAGTCGGTCCTGAGAGATGGGCGAGCGCCGTTC | >hg38_21_+_8433172_8446622_RNA45SN1_45S_50ntFlanks | 10411 | 10443 | 33 | CAGTCGGTCCTGAGAGATGGGCGAGCGCCGTTC | 4198.7 | -5.3 | 1.1 | -4.6 | 3.4E-06 | 1.6E-05 | BT20/MB468 | MP |
| >MI0000746\|hsa-mir-99b&WithFlank&19\|+\|51692606\|51692687@13.34.22(+1C)&[MIMAT0000689&hsa-miR-99b-5p&offsets\|0\|0(+1C);m-10&19\|+\|51692618\|51692639&offsets\|0\|0(+1C)]\|CACCCGTAGAACCGACCTTGCGC | >MI0000746\|hsa-mir-99b&WithFlank&19\|+\|51692606\|51692687 | 13 | 34 | 22(+1C) | CACCCGTAGAACCGACCTTGCGC | 239.9 | 7.4 | 1.7 | 4.4 | 1.1E-05 | 1.6E-05 | MB231/MB468 | MITO |
| >hg38_21_+_8433172_8446622_RNA45SN1_45S_50ntFlanks@10806.10827.22\|AACAATGTAGGTAAGGGAAGTC | >hg38_21_+_8433172_8446622_RNA45SN1_45S_50ntFlanks | 10806 | 10827 | 22 | AACAATGTAGGTAAGGGAAGTC | 184.0 | -3.8 | 0.9 | -4.4 | 1.0E-05 | 1.6E-05 | MB231/MB468 | CYTO |
| >hg38_21_+_8433172_8446622_RNA45SN1_45S_50ntFlanks@12416.12448.33\|TTCGATGTCGGCTCTTCCTATCATTGTGAAGCA | >hg38_21_+_8433172_8446622_RNA45SN1_45S_50ntFlanks | 12416 | 12448 | 33 | TTCGATGTCGGCTCTTCCTATCATTGTGAAGCA | 658.7 | 2.9 | 0.7 | 4.4 | 1.1E-05 | 1.6E-05 | MB231/MB468 | MITO |
| >hg38_21_+_8433172_8446622_RNA45SN1_45S_50ntFlanks@5485.5516.32\|AGCGCTGAGAAGACGGTCGAACTTGACTATCT | >hg38_21_+_8433172_8446622_RNA45SN1_45S_50ntFlanks | 5485 | 5516 | 32 | AGCGCTGAGAAGACGGTCGAACTTGACTATCT | 10162.3 | 3.9 | 0.9 | 4.4 | 9.3E-06 | 1.6E-05 | BT20/MB468 | CYTO |
| >hg38_21_+_8433172_8446622_RNA45SN1_45S_50ntFlanks@3815.3842.28\|ATGGTTCCTTTGGTCGCTCGCTCCTCTC | >hg38_21_+_8433172_8446622_RNA45SN1_45S_50ntFlanks | 3815 | 3842 | 28 | ATGGTTCCTTTGGTCGCTCGCTCCTCTC | 276.3 | -4.9 | 1.1 | -4.4 | 1.1E-05 | 1.6E-05 | MB231/MB468 | NUC |
| >hg38_21_+_8433172_8446622_RNA45SN1_45S_50ntFlanks@12646.12669.24\|TGTGGGATTATGACTGAACGCCTC | >hg38_21_+_8433172_8446622_RNA45SN1_45S_50ntFlanks | 12646 | 12669 | 24 | TGTGGGATTATGACTGAACGCCTC | 82.9 | 4.7 | 1.1 | 4.4 | 1.1E-05 | 1.6E-05 | BT20/MB468 | NUC |
| >trna134_GluTTC_1_-_16861774_16861845@1.35.35,trna5_GluTTC_1_+_17199078_17199149@1.35.35,trna84_GluTTC_1_-_161391883_161391954@1.35.35,trna94_GluTTC_1_-_149664355_149664427@1.35.35\|TCCCTGGTGGTCTAGTGGCTAGGATTCGGCGCTTT | >trna134_GluTTC_1_-_16861774_16861845 | 1 | 35 | 35 | TCCCTGGTGGTCTAGTGGCTAGGATTCGGCGCTTT | 193.5 | 4.7 | 1.0 | 4.9 | 1.0E-06 | 1.6E-05 | BT20/MB231 | CYTO |
| >trnaMT_GluTTC_MT_-_14674_14742@31.72.42,trnalookalike8_GluTTC_5_-_93905172_93905240@31.72.42\|TTCATATCATTGGTCGTGGTTGTAGTCCGTGCGAGAATACCA | >trnaMT_GluTTC_MT_-_14674_14742 | 31 | 72 | 42 | TTCATATCATTGGTCGTGGTTGTAGTCCGTGCGAGAATACCA | 2063.6 | -4.2 | 0.9 | -4.9 | 1.0E-06 | 1.6E-05 | BT20/MB231 | CYTO |
| >hg38_21_+_8433172_8446622_RNA45SN1_45S_50ntFlanks@10024.10042.19\|GCGCTGGAGCGTCGGGCCC | >hg38_21_+_8433172_8446622_RNA45SN1_45S_50ntFlanks | 10024 | 10042 | 19 | GCGCTGGAGCGTCGGGCCC | 259.6 | -4.9 | 1.0 | -4.9 | 1.0E-06 | 1.6E-05 | BT20/MB231 | CYTO |
| >hg38_21_+_8433172_8446622_RNA45SN1_45S_50ntFlanks@12388.12433.46\|CGTTCATAGCGACGTCGCTTTTTGATCCTTCGATGTCGGCTCTTCC | >hg38_21_+_8433172_8446622_RNA45SN1_45S_50ntFlanks | 12388 | 12433 | 46 | CGTTCATAGCGACGTCGCTTTTTGATCCTTCGATGTCGGCTCTTCC | 2017.6 | -4.7 | 1.0 | -4.9 | 1.0E-06 | 1.6E-05 | BT20/MB231 | CYTO |
| >trnaMT_PheGAA_MT_+_577_647@35.56.22\|GAAAATGTTTAGACGGGCTCAC | >trnaMT_PheGAA_MT_+_577_647 | 35 | 56 | 22 | GAAAATGTTTAGACGGGCTCAC | 473.6 | 4.3 | 1.0 | 4.4 | 1.1E-05 | 1.6E-05 | MB231/MB468 | MITO |
| >hg38_21_+_8433172_8446622_RNA45SN1_45S_50ntFlanks@4724.4758.35\|ATTAATCAAGAACGAAAGTCGGAGGTTCGAAGACG | >hg38_21_+_8433172_8446622_RNA45SN1_45S_50ntFlanks | 4724 | 4758 | 35 | ATTAATCAAGAACGAAAGTCGGAGGTTCGAAGACG | 6554.4 | -3.3 | 0.7 | -4.4 | 1.0E-05 | 1.6E-05 | MB231/MB468 | CYTO |
| >hg38_21_+_8433172_8446622_RNA45SN1_45S_50ntFlanks@10771.10817.47\|CAGGTCTCCAAGGTGAACAGCCTCTGGCATGTTGGAACAATGTAGGT | >hg38_21_+_8433172_8446622_RNA45SN1_45S_50ntFlanks | 10771 | 10817 | 47 | CAGGTCTCCAAGGTGAACAGCCTCTGGCATGTTGGAACAATGTAGGT | 267.8 | -6.7 | 1.5 | -4.4 | 1.1E-05 | 1.6E-05 | MB231/MB468 | MITO |
| >hg38_21_+_8433172_8446622_RNA45SN1_45S_50ntFlanks@12892.12932.41\|CCCTCGCCCGTCACGCACCGCACGTTCGTGGGGAACCTGGC | >hg38_21_+_8433172_8446622_RNA45SN1_45S_50ntFlanks | 12892 | 12932 | 41 | CCCTCGCCCGTCACGCACCGCACGTTCGTGGGGAACCTGGC | 180.2 | -3.8 | 0.9 | -4.4 | 1.0E-05 | 1.6E-05 | MB231/MB468 | CYTO |
| >trna11_GluTTC_15_-_26327381_26327452@41.73.33,trna3_GluTTC_13_-_45492062_45492133@41.73.33\|AGGCGGCCCGGGTTCGACTCCCGGTGTGGGAAC | >trna11_GluTTC_15_-_26327381_26327452 | 41 | 73 | 33 | AGGCGGCCCGGGTTCGACTCCCGGTGTGGGAAC | 251.5 | 7.6 | 1.7 | 4.4 | 1.1E-05 | 1.6E-05 | MB231/MB468 | MITO |
| >hg38_21_+_8433172_8446622_RNA45SN1_45S_50ntFlanks@10869.10891.23\|AGGGCTGGGTCGGTCGGGCTGGG | >hg38_21_+_8433172_8446622_RNA45SN1_45S_50ntFlanks | 10869 | 10891 | 23 | AGGGCTGGGTCGGTCGGGCTGGG | 1090.8 | -3.9 | 0.8 | -4.6 | 3.4E-06 | 1.6E-05 | BT20/MB468 | MP |
| >hg38_21_+_8433172_8446622_RNA45SN1_45S_50ntFlanks@6773.6807.35\|TTCCTCCCGGGGCTACGCCTGTCTGAGCGTCGCTT | >hg38_21_+_8433172_8446622_RNA45SN1_45S_50ntFlanks | 6773 | 6807 | 35 | TTCCTCCCGGGGCTACGCCTGTCTGAGCGTCGCTT | 275.3 | -5.2 | 1.2 | -4.4 | 9.1E-06 | 1.6E-05 | MB231/MB468 | MP |
| >hg38_21_+_8433172_8446622_RNA45SN1_45S_50ntFlanks@8194.8215.22\|CCCGTGGACGGTGTGAGGCCGG | >hg38_21_+_8433172_8446622_RNA45SN1_45S_50ntFlanks | 8194 | 8215 | 22 | CCCGTGGACGGTGTGAGGCCGG | 1721.3 | -7.7 | 1.6 | -4.6 | 3.4E-06 | 1.6E-05 | BT20/MB468 | MP |
| >hg38_21_+_8433172_8446622_RNA45SN1_45S_50ntFlanks@12416.12456.41\|TTCGATGTCGGCTCTTCCTATCATTGTGAAGCAGAATTCAC | >hg38_21_+_8433172_8446622_RNA45SN1_45S_50ntFlanks | 12416 | 12456 | 41 | TTCGATGTCGGCTCTTCCTATCATTGTGAAGCAGAATTCAC | 378.6 | -4.1 | 0.8 | -4.9 | 1.0E-06 | 1.6E-05 | BT20/MB231 | CYTO |
| >trnaMT_GluTTC_MT_-_14674_14742@27.61.35,trnalookalike8_GluTTC_5_-_93905172_93905240@27.61.35\|GTTTTTCATATCATTGGTCGTGGTTGTAGTCCGTG | >trnaMT_GluTTC_MT_-_14674_14742 | 27 | 61 | 35 | GTTTTTCATATCATTGGTCGTGGTTGTAGTCCGTG | 1243.4 | -8.1 | 1.8 | -4.4 | 1.2E-05 | 1.6E-05 | BT20/MB468 | NUC |
| >hg38_21_+_8433172_8446622_RNA45SN1_45S_50ntFlanks@12416.12454.39\|TTCGATGTCGGCTCTTCCTATCATTGTGAAGCAGAATTC | >hg38_21_+_8433172_8446622_RNA45SN1_45S_50ntFlanks | 12416 | 12454 | 39 | TTCGATGTCGGCTCTTCCTATCATTGTGAAGCAGAATTC | 261.6 | 4.1 | 0.9 | 4.4 | 1.1E-05 | 1.6E-05 | MB231/MB468 | NUC |
| >hg38_21_+_8433172_8446622_RNA45SN1_45S_50ntFlanks@6648.6672.25\|GTACGACTCTTAGCGGTGGATCACT | >hg38_21_+_8433172_8446622_RNA45SN1_45S_50ntFlanks | 6648 | 6672 | 25 | GTACGACTCTTAGCGGTGGATCACT | 197.8 | 2.2 | 0.5 | 4.4 | 1.2E-05 | 1.6E-05 | BT20/MB468 | NUC |
| >hg38_21_+_8433172_8446622_RNA45SN1_45S_50ntFlanks@9504.9521.18\|GTGAACTATGCCTGGGCA | >hg38_21_+_8433172_8446622_RNA45SN1_45S_50ntFlanks | 9504 | 9521 | 18 | GTGAACTATGCCTGGGCA | 570.4 | -2.6 | 0.6 | -4.5 | 7.9E-06 | 1.6E-05 | BT20/MB468 | MITO |
| >hg38_21_+_8433172_8446622_RNA45SN1_45S_50ntFlanks@4046.4069.24\|CATTCGAACGTCTGCCCTATCAAC | >hg38_21_+_8433172_8446622_RNA45SN1_45S_50ntFlanks | 4046 | 4069 | 24 | CATTCGAACGTCTGCCCTATCAAC | 627.2 | -4.4 | 1.0 | -4.4 | 1.1E-05 | 1.6E-05 | MB231/MB468 | MITO |
| >hg38_21_+_8433172_8446622_RNA45SN1_45S_50ntFlanks@4090.4111.22\|CCTACCATGGTGACCACGGGTG | >hg38_21_+_8433172_8446622_RNA45SN1_45S_50ntFlanks | 4090 | 4111 | 22 | CCTACCATGGTGACCACGGGTG | 92.3 | -10.4 | 2.3 | -4.4 | 9.2E-06 | 1.6E-05 | MB231/MB468 | MP |
| >hg38_21_+_8433172_8446622_RNA45SN1_45S_50ntFlanks@4558.4587.30\|AGCTAGGAATAATGGAATAGGACCGCGGTT | >hg38_21_+_8433172_8446622_RNA45SN1_45S_50ntFlanks | 4558 | 4587 | 30 | AGCTAGGAATAATGGAATAGGACCGCGGTT | 140.3 | -2.9 | 0.7 | -4.4 | 1.0E-05 | 1.6E-05 | MB231/MB468 | CYTO |
| >hg38_21_+_8433172_8446622_RNA45SN1_45S_50ntFlanks@11595.11626.32\|AGGGGAATCCGACTGTTTAATTAAAACAAAGC | >hg38_21_+_8433172_8446622_RNA45SN1_45S_50ntFlanks | 11595 | 11626 | 32 | AGGGGAATCCGACTGTTTAATTAAAACAAAGC | 147.9 | 10.5 | 2.2 | 4.9 | 1.0E-06 | 1.6E-05 | BT20/MB231 | MP |
| >hg38_21_+_8433172_8446622_RNA45SN1_45S_50ntFlanks@6645.6680.36\|CTCGTACGACTCTTAGCGGTGGATCACTCGGCTCGT | >hg38_21_+_8433172_8446622_RNA45SN1_45S_50ntFlanks | 6645 | 6680 | 36 | CTCGTACGACTCTTAGCGGTGGATCACTCGGCTCGT | 492.4 | 2.5 | 0.6 | 4.4 | 1.0E-05 | 1.6E-05 | MB231/MB468 | CYTO |
| >hg38_21_+_8433172_8446622_RNA45SN1_45S_50ntFlanks@11540.11582.43\|CGCGGCGCCTCGCCTCGGCCGGCGCCTAGCAGCCGACTTAGAA | >hg38_21_+_8433172_8446622_RNA45SN1_45S_50ntFlanks | 11540 | 11582 | 43 | CGCGGCGCCTCGCCTCGGCCGGCGCCTAGCAGCCGACTTAGAA | 896.1 | -4.3 | 0.9 | -4.9 | 1.0E-06 | 1.6E-05 | BT20/MB231 | CYTO |
| >hg38_21_+_8433172_8446622_RNA45SN1_45S_50ntFlanks@11915.11943.29\|GAGACATGAGAGGTGTAGAATAAGTGGGA | >hg38_21_+_8433172_8446622_RNA45SN1_45S_50ntFlanks | 11915 | 11943 | 29 | GAGACATGAGAGGTGTAGAATAAGTGGGA | 41.9 | 9.1 | 2.0 | 4.6 | 3.4E-06 | 1.6E-05 | BT20/MB468 | MP |
| >hg38_21_+_8433172_8446622_RNA45SN1_45S_50ntFlanks@12720.12742.23\|CGGTTGGCCTCGGATAGCCGGTC | >hg38_21_+_8433172_8446622_RNA45SN1_45S_50ntFlanks | 12720 | 12742 | 23 | CGGTTGGCCTCGGATAGCCGGTC | 65.1 | -9.1 | 2.0 | -4.6 | 3.5E-06 | 1.6E-05 | BT20/MB468 | MP |
| >hg38_21_+_8433172_8446622_RNA45SN1_45S_50ntFlanks@4729.4752.24\|TCAAGAACGAAAGTCGGAGGTTCG | >hg38_21_+_8433172_8446622_RNA45SN1_45S_50ntFlanks | 4729 | 4752 | 24 | TCAAGAACGAAAGTCGGAGGTTCG | 175.0 | -3.0 | 0.7 | -4.4 | 9.4E-06 | 1.6E-05 | BT20/MB468 | CYTO |
| >MI0000487\|hsa-mir-193a&WithFlank&17\|+\|31559990\|31560089@27.47.21(+1U)&[MIMAT0004614&hsa-miR-193a-5p&offsets\|0\|-1(+1U);m-123&17\|+\|31560016\|31560037&offsets\|0\|-1(+1U)]\|TGGGTCTTTGCGGGCGAGATGT | >MI0000487\|hsa-mir-193a&WithFlank&17\|+\|31559990\|31560089 | 27 | 47 | 21(+1U) | TGGGTCTTTGCGGGCGAGATGT | 420.1 | 7.9 | 1.7 | 4.6 | 3.5E-06 | 1.6E-05 | BT20/MB468 | MP |
| >hg38_21_+_8433172_8446622_RNA45SN1_45S_50ntFlanks@8182.8223.42\|ATCGAGGCCCAGCCCGTGGACGGTGTGAGGCCGGTAGCGGCC | >hg38_21_+_8433172_8446622_RNA45SN1_45S_50ntFlanks | 8182 | 8223 | 42 | ATCGAGGCCCAGCCCGTGGACGGTGTGAGGCCGGTAGCGGCC | 68.6 | -9.2 | 2.0 | -4.6 | 3.5E-06 | 1.6E-05 | BT20/MB468 | MP |
| >hg38_MT_+_1621_3279_RNR2_16S_50ntFlanks@1000.1034.35\|GGGACCTGTATGAATGGCTCCACGAGGGTTCAGCT | >hg38_MT_+_1621_3279_RNR2_16S_50ntFlanks | 1000 | 1034 | 35 | GGGACCTGTATGAATGGCTCCACGAGGGTTCAGCT | 32.3 | 8.2 | 1.9 | 4.4 | 9.2E-06 | 1.6E-05 | MB231/MB468 | MP |

| | | | | | | | | | | | | | |
|---|---|---|---|---|---|---|---|---|---|---|---|---|---|
| >MI0002468\|hsa-mir-484&WithFlank&16\|+\|15643288\|15643378@14.34.21(+1A)&[MIMAT0002174&hsa-miR-484&offsets\|0\|-1(+1A);m-127&16\|+\|15643301\|15643321&offsets\|0\|0(+1A)]\|TCAGGCTCAGTCCCCTCCCGAA | >MI0002468\|hsa-mir-484&WithFlank&16\|+\|15643288\|15643378 | 14 | 34 | 21(+1A) | TCAGGCTCAGTCCCCTCCCGAA | 210.4 | 4.7 | 1.1 | 4.4 | 1.1E-05 | 1.6E-05 | MB231/MB468 | MITO |
| >hg38_21_+_8433172_8446622_RNA45SN1_45S_50ntFlanks@10410.10432.23\|TCAGTCGGTCCTGAGAGATGGGC | >hg38_21_+_8433172_8446622_RNA45SN1_45S_50ntFlanks | 10410 | 10432 | 23 | TCAGTCGGTCCTGAGAGATGGGC | 140.7 | 10.5 | 2.1 | 4.9 | 1.0E-06 | 1.6E-05 | BT20/MB231 | MP |
| >hg38_21_+_8433172_8446622_RNA45SN1_45S_50ntFlanks@11581.11602.22\|AACTGGTGCGGACCAGGGGAAT | >hg38_21_+_8433172_8446622_RNA45SN1_45S_50ntFlanks | 11581 | 11602 | 22 | AACTGGTGCGGACCAGGGGAAT | 1028.9 | -6.2 | 1.3 | -4.6 | 3.5E-06 | 1.6E-05 | BT20/MB468 | MP |
| >hg38_21_+_8433172_8446622_RNA45SN1_45S_50ntFlanks@3840.3868.29\|CTCCTACTTGGATAACTGTGGTAATTCTA | >hg38_21_+_8433172_8446622_RNA45SN1_45S_50ntFlanks | 3840 | 3868 | 29 | CTCCTACTTGGATAACTGTGGTAATTCTA | 979.3 | 3.3 | 0.7 | 4.5 | 8.0E-06 | 1.6E-05 | BT20/MB468 | MITO |
| >hg38_21_+_8433172_8446622_RNA45SN1_45S_50ntFlanks@6651.6677.27\|CGACTCTTAGCGGTGGATCACTCGGCT | >hg38_21_+_8433172_8446622_RNA45SN1_45S_50ntFlanks | 6651 | 6677 | 27 | CGACTCTTAGCGGTGGATCACTCGGCT | 3469.5 | 2.9 | 0.7 | 4.4 | 1.0E-05 | 1.6E-05 | MB231/MB468 | CYTO |
| >trnaMT_SerGCT_MT_+_12207_12265@1.36.36\|GAGAAAGCTCACAAGAACTGCTAACTCATGCCCCCA | >trnaMT_SerGCT_MT_+_12207_12265 | 1 | 36 | 36 | GAGAAAGCTCACAAGAACTGCTAACTCATGCCCCCA | 36.4 | 8.4 | 1.9 | 4.4 | 9.2E-06 | 1.6E-05 | MB231/MB468 | MP |
| >hg38_21_+_8433172_8446622_RNA45SN1_45S_50ntFlanks@10475.10493.19\|CGGCCGATCGAAAGGGAGT | >hg38_21_+_8433172_8446622_RNA45SN1_45S_50ntFlanks | 10475 | 10493 | 19 | CGGCCGATCGAAAGGGAGT | 182.0 | -6.5 | 1.5 | -4.4 | 9.2E-06 | 1.6E-05 | MB231/MB468 | MP |
| >MI0000434\|hsa-let-7i&WithFlank&12\|+\|62603680\|62603775@12.33.22(+1A)&[MIMAT0000415&hsa-let-7i-5p&offsets\|0\|0(+1A);m-49&12\|+\|62603691\|62603712&offsets\|0\|0(+1A)]\|TGAGGTAGTAGTTTGTGCTGTTA | >MI0000434\|hsa-let-7i&WithFlank&12\|+\|62603680\|62603775 | 12 | 33 | 22(+1A) | TGAGGTAGTAGTTTGTGCTGTTA | 1557.4 | 4.1 | 0.9 | 4.4 | 1.1E-05 | 1.6E-05 | MB231/MB468 | MITO |
| >hg38_21_+_8433172_8446622_RNA45SN1_45S_50ntFlanks@12397.12415.19\|CGACGTCGCTTTTTGATCC | >hg38_21_+_8433172_8446622_RNA45SN1_45S_50ntFlanks | 12397 | 12415 | 19 | CGACGTCGCTTTTTGATCC | 250.9 | 2.4 | 0.6 | 4.4 | 1.1E-05 | 1.6E-05 | MB231/MB468 | CYTO |
| >hg38_21_+_8433172_8446622_RNA45SN1_45S_50ntFlanks@11207.11254.48\|GGCGGCGACTCTGGACGCGAGCCGGGCCCTTCCCGTGGATCGCCCCAG | >hg38_21_+_8433172_8446622_RNA45SN1_45S_50ntFlanks | 11207 | 11254 | 48 | GGCGGCGACTCTGGACGCGAGCCGGGCCCTTCCCGTGGATCGCCCCAG | 362.0 | -4.1 | 0.9 | -4.4 | 1.1E-05 | 1.6E-05 | MB231/MB468 | CYTO |
| >am_trnaMT_ValTAC_MT_+_1602_1670@44.68.25\|ATTTCAACTTAACTTGACCGCTCTG | >am_trnaMT_ValTAC_MT_+_1602_1670 | 44 | 68 | 25 | ATTTCAACTTAACTTGACCGCTCTG | 2012.7 | -8.8 | 2.0 | -4.4 | 1.2E-05 | 1.6E-05 | BT20/MB468 | NUC |
| >hg38_21_+_8433172_8446622_RNA45SN1_45S_50ntFlanks@9566.9592.27\|GACGTGCAAATCGGTCGTCCGACCTGG | >hg38_21_+_8433172_8446622_RNA45SN1_45S_50ntFlanks | 9566 | 9592 | 27 | GACGTGCAAATCGGTCGTCCGACCTGG | 308.8 | -5.3 | 1.2 | -4.4 | 1.2E-05 | 1.6E-05 | BT20/MB468 | NUC |
| >MI0000102\|hsa-mir-100&WithFlank&11\|-\|122152223\|122152314@19.40.22(+1A)&[MIMAT0000098&hsa-miR-100-5p&offsets\|0\|0(+1A);m-27&11\|-\|122152276\|122152296&offsets\|0\|+1(+1A)]\|AACCCGTAGATCCGAACTTGTGA | >MI0000102\|hsa-mir-100&WithFlank&11\|-\|122152223\|122152314 | 19 | 40 | 22(+1A) | AACCCGTAGATCCGAACTTGTGA | 741.6 | 8.7 | 2.0 | 4.5 | 8.0E-06 | 1.6E-05 | BT20/MB468 | MITO |
| >hg38_21_+_8433172_8446622_RNA45SN1_45S_50ntFlanks@12617.12654.38\|TGAGGAGCCAATGGGGCGAAGCTACCATCTGTGGGATT | >hg38_21_+_8433172_8446622_RNA45SN1_45S_50ntFlanks | 12617 | 12654 | 38 | TGAGGAGCCAATGGGGCGAAGCTACCATCTGTGGGATT | 1573.6 | -3.7 | 0.8 | -4.5 | 8.0E-06 | 1.6E-05 | BT20/MB468 | MITO |
| >hg38_21_+_8433172_8446622_RNA45SN1_45S_50ntFlanks@11584.11607.24\|TGGTGCGGACCAGGGGAATCCGAC | >hg38_21_+_8433172_8446622_RNA45SN1_45S_50ntFlanks | 11584 | 11607 | 24 | TGGTGCGGACCAGGGGAATCCGAC | 213.7 | 4.8 | 1.0 | 4.9 | 1.2E-06 | 1.6E-05 | BT20/MB231 | MITO |
| >hg38_21_+_8433172_8446622_RNA45SN1_45S_50ntFlanks@4574.4611.38\|ATAGGACCGCGGTTCTATTTTGTTGGTTTTCGGAACTG | >hg38_21_+_8433172_8446622_RNA45SN1_45S_50ntFlanks | 4574 | 4611 | 38 | ATAGGACCGCGGTTCTATTTTGTTGGTTTTCGGAACTG | 164.0 | -6.5 | 1.5 | -4.4 | 9.5E-06 | 1.6E-05 | BT20/MB468 | CYTO |
| >MI0000783\|hsa-mir-375&WithFlank&2\|-\|219001639\|219001714@46.67.22&[MIMAT0000728&hsa-miR-375-3p&offsets\|0\|0;m-92&2\|-\|219001648\|219001669&offsets\|0\|0]\|TTTGTTCGTTCGGCTCGCGTGA | >MI0000783\|hsa-mir-375&WithFlank&2\|-\|219001639\|219001714 | 46 | 67 | 22 | TTTGTTCGTTCGGCTCGCGTGA | 85.0 | 9.7 | 2.0 | 4.9 | 1.0E-06 | 1.6E-05 | BT20/MB231 | MP |
| >MI0000087\|hsa-mir-29a&WithFlank&7\|-\|130876741\|130876816@48.68.21&[MIMAT0000086&hsa-miR-29a-3p&offsets\|0\|-1;m-26&7\|-\|130876749\|130876769&offsets\|0\|0]\|TAGCACCATCTGAAATCGGTT | >MI0000087\|hsa-mir-29a&WithFlank&7\|-\|130876741\|130876816 | 48 | 68 | 21 | TAGCACCATCTGAAATCGGTT | 18399.7 | 6.2 | 1.3 | 4.6 | 3.5E-06 | 1.6E-05 | BT20/MB468 | MP |
| >hg38_21_+_8433172_8446622_RNA45SN1_45S_50ntFlanks@10766.10785.20\|CGCAGCAGGTCTCCAAGGTG | >hg38_21_+_8433172_8446622_RNA45SN1_45S_50ntFlanks | 10766 | 10785 | 20 | CGCAGCAGGTCTCCAAGGTG | 153.8 | -2.5 | 0.6 | -4.5 | 8.1E-06 | 1.6E-05 | BT20/MB468 | MITO |
| >hg38_21_+_8433172_8446622_RNA45SN1_45S_50ntFlanks@12457.12480.24\|CAAGCGTTGGATTGTTCACCCACT | >hg38_21_+_8433172_8446622_RNA45SN1_45S_50ntFlanks | 12457 | 12480 | 24 | CAAGCGTTGGATTGTTCACCCACT | 268.8 | -6.5 | 1.5 | -4.4 | 9.3E-06 | 1.6E-05 | MB231/MB468 | MP |
| >hg38_21_+_8433172_8446622_RNA45SN1_45S_50ntFlanks@10669.10706.38\|AAAGCGTCGCGGTTCCGGCGGCGTCCGGTGAGCTCTCG | >hg38_21_+_8433172_8446622_RNA45SN1_45S_50ntFlanks | 10669 | 10706 | 38 | AAAGCGTCGCGGTTCCGGCGGCGTCCGGTGAGCTCTCG | 1422.1 | -7.5 | 1.7 | -4.4 | 1.1E-05 | 1.6E-05 | MB231/MB468 | NUC |
| >MI0000285\|hsa-mir-205&WithFlank&1\|+\|209432127\|209432248@40.60.21(+1A)&[MIMAT0000266&hsa-miR-205-5p&offsets\|0\|-1(+1A);m-351&1\|+\|209432166\|209432187&offsets\|0\|-1(+1A)]\|TCCTTCATTCCACCGGAGTCTA | >MI0000285\|hsa-mir-205&WithFlank&1\|+\|209432127\|209432248 | 40 | 60 | 21(+1A) | TCCTTCATTCCACCGGAGTCTA | 1535.3 | 3.5 | 0.8 | 4.4 | 1.2E-05 | 1.6E-05 | BT20/MB468 | NUC |
| >hg38_21_+_8433172_8446622_RNA45SN1_45S_50ntFlanks@9533.9563.31\|GAGGAAACTCTGGTGGAGGTCCGTAGCGGTC | >hg38_21_+_8433172_8446622_RNA45SN1_45S_50ntFlanks | 9533 | 9563 | 31 | GAGGAAACTCTGGTGGAGGTCCGTAGCGGTC | 1386.1 | -4.2 | 0.9 | -4.6 | 3.5E-06 | 1.6E-05 | BT20/MB468 | MP |
| >MI0000298\|hsa-mir-221&WithFlank&X\|-\|45746151\|45746272@71.93.23(+2U)&[MIMAT0000278&hsa-miR-221-3p&offsets\|0\|0(+2U);m-91&X\|-\|45746181\|45746202&offsets\|0\|+1(+2U)]\|AGCTACATTGTCTGCTGGGTTTCTT | >MI0000298\|hsa-mir-221&WithFlank&X\|-\|45746151\|45746272 | 71 | 93 | 23(+2U) | AGCTACATTGTCTGCTGGGTTTCTT | 148.1 | 7.2 | 1.6 | 4.4 | 1.1E-05 | 1.6E-05 | MB231/MB468 | MITO |
| >hg38_21_+_8433172_8446622_RNA45SN1_45S_50ntFlanks@9535.9563.29\|GGAAACTCTGGTGGAGGTCCGTAGCGGTC | >hg38_21_+_8433172_8446622_RNA45SN1_45S_50ntFlanks | 9535 | 9563 | 29 | GGAAACTCTGGTGGAGGTCCGTAGCGGTC | 181.4 | -6.4 | 1.5 | -4.4 | 9.4E-06 | 1.6E-05 | MB231/MB468 | MP |
| >MI0000342\|hsa-mir-200b&WithFlank&1\|+\|1167098\|1167204@63.85.23(+1A)&[MIMAT0000318&hsa-miR-200b-3p&offsets\|0\|+1(+1A);m-93&1\|+\|1167160\|1167182&offsets\|0\|0(+1A)]\|TAATACTGCCTGGTAATGATGACA | >MI0000342\|hsa-mir-200b&WithFlank&1\|+\|1167098\|1167204 | 63 | 85 | 23(+1A) | TAATACTGCCTGGTAATGATGACA | 250.5 | 4.2 | 0.9 | 4.9 | 1.0E-06 | 1.6E-05 | BT20/MB231 | CYTO |
| >hg38_21_+_8433172_8446622_RNA45SN1_45S_50ntFlanks@6734.6774.41\|ATTGATCATCGACACTTCGAACGCACTTGCGGCCCCGGGTT | >hg38_21_+_8433172_8446622_RNA45SN1_45S_50ntFlanks | 6734 | 6774 | 41 | ATTGATCATCGACACTTCGAACGCACTTGCGGCCCCGGGTT | 189.6 | 3.1 | 0.7 | 4.4 | 1.2E-05 | 1.6E-05 | BT20/MB468 | NUC |
| >MI0005559\|hsa-mir-744&WithFlank&17\|+\|12081893\|12082002@17.37.21&[MIMAT0004945&hsa-miR-744-5p&offsets\|0\|-1;m-172&17\|+\|12081909\|12081929&offsets\|0\|0]\|TGCGGGGCTAGGGCTAACAGC | >MI0005559\|hsa-mir-744&WithFlank&17\|+\|12081893\|12082002 | 17 | 37 | 21 | TGCGGGGCTAGGGCTAACAGC | 374.7 | 4.2 | 1.0 | 4.4 | 1.1E-05 | 1.6E-05 | MB231/MB468 | MITO |
| >hg38_21_+_8433172_8446622_RNA45SN1_45S_50ntFlanks@5473.5513.41\|CGGCCCTGGCGGAGCGCTGAGAAGACGGTCGAACTTGACTA | >hg38_21_+_8433172_8446622_RNA45SN1_45S_50ntFlanks | 5473 | 5513 | 41 | CGGCCCTGGCGGAGCGCTGAGAAGACGGTCGAACTTGACTA | 240.2 | 3.4 | 0.8 | 4.4 | 1.1E-05 | 1.6E-05 | MB231/MB468 | CYTO |
| >hg38_21_+_8433172_8446622_RNA45SN1_45S_50ntFlanks@4005.4025.21\|AACCTCGGGCCGATCGCACGC | >hg38_21_+_8433172_8446622_RNA45SN1_45S_50ntFlanks | 4005 | 4025 | 21 | AACCTCGGGCCGATCGCACGC | 296.5 | -3.5 | 0.8 | -4.5 | 8.2E-06 | 1.6E-05 | BT20/MB468 | MITO |
| >hg38_21_+_8433172_8446622_RNA45SN1_45S_50ntFlanks@6779.6803.25\|CCGGGGCTACGCCTGTCTGAGCGTC | >hg38_21_+_8433172_8446622_RNA45SN1_45S_50ntFlanks | 6779 | 6803 | 25 | CCGGGGCTACGCCTGTCTGAGCGTC | 797.2 | -4.1 | 0.9 | -4.6 | 3.5E-06 | 1.6E-05 | BT20/MB468 | MP |
| >hg38_21_+_8433172_8446622_RNA45SN1_45S_50ntFlanks@11923.11949.27\|AGAGGTGTAGAATAAGTGGGAGGCCCC | >hg38_21_+_8433172_8446622_RNA45SN1_45S_50ntFlanks | 11923 | 11949 | 27 | AGAGGTGTAGAATAAGTGGGAGGCCCC | 160.5 | -6.3 | 1.4 | -4.4 | 9.4E-06 | 1.6E-05 | MB231/MB468 | MP |
| >MI0000342\|hsa-mir-200b&WithFlank&1\|+\|1167098\|1167204@64.85.22(+1U)&[MIMAT0000318&hsa-miR-200b-3p&offsets\|+1\|+1(+1U);m-93&1\|+\|1167160\|1167182&offsets\|+1\|0(+1U)]\|AATACTGCCTGGTAATGATGACT | >MI0000342\|hsa-mir-200b&WithFlank&1\|+\|1167098\|1167204 | 64 | 85 | 22(+1U) | AATACTGCCTGGTAATGATGACT | 453.7 | -5.1 | 1.2 | -4.4 | 1.1E-05 | 1.6E-05 | MB231/MB468 | NUC |
| >MI0000298\|hsa-mir-221&WithFlank&X\|-\|45746151\|45746272@71.90.20&[MIMAT0000278&hsa-miR-221-3p&offsets\|0\|-3;m-91&X\|-\|45746181\|45746202&offsets\|0\|-2]\|AGCTACATTGTCTGCTGGGT | >MI0000298\|hsa-mir-221&WithFlank&X\|-\|45746151\|45746272 | 71 | 90 | 20 | AGCTACATTGTCTGCTGGGT | 1000.2 | 3.9 | 0.9 | 4.4 | 9.4E-06 | 1.6E-05 | MB231/MB468 | MP |
| >hg38_21_+_8433172_8446622_RNA45SN1_45S_50ntFlanks@10350.10388.39\|ACGAGAACTTTGAAGGCCGAAGTGGAGAAGGGTTCCATG | >hg38_21_+_8433172_8446622_RNA45SN1_45S_50ntFlanks | 10350 | 10388 | 39 | ACGAGAACTTTGAAGGCCGAAGTGGAGAAGGGTTCCATG | 1941.8 | -4.0 | 0.9 | -4.6 | 3.6E-06 | 1.6E-05 | BT20/MB468 | MP |
| >trnaMT_ProTGG_MT_-_15956_16023@41.67.27\|ATGGTGGAGTTAAAGACTTTTTCTCTG | >trnaMT_ProTGG_MT_-_15956_16023 | 41 | 67 | 27 | ATGGTGGAGTTAAAGACTTTTTCTCTG | 114.9 | -6.9 | 1.6 | -4.4 | 9.6E-06 | 1.6E-05 | BT20/MB468 | CYTO |
| >hg38_21_+_8433172_8446622_RNA45SN1_45S_50ntFlanks@3900.3936.37\|CTTCGCGGGGGGGATGCGTGCATTTATCAGATCAAAA | >hg38_21_+_8433172_8446622_RNA45SN1_45S_50ntFlanks | 3900 | 3936 | 37 | CTTCGCGGGGGGGATGCGTGCATTTATCAGATCAAAA | 76.4 | -9.3 | 2.0 | -4.6 | 3.6E-06 | 1.6E-05 | BT20/MB468 | MP |
| >hg38_21_+_8433172_8446622_RNA45SN1_45S_50ntFlanks@11576.11601.26\|CTTAGAACTGGTGCGGACCAGGGGAA | >hg38_21_+_8433172_8446622_RNA45SN1_45S_50ntFlanks | 11576 | 11601 | 26 | CTTAGAACTGGTGCGGACCAGGGGAA | 1521.0 | -5.9 | 1.3 | -4.4 | 9.6E-06 | 1.6E-05 | BT20/MB468 | CYTO |
| >hg38_21_+_8433172_8446622_RNA45SN1_45S_50ntFlanks@10248.10283.36\|CGACGAGTAGGAGGGCCGCTGCGGTGAGCCTTGAAG | >hg38_21_+_8433172_8446622_RNA45SN1_45S_50ntFlanks | 10248 | 10283 | 36 | CGACGAGTAGGAGGGCCGCTGCGGTGAGCCTTGAAG | 134.4 | 10.4 | 2.1 | 4.9 | 1.0E-06 | 1.6E-05 | BT20/MB231 | MP |
| >hg38_21_+_8433172_8446622_RNA45SN1_45S_50ntFlanks@8364.8395.32\|CTTTGAAGAGAGAGTTCAAGAGGGCGTGAAAC | >hg38_21_+_8433172_8446622_RNA45SN1_45S_50ntFlanks | 8364 | 8395 | 32 | CTTTGAAGAGAGAGTTCAAGAGGGCGTGAAAC | 129.2 | 10.3 | 2.1 | 4.9 | 1.0E-06 | 1.6E-05 | BT20/MB231 | MP |
| >am_trna119_LysCTT_1_-_145395522_145395594@7.34.28,trna11_LysCTT_5_-_180648979_180649051@7.34.28,trna128_LysCTT_1_-_55423542_55423614@7.34.28,trna13_LysCTT_6_+_26556774_26556846@7.34.28,trna6_LysCTT_19_-_52425396_52425468@7.34.28,trna7_LysCTT_16_+_3225692_3225764@7.34.28,trna9_LysCTT_5_+_180634755_180634827@7.34.28\|CTAGCTCAGTCGGTAGAGCATGAGACTC | >am_trna119_LysCTT_1_-_145395522_145395594 | 7 | 34 | 28 | CTAGCTCAGTCGGTAGAGCATGAGACTC | 91.2 | -9.9 | 2.0 | -4.9 | 1.0E-06 | 1.6E-05 | BT20/MB231 | MP |
| >hg38_21_+_8433172_8446622_RNA45SN1_45S_50ntFlanks@12478.12496.19\|ACTAATAGGGAACGTGAGC | >hg38_21_+_8433172_8446622_RNA45SN1_45S_50ntFlanks | 12478 | 12496 | 19 | ACTAATAGGGAACGTGAGC | 1487.9 | 2.4 | 0.5 | 4.4 | 1.1E-05 | 1.6E-05 | MB231/MB468 | CYTO |
| >trna10_AspGTC_12_-_125424193_125424264@1.36.36,trna12_AspGTC_12_-_125411891_125411962@1.36.36,trna38_AspGTC_17_-_8125556_8125627@1.36.36,trna45_AspGTC_6_+_27447453_27447524@1.36.36,trna48_AspGTC_6_+_27471523_27471594@1.36.36,trna4_AspGTC_12_+_96429799_96429870@1.36.36,trna69_AspGTC_1_-_161440205_161440276@1.36.36,trna72_AspGTC_1_-_161432824_161432895@1.36.36,trna75_AspGTC_1_-_161425414_161425485@1.36.36,trna78_AspGTC_1_-_161418033_161418104@1.36.36,trna81_AspGTC_1_-_161410615_161410686@1.36.36\|TCCTCGTTAGTATAGTGGTGAGTATCCCCGCCTGTC | >trna10_AspGTC_12_-_125424193_125424264 | 1 | 36 | 36 | TCCTCGTTAGTATAGTGGTGAGTATCCCCGCCTGTC | 1442.6 | 11.1 | 2.5 | 4.4 | 1.1E-05 | 1.6E-05 | MB231/MB468 | CYTO |

| | | | | | | | | | | | | | |
|---|---|---|---|---|---|---|---|---|---|---|---|---|---|
| >MI0003560\|hsa-mir-92b&WithFlank&1\|+\|155195171\|155195278@67.86.20(+2U)&[MIMAT0003218&hsa-miR-92b-3p&offsets\|0\|-2(+2U);m-115&1\|+\|155195237\|155195258&offsets\|0\|-2(+2U)]\|TATTGCACTCGTCCCGGCCTTT | >MI0003560\|hsa-mir-92b&WithFlank&1\|+\|155195171\|155195278 | 67 | 86 | 20(+2U) | TATTGCACTCGTCCCGGCCTTT | 45.9 | 8.7 | 2.0 | 4.4 | 9.4E-06 | 1.6E-05 | MB231/MB468 | MP |
| >hg38_21_+_8433172_8446622_RNA45SN1_45S_50ntFlanks@8081.8101.21\|GAGCCCAGCGCCGAATCCCCG | >hg38_21_+_8433172_8446622_RNA45SN1_45S_50ntFlanks | 8081 | 8101 | 21 | GAGCCCAGCGCCGAATCCCCG | 249.2 | 5.4 | 1.1 | 4.9 | 1.2E-06 | 1.6E-05 | BT20/MB231 | MITO |
| >hg38_21_+_8433172_8446622_RNA45SN1_45S_50ntFlanks@12638.12661.24\|CTACCATCTGTGGGATTATGACTG | >hg38_21_+_8433172_8446622_RNA45SN1_45S_50ntFlanks | 12638 | 12661 | 24 | CTACCATCTGTGGGATTATGACTG | 3840.5 | -3.8 | 0.9 | -4.5 | 8.2E-06 | 1.6E-05 | BT20/MB468 | MITO |
| >hg38_21_+_8433172_8446622_RNA45SN1_45S_50ntFlanks@10797.10817.21\|GCATGTTGGAACAATGTAGGT | >hg38_21_+_8433172_8446622_RNA45SN1_45S_50ntFlanks | 10797 | 10817 | 21 | GCATGTTGGAACAATGTAGGT | 360.0 | -5.0 | 1.1 | -4.4 | 1.1E-05 | 1.6E-05 | MB231/MB468 | MITO |
| >hg38_21_+_8433172_8446622_RNA45SN1_45S_50ntFlanks@9459.9503.45\|CCGGGGAGGTGGAGCACGAGCGCACGTGTTAGGACCCGAAAGATG | >hg38_21_+_8433172_8446622_RNA45SN1_45S_50ntFlanks | 9459 | 9503 | 45 | CCGGGGAGGTGGAGCACGAGCGCACGTGTTAGGACCCGAAAGATG | 825.6 | -8.1 | 1.7 | -4.6 | 3.6E-06 | 1.6E-05 | BT20/MB468 | MP |
| >hg38_21_+_8433172_8446622_RNA45SN1_45S_50ntFlanks@12183.12200.18\|AAACGGTAACGCAGGTGT | >hg38_21_+_8433172_8446622_RNA45SN1_45S_50ntFlanks | 12183 | 12200 | 18 | AAACGGTAACGCAGGTGT | 58.3 | -8.9 | 1.9 | -4.6 | 3.6E-06 | 1.6E-05 | BT20/MB468 | MP |
| >hg38_21_+_8433172_8446622_RNA45SN1_45S_50ntFlanks@10391.10429.39\|AACAGCAGTTGAACATGGGTCAGTCGGTCCTGAGAGATG | >hg38_21_+_8433172_8446622_RNA45SN1_45S_50ntFlanks | 10391 | 10429 | 39 | AACAGCAGTTGAACATGGGTCAGTCGGTCCTGAGAGATG | 2022.0 | -3.0 | 0.7 | -4.4 | 1.1E-05 | 1.6E-05 | MB231/MB468 | CYTO |
| >MI0000085\|hsa-mir-27a&WithFlank&19\|-\|13836434\|13836523@57.77.21(+1U)&[MIMAT0000084&hsa-miR-27a-3p&offsets\|0\|0(+1U);m-43&19\|-\|13836447\|13836467&offsets\|0\|0(+1U)]\|TTCACAGTGGCTAAGTTCCGCT | >MI0000085\|hsa-mir-27a&WithFlank&19\|-\|13836434\|13836523 | 57 | 77 | 21(+1U) | TTCACAGTGGCTAAGTTCCGCT | 519.4 | 6.2 | 1.4 | 4.4 | 1.2E-05 | 1.6E-05 | MB231/MB468 | MITO |
| >hg38_MT_+_1621_3279_RNR2_16S_50ntFlanks@625.664.40\|AACTGAACTCCTCACACCCAATTGGACCAATCTATCACCC | >hg38_MT_+_1621_3279_RNR2_16S_50ntFlanks | 625 | 664 | 40 | AACTGAACTCCTCACACCCAATTGGACCAATCTATCACCC | 58.2 | 9.6 | 2.1 | 4.6 | 3.6E-06 | 1.6E-05 | BT20/MB468 | MP |
| >hg38_21_+_8433172_8446622_RNA45SN1_45S_50ntFlanks@9129.9152.24\|CGGGTCGGGGGGCGGGGCGGACTG | >hg38_21_+_8433172_8446622_RNA45SN1_45S_50ntFlanks | 9129 | 9152 | 24 | CGGGTCGGGGGGCGGGGCGGACTG | 1497.0 | -7.6 | 1.6 | -4.6 | 3.6E-06 | 1.6E-05 | BT20/MB468 | MP |
| >hg38_21_+_8433172_8446622_RNA45SN1_45S_50ntFlanks@12938.12955.18\|ACCATTCGTAGACGACCT | >hg38_21_+_8433172_8446622_RNA45SN1_45S_50ntFlanks | 12938 | 12955 | 18 | ACCATTCGTAGACGACCT | 173.0 | -2.8 | 0.6 | -4.4 | 1.1E-05 | 1.7E-05 | MB231/MB468 | NUC |
| >hg38_21_+_8433172_8446622_RNA45SN1_45S_50ntFlanks@3843.3872.30\|CTACTTGGATAACTGTGGTAATTCTAGAGC | >hg38_21_+_8433172_8446622_RNA45SN1_45S_50ntFlanks | 3843 | 3872 | 30 | CTACTTGGATAACTGTGGTAATTCTAGAGC | 428.9 | -5.4 | 1.2 | -4.4 | 1.2E-05 | 1.7E-05 | MB231/MB468 | MITO |
| >hg38_21_+_8433172_8446622_RNA45SN1_45S_50ntFlanks@7975.8010.36\|CGCGACCTCAGATCAGACGTGGCGACCCGCTGAATT | >hg38_21_+_8433172_8446622_RNA45SN1_45S_50ntFlanks | 7975 | 8010 | 36 | CGCGACCTCAGATCAGACGTGGCGACCCGCTGAATT | 43921.3 | 5.4 | 1.2 | 4.6 | 3.6E-06 | 1.7E-05 | BT20/MB468 | MP |
| >MI0000476\|hsa-mir-138-1&WithFlank&3\|+\|44114206\|44114316@29.52.24(+1A)&[MIMAT0000430_1&hsa-miR-138-5p&offsets\|0\|+1(+1A);m-497&3\|+\|44114234\|44114256&offsets\|0\|+1(+1A)]\|AGCTGGTGTTGTGAATCAGGCCGTA | >MI0000476\|hsa-mir-138-1&WithFlank&3\|+\|44114206\|44114316 | 29 | 52 | 24(+1A) | AGCTGGTGTTGTGAATCAGGCCGTA | 33.2 | 8.3 | 1.9 | 4.4 | 9.6E-06 | 1.7E-05 | MB231/MB468 | MP |
| >hg38_21_+_8433172_8446622_RNA45SN1_45S_50ntFlanks@10409.10432.24\|GTCAGTCGGTCCTGAGAGATGGGC | >hg38_21_+_8433172_8446622_RNA45SN1_45S_50ntFlanks | 10409 | 10432 | 24 | GTCAGTCGGTCCTGAGAGATGGGC | 242.8 | -1.9 | 0.4 | -4.4 | 1.1E-05 | 1.7E-05 | MB231/MB468 | CYTO |
| >MI0000100\|hsa-mir-98&WithFlank&X\|-\|53556217\|53556347@28.48.21&[MIMAT0000096&hsa-miR-98-5p&offsets\|0\|-1;m-143&X\|-\|53556299\|53556320&offsets\|0\|-1]\|TGAGGTAGTAAGTTGTATTGT | >MI0000100\|hsa-mir-98&WithFlank&X\|-\|53556217\|53556347 | 28 | 48 | 21 | TGAGGTAGTAAGTTGTATTGT | 197.2 | 4.2 | 1.0 | 4.4 | 1.2E-05 | 1.7E-05 | MB231/MB468 | MITO |
| >hg38_21_+_8433172_8446622_RNA45SN1_45S_50ntFlanks@12140.12180.41\|GACAGTGCCAGGTGGGGAGTTTGACTGGGGCGGTACACCTG | >hg38_21_+_8433172_8446622_RNA45SN1_45S_50ntFlanks | 12140 | 12180 | 41 | GACAGTGCCAGGTGGGGAGTTTGACTGGGGCGGTACACCTG | 86.1 | -10.3 | 2.3 | -4.4 | 9.6E-06 | 1.7E-05 | MB231/MB468 | MP |
| >am_trnaMT_HisGTG_MT_+_12138_12206@26.44.19\|AGATTGTGAATCTGACAAC | >am_trnaMT_HisGTG_MT_+_12138_12206 | 26 | 44 | 19 | AGATTGTGAATCTGACAAC | 1377.5 | 4.4 | 1.0 | 4.5 | 8.3E-06 | 1.7E-05 | BT20/MB468 | MITO |
| >am_trnaMT_ValTAC_MT_+_1602_1670@50.67.18\|ACTTAACTTGACCGCTCT | >am_trnaMT_ValTAC_MT_+_1602_1670 | 50 | 67 | 18 | ACTTAACTTGACCGCTCT | 9952.0 | 5.6 | 1.3 | 4.4 | 9.8E-06 | 1.7E-05 | BT20/MB468 | CYTO |
| >hg38_21_+_8433172_8446622_RNA45SN1_45S_50ntFlanks@9563.9597.35\|CCTGACGTGCAAATCGGTCGTCCGACCTGGGTATA | >hg38_21_+_8433172_8446622_RNA45SN1_45S_50ntFlanks | 9563 | 9597 | 35 | CCTGACGTGCAAATCGGTCGTCCGACCTGGGTATA | 444.1 | -6.2 | 1.4 | -4.4 | 9.8E-06 | 1.7E-05 | BT20/MB468 | CYTO |
| >hg38_21_+_8433172_8446622_RNA45SN1_45S_50ntFlanks@12447.12474.28\|CAGAATTCACCAAGCGTTGGATTGTTCA | >hg38_21_+_8433172_8446622_RNA45SN1_45S_50ntFlanks | 12447 | 12474 | 28 | CAGAATTCACCAAGCGTTGGATTGTTCA | 78.6 | -9.4 | 2.0 | -4.6 | 3.6E-06 | 1.7E-05 | BT20/MB468 | MP |
| >hg38_21_+_8433172_8446622_RNA45SN1_45S_50ntFlanks@4725.4759.35\|TTAATCAAGAACGAAAGTCGGAGGTTCGAAGACGA | >hg38_21_+_8433172_8446622_RNA45SN1_45S_50ntFlanks | 4725 | 4759 | 35 | TTAATCAAGAACGAAAGTCGGAGGTTCGAAGACGA | 301.7 | -2.8 | 0.6 | -4.5 | 8.4E-06 | 1.7E-05 | BT20/MB468 | MITO |
| >MI0003776\|hsa-mir-320b-1&WithFlank&1\|+\|116671740\|116671823@48.67.20(+1C);MI0003839\|hsa-mir-320b-2&WithFlank&1\|-\|224257034\|224257116@47.66.20(+1C);MI0000542\|hsa-mir-320a&WithFlank&8\|-\|22244960\|22245043@48.67.20(+1C)&[MIMAT0005792&hsa-miR-320b&offsets\|0\|-2(+1C);m-183&1\|+\|116671787\|116671806&offsets\|0\|0(+1C)];[MIMAT0005792_1&hsa-miR-320b&offsets\|0\|-2(+1C);m-183&1\|-\|224257051\|224257070&offsets\|0\|0(+1C)];[MIMAT0000510&hsa-miR-320a-3p&offsets\|0\|-2(+1C);m-60&8\|-\|22244975\|22244996&offsets\|0\|-2(+1C)]\|AAAAGCTGGGTTGAGAGGGCC | >MI0003776\|hsa-mir-320b-1&WithFlank&1\|+\|116671740\|116671823 | 48 | 67 | 20(+1C) | AAAAGCTGGGTTGAGAGGGCC | 228.6 | 4.1 | 0.8 | 4.9 | 1.1E-06 | 1.7E-05 | BT20/MB231 | CYTO |
| >hg38_21_+_8433172_8446622_RNA45SN1_45S_50ntFlanks@12656.12674.19\|TGACTGAACGCCTCTAAGT | >hg38_21_+_8433172_8446622_RNA45SN1_45S_50ntFlanks | 12656 | 12674 | 19 | TGACTGAACGCCTCTAAGT | 236.3 | -2.1 | 0.5 | -4.4 | 1.1E-05 | 1.7E-05 | MB231/MB468 | CYTO |
| >hg38_21_+_8433172_8446622_RNA45SN1_45S_50ntFlanks@12004.12033.30\|CGGGCCGCCGGTGAAATACCACTACTCTGA | >hg38_21_+_8433172_8446622_RNA45SN1_45S_50ntFlanks | 12004 | 12033 | 30 | CGGGCCGCCGGTGAAATACCACTACTCTGA | 134.7 | 4.0 | 0.9 | 4.4 | 1.2E-05 | 1.7E-05 | BT20/MB468 | NUC |
| >trna116_GluCTC_1_-_145399233_145399304@1.34.34,trna59_GluCTC_1_+_249168447_249168518@1.34.34,trna71_GluCTC_1_-_161439189_161439260@1.34.34,trna74_GluCTC_1_-_161431809_161431880@1.34.34,trna77_GluCTC_1_-_161424398_161424469@1.34.34,trna77_GluCTC_6_+_28949976_28950047@1.34.34,trna80_GluCTC_1_-_161417018_161417089@1.34.34,trna87_GluCTC_6_-_126101393_126101464@1.34.34\|TCCCTGGTGGTCTAGTGGTTAGGATTCGGCGCTC | >trna116_GluCTC_1_-_145399233_145399304 | 1 | 34 | 34 | TCCCTGGTGGTCTAGTGGTTAGGATTCGGCGCTC | 2685.4 | 4.5 | 1.0 | 4.4 | 1.1E-05 | 1.7E-05 | MB231/MB468 | NUC |
| >MI0000746\|hsa-mir-99b&WithFlank&19\|+\|51692606\|51692687@13.31.19(+1A)&[MIMAT0000689&hsa-miR-99b-5p&offsets\|0\|-3(+1A);m-10&19\|+\|51692618\|51692639&offsets\|0\|-3(+1A)]\|CACCCGTAGAACCGACCTTA | >MI0000746\|hsa-mir-99b&WithFlank&19\|+\|51692606\|51692687 | 13 | 31 | 19(+1A) | CACCCGTAGAACCGACCTTA | 226.3 | 8.2 | 1.8 | 4.6 | 3.6E-06 | 1.7E-05 | BT20/MB468 | MP |
| >hg38_21_+_8433172_8446622_RNA45SN1_45S_50ntFlanks@12437.12476.40\|CATTGTGAAGCAGAATTCACCAAGCGTTGGATTGTTCACC | >hg38_21_+_8433172_8446622_RNA45SN1_45S_50ntFlanks | 12437 | 12476 | 40 | CATTGTGAAGCAGAATTCACCAAGCGTTGGATTGTTCACC | 80.4 | -9.4 | 2.0 | -4.6 | 3.7E-06 | 1.7E-05 | BT20/MB468 | MP |
| >hg38_21_+_8433172_8446622_RNA45SN1_45S_50ntFlanks@4554.4572.19\|CCGCAGCTAGGAATAATGG | >hg38_21_+_8433172_8446622_RNA45SN1_45S_50ntFlanks | 4554 | 4572 | 19 | CCGCAGCTAGGAATAATGG | 174.8 | -5.9 | 1.3 | -4.4 | 1.1E-05 | 1.7E-05 | MB231/MB468 | NUC |
| >hg38_21_+_8433172_8446622_RNA45SN1_45S_50ntFlanks@8192.8215.24\|AGCCCGTGGACGGTGTGAGGCCGG | >hg38_21_+_8433172_8446622_RNA45SN1_45S_50ntFlanks | 8192 | 8215 | 24 | AGCCCGTGGACGGTGTGAGGCCGG | 664.6 | -7.3 | 1.7 | -4.4 | 1.2E-05 | 1.7E-05 | BT20/MB468 | NUC |
| >trnaMT_MetCAT_MT_+_4402_4469@36.71.36\|CCCGAAAATGTTGGTTATACCCTTCCCGTACTACCA | >trnaMT_MetCAT_MT_+_4402_4469 | 36 | 71 | 36 | CCCGAAAATGTTGGTTATACCCTTCCCGTACTACCA | 499.4 | -3.9 | 0.8 | -4.9 | 1.1E-06 | 1.7E-05 | BT20/MB231 | CYTO |
| >hg38_21_+_8433172_8446622_RNA45SN1_45S_50ntFlanks@10808.10831.24\|CAATGTAGGTAAGGGAAGTCGGCA | >hg38_21_+_8433172_8446622_RNA45SN1_45S_50ntFlanks | 10808 | 10831 | 24 | CAATGTAGGTAAGGGAAGTCGGCA | 197.3 | -3.9 | 0.9 | -4.4 | 9.9E-06 | 1.7E-05 | BT20/MB468 | CYTO |
| >hg38_21_+_8433172_8446622_RNA45SN1_45S_50ntFlanks@12635.12657.23\|AAGCTACCATCTGTGGGATTATG | >hg38_21_+_8433172_8446622_RNA45SN1_45S_50ntFlanks | 12635 | 12657 | 23 | AAGCTACCATCTGTGGGATTATG | 1179.7 | -5.3 | 1.2 | -4.5 | 8.4E-06 | 1.7E-05 | BT20/MB468 | MITO |
| >hg38_1_-_228634819_228635039_RNA5S12_5S_50ntFlanks@134.171.38\|TGGGAGACCGCCTGGGAATACCGGGTGCTGTAGGCTTT | >hg38_1_-_228634819_228635039_RNA5S12_5S_50ntFlanks | 134 | 171 | 38 | TGGGAGACCGCCTGGGAATACCGGGTGCTGTAGGCTTT | 9111.0 | 4.6 | 1.0 | 4.4 | 9.7E-06 | 1.7E-05 | MB231/MB468 | MP |
| >hg38_21_+_8433172_8446622_RNA45SN1_45S_50ntFlanks@12457.12487.31\|CAAGCGTTGGATTGTTCACCCACTAATAGGG | >hg38_21_+_8433172_8446622_RNA45SN1_45S_50ntFlanks | 12457 | 12487 | 31 | CAAGCGTTGGATTGTTCACCCACTAATAGGG | 1586.4 | -6.9 | 1.6 | -4.4 | 9.9E-06 | 1.7E-05 | BT20/MB468 | CYTO |
| >hg38_21_+_8433172_8446622_RNA45SN1_45S_50ntFlanks@4445.4472.28\|CTTGCCTCTCGGCGCCCCCTCGATGCTC | >hg38_21_+_8433172_8446622_RNA45SN1_45S_50ntFlanks | 4445 | 4472 | 28 | CTTGCCTCTCGGCGCCCCCTCGATGCTC | 275.6 | -5.7 | 1.3 | -4.4 | 9.9E-06 | 1.7E-05 | BT20/MB468 | CYTO |
| >hg38_21_+_8433172_8446622_RNA45SN1_45S_50ntFlanks@4046.4065.20\|CATTCGAACGTCTGCCCTAT | >hg38_21_+_8433172_8446622_RNA45SN1_45S_50ntFlanks | 4046 | 4065 | 20 | CATTCGAACGTCTGCCCTAT | 43.1 | -8.5 | 1.8 | -4.6 | 3.7E-06 | 1.7E-05 | BT20/MB468 | MP |
| >MI0000088\|hsa-mir-30a&WithFlank&6\|-\|71403545\|71403627@54.73.20&[MIMAT0000088&hsa-miR-30a-3p&offsets\|+1\|-1;m-38&6\|-\|71403554\|71403575&offsets\|+1\|-1]\|TTTCAGTCGGATGTTTGCAG | >MI0000088\|hsa-mir-30a&WithFlank&6\|-\|71403545\|71403627 | 54 | 73 | 20 | TTTCAGTCGGATGTTTGCAG | 422.7 | 5.3 | 1.2 | 4.4 | 9.7E-06 | 1.7E-05 | MB231/MB468 | MP |
| >hg38_21_+_8433172_8446622_RNA45SN1_45S_50ntFlanks@9486.9510.25\|GTTAGGACCCGAAAGATGGTGAACT | >hg38_21_+_8433172_8446622_RNA45SN1_45S_50ntFlanks | 9486 | 9510 | 25 | GTTAGGACCCGAAAGATGGTGAACT | 45.3 | 9.2 | 2.0 | 4.6 | 3.7E-06 | 1.7E-05 | BT20/MB468 | MP |
| >hg38_21_+_8433172_8446622_RNA45SN1_45S_50ntFlanks@11778.11803.26\|ACGCGCATGAATGGATGAACGAGATT | >hg38_21_+_8433172_8446622_RNA45SN1_45S_50ntFlanks | 11778 | 11803 | 26 | ACGCGCATGAATGGATGAACGAGATT | 320.4 | -3.1 | 0.7 | -4.4 | 1.2E-05 | 1.7E-05 | MB231/MB468 | NUC |
| >MI0000433\|hsa-let-7g&WithFlank&3\|-\|52268272\|52268367@11.28.18&[MIMAT0000414&hsa-let-7g-5p&offsets\|0\|-4;m-58&3\|-\|52268336\|52268357&offsets\|0\|-4]\|TGAGGTAGTAGTTTGTAC | >MI0000433\|hsa-let-7g&WithFlank&3\|-\|52268272\|52268367 | 11 | 28 | 18 | TGAGGTAGTAGTTTGTAC | 1005.3 | 4.5 | 1.0 | 4.4 | 1.2E-05 | 1.7E-05 | MB231/MB468 | MITO |
| >hg38_21_+_8433172_8446622_RNA45SN1_45S_50ntFlanks@11572.11597.26\|CCGACTTAGAACTGGTGCGGACCAGG | >hg38_21_+_8433172_8446622_RNA45SN1_45S_50ntFlanks | 11572 | 11597 | 26 | CCGACTTAGAACTGGTGCGGACCAGG | 601.7 | -3.6 | 0.8 | -4.4 | 1.0E-05 | 1.7E-05 | BT20/MB468 | CYTO |
| >hg38_21_+_8433172_8446622_RNA45SN1_45S_50ntFlanks@7982.8004.23\|TCAGATCAGACGTGGCGACCCGC | >hg38_21_+_8433172_8446622_RNA45SN1_45S_50ntFlanks | 7982 | 8004 | 23 | TCAGATCAGACGTGGCGACCCGC | 234.2 | -3.2 | 0.7 | -4.4 | 1.2E-05 | 1.7E-05 | MB231/MB468 | NUC |
| >hg38_21_+_8433172_8446622_RNA45SN1_45S_50ntFlanks@10351.10385.35\|CGAGAACTTTGAAGGCCGAAGTGGAGAAGGGTTCC | >hg38_21_+_8433172_8446622_RNA45SN1_45S_50ntFlanks | 10351 | 10385 | 35 | CGAGAACTTTGAAGGCCGAAGTGGAGAAGGGTTCC | 205.4 | -2.8 | 0.6 | -4.4 | 1.0E-05 | 1.7E-05 | BT20/MB468 | CYTO |
| >hg38_21_+_8433172_8446622_RNA45SN1_45S_50ntFlanks@12999.13029.31\|CTGCGATCTATTGAAAGTCAGCCCTCGACAC | >hg38_21_+_8433172_8446622_RNA45SN1_45S_50ntFlanks | 12999 | 13029 | 31 | CTGCGATCTATTGAAAGTCAGCCCTCGACAC | 366.5 | -7.1 | 1.6 | -4.4 | 1.2E-05 | 1.7E-05 | BT20/MB468 | NUC |
| >MI0000542\|hsa-mir-320a&WithFlank&8\|-\|22244960\|22245043@48.68.21(+1G)&[MIMAT0000510&hsa-miR-320a-3p&offsets\|0\|-1(+1G);m-60&8\|-\|22244975\|22244996&offsets\|0\|-1(+1G)]\|AAAAGCTGGGTTGAGAGGGCGG | >MI0000542\|hsa-mir-320a&WithFlank&8\|-\|22244960\|22245043 | 48 | 68 | 21(+1G) | AAAAGCTGGGTTGAGAGGGCGG | 118.9 | 5.1 | 1.2 | 4.4 | 1.2E-05 | 1.7E-05 | BT20/MB468 | NUC |

| | | | | | | | | | | | | | |
|---|---|---|---|---|---|---|---|---|---|---|---|---|---|
| >hg38_21_+_8433172_8446622_RNA45SN1_45S_50ntFlanks@11569.11594.26\|CAGCCGACTTAGAACTGGTGCGGACC | >hg38_21_+_8433172_8446622_RNA45SN1_45S_50ntFlanks | 11569 | 11594 | 26 | CAGCCGACTTAGAACTGGTGCGGACC | 383.6 | -7.1 | 1.6 | -4.4 | 1.2E-05 | 1.7E-05 | BT20/MB468 | NUC |
| >hg38_21_+_8433172_8446622_RNA45SN1_45S_50ntFlanks@8194.8216.23\|CCCGTGGACGGTGTGAGGCCGGT | >hg38_21_+_8433172_8446622_RNA45SN1_45S_50ntFlanks | 8194 | 8216 | 23 | CCCGTGGACGGTGTGAGGCCGGT | 258.5 | -3.4 | 0.7 | -4.6 | 3.7E-06 | 1.7E-05 | BT20/MB468 | MP |
| >hg38_21_+_8433172_8446622_RNA45SN1_45S_50ntFlanks@11873.11901.29\|AAGAAGACCCTGTTGAGCTTGACTCTAGT | >hg38_21_+_8433172_8446622_RNA45SN1_45S_50ntFlanks | 11873 | 11901 | 29 | AAGAAGACCCTGTTGAGCTTGACTCTAGT | 393.6 | -7.0 | 1.6 | -4.4 | 1.2E-05 | 1.7E-05 | BT20/MB468 | NUC |
| >hg38_21_+_8433172_8446622_RNA45SN1_45S_50ntFlanks@11875.11912.38\|GAAGACCCTGTTGAGCTTGACTCTAGTCTGGCACGGTG | >hg38_21_+_8433172_8446622_RNA45SN1_45S_50ntFlanks | 11875 | 11912 | 38 | GAAGACCCTGTTGAGCTTGACTCTAGTCTGGCACGGTG | 50.7 | -8.7 | 1.9 | -4.6 | 3.7E-06 | 1.7E-05 | BT20/MB468 | MP |
| >hg38_21_+_8433172_8446622_RNA45SN1_45S_50ntFlanks@5084.5108.25\|CGAACGAGACTCTGGCATGCTAACT | >hg38_21_+_8433172_8446622_RNA45SN1_45S_50ntFlanks | 5084 | 5108 | 25 | CGAACGAGACTCTGGCATGCTAACT | 612.8 | -6.1 | 1.4 | -4.4 | 1.2E-05 | 1.7E-05 | BT20/MB468 | NUC |
| >MI0000078\|hsa-mir-22&WithFlank&17\|-\|1713897\|1713993@59.77.19&[MIMAT0000077&hsa-miR-22-3p&offsets\|0\|-3;m-2&17\|-\|1713914\|1713935&offsets\|0\|-3]\|AAGCTGCCAGTTGAAGAAC | >MI0000078\|hsa-mir-22&WithFlank&17\|-\|1713897\|1713993 | 59 | 77 | 19 | AAGCTGCCAGTTGAAGAAC | 147.6 | 7.1 | 1.6 | 4.4 | 9.8E-06 | 1.7E-05 | MB231/MB468 | MP |
| >trna10_AspGTC_12_-_125424193_125424264@37.70.34,trna12_AspGTC_12_-_125411891_125411962@37.70.34,trna144_AspGTC_6_-_27551236_27551307@37.70.34,trna38_AspGTC_17_-_8125556_8125627@37.70.34,trna45_AspGTC_6_+_27447453_27447524@37.70.34,trna48_AspGTC_6_+_27471523_27471594@37.70.34,trna4_AspGTC_12_+_96429799_96429870@37.70.34,trna69_AspGTC_1_-_161440205_161440276@37.70.34,trna72_AspGTC_1_-_161432824_161432895@37.70.34,trna75_AspGTC_1_-_161425414_161425485@37.70.34,trna78_AspGTC_1_-_161418033_161418104@37.70.34,trna81_AspGTC_1_-_161410615_161410686@37.70.34\|ACGCGGGAGACCGGGGTTCGATTCCCCGACGGGG | >trna10_AspGTC_12_-_125424193_125424264 | 37 | 70 | 34 | ACGCGGGAGACCGGGGTTCGATTCCCCGACGGGG | 3282.3 | -3.8 | 0.9 | -4.4 | 9.8E-06 | 1.7E-05 | MB231/MB468 | MP |
| >hg38_21_+_8433172_8446622_RNA45SN1_45S_50ntFlanks@10791.10823.33\|CCTCTGGCATGTTGGAACAATGTAGGTAAGGGA | >hg38_21_+_8433172_8446622_RNA45SN1_45S_50ntFlanks | 10791 | 10823 | 33 | CCTCTGGCATGTTGGAACAATGTAGGTAAGGGA | 436.7 | 5.1 | 1.1 | 4.8 | 1.3E-06 | 1.7E-05 | BT20/MB231 | MITO |
| >MI0000285\|hsa-mir-205&WithFlank&1\|+\|209432127\|209432248@40.59.20&[MIMAT0000266&hsa-miR-205-5p&offsets\|0\|-2;m-351&1\|+\|209432166\|209432187&offsets\|0\|-2]\|TCCTTCATTCCACCGGAGTC | >MI0000285\|hsa-mir-205&WithFlank&1\|+\|209432127\|209432248 | 40 | 59 | 20 | TCCTTCATTCCACCGGAGTC | 170.9 | 3.2 | 0.7 | 4.4 | 1.2E-05 | 1.7E-05 | BT20/MB468 | NUC |
| >hg38_21_+_8433172_8446622_RNA45SN1_45S_50ntFlanks@4731.4752.22\|AAGAACGAAAGTCGGAGGTTCG | >hg38_21_+_8433172_8446622_RNA45SN1_45S_50ntFlanks | 4731 | 4752 | 22 | AAGAACGAAAGTCGGAGGTTCG | 445.8 | -6.5 | 1.5 | -4.4 | 9.8E-06 | 1.7E-05 | MB231/MB468 | MP |
| >hg38_21_+_8433172_8446622_RNA45SN1_45S_50ntFlanks@8191.8219.29\|CAGCCCGTGGACGGTGTGAGGCCGGTAGC | >hg38_21_+_8433172_8446622_RNA45SN1_45S_50ntFlanks | 8191 | 8219 | 29 | CAGCCCGTGGACGGTGTGAGGCCGGTAGC | 265.4 | -3.1 | 0.7 | -4.4 | 1.2E-05 | 1.7E-05 | MB231/MB468 | NUC |
| >hg38_21_+_8433172_8446622_RNA45SN1_45S_50ntFlanks@9544.9563.20\|GGTGGAGGTCCGTAGCGGTC | >hg38_21_+_8433172_8446622_RNA45SN1_45S_50ntFlanks | 9544 | 9563 | 20 | GGTGGAGGTCCGTAGCGGTC | 152.9 | -6.7 | 1.5 | -4.4 | 9.9E-06 | 1.7E-05 | MB231/MB468 | MP |
| >am_trnaMT_HisGTG_MT_+_12138_12206@18.50.33\|AAAACATCAGATTGTGAATCTGACAACAGAGGC | >am_trnaMT_HisGTG_MT_+_12138_12206 | 18 | 50 | 33 | AAAACATCAGATTGTGAATCTGACAACAGAGGC | 33.3 | 8.3 | 1.9 | 4.4 | 9.9E-06 | 1.7E-05 | MB231/MB468 | MP |
| >hg38_21_+_8433172_8446622_RNA45SN1_45S_50ntFlanks@9507.9527.21\|AACTATGCCTGGGCAGGGCGA | >hg38_21_+_8433172_8446622_RNA45SN1_45S_50ntFlanks | 9507 | 9527 | 21 | AACTATGCCTGGGCAGGGCGA | 103.0 | -10.6 | 2.4 | -4.4 | 9.9E-06 | 1.7E-05 | MB231/MB468 | MP |
| >trnaMT_PheGAA_MT_+_577_647@35.54.20\|GAAAATGTTTAGACGGGCTC | >trnaMT_PheGAA_MT_+_577_647 | 35 | 54 | 20 | GAAAATGTTTAGACGGGCTC | 462.7 | 3.7 | 0.8 | 4.4 | 9.9E-06 | 1.7E-05 | MB231/MB468 | MP |
| >hg38_21_+_8433172_8446622_RNA45SN1_45S_50ntFlanks@6721.6739.19\|AATTGCAGGACACATTGAT | >hg38_21_+_8433172_8446622_RNA45SN1_45S_50ntFlanks | 6721 | 6739 | 19 | AATTGCAGGACACATTGAT | 420.9 | -3.2 | 0.7 | -4.5 | 8.6E-06 | 1.7E-05 | BT20/MB468 | MITO |
| >hg38_21_+_8433172_8446622_RNA45SN1_45S_50ntFlanks@11552.11569.18\|CCTCGGCCGGCGCCTAGC | >hg38_21_+_8433172_8446622_RNA45SN1_45S_50ntFlanks | 11552 | 11569 | 18 | CCTCGGCCGGCGCCTAGC | 1236.3 | -3.2 | 0.7 | -4.4 | 1.2E-05 | 1.7E-05 | MB231/MB468 | MITO |
| >hg38_1_-_228634819_228635039_RNA5S12_5S_50ntFlanks@128.169.42\|CTTGGATGGGAGACCGCCTGGGAATACCGGGTGCTGTAGGCT | >hg38_1_-_228634819_228635039_RNA5S12_5S_50ntFlanks | 128 | 169 | 42 | CTTGGATGGGAGACCGCCTGGGAATACCGGGTGCTGTAGGCT | 369.3 | 6.8 | 1.5 | 4.4 | 8.6E-06 | 1.7E-05 | BT20/MB468 | MITO |
| >hg38_21_+_8433172_8446622_RNA45SN1_45S_50ntFlanks@12946.12988.43\|TAGACGACCTGCTTCTGGGTCGGGGTTTCGTACGTAGCAGAGC | >hg38_21_+_8433172_8446622_RNA45SN1_45S_50ntFlanks | 12946 | 12988 | 43 | TAGACGACCTGCTTCTGGGTCGGGGTTTCGTACGTAGCAGAGC | 866.0 | -4.4 | 0.9 | -4.9 | 1.1E-06 | 1.7E-05 | BT20/MB231 | CYTO |
| >hg38_21_+_8433172_8446622_RNA45SN1_45S_50ntFlanks@3840.3868.29\|CTCCTACTTGGATAACTGTGGTAATTCTA | >hg38_21_+_8433172_8446622_RNA45SN1_45S_50ntFlanks | 3840 | 3868 | 29 | CTCCTACTTGGATAACTGTGGTAATTCTA | 116.1 | -10.7 | 2.4 | -4.4 | 9.9E-06 | 1.7E-05 | MB231/MB468 | MP |
| >hg38_21_+_8433172_8446622_RNA45SN1_45S_50ntFlanks@12415.12449.35\|CTTCGATGTCGGCTCTTCCTATCATTGTGAAGCAG | >hg38_21_+_8433172_8446622_RNA45SN1_45S_50ntFlanks | 12415 | 12449 | 35 | CTTCGATGTCGGCTCTTCCTATCATTGTGAAGCAG | 1052.2 | 2.7 | 0.6 | 4.4 | 1.1E-05 | 1.7E-05 | MB231/MB468 | CYTO |
| >MI0000078\|hsa-mir-22&WithFlank&17\|-\|1713897\|1713993@21.41.21&[MIMAT0004495&hsa-miR-22-5p&offsets\|0\|-1;m-217&17\|-\|1713953\|1713973&offsets\|0\|0]\|AGTTCTTCAGTGGCAAGCTTT | >MI0000078\|hsa-mir-22&WithFlank&17\|-\|1713897\|1713993 | 21 | 41 | 21 | AGTTCTTCAGTGGCAAGCTTT | 906.2 | 4.5 | 1.0 | 4.6 | 3.7E-06 | 1.7E-05 | BT20/MB468 | MP |
| >hg38_21_+_8433172_8446622_RNA45SN1_45S_50ntFlanks@11733.11751.19\|TATGACTCTCTTAAGGTAG | >hg38_21_+_8433172_8446622_RNA45SN1_45S_50ntFlanks | 11733 | 11751 | 19 | TATGACTCTCTTAAGGTAG | 203.8 | -5.5 | 1.2 | -4.4 | 1.0E-05 | 1.7E-05 | BT20/MB468 | CYTO |
| >hg38_21_+_8433172_8446622_RNA45SN1_45S_50ntFlanks@12002.12031.30\|TGCGGGCCGCCGGTGAAATACCACTACTCT | >hg38_21_+_8433172_8446622_RNA45SN1_45S_50ntFlanks | 12002 | 12031 | 30 | TGCGGGCCGCCGGTGAAATACCACTACTCT | 894.0 | -4.9 | 1.1 | -4.4 | 1.0E-05 | 1.7E-05 | BT20/MB468 | CYTO |
| >hg38_21_+_8433172_8446622_RNA45SN1_45S_50ntFlanks@10686.10706.21\|GCGGCGTCCGGTGAGCTCTCG | >hg38_21_+_8433172_8446622_RNA45SN1_45S_50ntFlanks | 10686 | 10706 | 21 | GCGGCGTCCGGTGAGCTCTCG | 813.9 | -3.7 | 0.8 | -4.6 | 3.7E-06 | 1.7E-05 | BT20/MB468 | MP |
| >hg38_21_+_8433172_8446622_RNA45SN1_45S_50ntFlanks@10806.10829.24\|AACAATGTAGGTAAGGGAAGTCGG | >hg38_21_+_8433172_8446622_RNA45SN1_45S_50ntFlanks | 10806 | 10829 | 24 | AACAATGTAGGTAAGGGAAGTCGG | 153.8 | -5.3 | 1.2 | -4.4 | 1.0E-05 | 1.7E-05 | BT20/MB468 | CYTO |
| >hg38_21_+_8433172_8446622_RNA45SN1_45S_50ntFlanks@9952.9981.30\|CATGGAAGTCGGAATCCGCTAAGGAGTGTG | >hg38_21_+_8433172_8446622_RNA45SN1_45S_50ntFlanks | 9952 | 9981 | 30 | CATGGAAGTCGGAATCCGCTAAGGAGTGTG | 40.5 | 9.1 | 2.0 | 4.6 | 3.8E-06 | 1.7E-05 | BT20/MB468 | MP |
| >hg38_21_+_8433172_8446622_RNA45SN1_45S_50ntFlanks@11562.11600.39\|CGCCTAGCAGCCGACTTAGAACTGGTGCGGACCAGGGGA | >hg38_21_+_8433172_8446622_RNA45SN1_45S_50ntFlanks | 11562 | 11600 | 39 | CGCCTAGCAGCCGACTTAGAACTGGTGCGGACCAGGGGA | 381.4 | -7.1 | 1.6 | -4.4 | 1.3E-05 | 1.7E-05 | BT20/MB468 | NUC |
| >trna10_AspGTC_12_-_125424193_125424264@37.72.36,trna12_AspGTC_12_-_125411891_125411962@37.72.36,trna144_AspGTC_6_-_27551236_27551307@37.72.36,trna38_AspGTC_17_-_8125556_8125627@37.72.36,trna45_AspGTC_6_+_27447453_27447524@37.72.36,trna48_AspGTC_6_+_27471523_27471594@37.72.36,trna4_AspGTC_12_+_96429799_96429870@37.72.36,trna69_AspGTC_1_-_161440205_161440276@37.72.36,trna72_AspGTC_1_-_161432824_161432895@37.72.36,trna75_AspGTC_1_-_161425414_161425485@37.72.36,trna78_AspGTC_1_-_161418033_161418104@37.72.36,trna81_AspGTC_1_-_161410615_161410686@37.72.36\|ACGCGGGAGACCGGGGTTCGATTCCCCGACGGGGAG | >trna10_AspGTC_12_-_125424193_125424264 | 37 | 72 | 36 | ACGCGGGAGACCGGGGTTCGATTCCCCGACGGGGAG | 742.5 | -1.6 | 0.4 | -4.4 | 1.0E-05 | 1.7E-05 | BT20/MB468 | CYTO |
| >hg38_21_+_8433172_8446622_RNA45SN1_45S_50ntFlanks@5080.5117.38\|ATAACGAACGAGACTCTGGCATGCTAACTAGTTACGCG | >hg38_21_+_8433172_8446622_RNA45SN1_45S_50ntFlanks | 5080 | 5117 | 38 | ATAACGAACGAGACTCTGGCATGCTAACTAGTTACGCG | 75.7 | -10.1 | 2.3 | -4.4 | 9.9E-06 | 1.7E-05 | MB231/MB468 | MP |
| >hg38_21_+_8433172_8446622_RNA45SN1_45S_50ntFlanks@9362.9380.19\|ATGAAGGTGAAGGCCGGCG | >hg38_21_+_8433172_8446622_RNA45SN1_45S_50ntFlanks | 9362 | 9380 | 19 | ATGAAGGTGAAGGCCGGCG | 59.8 | -9.0 | 1.9 | -4.6 | 3.8E-06 | 1.7E-05 | BT20/MB468 | MP |
| >hg38_21_+_8433172_8446622_RNA45SN1_45S_50ntFlanks@3840.3861.22\|CTCCTACTTGGATAACTGTGGT | >hg38_21_+_8433172_8446622_RNA45SN1_45S_50ntFlanks | 3840 | 3861 | 22 | CTCCTACTTGGATAACTGTGGT | 402.7 | 2.6 | 0.6 | 4.4 | 1.1E-05 | 1.7E-05 | MB231/MB468 | CYTO |
| >hg38_21_+_8433172_8446622_RNA45SN1_45S_50ntFlanks@3771.3791.21\|CACGGCCGGTACAGTGAAACT | >hg38_21_+_8433172_8446622_RNA45SN1_45S_50ntFlanks | 3771 | 3791 | 21 | CACGGCCGGTACAGTGAAACT | 438.4 | -5.4 | 1.2 | -4.4 | 1.2E-05 | 1.7E-05 | MB231/MB468 | MITO |
| >hg38_21_+_8433172_8446622_RNA45SN1_45S_50ntFlanks@8153.8181.29\|GCCGCTCGTGGGGGGCCCAAGTCCTTCTG | >hg38_21_+_8433172_8446622_RNA45SN1_45S_50ntFlanks | 8153 | 8181 | 29 | GCCGCTCGTGGGGGGCCCAAGTCCTTCTG | 1059.3 | -3.4 | 0.8 | -4.4 | 8.7E-06 | 1.7E-05 | BT20/MB468 | MITO |
| >hg38_21_+_8433172_8446622_RNA45SN1_45S_50ntFlanks@10830.10852.23\|CAAGCCGGATCCGTAACTTCGGG | >hg38_21_+_8433172_8446622_RNA45SN1_45S_50ntFlanks | 10830 | 10852 | 23 | CAAGCCGGATCCGTAACTTCGGG | 283.9 | -6.1 | 1.4 | -4.4 | 1.0E-05 | 1.7E-05 | BT20/MB468 | CYTO |
| >MI0000457\|hsa-mir-141&WithFlank&12\|+\|6964091\|6964197@64.85.22&[MIMAT0000432&hsa-miR-141-3p&offsets\|-1\|-1;m-134&12\|+\|6964155\|6964175&offsets\|-1\|0]\|CTAACACTGTCTGGTAAAGATG | >MI0000457\|hsa-mir-141&WithFlank&12\|+\|6964091\|6964197 | 64 | 85 | 22 | CTAACACTGTCTGGTAAAGATG | 84.0 | 9.7 | 2.0 | 4.9 | 1.1E-06 | 1.7E-05 | BT20/MB231 | MP |
| >hg38_21_+_8433172_8446622_RNA45SN1_45S_50ntFlanks@10411.10430.20\|CAGTCGGTCCTGAGAGATGG | >hg38_21_+_8433172_8446622_RNA45SN1_45S_50ntFlanks | 10411 | 10430 | 20 | CAGTCGGTCCTGAGAGATGG | 155.2 | 10.6 | 2.2 | 4.9 | 1.1E-06 | 1.7E-05 | BT20/MB231 | MP |
| >hg38_21_+_8433172_8446622_RNA45SN1_45S_50ntFlanks@11600.11625.26\|AATCCGACTGTTTAATTAAAACAAAG | >hg38_21_+_8433172_8446622_RNA45SN1_45S_50ntFlanks | 11600 | 11625 | 26 | AATCCGACTGTTTAATTAAAACAAAG | 149.2 | 10.6 | 2.2 | 4.9 | 1.1E-06 | 1.7E-05 | BT20/MB231 | MP |
| >hg38_21_+_8433172_8446622_RNA45SN1_45S_50ntFlanks@11553.11591.39\|CTCGGCCGGCGCCTAGCAGCCGACTTAGAACTGGTGCGG | >hg38_21_+_8433172_8446622_RNA45SN1_45S_50ntFlanks | 11553 | 11591 | 39 | CTCGGCCGGCGCCTAGCAGCCGACTTAGAACTGGTGCGG | 518.5 | -7.6 | 1.7 | -4.4 | 1.3E-05 | 1.7E-05 | BT20/MB468 | NUC |
| >am_trnaMT_PheGAA_MT_+_577_647@44.62.19\|TAGACGGGCTCACATCACC | >am_trnaMT_PheGAA_MT_+_577_647 | 44 | 62 | 19 | TAGACGGGCTCACATCACC | 1488.4 | 6.0 | 1.4 | 4.4 | 1.0E-05 | 1.7E-05 | BT20/MB468 | CYTO |
| >trnaMT_GluTTC_MT_-_14674_14742@39.57.19,trnalookalike8_GluTTC_5_-_93905172_93905240@39.57.19\|ATTGGTCGTGGTTGTAGTC | >trnaMT_GluTTC_MT_-_14674_14742 | 39 | 57 | 19 | ATTGGTCGTGGTTGTAGTC | 314.6 | 5.8 | 1.3 | 4.4 | 1.1E-05 | 1.7E-05 | MB231/MB468 | CYTO |
| >hg38_21_+_8433172_8446622_RNA45SN1_45S_50ntFlanks@7990.8009.20\|GACGTGGCGACCCGCTGAAT | >hg38_21_+_8433172_8446622_RNA45SN1_45S_50ntFlanks | 7990 | 8009 | 20 | GACGTGGCGACCCGCTGAAT | 1422.3 | -3.0 | 0.7 | -4.4 | 1.0E-05 | 1.7E-05 | MB231/MB468 | MP |
| >hg38_21_+_8433172_8446622_RNA45SN1_45S_50ntFlanks@10349.10390.42\|AACGAGAACTTTGAAGGCCGAAGTGGAGAAGGGTTCCATGTG | >hg38_21_+_8433172_8446622_RNA45SN1_45S_50ntFlanks | 10349 | 10390 | 42 | AACGAGAACTTTGAAGGCCGAAGTGGAGAAGGGTTCCATGTG | 710.4 | -3.6 | 0.8 | -4.4 | 1.0E-05 | 1.7E-05 | MB231/MB468 | MP |
| >hg38_21_+_8433172_8446622_RNA45SN1_45S_50ntFlanks@11566.11586.21\|TAGCAGCCGACTTAGAACTGG | >hg38_21_+_8433172_8446622_RNA45SN1_45S_50ntFlanks | 11566 | 11586 | 21 | TAGCAGCCGACTTAGAACTGG | 271.7 | 3.0 | 0.7 | 4.4 | 1.1E-05 | 1.7E-05 | MB231/MB468 | CYTO |
| >hg38_21_+_8433172_8446622_RNA45SN1_45S_50ntFlanks@7972.8011.40\|AGACGCGACCTCAGATCAGACGTGGCGACCCGCTGAATTT | >hg38_21_+_8433172_8446622_RNA45SN1_45S_50ntFlanks | 7972 | 8011 | 40 | AGACGCGACCTCAGATCAGACGTGGCGACCCGCTGAATTT | 241.9 | 3.2 | 0.7 | 4.4 | 1.3E-05 | 1.7E-05 | BT20/MB468 | NUC |
| >hg38_MT_+_598_1651_RNR1_12S_50ntFlanks@978.1002.25\|TGTACTGGAAAGTGCACTTGGACGA | >hg38_MT_+_598_1651_RNR1_12S_50ntFlanks | 978 | 1002 | 25 | TGTACTGGAAAGTGCACTTGGACGA | 34.8 | 8.3 | 1.9 | 4.4 | 1.0E-05 | 1.7E-05 | MB231/MB468 | MP |

<table>
<tr><td>>hg38_21_+_8433172_8446622_RNA45SN1_45S_50ntFlanks@11212.11232.21|CGACTCTGGACGCGAGCCGGG</td><td>>hg38_21_+_8433172_8446622_RNA45SN1_45S_50ntFlanks</td><td>11212</td><td>11232</td><td>21</td><td>CGACTCTGGACGCGAGCCGGG</td><td>463.3</td><td>-3.5</td><td>0.8</td><td>-4.4</td><td>8.7E-06</td><td>1.7E-05</td><td>BT20/MB468</td><td>MITO</td></tr>
<tr><td>>hg38_21_+_8433172_8446622_RNA45SN1_45S_50ntFlanks@11703.11738.36|AATGAAGCGCGGGTAAACGGCGGGAGTAACTATGAC</td><td>>hg38_21_+_8433172_8446622_RNA45SN1_45S_50ntFlanks</td><td>11703</td><td>11738</td><td>36</td><td>AATGAAGCGCGGGTAAACGGCGGGAGTAACTATGAC</td><td>204.6</td><td>-4.9</td><td>1.1</td><td>-4.4</td><td>1.2E-05</td><td>1.7E-05</td><td>MB231/MB468</td><td>NUC</td></tr>
<tr><td>>hg38_21_+_8433172_8446622_RNA45SN1_45S_50ntFlanks@10686.10709.24|GCGGCGTCCGGTGAGCTCTCGCTG</td><td>>hg38_21_+_8433172_8446622_RNA45SN1_45S_50ntFlanks</td><td>10686</td><td>10709</td><td>24</td><td>GCGGCGTCCGGTGAGCTCTCGCTG</td><td>1002.3</td><td>-2.8</td><td>0.6</td><td>-4.4</td><td>1.2E-05</td><td>1.7E-05</td><td>MB231/MB468</td><td>NUC</td></tr>
<tr><td>>hg38_21_+_8433172_8446622_RNA45SN1_45S_50ntFlanks@8176.8213.38|CTTCTGATCGAGGCCCAGCCCGTGGACGGTGTGAGGCC</td><td>>hg38_21_+_8433172_8446622_RNA45SN1_45S_50ntFlanks</td><td>8176</td><td>8213</td><td>38</td><td>CTTCTGATCGAGGCCCAGCCCGTGGACGGTGTGAGGCC</td><td>319.5</td><td>-3.4</td><td>0.8</td><td>-4.4</td><td>1.2E-05</td><td>1.7E-05</td><td>MB231/MB468</td><td>MITO</td></tr>
<tr><td>>hg38_21_+_8433172_8446622_RNA45SN1_45S_50ntFlanks@5444.5465.22|CTCGGATCGGCCCCGCCGGGGT</td><td>>hg38_21_+_8433172_8446622_RNA45SN1_45S_50ntFlanks</td><td>5444</td><td>5465</td><td>22</td><td>CTCGGATCGGCCCCGCCGGGGT</td><td>360.8</td><td>-4.9</td><td>1.1</td><td>-4.4</td><td>1.2E-05</td><td>1.7E-05</td><td>MB231/MB468</td><td>NUC</td></tr>
<tr><td>>MI0000239|hsa-mir-197&WithFlank&1|+|109598887|109598973@54.76.23&[MIMAT0000227&hsa-miR-197-3p&offsets|0|+1;m-99&1|+|109598940|109598961&offsets|0|+1]|TTCACCACCTTCTCCACCCAGCA</td><td>>MI0000239|hsa-mir-197&WithFlank&1|+|109598887|109598973</td><td>54</td><td>76</td><td>23</td><td>TTCACCACCTTCTCCACCCAGCA</td><td>407.4</td><td>3.7</td><td>0.8</td><td>4.9</td><td>1.1E-06</td><td>1.7E-05</td><td>BT20/MB231</td><td>CYTO</td></tr>
<tr><td>>hg38_21_+_8433172_8446622_RNA45SN1_45S_50ntFlanks@10352.10369.18|GAGAACTTTGAAGGCCGA</td><td>>hg38_21_+_8433172_8446622_RNA45SN1_45S_50ntFlanks</td><td>10352</td><td>10369</td><td>18</td><td>GAGAACTTTGAAGGCCGA</td><td>414.5</td><td>-2.8</td><td>0.6</td><td>-4.8</td><td>1.4E-06</td><td>1.7E-05</td><td>BT20/MB231</td><td>NUC</td></tr>
<tr><td>>MI0000093|hsa-mir-92a-1&WithFlank&13|+|91351308|91351397@54.75.22(+1A);MI0000094|hsa-mir-92a-2&WithFlank&X|-|134169532|134169618@54.75.22(+1A)&[MIMAT0000092&hsa-miR-92a-3p&offsets|0|0(+1A);m-9&13|+|91351361|91351382&offsets|0|0(+1A)];[MIMAT0000092_1&hsa-miR-92a-3p&offsets|0|0(+1A);m-11&X|-|134169544|134169565&offsets|0|0(+1A)]|TATTGCACTTGTCCCGGCCTGTA</td><td>>MI0000093|hsa-mir-92a-1&WithFlank&13|+|91351308|91351397</td><td>54</td><td>75</td><td>22(+1A)</td><td>TATTGCACTTGTCCCGGCCTGTA</td><td>9311.9</td><td>2.8</td><td>0.6</td><td>4.4</td><td>1.2E-05</td><td>1.7E-05</td><td>MB231/MB468</td><td>NUC</td></tr>
<tr><td>>am_MI0000748|hsa-mir-130b&WithFlank&22|+|21653298|21653391@19.39.21&[MIMAT0004680&hsa-miR-130b-5p&offsets|0|0;m-234&22|+|21653316|21653337&offsets|0|-1]|ACTCTTTCCCTGTTGCACTAC</td><td>>am_MI0000748|hsa-mir-130b&WithFlank&22|+|21653298|21653391</td><td>19</td><td>39</td><td>21</td><td>ACTCTTTCCCTGTTGCACTAC</td><td>717.6</td><td>4.5</td><td>1.0</td><td>4.4</td><td>1.2E-05</td><td>1.7E-05</td><td>MB231/MB468</td><td>NUC</td></tr>
<tr><td>>hg38_21_+_8433172_8446622_RNA45SN1_45S_50ntFlanks@6645.6668.24|CTCGTACGACTCTTAGCGGTGGAT</td><td>>hg38_21_+_8433172_8446622_RNA45SN1_45S_50ntFlanks</td><td>6645</td><td>6668</td><td>24</td><td>CTCGTACGACTCTTAGCGGTGGAT</td><td>1234.3</td><td>3.0</td><td>0.7</td><td>4.6</td><td>3.8E-06</td><td>1.7E-05</td><td>BT20/MB468</td><td>MP</td></tr>
<tr><td>>hg38_21_+_8433172_8446622_RNA45SN1_45S_50ntFlanks@12415.12434.20|CTTCGATGTCGGCTCTTCCT</td><td>>hg38_21_+_8433172_8446622_RNA45SN1_45S_50ntFlanks</td><td>12415</td><td>12434</td><td>20</td><td>CTTCGATGTCGGCTCTTCCT</td><td>4154.1</td><td>-3.8</td><td>0.9</td><td>-4.4</td><td>1.2E-05</td><td>1.7E-05</td><td>MB231/MB468</td><td>MITO</td></tr>
<tr><td>>hg38_21_+_8433172_8446622_RNA45SN1_45S_50ntFlanks@11197.11225.29|GGCCGGCGGCGGCGGCGACTCTGGACGCG</td><td>>hg38_21_+_8433172_8446622_RNA45SN1_45S_50ntFlanks</td><td>11197</td><td>11225</td><td>29</td><td>GGCCGGCGGCGGCGGCGACTCTGGACGCG</td><td>155.7</td><td>-5.4</td><td>1.2</td><td>-4.4</td><td>1.0E-05</td><td>1.7E-05</td><td>BT20/MB468</td><td>CYTO</td></tr>
<tr><td>>hg38_21_+_8433172_8446622_RNA45SN1_45S_50ntFlanks@9546.9574.29|TGGAGGTCCGTAGCGGTCCTGACGTGCAA</td><td>>hg38_21_+_8433172_8446622_RNA45SN1_45S_50ntFlanks</td><td>9546</td><td>9574</td><td>29</td><td>TGGAGGTCCGTAGCGGTCCTGACGTGCAA</td><td>444.4</td><td>-7.1</td><td>1.6</td><td>-4.4</td><td>1.3E-05</td><td>1.7E-05</td><td>BT20/MB468</td><td>NUC</td></tr>
<tr><td>>MI0002470|hsa-mir-486-1&WithFlank&8|-|41660435|41660514@10.29.20;MI0023622|hsa-mir-486-2&WithFlank&8|+|41660438|41660513@7.26.20&[MIMAT0002177&hsa-miR-486-5p&offsets|0|-2;m-107&8|-|41660484|41660505&offsets|0|-2];[MIMAT0002177_1&hsa-miR-486-5p&offsets|0|-2;TJU_CMC_MD2.ID03138.5p-miR&offsets|0|-4;m-106&8|+|41660444|41660465&offsets|0|-2]|TCCTGTACTGAGCTGCCCCG</td><td>>MI0002470|hsa-mir-486-1&WithFlank&8|-|41660435|41660514</td><td>10</td><td>29</td><td>20</td><td>TCCTGTACTGAGCTGCCCCG</td><td>79.9</td><td>-9.7</td><td>2.0</td><td>-4.9</td><td>1.1E-06</td><td>1.7E-05</td><td>BT20/MB231</td><td>MP</td></tr>
<tr><td>>am_trna116_GluCTC_1_-_145399233_145399304@1.29.29,trna59_GluCTC_1_+_249168447_249168518@1.29.29,trna71_GluCTC_1_-_161439189_161439260@1.29.29,trna74_GluCTC_1_-_161431809_161431880@1.29.29,trna77_GluCTC_1_-_161424398_161424469@1.29.29,trna77_GluCTC_6_+_28949976_28950047@1.29.29,trna80_GluCTC_1_-_161417018_161417089@1.29.29,trna87_GluCTC_6_-_126101393_126101464@1.29.29|TCCCTGGTGGTCTAGTGGTTAGGATTCGG</td><td>>am_trna116_GluCTC_1_-_145399233_145399304</td><td>1</td><td>29</td><td>29</td><td>TCCCTGGTGGTCTAGTGGTTAGGATTCGG</td><td>76240.2</td><td>-3.5</td><td>0.8</td><td>-4.4</td><td>1.1E-05</td><td>1.7E-05</td><td>MB231/MB468</td><td>CYTO</td></tr>
<tr><td>>hg38_21_+_8433172_8446622_RNA45SN1_45S_50ntFlanks@11570.11600.31|AGCCGACTTAGAACTGGTGCGGACCAGGGGA</td><td>>hg38_21_+_8433172_8446622_RNA45SN1_45S_50ntFlanks</td><td>11570</td><td>11600</td><td>31</td><td>AGCCGACTTAGAACTGGTGCGGACCAGGGGA</td><td>369.0</td><td>-4.9</td><td>1.1</td><td>-4.4</td><td>1.2E-05</td><td>1.7E-05</td><td>MB231/MB468</td><td>MITO</td></tr>
<tr><td>>hg38_21_+_8433172_8446622_RNA45SN1_45S_50ntFlanks@9568.9603.36|CGTGCAAATCGGTCGTCCGACCTGGGTATAGGGGCG</td><td>>hg38_21_+_8433172_8446622_RNA45SN1_45S_50ntFlanks</td><td>9568</td><td>9603</td><td>36</td><td>CGTGCAAATCGGTCGTCCGACCTGGGTATAGGGGCG</td><td>1354.1</td><td>-7.8</td><td>1.8</td><td>-4.4</td><td>1.3E-05</td><td>1.7E-05</td><td>BT20/MB468</td><td>NUC</td></tr>
<tr><td>>hg38_21_+_8433172_8446622_RNA45SN1_45S_50ntFlanks@10766.10785.20|CGCAGCAGGTCTCCAAGGTG</td><td>>hg38_21_+_8433172_8446622_RNA45SN1_45S_50ntFlanks</td><td>10766</td><td>10785</td><td>20</td><td>CGCAGCAGGTCTCCAAGGTG</td><td>223.7</td><td>-2.1</td><td>0.5</td><td>-4.4</td><td>1.2E-05</td><td>1.7E-05</td><td>MB231/MB468</td><td>CYTO</td></tr>
<tr><td>>MI0000293|hsa-mir-217&WithFlank&2|-|55982961|55983082@41.63.23&[MIMAT0000274&hsa-miR-217-5p&offsets|0|0;m-311&2|-|55983020|55983042&offsets|0|0]|TACTGCATCAGGAACTGATTGGA</td><td>>MI0000293|hsa-mir-217&WithFlank&2|-|55982961|55983082</td><td>41</td><td>63</td><td>23</td><td>TACTGCATCAGGAACTGATTGGA</td><td>32.9</td><td>8.3</td><td>1.9</td><td>4.4</td><td>1.0E-05</td><td>1.7E-05</td><td>MB231/MB468</td><td>MP</td></tr>
<tr><td>>hg38_21_+_8433172_8446622_RNA45SN1_45S_50ntFlanks@4373.4391.19|AAAAGCTCGTAGTTGGATC</td><td>>hg38_21_+_8433172_8446622_RNA45SN1_45S_50ntFlanks</td><td>4373</td><td>4391</td><td>19</td><td>AAAAGCTCGTAGTTGGATC</td><td>112.6</td><td>-4.9</td><td>1.1</td><td>-4.4</td><td>1.0E-05</td><td>1.7E-05</td><td>MB231/MB468</td><td>MP</td></tr>
<tr><td>>MI0000433|hsa-let-7g&WithFlank&3|-|52268272|52268367@11.32.22(+1A)&[MIMAT0000414&hsa-let-7g-5p&offsets|0|0(+1A);m-58&3|-|52268336|52268357&offsets|0|0(+1A)]|TGAGGTAGTAGTTTGTACAGTTA</td><td>>MI0000433|hsa-let-7g&WithFlank&3|-|52268272|52268367</td><td>11</td><td>32</td><td>22(+1A)</td><td>TGAGGTAGTAGTTTGTACAGTTA</td><td>375.6</td><td>3.9</td><td>0.9</td><td>4.4</td><td>1.2E-05</td><td>1.7E-05</td><td>MB231/MB468</td><td>MITO</td></tr>
<tr><td>>hg38_21_+_8433172_8446622_RNA45SN1_45S_50ntFlanks@6646.6689.44|TCGTACGACTCTTAGCGGTGGATCACTCGGCTCGTGCGTCGATG</td><td>>hg38_21_+_8433172_8446622_RNA45SN1_45S_50ntFlanks</td><td>6646</td><td>6689</td><td>44</td><td>TCGTACGACTCTTAGCGGTGGATCACTCGGCTCGTGCGTCGATG</td><td>56829.8</td><td>-4.1</td><td>0.9</td><td>-4.4</td><td>1.2E-05</td><td>1.7E-05</td><td>MB231/MB468</td><td>MITO</td></tr>
<tr><td>>hg38_21_+_8433172_8446622_RNA45SN1_45S_50ntFlanks@4976.5001.26|CCGGACACGGACAGGATTGACAGATT</td><td>>hg38_21_+_8433172_8446622_RNA45SN1_45S_50ntFlanks</td><td>4976</td><td>5001</td><td>26</td><td>CCGGACACGGACAGGATTGACAGATT</td><td>116.5</td><td>2.1</td><td>0.5</td><td>4.4</td><td>1.3E-05</td><td>1.8E-05</td><td>BT20/MB468</td><td>NUC</td></tr>
<tr><td>>hg38_21_+_8433172_8446622_RNA45SN1_45S_50ntFlanks@10794.10823.30|CTGGCATGTTGGAACAATGTAGGTAAGGGA</td><td>>hg38_21_+_8433172_8446622_RNA45SN1_45S_50ntFlanks</td><td>10794</td><td>10823</td><td>30</td><td>CTGGCATGTTGGAACAATGTAGGTAAGGGA</td><td>472.7</td><td>-3.2</td><td>0.7</td><td>-4.6</td><td>3.8E-06</td><td>1.8E-05</td><td>BT20/MB468</td><td>MP</td></tr>
<tr><td>>hg38_21_+_8433172_8446622_RNA45SN1_45S_50ntFlanks@12139.12158.20|GGACAGTGCCAGGTGGGGAG</td><td>>hg38_21_+_8433172_8446622_RNA45SN1_45S_50ntFlanks</td><td>12139</td><td>12158</td><td>20</td><td>GGACAGTGCCAGGTGGGGAG</td><td>45.8</td><td>-8.6</td><td>1.9</td><td>-4.6</td><td>3.8E-06</td><td>1.8E-05</td><td>BT20/MB468</td><td>MP</td></tr>
<tr><td>>MI0003776|hsa-mir-320b-1&WithFlank&1|+|116671740|116671823@48.69.22;MI0003839|hsa-mir-320b-2&WithFlank&1|-|224257034|224257116@47.68.22&[MIMAT0005792&hsa-miR-320b&offsets|0|0;m-183&1|+|116671787|116671806&offsets|0|+2];[MIMAT0005792_1&hsa-miR-320b&offsets|0|0;m-183&1|-|224257051|224257070&offsets|0|+2]|AAAAGCTGGGTTGAGAGGGCAA</td><td>>MI0003776|hsa-mir-320b-1&WithFlank&1|+|116671740|116671823</td><td>48</td><td>69</td><td>22</td><td>AAAAGCTGGGTTGAGAGGGCAA</td><td>164.4</td><td>4.2</td><td>0.9</td><td>4.9</td><td>1.2E-06</td><td>1.8E-05</td><td>BT20/MB231</td><td>CYTO</td></tr>
<tr><td>>am_trna111_HisGTG_1_-_147774845_147774916@-1G.23.24,trna118_HisGTG_1_-_145396881_145396952@-1G.23.24,trna16_HisGTG_1_+_146544773_146544844@-1G.23.24,trna1_HisGTG_15_+_45493349_45493420@-1G.23.24,trna21_HisGTG_1_+_147753471_147753542@-1G.23.24,trna33_HisGTG_6_+_27125906_27125977@-1G.23.24,trna7_HisGTG_9_-_14433938_14434009@-1G.23.24,trna8_HisGTG_15_-_45492611_45492682@-1G.23.24,trna9_HisGTG_15_-_45490804_45490875@-1G.23.24|GGCCGTGATCGTATAGTGGTTAGT</td><td>>am_trna111_HisGTG_1_-_147774845_147774916</td><td>-1G</td><td>23</td><td>24</td><td>GGCCGTGATCGTATAGTGGTTAGT</td><td>635.5</td><td>-4.3</td><td>0.9</td><td>-4.9</td><td>1.2E-06</td><td>1.8E-05</td><td>BT20/MB231</td><td>CYTO</td></tr>
<tr><td>>hg38_21_+_8433172_8446622_RNA45SN1_45S_50ntFlanks@12419.12461.43|GATGTCGGCTCTTCCTATCATTGTGAAGCAGAATTCACCAAGC</td><td>>hg38_21_+_8433172_8446622_RNA45SN1_45S_50ntFlanks</td><td>12419</td><td>12461</td><td>43</td><td>GATGTCGGCTCTTCCTATCATTGTGAAGCAGAATTCACCAAGC</td><td>313.2</td><td>-4.3</td><td>0.9</td><td>-4.9</td><td>1.2E-06</td><td>1.8E-05</td><td>BT20/MB231</td><td>CYTO</td></tr>
<tr><td>>hg38_21_+_8433172_8446622_RNA45SN1_45S_50ntFlanks@11781.11806.26|CGCATGAATGGATGAACGAGATTCCC</td><td>>hg38_21_+_8433172_8446622_RNA45SN1_45S_50ntFlanks</td><td>11781</td><td>11806</td><td>26</td><td>CGCATGAATGGATGAACGAGATTCCC</td><td>522.8</td><td>-2.8</td><td>0.6</td><td>-4.4</td><td>8.8E-06</td><td>1.8E-05</td><td>BT20/MB468</td><td>MITO</td></tr>
<tr><td>>MI0000465|hsa-mir-191&WithFlank&3|-|49020612|49020715@22.42.21&[MIMAT0000440&hsa-miR-191-5p&offsets|0|-2;m-48&3|-|49020672|49020694&offsets|0|-2]|CAACGGAATCCCAAAAGCAGC</td><td>>MI0000465|hsa-mir-191&WithFlank&3|-|49020612|49020715</td><td>22</td><td>42</td><td>21</td><td>CAACGGAATCCCAAAAGCAGC</td><td>280.0</td><td>1.8</td><td>0.4</td><td>4.4</td><td>1.3E-05</td><td>1.8E-05</td><td>BT20/MB468</td><td>NUC</td></tr>
<tr><td>>hg38_21_+_8433172_8446622_RNA45SN1_45S_50ntFlanks@11199.11220.22|CCGGCGGCGGCGGCGACTCTGG</td><td>>hg38_21_+_8433172_8446622_RNA45SN1_45S_50ntFlanks</td><td>11199</td><td>11220</td><td>22</td><td>CCGGCGGCGGCGGCGACTCTGG</td><td>758.1</td><td>-6.8</td><td>1.6</td><td>-4.4</td><td>1.2E-05</td><td>1.8E-05</td><td>MB231/MB468</td><td>NUC</td></tr>
<tr><td>>hg38_21_+_8433172_8446622_RNA45SN1_45S_50ntFlanks@3840.3860.21|CTCCTACTTGGATAACTGTGG</td><td>>hg38_21_+_8433172_8446622_RNA45SN1_45S_50ntFlanks</td><td>3840</td><td>3860</td><td>21</td><td>CTCCTACTTGGATAACTGTGG</td><td>2061.3</td><td>2.4</td><td>0.5</td><td>4.4</td><td>1.0E-05</td><td>1.8E-05</td><td>BT20/MB468</td><td>CYTO</td></tr>
<tr><td>>hg38_21_+_8433172_8446622_RNA45SN1_45S_50ntFlanks@10839.10868.30|TCCGTAACTTCGGGATAAGGATTGGCTCTA</td><td>>hg38_21_+_8433172_8446622_RNA45SN1_45S_50ntFlanks</td><td>10839</td><td>10868</td><td>30</td><td>TCCGTAACTTCGGGATAAGGATTGGCTCTA</td><td>380.3</td><td>-4.1</td><td>0.9</td><td>-4.4</td><td>1.0E-05</td><td>1.8E-05</td><td>BT20/MB468</td><td>CYTO</td></tr>
<tr><td>>MI0000803|hsa-mir-330&WithFlank&19|-|45638988|45639093@64.86.23(+1U)&[MIMAT0000751&hsa-miR-330-3p&offsets|+1|+1(+1U);m-347&19|-|45639009|45639030&offsets|0|+1(+1U)]|CAAAGCACACGGCCTGCAGAGAGT</td><td>>MI0000803|hsa-mir-330&WithFlank&19|-|45638988|45639093</td><td>64</td><td>86</td><td>23(+1U)</td><td>CAAAGCACACGGCCTGCAGAGAGT</td><td>185.5</td><td>7.3</td><td>1.7</td><td>4.4</td><td>1.2E-05</td><td>1.8E-05</td><td>MB231/MB468</td><td>MITO</td></tr>
<tr><td>>MI0000102|hsa-mir-100&WithFlank&11|-|122152223|122152314@19.38.20(+1G)&[MIMAT0000098&hsa-miR-100-5p&offsets|0|-2(+1G);m-27&11|-|122152276|122152296&offsets|0|-1(+1G)]|AACCCGTAGATCCGAACTTGG</td><td>>MI0000102|hsa-mir-100&WithFlank&11|-|122152223|122152314</td><td>19</td><td>38</td><td>20(+1G)</td><td>AACCCGTAGATCCGAACTTGG</td><td>2490.7</td><td>-6.0</td><td>1.2</td><td>-4.8</td><td>1.4E-06</td><td>1.8E-05</td><td>BT20/MB231</td><td>NUC</td></tr>
<tr><td>>hg38_21_+_8433172_8446622_RNA45SN1_45S_50ntFlanks@10466.10497.32|CGTTGCCCTCGGCCGATCGAAAGGGAGTCGGG</td><td>>hg38_21_+_8433172_8446622_RNA45SN1_45S_50ntFlanks</td><td>10466</td><td>10497</td><td>32</td><td>CGTTGCCCTCGGCCGATCGAAAGGGAGTCGGG</td><td>947.1</td><td>-6.7</td><td>1.5</td><td>-4.4</td><td>1.2E-05</td><td>1.8E-05</td><td>MB231/MB468</td><td>NUC</td></tr>
<tr><td>>hg38_21_+_8433172_8446622_RNA45SN1_45S_50ntFlanks@5486.5516.31|GCGCTGAGAAGACGGTCGAACTTGACTATCT</td><td>>hg38_21_+_8433172_8446622_RNA45SN1_45S_50ntFlanks</td><td>5486</td><td>5516</td><td>31</td><td>GCGCTGAGAAGACGGTCGAACTTGACTATCT</td><td>768.4</td><td>4.4</td><td>0.9</td><td>4.8</td><td>1.3E-06</td><td>1.8E-05</td><td>BT20/MB231</td><td>MITO</td></tr>
<tr><td>>hg38_21_+_8433172_8446622_RNA45SN1_45S_50ntFlanks@9565.9595.31|TGACGTGCAAATCGGTCGTCCGACCTGGGTA</td><td>>hg38_21_+_8433172_8446622_RNA45SN1_45S_50ntFlanks</td><td>9565</td><td>9595</td><td>31</td><td>TGACGTGCAAATCGGTCGTCCGACCTGGGTA</td><td>2845.1</td><td>-8.4</td><td>1.9</td><td>-4.4</td><td>1.3E-05</td><td>1.8E-05</td><td>BT20/MB468</td><td>NUC</td></tr>
<tr><td>>hg38_21_+_8433172_8446622_RNA45SN1_45S_50ntFlanks@7986.8017.32|ATCAGACGTGGCGACCCGCTGAATTTAAGCAT</td><td>>hg38_21_+_8433172_8446622_RNA45SN1_45S_50ntFlanks</td><td>7986</td><td>8017</td><td>32</td><td>ATCAGACGTGGCGACCCGCTGAATTTAAGCAT</td><td>1987.7</td><td>-7.0</td><td>1.6</td><td>-4.4</td><td>1.2E-05</td><td>1.8E-05</td><td>MB231/MB468</td><td>MITO</td></tr>
<tr><td>>hg38_21_+_8433172_8446622_RNA45SN1_45S_50ntFlanks@11561.11605.45|GCGCCTAGCAGCCGACTTAGAACTGGTGCGGACCAGGGGAATCCG</td><td>>hg38_21_+_8433172_8446622_RNA45SN1_45S_50ntFlanks</td><td>11561</td><td>11605</td><td>45</td><td>GCGCCTAGCAGCCGACTTAGAACTGGTGCGGACCAGGGGAATCCG</td><td>94.4</td><td>-10.4</td><td>2.4</td><td>-4.4</td><td>1.0E-05</td><td>1.8E-05</td><td>MB231/MB468</td><td>MP</td></tr>
<tr><td>>hg38_21_+_8433172_8446622_RNA45SN1_45S_50ntFlanks@4128.4162.35|CGATTCCGGAGAGGGAGCCTGAGAAACGGCTACCA</td><td>>hg38_21_+_8433172_8446622_RNA45SN1_45S_50ntFlanks</td><td>4128</td><td>4162</td><td>35</td><td>CGATTCCGGAGAGGGAGCCTGAGAAACGGCTACCA</td><td>454.2</td><td>-6.1</td><td>1.4</td><td>-4.4</td><td>1.0E-05</td><td>1.8E-05</td><td>BT20/MB468</td><td>CYTO</td></tr>
<tr><td>>MI0000077|hsa-mir-21&WithFlank&17|+|59841260|59841343@52.73.22&[MIMAT0004494&hsa-miR-21-3p&offsets|0|+1;m-52&17|+|59841311|59841332&offsets|0|0]|CAACACCAGTCGATGGGCTGTC</td><td>>MI0000077|hsa-mir-21&WithFlank&17|+|59841260|59841343</td><td>52</td><td>73</td><td>22</td><td>CAACACCAGTCGATGGGCTGTC</td><td>1229.9</td><td>4.0</td><td>0.9</td><td>4.4</td><td>1.0E-05</td><td>1.8E-05</td><td>MB231/MB468</td><td>MP</td></tr>
</table>

<table>
<tr><td>>hg38_21_+_8433172_8446622_RNA45SN1_45S_50ntFlanks@6655.6686.32|TCTTAGCGGTGGATCACTCGGCTCGTGCGTCG</td><td>>hg38_21_+_8433172_8446622_RNA45SN1_45S_50ntFlanks</td><td>6655</td><td>6686</td><td>32</td><td>TCTTAGCGGTGGATCACTCGGCTCGTGCGTCG</td><td>543.3</td><td>-3.3</td><td>0.8</td><td>-4.4</td><td>1.2E-05</td><td>1.8E-05</td><td>MB231/MB468</td><td>MITO</td></tr>
<tr><td>>hg38_21_+_8433172_8446622_RNA45SN1_45S_50ntFlanks@10514.10533.20|CGGAGTGGCGGAGATGGGCG</td><td>>hg38_21_+_8433172_8446622_RNA45SN1_45S_50ntFlanks</td><td>10514</td><td>10533</td><td>20</td><td>CGGAGTGGCGGAGATGGGCG</td><td>1002.7</td><td>-7.9</td><td>1.8</td><td>-4.4</td><td>1.3E-05</td><td>1.8E-05</td><td>BT20/MB468</td><td>NUC</td></tr>
<tr><td>>hg38_MT_+_1621_3279_RNR2_16S_50ntFlanks@1095.1117.23|AGACGAGAAGACCCTATGGAGCT</td><td>>hg38_MT_+_1621_3279_RNR2_16S_50ntFlanks</td><td>1095</td><td>1117</td><td>23</td><td>AGACGAGAAGACCCTATGGAGCT</td><td>403.8</td><td>4.2</td><td>0.9</td><td>4.6</td><td>3.9E-06</td><td>1.8E-05</td><td>BT20/MB468</td><td>MP</td></tr>
<tr><td>>hg38_21_+_8433172_8446622_RNA45SN1_45S_50ntFlanks@8123.8142.20|ATGTGGCGTACGGAAGACCC</td><td>>hg38_21_+_8433172_8446622_RNA45SN1_45S_50ntFlanks</td><td>8123</td><td>8142</td><td>20</td><td>ATGTGGCGTACGGAAGACCC</td><td>83.1</td><td>9.7</td><td>2.0</td><td>4.9</td><td>1.1E-06</td><td>1.8E-05</td><td>BT20/MB231</td><td>MP</td></tr>
<tr><td>>MI0002470|hsa-mir-486-1&WithFlank&8|-|41660435|41660514@10.31.22(+1A);MI0023622|hsa-mir-486-2&WithFlank&8|+|41660438|41660513@7.28.22(+1A)&[MIMAT0002177&hsa-miR-486-5p&offsets|0|0(+1A);m-107&8|-|41660484|41660505&offsets|0|0(+1A)];[MIMAT0002177_1&hsa-miR-486-5p&offsets|0|0(+1A);TJU_CMC_MD2.ID03138.5p-miR&offsets|0|-2(+1A);m-106&8|+|41660444|41660465&offsets|0|0(+1A)]|TCCTGTACTGAGCTGCCCCGAGA</td><td>>MI0002470|hsa-mir-486-1&WithFlank&8|-|41660435|41660514</td><td>10</td><td>31</td><td>22(+1A)</td><td>TCCTGTACTGAGCTGCCCCGAGA</td><td>77.9</td><td>-9.7</td><td>2.0</td><td>-4.9</td><td>1.2E-06</td><td>1.8E-05</td><td>BT20/MB231</td><td>MP</td></tr>
<tr><td>>hg38_21_+_8433172_8446622_RNA45SN1_45S_50ntFlanks@5423.5442.20|ATTGGATGGTTTAGTGAGGC</td><td>>hg38_21_+_8433172_8446622_RNA45SN1_45S_50ntFlanks</td><td>5423</td><td>5442</td><td>20</td><td>ATTGGATGGTTTAGTGAGGC</td><td>75.0</td><td>-10.1</td><td>2.3</td><td>-4.4</td><td>1.0E-05</td><td>1.8E-05</td><td>MB231/MB468</td><td>MP</td></tr>
<tr><td>>hg38_21_+_8433172_8446622_RNA45SN1_45S_50ntFlanks@4380.4404.25|CGTAGTTGGATCTTGGGAGCGGGCG</td><td>>hg38_21_+_8433172_8446622_RNA45SN1_45S_50ntFlanks</td><td>4380</td><td>4404</td><td>25</td><td>CGTAGTTGGATCTTGGGAGCGGGCG</td><td>108.6</td><td>-6.5</td><td>1.5</td><td>-4.4</td><td>1.0E-05</td><td>1.8E-05</td><td>MB231/MB468</td><td>MP</td></tr>
<tr><td>>MI0000289|hsa-mir-181a-1&WithFlank&1|-|198859038|198859159@30.51.22(+1A);MI0000269|hsa-mir-181a-2&WithFlank&9|+|124692436|124692557@45.66.22(+1A)&[MIMAT0000256&hsa-miR-181a-5p&offsets|0|-1(+1A);m-32&1|-|198859108|198859130&offsets|0|-1(+1A)];[MIMAT0000256_1&hsa-miR-181a-5p&offsets|0|-1(+1A);m-32&9|+|124692480|124692502&offsets|0|-1(+1A)]|AACATTCAACGCTGTCGGTGAGA</td><td>>MI0000289|hsa-mir-181a-1&WithFlank&1|-|198859038|198859159</td><td>30</td><td>51</td><td>22(+1A)</td><td>AACATTCAACGCTGTCGGTGAGA</td><td>311.7</td><td>4.3</td><td>1.0</td><td>4.4</td><td>1.3E-05</td><td>1.8E-05</td><td>MB231/MB468</td><td>MITO</td></tr>
<tr><td>>hg38_21_+_8433172_8446622_RNA45SN1_45S_50ntFlanks@5525.5550.26|TAAAAGTCGTAACAAGGTTTCCGTAG</td><td>>hg38_21_+_8433172_8446622_RNA45SN1_45S_50ntFlanks</td><td>5525</td><td>5550</td><td>26</td><td>TAAAAGTCGTAACAAGGTTTCCGTAG</td><td>242.9</td><td>-5.7</td><td>1.3</td><td>-4.4</td><td>1.1E-05</td><td>1.8E-05</td><td>BT20/MB468</td><td>CYTO</td></tr>
<tr><td>>MI0003591|hsa-mir-584&WithFlank&5|-|149062307|149062415@61.82.22&[MIMAT0022708&hsa-miR-584-3p&offsets|0|0;m-792&5|-|149062334|149062355&offsets|0|0]|TCAGTTCCAGGCCAACCAGGCT</td><td>>MI0003591|hsa-mir-584&WithFlank&5|-|149062307|149062415</td><td>61</td><td>82</td><td>22</td><td>TCAGTTCCAGGCCAACCAGGCT</td><td>32.9</td><td>8.3</td><td>1.9</td><td>4.4</td><td>1.0E-05</td><td>1.8E-05</td><td>MB231/MB468</td><td>MP</td></tr>
<tr><td>>MI0000750|hsa-mir-26a-2&WithFlank&12|-|57824603|57824698@20.40.21(+1C);MI0000083|hsa-mir-26a-1&WithFlank&3|+|37969398|37969486@16.36.21(+1C)&[MIMAT0000082&hsa-miR-26a-5p&offsets|0|-1(+1C);m-33&12|-|57824658|57824679&offsets|0|-1(+1C)];[MIMAT0000082_1&hsa-miR-26a-5p&offsets|0|-1(+1C);m-34&3|+|37969413|37969434&offsets|0|-1(+1C)]|TTCAAGTAATCCAGGATAGGCC</td><td>>MI0000750|hsa-mir-26a-2&WithFlank&12|-|57824603|57824698</td><td>20</td><td>40</td><td>21(+1C)</td><td>TTCAAGTAATCCAGGATAGGCC</td><td>12279.0</td><td>4.1</td><td>0.9</td><td>4.4</td><td>1.0E-05</td><td>1.8E-05</td><td>MB231/MB468</td><td>MP</td></tr>
<tr><td>>MI0005756|hsa-mir-934&WithFlank&X|+|136550872|136550966@21.41.21(+1A)&[MIMAT0004977&hsa-miR-934&offsets|0|-1(+1A);m-943&X|+|136550892|136550913&offsets|0|-1(+1A)]|TGTCTACTACTGGAGACACTGA</td><td>>MI0005756|hsa-mir-934&WithFlank&X|+|136550872|136550966</td><td>21</td><td>41</td><td>21(+1A)</td><td>TGTCTACTACTGGAGACACTGA</td><td>128.7</td><td>5.4</td><td>1.2</td><td>4.4</td><td>1.3E-05</td><td>1.8E-05</td><td>BT20/MB468</td><td>NUC</td></tr>
<tr><td>>hg38_21_+_8433172_8446622_RNA45SN1_45S_50ntFlanks@7988.8012.25|CAGACGTGGCGACCCGCTGAATTTA</td><td>>hg38_21_+_8433172_8446622_RNA45SN1_45S_50ntFlanks</td><td>7988</td><td>8012</td><td>25</td><td>CAGACGTGGCGACCCGCTGAATTTA</td><td>953.2</td><td>-2.8</td><td>0.6</td><td>-4.4</td><td>1.2E-05</td><td>1.8E-05</td><td>MB231/MB468</td><td>NUC</td></tr>
<tr><td>>hg38_21_+_8433172_8446622_RNA45SN1_45S_50ntFlanks@10823.10865.43|AAGTCGGCAAGCCGGATCCGTAACTTCGGGATAAGGATTGGCT</td><td>>hg38_21_+_8433172_8446622_RNA45SN1_45S_50ntFlanks</td><td>10823</td><td>10865</td><td>43</td><td>AAGTCGGCAAGCCGGATCCGTAACTTCGGGATAAGGATTGGCT</td><td>75.9</td><td>-10.1</td><td>2.3</td><td>-4.4</td><td>1.0E-05</td><td>1.8E-05</td><td>MB231/MB468</td><td>MP</td></tr>
<tr><td>>hg38_21_+_8433172_8446622_RNA45SN1_45S_50ntFlanks@8357.8387.31|AAAAGAACTTTGAAGAGAGAGTTCAAGAGGG</td><td>>hg38_21_+_8433172_8446622_RNA45SN1_45S_50ntFlanks</td><td>8357</td><td>8387</td><td>31</td><td>AAAAGAACTTTGAAGAGAGAGTTCAAGAGGG</td><td>292.3</td><td>-5.8</td><td>1.3</td><td>-4.4</td><td>1.1E-05</td><td>1.8E-05</td><td>BT20/MB468</td><td>CYTO</td></tr>
<tr><td>>am_trna10_ValCAC_5_-_180649395_180649467@1.36.36,trna18_ValCAC_5_-_180529253_180529325@1.36.36,trna2_ValCAC_5_+_180524070_180524142@1.36.36,trna6_ValCAC_5_+_180600650_180600722@1.36.36,trna85_ValCAC_1_-_161369490_161369562@1.36.36,trna90_ValCAC_1_-_149684088_149684161@1.36.36,trna98_ValCAC_1_-_149298555_149298627@1.36.36,trna9_ValCAC_6_+_26538282_26538354@1.36.36|GTTTCCGTAGTGTAGTGGTTATCACGTTCGCCTCAC</td><td>>am_trna10_ValCAC_5_-_180649395_180649467</td><td>1</td><td>36</td><td>36</td><td>GTTTCCGTAGTGTAGTGGTTATCACGTTCGCCTCAC</td><td>55.4</td><td>9.0</td><td>2.0</td><td>4.4</td><td>1.0E-05</td><td>1.8E-05</td><td>MB231/MB468</td><td>MP</td></tr>
<tr><td>>MI0000061|hsa-let-7a-2&WithFlank&11|-|122146516|122146599@56.76.21&[MIMAT0010195&hsa-let-7a-2-3p&offsets|0|-1;m-426&11|-|122146523|122146544&offsets|0|-1]|CTGTACAGCCTCCTAGCTTTC</td><td>>MI0000061|hsa-let-7a-2&WithFlank&11|-|122146516|122146599</td><td>56</td><td>76</td><td>21</td><td>CTGTACAGCCTCCTAGCTTTC</td><td>36.1</td><td>8.4</td><td>1.9</td><td>4.4</td><td>1.0E-05</td><td>1.8E-05</td><td>MB231/MB468</td><td>MP</td></tr>
<tr><td>>hg38_21_+_8433172_8446622_RNA45SN1_45S_50ntFlanks@10820.10838.19|GGGAAGTCGGCAAGCCGGA</td><td>>hg38_21_+_8433172_8446622_RNA45SN1_45S_50ntFlanks</td><td>10820</td><td>10838</td><td>19</td><td>GGGAAGTCGGCAAGCCGGA</td><td>255.3</td><td>-6.5</td><td>1.5</td><td>-4.4</td><td>1.0E-05</td><td>1.8E-05</td><td>MB231/MB468</td><td>MP</td></tr>
<tr><td>>hg38_21_+_8433172_8446622_RNA45SN1_45S_50ntFlanks@474.511.38|TCTTCGTGATCGATGTGGTGACGTCGTGCTCTCCCGGG</td><td>>hg38_21_+_8433172_8446622_RNA45SN1_45S_50ntFlanks</td><td>474</td><td>511</td><td>38</td><td>TCTTCGTGATCGATGTGGTGACGTCGTGCTCTCCCGGG</td><td>238.3</td><td>3.5</td><td>0.8</td><td>4.4</td><td>1.2E-05</td><td>1.8E-05</td><td>MB231/MB468</td><td>NUC</td></tr>
<tr><td>>hg38_21_+_8433172_8446622_RNA45SN1_45S_50ntFlanks@11918.11948.31|ACATGAGAGGTGTAGAATAAGTGGGAGGCCC</td><td>>hg38_21_+_8433172_8446622_RNA45SN1_45S_50ntFlanks</td><td>11918</td><td>11948</td><td>31</td><td>ACATGAGAGGTGTAGAATAAGTGGGAGGCCC</td><td>1329.7</td><td>-4.3</td><td>0.9</td><td>-4.6</td><td>3.9E-06</td><td>1.8E-05</td><td>BT20/MB468</td><td>MP</td></tr>
<tr><td>>hg38_21_+_8433172_8446622_RNA45SN1_45S_50ntFlanks@5236.5254.19|CACTGACTGGCTCAGCGTG</td><td>>hg38_21_+_8433172_8446622_RNA45SN1_45S_50ntFlanks</td><td>5236</td><td>5254</td><td>19</td><td>CACTGACTGGCTCAGCGTG</td><td>2445.6</td><td>-7.8</td><td>1.8</td><td>-4.4</td><td>1.3E-05</td><td>1.8E-05</td><td>MB231/MB468</td><td>MITO</td></tr>
<tr><td>>hg38_21_+_8433172_8446622_RNA45SN1_45S_50ntFlanks@12991.13027.37|CTCCCTCGCTGCGATCTATTGAAAGTCAGCCCTCGAC</td><td>>hg38_21_+_8433172_8446622_RNA45SN1_45S_50ntFlanks</td><td>12991</td><td>13027</td><td>37</td><td>CTCCCTCGCTGCGATCTATTGAAAGTCAGCCCTCGAC</td><td>61.3</td><td>-9.0</td><td>2.0</td><td>-4.6</td><td>3.9E-06</td><td>1.8E-05</td><td>BT20/MB468</td><td>MP</td></tr>
<tr><td>>hg38_21_+_8433172_8446622_RNA45SN1_45S_50ntFlanks@9574.9595.22|AATCGGTCGTCCGACCTGGGTA</td><td>>hg38_21_+_8433172_8446622_RNA45SN1_45S_50ntFlanks</td><td>9574</td><td>9595</td><td>22</td><td>AATCGGTCGTCCGACCTGGGTA</td><td>833.0</td><td>-6.1</td><td>1.4</td><td>-4.4</td><td>1.3E-05</td><td>1.8E-05</td><td>BT20/MB468</td><td>NUC</td></tr>
<tr><td>>hg38_21_+_8433172_8446622_RNA45SN1_45S_50ntFlanks@6648.6690.43|GTACGACTCTTAGCGGTGGATCACTCGGCTCGTGCGTCGATGA</td><td>>hg38_21_+_8433172_8446622_RNA45SN1_45S_50ntFlanks</td><td>6648</td><td>6690</td><td>43</td><td>GTACGACTCTTAGCGGTGGATCACTCGGCTCGTGCGTCGATGA</td><td>860.8</td><td>2.6</td><td>0.6</td><td>4.4</td><td>9.0E-06</td><td>1.8E-05</td><td>BT20/MB468</td><td>MITO</td></tr>
<tr><td>>hg38_21_+_8433172_8446622_RNA45SN1_45S_50ntFlanks@6646.6665.20|TCGTACGACTCTTAGCGGTG</td><td>>hg38_21_+_8433172_8446622_RNA45SN1_45S_50ntFlanks</td><td>6646</td><td>6665</td><td>20</td><td>TCGTACGACTCTTAGCGGTG</td><td>13240.1</td><td>3.3</td><td>0.7</td><td>4.4</td><td>1.2E-05</td><td>1.8E-05</td><td>MB231/MB468</td><td>NUC</td></tr>
<tr><td>>MI0000285|hsa-mir-205&WithFlank&1|+|209432127|209432248@40.62.23(+1A)&[MIMAT0000266&hsa-miR-205-5p&offsets|0|+1(+1A);m-351&1|+|209432166|209432187&offsets|0|+1(+1A)]|TCCTTCATTCCACCGGAGTCTGTA</td><td>>MI0000285|hsa-mir-205&WithFlank&1|+|209432127|209432248</td><td>40</td><td>62</td><td>23(+1A)</td><td>TCCTTCATTCCACCGGAGTCTGTA</td><td>3546.2</td><td>10.1</td><td>2.1</td><td>4.9</td><td>1.2E-06</td><td>1.8E-05</td><td>BT20/MB231</td><td>CYTO</td></tr>
<tr><td>>MI0000487|hsa-mir-193a&WithFlank&17|+|31559990|31560089@27.47.21&[MIMAT0004614&hsa-miR-193a-5p&offsets|0|-1;m-123&17|+|31560016|31560037&offsets|0|-1]|TGGGTCTTTGCGGGCGAGATG</td><td>>MI0000487|hsa-mir-193a&WithFlank&17|+|31559990|31560089</td><td>27</td><td>47</td><td>21</td><td>TGGGTCTTTGCGGGCGAGATG</td><td>416.6</td><td>4.0</td><td>0.8</td><td>4.9</td><td>1.2E-06</td><td>1.8E-05</td><td>BT20/MB231</td><td>CYTO</td></tr>
<tr><td>>MI0000736|hsa-mir-30c-1&WithFlank&1|+|40757278|40757378@23.45.23(+1G);MI0000254|hsa-mir-30c-2&WithFlank&6|-|71376954|71377037@13.35.23(+1G)&[MIMAT0000244&hsa-miR-30c-5p&offsets|0|0(+1G);m-57&1|+|40757300|40757323&offsets|0|-1(+1G)];[MIMAT0000244_1&hsa-miR-30c-5p&offsets|0|0(+1G);m-56&6|-|71377002|71377025&offsets|0|-1(+1G)]|TGTAAACATCCTACACTCTCAGCG</td><td>>MI0000736|hsa-mir-30c-1&WithFlank&1|+|40757278|40757378</td><td>23</td><td>45</td><td>23(+1G)</td><td>TGTAAACATCCTACACTCTCAGCG</td><td>1036.8</td><td>4.9</td><td>1.1</td><td>4.4</td><td>1.3E-05</td><td>1.8E-05</td><td>MB231/MB468</td><td>MITO</td></tr>
<tr><td>>hg38_21_+_8433172_8446622_RNA45SN1_45S_50ntFlanks@11567.11588.22|AGCAGCCGACTTAGAACTGGTG</td><td>>hg38_21_+_8433172_8446622_RNA45SN1_45S_50ntFlanks</td><td>11567</td><td>11588</td><td>22</td><td>AGCAGCCGACTTAGAACTGGTG</td><td>962.7</td><td>-3.5</td><td>0.8</td><td>-4.4</td><td>1.0E-05</td><td>1.8E-05</td><td>MB231/MB468</td><td>MP</td></tr>
<tr><td>>hg38_21_+_8433172_8446622_RNA45SN1_45S_50ntFlanks@11209.11232.24|CGGCGACTCTGGACGCGAGCCGGG</td><td>>hg38_21_+_8433172_8446622_RNA45SN1_45S_50ntFlanks</td><td>11209</td><td>11232</td><td>24</td><td>CGGCGACTCTGGACGCGAGCCGGG</td><td>522.5</td><td>4.5</td><td>0.9</td><td>4.8</td><td>1.4E-06</td><td>1.8E-05</td><td>BT20/MB231</td><td>MITO</td></tr>
<tr><td>>hg38_21_+_8433172_8446622_RNA45SN1_45S_50ntFlanks@11908.11929.22|CGGTGAAGAGACATGAGAGGTG</td><td>>hg38_21_+_8433172_8446622_RNA45SN1_45S_50ntFlanks</td><td>11908</td><td>11929</td><td>22</td><td>CGGTGAAGAGACATGAGAGGTG</td><td>131.1</td><td>10.4</td><td>2.1</td><td>4.9</td><td>1.2E-06</td><td>1.8E-05</td><td>BT20/MB231</td><td>MP</td></tr>
<tr><td>>hg38_21_+_8433172_8446622_RNA45SN1_45S_50ntFlanks@11922.11953.32|GAGAGGTGTAGAATAAGTGGGAGGCCCCCGGC</td><td>>hg38_21_+_8433172_8446622_RNA45SN1_45S_50ntFlanks</td><td>11922</td><td>11953</td><td>32</td><td>GAGAGGTGTAGAATAAGTGGGAGGCCCCCGGC</td><td>331.2</td><td>-5.3</td><td>1.1</td><td>-4.9</td><td>1.2E-06</td><td>1.8E-05</td><td>BT20/MB231</td><td>CYTO</td></tr>
<tr><td>>MI0000088|hsa-mir-30a&WithFlank&6|-|71403545|71403627@12.34.23&[MIMAT0000087&hsa-miR-30a-5p&offsets|0|+1;m-15&6|-|71403595|71403616&offsets|0|+1]|TGTAAACATCCTCGACTGGAAGC</td><td>>MI0000088|hsa-mir-30a&WithFlank&6|-|71403545|71403627</td><td>12</td><td>34</td><td>23</td><td>TGTAAACATCCTCGACTGGAAGC</td><td>4833.0</td><td>3.4</td><td>0.8</td><td>4.4</td><td>1.3E-05</td><td>1.8E-05</td><td>BT20/MB468</td><td>NUC</td></tr>
<tr><td>>hg38_21_+_8433172_8446622_RNA45SN1_45S_50ntFlanks@10885.10902.18|GGCTGGGGCGCGAAGCGG</td><td>>hg38_21_+_8433172_8446622_RNA45SN1_45S_50ntFlanks</td><td>10885</td><td>10902</td><td>18</td><td>GGCTGGGGCGCGAAGCGG</td><td>108.3</td><td>-5.1</td><td>1.2</td><td>-4.4</td><td>1.1E-05</td><td>1.8E-05</td><td>BT20/MB468</td><td>CYTO</td></tr>
<tr><td>>hg38_21_+_8433172_8446622_RNA45SN1_45S_50ntFlanks@10349.10374.26|AACGAGAACTTTGAAGGCCGAAGTGG</td><td>>hg38_21_+_8433172_8446622_RNA45SN1_45S_50ntFlanks</td><td>10349</td><td>10374</td><td>26</td><td>AACGAGAACTTTGAAGGCCGAAGTGG</td><td>562.5</td><td>-2.4</td><td>0.5</td><td>-4.4</td><td>9.1E-06</td><td>1.8E-05</td><td>BT20/MB468</td><td>MITO</td></tr>
<tr><td>>hg38_21_+_8433172_8446622_RNA45SN1_45S_50ntFlanks@3996.4022.27|ACTCTAGATAACCTCGGGCCGATCGCA</td><td>>hg38_21_+_8433172_8446622_RNA45SN1_45S_50ntFlanks</td><td>3996</td><td>4022</td><td>27</td><td>ACTCTAGATAACCTCGGGCCGATCGCA</td><td>452.0</td><td>-5.3</td><td>1.2</td><td>-4.4</td><td>1.2E-05</td><td>1.8E-05</td><td>MB231/MB468</td><td>NUC</td></tr>
<tr><td>>MI0005756|hsa-mir-934&WithFlank&X|+|136550872|136550966@21.42.22(+1C)&[MIMAT0004977&hsa-miR-934&offsets|0|0(+1C);m-943&X|+|136550892|136550913&offsets|0|0(+1C)]|TGTCTACTACTGGAGACACTGGC</td><td>>MI0005756|hsa-mir-934&WithFlank&X|+|136550872|136550966</td><td>21</td><td>42</td><td>22(+1C)</td><td>TGTCTACTACTGGAGACACTGGC</td><td>86.5</td><td>9.8</td><td>2.0</td><td>4.9</td><td>1.2E-06</td><td>1.8E-05</td><td>BT20/MB231</td><td>MP</td></tr>
<tr><td>>hg38_21_+_8433172_8446622_RNA45SN1_45S_50ntFlanks@12636.12659.24|AGCTACCATCTGTGGGATTATGAC</td><td>>hg38_21_+_8433172_8446622_RNA45SN1_45S_50ntFlanks</td><td>12636</td><td>12659</td><td>24</td><td>AGCTACCATCTGTGGGATTATGAC</td><td>195.6</td><td>-4.2</td><td>1.0</td><td>-4.4</td><td>1.1E-05</td><td>1.8E-05</td><td>BT20/MB468</td><td>CYTO</td></tr>
<tr><td>>MI0003188|hsa-mir-503&WithFlank&X|-|134546322|134546404@12.31.20(+1A)&[MIMAT0002874&hsa-miR-503-5p&offsets|0|-3(+1A);m-297&X|-|134546371|134546393&offsets|0|-3(+1A)]|TAGCAGCGGGAACAGTTCTGA</td><td>>MI0003188|hsa-mir-503&WithFlank&X|-|134546322|134546404</td><td>12</td><td>31</td><td>20(+1A)</td><td>TAGCAGCGGGAACAGTTCTGA</td><td>144.9</td><td>7.1</td><td>1.6</td><td>4.4</td><td>1.0E-05</td><td>1.8E-05</td><td>MB231/MB468</td><td>MP</td></tr>
<tr><td>>hg38_21_+_8433172_8446622_RNA45SN1_45S_50ntFlanks@12405.12435.31|CTTTTTGATCCTTCGATGTCGGCTCTTCCTA</td><td>>hg38_21_+_8433172_8446622_RNA45SN1_45S_50ntFlanks</td><td>12405</td><td>12435</td><td>31</td><td>CTTTTTGATCCTTCGATGTCGGCTCTTCCTA</td><td>4867.0</td><td>-6.0</td><td>1.4</td><td>-4.4</td><td>1.2E-05</td><td>1.8E-05</td><td>MB231/MB468</td><td>CYTO</td></tr>
<tr><td>>hg38_21_+_8433172_8446622_RNA45SN1_45S_50ntFlanks@9718.9735.18|TGATTAGAGGTCTTGGGG</td><td>>hg38_21_+_8433172_8446622_RNA45SN1_45S_50ntFlanks</td><td>9718</td><td>9735</td><td>18</td><td>TGATTAGAGGTCTTGGGG</td><td>46.1</td><td>-8.6</td><td>1.9</td><td>-4.6</td><td>4.0E-06</td><td>1.8E-05</td><td>BT20/MB468</td><td>MP</td></tr>
</table>

<table>
<tr><th></th><th></th><th></th><th></th><th></th><th></th><th></th><th></th><th></th><th></th><th></th><th></th><th></th><th></th></tr>
<tr><td>>am_trna128_GlyGCC_6_-_27870686_27870756@8.35.28,trna18_GlyGCC_16_+_70822597_70822667@8.35.28,trna19_GlyGCC_16_+_70823410_70823480@8.35.28,trna19_GlyGCC_2_-_157257659_157257729@8.35.28,trna24_GlyGCC_16_-_70812942_70813012@8.35.28,trna25_GlyGCC_16_-_70812114_70812184@8.35.28,trna2_GlyGCC_21_-_18827107_18827177@8.35.28,trna35_GlyGCC_1_+_161413094_161413164@8.35.28,trna37_GlyGCC_1_+_161420467_161420537@8.35.28,trna39_GlyGCC_1_+_161427898_161427968@8.35.28,trna41_GlyGCC_1_+_161435258_161435328@8.35.28,trna5_GlyGCC_17_+_8029064_8029134@8.35.28,trna68_GlyGCC_1_-_161493637_161493707@8.35.28|TGGTTCAGTGGTAGAATTCTCGCCTGCC</td><td>>am_trna128_GlyGCC_6_-_27870686_27870756</td><td>8</td><td>35</td><td>28</td><td>TGGTTCAGTGGTAGAATTCTCGCCTGCC</td><td>219.9</td><td>5.2</td><td>1.2</td><td>4.4</td><td>1.3E-05</td><td>1.8E-05</td><td>MB231/MB468</td><td>MITO</td></tr>
<tr><td>>hg38_21_+_8433172_8446622_RNA45SN1_45S_50ntFlanks@11233.11252.20|CCCTTCCCGTGGATCGCCCC</td><td>>hg38_21_+_8433172_8446622_RNA45SN1_45S_50ntFlanks</td><td>11233</td><td>11252</td><td>20</td><td>CCCTTCCCGTGGATCGCCCC</td><td>424.6</td><td>-4.9</td><td>1.1</td><td>-4.4</td><td>9.1E-06</td><td>1.8E-05</td><td>BT20/MB468</td><td>MITO</td></tr>
<tr><td>>MI0000088|hsa-mir-30a&WithFlank&6|-|71403545|71403627@54.75.22(+1U)&[MIMAT0000088&hsa-miR-30a-3p&offsets|+1|+1(+1U);m-38&6|-|71403554|71403575&offsets|+1|+1(+1U)]|TTTCAGTCGGATGTTTGCAGCTT</td><td>>MI0000088|hsa-mir-30a&WithFlank&6|-|71403545|71403627</td><td>54</td><td>75</td><td>22(+1U)</td><td>TTTCAGTCGGATGTTTGCAGCTT</td><td>82.8</td><td>-9.8</td><td>2.0</td><td>-4.9</td><td>1.2E-06</td><td>1.8E-05</td><td>BT20/MB231</td><td>MP</td></tr>
<tr><td>>hg38_21_+_8433172_8446622_RNA45SN1_45S_50ntFlanks@12457.12487.31|CAAGCGTTGGATTGTTCACCCACTAATAGGG</td><td>>hg38_21_+_8433172_8446622_RNA45SN1_45S_50ntFlanks</td><td>12457</td><td>12487</td><td>31</td><td>CAAGCGTTGGATTGTTCACCCACTAATAGGG</td><td>751.2</td><td>-5.3</td><td>1.2</td><td>-4.4</td><td>9.1E-06</td><td>1.8E-05</td><td>BT20/MB468</td><td>MITO</td></tr>
<tr><td>>hg38_21_+_8433172_8446622_RNA45SN1_45S_50ntFlanks@12677.12704.28|GAATCCCGCCCAGGCGGAACGATACGGC</td><td>>hg38_21_+_8433172_8446622_RNA45SN1_45S_50ntFlanks</td><td>12677</td><td>12704</td><td>28</td><td>GAATCCCGCCCAGGCGGAACGATACGGC</td><td>52.2</td><td>-8.8</td><td>1.9</td><td>-4.6</td><td>4.0E-06</td><td>1.8E-05</td><td>BT20/MB468</td><td>MP</td></tr>
<tr><td>>hg38_21_+_8433172_8446622_RNA45SN1_45S_50ntFlanks@3930.3952.23|ATCAAAACCAACCCGGTCAGCCC</td><td>>hg38_21_+_8433172_8446622_RNA45SN1_45S_50ntFlanks</td><td>3930</td><td>3952</td><td>23</td><td>ATCAAAACCAACCCGGTCAGCCC</td><td>43.4</td><td>-8.5</td><td>1.8</td><td>-4.6</td><td>4.0E-06</td><td>1.8E-05</td><td>BT20/MB468</td><td>MP</td></tr>
<tr><td>>trna111_HisGTG_1_-_147774845_147774916@2.33.32,trna118_HisGTG_1_-_145396881_145396952@2.33.32,trna16_HisGTG_1_+_146544773_146544844@2.33.32,trna1_HisGTG_15_+_45493349_45493420@2.33.32,trna21_HisGTG_1_+_147753471_147753542@2.33.32,trna33_HisGTG_6_+_27125906_27125977@2.33.32,trna7_HisGTG_9_-_14433938_14434009@2.33.32,trna8_HisGTG_15_-_45492611_45492682@2.33.32,trna9_HisGTG_15_-_45490804_45490875@2.33.32|CCGTGATCGTATAGTGGTTAGTACTCTGCGTT</td><td>>trna111_HisGTG_1_-_147774845_147774916</td><td>2</td><td>33</td><td>32</td><td>CCGTGATCGTATAGTGGTTAGTACTCTGCGTT</td><td>65.6</td><td>9.8</td><td>2.1</td><td>4.6</td><td>4.0E-06</td><td>1.8E-05</td><td>BT20/MB468</td><td>MP</td></tr>
<tr><td>>MI0000767|hsa-mir-365a&WithFlank&16|+|14309279|14309377@22.43.22&[MIMAT0009199&hsa-miR-365a-5p&offsets|0|-1;m-595&16|+|14309300|14309322&offsets|0|-1]|AGGGACTTTTGGGGGCAGATGT</td><td>>MI0000767|hsa-mir-365a&WithFlank&16|+|14309279|14309377</td><td>22</td><td>43</td><td>22</td><td>AGGGACTTTTGGGGGCAGATGT</td><td>368.3</td><td>8.3</td><td>1.8</td><td>4.6</td><td>4.0E-06</td><td>1.8E-05</td><td>BT20/MB468</td><td>MP</td></tr>
<tr><td>>MI0000478|hsa-mir-149&WithFlank&2|+|240455995|240456095@21.43.23(+1A)&[MIMAT0000450&hsa-miR-149-5p&offsets|0|0(+1A);m-195&2|+|240456015|240456037&offsets|0|0(+1A)]|TCTGGCTCCGTGTCTTCACTCCCA</td><td>>MI0000478|hsa-mir-149&WithFlank&2|+|240455995|240456095</td><td>21</td><td>43</td><td>23(+1A)</td><td>TCTGGCTCCGTGTCTTCACTCCCA</td><td>272.4</td><td>5.1</td><td>1.1</td><td>4.9</td><td>1.2E-06</td><td>1.8E-05</td><td>BT20/MB231</td><td>CYTO</td></tr>
<tr><td>>trnaMT_GluTTC_MT_-_14674_14742@40.72.33,trnalookalike8_GluTTC_5_-_93905172_93905240@40.72.33|TTGGTCGTGGTTGTAGTCCGTGCGAGAATACCA</td><td>>trnaMT_GluTTC_MT_-_14674_14742</td><td>40</td><td>72</td><td>33</td><td>TTGGTCGTGGTTGTAGTCCGTGCGAGAATACCA</td><td>10154.1</td><td>3.2</td><td>0.7</td><td>4.4</td><td>1.3E-05</td><td>1.8E-05</td><td>MB231/MB468</td><td>MITO</td></tr>
<tr><td>>hg38_21_+_8433172_8446622_RNA45SN1_45S_50ntFlanks@12357.12375.19|CCACAGGGATAACTGGCTT</td><td>>hg38_21_+_8433172_8446622_RNA45SN1_45S_50ntFlanks</td><td>12357</td><td>12375</td><td>19</td><td>CCACAGGGATAACTGGCTT</td><td>543.6</td><td>-7.3</td><td>1.7</td><td>-4.4</td><td>1.3E-05</td><td>1.8E-05</td><td>BT20/MB468</td><td>NUC</td></tr>
<tr><td>>trnaMT_HisGTG_MT_+_12138_12206@19.59.41|AAACATCAGATTGTGAATCTGACAACAGAGGCTTACGACCC</td><td>>trnaMT_HisGTG_MT_+_12138_12206</td><td>19</td><td>59</td><td>41</td><td>AAACATCAGATTGTGAATCTGACAACAGAGGCTTACGACCC</td><td>34.3</td><td>8.3</td><td>1.9</td><td>4.4</td><td>1.1E-05</td><td>1.8E-05</td><td>MB231/MB468</td><td>MP</td></tr>
<tr><td>>hg38_21_+_8433172_8446622_RNA45SN1_45S_50ntFlanks@8170.8199.30|CAAGTCCTTCTGATCGAGGCCCAGCCCGTG</td><td>>hg38_21_+_8433172_8446622_RNA45SN1_45S_50ntFlanks</td><td>8170</td><td>8199</td><td>30</td><td>CAAGTCCTTCTGATCGAGGCCCAGCCCGTG</td><td>290.4</td><td>-2.7</td><td>0.6</td><td>-4.4</td><td>9.2E-06</td><td>1.8E-05</td><td>BT20/MB468</td><td>MITO</td></tr>
<tr><td>>hg38_21_+_8433172_8446622_RNA45SN1_45S_50ntFlanks@9922.9949.28|GTTGGTTGATATAGACAGCAGGACGGTG</td><td>>hg38_21_+_8433172_8446622_RNA45SN1_45S_50ntFlanks</td><td>9922</td><td>9949</td><td>28</td><td>GTTGGTTGATATAGACAGCAGGACGGTG</td><td>89.2</td><td>9.8</td><td>2.0</td><td>4.9</td><td>1.2E-06</td><td>1.8E-05</td><td>BT20/MB231</td><td>MP</td></tr>
<tr><td>>hg38_21_+_8433172_8446622_RNA45SN1_45S_50ntFlanks@11315.11336.22|CCACCCCACGTCTCGTCGCGCG</td><td>>hg38_21_+_8433172_8446622_RNA45SN1_45S_50ntFlanks</td><td>11315</td><td>11336</td><td>22</td><td>CCACCCCACGTCTCGTCGCGCG</td><td>532.1</td><td>-4.9</td><td>1.1</td><td>-4.4</td><td>1.1E-05</td><td>1.8E-05</td><td>BT20/MB468</td><td>CYTO</td></tr>
<tr><td>>hg38_21_+_8433172_8446622_RNA45SN1_45S_50ntFlanks@10348.10374.27|AAACGAGAACTTTGAAGGCCGAAGTGG</td><td>>hg38_21_+_8433172_8446622_RNA45SN1_45S_50ntFlanks</td><td>10348</td><td>10374</td><td>27</td><td>AAACGAGAACTTTGAAGGCCGAAGTGG</td><td>1862.4</td><td>-4.9</td><td>1.1</td><td>-4.6</td><td>4.0E-06</td><td>1.8E-05</td><td>BT20/MB468</td><td>MP</td></tr>
<tr><td>>MI0000456|hsa-mir-140&WithFlank&16|+|69933075|69933186@69.90.22&[MIMAT0004597&hsa-miR-140-3p&offsets|+1|+2;m-45&16|+|69933143|69933164&offsets|0|0]|ACCACAGGGTAGAACCACGGAC</td><td>>MI0000456|hsa-mir-140&WithFlank&16|+|69933075|69933186</td><td>69</td><td>90</td><td>22</td><td>ACCACAGGGTAGAACCACGGAC</td><td>1427.3</td><td>5.8</td><td>1.3</td><td>4.4</td><td>1.1E-05</td><td>1.8E-05</td><td>MB231/MB468</td><td>MP</td></tr>
<tr><td>>hg38_1_-_228634819_228635039_RNA5S12_5S_50ntFlanks@72.92.21|AACGCGCCCGATCTCGTCTGA</td><td>>hg38_1_-_228634819_228635039_RNA5S12_5S_50ntFlanks</td><td>72</td><td>92</td><td>21</td><td>AACGCGCCCGATCTCGTCTGA</td><td>306.6</td><td>-3.5</td><td>0.8</td><td>-4.4</td><td>1.3E-05</td><td>1.8E-05</td><td>MB231/MB468</td><td>NUC</td></tr>
<tr><td>>hg38_21_+_8433172_8446622_RNA45SN1_45S_50ntFlanks@10458.10495.38|ATGGCCTCCGTTGCCCTCGGCCGATCGAAAGGGAGTCG</td><td>>hg38_21_+_8433172_8446622_RNA45SN1_45S_50ntFlanks</td><td>10458</td><td>10495</td><td>38</td><td>ATGGCCTCCGTTGCCCTCGGCCGATCGAAAGGGAGTCG</td><td>188.5</td><td>-5.8</td><td>1.3</td><td>-4.4</td><td>1.1E-05</td><td>1.8E-05</td><td>BT20/MB468</td><td>CYTO</td></tr>
<tr><td>>MI0001641|hsa-mir-429&WithFlank&1|+|1168999|1169093@57.77.21(+1C)&[MIMAT0001536&hsa-miR-429&offsets|0|-1(+1C);m-207&1|+|1169055|1169076&offsets|0|-1(+1C)]|TAATACTGTCTGGTAAAACCGC</td><td>>MI0001641|hsa-mir-429&WithFlank&1|+|1168999|1169093</td><td>57</td><td>77</td><td>21(+1C)</td><td>TAATACTGTCTGGTAAAACCGC</td><td>314.7</td><td>-5.1</td><td>1.2</td><td>-4.4</td><td>1.1E-05</td><td>1.8E-05</td><td>MB231/MB468</td><td>MP</td></tr>
<tr><td>>hg38_21_+_8433172_8446622_RNA45SN1_45S_50ntFlanks@12997.13015.19|CGCTGCGATCTATTGAAAG</td><td>>hg38_21_+_8433172_8446622_RNA45SN1_45S_50ntFlanks</td><td>12997</td><td>13015</td><td>19</td><td>CGCTGCGATCTATTGAAAG</td><td>1066.8</td><td>3.2</td><td>0.7</td><td>4.4</td><td>1.2E-05</td><td>1.8E-05</td><td>MB231/MB468</td><td>CYTO</td></tr>
<tr><td>>hg38_21_+_8433172_8446622_RNA45SN1_45S_50ntFlanks@3780.3801.22|TACAGTGAAACTGCGAATGGCT</td><td>>hg38_21_+_8433172_8446622_RNA45SN1_45S_50ntFlanks</td><td>3780</td><td>3801</td><td>22</td><td>TACAGTGAAACTGCGAATGGCT</td><td>380.2</td><td>-5.5</td><td>1.3</td><td>-4.4</td><td>1.1E-05</td><td>1.8E-05</td><td>MB231/MB468</td><td>MP</td></tr>
<tr><td>>MI0000269|hsa-mir-181a-2&WithFlank&9|+|124692436|124692557@83.102.20&[MIMAT0004558&hsa-miR-181a-2-3p&offsets|0|-2;m-71&9|+|124692518|124692539&offsets|0|-2]|ACCACTGACCGTTGACTGTA</td><td>>MI0000269|hsa-mir-181a-2&WithFlank&9|+|124692436|124692557</td><td>83</td><td>102</td><td>20</td><td>ACCACTGACCGTTGACTGTA</td><td>32.4</td><td>8.2</td><td>1.9</td><td>4.4</td><td>1.1E-05</td><td>1.8E-05</td><td>MB231/MB468</td><td>MP</td></tr>
<tr><td>>hg38_21_+_8433172_8446622_RNA45SN1_45S_50ntFlanks@11209.11228.20|CGGCGACTCTGGACGCGAGC</td><td>>hg38_21_+_8433172_8446622_RNA45SN1_45S_50ntFlanks</td><td>11209</td><td>11228</td><td>20</td><td>CGGCGACTCTGGACGCGAGC</td><td>186.5</td><td>-3.2</td><td>0.7</td><td>-4.4</td><td>1.2E-05</td><td>1.8E-05</td><td>MB231/MB468</td><td>CYTO</td></tr>
<tr><td>>hg38_21_+_8433172_8446622_RNA45SN1_45S_50ntFlanks@5078.5100.23|CGATAACGAACGAGACTCTGGCA</td><td>>hg38_21_+_8433172_8446622_RNA45SN1_45S_50ntFlanks</td><td>5078</td><td>5100</td><td>23</td><td>CGATAACGAACGAGACTCTGGCA</td><td>143.2</td><td>-6.9</td><td>1.6</td><td>-4.4</td><td>1.1E-05</td><td>1.8E-05</td><td>MB231/MB468</td><td>MP</td></tr>
<tr><td>>hg38_21_+_8433172_8446622_RNA45SN1_45S_50ntFlanks@4728.4753.26|ATCAAGAACGAAAGTCGGAGGTTCGA</td><td>>hg38_21_+_8433172_8446622_RNA45SN1_45S_50ntFlanks</td><td>4728</td><td>4753</td><td>26</td><td>ATCAAGAACGAAAGTCGGAGGTTCGA</td><td>214.0</td><td>-6.4</td><td>1.4</td><td>-4.4</td><td>1.1E-05</td><td>1.8E-05</td><td>MB231/MB468</td><td>MP</td></tr>
<tr><td>>hg38_21_+_8433172_8446622_RNA45SN1_45S_50ntFlanks@5472.5514.43|ACGGCCCTGGCGGAGCGCTGAGAAGACGGTCGAACTTGACTAT</td><td>>hg38_21_+_8433172_8446622_RNA45SN1_45S_50ntFlanks</td><td>5472</td><td>5514</td><td>43</td><td>ACGGCCCTGGCGGAGCGCTGAGAAGACGGTCGAACTTGACTAT</td><td>4442.8</td><td>-3.9</td><td>0.8</td><td>-4.8</td><td>1.3E-06</td><td>1.8E-05</td><td>BT20/MB231</td><td>CYTO</td></tr>
<tr><td>>hg38_21_+_8433172_8446622_RNA45SN1_45S_50ntFlanks@9739.9766.28|AAACGATCTCAACCTATTCTCAAACTTT</td><td>>hg38_21_+_8433172_8446622_RNA45SN1_45S_50ntFlanks</td><td>9739</td><td>9766</td><td>28</td><td>AAACGATCTCAACCTATTCTCAAACTTT</td><td>476.6</td><td>-5.5</td><td>1.3</td><td>-4.4</td><td>1.4E-05</td><td>1.8E-05</td><td>BT20/MB468</td><td>NUC</td></tr>
<tr><td>>MI0000102|hsa-mir-100&WithFlank&11|-|122152223|122152314@19.39.21&[MIMAT0000098&hsa-miR-100-5p&offsets|0|-1;m-27&11|-|122152276|122152296&offsets|0|0]|AACCCGTAGATCCGAACTTGT</td><td>>MI0000102|hsa-mir-100&WithFlank&11|-|122152223|122152314</td><td>19</td><td>39</td><td>21</td><td>AACCCGTAGATCCGAACTTGT</td><td>139110.3</td><td>13.1</td><td>3.0</td><td>4.4</td><td>1.3E-05</td><td>1.8E-05</td><td>MB231/MB468</td><td>MITO</td></tr>
<tr><td>>hg38_21_+_8433172_8446622_RNA45SN1_45S_50ntFlanks@6659.6686.28|AGCGGTGGATCACTCGGCTCGTGCGTCG</td><td>>hg38_21_+_8433172_8446622_RNA45SN1_45S_50ntFlanks</td><td>6659</td><td>6686</td><td>28</td><td>AGCGGTGGATCACTCGGCTCGTGCGTCG</td><td>135.8</td><td>-6.4</td><td>1.5</td><td>-4.4</td><td>1.1E-05</td><td>1.8E-05</td><td>MB231/MB468</td><td>MP</td></tr>
<tr><td>>hg38_21_+_8433172_8446622_RNA45SN1_45S_50ntFlanks@11873.11902.30|AAGAAGACCCTGTTGAGCTTGACTCTAGTC</td><td>>hg38_21_+_8433172_8446622_RNA45SN1_45S_50ntFlanks</td><td>11873</td><td>11902</td><td>30</td><td>AAGAAGACCCTGTTGAGCTTGACTCTAGTC</td><td>360.9</td><td>-3.3</td><td>0.7</td><td>-4.4</td><td>1.2E-05</td><td>1.8E-05</td><td>MB231/MB468</td><td>CYTO</td></tr>
<tr><td>>am_trna2_GlyGCC_21_-_18827107_18827177@1.27.27,trna35_GlyGCC_1_+_161413094_161413164@1.27.27,trna37_GlyGCC_1_+_161420467_161420537@1.27.27,trna39_GlyGCC_1_+_161427898_161427968@1.27.27,trna41_GlyGCC_1_+_161435258_161435328@1.27.27|GCATGGGTGGTTCAGTGGTAGAATTCT</td><td>>am_trna2_GlyGCC_21_-_18827107_18827177</td><td>1</td><td>27</td><td>27</td><td>GCATGGGTGGTTCAGTGGTAGAATTCT</td><td>2057.5</td><td>7.2</td><td>1.5</td><td>4.9</td><td>1.2E-06</td><td>1.8E-05</td><td>BT20/MB231</td><td>MP</td></tr>
<tr><td>>hg38_21_+_8433172_8446622_RNA45SN1_45S_50ntFlanks@11853.11886.34|CTTGGCGGAATCAGCGGGGAAAGAAGACCCTGTT</td><td>>hg38_21_+_8433172_8446622_RNA45SN1_45S_50ntFlanks</td><td>11853</td><td>11886</td><td>34</td><td>CTTGGCGGAATCAGCGGGGAAAGAAGACCCTGTT</td><td>130.9</td><td>10.4</td><td>2.1</td><td>4.9</td><td>1.2E-06</td><td>1.8E-05</td><td>BT20/MB231</td><td>MP</td></tr>
<tr><td>>MI0000298|hsa-mir-221&WithFlank&X|-|45746151|45746272@71.88.18(+4U)&[MIMAT0000278&hsa-miR-221-3p&offsets|0|-5(+4U);m-91&X|-|45746181|45746202&offsets|0|-4(+4U)]|AGCTACATTGTCTGCTGGTTTT</td><td>>MI0000298|hsa-mir-221&WithFlank&X|-|45746151|45746272</td><td>71</td><td>88</td><td>18(+4U)</td><td>AGCTACATTGTCTGCTGGTTTT</td><td>91.8</td><td>-9.9</td><td>2.0</td><td>-4.9</td><td>1.2E-06</td><td>1.8E-05</td><td>BT20/MB231</td><td>MP</td></tr>
<tr><td>>MI0000749|hsa-mir-30e&WithFlank&1|+|40754349|40754452@24.46.23(+2U)&[MIMAT0000692&hsa-miR-30e-5p&offsets|+1|+2(+2U);m-24&1|+|40754371|40754394&offsets|+1|0(+2U)]|GTAAACATCCTTGACTGGAAGCTTT</td><td>>MI0000749|hsa-mir-30e&WithFlank&1|+|40754349|40754452</td><td>24</td><td>46</td><td>23(+2U)</td><td>GTAAACATCCTTGACTGGAAGCTTT</td><td>81.7</td><td>-9.8</td><td>2.0</td><td>-4.9</td><td>1.2E-06</td><td>1.8E-05</td><td>BT20/MB231</td><td>MP</td></tr>
<tr><td>>hg38_21_+_8433172_8446622_RNA45SN1_45S_50ntFlanks@5469.5499.31|CCCACGGCCCTGGCGGAGCGCTGAGAAGACG</td><td>>hg38_21_+_8433172_8446622_RNA45SN1_45S_50ntFlanks</td><td>5469</td><td>5499</td><td>31</td><td>CCCACGGCCCTGGCGGAGCGCTGAGAAGACG</td><td>1413.7</td><td>-7.8</td><td>1.8</td><td>-4.4</td><td>1.4E-05</td><td>1.9E-05</td><td>BT20/MB468</td><td>NUC</td></tr>
<tr><td>>hg38_21_+_8433172_8446622_RNA45SN1_45S_50ntFlanks@11594.11611.18|CAGGGGAATCCGACTGTT</td><td>>hg38_21_+_8433172_8446622_RNA45SN1_45S_50ntFlanks</td><td>11594</td><td>11611</td><td>18</td><td>CAGGGGAATCCGACTGTT</td><td>1392.6</td><td>-4.4</td><td>0.9</td><td>-4.6</td><td>4.1E-06</td><td>1.9E-05</td><td>BT20/MB468</td><td>MP</td></tr>
<tr><td>>hg38_21_+_8433172_8446622_RNA45SN1_45S_50ntFlanks@12004.12043.40|CGGGCCGCCGGTGAAATACCACTACTCTGATCGTTTTTTC</td><td>>hg38_21_+_8433172_8446622_RNA45SN1_45S_50ntFlanks</td><td>12004</td><td>12043</td><td>40</td><td>CGGGCCGCCGGTGAAATACCACTACTCTGATCGTTTTTTC</td><td>99.1</td><td>-9.7</td><td>2.1</td><td>-4.6</td><td>4.1E-06</td><td>1.9E-05</td><td>BT20/MB468</td><td>MP</td></tr>
<tr><td>>hg38_21_+_8433172_8446622_RNA45SN1_45S_50ntFlanks@8176.8201.26|CTTCTGATCGAGGCCCAGCCCGTGGA</td><td>>hg38_21_+_8433172_8446622_RNA45SN1_45S_50ntFlanks</td><td>8176</td><td>8201</td><td>26</td><td>CTTCTGATCGAGGCCCAGCCCGTGGA</td><td>487.0</td><td>-4.9</td><td>1.1</td><td>-4.4</td><td>1.3E-05</td><td>1.9E-05</td><td>MB231/MB468</td><td>MITO</td></tr>
<tr><td>>hg38_21_+_8433172_8446622_RNA45SN1_45S_50ntFlanks@8380.8399.20|CAAGAGGGCGTGAAACCGTT</td><td>>hg38_21_+_8433172_8446622_RNA45SN1_45S_50ntFlanks</td><td>8380</td><td>8399</td><td>20</td><td>CAAGAGGGCGTGAAACCGTT</td><td>158.4</td><td>-5.7</td><td>1.3</td><td>-4.4</td><td>1.3E-05</td><td>1.9E-05</td><td>MB231/MB468</td><td>NUC</td></tr>
<tr><td>>hg38_21_+_8433172_8446622_RNA45SN1_45S_50ntFlanks@9534.9569.36|AGGAAACTCTGGTGGAGGTCCGTAGCGGTCCTGACG</td><td>>hg38_21_+_8433172_8446622_RNA45SN1_45S_50ntFlanks</td><td>9534</td><td>9569</td><td>36</td><td>AGGAAACTCTGGTGGAGGTCCGTAGCGGTCCTGACG</td><td>5333.2</td><td>-3.4</td><td>0.8</td><td>-4.4</td><td>1.2E-05</td><td>1.9E-05</td><td>MB231/MB468</td><td>CYTO</td></tr>
<tr><td>>hg38_21_+_8433172_8446622_RNA45SN1_45S_50ntFlanks@10358.10383.26|TTTGAAGGCCGAAGTGGAGAAGGGTT</td><td>>hg38_21_+_8433172_8446622_RNA45SN1_45S_50ntFlanks</td><td>10358</td><td>10383</td><td>26</td><td>TTTGAAGGCCGAAGTGGAGAAGGGTT</td><td>104.6</td><td>-5.0</td><td>1.1</td><td>-4.4</td><td>1.1E-05</td><td>1.9E-05</td><td>MB231/MB468</td><td>MP</td></tr>
<tr><td>>hg38_21_+_8433172_8446622_RNA45SN1_45S_50ntFlanks@11670.11704.35|CCCAGTGCTCTGAATGTCAAAGTGAAGAAATTCAA</td><td>>hg38_21_+_8433172_8446622_RNA45SN1_45S_50ntFlanks</td><td>11670</td><td>11704</td><td>35</td><td>CCCAGTGCTCTGAATGTCAAAGTGAAGAAATTCAA</td><td>296.6</td><td>-4.8</td><td>1.1</td><td>-4.4</td><td>1.3E-05</td><td>1.9E-05</td><td>MB231/MB468</td><td>MITO</td></tr>
<tr><td>>hg38_21_+_8433172_8446622_RNA45SN1_45S_50ntFlanks@8354.8392.39|TTGAAAAGAACTTTGAAGAGAGAGTTCAAGAGGGCGTGA</td><td>>hg38_21_+_8433172_8446622_RNA45SN1_45S_50ntFlanks</td><td>8354</td><td>8392</td><td>39</td><td>TTGAAAAGAACTTTGAAGAGAGAGTTCAAGAGGGCGTGA</td><td>103.5</td><td>10.0</td><td>2.1</td><td>4.9</td><td>1.2E-06</td><td>1.9E-05</td><td>BT20/MB231</td><td>MP</td></tr>
</table>

| | | | | | | | | | | | | | |
|---|---|---|---|---|---|---|---|---|---|---|---|---|---|
| >hg38_21_+_8433172_8446622_RNA45SN1_45S_50ntFlanks@9506.9526.21\|GAACTATGCCTGGGCAGGGCG | >hg38_21_+_8433172_8446622_RNA45SN1_45S_50ntFlanks | 9506 | 9526 | 21 | GAACTATGCCTGGGCAGGGCG | 88.8 | -10.3 | 2.3 | -4.4 | 1.1E-05 | 1.9E-05 | MB231/MB468 | MP |
| >hg38_21_+_8433172_8446622_RNA45SN1_45S_50ntFlanks@8156.8181.26\|GCTCGTGGGGGGCCCAAGTCCTTCTG | >hg38_21_+_8433172_8446622_RNA45SN1_45S_50ntFlanks | 8156 | 8181 | 26 | GCTCGTGGGGGGCCCAAGTCCTTCTG | 532.8 | -5.1 | 1.2 | -4.4 | 1.3E-05 | 1.9E-05 | MB231/MB468 | MITO |
| >trnaMT_PheGAA_MT_+_577_647@36.67.32\|AAAATGTTTAGACGGGCTCACATCACCCCATA | >trnaMT_PheGAA_MT_+_577_647 | 36 | 67 | 32 | AAAATGTTTAGACGGGCTCACATCACCCCATA | 251.9 | 11.7 | 2.5 | 4.6 | 4.1E-06 | 1.9E-05 | BT20/MB468 | MP |
| >hg38_21_+_8433172_8446622_RNA45SN1_45S_50ntFlanks@4562.4579.18\|AGGAATAATGGAATAGGA | >hg38_21_+_8433172_8446622_RNA45SN1_45S_50ntFlanks | 4562 | 4579 | 18 | AGGAATAATGGAATAGGA | 142.6 | -6.4 | 1.5 | -4.4 | 1.1E-05 | 1.9E-05 | MB231/MB468 | MP |
| >hg38_21_+_8433172_8446622_RNA45SN1_45S_50ntFlanks@11875.11897.23\|GAAGACCCTGTTGAGCTTGACTC | >hg38_21_+_8433172_8446622_RNA45SN1_45S_50ntFlanks | 11875 | 11897 | 23 | GAAGACCCTGTTGAGCTTGACTC | 114.8 | -6.2 | 1.4 | -4.4 | 1.1E-05 | 1.9E-05 | MB231/MB468 | MP |
| >hg38_21_+_8433172_8446622_RNA45SN1_45S_50ntFlanks@12419.12458.40\|GATGTCGGCTCTTCCTATCATTGTGAAGCAGAATTCACCA | >hg38_21_+_8433172_8446622_RNA45SN1_45S_50ntFlanks | 12419 | 12458 | 40 | GATGTCGGCTCTTCCTATCATTGTGAAGCAGAATTCACCA | 1315.9 | -3.6 | 0.8 | -4.8 | 1.3E-06 | 1.9E-05 | BT20/MB231 | CYTO |
| >trna119_LysCTT_1_-_145395522_145395594@3.33.31,trna11_LysCTT_5_-_180648979_180649051@3.33.31,trna13_LysCTT_6_+_26556774_26556846@3.33.31,trna7_LysCTT_16_+_3225692_3225764@3.33.31,trna9_LysCTT_5_+_180634755_180634827@3.33.31\|CCGGCTAGCTCAGTCGGTAGAGCATGAGACT | >trna119_LysCTT_1_-_145395522_145395594 | 3 | 33 | 31 | CCGGCTAGCTCAGTCGGTAGAGCATGAGACT | 739.9 | -4.5 | 0.9 | -4.8 | 1.3E-06 | 1.9E-05 | BT20/MB231 | CYTO |
| >MI0000061\|hsa-let-7a-2&WithFlank&11\|-\|122146516\|122146599@11.33.23;MI0000062\|hsa-let-7a-3&WithFlank&22\|+\|46112743\|46112828@10.32.23;MI0000060\|hsa-let-7a-1&WithFlank&9\|+\|94175951\|94176042@12.34.23&[MIMAT0000062&hsa-let-7a-5p&offsets\|0\|+1;m-6&11\|-\|122146568\|122146589&offsets\|0\|+1];[MIMAT0000062_1&hsa-let-7a-5p&offsets\|0\|+1;m-5&22\|+\|46112752\|46112773&offsets\|0\|+1];[MIMAT0000062_2&hsa-let-7a-5p&offsets\|0\|+1;m-4&9\|+\|94175962\|94175983&offsets\|0\|+1]\|TGAGGTAGTAGGTTGTATAGTTT | >MI0000061\|hsa-let-7a-2&WithFlank&11\|-\|122146516\|122146599 | 11 | 33 | 23 | TGAGGTAGTAGGTTGTATAGTTT | 725.8 | 1.9 | 0.4 | 4.3 | 1.4E-05 | 1.9E-05 | BT20/MB468 | NUC |
| >hg38_21_+_8433172_8446622_RNA45SN1_45S_50ntFlanks@10679.10712.34\|GGTTCCGGCGGCGTCCGGTGAGCTCTCGCTGGCC | >hg38_21_+_8433172_8446622_RNA45SN1_45S_50ntFlanks | 10679 | 10712 | 34 | GGTTCCGGCGGCGTCCGGTGAGCTCTCGCTGGCC | 562.4 | -6.3 | 1.4 | -4.4 | 9.4E-06 | 1.9E-05 | BT20/MB468 | MITO |
| >hg38_21_+_8433172_8446622_RNA45SN1_45S_50ntFlanks@8176.8205.30\|CTTCTGATCGAGGCCCAGCCCGTGGACGGT | >hg38_21_+_8433172_8446622_RNA45SN1_45S_50ntFlanks | 8176 | 8205 | 30 | CTTCTGATCGAGGCCCAGCCCGTGGACGGT | 424.9 | -1.9 | 0.4 | -4.4 | 1.3E-05 | 1.9E-05 | MB231/MB468 | NUC |
| >hg38_21_+_8433172_8446622_RNA45SN1_45S_50ntFlanks@11907.11931.25\|ACGGTGAAGAGACATGAGAGGTGTA | >hg38_21_+_8433172_8446622_RNA45SN1_45S_50ntFlanks | 11907 | 11931 | 25 | ACGGTGAAGAGACATGAGAGGTGTA | 130.5 | 10.4 | 2.1 | 4.8 | 1.2E-06 | 1.9E-05 | BT20/MB231 | MP |
| >hg38_21_+_8433172_8446622_RNA45SN1_45S_50ntFlanks@9926.9951.26\|GTTGATATAGACAGCAGGACGGTGGC | >hg38_21_+_8433172_8446622_RNA45SN1_45S_50ntFlanks | 9926 | 9951 | 26 | GTTGATATAGACAGCAGGACGGTGGC | 99.3 | 10.0 | 2.1 | 4.8 | 1.2E-06 | 1.9E-05 | BT20/MB231 | MP |
| >MI0000273\|hsa-mir-183&WithFlank&7\|-\|129774899\|129775020@33.55.23&[MIMAT0000261&hsa-miR-183-5p&offsets\|0\|+1;m-41&7\|-\|129774966\|129774987&offsets\|-1\|0]\|TATGGCACTGGTAGAATTCACTG | >MI0000273\|hsa-mir-183&WithFlank&7\|-\|129774899\|129775020 | 33 | 55 | 23 | TATGGCACTGGTAGAATTCACTG | 454.7 | 3.1 | 0.7 | 4.4 | 1.2E-05 | 1.9E-05 | MB231/MB468 | CYTO |
| >hg38_21_+_8433172_8446622_RNA45SN1_45S_50ntFlanks@6773.6799.27\|TTCCTCCCGGGGCTACGCCTGTCTGAG | >hg38_21_+_8433172_8446622_RNA45SN1_45S_50ntFlanks | 6773 | 6799 | 27 | TTCCTCCCGGGGCTACGCCTGTCTGAG | 278.5 | -6.7 | 1.5 | -4.6 | 4.1E-06 | 1.9E-05 | BT20/MB468 | MP |
| >hg38_21_+_8433172_8446622_RNA45SN1_45S_50ntFlanks@12996.13038.43\|TCGCTGCGATCTATTGAAAGTCAGCCCTCGACACAAGGGTTTG | >hg38_21_+_8433172_8446622_RNA45SN1_45S_50ntFlanks | 12996 | 13038 | 43 | TCGCTGCGATCTATTGAAAGTCAGCCCTCGACACAAGGGTTTG | 35.3 | 8.4 | 1.9 | 4.4 | 1.1E-05 | 1.9E-05 | MB231/MB468 | MP |
| >hg38_21_+_8433172_8446622_RNA45SN1_45S_50ntFlanks@9534.9567.34\|AGGAAACTCTGGTGGAGGTCCGTAGCGGTCCTGA | >hg38_21_+_8433172_8446622_RNA45SN1_45S_50ntFlanks | 9534 | 9567 | 34 | AGGAAACTCTGGTGGAGGTCCGTAGCGGTCCTGA | 1176.0 | -5.8 | 1.3 | -4.4 | 1.1E-05 | 1.9E-05 | BT20/MB468 | CYTO |
| >hg38_21_+_8433172_8446622_RNA45SN1_45S_50ntFlanks@10678.10712.35\|CGGTTCCGGCGGCGTCCGGTGAGCTCTCGCTGGCC | >hg38_21_+_8433172_8446622_RNA45SN1_45S_50ntFlanks | 10678 | 10712 | 35 | CGGTTCCGGCGGCGTCCGGTGAGCTCTCGCTGGCC | 7640.0 | -4.4 | 1.0 | -4.4 | 1.1E-05 | 1.9E-05 | BT20/MB468 | CYTO |
| >hg38_21_+_8433172_8446622_RNA45SN1_45S_50ntFlanks@12457.12488.32\|CAAGCGTTGGATTGTTCACCCACTAATAGGGA | >hg38_21_+_8433172_8446622_RNA45SN1_45S_50ntFlanks | 12457 | 12488 | 32 | CAAGCGTTGGATTGTTCACCCACTAATAGGGA | 555.0 | -6.5 | 1.5 | -4.4 | 1.1E-05 | 1.9E-05 | BT20/MB468 | CYTO |
| >MI0000073\|hsa-mir-19a&WithFlank&13\|+\|91350885\|91350978@55.76.22(+1U)&[MIMAT0000073&hsa-miR-19a-3p&offsets\|0\|-1(+1U);m-201&13\|+\|91350939\|91350961&offsets\|0\|-1(+1U)]\|TGTGCAAATCTATGCAAAACTGT | >MI0000073\|hsa-mir-19a&WithFlank&13\|+\|91350885\|91350978 | 55 | 76 | 22(+1U) | TGTGCAAATCTATGCAAAACTGT | 323.3 | 4.9 | 1.1 | 4.4 | 1.3E-05 | 1.9E-05 | MB231/MB468 | MITO |
| >trna2_GlyGCC_21_-_18827107_18827177@3.28.26,trna35_GlyGCC_1_+_161413094_161413164@3.28.26,trna37_GlyGCC_1_+_161420467_161420537@3.28.26,trna39_GlyGCC_1_+_161427898_161427968@3.28.26,trna41_GlyGCC_1_+_161435258_161435328@3.28.26\|ATGGGTGGTTCAGTGGTAGAATTCTC | >trna2_GlyGCC_21_-_18827107_18827177 | 3 | 28 | 26 | ATGGGTGGTTCAGTGGTAGAATTCTC | 107.5 | 10.1 | 2.1 | 4.8 | 1.2E-06 | 1.9E-05 | BT20/MB231 | MP |
| >trna10_AspGTC_12_-_125424193_125424264@39.75.37,trna12_AspGTC_12_-_125411891_125411962@39.75.37,trna144_AspGTC_6_-_27551236_27551307@39.75.37,trna38_AspGTC_17_-_8125556_8125627@39.75.37,trna45_AspGTC_6_+_27447453_27447524@39.75.37,trna48_AspGTC_6_+_27471523_27471594@39.75.37,trna4_AspGTC_12_+_96429799_96429870@39.75.37,trna69_AspGTC_1_-_161440205_161440276@39.75.37,trna72_AspGTC_1_-_161432824_161432895@39.75.37,trna75_AspGTC_1_-_161425414_161425485@39.75.37,trna78_AspGTC_1_-_161418033_161418104@39.75.37,trna81_AspGTC_1_-_161410615_161410686@39.75.37\|GCGGGAGACCGGGGTTCGATTCCCCGACGGGGAGCCA | >trna10_AspGTC_12_-_125424193_125424264 | 39 | 75 | 37 | GCGGGAGACCGGGGTTCGATTCCCCGACGGGGAGCCA | 7457.8 | 4.8 | 1.1 | 4.4 | 1.2E-05 | 1.9E-05 | MB231/MB468 | CYTO |
| >hg38_21_+_8433172_8446622_RNA45SN1_45S_50ntFlanks@9569.9597.29\|GTGCAAATCGGTCGTCCGACCTGGGTATA | >hg38_21_+_8433172_8446622_RNA45SN1_45S_50ntFlanks | 9569 | 9597 | 29 | GTGCAAATCGGTCGTCCGACCTGGGTATA | 663.0 | -4.6 | 1.1 | -4.4 | 1.2E-05 | 1.9E-05 | MB231/MB468 | CYTO |
| >am_trnaMT_ValTAC_MT_+_1602_1670@29.66.38\|ACTTACACTTAGGAGATTTCAACTTAACTTGACCGCTC | >am_trnaMT_ValTAC_MT_+_1602_1670 | 29 | 66 | 38 | ACTTACACTTAGGAGATTTCAACTTAACTTGACCGCTC | 319.0 | 12.1 | 2.6 | 4.6 | 4.1E-06 | 1.9E-05 | BT20/MB468 | MP |
| >hg38_21_+_8433172_8446622_RNA45SN1_45S_50ntFlanks@3882.3913.32\|CCGACGGGCGCTGACCCCCTTCGCGGGGGGGA | >hg38_21_+_8433172_8446622_RNA45SN1_45S_50ntFlanks | 3882 | 3913 | 32 | CCGACGGGCGCTGACCCCCTTCGCGGGGGGGA | 330.1 | 4.9 | 1.1 | 4.4 | 1.2E-05 | 1.9E-05 | MB231/MB468 | CYTO |
| >MI0000542\|hsa-mir-320a&WithFlank&8\|-\|22244960\|22245043@47.68.22(+1U)&[MIMAT0000510&hsa-miR-320a-3p&offsets\|-1\|-1(+1U);m-60&8\|-\|22244975\|22244996&offsets\|-1\|-1(+1U)]\|GAAAAGCTGGGTTGAGAGGGCGT | >MI0000542\|hsa-mir-320a&WithFlank&8\|-\|22244960\|22245043 | 47 | 68 | 22(+1U) | GAAAAGCTGGGTTGAGAGGGCGT | 546.7 | 7.5 | 1.6 | 4.6 | 4.2E-06 | 1.9E-05 | BT20/MB468 | MP |
| >MI0000273\|hsa-mir-183&WithFlank&7\|-\|129774899\|129775020@33.53.21(+1A)&[MIMAT0000261&hsa-miR-183-5p&offsets\|0\|-1(+1A);m-41&7\|-\|129774966\|129774987&offsets\|-1\|-2(+1A)]\|TATGGCACTGGTAGAATTCACA | >MI0000273\|hsa-mir-183&WithFlank&7\|-\|129774899\|129775020 | 33 | 53 | 21(+1A) | TATGGCACTGGTAGAATTCACA | 610.3 | 2.6 | 0.5 | 4.8 | 1.5E-06 | 1.9E-05 | BT20/MB231 | NUC |
| >hg38_21_+_8433172_8446622_RNA45SN1_45S_50ntFlanks@3851.3872.22\|ATAACTGTGGTAATTCTAGAGC | >hg38_21_+_8433172_8446622_RNA45SN1_45S_50ntFlanks | 3851 | 3872 | 22 | ATAACTGTGGTAATTCTAGAGC | 162.3 | -4.8 | 1.1 | -4.4 | 1.1E-05 | 1.9E-05 | MB231/MB468 | MP |
| >trnaMT_LysTTT_MT_+_8295_8364@38.72.35,trnalookalike7_LysTTT_1_+_568844_568913@38.72.35\|AAGATTAAGAGAACCAACACCTCTTTACAGTGACC | >trnaMT_LysTTT_MT_+_8295_8364 | 38 | 72 | 35 | AAGATTAAGAGAACCAACACCTCTTTACAGTGACC | 653.6 | 4.2 | 1.0 | 4.4 | 1.1E-05 | 1.9E-05 | MB231/MB468 | MP |
| >MI0000063\|hsa-let-7b&WithFlank&22\|+\|46113680\|46113774@66.86.21(+1U)&[MIMAT0004482&hsa-let-7b-3p&offsets\|0\|-1(+1U);m-171&22\|+\|46113745\|46113766&offsets\|0\|-1(+1U)]\|CTATACAACCTACTGCCTTCCT | >MI0000063\|hsa-let-7b&WithFlank&22\|+\|46113680\|46113774 | 66 | 86 | 21(+1U) | CTATACAACCTACTGCCTTCCT | 257.2 | 3.1 | 0.7 | 4.4 | 1.3E-05 | 1.9E-05 | MB231/MB468 | NUC |
| >trna10_AspGTC_12_-_125424193_125424264@39.74.36,trna12_AspGTC_12_-_125411891_125411962@39.74.36,trna144_AspGTC_6_-_27551236_27551307@39.74.36,trna38_AspGTC_17_-_8125556_8125627@39.74.36,trna45_AspGTC_6_+_27447453_27447524@39.74.36,trna48_AspGTC_6_+_27471523_27471594@39.74.36,trna4_AspGTC_12_+_96429799_96429870@39.74.36,trna69_AspGTC_1_-_161440205_161440276@39.74.36,trna72_AspGTC_1_-_161432824_161432895@39.74.36,trna75_AspGTC_1_-_161425414_161425485@39.74.36,trna78_AspGTC_1_-_161418033_161418104@39.74.36,trna81_AspGTC_1_-_161410615_161410686@39.74.36\|GCGGGAGACCGGGGTTCGATTCCCCGACGGGGAGCC | >trna10_AspGTC_12_-_125424193_125424264 | 39 | 74 | 36 | GCGGGAGACCGGGGTTCGATTCCCCGACGGGGAGCC | 985.8 | 4.4 | 1.0 | 4.4 | 1.1E-05 | 1.9E-05 | BT20/MB468 | CYTO |
| >hg38_21_+_8433172_8446622_RNA45SN1_45S_50ntFlanks@6698.6717.20\|AGCTAGCTGCGAGAATTAAT | >hg38_21_+_8433172_8446622_RNA45SN1_45S_50ntFlanks | 6698 | 6717 | 20 | AGCTAGCTGCGAGAATTAAT | 378.4 | -7.0 | 1.5 | -4.6 | 4.2E-06 | 1.9E-05 | BT20/MB468 | MP |
| >hg38_21_+_8433172_8446622_RNA45SN1_45S_50ntFlanks@6721.6770.50\|AATTGCAGGACACATTGATCATCGACACTTCGAACGCACTTGCGGCCCCG | >hg38_21_+_8433172_8446622_RNA45SN1_45S_50ntFlanks | 6721 | 6770 | 50 | AATTGCAGGACACATTGATCATCGACACTTCGAACGCACTTGCGGCCCCG | 1320.9 | -5.9 | 1.3 | -4.4 | 9.5E-06 | 1.9E-05 | BT20/MB468 | MITO |
| >hg38_21_+_8433172_8446622_RNA45SN1_45S_50ntFlanks@12748.12774.27\|CCTGTCCCCGCCGGCGGGCCGCCCCCC | >hg38_21_+_8433172_8446622_RNA45SN1_45S_50ntFlanks | 12748 | 12774 | 27 | CCTGTCCCCGCCGGCGGGCCGCCCCCC | 571.7 | -7.8 | 1.8 | -4.3 | 1.4E-05 | 1.9E-05 | BT20/MB468 | NUC |
| >hg38_21_+_8433172_8446622_RNA45SN1_45S_50ntFlanks@10815.10840.26\|GGTAAGGGAAGTCGGCAAGCCGGATC | >hg38_21_+_8433172_8446622_RNA45SN1_45S_50ntFlanks | 10815 | 10840 | 26 | GGTAAGGGAAGTCGGCAAGCCGGATC | 121.9 | -6.4 | 1.5 | -4.4 | 1.1E-05 | 1.9E-05 | MB231/MB468 | MP |
| >hg38_21_+_8433172_8446622_RNA45SN1_45S_50ntFlanks@6652.6696.45\|GACTCTTAGCGGTGGATCACTCGGCTCGTGCGTCGATGAAGAACG | >hg38_21_+_8433172_8446622_RNA45SN1_45S_50ntFlanks | 6652 | 6696 | 45 | GACTCTTAGCGGTGGATCACTCGGCTCGTGCGTCGATGAAGAACG | 378.2 | 3.8 | 0.9 | 4.4 | 1.1E-05 | 1.9E-05 | BT20/MB468 | CYTO |
| >hg38_21_+_8433172_8446622_RNA45SN1_45S_50ntFlanks@6651.6682.32\|CGACTCTTAGCGGTGGATCACTCGGCTCGTGC | >hg38_21_+_8433172_8446622_RNA45SN1_45S_50ntFlanks | 6651 | 6682 | 32 | CGACTCTTAGCGGTGGATCACTCGGCTCGTGC | 2023.3 | 4.6 | 1.0 | 4.4 | 1.3E-05 | 1.9E-05 | MB231/MB468 | CYTO |
| >hg38_21_+_8433172_8446622_RNA45SN1_45S_50ntFlanks@7981.7999.19\|CTCAGATCAGACGTGGCGA | >hg38_21_+_8433172_8446622_RNA45SN1_45S_50ntFlanks | 7981 | 7999 | 19 | CTCAGATCAGACGTGGCGA | 149.3 | -3.6 | 0.8 | -4.4 | 1.3E-05 | 1.9E-05 | MB231/MB468 | CYTO |
| >hg38_1_-_228634819_228635039_RNA5S12_5S_50ntFlanks@51.81.31\|GTCTACGGCCATACCACCCTGAACGCGCCCG | >hg38_1_-_228634819_228635039_RNA5S12_5S_50ntFlanks | 51 | 81 | 31 | GTCTACGGCCATACCACCCTGAACGCGCCCG | 2055.9 | 4.3 | 1.0 | 4.4 | 1.1E-05 | 1.9E-05 | BT20/MB468 | CYTO |
| >hg38_21_+_8433172_8446622_RNA45SN1_45S_50ntFlanks@10809.10830.22\|AATGTAGGTAAGGGAAGTCGGC | >hg38_21_+_8433172_8446622_RNA45SN1_45S_50ntFlanks | 10809 | 10830 | 22 | AATGTAGGTAAGGGAAGTCGGC | 1631.6 | 4.9 | 1.1 | 4.4 | 1.3E-05 | 1.9E-05 | MB231/MB468 | CYTO |
| >hg38_21_+_8433172_8446622_RNA45SN1_45S_50ntFlanks@6646.6665.20\|TCGTACGACTCTTAGCGGTG | >hg38_21_+_8433172_8446622_RNA45SN1_45S_50ntFlanks | 6646 | 6665 | 20 | TCGTACGACTCTTAGCGGTG | 8815.4 | 2.9 | 0.7 | 4.4 | 1.3E-05 | 1.9E-05 | MB231/MB468 | CYTO |

<table>
<tr><th></th><th></th><th></th><th></th><th></th><th></th><th></th><th></th><th></th><th></th><th></th><th></th><th></th><th></th></tr>
<tr><td>>hg38_21_+_8433172_8446622_RNA45SN1_45S_50ntFlanks@11580.11611.32|GAACTGGTGCGGACCAGGGGAATCCGACTGTT</td><td>>hg38_21_+_8433172_8446622_RNA45SN1_45S_50ntFlanks</td><td>11580</td><td>11611</td><td>32</td><td>GAACTGGTGCGGACCAGGGGAATCCGACTGTT</td><td>2429.4</td><td>-4.2</td><td>0.9</td><td>-4.6</td><td>4.2E-06</td><td>1.9E-05</td><td>BT20/MB468</td><td>MP</td></tr>
<tr><td>>am_trnaMT_PheGAA_MT_+_577_647@43.62.20|TTAGACGGGCTCACATCACC</td><td>>am_trnaMT_PheGAA_MT_+_577_647</td><td>43</td><td>62</td><td>20</td><td>TTAGACGGGCTCACATCACC</td><td>478.4</td><td>8.1</td><td>1.9</td><td>4.4</td><td>1.1E-05</td><td>1.9E-05</td><td>BT20/MB468</td><td>CYTO</td></tr>
<tr><td>>hg38_21_+_8433172_8446622_RNA45SN1_45S_50ntFlanks@3993.4020.28|GTGACTCTAGATAACCTCGGGCCGATCG</td><td>>hg38_21_+_8433172_8446622_RNA45SN1_45S_50ntFlanks</td><td>3993</td><td>4020</td><td>28</td><td>GTGACTCTAGATAACCTCGGGCCGATCG</td><td>737.0</td><td>-5.6</td><td>1.3</td><td>-4.4</td><td>1.3E-05</td><td>1.9E-05</td><td>MB231/MB468</td><td>MITO</td></tr>
<tr><td>>trnaMT_SerGCT_MT_+_12207_12265@1.21.21|GAGAAAGCTCACAAGAACTGC</td><td>>trnaMT_SerGCT_MT_+_12207_12265</td><td>1</td><td>21</td><td>21</td><td>GAGAAAGCTCACAAGAACTGC</td><td>8030.4</td><td>5.5</td><td>1.3</td><td>4.4</td><td>9.6E-06</td><td>1.9E-05</td><td>BT20/MB468</td><td>MITO</td></tr>
<tr><td>>MI0005566|hsa-mir-374b&WithFlank&X|-|74218541|74218624@17.38.22&[MIMAT0004955&hsa-miR-374b-5p&offsets|0|0;m-153&X|-|74218587|74218608&offsets|0|0]|ATATAATACAACCTGCTAAGTG</td><td>>MI0005566|hsa-mir-374b&WithFlank&X|-|74218541|74218624</td><td>17</td><td>38</td><td>22</td><td>ATATAATACAACCTGCTAAGTG</td><td>240.4</td><td>2.0</td><td>0.5</td><td>4.3</td><td>1.4E-05</td><td>1.9E-05</td><td>BT20/MB468</td><td>NUC</td></tr>
<tr><td>>hg38_21_+_8433172_8446622_RNA45SN1_45S_50ntFlanks@10409.10430.22|GTCAGTCGGTCCTGAGAGATGG</td><td>>hg38_21_+_8433172_8446622_RNA45SN1_45S_50ntFlanks</td><td>10409</td><td>10430</td><td>22</td><td>GTCAGTCGGTCCTGAGAGATGG</td><td>850.0</td><td>-4.5</td><td>1.0</td><td>-4.4</td><td>1.3E-05</td><td>1.9E-05</td><td>MB231/MB468</td><td>CYTO</td></tr>
<tr><td>>hg38_21_+_8433172_8446622_RNA45SN1_45S_50ntFlanks@10392.10422.31|ACAGCAGTTGAACATGGGTCAGTCGGTCCTG</td><td>>hg38_21_+_8433172_8446622_RNA45SN1_45S_50ntFlanks</td><td>10392</td><td>10422</td><td>31</td><td>ACAGCAGTTGAACATGGGTCAGTCGGTCCTG</td><td>710.0</td><td>-4.7</td><td>1.1</td><td>-4.4</td><td>1.3E-05</td><td>1.9E-05</td><td>MB231/MB468</td><td>CYTO</td></tr>
<tr><td>>MI0000102|hsa-mir-100&WithFlank&11|-|122152223|122152314@19.40.22&[MIMAT0000098&hsa-miR-100-5p&offsets|0|0;m-27&11|-|122152276|122152296&offsets|0|+1]|AACCCGTAGATCCGAACTTGTG</td><td>>MI0000102|hsa-mir-100&WithFlank&11|-|122152223|122152314</td><td>19</td><td>40</td><td>22</td><td>AACCCGTAGATCCGAACTTGTG</td><td>8705.8</td><td>7.9</td><td>1.8</td><td>4.4</td><td>9.6E-06</td><td>1.9E-05</td><td>BT20/MB468</td><td>MITO</td></tr>
<tr><td>>MI0000255|hsa-mir-30d&WithFlank&8|-|134804870|134804951@12.35.24(+1A)&[MIMAT0000245&hsa-miR-30d-5p&offsets|0|+2(+1A);m-21&8|-|134804919|134804940&offsets|0|+2(+1A)]|TGTAAACATCCCCGACTGGAAGCTA</td><td>>MI0000255|hsa-mir-30d&WithFlank&8|-|134804870|134804951</td><td>12</td><td>35</td><td>24(+1A)</td><td>TGTAAACATCCCCGACTGGAAGCTA</td><td>70.7</td><td>3.2</td><td>0.7</td><td>4.3</td><td>1.4E-05</td><td>1.9E-05</td><td>BT20/MB468</td><td>NUC</td></tr>
<tr><td>>hg38_21_+_8433172_8446622_RNA45SN1_45S_50ntFlanks@10361.10391.31|GAAGGCCGAAGTGGAGAAGGGTTCCATGTGA</td><td>>hg38_21_+_8433172_8446622_RNA45SN1_45S_50ntFlanks</td><td>10361</td><td>10391</td><td>31</td><td>GAAGGCCGAAGTGGAGAAGGGTTCCATGTGA</td><td>42.7</td><td>-8.5</td><td>1.8</td><td>-4.6</td><td>4.2E-06</td><td>1.9E-05</td><td>BT20/MB468</td><td>MP</td></tr>
<tr><td>>trnaMT_GluTTC_MT_-_14674_14742@27.61.35,trnalookalike8_GluTTC_5_-_93905172_93905240@27.61.35|GTTTTTCATATCATTGGTCGTGGTTGTAGTCCGTG</td><td>>trnaMT_GluTTC_MT_-_14674_14742</td><td>27</td><td>61</td><td>35</td><td>GTTTTTCATATCATTGGTCGTGGTTGTAGTCCGTG</td><td>604.5</td><td>-3.6</td><td>0.8</td><td>-4.4</td><td>1.3E-05</td><td>1.9E-05</td><td>MB231/MB468</td><td>NUC</td></tr>
<tr><td>>hg38_21_+_8433172_8446622_RNA45SN1_45S_50ntFlanks@6663.6681.19|GTGGATCACTCGGCTCGTG</td><td>>hg38_21_+_8433172_8446622_RNA45SN1_45S_50ntFlanks</td><td>6663</td><td>6681</td><td>19</td><td>GTGGATCACTCGGCTCGTG</td><td>904.9</td><td>-5.3</td><td>1.2</td><td>-4.4</td><td>1.3E-05</td><td>1.9E-05</td><td>MB231/MB468</td><td>CYTO</td></tr>
<tr><td>>MI0000299|hsa-mir-222&WithFlank&X|-|45747009|45747130@75.99.25(+3U)&[MIMAT0000279&hsa-miR-222-3p&offsets|0|+4(+3U);m-140&X|-|45747033|45747056&offsets|0|+1(+3U)]|AGCTACATCTGGCTACTGGGTCTCTTTT</td><td>>MI0000299|hsa-mir-222&WithFlank&X|-|45747009|45747130</td><td>75</td><td>99</td><td>25(+3U)</td><td>AGCTACATCTGGCTACTGGGTCTCTTTT</td><td>2892.1</td><td>-3.9</td><td>0.8</td><td>-4.8</td><td>1.5E-06</td><td>1.9E-05</td><td>BT20/MB231</td><td>MITO</td></tr>
<tr><td>>hg38_21_+_8433172_8446622_RNA45SN1_45S_50ntFlanks@12499.12520.22|GGTTTAGACCGTCGTGAGACAG</td><td>>hg38_21_+_8433172_8446622_RNA45SN1_45S_50ntFlanks</td><td>12499</td><td>12520</td><td>22</td><td>GGTTTAGACCGTCGTGAGACAG</td><td>101.5</td><td>-10.5</td><td>2.4</td><td>-4.4</td><td>1.1E-05</td><td>1.9E-05</td><td>MB231/MB468</td><td>MP</td></tr>
<tr><td>>hg38_21_+_8433172_8446622_RNA45SN1_45S_50ntFlanks@6776.6806.31|CTCCCGGGGCTACGCCTGTCTGAGCGTCGCT</td><td>>hg38_21_+_8433172_8446622_RNA45SN1_45S_50ntFlanks</td><td>6776</td><td>6806</td><td>31</td><td>CTCCCGGGGCTACGCCTGTCTGAGCGTCGCT</td><td>1669.1</td><td>2.8</td><td>0.6</td><td>4.4</td><td>1.3E-05</td><td>1.9E-05</td><td>MB231/MB468</td><td>NUC</td></tr>
<tr><td>>hg38_21_+_8433172_8446622_RNA45SN1_45S_50ntFlanks@9300.9332.33|ACGGACCAAGGAGTCTAACACGTGCGCGAGTCG</td><td>>hg38_21_+_8433172_8446622_RNA45SN1_45S_50ntFlanks</td><td>9300</td><td>9332</td><td>33</td><td>ACGGACCAAGGAGTCTAACACGTGCGCGAGTCG</td><td>39.0</td><td>9.0</td><td>2.0</td><td>4.6</td><td>4.2E-06</td><td>1.9E-05</td><td>BT20/MB468</td><td>MP</td></tr>
<tr><td>>hg38_21_+_8433172_8446622_RNA45SN1_45S_50ntFlanks@8106.8140.35|GCGGCGGGGCGCGGGACATGTGGCGTACGGAAGAC</td><td>>hg38_21_+_8433172_8446622_RNA45SN1_45S_50ntFlanks</td><td>8106</td><td>8140</td><td>35</td><td>GCGGCGGGGCGCGGGACATGTGGCGTACGGAAGAC</td><td>847.5</td><td>-3.7</td><td>0.8</td><td>-4.6</td><td>4.2E-06</td><td>1.9E-05</td><td>BT20/MB468</td><td>MP</td></tr>
<tr><td>>hg38_21_+_8433172_8446622_RNA45SN1_45S_50ntFlanks@5420.5466.47|CCGATTGGATGGTTTAGTGAGGCCCTCGGATCGGCCCCGCCGGGGTC</td><td>>hg38_21_+_8433172_8446622_RNA45SN1_45S_50ntFlanks</td><td>5420</td><td>5466</td><td>47</td><td>CCGATTGGATGGTTTAGTGAGGCCCTCGGATCGGCCCCGCCGGGGTC</td><td>69.8</td><td>-9.2</td><td>2.0</td><td>-4.6</td><td>4.2E-06</td><td>1.9E-05</td><td>BT20/MB468</td><td>MP</td></tr>
<tr><td>>hg38_21_+_8433172_8446622_RNA45SN1_45S_50ntFlanks@11233.11252.20|CCCTTCCCGTGGATCGCCCC</td><td>>hg38_21_+_8433172_8446622_RNA45SN1_45S_50ntFlanks</td><td>11233</td><td>11252</td><td>20</td><td>CCCTTCCCGTGGATCGCCCC</td><td>437.8</td><td>-6.2</td><td>1.4</td><td>-4.4</td><td>1.1E-05</td><td>1.9E-05</td><td>BT20/MB468</td><td>CYTO</td></tr>
<tr><td>>hg38_21_+_8433172_8446622_RNA45SN1_45S_50ntFlanks@3981.4001.21|CCGGCGGCTTTGGTGACTCTA</td><td>>hg38_21_+_8433172_8446622_RNA45SN1_45S_50ntFlanks</td><td>3981</td><td>4001</td><td>21</td><td>CCGGCGGCTTTGGTGACTCTA</td><td>92.9</td><td>-10.4</td><td>2.4</td><td>-4.4</td><td>1.1E-05</td><td>1.9E-05</td><td>MB231/MB468</td><td>MP</td></tr>
<tr><td>>hg38_21_+_8433172_8446622_RNA45SN1_45S_50ntFlanks@11568.11601.34|GCAGCCGACTTAGAACTGGTGCGGACCAGGGGAA</td><td>>hg38_21_+_8433172_8446622_RNA45SN1_45S_50ntFlanks</td><td>11568</td><td>11601</td><td>34</td><td>GCAGCCGACTTAGAACTGGTGCGGACCAGGGGAA</td><td>1119.7</td><td>-5.3</td><td>1.2</td><td>-4.4</td><td>1.4E-05</td><td>1.9E-05</td><td>MB231/MB468</td><td>MITO</td></tr>
<tr><td>>hg38_21_+_8433172_8446622_RNA45SN1_45S_50ntFlanks@10133.10157.25|GCGGCGGCGGCGGCGGGGGTGTGGG</td><td>>hg38_21_+_8433172_8446622_RNA45SN1_45S_50ntFlanks</td><td>10133</td><td>10157</td><td>25</td><td>GCGGCGGCGGCGGCGGGGGTGTGGG</td><td>152.3</td><td>-6.8</td><td>1.5</td><td>-4.4</td><td>1.1E-05</td><td>1.9E-05</td><td>MB231/MB468</td><td>MP</td></tr>
<tr><td>>MI0000081|hsa-mir-24-2&WithFlank&19|-|13836281|13836365@56.75.20;MI0000080|hsa-mir-24-1&WithFlank&9|+|95086015|95086094@50.69.20&[MIMAT0000080&hsa-miR-24-3p&offsets|0|-2;m-36&19|-|13836288|13836310&offsets|0|-3];[MIMAT0000080_1&hsa-miR-24-3p&offsets|0|-2;m-35&9|+|95086064|95086086&offsets|0|-3]|TGGCTCAGTTCAGCAGGAAC</td><td>>MI0000081|hsa-mir-24-2&WithFlank&19|-|13836281|13836365</td><td>56</td><td>75</td><td>20</td><td>TGGCTCAGTTCAGCAGGAAC</td><td>3794.7</td><td>1.8</td><td>0.4</td><td>4.3</td><td>1.4E-05</td><td>1.9E-05</td><td>BT20/MB468</td><td>NUC</td></tr>
<tr><td>>trna119_LysCTT_1_-_145395522_145395594@1.29.29,trna11_LysCTT_5_-_180648979_180649051@1.29.29,trna13_LysCTT_6_+_26556774_26556846@1.29.29,trna32_LysCTT_16_-_3207406_3207478@1.29.29,trna7_LysCTT_16_+_3225692_3225764@1.29.29,trna9_LysCTT_5_+_180634755_180634827@1.29.29|GCCCGGCTAGCTCAGTCGGTAGAGCATGA</td><td>>trna119_LysCTT_1_-_145395522_145395594</td><td>1</td><td>29</td><td>29</td><td>GCCCGGCTAGCTCAGTCGGTAGAGCATGA</td><td>2332.0</td><td>-4.1</td><td>0.9</td><td>-4.4</td><td>1.1E-05</td><td>1.9E-05</td><td>BT20/MB468</td><td>CYTO</td></tr>
<tr><td>>trna111_HisGTG_1_-_147774845_147774916@-1G.34.35,trna118_HisGTG_1_-_145396881_145396952@-1G.34.35,trna16_HisGTG_1_+_146544773_146544844@-1G.34.35,trna1_HisGTG_15_+_45493349_45493420@-1G.34.35,trna21_HisGTG_1_+_147753471_147753542@-1G.34.35,trna33_HisGTG_6_+_27125906_27125977@-1G.34.35,trna7_HisGTG_9_-_14433938_14434009@-1G.34.35,trna8_HisGTG_15_-_45492611_45492682@-1G.34.35,trna9_HisGTG_15_-_45490804_45490875@-1G.34.35|GGCCGTGATCGTATAGTGGTTAGTACTCTGCGTTG</td><td>>trna111_HisGTG_1_-_147774845_147774916</td><td>-1G</td><td>34</td><td>35</td><td>GGCCGTGATCGTATAGTGGTTAGTACTCTGCGTTG</td><td>11259.8</td><td>5.8</td><td>1.3</td><td>4.4</td><td>1.3E-05</td><td>1.9E-05</td><td>MB231/MB468</td><td>NUC</td></tr>
<tr><td>>hg38_21_+_8433172_8446622_RNA45SN1_45S_50ntFlanks@10350.10381.32|ACGAGAACTTTGAAGGCCGAAGTGGAGAAGGG</td><td>>hg38_21_+_8433172_8446622_RNA45SN1_45S_50ntFlanks</td><td>10350</td><td>10381</td><td>32</td><td>ACGAGAACTTTGAAGGCCGAAGTGGAGAAGGG</td><td>276.7</td><td>-2.4</td><td>0.5</td><td>-4.4</td><td>1.1E-05</td><td>1.9E-05</td><td>BT20/MB468</td><td>CYTO</td></tr>
<tr><td>>hg38_21_+_8433172_8446622_RNA45SN1_45S_50ntFlanks@5481.5501.21|GCGGAGCGCTGAGAAGACGGT</td><td>>hg38_21_+_8433172_8446622_RNA45SN1_45S_50ntFlanks</td><td>5481</td><td>5501</td><td>21</td><td>GCGGAGCGCTGAGAAGACGGT</td><td>148.1</td><td>10.5</td><td>2.2</td><td>4.8</td><td>1.3E-06</td><td>1.9E-05</td><td>BT20/MB231</td><td>MP</td></tr>
<tr><td>>hg38_21_+_8433172_8446622_RNA45SN1_45S_50ntFlanks@11661.11680.20|TGATTTCTGCCCAGTGCTCT</td><td>>hg38_21_+_8433172_8446622_RNA45SN1_45S_50ntFlanks</td><td>11661</td><td>11680</td><td>20</td><td>TGATTTCTGCCCAGTGCTCT</td><td>716.0</td><td>-5.8</td><td>1.3</td><td>-4.3</td><td>1.4E-05</td><td>1.9E-05</td><td>BT20/MB468</td><td>NUC</td></tr>
<tr><td>>hg38_21_+_8433172_8446622_RNA45SN1_45S_50ntFlanks@11582.11603.22|ACTGGTGCGGACCAGGGGAATC</td><td>>hg38_21_+_8433172_8446622_RNA45SN1_45S_50ntFlanks</td><td>11582</td><td>11603</td><td>22</td><td>ACTGGTGCGGACCAGGGGAATC</td><td>1518.0</td><td>-2.6</td><td>0.6</td><td>-4.4</td><td>1.3E-05</td><td>1.9E-05</td><td>MB231/MB468</td><td>CYTO</td></tr>
<tr><td>>hg38_21_+_8433172_8446622_RNA45SN1_45S_50ntFlanks@10846.10867.22|CTTCGGGATAAGGATTGGCTCT</td><td>>hg38_21_+_8433172_8446622_RNA45SN1_45S_50ntFlanks</td><td>10846</td><td>10867</td><td>22</td><td>CTTCGGGATAAGGATTGGCTCT</td><td>402.8</td><td>-5.1</td><td>1.2</td><td>-4.4</td><td>1.3E-05</td><td>1.9E-05</td><td>MB231/MB468</td><td>NUC</td></tr>
<tr><td>>MI0000088|hsa-mir-30a&WithFlank&6|-|71403545|71403627@13.35.23(+1U)&[MIMAT0000087&hsa-miR-30a-5p&offsets|+1|+2(+1U);m-15&6|-|71403595|71403616&offsets|+1|+2(+1U)]|GTAAACATCCTCGACTGGAAGCTT</td><td>>MI0000088|hsa-mir-30a&WithFlank&6|-|71403545|71403627</td><td>13</td><td>35</td><td>23(+1U)</td><td>GTAAACATCCTCGACTGGAAGCTT</td><td>299.2</td><td>-3.9</td><td>0.8</td><td>-4.8</td><td>1.5E-06</td><td>1.9E-05</td><td>BT20/MB231</td><td>MITO</td></tr>
<tr><td>>hg38_21_+_8433172_8446622_RNA45SN1_45S_50ntFlanks@8157.8181.25|CTCGTGGGGGGCCCAAGTCCTTCTG</td><td>>hg38_21_+_8433172_8446622_RNA45SN1_45S_50ntFlanks</td><td>8157</td><td>8181</td><td>25</td><td>CTCGTGGGGGGCCCAAGTCCTTCTG</td><td>1210.0</td><td>-6.0</td><td>1.3</td><td>-4.6</td><td>4.3E-06</td><td>1.9E-05</td><td>BT20/MB468</td><td>MP</td></tr>
<tr><td>>hg38_21_+_8433172_8446622_RNA45SN1_45S_50ntFlanks@7975.8009.35|CGCGACCTCAGATCAGACGTGGCGACCCGCTGAAT</td><td>>hg38_21_+_8433172_8446622_RNA45SN1_45S_50ntFlanks</td><td>7975</td><td>8009</td><td>35</td><td>CGCGACCTCAGATCAGACGTGGCGACCCGCTGAAT</td><td>106217.9</td><td>3.1</td><td>0.7</td><td>4.3</td><td>1.4E-05</td><td>1.9E-05</td><td>BT20/MB468</td><td>NUC</td></tr>
<tr><td>>hg38_21_+_8433172_8446622_RNA45SN1_45S_50ntFlanks@12458.12475.18|AAGCGTTGGATTGTTCAC</td><td>>hg38_21_+_8433172_8446622_RNA45SN1_45S_50ntFlanks</td><td>12458</td><td>12475</td><td>18</td><td>AAGCGTTGGATTGTTCAC</td><td>223.8</td><td>-6.5</td><td>1.5</td><td>-4.4</td><td>1.1E-05</td><td>1.9E-05</td><td>MB231/MB468</td><td>MP</td></tr>
<tr><td>>hg38_21_+_8433172_8446622_RNA45SN1_45S_50ntFlanks@12418.12457.40|CGATGTCGGCTCTTCCTATCATTGTGAAGCAGAATTCACC</td><td>>hg38_21_+_8433172_8446622_RNA45SN1_45S_50ntFlanks</td><td>12418</td><td>12457</td><td>40</td><td>CGATGTCGGCTCTTCCTATCATTGTGAAGCAGAATTCACC</td><td>358.5</td><td>-6.0</td><td>1.4</td><td>-4.3</td><td>1.4E-05</td><td>1.9E-05</td><td>BT20/MB468</td><td>NUC</td></tr>
<tr><td>>hg38_21_+_8433172_8446622_RNA45SN1_45S_50ntFlanks@4449.4472.24|CCTCTCGGCGCCCCCTCGATGCTC</td><td>>hg38_21_+_8433172_8446622_RNA45SN1_45S_50ntFlanks</td><td>4449</td><td>4472</td><td>24</td><td>CCTCTCGGCGCCCCCTCGATGCTC</td><td>528.3</td><td>-6.0</td><td>1.4</td><td>-4.4</td><td>1.2E-05</td><td>1.9E-05</td><td>BT20/MB468</td><td>CYTO</td></tr>
<tr><td>>hg38_21_+_8433172_8446622_RNA45SN1_45S_50ntFlanks@4724.4751.28|ATTAATCAAGAACGAAAGTCGGAGGTTC</td><td>>hg38_21_+_8433172_8446622_RNA45SN1_45S_50ntFlanks</td><td>4724</td><td>4751</td><td>28</td><td>ATTAATCAAGAACGAAAGTCGGAGGTTC</td><td>467.6</td><td>-6.3</td><td>1.4</td><td>-4.4</td><td>9.8E-06</td><td>1.9E-05</td><td>BT20/MB468</td><td>MITO</td></tr>
<tr><td>>hg38_21_+_8433172_8446622_RNA45SN1_45S_50ntFlanks@9554.9579.26|CGTAGCGGTCCTGACGTGCAAATCGG</td><td>>hg38_21_+_8433172_8446622_RNA45SN1_45S_50ntFlanks</td><td>9554</td><td>9579</td><td>26</td><td>CGTAGCGGTCCTGACGTGCAAATCGG</td><td>824.6</td><td>-7.6</td><td>1.7</td><td>-4.3</td><td>1.4E-05</td><td>1.9E-05</td><td>BT20/MB468</td><td>NUC</td></tr>
<tr><td>>hg38_21_+_8433172_8446622_RNA45SN1_45S_50ntFlanks@3858.3881.24|TGGTAATTCTAGAGCTAATACATG</td><td>>hg38_21_+_8433172_8446622_RNA45SN1_45S_50ntFlanks</td><td>3858</td><td>3881</td><td>24</td><td>TGGTAATTCTAGAGCTAATACATG</td><td>46.7</td><td>-8.6</td><td>1.9</td><td>-4.6</td><td>4.3E-06</td><td>1.9E-05</td><td>BT20/MB468</td><td>MP</td></tr>
<tr><td>>MI0000085|hsa-mir-27a&WithFlank&19|-|13836434|13836523@57.76.20(+1A)&[MIMAT0000084&hsa-miR-27a-3p&offsets|0|-1(+1A);m-43&19|-|13836447|13836467&offsets|0|-1(+1A)]|TTCACAGTGGCTAAGTTCCGA</td><td>>MI0000085|hsa-mir-27a&WithFlank&19|-|13836434|13836523</td><td>57</td><td>76</td><td>20(+1A)</td><td>TTCACAGTGGCTAAGTTCCGA</td><td>2919.9</td><td>3.9</td><td>0.9</td><td>4.3</td><td>1.4E-05</td><td>2.0E-05</td><td>MB231/MB468</td><td>MITO</td></tr>
<tr><td>>hg38_21_+_8433172_8446622_RNA45SN1_45S_50ntFlanks@6731.6758.28|CACATTGATCATCGACACTTCGAACGCA</td><td>>hg38_21_+_8433172_8446622_RNA45SN1_45S_50ntFlanks</td><td>6731</td><td>6758</td><td>28</td><td>CACATTGATCATCGACACTTCGAACGCA</td><td>220.6</td><td>-4.6</td><td>1.1</td><td>-4.4</td><td>1.3E-05</td><td>2.0E-05</td><td>MB231/MB468</td><td>NUC</td></tr>
<tr><td>>MI0000273|hsa-mir-183&WithFlank&7|-|129774899|129775020@34.54.21&[MIMAT0000261&hsa-miR-183-5p&offsets|+1|0;m-41&7|-|129774966|129774987&offsets|0|-1]|ATGGCACTGGTAGAATTCACT</td><td>>MI0000273|hsa-mir-183&WithFlank&7|-|129774899|129775020</td><td>34</td><td>54</td><td>21</td><td>ATGGCACTGGTAGAATTCACT</td><td>4577.7</td><td>3.7</td><td>0.8</td><td>4.8</td><td>1.3E-06</td><td>2.0E-05</td><td>BT20/MB231</td><td>CYTO</td></tr>
<tr><td>>MI0000071|hsa-mir-17&WithFlank&13|+|91350599|91350694@20.41.22&[MIMAT0000070&hsa-miR-17-5p&offsets|0|-1;m-88&13|+|91350618|91350640&offsets|0|-1]|CAAAGTGCTTACAGTGCAGGTA</td><td>>MI0000071|hsa-mir-17&WithFlank&13|+|91350599|91350694</td><td>20</td><td>41</td><td>22</td><td>CAAAGTGCTTACAGTGCAGGTA</td><td>395.3</td><td>4.4</td><td>1.0</td><td>4.4</td><td>1.3E-05</td><td>2.0E-05</td><td>MB231/MB468</td><td>CYTO</td></tr>
<tr><td>>hg38_21_+_8433172_8446622_RNA45SN1_45S_50ntFlanks@8168.8186.19|CCCAAGTCCTTCTGATCGA</td><td>>hg38_21_+_8433172_8446622_RNA45SN1_45S_50ntFlanks</td><td>8168</td><td>8186</td><td>19</td><td>CCCAAGTCCTTCTGATCGA</td><td>1136.8</td><td>-5.1</td><td>1.2</td><td>-4.4</td><td>1.3E-05</td><td>2.0E-05</td><td>MB231/MB468</td><td>CYTO</td></tr>
<tr><td>>hg38_1_-_228634819_228635039_RNA5S12_5S_50ntFlanks@152.171.20|TACCGGGTGCTGTAGGCTTT</td><td>>hg38_1_-_228634819_228635039_RNA5S12_5S_50ntFlanks</td><td>152</td><td>171</td><td>20</td><td>TACCGGGTGCTGTAGGCTTT</td><td>1657.1</td><td>4.0</td><td>0.9</td><td>4.4</td><td>1.3E-05</td><td>2.0E-05</td><td>MB231/MB468</td><td>NUC</td></tr>
<tr><td>>hg38_21_+_8433172_8446622_RNA45SN1_45S_50ntFlanks@4739.4758.20|AAGTCGGAGGTTCGAAGACG</td><td>>hg38_21_+_8433172_8446622_RNA45SN1_45S_50ntFlanks</td><td>4739</td><td>4758</td><td>20</td><td>AAGTCGGAGGTTCGAAGACG</td><td>153.4</td><td>-4.5</td><td>1.0</td><td>-4.4</td><td>1.3E-05</td><td>2.0E-05</td><td>MB231/MB468</td><td>NUC</td></tr>
<tr><td>>MI0000746|hsa-mir-99b&WithFlank&19|+|51692606|51692687@51.73.23&[MIMAT0004678&hsa-miR-99b-3p&offsets|0|+1;m-151&19|+|51692656|51692678&offsets|0|0]|CAAGCTCGTGTCTGTGGGTCCGT</td><td>>MI0000746|hsa-mir-99b&WithFlank&19|+|51692606|51692687</td><td>51</td><td>73</td><td>23</td><td>CAAGCTCGTGTCTGTGGGTCCGT</td><td>278.7</td><td>4.5</td><td>1.0</td><td>4.3</td><td>1.4E-05</td><td>2.0E-05</td><td>MB231/MB468</td><td>MITO</td></tr>
</table>

| | | | | | | | | | | | | | |
|---|---|---|---|---|---|---|---|---|---|---|---|---|---|
| >hg38_21_+_8433172_8446622_RNA45SN1_45S_50ntFlanks@11315.11344.30\|CCACCCCACGTCTCGTCGCGCGCGCGTCCG | >hg38_21_+_8433172_8446622_RNA45SN1_45S_50ntFlanks | 11315 | 11344 | 30 | CCACCCCACGTCTCGTCGCGCGCGCGTCCG | 288.1 | -6.7 | 1.5 | -4.6 | 4.3E-06 | 2.0E-05 | BT20/MB468 | MP |
| >hg38_21_+_8433172_8446622_RNA45SN1_45S_50ntFlanks@6648.6676.29\|GTACGACTCTTAGCGGTGGATCACTCGGC | >hg38_21_+_8433172_8446622_RNA45SN1_45S_50ntFlanks | 6648 | 6676 | 29 | GTACGACTCTTAGCGGTGGATCACTCGGC | 1244.9 | 4.3 | 1.0 | 4.4 | 1.4E-05 | 2.0E-05 | MB231/MB468 | NUC |
| >hg38_21_+_8433172_8446622_RNA45SN1_45S_50ntFlanks@6655.6686.32\|TCTTAGCGGTGGATCACTCGGCTCGTGCGTCG | >hg38_21_+_8433172_8446622_RNA45SN1_45S_50ntFlanks | 6655 | 6686 | 32 | TCTTAGCGGTGGATCACTCGGCTCGTGCGTCG | 391.9 | -2.6 | 0.6 | -4.3 | 1.4E-05 | 2.0E-05 | BT20/MB468 | NUC |
| >hg38_21_+_8433172_8446622_RNA45SN1_45S_50ntFlanks@11572.11602.31\|CCGACTTAGAACTGGTGCGGACCAGGGGAAT | >hg38_21_+_8433172_8446622_RNA45SN1_45S_50ntFlanks | 11572 | 11602 | 31 | CCGACTTAGAACTGGTGCGGACCAGGGGAAT | 3151.3 | -2.6 | 0.6 | -4.4 | 9.9E-06 | 2.0E-05 | BT20/MB468 | MITO |
| >hg38_21_+_8433172_8446622_RNA45SN1_45S_50ntFlanks@10782.10799.18\|GGTGAACAGCCTCTGGCA | >hg38_21_+_8433172_8446622_RNA45SN1_45S_50ntFlanks | 10782 | 10799 | 18 | GGTGAACAGCCTCTGGCA | 150.4 | -6.2 | 1.4 | -4.4 | 1.1E-05 | 2.0E-05 | MB231/MB468 | MP |
| >hg38_21_+_8433172_8446622_RNA45SN1_45S_50ntFlanks@12420.12447.28\|ATGTCGGCTCTTCCTATCATTGTGAAGC | >hg38_21_+_8433172_8446622_RNA45SN1_45S_50ntFlanks | 12420 | 12447 | 28 | ATGTCGGCTCTTCCTATCATTGTGAAGC | 1255.0 | 2.4 | 0.6 | 4.4 | 1.3E-05 | 2.0E-05 | MB231/MB468 | CYTO |
| >MI0000446\|hsa-mir-125b-1&WithFlank&11\|-\|122099751\|122099850@61.80.20(+1A)&[MIMAT0004592&hsa-miR-125b-1-3p&offsets\|0\|-2(+1A);m-485&11\|-\|122099769\|122099790&offsets\|0\|-2(+1A)]\|ACGGGTTAGGCTCTTGGGAGA | >MI0000446\|hsa-mir-125b-1&WithFlank&11\|-\|122099751\|122099850 | 61 | 80 | 20(+1A) | ACGGGTTAGGCTCTTGGGAGA | 38.9 | 8.5 | 1.9 | 4.4 | 1.1E-05 | 2.0E-05 | MB231/MB468 | MP |
| >hg38_21_+_8433172_8446622_RNA45SN1_45S_50ntFlanks@7985.8006.22\|GATCAGACGTGGCGACCCGCTG | >hg38_21_+_8433172_8446622_RNA45SN1_45S_50ntFlanks | 7985 | 8006 | 22 | GATCAGACGTGGCGACCCGCTG | 994.7 | -6.0 | 1.4 | -4.4 | 1.3E-05 | 2.0E-05 | MB231/MB468 | CYTO |
| >hg38_21_+_8433172_8446622_RNA45SN1_45S_50ntFlanks@12458.12478.21\|AAGCGTTGGATTGTTCACCCA | >hg38_21_+_8433172_8446622_RNA45SN1_45S_50ntFlanks | 12458 | 12478 | 21 | AAGCGTTGGATTGTTCACCCA | 536.6 | -5.6 | 1.3 | -4.4 | 1.1E-05 | 2.0E-05 | MB231/MB468 | MP |
| >hg38_21_+_8433172_8446622_RNA45SN1_45S_50ntFlanks@8180.8211.32\|TGATCGAGGCCCAGCCCGTGGACGGTGTGAGG | >hg38_21_+_8433172_8446622_RNA45SN1_45S_50ntFlanks | 8180 | 8211 | 32 | TGATCGAGGCCCAGCCCGTGGACGGTGTGAGG | 233.4 | -6.3 | 1.4 | -4.4 | 1.1E-05 | 2.0E-05 | MB231/MB468 | MP |
| >hg38_1_-_228634819_228635039_RNA5S12_5S_50ntFlanks@145.170.26\|CTGGGAATACCGGGTGCTGTAGGCTT | >hg38_1_-_228634819_228635039_RNA5S12_5S_50ntFlanks | 145 | 170 | 26 | CTGGGAATACCGGGTGCTGTAGGCTT | 1027.0 | 3.4 | 0.8 | 4.4 | 9.9E-06 | 2.0E-05 | BT20/MB468 | MITO |
| >hg38_21_+_8433172_8446622_RNA45SN1_45S_50ntFlanks@6699.6719.21\|GCTAGCTGCGAGAATTAATGT | >hg38_21_+_8433172_8446622_RNA45SN1_45S_50ntFlanks | 6699 | 6719 | 21 | GCTAGCTGCGAGAATTAATGT | 714.5 | -5.0 | 1.1 | -4.4 | 9.9E-06 | 2.0E-05 | BT20/MB468 | MITO |
| >hg38_21_+_8433172_8446622_RNA45SN1_45S_50ntFlanks@11783.11808.26\|CATGAATGGATGAACGAGATTCCCAC | >hg38_21_+_8433172_8446622_RNA45SN1_45S_50ntFlanks | 11783 | 11808 | 26 | CATGAATGGATGAACGAGATTCCCAC | 57.0 | -8.9 | 1.9 | -4.6 | 4.3E-06 | 2.0E-05 | BT20/MB468 | MP |
| >hg38_MT_+_1621_3279_RNR2_16S_50ntFlanks@1367.1407.41\|TCGATGTTGGATCAGGACATCCCGATGGTGCAGCCGCTATT | >hg38_MT_+_1621_3279_RNR2_16S_50ntFlanks | 1367 | 1407 | 41 | TCGATGTTGGATCAGGACATCCCGATGGTGCAGCCGCTATT | 51.6 | 9.4 | 2.1 | 4.6 | 4.4E-06 | 2.0E-05 | BT20/MB468 | MP |
| >hg38_21_+_8433172_8446622_RNA45SN1_45S_50ntFlanks@9806.9830.25\|CGTGGAATGCGAGTGCCTAGTGGGC | >hg38_21_+_8433172_8446622_RNA45SN1_45S_50ntFlanks | 9806 | 9830 | 25 | CGTGGAATGCGAGTGCCTAGTGGGC | 304.4 | -5.4 | 1.2 | -4.3 | 1.5E-05 | 2.0E-05 | BT20/MB468 | NUC |
| >hg38_21_+_8433172_8446622_RNA45SN1_45S_50ntFlanks@5487.5516.30\|CGCTGAGAAGACGGTCGAACTTGACTATCT | >hg38_21_+_8433172_8446622_RNA45SN1_45S_50ntFlanks | 5487 | 5516 | 30 | CGCTGAGAAGACGGTCGAACTTGACTATCT | 2655.7 | 2.7 | 0.6 | 4.4 | 1.0E-05 | 2.0E-05 | BT20/MB468 | MITO |
| >hg38_21_+_8433172_8446622_RNA45SN1_45S_50ntFlanks@5473.5499.27\|CGGCCCTGGCGGAGCGCTGAGAAGACG | >hg38_21_+_8433172_8446622_RNA45SN1_45S_50ntFlanks | 5473 | 5499 | 27 | CGGCCCTGGCGGAGCGCTGAGAAGACG | 2375.1 | -3.5 | 0.8 | -4.3 | 1.4E-05 | 2.0E-05 | MB231/MB468 | MITO |
| >hg38_21_+_8433172_8446622_RNA45SN1_45S_50ntFlanks@12436.12473.38\|TCATTGTGAAGCAGAATTCACCAAGCGTTGGATTGTTC | >hg38_21_+_8433172_8446622_RNA45SN1_45S_50ntFlanks | 12436 | 12473 | 38 | TCATTGTGAAGCAGAATTCACCAAGCGTTGGATTGTTC | 238.2 | -5.1 | 1.2 | -4.3 | 1.4E-05 | 2.0E-05 | MB231/MB468 | MITO |
| >hg38_21_+_8433172_8446622_RNA45SN1_45S_50ntFlanks@12615.12661.47\|GCTGAGGAGCCAATGGGGCGAAGCTACCATCTGTGGGATTATGACTG | >hg38_21_+_8433172_8446622_RNA45SN1_45S_50ntFlanks | 12615 | 12661 | 47 | GCTGAGGAGCCAATGGGGCGAAGCTACCATCTGTGGGATTATGACTG | 927.1 | -6.3 | 1.4 | -4.3 | 1.4E-05 | 2.0E-05 | MB231/MB468 | NUC |
| >hg38_21_+_8433172_8446622_RNA45SN1_45S_50ntFlanks@8183.8211.29\|TCGAGGCCCAGCCCGTGGACGGTGTGAGG | >hg38_21_+_8433172_8446622_RNA45SN1_45S_50ntFlanks | 8183 | 8211 | 29 | TCGAGGCCCAGCCCGTGGACGGTGTGAGG | 150.0 | -6.4 | 1.5 | -4.4 | 1.1E-05 | 2.0E-05 | MB231/MB468 | MP |
| >hg38_21_+_8433172_8446622_RNA45SN1_45S_50ntFlanks@11781.11805.25\|CGCATGAATGGATGAACGAGATTCC | >hg38_21_+_8433172_8446622_RNA45SN1_45S_50ntFlanks | 11781 | 11805 | 25 | CGCATGAATGGATGAACGAGATTCC | 167.5 | 10.7 | 2.2 | 4.8 | 1.3E-06 | 2.0E-05 | BT20/MB231 | MP |
| >hg38_21_+_8433172_8446622_RNA45SN1_45S_50ntFlanks@3862.3898.37\|AATTCTAGAGCTAATACATGCCGACGGGCGCTGACCC | >hg38_21_+_8433172_8446622_RNA45SN1_45S_50ntFlanks | 3862 | 3898 | 37 | AATTCTAGAGCTAATACATGCCGACGGGCGCTGACCC | 176.7 | -4.3 | 1.0 | -4.4 | 1.3E-05 | 2.0E-05 | MB231/MB468 | CYTO |
| >hg38_21_+_8433172_8446622_RNA45SN1_45S_50ntFlanks@11552.11591.40\|CCTCGGCCGGCGCCTAGCAGCCGACTTAGAACTGGTGCGG | >hg38_21_+_8433172_8446622_RNA45SN1_45S_50ntFlanks | 11552 | 11591 | 40 | CCTCGGCCGGCGCCTAGCAGCCGACTTAGAACTGGTGCGG | 70.1 | -10.0 | 2.3 | -4.4 | 1.2E-05 | 2.0E-05 | MB231/MB468 | MP |
| >hg38_21_+_8433172_8446622_RNA45SN1_45S_50ntFlanks@6650.6694.45\|ACGACTCTTAGCGGTGGATCACTCGGCTCGTGCGTCGATGAAGAA | >hg38_21_+_8433172_8446622_RNA45SN1_45S_50ntFlanks | 6650 | 6694 | 45 | ACGACTCTTAGCGGTGGATCACTCGGCTCGTGCGTCGATGAAGAA | 401.7 | 4.1 | 0.9 | 4.4 | 1.0E-05 | 2.0E-05 | BT20/MB468 | MITO |
| >hg38_21_+_8433172_8446622_RNA45SN1_45S_50ntFlanks@11768.11794.27\|CTAATTAGTGACGCGCATGAATGGATG | >hg38_21_+_8433172_8446622_RNA45SN1_45S_50ntFlanks | 11768 | 11794 | 27 | CTAATTAGTGACGCGCATGAATGGATG | 685.6 | -2.2 | 0.5 | -4.4 | 1.0E-05 | 2.0E-05 | BT20/MB468 | MITO |
| >hg38_21_+_8433172_8446622_RNA45SN1_45S_50ntFlanks@6652.6676.25\|GACTCTTAGCGGTGGATCACTCGGC | >hg38_21_+_8433172_8446622_RNA45SN1_45S_50ntFlanks | 6652 | 6676 | 25 | GACTCTTAGCGGTGGATCACTCGGC | 2922.9 | 4.4 | 1.0 | 4.6 | 4.4E-06 | 2.0E-05 | BT20/MB468 | MP |
| >trna111_HisGTG_1_-_147774845_147774916@1.33.33,trna118_HisGTG_1_-_145396881_145396952@1.33.33,trna16_HisGTG_1_+_146544773_146544844@1.33.33,trna1_HisGTG_15_+_45493349_45493420@1.33.33,trna21_HisGTG_1_+_147753471_147753542@1.33.33,trna33_HisGTG_6_+_27125906_27125977@1.33.33,trna7_HisGTG_9_-_14433938_14434009@1.33.33,trna8_HisGTG_15_-_45492611_45492682@1.33.33,trna9_HisGTG_15_-_45490804_45490875@1.33.33\|GCCGTGATCGTATAGTGGTTAGTACTCTGCGTT | >trna111_HisGTG_1_-_147774845_147774916 | 1 | 33 | 33 | GCCGTGATCGTATAGTGGTTAGTACTCTGCGTT | 2812.7 | 6.2 | 1.4 | 4.6 | 4.4E-06 | 2.0E-05 | BT20/MB468 | MP |
| >hg38_21_+_8433172_8446622_RNA45SN1_45S_50ntFlanks@12482.12499.18\|ATAGGGAACGTGAGCTGG | >hg38_21_+_8433172_8446622_RNA45SN1_45S_50ntFlanks | 12482 | 12499 | 18 | ATAGGGAACGTGAGCTGG | 200.3 | -4.7 | 1.1 | -4.3 | 1.4E-05 | 2.0E-05 | MB231/MB468 | NUC |
| >hg38_21_+_8433172_8446622_RNA45SN1_45S_50ntFlanks@5073.5112.40\|AATTCCGATAACGAACGAGACTCTGGCATGCTAACTAGTT | >hg38_21_+_8433172_8446622_RNA45SN1_45S_50ntFlanks | 5073 | 5112 | 40 | AATTCCGATAACGAACGAGACTCTGGCATGCTAACTAGTT | 562.1 | -6.1 | 1.4 | -4.3 | 1.5E-05 | 2.0E-05 | BT20/MB468 | NUC |
| >hg38_1_-_228634819_228635039_RNA5S12_5S_50ntFlanks@132.169.38\|GATGGGAGACCGCCTGGGAATACCGGGTGCTGTAGGCT | >hg38_1_-_228634819_228635039_RNA5S12_5S_50ntFlanks | 132 | 169 | 38 | GATGGGAGACCGCCTGGGAATACCGGGTGCTGTAGGCT | 4008.3 | 4.3 | 0.9 | 4.8 | 1.5E-06 | 2.0E-05 | BT20/MB231 | MITO |
| >hg38_21_+_8433172_8446622_RNA45SN1_45S_50ntFlanks@10402.10435.34\|AACATGGGTCAGTCGGTCCTGAGAGATGGGCGAG | >hg38_21_+_8433172_8446622_RNA45SN1_45S_50ntFlanks | 10402 | 10435 | 34 | AACATGGGTCAGTCGGTCCTGAGAGATGGGCGAG | 580.6 | -4.6 | 1.0 | -4.4 | 1.0E-05 | 2.0E-05 | BT20/MB468 | MITO |
| >MI0000746\|hsa-mir-99b&WithFlank&19\|+\|51692606\|51692687@51.70.20(+1U)&[MIMAT0004678&hsa-miR-99b-3p&offsets\|0\|-2(+1U);m-151&19\|+\|51692656\|51692678&offsets\|0\|-3(+1U)]\|CAAGCTCGTGTCTGTGGGTCT | >MI0000746\|hsa-mir-99b&WithFlank&19\|+\|51692606\|51692687 | 51 | 70 | 20(+1U) | CAAGCTCGTGTCTGTGGGTCT | 444.2 | 6.5 | 1.4 | 4.6 | 4.4E-06 | 2.0E-05 | BT20/MB468 | MP |
| >hg38_1_-_228634819_228635039_RNA5S12_5S_50ntFlanks@124.171.48\|AGTACTTGGATGGGAGACCGCCTGGGAATACCGGGTGCTGTAGGCTTT | >hg38_1_-_228634819_228635039_RNA5S12_5S_50ntFlanks | 124 | 171 | 48 | AGTACTTGGATGGGAGACCGCCTGGGAATACCGGGTGCTGTAGGCTTT | 680.3 | 4.3 | 1.0 | 4.4 | 1.3E-05 | 2.0E-05 | MB231/MB468 | CYTO |
| >hg38_21_+_8433172_8446622_RNA45SN1_45S_50ntFlanks@6664.6689.26\|TGGATCACTCGGCTCGTGCGTCGATG | >hg38_21_+_8433172_8446622_RNA45SN1_45S_50ntFlanks | 6664 | 6689 | 26 | TGGATCACTCGGCTCGTGCGTCGATG | 1513.2 | -4.0 | 0.9 | -4.6 | 4.4E-06 | 2.0E-05 | BT20/MB468 | MP |
| >MI0000456\|hsa-mir-140&WithFlank&16\|+\|69933075\|69933186@29.50.22&[MIMAT0000431&hsa-miR-140-5p&offsets\|0\|0;m-173&16\|+\|69933103\|69933124&offsets\|0\|0]\|CAGTGGTTTTACCCTATGGTAG | >MI0000456\|hsa-mir-140&WithFlank&16\|+\|69933075\|69933186 | 29 | 50 | 22 | CAGTGGTTTTACCCTATGGTAG | 765.9 | 5.3 | 1.1 | 4.6 | 4.4E-06 | 2.0E-05 | BT20/MB468 | MP |
| >hg38_21_+_8433172_8446622_RNA45SN1_45S_50ntFlanks@3838.3872.35\|CTCTCCTACTTGGATAACTGTGGTAATTCTAGAGC | >hg38_21_+_8433172_8446622_RNA45SN1_45S_50ntFlanks | 3838 | 3872 | 35 | CTCTCCTACTTGGATAACTGTGGTAATTCTAGAGC | 69.0 | -9.2 | 2.0 | -4.6 | 4.4E-06 | 2.0E-05 | BT20/MB468 | MP |
| >hg38_21_+_8433172_8446622_RNA45SN1_45S_50ntFlanks@9535.9569.35\|GGAAACTCTGGTGGAGGTCCGTAGCGGTCCTGACG | >hg38_21_+_8433172_8446622_RNA45SN1_45S_50ntFlanks | 9535 | 9569 | 35 | GGAAACTCTGGTGGAGGTCCGTAGCGGTCCTGACG | 444.0 | -7.3 | 1.7 | -4.3 | 1.5E-05 | 2.0E-05 | BT20/MB468 | NUC |
| >hg38_21_+_8433172_8446622_RNA45SN1_45S_50ntFlanks@7983.8007.25\|CAGATCAGACGTGGCGACCCGCTGA | >hg38_21_+_8433172_8446622_RNA45SN1_45S_50ntFlanks | 7983 | 8007 | 25 | CAGATCAGACGTGGCGACCCGCTGA | 208.2 | -6.0 | 1.4 | -4.3 | 1.4E-05 | 2.0E-05 | MB231/MB468 | NUC |
| >hg38_21_+_8433172_8446622_RNA45SN1_45S_50ntFlanks@10794.10817.24\|CTGGCATGTTGGAACAATGTAGGT | >hg38_21_+_8433172_8446622_RNA45SN1_45S_50ntFlanks | 10794 | 10817 | 24 | CTGGCATGTTGGAACAATGTAGGT | 737.3 | -6.0 | 1.3 | -4.6 | 4.4E-06 | 2.0E-05 | BT20/MB468 | MP |
| >hg38_21_+_8433172_8446622_RNA45SN1_45S_50ntFlanks@4660.4686.27\|GAGGTGAAATTCTTGGACCGGCGCAAG | >hg38_21_+_8433172_8446622_RNA45SN1_45S_50ntFlanks | 4660 | 4686 | 27 | GAGGTGAAATTCTTGGACCGGCGCAAG | 72.8 | -10.0 | 2.3 | -4.4 | 1.2E-05 | 2.0E-05 | MB231/MB468 | MP |
| >MI0001445\|hsa-mir-423&WithFlank&17\|+\|30117073\|30117178@23.44.22(+1C)&[MIMAT0004748&hsa-miR-423-5p&offsets\|0\|-1(+1C);m-141&17\|+\|30117095\|30117117&offsets\|0\|-1(+1C)]\|TGAGGGGCAGAGAGCGAGACTTC | >MI0001445\|hsa-mir-423&WithFlank&17\|+\|30117073\|30117178 | 23 | 44 | 22(+1C) | TGAGGGGCAGAGAGCGAGACTTC | 121.8 | 2.6 | 0.6 | 4.3 | 1.5E-05 | 2.0E-05 | BT20/MB468 | NUC |
| >hg38_21_+_8433172_8446622_RNA45SN1_45S_50ntFlanks@12893.12928.36\|CCTCGCCCGTCACGCACCGCACGTTCGTGGGGAACC | >hg38_21_+_8433172_8446622_RNA45SN1_45S_50ntFlanks | 12893 | 12928 | 36 | CCTCGCCCGTCACGCACCGCACGTTCGTGGGGAACC | 179.2 | -6.6 | 1.4 | -4.6 | 4.4E-06 | 2.0E-05 | BT20/MB468 | MP |
| >hg38_21_+_8433172_8446622_RNA45SN1_45S_50ntFlanks@4737.4758.22\|GAAAGTCGGAGGTTCGAAGACG | >hg38_21_+_8433172_8446622_RNA45SN1_45S_50ntFlanks | 4737 | 4758 | 22 | GAAAGTCGGAGGTTCGAAGACG | 276.7 | -2.6 | 0.6 | -4.4 | 1.2E-05 | 2.0E-05 | MB231/MB468 | MP |
| >hg38_21_+_8433172_8446622_RNA45SN1_45S_50ntFlanks@10866.10900.35\|CTAAGGGCTGGGTCGGTCGGGCTGGGGCGCGAAGC | >hg38_21_+_8433172_8446622_RNA45SN1_45S_50ntFlanks | 10866 | 10900 | 35 | CTAAGGGCTGGGTCGGTCGGGCTGGGGCGCGAAGC | 381.2 | -7.1 | 1.6 | -4.3 | 1.5E-05 | 2.0E-05 | BT20/MB468 | NUC |
| >trna101_AlaAGC_6_-_28831462_28831533@58.75.18,trna102_AlaAGC_6_-_28806221_28806292@58.75.18,trna110_AlaTGC_6_-_28757547_28757618@58.75.18,trna113_AlaTGC_6_-_28726141_28726212@58.75.18,trna65_AlaAGC_6_+_28574933_28575004@58.75.18\|TCCCCGGCACCTCCACCA | >trna101_AlaAGC_6_-_28831462_28831533 | 58 | 75 | 18 | TCCCCGGCACCTCCACCA | 1226.6 | 4.2 | 1.0 | 4.3 | 1.4E-05 | 2.0E-05 | MB231/MB468 | NUC |
| >hg38_21_+_8433172_8446622_RNA45SN1_45S_50ntFlanks@6661.6685.25\|CGGTGGATCACTCGGCTCGTGCGTC | >hg38_21_+_8433172_8446622_RNA45SN1_45S_50ntFlanks | 6661 | 6685 | 25 | CGGTGGATCACTCGGCTCGTGCGTC | 5326.5 | 3.9 | 0.8 | 4.8 | 1.6E-06 | 2.0E-05 | BT20/MB231 | MITO |

<table>
<tr><td>>MI0000088|hsa-mir-30a&WithFlank&6|-|71403545|71403627@12.33.22(+1A)&[MIMAT0000087&hsa-miR-30a-5p&offsets|0|0(+1A);m-15&6|-|71403595|71403616&offsets|0|0(+1A)]|TGTAAACATCCTCGACTGGAAGA</td><td>>MI0000088|hsa-mir-30a&WithFlank&6|-|71403545|71403627</td><td>12</td><td>33</td><td>22(+1A)</td><td>TGTAAACATCCTCGACTGGAAGA</td><td>431.9</td><td>5.6</td><td>1.3</td><td>4.4</td><td>1.3E-05</td><td>2.0E-05</td><td>MB231/MB468</td><td>CYTO</td></tr>
<tr><td>>hg38_21_+_8433172_8446622_RNA45SN1_45S_50ntFlanks@10786.10820.35|AACAGCCTCTGGCATGTTGGAACAATGTAGGTAAG</td><td>>hg38_21_+_8433172_8446622_RNA45SN1_45S_50ntFlanks</td><td>10786</td><td>10820</td><td>35</td><td>AACAGCCTCTGGCATGTTGGAACAATGTAGGTAAG</td><td>361.0</td><td>-3.7</td><td>0.8</td><td>-4.6</td><td>4.5E-06</td><td>2.0E-05</td><td>BT20/MB468</td><td>MP</td></tr>
<tr><td>>hg38_21_+_8433172_8446622_RNA45SN1_45S_50ntFlanks@12528.12546.19|TTACCCTACTGATGATGTG</td><td>>hg38_21_+_8433172_8446622_RNA45SN1_45S_50ntFlanks</td><td>12528</td><td>12546</td><td>19</td><td>TTACCCTACTGATGATGTG</td><td>52.9</td><td>-8.8</td><td>1.9</td><td>-4.6</td><td>4.5E-06</td><td>2.0E-05</td><td>BT20/MB468</td><td>MP</td></tr>
<tr><td>>hg38_21_+_8433172_8446622_RNA45SN1_45S_50ntFlanks@7974.7993.20|ACGCGACCTCAGATCAGACG</td><td>>hg38_21_+_8433172_8446622_RNA45SN1_45S_50ntFlanks</td><td>7974</td><td>7993</td><td>20</td><td>ACGCGACCTCAGATCAGACG</td><td>8947.6</td><td>3.8</td><td>0.8</td><td>4.6</td><td>4.5E-06</td><td>2.0E-05</td><td>BT20/MB468</td><td>MP</td></tr>
<tr><td>>hg38_21_+_8433172_8446622_RNA45SN1_45S_50ntFlanks@3899.3938.40|CCTTCGCGGGGGGGATGCGTGCATTTATCAGATCAAAACC</td><td>>hg38_21_+_8433172_8446622_RNA45SN1_45S_50ntFlanks</td><td>3899</td><td>3938</td><td>40</td><td>CCTTCGCGGGGGGGATGCGTGCATTTATCAGATCAAAACC</td><td>563.0</td><td>-6.2</td><td>1.4</td><td>-4.4</td><td>1.3E-05</td><td>2.0E-05</td><td>MB231/MB468</td><td>CYTO</td></tr>
<tr><td>>hg38_21_+_8433172_8446622_RNA45SN1_45S_50ntFlanks@5084.5102.19|CGAACGAGACTCTGGCATG</td><td>>hg38_21_+_8433172_8446622_RNA45SN1_45S_50ntFlanks</td><td>5084</td><td>5102</td><td>19</td><td>CGAACGAGACTCTGGCATG</td><td>836.0</td><td>-4.7</td><td>1.1</td><td>-4.4</td><td>1.0E-05</td><td>2.0E-05</td><td>BT20/MB468</td><td>MITO</td></tr>
<tr><td>>hg38_21_+_8433172_8446622_RNA45SN1_45S_50ntFlanks@12415.12441.27|CTTCGATGTCGGCTCTTCCTATCATTG</td><td>>hg38_21_+_8433172_8446622_RNA45SN1_45S_50ntFlanks</td><td>12415</td><td>12441</td><td>27</td><td>CTTCGATGTCGGCTCTTCCTATCATTG</td><td>1753.5</td><td>-1.7</td><td>0.4</td><td>-4.4</td><td>1.2E-05</td><td>2.0E-05</td><td>BT20/MB468</td><td>CYTO</td></tr>
<tr><td>>hg38_21_+_8433172_8446622_RNA45SN1_45S_50ntFlanks@10682.10710.29|TCCGGCGGCGTCCGGTGAGCTCTCGCTGG</td><td>>hg38_21_+_8433172_8446622_RNA45SN1_45S_50ntFlanks</td><td>10682</td><td>10710</td><td>29</td><td>TCCGGCGGCGTCCGGTGAGCTCTCGCTGG</td><td>134.6</td><td>-5.5</td><td>1.2</td><td>-4.4</td><td>1.0E-05</td><td>2.0E-05</td><td>BT20/MB468</td><td>MITO</td></tr>
<tr><td>>hg38_21_+_8433172_8446622_RNA45SN1_45S_50ntFlanks@12163.12187.25|ACTGGGGCGGTACACCTGTCAAACG</td><td>>hg38_21_+_8433172_8446622_RNA45SN1_45S_50ntFlanks</td><td>12163</td><td>12187</td><td>25</td><td>ACTGGGGCGGTACACCTGTCAAACG</td><td>107.7</td><td>-6.2</td><td>1.4</td><td>-4.4</td><td>1.2E-05</td><td>2.0E-05</td><td>MB231/MB468</td><td>MP</td></tr>
<tr><td>>hg38_21_+_8433172_8446622_RNA45SN1_45S_50ntFlanks@11562.11592.31|CGCCTAGCAGCCGACTTAGAACTGGTGCGGA</td><td>>hg38_21_+_8433172_8446622_RNA45SN1_45S_50ntFlanks</td><td>11562</td><td>11592</td><td>31</td><td>CGCCTAGCAGCCGACTTAGAACTGGTGCGGA</td><td>2848.0</td><td>-4.8</td><td>1.0</td><td>-4.6</td><td>4.5E-06</td><td>2.0E-05</td><td>BT20/MB468</td><td>MP</td></tr>
<tr><td>>hg38_MT_+_1621_3279_RNR2_16S_50ntFlanks@728.749.22|AGATTAAAACACTGAACTGACA</td><td>>hg38_MT_+_1621_3279_RNR2_16S_50ntFlanks</td><td>728</td><td>749</td><td>22</td><td>AGATTAAAACACTGAACTGACA</td><td>37.7</td><td>8.5</td><td>1.9</td><td>4.4</td><td>1.2E-05</td><td>2.0E-05</td><td>MB231/MB468</td><td>MP</td></tr>
<tr><td>>trna14_LysTTT_11_-_59327808_59327880@4.36.33,trna2_LysTTT_17_+_8022473_8022545@4.36.33,trna54_LysTTT_1_+_204475655_204475727@4.36.33,trna5_LysTTT_11_+_59323902_59323974@4.36.33,trna62_LysTTT_1_-_204476158_204476230@4.36.33,trna76_LysTTT_6_+_28918806_28918878@4.36.33|CGGATAGCTCAGTCGGTAGAGCATCAGACTTTT</td><td>>trna14_LysTTT_11_-_59327808_59327880</td><td>4</td><td>36</td><td>33</td><td>CGGATAGCTCAGTCGGTAGAGCATCAGACTTTT</td><td>87.9</td><td>-9.9</td><td>2.0</td><td>-4.8</td><td>1.4E-06</td><td>2.0E-05</td><td>BT20/MB231</td><td>MP</td></tr>
<tr><td>>hg38_21_+_8433172_8446622_RNA45SN1_45S_50ntFlanks@10391.10424.34|AACAGCAGTTGAACATGGGTCAGTCGGTCCTGAG</td><td>>hg38_21_+_8433172_8446622_RNA45SN1_45S_50ntFlanks</td><td>10391</td><td>10424</td><td>34</td><td>AACAGCAGTTGAACATGGGTCAGTCGGTCCTGAG</td><td>20144.3</td><td>4.3</td><td>0.9</td><td>4.8</td><td>1.6E-06</td><td>2.0E-05</td><td>BT20/MB231</td><td>MITO</td></tr>
<tr><td>>hg38_21_+_8433172_8446622_RNA45SN1_45S_50ntFlanks@10478.10512.35|CCGATCGAAAGGGAGTCGGGTTCAGATCCCCGAAT</td><td>>hg38_21_+_8433172_8446622_RNA45SN1_45S_50ntFlanks</td><td>10478</td><td>10512</td><td>35</td><td>CCGATCGAAAGGGAGTCGGGTTCAGATCCCCGAAT</td><td>264.7</td><td>-5.7</td><td>1.3</td><td>-4.4</td><td>1.2E-05</td><td>2.0E-05</td><td>BT20/MB468</td><td>CYTO</td></tr>
<tr><td>>hg38_21_+_8433172_8446622_RNA45SN1_45S_50ntFlanks@12409.12436.28|TTGATCCTTCGATGTCGGCTCTTCCTAT</td><td>>hg38_21_+_8433172_8446622_RNA45SN1_45S_50ntFlanks</td><td>12409</td><td>12436</td><td>28</td><td>TTGATCCTTCGATGTCGGCTCTTCCTAT</td><td>303.0</td><td>-2.9</td><td>0.7</td><td>-4.3</td><td>1.4E-05</td><td>2.0E-05</td><td>MB231/MB468</td><td>NUC</td></tr>
<tr><td>>hg38_21_+_8433172_8446622_RNA45SN1_45S_50ntFlanks@8104.8142.39|CCGCGGCGGGGCGCGGGACATGTGGCGTACGGAAGACCC</td><td>>hg38_21_+_8433172_8446622_RNA45SN1_45S_50ntFlanks</td><td>8104</td><td>8142</td><td>39</td><td>CCGCGGCGGGGCGCGGGACATGTGGCGTACGGAAGACCC</td><td>219.8</td><td>-6.0</td><td>1.4</td><td>-4.3</td><td>1.4E-05</td><td>2.0E-05</td><td>MB231/MB468</td><td>NUC</td></tr>
<tr><td>>MI0000078|hsa-mir-22&WithFlank&17|-|1713897|1713993@59.81.23&[MIMAT0000077&hsa-miR-22-3p&offsets|0|+1;m-2&17|-|1713914|1713935&offsets|0|+1]|AAGCTGCCAGTTGAAGAACTGTT</td><td>>MI0000078|hsa-mir-22&WithFlank&17|-|1713897|1713993</td><td>59</td><td>81</td><td>23</td><td>AAGCTGCCAGTTGAAGAACTGTT</td><td>874.4</td><td>5.8</td><td>1.3</td><td>4.4</td><td>1.2E-05</td><td>2.0E-05</td><td>MB231/MB468</td><td>MP</td></tr>
<tr><td>>hg38_21_+_8433172_8446622_RNA45SN1_45S_50ntFlanks@4554.4573.20|CCGCAGCTAGGAATAATGGA</td><td>>hg38_21_+_8433172_8446622_RNA45SN1_45S_50ntFlanks</td><td>4554</td><td>4573</td><td>20</td><td>CCGCAGCTAGGAATAATGGA</td><td>230.5</td><td>-4.7</td><td>1.1</td><td>-4.3</td><td>1.4E-05</td><td>2.0E-05</td><td>MB231/MB468</td><td>NUC</td></tr>
<tr><td>>MI0000273|hsa-mir-183&WithFlank&7|-|129774899|129775020@33.53.21(+1C)&[MIMAT0000261&hsa-miR-183-5p&offsets|0|-1(+1C);m-41&7|-|129774966|129774987&offsets|-1|-2(+1C)]|TATGGCACTGGTAGAATTCACC</td><td>>MI0000273|hsa-mir-183&WithFlank&7|-|129774899|129775020</td><td>33</td><td>53</td><td>21(+1C)</td><td>TATGGCACTGGTAGAATTCACC</td><td>2095.6</td><td>3.6</td><td>0.7</td><td>4.8</td><td>1.4E-06</td><td>2.0E-05</td><td>BT20/MB231</td><td>CYTO</td></tr>
<tr><td>>hg38_21_+_8433172_8446622_RNA45SN1_45S_50ntFlanks@6718.6743.26|GTGAATTGCAGGACACATTGATCATC</td><td>>hg38_21_+_8433172_8446622_RNA45SN1_45S_50ntFlanks</td><td>6718</td><td>6743</td><td>26</td><td>GTGAATTGCAGGACACATTGATCATC</td><td>385.2</td><td>-5.4</td><td>1.2</td><td>-4.3</td><td>1.5E-05</td><td>2.0E-05</td><td>BT20/MB468</td><td>NUC</td></tr>
<tr><td>>hg38_21_+_8433172_8446622_RNA45SN1_45S_50ntFlanks@6661.6692.32|CGGTGGATCACTCGGCTCGTGCGTCGATGAAG</td><td>>hg38_21_+_8433172_8446622_RNA45SN1_45S_50ntFlanks</td><td>6661</td><td>6692</td><td>32</td><td>CGGTGGATCACTCGGCTCGTGCGTCGATGAAG</td><td>347.9</td><td>-1.4</td><td>0.3</td><td>-4.3</td><td>1.5E-05</td><td>2.0E-05</td><td>BT20/MB468</td><td>NUC</td></tr>
<tr><td>>hg38_21_+_8433172_8446622_RNA45SN1_45S_50ntFlanks@10679.10710.32|GGTTCCGGCGGCGTCCGGTGAGCTCTCGCTGG</td><td>>hg38_21_+_8433172_8446622_RNA45SN1_45S_50ntFlanks</td><td>10679</td><td>10710</td><td>32</td><td>GGTTCCGGCGGCGTCCGGTGAGCTCTCGCTGG</td><td>24871.3</td><td>-4.7</td><td>1.1</td><td>-4.4</td><td>1.2E-05</td><td>2.0E-05</td><td>BT20/MB468</td><td>CYTO</td></tr>
<tr><td>>hg38_21_+_8433172_8446622_RNA45SN1_45S_50ntFlanks@8170.8195.26|CAAGTCCTTCTGATCGAGGCCCAGCC</td><td>>hg38_21_+_8433172_8446622_RNA45SN1_45S_50ntFlanks</td><td>8170</td><td>8195</td><td>26</td><td>CAAGTCCTTCTGATCGAGGCCCAGCC</td><td>1067.3</td><td>-5.3</td><td>1.2</td><td>-4.4</td><td>1.2E-05</td><td>2.0E-05</td><td>BT20/MB468</td><td>CYTO</td></tr>
<tr><td>>hg38_21_+_8433172_8446622_RNA45SN1_45S_50ntFlanks@4201.4222.22|CCGACCCGGGGAGGTAGTGACG</td><td>>hg38_21_+_8433172_8446622_RNA45SN1_45S_50ntFlanks</td><td>4201</td><td>4222</td><td>22</td><td>CCGACCCGGGGAGGTAGTGACG</td><td>460.3</td><td>-5.7</td><td>1.3</td><td>-4.4</td><td>1.2E-05</td><td>2.1E-05</td><td>MB231/MB468</td><td>MP</td></tr>
<tr><td>>MI0000542|hsa-mir-320a&WithFlank&8|-|22244960|22245043@49.71.23&[MIMAT0000510&hsa-miR-320a-3p&offsets|+1|+2;m-60&8|-|22244975|22244996&offsets|+1|+2]|AAAGCTGGGTTGAGAGGGCGAAA</td><td>>MI0000542|hsa-mir-320a&WithFlank&8|-|22244960|22245043</td><td>49</td><td>71</td><td>23</td><td>AAAGCTGGGTTGAGAGGGCGAAA</td><td>368.9</td><td>5.9</td><td>1.3</td><td>4.6</td><td>4.5E-06</td><td>2.1E-05</td><td>BT20/MB468</td><td>MP</td></tr>
<tr><td>>hg38_21_+_8433172_8446622_RNA45SN1_45S_50ntFlanks@12626.12653.28|AATGGGGCGAAGCTACCATCTGTGGGAT</td><td>>hg38_21_+_8433172_8446622_RNA45SN1_45S_50ntFlanks</td><td>12626</td><td>12653</td><td>28</td><td>AATGGGGCGAAGCTACCATCTGTGGGAT</td><td>2792.9</td><td>-5.3</td><td>1.2</td><td>-4.4</td><td>1.2E-05</td><td>2.1E-05</td><td>BT20/MB468</td><td>CYTO</td></tr>
<tr><td>>hg38_21_+_8433172_8446622_RNA45SN1_45S_50ntFlanks@11657.11687.31|GATGTGATTTCTGCCCAGTGCTCTGAATGTC</td><td>>hg38_21_+_8433172_8446622_RNA45SN1_45S_50ntFlanks</td><td>11657</td><td>11687</td><td>31</td><td>GATGTGATTTCTGCCCAGTGCTCTGAATGTC</td><td>2921.5</td><td>10.3</td><td>2.1</td><td>4.8</td><td>1.4E-06</td><td>2.1E-05</td><td>BT20/MB231</td><td>MP</td></tr>
<tr><td>>hg38_21_+_8433172_8446622_RNA45SN1_45S_50ntFlanks@4129.4162.34|GATTCCGGAGAGGGAGCCTGAGAAACGGCTACCA</td><td>>hg38_21_+_8433172_8446622_RNA45SN1_45S_50ntFlanks</td><td>4129</td><td>4162</td><td>34</td><td>GATTCCGGAGAGGGAGCCTGAGAAACGGCTACCA</td><td>433.0</td><td>-4.8</td><td>1.1</td><td>-4.4</td><td>1.2E-05</td><td>2.1E-05</td><td>BT20/MB468</td><td>CYTO</td></tr>
<tr><td>>hg38_21_+_8433172_8446622_RNA45SN1_45S_50ntFlanks@12892.12932.41|CCCTCGCCCGTCACGCACCGCACGTTCGTGGGGAACCTGGC</td><td>>hg38_21_+_8433172_8446622_RNA45SN1_45S_50ntFlanks</td><td>12892</td><td>12932</td><td>41</td><td>CCCTCGCCCGTCACGCACCGCACGTTCGTGGGGAACCTGGC</td><td>676.8</td><td>-5.3</td><td>1.2</td><td>-4.3</td><td>1.4E-05</td><td>2.1E-05</td><td>MB231/MB468</td><td>NUC</td></tr>
<tr><td>>hg38_21_+_8433172_8446622_RNA45SN1_45S_50ntFlanks@11920.11967.48|ATGAGAGGTGTAGAATAAGTGGGAGGCCCCCGGCGCCCCCCCGGTGTC</td><td>>hg38_21_+_8433172_8446622_RNA45SN1_45S_50ntFlanks</td><td>11920</td><td>11967</td><td>48</td><td>ATGAGAGGTGTAGAATAAGTGGGAGGCCCCCGGCGCCCCCCCGGTGTC</td><td>86.8</td><td>-10.3</td><td>2.4</td><td>-4.4</td><td>1.2E-05</td><td>2.1E-05</td><td>MB231/MB468</td><td>MP</td></tr>
<tr><td>>hg38_21_+_8433172_8446622_RNA45SN1_45S_50ntFlanks@10400.10424.25|TGAACATGGGTCAGTCGGTCCTGAG</td><td>>hg38_21_+_8433172_8446622_RNA45SN1_45S_50ntFlanks</td><td>10400</td><td>10424</td><td>25</td><td>TGAACATGGGTCAGTCGGTCCTGAG</td><td>470.1</td><td>3.4</td><td>0.8</td><td>4.3</td><td>1.5E-05</td><td>2.1E-05</td><td>MB231/MB468</td><td>MITO</td></tr>
<tr><td>>trna10_ValCAC_5_-_180649395_180649467@32.54.23,trna13_ValCAC_19_-_4724647_4724719@32.54.23,trna152_ValCAC_6_-_27248049_27248121@32.54.23,trna157_ValCAC_6_-_27173867_27173939@32.54.23,trna18_ValCAC_5_-_180529253_180529325@32.54.23,trna2_ValCAC_5_+_180524070_180524142@32.54.23,trna6_ValCAC_5_+_180600650_180600722@32.54.23,trna85_ValCAC_1_-_161369490_161369562@32.54.23,trna98_ValCAC_1_-_149298555_149298627@32.54.23,trna9_ValCAC_6_+_26538282_26538354@32.54.23|CTCACACGCGAAAGGTCCCCGGT</td><td>>trna10_ValCAC_5_-_180649395_180649467</td><td>32</td><td>54</td><td>23</td><td>CTCACACGCGAAAGGTCCCCGGT</td><td>92.7</td><td>-10.4</td><td>2.4</td><td>-4.4</td><td>1.2E-05</td><td>2.1E-05</td><td>MB231/MB468</td><td>MP</td></tr>
<tr><td>>hg38_21_+_8433172_8446622_RNA45SN1_45S_50ntFlanks@11589.11607.19|CGGACCAGGGGAATCCGAC</td><td>>hg38_21_+_8433172_8446622_RNA45SN1_45S_50ntFlanks</td><td>11589</td><td>11607</td><td>19</td><td>CGGACCAGGGGAATCCGAC</td><td>122.2</td><td>-6.3</td><td>1.4</td><td>-4.4</td><td>1.2E-05</td><td>2.1E-05</td><td>MB231/MB468</td><td>MP</td></tr>
<tr><td>>hg38_21_+_8433172_8446622_RNA45SN1_45S_50ntFlanks@11594.11612.19|CAGGGGAATCCGACTGTTT</td><td>>hg38_21_+_8433172_8446622_RNA45SN1_45S_50ntFlanks</td><td>11594</td><td>11612</td><td>19</td><td>CAGGGGAATCCGACTGTTT</td><td>808.5</td><td>-3.0</td><td>0.7</td><td>-4.4</td><td>1.2E-05</td><td>2.1E-05</td><td>BT20/MB468</td><td>CYTO</td></tr>
<tr><td>>trna10_LysCTT_16_+_3241501_3241573@1.36.36,trna13_LysCTT_14_-_58706613_58706685@1.36.36,trna2_LysCTT_15_+_79152904_79152976@1.36.36|GCCCGGCTAGCTCAGTCGGTAGAGCATGGGACTCTT</td><td>>trna10_LysCTT_16_+_3241501_3241573</td><td>1</td><td>36</td><td>36</td><td>GCCCGGCTAGCTCAGTCGGTAGAGCATGGGACTCTT</td><td>105.5</td><td>3.4</td><td>0.8</td><td>4.3</td><td>1.5E-05</td><td>2.1E-05</td><td>BT20/MB468</td><td>NUC</td></tr>
<tr><td>>hg38_21_+_8433172_8446622_RNA45SN1_45S_50ntFlanks@6776.6806.31|CTCCCGGGGCTACGCCTGTCTGAGCGTCGCT</td><td>>hg38_21_+_8433172_8446622_RNA45SN1_45S_50ntFlanks</td><td>6776</td><td>6806</td><td>31</td><td>CTCCCGGGGCTACGCCTGTCTGAGCGTCGCT</td><td>10098.0</td><td>3.6</td><td>0.8</td><td>4.4</td><td>1.0E-05</td><td>2.1E-05</td><td>BT20/MB468</td><td>MITO</td></tr>
<tr><td>>hg38_21_+_8433172_8446622_RNA45SN1_45S_50ntFlanks@10349.10378.30|AACGAGAACTTTGAAGGCCGAAGTGGAGAA</td><td>>hg38_21_+_8433172_8446622_RNA45SN1_45S_50ntFlanks</td><td>10349</td><td>10378</td><td>30</td><td>AACGAGAACTTTGAAGGCCGAAGTGGAGAA</td><td>392.3</td><td>-5.5</td><td>1.3</td><td>-4.4</td><td>1.2E-05</td><td>2.1E-05</td><td>MB231/MB468</td><td>MP</td></tr>
<tr><td>>hg38_21_+_8433172_8446622_RNA45SN1_45S_50ntFlanks@12072.12093.22|CCCGAGGGGCTCTCGCTTCTGG</td><td>>hg38_21_+_8433172_8446622_RNA45SN1_45S_50ntFlanks</td><td>12072</td><td>12093</td><td>22</td><td>CCCGAGGGGCTCTCGCTTCTGG</td><td>241.2</td><td>-5.7</td><td>1.3</td><td>-4.4</td><td>1.2E-05</td><td>2.1E-05</td><td>BT20/MB468</td><td>CYTO</td></tr>
<tr><td>>hg38_21_+_8433172_8446622_RNA45SN1_45S_50ntFlanks@12073.12093.21|CCGAGGGGCTCTCGCTTCTGG</td><td>>hg38_21_+_8433172_8446622_RNA45SN1_45S_50ntFlanks</td><td>12073</td><td>12093</td><td>21</td><td>CCGAGGGGCTCTCGCTTCTGG</td><td>216.0</td><td>-5.7</td><td>1.3</td><td>-4.4</td><td>1.2E-05</td><td>2.1E-05</td><td>BT20/MB468</td><td>CYTO</td></tr>
<tr><td>>am_MI0000786|hsa-mir-378a&WithFlank&5|+|149732819|149732896@49.69.21&[MIMAT0000732&hsa-miR-378a-3p&offsets|0|-1;m-44&5|+|149732867|149732888&offsets|0|-1]|ACTGGACTTGGAGTCAGAAGG</td><td>>am_MI0000786|hsa-mir-378a&WithFlank&5|+|149732819|149732896</td><td>49</td><td>69</td><td>21</td><td>ACTGGACTTGGAGTCAGAAGG</td><td>1464.1</td><td>3.6</td><td>0.8</td><td>4.3</td><td>1.5E-05</td><td>2.1E-05</td><td>MB231/MB468</td><td>MITO</td></tr>
<tr><td>>hg38_21_+_8433172_8446622_RNA45SN1_45S_50ntFlanks@8106.8143.38|GCGGCGGGGCGCGGGACATGTGGCGTACGGAAGACCCG</td><td>>hg38_21_+_8433172_8446622_RNA45SN1_45S_50ntFlanks</td><td>8106</td><td>8143</td><td>38</td><td>GCGGCGGGGCGCGGGACATGTGGCGTACGGAAGACCCG</td><td>1443.1</td><td>-6.4</td><td>1.4</td><td>-4.6</td><td>4.6E-06</td><td>2.1E-05</td><td>BT20/MB468</td><td>MP</td></tr>
<tr><td>>hg38_21_+_8433172_8446622_RNA45SN1_45S_50ntFlanks@4368.4391.24|AGTTAAAAAGCTCGTAGTTGGATC</td><td>>hg38_21_+_8433172_8446622_RNA45SN1_45S_50ntFlanks</td><td>4368</td><td>4391</td><td>24</td><td>AGTTAAAAAGCTCGTAGTTGGATC</td><td>159.2</td><td>-6.3</td><td>1.4</td><td>-4.4</td><td>1.2E-05</td><td>2.1E-05</td><td>MB231/MB468</td><td>MP</td></tr>
<tr><td>>hg38_21_+_8433172_8446622_RNA45SN1_45S_50ntFlanks@4763.4782.20|GATACCGTCGTAGTTCCGAC</td><td>>hg38_21_+_8433172_8446622_RNA45SN1_45S_50ntFlanks</td><td>4763</td><td>4782</td><td>20</td><td>GATACCGTCGTAGTTCCGAC</td><td>189.1</td><td>-6.5</td><td>1.4</td><td>-4.6</td><td>4.6E-06</td><td>2.1E-05</td><td>BT20/MB468</td><td>MP</td></tr>
<tr><td>>MI0000746|hsa-mir-99b&WithFlank&19|+|51692606|51692687@14.34.21&[MIMAT0000689&hsa-miR-99b-5p&offsets|+1|0;m-10&19|+|51692618|51692639&offsets|+1|0]|ACCCGTAGAACCGACCTTGCG</td><td>>MI0000746|hsa-mir-99b&WithFlank&19|+|51692606|51692687</td><td>14</td><td>34</td><td>21</td><td>ACCCGTAGAACCGACCTTGCG</td><td>78.7</td><td>1.8</td><td>0.4</td><td>4.3</td><td>1.5E-05</td><td>2.1E-05</td><td>BT20/MB468</td><td>NUC</td></tr>
<tr><td>>hg38_21_+_8433172_8446622_RNA45SN1_45S_50ntFlanks@12918.12935.18|CGTGGGGAACCTGGCGCT</td><td>>hg38_21_+_8433172_8446622_RNA45SN1_45S_50ntFlanks</td><td>12918</td><td>12935</td><td>18</td><td>CGTGGGGAACCTGGCGCT</td><td>113.4</td><td>-6.3</td><td>1.4</td><td>-4.4</td><td>1.2E-05</td><td>2.1E-05</td><td>MB231/MB468</td><td>MP</td></tr>
<tr><td>>hg38_21_+_8433172_8446622_RNA45SN1_45S_50ntFlanks@12622.12654.33|AGCCAATGGGGCGAAGCTACCATCTGTGGGATT</td><td>>hg38_21_+_8433172_8446622_RNA45SN1_45S_50ntFlanks</td><td>12622</td><td>12654</td><td>33</td><td>AGCCAATGGGGCGAAGCTACCATCTGTGGGATT</td><td>208.1</td><td>-6.0</td><td>1.4</td><td>-4.3</td><td>1.4E-05</td><td>2.1E-05</td><td>MB231/MB468</td><td>NUC</td></tr>
<tr><td>>hg38_21_+_8433172_8446622_RNA45SN1_45S_50ntFlanks@9454.9500.47|CCGCGCCGGGGAGGTGGAGCACGAGCGCACGTGTTAGGACCCGAAAG</td><td>>hg38_21_+_8433172_8446622_RNA45SN1_45S_50ntFlanks</td><td>9454</td><td>9500</td><td>47</td><td>CCGCGCCGGGGAGGTGGAGCACGAGCGCACGTGTTAGGACCCGAAAG</td><td>229.0</td><td>-5.7</td><td>1.3</td><td>-4.4</td><td>1.1E-05</td><td>2.1E-05</td><td>BT20/MB468</td><td>MITO</td></tr>
</table>

<table>
<tr><th></th><th></th><th></th><th></th><th></th><th></th><th></th><th></th><th></th><th></th><th></th><th></th><th></th><th></th></tr>
<tr><td>>hg38_21_+_8433172_8446622_RNA45SN1_45S_50ntFlanks@11583.11605.23|CTGGTGCGGACCAGGGGAATCCG</td><td>>hg38_21_+_8433172_8446622_RNA45SN1_45S_50ntFlanks</td><td>11583</td><td>11605</td><td>23</td><td>CTGGTGCGGACCAGGGGAATCCG</td><td>909.9</td><td>-5.0</td><td>1.1</td><td>-4.4</td><td>1.2E-05</td><td>2.1E-05</td><td>BT20/MB468</td><td>CYTO</td></tr>
<tr><td>>hg38_21_+_8433172_8446622_RNA45SN1_45S_50ntFlanks@11923.11943.21|AGAGGTGTAGAATAAGTGGGA</td><td>>hg38_21_+_8433172_8446622_RNA45SN1_45S_50ntFlanks</td><td>11923</td><td>11943</td><td>21</td><td>AGAGGTGTAGAATAAGTGGGA</td><td>41.5</td><td>9.1</td><td>2.0</td><td>4.6</td><td>4.6E-06</td><td>2.1E-05</td><td>BT20/MB468</td><td>MP</td></tr>
<tr><td>>am_MI0008190|hsa-mir-320d-1&WithFlank&13|-|40727810|40727893@48.66.19;MI0008192|hsa-mir-320d-2&WithFlank&X|-|140926154|140926237@48.66.19&[MIMAT0006764&hsa-miR-320d&offsets|0|0;m-246&13|-|40727829|40727846&offsets|0|+1];[MIMAT0006764_1&hsa-miR-320d&offsets|0|0;m-253&X|-|140926173|140926190&offsets|0|+1]|AAAAGCTGGGTTGAGAGGA</td><td>>am_MI0008190|hsa-mir-320d-1&WithFlank&13|-|40727810|40727893</td><td>48</td><td>66</td><td>19</td><td>AAAAGCTGGGTTGAGAGGA</td><td>519.3</td><td>7.5</td><td>1.6</td><td>4.6</td><td>4.6E-06</td><td>2.1E-05</td><td>BT20/MB468</td><td>MP</td></tr>
<tr><td>>am_trna162_MetCAT_6_-_26758550_26758622@36.54.19,trna164_MetCAT_6_-_26735574_26735646@36.54.19,trna20_MetCAT_16_+_71460396_71460468@36.54.19,trna21_MetCAT_6_+_26701712_26701784@36.54.19,trna27_MetCAT_6_+_26766444_26766516@36.54.19,trna75_MetCAT_6_+_28912352_28912424@36.54.19,trna97_MetCAT_6_-_28921042_28921114@36.54.19|TAATCTGAAGGTCCTGAGT</td><td>>am_trna162_MetCAT_6_-_26758550_26758622</td><td>36</td><td>54</td><td>19</td><td>TAATCTGAAGGTCCTGAGT</td><td>420.9</td><td>3.1</td><td>0.7</td><td>4.4</td><td>1.2E-05</td><td>2.1E-05</td><td>BT20/MB468</td><td>CYTO</td></tr>
<tr><td>>hg38_21_+_8433172_8446622_RNA45SN1_45S_50ntFlanks@10101.10126.26|CGCGCGCGTGTGGTGTGCGTCGGAGG</td><td>>hg38_21_+_8433172_8446622_RNA45SN1_45S_50ntFlanks</td><td>10101</td><td>10126</td><td>26</td><td>CGCGCGCGTGTGGTGTGCGTCGGAGG</td><td>471.8</td><td>-5.7</td><td>1.3</td><td>-4.6</td><td>4.6E-06</td><td>2.1E-05</td><td>BT20/MB468</td><td>MP</td></tr>
<tr><td>>hg38_21_+_8433172_8446622_RNA45SN1_45S_50ntFlanks@12630.12654.25|GGGCGAAGCTACCATCTGTGGGATT</td><td>>hg38_21_+_8433172_8446622_RNA45SN1_45S_50ntFlanks</td><td>12630</td><td>12654</td><td>25</td><td>GGGCGAAGCTACCATCTGTGGGATT</td><td>130.7</td><td>-5.2</td><td>1.2</td><td>-4.4</td><td>1.2E-05</td><td>2.1E-05</td><td>BT20/MB468</td><td>CYTO</td></tr>
<tr><td>>hg38_21_+_8433172_8446622_RNA45SN1_45S_50ntFlanks@11211.11251.41|GCGACTCTGGACGCGAGCCGGGCCCTTCCCGTGGATCGCCC</td><td>>hg38_21_+_8433172_8446622_RNA45SN1_45S_50ntFlanks</td><td>11211</td><td>11251</td><td>41</td><td>GCGACTCTGGACGCGAGCCGGGCCCTTCCCGTGGATCGCCC</td><td>279.5</td><td>4.2</td><td>0.9</td><td>4.8</td><td>1.6E-06</td><td>2.1E-05</td><td>BT20/MB231</td><td>MITO</td></tr>
<tr><td>>hg38_21_+_8433172_8446622_RNA45SN1_45S_50ntFlanks@9739.9762.24|AAACGATCTCAACCTATTCTCAAA</td><td>>hg38_21_+_8433172_8446622_RNA45SN1_45S_50ntFlanks</td><td>9739</td><td>9762</td><td>24</td><td>AAACGATCTCAACCTATTCTCAAA</td><td>204.3</td><td>-2.6</td><td>0.6</td><td>-4.3</td><td>1.4E-05</td><td>2.1E-05</td><td>MB231/MB468</td><td>NUC</td></tr>
<tr><td>>MI0005566|hsa-mir-374b&WithFlank&X|-|74218541|74218624@17.39.23&[MIMAT0004955&hsa-miR-374b-5p&offsets|0|+1;m-153&X|-|74218587|74218608&offsets|0|+1]|ATATAATACAACCTGCTAAGTGT</td><td>>MI0005566|hsa-mir-374b&WithFlank&X|-|74218541|74218624</td><td>17</td><td>39</td><td>23</td><td>ATATAATACAACCTGCTAAGTGT</td><td>688.1</td><td>-5.1</td><td>1.0</td><td>-4.8</td><td>1.4E-06</td><td>2.1E-05</td><td>BT20/MB231</td><td>MP</td></tr>
<tr><td>>MI0000061|hsa-let-7a-2&WithFlank&11|-|122146516|122146599@11.32.22(+1A);MI0000062|hsa-let-7a-3&WithFlank&22|+|46112743|46112828@10.31.22(+1A);MI0000060|hsa-let-7a-1&WithFlank&9|+|94175951|94176042@12.33.22(+1A)&[MIMAT0000062&hsa-let-7a-5p&offsets|0|0(+1A);m-6&11|-|122146568|122146589&offsets|0|0(+1A)];[MIMAT0000062_1&hsa-let-7a-5p&offsets|0|0(+1A);m-5&22|+|46112752|46112773&offsets|0|0(+1A)];[MIMAT0000062_2&hsa-let-7a-5p&offsets|0|0(+1A);m-4&9|+|94175962|94175983&offsets|0|0(+1A)]|TGAGGTAGTAGGTTGTATAGTTA</td><td>>MI0000061|hsa-let-7a-2&WithFlank&11|-|122146516|122146599</td><td>11</td><td>32</td><td>22(+1A)</td><td>TGAGGTAGTAGGTTGTATAGTTA</td><td>1230.9</td><td>4.6</td><td>1.0</td><td>4.6</td><td>4.6E-06</td><td>2.1E-05</td><td>BT20/MB468</td><td>MP</td></tr>
<tr><td>>hg38_21_+_8433172_8446622_RNA45SN1_45S_50ntFlanks@4498.4517.20|AAGCGTTTACTTTGAAAAAA</td><td>>hg38_21_+_8433172_8446622_RNA45SN1_45S_50ntFlanks</td><td>4498</td><td>4517</td><td>20</td><td>AAGCGTTTACTTTGAAAAAA</td><td>112.5</td><td>-10.7</td><td>2.4</td><td>-4.4</td><td>1.2E-05</td><td>2.1E-05</td><td>MB231/MB468</td><td>MP</td></tr>
<tr><td>>hg38_21_+_8433172_8446622_RNA45SN1_45S_50ntFlanks@10798.10822.25|CATGTTGGAACAATGTAGGTAAGGG</td><td>>hg38_21_+_8433172_8446622_RNA45SN1_45S_50ntFlanks</td><td>10798</td><td>10822</td><td>25</td><td>CATGTTGGAACAATGTAGGTAAGGG</td><td>568.8</td><td>-3.3</td><td>0.7</td><td>-4.6</td><td>4.7E-06</td><td>2.1E-05</td><td>BT20/MB468</td><td>MP</td></tr>
<tr><td>>MI0000272|hsa-mir-182&WithFlank&7|-|129770377|129770498@29.54.26&[MIMAT0000259&hsa-miR-182-5p&offsets|0|+2;m-22&7|-|129770449|129770470&offsets|0|+4]|TTTGGCAATGGTAGAACTCACACTGG</td><td>>MI0000272|hsa-mir-182&WithFlank&7|-|129770377|129770498</td><td>29</td><td>54</td><td>26</td><td>TTTGGCAATGGTAGAACTCACACTGG</td><td>5322.0</td><td>3.6</td><td>0.7</td><td>4.8</td><td>1.5E-06</td><td>2.1E-05</td><td>BT20/MB231</td><td>CYTO</td></tr>
<tr><td>>hg38_21_+_8433172_8446622_RNA45SN1_45S_50ntFlanks@3850.3869.20|GATAACTGTGGTAATTCTAG</td><td>>hg38_21_+_8433172_8446622_RNA45SN1_45S_50ntFlanks</td><td>3850</td><td>3869</td><td>20</td><td>GATAACTGTGGTAATTCTAG</td><td>213.3</td><td>-2.2</td><td>0.5</td><td>-4.4</td><td>1.3E-05</td><td>2.1E-05</td><td>BT20/MB468</td><td>CYTO</td></tr>
<tr><td>>hg38_21_+_8433172_8446622_RNA45SN1_45S_50ntFlanks@11581.11601.21|AACTGGTGCGGACCAGGGGAA</td><td>>hg38_21_+_8433172_8446622_RNA45SN1_45S_50ntFlanks</td><td>11581</td><td>11601</td><td>21</td><td>AACTGGTGCGGACCAGGGGAA</td><td>394.9</td><td>-2.7</td><td>0.6</td><td>-4.4</td><td>1.1E-05</td><td>2.1E-05</td><td>BT20/MB468</td><td>MITO</td></tr>
<tr><td>>MI0000063|hsa-let-7b&WithFlank&22|+|46113680|46113774@12.33.22(+1G)&[MIMAT0000063&hsa-let-7b-5p&offsets|0|0(+1G);m-14&22|+|46113691|46113712&offsets|0|0(+1G)]|TGAGGTAGTAGGTTGTGTGGTTG</td><td>>MI0000063|hsa-let-7b&WithFlank&22|+|46113680|46113774</td><td>12</td><td>33</td><td>22(+1G)</td><td>TGAGGTAGTAGGTTGTGTGGTTG</td><td>342.1</td><td>2.9</td><td>0.6</td><td>4.8</td><td>1.7E-06</td><td>2.1E-05</td><td>BT20/MB231</td><td>NUC</td></tr>
<tr><td>>hg38_21_+_8433172_8446622_RNA45SN1_45S_50ntFlanks@12633.12657.25|CGAAGCTACCATCTGTGGGATTATG</td><td>>hg38_21_+_8433172_8446622_RNA45SN1_45S_50ntFlanks</td><td>12633</td><td>12657</td><td>25</td><td>CGAAGCTACCATCTGTGGGATTATG</td><td>1761.8</td><td>-5.8</td><td>1.3</td><td>-4.6</td><td>4.7E-06</td><td>2.1E-05</td><td>BT20/MB468</td><td>MP</td></tr>
<tr><td>>hg38_21_+_8433172_8446622_RNA45SN1_45S_50ntFlanks@9712.9737.26|AGCGAATGATTAGAGGTCTTGGGGCC</td><td>>hg38_21_+_8433172_8446622_RNA45SN1_45S_50ntFlanks</td><td>9712</td><td>9737</td><td>26</td><td>AGCGAATGATTAGAGGTCTTGGGGCC</td><td>758.4</td><td>-5.1</td><td>1.2</td><td>-4.4</td><td>1.1E-05</td><td>2.1E-05</td><td>BT20/MB468</td><td>MITO</td></tr>
<tr><td>>hg38_21_+_8433172_8446622_RNA45SN1_45S_50ntFlanks@9738.9763.26|GAAACGATCTCAACCTATTCTCAAAC</td><td>>hg38_21_+_8433172_8446622_RNA45SN1_45S_50ntFlanks</td><td>9738</td><td>9763</td><td>26</td><td>GAAACGATCTCAACCTATTCTCAAAC</td><td>575.0</td><td>-2.6</td><td>0.6</td><td>-4.3</td><td>1.5E-05</td><td>2.1E-05</td><td>MB231/MB468</td><td>NUC</td></tr>
<tr><td>>am_MI0003639|hsa-mir-625&WithFlank&14|+|65471096|65471192@58.78.21(+1U)&[MIMAT0004808&hsa-miR-625-3p&offsets|0|-1(+1U);m-137&14|+|65471153|65471174&offsets|0|-1(+1U)]|GACTATAGAACTTTCCCCCTCT</td><td>>am_MI0003639|hsa-mir-625&WithFlank&14|+|65471096|65471192</td><td>58</td><td>78</td><td>21(+1U)</td><td>GACTATAGAACTTTCCCCCTCT</td><td>186.0</td><td>-3.4</td><td>0.7</td><td>-4.8</td><td>1.6E-06</td><td>2.1E-05</td><td>BT20/MB231</td><td>MITO</td></tr>
<tr><td>>MI0000078|hsa-mir-22&WithFlank&17|-|1713897|1713993@59.78.20&[MIMAT0000077&hsa-miR-22-3p&offsets|0|-2;m-2&17|-|1713914|1713935&offsets|0|-2]|AAGCTGCCAGTTGAAGAACT</td><td>>MI0000078|hsa-mir-22&WithFlank&17|-|1713897|1713993</td><td>59</td><td>78</td><td>20</td><td>AAGCTGCCAGTTGAAGAACT</td><td>411.2</td><td>5.4</td><td>1.2</td><td>4.4</td><td>1.2E-05</td><td>2.1E-05</td><td>MB231/MB468</td><td>MP</td></tr>
<tr><td>>hg38_21_+_8433172_8446622_RNA45SN1_45S_50ntFlanks@7992.8011.20|CGTGGCGACCCGCTGAATTT</td><td>>hg38_21_+_8433172_8446622_RNA45SN1_45S_50ntFlanks</td><td>7992</td><td>8011</td><td>20</td><td>CGTGGCGACCCGCTGAATTT</td><td>358.0</td><td>3.7</td><td>0.8</td><td>4.3</td><td>1.4E-05</td><td>2.1E-05</td><td>MB231/MB468</td><td>CYTO</td></tr>
<tr><td>>hg38_21_+_8433172_8446622_RNA45SN1_45S_50ntFlanks@11666.11702.37|TCTGCCCAGTGCTCTGAATGTCAAAGTGAAGAAATTC</td><td>>hg38_21_+_8433172_8446622_RNA45SN1_45S_50ntFlanks</td><td>11666</td><td>11702</td><td>37</td><td>TCTGCCCAGTGCTCTGAATGTCAAAGTGAAGAAATTC</td><td>794.2</td><td>2.4</td><td>0.5</td><td>4.4</td><td>1.3E-05</td><td>2.1E-05</td><td>BT20/MB468</td><td>CYTO</td></tr>
<tr><td>>hg38_21_+_8433172_8446622_RNA45SN1_45S_50ntFlanks@11581.11611.31|AACTGGTGCGGACCAGGGGAATCCGACTGTT</td><td>>hg38_21_+_8433172_8446622_RNA45SN1_45S_50ntFlanks</td><td>11581</td><td>11611</td><td>31</td><td>AACTGGTGCGGACCAGGGGAATCCGACTGTT</td><td>1277.4</td><td>-2.9</td><td>0.7</td><td>-4.3</td><td>1.4E-05</td><td>2.1E-05</td><td>MB231/MB468</td><td>CYTO</td></tr>
<tr><td>>MI0000471|hsa-mir-126&WithFlank&9|+|136670596|136670692@58.77.20(+1U)&[MIMAT0000445&hsa-miR-126-3p&offsets|0|-2(+1U);m-40&9|+|136670653|136670674&offsets|0|-2(+1U)]|TCGTACCGTGAGTAATAATGT</td><td>>MI0000471|hsa-mir-126&WithFlank&9|+|136670596|136670692</td><td>58</td><td>77</td><td>20(+1U)</td><td>TCGTACCGTGAGTAATAATGT</td><td>1344.5</td><td>8.2</td><td>1.9</td><td>4.4</td><td>1.2E-05</td><td>2.1E-05</td><td>MB231/MB468</td><td>MP</td></tr>
<tr><td>>hg38_21_+_8433172_8446622_RNA45SN1_45S_50ntFlanks@9484.9509.26|GTGTTAGGACCCGAAAGATGGTGAAC</td><td>>hg38_21_+_8433172_8446622_RNA45SN1_45S_50ntFlanks</td><td>9484</td><td>9509</td><td>26</td><td>GTGTTAGGACCCGAAAGATGGTGAAC</td><td>114.2</td><td>10.2</td><td>2.1</td><td>4.8</td><td>1.4E-06</td><td>2.1E-05</td><td>BT20/MB231</td><td>MP</td></tr>
<tr><td>>hg38_21_+_8433172_8446622_RNA45SN1_45S_50ntFlanks@11917.11939.23|GACATGAGAGGTGTAGAATAAGT</td><td>>hg38_21_+_8433172_8446622_RNA45SN1_45S_50ntFlanks</td><td>11917</td><td>11939</td><td>23</td><td>GACATGAGAGGTGTAGAATAAGT</td><td>40.9</td><td>9.1</td><td>2.0</td><td>4.6</td><td>4.7E-06</td><td>2.1E-05</td><td>BT20/MB468</td><td>MP</td></tr>
<tr><td>>hg38_21_+_8433172_8446622_RNA45SN1_45S_50ntFlanks@11846.11886.41|GAACGGGCTTGGCGGAATCAGCGGGGAAAGAAGACCCTGTT</td><td>>hg38_21_+_8433172_8446622_RNA45SN1_45S_50ntFlanks</td><td>11846</td><td>11886</td><td>41</td><td>GAACGGGCTTGGCGGAATCAGCGGGGAAAGAAGACCCTGTT</td><td>114.6</td><td>10.2</td><td>2.1</td><td>4.8</td><td>1.4E-06</td><td>2.1E-05</td><td>BT20/MB231</td><td>MP</td></tr>
<tr><td>>hg38_21_+_8433172_8446622_RNA45SN1_45S_50ntFlanks@8213.8235.23|CGGTAGCGGCCCCCGGCGCGCCG</td><td>>hg38_21_+_8433172_8446622_RNA45SN1_45S_50ntFlanks</td><td>8213</td><td>8235</td><td>23</td><td>CGGTAGCGGCCCCCGGCGCGCCG</td><td>259.3</td><td>-6.6</td><td>1.4</td><td>-4.6</td><td>4.7E-06</td><td>2.1E-05</td><td>BT20/MB468</td><td>MP</td></tr>
<tr><td>>MI0000483|hsa-mir-186&WithFlank&1|-|71067625|71067722@21.42.22&[MIMAT0000456&hsa-miR-186-5p&offsets|0|0;m-84&1|-|71067681|71067702&offsets|0|0]|CAAAGAATTCTCCTTTTGGGCT</td><td>>MI0000483|hsa-mir-186&WithFlank&1|-|71067625|71067722</td><td>21</td><td>42</td><td>22</td><td>CAAAGAATTCTCCTTTTGGGCT</td><td>370.3</td><td>3.9</td><td>0.9</td><td>4.3</td><td>1.4E-05</td><td>2.1E-05</td><td>MB231/MB468</td><td>CYTO</td></tr>
<tr><td>>hg38_21_+_8433172_8446622_RNA45SN1_45S_50ntFlanks@9837.9868.32|TGGTAAGCAGAACTGGCGCTGCGGGATGAACC</td><td>>hg38_21_+_8433172_8446622_RNA45SN1_45S_50ntFlanks</td><td>9837</td><td>9868</td><td>32</td><td>TGGTAAGCAGAACTGGCGCTGCGGGATGAACC</td><td>38.7</td><td>9.0</td><td>2.0</td><td>4.6</td><td>4.7E-06</td><td>2.1E-05</td><td>BT20/MB468</td><td>MP</td></tr>
<tr><td>>trna116_GluCTC_1_-_145399233_145399304@1.31.31,trna59_GluCTC_1_+_249168447_249168518@1.31.31,trna71_GluCTC_1_-_161439189_161439260@1.31.31,trna74_GluCTC_1_-_161431809_161431880@1.31.31,trna77_GluCTC_1_-_161424398_161424469@1.31.31,trna77_GluCTC_6_+_28949976_28950047@1.31.31,trna80_GluCTC_1_-_161417018_161417089@1.31.31,trna87_GluCTC_6_-_126101393_126101464@1.31.31|TCCCTGGTGGTCTAGTGGTTAGGATTCGGCG</td><td>>trna116_GluCTC_1_-_145399233_145399304</td><td>1</td><td>31</td><td>31</td><td>TCCCTGGTGGTCTAGTGGTTAGGATTCGGCG</td><td>25049.5</td><td>-3.9</td><td>0.9</td><td>-4.4</td><td>1.2E-05</td><td>2.1E-05</td><td>MB231/MB468</td><td>MP</td></tr>
<tr><td>>MI0000272|hsa-mir-182&WithFlank&7|-|129770377|129770498@29.50.22&[MIMAT0000259&hsa-miR-182-5p&offsets|0|-2;m-22&7|-|129770449|129770470&offsets|0|0]|TTTGGCAATGGTAGAACTCACA</td><td>>MI0000272|hsa-mir-182&WithFlank&7|-|129770377|129770498</td><td>29</td><td>50</td><td>22</td><td>TTTGGCAATGGTAGAACTCACA</td><td>378.3</td><td>3.5</td><td>0.8</td><td>4.3</td><td>1.4E-05</td><td>2.1E-05</td><td>MB231/MB468</td><td>CYTO</td></tr>
<tr><td>>trna124_ArgTCG_6_-_28510891_28510963@31.52.22,trna19_ArgTCG_17_+_73031208_73031280@31.52.22,trna3_ArgTCG_6_+_26299905_26299977@31.52.22,trna4_ArgTCG_6_+_26323046_26323118@31.52.22|ACTTCGGATCAGAAGATTGAGG</td><td>>trna124_ArgTCG_6_-_28510891_28510963</td><td>31</td><td>52</td><td>22</td><td>ACTTCGGATCAGAAGATTGAGG</td><td>65.0</td><td>-9.1</td><td>2.0</td><td>-4.6</td><td>4.7E-06</td><td>2.1E-05</td><td>BT20/MB468</td><td>MP</td></tr>
<tr><td>>MI0000477|hsa-mir-146a&WithFlank&5|+|160485346|160485456@27.47.21(+1C)&[MIMAT0000449&hsa-miR-146a-5p&offsets|0|-1(+1C);m-129&5|+|160485372|160485393&offsets|0|-1(+1C)]|TGAGAACTGAATTCCATGGGTC</td><td>>MI0000477|hsa-mir-146a&WithFlank&5|+|160485346|160485456</td><td>27</td><td>47</td><td>21(+1C)</td><td>TGAGAACTGAATTCCATGGGTC</td><td>3327.1</td><td>-10.6</td><td>2.2</td><td>-4.8</td><td>1.7E-06</td><td>2.1E-05</td><td>BT20/MB231</td><td>MITO</td></tr>
<tr><td>>hg38_21_+_8433172_8446622_RNA45SN1_45S_50ntFlanks@12263.12284.22|ATCTTGATTTTCAGTACGAATA</td><td>>hg38_21_+_8433172_8446622_RNA45SN1_45S_50ntFlanks</td><td>12263</td><td>12284</td><td>22</td><td>ATCTTGATTTTCAGTACGAATA</td><td>54.9</td><td>-8.9</td><td>1.9</td><td>-4.6</td><td>4.7E-06</td><td>2.1E-05</td><td>BT20/MB468</td><td>MP</td></tr>
<tr><td>>hg38_21_+_8433172_8446622_RNA45SN1_45S_50ntFlanks@12415.12439.25|CTTCGATGTCGGCTCTTCCTATCAT</td><td>>hg38_21_+_8433172_8446622_RNA45SN1_45S_50ntFlanks</td><td>12415</td><td>12439</td><td>25</td><td>CTTCGATGTCGGCTCTTCCTATCAT</td><td>715.2</td><td>2.8</td><td>0.6</td><td>4.4</td><td>1.3E-05</td><td>2.1E-05</td><td>BT20/MB468</td><td>CYTO</td></tr>
<tr><td>>hg38_21_+_8433172_8446622_RNA45SN1_45S_50ntFlanks@11847.11871.25|AACGGGCTTGGCGGAATCAGCGGGG</td><td>>hg38_21_+_8433172_8446622_RNA45SN1_45S_50ntFlanks</td><td>11847</td><td>11871</td><td>25</td><td>AACGGGCTTGGCGGAATCAGCGGGG</td><td>98.4</td><td>10.0</td><td>2.1</td><td>4.8</td><td>1.4E-06</td><td>2.1E-05</td><td>BT20/MB231</td><td>MP</td></tr>
<tr><td>>hg38_21_+_8433172_8446622_RNA45SN1_45S_50ntFlanks@476.523.48|TTCGTGATCGATGTGGTGACGTCGTGCTCTCCCGGGCCGGGTCCGAGC</td><td>>hg38_21_+_8433172_8446622_RNA45SN1_45S_50ntFlanks</td><td>476</td><td>523</td><td>48</td><td>TTCGTGATCGATGTGGTGACGTCGTGCTCTCCCGGGCCGGGTCCGAGC</td><td>481.9</td><td>6.9</td><td>1.6</td><td>4.4</td><td>1.1E-05</td><td>2.1E-05</td><td>BT20/MB468</td><td>MITO</td></tr>
<tr><td>>MI0000234|hsa-mir-192&WithFlank&11|-|64891131|64891252@30.50.21&[MIMAT0000222&hsa-miR-192-5p&offsets|0|0;m-59&11|-|64891203|64891223&offsets|0|0]|CTGACCTATGAATTGACAGCC</td><td>>MI0000234|hsa-mir-192&WithFlank&11|-|64891131|64891252</td><td>30</td><td>50</td><td>21</td><td>CTGACCTATGAATTGACAGCC</td><td>915.0</td><td>4.2</td><td>1.0</td><td>4.3</td><td>1.5E-05</td><td>2.1E-05</td><td>MB231/MB468</td><td>MITO</td></tr>
</table>

<table>
<tr><th></th><th></th><th></th><th></th><th></th><th></th><th></th><th></th><th></th><th></th><th></th><th></th><th></th><th></th></tr>
<tr><td>>MI0000061|hsa-let-7a-2&WithFlank&11|-|122146516|122146599@11.32.22;MI0000062|hsa-let-7a-3&WithFlank&22|+|46112743|46112828@10.31.22;MI0000060|hsa-let-7a-1&WithFlank&9|+|94175951|94176042@12.33.22&[MIMAT0000062&hsa-let-7a-5p&offsets|0|0;m-6&11|-|122146568|122146589&offsets|0|0];[MIMAT0000062_1&hsa-let-7a-5p&offsets|0|0;m-5&22|+|46112752|46112773&offsets|0|0];[MIMAT0000062_2&hsa-let-7a-5p&offsets|0|0;m-4&9|+|94175962|94175983&offsets|0|0]|TGAGGTAGTAGGTTGTATAGTT</td><td>>MI0000061|hsa-let-7a-2&WithFlank&11|-|122146516|122146599</td><td>11</td><td>32</td><td>22</td><td>TGAGGTAGTAGGTTGTATAGTT</td><td>32142.0</td><td>1.8</td><td>0.4</td><td>4.3</td><td>1.6E-05</td><td>2.1E-05</td><td>BT20/MB468</td><td>NUC</td></tr>
<tr><td>>MI0000746|hsa-mir-99b&WithFlank&19|+|51692606|51692687@13.32.20(+1U)&[MIMAT0000689&hsa-miR-99b-5p&offsets|0|-2(+1U);m-10&19|+|51692618|51692639&offsets|0|-2(+1U)]|CACCCGTAGAACCGACCTTGT</td><td>>MI0000746|hsa-mir-99b&WithFlank&19|+|51692606|51692687</td><td>13</td><td>32</td><td>20(+1U)</td><td>CACCCGTAGAACCGACCTTGT</td><td>865.4</td><td>3.2</td><td>0.7</td><td>4.4</td><td>1.2E-05</td><td>2.1E-05</td><td>MB231/MB468</td><td>MP</td></tr>
<tr><td>>hg38_21_+_8433172_8446622_RNA45SN1_45S_50ntFlanks@3882.3899.18|CCGACGGGCGCTGACCCC</td><td>>hg38_21_+_8433172_8446622_RNA45SN1_45S_50ntFlanks</td><td>3882</td><td>3899</td><td>18</td><td>CCGACGGGCGCTGACCCC</td><td>459.1</td><td>-6.5</td><td>1.5</td><td>-4.3</td><td>1.5E-05</td><td>2.1E-05</td><td>MB231/MB468</td><td>NUC</td></tr>
<tr><td>>MI0000762|hsa-mir-362&WithFlank&X|+|50008958|50009034@11.34.24&[MIMAT0000705&hsa-miR-362-5p&offsets|0|0;m-267&X|+|50008968|50008991&offsets|0|0]|AATCCTTGGAACCTAGGTGTGAGT</td><td>>MI0000762|hsa-mir-362&WithFlank&X|+|50008958|50009034</td><td>11</td><td>34</td><td>24</td><td>AATCCTTGGAACCTAGGTGTGAGT</td><td>327.6</td><td>4.8</td><td>1.1</td><td>4.3</td><td>1.5E-05</td><td>2.1E-05</td><td>MB231/MB468</td><td>MITO</td></tr>
<tr><td>>trna111_HisGTG_1_-_147774845_147774916@-1A.29.30,trna118_HisGTG_1_-_145396881_145396952@-1A.29.30,trna16_HisGTG_1_+_146544773_146544844@-1A.29.30,trna1_HisGTG_15_+_45493349_45493420@-1A.29.30,trna21_HisGTG_1_+_147753471_147753542@-1A.29.30,trna33_HisGTG_6_+_27125906_27125977@-1A.29.30,trna7_HisGTG_9_-_14433938_14434009@-1A.29.30,trna8_HisGTG_15_-_45492611_45492682@-1A.29.30,trna9_HisGTG_15_-_45490804_45490875@-1A.29.30|AGCCGTGATCGTATAGTGGTTAGTACTCTG</td><td>>trna111_HisGTG_1_-_147774845_147774916</td><td>-1A</td><td>29</td><td>30</td><td>AGCCGTGATCGTATAGTGGTTAGTACTCTG</td><td>309.9</td><td>-5.3</td><td>1.2</td><td>-4.3</td><td>1.5E-05</td><td>2.1E-05</td><td>MB231/MB468</td><td>MITO</td></tr>
<tr><td>>hg38_21_+_8433172_8446622_RNA45SN1_45S_50ntFlanks@10351.10373.23|CGAGAACTTTGAAGGCCGAAGTG</td><td>>hg38_21_+_8433172_8446622_RNA45SN1_45S_50ntFlanks</td><td>10351</td><td>10373</td><td>23</td><td>CGAGAACTTTGAAGGCCGAAGTG</td><td>633.0</td><td>-2.9</td><td>0.7</td><td>-4.3</td><td>1.5E-05</td><td>2.1E-05</td><td>MB231/MB468</td><td>NUC</td></tr>
<tr><td>>hg38_21_+_8433172_8446622_RNA45SN1_45S_50ntFlanks@9861.9893.33|GATGAACCGAACGCCGGGTTAAGGCGCCCGATG</td><td>>hg38_21_+_8433172_8446622_RNA45SN1_45S_50ntFlanks</td><td>9861</td><td>9893</td><td>33</td><td>GATGAACCGAACGCCGGGTTAAGGCGCCCGATG</td><td>171.6</td><td>-6.6</td><td>1.5</td><td>-4.4</td><td>1.2E-05</td><td>2.1E-05</td><td>MB231/MB468</td><td>MP</td></tr>
<tr><td>>hg38_21_+_8433172_8446622_RNA45SN1_45S_50ntFlanks@11200.11232.33|CGGCGGCGGCGGCGACTCTGGACGCGAGCCGGG</td><td>>hg38_21_+_8433172_8446622_RNA45SN1_45S_50ntFlanks</td><td>11200</td><td>11232</td><td>33</td><td>CGGCGGCGGCGGCGACTCTGGACGCGAGCCGGG</td><td>4555.4</td><td>-8.4</td><td>2.0</td><td>-4.3</td><td>1.6E-05</td><td>2.1E-05</td><td>BT20/MB468</td><td>NUC</td></tr>
<tr><td>>hg38_21_+_8433172_8446622_RNA45SN1_45S_50ntFlanks@9790.9819.30|CTGGCGTGGAGCCGGGCGTGGAATGCGAGT</td><td>>hg38_21_+_8433172_8446622_RNA45SN1_45S_50ntFlanks</td><td>9790</td><td>9819</td><td>30</td><td>CTGGCGTGGAGCCGGGCGTGGAATGCGAGT</td><td>130.2</td><td>10.4</td><td>2.1</td><td>4.8</td><td>1.4E-06</td><td>2.1E-05</td><td>BT20/MB231</td><td>MP</td></tr>
<tr><td>>hg38_21_+_8433172_8446622_RNA45SN1_45S_50ntFlanks@6654.6681.28|CTCTTAGCGGTGGATCACTCGGCTCGTG</td><td>>hg38_21_+_8433172_8446622_RNA45SN1_45S_50ntFlanks</td><td>6654</td><td>6681</td><td>28</td><td>CTCTTAGCGGTGGATCACTCGGCTCGTG</td><td>1416.6</td><td>-2.1</td><td>0.5</td><td>-4.4</td><td>1.1E-05</td><td>2.1E-05</td><td>BT20/MB468</td><td>MITO</td></tr>
<tr><td>>hg38_21_+_8433172_8446622_RNA45SN1_45S_50ntFlanks@11567.11592.26|AGCAGCCGACTTAGAACTGGTGCGGA</td><td>>hg38_21_+_8433172_8446622_RNA45SN1_45S_50ntFlanks</td><td>11567</td><td>11592</td><td>26</td><td>AGCAGCCGACTTAGAACTGGTGCGGA</td><td>213.5</td><td>-3.0</td><td>0.7</td><td>-4.3</td><td>1.4E-05</td><td>2.1E-05</td><td>MB231/MB468</td><td>CYTO</td></tr>
<tr><td>>hg38_21_+_8433172_8446622_RNA45SN1_45S_50ntFlanks@12405.12445.41|CTTTTTGATCCTTCGATGTCGGCTCTTCCTATCATTGTGAA</td><td>>hg38_21_+_8433172_8446622_RNA45SN1_45S_50ntFlanks</td><td>12405</td><td>12445</td><td>41</td><td>CTTTTTGATCCTTCGATGTCGGCTCTTCCTATCATTGTGAA</td><td>140.8</td><td>-5.5</td><td>1.3</td><td>-4.4</td><td>1.3E-05</td><td>2.1E-05</td><td>BT20/MB468</td><td>CYTO</td></tr>
<tr><td>>trnaMT_SerGCT_MT_+_12207_12265@1.25.25|GAGAAAGCTCACAAGAACTGCTAAC</td><td>>trnaMT_SerGCT_MT_+_12207_12265</td><td>1</td><td>25</td><td>25</td><td>GAGAAAGCTCACAAGAACTGCTAAC</td><td>903.2</td><td>5.4</td><td>1.3</td><td>4.3</td><td>1.4E-05</td><td>2.1E-05</td><td>MB231/MB468</td><td>CYTO</td></tr>
<tr><td>>hg38_MT_+_1621_3279_RNR2_16S_50ntFlanks@51.88.38|GCTAAACCTAGCCCCAAACCCACTCCACCTTACTACCA</td><td>>hg38_MT_+_1621_3279_RNR2_16S_50ntFlanks</td><td>51</td><td>88</td><td>38</td><td>GCTAAACCTAGCCCCAAACCCACTCCACCTTACTACCA</td><td>7719.5</td><td>4.6</td><td>1.1</td><td>4.3</td><td>1.5E-05</td><td>2.1E-05</td><td>MB231/MB468</td><td>MITO</td></tr>
<tr><td>>hg38_21_+_8433172_8446622_RNA45SN1_45S_50ntFlanks@12396.12414.19|GCGACGTCGCTTTTTGATC</td><td>>hg38_21_+_8433172_8446622_RNA45SN1_45S_50ntFlanks</td><td>12396</td><td>12414</td><td>19</td><td>GCGACGTCGCTTTTTGATC</td><td>934.5</td><td>-4.9</td><td>1.1</td><td>-4.4</td><td>1.2E-05</td><td>2.1E-05</td><td>MB231/MB468</td><td>MP</td></tr>
<tr><td>>hg38_21_+_8433172_8446622_RNA45SN1_45S_50ntFlanks@12638.12657.20|CTACCATCTGTGGGATTATG</td><td>>hg38_21_+_8433172_8446622_RNA45SN1_45S_50ntFlanks</td><td>12638</td><td>12657</td><td>20</td><td>CTACCATCTGTGGGATTATG</td><td>84.0</td><td>-6.0</td><td>1.4</td><td>-4.4</td><td>1.3E-05</td><td>2.1E-05</td><td>BT20/MB468</td><td>CYTO</td></tr>
<tr><td>>trnaMT_MetCAT_MT_+_4402_4469@36.71.36|CCCGAAAATGTTGGTTATACCCTTCCCGTACTACCA</td><td>>trnaMT_MetCAT_MT_+_4402_4469</td><td>36</td><td>71</td><td>36</td><td>CCCGAAAATGTTGGTTATACCCTTCCCGTACTACCA</td><td>526.5</td><td>-4.2</td><td>0.9</td><td>-4.8</td><td>1.8E-06</td><td>2.1E-05</td><td>BT20/MB231</td><td>NUC</td></tr>
<tr><td>>hg38_21_+_8433172_8446622_RNA45SN1_45S_50ntFlanks@4732.4768.37|AGAACGAAAGTCGGAGGTTCGAAGACGATCAGATACC</td><td>>hg38_21_+_8433172_8446622_RNA45SN1_45S_50ntFlanks</td><td>4732</td><td>4768</td><td>37</td><td>AGAACGAAAGTCGGAGGTTCGAAGACGATCAGATACC</td><td>125.1</td><td>7.8</td><td>1.7</td><td>4.6</td><td>4.8E-06</td><td>2.1E-05</td><td>BT20/MB468</td><td>MP</td></tr>
<tr><td>>hg38_21_+_8433172_8446622_RNA45SN1_45S_50ntFlanks@11579.11604.26|AGAACTGGTGCGGACCAGGGGAATCC</td><td>>hg38_21_+_8433172_8446622_RNA45SN1_45S_50ntFlanks</td><td>11579</td><td>11604</td><td>26</td><td>AGAACTGGTGCGGACCAGGGGAATCC</td><td>134.0</td><td>-6.5</td><td>1.5</td><td>-4.4</td><td>1.2E-05</td><td>2.1E-05</td><td>MB231/MB468</td><td>MP</td></tr>
<tr><td>>hg38_21_+_8433172_8446622_RNA45SN1_45S_50ntFlanks@5083.5120.38|ACGAACGAGACTCTGGCATGCTAACTAGTTACGCGACC</td><td>>hg38_21_+_8433172_8446622_RNA45SN1_45S_50ntFlanks</td><td>5083</td><td>5120</td><td>38</td><td>ACGAACGAGACTCTGGCATGCTAACTAGTTACGCGACC</td><td>671.0</td><td>-7.9</td><td>1.8</td><td>-4.3</td><td>1.6E-05</td><td>2.1E-05</td><td>BT20/MB468</td><td>NUC</td></tr>
<tr><td>>hg38_1_-_228634819_228635039_RNA5S12_5S_50ntFlanks@135.172.38|GGGAGACCGCCTGGGAATACCGGGTGCTGTAGGCTTTT</td><td>>hg38_1_-_228634819_228635039_RNA5S12_5S_50ntFlanks</td><td>135</td><td>172</td><td>38</td><td>GGGAGACCGCCTGGGAATACCGGGTGCTGTAGGCTTTT</td><td>813.6</td><td>5.4</td><td>1.2</td><td>4.4</td><td>1.3E-05</td><td>2.1E-05</td><td>BT20/MB468</td><td>CYTO</td></tr>
<tr><td>>hg38_21_+_8433172_8446622_RNA45SN1_45S_50ntFlanks@10458.10483.26|ATGGCCTCCGTTGCCCTCGGCCGATC</td><td>>hg38_21_+_8433172_8446622_RNA45SN1_45S_50ntFlanks</td><td>10458</td><td>10483</td><td>26</td><td>ATGGCCTCCGTTGCCCTCGGCCGATC</td><td>184.5</td><td>-5.9</td><td>1.4</td><td>-4.4</td><td>1.3E-05</td><td>2.1E-05</td><td>BT20/MB468</td><td>CYTO</td></tr>
<tr><td>>hg38_21_+_8433172_8446622_RNA45SN1_45S_50ntFlanks@10461.10488.28|GCCTCCGTTGCCCTCGGCCGATCGAAAG</td><td>>hg38_21_+_8433172_8446622_RNA45SN1_45S_50ntFlanks</td><td>10461</td><td>10488</td><td>28</td><td>GCCTCCGTTGCCCTCGGCCGATCGAAAG</td><td>507.1</td><td>-7.6</td><td>1.8</td><td>-4.3</td><td>1.6E-05</td><td>2.1E-05</td><td>BT20/MB468</td><td>NUC</td></tr>
<tr><td>>hg38_21_+_8433172_8446622_RNA45SN1_45S_50ntFlanks@10361.10384.24|GAAGGCCGAAGTGGAGAAGGGTTC</td><td>>hg38_21_+_8433172_8446622_RNA45SN1_45S_50ntFlanks</td><td>10361</td><td>10384</td><td>24</td><td>GAAGGCCGAAGTGGAGAAGGGTTC</td><td>234.3</td><td>-6.5</td><td>1.5</td><td>-4.4</td><td>1.3E-05</td><td>2.1E-05</td><td>MB231/MB468</td><td>MP</td></tr>
<tr><td>>hg38_21_+_8433172_8446622_RNA45SN1_45S_50ntFlanks@12476.12498.23|CCACTAATAGGGAACGTGAGCTG</td><td>>hg38_21_+_8433172_8446622_RNA45SN1_45S_50ntFlanks</td><td>12476</td><td>12498</td><td>23</td><td>CCACTAATAGGGAACGTGAGCTG</td><td>1921.0</td><td>-5.2</td><td>1.2</td><td>-4.4</td><td>1.3E-05</td><td>2.1E-05</td><td>BT20/MB468</td><td>CYTO</td></tr>
<tr><td>>hg38_21_+_8433172_8446622_RNA45SN1_45S_50ntFlanks@4000.4023.24|TAGATAACCTCGGGCCGATCGCAC</td><td>>hg38_21_+_8433172_8446622_RNA45SN1_45S_50ntFlanks</td><td>4000</td><td>4023</td><td>24</td><td>TAGATAACCTCGGGCCGATCGCAC</td><td>525.3</td><td>3.0</td><td>0.7</td><td>4.3</td><td>1.4E-05</td><td>2.1E-05</td><td>MB231/MB468</td><td>CYTO</td></tr>
<tr><td>>hg38_21_+_8433172_8446622_RNA45SN1_45S_50ntFlanks@10808.10829.22|CAATGTAGGTAAGGGAAGTCGG</td><td>>hg38_21_+_8433172_8446622_RNA45SN1_45S_50ntFlanks</td><td>10808</td><td>10829</td><td>22</td><td>CAATGTAGGTAAGGGAAGTCGG</td><td>400.8</td><td>-5.8</td><td>1.3</td><td>-4.4</td><td>1.3E-05</td><td>2.1E-05</td><td>BT20/MB468</td><td>CYTO</td></tr>
<tr><td>>hg38_21_+_8433172_8446622_RNA45SN1_45S_50ntFlanks@9714.9737.24|CGAATGATTAGAGGTCTTGGGGCC</td><td>>hg38_21_+_8433172_8446622_RNA45SN1_45S_50ntFlanks</td><td>9714</td><td>9737</td><td>24</td><td>CGAATGATTAGAGGTCTTGGGGCC</td><td>688.9</td><td>-4.7</td><td>1.1</td><td>-4.4</td><td>1.3E-05</td><td>2.1E-05</td><td>BT20/MB468</td><td>CYTO</td></tr>
<tr><td>>hg38_21_+_8433172_8446622_RNA45SN1_45S_50ntFlanks@11592.11621.30|ACCAGGGGAATCCGACTGTTTAATTAAAAC</td><td>>hg38_21_+_8433172_8446622_RNA45SN1_45S_50ntFlanks</td><td>11592</td><td>11621</td><td>30</td><td>ACCAGGGGAATCCGACTGTTTAATTAAAAC</td><td>121.2</td><td>10.3</td><td>2.1</td><td>4.8</td><td>1.4E-06</td><td>2.2E-05</td><td>BT20/MB231</td><td>MP</td></tr>
<tr><td>>hg38_21_+_8433172_8446622_RNA45SN1_45S_50ntFlanks@485.517.33|GATGTGGTGACGTCGTGCTCTCCCGGGCCGGGT</td><td>>hg38_21_+_8433172_8446622_RNA45SN1_45S_50ntFlanks</td><td>485</td><td>517</td><td>33</td><td>GATGTGGTGACGTCGTGCTCTCCCGGGCCGGGT</td><td>179.9</td><td>10.7</td><td>2.5</td><td>4.4</td><td>1.3E-05</td><td>2.2E-05</td><td>MB231/MB468</td><td>MP</td></tr>
<tr><td>>hg38_21_+_8433172_8446622_RNA45SN1_45S_50ntFlanks@12408.12432.25|TTTGATCCTTCGATGTCGGCTCTTC</td><td>>hg38_21_+_8433172_8446622_RNA45SN1_45S_50ntFlanks</td><td>12408</td><td>12432</td><td>25</td><td>TTTGATCCTTCGATGTCGGCTCTTC</td><td>198.8</td><td>-6.4</td><td>1.5</td><td>-4.4</td><td>1.3E-05</td><td>2.2E-05</td><td>MB231/MB468</td><td>MP</td></tr>
<tr><td>>hg38_21_+_8433172_8446622_RNA45SN1_45S_50ntFlanks@10786.10822.37|AACAGCCTCTGGCATGTTGGAACAATGTAGGTAAGGG</td><td>>hg38_21_+_8433172_8446622_RNA45SN1_45S_50ntFlanks</td><td>10786</td><td>10822</td><td>37</td><td>AACAGCCTCTGGCATGTTGGAACAATGTAGGTAAGGG</td><td>839.5</td><td>-4.3</td><td>0.9</td><td>-4.6</td><td>4.8E-06</td><td>2.2E-05</td><td>BT20/MB468</td><td>MP</td></tr>
<tr><td>>hg38_1_-_228634819_228635039_RNA5S12_5S_50ntFlanks@104.131.28|AAGCAGGGTCGGGCCTGGTTAGTACTTG</td><td>>hg38_1_-_228634819_228635039_RNA5S12_5S_50ntFlanks</td><td>104</td><td>131</td><td>28</td><td>AAGCAGGGTCGGGCCTGGTTAGTACTTG</td><td>175.6</td><td>-4.0</td><td>0.9</td><td>-4.3</td><td>1.4E-05</td><td>2.2E-05</td><td>MB231/MB468</td><td>CYTO</td></tr>
<tr><td>>hg38_21_+_8433172_8446622_RNA45SN1_45S_50ntFlanks@12570.12588.19|AGTACGAGAGGAACCGCAG</td><td>>hg38_21_+_8433172_8446622_RNA45SN1_45S_50ntFlanks</td><td>12570</td><td>12588</td><td>19</td><td>AGTACGAGAGGAACCGCAG</td><td>191.5</td><td>-6.4</td><td>1.5</td><td>-4.4</td><td>1.3E-05</td><td>2.2E-05</td><td>MB231/MB468</td><td>MP</td></tr>
<tr><td>>hg38_21_+_8433172_8446622_RNA45SN1_45S_50ntFlanks@6651.6693.43|CGACTCTTAGCGGTGGATCACTCGGCTCGTGCGTCGATGAAGA</td><td>>hg38_21_+_8433172_8446622_RNA45SN1_45S_50ntFlanks</td><td>6651</td><td>6693</td><td>43</td><td>CGACTCTTAGCGGTGGATCACTCGGCTCGTGCGTCGATGAAGA</td><td>4044.7</td><td>4.2</td><td>1.0</td><td>4.3</td><td>1.5E-05</td><td>2.2E-05</td><td>MB231/MB468</td><td>NUC</td></tr>
<tr><td>>hg38_21_+_8433172_8446622_RNA45SN1_45S_50ntFlanks@5486.5526.41|GCGCTGAGAAGACGGTCGAACTTGACTATCTAGAGGAAGTA</td><td>>hg38_21_+_8433172_8446622_RNA45SN1_45S_50ntFlanks</td><td>5486</td><td>5526</td><td>41</td><td>GCGCTGAGAAGACGGTCGAACTTGACTATCTAGAGGAAGTA</td><td>517.9</td><td>-6.0</td><td>1.4</td><td>-4.3</td><td>1.6E-05</td><td>2.2E-05</td><td>BT20/MB468</td><td>NUC</td></tr>
<tr><td>>MI0000273|hsa-mir-183&WithFlank&7|-|129774899|129775020@34.55.22&[MIMAT0000261&hsa-miR-183-5p&offsets|+1|+1;m-41&7|-|129774966|129774987&offsets|0|0]|ATGGCACTGGTAGAATTCACTG</td><td>>MI0000273|hsa-mir-183&WithFlank&7|-|129774899|129775020</td><td>34</td><td>55</td><td>22</td><td>ATGGCACTGGTAGAATTCACTG</td><td>3474.7</td><td>3.2</td><td>0.7</td><td>4.8</td><td>1.7E-06</td><td>2.2E-05</td><td>BT20/MB231</td><td>MITO</td></tr>
<tr><td>>hg38_21_+_8433172_8446622_RNA45SN1_45S_50ntFlanks@9305.9323.19|CCAAGGAGTCTAACACGTG</td><td>>hg38_21_+_8433172_8446622_RNA45SN1_45S_50ntFlanks</td><td>9305</td><td>9323</td><td>19</td><td>CCAAGGAGTCTAACACGTG</td><td>381.5</td><td>-3.3</td><td>0.7</td><td>-4.4</td><td>1.1E-05</td><td>2.2E-05</td><td>BT20/MB468</td><td>MITO</td></tr>
<tr><td>>hg38_21_+_8433172_8446622_RNA45SN1_45S_50ntFlanks@5485.5512.28|AGCGCTGAGAAGACGGTCGAACTTGACT</td><td>>hg38_21_+_8433172_8446622_RNA45SN1_45S_50ntFlanks</td><td>5485</td><td>5512</td><td>28</td><td>AGCGCTGAGAAGACGGTCGAACTTGACT</td><td>3336.7</td><td>5.3</td><td>1.2</td><td>4.4</td><td>1.3E-05</td><td>2.2E-05</td><td>BT20/MB468</td><td>CYTO</td></tr>
<tr><td>>hg38_21_+_8433172_8446622_RNA45SN1_45S_50ntFlanks@9201.9222.22|CTCTCGGGGCCACGCGCGCGTC</td><td>>hg38_21_+_8433172_8446622_RNA45SN1_45S_50ntFlanks</td><td>9201</td><td>9222</td><td>22</td><td>CTCTCGGGGCCACGCGCGCGTC</td><td>749.3</td><td>-7.2</td><td>1.6</td><td>-4.6</td><td>4.8E-06</td><td>2.2E-05</td><td>BT20/MB468</td><td>MP</td></tr>
<tr><td>>hg38_21_+_8433172_8446622_RNA45SN1_45S_50ntFlanks@11876.11900.25|AAGACCCTGTTGAGCTTGACTCTAG</td><td>>hg38_21_+_8433172_8446622_RNA45SN1_45S_50ntFlanks</td><td>11876</td><td>11900</td><td>25</td><td>AAGACCCTGTTGAGCTTGACTCTAG</td><td>493.9</td><td>-3.6</td><td>0.8</td><td>-4.4</td><td>1.3E-05</td><td>2.2E-05</td><td>BT20/MB468</td><td>CYTO</td></tr>
<tr><td>>MI0000102|hsa-mir-100&WithFlank&11|-|122152223|122152314@19.39.21(+1C)&[MIMAT0000098&hsa-miR-100-5p&offsets|0|-1(+1C);m-27&11|-|122152276|122152296&offsets|0|0(+1C)]|AACCCGTAGATCCGAACTTGTC</td><td>>MI0000102|hsa-mir-100&WithFlank&11|-|122152223|122152314</td><td>19</td><td>39</td><td>21(+1C)</td><td>AACCCGTAGATCCGAACTTGTC</td><td>1497.6</td><td>-5.6</td><td>1.2</td><td>-4.8</td><td>1.8E-06</td><td>2.2E-05</td><td>BT20/MB231</td><td>NUC</td></tr>
<tr><td>>hg38_21_+_8433172_8446622_RNA45SN1_45S_50ntFlanks@10352.10374.23|GAGAACTTTGAAGGCCGAAGTGG</td><td>>hg38_21_+_8433172_8446622_RNA45SN1_45S_50ntFlanks</td><td>10352</td><td>10374</td><td>23</td><td>GAGAACTTTGAAGGCCGAAGTGG</td><td>238.5</td><td>-5.3</td><td>1.2</td><td>-4.4</td><td>1.3E-05</td><td>2.2E-05</td><td>MB231/MB468</td><td>MP</td></tr>
<tr><td>>hg38_21_+_8433172_8446622_RNA45SN1_45S_50ntFlanks@10444.10462.19|CGAAGGGACGGGCGATGGC</td><td>>hg38_21_+_8433172_8446622_RNA45SN1_45S_50ntFlanks</td><td>10444</td><td>10462</td><td>19</td><td>CGAAGGGACGGGCGATGGC</td><td>324.5</td><td>-4.6</td><td>1.1</td><td>-4.4</td><td>1.3E-05</td><td>2.2E-05</td><td>BT20/MB468</td><td>CYTO</td></tr>
<tr><td>>hg38_21_+_8433172_8446622_RNA45SN1_45S_50ntFlanks@4203.4223.21|GACCCGGGGAGGTAGTGACGA</td><td>>hg38_21_+_8433172_8446622_RNA45SN1_45S_50ntFlanks</td><td>4203</td><td>4223</td><td>21</td><td>GACCCGGGGAGGTAGTGACGA</td><td>406.1</td><td>-6.8</td><td>1.6</td><td>-4.4</td><td>1.3E-05</td><td>2.2E-05</td><td>MB231/MB468</td><td>MP</td></tr>
<tr><td>>hg38_21_+_8433172_8446622_RNA45SN1_45S_50ntFlanks@10348.10376.29|AAACGAGAACTTTGAAGGCCGAAGTGGAG</td><td>>hg38_21_+_8433172_8446622_RNA45SN1_45S_50ntFlanks</td><td>10348</td><td>10376</td><td>29</td><td>AAACGAGAACTTTGAAGGCCGAAGTGGAG</td><td>499.9</td><td>-5.9</td><td>1.4</td><td>-4.4</td><td>1.3E-05</td><td>2.2E-05</td><td>MB231/MB468</td><td>MP</td></tr>
<tr><td>>hg38_21_+_8433172_8446622_RNA45SN1_45S_50ntFlanks@9938.9975.38|AGCAGGACGGTGGCCATGGAAGTCGGAATCCGCTAAGG</td><td>>hg38_21_+_8433172_8446622_RNA45SN1_45S_50ntFlanks</td><td>9938</td><td>9975</td><td>38</td><td>AGCAGGACGGTGGCCATGGAAGTCGGAATCCGCTAAGG</td><td>50.6</td><td>9.4</td><td>2.1</td><td>4.6</td><td>4.8E-06</td><td>2.2E-05</td><td>BT20/MB468</td><td>MP</td></tr>
</table>

<table>
<tr><td>>hg38_21_+_8433172_8446622_RNA45SN1_45S_50ntFlanks@10827.10865.39|CGGCAAGCCGGATCCGTAACTTCGGGATAAGGATTGGCT</td><td>>hg38_21_+_8433172_8446622_RNA45SN1_45S_50ntFlanks</td><td>10827</td><td>10865</td><td>39</td><td>CGGCAAGCCGGATCCGTAACTTCGGGATAAGGATTGGCT</td><td>51.3</td><td>-8.8</td><td>1.9</td><td>-4.6</td><td>4.8E-06</td><td>2.2E-05</td><td>BT20/MB468</td><td>MP</td></tr>
<tr><td>>hg38_21_+_8433172_8446622_RNA45SN1_45S_50ntFlanks@4760.4783.24|TCAGATACCGTCGTAGTTCCGACC</td><td>>hg38_21_+_8433172_8446622_RNA45SN1_45S_50ntFlanks</td><td>4760</td><td>4783</td><td>24</td><td>TCAGATACCGTCGTAGTTCCGACC</td><td>395.1</td><td>-3.0</td><td>0.7</td><td>-4.3</td><td>1.5E-05</td><td>2.2E-05</td><td>MB231/MB468</td><td>NUC</td></tr>
<tr><td>>hg38_21_+_8433172_8446622_RNA45SN1_45S_50ntFlanks@6652.6678.27|GACTCTTAGCGGTGGATCACTCGGCTC</td><td>>hg38_21_+_8433172_8446622_RNA45SN1_45S_50ntFlanks</td><td>6652</td><td>6678</td><td>27</td><td>GACTCTTAGCGGTGGATCACTCGGCTC</td><td>4381.7</td><td>5.2</td><td>1.1</td><td>4.6</td><td>4.8E-06</td><td>2.2E-05</td><td>BT20/MB468</td><td>MP</td></tr>
<tr><td>>hg38_21_+_8433172_8446622_RNA45SN1_45S_50ntFlanks@11845.11872.28|GGAACGGGCTTGGCGGAATCAGCGGGGA</td><td>>hg38_21_+_8433172_8446622_RNA45SN1_45S_50ntFlanks</td><td>11845</td><td>11872</td><td>28</td><td>GGAACGGGCTTGGCGGAATCAGCGGGGA</td><td>498.9</td><td>-4.6</td><td>1.0</td><td>-4.4</td><td>1.1E-05</td><td>2.2E-05</td><td>BT20/MB468</td><td>MITO</td></tr>
<tr><td>>MI0000067|hsa-let-7f-1&WithFlank&9|+|94176341|94176439@69.89.21&[MIMAT0004486&hsa-let-7f-1-3p&offsets|0|-1;m-275&9|+|94176409|94176430&offsets|0|-1]|CTATACAATCTATTGCCTTCC</td><td>>MI0000067|hsa-let-7f-1&WithFlank&9|+|94176341|94176439</td><td>69</td><td>89</td><td>21</td><td>CTATACAATCTATTGCCTTCC</td><td>36.8</td><td>8.4</td><td>1.9</td><td>4.4</td><td>1.3E-05</td><td>2.2E-05</td><td>MB231/MB468</td><td>MP</td></tr>
<tr><td>>hg38_21_+_8433172_8446622_RNA45SN1_45S_50ntFlanks@11854.11886.33|TTGGCGGAATCAGCGGGGAAAGAAGACCCTGTT</td><td>>hg38_21_+_8433172_8446622_RNA45SN1_45S_50ntFlanks</td><td>11854</td><td>11886</td><td>33</td><td>TTGGCGGAATCAGCGGGGAAAGAAGACCCTGTT</td><td>108.1</td><td>10.1</td><td>2.1</td><td>4.8</td><td>1.5E-06</td><td>2.2E-05</td><td>BT20/MB231</td><td>MP</td></tr>
<tr><td>>trnaMT_PheGAA_MT_+_577_647@26.57.32|CAATACACTGAAAATGTTTAGACGGGCTCACA</td><td>>trnaMT_PheGAA_MT_+_577_647</td><td>26</td><td>57</td><td>32</td><td>CAATACACTGAAAATGTTTAGACGGGCTCACA</td><td>53.8</td><td>9.5</td><td>2.1</td><td>4.6</td><td>4.9E-06</td><td>2.2E-05</td><td>BT20/MB468</td><td>MP</td></tr>
<tr><td>>hg38_21_+_8433172_8446622_RNA45SN1_45S_50ntFlanks@12993.13013.21|CCCTCGCTGCGATCTATTGAA</td><td>>hg38_21_+_8433172_8446622_RNA45SN1_45S_50ntFlanks</td><td>12993</td><td>13013</td><td>21</td><td>CCCTCGCTGCGATCTATTGAA</td><td>250.1</td><td>-2.8</td><td>0.6</td><td>-4.4</td><td>1.1E-05</td><td>2.2E-05</td><td>BT20/MB468</td><td>MITO</td></tr>
<tr><td>>hg38_21_+_8433172_8446622_RNA45SN1_45S_50ntFlanks@8184.8223.40|CGAGGCCCAGCCCGTGGACGGTGTGAGGCCGGTAGCGGCC</td><td>>hg38_21_+_8433172_8446622_RNA45SN1_45S_50ntFlanks</td><td>8184</td><td>8223</td><td>40</td><td>CGAGGCCCAGCCCGTGGACGGTGTGAGGCCGGTAGCGGCC</td><td>216.9</td><td>-5.8</td><td>1.3</td><td>-4.4</td><td>1.3E-05</td><td>2.2E-05</td><td>BT20/MB468</td><td>CYTO</td></tr>
<tr><td>>hg38_21_+_8433172_8446622_RNA45SN1_45S_50ntFlanks@4730.4755.26|CAAGAACGAAAGTCGGAGGTTCGAAG</td><td>>hg38_21_+_8433172_8446622_RNA45SN1_45S_50ntFlanks</td><td>4730</td><td>4755</td><td>26</td><td>CAAGAACGAAAGTCGGAGGTTCGAAG</td><td>107.5</td><td>-4.7</td><td>1.1</td><td>-4.4</td><td>1.3E-05</td><td>2.2E-05</td><td>MB231/MB468</td><td>MP</td></tr>
<tr><td>>hg38_21_+_8433172_8446622_RNA45SN1_45S_50ntFlanks@6651.6669.19|CGACTCTTAGCGGTGGATC</td><td>>hg38_21_+_8433172_8446622_RNA45SN1_45S_50ntFlanks</td><td>6651</td><td>6669</td><td>19</td><td>CGACTCTTAGCGGTGGATC</td><td>15656.1</td><td>3.2</td><td>0.7</td><td>4.8</td><td>1.8E-06</td><td>2.2E-05</td><td>BT20/MB231</td><td>NUC</td></tr>
<tr><td>>hg38_21_+_8433172_8446622_RNA45SN1_45S_50ntFlanks@4047.4064.18|ATTCGAACGTCTGCCCTA</td><td>>hg38_21_+_8433172_8446622_RNA45SN1_45S_50ntFlanks</td><td>4047</td><td>4064</td><td>18</td><td>ATTCGAACGTCTGCCCTA</td><td>405.1</td><td>-4.1</td><td>1.0</td><td>-4.3</td><td>1.4E-05</td><td>2.2E-05</td><td>MB231/MB468</td><td>CYTO</td></tr>
<tr><td>>MI0000441|hsa-mir-30b&WithFlank&8|-|134800514|134800613@23.44.22(+1U)&[MIMAT0000420&hsa-miR-30b-5p&offsets|0|0(+1U);m-76&8|-|134800570|134800591&offsets|0|0(+1U)]|TGTAAACATCCTACACTCAGCTT</td><td>>MI0000441|hsa-mir-30b&WithFlank&8|-|134800514|134800613</td><td>23</td><td>44</td><td>22(+1U)</td><td>TGTAAACATCCTACACTCAGCTT</td><td>76.2</td><td>-9.7</td><td>2.0</td><td>-4.8</td><td>1.5E-06</td><td>2.2E-05</td><td>BT20/MB231</td><td>MP</td></tr>
<tr><td>>hg38_21_+_8433172_8446622_RNA45SN1_45S_50ntFlanks@9574.9594.21|AATCGGTCGTCCGACCTGGGT</td><td>>hg38_21_+_8433172_8446622_RNA45SN1_45S_50ntFlanks</td><td>9574</td><td>9594</td><td>21</td><td>AATCGGTCGTCCGACCTGGGT</td><td>3972.1</td><td>-5.0</td><td>1.1</td><td>-4.6</td><td>4.9E-06</td><td>2.2E-05</td><td>BT20/MB468</td><td>MP</td></tr>
<tr><td>>MI0003560|hsa-mir-92b&WithFlank&1|+|155195171|155195278@67.88.22(+1U)&[MIMAT0003218&hsa-miR-92b-3p&offsets|0|0(+1U);m-115&1|+|155195237|155195258&offsets|0|0(+1U)]|TATTGCACTCGTCCCGGCCTCCT</td><td>>MI0003560|hsa-mir-92b&WithFlank&1|+|155195171|155195278</td><td>67</td><td>88</td><td>22(+1U)</td><td>TATTGCACTCGTCCCGGCCTCCT</td><td>343.5</td><td>6.8</td><td>1.5</td><td>4.6</td><td>4.9E-06</td><td>2.2E-05</td><td>BT20/MB468</td><td>MP</td></tr>
<tr><td>>hg38_21_+_8433172_8446622_RNA45SN1_45S_50ntFlanks@12941.12964.24|ATTCGTAGACGACCTGCTTCTGGG</td><td>>hg38_21_+_8433172_8446622_RNA45SN1_45S_50ntFlanks</td><td>12941</td><td>12964</td><td>24</td><td>ATTCGTAGACGACCTGCTTCTGGG</td><td>144.1</td><td>-3.3</td><td>0.8</td><td>-4.3</td><td>1.5E-05</td><td>2.2E-05</td><td>MB231/MB468</td><td>CYTO</td></tr>
<tr><td>>hg38_21_+_8433172_8446622_RNA45SN1_45S_50ntFlanks@6650.6685.36|ACGACTCTTAGCGGTGGATCACTCGGCTCGTGCGTC</td><td>>hg38_21_+_8433172_8446622_RNA45SN1_45S_50ntFlanks</td><td>6650</td><td>6685</td><td>36</td><td>ACGACTCTTAGCGGTGGATCACTCGGCTCGTGCGTC</td><td>893.7</td><td>4.2</td><td>0.9</td><td>4.6</td><td>4.9E-06</td><td>2.2E-05</td><td>BT20/MB468</td><td>MP</td></tr>
<tr><td>>hg38_21_+_8433172_8446622_RNA45SN1_45S_50ntFlanks@6697.6718.22|CAGCTAGCTGCGAGAATTAATG</td><td>>hg38_21_+_8433172_8446622_RNA45SN1_45S_50ntFlanks</td><td>6697</td><td>6718</td><td>22</td><td>CAGCTAGCTGCGAGAATTAATG</td><td>1216.4</td><td>-7.3</td><td>1.6</td><td>-4.6</td><td>4.9E-06</td><td>2.2E-05</td><td>BT20/MB468</td><td>MP</td></tr>
<tr><td>>hg38_21_+_8433172_8446622_RNA45SN1_45S_50ntFlanks@12413.12434.22|TCCTTCGATGTCGGCTCTTCCT</td><td>>hg38_21_+_8433172_8446622_RNA45SN1_45S_50ntFlanks</td><td>12413</td><td>12434</td><td>22</td><td>TCCTTCGATGTCGGCTCTTCCT</td><td>280.5</td><td>-3.1</td><td>0.7</td><td>-4.3</td><td>1.5E-05</td><td>2.2E-05</td><td>MB231/MB468</td><td>MITO</td></tr>
<tr><td>>hg38_21_+_8433172_8446622_RNA45SN1_45S_50ntFlanks@3845.3874.30|ACTTGGATAACTGTGGTAATTCTAGAGCTA</td><td>>hg38_21_+_8433172_8446622_RNA45SN1_45S_50ntFlanks</td><td>3845</td><td>3874</td><td>30</td><td>ACTTGGATAACTGTGGTAATTCTAGAGCTA</td><td>153.2</td><td>-5.4</td><td>1.2</td><td>-4.4</td><td>1.3E-05</td><td>2.2E-05</td><td>MB231/MB468</td><td>MP</td></tr>
<tr><td>>am_trna119_LysCTT_1_-_145395522_145395594@14.31.18,trna11_LysCTT_5_-_180648979_180649051@14.31.18,trna128_LysCTT_1_-_55423542_55423614@14.31.18,trna13_LysCTT_6_+_26556774_26556846@14.31.18,trna32_LysCTT_16_-_3207406_3207478@14.31.18,trna6_LysCTT_19_-_52425396_52425468@14.31.18,trna7_LysCTT_16_+_3225692_3225764@14.31.18,trna9_LysCTT_5_+_180634755_180634827@14.31.18|AGTCGGTAGAGCATGAGA</td><td>>am_trna119_LysCTT_1_-_145395522_145395594</td><td>14</td><td>31</td><td>18</td><td>AGTCGGTAGAGCATGAGA</td><td>292.3</td><td>-3.0</td><td>0.7</td><td>-4.4</td><td>1.3E-05</td><td>2.2E-05</td><td>MB231/MB468</td><td>MP</td></tr>
<tr><td>>hg38_21_+_8433172_8446622_RNA45SN1_45S_50ntFlanks@10349.10382.34|AACGAGAACTTTGAAGGCCGAAGTGGAGAAGGGT</td><td>>hg38_21_+_8433172_8446622_RNA45SN1_45S_50ntFlanks</td><td>10349</td><td>10382</td><td>34</td><td>AACGAGAACTTTGAAGGCCGAAGTGGAGAAGGGT</td><td>2011.8</td><td>3.0</td><td>0.7</td><td>4.3</td><td>1.5E-05</td><td>2.2E-05</td><td>MB231/MB468</td><td>CYTO</td></tr>
<tr><td>>hg38_21_+_8433172_8446622_RNA45SN1_45S_50ntFlanks@12417.12441.25|TCGATGTCGGCTCTTCCTATCATTG</td><td>>hg38_21_+_8433172_8446622_RNA45SN1_45S_50ntFlanks</td><td>12417</td><td>12441</td><td>25</td><td>TCGATGTCGGCTCTTCCTATCATTG</td><td>352.3</td><td>-6.6</td><td>1.5</td><td>-4.4</td><td>1.3E-05</td><td>2.2E-05</td><td>MB231/MB468</td><td>MP</td></tr>
<tr><td>>hg38_21_+_8433172_8446622_RNA45SN1_45S_50ntFlanks@9761.9787.27|AACTTTAAATGGGTAAGAAGCCCGGCT</td><td>>hg38_21_+_8433172_8446622_RNA45SN1_45S_50ntFlanks</td><td>9761</td><td>9787</td><td>27</td><td>AACTTTAAATGGGTAAGAAGCCCGGCT</td><td>103.1</td><td>-10.6</td><td>2.4</td><td>-4.4</td><td>1.3E-05</td><td>2.2E-05</td><td>MB231/MB468</td><td>MP</td></tr>
<tr><td>>hg38_21_+_8433172_8446622_RNA45SN1_45S_50ntFlanks@4130.4155.26|ATTCCGGAGAGGGAGCCTGAGAAACG</td><td>>hg38_21_+_8433172_8446622_RNA45SN1_45S_50ntFlanks</td><td>4130</td><td>4155</td><td>26</td><td>ATTCCGGAGAGGGAGCCTGAGAAACG</td><td>163.8</td><td>4.2</td><td>1.0</td><td>4.3</td><td>1.5E-05</td><td>2.2E-05</td><td>MB231/MB468</td><td>CYTO</td></tr>
<tr><td>>hg38_21_+_8433172_8446622_RNA45SN1_45S_50ntFlanks@4240.4258.19|ACTCTTTCGAGGCCCTGTA</td><td>>hg38_21_+_8433172_8446622_RNA45SN1_45S_50ntFlanks</td><td>4240</td><td>4258</td><td>19</td><td>ACTCTTTCGAGGCCCTGTA</td><td>168.0</td><td>-6.5</td><td>1.4</td><td>-4.6</td><td>4.9E-06</td><td>2.2E-05</td><td>BT20/MB468</td><td>MP</td></tr>
<tr><td>>hg38_21_+_8433172_8446622_RNA45SN1_45S_50ntFlanks@11921.11967.47|TGAGAGGTGTAGAATAAGTGGGAGGCCCCCGGCGCCCCCCCGGTGTC</td><td>>hg38_21_+_8433172_8446622_RNA45SN1_45S_50ntFlanks</td><td>11921</td><td>11967</td><td>47</td><td>TGAGAGGTGTAGAATAAGTGGGAGGCCCCCGGCGCCCCCCCGGTGTC</td><td>85.2</td><td>-10.3</td><td>2.4</td><td>-4.4</td><td>1.3E-05</td><td>2.2E-05</td><td>MB231/MB468</td><td>MP</td></tr>
<tr><td>>hg38_21_+_8433172_8446622_RNA45SN1_45S_50ntFlanks@10351.10377.27|CGAGAACTTTGAAGGCCGAAGTGGAGA</td><td>>hg38_21_+_8433172_8446622_RNA45SN1_45S_50ntFlanks</td><td>10351</td><td>10377</td><td>27</td><td>CGAGAACTTTGAAGGCCGAAGTGGAGA</td><td>23475.7</td><td>-5.6</td><td>1.2</td><td>-4.6</td><td>4.9E-06</td><td>2.2E-05</td><td>BT20/MB468</td><td>MP</td></tr>
<tr><td>>hg38_21_+_8433172_8446622_RNA45SN1_45S_50ntFlanks@4752.4769.18|GAAGACGATCAGATACCG</td><td>>hg38_21_+_8433172_8446622_RNA45SN1_45S_50ntFlanks</td><td>4752</td><td>4769</td><td>18</td><td>GAAGACGATCAGATACCG</td><td>231.4</td><td>-6.6</td><td>1.5</td><td>-4.4</td><td>1.3E-05</td><td>2.2E-05</td><td>MB231/MB468</td><td>MP</td></tr>
<tr><td>>hg38_21_+_8433172_8446622_RNA45SN1_45S_50ntFlanks@8197.8214.18|GTGGACGGTGTGAGGCCG</td><td>>hg38_21_+_8433172_8446622_RNA45SN1_45S_50ntFlanks</td><td>8197</td><td>8214</td><td>18</td><td>GTGGACGGTGTGAGGCCG</td><td>544.6</td><td>-3.9</td><td>0.9</td><td>-4.4</td><td>1.3E-05</td><td>2.2E-05</td><td>MB231/MB468</td><td>MP</td></tr>
<tr><td>>trnaMT_GluTTC_MT_-_14674_14742@31.71.41,trnalookalike8_GluTTC_5_-_93905172_93905240@31.71.41|TTCATATCATTGGTCGTGGTTGTAGTCCGTGCGAGAATACC</td><td>>trnaMT_GluTTC_MT_-_14674_14742</td><td>31</td><td>71</td><td>41</td><td>TTCATATCATTGGTCGTGGTTGTAGTCCGTGCGAGAATACC</td><td>280.8</td><td>-4.3</td><td>0.9</td><td>-4.8</td><td>1.5E-06</td><td>2.2E-05</td><td>BT20/MB231</td><td>CYTO</td></tr>
<tr><td>>hg38_21_+_8433172_8446622_RNA45SN1_45S_50ntFlanks@4983.5002.20|CGGACAGGATTGACAGATTG</td><td>>hg38_21_+_8433172_8446622_RNA45SN1_45S_50ntFlanks</td><td>4983</td><td>5002</td><td>20</td><td>CGGACAGGATTGACAGATTG</td><td>1041.9</td><td>-7.7</td><td>1.7</td><td>-4.4</td><td>1.1E-05</td><td>2.2E-05</td><td>BT20/MB468</td><td>MITO</td></tr>
<tr><td>>hg38_21_+_8433172_8446622_RNA45SN1_45S_50ntFlanks@10829.10852.24|GCAAGCCGGATCCGTAACTTCGGG</td><td>>hg38_21_+_8433172_8446622_RNA45SN1_45S_50ntFlanks</td><td>10829</td><td>10852</td><td>24</td><td>GCAAGCCGGATCCGTAACTTCGGG</td><td>205.9</td><td>-4.7</td><td>1.1</td><td>-4.3</td><td>1.5E-05</td><td>2.2E-05</td><td>MB231/MB468</td><td>NUC</td></tr>
<tr><td>>hg38_21_+_8433172_8446622_RNA45SN1_45S_50ntFlanks@3939.3958.20|AACCCGGTCAGCCCCTCTCC</td><td>>hg38_21_+_8433172_8446622_RNA45SN1_45S_50ntFlanks</td><td>3939</td><td>3958</td><td>20</td><td>AACCCGGTCAGCCCCTCTCC</td><td>245.4</td><td>-4.1</td><td>0.9</td><td>-4.3</td><td>1.5E-05</td><td>2.2E-05</td><td>MB231/MB468</td><td>CYTO</td></tr>
<tr><td>>hg38_21_+_8433172_8446622_RNA45SN1_45S_50ntFlanks@5490.5516.27|TGAGAAGACGGTCGAACTTGACTATCT</td><td>>hg38_21_+_8433172_8446622_RNA45SN1_45S_50ntFlanks</td><td>5490</td><td>5516</td><td>27</td><td>TGAGAAGACGGTCGAACTTGACTATCT</td><td>1147.1</td><td>8.7</td><td>1.9</td><td>4.6</td><td>5.0E-06</td><td>2.2E-05</td><td>BT20/MB468</td><td>MP</td></tr>
<tr><td>>hg38_21_+_8433172_8446622_RNA45SN1_45S_50ntFlanks@11661.11702.42|TGATTTCTGCCCAGTGCTCTGAATGTCAAAGTGAAGAAATTC</td><td>>hg38_21_+_8433172_8446622_RNA45SN1_45S_50ntFlanks</td><td>11661</td><td>11702</td><td>42</td><td>TGATTTCTGCCCAGTGCTCTGAATGTCAAAGTGAAGAAATTC</td><td>265.9</td><td>-4.1</td><td>1.0</td><td>-4.3</td><td>1.6E-05</td><td>2.2E-05</td><td>MB231/MB468</td><td>MITO</td></tr>
<tr><td>>hg38_21_+_8433172_8446622_RNA45SN1_45S_50ntFlanks@10786.10815.30|AACAGCCTCTGGCATGTTGGAACAATGTAG</td><td>>hg38_21_+_8433172_8446622_RNA45SN1_45S_50ntFlanks</td><td>10786</td><td>10815</td><td>30</td><td>AACAGCCTCTGGCATGTTGGAACAATGTAG</td><td>374.9</td><td>-4.9</td><td>1.1</td><td>-4.3</td><td>1.5E-05</td><td>2.2E-05</td><td>MB231/MB468</td><td>NUC</td></tr>
<tr><td>>hg38_21_+_8433172_8446622_RNA45SN1_45S_50ntFlanks@12331.12365.35|TTAAGCAGGAGGTGTCAGAAAAGTTACCACAGGGA</td><td>>hg38_21_+_8433172_8446622_RNA45SN1_45S_50ntFlanks</td><td>12331</td><td>12365</td><td>35</td><td>TTAAGCAGGAGGTGTCAGAAAAGTTACCACAGGGA</td><td>63.3</td><td>-9.8</td><td>2.3</td><td>-4.4</td><td>1.3E-05</td><td>2.2E-05</td><td>MB231/MB468</td><td>MP</td></tr>
<tr><td>>hg38_21_+_8433172_8446622_RNA45SN1_45S_50ntFlanks@12432.12454.23|CCTATCATTGTGAAGCAGAATTC</td><td>>hg38_21_+_8433172_8446622_RNA45SN1_45S_50ntFlanks</td><td>12432</td><td>12454</td><td>23</td><td>CCTATCATTGTGAAGCAGAATTC</td><td>140.4</td><td>-4.4</td><td>1.0</td><td>-4.3</td><td>1.5E-05</td><td>2.2E-05</td><td>MB231/MB468</td><td>NUC</td></tr>
<tr><td>>hg38_21_+_8433172_8446622_RNA45SN1_45S_50ntFlanks@12996.13014.19|TCGCTGCGATCTATTGAAA</td><td>>hg38_21_+_8433172_8446622_RNA45SN1_45S_50ntFlanks</td><td>12996</td><td>13014</td><td>19</td><td>TCGCTGCGATCTATTGAAA</td><td>1720.0</td><td>2.7</td><td>0.6</td><td>4.3</td><td>1.5E-05</td><td>2.2E-05</td><td>MB231/MB468</td><td>NUC</td></tr>
<tr><td>>hg38_21_+_8433172_8446622_RNA45SN1_45S_50ntFlanks@12996.13015.20|TCGCTGCGATCTATTGAAAG</td><td>>hg38_21_+_8433172_8446622_RNA45SN1_45S_50ntFlanks</td><td>12996</td><td>13015</td><td>20</td><td>TCGCTGCGATCTATTGAAAG</td><td>698.4</td><td>2.3</td><td>0.5</td><td>4.3</td><td>1.5E-05</td><td>2.2E-05</td><td>MB231/MB468</td><td>NUC</td></tr>
<tr><td>>hg38_21_+_8433172_8446622_RNA45SN1_45S_50ntFlanks@9143.9163.21|GGGCGGACTGTCCCCAGTGCG</td><td>>hg38_21_+_8433172_8446622_RNA45SN1_45S_50ntFlanks</td><td>9143</td><td>9163</td><td>21</td><td>GGGCGGACTGTCCCCAGTGCG</td><td>199.6</td><td>-6.7</td><td>1.5</td><td>-4.6</td><td>5.0E-06</td><td>2.2E-05</td><td>BT20/MB468</td><td>MP</td></tr>
<tr><td>>hg38_MT_+_1621_3279_RNR2_16S_50ntFlanks@51.95.45|GCTAAACCTAGCCCCAAACCCACTCCACCTTACTACCAGACAACC</td><td>>hg38_MT_+_1621_3279_RNR2_16S_50ntFlanks</td><td>51</td><td>95</td><td>45</td><td>GCTAAACCTAGCCCCAAACCCACTCCACCTTACTACCAGACAACC</td><td>12365.8</td><td>5.4</td><td>1.3</td><td>4.3</td><td>1.6E-05</td><td>2.2E-05</td><td>MB231/MB468</td><td>MITO</td></tr>
<tr><td>>hg38_21_+_8433172_8446622_RNA45SN1_45S_50ntFlanks@11894.11916.23|ACTCTAGTCTGGCACGGTGAAGA</td><td>>hg38_21_+_8433172_8446622_RNA45SN1_45S_50ntFlanks</td><td>11894</td><td>11916</td><td>23</td><td>ACTCTAGTCTGGCACGGTGAAGA</td><td>241.9</td><td>-3.6</td><td>0.8</td><td>-4.3</td><td>1.5E-05</td><td>2.2E-05</td><td>MB231/MB468</td><td>CYTO</td></tr>
<tr><td>>trna32_LysCTT_16_-_3207406_3207478@1.33.33|GCCCGGCTAGCTCAGTCGGTAGAGCATGAGACC</td><td>>trna32_LysCTT_16_-_3207406_3207478</td><td>1</td><td>33</td><td>33</td><td>GCCCGGCTAGCTCAGTCGGTAGAGCATGAGACC</td><td>472.0</td><td>4.5</td><td>1.0</td><td>4.4</td><td>1.3E-05</td><td>2.2E-05</td><td>MB231/MB468</td><td>MP</td></tr>
<tr><td>>hg38_21_+_8433172_8446622_RNA45SN1_45S_50ntFlanks@9198.9224.27|GTTCTCTCGGGGCCACGCGCGCGTCCC</td><td>>hg38_21_+_8433172_8446622_RNA45SN1_45S_50ntFlanks</td><td>9198</td><td>9224</td><td>27</td><td>GTTCTCTCGGGGCCACGCGCGCGTCCC</td><td>326.1</td><td>-5.7</td><td>1.3</td><td>-4.4</td><td>1.1E-05</td><td>2.2E-05</td><td>BT20/MB468</td><td>MITO</td></tr>
<tr><td>>am_MI0000086|hsa-mir-28&WithFlank&3|+|188688775|188688872@60.77.18(+1C)&[MIMAT0004502&hsa-miR-28-3p&offsets|0|-4(+1C);m-28&3|+|188688834|188688855&offsets|0|-4(+1C)]|CACTAGATTGTGAGCTCCC</td><td>>am_MI0000086|hsa-mir-28&WithFlank&3|+|188688775|188688872</td><td>60</td><td>77</td><td>18(+1C)</td><td>CACTAGATTGTGAGCTCCC</td><td>47.0</td><td>9.3</td><td>2.0</td><td>4.6</td><td>5.0E-06</td><td>2.2E-05</td><td>BT20/MB468</td><td>MP</td></tr>
<tr><td>>trna37_ValAAC_6_+_27203288_27203360@1.29.29|GTTTCCGTAGTGTAGTGGTTATCACGTTT</td><td>>trna37_ValAAC_6_+_27203288_27203360</td><td>1</td><td>29</td><td>29</td><td>GTTTCCGTAGTGTAGTGGTTATCACGTTT</td><td>589.2</td><td>-4.3</td><td>1.0</td><td>-4.3</td><td>1.6E-05</td><td>2.2E-05</td><td>MB231/MB468</td><td>MITO</td></tr>
<tr><td>>hg38_1_-_228634819_228635039_RNA5S12_5S_50ntFlanks@51.76.26|GTCTACGGCCATACCACCCTGAACGC</td><td>>hg38_1_-_228634819_228635039_RNA5S12_5S_50ntFlanks</td><td>51</td><td>76</td><td>26</td><td>GTCTACGGCCATACCACCCTGAACGC</td><td>495.7</td><td>3.5</td><td>0.8</td><td>4.3</td><td>1.5E-05</td><td>2.2E-05</td><td>MB231/MB468</td><td>NUC</td></tr>
<tr><td>>hg38_21_+_8433172_8446622_RNA45SN1_45S_50ntFlanks@12717.12753.37|CCTCGGTTGGCCTCGGATAGCCGGTCCCCCGCCTGTC</td><td>>hg38_21_+_8433172_8446622_RNA45SN1_45S_50ntFlanks</td><td>12717</td><td>12753</td><td>37</td><td>CCTCGGTTGGCCTCGGATAGCCGGTCCCCCGCCTGTC</td><td>170.4</td><td>-2.2</td><td>0.5</td><td>-4.4</td><td>1.1E-05</td><td>2.2E-05</td><td>BT20/MB468</td><td>MITO</td></tr>
<tr><td>>hg38_21_+_8433172_8446622_RNA45SN1_45S_50ntFlanks@10783.10800.18|GTGAACAGCCTCTGGCAT</td><td>>hg38_21_+_8433172_8446622_RNA45SN1_45S_50ntFlanks</td><td>10783</td><td>10800</td><td>18</td><td>GTGAACAGCCTCTGGCAT</td><td>46.0</td><td>-8.6</td><td>1.9</td><td>-4.6</td><td>5.0E-06</td><td>2.2E-05</td><td>BT20/MB468</td><td>MP</td></tr>
<tr><td>>hg38_21_+_8433172_8446622_RNA45SN1_45S_50ntFlanks@11729.11751.23|TAACTATGACTCTCTTAAGGTAG</td><td>>hg38_21_+_8433172_8446622_RNA45SN1_45S_50ntFlanks</td><td>11729</td><td>11751</td><td>23</td><td>TAACTATGACTCTCTTAAGGTAG</td><td>5435.9</td><td>-8.0</td><td>1.8</td><td>-4.6</td><td>5.0E-06</td><td>2.2E-05</td><td>BT20/MB468</td><td>MP</td></tr>
</table>

<table>
<tr><td>>hg38_21_+_8433172_8446622_RNA45SN1_45S_50ntFlanks@5472.5501.30|ACGGCCCTGGCGGAGCGCTGAGAAGACGGT</td><td>>hg38_21_+_8433172_8446622_RNA45SN1_45S_50ntFlanks</td><td>5472</td><td>5501</td><td>30</td><td>ACGGCCCTGGCGGAGCGCTGAGAAGACGGT</td><td>567.9</td><td>4.5</td><td>1.0</td><td>4.6</td><td>5.0E-06</td><td>2.2E-05</td><td>BT20/MB468</td><td>MP</td></tr>
<tr><td>>hg38_21_+_8433172_8446622_RNA45SN1_45S_50ntFlanks@9915.9937.23|AAAAGGTGTTGGTTGATATAGAC</td><td>>hg38_21_+_8433172_8446622_RNA45SN1_45S_50ntFlanks</td><td>9915</td><td>9937</td><td>23</td><td>AAAAGGTGTTGGTTGATATAGAC</td><td>37.8</td><td>9.0</td><td>2.0</td><td>4.6</td><td>5.0E-06</td><td>2.2E-05</td><td>BT20/MB468</td><td>MP</td></tr>
<tr><td>>hg38_21_+_8433172_8446622_RNA45SN1_45S_50ntFlanks@12241.12268.28|CAGAAGGGCAAAAGCTCGCTTGATCTTG</td><td>>hg38_21_+_8433172_8446622_RNA45SN1_45S_50ntFlanks</td><td>12241</td><td>12268</td><td>28</td><td>CAGAAGGGCAAAAGCTCGCTTGATCTTG</td><td>186.1</td><td>-5.7</td><td>1.3</td><td>-4.4</td><td>1.1E-05</td><td>2.2E-05</td><td>BT20/MB468</td><td>MITO</td></tr>
<tr><td>>hg38_21_+_8433172_8446622_RNA45SN1_45S_50ntFlanks@12678.12703.26|AATCCCGCCCAGGCGGAACGATACGG</td><td>>hg38_21_+_8433172_8446622_RNA45SN1_45S_50ntFlanks</td><td>12678</td><td>12703</td><td>26</td><td>AATCCCGCCCAGGCGGAACGATACGG</td><td>305.5</td><td>-4.6</td><td>1.0</td><td>-4.4</td><td>1.1E-05</td><td>2.2E-05</td><td>BT20/MB468</td><td>MITO</td></tr>
<tr><td>>MI0000076|hsa-mir-20a&WithFlank&13|+|91351059|91351141@14.36.23&[MIMAT0000075&hsa-miR-20a-5p&offsets|0|0;m-96&13|+|91351072|91351094&offsets|0|0]|TAAAGTGCTTATAGTGCAGGTAG</td><td>>MI0000076|hsa-mir-20a&WithFlank&13|+|91351059|91351141</td><td>14</td><td>36</td><td>23</td><td>TAAAGTGCTTATAGTGCAGGTAG</td><td>20246.3</td><td>2.7</td><td>0.6</td><td>4.4</td><td>1.3E-05</td><td>2.2E-05</td><td>MB231/MB468</td><td>MP</td></tr>
<tr><td>>hg38_21_+_8433172_8446622_RNA45SN1_45S_50ntFlanks@12416.12435.20|TTCGATGTCGGCTCTTCCTA</td><td>>hg38_21_+_8433172_8446622_RNA45SN1_45S_50ntFlanks</td><td>12416</td><td>12435</td><td>20</td><td>TTCGATGTCGGCTCTTCCTA</td><td>184.7</td><td>-6.2</td><td>1.4</td><td>-4.4</td><td>1.3E-05</td><td>2.2E-05</td><td>MB231/MB468</td><td>MP</td></tr>
<tr><td>>hg38_21_+_8433172_8446622_RNA45SN1_45S_50ntFlanks@11573.11593.21|CGACTTAGAACTGGTGCGGAC</td><td>>hg38_21_+_8433172_8446622_RNA45SN1_45S_50ntFlanks</td><td>11573</td><td>11593</td><td>21</td><td>CGACTTAGAACTGGTGCGGAC</td><td>573.2</td><td>-5.5</td><td>1.3</td><td>-4.4</td><td>1.3E-05</td><td>2.2E-05</td><td>MB231/MB468</td><td>MP</td></tr>
<tr><td>>hg38_21_+_8433172_8446622_RNA45SN1_45S_50ntFlanks@10868.10886.19|AAGGGCTGGGTCGGTCGGG</td><td>>hg38_21_+_8433172_8446622_RNA45SN1_45S_50ntFlanks</td><td>10868</td><td>10886</td><td>19</td><td>AAGGGCTGGGTCGGTCGGG</td><td>116.4</td><td>-6.4</td><td>1.5</td><td>-4.4</td><td>1.3E-05</td><td>2.2E-05</td><td>MB231/MB468</td><td>MP</td></tr>
<tr><td>>trna10_GlyTCC_17_+_8124866_8124937@1.30.30|GCGTTGGTGGTATAGTGGTAAGCATAGCTG</td><td>>trna10_GlyTCC_17_+_8124866_8124937</td><td>1</td><td>30</td><td>30</td><td>GCGTTGGTGGTATAGTGGTAAGCATAGCTG</td><td>342.0</td><td>-6.8</td><td>1.5</td><td>-4.6</td><td>5.0E-06</td><td>2.2E-05</td><td>BT20/MB468</td><td>MP</td></tr>
<tr><td>>hg38_21_+_8433172_8446622_RNA45SN1_45S_50ntFlanks@10401.10431.31|GAACATGGGTCAGTCGGTCCTGAGAGATGGG</td><td>>hg38_21_+_8433172_8446622_RNA45SN1_45S_50ntFlanks</td><td>10401</td><td>10431</td><td>31</td><td>GAACATGGGTCAGTCGGTCCTGAGAGATGGG</td><td>331.6</td><td>-3.6</td><td>0.8</td><td>-4.3</td><td>1.6E-05</td><td>2.2E-05</td><td>MB231/MB468</td><td>NUC</td></tr>
<tr><td>>trna111_HisGTG_1_-_147774845_147774916@-1T.23.24,trna118_HisGTG_1_-_145396881_145396952@-1T.23.24,trna16_HisGTG_1_+_146544773_146544844@-1T.23.24,trna1_HisGTG_15_+_45493349_45493420@-1T.23.24,trna21_HisGTG_1_+_147753471_147753542@-1T.23.24,trna33_HisGTG_6_+_27125906_27125977@-1T.23.24,trna7_HisGTG_9_-_14433938_14434009@-1T.23.24,trna8_HisGTG_15_-_45492611_45492682@-1T.23.24,trna9_HisGTG_15_-_45490804_45490875@-1T.23.24|TGCCGTGATCGTATAGTGGTTAGT</td><td>>trna111_HisGTG_1_-_147774845_147774916</td><td>-1T</td><td>23</td><td>24</td><td>TGCCGTGATCGTATAGTGGTTAGT</td><td>923.6</td><td>8.0</td><td>1.8</td><td>4.4</td><td>1.3E-05</td><td>2.2E-05</td><td>MB231/MB468</td><td>MP</td></tr>
<tr><td>>hg38_21_+_8433172_8446622_RNA45SN1_45S_50ntFlanks@6663.6689.27|GTGGATCACTCGGCTCGTGCGTCGATG</td><td>>hg38_21_+_8433172_8446622_RNA45SN1_45S_50ntFlanks</td><td>6663</td><td>6689</td><td>27</td><td>GTGGATCACTCGGCTCGTGCGTCGATG</td><td>232.1</td><td>3.3</td><td>0.7</td><td>4.8</td><td>1.8E-06</td><td>2.3E-05</td><td>BT20/MB231</td><td>MITO</td></tr>
<tr><td>>MI0000272|hsa-mir-182&WithFlank&7|-|129770377|129770498@29.48.20&[MIMAT0000259&hsa-miR-182-5p&offsets|0|-4;m-22&7|-|129770449|129770470&offsets|0|-2]|TTTGGCAATGGTAGAACTCA</td><td>>MI0000272|hsa-mir-182&WithFlank&7|-|129770377|129770498</td><td>29</td><td>48</td><td>20</td><td>TTTGGCAATGGTAGAACTCA</td><td>215.4</td><td>5.0</td><td>1.0</td><td>4.8</td><td>1.6E-06</td><td>2.3E-05</td><td>BT20/MB231</td><td>CYTO</td></tr>
<tr><td>>MI0000272|hsa-mir-182&WithFlank&7|-|129770377|129770498@29.50.22(+2U)&[MIMAT0000259&hsa-miR-182-5p&offsets|0|-2(+2U);m-22&7|-|129770449|129770470&offsets|0|0(+2U)]|TTTGGCAATGGTAGAACTCACATT</td><td>>MI0000272|hsa-mir-182&WithFlank&7|-|129770377|129770498</td><td>29</td><td>50</td><td>22(+2U)</td><td>TTTGGCAATGGTAGAACTCACATT</td><td>303.2</td><td>5.3</td><td>1.1</td><td>4.8</td><td>1.6E-06</td><td>2.3E-05</td><td>BT20/MB231</td><td>CYTO</td></tr>
<tr><td>>hg38_21_+_8433172_8446622_RNA45SN1_45S_50ntFlanks@11578.11613.36|TAGAACTGGTGCGGACCAGGGGAATCCGACTGTTTA</td><td>>hg38_21_+_8433172_8446622_RNA45SN1_45S_50ntFlanks</td><td>11578</td><td>11613</td><td>36</td><td>TAGAACTGGTGCGGACCAGGGGAATCCGACTGTTTA</td><td>1259.5</td><td>5.2</td><td>1.1</td><td>4.8</td><td>1.8E-06</td><td>2.3E-05</td><td>BT20/MB231</td><td>MITO</td></tr>
<tr><td>>hg38_21_+_8433172_8446622_RNA45SN1_45S_50ntFlanks@4730.4767.38|CAAGAACGAAAGTCGGAGGTTCGAAGACGATCAGATAC</td><td>>hg38_21_+_8433172_8446622_RNA45SN1_45S_50ntFlanks</td><td>4730</td><td>4767</td><td>38</td><td>CAAGAACGAAAGTCGGAGGTTCGAAGACGATCAGATAC</td><td>140.1</td><td>5.2</td><td>1.2</td><td>4.3</td><td>1.6E-05</td><td>2.3E-05</td><td>MB231/MB468</td><td>NUC</td></tr>
<tr><td>>hg38_MT_+_1621_3279_RNR2_16S_50ntFlanks@51.91.41|GCTAAACCTAGCCCCAAACCCACTCCACCTTACTACCAGAC</td><td>>hg38_MT_+_1621_3279_RNR2_16S_50ntFlanks</td><td>51</td><td>91</td><td>41</td><td>GCTAAACCTAGCCCCAAACCCACTCCACCTTACTACCAGAC</td><td>1424.9</td><td>4.6</td><td>1.1</td><td>4.3</td><td>1.6E-05</td><td>2.3E-05</td><td>MB231/MB468</td><td>MITO</td></tr>
<tr><td>>hg38_21_+_8433172_8446622_RNA45SN1_45S_50ntFlanks@3899.3919.21|CCTTCGCGGGGGGGATGCGTG</td><td>>hg38_21_+_8433172_8446622_RNA45SN1_45S_50ntFlanks</td><td>3899</td><td>3919</td><td>21</td><td>CCTTCGCGGGGGGGATGCGTG</td><td>4760.2</td><td>-8.0</td><td>1.8</td><td>-4.6</td><td>5.1E-06</td><td>2.3E-05</td><td>BT20/MB468</td><td>MP</td></tr>
<tr><td>>hg38_21_+_8433172_8446622_RNA45SN1_45S_50ntFlanks@5071.5105.35|TTAATTCCGATAACGAACGAGACTCTGGCATGCTA</td><td>>hg38_21_+_8433172_8446622_RNA45SN1_45S_50ntFlanks</td><td>5071</td><td>5105</td><td>35</td><td>TTAATTCCGATAACGAACGAGACTCTGGCATGCTA</td><td>80.0</td><td>-10.2</td><td>2.3</td><td>-4.4</td><td>1.3E-05</td><td>2.3E-05</td><td>MB231/MB468</td><td>MP</td></tr>
<tr><td>>hg38_21_+_8433172_8446622_RNA45SN1_45S_50ntFlanks@10806.10840.35|AACAATGTAGGTAAGGGAAGTCGGCAAGCCGGATC</td><td>>hg38_21_+_8433172_8446622_RNA45SN1_45S_50ntFlanks</td><td>10806</td><td>10840</td><td>35</td><td>AACAATGTAGGTAAGGGAAGTCGGCAAGCCGGATC</td><td>478.2</td><td>-4.0</td><td>0.9</td><td>-4.3</td><td>1.5E-05</td><td>2.3E-05</td><td>MB231/MB468</td><td>CYTO</td></tr>
<tr><td>>hg38_21_+_8433172_8446622_RNA45SN1_45S_50ntFlanks@12644.12661.18|TCTGTGGGATTATGACTG</td><td>>hg38_21_+_8433172_8446622_RNA45SN1_45S_50ntFlanks</td><td>12644</td><td>12661</td><td>18</td><td>TCTGTGGGATTATGACTG</td><td>1263.5</td><td>-3.4</td><td>0.8</td><td>-4.4</td><td>1.1E-05</td><td>2.3E-05</td><td>BT20/MB468</td><td>MITO</td></tr>
<tr><td>>hg38_21_+_8433172_8446622_RNA45SN1_45S_50ntFlanks@11569.11604.36|CAGCCGACTTAGAACTGGTGCGGACCAGGGGAATCC</td><td>>hg38_21_+_8433172_8446622_RNA45SN1_45S_50ntFlanks</td><td>11569</td><td>11604</td><td>36</td><td>CAGCCGACTTAGAACTGGTGCGGACCAGGGGAATCC</td><td>211.2</td><td>-2.8</td><td>0.7</td><td>-4.3</td><td>1.6E-05</td><td>2.3E-05</td><td>MB231/MB468</td><td>NUC</td></tr>
<tr><td>>MI0000742|hsa-mir-34b&WithFlank&11|+|111512932|111513027@57.78.22&[MIMAT0004676&hsa-miR-34b-3p&offsets|+1|+1;m-585&11|+|111512988|111513009&offsets|0|0]|AATCACTAACTCCACTGCCATC</td><td>>MI0000742|hsa-mir-34b&WithFlank&11|+|111512932|111513027</td><td>57</td><td>78</td><td>22</td><td>AATCACTAACTCCACTGCCATC</td><td>33.0</td><td>8.3</td><td>1.9</td><td>4.4</td><td>1.3E-05</td><td>2.3E-05</td><td>MB231/MB468</td><td>MP</td></tr>
<tr><td>>hg38_21_+_8433172_8446622_RNA45SN1_45S_50ntFlanks@12626.12667.42|AATGGGGCGAAGCTACCATCTGTGGGATTATGACTGAACGCC</td><td>>hg38_21_+_8433172_8446622_RNA45SN1_45S_50ntFlanks</td><td>12626</td><td>12667</td><td>42</td><td>AATGGGGCGAAGCTACCATCTGTGGGATTATGACTGAACGCC</td><td>487.6</td><td>4.8</td><td>1.1</td><td>4.3</td><td>1.6E-05</td><td>2.3E-05</td><td>MB231/MB468</td><td>NUC</td></tr>
<tr><td>>hg38_21_+_8433172_8446622_RNA45SN1_45S_50ntFlanks@5473.5491.19|CGGCCCTGGCGGAGCGCTG</td><td>>hg38_21_+_8433172_8446622_RNA45SN1_45S_50ntFlanks</td><td>5473</td><td>5491</td><td>19</td><td>CGGCCCTGGCGGAGCGCTG</td><td>1276.8</td><td>4.5</td><td>1.0</td><td>4.4</td><td>1.4E-05</td><td>2.3E-05</td><td>BT20/MB468</td><td>CYTO</td></tr>
<tr><td>>hg38_21_+_8433172_8446622_RNA45SN1_45S_50ntFlanks@9604.9621.18|AAAGACTAATCGAACCAT</td><td>>hg38_21_+_8433172_8446622_RNA45SN1_45S_50ntFlanks</td><td>9604</td><td>9621</td><td>18</td><td>AAAGACTAATCGAACCAT</td><td>652.6</td><td>-6.7</td><td>1.5</td><td>-4.4</td><td>1.3E-05</td><td>2.3E-05</td><td>MB231/MB468</td><td>MP</td></tr>
<tr><td>>hg38_21_+_8433172_8446622_RNA45SN1_45S_50ntFlanks@4243.4263.21|CTTTCGAGGCCCTGTAATTGG</td><td>>hg38_21_+_8433172_8446622_RNA45SN1_45S_50ntFlanks</td><td>4243</td><td>4263</td><td>21</td><td>CTTTCGAGGCCCTGTAATTGG</td><td>1035.5</td><td>-6.1</td><td>1.4</td><td>-4.3</td><td>1.7E-05</td><td>2.3E-05</td><td>BT20/MB468</td><td>NUC</td></tr>
<tr><td>>MI0000068|hsa-let-7f-2&WithFlank&X|-|53557186|53557280@14.37.24&[MIMAT0000067_1&hsa-let-7f-5p&offsets|0|+2;m-19&X|-|53557246|53557267&offsets|0|+2]|TGAGGTAGTAGATTGTATAGTTTT</td><td>>MI0000068|hsa-let-7f-2&WithFlank&X|-|53557186|53557280</td><td>14</td><td>37</td><td>24</td><td>TGAGGTAGTAGATTGTATAGTTTT</td><td>43.8</td><td>8.7</td><td>2.0</td><td>4.4</td><td>1.3E-05</td><td>2.3E-05</td><td>MB231/MB468</td><td>MP</td></tr>
<tr><td>>hg38_21_+_8433172_8446622_RNA45SN1_45S_50ntFlanks@12995.13039.45|CTCGCTGCGATCTATTGAAAGTCAGCCCTCGACACAAGGGTTTGT</td><td>>hg38_21_+_8433172_8446622_RNA45SN1_45S_50ntFlanks</td><td>12995</td><td>13039</td><td>45</td><td>CTCGCTGCGATCTATTGAAAGTCAGCCCTCGACACAAGGGTTTGT</td><td>240464.0</td><td>5.4</td><td>1.2</td><td>4.3</td><td>1.6E-05</td><td>2.3E-05</td><td>MB231/MB468</td><td>NUC</td></tr>
<tr><td>>hg38_21_+_8433172_8446622_RNA45SN1_45S_50ntFlanks@7982.8008.27|TCAGATCAGACGTGGCGACCCGCTGAA</td><td>>hg38_21_+_8433172_8446622_RNA45SN1_45S_50ntFlanks</td><td>7982</td><td>8008</td><td>27</td><td>TCAGATCAGACGTGGCGACCCGCTGAA</td><td>360.4</td><td>-2.8</td><td>0.6</td><td>-4.3</td><td>1.6E-05</td><td>2.3E-05</td><td>MB231/MB468</td><td>NUC</td></tr>
<tr><td>>hg38_21_+_8433172_8446622_RNA45SN1_45S_50ntFlanks@11540.11584.45|CGCGGCGCCTCGCCTCGGCCGGCGCCTAGCAGCCGACTTAGAACT</td><td>>hg38_21_+_8433172_8446622_RNA45SN1_45S_50ntFlanks</td><td>11540</td><td>11584</td><td>45</td><td>CGCGGCGCCTCGCCTCGGCCGGCGCCTAGCAGCCGACTTAGAACT</td><td>1219.0</td><td>5.0</td><td>1.1</td><td>4.4</td><td>1.2E-05</td><td>2.3E-05</td><td>BT20/MB468</td><td>MITO</td></tr>
<tr><td>>hg38_21_+_8433172_8446622_RNA45SN1_45S_50ntFlanks@4002.4022.21|GATAACCTCGGGCCGATCGCA</td><td>>hg38_21_+_8433172_8446622_RNA45SN1_45S_50ntFlanks</td><td>4002</td><td>4022</td><td>21</td><td>GATAACCTCGGGCCGATCGCA</td><td>529.4</td><td>-2.5</td><td>0.6</td><td>-4.3</td><td>1.5E-05</td><td>2.3E-05</td><td>MB231/MB468</td><td>CYTO</td></tr>
<tr><td>>hg38_21_+_8433172_8446622_RNA45SN1_45S_50ntFlanks@7968.8009.42|TCCGAGACGCGACCTCAGATCAGACGTGGCGACCCGCTGAAT</td><td>>hg38_21_+_8433172_8446622_RNA45SN1_45S_50ntFlanks</td><td>7968</td><td>8009</td><td>42</td><td>TCCGAGACGCGACCTCAGATCAGACGTGGCGACCCGCTGAAT</td><td>313.8</td><td>4.0</td><td>0.9</td><td>4.3</td><td>1.7E-05</td><td>2.3E-05</td><td>BT20/MB468</td><td>NUC</td></tr>
<tr><td>>hg38_21_+_8433172_8446622_RNA45SN1_45S_50ntFlanks@6729.6770.42|GACACATTGATCATCGACACTTCGAACGCACTTGCGGCCCCG</td><td>>hg38_21_+_8433172_8446622_RNA45SN1_45S_50ntFlanks</td><td>6729</td><td>6770</td><td>42</td><td>GACACATTGATCATCGACACTTCGAACGCACTTGCGGCCCCG</td><td>512.0</td><td>-3.7</td><td>0.9</td><td>-4.3</td><td>1.6E-05</td><td>2.3E-05</td><td>MB231/MB468</td><td>NUC</td></tr>
<tr><td>>hg38_21_+_8433172_8446622_RNA45SN1_45S_50ntFlanks@9300.9324.25|ACGGACCAAGGAGTCTAACACGTGC</td><td>>hg38_21_+_8433172_8446622_RNA45SN1_45S_50ntFlanks</td><td>9300</td><td>9324</td><td>25</td><td>ACGGACCAAGGAGTCTAACACGTGC</td><td>87.9</td><td>9.8</td><td>2.0</td><td>4.8</td><td>1.5E-06</td><td>2.3E-05</td><td>BT20/MB231</td><td>MP</td></tr>
<tr><td>>MI0000072|hsa-mir-18a&WithFlank&13|+|91350745|91350827@12.34.23&[MIMAT0000072&hsa-miR-18a-5p&offsets|0|0;m-262&13|+|91350756|91350778&offsets|0|0]|TAAGGTGCATCTAGTGCAGATAG</td><td>>MI0000072|hsa-mir-18a&WithFlank&13|+|91350745|91350827</td><td>12</td><td>34</td><td>23</td><td>TAAGGTGCATCTAGTGCAGATAG</td><td>556.6</td><td>-4.0</td><td>0.8</td><td>-4.8</td><td>1.8E-06</td><td>2.3E-05</td><td>BT20/MB231</td><td>MITO</td></tr>
<tr><td>>hg38_21_+_8433172_8446622_RNA45SN1_45S_50ntFlanks@12416.12447.32|TTCGATGTCGGCTCTTCCTATCATTGTGAAGC</td><td>>hg38_21_+_8433172_8446622_RNA45SN1_45S_50ntFlanks</td><td>12416</td><td>12447</td><td>32</td><td>TTCGATGTCGGCTCTTCCTATCATTGTGAAGC</td><td>499.5</td><td>5.3</td><td>1.2</td><td>4.3</td><td>1.6E-05</td><td>2.3E-05</td><td>MB231/MB468</td><td>NUC</td></tr>
<tr><td>>MI0000254|hsa-mir-30c-2&WithFlank&6|-|71376954|71377037@53.74.22&[MIMAT0004550&hsa-miR-30c-2-3p&offsets|0|0;m-206&6|-|71376964|71376985&offsets|0|0]|CTGGGAGAAGGCTGTTTACTCT</td><td>>MI0000254|hsa-mir-30c-2&WithFlank&6|-|71376954|71377037</td><td>53</td><td>74</td><td>22</td><td>CTGGGAGAAGGCTGTTTACTCT</td><td>270.9</td><td>1.7</td><td>0.4</td><td>4.3</td><td>1.7E-05</td><td>2.3E-05</td><td>BT20/MB468</td><td>NUC</td></tr>
<tr><td>>hg38_21_+_8433172_8446622_RNA45SN1_45S_50ntFlanks@4005.4023.19|AACCTCGGGCCGATCGCAC</td><td>>hg38_21_+_8433172_8446622_RNA45SN1_45S_50ntFlanks</td><td>4005</td><td>4023</td><td>19</td><td>AACCTCGGGCCGATCGCAC</td><td>1500.6</td><td>-4.0</td><td>0.9</td><td>-4.3</td><td>1.7E-05</td><td>2.3E-05</td><td>BT20/MB468</td><td>NUC</td></tr>
<tr><td>>hg38_21_+_8433172_8446622_RNA45SN1_45S_50ntFlanks@11578.11603.26|TAGAACTGGTGCGGACCAGGGGAATC</td><td>>hg38_21_+_8433172_8446622_RNA45SN1_45S_50ntFlanks</td><td>11578</td><td>11603</td><td>26</td><td>TAGAACTGGTGCGGACCAGGGGAATC</td><td>809.5</td><td>-4.3</td><td>0.9</td><td>-4.6</td><td>5.1E-06</td><td>2.3E-05</td><td>BT20/MB468</td><td>MP</td></tr>
<tr><td>>hg38_21_+_8433172_8446622_RNA45SN1_45S_50ntFlanks@10389.10424.36|TGAACAGCAGTTGAACATGGGTCAGTCGGTCCTGAG</td><td>>hg38_21_+_8433172_8446622_RNA45SN1_45S_50ntFlanks</td><td>10389</td><td>10424</td><td>36</td><td>TGAACAGCAGTTGAACATGGGTCAGTCGGTCCTGAG</td><td>17898.3</td><td>4.2</td><td>0.9</td><td>4.8</td><td>1.8E-06</td><td>2.3E-05</td><td>BT20/MB231</td><td>MITO</td></tr>
<tr><td>>MI0000071|hsa-mir-17&WithFlank&13|+|91350599|91350694@20.40.21&[MIMAT0000070&hsa-miR-17-5p&offsets|0|-2;m-88&13|+|91350618|91350640&offsets|0|-2]|CAAAGTGCTTACAGTGCAGGT</td><td>>MI0000071|hsa-mir-17&WithFlank&13|+|91350599|91350694</td><td>20</td><td>40</td><td>21</td><td>CAAAGTGCTTACAGTGCAGGT</td><td>374.2</td><td>4.5</td><td>1.0</td><td>4.3</td><td>1.5E-05</td><td>2.3E-05</td><td>MB231/MB468</td><td>CYTO</td></tr>
<tr><td>>hg38_21_+_8433172_8446622_RNA45SN1_45S_50ntFlanks@11729.11766.38|TAACTATGACTCTCTTAAGGTAGCCAAATGCCTCGTCA</td><td>>hg38_21_+_8433172_8446622_RNA45SN1_45S_50ntFlanks</td><td>11729</td><td>11766</td><td>38</td><td>TAACTATGACTCTCTTAAGGTAGCCAAATGCCTCGTCA</td><td>575.2</td><td>-6.5</td><td>1.5</td><td>-4.3</td><td>1.4E-05</td><td>2.3E-05</td><td>BT20/MB468</td><td>CYTO</td></tr>
<tr><td>>hg38_21_+_8433172_8446622_RNA45SN1_45S_50ntFlanks@8170.8199.30|CAAGTCCTTCTGATCGAGGCCCAGCCCGTG</td><td>>hg38_21_+_8433172_8446622_RNA45SN1_45S_50ntFlanks</td><td>8170</td><td>8199</td><td>30</td><td>CAAGTCCTTCTGATCGAGGCCCAGCCCGTG</td><td>299.5</td><td>-6.5</td><td>1.5</td><td>-4.3</td><td>1.6E-05</td><td>2.3E-05</td><td>MB231/MB468</td><td>NUC</td></tr>
<tr><td>>hg38_21_+_8433172_8446622_RNA45SN1_45S_50ntFlanks@7974.8005.32|ACGCGACCTCAGATCAGACGTGGCGACCCGCT</td><td>>hg38_21_+_8433172_8446622_RNA45SN1_45S_50ntFlanks</td><td>7974</td><td>8005</td><td>32</td><td>ACGCGACCTCAGATCAGACGTGGCGACCCGCT</td><td>1747.7</td><td>2.8</td><td>0.7</td><td>4.3</td><td>1.6E-05</td><td>2.3E-05</td><td>MB231/MB468</td><td>NUC</td></tr>
<tr><td>>hg38_21_+_8433172_8446622_RNA45SN1_45S_50ntFlanks@12241.12265.25|CAGAAGGGCAAAAGCTCGCTTGATC</td><td>>hg38_21_+_8433172_8446622_RNA45SN1_45S_50ntFlanks</td><td>12241</td><td>12265</td><td>25</td><td>CAGAAGGGCAAAAGCTCGCTTGATC</td><td>393.0</td><td>-5.4</td><td>1.2</td><td>-4.3</td><td>1.6E-05</td><td>2.3E-05</td><td>MB231/MB468</td><td>MITO</td></tr>
<tr><td>>hg38_21_+_8433172_8446622_RNA45SN1_45S_50ntFlanks@12008.12035.28|CCGCCGGTGAAATACCACTACTCTGATC</td><td>>hg38_21_+_8433172_8446622_RNA45SN1_45S_50ntFlanks</td><td>12008</td><td>12035</td><td>28</td><td>CCGCCGGTGAAATACCACTACTCTGATC</td><td>237.0</td><td>-4.5</td><td>1.1</td><td>-4.3</td><td>1.6E-05</td><td>2.3E-05</td><td>MB231/MB468</td><td>NUC</td></tr>
<tr><td>>hg38_21_+_8433172_8446622_RNA45SN1_45S_50ntFlanks@10683.10710.28|CCGGCGGCGTCCGGTGAGCTCTCGCTGG</td><td>>hg38_21_+_8433172_8446622_RNA45SN1_45S_50ntFlanks</td><td>10683</td><td>10710</td><td>28</td><td>CCGGCGGCGTCCGGTGAGCTCTCGCTGG</td><td>321.1</td><td>-6.7</td><td>1.5</td><td>-4.6</td><td>5.2E-06</td><td>2.3E-05</td><td>BT20/MB468</td><td>MP</td></tr>
<tr><td>>hg38_21_+_8433172_8446622_RNA45SN1_45S_50ntFlanks@12073.12097.25|CCGAGGGGCTCTCGCTTCTGGCGCC</td><td>>hg38_21_+_8433172_8446622_RNA45SN1_45S_50ntFlanks</td><td>12073</td><td>12097</td><td>25</td><td>CCGAGGGGCTCTCGCTTCTGGCGCC</td><td>2214.7</td><td>-3.5</td><td>0.8</td><td>-4.3</td><td>1.4E-05</td><td>2.3E-05</td><td>BT20/MB468</td><td>CYTO</td></tr>
<tr><td>>hg38_21_+_8433172_8446622_RNA45SN1_45S_50ntFlanks@11569.11612.44|CAGCCGACTTAGAACTGGTGCGGACCAGGGGAATCCGACTGTTT</td><td>>hg38_21_+_8433172_8446622_RNA45SN1_45S_50ntFlanks</td><td>11569</td><td>11612</td><td>44</td><td>CAGCCGACTTAGAACTGGTGCGGACCAGGGGAATCCGACTGTTT</td><td>850.1</td><td>-7.7</td><td>1.8</td><td>-4.4</td><td>1.3E-05</td><td>2.3E-05</td><td>MB231/MB468</td><td>MP</td></tr>
</table>

| | | | | | | | | | | | | | |
|---|---|---|---|---|---|---|---|---|---|---|---|---|---|
| >hg38_21_+_8433172_8446622_RNA45SN1_45S_50ntFlanks@12412.12435.24\|ATCCTTCGATGTCGGCTCTTCCTA | >hg38_21_+_8433172_8446622_RNA45SN1_45S_50ntFlanks | 12412 | 12435 | 24 | ATCCTTCGATGTCGGCTCTTCCTA | 1070.2 | -6.8 | 1.6 | -4.3 | 1.6E-05 | 2.3E-05 | MB231/MB468 | NUC |
| >hg38_21_+_8433172_8446622_RNA45SN1_45S_50ntFlanks@5473.5514.42\|CGGCCCTGGCGGAGCGCTGAGAAGACGGTCGAACTTGACTAT | >hg38_21_+_8433172_8446622_RNA45SN1_45S_50ntFlanks | 5473 | 5514 | 42 | CGGCCCTGGCGGAGCGCTGAGAAGACGGTCGAACTTGACTAT | 1213.0 | -6.3 | 1.5 | -4.3 | 1.7E-05 | 2.3E-05 | BT20/MB468 | NUC |
| >hg38_21_+_8433172_8446622_RNA45SN1_45S_50ntFlanks@9029.9058.30\|TCTCCTCGTGGGGGGGCCGGGCCACCCCTC | >hg38_21_+_8433172_8446622_RNA45SN1_45S_50ntFlanks | 9029 | 9058 | 30 | TCTCCTCGTGGGGGGGCCGGGCCACCCCTC | 42.5 | -8.5 | 1.9 | -4.6 | 5.2E-06 | 2.3E-05 | BT20/MB468 | MP |
| >MI0000273\|hsa-mir-183&WithFlank&7\|-\|129774899\|129775020@34.55.22&[MIMAT0000261&hsa-miR-183-5p&offsets\|+1\|+1;m-41&7\|-\|129774966\|129774987&offsets\|0\|0]\|ATGGCACTGGTAGAATTCACTG | >MI0000273\|hsa-mir-183&WithFlank&7\|-\|129774899\|129775020 | 34 | 55 | 22 | ATGGCACTGGTAGAATTCACTG | 379.1 | 3.4 | 0.8 | 4.3 | 1.5E-05 | 2.3E-05 | MB231/MB468 | CYTO |
| >hg38_21_+_8433172_8446622_RNA45SN1_45S_50ntFlanks@12396.12414.19\|GCGACGTCGCTTTTTGATC | >hg38_21_+_8433172_8446622_RNA45SN1_45S_50ntFlanks | 12396 | 12414 | 19 | GCGACGTCGCTTTTTGATC | 1879.1 | -3.3 | 0.8 | -4.3 | 1.6E-05 | 2.3E-05 | MB231/MB468 | MITO |
| >MI0003560\|hsa-mir-92b&WithFlank&1\|+\|155195171\|155195278@67.87.21(+1A)&[MIMAT0003218&hsa-miR-92b-3p&offsets\|0\|-1(+1A);m-115&1\|+\|155195237\|155195258&offsets\|0\|-1(+1A)]\|TATTGCACTCGTCCCGGCCTCA | >MI0003560\|hsa-mir-92b&WithFlank&1\|+\|155195171\|155195278 | 67 | 87 | 21(+1A) | TATTGCACTCGTCCCGGCCTCA | 609.5 | 5.9 | 1.4 | 4.4 | 1.4E-05 | 2.3E-05 | MB231/MB468 | MP |
| >hg38_21_+_8433172_8446622_RNA45SN1_45S_50ntFlanks@5186.5206.21\|CAATAACAGGTCTGTGATGCC | >hg38_21_+_8433172_8446622_RNA45SN1_45S_50ntFlanks | 5186 | 5206 | 21 | CAATAACAGGTCTGTGATGCC | 394.8 | -7.5 | 1.6 | -4.6 | 5.2E-06 | 2.3E-05 | BT20/MB468 | MP |
| >hg38_21_+_8433172_8446622_RNA45SN1_45S_50ntFlanks@6653.6692.40\|ACTCTTAGCGGTGGATCACTCGGCTCGTGCGTCGATGAAG | >hg38_21_+_8433172_8446622_RNA45SN1_45S_50ntFlanks | 6653 | 6692 | 40 | ACTCTTAGCGGTGGATCACTCGGCTCGTGCGTCGATGAAG | 193.6 | 2.6 | 0.6 | 4.3 | 1.6E-05 | 2.3E-05 | MB231/MB468 | NUC |
| >hg38_21_+_8433172_8446622_RNA45SN1_45S_50ntFlanks@9932.9949.18\|ATAGACAGCAGGACGGTG | >hg38_21_+_8433172_8446622_RNA45SN1_45S_50ntFlanks | 9932 | 9949 | 18 | ATAGACAGCAGGACGGTG | 319.1 | 3.5 | 0.8 | 4.6 | 5.2E-06 | 2.3E-05 | BT20/MB468 | MP |
| >hg38_21_+_8433172_8446622_RNA45SN1_45S_50ntFlanks@8556.8581.26\|CCGGAGGGGGCGGGCTCCGGCGGGTG | >hg38_21_+_8433172_8446622_RNA45SN1_45S_50ntFlanks | 8556 | 8581 | 26 | CCGGAGGGGGCGGGCTCCGGCGGGTG | 45.5 | -8.6 | 1.9 | -4.6 | 5.2E-06 | 2.3E-05 | BT20/MB468 | MP |
| >hg38_21_+_8433172_8446622_RNA45SN1_45S_50ntFlanks@8290.8308.19\|TAAACTCCATCTAAGGCTA | >hg38_21_+_8433172_8446622_RNA45SN1_45S_50ntFlanks | 8290 | 8308 | 19 | TAAACTCCATCTAAGGCTA | 78.9 | -10.2 | 2.3 | -4.4 | 1.4E-05 | 2.3E-05 | MB231/MB468 | MP |
| >hg38_MT_+_1621_3279_RNR2_16S_50ntFlanks@1383.1415.33\|ACATCCCGATGGTGCAGCCGCTATTAAAGGTTC | >hg38_MT_+_1621_3279_RNR2_16S_50ntFlanks | 1383 | 1415 | 33 | ACATCCCGATGGTGCAGCCGCTATTAAAGGTTC | 35.1 | 8.4 | 1.9 | 4.4 | 1.4E-05 | 2.3E-05 | MB231/MB468 | MP |
| >hg38_21_+_8433172_8446622_RNA45SN1_45S_50ntFlanks@7989.8007.19\|AGACGTGGCGACCCGCTGA | >hg38_21_+_8433172_8446622_RNA45SN1_45S_50ntFlanks | 7989 | 8007 | 19 | AGACGTGGCGACCCGCTGA | 784.0 | -3.1 | 0.7 | -4.3 | 1.7E-05 | 2.3E-05 | BT20/MB468 | NUC |
| >hg38_21_+_8433172_8446622_RNA45SN1_45S_50ntFlanks@7994.8011.18\|TGGCGACCCGCTGAATTT | >hg38_21_+_8433172_8446622_RNA45SN1_45S_50ntFlanks | 7994 | 8011 | 18 | TGGCGACCCGCTGAATTT | 275.8 | 4.7 | 1.0 | 4.8 | 1.9E-06 | 2.3E-05 | BT20/MB231 | MITO |
| >hg38_21_+_8433172_8446622_RNA45SN1_45S_50ntFlanks@4738.4759.22\|AAAGTCGGAGGTTCGAAGACGA | >hg38_21_+_8433172_8446622_RNA45SN1_45S_50ntFlanks | 4738 | 4759 | 22 | AAAGTCGGAGGTTCGAAGACGA | 1938.6 | -3.7 | 0.8 | -4.6 | 5.2E-06 | 2.3E-05 | BT20/MB468 | MP |
| >hg38_21_+_8433172_8446622_RNA45SN1_45S_50ntFlanks@12936.12970.35\|AAACCATTCGTAGACGACCTGCTTCTGGGTCGGGG | >hg38_21_+_8433172_8446622_RNA45SN1_45S_50ntFlanks | 12936 | 12970 | 35 | AAACCATTCGTAGACGACCTGCTTCTGGGTCGGGG | 43.3 | -8.5 | 1.9 | -4.6 | 5.2E-06 | 2.3E-05 | BT20/MB468 | MP |
| >hg38_21_+_8433172_8446622_RNA45SN1_45S_50ntFlanks@6645.6665.21\|CTCGTACGACTCTTAGCGGTG | >hg38_21_+_8433172_8446622_RNA45SN1_45S_50ntFlanks | 6645 | 6665 | 21 | CTCGTACGACTCTTAGCGGTG | 4784.9 | 3.3 | 0.8 | 4.3 | 1.6E-05 | 2.3E-05 | MB231/MB468 | NUC |
| >hg38_21_+_8433172_8446622_RNA45SN1_45S_50ntFlanks@6730.6752.23\|ACACATTGATCATCGACACTTCG | >hg38_21_+_8433172_8446622_RNA45SN1_45S_50ntFlanks | 6730 | 6752 | 23 | ACACATTGATCATCGACACTTCG | 197.3 | -2.4 | 0.6 | -4.3 | 1.6E-05 | 2.3E-05 | MB231/MB468 | NUC |
| >hg38_21_+_8433172_8446622_RNA45SN1_45S_50ntFlanks@11767.11803.37\|TCTAATTAGTGACGCGCATGAATGGATGAACGAGATT | >hg38_21_+_8433172_8446622_RNA45SN1_45S_50ntFlanks | 11767 | 11803 | 37 | TCTAATTAGTGACGCGCATGAATGGATGAACGAGATT | 293.3 | -6.7 | 1.5 | -4.4 | 1.4E-05 | 2.3E-05 | MB231/MB468 | MP |
| >hg38_21_+_8433172_8446622_RNA45SN1_45S_50ntFlanks@3837.3871.35\|CCTCTCCTACTTGGATAACTGTGGTAATTCTAGAG | >hg38_21_+_8433172_8446622_RNA45SN1_45S_50ntFlanks | 3837 | 3871 | 35 | CCTCTCCTACTTGGATAACTGTGGTAATTCTAGAG | 245.1 | -2.4 | 0.5 | -4.4 | 1.2E-05 | 2.3E-05 | BT20/MB468 | MITO |
| >hg38_21_+_8433172_8446622_RNA45SN1_45S_50ntFlanks@10329.10362.34\|GTGGTAGTAGCAAATATTCAAACGAGAACTTTGA | >hg38_21_+_8433172_8446622_RNA45SN1_45S_50ntFlanks | 10329 | 10362 | 34 | GTGGTAGTAGCAAATATTCAAACGAGAACTTTGA | 41.0 | 9.1 | 2.0 | 4.6 | 5.2E-06 | 2.3E-05 | BT20/MB468 | MP |
| >trnaMT_GluTTC_MT_-_14674_14742@39.57.19,trnalookalike8_GluTTC_5_-_93905172_93905240@39.57.19\|ATTGGTCGTGGTTGTAGTC | >trnaMT_GluTTC_MT_-_14674_14742 | 39 | 57 | 19 | ATTGGTCGTGGTTGTAGTC | 782.8 | 4.7 | 1.0 | 4.6 | 5.2E-06 | 2.3E-05 | BT20/MB468 | MP |
| >hg38_21_+_8433172_8446622_RNA45SN1_45S_50ntFlanks@5492.5516.25\|AGAAGACGGTCGAACTTGACTATCT | >hg38_21_+_8433172_8446622_RNA45SN1_45S_50ntFlanks | 5492 | 5516 | 25 | AGAAGACGGTCGAACTTGACTATCT | 15681.3 | 4.6 | 1.0 | 4.8 | 1.9E-06 | 2.3E-05 | BT20/MB231 | MITO |
| >hg38_21_+_8433172_8446622_RNA45SN1_45S_50ntFlanks@12936.12953.18\|AAACCATTCGTAGACGAC | >hg38_21_+_8433172_8446622_RNA45SN1_45S_50ntFlanks | 12936 | 12953 | 18 | AAACCATTCGTAGACGAC | 603.9 | -5.1 | 1.2 | -4.3 | 1.6E-05 | 2.3E-05 | MB231/MB468 | NUC |
| >MI0000085\|hsa-mir-27a&WithFlank&19\|-\|13836434\|13836523@57.76.20(+3U)&[MIMAT0000084&hsa-miR-27a-3p&offsets\|0\|-1(+3U);m-43&19\|-\|13836447\|13836467&offsets\|0\|-1(+3U)]\|TTCACAGTGGCTAAGTTCCGTTT | >MI0000085\|hsa-mir-27a&WithFlank&19\|-\|13836434\|13836523 | 57 | 76 | 20(+3U) | TTCACAGTGGCTAAGTTCCGTTT | 3382.1 | -5.4 | 1.1 | -4.8 | 1.6E-06 | 2.3E-05 | BT20/MB231 | MP |
| >hg38_21_+_8433172_8446622_RNA45SN1_45S_50ntFlanks@6648.6690.43\|GTACGACTCTTAGCGGTGGATCACTCGGCTCGTGCGTCGATGA | >hg38_21_+_8433172_8446622_RNA45SN1_45S_50ntFlanks | 6648 | 6690 | 43 | GTACGACTCTTAGCGGTGGATCACTCGGCTCGTGCGTCGATGA | 458.6 | 3.3 | 0.8 | 4.3 | 1.6E-05 | 2.3E-05 | MB231/MB468 | NUC |
| >hg38_21_+_8433172_8446622_RNA45SN1_45S_50ntFlanks@9602.9622.21\|CGAAAGACTAATCGAACCATC | >hg38_21_+_8433172_8446622_RNA45SN1_45S_50ntFlanks | 9602 | 9622 | 21 | CGAAAGACTAATCGAACCATC | 186.0 | -4.6 | 1.1 | -4.3 | 1.6E-05 | 2.3E-05 | MB231/MB468 | NUC |
| >am_trna128_GlyGCC_6_-_27870686_27870756@4.35.32,trna18_GlyGCC_16_+_70822597_70822667@4.35.32,trna19_GlyGCC_16_+_70823410_70823480@4.35.32,trna19_GlyGCC_2_-_157257659_157257729@4.35.32,trna24_GlyGCC_16_-_70812942_70813012@4.35.32,trna25_GlyGCC_16_-_70812114_70812184@4.35.32,trna5_GlyGCC_17_+_8029064_8029134@4.35.32,trna68_GlyGCC_1_-_161493637_161493707@4.35.32\|TTGGTGGTTCAGTGGTAGAATTCTCGCCTGCC | >am_trna128_GlyGCC_6_-_27870686_27870756 | 4 | 35 | 32 | TTGGTGGTTCAGTGGTAGAATTCTCGCCTGCC | 267.8 | 5.3 | 1.2 | 4.3 | 1.6E-05 | 2.3E-05 | MB231/MB468 | MITO |
| >hg38_21_+_8433172_8446622_RNA45SN1_45S_50ntFlanks@8356.8391.36\|GAAAAGAACTTTGAAGAGAGAGTTCAAGAGGGCGTG | >hg38_21_+_8433172_8446622_RNA45SN1_45S_50ntFlanks | 8356 | 8391 | 36 | GAAAAGAACTTTGAAGAGAGAGTTCAAGAGGGCGTG | 240.4 | -3.6 | 0.8 | -4.3 | 1.7E-05 | 2.3E-05 | MB231/MB468 | MITO |
| >hg38_21_+_8433172_8446622_RNA45SN1_45S_50ntFlanks@4002.4025.24\|GATAACCTCGGGCCGATCGCACGC | >hg38_21_+_8433172_8446622_RNA45SN1_45S_50ntFlanks | 4002 | 4025 | 24 | GATAACCTCGGGCCGATCGCACGC | 387.2 | -5.8 | 1.3 | -4.3 | 1.4E-05 | 2.3E-05 | BT20/MB468 | CYTO |
| >hg38_21_+_8433172_8446622_RNA45SN1_45S_50ntFlanks@10683.10709.27\|CCGGCGGCGTCCGGTGAGCTCTCGCTG | >hg38_21_+_8433172_8446622_RNA45SN1_45S_50ntFlanks | 10683 | 10709 | 27 | CCGGCGGCGTCCGGTGAGCTCTCGCTG | 655.5 | -7.4 | 1.7 | -4.3 | 1.7E-05 | 2.3E-05 | BT20/MB468 | NUC |
| >hg38_21_+_8433172_8446622_RNA45SN1_45S_50ntFlanks@9528.9570.43\|AGCCAGAGGAAACTCTGGTGGAGGTCCGTAGCGGTCCTGACGT | >hg38_21_+_8433172_8446622_RNA45SN1_45S_50ntFlanks | 9528 | 9570 | 43 | AGCCAGAGGAAACTCTGGTGGAGGTCCGTAGCGGTCCTGACGT | 38.2 | 9.0 | 2.0 | 4.6 | 5.3E-06 | 2.3E-05 | BT20/MB468 | MP |
| >hg38_21_+_8433172_8446622_RNA45SN1_45S_50ntFlanks@10873.10890.18\|CTGGGTCGGTCGGGCTGG | >hg38_21_+_8433172_8446622_RNA45SN1_45S_50ntFlanks | 10873 | 10890 | 18 | CTGGGTCGGTCGGGCTGG | 667.2 | -3.0 | 0.7 | -4.3 | 1.4E-05 | 2.3E-05 | BT20/MB468 | CYTO |
| >hg38_21_+_8433172_8446622_RNA45SN1_45S_50ntFlanks@10127.10149.23\|GCGGCGGCGGCGGCGGCGGCGGG | >hg38_21_+_8433172_8446622_RNA45SN1_45S_50ntFlanks | 10127 | 10149 | 23 | GCGGCGGCGGCGGCGGCGGCGGG | 224.9 | -6.8 | 1.5 | -4.6 | 5.3E-06 | 2.3E-05 | BT20/MB468 | MP |
| >hg38_21_+_8433172_8446622_RNA45SN1_45S_50ntFlanks@8844.8866.23\|GCGGCGGCGGCGGCGGCGGCGGG | >hg38_21_+_8433172_8446622_RNA45SN1_45S_50ntFlanks | 8844 | 8866 | 23 | GCGGCGGCGGCGGCGGCGGCGGG | 224.9 | -6.8 | 1.5 | -4.6 | 5.3E-06 | 2.3E-05 | BT20/MB468 | MP |
| >hg38_21_+_8433172_8446622_RNA45SN1_45S_50ntFlanks@3985.4026.42\|CGGCTTTGGTGACTCTAGATAACCTCGGGCCGATCGCACGCC | >hg38_21_+_8433172_8446622_RNA45SN1_45S_50ntFlanks | 3985 | 4026 | 42 | CGGCTTTGGTGACTCTAGATAACCTCGGGCCGATCGCACGCC | 1502.3 | -7.3 | 1.7 | -4.3 | 1.4E-05 | 2.3E-05 | BT20/MB468 | CYTO |
| >hg38_21_+_8433172_8446622_RNA45SN1_45S_50ntFlanks@11581.11609.29\|AACTGGTGCGGACCAGGGGAATCCGACTG | >hg38_21_+_8433172_8446622_RNA45SN1_45S_50ntFlanks | 11581 | 11609 | 29 | AACTGGTGCGGACCAGGGGAATCCGACTG | 2436.2 | -3.4 | 0.7 | -4.6 | 5.3E-06 | 2.3E-05 | BT20/MB468 | MP |
| >hg38_21_+_8433172_8446622_RNA45SN1_45S_50ntFlanks@12634.12653.20\|GAAGCTACCATCTGTGGGAT | >hg38_21_+_8433172_8446622_RNA45SN1_45S_50ntFlanks | 12634 | 12653 | 20 | GAAGCTACCATCTGTGGGAT | 482.9 | -6.8 | 1.5 | -4.6 | 5.3E-06 | 2.3E-05 | BT20/MB468 | MP |
| >hg38_21_+_8433172_8446622_RNA45SN1_45S_50ntFlanks@6644.6664.21\|CCTCGTACGACTCTTAGCGGT | >hg38_21_+_8433172_8446622_RNA45SN1_45S_50ntFlanks | 6644 | 6664 | 21 | CCTCGTACGACTCTTAGCGGT | 336.5 | 3.6 | 0.8 | 4.3 | 1.7E-05 | 2.3E-05 | MB231/MB468 | MITO |
| >hg38_21_+_8433172_8446622_RNA45SN1_45S_50ntFlanks@12416.12457.42\|TTCGATGTCGGCTCTTCCTATCATTGTGAAGCAGAATTCACC | >hg38_21_+_8433172_8446622_RNA45SN1_45S_50ntFlanks | 12416 | 12457 | 42 | TTCGATGTCGGCTCTTCCTATCATTGTGAAGCAGAATTCACC | 511.5 | 6.7 | 1.6 | 4.3 | 1.6E-05 | 2.3E-05 | MB231/MB468 | NUC |
| >hg38_21_+_8433172_8446622_RNA45SN1_45S_50ntFlanks@9554.9582.29\|CGTAGCGGTCCTGACGTGCAAATCGGTCG | >hg38_21_+_8433172_8446622_RNA45SN1_45S_50ntFlanks | 9554 | 9582 | 29 | CGTAGCGGTCCTGACGTGCAAATCGGTCG | 551.3 | -6.3 | 1.5 | -4.3 | 1.7E-05 | 2.3E-05 | MB231/MB468 | MITO |
| >hg38_21_+_8433172_8446622_RNA45SN1_45S_50ntFlanks@11577.11613.37\|TTAGAACTGGTGCGGACCAGGGGAATCCGACTGTTTA | >hg38_21_+_8433172_8446622_RNA45SN1_45S_50ntFlanks | 11577 | 11613 | 37 | TTAGAACTGGTGCGGACCAGGGGAATCCGACTGTTTA | 521.7 | 4.0 | 0.8 | 4.8 | 1.9E-06 | 2.4E-05 | BT20/MB231 | MITO |
| >MI0000298\|hsa-mir-221&WithFlank&X\|-\|45746151\|45746272@31.55.25(+1C)&[MIMAT0004568&hsa-miR-221-5p&offsets\|0\|+3(+1C);m-405&X\|-\|45746220\|45746242&offsets\|0\|+2(+1C)]\|ACCTGGCATACAATGTAGATTTCTGC | >MI0000298\|hsa-mir-221&WithFlank&X\|-\|45746151\|45746272 | 31 | 55 | 25(+1C) | ACCTGGCATACAATGTAGATTTCTGC | 241.9 | -3.6 | 0.8 | -4.8 | 1.9E-06 | 2.4E-05 | BT20/MB231 | MITO |
| >hg38_21_+_8433172_8446622_RNA45SN1_45S_50ntFlanks@10547.10573.27\|AGTGCGGTAACGCGACCGATCCCGGAG | >hg38_21_+_8433172_8446622_RNA45SN1_45S_50ntFlanks | 10547 | 10573 | 27 | AGTGCGGTAACGCGACCGATCCCGGAG | 91.6 | -6.2 | 1.4 | -4.3 | 1.4E-05 | 2.4E-05 | MB231/MB468 | MP |
| >trnaMT_PheGAA_MT_+_577_647@35.63.29\|GAAAATGTTTAGACGGGCTCACATCACCC | >trnaMT_PheGAA_MT_+_577_647 | 35 | 63 | 29 | GAAAATGTTTAGACGGGCTCACATCACCC | 225.2 | 11.6 | 2.5 | 4.6 | 5.3E-06 | 2.4E-05 | BT20/MB468 | MP |
| >hg38_21_+_8433172_8446622_RNA45SN1_45S_50ntFlanks@11666.11687.22\|TCTGCCCAGTGCTCTGAATGTC | >hg38_21_+_8433172_8446622_RNA45SN1_45S_50ntFlanks | 11666 | 11687 | 22 | TCTGCCCAGTGCTCTGAATGTC | 696.6 | 2.5 | 0.6 | 4.3 | 1.6E-05 | 2.4E-05 | MB231/MB468 | NUC |
| >hg38_1_-_228634819_228635039_RNA5S12_5S_50ntFlanks@128.171.44\|CTTGGATGGGAGACCGCCTGGGAATACCGGGTGCTGTAGGCTTT | >hg38_1_-_228634819_228635039_RNA5S12_5S_50ntFlanks | 128 | 171 | 44 | CTTGGATGGGAGACCGCCTGGGAATACCGGGTGCTGTAGGCTTT | 688.9 | 5.1 | 1.2 | 4.3 | 1.6E-05 | 2.4E-05 | MB231/MB468 | NUC |
| >hg38_21_+_8433172_8446622_RNA45SN1_45S_50ntFlanks@8176.8199.24\|CTTCTGATCGAGGCCCAGCCCGTG | >hg38_21_+_8433172_8446622_RNA45SN1_45S_50ntFlanks | 8176 | 8199 | 24 | CTTCTGATCGAGGCCCAGCCCGTG | 396.9 | -3.9 | 0.8 | -4.6 | 5.3E-06 | 2.4E-05 | BT20/MB468 | MP |
| >hg38_21_+_8433172_8446622_RNA45SN1_45S_50ntFlanks@9571.9596.26\|GCAAATCGGTCGTCCGACCTGGGTAT | >hg38_21_+_8433172_8446622_RNA45SN1_45S_50ntFlanks | 9571 | 9596 | 26 | GCAAATCGGTCGTCCGACCTGGGTAT | 179.7 | -5.5 | 1.3 | -4.3 | 1.4E-05 | 2.4E-05 | BT20/MB468 | CYTO |
| >hg38_21_+_8433172_8446622_RNA45SN1_45S_50ntFlanks@12405.12422.18\|CTTTTTGATCCTTCGATG | >hg38_21_+_8433172_8446622_RNA45SN1_45S_50ntFlanks | 12405 | 12422 | 18 | CTTTTTGATCCTTCGATG | 1758.9 | -7.0 | 1.6 | -4.3 | 1.4E-05 | 2.4E-05 | BT20/MB468 | CYTO |
| >hg38_MT_+_598_1651_RNR1_12S_50ntFlanks@628.657.30\|CCGATCAACCTCACCACCTCTTGCTCAGCC | >hg38_MT_+_598_1651_RNR1_12S_50ntFlanks | 628 | 657 | 30 | CCGATCAACCTCACCACCTCTTGCTCAGCC | 34.1 | 8.3 | 1.9 | 4.3 | 1.4E-05 | 2.4E-05 | MB231/MB468 | MP |
| >hg38_21_+_8433172_8446622_RNA45SN1_45S_50ntFlanks@3999.4024.26\|CTAGATAACCTCGGGCCGATCGCACG | >hg38_21_+_8433172_8446622_RNA45SN1_45S_50ntFlanks | 3999 | 4024 | 26 | CTAGATAACCTCGGGCCGATCGCACG | 370.9 | -4.9 | 1.1 | -4.3 | 1.6E-05 | 2.4E-05 | MB231/MB468 | NUC |

<table>
<tr><td>>MI0000477|hsa-mir-146a&WithFlank&5|+|160485346|160485456@26.47.22(+1C)&[MIMAT0000449&hsa-miR-146a-5p&offsets|-1|-1(+1C);m-129&5|+|160485372|160485393&offsets|-1|-1(+1C)]|TTGAGAACTGAATTCCATGGGTC</td><td>>MI0000477|hsa-mir-146a&WithFlank&5|+|160485346|160485456</td><td>26</td><td>47</td><td>22(+1C)</td><td>TTGAGAACTGAATTCCATGGGTC</td><td>95.9</td><td>-10.0</td><td>2.1</td><td>-4.8</td><td>1.6E-06</td><td>2.4E-05</td><td>BT20/MB231</td><td>MP</td></tr>
<tr><td>>hg38_21_+_8433172_8446622_RNA45SN1_45S_50ntFlanks@4722.4758.37|TCATTAATCAAGAACGAAAGTCGGAGGTTCGAAGACG</td><td>>hg38_21_+_8433172_8446622_RNA45SN1_45S_50ntFlanks</td><td>4722</td><td>4758</td><td>37</td><td>TCATTAATCAAGAACGAAAGTCGGAGGTTCGAAGACG</td><td>6725.0</td><td>-3.8</td><td>0.9</td><td>-4.3</td><td>1.6E-05</td><td>2.4E-05</td><td>MB231/MB468</td><td>CYTO</td></tr>
<tr><td>>hg38_21_+_8433172_8446622_RNA45SN1_45S_50ntFlanks@11576.11605.30|CTTAGAACTGGTGCGGACCAGGGGAATCCG</td><td>>hg38_21_+_8433172_8446622_RNA45SN1_45S_50ntFlanks</td><td>11576</td><td>11605</td><td>30</td><td>CTTAGAACTGGTGCGGACCAGGGGAATCCG</td><td>678.8</td><td>-5.6</td><td>1.2</td><td>-4.6</td><td>5.3E-06</td><td>2.4E-05</td><td>BT20/MB468</td><td>MP</td></tr>
<tr><td>>hg38_21_+_8433172_8446622_RNA45SN1_45S_50ntFlanks@10681.10705.25|TTCCGGCGGCGTCCGGTGAGCTCTC</td><td>>hg38_21_+_8433172_8446622_RNA45SN1_45S_50ntFlanks</td><td>10681</td><td>10705</td><td>25</td><td>TTCCGGCGGCGTCCGGTGAGCTCTC</td><td>227.8</td><td>-3.2</td><td>0.7</td><td>-4.3</td><td>1.6E-05</td><td>2.4E-05</td><td>MB231/MB468</td><td>CYTO</td></tr>
<tr><td>>hg38_21_+_8433172_8446622_RNA45SN1_45S_50ntFlanks@9556.9584.29|TAGCGGTCCTGACGTGCAAATCGGTCGTC</td><td>>hg38_21_+_8433172_8446622_RNA45SN1_45S_50ntFlanks</td><td>9556</td><td>9584</td><td>29</td><td>TAGCGGTCCTGACGTGCAAATCGGTCGTC</td><td>236.7</td><td>-6.2</td><td>1.4</td><td>-4.3</td><td>1.4E-05</td><td>2.4E-05</td><td>BT20/MB468</td><td>CYTO</td></tr>
<tr><td>>hg38_21_+_8433172_8446622_RNA45SN1_45S_50ntFlanks@5491.5516.26|GAGAAGACGGTCGAACTTGACTATCT</td><td>>hg38_21_+_8433172_8446622_RNA45SN1_45S_50ntFlanks</td><td>5491</td><td>5516</td><td>26</td><td>GAGAAGACGGTCGAACTTGACTATCT</td><td>4952.5</td><td>6.0</td><td>1.4</td><td>4.3</td><td>1.8E-05</td><td>2.4E-05</td><td>BT20/MB468</td><td>NUC</td></tr>
<tr><td>>hg38_21_+_8433172_8446622_RNA45SN1_45S_50ntFlanks@6650.6693.44|ACGACTCTTAGCGGTGGATCACTCGGCTCGTGCGTCGATGAAGA</td><td>>hg38_21_+_8433172_8446622_RNA45SN1_45S_50ntFlanks</td><td>6650</td><td>6693</td><td>44</td><td>ACGACTCTTAGCGGTGGATCACTCGGCTCGTGCGTCGATGAAGA</td><td>361.1</td><td>4.7</td><td>1.1</td><td>4.3</td><td>1.7E-05</td><td>2.4E-05</td><td>MB231/MB468</td><td>NUC</td></tr>
<tr><td>>hg38_21_+_8433172_8446622_RNA45SN1_45S_50ntFlanks@9570.9594.25|TGCAAATCGGTCGTCCGACCTGGGT</td><td>>hg38_21_+_8433172_8446622_RNA45SN1_45S_50ntFlanks</td><td>9570</td><td>9594</td><td>25</td><td>TGCAAATCGGTCGTCCGACCTGGGT</td><td>854.7</td><td>-4.5</td><td>1.0</td><td>-4.5</td><td>5.4E-06</td><td>2.4E-05</td><td>BT20/MB468</td><td>MP</td></tr>
<tr><td>>MI0000542|hsa-mir-320a&WithFlank&8|-|22244960|22245043@50.70.21&[MIMAT0000510&hsa-miR-320a-3p&offsets|+2|+1;m-60&8|-|22244975|22244996&offsets|+2|+1]|AAGCTGGGTTGAGAGGGCGAA</td><td>>MI0000542|hsa-mir-320a&WithFlank&8|-|22244960|22245043</td><td>50</td><td>70</td><td>21</td><td>AAGCTGGGTTGAGAGGGCGAA</td><td>426.1</td><td>7.2</td><td>1.6</td><td>4.5</td><td>5.4E-06</td><td>2.4E-05</td><td>BT20/MB468</td><td>MP</td></tr>
<tr><td>>hg38_21_+_8433172_8446622_RNA45SN1_45S_50ntFlanks@12526.12546.21|TTTTACCCTACTGATGATGTG</td><td>>hg38_21_+_8433172_8446622_RNA45SN1_45S_50ntFlanks</td><td>12526</td><td>12546</td><td>21</td><td>TTTTACCCTACTGATGATGTG</td><td>155.3</td><td>-5.8</td><td>1.3</td><td>-4.3</td><td>1.4E-05</td><td>2.4E-05</td><td>BT20/MB468</td><td>CYTO</td></tr>
<tr><td>>hg38_21_+_8433172_8446622_RNA45SN1_45S_50ntFlanks@10467.10487.21|GTTGCCCTCGGCCGATCGAAA</td><td>>hg38_21_+_8433172_8446622_RNA45SN1_45S_50ntFlanks</td><td>10467</td><td>10487</td><td>21</td><td>GTTGCCCTCGGCCGATCGAAA</td><td>243.8</td><td>3.3</td><td>0.8</td><td>4.3</td><td>1.6E-05</td><td>2.4E-05</td><td>MB231/MB468</td><td>CYTO</td></tr>
<tr><td>>hg38_21_+_8433172_8446622_RNA45SN1_45S_50ntFlanks@8357.8377.21|AAAAGAACTTTGAAGAGAGAG</td><td>>hg38_21_+_8433172_8446622_RNA45SN1_45S_50ntFlanks</td><td>8357</td><td>8377</td><td>21</td><td>AAAAGAACTTTGAAGAGAGAG</td><td>1833.7</td><td>-3.2</td><td>0.7</td><td>-4.4</td><td>1.2E-05</td><td>2.4E-05</td><td>BT20/MB468</td><td>MITO</td></tr>
<tr><td>>hg38_21_+_8433172_8446622_RNA45SN1_45S_50ntFlanks@9766.9788.23|TAAATGGGTAAGAAGCCCGGCTC</td><td>>hg38_21_+_8433172_8446622_RNA45SN1_45S_50ntFlanks</td><td>9766</td><td>9788</td><td>23</td><td>TAAATGGGTAAGAAGCCCGGCTC</td><td>55.2</td><td>-8.9</td><td>1.9</td><td>-4.5</td><td>5.4E-06</td><td>2.4E-05</td><td>BT20/MB468</td><td>MP</td></tr>
<tr><td>>MI0000086|hsa-mir-28&WithFlank&3|+|188688775|188688872@60.79.20(+1A)&[MIMAT0004502&hsa-miR-28-3p&offsets|0|-2(+1A);m-28&3|+|188688834|188688855&offsets|0|-2(+1A)]|CACTAGATTGTGAGCTCCTGA</td><td>>MI0000086|hsa-mir-28&WithFlank&3|+|188688775|188688872</td><td>60</td><td>79</td><td>20(+1A)</td><td>CACTAGATTGTGAGCTCCTGA</td><td>204.4</td><td>8.1</td><td>1.8</td><td>4.5</td><td>5.4E-06</td><td>2.4E-05</td><td>BT20/MB468</td><td>MP</td></tr>
<tr><td>>hg38_21_+_8433172_8446622_RNA45SN1_45S_50ntFlanks@11573.11597.25|CGACTTAGAACTGGTGCGGACCAGG</td><td>>hg38_21_+_8433172_8446622_RNA45SN1_45S_50ntFlanks</td><td>11573</td><td>11597</td><td>25</td><td>CGACTTAGAACTGGTGCGGACCAGG</td><td>105.1</td><td>10.0</td><td>2.1</td><td>4.8</td><td>1.6E-06</td><td>2.4E-05</td><td>BT20/MB231</td><td>MP</td></tr>
<tr><td>>hg38_21_+_8433172_8446622_RNA45SN1_45S_50ntFlanks@10845.10867.23|ACTTCGGGATAAGGATTGGCTCT</td><td>>hg38_21_+_8433172_8446622_RNA45SN1_45S_50ntFlanks</td><td>10845</td><td>10867</td><td>23</td><td>ACTTCGGGATAAGGATTGGCTCT</td><td>500.8</td><td>-3.0</td><td>0.7</td><td>-4.3</td><td>1.4E-05</td><td>2.4E-05</td><td>BT20/MB468</td><td>CYTO</td></tr>
<tr><td>>hg38_21_+_8433172_8446622_RNA45SN1_45S_50ntFlanks@12636.12660.25|AGCTACCATCTGTGGGATTATGACT</td><td>>hg38_21_+_8433172_8446622_RNA45SN1_45S_50ntFlanks</td><td>12636</td><td>12660</td><td>25</td><td>AGCTACCATCTGTGGGATTATGACT</td><td>356.1</td><td>-5.4</td><td>1.2</td><td>-4.3</td><td>1.7E-05</td><td>2.4E-05</td><td>MB231/MB468</td><td>MITO</td></tr>
<tr><td>>hg38_21_+_8433172_8446622_RNA45SN1_45S_50ntFlanks@3785.3802.18|TGAAACTGCGAATGGCTC</td><td>>hg38_21_+_8433172_8446622_RNA45SN1_45S_50ntFlanks</td><td>3785</td><td>3802</td><td>18</td><td>TGAAACTGCGAATGGCTC</td><td>357.8</td><td>-6.9</td><td>1.6</td><td>-4.3</td><td>1.4E-05</td><td>2.4E-05</td><td>MB231/MB468</td><td>MP</td></tr>
<tr><td>>hg38_21_+_8433172_8446622_RNA45SN1_45S_50ntFlanks@10919.10936.18|CGGCTGGACGAGGCGCCG</td><td>>hg38_21_+_8433172_8446622_RNA45SN1_45S_50ntFlanks</td><td>10919</td><td>10936</td><td>18</td><td>CGGCTGGACGAGGCGCCG</td><td>41.6</td><td>-8.5</td><td>1.9</td><td>-4.5</td><td>5.4E-06</td><td>2.4E-05</td><td>BT20/MB468</td><td>MP</td></tr>
<tr><td>>hg38_21_+_8433172_8446622_RNA45SN1_45S_50ntFlanks@5083.5112.30|ACGAACGAGACTCTGGCATGCTAACTAGTT</td><td>>hg38_21_+_8433172_8446622_RNA45SN1_45S_50ntFlanks</td><td>5083</td><td>5112</td><td>30</td><td>ACGAACGAGACTCTGGCATGCTAACTAGTT</td><td>377.4</td><td>-6.8</td><td>1.6</td><td>-4.3</td><td>1.7E-05</td><td>2.4E-05</td><td>MB231/MB468</td><td>MITO</td></tr>
<tr><td>>hg38_21_+_8433172_8446622_RNA45SN1_45S_50ntFlanks@12418.12457.40|CGATGTCGGCTCTTCCTATCATTGTGAAGCAGAATTCACC</td><td>>hg38_21_+_8433172_8446622_RNA45SN1_45S_50ntFlanks</td><td>12418</td><td>12457</td><td>40</td><td>CGATGTCGGCTCTTCCTATCATTGTGAAGCAGAATTCACC</td><td>896.8</td><td>-3.7</td><td>0.8</td><td>-4.8</td><td>1.7E-06</td><td>2.4E-05</td><td>BT20/MB231</td><td>CYTO</td></tr>
<tr><td>>hg38_21_+_8433172_8446622_RNA45SN1_45S_50ntFlanks@3780.3801.22|TACAGTGAAACTGCGAATGGCT</td><td>>hg38_21_+_8433172_8446622_RNA45SN1_45S_50ntFlanks</td><td>3780</td><td>3801</td><td>22</td><td>TACAGTGAAACTGCGAATGGCT</td><td>189.5</td><td>-2.3</td><td>0.5</td><td>-4.3</td><td>1.4E-05</td><td>2.4E-05</td><td>BT20/MB468</td><td>CYTO</td></tr>
<tr><td>>hg38_21_+_8433172_8446622_RNA45SN1_45S_50ntFlanks@11641.11660.20|CGGCGGGTGTTGACGCGATG</td><td>>hg38_21_+_8433172_8446622_RNA45SN1_45S_50ntFlanks</td><td>11641</td><td>11660</td><td>20</td><td>CGGCGGGTGTTGACGCGATG</td><td>1557.3</td><td>-4.1</td><td>0.9</td><td>-4.3</td><td>1.4E-05</td><td>2.4E-05</td><td>BT20/MB468</td><td>CYTO</td></tr>
<tr><td>>MI0000447|hsa-mir-128-1&WithFlank&2|+|135665391|135665484@56.75.20(+1C);MI0000727|hsa-mir-128-2&WithFlank&3|+|35744470|35744565@58.77.20(+1C)&[MIMAT0000424&hsa-miR-128-3p&offsets|0|-1(+1C);m-108&2|+|135665446|135665466&offsets|0|-1(+1C)];[MIMAT0000424_1&hsa-miR-128-3p&offsets|0|-1(+1C);m-117&3|+|35744527|35744547&offsets|0|-1(+1C)]|TCACAGTGAACCGGTCTCTTC</td><td>>MI0000447|hsa-mir-128-1&WithFlank&2|+|135665391|135665484</td><td>56</td><td>75</td><td>20(+1C)</td><td>TCACAGTGAACCGGTCTCTTC</td><td>279.3</td><td>3.3</td><td>0.8</td><td>4.3</td><td>1.6E-05</td><td>2.4E-05</td><td>MB231/MB468</td><td>CYTO</td></tr>
<tr><td>>MI0000750|hsa-mir-26a-2&WithFlank&12|-|57824603|57824698@20.38.19(+1A);MI0000083|hsa-mir-26a-1&WithFlank&3|+|37969398|37969486@16.34.19(+1A)&[MIMAT0000082&hsa-miR-26a-5p&offsets|0|-3(+1A);m-33&12|-|57824658|57824679&offsets|0|-3(+1A)];[MIMAT0000082_1&hsa-miR-26a-5p&offsets|0|-3(+1A);m-34&3|+|37969413|37969434&offsets|0|-3(+1A)]|TTCAAGTAATCCAGGATAGA</td><td>>MI0000750|hsa-mir-26a-2&WithFlank&12|-|57824603|57824698</td><td>20</td><td>38</td><td>19(+1A)</td><td>TTCAAGTAATCCAGGATAGA</td><td>86.2</td><td>-9.8</td><td>2.1</td><td>-4.8</td><td>1.6E-06</td><td>2.4E-05</td><td>BT20/MB231</td><td>MP</td></tr>
<tr><td>>MI0000289|hsa-mir-181a-1&WithFlank&1|-|198859038|198859159@30.53.24(+1G);MI0000269|hsa-mir-181a-2&WithFlank&9|+|124692436|124692557@45.68.24(+1G)&[MIMAT0000256&hsa-miR-181a-5p&offsets|0|+1(+1G);m-32&1|-|198859108|198859130&offsets|0|+1(+1G)];[MIMAT0000256_1&hsa-miR-181a-5p&offsets|0|+1(+1G);m-32&9|+|124692480|124692502&offsets|0|+1(+1G)]|AACATTCAACGCTGTCGGTGAGTTG</td><td>>MI0000289|hsa-mir-181a-1&WithFlank&1|-|198859038|198859159</td><td>30</td><td>53</td><td>24(+1G)</td><td>AACATTCAACGCTGTCGGTGAGTTG</td><td>80.1</td><td>-9.7</td><td>2.0</td><td>-4.8</td><td>1.6E-06</td><td>2.4E-05</td><td>BT20/MB231</td><td>MP</td></tr>
<tr><td>>hg38_21_+_8433172_8446622_RNA45SN1_45S_50ntFlanks@11209.11254.46|CGGCGACTCTGGACGCGAGCCGGGCCCTTCCCGTGGATCGCCCCAG</td><td>>hg38_21_+_8433172_8446622_RNA45SN1_45S_50ntFlanks</td><td>11209</td><td>11254</td><td>46</td><td>CGGCGACTCTGGACGCGAGCCGGGCCCTTCCCGTGGATCGCCCCAG</td><td>142.4</td><td>10.5</td><td>2.2</td><td>4.8</td><td>1.7E-06</td><td>2.4E-05</td><td>BT20/MB231</td><td>MP</td></tr>
<tr><td>>MI0003643|hsa-mir-629&WithFlank&15|-|70079366|70079474@28.50.23&[MIMAT0004810&hsa-miR-629-5p&offsets|0|+2;m-150&15|-|70079427|70079447&offsets|0|+2]|TGGGTTTACGTTGGGAGAACTTT</td><td>>MI0003643|hsa-mir-629&WithFlank&15|-|70079366|70079474</td><td>28</td><td>50</td><td>23</td><td>TGGGTTTACGTTGGGAGAACTTT</td><td>232.9</td><td>11.6</td><td>2.6</td><td>4.5</td><td>5.5E-06</td><td>2.4E-05</td><td>BT20/MB468</td><td>MP</td></tr>
<tr><td>>hg38_21_+_8433172_8446622_RNA45SN1_45S_50ntFlanks@3813.3841.29|TTATGGTTCCTTTGGTCGCTCGCTCCTCT</td><td>>hg38_21_+_8433172_8446622_RNA45SN1_45S_50ntFlanks</td><td>3813</td><td>3841</td><td>29</td><td>TTATGGTTCCTTTGGTCGCTCGCTCCTCT</td><td>267.3</td><td>-5.0</td><td>1.2</td><td>-4.3</td><td>1.7E-05</td><td>2.4E-05</td><td>MB231/MB468</td><td>NUC</td></tr>
<tr><td>>hg38_21_+_8433172_8446622_RNA45SN1_45S_50ntFlanks@12423.12446.24|TCGGCTCTTCCTATCATTGTGAAG</td><td>>hg38_21_+_8433172_8446622_RNA45SN1_45S_50ntFlanks</td><td>12423</td><td>12446</td><td>24</td><td>TCGGCTCTTCCTATCATTGTGAAG</td><td>139.9</td><td>-3.6</td><td>0.8</td><td>-4.3</td><td>1.6E-05</td><td>2.4E-05</td><td>MB231/MB468</td><td>CYTO</td></tr>
<tr><td>>trna13_LysCTT_14_-_58706613_58706685@1.41.41,trna2_LysCTT_15_+_79152904_79152976@1.41.41|GCCCGGCTAGCTCAGTCGGTAGAGCATGGGACTCTTAATCC</td><td>>trna13_LysCTT_14_-_58706613_58706685</td><td>1</td><td>41</td><td>41</td><td>GCCCGGCTAGCTCAGTCGGTAGAGCATGGGACTCTTAATCC</td><td>467.4</td><td>6.4</td><td>1.5</td><td>4.3</td><td>1.5E-05</td><td>2.4E-05</td><td>BT20/MB468</td><td>CYTO</td></tr>
<tr><td>>MI0016887|hsa-mir-4521&WithFlank&17|+|8186939|8187010@10.34.25&[MIMAT0019058&hsa-miR-4521&offsets|0|+3;m-765&17|+|8186948|8186969&offsets|0|+3]|GCTAAGGAAGTCCTGTGCTCAGTTT</td><td>>MI0016887|hsa-mir-4521&WithFlank&17|+|8186939|8187010</td><td>10</td><td>34</td><td>25</td><td>GCTAAGGAAGTCCTGTGCTCAGTTT</td><td>1029.2</td><td>4.7</td><td>1.1</td><td>4.3</td><td>1.6E-05</td><td>2.4E-05</td><td>MB231/MB468</td><td>CYTO</td></tr>
<tr><td>>hg38_21_+_8433172_8446622_RNA45SN1_45S_50ntFlanks@11786.11803.18|GAATGGATGAACGAGATT</td><td>>hg38_21_+_8433172_8446622_RNA45SN1_45S_50ntFlanks</td><td>11786</td><td>11803</td><td>18</td><td>GAATGGATGAACGAGATT</td><td>939.5</td><td>-3.7</td><td>0.8</td><td>-4.5</td><td>5.5E-06</td><td>2.4E-05</td><td>BT20/MB468</td><td>MP</td></tr>
<tr><td>>hg38_21_+_8433172_8446622_RNA45SN1_45S_50ntFlanks@9556.9584.29|TAGCGGTCCTGACGTGCAAATCGGTCGTC</td><td>>hg38_21_+_8433172_8446622_RNA45SN1_45S_50ntFlanks</td><td>9556</td><td>9584</td><td>29</td><td>TAGCGGTCCTGACGTGCAAATCGGTCGTC</td><td>115.8</td><td>-6.3</td><td>1.4</td><td>-4.3</td><td>1.4E-05</td><td>2.4E-05</td><td>MB231/MB468</td><td>MP</td></tr>
<tr><td>>trna20_GluTTC_2_-_131094701_131094772@1.30.30,trna5_GluTTC_13_-_41634874_41634945@1.30.30|TCCCATATGGTCTAGCGGTTAGGATTCCTG</td><td>>trna20_GluTTC_2_-_131094701_131094772</td><td>1</td><td>30</td><td>30</td><td>TCCCATATGGTCTAGCGGTTAGGATTCCTG</td><td>310.0</td><td>-2.9</td><td>0.7</td><td>-4.3</td><td>1.5E-05</td><td>2.4E-05</td><td>BT20/MB468</td><td>CYTO</td></tr>
<tr><td>>hg38_21_+_8433172_8446622_RNA45SN1_45S_50ntFlanks@11125.11145.21|CGGCGGCGGCCCGCCGCGGGG</td><td>>hg38_21_+_8433172_8446622_RNA45SN1_45S_50ntFlanks</td><td>11125</td><td>11145</td><td>21</td><td>CGGCGGCGGCCCGCCGCGGGG</td><td>76.9</td><td>-9.3</td><td>2.1</td><td>-4.5</td><td>5.5E-06</td><td>2.4E-05</td><td>BT20/MB468</td><td>MP</td></tr>
<tr><td>>hg38_21_+_8433172_8446622_RNA45SN1_45S_50ntFlanks@10388.10409.22|GTGAACAGCAGTTGAACATGGG</td><td>>hg38_21_+_8433172_8446622_RNA45SN1_45S_50ntFlanks</td><td>10388</td><td>10409</td><td>22</td><td>GTGAACAGCAGTTGAACATGGG</td><td>155.7</td><td>-5.4</td><td>1.2</td><td>-4.3</td><td>1.5E-05</td><td>2.4E-05</td><td>BT20/MB468</td><td>CYTO</td></tr>
<tr><td>>trnaMT_PheGAA_MT_+_577_647@39.59.21|ATGTTTAGACGGGCTCACATC</td><td>>trnaMT_PheGAA_MT_+_577_647</td><td>39</td><td>59</td><td>21</td><td>ATGTTTAGACGGGCTCACATC</td><td>279.5</td><td>3.5</td><td>0.8</td><td>4.3</td><td>1.7E-05</td><td>2.4E-05</td><td>MB231/MB468</td><td>MITO</td></tr>
<tr><td>>MI0000465|hsa-mir-191&WithFlank&3|-|49020612|49020715@22.45.24&[MIMAT0000440&hsa-miR-191-5p&offsets|0|+1;m-48&3|-|49020672|49020694&offsets|0|+1]|CAACGGAATCCCAAAAGCAGCTGT</td><td>>MI0000465|hsa-mir-191&WithFlank&3|-|49020612|49020715</td><td>22</td><td>45</td><td>24</td><td>CAACGGAATCCCAAAAGCAGCTGT</td><td>6634.2</td><td>4.2</td><td>1.0</td><td>4.3</td><td>1.4E-05</td><td>2.4E-05</td><td>MB231/MB468</td><td>MP</td></tr>
<tr><td>>hg38_21_+_8433172_8446622_RNA45SN1_45S_50ntFlanks@4673.4691.19|TGGACCGGCGCAAGACGGA</td><td>>hg38_21_+_8433172_8446622_RNA45SN1_45S_50ntFlanks</td><td>4673</td><td>4691</td><td>19</td><td>TGGACCGGCGCAAGACGGA</td><td>85.7</td><td>-10.3</td><td>2.4</td><td>-4.3</td><td>1.4E-05</td><td>2.4E-05</td><td>MB231/MB468</td><td>MP</td></tr>
<tr><td>>MI0000077|hsa-mir-21&WithFlank&17|+|59841260|59841343@14.33.20(+1U)&[MIMAT0000076&hsa-miR-21-5p&offsets|0|-2(+1U);m-1&17|+|59841273|59841294&offsets|0|-2(+1U)]|TAGCTTATCAGACTGATGTTT</td><td>>MI0000077|hsa-mir-21&WithFlank&17|+|59841260|59841343</td><td>14</td><td>33</td><td>20(+1U)</td><td>TAGCTTATCAGACTGATGTTT</td><td>721.2</td><td>8.6</td><td>2.0</td><td>4.3</td><td>1.5E-05</td><td>2.4E-05</td><td>BT20/MB468</td><td>CYTO</td></tr>
<tr><td>>hg38_21_+_8433172_8446622_RNA45SN1_45S_50ntFlanks@4759.4786.28|ATCAGATACCGTCGTAGTTCCGACCATA</td><td>>hg38_21_+_8433172_8446622_RNA45SN1_45S_50ntFlanks</td><td>4759</td><td>4786</td><td>28</td><td>ATCAGATACCGTCGTAGTTCCGACCATA</td><td>619.9</td><td>-5.9</td><td>1.4</td><td>-4.3</td><td>1.6E-05</td><td>2.4E-05</td><td>MB231/MB468</td><td>CYTO</td></tr>
<tr><td>>hg38_21_+_8433172_8446622_RNA45SN1_45S_50ntFlanks@11234.11254.21|CCTTCCCGTGGATCGCCCCAG</td><td>>hg38_21_+_8433172_8446622_RNA45SN1_45S_50ntFlanks</td><td>11234</td><td>11254</td><td>21</td><td>CCTTCCCGTGGATCGCCCCAG</td><td>349.0</td><td>-6.0</td><td>1.4</td><td>-4.3</td><td>1.5E-05</td><td>2.4E-05</td><td>BT20/MB468</td><td>CYTO</td></tr>
<tr><td>>hg38_21_+_8433172_8446622_RNA45SN1_45S_50ntFlanks@11563.11584.22|GCCTAGCAGCCGACTTAGAACT</td><td>>hg38_21_+_8433172_8446622_RNA45SN1_45S_50ntFlanks</td><td>11563</td><td>11584</td><td>22</td><td>GCCTAGCAGCCGACTTAGAACT</td><td>112.4</td><td>7.6</td><td>1.7</td><td>4.5</td><td>5.5E-06</td><td>2.4E-05</td><td>BT20/MB468</td><td>MP</td></tr>
<tr><td>>MI0000286|hsa-mir-210&WithFlank&11|-|568083|568204@72.93.22(+2A)&[MIMAT0000267&hsa-miR-210-3p&offsets|0|0(+2A);m-102&11|-|568112|568133&offsets|0|0(+2A)]|CTGTGCGTGTGACAGCGGCTGAAA</td><td>>MI0000286|hsa-mir-210&WithFlank&11|-|568083|568204</td><td>72</td><td>93</td><td>22(+2A)</td><td>CTGTGCGTGTGACAGCGGCTGAAA</td><td>1289.7</td><td>4.0</td><td>0.9</td><td>4.3</td><td>1.4E-05</td><td>2.4E-05</td><td>MB231/MB468</td><td>MP</td></tr>
<tr><td>>MI0001641|hsa-mir-429&WithFlank&1|+|1168999|1169093@57.77.21&[MIMAT0001536&hsa-miR-429&offsets|0|-1;m-207&1|+|1169055|1169076&offsets|0|-1]|TAATACTGTCTGGTAAAACCG</td><td>>MI0001641|hsa-mir-429&WithFlank&1|+|1168999|1169093</td><td>57</td><td>77</td><td>21</td><td>TAATACTGTCTGGTAAAACCG</td><td>93.4</td><td>-6.0</td><td>1.4</td><td>-4.3</td><td>1.4E-05</td><td>2.4E-05</td><td>MB231/MB468</td><td>MP</td></tr>
</table>

| | | | | | | | | | | | | | |
|---|---|---|---|---|---|---|---|---|---|---|---|---|---|
| >hg38_MT_+_598_1651_RNR1_12S_50ntFlanks@428.470.43\|AAAGTGGCTTTAACATATCTGAACACACAATAGCTAAGACCCA | >hg38_MT_+_598_1651_RNR1_12S_50ntFlanks | 428 | 470 | 43 | AAAGTGGCTTTAACATATCTGAACACACAATAGCTAAGACCCA | 32.7 | 8.3 | 1.9 | 4.3 | 1.4E-05 | 2.4E-05 | MB231/MB468 | MP |
| >hg38_21_+_8433172_8446622_RNA45SN1_45S_50ntFlanks@11874.11908.35\|AGAAGACCCTGTTGAGCTTGACTCTAGTCTGGCAC | >hg38_21_+_8433172_8446622_RNA45SN1_45S_50ntFlanks | 11874 | 11908 | 35 | AGAAGACCCTGTTGAGCTTGACTCTAGTCTGGCAC | 377.4 | -5.8 | 1.3 | -4.3 | 1.5E-05 | 2.4E-05 | BT20/MB468 | CYTO |
| >hg38_21_+_8433172_8446622_RNA45SN1_45S_50ntFlanks@4759.4790.32\|ATCAGATACCGTCGTAGTTCCGACCATAAACG | >hg38_21_+_8433172_8446622_RNA45SN1_45S_50ntFlanks | 4759 | 4790 | 32 | ATCAGATACCGTCGTAGTTCCGACCATAAACG | 41.1 | -8.4 | 1.9 | -4.5 | 5.5E-06 | 2.4E-05 | BT20/MB468 | MP |
| >hg38_21_+_8433172_8446622_RNA45SN1_45S_50ntFlanks@9554.9586.33\|CGTAGCGGTCCTGACGTGCAAATCGGTCGTCCG | >hg38_21_+_8433172_8446622_RNA45SN1_45S_50ntFlanks | 9554 | 9586 | 33 | CGTAGCGGTCCTGACGTGCAAATCGGTCGTCCG | 43.4 | -8.5 | 1.9 | -4.5 | 5.5E-06 | 2.4E-05 | BT20/MB468 | MP |
| >hg38_21_+_8433172_8446622_RNA45SN1_45S_50ntFlanks@4759.4780.22\|ATCAGATACCGTCGTAGTTCCG | >hg38_21_+_8433172_8446622_RNA45SN1_45S_50ntFlanks | 4759 | 4780 | 22 | ATCAGATACCGTCGTAGTTCCG | 9753.1 | -7.0 | 1.6 | -4.3 | 1.6E-05 | 2.4E-05 | MB231/MB468 | CYTO |
| >hg38_21_+_8433172_8446622_RNA45SN1_45S_50ntFlanks@11785.11805.21\|TGAATGGATGAACGAGATTCC | >hg38_21_+_8433172_8446622_RNA45SN1_45S_50ntFlanks | 11785 | 11805 | 21 | TGAATGGATGAACGAGATTCC | 930.9 | -6.0 | 1.4 | -4.3 | 1.5E-05 | 2.4E-05 | BT20/MB468 | CYTO |
| >hg38_1_-_228634819_228635039_RNA5S12_5S_50ntFlanks@51.81.31\|GTCTACGGCCATACCACCCTGAACGCGCCCG | >hg38_1_-_228634819_228635039_RNA5S12_5S_50ntFlanks | 51 | 81 | 31 | GTCTACGGCCATACCACCCTGAACGCGCCCG | 1693.2 | -3.0 | 0.7 | -4.3 | 1.8E-05 | 2.4E-05 | BT20/MB468 | NUC |
| >MI0016746\|hsa-mir-548o-2&WithFlank&20\|+\|38516557\|38516638@51.72.22;MI0006402\|hsa-mir-548o&WithFlank&7\|-\|102405736\|102405861@93.114.22&[MIMAT0005919&hsa-miR-548o-3p&offsets\|0\|0;m-431&20\|+\|38516607\|38516628&offsets\|0\|0];[MIMAT0005919_1&hsa-miR-548o-3p&offsets\|0\|0;m-430&7\|-\|102405748\|102405769&offsets\|0\|0]\|CCAAAACTGCAGTTACTTTTGC | >MI0016746\|hsa-mir-548o-2&WithFlank&20\|+\|38516557\|38516638 | 51 | 72 | 22 | CCAAAACTGCAGTTACTTTTGC | 504.2 | 4.3 | 1.0 | 4.5 | 5.5E-06 | 2.4E-05 | BT20/MB468 | MP |
| >hg38_MT_+_1621_3279_RNR2_16S_50ntFlanks@1427.1453.27\|GATTAAAGTCCTACGTGATCTGAGTTC | >hg38_MT_+_1621_3279_RNR2_16S_50ntFlanks | 1427 | 1453 | 27 | GATTAAAGTCCTACGTGATCTGAGTTC | 270.1 | 3.9 | 0.9 | 4.3 | 1.4E-05 | 2.5E-05 | MB231/MB468 | MP |
| >hg38_21_+_8433172_8446622_RNA45SN1_45S_50ntFlanks@9936.9960.25\|ACAGCAGGACGGTGGCCATGGAAGT | >hg38_21_+_8433172_8446622_RNA45SN1_45S_50ntFlanks | 9936 | 9960 | 25 | ACAGCAGGACGGTGGCCATGGAAGT | 106.8 | 10.1 | 2.1 | 4.8 | 1.7E-06 | 2.5E-05 | BT20/MB231 | MP |
| >hg38_21_+_8433172_8446622_RNA45SN1_45S_50ntFlanks@4723.4748.26\|CATTAATCAAGAACGAAAGTCGGAGG | >hg38_21_+_8433172_8446622_RNA45SN1_45S_50ntFlanks | 4723 | 4748 | 26 | CATTAATCAAGAACGAAAGTCGGAGG | 391.7 | -4.7 | 1.1 | -4.4 | 1.2E-05 | 2.5E-05 | BT20/MB468 | MITO |
| >MI0000254\|hsa-mir-30c-2&WithFlank&6\|-\|71376954\|71377037@53.75.23&[MIMAT0004550&hsa-miR-30c-2-3p&offsets\|0\|+1;m-206&6\|-\|71376964\|71376985&offsets\|0\|+1]\|CTGGGAGAAGGCTGTTTACTCTT | >MI0000254\|hsa-mir-30c-2&WithFlank&6\|-\|71376954\|71377037 | 53 | 75 | 23 | CTGGGAGAAGGCTGTTTACTCTT | 189.4 | 6.2 | 1.4 | 4.3 | 1.7E-05 | 2.5E-05 | MB231/MB468 | MITO |
| >hg38_21_+_8433172_8446622_RNA45SN1_45S_50ntFlanks@9714.9735.22\|CGAATGATTAGAGGTCTTGGGG | >hg38_21_+_8433172_8446622_RNA45SN1_45S_50ntFlanks | 9714 | 9735 | 22 | CGAATGATTAGAGGTCTTGGGG | 256.7 | -4.7 | 1.1 | -4.3 | 1.6E-05 | 2.5E-05 | MB231/MB468 | CYTO |
| >hg38_21_+_8433172_8446622_RNA45SN1_45S_50ntFlanks@11104.11123.20\|AAGGGTCGGGGCGGCAGGGG | >hg38_21_+_8433172_8446622_RNA45SN1_45S_50ntFlanks | 11104 | 11123 | 20 | AAGGGTCGGGGCGGCAGGGG | 2266.7 | -8.0 | 1.9 | -4.3 | 1.8E-05 | 2.5E-05 | BT20/MB468 | NUC |
| >hg38_21_+_8433172_8446622_RNA45SN1_45S_50ntFlanks@12490.12521.32\|CGTGAGCTGGGTTTAGACCGTCGTGAGACAGG | >hg38_21_+_8433172_8446622_RNA45SN1_45S_50ntFlanks | 12490 | 12521 | 32 | CGTGAGCTGGGTTTAGACCGTCGTGAGACAGG | 102.6 | -5.0 | 1.2 | -4.3 | 1.5E-05 | 2.5E-05 | BT20/MB468 | CYTO |
| >hg38_21_+_8433172_8446622_RNA45SN1_45S_50ntFlanks@11671.11701.31\|CCAGTGCTCTGAATGTCAAAGTGAAGAAATT | >hg38_21_+_8433172_8446622_RNA45SN1_45S_50ntFlanks | 11671 | 11701 | 31 | CCAGTGCTCTGAATGTCAAAGTGAAGAAATT | 338.0 | -2.3 | 0.5 | -4.3 | 1.5E-05 | 2.5E-05 | BT20/MB468 | CYTO |
| >MI0000342\|hsa-mir-200b&WithFlank&1\|+\|1167098\|1167204@63.82.20(+1A)&[MIMAT0000318&hsa-miR-200b-3p&offsets\|0\|-2(+1A);m-93&1\|+\|1167160\|1167182&offsets\|0\|-3(+1A)]\|TAATACTGCCTGGTAATGATA | >MI0000342\|hsa-mir-200b&WithFlank&1\|+\|1167098\|1167204 | 63 | 82 | 20(+1A) | TAATACTGCCTGGTAATGATA | 173.1 | -6.2 | 1.4 | -4.3 | 1.4E-05 | 2.5E-05 | MB231/MB468 | MP |
| >hg38_21_+_8433172_8446622_RNA45SN1_45S_50ntFlanks@11885.11902.18\|TTGAGCTTGACTCTAGTC | >hg38_21_+_8433172_8446622_RNA45SN1_45S_50ntFlanks | 11885 | 11902 | 18 | TTGAGCTTGACTCTAGTC | 47.0 | -8.6 | 1.9 | -4.5 | 5.6E-06 | 2.5E-05 | BT20/MB468 | MP |
| >hg38_21_+_8433172_8446622_RNA45SN1_45S_50ntFlanks@10391.10408.18\|AACAGCAGTTGAACATGG | >hg38_21_+_8433172_8446622_RNA45SN1_45S_50ntFlanks | 10391 | 10408 | 18 | AACAGCAGTTGAACATGG | 443.5 | -4.8 | 1.1 | -4.3 | 1.7E-05 | 2.5E-05 | MB231/MB468 | NUC |
| >hg38_MT_+_1621_3279_RNR2_16S_50ntFlanks@1438.1477.40\|TACGTGATCTGAGTTCAGACCGGAGTAATCCAGGTCGGTT | >hg38_MT_+_1621_3279_RNR2_16S_50ntFlanks | 1438 | 1477 | 40 | TACGTGATCTGAGTTCAGACCGGAGTAATCCAGGTCGGTT | 226.1 | 3.2 | 0.7 | 4.3 | 1.7E-05 | 2.5E-05 | MB231/MB468 | MITO |
| >hg38_21_+_8433172_8446622_RNA45SN1_45S_50ntFlanks@10458.10489.32\|ATGGCCTCCGTTGCCCTCGGCCGATCGAAAGG | >hg38_21_+_8433172_8446622_RNA45SN1_45S_50ntFlanks | 10458 | 10489 | 32 | ATGGCCTCCGTTGCCCTCGGCCGATCGAAAGG | 211.7 | -4.8 | 1.1 | -4.3 | 1.7E-05 | 2.5E-05 | MB231/MB468 | NUC |
| >hg38_21_+_8433172_8446622_RNA45SN1_45S_50ntFlanks@11886.11904.19\|TGAGCTTGACTCTAGTCTG | >hg38_21_+_8433172_8446622_RNA45SN1_45S_50ntFlanks | 11886 | 11904 | 19 | TGAGCTTGACTCTAGTCTG | 944.2 | -7.1 | 1.6 | -4.5 | 5.6E-06 | 2.5E-05 | BT20/MB468 | MP |
| >hg38_21_+_8433172_8446622_RNA45SN1_45S_50ntFlanks@12627.12657.31\|ATGGGGCGAAGCTACCATCTGTGGGATTATG | >hg38_21_+_8433172_8446622_RNA45SN1_45S_50ntFlanks | 12627 | 12657 | 31 | ATGGGGCGAAGCTACCATCTGTGGGATTATG | 277.4 | -5.8 | 1.3 | -4.3 | 1.5E-05 | 2.5E-05 | BT20/MB468 | CYTO |
| >hg38_21_+_8433172_8446622_RNA45SN1_45S_50ntFlanks@11703.11730.28\|AATGAAGCGCGGGTAAACGGCGGGAGTA | >hg38_21_+_8433172_8446622_RNA45SN1_45S_50ntFlanks | 11703 | 11730 | 28 | AATGAAGCGCGGGTAAACGGCGGGAGTA | 89.2 | -5.0 | 1.1 | -4.3 | 1.4E-05 | 2.5E-05 | MB231/MB468 | MP |
| >MI0000087\|hsa-mir-29a&WithFlank&7\|-\|130876741\|130876816@48.68.21(+1C)&[MIMAT0000086&hsa-miR-29a-3p&offsets\|0\|-1(+1C);m-26&7\|-\|130876749\|130876769&offsets\|0\|0(+1C)]\|TAGCACCATCTGAAATCGGTTC | >MI0000087\|hsa-mir-29a&WithFlank&7\|-\|130876741\|130876816 | 48 | 68 | 21(+1C) | TAGCACCATCTGAAATCGGTTC | 225.2 | 11.6 | 2.5 | 4.5 | 5.6E-06 | 2.5E-05 | BT20/MB468 | MP |
| >hg38_21_+_8433172_8446622_RNA45SN1_45S_50ntFlanks@8369.8387.19\|AAGAGAGAGTTCAAGAGGG | >hg38_21_+_8433172_8446622_RNA45SN1_45S_50ntFlanks | 8369 | 8387 | 19 | AAGAGAGAGTTCAAGAGGG | 91.0 | -6.1 | 1.4 | -4.3 | 1.4E-05 | 2.5E-05 | MB231/MB468 | MP |
| >hg38_21_+_8433172_8446622_RNA45SN1_45S_50ntFlanks@12991.13039.49\|CTCCCTCGCTGCGATCTATTGAAAGTCAGCCCTCGACACAAGGGTTTGT | >hg38_21_+_8433172_8446622_RNA45SN1_45S_50ntFlanks | 12991 | 13039 | 49 | CTCCCTCGCTGCGATCTATTGAAAGTCAGCCCTCGACACAAGGGTTTGT | 14580.3 | -5.3 | 1.2 | -4.3 | 1.8E-05 | 2.5E-05 | MB231/MB468 | MITO |
| >hg38_21_+_8433172_8446622_RNA45SN1_45S_50ntFlanks@4923.4944.22\|CAGGAGTGGAGCCTGCGGCTTA | >hg38_21_+_8433172_8446622_RNA45SN1_45S_50ntFlanks | 4923 | 4944 | 22 | CAGGAGTGGAGCCTGCGGCTTA | 198.3 | -7.0 | 1.6 | -4.3 | 1.5E-05 | 2.5E-05 | MB231/MB468 | MP |
| >trna111_HisGTG_1_-_147774845_147774916@-1T.35.36,trna118_HisGTG_1_-_145396881_145396952@-1T.35.36,trna16_HisGTG_1_+_146544773_146544844@-1T.35.36,trna1_HisGTG_15_+_45493349_45493420@-1T.35.36,trna21_HisGTG_1_+_147753471_147753542@-1T.35.36,trna33_HisGTG_6_+_27125906_27125977@-1T.35.36,trna7_HisGTG_9_-_14433938_14434009@-1T.35.36,trna8_HisGTG_15_-_45492611_45492682@-1T.35.36,trna9_HisGTG_15_-_45490804_45490875@-1T.35.36\|TGCCGTGATCGTATAGTGGTTAGTACTCTGCGTTGT | >trna111_HisGTG_1_-_147774845_147774916 | -1T | 35 | 36 | TGCCGTGATCGTATAGTGGTTAGTACTCTGCGTTGT | 43.2 | 8.7 | 2.0 | 4.3 | 1.5E-05 | 2.5E-05 | MB231/MB468 | MP |
| >hg38_21_+_8433172_8446622_RNA45SN1_45S_50ntFlanks@11766.11790.25\|ATCTAATTAGTGACGCGCATGAATG | >hg38_21_+_8433172_8446622_RNA45SN1_45S_50ntFlanks | 11766 | 11790 | 25 | ATCTAATTAGTGACGCGCATGAATG | 476.6 | -6.1 | 1.4 | -4.3 | 1.7E-05 | 2.5E-05 | MB231/MB468 | NUC |
| >hg38_21_+_8433172_8446622_RNA45SN1_45S_50ntFlanks@12204.12221.18\|AAGGCGAGCTCAGGGAGG | >hg38_21_+_8433172_8446622_RNA45SN1_45S_50ntFlanks | 12204 | 12221 | 18 | AAGGCGAGCTCAGGGAGG | 619.4 | -6.2 | 1.4 | -4.3 | 1.8E-05 | 2.5E-05 | MB231/MB468 | MITO |
| >hg38_21_+_8433172_8446622_RNA45SN1_45S_50ntFlanks@12073.12110.38\|CCGAGGGGCTCTCGCTTCTGGCGCCAAGCGCCCGGCCG | >hg38_21_+_8433172_8446622_RNA45SN1_45S_50ntFlanks | 12073 | 12110 | 38 | CCGAGGGGCTCTCGCTTCTGGCGCCAAGCGCCCGGCCG | 599.3 | -7.6 | 1.8 | -4.3 | 1.8E-05 | 2.5E-05 | BT20/MB468 | NUC |
| >MI0001445\|hsa-mir-423&WithFlank&17\|+\|30117073\|30117178@59.78.20(+1U)&[MIMAT0001340&hsa-miR-423-3p&offsets\|0\|-3(+1U);m-100&17\|+\|30117131\|30117153&offsets\|0\|-3(+1U)]\|AGCTCGGTCTGAGGCCCCTCT | >MI0001445\|hsa-mir-423&WithFlank&17\|+\|30117073\|30117178 | 59 | 78 | 20(+1U) | AGCTCGGTCTGAGGCCCCTCT | 138.7 | 7.9 | 1.7 | 4.5 | 5.6E-06 | 2.5E-05 | BT20/MB468 | MP |
| >hg38_21_+_8433172_8446622_RNA45SN1_45S_50ntFlanks@8117.8134.18\|CGGGACATGTGGCGTACG | >hg38_21_+_8433172_8446622_RNA45SN1_45S_50ntFlanks | 8117 | 8134 | 18 | CGGGACATGTGGCGTACG | 114.3 | -6.1 | 1.4 | -4.3 | 1.5E-05 | 2.5E-05 | MB231/MB468 | MP |
| >hg38_21_+_8433172_8446622_RNA45SN1_45S_50ntFlanks@4657.4688.32\|CTAGAGGTGAAATTCTTGGACCGGCGCAAGAC | >hg38_21_+_8433172_8446622_RNA45SN1_45S_50ntFlanks | 4657 | 4688 | 32 | CTAGAGGTGAAATTCTTGGACCGGCGCAAGAC | 452.6 | -6.2 | 1.4 | -4.3 | 1.7E-05 | 2.5E-05 | MB231/MB468 | CYTO |
| >hg38_21_+_8433172_8446622_RNA45SN1_45S_50ntFlanks@10841.10872.32\|CGTAACTTCGGGATAAGGATTGGCTCTAAGGG | >hg38_21_+_8433172_8446622_RNA45SN1_45S_50ntFlanks | 10841 | 10872 | 32 | CGTAACTTCGGGATAAGGATTGGCTCTAAGGG | 1170.8 | -5.2 | 1.2 | -4.3 | 1.7E-05 | 2.5E-05 | MB231/MB468 | NUC |
| >hg38_21_+_8433172_8446622_RNA45SN1_45S_50ntFlanks@8183.8210.28\|TCGAGGCCCAGCCCGTGGACGGTGTGAG | >hg38_21_+_8433172_8446622_RNA45SN1_45S_50ntFlanks | 8183 | 8210 | 28 | TCGAGGCCCAGCCCGTGGACGGTGTGAG | 568.7 | -7.3 | 1.7 | -4.3 | 1.8E-05 | 2.5E-05 | BT20/MB468 | NUC |
| >MI0000111\|hsa-mir-105-1&WithFlank&X\|-\|152392213\|152392305@19.41.23;MI0000112\|hsa-mir-105-2&WithFlank&X\|-\|152394406\|152394498@19.41.23&[MIMAT0000102&hsa-miR-105-5p&offsets\|0\|0;m-735&X\|-\|152392266\|152392287&offsets\|0\|+1];[MIMAT0000102_1&hsa-miR-105-5p&offsets\|0\|0;m-735&X\|-\|152394459\|152394480&offsets\|0\|+1]\|TCAAATGCTCAGACTCCTGTGGT | >MI0000111\|hsa-mir-105-1&WithFlank&X\|-\|152392213\|152392305 | 19 | 41 | 23 | TCAAATGCTCAGACTCCTGTGGT | 82.8 | 9.7 | 2.0 | 4.8 | 1.7E-06 | 2.5E-05 | BT20/MB231 | MP |
| >hg38_21_+_8433172_8446622_RNA45SN1_45S_50ntFlanks@9925.9955.31\|GGTTGATATAGACAGCAGGACGGTGGCCATG | >hg38_21_+_8433172_8446622_RNA45SN1_45S_50ntFlanks | 9925 | 9955 | 31 | GGTTGATATAGACAGCAGGACGGTGGCCATG | 73.9 | 9.5 | 2.0 | 4.8 | 1.7E-06 | 2.5E-05 | BT20/MB231 | MP |
| >MI0000102\|hsa-mir-100&WithFlank&11\|-\|122152223\|122152314@20.38.19&[MIMAT0000098&hsa-miR-100-5p&offsets\|+1\|-2;m-27&11\|-\|122152276\|122152296&offsets\|+1\|-1]\|ACCCGTAGATCCGAACTTG | >MI0000102\|hsa-mir-100&WithFlank&11\|-\|122152223\|122152314 | 20 | 38 | 19 | ACCCGTAGATCCGAACTTG | 102.7 | -10.1 | 2.1 | -4.8 | 1.7E-06 | 2.5E-05 | BT20/MB231 | MP |
| >hg38_21_+_8433172_8446622_RNA45SN1_45S_50ntFlanks@12996.13036.41\|TCGCTGCGATCTATTGAAAGTCAGCCCTCGACACAAGGGTT | >hg38_21_+_8433172_8446622_RNA45SN1_45S_50ntFlanks | 12996 | 13036 | 41 | TCGCTGCGATCTATTGAAAGTCAGCCCTCGACACAAGGGTT | 1525.5 | 4.9 | 1.2 | 4.3 | 1.7E-05 | 2.5E-05 | MB231/MB468 | NUC |
| >hg38_21_+_8433172_8446622_RNA45SN1_45S_50ntFlanks@5227.5246.20\|CGCGCGCTACACTGACTGGC | >hg38_21_+_8433172_8446622_RNA45SN1_45S_50ntFlanks | 5227 | 5246 | 20 | CGCGCGCTACACTGACTGGC | 87.5 | -10.3 | 2.4 | -4.3 | 1.5E-05 | 2.5E-05 | MB231/MB468 | MP |
| >hg38_21_+_8433172_8446622_RNA45SN1_45S_50ntFlanks@11786.11805.20\|GAATGGATGAACGAGATTCC | >hg38_21_+_8433172_8446622_RNA45SN1_45S_50ntFlanks | 11786 | 11805 | 20 | GAATGGATGAACGAGATTCC | 139.3 | -6.2 | 1.4 | -4.3 | 1.5E-05 | 2.5E-05 | MB231/MB468 | MP |
| >MI0000477\|hsa-mir-146a&WithFlank&5\|+\|160485346\|160485456@27.46.20(+2C)&[MIMAT0000449&hsa-miR-146a-5p&offsets\|0\|-2(+2C);m-129&5\|+\|160485372\|160485393&offsets\|0\|-2(+2C)]\|TGAGAACTGAATTCCATGGGCC | >MI0000477\|hsa-mir-146a&WithFlank&5\|+\|160485346\|160485456 | 27 | 46 | 20(+2C) | TGAGAACTGAATTCCATGGGCC | 94.0 | -10.0 | 2.1 | -4.8 | 1.7E-06 | 2.5E-05 | BT20/MB231 | MP |
| >hg38_21_+_8433172_8446622_RNA45SN1_45S_50ntFlanks@4047.4069.23\|ATTCGAACGTCTGCCCTATCAAC | >hg38_21_+_8433172_8446622_RNA45SN1_45S_50ntFlanks | 4047 | 4069 | 23 | ATTCGAACGTCTGCCCTATCAAC | 143.0 | -6.7 | 1.5 | -4.3 | 1.5E-05 | 2.5E-05 | MB231/MB468 | MP |
| >hg38_21_+_8433172_8446622_RNA45SN1_45S_50ntFlanks@5236.5254.19\|CACTGACTGGCTCAGCGTG | >hg38_21_+_8433172_8446622_RNA45SN1_45S_50ntFlanks | 5236 | 5254 | 19 | CACTGACTGGCTCAGCGTG | 348.1 | -4.0 | 0.9 | -4.3 | 1.5E-05 | 2.5E-05 | MB231/MB468 | MP |
| >hg38_21_+_8433172_8446622_RNA45SN1_45S_50ntFlanks@8169.8215.47\|CCAAGTCCTTCTGATCGAGGCCCAGCCCGTGGACGGTGTGAGGCCGG | >hg38_21_+_8433172_8446622_RNA45SN1_45S_50ntFlanks | 8169 | 8215 | 47 | CCAAGTCCTTCTGATCGAGGCCCAGCCCGTGGACGGTGTGAGGCCGG | 416.4 | -5.5 | 1.3 | -4.3 | 1.7E-05 | 2.5E-05 | MB231/MB468 | NUC |

<table>
<tr><td></td><td></td><td></td><td></td><td></td><td></td><td></td><td></td><td></td><td></td><td></td><td></td><td></td><td></td></tr>
<tr><td>>MI0000082|hsa-mir-25&WithFlank&7|-|100093554|100093649@58.78.21(+1U)&[MIMAT0000081&hsa-miR-25-3p&offsets|0|-1(+1U);m-23&7|-|100093571|100093592&offsets|0|-1(+1U)]|CATTGCACTTGTCTCGGTCTGT</td><td>>MI0000082|hsa-mir-25&WithFlank&7|-|100093554|100093649</td><td>58</td><td>78</td><td>21(+1U)</td><td>CATTGCACTTGTCTCGGTCTGT</td><td>1808.0</td><td>3.6</td><td>0.8</td><td>4.3</td><td>1.5E-05</td><td>2.5E-05</td><td>MB231/MB468</td><td>MP</td></tr>
<tr><td>>hg38_21_+_8433172_8446622_RNA45SN1_45S_50ntFlanks@3860.3898.39|GTAATTCTAGAGCTAATACATGCCGACGGGCGCTGACCC</td><td>>hg38_21_+_8433172_8446622_RNA45SN1_45S_50ntFlanks</td><td>3860</td><td>3898</td><td>39</td><td>GTAATTCTAGAGCTAATACATGCCGACGGGCGCTGACCC</td><td>78.9</td><td>-10.2</td><td>2.3</td><td>-4.3</td><td>1.5E-05</td><td>2.5E-05</td><td>MB231/MB468</td><td>MP</td></tr>
<tr><td>>hg38_21_+_8433172_8446622_RNA45SN1_45S_50ntFlanks@11572.11599.28|CCGACTTAGAACTGGTGCGGACCAGGGG</td><td>>hg38_21_+_8433172_8446622_RNA45SN1_45S_50ntFlanks</td><td>11572</td><td>11599</td><td>28</td><td>CCGACTTAGAACTGGTGCGGACCAGGGG</td><td>776.8</td><td>-7.4</td><td>1.7</td><td>-4.3</td><td>1.9E-05</td><td>2.5E-05</td><td>BT20/MB468</td><td>NUC</td></tr>
<tr><td>>hg38_21_+_8433172_8446622_RNA45SN1_45S_50ntFlanks@3922.3959.38|TTTATCAGATCAAAACCAACCCGGTCAGCCCCTCTCCG</td><td>>hg38_21_+_8433172_8446622_RNA45SN1_45S_50ntFlanks</td><td>3922</td><td>3959</td><td>38</td><td>TTTATCAGATCAAAACCAACCCGGTCAGCCCCTCTCCG</td><td>236.9</td><td>-5.7</td><td>1.3</td><td>-4.3</td><td>1.5E-05</td><td>2.5E-05</td><td>BT20/MB468</td><td>CYTO</td></tr>
<tr><td>>hg38_21_+_8433172_8446622_RNA45SN1_45S_50ntFlanks@12457.12474.18|CAAGCGTTGGATTGTTCA</td><td>>hg38_21_+_8433172_8446622_RNA45SN1_45S_50ntFlanks</td><td>12457</td><td>12474</td><td>18</td><td>CAAGCGTTGGATTGTTCA</td><td>123.6</td><td>-2.3</td><td>0.5</td><td>-4.4</td><td>1.3E-05</td><td>2.5E-05</td><td>BT20/MB468</td><td>MITO</td></tr>
<tr><td>>hg38_21_+_8433172_8446622_RNA45SN1_45S_50ntFlanks@475.516.42|CTTCGTGATCGATGTGGTGACGTCGTGCTCTCCCGGGCCGGG</td><td>>hg38_21_+_8433172_8446622_RNA45SN1_45S_50ntFlanks</td><td>475</td><td>516</td><td>42</td><td>CTTCGTGATCGATGTGGTGACGTCGTGCTCTCCCGGGCCGGG</td><td>1581.1</td><td>5.5</td><td>1.3</td><td>4.3</td><td>1.8E-05</td><td>2.5E-05</td><td>MB231/MB468</td><td>MITO</td></tr>
<tr><td>>hg38_21_+_8433172_8446622_RNA45SN1_45S_50ntFlanks@8358.8377.20|AAAGAACTTTGAAGAGAGAG</td><td>>hg38_21_+_8433172_8446622_RNA45SN1_45S_50ntFlanks</td><td>8358</td><td>8377</td><td>20</td><td>AAAGAACTTTGAAGAGAGAG</td><td>333.9</td><td>-5.9</td><td>1.4</td><td>-4.3</td><td>1.5E-05</td><td>2.5E-05</td><td>MB231/MB468</td><td>MP</td></tr>
<tr><td>>hg38_21_+_8433172_8446622_RNA45SN1_45S_50ntFlanks@12973.12994.22|TCGTACGTAGCAGAGCAGCTCC</td><td>>hg38_21_+_8433172_8446622_RNA45SN1_45S_50ntFlanks</td><td>12973</td><td>12994</td><td>22</td><td>TCGTACGTAGCAGAGCAGCTCC</td><td>152.5</td><td>-2.0</td><td>0.5</td><td>-4.3</td><td>1.8E-05</td><td>2.5E-05</td><td>MB231/MB468</td><td>NUC</td></tr>
<tr><td>>hg38_21_+_8433172_8446622_RNA45SN1_45S_50ntFlanks@4661.4689.29|AGGTGAAATTCTTGGACCGGCGCAAGACG</td><td>>hg38_21_+_8433172_8446622_RNA45SN1_45S_50ntFlanks</td><td>4661</td><td>4689</td><td>29</td><td>AGGTGAAATTCTTGGACCGGCGCAAGACG</td><td>288.9</td><td>-6.2</td><td>1.4</td><td>-4.3</td><td>1.8E-05</td><td>2.5E-05</td><td>MB231/MB468</td><td>NUC</td></tr>
<tr><td>>hg38_21_+_8433172_8446622_RNA45SN1_45S_50ntFlanks@4003.4022.20|ATAACCTCGGGCCGATCGCA</td><td>>hg38_21_+_8433172_8446622_RNA45SN1_45S_50ntFlanks</td><td>4003</td><td>4022</td><td>20</td><td>ATAACCTCGGGCCGATCGCA</td><td>585.4</td><td>-2.6</td><td>0.6</td><td>-4.3</td><td>1.5E-05</td><td>2.5E-05</td><td>BT20/MB468</td><td>CYTO</td></tr>
<tr><td>>hg38_21_+_8433172_8446622_RNA45SN1_45S_50ntFlanks@8153.8182.30|GCCGCTCGTGGGGGGCCCAAGTCCTTCTGA</td><td>>hg38_21_+_8433172_8446622_RNA45SN1_45S_50ntFlanks</td><td>8153</td><td>8182</td><td>30</td><td>GCCGCTCGTGGGGGGCCCAAGTCCTTCTGA</td><td>597.8</td><td>-4.1</td><td>0.9</td><td>-4.4</td><td>1.3E-05</td><td>2.5E-05</td><td>BT20/MB468</td><td>MITO</td></tr>
<tr><td>>hg38_21_+_8433172_8446622_RNA45SN1_45S_50ntFlanks@6647.6673.27|CGTACGACTCTTAGCGGTGGATCACTC</td><td>>hg38_21_+_8433172_8446622_RNA45SN1_45S_50ntFlanks</td><td>6647</td><td>6673</td><td>27</td><td>CGTACGACTCTTAGCGGTGGATCACTC</td><td>262.5</td><td>3.8</td><td>0.8</td><td>4.5</td><td>5.7E-06</td><td>2.5E-05</td><td>BT20/MB468</td><td>MP</td></tr>
<tr><td>>hg38_21_+_8433172_8446622_RNA45SN1_45S_50ntFlanks@9507.9524.18|AACTATGCCTGGGCAGGG</td><td>>hg38_21_+_8433172_8446622_RNA45SN1_45S_50ntFlanks</td><td>9507</td><td>9524</td><td>18</td><td>AACTATGCCTGGGCAGGG</td><td>80.3</td><td>-10.2</td><td>2.4</td><td>-4.3</td><td>1.5E-05</td><td>2.5E-05</td><td>MB231/MB468</td><td>MP</td></tr>
<tr><td>>hg38_21_+_8433172_8446622_RNA45SN1_45S_50ntFlanks@10236.10264.29|GACGCTACGCCGCGACGAGTAGGAGGGCC</td><td>>hg38_21_+_8433172_8446622_RNA45SN1_45S_50ntFlanks</td><td>10236</td><td>10264</td><td>29</td><td>GACGCTACGCCGCGACGAGTAGGAGGGCC</td><td>111.8</td><td>10.1</td><td>2.1</td><td>4.8</td><td>1.7E-06</td><td>2.5E-05</td><td>BT20/MB231</td><td>MP</td></tr>
<tr><td>>hg38_21_+_8433172_8446622_RNA45SN1_45S_50ntFlanks@9533.9561.29|GAGGAAACTCTGGTGGAGGTCCGTAGCGG</td><td>>hg38_21_+_8433172_8446622_RNA45SN1_45S_50ntFlanks</td><td>9533</td><td>9561</td><td>29</td><td>GAGGAAACTCTGGTGGAGGTCCGTAGCGG</td><td>513.0</td><td>-4.6</td><td>1.1</td><td>-4.3</td><td>1.7E-05</td><td>2.5E-05</td><td>MB231/MB468</td><td>CYTO</td></tr>
<tr><td>>hg38_21_+_8433172_8446622_RNA45SN1_45S_50ntFlanks@10398.10424.27|GTTGAACATGGGTCAGTCGGTCCTGAG</td><td>>hg38_21_+_8433172_8446622_RNA45SN1_45S_50ntFlanks</td><td>10398</td><td>10424</td><td>27</td><td>GTTGAACATGGGTCAGTCGGTCCTGAG</td><td>13789.6</td><td>4.2</td><td>0.9</td><td>4.7</td><td>2.1E-06</td><td>2.5E-05</td><td>BT20/MB231</td><td>MITO</td></tr>
<tr><td>>hg38_21_+_8433172_8446622_RNA45SN1_45S_50ntFlanks@11575.11612.38|ACTTAGAACTGGTGCGGACCAGGGGAATCCGACTGTTT</td><td>>hg38_21_+_8433172_8446622_RNA45SN1_45S_50ntFlanks</td><td>11575</td><td>11612</td><td>38</td><td>ACTTAGAACTGGTGCGGACCAGGGGAATCCGACTGTTT</td><td>182.3</td><td>-5.2</td><td>1.2</td><td>-4.3</td><td>1.5E-05</td><td>2.5E-05</td><td>MB231/MB468</td><td>MP</td></tr>
<tr><td>>hg38_21_+_8433172_8446622_RNA45SN1_45S_50ntFlanks@11879.11905.27|ACCCTGTTGAGCTTGACTCTAGTCTGG</td><td>>hg38_21_+_8433172_8446622_RNA45SN1_45S_50ntFlanks</td><td>11879</td><td>11905</td><td>27</td><td>ACCCTGTTGAGCTTGACTCTAGTCTGG</td><td>85.0</td><td>-6.0</td><td>1.4</td><td>-4.3</td><td>1.5E-05</td><td>2.5E-05</td><td>MB231/MB468</td><td>MP</td></tr>
<tr><td>>hg38_21_+_8433172_8446622_RNA45SN1_45S_50ntFlanks@12729.12747.19|TCGGATAGCCGGTCCCCCG</td><td>>hg38_21_+_8433172_8446622_RNA45SN1_45S_50ntFlanks</td><td>12729</td><td>12747</td><td>19</td><td>TCGGATAGCCGGTCCCCCG</td><td>82.0</td><td>-10.2</td><td>2.4</td><td>-4.3</td><td>1.5E-05</td><td>2.5E-05</td><td>MB231/MB468</td><td>MP</td></tr>
<tr><td>>hg38_21_+_8433172_8446622_RNA45SN1_45S_50ntFlanks@9561.9597.37|GTCCTGACGTGCAAATCGGTCGTCCGACCTGGGTATA</td><td>>hg38_21_+_8433172_8446622_RNA45SN1_45S_50ntFlanks</td><td>9561</td><td>9597</td><td>37</td><td>GTCCTGACGTGCAAATCGGTCGTCCGACCTGGGTATA</td><td>71.3</td><td>-10.0</td><td>2.3</td><td>-4.3</td><td>1.5E-05</td><td>2.5E-05</td><td>MB231/MB468</td><td>MP</td></tr>
<tr><td>>hg38_21_+_8433172_8446622_RNA45SN1_45S_50ntFlanks@11805.11823.19|CCACTGTCCCTACCTACTA</td><td>>hg38_21_+_8433172_8446622_RNA45SN1_45S_50ntFlanks</td><td>11805</td><td>11823</td><td>19</td><td>CCACTGTCCCTACCTACTA</td><td>118.1</td><td>-3.8</td><td>0.9</td><td>-4.3</td><td>1.5E-05</td><td>2.5E-05</td><td>BT20/MB468</td><td>CYTO</td></tr>
<tr><td>>hg38_21_+_8433172_8446622_RNA45SN1_45S_50ntFlanks@7973.8005.33|GACGCGACCTCAGATCAGACGTGGCGACCCGCT</td><td>>hg38_21_+_8433172_8446622_RNA45SN1_45S_50ntFlanks</td><td>7973</td><td>8005</td><td>33</td><td>GACGCGACCTCAGATCAGACGTGGCGACCCGCT</td><td>455.3</td><td>2.9</td><td>0.7</td><td>4.3</td><td>1.8E-05</td><td>2.5E-05</td><td>MB231/MB468</td><td>NUC</td></tr>
<tr><td>>hg38_21_+_8433172_8446622_RNA45SN1_45S_50ntFlanks@9545.9566.22|GTGGAGGTCCGTAGCGGTCCTG</td><td>>hg38_21_+_8433172_8446622_RNA45SN1_45S_50ntFlanks</td><td>9545</td><td>9566</td><td>22</td><td>GTGGAGGTCCGTAGCGGTCCTG</td><td>827.7</td><td>-7.0</td><td>1.5</td><td>-4.5</td><td>5.7E-06</td><td>2.5E-05</td><td>BT20/MB468</td><td>MP</td></tr>
<tr><td>>hg38_21_+_8433172_8446622_RNA45SN1_45S_50ntFlanks@5186.5225.40|CAATAACAGGTCTGTGATGCCCTTAGATGTCCGGGGCTGC</td><td>>hg38_21_+_8433172_8446622_RNA45SN1_45S_50ntFlanks</td><td>5186</td><td>5225</td><td>40</td><td>CAATAACAGGTCTGTGATGCCCTTAGATGTCCGGGGCTGC</td><td>78.8</td><td>-10.2</td><td>2.3</td><td>-4.3</td><td>1.5E-05</td><td>2.5E-05</td><td>MB231/MB468</td><td>MP</td></tr>
<tr><td>>MI0000273|hsa-mir-183&WithFlank&7|-|129774899|129775020@33.53.21(+1G)&[MIMAT0000261&hsa-miR-183-5p&offsets|0|-1(+1G);m-41&7|-|129774966|129774987&offsets|-1|-2(+1G)]|TATGGCACTGGTAGAATTCACG</td><td>>MI0000273|hsa-mir-183&WithFlank&7|-|129774899|129775020</td><td>33</td><td>53</td><td>21(+1G)</td><td>TATGGCACTGGTAGAATTCACG</td><td>323.3</td><td>2.8</td><td>0.6</td><td>4.7</td><td>2.1E-06</td><td>2.5E-05</td><td>BT20/MB231</td><td>NUC</td></tr>
<tr><td>>hg38_21_+_8433172_8446622_RNA45SN1_45S_50ntFlanks@12409.12431.23|TTGATCCTTCGATGTCGGCTCTT</td><td>>hg38_21_+_8433172_8446622_RNA45SN1_45S_50ntFlanks</td><td>12409</td><td>12431</td><td>23</td><td>TTGATCCTTCGATGTCGGCTCTT</td><td>104.9</td><td>-6.2</td><td>1.4</td><td>-4.3</td><td>1.5E-05</td><td>2.5E-05</td><td>MB231/MB468</td><td>MP</td></tr>
<tr><td>>trna10_AspGTC_12_-_125424193_125424264@37.73.37,trna12_AspGTC_12_-_125411891_125411962@37.73.37,trna144_AspGTC_6_-_27551236_27551307@37.73.37,trna38_AspGTC_17_-_8125556_8125627@37.73.37,trna45_AspGTC_6_+_27447453_27447524@37.73.37,trna48_AspGTC_6_+_27471523_27471594@37.73.37,trna4_AspGTC_12_+_96429799_96429870@37.73.37,trna69_AspGTC_1_-_161440205_161440276@37.73.37,trna72_AspGTC_1_-_161432824_161432895@37.73.37,trna75_AspGTC_1_-_161425414_161425485@37.73.37,trna78_AspGTC_1_-_161418033_161418104@37.73.37,trna81_AspGTC_1_-_161410615_161410686@37.73.37|ACGCGGGAGACCGGGGTTCGATTCCCCGACGGGGAGC</td><td>>trna10_AspGTC_12_-_125424193_125424264</td><td>37</td><td>73</td><td>37</td><td>ACGCGGGAGACCGGGGTTCGATTCCCCGACGGGGAGC</td><td>1713.1</td><td>5.4</td><td>1.3</td><td>4.3</td><td>1.8E-05</td><td>2.5E-05</td><td>MB231/MB468</td><td>NUC</td></tr>
<tr><td>>MI0003776|hsa-mir-320b-1&WithFlank&1|+|116671740|116671823@48.67.20;MI0003839|hsa-mir-320b-2&WithFlank&1|-|224257034|224257116@47.66.20;MI0000542|hsa-mir-320a&WithFlank&8|-|22244960|22245043@48.67.20&[MIMAT0005792&hsa-miR-320b&offsets|0|-2;m-183&1|+|116671787|116671806&offsets|0|0];[MIMAT0005792_1&hsa-miR-320b&offsets|0|-2;m-183&1|-|224257051|224257070&offsets|0|0];[MIMAT0000510&hsa-miR-320a-3p&offsets|0|-2;m-60&8|-|22244975|22244996&offsets|0|-2]|AAAAGCTGGGTTGAGAGGGC</td><td>>MI0003776|hsa-mir-320b-1&WithFlank&1|+|116671740|116671823</td><td>48</td><td>67</td><td>20</td><td>AAAAGCTGGGTTGAGAGGGC</td><td>261.4</td><td>3.4</td><td>0.8</td><td>4.3</td><td>1.8E-05</td><td>2.5E-05</td><td>MB231/MB468</td><td>NUC</td></tr>
<tr><td>>hg38_21_+_8433172_8446622_RNA45SN1_45S_50ntFlanks@4760.4777.18|TCAGATACCGTCGTAGTT</td><td>>hg38_21_+_8433172_8446622_RNA45SN1_45S_50ntFlanks</td><td>4760</td><td>4777</td><td>18</td><td>TCAGATACCGTCGTAGTT</td><td>136.6</td><td>-5.5</td><td>1.3</td><td>-4.3</td><td>1.5E-05</td><td>2.5E-05</td><td>BT20/MB468</td><td>CYTO</td></tr>
<tr><td>>hg38_21_+_8433172_8446622_RNA45SN1_45S_50ntFlanks@11567.11592.26|AGCAGCCGACTTAGAACTGGTGCGGA</td><td>>hg38_21_+_8433172_8446622_RNA45SN1_45S_50ntFlanks</td><td>11567</td><td>11592</td><td>26</td><td>AGCAGCCGACTTAGAACTGGTGCGGA</td><td>782.7</td><td>-5.2</td><td>1.2</td><td>-4.5</td><td>5.8E-06</td><td>2.5E-05</td><td>BT20/MB468</td><td>MP</td></tr>
<tr><td>>hg38_21_+_8433172_8446622_RNA45SN1_45S_50ntFlanks@5073.5092.20|AATTCCGATAACGAACGAGA</td><td>>hg38_21_+_8433172_8446622_RNA45SN1_45S_50ntFlanks</td><td>5073</td><td>5092</td><td>20</td><td>AATTCCGATAACGAACGAGA</td><td>222.0</td><td>-7.0</td><td>1.6</td><td>-4.3</td><td>1.5E-05</td><td>2.5E-05</td><td>MB231/MB468</td><td>MP</td></tr>
<tr><td>>hg38_21_+_8433172_8446622_RNA45SN1_45S_50ntFlanks@6651.6685.35|CGACTCTTAGCGGTGGATCACTCGGCTCGTGCGTC</td><td>>hg38_21_+_8433172_8446622_RNA45SN1_45S_50ntFlanks</td><td>6651</td><td>6685</td><td>35</td><td>CGACTCTTAGCGGTGGATCACTCGGCTCGTGCGTC</td><td>412.9</td><td>2.6</td><td>0.6</td><td>4.3</td><td>1.7E-05</td><td>2.5E-05</td><td>MB231/MB468</td><td>CYTO</td></tr>
<tr><td>>hg38_MT_+_1621_3279_RNR2_16S_50ntFlanks@1381.1414.34|GGACATCCCGATGGTGCAGCCGCTATTAAAGGTT</td><td>>hg38_MT_+_1621_3279_RNR2_16S_50ntFlanks</td><td>1381</td><td>1414</td><td>34</td><td>GGACATCCCGATGGTGCAGCCGCTATTAAAGGTT</td><td>31.6</td><td>8.2</td><td>1.9</td><td>4.3</td><td>1.5E-05</td><td>2.5E-05</td><td>MB231/MB468</td><td>MP</td></tr>
<tr><td>>hg38_21_+_8433172_8446622_RNA45SN1_45S_50ntFlanks@11875.11892.18|GAAGACCCTGTTGAGCTT</td><td>>hg38_21_+_8433172_8446622_RNA45SN1_45S_50ntFlanks</td><td>11875</td><td>11892</td><td>18</td><td>GAAGACCCTGTTGAGCTT</td><td>151.0</td><td>-6.2</td><td>1.4</td><td>-4.3</td><td>1.5E-05</td><td>2.5E-05</td><td>MB231/MB468</td><td>MP</td></tr>
<tr><td>>MI0000456|hsa-mir-140&WithFlank&16|+|69933075|69933186@69.88.20&[MIMAT0004597&hsa-miR-140-3p&offsets|+1|0;m-45&16|+|69933143|69933164&offsets|0|-2]|ACCACAGGGTAGAACCACGG</td><td>>MI0000456|hsa-mir-140&WithFlank&16|+|69933075|69933186</td><td>69</td><td>88</td><td>20</td><td>ACCACAGGGTAGAACCACGG</td><td>426.9</td><td>6.4</td><td>1.4</td><td>4.5</td><td>5.8E-06</td><td>2.5E-05</td><td>BT20/MB468</td><td>MP</td></tr>
<tr><td>>hg38_21_+_8433172_8446622_RNA45SN1_45S_50ntFlanks@9738.9765.28|GAAACGATCTCAACCTATTCTCAAACTT</td><td>>hg38_21_+_8433172_8446622_RNA45SN1_45S_50ntFlanks</td><td>9738</td><td>9765</td><td>28</td><td>GAAACGATCTCAACCTATTCTCAAACTT</td><td>3499.4</td><td>-3.8</td><td>0.9</td><td>-4.3</td><td>1.8E-05</td><td>2.5E-05</td><td>MB231/MB468</td><td>NUC</td></tr>
<tr><td>>hg38_MT_+_1621_3279_RNR2_16S_50ntFlanks@51.72.22|GCTAAACCTAGCCCCAAACCCA</td><td>>hg38_MT_+_1621_3279_RNR2_16S_50ntFlanks</td><td>51</td><td>72</td><td>22</td><td>GCTAAACCTAGCCCCAAACCCA</td><td>4784.9</td><td>4.5</td><td>1.1</td><td>4.3</td><td>1.8E-05</td><td>2.5E-05</td><td>MB231/MB468</td><td>MITO</td></tr>
<tr><td>>hg38_21_+_8433172_8446622_RNA45SN1_45S_50ntFlanks@9572.9601.30|CAAATCGGTCGTCCGACCTGGGTATAGGGG</td><td>>hg38_21_+_8433172_8446622_RNA45SN1_45S_50ntFlanks</td><td>9572</td><td>9601</td><td>30</td><td>CAAATCGGTCGTCCGACCTGGGTATAGGGG</td><td>400.2</td><td>-6.9</td><td>1.6</td><td>-4.3</td><td>1.9E-05</td><td>2.6E-05</td><td>BT20/MB468</td><td>NUC</td></tr>
<tr><td>>trnaMT_MetCAT_MT_+_4402_4469@51.71.21|TATACCCTTCCCGTACTACCA</td><td>>trnaMT_MetCAT_MT_+_4402_4469</td><td>51</td><td>71</td><td>21</td><td>TATACCCTTCCCGTACTACCA</td><td>212.5</td><td>5.6</td><td>1.3</td><td>4.3</td><td>1.8E-05</td><td>2.6E-05</td><td>MB231/MB468</td><td>NUC</td></tr>
<tr><td>>hg38_21_+_8433172_8446622_RNA45SN1_45S_50ntFlanks@8194.8213.20|CCCGTGGACGGTGTGAGGCC</td><td>>hg38_21_+_8433172_8446622_RNA45SN1_45S_50ntFlanks</td><td>8194</td><td>8213</td><td>20</td><td>CCCGTGGACGGTGTGAGGCC</td><td>651.2</td><td>-1.9</td><td>0.4</td><td>-4.4</td><td>1.3E-05</td><td>2.6E-05</td><td>BT20/MB468</td><td>MITO</td></tr>
<tr><td>>hg38_21_+_8433172_8446622_RNA45SN1_45S_50ntFlanks@3773.3791.19|CGGCCGGTACAGTGAAACT</td><td>>hg38_21_+_8433172_8446622_RNA45SN1_45S_50ntFlanks</td><td>3773</td><td>3791</td><td>19</td><td>CGGCCGGTACAGTGAAACT</td><td>291.1</td><td>-5.1</td><td>1.2</td><td>-4.3</td><td>1.9E-05</td><td>2.6E-05</td><td>BT20/MB468</td><td>NUC</td></tr>
<tr><td>>hg38_21_+_8433172_8446622_RNA45SN1_45S_50ntFlanks@10387.10424.38|TGTGAACAGCAGTTGAACATGGGTCAGTCGGTCCTGAG</td><td>>hg38_21_+_8433172_8446622_RNA45SN1_45S_50ntFlanks</td><td>10387</td><td>10424</td><td>38</td><td>TGTGAACAGCAGTTGAACATGGGTCAGTCGGTCCTGAG</td><td>14208.5</td><td>4.1</td><td>0.9</td><td>4.7</td><td>2.1E-06</td><td>2.6E-05</td><td>BT20/MB231</td><td>MITO</td></tr>
<tr><td>>trnaMT_ValTAC_MT_+_1602_1670@50.72.23|ACTTAACTTGACCGCTCTGACCA</td><td>>trnaMT_ValTAC_MT_+_1602_1670</td><td>50</td><td>72</td><td>23</td><td>ACTTAACTTGACCGCTCTGACCA</td><td>514.7</td><td>3.9</td><td>0.9</td><td>4.3</td><td>1.7E-05</td><td>2.6E-05</td><td>MB231/MB468</td><td>CYTO</td></tr>
<tr><td>>hg38_21_+_8433172_8446622_RNA45SN1_45S_50ntFlanks@6652.6696.45|GACTCTTAGCGGTGGATCACTCGGCTCGTGCGTCGATGAAGAACG</td><td>>hg38_21_+_8433172_8446622_RNA45SN1_45S_50ntFlanks</td><td>6652</td><td>6696</td><td>45</td><td>GACTCTTAGCGGTGGATCACTCGGCTCGTGCGTCGATGAAGAACG</td><td>5200.1</td><td>3.9</td><td>0.9</td><td>4.3</td><td>1.8E-05</td><td>2.6E-05</td><td>MB231/MB468</td><td>NUC</td></tr>
<tr><td>>hg38_21_+_8433172_8446622_RNA45SN1_45S_50ntFlanks@9454.9503.50|CCGCGCCGGGGAGGTGGAGCACGAGCGCACGTGTTAGGACCCGAAAGATG</td><td>>hg38_21_+_8433172_8446622_RNA45SN1_45S_50ntFlanks</td><td>9454</td><td>9503</td><td>50</td><td>CCGCGCCGGGGAGGTGGAGCACGAGCGCACGTGTTAGGACCCGAAAGATG</td><td>256.4</td><td>-6.0</td><td>1.4</td><td>-4.3</td><td>1.5E-05</td><td>2.6E-05</td><td>BT20/MB468</td><td>CYTO</td></tr>
<tr><td>>hg38_21_+_8433172_8446622_RNA45SN1_45S_50ntFlanks@475.509.35|CTTCGTGATCGATGTGGTGACGTCGTGCTCTCCCG</td><td>>hg38_21_+_8433172_8446622_RNA45SN1_45S_50ntFlanks</td><td>475</td><td>509</td><td>35</td><td>CTTCGTGATCGATGTGGTGACGTCGTGCTCTCCCG</td><td>943.9</td><td>4.6</td><td>1.1</td><td>4.3</td><td>1.8E-05</td><td>2.6E-05</td><td>MB231/MB468</td><td>MITO</td></tr>
<tr><td>>hg38_21_+_8433172_8446622_RNA45SN1_45S_50ntFlanks@4995.5013.19|ACAGATTGATAGCTCTTTC</td><td>>hg38_21_+_8433172_8446622_RNA45SN1_45S_50ntFlanks</td><td>4995</td><td>5013</td><td>19</td><td>ACAGATTGATAGCTCTTTC</td><td>1529.6</td><td>-7.6</td><td>1.7</td><td>-4.5</td><td>5.8E-06</td><td>2.6E-05</td><td>BT20/MB468</td><td>MP</td></tr>
<tr><td>>hg38_21_+_8433172_8446622_RNA45SN1_45S_50ntFlanks@11540.11560.21|CGCGGCGCCTCGCCTCGGCCG</td><td>>hg38_21_+_8433172_8446622_RNA45SN1_45S_50ntFlanks</td><td>11540</td><td>11560</td><td>21</td><td>CGCGGCGCCTCGCCTCGGCCG</td><td>260.9</td><td>-6.0</td><td>1.4</td><td>-4.3</td><td>1.5E-05</td><td>2.6E-05</td><td>BT20/MB468</td><td>CYTO</td></tr>
<tr><td>>am_trna119_LysCTT_1_-_145395522_145395594@13.34.22,trna11_LysCTT_5_-_180648979_180649051@13.34.22,trna128_LysCTT_1_-_55423542_55423614@13.34.22,trna13_LysCTT_6_+_26556774_26556846@13.34.22,trna6_LysCTT_19_-_52425396_52425468@13.34.22,trna7_LysCTT_16_+_3225692_3225764@13.34.22,trna9_LysCTT_5_+_180634755_180634827@13.34.22|CAGTCGGTAGAGCATGAGACTC</td><td>>am_trna119_LysCTT_1_-_145395522_145395594</td><td>13</td><td>34</td><td>22</td><td>CAGTCGGTAGAGCATGAGACTC</td><td>384.9</td><td>-4.8</td><td>1.0</td><td>-4.7</td><td>2.1E-06</td><td>2.6E-05</td><td>BT20/MB231</td><td>MITO</td></tr>
</table>

<table>
<tr><td>>hg38_21_+_8433172_8446622_RNA45SN1_45S_50ntFlanks@11914.11948.35|AGAGACATGAGAGGTGTAGAATAAGTGGGAGGCCC</td><td>>hg38_21_+_8433172_8446622_RNA45SN1_45S_50ntFlanks</td><td>11914</td><td>11948</td><td>35</td><td>AGAGACATGAGAGGTGTAGAATAAGTGGGAGGCCC</td><td>146.8</td><td>10.5</td><td>2.2</td><td>4.8</td><td>1.8E-06</td><td>2.6E-05</td><td>BT20/MB231</td><td>MP</td></tr>
<tr><td>>hg38_21_+_8433172_8446622_RNA45SN1_45S_50ntFlanks@12714.12747.34|GAGCCTCGGTTGGCCTCGGATAGCCGGTCCCCCG</td><td>>hg38_21_+_8433172_8446622_RNA45SN1_45S_50ntFlanks</td><td>12714</td><td>12747</td><td>34</td><td>GAGCCTCGGTTGGCCTCGGATAGCCGGTCCCCCG</td><td>836.4</td><td>-7.5</td><td>1.6</td><td>-4.5</td><td>5.8E-06</td><td>2.6E-05</td><td>BT20/MB468</td><td>MP</td></tr>
<tr><td>>hg38_1_-_228634819_228635039_RNA5S12_5S_50ntFlanks@151.170.20|ATACCGGGTGCTGTAGGCTT</td><td>>hg38_1_-_228634819_228635039_RNA5S12_5S_50ntFlanks</td><td>151</td><td>170</td><td>20</td><td>ATACCGGGTGCTGTAGGCTT</td><td>2285.8</td><td>-2.9</td><td>0.7</td><td>-4.3</td><td>1.8E-05</td><td>2.6E-05</td><td>MB231/MB468</td><td>MITO</td></tr>
<tr><td>>hg38_21_+_8433172_8446622_RNA45SN1_45S_50ntFlanks@6697.6718.22|CAGCTAGCTGCGAGAATTAATG</td><td>>hg38_21_+_8433172_8446622_RNA45SN1_45S_50ntFlanks</td><td>6697</td><td>6718</td><td>22</td><td>CAGCTAGCTGCGAGAATTAATG</td><td>1145.7</td><td>-7.6</td><td>1.8</td><td>-4.3</td><td>1.9E-05</td><td>2.6E-05</td><td>BT20/MB468</td><td>NUC</td></tr>
<tr><td>>hg38_21_+_8433172_8446622_RNA45SN1_45S_50ntFlanks@8112.8139.28|GGGCGCGGGACATGTGGCGTACGGAAGA</td><td>>hg38_21_+_8433172_8446622_RNA45SN1_45S_50ntFlanks</td><td>8112</td><td>8139</td><td>28</td><td>GGGCGCGGGACATGTGGCGTACGGAAGA</td><td>500.2</td><td>-6.9</td><td>1.6</td><td>-4.3</td><td>1.5E-05</td><td>2.6E-05</td><td>MB231/MB468</td><td>MP</td></tr>
<tr><td>>hg38_21_+_8433172_8446622_RNA45SN1_45S_50ntFlanks@11686.11728.43|TCAAAGTGAAGAAATTCAATGAAGCGCGGGTAAACGGCGGGAG</td><td>>hg38_21_+_8433172_8446622_RNA45SN1_45S_50ntFlanks</td><td>11686</td><td>11728</td><td>43</td><td>TCAAAGTGAAGAAATTCAATGAAGCGCGGGTAAACGGCGGGAG</td><td>72.3</td><td>-8.3</td><td>1.9</td><td>-4.4</td><td>1.3E-05</td><td>2.6E-05</td><td>BT20/MB468</td><td>MITO</td></tr>
<tr><td>>hg38_21_+_8433172_8446622_RNA45SN1_45S_50ntFlanks@5524.5550.27|GTAAAAGTCGTAACAAGGTTTCCGTAG</td><td>>hg38_21_+_8433172_8446622_RNA45SN1_45S_50ntFlanks</td><td>5524</td><td>5550</td><td>27</td><td>GTAAAAGTCGTAACAAGGTTTCCGTAG</td><td>245.4</td><td>-5.6</td><td>1.3</td><td>-4.3</td><td>1.6E-05</td><td>2.6E-05</td><td>BT20/MB468</td><td>CYTO</td></tr>
<tr><td>>am_MI0015825|hsa-mir-378c&WithFlank&10|-|130962582|130962674@17.36.20;MI0000786|hsa-mir-378a&WithFlank&5|+|149732819|149732896@49.68.20&[MIMAT0016847&hsa-miR-378c&offsets|0|-5;m-130&10|-|130962639|130962658&offsets|0|0];[MIMAT0000732&hsa-miR-378a-3p&offsets|0|-2;m-44&5|+|149732867|149732888&offsets|0|-2]|ACTGGACTTGGAGTCAGAAG</td><td>>am_MI0015825|hsa-mir-378c&WithFlank&10|-|130962582|130962674</td><td>17</td><td>36</td><td>20</td><td>ACTGGACTTGGAGTCAGAAG</td><td>277.1</td><td>4.4</td><td>1.0</td><td>4.3</td><td>1.8E-05</td><td>2.6E-05</td><td>MB231/MB468</td><td>MITO</td></tr>
<tr><td>>hg38_21_+_8433172_8446622_RNA45SN1_45S_50ntFlanks@8799.8823.25|CCTCGGGAGGGCGCGCGGGTCGGGG</td><td>>hg38_21_+_8433172_8446622_RNA45SN1_45S_50ntFlanks</td><td>8799</td><td>8823</td><td>25</td><td>CCTCGGGAGGGCGCGCGGGTCGGGG</td><td>803.9</td><td>-7.6</td><td>1.8</td><td>-4.3</td><td>1.9E-05</td><td>2.6E-05</td><td>BT20/MB468</td><td>NUC</td></tr>
<tr><td>>hg38_21_+_8433172_8446622_RNA45SN1_45S_50ntFlanks@11575.11609.35|ACTTAGAACTGGTGCGGACCAGGGGAATCCGACTG</td><td>>hg38_21_+_8433172_8446622_RNA45SN1_45S_50ntFlanks</td><td>11575</td><td>11609</td><td>35</td><td>ACTTAGAACTGGTGCGGACCAGGGGAATCCGACTG</td><td>494.5</td><td>-4.1</td><td>1.0</td><td>-4.3</td><td>1.7E-05</td><td>2.6E-05</td><td>MB231/MB468</td><td>CYTO</td></tr>
<tr><td>>hg38_21_+_8433172_8446622_RNA45SN1_45S_50ntFlanks@6719.6752.34|TGAATTGCAGGACACATTGATCATCGACACTTCG</td><td>>hg38_21_+_8433172_8446622_RNA45SN1_45S_50ntFlanks</td><td>6719</td><td>6752</td><td>34</td><td>TGAATTGCAGGACACATTGATCATCGACACTTCG</td><td>180.7</td><td>-5.8</td><td>1.3</td><td>-4.3</td><td>1.6E-05</td><td>2.6E-05</td><td>BT20/MB468</td><td>CYTO</td></tr>
<tr><td>>hg38_1_-_228634819_228635039_RNA5S12_5S_50ntFlanks@133.170.38|ATGGGAGACCGCCTGGGAATACCGGGTGCTGTAGGCTT</td><td>>hg38_1_-_228634819_228635039_RNA5S12_5S_50ntFlanks</td><td>133</td><td>170</td><td>38</td><td>ATGGGAGACCGCCTGGGAATACCGGGTGCTGTAGGCTT</td><td>7710.8</td><td>3.1</td><td>0.7</td><td>4.3</td><td>1.8E-05</td><td>2.6E-05</td><td>MB231/MB468</td><td>NUC</td></tr>
<tr><td>>hg38_21_+_8433172_8446622_RNA45SN1_45S_50ntFlanks@10351.10370.20|CGAGAACTTTGAAGGCCGAA</td><td>>hg38_21_+_8433172_8446622_RNA45SN1_45S_50ntFlanks</td><td>10351</td><td>10370</td><td>20</td><td>CGAGAACTTTGAAGGCCGAA</td><td>374.1</td><td>-4.8</td><td>1.1</td><td>-4.3</td><td>1.8E-05</td><td>2.6E-05</td><td>MB231/MB468</td><td>MITO</td></tr>
<tr><td>>MI0000114|hsa-mir-107&WithFlank&10|-|89592741|89592833@56.74.19(+2U);MI0000108|hsa-mir-103a-2&WithFlank&20|+|3917488|3917577@54.72.19(+2U);MI0000109|hsa-mir-103a-1&WithFlank&5|-|168560890|168560979@54.72.19(+2U)&[MIMAT0000104&hsa-miR-107&offsets|0|-4(+2U);m-78&10|-|89592758|89592778&offsets|0|-2(+2U)];[MIMAT0000101&hsa-miR-103a-3p&offsets|0|-4(+2U);m-12&20|+|3917541|3917563&offsets|0|-4(+2U)];[MIMAT0000101_1&hsa-miR-103a-3p&offsets|0|-4(+2U);m-13&5|-|168560904|168560926&offsets|0|-4(+2U);m-44220&5|-|168560908|168560926&offsets|0|0(+2U)]|AGCAGCATTGTACAGGGCTTT</td><td>>MI0000114|hsa-mir-107&WithFlank&10|-|89592741|89592833</td><td>56</td><td>74</td><td>19(+2U)</td><td>AGCAGCATTGTACAGGGCTTT</td><td>34.3</td><td>8.3</td><td>1.9</td><td>4.3</td><td>1.5E-05</td><td>2.6E-05</td><td>MB231/MB468</td><td>MP</td></tr>
<tr><td>>hg38_21_+_8433172_8446622_RNA45SN1_45S_50ntFlanks@4921.4941.21|ACCAGGAGTGGAGCCTGCGGC</td><td>>hg38_21_+_8433172_8446622_RNA45SN1_45S_50ntFlanks</td><td>4921</td><td>4941</td><td>21</td><td>ACCAGGAGTGGAGCCTGCGGC</td><td>108.1</td><td>-3.1</td><td>0.7</td><td>-4.3</td><td>1.5E-05</td><td>2.6E-05</td><td>MB231/MB468</td><td>MP</td></tr>
<tr><td>>hg38_21_+_8433172_8446622_RNA45SN1_45S_50ntFlanks@8105.8129.25|CGCGGCGGGGCGCGGGACATGTGGC</td><td>>hg38_21_+_8433172_8446622_RNA45SN1_45S_50ntFlanks</td><td>8105</td><td>8129</td><td>25</td><td>CGCGGCGGGGCGCGGGACATGTGGC</td><td>114.1</td><td>10.2</td><td>2.1</td><td>4.8</td><td>1.8E-06</td><td>2.6E-05</td><td>BT20/MB231</td><td>MP</td></tr>
<tr><td>>hg38_21_+_8433172_8446622_RNA45SN1_45S_50ntFlanks@4006.4026.21|ACCTCGGGCCGATCGCACGCC</td><td>>hg38_21_+_8433172_8446622_RNA45SN1_45S_50ntFlanks</td><td>4006</td><td>4026</td><td>21</td><td>ACCTCGGGCCGATCGCACGCC</td><td>294.5</td><td>-5.7</td><td>1.3</td><td>-4.3</td><td>1.6E-05</td><td>2.6E-05</td><td>BT20/MB468</td><td>CYTO</td></tr>
<tr><td>>MI0000272|hsa-mir-182&WithFlank&7|-|129770377|129770498@29.51.23&[MIMAT0000259&hsa-miR-182-5p&offsets|0|-1;m-22&7|-|129770449|129770470&offsets|0|+1]|TTTGGCAATGGTAGAACTCACAC</td><td>>MI0000272|hsa-mir-182&WithFlank&7|-|129770377|129770498</td><td>29</td><td>51</td><td>23</td><td>TTTGGCAATGGTAGAACTCACAC</td><td>354.7</td><td>2.6</td><td>0.6</td><td>4.3</td><td>1.8E-05</td><td>2.6E-05</td><td>MB231/MB468</td><td>NUC</td></tr>
<tr><td>>hg38_21_+_8433172_8446622_RNA45SN1_45S_50ntFlanks@12641.12659.19|CCATCTGTGGGATTATGAC</td><td>>hg38_21_+_8433172_8446622_RNA45SN1_45S_50ntFlanks</td><td>12641</td><td>12659</td><td>19</td><td>CCATCTGTGGGATTATGAC</td><td>798.7</td><td>-3.9</td><td>0.9</td><td>-4.3</td><td>1.7E-05</td><td>2.6E-05</td><td>MB231/MB468</td><td>CYTO</td></tr>
<tr><td>>hg38_21_+_8433172_8446622_RNA45SN1_45S_50ntFlanks@3811.3837.27|AGTTATGGTTCCTTTGGTCGCTCGCTC</td><td>>hg38_21_+_8433172_8446622_RNA45SN1_45S_50ntFlanks</td><td>3811</td><td>3837</td><td>27</td><td>AGTTATGGTTCCTTTGGTCGCTCGCTC</td><td>211.6</td><td>-4.6</td><td>1.1</td><td>-4.3</td><td>1.8E-05</td><td>2.6E-05</td><td>MB231/MB468</td><td>NUC</td></tr>
<tr><td>>hg38_21_+_8433172_8446622_RNA45SN1_45S_50ntFlanks@5467.5498.32|GGCCCACGGCCCTGGCGGAGCGCTGAGAAGAC</td><td>>hg38_21_+_8433172_8446622_RNA45SN1_45S_50ntFlanks</td><td>5467</td><td>5498</td><td>32</td><td>GGCCCACGGCCCTGGCGGAGCGCTGAGAAGAC</td><td>793.0</td><td>-5.9</td><td>1.4</td><td>-4.3</td><td>1.9E-05</td><td>2.6E-05</td><td>BT20/MB468</td><td>NUC</td></tr>
<tr><td>>hg38_21_+_8433172_8446622_RNA45SN1_45S_50ntFlanks@3705.3722.18|TACCTGGTTGATCCTGCC</td><td>>hg38_21_+_8433172_8446622_RNA45SN1_45S_50ntFlanks</td><td>3705</td><td>3722</td><td>18</td><td>TACCTGGTTGATCCTGCC</td><td>4043.5</td><td>2.7</td><td>0.6</td><td>4.3</td><td>1.8E-05</td><td>2.6E-05</td><td>MB231/MB468</td><td>NUC</td></tr>
<tr><td>>hg38_21_+_8433172_8446622_RNA45SN1_45S_50ntFlanks@11540.11582.43|CGCGGCGCCTCGCCTCGGCCGGCGCCTAGCAGCCGACTTAGAA</td><td>>hg38_21_+_8433172_8446622_RNA45SN1_45S_50ntFlanks</td><td>11540</td><td>11582</td><td>43</td><td>CGCGGCGCCTCGCCTCGGCCGGCGCCTAGCAGCCGACTTAGAA</td><td>1660.5</td><td>-5.4</td><td>1.3</td><td>-4.3</td><td>1.9E-05</td><td>2.6E-05</td><td>BT20/MB468</td><td>NUC</td></tr>
<tr><td>>hg38_21_+_8433172_8446622_RNA45SN1_45S_50ntFlanks@10685.10706.22|GGCGGCGTCCGGTGAGCTCTCG</td><td>>hg38_21_+_8433172_8446622_RNA45SN1_45S_50ntFlanks</td><td>10685</td><td>10706</td><td>22</td><td>GGCGGCGTCCGGTGAGCTCTCG</td><td>519.9</td><td>-5.3</td><td>1.2</td><td>-4.3</td><td>1.8E-05</td><td>2.6E-05</td><td>MB231/MB468</td><td>NUC</td></tr>
<tr><td>>hg38_21_+_8433172_8446622_RNA45SN1_45S_50ntFlanks@10316.10352.37|GGTGCAGATCTTGGTGGTAGTAGCAAATATTCAAACG</td><td>>hg38_21_+_8433172_8446622_RNA45SN1_45S_50ntFlanks</td><td>10316</td><td>10352</td><td>37</td><td>GGTGCAGATCTTGGTGGTAGTAGCAAATATTCAAACG</td><td>37.8</td><td>9.0</td><td>2.0</td><td>4.5</td><td>5.9E-06</td><td>2.6E-05</td><td>BT20/MB468</td><td>MP</td></tr>
<tr><td>>hg38_21_+_8433172_8446622_RNA45SN1_45S_50ntFlanks@9808.9838.31|TGGAATGCGAGTGCCTAGTGGGCCACTTTTG</td><td>>hg38_21_+_8433172_8446622_RNA45SN1_45S_50ntFlanks</td><td>9808</td><td>9838</td><td>31</td><td>TGGAATGCGAGTGCCTAGTGGGCCACTTTTG</td><td>64.4</td><td>-9.1</td><td>2.0</td><td>-4.5</td><td>5.9E-06</td><td>2.6E-05</td><td>BT20/MB468</td><td>MP</td></tr>
<tr><td>>hg38_21_+_8433172_8446622_RNA45SN1_45S_50ntFlanks@12003.12033.31|GCGGGCCGCCGGTGAAATACCACTACTCTGA</td><td>>hg38_21_+_8433172_8446622_RNA45SN1_45S_50ntFlanks</td><td>12003</td><td>12033</td><td>31</td><td>GCGGGCCGCCGGTGAAATACCACTACTCTGA</td><td>964.5</td><td>4.0</td><td>0.9</td><td>4.3</td><td>1.8E-05</td><td>2.6E-05</td><td>MB231/MB468</td><td>NUC</td></tr>
<tr><td>>MI0000440|hsa-mir-27b&WithFlank&9|+|95085439|95085547@67.85.19(+1A)&[MIMAT0000419&hsa-miR-27b-3p&offsets|0|-2(+1A);m-39&9|+|95085505|95085525&offsets|0|-2(+1A)]|TTCACAGTGGCTAAGTTCTA</td><td>>MI0000440|hsa-mir-27b&WithFlank&9|+|95085439|95085547</td><td>67</td><td>85</td><td>19(+1A)</td><td>TTCACAGTGGCTAAGTTCTA</td><td>598.5</td><td>4.7</td><td>1.0</td><td>4.5</td><td>5.9E-06</td><td>2.6E-05</td><td>BT20/MB468</td><td>MP</td></tr>
<tr><td>>hg38_21_+_8433172_8446622_RNA45SN1_45S_50ntFlanks@11768.11804.37|CTAATTAGTGACGCGCATGAATGGATGAACGAGATTC</td><td>>hg38_21_+_8433172_8446622_RNA45SN1_45S_50ntFlanks</td><td>11768</td><td>11804</td><td>37</td><td>CTAATTAGTGACGCGCATGAATGGATGAACGAGATTC</td><td>390.5</td><td>-5.9</td><td>1.4</td><td>-4.3</td><td>1.6E-05</td><td>2.6E-05</td><td>BT20/MB468</td><td>CYTO</td></tr>
<tr><td>>hg38_21_+_8433172_8446622_RNA45SN1_45S_50ntFlanks@3842.3884.43|CCTACTTGGATAACTGTGGTAATTCTAGAGCTAATACATGCCG</td><td>>hg38_21_+_8433172_8446622_RNA45SN1_45S_50ntFlanks</td><td>3842</td><td>3884</td><td>43</td><td>CCTACTTGGATAACTGTGGTAATTCTAGAGCTAATACATGCCG</td><td>520.5</td><td>6.6</td><td>1.5</td><td>4.3</td><td>1.7E-05</td><td>2.6E-05</td><td>MB231/MB468</td><td>CYTO</td></tr>
<tr><td>>am_trna128_GlyGCC_6_-_27870686_27870756@1.29.29,trna133_GlyCCC_1_-_16872434_16872504@1.29.29,trna18_GlyGCC_16_+_70822597_70822667@1.29.29,trna19_GlyGCC_16_+_70823410_70823480@1.29.29,trna19_GlyGCC_2_-_157257659_157257729@1.29.29,trna24_GlyGCC_16_-_70812942_70813012@1.29.29,trna25_GlyGCC_16_-_70812114_70812184@1.29.29,trna4_GlyCCC_1_+_17188416_17188486@1.29.29,trna5_GlyGCC_17_+_8029064_8029134@1.29.29,trna68_GlyGCC_1_-_161493637_161493707@1.29.29|GCATTGGTGGTTCAGTGGTAGAATTCTCG</td><td>>am_trna128_GlyGCC_6_-_27870686_27870756</td><td>1</td><td>29</td><td>29</td><td>GCATTGGTGGTTCAGTGGTAGAATTCTCG</td><td>49049.2</td><td>-3.4</td><td>0.8</td><td>-4.3</td><td>1.7E-05</td><td>2.6E-05</td><td>MB231/MB468</td><td>CYTO</td></tr>
<tr><td>>MI0016005|hsa-mir-3615&WithFlank&17|+|74748607|74748705@57.79.23&[MIMAT0017994&hsa-miR-3615&offsets|0|+2;m-271&17|+|74748663|74748684&offsets|0|+1]|TCTCTCGGCTCCTCGCGGCTCGC</td><td>>MI0016005|hsa-mir-3615&WithFlank&17|+|74748607|74748705</td><td>57</td><td>79</td><td>23</td><td>TCTCTCGGCTCCTCGCGGCTCGC</td><td>300.9</td><td>3.4</td><td>0.8</td><td>4.3</td><td>1.8E-05</td><td>2.6E-05</td><td>MB231/MB468</td><td>NUC</td></tr>
<tr><td>>hg38_21_+_8433172_8446622_RNA45SN1_45S_50ntFlanks@10357.10390.34|CTTTGAAGGCCGAAGTGGAGAAGGGTTCCATGTG</td><td>>hg38_21_+_8433172_8446622_RNA45SN1_45S_50ntFlanks</td><td>10357</td><td>10390</td><td>34</td><td>CTTTGAAGGCCGAAGTGGAGAAGGGTTCCATGTG</td><td>285.8</td><td>-4.7</td><td>1.1</td><td>-4.3</td><td>1.9E-05</td><td>2.6E-05</td><td>MB231/MB468</td><td>MITO</td></tr>
<tr><td>>MI0001445|hsa-mir-423&WithFlank&17|+|30117073|30117178@58.80.23(+1C)&[MIMAT0001340&hsa-miR-423-3p&offsets|-1|-1(+1C);m-100&17|+|30117131|30117153&offsets|-1|-1(+1C)]|AAGCTCGGTCTGAGGCCCCTCAGC</td><td>>MI0001445|hsa-mir-423&WithFlank&17|+|30117073|30117178</td><td>58</td><td>80</td><td>23(+1C)</td><td>AAGCTCGGTCTGAGGCCCCTCAGC</td><td>247.1</td><td>7.0</td><td>1.6</td><td>4.5</td><td>6.0E-06</td><td>2.6E-05</td><td>BT20/MB468</td><td>MP</td></tr>
<tr><td>>hg38_21_+_8433172_8446622_RNA45SN1_45S_50ntFlanks@11322.11342.21|ACGTCTCGTCGCGCGCGCGTC</td><td>>hg38_21_+_8433172_8446622_RNA45SN1_45S_50ntFlanks</td><td>11322</td><td>11342</td><td>21</td><td>ACGTCTCGTCGCGCGCGCGTC</td><td>680.3</td><td>-3.8</td><td>0.8</td><td>-4.5</td><td>6.0E-06</td><td>2.6E-05</td><td>BT20/MB468</td><td>MP</td></tr>
<tr><td>>hg38_21_+_8433172_8446622_RNA45SN1_45S_50ntFlanks@3838.3873.36|CTCTCCTACTTGGATAACTGTGGTAATTCTAGAGCT</td><td>>hg38_21_+_8433172_8446622_RNA45SN1_45S_50ntFlanks</td><td>3838</td><td>3873</td><td>36</td><td>CTCTCCTACTTGGATAACTGTGGTAATTCTAGAGCT</td><td>848.1</td><td>-5.7</td><td>1.3</td><td>-4.3</td><td>1.9E-05</td><td>2.6E-05</td><td>BT20/MB468</td><td>NUC</td></tr>
<tr><td>>hg38_21_+_8433172_8446622_RNA45SN1_45S_50ntFlanks@11586.11612.27|GTGCGGACCAGGGGAATCCGACTGTTT</td><td>>hg38_21_+_8433172_8446622_RNA45SN1_45S_50ntFlanks</td><td>11586</td><td>11612</td><td>27</td><td>GTGCGGACCAGGGGAATCCGACTGTTT</td><td>1086.5</td><td>-3.1</td><td>0.7</td><td>-4.3</td><td>1.6E-05</td><td>2.6E-05</td><td>BT20/MB468</td><td>CYTO</td></tr>
<tr><td>>hg38_21_+_8433172_8446622_RNA45SN1_45S_50ntFlanks@12423.12443.21|TCGGCTCTTCCTATCATTGTG</td><td>>hg38_21_+_8433172_8446622_RNA45SN1_45S_50ntFlanks</td><td>12423</td><td>12443</td><td>21</td><td>TCGGCTCTTCCTATCATTGTG</td><td>407.2</td><td>-4.4</td><td>1.0</td><td>-4.3</td><td>1.6E-05</td><td>2.6E-05</td><td>BT20/MB468</td><td>CYTO</td></tr>
<tr><td>>hg38_21_+_8433172_8446622_RNA45SN1_45S_50ntFlanks@12999.13039.41|CTGCGATCTATTGAAAGTCAGCCCTCGACACAAGGGTTTGT</td><td>>hg38_21_+_8433172_8446622_RNA45SN1_45S_50ntFlanks</td><td>12999</td><td>13039</td><td>41</td><td>CTGCGATCTATTGAAAGTCAGCCCTCGACACAAGGGTTTGT</td><td>6658.0</td><td>4.1</td><td>0.9</td><td>4.4</td><td>1.3E-05</td><td>2.6E-05</td><td>BT20/MB468</td><td>MITO</td></tr>
<tr><td>>hg38_21_+_8433172_8446622_RNA45SN1_45S_50ntFlanks@12473.12497.25|CACCCACTAATAGGGAACGTGAGCT</td><td>>hg38_21_+_8433172_8446622_RNA45SN1_45S_50ntFlanks</td><td>12473</td><td>12497</td><td>25</td><td>CACCCACTAATAGGGAACGTGAGCT</td><td>222.7</td><td>2.7</td><td>0.6</td><td>4.3</td><td>1.8E-05</td><td>2.6E-05</td><td>MB231/MB468</td><td>CYTO</td></tr>
<tr><td>>hg38_1_-_228634819_228635039_RNA5S12_5S_50ntFlanks@142.170.29|CGCCTGGGAATACCGGGTGCTGTAGGCTT</td><td>>hg38_1_-_228634819_228635039_RNA5S12_5S_50ntFlanks</td><td>142</td><td>170</td><td>29</td><td>CGCCTGGGAATACCGGGTGCTGTAGGCTT</td><td>1424.3</td><td>-2.5</td><td>0.6</td><td>-4.3</td><td>1.5E-05</td><td>2.6E-05</td><td>MB231/MB468</td><td>MP</td></tr>
<tr><td>>hg38_21_+_8433172_8446622_RNA45SN1_45S_50ntFlanks@9508.9526.19|ACTATGCCTGGGCAGGGCG</td><td>>hg38_21_+_8433172_8446622_RNA45SN1_45S_50ntFlanks</td><td>9508</td><td>9526</td><td>19</td><td>ACTATGCCTGGGCAGGGCG</td><td>103.9</td><td>-10.6</td><td>2.4</td><td>-4.3</td><td>1.5E-05</td><td>2.6E-05</td><td>MB231/MB468</td><td>MP</td></tr>
<tr><td>>hg38_21_+_8433172_8446622_RNA45SN1_45S_50ntFlanks@9307.9337.31|AAGGAGTCTAACACGTGCGCGAGTCGGGGGC</td><td>>hg38_21_+_8433172_8446622_RNA45SN1_45S_50ntFlanks</td><td>9307</td><td>9337</td><td>31</td><td>AAGGAGTCTAACACGTGCGCGAGTCGGGGGC</td><td>631.8</td><td>-5.5</td><td>1.3</td><td>-4.3</td><td>1.5E-05</td><td>2.6E-05</td><td>MB231/MB468</td><td>MP</td></tr>
<tr><td>>hg38_MT_+_598_1651_RNR1_12S_50ntFlanks@634.657.24|AACCTCACCACCTCTTGCTCAGCC</td><td>>hg38_MT_+_598_1651_RNR1_12S_50ntFlanks</td><td>634</td><td>657</td><td>24</td><td>AACCTCACCACCTCTTGCTCAGCC</td><td>531.1</td><td>8.3</td><td>1.9</td><td>4.4</td><td>1.3E-05</td><td>2.6E-05</td><td>BT20/MB468</td><td>MITO</td></tr>
<tr><td>>hg38_21_+_8433172_8446622_RNA45SN1_45S_50ntFlanks@10065.10082.18|CGAGAGTGGACGGGAGCG</td><td>>hg38_21_+_8433172_8446622_RNA45SN1_45S_50ntFlanks</td><td>10065</td><td>10082</td><td>18</td><td>CGAGAGTGGACGGGAGCG</td><td>85.4</td><td>9.7</td><td>2.0</td><td>4.8</td><td>1.8E-06</td><td>2.6E-05</td><td>BT20/MB231</td><td>MP</td></tr>
</table>

<table>
<tr><td>>hg38_21_+_8433172_8446622_RNA45SN1_45S_50ntFlanks@10237.10260.24|ACGCTACGCCGCGACGAGTAGGAG</td><td>>hg38_21_+_8433172_8446622_RNA45SN1_45S_50ntFlanks</td><td>10237</td><td>10260</td><td>24</td><td>ACGCTACGCCGCGACGAGTAGGAG</td><td>102.2</td><td>10.0</td><td>2.1</td><td>4.8</td><td>1.8E-06</td><td>2.6E-05</td><td>BT20/MB231</td><td>MP</td></tr>
<tr><td>>hg38_21_+_8433172_8446622_RNA45SN1_45S_50ntFlanks@11848.11886.39|ACGGGCTTGGCGGAATCAGCGGGGAAAGAAGACCCTGTT</td><td>>hg38_21_+_8433172_8446622_RNA45SN1_45S_50ntFlanks</td><td>11848</td><td>11886</td><td>39</td><td>ACGGGCTTGGCGGAATCAGCGGGGAAAGAAGACCCTGTT</td><td>98.3</td><td>10.0</td><td>2.1</td><td>4.8</td><td>1.8E-06</td><td>2.6E-05</td><td>BT20/MB231</td><td>MP</td></tr>
<tr><td>>hg38_21_+_8433172_8446622_RNA45SN1_45S_50ntFlanks@5468.5514.47|GCCCACGGCCCTGGCGGAGCGCTGAGAAGACGGTCGAACTTGACTAT</td><td>>hg38_21_+_8433172_8446622_RNA45SN1_45S_50ntFlanks</td><td>5468</td><td>5514</td><td>47</td><td>GCCCACGGCCCTGGCGGAGCGCTGAGAAGACGGTCGAACTTGACTAT</td><td>100.7</td><td>-6.3</td><td>1.5</td><td>-4.3</td><td>1.6E-05</td><td>2.6E-05</td><td>BT20/MB468</td><td>CYTO</td></tr>
<tr><td>>trna10_AspGTC_12_-_125424193_125424264@36.71.36,trna12_AspGTC_12_-_125411891_125411962@36.71.36,trna144_AspGTC_6_-_27551236_27551307@36.71.36,trna38_AspGTC_17_-_8125556_8125627@36.71.36,trna45_AspGTC_6_+_27447453_27447524@36.71.36,trna48_AspGTC_6_+_27471523_27471594@36.71.36,trna4_AspGTC_12_+_96429799_96429870@36.71.36,trna69_AspGTC_1_-_161440205_161440276@36.71.36,trna72_AspGTC_1_-_161432824_161432895@36.71.36,trna75_AspGTC_1_-_161425414_161425485@36.71.36,trna78_AspGTC_1_-_161418033_161418104@36.71.36,trna81_AspGTC_1_-_161410615_161410686@36.71.36|CACGCGGGAGACCGGGGTTCGATTCCCCGACGGGGA</td><td>>trna10_AspGTC_12_-_125424193_125424264</td><td>36</td><td>71</td><td>36</td><td>CACGCGGGAGACCGGGGTTCGATTCCCCGACGGGGA</td><td>473.7</td><td>4.1</td><td>1.0</td><td>4.3</td><td>1.8E-05</td><td>2.6E-05</td><td>MB231/MB468</td><td>CYTO</td></tr>
<tr><td>>MI0000084|hsa-mir-26b&WithFlank&2|+|218402640|218402728@18.39.22(+2U)&[MIMAT0000083&hsa-miR-26b-5p&offsets|0|+1(+2U);m-53&2|+|218402657|218402678&offsets|0|0(+2U)]|TTCAAGTAATTCAGGATAGGTTTT</td><td>>MI0000084|hsa-mir-26b&WithFlank&2|+|218402640|218402728</td><td>18</td><td>39</td><td>22(+2U)</td><td>TTCAAGTAATTCAGGATAGGTTTT</td><td>38.1</td><td>8.5</td><td>2.0</td><td>4.3</td><td>1.5E-05</td><td>2.6E-05</td><td>MB231/MB468</td><td>MP</td></tr>
<tr><td>>hg38_21_+_8433172_8446622_RNA45SN1_45S_50ntFlanks@5472.5515.44|ACGGCCCTGGCGGAGCGCTGAGAAGACGGTCGAACTTGACTATC</td><td>>hg38_21_+_8433172_8446622_RNA45SN1_45S_50ntFlanks</td><td>5472</td><td>5515</td><td>44</td><td>ACGGCCCTGGCGGAGCGCTGAGAAGACGGTCGAACTTGACTATC</td><td>369.8</td><td>4.5</td><td>1.0</td><td>4.3</td><td>1.6E-05</td><td>2.6E-05</td><td>BT20/MB468</td><td>CYTO</td></tr>
<tr><td>>hg38_21_+_8433172_8446622_RNA45SN1_45S_50ntFlanks@12607.12626.20|TGTGCTTGGCTGAGGAGCCA</td><td>>hg38_21_+_8433172_8446622_RNA45SN1_45S_50ntFlanks</td><td>12607</td><td>12626</td><td>20</td><td>TGTGCTTGGCTGAGGAGCCA</td><td>157.8</td><td>-4.5</td><td>1.1</td><td>-4.3</td><td>1.8E-05</td><td>2.6E-05</td><td>MB231/MB468</td><td>NUC</td></tr>
<tr><td>>hg38_21_+_8433172_8446622_RNA45SN1_45S_50ntFlanks@8184.8223.40|CGAGGCCCAGCCCGTGGACGGTGTGAGGCCGGTAGCGGCC</td><td>>hg38_21_+_8433172_8446622_RNA45SN1_45S_50ntFlanks</td><td>8184</td><td>8223</td><td>40</td><td>CGAGGCCCAGCCCGTGGACGGTGTGAGGCCGGTAGCGGCC</td><td>46.2</td><td>-8.6</td><td>1.9</td><td>-4.5</td><td>6.0E-06</td><td>2.6E-05</td><td>BT20/MB468</td><td>MP</td></tr>
<tr><td>>hg38_MT_+_1621_3279_RNR2_16S_50ntFlanks@496.526.31|CTTTGGACACTAGGAAAAAACCTTGTAGAGA</td><td>>hg38_MT_+_1621_3279_RNR2_16S_50ntFlanks</td><td>496</td><td>526</td><td>31</td><td>CTTTGGACACTAGGAAAAAACCTTGTAGAGA</td><td>443.6</td><td>7.2</td><td>1.6</td><td>4.5</td><td>6.0E-06</td><td>2.6E-05</td><td>BT20/MB468</td><td>MP</td></tr>
<tr><td>>hg38_21_+_8433172_8446622_RNA45SN1_45S_50ntFlanks@10669.10706.38|AAAGCGTCGCGGTTCCGGCGGCGTCCGGTGAGCTCTCG</td><td>>hg38_21_+_8433172_8446622_RNA45SN1_45S_50ntFlanks</td><td>10669</td><td>10706</td><td>38</td><td>AAAGCGTCGCGGTTCCGGCGGCGTCCGGTGAGCTCTCG</td><td>527.6</td><td>-5.5</td><td>1.3</td><td>-4.3</td><td>1.6E-05</td><td>2.6E-05</td><td>BT20/MB468</td><td>CYTO</td></tr>
<tr><td>>hg38_21_+_8433172_8446622_RNA45SN1_45S_50ntFlanks@11207.11233.27|GGCGGCGACTCTGGACGCGAGCCGGGC</td><td>>hg38_21_+_8433172_8446622_RNA45SN1_45S_50ntFlanks</td><td>11207</td><td>11233</td><td>27</td><td>GGCGGCGACTCTGGACGCGAGCCGGGC</td><td>213.8</td><td>-2.4</td><td>0.6</td><td>-4.3</td><td>1.6E-05</td><td>2.6E-05</td><td>BT20/MB468</td><td>CYTO</td></tr>
<tr><td>>hg38_21_+_8433172_8446622_RNA45SN1_45S_50ntFlanks@12686.12711.26|CCAGGCGGAACGATACGGCAGCGCCG</td><td>>hg38_21_+_8433172_8446622_RNA45SN1_45S_50ntFlanks</td><td>12686</td><td>12711</td><td>26</td><td>CCAGGCGGAACGATACGGCAGCGCCG</td><td>324.7</td><td>-3.5</td><td>0.8</td><td>-4.4</td><td>1.3E-05</td><td>2.6E-05</td><td>BT20/MB468</td><td>MITO</td></tr>
<tr><td>>hg38_21_+_8433172_8446622_RNA45SN1_45S_50ntFlanks@10482.10511.30|TCGAAAGGGAGTCGGGTTCAGATCCCCGAA</td><td>>hg38_21_+_8433172_8446622_RNA45SN1_45S_50ntFlanks</td><td>10482</td><td>10511</td><td>30</td><td>TCGAAAGGGAGTCGGGTTCAGATCCCCGAA</td><td>417.5</td><td>-6.2</td><td>1.4</td><td>-4.3</td><td>1.6E-05</td><td>2.6E-05</td><td>BT20/MB468</td><td>CYTO</td></tr>
<tr><td>>hg38_21_+_8433172_8446622_RNA45SN1_45S_50ntFlanks@5490.5525.36|TGAGAAGACGGTCGAACTTGACTATCTAGAGGAAGT</td><td>>hg38_21_+_8433172_8446622_RNA45SN1_45S_50ntFlanks</td><td>5490</td><td>5525</td><td>36</td><td>TGAGAAGACGGTCGAACTTGACTATCTAGAGGAAGT</td><td>31.0</td><td>8.2</td><td>1.9</td><td>4.3</td><td>1.6E-05</td><td>2.6E-05</td><td>MB231/MB468</td><td>MP</td></tr>
<tr><td>>hg38_21_+_8433172_8446622_RNA45SN1_45S_50ntFlanks@10351.10392.42|CGAGAACTTTGAAGGCCGAAGTGGAGAAGGGTTCCATGTGAA</td><td>>hg38_21_+_8433172_8446622_RNA45SN1_45S_50ntFlanks</td><td>10351</td><td>10392</td><td>42</td><td>CGAGAACTTTGAAGGCCGAAGTGGAGAAGGGTTCCATGTGAA</td><td>473.5</td><td>-7.3</td><td>1.7</td><td>-4.3</td><td>2.0E-05</td><td>2.6E-05</td><td>BT20/MB468</td><td>NUC</td></tr>
<tr><td>>hg38_21_+_8433172_8446622_RNA45SN1_45S_50ntFlanks@12416.12444.29|TTCGATGTCGGCTCTTCCTATCATTGTGA</td><td>>hg38_21_+_8433172_8446622_RNA45SN1_45S_50ntFlanks</td><td>12416</td><td>12444</td><td>29</td><td>TTCGATGTCGGCTCTTCCTATCATTGTGA</td><td>699.4</td><td>2.9</td><td>0.7</td><td>4.4</td><td>1.3E-05</td><td>2.6E-05</td><td>BT20/MB468</td><td>MITO</td></tr>
<tr><td>>hg38_21_+_8433172_8446622_RNA45SN1_45S_50ntFlanks@4567.4589.23|TAATGGAATAGGACCGCGGTTCT</td><td>>hg38_21_+_8433172_8446622_RNA45SN1_45S_50ntFlanks</td><td>4567</td><td>4589</td><td>23</td><td>TAATGGAATAGGACCGCGGTTCT</td><td>338.0</td><td>-2.8</td><td>0.7</td><td>-4.3</td><td>1.8E-05</td><td>2.6E-05</td><td>MB231/MB468</td><td>CYTO</td></tr>
<tr><td>>hg38_21_+_8433172_8446622_RNA45SN1_45S_50ntFlanks@8356.8378.23|GAAAAGAACTTTGAAGAGAGAGT</td><td>>hg38_21_+_8433172_8446622_RNA45SN1_45S_50ntFlanks</td><td>8356</td><td>8378</td><td>23</td><td>GAAAAGAACTTTGAAGAGAGAGT</td><td>688.1</td><td>-3.0</td><td>0.7</td><td>-4.3</td><td>1.6E-05</td><td>2.6E-05</td><td>BT20/MB468</td><td>CYTO</td></tr>
<tr><td>>hg38_21_+_8433172_8446622_RNA45SN1_45S_50ntFlanks@6734.6758.25|ATTGATCATCGACACTTCGAACGCA</td><td>>hg38_21_+_8433172_8446622_RNA45SN1_45S_50ntFlanks</td><td>6734</td><td>6758</td><td>25</td><td>ATTGATCATCGACACTTCGAACGCA</td><td>120.2</td><td>-4.7</td><td>1.1</td><td>-4.3</td><td>1.6E-05</td><td>2.6E-05</td><td>MB231/MB468</td><td>MP</td></tr>
<tr><td>>hg38_21_+_8433172_8446622_RNA45SN1_45S_50ntFlanks@11569.11592.24|CAGCCGACTTAGAACTGGTGCGGA</td><td>>hg38_21_+_8433172_8446622_RNA45SN1_45S_50ntFlanks</td><td>11569</td><td>11592</td><td>24</td><td>CAGCCGACTTAGAACTGGTGCGGA</td><td>361.7</td><td>-4.6</td><td>1.1</td><td>-4.3</td><td>1.6E-05</td><td>2.7E-05</td><td>BT20/MB468</td><td>CYTO</td></tr>
<tr><td>>hg38_21_+_8433172_8446622_RNA45SN1_45S_50ntFlanks@5084.5121.38|CGAACGAGACTCTGGCATGCTAACTAGTTACGCGACCC</td><td>>hg38_21_+_8433172_8446622_RNA45SN1_45S_50ntFlanks</td><td>5084</td><td>5121</td><td>38</td><td>CGAACGAGACTCTGGCATGCTAACTAGTTACGCGACCC</td><td>664.0</td><td>-4.5</td><td>1.1</td><td>-4.3</td><td>1.9E-05</td><td>2.7E-05</td><td>MB231/MB468</td><td>NUC</td></tr>
<tr><td>>hg38_MT_+_598_1651_RNR1_12S_50ntFlanks@428.469.42|AAAGTGGCTTTAACATATCTGAACACACAATAGCTAAGACCC</td><td>>hg38_MT_+_598_1651_RNR1_12S_50ntFlanks</td><td>428</td><td>469</td><td>42</td><td>AAAGTGGCTTTAACATATCTGAACACACAATAGCTAAGACCC</td><td>579.0</td><td>7.2</td><td>1.7</td><td>4.3</td><td>1.9E-05</td><td>2.7E-05</td><td>MB231/MB468</td><td>MITO</td></tr>
<tr><td>>MI0000086|hsa-mir-28&WithFlank&3|+|188688775|188688872@60.81.22(+1U)&[MIMAT0004502&hsa-miR-28-3p&offsets|0|0(+1U);m-28&3|+|188688834|188688855&offsets|0|0(+1U)]|CACTAGATTGTGAGCTCCTGGAT</td><td>>MI0000086|hsa-mir-28&WithFlank&3|+|188688775|188688872</td><td>60</td><td>81</td><td>22(+1U)</td><td>CACTAGATTGTGAGCTCCTGGAT</td><td>320.9</td><td>5.2</td><td>1.2</td><td>4.3</td><td>1.6E-05</td><td>2.7E-05</td><td>MB231/MB468</td><td>MP</td></tr>
<tr><td>>trna10_LysCTT_16_+_3241501_3241573@8.33.26,trna13_LysCTT_14_-_58706613_58706685@8.33.26,trna2_LysCTT_15_+_79152904_79152976@8.33.26,trna4_LysCTT_18_-_43669271_43669343@8.33.26|TAGCTCAGTCGGTAGAGCATGGGACT</td><td>>trna10_LysCTT_16_+_3241501_3241573</td><td>8</td><td>33</td><td>26</td><td>TAGCTCAGTCGGTAGAGCATGGGACT</td><td>31.6</td><td>8.2</td><td>1.9</td><td>4.3</td><td>1.6E-05</td><td>2.7E-05</td><td>MB231/MB468</td><td>MP</td></tr>
<tr><td>>hg38_MT_+_1621_3279_RNR2_16S_50ntFlanks@1175.1215.41|TGCATTAAAAATTTCGGTTGGGGCGACCTCGGAGCAGAACC</td><td>>hg38_MT_+_1621_3279_RNR2_16S_50ntFlanks</td><td>1175</td><td>1215</td><td>41</td><td>TGCATTAAAAATTTCGGTTGGGGCGACCTCGGAGCAGAACC</td><td>460.6</td><td>8.0</td><td>1.8</td><td>4.4</td><td>1.4E-05</td><td>2.7E-05</td><td>BT20/MB468</td><td>MITO</td></tr>
<tr><td>>MI0000478|hsa-mir-149&WithFlank&2|+|240455995|240456095@21.43.23&[MIMAT0000450&hsa-miR-149-5p&offsets|0|0;m-195&2|+|240456015|240456037&offsets|0|0]|TCTGGCTCCGTGTCTTCACTCCC</td><td>>MI0000478|hsa-mir-149&WithFlank&2|+|240455995|240456095</td><td>21</td><td>43</td><td>23</td><td>TCTGGCTCCGTGTCTTCACTCCC</td><td>2106.5</td><td>3.6</td><td>0.8</td><td>4.7</td><td>2.2E-06</td><td>2.7E-05</td><td>BT20/MB231</td><td>MITO</td></tr>
<tr><td>>MI0000255|hsa-mir-30d&WithFlank&8|-|134804870|134804951@12.35.24(+4U)&[MIMAT0000245&hsa-miR-30d-5p&offsets|0|+2(+4U);m-21&8|-|134804919|134804940&offsets|0|+2(+4U)]|TGTAAACATCCCCGACTGGAAGCTTTTT</td><td>>MI0000255|hsa-mir-30d&WithFlank&8|-|134804870|134804951</td><td>12</td><td>35</td><td>24(+4U)</td><td>TGTAAACATCCCCGACTGGAAGCTTTTT</td><td>213.7</td><td>-4.7</td><td>1.0</td><td>-4.7</td><td>2.2E-06</td><td>2.7E-05</td><td>BT20/MB231</td><td>MITO</td></tr>
<tr><td>>hg38_21_+_8433172_8446622_RNA45SN1_45S_50ntFlanks@7989.8006.18|AGACGTGGCGACCCGCTG</td><td>>hg38_21_+_8433172_8446622_RNA45SN1_45S_50ntFlanks</td><td>7989</td><td>8006</td><td>18</td><td>AGACGTGGCGACCCGCTG</td><td>18064.9</td><td>-5.2</td><td>1.2</td><td>-4.5</td><td>6.1E-06</td><td>2.7E-05</td><td>BT20/MB468</td><td>MP</td></tr>
<tr><td>>hg38_21_+_8433172_8446622_RNA45SN1_45S_50ntFlanks@10731.10757.27|AGGGTGTAAATCTCGCGCCGGGCCGTA</td><td>>hg38_21_+_8433172_8446622_RNA45SN1_45S_50ntFlanks</td><td>10731</td><td>10757</td><td>27</td><td>AGGGTGTAAATCTCGCGCCGGGCCGTA</td><td>123.5</td><td>-3.8</td><td>0.9</td><td>-4.3</td><td>1.6E-05</td><td>2.7E-05</td><td>BT20/MB468</td><td>CYTO</td></tr>
<tr><td>>MI0003763|hsa-mir-767&WithFlank&X|-|152393415|152393535@33.56.24&[MIMAT0003882&hsa-miR-767-5p&offsets|0|+1;m-788&X|-|152393481|152393503&offsets|0|+1]|TGCACCATGGTTGTCTGAGCATGC</td><td>>MI0003763|hsa-mir-767&WithFlank&X|-|152393415|152393535</td><td>33</td><td>56</td><td>24</td><td>TGCACCATGGTTGTCTGAGCATGC</td><td>73.9</td><td>9.5</td><td>2.0</td><td>4.8</td><td>1.9E-06</td><td>2.7E-05</td><td>BT20/MB231</td><td>MP</td></tr>
<tr><td>>hg38_21_+_8433172_8446622_RNA45SN1_45S_50ntFlanks@10466.10488.23|CGTTGCCCTCGGCCGATCGAAAG</td><td>>hg38_21_+_8433172_8446622_RNA45SN1_45S_50ntFlanks</td><td>10466</td><td>10488</td><td>23</td><td>CGTTGCCCTCGGCCGATCGAAAG</td><td>401.3</td><td>-3.7</td><td>0.9</td><td>-4.3</td><td>1.9E-05</td><td>2.7E-05</td><td>MB231/MB468</td><td>NUC</td></tr>
<tr><td>>hg38_21_+_8433172_8446622_RNA45SN1_45S_50ntFlanks@11567.11587.21|AGCAGCCGACTTAGAACTGGT</td><td>>hg38_21_+_8433172_8446622_RNA45SN1_45S_50ntFlanks</td><td>11567</td><td>11587</td><td>21</td><td>AGCAGCCGACTTAGAACTGGT</td><td>387.9</td><td>3.1</td><td>0.7</td><td>4.3</td><td>1.9E-05</td><td>2.7E-05</td><td>MB231/MB468</td><td>NUC</td></tr>
<tr><td>>hg38_21_+_8433172_8446622_RNA45SN1_45S_50ntFlanks@12408.12433.26|TTTGATCCTTCGATGTCGGCTCTTCC</td><td>>hg38_21_+_8433172_8446622_RNA45SN1_45S_50ntFlanks</td><td>12408</td><td>12433</td><td>26</td><td>TTTGATCCTTCGATGTCGGCTCTTCC</td><td>441.2</td><td>-3.4</td><td>0.8</td><td>-4.3</td><td>2.0E-05</td><td>2.7E-05</td><td>BT20/MB468</td><td>NUC</td></tr>
<tr><td>>hg38_21_+_8433172_8446622_RNA45SN1_45S_50ntFlanks@9471.9503.33|AGCACGAGCGCACGTGTTAGGACCCGAAAGATG</td><td>>hg38_21_+_8433172_8446622_RNA45SN1_45S_50ntFlanks</td><td>9471</td><td>9503</td><td>33</td><td>AGCACGAGCGCACGTGTTAGGACCCGAAAGATG</td><td>328.8</td><td>-6.8</td><td>1.6</td><td>-4.3</td><td>2.0E-05</td><td>2.7E-05</td><td>BT20/MB468</td><td>NUC</td></tr>
<tr><td>>hg38_21_+_8433172_8446622_RNA45SN1_45S_50ntFlanks@5471.5501.31|CACGGCCCTGGCGGAGCGCTGAGAAGACGGT</td><td>>hg38_21_+_8433172_8446622_RNA45SN1_45S_50ntFlanks</td><td>5471</td><td>5501</td><td>31</td><td>CACGGCCCTGGCGGAGCGCTGAGAAGACGGT</td><td>2457.6</td><td>5.1</td><td>1.2</td><td>4.3</td><td>1.6E-05</td><td>2.7E-05</td><td>BT20/MB468</td><td>CYTO</td></tr>
<tr><td>>hg38_21_+_8433172_8446622_RNA45SN1_45S_50ntFlanks@12949.12972.24|ACGACCTGCTTCTGGGTCGGGGTT</td><td>>hg38_21_+_8433172_8446622_RNA45SN1_45S_50ntFlanks</td><td>12949</td><td>12972</td><td>24</td><td>ACGACCTGCTTCTGGGTCGGGGTT</td><td>56.9</td><td>-8.9</td><td>2.0</td><td>-4.5</td><td>6.1E-06</td><td>2.7E-05</td><td>BT20/MB468</td><td>MP</td></tr>
<tr><td>>hg38_21_+_8433172_8446622_RNA45SN1_45S_50ntFlanks@12669.12692.24|CTAAGTCAGAATCCCGCCCAGGCG</td><td>>hg38_21_+_8433172_8446622_RNA45SN1_45S_50ntFlanks</td><td>12669</td><td>12692</td><td>24</td><td>CTAAGTCAGAATCCCGCCCAGGCG</td><td>126.3</td><td>-6.3</td><td>1.5</td><td>-4.3</td><td>1.6E-05</td><td>2.7E-05</td><td>MB231/MB468</td><td>MP</td></tr>
<tr><td>>hg38_21_+_8433172_8446622_RNA45SN1_45S_50ntFlanks@6772.6807.36|GTTCCTCCCGGGGCTACGCCTGTCTGAGCGTCGCTT</td><td>>hg38_21_+_8433172_8446622_RNA45SN1_45S_50ntFlanks</td><td>6772</td><td>6807</td><td>36</td><td>GTTCCTCCCGGGGCTACGCCTGTCTGAGCGTCGCTT</td><td>2262.2</td><td>-1.9</td><td>0.4</td><td>-4.4</td><td>1.4E-05</td><td>2.7E-05</td><td>BT20/MB468</td><td>MITO</td></tr>
<tr><td>>hg38_21_+_8433172_8446622_RNA45SN1_45S_50ntFlanks@7968.8010.43|TCCGAGACGCGACCTCAGATCAGACGTGGCGACCCGCTGAATT</td><td>>hg38_21_+_8433172_8446622_RNA45SN1_45S_50ntFlanks</td><td>7968</td><td>8010</td><td>43</td><td>TCCGAGACGCGACCTCAGATCAGACGTGGCGACCCGCTGAATT</td><td>484.3</td><td>4.9</td><td>1.1</td><td>4.3</td><td>1.9E-05</td><td>2.7E-05</td><td>MB231/MB468</td><td>NUC</td></tr>
<tr><td>>hg38_21_+_8433172_8446622_RNA45SN1_45S_50ntFlanks@9574.9610.37|AATCGGTCGTCCGACCTGGGTATAGGGGCGAAAGACT</td><td>>hg38_21_+_8433172_8446622_RNA45SN1_45S_50ntFlanks</td><td>9574</td><td>9610</td><td>37</td><td>AATCGGTCGTCCGACCTGGGTATAGGGGCGAAAGACT</td><td>681.2</td><td>4.4</td><td>1.0</td><td>4.3</td><td>1.4E-05</td><td>2.7E-05</td><td>BT20/MB468</td><td>MITO</td></tr>
<tr><td>>hg38_21_+_8433172_8446622_RNA45SN1_45S_50ntFlanks@11769.11804.36|TAATTAGTGACGCGCATGAATGGATGAACGAGATTC</td><td>>hg38_21_+_8433172_8446622_RNA45SN1_45S_50ntFlanks</td><td>11769</td><td>11804</td><td>36</td><td>TAATTAGTGACGCGCATGAATGGATGAACGAGATTC</td><td>706.2</td><td>-4.3</td><td>1.0</td><td>-4.3</td><td>1.8E-05</td><td>2.7E-05</td><td>MB231/MB468</td><td>CYTO</td></tr>
<tr><td>>hg38_21_+_8433172_8446622_RNA45SN1_45S_50ntFlanks@10927.10946.20|CGAGGCGCCGCCGCCCCCCC</td><td>>hg38_21_+_8433172_8446622_RNA45SN1_45S_50ntFlanks</td><td>10927</td><td>10946</td><td>20</td><td>CGAGGCGCCGCCGCCCCCCC</td><td>873.3</td><td>-8.2</td><td>1.8</td><td>-4.5</td><td>6.1E-06</td><td>2.7E-05</td><td>BT20/MB468</td><td>MP</td></tr>
<tr><td>>hg38_21_+_8433172_8446622_RNA45SN1_45S_50ntFlanks@12324.12366.43|TTGGGTTTTAAGCAGGAGGTGTCAGAAAAGTTACCACAGGGAT</td><td>>hg38_21_+_8433172_8446622_RNA45SN1_45S_50ntFlanks</td><td>12324</td><td>12366</td><td>43</td><td>TTGGGTTTTAAGCAGGAGGTGTCAGAAAAGTTACCACAGGGAT</td><td>66.4</td><td>-9.9</td><td>2.3</td><td>-4.3</td><td>1.6E-05</td><td>2.7E-05</td><td>MB231/MB468</td><td>MP</td></tr>
<tr><td>>hg38_21_+_8433172_8446622_RNA45SN1_45S_50ntFlanks@10392.10431.40|ACAGCAGTTGAACATGGGTCAGTCGGTCCTGAGAGATGGG</td><td>>hg38_21_+_8433172_8446622_RNA45SN1_45S_50ntFlanks</td><td>10392</td><td>10431</td><td>40</td><td>ACAGCAGTTGAACATGGGTCAGTCGGTCCTGAGAGATGGG</td><td>325.3</td><td>-4.3</td><td>1.0</td><td>-4.3</td><td>1.8E-05</td><td>2.7E-05</td><td>MB231/MB468</td><td>CYTO</td></tr>
</table>

| | | | | | | | | | | | | | |
|---|---|---|---|---|---|---|---|---|---|---|---|---|---|
| >hg38_21_+_8433172_8446622_RNA45SN1_45S_50ntFlanks@3989.4024.36\|TTTGGTGACTCTAGATAACCTCGGGCCGATCGCACG | >hg38_21_+_8433172_8446622_RNA45SN1_45S_50ntFlanks | 3989 | 4024 | 36 | TTTGGTGACTCTAGATAACCTCGGGCCGATCGCACG | 142.6 | -5.3 | 1.2 | -4.3 | 1.6E-05 | 2.7E-05 | MB231/MB468 | MP |
| >hg38_21_+_8433172_8446622_RNA45SN1_45S_50ntFlanks@9534.9572.39\|AGGAAACTCTGGTGGAGGTCCGTAGCGGTCCTGACGTGC | >hg38_21_+_8433172_8446622_RNA45SN1_45S_50ntFlanks | 9534 | 9572 | 39 | AGGAAACTCTGGTGGAGGTCCGTAGCGGTCCTGACGTGC | 3774.4 | -2.6 | 0.6 | -4.3 | 1.8E-05 | 2.7E-05 | MB231/MB468 | CYTO |
| >hg38_21_+_8433172_8446622_RNA45SN1_45S_50ntFlanks@9917.9947.31\|AAGGTGTTGGTTGATATAGACAGCAGGACGG | >hg38_21_+_8433172_8446622_RNA45SN1_45S_50ntFlanks | 9917 | 9947 | 31 | AAGGTGTTGGTTGATATAGACAGCAGGACGG | 94.4 | 9.9 | 2.1 | 4.8 | 1.9E-06 | 2.7E-05 | BT20/MB231 | MP |
| >MI0000102\|hsa-mir-100&WithFlank&11\|-\|122152223\|122152314@22.39.18(+1A)&[MIMAT0000098&hsa-miR-100-5p&offsets\|+3\|-1(+1A);m-27&11\|-\|122152276\|122152296&offsets\|+3\|0(+1A)]\|CCGTAGATCCGAACTTGTA | >MI0000102\|hsa-mir-100&WithFlank&11\|-\|122152223\|122152314 | 22 | 39 | 18(+1A) | CCGTAGATCCGAACTTGTA | 108.5 | -10.2 | 2.1 | -4.8 | 1.9E-06 | 2.7E-05 | BT20/MB231 | MP |
| >hg38_21_+_8433172_8446622_RNA45SN1_45S_50ntFlanks@10348.10383.36\|AAACGAGAACTTTGAAGGCCGAAGTGGAGAAGGGTT | >hg38_21_+_8433172_8446622_RNA45SN1_45S_50ntFlanks | 10348 | 10383 | 36 | AAACGAGAACTTTGAAGGCCGAAGTGGAGAAGGGTT | 1289.0 | -3.3 | 0.8 | -4.3 | 1.9E-05 | 2.7E-05 | MB231/MB468 | MITO |
| >hg38_21_+_8433172_8446622_RNA45SN1_45S_50ntFlanks@5083.5108.26\|ACGAACGAGACTCTGGCATGCTAACT | >hg38_21_+_8433172_8446622_RNA45SN1_45S_50ntFlanks | 5083 | 5108 | 26 | ACGAACGAGACTCTGGCATGCTAACT | 449.0 | -5.0 | 1.2 | -4.3 | 1.9E-05 | 2.7E-05 | MB231/MB468 | MITO |
| >hg38_21_+_8433172_8446622_RNA45SN1_45S_50ntFlanks@12942.12973.32\|TTCGTAGACGACCTGCTTCTGGGTCGGGGTTT | >hg38_21_+_8433172_8446622_RNA45SN1_45S_50ntFlanks | 12942 | 12973 | 32 | TTCGTAGACGACCTGCTTCTGGGTCGGGGTTT | 312.5 | -6.0 | 1.4 | -4.3 | 1.6E-05 | 2.7E-05 | BT20/MB468 | CYTO |
| >hg38_21_+_8433172_8446622_RNA45SN1_45S_50ntFlanks@6651.6678.28\|CGACTCTTAGCGGTGGATCACTCGGCTC | >hg38_21_+_8433172_8446622_RNA45SN1_45S_50ntFlanks | 6651 | 6678 | 28 | CGACTCTTAGCGGTGGATCACTCGGCTC | 13913.8 | 4.5 | 1.0 | 4.5 | 6.1E-06 | 2.7E-05 | BT20/MB468 | MP |
| >hg38_21_+_8433172_8446622_RNA45SN1_45S_50ntFlanks@12154.12181.28\|GGGAGTTTGACTGGGGCGGTACACCTGT | >hg38_21_+_8433172_8446622_RNA45SN1_45S_50ntFlanks | 12154 | 12181 | 28 | GGGAGTTTGACTGGGGCGGTACACCTGT | 260.3 | -6.5 | 1.4 | -4.5 | 6.1E-06 | 2.7E-05 | BT20/MB468 | MP |
| >hg38_21_+_8433172_8446622_RNA45SN1_45S_50ntFlanks@6773.6806.34\|TTCCTCCCGGGGCTACGCCTGTCTGAGCGTCGCT | >hg38_21_+_8433172_8446622_RNA45SN1_45S_50ntFlanks | 6773 | 6806 | 34 | TTCCTCCCGGGGCTACGCCTGTCTGAGCGTCGCT | 1275.9 | -3.6 | 0.8 | -4.3 | 1.6E-05 | 2.7E-05 | MB231/MB468 | MP |
| >MI0000469\|hsa-mir-125a&WithFlank&19\|+\|51693248\|51693345@59.78.20(+1U)&[MIMAT0004602&hsa-miR-125a-3p&offsets\|0\|-2(+1U);m-333&19\|+\|51693306\|51693326&offsets\|0\|-1(+1U)]\|ACAGGTGAGGTTCTTGGGAGT | >MI0000469\|hsa-mir-125a&WithFlank&19\|+\|51693248\|51693345 | 59 | 78 | 20(+1U) | ACAGGTGAGGTTCTTGGGAGT | 37.2 | 8.4 | 2.0 | 4.3 | 1.6E-05 | 2.7E-05 | MB231/MB468 | MP |
| >hg38_21_+_8433172_8446622_RNA45SN1_45S_50ntFlanks@6652.6699.48\|GACTCTTAGCGGTGGATCACTCGGCTCGTGCGTCGATGAAGAACGCAG | >hg38_21_+_8433172_8446622_RNA45SN1_45S_50ntFlanks | 6652 | 6699 | 48 | GACTCTTAGCGGTGGATCACTCGGCTCGTGCGTCGATGAAGAACGCAG | 15249.6 | -3.6 | 0.8 | -4.3 | 1.9E-05 | 2.7E-05 | MB231/MB468 | MITO |
| >hg38_21_+_8433172_8446622_RNA45SN1_45S_50ntFlanks@3866.3899.34\|CTAGAGCTAATACATGCCGACGGGCGCTGACCCC | >hg38_21_+_8433172_8446622_RNA45SN1_45S_50ntFlanks | 3866 | 3899 | 34 | CTAGAGCTAATACATGCCGACGGGCGCTGACCCC | 78.5 | -10.2 | 2.4 | -4.3 | 1.6E-05 | 2.7E-05 | MB231/MB468 | MP |
| >hg38_21_+_8433172_8446622_RNA45SN1_45S_50ntFlanks@11569.11602.34\|CAGCCGACTTAGAACTGGTGCGGACCAGGGGAAT | >hg38_21_+_8433172_8446622_RNA45SN1_45S_50ntFlanks | 11569 | 11602 | 34 | CAGCCGACTTAGAACTGGTGCGGACCAGGGGAAT | 1985.8 | -3.5 | 0.8 | -4.3 | 1.8E-05 | 2.7E-05 | MB231/MB468 | CYTO |
| >trnaMT_GluTTC_MT_-_14674_14742@46.72.27,trnalookalike8_GluTTC_5_-_93905172_93905240@46.72.27\|GTGGTTGTAGTCCGTGCGAGAATACCA | >trnaMT_GluTTC_MT_-_14674_14742 | 46 | 72 | 27 | GTGGTTGTAGTCCGTGCGAGAATACCA | 670.4 | 3.5 | 0.8 | 4.3 | 1.9E-05 | 2.7E-05 | MB231/MB468 | MITO |
| >hg38_21_+_8433172_8446622_RNA45SN1_45S_50ntFlanks@11580.11611.32\|GAACTGGTGCGGACCAGGGGAATCCGACTGTT | >hg38_21_+_8433172_8446622_RNA45SN1_45S_50ntFlanks | 11580 | 11611 | 32 | GAACTGGTGCGGACCAGGGGAATCCGACTGTT | 941.9 | -2.3 | 0.5 | -4.3 | 1.9E-05 | 2.7E-05 | MB231/MB468 | NUC |
| >MI0000750\|hsa-mir-26a-2&WithFlank&12\|-\|57824603\|57824698@20.41.22;MI0000083\|hsa-mir-26a-1&WithFlank&3\|+\|37969398\|37969486@16.37.22&[MIMAT0000082&hsa-miR-26a-5p&offsets\|0\|0;m-33&12\|-\|57824658\|57824679&offsets\|0\|0];[MIMAT0000082_1&hsa-miR-26a-5p&offsets\|0\|0;m-34&3\|+\|37969413\|37969434&offsets\|0\|0]\|TTCAAGTAATCCAGGATAGGCT | >MI0000750\|hsa-mir-26a-2&WithFlank&12\|-\|57824603\|57824698 | 20 | 41 | 22 | TTCAAGTAATCCAGGATAGGCT | 72660.6 | 3.8 | 0.9 | 4.3 | 1.6E-05 | 2.7E-05 | MB231/MB468 | MP |
| >am_MI0000809\|hsa-mir-151a&WithFlank&8\|-\|140732558\|140732659@52.73.22&[MIMAT0000757&hsa-miR-151a-3p&offsets\|-1\|0;m-30&8\|-\|140732587\|140732607&offsets\|-1\|0]\|ACTAGACTGAAGCTCCTTGAGG | >am_MI0000809\|hsa-mir-151a&WithFlank&8\|-\|140732558\|140732659 | 52 | 73 | 22 | ACTAGACTGAAGCTCCTTGAGG | 983.5 | 4.2 | 0.9 | 4.5 | 6.2E-06 | 2.7E-05 | BT20/MB468 | MP |
| >hg38_21_+_8433172_8446622_RNA45SN1_45S_50ntFlanks@3853.3881.29\|AACTGTGGTAATTCTAGAGCTAATACATG | >hg38_21_+_8433172_8446622_RNA45SN1_45S_50ntFlanks | 3853 | 3881 | 29 | AACTGTGGTAATTCTAGAGCTAATACATG | 485.4 | -4.8 | 1.1 | -4.3 | 1.9E-05 | 2.7E-05 | MB231/MB468 | NUC |
| >MI0000086\|hsa-mir-28&WithFlank&3\|+\|188688775\|188688872@60.82.23&[MIMAT0004502&hsa-miR-28-3p&offsets\|0\|+1;m-28&3\|+\|188688834\|188688855&offsets\|0\|+1]\|CACTAGATTGTGAGCTCCTGGAG | >MI0000086\|hsa-mir-28&WithFlank&3\|+\|188688775\|188688872 | 60 | 82 | 23 | CACTAGATTGTGAGCTCCTGGAG | 472.3 | 7.2 | 1.6 | 4.5 | 6.2E-06 | 2.7E-05 | BT20/MB468 | MP |
| >hg38_21_+_8433172_8446622_RNA45SN1_45S_50ntFlanks@6676.6696.21\|CTCGTGCGTCGATGAAGAACG | >hg38_21_+_8433172_8446622_RNA45SN1_45S_50ntFlanks | 6676 | 6696 | 21 | CTCGTGCGTCGATGAAGAACG | 320.4 | -5.6 | 1.3 | -4.3 | 2.0E-05 | 2.7E-05 | BT20/MB468 | NUC |
| >hg38_21_+_8433172_8446622_RNA45SN1_45S_50ntFlanks@12416.12459.44\|TTCGATGTCGGCTCTTCCTATCATTGTGAAGCAGAATTCACCAA | >hg38_21_+_8433172_8446622_RNA45SN1_45S_50ntFlanks | 12416 | 12459 | 44 | TTCGATGTCGGCTCTTCCTATCATTGTGAAGCAGAATTCACCAA | 255.5 | 7.3 | 1.7 | 4.3 | 1.9E-05 | 2.7E-05 | MB231/MB468 | MITO |
| >MI0003188\|hsa-mir-503&WithFlank&X\|-\|134546322\|134546404@12.31.20(+1U)&[MIMAT0002874&hsa-miR-503-5p&offsets\|0\|-3(+1U);m-297&X\|-\|134546371\|134546393&offsets\|0\|-3(+1U)]\|TAGCAGCGGGAACAGTTCTGT | >MI0003188\|hsa-mir-503&WithFlank&X\|-\|134546322\|134546404 | 12 | 31 | 20(+1U) | TAGCAGCGGGAACAGTTCTGT | 33.8 | 8.3 | 1.9 | 4.3 | 1.6E-05 | 2.7E-05 | MB231/MB468 | MP |
| >am_trna10_ValCAC_5_-_180649395_180649467@1.23.23,trna12_ValAAC_5_-_180645270_180645342@1.23.23,trna132_ValAAC_6_-_27721179_27721251@1.23.23,trna136_ValAAC_6_-_27648885_27648957@1.23.23,trna139_ValAAC_6_-_27618707_27618779@1.23.23,trna18_ValCAC_5_-_180529253_180529325@1.23.23,trna2_ValAAC_3_+_169490018_169490090@1.23.23,trna2_ValCAC_5_+_180524070_180524142@1.23.23,trna37_ValAAC_6_+_27203288_27203360@1.23.23,trna4_ValAAC_5_+_180591154_180591226@1.23.23,trna5_ValAAC_5_+_180596610_180596682@1.23.23,trna6_ValCAC_5_+_180600650_180600722@1.23.23,trna85_ValCAC_1_-_161369490_161369562@1.23.23,trna90_ValCAC_1_-_149684088_149684161@1.23.23,trna98_ValCAC_1_-_149298555_149298627@1.23.23,trna9_ValCAC_6_+_26538282_26538354@1.23.23\|GTTTCCGTAGTGTAGTGGTTATC | >am_trna10_ValCAC_5_-_180649395_180649467 | 1 | 23 | 23 | GTTTCCGTAGTGTAGTGGTTATC | 6054.6 | 4.9 | 1.1 | 4.3 | 1.4E-05 | 2.7E-05 | BT20/MB468 | MITO |
| >hg38_21_+_8433172_8446622_RNA45SN1_45S_50ntFlanks@12405.12425.21\|CTTTTTGATCCTTCGATGTCG | >hg38_21_+_8433172_8446622_RNA45SN1_45S_50ntFlanks | 12405 | 12425 | 21 | CTTTTTGATCCTTCGATGTCG | 365.8 | -5.9 | 1.4 | -4.3 | 1.4E-05 | 2.7E-05 | BT20/MB468 | MITO |
| >MI0000542\|hsa-mir-320a&WithFlank&8\|-\|22244960\|22245043@47.69.23&[MIMAT0000510&hsa-miR-320a-3p&offsets\|-1\|0;m-60&8\|-\|22244975\|22244996&offsets\|-1\|0]\|GAAAAGCTGGGTTGAGAGGGCGA | >MI0000542\|hsa-mir-320a&WithFlank&8\|-\|22244960\|22245043 | 47 | 69 | 23 | GAAAAGCTGGGTTGAGAGGGCGA | 319.2 | 3.9 | 0.8 | 4.8 | 1.9E-06 | 2.7E-05 | BT20/MB231 | CYTO |
| >hg38_21_+_8433172_8446622_RNA45SN1_45S_50ntFlanks@3771.3792.22\|CACGGCCGGTACAGTGAAACTG | >hg38_21_+_8433172_8446622_RNA45SN1_45S_50ntFlanks | 3771 | 3792 | 22 | CACGGCCGGTACAGTGAAACTG | 325.0 | -4.6 | 1.1 | -4.3 | 1.6E-05 | 2.7E-05 | BT20/MB468 | CYTO |
| >hg38_21_+_8433172_8446622_RNA45SN1_45S_50ntFlanks@12348.12376.29\|GAAAAGTTACCACAGGGATAACTGGCTTG | >hg38_21_+_8433172_8446622_RNA45SN1_45S_50ntFlanks | 12348 | 12376 | 29 | GAAAAGTTACCACAGGGATAACTGGCTTG | 46.0 | -8.6 | 1.9 | -4.5 | 6.2E-06 | 2.7E-05 | BT20/MB468 | MP |
| >hg38_21_+_8433172_8446622_RNA45SN1_45S_50ntFlanks@4785.4812.28\|TAAACGATGCCGACCGGCGATGCGGCGG | >hg38_21_+_8433172_8446622_RNA45SN1_45S_50ntFlanks | 4785 | 4812 | 28 | TAAACGATGCCGACCGGCGATGCGGCGG | 45.2 | -8.6 | 1.9 | -4.5 | 6.2E-06 | 2.7E-05 | BT20/MB468 | MP |
| >hg38_21_+_8433172_8446622_RNA45SN1_45S_50ntFlanks@4685.4707.23\|AGACGGACCAGAGCGAAAGCATT | >hg38_21_+_8433172_8446622_RNA45SN1_45S_50ntFlanks | 4685 | 4707 | 23 | AGACGGACCAGAGCGAAAGCATT | 410.5 | -3.7 | 0.9 | -4.3 | 1.9E-05 | 2.7E-05 | MB231/MB468 | NUC |
| >hg38_21_+_8433172_8446622_RNA45SN1_45S_50ntFlanks@9545.9572.28\|GTGGAGGTCCGTAGCGGTCCTGACGTGC | >hg38_21_+_8433172_8446622_RNA45SN1_45S_50ntFlanks | 9545 | 9572 | 28 | GTGGAGGTCCGTAGCGGTCCTGACGTGC | 576.4 | 4.2 | 0.9 | 4.7 | 2.3E-06 | 2.7E-05 | BT20/MB231 | MITO |
| >MI0000065\|hsa-let-7d&WithFlank&9\|+\|94178828\|94178926@68.88.21(+1C)&[MIMAT0004484&hsa-let-7d-3p&offsets\|0\|-1(+1C);m-98&9\|+\|94178895\|94178916&offsets\|0\|-1(+1C)]\|CTATACGACCTGCTGCCTTTCC | >MI0000065\|hsa-let-7d&WithFlank&9\|+\|94178828\|94178926 | 68 | 88 | 21(+1C) | CTATACGACCTGCTGCCTTTCC | 666.6 | 5.0 | 1.1 | 4.5 | 6.2E-06 | 2.7E-05 | BT20/MB468 | MP |
| >hg38_21_+_8433172_8446622_RNA45SN1_45S_50ntFlanks@10246.10272.27\|CGCGACGAGTAGGAGGGCCGCTGCGGT | >hg38_21_+_8433172_8446622_RNA45SN1_45S_50ntFlanks | 10246 | 10272 | 27 | CGCGACGAGTAGGAGGGCCGCTGCGGT | 38.3 | 9.0 | 2.0 | 4.5 | 6.2E-06 | 2.7E-05 | BT20/MB468 | MP |
| >MI0000255\|hsa-mir-30d&WithFlank&8\|-\|134804870\|134804951@12.34.23(+1A)&[MIMAT0000245&hsa-miR-30d-5p&offsets\|0\|+1(+1A);m-21&8\|-\|134804919\|134804940&offsets\|0\|+1(+1A)]\|TGTAAACATCCCCGACTGGAAGCA | >MI0000255\|hsa-mir-30d&WithFlank&8\|-\|134804870\|134804951 | 12 | 34 | 23(+1A) | TGTAAACATCCCCGACTGGAAGCA | 1768.6 | 3.6 | 0.8 | 4.3 | 1.6E-05 | 2.7E-05 | MB231/MB468 | MP |
| >hg38_21_+_8433172_8446622_RNA45SN1_45S_50ntFlanks@7972.8005.34\|AGACGCGACCTCAGATCAGACGTGGCGACCCGCT | >hg38_21_+_8433172_8446622_RNA45SN1_45S_50ntFlanks | 7972 | 8005 | 34 | AGACGCGACCTCAGATCAGACGTGGCGACCCGCT | 918.6 | 5.5 | 1.2 | 4.7 | 2.3E-06 | 2.7E-05 | BT20/MB231 | NUC |
| >hg38_21_+_8433172_8446622_RNA45SN1_45S_50ntFlanks@10391.10422.32\|AACAGCAGTTGAACATGGGTCAGTCGGTCCTG | >hg38_21_+_8433172_8446622_RNA45SN1_45S_50ntFlanks | 10391 | 10422 | 32 | AACAGCAGTTGAACATGGGTCAGTCGGTCCTG | 469.0 | -4.1 | 1.0 | -4.3 | 1.8E-05 | 2.7E-05 | MB231/MB468 | CYTO |
| >hg38_21_+_8433172_8446622_RNA45SN1_45S_50ntFlanks@12435.12454.20\|ATCATTGTGAAGCAGAATTC | >hg38_21_+_8433172_8446622_RNA45SN1_45S_50ntFlanks | 12435 | 12454 | 20 | ATCATTGTGAAGCAGAATTC | 428.0 | -6.3 | 1.5 | -4.3 | 1.8E-05 | 2.7E-05 | MB231/MB468 | CYTO |
| >hg38_21_+_8433172_8446622_RNA45SN1_45S_50ntFlanks@10777.10818.42\|TCCAAGGTGAACAGCCTCTGGCATGTTGGAACAATGTAGGTA | >hg38_21_+_8433172_8446622_RNA45SN1_45S_50ntFlanks | 10777 | 10818 | 42 | TCCAAGGTGAACAGCCTCTGGCATGTTGGAACAATGTAGGTA | 452.5 | -3.4 | 0.8 | -4.3 | 1.8E-05 | 2.7E-05 | MB231/MB468 | CYTO |
| >hg38_21_+_8433172_8446622_RNA45SN1_45S_50ntFlanks@4995.5013.19\|ACAGATTGATAGCTCTTTC | >hg38_21_+_8433172_8446622_RNA45SN1_45S_50ntFlanks | 4995 | 5013 | 19 | ACAGATTGATAGCTCTTTC | 587.1 | -5.9 | 1.4 | -4.3 | 1.4E-05 | 2.7E-05 | BT20/MB468 | MITO |
| >trna16_ValTAC_11_-_59318460_59318532@31.53.23,trna17_ValTAC_11_-_59318102_59318174@31.53.23,trna4_ValTAC_X_-_18693029_18693101@31.53.23,trna6_ValTAC_10_-_5895674_5895746@31.53.23\|CTTTACACGCAGAAGGTCCTGGG | >trna16_ValTAC_11_-_59318460_59318532 | 31 | 53 | 23 | CTTTACACGCAGAAGGTCCTGGG | 45.0 | -8.6 | 1.9 | -4.5 | 6.2E-06 | 2.7E-05 | BT20/MB468 | MP |
| >hg38_21_+_8433172_8446622_RNA45SN1_45S_50ntFlanks@9465.9500.36\|AGGTGGAGCACGAGCGCACGTGTTAGGACCCGAAAG | >hg38_21_+_8433172_8446622_RNA45SN1_45S_50ntFlanks | 9465 | 9500 | 36 | AGGTGGAGCACGAGCGCACGTGTTAGGACCCGAAAG | 39.0 | -8.4 | 1.9 | -4.5 | 6.2E-06 | 2.7E-05 | BT20/MB468 | MP |

<table>
<tr><td>>trna10_LysCTT_16_+_3241501_3241573@1.36.36,trna13_LysCTT_14_-_58706613_58706685@1.36.36,trna2_LysCTT_15_+_79152904_79152976@1.36.36|GCCCGGCTAGCTCAGTCGGTAGAGCATGGGACTCTT</td><td>>trna10_LysCTT_16_+_3241501_3241573</td><td>1</td><td>36</td><td>36</td><td>GCCCGGCTAGCTCAGTCGGTAGAGCATGGGACTCTT</td><td>8173.6</td><td>6.5</td><td>1.4</td><td>4.5</td><td>6.2E-06</td><td>2.7E-05</td><td>BT20/MB468</td><td>MP</td></tr>
<tr><td>>hg38_21_+_8433172_8446622_RNA45SN1_45S_50ntFlanks@5237.5284.48|ACTGACTGGCTCAGCGTGTGCCTACCCTACGCCGGCAGGCGCGGGTAA</td><td>>hg38_21_+_8433172_8446622_RNA45SN1_45S_50ntFlanks</td><td>5237</td><td>5284</td><td>48</td><td>ACTGACTGGCTCAGCGTGTGCCTACCCTACGCCGGCAGGCGCGGGTAA</td><td>213.2</td><td>11.5</td><td>2.5</td><td>4.5</td><td>6.2E-06</td><td>2.7E-05</td><td>BT20/MB468</td><td>MP</td></tr>
<tr><td>>hg38_21_+_8433172_8446622_RNA45SN1_45S_50ntFlanks@11565.11585.21|CTAGCAGCCGACTTAGAACTG</td><td>>hg38_21_+_8433172_8446622_RNA45SN1_45S_50ntFlanks</td><td>11565</td><td>11585</td><td>21</td><td>CTAGCAGCCGACTTAGAACTG</td><td>206.4</td><td>-6.2</td><td>1.4</td><td>-4.3</td><td>1.6E-05</td><td>2.7E-05</td><td>MB231/MB468</td><td>MP</td></tr>
<tr><td>>hg38_21_+_8433172_8446622_RNA45SN1_45S_50ntFlanks@12394.12416.23|TAGCGACGTCGCTTTTTGATCCT</td><td>>hg38_21_+_8433172_8446622_RNA45SN1_45S_50ntFlanks</td><td>12394</td><td>12416</td><td>23</td><td>TAGCGACGTCGCTTTTTGATCCT</td><td>163.9</td><td>-6.6</td><td>1.5</td><td>-4.3</td><td>1.6E-05</td><td>2.7E-05</td><td>MB231/MB468</td><td>MP</td></tr>
<tr><td>>hg38_21_+_8433172_8446622_RNA45SN1_45S_50ntFlanks@4445.4472.28|CTTGCCTCTCGGCGCCCCCTCGATGCTC</td><td>>hg38_21_+_8433172_8446622_RNA45SN1_45S_50ntFlanks</td><td>4445</td><td>4472</td><td>28</td><td>CTTGCCTCTCGGCGCCCCCTCGATGCTC</td><td>327.4</td><td>-5.3</td><td>1.2</td><td>-4.3</td><td>1.8E-05</td><td>2.7E-05</td><td>MB231/MB468</td><td>CYTO</td></tr>
<tr><td>>hg38_21_+_8433172_8446622_RNA45SN1_45S_50ntFlanks@10726.10748.23|GGGAGAGGGTGTAAATCTCGCGC</td><td>>hg38_21_+_8433172_8446622_RNA45SN1_45S_50ntFlanks</td><td>10726</td><td>10748</td><td>23</td><td>GGGAGAGGGTGTAAATCTCGCGC</td><td>50.2</td><td>-8.7</td><td>1.9</td><td>-4.5</td><td>6.3E-06</td><td>2.7E-05</td><td>BT20/MB468</td><td>MP</td></tr>
<tr><td>>hg38_21_+_8433172_8446622_RNA45SN1_45S_50ntFlanks@8159.8191.33|CGTGGGGGGCCCAAGTCCTTCTGATCGAGGCCC</td><td>>hg38_21_+_8433172_8446622_RNA45SN1_45S_50ntFlanks</td><td>8159</td><td>8191</td><td>33</td><td>CGTGGGGGGCCCAAGTCCTTCTGATCGAGGCCC</td><td>218.9</td><td>3.8</td><td>0.9</td><td>4.3</td><td>1.9E-05</td><td>2.7E-05</td><td>MB231/MB468</td><td>MITO</td></tr>
<tr><td>>hg38_21_+_8433172_8446622_RNA45SN1_45S_50ntFlanks@3843.3861.19|CTACTTGGATAACTGTGGT</td><td>>hg38_21_+_8433172_8446622_RNA45SN1_45S_50ntFlanks</td><td>3843</td><td>3861</td><td>19</td><td>CTACTTGGATAACTGTGGT</td><td>1355.8</td><td>-2.1</td><td>0.5</td><td>-4.3</td><td>1.8E-05</td><td>2.7E-05</td><td>MB231/MB468</td><td>CYTO</td></tr>
<tr><td>>hg38_21_+_8433172_8446622_RNA45SN1_45S_50ntFlanks@4445.4471.27|CTTGCCTCTCGGCGCCCCCTCGATGCT</td><td>>hg38_21_+_8433172_8446622_RNA45SN1_45S_50ntFlanks</td><td>4445</td><td>4471</td><td>27</td><td>CTTGCCTCTCGGCGCCCCCTCGATGCT</td><td>1472.4</td><td>-7.6</td><td>1.8</td><td>-4.3</td><td>2.0E-05</td><td>2.7E-05</td><td>BT20/MB468</td><td>NUC</td></tr>
<tr><td>>hg38_21_+_8433172_8446622_RNA45SN1_45S_50ntFlanks@10788.10815.28|CAGCCTCTGGCATGTTGGAACAATGTAG</td><td>>hg38_21_+_8433172_8446622_RNA45SN1_45S_50ntFlanks</td><td>10788</td><td>10815</td><td>28</td><td>CAGCCTCTGGCATGTTGGAACAATGTAG</td><td>758.1</td><td>-5.2</td><td>1.2</td><td>-4.3</td><td>1.9E-05</td><td>2.7E-05</td><td>MB231/MB468</td><td>NUC</td></tr>
<tr><td>>MI0000102|hsa-mir-100&WithFlank&11|-|122152223|122152314@19.37.19(+1U)&[MIMAT0000098&hsa-miR-100-5p&offsets|0|-3(+1U);m-27&11|-|122152276|122152296&offsets|0|-2(+1U)]|AACCCGTAGATCCGAACTTT</td><td>>MI0000102|hsa-mir-100&WithFlank&11|-|122152223|122152314</td><td>19</td><td>37</td><td>19(+1U)</td><td>AACCCGTAGATCCGAACTTT</td><td>711.0</td><td>-8.4</td><td>1.8</td><td>-4.8</td><td>1.9E-06</td><td>2.7E-05</td><td>BT20/MB231</td><td>MP</td></tr>
<tr><td>>hg38_21_+_8433172_8446622_RNA45SN1_45S_50ntFlanks@9791.9811.21|TGGCGTGGAGCCGGGCGTGGA</td><td>>hg38_21_+_8433172_8446622_RNA45SN1_45S_50ntFlanks</td><td>9791</td><td>9811</td><td>21</td><td>TGGCGTGGAGCCGGGCGTGGA</td><td>141.1</td><td>10.5</td><td>2.2</td><td>4.8</td><td>1.9E-06</td><td>2.7E-05</td><td>BT20/MB231</td><td>MP</td></tr>
<tr><td>>hg38_21_+_8433172_8446622_RNA45SN1_45S_50ntFlanks@12163.12183.21|ACTGGGGCGGTACACCTGTCA</td><td>>hg38_21_+_8433172_8446622_RNA45SN1_45S_50ntFlanks</td><td>12163</td><td>12183</td><td>21</td><td>ACTGGGGCGGTACACCTGTCA</td><td>553.8</td><td>-6.1</td><td>1.4</td><td>-4.3</td><td>1.4E-05</td><td>2.7E-05</td><td>BT20/MB468</td><td>MITO</td></tr>
<tr><td>>hg38_21_+_8433172_8446622_RNA45SN1_45S_50ntFlanks@8465.8497.33|GTGTCGGCGGCCCGGCGGATCTTTCCCGCCCCC</td><td>>hg38_21_+_8433172_8446622_RNA45SN1_45S_50ntFlanks</td><td>8465</td><td>8497</td><td>33</td><td>GTGTCGGCGGCCCGGCGGATCTTTCCCGCCCCC</td><td>608.1</td><td>-7.4</td><td>1.7</td><td>-4.3</td><td>2.0E-05</td><td>2.7E-05</td><td>BT20/MB468</td><td>NUC</td></tr>
<tr><td>>hg38_21_+_8433172_8446622_RNA45SN1_45S_50ntFlanks@11582.11609.28|ACTGGTGCGGACCAGGGGAATCCGACTG</td><td>>hg38_21_+_8433172_8446622_RNA45SN1_45S_50ntFlanks</td><td>11582</td><td>11609</td><td>28</td><td>ACTGGTGCGGACCAGGGGAATCCGACTG</td><td>352.8</td><td>-3.5</td><td>0.8</td><td>-4.3</td><td>2.0E-05</td><td>2.8E-05</td><td>MB231/MB468</td><td>MITO</td></tr>
<tr><td>>hg38_21_+_8433172_8446622_RNA45SN1_45S_50ntFlanks@4380.4406.27|CGTAGTTGGATCTTGGGAGCGGGCGGG</td><td>>hg38_21_+_8433172_8446622_RNA45SN1_45S_50ntFlanks</td><td>4380</td><td>4406</td><td>27</td><td>CGTAGTTGGATCTTGGGAGCGGGCGGG</td><td>415.5</td><td>-5.3</td><td>1.2</td><td>-4.3</td><td>2.0E-05</td><td>2.8E-05</td><td>BT20/MB468</td><td>NUC</td></tr>
<tr><td>>hg38_21_+_8433172_8446622_RNA45SN1_45S_50ntFlanks@12157.12182.26|AGTTTGACTGGGGCGGTACACCTGTC</td><td>>hg38_21_+_8433172_8446622_RNA45SN1_45S_50ntFlanks</td><td>12157</td><td>12182</td><td>26</td><td>AGTTTGACTGGGGCGGTACACCTGTC</td><td>123.7</td><td>-5.5</td><td>1.3</td><td>-4.3</td><td>1.7E-05</td><td>2.8E-05</td><td>BT20/MB468</td><td>CYTO</td></tr>
<tr><td>>hg38_21_+_8433172_8446622_RNA45SN1_45S_50ntFlanks@6645.6670.26|CTCGTACGACTCTTAGCGGTGGATCA</td><td>>hg38_21_+_8433172_8446622_RNA45SN1_45S_50ntFlanks</td><td>6645</td><td>6670</td><td>26</td><td>CTCGTACGACTCTTAGCGGTGGATCA</td><td>4863.1</td><td>-3.4</td><td>0.8</td><td>-4.3</td><td>2.0E-05</td><td>2.8E-05</td><td>MB231/MB468</td><td>MITO</td></tr>
<tr><td>>MI0000067|hsa-let-7f-1&WithFlank&9|+|94176341|94176439@69.89.21(+2U)&[MIMAT0004486&hsa-let-7f-1-3p&offsets|0|-1(+2U);m-275&9|+|94176409|94176430&offsets|0|-1(+2U)]|CTATACAATCTATTGCCTTCCTT</td><td>>MI0000067|hsa-let-7f-1&WithFlank&9|+|94176341|94176439</td><td>69</td><td>89</td><td>21(+2U)</td><td>CTATACAATCTATTGCCTTCCTT</td><td>35.5</td><td>8.4</td><td>1.9</td><td>4.3</td><td>1.6E-05</td><td>2.8E-05</td><td>MB231/MB468</td><td>MP</td></tr>
<tr><td>>hg38_21_+_8433172_8446622_RNA45SN1_45S_50ntFlanks@9914.9946.33|GAAAAGGTGTTGGTTGATATAGACAGCAGGACG</td><td>>hg38_21_+_8433172_8446622_RNA45SN1_45S_50ntFlanks</td><td>9914</td><td>9946</td><td>33</td><td>GAAAAGGTGTTGGTTGATATAGACAGCAGGACG</td><td>116.2</td><td>10.2</td><td>2.1</td><td>4.8</td><td>1.9E-06</td><td>2.8E-05</td><td>BT20/MB231</td><td>MP</td></tr>
<tr><td>>hg38_21_+_8433172_8446622_RNA45SN1_45S_50ntFlanks@5526.5550.25|AAAAGTCGTAACAAGGTTTCCGTAG</td><td>>hg38_21_+_8433172_8446622_RNA45SN1_45S_50ntFlanks</td><td>5526</td><td>5550</td><td>25</td><td>AAAAGTCGTAACAAGGTTTCCGTAG</td><td>166.3</td><td>-3.9</td><td>0.9</td><td>-4.3</td><td>1.9E-05</td><td>2.8E-05</td><td>MB231/MB468</td><td>CYTO</td></tr>
<tr><td>>hg38_21_+_8433172_8446622_RNA45SN1_45S_50ntFlanks@8183.8208.26|TCGAGGCCCAGCCCGTGGACGGTGTG</td><td>>hg38_21_+_8433172_8446622_RNA45SN1_45S_50ntFlanks</td><td>8183</td><td>8208</td><td>26</td><td>TCGAGGCCCAGCCCGTGGACGGTGTG</td><td>117.5</td><td>-6.3</td><td>1.5</td><td>-4.3</td><td>1.6E-05</td><td>2.8E-05</td><td>MB231/MB468</td><td>MP</td></tr>
<tr><td>>hg38_21_+_8433172_8446622_RNA45SN1_45S_50ntFlanks@3996.4026.31|ACTCTAGATAACCTCGGGCCGATCGCACGCC</td><td>>hg38_21_+_8433172_8446622_RNA45SN1_45S_50ntFlanks</td><td>3996</td><td>4026</td><td>31</td><td>ACTCTAGATAACCTCGGGCCGATCGCACGCC</td><td>322.3</td><td>-6.6</td><td>1.5</td><td>-4.3</td><td>1.9E-05</td><td>2.8E-05</td><td>MB231/MB468</td><td>NUC</td></tr>
<tr><td>>MI0000477|hsa-mir-146a&WithFlank&5|+|160485346|160485456@27.48.22(+2A)&[MIMAT0000449&hsa-miR-146a-5p&offsets|0|0(+2A);m-129&5|+|160485372|160485393&offsets|0|0(+2A)]|TGAGAACTGAATTCCATGGGTTAA</td><td>>MI0000477|hsa-mir-146a&WithFlank&5|+|160485346|160485456</td><td>27</td><td>48</td><td>22(+2A)</td><td>TGAGAACTGAATTCCATGGGTTAA</td><td>80.2</td><td>-9.7</td><td>2.0</td><td>-4.8</td><td>2.0E-06</td><td>2.8E-05</td><td>BT20/MB231</td><td>MP</td></tr>
<tr><td>>MI0000253|hsa-mir-148a&WithFlank&7|-|25949913|25949992@50.71.22(+1U)&[MIMAT0000243&hsa-miR-148a-3p&offsets|0|0(+1U);m-7&7|-|25949922|25949943&offsets|0|0(+1U)]|TCAGTGCACTACAGAACTTTGTT</td><td>>MI0000253|hsa-mir-148a&WithFlank&7|-|25949913|25949992</td><td>50</td><td>71</td><td>22(+1U)</td><td>TCAGTGCACTACAGAACTTTGTT</td><td>650.5</td><td>5.3</td><td>1.1</td><td>4.8</td><td>2.0E-06</td><td>2.8E-05</td><td>BT20/MB231</td><td>CYTO</td></tr>
<tr><td>>hg38_21_+_8433172_8446622_RNA45SN1_45S_50ntFlanks@10408.10428.21|GGTCAGTCGGTCCTGAGAGAT</td><td>>hg38_21_+_8433172_8446622_RNA45SN1_45S_50ntFlanks</td><td>10408</td><td>10428</td><td>21</td><td>GGTCAGTCGGTCCTGAGAGAT</td><td>257.6</td><td>-5.7</td><td>1.3</td><td>-4.3</td><td>1.7E-05</td><td>2.8E-05</td><td>BT20/MB468</td><td>CYTO</td></tr>
<tr><td>>hg38_21_+_8433172_8446622_RNA45SN1_45S_50ntFlanks@8170.8215.46|CAAGTCCTTCTGATCGAGGCCCAGCCCGTGGACGGTGTGAGGCCGG</td><td>>hg38_21_+_8433172_8446622_RNA45SN1_45S_50ntFlanks</td><td>8170</td><td>8215</td><td>46</td><td>CAAGTCCTTCTGATCGAGGCCCAGCCCGTGGACGGTGTGAGGCCGG</td><td>997.3</td><td>-5.3</td><td>1.2</td><td>-4.3</td><td>1.9E-05</td><td>2.8E-05</td><td>MB231/MB468</td><td>CYTO</td></tr>
<tr><td>>trnaMT_GluTTC_MT_-_14674_14742@35.72.38,trnalookalike8_GluTTC_5_-_93905172_93905240@35.72.38|TATCATTGGTCGTGGTTGTAGTCCGTGCGAGAATACCA</td><td>>trnaMT_GluTTC_MT_-_14674_14742</td><td>35</td><td>72</td><td>38</td><td>TATCATTGGTCGTGGTTGTAGTCCGTGCGAGAATACCA</td><td>7407.7</td><td>4.7</td><td>1.1</td><td>4.3</td><td>1.9E-05</td><td>2.8E-05</td><td>MB231/MB468</td><td>NUC</td></tr>
<tr><td>>hg38_21_+_8433172_8446622_RNA45SN1_45S_50ntFlanks@11027.11053.27|CTCTCTCTCTCTCTCCCCCGCTCCCCG</td><td>>hg38_21_+_8433172_8446622_RNA45SN1_45S_50ntFlanks</td><td>11027</td><td>11053</td><td>27</td><td>CTCTCTCTCTCTCTCCCCCGCTCCCCG</td><td>4388.8</td><td>-8.7</td><td>2.0</td><td>-4.3</td><td>2.1E-05</td><td>2.8E-05</td><td>BT20/MB468</td><td>NUC</td></tr>
<tr><td>>hg38_21_+_8433172_8446622_RNA45SN1_45S_50ntFlanks@8179.8203.25|CTGATCGAGGCCCAGCCCGTGGACG</td><td>>hg38_21_+_8433172_8446622_RNA45SN1_45S_50ntFlanks</td><td>8179</td><td>8203</td><td>25</td><td>CTGATCGAGGCCCAGCCCGTGGACG</td><td>221.0</td><td>-4.9</td><td>1.1</td><td>-4.3</td><td>1.9E-05</td><td>2.8E-05</td><td>MB231/MB468</td><td>NUC</td></tr>
<tr><td>>hg38_21_+_8433172_8446622_RNA45SN1_45S_50ntFlanks@10410.10440.31|TCAGTCGGTCCTGAGAGATGGGCGAGCGCCG</td><td>>hg38_21_+_8433172_8446622_RNA45SN1_45S_50ntFlanks</td><td>10410</td><td>10440</td><td>31</td><td>TCAGTCGGTCCTGAGAGATGGGCGAGCGCCG</td><td>196.6</td><td>-4.0</td><td>0.9</td><td>-4.3</td><td>1.7E-05</td><td>2.8E-05</td><td>BT20/MB468</td><td>CYTO</td></tr>
<tr><td>>hg38_21_+_8433172_8446622_RNA45SN1_45S_50ntFlanks@6775.6807.33|CCTCCCGGGGCTACGCCTGTCTGAGCGTCGCTT</td><td>>hg38_21_+_8433172_8446622_RNA45SN1_45S_50ntFlanks</td><td>6775</td><td>6807</td><td>33</td><td>CCTCCCGGGGCTACGCCTGTCTGAGCGTCGCTT</td><td>699.5</td><td>3.4</td><td>0.8</td><td>4.3</td><td>1.4E-05</td><td>2.8E-05</td><td>BT20/MB468</td><td>MITO</td></tr>
<tr><td>>hg38_21_+_8433172_8446622_RNA45SN1_45S_50ntFlanks@11534.11552.19|GTTCCGCGCGGCGCCTCGC</td><td>>hg38_21_+_8433172_8446622_RNA45SN1_45S_50ntFlanks</td><td>11534</td><td>11552</td><td>19</td><td>GTTCCGCGCGGCGCCTCGC</td><td>624.0</td><td>-6.2</td><td>1.4</td><td>-4.3</td><td>1.4E-05</td><td>2.8E-05</td><td>BT20/MB468</td><td>MITO</td></tr>
<tr><td>>am_trna119_LysCTT_1_-_145395522_145395594@8.35.28,trna11_LysCTT_5_-_180648979_180649051@8.35.28,trna128_LysCTT_1_-_55423542_55423614@8.35.28,trna13_LysCTT_6_+_26556774_26556846@8.35.28,trna6_LysCTT_19_-_52425396_52425468@8.35.28,trna7_LysCTT_16_+_3225692_3225764@8.35.28,trna9_LysCTT_5_+_180634755_180634827@8.35.28|TAGCTCAGTCGGTAGAGCATGAGACTCT</td><td>>am_trna119_LysCTT_1_-_145395522_145395594</td><td>8</td><td>35</td><td>28</td><td>TAGCTCAGTCGGTAGAGCATGAGACTCT</td><td>183.2</td><td>7.2</td><td>1.7</td><td>4.3</td><td>2.0E-05</td><td>2.8E-05</td><td>MB231/MB468</td><td>MITO</td></tr>
<tr><td>>hg38_21_+_8433172_8446622_RNA45SN1_45S_50ntFlanks@9574.9595.22|AATCGGTCGTCCGACCTGGGTA</td><td>>hg38_21_+_8433172_8446622_RNA45SN1_45S_50ntFlanks</td><td>9574</td><td>9595</td><td>22</td><td>AATCGGTCGTCCGACCTGGGTA</td><td>407.3</td><td>-3.0</td><td>0.7</td><td>-4.3</td><td>2.0E-05</td><td>2.8E-05</td><td>MB231/MB468</td><td>NUC</td></tr>
<tr><td>>MI0000084|hsa-mir-26b&WithFlank&2|+|218402640|218402728@18.39.22&[MIMAT0000083&hsa-miR-26b-5p&offsets|0|+1;m-53&2|+|218402657|218402678&offsets|0|0]|TTCAAGTAATTCAGGATAGGTT</td><td>>MI0000084|hsa-mir-26b&WithFlank&2|+|218402640|218402728</td><td>18</td><td>39</td><td>22</td><td>TTCAAGTAATTCAGGATAGGTT</td><td>2496.4</td><td>3.7</td><td>0.9</td><td>4.3</td><td>1.9E-05</td><td>2.8E-05</td><td>MB231/MB468</td><td>CYTO</td></tr>
<tr><td>>hg38_21_+_8433172_8446622_RNA45SN1_45S_50ntFlanks@9545.9574.30|GTGGAGGTCCGTAGCGGTCCTGACGTGCAA</td><td>>hg38_21_+_8433172_8446622_RNA45SN1_45S_50ntFlanks</td><td>9545</td><td>9574</td><td>30</td><td>GTGGAGGTCCGTAGCGGTCCTGACGTGCAA</td><td>441.4</td><td>-6.6</td><td>1.5</td><td>-4.3</td><td>1.6E-05</td><td>2.8E-05</td><td>MB231/MB468</td><td>MP</td></tr>
<tr><td>>hg38_21_+_8433172_8446622_RNA45SN1_45S_50ntFlanks@4160.4183.24|CCACATCCAAGGAAGGCAGCAGGC</td><td>>hg38_21_+_8433172_8446622_RNA45SN1_45S_50ntFlanks</td><td>4160</td><td>4183</td><td>24</td><td>CCACATCCAAGGAAGGCAGCAGGC</td><td>67.0</td><td>-9.9</td><td>2.3</td><td>-4.3</td><td>1.6E-05</td><td>2.8E-05</td><td>MB231/MB468</td><td>MP</td></tr>
<tr><td>>hg38_21_+_8433172_8446622_RNA45SN1_45S_50ntFlanks@10238.10273.36|CGCTACGCCGCGACGAGTAGGAGGGCCGCTGCGGTG</td><td>>hg38_21_+_8433172_8446622_RNA45SN1_45S_50ntFlanks</td><td>10238</td><td>10273</td><td>36</td><td>CGCTACGCCGCGACGAGTAGGAGGGCCGCTGCGGTG</td><td>1629.5</td><td>5.2</td><td>1.2</td><td>4.5</td><td>6.4E-06</td><td>2.8E-05</td><td>BT20/MB468</td><td>MP</td></tr>
<tr><td>>hg38_21_+_8433172_8446622_RNA45SN1_45S_50ntFlanks@6661.6686.26|CGGTGGATCACTCGGCTCGTGCGTCG</td><td>>hg38_21_+_8433172_8446622_RNA45SN1_45S_50ntFlanks</td><td>6661</td><td>6686</td><td>26</td><td>CGGTGGATCACTCGGCTCGTGCGTCG</td><td>3517.0</td><td>-3.4</td><td>0.8</td><td>-4.3</td><td>2.0E-05</td><td>2.8E-05</td><td>MB231/MB468</td><td>MITO</td></tr>
<tr><td>>hg38_21_+_8433172_8446622_RNA45SN1_45S_50ntFlanks@12631.12661.31|GGCGAAGCTACCATCTGTGGGATTATGACTG</td><td>>hg38_21_+_8433172_8446622_RNA45SN1_45S_50ntFlanks</td><td>12631</td><td>12661</td><td>31</td><td>GGCGAAGCTACCATCTGTGGGATTATGACTG</td><td>1342.8</td><td>-5.7</td><td>1.3</td><td>-4.3</td><td>2.0E-05</td><td>2.8E-05</td><td>MB231/MB468</td><td>MITO</td></tr>
<tr><td>>hg38_21_+_8433172_8446622_RNA45SN1_45S_50ntFlanks@6645.6689.45|CTCGTACGACTCTTAGCGGTGGATCACTCGGCTCGTGCGTCGATG</td><td>>hg38_21_+_8433172_8446622_RNA45SN1_45S_50ntFlanks</td><td>6645</td><td>6689</td><td>45</td><td>CTCGTACGACTCTTAGCGGTGGATCACTCGGCTCGTGCGTCGATG</td><td>2458.5</td><td>-4.0</td><td>0.9</td><td>-4.3</td><td>2.0E-05</td><td>2.8E-05</td><td>MB231/MB468</td><td>MITO</td></tr>
<tr><td>>hg38_21_+_8433172_8446622_RNA45SN1_45S_50ntFlanks@10391.10411.21|AACAGCAGTTGAACATGGGTC</td><td>>hg38_21_+_8433172_8446622_RNA45SN1_45S_50ntFlanks</td><td>10391</td><td>10411</td><td>21</td><td>AACAGCAGTTGAACATGGGTC</td><td>175.6</td><td>-6.3</td><td>1.5</td><td>-4.3</td><td>1.6E-05</td><td>2.8E-05</td><td>MB231/MB468</td><td>MP</td></tr>
<tr><td>>hg38_21_+_8433172_8446622_RNA45SN1_45S_50ntFlanks@8182.8223.42|ATCGAGGCCCAGCCCGTGGACGGTGTGAGGCCGGTAGCGGCC</td><td>>hg38_21_+_8433172_8446622_RNA45SN1_45S_50ntFlanks</td><td>8182</td><td>8223</td><td>42</td><td>ATCGAGGCCCAGCCCGTGGACGGTGTGAGGCCGGTAGCGGCC</td><td>790.3</td><td>-6.5</td><td>1.5</td><td>-4.3</td><td>2.0E-05</td><td>2.8E-05</td><td>MB231/MB468</td><td>NUC</td></tr>
<tr><td>>am_trnaMT_ArgTCG_MT_+_10405_10469@28.48.21|ATTTCGACTCATTAAATTATG</td><td>>am_trnaMT_ArgTCG_MT_+_10405_10469</td><td>28</td><td>48</td><td>21</td><td>ATTTCGACTCATTAAATTATG</td><td>667.0</td><td>-6.6</td><td>1.5</td><td>-4.3</td><td>1.4E-05</td><td>2.8E-05</td><td>BT20/MB468</td><td>MITO</td></tr>
<tr><td>>hg38_21_+_8433172_8446622_RNA45SN1_45S_50ntFlanks@476.520.45|TTCGTGATCGATGTGGTGACGTCGTGCTCTCCCGGGCCGGGTCCG</td><td>>hg38_21_+_8433172_8446622_RNA45SN1_45S_50ntFlanks</td><td>476</td><td>520</td><td>45</td><td>TTCGTGATCGATGTGGTGACGTCGTGCTCTCCCGGGCCGGGTCCG</td><td>4017.0</td><td>4.4</td><td>1.0</td><td>4.3</td><td>2.0E-05</td><td>2.8E-05</td><td>MB231/MB468</td><td>NUC</td></tr>
<tr><td>>hg38_21_+_8433172_8446622_RNA45SN1_45S_50ntFlanks@3985.4002.18|CGGCTTTGGTGACTCTAG</td><td>>hg38_21_+_8433172_8446622_RNA45SN1_45S_50ntFlanks</td><td>3985</td><td>4002</td><td>18</td><td>CGGCTTTGGTGACTCTAG</td><td>2471.4</td><td>-5.4</td><td>1.3</td><td>-4.3</td><td>1.9E-05</td><td>2.8E-05</td><td>MB231/MB468</td><td>CYTO</td></tr>
<tr><td>>hg38_21_+_8433172_8446622_RNA45SN1_45S_50ntFlanks@8195.8214.20|CCGTGGACGGTGTGAGGCCG</td><td>>hg38_21_+_8433172_8446622_RNA45SN1_45S_50ntFlanks</td><td>8195</td><td>8214</td><td>20</td><td>CCGTGGACGGTGTGAGGCCG</td><td>228.5</td><td>-5.3</td><td>1.2</td><td>-4.3</td><td>1.6E-05</td><td>2.8E-05</td><td>MB231/MB468</td><td>MP</td></tr>
</table>

<table>
<tr><th></th><th></th><th></th><th></th><th></th><th></th><th></th><th></th><th></th><th></th><th></th><th></th><th></th><th></th></tr>
<tr><td>>hg38_21_+_8433172_8446622_RNA45SN1_45S_50ntFlanks@11569.11611.43|CAGCCGACTTAGAACTGGTGCGGACCAGGGGAATCCGACTGTT</td><td>>hg38_21_+_8433172_8446622_RNA45SN1_45S_50ntFlanks</td><td>11569</td><td>11611</td><td>43</td><td>CAGCCGACTTAGAACTGGTGCGGACCAGGGGAATCCGACTGTT</td><td>259.4</td><td>-6.8</td><td>1.6</td><td>-4.3</td><td>1.6E-05</td><td>2.8E-05</td><td>MB231/MB468</td><td>MP</td></tr>
<tr><td>>MI0003188|hsa-mir-503&WithFlank&X|-|134546322|134546404@12.31.20(+3A)&[MIMAT0002874&hsa-miR-503-5p&offsets|0|-3(+3A);m-297&X|-|134546371|134546393&offsets|0|-3(+3A)]|TAGCAGCGGGAACAGTTCTGAAA</td><td>>MI0003188|hsa-mir-503&WithFlank&X|-|134546322|134546404</td><td>12</td><td>31</td><td>20(+3A)</td><td>TAGCAGCGGGAACAGTTCTGAAA</td><td>31.0</td><td>8.2</td><td>1.9</td><td>4.3</td><td>1.6E-05</td><td>2.8E-05</td><td>MB231/MB468</td><td>MP</td></tr>
<tr><td>>hg38_21_+_8433172_8446622_RNA45SN1_45S_50ntFlanks@11580.11607.28|GAACTGGTGCGGACCAGGGGAATCCGAC</td><td>>hg38_21_+_8433172_8446622_RNA45SN1_45S_50ntFlanks</td><td>11580</td><td>11607</td><td>28</td><td>GAACTGGTGCGGACCAGGGGAATCCGAC</td><td>636.3</td><td>-5.5</td><td>1.3</td><td>-4.3</td><td>1.6E-05</td><td>2.8E-05</td><td>MB231/MB468</td><td>MP</td></tr>
<tr><td>>hg38_21_+_8433172_8446622_RNA45SN1_45S_50ntFlanks@11581.11623.43|AACTGGTGCGGACCAGGGGAATCCGACTGTTTAATTAAAACAA</td><td>>hg38_21_+_8433172_8446622_RNA45SN1_45S_50ntFlanks</td><td>11581</td><td>11623</td><td>43</td><td>AACTGGTGCGGACCAGGGGAATCCGACTGTTTAATTAAAACAA</td><td>555.0</td><td>4.7</td><td>1.0</td><td>4.7</td><td>2.3E-06</td><td>2.8E-05</td><td>BT20/MB231</td><td>MITO</td></tr>
<tr><td>>hg38_21_+_8433172_8446622_RNA45SN1_45S_50ntFlanks@8170.8211.42|CAAGTCCTTCTGATCGAGGCCCAGCCCGTGGACGGTGTGAGG</td><td>>hg38_21_+_8433172_8446622_RNA45SN1_45S_50ntFlanks</td><td>8170</td><td>8211</td><td>42</td><td>CAAGTCCTTCTGATCGAGGCCCAGCCCGTGGACGGTGTGAGG</td><td>251.1</td><td>-6.2</td><td>1.4</td><td>-4.3</td><td>1.4E-05</td><td>2.8E-05</td><td>BT20/MB468</td><td>MITO</td></tr>
<tr><td>>hg38_21_+_8433172_8446622_RNA45SN1_45S_50ntFlanks@3842.3868.27|CCTACTTGGATAACTGTGGTAATTCTA</td><td>>hg38_21_+_8433172_8446622_RNA45SN1_45S_50ntFlanks</td><td>3842</td><td>3868</td><td>27</td><td>CCTACTTGGATAACTGTGGTAATTCTA</td><td>306.9</td><td>-6.0</td><td>1.4</td><td>-4.3</td><td>1.7E-05</td><td>2.8E-05</td><td>MB231/MB468</td><td>MP</td></tr>
<tr><td>>hg38_21_+_8433172_8446622_RNA45SN1_45S_50ntFlanks@8170.8214.45|CAAGTCCTTCTGATCGAGGCCCAGCCCGTGGACGGTGTGAGGCCG</td><td>>hg38_21_+_8433172_8446622_RNA45SN1_45S_50ntFlanks</td><td>8170</td><td>8214</td><td>45</td><td>CAAGTCCTTCTGATCGAGGCCCAGCCCGTGGACGGTGTGAGGCCG</td><td>3907.3</td><td>-4.5</td><td>1.0</td><td>-4.3</td><td>1.4E-05</td><td>2.8E-05</td><td>BT20/MB468</td><td>MITO</td></tr>
<tr><td>>hg38_21_+_8433172_8446622_RNA45SN1_45S_50ntFlanks@5080.5110.31|ATAACGAACGAGACTCTGGCATGCTAACTAG</td><td>>hg38_21_+_8433172_8446622_RNA45SN1_45S_50ntFlanks</td><td>5080</td><td>5110</td><td>31</td><td>ATAACGAACGAGACTCTGGCATGCTAACTAG</td><td>344.3</td><td>-7.3</td><td>1.7</td><td>-4.3</td><td>1.7E-05</td><td>2.8E-05</td><td>MB231/MB468</td><td>MP</td></tr>
<tr><td>>am_trnaMT_PheGAA_MT_+_577_647@40.62.23|TGTTTAGACGGGCTCACATCACC</td><td>>am_trnaMT_PheGAA_MT_+_577_647</td><td>40</td><td>62</td><td>23</td><td>TGTTTAGACGGGCTCACATCACC</td><td>410.0</td><td>5.3</td><td>1.2</td><td>4.5</td><td>6.4E-06</td><td>2.8E-05</td><td>BT20/MB468</td><td>MP</td></tr>
<tr><td>>hg38_21_+_8433172_8446622_RNA45SN1_45S_50ntFlanks@11833.11872.40|ACCACAGCCAAGGGAACGGGCTTGGCGGAATCAGCGGGGA</td><td>>hg38_21_+_8433172_8446622_RNA45SN1_45S_50ntFlanks</td><td>11833</td><td>11872</td><td>40</td><td>ACCACAGCCAAGGGAACGGGCTTGGCGGAATCAGCGGGGA</td><td>45.5</td><td>-8.6</td><td>1.9</td><td>-4.5</td><td>6.4E-06</td><td>2.8E-05</td><td>BT20/MB468</td><td>MP</td></tr>
<tr><td>>hg38_21_+_8433172_8446622_RNA45SN1_45S_50ntFlanks@11210.11233.24|GGCGACTCTGGACGCGAGCCGGGC</td><td>>hg38_21_+_8433172_8446622_RNA45SN1_45S_50ntFlanks</td><td>11210</td><td>11233</td><td>24</td><td>GGCGACTCTGGACGCGAGCCGGGC</td><td>708.3</td><td>-5.9</td><td>1.4</td><td>-4.3</td><td>2.1E-05</td><td>2.8E-05</td><td>BT20/MB468</td><td>NUC</td></tr>
<tr><td>>hg38_MT_+_598_1651_RNR1_12S_50ntFlanks@628.665.38|CCGATCAACCTCACCACCTCTTGCTCAGCCTATATACC</td><td>>hg38_MT_+_598_1651_RNR1_12S_50ntFlanks</td><td>628</td><td>665</td><td>38</td><td>CCGATCAACCTCACCACCTCTTGCTCAGCCTATATACC</td><td>31.3</td><td>8.2</td><td>1.9</td><td>4.3</td><td>1.7E-05</td><td>2.8E-05</td><td>MB231/MB468</td><td>MP</td></tr>
<tr><td>>hg38_21_+_8433172_8446622_RNA45SN1_45S_50ntFlanks@10392.10427.36|ACAGCAGTTGAACATGGGTCAGTCGGTCCTGAGAGA</td><td>>hg38_21_+_8433172_8446622_RNA45SN1_45S_50ntFlanks</td><td>10392</td><td>10427</td><td>36</td><td>ACAGCAGTTGAACATGGGTCAGTCGGTCCTGAGAGA</td><td>122.9</td><td>-6.4</td><td>1.5</td><td>-4.3</td><td>1.7E-05</td><td>2.8E-05</td><td>MB231/MB468</td><td>MP</td></tr>
<tr><td>>hg38_21_+_8433172_8446622_RNA45SN1_45S_50ntFlanks@11580.11602.23|GAACTGGTGCGGACCAGGGGAAT</td><td>>hg38_21_+_8433172_8446622_RNA45SN1_45S_50ntFlanks</td><td>11580</td><td>11602</td><td>23</td><td>GAACTGGTGCGGACCAGGGGAAT</td><td>1130.6</td><td>-3.2</td><td>0.8</td><td>-4.3</td><td>1.7E-05</td><td>2.8E-05</td><td>BT20/MB468</td><td>CYTO</td></tr>
<tr><td>>hg38_21_+_8433172_8446622_RNA45SN1_45S_50ntFlanks@8184.8208.25|CGAGGCCCAGCCCGTGGACGGTGTG</td><td>>hg38_21_+_8433172_8446622_RNA45SN1_45S_50ntFlanks</td><td>8184</td><td>8208</td><td>25</td><td>CGAGGCCCAGCCCGTGGACGGTGTG</td><td>289.8</td><td>-5.4</td><td>1.2</td><td>-4.3</td><td>1.7E-05</td><td>2.8E-05</td><td>MB231/MB468</td><td>MP</td></tr>
<tr><td>>hg38_21_+_8433172_8446622_RNA45SN1_45S_50ntFlanks@10797.10821.25|GCATGTTGGAACAATGTAGGTAAGG</td><td>>hg38_21_+_8433172_8446622_RNA45SN1_45S_50ntFlanks</td><td>10797</td><td>10821</td><td>25</td><td>GCATGTTGGAACAATGTAGGTAAGG</td><td>296.0</td><td>-3.5</td><td>0.8</td><td>-4.3</td><td>1.7E-05</td><td>2.8E-05</td><td>MB231/MB468</td><td>MP</td></tr>
<tr><td>>hg38_MT_+_1621_3279_RNR2_16S_50ntFlanks@286.310.25|GAAACCAGACGAGCTACCTAAGAAC</td><td>>hg38_MT_+_1621_3279_RNR2_16S_50ntFlanks</td><td>286</td><td>310</td><td>25</td><td>GAAACCAGACGAGCTACCTAAGAAC</td><td>35.6</td><td>8.4</td><td>1.9</td><td>4.3</td><td>1.7E-05</td><td>2.8E-05</td><td>MB231/MB468</td><td>MP</td></tr>
<tr><td>>hg38_21_+_8433172_8446622_RNA45SN1_45S_50ntFlanks@12432.12454.23|CCTATCATTGTGAAGCAGAATTC</td><td>>hg38_21_+_8433172_8446622_RNA45SN1_45S_50ntFlanks</td><td>12432</td><td>12454</td><td>23</td><td>CCTATCATTGTGAAGCAGAATTC</td><td>212.6</td><td>-6.7</td><td>1.5</td><td>-4.5</td><td>6.5E-06</td><td>2.8E-05</td><td>BT20/MB468</td><td>MP</td></tr>
<tr><td>>hg38_21_+_8433172_8446622_RNA45SN1_45S_50ntFlanks@12004.12033.30|CGGGCCGCCGGTGAAATACCACTACTCTGA</td><td>>hg38_21_+_8433172_8446622_RNA45SN1_45S_50ntFlanks</td><td>12004</td><td>12033</td><td>30</td><td>CGGGCCGCCGGTGAAATACCACTACTCTGA</td><td>1648.7</td><td>4.0</td><td>0.9</td><td>4.7</td><td>2.4E-06</td><td>2.8E-05</td><td>BT20/MB231</td><td>MITO</td></tr>
<tr><td>>trna14_LysTTT_11_-_59327808_59327880@3.36.34,trna2_LysTTT_17_+_8022473_8022545@3.36.34,trna54_LysTTT_1_+_204475655_204475727@3.36.34,trna5_LysTTT_11_+_59323902_59323974@3.36.34,trna62_LysTTT_1_-_204476158_204476230@3.36.34,trna76_LysTTT_6_+_28918806_28918878@3.36.34|CCGGATAGCTCAGTCGGTAGAGCATCAGACTTTT</td><td>>trna14_LysTTT_11_-_59327808_59327880</td><td>3</td><td>36</td><td>34</td><td>CCGGATAGCTCAGTCGGTAGAGCATCAGACTTTT</td><td>311.3</td><td>-5.1</td><td>1.1</td><td>-4.7</td><td>2.4E-06</td><td>2.8E-05</td><td>BT20/MB231</td><td>MITO</td></tr>
<tr><td>>MI0000061|hsa-let-7a-2&WithFlank&11|-|122146516|122146599@11.32.22(+1C);MI0000062|hsa-let-7a-3&WithFlank&22|+|46112743|46112828@10.31.22(+1C);MI0000060|hsa-let-7a-1&WithFlank&9|+|94175951|94176042@12.33.22(+1C)&[MIMAT0000062&hsa-let-7a-5p&offsets|0|0(+1C);m-6&11|-|122146568|122146589&offsets|0|0(+1C)];[MIMAT0000062_1&hsa-let-7a-5p&offsets|0|0(+1C);m-5&22|+|46112752|46112773&offsets|0|0(+1C)];[MIMAT0000062_2&hsa-let-7a-5p&offsets|0|0(+1C);m-4&9|+|94175962|94175983&offsets|0|0(+1C)]|TGAGGTAGTAGGTTGTATAGTTC</td><td>>MI0000061|hsa-let-7a-2&WithFlank&11|-|122146516|122146599</td><td>11</td><td>32</td><td>22(+1C)</td><td>TGAGGTAGTAGGTTGTATAGTTC</td><td>657.7</td><td>5.4</td><td>1.3</td><td>4.3</td><td>1.7E-05</td><td>2.8E-05</td><td>MB231/MB468</td><td>MP</td></tr>
<tr><td>>hg38_21_+_8433172_8446622_RNA45SN1_45S_50ntFlanks@12003.12033.31|GCGGGCCGCCGGTGAAATACCACTACTCTGA</td><td>>hg38_21_+_8433172_8446622_RNA45SN1_45S_50ntFlanks</td><td>12003</td><td>12033</td><td>31</td><td>GCGGGCCGCCGGTGAAATACCACTACTCTGA</td><td>558.0</td><td>3.3</td><td>0.8</td><td>4.3</td><td>1.7E-05</td><td>2.8E-05</td><td>BT20/MB468</td><td>CYTO</td></tr>
<tr><td>>hg38_21_+_8433172_8446622_RNA45SN1_45S_50ntFlanks@5417.5445.29|CTACCGATTGGATGGTTTAGTGAGGCCCT</td><td>>hg38_21_+_8433172_8446622_RNA45SN1_45S_50ntFlanks</td><td>5417</td><td>5445</td><td>29</td><td>CTACCGATTGGATGGTTTAGTGAGGCCCT</td><td>261.6</td><td>-5.9</td><td>1.4</td><td>-4.3</td><td>1.5E-05</td><td>2.8E-05</td><td>BT20/MB468</td><td>MITO</td></tr>
<tr><td>>hg38_21_+_8433172_8446622_RNA45SN1_45S_50ntFlanks@11581.11599.19|AACTGGTGCGGACCAGGGG</td><td>>hg38_21_+_8433172_8446622_RNA45SN1_45S_50ntFlanks</td><td>11581</td><td>11599</td><td>19</td><td>AACTGGTGCGGACCAGGGG</td><td>193.4</td><td>-5.6</td><td>1.3</td><td>-4.3</td><td>1.7E-05</td><td>2.8E-05</td><td>BT20/MB468</td><td>CYTO</td></tr>
<tr><td>>hg38_21_+_8433172_8446622_RNA45SN1_45S_50ntFlanks@3775.3797.23|GCCGGTACAGTGAAACTGCGAAT</td><td>>hg38_21_+_8433172_8446622_RNA45SN1_45S_50ntFlanks</td><td>3775</td><td>3797</td><td>23</td><td>GCCGGTACAGTGAAACTGCGAAT</td><td>245.9</td><td>-7.3</td><td>1.7</td><td>-4.3</td><td>1.7E-05</td><td>2.8E-05</td><td>MB231/MB468</td><td>MP</td></tr>
<tr><td>>MI0003560|hsa-mir-92b&WithFlank&1|+|155195171|155195278@67.88.22(+1U)&[MIMAT0003218&hsa-miR-92b-3p&offsets|0|0(+1U);m-115&1|+|155195237|155195258&offsets|0|0(+1U)]|TATTGCACTCGTCCCGGCCTCCT</td><td>>MI0003560|hsa-mir-92b&WithFlank&1|+|155195171|155195278</td><td>67</td><td>88</td><td>22(+1U)</td><td>TATTGCACTCGTCCCGGCCTCCT</td><td>210.6</td><td>5.5</td><td>1.3</td><td>4.3</td><td>1.7E-05</td><td>2.8E-05</td><td>MB231/MB468</td><td>MP</td></tr>
<tr><td>>hg38_21_+_8433172_8446622_RNA45SN1_45S_50ntFlanks@9567.9585.19|ACGTGCAAATCGGTCGTCC</td><td>>hg38_21_+_8433172_8446622_RNA45SN1_45S_50ntFlanks</td><td>9567</td><td>9585</td><td>19</td><td>ACGTGCAAATCGGTCGTCC</td><td>387.0</td><td>-4.6</td><td>1.1</td><td>-4.3</td><td>1.9E-05</td><td>2.8E-05</td><td>MB231/MB468</td><td>CYTO</td></tr>
<tr><td>>hg38_21_+_8433172_8446622_RNA45SN1_45S_50ntFlanks@10351.10379.29|CGAGAACTTTGAAGGCCGAAGTGGAGAAG</td><td>>hg38_21_+_8433172_8446622_RNA45SN1_45S_50ntFlanks</td><td>10351</td><td>10379</td><td>29</td><td>CGAGAACTTTGAAGGCCGAAGTGGAGAAG</td><td>596.6</td><td>-2.4</td><td>0.6</td><td>-4.3</td><td>1.7E-05</td><td>2.8E-05</td><td>BT20/MB468</td><td>CYTO</td></tr>
<tr><td>>hg38_21_+_8433172_8446622_RNA45SN1_45S_50ntFlanks@11874.11900.27|AGAAGACCCTGTTGAGCTTGACTCTAG</td><td>>hg38_21_+_8433172_8446622_RNA45SN1_45S_50ntFlanks</td><td>11874</td><td>11900</td><td>27</td><td>AGAAGACCCTGTTGAGCTTGACTCTAG</td><td>908.2</td><td>-6.4</td><td>1.5</td><td>-4.3</td><td>1.7E-05</td><td>2.8E-05</td><td>BT20/MB468</td><td>CYTO</td></tr>
<tr><td>>hg38_21_+_8433172_8446622_RNA45SN1_45S_50ntFlanks@9573.9603.31|AAATCGGTCGTCCGACCTGGGTATAGGGGCG</td><td>>hg38_21_+_8433172_8446622_RNA45SN1_45S_50ntFlanks</td><td>9573</td><td>9603</td><td>31</td><td>AAATCGGTCGTCCGACCTGGGTATAGGGGCG</td><td>3740.5</td><td>-3.9</td><td>0.9</td><td>-4.3</td><td>2.0E-05</td><td>2.8E-05</td><td>MB231/MB468</td><td>MITO</td></tr>
<tr><td>>hg38_21_+_8433172_8446622_RNA45SN1_45S_50ntFlanks@4731.4759.29|AAGAACGAAAGTCGGAGGTTCGAAGACGA</td><td>>hg38_21_+_8433172_8446622_RNA45SN1_45S_50ntFlanks</td><td>4731</td><td>4759</td><td>29</td><td>AAGAACGAAAGTCGGAGGTTCGAAGACGA</td><td>2378.4</td><td>-3.2</td><td>0.7</td><td>-4.3</td><td>2.0E-05</td><td>2.8E-05</td><td>MB231/MB468</td><td>NUC</td></tr>
<tr><td>>MI0000103|hsa-mir-101-1&WithFlank&1|-|65058428|65058514@53.74.22(+1U);MI0000739|hsa-mir-101-2&WithFlank&9|+|4850291|4850381@55.76.22(+1U)&[MIMAT0000099&hsa-miR-101-3p&offsets|0|+1(+1U);m-17&1|-|65058443|65058462&offsets|0|+2(+1U)];[MIMAT0000099_1&hsa-miR-101-3p&offsets|0|+1(+1U);m-16&9|+|4850345|4850364&offsets|0|+2(+1U)]|TACAGTACTGTGATAACTGAAGT</td><td>>MI0000103|hsa-mir-101-1&WithFlank&1|-|65058428|65058514</td><td>53</td><td>74</td><td>22(+1U)</td><td>TACAGTACTGTGATAACTGAAGT</td><td>564.4</td><td>6.1</td><td>1.4</td><td>4.3</td><td>2.0E-05</td><td>2.8E-05</td><td>MB231/MB468</td><td>MITO</td></tr>
<tr><td>>MI0003820|hsa-mir-454&WithFlank&17|-|59137752|59137878@70.93.24&[MIMAT0003885&hsa-miR-454-3p&offsets|0|+1;m-272&17|-|59137787|59137809&offsets|0|+1]|TAGTGCAATATTGCTTATAGGGTT</td><td>>MI0003820|hsa-mir-454&WithFlank&17|-|59137752|59137878</td><td>70</td><td>93</td><td>24</td><td>TAGTGCAATATTGCTTATAGGGTT</td><td>197.1</td><td>2.7</td><td>0.6</td><td>4.3</td><td>2.1E-05</td><td>2.8E-05</td><td>BT20/MB468</td><td>NUC</td></tr>
<tr><td>>MI0000456|hsa-mir-140&WithFlank&16|+|69933075|69933186@68.90.23&[MIMAT0004597&hsa-miR-140-3p&offsets|0|+2;m-45&16|+|69933143|69933164&offsets|-1|0]|TACCACAGGGTAGAACCACGGAC</td><td>>MI0000456|hsa-mir-140&WithFlank&16|+|69933075|69933186</td><td>68</td><td>90</td><td>23</td><td>TACCACAGGGTAGAACCACGGAC</td><td>390.7</td><td>5.3</td><td>1.2</td><td>4.3</td><td>1.7E-05</td><td>2.9E-05</td><td>MB231/MB468</td><td>MP</td></tr>
<tr><td>>hg38_21_+_8433172_8446622_RNA45SN1_45S_50ntFlanks@3852.3872.21|TAACTGTGGTAATTCTAGAGC</td><td>>hg38_21_+_8433172_8446622_RNA45SN1_45S_50ntFlanks</td><td>3852</td><td>3872</td><td>21</td><td>TAACTGTGGTAATTCTAGAGC</td><td>143.8</td><td>-6.1</td><td>1.4</td><td>-4.3</td><td>1.7E-05</td><td>2.9E-05</td><td>MB231/MB468</td><td>MP</td></tr>
<tr><td>>hg38_21_+_8433172_8446622_RNA45SN1_45S_50ntFlanks@12638.12659.22|CTACCATCTGTGGGATTATGAC</td><td>>hg38_21_+_8433172_8446622_RNA45SN1_45S_50ntFlanks</td><td>12638</td><td>12659</td><td>22</td><td>CTACCATCTGTGGGATTATGAC</td><td>202.4</td><td>-5.6</td><td>1.3</td><td>-4.3</td><td>1.7E-05</td><td>2.9E-05</td><td>BT20/MB468</td><td>CYTO</td></tr>
<tr><td>>hg38_21_+_8433172_8446622_RNA45SN1_45S_50ntFlanks@9917.9948.32|AAGGTGTTGGTTGATATAGACAGCAGGACGGT</td><td>>hg38_21_+_8433172_8446622_RNA45SN1_45S_50ntFlanks</td><td>9917</td><td>9948</td><td>32</td><td>AAGGTGTTGGTTGATATAGACAGCAGGACGGT</td><td>88.0</td><td>9.8</td><td>2.1</td><td>4.8</td><td>2.0E-06</td><td>2.9E-05</td><td>BT20/MB231</td><td>MP</td></tr>
<tr><td>>hg38_21_+_8433172_8446622_RNA45SN1_45S_50ntFlanks@3899.3920.22|CCTTCGCGGGGGGGATGCGTGC</td><td>>hg38_21_+_8433172_8446622_RNA45SN1_45S_50ntFlanks</td><td>3899</td><td>3920</td><td>22</td><td>CCTTCGCGGGGGGGATGCGTGC</td><td>1036.0</td><td>-7.2</td><td>1.6</td><td>-4.5</td><td>6.5E-06</td><td>2.9E-05</td><td>BT20/MB468</td><td>MP</td></tr>
<tr><td>>hg38_21_+_8433172_8446622_RNA45SN1_45S_50ntFlanks@4726.4766.41|TAATCAAGAACGAAAGTCGGAGGTTCGAAGACGATCAGATA</td><td>>hg38_21_+_8433172_8446622_RNA45SN1_45S_50ntFlanks</td><td>4726</td><td>4766</td><td>41</td><td>TAATCAAGAACGAAAGTCGGAGGTTCGAAGACGATCAGATA</td><td>254.4</td><td>4.4</td><td>1.0</td><td>4.3</td><td>1.9E-05</td><td>2.9E-05</td><td>MB231/MB468</td><td>CYTO</td></tr>
<tr><td>>hg38_MT_+_598_1651_RNR1_12S_50ntFlanks@930.950.21|AAAACCCCTACGCATTTATAT</td><td>>hg38_MT_+_598_1651_RNR1_12S_50ntFlanks</td><td>930</td><td>950</td><td>21</td><td>AAAACCCCTACGCATTTATAT</td><td>37.0</td><td>8.4</td><td>2.0</td><td>4.3</td><td>1.7E-05</td><td>2.9E-05</td><td>MB231/MB468</td><td>MP</td></tr>
<tr><td>>MI0000077|hsa-mir-21&WithFlank&17|+|59841260|59841343@14.34.21(+2C)&[MIMAT0000076&hsa-miR-21-5p&offsets|0|-1(+2C);m-1&17|+|59841273|59841294&offsets|0|-1(+2C)]|TAGCTTATCAGACTGATGTTGCC</td><td>>MI0000077|hsa-mir-21&WithFlank&17|+|59841260|59841343</td><td>14</td><td>34</td><td>21(+2C)</td><td>TAGCTTATCAGACTGATGTTGCC</td><td>1659.4</td><td>-5.4</td><td>1.1</td><td>-4.8</td><td>2.0E-06</td><td>2.9E-05</td><td>BT20/MB231</td><td>MP</td></tr>
<tr><td>>hg38_21_+_8433172_8446622_RNA45SN1_45S_50ntFlanks@10802.10821.20|TTGGAACAATGTAGGTAAGG</td><td>>hg38_21_+_8433172_8446622_RNA45SN1_45S_50ntFlanks</td><td>10802</td><td>10821</td><td>20</td><td>TTGGAACAATGTAGGTAAGG</td><td>187.9</td><td>-6.2</td><td>1.5</td><td>-4.3</td><td>1.7E-05</td><td>2.9E-05</td><td>MB231/MB468</td><td>MP</td></tr>
<tr><td>>hg38_21_+_8433172_8446622_RNA45SN1_45S_50ntFlanks@9572.9597.26|CAAATCGGTCGTCCGACCTGGGTATA</td><td>>hg38_21_+_8433172_8446622_RNA45SN1_45S_50ntFlanks</td><td>9572</td><td>9597</td><td>26</td><td>CAAATCGGTCGTCCGACCTGGGTATA</td><td>4519.6</td><td>-7.3</td><td>1.7</td><td>-4.3</td><td>1.7E-05</td><td>2.9E-05</td><td>BT20/MB468</td><td>CYTO</td></tr>
<tr><td>>hg38_21_+_8433172_8446622_RNA45SN1_45S_50ntFlanks@12927.12960.34|CCTGGCGCTAAACCATTCGTAGACGACCTGCTTC</td><td>>hg38_21_+_8433172_8446622_RNA45SN1_45S_50ntFlanks</td><td>12927</td><td>12960</td><td>34</td><td>CCTGGCGCTAAACCATTCGTAGACGACCTGCTTC</td><td>244.2</td><td>-4.7</td><td>1.1</td><td>-4.3</td><td>2.0E-05</td><td>2.9E-05</td><td>MB231/MB468</td><td>MITO</td></tr>
<tr><td>>hg38_21_+_8433172_8446622_RNA45SN1_45S_50ntFlanks@6650.6672.23|ACGACTCTTAGCGGTGGATCACT</td><td>>hg38_21_+_8433172_8446622_RNA45SN1_45S_50ntFlanks</td><td>6650</td><td>6672</td><td>23</td><td>ACGACTCTTAGCGGTGGATCACT</td><td>340.9</td><td>3.1</td><td>0.7</td><td>4.3</td><td>2.1E-05</td><td>2.9E-05</td><td>BT20/MB468</td><td>NUC</td></tr>
</table>

| | | | | | | | | | | | | | |
|---|---|---|---|---|---|---|---|---|---|---|---|---|---|
| >MI0000234\|hsa-mir-192&WithFlank&11\|-\|64891131\|64891252@31.53.23&[MIMAT0000222&hsa-miR-192-5p&offsets\|+1\|+3;m-59&11\|-\|64891203\|64891223&offsets\|+1\|+3]\|TGACCTATGAATTGACAGCCAGT | >MI0000234\|hsa-mir-192&WithFlank&11\|-\|64891131\|64891252 | 31 | 53 | 23 | TGACCTATGAATTGACAGCCAGT | 408.2 | 4.3 | 1.0 | 4.3 | 2.0E-05 | 2.9E-05 | MB231/MB468 | MITO |
| >hg38_21_+_8433172_8446622_RNA45SN1_45S_50ntFlanks@4884.4906.23\|CAAAGCTGAAACTTAAAGGAATT | >hg38_21_+_8433172_8446622_RNA45SN1_45S_50ntFlanks | 4884 | 4906 | 23 | CAAAGCTGAAACTTAAAGGAATT | 201.9 | -6.4 | 1.4 | -4.5 | 6.6E-06 | 2.9E-05 | BT20/MB468 | MP |
| >hg38_21_+_8433172_8446622_RNA45SN1_45S_50ntFlanks@5188.5205.18\|ATAACAGGTCTGTGATGC | >hg38_21_+_8433172_8446622_RNA45SN1_45S_50ntFlanks | 5188 | 5205 | 18 | ATAACAGGTCTGTGATGC | 284.2 | -7.3 | 1.7 | -4.3 | 1.7E-05 | 2.9E-05 | MB231/MB468 | MP |
| >hg38_21_+_8433172_8446622_RNA45SN1_45S_50ntFlanks@12249.12272.24\|CAAAAGCTCGCTTGATCTTGATTT | >hg38_21_+_8433172_8446622_RNA45SN1_45S_50ntFlanks | 12249 | 12272 | 24 | CAAAAGCTCGCTTGATCTTGATTT | 1370.2 | -4.3 | 1.0 | -4.3 | 1.5E-05 | 2.9E-05 | BT20/MB468 | MITO |
| >hg38_21_+_8433172_8446622_RNA45SN1_45S_50ntFlanks@11316.11343.28\|CACCCCACGTCTCGTCGCGCGCGCGTCC | >hg38_21_+_8433172_8446622_RNA45SN1_45S_50ntFlanks | 11316 | 11343 | 28 | CACCCCACGTCTCGTCGCGCGCGCGTCC | 7366.0 | -8.7 | 2.0 | -4.3 | 2.1E-05 | 2.9E-05 | BT20/MB468 | NUC |
| >hg38_21_+_8433172_8446622_RNA45SN1_45S_50ntFlanks@5090.5110.21\|AGACTCTGGCATGCTAACTAG | >hg38_21_+_8433172_8446622_RNA45SN1_45S_50ntFlanks | 5090 | 5110 | 21 | AGACTCTGGCATGCTAACTAG | 86.6 | -10.3 | 2.4 | -4.3 | 1.7E-05 | 2.9E-05 | MB231/MB468 | MP |
| >MI0003778\|hsa-mir-320c-1&WithFlank&18\|+\|21683512\|21683595@48.67.20;MI0008191\|hsa-mir-320c-2&WithFlank&18\|+\|24321669\|24321752@48.67.20&[MIMAT0005793&hsa-miR-320c&offsets\|0\|0;m-222&18\|+\|21683559\|21683578&offsets\|0\|0];[MIMAT0005793_1&hsa-miR-320c&offsets\|0\|0;m-219&18\|+\|24321716\|24321735&offsets\|0\|0]\|AAAAGCTGGGTTGAGAGGGT | >MI0003778\|hsa-mir-320c-1&WithFlank&18\|+\|21683512\|21683595 | 48 | 67 | 20 | AAAAGCTGGGTTGAGAGGGT | 508.8 | 2.9 | 0.6 | 4.7 | 2.4E-06 | 2.9E-05 | BT20/MB231 | NUC |
| >hg38_21_+_8433172_8446622_RNA45SN1_45S_50ntFlanks@10789.10812.24\|AGCCTCTGGCATGTTGGAACAATG | >hg38_21_+_8433172_8446622_RNA45SN1_45S_50ntFlanks | 10789 | 10812 | 24 | AGCCTCTGGCATGTTGGAACAATG | 92.8 | 9.9 | 2.1 | 4.7 | 2.0E-06 | 2.9E-05 | BT20/MB231 | MP |
| >am_trna2_GlyGCC_21_-_18827107_18827177@1.25.25,trna35_GlyGCC_1_+_161413094_161413164@1.25.25,trna37_GlyGCC_1_+_161420467_161420537@1.25.25,trna39_GlyGCC_1_+_161427898_161427968@1.25.25,trna41_GlyGCC_1_+_161435258_161435328@1.25.25\|GCATGGGTGGTTCAGTGGTAGAATT | >am_trna2_GlyGCC_21_-_18827107_18827177 | 1 | 25 | 25 | GCATGGGTGGTTCAGTGGTAGAATT | 280.1 | -3.1 | 0.7 | -4.3 | 1.9E-05 | 2.9E-05 | MB231/MB468 | CYTO |
| >MI0003137\|hsa-mir-193b&WithFlank&16\|+\|14303961\|14304055@57.79.23&[MIMAT0002819&hsa-miR-193b-3p&offsets\|0\|+1;m-166&16\|+\|14304017\|14304038&offsets\|0\|+1]\|AACTGGCCCTCAAAGTCCCGCTT | >MI0003137\|hsa-mir-193b&WithFlank&16\|+\|14303961\|14304055 | 57 | 79 | 23 | AACTGGCCCTCAAAGTCCCGCTT | 1609.5 | 7.1 | 1.6 | 4.5 | 6.6E-06 | 2.9E-05 | BT20/MB468 | MP |
| >hg38_21_+_8433172_8446622_RNA45SN1_45S_50ntFlanks@12627.12661.35\|ATGGGGCGAAGCTACCATCTGTGGGATTATGACTG | >hg38_21_+_8433172_8446622_RNA45SN1_45S_50ntFlanks | 12627 | 12661 | 35 | ATGGGGCGAAGCTACCATCTGTGGGATTATGACTG | 385.4 | -4.1 | 1.0 | -4.3 | 1.9E-05 | 2.9E-05 | MB231/MB468 | CYTO |
| >hg38_21_+_8433172_8446622_RNA45SN1_45S_50ntFlanks@11661.11684.24\|TGATTTCTGCCCAGTGCTCTGAAT | >hg38_21_+_8433172_8446622_RNA45SN1_45S_50ntFlanks | 11661 | 11684 | 24 | TGATTTCTGCCCAGTGCTCTGAAT | 215.8 | -4.1 | 1.0 | -4.3 | 1.9E-05 | 2.9E-05 | MB231/MB468 | CYTO |
| >hg38_21_+_8433172_8446622_RNA45SN1_45S_50ntFlanks@6786.6806.21\|TACGCCTGTCTGAGCGTCGCT | >hg38_21_+_8433172_8446622_RNA45SN1_45S_50ntFlanks | 6786 | 6806 | 21 | TACGCCTGTCTGAGCGTCGCT | 3273.4 | 3.9 | 0.9 | 4.3 | 2.0E-05 | 2.9E-05 | MB231/MB468 | MITO |
| >hg38_21_+_8433172_8446622_RNA45SN1_45S_50ntFlanks@11586.11611.26\|GTGCGGACCAGGGGAATCCGACTGTT | >hg38_21_+_8433172_8446622_RNA45SN1_45S_50ntFlanks | 11586 | 11611 | 26 | GTGCGGACCAGGGGAATCCGACTGTT | 2263.2 | -3.9 | 0.9 | -4.5 | 6.6E-06 | 2.9E-05 | BT20/MB468 | MP |
| >hg38_21_+_8433172_8446622_RNA45SN1_45S_50ntFlanks@6650.6678.29\|ACGACTCTTAGCGGTGGATCACTCGGCTC | >hg38_21_+_8433172_8446622_RNA45SN1_45S_50ntFlanks | 6650 | 6678 | 29 | ACGACTCTTAGCGGTGGATCACTCGGCTC | 1074.7 | 4.4 | 1.0 | 4.5 | 6.6E-06 | 2.9E-05 | BT20/MB468 | MP |
| >trnaMT_GluTTC_MT_-_14674_14742@36.72.37,trnalookalike8_GluTTC_5_-_93905172_93905240@36.72.37\|ATCATTGGTCGTGGTTGTAGTCCGTGCGAGAATACCA | >trnaMT_GluTTC_MT_-_14674_14742 | 36 | 72 | 37 | ATCATTGGTCGTGGTTGTAGTCCGTGCGAGAATACCA | 181.8 | -10.6 | 2.3 | -4.5 | 6.6E-06 | 2.9E-05 | BT20/MB468 | MP |
| >hg38_21_+_8433172_8446622_RNA45SN1_45S_50ntFlanks@11199.11234.36\|CCGGCGGCGGCGGCGACTCTGGACGCGAGCCGGGCC | >hg38_21_+_8433172_8446622_RNA45SN1_45S_50ntFlanks | 11199 | 11234 | 36 | CCGGCGGCGGCGGCGACTCTGGACGCGAGCCGGGCC | 44.3 | -8.6 | 1.9 | -4.5 | 6.6E-06 | 2.9E-05 | BT20/MB468 | MP |
| >hg38_21_+_8433172_8446622_RNA45SN1_45S_50ntFlanks@12044.12065.22\|ACTGACCCGGTGAGGCGGGGGG | >hg38_21_+_8433172_8446622_RNA45SN1_45S_50ntFlanks | 12044 | 12065 | 22 | ACTGACCCGGTGAGGCGGGGGG | 63.0 | -9.1 | 2.0 | -4.5 | 6.6E-06 | 2.9E-05 | BT20/MB468 | MP |
| >hg38_21_+_8433172_8446622_RNA45SN1_45S_50ntFlanks@9577.9595.19\|CGGTCGTCCGACCTGGGTA | >hg38_21_+_8433172_8446622_RNA45SN1_45S_50ntFlanks | 9577 | 9595 | 19 | CGGTCGTCCGACCTGGGTA | 75.7 | -10.1 | 2.3 | -4.3 | 1.7E-05 | 2.9E-05 | MB231/MB468 | MP |
| >hg38_21_+_8433172_8446622_RNA45SN1_45S_50ntFlanks@10841.10866.26\|CGTAACTTCGGGATAAGGATTGGCTC | >hg38_21_+_8433172_8446622_RNA45SN1_45S_50ntFlanks | 10841 | 10866 | 26 | CGTAACTTCGGGATAAGGATTGGCTC | 722.1 | -5.5 | 1.3 | -4.3 | 1.7E-05 | 2.9E-05 | MB231/MB468 | MP |
| >hg38_21_+_8433172_8446622_RNA45SN1_45S_50ntFlanks@6722.6740.19\|ATTGCAGGACACATTGATC | >hg38_21_+_8433172_8446622_RNA45SN1_45S_50ntFlanks | 6722 | 6740 | 19 | ATTGCAGGACACATTGATC | 344.6 | -4.8 | 1.1 | -4.3 | 2.1E-05 | 2.9E-05 | MB231/MB468 | MITO |
| >hg38_21_+_8433172_8446622_RNA45SN1_45S_50ntFlanks@4590.4614.25\|ATTTTGTTGGTTTTCGGAACTGAGG | >hg38_21_+_8433172_8446622_RNA45SN1_45S_50ntFlanks | 4590 | 4614 | 25 | ATTTTGTTGGTTTTCGGAACTGAGG | 175.2 | -5.7 | 1.3 | -4.3 | 2.0E-05 | 2.9E-05 | MB231/MB468 | NUC |
| >hg38_21_+_8433172_8446622_RNA45SN1_45S_50ntFlanks@4734.4767.34\|AACGAAAGTCGGAGGTTCGAAGACGATCAGATAC | >hg38_21_+_8433172_8446622_RNA45SN1_45S_50ntFlanks | 4734 | 4767 | 34 | AACGAAAGTCGGAGGTTCGAAGACGATCAGATAC | 150.4 | 6.2 | 1.5 | 4.3 | 2.0E-05 | 2.9E-05 | MB231/MB468 | NUC |
| >hg38_21_+_8433172_8446622_RNA45SN1_45S_50ntFlanks@8183.8210.28\|TCGAGGCCCAGCCCGTGGACGGTGTGAG | >hg38_21_+_8433172_8446622_RNA45SN1_45S_50ntFlanks | 8183 | 8210 | 28 | TCGAGGCCCAGCCCGTGGACGGTGTGAG | 301.7 | -3.7 | 0.9 | -4.3 | 2.0E-05 | 2.9E-05 | MB231/MB468 | NUC |
| >trna10_AspGTC_12_-_125424193_125424264@39.74.36,trna12_AspGTC_12_-_125411891_125411962@39.74.36,trna144_AspGTC_6_-_27551236_27551307@39.74.36,trna38_AspGTC_17_-_8125556_8125627@39.74.36,trna45_AspGTC_6_+_27447453_27447524@39.74.36,trna48_AspGTC_6_+_27471523_27471594@39.74.36,trna4_AspGTC_12_+_96429799_96429870@39.74.36,trna69_AspGTC_1_-_161440205_161440276@39.74.36,trna72_AspGTC_1_-_161432824_161432895@39.74.36,trna75_AspGTC_1_-_161425414_161425485@39.74.36,trna78_AspGTC_1_-_161418033_161418104@39.74.36,trna81_AspGTC_1_-_161410615_161410686@39.74.36\|GCGGGAGACCGGGGTTCGATTCCCCGACGGGGAGCC | >trna10_AspGTC_12_-_125424193_125424264 | 39 | 74 | 36 | GCGGGAGACCGGGGTTCGATTCCCCGACGGGGAGCC | 34.1 | 8.3 | 1.9 | 4.3 | 1.7E-05 | 2.9E-05 | MB231/MB468 | MP |
| >hg38_21_+_8433172_8446622_RNA45SN1_45S_50ntFlanks@9922.9951.30\|GTTGGTTGATATAGACAGCAGGACGGTGGC | >hg38_21_+_8433172_8446622_RNA45SN1_45S_50ntFlanks | 9922 | 9951 | 30 | GTTGGTTGATATAGACAGCAGGACGGTGGC | 81.3 | 9.7 | 2.0 | 4.7 | 2.1E-06 | 2.9E-05 | BT20/MB231 | MP |
| >MI0000101\|hsa-mir-99a&WithFlank&21\|+\|16539083\|16539175@19.38.20&[MIMAT0000097&hsa-miR-99a-5p&offsets\|0\|-2;m-64&21\|+\|16539101\|16539121&offsets\|0\|-1]\|AACCCGTAGATCCGATCTTG | >MI0000101\|hsa-mir-99a&WithFlank&21\|+\|16539083\|16539175 | 19 | 38 | 20 | AACCCGTAGATCCGATCTTG | 99.8 | -10.0 | 2.1 | -4.7 | 2.1E-06 | 2.9E-05 | BT20/MB231 | MP |
| >hg38_21_+_8433172_8446622_RNA45SN1_45S_50ntFlanks@9914.9947.34\|GAAAAGGTGTTGGTTGATATAGACAGCAGGACGG | >hg38_21_+_8433172_8446622_RNA45SN1_45S_50ntFlanks | 9914 | 9947 | 34 | GAAAAGGTGTTGGTTGATATAGACAGCAGGACGG | 111.0 | 10.1 | 2.1 | 4.7 | 2.1E-06 | 2.9E-05 | BT20/MB231 | MP |
| >hg38_21_+_8433172_8446622_RNA45SN1_45S_50ntFlanks@4257.4282.26\|TAATTGGAATGAGTCCACTTTAAATC | >hg38_21_+_8433172_8446622_RNA45SN1_45S_50ntFlanks | 4257 | 4282 | 26 | TAATTGGAATGAGTCCACTTTAAATC | 45.4 | -8.6 | 1.9 | -4.5 | 6.7E-06 | 2.9E-05 | BT20/MB468 | MP |
| >hg38_1_-_228634819_228635039_RNA5S12_5S_50ntFlanks@152.170.19\|TACCGGGTGCTGTAGGCTT | >hg38_1_-_228634819_228635039_RNA5S12_5S_50ntFlanks | 152 | 170 | 19 | TACCGGGTGCTGTAGGCTT | 5758.2 | -3.0 | 0.7 | -4.3 | 1.7E-05 | 2.9E-05 | MB231/MB468 | MP |
| >hg38_21_+_8433172_8446622_RNA45SN1_45S_50ntFlanks@8198.8220.23\|TGGACGGTGTGAGGCCGGTAGCG | >hg38_21_+_8433172_8446622_RNA45SN1_45S_50ntFlanks | 8198 | 8220 | 23 | TGGACGGTGTGAGGCCGGTAGCG | 188.9 | -4.9 | 1.1 | -4.3 | 1.7E-05 | 2.9E-05 | MB231/MB468 | MP |
| >hg38_21_+_8433172_8446622_RNA45SN1_45S_50ntFlanks@10547.10576.30\|AGTGCGGTAACGCGACCGATCCCGGAGAAG | >hg38_21_+_8433172_8446622_RNA45SN1_45S_50ntFlanks | 10547 | 10576 | 30 | AGTGCGGTAACGCGACCGATCCCGGAGAAG | 386.0 | 3.3 | 0.8 | 4.3 | 1.5E-05 | 2.9E-05 | BT20/MB468 | MITO |
| >hg38_21_+_8433172_8446622_RNA45SN1_45S_50ntFlanks@10853.10873.21\|ATAAGGATTGGCTCTAAGGGC | >hg38_21_+_8433172_8446622_RNA45SN1_45S_50ntFlanks | 10853 | 10873 | 21 | ATAAGGATTGGCTCTAAGGGC | 231.2 | -3.1 | 0.7 | -4.3 | 2.0E-05 | 2.9E-05 | MB231/MB468 | NUC |
| >hg38_21_+_8433172_8446622_RNA45SN1_45S_50ntFlanks@11783.11800.18\|CATGAATGGATGAACGAG | >hg38_21_+_8433172_8446622_RNA45SN1_45S_50ntFlanks | 11783 | 11800 | 18 | CATGAATGGATGAACGAG | 724.9 | -5.4 | 1.2 | -4.5 | 6.7E-06 | 2.9E-05 | BT20/MB468 | MP |
| >hg38_21_+_8433172_8446622_RNA45SN1_45S_50ntFlanks@10476.10497.22\|GGCCGATCGAAAGGGAGTCGGG | >hg38_21_+_8433172_8446622_RNA45SN1_45S_50ntFlanks | 10476 | 10497 | 22 | GGCCGATCGAAAGGGAGTCGGG | 55.9 | -8.9 | 2.0 | -4.5 | 6.7E-06 | 2.9E-05 | BT20/MB468 | MP |
| >hg38_21_+_8433172_8446622_RNA45SN1_45S_50ntFlanks@3912.3937.26\|GATGCGTGCATTTATCAGATCAAAAC | >hg38_21_+_8433172_8446622_RNA45SN1_45S_50ntFlanks | 3912 | 3937 | 26 | GATGCGTGCATTTATCAGATCAAAAC | 54.7 | -8.9 | 2.0 | -4.5 | 6.7E-06 | 2.9E-05 | BT20/MB468 | MP |
| >trna37_ValAAC_6_+_27203288_27203360@1.29.29\|GTTTCCGTAGTGTAGTGGTTATCACGTTT | >trna37_ValAAC_6_+_27203288_27203360 | 1 | 29 | 29 | GTTTCCGTAGTGTAGTGGTTATCACGTTT | 1525.3 | -4.1 | 1.0 | -4.3 | 1.7E-05 | 2.9E-05 | MB231/MB468 | MP |
| >hg38_21_+_8433172_8446622_RNA45SN1_45S_50ntFlanks@3842.3881.40\|CCTACTTGGATAACTGTGGTAATTCTAGAGCTAATACATG | >hg38_21_+_8433172_8446622_RNA45SN1_45S_50ntFlanks | 3842 | 3881 | 40 | CCTACTTGGATAACTGTGGTAATTCTAGAGCTAATACATG | 69.6 | -10.0 | 2.3 | -4.3 | 1.7E-05 | 2.9E-05 | MB231/MB468 | MP |
| >trnaMT_PheGAA_MT_+_577_647@35.53.19\|GAAAATGTTTAGACGGGCT | >trnaMT_PheGAA_MT_+_577_647 | 35 | 53 | 19 | GAAAATGTTTAGACGGGCT | 870.4 | 4.9 | 1.1 | 4.5 | 6.7E-06 | 2.9E-05 | BT20/MB468 | MP |
| >hg38_21_+_8433172_8446622_RNA45SN1_45S_50ntFlanks@8191.8212.22\|CAGCCCGTGGACGGTGTGAGGC | >hg38_21_+_8433172_8446622_RNA45SN1_45S_50ntFlanks | 8191 | 8212 | 22 | CAGCCCGTGGACGGTGTGAGGC | 1686.8 | -6.2 | 1.4 | -4.5 | 6.7E-06 | 2.9E-05 | BT20/MB468 | MP |
| >trna10_LysCTT_16_+_3241501_3241573@13.33.21,trna13_LysCTT_14_-_58706613_58706685@13.33.21,trna2_LysCTT_15_+_79152904_79152976@13.33.21,trna4_LysCTT_18_-_43669271_43669343@13.33.21\|CAGTCGGTAGAGCATGGGACT | >trna10_LysCTT_16_+_3241501_3241573 | 13 | 33 | 21 | CAGTCGGTAGAGCATGGGACT | 176.1 | 7.5 | 1.8 | 4.3 | 2.1E-05 | 2.9E-05 | MB231/MB468 | MITO |
| >hg38_21_+_8433172_8446622_RNA45SN1_45S_50ntFlanks@9313.9334.22\|TCTAACACGTGCGCGAGTCGGG | >hg38_21_+_8433172_8446622_RNA45SN1_45S_50ntFlanks | 9313 | 9334 | 22 | TCTAACACGTGCGCGAGTCGGG | 354.5 | -4.6 | 1.0 | -4.5 | 6.7E-06 | 2.9E-05 | BT20/MB468 | MP |
| >MI0000085\|hsa-mir-27a&WithFlank&19\|-\|13836434\|13836523@57.77.21&[MIMAT0000084&hsa-miR-27a-3p&offsets\|0\|0;m-43&19\|-\|13836447\|13836467&offsets\|0\|0]\|TTCACAGTGGCTAAGTTCCGC | >MI0000085\|hsa-mir-27a&WithFlank&19\|-\|13836434\|13836523 | 57 | 77 | 21 | TTCACAGTGGCTAAGTTCCGC | 637.7 | 4.3 | 1.0 | 4.3 | 2.0E-05 | 2.9E-05 | MB231/MB468 | CYTO |
| >hg38_21_+_8433172_8446622_RNA45SN1_45S_50ntFlanks@12957.12979.23\|CTTCTGGGTCGGGGTTTCGTACG | >hg38_21_+_8433172_8446622_RNA45SN1_45S_50ntFlanks | 12957 | 12979 | 23 | CTTCTGGGTCGGGGTTTCGTACG | 733.0 | -4.7 | 1.1 | -4.3 | 2.0E-05 | 2.9E-05 | MB231/MB468 | CYTO |
| >hg38_MT_+_1621_3279_RNR2_16S_50ntFlanks@722.749.28\|TGCGTCAGATTAAAACACTGAACTGACA | >hg38_MT_+_1621_3279_RNR2_16S_50ntFlanks | 722 | 749 | 28 | TGCGTCAGATTAAAACACTGAACTGACA | 31.3 | 8.2 | 1.9 | 4.3 | 1.7E-05 | 2.9E-05 | MB231/MB468 | MP |
| >hg38_21_+_8433172_8446622_RNA45SN1_45S_50ntFlanks@11212.11251.40\|CGACTCTGGACGCGAGCCGGGCCCTTCCCGTGGATCGCCC | >hg38_21_+_8433172_8446622_RNA45SN1_45S_50ntFlanks | 11212 | 11251 | 40 | CGACTCTGGACGCGAGCCGGGCCCTTCCCGTGGATCGCCC | 784.1 | -6.0 | 1.4 | -4.2 | 2.2E-05 | 2.9E-05 | BT20/MB468 | NUC |
| >hg38_21_+_8433172_8446622_RNA45SN1_45S_50ntFlanks@12615.12654.40\|GCTGAGGAGCCAATGGGGCGAAGCTACCATCTGTGGGATT | >hg38_21_+_8433172_8446622_RNA45SN1_45S_50ntFlanks | 12615 | 12654 | 40 | GCTGAGGAGCCAATGGGGCGAAGCTACCATCTGTGGGATT | 3815.6 | -3.8 | 0.9 | -4.3 | 2.0E-05 | 2.9E-05 | MB231/MB468 | CYTO |
| >hg38_21_+_8433172_8446622_RNA45SN1_45S_50ntFlanks@4727.4767.41\|AATCAAGAACGAAAGTCGGAGGTTCGAAGACGATCAGATAC | >hg38_21_+_8433172_8446622_RNA45SN1_45S_50ntFlanks | 4727 | 4767 | 41 | AATCAAGAACGAAAGTCGGAGGTTCGAAGACGATCAGATAC | 205.8 | 5.8 | 1.4 | 4.3 | 2.1E-05 | 2.9E-05 | MB231/MB468 | NUC |

<table>
<tr><td></td><td></td><td></td><td></td><td></td><td></td><td></td><td></td><td></td><td></td><td></td><td></td><td></td><td></td><td></td></tr>
<tr><td>>hg38_21_+_8433172_8446622_RNA45SN1_45S_50ntFlanks@11208.11231.24|GCGGCGACTCTGGACGCGAGCCGG</td><td>>hg38_21_+_8433172_8446622_RNA45SN1_45S_50ntFlanks</td><td>11208</td><td>11231</td><td>24</td><td>GCGGCGACTCTGGACGCGAGCCGG</td><td>4417.4</td><td>-6.7</td><td>1.6</td><td>-4.2</td><td>2.2E-05</td><td>2.9E-05</td><td>BT20/MB468</td><td>NUC</td></tr>
<tr><td>>MI0006273|hsa-mir-1180&WithFlank&17|-|19344500|19344580@47.67.21(+1A)&[MIMAT0005825&hsa-miR-1180-3p&offsets|0|-1(+1A);m-214&17|-|19344513|19344534&offsets|0|-1(+1A)]|TTTCCGGCTCGCGTGGGTGTGA</td><td>>MI0006273|hsa-mir-1180&WithFlank&17|-|19344500|19344580</td><td>47</td><td>67</td><td>21(+1A)</td><td>TTTCCGGCTCGCGTGGGTGTGA</td><td>264.4</td><td>3.7</td><td>0.8</td><td>4.7</td><td>2.1E-06</td><td>2.9E-05</td><td>BT20/MB231</td><td>CYTO</td></tr>
<tr><td>>hg38_21_+_8433172_8446622_RNA45SN1_45S_50ntFlanks@8191.8216.26|CAGCCCGTGGACGGTGTGAGGCCGGT</td><td>>hg38_21_+_8433172_8446622_RNA45SN1_45S_50ntFlanks</td><td>8191</td><td>8216</td><td>26</td><td>CAGCCCGTGGACGGTGTGAGGCCGGT</td><td>301.2</td><td>-5.4</td><td>1.3</td><td>-4.3</td><td>1.7E-05</td><td>2.9E-05</td><td>MB231/MB468</td><td>MP</td></tr>
<tr><td>>MI0016003|hsa-mir-3613&WithFlank&13|-|49996409|49996507@22.43.22&[MIMAT0017990&hsa-miR-3613-5p&offsets|0|0;m-259&13|-|49996465|49996486&offsets|0|0]|TGTTGTACTTTTTTTTTTGTTC</td><td>>MI0016003|hsa-mir-3613&WithFlank&13|-|49996409|49996507</td><td>22</td><td>43</td><td>22</td><td>TGTTGTACTTTTTTTTTTGTTC</td><td>426.3</td><td>6.1</td><td>1.4</td><td>4.3</td><td>1.8E-05</td><td>2.9E-05</td><td>BT20/MB468</td><td>CYTO</td></tr>
<tr><td>>hg38_21_+_8433172_8446622_RNA45SN1_45S_50ntFlanks@12183.12201.19|AAACGGTAACGCAGGTGTC</td><td>>hg38_21_+_8433172_8446622_RNA45SN1_45S_50ntFlanks</td><td>12183</td><td>12201</td><td>19</td><td>AAACGGTAACGCAGGTGTC</td><td>50.2</td><td>-8.7</td><td>1.9</td><td>-4.5</td><td>6.8E-06</td><td>2.9E-05</td><td>BT20/MB468</td><td>MP</td></tr>
<tr><td>>hg38_21_+_8433172_8446622_RNA45SN1_45S_50ntFlanks@11805.11827.23|CCACTGTCCCTACCTACTATCCA</td><td>>hg38_21_+_8433172_8446622_RNA45SN1_45S_50ntFlanks</td><td>11805</td><td>11827</td><td>23</td><td>CCACTGTCCCTACCTACTATCCA</td><td>230.7</td><td>-6.2</td><td>1.5</td><td>-4.3</td><td>2.1E-05</td><td>2.9E-05</td><td>MB231/MB468</td><td>MITO</td></tr>
<tr><td>>hg38_21_+_8433172_8446622_RNA45SN1_45S_50ntFlanks@11784.11806.23|ATGAATGGATGAACGAGATTCCC</td><td>>hg38_21_+_8433172_8446622_RNA45SN1_45S_50ntFlanks</td><td>11784</td><td>11806</td><td>23</td><td>ATGAATGGATGAACGAGATTCCC</td><td>560.1</td><td>-5.4</td><td>1.2</td><td>-4.3</td><td>1.8E-05</td><td>2.9E-05</td><td>BT20/MB468</td><td>CYTO</td></tr>
<tr><td>>hg38_21_+_8433172_8446622_RNA45SN1_45S_50ntFlanks@10806.10825.20|AACAATGTAGGTAAGGGAAG</td><td>>hg38_21_+_8433172_8446622_RNA45SN1_45S_50ntFlanks</td><td>10806</td><td>10825</td><td>20</td><td>AACAATGTAGGTAAGGGAAG</td><td>393.5</td><td>-4.6</td><td>1.1</td><td>-4.3</td><td>1.5E-05</td><td>2.9E-05</td><td>BT20/MB468</td><td>MITO</td></tr>
<tr><td>>hg38_21_+_8433172_8446622_RNA45SN1_45S_50ntFlanks@8116.8135.20|GCGGGACATGTGGCGTACGG</td><td>>hg38_21_+_8433172_8446622_RNA45SN1_45S_50ntFlanks</td><td>8116</td><td>8135</td><td>20</td><td>GCGGGACATGTGGCGTACGG</td><td>309.6</td><td>8.4</td><td>1.9</td><td>4.5</td><td>6.8E-06</td><td>2.9E-05</td><td>BT20/MB468</td><td>MP</td></tr>
<tr><td>>hg38_21_+_8433172_8446622_RNA45SN1_45S_50ntFlanks@12993.13013.21|CCCTCGCTGCGATCTATTGAA</td><td>>hg38_21_+_8433172_8446622_RNA45SN1_45S_50ntFlanks</td><td>12993</td><td>13013</td><td>21</td><td>CCCTCGCTGCGATCTATTGAA</td><td>122.0</td><td>-6.2</td><td>1.4</td><td>-4.3</td><td>1.7E-05</td><td>2.9E-05</td><td>MB231/MB468</td><td>MP</td></tr>
<tr><td>>hg38_21_+_8433172_8446622_RNA45SN1_45S_50ntFlanks@12324.12357.34|TTGGGTTTTAAGCAGGAGGTGTCAGAAAAGTTAC</td><td>>hg38_21_+_8433172_8446622_RNA45SN1_45S_50ntFlanks</td><td>12324</td><td>12357</td><td>34</td><td>TTGGGTTTTAAGCAGGAGGTGTCAGAAAAGTTAC</td><td>49.4</td><td>-8.7</td><td>1.9</td><td>-4.5</td><td>6.8E-06</td><td>2.9E-05</td><td>BT20/MB468</td><td>MP</td></tr>
<tr><td>>trna111_HisGTG_1_-_147774845_147774916@-1G.31.32,trna118_HisGTG_1_-_145396881_145396952@-1G.31.32,trna16_HisGTG_1_+_146544773_146544844@-1G.31.32,trna1_HisGTG_15_+_45493349_45493420@-1G.31.32,trna21_HisGTG_1_+_147753471_147753542@-1G.31.32,trna33_HisGTG_6_+_27125906_27125977@-1G.31.32,trna7_HisGTG_9_-_14433938_14434009@-1G.31.32,trna8_HisGTG_15_-_45492611_45492682@-1G.31.32,trna9_HisGTG_15_-_45490804_45490875@-1G.31.32|GGCCGTGATCGTATAGTGGTTAGTACTCTGCG</td><td>>trna111_HisGTG_1_-_147774845_147774916</td><td>-1G</td><td>31</td><td>32</td><td>GGCCGTGATCGTATAGTGGTTAGTACTCTGCG</td><td>6561.4</td><td>-4.9</td><td>1.0</td><td>-4.7</td><td>2.5E-06</td><td>2.9E-05</td><td>BT20/MB231</td><td>MITO</td></tr>
<tr><td>>am_trnaMT_ValTAC_MT_+_1602_1670@48.68.21|CAACTTAACTTGACCGCTCTG</td><td>>am_trnaMT_ValTAC_MT_+_1602_1670</td><td>48</td><td>68</td><td>21</td><td>CAACTTAACTTGACCGCTCTG</td><td>1654.8</td><td>-4.7</td><td>1.1</td><td>-4.2</td><td>2.2E-05</td><td>2.9E-05</td><td>BT20/MB468</td><td>NUC</td></tr>
<tr><td>>hg38_21_+_8433172_8446622_RNA45SN1_45S_50ntFlanks@4555.4575.21|CGCAGCTAGGAATAATGGAAT</td><td>>hg38_21_+_8433172_8446622_RNA45SN1_45S_50ntFlanks</td><td>4555</td><td>4575</td><td>21</td><td>CGCAGCTAGGAATAATGGAAT</td><td>262.0</td><td>-4.8</td><td>1.1</td><td>-4.3</td><td>2.1E-05</td><td>2.9E-05</td><td>MB231/MB468</td><td>NUC</td></tr>
<tr><td>>hg38_21_+_8433172_8446622_RNA45SN1_45S_50ntFlanks@6669.6686.18|CACTCGGCTCGTGCGTCG</td><td>>hg38_21_+_8433172_8446622_RNA45SN1_45S_50ntFlanks</td><td>6669</td><td>6686</td><td>18</td><td>CACTCGGCTCGTGCGTCG</td><td>339.0</td><td>-2.8</td><td>0.6</td><td>-4.3</td><td>1.5E-05</td><td>2.9E-05</td><td>BT20/MB468</td><td>MITO</td></tr>
<tr><td>>MI0001445|hsa-mir-423&WithFlank&17|+|30117073|30117178@23.43.21(+1A)&[MIMAT0004748&hsa-miR-423-5p&offsets|0|-2(+1A);m-141&17|+|30117095|30117117&offsets|0|-2(+1A)]|TGAGGGGCAGAGAGCGAGACTA</td><td>>MI0001445|hsa-mir-423&WithFlank&17|+|30117073|30117178</td><td>23</td><td>43</td><td>21(+1A)</td><td>TGAGGGGCAGAGAGCGAGACTA</td><td>230.6</td><td>7.2</td><td>1.6</td><td>4.5</td><td>6.8E-06</td><td>3.0E-05</td><td>BT20/MB468</td><td>MP</td></tr>
<tr><td>>hg38_21_+_8433172_8446622_RNA45SN1_45S_50ntFlanks@11682.11706.25|AATGTCAAAGTGAAGAAATTCAATG</td><td>>hg38_21_+_8433172_8446622_RNA45SN1_45S_50ntFlanks</td><td>11682</td><td>11706</td><td>25</td><td>AATGTCAAAGTGAAGAAATTCAATG</td><td>184.8</td><td>-6.5</td><td>1.4</td><td>-4.5</td><td>6.8E-06</td><td>3.0E-05</td><td>BT20/MB468</td><td>MP</td></tr>
<tr><td>>hg38_21_+_8433172_8446622_RNA45SN1_45S_50ntFlanks@7980.8010.31|CCTCAGATCAGACGTGGCGACCCGCTGAATT</td><td>>hg38_21_+_8433172_8446622_RNA45SN1_45S_50ntFlanks</td><td>7980</td><td>8010</td><td>31</td><td>CCTCAGATCAGACGTGGCGACCCGCTGAATT</td><td>129.7</td><td>-4.6</td><td>1.1</td><td>-4.3</td><td>1.7E-05</td><td>3.0E-05</td><td>MB231/MB468</td><td>MP</td></tr>
<tr><td>>hg38_21_+_8433172_8446622_RNA45SN1_45S_50ntFlanks@11587.11609.23|TGCGGACCAGGGGAATCCGACTG</td><td>>hg38_21_+_8433172_8446622_RNA45SN1_45S_50ntFlanks</td><td>11587</td><td>11609</td><td>23</td><td>TGCGGACCAGGGGAATCCGACTG</td><td>297.5</td><td>-5.2</td><td>1.2</td><td>-4.3</td><td>1.8E-05</td><td>3.0E-05</td><td>MB231/MB468</td><td>MP</td></tr>
<tr><td>>hg38_21_+_8433172_8446622_RNA45SN1_45S_50ntFlanks@6651.6692.42|CGACTCTTAGCGGTGGATCACTCGGCTCGTGCGTCGATGAAG</td><td>>hg38_21_+_8433172_8446622_RNA45SN1_45S_50ntFlanks</td><td>6651</td><td>6692</td><td>42</td><td>CGACTCTTAGCGGTGGATCACTCGGCTCGTGCGTCGATGAAG</td><td>4462.7</td><td>4.3</td><td>1.0</td><td>4.3</td><td>1.8E-05</td><td>3.0E-05</td><td>BT20/MB468</td><td>CYTO</td></tr>
<tr><td>>hg38_21_+_8433172_8446622_RNA45SN1_45S_50ntFlanks@12006.12026.21|GGCCGCCGGTGAAATACCACT</td><td>>hg38_21_+_8433172_8446622_RNA45SN1_45S_50ntFlanks</td><td>12006</td><td>12026</td><td>21</td><td>GGCCGCCGGTGAAATACCACT</td><td>393.6</td><td>8.4</td><td>1.9</td><td>4.5</td><td>6.8E-06</td><td>3.0E-05</td><td>BT20/MB468</td><td>MP</td></tr>
<tr><td>>hg38_21_+_8433172_8446622_RNA45SN1_45S_50ntFlanks@5423.5455.33|ATTGGATGGTTTAGTGAGGCCCTCGGATCGGCC</td><td>>hg38_21_+_8433172_8446622_RNA45SN1_45S_50ntFlanks</td><td>5423</td><td>5455</td><td>33</td><td>ATTGGATGGTTTAGTGAGGCCCTCGGATCGGCC</td><td>45.3</td><td>-8.6</td><td>1.9</td><td>-4.5</td><td>6.8E-06</td><td>3.0E-05</td><td>BT20/MB468</td><td>MP</td></tr>
<tr><td>>hg38_21_+_8433172_8446622_RNA45SN1_45S_50ntFlanks@6745.6770.26|ACACTTCGAACGCACTTGCGGCCCCG</td><td>>hg38_21_+_8433172_8446622_RNA45SN1_45S_50ntFlanks</td><td>6745</td><td>6770</td><td>26</td><td>ACACTTCGAACGCACTTGCGGCCCCG</td><td>8076.6</td><td>-8.0</td><td>1.8</td><td>-4.5</td><td>6.8E-06</td><td>3.0E-05</td><td>BT20/MB468</td><td>MP</td></tr>
<tr><td>>hg38_21_+_8433172_8446622_RNA45SN1_45S_50ntFlanks@4721.4758.38|TTCATTAATCAAGAACGAAAGTCGGAGGTTCGAAGACG</td><td>>hg38_21_+_8433172_8446622_RNA45SN1_45S_50ntFlanks</td><td>4721</td><td>4758</td><td>38</td><td>TTCATTAATCAAGAACGAAAGTCGGAGGTTCGAAGACG</td><td>260.1</td><td>-2.7</td><td>0.6</td><td>-4.3</td><td>2.1E-05</td><td>3.0E-05</td><td>MB231/MB468</td><td>NUC</td></tr>
<tr><td>>MI0003180|hsa-mir-516a-1&WithFlank&19|+|53756735|53756836@22.43.22;MI0003181|hsa-mir-516a-2&WithFlank&19|+|53761127|53761228@22.43.22&[MIMAT0004770&hsa-miR-516a-5p&offsets|0|-1;m-852&19|+|53756756|53756777&offsets|0|0];[MIMAT0004770_1&hsa-miR-516a-5p&offsets|0|-1;m-852&19|+|53761148|53761169&offsets|0|0]|TTCTCGAGGAAAGAAGCACTTT</td><td>>MI0003180|hsa-mir-516a-1&WithFlank&19|+|53756735|53756836</td><td>22</td><td>43</td><td>22</td><td>TTCTCGAGGAAAGAAGCACTTT</td><td>108.7</td><td>6.9</td><td>1.6</td><td>4.3</td><td>1.8E-05</td><td>3.0E-05</td><td>MB231/MB468</td><td>MP</td></tr>
<tr><td>>hg38_21_+_8433172_8446622_RNA45SN1_45S_50ntFlanks@9623.9658.36|TAGTAGCTGGTTCCCTCCGAAGTTTCCCTCAGGATA</td><td>>hg38_21_+_8433172_8446622_RNA45SN1_45S_50ntFlanks</td><td>9623</td><td>9658</td><td>36</td><td>TAGTAGCTGGTTCCCTCCGAAGTTTCCCTCAGGATA</td><td>1054.8</td><td>-2.4</td><td>0.6</td><td>-4.3</td><td>1.5E-05</td><td>3.0E-05</td><td>BT20/MB468</td><td>MITO</td></tr>
<tr><td>>hg38_21_+_8433172_8446622_RNA45SN1_45S_50ntFlanks@7991.8016.26|ACGTGGCGACCCGCTGAATTTAAGCA</td><td>>hg38_21_+_8433172_8446622_RNA45SN1_45S_50ntFlanks</td><td>7991</td><td>8016</td><td>26</td><td>ACGTGGCGACCCGCTGAATTTAAGCA</td><td>571.8</td><td>-3.5</td><td>0.8</td><td>-4.5</td><td>6.9E-06</td><td>3.0E-05</td><td>BT20/MB468</td><td>MP</td></tr>
<tr><td>>hg38_21_+_8433172_8446622_RNA45SN1_45S_50ntFlanks@11786.11804.19|GAATGGATGAACGAGATTC</td><td>>hg38_21_+_8433172_8446622_RNA45SN1_45S_50ntFlanks</td><td>11786</td><td>11804</td><td>19</td><td>GAATGGATGAACGAGATTC</td><td>961.0</td><td>-5.9</td><td>1.4</td><td>-4.3</td><td>1.8E-05</td><td>3.0E-05</td><td>BT20/MB468</td><td>CYTO</td></tr>
<tr><td>>MI0000088|hsa-mir-30a&WithFlank&6|-|71403545|71403627@12.32.21&[MIMAT0000087&hsa-miR-30a-5p&offsets|0|-1;m-15&6|-|71403595|71403616&offsets|0|-1]|TGTAAACATCCTCGACTGGAA</td><td>>MI0000088|hsa-mir-30a&WithFlank&6|-|71403545|71403627</td><td>12</td><td>32</td><td>21</td><td>TGTAAACATCCTCGACTGGAA</td><td>1718.2</td><td>2.8</td><td>0.7</td><td>4.3</td><td>1.8E-05</td><td>3.0E-05</td><td>MB231/MB468</td><td>MP</td></tr>
<tr><td>>hg38_21_+_8433172_8446622_RNA45SN1_45S_50ntFlanks@12627.12667.41|ATGGGGCGAAGCTACCATCTGTGGGATTATGACTGAACGCC</td><td>>hg38_21_+_8433172_8446622_RNA45SN1_45S_50ntFlanks</td><td>12627</td><td>12667</td><td>41</td><td>ATGGGGCGAAGCTACCATCTGTGGGATTATGACTGAACGCC</td><td>207.1</td><td>4.2</td><td>1.0</td><td>4.3</td><td>2.1E-05</td><td>3.0E-05</td><td>MB231/MB468</td><td>NUC</td></tr>
<tr><td>>hg38_21_+_8433172_8446622_RNA45SN1_45S_50ntFlanks@10808.10825.18|CAATGTAGGTAAGGGAAG</td><td>>hg38_21_+_8433172_8446622_RNA45SN1_45S_50ntFlanks</td><td>10808</td><td>10825</td><td>18</td><td>CAATGTAGGTAAGGGAAG</td><td>1453.1</td><td>-6.8</td><td>1.6</td><td>-4.3</td><td>1.8E-05</td><td>3.0E-05</td><td>BT20/MB468</td><td>CYTO</td></tr>
<tr><td>>hg38_21_+_8433172_8446622_RNA45SN1_45S_50ntFlanks@10841.10859.19|CGTAACTTCGGGATAAGGA</td><td>>hg38_21_+_8433172_8446622_RNA45SN1_45S_50ntFlanks</td><td>10841</td><td>10859</td><td>19</td><td>CGTAACTTCGGGATAAGGA</td><td>1039.0</td><td>-5.8</td><td>1.3</td><td>-4.3</td><td>1.8E-05</td><td>3.0E-05</td><td>BT20/MB468</td><td>CYTO</td></tr>
<tr><td>>hg38_21_+_8433172_8446622_RNA45SN1_45S_50ntFlanks@6699.6721.23|GCTAGCTGCGAGAATTAATGTGA</td><td>>hg38_21_+_8433172_8446622_RNA45SN1_45S_50ntFlanks</td><td>6699</td><td>6721</td><td>23</td><td>GCTAGCTGCGAGAATTAATGTGA</td><td>289.8</td><td>-4.7</td><td>1.1</td><td>-4.3</td><td>2.1E-05</td><td>3.0E-05</td><td>MB231/MB468</td><td>MITO</td></tr>
<tr><td>>hg38_21_+_8433172_8446622_RNA45SN1_45S_50ntFlanks@8683.8715.33|CGACCGGCTCCGGGACGGCTGGGAAGGCCCGGC</td><td>>hg38_21_+_8433172_8446622_RNA45SN1_45S_50ntFlanks</td><td>8683</td><td>8715</td><td>33</td><td>CGACCGGCTCCGGGACGGCTGGGAAGGCCCGGC</td><td>44.1</td><td>9.2</td><td>2.0</td><td>4.5</td><td>6.9E-06</td><td>3.0E-05</td><td>BT20/MB468</td><td>MP</td></tr>
<tr><td>>hg38_21_+_8433172_8446622_RNA45SN1_45S_50ntFlanks@11661.11684.24|TGATTTCTGCCCAGTGCTCTGAAT</td><td>>hg38_21_+_8433172_8446622_RNA45SN1_45S_50ntFlanks</td><td>11661</td><td>11684</td><td>24</td><td>TGATTTCTGCCCAGTGCTCTGAAT</td><td>191.1</td><td>-3.1</td><td>0.7</td><td>-4.3</td><td>1.8E-05</td><td>3.0E-05</td><td>BT20/MB468</td><td>CYTO</td></tr>
<tr><td>>MI0003643|hsa-mir-629&WithFlank&15|-|70079366|70079474@28.49.22&[MIMAT0004810&hsa-miR-629-5p&offsets|0|+1;m-150&15|-|70079427|70079447&offsets|0|+1]|TGGGTTTACGTTGGGAGAACTT</td><td>>MI0003643|hsa-mir-629&WithFlank&15|-|70079366|70079474</td><td>28</td><td>49</td><td>22</td><td>TGGGTTTACGTTGGGAGAACTT</td><td>1850.5</td><td>2.8</td><td>0.6</td><td>4.7</td><td>2.5E-06</td><td>3.0E-05</td><td>BT20/MB231</td><td>NUC</td></tr>
<tr><td>>MI0003776|hsa-mir-320b-1&WithFlank&1|+|116671740|116671823@48.66.19;MI0003839|hsa-mir-320b-2&WithFlank&1|-|224257034|224257116@47.65.19;MI0003778|hsa-mir-320c-1&WithFlank&18|+|21683512|21683595@48.66.19;MI0008191|hsa-mir-320c-2&WithFlank&18|+|24321669|24321752@48.66.19;MI0000542|hsa-mir-320a&WithFlank&8|-|22244960|22245043@48.66.19&[MIMAT0005792&hsa-miR-320b&offsets|0|-3;m-183&1|+|116671787|116671806&offsets|0|-1];[MIMAT0005792_1&hsa-miR-320b&offsets|0|-3;m-183&1|-|224257051|224257070&offsets|0|-1];[MIMAT0005793&hsa-miR-320c&offsets|0|-1;m-222&18|+|21683559|21683578&offsets|0|-1];[MIMAT0005793_1&hsa-miR-320c&offsets|0|-1;m-219&18|+|24321716|24321735&offsets|0|-1];[MIMAT0000510&hsa-miR-320a-3p&offsets|0|-3;m-60&8|-|22244975|22244996&offsets|0|-3]|AAAAGCTGGGTTGAGAGGG</td><td>>MI0003776|hsa-mir-320b-1&WithFlank&1|+|116671740|116671823</td><td>48</td><td>66</td><td>19</td><td>AAAAGCTGGGTTGAGAGGG</td><td>349.5</td><td>3.0</td><td>0.6</td><td>4.7</td><td>2.5E-06</td><td>3.0E-05</td><td>BT20/MB231</td><td>NUC</td></tr>
<tr><td>>hg38_21_+_8433172_8446622_RNA45SN1_45S_50ntFlanks@6655.6686.32|TCTTAGCGGTGGATCACTCGGCTCGTGCGTCG</td><td>>hg38_21_+_8433172_8446622_RNA45SN1_45S_50ntFlanks</td><td>6655</td><td>6686</td><td>32</td><td>TCTTAGCGGTGGATCACTCGGCTCGTGCGTCG</td><td>117.3</td><td>-6.4</td><td>1.5</td><td>-4.3</td><td>1.8E-05</td><td>3.0E-05</td><td>MB231/MB468</td><td>MP</td></tr>
<tr><td>>hg38_21_+_8433172_8446622_RNA45SN1_45S_50ntFlanks@12416.12437.22|TTCGATGTCGGCTCTTCCTATC</td><td>>hg38_21_+_8433172_8446622_RNA45SN1_45S_50ntFlanks</td><td>12416</td><td>12437</td><td>22</td><td>TTCGATGTCGGCTCTTCCTATC</td><td>803.0</td><td>2.0</td><td>0.5</td><td>4.3</td><td>1.5E-05</td><td>3.0E-05</td><td>BT20/MB468</td><td>MITO</td></tr>
<tr><td>>hg38_21_+_8433172_8446622_RNA45SN1_45S_50ntFlanks@10350.10369.20|ACGAGAACTTTGAAGGCCGA</td><td>>hg38_21_+_8433172_8446622_RNA45SN1_45S_50ntFlanks</td><td>10350</td><td>10369</td><td>20</td><td>ACGAGAACTTTGAAGGCCGA</td><td>1272.4</td><td>-3.2</td><td>0.8</td><td>-4.3</td><td>1.8E-05</td><td>3.0E-05</td><td>MB231/MB468</td><td>MP</td></tr>
<tr><td>>hg38_21_+_8433172_8446622_RNA45SN1_45S_50ntFlanks@11827.11844.18|AGCGAAACCACAGCCAAG</td><td>>hg38_21_+_8433172_8446622_RNA45SN1_45S_50ntFlanks</td><td>11827</td><td>11844</td><td>18</td><td>AGCGAAACCACAGCCAAG</td><td>51.8</td><td>-8.8</td><td>2.0</td><td>-4.5</td><td>6.9E-06</td><td>3.0E-05</td><td>BT20/MB468</td><td>MP</td></tr>
<tr><td>>hg38_21_+_8433172_8446622_RNA45SN1_45S_50ntFlanks@13007.13028.22|TATTGAAAGTCAGCCCTCGACA</td><td>>hg38_21_+_8433172_8446622_RNA45SN1_45S_50ntFlanks</td><td>13007</td><td>13028</td><td>22</td><td>TATTGAAAGTCAGCCCTCGACA</td><td>872.4</td><td>-3.9</td><td>0.9</td><td>-4.3</td><td>1.8E-05</td><td>3.0E-05</td><td>BT20/MB468</td><td>CYTO</td></tr>
<tr><td>>hg38_21_+_8433172_8446622_RNA45SN1_45S_50ntFlanks@10350.10373.24|ACGAGAACTTTGAAGGCCGAAGTG</td><td>>hg38_21_+_8433172_8446622_RNA45SN1_45S_50ntFlanks</td><td>10350</td><td>10373</td><td>24</td><td>ACGAGAACTTTGAAGGCCGAAGTG</td><td>499.2</td><td>-1.9</td><td>0.5</td><td>-4.3</td><td>1.8E-05</td><td>3.0E-05</td><td>BT20/MB468</td><td>CYTO</td></tr>
</table>

<table>
<tr><td>>hg38_21_+_8433172_8446622_RNA45SN1_45S_50ntFlanks@12007.12043.37|GCCGCCGGTGAAATACCACTACTCTGATCGTTTTTTC</td><td>>hg38_21_+_8433172_8446622_RNA45SN1_45S_50ntFlanks</td><td>12007</td><td>12043</td><td>37</td><td>GCCGCCGGTGAAATACCACTACTCTGATCGTTTTTTC</td><td>4547.7</td><td>-4.5</td><td>1.1</td><td>-4.3</td><td>1.5E-05</td><td>3.0E-05</td><td>BT20/MB468</td><td>MITO</td></tr>
<tr><td>>hg38_21_+_8433172_8446622_RNA45SN1_45S_50ntFlanks@10574.10596.23|AAGCCGGCGGGAGCCCCGGGGAG</td><td>>hg38_21_+_8433172_8446622_RNA45SN1_45S_50ntFlanks</td><td>10574</td><td>10596</td><td>23</td><td>AAGCCGGCGGGAGCCCCGGGGAG</td><td>120.9</td><td>-5.3</td><td>1.2</td><td>-4.3</td><td>1.5E-05</td><td>3.0E-05</td><td>BT20/MB468</td><td>MITO</td></tr>
<tr><td>>hg38_21_+_8433172_8446622_RNA45SN1_45S_50ntFlanks@12457.12486.30|CAAGCGTTGGATTGTTCACCCACTAATAGG</td><td>>hg38_21_+_8433172_8446622_RNA45SN1_45S_50ntFlanks</td><td>12457</td><td>12486</td><td>30</td><td>CAAGCGTTGGATTGTTCACCCACTAATAGG</td><td>411.4</td><td>-5.2</td><td>1.2</td><td>-4.2</td><td>2.1E-05</td><td>3.0E-05</td><td>MB231/MB468</td><td>MITO</td></tr>
<tr><td>>hg38_21_+_8433172_8446622_RNA45SN1_45S_50ntFlanks@12513.12541.29|TGAGACAGGTTAGTTTTACCCTACTGATG</td><td>>hg38_21_+_8433172_8446622_RNA45SN1_45S_50ntFlanks</td><td>12513</td><td>12541</td><td>29</td><td>TGAGACAGGTTAGTTTTACCCTACTGATG</td><td>128.9</td><td>-6.0</td><td>1.4</td><td>-4.3</td><td>1.8E-05</td><td>3.0E-05</td><td>MB231/MB468</td><td>MP</td></tr>
<tr><td>>hg38_21_+_8433172_8446622_RNA45SN1_45S_50ntFlanks@11594.11612.19|CAGGGGAATCCGACTGTTT</td><td>>hg38_21_+_8433172_8446622_RNA45SN1_45S_50ntFlanks</td><td>11594</td><td>11612</td><td>19</td><td>CAGGGGAATCCGACTGTTT</td><td>827.2</td><td>-2.9</td><td>0.7</td><td>-4.3</td><td>1.5E-05</td><td>3.0E-05</td><td>BT20/MB468</td><td>MITO</td></tr>
<tr><td>>hg38_21_+_8433172_8446622_RNA45SN1_45S_50ntFlanks@3899.3927.29|CCTTCGCGGGGGGGATGCGTGCATTTATC</td><td>>hg38_21_+_8433172_8446622_RNA45SN1_45S_50ntFlanks</td><td>3899</td><td>3927</td><td>29</td><td>CCTTCGCGGGGGGGATGCGTGCATTTATC</td><td>460.9</td><td>-5.8</td><td>1.3</td><td>-4.5</td><td>7.0E-06</td><td>3.0E-05</td><td>BT20/MB468</td><td>MP</td></tr>
<tr><td>>hg38_21_+_8433172_8446622_RNA45SN1_45S_50ntFlanks@11604.11622.19|CGACTGTTTAATTAAAACA</td><td>>hg38_21_+_8433172_8446622_RNA45SN1_45S_50ntFlanks</td><td>11604</td><td>11622</td><td>19</td><td>CGACTGTTTAATTAAAACA</td><td>529.4</td><td>3.5</td><td>0.8</td><td>4.3</td><td>1.6E-05</td><td>3.0E-05</td><td>BT20/MB468</td><td>MITO</td></tr>
<tr><td>>MI0000085|hsa-mir-27a&WithFlank&19|-|13836434|13836523@57.74.18;MI0000440|hsa-mir-27b&WithFlank&9|+|95085439|95085547@67.84.18&[MIMAT0000084&hsa-miR-27a-3p&offsets|0|-3;m-43&19|-|13836447|13836467&offsets|0|-3];[MIMAT0000419&hsa-miR-27b-3p&offsets|0|-3;m-39&9|+|95085505|95085525&offsets|0|-3]|TTCACAGTGGCTAAGTTC</td><td>>MI0000085|hsa-mir-27a&WithFlank&19|-|13836434|13836523</td><td>57</td><td>74</td><td>18</td><td>TTCACAGTGGCTAAGTTC</td><td>1331.5</td><td>3.5</td><td>0.8</td><td>4.3</td><td>1.8E-05</td><td>3.0E-05</td><td>MB231/MB468</td><td>MP</td></tr>
<tr><td>>trnaMT_HisGTG_MT_+_12138_12206@25.72.48|CAGATTGTGAATCTGACAACAGAGGCTTACGACCCCTTATTTACCCCA</td><td>>trnaMT_HisGTG_MT_+_12138_12206</td><td>25</td><td>72</td><td>48</td><td>CAGATTGTGAATCTGACAACAGAGGCTTACGACCCCTTATTTACCCCA</td><td>249.9</td><td>11.7</td><td>2.6</td><td>4.5</td><td>7.0E-06</td><td>3.0E-05</td><td>BT20/MB468</td><td>MP</td></tr>
<tr><td>>hg38_21_+_8433172_8446622_RNA45SN1_45S_50ntFlanks@10797.10816.20|GCATGTTGGAACAATGTAGG</td><td>>hg38_21_+_8433172_8446622_RNA45SN1_45S_50ntFlanks</td><td>10797</td><td>10816</td><td>20</td><td>GCATGTTGGAACAATGTAGG</td><td>246.4</td><td>-4.2</td><td>1.0</td><td>-4.3</td><td>2.0E-05</td><td>3.0E-05</td><td>MB231/MB468</td><td>CYTO</td></tr>
<tr><td>>hg38_21_+_8433172_8446622_RNA45SN1_45S_50ntFlanks@8379.8399.21|TCAAGAGGGCGTGAAACCGTT</td><td>>hg38_21_+_8433172_8446622_RNA45SN1_45S_50ntFlanks</td><td>8379</td><td>8399</td><td>21</td><td>TCAAGAGGGCGTGAAACCGTT</td><td>191.9</td><td>-4.1</td><td>1.0</td><td>-4.3</td><td>1.8E-05</td><td>3.0E-05</td><td>MB231/MB468</td><td>MP</td></tr>
<tr><td>>hg38_21_+_8433172_8446622_RNA45SN1_45S_50ntFlanks@12436.12473.38|TCATTGTGAAGCAGAATTCACCAAGCGTTGGATTGTTC</td><td>>hg38_21_+_8433172_8446622_RNA45SN1_45S_50ntFlanks</td><td>12436</td><td>12473</td><td>38</td><td>TCATTGTGAAGCAGAATTCACCAAGCGTTGGATTGTTC</td><td>978.8</td><td>-5.3</td><td>1.2</td><td>-4.3</td><td>1.8E-05</td><td>3.0E-05</td><td>BT20/MB468</td><td>CYTO</td></tr>
<tr><td>>hg38_21_+_8433172_8446622_RNA45SN1_45S_50ntFlanks@6661.6703.43|CGGTGGATCACTCGGCTCGTGCGTCGATGAAGAACGCAGCTAG</td><td>>hg38_21_+_8433172_8446622_RNA45SN1_45S_50ntFlanks</td><td>6661</td><td>6703</td><td>43</td><td>CGGTGGATCACTCGGCTCGTGCGTCGATGAAGAACGCAGCTAG</td><td>76.1</td><td>-10.1</td><td>2.4</td><td>-4.3</td><td>1.8E-05</td><td>3.0E-05</td><td>MB231/MB468</td><td>MP</td></tr>
<tr><td>>hg38_21_+_8433172_8446622_RNA45SN1_45S_50ntFlanks@7991.8018.28|ACGTGGCGACCCGCTGAATTTAAGCATA</td><td>>hg38_21_+_8433172_8446622_RNA45SN1_45S_50ntFlanks</td><td>7991</td><td>8018</td><td>28</td><td>ACGTGGCGACCCGCTGAATTTAAGCATA</td><td>274.5</td><td>-5.2</td><td>1.2</td><td>-4.2</td><td>2.3E-05</td><td>3.0E-05</td><td>BT20/MB468</td><td>NUC</td></tr>
<tr><td>>hg38_21_+_8433172_8446622_RNA45SN1_45S_50ntFlanks@12073.12098.26|CCGAGGGGCTCTCGCTTCTGGCGCCA</td><td>>hg38_21_+_8433172_8446622_RNA45SN1_45S_50ntFlanks</td><td>12073</td><td>12098</td><td>26</td><td>CCGAGGGGCTCTCGCTTCTGGCGCCA</td><td>470.1</td><td>-7.1</td><td>1.7</td><td>-4.2</td><td>2.3E-05</td><td>3.0E-05</td><td>BT20/MB468</td><td>NUC</td></tr>
<tr><td>>hg38_21_+_8433172_8446622_RNA45SN1_45S_50ntFlanks@4050.4067.18|CGAACGTCTGCCCTATCA</td><td>>hg38_21_+_8433172_8446622_RNA45SN1_45S_50ntFlanks</td><td>4050</td><td>4067</td><td>18</td><td>CGAACGTCTGCCCTATCA</td><td>208.1</td><td>2.5</td><td>0.6</td><td>4.3</td><td>2.0E-05</td><td>3.0E-05</td><td>MB231/MB468</td><td>CYTO</td></tr>
<tr><td>>hg38_21_+_8433172_8446622_RNA45SN1_45S_50ntFlanks@12996.13039.44|TCGCTGCGATCTATTGAAAGTCAGCCCTCGACACAAGGGTTTGT</td><td>>hg38_21_+_8433172_8446622_RNA45SN1_45S_50ntFlanks</td><td>12996</td><td>13039</td><td>44</td><td>TCGCTGCGATCTATTGAAAGTCAGCCCTCGACACAAGGGTTTGT</td><td>103229.4</td><td>4.8</td><td>1.1</td><td>4.2</td><td>2.2E-05</td><td>3.0E-05</td><td>MB231/MB468</td><td>MITO</td></tr>
<tr><td>>MI0000434|hsa-let-7i&WithFlank&12|+|62603680|62603775@12.29.18(+1C)&[MIMAT0000415&hsa-let-7i-5p&offsets|0|-4(+1C);m-49&12|+|62603691|62603712&offsets|0|-4(+1C)]|TGAGGTAGTAGTTTGTGCC</td><td>>MI0000434|hsa-let-7i&WithFlank&12|+|62603680|62603775</td><td>12</td><td>29</td><td>18(+1C)</td><td>TGAGGTAGTAGTTTGTGCC</td><td>43.5</td><td>8.7</td><td>2.0</td><td>4.3</td><td>1.8E-05</td><td>3.0E-05</td><td>MB231/MB468</td><td>MP</td></tr>
<tr><td>>hg38_21_+_8433172_8446622_RNA45SN1_45S_50ntFlanks@10402.10426.25|AACATGGGTCAGTCGGTCCTGAGAG</td><td>>hg38_21_+_8433172_8446622_RNA45SN1_45S_50ntFlanks</td><td>10402</td><td>10426</td><td>25</td><td>AACATGGGTCAGTCGGTCCTGAGAG</td><td>433.2</td><td>-6.1</td><td>1.4</td><td>-4.3</td><td>1.9E-05</td><td>3.0E-05</td><td>BT20/MB468</td><td>CYTO</td></tr>
<tr><td>>hg38_21_+_8433172_8446622_RNA45SN1_45S_50ntFlanks@11666.11685.20|TCTGCCCAGTGCTCTGAATG</td><td>>hg38_21_+_8433172_8446622_RNA45SN1_45S_50ntFlanks</td><td>11666</td><td>11685</td><td>20</td><td>TCTGCCCAGTGCTCTGAATG</td><td>1865.4</td><td>-2.8</td><td>0.7</td><td>-4.3</td><td>1.6E-05</td><td>3.0E-05</td><td>BT20/MB468</td><td>MITO</td></tr>
<tr><td>>MI0000098|hsa-mir-96&WithFlank&7|-|129774686|129774775@15.36.22(+1C)&[MIMAT0000095&hsa-miR-96-5p&offsets|0|-1(+1C);m-248&7|-|129774739|129774761&offsets|0|-1(+1C)]|TTTGGCACTAGCACATTTTTGCC</td><td>>MI0000098|hsa-mir-96&WithFlank&7|-|129774686|129774775</td><td>15</td><td>36</td><td>22(+1C)</td><td>TTTGGCACTAGCACATTTTTGCC</td><td>1065.4</td><td>3.6</td><td>0.8</td><td>4.7</td><td>2.6E-06</td><td>3.0E-05</td><td>BT20/MB231</td><td>NUC</td></tr>
<tr><td>>hg38_21_+_8433172_8446622_RNA45SN1_45S_50ntFlanks@11564.11583.20|CCTAGCAGCCGACTTAGAAC</td><td>>hg38_21_+_8433172_8446622_RNA45SN1_45S_50ntFlanks</td><td>11564</td><td>11583</td><td>20</td><td>CCTAGCAGCCGACTTAGAAC</td><td>238.9</td><td>-6.3</td><td>1.5</td><td>-4.3</td><td>1.8E-05</td><td>3.0E-05</td><td>MB231/MB468</td><td>MP</td></tr>
<tr><td>>hg38_21_+_8433172_8446622_RNA45SN1_45S_50ntFlanks@12404.12432.29|GCTTTTTGATCCTTCGATGTCGGCTCTTC</td><td>>hg38_21_+_8433172_8446622_RNA45SN1_45S_50ntFlanks</td><td>12404</td><td>12432</td><td>29</td><td>GCTTTTTGATCCTTCGATGTCGGCTCTTC</td><td>321.2</td><td>-2.6</td><td>0.6</td><td>-4.2</td><td>2.2E-05</td><td>3.0E-05</td><td>MB231/MB468</td><td>NUC</td></tr>
<tr><td>>trna10_GluCTC_3_-_103879524_103879595@16.42.27,trna116_GluCTC_1_-_145399233_145399304@16.42.27,trna59_GluCTC_1_+_249168447_249168518@16.42.27,trna71_GluCTC_1_-_161439189_161439260@16.42.27,trna74_GluCTC_1_-_161431809_161431880@16.42.27,trna77_GluCTC_1_-_161424398_161424469@16.42.27,trna77_GluCTC_6_+_28949976_28950047@16.42.27,trna80_GluCTC_1_-_161417018_161417089@16.42.27,trna87_GluCTC_6_-_126101393_126101464@16.42.27|TGGTTAGGATTCGGCGCTCTCACCGCC</td><td>>trna10_GluCTC_3_-_103879524_103879595</td><td>16</td><td>42</td><td>27</td><td>TGGTTAGGATTCGGCGCTCTCACCGCC</td><td>2308.5</td><td>4.2</td><td>1.0</td><td>4.3</td><td>1.9E-05</td><td>3.0E-05</td><td>BT20/MB468</td><td>CYTO</td></tr>
<tr><td>>hg38_21_+_8433172_8446622_RNA45SN1_45S_50ntFlanks@12001.12031.31|CTGCGGGCCGCCGGTGAAATACCACTACTCT</td><td>>hg38_21_+_8433172_8446622_RNA45SN1_45S_50ntFlanks</td><td>12001</td><td>12031</td><td>31</td><td>CTGCGGGCCGCCGGTGAAATACCACTACTCT</td><td>692.2</td><td>-5.1</td><td>1.2</td><td>-4.2</td><td>2.2E-05</td><td>3.1E-05</td><td>MB231/MB468</td><td>NUC</td></tr>
<tr><td>>MI0000449|hsa-mir-132&WithFlank&17|-|2049902|2050014@65.86.22(+1U)&[MIMAT0000426&hsa-miR-132-3p&offsets|0|0(+1U);m-138&17|-|2049929|2049950&offsets|0|0(+1U)]|TAACAGTCTACAGCCATGGTCGT</td><td>>MI0000449|hsa-mir-132&WithFlank&17|-|2049902|2050014</td><td>65</td><td>86</td><td>22(+1U)</td><td>TAACAGTCTACAGCCATGGTCGT</td><td>303.6</td><td>8.3</td><td>1.9</td><td>4.5</td><td>7.1E-06</td><td>3.1E-05</td><td>BT20/MB468</td><td>MP</td></tr>
<tr><td>>hg38_21_+_8433172_8446622_RNA45SN1_45S_50ntFlanks@11784.11805.22|ATGAATGGATGAACGAGATTCC</td><td>>hg38_21_+_8433172_8446622_RNA45SN1_45S_50ntFlanks</td><td>11784</td><td>11805</td><td>22</td><td>ATGAATGGATGAACGAGATTCC</td><td>708.1</td><td>-4.1</td><td>1.0</td><td>-4.3</td><td>2.1E-05</td><td>3.1E-05</td><td>MB231/MB468</td><td>CYTO</td></tr>
<tr><td>>hg38_21_+_8433172_8446622_RNA45SN1_45S_50ntFlanks@12913.12933.21|ACGTTCGTGGGGAACCTGGCG</td><td>>hg38_21_+_8433172_8446622_RNA45SN1_45S_50ntFlanks</td><td>12913</td><td>12933</td><td>21</td><td>ACGTTCGTGGGGAACCTGGCG</td><td>342.4</td><td>-5.5</td><td>1.3</td><td>-4.2</td><td>2.2E-05</td><td>3.1E-05</td><td>MB231/MB468</td><td>MITO</td></tr>
<tr><td>>hg38_21_+_8433172_8446622_RNA45SN1_45S_50ntFlanks@11604.11623.20|CGACTGTTTAATTAAAACAA</td><td>>hg38_21_+_8433172_8446622_RNA45SN1_45S_50ntFlanks</td><td>11604</td><td>11623</td><td>20</td><td>CGACTGTTTAATTAAAACAA</td><td>3776.6</td><td>4.1</td><td>0.9</td><td>4.7</td><td>2.6E-06</td><td>3.1E-05</td><td>BT20/MB231</td><td>MITO</td></tr>
<tr><td>>hg38_21_+_8433172_8446622_RNA45SN1_45S_50ntFlanks@9859.9885.27|GGGATGAACCGAACGCCGGGTTAAGGC</td><td>>hg38_21_+_8433172_8446622_RNA45SN1_45S_50ntFlanks</td><td>9859</td><td>9885</td><td>27</td><td>GGGATGAACCGAACGCCGGGTTAAGGC</td><td>97.5</td><td>9.9</td><td>2.1</td><td>4.7</td><td>2.2E-06</td><td>3.1E-05</td><td>BT20/MB231</td><td>MP</td></tr>
<tr><td>>hg38_21_+_8433172_8446622_RNA45SN1_45S_50ntFlanks@11583.11601.19|CTGGTGCGGACCAGGGGAA</td><td>>hg38_21_+_8433172_8446622_RNA45SN1_45S_50ntFlanks</td><td>11583</td><td>11601</td><td>19</td><td>CTGGTGCGGACCAGGGGAA</td><td>345.2</td><td>-2.4</td><td>0.6</td><td>-4.3</td><td>1.6E-05</td><td>3.1E-05</td><td>BT20/MB468</td><td>MITO</td></tr>
<tr><td>>hg38_21_+_8433172_8446622_RNA45SN1_45S_50ntFlanks@10393.10430.38|CAGCAGTTGAACATGGGTCAGTCGGTCCTGAGAGATGG</td><td>>hg38_21_+_8433172_8446622_RNA45SN1_45S_50ntFlanks</td><td>10393</td><td>10430</td><td>38</td><td>CAGCAGTTGAACATGGGTCAGTCGGTCCTGAGAGATGG</td><td>567.6</td><td>-4.5</td><td>1.1</td><td>-4.3</td><td>2.1E-05</td><td>3.1E-05</td><td>MB231/MB468</td><td>CYTO</td></tr>
<tr><td>>hg38_21_+_8433172_8446622_RNA45SN1_45S_50ntFlanks@13007.13028.22|TATTGAAAGTCAGCCCTCGACA</td><td>>hg38_21_+_8433172_8446622_RNA45SN1_45S_50ntFlanks</td><td>13007</td><td>13028</td><td>22</td><td>TATTGAAAGTCAGCCCTCGACA</td><td>108.4</td><td>-6.2</td><td>1.4</td><td>-4.3</td><td>1.8E-05</td><td>3.1E-05</td><td>MB231/MB468</td><td>MP</td></tr>
<tr><td>>hg38_21_+_8433172_8446622_RNA45SN1_45S_50ntFlanks@5478.5516.39|CTGGCGGAGCGCTGAGAAGACGGTCGAACTTGACTATCT</td><td>>hg38_21_+_8433172_8446622_RNA45SN1_45S_50ntFlanks</td><td>5478</td><td>5516</td><td>39</td><td>CTGGCGGAGCGCTGAGAAGACGGTCGAACTTGACTATCT</td><td>193.5</td><td>7.8</td><td>1.7</td><td>4.5</td><td>7.1E-06</td><td>3.1E-05</td><td>BT20/MB468</td><td>MP</td></tr>
<tr><td>>hg38_21_+_8433172_8446622_RNA45SN1_45S_50ntFlanks@10457.10481.25|GATGGCCTCCGTTGCCCTCGGCCGA</td><td>>hg38_21_+_8433172_8446622_RNA45SN1_45S_50ntFlanks</td><td>10457</td><td>10481</td><td>25</td><td>GATGGCCTCCGTTGCCCTCGGCCGA</td><td>48.8</td><td>-8.7</td><td>1.9</td><td>-4.5</td><td>7.1E-06</td><td>3.1E-05</td><td>BT20/MB468</td><td>MP</td></tr>
<tr><td>>hg38_21_+_8433172_8446622_RNA45SN1_45S_50ntFlanks@12949.12973.25|ACGACCTGCTTCTGGGTCGGGGTTT</td><td>>hg38_21_+_8433172_8446622_RNA45SN1_45S_50ntFlanks</td><td>12949</td><td>12973</td><td>25</td><td>ACGACCTGCTTCTGGGTCGGGGTTT</td><td>58.8</td><td>-9.0</td><td>2.0</td><td>-4.5</td><td>7.1E-06</td><td>3.1E-05</td><td>BT20/MB468</td><td>MP</td></tr>
<tr><td>>MI0000084|hsa-mir-26b&WithFlank&2|+|218402640|218402728@18.39.22(+1U)&[MIMAT0000083&hsa-miR-26b-5p&offsets|0|+1(+1U);m-53&2|+|218402657|218402678&offsets|0|0(+1U)]|TTCAAGTAATTCAGGATAGGTTT</td><td>>MI0000084|hsa-mir-26b&WithFlank&2|+|218402640|218402728</td><td>18</td><td>39</td><td>22(+1U)</td><td>TTCAAGTAATTCAGGATAGGTTT</td><td>92.6</td><td>2.4</td><td>0.6</td><td>4.2</td><td>2.3E-05</td><td>3.1E-05</td><td>BT20/MB468</td><td>NUC</td></tr>
<tr><td>>hg38_21_+_8433172_8446622_RNA45SN1_45S_50ntFlanks@9736.9755.20|CCGAAACGATCTCAACCTAT</td><td>>hg38_21_+_8433172_8446622_RNA45SN1_45S_50ntFlanks</td><td>9736</td><td>9755</td><td>20</td><td>CCGAAACGATCTCAACCTAT</td><td>335.5</td><td>-6.0</td><td>1.4</td><td>-4.2</td><td>2.2E-05</td><td>3.1E-05</td><td>MB231/MB468</td><td>NUC</td></tr>
<tr><td>>hg38_21_+_8433172_8446622_RNA45SN1_45S_50ntFlanks@12626.12652.27|AATGGGGCGAAGCTACCATCTGTGGGA</td><td>>hg38_21_+_8433172_8446622_RNA45SN1_45S_50ntFlanks</td><td>12626</td><td>12652</td><td>27</td><td>AATGGGGCGAAGCTACCATCTGTGGGA</td><td>1239.0</td><td>-7.5</td><td>1.7</td><td>-4.5</td><td>7.2E-06</td><td>3.1E-05</td><td>BT20/MB468</td><td>MP</td></tr>
<tr><td>>hg38_21_+_8433172_8446622_RNA45SN1_45S_50ntFlanks@10679.10706.28|GGTTCCGGCGGCGTCCGGTGAGCTCTCG</td><td>>hg38_21_+_8433172_8446622_RNA45SN1_45S_50ntFlanks</td><td>10679</td><td>10706</td><td>28</td><td>GGTTCCGGCGGCGTCCGGTGAGCTCTCG</td><td>4210.2</td><td>-5.4</td><td>1.3</td><td>-4.3</td><td>1.9E-05</td><td>3.1E-05</td><td>BT20/MB468</td><td>CYTO</td></tr>
<tr><td>>hg38_21_+_8433172_8446622_RNA45SN1_45S_50ntFlanks@4220.4243.24|ACGAAAAATAACAATACAGGACTC</td><td>>hg38_21_+_8433172_8446622_RNA45SN1_45S_50ntFlanks</td><td>4220</td><td>4243</td><td>24</td><td>ACGAAAAATAACAATACAGGACTC</td><td>258.4</td><td>-2.5</td><td>0.6</td><td>-4.3</td><td>2.1E-05</td><td>3.1E-05</td><td>MB231/MB468</td><td>CYTO</td></tr>
<tr><td>>hg38_21_+_8433172_8446622_RNA45SN1_45S_50ntFlanks@11767.11790.24|TCTAATTAGTGACGCGCATGAATG</td><td>>hg38_21_+_8433172_8446622_RNA45SN1_45S_50ntFlanks</td><td>11767</td><td>11790</td><td>24</td><td>TCTAATTAGTGACGCGCATGAATG</td><td>125.7</td><td>-6.2</td><td>1.4</td><td>-4.3</td><td>1.8E-05</td><td>3.1E-05</td><td>MB231/MB468</td><td>MP</td></tr>
</table>

<table>
<tr><td></td><td></td><td></td><td></td><td></td><td></td><td></td><td></td><td></td><td></td><td></td><td></td><td></td><td></td></tr>
<tr><td>>trna10_AspGTC_12_-_125424193_125424264@37.75.39,trna12_AspGTC_12_-_125411891_125411962@37.75.39,trna144_AspGTC_6_-_27551236_27551307@37.75.39,trna38_AspGTC_17_-_8125556_8125627@37.75.39,trna45_AspGTC_6_+_27447453_27447524@37.75.39,trna48_AspGTC_6_+_27471523_27471594@37.75.39,trna4_AspGTC_12_+_96429799_96429870@37.75.39,trna69_AspGTC_1_-_161440205_161440276@37.75.39,trna72_AspGTC_1_-_161432824_161432895@37.75.39,trna75_AspGTC_1_-_161425414_161425485@37.75.39,trna78_AspGTC_1_-_161418033_161418104@37.75.39,trna81_AspGTC_1_-_161410615_161410686@37.75.39|ACGCGGGAGACCGGGGTTCGATTCCCCGACGGGGAGCCA</td><td>>trna10_AspGTC_12_-_125424193_125424264</td><td>37</td><td>75</td><td>39</td><td>ACGCGGGAGACCGGGGTTCGATTCCCCGACGGGGAGCCA</td><td>1412.0</td><td>4.4</td><td>1.0</td><td>4.3</td><td>1.9E-05</td><td>3.1E-05</td><td>BT20/MB468</td><td>CYTO</td></tr>
<tr><td>>hg38_21_+_8433172_8446622_RNA45SN1_45S_50ntFlanks@3783.3801.19|AGTGAAACTGCGAATGGCT</td><td>>hg38_21_+_8433172_8446622_RNA45SN1_45S_50ntFlanks</td><td>3783</td><td>3801</td><td>19</td><td>AGTGAAACTGCGAATGGCT</td><td>598.3</td><td>-4.1</td><td>0.9</td><td>-4.3</td><td>1.6E-05</td><td>3.1E-05</td><td>BT20/MB468</td><td>MITO</td></tr>
<tr><td>>hg38_21_+_8433172_8446622_RNA45SN1_45S_50ntFlanks@11566.11599.34|TAGCAGCCGACTTAGAACTGGTGCGGACCAGGGG</td><td>>hg38_21_+_8433172_8446622_RNA45SN1_45S_50ntFlanks</td><td>11566</td><td>11599</td><td>34</td><td>TAGCAGCCGACTTAGAACTGGTGCGGACCAGGGG</td><td>117.7</td><td>10.2</td><td>2.2</td><td>4.7</td><td>2.2E-06</td><td>3.1E-05</td><td>BT20/MB231</td><td>MP</td></tr>
<tr><td>>hg38_21_+_8433172_8446622_RNA45SN1_45S_50ntFlanks@11785.11804.20|TGAATGGATGAACGAGATTC</td><td>>hg38_21_+_8433172_8446622_RNA45SN1_45S_50ntFlanks</td><td>11785</td><td>11804</td><td>20</td><td>TGAATGGATGAACGAGATTC</td><td>1235.3</td><td>-2.7</td><td>0.6</td><td>-4.3</td><td>1.6E-05</td><td>3.1E-05</td><td>BT20/MB468</td><td>MITO</td></tr>
<tr><td>>hg38_21_+_8433172_8446622_RNA45SN1_45S_50ntFlanks@11587.11612.26|TGCGGACCAGGGGAATCCGACTGTTT</td><td>>hg38_21_+_8433172_8446622_RNA45SN1_45S_50ntFlanks</td><td>11587</td><td>11612</td><td>26</td><td>TGCGGACCAGGGGAATCCGACTGTTT</td><td>387.4</td><td>-3.0</td><td>0.7</td><td>-4.2</td><td>2.2E-05</td><td>3.1E-05</td><td>MB231/MB468</td><td>MITO</td></tr>
<tr><td>>hg38_21_+_8433172_8446622_RNA45SN1_45S_50ntFlanks@3787.3805.19|AAACTGCGAATGGCTCATT</td><td>>hg38_21_+_8433172_8446622_RNA45SN1_45S_50ntFlanks</td><td>3787</td><td>3805</td><td>19</td><td>AAACTGCGAATGGCTCATT</td><td>585.5</td><td>-4.0</td><td>0.9</td><td>-4.2</td><td>2.2E-05</td><td>3.1E-05</td><td>MB231/MB468</td><td>MITO</td></tr>
<tr><td>>MI0003581|hsa-mir-574&WithFlank&4|+|38868026|38868133@67.87.21&[MIMAT0003239&hsa-miR-574-3p&offsets|0|-1;m-149&4|+|38868092|38868113&offsets|0|-1]|CACGCTCATGCACACACCCAC</td><td>>MI0003581|hsa-mir-574&WithFlank&4|+|38868026|38868133</td><td>67</td><td>87</td><td>21</td><td>CACGCTCATGCACACACCCAC</td><td>385.3</td><td>7.9</td><td>1.8</td><td>4.3</td><td>1.9E-05</td><td>3.1E-05</td><td>BT20/MB468</td><td>CYTO</td></tr>
<tr><td>>hg38_21_+_8433172_8446622_RNA45SN1_45S_50ntFlanks@9862.9893.32|ATGAACCGAACGCCGGGTTAAGGCGCCCGATG</td><td>>hg38_21_+_8433172_8446622_RNA45SN1_45S_50ntFlanks</td><td>9862</td><td>9893</td><td>32</td><td>ATGAACCGAACGCCGGGTTAAGGCGCCCGATG</td><td>99.8</td><td>10.0</td><td>2.1</td><td>4.7</td><td>2.2E-06</td><td>3.1E-05</td><td>BT20/MB231</td><td>MP</td></tr>
<tr><td>>hg38_21_+_8433172_8446622_RNA45SN1_45S_50ntFlanks@9740.9758.19|AACGATCTCAACCTATTCT</td><td>>hg38_21_+_8433172_8446622_RNA45SN1_45S_50ntFlanks</td><td>9740</td><td>9758</td><td>19</td><td>AACGATCTCAACCTATTCT</td><td>335.9</td><td>-2.8</td><td>0.6</td><td>-4.3</td><td>1.9E-05</td><td>3.1E-05</td><td>BT20/MB468</td><td>CYTO</td></tr>
<tr><td>>hg38_21_+_8433172_8446622_RNA45SN1_45S_50ntFlanks@3784.3801.18|GTGAAACTGCGAATGGCT</td><td>>hg38_21_+_8433172_8446622_RNA45SN1_45S_50ntFlanks</td><td>3784</td><td>3801</td><td>18</td><td>GTGAAACTGCGAATGGCT</td><td>473.5</td><td>-5.1</td><td>1.2</td><td>-4.2</td><td>2.2E-05</td><td>3.1E-05</td><td>MB231/MB468</td><td>MITO</td></tr>
<tr><td>>hg38_MT_+_598_1651_RNR1_12S_50ntFlanks@853.873.21|CTTAGTTGAACAGGGCCCTGA</td><td>>hg38_MT_+_598_1651_RNR1_12S_50ntFlanks</td><td>853</td><td>873</td><td>21</td><td>CTTAGTTGAACAGGGCCCTGA</td><td>52.2</td><td>9.5</td><td>2.1</td><td>4.5</td><td>7.2E-06</td><td>3.1E-05</td><td>BT20/MB468</td><td>MP</td></tr>
<tr><td>>hg38_21_+_8433172_8446622_RNA45SN1_45S_50ntFlanks@11672.11708.37|CAGTGCTCTGAATGTCAAAGTGAAGAAATTCAATGAA</td><td>>hg38_21_+_8433172_8446622_RNA45SN1_45S_50ntFlanks</td><td>11672</td><td>11708</td><td>37</td><td>CAGTGCTCTGAATGTCAAAGTGAAGAAATTCAATGAA</td><td>249.8</td><td>3.8</td><td>0.9</td><td>4.3</td><td>2.1E-05</td><td>3.1E-05</td><td>MB231/MB468</td><td>CYTO</td></tr>
<tr><td>>hg38_21_+_8433172_8446622_RNA45SN1_45S_50ntFlanks@10846.10866.21|CTTCGGGATAAGGATTGGCTC</td><td>>hg38_21_+_8433172_8446622_RNA45SN1_45S_50ntFlanks</td><td>10846</td><td>10866</td><td>21</td><td>CTTCGGGATAAGGATTGGCTC</td><td>302.3</td><td>-4.1</td><td>1.0</td><td>-4.3</td><td>2.1E-05</td><td>3.1E-05</td><td>MB231/MB468</td><td>CYTO</td></tr>
<tr><td>>hg38_21_+_8433172_8446622_RNA45SN1_45S_50ntFlanks@10461.10484.24|GCCTCCGTTGCCCTCGGCCGATCG</td><td>>hg38_21_+_8433172_8446622_RNA45SN1_45S_50ntFlanks</td><td>10461</td><td>10484</td><td>24</td><td>GCCTCCGTTGCCCTCGGCCGATCG</td><td>473.0</td><td>-7.2</td><td>1.7</td><td>-4.2</td><td>2.3E-05</td><td>3.1E-05</td><td>BT20/MB468</td><td>NUC</td></tr>
<tr><td>>hg38_21_+_8433172_8446622_RNA45SN1_45S_50ntFlanks@7975.8019.45|CGCGACCTCAGATCAGACGTGGCGACCCGCTGAATTTAAGCATAT</td><td>>hg38_21_+_8433172_8446622_RNA45SN1_45S_50ntFlanks</td><td>7975</td><td>8019</td><td>45</td><td>CGCGACCTCAGATCAGACGTGGCGACCCGCTGAATTTAAGCATAT</td><td>817.2</td><td>4.4</td><td>1.0</td><td>4.2</td><td>2.2E-05</td><td>3.1E-05</td><td>MB231/MB468</td><td>MITO</td></tr>
<tr><td>>trna119_LysCTT_1_-_145395522_145395594@3.36.34,trna11_LysCTT_5_-_180648979_180649051@3.36.34,trna13_LysCTT_6_+_26556774_26556846@3.36.34,trna7_LysCTT_16_+_3225692_3225764@3.36.34,trna9_LysCTT_5_+_180634755_180634827@3.36.34|CCGGCTAGCTCAGTCGGTAGAGCATGAGACTCTT</td><td>>trna119_LysCTT_1_-_145395522_145395594</td><td>3</td><td>36</td><td>34</td><td>CCGGCTAGCTCAGTCGGTAGAGCATGAGACTCTT</td><td>307.5</td><td>6.6</td><td>1.5</td><td>4.2</td><td>2.2E-05</td><td>3.1E-05</td><td>MB231/MB468</td><td>MITO</td></tr>
<tr><td>>hg38_21_+_8433172_8446622_RNA45SN1_45S_50ntFlanks@11570.11609.40|AGCCGACTTAGAACTGGTGCGGACCAGGGGAATCCGACTG</td><td>>hg38_21_+_8433172_8446622_RNA45SN1_45S_50ntFlanks</td><td>11570</td><td>11609</td><td>40</td><td>AGCCGACTTAGAACTGGTGCGGACCAGGGGAATCCGACTG</td><td>313.5</td><td>-3.7</td><td>0.9</td><td>-4.2</td><td>2.2E-05</td><td>3.1E-05</td><td>MB231/MB468</td><td>MITO</td></tr>
<tr><td>>hg38_21_+_8433172_8446622_RNA45SN1_45S_50ntFlanks@12396.12417.22|GCGACGTCGCTTTTTGATCCTT</td><td>>hg38_21_+_8433172_8446622_RNA45SN1_45S_50ntFlanks</td><td>12396</td><td>12417</td><td>22</td><td>GCGACGTCGCTTTTTGATCCTT</td><td>973.1</td><td>-3.9</td><td>0.9</td><td>-4.3</td><td>1.6E-05</td><td>3.1E-05</td><td>BT20/MB468</td><td>MITO</td></tr>
<tr><td>>hg38_21_+_8433172_8446622_RNA45SN1_45S_50ntFlanks@10853.10873.21|ATAAGGATTGGCTCTAAGGGC</td><td>>hg38_21_+_8433172_8446622_RNA45SN1_45S_50ntFlanks</td><td>10853</td><td>10873</td><td>21</td><td>ATAAGGATTGGCTCTAAGGGC</td><td>375.5</td><td>-2.9</td><td>0.7</td><td>-4.3</td><td>2.1E-05</td><td>3.1E-05</td><td>MB231/MB468</td><td>CYTO</td></tr>
<tr><td>>hg38_21_+_8433172_8446622_RNA45SN1_45S_50ntFlanks@9545.9575.31|GTGGAGGTCCGTAGCGGTCCTGACGTGCAAA</td><td>>hg38_21_+_8433172_8446622_RNA45SN1_45S_50ntFlanks</td><td>9545</td><td>9575</td><td>31</td><td>GTGGAGGTCCGTAGCGGTCCTGACGTGCAAA</td><td>144.5</td><td>-6.3</td><td>1.5</td><td>-4.3</td><td>1.9E-05</td><td>3.1E-05</td><td>MB231/MB468</td><td>MP</td></tr>
<tr><td>>hg38_21_+_8433172_8446622_RNA45SN1_45S_50ntFlanks@7990.8011.22|GACGTGGCGACCCGCTGAATTT</td><td>>hg38_21_+_8433172_8446622_RNA45SN1_45S_50ntFlanks</td><td>7990</td><td>8011</td><td>22</td><td>GACGTGGCGACCCGCTGAATTT</td><td>1754.5</td><td>3.3</td><td>0.8</td><td>4.3</td><td>1.6E-05</td><td>3.1E-05</td><td>BT20/MB468</td><td>MITO</td></tr>
<tr><td>>hg38_21_+_8433172_8446622_RNA45SN1_45S_50ntFlanks@11622.11660.39|AAAGCATCGCGAAGGCCCGCGGCGGGTGTTGACGCGATG</td><td>>hg38_21_+_8433172_8446622_RNA45SN1_45S_50ntFlanks</td><td>11622</td><td>11660</td><td>39</td><td>AAAGCATCGCGAAGGCCCGCGGCGGGTGTTGACGCGATG</td><td>44.2</td><td>-8.5</td><td>1.9</td><td>-4.5</td><td>7.3E-06</td><td>3.1E-05</td><td>BT20/MB468</td><td>MP</td></tr>
<tr><td>>hg38_21_+_8433172_8446622_RNA45SN1_45S_50ntFlanks@11729.11761.33|TAACTATGACTCTCTTAAGGTAGCCAAATGCCT</td><td>>hg38_21_+_8433172_8446622_RNA45SN1_45S_50ntFlanks</td><td>11729</td><td>11761</td><td>33</td><td>TAACTATGACTCTCTTAAGGTAGCCAAATGCCT</td><td>408.0</td><td>-4.3</td><td>1.0</td><td>-4.3</td><td>1.9E-05</td><td>3.1E-05</td><td>BT20/MB468</td><td>CYTO</td></tr>
<tr><td>>hg38_21_+_8433172_8446622_RNA45SN1_45S_50ntFlanks@4048.4066.19|TTCGAACGTCTGCCCTATC</td><td>>hg38_21_+_8433172_8446622_RNA45SN1_45S_50ntFlanks</td><td>4048</td><td>4066</td><td>19</td><td>TTCGAACGTCTGCCCTATC</td><td>440.4</td><td>-3.9</td><td>0.9</td><td>-4.3</td><td>1.6E-05</td><td>3.1E-05</td><td>BT20/MB468</td><td>MITO</td></tr>
<tr><td>>hg38_21_+_8433172_8446622_RNA45SN1_45S_50ntFlanks@6646.6693.48|TCGTACGACTCTTAGCGGTGGATCACTCGGCTCGTGCGTCGATGAAGA</td><td>>hg38_21_+_8433172_8446622_RNA45SN1_45S_50ntFlanks</td><td>6646</td><td>6693</td><td>48</td><td>TCGTACGACTCTTAGCGGTGGATCACTCGGCTCGTGCGTCGATGAAGA</td><td>1818.6</td><td>4.5</td><td>1.1</td><td>4.3</td><td>1.6E-05</td><td>3.2E-05</td><td>BT20/MB468</td><td>MITO</td></tr>
<tr><td>>hg38_21_+_8433172_8446622_RNA45SN1_45S_50ntFlanks@11576.11606.31|CTTAGAACTGGTGCGGACCAGGGGAATCCGA</td><td>>hg38_21_+_8433172_8446622_RNA45SN1_45S_50ntFlanks</td><td>11576</td><td>11606</td><td>31</td><td>CTTAGAACTGGTGCGGACCAGGGGAATCCGA</td><td>185.2</td><td>-3.3</td><td>0.8</td><td>-4.3</td><td>1.9E-05</td><td>3.2E-05</td><td>BT20/MB468</td><td>CYTO</td></tr>
<tr><td>>hg38_21_+_8433172_8446622_RNA45SN1_45S_50ntFlanks@11703.11724.22|AATGAAGCGCGGGTAAACGGCG</td><td>>hg38_21_+_8433172_8446622_RNA45SN1_45S_50ntFlanks</td><td>11703</td><td>11724</td><td>22</td><td>AATGAAGCGCGGGTAAACGGCG</td><td>244.1</td><td>-6.7</td><td>1.6</td><td>-4.3</td><td>1.9E-05</td><td>3.2E-05</td><td>MB231/MB468</td><td>MP</td></tr>
<tr><td>>MI0000103|hsa-mir-101-1&WithFlank&1|-|65058428|65058514@53.73.21(+1U);MI0000739|hsa-mir-101-2&WithFlank&9|+|4850291|4850381@55.75.21(+1U)&[MIMAT0000099&hsa-miR-101-3p&offsets|0|0(+1U);m-17&1|-|65058443|65058462&offsets|0|+1(+1U)];[MIMAT0000099_1&hsa-miR-101-3p&offsets|0|0(+1U);m-16&9|+|4850345|4850364&offsets|0|+1(+1U)]|TACAGTACTGTGATAACTGAAT</td><td>>MI0000103|hsa-mir-101-1&WithFlank&1|-|65058428|65058514</td><td>53</td><td>73</td><td>21(+1U)</td><td>TACAGTACTGTGATAACTGAAT</td><td>556.4</td><td>-4.7</td><td>1.0</td><td>-4.7</td><td>2.7E-06</td><td>3.2E-05</td><td>BT20/MB231</td><td>MITO</td></tr>
<tr><td>>hg38_21_+_8433172_8446622_RNA45SN1_45S_50ntFlanks@9460.9488.29|CGGGGAGGTGGAGCACGAGCGCACGTGTT</td><td>>hg38_21_+_8433172_8446622_RNA45SN1_45S_50ntFlanks</td><td>9460</td><td>9488</td><td>29</td><td>CGGGGAGGTGGAGCACGAGCGCACGTGTT</td><td>500.1</td><td>6.5</td><td>1.5</td><td>4.3</td><td>2.1E-05</td><td>3.2E-05</td><td>MB231/MB468</td><td>CYTO</td></tr>
<tr><td>>hg38_21_+_8433172_8446622_RNA45SN1_45S_50ntFlanks@4169.4186.18|AGGAAGGCAGCAGGCGCG</td><td>>hg38_21_+_8433172_8446622_RNA45SN1_45S_50ntFlanks</td><td>4169</td><td>4186</td><td>18</td><td>AGGAAGGCAGCAGGCGCG</td><td>111.5</td><td>-6.1</td><td>1.4</td><td>-4.3</td><td>1.9E-05</td><td>3.2E-05</td><td>MB231/MB468</td><td>MP</td></tr>
<tr><td>>hg38_21_+_8433172_8446622_RNA45SN1_45S_50ntFlanks@10546.10566.21|CAGTGCGGTAACGCGACCGAT</td><td>>hg38_21_+_8433172_8446622_RNA45SN1_45S_50ntFlanks</td><td>10546</td><td>10566</td><td>21</td><td>CAGTGCGGTAACGCGACCGAT</td><td>43.0</td><td>-8.5</td><td>1.9</td><td>-4.5</td><td>7.3E-06</td><td>3.2E-05</td><td>BT20/MB468</td><td>MP</td></tr>
<tr><td>>hg38_21_+_8433172_8446622_RNA45SN1_45S_50ntFlanks@11767.11803.37|TCTAATTAGTGACGCGCATGAATGGATGAACGAGATT</td><td>>hg38_21_+_8433172_8446622_RNA45SN1_45S_50ntFlanks</td><td>11767</td><td>11803</td><td>37</td><td>TCTAATTAGTGACGCGCATGAATGGATGAACGAGATT</td><td>266.7</td><td>-4.7</td><td>1.1</td><td>-4.2</td><td>2.3E-05</td><td>3.2E-05</td><td>MB231/MB468</td><td>MITO</td></tr>
<tr><td>>MI0000061|hsa-let-7a-2&WithFlank&11|-|122146516|122146599@11.31.21(+1A);MI0000062|hsa-let-7a-3&WithFlank&22|+|46112743|46112828@10.30.21(+1A);MI0000060|hsa-let-7a-1&WithFlank&9|+|94175951|94176042@12.32.21(+1A)&[MIMAT0000062&hsa-let-7a-5p&offsets|0|-1(+1A);m-6&11|-|122146568|122146589&offsets|0|-1(+1A)];[MIMAT0000062_1&hsa-let-7a-5p&offsets|0|-1(+1A);m-5&22|+|46112752|46112773&offsets|0|-1(+1A)];[MIMAT0000062_2&hsa-let-7a-5p&offsets|0|-1(+1A);m-4&9|+|94175962|94175983&offsets|0|-1(+1A)]|TGAGGTAGTAGGTTGTATAGTA</td><td>>MI0000061|hsa-let-7a-2&WithFlank&11|-|122146516|122146599</td><td>11</td><td>31</td><td>21(+1A)</td><td>TGAGGTAGTAGGTTGTATAGTA</td><td>1623.5</td><td>4.3</td><td>1.0</td><td>4.3</td><td>1.9E-05</td><td>3.2E-05</td><td>MB231/MB468</td><td>MP</td></tr>
<tr><td>>hg38_21_+_8433172_8446622_RNA45SN1_45S_50ntFlanks@8403.8442.40|AGGTAAACGGGTGGGGTCCGCGCAGTCCGCCCGGAGGATT</td><td>>hg38_21_+_8433172_8446622_RNA45SN1_45S_50ntFlanks</td><td>8403</td><td>8442</td><td>40</td><td>AGGTAAACGGGTGGGGTCCGCGCAGTCCGCCCGGAGGATT</td><td>186.3</td><td>-5.4</td><td>1.3</td><td>-4.3</td><td>1.6E-05</td><td>3.2E-05</td><td>BT20/MB468</td><td>MITO</td></tr>
<tr><td>>hg38_21_+_8433172_8446622_RNA45SN1_45S_50ntFlanks@10391.10410.20|AACAGCAGTTGAACATGGGT</td><td>>hg38_21_+_8433172_8446622_RNA45SN1_45S_50ntFlanks</td><td>10391</td><td>10410</td><td>20</td><td>AACAGCAGTTGAACATGGGT</td><td>712.0</td><td>-2.6</td><td>0.6</td><td>-4.3</td><td>1.6E-05</td><td>3.2E-05</td><td>BT20/MB468</td><td>MITO</td></tr>
<tr><td>>hg38_21_+_8433172_8446622_RNA45SN1_45S_50ntFlanks@5077.5094.18|CCGATAACGAACGAGACT</td><td>>hg38_21_+_8433172_8446622_RNA45SN1_45S_50ntFlanks</td><td>5077</td><td>5094</td><td>18</td><td>CCGATAACGAACGAGACT</td><td>135.8</td><td>-5.5</td><td>1.3</td><td>-4.3</td><td>1.9E-05</td><td>3.2E-05</td><td>MB231/MB468</td><td>MP</td></tr>
<tr><td>>hg38_21_+_8433172_8446622_RNA45SN1_45S_50ntFlanks@11876.11900.25|AAGACCCTGTTGAGCTTGACTCTAG</td><td>>hg38_21_+_8433172_8446622_RNA45SN1_45S_50ntFlanks</td><td>11876</td><td>11900</td><td>25</td><td>AAGACCCTGTTGAGCTTGACTCTAG</td><td>500.1</td><td>-4.9</td><td>1.2</td><td>-4.2</td><td>2.2E-05</td><td>3.2E-05</td><td>MB231/MB468</td><td>NUC</td></tr>
<tr><td>>hg38_21_+_8433172_8446622_RNA45SN1_45S_50ntFlanks@9339.9369.31|CGCACGAAAGCCGCCGTGGCGCAATGAAGGT</td><td>>hg38_21_+_8433172_8446622_RNA45SN1_45S_50ntFlanks</td><td>9339</td><td>9369</td><td>31</td><td>CGCACGAAAGCCGCCGTGGCGCAATGAAGGT</td><td>477.6</td><td>-7.0</td><td>1.6</td><td>-4.5</td><td>7.4E-06</td><td>3.2E-05</td><td>BT20/MB468</td><td>MP</td></tr>
<tr><td>>am_trna143_LysTTT_6_-_27559593_27559665@7.26.20,trna14_LysTTT_11_-_59327808_59327880@7.26.20,trna2_LysTTT_17_+_8022473_8022545@7.26.20,trna54_LysTTT_1_+_204475655_204475727@7.26.20,trna5_LysTTT_11_+_59323902_59323974@7.26.20,trna62_LysTTT_1_-_204476158_204476230@7.26.20,trna76_LysTTT_6_+_28918806_28918878@7.26.20|ATAGCTCAGTCGGTAGAGCA</td><td>>am_trna143_LysTTT_6_-_27559593_27559665</td><td>7</td><td>26</td><td>20</td><td>ATAGCTCAGTCGGTAGAGCA</td><td>38.3</td><td>-8.3</td><td>1.9</td><td>-4.5</td><td>7.4E-06</td><td>3.2E-05</td><td>BT20/MB468</td><td>MP</td></tr>
<tr><td>>hg38_21_+_8433172_8446622_RNA45SN1_45S_50ntFlanks@8182.8209.28|ATCGAGGCCCAGCCCGTGGACGGTGTGA</td><td>>hg38_21_+_8433172_8446622_RNA45SN1_45S_50ntFlanks</td><td>8182</td><td>8209</td><td>28</td><td>ATCGAGGCCCAGCCCGTGGACGGTGTGA</td><td>165.9</td><td>-6.5</td><td>1.5</td><td>-4.3</td><td>1.9E-05</td><td>3.2E-05</td><td>MB231/MB468</td><td>MP</td></tr>
<tr><td>>hg38_21_+_8433172_8446622_RNA45SN1_45S_50ntFlanks@10238.10262.25|CGCTACGCCGCGACGAGTAGGAGGG</td><td>>hg38_21_+_8433172_8446622_RNA45SN1_45S_50ntFlanks</td><td>10238</td><td>10262</td><td>25</td><td>CGCTACGCCGCGACGAGTAGGAGGG</td><td>198.5</td><td>-6.3</td><td>1.5</td><td>-4.3</td><td>1.9E-05</td><td>3.2E-05</td><td>MB231/MB468</td><td>MP</td></tr>
<tr><td>>hg38_21_+_8433172_8446622_RNA45SN1_45S_50ntFlanks@10843.10861.19|TAACTTCGGGATAAGGATT</td><td>>hg38_21_+_8433172_8446622_RNA45SN1_45S_50ntFlanks</td><td>10843</td><td>10861</td><td>19</td><td>TAACTTCGGGATAAGGATT</td><td>350.3</td><td>-3.1</td><td>0.7</td><td>-4.3</td><td>1.9E-05</td><td>3.2E-05</td><td>BT20/MB468</td><td>CYTO</td></tr>
<tr><td>>hg38_21_+_8433172_8446622_RNA45SN1_45S_50ntFlanks@10391.10430.40|AACAGCAGTTGAACATGGGTCAGTCGGTCCTGAGAGATGG</td><td>>hg38_21_+_8433172_8446622_RNA45SN1_45S_50ntFlanks</td><td>10391</td><td>10430</td><td>40</td><td>AACAGCAGTTGAACATGGGTCAGTCGGTCCTGAGAGATGG</td><td>605.7</td><td>-3.8</td><td>0.9</td><td>-4.3</td><td>1.6E-05</td><td>3.2E-05</td><td>BT20/MB468</td><td>MITO</td></tr>
<tr><td>>hg38_21_+_8433172_8446622_RNA45SN1_45S_50ntFlanks@11874.11908.35|AGAAGACCCTGTTGAGCTTGACTCTAGTCTGGCAC</td><td>>hg38_21_+_8433172_8446622_RNA45SN1_45S_50ntFlanks</td><td>11874</td><td>11908</td><td>35</td><td>AGAAGACCCTGTTGAGCTTGACTCTAGTCTGGCAC</td><td>823.0</td><td>-2.6</td><td>0.6</td><td>-4.3</td><td>1.6E-05</td><td>3.2E-05</td><td>BT20/MB468</td><td>MITO</td></tr>
</table>

| | | | | | | | | | | | | | |
|---|---|---|---|---|---|---|---|---|---|---|---|---|---|
| >hg38_21_+_8433172_8446622_RNA45SN1_45S_50ntFlanks@475.510.36\|CTTCGTGATCGATGTGGTGACGTCGTGCTCTCCCGG | >hg38_21_+_8433172_8446622_RNA45SN1_45S_50ntFlanks | 475 | 510 | 36 | CTTCGTGATCGATGTGGTGACGTCGTGCTCTCCCGG | 1286.4 | 4.5 | 1.1 | 4.2 | 2.3E-05 | 3.2E-05 | MB231/MB468 | MITO |
| >hg38_21_+_8433172_8446622_RNA45SN1_45S_50ntFlanks@6735.6771.37\|TTGATCATCGACACTTCGAACGCACTTGCGGCCCCGG | >hg38_21_+_8433172_8446622_RNA45SN1_45S_50ntFlanks | 6735 | 6771 | 37 | TTGATCATCGACACTTCGAACGCACTTGCGGCCCCGG | 941.5 | -7.6 | 1.8 | -4.2 | 2.4E-05 | 3.2E-05 | BT20/MB468 | NUC |
| >hg38_21_+_8433172_8446622_RNA45SN1_45S_50ntFlanks@6647.6672.26\|CGTACGACTCTTAGCGGTGGATCACT | >hg38_21_+_8433172_8446622_RNA45SN1_45S_50ntFlanks | 6647 | 6672 | 26 | CGTACGACTCTTAGCGGTGGATCACT | 334.7 | 3.7 | 0.8 | 4.5 | 7.4E-06 | 3.2E-05 | BT20/MB468 | MP |
| >MI0000103\|hsa-mir-101-1&WithFlank&1\|-\|65058428\|65058514@53.73.21;MI0000739\|hsa-mir-101-2&WithFlank&9\|+\|4850291\|4850381@55.75.21&[MIMAT0000099&hsa-miR-101-3p&offsets\|0\|0;m-17&1\|-\|65058443\|65058462&offsets\|0\|+1];[MIMAT0000099_1&hsa-miR-101-3p&offsets\|0\|0;m-16&9\|+\|4850345\|4850364&offsets\|0\|+1]\|TACAGTACTGTGATAACTGAA | >MI0000103\|hsa-mir-101-1&WithFlank&1\|-\|65058428\|65058514 | 53 | 73 | 21 | TACAGTACTGTGATAACTGAA | 407.9 | 4.7 | 1.1 | 4.2 | 2.3E-05 | 3.2E-05 | MB231/MB468 | MITO |
| >hg38_21_+_8433172_8446622_RNA45SN1_45S_50ntFlanks@10692.10711.20\|TCCGGTGAGCTCTCGCTGGC | >hg38_21_+_8433172_8446622_RNA45SN1_45S_50ntFlanks | 10692 | 10711 | 20 | TCCGGTGAGCTCTCGCTGGC | 45.4 | -8.6 | 1.9 | -4.5 | 7.4E-06 | 3.2E-05 | BT20/MB468 | MP |
| >hg38_21_+_8433172_8446622_RNA45SN1_45S_50ntFlanks@12658.12677.20\|ACTGAACGCCTCTAAGTCAG | >hg38_21_+_8433172_8446622_RNA45SN1_45S_50ntFlanks | 12658 | 12677 | 20 | ACTGAACGCCTCTAAGTCAG | 70.3 | -10.0 | 2.3 | -4.3 | 1.9E-05 | 3.2E-05 | MB231/MB468 | MP |
| >am_trnaMT_MetCAT_MT_+_4402_4469@1.45.45\|AGTAAGGTCAGCTAAATAAGCTATCGGGCCCATACCCCGAAAATG | >am_trnaMT_MetCAT_MT_+_4402_4469 | 1 | 45 | 45 | AGTAAGGTCAGCTAAATAAGCTATCGGGCCCATACCCCGAAAATG | 232.0 | 11.6 | 2.6 | 4.5 | 7.4E-06 | 3.2E-05 | BT20/MB468 | MP |
| >trnaMT_HisGTG_MT_+_12138_12206@25.53.29\|CAGATTGTGAATCTGACAACAGAGGCTTA | >trnaMT_HisGTG_MT_+_12138_12206 | 25 | 53 | 29 | CAGATTGTGAATCTGACAACAGAGGCTTA | 57.0 | 9.6 | 2.1 | 4.5 | 7.4E-06 | 3.2E-05 | BT20/MB468 | MP |
| >MI0005559\|hsa-mir-744&WithFlank&17\|+\|12081893\|12082002@17.34.18&[MIMAT0004945&hsa-miR-744-5p&offsets\|0\|-4;m-172&17\|+\|12081909\|12081929&offsets\|0\|-3]\|TGCGGGGCTAGGGCTAAC | >MI0005559\|hsa-mir-744&WithFlank&17\|+\|12081893\|12082002 | 17 | 34 | 18 | TGCGGGGCTAGGGCTAAC | 295.8 | 3.5 | 0.7 | 4.7 | 2.7E-06 | 3.2E-05 | BT20/MB231 | MITO |
| >hg38_21_+_8433172_8446622_RNA45SN1_45S_50ntFlanks@9913.9947.35\|AGAAAAGGTGTTGGTTGATATAGACAGCAGGACGG | >hg38_21_+_8433172_8446622_RNA45SN1_45S_50ntFlanks | 9913 | 9947 | 35 | AGAAAAGGTGTTGGTTGATATAGACAGCAGGACGG | 135.0 | 10.4 | 2.2 | 4.7 | 2.3E-06 | 3.2E-05 | BT20/MB231 | MP |
| >hg38_21_+_8433172_8446622_RNA45SN1_45S_50ntFlanks@3854.3881.28\|ACTGTGGTAATTCTAGAGCTAATACATG | >hg38_21_+_8433172_8446622_RNA45SN1_45S_50ntFlanks | 3854 | 3881 | 28 | ACTGTGGTAATTCTAGAGCTAATACATG | 1043.5 | -7.3 | 1.6 | -4.5 | 7.4E-06 | 3.2E-05 | BT20/MB468 | MP |
| >MI0000062\|hsa-let-7a-3&WithFlank&22\|+\|46112743\|46112828@58.78.21(+1U);MI0000060\|hsa-let-7a-1&WithFlank&9\|+\|94175951\|94176042@63.83.21(+1U)&[MIMAT0004481&hsa-let-7a-3p&offsets\|0\|0(+1U);m-180&22\|+\|46112800\|46112821&offsets\|0\|-1(+1U)];[MIMAT0004481_1&hsa-let-7a-3p&offsets\|0\|0(+1U);m-181&9\|+\|94176013\|94176034&offsets\|0\|-1(+1U)]\|CTATACAATCTACTGTCTTTCT | >MI0000062\|hsa-let-7a-3&WithFlank&22\|+\|46112743\|46112828 | 58 | 78 | 21(+1U) | CTATACAATCTACTGTCTTTCT | 833.9 | 3.2 | 0.7 | 4.3 | 1.9E-05 | 3.2E-05 | MB231/MB468 | MP |
| >hg38_21_+_8433172_8446622_RNA45SN1_45S_50ntFlanks@11572.11589.18\|CCGACTTAGAACTGGTGC | >hg38_21_+_8433172_8446622_RNA45SN1_45S_50ntFlanks | 11572 | 11589 | 18 | CCGACTTAGAACTGGTGC | 278.1 | -2.2 | 0.5 | -4.2 | 2.3E-05 | 3.2E-05 | MB231/MB468 | NUC |
| >MI0000456\|hsa-mir-140&WithFlank&16\|+\|69933075\|69933186@69.87.19&[MIMAT0004597&hsa-miR-140-3p&offsets\|+1\|-1;m-45&16\|+\|69933143\|69933164&offsets\|0\|-3]\|ACCACAGGGTAGAACCACG | >MI0000456\|hsa-mir-140&WithFlank&16\|+\|69933075\|69933186 | 69 | 87 | 19 | ACCACAGGGTAGAACCACG | 46.6 | 9.3 | 2.1 | 4.5 | 7.4E-06 | 3.2E-05 | BT20/MB468 | MP |
| >hg38_21_+_8433172_8446622_RNA45SN1_45S_50ntFlanks@7977.8009.33\|CGACCTCAGATCAGACGTGGCGACCCGCTGAAT | >hg38_21_+_8433172_8446622_RNA45SN1_45S_50ntFlanks | 7977 | 8009 | 33 | CGACCTCAGATCAGACGTGGCGACCCGCTGAAT | 340.6 | 1.7 | 0.4 | 4.2 | 2.3E-05 | 3.2E-05 | MB231/MB468 | NUC |
| >am_trnaMT_ArgTCG_MT_+_10405_10469@29.48.20\|TTTCGACTCATTAAATTATG | >am_trnaMT_ArgTCG_MT_+_10405_10469 | 29 | 48 | 20 | TTTCGACTCATTAAATTATG | 662.1 | -6.1 | 1.5 | -4.2 | 2.3E-05 | 3.2E-05 | MB231/MB468 | NUC |
| >hg38_21_+_8433172_8446622_RNA45SN1_45S_50ntFlanks@6645.6678.34\|CTCGTACGACTCTTAGCGGTGGATCACTCGGCTC | >hg38_21_+_8433172_8446622_RNA45SN1_45S_50ntFlanks | 6645 | 6678 | 34 | CTCGTACGACTCTTAGCGGTGGATCACTCGGCTC | 658.4 | 3.8 | 0.9 | 4.5 | 7.5E-06 | 3.2E-05 | BT20/MB468 | MP |
| >MI0000542\|hsa-mir-320a&WithFlank&8\|-\|22244960\|22245043@50.69.20&[MIMAT0000510&hsa-miR-320a-3p&offsets\|+2\|0;m-60&8\|-\|22244975\|22244996&offsets\|+2\|0]\|AAGCTGGGTTGAGAGGGCGA | >MI0000542\|hsa-mir-320a&WithFlank&8\|-\|22244960\|22245043 | 50 | 69 | 20 | AAGCTGGGTTGAGAGGGCGA | 506.3 | 7.3 | 1.6 | 4.5 | 7.5E-06 | 3.2E-05 | BT20/MB468 | MP |
| >hg38_21_+_8433172_8446622_RNA45SN1_45S_50ntFlanks@10737.10757.21\|TAAATCTCGCGCCGGGCCGTA | >hg38_21_+_8433172_8446622_RNA45SN1_45S_50ntFlanks | 10737 | 10757 | 21 | TAAATCTCGCGCCGGGCCGTA | 223.0 | -4.1 | 1.0 | -4.2 | 2.2E-05 | 3.2E-05 | MB231/MB468 | CYTO |
| >hg38_21_+_8433172_8446622_RNA45SN1_45S_50ntFlanks@10386.10410.25\|ATGTGAACAGCAGTTGAACATGGGT | >hg38_21_+_8433172_8446622_RNA45SN1_45S_50ntFlanks | 10386 | 10410 | 25 | ATGTGAACAGCAGTTGAACATGGGT | 245.0 | -3.0 | 0.7 | -4.3 | 2.0E-05 | 3.2E-05 | BT20/MB468 | CYTO |
| >hg38_21_+_8433172_8446622_RNA45SN1_45S_50ntFlanks@7977.8016.40\|CGACCTCAGATCAGACGTGGCGACCCGCTGAATTTAAGCA | >hg38_21_+_8433172_8446622_RNA45SN1_45S_50ntFlanks | 7977 | 8016 | 40 | CGACCTCAGATCAGACGTGGCGACCCGCTGAATTTAAGCA | 686.8 | 4.1 | 0.9 | 4.3 | 1.7E-05 | 3.2E-05 | BT20/MB468 | MITO |
| >hg38_21_+_8433172_8446622_RNA45SN1_45S_50ntFlanks@3844.3868.25\|TACTTGGATAACTGTGGTAATTCTA | >hg38_21_+_8433172_8446622_RNA45SN1_45S_50ntFlanks | 3844 | 3868 | 25 | TACTTGGATAACTGTGGTAATTCTA | 226.0 | -5.7 | 1.3 | -4.3 | 1.9E-05 | 3.2E-05 | MB231/MB468 | MP |
| >MI0000102\|hsa-mir-100&WithFlank&11\|-\|122152223\|122152314@19.40.22(+3A)&[MIMAT0000098&hsa-miR-100-5p&offsets\|0\|0(+3A);m-27&11\|-\|122152276\|122152296&offsets\|0\|+1(+3A)]\|AACCCGTAGATCCGAACTTGTGAAA | >MI0000102\|hsa-mir-100&WithFlank&11\|-\|122152223\|122152314 | 19 | 40 | 22(+3A) | AACCCGTAGATCCGAACTTGTGAAA | 34.4 | 8.3 | 1.9 | 4.3 | 1.9E-05 | 3.2E-05 | MB231/MB468 | MP |
| >hg38_21_+_8433172_8446622_RNA45SN1_45S_50ntFlanks@11577.11613.37\|TTAGAACTGGTGCGGACCAGGGGAATCCGACTGTTTA | >hg38_21_+_8433172_8446622_RNA45SN1_45S_50ntFlanks | 11577 | 11613 | 37 | TTAGAACTGGTGCGGACCAGGGGAATCCGACTGTTTA | 833.2 | 3.9 | 0.9 | 4.3 | 1.7E-05 | 3.2E-05 | BT20/MB468 | MITO |
| >MI0000070\|hsa-mir-16-1&WithFlank&13\|-\|50048967\|50049067@20.41.22(+1C);MI0000115\|hsa-mir-16-2&WithFlank&3\|+\|160404739\|160404831@16.37.22(+1C)&[MIMAT0000069&hsa-miR-16-5p&offsets\|0\|0(+1C);m-62&13\|-\|50049027\|50049048&offsets\|0\|0(+1C)];[MIMAT0000069_1&hsa-miR-16-5p&offsets\|0\|0(+1C);m-61&3\|+\|160404754\|160404775&offsets\|0\|0(+1C)]\|TAGCAGCACGTAAATATTGGCGC | >MI0000070\|hsa-mir-16-1&WithFlank&13\|-\|50048967\|50049067 | 20 | 41 | 22(+1C) | TAGCAGCACGTAAATATTGGCGC | 158.7 | 4.6 | 1.1 | 4.2 | 2.4E-05 | 3.2E-05 | BT20/MB468 | NUC |
| >hg38_21_+_8433172_8446622_RNA45SN1_45S_50ntFlanks@9554.9575.22\|CGTAGCGGTCCTGACGTGCAAA | >hg38_21_+_8433172_8446622_RNA45SN1_45S_50ntFlanks | 9554 | 9575 | 22 | CGTAGCGGTCCTGACGTGCAAA | 589.6 | -2.4 | 0.6 | -4.3 | 1.7E-05 | 3.2E-05 | BT20/MB468 | MITO |
| >am_MI0015825\|hsa-mir-378c&WithFlank&10\|-\|130962582\|130962674@17.34.18;MI0016749\|hsa-mir-378d-1&WithFlank&4\|-\|5923269\|5923334@37.54.18;MI0000786\|hsa-mir-378a&WithFlank&5\|+\|149732819\|149732896@49.66.18;MI0003840\|hsa-mir-378d-2&WithFlank&8\|-\|93916016\|93916125@28.45.18&[MIMAT0016847&hsa-miR-378c&offsets\|0\|-7;m-130&10\|-\|130962639\|130962658&offsets\|0\|-2];[MIMAT0018926&hsa-miR-378d&offsets\|0\|-2;m-228&4\|-\|5923281\|5923298&offsets\|0\|0];[MIMAT0000732&hsa-miR-378a-3p&offsets\|0\|-4;m-44&5\|+\|149732867\|149732888&offsets\|0\|-4];[MIMAT0018926_1&hsa-miR-378d&offsets\|0\|-2;m-227&8\|-\|93916081\|93916098&offsets\|0\|0]\|ACTGGACTTGGAGTCAGA | >am_MI0015825\|hsa-mir-378c&WithFlank&10\|-\|130962582\|130962674 | 17 | 34 | 18 | ACTGGACTTGGAGTCAGA | 228.8 | 3.7 | 0.9 | 4.2 | 2.3E-05 | 3.2E-05 | MB231/MB468 | NUC |
| >hg38_21_+_8433172_8446622_RNA45SN1_45S_50ntFlanks@5471.5501.31\|CACGGCCCTGGCGGAGCGCTGAGAAGACGGT | >hg38_21_+_8433172_8446622_RNA45SN1_45S_50ntFlanks | 5471 | 5501 | 31 | CACGGCCCTGGCGGAGCGCTGAGAAGACGGT | 113.4 | 10.2 | 2.2 | 4.7 | 2.3E-06 | 3.2E-05 | BT20/MB231 | MP |
| >hg38_21_+_8433172_8446622_RNA45SN1_45S_50ntFlanks@11205.11225.21\|GCGGCGGCGACTCTGGACGCG | >hg38_21_+_8433172_8446622_RNA45SN1_45S_50ntFlanks | 11205 | 11225 | 21 | GCGGCGGCGACTCTGGACGCG | 542.1 | -4.7 | 1.1 | -4.3 | 2.0E-05 | 3.2E-05 | BT20/MB468 | CYTO |
| >hg38_21_+_8433172_8446622_RNA45SN1_45S_50ntFlanks@9565.9594.30\|TGACGTGCAAATCGGTCGTCCGACCTGGGT | >hg38_21_+_8433172_8446622_RNA45SN1_45S_50ntFlanks | 9565 | 9594 | 30 | TGACGTGCAAATCGGTCGTCCGACCTGGGT | 393.9 | -2.5 | 0.6 | -4.2 | 2.3E-05 | 3.2E-05 | MB231/MB468 | NUC |
| >MI0000078\|hsa-mir-22&WithFlank&17\|-\|1713897\|1713993@59.79.21(+1A)&[MIMAT0000077&hsa-miR-22-3p&offsets\|0\|-1(+1A);m-2&17\|-\|1713914\|1713935&offsets\|0\|-1(+1A)]\|AAGCTGCCAGTTGAAGAACTGA | >MI0000078\|hsa-mir-22&WithFlank&17\|-\|1713897\|1713993 | 59 | 79 | 21(+1A) | AAGCTGCCAGTTGAAGAACTGA | 320.3 | 6.5 | 1.5 | 4.2 | 2.3E-05 | 3.2E-05 | MB231/MB468 | MITO |
| >hg38_21_+_8433172_8446622_RNA45SN1_45S_50ntFlanks@9202.9223.22\|TCTCGGGGCCACGCGCGCGTCC | >hg38_21_+_8433172_8446622_RNA45SN1_45S_50ntFlanks | 9202 | 9223 | 22 | TCTCGGGGCCACGCGCGCGTCC | 317.0 | -5.5 | 1.2 | -4.5 | 7.5E-06 | 3.2E-05 | BT20/MB468 | MP |
| >hg38_21_+_8433172_8446622_RNA45SN1_45S_50ntFlanks@10786.10828.43\|AACAGCCTCTGGCATGTTGGAACAATGTAGGTAAGGGAAGTCG | >hg38_21_+_8433172_8446622_RNA45SN1_45S_50ntFlanks | 10786 | 10828 | 43 | AACAGCCTCTGGCATGTTGGAACAATGTAGGTAAGGGAAGTCG | 262.3 | -6.4 | 1.5 | -4.2 | 2.3E-05 | 3.2E-05 | MB231/MB468 | NUC |
| >hg38_21_+_8433172_8446622_RNA45SN1_45S_50ntFlanks@11567.11605.39\|AGCAGCCGACTTAGAACTGGTGCGGACCAGGGGAATCCG | >hg38_21_+_8433172_8446622_RNA45SN1_45S_50ntFlanks | 11567 | 11605 | 39 | AGCAGCCGACTTAGAACTGGTGCGGACCAGGGGAATCCG | 218.2 | -4.6 | 1.1 | -4.2 | 2.3E-05 | 3.2E-05 | MB231/MB468 | NUC |
| >hg38_21_+_8433172_8446622_RNA45SN1_45S_50ntFlanks@11807.11828.22\|ACTGTCCCTACCTACTATCCAG | >hg38_21_+_8433172_8446622_RNA45SN1_45S_50ntFlanks | 11807 | 11828 | 22 | ACTGTCCCTACCTACTATCCAG | 49.6 | -8.7 | 1.9 | -4.5 | 7.5E-06 | 3.2E-05 | BT20/MB468 | MP |
| >trnaMT_PheGAA_MT_+_577_647@35.56.22\|GAAAATGTTTAGACGGGCTCAC | >trnaMT_PheGAA_MT_+_577_647 | 35 | 56 | 22 | GAAAATGTTTAGACGGGCTCAC | 1398.9 | 5.4 | 1.2 | 4.5 | 7.5E-06 | 3.2E-05 | BT20/MB468 | MP |
| >hg38_21_+_8433172_8446622_RNA45SN1_45S_50ntFlanks@10314.10334.21\|CAGGTGCAGATCTTGGTGGTA | >hg38_21_+_8433172_8446622_RNA45SN1_45S_50ntFlanks | 10314 | 10334 | 21 | CAGGTGCAGATCTTGGTGGTA | 109.3 | 10.1 | 2.1 | 4.7 | 2.3E-06 | 3.2E-05 | BT20/MB231 | MP |
| >hg38_21_+_8433172_8446622_RNA45SN1_45S_50ntFlanks@10410.10453.44\|TCAGTCGGTCCTGAGAGATGGGCGAGCGCCGTTCCGAAGGGACG | >hg38_21_+_8433172_8446622_RNA45SN1_45S_50ntFlanks | 10410 | 10453 | 44 | TCAGTCGGTCCTGAGAGATGGGCGAGCGCCGTTCCGAAGGGACG | 792.7 | -4.1 | 1.0 | -4.2 | 2.3E-05 | 3.2E-05 | MB231/MB468 | NUC |
| >hg38_21_+_8433172_8446622_RNA45SN1_45S_50ntFlanks@6659.6685.27\|AGCGGTGGATCACTCGGCTCGTGCGTC | >hg38_21_+_8433172_8446622_RNA45SN1_45S_50ntFlanks | 6659 | 6685 | 27 | AGCGGTGGATCACTCGGCTCGTGCGTC | 313.3 | 3.9 | 0.9 | 4.5 | 7.6E-06 | 3.2E-05 | BT20/MB468 | MP |
| >MI0000264\|hsa-mir-7-2&WithFlank&15\|+\|88611819\|88611940@38.60.23(+1C);MI0000265\|hsa-mir-7-3&WithFlank&19\|+\|4770664\|4770785@37.59.23(+1C);MI0000263\|hsa-mir-7-1&WithFlank&9\|-\|83969742\|83969863@30.52.23(+1C)&[MIMAT0000252&hsa-miR-7-5p&offsets\|0\|-1(+1C);m-400&15\|+\|88611856\|88611879&offsets\|0\|-1(+1C)];[MIMAT0000252_1&hsa-miR-7-5p&offsets\|0\|-1(+1C);m-402&19\|+\|4770700\|4770723&offsets\|0\|-1(+1C)];[MIMAT0000252_2&hsa-miR-7-5p&offsets\|0\|-1(+1C);m-401&9\|-\|83969811\|83969834&offsets\|0\|-1(+1C)]\|TGGAAGACTAGTGATTTTGTTGTC | >MI0000264\|hsa-mir-7-2&WithFlank&15\|+\|88611819\|88611940 | 38 | 60 | 23(+1C) | TGGAAGACTAGTGATTTTGTTGTC | 5271.7 | 3.6 | 0.8 | 4.3 | 1.9E-05 | 3.2E-05 | MB231/MB468 | MP |
| >hg38_21_+_8433172_8446622_RNA45SN1_45S_50ntFlanks@10677.10707.31\|GCGGTTCCGGCGGCGTCCGGTGAGCTCTCGC | >hg38_21_+_8433172_8446622_RNA45SN1_45S_50ntFlanks | 10677 | 10707 | 31 | GCGGTTCCGGCGGCGTCCGGTGAGCTCTCGC | 378.7 | 6.3 | 1.4 | 4.5 | 7.6E-06 | 3.2E-05 | BT20/MB468 | MP |
| >hg38_21_+_8433172_8446622_RNA45SN1_45S_50ntFlanks@11710.11736.27\|CGCGGGTAAACGGCGGGAGTAACTATG | >hg38_21_+_8433172_8446622_RNA45SN1_45S_50ntFlanks | 11710 | 11736 | 27 | CGCGGGTAAACGGCGGGAGTAACTATG | 367.7 | -3.9 | 0.9 | -4.3 | 1.9E-05 | 3.3E-05 | MB231/MB468 | MP |

| | | | | | | | | | | | | | |
|---|---|---|---|---|---|---|---|---|---|---|---|---|---|
| >hg38_21_+_8433172_8446622_RNA45SN1_45S_50ntFlanks@12077.12095.19\|GGGGCTCTCGCTTCTGGCG | >hg38_21_+_8433172_8446622_RNA45SN1_45S_50ntFlanks | 12077 | 12095 | 19 | GGGGCTCTCGCTTCTGGCG | 1456.0 | -7.7 | 1.8 | -4.2 | 2.4E-05 | 3.3E-05 | BT20/MB468 | NUC |
| >hg38_21_+_8433172_8446622_RNA45SN1_45S_50ntFlanks@6646.6672.27\|TCGTACGACTCTTAGCGGTGGATCACT | >hg38_21_+_8433172_8446622_RNA45SN1_45S_50ntFlanks | 6646 | 6672 | 27 | TCGTACGACTCTTAGCGGTGGATCACT | 1917.3 | 3.8 | 0.9 | 4.5 | 7.6E-06 | 3.3E-05 | BT20/MB468 | MP |
| >hg38_21_+_8433172_8446622_RNA45SN1_45S_50ntFlanks@6651.6699.49\|CGACTCTTAGCGGTGGATCACTCGGCTCGTGCGTCGATGAAGAACGCAG | >hg38_21_+_8433172_8446622_RNA45SN1_45S_50ntFlanks | 6651 | 6699 | 49 | CGACTCTTAGCGGTGGATCACTCGGCTCGTGCGTCGATGAAGAACGCAG | 85950.7 | -3.8 | 0.9 | -4.2 | 2.3E-05 | 3.3E-05 | MB231/MB468 | MITO |
| >hg38_1_-_228634819_228635039_RNA5S12_5S_50ntFlanks@147.171.25\|GGGAATACCGGGTGCTGTAGGCTTT | >hg38_1_-_228634819_228635039_RNA5S12_5S_50ntFlanks | 147 | 171 | 25 | GGGAATACCGGGTGCTGTAGGCTTT | 1384.7 | 3.1 | 0.7 | 4.2 | 2.3E-05 | 3.3E-05 | MB231/MB468 | NUC |
| >hg38_21_+_8433172_8446622_RNA45SN1_45S_50ntFlanks@5417.5443.27\|CTACCGATTGGATGGTTTAGTGAGGCC | >hg38_21_+_8433172_8446622_RNA45SN1_45S_50ntFlanks | 5417 | 5443 | 27 | CTACCGATTGGATGGTTTAGTGAGGCC | 233.0 | -6.4 | 1.4 | -4.5 | 7.6E-06 | 3.3E-05 | BT20/MB468 | MP |
| >hg38_21_+_8433172_8446622_RNA45SN1_45S_50ntFlanks@11561.11592.32\|GCGCCTAGCAGCCGACTTAGAACTGGTGCGGA | >hg38_21_+_8433172_8446622_RNA45SN1_45S_50ntFlanks | 11561 | 11592 | 32 | GCGCCTAGCAGCCGACTTAGAACTGGTGCGGA | 449.5 | -6.2 | 1.5 | -4.2 | 2.3E-05 | 3.3E-05 | MB231/MB468 | NUC |
| >hg38_21_+_8433172_8446622_RNA45SN1_45S_50ntFlanks@3844.3873.30\|TACTTGGATAACTGTGGTAATTCTAGAGCT | >hg38_21_+_8433172_8446622_RNA45SN1_45S_50ntFlanks | 3844 | 3873 | 30 | TACTTGGATAACTGTGGTAATTCTAGAGCT | 356.1 | 3.5 | 0.8 | 4.3 | 2.0E-05 | 3.3E-05 | BT20/MB468 | CYTO |
| >hg38_21_+_8433172_8446622_RNA45SN1_45S_50ntFlanks@11573.11592.20\|CGACTTAGAACTGGTGCGGA | >hg38_21_+_8433172_8446622_RNA45SN1_45S_50ntFlanks | 11573 | 11592 | 20 | CGACTTAGAACTGGTGCGGA | 403.4 | -4.7 | 1.1 | -4.2 | 2.3E-05 | 3.3E-05 | MB231/MB468 | NUC |
| >hg38_21_+_8433172_8446622_RNA45SN1_45S_50ntFlanks@9533.9561.29\|GAGGAAACTCTGGTGGAGGTCCGTAGCGG | >hg38_21_+_8433172_8446622_RNA45SN1_45S_50ntFlanks | 9533 | 9561 | 29 | GAGGAAACTCTGGTGGAGGTCCGTAGCGG | 2047.6 | -8.2 | 1.9 | -4.2 | 2.4E-05 | 3.3E-05 | BT20/MB468 | NUC |
| >MI0001448\|hsa-mir-425&WithFlank&3\|-\|49020142\|49020240@21.44.24&[MIMAT0003393&hsa-miR-425-5p&offsets\|+1\|+2;m-95&3\|-\|49020200\|49020221&offsets\|+1\|+3]\|ATGACACGATCACTCCCGTTGAGT | >MI0001448\|hsa-mir-425&WithFlank&3\|-\|49020142\|49020240 | 21 | 44 | 24 | ATGACACGATCACTCCCGTTGAGT | 255.0 | 6.4 | 1.5 | 4.2 | 2.3E-05 | 3.3E-05 | MB231/MB468 | MITO |
| >hg38_21_+_8433172_8446622_RNA45SN1_45S_50ntFlanks@5073.5110.38\|AATTCCGATAACGAACGAGACTCTGGCATGCTAACTAG | >hg38_21_+_8433172_8446622_RNA45SN1_45S_50ntFlanks | 5073 | 5110 | 38 | AATTCCGATAACGAACGAGACTCTGGCATGCTAACTAG | 636.5 | -4.9 | 1.1 | -4.3 | 1.7E-05 | 3.3E-05 | BT20/MB468 | MITO |
| >am_trnaMT_GluTTC_MT_-_14674_14742@27.48.22,trnalookalike8_GluTTC_5_-_93905172_93905240@27.48.22\|GTTTTTCATATCATTGGTCGTG | >am_trnaMT_GluTTC_MT_-_14674_14742 | 27 | 48 | 22 | GTTTTTCATATCATTGGTCGTG | 137.1 | -5.4 | 1.3 | -4.3 | 2.0E-05 | 3.3E-05 | BT20/MB468 | CYTO |
| >hg38_21_+_8433172_8446622_RNA45SN1_45S_50ntFlanks@6722.6740.19\|ATTGCAGGACACATTGATC | >hg38_21_+_8433172_8446622_RNA45SN1_45S_50ntFlanks | 6722 | 6740 | 19 | ATTGCAGGACACATTGATC | 894.6 | -4.7 | 1.1 | -4.2 | 2.2E-05 | 3.3E-05 | MB231/MB468 | CYTO |
| >hg38_21_+_8433172_8446622_RNA45SN1_45S_50ntFlanks@9621.9658.38\|TCTAGTAGCTGGTTCCCTCCGAAGTTTCCCTCAGGATA | >hg38_21_+_8433172_8446622_RNA45SN1_45S_50ntFlanks | 9621 | 9658 | 38 | TCTAGTAGCTGGTTCCCTCCGAAGTTTCCCTCAGGATA | 499.6 | -2.8 | 0.6 | -4.3 | 1.7E-05 | 3.3E-05 | BT20/MB468 | MITO |
| >hg38_21_+_8433172_8446622_RNA45SN1_45S_50ntFlanks@11785.11802.18\|TGAATGGATGAACGAGAT | >hg38_21_+_8433172_8446622_RNA45SN1_45S_50ntFlanks | 11785 | 11802 | 18 | TGAATGGATGAACGAGAT | 201.6 | -4.6 | 1.1 | -4.2 | 2.3E-05 | 3.3E-05 | MB231/MB468 | NUC |
| >hg38_21_+_8433172_8446622_RNA45SN1_45S_50ntFlanks@12449.12486.38\|GAATTCACCAAGCGTTGGATTGTTCACCCACTAATAGG | >hg38_21_+_8433172_8446622_RNA45SN1_45S_50ntFlanks | 12449 | 12486 | 38 | GAATTCACCAAGCGTTGGATTGTTCACCCACTAATAGG | 149.9 | 4.8 | 1.1 | 4.2 | 2.3E-05 | 3.3E-05 | MB231/MB468 | NUC |
| >hg38_MT_+_1621_3279_RNR2_16S_50ntFlanks@622.663.42\|TATAACTGAACTCCTCACACCCAATTGGACCAATCTATCACC | >hg38_MT_+_1621_3279_RNR2_16S_50ntFlanks | 622 | 663 | 42 | TATAACTGAACTCCTCACACCCAATTGGACCAATCTATCACC | 219.2 | 11.5 | 2.6 | 4.5 | 7.6E-06 | 3.3E-05 | BT20/MB468 | MP |
| >hg38_21_+_8433172_8446622_RNA45SN1_45S_50ntFlanks@5423.5466.44\|ATTGGATGGTTTAGTGAGGCCCTCGGATCGGCCCCGCCGGGGTC | >hg38_21_+_8433172_8446622_RNA45SN1_45S_50ntFlanks | 5423 | 5466 | 44 | ATTGGATGGTTTAGTGAGGCCCTCGGATCGGCCCCGCCGGGGTC | 988.7 | -4.2 | 1.0 | -4.3 | 2.0E-05 | 3.3E-05 | BT20/MB468 | CYTO |
| >hg38_21_+_8433172_8446622_RNA45SN1_45S_50ntFlanks@4588.4613.26\|CTATTTTGTTGGTTTTCGGAACTGAG | >hg38_21_+_8433172_8446622_RNA45SN1_45S_50ntFlanks | 4588 | 4613 | 26 | CTATTTTGTTGGTTTTCGGAACTGAG | 220.6 | -5.0 | 1.2 | -4.2 | 2.2E-05 | 3.3E-05 | MB231/MB468 | CYTO |
| >hg38_21_+_8433172_8446622_RNA45SN1_45S_50ntFlanks@4243.4262.20\|CTTTCGAGGCCCTGTAATTG | >hg38_21_+_8433172_8446622_RNA45SN1_45S_50ntFlanks | 4243 | 4262 | 20 | CTTTCGAGGCCCTGTAATTG | 486.9 | -6.9 | 1.6 | -4.2 | 2.4E-05 | 3.3E-05 | BT20/MB468 | NUC |
| >hg38_21_+_8433172_8446622_RNA45SN1_45S_50ntFlanks@11586.11604.19\|GTGCGGACCAGGGGAATCC | >hg38_21_+_8433172_8446622_RNA45SN1_45S_50ntFlanks | 11586 | 11604 | 19 | GTGCGGACCAGGGGAATCC | 130.8 | -6.4 | 1.5 | -4.3 | 1.9E-05 | 3.3E-05 | MB231/MB468 | MP |
| >MI0000255\|hsa-mir-30d&WithFlank&8\|-\|134804870\|134804951@12.32.21(+1A)&[MIMAT0000245&hsa-miR-30d-5p&offsets\|0\|-1(+1A);m-21&8\|-\|134804919\|134804940&offsets\|0\|-1(+1A)]\|TGTAAACATCCCCGACTGGAAA | >MI0000255\|hsa-mir-30d&WithFlank&8\|-\|134804870\|134804951 | 12 | 32 | 21(+1A) | TGTAAACATCCCCGACTGGAAA | 887.5 | 4.2 | 0.9 | 4.7 | 2.3E-06 | 3.3E-05 | BT20/MB231 | CYTO |
| >hg38_21_+_8433172_8446622_RNA45SN1_45S_50ntFlanks@3803.3829.27\|ATTAAATCAGTTATGGTTCCTTTGGTC | >hg38_21_+_8433172_8446622_RNA45SN1_45S_50ntFlanks | 3803 | 3829 | 27 | ATTAAATCAGTTATGGTTCCTTTGGTC | 176.1 | -4.5 | 1.1 | -4.2 | 2.3E-05 | 3.3E-05 | MB231/MB468 | NUC |
| >hg38_21_+_8433172_8446622_RNA45SN1_45S_50ntFlanks@11821.11850.30\|CTATCCAGCGAAACCACAGCCAAGGGAACG | >hg38_21_+_8433172_8446622_RNA45SN1_45S_50ntFlanks | 11821 | 11850 | 30 | CTATCCAGCGAAACCACAGCCAAGGGAACG | 46.4 | -8.6 | 1.9 | -4.5 | 7.7E-06 | 3.3E-05 | BT20/MB468 | MP |
| >hg38_21_+_8433172_8446622_RNA45SN1_45S_50ntFlanks@9629.9648.20\|CTGGTTCCCTCCGAAGTTTC | >hg38_21_+_8433172_8446622_RNA45SN1_45S_50ntFlanks | 9629 | 9648 | 20 | CTGGTTCCCTCCGAAGTTTC | 79.1 | -9.4 | 2.1 | -4.5 | 7.7E-06 | 3.3E-05 | BT20/MB468 | MP |
| >hg38_21_+_8433172_8446622_RNA45SN1_45S_50ntFlanks@6775.6803.29\|CCTCCCGGGGCTACGCCTGTCTGAGCGTC | >hg38_21_+_8433172_8446622_RNA45SN1_45S_50ntFlanks | 6775 | 6803 | 29 | CCTCCCGGGGCTACGCCTGTCTGAGCGTC | 195.3 | -2.2 | 0.5 | -4.2 | 2.3E-05 | 3.3E-05 | MB231/MB468 | NUC |
| >hg38_21_+_8433172_8446622_RNA45SN1_45S_50ntFlanks@12941.12958.18\|ATTCGTAGACGACCTGCT | >hg38_21_+_8433172_8446622_RNA45SN1_45S_50ntFlanks | 12941 | 12958 | 18 | ATTCGTAGACGACCTGCT | 459.7 | 2.0 | 0.5 | 4.3 | 2.0E-05 | 3.3E-05 | BT20/MB468 | CYTO |
| >MI0000077\|hsa-mir-21&WithFlank&17\|+\|59841260\|59841343@14.35.22(+1U)&[MIMAT0000076&hsa-miR-21-5p&offsets\|0\|0(+1U);m-1&17\|+\|59841273\|59841294&offsets\|0\|0(+1U)]\|TAGCTTATCAGACTGATGTTGAT | >MI0000077\|hsa-mir-21&WithFlank&17\|+\|59841260\|59841343 | 14 | 35 | 22(+1U) | TAGCTTATCAGACTGATGTTGAT | 72514.8 | 4.0 | 0.9 | 4.3 | 2.0E-05 | 3.3E-05 | MB231/MB468 | MP |
| >hg38_21_+_8433172_8446622_RNA45SN1_45S_50ntFlanks@11919.11957.39\|CATGAGAGGTGTAGAATAAGTGGGAGGCCCCCGGCGCCC | >hg38_21_+_8433172_8446622_RNA45SN1_45S_50ntFlanks | 11919 | 11957 | 39 | CATGAGAGGTGTAGAATAAGTGGGAGGCCCCCGGCGCCC | 68.0 | -10.0 | 2.3 | -4.3 | 2.0E-05 | 3.3E-05 | MB231/MB468 | MP |
| >hg38_21_+_8433172_8446622_RNA45SN1_45S_50ntFlanks@6721.6747.27\|AATTGCAGGACACATTGATCATCGACA | >hg38_21_+_8433172_8446622_RNA45SN1_45S_50ntFlanks | 6721 | 6747 | 27 | AATTGCAGGACACATTGATCATCGACA | 2115.7 | -3.4 | 0.8 | -4.2 | 2.2E-05 | 3.3E-05 | MB231/MB468 | CYTO |
| >hg38_1_-_228634819_228635039_RNA5S12_5S_50ntFlanks@130.169.40\|TGGATGGGAGACCGCCTGGGAATACCGGGTGCTGTAGGCT | >hg38_1_-_228634819_228635039_RNA5S12_5S_50ntFlanks | 130 | 169 | 40 | TGGATGGGAGACCGCCTGGGAATACCGGGTGCTGTAGGCT | 505.5 | 3.8 | 0.8 | 4.7 | 2.8E-06 | 3.3E-05 | BT20/MB231 | MITO |
| >hg38_1_-_228634819_228635039_RNA5S12_5S_50ntFlanks@147.169.23\|GGGAATACCGGGTGCTGTAGGCT | >hg38_1_-_228634819_228635039_RNA5S12_5S_50ntFlanks | 147 | 169 | 23 | GGGAATACCGGGTGCTGTAGGCT | 675.5 | 3.7 | 0.8 | 4.7 | 2.8E-06 | 3.3E-05 | BT20/MB231 | MITO |
| >hg38_21_+_8433172_8446622_RNA45SN1_45S_50ntFlanks@10399.10424.26\|TTGAACATGGGTCAGTCGGTCCTGAG | >hg38_21_+_8433172_8446622_RNA45SN1_45S_50ntFlanks | 10399 | 10424 | 26 | TTGAACATGGGTCAGTCGGTCCTGAG | 2616.1 | 3.8 | 0.8 | 4.7 | 2.9E-06 | 3.3E-05 | BT20/MB231 | MITO |
| >MI0000289\|hsa-mir-181a-1&WithFlank&1\|-\|198859038\|198859159@30.53.24(+1C);MI0000269\|hsa-mir-181a-2&WithFlank&9\|+\|124692436\|124692557@45.68.24(+1C)&[MIMAT0000256&hsa-miR-181a-5p&offsets\|0\|+1(+1C);m-32&1\|-\|198859108\|198859130&offsets\|0\|+1(+1C)];[MIMAT0000256_1&hsa-miR-181a-5p&offsets\|0\|+1(+1C);m-32&9\|+\|124692480\|124692502&offsets\|0\|+1(+1C)]\|AACATTCAACGCTGTCGGTGAGTTC | >MI0000289\|hsa-mir-181a-1&WithFlank&1\|-\|198859038\|198859159 | 30 | 53 | 24(+1C) | AACATTCAACGCTGTCGGTGAGTTC | 510.8 | -5.4 | 1.2 | -4.7 | 2.9E-06 | 3.3E-05 | BT20/MB231 | MITO |
| >hg38_21_+_8433172_8446622_RNA45SN1_45S_50ntFlanks@3773.3792.20\|CGGCCGGTACAGTGAAACTG | >hg38_21_+_8433172_8446622_RNA45SN1_45S_50ntFlanks | 3773 | 3792 | 20 | CGGCCGGTACAGTGAAACTG | 827.3 | -7.0 | 1.6 | -4.3 | 1.7E-05 | 3.3E-05 | BT20/MB468 | MITO |
| >hg38_1_-_228634819_228635039_RNA5S12_5S_50ntFlanks@89.123.35\|CTGATCTCGGAAGCTAAGCAGGGTCGGGCCTGGTT | >hg38_1_-_228634819_228635039_RNA5S12_5S_50ntFlanks | 89 | 123 | 35 | CTGATCTCGGAAGCTAAGCAGGGTCGGGCCTGGTT | 236.3 | 4.1 | 1.0 | 4.2 | 2.3E-05 | 3.3E-05 | MB231/MB468 | NUC |
| >hg38_21_+_8433172_8446622_RNA45SN1_45S_50ntFlanks@6780.6799.20\|CGGGGCTACGCCTGTCTGAG | >hg38_21_+_8433172_8446622_RNA45SN1_45S_50ntFlanks | 6780 | 6799 | 20 | CGGGGCTACGCCTGTCTGAG | 159.4 | -6.5 | 1.5 | -4.5 | 7.7E-06 | 3.3E-05 | BT20/MB468 | MP |
| >hg38_21_+_8433172_8446622_RNA45SN1_45S_50ntFlanks@8380.8404.25\|CAAGAGGGCGTGAAACCGTTAAGAG | >hg38_21_+_8433172_8446622_RNA45SN1_45S_50ntFlanks | 8380 | 8404 | 25 | CAAGAGGGCGTGAAACCGTTAAGAG | 645.5 | -6.8 | 1.5 | -4.5 | 7.7E-06 | 3.3E-05 | BT20/MB468 | MP |
| >hg38_21_+_8433172_8446622_RNA45SN1_45S_50ntFlanks@11190.11225.36\|CCCGGGGGGCCGGCGGCGGCGGCGACTCTGGACGCG | >hg38_21_+_8433172_8446622_RNA45SN1_45S_50ntFlanks | 11190 | 11225 | 36 | CCCGGGGGGCCGGCGGCGGCGGCGACTCTGGACGCG | 2447.0 | -8.3 | 2.0 | -4.2 | 2.5E-05 | 3.3E-05 | BT20/MB468 | NUC |
| >hg38_21_+_8433172_8446622_RNA45SN1_45S_50ntFlanks@11593.11612.20\|CCAGGGGAATCCGACTGTTT | >hg38_21_+_8433172_8446622_RNA45SN1_45S_50ntFlanks | 11593 | 11612 | 20 | CCAGGGGAATCCGACTGTTT | 1093.7 | -5.7 | 1.3 | -4.5 | 7.7E-06 | 3.3E-05 | BT20/MB468 | MP |
| >MI0001733\|hsa-mir-452&WithFlank&X\|-\|151959622\|151959718@20.42.23&[MIMAT0001635&hsa-miR-452-5p&offsets\|0\|+1;m-188&X\|-\|151959677\|151959699&offsets\|0\|0]\|AACTGTTTGCAGAGGAAACTGAG | >MI0001733\|hsa-mir-452&WithFlank&X\|-\|151959622\|151959718 | 20 | 42 | 23 | AACTGTTTGCAGAGGAAACTGAG | 104.6 | 6.9 | 1.6 | 4.3 | 2.0E-05 | 3.3E-05 | MB231/MB468 | MP |
| >MI0000077\|hsa-mir-21&WithFlank&17\|+\|59841260\|59841343@14.34.21(+3C)&[MIMAT0000076&hsa-miR-21-5p&offsets\|0\|-1(+3C);m-1&17\|+\|59841273\|59841294&offsets\|0\|-1(+3C)]\|TAGCTTATCAGACTGATGTTGCCC | >MI0000077\|hsa-mir-21&WithFlank&17\|+\|59841260\|59841343 | 14 | 34 | 21(+3C) | TAGCTTATCAGACTGATGTTGCCC | 76.5 | -9.7 | 2.0 | -4.7 | 2.4E-06 | 3.3E-05 | BT20/MB231 | MP |
| >hg38_21_+_8433172_8446622_RNA45SN1_45S_50ntFlanks@8194.8212.19\|CCCGTGGACGGTGTGAGGC | >hg38_21_+_8433172_8446622_RNA45SN1_45S_50ntFlanks | 8194 | 8212 | 19 | CCCGTGGACGGTGTGAGGC | 368.7 | -2.8 | 0.7 | -4.3 | 1.7E-05 | 3.3E-05 | BT20/MB468 | MITO |
| >hg38_21_+_8433172_8446622_RNA45SN1_45S_50ntFlanks@9504.9532.29\|GTGAACTATGCCTGGGCAGGGCGAAGCCA | >hg38_21_+_8433172_8446622_RNA45SN1_45S_50ntFlanks | 9504 | 9532 | 29 | GTGAACTATGCCTGGGCAGGGCGAAGCCA | 364.2 | -7.1 | 1.6 | -4.5 | 7.7E-06 | 3.3E-05 | BT20/MB468 | MP |
| >hg38_21_+_8433172_8446622_RNA45SN1_45S_50ntFlanks@6771.6801.31\|GGTTCCTCCCGGGGCTACGCCTGTCTGAGCG | >hg38_21_+_8433172_8446622_RNA45SN1_45S_50ntFlanks | 6771 | 6801 | 31 | GGTTCCTCCCGGGGCTACGCCTGTCTGAGCG | 86.5 | -6.0 | 1.4 | -4.3 | 2.0E-05 | 3.3E-05 | MB231/MB468 | MP |
| >hg38_21_+_8433172_8446622_RNA45SN1_45S_50ntFlanks@12938.12957.20\|ACCATTCGTAGACGACCTGC | >hg38_21_+_8433172_8446622_RNA45SN1_45S_50ntFlanks | 12938 | 12957 | 20 | ACCATTCGTAGACGACCTGC | 495.2 | 3.2 | 0.8 | 4.2 | 2.2E-05 | 3.3E-05 | MB231/MB468 | CYTO |
| >hg38_21_+_8433172_8446622_RNA45SN1_45S_50ntFlanks@3985.4024.40\|CGGCTTTGGTGACTCTAGATAACCTCGGGCCGATCGCACG | >hg38_21_+_8433172_8446622_RNA45SN1_45S_50ntFlanks | 3985 | 4024 | 40 | CGGCTTTGGTGACTCTAGATAACCTCGGGCCGATCGCACG | 540.9 | -7.1 | 1.7 | -4.2 | 2.5E-05 | 3.3E-05 | BT20/MB468 | NUC |
| >hg38_21_+_8433172_8446622_RNA45SN1_45S_50ntFlanks@12459.12486.28\|AGCGTTGGATTGTTCACCCACTAATAGG | >hg38_21_+_8433172_8446622_RNA45SN1_45S_50ntFlanks | 12459 | 12486 | 28 | AGCGTTGGATTGTTCACCCACTAATAGG | 130.0 | -4.7 | 1.1 | -4.2 | 2.3E-05 | 3.3E-05 | MB231/MB468 | NUC |
| >hg38_21_+_8433172_8446622_RNA45SN1_45S_50ntFlanks@3940.3959.20\|ACCCGGTCAGCCCCTCTCCG | >hg38_21_+_8433172_8446622_RNA45SN1_45S_50ntFlanks | 3940 | 3959 | 20 | ACCCGGTCAGCCCCTCTCCG | 468.4 | -7.1 | 1.7 | -4.2 | 2.5E-05 | 3.3E-05 | BT20/MB468 | NUC |
| >MI0000816\|hsa-mir-335&WithFlank&7\|+\|130496105\|130496210@58.79.22&[MIMAT0004703&hsa-miR-335-3p&offsets\|0\|0;m-159&7\|+\|130496162\|130496183&offsets\|0\|0]\|TTTTTCATTATTGCTCCTGACC | >MI0000816\|hsa-mir-335&WithFlank&7\|+\|130496105\|130496210 | 58 | 79 | 22 | TTTTTCATTATTGCTCCTGACC | 605.7 | 3.3 | 0.8 | 4.3 | 2.0E-05 | 3.3E-05 | MB231/MB468 | MP |
| >hg38_21_+_8433172_8446622_RNA45SN1_45S_50ntFlanks@6739.6770.32\|TCATCGACACTTCGAACGCACTTGCGGCCCCG | >hg38_21_+_8433172_8446622_RNA45SN1_45S_50ntFlanks | 6739 | 6770 | 32 | TCATCGACACTTCGAACGCACTTGCGGCCCCG | 469.6 | -6.9 | 1.5 | -4.5 | 7.7E-06 | 3.3E-05 | BT20/MB468 | MP |

<table>
<tr><td>>MI0000273|hsa-mir-183&WithFlank&7|-|129774899|129775020@33.53.21(+1C)&[MIMAT0000261&hsa-miR-183-5p&offsets|0|-1(+1C);m-41&7|-|129774966|129774987&offsets|-1|-2(+1C)]|TATGGCACTGGTAGAATTCACC</td><td>>MI0000273|hsa-mir-183&WithFlank&7|-|129774899|129775020</td><td>33</td><td>53</td><td>21(+1C)</td><td>TATGGCACTGGTAGAATTCACC</td><td>1736.8</td><td>2.4</td><td>0.5</td><td>4.7</td><td>2.9E-06</td><td>3.3E-05</td><td>BT20/MB231</td><td>NUC</td></tr>
<tr><td>>hg38_21_+_8433172_8446622_RNA45SN1_45S_50ntFlanks@11672.11706.35|CAGTGCTCTGAATGTCAAAGTGAAGAAATTCAATG</td><td>>hg38_21_+_8433172_8446622_RNA45SN1_45S_50ntFlanks</td><td>11672</td><td>11706</td><td>35</td><td>CAGTGCTCTGAATGTCAAAGTGAAGAAATTCAATG</td><td>337.3</td><td>3.8</td><td>0.9</td><td>4.3</td><td>1.7E-05</td><td>3.3E-05</td><td>BT20/MB468</td><td>MITO</td></tr>
<tr><td>>hg38_21_+_8433172_8446622_RNA45SN1_45S_50ntFlanks@10101.10125.25|CGCGCGCGTGTGGTGTGCGTCGGAG</td><td>>hg38_21_+_8433172_8446622_RNA45SN1_45S_50ntFlanks</td><td>10101</td><td>10125</td><td>25</td><td>CGCGCGCGTGTGGTGTGCGTCGGAG</td><td>246.5</td><td>-6.1</td><td>1.4</td><td>-4.3</td><td>2.0E-05</td><td>3.3E-05</td><td>BT20/MB468</td><td>CYTO</td></tr>
<tr><td>>hg38_21_+_8433172_8446622_RNA45SN1_45S_50ntFlanks@4050.4081.32|CGAACGTCTGCCCTATCAACTTTCGATGGTAG</td><td>>hg38_21_+_8433172_8446622_RNA45SN1_45S_50ntFlanks</td><td>4050</td><td>4081</td><td>32</td><td>CGAACGTCTGCCCTATCAACTTTCGATGGTAG</td><td>183.2</td><td>-2.6</td><td>0.6</td><td>-4.2</td><td>2.2E-05</td><td>3.3E-05</td><td>MB231/MB468</td><td>CYTO</td></tr>
<tr><td>>hg38_21_+_8433172_8446622_RNA45SN1_45S_50ntFlanks@6732.6756.25|ACATTGATCATCGACACTTCGAACG</td><td>>hg38_21_+_8433172_8446622_RNA45SN1_45S_50ntFlanks</td><td>6732</td><td>6756</td><td>25</td><td>ACATTGATCATCGACACTTCGAACG</td><td>765.8</td><td>-5.1</td><td>1.2</td><td>-4.3</td><td>2.0E-05</td><td>3.3E-05</td><td>MB231/MB468</td><td>MP</td></tr>
<tr><td>>hg38_21_+_8433172_8446622_RNA45SN1_45S_50ntFlanks@4731.4751.21|AAGAACGAAAGTCGGAGGTTC</td><td>>hg38_21_+_8433172_8446622_RNA45SN1_45S_50ntFlanks</td><td>4731</td><td>4751</td><td>21</td><td>AAGAACGAAAGTCGGAGGTTC</td><td>1235.6</td><td>-2.9</td><td>0.7</td><td>-4.2</td><td>2.4E-05</td><td>3.3E-05</td><td>MB231/MB468</td><td>NUC</td></tr>
<tr><td>>hg38_MT_+_1621_3279_RNR2_16S_50ntFlanks@825.860.36|TAAGGAAAGGTTAAAAAAAGTAAAAGGAACTCGGCA</td><td>>hg38_MT_+_1621_3279_RNR2_16S_50ntFlanks</td><td>825</td><td>860</td><td>36</td><td>TAAGGAAAGGTTAAAAAAAGTAAAAGGAACTCGGCA</td><td>29.6</td><td>8.1</td><td>1.9</td><td>4.3</td><td>2.0E-05</td><td>3.3E-05</td><td>MB231/MB468</td><td>MP</td></tr>
<tr><td>>hg38_21_+_8433172_8446622_RNA45SN1_45S_50ntFlanks@4050.4081.32|CGAACGTCTGCCCTATCAACTTTCGATGGTAG</td><td>>hg38_21_+_8433172_8446622_RNA45SN1_45S_50ntFlanks</td><td>4050</td><td>4081</td><td>32</td><td>CGAACGTCTGCCCTATCAACTTTCGATGGTAG</td><td>184.0</td><td>-5.8</td><td>1.4</td><td>-4.2</td><td>2.4E-05</td><td>3.3E-05</td><td>MB231/MB468</td><td>NUC</td></tr>
<tr><td>>hg38_21_+_8433172_8446622_RNA45SN1_45S_50ntFlanks@12447.12476.30|CAGAATTCACCAAGCGTTGGATTGTTCACC</td><td>>hg38_21_+_8433172_8446622_RNA45SN1_45S_50ntFlanks</td><td>12447</td><td>12476</td><td>30</td><td>CAGAATTCACCAAGCGTTGGATTGTTCACC</td><td>222.5</td><td>-4.8</td><td>1.1</td><td>-4.2</td><td>2.4E-05</td><td>3.3E-05</td><td>MB231/MB468</td><td>MITO</td></tr>
<tr><td>>am_trnaMT_ValTAC_MT_+_1602_1670@42.69.28|AGATTTCAACTTAACTTGACCGCTCTGA</td><td>>am_trnaMT_ValTAC_MT_+_1602_1670</td><td>42</td><td>69</td><td>28</td><td>AGATTTCAACTTAACTTGACCGCTCTGA</td><td>422.1</td><td>4.3</td><td>1.0</td><td>4.2</td><td>2.4E-05</td><td>3.3E-05</td><td>MB231/MB468</td><td>MITO</td></tr>
<tr><td>>hg38_21_+_8433172_8446622_RNA45SN1_45S_50ntFlanks@4564.4587.24|GAATAATGGAATAGGACCGCGGTT</td><td>>hg38_21_+_8433172_8446622_RNA45SN1_45S_50ntFlanks</td><td>4564</td><td>4587</td><td>24</td><td>GAATAATGGAATAGGACCGCGGTT</td><td>199.0</td><td>-4.0</td><td>0.9</td><td>-4.2</td><td>2.2E-05</td><td>3.3E-05</td><td>MB231/MB468</td><td>CYTO</td></tr>
<tr><td>>hg38_21_+_8433172_8446622_RNA45SN1_45S_50ntFlanks@4102.4119.18|ACCACGGGTGACGGGGAA</td><td>>hg38_21_+_8433172_8446622_RNA45SN1_45S_50ntFlanks</td><td>4102</td><td>4119</td><td>18</td><td>ACCACGGGTGACGGGGAA</td><td>140.2</td><td>-6.1</td><td>1.4</td><td>-4.3</td><td>2.0E-05</td><td>3.3E-05</td><td>MB231/MB468</td><td>MP</td></tr>
<tr><td>>hg38_21_+_8433172_8446622_RNA45SN1_45S_50ntFlanks@12710.12742.33|CGCGGAGCCTCGGTTGGCCTCGGATAGCCGGTC</td><td>>hg38_21_+_8433172_8446622_RNA45SN1_45S_50ntFlanks</td><td>12710</td><td>12742</td><td>33</td><td>CGCGGAGCCTCGGTTGGCCTCGGATAGCCGGTC</td><td>170.3</td><td>-5.4</td><td>1.3</td><td>-4.3</td><td>1.7E-05</td><td>3.3E-05</td><td>BT20/MB468</td><td>MITO</td></tr>
<tr><td>>trnaMT_MetCAT_MT_+_4402_4469@52.71.20|ATACCCTTCCCGTACTACCA</td><td>>trnaMT_MetCAT_MT_+_4402_4469</td><td>52</td><td>71</td><td>20</td><td>ATACCCTTCCCGTACTACCA</td><td>3473.7</td><td>11.1</td><td>2.6</td><td>4.2</td><td>2.3E-05</td><td>3.3E-05</td><td>MB231/MB468</td><td>CYTO</td></tr>
<tr><td>>MI0000462|hsa-mir-152&WithFlank&17|-|48037155|48037253@60.79.20(+1U)&[MIMAT0000438&hsa-miR-152-3p&offsets|0|-1(+1U);m-97&17|-|48037174|48037194&offsets|0|-1(+1U)]|TCAGTGCATGACAGAACTTGT</td><td>>MI0000462|hsa-mir-152&WithFlank&17|-|48037155|48037253</td><td>60</td><td>79</td><td>20(+1U)</td><td>TCAGTGCATGACAGAACTTGT</td><td>32.0</td><td>8.2</td><td>1.9</td><td>4.3</td><td>2.0E-05</td><td>3.3E-05</td><td>MB231/MB468</td><td>MP</td></tr>
<tr><td>>hg38_21_+_8433172_8446622_RNA45SN1_45S_50ntFlanks@9802.9820.19|CGGGCGTGGAATGCGAGTG</td><td>>hg38_21_+_8433172_8446622_RNA45SN1_45S_50ntFlanks</td><td>9802</td><td>9820</td><td>19</td><td>CGGGCGTGGAATGCGAGTG</td><td>106.1</td><td>10.1</td><td>2.1</td><td>4.7</td><td>2.4E-06</td><td>3.3E-05</td><td>BT20/MB231</td><td>MP</td></tr>
<tr><td>>hg38_21_+_8433172_8446622_RNA45SN1_45S_50ntFlanks@10807.10826.20|ACAATGTAGGTAAGGGAAGT</td><td>>hg38_21_+_8433172_8446622_RNA45SN1_45S_50ntFlanks</td><td>10807</td><td>10826</td><td>20</td><td>ACAATGTAGGTAAGGGAAGT</td><td>168.7</td><td>-6.4</td><td>1.4</td><td>-4.5</td><td>7.8E-06</td><td>3.3E-05</td><td>BT20/MB468</td><td>MP</td></tr>
<tr><td>>hg38_21_+_8433172_8446622_RNA45SN1_45S_50ntFlanks@9554.9575.22|CGTAGCGGTCCTGACGTGCAAA</td><td>>hg38_21_+_8433172_8446622_RNA45SN1_45S_50ntFlanks</td><td>9554</td><td>9575</td><td>22</td><td>CGTAGCGGTCCTGACGTGCAAA</td><td>782.2</td><td>-3.9</td><td>0.9</td><td>-4.5</td><td>7.8E-06</td><td>3.3E-05</td><td>BT20/MB468</td><td>MP</td></tr>
<tr><td>>MI0003560|hsa-mir-92b&WithFlank&1|+|155195171|155195278@67.86.20(+1U)&[MIMAT0003218&hsa-miR-92b-3p&offsets|0|-2(+1U);m-115&1|+|155195237|155195258&offsets|0|-2(+1U)]|TATTGCACTCGTCCCGGCCTT</td><td>>MI0003560|hsa-mir-92b&WithFlank&1|+|155195171|155195278</td><td>67</td><td>86</td><td>20(+1U)</td><td>TATTGCACTCGTCCCGGCCTT</td><td>87.2</td><td>3.7</td><td>0.9</td><td>4.2</td><td>2.5E-05</td><td>3.3E-05</td><td>BT20/MB468</td><td>NUC</td></tr>
<tr><td>>hg38_21_+_8433172_8446622_RNA45SN1_45S_50ntFlanks@7973.7991.19|GACGCGACCTCAGATCAGA</td><td>>hg38_21_+_8433172_8446622_RNA45SN1_45S_50ntFlanks</td><td>7973</td><td>7991</td><td>19</td><td>GACGCGACCTCAGATCAGA</td><td>283.1</td><td>3.2</td><td>0.8</td><td>4.3</td><td>2.0E-05</td><td>3.3E-05</td><td>BT20/MB468</td><td>CYTO</td></tr>
<tr><td>>MI0000542|hsa-mir-320a&WithFlank&8|-|22244960|22245043@49.69.21&[MIMAT0000510&hsa-miR-320a-3p&offsets|+1|0;m-60&8|-|22244975|22244996&offsets|+1|0]|AAAGCTGGGTTGAGAGGGCGA</td><td>>MI0000542|hsa-mir-320a&WithFlank&8|-|22244960|22245043</td><td>49</td><td>69</td><td>21</td><td>AAAGCTGGGTTGAGAGGGCGA</td><td>1287.6</td><td>4.0</td><td>0.8</td><td>4.7</td><td>2.4E-06</td><td>3.3E-05</td><td>BT20/MB231</td><td>CYTO</td></tr>
<tr><td>>hg38_21_+_8433172_8446622_RNA45SN1_45S_50ntFlanks@8359.8390.32|AAGAACTTTGAAGAGAGAGTTCAAGAGGGCGT</td><td>>hg38_21_+_8433172_8446622_RNA45SN1_45S_50ntFlanks</td><td>8359</td><td>8390</td><td>32</td><td>AAGAACTTTGAAGAGAGAGTTCAAGAGGGCGT</td><td>96.9</td><td>9.9</td><td>2.1</td><td>4.7</td><td>2.4E-06</td><td>3.3E-05</td><td>BT20/MB231</td><td>MP</td></tr>
<tr><td>>hg38_21_+_8433172_8446622_RNA45SN1_45S_50ntFlanks@9552.9571.20|TCCGTAGCGGTCCTGACGTG</td><td>>hg38_21_+_8433172_8446622_RNA45SN1_45S_50ntFlanks</td><td>9552</td><td>9571</td><td>20</td><td>TCCGTAGCGGTCCTGACGTG</td><td>1155.9</td><td>-7.5</td><td>1.8</td><td>-4.2</td><td>2.5E-05</td><td>3.3E-05</td><td>BT20/MB468</td><td>NUC</td></tr>
<tr><td>>MI0000737|hsa-mir-200a&WithFlank&1|+|1167857|1167958@60.82.23(+1U)&[MIMAT0000682&hsa-miR-200a-3p&offsets|0|+1(+1U);m-142&1|+|1167916|1167938&offsets|0|0(+1U)]|TAACACTGTCTGGTAACGATGTTT</td><td>>MI0000737|hsa-mir-200a&WithFlank&1|+|1167857|1167958</td><td>60</td><td>82</td><td>23(+1U)</td><td>TAACACTGTCTGGTAACGATGTTT</td><td>892.9</td><td>-6.0</td><td>1.3</td><td>-4.5</td><td>7.8E-06</td><td>3.3E-05</td><td>BT20/MB468</td><td>MP</td></tr>
<tr><td>>MI0000466|hsa-mir-9-1&WithFlank&1|-|156420335|156420435@22.44.23;MI0000468|hsa-mir-9-3&WithFlank&15|+|89368011|89368112@22.44.23;MI0000467|hsa-mir-9-2&WithFlank&5|-|88666847|88666945@22.44.23&[MIMAT0000441&hsa-miR-9-5p&offsets|0|0;m-104&1|-|156420392|156420414&offsets|0|0];[MIMAT0000441_1&hsa-miR-9-5p&offsets|0|0;m-105&15|+|89368032|89368054&offsets|0|0];[MIMAT0000441_2&hsa-miR-9-5p&offsets|0|0;m-104&5|-|88666902|88666924&offsets|0|0]|TCTTTGGTTATCTAGCTGTATGA</td><td>>MI0000466|hsa-mir-9-1&WithFlank&1|-|156420335|156420435</td><td>22</td><td>44</td><td>23</td><td>TCTTTGGTTATCTAGCTGTATGA</td><td>300.7</td><td>5.5</td><td>1.3</td><td>4.2</td><td>2.4E-05</td><td>3.3E-05</td><td>MB231/MB468</td><td>MITO</td></tr>
<tr><td>>MI0000102|hsa-mir-100&WithFlank&11|-|122152223|122152314@19.39.21(+2A)&[MIMAT0000098&hsa-miR-100-5p&offsets|0|-1(+2A);m-27&11|-|122152276|122152296&offsets|0|0(+2A)]|AACCCGTAGATCCGAACTTGTAA</td><td>>MI0000102|hsa-mir-100&WithFlank&11|-|122152223|122152314</td><td>19</td><td>39</td><td>21(+2A)</td><td>AACCCGTAGATCCGAACTTGTAA</td><td>30.3</td><td>8.1</td><td>1.9</td><td>4.3</td><td>2.0E-05</td><td>3.3E-05</td><td>MB231/MB468</td><td>MP</td></tr>
<tr><td>>hg38_21_+_8433172_8446622_RNA45SN1_45S_50ntFlanks@10788.10828.41|CAGCCTCTGGCATGTTGGAACAATGTAGGTAAGGGAAGTCG</td><td>>hg38_21_+_8433172_8446622_RNA45SN1_45S_50ntFlanks</td><td>10788</td><td>10828</td><td>41</td><td>CAGCCTCTGGCATGTTGGAACAATGTAGGTAAGGGAAGTCG</td><td>58.6</td><td>-9.7</td><td>2.3</td><td>-4.3</td><td>2.0E-05</td><td>3.3E-05</td><td>MB231/MB468</td><td>MP</td></tr>
<tr><td>>hg38_21_+_8433172_8446622_RNA45SN1_45S_50ntFlanks@12420.12441.22|ATGTCGGCTCTTCCTATCATTG</td><td>>hg38_21_+_8433172_8446622_RNA45SN1_45S_50ntFlanks</td><td>12420</td><td>12441</td><td>22</td><td>ATGTCGGCTCTTCCTATCATTG</td><td>2237.9</td><td>-3.9</td><td>0.9</td><td>-4.2</td><td>2.4E-05</td><td>3.3E-05</td><td>MB231/MB468</td><td>MITO</td></tr>
<tr><td>>trna10_AspGTC_12_-_125424193_125424264@32.73.42,trna12_AspGTC_12_-_125411891_125411962@32.73.42,trna144_AspGTC_6_-_27551236_27551307@32.73.42,trna38_AspGTC_17_-_8125556_8125627@32.73.42,trna45_AspGTC_6_+_27447453_27447524@32.73.42,trna48_AspGTC_6_+_27471523_27471594@32.73.42,trna4_AspGTC_12_+_96429799_96429870@32.73.42,trna69_AspGTC_1_-_161440205_161440276@32.73.42,trna72_AspGTC_1_-_161432824_161432895@32.73.42,trna75_AspGTC_1_-_161425414_161425485@32.73.42,trna78_AspGTC_1_-_161418033_161418104@32.73.42,trna81_AspGTC_1_-_161410615_161410686@32.73.42|CTGTCACGCGGGAGACCGGGGTTCGATTCCCCGACGGGGAGC</td><td>>trna10_AspGTC_12_-_125424193_125424264</td><td>32</td><td>73</td><td>42</td><td>CTGTCACGCGGGAGACCGGGGTTCGATTCCCCGACGGGGAGC</td><td>709.2</td><td>-7.4</td><td>1.8</td><td>-4.2</td><td>2.5E-05</td><td>3.4E-05</td><td>BT20/MB468</td><td>NUC</td></tr>
<tr><td>>hg38_21_+_8433172_8446622_RNA45SN1_45S_50ntFlanks@8439.8471.33|GATTCAACCCGGCGGCGGGTCCGGCCGTGTCGG</td><td>>hg38_21_+_8433172_8446622_RNA45SN1_45S_50ntFlanks</td><td>8439</td><td>8471</td><td>33</td><td>GATTCAACCCGGCGGCGGGTCCGGCCGTGTCGG</td><td>155.3</td><td>-5.5</td><td>1.3</td><td>-4.3</td><td>1.7E-05</td><td>3.4E-05</td><td>BT20/MB468</td><td>MITO</td></tr>
<tr><td>>hg38_21_+_8433172_8446622_RNA45SN1_45S_50ntFlanks@11208.11250.43|GCGGCGACTCTGGACGCGAGCCGGGCCCTTCCCGTGGATCGCC</td><td>>hg38_21_+_8433172_8446622_RNA45SN1_45S_50ntFlanks</td><td>11208</td><td>11250</td><td>43</td><td>GCGGCGACTCTGGACGCGAGCCGGGCCCTTCCCGTGGATCGCC</td><td>719.8</td><td>-6.5</td><td>1.5</td><td>-4.2</td><td>2.4E-05</td><td>3.4E-05</td><td>MB231/MB468</td><td>NUC</td></tr>
<tr><td>>hg38_21_+_8433172_8446622_RNA45SN1_45S_50ntFlanks@8381.8399.19|AAGAGGGCGTGAAACCGTT</td><td>>hg38_21_+_8433172_8446622_RNA45SN1_45S_50ntFlanks</td><td>8381</td><td>8399</td><td>19</td><td>AAGAGGGCGTGAAACCGTT</td><td>909.7</td><td>-4.4</td><td>1.0</td><td>-4.5</td><td>7.8E-06</td><td>3.4E-05</td><td>BT20/MB468</td><td>MP</td></tr>
<tr><td>>hg38_21_+_8433172_8446622_RNA45SN1_45S_50ntFlanks@10802.10819.18|TTGGAACAATGTAGGTAA</td><td>>hg38_21_+_8433172_8446622_RNA45SN1_45S_50ntFlanks</td><td>10802</td><td>10819</td><td>18</td><td>TTGGAACAATGTAGGTAA</td><td>78.0</td><td>-6.0</td><td>1.4</td><td>-4.3</td><td>2.0E-05</td><td>3.4E-05</td><td>MB231/MB468</td><td>MP</td></tr>
<tr><td>>hg38_21_+_8433172_8446622_RNA45SN1_45S_50ntFlanks@4884.4903.20|CAAAGCTGAAACTTAAAGGA</td><td>>hg38_21_+_8433172_8446622_RNA45SN1_45S_50ntFlanks</td><td>4884</td><td>4903</td><td>20</td><td>CAAAGCTGAAACTTAAAGGA</td><td>281.6</td><td>-3.7</td><td>0.9</td><td>-4.3</td><td>2.0E-05</td><td>3.4E-05</td><td>MB231/MB468</td><td>MP</td></tr>
<tr><td>>hg38_21_+_8433172_8446622_RNA45SN1_45S_50ntFlanks@5083.5104.22|ACGAACGAGACTCTGGCATGCT</td><td>>hg38_21_+_8433172_8446622_RNA45SN1_45S_50ntFlanks</td><td>5083</td><td>5104</td><td>22</td><td>ACGAACGAGACTCTGGCATGCT</td><td>421.1</td><td>-4.3</td><td>1.0</td><td>-4.3</td><td>2.0E-05</td><td>3.4E-05</td><td>MB231/MB468</td><td>MP</td></tr>
<tr><td>>trna10_LysCTT_16_+_3241501_3241573@1.40.40,trna13_LysCTT_14_-_58706613_58706685@1.40.40,trna2_LysCTT_15_+_79152904_79152976@1.40.40|GCCCGGCTAGCTCAGTCGGTAGAGCATGGGACTCTTAATC</td><td>>trna10_LysCTT_16_+_3241501_3241573</td><td>1</td><td>40</td><td>40</td><td>GCCCGGCTAGCTCAGTCGGTAGAGCATGGGACTCTTAATC</td><td>1366.9</td><td>6.7</td><td>1.6</td><td>4.3</td><td>2.1E-05</td><td>3.4E-05</td><td>BT20/MB468</td><td>CYTO</td></tr>
<tr><td>>hg38_21_+_8433172_8446622_RNA45SN1_45S_50ntFlanks@10361.10390.30|GAAGGCCGAAGTGGAGAAGGGTTCCATGTG</td><td>>hg38_21_+_8433172_8446622_RNA45SN1_45S_50ntFlanks</td><td>10361</td><td>10390</td><td>30</td><td>GAAGGCCGAAGTGGAGAAGGGTTCCATGTG</td><td>134.2</td><td>-5.3</td><td>1.2</td><td>-4.3</td><td>1.7E-05</td><td>3.4E-05</td><td>BT20/MB468</td><td>MITO</td></tr>
<tr><td>>MI0000285|hsa-mir-205&WithFlank&1|+|209432127|209432248@40.62.23(+1A)&[MIMAT0000266&hsa-miR-205-5p&offsets|0|+1(+1A);m-351&1|+|209432166|209432187&offsets|0|+1(+1A)]|TCCTTCATTCCACCGGAGTCTGTA</td><td>>MI0000285|hsa-mir-205&WithFlank&1|+|209432127|209432248</td><td>40</td><td>62</td><td>23(+1A)</td><td>TCCTTCATTCCACCGGAGTCTGTA</td><td>75.8</td><td>9.6</td><td>2.0</td><td>4.7</td><td>2.4E-06</td><td>3.4E-05</td><td>BT20/MB231</td><td>MP</td></tr>
<tr><td>>MI0000279|hsa-mir-196a-2&WithFlank&12|+|53991732|53991853@31.53.23&[MIMAT0000226&hsa-miR-196a-5p&offsets|0|+1;m-261&12|+|53991762|53991783&offsets|0|+1]|TAGGTAGTTTCATGTTGTTGGGA</td><td>>MI0000279|hsa-mir-196a-2&WithFlank&12|+|53991732|53991853</td><td>31</td><td>53</td><td>23</td><td>TAGGTAGTTTCATGTTGTTGGGA</td><td>1137.2</td><td>4.7</td><td>1.1</td><td>4.5</td><td>7.9E-06</td><td>3.4E-05</td><td>BT20/MB468</td><td>MP</td></tr>
<tr><td>>hg38_21_+_8433172_8446622_RNA45SN1_45S_50ntFlanks@10349.10368.20|AACGAGAACTTTGAAGGCCG</td><td>>hg38_21_+_8433172_8446622_RNA45SN1_45S_50ntFlanks</td><td>10349</td><td>10368</td><td>20</td><td>AACGAGAACTTTGAAGGCCG</td><td>850.1</td><td>-2.4</td><td>0.6</td><td>-4.3</td><td>1.7E-05</td><td>3.4E-05</td><td>BT20/MB468</td><td>MITO</td></tr>
<tr><td>>hg38_21_+_8433172_8446622_RNA45SN1_45S_50ntFlanks@9554.9579.26|CGTAGCGGTCCTGACGTGCAAATCGG</td><td>>hg38_21_+_8433172_8446622_RNA45SN1_45S_50ntFlanks</td><td>9554</td><td>9579</td><td>26</td><td>CGTAGCGGTCCTGACGTGCAAATCGG</td><td>354.1</td><td>-2.8</td><td>0.6</td><td>-4.2</td><td>2.3E-05</td><td>3.4E-05</td><td>MB231/MB468</td><td>CYTO</td></tr>
<tr><td>>hg38_21_+_8433172_8446622_RNA45SN1_45S_50ntFlanks@5423.5466.44|ATTGGATGGTTTAGTGAGGCCCTCGGATCGGCCCCGCCGGGGTC</td><td>>hg38_21_+_8433172_8446622_RNA45SN1_45S_50ntFlanks</td><td>5423</td><td>5466</td><td>44</td><td>ATTGGATGGTTTAGTGAGGCCCTCGGATCGGCCCCGCCGGGGTC</td><td>1166.6</td><td>-3.8</td><td>0.9</td><td>-4.2</td><td>2.3E-05</td><td>3.4E-05</td><td>MB231/MB468</td><td>CYTO</td></tr>
<tr><td>>MI0000266|hsa-mir-10a&WithFlank&17|-|48579832|48579953@29.48.20(+1A)&[MIMAT0000253&hsa-miR-10a-5p&offsets|+1|-2(+1A);m-18&17|-|48579905|48579926&offsets|+1|-1(+1A)]|ACCCTGTAGATCCGAATTTGA</td><td>>MI0000266|hsa-mir-10a&WithFlank&17|-|48579832|48579953</td><td>29</td><td>48</td><td>20(+1A)</td><td>ACCCTGTAGATCCGAATTTGA</td><td>30.0</td><td>8.1</td><td>1.9</td><td>4.3</td><td>2.0E-05</td><td>3.4E-05</td><td>MB231/MB468</td><td>MP</td></tr>
<tr><td>>MI0005566|hsa-mir-374b&WithFlank&X|-|74218541|74218624@17.34.18&[MIMAT0004955&hsa-miR-374b-5p&offsets|0|-4;m-153&X|-|74218587|74218608&offsets|0|-4]|ATATAATACAACCTGCTA</td><td>>MI0005566|hsa-mir-374b&WithFlank&X|-|74218541|74218624</td><td>17</td><td>34</td><td>18</td><td>ATATAATACAACCTGCTA</td><td>218.2</td><td>6.2</td><td>1.5</td><td>4.2</td><td>2.4E-05</td><td>3.4E-05</td><td>MB231/MB468</td><td>MITO</td></tr>
</table>

| | | | | | | | | | | | | | |
|---|---|---|---|---|---|---|---|---|---|---|---|---|---|
| >hg38_21_+_8433172_8446622_RNA45SN1_45S_50ntFlanks@10686.10710.25\|GCGGCGTCCGGTGAGCTCTCGCTGG | >hg38_21_+_8433172_8446622_RNA45SN1_45S_50ntFlanks | 10686 | 10710 | 25 | GCGGCGTCCGGTGAGCTCTCGCTGG | 1309.0 | -5.8 | 1.4 | -4.3 | 2.0E-05 | 3.4E-05 | MB231/MB468 | MP |
| >hg38_21_+_8433172_8446622_RNA45SN1_45S_50ntFlanks@9790.9831.42\|CTGGCGTGGAGCCGGGCGTGGAATGCGAGTGCCTAGTGGGCC | >hg38_21_+_8433172_8446622_RNA45SN1_45S_50ntFlanks | 9790 | 9831 | 42 | CTGGCGTGGAGCCGGGCGTGGAATGCGAGTGCCTAGTGGGCC | 116.3 | 10.2 | 2.2 | 4.7 | 2.5E-06 | 3.4E-05 | BT20/MB231 | MP |
| >hg38_21_+_8433172_8446622_RNA45SN1_45S_50ntFlanks@12637.12661.25\|GCTACCATCTGTGGGATTATGACTG | >hg38_21_+_8433172_8446622_RNA45SN1_45S_50ntFlanks | 12637 | 12661 | 25 | GCTACCATCTGTGGGATTATGACTG | 1866.4 | -3.3 | 0.8 | -4.3 | 1.7E-05 | 3.4E-05 | BT20/MB468 | MITO |
| >hg38_21_+_8433172_8446622_RNA45SN1_45S_50ntFlanks@13007.13025.19\|TATTGAAAGTCAGCCCTCG | >hg38_21_+_8433172_8446622_RNA45SN1_45S_50ntFlanks | 13007 | 13025 | 19 | TATTGAAAGTCAGCCCTCG | 1026.3 | -6.6 | 1.5 | -4.3 | 2.1E-05 | 3.4E-05 | BT20/MB468 | CYTO |
| >hg38_21_+_8433172_8446622_RNA45SN1_45S_50ntFlanks@11874.11901.28\|AGAAGACCCTGTTGAGCTTGACTCTAGT | >hg38_21_+_8433172_8446622_RNA45SN1_45S_50ntFlanks | 11874 | 11901 | 28 | AGAAGACCCTGTTGAGCTTGACTCTAGT | 153.9 | -1.8 | 0.4 | -4.3 | 2.1E-05 | 3.4E-05 | BT20/MB468 | CYTO |
| >MI0000440\|hsa-mir-27b&WithFlank&9\|+\|95085439\|95085547@67.86.20&[MIMAT0000419&hsa-miR-27b-3p&offsets\|0\|-1;m-39&9\|+\|95085505\|95085525&offsets\|0\|-1]\|TTCACAGTGGCTAAGTTCTG | >MI0000440\|hsa-mir-27b&WithFlank&9\|+\|95085439\|95085547 | 67 | 86 | 20 | TTCACAGTGGCTAAGTTCTG | 5045.9 | 4.0 | 0.9 | 4.5 | 7.9E-06 | 3.4E-05 | BT20/MB468 | MP |
| >hg38_1_-_228634819_228635039_RNA5S12_5S_50ntFlanks@96.133.38\|CGGAAGCTAAGCAGGGTCGGGCCTGGTTAGTACTTGGA | >hg38_1_-_228634819_228635039_RNA5S12_5S_50ntFlanks | 96 | 133 | 38 | CGGAAGCTAAGCAGGGTCGGGCCTGGTTAGTACTTGGA | 35.6 | 8.9 | 2.0 | 4.5 | 7.9E-06 | 3.4E-05 | BT20/MB468 | MP |
| >hg38_21_+_8433172_8446622_RNA45SN1_45S_50ntFlanks@12456.12476.21\|CCAAGCGTTGGATTGTTCACC | >hg38_21_+_8433172_8446622_RNA45SN1_45S_50ntFlanks | 12456 | 12476 | 21 | CCAAGCGTTGGATTGTTCACC | 35355.2 | -6.1 | 1.4 | -4.5 | 7.9E-06 | 3.4E-05 | BT20/MB468 | MP |
| >hg38_21_+_8433172_8446622_RNA45SN1_45S_50ntFlanks@9573.9591.19\|AAATCGGTCGTCCGACCTG | >hg38_21_+_8433172_8446622_RNA45SN1_45S_50ntFlanks | 9573 | 9591 | 19 | AAATCGGTCGTCCGACCTG | 69.2 | -10.0 | 2.3 | -4.3 | 2.0E-05 | 3.4E-05 | MB231/MB468 | MP |
| >hg38_21_+_8433172_8446622_RNA45SN1_45S_50ntFlanks@4557.4575.19\|CAGCTAGGAATAATGGAAT | >hg38_21_+_8433172_8446622_RNA45SN1_45S_50ntFlanks | 4557 | 4575 | 19 | CAGCTAGGAATAATGGAAT | 251.7 | -5.1 | 1.2 | -4.2 | 2.5E-05 | 3.4E-05 | BT20/MB468 | NUC |
| >hg38_21_+_8433172_8446622_RNA45SN1_45S_50ntFlanks@12073.12092.20\|CCGAGGGGCTCTCGCTTCTG | >hg38_21_+_8433172_8446622_RNA45SN1_45S_50ntFlanks | 12073 | 12092 | 20 | CCGAGGGGCTCTCGCTTCTG | 282.7 | -6.5 | 1.5 | -4.5 | 7.9E-06 | 3.4E-05 | BT20/MB468 | MP |
| >hg38_21_+_8433172_8446622_RNA45SN1_45S_50ntFlanks@12578.12614.37\|AGGAACCGCAGGTTCAGACATTTGGTGTATGTGCTTG | >hg38_21_+_8433172_8446622_RNA45SN1_45S_50ntFlanks | 12578 | 12614 | 37 | AGGAACCGCAGGTTCAGACATTTGGTGTATGTGCTTG | 45.1 | -8.6 | 1.9 | -4.5 | 7.9E-06 | 3.4E-05 | BT20/MB468 | MP |
| >hg38_21_+_8433172_8446622_RNA45SN1_45S_50ntFlanks@12419.12449.31\|GATGTCGGCTCTTCCTATCATTGTGAAGCAG | >hg38_21_+_8433172_8446622_RNA45SN1_45S_50ntFlanks | 12419 | 12449 | 31 | GATGTCGGCTCTTCCTATCATTGTGAAGCAG | 311.4 | -3.3 | 0.8 | -4.2 | 2.5E-05 | 3.4E-05 | BT20/MB468 | NUC |
| >hg38_21_+_8433172_8446622_RNA45SN1_45S_50ntFlanks@12478.12500.23\|ACTAATAGGGAACGTGAGCTGGG | >hg38_21_+_8433172_8446622_RNA45SN1_45S_50ntFlanks | 12478 | 12500 | 23 | ACTAATAGGGAACGTGAGCTGGG | 586.4 | -7.2 | 1.7 | -4.2 | 2.5E-05 | 3.4E-05 | BT20/MB468 | NUC |
| >hg38_21_+_8433172_8446622_RNA45SN1_45S_50ntFlanks@3777.3814.38\|CGGTACAGTGAAACTGCGAATGGCTCATTAAATCAGTT | >hg38_21_+_8433172_8446622_RNA45SN1_45S_50ntFlanks | 3777 | 3814 | 38 | CGGTACAGTGAAACTGCGAATGGCTCATTAAATCAGTT | 70.1 | -10.0 | 2.3 | -4.3 | 2.0E-05 | 3.4E-05 | MB231/MB468 | MP |
| >hg38_21_+_8433172_8446622_RNA45SN1_45S_50ntFlanks@11641.11668.28\|CGGCGGGTGTTGACGCGATGTGATTTCT | >hg38_21_+_8433172_8446622_RNA45SN1_45S_50ntFlanks | 11641 | 11668 | 28 | CGGCGGGTGTTGACGCGATGTGATTTCT | 174.2 | -3.6 | 0.8 | -4.2 | 2.3E-05 | 3.4E-05 | MB231/MB468 | CYTO |
| >hg38_21_+_8433172_8446622_RNA45SN1_45S_50ntFlanks@10773.10796.24\|GGTCTCCAAGGTGAACAGCCTCTG | >hg38_21_+_8433172_8446622_RNA45SN1_45S_50ntFlanks | 10773 | 10796 | 24 | GGTCTCCAAGGTGAACAGCCTCTG | 305.2 | -6.8 | 1.6 | -4.2 | 2.5E-05 | 3.4E-05 | BT20/MB468 | NUC |
| >hg38_21_+_8433172_8446622_RNA45SN1_45S_50ntFlanks@9568.9585.18\|CGTGCAAATCGGTCGTCC | >hg38_21_+_8433172_8446622_RNA45SN1_45S_50ntFlanks | 9568 | 9585 | 18 | CGTGCAAATCGGTCGTCC | 229.0 | -6.1 | 1.4 | -4.3 | 2.1E-05 | 3.4E-05 | BT20/MB468 | CYTO |
| >hg38_21_+_8433172_8446622_RNA45SN1_45S_50ntFlanks@3769.3794.26\|CGCACGGCCGGTACAGTGAAACTGCG | >hg38_21_+_8433172_8446622_RNA45SN1_45S_50ntFlanks | 3769 | 3794 | 26 | CGCACGGCCGGTACAGTGAAACTGCG | 226.4 | -5.1 | 1.2 | -4.3 | 2.0E-05 | 3.4E-05 | MB231/MB468 | MP |
| >trna143_LysTTT_6_-_27559593_27559665@13.33.21,trna149_LysTTT_6_-_27302769_27302841@13.33.21,trna14_LysTTT_11_-_59327808_59327880@13.33.21,trna2_LysTTT_17_+_8022473_8022545@13.33.21,trna54_LysTTT_1_+_204475655_204475727@13.33.21,trna5_LysTTT_11_+_59323902_59323974@13.33.21,trna62_LysTTT_1_-_204476158_204476230@13.33.21,trna76_LysTTT_6_+_28918806_28918878@13.33.21\|CAGTCGGTAGAGCATCAGACT | >trna143_LysTTT_6_-_27559593_27559665 | 13 | 33 | 21 | CAGTCGGTAGAGCATCAGACT | 70.2 | -9.5 | 2.0 | -4.7 | 2.5E-06 | 3.4E-05 | BT20/MB231 | MP |
| >hg38_MT_+_1621_3279_RNR2_16S_50ntFlanks@829.864.36\|GAAAGGTTAAAAAAAGTAAAAGGAACTCGGCAAATC | >hg38_MT_+_1621_3279_RNR2_16S_50ntFlanks | 829 | 864 | 36 | GAAAGGTTAAAAAAAGTAAAAGGAACTCGGCAAATC | 36.3 | 8.4 | 2.0 | 4.3 | 2.0E-05 | 3.4E-05 | MB231/MB468 | MP |
| >hg38_21_+_8433172_8446622_RNA45SN1_45S_50ntFlanks@11771.11791.21\|ATTAGTGACGCGCATGAATGG | >hg38_21_+_8433172_8446622_RNA45SN1_45S_50ntFlanks | 11771 | 11791 | 21 | ATTAGTGACGCGCATGAATGG | 386.2 | -4.9 | 1.2 | -4.2 | 2.4E-05 | 3.4E-05 | MB231/MB468 | NUC |
| >hg38_21_+_8433172_8446622_RNA45SN1_45S_50ntFlanks@12627.12659.33\|ATGGGGCGAAGCTACCATCTGTGGGATTATGAC | >hg38_21_+_8433172_8446622_RNA45SN1_45S_50ntFlanks | 12627 | 12659 | 33 | ATGGGGCGAAGCTACCATCTGTGGGATTATGAC | 462.4 | -4.3 | 1.0 | -4.3 | 2.1E-05 | 3.4E-05 | BT20/MB468 | CYTO |
| >hg38_21_+_8433172_8446622_RNA45SN1_45S_50ntFlanks@7980.8008.29\|CCTCAGATCAGACGTGGCGACCCGCTGAA | >hg38_21_+_8433172_8446622_RNA45SN1_45S_50ntFlanks | 7980 | 8008 | 29 | CCTCAGATCAGACGTGGCGACCCGCTGAA | 1589.5 | -4.5 | 1.0 | -4.5 | 8.0E-06 | 3.4E-05 | BT20/MB468 | MP |
| >MI0000542\|hsa-mir-320a&WithFlank&8\|-\|22244960\|22245043@49.70.22(+1U)&[MIMAT0000510&hsa-miR-320a-3p&offsets\|+1\|+1(+1U);m-60&8\|-\|22244975\|22244996&offsets\|+1\|+1(+1U)]\|AAAGCTGGGTTGAGAGGGCGAAT | >MI0000542\|hsa-mir-320a&WithFlank&8\|-\|22244960\|22245043 | 49 | 70 | 22(+1U) | AAAGCTGGGTTGAGAGGGCGAAT | 1162.7 | 6.2 | 1.4 | 4.5 | 8.0E-06 | 3.4E-05 | BT20/MB468 | MP |
| >hg38_21_+_8433172_8446622_RNA45SN1_45S_50ntFlanks@11452.11469.18\|CTCCTCGGCGCGCGGCGG | >hg38_21_+_8433172_8446622_RNA45SN1_45S_50ntFlanks | 11452 | 11469 | 18 | CTCCTCGGCGCGCGGCGG | 138.6 | -3.9 | 0.9 | -4.3 | 2.1E-05 | 3.4E-05 | BT20/MB468 | CYTO |
| >hg38_21_+_8433172_8446622_RNA45SN1_45S_50ntFlanks@10687.10709.23\|CGGCGTCCGGTGAGCTCTCGCTG | >hg38_21_+_8433172_8446622_RNA45SN1_45S_50ntFlanks | 10687 | 10709 | 23 | CGGCGTCCGGTGAGCTCTCGCTG | 4940.6 | -8.4 | 2.0 | -4.2 | 2.5E-05 | 3.4E-05 | BT20/MB468 | NUC |
| >hg38_21_+_8433172_8446622_RNA45SN1_45S_50ntFlanks@10845.10871.27\|ACTTCGGGATAAGGATTGGCTCTAAGG | >hg38_21_+_8433172_8446622_RNA45SN1_45S_50ntFlanks | 10845 | 10871 | 27 | ACTTCGGGATAAGGATTGGCTCTAAGG | 295.2 | -6.6 | 1.6 | -4.2 | 2.6E-05 | 3.4E-05 | BT20/MB468 | NUC |
| >hg38_21_+_8433172_8446622_RNA45SN1_45S_50ntFlanks@10069.10086.18\|AGTGGACGGGAGCGGCGG | >hg38_21_+_8433172_8446622_RNA45SN1_45S_50ntFlanks | 10069 | 10086 | 18 | AGTGGACGGGAGCGGCGG | 105.1 | 10.0 | 2.1 | 4.7 | 2.5E-06 | 3.4E-05 | BT20/MB231 | MP |
| >hg38_21_+_8433172_8446622_RNA45SN1_45S_50ntFlanks@12404.12432.29\|GCTTTTTGATCCTTCGATGTCGGCTCTTC | >hg38_21_+_8433172_8446622_RNA45SN1_45S_50ntFlanks | 12404 | 12432 | 29 | GCTTTTTGATCCTTCGATGTCGGCTCTTC | 377.1 | -2.8 | 0.6 | -4.3 | 2.1E-05 | 3.4E-05 | BT20/MB468 | CYTO |
| >hg38_21_+_8433172_8446622_RNA45SN1_45S_50ntFlanks@8194.8211.18\|CCCGTGGACGGTGTGAGG | >hg38_21_+_8433172_8446622_RNA45SN1_45S_50ntFlanks | 8194 | 8211 | 18 | CCCGTGGACGGTGTGAGG | 643.0 | -4.6 | 1.1 | -4.3 | 2.1E-05 | 3.4E-05 | BT20/MB468 | CYTO |
| >hg38_21_+_8433172_8446622_RNA45SN1_45S_50ntFlanks@6779.6802.24\|CCGGGGCTACGCCTGTCTGAGCGT | >hg38_21_+_8433172_8446622_RNA45SN1_45S_50ntFlanks | 6779 | 6802 | 24 | CCGGGGCTACGCCTGTCTGAGCGT | 563.2 | 3.2 | 0.7 | 4.2 | 2.3E-05 | 3.4E-05 | MB231/MB468 | CYTO |
| >MI0000299\|hsa-mir-222&WithFlank&X\|-\|45747009\|45747130@75.96.22&[MIMAT0000279&hsa-miR-222-3p&offsets\|0\|+1;m-140&X\|-\|45747033\|45747056&offsets\|0\|-2]\|AGCTACATCTGGCTACTGGGTC | >MI0000299\|hsa-mir-222&WithFlank&X\|-\|45747009\|45747130 | 75 | 96 | 22 | AGCTACATCTGGCTACTGGGTC | 1648.8 | 3.9 | 0.9 | 4.3 | 2.0E-05 | 3.4E-05 | MB231/MB468 | MP |
| >hg38_MT_+_1621_3279_RNR2_16S_50ntFlanks@51.86.36\|GCTAAACCTAGCCCCAAACCCACTCCACCTTACTAC | >hg38_MT_+_1621_3279_RNR2_16S_50ntFlanks | 51 | 86 | 36 | GCTAAACCTAGCCCCAAACCCACTCCACCTTACTAC | 256.3 | 4.0 | 0.9 | 4.2 | 2.4E-05 | 3.4E-05 | MB231/MB468 | MITO |
| >hg38_21_+_8433172_8446622_RNA45SN1_45S_50ntFlanks@4004.4024.21\|TAACCTCGGGCCGATCGCACG | >hg38_21_+_8433172_8446622_RNA45SN1_45S_50ntFlanks | 4004 | 4024 | 21 | TAACCTCGGGCCGATCGCACG | 191.2 | -3.7 | 0.9 | -4.3 | 2.1E-05 | 3.4E-05 | BT20/MB468 | CYTO |
| >MI0000078\|hsa-mir-22&WithFlank&17\|-\|1713897\|1713993@59.80.22(+1C)&[MIMAT0000077&hsa-miR-22-3p&offsets\|0\|0(+1C);m-2&17\|-\|1713914\|1713935&offsets\|0\|0(+1C)]\|AAGCTGCCAGTTGAAGAACTGTC | >MI0000078\|hsa-mir-22&WithFlank&17\|-\|1713897\|1713993 | 59 | 80 | 22(+1C) | AAGCTGCCAGTTGAAGAACTGTC | 150.9 | 7.4 | 1.7 | 4.3 | 2.0E-05 | 3.4E-05 | MB231/MB468 | MP |
| >am_trnaMT_ValTAC_MT_+_1602_1670@44.68.25\|ATTTCAACTTAACTTGACCGCTCTG | >am_trnaMT_ValTAC_MT_+_1602_1670 | 44 | 68 | 25 | ATTTCAACTTAACTTGACCGCTCTG | 94.0 | -6.1 | 1.4 | -4.3 | 2.0E-05 | 3.4E-05 | MB231/MB468 | MP |
| >hg38_21_+_8433172_8446622_RNA45SN1_45S_50ntFlanks@12007.12043.37\|GCCGCCGGTGAAATACCACTACTCTGATCGTTTTTTC | >hg38_21_+_8433172_8446622_RNA45SN1_45S_50ntFlanks | 12007 | 12043 | 37 | GCCGCCGGTGAAATACCACTACTCTGATCGTTTTTTC | 767.4 | -6.7 | 1.6 | -4.3 | 2.1E-05 | 3.4E-05 | BT20/MB468 | CYTO |
| >hg38_21_+_8433172_8446622_RNA45SN1_45S_50ntFlanks@12073.12106.34\|CCGAGGGGCTCTCGCTTCTGGCGCCAAGCGCCCG | >hg38_21_+_8433172_8446622_RNA45SN1_45S_50ntFlanks | 12073 | 12106 | 34 | CCGAGGGGCTCTCGCTTCTGGCGCCAAGCGCCCG | 227.5 | -5.8 | 1.4 | -4.3 | 2.1E-05 | 3.4E-05 | BT20/MB468 | CYTO |
| >hg38_21_+_8433172_8446622_RNA45SN1_45S_50ntFlanks@12413.12432.20\|TCCTTCGATGTCGGCTCTTC | >hg38_21_+_8433172_8446622_RNA45SN1_45S_50ntFlanks | 12413 | 12432 | 20 | TCCTTCGATGTCGGCTCTTC | 679.4 | -3.2 | 0.8 | -4.3 | 2.1E-05 | 3.4E-05 | BT20/MB468 | CYTO |
| >MI0000067\|hsa-let-7f-1&WithFlank&9\|+\|94176341\|94176439@13.33.21;MI0000068\|hsa-let-7f-2&WithFlank&X\|-\|53557186\|53557280@14.34.21&[MIMAT0000067&hsa-let-7f-5p&offsets\|0\|-1;m-20&9\|+\|94176353\|94176374&offsets\|0\|-1];[MIMAT0000067_1&hsa-let-7f-5p&offsets\|0\|-1;m-19&X\|-\|53557246\|53557267&offsets\|0\|-1]\|TGAGGTAGTAGATTGTATAGT | >MI0000067\|hsa-let-7f-1&WithFlank&9\|+\|94176341\|94176439 | 13 | 33 | 21 | TGAGGTAGTAGATTGTATAGT | 5548.3 | 3.6 | 0.8 | 4.3 | 2.0E-05 | 3.4E-05 | MB231/MB468 | MP |
| >MI0000465\|hsa-mir-191&WithFlank&3\|-\|49020612\|49020715@23.44.22&[MIMAT0000440&hsa-miR-191-5p&offsets\|+1\|0;m-48&3\|-\|49020672\|49020694&offsets\|+1\|0]\|AACGGAATCCCAAAAGCAGCTG | >MI0000465\|hsa-mir-191&WithFlank&3\|-\|49020612\|49020715 | 23 | 44 | 22 | AACGGAATCCCAAAAGCAGCTG | 387.6 | 3.3 | 0.8 | 4.3 | 2.0E-05 | 3.4E-05 | MB231/MB468 | MP |
| >hg38_21_+_8433172_8446622_RNA45SN1_45S_50ntFlanks@4046.4064.19\|CATTCGAACGTCTGCCCTA | >hg38_21_+_8433172_8446622_RNA45SN1_45S_50ntFlanks | 4046 | 4064 | 19 | CATTCGAACGTCTGCCCTA | 323.6 | -5.2 | 1.2 | -4.2 | 2.6E-05 | 3.4E-05 | BT20/MB468 | NUC |
| >MI0000065\|hsa-let-7d&WithFlank&9\|+\|94178828\|94178926@14.31.18&[MIMAT0000065&hsa-let-7d-5p&offsets\|0\|-4;m-90&9\|+\|94178841\|94178862&offsets\|0\|-4]\|AGAGGTAGTAGGTTGCAT | >MI0000065\|hsa-let-7d&WithFlank&9\|+\|94178828\|94178926 | 14 | 31 | 18 | AGAGGTAGTAGGTTGCAT | 253.3 | 4.3 | 1.0 | 4.2 | 2.4E-05 | 3.4E-05 | MB231/MB468 | NUC |
| >hg38_21_+_8433172_8446622_RNA45SN1_45S_50ntFlanks@11314.11342.29\|CCCACCCCACGTCTCGTCGCGCGCGCGTC | >hg38_21_+_8433172_8446622_RNA45SN1_45S_50ntFlanks | 11314 | 11342 | 29 | CCCACCCCACGTCTCGTCGCGCGCGCGTC | 52.6 | -8.8 | 2.0 | -4.5 | 8.1E-06 | 3.4E-05 | BT20/MB468 | MP |
| >hg38_21_+_8433172_8446622_RNA45SN1_45S_50ntFlanks@10355.10388.34\|AACTTTGAAGGCCGAAGTGGAGAAGGGTTCCATG | >hg38_21_+_8433172_8446622_RNA45SN1_45S_50ntFlanks | 10355 | 10388 | 34 | AACTTTGAAGGCCGAAGTGGAGAAGGGTTCCATG | 657.8 | -5.5 | 1.3 | -4.3 | 2.1E-05 | 3.4E-05 | BT20/MB468 | CYTO |
| >MI0000542\|hsa-mir-320a&WithFlank&8\|-\|22244960\|22245043@48.69.22(+1U)&[MIMAT0000510&hsa-miR-320a-3p&offsets\|0\|0(+1U);m-60&8\|-\|22244975\|22244996&offsets\|0\|0(+1U)]\|AAAAGCTGGGTTGAGAGGGCGAT | >MI0000542\|hsa-mir-320a&WithFlank&8\|-\|22244960\|22245043 | 48 | 69 | 22(+1U) | AAAAGCTGGGTTGAGAGGGCGAT | 30170.5 | 3.9 | 0.8 | 4.7 | 2.5E-06 | 3.4E-05 | BT20/MB231 | CYTO |
| >hg38_21_+_8433172_8446622_RNA45SN1_45S_50ntFlanks@6655.6680.26\|TCTTAGCGGTGGATCACTCGGCTCGT | >hg38_21_+_8433172_8446622_RNA45SN1_45S_50ntFlanks | 6655 | 6680 | 26 | TCTTAGCGGTGGATCACTCGGCTCGT | 134.9 | -6.3 | 1.5 | -4.3 | 2.1E-05 | 3.4E-05 | MB231/MB468 | MP |
| >MI0003591\|hsa-mir-584&WithFlank&5\|-\|149062307\|149062415@22.41.20(+1C)&[MIMAT0003249&hsa-miR-584-5p&offsets\|0\|-2(+1C);m-167&5\|-\|149062374\|149062394&offsets\|0\|-1(+1C)]\|TTATGGTTTGCCTGGGACTGC | >MI0003591\|hsa-mir-584&WithFlank&5\|-\|149062307\|149062415 | 22 | 41 | 20(+1C) | TTATGGTTTGCCTGGGACTGC | 237.4 | 8.3 | 1.9 | 4.5 | 8.1E-06 | 3.4E-05 | BT20/MB468 | MP |
| >hg38_21_+_8433172_8446622_RNA45SN1_45S_50ntFlanks@12071.12096.26\|CCCCGAGGGGCTCTCGCTTCTGGCGC | >hg38_21_+_8433172_8446622_RNA45SN1_45S_50ntFlanks | 12071 | 12096 | 26 | CCCCGAGGGGCTCTCGCTTCTGGCGC | 254.8 | -5.6 | 1.3 | -4.3 | 1.8E-05 | 3.4E-05 | BT20/MB468 | MITO |

| | | | | | | | | | | | | | |
|---|---|---|---|---|---|---|---|---|---|---|---|---|---|
| >hg38_21_+_8433172_8446622_RNA45SN1_45S_50ntFlanks@10411.10443.33\|CAGTCGGTCCTGAGAGATGGGCGAGCGCCGTTC | >hg38_21_+_8433172_8446622_RNA45SN1_45S_50ntFlanks | 10411 | 10443 | 33 | CAGTCGGTCCTGAGAGATGGGCGAGCGCCGTTC | 1206.8 | -5.4 | 1.2 | -4.3 | 1.8E-05 | 3.4E-05 | BT20/MB468 | MITO |
| >hg38_21_+_8433172_8446622_RNA45SN1_45S_50ntFlanks@12932.12973.42\|CGCTAAACCATTCGTAGACGACCTGCTTCTGGGTCGGGGTTT | >hg38_21_+_8433172_8446622_RNA45SN1_45S_50ntFlanks | 12932 | 12973 | 42 | CGCTAAACCATTCGTAGACGACCTGCTTCTGGGTCGGGGTTT | 70.4 | -9.2 | 2.1 | -4.5 | 8.1E-06 | 3.5E-05 | BT20/MB468 | MP |
| >hg38_21_+_8433172_8446622_RNA45SN1_45S_50ntFlanks@9534.9571.38\|AGGAAACTCTGGTGGAGGTCCGTAGCGGTCCTGACGTG | >hg38_21_+_8433172_8446622_RNA45SN1_45S_50ntFlanks | 9534 | 9571 | 38 | AGGAAACTCTGGTGGAGGTCCGTAGCGGTCCTGACGTG | 30585.0 | -3.6 | 0.8 | -4.2 | 2.3E-05 | 3.5E-05 | MB231/MB468 | CYTO |
| >MI0000265\|hsa-mir-7-3&WithFlank&19\|+\|4770664\|4770785@37.61.25&[MIMAT0000252_1&hsa-miR-7-5p&offsets\|0\|+1;m-402&19\|+\|4770700\|4770723&offsets\|0\|+1]\|TGGAAGACTAGTGATTTTGTTGTTC | >MI0000265\|hsa-mir-7-3&WithFlank&19\|+\|4770664\|4770785 | 37 | 61 | 25 | TGGAAGACTAGTGATTTTGTTGTTC | 226.2 | 6.5 | 1.5 | 4.2 | 2.5E-05 | 3.5E-05 | MB231/MB468 | MITO |
| >MI0000088\|hsa-mir-30a&WithFlank&6\|-\|71403545\|71403627@53.72.20(+1A)&[MIMAT0000088&hsa-miR-30a-3p&offsets\|0\|-2(+1A);m-38&6\|-\|71403554\|71403575&offsets\|0\|-2(+1A)]\|CTTTCAGTCGGATGTTTGCAA | >MI0000088\|hsa-mir-30a&WithFlank&6\|-\|71403545\|71403627 | 53 | 72 | 20(+1A) | CTTTCAGTCGGATGTTTGCAA | 224.6 | 5.1 | 1.2 | 4.3 | 2.1E-05 | 3.5E-05 | MB231/MB468 | MP |
| >hg38_21_+_8433172_8446622_RNA45SN1_45S_50ntFlanks@6652.6700.49\|GACTCTTAGCGGTGGATCACTCGGCTCGTGCGTCGATGAAGAACGCAGC | >hg38_21_+_8433172_8446622_RNA45SN1_45S_50ntFlanks | 6652 | 6700 | 49 | GACTCTTAGCGGTGGATCACTCGGCTCGTGCGTCGATGAAGAACGCAGC | 1107.2 | 5.1 | 1.2 | 4.2 | 2.5E-05 | 3.5E-05 | MB231/MB468 | NUC |
| >hg38_21_+_8433172_8446622_RNA45SN1_45S_50ntFlanks@11920.11959.40\|ATGAGAGGTGTAGAATAAGTGGGAGGCCCCCGGCGCCCCC | >hg38_21_+_8433172_8446622_RNA45SN1_45S_50ntFlanks | 11920 | 11959 | 40 | ATGAGAGGTGTAGAATAAGTGGGAGGCCCCCGGCGCCCCC | 69.0 | -10.0 | 2.3 | -4.3 | 2.1E-05 | 3.5E-05 | MB231/MB468 | MP |
| >hg38_21_+_8433172_8446622_RNA45SN1_45S_50ntFlanks@12241.12258.18\|CAGAAGGGCAAAAGCTCG | >hg38_21_+_8433172_8446622_RNA45SN1_45S_50ntFlanks | 12241 | 12258 | 18 | CAGAAGGGCAAAAGCTCG | 51.5 | -8.8 | 2.0 | -4.5 | 8.2E-06 | 3.5E-05 | BT20/MB468 | MP |
| >hg38_21_+_8433172_8446622_RNA45SN1_45S_50ntFlanks@12457.12485.29\|CAAGCGTTGGATTGTTCACCCACTAATAG | >hg38_21_+_8433172_8446622_RNA45SN1_45S_50ntFlanks | 12457 | 12485 | 29 | CAAGCGTTGGATTGTTCACCCACTAATAG | 259.3 | -5.0 | 1.2 | -4.2 | 2.5E-05 | 3.5E-05 | MB231/MB468 | NUC |
| >am_trna128_GlyGCC_6_-_27870686_27870756@1.31.31,trna133_GlyCCC_1_-_16872434_16872504@1.31.31,trna18_GlyGCC_16_+_70822597_70822667@1.31.31,trna19_GlyGCC_16_+_70823410_70823480@1.31.31,trna19_GlyGCC_2_-_157257659_157257729@1.31.31,trna24_GlyGCC_16_-_70812942_70813012@1.31.31,trna25_GlyGCC_16_-_70812114_70812184@1.31.31,trna4_GlyCCC_1_+_17188416_17188486@1.31.31,trna5_GlyGCC_17_+_8029064_8029134@1.31.31,trna68_GlyGCC_1_-_161493637_161493707@1.31.31\|GCATTGGTGGTTCAGTGGTAGAATTCTCGCC | >am_trna128_GlyGCC_6_-_27870686_27870756 | 1 | 31 | 31 | GCATTGGTGGTTCAGTGGTAGAATTCTCGCC | 23893.3 | 5.4 | 1.2 | 4.5 | 8.2E-06 | 3.5E-05 | BT20/MB468 | MP |
| >hg38_21_+_8433172_8446622_RNA45SN1_45S_50ntFlanks@12539.12557.19\|ATGATGTGTTGTTGCCATG | >hg38_21_+_8433172_8446622_RNA45SN1_45S_50ntFlanks | 12539 | 12557 | 19 | ATGATGTGTTGTTGCCATG | 542.0 | -6.1 | 1.4 | -4.5 | 8.2E-06 | 3.5E-05 | BT20/MB468 | MP |
| >hg38_21_+_8433172_8446622_RNA45SN1_45S_50ntFlanks@9930.9955.26\|ATATAGACAGCAGGACGGTGGCCATG | >hg38_21_+_8433172_8446622_RNA45SN1_45S_50ntFlanks | 9930 | 9955 | 26 | ATATAGACAGCAGGACGGTGGCCATG | 215.8 | -6.5 | 1.5 | -4.3 | 2.1E-05 | 3.5E-05 | MB231/MB468 | MP |
| >hg38_21_+_8433172_8446622_RNA45SN1_45S_50ntFlanks@10797.10826.30\|GCATGTTGGAACAATGTAGGTAAGGGAAGT | >hg38_21_+_8433172_8446622_RNA45SN1_45S_50ntFlanks | 10797 | 10826 | 30 | GCATGTTGGAACAATGTAGGTAAGGGAAGT | 245.6 | -1.8 | 0.4 | -4.3 | 1.8E-05 | 3.5E-05 | BT20/MB468 | MITO |
| >hg38_21_+_8433172_8446622_RNA45SN1_45S_50ntFlanks@12686.12711.26\|CCAGGCGGAACGATACGGCAGCGCCG | >hg38_21_+_8433172_8446622_RNA45SN1_45S_50ntFlanks | 12686 | 12711 | 26 | CCAGGCGGAACGATACGGCAGCGCCG | 214.9 | -6.4 | 1.4 | -4.5 | 8.2E-06 | 3.5E-05 | BT20/MB468 | MP |
| >hg38_21_+_8433172_8446622_RNA45SN1_45S_50ntFlanks@8355.8391.37\|TGAAAAGAACTTTGAAGAGAGAGTTCAAGAGGGCGTG | >hg38_21_+_8433172_8446622_RNA45SN1_45S_50ntFlanks | 8355 | 8391 | 37 | TGAAAAGAACTTTGAAGAGAGAGTTCAAGAGGGCGTG | 139.4 | 7.5 | 1.7 | 4.5 | 8.2E-06 | 3.5E-05 | BT20/MB468 | MP |
| >hg38_21_+_8433172_8446622_RNA45SN1_45S_50ntFlanks@8357.8377.21\|AAAAGAACTTTGAAGAGAGAG | >hg38_21_+_8433172_8446622_RNA45SN1_45S_50ntFlanks | 8357 | 8377 | 21 | AAAAGAACTTTGAAGAGAGAG | 9255.0 | -8.4 | 1.9 | -4.5 | 8.2E-06 | 3.5E-05 | BT20/MB468 | MP |
| >hg38_21_+_8433172_8446622_RNA45SN1_45S_50ntFlanks@12938.12955.18\|ACCATTCGTAGACGACCT | >hg38_21_+_8433172_8446622_RNA45SN1_45S_50ntFlanks | 12938 | 12955 | 18 | ACCATTCGTAGACGACCT | 175.1 | -2.0 | 0.5 | -4.2 | 2.4E-05 | 3.5E-05 | MB231/MB468 | CYTO |
| >hg38_21_+_8433172_8446622_RNA45SN1_45S_50ntFlanks@11783.11820.38\|CATGAATGGATGAACGAGATTCCCACTGTCCCTACCTA | >hg38_21_+_8433172_8446622_RNA45SN1_45S_50ntFlanks | 11783 | 11820 | 38 | CATGAATGGATGAACGAGATTCCCACTGTCCCTACCTA | 486.1 | -6.2 | 1.5 | -4.3 | 1.8E-05 | 3.5E-05 | BT20/MB468 | MITO |
| >hg38_21_+_8433172_8446622_RNA45SN1_45S_50ntFlanks@11569.11594.26\|CAGCCGACTTAGAACTGGTGCGGACC | >hg38_21_+_8433172_8446622_RNA45SN1_45S_50ntFlanks | 11569 | 11594 | 26 | CAGCCGACTTAGAACTGGTGCGGACC | 440.2 | -3.4 | 0.8 | -4.2 | 2.2E-05 | 3.5E-05 | BT20/MB468 | CYTO |
| >hg38_21_+_8433172_8446622_RNA45SN1_45S_50ntFlanks@4417.4438.22\|CGAGGCGAGCCACCGCCCGTCC | >hg38_21_+_8433172_8446622_RNA45SN1_45S_50ntFlanks | 4417 | 4438 | 22 | CGAGGCGAGCCACCGCCCGTCC | 75.2 | -10.1 | 2.4 | -4.3 | 2.1E-05 | 3.5E-05 | MB231/MB468 | MP |
| >hg38_21_+_8433172_8446622_RNA45SN1_45S_50ntFlanks@11666.11693.28\|TCTGCCCAGTGCTCTGAATGTCAAAGTG | >hg38_21_+_8433172_8446622_RNA45SN1_45S_50ntFlanks | 11666 | 11693 | 28 | TCTGCCCAGTGCTCTGAATGTCAAAGTG | 405.4 | 3.6 | 0.9 | 4.2 | 2.4E-05 | 3.5E-05 | MB231/MB468 | CYTO |
| >hg38_21_+_8433172_8446622_RNA45SN1_45S_50ntFlanks@8357.8378.22\|AAAAGAACTTTGAAGAGAGAGT | >hg38_21_+_8433172_8446622_RNA45SN1_45S_50ntFlanks | 8357 | 8378 | 22 | AAAAGAACTTTGAAGAGAGAGT | 2442.8 | -3.0 | 0.7 | -4.3 | 1.8E-05 | 3.5E-05 | BT20/MB468 | MITO |
| >hg38_21_+_8433172_8446622_RNA45SN1_45S_50ntFlanks@12349.12375.27\|AAAAGTTACCACAGGGATAACTGGCTT | >hg38_21_+_8433172_8446622_RNA45SN1_45S_50ntFlanks | 12349 | 12375 | 27 | AAAAGTTACCACAGGGATAACTGGCTT | 85.6 | -10.3 | 2.4 | -4.3 | 2.1E-05 | 3.5E-05 | MB231/MB468 | MP |
| >hg38_21_+_8433172_8446622_RNA45SN1_45S_50ntFlanks@3789.3814.26\|ACTGCGAATGGCTCATTAAATCAGTT | >hg38_21_+_8433172_8446622_RNA45SN1_45S_50ntFlanks | 3789 | 3814 | 26 | ACTGCGAATGGCTCATTAAATCAGTT | 40.3 | -8.4 | 1.9 | -4.5 | 8.3E-06 | 3.5E-05 | BT20/MB468 | MP |
| >MI0000067\|hsa-let-7f-1&WithFlank&9\|+\|94176341\|94176439@13.32.20(+1C);MI0000068\|hsa-let-7f-2&WithFlank&X\|-\|53557186\|53557280@14.33.20(+1C)&[MIMAT0000067&hsa-let-7f-5p&offsets\|0\|-2(+1C);m-20&9\|+\|94176353\|94176374&offsets\|0\|-2(+1C)];[MIMAT0000067_1&hsa-let-7f-5p&offsets\|0\|-2(+1C);m-19&X\|-\|53557246\|53557267&offsets\|0\|-2(+1C)]\|TGAGGTAGTAGATTGTATAGC | >MI0000067\|hsa-let-7f-1&WithFlank&9\|+\|94176341\|94176439 | 13 | 32 | 20(+1C) | TGAGGTAGTAGATTGTATAGC | 799.2 | 3.9 | 0.9 | 4.3 | 2.1E-05 | 3.5E-05 | MB231/MB468 | MP |
| >hg38_21_+_8433172_8446622_RNA45SN1_45S_50ntFlanks@12244.12272.29\|AAGGGCAAAAGCTCGCTTGATCTTGATTT | >hg38_21_+_8433172_8446622_RNA45SN1_45S_50ntFlanks | 12244 | 12272 | 29 | AAGGGCAAAAGCTCGCTTGATCTTGATTT | 264.4 | -6.3 | 1.5 | -4.3 | 2.1E-05 | 3.5E-05 | MB231/MB468 | MP |
| >hg38_21_+_8433172_8446622_RNA45SN1_45S_50ntFlanks@10738.10755.18\|AAATCTCGCGCCGGGCCG | >hg38_21_+_8433172_8446622_RNA45SN1_45S_50ntFlanks | 10738 | 10755 | 18 | AAATCTCGCGCCGGGCCG | 2570.0 | -3.6 | 0.9 | -4.2 | 2.5E-05 | 3.5E-05 | MB231/MB468 | NUC |
| >hg38_21_+_8433172_8446622_RNA45SN1_45S_50ntFlanks@12154.12182.29\|GGGAGTTTGACTGGGGCGGTACACCTGTC | >hg38_21_+_8433172_8446622_RNA45SN1_45S_50ntFlanks | 12154 | 12182 | 29 | GGGAGTTTGACTGGGGCGGTACACCTGTC | 745.7 | -5.6 | 1.3 | -4.5 | 8.3E-06 | 3.5E-05 | BT20/MB468 | MP |
| >hg38_21_+_8433172_8446622_RNA45SN1_45S_50ntFlanks@5471.5509.39\|CACGGCCCTGGCGGAGCGCTGAGAAGACGGTCGAACTTG | >hg38_21_+_8433172_8446622_RNA45SN1_45S_50ntFlanks | 5471 | 5509 | 39 | CACGGCCCTGGCGGAGCGCTGAGAAGACGGTCGAACTTG | 134.1 | -6.6 | 1.5 | -4.3 | 2.1E-05 | 3.5E-05 | MB231/MB468 | MP |
| >hg38_21_+_8433172_8446622_RNA45SN1_45S_50ntFlanks@10511.10532.22\|ATCCGGAGTGGCGGAGATGGGC | >hg38_21_+_8433172_8446622_RNA45SN1_45S_50ntFlanks | 10511 | 10532 | 22 | ATCCGGAGTGGCGGAGATGGGC | 230.5 | -6.5 | 1.5 | -4.3 | 2.1E-05 | 3.5E-05 | MB231/MB468 | MP |
| >hg38_21_+_8433172_8446622_RNA45SN1_45S_50ntFlanks@9027.9060.34\|CGTCTCCTCGTGGGGGGGCCGGGCCACCCCTCCC | >hg38_21_+_8433172_8446622_RNA45SN1_45S_50ntFlanks | 9027 | 9060 | 34 | CGTCTCCTCGTGGGGGGGCCGGGCCACCCCTCCC | 133.0 | -5.4 | 1.3 | -4.3 | 1.8E-05 | 3.5E-05 | BT20/MB468 | MITO |
| >hg38_21_+_8433172_8446622_RNA45SN1_45S_50ntFlanks@3853.3882.30\|AACTGTGGTAATTCTAGAGCTAATACATGC | >hg38_21_+_8433172_8446622_RNA45SN1_45S_50ntFlanks | 3853 | 3882 | 30 | AACTGTGGTAATTCTAGAGCTAATACATGC | 107.1 | -6.4 | 1.5 | -4.3 | 2.1E-05 | 3.5E-05 | MB231/MB468 | MP |
| >MI0003776\|hsa-mir-320b-1&WithFlank&1\|+\|116671740\|116671823@48.66.19(+2A);MI0003839\|hsa-mir-320b-2&WithFlank&1\|-\|224257034\|224257116@47.65.19(+2A);MI0003778\|hsa-mir-320c-1&WithFlank&18\|+\|21683512\|21683595@48.66.19(+2A);MI0008191\|hsa-mir-320c-2&WithFlank&18\|+\|24321669\|24321752@48.66.19(+2A);MI0000542\|hsa-mir-320a&WithFlank&8\|-\|22244960\|22245043@48.66.19(+2A)&[MIMAT0005792&hsa-miR-320b&offsets\|0\|-3(+2A);m-183&1\|+\|116671787\|116671806&offsets\|0\|-1(+2A)];[MIMAT0005792_1&hsa-miR-320b&offsets\|0\|-3(+2A);m-183&1\|-\|224257051\|224257070&offsets\|0\|-1(+2A)];[MIMAT0005793&hsa-miR-320c&offsets\|0\|-1(+2A);m-222&18\|+\|21683559\|21683578&offsets\|0\|-1(+2A)];[MIMAT0005793_1&hsa-miR-320c&offsets\|0\|-1(+2A);m-219&18\|+\|24321716\|24321735&offsets\|0\|-1(+2A)];[MIMAT0000510&hsa-miR-320a-3p&offsets\|0\|-3(+2A);m-60&8\|-\|22244975\|22244996&offsets\|0\|-3(+2A)]\|AAAAGCTGGGTTGAGAGGGAA | >MI0003776\|hsa-mir-320b-1&WithFlank&1\|+\|116671740\|116671823 | 48 | 66 | 19(+2A) | AAAAGCTGGGTTGAGAGGGAA | 172.4 | 8.0 | 1.8 | 4.5 | 8.3E-06 | 3.5E-05 | BT20/MB468 | MP |
| >hg38_1_-_228634819_228635039_RNA5S12_5S_50ntFlanks@51.92.42\|GTCTACGGCCATACCACCCTGAACGCGCCCGATCTCGTCTGA | >hg38_1_-_228634819_228635039_RNA5S12_5S_50ntFlanks | 51 | 92 | 42 | GTCTACGGCCATACCACCCTGAACGCGCCCGATCTCGTCTGA | 984.2 | 4.0 | 0.9 | 4.3 | 1.8E-05 | 3.5E-05 | BT20/MB468 | MITO |
| >MI0000253\|hsa-mir-148a&WithFlank&7\|-\|25949913\|25949992@50.70.21&[MIMAT0000243&hsa-miR-148a-3p&offsets\|0\|-1;m-7&7\|-\|25949922\|25949943&offsets\|0\|-1]\|TCAGTGCACTACAGAACTTTG | >MI0000253\|hsa-mir-148a&WithFlank&7\|-\|25949913\|25949992 | 50 | 70 | 21 | TCAGTGCACTACAGAACTTTG | 378.2 | 5.5 | 1.2 | 4.7 | 3.1E-06 | 3.5E-05 | BT20/MB231 | NUC |
| >hg38_21_+_8433172_8446622_RNA45SN1_45S_50ntFlanks@11707.11740.34\|AAGCGCGGGTAAACGGCGGGAGTAACTATGACTC | >hg38_21_+_8433172_8446622_RNA45SN1_45S_50ntFlanks | 11707 | 11740 | 34 | AAGCGCGGGTAAACGGCGGGAGTAACTATGACTC | 386.5 | -5.1 | 1.2 | -4.2 | 2.5E-05 | 3.5E-05 | MB231/MB468 | NUC |
| >hg38_21_+_8433172_8446622_RNA45SN1_45S_50ntFlanks@10423.10455.33\|AGAGATGGGCGAGCGCCGTTCCGAAGGGACGGG | >hg38_21_+_8433172_8446622_RNA45SN1_45S_50ntFlanks | 10423 | 10455 | 33 | AGAGATGGGCGAGCGCCGTTCCGAAGGGACGGG | 150.8 | -2.0 | 0.5 | -4.2 | 2.2E-05 | 3.5E-05 | BT20/MB468 | CYTO |
| >MI0000102\|hsa-mir-100&WithFlank&11\|-\|122152223\|122152314@20.38.19(+1C)&[MIMAT0000098&hsa-miR-100-5p&offsets\|+1\|-2(+1C);m-27&11\|-\|122152276\|122152296&offsets\|+1\|-1(+1C)]\|ACCCGTAGATCCGAACTTGC | >MI0000102\|hsa-mir-100&WithFlank&11\|-\|122152223\|122152314 | 20 | 38 | 19(+1C) | ACCCGTAGATCCGAACTTGC | 78.8 | -9.7 | 2.1 | -4.7 | 2.6E-06 | 3.5E-05 | BT20/MB231 | MP |
| >hg38_21_+_8433172_8446622_RNA45SN1_45S_50ntFlanks@4728.4755.28\|ATCAAGAACGAAAGTCGGAGGTTCGAAG | >hg38_21_+_8433172_8446622_RNA45SN1_45S_50ntFlanks | 4728 | 4755 | 28 | ATCAAGAACGAAAGTCGGAGGTTCGAAG | 321.3 | -6.3 | 1.5 | -4.3 | 2.1E-05 | 3.5E-05 | MB231/MB468 | MP |
| >MI0000750\|hsa-mir-26a-2&WithFlank&12\|-\|57824603\|57824698@20.39.20(+2U);MI0000083\|hsa-mir-26a-1&WithFlank&3\|+\|37969398\|37969486@16.35.20(+2U)&[MIMAT0000082&hsa-miR-26a-5p&offsets\|0\|-2(+2U);m-33&12\|-\|57824658\|57824679&offsets\|0\|-2(+2U)];[MIMAT0000082_1&hsa-miR-26a-5p&offsets\|0\|-2(+2U);m-34&3\|+\|37969413\|37969434&offsets\|0\|-2(+2U)]\|TTCAAGTAATCCAGGATAGGTT | >MI0000750\|hsa-mir-26a-2&WithFlank&12\|-\|57824603\|57824698 | 20 | 39 | 20(+2U) | TTCAAGTAATCCAGGATAGGTT | 334.3 | 4.7 | 1.1 | 4.2 | 2.5E-05 | 3.6E-05 | MB231/MB468 | MITO |
| >hg38_21_+_8433172_8446622_RNA45SN1_45S_50ntFlanks@7975.8002.28\|CGCGACCTCAGATCAGACGTGGCGACCC | >hg38_21_+_8433172_8446622_RNA45SN1_45S_50ntFlanks | 7975 | 8002 | 28 | CGCGACCTCAGATCAGACGTGGCGACCC | 9582.5 | 3.7 | 0.8 | 4.5 | 8.4E-06 | 3.6E-05 | BT20/MB468 | MP |

<table>
<tr><td>>hg38_21_+_8433172_8446622_RNA45SN1_45S_50ntFlanks@3782.3812.31|CAGTGAAACTGCGAATGGCTCATTAAATCAG</td><td>>hg38_21_+_8433172_8446622_RNA45SN1_45S_50ntFlanks</td><td>3782</td><td>3812</td><td>31</td><td>CAGTGAAACTGCGAATGGCTCATTAAATCAG</td><td>178.6</td><td>-4.8</td><td>1.1</td><td>-4.2</td><td>2.4E-05</td><td>3.6E-05</td><td>MB231/MB468</td><td>CYTO</td></tr>
<tr><td>>MI0000100|hsa-mir-98&WithFlank&X|-|53556217|53556347@28.48.21(+1C)&[MIMAT0000096&hsa-miR-98-5p&offsets|0|-1(+1C);m-143&X|-|53556299|53556320&offsets|0|-1(+1C)]|TGAGGTAGTAAGTTGTATTGTC</td><td>>MI0000100|hsa-mir-98&WithFlank&X|-|53556217|53556347</td><td>28</td><td>48</td><td>21(+1C)</td><td>TGAGGTAGTAAGTTGTATTGTC</td><td>400.3</td><td>4.1</td><td>1.0</td><td>4.2</td><td>2.5E-05</td><td>3.6E-05</td><td>MB231/MB468</td><td>MITO</td></tr>
<tr><td>>hg38_21_+_8433172_8446622_RNA45SN1_45S_50ntFlanks@12633.12651.19|CGAAGCTACCATCTGTGGG</td><td>>hg38_21_+_8433172_8446622_RNA45SN1_45S_50ntFlanks</td><td>12633</td><td>12651</td><td>19</td><td>CGAAGCTACCATCTGTGGG</td><td>617.6</td><td>-4.7</td><td>1.1</td><td>-4.2</td><td>2.4E-05</td><td>3.6E-05</td><td>MB231/MB468</td><td>CYTO</td></tr>
<tr><td>>hg38_21_+_8433172_8446622_RNA45SN1_45S_50ntFlanks@5189.5206.18|TAACAGGTCTGTGATGCC</td><td>>hg38_21_+_8433172_8446622_RNA45SN1_45S_50ntFlanks</td><td>5189</td><td>5206</td><td>18</td><td>TAACAGGTCTGTGATGCC</td><td>54.5</td><td>-8.8</td><td>2.0</td><td>-4.5</td><td>8.4E-06</td><td>3.6E-05</td><td>BT20/MB468</td><td>MP</td></tr>
<tr><td>>hg38_21_+_8433172_8446622_RNA45SN1_45S_50ntFlanks@9939.9961.23|GCAGGACGGTGGCCATGGAAGTC</td><td>>hg38_21_+_8433172_8446622_RNA45SN1_45S_50ntFlanks</td><td>9939</td><td>9961</td><td>23</td><td>GCAGGACGGTGGCCATGGAAGTC</td><td>251.3</td><td>8.1</td><td>1.8</td><td>4.5</td><td>8.4E-06</td><td>3.6E-05</td><td>BT20/MB468</td><td>MP</td></tr>
<tr><td>>MI0000076|hsa-mir-20a&WithFlank&13|+|91351059|91351141@14.34.21&[MIMAT0000075&hsa-miR-20a-5p&offsets|0|-2;m-96&13|+|91351072|91351094&offsets|0|-2]|TAAAGTGCTTATAGTGCAGGT</td><td>>MI0000076|hsa-mir-20a&WithFlank&13|+|91351059|91351141</td><td>14</td><td>34</td><td>21</td><td>TAAAGTGCTTATAGTGCAGGT</td><td>2022.6</td><td>3.2</td><td>0.8</td><td>4.3</td><td>2.1E-05</td><td>3.6E-05</td><td>MB231/MB468</td><td>MP</td></tr>
<tr><td>>trna13_LysCTT_14_-_58706613_58706685@1.41.41,trna2_LysCTT_15_+_79152904_79152976@1.41.41|GCCCGGCTAGCTCAGTCGGTAGAGCATGGGACTCTTAATCC</td><td>>trna13_LysCTT_14_-_58706613_58706685</td><td>1</td><td>41</td><td>41</td><td>GCCCGGCTAGCTCAGTCGGTAGAGCATGGGACTCTTAATCC</td><td>228.4</td><td>5.1</td><td>1.2</td><td>4.2</td><td>2.4E-05</td><td>3.6E-05</td><td>MB231/MB468</td><td>CYTO</td></tr>
<tr><td>>hg38_MT_+_598_1651_RNR1_12S_50ntFlanks@370.404.35|ATAAAGCTAAAACTCACCTGAGTTGTAAAAAACTC</td><td>>hg38_MT_+_598_1651_RNR1_12S_50ntFlanks</td><td>370</td><td>404</td><td>35</td><td>ATAAAGCTAAAACTCACCTGAGTTGTAAAAAACTC</td><td>918.6</td><td>3.8</td><td>0.9</td><td>4.2</td><td>2.6E-05</td><td>3.6E-05</td><td>MB231/MB468</td><td>MITO</td></tr>
<tr><td>>trna10_LysCTT_16_+_3241501_3241573@3.35.33,trna13_LysCTT_14_-_58706613_58706685@3.35.33,trna2_LysCTT_15_+_79152904_79152976@3.35.33|CCGGCTAGCTCAGTCGGTAGAGCATGGGACTCT</td><td>>trna10_LysCTT_16_+_3241501_3241573</td><td>3</td><td>35</td><td>33</td><td>CCGGCTAGCTCAGTCGGTAGAGCATGGGACTCT</td><td>2335.9</td><td>-4.0</td><td>0.8</td><td>-4.7</td><td>3.1E-06</td><td>3.6E-05</td><td>BT20/MB231</td><td>MITO</td></tr>
<tr><td>>hg38_21_+_8433172_8446622_RNA45SN1_45S_50ntFlanks@9503.9527.25|GGTGAACTATGCCTGGGCAGGGCGA</td><td>>hg38_21_+_8433172_8446622_RNA45SN1_45S_50ntFlanks</td><td>9503</td><td>9527</td><td>25</td><td>GGTGAACTATGCCTGGGCAGGGCGA</td><td>72.1</td><td>-10.0</td><td>2.4</td><td>-4.3</td><td>2.1E-05</td><td>3.6E-05</td><td>MB231/MB468</td><td>MP</td></tr>
<tr><td>>trnaMT_ThrTGT_MT_+_15888_15953@36.68.33|AACCGGAGATGAAAACCTTTTTCCAAGGACACC</td><td>>trnaMT_ThrTGT_MT_+_15888_15953</td><td>36</td><td>68</td><td>33</td><td>AACCGGAGATGAAAACCTTTTTCCAAGGACACC</td><td>28.2</td><td>8.0</td><td>1.9</td><td>4.3</td><td>2.1E-05</td><td>3.6E-05</td><td>MB231/MB468</td><td>MP</td></tr>
<tr><td>>hg38_21_+_8433172_8446622_RNA45SN1_45S_50ntFlanks@10248.10282.35|CGACGAGTAGGAGGGCCGCTGCGGTGAGCCTTGAA</td><td>>hg38_21_+_8433172_8446622_RNA45SN1_45S_50ntFlanks</td><td>10248</td><td>10282</td><td>35</td><td>CGACGAGTAGGAGGGCCGCTGCGGTGAGCCTTGAA</td><td>113.9</td><td>10.2</td><td>2.2</td><td>4.7</td><td>2.6E-06</td><td>3.6E-05</td><td>BT20/MB231</td><td>MP</td></tr>
<tr><td>>hg38_21_+_8433172_8446622_RNA45SN1_45S_50ntFlanks@12405.12437.33|CTTTTTGATCCTTCGATGTCGGCTCTTCCTATC</td><td>>hg38_21_+_8433172_8446622_RNA45SN1_45S_50ntFlanks</td><td>12405</td><td>12437</td><td>33</td><td>CTTTTTGATCCTTCGATGTCGGCTCTTCCTATC</td><td>2886.4</td><td>-5.5</td><td>1.3</td><td>-4.2</td><td>2.2E-05</td><td>3.6E-05</td><td>BT20/MB468</td><td>CYTO</td></tr>
<tr><td>>hg38_21_+_8433172_8446622_RNA45SN1_45S_50ntFlanks@12073.12110.38|CCGAGGGGCTCTCGCTTCTGGCGCCAAGCGCCCGGCCG</td><td>>hg38_21_+_8433172_8446622_RNA45SN1_45S_50ntFlanks</td><td>12073</td><td>12110</td><td>38</td><td>CCGAGGGGCTCTCGCTTCTGGCGCCAAGCGCCCGGCCG</td><td>765.0</td><td>-6.1</td><td>1.4</td><td>-4.3</td><td>1.9E-05</td><td>3.6E-05</td><td>BT20/MB468</td><td>MITO</td></tr>
<tr><td>>hg38_21_+_8433172_8446622_RNA45SN1_45S_50ntFlanks@11778.11804.27|ACGCGCATGAATGGATGAACGAGATTC</td><td>>hg38_21_+_8433172_8446622_RNA45SN1_45S_50ntFlanks</td><td>11778</td><td>11804</td><td>27</td><td>ACGCGCATGAATGGATGAACGAGATTC</td><td>396.1</td><td>-6.2</td><td>1.5</td><td>-4.2</td><td>2.6E-05</td><td>3.6E-05</td><td>MB231/MB468</td><td>MITO</td></tr>
<tr><td>>trna116_GluCTC_1_-_145399233_145399304@37.70.34,trna134_GluTTC_1_-_16861774_16861845@37.70.34,trna71_GluCTC_1_-_161439189_161439260@37.70.34,trna74_GluCTC_1_-_161431809_161431880@37.70.34,trna77_GluCTC_1_-_161424398_161424469@37.70.34,trna77_GluCTC_6_+_28949976_28950047@37.70.34,trna80_GluCTC_1_-_161417018_161417089@37.70.34,trna84_GluTTC_1_-_161391883_161391954@37.70.34,trna87_GluCTC_6_-_126101393_126101464@37.70.34|ACCGCCGCGGCCCGGGTTCGATTCCCGGTCAGGG</td><td>>trna116_GluCTC_1_-_145399233_145399304</td><td>37</td><td>70</td><td>34</td><td>ACCGCCGCGGCCCGGGTTCGATTCCCGGTCAGGG</td><td>148.3</td><td>-3.7</td><td>0.9</td><td>-4.2</td><td>2.2E-05</td><td>3.6E-05</td><td>BT20/MB468</td><td>CYTO</td></tr>
<tr><td>>hg38_21_+_8433172_8446622_RNA45SN1_45S_50ntFlanks@10458.10486.29|ATGGCCTCCGTTGCCCTCGGCCGATCGAA</td><td>>hg38_21_+_8433172_8446622_RNA45SN1_45S_50ntFlanks</td><td>10458</td><td>10486</td><td>29</td><td>ATGGCCTCCGTTGCCCTCGGCCGATCGAA</td><td>237.8</td><td>-2.9</td><td>0.7</td><td>-4.3</td><td>1.9E-05</td><td>3.6E-05</td><td>BT20/MB468</td><td>MITO</td></tr>
<tr><td>>hg38_21_+_8433172_8446622_RNA45SN1_45S_50ntFlanks@4685.4707.23|AGACGGACCAGAGCGAAAGCATT</td><td>>hg38_21_+_8433172_8446622_RNA45SN1_45S_50ntFlanks</td><td>4685</td><td>4707</td><td>23</td><td>AGACGGACCAGAGCGAAAGCATT</td><td>876.8</td><td>-6.2</td><td>1.5</td><td>-4.2</td><td>2.7E-05</td><td>3.6E-05</td><td>BT20/MB468</td><td>NUC</td></tr>
<tr><td>>hg38_21_+_8433172_8446622_RNA45SN1_45S_50ntFlanks@9570.9601.32|TGCAAATCGGTCGTCCGACCTGGGTATAGGGG</td><td>>hg38_21_+_8433172_8446622_RNA45SN1_45S_50ntFlanks</td><td>9570</td><td>9601</td><td>32</td><td>TGCAAATCGGTCGTCCGACCTGGGTATAGGGG</td><td>191.9</td><td>-6.3</td><td>1.5</td><td>-4.2</td><td>2.1E-05</td><td>3.6E-05</td><td>MB231/MB468</td><td>MP</td></tr>
<tr><td>>MI0000298|hsa-mir-221&WithFlank&X|-|45746151|45746272@71.88.18(+3U)&[MIMAT0000278&hsa-miR-221-3p&offsets|0|-5(+3U);m-91&X|-|45746181|45746202&offsets|0|-4(+3U)]|AGCTACATTGTCTGCTGGTTT</td><td>>MI0000298|hsa-mir-221&WithFlank&X|-|45746151|45746272</td><td>71</td><td>88</td><td>18(+3U)</td><td>AGCTACATTGTCTGCTGGTTT</td><td>31.4</td><td>8.2</td><td>1.9</td><td>4.2</td><td>2.1E-05</td><td>3.6E-05</td><td>MB231/MB468</td><td>MP</td></tr>
<tr><td>>hg38_21_+_8433172_8446622_RNA45SN1_45S_50ntFlanks@11537.11560.24|CCGCGCGGCGCCTCGCCTCGGCCG</td><td>>hg38_21_+_8433172_8446622_RNA45SN1_45S_50ntFlanks</td><td>11537</td><td>11560</td><td>24</td><td>CCGCGCGGCGCCTCGCCTCGGCCG</td><td>220.2</td><td>-5.8</td><td>1.4</td><td>-4.3</td><td>1.9E-05</td><td>3.6E-05</td><td>BT20/MB468</td><td>MITO</td></tr>
<tr><td>>hg38_21_+_8433172_8446622_RNA45SN1_45S_50ntFlanks@11581.11608.28|AACTGGTGCGGACCAGGGGAATCCGACT</td><td>>hg38_21_+_8433172_8446622_RNA45SN1_45S_50ntFlanks</td><td>11581</td><td>11608</td><td>28</td><td>AACTGGTGCGGACCAGGGGAATCCGACT</td><td>192.4</td><td>-6.2</td><td>1.5</td><td>-4.2</td><td>2.1E-05</td><td>3.6E-05</td><td>MB231/MB468</td><td>MP</td></tr>
<tr><td>>hg38_21_+_8433172_8446622_RNA45SN1_45S_50ntFlanks@12404.12431.28|GCTTTTTGATCCTTCGATGTCGGCTCTT</td><td>>hg38_21_+_8433172_8446622_RNA45SN1_45S_50ntFlanks</td><td>12404</td><td>12431</td><td>28</td><td>GCTTTTTGATCCTTCGATGTCGGCTCTT</td><td>250.8</td><td>-4.7</td><td>1.1</td><td>-4.2</td><td>2.6E-05</td><td>3.6E-05</td><td>MB231/MB468</td><td>NUC</td></tr>
<tr><td>>hg38_21_+_8433172_8446622_RNA45SN1_45S_50ntFlanks@10679.10709.31|GGTTCCGGCGGCGTCCGGTGAGCTCTCGCTG</td><td>>hg38_21_+_8433172_8446622_RNA45SN1_45S_50ntFlanks</td><td>10679</td><td>10709</td><td>31</td><td>GGTTCCGGCGGCGTCCGGTGAGCTCTCGCTG</td><td>22659.1</td><td>-9.1</td><td>2.2</td><td>-4.2</td><td>2.7E-05</td><td>3.6E-05</td><td>BT20/MB468</td><td>NUC</td></tr>
<tr><td>>trna116_GluCTC_1_-_145399233_145399304@4.35.32,trna59_GluCTC_1_+_249168447_249168518@4.35.32,trna71_GluCTC_1_-_161439189_161439260@4.35.32,trna74_GluCTC_1_-_161431809_161431880@4.35.32,trna77_GluCTC_1_-_161424398_161424469@4.35.32,trna77_GluCTC_6_+_28949976_28950047@4.35.32,trna80_GluCTC_1_-_161417018_161417089@4.35.32,trna87_GluCTC_6_-_126101393_126101464@4.35.32|CTGGTGGTCTAGTGGTTAGGATTCGGCGCTCT</td><td>>trna116_GluCTC_1_-_145399233_145399304</td><td>4</td><td>35</td><td>32</td><td>CTGGTGGTCTAGTGGTTAGGATTCGGCGCTCT</td><td>39.2</td><td>8.5</td><td>2.0</td><td>4.2</td><td>2.2E-05</td><td>3.6E-05</td><td>MB231/MB468</td><td>MP</td></tr>
<tr><td>>hg38_21_+_8433172_8446622_RNA45SN1_45S_50ntFlanks@7979.8005.27|ACCTCAGATCAGACGTGGCGACCCGCT</td><td>>hg38_21_+_8433172_8446622_RNA45SN1_45S_50ntFlanks</td><td>7979</td><td>8005</td><td>27</td><td>ACCTCAGATCAGACGTGGCGACCCGCT</td><td>351.8</td><td>-2.7</td><td>0.6</td><td>-4.2</td><td>2.6E-05</td><td>3.6E-05</td><td>MB231/MB468</td><td>NUC</td></tr>
<tr><td>>hg38_21_+_8433172_8446622_RNA45SN1_45S_50ntFlanks@12416.12434.19|TTCGATGTCGGCTCTTCCT</td><td>>hg38_21_+_8433172_8446622_RNA45SN1_45S_50ntFlanks</td><td>12416</td><td>12434</td><td>19</td><td>TTCGATGTCGGCTCTTCCT</td><td>7756.2</td><td>3.3</td><td>0.8</td><td>4.2</td><td>2.4E-05</td><td>3.6E-05</td><td>MB231/MB468</td><td>CYTO</td></tr>
<tr><td>>hg38_21_+_8433172_8446622_RNA45SN1_45S_50ntFlanks@6745.6772.28|ACACTTCGAACGCACTTGCGGCCCCGGG</td><td>>hg38_21_+_8433172_8446622_RNA45SN1_45S_50ntFlanks</td><td>6745</td><td>6772</td><td>28</td><td>ACACTTCGAACGCACTTGCGGCCCCGGG</td><td>264.9</td><td>-5.6</td><td>1.3</td><td>-4.2</td><td>2.2E-05</td><td>3.6E-05</td><td>BT20/MB468</td><td>CYTO</td></tr>
<tr><td>>hg38_21_+_8433172_8446622_RNA45SN1_45S_50ntFlanks@9938.9981.44|AGCAGGACGGTGGCCATGGAAGTCGGAATCCGCTAAGGAGTGTG</td><td>>hg38_21_+_8433172_8446622_RNA45SN1_45S_50ntFlanks</td><td>9938</td><td>9981</td><td>44</td><td>AGCAGGACGGTGGCCATGGAAGTCGGAATCCGCTAAGGAGTGTG</td><td>267.2</td><td>8.5</td><td>1.9</td><td>4.5</td><td>8.5E-06</td><td>3.6E-05</td><td>BT20/MB468</td><td>MP</td></tr>
<tr><td>>hg38_21_+_8433172_8446622_RNA45SN1_45S_50ntFlanks@11027.11055.29|CTCTCTCTCTCTCTCCCCCGCTCCCCGTC</td><td>>hg38_21_+_8433172_8446622_RNA45SN1_45S_50ntFlanks</td><td>11027</td><td>11055</td><td>29</td><td>CTCTCTCTCTCTCTCCCCCGCTCCCCGTC</td><td>335.5</td><td>-4.0</td><td>0.9</td><td>-4.2</td><td>2.4E-05</td><td>3.6E-05</td><td>MB231/MB468</td><td>CYTO</td></tr>
<tr><td>>hg38_21_+_8433172_8446622_RNA45SN1_45S_50ntFlanks@11319.11344.26|CCCACGTCTCGTCGCGCGCGCGTCCG</td><td>>hg38_21_+_8433172_8446622_RNA45SN1_45S_50ntFlanks</td><td>11319</td><td>11344</td><td>26</td><td>CCCACGTCTCGTCGCGCGCGCGTCCG</td><td>5277.6</td><td>-7.3</td><td>1.7</td><td>-4.2</td><td>2.6E-05</td><td>3.6E-05</td><td>MB231/MB468</td><td>NUC</td></tr>
<tr><td>>MI0000298|hsa-mir-221&WithFlank&X|-|45746151|45746272@71.92.22(+2U)&[MIMAT0000278&hsa-miR-221-3p&offsets|0|-1(+2U);m-91&X|-|45746181|45746202&offsets|0|0(+2U)]|AGCTACATTGTCTGCTGGGTTTTT</td><td>>MI0000298|hsa-mir-221&WithFlank&X|-|45746151|45746272</td><td>71</td><td>92</td><td>22(+2U)</td><td>AGCTACATTGTCTGCTGGGTTTTT</td><td>536.6</td><td>5.1</td><td>1.2</td><td>4.2</td><td>2.2E-05</td><td>3.6E-05</td><td>MB231/MB468</td><td>MP</td></tr>
<tr><td>>MI0000734|hsa-mir-106b&WithFlank&7|-|100093987|100094080@58.78.21(+1A)&[MIMAT0004672&hsa-miR-106b-3p&offsets|0|-1(+1A);m-87&7|-|100094003|100094023&offsets|0|0(+1A)]|CCGCACTGTGGGTACTTGCTGA</td><td>>MI0000734|hsa-mir-106b&WithFlank&7|-|100093987|100094080</td><td>58</td><td>78</td><td>21(+1A)</td><td>CCGCACTGTGGGTACTTGCTGA</td><td>134.4</td><td>5.1</td><td>1.2</td><td>4.2</td><td>2.6E-05</td><td>3.6E-05</td><td>MB231/MB468</td><td>MITO</td></tr>
<tr><td>>hg38_21_+_8433172_8446622_RNA45SN1_45S_50ntFlanks@11770.11792.23|AATTAGTGACGCGCATGAATGGA</td><td>>hg38_21_+_8433172_8446622_RNA45SN1_45S_50ntFlanks</td><td>11770</td><td>11792</td><td>23</td><td>AATTAGTGACGCGCATGAATGGA</td><td>321.6</td><td>-6.7</td><td>1.6</td><td>-4.2</td><td>2.7E-05</td><td>3.6E-05</td><td>BT20/MB468</td><td>NUC</td></tr>
<tr><td>>hg38_21_+_8433172_8446622_RNA45SN1_45S_50ntFlanks@12479.12500.22|CTAATAGGGAACGTGAGCTGGG</td><td>>hg38_21_+_8433172_8446622_RNA45SN1_45S_50ntFlanks</td><td>12479</td><td>12500</td><td>22</td><td>CTAATAGGGAACGTGAGCTGGG</td><td>317.3</td><td>-5.0</td><td>1.2</td><td>-4.2</td><td>2.2E-05</td><td>3.6E-05</td><td>MB231/MB468</td><td>MP</td></tr>
<tr><td>>hg38_21_+_8433172_8446622_RNA45SN1_45S_50ntFlanks@12945.12965.21|GTAGACGACCTGCTTCTGGGT</td><td>>hg38_21_+_8433172_8446622_RNA45SN1_45S_50ntFlanks</td><td>12945</td><td>12965</td><td>21</td><td>GTAGACGACCTGCTTCTGGGT</td><td>95.2</td><td>-6.2</td><td>1.5</td><td>-4.2</td><td>2.2E-05</td><td>3.6E-05</td><td>MB231/MB468</td><td>MP</td></tr>
<tr><td>>hg38_21_+_8433172_8446622_RNA45SN1_45S_50ntFlanks@3838.3872.35|CTCTCCTACTTGGATAACTGTGGTAATTCTAGAGC</td><td>>hg38_21_+_8433172_8446622_RNA45SN1_45S_50ntFlanks</td><td>3838</td><td>3872</td><td>35</td><td>CTCTCCTACTTGGATAACTGTGGTAATTCTAGAGC</td><td>647.4</td><td>-7.2</td><td>1.7</td><td>-4.2</td><td>2.7E-05</td><td>3.6E-05</td><td>BT20/MB468</td><td>NUC</td></tr>
<tr><td>>hg38_21_+_8433172_8446622_RNA45SN1_45S_50ntFlanks@11874.11902.29|AGAAGACCCTGTTGAGCTTGACTCTAGTC</td><td>>hg38_21_+_8433172_8446622_RNA45SN1_45S_50ntFlanks</td><td>11874</td><td>11902</td><td>29</td><td>AGAAGACCCTGTTGAGCTTGACTCTAGTC</td><td>370.1</td><td>-4.9</td><td>1.2</td><td>-4.2</td><td>2.6E-05</td><td>3.6E-05</td><td>MB231/MB468</td><td>NUC</td></tr>
<tr><td>>hg38_21_+_8433172_8446622_RNA45SN1_45S_50ntFlanks@10291.10321.31|CGCGGGCCCGGGTGGAGCCGCCGCAGGTGCA</td><td>>hg38_21_+_8433172_8446622_RNA45SN1_45S_50ntFlanks</td><td>10291</td><td>10321</td><td>31</td><td>CGCGGGCCCGGGTGGAGCCGCCGCAGGTGCA</td><td>68.6</td><td>-10.0</td><td>2.3</td><td>-4.2</td><td>2.2E-05</td><td>3.6E-05</td><td>MB231/MB468</td><td>MP</td></tr>
<tr><td>>hg38_21_+_8433172_8446622_RNA45SN1_45S_50ntFlanks@9574.9602.29|AATCGGTCGTCCGACCTGGGTATAGGGGC</td><td>>hg38_21_+_8433172_8446622_RNA45SN1_45S_50ntFlanks</td><td>9574</td><td>9602</td><td>29</td><td>AATCGGTCGTCCGACCTGGGTATAGGGGC</td><td>470.8</td><td>2.8</td><td>0.7</td><td>4.2</td><td>2.5E-05</td><td>3.6E-05</td><td>MB231/MB468</td><td>CYTO</td></tr>
<tr><td>>hg38_1_-_228634819_228635039_RNA5S12_5S_50ntFlanks@148.170.23|GGAATACCGGGTGCTGTAGGCTT</td><td>>hg38_1_-_228634819_228635039_RNA5S12_5S_50ntFlanks</td><td>148</td><td>170</td><td>23</td><td>GGAATACCGGGTGCTGTAGGCTT</td><td>139853.0</td><td>7.9</td><td>1.7</td><td>4.7</td><td>2.7E-06</td><td>3.6E-05</td><td>BT20/MB231</td><td>MP</td></tr>
<tr><td>>hg38_21_+_8433172_8446622_RNA45SN1_45S_50ntFlanks@11876.11897.22|AAGACCCTGTTGAGCTTGACTC</td><td>>hg38_21_+_8433172_8446622_RNA45SN1_45S_50ntFlanks</td><td>11876</td><td>11897</td><td>22</td><td>AAGACCCTGTTGAGCTTGACTC</td><td>657.6</td><td>-2.2</td><td>0.5</td><td>-4.2</td><td>2.2E-05</td><td>3.6E-05</td><td>BT20/MB468</td><td>CYTO</td></tr>
<tr><td>>hg38_21_+_8433172_8446622_RNA45SN1_45S_50ntFlanks@11561.11592.32|GCGCCTAGCAGCCGACTTAGAACTGGTGCGGA</td><td>>hg38_21_+_8433172_8446622_RNA45SN1_45S_50ntFlanks</td><td>11561</td><td>11592</td><td>32</td><td>GCGCCTAGCAGCCGACTTAGAACTGGTGCGGA</td><td>1126.1</td><td>-6.4</td><td>1.5</td><td>-4.2</td><td>2.2E-05</td><td>3.6E-05</td><td>BT20/MB468</td><td>CYTO</td></tr>
</table>

| | | | | | | | | | | | | | |
|---|---|---|---|---|---|---|---|---|---|---|---|---|---|
| >MI0000481\|hsa-mir-184&WithFlank&15\|+\|79209782\|79209877@59.80.22(+1U)&[MIMAT0000454&hsa-miR-184&offsets\|0\|0(+1U);m-460&15\|+\|79209840\|79209861&offsets\|0\|0(+1U)]\|TGGACGGAGAACTGATAAGGGTT | >MI0000481\|hsa-mir-184&WithFlank&15\|+\|79209782\|79209877 | 59 | 80 | 22(+1U) | TGGACGGAGAACTGATAAGGGTT | 76.5 | 9.6 | 2.0 | 4.7 | 2.7E-06 | 3.6E-05 | BT20/MB231 | MP |
| >hg38_21_+_8433172_8446622_RNA45SN1_45S_50ntFlanks@7981.8002.22\|CTCAGATCAGACGTGGCGACCC | >hg38_21_+_8433172_8446622_RNA45SN1_45S_50ntFlanks | 7981 | 8002 | 22 | CTCAGATCAGACGTGGCGACCC | 90.5 | 9.8 | 2.1 | 4.7 | 2.7E-06 | 3.6E-05 | BT20/MB231 | MP |
| >trnaMT_GluTTC_MT_-_14674_14742@39.57.19,trnalookalike8_GluTTC_5_-_93905172_93905240@39.57.19\|ATTGGTCGTGGTTGTAGTC | >trnaMT_GluTTC_MT_-_14674_14742 | 39 | 57 | 19 | ATTGGTCGTGGTTGTAGTC | 389.3 | 6.4 | 1.5 | 4.2 | 2.2E-05 | 3.6E-05 | BT20/MB468 | CYTO |
| >hg38_21_+_8433172_8446622_RNA45SN1_45S_50ntFlanks@6652.6685.34\|GACTCTTAGCGGTGGATCACTCGGCTCGTGCGTC | >hg38_21_+_8433172_8446622_RNA45SN1_45S_50ntFlanks | 6652 | 6685 | 34 | GACTCTTAGCGGTGGATCACTCGGCTCGTGCGTC | 2828.3 | 4.8 | 1.1 | 4.4 | 8.6E-06 | 3.6E-05 | BT20/MB468 | MP |
| >hg38_21_+_8433172_8446622_RNA45SN1_45S_50ntFlanks@11317.11343.27\|ACCCCACGTCTCGTCGCGCGCGCGTCC | >hg38_21_+_8433172_8446622_RNA45SN1_45S_50ntFlanks | 11317 | 11343 | 27 | ACCCCACGTCTCGTCGCGCGCGCGTCC | 4953.1 | -8.4 | 2.0 | -4.2 | 2.7E-05 | 3.6E-05 | BT20/MB468 | NUC |
| >am_trnaMT_ValTAC_MT_+_1602_1670@29.68.40\|ACTTACACTTAGGAGATTTCAACTTAACTTGACCGCTCTG | >am_trnaMT_ValTAC_MT_+_1602_1670 | 29 | 68 | 40 | ACTTACACTTAGGAGATTTCAACTTAACTTGACCGCTCTG | 1332.8 | -8.3 | 2.0 | -4.2 | 2.7E-05 | 3.6E-05 | BT20/MB468 | NUC |
| >MI0000077\|hsa-mir-21&WithFlank&17\|+\|59841260\|59841343@14.32.19(+1C)&[MIMAT0000076&hsa-miR-21-5p&offsets\|0\|-3(+1C);m-1&17\|+\|59841273\|59841294&offsets\|0\|-3(+1C)]\|TAGCTTATCAGACTGATGTC | >MI0000077\|hsa-mir-21&WithFlank&17\|+\|59841260\|59841343 | 14 | 32 | 19(+1C) | TAGCTTATCAGACTGATGTC | 382.7 | 4.0 | 0.9 | 4.2 | 2.2E-05 | 3.6E-05 | MB231/MB468 | MP |
| >hg38_21_+_8433172_8446622_RNA45SN1_45S_50ntFlanks@11694.11717.24\|AAGAAATTCAATGAAGCGCGGGTA | >hg38_21_+_8433172_8446622_RNA45SN1_45S_50ntFlanks | 11694 | 11717 | 24 | AAGAAATTCAATGAAGCGCGGGTA | 336.6 | -6.0 | 1.4 | -4.2 | 2.6E-05 | 3.7E-05 | MB231/MB468 | NUC |
| >hg38_21_+_8433172_8446622_RNA45SN1_45S_50ntFlanks@10350.10367.18\|ACGAGAACTTTGAAGGCC | >hg38_21_+_8433172_8446622_RNA45SN1_45S_50ntFlanks | 10350 | 10367 | 18 | ACGAGAACTTTGAAGGCC | 1393.5 | -2.5 | 0.6 | -4.2 | 2.5E-05 | 3.7E-05 | MB231/MB468 | CYTO |
| >hg38_21_+_8433172_8446622_RNA45SN1_45S_50ntFlanks@9923.9946.24\|TTGGTTGATATAGACAGCAGGACG | >hg38_21_+_8433172_8446622_RNA45SN1_45S_50ntFlanks | 9923 | 9946 | 24 | TTGGTTGATATAGACAGCAGGACG | 96.8 | 9.9 | 2.1 | 4.7 | 2.7E-06 | 3.7E-05 | BT20/MB231 | MP |
| >hg38_21_+_8433172_8446622_RNA45SN1_45S_50ntFlanks@10059.10081.23\|GGCAGTCGAGAGTGGACGGGAGC | >hg38_21_+_8433172_8446622_RNA45SN1_45S_50ntFlanks | 10059 | 10081 | 23 | GGCAGTCGAGAGTGGACGGGAGC | 149.7 | 7.6 | 1.7 | 4.4 | 8.6E-06 | 3.7E-05 | BT20/MB468 | MP |
| >trna152_ValCAC_6_-_27248049_27248121@1.33.33\|GCTTCTGTAGTGTAGTGGTTATCACGTTCGCCT | >trna152_ValCAC_6_-_27248049_27248121 | 1 | 33 | 33 | GCTTCTGTAGTGTAGTGGTTATCACGTTCGCCT | 644.2 | 8.0 | 1.8 | 4.4 | 8.6E-06 | 3.7E-05 | BT20/MB468 | MP |
| >hg38_21_+_8433172_8446622_RNA45SN1_45S_50ntFlanks@4983.5015.33\|CGGACAGGATTGACAGATTGATAGCTCTTTCTC | >hg38_21_+_8433172_8446622_RNA45SN1_45S_50ntFlanks | 4983 | 5015 | 33 | CGGACAGGATTGACAGATTGATAGCTCTTTCTC | 709.7 | -6.1 | 1.4 | -4.3 | 1.9E-05 | 3.7E-05 | BT20/MB468 | MITO |
| >hg38_21_+_8433172_8446622_RNA45SN1_45S_50ntFlanks@11564.11587.24\|CCTAGCAGCCGACTTAGAACTGGT | >hg38_21_+_8433172_8446622_RNA45SN1_45S_50ntFlanks | 11564 | 11587 | 24 | CCTAGCAGCCGACTTAGAACTGGT | 228.7 | -2.5 | 0.6 | -4.2 | 2.5E-05 | 3.7E-05 | MB231/MB468 | CYTO |
| >hg38_21_+_8433172_8446622_RNA45SN1_45S_50ntFlanks@9178.9197.20\|CGCCGTCGGGCCCGGGGGAG | >hg38_21_+_8433172_8446622_RNA45SN1_45S_50ntFlanks | 9178 | 9197 | 20 | CGCCGTCGGGCCCGGGGGAG | 289.2 | -6.4 | 1.5 | -4.2 | 2.6E-05 | 3.7E-05 | MB231/MB468 | NUC |
| >hg38_21_+_8433172_8446622_RNA45SN1_45S_50ntFlanks@11718.11745.28\|AACGGCGGGAGTAACTATGACTCTCTTA | >hg38_21_+_8433172_8446622_RNA45SN1_45S_50ntFlanks | 11718 | 11745 | 28 | AACGGCGGGAGTAACTATGACTCTCTTA | 117.0 | -5.0 | 1.2 | -4.2 | 2.2E-05 | 3.7E-05 | MB231/MB468 | MP |
| >MI0000650\|hsa-mir-200c&WithFlank&12\|+\|6963693\|6963772@50.70.21(+1U)&[MIMAT0000617&hsa-miR-200c-3p&offsets\|0\|-2(+1U);m-54&12\|+\|6963742\|6963764&offsets\|0\|-2(+1U)]\|TAATACTGCCGGGTAATGATGT | >MI0000650\|hsa-mir-200c&WithFlank&12\|+\|6963693\|6963772 | 50 | 70 | 21(+1U) | TAATACTGCCGGGTAATGATGT | 608.3 | 6.4 | 1.4 | 4.4 | 8.6E-06 | 3.7E-05 | BT20/MB468 | MP |
| >hg38_21_+_8433172_8446622_RNA45SN1_45S_50ntFlanks@10797.10826.30\|GCATGTTGGAACAATGTAGGTAAGGGAAGT | >hg38_21_+_8433172_8446622_RNA45SN1_45S_50ntFlanks | 10797 | 10826 | 30 | GCATGTTGGAACAATGTAGGTAAGGGAAGT | 482.0 | -6.1 | 1.5 | -4.2 | 2.6E-05 | 3.7E-05 | MB231/MB468 | MITO |
| >am_trnaMT_ValTAC_MT_+_1602_1670@1.41.41\|CAGAGTGTAGCTTAACACAAAGCACCCAACTTACACTTAGG | >am_trnaMT_ValTAC_MT_+_1602_1670 | 1 | 41 | 41 | CAGAGTGTAGCTTAACACAAAGCACCCAACTTACACTTAGG | 36.7 | 8.4 | 2.0 | 4.2 | 2.2E-05 | 3.7E-05 | MB231/MB468 | MP |
| >trna111_HisGTG_1_-_147774845_147774916@-1A.29.30,trna118_HisGTG_1_-_145396881_145396952@-1A.29.30,trna16_HisGTG_1_+_146544773_146544844@-1A.29.30,trna1_HisGTG_15_+_45493349_45493420@-1A.29.30,trna21_HisGTG_1_+_147753471_147753542@-1A.29.30,trna33_HisGTG_6_+_27125906_27125977@-1A.29.30,trna7_HisGTG_9_-_14433938_14434009@-1A.29.30,trna8_HisGTG_15_-_45492611_45492682@-1A.29.30,trna9_HisGTG_15_-_45490804_45490875@-1A.29.30\|AGCCGTGATCGTATAGTGGTTAGTACTCTG | >trna111_HisGTG_1_-_147774845_147774916 | -1A | 29 | 30 | AGCCGTGATCGTATAGTGGTTAGTACTCTG | 131.8 | -5.4 | 1.3 | -4.3 | 1.9E-05 | 3.7E-05 | BT20/MB468 | MITO |
| >hg38_21_+_8433172_8446622_RNA45SN1_45S_50ntFlanks@4735.4758.24\|ACGAAAGTCGGAGGTTCGAAGACG | >hg38_21_+_8433172_8446622_RNA45SN1_45S_50ntFlanks | 4735 | 4758 | 24 | ACGAAAGTCGGAGGTTCGAAGACG | 1426.7 | -3.2 | 0.7 | -4.2 | 2.2E-05 | 3.7E-05 | MB231/MB468 | MP |
| >hg38_21_+_8433172_8446622_RNA45SN1_45S_50ntFlanks@6722.6758.37\|ATTGCAGGACACATTGATCATCGACACTTCGAACGCA | >hg38_21_+_8433172_8446622_RNA45SN1_45S_50ntFlanks | 6722 | 6758 | 37 | ATTGCAGGACACATTGATCATCGACACTTCGAACGCA | 196.5 | -4.9 | 1.2 | -4.2 | 2.6E-05 | 3.7E-05 | MB231/MB468 | MITO |
| >hg38_21_+_8433172_8446622_RNA45SN1_45S_50ntFlanks@4499.4517.19\|AGCGTTTACTTTGAAAAAA | >hg38_21_+_8433172_8446622_RNA45SN1_45S_50ntFlanks | 4499 | 4517 | 19 | AGCGTTTACTTTGAAAAAA | 309.9 | -6.7 | 1.6 | -4.2 | 2.6E-05 | 3.7E-05 | MB231/MB468 | MITO |
| >hg38_21_+_8433172_8446622_RNA45SN1_45S_50ntFlanks@9507.9529.23\|AACTATGCCTGGGCAGGGCGAAG | >hg38_21_+_8433172_8446622_RNA45SN1_45S_50ntFlanks | 9507 | 9529 | 23 | AACTATGCCTGGGCAGGGCGAAG | 69.6 | -10.0 | 2.4 | -4.2 | 2.2E-05 | 3.7E-05 | MB231/MB468 | MP |
| >hg38_21_+_8433172_8446622_RNA45SN1_45S_50ntFlanks@476.521.46\|TTCGTGATCGATGTGGTGACGTCGTGCTCTCCCGGGCCGGGTCCGA | >hg38_21_+_8433172_8446622_RNA45SN1_45S_50ntFlanks | 476 | 521 | 46 | TTCGTGATCGATGTGGTGACGTCGTGCTCTCCCGGGCCGGGTCCGA | 414.6 | 6.6 | 1.5 | 4.3 | 1.9E-05 | 3.7E-05 | BT20/MB468 | MITO |
| >hg38_21_+_8433172_8446622_RNA45SN1_45S_50ntFlanks@7973.8005.33\|GACGCGACCTCAGATCAGACGTGGCGACCCGCT | >hg38_21_+_8433172_8446622_RNA45SN1_45S_50ntFlanks | 7973 | 8005 | 33 | GACGCGACCTCAGATCAGACGTGGCGACCCGCT | 10833.0 | 5.0 | 1.1 | 4.7 | 3.2E-06 | 3.7E-05 | BT20/MB231 | NUC |
| >am_MI0003639\|hsa-mir-625&WithFlank&14\|+\|65471096\|65471192@58.78.21(+1C)&[MIMAT0004808&hsa-miR-625-3p&offsets\|0\|-1(+1C);m-137&14\|+\|65471153\|65471174&offsets\|0\|-1(+1C)]\|GACTATAGAACTTTCCCCCTCC | >am_MI0003639\|hsa-mir-625&WithFlank&14\|+\|65471096\|65471192 | 58 | 78 | 21(+1C) | GACTATAGAACTTTCCCCCTCC | 32.0 | 8.2 | 1.9 | 4.2 | 2.2E-05 | 3.7E-05 | MB231/MB468 | MP |
| >trna152_ValCAC_6_-_27248049_27248121@3.36.34\|TTCTGTAGTGTAGTGGTTATCACGTTCGCCTCAC | >trna152_ValCAC_6_-_27248049_27248121 | 3 | 36 | 34 | TTCTGTAGTGTAGTGGTTATCACGTTCGCCTCAC | 149.7 | 6.7 | 1.6 | 4.2 | 2.6E-05 | 3.7E-05 | MB231/MB468 | MITO |
| >hg38_21_+_8433172_8446622_RNA45SN1_45S_50ntFlanks@11599.11623.25\|GAATCCGACTGTTTAATTAAAACAA | >hg38_21_+_8433172_8446622_RNA45SN1_45S_50ntFlanks | 11599 | 11623 | 25 | GAATCCGACTGTTTAATTAAAACAA | 372.2 | 3.8 | 0.8 | 4.7 | 3.2E-06 | 3.7E-05 | BT20/MB231 | MITO |
| >hg38_21_+_8433172_8446622_RNA45SN1_45S_50ntFlanks@475.513.39\|CTTCGTGATCGATGTGGTGACGTCGTGCTCTCCCGGGCC | >hg38_21_+_8433172_8446622_RNA45SN1_45S_50ntFlanks | 475 | 513 | 39 | CTTCGTGATCGATGTGGTGACGTCGTGCTCTCCCGGGCC | 1555.9 | 3.6 | 0.8 | 4.2 | 2.6E-05 | 3.7E-05 | MB231/MB468 | NUC |
| >hg38_21_+_8433172_8446622_RNA45SN1_45S_50ntFlanks@8123.8140.18\|ATGTGGCGTACGGAAGAC | >hg38_21_+_8433172_8446622_RNA45SN1_45S_50ntFlanks | 8123 | 8140 | 18 | ATGTGGCGTACGGAAGAC | 83.9 | 9.7 | 2.1 | 4.7 | 2.7E-06 | 3.7E-05 | BT20/MB231 | MP |
| >hg38_21_+_8433172_8446622_RNA45SN1_45S_50ntFlanks@6644.6663.20\|CCTCGTACGACTCTTAGCGG | >hg38_21_+_8433172_8446622_RNA45SN1_45S_50ntFlanks | 6644 | 6663 | 20 | CCTCGTACGACTCTTAGCGG | 696.4 | 3.8 | 0.9 | 4.2 | 2.6E-05 | 3.7E-05 | MB231/MB468 | NUC |
| >hg38_21_+_8433172_8446622_RNA45SN1_45S_50ntFlanks@12635.12662.28\|AAGCTACCATCTGTGGGATTATGACTGA | >hg38_21_+_8433172_8446622_RNA45SN1_45S_50ntFlanks | 12635 | 12662 | 28 | AAGCTACCATCTGTGGGATTATGACTGA | 403.7 | -6.3 | 1.5 | -4.2 | 2.6E-05 | 3.7E-05 | MB231/MB468 | MITO |
| >hg38_21_+_8433172_8446622_RNA45SN1_45S_50ntFlanks@10871.10890.20\|GGCTGGGTCGGTCGGGCTGG | >hg38_21_+_8433172_8446622_RNA45SN1_45S_50ntFlanks | 10871 | 10890 | 20 | GGCTGGGTCGGTCGGGCTGG | 108.9 | -4.9 | 1.2 | -4.2 | 2.2E-05 | 3.7E-05 | MB231/MB468 | MP |
| >hg38_21_+_8433172_8446622_RNA45SN1_45S_50ntFlanks@10186.10207.22\|TCCTCCCCTCCTCCCGCCCACG | >hg38_21_+_8433172_8446622_RNA45SN1_45S_50ntFlanks | 10186 | 10207 | 22 | TCCTCCCCTCCTCCCGCCCACG | 378.6 | -4.5 | 1.1 | -4.2 | 2.3E-05 | 3.7E-05 | BT20/MB468 | CYTO |
| >trnaMT_TyrGTA_MT_-_5826_5891@38.68.31\|AAAGACAGGGGTTAGGCCTCTTTTTACCACC | >trnaMT_TyrGTA_MT_-_5826_5891 | 38 | 68 | 31 | AAAGACAGGGGTTAGGCCTCTTTTTACCACC | 31.3 | 8.2 | 1.9 | 4.2 | 2.2E-05 | 3.7E-05 | MB231/MB468 | MP |
| >hg38_21_+_8433172_8446622_RNA45SN1_45S_50ntFlanks@9928.9955.28\|TGATATAGACAGCAGGACGGTGGCCATG | >hg38_21_+_8433172_8446622_RNA45SN1_45S_50ntFlanks | 9928 | 9955 | 28 | TGATATAGACAGCAGGACGGTGGCCATG | 201.6 | 7.9 | 1.8 | 4.4 | 8.7E-06 | 3.7E-05 | BT20/MB468 | MP |
| >hg38_21_+_8433172_8446622_RNA45SN1_45S_50ntFlanks@9306.9330.25\|CAAGGAGTCTAACACGTGCGCGAGT | >hg38_21_+_8433172_8446622_RNA45SN1_45S_50ntFlanks | 9306 | 9330 | 25 | CAAGGAGTCTAACACGTGCGCGAGT | 71.7 | 9.5 | 2.0 | 4.7 | 2.7E-06 | 3.7E-05 | BT20/MB231 | MP |
| >hg38_21_+_8433172_8446622_RNA45SN1_45S_50ntFlanks@6734.6770.37\|ATTGATCATCGACACTTCGAACGCACTTGCGGCCCCG | >hg38_21_+_8433172_8446622_RNA45SN1_45S_50ntFlanks | 6734 | 6770 | 37 | ATTGATCATCGACACTTCGAACGCACTTGCGGCCCCG | 1302.7 | -1.8 | 0.4 | -4.3 | 1.9E-05 | 3.7E-05 | BT20/MB468 | MITO |
| >hg38_21_+_8433172_8446622_RNA45SN1_45S_50ntFlanks@3840.3881.42\|CTCCTACTTGGATAACTGTGGTAATTCTAGAGCTAATACATG | >hg38_21_+_8433172_8446622_RNA45SN1_45S_50ntFlanks | 3840 | 3881 | 42 | CTCCTACTTGGATAACTGTGGTAATTCTAGAGCTAATACATG | 220.5 | -3.7 | 0.9 | -4.2 | 2.6E-05 | 3.7E-05 | MB231/MB468 | NUC |
| >am_trnaMT_SerGCT_MT_+_12207_12265@12.32.21\|CAAGAACTGCTAACTCATGCC | >am_trnaMT_SerGCT_MT_+_12207_12265 | 12 | 32 | 21 | CAAGAACTGCTAACTCATGCC | 486.6 | 5.2 | 1.2 | 4.2 | 2.3E-05 | 3.7E-05 | BT20/MB468 | CYTO |
| >hg38_21_+_8433172_8446622_RNA45SN1_45S_50ntFlanks@11675.11709.35\|TGCTCTGAATGTCAAAGTGAAGAAATTCAATGAAG | >hg38_21_+_8433172_8446622_RNA45SN1_45S_50ntFlanks | 11675 | 11709 | 35 | TGCTCTGAATGTCAAAGTGAAGAAATTCAATGAAG | 359.6 | 3.7 | 0.9 | 4.2 | 2.3E-05 | 3.7E-05 | BT20/MB468 | CYTO |
| >hg38_21_+_8433172_8446622_RNA45SN1_45S_50ntFlanks@3845.3873.29\|ACTTGGATAACTGTGGTAATTCTAGAGCT | >hg38_21_+_8433172_8446622_RNA45SN1_45S_50ntFlanks | 3845 | 3873 | 29 | ACTTGGATAACTGTGGTAATTCTAGAGCT | 349.5 | 3.3 | 0.8 | 4.2 | 2.3E-05 | 3.7E-05 | BT20/MB468 | CYTO |
| >hg38_21_+_8433172_8446622_RNA45SN1_45S_50ntFlanks@12307.12331.25\|ACGATCCTTCTGACCTTTTGGGTTT | >hg38_21_+_8433172_8446622_RNA45SN1_45S_50ntFlanks | 12307 | 12331 | 25 | ACGATCCTTCTGACCTTTTGGGTTT | 66.6 | -9.9 | 2.3 | -4.2 | 2.2E-05 | 3.7E-05 | MB231/MB468 | MP |
| >hg38_21_+_8433172_8446622_RNA45SN1_45S_50ntFlanks@12488.12517.30\|AACGTGAGCTGGGTTTAGACCGTCGTGAGA | >hg38_21_+_8433172_8446622_RNA45SN1_45S_50ntFlanks | 12488 | 12517 | 30 | AACGTGAGCTGGGTTTAGACCGTCGTGAGA | 46.9 | -8.6 | 1.9 | -4.4 | 8.8E-06 | 3.7E-05 | BT20/MB468 | MP |
| >hg38_21_+_8433172_8446622_RNA45SN1_45S_50ntFlanks@11668.11700.33\|TGCCCAGTGCTCTGAATGTCAAAGTGAAGAAAT | >hg38_21_+_8433172_8446622_RNA45SN1_45S_50ntFlanks | 11668 | 11700 | 33 | TGCCCAGTGCTCTGAATGTCAAAGTGAAGAAAT | 432.0 | -6.9 | 1.7 | -4.2 | 2.8E-05 | 3.7E-05 | BT20/MB468 | NUC |
| >MI0000434\|hsa-let-7i&WithFlank&12\|+\|62603680\|62603775@12.33.22(+1U)&[MIMAT0000415&hsa-let-7i-5p&offsets\|0\|0(+1U);m-49&12\|+\|62603691\|62603712&offsets\|0\|0(+1U)]\|TGAGGTAGTAGTTTGTGCTGTTT | >MI0000434\|hsa-let-7i&WithFlank&12\|+\|62603680\|62603775 | 12 | 33 | 22(+1U) | TGAGGTAGTAGTTTGTGCTGTTT | 6510.0 | 5.0 | 1.1 | 4.4 | 8.8E-06 | 3.7E-05 | BT20/MB468 | MP |
| >hg38_21_+_8433172_8446622_RNA45SN1_45S_50ntFlanks@11602.11640.39\|TCCGACTGTTTAATTAAAACAAAGCATCGCGAAGGCCCG | >hg38_21_+_8433172_8446622_RNA45SN1_45S_50ntFlanks | 11602 | 11640 | 39 | TCCGACTGTTTAATTAAAACAAAGCATCGCGAAGGCCCG | 54.7 | -8.9 | 2.0 | -4.4 | 8.8E-06 | 3.7E-05 | BT20/MB468 | MP |
| >hg38_21_+_8433172_8446622_RNA45SN1_45S_50ntFlanks@9571.9596.26\|GCAAATCGGTCGTCCGACCTGGGTAT | >hg38_21_+_8433172_8446622_RNA45SN1_45S_50ntFlanks | 9571 | 9596 | 26 | GCAAATCGGTCGTCCGACCTGGGTAT | 338.0 | -4.7 | 1.1 | -4.3 | 1.9E-05 | 3.7E-05 | BT20/MB468 | MITO |
| >hg38_21_+_8433172_8446622_RNA45SN1_45S_50ntFlanks@4759.4779.21\|ATCAGATACCGTCGTAGTTCC | >hg38_21_+_8433172_8446622_RNA45SN1_45S_50ntFlanks | 4759 | 4779 | 21 | ATCAGATACCGTCGTAGTTCC | 804.3 | -5.7 | 1.4 | -4.2 | 2.5E-05 | 3.7E-05 | MB231/MB468 | CYTO |

| | | | | | | | | | | | | | |
|---|---|---|---|---|---|---|---|---|---|---|---|---|---|
| >MI0000087\|hsa-mir-29a&WithFlank&7\|-\|130876741\|130876816@48.65.18(+3U)&[MIMAT0000086&hsa-miR-29a-3p&offsets\|0\|-4(+3U);m-26&7\|-\|130876749\|130876769&offsets\|0\|-3(+3U)]\|TAGCACCATCTGAAATCGTTT | >MI0000087\|hsa-mir-29a&WithFlank&7\|-\|130876741\|130876816 | 48 | 65 | 18(+3U) | TAGCACCATCTGAAATCGTTT | 37.4 | 8.4 | 2.0 | 4.2 | 2.2E-05 | 3.7E-05 | MB231/MB468 | MP |
| >hg38_21_+_8433172_8446622_RNA45SN1_45S_50ntFlanks@8172.8212.41\|AGTCCTTCTGATCGAGGCCCAGCCCGTGGACGGTGTGAGGC | >hg38_21_+_8433172_8446622_RNA45SN1_45S_50ntFlanks | 8172 | 8212 | 41 | AGTCCTTCTGATCGAGGCCCAGCCCGTGGACGGTGTGAGGC | 286.4 | -7.2 | 1.7 | -4.2 | 2.2E-05 | 3.7E-05 | MB231/MB468 | MP |
| >hg38_21_+_8433172_8446622_RNA45SN1_45S_50ntFlanks@12366.12392.27\|TAACTGGCTTGTGGCGGCCAAGCGTTC | >hg38_21_+_8433172_8446622_RNA45SN1_45S_50ntFlanks | 12366 | 12392 | 27 | TAACTGGCTTGTGGCGGCCAAGCGTTC | 40.4 | -8.4 | 1.9 | -4.4 | 8.8E-06 | 3.7E-05 | BT20/MB468 | MP |
| >trnaMT_IleGAT_MT_+_4263_4331@47.71.25\|GAGCTTAAACCCCCTTATTTCTACC | >trnaMT_IleGAT_MT_+_4263_4331 | 47 | 71 | 25 | GAGCTTAAACCCCCTTATTTCTACC | 30.9 | 8.2 | 1.9 | 4.2 | 2.2E-05 | 3.7E-05 | MB231/MB468 | MP |
| >hg38_21_+_8433172_8446622_RNA45SN1_45S_50ntFlanks@11564.11603.40\|CCTAGCAGCCGACTTAGAACTGGTGCGGACCAGGGGAATC | >hg38_21_+_8433172_8446622_RNA45SN1_45S_50ntFlanks | 11564 | 11603 | 40 | CCTAGCAGCCGACTTAGAACTGGTGCGGACCAGGGGAATC | 101.1 | 10.0 | 2.1 | 4.7 | 2.8E-06 | 3.7E-05 | BT20/MB231 | MP |
| >hg38_MT_+_598_1651_RNR1_12S_50ntFlanks@819.840.22\|AAGGTGGATTTAGCAGTAAACT | >hg38_MT_+_598_1651_RNR1_12S_50ntFlanks | 819 | 840 | 22 | AAGGTGGATTTAGCAGTAAACT | 458.0 | 7.5 | 1.7 | 4.4 | 8.8E-06 | 3.7E-05 | BT20/MB468 | MP |
| >hg38_21_+_8433172_8446622_RNA45SN1_45S_50ntFlanks@6648.6676.29\|GTACGACTCTTAGCGGTGGATCACTCGGC | >hg38_21_+_8433172_8446622_RNA45SN1_45S_50ntFlanks | 6648 | 6676 | 29 | GTACGACTCTTAGCGGTGGATCACTCGGC | 955.6 | 4.4 | 1.0 | 4.4 | 8.8E-06 | 3.7E-05 | BT20/MB468 | MP |
| >hg38_21_+_8433172_8446622_RNA45SN1_45S_50ntFlanks@9759.9787.29\|CAAACTTTAAATGGGTAAGAAGCCCGGCT | >hg38_21_+_8433172_8446622_RNA45SN1_45S_50ntFlanks | 9759 | 9787 | 29 | CAAACTTTAAATGGGTAAGAAGCCCGGCT | 295.1 | -7.3 | 1.6 | -4.4 | 8.8E-06 | 3.7E-05 | BT20/MB468 | MP |
| >hg38_21_+_8433172_8446622_RNA45SN1_45S_50ntFlanks@6660.6688.29\|GCGGTGGATCACTCGGCTCGTGCGTCGAT | >hg38_21_+_8433172_8446622_RNA45SN1_45S_50ntFlanks | 6660 | 6688 | 29 | GCGGTGGATCACTCGGCTCGTGCGTCGAT | 915.9 | 2.8 | 0.6 | 4.3 | 1.9E-05 | 3.7E-05 | BT20/MB468 | MITO |
| >hg38_21_+_8433172_8446622_RNA45SN1_45S_50ntFlanks@10402.10437.36\|AACATGGGTCAGTCGGTCCTGAGAGATGGGCGAGCG | >hg38_21_+_8433172_8446622_RNA45SN1_45S_50ntFlanks | 10402 | 10437 | 36 | AACATGGGTCAGTCGGTCCTGAGAGATGGGCGAGCG | 125.4 | -6.2 | 1.5 | -4.2 | 2.2E-05 | 3.7E-05 | MB231/MB468 | MP |
| >hg38_21_+_8433172_8446622_RNA45SN1_45S_50ntFlanks@12912.12929.18\|CACGTTCGTGGGGAACCT | >hg38_21_+_8433172_8446622_RNA45SN1_45S_50ntFlanks | 12912 | 12929 | 18 | CACGTTCGTGGGGAACCT | 97.9 | -6.2 | 1.5 | -4.2 | 2.2E-05 | 3.7E-05 | MB231/MB468 | MP |
| >am_trnaMT_ArgTCG_MT_+_10405_10469@18.49.32\|AAAACGAATGATTTCGACTCATTAAATTATGA | >am_trnaMT_ArgTCG_MT_+_10405_10469 | 18 | 49 | 32 | AAAACGAATGATTTCGACTCATTAAATTATGA | 28.3 | 8.0 | 1.9 | 4.2 | 2.2E-05 | 3.7E-05 | MB231/MB468 | MP |
| >hg38_21_+_8433172_8446622_RNA45SN1_45S_50ntFlanks@4258.4282.25\|AATTGGAATGAGTCCACTTTAAATC | >hg38_21_+_8433172_8446622_RNA45SN1_45S_50ntFlanks | 4258 | 4282 | 25 | AATTGGAATGAGTCCACTTTAAATC | 74.6 | -10.1 | 2.4 | -4.2 | 2.2E-05 | 3.7E-05 | MB231/MB468 | MP |
| >hg38_21_+_8433172_8446622_RNA45SN1_45S_50ntFlanks@7984.8012.29\|AGATCAGACGTGGCGACCCGCTGAATTTA | >hg38_21_+_8433172_8446622_RNA45SN1_45S_50ntFlanks | 7984 | 8012 | 29 | AGATCAGACGTGGCGACCCGCTGAATTTA | 427.4 | -4.8 | 1.1 | -4.3 | 1.9E-05 | 3.7E-05 | BT20/MB468 | MITO |
| >hg38_21_+_8433172_8446622_RNA45SN1_45S_50ntFlanks@3996.4025.30\|ACTCTAGATAACCTCGGGCCGATCGCACGC | >hg38_21_+_8433172_8446622_RNA45SN1_45S_50ntFlanks | 3996 | 4025 | 30 | ACTCTAGATAACCTCGGGCCGATCGCACGC | 290.6 | -6.3 | 1.5 | -4.2 | 2.7E-05 | 3.7E-05 | MB231/MB468 | NUC |
| >hg38_21_+_8433172_8446622_RNA45SN1_45S_50ntFlanks@3823.3850.28\|TTTGGTCGCTCGCTCCTCTCCTACTTGG | >hg38_21_+_8433172_8446622_RNA45SN1_45S_50ntFlanks | 3823 | 3850 | 28 | TTTGGTCGCTCGCTCCTCTCCTACTTGG | 333.2 | -2.6 | 0.6 | -4.2 | 2.7E-05 | 3.7E-05 | MB231/MB468 | NUC |
| >MI0003581\|hsa-mir-574&WithFlank&4\|+\|38868026\|38868133@67.87.21&[MIMAT0003239&hsa-miR-574-3p&offsets\|0\|-1;m-149&4\|+\|38868092\|38868113&offsets\|0\|-1]\|CACGCTCATGCACACACCCAC | >MI0003581\|hsa-mir-574&WithFlank&4\|+\|38868026\|38868133 | 67 | 87 | 21 | CACGCTCATGCACACACCCAC | 183.6 | 5.2 | 1.2 | 4.2 | 2.7E-05 | 3.7E-05 | MB231/MB468 | MITO |
| >hg38_21_+_8433172_8446622_RNA45SN1_45S_50ntFlanks@11320.11343.24\|CCACGTCTCGTCGCGCGCGCGTCC | >hg38_21_+_8433172_8446622_RNA45SN1_45S_50ntFlanks | 11320 | 11343 | 24 | CCACGTCTCGTCGCGCGCGCGTCC | 964.4 | -5.5 | 1.3 | -4.2 | 2.7E-05 | 3.7E-05 | MB231/MB468 | NUC |
| >hg38_21_+_8433172_8446622_RNA45SN1_45S_50ntFlanks@12931.12965.35\|GCGCTAAACCATTCGTAGACGACCTGCTTCTGGGT | >hg38_21_+_8433172_8446622_RNA45SN1_45S_50ntFlanks | 12931 | 12965 | 35 | GCGCTAAACCATTCGTAGACGACCTGCTTCTGGGT | 59.5 | -9.8 | 2.3 | -4.2 | 2.2E-05 | 3.7E-05 | MB231/MB468 | MP |
| >MI0000067\|hsa-let-7f-1&WithFlank&9\|+\|94176341\|94176439@13.32.20(+1C);MI0000068\|hsa-let-7f-2&WithFlank&X\|-\|53557186\|53557280@14.33.20(+1C)&[MIMAT0000067&hsa-let-7f-5p&offsets\|0\|-2(+1C);m-20&9\|+\|94176353\|94176374&offsets\|0\|-2(+1C)];[MIMAT0000067_1&hsa-let-7f-5p&offsets\|0\|-2(+1C);m-19&X\|-\|53557246\|53557267&offsets\|0\|-2(+1C)]\|TGAGGTAGTAGATTGTATAGC | >MI0000067\|hsa-let-7f-1&WithFlank&9\|+\|94176341\|94176439 | 13 | 32 | 20(+1C) | TGAGGTAGTAGATTGTATAGC | 461.8 | 2.1 | 0.5 | 4.2 | 2.8E-05 | 3.7E-05 | BT20/MB468 | NUC |
| >hg38_21_+_8433172_8446622_RNA45SN1_45S_50ntFlanks@12456.12491.36\|CCAAGCGTTGGATTGTTCACCCACTAATAGGGAACG | >hg38_21_+_8433172_8446622_RNA45SN1_45S_50ntFlanks | 12456 | 12491 | 36 | CCAAGCGTTGGATTGTTCACCCACTAATAGGGAACG | 187.1 | -5.8 | 1.4 | -4.2 | 2.3E-05 | 3.8E-05 | BT20/MB468 | CYTO |
| >am_trnaMT_PheGAA_MT_+_577_647@42.62.21\|TTTAGACGGGCTCACATCACC | >am_trnaMT_PheGAA_MT_+_577_647 | 42 | 62 | 21 | TTTAGACGGGCTCACATCACC | 505.2 | 5.4 | 1.2 | 4.4 | 8.9E-06 | 3.8E-05 | BT20/MB468 | MP |
| >MI0000095\|hsa-mir-93&WithFlank&7\|-\|100093762\|100093853@17.38.22(+1U)&[MIMAT0000093&hsa-miR-93-5p&offsets\|0\|-1(+1U);m-29&7\|-\|100093815\|100093837&offsets\|0\|-1(+1U)]\|CAAAGTGCTGTTCGTGCAGGTAT | >MI0000095\|hsa-mir-93&WithFlank&7\|-\|100093762\|100093853 | 17 | 38 | 22(+1U) | CAAAGTGCTGTTCGTGCAGGTAT | 525.4 | 3.7 | 0.9 | 4.2 | 2.3E-05 | 3.8E-05 | MB231/MB468 | MP |
| >hg38_21_+_8433172_8446622_RNA45SN1_45S_50ntFlanks@12991.13028.38\|CTCCCTCGCTGCGATCTATTGAAAGTCAGCCCTCGACA | >hg38_21_+_8433172_8446622_RNA45SN1_45S_50ntFlanks | 12991 | 13028 | 38 | CTCCCTCGCTGCGATCTATTGAAAGTCAGCCCTCGACA | 49.2 | -8.7 | 2.0 | -4.4 | 8.9E-06 | 3.8E-05 | BT20/MB468 | MP |
| >hg38_21_+_8433172_8446622_RNA45SN1_45S_50ntFlanks@5468.5516.49\|GCCCACGGCCCTGGCGGAGCGCTGAGAAGACGGTCGAACTTGACTATCT | >hg38_21_+_8433172_8446622_RNA45SN1_45S_50ntFlanks | 5468 | 5516 | 49 | GCCCACGGCCCTGGCGGAGCGCTGAGAAGACGGTCGAACTTGACTATCT | 885.3 | 8.0 | 1.9 | 4.3 | 2.0E-05 | 3.8E-05 | BT20/MB468 | MITO |
| >hg38_21_+_8433172_8446622_RNA45SN1_45S_50ntFlanks@7980.8003.24\|CCTCAGATCAGACGTGGCGACCCG | >hg38_21_+_8433172_8446622_RNA45SN1_45S_50ntFlanks | 7980 | 8003 | 24 | CCTCAGATCAGACGTGGCGACCCG | 769.3 | -3.9 | 0.9 | -4.2 | 2.7E-05 | 3.8E-05 | MB231/MB468 | NUC |
| >am_trna119_LysCTT_1_-_145395522_145395594@6.35.30,trna11_LysCTT_5_-_180648979_180649051@6.35.30,trna128_LysCTT_1_-_55423542_55423614@6.35.30,trna13_LysCTT_6_+_26556774_26556846@6.35.30,trna6_LysCTT_19_-_52425396_52425468@6.35.30,trna7_LysCTT_16_+_3225692_3225764@6.35.30,trna9_LysCTT_5_+_180634755_180634827@6.35.30\|GCTAGCTCAGTCGGTAGAGCATGAGACTCT | >am_trna119_LysCTT_1_-_145395522_145395594 | 6 | 35 | 30 | GCTAGCTCAGTCGGTAGAGCATGAGACTCT | 222.0 | 5.5 | 1.3 | 4.2 | 2.7E-05 | 3.8E-05 | MB231/MB468 | MITO |
| >trnaMT_PheGAA_MT_+_577_647@1.35.35\|GTTTATGTAGCTTACCTCCTCAAAGCAATACACTG | >trnaMT_PheGAA_MT_+_577_647 | 1 | 35 | 35 | GTTTATGTAGCTTACCTCCTCAAAGCAATACACTG | 1153.5 | 4.7 | 1.1 | 4.2 | 2.7E-05 | 3.8E-05 | MB231/MB468 | MITO |
| >hg38_21_+_8433172_8446622_RNA45SN1_45S_50ntFlanks@9483.9510.28\|CGTGTTAGGACCCGAAAGATGGTGAACT | >hg38_21_+_8433172_8446622_RNA45SN1_45S_50ntFlanks | 9483 | 9510 | 28 | CGTGTTAGGACCCGAAAGATGGTGAACT | 172.4 | 7.8 | 1.8 | 4.4 | 9.0E-06 | 3.8E-05 | BT20/MB468 | MP |
| >hg38_21_+_8433172_8446622_RNA45SN1_45S_50ntFlanks@10846.10870.25\|CTTCGGGATAAGGATTGGCTCTAAG | >hg38_21_+_8433172_8446622_RNA45SN1_45S_50ntFlanks | 10846 | 10870 | 25 | CTTCGGGATAAGGATTGGCTCTAAG | 353.7 | -6.8 | 1.6 | -4.2 | 2.8E-05 | 3.8E-05 | BT20/MB468 | NUC |
| >hg38_21_+_8433172_8446622_RNA45SN1_45S_50ntFlanks@6654.6689.36\|CTCTTAGCGGTGGATCACTCGGCTCGTGCGTCGATG | >hg38_21_+_8433172_8446622_RNA45SN1_45S_50ntFlanks | 6654 | 6689 | 36 | CTCTTAGCGGTGGATCACTCGGCTCGTGCGTCGATG | 842.1 | -2.6 | 0.6 | -4.2 | 2.7E-05 | 3.8E-05 | MB231/MB468 | NUC |
| >hg38_21_+_8433172_8446622_RNA45SN1_45S_50ntFlanks@8169.8212.44\|CCAAGTCCTTCTGATCGAGGCCCAGCCCGTGGACGGTGTGAGGC | >hg38_21_+_8433172_8446622_RNA45SN1_45S_50ntFlanks | 8169 | 8212 | 44 | CCAAGTCCTTCTGATCGAGGCCCAGCCCGTGGACGGTGTGAGGC | 106.0 | -5.3 | 1.2 | -4.2 | 2.3E-05 | 3.8E-05 | BT20/MB468 | CYTO |
| >hg38_21_+_8433172_8446622_RNA45SN1_45S_50ntFlanks@12625.12653.29\|CAATGGGGCGAAGCTACCATCTGTGGGAT | >hg38_21_+_8433172_8446622_RNA45SN1_45S_50ntFlanks | 12625 | 12653 | 29 | CAATGGGGCGAAGCTACCATCTGTGGGAT | 1750.2 | -7.3 | 1.6 | -4.4 | 9.0E-06 | 3.8E-05 | BT20/MB468 | MP |
| >trna10_AspGTC_12_-_125424193_125424264@38.69.32,trna12_AspGTC_12_-_125411891_125411962@38.69.32,trna144_AspGTC_6_-_27551236_27551307@38.69.32,trna38_AspGTC_17_-_8125556_8125627@38.69.32,trna45_AspGTC_6_+_27447453_27447524@38.69.32,trna48_AspGTC_6_+_27471523_27471594@38.69.32,trna4_AspGTC_12_+_96429799_96429870@38.69.32,trna69_AspGTC_1_-_161440205_161440276@38.69.32,trna72_AspGTC_1_-_161432824_161432895@38.69.32,trna75_AspGTC_1_-_161425414_161425485@38.69.32,trna78_AspGTC_1_-_161418033_161418104@38.69.32,trna81_AspGTC_1_-_161410615_161410686@38.69.32\|CGCGGGAGACCGGGGTTCGATTCCCCGACGGG | >trna10_AspGTC_12_-_125424193_125424264 | 38 | 69 | 32 | CGCGGGAGACCGGGGTTCGATTCCCCGACGGG | 973.6 | 4.9 | 1.2 | 4.2 | 2.3E-05 | 3.8E-05 | BT20/MB468 | CYTO |
| >hg38_21_+_8433172_8446622_RNA45SN1_45S_50ntFlanks@11586.11609.24\|GTGCGGACCAGGGGAATCCGACTG | >hg38_21_+_8433172_8446622_RNA45SN1_45S_50ntFlanks | 11586 | 11609 | 24 | GTGCGGACCAGGGGAATCCGACTG | 361.2 | -2.8 | 0.7 | -4.2 | 2.7E-05 | 3.8E-05 | MB231/MB468 | NUC |
| >hg38_21_+_8433172_8446622_RNA45SN1_45S_50ntFlanks@6644.6661.18\|CCTCGTACGACTCTTAGC | >hg38_21_+_8433172_8446622_RNA45SN1_45S_50ntFlanks | 6644 | 6661 | 18 | CCTCGTACGACTCTTAGC | 423.3 | 4.3 | 1.0 | 4.2 | 2.7E-05 | 3.8E-05 | MB231/MB468 | MITO |
| >hg38_21_+_8433172_8446622_RNA45SN1_45S_50ntFlanks@9921.9951.31\|TGTTGGTTGATATAGACAGCAGGACGGTGGC | >hg38_21_+_8433172_8446622_RNA45SN1_45S_50ntFlanks | 9921 | 9951 | 31 | TGTTGGTTGATATAGACAGCAGGACGGTGGC | 83.5 | 9.7 | 2.1 | 4.7 | 2.8E-06 | 3.8E-05 | BT20/MB231 | MP |
| >trnaMT_GluTTC_MT_-_14674_14742@24.72.49,trnalookalike8_GluTTC_5_-_93905172_93905240@24.72.49\|ATGGTTTTTCATATCATTGGTCGTGGTTGTAGTCCGTGCGAGAATACCA | >trnaMT_GluTTC_MT_-_14674_14742 | 24 | 72 | 49 | ATGGTTTTTCATATCATTGGTCGTGGTTGTAGTCCGTGCGAGAATACCA | 207.3 | 11.4 | 2.6 | 4.4 | 9.0E-06 | 3.8E-05 | BT20/MB468 | MP |
| >hg38_21_+_8433172_8446622_RNA45SN1_45S_50ntFlanks@9554.9574.21\|CGTAGCGGTCCTGACGTGCAA | >hg38_21_+_8433172_8446622_RNA45SN1_45S_50ntFlanks | 9554 | 9574 | 21 | CGTAGCGGTCCTGACGTGCAA | 1844.4 | -4.2 | 0.9 | -4.4 | 9.0E-06 | 3.8E-05 | BT20/MB468 | MP |
| >hg38_21_+_8433172_8446622_RNA45SN1_45S_50ntFlanks@10685.10708.24\|GGCGGCGTCCGGTGAGCTCTCGCT | >hg38_21_+_8433172_8446622_RNA45SN1_45S_50ntFlanks | 10685 | 10708 | 24 | GGCGGCGTCCGGTGAGCTCTCGCT | 963.8 | 4.0 | 0.9 | 4.3 | 2.0E-05 | 3.8E-05 | BT20/MB468 | MITO |
| >am_trnaMT_MetCAT_MT_+_4402_4469@1.46.46\|AGTAAGGTCAGCTAAATAAGCTATCGGGCCCATACCCCGAAAATGT | >am_trnaMT_MetCAT_MT_+_4402_4469 | 1 | 46 | 46 | AGTAAGGTCAGCTAAATAAGCTATCGGGCCCATACCCCGAAAATGT | 201.9 | 11.4 | 2.6 | 4.4 | 9.0E-06 | 3.8E-05 | BT20/MB468 | MP |
| >hg38_1_-_228634819_228635039_RNA5S12_5S_50ntFlanks@131.171.41\|GGATGGGAGACCGCCTGGGAATACCGGGTGCTGTAGGCTTT | >hg38_1_-_228634819_228635039_RNA5S12_5S_50ntFlanks | 131 | 171 | 41 | GGATGGGAGACCGCCTGGGAATACCGGGTGCTGTAGGCTTT | 744.0 | 5.8 | 1.3 | 4.4 | 9.0E-06 | 3.8E-05 | BT20/MB468 | MP |
| >hg38_21_+_8433172_8446622_RNA45SN1_45S_50ntFlanks@11729.11749.21\|TAACTATGACTCTCTTAAGGT | >hg38_21_+_8433172_8446622_RNA45SN1_45S_50ntFlanks | 11729 | 11749 | 21 | TAACTATGACTCTCTTAAGGT | 1064.9 | -3.9 | 0.9 | -4.4 | 9.0E-06 | 3.8E-05 | BT20/MB468 | MP |
| >hg38_21_+_8433172_8446622_RNA45SN1_45S_50ntFlanks@8190.8210.21\|CCAGCCCGTGGACGGTGTGAG | >hg38_21_+_8433172_8446622_RNA45SN1_45S_50ntFlanks | 8190 | 8210 | 21 | CCAGCCCGTGGACGGTGTGAG | 40.8 | -8.4 | 1.9 | -4.4 | 9.0E-06 | 3.8E-05 | BT20/MB468 | MP |
| >MI0003820\|hsa-mir-454&WithFlank&17\|-\|59137752\|59137878@70.93.24&[MIMAT0003885&hsa-miR-454-3p&offsets\|0\|+1;m-272&17\|-\|59137787\|59137809&offsets\|0\|+1]\|TAGTGCAATATTGCTTATAGGGTT | >MI0003820\|hsa-mir-454&WithFlank&17\|-\|59137752\|59137878 | 70 | 93 | 24 | TAGTGCAATATTGCTTATAGGGTT | 714.0 | 3.7 | 0.9 | 4.2 | 2.3E-05 | 3.8E-05 | MB231/MB468 | MP |

<table>
<tr><th></th><th></th><th></th><th></th><th></th><th></th><th></th><th></th><th></th><th></th><th></th><th></th><th></th><th></th></tr>
<tr><td>>MI0003560|hsa-mir-92b&WithFlank&1|+|155195171|155195278@67.88.22(+1U)&[MIMAT0003218&hsa-miR-92b-3p&offsets|0|0(+1U);m-115&1|+|155195237|155195258&offsets|0|0(+1U)]|TATTGCACTCGTCCCGGCCTCCT</td><td>>MI0003560|hsa-mir-92b&WithFlank&1|+|155195171|155195278</td><td>67</td><td>88</td><td>22(+1U)</td><td>TATTGCACTCGTCCCGGCCTCCT</td><td>138.2</td><td>3.1</td><td>0.7</td><td>4.2</td><td>2.9E-05</td><td>3.8E-05</td><td>BT20/MB468</td><td>NUC</td></tr>
<tr><td>>hg38_21_+_8433172_8446622_RNA45SN1_45S_50ntFlanks@9352.9377.26|CCGTGGCGCAATGAAGGTGAAGGCCG</td><td>>hg38_21_+_8433172_8446622_RNA45SN1_45S_50ntFlanks</td><td>9352</td><td>9377</td><td>26</td><td>CCGTGGCGCAATGAAGGTGAAGGCCG</td><td>297.7</td><td>-7.0</td><td>1.6</td><td>-4.4</td><td>9.1E-06</td><td>3.8E-05</td><td>BT20/MB468</td><td>MP</td></tr>
<tr><td>>hg38_21_+_8433172_8446622_RNA45SN1_45S_50ntFlanks@10921.10939.19|GCTGGACGAGGCGCCGCCG</td><td>>hg38_21_+_8433172_8446622_RNA45SN1_45S_50ntFlanks</td><td>10921</td><td>10939</td><td>19</td><td>GCTGGACGAGGCGCCGCCG</td><td>357.3</td><td>-6.6</td><td>1.6</td><td>-4.2</td><td>2.7E-05</td><td>3.8E-05</td><td>MB231/MB468</td><td>NUC</td></tr>
<tr><td>>hg38_21_+_8433172_8446622_RNA45SN1_45S_50ntFlanks@4673.4690.18|TGGACCGGCGCAAGACGG</td><td>>hg38_21_+_8433172_8446622_RNA45SN1_45S_50ntFlanks</td><td>4673</td><td>4690</td><td>18</td><td>TGGACCGGCGCAAGACGG</td><td>74.1</td><td>-10.1</td><td>2.4</td><td>-4.2</td><td>2.3E-05</td><td>3.8E-05</td><td>MB231/MB468</td><td>MP</td></tr>
<tr><td>>hg38_21_+_8433172_8446622_RNA45SN1_45S_50ntFlanks@5090.5112.23|AGACTCTGGCATGCTAACTAGTT</td><td>>hg38_21_+_8433172_8446622_RNA45SN1_45S_50ntFlanks</td><td>5090</td><td>5112</td><td>23</td><td>AGACTCTGGCATGCTAACTAGTT</td><td>80.1</td><td>-10.2</td><td>2.4</td><td>-4.2</td><td>2.3E-05</td><td>3.8E-05</td><td>MB231/MB468</td><td>MP</td></tr>
<tr><td>>hg38_21_+_8433172_8446622_RNA45SN1_45S_50ntFlanks@12457.12486.30|CAAGCGTTGGATTGTTCACCCACTAATAGG</td><td>>hg38_21_+_8433172_8446622_RNA45SN1_45S_50ntFlanks</td><td>12457</td><td>12486</td><td>30</td><td>CAAGCGTTGGATTGTTCACCCACTAATAGG</td><td>278.9</td><td>-2.9</td><td>0.7</td><td>-4.2</td><td>2.6E-05</td><td>3.8E-05</td><td>MB231/MB468</td><td>CYTO</td></tr>
<tr><td>>MI0000736|hsa-mir-30c-1&WithFlank&1|+|40757278|40757378@23.44.22;MI0000254|hsa-mir-30c-2&WithFlank&6|-|71376954|71377037@13.34.22&[MIMAT0000244&hsa-miR-30c-5p&offsets|0|-1;m-57&1|+|40757300|40757323&offsets|0|-2];[MIMAT0000244_1&hsa-miR-30c-5p&offsets|0|-1;m-56&6|-|71377002|71377025&offsets|0|-2]|TGTAAACATCCTACACTCTCAG</td><td>>MI0000736|hsa-mir-30c-1&WithFlank&1|+|40757278|40757378</td><td>23</td><td>44</td><td>22</td><td>TGTAAACATCCTACACTCTCAG</td><td>374.5</td><td>3.8</td><td>0.9</td><td>4.2</td><td>2.3E-05</td><td>3.8E-05</td><td>MB231/MB468</td><td>MP</td></tr>
<tr><td>>hg38_21_+_8433172_8446622_RNA45SN1_45S_50ntFlanks@6775.6803.29|CCTCCCGGGGCTACGCCTGTCTGAGCGTC</td><td>>hg38_21_+_8433172_8446622_RNA45SN1_45S_50ntFlanks</td><td>6775</td><td>6803</td><td>29</td><td>CCTCCCGGGGCTACGCCTGTCTGAGCGTC</td><td>224.3</td><td>-2.5</td><td>0.6</td><td>-4.2</td><td>2.6E-05</td><td>3.8E-05</td><td>MB231/MB468</td><td>CYTO</td></tr>
<tr><td>>hg38_21_+_8433172_8446622_RNA45SN1_45S_50ntFlanks@10382.10424.43|TTCCATGTGAACAGCAGTTGAACATGGGTCAGTCGGTCCTGAG</td><td>>hg38_21_+_8433172_8446622_RNA45SN1_45S_50ntFlanks</td><td>10382</td><td>10424</td><td>43</td><td>TTCCATGTGAACAGCAGTTGAACATGGGTCAGTCGGTCCTGAG</td><td>8198.5</td><td>4.1</td><td>0.9</td><td>4.6</td><td>3.4E-06</td><td>3.8E-05</td><td>BT20/MB231</td><td>MITO</td></tr>
<tr><td>>hg38_21_+_8433172_8446622_RNA45SN1_45S_50ntFlanks@11718.11737.20|AACGGCGGGAGTAACTATGA</td><td>>hg38_21_+_8433172_8446622_RNA45SN1_45S_50ntFlanks</td><td>11718</td><td>11737</td><td>20</td><td>AACGGCGGGAGTAACTATGA</td><td>725.2</td><td>-7.3</td><td>1.8</td><td>-4.2</td><td>2.9E-05</td><td>3.8E-05</td><td>BT20/MB468</td><td>NUC</td></tr>
<tr><td>>MI0000477|hsa-mir-146a&WithFlank&5|+|160485346|160485456@26.48.23(+1U)&[MIMAT0000449&hsa-miR-146a-5p&offsets|-1|0(+1U);m-129&5|+|160485372|160485393&offsets|-1|0(+1U)]|TTGAGAACTGAATTCCATGGGTTT</td><td>>MI0000477|hsa-mir-146a&WithFlank&5|+|160485346|160485456</td><td>26</td><td>48</td><td>23(+1U)</td><td>TTGAGAACTGAATTCCATGGGTTT</td><td>32.9</td><td>8.3</td><td>2.0</td><td>4.2</td><td>2.3E-05</td><td>3.8E-05</td><td>MB231/MB468</td><td>MP</td></tr>
<tr><td>>hg38_21_+_8433172_8446622_RNA45SN1_45S_50ntFlanks@9533.9557.25|GAGGAAACTCTGGTGGAGGTCCGTA</td><td>>hg38_21_+_8433172_8446622_RNA45SN1_45S_50ntFlanks</td><td>9533</td><td>9557</td><td>25</td><td>GAGGAAACTCTGGTGGAGGTCCGTA</td><td>60.8</td><td>-9.0</td><td>2.0</td><td>-4.4</td><td>9.1E-06</td><td>3.8E-05</td><td>BT20/MB468</td><td>MP</td></tr>
<tr><td>>hg38_21_+_8433172_8446622_RNA45SN1_45S_50ntFlanks@5471.5499.29|CACGGCCCTGGCGGAGCGCTGAGAAGACG</td><td>>hg38_21_+_8433172_8446622_RNA45SN1_45S_50ntFlanks</td><td>5471</td><td>5499</td><td>29</td><td>CACGGCCCTGGCGGAGCGCTGAGAAGACG</td><td>3491.7</td><td>-2.5</td><td>0.6</td><td>-4.2</td><td>2.6E-05</td><td>3.8E-05</td><td>MB231/MB468</td><td>CYTO</td></tr>
<tr><td>>hg38_21_+_8433172_8446622_RNA45SN1_45S_50ntFlanks@8246.8269.24|TTCCCGGAGTCGGGTTGCTTGGGA</td><td>>hg38_21_+_8433172_8446622_RNA45SN1_45S_50ntFlanks</td><td>8246</td><td>8269</td><td>24</td><td>TTCCCGGAGTCGGGTTGCTTGGGA</td><td>81.7</td><td>-6.1</td><td>1.4</td><td>-4.2</td><td>2.3E-05</td><td>3.8E-05</td><td>MB231/MB468</td><td>MP</td></tr>
<tr><td>>hg38_21_+_8433172_8446622_RNA45SN1_45S_50ntFlanks@6651.6676.26|CGACTCTTAGCGGTGGATCACTCGGC</td><td>>hg38_21_+_8433172_8446622_RNA45SN1_45S_50ntFlanks</td><td>6651</td><td>6676</td><td>26</td><td>CGACTCTTAGCGGTGGATCACTCGGC</td><td>8651.4</td><td>4.5</td><td>1.0</td><td>4.4</td><td>9.2E-06</td><td>3.9E-05</td><td>BT20/MB468</td><td>MP</td></tr>
<tr><td>>MI0000811|hsa-mir-148b&WithFlank&12|+|54337210|54337320@69.90.22(+3U)&[MIMAT0000759&hsa-miR-148b-3p&offsets|0|0(+3U);m-77&12|+|54337278|54337299&offsets|0|0(+3U)]|TCAGTGCATCACAGAACTTTGTTTT</td><td>>MI0000811|hsa-mir-148b&WithFlank&12|+|54337210|54337320</td><td>69</td><td>90</td><td>22(+3U)</td><td>TCAGTGCATCACAGAACTTTGTTTT</td><td>267.6</td><td>6.2</td><td>1.5</td><td>4.2</td><td>2.8E-05</td><td>3.9E-05</td><td>MB231/MB468</td><td>MITO</td></tr>
<tr><td>>hg38_21_+_8433172_8446622_RNA45SN1_45S_50ntFlanks@11772.11794.23|TTAGTGACGCGCATGAATGGATG</td><td>>hg38_21_+_8433172_8446622_RNA45SN1_45S_50ntFlanks</td><td>11772</td><td>11794</td><td>23</td><td>TTAGTGACGCGCATGAATGGATG</td><td>306.6</td><td>-3.4</td><td>0.8</td><td>-4.2</td><td>2.3E-05</td><td>3.9E-05</td><td>MB231/MB468</td><td>MP</td></tr>
<tr><td>>hg38_21_+_8433172_8446622_RNA45SN1_45S_50ntFlanks@6734.6773.40|ATTGATCATCGACACTTCGAACGCACTTGCGGCCCCGGGT</td><td>>hg38_21_+_8433172_8446622_RNA45SN1_45S_50ntFlanks</td><td>6734</td><td>6773</td><td>40</td><td>ATTGATCATCGACACTTCGAACGCACTTGCGGCCCCGGGT</td><td>187.5</td><td>7.8</td><td>1.8</td><td>4.4</td><td>9.2E-06</td><td>3.9E-05</td><td>BT20/MB468</td><td>MP</td></tr>
<tr><td>>hg38_21_+_8433172_8446622_RNA45SN1_45S_50ntFlanks@10822.10840.19|GAAGTCGGCAAGCCGGATC</td><td>>hg38_21_+_8433172_8446622_RNA45SN1_45S_50ntFlanks</td><td>10822</td><td>10840</td><td>19</td><td>GAAGTCGGCAAGCCGGATC</td><td>453.8</td><td>-5.2</td><td>1.2</td><td>-4.2</td><td>2.4E-05</td><td>3.9E-05</td><td>BT20/MB468</td><td>CYTO</td></tr>
<tr><td>>hg38_21_+_8433172_8446622_RNA45SN1_45S_50ntFlanks@10425.10455.31|AGATGGGCGAGCGCCGTTCCGAAGGGACGGG</td><td>>hg38_21_+_8433172_8446622_RNA45SN1_45S_50ntFlanks</td><td>10425</td><td>10455</td><td>31</td><td>AGATGGGCGAGCGCCGTTCCGAAGGGACGGG</td><td>173.7</td><td>-2.2</td><td>0.5</td><td>-4.2</td><td>2.6E-05</td><td>3.9E-05</td><td>MB231/MB468</td><td>CYTO</td></tr>
<tr><td>>hg38_21_+_8433172_8446622_RNA45SN1_45S_50ntFlanks@5473.5516.44|CGGCCCTGGCGGAGCGCTGAGAAGACGGTCGAACTTGACTATCT</td><td>>hg38_21_+_8433172_8446622_RNA45SN1_45S_50ntFlanks</td><td>5473</td><td>5516</td><td>44</td><td>CGGCCCTGGCGGAGCGCTGAGAAGACGGTCGAACTTGACTATCT</td><td>30580.7</td><td>-4.0</td><td>0.8</td><td>-4.7</td><td>2.8E-06</td><td>3.9E-05</td><td>BT20/MB231</td><td>CYTO</td></tr>
<tr><td>>MI0010635|hsa-mir-2116&WithFlank&15|-|59171177|59171268@56.76.21&[MIMAT0011161&hsa-miR-2116-3p&offsets|-1|-1;m-656&15|-|59171193|59171213&offsets|0|0]|TCCTCCCATGCCAAGAACTCC</td><td>>MI0010635|hsa-mir-2116&WithFlank&15|-|59171177|59171268</td><td>56</td><td>76</td><td>21</td><td>TCCTCCCATGCCAAGAACTCC</td><td>182.1</td><td>4.2</td><td>0.9</td><td>4.7</td><td>2.8E-06</td><td>3.9E-05</td><td>BT20/MB231</td><td>CYTO</td></tr>
<tr><td>>hg38_21_+_8433172_8446622_RNA45SN1_45S_50ntFlanks@10317.10334.18|GTGCAGATCTTGGTGGTA</td><td>>hg38_21_+_8433172_8446622_RNA45SN1_45S_50ntFlanks</td><td>10317</td><td>10334</td><td>18</td><td>GTGCAGATCTTGGTGGTA</td><td>100.6</td><td>10.0</td><td>2.1</td><td>4.7</td><td>2.9E-06</td><td>3.9E-05</td><td>BT20/MB231</td><td>MP</td></tr>
<tr><td>>hg38_21_+_8433172_8446622_RNA45SN1_45S_50ntFlanks@8157.8184.28|CTCGTGGGGGGCCCAAGTCCTTCTGATC</td><td>>hg38_21_+_8433172_8446622_RNA45SN1_45S_50ntFlanks</td><td>8157</td><td>8184</td><td>28</td><td>CTCGTGGGGGGCCCAAGTCCTTCTGATC</td><td>104.5</td><td>-5.9</td><td>1.4</td><td>-4.2</td><td>2.3E-05</td><td>3.9E-05</td><td>MB231/MB468</td><td>MP</td></tr>
<tr><td>>hg38_21_+_8433172_8446622_RNA45SN1_45S_50ntFlanks@4104.4127.24|CACGGGTGACGGGGAATCAGGGTT</td><td>>hg38_21_+_8433172_8446622_RNA45SN1_45S_50ntFlanks</td><td>4104</td><td>4127</td><td>24</td><td>CACGGGTGACGGGGAATCAGGGTT</td><td>188.6</td><td>-5.0</td><td>1.2</td><td>-4.2</td><td>2.3E-05</td><td>3.9E-05</td><td>MB231/MB468</td><td>MP</td></tr>
<tr><td>>hg38_21_+_8433172_8446622_RNA45SN1_45S_50ntFlanks@9938.9961.24|AGCAGGACGGTGGCCATGGAAGTC</td><td>>hg38_21_+_8433172_8446622_RNA45SN1_45S_50ntFlanks</td><td>9938</td><td>9961</td><td>24</td><td>AGCAGGACGGTGGCCATGGAAGTC</td><td>379.2</td><td>6.9</td><td>1.6</td><td>4.4</td><td>9.2E-06</td><td>3.9E-05</td><td>BT20/MB468</td><td>MP</td></tr>
<tr><td>>MI0000102|hsa-mir-100&WithFlank&11|-|122152223|122152314@22.39.18&[MIMAT0000098&hsa-miR-100-5p&offsets|+3|-1;m-27&11|-|122152276|122152296&offsets|+3|0]|CCGTAGATCCGAACTTGT</td><td>>MI0000102|hsa-mir-100&WithFlank&11|-|122152223|122152314</td><td>22</td><td>39</td><td>18</td><td>CCGTAGATCCGAACTTGT</td><td>153.2</td><td>10.5</td><td>2.5</td><td>4.2</td><td>2.3E-05</td><td>3.9E-05</td><td>MB231/MB468</td><td>MP</td></tr>
<tr><td>>hg38_21_+_8433172_8446622_RNA45SN1_45S_50ntFlanks@6650.6670.21|ACGACTCTTAGCGGTGGATCA</td><td>>hg38_21_+_8433172_8446622_RNA45SN1_45S_50ntFlanks</td><td>6650</td><td>6670</td><td>21</td><td>ACGACTCTTAGCGGTGGATCA</td><td>804.8</td><td>-3.5</td><td>0.8</td><td>-4.2</td><td>2.8E-05</td><td>3.9E-05</td><td>MB231/MB468</td><td>MITO</td></tr>
<tr><td>>MI0006444|hsa-mir-1307&WithFlank&10|-|103394247|103394407@86.107.22(+1A)&[MIMAT0005951&hsa-miR-1307-3p&offsets|0|0(+1A);m-66&10|-|103394300|103394322&offsets|0|-1(+1A)]|ACTCGGCGTGGCGTCGGTCGTGA</td><td>>MI0006444|hsa-mir-1307&WithFlank&10|-|103394247|103394407</td><td>86</td><td>107</td><td>22(+1A)</td><td>ACTCGGCGTGGCGTCGGTCGTGA</td><td>124.0</td><td>1.9</td><td>0.4</td><td>4.2</td><td>2.9E-05</td><td>3.9E-05</td><td>BT20/MB468</td><td>NUC</td></tr>
<tr><td>>hg38_21_+_8433172_8446622_RNA45SN1_45S_50ntFlanks@10463.10484.22|CTCCGTTGCCCTCGGCCGATCG</td><td>>hg38_21_+_8433172_8446622_RNA45SN1_45S_50ntFlanks</td><td>10463</td><td>10484</td><td>22</td><td>CTCCGTTGCCCTCGGCCGATCG</td><td>149.3</td><td>-5.2</td><td>1.2</td><td>-4.2</td><td>2.4E-05</td><td>3.9E-05</td><td>BT20/MB468</td><td>CYTO</td></tr>
<tr><td>>hg38_21_+_8433172_8446622_RNA45SN1_45S_50ntFlanks@485.508.24|GATGTGGTGACGTCGTGCTCTCCC</td><td>>hg38_21_+_8433172_8446622_RNA45SN1_45S_50ntFlanks</td><td>485</td><td>508</td><td>24</td><td>GATGTGGTGACGTCGTGCTCTCCC</td><td>219.2</td><td>11.5</td><td>2.6</td><td>4.4</td><td>9.3E-06</td><td>3.9E-05</td><td>BT20/MB468</td><td>MP</td></tr>
<tr><td>>hg38_21_+_8433172_8446622_RNA45SN1_45S_50ntFlanks@6749.6770.22|TTCGAACGCACTTGCGGCCCCG</td><td>>hg38_21_+_8433172_8446622_RNA45SN1_45S_50ntFlanks</td><td>6749</td><td>6770</td><td>22</td><td>TTCGAACGCACTTGCGGCCCCG</td><td>143.8</td><td>-6.4</td><td>1.4</td><td>-4.4</td><td>9.3E-06</td><td>3.9E-05</td><td>BT20/MB468</td><td>MP</td></tr>
<tr><td>>hg38_21_+_8433172_8446622_RNA45SN1_45S_50ntFlanks@11714.11740.27|GGTAAACGGCGGGAGTAACTATGACTC</td><td>>hg38_21_+_8433172_8446622_RNA45SN1_45S_50ntFlanks</td><td>11714</td><td>11740</td><td>27</td><td>GGTAAACGGCGGGAGTAACTATGACTC</td><td>579.3</td><td>-7.3</td><td>1.7</td><td>-4.2</td><td>2.9E-05</td><td>3.9E-05</td><td>BT20/MB468</td><td>NUC</td></tr>
<tr><td>>hg38_MT_+_598_1651_RNR1_12S_50ntFlanks@429.468.40|AAGTGGCTTTAACATATCTGAACACACAATAGCTAAGACC</td><td>>hg38_MT_+_598_1651_RNR1_12S_50ntFlanks</td><td>429</td><td>468</td><td>40</td><td>AAGTGGCTTTAACATATCTGAACACACAATAGCTAAGACC</td><td>121.7</td><td>6.8</td><td>1.6</td><td>4.2</td><td>2.8E-05</td><td>3.9E-05</td><td>MB231/MB468</td><td>MITO</td></tr>
<tr><td>>hg38_21_+_8433172_8446622_RNA45SN1_45S_50ntFlanks@10737.10757.21|TAAATCTCGCGCCGGGCCGTA</td><td>>hg38_21_+_8433172_8446622_RNA45SN1_45S_50ntFlanks</td><td>10737</td><td>10757</td><td>21</td><td>TAAATCTCGCGCCGGGCCGTA</td><td>187.1</td><td>-3.8</td><td>0.9</td><td>-4.2</td><td>2.4E-05</td><td>3.9E-05</td><td>BT20/MB468</td><td>CYTO</td></tr>
<tr><td>>hg38_21_+_8433172_8446622_RNA45SN1_45S_50ntFlanks@12641.12661.21|CCATCTGTGGGATTATGACTG</td><td>>hg38_21_+_8433172_8446622_RNA45SN1_45S_50ntFlanks</td><td>12641</td><td>12661</td><td>21</td><td>CCATCTGTGGGATTATGACTG</td><td>2207.1</td><td>-3.2</td><td>0.7</td><td>-4.3</td><td>2.0E-05</td><td>3.9E-05</td><td>BT20/MB468</td><td>MITO</td></tr>
<tr><td>>hg38_21_+_8433172_8446622_RNA45SN1_45S_50ntFlanks@6654.6680.27|CTCTTAGCGGTGGATCACTCGGCTCGT</td><td>>hg38_21_+_8433172_8446622_RNA45SN1_45S_50ntFlanks</td><td>6654</td><td>6680</td><td>27</td><td>CTCTTAGCGGTGGATCACTCGGCTCGT</td><td>117.1</td><td>-6.2</td><td>1.5</td><td>-4.2</td><td>2.3E-05</td><td>3.9E-05</td><td>MB231/MB468</td><td>MP</td></tr>
<tr><td>>MI0000088|hsa-mir-30a&WithFlank&6|-|71403545|71403627@53.73.21(+3U)&[MIMAT0000088&hsa-miR-30a-3p&offsets|0|-1(+3U);m-38&6|-|71403554|71403575&offsets|0|-1(+3U)]|CTTTCAGTCGGATGTTTGCAGTTT</td><td>>MI0000088|hsa-mir-30a&WithFlank&6|-|71403545|71403627</td><td>53</td><td>73</td><td>21(+3U)</td><td>CTTTCAGTCGGATGTTTGCAGTTT</td><td>29.1</td><td>8.1</td><td>1.9</td><td>4.2</td><td>2.3E-05</td><td>3.9E-05</td><td>MB231/MB468</td><td>MP</td></tr>
<tr><td>>hg38_21_+_8433172_8446622_RNA45SN1_45S_50ntFlanks@7988.8015.28|CAGACGTGGCGACCCGCTGAATTTAAGC</td><td>>hg38_21_+_8433172_8446622_RNA45SN1_45S_50ntFlanks</td><td>7988</td><td>8015</td><td>28</td><td>CAGACGTGGCGACCCGCTGAATTTAAGC</td><td>383.9</td><td>-3.1</td><td>0.7</td><td>-4.2</td><td>2.8E-05</td><td>3.9E-05</td><td>MB231/MB468</td><td>MITO</td></tr>
<tr><td>>hg38_21_+_8433172_8446622_RNA45SN1_45S_50ntFlanks@475.505.31|CTTCGTGATCGATGTGGTGACGTCGTGCTCT</td><td>>hg38_21_+_8433172_8446622_RNA45SN1_45S_50ntFlanks</td><td>475</td><td>505</td><td>31</td><td>CTTCGTGATCGATGTGGTGACGTCGTGCTCT</td><td>808.4</td><td>3.6</td><td>0.9</td><td>4.2</td><td>2.4E-05</td><td>3.9E-05</td><td>MB231/MB468</td><td>MP</td></tr>
<tr><td>>hg38_21_+_8433172_8446622_RNA45SN1_45S_50ntFlanks@11781.11798.18|CGCATGAATGGATGAACG</td><td>>hg38_21_+_8433172_8446622_RNA45SN1_45S_50ntFlanks</td><td>11781</td><td>11798</td><td>18</td><td>CGCATGAATGGATGAACG</td><td>671.1</td><td>-4.1</td><td>0.9</td><td>-4.4</td><td>9.3E-06</td><td>3.9E-05</td><td>BT20/MB468</td><td>MP</td></tr>
<tr><td>>hg38_21_+_8433172_8446622_RNA45SN1_45S_50ntFlanks@12995.13040.46|CTCGCTGCGATCTATTGAAAGTCAGCCCTCGACACAAGGGTTTGTC</td><td>>hg38_21_+_8433172_8446622_RNA45SN1_45S_50ntFlanks</td><td>12995</td><td>13040</td><td>46</td><td>CTCGCTGCGATCTATTGAAAGTCAGCCCTCGACACAAGGGTTTGTC</td><td>735.0</td><td>6.9</td><td>1.7</td><td>4.2</td><td>2.8E-05</td><td>3.9E-05</td><td>MB231/MB468</td><td>NUC</td></tr>
<tr><td>>hg38_21_+_8433172_8446622_RNA45SN1_45S_50ntFlanks@8171.8197.27|AAGTCCTTCTGATCGAGGCCCAGCCCG</td><td>>hg38_21_+_8433172_8446622_RNA45SN1_45S_50ntFlanks</td><td>8171</td><td>8197</td><td>27</td><td>AAGTCCTTCTGATCGAGGCCCAGCCCG</td><td>696.5</td><td>-4.0</td><td>0.9</td><td>-4.3</td><td>2.0E-05</td><td>3.9E-05</td><td>BT20/MB468</td><td>MITO</td></tr>
<tr><td>>hg38_21_+_8433172_8446622_RNA45SN1_45S_50ntFlanks@3899.3937.39|CCTTCGCGGGGGGGATGCGTGCATTTATCAGATCAAAAC</td><td>>hg38_21_+_8433172_8446622_RNA45SN1_45S_50ntFlanks</td><td>3899</td><td>3937</td><td>39</td><td>CCTTCGCGGGGGGGATGCGTGCATTTATCAGATCAAAAC</td><td>422.0</td><td>-6.6</td><td>1.6</td><td>-4.2</td><td>2.4E-05</td><td>3.9E-05</td><td>BT20/MB468</td><td>CYTO</td></tr>
<tr><td>>am_trnaMT_PheGAA_MT_+_577_647@41.58.18|GTTTAGACGGGCTCACAT</td><td>>am_trnaMT_PheGAA_MT_+_577_647</td><td>41</td><td>58</td><td>18</td><td>GTTTAGACGGGCTCACAT</td><td>347.5</td><td>3.3</td><td>0.8</td><td>4.2</td><td>2.4E-05</td><td>3.9E-05</td><td>MB231/MB468</td><td>MP</td></tr>
<tr><td>>hg38_21_+_8433172_8446622_RNA45SN1_45S_50ntFlanks@12159.12180.22|TTTGACTGGGGCGGTACACCTG</td><td>>hg38_21_+_8433172_8446622_RNA45SN1_45S_50ntFlanks</td><td>12159</td><td>12180</td><td>22</td><td>TTTGACTGGGGCGGTACACCTG</td><td>606.1</td><td>-6.0</td><td>1.4</td><td>-4.2</td><td>2.4E-05</td><td>3.9E-05</td><td>BT20/MB468</td><td>CYTO</td></tr>
<tr><td>>hg38_21_+_8433172_8446622_RNA45SN1_45S_50ntFlanks@4007.4024.18|CCTCGGGCCGATCGCACG</td><td>>hg38_21_+_8433172_8446622_RNA45SN1_45S_50ntFlanks</td><td>4007</td><td>4024</td><td>18</td><td>CCTCGGGCCGATCGCACG</td><td>374.6</td><td>-2.9</td><td>0.7</td><td>-4.2</td><td>2.8E-05</td><td>3.9E-05</td><td>MB231/MB468</td><td>NUC</td></tr>
<tr><td>>hg38_21_+_8433172_8446622_RNA45SN1_45S_50ntFlanks@8194.8211.18|CCCGTGGACGGTGTGAGG</td><td>>hg38_21_+_8433172_8446622_RNA45SN1_45S_50ntFlanks</td><td>8194</td><td>8211</td><td>18</td><td>CCCGTGGACGGTGTGAGG</td><td>3291.4</td><td>-4.6</td><td>1.0</td><td>-4.4</td><td>9.3E-06</td><td>3.9E-05</td><td>BT20/MB468</td><td>MP</td></tr>
</table>

<table>
<tr><td>>hg38_21_+_8433172_8446622_RNA45SN1_45S_50ntFlanks@10808.10826.19|CAATGTAGGTAAGGGAAGT</td><td>>hg38_21_+_8433172_8446622_RNA45SN1_45S_50ntFlanks</td><td>10808</td><td>10826</td><td>19</td><td>CAATGTAGGTAAGGGAAGT</td><td>1176.8</td><td>-6.4</td><td>1.5</td><td>-4.2</td><td>2.4E-05</td><td>3.9E-05</td><td>BT20/MB468</td><td>CYTO</td></tr>
<tr><td>>hg38_MT_+_1621_3279_RNR2_16S_50ntFlanks@1092.1117.26|GCAAGACGAGAAGACCCTATGGAGCT</td><td>>hg38_MT_+_1621_3279_RNR2_16S_50ntFlanks</td><td>1092</td><td>1117</td><td>26</td><td>GCAAGACGAGAAGACCCTATGGAGCT</td><td>214.0</td><td>4.6</td><td>1.0</td><td>4.4</td><td>9.4E-06</td><td>3.9E-05</td><td>BT20/MB468</td><td>MP</td></tr>
<tr><td>>hg38_21_+_8433172_8446622_RNA45SN1_45S_50ntFlanks@9573.9594.22|AAATCGGTCGTCCGACCTGGGT</td><td>>hg38_21_+_8433172_8446622_RNA45SN1_45S_50ntFlanks</td><td>9573</td><td>9594</td><td>22</td><td>AAATCGGTCGTCCGACCTGGGT</td><td>1884.7</td><td>-4.4</td><td>1.0</td><td>-4.4</td><td>9.4E-06</td><td>3.9E-05</td><td>BT20/MB468</td><td>MP</td></tr>
<tr><td>>MI0001446|hsa-mir-424&WithFlank&X|-|134546608|134546717@54.72.19&[MIMAT0004749&hsa-miR-424-3p&offsets|0|-2;m-305&X|-|134546644|134546664&offsets|0|-2]|CAAAACGTGAGGCGCTGCT</td><td>>MI0001446|hsa-mir-424&WithFlank&X|-|134546608|134546717</td><td>54</td><td>72</td><td>19</td><td>CAAAACGTGAGGCGCTGCT</td><td>39.5</td><td>8.5</td><td>2.0</td><td>4.2</td><td>2.4E-05</td><td>3.9E-05</td><td>MB231/MB468</td><td>MP</td></tr>
<tr><td>>hg38_21_+_8433172_8446622_RNA45SN1_45S_50ntFlanks@11561.11594.34|GCGCCTAGCAGCCGACTTAGAACTGGTGCGGACC</td><td>>hg38_21_+_8433172_8446622_RNA45SN1_45S_50ntFlanks</td><td>11561</td><td>11594</td><td>34</td><td>GCGCCTAGCAGCCGACTTAGAACTGGTGCGGACC</td><td>425.9</td><td>-3.2</td><td>0.8</td><td>-4.2</td><td>2.4E-05</td><td>3.9E-05</td><td>BT20/MB468</td><td>CYTO</td></tr>
<tr><td>>hg38_21_+_8433172_8446622_RNA45SN1_45S_50ntFlanks@6661.6699.39|CGGTGGATCACTCGGCTCGTGCGTCGATGAAGAACGCAG</td><td>>hg38_21_+_8433172_8446622_RNA45SN1_45S_50ntFlanks</td><td>6661</td><td>6699</td><td>39</td><td>CGGTGGATCACTCGGCTCGTGCGTCGATGAAGAACGCAG</td><td>8180.5</td><td>-3.9</td><td>0.9</td><td>-4.2</td><td>2.8E-05</td><td>3.9E-05</td><td>MB231/MB468</td><td>MITO</td></tr>
<tr><td>>hg38_21_+_8433172_8446622_RNA45SN1_45S_50ntFlanks@12154.12183.30|GGGAGTTTGACTGGGGCGGTACACCTGTCA</td><td>>hg38_21_+_8433172_8446622_RNA45SN1_45S_50ntFlanks</td><td>12154</td><td>12183</td><td>30</td><td>GGGAGTTTGACTGGGGCGGTACACCTGTCA</td><td>331.8</td><td>-6.3</td><td>1.5</td><td>-4.2</td><td>2.4E-05</td><td>3.9E-05</td><td>MB231/MB468</td><td>MP</td></tr>
<tr><td>>hg38_21_+_8433172_8446622_RNA45SN1_45S_50ntFlanks@5527.5549.23|AAAGTCGTAACAAGGTTTCCGTA</td><td>>hg38_21_+_8433172_8446622_RNA45SN1_45S_50ntFlanks</td><td>5527</td><td>5549</td><td>23</td><td>AAAGTCGTAACAAGGTTTCCGTA</td><td>202.9</td><td>-5.6</td><td>1.3</td><td>-4.2</td><td>2.4E-05</td><td>3.9E-05</td><td>BT20/MB468</td><td>CYTO</td></tr>
<tr><td>>hg38_21_+_8433172_8446622_RNA45SN1_45S_50ntFlanks@3846.3881.36|CTTGGATAACTGTGGTAATTCTAGAGCTAATACATG</td><td>>hg38_21_+_8433172_8446622_RNA45SN1_45S_50ntFlanks</td><td>3846</td><td>3881</td><td>36</td><td>CTTGGATAACTGTGGTAATTCTAGAGCTAATACATG</td><td>247.6</td><td>-7.1</td><td>1.6</td><td>-4.4</td><td>9.4E-06</td><td>4.0E-05</td><td>BT20/MB468</td><td>MP</td></tr>
<tr><td>>hg38_21_+_8433172_8446622_RNA45SN1_45S_50ntFlanks@6652.6701.50|GACTCTTAGCGGTGGATCACTCGGCTCGTGCGTCGATGAAGAACGCAGCT</td><td>>hg38_21_+_8433172_8446622_RNA45SN1_45S_50ntFlanks</td><td>6652</td><td>6701</td><td>50</td><td>GACTCTTAGCGGTGGATCACTCGGCTCGTGCGTCGATGAAGAACGCAGCT</td><td>1083.2</td><td>4.5</td><td>1.1</td><td>4.2</td><td>2.8E-05</td><td>4.0E-05</td><td>MB231/MB468</td><td>MITO</td></tr>
<tr><td>>MI0000102|hsa-mir-100&WithFlank&11|-|122152223|122152314@19.38.20(+1G)&[MIMAT0000098&hsa-miR-100-5p&offsets|0|-2(+1G);m-27&11|-|122152276|122152296&offsets|0|-1(+1G)]|AACCCGTAGATCCGAACTTGG</td><td>>MI0000102|hsa-mir-100&WithFlank&11|-|122152223|122152314</td><td>19</td><td>38</td><td>20(+1G)</td><td>AACCCGTAGATCCGAACTTGG</td><td>1767.8</td><td>-9.3</td><td>2.0</td><td>-4.6</td><td>3.5E-06</td><td>4.0E-05</td><td>BT20/MB231</td><td>MITO</td></tr>
<tr><td>>hg38_21_+_8433172_8446622_RNA45SN1_45S_50ntFlanks@5492.5510.19|AGAAGACGGTCGAACTTGA</td><td>>hg38_21_+_8433172_8446622_RNA45SN1_45S_50ntFlanks</td><td>5492</td><td>5510</td><td>19</td><td>AGAAGACGGTCGAACTTGA</td><td>186.7</td><td>-4.4</td><td>1.1</td><td>-4.2</td><td>2.8E-05</td><td>4.0E-05</td><td>MB231/MB468</td><td>NUC</td></tr>
<tr><td>>hg38_1_-_228634819_228635039_RNA5S12_5S_50ntFlanks@133.171.39|ATGGGAGACCGCCTGGGAATACCGGGTGCTGTAGGCTTT</td><td>>hg38_1_-_228634819_228635039_RNA5S12_5S_50ntFlanks</td><td>133</td><td>171</td><td>39</td><td>ATGGGAGACCGCCTGGGAATACCGGGTGCTGTAGGCTTT</td><td>8682.3</td><td>4.1</td><td>1.0</td><td>4.2</td><td>2.7E-05</td><td>4.0E-05</td><td>MB231/MB468</td><td>CYTO</td></tr>
<tr><td>>hg38_21_+_8433172_8446622_RNA45SN1_45S_50ntFlanks@10510.10532.23|AATCCGGAGTGGCGGAGATGGGC</td><td>>hg38_21_+_8433172_8446622_RNA45SN1_45S_50ntFlanks</td><td>10510</td><td>10532</td><td>23</td><td>AATCCGGAGTGGCGGAGATGGGC</td><td>841.1</td><td>-5.4</td><td>1.3</td><td>-4.2</td><td>2.4E-05</td><td>4.0E-05</td><td>MB231/MB468</td><td>MP</td></tr>
<tr><td>>trnaMT_ProTGG_MT_-_15956_16023@43.60.18|GGTGGAGTTAAAGACTTT</td><td>>trnaMT_ProTGG_MT_-_15956_16023</td><td>43</td><td>60</td><td>18</td><td>GGTGGAGTTAAAGACTTT</td><td>32.7</td><td>8.2</td><td>2.0</td><td>4.2</td><td>2.4E-05</td><td>4.0E-05</td><td>MB231/MB468</td><td>MP</td></tr>
<tr><td>>MI0000433|hsa-let-7g&WithFlank&3|-|52268272|52268367@11.31.21&[MIMAT0000414&hsa-let-7g-5p&offsets|0|-1;m-58&3|-|52268336|52268357&offsets|0|-1]|TGAGGTAGTAGTTTGTACAGT</td><td>>MI0000433|hsa-let-7g&WithFlank&3|-|52268272|52268367</td><td>11</td><td>31</td><td>21</td><td>TGAGGTAGTAGTTTGTACAGT</td><td>3029.6</td><td>4.4</td><td>1.0</td><td>4.2</td><td>2.4E-05</td><td>4.0E-05</td><td>MB231/MB468</td><td>MP</td></tr>
<tr><td>>hg38_21_+_8433172_8446622_RNA45SN1_45S_50ntFlanks@8190.8212.23|CCAGCCCGTGGACGGTGTGAGGC</td><td>>hg38_21_+_8433172_8446622_RNA45SN1_45S_50ntFlanks</td><td>8190</td><td>8212</td><td>23</td><td>CCAGCCCGTGGACGGTGTGAGGC</td><td>398.0</td><td>-4.0</td><td>0.9</td><td>-4.2</td><td>2.7E-05</td><td>4.0E-05</td><td>MB231/MB468</td><td>CYTO</td></tr>
<tr><td>>MI0000088|hsa-mir-30a&WithFlank&6|-|71403545|71403627@12.35.24(+1A)&[MIMAT0000087&hsa-miR-30a-5p&offsets|0|+2(+1A);m-15&6|-|71403595|71403616&offsets|0|+2(+1A)]|TGTAAACATCCTCGACTGGAAGCTA</td><td>>MI0000088|hsa-mir-30a&WithFlank&6|-|71403545|71403627</td><td>12</td><td>35</td><td>24(+1A)</td><td>TGTAAACATCCTCGACTGGAAGCTA</td><td>1084.3</td><td>-4.1</td><td>0.9</td><td>-4.6</td><td>3.5E-06</td><td>4.0E-05</td><td>BT20/MB231</td><td>MITO</td></tr>
<tr><td>>hg38_21_+_8433172_8446622_RNA45SN1_45S_50ntFlanks@10388.10408.21|GTGAACAGCAGTTGAACATGG</td><td>>hg38_21_+_8433172_8446622_RNA45SN1_45S_50ntFlanks</td><td>10388</td><td>10408</td><td>21</td><td>GTGAACAGCAGTTGAACATGG</td><td>65.2</td><td>-9.9</td><td>2.3</td><td>-4.2</td><td>2.4E-05</td><td>4.0E-05</td><td>MB231/MB468</td><td>MP</td></tr>
<tr><td>>hg38_21_+_8433172_8446622_RNA45SN1_45S_50ntFlanks@12447.12486.40|CAGAATTCACCAAGCGTTGGATTGTTCACCCACTAATAGG</td><td>>hg38_21_+_8433172_8446622_RNA45SN1_45S_50ntFlanks</td><td>12447</td><td>12486</td><td>40</td><td>CAGAATTCACCAAGCGTTGGATTGTTCACCCACTAATAGG</td><td>371.2</td><td>-3.8</td><td>0.9</td><td>-4.2</td><td>2.8E-05</td><td>4.0E-05</td><td>MB231/MB468</td><td>NUC</td></tr>
<tr><td>>hg38_21_+_8433172_8446622_RNA45SN1_45S_50ntFlanks@11911.11929.19|TGAAGAGACATGAGAGGTG</td><td>>hg38_21_+_8433172_8446622_RNA45SN1_45S_50ntFlanks</td><td>11911</td><td>11929</td><td>19</td><td>TGAAGAGACATGAGAGGTG</td><td>107.5</td><td>10.1</td><td>2.2</td><td>4.7</td><td>3.0E-06</td><td>4.0E-05</td><td>BT20/MB231</td><td>MP</td></tr>
<tr><td>>hg38_21_+_8433172_8446622_RNA45SN1_45S_50ntFlanks@9554.9581.28|CGTAGCGGTCCTGACGTGCAAATCGGTC</td><td>>hg38_21_+_8433172_8446622_RNA45SN1_45S_50ntFlanks</td><td>9554</td><td>9581</td><td>28</td><td>CGTAGCGGTCCTGACGTGCAAATCGGTC</td><td>1948.6</td><td>2.6</td><td>0.6</td><td>4.2</td><td>2.7E-05</td><td>4.0E-05</td><td>MB231/MB468</td><td>CYTO</td></tr>
<tr><td>>hg38_21_+_8433172_8446622_RNA45SN1_45S_50ntFlanks@6787.6806.20|ACGCCTGTCTGAGCGTCGCT</td><td>>hg38_21_+_8433172_8446622_RNA45SN1_45S_50ntFlanks</td><td>6787</td><td>6806</td><td>20</td><td>ACGCCTGTCTGAGCGTCGCT</td><td>409.7</td><td>2.2</td><td>0.5</td><td>4.2</td><td>3.0E-05</td><td>4.0E-05</td><td>BT20/MB468</td><td>NUC</td></tr>
<tr><td>>hg38_21_+_8433172_8446622_RNA45SN1_45S_50ntFlanks@11567.11588.22|AGCAGCCGACTTAGAACTGGTG</td><td>>hg38_21_+_8433172_8446622_RNA45SN1_45S_50ntFlanks</td><td>11567</td><td>11588</td><td>22</td><td>AGCAGCCGACTTAGAACTGGTG</td><td>328.8</td><td>-2.2</td><td>0.5</td><td>-4.2</td><td>2.5E-05</td><td>4.0E-05</td><td>BT20/MB468</td><td>CYTO</td></tr>
<tr><td>>MI0000542|hsa-mir-320a&WithFlank&8|-|22244960|22245043@47.68.22&[MIMAT0000510&hsa-miR-320a-3p&offsets|-1|-1;m-60&8|-|22244975|22244996&offsets|-1|-1]|GAAAAGCTGGGTTGAGAGGGCG</td><td>>MI0000542|hsa-mir-320a&WithFlank&8|-|22244960|22245043</td><td>47</td><td>68</td><td>22</td><td>GAAAAGCTGGGTTGAGAGGGCG</td><td>345.8</td><td>8.2</td><td>1.8</td><td>4.4</td><td>9.5E-06</td><td>4.0E-05</td><td>BT20/MB468</td><td>MP</td></tr>
<tr><td>>MI0000471|hsa-mir-126&WithFlank&9|+|136670596|136670692@58.76.19(+1A)&[MIMAT0000445&hsa-miR-126-3p&offsets|0|-3(+1A);m-40&9|+|136670653|136670674&offsets|0|-3(+1A)]|TCGTACCGTGAGTAATAATA</td><td>>MI0000471|hsa-mir-126&WithFlank&9|+|136670596|136670692</td><td>58</td><td>76</td><td>19(+1A)</td><td>TCGTACCGTGAGTAATAATA</td><td>42.5</td><td>8.6</td><td>2.0</td><td>4.2</td><td>2.4E-05</td><td>4.0E-05</td><td>MB231/MB468</td><td>MP</td></tr>
<tr><td>>hg38_21_+_8433172_8446622_RNA45SN1_45S_50ntFlanks@11602.11619.18|TCCGACTGTTTAATTAAA</td><td>>hg38_21_+_8433172_8446622_RNA45SN1_45S_50ntFlanks</td><td>11602</td><td>11619</td><td>18</td><td>TCCGACTGTTTAATTAAA</td><td>1365.0</td><td>-3.2</td><td>0.8</td><td>-4.2</td><td>2.9E-05</td><td>4.0E-05</td><td>MB231/MB468</td><td>NUC</td></tr>
<tr><td>>hg38_21_+_8433172_8446622_RNA45SN1_45S_50ntFlanks@6647.6674.28|CGTACGACTCTTAGCGGTGGATCACTCG</td><td>>hg38_21_+_8433172_8446622_RNA45SN1_45S_50ntFlanks</td><td>6647</td><td>6674</td><td>28</td><td>CGTACGACTCTTAGCGGTGGATCACTCG</td><td>476.1</td><td>4.1</td><td>1.0</td><td>4.2</td><td>2.9E-05</td><td>4.0E-05</td><td>MB231/MB468</td><td>NUC</td></tr>
<tr><td>>hg38_21_+_8433172_8446622_RNA45SN1_45S_50ntFlanks@4385.4423.39|TTGGATCTTGGGAGCGGGCGGGCGGTCCGCCGCGAGGCG</td><td>>hg38_21_+_8433172_8446622_RNA45SN1_45S_50ntFlanks</td><td>4385</td><td>4423</td><td>39</td><td>TTGGATCTTGGGAGCGGGCGGGCGGTCCGCCGCGAGGCG</td><td>298.8</td><td>-6.0</td><td>1.4</td><td>-4.2</td><td>2.9E-05</td><td>4.0E-05</td><td>MB231/MB468</td><td>NUC</td></tr>
<tr><td>>trnaMT_LeuTAA_MT_+_3230_3304@1.30.30|GTTAAGATGGCAGAGCCCGGTAATCGCATA</td><td>>trnaMT_LeuTAA_MT_+_3230_3304</td><td>1</td><td>30</td><td>30</td><td>GTTAAGATGGCAGAGCCCGGTAATCGCATA</td><td>6099.6</td><td>6.3</td><td>1.4</td><td>4.4</td><td>9.5E-06</td><td>4.0E-05</td><td>BT20/MB468</td><td>MP</td></tr>
<tr><td>>hg38_21_+_8433172_8446622_RNA45SN1_45S_50ntFlanks@12074.12095.22|CGAGGGGCTCTCGCTTCTGGCG</td><td>>hg38_21_+_8433172_8446622_RNA45SN1_45S_50ntFlanks</td><td>12074</td><td>12095</td><td>22</td><td>CGAGGGGCTCTCGCTTCTGGCG</td><td>1081.3</td><td>-4.8</td><td>1.1</td><td>-4.2</td><td>2.5E-05</td><td>4.0E-05</td><td>BT20/MB468</td><td>CYTO</td></tr>
<tr><td>>trna128_GlyGCC_6_-_27870686_27870756@16.52.37,trna19_GlyGCC_16_+_70823410_70823480@16.52.37,trna19_GlyGCC_2_-_157257659_157257729@16.52.37,trna24_GlyGCC_16_-_70812942_70813012@16.52.37,trna25_GlyGCC_16_-_70812114_70812184@16.52.37,trna2_GlyGCC_21_-_18827107_18827177@16.52.37,trna35_GlyGCC_1_+_161413094_161413164@16.52.37,trna37_GlyGCC_1_+_161420467_161420537@16.52.37,trna39_GlyGCC_1_+_161427898_161427968@16.52.37,trna41_GlyGCC_1_+_161435258_161435328@16.52.37,trna5_GlyGCC_17_+_8029064_8029134@16.52.37,trna68_GlyGCC_1_-_161493637_161493707@16.52.37|TGGTAGAATTCTCGCCTGCCACGCGGGAGGCCCGGGT</td><td>>trna128_GlyGCC_6_-_27870686_27870756</td><td>16</td><td>52</td><td>37</td><td>TGGTAGAATTCTCGCCTGCCACGCGGGAGGCCCGGGT</td><td>592.3</td><td>3.7</td><td>0.8</td><td>4.7</td><td>2.9E-06</td><td>4.0E-05</td><td>BT20/MB231</td><td>CYTO</td></tr>
<tr><td>>MI0000086|hsa-mir-28&WithFlank&3|+|188688775|188688872@62.82.21(+1U)&[MIMAT0004502&hsa-miR-28-3p&offsets|+2|+1(+1U);m-28&3|+|188688834|188688855&offsets|+2|+1(+1U)]|CTAGATTGTGAGCTCCTGGAGT</td><td>>MI0000086|hsa-mir-28&WithFlank&3|+|188688775|188688872</td><td>62</td><td>82</td><td>21(+1U)</td><td>CTAGATTGTGAGCTCCTGGAGT</td><td>909.9</td><td>7.5</td><td>1.7</td><td>4.4</td><td>9.6E-06</td><td>4.0E-05</td><td>BT20/MB468</td><td>MP</td></tr>
<tr><td>>hg38_21_+_8433172_8446622_RNA45SN1_45S_50ntFlanks@8184.8221.38|CGAGGCCCAGCCCGTGGACGGTGTGAGGCCGGTAGCGG</td><td>>hg38_21_+_8433172_8446622_RNA45SN1_45S_50ntFlanks</td><td>8184</td><td>8221</td><td>38</td><td>CGAGGCCCAGCCCGTGGACGGTGTGAGGCCGGTAGCGG</td><td>45.5</td><td>-8.6</td><td>1.9</td><td>-4.4</td><td>9.6E-06</td><td>4.0E-05</td><td>BT20/MB468</td><td>MP</td></tr>
<tr><td>>hg38_21_+_8433172_8446622_RNA45SN1_45S_50ntFlanks@9556.9584.29|TAGCGGTCCTGACGTGCAAATCGGTCGTC</td><td>>hg38_21_+_8433172_8446622_RNA45SN1_45S_50ntFlanks</td><td>9556</td><td>9584</td><td>29</td><td>TAGCGGTCCTGACGTGCAAATCGGTCGTC</td><td>284.9</td><td>-4.3</td><td>1.0</td><td>-4.2</td><td>2.7E-05</td><td>4.0E-05</td><td>MB231/MB468</td><td>CYTO</td></tr>
<tr><td>>hg38_21_+_8433172_8446622_RNA45SN1_45S_50ntFlanks@10680.10712.33|GTTCCGGCGGCGTCCGGTGAGCTCTCGCTGGCC</td><td>>hg38_21_+_8433172_8446622_RNA45SN1_45S_50ntFlanks</td><td>10680</td><td>10712</td><td>33</td><td>GTTCCGGCGGCGTCCGGTGAGCTCTCGCTGGCC</td><td>2085.2</td><td>-4.3</td><td>1.0</td><td>-4.4</td><td>9.6E-06</td><td>4.0E-05</td><td>BT20/MB468</td><td>MP</td></tr>
<tr><td>>hg38_21_+_8433172_8446622_RNA45SN1_45S_50ntFlanks@12444.12481.38|AAGCAGAATTCACCAAGCGTTGGATTGTTCACCCACTA</td><td>>hg38_21_+_8433172_8446622_RNA45SN1_45S_50ntFlanks</td><td>12444</td><td>12481</td><td>38</td><td>AAGCAGAATTCACCAAGCGTTGGATTGTTCACCCACTA</td><td>42.7</td><td>-8.5</td><td>1.9</td><td>-4.4</td><td>9.6E-06</td><td>4.0E-05</td><td>BT20/MB468</td><td>MP</td></tr>
<tr><td>>hg38_21_+_8433172_8446622_RNA45SN1_45S_50ntFlanks@11568.11599.32|GCAGCCGACTTAGAACTGGTGCGGACCAGGGG</td><td>>hg38_21_+_8433172_8446622_RNA45SN1_45S_50ntFlanks</td><td>11568</td><td>11599</td><td>32</td><td>GCAGCCGACTTAGAACTGGTGCGGACCAGGGG</td><td>112.2</td><td>-4.8</td><td>1.1</td><td>-4.2</td><td>2.4E-05</td><td>4.0E-05</td><td>MB231/MB468</td><td>MP</td></tr>
<tr><td>>hg38_21_+_8433172_8446622_RNA45SN1_45S_50ntFlanks@12406.12434.29|TTTTTGATCCTTCGATGTCGGCTCTTCCT</td><td>>hg38_21_+_8433172_8446622_RNA45SN1_45S_50ntFlanks</td><td>12406</td><td>12434</td><td>29</td><td>TTTTTGATCCTTCGATGTCGGCTCTTCCT</td><td>2325.0</td><td>-2.9</td><td>0.7</td><td>-4.2</td><td>2.9E-05</td><td>4.0E-05</td><td>MB231/MB468</td><td>NUC</td></tr>
<tr><td>>hg38_21_+_8433172_8446622_RNA45SN1_45S_50ntFlanks@12285.12323.39|CAGACCGTGAAAGCGGGGCCTCACGATCCTTCTGACCTT</td><td>>hg38_21_+_8433172_8446622_RNA45SN1_45S_50ntFlanks</td><td>12285</td><td>12323</td><td>39</td><td>CAGACCGTGAAAGCGGGGCCTCACGATCCTTCTGACCTT</td><td>60.0</td><td>-9.8</td><td>2.3</td><td>-4.2</td><td>2.4E-05</td><td>4.0E-05</td><td>MB231/MB468</td><td>MP</td></tr>
<tr><td>>hg38_21_+_8433172_8446622_RNA45SN1_45S_50ntFlanks@12934.12958.25|CTAAACCATTCGTAGACGACCTGCT</td><td>>hg38_21_+_8433172_8446622_RNA45SN1_45S_50ntFlanks</td><td>12934</td><td>12958</td><td>25</td><td>CTAAACCATTCGTAGACGACCTGCT</td><td>264.7</td><td>-2.0</td><td>0.5</td><td>-4.2</td><td>2.5E-05</td><td>4.0E-05</td><td>BT20/MB468</td><td>CYTO</td></tr>
<tr><td>>hg38_21_+_8433172_8446622_RNA45SN1_45S_50ntFlanks@6650.6682.33|ACGACTCTTAGCGGTGGATCACTCGGCTCGTGC</td><td>>hg38_21_+_8433172_8446622_RNA45SN1_45S_50ntFlanks</td><td>6650</td><td>6682</td><td>33</td><td>ACGACTCTTAGCGGTGGATCACTCGGCTCGTGC</td><td>587.6</td><td>5.2</td><td>1.2</td><td>4.2</td><td>2.9E-05</td><td>4.0E-05</td><td>MB231/MB468</td><td>NUC</td></tr>
<tr><td>>hg38_21_+_8433172_8446622_RNA45SN1_45S_50ntFlanks@12914.12933.20|CGTTCGTGGGGAACCTGGCG</td><td>>hg38_21_+_8433172_8446622_RNA45SN1_45S_50ntFlanks</td><td>12914</td><td>12933</td><td>20</td><td>CGTTCGTGGGGAACCTGGCG</td><td>167.4</td><td>-3.0</td><td>0.7</td><td>-4.2</td><td>2.9E-05</td><td>4.0E-05</td><td>MB231/MB468</td><td>NUC</td></tr>
<tr><td>>hg38_21_+_8433172_8446622_RNA45SN1_45S_50ntFlanks@12150.12183.34|GGTGGGGAGTTTGACTGGGGCGGTACACCTGTCA</td><td>>hg38_21_+_8433172_8446622_RNA45SN1_45S_50ntFlanks</td><td>12150</td><td>12183</td><td>34</td><td>GGTGGGGAGTTTGACTGGGGCGGTACACCTGTCA</td><td>42.1</td><td>-8.5</td><td>1.9</td><td>-4.4</td><td>9.6E-06</td><td>4.0E-05</td><td>BT20/MB468</td><td>MP</td></tr>
<tr><td>>hg38_21_+_8433172_8446622_RNA45SN1_45S_50ntFlanks@10692.10709.18|TCCGGTGAGCTCTCGCTG</td><td>>hg38_21_+_8433172_8446622_RNA45SN1_45S_50ntFlanks</td><td>10692</td><td>10709</td><td>18</td><td>TCCGGTGAGCTCTCGCTG</td><td>267.1</td><td>-6.6</td><td>1.5</td><td>-4.4</td><td>9.6E-06</td><td>4.0E-05</td><td>BT20/MB468</td><td>MP</td></tr>
<tr><td>>hg38_21_+_8433172_8446622_RNA45SN1_45S_50ntFlanks@12909.12935.27|CCGCACGTTCGTGGGGAACCTGGCGCT</td><td>>hg38_21_+_8433172_8446622_RNA45SN1_45S_50ntFlanks</td><td>12909</td><td>12935</td><td>27</td><td>CCGCACGTTCGTGGGGAACCTGGCGCT</td><td>84.6</td><td>-6.1</td><td>1.4</td><td>-4.2</td><td>2.4E-05</td><td>4.0E-05</td><td>MB231/MB468</td><td>MP</td></tr>
<tr><td>>hg38_21_+_8433172_8446622_RNA45SN1_45S_50ntFlanks@12957.12980.24|CTTCTGGGTCGGGGTTTCGTACGT</td><td>>hg38_21_+_8433172_8446622_RNA45SN1_45S_50ntFlanks</td><td>12957</td><td>12980</td><td>24</td><td>CTTCTGGGTCGGGGTTTCGTACGT</td><td>149.9</td><td>-5.4</td><td>1.3</td><td>-4.3</td><td>2.1E-05</td><td>4.0E-05</td><td>BT20/MB468</td><td>MITO</td></tr>
<tr><td>>hg38_21_+_8433172_8446622_RNA45SN1_45S_50ntFlanks@4657.4682.26|CTAGAGGTGAAATTCTTGGACCGGCG</td><td>>hg38_21_+_8433172_8446622_RNA45SN1_45S_50ntFlanks</td><td>4657</td><td>4682</td><td>26</td><td>CTAGAGGTGAAATTCTTGGACCGGCG</td><td>700.7</td><td>-4.9</td><td>1.2</td><td>-4.2</td><td>2.7E-05</td><td>4.0E-05</td><td>MB231/MB468</td><td>CYTO</td></tr>
</table>

| | | | | | | | | | | | | | |
|---|---|---|---|---|---|---|---|---|---|---|---|---|---|
| >MI0000081\|hsa-mir-24-2&WithFlank&19\|-\|13836281\|13836365@18.41.24&[MIMAT0004497&hsa-miR-24-2-5p&offsets\|-1\|+1;m-226&19\|-\|13836326\|13836348&offsets\|0\|+1]\|GTGCCTACTGAGCTGAAACACAGT | >MI0000081\|hsa-mir-24-2&WithFlank&19\|-\|13836281\|13836365 | 18 | 41 | 24 | GTGCCTACTGAGCTGAAACACAGT | 392.1 | 4.2 | 1.0 | 4.2 | 2.9E-05 | 4.0E-05 | MB231/MB468 | MITO |
| >hg38_21_+_8433172_8446622_RNA45SN1_45S_50ntFlanks@10689.10712.24\|GCGTCCGGTGAGCTCTCGCTGGCC | >hg38_21_+_8433172_8446622_RNA45SN1_45S_50ntFlanks | 10689 | 10712 | 24 | GCGTCCGGTGAGCTCTCGCTGGCC | 79.5 | -10.2 | 2.4 | -4.2 | 2.4E-05 | 4.0E-05 | MB231/MB468 | MP |
| >hg38_21_+_8433172_8446622_RNA45SN1_45S_50ntFlanks@4735.4758.24\|ACGAAAGTCGGAGGTTCGAAGACG | >hg38_21_+_8433172_8446622_RNA45SN1_45S_50ntFlanks | 4735 | 4758 | 24 | ACGAAAGTCGGAGGTTCGAAGACG | 535.3 | -2.8 | 0.7 | -4.2 | 2.9E-05 | 4.0E-05 | MB231/MB468 | MITO |
| >hg38_21_+_8433172_8446622_RNA45SN1_45S_50ntFlanks@5073.5097.25\|AATTCCGATAACGAACGAGACTCTG | >hg38_21_+_8433172_8446622_RNA45SN1_45S_50ntFlanks | 5073 | 5097 | 25 | AATTCCGATAACGAACGAGACTCTG | 868.6 | -4.0 | 1.0 | -4.2 | 2.9E-05 | 4.0E-05 | MB231/MB468 | NUC |
| >hg38_21_+_8433172_8446622_RNA45SN1_45S_50ntFlanks@10351.10372.22\|CGAGAACTTTGAAGGCCGAAGT | >hg38_21_+_8433172_8446622_RNA45SN1_45S_50ntFlanks | 10351 | 10372 | 22 | CGAGAACTTTGAAGGCCGAAGT | 593.9 | -3.1 | 0.7 | -4.2 | 2.9E-05 | 4.0E-05 | MB231/MB468 | MITO |
| >MI0000070\|hsa-mir-16-1&WithFlank&13\|-\|50048967\|50049067@20.40.21;MI0000115\|hsa-mir-16-2&WithFlank&3\|+\|160404739\|160404831@16.36.21&[MIMAT0000069&hsa-miR-16-5p&offsets\|0\|-1;m-62&13\|-\|50049027\|50049048&offsets\|0\|-1];[MIMAT0000069_1&hsa-miR-16-5p&offsets\|0\|-1;m-61&3\|+\|160404754\|160404775&offsets\|0\|-1]\|TAGCAGCACGTAAATATTGGC | >MI0000070\|hsa-mir-16-1&WithFlank&13\|-\|50048967\|50049067 | 20 | 40 | 21 | TAGCAGCACGTAAATATTGGC | 519.3 | 2.4 | 0.6 | 4.2 | 3.0E-05 | 4.0E-05 | BT20/MB468 | NUC |
| >hg38_21_+_8433172_8446622_RNA45SN1_45S_50ntFlanks@10922.10944.23\|CTGGACGAGGCGCCGCCGCCCCC | >hg38_21_+_8433172_8446622_RNA45SN1_45S_50ntFlanks | 10922 | 10944 | 23 | CTGGACGAGGCGCCGCCGCCCCC | 159.0 | -5.6 | 1.3 | -4.2 | 2.9E-05 | 4.0E-05 | MB231/MB468 | NUC |
| >hg38_21_+_8433172_8446622_RNA45SN1_45S_50ntFlanks@12481.12500.20\|AATAGGGAACGTGAGCTGGG | >hg38_21_+_8433172_8446622_RNA45SN1_45S_50ntFlanks | 12481 | 12500 | 20 | AATAGGGAACGTGAGCTGGG | 233.6 | -2.3 | 0.6 | -4.2 | 2.9E-05 | 4.0E-05 | MB231/MB468 | NUC |
| >trnaMT_HisGTG_MT_+_12138_12206@28.72.45\|ATTGTGAATCTGACAACAGAGGCTTACGACCCCTTATTTACCCCA | >trnaMT_HisGTG_MT_+_12138_12206 | 28 | 72 | 45 | ATTGTGAATCTGACAACAGAGGCTTACGACCCCTTATTTACCCCA | 888.8 | -8.3 | 2.0 | -4.2 | 3.0E-05 | 4.0E-05 | BT20/MB468 | NUC |
| >hg38_21_+_8433172_8446622_RNA45SN1_45S_50ntFlanks@6646.6677.32\|TCGTACGACTCTTAGCGGTGGATCACTCGGCT | >hg38_21_+_8433172_8446622_RNA45SN1_45S_50ntFlanks | 6646 | 6677 | 32 | TCGTACGACTCTTAGCGGTGGATCACTCGGCT | 3766.9 | 3.4 | 0.8 | 4.4 | 9.7E-06 | 4.0E-05 | BT20/MB468 | MP |
| >hg38_21_+_8433172_8446622_RNA45SN1_45S_50ntFlanks@8365.8395.31\|TTTGAAGAGAGAGTTCAAGAGGGCGTGAAAC | >hg38_21_+_8433172_8446622_RNA45SN1_45S_50ntFlanks | 8365 | 8395 | 31 | TTTGAAGAGAGAGTTCAAGAGGGCGTGAAAC | 110.9 | 10.1 | 2.2 | 4.7 | 3.0E-06 | 4.0E-05 | BT20/MB231 | MP |
| >trna116_GluCTC_1_-_145399233_145399304@30.70.41,trna71_GluCTC_1_-_161439189_161439260@30.70.41,trna74_GluCTC_1_-_161431809_161431880@30.70.41,trna77_GluCTC_1_-_161424398_161424469@30.70.41,trna77_GluCTC_6_+_28949976_28950047@30.70.41,trna80_GluCTC_1_-_161417018_161417089@30.70.41,trna87_GluCTC_6_-_126101393_126101464@30.70.41\|CGCTCTCACCGCCGCGGCCCGGGTTCGATTCCCGGTCAGGG | >trna116_GluCTC_1_-_145399233_145399304 | 30 | 70 | 41 | CGCTCTCACCGCCGCGGCCCGGGTTCGATTCCCGGTCAGGG | 346.0 | -4.4 | 1.0 | -4.2 | 2.5E-05 | 4.0E-05 | BT20/MB468 | CYTO |
| >hg38_21_+_8433172_8446622_RNA45SN1_45S_50ntFlanks@10841.10866.26\|CGTAACTTCGGGATAAGGATTGGCTC | >hg38_21_+_8433172_8446622_RNA45SN1_45S_50ntFlanks | 10841 | 10866 | 26 | CGTAACTTCGGGATAAGGATTGGCTC | 1931.8 | -5.4 | 1.3 | -4.2 | 2.9E-05 | 4.0E-05 | MB231/MB468 | NUC |
| >hg38_21_+_8433172_8446622_RNA45SN1_45S_50ntFlanks@10351.10373.23\|CGAGAACTTTGAAGGCCGAAGTG | >hg38_21_+_8433172_8446622_RNA45SN1_45S_50ntFlanks | 10351 | 10373 | 23 | CGAGAACTTTGAAGGCCGAAGTG | 335.0 | -2.7 | 0.6 | -4.3 | 2.1E-05 | 4.0E-05 | BT20/MB468 | MITO |
| >am_trna127_CysGCA_1_-_93981834_93981906@17.35.19,trna15_CysGCA_17_+_37023898_37023969@16.34.19,trna26_CysGCA_17_-_37310744_37310815@16.34.19,trna27_CysGCA_17_-_37309987_37310058@16.34.19,trna28_CysGCA_17_-_37025545_37025616@16.34.19,trna3_CysGCA_4_-_124430005_124430076@16.34.19,trna5_CysGCA_7_+_149007281_149007352@16.34.19,trna7_CysGCA_3_-_131947944_131948015@16.34.19\|TGGTAGAGCATTTGACTGC | >am_trna127_CysGCA_1_-_93981834_93981906 | 17 | 35 | 19 | TGGTAGAGCATTTGACTGC | 28.1 | 8.0 | 1.9 | 4.2 | 2.4E-05 | 4.1E-05 | MB231/MB468 | MP |
| >hg38_21_+_8433172_8446622_RNA45SN1_45S_50ntFlanks@9484.9513.30\|GTGTTAGGACCCGAAAGATGGTGAACTATG | >hg38_21_+_8433172_8446622_RNA45SN1_45S_50ntFlanks | 9484 | 9513 | 30 | GTGTTAGGACCCGAAAGATGGTGAACTATG | 38.5 | 9.0 | 2.0 | 4.4 | 9.7E-06 | 4.1E-05 | BT20/MB468 | MP |
| >hg38_21_+_8433172_8446622_RNA45SN1_45S_50ntFlanks@11821.11847.27\|CTATCCAGCGAAACCACAGCCAAGGGA | >hg38_21_+_8433172_8446622_RNA45SN1_45S_50ntFlanks | 11821 | 11847 | 27 | CTATCCAGCGAAACCACAGCCAAGGGA | 43.5 | -8.5 | 1.9 | -4.4 | 9.7E-06 | 4.1E-05 | BT20/MB468 | MP |
| >hg38_21_+_8433172_8446622_RNA45SN1_45S_50ntFlanks@9597.9622.26\|AGGGGCGAAAGACTAATCGAACCATC | >hg38_21_+_8433172_8446622_RNA45SN1_45S_50ntFlanks | 9597 | 9622 | 26 | AGGGGCGAAAGACTAATCGAACCATC | 530.6 | -7.1 | 1.7 | -4.2 | 2.4E-05 | 4.1E-05 | MB231/MB468 | MP |
| >MI0000095\|hsa-mir-93&WithFlank&7\|-\|100093762\|100093853@17.39.23&[MIMAT0000093&hsa-miR-93-5p&offsets\|0\|0;m-29&7\|-\|100093815\|100093837&offsets\|0\|0]\|CAAAGTGCTGTTCGTGCAGGTAG | >MI0000095\|hsa-mir-93&WithFlank&7\|-\|100093762\|100093853 | 17 | 39 | 23 | CAAAGTGCTGTTCGTGCAGGTAG | 25315.8 | 3.2 | 0.8 | 4.2 | 2.4E-05 | 4.1E-05 | MB231/MB468 | MP |
| >trnaMT_PheGAA_MT_+_577_647@39.56.18\|ATGTTTAGACGGGCTCAC | >trnaMT_PheGAA_MT_+_577_647 | 39 | 56 | 18 | ATGTTTAGACGGGCTCAC | 1194.6 | 2.8 | 0.7 | 4.2 | 2.4E-05 | 4.1E-05 | MB231/MB468 | MP |
| >hg38_21_+_8433172_8446622_RNA45SN1_45S_50ntFlanks@9572.9589.18\|CAAATCGGTCGTCCGACC | >hg38_21_+_8433172_8446622_RNA45SN1_45S_50ntFlanks | 9572 | 9589 | 18 | CAAATCGGTCGTCCGACC | 73.5 | -10.1 | 2.4 | -4.2 | 2.4E-05 | 4.1E-05 | MB231/MB468 | MP |
| >hg38_21_+_8433172_8446622_RNA45SN1_45S_50ntFlanks@10411.10445.35\|CAGTCGGTCCTGAGAGATGGGCGAGCGCCGTTCCG | >hg38_21_+_8433172_8446622_RNA45SN1_45S_50ntFlanks | 10411 | 10445 | 35 | CAGTCGGTCCTGAGAGATGGGCGAGCGCCGTTCCG | 299.7 | -6.0 | 1.4 | -4.2 | 2.9E-05 | 4.1E-05 | MB231/MB468 | NUC |
| >hg38_21_+_8433172_8446622_RNA45SN1_45S_50ntFlanks@6733.6773.41\|CATTGATCATCGACACTTCGAACGCACTTGCGGCCCCGGGT | >hg38_21_+_8433172_8446622_RNA45SN1_45S_50ntFlanks | 6733 | 6773 | 41 | CATTGATCATCGACACTTCGAACGCACTTGCGGCCCCGGGT | 548.0 | 3.5 | 0.8 | 4.2 | 2.9E-05 | 4.1E-05 | MB231/MB468 | MITO |
| >hg38_21_+_8433172_8446622_RNA45SN1_45S_50ntFlanks@12996.13037.42\|TCGCTGCGATCTATTGAAAGTCAGCCCTCGACACAAGGGTTT | >hg38_21_+_8433172_8446622_RNA45SN1_45S_50ntFlanks | 12996 | 13037 | 42 | TCGCTGCGATCTATTGAAAGTCAGCCCTCGACACAAGGGTTT | 143.4 | 5.7 | 1.4 | 4.2 | 2.9E-05 | 4.1E-05 | MB231/MB468 | MITO |
| >hg38_21_+_8433172_8446622_RNA45SN1_45S_50ntFlanks@10797.10827.31\|GCATGTTGGAACAATGTAGGTAAGGGAAGTC | >hg38_21_+_8433172_8446622_RNA45SN1_45S_50ntFlanks | 10797 | 10827 | 31 | GCATGTTGGAACAATGTAGGTAAGGGAAGTC | 392.4 | -5.0 | 1.2 | -4.2 | 2.4E-05 | 4.1E-05 | MB231/MB468 | MP |
| >hg38_21_+_8433172_8446622_RNA45SN1_45S_50ntFlanks@6653.6687.35\|ACTCTTAGCGGTGGATCACTCGGCTCGTGCGTCGA | >hg38_21_+_8433172_8446622_RNA45SN1_45S_50ntFlanks | 6653 | 6687 | 35 | ACTCTTAGCGGTGGATCACTCGGCTCGTGCGTCGA | 620.6 | 3.3 | 0.8 | 4.3 | 2.1E-05 | 4.1E-05 | BT20/MB468 | MITO |
| >MI0000342\|hsa-mir-200b&WithFlank&1\|+\|1167098\|1167204@63.85.23(+1C)&[MIMAT0000318&hsa-miR-200b-3p&offsets\|0\|+1(+1C);m-93&1\|+\|1167160\|1167182&offsets\|0\|0(+1C)]\|TAATACTGCCTGGTAATGATGACC | >MI0000342\|hsa-mir-200b&WithFlank&1\|+\|1167098\|1167204 | 63 | 85 | 23(+1C) | TAATACTGCCTGGTAATGATGACC | 306.4 | -4.6 | 1.1 | -4.2 | 2.9E-05 | 4.1E-05 | MB231/MB468 | NUC |
| >hg38_21_+_8433172_8446622_RNA45SN1_45S_50ntFlanks@11891.11910.20\|TTGACTCTAGTCTGGCACGG | >hg38_21_+_8433172_8446622_RNA45SN1_45S_50ntFlanks | 11891 | 11910 | 20 | TTGACTCTAGTCTGGCACGG | 439.4 | -2.5 | 0.6 | -4.2 | 2.5E-05 | 4.1E-05 | BT20/MB468 | CYTO |
| >MI0003591\|hsa-mir-584&WithFlank&5\|-\|149062307\|149062415@22.39.18(+1A)&[MIMAT0003249&hsa-miR-584-5p&offsets\|0\|-4(+1A);m-167&5\|-\|149062374\|149062394&offsets\|0\|-3(+1A)]\|TTATGGTTTGCCTGGGACA | >MI0003591\|hsa-mir-584&WithFlank&5\|-\|149062307\|149062415 | 22 | 39 | 18(+1A) | TTATGGTTTGCCTGGGACA | 165.8 | 8.0 | 1.8 | 4.4 | 9.7E-06 | 4.1E-05 | BT20/MB468 | MP |
| >MI0000088\|hsa-mir-30a&WithFlank&6\|-\|71403545\|71403627@53.74.22&[MIMAT0000088&hsa-miR-30a-3p&offsets\|0\|0;m-38&6\|-\|71403554\|71403575&offsets\|0\|0]\|CTTTCAGTCGGATGTTTGCAGC | >MI0000088\|hsa-mir-30a&WithFlank&6\|-\|71403545\|71403627 | 53 | 74 | 22 | CTTTCAGTCGGATGTTTGCAGC | 12502.0 | -4.0 | 0.9 | -4.7 | 3.0E-06 | 4.1E-05 | BT20/MB231 | MP |
| >hg38_21_+_8433172_8446622_RNA45SN1_45S_50ntFlanks@6648.6666.19\|GTACGACTCTTAGCGGTGG | >hg38_21_+_8433172_8446622_RNA45SN1_45S_50ntFlanks | 6648 | 6666 | 19 | GTACGACTCTTAGCGGTGG | 1607.3 | 3.7 | 0.9 | 4.3 | 2.1E-05 | 4.1E-05 | BT20/MB468 | MITO |
| >hg38_21_+_8433172_8446622_RNA45SN1_45S_50ntFlanks@11896.11935.40\|TCTAGTCTGGCACGGTGAAGAGACATGAGAGGTGTAGAAT | >hg38_21_+_8433172_8446622_RNA45SN1_45S_50ntFlanks | 11896 | 11935 | 40 | TCTAGTCTGGCACGGTGAAGAGACATGAGAGGTGTAGAAT | 38.3 | 9.0 | 2.0 | 4.4 | 9.8E-06 | 4.1E-05 | BT20/MB468 | MP |
| >trna111_HisGTG_1_-_147774845_147774916@-1T.20.21,trna118_HisGTG_1_-_145396881_145396952@-1T.20.21,trna16_HisGTG_1_+_146544773_146544844@-1T.20.21,trna1_HisGTG_15_+_45493349_45493420@-1T.20.21,trna21_HisGTG_1_+_147753471_147753542@-1T.20.21,trna33_HisGTG_6_+_27125906_27125977@-1T.20.21,trna7_HisGTG_9_-_14433938_14434009@-1T.20.21,trna8_HisGTG_15_-_45492611_45492682@-1T.20.21,trna9_HisGTG_15_-_45490804_45490875@-1T.20.21\|TGCCGTGATCGTATAGTGGTT | >trna111_HisGTG_1_-_147774845_147774916 | -1T | 20 | 21 | TGCCGTGATCGTATAGTGGTT | 171.6 | 7.2 | 1.7 | 4.2 | 2.9E-05 | 4.1E-05 | MB231/MB468 | MITO |
| >MI0000283\|hsa-mir-203a&WithFlank&14\|+\|104117399\|104117520@71.91.21(+1A)&[MIMAT0000264&hsa-miR-203a-3p&offsets\|0\|-1(+1A);m-79&14\|+\|104117469\|104117490&offsets\|0\|-1(+1A)]\|GTGAAATGTTTAGGACCACTAA | >MI0000283\|hsa-mir-203a&WithFlank&14\|+\|104117399\|104117520 | 71 | 91 | 21(+1A) | GTGAAATGTTTAGGACCACTAA | 78.4 | 9.6 | 2.1 | 4.7 | 3.1E-06 | 4.1E-05 | BT20/MB231 | MP |
| >MI0000102\|hsa-mir-100&WithFlank&11\|-\|122152223\|122152314@19.41.23&[MIMAT0000098&hsa-miR-100-5p&offsets\|0\|+1;m-27&11\|-\|122152276\|122152296&offsets\|0\|+2]\|AACCCGTAGATCCGAACTTGTGG | >MI0000102\|hsa-mir-100&WithFlank&11\|-\|122152223\|122152314 | 19 | 41 | 23 | AACCCGTAGATCCGAACTTGTGG | 2375.7 | -5.7 | 1.2 | -4.6 | 3.6E-06 | 4.1E-05 | BT20/MB231 | NUC |
| >hg38_21_+_8433172_8446622_RNA45SN1_45S_50ntFlanks@12658.12681.24\|ACTGAACGCCTCTAAGTCAGAATC | >hg38_21_+_8433172_8446622_RNA45SN1_45S_50ntFlanks | 12658 | 12681 | 24 | ACTGAACGCCTCTAAGTCAGAATC | 235.5 | -7.1 | 1.6 | -4.4 | 9.8E-06 | 4.1E-05 | BT20/MB468 | MP |
| >hg38_21_+_8433172_8446622_RNA45SN1_45S_50ntFlanks@9564.9591.28\|CTGACGTGCAAATCGGTCGTCCGACCTG | >hg38_21_+_8433172_8446622_RNA45SN1_45S_50ntFlanks | 9564 | 9591 | 28 | CTGACGTGCAAATCGGTCGTCCGACCTG | 164.2 | -4.0 | 0.9 | -4.2 | 2.5E-05 | 4.1E-05 | BT20/MB468 | CYTO |
| >hg38_21_+_8433172_8446622_RNA45SN1_45S_50ntFlanks@12615.12654.40\|GCTGAGGAGCCAATGGGGCGAAGCTACCATCTGTGGGATT | >hg38_21_+_8433172_8446622_RNA45SN1_45S_50ntFlanks | 12615 | 12654 | 40 | GCTGAGGAGCCAATGGGGCGAAGCTACCATCTGTGGGATT | 3177.4 | -4.0 | 0.9 | -4.2 | 2.5E-05 | 4.1E-05 | BT20/MB468 | CYTO |
| >hg38_21_+_8433172_8446622_RNA45SN1_45S_50ntFlanks@6651.6673.23\|CGACTCTTAGCGGTGGATCACTC | >hg38_21_+_8433172_8446622_RNA45SN1_45S_50ntFlanks | 6651 | 6673 | 23 | CGACTCTTAGCGGTGGATCACTC | 1593.5 | 3.6 | 0.8 | 4.4 | 9.8E-06 | 4.1E-05 | BT20/MB468 | MP |
| >hg38_21_+_8433172_8446622_RNA45SN1_45S_50ntFlanks@10391.10431.41\|AACAGCAGTTGAACATGGGTCAGTCGGTCCTGAGAGATGGG | >hg38_21_+_8433172_8446622_RNA45SN1_45S_50ntFlanks | 10391 | 10431 | 41 | AACAGCAGTTGAACATGGGTCAGTCGGTCCTGAGAGATGGG | 180.5 | -5.6 | 1.3 | -4.2 | 2.5E-05 | 4.1E-05 | BT20/MB468 | CYTO |
| >hg38_21_+_8433172_8446622_RNA45SN1_45S_50ntFlanks@10512.10533.22\|TCCGGAGTGGCGGAGATGGGCG | >hg38_21_+_8433172_8446622_RNA45SN1_45S_50ntFlanks | 10512 | 10533 | 22 | TCCGGAGTGGCGGAGATGGGCG | 994.7 | -5.3 | 1.3 | -4.2 | 2.5E-05 | 4.1E-05 | BT20/MB468 | CYTO |
| >hg38_21_+_8433172_8446622_RNA45SN1_45S_50ntFlanks@12488.12509.22\|AACGTGAGCTGGGTTTAGACCG | >hg38_21_+_8433172_8446622_RNA45SN1_45S_50ntFlanks | 12488 | 12509 | 22 | AACGTGAGCTGGGTTTAGACCG | 3352.4 | -5.5 | 1.3 | -4.2 | 2.5E-05 | 4.1E-05 | BT20/MB468 | CYTO |

<table>
<tr><td>>MI0000108|hsa-mir-103a-2&WithFlank&20|+|3917488|3917577@56.76.21;MI0000109|hsa-mir-103a-1&WithFlank&5|-|168560890|168560979@56.76.21&[MIMAT0000101&hsa-miR-103a-3p&offsets|+2|0;m-12&20|+|3917541|3917563&offsets|+2|0];[MIMAT0000101_1&hsa-miR-103a-3p&offsets|+2|0;m-13&5|-|168560904|168560926&offsets|+2|0;m-44220&5|-|168560908|168560926&offsets|+2|+4]|CAGCATTGTACAGGGCTATGA</td><td>>MI0000108|hsa-mir-103a-2&WithFlank&20|+|3917488|3917577</td><td>56</td><td>76</td><td>21</td><td>CAGCATTGTACAGGGCTATGA</td><td>314.4</td><td>4.5</td><td>1.1</td><td>4.2</td><td>2.9E-05</td><td>4.1E-05</td><td>MB231/MB468</td><td>MITO</td></tr>
<tr><td>>hg38_21_+_8433172_8446622_RNA45SN1_45S_50ntFlanks@8176.8206.31|CTTCTGATCGAGGCCCAGCCCGTGGACGGTG</td><td>>hg38_21_+_8433172_8446622_RNA45SN1_45S_50ntFlanks</td><td>8176</td><td>8206</td><td>31</td><td>CTTCTGATCGAGGCCCAGCCCGTGGACGGTG</td><td>811.1</td><td>-2.5</td><td>0.6</td><td>-4.2</td><td>2.5E-05</td><td>4.1E-05</td><td>BT20/MB468</td><td>CYTO</td></tr>
<tr><td>>hg38_21_+_8433172_8446622_RNA45SN1_45S_50ntFlanks@11683.11701.19|ATGTCAAAGTGAAGAAATT</td><td>>hg38_21_+_8433172_8446622_RNA45SN1_45S_50ntFlanks</td><td>11683</td><td>11701</td><td>19</td><td>ATGTCAAAGTGAAGAAATT</td><td>44.4</td><td>-8.6</td><td>1.9</td><td>-4.4</td><td>9.8E-06</td><td>4.1E-05</td><td>BT20/MB468</td><td>MP</td></tr>
<tr><td>>hg38_21_+_8433172_8446622_RNA45SN1_45S_50ntFlanks@4566.4587.22|ATAATGGAATAGGACCGCGGTT</td><td>>hg38_21_+_8433172_8446622_RNA45SN1_45S_50ntFlanks</td><td>4566</td><td>4587</td><td>22</td><td>ATAATGGAATAGGACCGCGGTT</td><td>358.5</td><td>-2.8</td><td>0.7</td><td>-4.2</td><td>2.8E-05</td><td>4.1E-05</td><td>MB231/MB468</td><td>CYTO</td></tr>
<tr><td>>MI0000114|hsa-mir-107&WithFlank&10|-|89592741|89592833@56.76.21;MI0000108|hsa-mir-103a-2&WithFlank&20|+|3917488|3917577@54.74.21;MI0000109|hsa-mir-103a-1&WithFlank&5|-|168560890|168560979@54.74.21&[MIMAT0000104&hsa-miR-107&offsets|0|-2;m-78&10|-|89592758|89592778&offsets|0|0];[MIMAT0000101&hsa-miR-103a-3p&offsets|0|-2;m-12&20|+|3917541|3917563&offsets|0|-2];[MIMAT0000101_1&hsa-miR-103a-3p&offsets|0|-2;m-13&5|-|168560904|168560926&offsets|0|-2;m-44220&5|-|168560908|168560926&offsets|0|+2]|AGCAGCATTGTACAGGGCTAT</td><td>>MI0000114|hsa-mir-107&WithFlank&10|-|89592741|89592833</td><td>56</td><td>76</td><td>21</td><td>AGCAGCATTGTACAGGGCTAT</td><td>2564.8</td><td>3.8</td><td>0.9</td><td>4.2</td><td>2.5E-05</td><td>4.1E-05</td><td>MB231/MB468</td><td>MP</td></tr>
<tr><td>>hg38_21_+_8433172_8446622_RNA45SN1_45S_50ntFlanks@8079.8105.27|AAGAGCCCAGCGCCGAATCCCCGCCCC</td><td>>hg38_21_+_8433172_8446622_RNA45SN1_45S_50ntFlanks</td><td>8079</td><td>8105</td><td>27</td><td>AAGAGCCCAGCGCCGAATCCCCGCCCC</td><td>640.6</td><td>-6.2</td><td>1.4</td><td>-4.4</td><td>9.8E-06</td><td>4.1E-05</td><td>BT20/MB468</td><td>MP</td></tr>
<tr><td>>hg38_21_+_8433172_8446622_RNA45SN1_45S_50ntFlanks@12416.12433.18|TTCGATGTCGGCTCTTCC</td><td>>hg38_21_+_8433172_8446622_RNA45SN1_45S_50ntFlanks</td><td>12416</td><td>12433</td><td>18</td><td>TTCGATGTCGGCTCTTCC</td><td>1090.9</td><td>2.9</td><td>0.7</td><td>4.2</td><td>2.9E-05</td><td>4.1E-05</td><td>MB231/MB468</td><td>NUC</td></tr>
<tr><td>>hg38_21_+_8433172_8446622_RNA45SN1_45S_50ntFlanks@4129.4148.20|GATTCCGGAGAGGGAGCCTG</td><td>>hg38_21_+_8433172_8446622_RNA45SN1_45S_50ntFlanks</td><td>4129</td><td>4148</td><td>20</td><td>GATTCCGGAGAGGGAGCCTG</td><td>140.6</td><td>-6.1</td><td>1.4</td><td>-4.2</td><td>2.5E-05</td><td>4.1E-05</td><td>MB231/MB468</td><td>MP</td></tr>
<tr><td>>hg38_21_+_8433172_8446622_RNA45SN1_45S_50ntFlanks@12018.12035.18|AATACCACTACTCTGATC</td><td>>hg38_21_+_8433172_8446622_RNA45SN1_45S_50ntFlanks</td><td>12018</td><td>12035</td><td>18</td><td>AATACCACTACTCTGATC</td><td>38.0</td><td>9.0</td><td>2.0</td><td>4.4</td><td>9.8E-06</td><td>4.1E-05</td><td>BT20/MB468</td><td>MP</td></tr>
<tr><td>>hg38_21_+_8433172_8446622_RNA45SN1_45S_50ntFlanks@9534.9573.40|AGGAAACTCTGGTGGAGGTCCGTAGCGGTCCTGACGTGCA</td><td>>hg38_21_+_8433172_8446622_RNA45SN1_45S_50ntFlanks</td><td>9534</td><td>9573</td><td>40</td><td>AGGAAACTCTGGTGGAGGTCCGTAGCGGTCCTGACGTGCA</td><td>993.3</td><td>-2.1</td><td>0.5</td><td>-4.2</td><td>2.6E-05</td><td>4.1E-05</td><td>BT20/MB468</td><td>CYTO</td></tr>
<tr><td>>hg38_21_+_8433172_8446622_RNA45SN1_45S_50ntFlanks@5083.5120.38|ACGAACGAGACTCTGGCATGCTAACTAGTTACGCGACC</td><td>>hg38_21_+_8433172_8446622_RNA45SN1_45S_50ntFlanks</td><td>5083</td><td>5120</td><td>38</td><td>ACGAACGAGACTCTGGCATGCTAACTAGTTACGCGACC</td><td>399.9</td><td>-3.6</td><td>0.9</td><td>-4.2</td><td>3.0E-05</td><td>4.1E-05</td><td>MB231/MB468</td><td>MITO</td></tr>
<tr><td>>hg38_21_+_8433172_8446622_RNA45SN1_45S_50ntFlanks@8190.8208.19|CCAGCCCGTGGACGGTGTG</td><td>>hg38_21_+_8433172_8446622_RNA45SN1_45S_50ntFlanks</td><td>8190</td><td>8208</td><td>19</td><td>CCAGCCCGTGGACGGTGTG</td><td>136.5</td><td>-6.2</td><td>1.5</td><td>-4.2</td><td>2.5E-05</td><td>4.1E-05</td><td>MB231/MB468</td><td>MP</td></tr>
<tr><td>>hg38_21_+_8433172_8446622_RNA45SN1_45S_50ntFlanks@11208.11232.25|GCGGCGACTCTGGACGCGAGCCGGG</td><td>>hg38_21_+_8433172_8446622_RNA45SN1_45S_50ntFlanks</td><td>11208</td><td>11232</td><td>25</td><td>GCGGCGACTCTGGACGCGAGCCGGG</td><td>37788.7</td><td>-9.2</td><td>2.2</td><td>-4.2</td><td>3.1E-05</td><td>4.1E-05</td><td>BT20/MB468</td><td>NUC</td></tr>
<tr><td>>am_MI0000786|hsa-mir-378a&WithFlank&5|+|149732819|149732896@49.69.21(+1U)&[MIMAT0000732&hsa-miR-378a-3p&offsets|0|-1(+1U);m-44&5|+|149732867|149732888&offsets|0|-1(+1U)]|ACTGGACTTGGAGTCAGAAGGT</td><td>>am_MI0000786|hsa-mir-378a&WithFlank&5|+|149732819|149732896</td><td>49</td><td>69</td><td>21(+1U)</td><td>ACTGGACTTGGAGTCAGAAGGT</td><td>2137.4</td><td>3.7</td><td>0.9</td><td>4.2</td><td>3.0E-05</td><td>4.1E-05</td><td>MB231/MB468</td><td>MITO</td></tr>
<tr><td>>hg38_21_+_8433172_8446622_RNA45SN1_45S_50ntFlanks@7991.8009.19|ACGTGGCGACCCGCTGAAT</td><td>>hg38_21_+_8433172_8446622_RNA45SN1_45S_50ntFlanks</td><td>7991</td><td>8009</td><td>19</td><td>ACGTGGCGACCCGCTGAAT</td><td>9037.5</td><td>-3.8</td><td>0.9</td><td>-4.2</td><td>3.0E-05</td><td>4.1E-05</td><td>MB231/MB468</td><td>MITO</td></tr>
<tr><td>>hg38_21_+_8433172_8446622_RNA45SN1_45S_50ntFlanks@7974.8004.31|ACGCGACCTCAGATCAGACGTGGCGACCCGC</td><td>>hg38_21_+_8433172_8446622_RNA45SN1_45S_50ntFlanks</td><td>7974</td><td>8004</td><td>31</td><td>ACGCGACCTCAGATCAGACGTGGCGACCCGC</td><td>1657.3</td><td>3.9</td><td>0.9</td><td>4.4</td><td>9.9E-06</td><td>4.1E-05</td><td>BT20/MB468</td><td>MP</td></tr>
<tr><td>>MI0033423|hsa-mir-10399&WithFlank&7|-|139044094|139044163@45.63.19(+1U)&[MIMAT0041630&hsa-miR-10399-3p&offsets|+1|-1(+1U);TJU_CMC_MD2.ID03061.3p-miR&offsets|+5|+2(+1U);m-2657&7|-|139044103|139044124&offsets|+5|+2(+1U)]|TCTCGGACAAGCTGTAGGTT</td><td>>MI0033423|hsa-mir-10399&WithFlank&7|-|139044094|139044163</td><td>45</td><td>63</td><td>19(+1U)</td><td>TCTCGGACAAGCTGTAGGTT</td><td>69.2</td><td>-9.5</td><td>2.0</td><td>-4.7</td><td>3.1E-06</td><td>4.1E-05</td><td>BT20/MB231</td><td>MP</td></tr>
<tr><td>>hg38_21_+_8433172_8446622_RNA45SN1_45S_50ntFlanks@12615.12653.39|GCTGAGGAGCCAATGGGGCGAAGCTACCATCTGTGGGAT</td><td>>hg38_21_+_8433172_8446622_RNA45SN1_45S_50ntFlanks</td><td>12615</td><td>12653</td><td>39</td><td>GCTGAGGAGCCAATGGGGCGAAGCTACCATCTGTGGGAT</td><td>1056.3</td><td>-7.3</td><td>1.7</td><td>-4.4</td><td>9.9E-06</td><td>4.1E-05</td><td>BT20/MB468</td><td>MP</td></tr>
<tr><td>>MI0000108|hsa-mir-103a-2&WithFlank&20|+|3917488|3917577@54.75.22(+2U);MI0000109|hsa-mir-103a-1&WithFlank&5|-|168560890|168560979@54.75.22(+2U)&[MIMAT0000101&hsa-miR-103a-3p&offsets|0|-1(+2U);m-12&20|+|3917541|3917563&offsets|0|-1(+2U)];[MIMAT0000101_1&hsa-miR-103a-3p&offsets|0|-1(+2U);m-13&5|-|168560904|168560926&offsets|0|-1(+2U);m-44220&5|-|168560908|168560926&offsets|0|+3(+2U)]|AGCAGCATTGTACAGGGCTATGTT</td><td>>MI0000108|hsa-mir-103a-2&WithFlank&20|+|3917488|3917577</td><td>54</td><td>75</td><td>22(+2U)</td><td>AGCAGCATTGTACAGGGCTATGTT</td><td>223.9</td><td>7.1</td><td>1.7</td><td>4.2</td><td>3.0E-05</td><td>4.1E-05</td><td>MB231/MB468</td><td>MITO</td></tr>
<tr><td>>hg38_21_+_8433172_8446622_RNA45SN1_45S_50ntFlanks@475.514.40|CTTCGTGATCGATGTGGTGACGTCGTGCTCTCCCGGGCCG</td><td>>hg38_21_+_8433172_8446622_RNA45SN1_45S_50ntFlanks</td><td>475</td><td>514</td><td>40</td><td>CTTCGTGATCGATGTGGTGACGTCGTGCTCTCCCGGGCCG</td><td>1248.4</td><td>7.1</td><td>1.7</td><td>4.2</td><td>2.2E-05</td><td>4.1E-05</td><td>BT20/MB468</td><td>MITO</td></tr>
<tr><td>>hg38_21_+_8433172_8446622_RNA45SN1_45S_50ntFlanks@12394.12419.26|TAGCGACGTCGCTTTTTGATCCTTCG</td><td>>hg38_21_+_8433172_8446622_RNA45SN1_45S_50ntFlanks</td><td>12394</td><td>12419</td><td>26</td><td>TAGCGACGTCGCTTTTTGATCCTTCG</td><td>597.5</td><td>-2.8</td><td>0.7</td><td>-4.2</td><td>2.6E-05</td><td>4.1E-05</td><td>BT20/MB468</td><td>CYTO</td></tr>
<tr><td>>hg38_21_+_8433172_8446622_RNA45SN1_45S_50ntFlanks@5515.5535.21|CTAGAGGAAGTAAAAGTCGTA</td><td>>hg38_21_+_8433172_8446622_RNA45SN1_45S_50ntFlanks</td><td>5515</td><td>5535</td><td>21</td><td>CTAGAGGAAGTAAAAGTCGTA</td><td>537.0</td><td>-3.2</td><td>0.8</td><td>-4.2</td><td>2.8E-05</td><td>4.1E-05</td><td>MB231/MB468</td><td>CYTO</td></tr>
<tr><td>>MI0003674|hsa-mir-653&WithFlank&7|-|93482754|93482861@22.42.21(+1U)&[MIMAT0003328&hsa-miR-653-5p&offsets|+3|+3(+1U);m-368&7|-|93482819|93482840&offsets|0|-1(+1U)]|TTGAAACAATCTCTACTGAACT</td><td>>MI0003674|hsa-mir-653&WithFlank&7|-|93482754|93482861</td><td>22</td><td>42</td><td>21(+1U)</td><td>TTGAAACAATCTCTACTGAACT</td><td>68.7</td><td>-9.5</td><td>2.0</td><td>-4.7</td><td>3.1E-06</td><td>4.1E-05</td><td>BT20/MB231</td><td>MP</td></tr>
<tr><td>>MI0006444|hsa-mir-1307&WithFlank&10|-|103394247|103394407@86.106.21&[MIMAT0005951&hsa-miR-1307-3p&offsets|0|-1;m-66&10|-|103394300|103394322&offsets|0|-2]|ACTCGGCGTGGCGTCGGTCGT</td><td>>MI0006444|hsa-mir-1307&WithFlank&10|-|103394247|103394407</td><td>86</td><td>106</td><td>21</td><td>ACTCGGCGTGGCGTCGGTCGT</td><td>176.9</td><td>7.2</td><td>1.7</td><td>4.2</td><td>3.0E-05</td><td>4.1E-05</td><td>MB231/MB468</td><td>MITO</td></tr>
<tr><td>>MI0000438|hsa-mir-15b&WithFlank&3|+|160404582|160404691@64.84.21&[MIMAT0004586&hsa-miR-15b-3p&offsets|0|-1;m-204&3|+|160404645|160404665&offsets|0|0]|CGAATCATTATTTGCTGCTCT</td><td>>MI0000438|hsa-mir-15b&WithFlank&3|+|160404582|160404691</td><td>64</td><td>84</td><td>21</td><td>CGAATCATTATTTGCTGCTCT</td><td>368.0</td><td>4.4</td><td>1.0</td><td>4.2</td><td>3.0E-05</td><td>4.1E-05</td><td>MB231/MB468</td><td>MITO</td></tr>
<tr><td>>hg38_21_+_8433172_8446622_RNA45SN1_45S_50ntFlanks@10914.10936.23|CGCCGCGGCTGGACGAGGCGCCG</td><td>>hg38_21_+_8433172_8446622_RNA45SN1_45S_50ntFlanks</td><td>10914</td><td>10936</td><td>23</td><td>CGCCGCGGCTGGACGAGGCGCCG</td><td>541.3</td><td>-6.2</td><td>1.5</td><td>-4.2</td><td>2.6E-05</td><td>4.1E-05</td><td>BT20/MB468</td><td>CYTO</td></tr>
<tr><td>>hg38_21_+_8433172_8446622_RNA45SN1_45S_50ntFlanks@9494.9520.27|CCGAAAGATGGTGAACTATGCCTGGGC</td><td>>hg38_21_+_8433172_8446622_RNA45SN1_45S_50ntFlanks</td><td>9494</td><td>9520</td><td>27</td><td>CCGAAAGATGGTGAACTATGCCTGGGC</td><td>124.1</td><td>10.3</td><td>2.2</td><td>4.7</td><td>3.1E-06</td><td>4.1E-05</td><td>BT20/MB231</td><td>MP</td></tr>
<tr><td>>trnaMT_SerGCT_MT_+_12207_12265@15.61.47|GAACTGCTAACTCATGCCCCCATGTCTAACAACATGGCTTTCTCACC</td><td>>trnaMT_SerGCT_MT_+_12207_12265</td><td>15</td><td>61</td><td>47</td><td>GAACTGCTAACTCATGCCCCCATGTCTAACAACATGGCTTTCTCACC</td><td>186.4</td><td>11.3</td><td>2.6</td><td>4.4</td><td>9.9E-06</td><td>4.1E-05</td><td>BT20/MB468</td><td>MP</td></tr>
<tr><td>>trnaMT_HisGTG_MT_+_12138_12206@27.72.46|GATTGTGAATCTGACAACAGAGGCTTACGACCCCTTATTTACCCCA</td><td>>trnaMT_HisGTG_MT_+_12138_12206</td><td>27</td><td>72</td><td>46</td><td>GATTGTGAATCTGACAACAGAGGCTTACGACCCCTTATTTACCCCA</td><td>184.3</td><td>11.3</td><td>2.6</td><td>4.4</td><td>1.0E-05</td><td>4.2E-05</td><td>BT20/MB468</td><td>MP</td></tr>
<tr><td>>hg38_21_+_8433172_8446622_RNA45SN1_45S_50ntFlanks@12397.12416.20|CGACGTCGCTTTTTGATCCT</td><td>>hg38_21_+_8433172_8446622_RNA45SN1_45S_50ntFlanks</td><td>12397</td><td>12416</td><td>20</td><td>CGACGTCGCTTTTTGATCCT</td><td>400.1</td><td>-5.3</td><td>1.3</td><td>-4.2</td><td>3.1E-05</td><td>4.2E-05</td><td>BT20/MB468</td><td>NUC</td></tr>
<tr><td>>hg38_21_+_8433172_8446622_RNA45SN1_45S_50ntFlanks@12008.12042.35|CCGCCGGTGAAATACCACTACTCTGATCGTTTTTT</td><td>>hg38_21_+_8433172_8446622_RNA45SN1_45S_50ntFlanks</td><td>12008</td><td>12042</td><td>35</td><td>CCGCCGGTGAAATACCACTACTCTGATCGTTTTTT</td><td>270.6</td><td>-7.3</td><td>1.7</td><td>-4.4</td><td>1.0E-05</td><td>4.2E-05</td><td>BT20/MB468</td><td>MP</td></tr>
<tr><td>>hg38_MT_+_1621_3279_RNR2_16S_50ntFlanks@623.642.20|ATAACTGAACTCCTCACACC</td><td>>hg38_MT_+_1621_3279_RNR2_16S_50ntFlanks</td><td>623</td><td>642</td><td>20</td><td>ATAACTGAACTCCTCACACC</td><td>29.9</td><td>8.1</td><td>1.9</td><td>4.2</td><td>2.5E-05</td><td>4.2E-05</td><td>MB231/MB468</td><td>MP</td></tr>
<tr><td>>hg38_21_+_8433172_8446622_RNA45SN1_45S_50ntFlanks@11881.11898.18|CCTGTTGAGCTTGACTCT</td><td>>hg38_21_+_8433172_8446622_RNA45SN1_45S_50ntFlanks</td><td>11881</td><td>11898</td><td>18</td><td>CCTGTTGAGCTTGACTCT</td><td>152.3</td><td>-2.3</td><td>0.5</td><td>-4.2</td><td>2.6E-05</td><td>4.2E-05</td><td>BT20/MB468</td><td>CYTO</td></tr>
<tr><td>>hg38_21_+_8433172_8446622_RNA45SN1_45S_50ntFlanks@8190.8212.23|CCAGCCCGTGGACGGTGTGAGGC</td><td>>hg38_21_+_8433172_8446622_RNA45SN1_45S_50ntFlanks</td><td>8190</td><td>8212</td><td>23</td><td>CCAGCCCGTGGACGGTGTGAGGC</td><td>349.7</td><td>-2.8</td><td>0.7</td><td>-4.2</td><td>2.2E-05</td><td>4.2E-05</td><td>BT20/MB468</td><td>MITO</td></tr>
<tr><td>>hg38_21_+_8433172_8446622_RNA45SN1_45S_50ntFlanks@10820.10839.20|GGGAAGTCGGCAAGCCGGAT</td><td>>hg38_21_+_8433172_8446622_RNA45SN1_45S_50ntFlanks</td><td>10820</td><td>10839</td><td>20</td><td>GGGAAGTCGGCAAGCCGGAT</td><td>290.5</td><td>-5.3</td><td>1.3</td><td>-4.2</td><td>2.5E-05</td><td>4.2E-05</td><td>MB231/MB468</td><td>MP</td></tr>
<tr><td>>hg38_21_+_8433172_8446622_RNA45SN1_45S_50ntFlanks@11561.11593.33|GCGCCTAGCAGCCGACTTAGAACTGGTGCGGAC</td><td>>hg38_21_+_8433172_8446622_RNA45SN1_45S_50ntFlanks</td><td>11561</td><td>11593</td><td>33</td><td>GCGCCTAGCAGCCGACTTAGAACTGGTGCGGAC</td><td>424.0</td><td>-5.9</td><td>1.4</td><td>-4.2</td><td>2.6E-05</td><td>4.2E-05</td><td>BT20/MB468</td><td>CYTO</td></tr>
<tr><td>>am_trnaMT_ValTAC_MT_+_1602_1670@33.67.35|ACACTTAGGAGATTTCAACTTAACTTGACCGCTCT</td><td>>am_trnaMT_ValTAC_MT_+_1602_1670</td><td>33</td><td>67</td><td>35</td><td>ACACTTAGGAGATTTCAACTTAACTTGACCGCTCT</td><td>200.0</td><td>11.4</td><td>2.6</td><td>4.4</td><td>1.0E-05</td><td>4.2E-05</td><td>BT20/MB468</td><td>MP</td></tr>
<tr><td>>hg38_21_+_8433172_8446622_RNA45SN1_45S_50ntFlanks@8107.8138.32|CGGCGGGGCGCGGGACATGTGGCGTACGGAAG</td><td>>hg38_21_+_8433172_8446622_RNA45SN1_45S_50ntFlanks</td><td>8107</td><td>8138</td><td>32</td><td>CGGCGGGGCGCGGGACATGTGGCGTACGGAAG</td><td>1030.9</td><td>-3.8</td><td>0.9</td><td>-4.4</td><td>1.0E-05</td><td>4.2E-05</td><td>BT20/MB468</td><td>MP</td></tr>
<tr><td>>hg38_21_+_8433172_8446622_RNA45SN1_45S_50ntFlanks@10727.10757.31|GGAGAGGGTGTAAATCTCGCGCCGGGCCGTA</td><td>>hg38_21_+_8433172_8446622_RNA45SN1_45S_50ntFlanks</td><td>10727</td><td>10757</td><td>31</td><td>GGAGAGGGTGTAAATCTCGCGCCGGGCCGTA</td><td>293.9</td><td>-5.6</td><td>1.3</td><td>-4.2</td><td>2.6E-05</td><td>4.2E-05</td><td>BT20/MB468</td><td>CYTO</td></tr>
<tr><td>>hg38_21_+_8433172_8446622_RNA45SN1_45S_50ntFlanks@11572.11589.18|CCGACTTAGAACTGGTGC</td><td>>hg38_21_+_8433172_8446622_RNA45SN1_45S_50ntFlanks</td><td>11572</td><td>11589</td><td>18</td><td>CCGACTTAGAACTGGTGC</td><td>300.2</td><td>-2.5</td><td>0.6</td><td>-4.2</td><td>2.6E-05</td><td>4.2E-05</td><td>BT20/MB468</td><td>CYTO</td></tr>
<tr><td>>hg38_21_+_8433172_8446622_RNA45SN1_45S_50ntFlanks@8174.8212.39|TCCTTCTGATCGAGGCCCAGCCCGTGGACGGTGTGAGGC</td><td>>hg38_21_+_8433172_8446622_RNA45SN1_45S_50ntFlanks</td><td>8174</td><td>8212</td><td>39</td><td>TCCTTCTGATCGAGGCCCAGCCCGTGGACGGTGTGAGGC</td><td>408.9</td><td>-5.4</td><td>1.3</td><td>-4.2</td><td>3.2E-05</td><td>4.2E-05</td><td>BT20/MB468</td><td>NUC</td></tr>
<tr><td>>hg38_21_+_8433172_8446622_RNA45SN1_45S_50ntFlanks@3782.3802.21|CAGTGAAACTGCGAATGGCTC</td><td>>hg38_21_+_8433172_8446622_RNA45SN1_45S_50ntFlanks</td><td>3782</td><td>3802</td><td>21</td><td>CAGTGAAACTGCGAATGGCTC</td><td>278.3</td><td>-4.9</td><td>1.2</td><td>-4.2</td><td>2.2E-05</td><td>4.2E-05</td><td>BT20/MB468</td><td>MITO</td></tr>
<tr><td>>hg38_21_+_8433172_8446622_RNA45SN1_45S_50ntFlanks@11593.11613.21|CCAGGGGAATCCGACTGTTTA</td><td>>hg38_21_+_8433172_8446622_RNA45SN1_45S_50ntFlanks</td><td>11593</td><td>11613</td><td>21</td><td>CCAGGGGAATCCGACTGTTTA</td><td>359.9</td><td>-3.3</td><td>0.7</td><td>-4.4</td><td>1.0E-05</td><td>4.2E-05</td><td>BT20/MB468</td><td>MP</td></tr>
<tr><td>>hg38_21_+_8433172_8446622_RNA45SN1_45S_50ntFlanks@11582.11613.32|ACTGGTGCGGACCAGGGGAATCCGACTGTTTA</td><td>>hg38_21_+_8433172_8446622_RNA45SN1_45S_50ntFlanks</td><td>11582</td><td>11613</td><td>32</td><td>ACTGGTGCGGACCAGGGGAATCCGACTGTTTA</td><td>296.6</td><td>-3.3</td><td>0.8</td><td>-4.2</td><td>3.0E-05</td><td>4.2E-05</td><td>MB231/MB468</td><td>MITO</td></tr>
<tr><td>>hg38_21_+_8433172_8446622_RNA45SN1_45S_50ntFlanks@10914.10931.18|CGCCGCGGCTGGACGAGG</td><td>>hg38_21_+_8433172_8446622_RNA45SN1_45S_50ntFlanks</td><td>10914</td><td>10931</td><td>18</td><td>CGCCGCGGCTGGACGAGG</td><td>2498.2</td><td>-8.3</td><td>2.0</td><td>-4.2</td><td>3.2E-05</td><td>4.2E-05</td><td>BT20/MB468</td><td>NUC</td></tr>
<tr><td>>hg38_21_+_8433172_8446622_RNA45SN1_45S_50ntFlanks@10677.10711.35|GCGGTTCCGGCGGCGTCCGGTGAGCTCTCGCTGGC</td><td>>hg38_21_+_8433172_8446622_RNA45SN1_45S_50ntFlanks</td><td>10677</td><td>10711</td><td>35</td><td>GCGGTTCCGGCGGCGTCCGGTGAGCTCTCGCTGGC</td><td>109.6</td><td>10.1</td><td>2.2</td><td>4.7</td><td>3.2E-06</td><td>4.2E-05</td><td>BT20/MB231</td><td>MP</td></tr>
<tr><td>>hg38_21_+_8433172_8446622_RNA45SN1_45S_50ntFlanks@12415.12443.29|CTTCGATGTCGGCTCTTCCTATCATTGTG</td><td>>hg38_21_+_8433172_8446622_RNA45SN1_45S_50ntFlanks</td><td>12415</td><td>12443</td><td>29</td><td>CTTCGATGTCGGCTCTTCCTATCATTGTG</td><td>5622.0</td><td>-2.7</td><td>0.6</td><td>-4.2</td><td>2.2E-05</td><td>4.2E-05</td><td>BT20/MB468</td><td>MITO</td></tr>
</table>

| | | | | | | | | | | | | | |
|---|---|---|---|---|---|---|---|---|---|---|---|---|---|
| >hg38_21_+_8433172_8446622_RNA45SN1_45S_50ntFlanks@6699.6721.23\|GCTAGCTGCGAGAATTAATGTGA | >hg38_21_+_8433172_8446622_RNA45SN1_45S_50ntFlanks | 6699 | 6721 | 23 | GCTAGCTGCGAGAATTAATGTGA | 618.9 | -5.0 | 1.2 | -4.2 | 2.5E-05 | 4.2E-05 | MB231/MB468 | MP |
| >MI0000477\|hsa-mir-146a&WithFlank&5\|+\|160485346\|160485456@27.45.19(+2U)&[MIMAT0000449&hsa-miR-146a-5p&offsets\|0\|-3(+2U);m-129&5\|+\|160485372\|160485393&offsets\|0\|-3(+2U)]\|TGAGAACTGAATTCCATGGTT | >MI0000477\|hsa-mir-146a&WithFlank&5\|+\|160485346\|160485456 | 27 | 45 | 19(+2U) | TGAGAACTGAATTCCATGGTT | 41.7 | 8.6 | 2.0 | 4.2 | 2.5E-05 | 4.2E-05 | MB231/MB468 | MP |
| >hg38_21_+_8433172_8446622_RNA45SN1_45S_50ntFlanks@9573.9593.21\|AAATCGGTCGTCCGACCTGGG | >hg38_21_+_8433172_8446622_RNA45SN1_45S_50ntFlanks | 9573 | 9593 | 21 | AAATCGGTCGTCCGACCTGGG | 157.9 | -3.7 | 0.9 | -4.2 | 2.5E-05 | 4.2E-05 | MB231/MB468 | MP |
| >hg38_21_+_8433172_8446622_RNA45SN1_45S_50ntFlanks@6652.6698.47\|GACTCTTAGCGGTGGATCACTCGGCTCGTGCGTCGATGAAGAACGCA | >hg38_21_+_8433172_8446622_RNA45SN1_45S_50ntFlanks | 6652 | 6698 | 47 | GACTCTTAGCGGTGGATCACTCGGCTCGTGCGTCGATGAAGAACGCA | 10499.6 | -4.2 | 1.0 | -4.2 | 3.2E-05 | 4.2E-05 | BT20/MB468 | NUC |
| >hg38_21_+_8433172_8446622_RNA45SN1_45S_50ntFlanks@5490.5516.27\|TGAGAAGACGGTCGAACTTGACTATCT | >hg38_21_+_8433172_8446622_RNA45SN1_45S_50ntFlanks | 5490 | 5516 | 27 | TGAGAAGACGGTCGAACTTGACTATCT | 762.1 | 5.1 | 1.2 | 4.2 | 3.2E-05 | 4.2E-05 | BT20/MB468 | NUC |
| >hg38_21_+_8433172_8446622_RNA45SN1_45S_50ntFlanks@3993.4011.19\|GTGACTCTAGATAACCTCG | >hg38_21_+_8433172_8446622_RNA45SN1_45S_50ntFlanks | 3993 | 4011 | 19 | GTGACTCTAGATAACCTCG | 279.5 | -4.9 | 1.2 | -4.2 | 3.2E-05 | 4.2E-05 | BT20/MB468 | NUC |
| >hg38_21_+_8433172_8446622_RNA45SN1_45S_50ntFlanks@10411.10431.21\|CAGTCGGTCCTGAGAGATGGG | >hg38_21_+_8433172_8446622_RNA45SN1_45S_50ntFlanks | 10411 | 10431 | 21 | CAGTCGGTCCTGAGAGATGGG | 2298.5 | -4.2 | 1.0 | -4.4 | 1.0E-05 | 4.2E-05 | BT20/MB468 | MP |
| >hg38_21_+_8433172_8446622_RNA45SN1_45S_50ntFlanks@12348.12365.18\|GAAAAGTTACCACAGGGA | >hg38_21_+_8433172_8446622_RNA45SN1_45S_50ntFlanks | 12348 | 12365 | 18 | GAAAAGTTACCACAGGGA | 44.6 | -8.6 | 1.9 | -4.4 | 1.0E-05 | 4.2E-05 | BT20/MB468 | MP |
| >MI0003591\|hsa-mir-584&WithFlank&5\|-\|149062307\|149062415@22.43.22(+1U)&[MIMAT0003249&hsa-miR-584-5p&offsets\|0\|0(+1U);m-167&5\|-\|149062374\|149062394&offsets\|0\|+1(+1U)]\|TTATGGTTTGCCTGGGACTGAGT | >MI0003591\|hsa-mir-584&WithFlank&5\|-\|149062307\|149062415 | 22 | 43 | 22(+1U) | TTATGGTTTGCCTGGGACTGAGT | 191.8 | 4.9 | 1.2 | 4.2 | 2.5E-05 | 4.2E-05 | MB231/MB468 | MP |
| >hg38_MT_+_1621_3279_RNR2_16S_50ntFlanks@617.656.40\|AAACATATAACTGAACTCCTCACACCCAATTGGACCAATC | >hg38_MT_+_1621_3279_RNR2_16S_50ntFlanks | 617 | 656 | 40 | AAACATATAACTGAACTCCTCACACCCAATTGGACCAATC | 27.8 | 8.0 | 1.9 | 4.2 | 2.6E-05 | 4.2E-05 | MB231/MB468 | MP |
| >hg38_21_+_8433172_8446622_RNA45SN1_45S_50ntFlanks@9545.9573.29\|GTGGAGGTCCGTAGCGGTCCTGACGTGCA | >hg38_21_+_8433172_8446622_RNA45SN1_45S_50ntFlanks | 9545 | 9573 | 29 | GTGGAGGTCCGTAGCGGTCCTGACGTGCA | 682.2 | -5.1 | 1.2 | -4.2 | 2.6E-05 | 4.2E-05 | MB231/MB468 | MP |
| >hg38_21_+_8433172_8446622_RNA45SN1_45S_50ntFlanks@9570.9597.28\|TGCAAATCGGTCGTCCGACCTGGGTATA | >hg38_21_+_8433172_8446622_RNA45SN1_45S_50ntFlanks | 9570 | 9597 | 28 | TGCAAATCGGTCGTCCGACCTGGGTATA | 1612.5 | -6.7 | 1.6 | -4.2 | 2.6E-05 | 4.2E-05 | BT20/MB468 | CYTO |
| >hg38_21_+_8433172_8446622_RNA45SN1_45S_50ntFlanks@10344.10388.45\|ATTCAAACGAGAACTTTGAAGGCCGAAGTGGAGAAGGGTTCCATG | >hg38_21_+_8433172_8446622_RNA45SN1_45S_50ntFlanks | 10344 | 10388 | 45 | ATTCAAACGAGAACTTTGAAGGCCGAAGTGGAGAAGGGTTCCATG | 501.3 | 5.9 | 1.4 | 4.2 | 2.2E-05 | 4.2E-05 | BT20/MB468 | MITO |
| >hg38_21_+_8433172_8446622_RNA45SN1_45S_50ntFlanks@4588.4622.35\|CTATTTTGTTGGTTTTCGGAACTGAGGCCATGATT | >hg38_21_+_8433172_8446622_RNA45SN1_45S_50ntFlanks | 4588 | 4622 | 35 | CTATTTTGTTGGTTTTCGGAACTGAGGCCATGATT | 238.0 | -5.4 | 1.3 | -4.2 | 2.6E-05 | 4.2E-05 | MB231/MB468 | MP |
| >hg38_21_+_8433172_8446622_RNA45SN1_45S_50ntFlanks@8176.8196.21\|CTTCTGATCGAGGCCCAGCCC | >hg38_21_+_8433172_8446622_RNA45SN1_45S_50ntFlanks | 8176 | 8196 | 21 | CTTCTGATCGAGGCCCAGCCC | 190.2 | -4.9 | 1.2 | -4.2 | 2.6E-05 | 4.2E-05 | MB231/MB468 | MP |
| >hg38_21_+_8433172_8446622_RNA45SN1_45S_50ntFlanks@4007.4026.20\|CCTCGGGCCGATCGCACGCC | >hg38_21_+_8433172_8446622_RNA45SN1_45S_50ntFlanks | 4007 | 4026 | 20 | CCTCGGGCCGATCGCACGCC | 230.0 | -5.9 | 1.4 | -4.2 | 2.2E-05 | 4.2E-05 | BT20/MB468 | MITO |
| >hg38_21_+_8433172_8446622_RNA45SN1_45S_50ntFlanks@7978.8008.31\|GACCTCAGATCAGACGTGGCGACCCGCTGAA | >hg38_21_+_8433172_8446622_RNA45SN1_45S_50ntFlanks | 7978 | 8008 | 31 | GACCTCAGATCAGACGTGGCGACCCGCTGAA | 443.7 | 2.1 | 0.5 | 4.2 | 2.2E-05 | 4.3E-05 | BT20/MB468 | MITO |
| >hg38_21_+_8433172_8446622_RNA45SN1_45S_50ntFlanks@11317.11342.26\|ACCCCACGTCTCGTCGCGCGCGTC | >hg38_21_+_8433172_8446622_RNA45SN1_45S_50ntFlanks | 11317 | 11342 | 26 | ACCCCACGTCTCGTCGCGCGCGTC | 8063.9 | -8.5 | 2.0 | -4.2 | 3.2E-05 | 4.3E-05 | BT20/MB468 | NUC |
| >hg38_MT_+_1621_3279_RNR2_16S_50ntFlanks@757.777.21\|GCCCAATATCTACAATCAACC | >hg38_MT_+_1621_3279_RNR2_16S_50ntFlanks | 757 | 777 | 21 | GCCCAATATCTACAATCAACC | 30.9 | 8.2 | 1.9 | 4.2 | 2.6E-05 | 4.3E-05 | MB231/MB468 | MP |
| >hg38_21_+_8433172_8446622_RNA45SN1_45S_50ntFlanks@11603.11621.19\|CCGACTGTTTAATTAAAAC | >hg38_21_+_8433172_8446622_RNA45SN1_45S_50ntFlanks | 11603 | 11621 | 19 | CCGACTGTTTAATTAAAAC | 288.4 | -5.2 | 1.2 | -4.2 | 3.2E-05 | 4.3E-05 | BT20/MB468 | NUC |
| >trna27_GlyCCC_2_-_70476123_70476193@1.35.35,trna34_GlyCCC_16_-_686736_686806@1.35.35\|GCGCCGCTGGTGTAGTGGTATCATGCAAGATTCCC | >trna27_GlyCCC_2_-_70476123_70476193 | 1 | 35 | 35 | GCGCCGCTGGTGTAGTGGTATCATGCAAGATTCCC | 212.8 | 11.5 | 2.6 | 4.4 | 1.0E-05 | 4.3E-05 | BT20/MB468 | MP |
| >hg38_21_+_8433172_8446622_RNA45SN1_45S_50ntFlanks@4132.4151.20\|TCCGGAGAGGGAGCCTGAGA | >hg38_21_+_8433172_8446622_RNA45SN1_45S_50ntFlanks | 4132 | 4151 | 20 | TCCGGAGAGGGAGCCTGAGA | 319.6 | 6.1 | 1.4 | 4.4 | 1.0E-05 | 4.3E-05 | BT20/MB468 | MP |
| >hg38_21_+_8433172_8446622_RNA45SN1_45S_50ntFlanks@4372.4391.20\|AAAAAGCTCGTAGTTGGATC | >hg38_21_+_8433172_8446622_RNA45SN1_45S_50ntFlanks | 4372 | 4391 | 20 | AAAAAGCTCGTAGTTGGATC | 292.8 | -3.6 | 0.8 | -4.4 | 1.0E-05 | 4.3E-05 | BT20/MB468 | MP |
| >MI0000255\|hsa-mir-30d&WithFlank&8\|-\|134804870\|134804951@12.34.23(+1A)&[MIMAT0000245&hsa-miR-30d-5p&offsets\|0\|+1(+1A);m-21&8\|-\|134804919\|134804940&offsets\|0\|+1(+1A)]\|TGTAAACATCCCCGACTGGAAGCA | >MI0000255\|hsa-mir-30d&WithFlank&8\|-\|134804870\|134804951 | 12 | 34 | 23(+1A) | TGTAAACATCCCCGACTGGAAGCA | 2012.3 | 4.4 | 1.0 | 4.4 | 1.0E-05 | 4.3E-05 | BT20/MB468 | MP |
| >hg38_21_+_8433172_8446622_RNA45SN1_45S_50ntFlanks@10793.10814.22\|TCTGGCATGTTGGAACAATGTA | >hg38_21_+_8433172_8446622_RNA45SN1_45S_50ntFlanks | 10793 | 10814 | 22 | TCTGGCATGTTGGAACAATGTA | 95.7 | 9.9 | 2.1 | 4.7 | 3.2E-06 | 4.3E-05 | BT20/MB231 | MP |
| >trna119_LysCTT_1_-_145395522_145395594@3.33.31,trna11_LysCTT_5_-_180648979_180649051@3.33.31,trna13_LysCTT_6_+_26556774_26556846@3.33.31,trna7_LysCTT_16_+_3225692_3225764@3.33.31,trna9_LysCTT_5_+_180634755_180634827@3.33.31\|CCGGCTAGCTCAGTCGGTAGAGCATGAGACT | >trna119_LysCTT_1_-_145395522_145395594 | 3 | 33 | 31 | CCGGCTAGCTCAGTCGGTAGAGCATGAGACT | 227.5 | 7.1 | 1.7 | 4.2 | 2.6E-05 | 4.3E-05 | MB231/MB468 | MP |
| >hg38_21_+_8433172_8446622_RNA45SN1_45S_50ntFlanks@5493.5526.34\|GAAGACGGTCGAACTTGACTATCTAGAGGAAGTA | >hg38_21_+_8433172_8446622_RNA45SN1_45S_50ntFlanks | 5493 | 5526 | 34 | GAAGACGGTCGAACTTGACTATCTAGAGGAAGTA | 117.5 | -6.1 | 1.4 | -4.2 | 2.6E-05 | 4.3E-05 | MB231/MB468 | MP |
| >hg38_21_+_8433172_8446622_RNA45SN1_45S_50ntFlanks@475.521.47\|CTTCGTGATCGATGTGGTGACGTCGTGCTCTCCCGGGCCGGGTCCGA | >hg38_21_+_8433172_8446622_RNA45SN1_45S_50ntFlanks | 475 | 521 | 47 | CTTCGTGATCGATGTGGTGACGTCGTGCTCTCCCGGGCCGGGTCCGA | 36.6 | 8.4 | 2.0 | 4.2 | 2.6E-05 | 4.3E-05 | MB231/MB468 | MP |
| >trna111_HisGTG_1_-_147774845_147774916@2.30.29,trna118_HisGTG_1_-_145396881_145396952@2.30.29,trna16_HisGTG_1_+_146544773_146544844@2.30.29,trna1_HisGTG_15_+_45493349_45493420@2.30.29,trna21_HisGTG_1_+_147753471_147753542@2.30.29,trna33_HisGTG_6_+_27125906_27125977@2.30.29,trna7_HisGTG_9_-_14433938_14434009@2.30.29,trna8_HisGTG_15_-_45492611_45492682@2.30.29,trna9_HisGTG_15_-_45490804_45490875@2.30.29\|CCGTGATCGTATAGTGGTTAGTACTCTGC | >trna111_HisGTG_1_-_147774845_147774916 | 2 | 30 | 29 | CCGTGATCGTATAGTGGTTAGTACTCTGC | 524.4 | 5.4 | 1.2 | 4.4 | 1.0E-05 | 4.3E-05 | BT20/MB468 | MP |
| >hg38_21_+_8433172_8446622_RNA45SN1_45S_50ntFlanks@10351.10380.30\|CGAGAACTTTGAAGGCCGAAGTGGAGAAGG | >hg38_21_+_8433172_8446622_RNA45SN1_45S_50ntFlanks | 10351 | 10380 | 30 | CGAGAACTTTGAAGGCCGAAGTGGAGAAGG | 474.9 | -2.3 | 0.5 | -4.2 | 2.7E-05 | 4.3E-05 | BT20/MB468 | CYTO |
| >hg38_21_+_8433172_8446622_RNA45SN1_45S_50ntFlanks@10687.10709.23\|CGGCGTCCGGTGAGCTCTCGCTG | >hg38_21_+_8433172_8446622_RNA45SN1_45S_50ntFlanks | 10687 | 10709 | 23 | CGGCGTCCGGTGAGCTCTCGCTG | 2889.1 | -5.1 | 1.2 | -4.2 | 2.7E-05 | 4.3E-05 | BT20/MB468 | CYTO |
| >hg38_21_+_8433172_8446622_RNA45SN1_45S_50ntFlanks@6680.6718.39\|TGCGTCGATGAAGAACGCAGCTAGCTGCGAGAATTAATG | >hg38_21_+_8433172_8446622_RNA45SN1_45S_50ntFlanks | 6680 | 6718 | 39 | TGCGTCGATGAAGAACGCAGCTAGCTGCGAGAATTAATG | 62.5 | -9.8 | 2.3 | -4.2 | 2.6E-05 | 4.3E-05 | MB231/MB468 | MP |
| >hg38_21_+_8433172_8446622_RNA45SN1_45S_50ntFlanks@10572.10608.37\|AGAAGCCGGCGGGAGCCCCGGGGAGAGTTCTCTTTTC | >hg38_21_+_8433172_8446622_RNA45SN1_45S_50ntFlanks | 10572 | 10608 | 37 | AGAAGCCGGCGGGAGCCCCGGGGAGAGTTCTCTTTTC | 309.5 | -3.0 | 0.7 | -4.2 | 2.2E-05 | 4.3E-05 | BT20/MB468 | MITO |
| >hg38_21_+_8433172_8446622_RNA45SN1_45S_50ntFlanks@6747.6771.25\|ACTTCGAACGCACTTGCGGCCCCGG | >hg38_21_+_8433172_8446622_RNA45SN1_45S_50ntFlanks | 6747 | 6771 | 25 | ACTTCGAACGCACTTGCGGCCCCGG | 387.1 | -5.7 | 1.3 | -4.2 | 2.2E-05 | 4.3E-05 | BT20/MB468 | MITO |
| >hg38_21_+_8433172_8446622_RNA45SN1_45S_50ntFlanks@12909.12932.24\|CCGCACGTTCGTGGGGAACCTGGC | >hg38_21_+_8433172_8446622_RNA45SN1_45S_50ntFlanks | 12909 | 12932 | 24 | CCGCACGTTCGTGGGGAACCTGGC | 98.5 | -6.2 | 1.5 | -4.2 | 2.6E-05 | 4.3E-05 | MB231/MB468 | MP |
| >hg38_21_+_8433172_8446622_RNA45SN1_45S_50ntFlanks@5515.5535.21\|CTAGAGGAAGTAAAAGTCGTA | >hg38_21_+_8433172_8446622_RNA45SN1_45S_50ntFlanks | 5515 | 5535 | 21 | CTAGAGGAAGTAAAAGTCGTA | 162.3 | -5.4 | 1.3 | -4.2 | 3.1E-05 | 4.3E-05 | MB231/MB468 | NUC |
| >trnaMT_PheGAA_MT_+_577_647@1.34.34\|GTTTATGTAGCTTACCTCCTCAAAGCAATACACT | >trnaMT_PheGAA_MT_+_577_647 | 1 | 34 | 34 | GTTTATGTAGCTTACCTCCTCAAAGCAATACACT | 2391.7 | 6.2 | 1.5 | 4.2 | 3.1E-05 | 4.3E-05 | MB231/MB468 | MITO |
| >hg38_21_+_8433172_8446622_RNA45SN1_45S_50ntFlanks@12633.12659.27\|CGAAGCTACCATCTGTGGGATTATGAC | >hg38_21_+_8433172_8446622_RNA45SN1_45S_50ntFlanks | 12633 | 12659 | 27 | CGAAGCTACCATCTGTGGGATTATGAC | 320.6 | -4.3 | 1.0 | -4.2 | 2.7E-05 | 4.3E-05 | BT20/MB468 | CYTO |
| >hg38_21_+_8433172_8446622_RNA45SN1_45S_50ntFlanks@12948.12990.43\|GACGACCTGCTTCTGGGTCGGGGTTTCGTACGTAGCAGAGCAG | >hg38_21_+_8433172_8446622_RNA45SN1_45S_50ntFlanks | 12948 | 12990 | 43 | GACGACCTGCTTCTGGGTCGGGGTTTCGTACGTAGCAGAGCAG | 94.7 | -5.7 | 1.3 | -4.2 | 2.7E-05 | 4.3E-05 | BT20/MB468 | CYTO |
| >hg38_21_+_8433172_8446622_RNA45SN1_45S_50ntFlanks@9913.9946.34\|AGAAAAGGTGTTGGTTGATATAGACAGCAGGACG | >hg38_21_+_8433172_8446622_RNA45SN1_45S_50ntFlanks | 9913 | 9946 | 34 | AGAAAAGGTGTTGGTTGATATAGACAGCAGGACG | 82.9 | 9.7 | 2.1 | 4.7 | 3.2E-06 | 4.3E-05 | BT20/MB231 | MP |
| >hg38_21_+_8433172_8446622_RNA45SN1_45S_50ntFlanks@12502.12520.19\|TTAGACCGTCGTGAGACAG | >hg38_21_+_8433172_8446622_RNA45SN1_45S_50ntFlanks | 12502 | 12520 | 19 | TTAGACCGTCGTGAGACAG | 54.4 | -8.8 | 2.0 | -4.4 | 1.0E-05 | 4.3E-05 | BT20/MB468 | MP |
| >hg38_21_+_8433172_8446622_RNA45SN1_45S_50ntFlanks@9568.9603.36\|CGTGCAAATCGGTCGTCCGACCTGGGTATAGGGGCG | >hg38_21_+_8433172_8446622_RNA45SN1_45S_50ntFlanks | 9568 | 9603 | 36 | CGTGCAAATCGGTCGTCCGACCTGGGTATAGGGGCG | 300.4 | -4.1 | 1.0 | -4.2 | 2.7E-05 | 4.3E-05 | BT20/MB468 | CYTO |
| >hg38_21_+_8433172_8446622_RNA45SN1_45S_50ntFlanks@6646.6674.29\|TCGTACGACTCTTAGCGGTGGATCACTCG | >hg38_21_+_8433172_8446622_RNA45SN1_45S_50ntFlanks | 6646 | 6674 | 29 | TCGTACGACTCTTAGCGGTGGATCACTCG | 1177.7 | 3.5 | 0.8 | 4.2 | 2.9E-05 | 4.3E-05 | MB231/MB468 | CYTO |
| >hg38_21_+_8433172_8446622_RNA45SN1_45S_50ntFlanks@11710.11740.31\|CGCGGGTAAACGGCGGGAGTAACTATGACTC | >hg38_21_+_8433172_8446622_RNA45SN1_45S_50ntFlanks | 11710 | 11740 | 31 | CGCGGGTAAACGGCGGGAGTAACTATGACTC | 212.4 | -5.6 | 1.3 | -4.2 | 2.6E-05 | 4.3E-05 | MB231/MB468 | MP |
| >hg38_21_+_8433172_8446622_RNA45SN1_45S_50ntFlanks@6787.6806.20\|ACGCCTGTCTGAGCGTCGCT | >hg38_21_+_8433172_8446622_RNA45SN1_45S_50ntFlanks | 6787 | 6806 | 20 | ACGCCTGTCTGAGCGTCGCT | 2627.1 | 3.5 | 0.8 | 4.2 | 3.1E-05 | 4.3E-05 | MB231/MB468 | MITO |
| >hg38_21_+_8433172_8446622_RNA45SN1_45S_50ntFlanks@12415.12446.32\|CTTCGATGTCGGCTCTTCCTATCATTGTGAAG | >hg38_21_+_8433172_8446622_RNA45SN1_45S_50ntFlanks | 12415 | 12446 | 32 | CTTCGATGTCGGCTCTTCCTATCATTGTGAAG | 2483.7 | -3.4 | 0.8 | -4.2 | 3.1E-05 | 4.3E-05 | MB231/MB468 | MITO |
| >hg38_21_+_8433172_8446622_RNA45SN1_45S_50ntFlanks@11580.11602.23\|GAACTGGTGCGGACCAGGGGAAT | >hg38_21_+_8433172_8446622_RNA45SN1_45S_50ntFlanks | 11580 | 11602 | 23 | GAACTGGTGCGGACCAGGGGAAT | 1332.3 | -3.2 | 0.8 | -4.2 | 2.9E-05 | 4.3E-05 | MB231/MB468 | CYTO |

| | | | | | | | | | | | | | |
|---|---|---|---|---|---|---|---|---|---|---|---|---|---|
| >trna116_GluCTC_1_-_145399233_145399304@37.74.38,trna134_GluTTC_1_-_16861774_16861845@37.74.38,trna71_GluCTC_1_-_161439189_161439260@37.74.38,trna74_GluCTC_1_-_161431809_161431880@37.74.38,trna77_GluCTC_1_-_161424398_161424469@37.74.38,trna77_GluCTC_6_+_28949976_28950047@37.74.38,trna80_GluCTC_1_-_161417018_161417089@37.74.38,trna84_GluTTC_1_-_161391883_161391954@37.74.38,trna87_GluCTC_6_-_126101393_126101464@37.74.38\|ACCGCCGCGGCCCGGGTTCGATTCCCGGTCAGGGAACC | >trna116_GluCTC_1_-_145399233_145399304 | 37 | 74 | 38 | ACCGCCGCGGCCCGGGTTCGATTCCCGGTCAGGGAACC | 14508.6 | -5.3 | 1.1 | -4.7 | 3.2E-06 | 4.3E-05 | BT20/MB231 | CYTO |
| >MI0000478\|hsa-mir-149&WithFlank&2\|+\|240455995\|240456095@21.41.21(+1U)&[MIMAT0000450&hsa-miR-149-5p&offsets\|0\|-2(+1U);m-195&2\|+\|240456015\|240456037&offsets\|0\|-2(+1U)]\|TCTGGCTCCGTGTCTTCACTCT | >MI0000478\|hsa-mir-149&WithFlank&2\|+\|240455995\|240456095 | 21 | 41 | 21(+1U) | TCTGGCTCCGTGTCTTCACTCT | 585.6 | 3.4 | 0.7 | 4.6 | 3.8E-06 | 4.3E-05 | BT20/MB231 | NUC |
| >MI0000811\|hsa-mir-148b&WithFlank&12\|+\|54337210\|54337320@69.89.21&[MIMAT0000759&hsa-miR-148b-3p&offsets\|0\|-1;m-77&12\|+\|54337278\|54337299&offsets\|0\|-1]\|TCAGTGCATCACAGAACTTTG | >MI0000811\|hsa-mir-148b&WithFlank&12\|+\|54337210\|54337320 | 69 | 89 | 21 | TCAGTGCATCACAGAACTTTG | 1021.8 | 3.5 | 0.8 | 4.2 | 2.6E-05 | 4.3E-05 | MB231/MB468 | MP |
| >hg38_21_+_8433172_8446622_RNA45SN1_45S_50ntFlanks@8176.8208.33\|CTTCTGATCGAGGCCCAGCCCGTGGACGGTGTG | >hg38_21_+_8433172_8446622_RNA45SN1_45S_50ntFlanks | 8176 | 8208 | 33 | CTTCTGATCGAGGCCCAGCCCGTGGACGGTGTG | 391.4 | -4.7 | 1.1 | -4.2 | 3.1E-05 | 4.3E-05 | MB231/MB468 | MITO |
| >hg38_21_+_8433172_8446622_RNA45SN1_45S_50ntFlanks@10826.10858.33\|TCGGCAAGCCGGATCCGTAACTTCGGGATAAGG | >hg38_21_+_8433172_8446622_RNA45SN1_45S_50ntFlanks | 10826 | 10858 | 33 | TCGGCAAGCCGGATCCGTAACTTCGGGATAAGG | 393.0 | -5.9 | 1.3 | -4.4 | 1.0E-05 | 4.3E-05 | BT20/MB468 | MP |
| >hg38_21_+_8433172_8446622_RNA45SN1_45S_50ntFlanks@3984.4027.44\|GCGGCTTTGGTGACTCTAGATAACCTCGGGCCGATCGCACGCCC | >hg38_21_+_8433172_8446622_RNA45SN1_45S_50ntFlanks | 3984 | 4027 | 44 | GCGGCTTTGGTGACTCTAGATAACCTCGGGCCGATCGCACGCCC | 68.2 | -10.0 | 2.4 | -4.2 | 2.6E-05 | 4.3E-05 | MB231/MB468 | MP |
| >hg38_21_+_8433172_8446622_RNA45SN1_45S_50ntFlanks@10665.10706.42\|TTGGAAAGCGTCGCGGTTCCGGCGGCGTCCGGTGAGCTCTCG | >hg38_21_+_8433172_8446622_RNA45SN1_45S_50ntFlanks | 10665 | 10706 | 42 | TTGGAAAGCGTCGCGGTTCCGGCGGCGTCCGGTGAGCTCTCG | 201.0 | -6.2 | 1.5 | -4.2 | 3.1E-05 | 4.3E-05 | MB231/MB468 | NUC |
| >MI0000650\|hsa-mir-200c&WithFlank&12\|+\|6963693\|6963772@50.71.22(+1U)&[MIMAT0000617&hsa-miR-200c-3p&offsets\|0\|-1(+1U);m-54&12\|+\|6963742\|6963764&offsets\|0\|-1(+1U)]\|TAATACTGCCGGGTAATGATGGT | >MI0000650\|hsa-mir-200c&WithFlank&12\|+\|6963693\|6963772 | 50 | 71 | 22(+1U) | TAATACTGCCGGGTAATGATGGT | 2343.2 | 4.9 | 1.1 | 4.4 | 1.0E-05 | 4.3E-05 | BT20/MB468 | MP |
| >trna10_AspGTC_12_-_125424193_125424264@35.74.40,trna12_AspGTC_12_-_125411891_125411962@35.74.40,trna144_AspGTC_6_-_27551236_27551307@35.74.40,trna38_AspGTC_17_-_8125556_8125627@35.74.40,trna45_AspGTC_6_+_27447453_27447524@35.74.40,trna48_AspGTC_6_+_27471523_27471594@35.74.40,trna4_AspGTC_12_+_96429799_96429870@35.74.40,trna69_AspGTC_1_-_161440205_161440276@35.74.40,trna72_AspGTC_1_-_161432824_161432895@35.74.40,trna75_AspGTC_1_-_161425414_161425485@35.74.40,trna78_AspGTC_1_-_161418033_161418104@35.74.40,trna81_AspGTC_1_-_161410615_161410686@35.74.40\|TCACGCGGGAGACCGGGGTTCGATTCCCCGACGGGGAGCC | >trna10_AspGTC_12_-_125424193_125424264 | 35 | 74 | 40 | TCACGCGGGAGACCGGGGTTCGATTCCCCGACGGGGAGCC | 613.5 | 5.4 | 1.3 | 4.2 | 3.1E-05 | 4.3E-05 | MB231/MB468 | NUC |
| >hg38_21_+_8433172_8446622_RNA45SN1_45S_50ntFlanks@11568.11603.36\|GCAGCCGACTTAGAACTGGTGCGGACCAGGGGAATC | >hg38_21_+_8433172_8446622_RNA45SN1_45S_50ntFlanks | 11568 | 11603 | 36 | GCAGCCGACTTAGAACTGGTGCGGACCAGGGGAATC | 366.9 | -2.3 | 0.5 | -4.2 | 2.7E-05 | 4.3E-05 | BT20/MB468 | CYTO |
| >hg38_21_+_8433172_8446622_RNA45SN1_45S_50ntFlanks@12456.12489.34\|CCAAGCGTTGGATTGTTCACCCACTAATAGGGAA | >hg38_21_+_8433172_8446622_RNA45SN1_45S_50ntFlanks | 12456 | 12489 | 34 | CCAAGCGTTGGATTGTTCACCCACTAATAGGGAA | 1239.4 | -7.6 | 1.7 | -4.4 | 1.0E-05 | 4.3E-05 | BT20/MB468 | MP |
| >am_trna128_GlyGCC_6_-_27870686_27870756@1.35.35,trna18_GlyGCC_16_+_70822597_70822667@1.35.35,trna19_GlyGCC_16_+_70823410_70823480@1.35.35,trna19_GlyGCC_2_-_157257659_157257729@1.35.35,trna24_GlyGCC_16_-_70812942_70813012@1.35.35,trna25_GlyGCC_16_-_70812114_70812184@1.35.35,trna5_GlyGCC_17_+_8029064_8029134@1.35.35,trna68_GlyGCC_1_-_161493637_161493707@1.35.35\|GCATTGGTGGTTCAGTGGTAGAATTCTCGCCTGCC | >am_trna128_GlyGCC_6_-_27870686_27870756 | 1 | 35 | 35 | GCATTGGTGGTTCAGTGGTAGAATTCTCGCCTGCC | 2424.8 | -4.3 | 0.9 | -4.6 | 3.8E-06 | 4.3E-05 | BT20/MB231 | NUC |
| >hg38_21_+_8433172_8446622_RNA45SN1_45S_50ntFlanks@9554.9574.21\|CGTAGCGGTCCTGACGTGCAA | >hg38_21_+_8433172_8446622_RNA45SN1_45S_50ntFlanks | 9554 | 9574 | 21 | CGTAGCGGTCCTGACGTGCAA | 2839.9 | -6.0 | 1.4 | -4.2 | 2.6E-05 | 4.3E-05 | MB231/MB468 | MP |
| >trnaMT_ValTAC_MT_+_1602_1670@44.71.28\|ATTTCAACTTAACTTGACCGCTCTGACC | >trnaMT_ValTAC_MT_+_1602_1670 | 44 | 71 | 28 | ATTTCAACTTAACTTGACCGCTCTGACC | 2860.5 | 4.5 | 1.1 | 4.2 | 2.3E-05 | 4.3E-05 | BT20/MB468 | MITO |
| >MI0000086\|hsa-mir-28&WithFlank&3\|+\|188688775\|188688872@61.82.22(+1A)&[MIMAT0004502&hsa-miR-28-3p&offsets\|+1\|+1(+1A);m-28&3\|+\|188688834\|188688855&offsets\|+1\|+1(+1A)]\|ACTAGATTGTGAGCTCCTGGAGA | >MI0000086\|hsa-mir-28&WithFlank&3\|+\|188688775\|188688872 | 61 | 82 | 22(+1A) | ACTAGATTGTGAGCTCCTGGAGA | 147.2 | 6.6 | 1.5 | 4.4 | 1.0E-05 | 4.3E-05 | BT20/MB468 | MP |
| >hg38_21_+_8433172_8446622_RNA45SN1_45S_50ntFlanks@9739.9763.25\|AAACGATCTCAACCTATTCTCAAAC | >hg38_21_+_8433172_8446622_RNA45SN1_45S_50ntFlanks | 9739 | 9763 | 25 | AAACGATCTCAACCTATTCTCAAAC | 325.6 | -4.8 | 1.2 | -4.2 | 3.1E-05 | 4.3E-05 | MB231/MB468 | MITO |
| >hg38_21_+_8433172_8446622_RNA45SN1_45S_50ntFlanks@4297.4318.22\|CATTGGAGGGCAAGTCTGGTGC | >hg38_21_+_8433172_8446622_RNA45SN1_45S_50ntFlanks | 4297 | 4318 | 22 | CATTGGAGGGCAAGTCTGGTGC | 196.5 | -4.3 | 1.0 | -4.2 | 3.1E-05 | 4.3E-05 | MB231/MB468 | NUC |
| >hg38_21_+_8433172_8446622_RNA45SN1_45S_50ntFlanks@10409.10426.18\|GTCAGTCGGTCCTGAGAG | >hg38_21_+_8433172_8446622_RNA45SN1_45S_50ntFlanks | 10409 | 10426 | 18 | GTCAGTCGGTCCTGAGAG | 172.6 | -4.4 | 1.1 | -4.2 | 3.1E-05 | 4.3E-05 | MB231/MB468 | NUC |
| >hg38_21_+_8433172_8446622_RNA45SN1_45S_50ntFlanks@12637.12659.23\|GCTACCATCTGTGGGATTATGAC | >hg38_21_+_8433172_8446622_RNA45SN1_45S_50ntFlanks | 12637 | 12659 | 23 | GCTACCATCTGTGGGATTATGAC | 276.0 | -3.1 | 0.7 | -4.2 | 3.1E-05 | 4.4E-05 | MB231/MB468 | NUC |
| >hg38_21_+_8433172_8446622_RNA45SN1_45S_50ntFlanks@10346.10377.32\|TCAAACGAGAACTTTGAAGGCCGAAGTGGAGA | >hg38_21_+_8433172_8446622_RNA45SN1_45S_50ntFlanks | 10346 | 10377 | 32 | TCAAACGAGAACTTTGAAGGCCGAAGTGGAGA | 197.7 | -3.4 | 0.8 | -4.2 | 2.6E-05 | 4.4E-05 | MB231/MB468 | MP |
| >hg38_21_+_8433172_8446622_RNA45SN1_45S_50ntFlanks@12416.12452.37\|TTCGATGTCGGCTCTTCCTATCATTGTGAAGCAGAAT | >hg38_21_+_8433172_8446622_RNA45SN1_45S_50ntFlanks | 12416 | 12452 | 37 | TTCGATGTCGGCTCTTCCTATCATTGTGAAGCAGAAT | 205.7 | 5.8 | 1.4 | 4.2 | 3.1E-05 | 4.4E-05 | MB231/MB468 | MITO |
| >trnaMT_PheGAA_MT_+_577_647@35.59.25\|GAAAATGTTTAGACGGGCTCACATC | >trnaMT_PheGAA_MT_+_577_647 | 35 | 59 | 25 | GAAAATGTTTAGACGGGCTCACATC | 769.4 | 5.4 | 1.2 | 4.4 | 1.1E-05 | 4.4E-05 | BT20/MB468 | MP |
| >hg38_21_+_8433172_8446622_RNA45SN1_45S_50ntFlanks@10843.10872.30\|TAACTTCGGGATAAGGATTGGCTCTAAGGG | >hg38_21_+_8433172_8446622_RNA45SN1_45S_50ntFlanks | 10843 | 10872 | 30 | TAACTTCGGGATAAGGATTGGCTCTAAGGG | 206.2 | -2.7 | 0.6 | -4.2 | 2.7E-05 | 4.4E-05 | BT20/MB468 | CYTO |
| >MI0000077\|hsa-mir-21&WithFlank&17\|+\|59841260\|59841343@14.39.26(+1U)&[MIMAT0000076&hsa-miR-21-5p&offsets\|0\|+4(+1U);m-1&17\|+\|59841273\|59841294&offsets\|0\|+4(+1U)]\|TAGCTTATCAGACTGATGTTGACTGTT | >MI0000077\|hsa-mir-21&WithFlank&17\|+\|59841260\|59841343 | 14 | 39 | 26(+1U) | TAGCTTATCAGACTGATGTTGACTGTT | 218.4 | 4.4 | 1.0 | 4.2 | 3.1E-05 | 4.4E-05 | MB231/MB468 | MITO |
| >MI0000081\|hsa-mir-24-2&WithFlank&19\|-\|13836281\|13836365@56.77.22(+1A);MI0000080\|hsa-mir-24-1&WithFlank&9\|+\|95086015\|95086094@50.71.22(+1A)&[MIMAT0000080&hsa-miR-24-3p&offsets\|0\|0(+1A);m-36&19\|-\|13836288\|13836310&offsets\|0\|-1(+1A)];[MIMAT0000080_1&hsa-miR-24-3p&offsets\|0\|0(+1A);m-35&9\|+\|95086064\|95086086&offsets\|0\|-1(+1A)]\|TGGCTCAGTTCAGCAGGAACAGA | >MI0000081\|hsa-mir-24-2&WithFlank&19\|-\|13836281\|13836365 | 56 | 77 | 22(+1A) | TGGCTCAGTTCAGCAGGAACAGA | 456.8 | 4.3 | 1.0 | 4.2 | 3.1E-05 | 4.4E-05 | MB231/MB468 | MITO |
| >hg38_21_+_8433172_8446622_RNA45SN1_45S_50ntFlanks@4764.4786.23\|ATACCGTCGTAGTTCCGACCATA | >hg38_21_+_8433172_8446622_RNA45SN1_45S_50ntFlanks | 4764 | 4786 | 23 | ATACCGTCGTAGTTCCGACCATA | 64.2 | -9.9 | 2.3 | -4.2 | 2.6E-05 | 4.4E-05 | MB231/MB468 | MP |
| >MI0001448\|hsa-mir-425&WithFlank&3\|-\|49020142\|49020240@60.81.22&[MIMAT0001343&hsa-miR-425-3p&offsets\|-1\|-1;m-258&3\|-\|49020160\|49020181&offsets\|0\|0]\|CATCGGGAATGTCGTGTCCGCC | >MI0001448\|hsa-mir-425&WithFlank&3\|-\|49020142\|49020240 | 60 | 81 | 22 | CATCGGGAATGTCGTGTCCGCC | 584.9 | 5.1 | 1.2 | 4.4 | 1.1E-05 | 4.4E-05 | BT20/MB468 | MP |
| >hg38_21_+_8433172_8446622_RNA45SN1_45S_50ntFlanks@7975.8000.26\|CGCGACCTCAGATCAGACGTGGCGAC | >hg38_21_+_8433172_8446622_RNA45SN1_45S_50ntFlanks | 7975 | 8000 | 26 | CGCGACCTCAGATCAGACGTGGCGAC | 1797.4 | 3.5 | 0.8 | 4.6 | 3.9E-06 | 4.4E-05 | BT20/MB231 | NUC |
| >trna111_HisGTG_1_-_147774845_147774916@1.33.33,trna118_HisGTG_1_-_145396881_145396952@1.33.33,trna16_HisGTG_1_+_146544773_146544844@1.33.33,trna1_HisGTG_15_+_45493349_45493420@1.33.33,trna21_HisGTG_1_+_147753471_147753542@1.33.33,trna33_HisGTG_6_+_27125906_27125977@1.33.33,trna7_HisGTG_9_-_14433938_14434009@1.33.33,trna8_HisGTG_15_-_45492611_45492682@1.33.33,trna9_HisGTG_15_-_45490804_45490875@1.33.33\|GCCGTGATCGTATAGTGGTTAGTACTCTGCGTT | >trna111_HisGTG_1_-_147774845_147774916 | 1 | 33 | 33 | GCCGTGATCGTATAGTGGTTAGTACTCTGCGTT | 723.0 | 3.7 | 0.9 | 4.2 | 2.6E-05 | 4.4E-05 | MB231/MB468 | MP |
| >hg38_1_-_228634819_228635039_RNA5S12_5S_50ntFlanks@51.99.49\|GTCTACGGCCATACCACCCTGAACGCGCCCGATCTCGTCTGATCTCGGA | >hg38_1_-_228634819_228635039_RNA5S12_5S_50ntFlanks | 51 | 99 | 49 | GTCTACGGCCATACCACCCTGAACGCGCCCGATCTCGTCTGATCTCGGA | 191.2 | 11.3 | 2.6 | 4.4 | 1.1E-05 | 4.4E-05 | BT20/MB468 | MP |
| >hg38_21_+_8433172_8446622_RNA45SN1_45S_50ntFlanks@10680.10709.30\|GTTCCGGCGGCGTCCGGTGAGCTCTCGCTG | >hg38_21_+_8433172_8446622_RNA45SN1_45S_50ntFlanks | 10680 | 10709 | 30 | GTTCCGGCGGCGTCCGGTGAGCTCTCGCTG | 3186.7 | -5.2 | 1.2 | -4.2 | 2.7E-05 | 4.4E-05 | BT20/MB468 | CYTO |
| >hg38_21_+_8433172_8446622_RNA45SN1_45S_50ntFlanks@4726.4765.40\|TAATCAAGAACGAAAGTCGGAGGTTCGAAGACGATCAGAT | >hg38_21_+_8433172_8446622_RNA45SN1_45S_50ntFlanks | 4726 | 4765 | 40 | TAATCAAGAACGAAAGTCGGAGGTTCGAAGACGATCAGAT | 309.8 | 5.4 | 1.3 | 4.2 | 2.7E-05 | 4.4E-05 | BT20/MB468 | CYTO |
| >MI0000749\|hsa-mir-30e&WithFlank&1\|+\|40754349\|40754452@23.46.24(+1U)&[MIMAT0000692&hsa-miR-30e-5p&offsets\|0\|+2(+1U);m-24&1\|+\|40754371\|40754394&offsets\|0\|0(+1U)]\|TGTAAACATCCTTGACTGGAAGCTT | >MI0000749\|hsa-mir-30e&WithFlank&1\|+\|40754349\|40754452 | 23 | 46 | 24(+1U) | TGTAAACATCCTTGACTGGAAGCTT | 649.2 | 4.7 | 1.1 | 4.2 | 3.2E-05 | 4.4E-05 | MB231/MB468 | MITO |
| >trnaMT_HisGTG_MT_+_12138_12206@26.72.47\|AGATTGTGAATCTGACAACAGAGGCTTACGACCCCTTATTTACCCCA | >trnaMT_HisGTG_MT_+_12138_12206 | 26 | 72 | 47 | AGATTGTGAATCTGACAACAGAGGCTTACGACCCCTTATTTACCCCA | 199.1 | 11.4 | 2.6 | 4.4 | 1.1E-05 | 4.4E-05 | BT20/MB468 | MP |
| >hg38_21_+_8433172_8446622_RNA45SN1_45S_50ntFlanks@12420.12437.18\|ATGTCGGCTCTTCCTATC | >hg38_21_+_8433172_8446622_RNA45SN1_45S_50ntFlanks | 12420 | 12437 | 18 | ATGTCGGCTCTTCCTATC | 987.7 | -3.7 | 0.8 | -4.4 | 1.1E-05 | 4.4E-05 | BT20/MB468 | MP |
| >hg38_21_+_8433172_8446622_RNA45SN1_45S_50ntFlanks@5055.5081.27\|GGAGCGATTTGTCTGGTTAATTCCGAT | >hg38_21_+_8433172_8446622_RNA45SN1_45S_50ntFlanks | 5055 | 5081 | 27 | GGAGCGATTTGTCTGGTTAATTCCGAT | 190.9 | 3.3 | 0.7 | 4.6 | 3.9E-06 | 4.4E-05 | BT20/MB231 | MITO |

| | | | | | | | | | | | | | |
|---|---|---|---|---|---|---|---|---|---|---|---|---|---|
| >MI0000298\|hsa-mir-221&WithFlank&X\|-\|45746151\|45746272@71.92.22(+2U)&[MIMAT0000278&hsa-miR-221-3p&offsets\|0\|-1(+2U);m-91&X\|-\|45746181\|45746202&offsets\|0\|0(+2U)]\|AGCTACATTGTCTGCTGGGTTTTT | >MI0000298\|hsa-mir-221&WithFlank&X\|-\|45746151\|45746272 | 71 | 92 | 22(+2U) | AGCTACATTGTCTGCTGGGTTTTT | 723.7 | -3.8 | 0.8 | -4.6 | 3.9E-06 | 4.4E-05 | BT20/MB231 | MITO |
| >trna111_HisGTG_1_-_147774845_147774916@-1G.36.37,trna118_HisGTG_1_-_145396881_145396952@-1G.36.37,trna16_HisGTG_1_+_146544773_146544844@-1G.36.37,trna1_HisGTG_15_+_45493349_45493420@-1G.36.37,trna21_HisGTG_1_+_147753471_147753542@-1G.36.37,trna33_HisGTG_6_+_27125906_27125977@-1G.36.37,trna7_HisGTG_9_-_14433938_14434009@-1G.36.37,trna8_HisGTG_15_-_45492611_45492682@-1G.36.37,trna9_HisGTG_15_-_45490804_45490875@-1G.36.37\|GGCCGTGATCGTATAGTGGTTAGTACTCTGCGTTGTG | >trna111_HisGTG_1_-_147774845_147774916 | -1G | 36 | 37 | GGCCGTGATCGTATAGTGGTTAGTACTCTGCGTTGTG | 668.3 | -3.4 | 0.7 | -4.6 | 3.9E-06 | 4.4E-05 | BT20/MB231 | MITO |
| >MI0003560\|hsa-mir-92b&WithFlank&1\|+\|155195171\|155195278@67.84.18(+1U)&[MIMAT0003218&hsa-miR-92b-3p&offsets\|0\|-4(+1U);m-115&1\|+\|155195237\|155195258&offsets\|0\|-4(+1U)]\|TATTGCACTCGTCCCGGCT | >MI0003560\|hsa-mir-92b&WithFlank&1\|+\|155195171\|155195278 | 67 | 84 | 18(+1U) | TATTGCACTCGTCCCGGCT | 39.3 | 8.5 | 2.0 | 4.2 | 2.6E-05 | 4.4E-05 | MB231/MB468 | MP |
| >hg38_21_+_8433172_8446622_RNA45SN1_45S_50ntFlanks@4727.4759.33\|AATCAAGAACGAAAGTCGGAGGTTCGAAGACGA | >hg38_21_+_8433172_8446622_RNA45SN1_45S_50ntFlanks | 4727 | 4759 | 33 | AATCAAGAACGAAAGTCGGAGGTTCGAAGACGA | 534.0 | -5.4 | 1.3 | -4.2 | 3.3E-05 | 4.4E-05 | BT20/MB468 | NUC |
| >hg38_21_+_8433172_8446622_RNA45SN1_45S_50ntFlanks@10392.10433.42\|ACAGCAGTTGAACATGGGTCAGTCGGTCCTGAGAGATGGGCG | >hg38_21_+_8433172_8446622_RNA45SN1_45S_50ntFlanks | 10392 | 10433 | 42 | ACAGCAGTTGAACATGGGTCAGTCGGTCCTGAGAGATGGGCG | 368.1 | -6.9 | 1.6 | -4.2 | 2.6E-05 | 4.4E-05 | MB231/MB468 | MP |
| >MI0000077\|hsa-mir-21&WithFlank&17\|+\|59841260\|59841343@14.36.23&[MIMAT0000076&hsa-miR-21-5p&offsets\|0\|+1;m-1&17\|+\|59841273\|59841294&offsets\|0\|+1]\|TAGCTTATCAGACTGATGTTGAC | >MI0000077\|hsa-mir-21&WithFlank&17\|+\|59841260\|59841343 | 14 | 36 | 23 | TAGCTTATCAGACTGATGTTGAC | 418749.2 | 3.3 | 0.8 | 4.2 | 2.7E-05 | 4.4E-05 | MB231/MB468 | MP |
| >hg38_21_+_8433172_8446622_RNA45SN1_45S_50ntFlanks@11873.11907.35\|AAGAAGACCCTGTTGAGCTTGACTCTAGTCTGGCA | >hg38_21_+_8433172_8446622_RNA45SN1_45S_50ntFlanks | 11873 | 11907 | 35 | AAGAAGACCCTGTTGAGCTTGACTCTAGTCTGGCA | 280.1 | -5.6 | 1.3 | -4.2 | 2.7E-05 | 4.4E-05 | BT20/MB468 | CYTO |
| >hg38_21_+_8433172_8446622_RNA45SN1_45S_50ntFlanks@10468.10495.28\|TTGCCCTCGGCCGATCGAAAGGGAGTCG | >hg38_21_+_8433172_8446622_RNA45SN1_45S_50ntFlanks | 10468 | 10495 | 28 | TTGCCCTCGGCCGATCGAAAGGGAGTCG | 101.7 | -4.8 | 1.1 | -4.2 | 2.7E-05 | 4.4E-05 | MB231/MB468 | MP |
| >hg38_21_+_8433172_8446622_RNA45SN1_45S_50ntFlanks@9940.9975.36\|CAGGACGGTGGCCATGGAAGTCGGAATCCGCTAAGG | >hg38_21_+_8433172_8446622_RNA45SN1_45S_50ntFlanks | 9940 | 9975 | 36 | CAGGACGGTGGCCATGGAAGTCGGAATCCGCTAAGG | 158.0 | 7.5 | 1.7 | 4.4 | 1.1E-05 | 4.4E-05 | BT20/MB468 | MP |
| >MI0000088\|hsa-mir-30a&WithFlank&6\|-\|71403545\|71403627@12.33.22&[MIMAT0000087&hsa-miR-30a-5p&offsets\|0\|0;m-15&6\|-\|71403595\|71403616&offsets\|0\|0]\|TGTAAACATCCTCGACTGGAAG | >MI0000088\|hsa-mir-30a&WithFlank&6\|-\|71403545\|71403627 | 12 | 33 | 22 | TGTAAACATCCTCGACTGGAAG | 842.2 | 3.5 | 0.8 | 4.2 | 3.3E-05 | 4.4E-05 | BT20/MB468 | NUC |
| >hg38_1_-_228634819_228635039_RNA5S12_5S_50ntFlanks@133.169.37\|ATGGGAGACCGCCTGGGAATACCGGGTGCTGTAGGCT | >hg38_1_-_228634819_228635039_RNA5S12_5S_50ntFlanks | 133 | 169 | 37 | ATGGGAGACCGCCTGGGAATACCGGGTGCTGTAGGCT | 2009.2 | 3.8 | 0.8 | 4.6 | 4.0E-06 | 4.4E-05 | BT20/MB231 | MITO |
| >hg38_21_+_8433172_8446622_RNA45SN1_45S_50ntFlanks@6668.6696.29\|TCACTCGGCTCGTGCGTCGATGAAGAACG | >hg38_21_+_8433172_8446622_RNA45SN1_45S_50ntFlanks | 6668 | 6696 | 29 | TCACTCGGCTCGTGCGTCGATGAAGAACG | 74.8 | -5.9 | 1.4 | -4.2 | 2.7E-05 | 4.4E-05 | MB231/MB468 | MP |
| >hg38_21_+_8433172_8446622_RNA45SN1_45S_50ntFlanks@5423.5466.44\|ATTGGATGGTTTAGTGAGGCCCTCGGATCGGCCCCGCCGGGGTC | >hg38_21_+_8433172_8446622_RNA45SN1_45S_50ntFlanks | 5423 | 5466 | 44 | ATTGGATGGTTTAGTGAGGCCCTCGGATCGGCCCCGCCGGGGTC | 119.1 | 10.2 | 2.2 | 4.6 | 3.3E-06 | 4.4E-05 | BT20/MB231 | MP |
| >MI0000298\|hsa-mir-221&WithFlank&X\|-\|45746151\|45746272@30.53.24&[MIMAT0004568&hsa-miR-221-5p&offsets\|-1\|+1;m-405&X\|-\|45746220\|45746242&offsets\|-1\|0]\|AACCTGGCATACAATGTAGATTTC | >MI0000298\|hsa-mir-221&WithFlank&X\|-\|45746151\|45746272 | 30 | 53 | 24 | AACCTGGCATACAATGTAGATTTC | 29.7 | 8.1 | 1.9 | 4.2 | 2.7E-05 | 4.4E-05 | MB231/MB468 | MP |
| >trnaMT_MetCAT_MT_+_4402_4469@39.70.32\|GAAAATGTTGGTTATACCCTTCCCGTACTACC | >trnaMT_MetCAT_MT_+_4402_4469 | 39 | 70 | 32 | GAAAATGTTGGTTATACCCTTCCCGTACTACC | 27.6 | 8.0 | 1.9 | 4.2 | 2.7E-05 | 4.4E-05 | MB231/MB468 | MP |
| >hg38_21_+_8433172_8446622_RNA45SN1_45S_50ntFlanks@10378.10401.24\|AGGGTTCCATGTGAACAGCAGTTG | >hg38_21_+_8433172_8446622_RNA45SN1_45S_50ntFlanks | 10378 | 10401 | 24 | AGGGTTCCATGTGAACAGCAGTTG | 51.5 | -8.8 | 2.0 | -4.4 | 1.1E-05 | 4.4E-05 | BT20/MB468 | MP |
| >MI0003776\|hsa-mir-320b-1&WithFlank&1\|+\|116671740\|116671823@48.67.20(+1C);MI0003839\|hsa-mir-320b-2&WithFlank&1\|-\|224257034\|224257116@47.66.20(+1C);MI0000542\|hsa-mir-320a&WithFlank&8\|-\|22244960\|22245043@48.67.20(+1C)&[MIMAT0005792&hsa-miR-320b&offsets\|0\|-2(+1C);m-183&1\|+\|116671787\|116671806&offsets\|0\|0(+1C)];[MIMAT0005792_1&hsa-miR-320b&offsets\|0\|-2(+1C);m-183&1\|-\|224257051\|224257070&offsets\|0\|0(+1C)];[MIMAT0000510&hsa-miR-320a-3p&offsets\|0\|-2(+1C);m-60&8\|-\|22244975\|22244996&offsets\|0\|-2(+1C)]\|AAAAGCTGGGTTGAGAGGGCC | >MI0003776\|hsa-mir-320b-1&WithFlank&1\|+\|116671740\|116671823 | 48 | 67 | 20(+1C) | AAAAGCTGGGTTGAGAGGGCC | 342.7 | 7.2 | 1.6 | 4.4 | 1.1E-05 | 4.4E-05 | BT20/MB468 | MP |
| >hg38_21_+_8433172_8446622_RNA45SN1_45S_50ntFlanks@10869.10886.18\|AGGGCTGGGTCGGTCGGG | >hg38_21_+_8433172_8446622_RNA45SN1_45S_50ntFlanks | 10869 | 10886 | 18 | AGGGCTGGGTCGGTCGGG | 644.4 | -5.0 | 1.2 | -4.2 | 2.8E-05 | 4.4E-05 | BT20/MB468 | CYTO |
| >hg38_MT_+_598_1651_RNR1_12S_50ntFlanks@427.469.43\|GAAAGTGGCTTTAACATATCTGAACACACAATAGCTAAGACCC | >hg38_MT_+_598_1651_RNR1_12S_50ntFlanks | 427 | 469 | 43 | GAAAGTGGCTTTAACATATCTGAACACACAATAGCTAAGACCC | 31.5 | 8.2 | 2.0 | 4.2 | 2.7E-05 | 4.4E-05 | MB231/MB468 | MP |
| >hg38_21_+_8433172_8446622_RNA45SN1_45S_50ntFlanks@10315.10336.22\|AGGTGCAGATCTTGGTGGTAGT | >hg38_21_+_8433172_8446622_RNA45SN1_45S_50ntFlanks | 10315 | 10336 | 22 | AGGTGCAGATCTTGGTGGTAGT | 117.9 | 10.2 | 2.2 | 4.6 | 3.4E-06 | 4.4E-05 | BT20/MB231 | MP |
| >MI0000433\|hsa-let-7g&WithFlank&3\|-\|52268272\|52268367@11.31.21&[MIMAT0000414&hsa-let-7g-5p&offsets\|0\|-1;m-58&3\|-\|52268336\|52268357&offsets\|0\|-1]\|TGAGGTAGTAGTTTGTACAGT | >MI0000433\|hsa-let-7g&WithFlank&3\|-\|52268272\|52268367 | 11 | 31 | 21 | TGAGGTAGTAGTTTGTACAGT | 1911.2 | 4.4 | 1.0 | 4.4 | 1.1E-05 | 4.4E-05 | BT20/MB468 | MP |
| >hg38_21_+_8433172_8446622_RNA45SN1_45S_50ntFlanks@4971.4999.29\|CCGGCCCGGACACGGACAGGATTGACAGA | >hg38_21_+_8433172_8446622_RNA45SN1_45S_50ntFlanks | 4971 | 4999 | 29 | CCGGCCCGGACACGGACAGGATTGACAGA | 469.1 | -3.7 | 0.9 | -4.2 | 2.3E-05 | 4.4E-05 | BT20/MB468 | MITO |
| >hg38_21_+_8433172_8446622_RNA45SN1_45S_50ntFlanks@12076.12095.20\|AGGGGCTCTCGCTTCTGGCG | >hg38_21_+_8433172_8446622_RNA45SN1_45S_50ntFlanks | 12076 | 12095 | 20 | AGGGGCTCTCGCTTCTGGCG | 8398.0 | -8.5 | 2.0 | -4.1 | 3.3E-05 | 4.4E-05 | BT20/MB468 | NUC |
| >hg38_21_+_8433172_8446622_RNA45SN1_45S_50ntFlanks@6697.6718.22\|CAGCTAGCTGCGAGAATTAATG | >hg38_21_+_8433172_8446622_RNA45SN1_45S_50ntFlanks | 6697 | 6718 | 22 | CAGCTAGCTGCGAGAATTAATG | 620.8 | -2.6 | 0.6 | -4.2 | 3.2E-05 | 4.4E-05 | MB231/MB468 | NUC |
| >hg38_21_+_8433172_8446622_RNA45SN1_45S_50ntFlanks@10848.10865.18\|TCGGGATAAGGATTGGCT | >hg38_21_+_8433172_8446622_RNA45SN1_45S_50ntFlanks | 10848 | 10865 | 18 | TCGGGATAAGGATTGGCT | 362.9 | -3.1 | 0.7 | -4.2 | 2.8E-05 | 4.4E-05 | BT20/MB468 | CYTO |
| >hg38_21_+_8433172_8446622_RNA45SN1_45S_50ntFlanks@6721.6747.27\|AATTGCAGGACACATTGATCATCGACA | >hg38_21_+_8433172_8446622_RNA45SN1_45S_50ntFlanks | 6721 | 6747 | 27 | AATTGCAGGACACATTGATCATCGACA | 1599.6 | -2.3 | 0.5 | -4.2 | 2.3E-05 | 4.4E-05 | BT20/MB468 | MITO |
| >hg38_MT_+_1621_3279_RNR2_16S_50ntFlanks@1372.1404.33\|GTTGGATCAGGACATCCCGATGGTGCAGCCGCT | >hg38_MT_+_1621_3279_RNR2_16S_50ntFlanks | 1372 | 1404 | 33 | GTTGGATCAGGACATCCCGATGGTGCAGCCGCT | 28.6 | 8.1 | 1.9 | 4.2 | 2.7E-05 | 4.4E-05 | MB231/MB468 | MP |
| >hg38_21_+_8433172_8446622_RNA45SN1_45S_50ntFlanks@11561.11602.42\|GCGCCTAGCAGCCGACTTAGAACTGGTGCGGACCAGGGGAAT | >hg38_21_+_8433172_8446622_RNA45SN1_45S_50ntFlanks | 11561 | 11602 | 42 | GCGCCTAGCAGCCGACTTAGAACTGGTGCGGACCAGGGGAAT | 1826.0 | -3.5 | 0.8 | -4.2 | 2.3E-05 | 4.4E-05 | BT20/MB468 | MITO |
| >hg38_21_+_8433172_8446622_RNA45SN1_45S_50ntFlanks@10839.10868.30\|TCCGTAACTTCGGGATAAGGATTGGCTCTA | >hg38_21_+_8433172_8446622_RNA45SN1_45S_50ntFlanks | 10839 | 10868 | 30 | TCCGTAACTTCGGGATAAGGATTGGCTCTA | 437.6 | -4.6 | 1.1 | -4.2 | 3.0E-05 | 4.4E-05 | MB231/MB468 | CYTO |
| >hg38_21_+_8433172_8446622_RNA45SN1_45S_50ntFlanks@11920.11939.20\|ATGAGAGGTGTAGAATAAGT | >hg38_21_+_8433172_8446622_RNA45SN1_45S_50ntFlanks | 11920 | 11939 | 20 | ATGAGAGGTGTAGAATAAGT | 105.6 | 10.1 | 2.2 | 4.6 | 3.4E-06 | 4.4E-05 | BT20/MB231 | MP |
| >hg38_21_+_8433172_8446622_RNA45SN1_45S_50ntFlanks@12463.12487.25\|TTGGATTGTTCACCCACTAATAGGG | >hg38_21_+_8433172_8446622_RNA45SN1_45S_50ntFlanks | 12463 | 12487 | 25 | TTGGATTGTTCACCCACTAATAGGG | 106.4 | -6.0 | 1.4 | -4.2 | 2.7E-05 | 4.5E-05 | MB231/MB468 | MP |
| >hg38_21_+_8433172_8446622_RNA45SN1_45S_50ntFlanks@4760.4781.22\|TCAGATACCGTCGTAGTTCCGA | >hg38_21_+_8433172_8446622_RNA45SN1_45S_50ntFlanks | 4760 | 4781 | 22 | TCAGATACCGTCGTAGTTCCGA | 98.2 | -6.0 | 1.4 | -4.2 | 2.7E-05 | 4.5E-05 | MB231/MB468 | MP |
| >MI0000283\|hsa-mir-203a&WithFlank&14\|+\|104117399\|104117520@71.93.23&[MIMAT0000264&hsa-miR-203a-3p&offsets\|0\|+1;m-79&14\|+\|104117469\|104117490&offsets\|0\|+1]\|GTGAAATGTTTAGGACCACTAGA | >MI0000283\|hsa-mir-203a&WithFlank&14\|+\|104117399\|104117520 | 71 | 93 | 23 | GTGAAATGTTTAGGACCACTAGA | 69.6 | 9.5 | 2.0 | 4.6 | 3.4E-06 | 4.5E-05 | BT20/MB231 | MP |
| >MI0003628\|hsa-mir-615&WithFlank&12\|+\|54033944\|54034051@67.87.21&[MIMAT0003283&hsa-miR-615-3p&offsets\|0\|-1;m-495&12\|+\|54034010\|54034030&offsets\|0\|0]\|TCCGAGCCTGGGTCTCCCTCT | >MI0003628\|hsa-mir-615&WithFlank&12\|+\|54033944\|54034051 | 67 | 87 | 21 | TCCGAGCCTGGGTCTCCCTCT | 290.5 | 5.6 | 1.3 | 4.4 | 1.1E-05 | 4.5E-05 | BT20/MB468 | MP |
| >hg38_21_+_8433172_8446622_RNA45SN1_45S_50ntFlanks@6652.6674.23\|GACTCTTAGCGGTGGATCACTCG | >hg38_21_+_8433172_8446622_RNA45SN1_45S_50ntFlanks | 6652 | 6674 | 23 | GACTCTTAGCGGTGGATCACTCG | 636.5 | 4.4 | 1.0 | 4.4 | 1.1E-05 | 4.5E-05 | BT20/MB468 | MP |
| >hg38_21_+_8433172_8446622_RNA45SN1_45S_50ntFlanks@10420.10456.37\|CTGAGAGATGGGCGAGCGCCGTTCCGAAGGGACGGGC | >hg38_21_+_8433172_8446622_RNA45SN1_45S_50ntFlanks | 10420 | 10456 | 37 | CTGAGAGATGGGCGAGCGCCGTTCCGAAGGGACGGGC | 332.1 | -2.5 | 0.6 | -4.2 | 2.3E-05 | 4.5E-05 | BT20/MB468 | MITO |
| >hg38_21_+_8433172_8446622_RNA45SN1_45S_50ntFlanks@12073.12090.18\|CCGAGGGGCTCTCGCTTC | >hg38_21_+_8433172_8446622_RNA45SN1_45S_50ntFlanks | 12073 | 12090 | 18 | CCGAGGGGCTCTCGCTTC | 281.3 | -5.6 | 1.3 | -4.2 | 2.3E-05 | 4.5E-05 | BT20/MB468 | MITO |
| >MI0000081\|hsa-mir-24-2&WithFlank&19\|-\|13836281\|13836365@56.77.22;MI0000080\|hsa-mir-24-1&WithFlank&9\|+\|95086015\|95086094@50.71.22&[MIMAT0000080&hsa-miR-24-3p&offsets\|0\|0;m-36&19\|-\|13836288\|13836310&offsets\|0\|-1];[MIMAT0000080_1&hsa-miR-24-3p&offsets\|0\|0;m-35&9\|+\|95086064\|95086086&offsets\|0\|-1]\|TGGCTCAGTTCAGCAGGAACAG | >MI0000081\|hsa-mir-24-2&WithFlank&19\|-\|13836281\|13836365 | 56 | 77 | 22 | TGGCTCAGTTCAGCAGGAACAG | 35311.1 | 3.2 | 0.8 | 4.2 | 2.7E-05 | 4.5E-05 | MB231/MB468 | MP |
| >hg38_21_+_8433172_8446622_RNA45SN1_45S_50ntFlanks@7979.8012.34\|ACCTCAGATCAGACGTGGCGACCCGCTGAATTTA | >hg38_21_+_8433172_8446622_RNA45SN1_45S_50ntFlanks | 7979 | 8012 | 34 | ACCTCAGATCAGACGTGGCGACCCGCTGAATTTA | 1012.3 | -2.8 | 0.7 | -4.2 | 2.3E-05 | 4.5E-05 | BT20/MB468 | MITO |
| >hg38_21_+_8433172_8446622_RNA45SN1_45S_50ntFlanks@12045.12068.24\|CTGACCCGGTGAGGCGGGGGGGCG | >hg38_21_+_8433172_8446622_RNA45SN1_45S_50ntFlanks | 12045 | 12068 | 24 | CTGACCCGGTGAGGCGGGGGGGCG | 325.3 | -6.0 | 1.4 | -4.2 | 2.3E-05 | 4.5E-05 | BT20/MB468 | MITO |
| >hg38_21_+_8433172_8446622_RNA45SN1_45S_50ntFlanks@12993.13015.23\|CCCTCGCTGCGATCTATTGAAAG | >hg38_21_+_8433172_8446622_RNA45SN1_45S_50ntFlanks | 12993 | 13015 | 23 | CCCTCGCTGCGATCTATTGAAAG | 350.2 | -4.8 | 1.2 | -4.2 | 3.2E-05 | 4.5E-05 | MB231/MB468 | MITO |
| >hg38_21_+_8433172_8446622_RNA45SN1_45S_50ntFlanks@476.520.45\|TTCGTGATCGATGTGGTGACGTCGTGCTCTCCCGGGCCGGGTCCG | >hg38_21_+_8433172_8446622_RNA45SN1_45S_50ntFlanks | 476 | 520 | 45 | TTCGTGATCGATGTGGTGACGTCGTGCTCTCCCGGGCCGGGTCCG | 317.7 | 6.3 | 1.5 | 4.2 | 2.3E-05 | 4.5E-05 | BT20/MB468 | MITO |
| >hg38_21_+_8433172_8446622_RNA45SN1_45S_50ntFlanks@10795.10812.18\|TGGCATGTTGGAACAATG | >hg38_21_+_8433172_8446622_RNA45SN1_45S_50ntFlanks | 10795 | 10812 | 18 | TGGCATGTTGGAACAATG | 105.5 | 10.1 | 2.2 | 4.6 | 3.4E-06 | 4.5E-05 | BT20/MB231 | MP |
| >hg38_21_+_8433172_8446622_RNA45SN1_45S_50ntFlanks@12933.12953.21\|GCTAAACCATTCGTAGACGAC | >hg38_21_+_8433172_8446622_RNA45SN1_45S_50ntFlanks | 12933 | 12953 | 21 | GCTAAACCATTCGTAGACGAC | 374.7 | -3.1 | 0.7 | -4.2 | 3.2E-05 | 4.5E-05 | MB231/MB468 | NUC |

| | | | | | | | | | | | | | |
|---|---|---|---|---|---|---|---|---|---|---|---|---|---|
| >MI0000086\|hsa-mir-28&WithFlank&3\|+\|188688775\|188688872@60.81.22(+2A)&[MIMAT0004502&hsa-miR-28-3p&offsets\|0\|0(+2A);m-28&3\|+\|188688834\|188688855&offsets\|0\|0(+2A)]\|CACTAGATTGTGAGCTCCTGGAAA | >MI0000086\|hsa-mir-28&WithFlank&3\|+\|188688775\|188688872 | 60 | 81 | 22(+2A) | CACTAGATTGTGAGCTCCTGGAAA | 132.1 | 7.7 | 1.8 | 4.4 | 1.1E-05 | 4.5E-05 | BT20/MB468 | MP |
| >trnaMT_PheGAA_MT_+_577_647@35.62.28\|GAAAATGTTTAGACGGGCTCACATCACC | >trnaMT_PheGAA_MT_+_577_647 | 35 | 62 | 28 | GAAAATGTTTAGACGGGCTCACATCACC | 688.3 | 6.3 | 1.5 | 4.2 | 3.2E-05 | 4.5E-05 | MB231/MB468 | MITO |
| >hg38_21_+_8433172_8446622_RNA45SN1_45S_50ntFlanks@3858.3898.41\|TGGTAATTCTAGAGCTAATACATGCCGACGGGCGCTGACCC | >hg38_21_+_8433172_8446622_RNA45SN1_45S_50ntFlanks | 3858 | 3898 | 41 | TGGTAATTCTAGAGCTAATACATGCCGACGGGCGCTGACCC | 245.8 | -6.5 | 1.6 | -4.2 | 3.2E-05 | 4.5E-05 | MB231/MB468 | NUC |
| >hg38_21_+_8433172_8446622_RNA45SN1_45S_50ntFlanks@3839.3860.22\|TCTCCTACTTGGATAACTGTGG | >hg38_21_+_8433172_8446622_RNA45SN1_45S_50ntFlanks | 3839 | 3860 | 22 | TCTCCTACTTGGATAACTGTGG | 1027.6 | 2.7 | 0.6 | 4.2 | 3.1E-05 | 4.5E-05 | MB231/MB468 | CYTO |
| >hg38_21_+_8433172_8446622_RNA45SN1_45S_50ntFlanks@5472.5518.47\|ACGGCCCTGGCGGAGCGCTGAGAAGACGGTCGAACTTGACTATCTAG | >hg38_21_+_8433172_8446622_RNA45SN1_45S_50ntFlanks | 5472 | 5518 | 47 | ACGGCCCTGGCGGAGCGCTGAGAAGACGGTCGAACTTGACTATCTAG | 422.7 | 4.2 | 1.0 | 4.2 | 3.1E-05 | 4.5E-05 | MB231/MB468 | CYTO |
| >hg38_21_+_8433172_8446622_RNA45SN1_45S_50ntFlanks@6646.6667.22\|TCGTACGACTCTTAGCGGTGGA | >hg38_21_+_8433172_8446622_RNA45SN1_45S_50ntFlanks | 6646 | 6667 | 22 | TCGTACGACTCTTAGCGGTGGA | 3343.4 | 2.5 | 0.6 | 4.1 | 3.4E-05 | 4.5E-05 | BT20/MB468 | NUC |
| >hg38_21_+_8433172_8446622_RNA45SN1_45S_50ntFlanks@10392.10432.41\|ACAGCAGTTGAACATGGGTCAGTCGGTCCTGAGAGATGGGC | >hg38_21_+_8433172_8446622_RNA45SN1_45S_50ntFlanks | 10392 | 10432 | 41 | ACAGCAGTTGAACATGGGTCAGTCGGTCCTGAGAGATGGGC | 280.0 | -5.0 | 1.2 | -4.2 | 3.2E-05 | 4.5E-05 | MB231/MB468 | MITO |
| >hg38_21_+_8433172_8446622_RNA45SN1_45S_50ntFlanks@4731.4755.25\|AAGAACGAAAGTCGGAGGTTCGAAG | >hg38_21_+_8433172_8446622_RNA45SN1_45S_50ntFlanks | 4731 | 4755 | 25 | AAGAACGAAAGTCGGAGGTTCGAAG | 202.8 | -4.8 | 1.1 | -4.2 | 2.7E-05 | 4.5E-05 | MB231/MB468 | MP |
| >hg38_21_+_8433172_8446622_RNA45SN1_45S_50ntFlanks@12633.12651.19\|CGAAGCTACCATCTGTGGG | >hg38_21_+_8433172_8446622_RNA45SN1_45S_50ntFlanks | 12633 | 12651 | 19 | CGAAGCTACCATCTGTGGG | 363.3 | -4.6 | 1.1 | -4.2 | 3.2E-05 | 4.5E-05 | MB231/MB468 | NUC |
| >MI0000812\|hsa-mir-331&WithFlank&12\|+\|95308414\|95308519@32.52.21(+1U)&[MIMAT0004700&hsa-miR-331-5p&offsets\|0\|-1(+1U);m-377&12\|+\|95308445\|95308465&offsets\|0\|0(+1U)]\|CTAGGTATGGTCCCAGGGATCT | >MI0000812\|hsa-mir-331&WithFlank&12\|+\|95308414\|95308519 | 32 | 52 | 21(+1U) | CTAGGTATGGTCCCAGGGATCT | 34.3 | 8.3 | 2.0 | 4.2 | 2.7E-05 | 4.5E-05 | MB231/MB468 | MP |
| >MI0000439\|hsa-mir-23b&WithFlank&9\|+\|95085202\|95085310@67.86.20&[MIMAT0000418&hsa-miR-23b-3p&offsets\|+3\|0;m-42&9\|+\|95085265\|95085285&offsets\|+3\|+2]\|ACATTGCCAGGGATTACCAC | >MI0000439\|hsa-mir-23b&WithFlank&9\|+\|95085202\|95085310 | 67 | 86 | 20 | ACATTGCCAGGGATTACCAC | 148.3 | 10.4 | 2.5 | 4.2 | 2.7E-05 | 4.5E-05 | MB231/MB468 | MP |
| >hg38_21_+_8433172_8446622_RNA45SN1_45S_50ntFlanks@11807.11827.21\|ACTGTCCCTACCTACTATCCA | >hg38_21_+_8433172_8446622_RNA45SN1_45S_50ntFlanks | 11807 | 11827 | 21 | ACTGTCCCTACCTACTATCCA | 78.6 | -10.2 | 2.4 | -4.2 | 2.7E-05 | 4.5E-05 | MB231/MB468 | MP |
| >hg38_21_+_8433172_8446622_RNA45SN1_45S_50ntFlanks@5487.5526.40\|CGCTGAGAAGACGGTCGAACTTGACTATCTAGAGGAAGTA | >hg38_21_+_8433172_8446622_RNA45SN1_45S_50ntFlanks | 5487 | 5526 | 40 | CGCTGAGAAGACGGTCGAACTTGACTATCTAGAGGAAGTA | 708.1 | -3.8 | 0.9 | -4.2 | 3.1E-05 | 4.5E-05 | MB231/MB468 | CYTO |
| >hg38_21_+_8433172_8446622_RNA45SN1_45S_50ntFlanks@10398.10424.27\|GTTGAACATGGGTCAGTCGGTCCTGAG | >hg38_21_+_8433172_8446622_RNA45SN1_45S_50ntFlanks | 10398 | 10424 | 27 | GTTGAACATGGGTCAGTCGGTCCTGAG | 450.5 | 3.5 | 0.8 | 4.2 | 3.3E-05 | 4.5E-05 | MB231/MB468 | MITO |
| >hg38_21_+_8433172_8446622_RNA45SN1_45S_50ntFlanks@10840.10863.24\|CCGTAACTTCGGGATAAGGATTGG | >hg38_21_+_8433172_8446622_RNA45SN1_45S_50ntFlanks | 10840 | 10863 | 24 | CCGTAACTTCGGGATAAGGATTGG | 162.7 | -6.6 | 1.5 | -4.4 | 1.1E-05 | 4.5E-05 | BT20/MB468 | MP |
| >trnaMT_GluTTC_MT_-_14674_14742@40.71.32,trnalookalike8_GluTTC_5_-_93905172_93905240@40.71.32\|TTGGTCGTGGTTGTAGTCCGTGCGAGAATACC | >trnaMT_GluTTC_MT_-_14674_14742 | 40 | 71 | 32 | TTGGTCGTGGTTGTAGTCCGTGCGAGAATACC | 1453.0 | -3.9 | 0.8 | -4.7 | 3.3E-06 | 4.5E-05 | BT20/MB231 | CYTO |
| >hg38_21_+_8433172_8446622_RNA45SN1_45S_50ntFlanks@4588.4621.34\|CTATTTTGTTGGTTTTCGGAACTGAGGCCATGAT | >hg38_21_+_8433172_8446622_RNA45SN1_45S_50ntFlanks | 4588 | 4621 | 34 | CTATTTTGTTGGTTTTCGGAACTGAGGCCATGAT | 275.9 | -2.5 | 0.6 | -4.2 | 2.4E-05 | 4.5E-05 | BT20/MB468 | MITO |
| >hg38_21_+_8433172_8446622_RNA45SN1_45S_50ntFlanks@12476.12493.18\|CCACTAATAGGGAACGTG | >hg38_21_+_8433172_8446622_RNA45SN1_45S_50ntFlanks | 12476 | 12493 | 18 | CCACTAATAGGGAACGTG | 378.7 | -5.7 | 1.4 | -4.2 | 2.8E-05 | 4.5E-05 | BT20/MB468 | CYTO |
| >hg38_21_+_8433172_8446622_RNA45SN1_45S_50ntFlanks@3988.4023.36\|CTTTGGTGACTCTAGATAACCTCGGGCCGATCGCAC | >hg38_21_+_8433172_8446622_RNA45SN1_45S_50ntFlanks | 3988 | 4023 | 36 | CTTTGGTGACTCTAGATAACCTCGGGCCGATCGCAC | 4295.9 | 7.3 | 1.6 | 4.6 | 3.4E-06 | 4.5E-05 | BT20/MB231 | MP |
| >trna27_GlyCCC_2_-_70476123_70476193@4.35.32,trna34_GlyCCC_16_-_686736_686806@4.35.32\|CCGCTGGTGTAGTGGTATCATGCAAGATTCCC | >trna27_GlyCCC_2_-_70476123_70476193 | 4 | 35 | 32 | CCGCTGGTGTAGTGGTATCATGCAAGATTCCC | 119.1 | 7.0 | 1.7 | 4.2 | 3.3E-05 | 4.5E-05 | MB231/MB468 | MITO |
| >hg38_21_+_8433172_8446622_RNA45SN1_45S_50ntFlanks@6648.6668.21\|GTACGACTCTTAGCGGTGGAT | >hg38_21_+_8433172_8446622_RNA45SN1_45S_50ntFlanks | 6648 | 6668 | 21 | GTACGACTCTTAGCGGTGGAT | 859.8 | 3.1 | 0.7 | 4.4 | 1.1E-05 | 4.5E-05 | BT20/MB468 | MP |
| >hg38_21_+_8433172_8446622_RNA45SN1_45S_50ntFlanks@9545.9562.18\|GTGGAGGTCCGTAGCGGT | >hg38_21_+_8433172_8446622_RNA45SN1_45S_50ntFlanks | 9545 | 9562 | 18 | GTGGAGGTCCGTAGCGGT | 269.8 | -4.0 | 0.9 | -4.2 | 2.8E-05 | 4.5E-05 | BT20/MB468 | CYTO |
| >am_MI0003581\|hsa-mir-574&WithFlank&4\|+\|38868026\|38868133@31.53.23&[MIMAT0004795&hsa-miR-574-5p&offsets\|0\|0;m-213&4\|+\|38868056\|38868078&offsets\|0\|0]\|TGAGTGTGTGTGTGTGAGTGTGT | >am_MI0003581\|hsa-mir-574&WithFlank&4\|+\|38868026\|38868133 | 31 | 53 | 23 | TGAGTGTGTGTGTGTGAGTGTGT | 150.6 | 1.8 | 0.4 | 4.1 | 3.4E-05 | 4.5E-05 | BT20/MB468 | NUC |
| >hg38_21_+_8433172_8446622_RNA45SN1_45S_50ntFlanks@6652.6682.31\|GACTCTTAGCGGTGGATCACTCGGCTCGTGC | >hg38_21_+_8433172_8446622_RNA45SN1_45S_50ntFlanks | 6652 | 6682 | 31 | GACTCTTAGCGGTGGATCACTCGGCTCGTGC | 1635.3 | 4.5 | 1.1 | 4.2 | 3.3E-05 | 4.5E-05 | MB231/MB468 | NUC |
| >hg38_21_+_8433172_8446622_RNA45SN1_45S_50ntFlanks@9120.9138.19\|TGCCGCGCGCGGGTCGGGG | >hg38_21_+_8433172_8446622_RNA45SN1_45S_50ntFlanks | 9120 | 9138 | 19 | TGCCGCGCGCGGGTCGGGG | 131.7 | -5.2 | 1.2 | -4.2 | 2.4E-05 | 4.5E-05 | BT20/MB468 | MITO |
| >hg38_21_+_8433172_8446622_RNA45SN1_45S_50ntFlanks@3855.3872.18\|CTGTGGTAATTCTAGAGC | >hg38_21_+_8433172_8446622_RNA45SN1_45S_50ntFlanks | 3855 | 3872 | 18 | CTGTGGTAATTCTAGAGC | 147.9 | -6.3 | 1.4 | -4.4 | 1.1E-05 | 4.5E-05 | BT20/MB468 | MP |
| >hg38_21_+_8433172_8446622_RNA45SN1_45S_50ntFlanks@9496.9524.29\|GAAAGATGGTGAACTATGCCTGGGCAGGG | >hg38_21_+_8433172_8446622_RNA45SN1_45S_50ntFlanks | 9496 | 9524 | 29 | GAAAGATGGTGAACTATGCCTGGGCAGGG | 106.7 | 10.1 | 2.2 | 4.6 | 3.5E-06 | 4.5E-05 | BT20/MB231 | MP |
| >hg38_21_+_8433172_8446622_RNA45SN1_45S_50ntFlanks@11707.11728.22\|AAGCGCGGGTAAACGGCGGGAG | >hg38_21_+_8433172_8446622_RNA45SN1_45S_50ntFlanks | 11707 | 11728 | 22 | AAGCGCGGGTAAACGGCGGGAG | 183.2 | -5.4 | 1.3 | -4.2 | 2.4E-05 | 4.5E-05 | BT20/MB468 | MITO |
| >hg38_21_+_8433172_8446622_RNA45SN1_45S_50ntFlanks@10128.10148.21\|CGGCGGCGGCGGCGGCGGCGG | >hg38_21_+_8433172_8446622_RNA45SN1_45S_50ntFlanks | 10128 | 10148 | 21 | CGGCGGCGGCGGCGGCGGCGG | 216.8 | -6.5 | 1.5 | -4.4 | 1.1E-05 | 4.5E-05 | BT20/MB468 | MP |
| >hg38_21_+_8433172_8446622_RNA45SN1_45S_50ntFlanks@8845.8865.21\|CGGCGGCGGCGGCGGCGGCGG | >hg38_21_+_8433172_8446622_RNA45SN1_45S_50ntFlanks | 8845 | 8865 | 21 | CGGCGGCGGCGGCGGCGGCGG | 216.8 | -6.5 | 1.5 | -4.4 | 1.1E-05 | 4.5E-05 | BT20/MB468 | MP |
| >hg38_21_+_8433172_8446622_RNA45SN1_45S_50ntFlanks@10350.10370.21\|ACGAGAACTTTGAAGGCCGAA | >hg38_21_+_8433172_8446622_RNA45SN1_45S_50ntFlanks | 10350 | 10370 | 21 | ACGAGAACTTTGAAGGCCGAA | 283.7 | -3.0 | 0.7 | -4.2 | 3.3E-05 | 4.5E-05 | MB231/MB468 | MITO |
| >hg38_21_+_8433172_8446622_RNA45SN1_45S_50ntFlanks@12990.13015.26\|GCTCCCTCGCTGCGATCTATTGAAAG | >hg38_21_+_8433172_8446622_RNA45SN1_45S_50ntFlanks | 12990 | 13015 | 26 | GCTCCCTCGCTGCGATCTATTGAAAG | 693.5 | -5.0 | 1.2 | -4.2 | 3.3E-05 | 4.5E-05 | MB231/MB468 | MITO |
| >hg38_21_+_8433172_8446622_RNA45SN1_45S_50ntFlanks@9460.9488.29\|CGGGGAGGTGGAGCACGAGCGCACGTGTT | >hg38_21_+_8433172_8446622_RNA45SN1_45S_50ntFlanks | 9460 | 9488 | 29 | CGGGGAGGTGGAGCACGAGCGCACGTGTT | 392.8 | -4.4 | 0.9 | -4.6 | 3.3E-06 | 4.5E-05 | BT20/MB231 | CYTO |
| >hg38_MT_+_598_1651_RNR1_12S_50ntFlanks@971.1004.34\|TGGTAAGTGTACTGGAAAGTGCACTTGGACGAAC | >hg38_MT_+_598_1651_RNR1_12S_50ntFlanks | 971 | 1004 | 34 | TGGTAAGTGTACTGGAAAGTGCACTTGGACGAAC | 242.3 | 5.0 | 1.2 | 4.2 | 3.3E-05 | 4.5E-05 | MB231/MB468 | MITO |
| >hg38_21_+_8433172_8446622_RNA45SN1_45S_50ntFlanks@11235.11254.20\|CTTCCCGTGGATCGCCCCAG | >hg38_21_+_8433172_8446622_RNA45SN1_45S_50ntFlanks | 11235 | 11254 | 20 | CTTCCCGTGGATCGCCCCAG | 3203.0 | -7.1 | 1.7 | -4.2 | 2.8E-05 | 4.5E-05 | BT20/MB468 | CYTO |
| >hg38_21_+_8433172_8446622_RNA45SN1_45S_50ntFlanks@9500.9525.26\|GATGGTGAACTATGCCTGGGCAGGGC | >hg38_21_+_8433172_8446622_RNA45SN1_45S_50ntFlanks | 9500 | 9525 | 26 | GATGGTGAACTATGCCTGGGCAGGGC | 146.8 | 8.0 | 1.8 | 4.4 | 1.1E-05 | 4.6E-05 | BT20/MB468 | MP |
| >hg38_21_+_8433172_8446622_RNA45SN1_45S_50ntFlanks@12397.12419.23\|CGACGTCGCTTTTTGATCCTTCG | >hg38_21_+_8433172_8446622_RNA45SN1_45S_50ntFlanks | 12397 | 12419 | 23 | CGACGTCGCTTTTTGATCCTTCG | 133.8 | -6.0 | 1.4 | -4.2 | 2.8E-05 | 4.6E-05 | MB231/MB468 | MP |
| >MI0000088\|hsa-mir-30a&WithFlank&6\|-\|71403545\|71403627@12.34.23&[MIMAT0000087&hsa-miR-30a-5p&offsets\|0\|+1;m-15&6\|-\|71403595\|71403616&offsets\|0\|+1]\|TGTAAACATCCTCGACTGGAAGC | >MI0000088\|hsa-mir-30a&WithFlank&6\|-\|71403545\|71403627 | 12 | 34 | 23 | TGTAAACATCCTCGACTGGAAGC | 41762.2 | 3.2 | 0.8 | 4.2 | 2.8E-05 | 4.6E-05 | MB231/MB468 | MP |
| >hg38_21_+_8433172_8446622_RNA45SN1_45S_50ntFlanks@12159.12178.20\|TTTGACTGGGGCGGTACACC | >hg38_21_+_8433172_8446622_RNA45SN1_45S_50ntFlanks | 12159 | 12178 | 20 | TTTGACTGGGGCGGTACACC | 497.1 | -6.2 | 1.5 | -4.2 | 3.3E-05 | 4.6E-05 | MB231/MB468 | NUC |
| >hg38_21_+_8433172_8446622_RNA45SN1_45S_50ntFlanks@9955.9994.40\|GGAAGTCGGAATCCGCTAAGGAGTGTGTAACAACTCACCT | >hg38_21_+_8433172_8446622_RNA45SN1_45S_50ntFlanks | 9955 | 9994 | 40 | GGAAGTCGGAATCCGCTAAGGAGTGTGTAACAACTCACCT | 36.9 | 9.0 | 2.0 | 4.4 | 1.1E-05 | 4.6E-05 | BT20/MB468 | MP |
| >hg38_21_+_8433172_8446622_RNA45SN1_45S_50ntFlanks@9957.9983.27\|AAGTCGGAATCCGCTAAGGAGTGTGTA | >hg38_21_+_8433172_8446622_RNA45SN1_45S_50ntFlanks | 9957 | 9983 | 27 | AAGTCGGAATCCGCTAAGGAGTGTGTA | 35.0 | 8.9 | 2.0 | 4.4 | 1.1E-05 | 4.6E-05 | BT20/MB468 | MP |
| >hg38_21_+_8433172_8446622_RNA45SN1_45S_50ntFlanks@11671.11693.23\|CCAGTGCTCTGAATGTCAAAGTG | >hg38_21_+_8433172_8446622_RNA45SN1_45S_50ntFlanks | 11671 | 11693 | 23 | CCAGTGCTCTGAATGTCAAAGTG | 1019.5 | -3.4 | 0.8 | -4.2 | 3.3E-05 | 4.6E-05 | MB231/MB468 | NUC |
| >am_MI0000809\|hsa-mir-151a&WithFlank&8\|-\|140732558\|140732659@53.72.20(+1C)&[MIMAT0000757&hsa-miR-151a-3p&offsets\|0\|-1(+1C);m-30&8\|-\|140732587\|140732607&offsets\|0\|-1(+1C)]\|CTAGACTGAAGCTCCTTGAGC | >am_MI0000809\|hsa-mir-151a&WithFlank&8\|-\|140732558\|140732659 | 53 | 72 | 20(+1C) | CTAGACTGAAGCTCCTTGAGC | 916.1 | 6.1 | 1.4 | 4.4 | 1.1E-05 | 4.6E-05 | BT20/MB468 | MP |
| >hg38_21_+_8433172_8446622_RNA45SN1_45S_50ntFlanks@12336.12370.35\|CAGGAGGTGTCAGAAAAGTTACCACAGGGATAACT | >hg38_21_+_8433172_8446622_RNA45SN1_45S_50ntFlanks | 12336 | 12370 | 35 | CAGGAGGTGTCAGAAAAGTTACCACAGGGATAACT | 111.2 | 10.1 | 2.2 | 4.6 | 3.5E-06 | 4.6E-05 | BT20/MB231 | MP |
| >hg38_21_+_8433172_8446622_RNA45SN1_45S_50ntFlanks@8107.8138.32\|CGGCGGGGCGCGGGACATGTGGCGTACGGAAG | >hg38_21_+_8433172_8446622_RNA45SN1_45S_50ntFlanks | 8107 | 8138 | 32 | CGGCGGGGCGCGGGACATGTGGCGTACGGAAG | 106.9 | 10.1 | 2.2 | 4.6 | 3.5E-06 | 4.6E-05 | BT20/MB231 | MP |
| >hg38_21_+_8433172_8446622_RNA45SN1_45S_50ntFlanks@9539.9563.25\|ACTCTGGTGGAGGTCCGTAGCGGTC | >hg38_21_+_8433172_8446622_RNA45SN1_45S_50ntFlanks | 9539 | 9563 | 25 | ACTCTGGTGGAGGTCCGTAGCGGTC | 266.8 | -4.6 | 1.1 | -4.2 | 2.9E-05 | 4.6E-05 | BT20/MB468 | CYTO |
| >hg38_21_+_8433172_8446622_RNA45SN1_45S_50ntFlanks@11321.11343.23\|CACGTCTCGTCGCGCGCGCGTCC | >hg38_21_+_8433172_8446622_RNA45SN1_45S_50ntFlanks | 11321 | 11343 | 23 | CACGTCTCGTCGCGCGCGCGTCC | 2061.2 | -5.6 | 1.4 | -4.2 | 3.3E-05 | 4.6E-05 | MB231/MB468 | NUC |
| >hg38_21_+_8433172_8446622_RNA45SN1_45S_50ntFlanks@10869.10888.20\|AGGGCTGGGTCGGTCGGGCT | >hg38_21_+_8433172_8446622_RNA45SN1_45S_50ntFlanks | 10869 | 10888 | 20 | AGGGCTGGGTCGGTCGGGCT | 136.2 | -6.2 | 1.5 | -4.2 | 2.8E-05 | 4.6E-05 | MB231/MB468 | MP |
| >hg38_21_+_8433172_8446622_RNA45SN1_45S_50ntFlanks@12957.12990.34\|CTTCTGGGTCGGGGTTTCGTACGTAGCAGAGCAG | >hg38_21_+_8433172_8446622_RNA45SN1_45S_50ntFlanks | 12957 | 12990 | 34 | CTTCTGGGTCGGGGTTTCGTACGTAGCAGAGCAG | 391.6 | -5.7 | 1.4 | -4.2 | 2.9E-05 | 4.6E-05 | BT20/MB468 | CYTO |
| >hg38_21_+_8433172_8446622_RNA45SN1_45S_50ntFlanks@12078.12095.18\|GGGCTCTCGCTTCTGGCG | >hg38_21_+_8433172_8446622_RNA45SN1_45S_50ntFlanks | 12078 | 12095 | 18 | GGGCTCTCGCTTCTGGCG | 628.9 | -4.6 | 1.1 | -4.2 | 3.1E-05 | 4.6E-05 | MB231/MB468 | CYTO |
| >hg38_21_+_8433172_8446622_RNA45SN1_45S_50ntFlanks@13002.13027.26\|CGATCTATTGAAAGTCAGCCCTCGAC | >hg38_21_+_8433172_8446622_RNA45SN1_45S_50ntFlanks | 13002 | 13027 | 26 | CGATCTATTGAAAGTCAGCCCTCGAC | 13238.7 | -7.0 | 1.7 | -4.2 | 3.1E-05 | 4.6E-05 | MB231/MB468 | CYTO |

| | | | | | | | | | | | | | |
|---|---|---|---|---|---|---|---|---|---|---|---|---|---|
| >hg38_21_+_8433172_8446622_RNA45SN1_45S_50ntFlanks@11845.11886.42\|GGAACGGGCTTGGCGGAATCAGCGGGGAAAGAAGACCCTGTT | >hg38_21_+_8433172_8446622_RNA45SN1_45S_50ntFlanks | 11845 | 11886 | 42 | GGAACGGGCTTGGCGGAATCAGCGGGGAAAGAAGACCCTGTT | 92.0 | 9.9 | 2.1 | 4.6 | 3.5E-06 | 4.6E-05 | BT20/MB231 | MP |
| >hg38_21_+_8433172_8446622_RNA45SN1_45S_50ntFlanks@10061.10078.18\|CAGTCGAGAGTGGACGGG | >hg38_21_+_8433172_8446622_RNA45SN1_45S_50ntFlanks | 10061 | 10078 | 18 | CAGTCGAGAGTGGACGGG | 162.0 | -5.4 | 1.3 | -4.2 | 2.8E-05 | 4.6E-05 | MB231/MB468 | MP |
| >am_trnaMT_ValTAC_MT_+_1602_1670@48.67.20\|CAACTTAACTTGACCGCTCT | >am_trnaMT_ValTAC_MT_+_1602_1670 | 48 | 67 | 20 | CAACTTAACTTGACCGCTCT | 732.0 | 3.5 | 0.8 | 4.2 | 3.3E-05 | 4.6E-05 | MB231/MB468 | MITO |
| >hg38_21_+_8433172_8446622_RNA45SN1_45S_50ntFlanks@475.511.37\|CTTCGTGATCGATGTGGTGACGTCGTGCTCTCCCGGG | >hg38_21_+_8433172_8446622_RNA45SN1_45S_50ntFlanks | 475 | 511 | 37 | CTTCGTGATCGATGTGGTGACGTCGTGCTCTCCCGGG | 6791.4 | 3.2 | 0.8 | 4.2 | 3.3E-05 | 4.6E-05 | MB231/MB468 | NUC |
| >hg38_21_+_8433172_8446622_RNA45SN1_45S_50ntFlanks@6650.6667.18\|ACGACTCTTAGCGGTGGA | >hg38_21_+_8433172_8446622_RNA45SN1_45S_50ntFlanks | 6650 | 6667 | 18 | ACGACTCTTAGCGGTGGA | 115.0 | 3.4 | 0.8 | 4.1 | 3.5E-05 | 4.6E-05 | BT20/MB468 | NUC |
| >hg38_21_+_8433172_8446622_RNA45SN1_45S_50ntFlanks@3874.3899.26\|AATACATGCCGACGGGCGCTGACCCC | >hg38_21_+_8433172_8446622_RNA45SN1_45S_50ntFlanks | 3874 | 3899 | 26 | AATACATGCCGACGGGCGCTGACCCC | 683.6 | -5.4 | 1.3 | -4.2 | 3.3E-05 | 4.6E-05 | MB231/MB468 | NUC |
| >hg38_21_+_8433172_8446622_RNA45SN1_45S_50ntFlanks@11323.11342.20\|CGTCTCGTCGCGCGCGCGTC | >hg38_21_+_8433172_8446622_RNA45SN1_45S_50ntFlanks | 11323 | 11342 | 20 | CGTCTCGTCGCGCGCGCGTC | 3639.6 | -5.9 | 1.4 | -4.4 | 1.1E-05 | 4.6E-05 | BT20/MB468 | MP |
| >hg38_MT_+_1621_3279_RNR2_16S_50ntFlanks@623.652.30\|ATAACTGAACTCCTCACACCCAATTGGACC | >hg38_MT_+_1621_3279_RNR2_16S_50ntFlanks | 623 | 652 | 30 | ATAACTGAACTCCTCACACCCAATTGGACC | 291.5 | 8.3 | 1.9 | 4.4 | 1.1E-05 | 4.6E-05 | BT20/MB468 | MP |
| >MI0000253\|hsa-mir-148a&WithFlank&7\|-\|25949913\|25949992@50.70.21&[MIMAT0000243&hsa-miR-148a-3p&offsets\|0\|-1;m-7&7\|-\|25949922\|25949943&offsets\|0\|-1]\|TCAGTGCACTACAGAACTTTG | >MI0000253\|hsa-mir-148a&WithFlank&7\|-\|25949913\|25949992 | 50 | 70 | 21 | TCAGTGCACTACAGAACTTTG | 115.7 | 4.8 | 1.2 | 4.1 | 3.5E-05 | 4.6E-05 | BT20/MB468 | NUC |
| >hg38_21_+_8433172_8446622_RNA45SN1_45S_50ntFlanks@10796.10818.23\|GGCATGTTGGAACAATGTAGGTA | >hg38_21_+_8433172_8446622_RNA45SN1_45S_50ntFlanks | 10796 | 10818 | 23 | GGCATGTTGGAACAATGTAGGTA | 530.6 | -7.1 | 1.7 | -4.1 | 3.5E-05 | 4.6E-05 | BT20/MB468 | NUC |
| >hg38_21_+_8433172_8446622_RNA45SN1_45S_50ntFlanks@7975.8014.40\|CGCGACCTCAGATCAGACGTGGCGACCCGCTGAATTTAAG | >hg38_21_+_8433172_8446622_RNA45SN1_45S_50ntFlanks | 7975 | 8014 | 40 | CGCGACCTCAGATCAGACGTGGCGACCCGCTGAATTTAAG | 8477.1 | 3.1 | 0.7 | 4.2 | 3.3E-05 | 4.6E-05 | MB231/MB468 | NUC |
| >MI0003599\|hsa-mir-589&WithFlank&7\|-\|5495813\|5495923@30.50.21&[MIMAT0004799&hsa-miR-589-5p&offsets\|0\|-1;m-162&7\|-\|5495874\|5495894&offsets\|0\|0]\|TGAGAACCACGTCTGCTCTGA | >MI0003599\|hsa-mir-589&WithFlank&7\|-\|5495813\|5495923 | 30 | 50 | 21 | TGAGAACCACGTCTGCTCTGA | 398.2 | 4.9 | 1.2 | 4.2 | 2.8E-05 | 4.6E-05 | MB231/MB468 | MP |
| >MI0000446\|hsa-mir-125b-1&WithFlank&11\|-\|122099751\|122099850@61.81.21&[MIMAT0004592&hsa-miR-125b-1-3p&offsets\|0\|-1;m-485&11\|-\|122099769\|122099790&offsets\|0\|-1]\|ACGGGTTAGGCTCTTGGGAGC | >MI0000446\|hsa-mir-125b-1&WithFlank&11\|-\|122099751\|122099850 | 61 | 81 | 21 | ACGGGTTAGGCTCTTGGGAGC | 143.6 | -5.2 | 1.1 | -4.6 | 4.2E-06 | 4.6E-05 | BT20/MB231 | MITO |
| >hg38_21_+_8433172_8446622_RNA45SN1_45S_50ntFlanks@11677.11702.26\|CTCTGAATGTCAAAGTGAAGAAATTC | >hg38_21_+_8433172_8446622_RNA45SN1_45S_50ntFlanks | 11677 | 11702 | 26 | CTCTGAATGTCAAAGTGAAGAAATTC | 533.3 | -5.1 | 1.2 | -4.4 | 1.1E-05 | 4.6E-05 | BT20/MB468 | MP |
| >hg38_21_+_8433172_8446622_RNA45SN1_45S_50ntFlanks@4595.4616.22\|GTTGGTTTTCGGAACTGAGGCC | >hg38_21_+_8433172_8446622_RNA45SN1_45S_50ntFlanks | 4595 | 4616 | 22 | GTTGGTTTTCGGAACTGAGGCC | 156.7 | -4.9 | 1.2 | -4.2 | 2.8E-05 | 4.6E-05 | MB231/MB468 | MP |
| >hg38_MT_+_598_1651_RNR1_12S_50ntFlanks@440.470.31\|ACATATCTGAACACACAATAGCTAAGACCCA | >hg38_MT_+_598_1651_RNR1_12S_50ntFlanks | 440 | 470 | 31 | ACATATCTGAACACACAATAGCTAAGACCCA | 31.7 | 8.2 | 2.0 | 4.2 | 2.8E-05 | 4.6E-05 | MB231/MB468 | MP |
| >hg38_21_+_8433172_8446622_RNA45SN1_45S_50ntFlanks@11318.11342.25\|CCCCACGTCTCGTCGCGCGCGCGTC | >hg38_21_+_8433172_8446622_RNA45SN1_45S_50ntFlanks | 11318 | 11342 | 25 | CCCCACGTCTCGTCGCGCGCGCGTC | 6029.3 | -8.3 | 2.0 | -4.1 | 3.5E-05 | 4.6E-05 | BT20/MB468 | NUC |
| >hg38_21_+_8433172_8446622_RNA45SN1_45S_50ntFlanks@11598.11627.30\|GGAATCCGACTGTTTAATTAAAACAAAGCA | >hg38_21_+_8433172_8446622_RNA45SN1_45S_50ntFlanks | 11598 | 11627 | 30 | GGAATCCGACTGTTTAATTAAAACAAAGCA | 104.7 | 10.0 | 2.2 | 4.6 | 3.6E-06 | 4.6E-05 | BT20/MB231 | MP |
| >hg38_21_+_8433172_8446622_RNA45SN1_45S_50ntFlanks@5125.5158.34\|AGCGGTCGGCGTCCCCCAACTTCTTAGAGGGACA | >hg38_21_+_8433172_8446622_RNA45SN1_45S_50ntFlanks | 5125 | 5158 | 34 | AGCGGTCGGCGTCCCCCAACTTCTTAGAGGGACA | 113.0 | 10.2 | 2.2 | 4.6 | 3.6E-06 | 4.6E-05 | BT20/MB231 | MP |
| >hg38_21_+_8433172_8446622_RNA45SN1_45S_50ntFlanks@12419.12459.41\|GATGTCGGCTCTTCCTATCATTGTGAAGCAGAATTCACCAA | >hg38_21_+_8433172_8446622_RNA45SN1_45S_50ntFlanks | 12419 | 12459 | 41 | GATGTCGGCTCTTCCTATCATTGTGAAGCAGAATTCACCAA | 371.2 | -3.4 | 0.7 | -4.6 | 3.4E-06 | 4.6E-05 | BT20/MB231 | CYTO |
| >hg38_21_+_8433172_8446622_RNA45SN1_45S_50ntFlanks@6652.6700.49\|GACTCTTAGCGGTGGATCACTCGGCTCGTGCGTCGATGAAGAACGCAGC | >hg38_21_+_8433172_8446622_RNA45SN1_45S_50ntFlanks | 6652 | 6700 | 49 | GACTCTTAGCGGTGGATCACTCGGCTCGTGCGTCGATGAAGAACGCAGC | 1611.7 | 3.8 | 0.9 | 4.1 | 3.4E-05 | 4.6E-05 | MB231/MB468 | MITO |
| >hg38_21_+_8433172_8446622_RNA45SN1_45S_50ntFlanks@6779.6803.25\|CCGGGGCTACGCCTGTCTGAGCGTC | >hg38_21_+_8433172_8446622_RNA45SN1_45S_50ntFlanks | 6779 | 6803 | 25 | CCGGGGCTACGCCTGTCTGAGCGTC | 521.6 | -5.4 | 1.3 | -4.1 | 3.5E-05 | 4.6E-05 | BT20/MB468 | NUC |
| >am_MI0000786\|hsa-mir-378a&WithFlank&5\|+\|149732819\|149732896@49.70.22&[MIMAT0000732&hsa-miR-378a-3p&offsets\|0\|0;m-44&5\|+\|149732867\|149732888&offsets\|0\|0]\|ACTGGACTTGGAGTCAGAAGGC | >am_MI0000786\|hsa-mir-378a&WithFlank&5\|+\|149732819\|149732896 | 49 | 70 | 22 | ACTGGACTTGGAGTCAGAAGGC | 9161.2 | 3.5 | 0.8 | 4.2 | 3.2E-05 | 4.6E-05 | MB231/MB468 | CYTO |
| >MI0000767\|hsa-mir-365a&WithFlank&16\|+\|14309279\|14309377@62.81.20(+2U);MI0000769\|hsa-mir-365b&WithFlank&17\|+\|31575405\|31575527@74.93.20(+2U)&[MIMAT0000710&hsa-miR-365a-3p&offsets\|0\|-2(+2U);m-176&16\|+\|14309340\|14309361&offsets\|0\|-2(+2U)];[MIMAT0022834&hsa-miR-365b-3p&offsets\|0\|-2(+2U);m-175&17\|+\|31575478\|31575499&offsets\|0\|-2(+2U)]\|TAATGCCCCTAAAAATCCTTTT | >MI0000767\|hsa-mir-365a&WithFlank&16\|+\|14309279\|14309377 | 62 | 81 | 20(+2U) | TAATGCCCCTAAAAATCCTTTT | 510.5 | 3.9 | 0.8 | 4.6 | 3.4E-06 | 4.6E-05 | BT20/MB231 | CYTO |
| >MI0000085\|hsa-mir-27a&WithFlank&19\|-\|13836434\|13836523@57.76.20(+2U)&[MIMAT0000084&hsa-miR-27a-3p&offsets\|0\|-1(+2U);m-43&19\|-\|13836447\|13836467&offsets\|0\|-1(+2U)]\|TTCACAGTGGCTAAGTTCCGTT | >MI0000085\|hsa-mir-27a&WithFlank&19\|-\|13836434\|13836523 | 57 | 76 | 20(+2U) | TTCACAGTGGCTAAGTTCCGTT | 2798.0 | 3.9 | 1.0 | 4.1 | 3.4E-05 | 4.7E-05 | MB231/MB468 | MITO |
| >hg38_21_+_8433172_8446622_RNA45SN1_45S_50ntFlanks@8104.8138.35\|CCGCGGCGGGGCGCGGGACATGTGGCGTACGGAAG | >hg38_21_+_8433172_8446622_RNA45SN1_45S_50ntFlanks | 8104 | 8138 | 35 | CCGCGGCGGGGCGCGGGACATGTGGCGTACGGAAG | 315.3 | -4.7 | 1.1 | -4.1 | 3.4E-05 | 4.7E-05 | MB231/MB468 | NUC |
| >hg38_21_+_8433172_8446622_RNA45SN1_45S_50ntFlanks@12973.12990.18\|TCGTACGTAGCAGAGCAG | >hg38_21_+_8433172_8446622_RNA45SN1_45S_50ntFlanks | 12973 | 12990 | 18 | TCGTACGTAGCAGAGCAG | 5340.4 | -7.1 | 1.7 | -4.2 | 2.9E-05 | 4.7E-05 | BT20/MB468 | CYTO |
| >am_MI0000809\|hsa-mir-151a&WithFlank&8\|-\|140732558\|140732659@53.73.21(+4U)&[MIMAT0000757&hsa-miR-151a-3p&offsets\|0\|0(+4U);m-30&8\|-\|140732587\|140732607&offsets\|0\|0(+4U)]\|CTAGACTGAAGCTCCTTGAGGTTTT | >am_MI0000809\|hsa-mir-151a&WithFlank&8\|-\|140732558\|140732659 | 53 | 73 | 21(+4U) | CTAGACTGAAGCTCCTTGAGGTTTT | 31.2 | 8.2 | 2.0 | 4.2 | 2.8E-05 | 4.7E-05 | MB231/MB468 | MP |
| >hg38_21_+_8433172_8446622_RNA45SN1_45S_50ntFlanks@10348.10368.21\|AAACGAGAACTTTGAAGGCCG | >hg38_21_+_8433172_8446622_RNA45SN1_45S_50ntFlanks | 10348 | 10368 | 21 | AAACGAGAACTTTGAAGGCCG | 1388.4 | -3.3 | 0.8 | -4.1 | 3.4E-05 | 4.7E-05 | MB231/MB468 | NUC |
| >hg38_21_+_8433172_8446622_RNA45SN1_45S_50ntFlanks@5445.5467.23\|TCGGATCGGCCCCGCCGGGGTCG | >hg38_21_+_8433172_8446622_RNA45SN1_45S_50ntFlanks | 5445 | 5467 | 23 | TCGGATCGGCCCCGCCGGGGTCG | 167.9 | -4.3 | 1.0 | -4.1 | 3.4E-05 | 4.7E-05 | MB231/MB468 | NUC |
| >hg38_21_+_8433172_8446622_RNA45SN1_45S_50ntFlanks@12935.12966.32\|TAAACCATTCGTAGACGACCTGCTTCTGGGTC | >hg38_21_+_8433172_8446622_RNA45SN1_45S_50ntFlanks | 12935 | 12966 | 32 | TAAACCATTCGTAGACGACCTGCTTCTGGGTC | 44.6 | -8.6 | 2.0 | -4.4 | 1.1E-05 | 4.7E-05 | BT20/MB468 | MP |
| >hg38_21_+_8433172_8446622_RNA45SN1_45S_50ntFlanks@10907.10930.24\|GGGCGCGCGCCGCGGCTGGACGAG | >hg38_21_+_8433172_8446622_RNA45SN1_45S_50ntFlanks | 10907 | 10930 | 24 | GGGCGCGCGCCGCGGCTGGACGAG | 3285.9 | 5.7 | 1.2 | 4.6 | 4.2E-06 | 4.7E-05 | BT20/MB231 | MITO |
| >hg38_21_+_8433172_8446622_RNA45SN1_45S_50ntFlanks@11567.11602.36\|AGCAGCCGACTTAGAACTGGTGCGGACCAGGGGAAT | >hg38_21_+_8433172_8446622_RNA45SN1_45S_50ntFlanks | 11567 | 11602 | 36 | AGCAGCCGACTTAGAACTGGTGCGGACCAGGGGAAT | 350.2 | -3.3 | 0.8 | -4.2 | 2.8E-05 | 4.7E-05 | MB231/MB468 | MP |
| >hg38_21_+_8433172_8446622_RNA45SN1_45S_50ntFlanks@9004.9025.22\|TCTCCCCCGCGGGGGCGCGCCG | >hg38_21_+_8433172_8446622_RNA45SN1_45S_50ntFlanks | 9004 | 9025 | 22 | TCTCCCCCGCGGGGGCGCGCCG | 123.6 | -5.3 | 1.3 | -4.2 | 2.5E-05 | 4.7E-05 | BT20/MB468 | MITO |
| >hg38_21_+_8433172_8446622_RNA45SN1_45S_50ntFlanks@4894.4918.25\|ACTTAAAGGAATTGACGGAAGGGCA | >hg38_21_+_8433172_8446622_RNA45SN1_45S_50ntFlanks | 4894 | 4918 | 25 | ACTTAAAGGAATTGACGGAAGGGCA | 49.3 | -8.7 | 2.0 | -4.4 | 1.1E-05 | 4.7E-05 | BT20/MB468 | MP |
| >MI0000811\|hsa-mir-148b&WithFlank&12\|+\|54337210\|54337320@69.89.21(+1A)&[MIMAT0000759&hsa-miR-148b-3p&offsets\|0\|-1(+1A);m-77&12\|+\|54337278\|54337299&offsets\|0\|-1(+1A)]\|TCAGTGCATCACAGAACTTTGA | >MI0000811\|hsa-mir-148b&WithFlank&12\|+\|54337210\|54337320 | 69 | 89 | 21(+1A) | TCAGTGCATCACAGAACTTTGA | 413.8 | 7.8 | 1.9 | 4.2 | 2.9E-05 | 4.7E-05 | BT20/MB468 | CYTO |
| >hg38_21_+_8433172_8446622_RNA45SN1_45S_50ntFlanks@11682.11699.18\|AATGTCAAAGTGAAGAAA | >hg38_21_+_8433172_8446622_RNA45SN1_45S_50ntFlanks | 11682 | 11699 | 18 | AATGTCAAAGTGAAGAAA | 43.8 | -8.5 | 1.9 | -4.4 | 1.1E-05 | 4.7E-05 | BT20/MB468 | MP |
| >hg38_21_+_8433172_8446622_RNA45SN1_45S_50ntFlanks@5472.5499.28\|ACGGCCCTGGCGGAGCGCTGAGAAGACG | >hg38_21_+_8433172_8446622_RNA45SN1_45S_50ntFlanks | 5472 | 5499 | 28 | ACGGCCCTGGCGGAGCGCTGAGAAGACG | 2344.8 | -3.9 | 0.9 | -4.2 | 2.8E-05 | 4.7E-05 | MB231/MB468 | MP |
| >MI0000542\|hsa-mir-320a&WithFlank&8\|-\|22244960\|22245043@49.70.22&[MIMAT0000510&hsa-miR-320a-3p&offsets\|+1\|+1;m-60&8\|-\|22244975\|22244996&offsets\|+1\|+1]\|AAAGCTGGGTTGAGAGGGCGAA | >MI0000542\|hsa-mir-320a&WithFlank&8\|-\|22244960\|22245043 | 49 | 70 | 22 | AAAGCTGGGTTGAGAGGGCGAA | 958.1 | 3.7 | 0.8 | 4.6 | 3.5E-06 | 4.7E-05 | BT20/MB231 | CYTO |
| >hg38_21_+_8433172_8446622_RNA45SN1_45S_50ntFlanks@7983.8011.29\|CAGATCAGACGTGGCGACCCGCTGAATTT | >hg38_21_+_8433172_8446622_RNA45SN1_45S_50ntFlanks | 7983 | 8011 | 29 | CAGATCAGACGTGGCGACCCGCTGAATTT | 387.1 | -3.5 | 0.8 | -4.1 | 3.4E-05 | 4.7E-05 | MB231/MB468 | MITO |
| >hg38_21_+_8433172_8446622_RNA45SN1_45S_50ntFlanks@6722.6744.23\|ATTGCAGGACACATTGATCATCG | >hg38_21_+_8433172_8446622_RNA45SN1_45S_50ntFlanks | 6722 | 6744 | 23 | ATTGCAGGACACATTGATCATCG | 300.5 | -3.0 | 0.7 | -4.2 | 2.5E-05 | 4.7E-05 | BT20/MB468 | MITO |
| >hg38_21_+_8433172_8446622_RNA45SN1_45S_50ntFlanks@5477.5501.25\|CCTGGCGGAGCGCTGAGAAGACGGT | >hg38_21_+_8433172_8446622_RNA45SN1_45S_50ntFlanks | 5477 | 5501 | 25 | CCTGGCGGAGCGCTGAGAAGACGGT | 97.3 | 9.9 | 2.1 | 4.6 | 3.6E-06 | 4.7E-05 | BT20/MB231 | MP |
| >hg38_21_+_8433172_8446622_RNA45SN1_45S_50ntFlanks@8113.8136.24\|GGCGCGGGACATGTGGCGTACGGA | >hg38_21_+_8433172_8446622_RNA45SN1_45S_50ntFlanks | 8113 | 8136 | 24 | GGCGCGGGACATGTGGCGTACGGA | 1370.2 | 8.3 | 1.8 | 4.6 | 3.6E-06 | 4.7E-05 | BT20/MB231 | MP |
| >hg38_21_+_8433172_8446622_RNA45SN1_45S_50ntFlanks@4668.4695.28\|ATTCTTGGACCGGCGCAAGACGGACCAG | >hg38_21_+_8433172_8446622_RNA45SN1_45S_50ntFlanks | 4668 | 4695 | 28 | ATTCTTGGACCGGCGCAAGACGGACCAG | 43.4 | -8.5 | 1.9 | -4.4 | 1.1E-05 | 4.7E-05 | BT20/MB468 | MP |
| >hg38_21_+_8433172_8446622_RNA45SN1_45S_50ntFlanks@8339.8378.40\|TACCGTAAGGGAAAGTTGAAAAGAACTTTGAAGAGAGAGT | >hg38_21_+_8433172_8446622_RNA45SN1_45S_50ntFlanks | 8339 | 8378 | 40 | TACCGTAAGGGAAAGTTGAAAAGAACTTTGAAGAGAGAGT | 109.1 | 7.6 | 1.7 | 4.4 | 1.1E-05 | 4.7E-05 | BT20/MB468 | MP |
| >hg38_21_+_8433172_8446622_RNA45SN1_45S_50ntFlanks@12243.12275.33\|GAAGGGCAAAAGCTCGCTTGATCTTGATTTTCA | >hg38_21_+_8433172_8446622_RNA45SN1_45S_50ntFlanks | 12243 | 12275 | 33 | GAAGGGCAAAAGCTCGCTTGATCTTGATTTTCA | 348.8 | 3.7 | 0.9 | 4.1 | 3.4E-05 | 4.7E-05 | MB231/MB468 | MITO |
| >hg38_21_+_8433172_8446622_RNA45SN1_45S_50ntFlanks@9536.9563.28\|GAAACTCTGGTGGAGGTCCGTAGCGGTC | >hg38_21_+_8433172_8446622_RNA45SN1_45S_50ntFlanks | 9536 | 9563 | 28 | GAAACTCTGGTGGAGGTCCGTAGCGGTC | 274.5 | -2.9 | 0.7 | -4.2 | 3.2E-05 | 4.7E-05 | MB231/MB468 | CYTO |
| >hg38_21_+_8433172_8446622_RNA45SN1_45S_50ntFlanks@12712.12752.41\|CGGAGCCTCGGTTGGCCTCGGATAGCCGGTCCCCCGCCTGT | >hg38_21_+_8433172_8446622_RNA45SN1_45S_50ntFlanks | 12712 | 12752 | 41 | CGGAGCCTCGGTTGGCCTCGGATAGCCGGTCCCCCGCCTGT | 62.6 | -9.8 | 2.3 | -4.2 | 2.8E-05 | 4.7E-05 | MB231/MB468 | MP |
| >hg38_21_+_8433172_8446622_RNA45SN1_45S_50ntFlanks@10410.10453.44\|TCAGTCGGTCCTGAGAGATGGGCGAGCGCCGTTCCGAAGGGACG | >hg38_21_+_8433172_8446622_RNA45SN1_45S_50ntFlanks | 10410 | 10453 | 44 | TCAGTCGGTCCTGAGAGATGGGCGAGCGCCGTTCCGAAGGGACG | 207.3 | -5.4 | 1.3 | -4.2 | 2.9E-05 | 4.7E-05 | MB231/MB468 | MP |

| | | | | | | | | | | | | | |
|---|---|---|---|---|---|---|---|---|---|---|---|---|---|
| >hg38_21_+_8433172_8446622_RNA45SN1_45S_50ntFlanks@7984.8017.34\|AGATCAGACGTGGCGACCCGCTGAATTTAAGCAT | >hg38_21_+_8433172_8446622_RNA45SN1_45S_50ntFlanks | 7984 | 8017 | 34 | AGATCAGACGTGGCGACCCGCTGAATTTAAGCAT | 131.8 | -6.2 | 1.5 | -4.2 | 2.9E-05 | 4.7E-05 | MB231/MB468 | MP |
| >hg38_21_+_8433172_8446622_RNA45SN1_45S_50ntFlanks@4661.4689.29\|AGGTGAAATTCTTGGACCGGCGCAAGACG | >hg38_21_+_8433172_8446622_RNA45SN1_45S_50ntFlanks | 4661 | 4689 | 29 | AGGTGAAATTCTTGGACCGGCGCAAGACG | 238.6 | -2.5 | 0.6 | -4.2 | 2.5E-05 | 4.7E-05 | BT20/MB468 | MITO |
| >hg38_21_+_8433172_8446622_RNA45SN1_45S_50ntFlanks@11583.11623.41\|CTGGTGCGGACCAGGGGAATCCGACTGTTTAATTAAAACAA | >hg38_21_+_8433172_8446622_RNA45SN1_45S_50ntFlanks | 11583 | 11623 | 41 | CTGGTGCGGACCAGGGGAATCCGACTGTTTAATTAAAACAA | 1550.0 | 4.5 | 1.0 | 4.6 | 4.3E-06 | 4.7E-05 | BT20/MB231 | MITO |
| >trna15_ValAAC_5_-_180615416_180615488@1.24.24\|GTTTCCGTAGTGTAGTGGTCATCA | >trna15_ValAAC_5_-_180615416_180615488 | 1 | 24 | 24 | GTTTCCGTAGTGTAGTGGTCATCA | 79.8 | -10.2 | 2.4 | -4.2 | 2.9E-05 | 4.7E-05 | MB231/MB468 | MP |
| >hg38_21_+_8433172_8446622_RNA45SN1_45S_50ntFlanks@11876.11899.24\|AAGACCCTGTTGAGCTTGACTCTA | >hg38_21_+_8433172_8446622_RNA45SN1_45S_50ntFlanks | 11876 | 11899 | 24 | AAGACCCTGTTGAGCTTGACTCTA | 383.2 | -2.3 | 0.6 | -4.2 | 2.9E-05 | 4.7E-05 | BT20/MB468 | CYTO |
| >hg38_21_+_8433172_8446622_RNA45SN1_45S_50ntFlanks@12072.12106.35\|CCCGAGGGGCTCTCGCTTCTGGCGCCAAGCGCCCG | >hg38_21_+_8433172_8446622_RNA45SN1_45S_50ntFlanks | 12072 | 12106 | 35 | CCCGAGGGGCTCTCGCTTCTGGCGCCAAGCGCCCG | 171.6 | -5.5 | 1.3 | -4.2 | 2.9E-05 | 4.7E-05 | BT20/MB468 | CYTO |
| >hg38_21_+_8433172_8446622_RNA45SN1_45S_50ntFlanks@12223.12247.25\|CAGAAACCTCCCGTGGAGCAGAAGG | >hg38_21_+_8433172_8446622_RNA45SN1_45S_50ntFlanks | 12223 | 12247 | 25 | CAGAAACCTCCCGTGGAGCAGAAGG | 61.9 | -9.8 | 2.3 | -4.2 | 2.9E-05 | 4.7E-05 | MB231/MB468 | MP |
| >hg38_MT_+_1621_3279_RNR2_16S_50ntFlanks@962.992.31\|ACCGTGCAAAGGTAGCATAATCACTTGTTCC | >hg38_MT_+_1621_3279_RNR2_16S_50ntFlanks | 962 | 992 | 31 | ACCGTGCAAAGGTAGCATAATCACTTGTTCC | 28.0 | 8.0 | 1.9 | 4.2 | 2.9E-05 | 4.7E-05 | MB231/MB468 | MP |
| >hg38_21_+_8433172_8446622_RNA45SN1_45S_50ntFlanks@8356.8389.34\|GAAAAGAACTTTGAAGAGAGAGTTCAAGAGGGCG | >hg38_21_+_8433172_8446622_RNA45SN1_45S_50ntFlanks | 8356 | 8389 | 34 | GAAAAGAACTTTGAAGAGAGAGTTCAAGAGGGCG | 109.4 | -6.1 | 1.5 | -4.2 | 2.9E-05 | 4.7E-05 | MB231/MB468 | MP |
| >MI0000074\|hsa-mir-19b-1&WithFlank&13\|+\|91351186\|91351284@60.82.23;MI0000075\|hsa-mir-19b-2&WithFlank&X\|-\|134169665\|134169772@68.90.23&[MIMAT0000074&hsa-miR-19b-3p&offsets\|0\|0;m-122&13\|+\|91351245\|91351267&offsets\|0\|0];[MIMAT0000074_1&hsa-miR-19b-3p&offsets\|0\|0;m-121&X\|-\|134169683\|134169705&offsets\|0\|0]\|TGTGCAAATCCATGCAAAACTGA | >MI0000074\|hsa-mir-19b-1&WithFlank&13\|+\|91351186\|91351284 | 60 | 82 | 23 | TGTGCAAATCCATGCAAAACTGA | 175.7 | 3.3 | 0.8 | 4.1 | 3.6E-05 | 4.7E-05 | BT20/MB468 | NUC |
| >MI0000076\|hsa-mir-20a&WithFlank&13\|+\|91351059\|91351141@14.36.23(+1A)&[MIMAT0000075&hsa-miR-20a-5p&offsets\|0\|0(+1A);m-96&13\|+\|91351072\|91351094&offsets\|0\|0(+1A)]\|TAAAGTGCTTATAGTGCAGGTAGA | >MI0000076\|hsa-mir-20a&WithFlank&13\|+\|91351059\|91351141 | 14 | 36 | 23(+1A) | TAAAGTGCTTATAGTGCAGGTAGA | 5393.1 | -4.3 | 0.9 | -4.6 | 4.3E-06 | 4.7E-05 | BT20/MB231 | MITO |
| >hg38_21_+_8433172_8446622_RNA45SN1_45S_50ntFlanks@6651.6682.32\|CGACTCTTAGCGGTGGATCACTCGGCTCGTGC | >hg38_21_+_8433172_8446622_RNA45SN1_45S_50ntFlanks | 6651 | 6682 | 32 | CGACTCTTAGCGGTGGATCACTCGGCTCGTGC | 1649.3 | 3.7 | 0.8 | 4.4 | 1.2E-05 | 4.7E-05 | BT20/MB468 | MP |
| >hg38_MT_+_1621_3279_RNR2_16S_50ntFlanks@51.80.30\|GCTAAACCTAGCCCCAAACCCACTCCACCT | >hg38_MT_+_1621_3279_RNR2_16S_50ntFlanks | 51 | 80 | 30 | GCTAAACCTAGCCCCAAACCCACTCCACCT | 1000.1 | 3.3 | 0.8 | 4.1 | 3.4E-05 | 4.7E-05 | MB231/MB468 | MITO |
| >hg38_21_+_8433172_8446622_RNA45SN1_45S_50ntFlanks@10099.10117.19\|CGCGCGCGCGTGTGGTGTG | >hg38_21_+_8433172_8446622_RNA45SN1_45S_50ntFlanks | 10099 | 10117 | 19 | CGCGCGCGCGTGTGGTGTG | 350.4 | -4.2 | 1.0 | -4.2 | 3.0E-05 | 4.7E-05 | BT20/MB468 | CYTO |
| >hg38_21_+_8433172_8446622_RNA45SN1_45S_50ntFlanks@9566.9595.30\|GACGTGCAAATCGGTCGTCCGACCTGGGTA | >hg38_21_+_8433172_8446622_RNA45SN1_45S_50ntFlanks | 9566 | 9595 | 30 | GACGTGCAAATCGGTCGTCCGACCTGGGTA | 474.7 | -2.4 | 0.6 | -4.1 | 3.4E-05 | 4.7E-05 | MB231/MB468 | NUC |
| >hg38_21_+_8433172_8446622_RNA45SN1_45S_50ntFlanks@5175.5206.32\|CCGAGATTGAGCAATAACAGGTCTGTGATGCC | >hg38_21_+_8433172_8446622_RNA45SN1_45S_50ntFlanks | 5175 | 5206 | 32 | CCGAGATTGAGCAATAACAGGTCTGTGATGCC | 65.8 | -9.9 | 2.4 | -4.2 | 2.9E-05 | 4.7E-05 | MB231/MB468 | MP |
| >am_trnaMT_ValTAC_MT_+_1602_1670@51.69.19\|CTTAACTTGACCGCTCTGA | >am_trnaMT_ValTAC_MT_+_1602_1670 | 51 | 69 | 19 | CTTAACTTGACCGCTCTGA | 195.0 | -4.6 | 1.1 | -4.2 | 3.2E-05 | 4.8E-05 | MB231/MB468 | CYTO |
| >MI0000736\|hsa-mir-30c-1&WithFlank&1\|+\|40757278\|40757378@23.41.19;MI0000254\|hsa-mir-30c-2&WithFlank&6\|-\|71376954\|71377037@13.31.19&[MIMAT0000244&hsa-miR-30c-5p&offsets\|0\|-4;m-57&1\|+\|40757300\|40757323&offsets\|0\|-5];[MIMAT0000244_1&hsa-miR-30c-5p&offsets\|0\|-4;m-56&6\|-\|71377002\|71377025&offsets\|0\|-5]\|TGTAAACATCCTACACTCT | >MI0000736\|hsa-mir-30c-1&WithFlank&1\|+\|40757278\|40757378 | 23 | 41 | 19 | TGTAAACATCCTACACTCT | 36.9 | 8.4 | 2.0 | 4.2 | 2.9E-05 | 4.8E-05 | MB231/MB468 | MP |
| >MI0000093\|hsa-mir-92a-1&WithFlank&13\|+\|91351308\|91351397@54.72.19(+1A);MI0000094\|hsa-mir-92a-2&WithFlank&X\|-\|134169532\|134169618@54.72.19(+1A)&[MIMAT0000092&hsa-miR-92a-3p&offsets\|0\|-3(+1A);m-9&13\|+\|91351361\|91351382&offsets\|0\|-3(+1A)];[MIMAT0000092_1&hsa-miR-92a-3p&offsets\|0\|-3(+1A);m-11&X\|-\|134169544\|134169565&offsets\|0\|-3(+1A)]\|TATTGCACTTGTCCCGGCCA | >MI0000093\|hsa-mir-92a-1&WithFlank&13\|+\|91351308\|91351397 | 54 | 72 | 19(+1A) | TATTGCACTTGTCCCGGCCA | 36.8 | 8.4 | 2.0 | 4.2 | 2.9E-05 | 4.8E-05 | MB231/MB468 | MP |
| >MI0000542\|hsa-mir-320a&WithFlank&8\|-\|22244960\|22245043@48.69.22(+2U)&[MIMAT0000510&hsa-miR-320a-3p&offsets\|0\|0(+2U);m-60&8\|-\|22244975\|22244996&offsets\|0\|0(+2U)]\|AAAAGCTGGGTTGAGAGGGCGATT | >MI0000542\|hsa-mir-320a&WithFlank&8\|-\|22244960\|22245043 | 48 | 69 | 22(+2U) | AAAAGCTGGGTTGAGAGGGCGATT | 17354.7 | 3.9 | 0.8 | 4.6 | 3.6E-06 | 4.8E-05 | BT20/MB231 | CYTO |
| >MI0000077\|hsa-mir-21&WithFlank&17\|+\|59841260\|59841343@14.39.26&[MIMAT0000076&hsa-miR-21-5p&offsets\|0\|+4;m-1&17\|+\|59841273\|59841294&offsets\|0\|+4]\|TAGCTTATCAGACTGATGTTGACTGT | >MI0000077\|hsa-mir-21&WithFlank&17\|+\|59841260\|59841343 | 14 | 39 | 26 | TAGCTTATCAGACTGATGTTGACTGT | 83.4 | 4.4 | 1.1 | 4.1 | 3.6E-05 | 4.8E-05 | BT20/MB468 | NUC |
| >hg38_21_+_8433172_8446622_RNA45SN1_45S_50ntFlanks@11876.11898.23\|AAGACCCTGTTGAGCTTGACTCT | >hg38_21_+_8433172_8446622_RNA45SN1_45S_50ntFlanks | 11876 | 11898 | 23 | AAGACCCTGTTGAGCTTGACTCT | 946.3 | -2.5 | 0.6 | -4.2 | 3.0E-05 | 4.8E-05 | BT20/MB468 | CYTO |
| >hg38_21_+_8433172_8446622_RNA45SN1_45S_50ntFlanks@9938.9955.18\|AGCAGGACGGTGGCCATG | >hg38_21_+_8433172_8446622_RNA45SN1_45S_50ntFlanks | 9938 | 9955 | 18 | AGCAGGACGGTGGCCATG | 2263.5 | 6.8 | 1.6 | 4.4 | 1.2E-05 | 4.8E-05 | BT20/MB468 | MP |
| >MI0003628\|hsa-mir-615&WithFlank&12\|+\|54033944\|54034051@67.85.19&[MIMAT0003283&hsa-miR-615-3p&offsets\|0\|-3;m-495&12\|+\|54034010\|54034030&offsets\|0\|-2]\|TCCGAGCCTGGGTCTCCCT | >MI0003628\|hsa-mir-615&WithFlank&12\|+\|54033944\|54034051 | 67 | 85 | 19 | TCCGAGCCTGGGTCTCCCT | 91.6 | 1.8 | 0.4 | 4.1 | 3.6E-05 | 4.8E-05 | BT20/MB468 | NUC |
| >hg38_21_+_8433172_8446622_RNA45SN1_45S_50ntFlanks@7975.7993.19\|CGCGACCTCAGATCAGACG | >hg38_21_+_8433172_8446622_RNA45SN1_45S_50ntFlanks | 7975 | 7993 | 19 | CGCGACCTCAGATCAGACG | 33265.7 | 3.3 | 0.8 | 4.4 | 1.2E-05 | 4.8E-05 | BT20/MB468 | MP |
| >hg38_21_+_8433172_8446622_RNA45SN1_45S_50ntFlanks@12457.12477.21\|CAAGCGTTGGATTGTTCACCC | >hg38_21_+_8433172_8446622_RNA45SN1_45S_50ntFlanks | 12457 | 12477 | 21 | CAAGCGTTGGATTGTTCACCC | 5537.1 | -3.6 | 0.9 | -4.2 | 3.3E-05 | 4.8E-05 | MB231/MB468 | CYTO |
| >MI0000093\|hsa-mir-92a-1&WithFlank&13\|+\|91351308\|91351397@54.76.23&[MIMAT0000092&hsa-miR-92a-3p&offsets\|0\|+1;m-9&13\|+\|91351361\|91351382&offsets\|0\|+1]\|TATTGCACTTGTCCCGGCCTGTT | >MI0000093\|hsa-mir-92a-1&WithFlank&13\|+\|91351308\|91351397 | 54 | 76 | 23 | TATTGCACTTGTCCCGGCCTGTT | 2426.4 | 3.3 | 0.8 | 4.2 | 2.9E-05 | 4.8E-05 | MB231/MB468 | MP |
| >hg38_21_+_8433172_8446622_RNA45SN1_45S_50ntFlanks@6656.6700.45\|CTTAGCGGTGGATCACTCGGCTCGTGCGTCGATGAAGAACGCAGC | >hg38_21_+_8433172_8446622_RNA45SN1_45S_50ntFlanks | 6656 | 6700 | 45 | CTTAGCGGTGGATCACTCGGCTCGTGCGTCGATGAAGAACGCAGC | 47.4 | 9.3 | 2.1 | 4.4 | 1.2E-05 | 4.8E-05 | BT20/MB468 | MP |
| >hg38_21_+_8433172_8446622_RNA45SN1_45S_50ntFlanks@10341.10373.33\|AATATTCAAACGAGAACTTTGAAGGCCGAAGTG | >hg38_21_+_8433172_8446622_RNA45SN1_45S_50ntFlanks | 10341 | 10373 | 33 | AATATTCAAACGAGAACTTTGAAGGCCGAAGTG | 229.5 | 4.3 | 1.0 | 4.4 | 1.2E-05 | 4.8E-05 | BT20/MB468 | MP |
| >hg38_21_+_8433172_8446622_RNA45SN1_45S_50ntFlanks@12627.12657.31\|ATGGGGCGAAGCTACCATCTGTGGGATTATG | >hg38_21_+_8433172_8446622_RNA45SN1_45S_50ntFlanks | 12627 | 12657 | 31 | ATGGGGCGAAGCTACCATCTGTGGGATTATG | 346.4 | -4.0 | 1.0 | -4.2 | 3.3E-05 | 4.8E-05 | MB231/MB468 | CYTO |
| >hg38_21_+_8433172_8446622_RNA45SN1_45S_50ntFlanks@8190.8212.23\|CCAGCCCGTGGACGGTGTGAGGC | >hg38_21_+_8433172_8446622_RNA45SN1_45S_50ntFlanks | 8190 | 8212 | 23 | CCAGCCCGTGGACGGTGTGAGGC | 579.0 | -5.1 | 1.2 | -4.1 | 3.5E-05 | 4.8E-05 | MB231/MB468 | NUC |
| >trnaMT_HisGTG_MT_+_12138_12206@18.58.41\|AAAACATCAGATTGTGAATCTGACAACAGAGGCTTACGACC | >trnaMT_HisGTG_MT_+_12138_12206 | 18 | 58 | 41 | AAAACATCAGATTGTGAATCTGACAACAGAGGCTTACGACC | 190.8 | 11.3 | 2.6 | 4.4 | 1.2E-05 | 4.8E-05 | BT20/MB468 | MP |
| >hg38_21_+_8433172_8446622_RNA45SN1_45S_50ntFlanks@51.72.22\|GCTGACACGCTGTCCTCTGGCG | >hg38_21_+_8433172_8446622_RNA45SN1_45S_50ntFlanks | 51 | 72 | 22 | GCTGACACGCTGTCCTCTGGCG | 174.0 | -2.3 | 0.6 | -4.2 | 3.3E-05 | 4.8E-05 | MB231/MB468 | CYTO |
| >hg38_21_+_8433172_8446622_RNA45SN1_45S_50ntFlanks@4499.4517.19\|AGCGTTTACTTTGAAAAAA | >hg38_21_+_8433172_8446622_RNA45SN1_45S_50ntFlanks | 4499 | 4517 | 19 | AGCGTTTACTTTGAAAAAA | 848.0 | 3.8 | 0.9 | 4.2 | 3.0E-05 | 4.8E-05 | BT20/MB468 | CYTO |
| >hg38_21_+_8433172_8446622_RNA45SN1_45S_50ntFlanks@3849.3866.18\|GGATAACTGTGGTAATTC | >hg38_21_+_8433172_8446622_RNA45SN1_45S_50ntFlanks | 3849 | 3866 | 18 | GGATAACTGTGGTAATTC | 139.4 | 4.9 | 1.2 | 4.1 | 3.5E-05 | 4.8E-05 | MB231/MB468 | NUC |
| >hg38_21_+_8433172_8446622_RNA45SN1_45S_50ntFlanks@11578.11609.32\|TAGAACTGGTGCGGACCAGGGGAATCCGACTG | >hg38_21_+_8433172_8446622_RNA45SN1_45S_50ntFlanks | 11578 | 11609 | 32 | TAGAACTGGTGCGGACCAGGGGAATCCGACTG | 424.6 | 2.5 | 0.6 | 4.2 | 2.5E-05 | 4.8E-05 | BT20/MB468 | MITO |
| >hg38_21_+_8433172_8446622_RNA45SN1_45S_50ntFlanks@3841.3871.31\|TCCTACTTGGATAACTGTGGTAATTCTAGAG | >hg38_21_+_8433172_8446622_RNA45SN1_45S_50ntFlanks | 3841 | 3871 | 31 | TCCTACTTGGATAACTGTGGTAATTCTAGAG | 371.2 | 3.7 | 0.9 | 4.2 | 3.0E-05 | 4.8E-05 | BT20/MB468 | CYTO |
| >hg38_21_+_8433172_8446622_RNA45SN1_45S_50ntFlanks@9572.9602.31\|CAAATCGGTCGTCCGACCTGGGTATAGGGGC | >hg38_21_+_8433172_8446622_RNA45SN1_45S_50ntFlanks | 9572 | 9602 | 31 | CAAATCGGTCGTCCGACCTGGGTATAGGGGC | 699.2 | -3.5 | 0.9 | -4.1 | 3.5E-05 | 4.8E-05 | MB231/MB468 | MITO |
| >hg38_21_+_8433172_8446622_RNA45SN1_45S_50ntFlanks@10402.10422.21\|AACATGGGTCAGTCGGTCCTG | >hg38_21_+_8433172_8446622_RNA45SN1_45S_50ntFlanks | 10402 | 10422 | 21 | AACATGGGTCAGTCGGTCCTG | 259.1 | -4.6 | 1.1 | -4.1 | 3.5E-05 | 4.8E-05 | MB231/MB468 | MITO |
| >hg38_1_-_228634819_228635039_RNA5S12_5S_50ntFlanks@149.171.23\|GAATACCGGGTGCTGTAGGCTTT | >hg38_1_-_228634819_228635039_RNA5S12_5S_50ntFlanks | 149 | 171 | 23 | GAATACCGGGTGCTGTAGGCTTT | 1742.9 | 3.8 | 0.9 | 4.1 | 3.5E-05 | 4.8E-05 | MB231/MB468 | NUC |
| >hg38_21_+_8433172_8446622_RNA45SN1_45S_50ntFlanks@12488.12524.37\|AACGTGAGCTGGGTTTAGACCGTCGTGAGACAGGTTA | >hg38_21_+_8433172_8446622_RNA45SN1_45S_50ntFlanks | 12488 | 12524 | 37 | AACGTGAGCTGGGTTTAGACCGTCGTGAGACAGGTTA | 945.0 | -7.6 | 1.8 | -4.1 | 3.6E-05 | 4.8E-05 | BT20/MB468 | NUC |
| >hg38_21_+_8433172_8446622_RNA45SN1_45S_50ntFlanks@10513.10531.19\|CCGGAGTGGCGGAGATGGG | >hg38_21_+_8433172_8446622_RNA45SN1_45S_50ntFlanks | 10513 | 10531 | 19 | CCGGAGTGGCGGAGATGGG | 97.4 | -6.3 | 1.5 | -4.2 | 2.9E-05 | 4.8E-05 | MB231/MB468 | MP |
| >hg38_21_+_8433172_8446622_RNA45SN1_45S_50ntFlanks@6651.6700.50\|CGACTCTTAGCGGTGGATCACTCGGCTCGTGCGTCGATGAAGAACGCAGC | >hg38_21_+_8433172_8446622_RNA45SN1_45S_50ntFlanks | 6651 | 6700 | 50 | CGACTCTTAGCGGTGGATCACTCGGCTCGTGCGTCGATGAAGAACGCAGC | 7016.7 | 4.0 | 1.0 | 4.1 | 3.5E-05 | 4.8E-05 | MB231/MB468 | MITO |
| >hg38_21_+_8433172_8446622_RNA45SN1_45S_50ntFlanks@5142.5162.21\|AACTTCTTAGAGGGACAAGTG | >hg38_21_+_8433172_8446622_RNA45SN1_45S_50ntFlanks | 5142 | 5162 | 21 | AACTTCTTAGAGGGACAAGTG | 154.1 | -7.0 | 1.7 | -4.2 | 2.9E-05 | 4.8E-05 | MB231/MB468 | MP |
| >hg38_21_+_8433172_8446622_RNA45SN1_45S_50ntFlanks@9569.9596.28\|GTGCAAATCGGTCGTCCGACCTGGGTAT | >hg38_21_+_8433172_8446622_RNA45SN1_45S_50ntFlanks | 9569 | 9596 | 28 | GTGCAAATCGGTCGTCCGACCTGGGTAT | 327.8 | -5.9 | 1.4 | -4.2 | 3.0E-05 | 4.8E-05 | BT20/MB468 | CYTO |
| >hg38_21_+_8433172_8446622_RNA45SN1_45S_50ntFlanks@5088.5110.23\|CGAGACTCTGGCATGCTAACTAG | >hg38_21_+_8433172_8446622_RNA45SN1_45S_50ntFlanks | 5088 | 5110 | 23 | CGAGACTCTGGCATGCTAACTAG | 79.2 | -10.2 | 2.4 | -4.2 | 2.9E-05 | 4.8E-05 | MB231/MB468 | MP |
| >hg38_21_+_8433172_8446622_RNA45SN1_45S_50ntFlanks@4048.4066.19\|TTCGAACGTCTGCCCTATC | >hg38_21_+_8433172_8446622_RNA45SN1_45S_50ntFlanks | 4048 | 4066 | 19 | TTCGAACGTCTGCCCTATC | 248.2 | -6.2 | 1.5 | -4.2 | 2.9E-05 | 4.8E-05 | MB231/MB468 | MP |
| >am_trna10_ValCAC_5_-_180649395_180649467@1.36.36,trna18_ValCAC_5_-_180529253_180529325@1.36.36,trna2_ValCAC_5_+_180524070_180524142@1.36.36,trna6_ValCAC_5_+_180600650_180600722@1.36.36,trna85_ValCAC_1_-_161369490_161369562@1.36.36,trna90_ValCAC_1_-_149684088_149684161@1.36.36,trna98_ValCAC_1_-_149298555_149298627@1.36.36,trna9_ValCAC_6_+_26538282_26538354@1.36.36\|GTTTCCGTAGTGTAGTGGTTATCACGTTCGCCTCAC | >am_trna10_ValCAC_5_-_180649395_180649467 | 1 | 36 | 36 | GTTTCCGTAGTGTAGTGGTTATCACGTTCGCCTCAC | 1668.8 | 10.7 | 2.6 | 4.2 | 3.3E-05 | 4.8E-05 | MB231/MB468 | CYTO |

<table>
<tr><th></th><th></th><th></th><th></th><th></th><th></th><th></th><th></th><th></th><th></th><th></th><th></th><th></th><th></th></tr>
<tr><td>>hg38_21_+_8433172_8446622_RNA45SN1_45S_50ntFlanks@11561.11591.31|GCGCCTAGCAGCCGACTTAGAACTGGTGCGG</td><td>>hg38_21_+_8433172_8446622_RNA45SN1_45S_50ntFlanks</td><td>11561</td><td>11591</td><td>31</td><td>GCGCCTAGCAGCCGACTTAGAACTGGTGCGG</td><td>805.8</td><td>-6.2</td><td>1.5</td><td>-4.2</td><td>3.0E-05</td><td>4.8E-05</td><td>BT20/MB468</td><td>CYTO</td></tr>
<tr><td>>hg38_21_+_8433172_8446622_RNA45SN1_45S_50ntFlanks@12996.13039.44|TCGCTGCGATCTATTGAAAGTCAGCCCTCGACACAAGGGTTTGT</td><td>>hg38_21_+_8433172_8446622_RNA45SN1_45S_50ntFlanks</td><td>12996</td><td>13039</td><td>44</td><td>TCGCTGCGATCTATTGAAAGTCAGCCCTCGACACAAGGGTTTGT</td><td>3032.7</td><td>4.1</td><td>1.0</td><td>4.2</td><td>2.9E-05</td><td>4.8E-05</td><td>MB231/MB468</td><td>MP</td></tr>
<tr><td>>hg38_21_+_8433172_8446622_RNA45SN1_45S_50ntFlanks@10771.10796.26|CAGGTCTCCAAGGTGAACAGCCTCTG</td><td>>hg38_21_+_8433172_8446622_RNA45SN1_45S_50ntFlanks</td><td>10771</td><td>10796</td><td>26</td><td>CAGGTCTCCAAGGTGAACAGCCTCTG</td><td>570.1</td><td>-4.2</td><td>1.0</td><td>-4.2</td><td>3.3E-05</td><td>4.8E-05</td><td>MB231/MB468</td><td>CYTO</td></tr>
<tr><td>>hg38_21_+_8433172_8446622_RNA45SN1_45S_50ntFlanks@4733.4758.26|GAACGAAAGTCGGAGGTTCGAAGACG</td><td>>hg38_21_+_8433172_8446622_RNA45SN1_45S_50ntFlanks</td><td>4733</td><td>4758</td><td>26</td><td>GAACGAAAGTCGGAGGTTCGAAGACG</td><td>1083.6</td><td>-2.3</td><td>0.5</td><td>-4.1</td><td>3.5E-05</td><td>4.8E-05</td><td>MB231/MB468</td><td>NUC</td></tr>
<tr><td>>hg38_21_+_8433172_8446622_RNA45SN1_45S_50ntFlanks@9538.9569.32|AACTCTGGTGGAGGTCCGTAGCGGTCCTGACG</td><td>>hg38_21_+_8433172_8446622_RNA45SN1_45S_50ntFlanks</td><td>9538</td><td>9569</td><td>32</td><td>AACTCTGGTGGAGGTCCGTAGCGGTCCTGACG</td><td>326.4</td><td>-3.7</td><td>0.9</td><td>-4.1</td><td>3.5E-05</td><td>4.9E-05</td><td>MB231/MB468</td><td>NUC</td></tr>
<tr><td>>hg38_21_+_8433172_8446622_RNA45SN1_45S_50ntFlanks@11767.11791.25|TCTAATTAGTGACGCGCATGAATGG</td><td>>hg38_21_+_8433172_8446622_RNA45SN1_45S_50ntFlanks</td><td>11767</td><td>11791</td><td>25</td><td>TCTAATTAGTGACGCGCATGAATGG</td><td>194.4</td><td>-5.4</td><td>1.3</td><td>-4.2</td><td>3.0E-05</td><td>4.9E-05</td><td>BT20/MB468</td><td>CYTO</td></tr>
<tr><td>>hg38_21_+_8433172_8446622_RNA45SN1_45S_50ntFlanks@11576.11602.27|CTTAGAACTGGTGCGGACCAGGGGAAT</td><td>>hg38_21_+_8433172_8446622_RNA45SN1_45S_50ntFlanks</td><td>11576</td><td>11602</td><td>27</td><td>CTTAGAACTGGTGCGGACCAGGGGAAT</td><td>249.4</td><td>-2.3</td><td>0.5</td><td>-4.2</td><td>2.6E-05</td><td>4.9E-05</td><td>BT20/MB468</td><td>MITO</td></tr>
<tr><td>>hg38_21_+_8433172_8446622_RNA45SN1_45S_50ntFlanks@12478.12499.22|ACTAATAGGGAACGTGAGCTGG</td><td>>hg38_21_+_8433172_8446622_RNA45SN1_45S_50ntFlanks</td><td>12478</td><td>12499</td><td>22</td><td>ACTAATAGGGAACGTGAGCTGG</td><td>168.3</td><td>-4.8</td><td>1.1</td><td>-4.2</td><td>2.9E-05</td><td>4.9E-05</td><td>MB231/MB468</td><td>MP</td></tr>
<tr><td>>hg38_21_+_8433172_8446622_RNA45SN1_45S_50ntFlanks@11209.11227.19|CGGCGACTCTGGACGCGAG</td><td>>hg38_21_+_8433172_8446622_RNA45SN1_45S_50ntFlanks</td><td>11209</td><td>11227</td><td>19</td><td>CGGCGACTCTGGACGCGAG</td><td>1981.5</td><td>-5.8</td><td>1.3</td><td>-4.4</td><td>1.2E-05</td><td>4.9E-05</td><td>BT20/MB468</td><td>MP</td></tr>
<tr><td>>hg38_21_+_8433172_8446622_RNA45SN1_45S_50ntFlanks@3984.4005.22|GCGGCTTTGGTGACTCTAGATA</td><td>>hg38_21_+_8433172_8446622_RNA45SN1_45S_50ntFlanks</td><td>3984</td><td>4005</td><td>22</td><td>GCGGCTTTGGTGACTCTAGATA</td><td>312.9</td><td>-6.0</td><td>1.4</td><td>-4.2</td><td>2.6E-05</td><td>4.9E-05</td><td>BT20/MB468</td><td>MITO</td></tr>
<tr><td>>hg38_21_+_8433172_8446622_RNA45SN1_45S_50ntFlanks@9622.9658.37|CTAGTAGCTGGTTCCCTCCGAAGTTTCCCTCAGGATA</td><td>>hg38_21_+_8433172_8446622_RNA45SN1_45S_50ntFlanks</td><td>9622</td><td>9658</td><td>37</td><td>CTAGTAGCTGGTTCCCTCCGAAGTTTCCCTCAGGATA</td><td>554.1</td><td>-2.5</td><td>0.6</td><td>-4.2</td><td>2.6E-05</td><td>4.9E-05</td><td>BT20/MB468</td><td>MITO</td></tr>
<tr><td>>MI0000434|hsa-let-7i&WithFlank&12|+|62603680|62603775@12.32.21(+1G)&[MIMAT0000415&hsa-let-7i-5p&offsets|0|-1(+1G);m-49&12|+|62603691|62603712&offsets|0|-1(+1G)]|TGAGGTAGTAGTTTGTGCTGTG</td><td>>MI0000434|hsa-let-7i&WithFlank&12|+|62603680|62603775</td><td>12</td><td>32</td><td>21(+1G)</td><td>TGAGGTAGTAGTTTGTGCTGTG</td><td>1786.8</td><td>3.7</td><td>0.9</td><td>4.2</td><td>2.9E-05</td><td>4.9E-05</td><td>MB231/MB468</td><td>MP</td></tr>
<tr><td>>hg38_21_+_8433172_8446622_RNA45SN1_45S_50ntFlanks@481.505.25|GATCGATGTGGTGACGTCGTGCTCT</td><td>>hg38_21_+_8433172_8446622_RNA45SN1_45S_50ntFlanks</td><td>481</td><td>505</td><td>25</td><td>GATCGATGTGGTGACGTCGTGCTCT</td><td>41.3</td><td>-8.5</td><td>1.9</td><td>-4.4</td><td>1.2E-05</td><td>4.9E-05</td><td>BT20/MB468</td><td>MP</td></tr>
<tr><td>>am_trnaMT_TyrGTA_MT_-_5826_5891@13.32.20|GAGTGAAGCATTGGACTGTA</td><td>>am_trnaMT_TyrGTA_MT_-_5826_5891</td><td>13</td><td>32</td><td>20</td><td>GAGTGAAGCATTGGACTGTA</td><td>223.7</td><td>3.8</td><td>0.9</td><td>4.4</td><td>1.2E-05</td><td>4.9E-05</td><td>BT20/MB468</td><td>MP</td></tr>
<tr><td>>hg38_21_+_8433172_8446622_RNA45SN1_45S_50ntFlanks@12704.12742.39|CAGCGCCGCGGAGCCTCGGTTGGCCTCGGATAGCCGGTC</td><td>>hg38_21_+_8433172_8446622_RNA45SN1_45S_50ntFlanks</td><td>12704</td><td>12742</td><td>39</td><td>CAGCGCCGCGGAGCCTCGGTTGGCCTCGGATAGCCGGTC</td><td>43.1</td><td>-8.5</td><td>1.9</td><td>-4.4</td><td>1.2E-05</td><td>4.9E-05</td><td>BT20/MB468</td><td>MP</td></tr>
<tr><td>>hg38_21_+_8433172_8446622_RNA45SN1_45S_50ntFlanks@11885.11912.28|TTGAGCTTGACTCTAGTCTGGCACGGTG</td><td>>hg38_21_+_8433172_8446622_RNA45SN1_45S_50ntFlanks</td><td>11885</td><td>11912</td><td>28</td><td>TTGAGCTTGACTCTAGTCTGGCACGGTG</td><td>484.5</td><td>-6.9</td><td>1.6</td><td>-4.4</td><td>1.2E-05</td><td>4.9E-05</td><td>BT20/MB468</td><td>MP</td></tr>
<tr><td>>hg38_21_+_8433172_8446622_RNA45SN1_45S_50ntFlanks@4739.4758.20|AAGTCGGAGGTTCGAAGACG</td><td>>hg38_21_+_8433172_8446622_RNA45SN1_45S_50ntFlanks</td><td>4739</td><td>4758</td><td>20</td><td>AAGTCGGAGGTTCGAAGACG</td><td>433.8</td><td>-4.0</td><td>1.0</td><td>-4.1</td><td>3.5E-05</td><td>4.9E-05</td><td>MB231/MB468</td><td>MITO</td></tr>
<tr><td>>hg38_21_+_8433172_8446622_RNA45SN1_45S_50ntFlanks@12476.12496.21|CCACTAATAGGGAACGTGAGC</td><td>>hg38_21_+_8433172_8446622_RNA45SN1_45S_50ntFlanks</td><td>12476</td><td>12496</td><td>21</td><td>CCACTAATAGGGAACGTGAGC</td><td>1803.6</td><td>-2.8</td><td>0.7</td><td>-4.2</td><td>3.0E-05</td><td>4.9E-05</td><td>BT20/MB468</td><td>CYTO</td></tr>
<tr><td>>hg38_21_+_8433172_8446622_RNA45SN1_45S_50ntFlanks@12455.12474.20|ACCAAGCGTTGGATTGTTCA</td><td>>hg38_21_+_8433172_8446622_RNA45SN1_45S_50ntFlanks</td><td>12455</td><td>12474</td><td>20</td><td>ACCAAGCGTTGGATTGTTCA</td><td>207.9</td><td>-5.1</td><td>1.2</td><td>-4.2</td><td>3.0E-05</td><td>4.9E-05</td><td>MB231/MB468</td><td>MP</td></tr>
<tr><td>>hg38_21_+_8433172_8446622_RNA45SN1_45S_50ntFlanks@8171.8214.44|AAGTCCTTCTGATCGAGGCCCAGCCCGTGGACGGTGTGAGGCCG</td><td>>hg38_21_+_8433172_8446622_RNA45SN1_45S_50ntFlanks</td><td>8171</td><td>8214</td><td>44</td><td>AAGTCCTTCTGATCGAGGCCCAGCCCGTGGACGGTGTGAGGCCG</td><td>1218.3</td><td>-3.3</td><td>0.8</td><td>-4.1</td><td>3.5E-05</td><td>4.9E-05</td><td>MB231/MB468</td><td>NUC</td></tr>
<tr><td>>hg38_21_+_8433172_8446622_RNA45SN1_45S_50ntFlanks@10572.10596.25|AGAAGCCGGCGGGAGCCCCGGGGAG</td><td>>hg38_21_+_8433172_8446622_RNA45SN1_45S_50ntFlanks</td><td>10572</td><td>10596</td><td>25</td><td>AGAAGCCGGCGGGAGCCCCGGGGAG</td><td>143.3</td><td>-5.2</td><td>1.2</td><td>-4.2</td><td>3.1E-05</td><td>4.9E-05</td><td>BT20/MB468</td><td>CYTO</td></tr>
<tr><td>>hg38_21_+_8433172_8446622_RNA45SN1_45S_50ntFlanks@7974.8016.43|ACGCGACCTCAGATCAGACGTGGCGACCCGCTGAATTTAAGCA</td><td>>hg38_21_+_8433172_8446622_RNA45SN1_45S_50ntFlanks</td><td>7974</td><td>8016</td><td>43</td><td>ACGCGACCTCAGATCAGACGTGGCGACCCGCTGAATTTAAGCA</td><td>2031.0</td><td>5.1</td><td>1.2</td><td>4.1</td><td>3.5E-05</td><td>4.9E-05</td><td>MB231/MB468</td><td>NUC</td></tr>
<tr><td>>hg38_21_+_8433172_8446622_RNA45SN1_45S_50ntFlanks@6774.6806.33|TCCTCCCGGGGCTACGCCTGTCTGAGCGTCGCT</td><td>>hg38_21_+_8433172_8446622_RNA45SN1_45S_50ntFlanks</td><td>6774</td><td>6806</td><td>33</td><td>TCCTCCCGGGGCTACGCCTGTCTGAGCGTCGCT</td><td>2697.4</td><td>2.5</td><td>0.5</td><td>4.6</td><td>4.3E-06</td><td>4.9E-05</td><td>BT20/MB231</td><td>NUC</td></tr>
<tr><td>>hg38_21_+_8433172_8446622_RNA45SN1_45S_50ntFlanks@12071.12108.38|CCCCGAGGGGCTCTCGCTTCTGGCGCCAAGCGCCCGGC</td><td>>hg38_21_+_8433172_8446622_RNA45SN1_45S_50ntFlanks</td><td>12071</td><td>12108</td><td>38</td><td>CCCCGAGGGGCTCTCGCTTCTGGCGCCAAGCGCCCGGC</td><td>58.0</td><td>-9.7</td><td>2.3</td><td>-4.2</td><td>3.0E-05</td><td>4.9E-05</td><td>MB231/MB468</td><td>MP</td></tr>
<tr><td>>hg38_21_+_8433172_8446622_RNA45SN1_45S_50ntFlanks@6648.6674.27|GTACGACTCTTAGCGGTGGATCACTCG</td><td>>hg38_21_+_8433172_8446622_RNA45SN1_45S_50ntFlanks</td><td>6648</td><td>6674</td><td>27</td><td>GTACGACTCTTAGCGGTGGATCACTCG</td><td>603.4</td><td>4.4</td><td>1.1</td><td>4.1</td><td>3.5E-05</td><td>4.9E-05</td><td>MB231/MB468</td><td>NUC</td></tr>
<tr><td>>hg38_21_+_8433172_8446622_RNA45SN1_45S_50ntFlanks@5059.5097.39|CGATTTGTCTGGTTAATTCCGATAACGAACGAGACTCTG</td><td>>hg38_21_+_8433172_8446622_RNA45SN1_45S_50ntFlanks</td><td>5059</td><td>5097</td><td>39</td><td>CGATTTGTCTGGTTAATTCCGATAACGAACGAGACTCTG</td><td>480.7</td><td>-7.8</td><td>1.9</td><td>-4.2</td><td>3.0E-05</td><td>4.9E-05</td><td>MB231/MB468</td><td>MP</td></tr>
<tr><td>>MI0000746|hsa-mir-99b&WithFlank&19|+|51692606|51692687@13.33.21(+1A)&[MIMAT0000689&hsa-miR-99b-5p&offsets|0|-1(+1A);m-10&19|+|51692618|51692639&offsets|0|-1(+1A)]|CACCCGTAGAACCGACCTTGCA</td><td>>MI0000746|hsa-mir-99b&WithFlank&19|+|51692606|51692687</td><td>13</td><td>33</td><td>21(+1A)</td><td>CACCCGTAGAACCGACCTTGCA</td><td>1699.4</td><td>1.7</td><td>0.4</td><td>4.1</td><td>3.7E-05</td><td>4.9E-05</td><td>BT20/MB468</td><td>NUC</td></tr>
<tr><td>>MI0000263|hsa-mir-7-1&WithFlank&9|-|83969742|83969863@30.55.26&[MIMAT0000252_2&hsa-miR-7-5p&offsets|0|+2;m-401&9|-|83969811|83969834&offsets|0|+2]|TGGAAGACTAGTGATTTTGTTGTTTT</td><td>>MI0000263|hsa-mir-7-1&WithFlank&9|-|83969742|83969863</td><td>30</td><td>55</td><td>26</td><td>TGGAAGACTAGTGATTTTGTTGTTTT</td><td>264.7</td><td>7.1</td><td>1.7</td><td>4.1</td><td>3.5E-05</td><td>4.9E-05</td><td>MB231/MB468</td><td>MITO</td></tr>
<tr><td>>am_trnaMT_SerGCT_MT_+_12207_12265@11.38.28|ACAAGAACTGCTAACTCATGCCCCCATG</td><td>>am_trnaMT_SerGCT_MT_+_12207_12265</td><td>11</td><td>38</td><td>28</td><td>ACAAGAACTGCTAACTCATGCCCCCATG</td><td>357.5</td><td>-3.3</td><td>0.8</td><td>-4.1</td><td>3.5E-05</td><td>4.9E-05</td><td>MB231/MB468</td><td>MITO</td></tr>
<tr><td>>trnaMT_PheGAA_MT_+_577_647@35.62.28|GAAAATGTTTAGACGGGCTCACATCACC</td><td>>trnaMT_PheGAA_MT_+_577_647</td><td>35</td><td>62</td><td>28</td><td>GAAAATGTTTAGACGGGCTCACATCACC</td><td>950.7</td><td>5.9</td><td>1.3</td><td>4.4</td><td>1.2E-05</td><td>4.9E-05</td><td>BT20/MB468</td><td>MP</td></tr>
<tr><td>>hg38_21_+_8433172_8446622_RNA45SN1_45S_50ntFlanks@12456.12473.18|CCAAGCGTTGGATTGTTC</td><td>>hg38_21_+_8433172_8446622_RNA45SN1_45S_50ntFlanks</td><td>12456</td><td>12473</td><td>18</td><td>CCAAGCGTTGGATTGTTC</td><td>992.1</td><td>-5.0</td><td>1.2</td><td>-4.2</td><td>2.6E-05</td><td>4.9E-05</td><td>BT20/MB468</td><td>MITO</td></tr>
<tr><td>>hg38_21_+_8433172_8446622_RNA45SN1_45S_50ntFlanks@12934.12962.29|CTAAACCATTCGTAGACGACCTGCTTCTG</td><td>>hg38_21_+_8433172_8446622_RNA45SN1_45S_50ntFlanks</td><td>12934</td><td>12962</td><td>29</td><td>CTAAACCATTCGTAGACGACCTGCTTCTG</td><td>330.4</td><td>-4.4</td><td>1.1</td><td>-4.2</td><td>3.1E-05</td><td>4.9E-05</td><td>BT20/MB468</td><td>CYTO</td></tr>
<tr><td>>hg38_21_+_8433172_8446622_RNA45SN1_45S_50ntFlanks@6731.6752.22|CACATTGATCATCGACACTTCG</td><td>>hg38_21_+_8433172_8446622_RNA45SN1_45S_50ntFlanks</td><td>6731</td><td>6752</td><td>22</td><td>CACATTGATCATCGACACTTCG</td><td>675.4</td><td>-4.1</td><td>1.0</td><td>-4.1</td><td>3.3E-05</td><td>4.9E-05</td><td>MB231/MB468</td><td>CYTO</td></tr>
<tr><td>>hg38_21_+_8433172_8446622_RNA45SN1_45S_50ntFlanks@9861.9880.20|GATGAACCGAACGCCGGGTT</td><td>>hg38_21_+_8433172_8446622_RNA45SN1_45S_50ntFlanks</td><td>9861</td><td>9880</td><td>20</td><td>GATGAACCGAACGCCGGGTT</td><td>83.1</td><td>-10.2</td><td>2.5</td><td>-4.2</td><td>3.0E-05</td><td>4.9E-05</td><td>MB231/MB468</td><td>MP</td></tr>
<tr><td>>hg38_21_+_8433172_8446622_RNA45SN1_45S_50ntFlanks@11217.11243.27|CTGGACGCGAGCCGGGCCCTTCCCGTG</td><td>>hg38_21_+_8433172_8446622_RNA45SN1_45S_50ntFlanks</td><td>11217</td><td>11243</td><td>27</td><td>CTGGACGCGAGCCGGGCCCTTCCCGTG</td><td>112.1</td><td>-6.1</td><td>1.5</td><td>-4.2</td><td>3.0E-05</td><td>4.9E-05</td><td>MB231/MB468</td><td>MP</td></tr>
<tr><td>>hg38_21_+_8433172_8446622_RNA45SN1_45S_50ntFlanks@3840.3868.29|CTCCTACTTGGATAACTGTGGTAATTCTA</td><td>>hg38_21_+_8433172_8446622_RNA45SN1_45S_50ntFlanks</td><td>3840</td><td>3868</td><td>29</td><td>CTCCTACTTGGATAACTGTGGTAATTCTA</td><td>964.6</td><td>2.6</td><td>0.6</td><td>4.1</td><td>3.4E-05</td><td>4.9E-05</td><td>MB231/MB468</td><td>CYTO</td></tr>
<tr><td>>hg38_21_+_8433172_8446622_RNA45SN1_45S_50ntFlanks@9504.9524.21|GTGAACTATGCCTGGGCAGGG</td><td>>hg38_21_+_8433172_8446622_RNA45SN1_45S_50ntFlanks</td><td>9504</td><td>9524</td><td>21</td><td>GTGAACTATGCCTGGGCAGGG</td><td>788.8</td><td>-5.6</td><td>1.3</td><td>-4.4</td><td>1.2E-05</td><td>4.9E-05</td><td>BT20/MB468</td><td>MP</td></tr>
<tr><td>>hg38_21_+_8433172_8446622_RNA45SN1_45S_50ntFlanks@11569.11600.32|CAGCCGACTTAGAACTGGTGCGGACCAGGGGA</td><td>>hg38_21_+_8433172_8446622_RNA45SN1_45S_50ntFlanks</td><td>11569</td><td>11600</td><td>32</td><td>CAGCCGACTTAGAACTGGTGCGGACCAGGGGA</td><td>2949.9</td><td>-3.5</td><td>0.8</td><td>-4.4</td><td>1.2E-05</td><td>4.9E-05</td><td>BT20/MB468</td><td>MP</td></tr>
<tr><td>>MI0000482|hsa-mir-185&WithFlank&22|+|20033133|20033226@21.42.22&[MIMAT0000455&hsa-miR-185-5p&offsets|0|0;m-145&22|+|20033153|20033174&offsets|0|0]|TGGAGAGAAAGGCAGTTCCTGA</td><td>>MI0000482|hsa-mir-185&WithFlank&22|+|20033133|20033226</td><td>21</td><td>42</td><td>22</td><td>TGGAGAGAAAGGCAGTTCCTGA</td><td>497.0</td><td>3.7</td><td>0.9</td><td>4.1</td><td>3.6E-05</td><td>4.9E-05</td><td>MB231/MB468</td><td>MITO</td></tr>
<tr><td>>trnaMT_GluTTC_MT_-_14674_14742@32.72.41,trnalookalike8_GluTTC_5_-_93905172_93905240@32.72.41|TCATATCATTGGTCGTGGTTGTAGTCCGTGCGAGAATACCA</td><td>>trnaMT_GluTTC_MT_-_14674_14742</td><td>32</td><td>72</td><td>41</td><td>TCATATCATTGGTCGTGGTTGTAGTCCGTGCGAGAATACCA</td><td>619.5</td><td>3.3</td><td>0.8</td><td>4.1</td><td>3.6E-05</td><td>4.9E-05</td><td>MB231/MB468</td><td>MITO</td></tr>
<tr><td>>MI0000804|hsa-mir-328&WithFlank&16|-|67202315|67202401@54.75.22&[MIMAT0000752&hsa-miR-328-3p&offsets|0|0;m-160&16|-|67202327|67202348&offsets|0|0]|CTGGCCCTCTCTGCCCTTCCGT</td><td>>MI0000804|hsa-mir-328&WithFlank&16|-|67202315|67202401</td><td>54</td><td>75</td><td>22</td><td>CTGGCCCTCTCTGCCCTTCCGT</td><td>208.3</td><td>2.7</td><td>0.6</td><td>4.1</td><td>3.7E-05</td><td>4.9E-05</td><td>BT20/MB468</td><td>NUC</td></tr>
<tr><td>>hg38_21_+_8433172_8446622_RNA45SN1_45S_50ntFlanks@4555.4573.19|CGCAGCTAGGAATAATGGA</td><td>>hg38_21_+_8433172_8446622_RNA45SN1_45S_50ntFlanks</td><td>4555</td><td>4573</td><td>19</td><td>CGCAGCTAGGAATAATGGA</td><td>127.8</td><td>-2.5</td><td>0.6</td><td>-4.1</td><td>3.4E-05</td><td>4.9E-05</td><td>MB231/MB468</td><td>CYTO</td></tr>
<tr><td>>am_MI0015825|hsa-mir-378c&WithFlank&10|-|130962582|130962674@17.37.21&[MIMAT0016847&hsa-miR-378c&offsets|0|-4;m-130&10|-|130962639|130962658&offsets|0|+1]|ACTGGACTTGGAGTCAGAAGA</td><td>>am_MI0015825|hsa-mir-378c&WithFlank&10|-|130962582|130962674</td><td>17</td><td>37</td><td>21</td><td>ACTGGACTTGGAGTCAGAAGA</td><td>216.6</td><td>4.2</td><td>1.0</td><td>4.1</td><td>3.6E-05</td><td>4.9E-05</td><td>MB231/MB468</td><td>MITO</td></tr>
<tr><td>>hg38_21_+_8433172_8446622_RNA45SN1_45S_50ntFlanks@8355.8395.41|TGAAAAGAACTTTGAAGAGAGAGTTCAAGAGGGCGTGAAAC</td><td>>hg38_21_+_8433172_8446622_RNA45SN1_45S_50ntFlanks</td><td>8355</td><td>8395</td><td>41</td><td>TGAAAAGAACTTTGAAGAGAGAGTTCAAGAGGGCGTGAAAC</td><td>34.8</td><td>8.9</td><td>2.0</td><td>4.4</td><td>1.2E-05</td><td>4.9E-05</td><td>BT20/MB468</td><td>MP</td></tr>
<tr><td>>hg38_21_+_8433172_8446622_RNA45SN1_45S_50ntFlanks@475.516.42|CTTCGTGATCGATGTGGTGACGTCGTGCTCTCCCGGGCCGGG</td><td>>hg38_21_+_8433172_8446622_RNA45SN1_45S_50ntFlanks</td><td>475</td><td>516</td><td>42</td><td>CTTCGTGATCGATGTGGTGACGTCGTGCTCTCCCGGGCCGGG</td><td>23427.9</td><td>3.8</td><td>0.9</td><td>4.1</td><td>3.6E-05</td><td>4.9E-05</td><td>MB231/MB468</td><td>NUC</td></tr>
<tr><td>>hg38_1_-_228634819_228635039_RNA5S12_5S_50ntFlanks@136.171.36|GGAGACCGCCTGGGAATACCGGGTGCTGTAGGCTTT</td><td>>hg38_1_-_228634819_228635039_RNA5S12_5S_50ntFlanks</td><td>136</td><td>171</td><td>36</td><td>GGAGACCGCCTGGGAATACCGGGTGCTGTAGGCTTT</td><td>4707.5</td><td>3.8</td><td>0.9</td><td>4.2</td><td>2.6E-05</td><td>4.9E-05</td><td>BT20/MB468</td><td>MITO</td></tr>
<tr><td>>hg38_21_+_8433172_8446622_RNA45SN1_45S_50ntFlanks@3816.3837.22|TGGTTCCTTTGGTCGCTCGCTC</td><td>>hg38_21_+_8433172_8446622_RNA45SN1_45S_50ntFlanks</td><td>3816</td><td>3837</td><td>22</td><td>TGGTTCCTTTGGTCGCTCGCTC</td><td>231.9</td><td>-4.5</td><td>1.1</td><td>-4.1</td><td>3.6E-05</td><td>4.9E-05</td><td>MB231/MB468</td><td>NUC</td></tr>
<tr><td>>am_trnaMT_ValTAC_MT_+_1602_1670@39.62.24|AGGAGATTTCAACTTAACTTGACC</td><td>>am_trnaMT_ValTAC_MT_+_1602_1670</td><td>39</td><td>62</td><td>24</td><td>AGGAGATTTCAACTTAACTTGACC</td><td>526.6</td><td>6.1</td><td>1.4</td><td>4.4</td><td>1.2E-05</td><td>4.9E-05</td><td>BT20/MB468</td><td>MP</td></tr>
<tr><td>>hg38_21_+_8433172_8446622_RNA45SN1_45S_50ntFlanks@10734.10755.22|GTGTAAATCTCGCGCCGGGCCG</td><td>>hg38_21_+_8433172_8446622_RNA45SN1_45S_50ntFlanks</td><td>10734</td><td>10755</td><td>22</td><td>GTGTAAATCTCGCGCCGGGCCG</td><td>170.8</td><td>-4.7</td><td>1.1</td><td>-4.1</td><td>3.4E-05</td><td>5.0E-05</td><td>MB231/MB468</td><td>CYTO</td></tr>
<tr><td>>hg38_MT_+_598_1651_RNR1_12S_50ntFlanks@852.872.21|GCTTAGTTGAACAGGGCCCTG</td><td>>hg38_MT_+_598_1651_RNR1_12S_50ntFlanks</td><td>852</td><td>872</td><td>21</td><td>GCTTAGTTGAACAGGGCCCTG</td><td>43.1</td><td>9.2</td><td>2.1</td><td>4.4</td><td>1.2E-05</td><td>5.0E-05</td><td>BT20/MB468</td><td>MP</td></tr>
<tr><td>>trna119_LysCTT_1_-_145395522_145395594@1.29.29,trna11_LysCTT_5_-_180648979_180649051@1.29.29,trna13_LysCTT_6_+_26556774_26556846@1.29.29,trna32_LysCTT_16_-_3207406_3207478@1.29.29,trna7_LysCTT_16_+_3225692_3225764@1.29.29,trna9_LysCTT_5_+_180634755_180634827@1.29.29|GCCCGGCTAGCTCAGTCGGTAGAGCATGA</td><td>>trna119_LysCTT_1_-_145395522_145395594</td><td>1</td><td>29</td><td>29</td><td>GCCCGGCTAGCTCAGTCGGTAGAGCATGA</td><td>1095.4</td><td>-3.1</td><td>0.7</td><td>-4.1</td><td>3.6E-05</td><td>5.0E-05</td><td>MB231/MB468</td><td>MITO</td></tr>
</table>

<table>
<tr><th></th><th></th><th></th><th></th><th></th><th></th><th></th><th></th><th></th><th></th><th></th><th></th><th></th><th></th><th></th><th></th></tr>
<tr><td>>hg38_21_+_8433172_8446622_RNA45SN1_45S_50ntFlanks@11572.11613.42|CCGACTTAGAACTGGTGCGGACCAGGGGAATCCGACTGTTTA</td><td>>hg38_21_+_8433172_8446622_RNA45SN1_45S_50ntFlanks</td><td>11572</td><td>11613</td><td>42</td><td>CCGACTTAGAACTGGTGCGGACCAGGGGAATCCGACTGTTTA</td><td>1706.2</td><td>-4.1</td><td>1.0</td><td>-4.1</td><td>3.6E-05</td><td>5.0E-05</td><td>MB231/MB468</td><td>MITO</td></tr>
<tr><td>>hg38_21_+_8433172_8446622_RNA45SN1_45S_50ntFlanks@12419.12445.27|GATGTCGGCTCTTCCTATCATTGTGAA</td><td>>hg38_21_+_8433172_8446622_RNA45SN1_45S_50ntFlanks</td><td>12419</td><td>12445</td><td>27</td><td>GATGTCGGCTCTTCCTATCATTGTGAA</td><td>369.7</td><td>2.5</td><td>0.6</td><td>4.2</td><td>3.1E-05</td><td>5.0E-05</td><td>BT20/MB468</td><td>CYTO</td></tr>
<tr><td>>hg38_21_+_8433172_8446622_RNA45SN1_45S_50ntFlanks@10348.10374.27|AAACGAGAACTTTGAAGGCCGAAGTGG</td><td>>hg38_21_+_8433172_8446622_RNA45SN1_45S_50ntFlanks</td><td>10348</td><td>10374</td><td>27</td><td>AAACGAGAACTTTGAAGGCCGAAGTGG</td><td>637.2</td><td>-2.4</td><td>0.6</td><td>-4.2</td><td>3.1E-05</td><td>5.0E-05</td><td>BT20/MB468</td><td>CYTO</td></tr>
<tr><td>>hg38_21_+_8433172_8446622_RNA45SN1_45S_50ntFlanks@9554.9579.26|CGTAGCGGTCCTGACGTGCAAATCGG</td><td>>hg38_21_+_8433172_8446622_RNA45SN1_45S_50ntFlanks</td><td>9554</td><td>9579</td><td>26</td><td>CGTAGCGGTCCTGACGTGCAAATCGG</td><td>281.4</td><td>-4.2</td><td>1.0</td><td>-4.2</td><td>3.1E-05</td><td>5.0E-05</td><td>BT20/MB468</td><td>CYTO</td></tr>
<tr><td>>MI0000542|hsa-mir-320a&WithFlank&8|-|22244960|22245043@48.69.22&[MIMAT0000510&hsa-miR-320a-3p&offsets|0|0;m-60&8|-|22244975|22244996&offsets|0|0]|AAAAGCTGGGTTGAGAGGGCGA</td><td>>MI0000542|hsa-mir-320a&WithFlank&8|-|22244960|22245043</td><td>48</td><td>69</td><td>22</td><td>AAAAGCTGGGTTGAGAGGGCGA</td><td>7395.6</td><td>3.9</td><td>0.9</td><td>4.2</td><td>3.0E-05</td><td>5.0E-05</td><td>MB231/MB468</td><td>MP</td></tr>
<tr><td>>hg38_21_+_8433172_8446622_RNA45SN1_45S_50ntFlanks@6649.6682.34|TACGACTCTTAGCGGTGGATCACTCGGCTCGTGC</td><td>>hg38_21_+_8433172_8446622_RNA45SN1_45S_50ntFlanks</td><td>6649</td><td>6682</td><td>34</td><td>TACGACTCTTAGCGGTGGATCACTCGGCTCGTGC</td><td>363.1</td><td>4.7</td><td>1.1</td><td>4.1</td><td>3.6E-05</td><td>5.0E-05</td><td>MB231/MB468</td><td>NUC</td></tr>
<tr><td>>hg38_21_+_8433172_8446622_RNA45SN1_45S_50ntFlanks@8381.8402.22|AAGAGGGCGTGAAACCGTTAAG</td><td>>hg38_21_+_8433172_8446622_RNA45SN1_45S_50ntFlanks</td><td>8381</td><td>8402</td><td>22</td><td>AAGAGGGCGTGAAACCGTTAAG</td><td>579.1</td><td>-4.8</td><td>1.2</td><td>-4.1</td><td>3.6E-05</td><td>5.0E-05</td><td>MB231/MB468</td><td>NUC</td></tr>
<tr><td>>trna27_GlyCCC_2_-_70476123_70476193@4.34.31,trna34_GlyCCC_16_-_686736_686806@4.34.31|CCGCTGGTGTAGTGGTATCATGCAAGATTCC</td><td>>trna27_GlyCCC_2_-_70476123_70476193</td><td>4</td><td>34</td><td>31</td><td>CCGCTGGTGTAGTGGTATCATGCAAGATTCC</td><td>124.5</td><td>6.7</td><td>1.6</td><td>4.1</td><td>3.6E-05</td><td>5.0E-05</td><td>MB231/MB468</td><td>MITO</td></tr>
<tr><td>>hg38_21_+_8433172_8446622_RNA45SN1_45S_50ntFlanks@6647.6677.31|CGTACGACTCTTAGCGGTGGATCACTCGGCT</td><td>>hg38_21_+_8433172_8446622_RNA45SN1_45S_50ntFlanks</td><td>6647</td><td>6677</td><td>31</td><td>CGTACGACTCTTAGCGGTGGATCACTCGGCT</td><td>722.8</td><td>3.0</td><td>0.7</td><td>4.1</td><td>3.6E-05</td><td>5.0E-05</td><td>MB231/MB468</td><td>NUC</td></tr>
<tr><td>>hg38_21_+_8433172_8446622_RNA45SN1_45S_50ntFlanks@3850.3873.24|GATAACTGTGGTAATTCTAGAGCT</td><td>>hg38_21_+_8433172_8446622_RNA45SN1_45S_50ntFlanks</td><td>3850</td><td>3873</td><td>24</td><td>GATAACTGTGGTAATTCTAGAGCT</td><td>399.3</td><td>-3.6</td><td>0.9</td><td>-4.1</td><td>3.8E-05</td><td>5.0E-05</td><td>BT20/MB468</td><td>NUC</td></tr>
<tr><td>>hg38_21_+_8433172_8446622_RNA45SN1_45S_50ntFlanks@4728.4759.32|ATCAAGAACGAAAGTCGGAGGTTCGAAGACGA</td><td>>hg38_21_+_8433172_8446622_RNA45SN1_45S_50ntFlanks</td><td>4728</td><td>4759</td><td>32</td><td>ATCAAGAACGAAAGTCGGAGGTTCGAAGACGA</td><td>444.6</td><td>-5.9</td><td>1.4</td><td>-4.1</td><td>3.6E-05</td><td>5.0E-05</td><td>MB231/MB468</td><td>NUC</td></tr>
<tr><td>>hg38_21_+_8433172_8446622_RNA45SN1_45S_50ntFlanks@5471.5512.42|CACGGCCCTGGCGGAGCGCTGAGAAGACGGTCGAACTTGACT</td><td>>hg38_21_+_8433172_8446622_RNA45SN1_45S_50ntFlanks</td><td>5471</td><td>5512</td><td>42</td><td>CACGGCCCTGGCGGAGCGCTGAGAAGACGGTCGAACTTGACT</td><td>638.2</td><td>3.3</td><td>0.8</td><td>4.1</td><td>3.4E-05</td><td>5.0E-05</td><td>MB231/MB468</td><td>CYTO</td></tr>
<tr><td>>hg38_21_+_8433172_8446622_RNA45SN1_45S_50ntFlanks@8814.8844.31|CGGGTCGGGGCGGCGGCGGCGGCGGCGGTGG</td><td>>hg38_21_+_8433172_8446622_RNA45SN1_45S_50ntFlanks</td><td>8814</td><td>8844</td><td>31</td><td>CGGGTCGGGGCGGCGGCGGCGGCGGCGGTGG</td><td>44.3</td><td>-8.5</td><td>2.0</td><td>-4.4</td><td>1.2E-05</td><td>5.0E-05</td><td>BT20/MB468</td><td>MP</td></tr>
<tr><td>>hg38_21_+_8433172_8446622_RNA45SN1_45S_50ntFlanks@8176.8213.38|CTTCTGATCGAGGCCCAGCCCGTGGACGGTGTGAGGCC</td><td>>hg38_21_+_8433172_8446622_RNA45SN1_45S_50ntFlanks</td><td>8176</td><td>8213</td><td>38</td><td>CTTCTGATCGAGGCCCAGCCCGTGGACGGTGTGAGGCC</td><td>930.7</td><td>-5.5</td><td>1.3</td><td>-4.2</td><td>3.0E-05</td><td>5.0E-05</td><td>MB231/MB468</td><td>MP</td></tr>
<tr><td>>hg38_21_+_8433172_8446622_RNA45SN1_45S_50ntFlanks@8107.8141.35|CGGCGGGGCGCGGGACATGTGGCGTACGGAAGACC</td><td>>hg38_21_+_8433172_8446622_RNA45SN1_45S_50ntFlanks</td><td>8107</td><td>8141</td><td>35</td><td>CGGCGGGGCGCGGGACATGTGGCGTACGGAAGACC</td><td>677.2</td><td>-6.7</td><td>1.6</td><td>-4.2</td><td>3.0E-05</td><td>5.0E-05</td><td>MB231/MB468</td><td>MP</td></tr>
<tr><td>>hg38_21_+_8433172_8446622_RNA45SN1_45S_50ntFlanks@5067.5098.32|CTGGTTAATTCCGATAACGAACGAGACTCTGG</td><td>>hg38_21_+_8433172_8446622_RNA45SN1_45S_50ntFlanks</td><td>5067</td><td>5098</td><td>32</td><td>CTGGTTAATTCCGATAACGAACGAGACTCTGG</td><td>56.9</td><td>-8.9</td><td>2.0</td><td>-4.4</td><td>1.2E-05</td><td>5.0E-05</td><td>BT20/MB468</td><td>MP</td></tr>
<tr><td>>hg38_21_+_8433172_8446622_RNA45SN1_45S_50ntFlanks@7968.7987.20|TCCGAGACGCGACCTCAGAT</td><td>>hg38_21_+_8433172_8446622_RNA45SN1_45S_50ntFlanks</td><td>7968</td><td>7987</td><td>20</td><td>TCCGAGACGCGACCTCAGAT</td><td>131.5</td><td>3.1</td><td>0.7</td><td>4.1</td><td>3.8E-05</td><td>5.0E-05</td><td>BT20/MB468</td><td>NUC</td></tr>
<tr><td>>hg38_21_+_8433172_8446622_RNA45SN1_45S_50ntFlanks@9535.9563.29|GGAAACTCTGGTGGAGGTCCGTAGCGGTC</td><td>>hg38_21_+_8433172_8446622_RNA45SN1_45S_50ntFlanks</td><td>9535</td><td>9563</td><td>29</td><td>GGAAACTCTGGTGGAGGTCCGTAGCGGTC</td><td>141.6</td><td>-2.7</td><td>0.7</td><td>-4.2</td><td>3.1E-05</td><td>5.0E-05</td><td>BT20/MB468</td><td>CYTO</td></tr>
<tr><td>>hg38_21_+_8433172_8446622_RNA45SN1_45S_50ntFlanks@4046.4065.20|CATTCGAACGTCTGCCCTAT</td><td>>hg38_21_+_8433172_8446622_RNA45SN1_45S_50ntFlanks</td><td>4046</td><td>4065</td><td>20</td><td>CATTCGAACGTCTGCCCTAT</td><td>219.9</td><td>-5.5</td><td>1.3</td><td>-4.1</td><td>3.6E-05</td><td>5.0E-05</td><td>MB231/MB468</td><td>NUC</td></tr>
<tr><td>>hg38_21_+_8433172_8446622_RNA45SN1_45S_50ntFlanks@11644.11662.19|CGGGTGTTGACGCGATGTG</td><td>>hg38_21_+_8433172_8446622_RNA45SN1_45S_50ntFlanks</td><td>11644</td><td>11662</td><td>19</td><td>CGGGTGTTGACGCGATGTG</td><td>109.8</td><td>10.1</td><td>2.2</td><td>4.6</td><td>3.9E-06</td><td>5.0E-05</td><td>BT20/MB231</td><td>MP</td></tr>
<tr><td>>hg38_21_+_8433172_8446622_RNA45SN1_45S_50ntFlanks@11579.11602.24|AGAACTGGTGCGGACCAGGGGAAT</td><td>>hg38_21_+_8433172_8446622_RNA45SN1_45S_50ntFlanks</td><td>11579</td><td>11602</td><td>24</td><td>AGAACTGGTGCGGACCAGGGGAAT</td><td>878.4</td><td>-3.7</td><td>0.9</td><td>-4.1</td><td>3.4E-05</td><td>5.0E-05</td><td>MB231/MB468</td><td>CYTO</td></tr>
<tr><td>>hg38_21_+_8433172_8446622_RNA45SN1_45S_50ntFlanks@8182.8204.23|ATCGAGGCCCAGCCCGTGGACGG</td><td>>hg38_21_+_8433172_8446622_RNA45SN1_45S_50ntFlanks</td><td>8182</td><td>8204</td><td>23</td><td>ATCGAGGCCCAGCCCGTGGACGG</td><td>493.5</td><td>-6.2</td><td>1.5</td><td>-4.1</td><td>3.6E-05</td><td>5.0E-05</td><td>MB231/MB468</td><td>MITO</td></tr>
<tr><td>>hg38_21_+_8433172_8446622_RNA45SN1_45S_50ntFlanks@8378.8399.22|TTCAAGAGGGCGTGAAACCGTT</td><td>>hg38_21_+_8433172_8446622_RNA45SN1_45S_50ntFlanks</td><td>8378</td><td>8399</td><td>22</td><td>TTCAAGAGGGCGTGAAACCGTT</td><td>82.8</td><td>-6.1</td><td>1.5</td><td>-4.2</td><td>3.0E-05</td><td>5.0E-05</td><td>MB231/MB468</td><td>MP</td></tr>
<tr><td>>hg38_21_+_8433172_8446622_RNA45SN1_45S_50ntFlanks@11564.11592.29|CCTAGCAGCCGACTTAGAACTGGTGCGGA</td><td>>hg38_21_+_8433172_8446622_RNA45SN1_45S_50ntFlanks</td><td>11564</td><td>11592</td><td>29</td><td>CCTAGCAGCCGACTTAGAACTGGTGCGGA</td><td>1859.7</td><td>-7.8</td><td>1.8</td><td>-4.4</td><td>1.2E-05</td><td>5.0E-05</td><td>BT20/MB468</td><td>MP</td></tr>
<tr><td>>hg38_21_+_8433172_8446622_RNA45SN1_45S_50ntFlanks@3777.3805.29|CGGTACAGTGAAACTGCGAATGGCTCATT</td><td>>hg38_21_+_8433172_8446622_RNA45SN1_45S_50ntFlanks</td><td>3777</td><td>3805</td><td>29</td><td>CGGTACAGTGAAACTGCGAATGGCTCATT</td><td>303.1</td><td>-6.6</td><td>1.6</td><td>-4.2</td><td>3.0E-05</td><td>5.0E-05</td><td>MB231/MB468</td><td>MP</td></tr>
<tr><td>>hg38_21_+_8433172_8446622_RNA45SN1_45S_50ntFlanks@7973.7992.20|GACGCGACCTCAGATCAGAC</td><td>>hg38_21_+_8433172_8446622_RNA45SN1_45S_50ntFlanks</td><td>7973</td><td>7992</td><td>20</td><td>GACGCGACCTCAGATCAGAC</td><td>861.0</td><td>4.2</td><td>1.0</td><td>4.4</td><td>1.2E-05</td><td>5.0E-05</td><td>BT20/MB468</td><td>MP</td></tr>
<tr><td>>hg38_21_+_8433172_8446622_RNA45SN1_45S_50ntFlanks@10395.10429.35|GCAGTTGAACATGGGTCAGTCGGTCCTGAGAGATG</td><td>>hg38_21_+_8433172_8446622_RNA45SN1_45S_50ntFlanks</td><td>10395</td><td>10429</td><td>35</td><td>GCAGTTGAACATGGGTCAGTCGGTCCTGAGAGATG</td><td>395.2</td><td>-4.0</td><td>1.0</td><td>-4.2</td><td>3.1E-05</td><td>5.0E-05</td><td>MB231/MB468</td><td>MP</td></tr>
<tr><td>>hg38_21_+_8433172_8446622_RNA45SN1_45S_50ntFlanks@10781.10825.45|AGGTGAACAGCCTCTGGCATGTTGGAACAATGTAGGTAAGGGAAG</td><td>>hg38_21_+_8433172_8446622_RNA45SN1_45S_50ntFlanks</td><td>10781</td><td>10825</td><td>45</td><td>AGGTGAACAGCCTCTGGCATGTTGGAACAATGTAGGTAAGGGAAG</td><td>54.8</td><td>-8.9</td><td>2.0</td><td>-4.4</td><td>1.2E-05</td><td>5.0E-05</td><td>BT20/MB468</td><td>MP</td></tr>
<tr><td>>hg38_21_+_8433172_8446622_RNA45SN1_45S_50ntFlanks@12008.12040.33|CCGCCGGTGAAATACCACTACTCTGATCGTTTT</td><td>>hg38_21_+_8433172_8446622_RNA45SN1_45S_50ntFlanks</td><td>12008</td><td>12040</td><td>33</td><td>CCGCCGGTGAAATACCACTACTCTGATCGTTTT</td><td>43.3</td><td>-8.5</td><td>1.9</td><td>-4.4</td><td>1.2E-05</td><td>5.0E-05</td><td>BT20/MB468</td><td>MP</td></tr>
<tr><td>>hg38_21_+_8433172_8446622_RNA45SN1_45S_50ntFlanks@9495.9526.32|CGAAAGATGGTGAACTATGCCTGGGCAGGGCG</td><td>>hg38_21_+_8433172_8446622_RNA45SN1_45S_50ntFlanks</td><td>9495</td><td>9526</td><td>32</td><td>CGAAAGATGGTGAACTATGCCTGGGCAGGGCG</td><td>87.5</td><td>-6.1</td><td>1.5</td><td>-4.2</td><td>3.1E-05</td><td>5.0E-05</td><td>MB231/MB468</td><td>MP</td></tr>
<tr><td>>hg38_21_+_8433172_8446622_RNA45SN1_45S_50ntFlanks@11886.11910.25|TGAGCTTGACTCTAGTCTGGCACGG</td><td>>hg38_21_+_8433172_8446622_RNA45SN1_45S_50ntFlanks</td><td>11886</td><td>11910</td><td>25</td><td>TGAGCTTGACTCTAGTCTGGCACGG</td><td>40.8</td><td>-8.4</td><td>1.9</td><td>-4.4</td><td>1.2E-05</td><td>5.0E-05</td><td>BT20/MB468</td><td>MP</td></tr>
<tr><td>>hg38_21_+_8433172_8446622_RNA45SN1_45S_50ntFlanks@5084.5120.37|CGAACGAGACTCTGGCATGCTAACTAGTTACGCGACC</td><td>>hg38_21_+_8433172_8446622_RNA45SN1_45S_50ntFlanks</td><td>5084</td><td>5120</td><td>37</td><td>CGAACGAGACTCTGGCATGCTAACTAGTTACGCGACC</td><td>414.0</td><td>-3.5</td><td>0.8</td><td>-4.2</td><td>2.7E-05</td><td>5.0E-05</td><td>BT20/MB468</td><td>MITO</td></tr>
<tr><td>>hg38_21_+_8433172_8446622_RNA45SN1_45S_50ntFlanks@6651.6699.49|CGACTCTTAGCGGTGGATCACTCGGCTCGTGCGTCGATGAAGAACGCAG</td><td>>hg38_21_+_8433172_8446622_RNA45SN1_45S_50ntFlanks</td><td>6651</td><td>6699</td><td>49</td><td>CGACTCTTAGCGGTGGATCACTCGGCTCGTGCGTCGATGAAGAACGCAG</td><td>31221.9</td><td>-4.5</td><td>1.1</td><td>-4.1</td><td>3.8E-05</td><td>5.0E-05</td><td>BT20/MB468</td><td>NUC</td></tr>
<tr><td>>trnaMT_GluTTC_MT_-_14674_14742@40.57.18,trnalookalike8_GluTTC_5_-_93905172_93905240@40.57.18|TTGGTCGTGGTTGTAGTC</td><td>>trnaMT_GluTTC_MT_-_14674_14742</td><td>40</td><td>57</td><td>18</td><td>TTGGTCGTGGTTGTAGTC</td><td>430.5</td><td>4.1</td><td>0.9</td><td>4.4</td><td>1.2E-05</td><td>5.0E-05</td><td>BT20/MB468</td><td>MP</td></tr>
<tr><td>>MI0000465|hsa-mir-191&WithFlank&3|-|49020612|49020715@23.44.22&[MIMAT0000440&hsa-miR-191-5p&offsets|+1|0;m-48&3|-|49020672|49020694&offsets|+1|0]|AACGGAATCCCAAAAGCAGCTG</td><td>>MI0000465|hsa-mir-191&WithFlank&3|-|49020612|49020715</td><td>23</td><td>44</td><td>22</td><td>AACGGAATCCCAAAAGCAGCTG</td><td>173.8</td><td>5.1</td><td>1.2</td><td>4.1</td><td>3.7E-05</td><td>5.0E-05</td><td>MB231/MB468</td><td>MITO</td></tr>
<tr><td>>hg38_21_+_8433172_8446622_RNA45SN1_45S_50ntFlanks@8177.8204.28|TTCTGATCGAGGCCCAGCCCGTGGACGG</td><td>>hg38_21_+_8433172_8446622_RNA45SN1_45S_50ntFlanks</td><td>8177</td><td>8204</td><td>28</td><td>TTCTGATCGAGGCCCAGCCCGTGGACGG</td><td>536.6</td><td>-2.6</td><td>0.6</td><td>-4.1</td><td>3.6E-05</td><td>5.0E-05</td><td>MB231/MB468</td><td>NUC</td></tr>
<tr><td>>trnaMT_HisGTG_MT_+_12138_12206@26.59.34|AGATTGTGAATCTGACAACAGAGGCTTACGACCC</td><td>>trnaMT_HisGTG_MT_+_12138_12206</td><td>26</td><td>59</td><td>34</td><td>AGATTGTGAATCTGACAACAGAGGCTTACGACCC</td><td>30.1</td><td>8.1</td><td>2.0</td><td>4.2</td><td>3.1E-05</td><td>5.0E-05</td><td>MB231/MB468</td><td>MP</td></tr>
<tr><td>>hg38_21_+_8433172_8446622_RNA45SN1_45S_50ntFlanks@11668.11693.26|TGCCCAGTGCTCTGAATGTCAAAGTG</td><td>>hg38_21_+_8433172_8446622_RNA45SN1_45S_50ntFlanks</td><td>11668</td><td>11693</td><td>26</td><td>TGCCCAGTGCTCTGAATGTCAAAGTG</td><td>113.4</td><td>3.9</td><td>1.0</td><td>4.1</td><td>3.7E-05</td><td>5.0E-05</td><td>MB231/MB468</td><td>MITO</td></tr>
<tr><td>>hg38_21_+_8433172_8446622_RNA45SN1_45S_50ntFlanks@9739.9765.27|AAACGATCTCAACCTATTCTCAAACTT</td><td>>hg38_21_+_8433172_8446622_RNA45SN1_45S_50ntFlanks</td><td>9739</td><td>9765</td><td>27</td><td>AAACGATCTCAACCTATTCTCAAACTT</td><td>2556.0</td><td>-3.3</td><td>0.8</td><td>-4.1</td><td>3.6E-05</td><td>5.0E-05</td><td>MB231/MB468</td><td>NUC</td></tr>
<tr><td>>hg38_21_+_8433172_8446622_RNA45SN1_45S_50ntFlanks@12624.12652.29|CCAATGGGGCGAAGCTACCATCTGTGGGA</td><td>>hg38_21_+_8433172_8446622_RNA45SN1_45S_50ntFlanks</td><td>12624</td><td>12652</td><td>29</td><td>CCAATGGGGCGAAGCTACCATCTGTGGGA</td><td>221.2</td><td>-5.4</td><td>1.3</td><td>-4.2</td><td>2.7E-05</td><td>5.1E-05</td><td>BT20/MB468</td><td>MITO</td></tr>
<tr><td>>hg38_21_+_8433172_8446622_RNA45SN1_45S_50ntFlanks@4364.4394.31|CTGCAGTTAAAAAGCTCGTAGTTGGATCTTG</td><td>>hg38_21_+_8433172_8446622_RNA45SN1_45S_50ntFlanks</td><td>4364</td><td>4394</td><td>31</td><td>CTGCAGTTAAAAAGCTCGTAGTTGGATCTTG</td><td>38.4</td><td>-8.3</td><td>1.9</td><td>-4.4</td><td>1.2E-05</td><td>5.1E-05</td><td>BT20/MB468</td><td>MP</td></tr>
<tr><td>>hg38_21_+_8433172_8446622_RNA45SN1_45S_50ntFlanks@8358.8396.39|AAAGAACTTTGAAGAGAGAGTTCAAGAGGGCGTGAAACC</td><td>>hg38_21_+_8433172_8446622_RNA45SN1_45S_50ntFlanks</td><td>8358</td><td>8396</td><td>39</td><td>AAAGAACTTTGAAGAGAGAGTTCAAGAGGGCGTGAAACC</td><td>104.1</td><td>10.0</td><td>2.2</td><td>4.6</td><td>3.9E-06</td><td>5.1E-05</td><td>BT20/MB231</td><td>MP</td></tr>
<tr><td>>hg38_21_+_8433172_8446622_RNA45SN1_45S_50ntFlanks@6700.6721.22|CTAGCTGCGAGAATTAATGTGA</td><td>>hg38_21_+_8433172_8446622_RNA45SN1_45S_50ntFlanks</td><td>6700</td><td>6721</td><td>22</td><td>CTAGCTGCGAGAATTAATGTGA</td><td>200.4</td><td>-4.6</td><td>1.1</td><td>-4.2</td><td>3.1E-05</td><td>5.1E-05</td><td>MB231/MB468</td><td>MP</td></tr>
<tr><td>>hg38_21_+_8433172_8446622_RNA45SN1_45S_50ntFlanks@11769.11804.36|TAATTAGTGACGCGCATGAATGGATGAACGAGATTC</td><td>>hg38_21_+_8433172_8446622_RNA45SN1_45S_50ntFlanks</td><td>11769</td><td>11804</td><td>36</td><td>TAATTAGTGACGCGCATGAATGGATGAACGAGATTC</td><td>592.4</td><td>-4.4</td><td>1.1</td><td>-4.2</td><td>3.2E-05</td><td>5.1E-05</td><td>BT20/MB468</td><td>CYTO</td></tr>
<tr><td>>hg38_21_+_8433172_8446622_RNA45SN1_45S_50ntFlanks@5492.5514.23|AGAAGACGGTCGAACTTGACTAT</td><td>>hg38_21_+_8433172_8446622_RNA45SN1_45S_50ntFlanks</td><td>5492</td><td>5514</td><td>23</td><td>AGAAGACGGTCGAACTTGACTAT</td><td>1027.8</td><td>-5.0</td><td>1.2</td><td>-4.2</td><td>2.7E-05</td><td>5.1E-05</td><td>BT20/MB468</td><td>MITO</td></tr>
<tr><td>>hg38_21_+_8433172_8446622_RNA45SN1_45S_50ntFlanks@7973.8004.32|GACGCGACCTCAGATCAGACGTGGCGACCCGC</td><td>>hg38_21_+_8433172_8446622_RNA45SN1_45S_50ntFlanks</td><td>7973</td><td>8004</td><td>32</td><td>GACGCGACCTCAGATCAGACGTGGCGACCCGC</td><td>409.4</td><td>3.8</td><td>0.9</td><td>4.1</td><td>3.8E-05</td><td>5.1E-05</td><td>BT20/MB468</td><td>NUC</td></tr>
<tr><td>>hg38_21_+_8433172_8446622_RNA45SN1_45S_50ntFlanks@3842.3871.30|CCTACTTGGATAACTGTGGTAATTCTAGAG</td><td>>hg38_21_+_8433172_8446622_RNA45SN1_45S_50ntFlanks</td><td>3842</td><td>3871</td><td>30</td><td>CCTACTTGGATAACTGTGGTAATTCTAGAG</td><td>115.8</td><td>-6.1</td><td>1.5</td><td>-4.2</td><td>3.1E-05</td><td>5.1E-05</td><td>MB231/MB468</td><td>MP</td></tr>
<tr><td>>am_trnaMT_ArgTCG_MT_+_10405_10469@29.48.20|TTTCGACTCATTAAATTATG</td><td>>am_trnaMT_ArgTCG_MT_+_10405_10469</td><td>29</td><td>48</td><td>20</td><td>TTTCGACTCATTAAATTATG</td><td>244.0</td><td>-4.7</td><td>1.1</td><td>-4.1</td><td>3.5E-05</td><td>5.1E-05</td><td>MB231/MB468</td><td>CYTO</td></tr>
<tr><td>>MI0000542|hsa-mir-320a&WithFlank&8|-|22244960|22245043@48.71.24(+1C)&[MIMAT0000510&hsa-miR-320a-3p&offsets|0|+2(+1C);m-60&8|-|22244975|22244996&offsets|0|+2(+1C)]|AAAAGCTGGGTTGAGAGGGCGAAAC</td><td>>MI0000542|hsa-mir-320a&WithFlank&8|-|22244960|22245043</td><td>48</td><td>71</td><td>24(+1C)</td><td>AAAAGCTGGGTTGAGAGGGCGAAAC</td><td>34.1</td><td>8.3</td><td>2.0</td><td>4.2</td><td>3.1E-05</td><td>5.1E-05</td><td>MB231/MB468</td><td>MP</td></tr>
<tr><td>>hg38_21_+_8433172_8446622_RNA45SN1_45S_50ntFlanks@8359.8379.21|AAGAACTTTGAAGAGAGAGTT</td><td>>hg38_21_+_8433172_8446622_RNA45SN1_45S_50ntFlanks</td><td>8359</td><td>8379</td><td>21</td><td>AAGAACTTTGAAGAGAGAGTT</td><td>731.5</td><td>-7.0</td><td>1.6</td><td>-4.4</td><td>1.2E-05</td><td>5.1E-05</td><td>BT20/MB468</td><td>MP</td></tr>
<tr><td>>hg38_21_+_8433172_8446622_RNA45SN1_45S_50ntFlanks@12155.12180.26|GGAGTTTGACTGGGGCGGTACACCTG</td><td>>hg38_21_+_8433172_8446622_RNA45SN1_45S_50ntFlanks</td><td>12155</td><td>12180</td><td>26</td><td>GGAGTTTGACTGGGGCGGTACACCTG</td><td>262.9</td><td>-6.5</td><td>1.5</td><td>-4.4</td><td>1.2E-05</td><td>5.1E-05</td><td>BT20/MB468</td><td>MP</td></tr>
<tr><td>>hg38_21_+_8433172_8446622_RNA45SN1_45S_50ntFlanks@9575.9603.29|ATCGGTCGTCCGACCTGGGTATAGGGGCG</td><td>>hg38_21_+_8433172_8446622_RNA45SN1_45S_50ntFlanks</td><td>9575</td><td>9603</td><td>29</td><td>ATCGGTCGTCCGACCTGGGTATAGGGGCG</td><td>3463.7</td><td>-7.4</td><td>1.8</td><td>-4.2</td><td>3.1E-05</td><td>5.1E-05</td><td>MB231/MB468</td><td>MP</td></tr>
<tr><td>>hg38_21_+_8433172_8446622_RNA45SN1_45S_50ntFlanks@4734.4752.19|AACGAAAGTCGGAGGTTCG</td><td>>hg38_21_+_8433172_8446622_RNA45SN1_45S_50ntFlanks</td><td>4734</td><td>4752</td><td>19</td><td>AACGAAAGTCGGAGGTTCG</td><td>442.1</td><td>-2.4</td><td>0.6</td><td>-4.1</td><td>3.5E-05</td><td>5.1E-05</td><td>MB231/MB468</td><td>CYTO</td></tr>
</table>

| | | | | | | | | | | | | | |
|---|---|---|---|---|---|---|---|---|---|---|---|---|---|
| >hg38_21_+_8433172_8446622_RNA45SN1_45S_50ntFlanks@12941.12962.22\|ATTCGTAGACGACCTGCTTCTG | >hg38_21_+_8433172_8446622_RNA45SN1_45S_50ntFlanks | 12941 | 12962 | 22 | ATTCGTAGACGACCTGCTTCTG | 106.6 | -6.0 | 1.4 | -4.2 | 3.1E-05 | 5.1E-05 | MB231/MB468 | MP |
| >hg38_21_+_8433172_8446622_RNA45SN1_45S_50ntFlanks@11189.11207.19\|ACCCGGGGGGCCGGCGGCG | >hg38_21_+_8433172_8446622_RNA45SN1_45S_50ntFlanks | 11189 | 11207 | 19 | ACCCGGGGGGCCGGCGGCG | 596.2 | -7.2 | 1.8 | -4.1 | 3.8E-05 | 5.1E-05 | BT20/MB468 | NUC |
| >hg38_21_+_8433172_8446622_RNA45SN1_45S_50ntFlanks@11586.11605.20\|GTGCGGACCAGGGGAATCCG | >hg38_21_+_8433172_8446622_RNA45SN1_45S_50ntFlanks | 11586 | 11605 | 20 | GTGCGGACCAGGGGAATCCG | 274.2 | -5.6 | 1.3 | -4.2 | 3.2E-05 | 5.1E-05 | BT20/MB468 | CYTO |
| >hg38_21_+_8433172_8446622_RNA45SN1_45S_50ntFlanks@9568.9594.27\|CGTGCAAATCGGTCGTCCGACCTGGGT | >hg38_21_+_8433172_8446622_RNA45SN1_45S_50ntFlanks | 9568 | 9594 | 27 | CGTGCAAATCGGTCGTCCGACCTGGGT | 341.5 | -4.6 | 1.1 | -4.1 | 3.7E-05 | 5.1E-05 | MB231/MB468 | NUC |
| >trna37_ValAAC_6_+_27203288_27203360@1.29.29\|GTTTCCGTAGTGTAGTGGTTATCACGTTT | >trna37_ValAAC_6_+_27203288_27203360 | 1 | 29 | 29 | GTTTCCGTAGTGTAGTGGTTATCACGTTT | 272.0 | -4.7 | 1.1 | -4.1 | 3.7E-05 | 5.1E-05 | MB231/MB468 | NUC |
| >hg38_MT_+_598_1651_RNR1_12S_50ntFlanks@994.1012.19\|CTTGGACGAACCAGAGTGT | >hg38_MT_+_598_1651_RNR1_12S_50ntFlanks | 994 | 1012 | 19 | CTTGGACGAACCAGAGTGT | 67.0 | -9.5 | 2.1 | -4.6 | 4.0E-06 | 5.1E-05 | BT20/MB231 | MP |
| >hg38_21_+_8433172_8446622_RNA45SN1_45S_50ntFlanks@12628.12652.25\|TGGGGCGAAGCTACCATCTGTGGGA | >hg38_21_+_8433172_8446622_RNA45SN1_45S_50ntFlanks | 12628 | 12652 | 25 | TGGGGCGAAGCTACCATCTGTGGGA | 213.9 | -6.6 | 1.5 | -4.4 | 1.3E-05 | 5.1E-05 | BT20/MB468 | MP |
| >hg38_21_+_8433172_8446622_RNA45SN1_45S_50ntFlanks@12003.12029.27\|GCGGGCCGCCGGTGAAATACCACTACT | >hg38_21_+_8433172_8446622_RNA45SN1_45S_50ntFlanks | 12003 | 12029 | 27 | GCGGGCCGCCGGTGAAATACCACTACT | 153.2 | -3.4 | 0.8 | -4.2 | 3.1E-05 | 5.1E-05 | MB231/MB468 | MP |
| >hg38_21_+_8433172_8446622_RNA45SN1_45S_50ntFlanks@7983.8012.30\|CAGATCAGACGTGGCGACCCGCTGAATTTA | >hg38_21_+_8433172_8446622_RNA45SN1_45S_50ntFlanks | 7983 | 8012 | 30 | CAGATCAGACGTGGCGACCCGCTGAATTTA | 117.7 | -4.4 | 1.1 | -4.1 | 3.7E-05 | 5.1E-05 | MB231/MB468 | NUC |
| >hg38_21_+_8433172_8446622_RNA45SN1_45S_50ntFlanks@10682.10712.31\|TCCGGCGGCGTCCGGTGAGCTCTCGCTGGCC | >hg38_21_+_8433172_8446622_RNA45SN1_45S_50ntFlanks | 10682 | 10712 | 31 | TCCGGCGGCGTCCGGTGAGCTCTCGCTGGCC | 155.2 | -6.0 | 1.4 | -4.2 | 3.1E-05 | 5.1E-05 | MB231/MB468 | MP |
| >hg38_MT_+_1621_3279_RNR2_16S_50ntFlanks@1367.1407.41\|TCGATGTTGGATCAGGACATCCCGATGGTGCAGCCGCTATT | >hg38_MT_+_1621_3279_RNR2_16S_50ntFlanks | 1367 | 1407 | 41 | TCGATGTTGGATCAGGACATCCCGATGGTGCAGCCGCTATT | 29.1 | 8.1 | 1.9 | 4.2 | 3.1E-05 | 5.1E-05 | MB231/MB468 | MP |
| >hg38_21_+_8433172_8446622_RNA45SN1_45S_50ntFlanks@11600.11622.23\|AATCCGACTGTTTAATTAAAACA | >hg38_21_+_8433172_8446622_RNA45SN1_45S_50ntFlanks | 11600 | 11622 | 23 | AATCCGACTGTTTAATTAAAACA | 109.7 | 7.6 | 1.7 | 4.4 | 1.3E-05 | 5.1E-05 | BT20/MB468 | MP |
| >hg38_21_+_8433172_8446622_RNA45SN1_45S_50ntFlanks@4297.4332.36\|CATTGGAGGGCAAGTCTGGTGCCAGCAGCCGCGGTA | >hg38_21_+_8433172_8446622_RNA45SN1_45S_50ntFlanks | 4297 | 4332 | 36 | CATTGGAGGGCAAGTCTGGTGCCAGCAGCCGCGGTA | 228.9 | -5.8 | 1.4 | -4.1 | 3.7E-05 | 5.1E-05 | MB231/MB468 | NUC |
| >hg38_21_+_8433172_8446622_RNA45SN1_45S_50ntFlanks@10684.10708.25\|CGGCGGCGTCCGGTGAGCTCTCGCT | >hg38_21_+_8433172_8446622_RNA45SN1_45S_50ntFlanks | 10684 | 10708 | 25 | CGGCGGCGTCCGGTGAGCTCTCGCT | 619.5 | -2.5 | 0.6 | -4.2 | 3.2E-05 | 5.1E-05 | BT20/MB468 | CYTO |
| >hg38_21_+_8433172_8446622_RNA45SN1_45S_50ntFlanks@11769.11786.18\|TAATTAGTGACGCGCATG | >hg38_21_+_8433172_8446622_RNA45SN1_45S_50ntFlanks | 11769 | 11786 | 18 | TAATTAGTGACGCGCATG | 665.6 | -4.8 | 1.1 | -4.1 | 3.5E-05 | 5.1E-05 | MB231/MB468 | CYTO |
| >hg38_21_+_8433172_8446622_RNA45SN1_45S_50ntFlanks@12435.12473.39\|ATCATTGTGAAGCAGAATTCACCAAGCGTTGGATTGTTC | >hg38_21_+_8433172_8446622_RNA45SN1_45S_50ntFlanks | 12435 | 12473 | 39 | ATCATTGTGAAGCAGAATTCACCAAGCGTTGGATTGTTC | 40.1 | -8.4 | 1.9 | -4.4 | 1.3E-05 | 5.1E-05 | BT20/MB468 | MP |
| >hg38_21_+_8433172_8446622_RNA45SN1_45S_50ntFlanks@9308.9325.18\|AGGAGTCTAACACGTGCG | >hg38_21_+_8433172_8446622_RNA45SN1_45S_50ntFlanks | 9308 | 9325 | 18 | AGGAGTCTAACACGTGCG | 63.5 | -9.9 | 2.4 | -4.2 | 3.1E-05 | 5.1E-05 | MB231/MB468 | MP |
| >MI0003643\|hsa-mir-629&WithFlank&15\|-\|70079366\|70079474@28.48.21(+1C)&[MIMAT0004810&hsa-miR-629-5p&offsets\|0\|0(+1C);m-150&15\|-\|70079427\|70079447&offsets\|0\|0(+1C)]\|TGGGTTTACGTTGGGAGAACTC | >MI0003643\|hsa-mir-629&WithFlank&15\|-\|70079366\|70079474 | 28 | 48 | 21(+1C) | TGGGTTTACGTTGGGAGAACTC | 237.2 | 3.2 | 0.7 | 4.6 | 4.6E-06 | 5.1E-05 | BT20/MB231 | NUC |
| >hg38_21_+_8433172_8446622_RNA45SN1_45S_50ntFlanks@10922.10939.18\|CTGGACGAGGCGCCGCCG | >hg38_21_+_8433172_8446622_RNA45SN1_45S_50ntFlanks | 10922 | 10939 | 18 | CTGGACGAGGCGCCGCCG | 1495.9 | -5.7 | 1.3 | -4.4 | 1.3E-05 | 5.1E-05 | BT20/MB468 | MP |
| >hg38_21_+_8433172_8446622_RNA45SN1_45S_50ntFlanks@10797.10822.26\|GCATGTTGGAACAATGTAGGTAAGGG | >hg38_21_+_8433172_8446622_RNA45SN1_45S_50ntFlanks | 10797 | 10822 | 26 | GCATGTTGGAACAATGTAGGTAAGGG | 715.8 | -3.3 | 0.8 | -4.4 | 1.3E-05 | 5.1E-05 | BT20/MB468 | MP |
| >am_trna134_GluTTC_1_-_16861774_16861845@1.29.29,trna5_GluTTC_1_+_17199078_17199149@1.29.29,trna84_GluTTC_1_-_161391883_161391954@1.29.29,trna94_GluTTC_1_-_149664355_149664427@1.29.29\|TCCCTGGTGGTCTAGTGGCTAGGATTCGG | >am_trna134_GluTTC_1_-_16861774_16861845 | 1 | 29 | 29 | TCCCTGGTGGTCTAGTGGCTAGGATTCGG | 5763.6 | -8.4 | 2.0 | -4.1 | 3.9E-05 | 5.1E-05 | BT20/MB468 | NUC |
| >hg38_21_+_8433172_8446622_RNA45SN1_45S_50ntFlanks@4588.4611.24\|CTATTTTGTTGGTTTTCGGAACTG | >hg38_21_+_8433172_8446622_RNA45SN1_45S_50ntFlanks | 4588 | 4611 | 24 | CTATTTTGTTGGTTTTCGGAACTG | 207.8 | -5.4 | 1.3 | -4.2 | 3.2E-05 | 5.1E-05 | BT20/MB468 | CYTO |
| >MI0000102\|hsa-mir-100&WithFlank&11\|-\|122152223\|122152314@21.40.20&[MIMAT0000098&hsa-miR-100-5p&offsets\|+2\|0;m-27&11\|-\|122152276\|122152296&offsets\|+2\|+1]\|CCCGTAGATCCGAACTTGTG | >MI0000102\|hsa-mir-100&WithFlank&11\|-\|122152223\|122152314 | 21 | 40 | 20 | CCCGTAGATCCGAACTTGTG | 67.8 | -9.5 | 2.1 | -4.6 | 4.0E-06 | 5.1E-05 | BT20/MB231 | MP |
| >MI0000446\|hsa-mir-125b-1&WithFlank&11\|-\|122099751\|122099850@61.82.22(+1C)&[MIMAT0004592&hsa-miR-125b-1-3p&offsets\|0\|0(+1C);m-485&11\|-\|122099769\|122099790&offsets\|0\|0(+1C)]\|ACGGGTTAGGCTCTTGGGAGCTC | >MI0000446\|hsa-mir-125b-1&WithFlank&11\|-\|122099751\|122099850 | 61 | 82 | 22(+1C) | ACGGGTTAGGCTCTTGGGAGCTC | 68.8 | -9.5 | 2.1 | -4.6 | 4.0E-06 | 5.1E-05 | BT20/MB231 | MP |
| >trna116_GluCTC_1_-_145399233_145399304@38.74.37,trna134_GluTTC_1_-_16861774_16861845@38.74.37,trna71_GluCTC_1_-_161439189_161439260@38.74.37,trna74_GluCTC_1_-_161431809_161431880@38.74.37,trna77_GluCTC_1_-_161424398_161424469@38.74.37,trna77_GluCTC_6_+_28949976_28950047@38.74.37,trna80_GluCTC_1_-_161417018_161417089@38.74.37,trna84_GluTTC_1_-_161391883_161391954@38.74.37,trna87_GluCTC_6_-_126101393_126101464@38.74.37\|CCGCCGCGGCCCGGGTTCGATTCCCGGTCAGGGAACC | >trna116_GluCTC_1_-_145399233_145399304 | 38 | 74 | 37 | CCGCCGCGGCCCGGGTTCGATTCCCGGTCAGGGAACC | 644.7 | 5.2 | 1.3 | 4.1 | 3.5E-05 | 5.2E-05 | MB231/MB468 | CYTO |
| >hg38_21_+_8433172_8446622_RNA45SN1_45S_50ntFlanks@12076.12096.21\|AGGGGCTCTCGCTTCTGGCGC | >hg38_21_+_8433172_8446622_RNA45SN1_45S_50ntFlanks | 12076 | 12096 | 21 | AGGGGCTCTCGCTTCTGGCGC | 805.8 | -7.3 | 1.8 | -4.1 | 3.9E-05 | 5.2E-05 | BT20/MB468 | NUC |
| >hg38_MT_+_1621_3279_RNR2_16S_50ntFlanks@717.749.33\|AAGCCTGCGTCAGATTAAAACACTGAACTGACA | >hg38_MT_+_1621_3279_RNR2_16S_50ntFlanks | 717 | 749 | 33 | AAGCCTGCGTCAGATTAAAACACTGAACTGACA | 28.6 | 8.1 | 1.9 | 4.2 | 3.1E-05 | 5.2E-05 | MB231/MB468 | MP |
| >trna116_GluCTC_1_-_145399233_145399304@37.73.37,trna134_GluTTC_1_-_16861774_16861845@37.73.37,trna71_GluCTC_1_-_161439189_161439260@37.73.37,trna74_GluCTC_1_-_161431809_161431880@37.73.37,trna77_GluCTC_1_-_161424398_161424469@37.73.37,trna77_GluCTC_6_+_28949976_28950047@37.73.37,trna80_GluCTC_1_-_161417018_161417089@37.73.37,trna84_GluTTC_1_-_161391883_161391954@37.73.37,trna87_GluCTC_6_-_126101393_126101464@37.73.37\|ACCGCCGCGGCCCGGGTTCGATTCCCGGTCAGGGAAC | >trna116_GluCTC_1_-_145399233_145399304 | 37 | 73 | 37 | ACCGCCGCGGCCCGGGTTCGATTCCCGGTCAGGGAAC | 4237.4 | 5.3 | 1.3 | 4.1 | 3.5E-05 | 5.2E-05 | MB231/MB468 | CYTO |
| >hg38_21_+_8433172_8446622_RNA45SN1_45S_50ntFlanks@10916.10933.18\|CCGCGGCTGGACGAGGCG | >hg38_21_+_8433172_8446622_RNA45SN1_45S_50ntFlanks | 10916 | 10933 | 18 | CCGCGGCTGGACGAGGCG | 1386.2 | -7.8 | 1.9 | -4.1 | 3.9E-05 | 5.2E-05 | BT20/MB468 | NUC |
| >hg38_21_+_8433172_8446622_RNA45SN1_45S_50ntFlanks@10686.10716.31\|GCGGCGTCCGGTGAGCTCTCGCTGGCCCTTG | >hg38_21_+_8433172_8446622_RNA45SN1_45S_50ntFlanks | 10686 | 10716 | 31 | GCGGCGTCCGGTGAGCTCTCGCTGGCCCTTG | 29.8 | 8.1 | 2.0 | 4.2 | 3.1E-05 | 5.2E-05 | MB231/MB468 | MP |
| >MI0005756\|hsa-mir-934&WithFlank&X\|+\|136550872\|136550966@21.42.22(+2A)&[MIMAT0004977&hsa-miR-934&offsets\|0\|0(+2A);m-943&X\|+\|136550892\|136550913&offsets\|0\|0(+2A)]\|TGTCTACTACTGGAGACACTGGAA | >MI0005756\|hsa-mir-934&WithFlank&X\|+\|136550872\|136550966 | 21 | 42 | 22(+2A) | TGTCTACTACTGGAGACACTGGAA | 72.6 | 9.5 | 2.1 | 4.6 | 4.0E-06 | 5.2E-05 | BT20/MB231 | MP |
| >hg38_21_+_8433172_8446622_RNA45SN1_45S_50ntFlanks@10378.10424.47\|AGGGTTCCATGTGAACAGCAGTTGAACATGGGTCAGTCGGTCCTGAG | >hg38_21_+_8433172_8446622_RNA45SN1_45S_50ntFlanks | 10378 | 10424 | 47 | AGGGTTCCATGTGAACAGCAGTTGAACATGGGTCAGTCGGTCCTGAG | 2761.7 | 4.6 | 1.0 | 4.6 | 4.7E-06 | 5.2E-05 | BT20/MB231 | MITO |
| >am_MI0000809\|hsa-mir-151a&WithFlank&8\|-\|140732558\|140732659@53.73.21(+1C)&[MIMAT0000757&hsa-miR-151a-3p&offsets\|0\|0(+1C);m-30&8\|-\|140732587\|140732607&offsets\|0\|0(+1C)]\|CTAGACTGAAGCTCCTTGAGGC | >am_MI0000809\|hsa-mir-151a&WithFlank&8\|-\|140732558\|140732659 | 53 | 73 | 21(+1C) | CTAGACTGAAGCTCCTTGAGGC | 1088.1 | 6.0 | 1.4 | 4.4 | 1.3E-05 | 5.2E-05 | BT20/MB468 | MP |
| >hg38_21_+_8433172_8446622_RNA45SN1_45S_50ntFlanks@10546.10571.26\|CAGTGCGGTAACGCGACCGATCCCGG | >hg38_21_+_8433172_8446622_RNA45SN1_45S_50ntFlanks | 10546 | 10571 | 26 | CAGTGCGGTAACGCGACCGATCCCGG | 326.9 | -5.3 | 1.3 | -4.2 | 3.2E-05 | 5.2E-05 | MB231/MB468 | MP |
| >hg38_21_+_8433172_8446622_RNA45SN1_45S_50ntFlanks@10821.10862.42\|GGAAGTCGGCAAGCCGGATCCGTAACTTCGGGATAAGGATTG | >hg38_21_+_8433172_8446622_RNA45SN1_45S_50ntFlanks | 10821 | 10862 | 42 | GGAAGTCGGCAAGCCGGATCCGTAACTTCGGGATAAGGATTG | 52.0 | -8.8 | 2.0 | -4.4 | 1.3E-05 | 5.2E-05 | BT20/MB468 | MP |
| >MI0000239\|hsa-mir-197&WithFlank&1\|+\|109598887\|109598973@54.74.21(+1A)&[MIMAT0000227&hsa-miR-197-3p&offsets\|0\|-1(+1A);m-99&1\|+\|109598940\|109598961&offsets\|0\|-1(+1A)]\|TTCACCACCTTCTCCACCCAGA | >MI0000239\|hsa-mir-197&WithFlank&1\|+\|109598887\|109598973 | 54 | 74 | 21(+1A) | TTCACCACCTTCTCCACCCAGA | 1701.6 | 3.9 | 0.8 | 4.6 | 3.9E-06 | 5.2E-05 | BT20/MB231 | CYTO |
| >am_trnaMT_PheGAA_MT_+_577_647@45.62.18\|AGACGGGCTCACATCACC | >am_trnaMT_PheGAA_MT_+_577_647 | 45 | 62 | 18 | AGACGGGCTCACATCACC | 1075.7 | 4.0 | 0.9 | 4.2 | 2.7E-05 | 5.2E-05 | BT20/MB468 | MITO |
| >hg38_21_+_8433172_8446622_RNA45SN1_45S_50ntFlanks@10678.10710.33\|CGGTTCCGGCGGCGTCCGGTGAGCTCTCGCTGG | >hg38_21_+_8433172_8446622_RNA45SN1_45S_50ntFlanks | 10678 | 10710 | 33 | CGGTTCCGGCGGCGTCCGGTGAGCTCTCGCTGG | 149299.4 | -4.7 | 1.1 | -4.2 | 3.3E-05 | 5.2E-05 | BT20/MB468 | CYTO |
| >MI0000298\|hsa-mir-221&WithFlank&X\|-\|45746151\|45746272@31.51.21&[MIMAT0004568&hsa-miR-221-5p&offsets\|0\|-1;m-405&X\|-\|45746220\|45746242&offsets\|0\|-2]\|ACCTGGCATACAATGTAGATT | >MI0000298\|hsa-mir-221&WithFlank&X\|-\|45746151\|45746272 | 31 | 51 | 21 | ACCTGGCATACAATGTAGATT | 35.2 | 8.4 | 2.0 | 4.2 | 3.2E-05 | 5.2E-05 | MB231/MB468 | MP |
| >MI0000272\|hsa-mir-182&WithFlank&7\|-\|129770377\|129770498@29.53.25&[MIMAT0000259&hsa-miR-182-5p&offsets\|0\|+1;m-22&7\|-\|129770449\|129770470&offsets\|0\|+3]\|TTTGGCAATGGTAGAACTCACACTG | >MI0000272\|hsa-mir-182&WithFlank&7\|-\|129770377\|129770498 | 29 | 53 | 25 | TTTGGCAATGGTAGAACTCACACTG | 12941.4 | 3.5 | 0.7 | 4.6 | 3.9E-06 | 5.2E-05 | BT20/MB231 | CYTO |

| | | | | | | | | | | | | | |
|---|---|---|---|---|---|---|---|---|---|---|---|---|---|
| >MI0003776\|hsa-mir-320b-1&WithFlank&1\|+\|116671740\|116671823@48.66.19;MI0003839\|hsa-mir-320b-2&WithFlank&1\|-\|224257034\|224257116@47.65.19;MI0003778\|hsa-mir-320c-1&WithFlank&18\|+\|21683512\|21683595@48.66.19;MI0008191\|hsa-mir-320c-2&WithFlank&18\|+\|24321669\|24321752@48.66.19;MI0000542\|hsa-mir-320a&WithFlank&8\|-\|22244960\|22245043@48.66.19&[MIMAT0005792&hsa-miR-320b&offsets\|0\|-3;m-183&1\|+\|116671787\|116671806&offsets\|0\|-1];[MIMAT0005792_1&hsa-miR-320b&offsets\|0\|-3;m-183&1\|-\|224257051\|224257070&offsets\|0\|-1];[MIMAT0005793&hsa-miR-320c&offsets\|0\|-1;m-222&18\|+\|21683559\|21683578&offsets\|0\|-1];[MIMAT0005793_1&hsa-miR-320c&offsets\|0\|-1;m-219&18\|+\|24321716\|24321735&offsets\|0\|-1];[MIMAT0000510&hsa-miR-320a-3p&offsets\|0\|-3;m-60&8\|-\|22244975\|22244996&offsets\|0\|-3]\|AAAAGCTGGGTTGAGAGGG | >MI0003776\|hsa-mir-320b-1&WithFlank&1\|+\|116671740\|116671823 | 48 | 66 | 19 | AAAAGCTGGGTTGAGAGGG | 566.1 | 4.0 | 0.9 | 4.6 | 3.9E-06 | 5.2E-05 | BT20/MB231 | CYTO |
| >hg38_21_+_8433172_8446622_RNA45SN1_45S_50ntFlanks@8356.8375.20\|GAAAAGAACTTTGAAGAGAG | >hg38_21_+_8433172_8446622_RNA45SN1_45S_50ntFlanks | 8356 | 8375 | 20 | GAAAAGAACTTTGAAGAGAG | 606.3 | 5.4 | 1.2 | 4.6 | 4.8E-06 | 5.2E-05 | BT20/MB231 | MITO |
| >hg38_21_+_8433172_8446622_RNA45SN1_45S_50ntFlanks@12360.12382.23\|CAGGGATAACTGGCTTGTGGCGG | >hg38_21_+_8433172_8446622_RNA45SN1_45S_50ntFlanks | 12360 | 12382 | 23 | CAGGGATAACTGGCTTGTGGCGG | 187.8 | -2.5 | 0.6 | -4.2 | 2.7E-05 | 5.2E-05 | BT20/MB468 | MITO |
| >MI0000273\|hsa-mir-183&WithFlank&7\|-\|129774899\|129775020@33.53.21(+1G)&[MIMAT0000261&hsa-miR-183-5p&offsets\|0\|-1(+1G);m-41&7\|-\|129774966\|129774987&offsets\|-1\|-2(+1G)]\|TATGGCACTGGTAGAATTCACG | >MI0000273\|hsa-mir-183&WithFlank&7\|-\|129774899\|129775020 | 33 | 53 | 21(+1G) | TATGGCACTGGTAGAATTCACG | 432.6 | 4.0 | 0.9 | 4.6 | 3.9E-06 | 5.2E-05 | BT20/MB231 | CYTO |
| >hg38_21_+_8433172_8446622_RNA45SN1_45S_50ntFlanks@7973.7993.21\|GACGCGACCTCAGATCAGACG | >hg38_21_+_8433172_8446622_RNA45SN1_45S_50ntFlanks | 7973 | 7993 | 21 | GACGCGACCTCAGATCAGACG | 1517.5 | 3.8 | 0.9 | 4.4 | 1.3E-05 | 5.2E-05 | BT20/MB468 | MP |
| >hg38_21_+_8433172_8446622_RNA45SN1_45S_50ntFlanks@11682.11702.21\|AATGTCAAAGTGAAGAAATTC | >hg38_21_+_8433172_8446622_RNA45SN1_45S_50ntFlanks | 11682 | 11702 | 21 | AATGTCAAAGTGAAGAAATTC | 730.2 | -7.1 | 1.6 | -4.4 | 1.3E-05 | 5.2E-05 | BT20/MB468 | MP |
| >hg38_21_+_8433172_8446622_RNA45SN1_45S_50ntFlanks@10772.10796.25\|AGGTCTCCAAGGTGAACAGCCTCTG | >hg38_21_+_8433172_8446622_RNA45SN1_45S_50ntFlanks | 10772 | 10796 | 25 | AGGTCTCCAAGGTGAACAGCCTCTG | 417.5 | -6.4 | 1.5 | -4.2 | 3.2E-05 | 5.2E-05 | MB231/MB468 | MP |
| >MI0000087\|hsa-mir-29a&WithFlank&7\|-\|130876741\|130876816@48.65.18(+1A)&[MIMAT0000086&hsa-miR-29a-3p&offsets\|0\|-4(+1A);m-26&7\|-\|130876749\|130876769&offsets\|0\|-3(+1A)]\|TAGCACCATCTGAAATCGA | >MI0000087\|hsa-mir-29a&WithFlank&7\|-\|130876741\|130876816 | 48 | 65 | 18(+1A) | TAGCACCATCTGAAATCGA | 40.6 | 8.6 | 2.1 | 4.2 | 3.2E-05 | 5.2E-05 | MB231/MB468 | MP |
| >hg38_21_+_8433172_8446622_RNA45SN1_45S_50ntFlanks@4772.4793.22\|GTAGTTCCGACCATAAACGATG | >hg38_21_+_8433172_8446622_RNA45SN1_45S_50ntFlanks | 4772 | 4793 | 22 | GTAGTTCCGACCATAAACGATG | 264.4 | -6.0 | 1.4 | -4.1 | 3.8E-05 | 5.2E-05 | MB231/MB468 | MITO |
| >hg38_21_+_8433172_8446622_RNA45SN1_45S_50ntFlanks@5237.5267.31\|ACTGACTGGCTCAGCGTGTGCCTACCCTACG | >hg38_21_+_8433172_8446622_RNA45SN1_45S_50ntFlanks | 5237 | 5267 | 31 | ACTGACTGGCTCAGCGTGTGCCTACCCTACG | 167.4 | -6.1 | 1.5 | -4.2 | 3.2E-05 | 5.2E-05 | MB231/MB468 | MP |
| >trnaMT_HisGTG_MT_+_12138_12206@18.58.41\|AAAACATCAGATTGTGAATCTGACAACAGAGGCTTACGACC | >trnaMT_HisGTG_MT_+_12138_12206 | 18 | 58 | 41 | AAAACATCAGATTGTGAATCTGACAACAGAGGCTTACGACC | 30.1 | 8.1 | 2.0 | 4.2 | 3.2E-05 | 5.2E-05 | MB231/MB468 | MP |
| >hg38_21_+_8433172_8446622_RNA45SN1_45S_50ntFlanks@12444.12478.35\|AAGCAGAATTCACCAAGCGTTGGATTGTTCACCCA | >hg38_21_+_8433172_8446622_RNA45SN1_45S_50ntFlanks | 12444 | 12478 | 35 | AAGCAGAATTCACCAAGCGTTGGATTGTTCACCCA | 246.9 | -5.4 | 1.3 | -4.2 | 3.2E-05 | 5.2E-05 | MB231/MB468 | MP |
| >MI0000298\|hsa-mir-221&WithFlank&X\|-\|45746151\|45746272@71.90.20(+1C)&[MIMAT0000278&hsa-miR-221-3p&offsets\|0\|-3(+1C);m-91&X\|-\|45746181\|45746202&offsets\|0\|-2(+1C)]\|AGCTACATTGTCTGCTGGGTC | >MI0000298\|hsa-mir-221&WithFlank&X\|-\|45746151\|45746272 | 71 | 90 | 20(+1C) | AGCTACATTGTCTGCTGGGTC | 845.4 | 5.3 | 1.3 | 4.2 | 3.2E-05 | 5.2E-05 | MB231/MB468 | MP |
| >hg38_21_+_8433172_8446622_RNA45SN1_45S_50ntFlanks@11317.11336.20\|ACCCCACGTCTCGTCGCGCG | >hg38_21_+_8433172_8446622_RNA45SN1_45S_50ntFlanks | 11317 | 11336 | 20 | ACCCCACGTCTCGTCGCGCG | 893.3 | -4.7 | 1.1 | -4.2 | 3.3E-05 | 5.2E-05 | BT20/MB468 | CYTO |
| >hg38_21_+_8433172_8446622_RNA45SN1_45S_50ntFlanks@4472.4493.22\|CTTAGCTGAGTGTCCCGCGGGG | >hg38_21_+_8433172_8446622_RNA45SN1_45S_50ntFlanks | 4472 | 4493 | 22 | CTTAGCTGAGTGTCCCGCGGGG | 167.7 | -5.6 | 1.3 | -4.2 | 2.8E-05 | 5.2E-05 | BT20/MB468 | MITO |
| >hg38_21_+_8433172_8446622_RNA45SN1_45S_50ntFlanks@10423.10456.34\|AGAGATGGGCGAGCGCCGTTCCGAAGGGACGGGC | >hg38_21_+_8433172_8446622_RNA45SN1_45S_50ntFlanks | 10423 | 10456 | 34 | AGAGATGGGCGAGCGCCGTTCCGAAGGGACGGGC | 122.0 | -6.0 | 1.4 | -4.2 | 3.2E-05 | 5.2E-05 | MB231/MB468 | MP |
| >hg38_21_+_8433172_8446622_RNA45SN1_45S_50ntFlanks@10829.10853.25\|GCAAGCCGGATCCGTAACTTCGGGA | >hg38_21_+_8433172_8446622_RNA45SN1_45S_50ntFlanks | 10829 | 10853 | 25 | GCAAGCCGGATCCGTAACTTCGGGA | 242.1 | -3.2 | 0.8 | -4.1 | 3.6E-05 | 5.2E-05 | MB231/MB468 | CYTO |
| >hg38_21_+_8433172_8446622_RNA45SN1_45S_50ntFlanks@4297.4319.23\|CATTGGAGGGCAAGTCTGGTGCC | >hg38_21_+_8433172_8446622_RNA45SN1_45S_50ntFlanks | 4297 | 4319 | 23 | CATTGGAGGGCAAGTCTGGTGCC | 285.1 | -3.8 | 0.9 | -4.1 | 3.6E-05 | 5.2E-05 | MB231/MB468 | CYTO |
| >MI0000077\|hsa-mir-21&WithFlank&17\|+\|59841260\|59841343@14.34.21(+1C)&[MIMAT0000076&hsa-miR-21-5p&offsets\|0\|-1(+1C);m-1&17\|+\|59841273\|59841294&offsets\|0\|-1(+1C)]\|TAGCTTATCAGACTGATGTTGC | >MI0000077\|hsa-mir-21&WithFlank&17\|+\|59841260\|59841343 | 14 | 34 | 21(+1C) | TAGCTTATCAGACTGATGTTGC | 462.2 | 8.1 | 2.0 | 4.2 | 3.3E-05 | 5.3E-05 | BT20/MB468 | CYTO |
| >am_trnaMT_LysTTT_MT_+_8295_8364@30.52.23,trnalookalike7_LysTTT_1_+_568844_568913@30.52.23\|TTAAGTTAAAGATTAAGAGAACC | >am_trnaMT_LysTTT_MT_+_8295_8364 | 30 | 52 | 23 | TTAAGTTAAAGATTAAGAGAACC | 527.7 | 3.8 | 0.8 | 4.6 | 4.0E-06 | 5.3E-05 | BT20/MB231 | CYTO |
| >am_trna128_GlyGCC_6_-_27870686_27870756@2.30.29,trna133_GlyCCC_1_-_16872434_16872504@2.30.29,trna18_GlyGCC_16_+_70822597_70822667@2.30.29,trna19_GlyGCC_16_+_70823410_70823480@2.30.29,trna19_GlyGCC_2_-_157257659_157257729@2.30.29,trna24_GlyGCC_16_-_70812942_70813012@2.30.29,trna25_GlyGCC_16_-_70812114_70812184@2.30.29,trna4_GlyCCC_1_+_17188416_17188486@2.30.29,trna5_GlyGCC_17_+_8029064_8029134@2.30.29,trna68_GlyGCC_1_-_161493637_161493707@2.30.29\|CATTGGTGGTTCAGTGGTAGAATTCTCGC | >am_trna128_GlyGCC_6_-_27870686_27870756 | 2 | 30 | 29 | CATTGGTGGTTCAGTGGTAGAATTCTCGC | 271.9 | 6.3 | 1.5 | 4.2 | 3.2E-05 | 5.3E-05 | MB231/MB468 | MP |
| >hg38_21_+_8433172_8446622_RNA45SN1_45S_50ntFlanks@9821.9838.18\|CCTAGTGGGCCACTTTTG | >hg38_21_+_8433172_8446622_RNA45SN1_45S_50ntFlanks | 9821 | 9838 | 18 | CCTAGTGGGCCACTTTTG | 316.3 | -6.9 | 1.6 | -4.4 | 1.3E-05 | 5.3E-05 | BT20/MB468 | MP |
| >MI0003137\|hsa-mir-193b&WithFlank&16\|+\|14303961\|14304055@57.78.22&[MIMAT0002819&hsa-miR-193b-3p&offsets\|0\|0;m-166&16\|+\|14304017\|14304038&offsets\|0\|0]\|AACTGGCCCTCAAAGTCCCGCT | >MI0003137\|hsa-mir-193b&WithFlank&16\|+\|14303961\|14304055 | 57 | 78 | 22 | AACTGGCCCTCAAAGTCCCGCT | 347.9 | 4.5 | 1.1 | 4.1 | 3.8E-05 | 5.3E-05 | MB231/MB468 | MITO |
| >hg38_21_+_8433172_8446622_RNA45SN1_45S_50ntFlanks@10351.10391.41\|CGAGAACTTTGAAGGCCGAAGTGGAGAAGGGTTCCATGTGA | >hg38_21_+_8433172_8446622_RNA45SN1_45S_50ntFlanks | 10351 | 10391 | 41 | CGAGAACTTTGAAGGCCGAAGTGGAGAAGGGTTCCATGTGA | 466.6 | -3.5 | 0.8 | -4.1 | 3.6E-05 | 5.3E-05 | MB231/MB468 | CYTO |
| >hg38_21_+_8433172_8446622_RNA45SN1_45S_50ntFlanks@12388.12437.50\|CGTTCATAGCGACGTCGCTTTTTGATCCTTCGATGTCGGCTCTTCCTATC | >hg38_21_+_8433172_8446622_RNA45SN1_45S_50ntFlanks | 12388 | 12437 | 50 | CGTTCATAGCGACGTCGCTTTTTGATCCTTCGATGTCGGCTCTTCCTATC | 194.8 | -5.1 | 1.1 | -4.6 | 4.0E-06 | 5.3E-05 | BT20/MB231 | CYTO |
| >hg38_21_+_8433172_8446622_RNA45SN1_45S_50ntFlanks@11235.11259.25\|CTTCCCGTGGATCGCCCCAGCTGCG | >hg38_21_+_8433172_8446622_RNA45SN1_45S_50ntFlanks | 11235 | 11259 | 25 | CTTCCCGTGGATCGCCCCAGCTGCG | 119.0 | -3.4 | 0.8 | -4.2 | 3.3E-05 | 5.3E-05 | BT20/MB468 | CYTO |
| >hg38_21_+_8433172_8446622_RNA45SN1_45S_50ntFlanks@4735.4752.18\|ACGAAAGTCGGAGGTTCG | >hg38_21_+_8433172_8446622_RNA45SN1_45S_50ntFlanks | 4735 | 4752 | 18 | ACGAAAGTCGGAGGTTCG | 130.1 | -6.0 | 1.4 | -4.2 | 3.2E-05 | 5.3E-05 | MB231/MB468 | MP |
| >hg38_21_+_8433172_8446622_RNA45SN1_45S_50ntFlanks@12458.12475.18\|AAGCGTTGGATTGTTCAC | >hg38_21_+_8433172_8446622_RNA45SN1_45S_50ntFlanks | 12458 | 12475 | 18 | AAGCGTTGGATTGTTCAC | 250.6 | -2.8 | 0.7 | -4.1 | 3.8E-05 | 5.3E-05 | MB231/MB468 | NUC |
| >hg38_21_+_8433172_8446622_RNA45SN1_45S_50ntFlanks@12927.12956.30\|CCTGGCGCTAAACCATTCGTAGACGACCTG | >hg38_21_+_8433172_8446622_RNA45SN1_45S_50ntFlanks | 12927 | 12956 | 30 | CCTGGCGCTAAACCATTCGTAGACGACCTG | 559.5 | -5.8 | 1.4 | -4.2 | 2.8E-05 | 5.3E-05 | BT20/MB468 | MITO |
| >hg38_21_+_8433172_8446622_RNA45SN1_45S_50ntFlanks@4760.4786.27\|TCAGATACCGTCGTAGTTCCGACCATA | >hg38_21_+_8433172_8446622_RNA45SN1_45S_50ntFlanks | 4760 | 4786 | 27 | TCAGATACCGTCGTAGTTCCGACCATA | 412.0 | -6.1 | 1.5 | -4.2 | 2.8E-05 | 5.3E-05 | BT20/MB468 | MITO |
| >hg38_21_+_8433172_8446622_RNA45SN1_45S_50ntFlanks@5379.5397.19\|ATTAAGTCCCTGCCCTTTG | >hg38_21_+_8433172_8446622_RNA45SN1_45S_50ntFlanks | 5379 | 5397 | 19 | ATTAAGTCCCTGCCCTTTG | 1200.1 | -4.6 | 1.1 | -4.2 | 2.8E-05 | 5.3E-05 | BT20/MB468 | MITO |
| >trnaMT_SerGCT_MT_+_12207_12265@13.48.36\|AAGAACTGCTAACTCATGCCCCCATGTCTAACAACA | >trnaMT_SerGCT_MT_+_12207_12265 | 13 | 48 | 36 | AAGAACTGCTAACTCATGCCCCCATGTCTAACAACA | 136.9 | 5.0 | 1.2 | 4.1 | 3.9E-05 | 5.3E-05 | MB231/MB468 | MITO |
| >trna116_GluCTC_1_-_145399233_145399304@1.32.32,trna59_GluCTC_1_+_249168447_249168518@1.32.32,trna71_GluCTC_1_-_161439189_161439260@1.32.32,trna74_GluCTC_1_-_161431809_161431880@1.32.32,trna77_GluCTC_1_-_161424398_161424469@1.32.32,trna77_GluCTC_6_+_28949976_28950047@1.32.32,trna80_GluCTC_1_-_161417018_161417089@1.32.32,trna87_GluCTC_6_-_126101393_126101464@1.32.32\|TCCCTGGTGGTCTAGTGGTTAGGATTCGGCGC | >trna116_GluCTC_1_-_145399233_145399304 | 1 | 32 | 32 | TCCCTGGTGGTCTAGTGGTTAGGATTCGGCGC | 2117.1 | 4.8 | 1.2 | 4.1 | 3.8E-05 | 5.3E-05 | MB231/MB468 | NUC |
| >trna116_GluCTC_1_-_145399233_145399304@1.32.32,trna59_GluCTC_1_+_249168447_249168518@1.32.32,trna71_GluCTC_1_-_161439189_161439260@1.32.32,trna74_GluCTC_1_-_161431809_161431880@1.32.32,trna77_GluCTC_1_-_161424398_161424469@1.32.32,trna77_GluCTC_6_+_28949976_28950047@1.32.32,trna80_GluCTC_1_-_161417018_161417089@1.32.32,trna87_GluCTC_6_-_126101393_126101464@1.32.32\|TCCCTGGTGGTCTAGTGGTTAGGATTCGGCGC | >trna116_GluCTC_1_-_145399233_145399304 | 1 | 32 | 32 | TCCCTGGTGGTCTAGTGGTTAGGATTCGGCGC | 1182.9 | 5.2 | 1.2 | 4.4 | 1.3E-05 | 5.3E-05 | BT20/MB468 | MP |
| >MI0000342\|hsa-mir-200b&WithFlank&1\|+\|1167098\|1167204@63.85.23(+1U)&[MIMAT0000318&hsa-miR-200b-3p&offsets\|0\|+1(+1U);m-93&1\|+\|1167160\|1167182&offsets\|0\|0(+1U)]\|TAATACTGCCTGGTAATGATGACT | >MI0000342\|hsa-mir-200b&WithFlank&1\|+\|1167098\|1167204 | 63 | 85 | 23(+1U) | TAATACTGCCTGGTAATGATGACT | 914.4 | -3.5 | 0.8 | -4.1 | 3.6E-05 | 5.3E-05 | MB231/MB468 | CYTO |
| >hg38_21_+_8433172_8446622_RNA45SN1_45S_50ntFlanks@9747.9767.21\|TCAACCTATTCTCAAACTTTA | >hg38_21_+_8433172_8446622_RNA45SN1_45S_50ntFlanks | 9747 | 9767 | 21 | TCAACCTATTCTCAAACTTTA | 277.7 | 4.6 | 1.0 | 4.6 | 4.9E-06 | 5.3E-05 | BT20/MB231 | MITO |
| >hg38_21_+_8433172_8446622_RNA45SN1_45S_50ntFlanks@10350.10390.41\|ACGAGAACTTTGAAGGCCGAAGTGGAGAAGGGTTCCATGTG | >hg38_21_+_8433172_8446622_RNA45SN1_45S_50ntFlanks | 10350 | 10390 | 41 | ACGAGAACTTTGAAGGCCGAAGTGGAGAAGGGTTCCATGTG | 2823.1 | -3.4 | 0.8 | -4.2 | 2.8E-05 | 5.3E-05 | BT20/MB468 | MITO |
| >hg38_21_+_8433172_8446622_RNA45SN1_45S_50ntFlanks@4046.4081.36\|CATTCGAACGTCTGCCCTATCAACTTTCGATGGTAG | >hg38_21_+_8433172_8446622_RNA45SN1_45S_50ntFlanks | 4046 | 4081 | 36 | CATTCGAACGTCTGCCCTATCAACTTTCGATGGTAG | 49.4 | -8.7 | 2.0 | -4.4 | 1.3E-05 | 5.3E-05 | BT20/MB468 | MP |
| >am_trnaMT_LysTTT_MT_+_8295_8364@35.52.18,trnalookalike7_LysTTT_1_+_568844_568913@35.52.18\|TTAAAGATTAAGAGAACC | >am_trnaMT_LysTTT_MT_+_8295_8364 | 35 | 52 | 18 | TTAAAGATTAAGAGAACC | 28.9 | 8.1 | 1.9 | 4.2 | 3.2E-05 | 5.3E-05 | MB231/MB468 | MP |

| | | | | | | | | | | | | | |
|---|---|---|---|---|---|---|---|---|---|---|---|---|---|
| >trna116_GluCTC_1_-_145399233_145399304@1.32.32,trna59_GluCTC_1_+_249168447_249168518@1.32.32,trna71_GluCTC_1_-_161439189_161439260@1.32.32,trna74_GluCTC_1_-_161431809_161431880@1.32.32,trna77_GluCTC_1_-_161424398_161424469@1.32.32,trna77_GluCTC_6_+_28949976_28950047@1.32.32,trna80_GluCTC_1_-_161417018_161417089@1.32.32,trna87_GluCTC_6_-_126101393_126101464@1.32.32\|TCCCTGGTGGTCTAGTGGTTAGGATTCGGCGC | >trna116_GluCTC_1_-_145399233_145399304 | 1 | 32 | 32 | TCCCTGGTGGTCTAGTGGTTAGGATTCGGCGC | 844.3 | 4.0 | 1.0 | 4.2 | 3.2E-05 | 5.3E-05 | MB231/MB468 | MP |
| >hg38_21_+_8433172_8446622_RNA45SN1_45S_50ntFlanks@6732.6753.22\|ACATTGATCATCGACACTTCGA | >hg38_21_+_8433172_8446622_RNA45SN1_45S_50ntFlanks | 6732 | 6753 | 22 | ACATTGATCATCGACACTTCGA | 1018.1 | 3.1 | 0.7 | 4.2 | 2.8E-05 | 5.3E-05 | BT20/MB468 | MITO |
| >hg38_21_+_8433172_8446622_RNA45SN1_45S_50ntFlanks@10351.10369.19\|CGAGAACTTTGAAGGCCGA | >hg38_21_+_8433172_8446622_RNA45SN1_45S_50ntFlanks | 10351 | 10369 | 19 | CGAGAACTTTGAAGGCCGA | 889.9 | -3.3 | 0.8 | -4.1 | 3.9E-05 | 5.3E-05 | MB231/MB468 | MITO |
| >hg38_21_+_8433172_8446622_RNA45SN1_45S_50ntFlanks@10830.10875.46\|CAAGCCGGATCCGTAACTTCGGGATAAGGATTGGCTCTAAGGGCTG | >hg38_21_+_8433172_8446622_RNA45SN1_45S_50ntFlanks | 10830 | 10875 | 46 | CAAGCCGGATCCGTAACTTCGGGATAAGGATTGGCTCTAAGGGCTG | 2003.5 | -6.4 | 1.5 | -4.1 | 3.9E-05 | 5.3E-05 | MB231/MB468 | NUC |
| >hg38_21_+_8433172_8446622_RNA45SN1_45S_50ntFlanks@11873.11909.37\|AAGAAGACCCTGTTGAGCTTGACTCTAGTCTGGCACG | >hg38_21_+_8433172_8446622_RNA45SN1_45S_50ntFlanks | 11873 | 11909 | 37 | AAGAAGACCCTGTTGAGCTTGACTCTAGTCTGGCACG | 227.2 | -4.2 | 1.0 | -4.1 | 3.7E-05 | 5.3E-05 | MB231/MB468 | CYTO |
| >hg38_21_+_8433172_8446622_RNA45SN1_45S_50ntFlanks@11567.11605.39\|AGCAGCCGACTTAGAACTGGTGCGGACCAGGGGAATCCG | >hg38_21_+_8433172_8446622_RNA45SN1_45S_50ntFlanks | 11567 | 11605 | 39 | AGCAGCCGACTTAGAACTGGTGCGGACCAGGGGAATCCG | 468.9 | -6.3 | 1.5 | -4.1 | 3.9E-05 | 5.3E-05 | MB231/MB468 | MITO |
| >hg38_1_-_228634819_228635039_RNA5S12_5S_50ntFlanks@140.170.31\|ACCGCCTGGGAATACCGGGTGCTGTAGGCTT | >hg38_1_-_228634819_228635039_RNA5S12_5S_50ntFlanks | 140 | 170 | 31 | ACCGCCTGGGAATACCGGGTGCTGTAGGCTT | 16807.3 | 3.5 | 0.9 | 4.1 | 3.7E-05 | 5.3E-05 | MB231/MB468 | CYTO |
| >hg38_21_+_8433172_8446622_RNA45SN1_45S_50ntFlanks@11537.11582.46\|CCGCGCGGCGCCTCGCCTCGGCCGGCGCCTAGCAGCCGACTTAGAA | >hg38_21_+_8433172_8446622_RNA45SN1_45S_50ntFlanks | 11537 | 11582 | 46 | CCGCGCGGCGCCTCGCCTCGGCCGGCGCCTAGCAGCCGACTTAGAA | 317.2 | -4.3 | 1.1 | -4.1 | 3.9E-05 | 5.3E-05 | MB231/MB468 | MITO |
| >trnaMT_PheGAA_MT_+_577_647@38.57.20\|AATGTTTAGACGGGCTCACA | >trnaMT_PheGAA_MT_+_577_647 | 38 | 57 | 20 | AATGTTTAGACGGGCTCACA | 241.1 | 4.4 | 1.0 | 4.4 | 1.3E-05 | 5.3E-05 | BT20/MB468 | MP |
| >hg38_21_+_8433172_8446622_RNA45SN1_45S_50ntFlanks@10548.10576.29\|GTGCGGTAACGCGACCGATCCCGGAGAAG | >hg38_21_+_8433172_8446622_RNA45SN1_45S_50ntFlanks | 10548 | 10576 | 29 | GTGCGGTAACGCGACCGATCCCGGAGAAG | 578.6 | 4.4 | 1.0 | 4.6 | 4.9E-06 | 5.3E-05 | BT20/MB231 | MITO |
| >hg38_21_+_8433172_8446622_RNA45SN1_45S_50ntFlanks@9311.9338.28\|AGTCTAACACGTGCGCGAGTCGGGGGCT | >hg38_21_+_8433172_8446622_RNA45SN1_45S_50ntFlanks | 9311 | 9338 | 28 | AGTCTAACACGTGCGCGAGTCGGGGGCT | 230.0 | -6.3 | 1.5 | -4.2 | 3.3E-05 | 5.4E-05 | MB231/MB468 | MP |
| >trnaMT_IleGAT_MT_+_4263_4331@25.63.39\|ACTTTGATAGAGTAAATAATAGGAGCTTAAACCCCCTTA | >trnaMT_IleGAT_MT_+_4263_4331 | 25 | 63 | 39 | ACTTTGATAGAGTAAATAATAGGAGCTTAAACCCCCTTA | 174.6 | 11.2 | 2.6 | 4.4 | 1.3E-05 | 5.4E-05 | BT20/MB468 | MP |
| >hg38_21_+_8433172_8446622_RNA45SN1_45S_50ntFlanks@10342.10361.20\|ATATTCAAACGAGAACTTTG | >hg38_21_+_8433172_8446622_RNA45SN1_45S_50ntFlanks | 10342 | 10361 | 20 | ATATTCAAACGAGAACTTTG | 694.1 | -3.2 | 0.8 | -4.1 | 3.4E-05 | 5.4E-05 | BT20/MB468 | CYTO |
| >hg38_21_+_8433172_8446622_RNA45SN1_45S_50ntFlanks@12457.12484.28\|CAAGCGTTGGATTGTTCACCCACTAATA | >hg38_21_+_8433172_8446622_RNA45SN1_45S_50ntFlanks | 12457 | 12484 | 28 | CAAGCGTTGGATTGTTCACCCACTAATA | 691.5 | -7.0 | 1.7 | -4.1 | 3.4E-05 | 5.4E-05 | BT20/MB468 | CYTO |
| >MI0000462\|hsa-mir-152&WithFlank&17\|-\|48037155\|48037253@60.80.21&[MIMAT0000438&hsa-miR-152-3p&offsets\|0\|0;m-97&17\|-\|48037174\|48037194&offsets\|0\|0]\|TCAGTGCATGACAGAACTTGG | >MI0000462\|hsa-mir-152&WithFlank&17\|-\|48037155\|48037253 | 60 | 80 | 21 | TCAGTGCATGACAGAACTTGG | 892.6 | 7.7 | 1.8 | 4.4 | 1.3E-05 | 5.4E-05 | BT20/MB468 | MP |
| >am_trnaMT_SerGCT_MT_+_12207_12265@11.38.28\|ACAAGAACTGCTAACTCATGCCCCCATG | >am_trnaMT_SerGCT_MT_+_12207_12265 | 11 | 38 | 28 | ACAAGAACTGCTAACTCATGCCCCCATG | 517.5 | -5.2 | 1.3 | -4.1 | 3.9E-05 | 5.4E-05 | MB231/MB468 | NUC |
| >hg38_21_+_8433172_8446622_RNA45SN1_45S_50ntFlanks@11769.11794.26\|TAATTAGTGACGCGCATGAATGGATG | >hg38_21_+_8433172_8446622_RNA45SN1_45S_50ntFlanks | 11769 | 11794 | 26 | TAATTAGTGACGCGCATGAATGGATG | 1127.8 | -2.1 | 0.5 | -4.1 | 3.4E-05 | 5.4E-05 | BT20/MB468 | CYTO |
| >hg38_21_+_8433172_8446622_RNA45SN1_45S_50ntFlanks@10664.10710.47\|CTTGGAAAGCGTCGCGGTTCCGGCGGCGTCCGGTGAGCTCTCGCTGG | >hg38_21_+_8433172_8446622_RNA45SN1_45S_50ntFlanks | 10664 | 10710 | 47 | CTTGGAAAGCGTCGCGGTTCCGGCGGCGTCCGGTGAGCTCTCGCTGG | 56.0 | -8.9 | 2.0 | -4.4 | 1.3E-05 | 5.4E-05 | BT20/MB468 | MP |
| >MI0000478\|hsa-mir-149&WithFlank&2\|+\|240455995\|240456095@21.40.20&[MIMAT0000450&hsa-miR-149-5p&offsets\|0\|-3;m-195&2\|+\|240456015\|240456037&offsets\|0\|-3]\|TCTGGCTCCGTGTCTTCACT | >MI0000478\|hsa-mir-149&WithFlank&2\|+\|240455995\|240456095 | 21 | 40 | 20 | TCTGGCTCCGTGTCTTCACT | 33.7 | 8.3 | 2.0 | 4.2 | 3.3E-05 | 5.4E-05 | MB231/MB468 | MP |
| >hg38_21_+_8433172_8446622_RNA45SN1_45S_50ntFlanks@12625.12661.37\|CAATGGGGCGAAGCTACCATCTGTGGGATTATGACTG | >hg38_21_+_8433172_8446622_RNA45SN1_45S_50ntFlanks | 12625 | 12661 | 37 | CAATGGGGCGAAGCTACCATCTGTGGGATTATGACTG | 678.8 | -7.1 | 1.7 | -4.1 | 4.1E-05 | 5.4E-05 | BT20/MB468 | NUC |
| >hg38_21_+_8433172_8446622_RNA45SN1_45S_50ntFlanks@6731.6753.23\|CACATTGATCATCGACACTTCGA | >hg38_21_+_8433172_8446622_RNA45SN1_45S_50ntFlanks | 6731 | 6753 | 23 | CACATTGATCATCGACACTTCGA | 381.4 | -3.0 | 0.7 | -4.1 | 3.7E-05 | 5.4E-05 | MB231/MB468 | CYTO |
| >hg38_21_+_8433172_8446622_RNA45SN1_45S_50ntFlanks@6649.6689.41\|TACGACTCTTAGCGGTGGATCACTCGGCTCGTGCGTCGATG | >hg38_21_+_8433172_8446622_RNA45SN1_45S_50ntFlanks | 6649 | 6689 | 41 | TACGACTCTTAGCGGTGGATCACTCGGCTCGTGCGTCGATG | 1373.7 | -2.9 | 0.7 | -4.1 | 4.1E-05 | 5.4E-05 | BT20/MB468 | NUC |
| >MI0005566\|hsa-mir-374b&WithFlank&X\|-\|74218541\|74218624@17.39.23&[MIMAT0004955&hsa-miR-374b-5p&offsets\|0\|+1;m-153&X\|-\|74218587\|74218608&offsets\|0\|+1]\|ATATAATACAACCTGCTAAGTGT | >MI0005566\|hsa-mir-374b&WithFlank&X\|-\|74218541\|74218624 | 17 | 39 | 23 | ATATAATACAACCTGCTAAGTGT | 542.6 | -4.0 | 0.9 | -4.6 | 5.0E-06 | 5.4E-05 | BT20/MB231 | MITO |
| >hg38_21_+_8433172_8446622_RNA45SN1_45S_50ntFlanks@10350.10385.36\|ACGAGAACTTTGAAGGCCGAAGTGGAGAAGGGTTCC | >hg38_21_+_8433172_8446622_RNA45SN1_45S_50ntFlanks | 10350 | 10385 | 36 | ACGAGAACTTTGAAGGCCGAAGTGGAGAAGGGTTCC | 189.5 | -5.1 | 1.2 | -4.2 | 3.3E-05 | 5.4E-05 | MB231/MB468 | MP |
| >hg38_21_+_8433172_8446622_RNA45SN1_45S_50ntFlanks@12332.12366.35\|TAAGCAGGAGGTGTCAGAAAAGTTACCACAGGGAT | >hg38_21_+_8433172_8446622_RNA45SN1_45S_50ntFlanks | 12332 | 12366 | 35 | TAAGCAGGAGGTGTCAGAAAAGTTACCACAGGGAT | 43.6 | -8.5 | 2.0 | -4.4 | 1.3E-05 | 5.4E-05 | BT20/MB468 | MP |
| >hg38_21_+_8433172_8446622_RNA45SN1_45S_50ntFlanks@12942.12960.19\|TTCGTAGACGACCTGCTTC | >hg38_21_+_8433172_8446622_RNA45SN1_45S_50ntFlanks | 12942 | 12960 | 19 | TTCGTAGACGACCTGCTTC | 297.8 | -3.1 | 0.7 | -4.4 | 1.3E-05 | 5.4E-05 | BT20/MB468 | MP |
| >hg38_21_+_8433172_8446622_RNA45SN1_45S_50ntFlanks@10356.10383.28\|ACTTTGAAGGCCGAAGTGGAGAAGGGTT | >hg38_21_+_8433172_8446622_RNA45SN1_45S_50ntFlanks | 10356 | 10383 | 28 | ACTTTGAAGGCCGAAGTGGAGAAGGGTT | 263.8 | -4.4 | 1.1 | -4.1 | 3.9E-05 | 5.4E-05 | MB231/MB468 | NUC |
| >MI0000088\|hsa-mir-30a&WithFlank&6\|-\|71403545\|71403627@54.71.18&[MIMAT0000088&hsa-miR-30a-3p&offsets\|+1\|-3;m-38&6\|-\|71403554\|71403575&offsets\|+1\|-3]\|TTTCAGTCGGATGTTTGC | >MI0000088\|hsa-mir-30a&WithFlank&6\|-\|71403545\|71403627 | 54 | 71 | 18 | TTTCAGTCGGATGTTTGC | 472.2 | 6.3 | 1.5 | 4.1 | 3.9E-05 | 5.4E-05 | MB231/MB468 | MITO |
| >hg38_21_+_8433172_8446622_RNA45SN1_45S_50ntFlanks@12994.13039.46\|CCTCGCTGCGATCTATTGAAAGTCAGCCCTCGACACAAGGGTTTGT | >hg38_21_+_8433172_8446622_RNA45SN1_45S_50ntFlanks | 12994 | 13039 | 46 | CCTCGCTGCGATCTATTGAAAGTCAGCCCTCGACACAAGGGTTTGT | 11054.2 | -4.7 | 1.0 | -4.6 | 4.8E-06 | 5.4E-05 | BT20/MB231 | NUC |
| >MI0000736\|hsa-mir-30c-1&WithFlank&1\|+\|40757278\|40757378@23.44.22(+1A);MI0000254\|hsa-mir-30c-2&WithFlank&6\|-\|71376954\|71377037@13.34.22(+1A)&[MIMAT0000244&hsa-miR-30c-5p&offsets\|0\|-1(+1A);m-57&1\|+\|40757300\|40757323&offsets\|0\|-2(+1A)];[MIMAT0000244_1&hsa-miR-30c-5p&offsets\|0\|-1(+1A);m-56&6\|-\|71377002\|71377025&offsets\|0\|-2(+1A)]\|TGTAAACATCCTACACTCTCAGA | >MI0000736\|hsa-mir-30c-1&WithFlank&1\|+\|40757278\|40757378 | 23 | 44 | 22(+1A) | TGTAAACATCCTACACTCTCAGA | 2783.2 | -3.8 | 0.8 | -4.6 | 5.0E-06 | 5.4E-05 | BT20/MB231 | MITO |
| >hg38_21_+_8433172_8446622_RNA45SN1_45S_50ntFlanks@4046.4074.29\|CATTCGAACGTCTGCCCTATCAACTTTCG | >hg38_21_+_8433172_8446622_RNA45SN1_45S_50ntFlanks | 4046 | 4074 | 29 | CATTCGAACGTCTGCCCTATCAACTTTCG | 81.2 | -6.1 | 1.5 | -4.2 | 3.3E-05 | 5.4E-05 | MB231/MB468 | MP |
| >MI0000750\|hsa-mir-26a-2&WithFlank&12\|-\|57824603\|57824698@20.40.21(+1A);MI0000083\|hsa-mir-26a-1&WithFlank&3\|+\|37969398\|37969486@16.36.21(+1A)&[MIMAT0000082&hsa-miR-26a-5p&offsets\|0\|-1(+1A);m-33&12\|-\|57824658\|57824679&offsets\|0\|-1(+1A)];[MIMAT0000082_1&hsa-miR-26a-5p&offsets\|0\|-1(+1A);m-34&3\|+\|37969413\|37969434&offsets\|0\|-1(+1A)]\|TTCAAGTAATCCAGGATAGGCA | >MI0000750\|hsa-mir-26a-2&WithFlank&12\|-\|57824603\|57824698 | 20 | 40 | 21(+1A) | TTCAAGTAATCCAGGATAGGCA | 1599.5 | 1.9 | 0.5 | 4.1 | 4.1E-05 | 5.4E-05 | BT20/MB468 | NUC |
| >hg38_21_+_8433172_8446622_RNA45SN1_45S_50ntFlanks@9737.9767.31\|CGAAACGATCTCAACCTATTCTCAAACTTTA | >hg38_21_+_8433172_8446622_RNA45SN1_45S_50ntFlanks | 9737 | 9767 | 31 | CGAAACGATCTCAACCTATTCTCAAACTTTA | 325.5 | -4.7 | 1.1 | -4.1 | 3.9E-05 | 5.4E-05 | MB231/MB468 | MITO |
| >hg38_21_+_8433172_8446622_RNA45SN1_45S_50ntFlanks@12602.12626.25\|GTGTATGTGCTTGGCTGAGGAGCCA | >hg38_21_+_8433172_8446622_RNA45SN1_45S_50ntFlanks | 12602 | 12626 | 25 | GTGTATGTGCTTGGCTGAGGAGCCA | 65.0 | -9.9 | 2.4 | -4.2 | 3.3E-05 | 5.4E-05 | MB231/MB468 | MP |
| >hg38_21_+_8433172_8446622_RNA45SN1_45S_50ntFlanks@12476.12500.25\|CCACTAATAGGGAACGTGAGCTGGG | >hg38_21_+_8433172_8446622_RNA45SN1_45S_50ntFlanks | 12476 | 12500 | 25 | CCACTAATAGGGAACGTGAGCTGGG | 150.4 | -2.5 | 0.6 | -4.2 | 2.9E-05 | 5.4E-05 | BT20/MB468 | MITO |
| >hg38_21_+_8433172_8446622_RNA45SN1_45S_50ntFlanks@6644.6681.38\|CCTCGTACGACTCTTAGCGGTGGATCACTCGGCTCGTG | >hg38_21_+_8433172_8446622_RNA45SN1_45S_50ntFlanks | 6644 | 6681 | 38 | CCTCGTACGACTCTTAGCGGTGGATCACTCGGCTCGTG | 999.9 | 2.7 | 0.7 | 4.1 | 3.9E-05 | 5.4E-05 | MB231/MB468 | NUC |
| >hg38_21_+_8433172_8446622_RNA45SN1_45S_50ntFlanks@11569.11602.34\|CAGCCGACTTAGAACTGGTGCGGACCAGGGGAAT | >hg38_21_+_8433172_8446622_RNA45SN1_45S_50ntFlanks | 11569 | 11602 | 34 | CAGCCGACTTAGAACTGGTGCGGACCAGGGGAAT | 2959.8 | -2.6 | 0.6 | -4.2 | 2.9E-05 | 5.4E-05 | BT20/MB468 | MITO |
| >MI0000103\|hsa-mir-101-1&WithFlank&1\|-\|65058428\|65058514@52.73.22;MI0000739\|hsa-mir-101-2&WithFlank&9\|+\|4850291\|4850381@54.75.22&[MIMAT0000099&hsa-miR-101-3p&offsets\|-1\|0;m-17&1\|-\|65058443\|65058462&offsets\|-1\|+1];[MIMAT0000099_1&hsa-miR-101-3p&offsets\|-1\|0;m-16&9\|+\|4850345\|4850364&offsets\|-1\|+1]\|GTACAGTACTGTGATAACTGAA | >MI0000103\|hsa-mir-101-1&WithFlank&1\|-\|65058428\|65058514 | 52 | 73 | 22 | GTACAGTACTGTGATAACTGAA | 408.0 | 2.7 | 0.7 | 4.1 | 4.1E-05 | 5.4E-05 | BT20/MB468 | NUC |
| >trna20_GluTTC_2_-_131094701_131094772@1.30.30,trna5_GluTTC_13_-_41634874_41634945@1.30.30\|TCCCATATGGTCTAGCGGTTAGGATTCCTG | >trna20_GluTTC_2_-_131094701_131094772 | 1 | 30 | 30 | TCCCATATGGTCTAGCGGTTAGGATTCCTG | 188.0 | -4.2 | 1.0 | -4.2 | 2.9E-05 | 5.4E-05 | BT20/MB468 | MITO |
| >hg38_21_+_8433172_8446622_RNA45SN1_45S_50ntFlanks@11564.11587.24\|CCTAGCAGCCGACTTAGAACTGGT | >hg38_21_+_8433172_8446622_RNA45SN1_45S_50ntFlanks | 11564 | 11587 | 24 | CCTAGCAGCCGACTTAGAACTGGT | 194.6 | -2.3 | 0.6 | -4.1 | 3.4E-05 | 5.4E-05 | BT20/MB468 | CYTO |
| >hg38_21_+_8433172_8446622_RNA45SN1_45S_50ntFlanks@4163.4182.20\|CATCCAAGGAAGGCAGCAGG | >hg38_21_+_8433172_8446622_RNA45SN1_45S_50ntFlanks | 4163 | 4182 | 20 | CATCCAAGGAAGGCAGCAGG | 716.3 | -7.0 | 1.6 | -4.4 | 1.3E-05 | 5.4E-05 | BT20/MB468 | MP |
| >MI0003628\|hsa-mir-615&WithFlank&12\|+\|54033944\|54034051@67.85.19&[MIMAT0003283&hsa-miR-615-3p&offsets\|0\|-3;m-495&12\|+\|54034010\|54034030&offsets\|0\|-2]\|TCCGAGCCTGGGTCTCCCT | >MI0003628\|hsa-mir-615&WithFlank&12\|+\|54033944\|54034051 | 67 | 85 | 19 | TCCGAGCCTGGGTCTCCCT | 76.9 | 9.6 | 2.1 | 4.6 | 4.2E-06 | 5.4E-05 | BT20/MB231 | MP |
| >hg38_21_+_8433172_8446622_RNA45SN1_45S_50ntFlanks@10393.10422.30\|CAGCAGTTGAACATGGGTCAGTCGGTCCTG | >hg38_21_+_8433172_8446622_RNA45SN1_45S_50ntFlanks | 10393 | 10422 | 30 | CAGCAGTTGAACATGGGTCAGTCGGTCCTG | 401.2 | -4.7 | 1.1 | -4.1 | 3.9E-05 | 5.4E-05 | MB231/MB468 | MITO |
| >am_trnaMT_ProTGG_MT_-_15956_16023@54.71.18\|AGACTTTTTCTCTGACCA | >am_trnaMT_ProTGG_MT_-_15956_16023 | 54 | 71 | 18 | AGACTTTTTCTCTGACCA | 1645.8 | 3.8 | 0.9 | 4.1 | 3.9E-05 | 5.4E-05 | MB231/MB468 | NUC |
| >hg38_21_+_8433172_8446622_RNA45SN1_45S_50ntFlanks@12419.12448.30\|GATGTCGGCTCTTCCTATCATTGTGAAGCA | >hg38_21_+_8433172_8446622_RNA45SN1_45S_50ntFlanks | 12419 | 12448 | 30 | GATGTCGGCTCTTCCTATCATTGTGAAGCA | 414.1 | 2.9 | 0.7 | 4.1 | 4.0E-05 | 5.4E-05 | MB231/MB468 | MITO |
| >hg38_21_+_8433172_8446622_RNA45SN1_45S_50ntFlanks@8196.8215.20\|CGTGGACGGTGTGAGGCCGG | >hg38_21_+_8433172_8446622_RNA45SN1_45S_50ntFlanks | 8196 | 8215 | 20 | CGTGGACGGTGTGAGGCCGG | 432.9 | -2.5 | 0.6 | -4.2 | 2.9E-05 | 5.4E-05 | BT20/MB468 | MITO |

| | | | | | | | | | | | | | |
|---|---|---|---|---|---|---|---|---|---|---|---|---|---|
| >hg38_21_+_8433172_8446622_RNA45SN1_45S_50ntFlanks@5492.5522.31\|AGAAGACGGTCGAACTTGACTATCTAGAGGA | >hg38_21_+_8433172_8446622_RNA45SN1_45S_50ntFlanks | 5492 | 5522 | 31 | AGAAGACGGTCGAACTTGACTATCTAGAGGA | 285.5 | -4.9 | 1.2 | -4.2 | 3.3E-05 | 5.4E-05 | MB231/MB468 | MP |
| >hg38_21_+_8433172_8446622_RNA45SN1_45S_50ntFlanks@5485.5512.28\|AGCGCTGAGAAGACGGTCGAACTTGACT | >hg38_21_+_8433172_8446622_RNA45SN1_45S_50ntFlanks | 5485 | 5512 | 28 | AGCGCTGAGAAGACGGTCGAACTTGACT | 2287.1 | 5.5 | 1.2 | 4.6 | 4.2E-06 | 5.4E-05 | BT20/MB231 | CYTO |
| >hg38_21_+_8433172_8446622_RNA45SN1_45S_50ntFlanks@8425.8469.45\|CAGTCCGCCCGGAGGATTCAACCCGGCGGCGGGTCCGGCCGTGTC | >hg38_21_+_8433172_8446622_RNA45SN1_45S_50ntFlanks | 8425 | 8469 | 45 | CAGTCCGCCCGGAGGATTCAACCCGGCGGCGGGTCCGGCCGTGTC | 71.5 | -9.2 | 2.1 | -4.4 | 1.3E-05 | 5.4E-05 | BT20/MB468 | MP |
| >MI0000809\|hsa-mir-151a&WithFlank&8\|-\|140732558\|140732659@53.74.22(+2A)&[MIMAT0000757&hsa-miR-151a-3p&offsets\|0\|+1(+2A);m-30&8\|-\|140732587\|140732607&offsets\|0\|+1(+2A)]\|CTAGACTGAAGCTCCTTGAGGAAA | >MI0000809\|hsa-mir-151a&WithFlank&8\|-\|140732558\|140732659 | 53 | 74 | 22(+2A) | CTAGACTGAAGCTCCTTGAGGAAA | 747.3 | 5.7 | 1.3 | 4.4 | 1.3E-05 | 5.4E-05 | BT20/MB468 | MP |
| >hg38_21_+_8433172_8446622_RNA45SN1_45S_50ntFlanks@4472.4490.19\|CTTAGCTGAGTGTCCCGCG | >hg38_21_+_8433172_8446622_RNA45SN1_45S_50ntFlanks | 4472 | 4490 | 19 | CTTAGCTGAGTGTCCCGCG | 58.0 | -9.7 | 2.3 | -4.2 | 3.3E-05 | 5.4E-05 | MB231/MB468 | MP |
| >hg38_21_+_8433172_8446622_RNA45SN1_45S_50ntFlanks@10468.10497.30\|TTGCCCTCGGCCGATCGAAAGGGAGTCGGG | >hg38_21_+_8433172_8446622_RNA45SN1_45S_50ntFlanks | 10468 | 10497 | 30 | TTGCCCTCGGCCGATCGAAAGGGAGTCGGG | 436.7 | -5.8 | 1.4 | -4.1 | 3.4E-05 | 5.4E-05 | BT20/MB468 | CYTO |
| >hg38_21_+_8433172_8446622_RNA45SN1_45S_50ntFlanks@12458.12488.31\|AAGCGTTGGATTGTTCACCCACTAATAGGGA | >hg38_21_+_8433172_8446622_RNA45SN1_45S_50ntFlanks | 12458 | 12488 | 31 | AAGCGTTGGATTGTTCACCCACTAATAGGGA | 176.0 | -4.3 | 1.0 | -4.1 | 3.9E-05 | 5.4E-05 | MB231/MB468 | NUC |
| >hg38_21_+_8433172_8446622_RNA45SN1_45S_50ntFlanks@475.518.44\|CTTCGTGATCGATGTGGTGACGTCGTGCTCTCCCGGGCCGGGTC | >hg38_21_+_8433172_8446622_RNA45SN1_45S_50ntFlanks | 475 | 518 | 44 | CTTCGTGATCGATGTGGTGACGTCGTGCTCTCCCGGGCCGGGTC | 915.7 | 5.2 | 1.3 | 4.1 | 4.0E-05 | 5.4E-05 | MB231/MB468 | MITO |
| >trnaMT_PheGAA_MT_+_577_647@56.74.19\|CATCACCCCATAAACACCA | >trnaMT_PheGAA_MT_+_577_647 | 56 | 74 | 19 | CATCACCCCATAAACACCA | 734.4 | 3.3 | 0.8 | 4.1 | 4.0E-05 | 5.5E-05 | MB231/MB468 | MITO |
| >hg38_21_+_8433172_8446622_RNA45SN1_45S_50ntFlanks@12542.12566.25\|ATGTGTTGTTGCCATGGTAATCCTG | >hg38_21_+_8433172_8446622_RNA45SN1_45S_50ntFlanks | 12542 | 12566 | 25 | ATGTGTTGTTGCCATGGTAATCCTG | 166.3 | -6.2 | 1.4 | -4.4 | 1.3E-05 | 5.5E-05 | BT20/MB468 | MP |
| >hg38_21_+_8433172_8446622_RNA45SN1_45S_50ntFlanks@9307.9338.32\|AAGGAGTCTAACACGTGCGCGAGTCGGGGGCT | >hg38_21_+_8433172_8446622_RNA45SN1_45S_50ntFlanks | 9307 | 9338 | 32 | AAGGAGTCTAACACGTGCGCGAGTCGGGGGCT | 1824.0 | -3.8 | 0.9 | -4.4 | 1.3E-05 | 5.5E-05 | BT20/MB468 | MP |
| >hg38_21_+_8433172_8446622_RNA45SN1_45S_50ntFlanks@9833.9851.19\|CTTTTGGTAAGCAGAACTG | >hg38_21_+_8433172_8446622_RNA45SN1_45S_50ntFlanks | 9833 | 9851 | 19 | CTTTTGGTAAGCAGAACTG | 48.1 | -8.7 | 2.0 | -4.4 | 1.3E-05 | 5.5E-05 | BT20/MB468 | MP |
| >hg38_21_+_8433172_8446622_RNA45SN1_45S_50ntFlanks@11672.11702.31\|CAGTGCTCTGAATGTCAAAGTGAAGAAATTC | >hg38_21_+_8433172_8446622_RNA45SN1_45S_50ntFlanks | 11672 | 11702 | 31 | CAGTGCTCTGAATGTCAAAGTGAAGAAATTC | 474.0 | -3.5 | 0.8 | -4.1 | 3.3E-05 | 5.5E-05 | MB231/MB468 | MP |
| >hg38_21_+_8433172_8446622_RNA45SN1_45S_50ntFlanks@11572.11600.29\|CCGACTTAGAACTGGTGCGGACCAGGGGA | >hg38_21_+_8433172_8446622_RNA45SN1_45S_50ntFlanks | 11572 | 11600 | 29 | CCGACTTAGAACTGGTGCGGACCAGGGGA | 5055.6 | -3.4 | 0.8 | -4.1 | 3.7E-05 | 5.5E-05 | MB231/MB468 | CYTO |
| >hg38_21_+_8433172_8446622_RNA45SN1_45S_50ntFlanks@12417.12452.36\|TCGATGTCGGCTCTTCCTATCATTGTGAAGCAGAAT | >hg38_21_+_8433172_8446622_RNA45SN1_45S_50ntFlanks | 12417 | 12452 | 36 | TCGATGTCGGCTCTTCCTATCATTGTGAAGCAGAAT | 182.6 | 2.9 | 0.7 | 4.1 | 4.0E-05 | 5.5E-05 | MB231/MB468 | NUC |
| >hg38_21_+_8433172_8446622_RNA45SN1_45S_50ntFlanks@11209.11252.44\|CGGCGACTCTGGACGCGAGCCGGGCCCTTCCCGTGGATCGCCCC | >hg38_21_+_8433172_8446622_RNA45SN1_45S_50ntFlanks | 11209 | 11252 | 44 | CGGCGACTCTGGACGCGAGCCGGGCCCTTCCCGTGGATCGCCCC | 664.4 | -3.6 | 0.9 | -4.1 | 3.8E-05 | 5.5E-05 | MB231/MB468 | CYTO |
| >hg38_21_+_8433172_8446622_RNA45SN1_45S_50ntFlanks@9913.9938.26\|AGAAAAGGTGTTGGTTGATATAGACA | >hg38_21_+_8433172_8446622_RNA45SN1_45S_50ntFlanks | 9913 | 9938 | 26 | AGAAAAGGTGTTGGTTGATATAGACA | 134.7 | 7.5 | 1.7 | 4.4 | 1.4E-05 | 5.5E-05 | BT20/MB468 | MP |
| >hg38_21_+_8433172_8446622_RNA45SN1_45S_50ntFlanks@12320.12346.27\|CCTTTTGGGTTTTAAGCAGGAGGTGTC | >hg38_21_+_8433172_8446622_RNA45SN1_45S_50ntFlanks | 12320 | 12346 | 27 | CCTTTTGGGTTTTAAGCAGGAGGTGTC | 47.4 | -8.6 | 2.0 | -4.4 | 1.4E-05 | 5.5E-05 | BT20/MB468 | MP |
| >hg38_21_+_8433172_8446622_RNA45SN1_45S_50ntFlanks@9201.9225.25\|CTCTCGGGGCCACGCGCGCGTCCCC | >hg38_21_+_8433172_8446622_RNA45SN1_45S_50ntFlanks | 9201 | 9225 | 25 | CTCTCGGGGCCACGCGCGCGTCCCC | 295.5 | -6.6 | 1.5 | -4.4 | 1.4E-05 | 5.5E-05 | BT20/MB468 | MP |
| >MI0000077\|hsa-mir-21&WithFlank&17\|+\|59841260\|59841343@13.33.21(+1A)&[MIMAT0000076&hsa-miR-21-5p&offsets\|-1\|-2(+1A);m-1&17\|+\|59841273\|59841294&offsets\|-1\|-2(+1A)]\|GTAGCTTATCAGACTGATGTTA | >MI0000077\|hsa-mir-21&WithFlank&17\|+\|59841260\|59841343 | 13 | 33 | 21(+1A) | GTAGCTTATCAGACTGATGTTA | 39.0 | 8.5 | 2.0 | 4.1 | 3.3E-05 | 5.5E-05 | MB231/MB468 | MP |
| >hg38_MT_+_1621_3279_RNR2_16S_50ntFlanks@51.96.46\|GCTAAACCTAGCCCCAAACCCACTCCACCTTACTACCAGACAACCT | >hg38_MT_+_1621_3279_RNR2_16S_50ntFlanks | 51 | 96 | 46 | GCTAAACCTAGCCCCAAACCCACTCCACCTTACTACCAGACAACCT | 2055.7 | 5.2 | 1.3 | 4.1 | 4.0E-05 | 5.5E-05 | MB231/MB468 | MITO |
| >hg38_21_+_8433172_8446622_RNA45SN1_45S_50ntFlanks@10678.10708.31\|CGGTTCCGGCGGCGTCCGGTGAGCTCTCGCT | >hg38_21_+_8433172_8446622_RNA45SN1_45S_50ntFlanks | 10678 | 10708 | 31 | CGGTTCCGGCGGCGTCCGGTGAGCTCTCGCT | 7051.1 | -4.2 | 1.0 | -4.4 | 1.4E-05 | 5.5E-05 | BT20/MB468 | MP |
| >trnaMT_GluTTC_MT_-_14674_14742@53.72.20,trnalookalike8_GluTTC_5_-_93905172_93905240@53.72.20\|TAGTCCGTGCGAGAATACCA | >trnaMT_GluTTC_MT_-_14674_14742 | 53 | 72 | 20 | TAGTCCGTGCGAGAATACCA | 1359.1 | -3.9 | 0.9 | -4.1 | 4.0E-05 | 5.5E-05 | MB231/MB468 | MITO |
| >hg38_21_+_8433172_8446622_RNA45SN1_45S_50ntFlanks@10568.10594.27\|CCGGAGAAGCCGGCGGGAGCCCCGGGG | >hg38_21_+_8433172_8446622_RNA45SN1_45S_50ntFlanks | 10568 | 10594 | 27 | CCGGAGAAGCCGGCGGGAGCCCCGGGG | 79.8 | -6.0 | 1.5 | -4.1 | 3.3E-05 | 5.5E-05 | MB231/MB468 | MP |
| >hg38_21_+_8433172_8446622_RNA45SN1_45S_50ntFlanks@7987.8017.31\|TCAGACGTGGCGACCCGCTGAATTTAAGCAT | >hg38_21_+_8433172_8446622_RNA45SN1_45S_50ntFlanks | 7987 | 8017 | 31 | TCAGACGTGGCGACCCGCTGAATTTAAGCAT | 76.3 | -6.0 | 1.4 | -4.1 | 3.3E-05 | 5.5E-05 | MB231/MB468 | MP |
| >hg38_21_+_8433172_8446622_RNA45SN1_45S_50ntFlanks@10843.10864.22\|TAACTTCGGGATAAGGATTGGC | >hg38_21_+_8433172_8446622_RNA45SN1_45S_50ntFlanks | 10843 | 10864 | 22 | TAACTTCGGGATAAGGATTGGC | 304.6 | -5.8 | 1.4 | -4.1 | 3.5E-05 | 5.5E-05 | BT20/MB468 | CYTO |
| >hg38_21_+_8433172_8446622_RNA45SN1_45S_50ntFlanks@475.495.21\|CTTCGTGATCGATGTGGTGAC | >hg38_21_+_8433172_8446622_RNA45SN1_45S_50ntFlanks | 475 | 495 | 21 | CTTCGTGATCGATGTGGTGAC | 758.0 | 6.9 | 1.7 | 4.2 | 2.9E-05 | 5.5E-05 | BT20/MB468 | MITO |
| >trnaMT_ProTGG_MT_-_15956_16023@51.71.21\|TAAAGACTTTTTCTCTGACCA | >trnaMT_ProTGG_MT_-_15956_16023 | 51 | 71 | 21 | TAAAGACTTTTTCTCTGACCA | 526.3 | -2.9 | 0.6 | -4.6 | 4.9E-06 | 5.5E-05 | BT20/MB231 | NUC |
| >hg38_21_+_8433172_8446622_RNA45SN1_45S_50ntFlanks@6648.6672.25\|GTACGACTCTTAGCGGTGGATCACT | >hg38_21_+_8433172_8446622_RNA45SN1_45S_50ntFlanks | 6648 | 6672 | 25 | GTACGACTCTTAGCGGTGGATCACT | 437.6 | 6.7 | 1.6 | 4.1 | 3.5E-05 | 5.5E-05 | BT20/MB468 | CYTO |
| >hg38_21_+_8433172_8446622_RNA45SN1_45S_50ntFlanks@6653.6688.36\|ACTCTTAGCGGTGGATCACTCGGCTCGTGCGTCGAT | >hg38_21_+_8433172_8446622_RNA45SN1_45S_50ntFlanks | 6653 | 6688 | 36 | ACTCTTAGCGGTGGATCACTCGGCTCGTGCGTCGAT | 233.4 | 1.9 | 0.5 | 4.1 | 3.8E-05 | 5.5E-05 | MB231/MB468 | CYTO |
| >hg38_21_+_8433172_8446622_RNA45SN1_45S_50ntFlanks@12836.12856.21\|CGGTGCGGAGTGCCCTTCGTC | >hg38_21_+_8433172_8446622_RNA45SN1_45S_50ntFlanks | 12836 | 12856 | 21 | CGGTGCGGAGTGCCCTTCGTC | 402.9 | -3.5 | 0.8 | -4.3 | 1.4E-05 | 5.5E-05 | BT20/MB468 | MP |
| >hg38_21_+_8433172_8446622_RNA45SN1_45S_50ntFlanks@7989.8019.31\|AGACGTGGCGACCCGCTGAATTTAAGCATAT | >hg38_21_+_8433172_8446622_RNA45SN1_45S_50ntFlanks | 7989 | 8019 | 31 | AGACGTGGCGACCCGCTGAATTTAAGCATAT | 294.6 | 4.6 | 1.1 | 4.1 | 4.0E-05 | 5.5E-05 | MB231/MB468 | NUC |
| >MI0000446\|hsa-mir-125b-1&WithFlank&11\|-\|122099751\|122099850@21.41.21;MI0000470\|hsa-mir-125b-2&WithFlank&21\|+\|16590231\|16590331@23.43.21&[MIMAT0000423&hsa-miR-125b-5p&offsets\|0\|-1;m-67&11\|-\|122099809\|122099830&offsets\|0\|-1];[MIMAT0000423_1&hsa-miR-125b-5p&offsets\|0\|-1;m-68&21\|+\|16590253\|16590274&offsets\|0\|-1]\|TCCCTGAGACCCTAACTTGTG | >MI0000446\|hsa-mir-125b-1&WithFlank&11\|-\|122099751\|122099850 | 21 | 41 | 21 | TCCCTGAGACCCTAACTTGTG | 9089.5 | -7.3 | 1.6 | -4.6 | 4.3E-06 | 5.5E-05 | BT20/MB231 | MP |
| >MI0000449\|hsa-mir-132&WithFlank&17\|-\|2049902\|2050014@65.87.23&[MIMAT0000426&hsa-miR-132-3p&offsets\|0\|+1;m-138&17\|-\|2049929\|2049950&offsets\|0\|+1]\|TAACAGTCTACAGCCATGGTCGC | >MI0000449\|hsa-mir-132&WithFlank&17\|-\|2049902\|2050014 | 65 | 87 | 23 | TAACAGTCTACAGCCATGGTCGC | 45.4 | 9.3 | 2.1 | 4.3 | 1.4E-05 | 5.5E-05 | BT20/MB468 | MP |
| >hg38_MT_+_1621_3279_RNR2_16S_50ntFlanks@51.71.21\|GCTAAACCTAGCCCCAAACCC | >hg38_MT_+_1621_3279_RNR2_16S_50ntFlanks | 51 | 71 | 21 | GCTAAACCTAGCCCCAAACCC | 5567.2 | 4.6 | 1.1 | 4.1 | 4.0E-05 | 5.5E-05 | MB231/MB468 | MITO |
| >MI0000081\|hsa-mir-24-2&WithFlank&19\|-\|13836281\|13836365@56.76.21;MI0000080\|hsa-mir-24-1&WithFlank&9\|+\|95086015\|95086094@50.70.21&[MIMAT0000080&hsa-miR-24-3p&offsets\|0\|-1;m-36&19\|-\|13836288\|13836310&offsets\|0\|-2];[MIMAT0000080_1&hsa-miR-24-3p&offsets\|0\|-1;m-35&9\|+\|95086064\|95086086&offsets\|0\|-2]\|TGGCTCAGTTCAGCAGGAACA | >MI0000081\|hsa-mir-24-2&WithFlank&19\|-\|13836281\|13836365 | 56 | 76 | 21 | TGGCTCAGTTCAGCAGGAACA | 2845.7 | 3.4 | 0.8 | 4.1 | 4.0E-05 | 5.5E-05 | MB231/MB468 | MITO |
| >hg38_21_+_8433172_8446622_RNA45SN1_45S_50ntFlanks@10841.10866.26\|CGTAACTTCGGGATAAGGATTGGCTC | >hg38_21_+_8433172_8446622_RNA45SN1_45S_50ntFlanks | 10841 | 10866 | 26 | CGTAACTTCGGGATAAGGATTGGCTC | 369.2 | -3.6 | 0.9 | -4.1 | 4.0E-05 | 5.5E-05 | MB231/MB468 | MITO |
| >hg38_21_+_8433172_8446622_RNA45SN1_45S_50ntFlanks@3840.3861.22\|CTCCTACTTGGATAACTGTGGT | >hg38_21_+_8433172_8446622_RNA45SN1_45S_50ntFlanks | 3840 | 3861 | 22 | CTCCTACTTGGATAACTGTGGT | 78.9 | -10.2 | 2.5 | -4.1 | 3.4E-05 | 5.5E-05 | MB231/MB468 | MP |
| >hg38_21_+_8433172_8446622_RNA45SN1_45S_50ntFlanks@9575.9604.30\|ATCGGTCGTCCGACCTGGGTATAGGGGCGA | >hg38_21_+_8433172_8446622_RNA45SN1_45S_50ntFlanks | 9575 | 9604 | 30 | ATCGGTCGTCCGACCTGGGTATAGGGGCGA | 184.2 | -6.3 | 1.5 | -4.1 | 3.4E-05 | 5.5E-05 | MB231/MB468 | MP |
| >hg38_21_+_8433172_8446622_RNA45SN1_45S_50ntFlanks@10351.10371.21\|CGAGAACTTTGAAGGCCGAAG | >hg38_21_+_8433172_8446622_RNA45SN1_45S_50ntFlanks | 10351 | 10371 | 21 | CGAGAACTTTGAAGGCCGAAG | 606.8 | -3.0 | 0.7 | -4.1 | 3.5E-05 | 5.5E-05 | BT20/MB468 | CYTO |
| >hg38_21_+_8433172_8446622_RNA45SN1_45S_50ntFlanks@10317.10353.37\|GTGCAGATCTTGGTGGTAGTAGCAAATATTCAAACGA | >hg38_21_+_8433172_8446622_RNA45SN1_45S_50ntFlanks | 10317 | 10353 | 37 | GTGCAGATCTTGGTGGTAGTAGCAAATATTCAAACGA | 86.5 | 9.8 | 2.1 | 4.6 | 4.3E-06 | 5.5E-05 | BT20/MB231 | MP |
| >hg38_21_+_8433172_8446622_RNA45SN1_45S_50ntFlanks@9252.9271.20\|CACGGGGTCGGCGGCGACGT | >hg38_21_+_8433172_8446622_RNA45SN1_45S_50ntFlanks | 9252 | 9271 | 20 | CACGGGGTCGGCGGCGACGT | 111.0 | 10.1 | 2.2 | 4.6 | 4.3E-06 | 5.5E-05 | BT20/MB231 | MP |
| >hg38_21_+_8433172_8446622_RNA45SN1_45S_50ntFlanks@9572.9605.34\|CAAATCGGTCGTCCGACCTGGGTATAGGGGCGAA | >hg38_21_+_8433172_8446622_RNA45SN1_45S_50ntFlanks | 9572 | 9605 | 34 | CAAATCGGTCGTCCGACCTGGGTATAGGGGCGAA | 880.7 | -2.5 | 0.6 | -4.2 | 2.9E-05 | 5.5E-05 | BT20/MB468 | MITO |
| >hg38_21_+_8433172_8446622_RNA45SN1_45S_50ntFlanks@8105.8135.31\|CGCGGCGGGGCGCGGGACATGTGGCGTACGG | >hg38_21_+_8433172_8446622_RNA45SN1_45S_50ntFlanks | 8105 | 8135 | 31 | CGCGGCGGGGCGCGGGACATGTGGCGTACGG | 119.4 | -6.4 | 1.5 | -4.1 | 3.4E-05 | 5.5E-05 | MB231/MB468 | MP |
| >hg38_21_+_8433172_8446622_RNA45SN1_45S_50ntFlanks@10349.10377.29\|AACGAGAACTTTGAAGGCCGAAGTGGAGA | >hg38_21_+_8433172_8446622_RNA45SN1_45S_50ntFlanks | 10349 | 10377 | 29 | AACGAGAACTTTGAAGGCCGAAGTGGAGA | 622.5 | -3.4 | 0.8 | -4.1 | 4.0E-05 | 5.5E-05 | MB231/MB468 | MITO |
| >hg38_21_+_8433172_8446622_RNA45SN1_45S_50ntFlanks@12423.12455.33\|TCGGCTCTTCCTATCATTGTGAAGCAGAATTCA | >hg38_21_+_8433172_8446622_RNA45SN1_45S_50ntFlanks | 12423 | 12455 | 33 | TCGGCTCTTCCTATCATTGTGAAGCAGAATTCA | 101.4 | -6.0 | 1.4 | -4.1 | 3.4E-05 | 5.5E-05 | MB231/MB468 | MP |
| >hg38_21_+_8433172_8446622_RNA45SN1_45S_50ntFlanks@10873.10901.29\|CTGGGTCGGTCGGGCTGGGGCGCGAAGCG | >hg38_21_+_8433172_8446622_RNA45SN1_45S_50ntFlanks | 10873 | 10901 | 29 | CTGGGTCGGTCGGGCTGGGGCGCGAAGCG | 233.0 | -4.0 | 1.0 | -4.1 | 3.5E-05 | 5.5E-05 | BT20/MB468 | CYTO |
| >hg38_21_+_8433172_8446622_RNA45SN1_45S_50ntFlanks@3705.3737.33\|TACCTGGTTGATCCTGCCAGTAGCATATGCTTG | >hg38_21_+_8433172_8446622_RNA45SN1_45S_50ntFlanks | 3705 | 3737 | 33 | TACCTGGTTGATCCTGCCAGTAGCATATGCTTG | 862.3 | 3.0 | 0.7 | 4.1 | 3.5E-05 | 5.5E-05 | BT20/MB468 | CYTO |
| >hg38_21_+_8433172_8446622_RNA45SN1_45S_50ntFlanks@8355.8388.34\|TGAAAAGAACTTTGAAGAGAGAGTTCAAGAGGGC | >hg38_21_+_8433172_8446622_RNA45SN1_45S_50ntFlanks | 8355 | 8388 | 34 | TGAAAAGAACTTTGAAGAGAGAGTTCAAGAGGGC | 95.9 | 9.9 | 2.2 | 4.6 | 4.4E-06 | 5.6E-05 | BT20/MB231 | MP |

<table>
<tr><th></th><th></th><th></th><th></th><th></th><th></th><th></th><th></th><th></th><th></th><th></th><th></th><th></th><th></th></tr>
<tr><td>>MI0000455|hsa-mir-138-2&WithFlank&16|+|56858512|56858607@16.36.21(+1U);MI0000476|hsa-mir-138-1&WithFlank&3|+|44114206|44114316@29.49.21(+1U)&[MIMAT0000430&hsa-miR-138-5p&offsets|0|-2(+1U);m-509&16|+|56858527|56858549&offsets|0|-2(+1U)];[MIMAT0000430_1&hsa-miR-138-5p&offsets|0|-2(+1U);m-497&3|+|44114234|44114256&offsets|0|-2(+1U)]|AGCTGGTGTTGTGAATCAGGCT</td><td>>MI0000455|hsa-mir-138-2&WithFlank&16|+|56858512|56858607</td><td>16</td><td>36</td><td>21(+1U)</td><td>AGCTGGTGTTGTGAATCAGGCT</td><td>68.7</td><td>-9.5</td><td>2.1</td><td>-4.6</td><td>4.4E-06</td><td>5.6E-05</td><td>BT20/MB231</td><td>MP</td></tr>
<tr><td>>hg38_21_+_8433172_8446622_RNA45SN1_45S_50ntFlanks@10808.10828.21|CAATGTAGGTAAGGGAAGTCG</td><td>>hg38_21_+_8433172_8446622_RNA45SN1_45S_50ntFlanks</td><td>10808</td><td>10828</td><td>21</td><td>CAATGTAGGTAAGGGAAGTCG</td><td>410.4</td><td>-6.5</td><td>1.5</td><td>-4.3</td><td>1.4E-05</td><td>5.6E-05</td><td>BT20/MB468</td><td>MP</td></tr>
<tr><td>>hg38_21_+_8433172_8446622_RNA45SN1_45S_50ntFlanks@12329.12346.18|TTTTAAGCAGGAGGTGTC</td><td>>hg38_21_+_8433172_8446622_RNA45SN1_45S_50ntFlanks</td><td>12329</td><td>12346</td><td>18</td><td>TTTTAAGCAGGAGGTGTC</td><td>207.1</td><td>-5.5</td><td>1.3</td><td>-4.1</td><td>3.5E-05</td><td>5.6E-05</td><td>BT20/MB468</td><td>CYTO</td></tr>
<tr><td>>hg38_21_+_8433172_8446622_RNA45SN1_45S_50ntFlanks@11664.11692.29|TTTCTGCCCAGTGCTCTGAATGTCAAAGT</td><td>>hg38_21_+_8433172_8446622_RNA45SN1_45S_50ntFlanks</td><td>11664</td><td>11692</td><td>29</td><td>TTTCTGCCCAGTGCTCTGAATGTCAAAGT</td><td>92.9</td><td>9.9</td><td>2.1</td><td>4.6</td><td>4.4E-06</td><td>5.6E-05</td><td>BT20/MB231</td><td>MP</td></tr>
<tr><td>>MI0000077|hsa-mir-21&WithFlank&17|+|59841260|59841343@14.33.20&[MIMAT0000076&hsa-miR-21-5p&offsets|0|-2;m-1&17|+|59841273|59841294&offsets|0|-2]|TAGCTTATCAGACTGATGTT</td><td>>MI0000077|hsa-mir-21&WithFlank&17|+|59841260|59841343</td><td>14</td><td>33</td><td>20</td><td>TAGCTTATCAGACTGATGTT</td><td>2240.4</td><td>3.4</td><td>0.8</td><td>4.1</td><td>3.4E-05</td><td>5.6E-05</td><td>MB231/MB468</td><td>MP</td></tr>
<tr><td>>MI0000438|hsa-mir-15b&WithFlank&3|+|160404582|160404691@26.46.21&[MIMAT0000417&hsa-miR-15b-5p&offsets|0|-1;m-110&3|+|160404607|160404628&offsets|0|-1]|TAGCAGCACATCATGGTTTAC</td><td>>MI0000438|hsa-mir-15b&WithFlank&3|+|160404582|160404691</td><td>26</td><td>46</td><td>21</td><td>TAGCAGCACATCATGGTTTAC</td><td>273.4</td><td>5.5</td><td>1.3</td><td>4.1</td><td>4.1E-05</td><td>5.6E-05</td><td>MB231/MB468</td><td>MITO</td></tr>
<tr><td>>hg38_21_+_8433172_8446622_RNA45SN1_45S_50ntFlanks@11582.11623.42|ACTGGTGCGGACCAGGGGAATCCGACTGTTTAATTAAAACAA</td><td>>hg38_21_+_8433172_8446622_RNA45SN1_45S_50ntFlanks</td><td>11582</td><td>11623</td><td>42</td><td>ACTGGTGCGGACCAGGGGAATCCGACTGTTTAATTAAAACAA</td><td>410.1</td><td>5.2</td><td>1.1</td><td>4.6</td><td>5.2E-06</td><td>5.6E-05</td><td>BT20/MB231</td><td>MITO</td></tr>
<tr><td>>hg38_21_+_8433172_8446622_RNA45SN1_45S_50ntFlanks@3835.3871.37|CTCCTCTCCTACTTGGATAACTGTGGTAATTCTAGAG</td><td>>hg38_21_+_8433172_8446622_RNA45SN1_45S_50ntFlanks</td><td>3835</td><td>3871</td><td>37</td><td>CTCCTCTCCTACTTGGATAACTGTGGTAATTCTAGAG</td><td>50.4</td><td>-9.5</td><td>2.3</td><td>-4.1</td><td>3.4E-05</td><td>5.6E-05</td><td>MB231/MB468</td><td>MP</td></tr>
<tr><td>>MI0003684|hsa-mir-660&WithFlank&X|+|50013235|50013343@22.44.23&[MIMAT0003338&hsa-miR-660-5p&offsets|0|+1;m-157&X|+|50013256|50013278&offsets|0|0]|TACCCATTGCATATCGGAGTTGT</td><td>>MI0003684|hsa-mir-660&WithFlank&X|+|50013235|50013343</td><td>22</td><td>44</td><td>23</td><td>TACCCATTGCATATCGGAGTTGT</td><td>329.6</td><td>4.2</td><td>1.0</td><td>4.1</td><td>4.1E-05</td><td>5.6E-05</td><td>MB231/MB468</td><td>MITO</td></tr>
<tr><td>>MI0000102|hsa-mir-100&WithFlank&11|-|122152223|122152314@19.41.23(+1A)&[MIMAT0000098&hsa-miR-100-5p&offsets|0|+1(+1A);m-27&11|-|122152276|122152296&offsets|0|+2(+1A)]|AACCCGTAGATCCGAACTTGTGGA</td><td>>MI0000102|hsa-mir-100&WithFlank&11|-|122152223|122152314</td><td>19</td><td>41</td><td>23(+1A)</td><td>AACCCGTAGATCCGAACTTGTGGA</td><td>66.6</td><td>-9.5</td><td>2.1</td><td>-4.6</td><td>4.4E-06</td><td>5.6E-05</td><td>BT20/MB231</td><td>MP</td></tr>
<tr><td>>hg38_MT_+_1621_3279_RNR2_16S_50ntFlanks@1350.1388.39|CAATAGGGTTTACGACCTCGATGTTGGATCAGGACATCC</td><td>>hg38_MT_+_1621_3279_RNR2_16S_50ntFlanks</td><td>1350</td><td>1388</td><td>39</td><td>CAATAGGGTTTACGACCTCGATGTTGGATCAGGACATCC</td><td>29.7</td><td>8.1</td><td>2.0</td><td>4.1</td><td>3.4E-05</td><td>5.6E-05</td><td>MB231/MB468</td><td>MP</td></tr>
<tr><td>>trnaMT_SerGCT_MT_+_12207_12265@1.22.22|GAGAAAGCTCACAAGAACTGCT</td><td>>trnaMT_SerGCT_MT_+_12207_12265</td><td>1</td><td>22</td><td>22</td><td>GAGAAAGCTCACAAGAACTGCT</td><td>761.7</td><td>3.1</td><td>0.8</td><td>4.1</td><td>4.2E-05</td><td>5.6E-05</td><td>BT20/MB468</td><td>NUC</td></tr>
<tr><td>>hg38_21_+_8433172_8446622_RNA45SN1_45S_50ntFlanks@10947.10981.35|CACGCCCGGGGCACCCCCCTCGCGGCCCTCCCCCG</td><td>>hg38_21_+_8433172_8446622_RNA45SN1_45S_50ntFlanks</td><td>10947</td><td>10981</td><td>35</td><td>CACGCCCGGGGCACCCCCCTCGCGGCCCTCCCCCG</td><td>106.7</td><td>-5.0</td><td>1.2</td><td>-4.1</td><td>3.5E-05</td><td>5.6E-05</td><td>BT20/MB468</td><td>CYTO</td></tr>
<tr><td>>MI0000086|hsa-mir-28&WithFlank&3|+|188688775|188688872@61.82.22(+1U)&[MIMAT0004502&hsa-miR-28-3p&offsets|+1|+1(+1U);m-28&3|+|188688834|188688855&offsets|+1|+1(+1U)]|ACTAGATTGTGAGCTCCTGGAGT</td><td>>MI0000086|hsa-mir-28&WithFlank&3|+|188688775|188688872</td><td>61</td><td>82</td><td>22(+1U)</td><td>ACTAGATTGTGAGCTCCTGGAGT</td><td>319.3</td><td>4.9</td><td>1.2</td><td>4.1</td><td>3.4E-05</td><td>5.6E-05</td><td>MB231/MB468</td><td>MP</td></tr>
<tr><td>>hg38_21_+_8433172_8446622_RNA45SN1_45S_50ntFlanks@9123.9140.18|CGCGCGCGGGTCGGGGGG</td><td>>hg38_21_+_8433172_8446622_RNA45SN1_45S_50ntFlanks</td><td>9123</td><td>9140</td><td>18</td><td>CGCGCGCGGGTCGGGGGG</td><td>60.1</td><td>-9.8</td><td>2.4</td><td>-4.1</td><td>3.4E-05</td><td>5.6E-05</td><td>MB231/MB468</td><td>MP</td></tr>
<tr><td>>trna11_GluTTC_15_-_26327381_26327452@1.32.32,trna3_GluTTC_13_-_45492062_45492133@1.32.32|TCCCACATGGTCTAGCGGTTAGGATTCCTGGT</td><td>>trna11_GluTTC_15_-_26327381_26327452</td><td>1</td><td>32</td><td>32</td><td>TCCCACATGGTCTAGCGGTTAGGATTCCTGGT</td><td>213.9</td><td>11.5</td><td>2.6</td><td>4.3</td><td>1.4E-05</td><td>5.6E-05</td><td>BT20/MB468</td><td>MP</td></tr>
<tr><td>>hg38_21_+_8433172_8446622_RNA45SN1_45S_50ntFlanks@9924.9949.26|TGGTTGATATAGACAGCAGGACGGTG</td><td>>hg38_21_+_8433172_8446622_RNA45SN1_45S_50ntFlanks</td><td>9924</td><td>9949</td><td>26</td><td>TGGTTGATATAGACAGCAGGACGGTG</td><td>75.3</td><td>9.6</td><td>2.1</td><td>4.6</td><td>4.4E-06</td><td>5.6E-05</td><td>BT20/MB231</td><td>MP</td></tr>
<tr><td>>MI0000289|hsa-mir-181a-1&WithFlank&1|-|198859038|198859159@30.54.25(+1A);MI0000269|hsa-mir-181a-2&WithFlank&9|+|124692436|124692557@45.69.25(+1A)&[MIMAT0000256&hsa-miR-181a-5p&offsets|0|+2(+1A);m-32&1|-|198859108|198859130&offsets|0|+2(+1A)];[MIMAT0000256_1&hsa-miR-181a-5p&offsets|0|+2(+1A);m-32&9|+|124692480|124692502&offsets|0|+2(+1A)]|AACATTCAACGCTGTCGGTGAGTTTA</td><td>>MI0000289|hsa-mir-181a-1&WithFlank&1|-|198859038|198859159</td><td>30</td><td>54</td><td>25(+1A)</td><td>AACATTCAACGCTGTCGGTGAGTTTA</td><td>69.3</td><td>-9.5</td><td>2.1</td><td>-4.6</td><td>4.4E-06</td><td>5.6E-05</td><td>BT20/MB231</td><td>MP</td></tr>
<tr><td>>MI0000446|hsa-mir-125b-1&WithFlank&11|-|122099751|122099850@21.40.20;MI0000470|hsa-mir-125b-2&WithFlank&21|+|16590231|16590331@23.42.20&[MIMAT0000423&hsa-miR-125b-5p&offsets|0|-2;m-67&11|-|122099809|122099830&offsets|0|-2];[MIMAT0000423_1&hsa-miR-125b-5p&offsets|0|-2;m-68&21|+|16590253|16590274&offsets|0|-2]|TCCCTGAGACCCTAACTTGT</td><td>>MI0000446|hsa-mir-125b-1&WithFlank&11|-|122099751|122099850</td><td>21</td><td>40</td><td>20</td><td>TCCCTGAGACCCTAACTTGT</td><td>1035.4</td><td>-8.0</td><td>1.7</td><td>-4.6</td><td>4.4E-06</td><td>5.6E-05</td><td>BT20/MB231</td><td>MP</td></tr>
<tr><td>>hg38_21_+_8433172_8446622_RNA45SN1_45S_50ntFlanks@475.515.41|CTTCGTGATCGATGTGGTGACGTCGTGCTCTCCCGGGCCGG</td><td>>hg38_21_+_8433172_8446622_RNA45SN1_45S_50ntFlanks</td><td>475</td><td>515</td><td>41</td><td>CTTCGTGATCGATGTGGTGACGTCGTGCTCTCCCGGGCCGG</td><td>4731.5</td><td>3.8</td><td>0.9</td><td>4.1</td><td>4.1E-05</td><td>5.6E-05</td><td>MB231/MB468</td><td>NUC</td></tr>
<tr><td>>trnaMT_PheGAA_MT_+_577_647@31.53.23|CACTGAAAATGTTTAGACGGGCT</td><td>>trnaMT_PheGAA_MT_+_577_647</td><td>31</td><td>53</td><td>23</td><td>CACTGAAAATGTTTAGACGGGCT</td><td>269.2</td><td>-5.2</td><td>1.3</td><td>-4.1</td><td>4.1E-05</td><td>5.6E-05</td><td>MB231/MB468</td><td>NUC</td></tr>
<tr><td>>hg38_21_+_8433172_8446622_RNA45SN1_45S_50ntFlanks@8352.8379.28|AGTTGAAAAGAACTTTGAAGAGAGAGTT</td><td>>hg38_21_+_8433172_8446622_RNA45SN1_45S_50ntFlanks</td><td>8352</td><td>8379</td><td>28</td><td>AGTTGAAAAGAACTTTGAAGAGAGAGTT</td><td>167.3</td><td>-4.8</td><td>1.2</td><td>-4.1</td><td>3.4E-05</td><td>5.6E-05</td><td>MB231/MB468</td><td>MP</td></tr>
<tr><td>>hg38_21_+_8433172_8446622_RNA45SN1_45S_50ntFlanks@11212.11250.39|CGACTCTGGACGCGAGCCGGGCCCTTCCCGTGGATCGCC</td><td>>hg38_21_+_8433172_8446622_RNA45SN1_45S_50ntFlanks</td><td>11212</td><td>11250</td><td>39</td><td>CGACTCTGGACGCGAGCCGGGCCCTTCCCGTGGATCGCC</td><td>891.5</td><td>-7.3</td><td>1.8</td><td>-4.1</td><td>4.3E-05</td><td>5.6E-05</td><td>BT20/MB468</td><td>NUC</td></tr>
<tr><td>>hg38_21_+_8433172_8446622_RNA45SN1_45S_50ntFlanks@3854.3880.27|ACTGTGGTAATTCTAGAGCTAATACAT</td><td>>hg38_21_+_8433172_8446622_RNA45SN1_45S_50ntFlanks</td><td>3854</td><td>3880</td><td>27</td><td>ACTGTGGTAATTCTAGAGCTAATACAT</td><td>547.5</td><td>-5.6</td><td>1.4</td><td>-4.1</td><td>4.3E-05</td><td>5.6E-05</td><td>BT20/MB468</td><td>NUC</td></tr>
<tr><td>>hg38_21_+_8433172_8446622_RNA45SN1_45S_50ntFlanks@9481.9509.29|CACGTGTTAGGACCCGAAAGATGGTGAAC</td><td>>hg38_21_+_8433172_8446622_RNA45SN1_45S_50ntFlanks</td><td>9481</td><td>9509</td><td>29</td><td>CACGTGTTAGGACCCGAAAGATGGTGAAC</td><td>116.2</td><td>10.2</td><td>2.2</td><td>4.6</td><td>4.5E-06</td><td>5.6E-05</td><td>BT20/MB231</td><td>MP</td></tr>
<tr><td>>hg38_21_+_8433172_8446622_RNA45SN1_45S_50ntFlanks@10683.10710.28|CCGGCGGCGTCCGGTGAGCTCTCGCTGG</td><td>>hg38_21_+_8433172_8446622_RNA45SN1_45S_50ntFlanks</td><td>10683</td><td>10710</td><td>28</td><td>CCGGCGGCGTCCGGTGAGCTCTCGCTGG</td><td>305.9</td><td>-6.1</td><td>1.5</td><td>-4.1</td><td>3.5E-05</td><td>5.6E-05</td><td>BT20/MB468</td><td>CYTO</td></tr>
<tr><td>>trnaMT_SerGCT_MT_+_12207_12265@14.48.35|AGAACTGCTAACTCATGCCCCCATGTCTAACAACA</td><td>>trnaMT_SerGCT_MT_+_12207_12265</td><td>14</td><td>48</td><td>35</td><td>AGAACTGCTAACTCATGCCCCCATGTCTAACAACA</td><td>300.1</td><td>4.9</td><td>1.2</td><td>4.1</td><td>4.1E-05</td><td>5.6E-05</td><td>MB231/MB468</td><td>MITO</td></tr>
<tr><td>>hg38_21_+_8433172_8446622_RNA45SN1_45S_50ntFlanks@11999.12035.37|CCCTGCGGGCCGCCGGTGAAATACCACTACTCTGATC</td><td>>hg38_21_+_8433172_8446622_RNA45SN1_45S_50ntFlanks</td><td>11999</td><td>12035</td><td>37</td><td>CCCTGCGGGCCGCCGGTGAAATACCACTACTCTGATC</td><td>982.6</td><td>-7.5</td><td>1.8</td><td>-4.1</td><td>4.3E-05</td><td>5.6E-05</td><td>BT20/MB468</td><td>NUC</td></tr>
<tr><td>>MI0000299|hsa-mir-222&WithFlank&X|-|45747009|45747130@75.94.20&[MIMAT0000279&hsa-miR-222-3p&offsets|0|-1;m-140&X|-|45747033|45747056&offsets|0|-4]|AGCTACATCTGGCTACTGGG</td><td>>MI0000299|hsa-mir-222&WithFlank&X|-|45747009|45747130</td><td>75</td><td>94</td><td>20</td><td>AGCTACATCTGGCTACTGGG</td><td>160.7</td><td>6.0</td><td>1.5</td><td>4.1</td><td>4.1E-05</td><td>5.6E-05</td><td>MB231/MB468</td><td>MITO</td></tr>
<tr><td>>hg38_21_+_8433172_8446622_RNA45SN1_45S_50ntFlanks@10351.10371.21|CGAGAACTTTGAAGGCCGAAG</td><td>>hg38_21_+_8433172_8446622_RNA45SN1_45S_50ntFlanks</td><td>10351</td><td>10371</td><td>21</td><td>CGAGAACTTTGAAGGCCGAAG</td><td>841.5</td><td>-2.6</td><td>0.6</td><td>-4.2</td><td>3.0E-05</td><td>5.6E-05</td><td>BT20/MB468</td><td>MITO</td></tr>
<tr><td>>hg38_21_+_8433172_8446622_RNA45SN1_45S_50ntFlanks@12415.12439.25|CTTCGATGTCGGCTCTTCCTATCAT</td><td>>hg38_21_+_8433172_8446622_RNA45SN1_45S_50ntFlanks</td><td>12415</td><td>12439</td><td>25</td><td>CTTCGATGTCGGCTCTTCCTATCAT</td><td>745.9</td><td>2.5</td><td>0.6</td><td>4.1</td><td>3.9E-05</td><td>5.6E-05</td><td>MB231/MB468</td><td>CYTO</td></tr>
<tr><td>>hg38_21_+_8433172_8446622_RNA45SN1_45S_50ntFlanks@9552.9570.19|TCCGTAGCGGTCCTGACGT</td><td>>hg38_21_+_8433172_8446622_RNA45SN1_45S_50ntFlanks</td><td>9552</td><td>9570</td><td>19</td><td>TCCGTAGCGGTCCTGACGT</td><td>307.1</td><td>-3.4</td><td>0.8</td><td>-4.1</td><td>4.3E-05</td><td>5.6E-05</td><td>BT20/MB468</td><td>NUC</td></tr>
<tr><td>>hg38_21_+_8433172_8446622_RNA45SN1_45S_50ntFlanks@9554.9579.26|CGTAGCGGTCCTGACGTGCAAATCGG</td><td>>hg38_21_+_8433172_8446622_RNA45SN1_45S_50ntFlanks</td><td>9554</td><td>9579</td><td>26</td><td>CGTAGCGGTCCTGACGTGCAAATCGG</td><td>421.4</td><td>-4.8</td><td>1.2</td><td>-4.1</td><td>4.1E-05</td><td>5.6E-05</td><td>MB231/MB468</td><td>NUC</td></tr>
<tr><td>>hg38_21_+_8433172_8446622_RNA45SN1_45S_50ntFlanks@10393.10420.28|CAGCAGTTGAACATGGGTCAGTCGGTCC</td><td>>hg38_21_+_8433172_8446622_RNA45SN1_45S_50ntFlanks</td><td>10393</td><td>10420</td><td>28</td><td>CAGCAGTTGAACATGGGTCAGTCGGTCC</td><td>178.0</td><td>-5.2</td><td>1.3</td><td>-4.1</td><td>3.6E-05</td><td>5.6E-05</td><td>BT20/MB468</td><td>CYTO</td></tr>
<tr><td>>hg38_21_+_8433172_8446622_RNA45SN1_45S_50ntFlanks@12934.12959.26|CTAAACCATTCGTAGACGACCTGCTT</td><td>>hg38_21_+_8433172_8446622_RNA45SN1_45S_50ntFlanks</td><td>12934</td><td>12959</td><td>26</td><td>CTAAACCATTCGTAGACGACCTGCTT</td><td>468.4</td><td>-4.3</td><td>1.0</td><td>-4.1</td><td>3.6E-05</td><td>5.6E-05</td><td>BT20/MB468</td><td>CYTO</td></tr>
<tr><td>>hg38_21_+_8433172_8446622_RNA45SN1_45S_50ntFlanks@10351.10374.24|CGAGAACTTTGAAGGCCGAAGTGG</td><td>>hg38_21_+_8433172_8446622_RNA45SN1_45S_50ntFlanks</td><td>10351</td><td>10374</td><td>24</td><td>CGAGAACTTTGAAGGCCGAAGTGG</td><td>7215.3</td><td>-6.6</td><td>1.5</td><td>-4.3</td><td>1.4E-05</td><td>5.6E-05</td><td>BT20/MB468</td><td>MP</td></tr>
<tr><td>>hg38_21_+_8433172_8446622_RNA45SN1_45S_50ntFlanks@12992.13015.24|TCCCTCGCTGCGATCTATTGAAAG</td><td>>hg38_21_+_8433172_8446622_RNA45SN1_45S_50ntFlanks</td><td>12992</td><td>13015</td><td>24</td><td>TCCCTCGCTGCGATCTATTGAAAG</td><td>359.3</td><td>-4.5</td><td>1.1</td><td>-4.2</td><td>3.0E-05</td><td>5.7E-05</td><td>BT20/MB468</td><td>MITO</td></tr>
<tr><td>>hg38_21_+_8433172_8446622_RNA45SN1_45S_50ntFlanks@10873.10894.22|CTGGGTCGGTCGGGCTGGGGCG</td><td>>hg38_21_+_8433172_8446622_RNA45SN1_45S_50ntFlanks</td><td>10873</td><td>10894</td><td>22</td><td>CTGGGTCGGTCGGGCTGGGGCG</td><td>756.4</td><td>-5.5</td><td>1.3</td><td>-4.3</td><td>1.4E-05</td><td>5.7E-05</td><td>BT20/MB468</td><td>MP</td></tr>
<tr><td>>MI0000273|hsa-mir-183&WithFlank&7|-|129774899|129775020@33.53.21(+1C)&[MIMAT0000261&hsa-miR-183-5p&offsets|0|-1(+1C);m-41&7|-|129774966|129774987&offsets|-1|-2(+1C)]|TATGGCACTGGTAGAATTCACC</td><td>>MI0000273|hsa-mir-183&WithFlank&7|-|129774899|129775020</td><td>33</td><td>53</td><td>21(+1C)</td><td>TATGGCACTGGTAGAATTCACC</td><td>470.2</td><td>2.6</td><td>0.6</td><td>4.1</td><td>4.1E-05</td><td>5.7E-05</td><td>MB231/MB468</td><td>NUC</td></tr>
<tr><td>>MI0000093|hsa-mir-92a-1&WithFlank&13|+|91351308|91351397@54.75.22(+2A);MI0000094|hsa-mir-92a-2&WithFlank&X|-|134169532|134169618@54.75.22(+2A)&[MIMAT0000092&hsa-miR-92a-3p&offsets|0|0(+2A);m-9&13|+|91351361|91351382&offsets|0|0(+2A)];[MIMAT0000092_1&hsa-miR-92a-3p&offsets|0|0(+2A);m-11&X|-|134169544|134169565&offsets|0|0(+2A)]|TATTGCACTTGTCCCGGCCTGTAA</td><td>>MI0000093|hsa-mir-92a-1&WithFlank&13|+|91351308|91351397</td><td>54</td><td>75</td><td>22(+2A)</td><td>TATTGCACTTGTCCCGGCCTGTAA</td><td>576.2</td><td>3.7</td><td>0.9</td><td>4.1</td><td>4.1E-05</td><td>5.7E-05</td><td>MB231/MB468</td><td>MITO</td></tr>
<tr><td>>hg38_21_+_8433172_8446622_RNA45SN1_45S_50ntFlanks@9737.9757.21|CGAAACGATCTCAACCTATTC</td><td>>hg38_21_+_8433172_8446622_RNA45SN1_45S_50ntFlanks</td><td>9737</td><td>9757</td><td>21</td><td>CGAAACGATCTCAACCTATTC</td><td>103.0</td><td>-6.2</td><td>1.5</td><td>-4.1</td><td>3.5E-05</td><td>5.7E-05</td><td>MB231/MB468</td><td>MP</td></tr>
<tr><td>>hg38_21_+_8433172_8446622_RNA45SN1_45S_50ntFlanks@11660.11685.26|GTGATTTCTGCCCAGTGCTCTGAATG</td><td>>hg38_21_+_8433172_8446622_RNA45SN1_45S_50ntFlanks</td><td>11660</td><td>11685</td><td>26</td><td>GTGATTTCTGCCCAGTGCTCTGAATG</td><td>404.1</td><td>-2.1</td><td>0.5</td><td>-4.1</td><td>3.6E-05</td><td>5.7E-05</td><td>BT20/MB468</td><td>CYTO</td></tr>
<tr><td>>MI0000085|hsa-mir-27a&WithFlank&19|-|13836434|13836523@57.76.20(+2U)&[MIMAT0000084&hsa-miR-27a-3p&offsets|0|-1(+2U);m-43&19|-|13836447|13836467&offsets|0|-1(+2U)]|TTCACAGTGGCTAAGTTCCGTT</td><td>>MI0000085|hsa-mir-27a&WithFlank&19|-|13836434|13836523</td><td>57</td><td>76</td><td>20(+2U)</td><td>TTCACAGTGGCTAAGTTCCGTT</td><td>804.6</td><td>3.8</td><td>0.9</td><td>4.1</td><td>3.9E-05</td><td>5.7E-05</td><td>MB231/MB468</td><td>CYTO</td></tr>
<tr><td>>hg38_21_+_8433172_8446622_RNA45SN1_45S_50ntFlanks@6644.6665.22|CCTCGTACGACTCTTAGCGGTG</td><td>>hg38_21_+_8433172_8446622_RNA45SN1_45S_50ntFlanks</td><td>6644</td><td>6665</td><td>22</td><td>CCTCGTACGACTCTTAGCGGTG</td><td>415.2</td><td>4.8</td><td>1.2</td><td>4.1</td><td>3.6E-05</td><td>5.7E-05</td><td>BT20/MB468</td><td>CYTO</td></tr>
<tr><td>>hg38_21_+_8433172_8446622_RNA45SN1_45S_50ntFlanks@9567.9596.30|ACGTGCAAATCGGTCGTCCGACCTGGGTAT</td><td>>hg38_21_+_8433172_8446622_RNA45SN1_45S_50ntFlanks</td><td>9567</td><td>9596</td><td>30</td><td>ACGTGCAAATCGGTCGTCCGACCTGGGTAT</td><td>4934.4</td><td>-7.0</td><td>1.7</td><td>-4.1</td><td>3.6E-05</td><td>5.7E-05</td><td>BT20/MB468</td><td>CYTO</td></tr>
<tr><td>>hg38_21_+_8433172_8446622_RNA45SN1_45S_50ntFlanks@5486.5516.31|GCGCTGAGAAGACGGTCGAACTTGACTATCT</td><td>>hg38_21_+_8433172_8446622_RNA45SN1_45S_50ntFlanks</td><td>5486</td><td>5516</td><td>31</td><td>GCGCTGAGAAGACGGTCGAACTTGACTATCT</td><td>794.0</td><td>3.5</td><td>0.9</td><td>4.1</td><td>3.9E-05</td><td>5.7E-05</td><td>MB231/MB468</td><td>CYTO</td></tr>
<tr><td>>hg38_21_+_8433172_8446622_RNA45SN1_45S_50ntFlanks@6646.6670.25|TCGTACGACTCTTAGCGGTGGATCA</td><td>>hg38_21_+_8433172_8446622_RNA45SN1_45S_50ntFlanks</td><td>6646</td><td>6670</td><td>25</td><td>TCGTACGACTCTTAGCGGTGGATCA</td><td>3591.2</td><td>-3.3</td><td>0.8</td><td>-4.3</td><td>1.4E-05</td><td>5.7E-05</td><td>BT20/MB468</td><td>MP</td></tr>
</table>

| | | | | | | | | | | | | | |
|---|---|---|---|---|---|---|---|---|---|---|---|---|---|
| >MI0000081\|hsa-mir-24-2&WithFlank&19\|-\|13836281\|13836365@56.76.21(+1U);MI0000080\|hsa-mir-24-1&WithFlank&9\|+\|95086015\|95086094@50.70.21(+1U)&[MIMAT0000080&hsa-miR-24-3p&offsets\|0\|-1(+1U);m-36&19\|-\|13836288\|13836310&offsets\|0\|-2(+1U)];[MIMAT0000080_1&hsa-miR-24-3p&offsets\|0\|-1(+1U);m-35&9\|+\|95086064\|95086086&offsets\|0\|-2(+1U)]\|TGGCTCAGTTCAGCAGGAACAT | >MI0000081\|hsa-mir-24-2&WithFlank&19\|-\|13836281\|13836365 | 56 | 76 | 21(+1U) | TGGCTCAGTTCAGCAGGAACAT | 622.7 | 4.6 | 1.1 | 4.3 | 1.4E-05 | 5.7E-05 | BT20/MB468 | MP |
| >hg38_21_+_8433172_8446622_RNA45SN1_45S_50ntFlanks@6645.6662.18\|CTCGTACGACTCTTAGCG | >hg38_21_+_8433172_8446622_RNA45SN1_45S_50ntFlanks | 6645 | 6662 | 18 | CTCGTACGACTCTTAGCG | 8079.5 | 3.3 | 0.8 | 4.3 | 1.4E-05 | 5.7E-05 | BT20/MB468 | MP |
| >hg38_21_+_8433172_8446622_RNA45SN1_45S_50ntFlanks@9623.9658.36\|TAGTAGCTGGTTCCCTCCGAAGTTTCCCTCAGGATA | >hg38_21_+_8433172_8446622_RNA45SN1_45S_50ntFlanks | 9623 | 9658 | 36 | TAGTAGCTGGTTCCCTCCGAAGTTTCCCTCAGGATA | 569.0 | -7.1 | 1.7 | -4.1 | 4.3E-05 | 5.7E-05 | BT20/MB468 | NUC |
| >hg38_21_+_8433172_8446622_RNA45SN1_45S_50ntFlanks@9787.9830.44\|TCGCTGGCGTGGAGCCGGGCGTGGAATGCGAGTGCCTAGTGGGC | >hg38_21_+_8433172_8446622_RNA45SN1_45S_50ntFlanks | 9787 | 9830 | 44 | TCGCTGGCGTGGAGCCGGGCGTGGAATGCGAGTGCCTAGTGGGC | 58.5 | -9.7 | 2.4 | -4.1 | 3.5E-05 | 5.7E-05 | MB231/MB468 | MP |
| >hg38_21_+_8433172_8446622_RNA45SN1_45S_50ntFlanks@10922.10939.18\|CTGGACGAGGCGCCGCCG | >hg38_21_+_8433172_8446622_RNA45SN1_45S_50ntFlanks | 10922 | 10939 | 18 | CTGGACGAGGCGCCGCCG | 2106.1 | -7.2 | 1.8 | -4.1 | 4.1E-05 | 5.7E-05 | MB231/MB468 | NUC |
| >hg38_21_+_8433172_8446622_RNA45SN1_45S_50ntFlanks@5472.5513.42\|ACGGCCCTGGCGGAGCGCTGAGAAGACGGTCGAACTTGACTA | >hg38_21_+_8433172_8446622_RNA45SN1_45S_50ntFlanks | 5472 | 5513 | 42 | ACGGCCCTGGCGGAGCGCTGAGAAGACGGTCGAACTTGACTA | 570.1 | 3.9 | 0.9 | 4.1 | 3.6E-05 | 5.7E-05 | BT20/MB468 | CYTO |
| >hg38_21_+_8433172_8446622_RNA45SN1_45S_50ntFlanks@11718.11744.27\|AACGGCGGGAGTAACTATGACTCTCTT | >hg38_21_+_8433172_8446622_RNA45SN1_45S_50ntFlanks | 11718 | 11744 | 27 | AACGGCGGGAGTAACTATGACTCTCTT | 382.3 | -6.4 | 1.6 | -4.1 | 4.1E-05 | 5.7E-05 | MB231/MB468 | MITO |
| >hg38_21_+_8433172_8446622_RNA45SN1_45S_50ntFlanks@12418.12455.38\|CGATGTCGGCTCTTCCTATCATTGTGAAGCAGAATTCA | >hg38_21_+_8433172_8446622_RNA45SN1_45S_50ntFlanks | 12418 | 12455 | 38 | CGATGTCGGCTCTTCCTATCATTGTGAAGCAGAATTCA | 4192.7 | -3.2 | 0.8 | -4.1 | 3.9E-05 | 5.7E-05 | MB231/MB468 | CYTO |
| >hg38_21_+_8433172_8446622_RNA45SN1_45S_50ntFlanks@5493.5510.18\|GAAGACGGTCGAACTTGA | >hg38_21_+_8433172_8446622_RNA45SN1_45S_50ntFlanks | 5493 | 5510 | 18 | GAAGACGGTCGAACTTGA | 79.9 | -6.0 | 1.5 | -4.1 | 3.5E-05 | 5.7E-05 | MB231/MB468 | MP |
| >hg38_21_+_8433172_8446622_RNA45SN1_45S_50ntFlanks@11572.11614.43\|CCGACTTAGAACTGGTGCGGACCAGGGGAATCCGACTGTTTAA | >hg38_21_+_8433172_8446622_RNA45SN1_45S_50ntFlanks | 11572 | 11614 | 43 | CCGACTTAGAACTGGTGCGGACCAGGGGAATCCGACTGTTTAA | 475.9 | -4.1 | 1.0 | -4.1 | 4.2E-05 | 5.7E-05 | MB231/MB468 | MITO |
| >hg38_21_+_8433172_8446622_RNA45SN1_45S_50ntFlanks@12249.12271.23\|CAAAAGCTCGCTTGATCTTGATT | >hg38_21_+_8433172_8446622_RNA45SN1_45S_50ntFlanks | 12249 | 12271 | 23 | CAAAAGCTCGCTTGATCTTGATT | 105.4 | -6.0 | 1.5 | -4.1 | 3.5E-05 | 5.7E-05 | MB231/MB468 | MP |
| >MI0000085\|hsa-mir-27a&WithFlank&19\|-\|13836434\|13836523@57.77.21(+1U)&[MIMAT0000084&hsa-miR-27a-3p&offsets\|0\|0(+1U);m-43&19\|-\|13836447\|13836467&offsets\|0\|0(+1U)]\|TTCACAGTGGCTAAGTTCCGCT | >MI0000085\|hsa-mir-27a&WithFlank&19\|-\|13836434\|13836523 | 57 | 77 | 21(+1U) | TTCACAGTGGCTAAGTTCCGCT | 501.1 | -5.3 | 1.2 | -4.6 | 5.3E-06 | 5.7E-05 | BT20/MB231 | MITO |
| >hg38_21_+_8433172_8446622_RNA45SN1_45S_50ntFlanks@12151.12180.30\|GTGGGGAGTTTGACTGGGGCGGTACACCTG | >hg38_21_+_8433172_8446622_RNA45SN1_45S_50ntFlanks | 12151 | 12180 | 30 | GTGGGGAGTTTGACTGGGGCGGTACACCTG | 264.5 | -5.0 | 1.2 | -4.1 | 3.9E-05 | 5.7E-05 | MB231/MB468 | CYTO |
| >hg38_21_+_8433172_8446622_RNA45SN1_45S_50ntFlanks@8021.8042.22\|AGTCAGCGGAGGAGAAGAAACT | >hg38_21_+_8433172_8446622_RNA45SN1_45S_50ntFlanks | 8021 | 8042 | 22 | AGTCAGCGGAGGAGAAGAAACT | 195.8 | -3.4 | 0.7 | -4.6 | 4.4E-06 | 5.7E-05 | BT20/MB231 | CYTO |
| >MI0000342\|hsa-mir-200b&WithFlank&1\|+\|1167098\|1167204@63.85.23(+1C)&[MIMAT0000318&hsa-miR-200b-3p&offsets\|0\|+1(+1C);m-93&1\|+\|1167160\|1167182&offsets\|0\|0(+1C)]\|TAATACTGCCTGGTAATGATGACC | >MI0000342\|hsa-mir-200b&WithFlank&1\|+\|1167098\|1167204 | 63 | 85 | 23(+1C) | TAATACTGCCTGGTAATGATGACC | 229.4 | -4.8 | 1.2 | -4.1 | 3.5E-05 | 5.7E-05 | MB231/MB468 | MP |
| >hg38_21_+_8433172_8446622_RNA45SN1_45S_50ntFlanks@12418.12446.29\|CGATGTCGGCTCTTCCTATCATTGTGAAG | >hg38_21_+_8433172_8446622_RNA45SN1_45S_50ntFlanks | 12418 | 12446 | 29 | CGATGTCGGCTCTTCCTATCATTGTGAAG | 3156.8 | -3.6 | 0.9 | -4.1 | 3.6E-05 | 5.7E-05 | BT20/MB468 | CYTO |
| >hg38_21_+_8433172_8446622_RNA45SN1_45S_50ntFlanks@10735.10755.21\|TGTAAATCTCGCGCCGGGCCG | >hg38_21_+_8433172_8446622_RNA45SN1_45S_50ntFlanks | 10735 | 10755 | 21 | TGTAAATCTCGCGCCGGGCCG | 1050.8 | -5.1 | 1.2 | -4.1 | 3.9E-05 | 5.7E-05 | MB231/MB468 | CYTO |
| >MI0000487\|hsa-mir-193a&WithFlank&17\|+\|31559990\|31560089@27.47.21&[MIMAT0004614&hsa-miR-193a-5p&offsets\|0\|-1;m-123&17\|+\|31560016\|31560037&offsets\|0\|-1]\|TGGGTCTTTGCGGGCGAGATG | >MI0000487\|hsa-mir-193a&WithFlank&17\|+\|31559990\|31560089 | 27 | 47 | 21 | TGGGTCTTTGCGGGCGAGATG | 823.5 | 4.7 | 1.0 | 4.6 | 4.5E-06 | 5.7E-05 | BT20/MB231 | MP |
| >hg38_21_+_8433172_8446622_RNA45SN1_45S_50ntFlanks@3840.3862.23\|CTCCTACTTGGATAACTGTGGTA | >hg38_21_+_8433172_8446622_RNA45SN1_45S_50ntFlanks | 3840 | 3862 | 23 | CTCCTACTTGGATAACTGTGGTA | 1991.2 | 3.6 | 0.8 | 4.6 | 4.4E-06 | 5.7E-05 | BT20/MB231 | CYTO |
| >trnaMT_PheGAA_MT_+_577_647@27.56.30\|AATACACTGAAAATGTTTAGACGGGCTCAC | >trnaMT_PheGAA_MT_+_577_647 | 27 | 56 | 30 | AATACACTGAAAATGTTTAGACGGGCTCAC | 46.3 | 9.3 | 2.1 | 4.3 | 1.4E-05 | 5.7E-05 | BT20/MB468 | MP |
| >hg38_21_+_8433172_8446622_RNA45SN1_45S_50ntFlanks@10869.10904.36\|AGGGCTGGGTCGGTCGGGCTGGGGCGCGAAGCGGGG | >hg38_21_+_8433172_8446622_RNA45SN1_45S_50ntFlanks | 10869 | 10904 | 36 | AGGGCTGGGTCGGTCGGGCTGGGGCGCGAAGCGGGG | 399.0 | -5.2 | 1.2 | -4.3 | 1.4E-05 | 5.7E-05 | BT20/MB468 | MP |
| >MI0000434\|hsa-let-7i&WithFlank&12\|+\|62603680\|62603775@12.32.21&[MIMAT0000415&hsa-let-7i-5p&offsets\|0\|-1;m-49&12\|+\|62603691\|62603712&offsets\|0\|-1]\|TGAGGTAGTAGTTTGTGCTGT | >MI0000434\|hsa-let-7i&WithFlank&12\|+\|62603680\|62603775 | 12 | 32 | 21 | TGAGGTAGTAGTTTGTGCTGT | 3659.0 | 4.6 | 1.1 | 4.3 | 1.4E-05 | 5.7E-05 | BT20/MB468 | MP |
| >hg38_21_+_8433172_8446622_RNA45SN1_45S_50ntFlanks@11568.11591.24\|GCAGCCGACTTAGAACTGGTGCGG | >hg38_21_+_8433172_8446622_RNA45SN1_45S_50ntFlanks | 11568 | 11591 | 24 | GCAGCCGACTTAGAACTGGTGCGG | 132.5 | -5.2 | 1.3 | -4.1 | 3.6E-05 | 5.7E-05 | BT20/MB468 | CYTO |
| >trna116_GluCTC_1_-_145399233_145399304@31.70.40,trna71_GluCTC_1_-_161439189_161439260@31.70.40,trna74_GluCTC_1_-_161431809_161431880@31.70.40,trna77_GluCTC_1_-_161424398_161424469@31.70.40,trna77_GluCTC_6_+_28949976_28950047@31.70.40,trna80_GluCTC_1_-_161417018_161417089@31.70.40,trna87_GluCTC_6_-_126101393_126101464@31.70.40\|GCTCTCACCGCCGCGGCCCGGGTTCGATTCCCGGTCAGGG | >trna116_GluCTC_1_-_145399233_145399304 | 31 | 70 | 40 | GCTCTCACCGCCGCGGCCCGGGTTCGATTCCCGGTCAGGG | 190.5 | -3.9 | 0.9 | -4.1 | 3.6E-05 | 5.7E-05 | BT20/MB468 | CYTO |
| >hg38_21_+_8433172_8446622_RNA45SN1_45S_50ntFlanks@11569.11589.21\|CAGCCGACTTAGAACTGGTGC | >hg38_21_+_8433172_8446622_RNA45SN1_45S_50ntFlanks | 11569 | 11589 | 21 | CAGCCGACTTAGAACTGGTGC | 158.7 | -6.6 | 1.6 | -4.1 | 3.5E-05 | 5.7E-05 | MB231/MB468 | MP |
| >hg38_21_+_8433172_8446622_RNA45SN1_45S_50ntFlanks@8176.8218.43\|CTTCTGATCGAGGCCCAGCCCGTGGACGGTGTGAGGCCGGTAG | >hg38_21_+_8433172_8446622_RNA45SN1_45S_50ntFlanks | 8176 | 8218 | 43 | CTTCTGATCGAGGCCCAGCCCGTGGACGGTGTGAGGCCGGTAG | 221.6 | -5.5 | 1.3 | -4.1 | 3.5E-05 | 5.7E-05 | MB231/MB468 | MP |
| >hg38_21_+_8433172_8446622_RNA45SN1_45S_50ntFlanks@10361.10388.28\|GAAGGCCGAAGTGGAGAAGGGTTCCATG | >hg38_21_+_8433172_8446622_RNA45SN1_45S_50ntFlanks | 10361 | 10388 | 28 | GAAGGCCGAAGTGGAGAAGGGTTCCATG | 171.1 | -3.1 | 0.7 | -4.2 | 3.0E-05 | 5.7E-05 | BT20/MB468 | MITO |
| >trna2_GlyGCC_21_-_18827107_18827177@1.30.30,trna35_GlyGCC_1_+_161413094_161413164@1.30.30,trna37_GlyGCC_1_+_161420467_161420537@1.30.30,trna39_GlyGCC_1_+_161427898_161427968@1.30.30,trna41_GlyGCC_1_+_161435258_161435328@1.30.30\|GCATGGGTGGTTCAGTGGTAGAATTCTCGC | >trna2_GlyGCC_21_-_18827107_18827177 | 1 | 30 | 30 | GCATGGGTGGTTCAGTGGTAGAATTCTCGC | 8934.9 | -4.0 | 1.0 | -4.1 | 3.5E-05 | 5.7E-05 | MB231/MB468 | MP |
| >hg38_21_+_8433172_8446622_RNA45SN1_45S_50ntFlanks@4590.4619.30\|ATTTTGTTGGTTTTCGGAACTGAGGCCATG | >hg38_21_+_8433172_8446622_RNA45SN1_45S_50ntFlanks | 4590 | 4619 | 30 | ATTTTGTTGGTTTTCGGAACTGAGGCCATG | 242.9 | -6.3 | 1.5 | -4.1 | 3.5E-05 | 5.7E-05 | MB231/MB468 | MP |
| >hg38_21_+_8433172_8446622_RNA45SN1_45S_50ntFlanks@4660.4679.20\|GAGGTGAAATTCTTGGACCG | >hg38_21_+_8433172_8446622_RNA45SN1_45S_50ntFlanks | 4660 | 4679 | 20 | GAGGTGAAATTCTTGGACCG | 59.0 | -9.7 | 2.4 | -4.1 | 3.5E-05 | 5.7E-05 | MB231/MB468 | MP |
| >hg38_21_+_8433172_8446622_RNA45SN1_45S_50ntFlanks@11580.11605.26\|GAACTGGTGCGGACCAGGGGAATCCG | >hg38_21_+_8433172_8446622_RNA45SN1_45S_50ntFlanks | 11580 | 11605 | 26 | GAACTGGTGCGGACCAGGGGAATCCG | 1597.8 | -5.8 | 1.3 | -4.3 | 1.4E-05 | 5.7E-05 | BT20/MB468 | MP |
| >hg38_21_+_8433172_8446622_RNA45SN1_45S_50ntFlanks@10679.10707.29\|GGTTCCGGCGGCGTCCGGTGAGCTCTCGC | >hg38_21_+_8433172_8446622_RNA45SN1_45S_50ntFlanks | 10679 | 10707 | 29 | GGTTCCGGCGGCGTCCGGTGAGCTCTCGC | 1927.3 | 9.9 | 2.2 | 4.6 | 4.6E-06 | 5.7E-05 | BT20/MB231 | MP |
| >MI0000283\|hsa-mir-203a&WithFlank&14\|+\|104117399\|104117520@72.92.21&[MIMAT0000264&hsa-miR-203a-3p&offsets\|+1\|0;m-79&14\|+\|104117469\|104117490&offsets\|+1\|0]\|TGAAATGTTTAGGACCACTAG | >MI0000283\|hsa-mir-203a&WithFlank&14\|+\|104117399\|104117520 | 72 | 92 | 21 | TGAAATGTTTAGGACCACTAG | 608.5 | 7.1 | 1.6 | 4.3 | 1.4E-05 | 5.7E-05 | BT20/MB468 | MP |
| >MI0000077\|hsa-mir-21&WithFlank&17\|+\|59841260\|59841343@52.72.21&[MIMAT0004494&hsa-miR-21-3p&offsets\|0\|0;m-52&17\|+\|59841311\|59841332&offsets\|0\|-1]\|CAACACCAGTCGATGGGCTGT | >MI0000077\|hsa-mir-21&WithFlank&17\|+\|59841260\|59841343 | 52 | 72 | 21 | CAACACCAGTCGATGGGCTGT | 1256.7 | 3.8 | 0.9 | 4.1 | 3.5E-05 | 5.7E-05 | MB231/MB468 | MP |
| >hg38_21_+_8433172_8446622_RNA45SN1_45S_50ntFlanks@10807.10826.20\|ACAATGTAGGTAAGGGAAGT | >hg38_21_+_8433172_8446622_RNA45SN1_45S_50ntFlanks | 10807 | 10826 | 20 | ACAATGTAGGTAAGGGAAGT | 248.0 | -5.5 | 1.3 | -4.1 | 3.6E-05 | 5.7E-05 | BT20/MB468 | CYTO |
| >hg38_21_+_8433172_8446622_RNA45SN1_45S_50ntFlanks@4000.4042.43\|TAGATAACCTCGGGCCGATCGCACGCCCCCCGTGGCGGCGACG | >hg38_21_+_8433172_8446622_RNA45SN1_45S_50ntFlanks | 4000 | 4042 | 43 | TAGATAACCTCGGGCCGATCGCACGCCCCCCGTGGCGGCGACG | 546.7 | -5.4 | 1.3 | -4.1 | 3.6E-05 | 5.7E-05 | BT20/MB468 | CYTO |
| >hg38_21_+_8433172_8446622_RNA45SN1_45S_50ntFlanks@10386.10424.39\|ATGTGAACAGCAGTTGAACATGGGTCAGTCGGTCCTGAG | >hg38_21_+_8433172_8446622_RNA45SN1_45S_50ntFlanks | 10386 | 10424 | 39 | ATGTGAACAGCAGTTGAACATGGGTCAGTCGGTCCTGAG | 14383.5 | 4.0 | 0.9 | 4.5 | 5.4E-06 | 5.7E-05 | BT20/MB231 | MITO |
| >hg38_21_+_8433172_8446622_RNA45SN1_45S_50ntFlanks@9535.9571.37\|GGAAACTCTGGTGGAGGTCCGTAGCGGTCCTGACGTG | >hg38_21_+_8433172_8446622_RNA45SN1_45S_50ntFlanks | 9535 | 9571 | 37 | GGAAACTCTGGTGGAGGTCCGTAGCGGTCCTGACGTG | 1548.6 | -3.0 | 0.7 | -4.1 | 4.2E-05 | 5.8E-05 | MB231/MB468 | NUC |
| >hg38_21_+_8433172_8446622_RNA45SN1_45S_50ntFlanks@7992.8010.19\|CGTGGCGACCCGCTGAATT | >hg38_21_+_8433172_8446622_RNA45SN1_45S_50ntFlanks | 7992 | 8010 | 19 | CGTGGCGACCCGCTGAATT | 159.6 | 1.8 | 0.4 | 4.1 | 4.4E-05 | 5.8E-05 | BT20/MB468 | NUC |
| >hg38_21_+_8433172_8446622_RNA45SN1_45S_50ntFlanks@10906.10925.20\|TGGGCGCGCGCCGCGGCTGG | >hg38_21_+_8433172_8446622_RNA45SN1_45S_50ntFlanks | 10906 | 10925 | 20 | TGGGCGCGCGCCGCGGCTGG | 204.2 | -4.5 | 1.1 | -4.1 | 3.6E-05 | 5.8E-05 | BT20/MB468 | CYTO |
| >hg38_21_+_8433172_8446622_RNA45SN1_45S_50ntFlanks@474.515.42\|TCTTCGTGATCGATGTGGTGACGTCGTGCTCTCCCGGGCCGG | >hg38_21_+_8433172_8446622_RNA45SN1_45S_50ntFlanks | 474 | 515 | 42 | TCTTCGTGATCGATGTGGTGACGTCGTGCTCTCCCGGGCCGG | 217.7 | 4.3 | 1.0 | 4.1 | 4.2E-05 | 5.8E-05 | MB231/MB468 | NUC |
| >hg38_21_+_8433172_8446622_RNA45SN1_45S_50ntFlanks@9955.9989.35\|GGAAGTCGGAATCCGCTAAGGAGTGTGTAACAACT | >hg38_21_+_8433172_8446622_RNA45SN1_45S_50ntFlanks | 9955 | 9989 | 35 | GGAAGTCGGAATCCGCTAAGGAGTGTGTAACAACT | 119.8 | 10.2 | 2.2 | 4.6 | 4.6E-06 | 5.8E-05 | BT20/MB231 | MP |
| >hg38_21_+_8433172_8446622_RNA45SN1_45S_50ntFlanks@9574.9601.28\|AATCGGTCGTCCGACCTGGGTATAGGGG | >hg38_21_+_8433172_8446622_RNA45SN1_45S_50ntFlanks | 9574 | 9601 | 28 | AATCGGTCGTCCGACCTGGGTATAGGGG | 1089.3 | -2.6 | 0.6 | -4.1 | 4.2E-05 | 5.8E-05 | MB231/MB468 | MITO |
| >hg38_21_+_8433172_8446622_RNA45SN1_45S_50ntFlanks@11561.11605.45\|GCGCCTAGCAGCCGACTTAGAACTGGTGCGGACCAGGGGAATCCG | >hg38_21_+_8433172_8446622_RNA45SN1_45S_50ntFlanks | 11561 | 11605 | 45 | GCGCCTAGCAGCCGACTTAGAACTGGTGCGGACCAGGGGAATCCG | 417.1 | -4.8 | 1.1 | -4.2 | 3.1E-05 | 5.8E-05 | BT20/MB468 | MITO |
| >trnaMT_HisGTG_MT_+_12138_12206@27.53.27\|GATTGTGAATCTGACAACAGAGGCTTA | >trnaMT_HisGTG_MT_+_12138_12206 | 27 | 53 | 27 | GATTGTGAATCTGACAACAGAGGCTTA | 156.0 | 8.1 | 1.9 | 4.3 | 1.4E-05 | 5.8E-05 | BT20/MB468 | MP |
| >trnaMT_ValTAC_MT_+_1602_1670@28.70.43\|AACTTACACTTAGGAGATTTCAACTTAACTTGACCGCTCTGAC | >trnaMT_ValTAC_MT_+_1602_1670 | 28 | 70 | 43 | AACTTACACTTAGGAGATTTCAACTTAACTTGACCGCTCTGAC | 294.6 | 8.8 | 2.0 | 4.3 | 1.4E-05 | 5.8E-05 | BT20/MB468 | MP |
| >hg38_21_+_8433172_8446622_RNA45SN1_45S_50ntFlanks@4006.4025.20\|ACCTCGGGCCGATCGCACGC | >hg38_21_+_8433172_8446622_RNA45SN1_45S_50ntFlanks | 4006 | 4025 | 20 | ACCTCGGGCCGATCGCACGC | 386.0 | -4.2 | 1.0 | -4.1 | 3.7E-05 | 5.8E-05 | BT20/MB468 | CYTO |
| >hg38_21_+_8433172_8446622_RNA45SN1_45S_50ntFlanks@9929.9955.27\|GATATAGACAGCAGGACGGTGGCCATG | >hg38_21_+_8433172_8446622_RNA45SN1_45S_50ntFlanks | 9929 | 9955 | 27 | GATATAGACAGCAGGACGGTGGCCATG | 1167.5 | -7.0 | 1.7 | -4.1 | 3.5E-05 | 5.8E-05 | MB231/MB468 | MP |
| >hg38_21_+_8433172_8446622_RNA45SN1_45S_50ntFlanks@9533.9551.19\|GAGGAAACTCTGGTGGAGG | >hg38_21_+_8433172_8446622_RNA45SN1_45S_50ntFlanks | 9533 | 9551 | 19 | GAGGAAACTCTGGTGGAGG | 673.5 | -2.5 | 0.6 | -4.1 | 4.0E-05 | 5.8E-05 | MB231/MB468 | CYTO |

| | | | | | | | | | | | | | |
|---|---|---|---|---|---|---|---|---|---|---|---|---|---|
| >hg38_21_+_8433172_8446622_RNA45SN1_45S_50ntFlanks@10792.10812.21\|CTCTGGCATGTTGGAACAATG | >hg38_21_+_8433172_8446622_RNA45SN1_45S_50ntFlanks | 10792 | 10812 | 21 | CTCTGGCATGTTGGAACAATG | 102.0 | 10.0 | 2.2 | 4.6 | 4.6E-06 | 5.8E-05 | BT20/MB231 | MP |
| >hg38_21_+_8433172_8446622_RNA45SN1_45S_50ntFlanks@4982.5003.22\|ACGGACAGGATTGACAGATTGA | >hg38_21_+_8433172_8446622_RNA45SN1_45S_50ntFlanks | 4982 | 5003 | 22 | ACGGACAGGATTGACAGATTGA | 65.5 | -9.9 | 2.4 | -4.1 | 3.5E-05 | 5.8E-05 | MB231/MB468 | MP |
| >hg38_21_+_8433172_8446622_RNA45SN1_45S_50ntFlanks@4995.5022.28\|ACAGATTGATAGCTCTTTCTCGATTCCG | >hg38_21_+_8433172_8446622_RNA45SN1_45S_50ntFlanks | 4995 | 5022 | 28 | ACAGATTGATAGCTCTTTCTCGATTCCG | 450.8 | -7.4 | 1.7 | -4.3 | 1.4E-05 | 5.8E-05 | BT20/MB468 | MP |
| >MI0001445\|hsa-mir-423&WithFlank&17\|+\|30117073\|30117178@23.43.21&[MIMAT0004748&hsa-miR-423-5p&offsets\|0\|-2;m-141&17\|+\|30117095\|30117117&offsets\|0\|-2]\|TGAGGGGCAGAGAGCGAGACT | >MI0001445\|hsa-mir-423&WithFlank&17\|+\|30117073\|30117178 | 23 | 43 | 21 | TGAGGGGCAGAGAGCGAGACT | 192.0 | 6.9 | 1.7 | 4.1 | 4.2E-05 | 5.8E-05 | MB231/MB468 | MITO |
| >hg38_21_+_8433172_8446622_RNA45SN1_45S_50ntFlanks@4374.4391.18\|AAAGCTCGTAGTTGGATC | >hg38_21_+_8433172_8446622_RNA45SN1_45S_50ntFlanks | 4374 | 4391 | 18 | AAAGCTCGTAGTTGGATC | 106.7 | -5.9 | 1.4 | -4.1 | 3.5E-05 | 5.8E-05 | MB231/MB468 | MP |
| >hg38_21_+_8433172_8446622_RNA45SN1_45S_50ntFlanks@12996.13039.44\|TCGCTGCGATCTATTGAAAGTCAGCCCTCGACACAAGGGTTTGT | >hg38_21_+_8433172_8446622_RNA45SN1_45S_50ntFlanks | 12996 | 13039 | 44 | TCGCTGCGATCTATTGAAAGTCAGCCCTCGACACAAGGGTTTGT | 161470.5 | 5.3 | 1.3 | 4.1 | 4.2E-05 | 5.8E-05 | MB231/MB468 | NUC |
| >am_trna111_HisGTG_1_-_147774845_147774916@-1G.26.27,trna118_HisGTG_1_-_145396881_145396952@-1G.26.27,trna16_HisGTG_1_+_146544773_146544844@-1G.26.27,trna1_HisGTG_15_+_45493349_45493420@-1G.26.27,trna21_HisGTG_1_+_147753471_147753542@-1G.26.27,trna33_HisGTG_6_+_27125906_27125977@-1G.26.27,trna7_HisGTG_9_-_14433938_14434009@-1G.26.27,trna8_HisGTG_15_-_45492611_45492682@-1G.26.27,trna9_HisGTG_15_-_45490804_45490875@-1G.26.27\|GGCCGTGATCGTATAGTGGTTAGTACT | >am_trna111_HisGTG_1_-_147774845_147774916 | -1G | 26 | 27 | GGCCGTGATCGTATAGTGGTTAGTACT | 2437.0 | 3.4 | 0.8 | 4.1 | 4.2E-05 | 5.8E-05 | MB231/MB468 | NUC |
| >hg38_21_+_8433172_8446622_RNA45SN1_45S_50ntFlanks@12004.12034.31\|CGGGCCGCCGGTGAAATACCACTACTCTGAT | >hg38_21_+_8433172_8446622_RNA45SN1_45S_50ntFlanks | 12004 | 12034 | 31 | CGGGCCGCCGGTGAAATACCACTACTCTGAT | 511.6 | 3.9 | 0.8 | 4.5 | 5.5E-06 | 5.8E-05 | BT20/MB231 | MITO |
| >hg38_21_+_8433172_8446622_RNA45SN1_45S_50ntFlanks@12489.12509.21\|ACGTGAGCTGGGTTTAGACCG | >hg38_21_+_8433172_8446622_RNA45SN1_45S_50ntFlanks | 12489 | 12509 | 21 | ACGTGAGCTGGGTTTAGACCG | 638.2 | -4.5 | 1.1 | -4.1 | 3.7E-05 | 5.8E-05 | BT20/MB468 | CYTO |
| >hg38_21_+_8433172_8446622_RNA45SN1_45S_50ntFlanks@9939.9968.30\|GCAGGACGGTGGCCATGGAAGTCGGAATCC | >hg38_21_+_8433172_8446622_RNA45SN1_45S_50ntFlanks | 9939 | 9968 | 30 | GCAGGACGGTGGCCATGGAAGTCGGAATCC | 555.8 | 4.9 | 1.1 | 4.3 | 1.4E-05 | 5.8E-05 | BT20/MB468 | MP |
| >trna10_LysCTT_16_+_3241501_3241573@5.36.32,trna13_LysCTT_14_-_58706613_58706685@5.36.32,trna2_LysCTT_15_+_79152904_79152976@5.36.32\|GGCTAGCTCAGTCGGTAGAGCATGGGACTCTT | >trna10_LysCTT_16_+_3241501_3241573 | 5 | 36 | 32 | GGCTAGCTCAGTCGGTAGAGCATGGGACTCTT | 532.2 | 6.3 | 1.5 | 4.1 | 4.2E-05 | 5.8E-05 | MB231/MB468 | MITO |
| >hg38_21_+_8433172_8446622_RNA45SN1_45S_50ntFlanks@8176.8208.33\|CTTCTGATCGAGGCCCAGCCCGTGGACGGTGTG | >hg38_21_+_8433172_8446622_RNA45SN1_45S_50ntFlanks | 8176 | 8208 | 33 | CTTCTGATCGAGGCCCAGCCCGTGGACGGTGTG | 699.6 | -6.5 | 1.6 | -4.1 | 3.6E-05 | 5.8E-05 | MB231/MB468 | MP |
| >hg38_21_+_8433172_8446622_RNA45SN1_45S_50ntFlanks@12844.12862.19\|AGTGCCCTTCGTCCTGGGA | >hg38_21_+_8433172_8446622_RNA45SN1_45S_50ntFlanks | 12844 | 12862 | 19 | AGTGCCCTTCGTCCTGGGA | 65.6 | -9.9 | 2.4 | -4.1 | 3.6E-05 | 5.8E-05 | MB231/MB468 | MP |
| >hg38_21_+_8433172_8446622_RNA45SN1_45S_50ntFlanks@12416.12452.37\|TTCGATGTCGGCTCTTCCTATCATTGTGAAGCAGAAT | >hg38_21_+_8433172_8446622_RNA45SN1_45S_50ntFlanks | 12416 | 12452 | 37 | TTCGATGTCGGCTCTTCCTATCATTGTGAAGCAGAAT | 273.6 | 5.9 | 1.4 | 4.1 | 4.2E-05 | 5.8E-05 | MB231/MB468 | NUC |
| >hg38_1_-_228634819_228635039_RNA5S12_5S_50ntFlanks@128.170.43\|CTTGGATGGGAGACCGCCTGGGAATACCGGGTGCTGTAGGCTT | >hg38_1_-_228634819_228635039_RNA5S12_5S_50ntFlanks | 128 | 170 | 43 | CTTGGATGGGAGACCGCCTGGGAATACCGGGTGCTGTAGGCTT | 741.5 | 4.6 | 1.1 | 4.1 | 4.2E-05 | 5.8E-05 | MB231/MB468 | NUC |
| >hg38_21_+_8433172_8446622_RNA45SN1_45S_50ntFlanks@12420.12459.40\|ATGTCGGCTCTTCCTATCATTGTGAAGCAGAATTCACCAA | >hg38_21_+_8433172_8446622_RNA45SN1_45S_50ntFlanks | 12420 | 12459 | 40 | ATGTCGGCTCTTCCTATCATTGTGAAGCAGAATTCACCAA | 1382.1 | 4.6 | 1.1 | 4.1 | 4.2E-05 | 5.8E-05 | MB231/MB468 | MITO |
| >hg38_21_+_8433172_8446622_RNA45SN1_45S_50ntFlanks@4243.4264.22\|CTTTCGAGGCCCTGTAATTGGA | >hg38_21_+_8433172_8446622_RNA45SN1_45S_50ntFlanks | 4243 | 4264 | 22 | CTTTCGAGGCCCTGTAATTGGA | 303.5 | -4.6 | 1.1 | -4.1 | 4.2E-05 | 5.8E-05 | MB231/MB468 | NUC |
| >hg38_21_+_8433172_8446622_RNA45SN1_45S_50ntFlanks@11235.11253.19\|CTTCCCGTGGATCGCCCCA | >hg38_21_+_8433172_8446622_RNA45SN1_45S_50ntFlanks | 11235 | 11253 | 19 | CTTCCCGTGGATCGCCCCA | 903.5 | -4.7 | 1.1 | -4.2 | 3.1E-05 | 5.8E-05 | BT20/MB468 | MITO |
| >MI0000088\|hsa-mir-30a&WithFlank&6\|-\|71403545\|71403627@54.75.22&[MIMAT0000088&hsa-miR-30a-3p&offsets\|+1\|+1;m-38&6\|-\|71403554\|71403575&offsets\|+1\|+1]\|TTTCAGTCGGATGTTTGCAGCT | >MI0000088\|hsa-mir-30a&WithFlank&6\|-\|71403545\|71403627 | 54 | 75 | 22 | TTTCAGTCGGATGTTTGCAGCT | 1118.2 | -4.8 | 1.1 | -4.6 | 4.7E-06 | 5.8E-05 | BT20/MB231 | MP |
| >MI0000446\|hsa-mir-125b-1&WithFlank&11\|-\|122099751\|122099850@21.41.21(+1U);MI0000470\|hsa-mir-125b-2&WithFlank&21\|+\|16590231\|16590331@23.43.21(+1U)&[MIMAT0000423&hsa-miR-125b-5p&offsets\|0\|-1(+1U);m-67&11\|-\|122099809\|122099830&offsets\|0\|-1(+1U)];[MIMAT0000423_1&hsa-miR-125b-5p&offsets\|0\|-1(+1U);m-68&21\|+\|16590253\|16590274&offsets\|0\|-1(+1U)]\|TCCCTGAGACCCTAACTTGTGT | >MI0000446\|hsa-mir-125b-1&WithFlank&11\|-\|122099751\|122099850 | 21 | 41 | 21(+1U) | TCCCTGAGACCCTAACTTGTGT | 825.9 | -7.8 | 1.7 | -4.6 | 4.7E-06 | 5.8E-05 | BT20/MB231 | MP |
| >hg38_21_+_8433172_8446622_RNA45SN1_45S_50ntFlanks@7983.8006.24\|CAGATCAGACGTGGCGACCCGCTG | >hg38_21_+_8433172_8446622_RNA45SN1_45S_50ntFlanks | 7983 | 8006 | 24 | CAGATCAGACGTGGCGACCCGCTG | 13933.0 | -5.2 | 1.2 | -4.3 | 1.5E-05 | 5.8E-05 | BT20/MB468 | MP |
| >MI0003513\|hsa-mir-455&WithFlank&9\|+\|114209428\|114209535@60.80.21(+1C)&[MIMAT0004784&hsa-miR-455-3p&offsets\|0\|0(+1C);m-135&9\|+\|114209487\|114209508&offsets\|0\|-1(+1C)]\|GCAGTCCATGGGCATATACACC | >MI0003513\|hsa-mir-455&WithFlank&9\|+\|114209428\|114209535 | 60 | 80 | 21(+1C) | GCAGTCCATGGGCATATACACC | 292.2 | -3.1 | 0.7 | -4.5 | 5.5E-06 | 5.8E-05 | BT20/MB231 | MITO |
| >hg38_21_+_8433172_8446622_RNA45SN1_45S_50ntFlanks@6661.6700.40\|CGGTGGATCACTCGGCTCGTGCGTCGATGAAGAACGCAGC | >hg38_21_+_8433172_8446622_RNA45SN1_45S_50ntFlanks | 6661 | 6700 | 40 | CGGTGGATCACTCGGCTCGTGCGTCGATGAAGAACGCAGC | 457.1 | 3.7 | 0.9 | 4.1 | 4.3E-05 | 5.8E-05 | MB231/MB468 | MITO |
| >hg38_21_+_8433172_8446622_RNA45SN1_45S_50ntFlanks@10067.10085.19\|AGAGTGGACGGGAGCGGCG | >hg38_21_+_8433172_8446622_RNA45SN1_45S_50ntFlanks | 10067 | 10085 | 19 | AGAGTGGACGGGAGCGGCG | 78.3 | 9.6 | 2.1 | 4.6 | 4.7E-06 | 5.8E-05 | BT20/MB231 | MP |
| >hg38_21_+_8433172_8446622_RNA45SN1_45S_50ntFlanks@9573.9605.33\|AAATCGGTCGTCCGACCTGGGTATAGGGGCGAA | >hg38_21_+_8433172_8446622_RNA45SN1_45S_50ntFlanks | 9573 | 9605 | 33 | AAATCGGTCGTCCGACCTGGGTATAGGGGCGAA | 772.3 | -3.3 | 0.8 | -4.1 | 4.3E-05 | 5.8E-05 | MB231/MB468 | MITO |
| >hg38_21_+_8433172_8446622_RNA45SN1_45S_50ntFlanks@6733.6758.26\|CATTGATCATCGACACTTCGAACGCA | >hg38_21_+_8433172_8446622_RNA45SN1_45S_50ntFlanks | 6733 | 6758 | 26 | CATTGATCATCGACACTTCGAACGCA | 268.0 | -1.9 | 0.5 | -4.1 | 3.7E-05 | 5.9E-05 | BT20/MB468 | CYTO |
| >hg38_21_+_8433172_8446622_RNA45SN1_45S_50ntFlanks@12405.12434.30\|CTTTTTGATCCTTCGATGTCGGCTCTTCCT | >hg38_21_+_8433172_8446622_RNA45SN1_45S_50ntFlanks | 12405 | 12434 | 30 | CTTTTTGATCCTTCGATGTCGGCTCTTCCT | 15513.7 | -4.5 | 1.1 | -4.1 | 4.0E-05 | 5.9E-05 | MB231/MB468 | CYTO |
| >hg38_21_+_8433172_8446622_RNA45SN1_45S_50ntFlanks@9567.9584.18\|ACGTGCAAATCGGTCGTC | >hg38_21_+_8433172_8446622_RNA45SN1_45S_50ntFlanks | 9567 | 9584 | 18 | ACGTGCAAATCGGTCGTC | 521.5 | -5.3 | 1.2 | -4.3 | 1.5E-05 | 5.9E-05 | BT20/MB468 | MP |
| >MI0000253\|hsa-mir-148a&WithFlank&7\|-\|25949913\|25949992@12.33.22&[MIMAT0004549&hsa-miR-148a-5p&offsets\|0\|0;m-189&7\|-\|25949960\|25949981&offsets\|0\|0]\|AAAGTTCTGAGACACTCCGACT | >MI0000253\|hsa-mir-148a&WithFlank&7\|-\|25949913\|25949992 | 12 | 33 | 22 | AAAGTTCTGAGACACTCCGACT | 425.0 | 4.1 | 0.9 | 4.3 | 1.5E-05 | 5.9E-05 | BT20/MB468 | MP |
| >hg38_21_+_8433172_8446622_RNA45SN1_45S_50ntFlanks@4761.4780.20\|CAGATACCGTCGTAGTTCCG | >hg38_21_+_8433172_8446622_RNA45SN1_45S_50ntFlanks | 4761 | 4780 | 20 | CAGATACCGTCGTAGTTCCG | 240.3 | -5.6 | 1.4 | -4.1 | 3.7E-05 | 5.9E-05 | BT20/MB468 | CYTO |
| >MI0001733\|hsa-mir-452&WithFlank&X\|-\|151959622\|151959718@20.43.24&[MIMAT0001635&hsa-miR-452-5p&offsets\|0\|+2;m-188&X\|-\|151959677\|151959699&offsets\|0\|+1]\|AACTGTTTGCAGAGGAAACTGAGA | >MI0001733\|hsa-mir-452&WithFlank&X\|-\|151959622\|151959718 | 20 | 43 | 24 | AACTGTTTGCAGAGGAAACTGAGA | 114.9 | 6.7 | 1.6 | 4.1 | 3.6E-05 | 5.9E-05 | MB231/MB468 | MP |
| >hg38_21_+_8433172_8446622_RNA45SN1_45S_50ntFlanks@7975.8004.30\|CGCGACCTCAGATCAGACGTGGCGACCCGC | >hg38_21_+_8433172_8446622_RNA45SN1_45S_50ntFlanks | 7975 | 8004 | 30 | CGCGACCTCAGATCAGACGTGGCGACCCGC | 4462.4 | 2.9 | 0.7 | 4.1 | 4.3E-05 | 5.9E-05 | MB231/MB468 | NUC |
| >MI0000081\|hsa-mir-24-2&WithFlank&19\|-\|13836281\|13836365@56.76.21;MI0000080\|hsa-mir-24-1&WithFlank&9\|+\|95086015\|95086094@50.70.21&[MIMAT0000080&hsa-miR-24-3p&offsets\|0\|-1;m-36&19\|-\|13836288\|13836310&offsets\|0\|-2];[MIMAT0000080_1&hsa-miR-24-3p&offsets\|0\|-1;m-35&9\|+\|95086064\|95086086&offsets\|0\|-2]\|TGGCTCAGTTCAGCAGGAACA | >MI0000081\|hsa-mir-24-2&WithFlank&19\|-\|13836281\|13836365 | 56 | 76 | 21 | TGGCTCAGTTCAGCAGGAACA | 1031.6 | 3.4 | 0.8 | 4.1 | 4.0E-05 | 5.9E-05 | MB231/MB468 | CYTO |
| >hg38_21_+_8433172_8446622_RNA45SN1_45S_50ntFlanks@5479.5516.38\|TGGCGGAGCGCTGAGAAGACGGTCGAACTTGACTATCT | >hg38_21_+_8433172_8446622_RNA45SN1_45S_50ntFlanks | 5479 | 5516 | 38 | TGGCGGAGCGCTGAGAAGACGGTCGAACTTGACTATCT | 35.7 | 8.4 | 2.0 | 4.1 | 3.6E-05 | 5.9E-05 | MB231/MB468 | MP |
| >hg38_21_+_8433172_8446622_RNA45SN1_45S_50ntFlanks@10248.10269.22\|CGACGAGTAGGAGGGCCGCTGC | >hg38_21_+_8433172_8446622_RNA45SN1_45S_50ntFlanks | 10248 | 10269 | 22 | CGACGAGTAGGAGGGCCGCTGC | 85.2 | 9.7 | 2.1 | 4.6 | 4.7E-06 | 5.9E-05 | BT20/MB231 | MP |
| >MI0000486\|hsa-mir-190a&WithFlank&15\|+\|62823951\|62824047@21.43.23&[MIMAT0000458&hsa-miR-190a-5p&offsets\|0\|+1;m-349&15\|+\|62823971\|62823993&offsets\|0\|0]\|TGATATGTTTGATATATTAGGTT | >MI0000486\|hsa-mir-190a&WithFlank&15\|+\|62823951\|62824047 | 21 | 43 | 23 | TGATATGTTTGATATATTAGGTT | 68.8 | -9.5 | 2.1 | -4.6 | 4.7E-06 | 5.9E-05 | BT20/MB231 | MP |
| >MI0000063\|hsa-let-7b&WithFlank&22\|+\|46113680\|46113774@12.33.22(+1G)&[MIMAT0000063&hsa-let-7b-5p&offsets\|0\|0(+1G);m-14&22\|+\|46113691\|46113712&offsets\|0\|0(+1G)]\|TGAGGTAGTAGGTTGTGTGGTTG | >MI0000063\|hsa-let-7b&WithFlank&22\|+\|46113680\|46113774 | 12 | 33 | 22(+1G) | TGAGGTAGTAGGTTGTGTGGTTG | 866.8 | 6.2 | 1.4 | 4.3 | 1.5E-05 | 5.9E-05 | BT20/MB468 | MP |
| >trnaMT_PheGAA_MT_+_577_647@27.65.39\|AATACACTGAAAATGTTTAGACGGGCTCACATCACCCCA | >trnaMT_PheGAA_MT_+_577_647 | 27 | 65 | 39 | AATACACTGAAAATGTTTAGACGGGCTCACATCACCCCA | 164.4 | 11.1 | 2.6 | 4.3 | 1.5E-05 | 5.9E-05 | BT20/MB468 | MP |
| >hg38_21_+_8433172_8446622_RNA45SN1_45S_50ntFlanks@12526.12546.21\|TTTTACCCTACTGATGATGTG | >hg38_21_+_8433172_8446622_RNA45SN1_45S_50ntFlanks | 12526 | 12546 | 21 | TTTTACCCTACTGATGATGTG | 291.9 | -6.7 | 1.5 | -4.3 | 1.5E-05 | 5.9E-05 | BT20/MB468 | MP |
| >hg38_21_+_8433172_8446622_RNA45SN1_45S_50ntFlanks@11568.11611.44\|GCAGCCGACTTAGAACTGGTGCGGACCAGGGGAATCCGACTGTT | >hg38_21_+_8433172_8446622_RNA45SN1_45S_50ntFlanks | 11568 | 11611 | 44 | GCAGCCGACTTAGAACTGGTGCGGACCAGGGGAATCCGACTGTT | 17144.1 | 4.9 | 1.2 | 4.2 | 3.1E-05 | 5.9E-05 | BT20/MB468 | MITO |
| >hg38_21_+_8433172_8446622_RNA45SN1_45S_50ntFlanks@3840.3869.30\|CTCCTACTTGGATAACTGTGGTAATTCTAG | >hg38_21_+_8433172_8446622_RNA45SN1_45S_50ntFlanks | 3840 | 3869 | 30 | CTCCTACTTGGATAACTGTGGTAATTCTAG | 602.2 | 3.8 | 0.9 | 4.1 | 4.0E-05 | 5.9E-05 | MB231/MB468 | CYTO |
| >hg38_1_-_228634819_228635039_RNA5S12_5S_50ntFlanks@152.170.19\|TACCGGGTGCTGTAGGCTT | >hg38_1_-_228634819_228635039_RNA5S12_5S_50ntFlanks | 152 | 170 | 19 | TACCGGGTGCTGTAGGCTT | 1525.1 | 3.5 | 0.9 | 4.1 | 4.3E-05 | 5.9E-05 | MB231/MB468 | NUC |
| >MI0000434\|hsa-let-7i&WithFlank&12\|+\|62603680\|62603775@12.33.22(+1C)&[MIMAT0000415&hsa-let-7i-5p&offsets\|0\|0(+1C);m-49&12\|+\|62603691\|62603712&offsets\|0\|0(+1C)]\|TGAGGTAGTAGTTTGTGCTGTTC | >MI0000434\|hsa-let-7i&WithFlank&12\|+\|62603680\|62603775 | 12 | 33 | 22(+1C) | TGAGGTAGTAGTTTGTGCTGTTC | 1588.0 | 5.5 | 1.3 | 4.1 | 3.6E-05 | 5.9E-05 | MB231/MB468 | MP |
| >hg38_21_+_8433172_8446622_RNA45SN1_45S_50ntFlanks@3705.3737.33\|TACCTGGTTGATCCTGCCAGTAGCATATGCTTG | >hg38_21_+_8433172_8446622_RNA45SN1_45S_50ntFlanks | 3705 | 3737 | 33 | TACCTGGTTGATCCTGCCAGTAGCATATGCTTG | 137.6 | -4.7 | 1.1 | -4.1 | 3.6E-05 | 5.9E-05 | MB231/MB468 | MP |
| >trnaMT_PheGAA_MT_+_577_647@35.55.21\|GAAAATGTTTAGACGGGCTCA | >trnaMT_PheGAA_MT_+_577_647 | 35 | 55 | 21 | GAAAATGTTTAGACGGGCTCA | 140.6 | 6.6 | 1.6 | 4.1 | 4.5E-05 | 5.9E-05 | BT20/MB468 | NUC |
| >hg38_21_+_8433172_8446622_RNA45SN1_45S_50ntFlanks@10458.10486.29\|ATGGCCTCCGTTGCCCTCGGCCGATCGAA | >hg38_21_+_8433172_8446622_RNA45SN1_45S_50ntFlanks | 10458 | 10486 | 29 | ATGGCCTCCGTTGCCCTCGGCCGATCGAA | 369.7 | -4.9 | 1.2 | -4.1 | 4.3E-05 | 5.9E-05 | MB231/MB468 | NUC |
| >am_trnaMT_MetCAT_MT_+_4402_4469@1.21.21\|AGTAAGGTCAGCTAAATAAGC | >am_trnaMT_MetCAT_MT_+_4402_4469 | 1 | 21 | 21 | AGTAAGGTCAGCTAAATAAGC | 3245.3 | 4.0 | 1.0 | 4.1 | 4.3E-05 | 5.9E-05 | MB231/MB468 | MITO |

<table>
<tr><td>>MI0000483|hsa-mir-186&WithFlank&1|-|71067625|71067722@21.40.20&[MIMAT0000456&hsa-miR-186-5p&offsets|0|-2;m-84&1|-|71067681|71067702&offsets|0|-2]|CAAAGAATTCTCCTTTTGGG</td><td>>MI0000483|hsa-mir-186&WithFlank&1|-|71067625|71067722</td><td>21</td><td>40</td><td>20</td><td>CAAAGAATTCTCCTTTTGGG</td><td>175.3</td><td>3.7</td><td>0.9</td><td>4.1</td><td>4.3E-05</td><td>5.9E-05</td><td>MB231/MB468</td><td>MITO</td></tr>
<tr><td>>hg38_21_+_8433172_8446622_RNA45SN1_45S_50ntFlanks@12633.12652.20|CGAAGCTACCATCTGTGGGA</td><td>>hg38_21_+_8433172_8446622_RNA45SN1_45S_50ntFlanks</td><td>12633</td><td>12652</td><td>20</td><td>CGAAGCTACCATCTGTGGGA</td><td>2182.4</td><td>-6.1</td><td>1.4</td><td>-4.3</td><td>1.5E-05</td><td>5.9E-05</td><td>BT20/MB468</td><td>MP</td></tr>
<tr><td>>MI0000683|hsa-mir-181b-2&WithFlank&9|+|124693704|124693804@22.46.25(+1U)&[MIMAT0000257_1&hsa-miR-181b-5p&offsets|0|+2(+1U);m-72&9|+|124693725|124693747&offsets|0|+2(+1U)]|AACATTCATTGCTGTCGGTGGGTTTT</td><td>>MI0000683|hsa-mir-181b-2&WithFlank&9|+|124693704|124693804</td><td>22</td><td>46</td><td>25(+1U)</td><td>AACATTCATTGCTGTCGGTGGGTTTT</td><td>453.0</td><td>5.1</td><td>1.2</td><td>4.1</td><td>3.6E-05</td><td>5.9E-05</td><td>MB231/MB468</td><td>MP</td></tr>
<tr><td>>hg38_21_+_8433172_8446622_RNA45SN1_45S_50ntFlanks@7978.8013.36|GACCTCAGATCAGACGTGGCGACCCGCTGAATTTAA</td><td>>hg38_21_+_8433172_8446622_RNA45SN1_45S_50ntFlanks</td><td>7978</td><td>8013</td><td>36</td><td>GACCTCAGATCAGACGTGGCGACCCGCTGAATTTAA</td><td>303.9</td><td>5.5</td><td>1.3</td><td>4.1</td><td>4.3E-05</td><td>5.9E-05</td><td>MB231/MB468</td><td>MITO</td></tr>
<tr><td>>hg38_21_+_8433172_8446622_RNA45SN1_45S_50ntFlanks@11641.11664.24|CGGCGGGTGTTGACGCGATGTGAT</td><td>>hg38_21_+_8433172_8446622_RNA45SN1_45S_50ntFlanks</td><td>11641</td><td>11664</td><td>24</td><td>CGGCGGGTGTTGACGCGATGTGAT</td><td>1436.6</td><td>-5.2</td><td>1.2</td><td>-4.2</td><td>3.1E-05</td><td>5.9E-05</td><td>BT20/MB468</td><td>MITO</td></tr>
<tr><td>>hg38_21_+_8433172_8446622_RNA45SN1_45S_50ntFlanks@12002.12043.42|TGCGGGCCGCCGGTGAAATACCACTACTCTGATCGTTTTTTC</td><td>>hg38_21_+_8433172_8446622_RNA45SN1_45S_50ntFlanks</td><td>12002</td><td>12043</td><td>42</td><td>TGCGGGCCGCCGGTGAAATACCACTACTCTGATCGTTTTTTC</td><td>1317.1</td><td>-6.5</td><td>1.6</td><td>-4.2</td><td>3.1E-05</td><td>5.9E-05</td><td>BT20/MB468</td><td>MITO</td></tr>
<tr><td>>MI0000085|hsa-mir-27a&WithFlank&19|-|13836434|13836523@57.75.19&[MIMAT0000084&hsa-miR-27a-3p&offsets|0|-2;m-43&19|-|13836447|13836467&offsets|0|-2]|TTCACAGTGGCTAAGTTCC</td><td>>MI0000085|hsa-mir-27a&WithFlank&19|-|13836434|13836523</td><td>57</td><td>75</td><td>19</td><td>TTCACAGTGGCTAAGTTCC</td><td>1066.4</td><td>3.4</td><td>0.8</td><td>4.1</td><td>3.6E-05</td><td>5.9E-05</td><td>MB231/MB468</td><td>MP</td></tr>
<tr><td>>MI0000285|hsa-mir-205&WithFlank&1|+|209432127|209432248@40.62.23&[MIMAT0000266&hsa-miR-205-5p&offsets|0|+1;m-351&1|+|209432166|209432187&offsets|0|+1]|TCCTTCATTCCACCGGAGTCTGT</td><td>>MI0000285|hsa-mir-205&WithFlank&1|+|209432127|209432248</td><td>40</td><td>62</td><td>23</td><td>TCCTTCATTCCACCGGAGTCTGT</td><td>14985.4</td><td>2.9</td><td>0.7</td><td>4.1</td><td>4.5E-05</td><td>5.9E-05</td><td>BT20/MB468</td><td>NUC</td></tr>
<tr><td>>hg38_21_+_8433172_8446622_RNA45SN1_45S_50ntFlanks@4729.4752.24|TCAAGAACGAAAGTCGGAGGTTCG</td><td>>hg38_21_+_8433172_8446622_RNA45SN1_45S_50ntFlanks</td><td>4729</td><td>4752</td><td>24</td><td>TCAAGAACGAAAGTCGGAGGTTCG</td><td>218.5</td><td>-6.3</td><td>1.4</td><td>-4.3</td><td>1.5E-05</td><td>5.9E-05</td><td>BT20/MB468</td><td>MP</td></tr>
<tr><td>>trna111_HisGTG_1_-_147774845_147774916@2.31.30,trna118_HisGTG_1_-_145396881_145396952@2.31.30,trna16_HisGTG_1_+_146544773_146544844@2.31.30,trna1_HisGTG_15_+_45493349_45493420@2.31.30,trna21_HisGTG_1_+_147753471_147753542@2.31.30,trna33_HisGTG_6_+_27125906_27125977@2.31.30,trna7_HisGTG_9_-_14433938_14434009@2.31.30,trna8_HisGTG_15_-_45492611_45492682@2.31.30,trna9_HisGTG_15_-_45490804_45490875@2.31.30|CCGTGATCGTATAGTGGTTAGTACTCTGCG</td><td>>trna111_HisGTG_1_-_147774845_147774916</td><td>2</td><td>31</td><td>30</td><td>CCGTGATCGTATAGTGGTTAGTACTCTGCG</td><td>1604.5</td><td>3.0</td><td>0.7</td><td>4.1</td><td>3.8E-05</td><td>5.9E-05</td><td>BT20/MB468</td><td>CYTO</td></tr>
<tr><td>>hg38_21_+_8433172_8446622_RNA45SN1_45S_50ntFlanks@11682.11701.20|AATGTCAAAGTGAAGAAATT</td><td>>hg38_21_+_8433172_8446622_RNA45SN1_45S_50ntFlanks</td><td>11682</td><td>11701</td><td>20</td><td>AATGTCAAAGTGAAGAAATT</td><td>38.8</td><td>-8.4</td><td>1.9</td><td>-4.3</td><td>1.5E-05</td><td>5.9E-05</td><td>BT20/MB468</td><td>MP</td></tr>
<tr><td>>hg38_21_+_8433172_8446622_RNA45SN1_45S_50ntFlanks@11540.11581.42|CGCGGCGCCTCGCCTCGGCCGGCGCCTAGCAGCCGACTTAGA</td><td>>hg38_21_+_8433172_8446622_RNA45SN1_45S_50ntFlanks</td><td>11540</td><td>11581</td><td>42</td><td>CGCGGCGCCTCGCCTCGGCCGGCGCCTAGCAGCCGACTTAGA</td><td>359.3</td><td>-3.6</td><td>0.8</td><td>-4.6</td><td>4.6E-06</td><td>5.9E-05</td><td>BT20/MB231</td><td>CYTO</td></tr>
<tr><td>>hg38_21_+_8433172_8446622_RNA45SN1_45S_50ntFlanks@10237.10259.23|ACGCTACGCCGCGACGAGTAGGA</td><td>>hg38_21_+_8433172_8446622_RNA45SN1_45S_50ntFlanks</td><td>10237</td><td>10259</td><td>23</td><td>ACGCTACGCCGCGACGAGTAGGA</td><td>302.4</td><td>4.2</td><td>1.0</td><td>4.3</td><td>1.5E-05</td><td>5.9E-05</td><td>BT20/MB468</td><td>MP</td></tr>
<tr><td>>hg38_21_+_8433172_8446622_RNA45SN1_45S_50ntFlanks@10572.10596.25|AGAAGCCGGCGGGAGCCCCGGGGAG</td><td>>hg38_21_+_8433172_8446622_RNA45SN1_45S_50ntFlanks</td><td>10572</td><td>10596</td><td>25</td><td>AGAAGCCGGCGGGAGCCCCGGGGAG</td><td>331.3</td><td>-5.9</td><td>1.4</td><td>-4.1</td><td>4.3E-05</td><td>6.0E-05</td><td>MB231/MB468</td><td>NUC</td></tr>
<tr><td>>hg38_21_+_8433172_8446622_RNA45SN1_45S_50ntFlanks@10781.10814.34|AGGTGAACAGCCTCTGGCATGTTGGAACAATGTA</td><td>>hg38_21_+_8433172_8446622_RNA45SN1_45S_50ntFlanks</td><td>10781</td><td>10814</td><td>34</td><td>AGGTGAACAGCCTCTGGCATGTTGGAACAATGTA</td><td>187.0</td><td>-3.2</td><td>0.8</td><td>-4.1</td><td>4.1E-05</td><td>6.0E-05</td><td>MB231/MB468</td><td>CYTO</td></tr>
<tr><td>>hg38_21_+_8433172_8446622_RNA45SN1_45S_50ntFlanks@11561.11603.43|GCGCCTAGCAGCCGACTTAGAACTGGTGCGGACCAGGGGAATC</td><td>>hg38_21_+_8433172_8446622_RNA45SN1_45S_50ntFlanks</td><td>11561</td><td>11603</td><td>43</td><td>GCGCCTAGCAGCCGACTTAGAACTGGTGCGGACCAGGGGAATC</td><td>440.2</td><td>-5.9</td><td>1.4</td><td>-4.1</td><td>3.8E-05</td><td>6.0E-05</td><td>BT20/MB468</td><td>CYTO</td></tr>
<tr><td>>hg38_MT_+_1621_3279_RNR2_16S_50ntFlanks@51.94.44|GCTAAACCTAGCCCCAAACCCACTCCACCTTACTACCAGACAAC</td><td>>hg38_MT_+_1621_3279_RNR2_16S_50ntFlanks</td><td>51</td><td>94</td><td>44</td><td>GCTAAACCTAGCCCCAAACCCACTCCACCTTACTACCAGACAAC</td><td>6235.6</td><td>4.9</td><td>1.2</td><td>4.1</td><td>4.4E-05</td><td>6.0E-05</td><td>MB231/MB468</td><td>MITO</td></tr>
<tr><td>>MI0000734|hsa-mir-106b&WithFlank&7|-|100093987|100094080@56.76.21(+1A)&[MIMAT0004672&hsa-miR-106b-3p&offsets|-2|-3(+1A);m-87&7|-|100094003|100094023&offsets|-2|-2(+1A)]|TACCGCACTGTGGGTACTTGCA</td><td>>MI0000734|hsa-mir-106b&WithFlank&7|-|100093987|100094080</td><td>56</td><td>76</td><td>21(+1A)</td><td>TACCGCACTGTGGGTACTTGCA</td><td>46.7</td><td>9.3</td><td>2.1</td><td>4.3</td><td>1.5E-05</td><td>6.0E-05</td><td>BT20/MB468</td><td>MP</td></tr>
<tr><td>>hg38_21_+_8433172_8446622_RNA45SN1_45S_50ntFlanks@9196.9222.27|AGGTTCTCTCGGGGCCACGCGCGCGTC</td><td>>hg38_21_+_8433172_8446622_RNA45SN1_45S_50ntFlanks</td><td>9196</td><td>9222</td><td>27</td><td>AGGTTCTCTCGGGGCCACGCGCGCGTC</td><td>178.6</td><td>-6.7</td><td>1.6</td><td>-4.3</td><td>1.5E-05</td><td>6.0E-05</td><td>BT20/MB468</td><td>MP</td></tr>
<tr><td>>am_trna11_ProAGG_16_+_3241989_3242060@1.29.29,trna12_ProAGG_6_+_26555498_26555569@1.29.29,trna12_ProTGG_11_-_75946869_75946940@1.29.29,trna14_ProTGG_5_-_180615854_180615925@1.29.29,trna22_ProAGG_14_-_21081560_21081631@1.29.29,trna23_ProAGG_14_-_21077495_21077566@1.29.29,trna28_ProTGG_16_-_3234133_3234204@1.29.29,trna29_ProAGG_16_-_3232635_3232706@1.29.29,trna2_ProAGG_7_+_128423504_128423575@1.29.29,trna30_ProCGG_6_+_27059521_27059592@1.29.29,trna37_ProCGG_17_-_8126151_8126222@1.29.29,trna3_ProTGG_16_+_3208923_3208994@1.29.29,trna52_ProCGG_1_+_167683962_167684033@1.29.29,trna65_ProAGG_1_-_167684725_167684796@1.29.29,trna6_ProCGG_16_+_3222049_3222120@1.29.29,trna6_ProTGG_14_+_21152175_21152246@1.29.29,trna8_ProTGG_16_+_3238094_3238165@1.29.29,trna9_ProAGG_11_+_75946557_75946628@1.29.29,trna9_ProAGG_16_+_3239634_3239705@1.29.29|GGCTCGTTGGTCTAGGGGTATGATTCTCG</td><td>>am_trna11_ProAGG_16_+_3241989_3242060</td><td>1</td><td>29</td><td>29</td><td>GGCTCGTTGGTCTAGGGGTATGATTCTCG</td><td>355.0</td><td>-5.8</td><td>1.4</td><td>-4.2</td><td>3.2E-05</td><td>6.0E-05</td><td>BT20/MB468</td><td>MITO</td></tr>
<tr><td>>hg38_MT_+_1621_3279_RNR2_16S_50ntFlanks@1380.1413.34|AGGACATCCCGATGGTGCAGCCGCTATTAAAGGT</td><td>>hg38_MT_+_1621_3279_RNR2_16S_50ntFlanks</td><td>1380</td><td>1413</td><td>34</td><td>AGGACATCCCGATGGTGCAGCCGCTATTAAAGGT</td><td>28.7</td><td>8.1</td><td>2.0</td><td>4.1</td><td>3.7E-05</td><td>6.0E-05</td><td>MB231/MB468</td><td>MP</td></tr>
<tr><td>>hg38_21_+_8433172_8446622_RNA45SN1_45S_50ntFlanks@8355.8389.35|TGAAAAGAACTTTGAAGAGAGAGTTCAAGAGGGCG</td><td>>hg38_21_+_8433172_8446622_RNA45SN1_45S_50ntFlanks</td><td>8355</td><td>8389</td><td>35</td><td>TGAAAAGAACTTTGAAGAGAGAGTTCAAGAGGGCG</td><td>90.3</td><td>9.8</td><td>2.1</td><td>4.6</td><td>4.8E-06</td><td>6.0E-05</td><td>BT20/MB231</td><td>MP</td></tr>
<tr><td>>hg38_21_+_8433172_8446622_RNA45SN1_45S_50ntFlanks@11564.11581.18|CCTAGCAGCCGACTTAGA</td><td>>hg38_21_+_8433172_8446622_RNA45SN1_45S_50ntFlanks</td><td>11564</td><td>11581</td><td>18</td><td>CCTAGCAGCCGACTTAGA</td><td>791.2</td><td>-6.1</td><td>1.5</td><td>-4.1</td><td>3.8E-05</td><td>6.0E-05</td><td>BT20/MB468</td><td>CYTO</td></tr>
<tr><td>>hg38_21_+_8433172_8446622_RNA45SN1_45S_50ntFlanks@12249.12272.24|CAAAAGCTCGCTTGATCTTGATTT</td><td>>hg38_21_+_8433172_8446622_RNA45SN1_45S_50ntFlanks</td><td>12249</td><td>12272</td><td>24</td><td>CAAAAGCTCGCTTGATCTTGATTT</td><td>216.3</td><td>-1.9</td><td>0.5</td><td>-4.1</td><td>3.8E-05</td><td>6.0E-05</td><td>BT20/MB468</td><td>CYTO</td></tr>
<tr><td>>hg38_21_+_8433172_8446622_RNA45SN1_45S_50ntFlanks@12487.12509.23|GAACGTGAGCTGGGTTTAGACCG</td><td>>hg38_21_+_8433172_8446622_RNA45SN1_45S_50ntFlanks</td><td>12487</td><td>12509</td><td>23</td><td>GAACGTGAGCTGGGTTTAGACCG</td><td>152.6</td><td>-6.3</td><td>1.5</td><td>-4.1</td><td>3.7E-05</td><td>6.0E-05</td><td>MB231/MB468</td><td>MP</td></tr>
<tr><td>>hg38_21_+_8433172_8446622_RNA45SN1_45S_50ntFlanks@9482.9512.31|ACGTGTTAGGACCCGAAAGATGGTGAACTAT</td><td>>hg38_21_+_8433172_8446622_RNA45SN1_45S_50ntFlanks</td><td>9482</td><td>9512</td><td>31</td><td>ACGTGTTAGGACCCGAAAGATGGTGAACTAT</td><td>113.4</td><td>10.2</td><td>2.2</td><td>4.6</td><td>4.8E-06</td><td>6.0E-05</td><td>BT20/MB231</td><td>MP</td></tr>
<tr><td>>hg38_21_+_8433172_8446622_RNA45SN1_45S_50ntFlanks@3902.3938.37|TCGCGGGGGGGATGCGTGCATTTATCAGATCAAAACC</td><td>>hg38_21_+_8433172_8446622_RNA45SN1_45S_50ntFlanks</td><td>3902</td><td>3938</td><td>37</td><td>TCGCGGGGGGGATGCGTGCATTTATCAGATCAAAACC</td><td>60.1</td><td>-9.8</td><td>2.4</td><td>-4.1</td><td>3.7E-05</td><td>6.0E-05</td><td>MB231/MB468</td><td>MP</td></tr>
<tr><td>>hg38_21_+_8433172_8446622_RNA45SN1_45S_50ntFlanks@3993.4013.21|GTGACTCTAGATAACCTCGGG</td><td>>hg38_21_+_8433172_8446622_RNA45SN1_45S_50ntFlanks</td><td>3993</td><td>4013</td><td>21</td><td>GTGACTCTAGATAACCTCGGG</td><td>623.3</td><td>-1.8</td><td>0.4</td><td>-4.1</td><td>3.8E-05</td><td>6.0E-05</td><td>BT20/MB468</td><td>CYTO</td></tr>
<tr><td>>hg38_21_+_8433172_8446622_RNA45SN1_45S_50ntFlanks@11670.11706.37|CCCAGTGCTCTGAATGTCAAAGTGAAGAAATTCAATG</td><td>>hg38_21_+_8433172_8446622_RNA45SN1_45S_50ntFlanks</td><td>11670</td><td>11706</td><td>37</td><td>CCCAGTGCTCTGAATGTCAAAGTGAAGAAATTCAATG</td><td>590.0</td><td>-2.7</td><td>0.7</td><td>-4.1</td><td>4.4E-05</td><td>6.0E-05</td><td>MB231/MB468</td><td>NUC</td></tr>
<tr><td>>am_trna117_GlyTCC_1_-_145397864_145397935@1.30.30,trna45_GlyTCC_1_+_161500903_161500974@1.30.30,trna70_GlyTCC_1_-_161439547_161439618@1.30.30,trna73_GlyTCC_1_-_161432166_161432237@1.30.30,trna76_GlyTCC_1_-_161424756_161424827@1.30.30,trna79_GlyTCC_1_-_161417375_161417446@1.30.30|GCGTTGGTGGTATAGTGGTGAGCATAGCTG</td><td>>am_trna117_GlyTCC_1_-_145397864_145397935</td><td>1</td><td>30</td><td>30</td><td>GCGTTGGTGGTATAGTGGTGAGCATAGCTG</td><td>2134.3</td><td>-6.0</td><td>1.4</td><td>-4.3</td><td>1.5E-05</td><td>6.0E-05</td><td>BT20/MB468</td><td>MP</td></tr>
<tr><td>>MI0000103|hsa-mir-101-1&WithFlank&1|-|65058428|65058514@53.73.21(+1U);MI0000739|hsa-mir-101-2&WithFlank&9|+|4850291|4850381@55.75.21(+1U)&[MIMAT0000099&hsa-miR-101-3p&offsets|0|0(+1U);m-17&1|-|65058443|65058462&offsets|0|+1(+1U)];[MIMAT0000099_1&hsa-miR-101-3p&offsets|0|0(+1U);m-16&9|+|4850345|4850364&offsets|0|+1(+1U)]|TACAGTACTGTGATAACTGAAT</td><td>>MI0000103|hsa-mir-101-1&WithFlank&1|-|65058428|65058514</td><td>53</td><td>73</td><td>21(+1U)</td><td>TACAGTACTGTGATAACTGAAT</td><td>458.6</td><td>3.4</td><td>0.8</td><td>4.1</td><td>3.7E-05</td><td>6.0E-05</td><td>MB231/MB468</td><td>MP</td></tr>
<tr><td>>hg38_21_+_8433172_8446622_RNA45SN1_45S_50ntFlanks@4726.4758.33|TAATCAAGAACGAAAGTCGGAGGTTCGAAGACG</td><td>>hg38_21_+_8433172_8446622_RNA45SN1_45S_50ntFlanks</td><td>4726</td><td>4758</td><td>33</td><td>TAATCAAGAACGAAAGTCGGAGGTTCGAAGACG</td><td>1992.6</td><td>-2.6</td><td>0.6</td><td>-4.1</td><td>3.8E-05</td><td>6.0E-05</td><td>BT20/MB468</td><td>CYTO</td></tr>
<tr><td>>hg38_21_+_8433172_8446622_RNA45SN1_45S_50ntFlanks@5156.5185.30|ACAAGTGGCGTTCAGCCACCCGAGATTGAG</td><td>>hg38_21_+_8433172_8446622_RNA45SN1_45S_50ntFlanks</td><td>5156</td><td>5185</td><td>30</td><td>ACAAGTGGCGTTCAGCCACCCGAGATTGAG</td><td>124.3</td><td>-6.4</td><td>1.5</td><td>-4.1</td><td>3.7E-05</td><td>6.0E-05</td><td>MB231/MB468</td><td>MP</td></tr>
<tr><td>>hg38_21_+_8433172_8446622_RNA45SN1_45S_50ntFlanks@12712.12749.38|CGGAGCCTCGGTTGGCCTCGGATAGCCGGTCCCCCGCC</td><td>>hg38_21_+_8433172_8446622_RNA45SN1_45S_50ntFlanks</td><td>12712</td><td>12749</td><td>38</td><td>CGGAGCCTCGGTTGGCCTCGGATAGCCGGTCCCCCGCC</td><td>57.1</td><td>-8.9</td><td>2.1</td><td>-4.3</td><td>1.5E-05</td><td>6.0E-05</td><td>BT20/MB468</td><td>MP</td></tr>
<tr><td>>hg38_1_-_228634819_228635039_RNA5S12_5S_50ntFlanks@137.169.33|GAGACCGCCTGGGAATACCGGGTGCTGTAGGCT</td><td>>hg38_1_-_228634819_228635039_RNA5S12_5S_50ntFlanks</td><td>137</td><td>169</td><td>33</td><td>GAGACCGCCTGGGAATACCGGGTGCTGTAGGCT</td><td>549.0</td><td>3.6</td><td>0.8</td><td>4.3</td><td>1.5E-05</td><td>6.0E-05</td><td>BT20/MB468</td><td>MP</td></tr>
<tr><td>>MI0000456|hsa-mir-140&WithFlank&16|+|69933075|69933186@68.90.23(+1U)&[MIMAT0004597&hsa-miR-140-3p&offsets|0|+2(+1U);m-45&16|+|69933143|69933164&offsets|-1|0(+1U)]|TACCACAGGGTAGAACCACGGACT</td><td>>MI0000456|hsa-mir-140&WithFlank&16|+|69933075|69933186</td><td>68</td><td>90</td><td>23(+1U)</td><td>TACCACAGGGTAGAACCACGGACT</td><td>318.3</td><td>6.2</td><td>1.5</td><td>4.1</td><td>3.7E-05</td><td>6.0E-05</td><td>MB231/MB468</td><td>MP</td></tr>
<tr><td>>trnaMT_HisGTG_MT_+_12138_12206@20.58.39|AACATCAGATTGTGAATCTGACAACAGAGGCTTACGACC</td><td>>trnaMT_HisGTG_MT_+_12138_12206</td><td>20</td><td>58</td><td>39</td><td>AACATCAGATTGTGAATCTGACAACAGAGGCTTACGACC</td><td>28.2</td><td>8.0</td><td>1.9</td><td>4.1</td><td>3.7E-05</td><td>6.0E-05</td><td>MB231/MB468</td><td>MP</td></tr>
<tr><td>>hg38_21_+_8433172_8446622_RNA45SN1_45S_50ntFlanks@11672.11691.20|CAGTGCTCTGAATGTCAAAG</td><td>>hg38_21_+_8433172_8446622_RNA45SN1_45S_50ntFlanks</td><td>11672</td><td>11691</td><td>20</td><td>CAGTGCTCTGAATGTCAAAG</td><td>251.3</td><td>-2.8</td><td>0.7</td><td>-4.1</td><td>3.7E-05</td><td>6.0E-05</td><td>MB231/MB468</td><td>MP</td></tr>
<tr><td>>hg38_21_+_8433172_8446622_RNA45SN1_45S_50ntFlanks@9739.9759.21|AAACGATCTCAACCTATTCTC</td><td>>hg38_21_+_8433172_8446622_RNA45SN1_45S_50ntFlanks</td><td>9739</td><td>9759</td><td>21</td><td>AAACGATCTCAACCTATTCTC</td><td>576.5</td><td>-2.5</td><td>0.6</td><td>-4.1</td><td>4.2E-05</td><td>6.0E-05</td><td>MB231/MB468</td><td>CYTO</td></tr>
</table>

| | | | | | | | | | | | | | |
|---|---|---|---|---|---|---|---|---|---|---|---|---|---|
| >hg38_1_-_228634819_228635039_RNA5S12_5S_50ntFlanks@136.171.36\|GGAGACCGCCTGGGAATACCGGGTGCTGTAGGCTTT | >hg38_1_-_228634819_228635039_RNA5S12_5S_50ntFlanks | 136 | 171 | 36 | GGAGACCGCCTGGGAATACCGGGTGCTGTAGGCTTT | 2958.3 | 4.2 | 1.0 | 4.1 | 4.2E-05 | 6.1E-05 | MB231/MB468 | CYTO |
| >hg38_21_+_8433172_8446622_RNA45SN1_45S_50ntFlanks@11733.11751.19\|TATGACTCTCTTAAGGTAG | >hg38_21_+_8433172_8446622_RNA45SN1_45S_50ntFlanks | 11733 | 11751 | 19 | TATGACTCTCTTAAGGTAG | 113.1 | -5.9 | 1.4 | -4.1 | 3.7E-05 | 6.1E-05 | MB231/MB468 | MP |
| >trnaMT_GluTTC_MT_-_14674_14742@39.61.23,trnalookalike8_GluTTC_5_-_93905172_93905240@39.61.23\|ATTGGTCGTGGTTGTAGTCCGTG | >trnaMT_GluTTC_MT_-_14674_14742 | 39 | 61 | 23 | ATTGGTCGTGGTTGTAGTCCGTG | 239.6 | 3.3 | 0.8 | 4.1 | 4.4E-05 | 6.1E-05 | MB231/MB468 | NUC |
| >hg38_21_+_8433172_8446622_RNA45SN1_45S_50ntFlanks@6645.6677.33\|CTCGTACGACTCTTAGCGGTGGATCACTCGGCT | >hg38_21_+_8433172_8446622_RNA45SN1_45S_50ntFlanks | 6645 | 6677 | 33 | CTCGTACGACTCTTAGCGGTGGATCACTCGGCT | 754.6 | 3.5 | 0.8 | 4.3 | 1.5E-05 | 6.1E-05 | BT20/MB468 | MP |
| >hg38_21_+_8433172_8446622_RNA45SN1_45S_50ntFlanks@11568.11605.38\|GCAGCCGACTTAGAACTGGTGCGGACCAGGGGAATCCG | >hg38_21_+_8433172_8446622_RNA45SN1_45S_50ntFlanks | 11568 | 11605 | 38 | GCAGCCGACTTAGAACTGGTGCGGACCAGGGGAATCCG | 462.5 | -6.1 | 1.5 | -4.1 | 4.4E-05 | 6.1E-05 | MB231/MB468 | NUC |
| >hg38_21_+_8433172_8446622_RNA45SN1_45S_50ntFlanks@11641.11667.27\|CGGCGGGTGTTGACGCGATGTGATTTC | >hg38_21_+_8433172_8446622_RNA45SN1_45S_50ntFlanks | 11641 | 11667 | 27 | CGGCGGGTGTTGACGCGATGTGATTTC | 966.8 | -5.3 | 1.3 | -4.2 | 3.2E-05 | 6.1E-05 | BT20/MB468 | MITO |
| >hg38_21_+_8433172_8446622_RNA45SN1_45S_50ntFlanks@10829.10875.47\|GCAAGCCGGATCCGTAACTTCGGGATAAGGATTGGCTCTAAGGGCTG | >hg38_21_+_8433172_8446622_RNA45SN1_45S_50ntFlanks | 10829 | 10875 | 47 | GCAAGCCGGATCCGTAACTTCGGGATAAGGATTGGCTCTAAGGGCTG | 1181.6 | -6.4 | 1.6 | -4.1 | 4.4E-05 | 6.1E-05 | MB231/MB468 | NUC |
| >MI0000076\|hsa-mir-20a&WithFlank&13\|+\|91351059\|91351141@14.35.22&[MIMAT0000075&hsa-miR-20a-5p&offsets\|0\|-1;m-96&13\|+\|91351072\|91351094&offsets\|0\|-1]\|TAAAGTGCTTATAGTGCAGGTA | >MI0000076\|hsa-mir-20a&WithFlank&13\|+\|91351059\|91351141 | 14 | 35 | 22 | TAAAGTGCTTATAGTGCAGGTA | 716.9 | -5.9 | 1.3 | -4.6 | 4.9E-06 | 6.1E-05 | BT20/MB231 | MP |
| >hg38_MT_+_598_1651_RNR1_12S_50ntFlanks@905.929.25\|AGTATACTTCAAAGGACATTTAACT | >hg38_MT_+_598_1651_RNR1_12S_50ntFlanks | 905 | 929 | 25 | AGTATACTTCAAAGGACATTTAACT | 2981.2 | 4.8 | 1.2 | 4.1 | 3.7E-05 | 6.1E-05 | MB231/MB468 | MP |
| >MI0000270\|hsa-mir-181b-1&WithFlank&1\|-\|198858867\|198858988@42.64.23(+1A);MI0000683\|hsa-mir-181b-2&WithFlank&9\|+\|124693704\|124693804@22.44.23(+1A)&[MIMAT0000257&hsa-miR-181b-5p&offsets\|0\|0(+1A);m-73&1\|-\|198858925\|198858947&offsets\|0\|0(+1A)];[MIMAT0000257_1&hsa-miR-181b-5p&offsets\|0\|0(+1A);m-72&9\|+\|124693725\|124693747&offsets\|0\|0(+1A)]\|AACATTCATTGCTGTCGGTGGGTA | >MI0000270\|hsa-mir-181b-1&WithFlank&1\|-\|198858867\|198858988 | 42 | 64 | 23(+1A) | AACATTCATTGCTGTCGGTGGGTA | 216.8 | 6.4 | 1.6 | 4.1 | 4.4E-05 | 6.1E-05 | MB231/MB468 | MITO |
| >trna116_GluCTC_1_-_145399233_145399304@4.36.33,trna59_GluCTC_1_+_249168447_249168518@4.36.33,trna71_GluCTC_1_-_161439189_161439260@4.36.33,trna74_GluCTC_1_-_161431809_161431880@4.36.33,trna77_GluCTC_1_-_161424398_161424469@4.36.33,trna77_GluCTC_6_+_28949976_28950047@4.36.33,trna80_GluCTC_1_-_161417018_161417089@4.36.33,trna87_GluCTC_6_-_126101393_126101464@4.36.33\|CTGGTGGTCTAGTGGTTAGGATTCGGCGCTCTC | >trna116_GluCTC_1_-_145399233_145399304 | 4 | 36 | 33 | CTGGTGGTCTAGTGGTTAGGATTCGGCGCTCTC | 207.7 | 4.9 | 1.2 | 4.1 | 4.5E-05 | 6.1E-05 | MB231/MB468 | MITO |
| >trna119_LysCTT_1_-_145395522_145395594@3.36.34,trna11_LysCTT_5_-_180648979_180649051@3.36.34,trna13_LysCTT_6_+_26556774_26556846@3.36.34,trna7_LysCTT_16_+_3225692_3225764@3.36.34,trna9_LysCTT_5_+_180634755_180634827@3.36.34\|CCGGCTAGCTCAGTCGGTAGAGCATGAGACTCTT | >trna119_LysCTT_1_-_145395522_145395594 | 3 | 36 | 34 | CCGGCTAGCTCAGTCGGTAGAGCATGAGACTCTT | 315.1 | -4.9 | 1.1 | -4.5 | 5.8E-06 | 6.1E-05 | BT20/MB231 | MITO |
| >hg38_21_+_8433172_8446622_RNA45SN1_45S_50ntFlanks@9954.9979.26\|TGGAAGTCGGAATCCGCTAAGGAGTG | >hg38_21_+_8433172_8446622_RNA45SN1_45S_50ntFlanks | 9954 | 9979 | 26 | TGGAAGTCGGAATCCGCTAAGGAGTG | 205.4 | -6.2 | 1.5 | -4.1 | 3.7E-05 | 6.1E-05 | MB231/MB468 | MP |
| >MI0000066\|hsa-let-7e&WithFlank&19\|+\|51692780\|51692870@14.34.21&[MIMAT0000066&hsa-let-7e-5p&offsets\|0\|-1;m-50&19\|+\|51692793\|51692814&offsets\|0\|-1]\|TGAGGTAGGAGGTTGTATAGT | >MI0000066\|hsa-let-7e&WithFlank&19\|+\|51692780\|51692870 | 14 | 34 | 21 | TGAGGTAGGAGGTTGTATAGT | 631.3 | 1.4 | 0.3 | 4.1 | 4.6E-05 | 6.1E-05 | BT20/MB468 | NUC |
| >hg38_21_+_8433172_8446622_RNA45SN1_45S_50ntFlanks@11534.11551.18\|GTTCCGCGCGGCGCCTCG | >hg38_21_+_8433172_8446622_RNA45SN1_45S_50ntFlanks | 11534 | 11551 | 18 | GTTCCGCGCGGCGCCTCG | 1065.4 | -4.9 | 1.2 | -4.2 | 3.2E-05 | 6.1E-05 | BT20/MB468 | MITO |
| >hg38_21_+_8433172_8446622_RNA45SN1_45S_50ntFlanks@9806.9832.27\|CGTGGAATGCGAGTGCCTAGTGGGCCA | >hg38_21_+_8433172_8446622_RNA45SN1_45S_50ntFlanks | 9806 | 9832 | 27 | CGTGGAATGCGAGTGCCTAGTGGGCCA | 222.4 | -6.9 | 1.6 | -4.3 | 1.5E-05 | 6.1E-05 | BT20/MB468 | MP |
| >hg38_21_+_8433172_8446622_RNA45SN1_45S_50ntFlanks@11323.11346.24\|CGTCTCGTCGCGCGCGCGTCCGCT | >hg38_21_+_8433172_8446622_RNA45SN1_45S_50ntFlanks | 11323 | 11346 | 24 | CGTCTCGTCGCGCGCGCGTCCGCT | 2100.4 | 5.0 | 1.1 | 4.5 | 5.8E-06 | 6.1E-05 | BT20/MB231 | MITO |
| >hg38_21_+_8433172_8446622_RNA45SN1_45S_50ntFlanks@12418.12447.30\|CGATGTCGGCTCTTCCTATCATTGTGAAGC | >hg38_21_+_8433172_8446622_RNA45SN1_45S_50ntFlanks | 12418 | 12447 | 30 | CGATGTCGGCTCTTCCTATCATTGTGAAGC | 600.8 | 2.6 | 0.6 | 4.1 | 4.2E-05 | 6.1E-05 | MB231/MB468 | CYTO |
| >hg38_21_+_8433172_8446622_RNA45SN1_45S_50ntFlanks@10766.10785.20\|CGCAGCAGGTCTCCAAGGTG | >hg38_21_+_8433172_8446622_RNA45SN1_45S_50ntFlanks | 10766 | 10785 | 20 | CGCAGCAGGTCTCCAAGGTG | 195.5 | -1.8 | 0.4 | -4.1 | 3.9E-05 | 6.1E-05 | BT20/MB468 | CYTO |
| >hg38_21_+_8433172_8446622_RNA45SN1_45S_50ntFlanks@4898.4915.18\|AAAGGAATTGACGGAAGG | >hg38_21_+_8433172_8446622_RNA45SN1_45S_50ntFlanks | 4898 | 4915 | 18 | AAAGGAATTGACGGAAGG | 160.5 | -6.2 | 1.5 | -4.1 | 3.8E-05 | 6.1E-05 | MB231/MB468 | MP |
| >hg38_21_+_8433172_8446622_RNA45SN1_45S_50ntFlanks@10792.10815.24\|CTCTGGCATGTTGGAACAATGTAG | >hg38_21_+_8433172_8446622_RNA45SN1_45S_50ntFlanks | 10792 | 10815 | 24 | CTCTGGCATGTTGGAACAATGTAG | 98.5 | 10.0 | 2.2 | 4.6 | 4.9E-06 | 6.1E-05 | BT20/MB231 | MP |
| >hg38_21_+_8433172_8446622_RNA45SN1_45S_50ntFlanks@6663.6699.37\|GTGGATCACTCGGCTCGTGCGTCGATGAAGAACGCAG | >hg38_21_+_8433172_8446622_RNA45SN1_45S_50ntFlanks | 6663 | 6699 | 37 | GTGGATCACTCGGCTCGTGCGTCGATGAAGAACGCAG | 450.3 | -2.4 | 0.6 | -4.2 | 3.3E-05 | 6.1E-05 | BT20/MB468 | MITO |
| >hg38_21_+_8433172_8446622_RNA45SN1_45S_50ntFlanks@3782.3802.21\|CAGTGAAACTGCGAATGGCTC | >hg38_21_+_8433172_8446622_RNA45SN1_45S_50ntFlanks | 3782 | 3802 | 21 | CAGTGAAACTGCGAATGGCTC | 198.8 | -2.0 | 0.5 | -4.1 | 4.2E-05 | 6.1E-05 | MB231/MB468 | CYTO |
| >am_trna146_GlnCTG_6_-_27515531_27515602@1.29.29,trna1_GlnCTG_6_+_18836402_18836473@1.29.29,trna3_GlnCTG_17_+_8023070_8023141@1.29.29,trna42_GlnCTG_6_+_27263212_27263283@1.29.29,trna49_GlnCTG_6_+_27487308_27487379@1.29.29,trna7_GlnCTG_15_-_66161400_66161471@1.29.29,trna99_GlnCTG_6_-_28909378_28909449@1.29.29\|GGTTCCATGGTGTAATGGTTAGCACTCTG | >am_trna146_GlnCTG_6_-_27515531_27515602 | 1 | 29 | 29 | GGTTCCATGGTGTAATGGTTAGCACTCTG | 5952.0 | -4.7 | 1.1 | -4.3 | 1.5E-05 | 6.1E-05 | BT20/MB468 | MP |
| >hg38_21_+_8433172_8446622_RNA45SN1_45S_50ntFlanks@6722.6740.19\|ATTGCAGGACACATTGATC | >hg38_21_+_8433172_8446622_RNA45SN1_45S_50ntFlanks | 6722 | 6740 | 19 | ATTGCAGGACACATTGATC | 792.9 | -3.5 | 0.8 | -4.1 | 3.9E-05 | 6.1E-05 | BT20/MB468 | CYTO |
| >hg38_21_+_8433172_8446622_RNA45SN1_45S_50ntFlanks@12995.13038.44\|CTCGCTGCGATCTATTGAAAGTCAGCCCTCGACACAAGGGTTTG | >hg38_21_+_8433172_8446622_RNA45SN1_45S_50ntFlanks | 12995 | 13038 | 44 | CTCGCTGCGATCTATTGAAAGTCAGCCCTCGACACAAGGGTTTG | 1037.4 | 6.8 | 1.6 | 4.2 | 3.3E-05 | 6.1E-05 | BT20/MB468 | MITO |
| >hg38_21_+_8433172_8446622_RNA45SN1_45S_50ntFlanks@9534.9551.18\|AGGAAACTCTGGTGGAGG | >hg38_21_+_8433172_8446622_RNA45SN1_45S_50ntFlanks | 9534 | 9551 | 18 | AGGAAACTCTGGTGGAGG | 541.3 | -2.4 | 0.6 | -4.2 | 3.3E-05 | 6.1E-05 | BT20/MB468 | MITO |
| >hg38_21_+_8433172_8446622_RNA45SN1_45S_50ntFlanks@11567.11585.19\|AGCAGCCGACTTAGAACTG | >hg38_21_+_8433172_8446622_RNA45SN1_45S_50ntFlanks | 11567 | 11585 | 19 | AGCAGCCGACTTAGAACTG | 1185.3 | 1.8 | 0.4 | 4.2 | 3.3E-05 | 6.1E-05 | BT20/MB468 | MITO |
| >hg38_21_+_8433172_8446622_RNA45SN1_45S_50ntFlanks@4003.4022.20\|ATAACCTCGGGCCGATCGCA | >hg38_21_+_8433172_8446622_RNA45SN1_45S_50ntFlanks | 4003 | 4022 | 20 | ATAACCTCGGGCCGATCGCA | 587.8 | -4.7 | 1.1 | -4.2 | 3.3E-05 | 6.1E-05 | BT20/MB468 | MITO |
| >hg38_21_+_8433172_8446622_RNA45SN1_45S_50ntFlanks@10785.10812.28\|GAACAGCCTCTGGCATGTTGGAACAATG | >hg38_21_+_8433172_8446622_RNA45SN1_45S_50ntFlanks | 10785 | 10812 | 28 | GAACAGCCTCTGGCATGTTGGAACAATG | 88.0 | 9.8 | 2.1 | 4.6 | 5.0E-06 | 6.1E-05 | BT20/MB231 | MP |
| >hg38_21_+_8433172_8446622_RNA45SN1_45S_50ntFlanks@10348.10382.35\|AAACGAGAACTTTGAAGGCCGAAGTGGAGAAGGGT | >hg38_21_+_8433172_8446622_RNA45SN1_45S_50ntFlanks | 10348 | 10382 | 35 | AAACGAGAACTTTGAAGGCCGAAGTGGAGAAGGGT | 555.3 | 3.1 | 0.7 | 4.2 | 3.3E-05 | 6.1E-05 | BT20/MB468 | MITO |
| >hg38_21_+_8433172_8446622_RNA45SN1_45S_50ntFlanks@8168.8215.48\|CCCAAGTCCTTCTGATCGAGGCCCAGCCCGTGGACGGTGTGAGGCCGG | >hg38_21_+_8433172_8446622_RNA45SN1_45S_50ntFlanks | 8168 | 8215 | 48 | CCCAAGTCCTTCTGATCGAGGCCCAGCCCGTGGACGGTGTGAGGCCGG | 243.5 | -4.7 | 1.1 | -4.1 | 4.2E-05 | 6.1E-05 | MB231/MB468 | CYTO |
| >hg38_21_+_8433172_8446622_RNA45SN1_45S_50ntFlanks@12454.12473.20\|CACCAAGCGTTGGATTGTTC | >hg38_21_+_8433172_8446622_RNA45SN1_45S_50ntFlanks | 12454 | 12473 | 20 | CACCAAGCGTTGGATTGTTC | 373.3 | -3.4 | 0.8 | -4.1 | 3.8E-05 | 6.1E-05 | MB231/MB468 | MP |
| >hg38_21_+_8433172_8446622_RNA45SN1_45S_50ntFlanks@10782.10818.37\|GGTGAACAGCCTCTGGCATGTTGGAACAATGTAGGTA | >hg38_21_+_8433172_8446622_RNA45SN1_45S_50ntFlanks | 10782 | 10818 | 37 | GGTGAACAGCCTCTGGCATGTTGGAACAATGTAGGTA | 269.1 | -5.5 | 1.3 | -4.2 | 3.3E-05 | 6.2E-05 | BT20/MB468 | MITO |
| >hg38_21_+_8433172_8446622_RNA45SN1_45S_50ntFlanks@10823.10842.20\|AAGTCGGCAAGCCGGATCCG | >hg38_21_+_8433172_8446622_RNA45SN1_45S_50ntFlanks | 10823 | 10842 | 20 | AAGTCGGCAAGCCGGATCCG | 137.1 | -6.2 | 1.4 | -4.3 | 1.5E-05 | 6.2E-05 | BT20/MB468 | MP |
| >trna10_AspGTC_12_-_125424193_125424264@38.69.32,trna12_AspGTC_12_-_125411891_125411962@38.69.32,trna144_AspGTC_6_-_27551236_27551307@38.69.32,trna38_AspGTC_17_-_8125556_8125627@38.69.32,trna45_AspGTC_6_+_27447453_27447524@38.69.32,trna48_AspGTC_6_+_27471523_27471594@38.69.32,trna4_AspGTC_12_+_96429799_96429870@38.69.32,trna69_AspGTC_1_-_161440205_161440276@38.69.32,trna72_AspGTC_1_-_161432824_161432895@38.69.32,trna75_AspGTC_1_-_161425414_161425485@38.69.32,trna78_AspGTC_1_-_161418033_161418104@38.69.32,trna81_AspGTC_1_-_161410615_161410686@38.69.32\|CGCGGGAGACCGGGGTTCGATTCCCCGACGGG | >trna10_AspGTC_12_-_125424193_125424264 | 38 | 69 | 32 | CGCGGGAGACCGGGGTTCGATTCCCCGACGGG | 448.0 | 4.2 | 1.0 | 4.2 | 3.3E-05 | 6.2E-05 | BT20/MB468 | MITO |
| >hg38_21_+_8433172_8446622_RNA45SN1_45S_50ntFlanks@10392.10426.35\|ACAGCAGTTGAACATGGGTCAGTCGGTCCTGAGAG | >hg38_21_+_8433172_8446622_RNA45SN1_45S_50ntFlanks | 10392 | 10426 | 35 | ACAGCAGTTGAACATGGGTCAGTCGGTCCTGAGAG | 287.7 | -2.7 | 0.7 | -4.2 | 3.3E-05 | 6.2E-05 | BT20/MB468 | MITO |
| >hg38_1_-_228634819_228635039_RNA5S12_5S_50ntFlanks@51.90.40\|GTCTACGGCCATACCACCCTGAACGCGCCCGATCTCGTCT | >hg38_1_-_228634819_228635039_RNA5S12_5S_50ntFlanks | 51 | 90 | 40 | GTCTACGGCCATACCACCCTGAACGCGCCCGATCTCGTCT | 659.3 | 3.7 | 0.9 | 4.1 | 4.5E-05 | 6.2E-05 | MB231/MB468 | NUC |
| >MI0000802\|hsa-mir-340&WithFlank&5\|-\|180015297\|180015403@22.42.21&[MIMAT0004692&hsa-miR-340-5p&offsets\|0\|-1;m-220&5\|-\|180015361\|180015382&offsets\|0\|-1]\|TTATAAAGCAATGAGACTGAT | >MI0000802\|hsa-mir-340&WithFlank&5\|-\|180015297\|180015403 | 22 | 42 | 21 | TTATAAAGCAATGAGACTGAT | 103.8 | 2.6 | 0.6 | 4.1 | 4.7E-05 | 6.2E-05 | BT20/MB468 | NUC |
| >MI0000465\|hsa-mir-191&WithFlank&3\|-\|49020612\|49020715@22.45.24&[MIMAT0000440&hsa-miR-191-5p&offsets\|0\|+1;m-48&3\|-\|49020672\|49020694&offsets\|0\|+1]\|CAACGGAATCCCAAAAGCAGCTGT | >MI0000465\|hsa-mir-191&WithFlank&3\|-\|49020612\|49020715 | 22 | 45 | 24 | CAACGGAATCCCAAAAGCAGCTGT | 935.7 | 2.6 | 0.6 | 4.1 | 4.7E-05 | 6.2E-05 | BT20/MB468 | NUC |
| >MI0000456\|hsa-mir-140&WithFlank&16\|+\|69933075\|69933186@29.48.20&[MIMAT0000431&hsa-miR-140-5p&offsets\|0\|-2;m-173&16\|+\|69933103\|69933124&offsets\|0\|-2]\|CAGTGGTTTTACCCTATGGT | >MI0000456\|hsa-mir-140&WithFlank&16\|+\|69933075\|69933186 | 29 | 48 | 20 | CAGTGGTTTTACCCTATGGT | 349.8 | 6.6 | 1.6 | 4.1 | 3.8E-05 | 6.2E-05 | MB231/MB468 | MP |
| >hg38_21_+_8433172_8446622_RNA45SN1_45S_50ntFlanks@12957.12990.34\|CTTCTGGGTCGGGGTTTCGTACGTAGCAGAGCAG | >hg38_21_+_8433172_8446622_RNA45SN1_45S_50ntFlanks | 12957 | 12990 | 34 | CTTCTGGGTCGGGGTTTCGTACGTAGCAGAGCAG | 292.3 | -5.5 | 1.3 | -4.2 | 3.3E-05 | 6.2E-05 | BT20/MB468 | MITO |
| >hg38_21_+_8433172_8446622_RNA45SN1_45S_50ntFlanks@11669.11709.41\|GCCCAGTGCTCTGAATGTCAAAGTGAAGAAATTCAATGAAG | >hg38_21_+_8433172_8446622_RNA45SN1_45S_50ntFlanks | 11669 | 11709 | 41 | GCCCAGTGCTCTGAATGTCAAAGTGAAGAAATTCAATGAAG | 1079.0 | -4.2 | 0.9 | -4.6 | 4.8E-06 | 6.2E-05 | BT20/MB231 | CYTO |

<table>
<tr><td>>hg38_21_+_8433172_8446622_RNA45SN1_45S_50ntFlanks@10407.10429.23|GGGTCAGTCGGTCCTGAGAGATG</td><td>>hg38_21_+_8433172_8446622_RNA45SN1_45S_50ntFlanks</td><td>10407</td><td>10429</td><td>23</td><td>GGGTCAGTCGGTCCTGAGAGATG</td><td>216.6</td><td>-4.1</td><td>1.0</td><td>-4.1</td><td>3.9E-05</td><td>6.2E-05</td><td>BT20/MB468</td><td>CYTO</td></tr>
<tr><td>>hg38_21_+_8433172_8446622_RNA45SN1_45S_50ntFlanks@11869.11893.25|GGGAAAGAAGACCCTGTTGAGCTTG</td><td>>hg38_21_+_8433172_8446622_RNA45SN1_45S_50ntFlanks</td><td>11869</td><td>11893</td><td>25</td><td>GGGAAAGAAGACCCTGTTGAGCTTG</td><td>97.2</td><td>9.9</td><td>2.2</td><td>4.6</td><td>5.0E-06</td><td>6.2E-05</td><td>BT20/MB231</td><td>MP</td></tr>
<tr><td>>hg38_21_+_8433172_8446622_RNA45SN1_45S_50ntFlanks@12539.12560.22|ATGATGTGTTGTTGCCATGGTA</td><td>>hg38_21_+_8433172_8446622_RNA45SN1_45S_50ntFlanks</td><td>12539</td><td>12560</td><td>22</td><td>ATGATGTGTTGTTGCCATGGTA</td><td>250.1</td><td>-6.6</td><td>1.5</td><td>-4.3</td><td>1.5E-05</td><td>6.2E-05</td><td>BT20/MB468</td><td>MP</td></tr>
<tr><td>>hg38_21_+_8433172_8446622_RNA45SN1_45S_50ntFlanks@10806.10825.20|AACAATGTAGGTAAGGGAAG</td><td>>hg38_21_+_8433172_8446622_RNA45SN1_45S_50ntFlanks</td><td>10806</td><td>10825</td><td>20</td><td>AACAATGTAGGTAAGGGAAG</td><td>366.1</td><td>-5.8</td><td>1.4</td><td>-4.1</td><td>3.9E-05</td><td>6.2E-05</td><td>BT20/MB468</td><td>CYTO</td></tr>
<tr><td>>hg38_21_+_8433172_8446622_RNA45SN1_45S_50ntFlanks@6660.6685.26|GCGGTGGATCACTCGGCTCGTGCGTC</td><td>>hg38_21_+_8433172_8446622_RNA45SN1_45S_50ntFlanks</td><td>6660</td><td>6685</td><td>26</td><td>GCGGTGGATCACTCGGCTCGTGCGTC</td><td>553.9</td><td>3.8</td><td>0.9</td><td>4.3</td><td>1.5E-05</td><td>6.2E-05</td><td>BT20/MB468</td><td>MP</td></tr>
<tr><td>>hg38_21_+_8433172_8446622_RNA45SN1_45S_50ntFlanks@6737.6770.34|GATCATCGACACTTCGAACGCACTTGCGGCCCCG</td><td>>hg38_21_+_8433172_8446622_RNA45SN1_45S_50ntFlanks</td><td>6737</td><td>6770</td><td>34</td><td>GATCATCGACACTTCGAACGCACTTGCGGCCCCG</td><td>463.3</td><td>-6.9</td><td>1.6</td><td>-4.3</td><td>1.5E-05</td><td>6.2E-05</td><td>BT20/MB468</td><td>MP</td></tr>
<tr><td>>hg38_21_+_8433172_8446622_RNA45SN1_45S_50ntFlanks@11210.11231.22|GGCGACTCTGGACGCGAGCCGG</td><td>>hg38_21_+_8433172_8446622_RNA45SN1_45S_50ntFlanks</td><td>11210</td><td>11231</td><td>22</td><td>GGCGACTCTGGACGCGAGCCGG</td><td>440.9</td><td>-5.2</td><td>1.3</td><td>-4.1</td><td>4.7E-05</td><td>6.2E-05</td><td>BT20/MB468</td><td>NUC</td></tr>
<tr><td>>hg38_21_+_8433172_8446622_RNA45SN1_45S_50ntFlanks@11836.11873.38|ACAGCCAAGGGAACGGGCTTGGCGGAATCAGCGGGGAA</td><td>>hg38_21_+_8433172_8446622_RNA45SN1_45S_50ntFlanks</td><td>11836</td><td>11873</td><td>38</td><td>ACAGCCAAGGGAACGGGCTTGGCGGAATCAGCGGGGAA</td><td>39.1</td><td>-8.4</td><td>1.9</td><td>-4.3</td><td>1.6E-05</td><td>6.2E-05</td><td>BT20/MB468</td><td>MP</td></tr>
<tr><td>>hg38_21_+_8433172_8446622_RNA45SN1_45S_50ntFlanks@11536.11553.18|TCCGCGCGGCGCCTCGCC</td><td>>hg38_21_+_8433172_8446622_RNA45SN1_45S_50ntFlanks</td><td>11536</td><td>11553</td><td>18</td><td>TCCGCGCGGCGCCTCGCC</td><td>214.0</td><td>-5.8</td><td>1.4</td><td>-4.1</td><td>4.5E-05</td><td>6.2E-05</td><td>MB231/MB468</td><td>NUC</td></tr>
<tr><td>>hg38_21_+_8433172_8446622_RNA45SN1_45S_50ntFlanks@3872.3899.28|CTAATACATGCCGACGGGCGCTGACCCC</td><td>>hg38_21_+_8433172_8446622_RNA45SN1_45S_50ntFlanks</td><td>3872</td><td>3899</td><td>28</td><td>CTAATACATGCCGACGGGCGCTGACCCC</td><td>321.2</td><td>-3.2</td><td>0.8</td><td>-4.1</td><td>4.3E-05</td><td>6.2E-05</td><td>MB231/MB468</td><td>CYTO</td></tr>
<tr><td>>hg38_21_+_8433172_8446622_RNA45SN1_45S_50ntFlanks@10350.10371.22|ACGAGAACTTTGAAGGCCGAAG</td><td>>hg38_21_+_8433172_8446622_RNA45SN1_45S_50ntFlanks</td><td>10350</td><td>10371</td><td>22</td><td>ACGAGAACTTTGAAGGCCGAAG</td><td>1761.6</td><td>-2.1</td><td>0.5</td><td>-4.1</td><td>4.5E-05</td><td>6.2E-05</td><td>MB231/MB468</td><td>NUC</td></tr>
<tr><td>>am_trnaMT_HisGTG_MT_+_12138_12206@19.51.33|AAACATCAGATTGTGAATCTGACAACAGAGGCT</td><td>>am_trnaMT_HisGTG_MT_+_12138_12206</td><td>19</td><td>51</td><td>33</td><td>AAACATCAGATTGTGAATCTGACAACAGAGGCT</td><td>1110.9</td><td>6.3</td><td>1.5</td><td>4.3</td><td>1.6E-05</td><td>6.2E-05</td><td>BT20/MB468</td><td>MP</td></tr>
<tr><td>>trnaMT_HisGTG_MT_+_12138_12206@27.62.36|GATTGTGAATCTGACAACAGAGGCTTACGACCCCTT</td><td>>trnaMT_HisGTG_MT_+_12138_12206</td><td>27</td><td>62</td><td>36</td><td>GATTGTGAATCTGACAACAGAGGCTTACGACCCCTT</td><td>26.7</td><td>8.0</td><td>1.9</td><td>4.1</td><td>3.8E-05</td><td>6.2E-05</td><td>MB231/MB468</td><td>MP</td></tr>
<tr><td>>hg38_21_+_8433172_8446622_RNA45SN1_45S_50ntFlanks@11672.11709.38|CAGTGCTCTGAATGTCAAAGTGAAGAAATTCAATGAAG</td><td>>hg38_21_+_8433172_8446622_RNA45SN1_45S_50ntFlanks</td><td>11672</td><td>11709</td><td>38</td><td>CAGTGCTCTGAATGTCAAAGTGAAGAAATTCAATGAAG</td><td>592.6</td><td>-3.7</td><td>0.8</td><td>-4.6</td><td>4.9E-06</td><td>6.2E-05</td><td>BT20/MB231</td><td>CYTO</td></tr>
<tr><td>>hg38_21_+_8433172_8446622_RNA45SN1_45S_50ntFlanks@11703.11740.38|AATGAAGCGCGGGTAAACGGCGGGAGTAACTATGACTC</td><td>>hg38_21_+_8433172_8446622_RNA45SN1_45S_50ntFlanks</td><td>11703</td><td>11740</td><td>38</td><td>AATGAAGCGCGGGTAAACGGCGGGAGTAACTATGACTC</td><td>375.9</td><td>-4.9</td><td>1.2</td><td>-4.1</td><td>4.6E-05</td><td>6.2E-05</td><td>MB231/MB468</td><td>MITO</td></tr>
<tr><td>>hg38_21_+_8433172_8446622_RNA45SN1_45S_50ntFlanks@4049.4066.18|TCGAACGTCTGCCCTATC</td><td>>hg38_21_+_8433172_8446622_RNA45SN1_45S_50ntFlanks</td><td>4049</td><td>4066</td><td>18</td><td>TCGAACGTCTGCCCTATC</td><td>497.5</td><td>-3.9</td><td>1.0</td><td>-4.1</td><td>4.6E-05</td><td>6.2E-05</td><td>MB231/MB468</td><td>MITO</td></tr>
<tr><td>>hg38_21_+_8433172_8446622_RNA45SN1_45S_50ntFlanks@11662.11681.20|GATTTCTGCCCAGTGCTCTG</td><td>>hg38_21_+_8433172_8446622_RNA45SN1_45S_50ntFlanks</td><td>11662</td><td>11681</td><td>20</td><td>GATTTCTGCCCAGTGCTCTG</td><td>285.1</td><td>-2.8</td><td>0.7</td><td>-4.1</td><td>4.5E-05</td><td>6.2E-05</td><td>MB231/MB468</td><td>NUC</td></tr>
<tr><td>>hg38_21_+_8433172_8446622_RNA45SN1_45S_50ntFlanks@3854.3883.30|ACTGTGGTAATTCTAGAGCTAATACATGCC</td><td>>hg38_21_+_8433172_8446622_RNA45SN1_45S_50ntFlanks</td><td>3854</td><td>3883</td><td>30</td><td>ACTGTGGTAATTCTAGAGCTAATACATGCC</td><td>150.1</td><td>-5.1</td><td>1.2</td><td>-4.1</td><td>3.8E-05</td><td>6.2E-05</td><td>MB231/MB468</td><td>MP</td></tr>
<tr><td>>hg38_21_+_8433172_8446622_RNA45SN1_45S_50ntFlanks@10389.10424.36|TGAACAGCAGTTGAACATGGGTCAGTCGGTCCTGAG</td><td>>hg38_21_+_8433172_8446622_RNA45SN1_45S_50ntFlanks</td><td>10389</td><td>10424</td><td>36</td><td>TGAACAGCAGTTGAACATGGGTCAGTCGGTCCTGAG</td><td>3322.3</td><td>2.3</td><td>0.6</td><td>4.1</td><td>4.0E-05</td><td>6.2E-05</td><td>BT20/MB468</td><td>CYTO</td></tr>
<tr><td>>hg38_21_+_8433172_8446622_RNA45SN1_45S_50ntFlanks@3853.3874.22|AACTGTGGTAATTCTAGAGCTA</td><td>>hg38_21_+_8433172_8446622_RNA45SN1_45S_50ntFlanks</td><td>3853</td><td>3874</td><td>22</td><td>AACTGTGGTAATTCTAGAGCTA</td><td>320.8</td><td>-5.1</td><td>1.2</td><td>-4.3</td><td>1.6E-05</td><td>6.2E-05</td><td>BT20/MB468</td><td>MP</td></tr>
<tr><td>>hg38_21_+_8433172_8446622_RNA45SN1_45S_50ntFlanks@8342.8365.24|CGTAAGGGAAAGTTGAAAAGAACT</td><td>>hg38_21_+_8433172_8446622_RNA45SN1_45S_50ntFlanks</td><td>8342</td><td>8365</td><td>24</td><td>CGTAAGGGAAAGTTGAAAAGAACT</td><td>706.8</td><td>5.6</td><td>1.4</td><td>4.1</td><td>4.6E-05</td><td>6.2E-05</td><td>MB231/MB468</td><td>NUC</td></tr>
<tr><td>>hg38_21_+_8433172_8446622_RNA45SN1_45S_50ntFlanks@8179.8204.26|CTGATCGAGGCCCAGCCCGTGGACGG</td><td>>hg38_21_+_8433172_8446622_RNA45SN1_45S_50ntFlanks</td><td>8179</td><td>8204</td><td>26</td><td>CTGATCGAGGCCCAGCCCGTGGACGG</td><td>180.9</td><td>-6.4</td><td>1.5</td><td>-4.3</td><td>1.6E-05</td><td>6.3E-05</td><td>BT20/MB468</td><td>MP</td></tr>
<tr><td>>hg38_21_+_8433172_8446622_RNA45SN1_45S_50ntFlanks@11535.11552.18|TTCCGCGCGGCGCCTCGC</td><td>>hg38_21_+_8433172_8446622_RNA45SN1_45S_50ntFlanks</td><td>11535</td><td>11552</td><td>18</td><td>TTCCGCGCGGCGCCTCGC</td><td>156.9</td><td>-5.3</td><td>1.3</td><td>-4.1</td><td>4.0E-05</td><td>6.3E-05</td><td>BT20/MB468</td><td>CYTO</td></tr>
<tr><td>>hg38_21_+_8433172_8446622_RNA45SN1_45S_50ntFlanks@10348.10366.19|AAACGAGAACTTTGAAGGC</td><td>>hg38_21_+_8433172_8446622_RNA45SN1_45S_50ntFlanks</td><td>10348</td><td>10366</td><td>19</td><td>AAACGAGAACTTTGAAGGC</td><td>1363.4</td><td>-3.9</td><td>0.9</td><td>-4.1</td><td>3.8E-05</td><td>6.3E-05</td><td>MB231/MB468</td><td>MP</td></tr>
<tr><td>>am_trnaMT_ValTAC_MT_+_1602_1670@48.67.20|CAACTTAACTTGACCGCTCT</td><td>>am_trnaMT_ValTAC_MT_+_1602_1670</td><td>48</td><td>67</td><td>20</td><td>CAACTTAACTTGACCGCTCT</td><td>1190.4</td><td>4.1</td><td>1.0</td><td>4.1</td><td>4.8E-05</td><td>6.3E-05</td><td>BT20/MB468</td><td>NUC</td></tr>
<tr><td>>hg38_21_+_8433172_8446622_RNA45SN1_45S_50ntFlanks@9571.9597.27|GCAAATCGGTCGTCCGACCTGGGTATA</td><td>>hg38_21_+_8433172_8446622_RNA45SN1_45S_50ntFlanks</td><td>9571</td><td>9597</td><td>27</td><td>GCAAATCGGTCGTCCGACCTGGGTATA</td><td>167.7</td><td>-5.4</td><td>1.3</td><td>-4.1</td><td>4.0E-05</td><td>6.3E-05</td><td>BT20/MB468</td><td>CYTO</td></tr>
<tr><td>>hg38_21_+_8433172_8446622_RNA45SN1_45S_50ntFlanks@9790.9813.24|CTGGCGTGGAGCCGGGCGTGGAAT</td><td>>hg38_21_+_8433172_8446622_RNA45SN1_45S_50ntFlanks</td><td>9790</td><td>9813</td><td>24</td><td>CTGGCGTGGAGCCGGGCGTGGAAT</td><td>110.0</td><td>10.1</td><td>2.2</td><td>4.6</td><td>5.1E-06</td><td>6.3E-05</td><td>BT20/MB231</td><td>MP</td></tr>
<tr><td>>hg38_21_+_8433172_8446622_RNA45SN1_45S_50ntFlanks@6661.6693.33|CGGTGGATCACTCGGCTCGTGCGTCGATGAAGA</td><td>>hg38_21_+_8433172_8446622_RNA45SN1_45S_50ntFlanks</td><td>6661</td><td>6693</td><td>33</td><td>CGGTGGATCACTCGGCTCGTGCGTCGATGAAGA</td><td>972.0</td><td>3.5</td><td>0.8</td><td>4.3</td><td>1.6E-05</td><td>6.3E-05</td><td>BT20/MB468</td><td>MP</td></tr>
<tr><td>>hg38_21_+_8433172_8446622_RNA45SN1_45S_50ntFlanks@9533.9563.31|GAGGAAACTCTGGTGGAGGTCCGTAGCGGTC</td><td>>hg38_21_+_8433172_8446622_RNA45SN1_45S_50ntFlanks</td><td>9533</td><td>9563</td><td>31</td><td>GAGGAAACTCTGGTGGAGGTCCGTAGCGGTC</td><td>660.7</td><td>-6.2</td><td>1.5</td><td>-4.1</td><td>4.6E-05</td><td>6.3E-05</td><td>MB231/MB468</td><td>MITO</td></tr>
<tr><td>>hg38_21_+_8433172_8446622_RNA45SN1_45S_50ntFlanks@10794.10826.33|CTGGCATGTTGGAACAATGTAGGTAAGGGAAGT</td><td>>hg38_21_+_8433172_8446622_RNA45SN1_45S_50ntFlanks</td><td>10794</td><td>10826</td><td>33</td><td>CTGGCATGTTGGAACAATGTAGGTAAGGGAAGT</td><td>195.6</td><td>-2.9</td><td>0.7</td><td>-4.1</td><td>4.0E-05</td><td>6.3E-05</td><td>BT20/MB468</td><td>CYTO</td></tr>
<tr><td>>MI0000433|hsa-let-7g&WithFlank&3|-|52268272|52268367@11.32.22(+1A)&[MIMAT0000414&hsa-let-7g-5p&offsets|0|0(+1A);m-58&3|-|52268336|52268357&offsets|0|0(+1A)]|TGAGGTAGTAGTTTGTACAGTTA</td><td>>MI0000433|hsa-let-7g&WithFlank&3|-|52268272|52268367</td><td>11</td><td>32</td><td>22(+1A)</td><td>TGAGGTAGTAGTTTGTACAGTTA</td><td>547.5</td><td>1.8</td><td>0.4</td><td>4.1</td><td>4.8E-05</td><td>6.3E-05</td><td>BT20/MB468</td><td>NUC</td></tr>
<tr><td>>hg38_21_+_8433172_8446622_RNA45SN1_45S_50ntFlanks@10509.10533.25|GAATCCGGAGTGGCGGAGATGGGCG</td><td>>hg38_21_+_8433172_8446622_RNA45SN1_45S_50ntFlanks</td><td>10509</td><td>10533</td><td>25</td><td>GAATCCGGAGTGGCGGAGATGGGCG</td><td>570.9</td><td>-7.3</td><td>1.8</td><td>-4.1</td><td>4.8E-05</td><td>6.3E-05</td><td>BT20/MB468</td><td>NUC</td></tr>
<tr><td>>hg38_21_+_8433172_8446622_RNA45SN1_45S_50ntFlanks@12419.12447.29|GATGTCGGCTCTTCCTATCATTGTGAAGC</td><td>>hg38_21_+_8433172_8446622_RNA45SN1_45S_50ntFlanks</td><td>12419</td><td>12447</td><td>29</td><td>GATGTCGGCTCTTCCTATCATTGTGAAGC</td><td>182.8</td><td>3.4</td><td>0.8</td><td>4.1</td><td>4.6E-05</td><td>6.3E-05</td><td>MB231/MB468</td><td>MITO</td></tr>
<tr><td>>hg38_21_+_8433172_8446622_RNA45SN1_45S_50ntFlanks@10406.10424.19|TGGGTCAGTCGGTCCTGAG</td><td>>hg38_21_+_8433172_8446622_RNA45SN1_45S_50ntFlanks</td><td>10406</td><td>10424</td><td>19</td><td>TGGGTCAGTCGGTCCTGAG</td><td>3087.9</td><td>3.9</td><td>0.9</td><td>4.5</td><td>6.0E-06</td><td>6.3E-05</td><td>BT20/MB231</td><td>MITO</td></tr>
<tr><td>>MI0000071|hsa-mir-17&WithFlank&13|+|91350599|91350694@20.41.22&[MIMAT0000070&hsa-miR-17-5p&offsets|0|-1;m-88&13|+|91350618|91350640&offsets|0|-1]|CAAAGTGCTTACAGTGCAGGTA</td><td>>MI0000071|hsa-mir-17&WithFlank&13|+|91350599|91350694</td><td>20</td><td>41</td><td>22</td><td>CAAAGTGCTTACAGTGCAGGTA</td><td>741.4</td><td>-4.0</td><td>0.9</td><td>-4.5</td><td>6.0E-06</td><td>6.3E-05</td><td>BT20/MB231</td><td>MITO</td></tr>
<tr><td>>hg38_21_+_8433172_8446622_RNA45SN1_45S_50ntFlanks@6788.6807.20|CGCCTGTCTGAGCGTCGCTT</td><td>>hg38_21_+_8433172_8446622_RNA45SN1_45S_50ntFlanks</td><td>6788</td><td>6807</td><td>20</td><td>CGCCTGTCTGAGCGTCGCTT</td><td>3580.2</td><td>3.6</td><td>0.9</td><td>4.1</td><td>3.4E-05</td><td>6.3E-05</td><td>BT20/MB468</td><td>MITO</td></tr>
<tr><td>>am_trna13_ThrTGT_5_-_180618687_180618758@1.29.29,trna20_ThrTGT_14_-_21099319_21099391@1.29.29|GGCTCCATAGCTCAGGGGTTAGAGCACTG</td><td>>am_trna13_ThrTGT_5_-_180618687_180618758</td><td>1</td><td>29</td><td>29</td><td>GGCTCCATAGCTCAGGGGTTAGAGCACTG</td><td>141.4</td><td>-2.8</td><td>0.7</td><td>-4.1</td><td>4.3E-05</td><td>6.3E-05</td><td>MB231/MB468</td><td>CYTO</td></tr>
<tr><td>>hg38_21_+_8433172_8446622_RNA45SN1_45S_50ntFlanks@10869.10890.22|AGGGCTGGGTCGGTCGGGCTGG</td><td>>hg38_21_+_8433172_8446622_RNA45SN1_45S_50ntFlanks</td><td>10869</td><td>10890</td><td>22</td><td>AGGGCTGGGTCGGTCGGGCTGG</td><td>1020.9</td><td>-3.3</td><td>0.8</td><td>-4.1</td><td>4.0E-05</td><td>6.3E-05</td><td>BT20/MB468</td><td>CYTO</td></tr>
<tr><td>>hg38_21_+_8433172_8446622_RNA45SN1_45S_50ntFlanks@5044.5081.38|TCTTAGTTGGTGGAGCGATTTGTCTGGTTAATTCCGAT</td><td>>hg38_21_+_8433172_8446622_RNA45SN1_45S_50ntFlanks</td><td>5044</td><td>5081</td><td>38</td><td>TCTTAGTTGGTGGAGCGATTTGTCTGGTTAATTCCGAT</td><td>83.3</td><td>9.7</td><td>2.1</td><td>4.6</td><td>5.1E-06</td><td>6.3E-05</td><td>BT20/MB231</td><td>MP</td></tr>
<tr><td>>hg38_21_+_8433172_8446622_RNA45SN1_45S_50ntFlanks@11890.11907.18|CTTGACTCTAGTCTGGCA</td><td>>hg38_21_+_8433172_8446622_RNA45SN1_45S_50ntFlanks</td><td>11890</td><td>11907</td><td>18</td><td>CTTGACTCTAGTCTGGCA</td><td>1218.9</td><td>-5.7</td><td>1.3</td><td>-4.3</td><td>1.6E-05</td><td>6.3E-05</td><td>BT20/MB468</td><td>MP</td></tr>
<tr><td>>hg38_21_+_8433172_8446622_RNA45SN1_45S_50ntFlanks@10841.10872.32|CGTAACTTCGGGATAAGGATTGGCTCTAAGGG</td><td>>hg38_21_+_8433172_8446622_RNA45SN1_45S_50ntFlanks</td><td>10841</td><td>10872</td><td>32</td><td>CGTAACTTCGGGATAAGGATTGGCTCTAAGGG</td><td>2348.3</td><td>-7.8</td><td>1.9</td><td>-4.1</td><td>4.8E-05</td><td>6.3E-05</td><td>BT20/MB468</td><td>NUC</td></tr>
<tr><td>>hg38_21_+_8433172_8446622_RNA45SN1_45S_50ntFlanks@11707.11736.30|AAGCGCGGGTAAACGGCGGGAGTAACTATG</td><td>>hg38_21_+_8433172_8446622_RNA45SN1_45S_50ntFlanks</td><td>11707</td><td>11736</td><td>30</td><td>AAGCGCGGGTAAACGGCGGGAGTAACTATG</td><td>467.9</td><td>-5.5</td><td>1.4</td><td>-4.1</td><td>4.8E-05</td><td>6.3E-05</td><td>BT20/MB468</td><td>NUC</td></tr>
<tr><td>>hg38_21_+_8433172_8446622_RNA45SN1_45S_50ntFlanks@12944.12963.20|CGTAGACGACCTGCTTCTGG</td><td>>hg38_21_+_8433172_8446622_RNA45SN1_45S_50ntFlanks</td><td>12944</td><td>12963</td><td>20</td><td>CGTAGACGACCTGCTTCTGG</td><td>72.8</td><td>-5.9</td><td>1.4</td><td>-4.1</td><td>3.9E-05</td><td>6.3E-05</td><td>MB231/MB468</td><td>MP</td></tr>
<tr><td>>hg38_21_+_8433172_8446622_RNA45SN1_45S_50ntFlanks@10821.10839.19|GGAAGTCGGCAAGCCGGAT</td><td>>hg38_21_+_8433172_8446622_RNA45SN1_45S_50ntFlanks</td><td>10821</td><td>10839</td><td>19</td><td>GGAAGTCGGCAAGCCGGAT</td><td>166.6</td><td>-2.8</td><td>0.7</td><td>-4.1</td><td>4.0E-05</td><td>6.3E-05</td><td>BT20/MB468</td><td>CYTO</td></tr>
<tr><td>>hg38_21_+_8433172_8446622_RNA45SN1_45S_50ntFlanks@10498.10516.19|TTCAGATCCCCGAATCCGG</td><td>>hg38_21_+_8433172_8446622_RNA45SN1_45S_50ntFlanks</td><td>10498</td><td>10516</td><td>19</td><td>TTCAGATCCCCGAATCCGG</td><td>187.7</td><td>-6.4</td><td>1.5</td><td>-4.3</td><td>1.6E-05</td><td>6.3E-05</td><td>BT20/MB468</td><td>MP</td></tr>
<tr><td>>hg38_21_+_8433172_8446622_RNA45SN1_45S_50ntFlanks@11784.11813.30|ATGAATGGATGAACGAGATTCCCACTGTCC</td><td>>hg38_21_+_8433172_8446622_RNA45SN1_45S_50ntFlanks</td><td>11784</td><td>11813</td><td>30</td><td>ATGAATGGATGAACGAGATTCCCACTGTCC</td><td>100.0</td><td>10.0</td><td>2.2</td><td>4.6</td><td>5.2E-06</td><td>6.3E-05</td><td>BT20/MB231</td><td>MP</td></tr>
<tr><td>>hg38_21_+_8433172_8446622_RNA45SN1_45S_50ntFlanks@5066.5097.32|TCTGGTTAATTCCGATAACGAACGAGACTCTG</td><td>>hg38_21_+_8433172_8446622_RNA45SN1_45S_50ntFlanks</td><td>5066</td><td>5097</td><td>32</td><td>TCTGGTTAATTCCGATAACGAACGAGACTCTG</td><td>67.0</td><td>-9.1</td><td>2.1</td><td>-4.3</td><td>1.6E-05</td><td>6.3E-05</td><td>BT20/MB468</td><td>MP</td></tr>
<tr><td>>hg38_21_+_8433172_8446622_RNA45SN1_45S_50ntFlanks@3776.3810.35|CCGGTACAGTGAAACTGCGAATGGCTCATTAAATC</td><td>>hg38_21_+_8433172_8446622_RNA45SN1_45S_50ntFlanks</td><td>3776</td><td>3810</td><td>35</td><td>CCGGTACAGTGAAACTGCGAATGGCTCATTAAATC</td><td>50.9</td><td>-9.5</td><td>2.3</td><td>-4.1</td><td>3.9E-05</td><td>6.3E-05</td><td>MB231/MB468</td><td>MP</td></tr>
<tr><td>>hg38_MT_+_598_1651_RNR1_12S_50ntFlanks@903.935.33|CAAGTATACTTCAAAGGACATTTAACTAAAACC</td><td>>hg38_MT_+_598_1651_RNR1_12S_50ntFlanks</td><td>903</td><td>935</td><td>33</td><td>CAAGTATACTTCAAAGGACATTTAACTAAAACC</td><td>28.1</td><td>8.0</td><td>2.0</td><td>4.1</td><td>3.9E-05</td><td>6.3E-05</td><td>MB231/MB468</td><td>MP</td></tr>
<tr><td>>hg38_21_+_8433172_8446622_RNA45SN1_45S_50ntFlanks@11874.11906.33|AGAAGACCCTGTTGAGCTTGACTCTAGTCTGGC</td><td>>hg38_21_+_8433172_8446622_RNA45SN1_45S_50ntFlanks</td><td>11874</td><td>11906</td><td>33</td><td>AGAAGACCCTGTTGAGCTTGACTCTAGTCTGGC</td><td>699.4</td><td>-6.3</td><td>1.5</td><td>-4.1</td><td>4.6E-05</td><td>6.3E-05</td><td>MB231/MB468</td><td>NUC</td></tr>
<tr><td>>hg38_21_+_8433172_8446622_RNA45SN1_45S_50ntFlanks@12942.12960.19|TTCGTAGACGACCTGCTTC</td><td>>hg38_21_+_8433172_8446622_RNA45SN1_45S_50ntFlanks</td><td>12942</td><td>12960</td><td>19</td><td>TTCGTAGACGACCTGCTTC</td><td>424.3</td><td>-4.8</td><td>1.2</td><td>-4.1</td><td>3.9E-05</td><td>6.3E-05</td><td>MB231/MB468</td><td>MP</td></tr>
<tr><td>>hg38_21_+_8433172_8446622_RNA45SN1_45S_50ntFlanks@9651.9668.18|TCAGGATAGCTGGCGCTC</td><td>>hg38_21_+_8433172_8446622_RNA45SN1_45S_50ntFlanks</td><td>9651</td><td>9668</td><td>18</td><td>TCAGGATAGCTGGCGCTC</td><td>158.9</td><td>-5.7</td><td>1.4</td><td>-4.1</td><td>4.6E-05</td><td>6.3E-05</td><td>MB231/MB468</td><td>NUC</td></tr>
<tr><td>>hg38_21_+_8433172_8446622_RNA45SN1_45S_50ntFlanks@11579.11602.24|AGAACTGGTGCGGACCAGGGGAAT</td><td>>hg38_21_+_8433172_8446622_RNA45SN1_45S_50ntFlanks</td><td>11579</td><td>11602</td><td>24</td><td>AGAACTGGTGCGGACCAGGGGAAT</td><td>632.9</td><td>-7.0</td><td>1.6</td><td>-4.3</td><td>1.6E-05</td><td>6.4E-05</td><td>BT20/MB468</td><td>MP</td></tr>
<tr><td>>hg38_21_+_8433172_8446622_RNA45SN1_45S_50ntFlanks@9738.9757.20|GAAACGATCTCAACCTATTC</td><td>>hg38_21_+_8433172_8446622_RNA45SN1_45S_50ntFlanks</td><td>9738</td><td>9757</td><td>20</td><td>GAAACGATCTCAACCTATTC</td><td>588.6</td><td>-7.0</td><td>1.6</td><td>-4.3</td><td>1.6E-05</td><td>6.4E-05</td><td>BT20/MB468</td><td>MP</td></tr>
<tr><td>>hg38_21_+_8433172_8446622_RNA45SN1_45S_50ntFlanks@10512.10533.22|TCCGGAGTGGCGGAGATGGGCG</td><td>>hg38_21_+_8433172_8446622_RNA45SN1_45S_50ntFlanks</td><td>10512</td><td>10533</td><td>22</td><td>TCCGGAGTGGCGGAGATGGGCG</td><td>10969.6</td><td>-6.7</td><td>1.5</td><td>-4.3</td><td>1.6E-05</td><td>6.4E-05</td><td>BT20/MB468</td><td>MP</td></tr>
<tr><td>>Antisense_hg38_21_-_8433172_8446622_RNA45SN1_45S_50ntFlanks@7674.7693.20|CTCTCTCTCTCTCTCCCCCG</td><td>>Antisense_hg38_21_-_8433172_8446622_RNA45SN1_45S_50ntFlanks</td><td>7674</td><td>7693</td><td>20</td><td>CTCTCTCTCTCTCTCCCCCG</td><td>888.3</td><td>-5.1</td><td>1.2</td><td>-4.1</td><td>4.0E-05</td><td>6.4E-05</td><td>BT20/MB468</td><td>CYTO</td></tr>
</table>

<table>
<tr><td>>hg38_21_+_8433172_8446622_RNA45SN1_45S_50ntFlanks@11027.11046.20|CTCTCTCTCTCTCTCCCCCG</td><td>>hg38_21_+_8433172_8446622_RNA45SN1_45S_50ntFlanks</td><td>11027</td><td>11046</td><td>20</td><td>CTCTCTCTCTCTCTCCCCCG</td><td>888.3</td><td>-5.1</td><td>1.2</td><td>-4.1</td><td>4.0E-05</td><td>6.4E-05</td><td>BT20/MB468</td><td>CYTO</td></tr>
<tr><td>>hg38_21_+_8433172_8446622_RNA45SN1_45S_50ntFlanks@10770.10818.49|GCAGGTCTCCAAGGTGAACAGCCTCTGGCATGTTGGAACAATGTAGGTA</td><td>>hg38_21_+_8433172_8446622_RNA45SN1_45S_50ntFlanks</td><td>10770</td><td>10818</td><td>49</td><td>GCAGGTCTCCAAGGTGAACAGCCTCTGGCATGTTGGAACAATGTAGGTA</td><td>355.1</td><td>-6.5</td><td>1.6</td><td>-4.1</td><td>4.7E-05</td><td>6.4E-05</td><td>MB231/MB468</td><td>MITO</td></tr>
<tr><td>>trna116_GluCTC_1_-_145399233_145399304@1.34.34,trna59_GluCTC_1_+_249168447_249168518@1.34.34,trna71_GluCTC_1_-_161439189_161439260@1.34.34,trna74_GluCTC_1_-_161431809_161431880@1.34.34,trna77_GluCTC_1_-_161424398_161424469@1.34.34,trna77_GluCTC_6_+_28949976_28950047@1.34.34,trna80_GluCTC_1_-_161417018_161417089@1.34.34,trna87_GluCTC_6_-_126101393_126101464@1.34.34|TCCCTGGTGGTCTAGTGGTTAGGATTCGGCGCTC</td><td>>trna116_GluCTC_1_-_145399233_145399304</td><td>1</td><td>34</td><td>34</td><td>TCCCTGGTGGTCTAGTGGTTAGGATTCGGCGCTC</td><td>1791.8</td><td>3.8</td><td>0.9</td><td>4.1</td><td>3.9E-05</td><td>6.4E-05</td><td>MB231/MB468</td><td>MP</td></tr>
<tr><td>>MI0000088|hsa-mir-30a&WithFlank&6|-|71403545|71403627@54.71.18&[MIMAT0000088&hsa-miR-30a-3p&offsets|+1|-3;m-38&6|-|71403554|71403575&offsets|+1|-3]|TTTCAGTCGGATGTTTGC</td><td>>MI0000088|hsa-mir-30a&WithFlank&6|-|71403545|71403627</td><td>54</td><td>71</td><td>18</td><td>TTTCAGTCGGATGTTTGC</td><td>870.7</td><td>5.5</td><td>1.3</td><td>4.1</td><td>3.9E-05</td><td>6.4E-05</td><td>MB231/MB468</td><td>MP</td></tr>
<tr><td>>hg38_21_+_8433172_8446622_RNA45SN1_45S_50ntFlanks@12907.12940.34|CACCGCACGTTCGTGGGGAACCTGGCGCTAAACC</td><td>>hg38_21_+_8433172_8446622_RNA45SN1_45S_50ntFlanks</td><td>12907</td><td>12940</td><td>34</td><td>CACCGCACGTTCGTGGGGAACCTGGCGCTAAACC</td><td>409.4</td><td>-6.1</td><td>1.5</td><td>-4.1</td><td>4.8E-05</td><td>6.4E-05</td><td>BT20/MB468</td><td>NUC</td></tr>
<tr><td>>trnaMT_SerGCT_MT_+_12207_12265@1.18.18|GAGAAAGCTCACAAGAAC</td><td>>trnaMT_SerGCT_MT_+_12207_12265</td><td>1</td><td>18</td><td>18</td><td>GAGAAAGCTCACAAGAAC</td><td>15464.3</td><td>4.5</td><td>1.1</td><td>4.1</td><td>3.4E-05</td><td>6.4E-05</td><td>BT20/MB468</td><td>MITO</td></tr>
<tr><td>>hg38_MT_+_1621_3279_RNR2_16S_50ntFlanks@621.652.32|ATATAACTGAACTCCTCACACCCAATTGGACC</td><td>>hg38_MT_+_1621_3279_RNR2_16S_50ntFlanks</td><td>621</td><td>652</td><td>32</td><td>ATATAACTGAACTCCTCACACCCAATTGGACC</td><td>339.6</td><td>7.4</td><td>1.7</td><td>4.3</td><td>1.6E-05</td><td>6.4E-05</td><td>BT20/MB468</td><td>MP</td></tr>
<tr><td>>hg38_21_+_8433172_8446622_RNA45SN1_45S_50ntFlanks@10404.10424.21|CATGGGTCAGTCGGTCCTGAG</td><td>>hg38_21_+_8433172_8446622_RNA45SN1_45S_50ntFlanks</td><td>10404</td><td>10424</td><td>21</td><td>CATGGGTCAGTCGGTCCTGAG</td><td>9932.7</td><td>3.8</td><td>0.8</td><td>4.5</td><td>6.1E-06</td><td>6.4E-05</td><td>BT20/MB231</td><td>MITO</td></tr>
<tr><td>>hg38_MT_+_598_1651_RNR1_12S_50ntFlanks@425.465.41|ACGAAAGTGGCTTTAACATATCTGAACACACAATAGCTAAG</td><td>>hg38_MT_+_598_1651_RNR1_12S_50ntFlanks</td><td>425</td><td>465</td><td>41</td><td>ACGAAAGTGGCTTTAACATATCTGAACACACAATAGCTAAG</td><td>30.2</td><td>8.1</td><td>2.0</td><td>4.1</td><td>3.9E-05</td><td>6.4E-05</td><td>MB231/MB468</td><td>MP</td></tr>
<tr><td>>am_MI0000809|hsa-mir-151a&WithFlank&8|-|140732558|140732659@54.73.20(+1U)&[MIMAT0000757&hsa-miR-151a-3p&offsets|+1|0(+1U);m-30&8|-|140732587|140732607&offsets|+1|0(+1U)]|TAGACTGAAGCTCCTTGAGGT</td><td>>am_MI0000809|hsa-mir-151a&WithFlank&8|-|140732558|140732659</td><td>54</td><td>73</td><td>20(+1U)</td><td>TAGACTGAAGCTCCTTGAGGT</td><td>30.0</td><td>8.1</td><td>2.0</td><td>4.1</td><td>3.9E-05</td><td>6.4E-05</td><td>MB231/MB468</td><td>MP</td></tr>
<tr><td>>trnaMT_GluTTC_MT_-_14674_14742@38.58.21,trnalookalike8_GluTTC_5_-_93905172_93905240@38.58.21|CATTGGTCGTGGTTGTAGTCC</td><td>>trnaMT_GluTTC_MT_-_14674_14742</td><td>38</td><td>58</td><td>21</td><td>CATTGGTCGTGGTTGTAGTCC</td><td>308.9</td><td>5.9</td><td>1.4</td><td>4.3</td><td>1.6E-05</td><td>6.4E-05</td><td>BT20/MB468</td><td>MP</td></tr>
<tr><td>>MI0000088|hsa-mir-30a&WithFlank&6|-|71403545|71403627@53.73.21(+1U)&[MIMAT0000088&hsa-miR-30a-3p&offsets|0|-1(+1U);m-38&6|-|71403554|71403575&offsets|0|-1(+1U)]|CTTTCAGTCGGATGTTTGCAGT</td><td>>MI0000088|hsa-mir-30a&WithFlank&6|-|71403545|71403627</td><td>53</td><td>73</td><td>21(+1U)</td><td>CTTTCAGTCGGATGTTTGCAGT</td><td>5478.7</td><td>3.5</td><td>0.9</td><td>4.1</td><td>3.9E-05</td><td>6.4E-05</td><td>MB231/MB468</td><td>MP</td></tr>
<tr><td>>hg38_21_+_8433172_8446622_RNA45SN1_45S_50ntFlanks@12973.13003.31|TCGTACGTAGCAGAGCAGCTCCCTCGCTGCG</td><td>>hg38_21_+_8433172_8446622_RNA45SN1_45S_50ntFlanks</td><td>12973</td><td>13003</td><td>31</td><td>TCGTACGTAGCAGAGCAGCTCCCTCGCTGCG</td><td>212.6</td><td>-5.4</td><td>1.3</td><td>-4.1</td><td>4.1E-05</td><td>6.4E-05</td><td>BT20/MB468</td><td>CYTO</td></tr>
<tr><td>>hg38_21_+_8433172_8446622_RNA45SN1_45S_50ntFlanks@9568.9596.29|CGTGCAAATCGGTCGTCCGACCTGGGTAT</td><td>>hg38_21_+_8433172_8446622_RNA45SN1_45S_50ntFlanks</td><td>9568</td><td>9596</td><td>29</td><td>CGTGCAAATCGGTCGTCCGACCTGGGTAT</td><td>3745.3</td><td>-7.5</td><td>1.7</td><td>-4.3</td><td>1.6E-05</td><td>6.4E-05</td><td>BT20/MB468</td><td>MP</td></tr>
<tr><td>>hg38_21_+_8433172_8446622_RNA45SN1_45S_50ntFlanks@12241.12272.32|CAGAAGGGCAAAAGCTCGCTTGATCTTGATTT</td><td>>hg38_21_+_8433172_8446622_RNA45SN1_45S_50ntFlanks</td><td>12241</td><td>12272</td><td>32</td><td>CAGAAGGGCAAAAGCTCGCTTGATCTTGATTT</td><td>492.6</td><td>-5.2</td><td>1.3</td><td>-4.1</td><td>3.9E-05</td><td>6.4E-05</td><td>MB231/MB468</td><td>MP</td></tr>
<tr><td>>hg38_21_+_8433172_8446622_RNA45SN1_45S_50ntFlanks@8104.8145.42|CCGCGGCGGGGCGCGGGACATGTGGCGTACGGAAGACCCGCT</td><td>>hg38_21_+_8433172_8446622_RNA45SN1_45S_50ntFlanks</td><td>8104</td><td>8145</td><td>42</td><td>CCGCGGCGGGGCGCGGGACATGTGGCGTACGGAAGACCCGCT</td><td>61.5</td><td>-9.8</td><td>2.4</td><td>-4.1</td><td>3.9E-05</td><td>6.4E-05</td><td>MB231/MB468</td><td>MP</td></tr>
<tr><td>>hg38_21_+_8433172_8446622_RNA45SN1_45S_50ntFlanks@6733.6756.24|CATTGATCATCGACACTTCGAACG</td><td>>hg38_21_+_8433172_8446622_RNA45SN1_45S_50ntFlanks</td><td>6733</td><td>6756</td><td>24</td><td>CATTGATCATCGACACTTCGAACG</td><td>630.8</td><td>-2.3</td><td>0.6</td><td>-4.1</td><td>4.1E-05</td><td>6.4E-05</td><td>BT20/MB468</td><td>CYTO</td></tr>
<tr><td>>trna116_GluCTC_1_-_145399233_145399304@35.75.41,trna134_GluTTC_1_-_16861774_16861845@35.75.41,trna71_GluCTC_1_-_161439189_161439260@35.75.41,trna74_GluCTC_1_-_161431809_161431880@35.75.41,trna77_GluCTC_1_-_161424398_161424469@35.75.41,trna77_GluCTC_6_+_28949976_28950047@35.75.41,trna80_GluCTC_1_-_161417018_161417089@35.75.41,trna84_GluTTC_1_-_161391883_161391954@35.75.41,trna87_GluCTC_6_-_126101393_126101464@35.75.41|TCACCGCCGCGGCCCGGGTTCGATTCCCGGTCAGGGAACCA</td><td>>trna116_GluCTC_1_-_145399233_145399304</td><td>35</td><td>75</td><td>41</td><td>TCACCGCCGCGGCCCGGGTTCGATTCCCGGTCAGGGAACCA</td><td>398.3</td><td>3.6</td><td>0.9</td><td>4.1</td><td>4.1E-05</td><td>6.4E-05</td><td>BT20/MB468</td><td>CYTO</td></tr>
<tr><td>>hg38_21_+_8433172_8446622_RNA45SN1_45S_50ntFlanks@10919.10946.28|CGGCTGGACGAGGCGCCGCCGCCCCCCC</td><td>>hg38_21_+_8433172_8446622_RNA45SN1_45S_50ntFlanks</td><td>10919</td><td>10946</td><td>28</td><td>CGGCTGGACGAGGCGCCGCCGCCCCCCC</td><td>325.6</td><td>-6.6</td><td>1.6</td><td>-4.1</td><td>4.9E-05</td><td>6.4E-05</td><td>BT20/MB468</td><td>NUC</td></tr>
<tr><td>>hg38_21_+_8433172_8446622_RNA45SN1_45S_50ntFlanks@7975.7993.19|CGCGACCTCAGATCAGACG</td><td>>hg38_21_+_8433172_8446622_RNA45SN1_45S_50ntFlanks</td><td>7975</td><td>7993</td><td>19</td><td>CGCGACCTCAGATCAGACG</td><td>10372.8</td><td>2.7</td><td>0.7</td><td>4.1</td><td>4.4E-05</td><td>6.4E-05</td><td>MB231/MB468</td><td>CYTO</td></tr>
<tr><td>>hg38_21_+_8433172_8446622_RNA45SN1_45S_50ntFlanks@4759.4786.28|ATCAGATACCGTCGTAGTTCCGACCATA</td><td>>hg38_21_+_8433172_8446622_RNA45SN1_45S_50ntFlanks</td><td>4759</td><td>4786</td><td>28</td><td>ATCAGATACCGTCGTAGTTCCGACCATA</td><td>2076.2</td><td>-6.8</td><td>1.7</td><td>-4.1</td><td>4.7E-05</td><td>6.4E-05</td><td>MB231/MB468</td><td>MITO</td></tr>
<tr><td>>hg38_21_+_8433172_8446622_RNA45SN1_45S_50ntFlanks@11570.11588.19|AGCCGACTTAGAACTGGTG</td><td>>hg38_21_+_8433172_8446622_RNA45SN1_45S_50ntFlanks</td><td>11570</td><td>11588</td><td>19</td><td>AGCCGACTTAGAACTGGTG</td><td>189.7</td><td>-2.7</td><td>0.7</td><td>-4.1</td><td>4.4E-05</td><td>6.4E-05</td><td>MB231/MB468</td><td>CYTO</td></tr>
<tr><td>>hg38_21_+_8433172_8446622_RNA45SN1_45S_50ntFlanks@6779.6802.24|CCGGGGCTACGCCTGTCTGAGCGT</td><td>>hg38_21_+_8433172_8446622_RNA45SN1_45S_50ntFlanks</td><td>6779</td><td>6802</td><td>24</td><td>CCGGGGCTACGCCTGTCTGAGCGT</td><td>415.0</td><td>3.0</td><td>0.7</td><td>4.1</td><td>4.1E-05</td><td>6.4E-05</td><td>BT20/MB468</td><td>CYTO</td></tr>
<tr><td>>hg38_21_+_8433172_8446622_RNA45SN1_45S_50ntFlanks@3985.4010.26|CGGCTTTGGTGACTCTAGATAACCTC</td><td>>hg38_21_+_8433172_8446622_RNA45SN1_45S_50ntFlanks</td><td>3985</td><td>4010</td><td>26</td><td>CGGCTTTGGTGACTCTAGATAACCTC</td><td>512.9</td><td>-4.7</td><td>1.2</td><td>-4.1</td><td>4.7E-05</td><td>6.4E-05</td><td>MB231/MB468</td><td>NUC</td></tr>
<tr><td>>hg38_21_+_8433172_8446622_RNA45SN1_45S_50ntFlanks@3852.3881.30|TAACTGTGGTAATTCTAGAGCTAATACATG</td><td>>hg38_21_+_8433172_8446622_RNA45SN1_45S_50ntFlanks</td><td>3852</td><td>3881</td><td>30</td><td>TAACTGTGGTAATTCTAGAGCTAATACATG</td><td>237.3</td><td>-2.9</td><td>0.7</td><td>-4.1</td><td>4.4E-05</td><td>6.4E-05</td><td>MB231/MB468</td><td>CYTO</td></tr>
<tr><td>>MI0000114|hsa-mir-107&WithFlank&10|-|89592741|89592833@56.77.22&[MIMAT0000104&hsa-miR-107&offsets|0|-1;m-78&10|-|89592758|89592778&offsets|0|+1]|AGCAGCATTGTACAGGGCTATC</td><td>>MI0000114|hsa-mir-107&WithFlank&10|-|89592741|89592833</td><td>56</td><td>77</td><td>22</td><td>AGCAGCATTGTACAGGGCTATC</td><td>2039.1</td><td>-4.6</td><td>1.0</td><td>-4.6</td><td>5.3E-06</td><td>6.4E-05</td><td>BT20/MB231</td><td>MP</td></tr>
<tr><td>>hg38_21_+_8433172_8446622_RNA45SN1_45S_50ntFlanks@6775.6803.29|CCTCCCGGGGCTACGCCTGTCTGAGCGTC</td><td>>hg38_21_+_8433172_8446622_RNA45SN1_45S_50ntFlanks</td><td>6775</td><td>6803</td><td>29</td><td>CCTCCCGGGGCTACGCCTGTCTGAGCGTC</td><td>355.5</td><td>-5.1</td><td>1.3</td><td>-4.1</td><td>4.9E-05</td><td>6.4E-05</td><td>BT20/MB468</td><td>NUC</td></tr>
<tr><td>>hg38_21_+_8433172_8446622_RNA45SN1_45S_50ntFlanks@5484.5518.35|GAGCGCTGAGAAGACGGTCGAACTTGACTATCTAG</td><td>>hg38_21_+_8433172_8446622_RNA45SN1_45S_50ntFlanks</td><td>5484</td><td>5518</td><td>35</td><td>GAGCGCTGAGAAGACGGTCGAACTTGACTATCTAG</td><td>448.5</td><td>-3.8</td><td>0.9</td><td>-4.1</td><td>4.4E-05</td><td>6.4E-05</td><td>MB231/MB468</td><td>CYTO</td></tr>
<tr><td>>MI0000077|hsa-mir-21&WithFlank&17|+|59841260|59841343@14.37.24(+1U)&[MIMAT0000076&hsa-miR-21-5p&offsets|0|+2(+1U);m-1&17|+|59841273|59841294&offsets|0|+2(+1U)]|TAGCTTATCAGACTGATGTTGACTT</td><td>>MI0000077|hsa-mir-21&WithFlank&17|+|59841260|59841343</td><td>14</td><td>37</td><td>24(+1U)</td><td>TAGCTTATCAGACTGATGTTGACTT</td><td>66.3</td><td>-9.4</td><td>2.1</td><td>-4.6</td><td>5.3E-06</td><td>6.4E-05</td><td>BT20/MB231</td><td>MP</td></tr>
<tr><td>>hg38_21_+_8433172_8446622_RNA45SN1_45S_50ntFlanks@8017.8042.26|TATTAGTCAGCGGAGGAGAAGAAACT</td><td>>hg38_21_+_8433172_8446622_RNA45SN1_45S_50ntFlanks</td><td>8017</td><td>8042</td><td>26</td><td>TATTAGTCAGCGGAGGAGAAGAAACT</td><td>393.2</td><td>-4.4</td><td>1.0</td><td>-4.6</td><td>5.0E-06</td><td>6.4E-05</td><td>BT20/MB231</td><td>CYTO</td></tr>
<tr><td>>hg38_21_+_8433172_8446622_RNA45SN1_45S_50ntFlanks@6664.6700.37|TGGATCACTCGGCTCGTGCGTCGATGAAGAACGCAGC</td><td>>hg38_21_+_8433172_8446622_RNA45SN1_45S_50ntFlanks</td><td>6664</td><td>6700</td><td>37</td><td>TGGATCACTCGGCTCGTGCGTCGATGAAGAACGCAGC</td><td>167.2</td><td>7.6</td><td>1.8</td><td>4.3</td><td>1.6E-05</td><td>6.5E-05</td><td>BT20/MB468</td><td>MP</td></tr>
<tr><td>>am_trnaMT_ValTAC_MT_+_1602_1670@32.67.36|TACACTTAGGAGATTTCAACTTAACTTGACCGCTCT</td><td>>am_trnaMT_ValTAC_MT_+_1602_1670</td><td>32</td><td>67</td><td>36</td><td>TACACTTAGGAGATTTCAACTTAACTTGACCGCTCT</td><td>174.2</td><td>11.2</td><td>2.6</td><td>4.3</td><td>1.6E-05</td><td>6.5E-05</td><td>BT20/MB468</td><td>MP</td></tr>
<tr><td>>MI0005756|hsa-mir-934&WithFlank&X|+|136550872|136550966@21.42.22(+1C)&[MIMAT0004977&hsa-miR-934&offsets|0|0(+1C);m-943&X|+|136550892|136550913&offsets|0|0(+1C)]|TGTCTACTACTGGAGACACTGGC</td><td>>MI0005756|hsa-mir-934&WithFlank&X|+|136550872|136550966</td><td>21</td><td>42</td><td>22(+1C)</td><td>TGTCTACTACTGGAGACACTGGC</td><td>63.8</td><td>-9.9</td><td>2.4</td><td>-4.1</td><td>4.0E-05</td><td>6.5E-05</td><td>MB231/MB468</td><td>MP</td></tr>
<tr><td>>hg38_21_+_8433172_8446622_RNA45SN1_45S_50ntFlanks@8177.8204.28|TTCTGATCGAGGCCCAGCCCGTGGACGG</td><td>>hg38_21_+_8433172_8446622_RNA45SN1_45S_50ntFlanks</td><td>8177</td><td>8204</td><td>28</td><td>TTCTGATCGAGGCCCAGCCCGTGGACGG</td><td>266.6</td><td>-5.0</td><td>1.2</td><td>-4.1</td><td>4.0E-05</td><td>6.5E-05</td><td>MB231/MB468</td><td>MP</td></tr>
<tr><td>>hg38_21_+_8433172_8446622_RNA45SN1_45S_50ntFlanks@10402.10431.30|AACATGGGTCAGTCGGTCCTGAGAGATGGG</td><td>>hg38_21_+_8433172_8446622_RNA45SN1_45S_50ntFlanks</td><td>10402</td><td>10431</td><td>30</td><td>AACATGGGTCAGTCGGTCCTGAGAGATGGG</td><td>418.3</td><td>-2.9</td><td>0.7</td><td>-4.1</td><td>3.5E-05</td><td>6.5E-05</td><td>BT20/MB468</td><td>MITO</td></tr>
<tr><td>>trnaMT_MetCAT_MT_+_4402_4469@53.71.19|TACCCTTCCCGTACTACCA</td><td>>trnaMT_MetCAT_MT_+_4402_4469</td><td>53</td><td>71</td><td>19</td><td>TACCCTTCCCGTACTACCA</td><td>242.1</td><td>5.3</td><td>1.3</td><td>4.1</td><td>4.7E-05</td><td>6.5E-05</td><td>MB231/MB468</td><td>NUC</td></tr>
<tr><td>>hg38_21_+_8433172_8446622_RNA45SN1_45S_50ntFlanks@9481.9515.35|CACGTGTTAGGACCCGAAAGATGGTGAACTATGCC</td><td>>hg38_21_+_8433172_8446622_RNA45SN1_45S_50ntFlanks</td><td>9481</td><td>9515</td><td>35</td><td>CACGTGTTAGGACCCGAAAGATGGTGAACTATGCC</td><td>106.0</td><td>10.1</td><td>2.2</td><td>4.6</td><td>5.3E-06</td><td>6.5E-05</td><td>BT20/MB231</td><td>MP</td></tr>
<tr><td>>am_MI0000809|hsa-mir-151a&WithFlank&8|-|140732558|140732659@53.73.21(+2U)&[MIMAT0000757&hsa-miR-151a-3p&offsets|0|0(+2U);m-30&8|-|140732587|140732607&offsets|0|0(+2U)]|CTAGACTGAAGCTCCTTGAGGTT</td><td>>am_MI0000809|hsa-mir-151a&WithFlank&8|-|140732558|140732659</td><td>53</td><td>73</td><td>21(+2U)</td><td>CTAGACTGAAGCTCCTTGAGGTT</td><td>634.1</td><td>6.1</td><td>1.4</td><td>4.3</td><td>1.6E-05</td><td>6.5E-05</td><td>BT20/MB468</td><td>MP</td></tr>
<tr><td>>hg38_21_+_8433172_8446622_RNA45SN1_45S_50ntFlanks@6645.6685.41|CTCGTACGACTCTTAGCGGTGGATCACTCGGCTCGTGCGTC</td><td>>hg38_21_+_8433172_8446622_RNA45SN1_45S_50ntFlanks</td><td>6645</td><td>6685</td><td>41</td><td>CTCGTACGACTCTTAGCGGTGGATCACTCGGCTCGTGCGTC</td><td>667.6</td><td>3.5</td><td>0.8</td><td>4.5</td><td>5.8E-06</td><td>6.5E-05</td><td>BT20/MB231</td><td>NUC</td></tr>
<tr><td>>hg38_21_+_8433172_8446622_RNA45SN1_45S_50ntFlanks@10391.10426.36|AACAGCAGTTGAACATGGGTCAGTCGGTCCTGAGAG</td><td>>hg38_21_+_8433172_8446622_RNA45SN1_45S_50ntFlanks</td><td>10391</td><td>10426</td><td>36</td><td>AACAGCAGTTGAACATGGGTCAGTCGGTCCTGAGAG</td><td>337.9</td><td>-4.0</td><td>1.0</td><td>-4.1</td><td>4.5E-05</td><td>6.5E-05</td><td>MB231/MB468</td><td>CYTO</td></tr>
<tr><td>>hg38_MT_+_1621_3279_RNR2_16S_50ntFlanks@498.531.34|TTGGACACTAGGAAAAAACCTTGTAGAGAGAGTA</td><td>>hg38_MT_+_1621_3279_RNR2_16S_50ntFlanks</td><td>498</td><td>531</td><td>34</td><td>TTGGACACTAGGAAAAAACCTTGTAGAGAGAGTA</td><td>193.4</td><td>11.3</td><td>2.6</td><td>4.3</td><td>1.6E-05</td><td>6.5E-05</td><td>BT20/MB468</td><td>MP</td></tr>
<tr><td>>am_trnaMT_PheGAA_MT_+_577_647@41.60.20|GTTTAGACGGGCTCACATCA</td><td>>am_trnaMT_PheGAA_MT_+_577_647</td><td>41</td><td>60</td><td>20</td><td>GTTTAGACGGGCTCACATCA</td><td>204.6</td><td>4.9</td><td>1.2</td><td>4.1</td><td>4.8E-05</td><td>6.5E-05</td><td>MB231/MB468</td><td>MITO</td></tr>
<tr><td>>hg38_21_+_8433172_8446622_RNA45SN1_45S_50ntFlanks@11316.11343.28|CACCCCACGTCTCGTCGCGCGCGCGTCC</td><td>>hg38_21_+_8433172_8446622_RNA45SN1_45S_50ntFlanks</td><td>11316</td><td>11343</td><td>28</td><td>CACCCCACGTCTCGTCGCGCGCGCGTCC</td><td>620.9</td><td>-4.4</td><td>1.1</td><td>-4.1</td><td>4.1E-05</td><td>6.5E-05</td><td>BT20/MB468</td><td>CYTO</td></tr>
<tr><td>>MI0000542|hsa-mir-320a&WithFlank&8|-|22244960|22245043@48.68.21(+1U)&[MIMAT0000510&hsa-miR-320a-3p&offsets|0|-1(+1U);m-60&8|-|22244975|22244996&offsets|0|-1(+1U)]|AAAAGCTGGGTTGAGAGGGCGT</td><td>>MI0000542|hsa-mir-320a&WithFlank&8|-|22244960|22245043</td><td>48</td><td>68</td><td>21(+1U)</td><td>AAAAGCTGGGTTGAGAGGGCGT</td><td>348.9</td><td>6.2</td><td>1.5</td><td>4.1</td><td>4.8E-05</td><td>6.5E-05</td><td>MB231/MB468</td><td>MITO</td></tr>
</table>

<table>
<tr><th></th><th></th><th></th><th></th><th></th><th></th><th></th><th></th><th></th><th></th><th></th><th></th><th></th><th></th></tr>
<tr><td>>MI0000289|hsa-mir-181a-1&WithFlank&1|-|198859038|198859159@30.51.22;MI0000269|hsa-mir-181a-2&WithFlank&9|+|124692436|124692557@45.66.22&[MIMAT0000256&hsa-miR-181a-5p&offsets|0|-1;m-32&1|-|198859108|198859130&offsets|0|-1];[MIMAT0000256_1&hsa-miR-181a-5p&offsets|0|-1;m-32&9|+|124692480|124692502&offsets|0|-1]|AACATTCAACGCTGTCGGTGAG</td><td>>MI0000289|hsa-mir-181a-1&WithFlank&1|-|198859038|198859159</td><td>30</td><td>51</td><td>22</td><td>AACATTCAACGCTGTCGGTGAG</td><td>476.5</td><td>4.1</td><td>1.0</td><td>4.1</td><td>4.5E-05</td><td>6.5E-05</td><td>MB231/MB468</td><td>CYTO</td></tr>
<tr><td>>MI0000767|hsa-mir-365a&WithFlank&16|+|14309279|14309377@62.82.21(+1C);MI0000769|hsa-mir-365b&WithFlank&17|+|31575405|31575527@74.94.21(+1C)&[MIMAT0000710&hsa-miR-365a-3p&offsets|0|-1(+1C);m-176&16|+|14309340|14309361&offsets|0|-1(+1C)];[MIMAT0022834&hsa-miR-365b-3p&offsets|0|-1(+1C);m-175&17|+|31575478|31575499&offsets|0|-1(+1C)]|TAATGCCCCTAAAAATCCTTAC</td><td>>MI0000767|hsa-mir-365a&WithFlank&16|+|14309279|14309377</td><td>62</td><td>82</td><td>21(+1C)</td><td>TAATGCCCCTAAAAATCCTTAC</td><td>207.6</td><td>5.2</td><td>1.2</td><td>4.5</td><td>5.9E-06</td><td>6.5E-05</td><td>BT20/MB231</td><td>NUC</td></tr>
<tr><td>>hg38_21_+_8433172_8446622_RNA45SN1_45S_50ntFlanks@11188.11207.20|CACCCGGGGGGCCGGCGGCG</td><td>>hg38_21_+_8433172_8446622_RNA45SN1_45S_50ntFlanks</td><td>11188</td><td>11207</td><td>20</td><td>CACCCGGGGGGCCGGCGGCG</td><td>452.2</td><td>-6.9</td><td>1.7</td><td>-4.1</td><td>5.0E-05</td><td>6.5E-05</td><td>BT20/MB468</td><td>NUC</td></tr>
<tr><td>>hg38_21_+_8433172_8446622_RNA45SN1_45S_50ntFlanks@9605.9624.20|AAGACTAATCGAACCATCTA</td><td>>hg38_21_+_8433172_8446622_RNA45SN1_45S_50ntFlanks</td><td>9605</td><td>9624</td><td>20</td><td>AAGACTAATCGAACCATCTA</td><td>58.0</td><td>-9.7</td><td>2.4</td><td>-4.1</td><td>4.0E-05</td><td>6.5E-05</td><td>MB231/MB468</td><td>MP</td></tr>
<tr><td>>hg38_21_+_8433172_8446622_RNA45SN1_45S_50ntFlanks@3988.4023.36|CTTTGGTGACTCTAGATAACCTCGGGCCGATCGCAC</td><td>>hg38_21_+_8433172_8446622_RNA45SN1_45S_50ntFlanks</td><td>3988</td><td>4023</td><td>36</td><td>CTTTGGTGACTCTAGATAACCTCGGGCCGATCGCAC</td><td>1528.1</td><td>5.6</td><td>1.3</td><td>4.3</td><td>1.6E-05</td><td>6.5E-05</td><td>BT20/MB468</td><td>MP</td></tr>
<tr><td>>hg38_21_+_8433172_8446622_RNA45SN1_45S_50ntFlanks@11783.11818.36|CATGAATGGATGAACGAGATTCCCACTGTCCCTACC</td><td>>hg38_21_+_8433172_8446622_RNA45SN1_45S_50ntFlanks</td><td>11783</td><td>11818</td><td>36</td><td>CATGAATGGATGAACGAGATTCCCACTGTCCCTACC</td><td>256.8</td><td>-4.6</td><td>1.1</td><td>-4.1</td><td>4.8E-05</td><td>6.5E-05</td><td>MB231/MB468</td><td>MITO</td></tr>
<tr><td>>MI0000264|hsa-mir-7-2&WithFlank&15|+|88611819|88611940@38.60.23(+1A);MI0000265|hsa-mir-7-3&WithFlank&19|+|4770664|4770785@37.59.23(+1A);MI0000263|hsa-mir-7-1&WithFlank&9|-|83969742|83969863@30.52.23(+1A)&[MIMAT0000252&hsa-miR-7-5p&offsets|0|-1(+1A);m-400&15|+|88611856|88611879&offsets|0|-1(+1A)];[MIMAT0000252_1&hsa-miR-7-5p&offsets|0|-1(+1A);m-402&19|+|4770700|4770723&offsets|0|-1(+1A)];[MIMAT0000252_2&hsa-miR-7-5p&offsets|0|-1(+1A);m-401&9|-|83969811|83969834&offsets|0|-1(+1A)]|TGGAAGACTAGTGATTTTGTTGTA</td><td>>MI0000264|hsa-mir-7-2&WithFlank&15|+|88611819|88611940</td><td>38</td><td>60</td><td>23(+1A)</td><td>TGGAAGACTAGTGATTTTGTTGTA</td><td>1273.9</td><td>3.6</td><td>0.9</td><td>4.1</td><td>4.0E-05</td><td>6.5E-05</td><td>MB231/MB468</td><td>MP</td></tr>
<tr><td>>trna116_GluCTC_1_-_145399233_145399304@41.74.34,trna134_GluTTC_1_-_16861774_16861845@41.74.34,trna71_GluCTC_1_-_161439189_161439260@41.74.34,trna74_GluCTC_1_-_161431809_161431880@41.74.34,trna77_GluCTC_1_-_161424398_161424469@41.74.34,trna77_GluCTC_6_+_28949976_28950047@41.74.34,trna80_GluCTC_1_-_161417018_161417089@41.74.34,trna84_GluTTC_1_-_161391883_161391954@41.74.34,trna87_GluCTC_6_-_126101393_126101464@41.74.34|CCGCGGCCCGGGTTCGATTCCCGGTCAGGGAACC</td><td>>trna116_GluCTC_1_-_145399233_145399304</td><td>41</td><td>74</td><td>34</td><td>CCGCGGCCCGGGTTCGATTCCCGGTCAGGGAACC</td><td>287.1</td><td>4.7</td><td>1.2</td><td>4.1</td><td>4.8E-05</td><td>6.5E-05</td><td>MB231/MB468</td><td>MITO</td></tr>
<tr><td>>hg38_21_+_8433172_8446622_RNA45SN1_45S_50ntFlanks@11027.11059.33|CTCTCTCTCTCTCTCCCCCGCTCCCCGTCCTCC</td><td>>hg38_21_+_8433172_8446622_RNA45SN1_45S_50ntFlanks</td><td>11027</td><td>11059</td><td>33</td><td>CTCTCTCTCTCTCTCCCCCGCTCCCCGTCCTCC</td><td>232.6</td><td>-5.1</td><td>1.2</td><td>-4.1</td><td>4.8E-05</td><td>6.5E-05</td><td>MB231/MB468</td><td>MITO</td></tr>
<tr><td>>MI0000439|hsa-mir-23b&WithFlank&9|+|95085202|95085310@67.86.20(+1A)&[MIMAT0000418&hsa-miR-23b-3p&offsets|+3|0(+1A);m-42&9|+|95085265|95085285&offsets|+3|+2(+1A)]|ACATTGCCAGGGATTACCACA</td><td>>MI0000439|hsa-mir-23b&WithFlank&9|+|95085202|95085310</td><td>67</td><td>86</td><td>20(+1A)</td><td>ACATTGCCAGGGATTACCACA</td><td>132.2</td><td>10.3</td><td>2.5</td><td>4.1</td><td>4.0E-05</td><td>6.5E-05</td><td>MB231/MB468</td><td>MP</td></tr>
<tr><td>>hg38_21_+_8433172_8446622_RNA45SN1_45S_50ntFlanks@4983.5001.19|CGGACAGGATTGACAGATT</td><td>>hg38_21_+_8433172_8446622_RNA45SN1_45S_50ntFlanks</td><td>4983</td><td>5001</td><td>19</td><td>CGGACAGGATTGACAGATT</td><td>163.1</td><td>-3.5</td><td>0.9</td><td>-4.1</td><td>4.0E-05</td><td>6.5E-05</td><td>MB231/MB468</td><td>MP</td></tr>
<tr><td>>hg38_21_+_8433172_8446622_RNA45SN1_45S_50ntFlanks@6661.6703.43|CGGTGGATCACTCGGCTCGTGCGTCGATGAAGAACGCAGCTAG</td><td>>hg38_21_+_8433172_8446622_RNA45SN1_45S_50ntFlanks</td><td>6661</td><td>6703</td><td>43</td><td>CGGTGGATCACTCGGCTCGTGCGTCGATGAAGAACGCAGCTAG</td><td>459.1</td><td>3.4</td><td>0.8</td><td>4.1</td><td>4.8E-05</td><td>6.5E-05</td><td>MB231/MB468</td><td>NUC</td></tr>
<tr><td>>hg38_21_+_8433172_8446622_RNA45SN1_45S_50ntFlanks@6721.6738.18|AATTGCAGGACACATTGA</td><td>>hg38_21_+_8433172_8446622_RNA45SN1_45S_50ntFlanks</td><td>6721</td><td>6738</td><td>18</td><td>AATTGCAGGACACATTGA</td><td>254.9</td><td>-6.5</td><td>1.5</td><td>-4.3</td><td>1.6E-05</td><td>6.5E-05</td><td>BT20/MB468</td><td>MP</td></tr>
<tr><td>>hg38_21_+_8433172_8446622_RNA45SN1_45S_50ntFlanks@7985.8009.25|GATCAGACGTGGCGACCCGCTGAAT</td><td>>hg38_21_+_8433172_8446622_RNA45SN1_45S_50ntFlanks</td><td>7985</td><td>8009</td><td>25</td><td>GATCAGACGTGGCGACCCGCTGAAT</td><td>461.4</td><td>-6.1</td><td>1.5</td><td>-4.1</td><td>4.8E-05</td><td>6.5E-05</td><td>MB231/MB468</td><td>MITO</td></tr>
<tr><td>>hg38_21_+_8433172_8446622_RNA45SN1_45S_50ntFlanks@12994.13039.46|CCTCGCTGCGATCTATTGAAAGTCAGCCCTCGACACAAGGGTTTGT</td><td>>hg38_21_+_8433172_8446622_RNA45SN1_45S_50ntFlanks</td><td>12994</td><td>13039</td><td>46</td><td>CCTCGCTGCGATCTATTGAAAGTCAGCCCTCGACACAAGGGTTTGT</td><td>7686.3</td><td>5.2</td><td>1.3</td><td>4.1</td><td>4.2E-05</td><td>6.5E-05</td><td>BT20/MB468</td><td>CYTO</td></tr>
<tr><td>>hg38_21_+_8433172_8446622_RNA45SN1_45S_50ntFlanks@7987.8009.23|TCAGACGTGGCGACCCGCTGAAT</td><td>>hg38_21_+_8433172_8446622_RNA45SN1_45S_50ntFlanks</td><td>7987</td><td>8009</td><td>23</td><td>TCAGACGTGGCGACCCGCTGAAT</td><td>201.2</td><td>-5.0</td><td>1.2</td><td>-4.1</td><td>4.0E-05</td><td>6.5E-05</td><td>MB231/MB468</td><td>MP</td></tr>
<tr><td>>hg38_21_+_8433172_8446622_RNA45SN1_45S_50ntFlanks@11318.11343.26|CCCCACGTCTCGTCGCGCGCGCGTCC</td><td>>hg38_21_+_8433172_8446622_RNA45SN1_45S_50ntFlanks</td><td>11318</td><td>11343</td><td>26</td><td>CCCCACGTCTCGTCGCGCGCGCGTCC</td><td>4576.1</td><td>-8.2</td><td>2.0</td><td>-4.1</td><td>5.0E-05</td><td>6.5E-05</td><td>BT20/MB468</td><td>NUC</td></tr>
<tr><td>>hg38_21_+_8433172_8446622_RNA45SN1_45S_50ntFlanks@12397.12418.22|CGACGTCGCTTTTTGATCCTTC</td><td>>hg38_21_+_8433172_8446622_RNA45SN1_45S_50ntFlanks</td><td>12397</td><td>12418</td><td>22</td><td>CGACGTCGCTTTTTGATCCTTC</td><td>2699.3</td><td>3.0</td><td>0.7</td><td>4.1</td><td>4.5E-05</td><td>6.5E-05</td><td>MB231/MB468</td><td>CYTO</td></tr>
<tr><td>>hg38_21_+_8433172_8446622_RNA45SN1_45S_50ntFlanks@11565.11600.36|CTAGCAGCCGACTTAGAACTGGTGCGGACCAGGGGA</td><td>>hg38_21_+_8433172_8446622_RNA45SN1_45S_50ntFlanks</td><td>11565</td><td>11600</td><td>36</td><td>CTAGCAGCCGACTTAGAACTGGTGCGGACCAGGGGA</td><td>514.7</td><td>-6.9</td><td>1.7</td><td>-4.1</td><td>5.0E-05</td><td>6.5E-05</td><td>BT20/MB468</td><td>NUC</td></tr>
<tr><td>>am_trna2_GlyGCC_21_-_18827107_18827177@1.25.25,trna35_GlyGCC_1_+_161413094_161413164@1.25.25,trna37_GlyGCC_1_+_161420467_161420537@1.25.25,trna39_GlyGCC_1_+_161427898_161427968@1.25.25,trna41_GlyGCC_1_+_161435258_161435328@1.25.25|GCATGGGTGGTTCAGTGGTAGAATT</td><td>>am_trna2_GlyGCC_21_-_18827107_18827177</td><td>1</td><td>25</td><td>25</td><td>GCATGGGTGGTTCAGTGGTAGAATT</td><td>4750.5</td><td>5.0</td><td>1.2</td><td>4.3</td><td>1.6E-05</td><td>6.6E-05</td><td>BT20/MB468</td><td>MP</td></tr>
<tr><td>>hg38_MT_+_598_1651_RNR1_12S_50ntFlanks@417.455.39|AATAGACTACGAAAGTGGCTTTAACATATCTGAACACAC</td><td>>hg38_MT_+_598_1651_RNR1_12S_50ntFlanks</td><td>417</td><td>455</td><td>39</td><td>AATAGACTACGAAAGTGGCTTTAACATATCTGAACACAC</td><td>26.6</td><td>8.0</td><td>1.9</td><td>4.1</td><td>4.0E-05</td><td>6.6E-05</td><td>MB231/MB468</td><td>MP</td></tr>
<tr><td>>trnaMT_GluTTC_MT_-_14674_14742@24.61.38,trnalookalike8_GluTTC_5_-_93905172_93905240@24.61.38|ATGGTTTTTCATATCATTGGTCGTGGTTGTAGTCCGTG</td><td>>trnaMT_GluTTC_MT_-_14674_14742</td><td>24</td><td>61</td><td>38</td><td>ATGGTTTTTCATATCATTGGTCGTGGTTGTAGTCCGTG</td><td>187.8</td><td>8.4</td><td>1.9</td><td>4.3</td><td>1.6E-05</td><td>6.6E-05</td><td>BT20/MB468</td><td>MP</td></tr>
<tr><td>>hg38_21_+_8433172_8446622_RNA45SN1_45S_50ntFlanks@11847.11873.27|AACGGGCTTGGCGGAATCAGCGGGGAA</td><td>>hg38_21_+_8433172_8446622_RNA45SN1_45S_50ntFlanks</td><td>11847</td><td>11873</td><td>27</td><td>AACGGGCTTGGCGGAATCAGCGGGGAA</td><td>459.4</td><td>-3.5</td><td>0.8</td><td>-4.3</td><td>1.7E-05</td><td>6.6E-05</td><td>BT20/MB468</td><td>MP</td></tr>
<tr><td>>hg38_21_+_8433172_8446622_RNA45SN1_45S_50ntFlanks@8115.8138.24|CGCGGGACATGTGGCGTACGGAAG</td><td>>hg38_21_+_8433172_8446622_RNA45SN1_45S_50ntFlanks</td><td>8115</td><td>8138</td><td>24</td><td>CGCGGGACATGTGGCGTACGGAAG</td><td>234.6</td><td>-5.8</td><td>1.4</td><td>-4.1</td><td>4.8E-05</td><td>6.6E-05</td><td>MB231/MB468</td><td>NUC</td></tr>
<tr><td>>MI0003513|hsa-mir-455&WithFlank&9|+|114209428|114209535@60.80.21&[MIMAT0004784&hsa-miR-455-3p&offsets|0|0;m-135&9|+|114209487|114209508&offsets|0|-1]|GCAGTCCATGGGCATATACAC</td><td>>MI0003513|hsa-mir-455&WithFlank&9|+|114209428|114209535</td><td>60</td><td>80</td><td>21</td><td>GCAGTCCATGGGCATATACAC</td><td>72.2</td><td>-9.6</td><td>2.1</td><td>-4.5</td><td>5.4E-06</td><td>6.6E-05</td><td>BT20/MB231</td><td>MP</td></tr>
<tr><td>>hg38_21_+_8433172_8446622_RNA45SN1_45S_50ntFlanks@12337.12370.34|AGGAGGTGTCAGAAAAGTTACCACAGGGATAACT</td><td>>hg38_21_+_8433172_8446622_RNA45SN1_45S_50ntFlanks</td><td>12337</td><td>12370</td><td>34</td><td>AGGAGGTGTCAGAAAAGTTACCACAGGGATAACT</td><td>96.0</td><td>9.9</td><td>2.2</td><td>4.5</td><td>5.4E-06</td><td>6.6E-05</td><td>BT20/MB231</td><td>MP</td></tr>
<tr><td>>hg38_21_+_8433172_8446622_RNA45SN1_45S_50ntFlanks@12936.12969.34|AAACCATTCGTAGACGACCTGCTTCTGGGTCGGG</td><td>>hg38_21_+_8433172_8446622_RNA45SN1_45S_50ntFlanks</td><td>12936</td><td>12969</td><td>34</td><td>AAACCATTCGTAGACGACCTGCTTCTGGGTCGGG</td><td>183.3</td><td>-5.7</td><td>1.4</td><td>-4.1</td><td>4.2E-05</td><td>6.6E-05</td><td>BT20/MB468</td><td>CYTO</td></tr>
<tr><td>>hg38_21_+_8433172_8446622_RNA45SN1_45S_50ntFlanks@7975.8020.46|CGCGACCTCAGATCAGACGTGGCGACCCGCTGAATTTAAGCATATT</td><td>>hg38_21_+_8433172_8446622_RNA45SN1_45S_50ntFlanks</td><td>7975</td><td>8020</td><td>46</td><td>CGCGACCTCAGATCAGACGTGGCGACCCGCTGAATTTAAGCATATT</td><td>631.3</td><td>4.3</td><td>1.1</td><td>4.1</td><td>4.8E-05</td><td>6.6E-05</td><td>MB231/MB468</td><td>MITO</td></tr>
<tr><td>>MI0000088|hsa-mir-30a&WithFlank&6|-|71403545|71403627@12.33.22(+1U)&[MIMAT0000087&hsa-miR-30a-5p&offsets|0|0(+1U);m-15&6|-|71403595|71403616&offsets|0|0(+1U)]|TGTAAACATCCTCGACTGGAAGT</td><td>>MI0000088|hsa-mir-30a&WithFlank&6|-|71403545|71403627</td><td>12</td><td>33</td><td>22(+1U)</td><td>TGTAAACATCCTCGACTGGAAGT</td><td>13578.3</td><td>3.6</td><td>0.9</td><td>4.1</td><td>4.0E-05</td><td>6.6E-05</td><td>MB231/MB468</td><td>MP</td></tr>
<tr><td>>hg38_21_+_8433172_8446622_RNA45SN1_45S_50ntFlanks@7977.7996.20|CGACCTCAGATCAGACGTGG</td><td>>hg38_21_+_8433172_8446622_RNA45SN1_45S_50ntFlanks</td><td>7977</td><td>7996</td><td>20</td><td>CGACCTCAGATCAGACGTGG</td><td>198.0</td><td>-6.1</td><td>1.5</td><td>-4.1</td><td>4.1E-05</td><td>6.6E-05</td><td>MB231/MB468</td><td>MP</td></tr>
<tr><td>>hg38_21_+_8433172_8446622_RNA45SN1_45S_50ntFlanks@6659.6706.48|AGCGGTGGATCACTCGGCTCGTGCGTCGATGAAGAACGCAGCTAGCTG</td><td>>hg38_21_+_8433172_8446622_RNA45SN1_45S_50ntFlanks</td><td>6659</td><td>6706</td><td>48</td><td>AGCGGTGGATCACTCGGCTCGTGCGTCGATGAAGAACGCAGCTAGCTG</td><td>329.0</td><td>5.1</td><td>1.2</td><td>4.1</td><td>3.5E-05</td><td>6.6E-05</td><td>BT20/MB468</td><td>MITO</td></tr>
<tr><td>>hg38_21_+_8433172_8446622_RNA45SN1_45S_50ntFlanks@8079.8104.26|AAGAGCCCAGCGCCGAATCCCCGCCC</td><td>>hg38_21_+_8433172_8446622_RNA45SN1_45S_50ntFlanks</td><td>8079</td><td>8104</td><td>26</td><td>AAGAGCCCAGCGCCGAATCCCCGCCC</td><td>904.9</td><td>-6.2</td><td>1.5</td><td>-4.1</td><td>3.5E-05</td><td>6.6E-05</td><td>BT20/MB468</td><td>MITO</td></tr>
<tr><td>>MI0000809|hsa-mir-151a&WithFlank&8|-|140732558|140732659@51.71.21&[MIMAT0000757&hsa-miR-151a-3p&offsets|-2|-2;m-30&8|-|140732587|140732607&offsets|-2|-2]|TACTAGACTGAAGCTCCTTGA</td><td>>MI0000809|hsa-mir-151a&WithFlank&8|-|140732558|140732659</td><td>51</td><td>71</td><td>21</td><td>TACTAGACTGAAGCTCCTTGA</td><td>208.7</td><td>7.0</td><td>1.6</td><td>4.3</td><td>1.7E-05</td><td>6.6E-05</td><td>BT20/MB468</td><td>MP</td></tr>
<tr><td>>hg38_21_+_8433172_8446622_RNA45SN1_45S_50ntFlanks@10783.10826.44|GTGAACAGCCTCTGGCATGTTGGAACAATGTAGGTAAGGGAAGT</td><td>>hg38_21_+_8433172_8446622_RNA45SN1_45S_50ntFlanks</td><td>10783</td><td>10826</td><td>44</td><td>GTGAACAGCCTCTGGCATGTTGGAACAATGTAGGTAAGGGAAGT</td><td>53.5</td><td>-9.6</td><td>2.3</td><td>-4.1</td><td>4.1E-05</td><td>6.6E-05</td><td>MB231/MB468</td><td>MP</td></tr>
<tr><td>>hg38_21_+_8433172_8446622_RNA45SN1_45S_50ntFlanks@6651.6682.32|CGACTCTTAGCGGTGGATCACTCGGCTCGTGC</td><td>>hg38_21_+_8433172_8446622_RNA45SN1_45S_50ntFlanks</td><td>6651</td><td>6682</td><td>32</td><td>CGACTCTTAGCGGTGGATCACTCGGCTCGTGC</td><td>1798.4</td><td>4.6</td><td>1.1</td><td>4.1</td><td>4.2E-05</td><td>6.6E-05</td><td>BT20/MB468</td><td>CYTO</td></tr>
<tr><td>>MI0003591|hsa-mir-584&WithFlank&5|-|149062307|149062415@22.40.19(+1A)&[MIMAT0003249&hsa-miR-584-5p&offsets|0|-3(+1A);m-167&5|-|149062374|149062394&offsets|0|-2(+1A)]|TTATGGTTTGCCTGGGACTA</td><td>>MI0003591|hsa-mir-584&WithFlank&5|-|149062307|149062415</td><td>22</td><td>40</td><td>19(+1A)</td><td>TTATGGTTTGCCTGGGACTA</td><td>172.1</td><td>7.8</td><td>1.8</td><td>4.3</td><td>1.7E-05</td><td>6.6E-05</td><td>BT20/MB468</td><td>MP</td></tr>
<tr><td>>hg38_21_+_8433172_8446622_RNA45SN1_45S_50ntFlanks@8472.8490.19|CGGCCCGGCGGATCTTTCC</td><td>>hg38_21_+_8433172_8446622_RNA45SN1_45S_50ntFlanks</td><td>8472</td><td>8490</td><td>19</td><td>CGGCCCGGCGGATCTTTCC</td><td>241.5</td><td>-5.8</td><td>1.4</td><td>-4.1</td><td>3.5E-05</td><td>6.6E-05</td><td>BT20/MB468</td><td>MITO</td></tr>
<tr><td>>trna112_GlnCTG_1_-_147737382_147737453@31.50.20,trna131_GlnCTG_6_-_27759135_27759206@31.50.20,trna146_GlnCTG_6_-_27515531_27515602@31.50.20,trna15_GlnCTG_1_+_145963304_145963375@31.50.20,trna19_GlnCTG_1_+_147505038_147505109@31.50.20,trna1_GlnCTG_6_+_18836402_18836473@31.50.20,trna28_GlnCTG_1_+_149186125_149186196@31.50.20,trna3_GlnCTG_17_+_8023070_8023141@31.50.20,trna49_GlnCTG_6_+_27487308_27487379@31.50.20,trna7_GlnCTG_15_-_66161400_66161471@31.50.20,trna99_GlnCTG_6_-_28909378_28909449@31.50.20|ACTCTGAATCCAGCGATCCG</td><td>>trna112_GlnCTG_1_-_147737382_147737453</td><td>31</td><td>50</td><td>20</td><td>ACTCTGAATCCAGCGATCCG</td><td>132.8</td><td>-5.1</td><td>1.2</td><td>-4.1</td><td>4.2E-05</td><td>6.6E-05</td><td>BT20/MB468</td><td>CYTO</td></tr>
<tr><td>>hg38_21_+_8433172_8446622_RNA45SN1_45S_50ntFlanks@9788.9819.32|CGCTGGCGTGGAGCCGGGCGTGGAATGCGAGT</td><td>>hg38_21_+_8433172_8446622_RNA45SN1_45S_50ntFlanks</td><td>9788</td><td>9819</td><td>32</td><td>CGCTGGCGTGGAGCCGGGCGTGGAATGCGAGT</td><td>845.6</td><td>9.0</td><td>2.0</td><td>4.5</td><td>5.5E-06</td><td>6.6E-05</td><td>BT20/MB231</td><td>MP</td></tr>
<tr><td>>am_trnaMT_ProTGG_MT_-_15956_16023@54.71.18|AGACTTTTTCTCTGACCA</td><td>>am_trnaMT_ProTGG_MT_-_15956_16023</td><td>54</td><td>71</td><td>18</td><td>AGACTTTTTCTCTGACCA</td><td>2291.3</td><td>4.0</td><td>1.0</td><td>4.1</td><td>4.6E-05</td><td>6.7E-05</td><td>MB231/MB468</td><td>CYTO</td></tr>
</table>

| | | | | | | | | | | | | | |
|---|---|---|---|---|---|---|---|---|---|---|---|---|---|
| >hg38_21_+_8433172_8446622_RNA45SN1_45S_50ntFlanks@12274.12299.26\|CAGTACGAATACAGACCGTGAAAGCG | >hg38_21_+_8433172_8446622_RNA45SN1_45S_50ntFlanks | 12274 | 12299 | 26 | CAGTACGAATACAGACCGTGAAAGCG | 586.4 | -4.8 | 1.2 | -4.1 | 3.6E-05 | 6.7E-05 | BT20/MB468 | MITO |
| >hg38_21_+_8433172_8446622_RNA45SN1_45S_50ntFlanks@5515.5535.21\|CTAGAGGAAGTAAAAGTCGTA | >hg38_21_+_8433172_8446622_RNA45SN1_45S_50ntFlanks | 5515 | 5535 | 21 | CTAGAGGAAGTAAAAGTCGTA | 297.3 | -3.2 | 0.8 | -4.1 | 3.6E-05 | 6.7E-05 | BT20/MB468 | MITO |
| >hg38_1_-_228634819_228635039_RNA5S12_5S_50ntFlanks@51.68.18\|GTCTACGGCCATACCACC | >hg38_1_-_228634819_228635039_RNA5S12_5S_50ntFlanks | 51 | 68 | 18 | GTCTACGGCCATACCACC | 1039.6 | 3.7 | 0.9 | 4.1 | 4.2E-05 | 6.7E-05 | BT20/MB468 | CYTO |
| >hg38_21_+_8433172_8446622_RNA45SN1_45S_50ntFlanks@9929.9949.21\|GATATAGACAGCAGGACGGTG | >hg38_21_+_8433172_8446622_RNA45SN1_45S_50ntFlanks | 9929 | 9949 | 21 | GATATAGACAGCAGGACGGTG | 498.9 | 4.6 | 1.0 | 4.5 | 5.5E-06 | 6.7E-05 | BT20/MB231 | MP |
| >MI0000446\|hsa-mir-125b-1&WithFlank&11\|-\|122099751\|122099850@21.42.22(+1A);MI0000470\|hsa-mir-125b-2&WithFlank&21\|+\|16590231\|16590331@23.44.22(+1A)&[MIMAT0000423&hsa-miR-125b-5p&offsets\|0\|0(+1A);m-67&11\|-\|122099809\|122099830&offsets\|0\|0(+1A)];[MIMAT0000423_1&hsa-miR-125b-5p&offsets\|0\|0(+1A);m-68&21\|+\|16590253\|16590274&offsets\|0\|0(+1A)]\|TCCCTGAGACCCTAACTTGTGAA | >MI0000446\|hsa-mir-125b-1&WithFlank&11\|-\|122099751\|122099850 | 21 | 42 | 22(+1A) | TCCCTGAGACCCTAACTTGTGAA | 1562.0 | -6.6 | 1.5 | -4.5 | 5.5E-06 | 6.7E-05 | BT20/MB231 | MP |
| >MI0005544\|hsa-mir-147b&WithFlank&15\|+\|45433044\|45433135@55.73.19(+1A)&[MIMAT0004928&hsa-miR-147b-3p&offsets\|0\|-2(+1A);m-575&15\|+\|45433098\|45433118&offsets\|0\|-2(+1A)]\|GTGTGCGGAAATGCTTCTGA | >MI0005544\|hsa-mir-147b&WithFlank&15\|+\|45433044\|45433135 | 55 | 73 | 19(+1A) | GTGTGCGGAAATGCTTCTGA | 66.2 | -9.4 | 2.1 | -4.5 | 5.5E-06 | 6.7E-05 | BT20/MB231 | MP |
| >hg38_MT_+_1621_3279_RNR2_16S_50ntFlanks@1216.1262.47\|CAACCTCCGAGCAGTACATGCTAAGACTTCACCAGTCAAAGCGAACT | >hg38_MT_+_1621_3279_RNR2_16S_50ntFlanks | 1216 | 1262 | 47 | CAACCTCCGAGCAGTACATGCTAAGACTTCACCAGTCAAAGCGAACT | 502.9 | -5.1 | 1.1 | -4.6 | 5.2E-06 | 6.7E-05 | BT20/MB231 | CYTO |
| >hg38_MT_+_1621_3279_RNR2_16S_50ntFlanks@51.89.39\|GCTAAACCTAGCCCCAAACCCACTCCACCTTACTACCAG | >hg38_MT_+_1621_3279_RNR2_16S_50ntFlanks | 51 | 89 | 39 | GCTAAACCTAGCCCCAAACCCACTCCACCTTACTACCAG | 615.3 | 4.2 | 1.0 | 4.1 | 4.9E-05 | 6.7E-05 | MB231/MB468 | MITO |
| >hg38_MT_+_598_1651_RNR1_12S_50ntFlanks@970.1004.35\|ATGGTAAGTGTACTGGAAAGTGCACTTGGACGAAC | >hg38_MT_+_598_1651_RNR1_12S_50ntFlanks | 970 | 1004 | 35 | ATGGTAAGTGTACTGGAAAGTGCACTTGGACGAAC | 380.3 | 6.0 | 1.5 | 4.1 | 4.9E-05 | 6.7E-05 | MB231/MB468 | MITO |
| >hg38_21_+_8433172_8446622_RNA45SN1_45S_50ntFlanks@12437.12474.38\|CATTGTGAAGCAGAATTCACCAAGCGTTGGATTGTTCA | >hg38_21_+_8433172_8446622_RNA45SN1_45S_50ntFlanks | 12437 | 12474 | 38 | CATTGTGAAGCAGAATTCACCAAGCGTTGGATTGTTCA | 375.3 | -5.1 | 1.3 | -4.1 | 4.9E-05 | 6.7E-05 | MB231/MB468 | NUC |
| >hg38_21_+_8433172_8446622_RNA45SN1_45S_50ntFlanks@10795.10817.23\|TGGCATGTTGGAACAATGTAGGT | >hg38_21_+_8433172_8446622_RNA45SN1_45S_50ntFlanks | 10795 | 10817 | 23 | TGGCATGTTGGAACAATGTAGGT | 172.4 | -2.5 | 0.6 | -4.1 | 4.6E-05 | 6.7E-05 | MB231/MB468 | CYTO |
| >trna10_AlaCGC_6_+_26553731_26553802@1.18.18,trna119_AlaCGC_6_-_28641613_28641684@1.18.18,trna120_AlaAGC_6_-_28626014_28626085@1.18.18,trna13_AlaCGC_2_+_157257281_157257352@1.18.18,trna13_AlaTGC_12_-_125406301_125406372@1.18.18,trna66_AlaTGC_6_+_28611222_28611293@1.18.18,trna8_AlaTGC_12_+_125424512_125424583@1.18.18,trna8_AlaTGC_5_+_180633868_180633939@1.18.18\|GGGGATGTAGCTCAGTGG | >trna10_AlaCGC_6_+_26553731_26553802 | 1 | 18 | 18 | GGGGATGTAGCTCAGTGG | 712.4 | 7.0 | 1.7 | 4.1 | 4.3E-05 | 6.7E-05 | BT20/MB468 | CYTO |
| >hg38_21_+_8433172_8446622_RNA45SN1_45S_50ntFlanks@3855.3881.27\|CTGTGGTAATTCTAGAGCTAATACATG | >hg38_21_+_8433172_8446622_RNA45SN1_45S_50ntFlanks | 3855 | 3881 | 27 | CTGTGGTAATTCTAGAGCTAATACATG | 206.2 | -4.4 | 1.1 | -4.1 | 4.9E-05 | 6.7E-05 | MB231/MB468 | NUC |
| >hg38_21_+_8433172_8446622_RNA45SN1_45S_50ntFlanks@12626.12654.29\|AATGGGGCGAAGCTACCATCTGTGGGATT | >hg38_21_+_8433172_8446622_RNA45SN1_45S_50ntFlanks | 12626 | 12654 | 29 | AATGGGGCGAAGCTACCATCTGTGGGATT | 10939.4 | -6.6 | 1.5 | -4.3 | 1.7E-05 | 6.7E-05 | BT20/MB468 | MP |
| >hg38_21_+_8433172_8446622_RNA45SN1_45S_50ntFlanks@7975.8009.35\|CGCGACCTCAGATCAGACGTGGCGACCCGCTGAAT | >hg38_21_+_8433172_8446622_RNA45SN1_45S_50ntFlanks | 7975 | 8009 | 35 | CGCGACCTCAGATCAGACGTGGCGACCCGCTGAAT | 93255.6 | 4.8 | 1.1 | 4.3 | 1.7E-05 | 6.7E-05 | BT20/MB468 | MP |
| >hg38_21_+_8433172_8446622_RNA45SN1_45S_50ntFlanks@12632.12653.22\|GCGAAGCTACCATCTGTGGGAT | >hg38_21_+_8433172_8446622_RNA45SN1_45S_50ntFlanks | 12632 | 12653 | 22 | GCGAAGCTACCATCTGTGGGAT | 408.6 | -5.7 | 1.4 | -4.1 | 4.3E-05 | 6.7E-05 | BT20/MB468 | CYTO |
| >hg38_1_-_228634819_228635039_RNA5S12_5S_50ntFlanks@136.169.34\|GGAGACCGCCTGGGAATACCGGGTGCTGTAGGCT | >hg38_1_-_228634819_228635039_RNA5S12_5S_50ntFlanks | 136 | 169 | 34 | GGAGACCGCCTGGGAATACCGGGTGCTGTAGGCT | 437.3 | 5.4 | 1.3 | 4.1 | 4.3E-05 | 6.7E-05 | BT20/MB468 | CYTO |
| >hg38_21_+_8433172_8446622_RNA45SN1_45S_50ntFlanks@8113.8138.26\|GGCGCGGGACATGTGGCGTACGGAAG | >hg38_21_+_8433172_8446622_RNA45SN1_45S_50ntFlanks | 8113 | 8138 | 26 | GGCGCGGGACATGTGGCGTACGGAAG | 6242.7 | -4.8 | 1.1 | -4.3 | 1.7E-05 | 6.7E-05 | BT20/MB468 | MP |
| >hg38_21_+_8433172_8446622_RNA45SN1_45S_50ntFlanks@10361.10390.30\|GAAGGCCGAAGTGGAGAAGGGTTCCATGTG | >hg38_21_+_8433172_8446622_RNA45SN1_45S_50ntFlanks | 10361 | 10390 | 30 | GAAGGCCGAAGTGGAGAAGGGTTCCATGTG | 445.9 | -6.1 | 1.5 | -4.1 | 4.3E-05 | 6.7E-05 | BT20/MB468 | CYTO |
| >hg38_21_+_8433172_8446622_RNA45SN1_45S_50ntFlanks@10403.10424.22\|ACATGGGTCAGTCGGTCCTGAG | >hg38_21_+_8433172_8446622_RNA45SN1_45S_50ntFlanks | 10403 | 10424 | 22 | ACATGGGTCAGTCGGTCCTGAG | 1535.7 | 2.9 | 0.7 | 4.1 | 4.9E-05 | 6.7E-05 | MB231/MB468 | NUC |
| >hg38_21_+_8433172_8446622_RNA45SN1_45S_50ntFlanks@6653.6692.40\|ACTCTTAGCGGTGGATCACTCGGCTCGTGCGTCGATGAAG | >hg38_21_+_8433172_8446622_RNA45SN1_45S_50ntFlanks | 6653 | 6692 | 40 | ACTCTTAGCGGTGGATCACTCGGCTCGTGCGTCGATGAAG | 445.2 | 2.7 | 0.7 | 4.1 | 3.6E-05 | 6.7E-05 | BT20/MB468 | MITO |
| >MI0000449\|hsa-mir-132&WithFlank&17\|-\|2049902\|2050014@65.86.22(+1U)&[MIMAT0000426&hsa-miR-132-3p&offsets\|0\|0(+1U);m-138&17\|-\|2049929\|2049950&offsets\|0\|0(+1U)]\|TAACAGTCTACAGCCATGGTCGT | >MI0000449\|hsa-mir-132&WithFlank&17\|-\|2049902\|2050014 | 65 | 86 | 22(+1U) | TAACAGTCTACAGCCATGGTCGT | 229.2 | 7.1 | 1.7 | 4.1 | 4.1E-05 | 6.7E-05 | MB231/MB468 | MP |
| >hg38_21_+_8433172_8446622_RNA45SN1_45S_50ntFlanks@4683.4723.41\|CAAGACGGACCAGAGCGAAAGCATTTGCCAAGAATGTTTTC | >hg38_21_+_8433172_8446622_RNA45SN1_45S_50ntFlanks | 4683 | 4723 | 41 | CAAGACGGACCAGAGCGAAAGCATTTGCCAAGAATGTTTTC | 510.5 | -3.0 | 0.7 | -4.1 | 3.6E-05 | 6.7E-05 | BT20/MB468 | MITO |
| >hg38_21_+_8433172_8446622_RNA45SN1_45S_50ntFlanks@4160.4182.23\|CCACATCCAAGGAAGGCAGCAGG | >hg38_21_+_8433172_8446622_RNA45SN1_45S_50ntFlanks | 4160 | 4182 | 23 | CCACATCCAAGGAAGGCAGCAGG | 129.4 | -5.0 | 1.2 | -4.1 | 4.3E-05 | 6.7E-05 | BT20/MB468 | CYTO |
| >hg38_21_+_8433172_8446622_RNA45SN1_45S_50ntFlanks@474.516.43\|TCTTCGTGATCGATGTGGTGACGTCGTGCTCTCCCGGGCCGGG | >hg38_21_+_8433172_8446622_RNA45SN1_45S_50ntFlanks | 474 | 516 | 43 | TCTTCGTGATCGATGTGGTGACGTCGTGCTCTCCCGGGCCGGG | 760.2 | 3.8 | 0.9 | 4.1 | 4.9E-05 | 6.7E-05 | MB231/MB468 | NUC |
| >hg38_21_+_8433172_8446622_RNA45SN1_45S_50ntFlanks@8380.8407.28\|CAAGAGGGCGTGAAACCGTTAAGAGGTA | >hg38_21_+_8433172_8446622_RNA45SN1_45S_50ntFlanks | 8380 | 8407 | 28 | CAAGAGGGCGTGAAACCGTTAAGAGGTA | 174.1 | -4.3 | 1.1 | -4.1 | 4.9E-05 | 6.7E-05 | MB231/MB468 | NUC |
| >hg38_21_+_8433172_8446622_RNA45SN1_45S_50ntFlanks@6647.6668.22\|CGTACGACTCTTAGCGGTGGAT | >hg38_21_+_8433172_8446622_RNA45SN1_45S_50ntFlanks | 6647 | 6668 | 22 | CGTACGACTCTTAGCGGTGGAT | 2139.7 | 3.1 | 0.7 | 4.3 | 1.7E-05 | 6.7E-05 | BT20/MB468 | MP |
| >hg38_21_+_8433172_8446622_RNA45SN1_45S_50ntFlanks@6651.6684.34\|CGACTCTTAGCGGTGGATCACTCGGCTCGTGCGT | >hg38_21_+_8433172_8446622_RNA45SN1_45S_50ntFlanks | 6651 | 6684 | 34 | CGACTCTTAGCGGTGGATCACTCGGCTCGTGCGT | 2390.2 | 1.6 | 0.4 | 4.1 | 5.1E-05 | 6.7E-05 | BT20/MB468 | NUC |
| >hg38_21_+_8433172_8446622_RNA45SN1_45S_50ntFlanks@11569.11604.36\|CAGCCGACTTAGAACTGGTGCGGACCAGGGGAATCC | >hg38_21_+_8433172_8446622_RNA45SN1_45S_50ntFlanks | 11569 | 11604 | 36 | CAGCCGACTTAGAACTGGTGCGGACCAGGGGAATCC | 404.4 | -5.2 | 1.3 | -4.1 | 5.1E-05 | 6.7E-05 | BT20/MB468 | NUC |
| >hg38_21_+_8433172_8446622_RNA45SN1_45S_50ntFlanks@6661.6707.47\|CGGTGGATCACTCGGCTCGTGCGTCGATGAAGAACGCAGCTAGCTGC | >hg38_21_+_8433172_8446622_RNA45SN1_45S_50ntFlanks | 6661 | 6707 | 47 | CGGTGGATCACTCGGCTCGTGCGTCGATGAAGAACGCAGCTAGCTGC | 12329.8 | 4.4 | 1.0 | 4.5 | 6.5E-06 | 6.7E-05 | BT20/MB231 | MITO |
| >MI0000261\|hsa-mir-139&WithFlank&11\|-\|72615057\|72615136@13.35.23&[MIMAT0000250&hsa-miR-139-5p&offsets\|0\|0;m-186&11\|-\|72615102\|72615124&offsets\|0\|0]\|TCTACAGTGCACGTGTCTCCAGT | >MI0000261\|hsa-mir-139&WithFlank&11\|-\|72615057\|72615136 | 13 | 35 | 23 | TCTACAGTGCACGTGTCTCCAGT | 2226.3 | -3.5 | 0.8 | -4.5 | 6.5E-06 | 6.7E-05 | BT20/MB231 | MITO |
| >MI0000764\|hsa-mir-363&WithFlank&X\|-\|134169372\|134169458@57.77.21&[MIMAT0000707&hsa-miR-363-3p&offsets\|+1\|0;m-178&X\|-\|134169381\|134169403&offsets\|+1\|-1]\|ATTGCACGGTATCCATCTGTA | >MI0000764\|hsa-mir-363&WithFlank&X\|-\|134169372\|134169458 | 57 | 77 | 21 | ATTGCACGGTATCCATCTGTA | 64.9 | 9.4 | 2.1 | 4.5 | 5.6E-06 | 6.7E-05 | BT20/MB231 | MP |
| >hg38_21_+_8433172_8446622_RNA45SN1_45S_50ntFlanks@7976.8006.31\|GCGACCTCAGATCAGACGTGGCGACCCGCTG | >hg38_21_+_8433172_8446622_RNA45SN1_45S_50ntFlanks | 7976 | 8006 | 31 | GCGACCTCAGATCAGACGTGGCGACCCGCTG | 926.9 | 8.9 | 2.0 | 4.5 | 5.6E-06 | 6.7E-05 | BT20/MB231 | MP |
| >hg38_21_+_8433172_8446622_RNA45SN1_45S_50ntFlanks@8356.8395.40\|GAAAAGAACTTTGAAGAGAGAGTTCAAGAGGGCGTGAAAC | >hg38_21_+_8433172_8446622_RNA45SN1_45S_50ntFlanks | 8356 | 8395 | 40 | GAAAAGAACTTTGAAGAGAGAGTTCAAGAGGGCGTGAAAC | 88.2 | 9.8 | 2.2 | 4.5 | 5.6E-06 | 6.7E-05 | BT20/MB231 | MP |
| >hg38_21_+_8433172_8446622_RNA45SN1_45S_50ntFlanks@8178.8203.26\|TCTGATCGAGGCCCAGCCCGTGGACG | >hg38_21_+_8433172_8446622_RNA45SN1_45S_50ntFlanks | 8178 | 8203 | 26 | TCTGATCGAGGCCCAGCCCGTGGACG | 329.2 | -5.1 | 1.2 | -4.1 | 4.1E-05 | 6.7E-05 | MB231/MB468 | MP |
| >MI0000809\|hsa-mir-151a&WithFlank&8\|-\|140732558\|140732659@51.73.23(+1U)&[MIMAT0000757&hsa-miR-151a-3p&offsets\|-2\|0(+1U);m-30&8\|-\|140732587\|140732607&offsets\|-2\|0(+1U)]\|TACTAGACTGAAGCTCCTTGAGGT | >MI0000809\|hsa-mir-151a&WithFlank&8\|-\|140732558\|140732659 | 51 | 73 | 23(+1U) | TACTAGACTGAAGCTCCTTGAGGT | 150.1 | 4.7 | 1.2 | 4.1 | 4.1E-05 | 6.7E-05 | MB231/MB468 | MP |
| >hg38_21_+_8433172_8446622_RNA45SN1_45S_50ntFlanks@11573.11600.28\|CGACTTAGAACTGGTGCGGACCAGGGGA | >hg38_21_+_8433172_8446622_RNA45SN1_45S_50ntFlanks | 11573 | 11600 | 28 | CGACTTAGAACTGGTGCGGACCAGGGGA | 281.8 | -4.5 | 1.1 | -4.1 | 4.9E-05 | 6.7E-05 | MB231/MB468 | MITO |
| >hg38_21_+_8433172_8446622_RNA45SN1_45S_50ntFlanks@5444.5466.23\|CTCGGATCGGCCCCGCCGGGGTC | >hg38_21_+_8433172_8446622_RNA45SN1_45S_50ntFlanks | 5444 | 5466 | 23 | CTCGGATCGGCCCCGCCGGGGTC | 21197.0 | -3.7 | 0.9 | -4.1 | 4.3E-05 | 6.7E-05 | BT20/MB468 | CYTO |
| >hg38_21_+_8433172_8446622_RNA45SN1_45S_50ntFlanks@7980.8011.32\|CCTCAGATCAGACGTGGCGACCCGCTGAATTT | >hg38_21_+_8433172_8446622_RNA45SN1_45S_50ntFlanks | 7980 | 8011 | 32 | CCTCAGATCAGACGTGGCGACCCGCTGAATTT | 440.2 | -3.2 | 0.8 | -4.1 | 4.9E-05 | 6.7E-05 | MB231/MB468 | MITO |
| >hg38_21_+_8433172_8446622_RNA45SN1_45S_50ntFlanks@9545.9573.29\|GTGGAGGTCCGTAGCGGTCCTGACGTGCA | >hg38_21_+_8433172_8446622_RNA45SN1_45S_50ntFlanks | 9545 | 9573 | 29 | GTGGAGGTCCGTAGCGGTCCTGACGTGCA | 1623.0 | -2.4 | 0.6 | -4.1 | 4.3E-05 | 6.7E-05 | BT20/MB468 | CYTO |
| >hg38_21_+_8433172_8446622_RNA45SN1_45S_50ntFlanks@10351.10391.41\|CGAGAACTTTGAAGGCCGAAGTGGAGAAGGGTTCCATGTGA | >hg38_21_+_8433172_8446622_RNA45SN1_45S_50ntFlanks | 10351 | 10391 | 41 | CGAGAACTTTGAAGGCCGAAGTGGAGAAGGGTTCCATGTGA | 393.8 | -3.8 | 0.9 | -4.1 | 4.3E-05 | 6.7E-05 | BT20/MB468 | CYTO |
| >hg38_21_+_8433172_8446622_RNA45SN1_45S_50ntFlanks@8114.8140.27\|GCGCGGGACATGTGGCGTACGGAAGAC | >hg38_21_+_8433172_8446622_RNA45SN1_45S_50ntFlanks | 8114 | 8140 | 27 | GCGCGGGACATGTGGCGTACGGAAGAC | 860.9 | -7.3 | 1.8 | -4.0 | 5.1E-05 | 6.7E-05 | BT20/MB468 | NUC |
| >hg38_21_+_8433172_8446622_RNA45SN1_45S_50ntFlanks@8168.8208.41\|CCCAAGTCCTTCTGATCGAGGCCCAGCCCGTGGACGGTGTG | >hg38_21_+_8433172_8446622_RNA45SN1_45S_50ntFlanks | 8168 | 8208 | 41 | CCCAAGTCCTTCTGATCGAGGCCCAGCCCGTGGACGGTGTG | 43.2 | -8.5 | 2.0 | -4.3 | 1.7E-05 | 6.7E-05 | BT20/MB468 | MP |
| >hg38_21_+_8433172_8446622_RNA45SN1_45S_50ntFlanks@10788.10818.31\|CAGCCTCTGGCATGTTGGAACAATGTAGGTA | >hg38_21_+_8433172_8446622_RNA45SN1_45S_50ntFlanks | 10788 | 10818 | 31 | CAGCCTCTGGCATGTTGGAACAATGTAGGTA | 1978.1 | -2.9 | 0.7 | -4.1 | 3.6E-05 | 6.7E-05 | BT20/MB468 | MITO |

<table>
<tr><th></th><th></th><th></th><th></th><th></th><th></th><th></th><th></th><th></th><th></th><th></th><th></th><th></th><th></th></tr>
<tr><td>>hg38_21_+_8433172_8446622_RNA45SN1_45S_50ntFlanks@10807.10825.19|ACAATGTAGGTAAGGGAAG</td><td>>hg38_21_+_8433172_8446622_RNA45SN1_45S_50ntFlanks</td><td>10807</td><td>10825</td><td>19</td><td>ACAATGTAGGTAAGGGAAG</td><td>313.8</td><td>-5.8</td><td>1.4</td><td>-4.1</td><td>4.3E-05</td><td>6.7E-05</td><td>BT20/MB468</td><td>CYTO</td></tr>
<tr><td>>hg38_21_+_8433172_8446622_RNA45SN1_45S_50ntFlanks@4006.4023.18|ACCTCGGGCCGATCGCAC</td><td>>hg38_21_+_8433172_8446622_RNA45SN1_45S_50ntFlanks</td><td>4006</td><td>4023</td><td>18</td><td>ACCTCGGGCCGATCGCAC</td><td>2212.2</td><td>-4.5</td><td>1.1</td><td>-4.0</td><td>5.1E-05</td><td>6.8E-05</td><td>BT20/MB468</td><td>NUC</td></tr>
<tr><td>>MI0003591|hsa-mir-584&WithFlank&5|-|149062307|149062415@22.40.19(+1U)&[MIMAT0003249&hsa-miR-584-5p&offsets|0|-3(+1U);m-167&5|-|149062374|149062394&offsets|0|-2(+1U)]|TTATGGTTTGCCTGGGACTT</td><td>>MI0003591|hsa-mir-584&WithFlank&5|-|149062307|149062415</td><td>22</td><td>40</td><td>19(+1U)</td><td>TTATGGTTTGCCTGGGACTT</td><td>32.9</td><td>8.3</td><td>2.0</td><td>4.1</td><td>4.2E-05</td><td>6.8E-05</td><td>MB231/MB468</td><td>MP</td></tr>
<tr><td>>hg38_21_+_8433172_8446622_RNA45SN1_45S_50ntFlanks@5073.5099.27|AATTCCGATAACGAACGAGACTCTGGC</td><td>>hg38_21_+_8433172_8446622_RNA45SN1_45S_50ntFlanks</td><td>5073</td><td>5099</td><td>27</td><td>AATTCCGATAACGAACGAGACTCTGGC</td><td>180.0</td><td>-7.0</td><td>1.7</td><td>-4.1</td><td>4.2E-05</td><td>6.8E-05</td><td>MB231/MB468</td><td>MP</td></tr>
<tr><td>>MI0000477|hsa-mir-146a&WithFlank&5|+|160485346|160485456@26.46.21(+1C)&[MIMAT0000449&hsa-miR-146a-5p&offsets|-1|-2(+1C);m-129&5|+|160485372|160485393&offsets|-1|-2(+1C)]|TTGAGAACTGAATTCCATGGGC</td><td>>MI0000477|hsa-mir-146a&WithFlank&5|+|160485346|160485456</td><td>26</td><td>46</td><td>21(+1C)</td><td>TTGAGAACTGAATTCCATGGGC</td><td>72.0</td><td>-9.6</td><td>2.1</td><td>-4.5</td><td>5.6E-06</td><td>6.8E-05</td><td>BT20/MB231</td><td>MP</td></tr>
<tr><td>>hg38_21_+_8433172_8446622_RNA45SN1_45S_50ntFlanks@11669.11693.25|GCCCAGTGCTCTGAATGTCAAAGTG</td><td>>hg38_21_+_8433172_8446622_RNA45SN1_45S_50ntFlanks</td><td>11669</td><td>11693</td><td>25</td><td>GCCCAGTGCTCTGAATGTCAAAGTG</td><td>316.3</td><td>3.1</td><td>0.8</td><td>4.1</td><td>4.9E-05</td><td>6.8E-05</td><td>MB231/MB468</td><td>NUC</td></tr>
<tr><td>>hg38_21_+_8433172_8446622_RNA45SN1_45S_50ntFlanks@11671.11699.29|CCAGTGCTCTGAATGTCAAAGTGAAGAAA</td><td>>hg38_21_+_8433172_8446622_RNA45SN1_45S_50ntFlanks</td><td>11671</td><td>11699</td><td>29</td><td>CCAGTGCTCTGAATGTCAAAGTGAAGAAA</td><td>364.3</td><td>-4.7</td><td>1.1</td><td>-4.1</td><td>4.9E-05</td><td>6.8E-05</td><td>MB231/MB468</td><td>NUC</td></tr>
<tr><td>>hg38_1_-_228634819_228635039_RNA5S12_5S_50ntFlanks@51.70.20|GTCTACGGCCATACCACCCT</td><td>>hg38_1_-_228634819_228635039_RNA5S12_5S_50ntFlanks</td><td>51</td><td>70</td><td>20</td><td>GTCTACGGCCATACCACCCT</td><td>1978.2</td><td>3.9</td><td>0.9</td><td>4.1</td><td>4.3E-05</td><td>6.8E-05</td><td>BT20/MB468</td><td>CYTO</td></tr>
<tr><td>>hg38_21_+_8433172_8446622_RNA45SN1_45S_50ntFlanks@10783.10825.43|GTGAACAGCCTCTGGCATGTTGGAACAATGTAGGTAAGGGAAG</td><td>>hg38_21_+_8433172_8446622_RNA45SN1_45S_50ntFlanks</td><td>10783</td><td>10825</td><td>43</td><td>GTGAACAGCCTCTGGCATGTTGGAACAATGTAGGTAAGGGAAG</td><td>51.6</td><td>-9.6</td><td>2.3</td><td>-4.1</td><td>4.2E-05</td><td>6.8E-05</td><td>MB231/MB468</td><td>MP</td></tr>
<tr><td>>hg38_21_+_8433172_8446622_RNA45SN1_45S_50ntFlanks@10786.10828.43|AACAGCCTCTGGCATGTTGGAACAATGTAGGTAAGGGAAGTCG</td><td>>hg38_21_+_8433172_8446622_RNA45SN1_45S_50ntFlanks</td><td>10786</td><td>10828</td><td>43</td><td>AACAGCCTCTGGCATGTTGGAACAATGTAGGTAAGGGAAGTCG</td><td>240.9</td><td>-4.1</td><td>1.0</td><td>-4.1</td><td>4.7E-05</td><td>6.8E-05</td><td>MB231/MB468</td><td>CYTO</td></tr>
<tr><td>>MI0000102|hsa-mir-100&WithFlank&11|-|122152223|122152314@22.39.18&[MIMAT0000098&hsa-miR-100-5p&offsets|+3|-1;m-27&11|-|122152276|122152296&offsets|+3|0]|CCGTAGATCCGAACTTGT</td><td>>MI0000102|hsa-mir-100&WithFlank&11|-|122152223|122152314</td><td>22</td><td>39</td><td>18</td><td>CCGTAGATCCGAACTTGT</td><td>263.1</td><td>-11.4</td><td>2.5</td><td>-4.5</td><td>5.6E-06</td><td>6.8E-05</td><td>BT20/MB231</td><td>MP</td></tr>
<tr><td>>trna116_GluCTC_1_-_145399233_145399304@1.43.43,trna59_GluCTC_1_+_249168447_249168518@1.43.43,trna71_GluCTC_1_-_161439189_161439260@1.43.43,trna74_GluCTC_1_-_161431809_161431880@1.43.43,trna77_GluCTC_1_-_161424398_161424469@1.43.43,trna77_GluCTC_6_+_28949976_28950047@1.43.43,trna80_GluCTC_1_-_161417018_161417089@1.43.43,trna87_GluCTC_6_-_126101393_126101464@1.43.43|TCCCTGGTGGTCTAGTGGTTAGGATTCGGCGCTCTCACCGCCG</td><td>>trna116_GluCTC_1_-_145399233_145399304</td><td>1</td><td>43</td><td>43</td><td>TCCCTGGTGGTCTAGTGGTTAGGATTCGGCGCTCTCACCGCCG</td><td>246.6</td><td>3.4</td><td>0.8</td><td>4.1</td><td>4.7E-05</td><td>6.8E-05</td><td>MB231/MB468</td><td>CYTO</td></tr>
<tr><td>>hg38_21_+_8433172_8446622_RNA45SN1_45S_50ntFlanks@10404.10435.32|CATGGGTCAGTCGGTCCTGAGAGATGGGCGAG</td><td>>hg38_21_+_8433172_8446622_RNA45SN1_45S_50ntFlanks</td><td>10404</td><td>10435</td><td>32</td><td>CATGGGTCAGTCGGTCCTGAGAGATGGGCGAG</td><td>175.0</td><td>-5.2</td><td>1.3</td><td>-4.1</td><td>4.3E-05</td><td>6.8E-05</td><td>BT20/MB468</td><td>CYTO</td></tr>
<tr><td>>hg38_21_+_8433172_8446622_RNA45SN1_45S_50ntFlanks@11661.11680.20|TGATTTCTGCCCAGTGCTCT</td><td>>hg38_21_+_8433172_8446622_RNA45SN1_45S_50ntFlanks</td><td>11661</td><td>11680</td><td>20</td><td>TGATTTCTGCCCAGTGCTCT</td><td>360.5</td><td>-4.6</td><td>1.1</td><td>-4.1</td><td>5.0E-05</td><td>6.8E-05</td><td>MB231/MB468</td><td>NUC</td></tr>
<tr><td>>hg38_21_+_8433172_8446622_RNA45SN1_45S_50ntFlanks@12419.12457.39|GATGTCGGCTCTTCCTATCATTGTGAAGCAGAATTCACC</td><td>>hg38_21_+_8433172_8446622_RNA45SN1_45S_50ntFlanks</td><td>12419</td><td>12457</td><td>39</td><td>GATGTCGGCTCTTCCTATCATTGTGAAGCAGAATTCACC</td><td>28.8</td><td>8.1</td><td>2.0</td><td>4.1</td><td>4.2E-05</td><td>6.8E-05</td><td>MB231/MB468</td><td>MP</td></tr>
<tr><td>>MI0000433|hsa-let-7g&WithFlank&3|-|52268272|52268367@11.33.23&[MIMAT0000414&hsa-let-7g-5p&offsets|0|+1;m-58&3|-|52268336|52268357&offsets|0|+1]|TGAGGTAGTAGTTTGTACAGTTT</td><td>>MI0000433|hsa-let-7g&WithFlank&3|-|52268272|52268367</td><td>11</td><td>33</td><td>23</td><td>TGAGGTAGTAGTTTGTACAGTTT</td><td>450.1</td><td>1.8</td><td>0.5</td><td>4.0</td><td>5.2E-05</td><td>6.8E-05</td><td>BT20/MB468</td><td>NUC</td></tr>
<tr><td>>hg38_21_+_8433172_8446622_RNA45SN1_45S_50ntFlanks@11849.11886.38|CGGGCTTGGCGGAATCAGCGGGGAAAGAAGACCCTGTT</td><td>>hg38_21_+_8433172_8446622_RNA45SN1_45S_50ntFlanks</td><td>11849</td><td>11886</td><td>38</td><td>CGGGCTTGGCGGAATCAGCGGGGAAAGAAGACCCTGTT</td><td>99.1</td><td>10.0</td><td>2.2</td><td>4.5</td><td>5.7E-06</td><td>6.8E-05</td><td>BT20/MB231</td><td>MP</td></tr>
<tr><td>>hg38_MT_+_1621_3279_RNR2_16S_50ntFlanks@253.282.30|AACTTTGCAAGGAGAGCCAAAGCTAAGACC</td><td>>hg38_MT_+_1621_3279_RNR2_16S_50ntFlanks</td><td>253</td><td>282</td><td>30</td><td>AACTTTGCAAGGAGAGCCAAAGCTAAGACC</td><td>163.2</td><td>11.1</td><td>2.6</td><td>4.3</td><td>1.7E-05</td><td>6.8E-05</td><td>BT20/MB468</td><td>MP</td></tr>
<tr><td>>hg38_21_+_8433172_8446622_RNA45SN1_45S_50ntFlanks@10802.10822.21|TTGGAACAATGTAGGTAAGGG</td><td>>hg38_21_+_8433172_8446622_RNA45SN1_45S_50ntFlanks</td><td>10802</td><td>10822</td><td>21</td><td>TTGGAACAATGTAGGTAAGGG</td><td>196.1</td><td>-5.3</td><td>1.3</td><td>-4.1</td><td>4.4E-05</td><td>6.8E-05</td><td>BT20/MB468</td><td>CYTO</td></tr>
<tr><td>>hg38_21_+_8433172_8446622_RNA45SN1_45S_50ntFlanks@11578.11613.36|TAGAACTGGTGCGGACCAGGGGAATCCGACTGTTTA</td><td>>hg38_21_+_8433172_8446622_RNA45SN1_45S_50ntFlanks</td><td>11578</td><td>11613</td><td>36</td><td>TAGAACTGGTGCGGACCAGGGGAATCCGACTGTTTA</td><td>896.9</td><td>4.6</td><td>1.0</td><td>4.5</td><td>6.2E-06</td><td>6.8E-05</td><td>BT20/MB231</td><td>NUC</td></tr>
<tr><td>>hg38_21_+_8433172_8446622_RNA45SN1_45S_50ntFlanks@9855.9874.20|CTGCGGGATGAACCGAACGC</td><td>>hg38_21_+_8433172_8446622_RNA45SN1_45S_50ntFlanks</td><td>9855</td><td>9874</td><td>20</td><td>CTGCGGGATGAACCGAACGC</td><td>298.4</td><td>3.4</td><td>0.8</td><td>4.3</td><td>1.7E-05</td><td>6.8E-05</td><td>BT20/MB468</td><td>MP</td></tr>
<tr><td>>hg38_21_+_8433172_8446622_RNA45SN1_45S_50ntFlanks@12714.12751.38|GAGCCTCGGTTGGCCTCGGATAGCCGGTCCCCCGCCTG</td><td>>hg38_21_+_8433172_8446622_RNA45SN1_45S_50ntFlanks</td><td>12714</td><td>12751</td><td>38</td><td>GAGCCTCGGTTGGCCTCGGATAGCCGGTCCCCCGCCTG</td><td>308.9</td><td>-7.4</td><td>1.7</td><td>-4.3</td><td>1.7E-05</td><td>6.8E-05</td><td>BT20/MB468</td><td>MP</td></tr>
<tr><td>>hg38_21_+_8433172_8446622_RNA45SN1_45S_50ntFlanks@9568.9592.25|CGTGCAAATCGGTCGTCCGACCTGG</td><td>>hg38_21_+_8433172_8446622_RNA45SN1_45S_50ntFlanks</td><td>9568</td><td>9592</td><td>25</td><td>CGTGCAAATCGGTCGTCCGACCTGG</td><td>213.4</td><td>-4.4</td><td>1.1</td><td>-4.1</td><td>5.0E-05</td><td>6.8E-05</td><td>MB231/MB468</td><td>NUC</td></tr>
<tr><td>>hg38_21_+_8433172_8446622_RNA45SN1_45S_50ntFlanks@6650.6669.20|ACGACTCTTAGCGGTGGATC</td><td>>hg38_21_+_8433172_8446622_RNA45SN1_45S_50ntFlanks</td><td>6650</td><td>6669</td><td>20</td><td>ACGACTCTTAGCGGTGGATC</td><td>2280.4</td><td>3.2</td><td>0.7</td><td>4.5</td><td>6.2E-06</td><td>6.8E-05</td><td>BT20/MB231</td><td>NUC</td></tr>
<tr><td>>hg38_21_+_8433172_8446622_RNA45SN1_45S_50ntFlanks@6661.6685.25|CGGTGGATCACTCGGCTCGTGCGTC</td><td>>hg38_21_+_8433172_8446622_RNA45SN1_45S_50ntFlanks</td><td>6661</td><td>6685</td><td>25</td><td>CGGTGGATCACTCGGCTCGTGCGTC</td><td>8042.5</td><td>4.5</td><td>1.0</td><td>4.3</td><td>1.7E-05</td><td>6.8E-05</td><td>BT20/MB468</td><td>MP</td></tr>
<tr><td>>hg38_21_+_8433172_8446622_RNA45SN1_45S_50ntFlanks@7977.8015.39|CGACCTCAGATCAGACGTGGCGACCCGCTGAATTTAAGC</td><td>>hg38_21_+_8433172_8446622_RNA45SN1_45S_50ntFlanks</td><td>7977</td><td>8015</td><td>39</td><td>CGACCTCAGATCAGACGTGGCGACCCGCTGAATTTAAGC</td><td>265.8</td><td>-3.4</td><td>0.8</td><td>-4.5</td><td>5.4E-06</td><td>6.8E-05</td><td>BT20/MB231</td><td>CYTO</td></tr>
<tr><td>>hg38_21_+_8433172_8446622_RNA45SN1_45S_50ntFlanks@4557.4576.20|CAGCTAGGAATAATGGAATA</td><td>>hg38_21_+_8433172_8446622_RNA45SN1_45S_50ntFlanks</td><td>4557</td><td>4576</td><td>20</td><td>CAGCTAGGAATAATGGAATA</td><td>101.4</td><td>-5.9</td><td>1.4</td><td>-4.1</td><td>4.2E-05</td><td>6.8E-05</td><td>MB231/MB468</td><td>MP</td></tr>
<tr><td>>hg38_21_+_8433172_8446622_RNA45SN1_45S_50ntFlanks@12416.12442.27|TTCGATGTCGGCTCTTCCTATCATTGT</td><td>>hg38_21_+_8433172_8446622_RNA45SN1_45S_50ntFlanks</td><td>12416</td><td>12442</td><td>27</td><td>TTCGATGTCGGCTCTTCCTATCATTGT</td><td>285.4</td><td>3.5</td><td>0.9</td><td>4.1</td><td>5.0E-05</td><td>6.8E-05</td><td>MB231/MB468</td><td>MITO</td></tr>
<tr><td>>trnaMT_GluTTC_MT_-_14674_14742@27.67.41,trnalookalike8_GluTTC_5_-_93905172_93905240@27.67.41|GTTTTTCATATCATTGGTCGTGGTTGTAGTCCGTGCGAGAA</td><td>>trnaMT_GluTTC_MT_-_14674_14742</td><td>27</td><td>67</td><td>41</td><td>GTTTTTCATATCATTGGTCGTGGTTGTAGTCCGTGCGAGAA</td><td>44.9</td><td>9.2</td><td>2.1</td><td>4.3</td><td>1.7E-05</td><td>6.8E-05</td><td>BT20/MB468</td><td>MP</td></tr>
<tr><td>>hg38_21_+_8433172_8446622_RNA45SN1_45S_50ntFlanks@7974.8004.31|ACGCGACCTCAGATCAGACGTGGCGACCCGC</td><td>>hg38_21_+_8433172_8446622_RNA45SN1_45S_50ntFlanks</td><td>7974</td><td>8004</td><td>31</td><td>ACGCGACCTCAGATCAGACGTGGCGACCCGC</td><td>5317.5</td><td>3.6</td><td>0.8</td><td>4.5</td><td>6.3E-06</td><td>6.8E-05</td><td>BT20/MB231</td><td>NUC</td></tr>
<tr><td>>hg38_1_-_228634819_228635039_RNA5S12_5S_50ntFlanks@150.171.22|AATACCGGGTGCTGTAGGCTTT</td><td>>hg38_1_-_228634819_228635039_RNA5S12_5S_50ntFlanks</td><td>150</td><td>171</td><td>22</td><td>AATACCGGGTGCTGTAGGCTTT</td><td>4761.3</td><td>-3.2</td><td>0.8</td><td>-4.1</td><td>5.0E-05</td><td>6.8E-05</td><td>MB231/MB468</td><td>MITO</td></tr>
<tr><td>>trna16_ValTAC_11_-_59318460_59318532@35.76.42,trna17_ValTAC_11_-_59318102_59318174@35.76.42,trna4_ValTAC_X_-_18693029_18693101@35.76.42|ACACGCAGAAGGTCCTGGGTTCGAGCCCCAGTGGAACCACCA</td><td>>trna16_ValTAC_11_-_59318460_59318532</td><td>35</td><td>76</td><td>42</td><td>ACACGCAGAAGGTCCTGGGTTCGAGCCCCAGTGGAACCACCA</td><td>683.8</td><td>-5.2</td><td>1.2</td><td>-4.5</td><td>5.4E-06</td><td>6.8E-05</td><td>BT20/MB231</td><td>CYTO</td></tr>
<tr><td>>hg38_21_+_8433172_8446622_RNA45SN1_45S_50ntFlanks@10394.10430.37|AGCAGTTGAACATGGGTCAGTCGGTCCTGAGAGATGG</td><td>>hg38_21_+_8433172_8446622_RNA45SN1_45S_50ntFlanks</td><td>10394</td><td>10430</td><td>37</td><td>AGCAGTTGAACATGGGTCAGTCGGTCCTGAGAGATGG</td><td>197.8</td><td>-4.4</td><td>1.1</td><td>-4.1</td><td>5.0E-05</td><td>6.8E-05</td><td>MB231/MB468</td><td>NUC</td></tr>
<tr><td>>hg38_21_+_8433172_8446622_RNA45SN1_45S_50ntFlanks@3899.3934.36|CCTTCGCGGGGGGGATGCGTGCATTTATCAGATCAA</td><td>>hg38_21_+_8433172_8446622_RNA45SN1_45S_50ntFlanks</td><td>3899</td><td>3934</td><td>36</td><td>CCTTCGCGGGGGGGATGCGTGCATTTATCAGATCAA</td><td>47.9</td><td>-8.7</td><td>2.0</td><td>-4.3</td><td>1.7E-05</td><td>6.8E-05</td><td>BT20/MB468</td><td>MP</td></tr>
<tr><td>>hg38_21_+_8433172_8446622_RNA45SN1_45S_50ntFlanks@8105.8142.38|CGCGGCGGGGCGCGGGACATGTGGCGTACGGAAGACCC</td><td>>hg38_21_+_8433172_8446622_RNA45SN1_45S_50ntFlanks</td><td>8105</td><td>8142</td><td>38</td><td>CGCGGCGGGGCGCGGGACATGTGGCGTACGGAAGACCC</td><td>425.8</td><td>-6.7</td><td>1.7</td><td>-4.0</td><td>5.2E-05</td><td>6.9E-05</td><td>BT20/MB468</td><td>NUC</td></tr>
<tr><td>>hg38_21_+_8433172_8446622_RNA45SN1_45S_50ntFlanks@3842.3873.32|CCTACTTGGATAACTGTGGTAATTCTAGAGCT</td><td>>hg38_21_+_8433172_8446622_RNA45SN1_45S_50ntFlanks</td><td>3842</td><td>3873</td><td>32</td><td>CCTACTTGGATAACTGTGGTAATTCTAGAGCT</td><td>474.1</td><td>3.4</td><td>0.8</td><td>4.1</td><td>4.4E-05</td><td>6.9E-05</td><td>BT20/MB468</td><td>CYTO</td></tr>
<tr><td>>trnaMT_IleGAT_MT_+_4263_4331@44.71.28|TAGGAGCTTAAACCCCCTTATTTCTACC</td><td>>trnaMT_IleGAT_MT_+_4263_4331</td><td>44</td><td>71</td><td>28</td><td>TAGGAGCTTAAACCCCCTTATTTCTACC</td><td>156.4</td><td>11.0</td><td>2.6</td><td>4.3</td><td>1.7E-05</td><td>6.9E-05</td><td>BT20/MB468</td><td>MP</td></tr>
<tr><td>>hg38_21_+_8433172_8446622_RNA45SN1_45S_50ntFlanks@10401.10433.33|GAACATGGGTCAGTCGGTCCTGAGAGATGGGCG</td><td>>hg38_21_+_8433172_8446622_RNA45SN1_45S_50ntFlanks</td><td>10401</td><td>10433</td><td>33</td><td>GAACATGGGTCAGTCGGTCCTGAGAGATGGGCG</td><td>197.0</td><td>-5.3</td><td>1.3</td><td>-4.1</td><td>4.2E-05</td><td>6.9E-05</td><td>MB231/MB468</td><td>MP</td></tr>
<tr><td>>trna116_GluCTC_1_-_145399233_145399304@2.33.32,trna59_GluCTC_1_+_249168447_249168518@2.33.32,trna71_GluCTC_1_-_161439189_161439260@2.33.32,trna74_GluCTC_1_-_161431809_161431880@2.33.32,trna77_GluCTC_1_-_161424398_161424469@2.33.32,trna77_GluCTC_6_+_28949976_28950047@2.33.32,trna80_GluCTC_1_-_161417018_161417089@2.33.32,trna87_GluCTC_6_-_126101393_126101464@2.33.32|CCCTGGTGGTCTAGTGGTTAGGATTCGGCGCT</td><td>>trna116_GluCTC_1_-_145399233_145399304</td><td>2</td><td>33</td><td>32</td><td>CCCTGGTGGTCTAGTGGTTAGGATTCGGCGCT</td><td>785.6</td><td>6.8</td><td>1.7</td><td>4.1</td><td>4.4E-05</td><td>6.9E-05</td><td>BT20/MB468</td><td>CYTO</td></tr>
<tr><td>>hg38_21_+_8433172_8446622_RNA45SN1_45S_50ntFlanks@4983.5020.38|CGGACAGGATTGACAGATTGATAGCTCTTTCTCGATTC</td><td>>hg38_21_+_8433172_8446622_RNA45SN1_45S_50ntFlanks</td><td>4983</td><td>5020</td><td>38</td><td>CGGACAGGATTGACAGATTGATAGCTCTTTCTCGATTC</td><td>179.4</td><td>-5.5</td><td>1.3</td><td>-4.1</td><td>4.2E-05</td><td>6.9E-05</td><td>MB231/MB468</td><td>MP</td></tr>
<tr><td>>hg38_MT_+_1621_3279_RNR2_16S_50ntFlanks@51.85.35|GCTAAACCTAGCCCCAAACCCACTCCACCTTACTA</td><td>>hg38_MT_+_1621_3279_RNR2_16S_50ntFlanks</td><td>51</td><td>85</td><td>35</td><td>GCTAAACCTAGCCCCAAACCCACTCCACCTTACTA</td><td>1226.5</td><td>4.0</td><td>1.0</td><td>4.1</td><td>5.1E-05</td><td>6.9E-05</td><td>MB231/MB468</td><td>MITO</td></tr>
<tr><td>>hg38_21_+_8433172_8446622_RNA45SN1_45S_50ntFlanks@11216.11234.19|TCTGGACGCGAGCCGGGCC</td><td>>hg38_21_+_8433172_8446622_RNA45SN1_45S_50ntFlanks</td><td>11216</td><td>11234</td><td>19</td><td>TCTGGACGCGAGCCGGGCC</td><td>159.6</td><td>-5.2</td><td>1.3</td><td>-4.1</td><td>4.4E-05</td><td>6.9E-05</td><td>BT20/MB468</td><td>CYTO</td></tr>
<tr><td>>hg38_21_+_8433172_8446622_RNA45SN1_45S_50ntFlanks@9563.9592.30|CCTGACGTGCAAATCGGTCGTCCGACCTGG</td><td>>hg38_21_+_8433172_8446622_RNA45SN1_45S_50ntFlanks</td><td>9563</td><td>9592</td><td>30</td><td>CCTGACGTGCAAATCGGTCGTCCGACCTGG</td><td>160.4</td><td>-5.2</td><td>1.3</td><td>-4.1</td><td>4.4E-05</td><td>6.9E-05</td><td>BT20/MB468</td><td>CYTO</td></tr>
<tr><td>>hg38_21_+_8433172_8446622_RNA45SN1_45S_50ntFlanks@6700.6736.37|CTAGCTGCGAGAATTAATGTGAATTGCAGGACACATT</td><td>>hg38_21_+_8433172_8446622_RNA45SN1_45S_50ntFlanks</td><td>6700</td><td>6736</td><td>37</td><td>CTAGCTGCGAGAATTAATGTGAATTGCAGGACACATT</td><td>48.3</td><td>-9.5</td><td>2.3</td><td>-4.1</td><td>4.2E-05</td><td>6.9E-05</td><td>MB231/MB468</td><td>MP</td></tr>
<tr><td>>trnaMT_IleGAT_MT_+_4263_4331@45.71.27|AGGAGCTTAAACCCCCTTATTTCTACC</td><td>>trnaMT_IleGAT_MT_+_4263_4331</td><td>45</td><td>71</td><td>27</td><td>AGGAGCTTAAACCCCCTTATTTCTACC</td><td>28.1</td><td>8.0</td><td>2.0</td><td>4.1</td><td>4.2E-05</td><td>6.9E-05</td><td>MB231/MB468</td><td>MP</td></tr>
<tr><td>>hg38_21_+_8433172_8446622_RNA45SN1_45S_50ntFlanks@11847.11864.18|AACGGGCTTGGCGGAATC</td><td>>hg38_21_+_8433172_8446622_RNA45SN1_45S_50ntFlanks</td><td>11847</td><td>11864</td><td>18</td><td>AACGGGCTTGGCGGAATC</td><td>109.7</td><td>-6.4</td><td>1.6</td><td>-4.1</td><td>4.2E-05</td><td>6.9E-05</td><td>MB231/MB468</td><td>MP</td></tr>
<tr><td>>hg38_21_+_8433172_8446622_RNA45SN1_45S_50ntFlanks@10664.10712.49|CTTGGAAAGCGTCGCGGTTCCGGCGGCGTCCGGTGAGCTCTCGCTGGCC</td><td>>hg38_21_+_8433172_8446622_RNA45SN1_45S_50ntFlanks</td><td>10664</td><td>10712</td><td>49</td><td>CTTGGAAAGCGTCGCGGTTCCGGCGGCGTCCGGTGAGCTCTCGCTGGCC</td><td>200.4</td><td>-5.7</td><td>1.4</td><td>-4.1</td><td>3.7E-05</td><td>6.9E-05</td><td>BT20/MB468</td><td>MITO</td></tr>
<tr><td>>hg38_21_+_8433172_8446622_RNA45SN1_45S_50ntFlanks@10808.10837.30|CAATGTAGGTAAGGGAAGTCGGCAAGCCGG</td><td>>hg38_21_+_8433172_8446622_RNA45SN1_45S_50ntFlanks</td><td>10808</td><td>10837</td><td>30</td><td>CAATGTAGGTAAGGGAAGTCGGCAAGCCGG</td><td>283.9</td><td>-4.3</td><td>1.0</td><td>-4.1</td><td>3.7E-05</td><td>6.9E-05</td><td>BT20/MB468</td><td>MITO</td></tr>
</table>

<table>
<tr><td>>hg38_21_+_8433172_8446622_RNA45SN1_45S_50ntFlanks@4657.4684.28|CTAGAGGTGAAATTCTTGGACCGGCGCA</td><td>>hg38_21_+_8433172_8446622_RNA45SN1_45S_50ntFlanks</td><td>4657</td><td>4684</td><td>28</td><td>CTAGAGGTGAAATTCTTGGACCGGCGCA</td><td>607.5</td><td>-5.6</td><td>1.4</td><td>-4.1</td><td>3.7E-05</td><td>6.9E-05</td><td>BT20/MB468</td><td>MITO</td></tr>
<tr><td>>hg38_21_+_8433172_8446622_RNA45SN1_45S_50ntFlanks@11778.11803.26|ACGCGCATGAATGGATGAACGAGATT</td><td>>hg38_21_+_8433172_8446622_RNA45SN1_45S_50ntFlanks</td><td>11778</td><td>11803</td><td>26</td><td>ACGCGCATGAATGGATGAACGAGATT</td><td>352.5</td><td>-3.9</td><td>0.9</td><td>-4.1</td><td>4.8E-05</td><td>6.9E-05</td><td>MB231/MB468</td><td>CYTO</td></tr>
<tr><td>>hg38_21_+_8433172_8446622_RNA45SN1_45S_50ntFlanks@3842.3862.21|CCTACTTGGATAACTGTGGTA</td><td>>hg38_21_+_8433172_8446622_RNA45SN1_45S_50ntFlanks</td><td>3842</td><td>3862</td><td>21</td><td>CCTACTTGGATAACTGTGGTA</td><td>130.8</td><td>-6.2</td><td>1.5</td><td>-4.1</td><td>4.3E-05</td><td>6.9E-05</td><td>MB231/MB468</td><td>MP</td></tr>
<tr><td>>hg38_21_+_8433172_8446622_RNA45SN1_45S_50ntFlanks@10808.10830.23|CAATGTAGGTAAGGGAAGTCGGC</td><td>>hg38_21_+_8433172_8446622_RNA45SN1_45S_50ntFlanks</td><td>10808</td><td>10830</td><td>23</td><td>CAATGTAGGTAAGGGAAGTCGGC</td><td>364.6</td><td>-4.4</td><td>1.1</td><td>-4.1</td><td>3.7E-05</td><td>6.9E-05</td><td>BT20/MB468</td><td>MITO</td></tr>
<tr><td>>hg38_MT_+_598_1651_RNR1_12S_50ntFlanks@430.468.39|AGTGGCTTTAACATATCTGAACACACAATAGCTAAGACC</td><td>>hg38_MT_+_598_1651_RNR1_12S_50ntFlanks</td><td>430</td><td>468</td><td>39</td><td>AGTGGCTTTAACATATCTGAACACACAATAGCTAAGACC</td><td>215.3</td><td>8.2</td><td>1.9</td><td>4.3</td><td>1.7E-05</td><td>6.9E-05</td><td>BT20/MB468</td><td>MP</td></tr>
<tr><td>>hg38_21_+_8433172_8446622_RNA45SN1_45S_50ntFlanks@11537.11580.44|CCGCGCGGCGCCTCGCCTCGGCCGGCGCCTAGCAGCCGACTTAG</td><td>>hg38_21_+_8433172_8446622_RNA45SN1_45S_50ntFlanks</td><td>11537</td><td>11580</td><td>44</td><td>CCGCGCGGCGCCTCGCCTCGGCCGGCGCCTAGCAGCCGACTTAG</td><td>1270.9</td><td>-5.0</td><td>1.2</td><td>-4.1</td><td>5.1E-05</td><td>6.9E-05</td><td>MB231/MB468</td><td>NUC</td></tr>
<tr><td>>am_MI0000809|hsa-mir-151a&WithFlank&8|-|140732558|140732659@53.71.19(+1A)&[MIMAT0000757&hsa-miR-151a-3p&offsets|0|-2(+1A);m-30&8|-|140732587|140732607&offsets|0|-2(+1A)]|CTAGACTGAAGCTCCTTGAA</td><td>>am_MI0000809|hsa-mir-151a&WithFlank&8|-|140732558|140732659</td><td>53</td><td>71</td><td>19(+1A)</td><td>CTAGACTGAAGCTCCTTGAA</td><td>622.8</td><td>6.0</td><td>1.4</td><td>4.3</td><td>1.8E-05</td><td>6.9E-05</td><td>BT20/MB468</td><td>MP</td></tr>
<tr><td>>MI0025242|hsa-mir-7706&WithFlank&15|+|85380590|85380668@50.72.23&[MIMAT0030021&hsa-miR-7706&offsets|0|-1;m-390&15|+|85380639|85380662&offsets|0|-1]|TGAAGCGCCTGTGCTCTGCCGAG</td><td>>MI0025242|hsa-mir-7706&WithFlank&15|+|85380590|85380668</td><td>50</td><td>72</td><td>23</td><td>TGAAGCGCCTGTGCTCTGCCGAG</td><td>332.0</td><td>3.5</td><td>0.9</td><td>4.1</td><td>5.1E-05</td><td>6.9E-05</td><td>MB231/MB468</td><td>NUC</td></tr>
<tr><td>>hg38_21_+_8433172_8446622_RNA45SN1_45S_50ntFlanks@11703.11736.34|AATGAAGCGCGGGTAAACGGCGGGAGTAACTATG</td><td>>hg38_21_+_8433172_8446622_RNA45SN1_45S_50ntFlanks</td><td>11703</td><td>11736</td><td>34</td><td>AATGAAGCGCGGGTAAACGGCGGGAGTAACTATG</td><td>255.1</td><td>-3.6</td><td>0.9</td><td>-4.1</td><td>4.3E-05</td><td>6.9E-05</td><td>MB231/MB468</td><td>MP</td></tr>
<tr><td>>hg38_21_+_8433172_8446622_RNA45SN1_45S_50ntFlanks@6733.6751.19|CATTGATCATCGACACTTC</td><td>>hg38_21_+_8433172_8446622_RNA45SN1_45S_50ntFlanks</td><td>6733</td><td>6751</td><td>19</td><td>CATTGATCATCGACACTTC</td><td>135.2</td><td>-4.8</td><td>1.2</td><td>-4.1</td><td>4.3E-05</td><td>6.9E-05</td><td>MB231/MB468</td><td>MP</td></tr>
<tr><td>>hg38_21_+_8433172_8446622_RNA45SN1_45S_50ntFlanks@9606.9624.19|AGACTAATCGAACCATCTA</td><td>>hg38_21_+_8433172_8446622_RNA45SN1_45S_50ntFlanks</td><td>9606</td><td>9624</td><td>19</td><td>AGACTAATCGAACCATCTA</td><td>59.0</td><td>-9.7</td><td>2.4</td><td>-4.1</td><td>4.3E-05</td><td>6.9E-05</td><td>MB231/MB468</td><td>MP</td></tr>
<tr><td>>MI0000077|hsa-mir-21&WithFlank&17|+|59841260|59841343@13.35.23(+1U)&[MIMAT0000076&hsa-miR-21-5p&offsets|-1|0(+1U);m-1&17|+|59841273|59841294&offsets|-1|0(+1U)]|GTAGCTTATCAGACTGATGTTGAT</td><td>>MI0000077|hsa-mir-21&WithFlank&17|+|59841260|59841343</td><td>13</td><td>35</td><td>23(+1U)</td><td>GTAGCTTATCAGACTGATGTTGAT</td><td>70.1</td><td>-9.5</td><td>2.1</td><td>-4.5</td><td>5.8E-06</td><td>6.9E-05</td><td>BT20/MB231</td><td>MP</td></tr>
<tr><td>>hg38_21_+_8433172_8446622_RNA45SN1_45S_50ntFlanks@12405.12439.35|CTTTTTGATCCTTCGATGTCGGCTCTTCCTATCAT</td><td>>hg38_21_+_8433172_8446622_RNA45SN1_45S_50ntFlanks</td><td>12405</td><td>12439</td><td>35</td><td>CTTTTTGATCCTTCGATGTCGGCTCTTCCTATCAT</td><td>278.1</td><td>-2.1</td><td>0.5</td><td>-4.1</td><td>3.7E-05</td><td>6.9E-05</td><td>BT20/MB468</td><td>MITO</td></tr>
<tr><td>>hg38_21_+_8433172_8446622_RNA45SN1_45S_50ntFlanks@11207.11234.28|GGCGGCGACTCTGGACGCGAGCCGGGCC</td><td>>hg38_21_+_8433172_8446622_RNA45SN1_45S_50ntFlanks</td><td>11207</td><td>11234</td><td>28</td><td>GGCGGCGACTCTGGACGCGAGCCGGGCC</td><td>272.0</td><td>-4.5</td><td>1.1</td><td>-4.1</td><td>4.5E-05</td><td>6.9E-05</td><td>BT20/MB468</td><td>CYTO</td></tr>
<tr><td>>hg38_21_+_8433172_8446622_RNA45SN1_45S_50ntFlanks@12932.12960.29|CGCTAAACCATTCGTAGACGACCTGCTTC</td><td>>hg38_21_+_8433172_8446622_RNA45SN1_45S_50ntFlanks</td><td>12932</td><td>12960</td><td>29</td><td>CGCTAAACCATTCGTAGACGACCTGCTTC</td><td>348.0</td><td>3.1</td><td>0.8</td><td>4.1</td><td>4.8E-05</td><td>6.9E-05</td><td>MB231/MB468</td><td>CYTO</td></tr>
<tr><td>>hg38_21_+_8433172_8446622_RNA45SN1_45S_50ntFlanks@4399.4423.25|CGGGCGGGCGGTCCGCCGCGAGGCG</td><td>>hg38_21_+_8433172_8446622_RNA45SN1_45S_50ntFlanks</td><td>4399</td><td>4423</td><td>25</td><td>CGGGCGGGCGGTCCGCCGCGAGGCG</td><td>326.8</td><td>-6.0</td><td>1.5</td><td>-4.1</td><td>5.1E-05</td><td>7.0E-05</td><td>MB231/MB468</td><td>NUC</td></tr>
<tr><td>>hg38_21_+_8433172_8446622_RNA45SN1_45S_50ntFlanks@9862.9900.39|ATGAACCGAACGCCGGGTTAAGGCGCCCGATGCCGACGC</td><td>>hg38_21_+_8433172_8446622_RNA45SN1_45S_50ntFlanks</td><td>9862</td><td>9900</td><td>39</td><td>ATGAACCGAACGCCGGGTTAAGGCGCCCGATGCCGACGC</td><td>74.0</td><td>-8.3</td><td>2.0</td><td>-4.1</td><td>3.7E-05</td><td>7.0E-05</td><td>BT20/MB468</td><td>MITO</td></tr>
<tr><td>>am_trna111_HisGTG_1_-_147774845_147774916@-1G.26.27,trna118_HisGTG_1_-_145396881_145396952@-1G.26.27,trna16_HisGTG_1_+_146544773_146544844@-1G.26.27,trna1_HisGTG_15_+_45493349_45493420@-1G.26.27,trna21_HisGTG_1_+_147753471_147753542@-1G.26.27,trna33_HisGTG_6_+_27125906_27125977@-1G.26.27,trna7_HisGTG_9_-_14433938_14434009@-1G.26.27,trna8_HisGTG_15_-_45492611_45492682@-1G.26.27,trna9_HisGTG_15_-_45490804_45490875@-1G.26.27|GGCCGTGATCGTATAGTGGTTAGTACT</td><td>>am_trna111_HisGTG_1_-_147774845_147774916</td><td>-1G</td><td>26</td><td>27</td><td>GGCCGTGATCGTATAGTGGTTAGTACT</td><td>526.3</td><td>3.5</td><td>0.8</td><td>4.1</td><td>4.3E-05</td><td>7.0E-05</td><td>MB231/MB468</td><td>MP</td></tr>
<tr><td>>hg38_21_+_8433172_8446622_RNA45SN1_45S_50ntFlanks@12399.12436.38|ACGTCGCTTTTTGATCCTTCGATGTCGGCTCTTCCTAT</td><td>>hg38_21_+_8433172_8446622_RNA45SN1_45S_50ntFlanks</td><td>12399</td><td>12436</td><td>38</td><td>ACGTCGCTTTTTGATCCTTCGATGTCGGCTCTTCCTAT</td><td>398.2</td><td>-4.2</td><td>1.0</td><td>-4.1</td><td>5.1E-05</td><td>7.0E-05</td><td>MB231/MB468</td><td>NUC</td></tr>
<tr><td>>hg38_21_+_8433172_8446622_RNA45SN1_45S_50ntFlanks@8170.8215.46|CAAGTCCTTCTGATCGAGGCCCAGCCCGTGGACGGTGTGAGGCCGG</td><td>>hg38_21_+_8433172_8446622_RNA45SN1_45S_50ntFlanks</td><td>8170</td><td>8215</td><td>46</td><td>CAAGTCCTTCTGATCGAGGCCCAGCCCGTGGACGGTGTGAGGCCGG</td><td>848.4</td><td>-5.4</td><td>1.3</td><td>-4.1</td><td>4.5E-05</td><td>7.0E-05</td><td>BT20/MB468</td><td>CYTO</td></tr>
<tr><td>>hg38_21_+_8433172_8446622_RNA45SN1_45S_50ntFlanks@12073.12096.24|CCGAGGGGCTCTCGCTTCTGGCGC</td><td>>hg38_21_+_8433172_8446622_RNA45SN1_45S_50ntFlanks</td><td>12073</td><td>12096</td><td>24</td><td>CCGAGGGGCTCTCGCTTCTGGCGC</td><td>537.3</td><td>-5.8</td><td>1.4</td><td>-4.1</td><td>3.7E-05</td><td>7.0E-05</td><td>BT20/MB468</td><td>MITO</td></tr>
<tr><td>>MI0000094|hsa-mir-92a-2&WithFlank&X|-|134169532|134169618@54.76.23&[MIMAT0000092_1&hsa-miR-92a-3p&offsets|0|+1;m-11&X|-|134169544|134169565&offsets|0|+1]|TATTGCACTTGTCCCGGCCTGTG</td><td>>MI0000094|hsa-mir-92a-2&WithFlank&X|-|134169532|134169618</td><td>54</td><td>76</td><td>23</td><td>TATTGCACTTGTCCCGGCCTGTG</td><td>291.1</td><td>4.7</td><td>1.2</td><td>4.1</td><td>4.8E-05</td><td>7.0E-05</td><td>MB231/MB468</td><td>CYTO</td></tr>
<tr><td>>MI0000068|hsa-let-7f-2&WithFlank&X|-|53557186|53557280@14.36.23&[MIMAT0000067_1&hsa-let-7f-5p&offsets|0|+1;m-19&X|-|53557246|53557267&offsets|0|+1]|TGAGGTAGTAGATTGTATAGTTT</td><td>>MI0000068|hsa-let-7f-2&WithFlank&X|-|53557186|53557280</td><td>14</td><td>36</td><td>23</td><td>TGAGGTAGTAGATTGTATAGTTT</td><td>2431.3</td><td>4.4</td><td>1.0</td><td>4.3</td><td>1.8E-05</td><td>7.0E-05</td><td>BT20/MB468</td><td>MP</td></tr>
<tr><td>>am_trnaMT_SerGCT_MT_+_12207_12265@13.35.23|AAGAACTGCTAACTCATGCCCCC</td><td>>am_trnaMT_SerGCT_MT_+_12207_12265</td><td>13</td><td>35</td><td>23</td><td>AAGAACTGCTAACTCATGCCCCC</td><td>272.1</td><td>7.3</td><td>1.7</td><td>4.3</td><td>1.8E-05</td><td>7.0E-05</td><td>BT20/MB468</td><td>MP</td></tr>
<tr><td>>hg38_21_+_8433172_8446622_RNA45SN1_45S_50ntFlanks@4666.4684.19|AAATTCTTGGACCGGCGCA</td><td>>hg38_21_+_8433172_8446622_RNA45SN1_45S_50ntFlanks</td><td>4666</td><td>4684</td><td>19</td><td>AAATTCTTGGACCGGCGCA</td><td>100.9</td><td>-4.8</td><td>1.2</td><td>-4.1</td><td>4.3E-05</td><td>7.0E-05</td><td>MB231/MB468</td><td>MP</td></tr>
<tr><td>>hg38_21_+_8433172_8446622_RNA45SN1_45S_50ntFlanks@7980.8006.27|CCTCAGATCAGACGTGGCGACCCGCTG</td><td>>hg38_21_+_8433172_8446622_RNA45SN1_45S_50ntFlanks</td><td>7980</td><td>8006</td><td>27</td><td>CCTCAGATCAGACGTGGCGACCCGCTG</td><td>6040.8</td><td>-4.9</td><td>1.1</td><td>-4.3</td><td>1.8E-05</td><td>7.0E-05</td><td>BT20/MB468</td><td>MP</td></tr>
<tr><td>>hg38_21_+_8433172_8446622_RNA45SN1_45S_50ntFlanks@10422.10456.35|GAGAGATGGGCGAGCGCCGTTCCGAAGGGACGGGC</td><td>>hg38_21_+_8433172_8446622_RNA45SN1_45S_50ntFlanks</td><td>10422</td><td>10456</td><td>35</td><td>GAGAGATGGGCGAGCGCCGTTCCGAAGGGACGGGC</td><td>830.7</td><td>-3.6</td><td>0.8</td><td>-4.5</td><td>5.6E-06</td><td>7.0E-05</td><td>BT20/MB231</td><td>CYTO</td></tr>
<tr><td>>trna16_GlnTTG_17_+_47269890_47269961@1.29.29,trna64_GlnTTG_6_+_28557156_28557227@1.29.29,trna84_GlnTTG_6_+_145503859_145503930@1.29.29|GGTCCCATGGTGTAATGGTTAGCACTCTG</td><td>>trna16_GlnTTG_17_+_47269890_47269961</td><td>1</td><td>29</td><td>29</td><td>GGTCCCATGGTGTAATGGTTAGCACTCTG</td><td>4267.9</td><td>-4.9</td><td>1.2</td><td>-4.1</td><td>4.5E-05</td><td>7.0E-05</td><td>BT20/MB468</td><td>CYTO</td></tr>
<tr><td>>hg38_21_+_8433172_8446622_RNA45SN1_45S_50ntFlanks@12539.12573.35|ATGATGTGTTGTTGCCATGGTAATCCTGCTCAGTA</td><td>>hg38_21_+_8433172_8446622_RNA45SN1_45S_50ntFlanks</td><td>12539</td><td>12573</td><td>35</td><td>ATGATGTGTTGTTGCCATGGTAATCCTGCTCAGTA</td><td>180.0</td><td>-5.2</td><td>1.3</td><td>-4.1</td><td>4.5E-05</td><td>7.0E-05</td><td>BT20/MB468</td><td>CYTO</td></tr>
<tr><td>>hg38_21_+_8433172_8446622_RNA45SN1_45S_50ntFlanks@9537.9571.35|AAACTCTGGTGGAGGTCCGTAGCGGTCCTGACGTG</td><td>>hg38_21_+_8433172_8446622_RNA45SN1_45S_50ntFlanks</td><td>9537</td><td>9571</td><td>35</td><td>AAACTCTGGTGGAGGTCCGTAGCGGTCCTGACGTG</td><td>116.7</td><td>-6.1</td><td>1.5</td><td>-4.1</td><td>4.3E-05</td><td>7.0E-05</td><td>MB231/MB468</td><td>MP</td></tr>
<tr><td>>hg38_21_+_8433172_8446622_RNA45SN1_45S_50ntFlanks@9787.9819.33|TCGCTGGCGTGGAGCCGGGCGTGGAATGCGAGT</td><td>>hg38_21_+_8433172_8446622_RNA45SN1_45S_50ntFlanks</td><td>9787</td><td>9819</td><td>33</td><td>TCGCTGGCGTGGAGCCGGGCGTGGAATGCGAGT</td><td>99.8</td><td>-5.9</td><td>1.5</td><td>-4.1</td><td>4.3E-05</td><td>7.0E-05</td><td>MB231/MB468</td><td>MP</td></tr>
<tr><td>>hg38_21_+_8433172_8446622_RNA45SN1_45S_50ntFlanks@9565.9594.30|TGACGTGCAAATCGGTCGTCCGACCTGGGT</td><td>>hg38_21_+_8433172_8446622_RNA45SN1_45S_50ntFlanks</td><td>9565</td><td>9594</td><td>30</td><td>TGACGTGCAAATCGGTCGTCCGACCTGGGT</td><td>759.5</td><td>-5.5</td><td>1.4</td><td>-4.0</td><td>5.3E-05</td><td>7.0E-05</td><td>BT20/MB468</td><td>NUC</td></tr>
<tr><td>>hg38_21_+_8433172_8446622_RNA45SN1_45S_50ntFlanks@10845.10868.24|ACTTCGGGATAAGGATTGGCTCTA</td><td>>hg38_21_+_8433172_8446622_RNA45SN1_45S_50ntFlanks</td><td>10845</td><td>10868</td><td>24</td><td>ACTTCGGGATAAGGATTGGCTCTA</td><td>1034.6</td><td>-5.3</td><td>1.3</td><td>-4.1</td><td>4.5E-05</td><td>7.0E-05</td><td>BT20/MB468</td><td>CYTO</td></tr>
<tr><td>>hg38_21_+_8433172_8446622_RNA45SN1_45S_50ntFlanks@11671.11703.33|CCAGTGCTCTGAATGTCAAAGTGAAGAAATTCA</td><td>>hg38_21_+_8433172_8446622_RNA45SN1_45S_50ntFlanks</td><td>11671</td><td>11703</td><td>33</td><td>CCAGTGCTCTGAATGTCAAAGTGAAGAAATTCA</td><td>568.0</td><td>-2.8</td><td>0.7</td><td>-4.1</td><td>4.5E-05</td><td>7.0E-05</td><td>BT20/MB468</td><td>CYTO</td></tr>
<tr><td>>MI0003628|hsa-mir-615&WithFlank&12|+|54033944|54034051@67.87.21&[MIMAT0003283&hsa-miR-615-3p&offsets|0|-1;m-495&12|+|54034010|54034030&offsets|0|0]|TCCGAGCCTGGGTCTCCCTCT</td><td>>MI0003628|hsa-mir-615&WithFlank&12|+|54033944|54034051</td><td>67</td><td>87</td><td>21</td><td>TCCGAGCCTGGGTCTCCCTCT</td><td>324.4</td><td>7.3</td><td>1.6</td><td>4.5</td><td>5.9E-06</td><td>7.0E-05</td><td>BT20/MB231</td><td>MP</td></tr>
<tr><td>>hg38_21_+_8433172_8446622_RNA45SN1_45S_50ntFlanks@11555.11588.34|CGGCCGGCGCCTAGCAGCCGACTTAGAACTGGTG</td><td>>hg38_21_+_8433172_8446622_RNA45SN1_45S_50ntFlanks</td><td>11555</td><td>11588</td><td>34</td><td>CGGCCGGCGCCTAGCAGCCGACTTAGAACTGGTG</td><td>105.6</td><td>10.1</td><td>2.2</td><td>4.5</td><td>5.9E-06</td><td>7.0E-05</td><td>BT20/MB231</td><td>MP</td></tr>
<tr><td>>hg38_21_+_8433172_8446622_RNA45SN1_45S_50ntFlanks@11923.11947.25|AGAGGTGTAGAATAAGTGGGAGGCC</td><td>>hg38_21_+_8433172_8446622_RNA45SN1_45S_50ntFlanks</td><td>11923</td><td>11947</td><td>25</td><td>AGAGGTGTAGAATAAGTGGGAGGCC</td><td>99.4</td><td>10.0</td><td>2.2</td><td>4.5</td><td>5.9E-06</td><td>7.0E-05</td><td>BT20/MB231</td><td>MP</td></tr>
<tr><td>>trnaMT_ValTAC_MT_+_1602_1670@27.72.46|CAACTTACACTTAGGAGATTTCAACTTAACTTGACCGCTCTGACCA</td><td>>trnaMT_ValTAC_MT_+_1602_1670</td><td>27</td><td>72</td><td>46</td><td>CAACTTACACTTAGGAGATTTCAACTTAACTTGACCGCTCTGACCA</td><td>966.7</td><td>5.0</td><td>1.2</td><td>4.0</td><td>5.2E-05</td><td>7.0E-05</td><td>MB231/MB468</td><td>MITO</td></tr>
<tr><td>>MI0003137|hsa-mir-193b&WithFlank&16|+|14303961|14304055@58.79.22&[MIMAT0002819&hsa-miR-193b-3p&offsets|+1|+1;m-166&16|+|14304017|14304038&offsets|+1|+1]|ACTGGCCCTCAAAGTCCCGCTT</td><td>>MI0003137|hsa-mir-193b&WithFlank&16|+|14303961|14304055</td><td>58</td><td>79</td><td>22</td><td>ACTGGCCCTCAAAGTCCCGCTT</td><td>184.2</td><td>8.2</td><td>1.9</td><td>4.3</td><td>1.8E-05</td><td>7.0E-05</td><td>BT20/MB468</td><td>MP</td></tr>
<tr><td>>hg38_21_+_8433172_8446622_RNA45SN1_45S_50ntFlanks@12015.12035.21|TGAAATACCACTACTCTGATC</td><td>>hg38_21_+_8433172_8446622_RNA45SN1_45S_50ntFlanks</td><td>12015</td><td>12035</td><td>21</td><td>TGAAATACCACTACTCTGATC</td><td>34.3</td><td>8.8</td><td>2.1</td><td>4.3</td><td>1.8E-05</td><td>7.0E-05</td><td>BT20/MB468</td><td>MP</td></tr>
<tr><td>>MI0000298|hsa-mir-221&WithFlank&X|-|45746151|45746272@71.93.23(+1U)&[MIMAT0000278&hsa-miR-221-3p&offsets|0|0(+1U);m-91&X|-|45746181|45746202&offsets|0|+1(+1U)]|AGCTACATTGTCTGCTGGGTTTCT</td><td>>MI0000298|hsa-mir-221&WithFlank&X|-|45746151|45746272</td><td>71</td><td>93</td><td>23(+1U)</td><td>AGCTACATTGTCTGCTGGGTTTCT</td><td>9437.9</td><td>-2.3</td><td>0.5</td><td>-4.5</td><td>6.5E-06</td><td>7.0E-05</td><td>BT20/MB231</td><td>NUC</td></tr>
<tr><td>>hg38_21_+_8433172_8446622_RNA45SN1_45S_50ntFlanks@5066.5102.37|TCTGGTTAATTCCGATAACGAACGAGACTCTGGCATG</td><td>>hg38_21_+_8433172_8446622_RNA45SN1_45S_50ntFlanks</td><td>5066</td><td>5102</td><td>37</td><td>TCTGGTTAATTCCGATAACGAACGAGACTCTGGCATG</td><td>55.6</td><td>-9.7</td><td>2.4</td><td>-4.1</td><td>4.3E-05</td><td>7.0E-05</td><td>MB231/MB468</td><td>MP</td></tr>
<tr><td>>trna116_GluCTC_1_-_145399233_145399304@1.39.39,trna59_GluCTC_1_+_249168447_249168518@1.39.39,trna71_GluCTC_1_-_161439189_161439260@1.39.39,trna74_GluCTC_1_-_161431809_161431880@1.39.39,trna77_GluCTC_1_-_161424398_161424469@1.39.39,trna77_GluCTC_6_+_28949976_28950047@1.39.39,trna80_GluCTC_1_-_161417018_161417089@1.39.39,trna87_GluCTC_6_-_126101393_126101464@1.39.39|TCCCTGGTGGTCTAGTGGTTAGGATTCGGCGCTCTCACC</td><td>>trna116_GluCTC_1_-_145399233_145399304</td><td>1</td><td>39</td><td>39</td><td>TCCCTGGTGGTCTAGTGGTTAGGATTCGGCGCTCTCACC</td><td>682.3</td><td>3.0</td><td>0.7</td><td>4.0</td><td>5.2E-05</td><td>7.0E-05</td><td>MB231/MB468</td><td>MITO</td></tr>
<tr><td>>hg38_21_+_8433172_8446622_RNA45SN1_45S_50ntFlanks@10793.10816.24|TCTGGCATGTTGGAACAATGTAGG</td><td>>hg38_21_+_8433172_8446622_RNA45SN1_45S_50ntFlanks</td><td>10793</td><td>10816</td><td>24</td><td>TCTGGCATGTTGGAACAATGTAGG</td><td>74.4</td><td>9.5</td><td>2.1</td><td>4.5</td><td>5.9E-06</td><td>7.0E-05</td><td>BT20/MB231</td><td>MP</td></tr>
<tr><td>>MI0000289|hsa-mir-181a-1&WithFlank&1|-|198859038|198859159@70.89.20&[MIMAT0000270&hsa-miR-181a-3p&offsets|0|-2;m-125&1|-|198859069|198859090&offsets|0|-2]|ACCATCGACCGTTGATTGTA</td><td>>MI0000289|hsa-mir-181a-1&WithFlank&1|-|198859038|198859159</td><td>70</td><td>89</td><td>20</td><td>ACCATCGACCGTTGATTGTA</td><td>62.4</td><td>-9.4</td><td>2.1</td><td>-4.5</td><td>5.9E-06</td><td>7.0E-05</td><td>BT20/MB231</td><td>MP</td></tr>
</table>

<table>
<tr><td>>hg38_21_+_8433172_8446622_RNA45SN1_45S_50ntFlanks@11582.11601.20|ACTGGTGCGGACCAGGGGAA</td><td>>hg38_21_+_8433172_8446622_RNA45SN1_45S_50ntFlanks</td><td>11582</td><td>11601</td><td>20</td><td>ACTGGTGCGGACCAGGGGAA</td><td>277.5</td><td>-2.5</td><td>0.6</td><td>-4.0</td><td>5.2E-05</td><td>7.0E-05</td><td>MB231/MB468</td><td>NUC</td></tr>
<tr><td>>hg38_21_+_8433172_8446622_RNA45SN1_45S_50ntFlanks@12942.12965.24|TTCGTAGACGACCTGCTTCTGGGT</td><td>>hg38_21_+_8433172_8446622_RNA45SN1_45S_50ntFlanks</td><td>12942</td><td>12965</td><td>24</td><td>TTCGTAGACGACCTGCTTCTGGGT</td><td>309.4</td><td>-4.5</td><td>1.1</td><td>-4.0</td><td>5.2E-05</td><td>7.0E-05</td><td>MB231/MB468</td><td>NUC</td></tr>
<tr><td>>hg38_21_+_8433172_8446622_RNA45SN1_45S_50ntFlanks@6733.6772.40|CATTGATCATCGACACTTCGAACGCACTTGCGGCCCCGGG</td><td>>hg38_21_+_8433172_8446622_RNA45SN1_45S_50ntFlanks</td><td>6733</td><td>6772</td><td>40</td><td>CATTGATCATCGACACTTCGAACGCACTTGCGGCCCCGGG</td><td>472.5</td><td>-3.2</td><td>0.8</td><td>-4.0</td><td>5.2E-05</td><td>7.1E-05</td><td>MB231/MB468</td><td>NUC</td></tr>
<tr><td>>hg38_21_+_8433172_8446622_RNA45SN1_45S_50ntFlanks@8171.8197.27|AAGTCCTTCTGATCGAGGCCCAGCCCG</td><td>>hg38_21_+_8433172_8446622_RNA45SN1_45S_50ntFlanks</td><td>8171</td><td>8197</td><td>27</td><td>AAGTCCTTCTGATCGAGGCCCAGCCCG</td><td>1132.8</td><td>-4.2</td><td>1.0</td><td>-4.0</td><td>5.2E-05</td><td>7.1E-05</td><td>MB231/MB468</td><td>NUC</td></tr>
<tr><td>>am_trnaMT_GluTTC_MT_-_14674_14742@27.52.26,trnalookalike8_GluTTC_5_-_93905172_93905240@27.52.26|GTTTTTCATATCATTGGTCGTGGTTG</td><td>>am_trnaMT_GluTTC_MT_-_14674_14742</td><td>27</td><td>52</td><td>26</td><td>GTTTTTCATATCATTGGTCGTGGTTG</td><td>254.0</td><td>-6.3</td><td>1.5</td><td>-4.3</td><td>1.8E-05</td><td>7.1E-05</td><td>BT20/MB468</td><td>MP</td></tr>
<tr><td>>hg38_21_+_8433172_8446622_RNA45SN1_45S_50ntFlanks@9738.9755.18|GAAACGATCTCAACCTAT</td><td>>hg38_21_+_8433172_8446622_RNA45SN1_45S_50ntFlanks</td><td>9738</td><td>9755</td><td>18</td><td>GAAACGATCTCAACCTAT</td><td>432.3</td><td>-4.7</td><td>1.1</td><td>-4.1</td><td>3.8E-05</td><td>7.1E-05</td><td>BT20/MB468</td><td>MITO</td></tr>
<tr><td>>hg38_21_+_8433172_8446622_RNA45SN1_45S_50ntFlanks@10041.10072.32|CCATACCCGGCCGTCGCCGGCAGTCGAGAGTG</td><td>>hg38_21_+_8433172_8446622_RNA45SN1_45S_50ntFlanks</td><td>10041</td><td>10072</td><td>32</td><td>CCATACCCGGCCGTCGCCGGCAGTCGAGAGTG</td><td>187.1</td><td>-2.9</td><td>0.7</td><td>-4.1</td><td>3.8E-05</td><td>7.1E-05</td><td>BT20/MB468</td><td>MITO</td></tr>
<tr><td>>hg38_21_+_8433172_8446622_RNA45SN1_45S_50ntFlanks@6776.6802.27|CTCCCGGGGCTACGCCTGTCTGAGCGT</td><td>>hg38_21_+_8433172_8446622_RNA45SN1_45S_50ntFlanks</td><td>6776</td><td>6802</td><td>27</td><td>CTCCCGGGGCTACGCCTGTCTGAGCGT</td><td>770.9</td><td>1.9</td><td>0.5</td><td>4.0</td><td>5.2E-05</td><td>7.1E-05</td><td>MB231/MB468</td><td>NUC</td></tr>
<tr><td>>hg38_21_+_8433172_8446622_RNA45SN1_45S_50ntFlanks@7988.8017.30|CAGACGTGGCGACCCGCTGAATTTAAGCAT</td><td>>hg38_21_+_8433172_8446622_RNA45SN1_45S_50ntFlanks</td><td>7988</td><td>8017</td><td>30</td><td>CAGACGTGGCGACCCGCTGAATTTAAGCAT</td><td>576.5</td><td>-3.5</td><td>0.9</td><td>-4.0</td><td>5.4E-05</td><td>7.1E-05</td><td>BT20/MB468</td><td>NUC</td></tr>
<tr><td>>hg38_MT_+_1621_3279_RNR2_16S_50ntFlanks@1184.1215.32|AATTTCGGTTGGGGCGACCTCGGAGCAGAACC</td><td>>hg38_MT_+_1621_3279_RNR2_16S_50ntFlanks</td><td>1184</td><td>1215</td><td>32</td><td>AATTTCGGTTGGGGCGACCTCGGAGCAGAACC</td><td>166.4</td><td>4.9</td><td>1.2</td><td>4.0</td><td>5.2E-05</td><td>7.1E-05</td><td>MB231/MB468</td><td>MITO</td></tr>
<tr><td>>hg38_1_-_228634819_228635039_RNA5S12_5S_50ntFlanks@84.112.29|CTCGTCTGATCTCGGAAGCTAAGCAGGGT</td><td>>hg38_1_-_228634819_228635039_RNA5S12_5S_50ntFlanks</td><td>84</td><td>112</td><td>29</td><td>CTCGTCTGATCTCGGAAGCTAAGCAGGGT</td><td>310.1</td><td>-5.2</td><td>1.3</td><td>-4.1</td><td>4.9E-05</td><td>7.1E-05</td><td>MB231/MB468</td><td>CYTO</td></tr>
<tr><td>>hg38_21_+_8433172_8446622_RNA45SN1_45S_50ntFlanks@10922.10939.18|CTGGACGAGGCGCCGCCG</td><td>>hg38_21_+_8433172_8446622_RNA45SN1_45S_50ntFlanks</td><td>10922</td><td>10939</td><td>18</td><td>CTGGACGAGGCGCCGCCG</td><td>4054.1</td><td>-8.3</td><td>2.1</td><td>-4.0</td><td>5.4E-05</td><td>7.1E-05</td><td>BT20/MB468</td><td>NUC</td></tr>
<tr><td>>hg38_21_+_8433172_8446622_RNA45SN1_45S_50ntFlanks@6661.6696.36|CGGTGGATCACTCGGCTCGTGCGTCGATGAAGAACG</td><td>>hg38_21_+_8433172_8446622_RNA45SN1_45S_50ntFlanks</td><td>6661</td><td>6696</td><td>36</td><td>CGGTGGATCACTCGGCTCGTGCGTCGATGAAGAACG</td><td>247.7</td><td>2.1</td><td>0.5</td><td>4.1</td><td>4.9E-05</td><td>7.1E-05</td><td>MB231/MB468</td><td>CYTO</td></tr>
<tr><td>>MI0000342|hsa-mir-200b&WithFlank&1|+|1167098|1167204@63.84.22&[MIMAT0000318&hsa-miR-200b-3p&offsets|0|0;m-93&1|+|1167160|1167182&offsets|0|-1]|TAATACTGCCTGGTAATGATGA</td><td>>MI0000342|hsa-mir-200b&WithFlank&1|+|1167098|1167204</td><td>63</td><td>84</td><td>22</td><td>TAATACTGCCTGGTAATGATGA</td><td>502.3</td><td>-3.4</td><td>0.8</td><td>-4.1</td><td>4.9E-05</td><td>7.1E-05</td><td>MB231/MB468</td><td>CYTO</td></tr>
<tr><td>>MI0000098|hsa-mir-96&WithFlank&7|-|129774686|129774775@15.37.23&[MIMAT0000095&hsa-miR-96-5p&offsets|0|0;m-248&7|-|129774739|129774761&offsets|0|0]|TTTGGCACTAGCACATTTTTGCT</td><td>>MI0000098|hsa-mir-96&WithFlank&7|-|129774686|129774775</td><td>15</td><td>37</td><td>23</td><td>TTTGGCACTAGCACATTTTTGCT</td><td>6515.9</td><td>3.1</td><td>0.7</td><td>4.5</td><td>6.6E-06</td><td>7.1E-05</td><td>BT20/MB231</td><td>NUC</td></tr>
<tr><td>>hg38_MT_+_598_1651_RNR1_12S_50ntFlanks@443.468.26|TATCTGAACACACAATAGCTAAGACC</td><td>>hg38_MT_+_598_1651_RNR1_12S_50ntFlanks</td><td>443</td><td>468</td><td>26</td><td>TATCTGAACACACAATAGCTAAGACC</td><td>281.7</td><td>4.4</td><td>1.1</td><td>4.1</td><td>4.4E-05</td><td>7.1E-05</td><td>MB231/MB468</td><td>MP</td></tr>
<tr><td>>hg38_21_+_8433172_8446622_RNA45SN1_45S_50ntFlanks@4223.4242.20|AAAAATAACAATACAGGACT</td><td>>hg38_21_+_8433172_8446622_RNA45SN1_45S_50ntFlanks</td><td>4223</td><td>4242</td><td>20</td><td>AAAAATAACAATACAGGACT</td><td>205.2</td><td>-2.4</td><td>0.6</td><td>-4.1</td><td>3.8E-05</td><td>7.1E-05</td><td>BT20/MB468</td><td>MITO</td></tr>
<tr><td>>hg38_21_+_8433172_8446622_RNA45SN1_45S_50ntFlanks@11783.11805.23|CATGAATGGATGAACGAGATTCC</td><td>>hg38_21_+_8433172_8446622_RNA45SN1_45S_50ntFlanks</td><td>11783</td><td>11805</td><td>23</td><td>CATGAATGGATGAACGAGATTCC</td><td>706.6</td><td>-2.4</td><td>0.6</td><td>-4.1</td><td>3.8E-05</td><td>7.1E-05</td><td>BT20/MB468</td><td>MITO</td></tr>
<tr><td>>MI0000067|hsa-let-7f-1&WithFlank&9|+|94176341|94176439@13.33.21(+1C);MI0000068|hsa-let-7f-2&WithFlank&X|-|53557186|53557280@14.34.21(+1C)&[MIMAT0000067&hsa-let-7f-5p&offsets|0|-1(+1C);m-20&9|+|94176353|94176374&offsets|0|-1(+1C)];[MIMAT0000067_1&hsa-let-7f-5p&offsets|0|-1(+1C);m-19&X|-|53557246|53557267&offsets|0|-1(+1C)]|TGAGGTAGTAGATTGTATAGTC</td><td>>MI0000067|hsa-let-7f-1&WithFlank&9|+|94176341|94176439</td><td>13</td><td>33</td><td>21(+1C)</td><td>TGAGGTAGTAGATTGTATAGTC</td><td>6944.4</td><td>2.0</td><td>0.5</td><td>4.0</td><td>5.4E-05</td><td>7.1E-05</td><td>BT20/MB468</td><td>NUC</td></tr>
<tr><td>>MI0000102|hsa-mir-100&WithFlank&11|-|122152223|122152314@19.38.20(+1A)&[MIMAT0000098&hsa-miR-100-5p&offsets|0|-2(+1A);m-27&11|-|122152276|122152296&offsets|0|-1(+1A)]|AACCCGTAGATCCGAACTTGA</td><td>>MI0000102|hsa-mir-100&WithFlank&11|-|122152223|122152314</td><td>19</td><td>38</td><td>20(+1A)</td><td>AACCCGTAGATCCGAACTTGA</td><td>2762.5</td><td>-5.6</td><td>1.2</td><td>-4.5</td><td>6.6E-06</td><td>7.1E-05</td><td>BT20/MB231</td><td>NUC</td></tr>
<tr><td>>hg38_21_+_8433172_8446622_RNA45SN1_45S_50ntFlanks@6661.6700.40|CGGTGGATCACTCGGCTCGTGCGTCGATGAAGAACGCAGC</td><td>>hg38_21_+_8433172_8446622_RNA45SN1_45S_50ntFlanks</td><td>6661</td><td>6700</td><td>40</td><td>CGGTGGATCACTCGGCTCGTGCGTCGATGAAGAACGCAGC</td><td>961.4</td><td>7.0</td><td>1.7</td><td>4.1</td><td>4.6E-05</td><td>7.1E-05</td><td>BT20/MB468</td><td>CYTO</td></tr>
<tr><td>>hg38_21_+_8433172_8446622_RNA45SN1_45S_50ntFlanks@476.524.49|TTCGTGATCGATGTGGTGACGTCGTGCTCTCCCGGGCCGGGTCCGAGCC</td><td>>hg38_21_+_8433172_8446622_RNA45SN1_45S_50ntFlanks</td><td>476</td><td>524</td><td>49</td><td>TTCGTGATCGATGTGGTGACGTCGTGCTCTCCCGGGCCGGGTCCGAGCC</td><td>175.0</td><td>7.0</td><td>1.7</td><td>4.0</td><td>5.2E-05</td><td>7.1E-05</td><td>MB231/MB468</td><td>MITO</td></tr>
<tr><td>>hg38_21_+_8433172_8446622_RNA45SN1_45S_50ntFlanks@9545.9572.28|GTGGAGGTCCGTAGCGGTCCTGACGTGC</td><td>>hg38_21_+_8433172_8446622_RNA45SN1_45S_50ntFlanks</td><td>9545</td><td>9572</td><td>28</td><td>GTGGAGGTCCGTAGCGGTCCTGACGTGC</td><td>3294.8</td><td>-2.4</td><td>0.6</td><td>-4.1</td><td>4.9E-05</td><td>7.1E-05</td><td>MB231/MB468</td><td>CYTO</td></tr>
<tr><td>>hg38_21_+_8433172_8446622_RNA45SN1_45S_50ntFlanks@10678.10709.32|CGGTTCCGGCGGCGTCCGGTGAGCTCTCGCTG</td><td>>hg38_21_+_8433172_8446622_RNA45SN1_45S_50ntFlanks</td><td>10678</td><td>10709</td><td>32</td><td>CGGTTCCGGCGGCGTCCGGTGAGCTCTCGCTG</td><td>19445.9</td><td>-5.9</td><td>1.4</td><td>-4.1</td><td>4.9E-05</td><td>7.1E-05</td><td>MB231/MB468</td><td>CYTO</td></tr>
<tr><td>>hg38_21_+_8433172_8446622_RNA45SN1_45S_50ntFlanks@10386.10410.25|ATGTGAACAGCAGTTGAACATGGGT</td><td>>hg38_21_+_8433172_8446622_RNA45SN1_45S_50ntFlanks</td><td>10386</td><td>10410</td><td>25</td><td>ATGTGAACAGCAGTTGAACATGGGT</td><td>823.8</td><td>-6.5</td><td>1.5</td><td>-4.3</td><td>1.8E-05</td><td>7.1E-05</td><td>BT20/MB468</td><td>MP</td></tr>
<tr><td>>hg38_21_+_8433172_8446622_RNA45SN1_45S_50ntFlanks@4069.4089.21|CTTTCGATGGTAGTCGCCGTG</td><td>>hg38_21_+_8433172_8446622_RNA45SN1_45S_50ntFlanks</td><td>4069</td><td>4089</td><td>21</td><td>CTTTCGATGGTAGTCGCCGTG</td><td>1105.8</td><td>-6.0</td><td>1.5</td><td>-4.1</td><td>3.8E-05</td><td>7.1E-05</td><td>BT20/MB468</td><td>MITO</td></tr>
<tr><td>>hg38_21_+_8433172_8446622_RNA45SN1_45S_50ntFlanks@5486.5522.37|GCGCTGAGAAGACGGTCGAACTTGACTATCTAGAGGA</td><td>>hg38_21_+_8433172_8446622_RNA45SN1_45S_50ntFlanks</td><td>5486</td><td>5522</td><td>37</td><td>GCGCTGAGAAGACGGTCGAACTTGACTATCTAGAGGA</td><td>133.7</td><td>-6.1</td><td>1.5</td><td>-4.1</td><td>4.4E-05</td><td>7.1E-05</td><td>MB231/MB468</td><td>MP</td></tr>
<tr><td>>hg38_21_+_8433172_8446622_RNA45SN1_45S_50ntFlanks@5236.5259.24|CACTGACTGGCTCAGCGTGTGCCT</td><td>>hg38_21_+_8433172_8446622_RNA45SN1_45S_50ntFlanks</td><td>5236</td><td>5259</td><td>24</td><td>CACTGACTGGCTCAGCGTGTGCCT</td><td>587.8</td><td>-6.4</td><td>1.6</td><td>-4.0</td><td>5.2E-05</td><td>7.1E-05</td><td>MB231/MB468</td><td>MITO</td></tr>
<tr><td>>hg38_21_+_8433172_8446622_RNA45SN1_45S_50ntFlanks@12455.12473.19|ACCAAGCGTTGGATTGTTC</td><td>>hg38_21_+_8433172_8446622_RNA45SN1_45S_50ntFlanks</td><td>12455</td><td>12473</td><td>19</td><td>ACCAAGCGTTGGATTGTTC</td><td>801.1</td><td>-5.5</td><td>1.3</td><td>-4.3</td><td>1.8E-05</td><td>7.1E-05</td><td>BT20/MB468</td><td>MP</td></tr>
<tr><td>>MI0000301|hsa-mir-224&WithFlank&X|-|151958572|151958664@14.36.23(+1A)&[MIMAT0000281&hsa-miR-224-5p&offsets|+1|-1(+1A);m-238&X|-|151958630|151958651&offsets|0|+1(+1A)]|CAAGTCACTAGTGGTTCCGTTTAA</td><td>>MI0000301|hsa-mir-224&WithFlank&X|-|151958572|151958664</td><td>14</td><td>36</td><td>23(+1A)</td><td>CAAGTCACTAGTGGTTCCGTTTAA</td><td>34.9</td><td>8.3</td><td>2.0</td><td>4.1</td><td>4.4E-05</td><td>7.1E-05</td><td>MB231/MB468</td><td>MP</td></tr>
<tr><td>>hg38_21_+_8433172_8446622_RNA45SN1_45S_50ntFlanks@8176.8195.20|CTTCTGATCGAGGCCCAGCC</td><td>>hg38_21_+_8433172_8446622_RNA45SN1_45S_50ntFlanks</td><td>8176</td><td>8195</td><td>20</td><td>CTTCTGATCGAGGCCCAGCC</td><td>1905.8</td><td>-2.8</td><td>0.7</td><td>-4.0</td><td>5.3E-05</td><td>7.1E-05</td><td>MB231/MB468</td><td>MITO</td></tr>
<tr><td>>hg38_21_+_8433172_8446622_RNA45SN1_45S_50ntFlanks@7979.8010.32|ACCTCAGATCAGACGTGGCGACCCGCTGAATT</td><td>>hg38_21_+_8433172_8446622_RNA45SN1_45S_50ntFlanks</td><td>7979</td><td>8010</td><td>32</td><td>ACCTCAGATCAGACGTGGCGACCCGCTGAATT</td><td>977.0</td><td>2.9</td><td>0.7</td><td>4.1</td><td>3.8E-05</td><td>7.1E-05</td><td>BT20/MB468</td><td>MITO</td></tr>
<tr><td>>trna111_HisGTG_1_-_147774845_147774916@-1A.22.23,trna118_HisGTG_1_-_145396881_145396952@-1A.22.23,trna16_HisGTG_1_+_146544773_146544844@-1A.22.23,trna1_HisGTG_15_+_45493349_45493420@-1A.22.23,trna21_HisGTG_1_+_147753471_147753542@-1A.22.23,trna33_HisGTG_6_+_27125906_27125977@-1A.22.23,trna7_HisGTG_9_-_14433938_14434009@-1A.22.23,trna8_HisGTG_15_-_45492611_45492682@-1A.22.23,trna9_HisGTG_15_-_45490804_45490875@-1A.22.23|AGCCGTGATCGTATAGTGGTTAG</td><td>>trna111_HisGTG_1_-_147774845_147774916</td><td>-1A</td><td>22</td><td>23</td><td>AGCCGTGATCGTATAGTGGTTAG</td><td>68.9</td><td>-9.5</td><td>2.1</td><td>-4.5</td><td>6.0E-06</td><td>7.1E-05</td><td>BT20/MB231</td><td>MP</td></tr>
<tr><td>>hg38_21_+_8433172_8446622_RNA45SN1_45S_50ntFlanks@9534.9560.27|AGGAAACTCTGGTGGAGGTCCGTAGCG</td><td>>hg38_21_+_8433172_8446622_RNA45SN1_45S_50ntFlanks</td><td>9534</td><td>9560</td><td>27</td><td>AGGAAACTCTGGTGGAGGTCCGTAGCG</td><td>713.0</td><td>-2.7</td><td>0.7</td><td>-4.1</td><td>4.6E-05</td><td>7.1E-05</td><td>BT20/MB468</td><td>CYTO</td></tr>
<tr><td>>hg38_21_+_8433172_8446622_RNA45SN1_45S_50ntFlanks@12624.12661.38|CCAATGGGGCGAAGCTACCATCTGTGGGATTATGACTG</td><td>>hg38_21_+_8433172_8446622_RNA45SN1_45S_50ntFlanks</td><td>12624</td><td>12661</td><td>38</td><td>CCAATGGGGCGAAGCTACCATCTGTGGGATTATGACTG</td><td>498.8</td><td>-1.9</td><td>0.5</td><td>-4.1</td><td>4.9E-05</td><td>7.1E-05</td><td>MB231/MB468</td><td>CYTO</td></tr>
<tr><td>>hg38_21_+_8433172_8446622_RNA45SN1_45S_50ntFlanks@11671.11688.18|CCAGTGCTCTGAATGTCA</td><td>>hg38_21_+_8433172_8446622_RNA45SN1_45S_50ntFlanks</td><td>11671</td><td>11688</td><td>18</td><td>CCAGTGCTCTGAATGTCA</td><td>470.8</td><td>-3.3</td><td>0.8</td><td>-4.0</td><td>5.2E-05</td><td>7.2E-05</td><td>MB231/MB468</td><td>NUC</td></tr>
<tr><td>>hg38_21_+_8433172_8446622_RNA45SN1_45S_50ntFlanks@8317.8356.40|CACGAGACCGATAGTCAACAAGTACCGTAAGGGAAAGTTG</td><td>>hg38_21_+_8433172_8446622_RNA45SN1_45S_50ntFlanks</td><td>8317</td><td>8356</td><td>40</td><td>CACGAGACCGATAGTCAACAAGTACCGTAAGGGAAAGTTG</td><td>513.9</td><td>-3.3</td><td>0.8</td><td>-4.1</td><td>5.0E-05</td><td>7.2E-05</td><td>MB231/MB468</td><td>CYTO</td></tr>
<tr><td>>MI0000293|hsa-mir-217&WithFlank&2|-|55982961|55983082@41.63.23&[MIMAT0000274&hsa-miR-217-5p&offsets|0|0;m-311&2|-|55983020|55983042&offsets|0|0]|TACTGCATCAGGAACTGATTGGA</td><td>>MI0000293|hsa-mir-217&WithFlank&2|-|55982961|55983082</td><td>41</td><td>63</td><td>23</td><td>TACTGCATCAGGAACTGATTGGA</td><td>66.0</td><td>-9.4</td><td>2.1</td><td>-4.5</td><td>6.0E-06</td><td>7.2E-05</td><td>BT20/MB231</td><td>MP</td></tr>
<tr><td>>hg38_21_+_8433172_8446622_RNA45SN1_45S_50ntFlanks@11728.11751.24|GTAACTATGACTCTCTTAAGGTAG</td><td>>hg38_21_+_8433172_8446622_RNA45SN1_45S_50ntFlanks</td><td>11728</td><td>11751</td><td>24</td><td>GTAACTATGACTCTCTTAAGGTAG</td><td>226.1</td><td>-3.4</td><td>0.8</td><td>-4.1</td><td>5.0E-05</td><td>7.2E-05</td><td>MB231/MB468</td><td>CYTO</td></tr>
<tr><td>>hg38_21_+_8433172_8446622_RNA45SN1_45S_50ntFlanks@11579.11607.29|AGAACTGGTGCGGACCAGGGGAATCCGAC</td><td>>hg38_21_+_8433172_8446622_RNA45SN1_45S_50ntFlanks</td><td>11579</td><td>11607</td><td>29</td><td>AGAACTGGTGCGGACCAGGGGAATCCGAC</td><td>352.3</td><td>-2.4</td><td>0.6</td><td>-4.0</td><td>5.3E-05</td><td>7.2E-05</td><td>MB231/MB468</td><td>NUC</td></tr>
<tr><td>>hg38_21_+_8433172_8446622_RNA45SN1_45S_50ntFlanks@5073.5110.38|AATTCCGATAACGAACGAGACTCTGGCATGCTAACTAG</td><td>>hg38_21_+_8433172_8446622_RNA45SN1_45S_50ntFlanks</td><td>5073</td><td>5110</td><td>38</td><td>AATTCCGATAACGAACGAGACTCTGGCATGCTAACTAG</td><td>587.8</td><td>-5.1</td><td>1.3</td><td>-4.0</td><td>5.3E-05</td><td>7.2E-05</td><td>MB231/MB468</td><td>NUC</td></tr>
<tr><td>>hg38_21_+_8433172_8446622_RNA45SN1_45S_50ntFlanks@3840.3874.35|CTCCTACTTGGATAACTGTGGTAATTCTAGAGCTA</td><td>>hg38_21_+_8433172_8446622_RNA45SN1_45S_50ntFlanks</td><td>3840</td><td>3874</td><td>35</td><td>CTCCTACTTGGATAACTGTGGTAATTCTAGAGCTA</td><td>59.2</td><td>-9.8</td><td>2.4</td><td>-4.1</td><td>4.4E-05</td><td>7.2E-05</td><td>MB231/MB468</td><td>MP</td></tr>
<tr><td>>hg38_21_+_8433172_8446622_RNA45SN1_45S_50ntFlanks@7991.8018.28|ACGTGGCGACCCGCTGAATTTAAGCATA</td><td>>hg38_21_+_8433172_8446622_RNA45SN1_45S_50ntFlanks</td><td>7991</td><td>8018</td><td>28</td><td>ACGTGGCGACCCGCTGAATTTAAGCATA</td><td>159.8</td><td>3.0</td><td>0.7</td><td>4.1</td><td>5.0E-05</td><td>7.2E-05</td><td>MB231/MB468</td><td>CYTO</td></tr>
<tr><td>>hg38_21_+_8433172_8446622_RNA45SN1_45S_50ntFlanks@10846.10866.21|CTTCGGGATAAGGATTGGCTC</td><td>>hg38_21_+_8433172_8446622_RNA45SN1_45S_50ntFlanks</td><td>10846</td><td>10866</td><td>21</td><td>CTTCGGGATAAGGATTGGCTC</td><td>269.9</td><td>-3.0</td><td>0.7</td><td>-4.1</td><td>4.6E-05</td><td>7.2E-05</td><td>BT20/MB468</td><td>CYTO</td></tr>
<tr><td>>MI0000469|hsa-mir-125a&WithFlank&19|+|51693248|51693345@21.42.22&[MIMAT0000443&hsa-miR-125a-5p&offsets|0|-2;m-63&19|+|51693268|51693289&offsets|0|0]|TCCCTGAGACCCTTTAACCTGT</td><td>>MI0000469|hsa-mir-125a&WithFlank&19|+|51693248|51693345</td><td>21</td><td>42</td><td>22</td><td>TCCCTGAGACCCTTTAACCTGT</td><td>1619.7</td><td>3.5</td><td>0.9</td><td>4.0</td><td>5.3E-05</td><td>7.2E-05</td><td>MB231/MB468</td><td>MITO</td></tr>
<tr><td>>MI0000736|hsa-mir-30c-1&WithFlank&1|+|40757278|40757378@23.45.23;MI0000254|hsa-mir-30c-2&WithFlank&6|-|71376954|71377037@13.35.23&[MIMAT0000244&hsa-miR-30c-5p&offsets|0|0;m-57&1|+|40757300|40757323&offsets|0|-1];[MIMAT0000244_1&hsa-miR-30c-5p&offsets|0|0;m-56&6|-|71377002|71377025&offsets|0|-1]|TGTAAACATCCTACACTCTCAGC</td><td>>MI0000736|hsa-mir-30c-1&WithFlank&1|+|40757278|40757378</td><td>23</td><td>45</td><td>23</td><td>TGTAAACATCCTACACTCTCAGC</td><td>5333.9</td><td>3.5</td><td>0.9</td><td>4.1</td><td>4.5E-05</td><td>7.2E-05</td><td>MB231/MB468</td><td>MP</td></tr>
<tr><td>>hg38_21_+_8433172_8446622_RNA45SN1_45S_50ntFlanks@5492.5513.22|AGAAGACGGTCGAACTTGACTA</td><td>>hg38_21_+_8433172_8446622_RNA45SN1_45S_50ntFlanks</td><td>5492</td><td>5513</td><td>22</td><td>AGAAGACGGTCGAACTTGACTA</td><td>991.4</td><td>-4.2</td><td>1.0</td><td>-4.3</td><td>1.8E-05</td><td>7.2E-05</td><td>BT20/MB468</td><td>MP</td></tr>
<tr><td>>hg38_MT_+_1621_3279_RNR2_16S_50ntFlanks@929.958.30|CAGTGACACATGTTTAACGGCCGCGGTACC</td><td>>hg38_MT_+_1621_3279_RNR2_16S_50ntFlanks</td><td>929</td><td>958</td><td>30</td><td>CAGTGACACATGTTTAACGGCCGCGGTACC</td><td>145.5</td><td>6.9</td><td>1.7</td><td>4.0</td><td>5.3E-05</td><td>7.2E-05</td><td>MB231/MB468</td><td>MITO</td></tr>
</table>

| | | | | | | | | | | | | | |
|---|---|---|---|---|---|---|---|---|---|---|---|---|---|
| >hg38_21_+_8433172_8446622_RNA45SN1_45S_50ntFlanks@12419.12453.35\|GATGTCGGCTCTTCCTATCATTGTGAAGCAGAATT | >hg38_21_+_8433172_8446622_RNA45SN1_45S_50ntFlanks | 12419 | 12453 | 35 | GATGTCGGCTCTTCCTATCATTGTGAAGCAGAATT | 406.6 | 4.0 | 1.0 | 4.0 | 5.3E-05 | 7.2E-05 | MB231/MB468 | NUC |
| >hg38_21_+_8433172_8446622_RNA45SN1_45S_50ntFlanks@8116.8136.21\|GCGGGACATGTGGCGTACGGA | >hg38_21_+_8433172_8446622_RNA45SN1_45S_50ntFlanks | 8116 | 8136 | 21 | GCGGGACATGTGGCGTACGGA | 938.3 | 5.5 | 1.3 | 4.3 | 1.8E-05 | 7.2E-05 | BT20/MB468 | MP |
| >MI0000076\|hsa-mir-20a&WithFlank&13\|+\|91351059\|91351141@14.33.20(+1C)&[MIMAT0000075&hsa-miR-20a-5p&offsets\|0\|-3(+1C);m-96&13\|+\|91351072\|91351094&offsets\|0\|-3(+1C)]\|TAAAGTGCTTATAGTGCAGGC | >MI0000076\|hsa-mir-20a&WithFlank&13\|+\|91351059\|91351141 | 14 | 33 | 20(+1C) | TAAAGTGCTTATAGTGCAGGC | 342.8 | 3.5 | 0.9 | 4.1 | 4.5E-05 | 7.2E-05 | MB231/MB468 | MP |
| >hg38_21_+_8433172_8446622_RNA45SN1_45S_50ntFlanks@4752.4783.32\|GAAGACGATCAGATACCGTCGTAGTTCCGACC | >hg38_21_+_8433172_8446622_RNA45SN1_45S_50ntFlanks | 4752 | 4783 | 32 | GAAGACGATCAGATACCGTCGTAGTTCCGACC | 619.6 | -5.3 | 1.3 | -4.0 | 5.3E-05 | 7.2E-05 | MB231/MB468 | MITO |
| >MI0000767\|hsa-mir-365a&WithFlank&16\|+\|14309279\|14309377@62.83.22;MI0000769\|hsa-mir-365b&WithFlank&17\|+\|31575405\|31575527@74.95.22&[MIMAT0000710&hsa-miR-365a-3p&offsets\|0\|0;m-176&16\|+\|14309340\|14309361&offsets\|0\|0];[MIMAT0022834&hsa-miR-365b-3p&offsets\|0\|0;m-175&17\|+\|31575478\|31575499&offsets\|0\|0]\|TAATGCCCCTAAAAATCCTTAT | >MI0000767\|hsa-mir-365a&WithFlank&16\|+\|14309279\|14309377 | 62 | 83 | 22 | TAATGCCCCTAAAAATCCTTAT | 1963.9 | 3.7 | 0.8 | 4.5 | 5.8E-06 | 7.2E-05 | BT20/MB231 | CYTO |
| >hg38_1_-_228634819_228635039_RNA5S12_5S_50ntFlanks@131.170.40\|GGATGGGAGACCGCCTGGGAATACCGGGTGCTGTAGGCTT | >hg38_1_-_228634819_228635039_RNA5S12_5S_50ntFlanks | 131 | 170 | 40 | GGATGGGAGACCGCCTGGGAATACCGGGTGCTGTAGGCTT | 1877.1 | 3.9 | 0.9 | 4.5 | 7.0E-06 | 7.2E-05 | BT20/MB231 | MITO |
| >hg38_21_+_8433172_8446622_RNA45SN1_45S_50ntFlanks@5074.5102.29\|ATTCCGATAACGAACGAGACTCTGGCATG | >hg38_21_+_8433172_8446622_RNA45SN1_45S_50ntFlanks | 5074 | 5102 | 29 | ATTCCGATAACGAACGAGACTCTGGCATG | 379.2 | -5.2 | 1.3 | -4.0 | 5.3E-05 | 7.2E-05 | MB231/MB468 | NUC |
| >hg38_21_+_8433172_8446622_RNA45SN1_45S_50ntFlanks@8356.8379.24\|GAAAAGAACTTTGAAGAGAGAGTT | >hg38_21_+_8433172_8446622_RNA45SN1_45S_50ntFlanks | 8356 | 8379 | 24 | GAAAAGAACTTTGAAGAGAGAGTT | 3214.8 | -3.4 | 0.8 | -4.1 | 3.9E-05 | 7.2E-05 | BT20/MB468 | MITO |
| >MI0000063\|hsa-let-7b&WithFlank&22\|+\|46113680\|46113774@12.33.22(+1C)&[MIMAT0000063&hsa-let-7b-5p&offsets\|0\|0(+1C);m-14&22\|+\|46113691\|46113712&offsets\|0\|0(+1C)]\|TGAGGTAGTAGGTTGTGTGGTTC | >MI0000063\|hsa-let-7b&WithFlank&22\|+\|46113680\|46113774 | 12 | 33 | 22(+1C) | TGAGGTAGTAGGTTGTGTGGTTC | 207.2 | 6.2 | 1.5 | 4.0 | 5.3E-05 | 7.2E-05 | MB231/MB468 | MITO |
| >hg38_21_+_8433172_8446622_RNA45SN1_45S_50ntFlanks@4367.4398.32\|CAGTTAAAAAGCTCGTAGTTGGATCTTGGGAG | >hg38_21_+_8433172_8446622_RNA45SN1_45S_50ntFlanks | 4367 | 4398 | 32 | CAGTTAAAAAGCTCGTAGTTGGATCTTGGGAG | 346.3 | -4.1 | 1.0 | -4.1 | 4.7E-05 | 7.2E-05 | BT20/MB468 | CYTO |
| >hg38_21_+_8433172_8446622_RNA45SN1_45S_50ntFlanks@8737.8756.20\|CCCGTCCGTCCGTCCGTCCG | >hg38_21_+_8433172_8446622_RNA45SN1_45S_50ntFlanks | 8737 | 8756 | 20 | CCCGTCCGTCCGTCCGTCCG | 156.0 | -5.3 | 1.3 | -4.1 | 3.9E-05 | 7.2E-05 | BT20/MB468 | MITO |
| >am_trna10_LysCTT_16_+_3241501_3241573@11.28.18,trna119_LysCTT_1_-_145395522_145395594@11.28.18,trna11_LysCTT_5_-_180648979_180649051@11.28.18,trna128_LysCTT_1_-_55423542_55423614@11.28.18,trna13_LysCTT_14_-_58706613_58706685@11.28.18,trna13_LysCTT_6_+_26556774_26556846@11.28.18,trna2_LysCTT_15_+_79152904_79152976@11.28.18,trna32_LysCTT_16_-_3207406_3207478@11.28.18,trna4_LysCTT_18_-_43669271_43669343@11.28.18,trna6_LysCTT_19_-_52425396_52425468@11.28.18,trna7_LysCTT_16_+_3225692_3225764@11.28.18,trna9_LysCTT_5_+_180634755_180634827@11.28.18\|CTCAGTCGGTAGAGCATG | >am_trna10_LysCTT_16_+_3241501_3241573 | 11 | 28 | 18 | CTCAGTCGGTAGAGCATG | 145.6 | -4.2 | 0.9 | -4.5 | 7.1E-06 | 7.2E-05 | BT20/MB231 | MITO |
| >hg38_21_+_8433172_8446622_RNA45SN1_45S_50ntFlanks@10849.10868.20\|CGGGATAAGGATTGGCTCTA | >hg38_21_+_8433172_8446622_RNA45SN1_45S_50ntFlanks | 10849 | 10868 | 20 | CGGGATAAGGATTGGCTCTA | 652.6 | -5.5 | 1.3 | -4.1 | 4.7E-05 | 7.2E-05 | BT20/MB468 | CYTO |
| >hg38_21_+_8433172_8446622_RNA45SN1_45S_50ntFlanks@11845.11871.27\|GGAACGGGCTTGGCGGAATCAGCGGGG | >hg38_21_+_8433172_8446622_RNA45SN1_45S_50ntFlanks | 11845 | 11871 | 27 | GGAACGGGCTTGGCGGAATCAGCGGGG | 596.6 | -3.5 | 0.8 | -4.3 | 1.8E-05 | 7.3E-05 | BT20/MB468 | MP |
| >hg38_21_+_8433172_8446622_RNA45SN1_45S_50ntFlanks@3802.3837.36\|CATTAAATCAGTTATGGTTCCTTTGGTCGCTCGCTC | >hg38_21_+_8433172_8446622_RNA45SN1_45S_50ntFlanks | 3802 | 3837 | 36 | CATTAAATCAGTTATGGTTCCTTTGGTCGCTCGCTC | 45.9 | -8.6 | 2.0 | -4.3 | 1.8E-05 | 7.3E-05 | BT20/MB468 | MP |
| >MI0000088\|hsa-mir-30a&WithFlank&6\|-\|71403545\|71403627@12.34.23(+1A)&[MIMAT0000087&hsa-miR-30a-5p&offsets\|0\|+1(+1A);m-15&6\|-\|71403595\|71403616&offsets\|0\|+1(+1A)]\|TGTAAACATCCTCGACTGGAAGCA | >MI0000088\|hsa-mir-30a&WithFlank&6\|-\|71403545\|71403627 | 12 | 34 | 23(+1A) | TGTAAACATCCTCGACTGGAAGCA | 1195.3 | 4.1 | 1.0 | 4.0 | 5.5E-05 | 7.3E-05 | BT20/MB468 | NUC |
| >hg38_21_+_8433172_8446622_RNA45SN1_45S_50ntFlanks@12995.13039.45\|CTCGCTGCGATCTATTGAAAGTCAGCCCTCGACACAAGGGTTTGT | >hg38_21_+_8433172_8446622_RNA45SN1_45S_50ntFlanks | 12995 | 13039 | 45 | CTCGCTGCGATCTATTGAAAGTCAGCCCTCGACACAAGGGTTTGT | 2253.6 | 3.6 | 0.9 | 4.1 | 4.5E-05 | 7.3E-05 | MB231/MB468 | MP |
| >hg38_21_+_8433172_8446622_RNA45SN1_45S_50ntFlanks@9941.9968.28\|AGGACGGTGGCCATGGAAGTCGGAATCC | >hg38_21_+_8433172_8446622_RNA45SN1_45S_50ntFlanks | 9941 | 9968 | 28 | AGGACGGTGGCCATGGAAGTCGGAATCC | 296.7 | 4.5 | 1.0 | 4.5 | 7.1E-06 | 7.3E-05 | BT20/MB231 | MITO |
| >MI0000446\|hsa-mir-125b-1&WithFlank&11\|-\|122099751\|122099850@21.42.22;MI0000470\|hsa-mir-125b-2&WithFlank&21\|+\|16590231\|16590331@23.44.22&[MIMAT0000423&hsa-miR-125b-5p&offsets\|0\|0;m-67&11\|-\|122099809\|122099830&offsets\|0\|0];[MIMAT0000423_1&hsa-miR-125b-5p&offsets\|0\|0;m-68&21\|+\|16590253\|16590274&offsets\|0\|0]\|TCCCTGAGACCCTAACTTGTGA | >MI0000446\|hsa-mir-125b-1&WithFlank&11\|-\|122099751\|122099850 | 21 | 42 | 22 | TCCCTGAGACCCTAACTTGTGA | 14585.6 | -5.3 | 1.2 | -4.5 | 6.8E-06 | 7.3E-05 | BT20/MB231 | NUC |
| >hg38_21_+_8433172_8446622_RNA45SN1_45S_50ntFlanks@11703.11738.36\|AATGAAGCGCGGGTAAACGGCGGGAGTAACTATGAC | >hg38_21_+_8433172_8446622_RNA45SN1_45S_50ntFlanks | 11703 | 11738 | 36 | AATGAAGCGCGGGTAAACGGCGGGAGTAACTATGAC | 240.7 | -4.7 | 1.2 | -4.0 | 5.4E-05 | 7.3E-05 | MB231/MB468 | MITO |
| >hg38_21_+_8433172_8446622_RNA45SN1_45S_50ntFlanks@10791.10823.33\|CCTCTGGCATGTTGGAACAATGTAGGTAAGGGA | >hg38_21_+_8433172_8446622_RNA45SN1_45S_50ntFlanks | 10791 | 10823 | 33 | CCTCTGGCATGTTGGAACAATGTAGGTAAGGGA | 299.2 | -4.9 | 1.2 | -4.1 | 4.5E-05 | 7.3E-05 | MB231/MB468 | MP |
| >hg38_21_+_8433172_8446622_RNA45SN1_45S_50ntFlanks@12635.12652.18\|AAGCTACCATCTGTGGGA | >hg38_21_+_8433172_8446622_RNA45SN1_45S_50ntFlanks | 12635 | 12652 | 18 | AAGCTACCATCTGTGGGA | 172.7 | -5.5 | 1.3 | -4.1 | 3.9E-05 | 7.3E-05 | BT20/MB468 | MITO |
| >hg38_21_+_8433172_8446622_RNA45SN1_45S_50ntFlanks@10396.10424.29\|CAGTTGAACATGGGTCAGTCGGTCCTGAG | >hg38_21_+_8433172_8446622_RNA45SN1_45S_50ntFlanks | 10396 | 10424 | 29 | CAGTTGAACATGGGTCAGTCGGTCCTGAG | 3968.1 | 2.8 | 0.7 | 4.0 | 5.3E-05 | 7.3E-05 | MB231/MB468 | NUC |
| >hg38_21_+_8433172_8446622_RNA45SN1_45S_50ntFlanks@5486.5519.34\|GCGCTGAGAAGACGGTCGAACTTGACTATCTAGA | >hg38_21_+_8433172_8446622_RNA45SN1_45S_50ntFlanks | 5486 | 5519 | 34 | GCGCTGAGAAGACGGTCGAACTTGACTATCTAGA | 126.8 | -6.0 | 1.5 | -4.1 | 4.5E-05 | 7.3E-05 | MB231/MB468 | MP |
| >trnaMT_LeuTAA_MT_+_3230_3304@18.44.27\|CGGTAATCGCATAAAACTTAAAACTTT | >trnaMT_LeuTAA_MT_+_3230_3304 | 18 | 44 | 27 | CGGTAATCGCATAAAACTTAAAACTTT | 1140.6 | -3.8 | 0.9 | -4.0 | 5.4E-05 | 7.3E-05 | MB231/MB468 | MITO |
| >hg38_21_+_8433172_8446622_RNA45SN1_45S_50ntFlanks@6739.6771.33\|TCATCGACACTTCGAACGCACTTGCGGCCCCGG | >hg38_21_+_8433172_8446622_RNA45SN1_45S_50ntFlanks | 6739 | 6771 | 33 | TCATCGACACTTCGAACGCACTTGCGGCCCCGG | 355.2 | -5.9 | 1.5 | -4.0 | 5.4E-05 | 7.3E-05 | MB231/MB468 | NUC |
| >MI0000063\|hsa-let-7b&WithFlank&22\|+\|46113680\|46113774@12.30.19(+1A)&[MIMAT0000063&hsa-let-7b-5p&offsets\|0\|-3(+1A);m-14&22\|+\|46113691\|46113712&offsets\|0\|-3(+1A)]\|TGAGGTAGTAGGTTGTGTGA | >MI0000063\|hsa-let-7b&WithFlank&22\|+\|46113680\|46113774 | 12 | 30 | 19(+1A) | TGAGGTAGTAGGTTGTGTGA | 255.5 | 4.0 | 0.9 | 4.5 | 5.8E-06 | 7.3E-05 | BT20/MB231 | CYTO |
| >hg38_21_+_8433172_8446622_RNA45SN1_45S_50ntFlanks@11603.11620.18\|CCGACTGTTTAATTAAAA | >hg38_21_+_8433172_8446622_RNA45SN1_45S_50ntFlanks | 11603 | 11620 | 18 | CCGACTGTTTAATTAAAA | 785.7 | -2.9 | 0.7 | -4.0 | 5.4E-05 | 7.3E-05 | MB231/MB468 | NUC |
| >hg38_21_+_8433172_8446622_RNA45SN1_45S_50ntFlanks@3993.4024.32\|GTGACTCTAGATAACCTCGGGCCGATCGCACG | >hg38_21_+_8433172_8446622_RNA45SN1_45S_50ntFlanks | 3993 | 4024 | 32 | GTGACTCTAGATAACCTCGGGCCGATCGCACG | 1300.9 | -3.0 | 0.7 | -4.1 | 3.9E-05 | 7.3E-05 | BT20/MB468 | MITO |
| >hg38_21_+_8433172_8446622_RNA45SN1_45S_50ntFlanks@12419.12441.23\|GATGTCGGCTCTTCCTATCATTG | >hg38_21_+_8433172_8446622_RNA45SN1_45S_50ntFlanks | 12419 | 12441 | 23 | GATGTCGGCTCTTCCTATCATTG | 2923.7 | 2.7 | 0.7 | 4.1 | 5.1E-05 | 7.3E-05 | MB231/MB468 | CYTO |
| >hg38_21_+_8433172_8446622_RNA45SN1_45S_50ntFlanks@9920.9951.32\|GTGTTGGTTGATATAGACAGCAGGACGGTGGC | >hg38_21_+_8433172_8446622_RNA45SN1_45S_50ntFlanks | 9920 | 9951 | 32 | GTGTTGGTTGATATAGACAGCAGGACGGTGGC | 73.1 | 9.5 | 2.1 | 4.5 | 6.2E-06 | 7.3E-05 | BT20/MB231 | MP |
| >hg38_21_+_8433172_8446622_RNA45SN1_45S_50ntFlanks@9574.9594.21\|AATCGGTCGTCCGACCTGGGT | >hg38_21_+_8433172_8446622_RNA45SN1_45S_50ntFlanks | 9574 | 9594 | 21 | AATCGGTCGTCCGACCTGGGT | 1104.5 | -4.5 | 1.1 | -4.0 | 5.4E-05 | 7.3E-05 | MB231/MB468 | MITO |
| >hg38_21_+_8433172_8446622_RNA45SN1_45S_50ntFlanks@3837.3860.24\|CCTCTCCTACTTGGATAACTGTGG | >hg38_21_+_8433172_8446622_RNA45SN1_45S_50ntFlanks | 3837 | 3860 | 24 | CCTCTCCTACTTGGATAACTGTGG | 359.5 | -4.5 | 1.1 | -4.0 | 5.4E-05 | 7.3E-05 | MB231/MB468 | NUC |
| >hg38_21_+_8433172_8446622_RNA45SN1_45S_50ntFlanks@12045.12066.22\|CTGACCCGGTGAGGCGGGGGGG | >hg38_21_+_8433172_8446622_RNA45SN1_45S_50ntFlanks | 12045 | 12066 | 22 | CTGACCCGGTGAGGCGGGGGGG | 45.3 | -8.6 | 2.0 | -4.3 | 1.9E-05 | 7.3E-05 | BT20/MB468 | MP |
| >hg38_21_+_8433172_8446622_RNA45SN1_45S_50ntFlanks@11581.11600.20\|AACTGGTGCGGACCAGGGGA | >hg38_21_+_8433172_8446622_RNA45SN1_45S_50ntFlanks | 11581 | 11600 | 20 | AACTGGTGCGGACCAGGGGA | 266.4 | -5.2 | 1.3 | -4.0 | 5.4E-05 | 7.3E-05 | MB231/MB468 | MITO |
| >hg38_21_+_8433172_8446622_RNA45SN1_45S_50ntFlanks@10238.10255.18\|CGCTACGCCGCGACGAGT | >hg38_21_+_8433172_8446622_RNA45SN1_45S_50ntFlanks | 10238 | 10255 | 18 | CGCTACGCCGCGACGAGT | 257.2 | 5.5 | 1.2 | 4.5 | 6.9E-06 | 7.3E-05 | BT20/MB231 | NUC |
| >hg38_21_+_8433172_8446622_RNA45SN1_45S_50ntFlanks@11804.11821.18\|CCCACTGTCCCTACCTAC | >hg38_21_+_8433172_8446622_RNA45SN1_45S_50ntFlanks | 11804 | 11821 | 18 | CCCACTGTCCCTACCTAC | 475.7 | -5.3 | 1.3 | -4.1 | 3.9E-05 | 7.3E-05 | BT20/MB468 | MITO |
| >hg38_21_+_8433172_8446622_RNA45SN1_45S_50ntFlanks@11876.11898.23\|AAGACCCTGTTGAGCTTGACTCT | >hg38_21_+_8433172_8446622_RNA45SN1_45S_50ntFlanks | 11876 | 11898 | 23 | AAGACCCTGTTGAGCTTGACTCT | 1370.8 | -3.3 | 0.8 | -4.0 | 5.4E-05 | 7.3E-05 | MB231/MB468 | NUC |
| >hg38_21_+_8433172_8446622_RNA45SN1_45S_50ntFlanks@3846.3879.34\|CTTGGATAACTGTGGTAATTCTAGAGCTAATACA | >hg38_21_+_8433172_8446622_RNA45SN1_45S_50ntFlanks | 3846 | 3879 | 34 | CTTGGATAACTGTGGTAATTCTAGAGCTAATACA | 402.8 | 3.5 | 0.9 | 4.1 | 5.1E-05 | 7.3E-05 | MB231/MB468 | CYTO |
| >hg38_21_+_8433172_8446622_RNA45SN1_45S_50ntFlanks@4046.4066.21\|CATTCGAACGTCTGCCCTATC | >hg38_21_+_8433172_8446622_RNA45SN1_45S_50ntFlanks | 4046 | 4066 | 21 | CATTCGAACGTCTGCCCTATC | 1841.1 | -4.0 | 1.0 | -4.1 | 3.9E-05 | 7.3E-05 | BT20/MB468 | MITO |
| >MI0000434\|hsa-let-7i&WithFlank&12\|+\|62603680\|62603775@12.33.22(+1C)&[MIMAT0000415&hsa-let-7i-5p&offsets\|0\|0(+1C);m-49&12\|+\|62603691\|62603712&offsets\|0\|0(+1C)]\|TGAGGTAGTAGTTTGTGCTGTTC | >MI0000434\|hsa-let-7i&WithFlank&12\|+\|62603680\|62603775 | 12 | 33 | 22(+1C) | TGAGGTAGTAGTTTGTGCTGTTC | 272.7 | 6.0 | 1.5 | 4.0 | 5.4E-05 | 7.3E-05 | MB231/MB468 | MITO |
| >hg38_21_+_8433172_8446622_RNA45SN1_45S_50ntFlanks@10391.10419.29\|AACAGCAGTTGAACATGGGTCAGTCGGTC | >hg38_21_+_8433172_8446622_RNA45SN1_45S_50ntFlanks | 10391 | 10419 | 29 | AACAGCAGTTGAACATGGGTCAGTCGGTC | 439.7 | -5.7 | 1.4 | -4.1 | 4.7E-05 | 7.3E-05 | BT20/MB468 | CYTO |
| >hg38_21_+_8433172_8446622_RNA45SN1_45S_50ntFlanks@10353.10388.36\|AGAACTTTGAAGGCCGAAGTGGAGAAGGGTTCCATG | >hg38_21_+_8433172_8446622_RNA45SN1_45S_50ntFlanks | 10353 | 10388 | 36 | AGAACTTTGAAGGCCGAAGTGGAGAAGGGTTCCATG | 381.7 | -6.5 | 1.6 | -4.1 | 4.5E-05 | 7.4E-05 | MB231/MB468 | MP |
| >hg38_21_+_8433172_8446622_RNA45SN1_45S_50ntFlanks@11593.11613.21\|CCAGGGGAATCCGACTGTTTA | >hg38_21_+_8433172_8446622_RNA45SN1_45S_50ntFlanks | 11593 | 11613 | 21 | CCAGGGGAATCCGACTGTTTA | 386.8 | -2.9 | 0.7 | -4.1 | 4.7E-05 | 7.4E-05 | BT20/MB468 | CYTO |
| >hg38_21_+_8433172_8446622_RNA45SN1_45S_50ntFlanks@4367.4400.34\|CAGTTAAAAAGCTCGTAGTTGGATCTTGGGAGCG | >hg38_21_+_8433172_8446622_RNA45SN1_45S_50ntFlanks | 4367 | 4400 | 34 | CAGTTAAAAAGCTCGTAGTTGGATCTTGGGAGCG | 163.5 | -2.9 | 0.7 | -4.1 | 5.1E-05 | 7.4E-05 | MB231/MB468 | CYTO |
| >hg38_21_+_8433172_8446622_RNA45SN1_45S_50ntFlanks@4722.4751.30\|TCATTAATCAAGAACGAAAGTCGGAGGTTC | >hg38_21_+_8433172_8446622_RNA45SN1_45S_50ntFlanks | 4722 | 4751 | 30 | TCATTAATCAAGAACGAAAGTCGGAGGTTC | 750.7 | -6.7 | 1.6 | -4.3 | 1.9E-05 | 7.4E-05 | BT20/MB468 | MP |

| | | | | | | | | | | | | | |
|---|---|---|---|---|---|---|---|---|---|---|---|---|---|
| >hg38_21_+_8433172_8446622_RNA45SN1_45S_50ntFlanks@9340.9370.31\|GCACGAAAGCCGCCGTGGCGCAATGAAGGTG | >hg38_21_+_8433172_8446622_RNA45SN1_45S_50ntFlanks | 9340 | 9370 | 31 | GCACGAAAGCCGCCGTGGCGCAATGAAGGTG | 42.5 | -8.5 | 2.0 | -4.3 | 1.9E-05 | 7.4E-05 | BT20/MB468 | MP |
| >hg38_21_+_8433172_8446622_RNA45SN1_45S_50ntFlanks@8170.8197.28\|CAAGTCCTTCTGATCGAGGCCCAGCCCG | >hg38_21_+_8433172_8446622_RNA45SN1_45S_50ntFlanks | 8170 | 8197 | 28 | CAAGTCCTTCTGATCGAGGCCCAGCCCG | 556.0 | -6.7 | 1.6 | -4.3 | 1.9E-05 | 7.4E-05 | BT20/MB468 | MP |
| >MI0006273\|hsa-mir-1180&WithFlank&17\|-\|19344500\|19344580@47.68.22&[MIMAT0005825&hsa-miR-1180-3p&offsets\|0\|0;m-214&17\|-\|19344513\|19344534&offsets\|0\|0]\|TTTCCGGCTCGCGTGGGTGTGT | >MI0006273\|hsa-mir-1180&WithFlank&17\|-\|19344500\|19344580 | 47 | 68 | 22 | TTTCCGGCTCGCGTGGGTGTGT | 336.4 | 3.7 | 0.9 | 4.0 | 5.4E-05 | 7.4E-05 | MB231/MB468 | MITO |
| >MI0000067\|hsa-let-7f-1&WithFlank&9\|+\|94176341\|94176439@13.31.19(+3U);MI0000068\|hsa-let-7f-2&WithFlank&X\|-\|53557186\|53557280@14.32.19(+3U)&[MIMAT0000067&hsa-let-7f-5p&offsets\|0\|-3(+3U);m-20&9\|+\|94176353\|94176374&offsets\|0\|-3(+3U)];[MIMAT0000067_1&hsa-let-7f-5p&offsets\|0\|-3(+3U);m-19&X\|-\|53557246\|53557267&offsets\|0\|-3(+3U)]\|TGAGGTAGTAGATTGTATATTT | >MI0000067\|hsa-let-7f-1&WithFlank&9\|+\|94176341\|94176439 | 13 | 31 | 19(+3U) | TGAGGTAGTAGATTGTATATTT | 31.8 | 8.2 | 2.0 | 4.1 | 4.6E-05 | 7.4E-05 | MB231/MB468 | MP |
| >hg38_21_+_8433172_8446622_RNA45SN1_45S_50ntFlanks@12407.12437.31\|TTTTGATCCTTCGATGTCGGCTCTTCCTATC | >hg38_21_+_8433172_8446622_RNA45SN1_45S_50ntFlanks | 12407 | 12437 | 31 | TTTTGATCCTTCGATGTCGGCTCTTCCTATC | 2767.3 | -2.7 | 0.7 | -4.0 | 5.4E-05 | 7.4E-05 | MB231/MB468 | NUC |
| >am_trna111_HisGTG_1_-_147774845_147774916@-1G.20.21,trna118_HisGTG_1_-_145396881_145396952@-1G.20.21,trna16_HisGTG_1_+_146544773_146544844@-1G.20.21,trna1_HisGTG_15_+_45493349_45493420@-1G.20.21,trna21_HisGTG_1_+_147753471_147753542@-1G.20.21,trna33_HisGTG_6_+_27125906_27125977@-1G.20.21,trna7_HisGTG_9_-_14433938_14434009@-1G.20.21,trna8_HisGTG_15_-_45492611_45492682@-1G.20.21,trna9_HisGTG_15_-_45490804_45490875@-1G.20.21\|GGCCGTGATCGTATAGTGGTT | >am_trna111_HisGTG_1_-_147774845_147774916 | -1G | 20 | 21 | GGCCGTGATCGTATAGTGGTT | 149.7 | 7.0 | 1.7 | 4.1 | 4.6E-05 | 7.4E-05 | MB231/MB468 | MP |
| >hg38_21_+_8433172_8446622_RNA45SN1_45S_50ntFlanks@3855.3873.19\|CTGTGGTAATTCTAGAGCT | >hg38_21_+_8433172_8446622_RNA45SN1_45S_50ntFlanks | 3855 | 3873 | 19 | CTGTGGTAATTCTAGAGCT | 144.1 | -6.1 | 1.4 | -4.3 | 1.9E-05 | 7.4E-05 | BT20/MB468 | MP |
| >hg38_21_+_8433172_8446622_RNA45SN1_45S_50ntFlanks@4565.4587.23\|AATAATGGAATAGGACCGCGGTT | >hg38_21_+_8433172_8446622_RNA45SN1_45S_50ntFlanks | 4565 | 4587 | 23 | AATAATGGAATAGGACCGCGGTT | 588.1 | -2.7 | 0.7 | -4.0 | 5.1E-05 | 7.4E-05 | MB231/MB468 | CYTO |
| >hg38_21_+_8433172_8446622_RNA45SN1_45S_50ntFlanks@10390.10424.35\|GAACAGCAGTTGAACATGGGTCAGTCGGTCCTGAG | >hg38_21_+_8433172_8446622_RNA45SN1_45S_50ntFlanks | 10390 | 10424 | 35 | GAACAGCAGTTGAACATGGGTCAGTCGGTCCTGAG | 280.7 | 3.9 | 1.0 | 4.0 | 5.5E-05 | 7.4E-05 | MB231/MB468 | MITO |
| >hg38_21_+_8433172_8446622_RNA45SN1_45S_50ntFlanks@12274.12309.36\|CAGTACGAATACAGACCGTGAAAGCGGGGCCTCACG | >hg38_21_+_8433172_8446622_RNA45SN1_45S_50ntFlanks | 12274 | 12309 | 36 | CAGTACGAATACAGACCGTGAAAGCGGGGCCTCACG | 125.4 | -6.1 | 1.5 | -4.1 | 4.6E-05 | 7.4E-05 | MB231/MB468 | MP |
| >hg38_21_+_8433172_8446622_RNA45SN1_45S_50ntFlanks@12457.12476.20\|CAAGCGTTGGATTGTTCACC | >hg38_21_+_8433172_8446622_RNA45SN1_45S_50ntFlanks | 12457 | 12476 | 20 | CAAGCGTTGGATTGTTCACC | 11325.7 | -3.5 | 0.9 | -4.0 | 5.1E-05 | 7.4E-05 | MB231/MB468 | CYTO |
| >hg38_21_+_8433172_8446622_RNA45SN1_45S_50ntFlanks@4043.4063.21\|ACCCATTCGAACGTCTGCCCT | >hg38_21_+_8433172_8446622_RNA45SN1_45S_50ntFlanks | 4043 | 4063 | 21 | ACCCATTCGAACGTCTGCCCT | 257.7 | -4.0 | 1.0 | -4.0 | 5.1E-05 | 7.4E-05 | MB231/MB468 | CYTO |
| >hg38_21_+_8433172_8446622_RNA45SN1_45S_50ntFlanks@12944.12962.19\|CGTAGACGACCTGCTTCTG | >hg38_21_+_8433172_8446622_RNA45SN1_45S_50ntFlanks | 12944 | 12962 | 19 | CGTAGACGACCTGCTTCTG | 441.0 | -6.8 | 1.7 | -4.0 | 5.7E-05 | 7.4E-05 | BT20/MB468 | NUC |
| >hg38_21_+_8433172_8446622_RNA45SN1_45S_50ntFlanks@13008.13027.20\|ATTGAAAGTCAGCCCTCGAC | >hg38_21_+_8433172_8446622_RNA45SN1_45S_50ntFlanks | 13008 | 13027 | 20 | ATTGAAAGTCAGCCCTCGAC | 846.3 | -5.6 | 1.4 | -4.1 | 4.8E-05 | 7.4E-05 | BT20/MB468 | CYTO |
| >MI0003780\|hsa-mir-1296&WithFlank&10\|-\|63372951\|63373054@22.43.22&[MIMAT0005794&hsa-miR-1296-5p&offsets\|0\|0;m-304&10\|-\|63373012\|63373033&offsets\|0\|0]\|TTAGGGCCCTGGCTCCATCTCC | >MI0003780\|hsa-mir-1296&WithFlank&10\|-\|63372951\|63373054 | 22 | 43 | 22 | TTAGGGCCCTGGCTCCATCTCC | 211.9 | 7.1 | 1.8 | 4.0 | 5.5E-05 | 7.4E-05 | MB231/MB468 | MITO |
| >hg38_1_-_228634819_228635039_RNA5S12_5S_50ntFlanks@148.168.21\|GGAATACCGGGTGCTGTAGGC | >hg38_1_-_228634819_228635039_RNA5S12_5S_50ntFlanks | 148 | 168 | 21 | GGAATACCGGGTGCTGTAGGC | 2860.4 | 10.0 | 2.2 | 4.5 | 6.3E-06 | 7.4E-05 | BT20/MB231 | MP |
| >MI0001448\|hsa-mir-425&WithFlank&3\|-\|49020142\|49020240@60.81.22&[MIMAT0001343&hsa-miR-425-3p&offsets\|-1\|-1;m-258&3\|-\|49020160\|49020181&offsets\|0\|0]\|CATCGGGAATGTCGTGTCCGCC | >MI0001448\|hsa-mir-425&WithFlank&3\|-\|49020142\|49020240 | 60 | 81 | 22 | CATCGGGAATGTCGTGTCCGCC | 322.5 | 3.6 | 0.9 | 4.1 | 4.6E-05 | 7.4E-05 | MB231/MB468 | MP |
| >hg38_21_+_8433172_8446622_RNA45SN1_45S_50ntFlanks@3822.3850.29\|CTTTGGTCGCTCGCTCCTCTCCTACTTGG | >hg38_21_+_8433172_8446622_RNA45SN1_45S_50ntFlanks | 3822 | 3850 | 29 | CTTTGGTCGCTCGCTCCTCTCCTACTTGG | 254.6 | -5.7 | 1.4 | -4.0 | 5.5E-05 | 7.4E-05 | MB231/MB468 | NUC |
| >hg38_21_+_8433172_8446622_RNA45SN1_45S_50ntFlanks@12717.12755.39\|CCTCGGTTGGCCTCGGATAGCCGGTCCCCCGCCTGTCCC | >hg38_21_+_8433172_8446622_RNA45SN1_45S_50ntFlanks | 12717 | 12755 | 39 | CCTCGGTTGGCCTCGGATAGCCGGTCCCCCGCCTGTCCC | 205.3 | -5.7 | 1.4 | -4.0 | 5.5E-05 | 7.4E-05 | MB231/MB468 | NUC |
| >hg38_21_+_8433172_8446622_RNA45SN1_45S_50ntFlanks@6650.6676.27\|ACGACTCTTAGCGGTGGATCACTCGGC | >hg38_21_+_8433172_8446622_RNA45SN1_45S_50ntFlanks | 6650 | 6676 | 27 | ACGACTCTTAGCGGTGGATCACTCGGC | 765.6 | 4.1 | 1.0 | 4.0 | 5.5E-05 | 7.4E-05 | MB231/MB468 | NUC |
| >hg38_21_+_8433172_8446622_RNA45SN1_45S_50ntFlanks@11874.11899.26\|AGAAGACCCTGTTGAGCTTGACTCTA | >hg38_21_+_8433172_8446622_RNA45SN1_45S_50ntFlanks | 11874 | 11899 | 26 | AGAAGACCCTGTTGAGCTTGACTCTA | 568.3 | -6.1 | 1.5 | -4.0 | 5.5E-05 | 7.4E-05 | MB231/MB468 | NUC |
| >hg38_21_+_8433172_8446622_RNA45SN1_45S_50ntFlanks@11581.11602.22\|AACTGGTGCGGACCAGGGGAAT | >hg38_21_+_8433172_8446622_RNA45SN1_45S_50ntFlanks | 11581 | 11602 | 22 | AACTGGTGCGGACCAGGGGAAT | 437.5 | -2.2 | 0.5 | -4.1 | 4.0E-05 | 7.4E-05 | BT20/MB468 | MITO |
| >hg38_21_+_8433172_8446622_RNA45SN1_45S_50ntFlanks@11774.11794.21\|AGTGACGCGCATGAATGGATG | >hg38_21_+_8433172_8446622_RNA45SN1_45S_50ntFlanks | 11774 | 11794 | 21 | AGTGACGCGCATGAATGGATG | 454.8 | 2.9 | 0.7 | 4.1 | 4.8E-05 | 7.4E-05 | BT20/MB468 | CYTO |
| >MI0003137\|hsa-mir-193b&WithFlank&16\|+\|14303961\|14304055@20.40.21&[MIMAT0004767&hsa-miR-193b-5p&offsets\|0\|-1;m-343&16\|+\|14303980\|14304001&offsets\|0\|-1]\|CGGGGTTTTGAGGGCGAGATG | >MI0003137\|hsa-mir-193b&WithFlank&16\|+\|14303961\|14304055 | 20 | 40 | 21 | CGGGGTTTTGAGGGCGAGATG | 297.7 | 7.2 | 1.6 | 4.5 | 6.3E-06 | 7.4E-05 | BT20/MB231 | MP |
| >hg38_MT_+_598_1651_RNR1_12S_50ntFlanks@427.468.42\|GAAAGTGGCTTTAACATATCTGAACACACAATAGCTAAGACC | >hg38_MT_+_598_1651_RNR1_12S_50ntFlanks | 427 | 468 | 42 | GAAAGTGGCTTTAACATATCTGAACACACAATAGCTAAGACC | 158.9 | 6.6 | 1.6 | 4.0 | 5.5E-05 | 7.4E-05 | MB231/MB468 | MITO |
| >trna2_GlyGCC_21_-_18827107_18827177@1.32.32,trna35_GlyGCC_1_+_161413094_161413164@1.32.32,trna37_GlyGCC_1_+_161420467_161420537@1.32.32,trna39_GlyGCC_1_+_161427898_161427968@1.32.32,trna41_GlyGCC_1_+_161435258_161435328@1.32.32\|GCATGGGTGGTTCAGTGGTAGAATTCTCGCCT | >trna2_GlyGCC_21_-_18827107_18827177 | 1 | 32 | 32 | GCATGGGTGGTTCAGTGGTAGAATTCTCGCCT | 8559.7 | 4.3 | 0.9 | 4.5 | 6.0E-06 | 7.4E-05 | BT20/MB231 | CYTO |
| >hg38_21_+_8433172_8446622_RNA45SN1_45S_50ntFlanks@9033.9068.36\|CTCGTGGGGGGGCCGGGCCACCCCTCCCACGGCGCG | >hg38_21_+_8433172_8446622_RNA45SN1_45S_50ntFlanks | 9033 | 9068 | 36 | CTCGTGGGGGGGCCGGGCCACCCCTCCCACGGCGCG | 170.0 | -5.4 | 1.3 | -4.1 | 4.0E-05 | 7.5E-05 | BT20/MB468 | MITO |
| >hg38_21_+_8433172_8446622_RNA45SN1_45S_50ntFlanks@3705.3743.39\|TACCTGGTTGATCCTGCCAGTAGCATATGCTTGTCTCAA | >hg38_21_+_8433172_8446622_RNA45SN1_45S_50ntFlanks | 3705 | 3743 | 39 | TACCTGGTTGATCCTGCCAGTAGCATATGCTTGTCTCAA | 558.8 | 2.7 | 0.7 | 4.0 | 5.5E-05 | 7.5E-05 | MB231/MB468 | NUC |
| >am_trna119_LysCTT_1_-_145395522_145395594@14.32.19,trna11_LysCTT_5_-_180648979_180649051@14.32.19,trna128_LysCTT_1_-_55423542_55423614@14.32.19,trna13_LysCTT_6_+_26556774_26556846@14.32.19,trna32_LysCTT_16_-_3207406_3207478@14.32.19,trna6_LysCTT_19_-_52425396_52425468@14.32.19,trna7_LysCTT_16_+_3225692_3225764@14.32.19,trna9_LysCTT_5_+_180634755_180634827@14.32.19\|AGTCGGTAGAGCATGAGAC | >am_trna119_LysCTT_1_-_145395522_145395594 | 14 | 32 | 19 | AGTCGGTAGAGCATGAGAC | 350.7 | 3.4 | 0.8 | 4.0 | 5.2E-05 | 7.5E-05 | MB231/MB468 | CYTO |
| >MI0000750\|hsa-mir-26a-2&WithFlank&12\|-\|57824603\|57824698@23.40.18(+1C);MI0000083\|hsa-mir-26a-1&WithFlank&3\|+\|37969398\|37969486@19.36.18(+1C)&[MIMAT0000082&hsa-miR-26a-5p&offsets\|+3\|-1(+1C);m-33&12\|-\|57824658\|57824679&offsets\|+3\|-1(+1C)];[MIMAT0000082_1&hsa-miR-26a-5p&offsets\|+3\|-1(+1C);m-34&3\|+\|37969413\|37969434&offsets\|+3\|-1(+1C)]\|AAGTAATCCAGGATAGGCC | >MI0000750\|hsa-mir-26a-2&WithFlank&12\|-\|57824603\|57824698 | 23 | 40 | 18(+1C) | AAGTAATCCAGGATAGGCC | 122.9 | 10.2 | 2.5 | 4.1 | 4.6E-05 | 7.5E-05 | MB231/MB468 | MP |
| >trnaMT_HisGTG_MT_+_12138_12206@32.72.41\|TGAATCTGACAACAGAGGCTTACGACCCCTTATTTACCCCA | >trnaMT_HisGTG_MT_+_12138_12206 | 32 | 72 | 41 | TGAATCTGACAACAGAGGCTTACGACCCCTTATTTACCCCA | 668.3 | 4.4 | 1.1 | 4.0 | 5.5E-05 | 7.5E-05 | MB231/MB468 | NUC |
| >hg38_21_+_8433172_8446622_RNA45SN1_45S_50ntFlanks@12417.12447.31\|TCGATGTCGGCTCTTCCTATCATTGTGAAGC | >hg38_21_+_8433172_8446622_RNA45SN1_45S_50ntFlanks | 12417 | 12447 | 31 | TCGATGTCGGCTCTTCCTATCATTGTGAAGC | 202.5 | 2.4 | 0.6 | 4.0 | 5.5E-05 | 7.5E-05 | MB231/MB468 | NUC |
| >hg38_21_+_8433172_8446622_RNA45SN1_45S_50ntFlanks@6664.6686.23\|TGGATCACTCGGCTCGTGCGTCG | >hg38_21_+_8433172_8446622_RNA45SN1_45S_50ntFlanks | 6664 | 6686 | 23 | TGGATCACTCGGCTCGTGCGTCG | 197.9 | -2.3 | 0.6 | -4.0 | 5.5E-05 | 7.5E-05 | MB231/MB468 | NUC |
| >hg38_21_+_8433172_8446622_RNA45SN1_45S_50ntFlanks@6723.6740.18\|TTGCAGGACACATTGATC | >hg38_21_+_8433172_8446622_RNA45SN1_45S_50ntFlanks | 6723 | 6740 | 18 | TTGCAGGACACATTGATC | 482.0 | -3.3 | 0.8 | -4.0 | 5.5E-05 | 7.5E-05 | MB231/MB468 | NUC |
| >hg38_21_+_8433172_8446622_RNA45SN1_45S_50ntFlanks@11576.11600.25\|CTTAGAACTGGTGCGGACCAGGGGA | >hg38_21_+_8433172_8446622_RNA45SN1_45S_50ntFlanks | 11576 | 11600 | 25 | CTTAGAACTGGTGCGGACCAGGGGA | 778.3 | -5.2 | 1.3 | -4.1 | 4.8E-05 | 7.5E-05 | BT20/MB468 | CYTO |
| >MI0000067\|hsa-let-7f-1&WithFlank&9\|+\|94176341\|94176439@13.34.22(+1C);MI0000068\|hsa-let-7f-2&WithFlank&X\|-\|53557186\|53557280@14.35.22(+1C)&[MIMAT0000067&hsa-let-7f-5p&offsets\|0\|0(+1C);m-20&9\|+\|94176353\|94176374&offsets\|0\|0(+1C)];[MIMAT0000067_1&hsa-let-7f-5p&offsets\|0\|0(+1C);m-19&X\|-\|53557246\|53557267&offsets\|0\|0(+1C)]\|TGAGGTAGTAGATTGTATAGTTC | >MI0000067\|hsa-let-7f-1&WithFlank&9\|+\|94176341\|94176439 | 13 | 34 | 22(+1C) | TGAGGTAGTAGATTGTATAGTTC | 692.6 | 5.4 | 1.3 | 4.1 | 4.6E-05 | 7.5E-05 | MB231/MB468 | MP |
| >hg38_21_+_8433172_8446622_RNA45SN1_45S_50ntFlanks@10352.10383.32\|GAGAACTTTGAAGGCCGAAGTGGAGAAGGGTT | >hg38_21_+_8433172_8446622_RNA45SN1_45S_50ntFlanks | 10352 | 10383 | 32 | GAGAACTTTGAAGGCCGAAGTGGAGAAGGGTT | 289.8 | -2.1 | 0.5 | -4.0 | 5.5E-05 | 7.5E-05 | MB231/MB468 | NUC |
| >trna11_GluTTC_15_-_26327381_26327452@41.74.34,trna3_GluTTC_13_-_45492062_45492133@41.74.34\|AGGCGGCCCGGGTTCGACTCCCGGTGTGGGAACC | >trna11_GluTTC_15_-_26327381_26327452 | 41 | 74 | 34 | AGGCGGCCCGGGTTCGACTCCCGGTGTGGGAACC | 195.3 | 6.2 | 1.5 | 4.0 | 5.5E-05 | 7.5E-05 | MB231/MB468 | MITO |
| >hg38_21_+_8433172_8446622_RNA45SN1_45S_50ntFlanks@9533.9563.31\|GAGGAAACTCTGGTGGAGGTCCGTAGCGGTC | >hg38_21_+_8433172_8446622_RNA45SN1_45S_50ntFlanks | 9533 | 9563 | 31 | GAGGAAACTCTGGTGGAGGTCCGTAGCGGTC | 3592.0 | -3.3 | 0.8 | -4.0 | 5.2E-05 | 7.5E-05 | MB231/MB468 | CYTO |
| >MI0000289\|hsa-mir-181a-1&WithFlank&1\|-\|198859038\|198859159@70.91.22&[MIMAT0000270&hsa-miR-181a-3p&offsets\|0\|0;m-125&1\|-\|198859069\|198859090&offsets\|0\|0]\|ACCATCGACCGTTGATTGTACC | >MI0000289\|hsa-mir-181a-1&WithFlank&1\|-\|198859038\|198859159 | 70 | 91 | 22 | ACCATCGACCGTTGATTGTACC | 528.0 | 3.2 | 0.8 | 4.1 | 4.6E-05 | 7.5E-05 | MB231/MB468 | MP |
| >trnaMT_GluTTC_MT_-_14674_14742@39.65.27,trnalookalike8_GluTTC_5_-_93905172_93905240@39.65.27\|ATTGGTCGTGGTTGTAGTCCGTGCGAG | >trnaMT_GluTTC_MT_-_14674_14742 | 39 | 65 | 27 | ATTGGTCGTGGTTGTAGTCCGTGCGAG | 559.3 | 4.5 | 1.0 | 4.3 | 1.9E-05 | 7.5E-05 | BT20/MB468 | MP |
| >MI0000077\|hsa-mir-21&WithFlank&17\|+\|59841260\|59841343@14.33.20(+1A)&[MIMAT0000076&hsa-miR-21-5p&offsets\|0\|-2(+1A);m-1&17\|+\|59841273\|59841294&offsets\|0\|-2(+1A)]\|TAGCTTATCAGACTGATGTTA | >MI0000077\|hsa-mir-21&WithFlank&17\|+\|59841260\|59841343 | 14 | 33 | 20(+1A) | TAGCTTATCAGACTGATGTTA | 504.5 | 5.0 | 1.2 | 4.0 | 5.2E-05 | 7.5E-05 | MB231/MB468 | CYTO |
| >hg38_21_+_8433172_8446622_RNA45SN1_45S_50ntFlanks@10789.10823.35\|AGCCTCTGGCATGTTGGAACAATGTAGGTAAGGGA | >hg38_21_+_8433172_8446622_RNA45SN1_45S_50ntFlanks | 10789 | 10823 | 35 | AGCCTCTGGCATGTTGGAACAATGTAGGTAAGGGA | 75.4 | 9.6 | 2.1 | 4.5 | 6.4E-06 | 7.5E-05 | BT20/MB231 | MP |

<table>
<tr><th></th><th></th><th></th><th></th><th></th><th></th><th></th><th></th><th></th><th></th><th></th><th></th><th></th><th></th><th></th></tr>
<tr><td>>hg38_21_+_8433172_8446622_RNA45SN1_45S_50ntFlanks@11564.11586.23|CCTAGCAGCCGACTTAGAACTGG</td><td>>hg38_21_+_8433172_8446622_RNA45SN1_45S_50ntFlanks</td><td>11564</td><td>11586</td><td>23</td><td>CCTAGCAGCCGACTTAGAACTGG</td><td>514.1</td><td>-3.9</td><td>1.0</td><td>-4.0</td><td>5.2E-05</td><td>7.5E-05</td><td>MB231/MB468</td><td>CYTO</td></tr>
<tr><td>>hg38_21_+_8433172_8446622_RNA45SN1_45S_50ntFlanks@11576.11613.38|CTTAGAACTGGTGCGGACCAGGGGAATCCGACTGTTTA</td><td>>hg38_21_+_8433172_8446622_RNA45SN1_45S_50ntFlanks</td><td>11576</td><td>11613</td><td>38</td><td>CTTAGAACTGGTGCGGACCAGGGGAATCCGACTGTTTA</td><td>929.0</td><td>-3.6</td><td>0.9</td><td>-4.0</td><td>5.5E-05</td><td>7.5E-05</td><td>MB231/MB468</td><td>MITO</td></tr>
<tr><td>>hg38_21_+_8433172_8446622_RNA45SN1_45S_50ntFlanks@3854.3883.30|ACTGTGGTAATTCTAGAGCTAATACATGCC</td><td>>hg38_21_+_8433172_8446622_RNA45SN1_45S_50ntFlanks</td><td>3854</td><td>3883</td><td>30</td><td>ACTGTGGTAATTCTAGAGCTAATACATGCC</td><td>463.0</td><td>-5.4</td><td>1.3</td><td>-4.0</td><td>5.7E-05</td><td>7.5E-05</td><td>BT20/MB468</td><td>NUC</td></tr>
<tr><td>>hg38_21_+_8433172_8446622_RNA45SN1_45S_50ntFlanks@10737.10769.33|TAAATCTCGCGCCGGGCCGTACCCATATCCGCA</td><td>>hg38_21_+_8433172_8446622_RNA45SN1_45S_50ntFlanks</td><td>10737</td><td>10769</td><td>33</td><td>TAAATCTCGCGCCGGGCCGTACCCATATCCGCA</td><td>271.4</td><td>-4.7</td><td>1.2</td><td>-4.0</td><td>5.5E-05</td><td>7.5E-05</td><td>MB231/MB468</td><td>NUC</td></tr>
<tr><td>>hg38_21_+_8433172_8446622_RNA45SN1_45S_50ntFlanks@3992.4013.22|GGTGACTCTAGATAACCTCGGG</td><td>>hg38_21_+_8433172_8446622_RNA45SN1_45S_50ntFlanks</td><td>3992</td><td>4013</td><td>22</td><td>GGTGACTCTAGATAACCTCGGG</td><td>165.2</td><td>-4.2</td><td>1.0</td><td>-4.0</td><td>5.5E-05</td><td>7.5E-05</td><td>MB231/MB468</td><td>NUC</td></tr>
<tr><td>>hg38_21_+_8433172_8446622_RNA45SN1_45S_50ntFlanks@11569.11602.34|CAGCCGACTTAGAACTGGTGCGGACCAGGGGAAT</td><td>>hg38_21_+_8433172_8446622_RNA45SN1_45S_50ntFlanks</td><td>11569</td><td>11602</td><td>34</td><td>CAGCCGACTTAGAACTGGTGCGGACCAGGGGAAT</td><td>3310.7</td><td>-3.6</td><td>0.9</td><td>-4.3</td><td>1.9E-05</td><td>7.5E-05</td><td>BT20/MB468</td><td>MP</td></tr>
<tr><td>>MI0000749|hsa-mir-30e&WithFlank&1|+|40754349|40754452@65.85.21(+1U)&[MIMAT0000693&hsa-miR-30e-3p&offsets|0|-1(+1U);m-25&1|+|40754413|40754434&offsets|0|-1(+1U)]|CTTTCAGTCGGATGTTTACAGT</td><td>>MI0000749|hsa-mir-30e&WithFlank&1|+|40754349|40754452</td><td>65</td><td>85</td><td>21(+1U)</td><td>CTTTCAGTCGGATGTTTACAGT</td><td>2100.4</td><td>3.6</td><td>0.9</td><td>4.0</td><td>5.6E-05</td><td>7.5E-05</td><td>MB231/MB468</td><td>MITO</td></tr>
<tr><td>>hg38_21_+_8433172_8446622_RNA45SN1_45S_50ntFlanks@12415.12445.31|CTTCGATGTCGGCTCTTCCTATCATTGTGAA</td><td>>hg38_21_+_8433172_8446622_RNA45SN1_45S_50ntFlanks</td><td>12415</td><td>12445</td><td>31</td><td>CTTCGATGTCGGCTCTTCCTATCATTGTGAA</td><td>983.6</td><td>-3.0</td><td>0.7</td><td>-4.0</td><td>5.6E-05</td><td>7.5E-05</td><td>MB231/MB468</td><td>MITO</td></tr>
<tr><td>>MI0000650|hsa-mir-200c&WithFlank&12|+|6963693|6963772@50.70.21(+1A)&[MIMAT0000617&hsa-miR-200c-3p&offsets|0|-2(+1A);m-54&12|+|6963742|6963764&offsets|0|-2(+1A)]|TAATACTGCCGGGTAATGATGA</td><td>>MI0000650|hsa-mir-200c&WithFlank&12|+|6963693|6963772</td><td>50</td><td>70</td><td>21(+1A)</td><td>TAATACTGCCGGGTAATGATGA</td><td>111.8</td><td>-6.0</td><td>1.5</td><td>-4.1</td><td>4.7E-05</td><td>7.5E-05</td><td>MB231/MB468</td><td>MP</td></tr>
<tr><td>>hg38_21_+_8433172_8446622_RNA45SN1_45S_50ntFlanks@12818.12840.23|CGCGCCGGGACCGGGGTCCGGTG</td><td>>hg38_21_+_8433172_8446622_RNA45SN1_45S_50ntFlanks</td><td>12818</td><td>12840</td><td>23</td><td>CGCGCCGGGACCGGGGTCCGGTG</td><td>226.3</td><td>-10.9</td><td>2.6</td><td>-4.3</td><td>1.9E-05</td><td>7.5E-05</td><td>BT20/MB468</td><td>MP</td></tr>
<tr><td>>MI0003591|hsa-mir-584&WithFlank&5|-|149062307|149062415@22.41.20(+1C)&[MIMAT0003249&hsa-miR-584-5p&offsets|0|-2(+1C);m-167&5|-|149062374|149062394&offsets|0|-1(+1C)]|TTATGGTTTGCCTGGGACTGC</td><td>>MI0003591|hsa-mir-584&WithFlank&5|-|149062307|149062415</td><td>22</td><td>41</td><td>20(+1C)</td><td>TTATGGTTTGCCTGGGACTGC</td><td>143.6</td><td>7.0</td><td>1.7</td><td>4.1</td><td>4.7E-05</td><td>7.5E-05</td><td>MB231/MB468</td><td>MP</td></tr>
<tr><td>>hg38_21_+_8433172_8446622_RNA45SN1_45S_50ntFlanks@10350.10367.18|ACGAGAACTTTGAAGGCC</td><td>>hg38_21_+_8433172_8446622_RNA45SN1_45S_50ntFlanks</td><td>10350</td><td>10367</td><td>18</td><td>ACGAGAACTTTGAAGGCC</td><td>1793.9</td><td>-5.4</td><td>1.3</td><td>-4.1</td><td>4.7E-05</td><td>7.5E-05</td><td>MB231/MB468</td><td>MP</td></tr>
<tr><td>>hg38_21_+_8433172_8446622_RNA45SN1_45S_50ntFlanks@5417.5443.27|CTACCGATTGGATGGTTTAGTGAGGCC</td><td>>hg38_21_+_8433172_8446622_RNA45SN1_45S_50ntFlanks</td><td>5417</td><td>5443</td><td>27</td><td>CTACCGATTGGATGGTTTAGTGAGGCC</td><td>225.2</td><td>-4.1</td><td>1.0</td><td>-4.1</td><td>4.9E-05</td><td>7.5E-05</td><td>BT20/MB468</td><td>CYTO</td></tr>
<tr><td>>hg38_21_+_8433172_8446622_RNA45SN1_45S_50ntFlanks@10822.10844.23|GAAGTCGGCAAGCCGGATCCGTA</td><td>>hg38_21_+_8433172_8446622_RNA45SN1_45S_50ntFlanks</td><td>10822</td><td>10844</td><td>23</td><td>GAAGTCGGCAAGCCGGATCCGTA</td><td>204.1</td><td>-2.8</td><td>0.7</td><td>-4.1</td><td>4.9E-05</td><td>7.6E-05</td><td>BT20/MB468</td><td>CYTO</td></tr>
<tr><td>>hg38_21_+_8433172_8446622_RNA45SN1_45S_50ntFlanks@10807.10830.24|ACAATGTAGGTAAGGGAAGTCGGC</td><td>>hg38_21_+_8433172_8446622_RNA45SN1_45S_50ntFlanks</td><td>10807</td><td>10830</td><td>24</td><td>ACAATGTAGGTAAGGGAAGTCGGC</td><td>139.1</td><td>-2.6</td><td>0.6</td><td>-4.0</td><td>5.2E-05</td><td>7.6E-05</td><td>MB231/MB468</td><td>CYTO</td></tr>
<tr><td>>hg38_21_+_8433172_8446622_RNA45SN1_45S_50ntFlanks@6669.6696.28|CACTCGGCTCGTGCGTCGATGAAGAACG</td><td>>hg38_21_+_8433172_8446622_RNA45SN1_45S_50ntFlanks</td><td>6669</td><td>6696</td><td>28</td><td>CACTCGGCTCGTGCGTCGATGAAGAACG</td><td>268.1</td><td>-4.6</td><td>1.1</td><td>-4.1</td><td>4.7E-05</td><td>7.6E-05</td><td>MB231/MB468</td><td>MP</td></tr>
<tr><td>>hg38_21_+_8433172_8446622_RNA45SN1_45S_50ntFlanks@4729.4751.23|TCAAGAACGAAAGTCGGAGGTTC</td><td>>hg38_21_+_8433172_8446622_RNA45SN1_45S_50ntFlanks</td><td>4729</td><td>4751</td><td>23</td><td>TCAAGAACGAAAGTCGGAGGTTC</td><td>318.2</td><td>-2.1</td><td>0.5</td><td>-4.0</td><td>5.3E-05</td><td>7.6E-05</td><td>MB231/MB468</td><td>CYTO</td></tr>
<tr><td>>hg38_21_+_8433172_8446622_RNA45SN1_45S_50ntFlanks@10806.10828.23|AACAATGTAGGTAAGGGAAGTCG</td><td>>hg38_21_+_8433172_8446622_RNA45SN1_45S_50ntFlanks</td><td>10806</td><td>10828</td><td>23</td><td>AACAATGTAGGTAAGGGAAGTCG</td><td>349.0</td><td>-5.5</td><td>1.3</td><td>-4.1</td><td>4.1E-05</td><td>7.6E-05</td><td>BT20/MB468</td><td>MITO</td></tr>
<tr><td>>hg38_21_+_8433172_8446622_RNA45SN1_45S_50ntFlanks@12942.12959.18|TTCGTAGACGACCTGCTT</td><td>>hg38_21_+_8433172_8446622_RNA45SN1_45S_50ntFlanks</td><td>12942</td><td>12959</td><td>18</td><td>TTCGTAGACGACCTGCTT</td><td>676.2</td><td>-7.1</td><td>1.8</td><td>-4.0</td><td>5.8E-05</td><td>7.6E-05</td><td>BT20/MB468</td><td>NUC</td></tr>
<tr><td>>hg38_21_+_8433172_8446622_RNA45SN1_45S_50ntFlanks@10351.10378.28|CGAGAACTTTGAAGGCCGAAGTGGAGAA</td><td>>hg38_21_+_8433172_8446622_RNA45SN1_45S_50ntFlanks</td><td>10351</td><td>10378</td><td>28</td><td>CGAGAACTTTGAAGGCCGAAGTGGAGAA</td><td>446.9</td><td>-2.6</td><td>0.6</td><td>-4.1</td><td>4.1E-05</td><td>7.6E-05</td><td>BT20/MB468</td><td>MITO</td></tr>
<tr><td>>MI0000438|hsa-mir-15b&WithFlank&3|+|160404582|160404691@64.84.21&[MIMAT0004586&hsa-miR-15b-3p&offsets|0|-1;m-204&3|+|160404645|160404665&offsets|0|0]|CGAATCATTATTTGCTGCTCT</td><td>>MI0000438|hsa-mir-15b&WithFlank&3|+|160404582|160404691</td><td>64</td><td>84</td><td>21</td><td>CGAATCATTATTTGCTGCTCT</td><td>518.6</td><td>3.4</td><td>0.8</td><td>4.1</td><td>4.7E-05</td><td>7.6E-05</td><td>MB231/MB468</td><td>MP</td></tr>
<tr><td>>MI0000476|hsa-mir-138-1&WithFlank&3|+|44114206|44114316@29.52.24(+1A)&[MIMAT0000430_1&hsa-miR-138-5p&offsets|0|+1(+1A);m-497&3|+|44114234|44114256&offsets|0|+1(+1A)]|AGCTGGTGTTGTGAATCAGGCCGTA</td><td>>MI0000476|hsa-mir-138-1&WithFlank&3|+|44114206|44114316</td><td>29</td><td>52</td><td>24(+1A)</td><td>AGCTGGTGTTGTGAATCAGGCCGTA</td><td>63.9</td><td>-9.4</td><td>2.1</td><td>-4.5</td><td>6.4E-06</td><td>7.6E-05</td><td>BT20/MB231</td><td>MP</td></tr>
<tr><td>>hg38_21_+_8433172_8446622_RNA45SN1_45S_50ntFlanks@11667.11685.19|CTGCCCAGTGCTCTGAATG</td><td>>hg38_21_+_8433172_8446622_RNA45SN1_45S_50ntFlanks</td><td>11667</td><td>11685</td><td>19</td><td>CTGCCCAGTGCTCTGAATG</td><td>1083.7</td><td>-3.9</td><td>1.0</td><td>-4.0</td><td>5.8E-05</td><td>7.6E-05</td><td>BT20/MB468</td><td>NUC</td></tr>
<tr><td>>hg38_21_+_8433172_8446622_RNA45SN1_45S_50ntFlanks@10411.10432.22|CAGTCGGTCCTGAGAGATGGGC</td><td>>hg38_21_+_8433172_8446622_RNA45SN1_45S_50ntFlanks</td><td>10411</td><td>10432</td><td>22</td><td>CAGTCGGTCCTGAGAGATGGGC</td><td>587.7</td><td>-2.9</td><td>0.7</td><td>-4.1</td><td>4.1E-05</td><td>7.6E-05</td><td>BT20/MB468</td><td>MITO</td></tr>
<tr><td>>hg38_21_+_8433172_8446622_RNA45SN1_45S_50ntFlanks@5488.5516.29|GCTGAGAAGACGGTCGAACTTGACTATCT</td><td>>hg38_21_+_8433172_8446622_RNA45SN1_45S_50ntFlanks</td><td>5488</td><td>5516</td><td>29</td><td>GCTGAGAAGACGGTCGAACTTGACTATCT</td><td>737.0</td><td>3.0</td><td>0.7</td><td>4.5</td><td>7.5E-06</td><td>7.6E-05</td><td>BT20/MB231</td><td>MITO</td></tr>
<tr><td>>am_MI0000809|hsa-mir-151a&WithFlank&8|-|140732558|140732659@53.73.21(+1C)&[MIMAT0000757&hsa-miR-151a-3p&offsets|0|0(+1C);m-30&8|-|140732587|140732607&offsets|0|0(+1C)]|CTAGACTGAAGCTCCTTGAGGC</td><td>>am_MI0000809|hsa-mir-151a&WithFlank&8|-|140732558|140732659</td><td>53</td><td>73</td><td>21(+1C)</td><td>CTAGACTGAAGCTCCTTGAGGC</td><td>927.8</td><td>5.2</td><td>1.3</td><td>4.1</td><td>4.7E-05</td><td>7.6E-05</td><td>MB231/MB468</td><td>MP</td></tr>
<tr><td>>hg38_21_+_8433172_8446622_RNA45SN1_45S_50ntFlanks@9306.9338.33|CAAGGAGTCTAACACGTGCGCGAGTCGGGGGCT</td><td>>hg38_21_+_8433172_8446622_RNA45SN1_45S_50ntFlanks</td><td>9306</td><td>9338</td><td>33</td><td>CAAGGAGTCTAACACGTGCGCGAGTCGGGGGCT</td><td>803.1</td><td>-4.8</td><td>1.2</td><td>-4.1</td><td>4.1E-05</td><td>7.6E-05</td><td>BT20/MB468</td><td>MITO</td></tr>
<tr><td>>MI0000081|hsa-mir-24-2&WithFlank&19|-|13836281|13836365@56.76.21(+1A);MI0000080|hsa-mir-24-1&WithFlank&9|+|95086015|95086094@50.70.21(+1A)&[MIMAT0000080&hsa-miR-24-3p&offsets|0|-1(+1A);m-36&19|-|13836288|13836310&offsets|0|-2(+1A)];[MIMAT0000080_1&hsa-miR-24-3p&offsets|0|-1(+1A);m-35&9|+|95086064|95086086&offsets|0|-2(+1A)]|TGGCTCAGTTCAGCAGGAACAA</td><td>>MI0000081|hsa-mir-24-2&WithFlank&19|-|13836281|13836365</td><td>56</td><td>76</td><td>21(+1A)</td><td>TGGCTCAGTTCAGCAGGAACAA</td><td>4643.2</td><td>4.2</td><td>1.0</td><td>4.3</td><td>1.9E-05</td><td>7.6E-05</td><td>BT20/MB468</td><td>MP</td></tr>
<tr><td>>hg38_21_+_8433172_8446622_RNA45SN1_45S_50ntFlanks@4733.4761.29|GAACGAAAGTCGGAGGTTCGAAGACGATC</td><td>>hg38_21_+_8433172_8446622_RNA45SN1_45S_50ntFlanks</td><td>4733</td><td>4761</td><td>29</td><td>GAACGAAAGTCGGAGGTTCGAAGACGATC</td><td>2059.4</td><td>5.1</td><td>1.2</td><td>4.3</td><td>1.9E-05</td><td>7.6E-05</td><td>BT20/MB468</td><td>MP</td></tr>
<tr><td>>hg38_21_+_8433172_8446622_RNA45SN1_45S_50ntFlanks@10069.10096.28|AGTGGACGGGAGCGGCGGGGGCGGCGCG</td><td>>hg38_21_+_8433172_8446622_RNA45SN1_45S_50ntFlanks</td><td>10069</td><td>10096</td><td>28</td><td>AGTGGACGGGAGCGGCGGGGGCGGCGCG</td><td>79.0</td><td>9.6</td><td>2.1</td><td>4.5</td><td>6.5E-06</td><td>7.6E-05</td><td>BT20/MB231</td><td>MP</td></tr>
<tr><td>>hg38_21_+_8433172_8446622_RNA45SN1_45S_50ntFlanks@10393.10435.43|CAGCAGTTGAACATGGGTCAGTCGGTCCTGAGAGATGGGCGAG</td><td>>hg38_21_+_8433172_8446622_RNA45SN1_45S_50ntFlanks</td><td>10393</td><td>10435</td><td>43</td><td>CAGCAGTTGAACATGGGTCAGTCGGTCCTGAGAGATGGGCGAG</td><td>391.6</td><td>-6.2</td><td>1.5</td><td>-4.1</td><td>4.9E-05</td><td>7.6E-05</td><td>BT20/MB468</td><td>CYTO</td></tr>
<tr><td>>hg38_MT_+_1621_3279_RNR2_16S_50ntFlanks@51.77.27|GCTAAACCTAGCCCCAAACCCACTCCA</td><td>>hg38_MT_+_1621_3279_RNR2_16S_50ntFlanks</td><td>51</td><td>77</td><td>27</td><td>GCTAAACCTAGCCCCAAACCCACTCCA</td><td>336.0</td><td>3.4</td><td>0.8</td><td>4.0</td><td>5.6E-05</td><td>7.6E-05</td><td>MB231/MB468</td><td>MITO</td></tr>
<tr><td>>hg38_21_+_8433172_8446622_RNA45SN1_45S_50ntFlanks@5467.5499.33|GGCCCACGGCCCTGGCGGAGCGCTGAGAAGACG</td><td>>hg38_21_+_8433172_8446622_RNA45SN1_45S_50ntFlanks</td><td>5467</td><td>5499</td><td>33</td><td>GGCCCACGGCCCTGGCGGAGCGCTGAGAAGACG</td><td>15115.7</td><td>-8.7</td><td>2.2</td><td>-4.0</td><td>5.8E-05</td><td>7.6E-05</td><td>BT20/MB468</td><td>NUC</td></tr>
<tr><td>>hg38_21_+_8433172_8446622_RNA45SN1_45S_50ntFlanks@6651.6700.50|CGACTCTTAGCGGTGGATCACTCGGCTCGTGCGTCGATGAAGAACGCAGC</td><td>>hg38_21_+_8433172_8446622_RNA45SN1_45S_50ntFlanks</td><td>6651</td><td>6700</td><td>50</td><td>CGACTCTTAGCGGTGGATCACTCGGCTCGTGCGTCGATGAAGAACGCAGC</td><td>4701.0</td><td>4.9</td><td>1.2</td><td>4.0</td><td>5.6E-05</td><td>7.6E-05</td><td>MB231/MB468</td><td>NUC</td></tr>
<tr><td>>MI0000434|hsa-let-7i&WithFlank&12|+|62603680|62603775@12.32.21(+1G)&[MIMAT0000415&hsa-let-7i-5p&offsets|0|-1(+1G);m-49&12|+|62603691|62603712&offsets|0|-1(+1G)]|TGAGGTAGTAGTTTGTGCTGTG</td><td>>MI0000434|hsa-let-7i&WithFlank&12|+|62603680|62603775</td><td>12</td><td>32</td><td>21(+1G)</td><td>TGAGGTAGTAGTTTGTGCTGTG</td><td>1024.1</td><td>3.8</td><td>0.9</td><td>4.0</td><td>5.3E-05</td><td>7.6E-05</td><td>MB231/MB468</td><td>CYTO</td></tr>
<tr><td>>MI0000063|hsa-let-7b&WithFlank&22|+|46113680|46113774@12.33.22(+1A)&[MIMAT0000063&hsa-let-7b-5p&offsets|0|0(+1A);m-14&22|+|46113691|46113712&offsets|0|0(+1A)]|TGAGGTAGTAGGTTGTGTGGTTA</td><td>>MI0000063|hsa-let-7b&WithFlank&22|+|46113680|46113774</td><td>12</td><td>33</td><td>22(+1A)</td><td>TGAGGTAGTAGGTTGTGTGGTTA</td><td>2543.4</td><td>2.6</td><td>0.6</td><td>4.5</td><td>7.2E-06</td><td>7.6E-05</td><td>BT20/MB231</td><td>NUC</td></tr>
<tr><td>>am_trna128_GlyGCC_6_-_27870686_27870756@12.29.18,trna133_GlyCCC_1_-_16872434_16872504@12.29.18,trna17_GluTTC_2_-_204229131_204229201@12.29.18,trna18_GlyGCC_16_+_70822597_70822667@12.29.18,trna19_GlyGCC_16_+_70823410_70823480@12.29.18,trna19_GlyGCC_2_-_157257659_157257729@12.29.18,trna24_GlyGCC_16_-_70812942_70813012@12.29.18,trna25_GlyGCC_16_-_70812114_70812184@12.29.18,trna2_GlyGCC_21_-_18827107_18827177@12.29.18,trna35_GlyGCC_1_+_161413094_161413164@12.29.18,trna37_GlyGCC_1_+_161420467_161420537@12.29.18,trna39_GlyGCC_1_+_161427898_161427968@12.29.18,trna41_GlyGCC_1_+_161435258_161435328@12.29.18,trna4_GlyCCC_1_+_17188416_17188486@12.29.18,trna5_GlyGCC_17_+_8029064_8029134@12.29.18,trna68_GlyGCC_1_-_161493637_161493707@12.29.18,trna91_GlyCCC_1_-_149680210_149680280@12.29.18,trna99_ValCAC_1_-_149294666_149294736@12.29.18|TCAGTGGTAGAATTCTCG</td><td>>am_trna128_GlyGCC_6_-_27870686_27870756</td><td>12</td><td>29</td><td>18</td><td>TCAGTGGTAGAATTCTCG</td><td>57.7</td><td>-9.7</td><td>2.4</td><td>-4.1</td><td>4.7E-05</td><td>7.7E-05</td><td>MB231/MB468</td><td>MP</td></tr>
<tr><td>>MI0000471|hsa-mir-126&WithFlank&9|+|136670596|136670692@58.80.23&[MIMAT0000445&hsa-miR-126-3p&offsets|0|+1;m-40&9|+|136670653|136670674&offsets|0|+1]|TCGTACCGTGAGTAATAATGCGC</td><td>>MI0000471|hsa-mir-126&WithFlank&9|+|136670596|136670692</td><td>58</td><td>80</td><td>23</td><td>TCGTACCGTGAGTAATAATGCGC</td><td>29.8</td><td>8.1</td><td>2.0</td><td>4.1</td><td>4.7E-05</td><td>7.7E-05</td><td>MB231/MB468</td><td>MP</td></tr>
<tr><td>>MI0000465|hsa-mir-191&WithFlank&3|-|49020612|49020715@22.44.23(+1C)&[MIMAT0000440&hsa-miR-191-5p&offsets|0|0(+1C);m-48&3|-|49020672|49020694&offsets|0|0(+1C)]|CAACGGAATCCCAAAAGCAGCTGC</td><td>>MI0000465|hsa-mir-191&WithFlank&3|-|49020612|49020715</td><td>22</td><td>44</td><td>23(+1C)</td><td>CAACGGAATCCCAAAAGCAGCTGC</td><td>137.6</td><td>3.0</td><td>0.7</td><td>4.0</td><td>5.9E-05</td><td>7.7E-05</td><td>BT20/MB468</td><td>NUC</td></tr>
<tr><td>>hg38_21_+_8433172_8446622_RNA45SN1_45S_50ntFlanks@8115.8141.27|CGCGGGACATGTGGCGTACGGAAGACC</td><td>>hg38_21_+_8433172_8446622_RNA45SN1_45S_50ntFlanks</td><td>8115</td><td>8141</td><td>27</td><td>CGCGGGACATGTGGCGTACGGAAGACC</td><td>1030.3</td><td>8.6</td><td>1.9</td><td>4.5</td><td>6.5E-06</td><td>7.7E-05</td><td>BT20/MB231</td><td>MP</td></tr>
<tr><td>>MI0000749|hsa-mir-30e&WithFlank&1|+|40754349|40754452@23.46.24(+2U)&[MIMAT0000692&hsa-miR-30e-5p&offsets|0|+2(+2U);m-24&1|+|40754371|40754394&offsets|0|0(+2U)]|TGTAAACATCCTTGACTGGAAGCTTT</td><td>>MI0000749|hsa-mir-30e&WithFlank&1|+|40754349|40754452</td><td>23</td><td>46</td><td>24(+2U)</td><td>TGTAAACATCCTTGACTGGAAGCTTT</td><td>324.3</td><td>4.8</td><td>1.2</td><td>4.0</td><td>5.7E-05</td><td>7.7E-05</td><td>MB231/MB468</td><td>MITO</td></tr>
</table>

<table>
<tr><td>>MI0003815|hsa-mir-1301&WithFlank&2|-|25328634|25328727@54.77.24&[MIMAT0005797&hsa-miR-1301-3p&offsets|0|0;m-256&2|-|25328650|25328674&offsets|0|-1]|TTGCAGCTGCCTGGGAGTGACTTC</td><td>>MI0003815|hsa-mir-1301&WithFlank&2|-|25328634|25328727</td><td>54</td><td>77</td><td>24</td><td>TTGCAGCTGCCTGGGAGTGACTTC</td><td>400.0</td><td>3.2</td><td>0.7</td><td>4.5</td><td>7.6E-06</td><td>7.7E-05</td><td>BT20/MB231</td><td>MITO</td></tr>
<tr><td>>MI0000301|hsa-mir-224&WithFlank&X|-|151958572|151958664@14.34.21(+1C)&[MIMAT0000281&hsa-miR-224-5p&offsets|+1|-3(+1C);m-238&X|-|151958630|151958651&offsets|0|-1(+1C)]|CAAGTCACTAGTGGTTCCGTTC</td><td>>MI0000301|hsa-mir-224&WithFlank&X|-|151958572|151958664</td><td>14</td><td>34</td><td>21(+1C)</td><td>CAAGTCACTAGTGGTTCCGTTC</td><td>1017.0</td><td>3.1</td><td>0.7</td><td>4.5</td><td>7.3E-06</td><td>7.7E-05</td><td>BT20/MB231</td><td>NUC</td></tr>
<tr><td>>hg38_21_+_8433172_8446622_RNA45SN1_45S_50ntFlanks@6648.6681.34|GTACGACTCTTAGCGGTGGATCACTCGGCTCGTG</td><td>>hg38_21_+_8433172_8446622_RNA45SN1_45S_50ntFlanks</td><td>6648</td><td>6681</td><td>34</td><td>GTACGACTCTTAGCGGTGGATCACTCGGCTCGTG</td><td>5564.1</td><td>-2.8</td><td>0.6</td><td>-4.5</td><td>7.3E-06</td><td>7.7E-05</td><td>BT20/MB231</td><td>NUC</td></tr>
<tr><td>>hg38_1_-_228634819_228635039_RNA5S12_5S_50ntFlanks@136.170.35|GGAGACCGCCTGGGAATACCGGGTGCTGTAGGCTT</td><td>>hg38_1_-_228634819_228635039_RNA5S12_5S_50ntFlanks</td><td>136</td><td>170</td><td>35</td><td>GGAGACCGCCTGGGAATACCGGGTGCTGTAGGCTT</td><td>1604.6</td><td>4.0</td><td>1.0</td><td>4.0</td><td>5.3E-05</td><td>7.7E-05</td><td>MB231/MB468</td><td>CYTO</td></tr>
<tr><td>>hg38_21_+_8433172_8446622_RNA45SN1_45S_50ntFlanks@12932.12969.38|CGCTAAACCATTCGTAGACGACCTGCTTCTGGGTCGGG</td><td>>hg38_21_+_8433172_8446622_RNA45SN1_45S_50ntFlanks</td><td>12932</td><td>12969</td><td>38</td><td>CGCTAAACCATTCGTAGACGACCTGCTTCTGGGTCGGG</td><td>283.0</td><td>-3.9</td><td>1.0</td><td>-4.0</td><td>5.3E-05</td><td>7.7E-05</td><td>MB231/MB468</td><td>CYTO</td></tr>
<tr><td>>hg38_21_+_8433172_8446622_RNA45SN1_45S_50ntFlanks@9567.9605.39|ACGTGCAAATCGGTCGTCCGACCTGGGTATAGGGGCGAA</td><td>>hg38_21_+_8433172_8446622_RNA45SN1_45S_50ntFlanks</td><td>9567</td><td>9605</td><td>39</td><td>ACGTGCAAATCGGTCGTCCGACCTGGGTATAGGGGCGAA</td><td>301.5</td><td>-4.8</td><td>1.2</td><td>-4.0</td><td>5.7E-05</td><td>7.7E-05</td><td>MB231/MB468</td><td>MITO</td></tr>
<tr><td>>hg38_21_+_8433172_8446622_RNA45SN1_45S_50ntFlanks@485.516.32|GATGTGGTGACGTCGTGCTCTCCCGGGCCGGG</td><td>>hg38_21_+_8433172_8446622_RNA45SN1_45S_50ntFlanks</td><td>485</td><td>516</td><td>32</td><td>GATGTGGTGACGTCGTGCTCTCCCGGGCCGGG</td><td>121.1</td><td>10.1</td><td>2.5</td><td>4.1</td><td>4.8E-05</td><td>7.7E-05</td><td>MB231/MB468</td><td>MP</td></tr>
<tr><td>>hg38_21_+_8433172_8446622_RNA45SN1_45S_50ntFlanks@10686.10712.27|GCGGCGTCCGGTGAGCTCTCGCTGGCC</td><td>>hg38_21_+_8433172_8446622_RNA45SN1_45S_50ntFlanks</td><td>10686</td><td>10712</td><td>27</td><td>GCGGCGTCCGGTGAGCTCTCGCTGGCC</td><td>660.7</td><td>-2.7</td><td>0.7</td><td>-4.0</td><td>5.7E-05</td><td>7.7E-05</td><td>MB231/MB468</td><td>MITO</td></tr>
<tr><td>>hg38_21_+_8433172_8446622_RNA45SN1_45S_50ntFlanks@11209.11252.44|CGGCGACTCTGGACGCGAGCCGGGCCCTTCCCGTGGATCGCCCC</td><td>>hg38_21_+_8433172_8446622_RNA45SN1_45S_50ntFlanks</td><td>11209</td><td>11252</td><td>44</td><td>CGGCGACTCTGGACGCGAGCCGGGCCCTTCCCGTGGATCGCCCC</td><td>569.6</td><td>-3.6</td><td>0.9</td><td>-4.1</td><td>5.0E-05</td><td>7.7E-05</td><td>BT20/MB468</td><td>CYTO</td></tr>
<tr><td>>hg38_21_+_8433172_8446622_RNA45SN1_45S_50ntFlanks@12417.12448.32|TCGATGTCGGCTCTTCCTATCATTGTGAAGCA</td><td>>hg38_21_+_8433172_8446622_RNA45SN1_45S_50ntFlanks</td><td>12417</td><td>12448</td><td>32</td><td>TCGATGTCGGCTCTTCCTATCATTGTGAAGCA</td><td>960.1</td><td>3.2</td><td>0.8</td><td>4.1</td><td>5.0E-05</td><td>7.7E-05</td><td>BT20/MB468</td><td>CYTO</td></tr>
<tr><td>>hg38_21_+_8433172_8446622_RNA45SN1_45S_50ntFlanks@3815.3852.38|ATGGTTCCTTTGGTCGCTCGCTCCTCTCCTACTTGGAT</td><td>>hg38_21_+_8433172_8446622_RNA45SN1_45S_50ntFlanks</td><td>3815</td><td>3852</td><td>38</td><td>ATGGTTCCTTTGGTCGCTCGCTCCTCTCCTACTTGGAT</td><td>426.2</td><td>-3.4</td><td>0.8</td><td>-4.0</td><td>5.7E-05</td><td>7.7E-05</td><td>MB231/MB468</td><td>NUC</td></tr>
<tr><td>>MI0000084|hsa-mir-26b&WithFlank&2|+|218402640|218402728@18.39.22(+1U)&[MIMAT0000083&hsa-miR-26b-5p&offsets|0|+1(+1U);m-53&2|+|218402657|218402678&offsets|0|0(+1U)]|TTCAAGTAATTCAGGATAGGTTT</td><td>>MI0000084|hsa-mir-26b&WithFlank&2|+|218402640|218402728</td><td>18</td><td>39</td><td>22(+1U)</td><td>TTCAAGTAATTCAGGATAGGTTT</td><td>899.5</td><td>3.3</td><td>0.8</td><td>4.1</td><td>4.8E-05</td><td>7.7E-05</td><td>MB231/MB468</td><td>MP</td></tr>
<tr><td>>hg38_21_+_8433172_8446622_RNA45SN1_45S_50ntFlanks@10534.10567.34|CCGCGAGGCGTCCAGTGCGGTAACGCGACCGATC</td><td>>hg38_21_+_8433172_8446622_RNA45SN1_45S_50ntFlanks</td><td>10534</td><td>10567</td><td>34</td><td>CCGCGAGGCGTCCAGTGCGGTAACGCGACCGATC</td><td>299.0</td><td>-5.6</td><td>1.4</td><td>-4.1</td><td>4.2E-05</td><td>7.7E-05</td><td>BT20/MB468</td><td>MITO</td></tr>
<tr><td>>hg38_21_+_8433172_8446622_RNA45SN1_45S_50ntFlanks@10355.10376.22|AACTTTGAAGGCCGAAGTGGAG</td><td>>hg38_21_+_8433172_8446622_RNA45SN1_45S_50ntFlanks</td><td>10355</td><td>10376</td><td>22</td><td>AACTTTGAAGGCCGAAGTGGAG</td><td>62.0</td><td>-9.8</td><td>2.4</td><td>-4.1</td><td>4.8E-05</td><td>7.7E-05</td><td>MB231/MB468</td><td>MP</td></tr>
<tr><td>>hg38_21_+_8433172_8446622_RNA45SN1_45S_50ntFlanks@11570.11603.34|AGCCGACTTAGAACTGGTGCGGACCAGGGGAATC</td><td>>hg38_21_+_8433172_8446622_RNA45SN1_45S_50ntFlanks</td><td>11570</td><td>11603</td><td>34</td><td>AGCCGACTTAGAACTGGTGCGGACCAGGGGAATC</td><td>1927.2</td><td>-2.6</td><td>0.6</td><td>-4.1</td><td>4.2E-05</td><td>7.7E-05</td><td>BT20/MB468</td><td>MITO</td></tr>
<tr><td>>hg38_21_+_8433172_8446622_RNA45SN1_45S_50ntFlanks@5467.5491.25|GGCCCACGGCCCTGGCGGAGCGCTG</td><td>>hg38_21_+_8433172_8446622_RNA45SN1_45S_50ntFlanks</td><td>5467</td><td>5491</td><td>25</td><td>GGCCCACGGCCCTGGCGGAGCGCTG</td><td>280.7</td><td>-5.2</td><td>1.3</td><td>-4.0</td><td>5.4E-05</td><td>7.7E-05</td><td>MB231/MB468</td><td>CYTO</td></tr>
<tr><td>>hg38_21_+_8433172_8446622_RNA45SN1_45S_50ntFlanks@12444.12478.35|AAGCAGAATTCACCAAGCGTTGGATTGTTCACCCA</td><td>>hg38_21_+_8433172_8446622_RNA45SN1_45S_50ntFlanks</td><td>12444</td><td>12478</td><td>35</td><td>AAGCAGAATTCACCAAGCGTTGGATTGTTCACCCA</td><td>286.2</td><td>-4.9</td><td>1.2</td><td>-4.0</td><td>5.7E-05</td><td>7.7E-05</td><td>MB231/MB468</td><td>MITO</td></tr>
<tr><td>>hg38_21_+_8433172_8446622_RNA45SN1_45S_50ntFlanks@4731.4757.27|AAGAACGAAAGTCGGAGGTTCGAAGAC</td><td>>hg38_21_+_8433172_8446622_RNA45SN1_45S_50ntFlanks</td><td>4731</td><td>4757</td><td>27</td><td>AAGAACGAAAGTCGGAGGTTCGAAGAC</td><td>529.2</td><td>3.2</td><td>0.8</td><td>4.0</td><td>5.4E-05</td><td>7.7E-05</td><td>MB231/MB468</td><td>CYTO</td></tr>
<tr><td>>hg38_21_+_8433172_8446622_RNA45SN1_45S_50ntFlanks@10775.10818.44|TCTCCAAGGTGAACAGCCTCTGGCATGTTGGAACAATGTAGGTA</td><td>>hg38_21_+_8433172_8446622_RNA45SN1_45S_50ntFlanks</td><td>10775</td><td>10818</td><td>44</td><td>TCTCCAAGGTGAACAGCCTCTGGCATGTTGGAACAATGTAGGTA</td><td>263.6</td><td>-6.4</td><td>1.6</td><td>-4.0</td><td>5.7E-05</td><td>7.7E-05</td><td>MB231/MB468</td><td>MITO</td></tr>
<tr><td>>hg38_21_+_8433172_8446622_RNA45SN1_45S_50ntFlanks@8380.8419.40|CAAGAGGGCGTGAAACCGTTAAGAGGTAAACGGGTGGGGT</td><td>>hg38_21_+_8433172_8446622_RNA45SN1_45S_50ntFlanks</td><td>8380</td><td>8419</td><td>40</td><td>CAAGAGGGCGTGAAACCGTTAAGAGGTAAACGGGTGGGGT</td><td>54.3</td><td>-9.6</td><td>2.4</td><td>-4.1</td><td>4.8E-05</td><td>7.7E-05</td><td>MB231/MB468</td><td>MP</td></tr>
<tr><td>>MI0000082|hsa-mir-25&WithFlank&7|-|100093554|100093649@58.78.21(+1C)&[MIMAT0000081&hsa-miR-25-3p&offsets|0|-1(+1C);m-23&7|-|100093571|100093592&offsets|0|-1(+1C)]|CATTGCACTTGTCTCGGTCTGC</td><td>>MI0000082|hsa-mir-25&WithFlank&7|-|100093554|100093649</td><td>58</td><td>78</td><td>21(+1C)</td><td>CATTGCACTTGTCTCGGTCTGC</td><td>95.7</td><td>3.2</td><td>0.8</td><td>4.0</td><td>5.9E-05</td><td>7.8E-05</td><td>BT20/MB468</td><td>NUC</td></tr>
<tr><td>>hg38_21_+_8433172_8446622_RNA45SN1_45S_50ntFlanks@12407.12435.29|TTTTGATCCTTCGATGTCGGCTCTTCCTA</td><td>>hg38_21_+_8433172_8446622_RNA45SN1_45S_50ntFlanks</td><td>12407</td><td>12435</td><td>29</td><td>TTTTGATCCTTCGATGTCGGCTCTTCCTA</td><td>499.6</td><td>-4.3</td><td>1.0</td><td>-4.1</td><td>5.0E-05</td><td>7.8E-05</td><td>BT20/MB468</td><td>CYTO</td></tr>
<tr><td>>MI0000085|hsa-mir-27a&WithFlank&19|-|13836434|13836523@57.75.19(+1A)&[MIMAT0000084&hsa-miR-27a-3p&offsets|0|-2(+1A);m-43&19|-|13836447|13836467&offsets|0|-2(+1A)]|TTCACAGTGGCTAAGTTCCA</td><td>>MI0000085|hsa-mir-27a&WithFlank&19|-|13836434|13836523</td><td>57</td><td>75</td><td>19(+1A)</td><td>TTCACAGTGGCTAAGTTCCA</td><td>3755.6</td><td>3.7</td><td>0.9</td><td>4.1</td><td>4.8E-05</td><td>7.8E-05</td><td>MB231/MB468</td><td>MP</td></tr>
<tr><td>>MI0000071|hsa-mir-17&WithFlank&13|+|91350599|91350694@20.42.23(+1A)&[MIMAT0000070&hsa-miR-17-5p&offsets|0|0(+1A);m-88&13|+|91350618|91350640&offsets|0|0(+1A)]|CAAAGTGCTTACAGTGCAGGTAGA</td><td>>MI0000071|hsa-mir-17&WithFlank&13|+|91350599|91350694</td><td>20</td><td>42</td><td>23(+1A)</td><td>CAAAGTGCTTACAGTGCAGGTAGA</td><td>2773.9</td><td>-4.2</td><td>0.9</td><td>-4.5</td><td>6.6E-06</td><td>7.8E-05</td><td>BT20/MB231</td><td>MP</td></tr>
<tr><td>>hg38_21_+_8433172_8446622_RNA45SN1_45S_50ntFlanks@10792.10814.23|CTCTGGCATGTTGGAACAATGTA</td><td>>hg38_21_+_8433172_8446622_RNA45SN1_45S_50ntFlanks</td><td>10792</td><td>10814</td><td>23</td><td>CTCTGGCATGTTGGAACAATGTA</td><td>90.6</td><td>9.8</td><td>2.2</td><td>4.5</td><td>6.6E-06</td><td>7.8E-05</td><td>BT20/MB231</td><td>MP</td></tr>
<tr><td>>MI0000088|hsa-mir-30a&WithFlank&6|-|71403545|71403627@53.70.18(+1A)&[MIMAT0000088&hsa-miR-30a-3p&offsets|0|-4(+1A);m-38&6|-|71403554|71403575&offsets|0|-4(+1A)]|CTTTCAGTCGGATGTTTGA</td><td>>MI0000088|hsa-mir-30a&WithFlank&6|-|71403545|71403627</td><td>53</td><td>70</td><td>18(+1A)</td><td>CTTTCAGTCGGATGTTTGA</td><td>190.0</td><td>5.1</td><td>1.3</td><td>4.1</td><td>4.8E-05</td><td>7.8E-05</td><td>MB231/MB468</td><td>MP</td></tr>
<tr><td>>hg38_21_+_8433172_8446622_RNA45SN1_45S_50ntFlanks@7978.8010.33|GACCTCAGATCAGACGTGGCGACCCGCTGAATT</td><td>>hg38_21_+_8433172_8446622_RNA45SN1_45S_50ntFlanks</td><td>7978</td><td>8010</td><td>33</td><td>GACCTCAGATCAGACGTGGCGACCCGCTGAATT</td><td>234.2</td><td>3.5</td><td>0.9</td><td>4.0</td><td>5.8E-05</td><td>7.8E-05</td><td>MB231/MB468</td><td>MITO</td></tr>
<tr><td>>MI0003628|hsa-mir-615&WithFlank&12|+|54033944|54034051@67.86.20&[MIMAT0003283&hsa-miR-615-3p&offsets|0|-2;m-495&12|+|54034010|54034030&offsets|0|-1]|TCCGAGCCTGGGTCTCCCTC</td><td>>MI0003628|hsa-mir-615&WithFlank&12|+|54033944|54034051</td><td>67</td><td>86</td><td>20</td><td>TCCGAGCCTGGGTCTCCCTC</td><td>217.7</td><td>1.6</td><td>0.4</td><td>4.0</td><td>6.0E-05</td><td>7.8E-05</td><td>BT20/MB468</td><td>NUC</td></tr>
<tr><td>>hg38_21_+_8433172_8446622_RNA45SN1_45S_50ntFlanks@9544.9571.28|GGTGGAGGTCCGTAGCGGTCCTGACGTG</td><td>>hg38_21_+_8433172_8446622_RNA45SN1_45S_50ntFlanks</td><td>9544</td><td>9571</td><td>28</td><td>GGTGGAGGTCCGTAGCGGTCCTGACGTG</td><td>1071.6</td><td>-4.9</td><td>1.2</td><td>-4.0</td><td>5.8E-05</td><td>7.8E-05</td><td>MB231/MB468</td><td>MITO</td></tr>
<tr><td>>MI0000456|hsa-mir-140&WithFlank&16|+|69933075|69933186@69.89.21&[MIMAT0004597&hsa-miR-140-3p&offsets|+1|+1;m-45&16|+|69933143|69933164&offsets|0|-1]|ACCACAGGGTAGAACCACGGA</td><td>>MI0000456|hsa-mir-140&WithFlank&16|+|69933075|69933186</td><td>69</td><td>89</td><td>21</td><td>ACCACAGGGTAGAACCACGGA</td><td>1388.8</td><td>5.3</td><td>1.3</td><td>4.1</td><td>4.8E-05</td><td>7.8E-05</td><td>MB231/MB468</td><td>MP</td></tr>
<tr><td>>hg38_1_-_228634819_228635039_RNA5S12_5S_50ntFlanks@124.169.46|AGTACTTGGATGGGAGACCGCCTGGGAATACCGGGTGCTGTAGGCT</td><td>>hg38_1_-_228634819_228635039_RNA5S12_5S_50ntFlanks</td><td>124</td><td>169</td><td>46</td><td>AGTACTTGGATGGGAGACCGCCTGGGAATACCGGGTGCTGTAGGCT</td><td>486.6</td><td>4.6</td><td>1.0</td><td>4.5</td><td>7.7E-06</td><td>7.8E-05</td><td>BT20/MB231</td><td>MITO</td></tr>
<tr><td>>trna10_LysCTT_16_+_3241501_3241573@13.35.23,trna13_LysCTT_14_-_58706613_58706685@13.35.23,trna2_LysCTT_15_+_79152904_79152976@13.35.23,trna4_LysCTT_18_-_43669271_43669343@13.35.23|CAGTCGGTAGAGCATGGGACTCT</td><td>>trna10_LysCTT_16_+_3241501_3241573</td><td>13</td><td>35</td><td>23</td><td>CAGTCGGTAGAGCATGGGACTCT</td><td>300.9</td><td>-4.6</td><td>1.0</td><td>-4.5</td><td>7.7E-06</td><td>7.8E-05</td><td>BT20/MB231</td><td>MITO</td></tr>
<tr><td>>hg38_21_+_8433172_8446622_RNA45SN1_45S_50ntFlanks@10849.10868.20|CGGGATAAGGATTGGCTCTA</td><td>>hg38_21_+_8433172_8446622_RNA45SN1_45S_50ntFlanks</td><td>10849</td><td>10868</td><td>20</td><td>CGGGATAAGGATTGGCTCTA</td><td>282.8</td><td>-3.5</td><td>0.8</td><td>-4.3</td><td>2.0E-05</td><td>7.8E-05</td><td>BT20/MB468</td><td>MP</td></tr>
<tr><td>>MI0000466|hsa-mir-9-1&WithFlank&1|-|156420335|156420435@22.44.23;MI0000468|hsa-mir-9-3&WithFlank&15|+|89368011|89368112@22.44.23;MI0000467|hsa-mir-9-2&WithFlank&5|-|88666847|88666945@22.44.23&[MIMAT0000441&hsa-miR-9-5p&offsets|0|0;m-104&1|-|156420392|156420414&offsets|0|0];[MIMAT0000441_1&hsa-miR-9-5p&offsets|0|0;m-105&15|+|89368032|89368054&offsets|0|0];[MIMAT0000441_2&hsa-miR-9-5p&offsets|0|0;m-104&5|-|88666902|88666924&offsets|0|0]|TCTTTGGTTATCTAGCTGTATGA</td><td>>MI0000466|hsa-mir-9-1&WithFlank&1|-|156420335|156420435</td><td>22</td><td>44</td><td>23</td><td>TCTTTGGTTATCTAGCTGTATGA</td><td>1106.6</td><td>4.0</td><td>0.9</td><td>4.5</td><td>6.3E-06</td><td>7.8E-05</td><td>BT20/MB231</td><td>CYTO</td></tr>
<tr><td>>hg38_21_+_8433172_8446622_RNA45SN1_45S_50ntFlanks@8687.8705.19|CGGCTCCGGGACGGCTGGG</td><td>>hg38_21_+_8433172_8446622_RNA45SN1_45S_50ntFlanks</td><td>8687</td><td>8705</td><td>19</td><td>CGGCTCCGGGACGGCTGGG</td><td>331.7</td><td>-3.5</td><td>0.8</td><td>-4.5</td><td>6.3E-06</td><td>7.8E-05</td><td>BT20/MB231</td><td>CYTO</td></tr>
<tr><td>>MI0000342|hsa-mir-200b&WithFlank&1|+|1167098|1167204@63.84.22(+1U)&[MIMAT0000318&hsa-miR-200b-3p&offsets|0|0(+1U);m-93&1|+|1167160|1167182&offsets|0|-1(+1U)]|TAATACTGCCTGGTAATGATGAT</td><td>>MI0000342|hsa-mir-200b&WithFlank&1|+|1167098|1167204</td><td>63</td><td>84</td><td>22(+1U)</td><td>TAATACTGCCTGGTAATGATGAT</td><td>181.5</td><td>3.7</td><td>0.8</td><td>4.5</td><td>6.3E-06</td><td>7.8E-05</td><td>BT20/MB231</td><td>CYTO</td></tr>
<tr><td>>hg38_21_+_8433172_8446622_RNA45SN1_45S_50ntFlanks@12995.13015.21|CTCGCTGCGATCTATTGAAAG</td><td>>hg38_21_+_8433172_8446622_RNA45SN1_45S_50ntFlanks</td><td>12995</td><td>13015</td><td>21</td><td>CTCGCTGCGATCTATTGAAAG</td><td>1183.4</td><td>3.1</td><td>0.8</td><td>4.0</td><td>5.4E-05</td><td>7.8E-05</td><td>MB231/MB468</td><td>CYTO</td></tr>
<tr><td>>hg38_21_+_8433172_8446622_RNA45SN1_45S_50ntFlanks@10350.10373.24|ACGAGAACTTTGAAGGCCGAAGTG</td><td>>hg38_21_+_8433172_8446622_RNA45SN1_45S_50ntFlanks</td><td>10350</td><td>10373</td><td>24</td><td>ACGAGAACTTTGAAGGCCGAAGTG</td><td>1808.6</td><td>-5.6</td><td>1.4</td><td>-4.1</td><td>4.9E-05</td><td>7.8E-05</td><td>MB231/MB468</td><td>MP</td></tr>
<tr><td>>hg38_21_+_8433172_8446622_RNA45SN1_45S_50ntFlanks@10393.10430.38|CAGCAGTTGAACATGGGTCAGTCGGTCCTGAGAGATGG</td><td>>hg38_21_+_8433172_8446622_RNA45SN1_45S_50ntFlanks</td><td>10393</td><td>10430</td><td>38</td><td>CAGCAGTTGAACATGGGTCAGTCGGTCCTGAGAGATGG</td><td>473.9</td><td>-6.0</td><td>1.5</td><td>-4.1</td><td>5.1E-05</td><td>7.8E-05</td><td>BT20/MB468</td><td>CYTO</td></tr>
<tr><td>>hg38_21_+_8433172_8446622_RNA45SN1_45S_50ntFlanks@12414.12448.35|CCTTCGATGTCGGCTCTTCCTATCATTGTGAAGCA</td><td>>hg38_21_+_8433172_8446622_RNA45SN1_45S_50ntFlanks</td><td>12414</td><td>12448</td><td>35</td><td>CCTTCGATGTCGGCTCTTCCTATCATTGTGAAGCA</td><td>178.6</td><td>3.3</td><td>0.8</td><td>4.0</td><td>5.8E-05</td><td>7.8E-05</td><td>MB231/MB468</td><td>MITO</td></tr>
<tr><td>>am_trna119_LysCTT_1_-_145395522_145395594@7.34.28,trna11_LysCTT_5_-_180648979_180649051@7.34.28,trna128_LysCTT_1_-_55423542_55423614@7.34.28,trna13_LysCTT_6_+_26556774_26556846@7.34.28,trna6_LysCTT_19_-_52425396_52425468@7.34.28,trna7_LysCTT_16_+_3225692_3225764@7.34.28,trna9_LysCTT_5_+_180634755_180634827@7.34.28|CTAGCTCAGTCGGTAGAGCATGAGACTC</td><td>>am_trna119_LysCTT_1_-_145395522_145395594</td><td>7</td><td>34</td><td>28</td><td>CTAGCTCAGTCGGTAGAGCATGAGACTC</td><td>268.7</td><td>6.9</td><td>1.7</td><td>4.0</td><td>5.8E-05</td><td>7.8E-05</td><td>MB231/MB468</td><td>MITO</td></tr>
<tr><td>>hg38_21_+_8433172_8446622_RNA45SN1_45S_50ntFlanks@12616.12654.39|CTGAGGAGCCAATGGGGCGAAGCTACCATCTGTGGGATT</td><td>>hg38_21_+_8433172_8446622_RNA45SN1_45S_50ntFlanks</td><td>12616</td><td>12654</td><td>39</td><td>CTGAGGAGCCAATGGGGCGAAGCTACCATCTGTGGGATT</td><td>784.2</td><td>-3.2</td><td>0.8</td><td>-4.0</td><td>5.5E-05</td><td>7.9E-05</td><td>MB231/MB468</td><td>CYTO</td></tr>
</table>

<table>
<tr><td>>MI0000802|hsa-mir-340&WithFlank&5|-|180015297|180015403@22.43.22&[MIMAT0004692&hsa-miR-340-5p&offsets|0|0;m-220&5|-|180015361|180015382&offsets|0|0]|TTATAAAGCAATGAGACTGATT</td><td>>MI0000802|hsa-mir-340&WithFlank&5|-|180015297|180015403</td><td>22</td><td>43</td><td>22</td><td>TTATAAAGCAATGAGACTGATT</td><td>1540.3</td><td>4.4</td><td>1.1</td><td>4.0</td><td>5.8E-05</td><td>7.9E-05</td><td>MB231/MB468</td><td>MITO</td></tr>
<tr><td>>hg38_21_+_8433172_8446622_RNA45SN1_45S_50ntFlanks@4439.4471.33|CCGCCCCTTGCCTCTCGGCGCCCCCTCGATGCT</td><td>>hg38_21_+_8433172_8446622_RNA45SN1_45S_50ntFlanks</td><td>4439</td><td>4471</td><td>33</td><td>CCGCCCCTTGCCTCTCGGCGCCCCCTCGATGCT</td><td>252.7</td><td>-5.4</td><td>1.3</td><td>-4.1</td><td>4.2E-05</td><td>7.9E-05</td><td>BT20/MB468</td><td>MITO</td></tr>
<tr><td>>hg38_21_+_8433172_8446622_RNA45SN1_45S_50ntFlanks@7988.8009.22|CAGACGTGGCGACCCGCTGAAT</td><td>>hg38_21_+_8433172_8446622_RNA45SN1_45S_50ntFlanks</td><td>7988</td><td>8009</td><td>22</td><td>CAGACGTGGCGACCCGCTGAAT</td><td>1716.9</td><td>-2.1</td><td>0.5</td><td>-4.1</td><td>4.2E-05</td><td>7.9E-05</td><td>BT20/MB468</td><td>MITO</td></tr>
<tr><td>>Antisense_hg38_21_-_8433172_8446622_RNA45SN1_45S_50ntFlanks@7674.7693.20|CTCTCTCTCTCTCTCCCCCG</td><td>>Antisense_hg38_21_-_8433172_8446622_RNA45SN1_45S_50ntFlanks</td><td>7674</td><td>7693</td><td>20</td><td>CTCTCTCTCTCTCTCCCCCG</td><td>3369.1</td><td>-8.3</td><td>2.1</td><td>-4.0</td><td>6.0E-05</td><td>7.9E-05</td><td>BT20/MB468</td><td>NUC</td></tr>
<tr><td>>hg38_21_+_8433172_8446622_RNA45SN1_45S_50ntFlanks@11027.11046.20|CTCTCTCTCTCTCTCCCCCG</td><td>>hg38_21_+_8433172_8446622_RNA45SN1_45S_50ntFlanks</td><td>11027</td><td>11046</td><td>20</td><td>CTCTCTCTCTCTCTCCCCCG</td><td>3369.1</td><td>-8.3</td><td>2.1</td><td>-4.0</td><td>6.0E-05</td><td>7.9E-05</td><td>BT20/MB468</td><td>NUC</td></tr>
<tr><td>>hg38_21_+_8433172_8446622_RNA45SN1_45S_50ntFlanks@11784.11806.23|ATGAATGGATGAACGAGATTCCC</td><td>>hg38_21_+_8433172_8446622_RNA45SN1_45S_50ntFlanks</td><td>11784</td><td>11806</td><td>23</td><td>ATGAATGGATGAACGAGATTCCC</td><td>1173.6</td><td>-4.0</td><td>0.9</td><td>-4.3</td><td>2.0E-05</td><td>7.9E-05</td><td>BT20/MB468</td><td>MP</td></tr>
<tr><td>>hg38_21_+_8433172_8446622_RNA45SN1_45S_50ntFlanks@51.72.22|GCTGACACGCTGTCCTCTGGCG</td><td>>hg38_21_+_8433172_8446622_RNA45SN1_45S_50ntFlanks</td><td>51</td><td>72</td><td>22</td><td>GCTGACACGCTGTCCTCTGGCG</td><td>991.9</td><td>-3.0</td><td>0.7</td><td>-4.0</td><td>5.8E-05</td><td>7.9E-05</td><td>MB231/MB468</td><td>NUC</td></tr>
<tr><td>>hg38_21_+_8433172_8446622_RNA45SN1_45S_50ntFlanks@11516.11539.24|TCCGCCCCCGGGGCCGCGGTTCCG</td><td>>hg38_21_+_8433172_8446622_RNA45SN1_45S_50ntFlanks</td><td>11516</td><td>11539</td><td>24</td><td>TCCGCCCCCGGGGCCGCGGTTCCG</td><td>236.6</td><td>-5.7</td><td>1.4</td><td>-4.0</td><td>5.8E-05</td><td>7.9E-05</td><td>MB231/MB468</td><td>NUC</td></tr>
<tr><td>>hg38_21_+_8433172_8446622_RNA45SN1_45S_50ntFlanks@10346.10371.26|TCAAACGAGAACTTTGAAGGCCGAAG</td><td>>hg38_21_+_8433172_8446622_RNA45SN1_45S_50ntFlanks</td><td>10346</td><td>10371</td><td>26</td><td>TCAAACGAGAACTTTGAAGGCCGAAG</td><td>596.5</td><td>-3.5</td><td>0.9</td><td>-4.0</td><td>5.8E-05</td><td>7.9E-05</td><td>MB231/MB468</td><td>MITO</td></tr>
<tr><td>>hg38_21_+_8433172_8446622_RNA45SN1_45S_50ntFlanks@10663.10711.49|CCTTGGAAAGCGTCGCGGTTCCGGCGGCGTCCGGTGAGCTCTCGCTGGC</td><td>>hg38_21_+_8433172_8446622_RNA45SN1_45S_50ntFlanks</td><td>10663</td><td>10711</td><td>49</td><td>CCTTGGAAAGCGTCGCGGTTCCGGCGGCGTCCGGTGAGCTCTCGCTGGC</td><td>394.8</td><td>-6.2</td><td>1.5</td><td>-4.1</td><td>4.3E-05</td><td>7.9E-05</td><td>BT20/MB468</td><td>MITO</td></tr>
<tr><td>>trnaMT_GluTTC_MT_-_14674_14742@39.62.24,trnalookalike8_GluTTC_5_-_93905172_93905240@39.62.24|ATTGGTCGTGGTTGTAGTCCGTGC</td><td>>trnaMT_GluTTC_MT_-_14674_14742</td><td>39</td><td>62</td><td>24</td><td>ATTGGTCGTGGTTGTAGTCCGTGC</td><td>357.2</td><td>5.8</td><td>1.4</td><td>4.3</td><td>2.0E-05</td><td>7.9E-05</td><td>BT20/MB468</td><td>MP</td></tr>
<tr><td>>MI0000478|hsa-mir-149&WithFlank&2|+|240455995|240456095@21.42.22&[MIMAT0000450&hsa-miR-149-5p&offsets|0|-1;m-195&2|+|240456015|240456037&offsets|0|-1]|TCTGGCTCCGTGTCTTCACTCC</td><td>>MI0000478|hsa-mir-149&WithFlank&2|+|240455995|240456095</td><td>21</td><td>42</td><td>22</td><td>TCTGGCTCCGTGTCTTCACTCC</td><td>1373.7</td><td>3.3</td><td>0.7</td><td>4.5</td><td>7.6E-06</td><td>7.9E-05</td><td>BT20/MB231</td><td>NUC</td></tr>
<tr><td>>hg38_21_+_8433172_8446622_RNA45SN1_45S_50ntFlanks@10411.10441.31|CAGTCGGTCCTGAGAGATGGGCGAGCGCCGT</td><td>>hg38_21_+_8433172_8446622_RNA45SN1_45S_50ntFlanks</td><td>10411</td><td>10441</td><td>31</td><td>CAGTCGGTCCTGAGAGATGGGCGAGCGCCGT</td><td>147.2</td><td>-5.0</td><td>1.2</td><td>-4.1</td><td>5.1E-05</td><td>7.9E-05</td><td>BT20/MB468</td><td>CYTO</td></tr>
<tr><td>>hg38_21_+_8433172_8446622_RNA45SN1_45S_50ntFlanks@6661.6682.22|CGGTGGATCACTCGGCTCGTGC</td><td>>hg38_21_+_8433172_8446622_RNA45SN1_45S_50ntFlanks</td><td>6661</td><td>6682</td><td>22</td><td>CGGTGGATCACTCGGCTCGTGC</td><td>431.6</td><td>4.4</td><td>1.0</td><td>4.3</td><td>2.0E-05</td><td>7.9E-05</td><td>BT20/MB468</td><td>MP</td></tr>
<tr><td>>hg38_21_+_8433172_8446622_RNA45SN1_45S_50ntFlanks@12242.12266.25|AGAAGGGCAAAAGCTCGCTTGATCT</td><td>>hg38_21_+_8433172_8446622_RNA45SN1_45S_50ntFlanks</td><td>12242</td><td>12266</td><td>25</td><td>AGAAGGGCAAAAGCTCGCTTGATCT</td><td>215.5</td><td>-6.0</td><td>1.5</td><td>-4.1</td><td>4.9E-05</td><td>7.9E-05</td><td>MB231/MB468</td><td>MP</td></tr>
<tr><td>>hg38_21_+_8433172_8446622_RNA45SN1_45S_50ntFlanks@9305.9336.32|CCAAGGAGTCTAACACGTGCGCGAGTCGGGGG</td><td>>hg38_21_+_8433172_8446622_RNA45SN1_45S_50ntFlanks</td><td>9305</td><td>9336</td><td>32</td><td>CCAAGGAGTCTAACACGTGCGCGAGTCGGGGG</td><td>1372.1</td><td>-5.7</td><td>1.3</td><td>-4.3</td><td>2.0E-05</td><td>7.9E-05</td><td>BT20/MB468</td><td>MP</td></tr>
<tr><td>>MI0000087|hsa-mir-29a&WithFlank&7|-|130876741|130876816@48.67.20(+1A)&[MIMAT0000086&hsa-miR-29a-3p&offsets|0|-2(+1A);m-26&7|-|130876749|130876769&offsets|0|-1(+1A)]|TAGCACCATCTGAAATCGGTA</td><td>>MI0000087|hsa-mir-29a&WithFlank&7|-|130876741|130876816</td><td>48</td><td>67</td><td>20(+1A)</td><td>TAGCACCATCTGAAATCGGTA</td><td>1107.3</td><td>6.6</td><td>1.6</td><td>4.1</td><td>4.9E-05</td><td>7.9E-05</td><td>MB231/MB468</td><td>MP</td></tr>
<tr><td>>hg38_21_+_8433172_8446622_RNA45SN1_45S_50ntFlanks@10842.10865.24|GTAACTTCGGGATAAGGATTGGCT</td><td>>hg38_21_+_8433172_8446622_RNA45SN1_45S_50ntFlanks</td><td>10842</td><td>10865</td><td>24</td><td>GTAACTTCGGGATAAGGATTGGCT</td><td>721.5</td><td>-2.8</td><td>0.7</td><td>-4.0</td><td>5.5E-05</td><td>8.0E-05</td><td>MB231/MB468</td><td>CYTO</td></tr>
<tr><td>>hg38_21_+_8433172_8446622_RNA45SN1_45S_50ntFlanks@10350.10376.27|ACGAGAACTTTGAAGGCCGAAGTGGAG</td><td>>hg38_21_+_8433172_8446622_RNA45SN1_45S_50ntFlanks</td><td>10350</td><td>10376</td><td>27</td><td>ACGAGAACTTTGAAGGCCGAAGTGGAG</td><td>613.8</td><td>-2.7</td><td>0.7</td><td>-4.1</td><td>4.3E-05</td><td>8.0E-05</td><td>BT20/MB468</td><td>MITO</td></tr>
<tr><td>>hg38_21_+_8433172_8446622_RNA45SN1_45S_50ntFlanks@10794.10812.19|CTGGCATGTTGGAACAATG</td><td>>hg38_21_+_8433172_8446622_RNA45SN1_45S_50ntFlanks</td><td>10794</td><td>10812</td><td>19</td><td>CTGGCATGTTGGAACAATG</td><td>86.2</td><td>9.8</td><td>2.2</td><td>4.5</td><td>6.8E-06</td><td>8.0E-05</td><td>BT20/MB231</td><td>MP</td></tr>
<tr><td>>hg38_21_+_8433172_8446622_RNA45SN1_45S_50ntFlanks@6651.6692.42|CGACTCTTAGCGGTGGATCACTCGGCTCGTGCGTCGATGAAG</td><td>>hg38_21_+_8433172_8446622_RNA45SN1_45S_50ntFlanks</td><td>6651</td><td>6692</td><td>42</td><td>CGACTCTTAGCGGTGGATCACTCGGCTCGTGCGTCGATGAAG</td><td>2000.1</td><td>2.7</td><td>0.7</td><td>4.0</td><td>5.5E-05</td><td>8.0E-05</td><td>MB231/MB468</td><td>CYTO</td></tr>
<tr><td>>hg38_21_+_8433172_8446622_RNA45SN1_45S_50ntFlanks@6646.6672.27|TCGTACGACTCTTAGCGGTGGATCACT</td><td>>hg38_21_+_8433172_8446622_RNA45SN1_45S_50ntFlanks</td><td>6646</td><td>6672</td><td>27</td><td>TCGTACGACTCTTAGCGGTGGATCACT</td><td>2980.1</td><td>2.3</td><td>0.6</td><td>4.0</td><td>6.1E-05</td><td>8.0E-05</td><td>BT20/MB468</td><td>NUC</td></tr>
<tr><td>>hg38_MT_+_1621_3279_RNR2_16S_50ntFlanks@51.79.29|GCTAAACCTAGCCCCAAACCCACTCCACC</td><td>>hg38_MT_+_1621_3279_RNR2_16S_50ntFlanks</td><td>51</td><td>79</td><td>29</td><td>GCTAAACCTAGCCCCAAACCCACTCCACC</td><td>2844.2</td><td>3.4</td><td>0.9</td><td>4.0</td><td>5.9E-05</td><td>8.0E-05</td><td>MB231/MB468</td><td>MITO</td></tr>
<tr><td>>hg38_1_-_228634819_228635039_RNA5S12_5S_50ntFlanks@84.102.19|CTCGTCTGATCTCGGAAGC</td><td>>hg38_1_-_228634819_228635039_RNA5S12_5S_50ntFlanks</td><td>84</td><td>102</td><td>19</td><td>CTCGTCTGATCTCGGAAGC</td><td>497.1</td><td>-3.3</td><td>0.8</td><td>-4.0</td><td>5.2E-05</td><td>8.0E-05</td><td>BT20/MB468</td><td>CYTO</td></tr>
<tr><td>>am_trna134_GluTTC_1_-_16861774_16861845@1.28.28,trna5_GluTTC_1_+_17199078_17199149@1.28.28,trna84_GluTTC_1_-_161391883_161391954@1.28.28,trna94_GluTTC_1_-_149664355_149664427@1.28.28|TCCCTGGTGGTCTAGTGGCTAGGATTCG</td><td>>am_trna134_GluTTC_1_-_16861774_16861845</td><td>1</td><td>28</td><td>28</td><td>TCCCTGGTGGTCTAGTGGCTAGGATTCG</td><td>1691.7</td><td>-7.6</td><td>1.9</td><td>-4.0</td><td>6.1E-05</td><td>8.0E-05</td><td>BT20/MB468</td><td>NUC</td></tr>
<tr><td>>hg38_21_+_8433172_8446622_RNA45SN1_45S_50ntFlanks@12907.12935.29|CACCGCACGTTCGTGGGGAACCTGGCGCT</td><td>>hg38_21_+_8433172_8446622_RNA45SN1_45S_50ntFlanks</td><td>12907</td><td>12935</td><td>29</td><td>CACCGCACGTTCGTGGGGAACCTGGCGCT</td><td>612.7</td><td>-7.8</td><td>1.9</td><td>-4.0</td><td>6.1E-05</td><td>8.0E-05</td><td>BT20/MB468</td><td>NUC</td></tr>
<tr><td>>hg38_21_+_8433172_8446622_RNA45SN1_45S_50ntFlanks@6648.6678.31|GTACGACTCTTAGCGGTGGATCACTCGGCTC</td><td>>hg38_21_+_8433172_8446622_RNA45SN1_45S_50ntFlanks</td><td>6648</td><td>6678</td><td>31</td><td>GTACGACTCTTAGCGGTGGATCACTCGGCTC</td><td>536.5</td><td>3.9</td><td>0.9</td><td>4.3</td><td>2.0E-05</td><td>8.0E-05</td><td>BT20/MB468</td><td>MP</td></tr>
<tr><td>>hg38_21_+_8433172_8446622_RNA45SN1_45S_50ntFlanks@6652.6670.19|GACTCTTAGCGGTGGATCA</td><td>>hg38_21_+_8433172_8446622_RNA45SN1_45S_50ntFlanks</td><td>6652</td><td>6670</td><td>19</td><td>GACTCTTAGCGGTGGATCA</td><td>380.0</td><td>-3.4</td><td>0.8</td><td>-4.0</td><td>5.9E-05</td><td>8.0E-05</td><td>MB231/MB468</td><td>MITO</td></tr>
<tr><td>>trna10_LysCTT_16_+_3241501_3241573@4.35.32,trna13_LysCTT_14_-_58706613_58706685@4.35.32,trna2_LysCTT_15_+_79152904_79152976@4.35.32|CGGCTAGCTCAGTCGGTAGAGCATGGGACTCT</td><td>>trna10_LysCTT_16_+_3241501_3241573</td><td>4</td><td>35</td><td>32</td><td>CGGCTAGCTCAGTCGGTAGAGCATGGGACTCT</td><td>250.6</td><td>5.0</td><td>1.2</td><td>4.1</td><td>5.0E-05</td><td>8.0E-05</td><td>MB231/MB468</td><td>MP</td></tr>
<tr><td>>MI0000478|hsa-mir-149&WithFlank&2|+|240455995|240456095@21.42.22(+1U)&[MIMAT0000450&hsa-miR-149-5p&offsets|0|-1(+1U);m-195&2|+|240456015|240456037&offsets|0|-1(+1U)]|TCTGGCTCCGTGTCTTCACTCCT</td><td>>MI0000478|hsa-mir-149&WithFlank&2|+|240455995|240456095</td><td>21</td><td>42</td><td>22(+1U)</td><td>TCTGGCTCCGTGTCTTCACTCCT</td><td>1028.0</td><td>3.6</td><td>0.8</td><td>4.5</td><td>8.0E-06</td><td>8.0E-05</td><td>BT20/MB231</td><td>MITO</td></tr>
<tr><td>>hg38_21_+_8433172_8446622_RNA45SN1_45S_50ntFlanks@6646.6680.35|TCGTACGACTCTTAGCGGTGGATCACTCGGCTCGT</td><td>>hg38_21_+_8433172_8446622_RNA45SN1_45S_50ntFlanks</td><td>6646</td><td>6680</td><td>35</td><td>TCGTACGACTCTTAGCGGTGGATCACTCGGCTCGT</td><td>2925.0</td><td>-3.4</td><td>0.8</td><td>-4.1</td><td>5.0E-05</td><td>8.0E-05</td><td>MB231/MB468</td><td>MP</td></tr>
<tr><td>>hg38_21_+_8433172_8446622_RNA45SN1_45S_50ntFlanks@10869.10891.23|AGGGCTGGGTCGGTCGGGCTGGG</td><td>>hg38_21_+_8433172_8446622_RNA45SN1_45S_50ntFlanks</td><td>10869</td><td>10891</td><td>23</td><td>AGGGCTGGGTCGGTCGGGCTGGG</td><td>712.7</td><td>-6.4</td><td>1.6</td><td>-4.0</td><td>5.9E-05</td><td>8.0E-05</td><td>MB231/MB468</td><td>NUC</td></tr>
<tr><td>>hg38_21_+_8433172_8446622_RNA45SN1_45S_50ntFlanks@12006.12032.27|GGCCGCCGGTGAAATACCACTACTCTG</td><td>>hg38_21_+_8433172_8446622_RNA45SN1_45S_50ntFlanks</td><td>12006</td><td>12032</td><td>27</td><td>GGCCGCCGGTGAAATACCACTACTCTG</td><td>603.9</td><td>-5.5</td><td>1.3</td><td>-4.3</td><td>2.1E-05</td><td>8.0E-05</td><td>BT20/MB468</td><td>MP</td></tr>
<tr><td>>hg38_21_+_8433172_8446622_RNA45SN1_45S_50ntFlanks@3839.3860.22|TCTCCTACTTGGATAACTGTGG</td><td>>hg38_21_+_8433172_8446622_RNA45SN1_45S_50ntFlanks</td><td>3839</td><td>3860</td><td>22</td><td>TCTCCTACTTGGATAACTGTGG</td><td>412.4</td><td>-7.3</td><td>1.8</td><td>-4.1</td><td>5.0E-05</td><td>8.0E-05</td><td>MB231/MB468</td><td>MP</td></tr>
<tr><td>>hg38_21_+_8433172_8446622_RNA45SN1_45S_50ntFlanks@9306.9323.18|CAAGGAGTCTAACACGTG</td><td>>hg38_21_+_8433172_8446622_RNA45SN1_45S_50ntFlanks</td><td>9306</td><td>9323</td><td>18</td><td>CAAGGAGTCTAACACGTG</td><td>120.3</td><td>-5.0</td><td>1.2</td><td>-4.0</td><td>5.2E-05</td><td>8.0E-05</td><td>BT20/MB468</td><td>CYTO</td></tr>
<tr><td>>hg38_1_-_228634819_228635039_RNA5S12_5S_50ntFlanks@134.172.39|TGGGAGACCGCCTGGGAATACCGGGTGCTGTAGGCTTTT</td><td>>hg38_1_-_228634819_228635039_RNA5S12_5S_50ntFlanks</td><td>134</td><td>172</td><td>39</td><td>TGGGAGACCGCCTGGGAATACCGGGTGCTGTAGGCTTTT</td><td>563.4</td><td>-4.4</td><td>1.0</td><td>-4.5</td><td>8.0E-06</td><td>8.0E-05</td><td>BT20/MB231</td><td>MITO</td></tr>
<tr><td>>hg38_21_+_8433172_8446622_RNA45SN1_45S_50ntFlanks@10409.10429.21|GTCAGTCGGTCCTGAGAGATG</td><td>>hg38_21_+_8433172_8446622_RNA45SN1_45S_50ntFlanks</td><td>10409</td><td>10429</td><td>21</td><td>GTCAGTCGGTCCTGAGAGATG</td><td>1363.5</td><td>-2.3</td><td>0.6</td><td>-4.0</td><td>5.6E-05</td><td>8.0E-05</td><td>MB231/MB468</td><td>CYTO</td></tr>
<tr><td>>MI0000294|hsa-mir-218-1&WithFlank&4|+|20528269|20528390@31.51.21(+1A);MI0000295|hsa-mir-218-2&WithFlank&5|-|168768140|168768261@31.51.21(+1A)&[MIMAT0000275&hsa-miR-218-5p&offsets|0|0(+1A);m-185&4|+|20528299|20528319&offsets|0|0(+1A)];[MIMAT0000275_1&hsa-miR-218-5p&offsets|0|0(+1A);m-187&5|-|168768211|168768231&offsets|0|0(+1A)]|TTGTGCTTGATCTAACCATGTA</td><td>>MI0000294|hsa-mir-218-1&WithFlank&4|+|20528269|20528390</td><td>31</td><td>51</td><td>21(+1A)</td><td>TTGTGCTTGATCTAACCATGTA</td><td>32.4</td><td>8.2</td><td>2.0</td><td>4.1</td><td>5.0E-05</td><td>8.0E-05</td><td>MB231/MB468</td><td>MP</td></tr>
<tr><td>>hg38_21_+_8433172_8446622_RNA45SN1_45S_50ntFlanks@11843.11860.18|AGGGAACGGGCTTGGCGG</td><td>>hg38_21_+_8433172_8446622_RNA45SN1_45S_50ntFlanks</td><td>11843</td><td>11860</td><td>18</td><td>AGGGAACGGGCTTGGCGG</td><td>85.2</td><td>9.7</td><td>2.2</td><td>4.5</td><td>6.9E-06</td><td>8.1E-05</td><td>BT20/MB231</td><td>MP</td></tr>
<tr><td>>hg38_21_+_8433172_8446622_RNA45SN1_45S_50ntFlanks@9485.9510.26|TGTTAGGACCCGAAAGATGGTGAACT</td><td>>hg38_21_+_8433172_8446622_RNA45SN1_45S_50ntFlanks</td><td>9485</td><td>9510</td><td>26</td><td>TGTTAGGACCCGAAAGATGGTGAACT</td><td>75.4</td><td>9.6</td><td>2.1</td><td>4.5</td><td>6.9E-06</td><td>8.1E-05</td><td>BT20/MB231</td><td>MP</td></tr>
<tr><td>>MI0000061|hsa-let-7a-2&WithFlank&11|-|122146516|122146599@56.76.21&[MIMAT0010195&hsa-let-7a-2-3p&offsets|0|-1;m-426&11|-|122146523|122146544&offsets|0|-1]|CTGTACAGCCTCCTAGCTTTC</td><td>>MI0000061|hsa-let-7a-2&WithFlank&11|-|122146516|122146599</td><td>56</td><td>76</td><td>21</td><td>CTGTACAGCCTCCTAGCTTTC</td><td>65.6</td><td>-9.4</td><td>2.1</td><td>-4.5</td><td>6.9E-06</td><td>8.1E-05</td><td>BT20/MB231</td><td>MP</td></tr>
<tr><td>>MI0000102|hsa-mir-100&WithFlank&11|-|122152223|122152314@19.37.19(+1A)&[MIMAT0000098&hsa-miR-100-5p&offsets|0|-3(+1A);m-27&11|-|122152276|122152296&offsets|0|-2(+1A)]|AACCCGTAGATCCGAACTTA</td><td>>MI0000102|hsa-mir-100&WithFlank&11|-|122152223|122152314</td><td>19</td><td>37</td><td>19(+1A)</td><td>AACCCGTAGATCCGAACTTA</td><td>9134.3</td><td>-7.6</td><td>1.7</td><td>-4.5</td><td>6.9E-06</td><td>8.1E-05</td><td>BT20/MB231</td><td>MP</td></tr>
<tr><td>>hg38_21_+_8433172_8446622_RNA45SN1_45S_50ntFlanks@10901.10930.30|GGGGCTGGGCGCGCGCCGCGGCTGGACGAG</td><td>>hg38_21_+_8433172_8446622_RNA45SN1_45S_50ntFlanks</td><td>10901</td><td>10930</td><td>30</td><td>GGGGCTGGGCGCGCGCCGCGGCTGGACGAG</td><td>34.6</td><td>8.9</td><td>2.1</td><td>4.3</td><td>2.1E-05</td><td>8.1E-05</td><td>BT20/MB468</td><td>MP</td></tr>
<tr><td>>hg38_21_+_8433172_8446622_RNA45SN1_45S_50ntFlanks@12995.13040.46|CTCGCTGCGATCTATTGAAAGTCAGCCCTCGACACAAGGGTTTGTC</td><td>>hg38_21_+_8433172_8446622_RNA45SN1_45S_50ntFlanks</td><td>12995</td><td>13040</td><td>46</td><td>CTCGCTGCGATCTATTGAAAGTCAGCCCTCGACACAAGGGTTTGTC</td><td>1129.6</td><td>7.5</td><td>1.9</td><td>4.0</td><td>6.0E-05</td><td>8.1E-05</td><td>MB231/MB468</td><td>MITO</td></tr>
<tr><td>>am_trnaMT_ValTAC_MT_+_1602_1670@27.67.41|CAACTTACACTTAGGAGATTTCAACTTAACTTGACCGCTCT</td><td>>am_trnaMT_ValTAC_MT_+_1602_1670</td><td>27</td><td>67</td><td>41</td><td>CAACTTACACTTAGGAGATTTCAACTTAACTTGACCGCTCT</td><td>161.7</td><td>11.1</td><td>2.6</td><td>4.3</td><td>2.1E-05</td><td>8.1E-05</td><td>BT20/MB468</td><td>MP</td></tr>
<tr><td>>hg38_21_+_8433172_8446622_RNA45SN1_45S_50ntFlanks@8176.8216.41|CTTCTGATCGAGGCCCAGCCCGTGGACGGTGTGAGGCCGGT</td><td>>hg38_21_+_8433172_8446622_RNA45SN1_45S_50ntFlanks</td><td>8176</td><td>8216</td><td>41</td><td>CTTCTGATCGAGGCCCAGCCCGTGGACGGTGTGAGGCCGGT</td><td>339.3</td><td>-3.4</td><td>0.8</td><td>-4.0</td><td>6.0E-05</td><td>8.1E-05</td><td>MB231/MB468</td><td>MITO</td></tr>
<tr><td>>MI0006353|hsa-mir-1291&WithFlank&12|-|48654438|48654536@19.40.22&[MIMAT0005881&hsa-miR-1291&offsets|-1|-3;m-616&12|-|48654494|48654517&offsets|-1|-3]|GTGGCCCTGACTGAAGACCAGC</td><td>>MI0006353|hsa-mir-1291&WithFlank&12|-|48654438|48654536</td><td>19</td><td>40</td><td>22</td><td>GTGGCCCTGACTGAAGACCAGC</td><td>248.8</td><td>3.7</td><td>0.8</td><td>4.5</td><td>7.7E-06</td><td>8.1E-05</td><td>BT20/MB231</td><td>NUC</td></tr>
<tr><td>>MI0000088|hsa-mir-30a&WithFlank&6|-|71403545|71403627@53.74.22(+1C)&[MIMAT0000088&hsa-miR-30a-3p&offsets|0|0(+1C);m-38&6|-|71403554|71403575&offsets|0|0(+1C)]|CTTTCAGTCGGATGTTTGCAGCC</td><td>>MI0000088|hsa-mir-30a&WithFlank&6|-|71403545|71403627</td><td>53</td><td>74</td><td>22(+1C)</td><td>CTTTCAGTCGGATGTTTGCAGCC</td><td>65.7</td><td>-9.4</td><td>2.1</td><td>-4.5</td><td>7.0E-06</td><td>8.1E-05</td><td>BT20/MB231</td><td>MP</td></tr>
</table>

<table>
<tr><td>>hg38_21_+_8433172_8446622_RNA45SN1_45S_50ntFlanks@10807.10828.22|ACAATGTAGGTAAGGGAAGTCG</td><td>>hg38_21_+_8433172_8446622_RNA45SN1_45S_50ntFlanks</td><td>10807</td><td>10828</td><td>22</td><td>ACAATGTAGGTAAGGGAAGTCG</td><td>208.7</td><td>-3.6</td><td>0.9</td><td>-4.1</td><td>5.0E-05</td><td>8.1E-05</td><td>MB231/MB468</td><td>MP</td></tr>
<tr><td>>hg38_21_+_8433172_8446622_RNA45SN1_45S_50ntFlanks@10788.10806.19|CAGCCTCTGGCATGTTGGA</td><td>>hg38_21_+_8433172_8446622_RNA45SN1_45S_50ntFlanks</td><td>10788</td><td>10806</td><td>19</td><td>CAGCCTCTGGCATGTTGGA</td><td>323.5</td><td>-5.2</td><td>1.3</td><td>-4.0</td><td>5.6E-05</td><td>8.1E-05</td><td>MB231/MB468</td><td>CYTO</td></tr>
<tr><td>>MI0000542|hsa-mir-320a&WithFlank&8|-|22244960|22245043@48.70.23&[MIMAT0000510&hsa-miR-320a-3p&offsets|0|+1;m-60&8|-|22244975|22244996&offsets|0|+1]|AAAAGCTGGGTTGAGAGGGCGAA</td><td>>MI0000542|hsa-mir-320a&WithFlank&8|-|22244960|22245043</td><td>48</td><td>70</td><td>23</td><td>AAAAGCTGGGTTGAGAGGGCGAA</td><td>1981.9</td><td>3.8</td><td>0.9</td><td>4.1</td><td>5.0E-05</td><td>8.1E-05</td><td>MB231/MB468</td><td>MP</td></tr>
<tr><td>>hg38_21_+_8433172_8446622_RNA45SN1_45S_50ntFlanks@8213.8237.25|CGGTAGCGGCCCCCGGCGCGCCGGG</td><td>>hg38_21_+_8433172_8446622_RNA45SN1_45S_50ntFlanks</td><td>8213</td><td>8237</td><td>25</td><td>CGGTAGCGGCCCCCGGCGCGCCGGG</td><td>46.3</td><td>-8.6</td><td>2.0</td><td>-4.3</td><td>2.1E-05</td><td>8.1E-05</td><td>BT20/MB468</td><td>MP</td></tr>
<tr><td>>hg38_21_+_8433172_8446622_RNA45SN1_45S_50ntFlanks@10389.10408.20|TGAACAGCAGTTGAACATGG</td><td>>hg38_21_+_8433172_8446622_RNA45SN1_45S_50ntFlanks</td><td>10389</td><td>10408</td><td>20</td><td>TGAACAGCAGTTGAACATGG</td><td>197.3</td><td>-3.8</td><td>0.9</td><td>-4.3</td><td>2.1E-05</td><td>8.1E-05</td><td>BT20/MB468</td><td>MP</td></tr>
<tr><td>>hg38_21_+_8433172_8446622_RNA45SN1_45S_50ntFlanks@6732.6772.41|ACATTGATCATCGACACTTCGAACGCACTTGCGGCCCCGGG</td><td>>hg38_21_+_8433172_8446622_RNA45SN1_45S_50ntFlanks</td><td>6732</td><td>6772</td><td>41</td><td>ACATTGATCATCGACACTTCGAACGCACTTGCGGCCCCGGG</td><td>1001.4</td><td>-2.9</td><td>0.7</td><td>-4.0</td><td>6.0E-05</td><td>8.1E-05</td><td>MB231/MB468</td><td>MITO</td></tr>
<tr><td>>MI0000289|hsa-mir-181a-1&WithFlank&1|-|198859038|198859159@30.52.23;MI0000269|hsa-mir-181a-2&WithFlank&9|+|124692436|124692557@45.67.23&[MIMAT0000256&hsa-miR-181a-5p&offsets|0|0;m-32&1|-|198859108|198859130&offsets|0|0];[MIMAT0000256_1&hsa-miR-181a-5p&offsets|0|0;m-32&9|+|124692480|124692502&offsets|0|0]|AACATTCAACGCTGTCGGTGAGT</td><td>>MI0000289|hsa-mir-181a-1&WithFlank&1|-|198859038|198859159</td><td>30</td><td>52</td><td>23</td><td>AACATTCAACGCTGTCGGTGAGT</td><td>20456.7</td><td>-4.4</td><td>1.0</td><td>-4.5</td><td>7.0E-06</td><td>8.1E-05</td><td>BT20/MB231</td><td>MP</td></tr>
<tr><td>>MI0003834|hsa-mir-769&WithFlank&19|+|46018926|46019055@36.56.21&[MIMAT0003886&hsa-miR-769-5p&offsets|0|-1;m-216&19|+|46018961|46018982&offsets|0|-1]|TGAGACCTCTGGGTTCTGAGC</td><td>>MI0003834|hsa-mir-769&WithFlank&19|+|46018926|46019055</td><td>36</td><td>56</td><td>21</td><td>TGAGACCTCTGGGTTCTGAGC</td><td>258.9</td><td>6.2</td><td>1.6</td><td>4.0</td><td>6.0E-05</td><td>8.1E-05</td><td>MB231/MB468</td><td>MITO</td></tr>
<tr><td>>MI0000286|hsa-mir-210&WithFlank&11|-|568083|568204@72.95.24&[MIMAT0000267&hsa-miR-210-3p&offsets|0|+2;m-102&11|-|568112|568133&offsets|0|+2]|CTGTGCGTGTGACAGCGGCTGATC</td><td>>MI0000286|hsa-mir-210&WithFlank&11|-|568083|568204</td><td>72</td><td>95</td><td>24</td><td>CTGTGCGTGTGACAGCGGCTGATC</td><td>31.8</td><td>8.2</td><td>2.0</td><td>4.1</td><td>5.1E-05</td><td>8.1E-05</td><td>MB231/MB468</td><td>MP</td></tr>
<tr><td>>hg38_21_+_8433172_8446622_RNA45SN1_45S_50ntFlanks@11857.11887.31|GCGGAATCAGCGGGGAAAGAAGACCCTGTTG</td><td>>hg38_21_+_8433172_8446622_RNA45SN1_45S_50ntFlanks</td><td>11857</td><td>11887</td><td>31</td><td>GCGGAATCAGCGGGGAAAGAAGACCCTGTTG</td><td>237.6</td><td>-3.6</td><td>0.9</td><td>-4.3</td><td>2.1E-05</td><td>8.1E-05</td><td>BT20/MB468</td><td>MP</td></tr>
<tr><td>>hg38_21_+_8433172_8446622_RNA45SN1_45S_50ntFlanks@11315.11332.18|CCACCCCACGTCTCGTCG</td><td>>hg38_21_+_8433172_8446622_RNA45SN1_45S_50ntFlanks</td><td>11315</td><td>11332</td><td>18</td><td>CCACCCCACGTCTCGTCG</td><td>551.2</td><td>-6.4</td><td>1.6</td><td>-4.0</td><td>5.3E-05</td><td>8.1E-05</td><td>BT20/MB468</td><td>CYTO</td></tr>
<tr><td>>hg38_21_+_8433172_8446622_RNA45SN1_45S_50ntFlanks@12274.12295.22|CAGTACGAATACAGACCGTGAA</td><td>>hg38_21_+_8433172_8446622_RNA45SN1_45S_50ntFlanks</td><td>12274</td><td>12295</td><td>22</td><td>CAGTACGAATACAGACCGTGAA</td><td>114.7</td><td>-5.8</td><td>1.4</td><td>-4.1</td><td>5.1E-05</td><td>8.2E-05</td><td>MB231/MB468</td><td>MP</td></tr>
<tr><td>>hg38_21_+_8433172_8446622_RNA45SN1_45S_50ntFlanks@11316.11342.27|CACCCCACGTCTCGTCGCGCGCGCGTC</td><td>>hg38_21_+_8433172_8446622_RNA45SN1_45S_50ntFlanks</td><td>11316</td><td>11342</td><td>27</td><td>CACCCCACGTCTCGTCGCGCGCGCGTC</td><td>5146.9</td><td>-4.4</td><td>1.1</td><td>-4.1</td><td>4.4E-05</td><td>8.2E-05</td><td>BT20/MB468</td><td>MITO</td></tr>
<tr><td>>hg38_21_+_8433172_8446622_RNA45SN1_45S_50ntFlanks@12044.12071.28|ACTGACCCGGTGAGGCGGGGGGGCGAGC</td><td>>hg38_21_+_8433172_8446622_RNA45SN1_45S_50ntFlanks</td><td>12044</td><td>12071</td><td>28</td><td>ACTGACCCGGTGAGGCGGGGGGGCGAGC</td><td>710.2</td><td>-6.9</td><td>1.6</td><td>-4.3</td><td>2.1E-05</td><td>8.2E-05</td><td>BT20/MB468</td><td>MP</td></tr>
<tr><td>>hg38_21_+_8433172_8446622_RNA45SN1_45S_50ntFlanks@10794.10825.32|CTGGCATGTTGGAACAATGTAGGTAAGGGAAG</td><td>>hg38_21_+_8433172_8446622_RNA45SN1_45S_50ntFlanks</td><td>10794</td><td>10825</td><td>32</td><td>CTGGCATGTTGGAACAATGTAGGTAAGGGAAG</td><td>337.0</td><td>-3.7</td><td>0.9</td><td>-4.0</td><td>5.7E-05</td><td>8.2E-05</td><td>MB231/MB468</td><td>CYTO</td></tr>
<tr><td>>MI0000272|hsa-mir-182&WithFlank&7|-|129770377|129770498@29.52.24&[MIMAT0000259&hsa-miR-182-5p&offsets|0|0;m-22&7|-|129770449|129770470&offsets|0|+2]|TTTGGCAATGGTAGAACTCACACT</td><td>>MI0000272|hsa-mir-182&WithFlank&7|-|129770377|129770498</td><td>29</td><td>52</td><td>24</td><td>TTTGGCAATGGTAGAACTCACACT</td><td>5709.0</td><td>3.0</td><td>0.7</td><td>4.0</td><td>5.7E-05</td><td>8.2E-05</td><td>MB231/MB468</td><td>CYTO</td></tr>
<tr><td>>hg38_21_+_8433172_8446622_RNA45SN1_45S_50ntFlanks@4297.4332.36|CATTGGAGGGCAAGTCTGGTGCCAGCAGCCGCGGTA</td><td>>hg38_21_+_8433172_8446622_RNA45SN1_45S_50ntFlanks</td><td>4297</td><td>4332</td><td>36</td><td>CATTGGAGGGCAAGTCTGGTGCCAGCAGCCGCGGTA</td><td>474.9</td><td>-5.4</td><td>1.3</td><td>-4.0</td><td>6.3E-05</td><td>8.2E-05</td><td>BT20/MB468</td><td>NUC</td></tr>
<tr><td>>hg38_21_+_8433172_8446622_RNA45SN1_45S_50ntFlanks@10868.10891.24|AAGGGCTGGGTCGGTCGGGCTGGG</td><td>>hg38_21_+_8433172_8446622_RNA45SN1_45S_50ntFlanks</td><td>10868</td><td>10891</td><td>24</td><td>AAGGGCTGGGTCGGTCGGGCTGGG</td><td>161.6</td><td>-3.5</td><td>0.9</td><td>-4.1</td><td>5.1E-05</td><td>8.2E-05</td><td>MB231/MB468</td><td>MP</td></tr>
<tr><td>>hg38_21_+_8433172_8446622_RNA45SN1_45S_50ntFlanks@9304.9336.33|ACCAAGGAGTCTAACACGTGCGCGAGTCGGGGG</td><td>>hg38_21_+_8433172_8446622_RNA45SN1_45S_50ntFlanks</td><td>9304</td><td>9336</td><td>33</td><td>ACCAAGGAGTCTAACACGTGCGCGAGTCGGGGG</td><td>478.0</td><td>-5.3</td><td>1.3</td><td>-4.3</td><td>2.1E-05</td><td>8.2E-05</td><td>BT20/MB468</td><td>MP</td></tr>
<tr><td>>hg38_21_+_8433172_8446622_RNA45SN1_45S_50ntFlanks@6732.6774.43|ACATTGATCATCGACACTTCGAACGCACTTGCGGCCCCGGGTT</td><td>>hg38_21_+_8433172_8446622_RNA45SN1_45S_50ntFlanks</td><td>6732</td><td>6774</td><td>43</td><td>ACATTGATCATCGACACTTCGAACGCACTTGCGGCCCCGGGTT</td><td>721.9</td><td>5.0</td><td>1.2</td><td>4.0</td><td>6.1E-05</td><td>8.2E-05</td><td>MB231/MB468</td><td>MITO</td></tr>
<tr><td>>hg38_21_+_8433172_8446622_RNA45SN1_45S_50ntFlanks@11582.11611.30|ACTGGTGCGGACCAGGGGAATCCGACTGTT</td><td>>hg38_21_+_8433172_8446622_RNA45SN1_45S_50ntFlanks</td><td>11582</td><td>11611</td><td>30</td><td>ACTGGTGCGGACCAGGGGAATCCGACTGTT</td><td>1152.4</td><td>-2.9</td><td>0.7</td><td>-4.0</td><td>6.3E-05</td><td>8.2E-05</td><td>BT20/MB468</td><td>NUC</td></tr>
<tr><td>>MI0000285|hsa-mir-205&WithFlank&1|+|209432127|209432248@40.61.22(+2A)&[MIMAT0000266&hsa-miR-205-5p&offsets|0|0(+2A);m-351&1|+|209432166|209432187&offsets|0|0(+2A)]|TCCTTCATTCCACCGGAGTCTGAA</td><td>>MI0000285|hsa-mir-205&WithFlank&1|+|209432127|209432248</td><td>40</td><td>61</td><td>22(+2A)</td><td>TCCTTCATTCCACCGGAGTCTGAA</td><td>76.0</td><td>9.6</td><td>2.1</td><td>4.5</td><td>7.1E-06</td><td>8.2E-05</td><td>BT20/MB231</td><td>MP</td></tr>
<tr><td>>hg38_21_+_8433172_8446622_RNA45SN1_45S_50ntFlanks@11565.11603.39|CTAGCAGCCGACTTAGAACTGGTGCGGACCAGGGGAATC</td><td>>hg38_21_+_8433172_8446622_RNA45SN1_45S_50ntFlanks</td><td>11565</td><td>11603</td><td>39</td><td>CTAGCAGCCGACTTAGAACTGGTGCGGACCAGGGGAATC</td><td>304.6</td><td>-4.8</td><td>1.2</td><td>-4.0</td><td>6.1E-05</td><td>8.2E-05</td><td>MB231/MB468</td><td>NUC</td></tr>
<tr><td>>hg38_21_+_8433172_8446622_RNA45SN1_45S_50ntFlanks@6728.6770.43|GGACACATTGATCATCGACACTTCGAACGCACTTGCGGCCCCG</td><td>>hg38_21_+_8433172_8446622_RNA45SN1_45S_50ntFlanks</td><td>6728</td><td>6770</td><td>43</td><td>GGACACATTGATCATCGACACTTCGAACGCACTTGCGGCCCCG</td><td>368.2</td><td>-6.9</td><td>1.7</td><td>-4.0</td><td>6.3E-05</td><td>8.2E-05</td><td>BT20/MB468</td><td>NUC</td></tr>
<tr><td>>hg38_21_+_8433172_8446622_RNA45SN1_45S_50ntFlanks@12539.12557.19|ATGATGTGTTGTTGCCATG</td><td>>hg38_21_+_8433172_8446622_RNA45SN1_45S_50ntFlanks</td><td>12539</td><td>12557</td><td>19</td><td>ATGATGTGTTGTTGCCATG</td><td>316.3</td><td>-4.7</td><td>1.1</td><td>-4.1</td><td>4.4E-05</td><td>8.2E-05</td><td>BT20/MB468</td><td>MITO</td></tr>
<tr><td>>hg38_21_+_8433172_8446622_RNA45SN1_45S_50ntFlanks@3854.3889.36|ACTGTGGTAATTCTAGAGCTAATACATGCCGACGGG</td><td>>hg38_21_+_8433172_8446622_RNA45SN1_45S_50ntFlanks</td><td>3854</td><td>3889</td><td>36</td><td>ACTGTGGTAATTCTAGAGCTAATACATGCCGACGGG</td><td>177.3</td><td>-6.8</td><td>1.7</td><td>-4.1</td><td>5.1E-05</td><td>8.2E-05</td><td>MB231/MB468</td><td>MP</td></tr>
<tr><td>>trna2_GlyGCC_21_-_18827107_18827177@1.30.30,trna35_GlyGCC_1_+_161413094_161413164@1.30.30,trna37_GlyGCC_1_+_161420467_161420537@1.30.30,trna39_GlyGCC_1_+_161427898_161427968@1.30.30,trna41_GlyGCC_1_+_161435258_161435328@1.30.30|GCATGGGTGGTTCAGTGGTAGAATTCTCGC</td><td>>trna2_GlyGCC_21_-_18827107_18827177</td><td>1</td><td>30</td><td>30</td><td>GCATGGGTGGTTCAGTGGTAGAATTCTCGC</td><td>1569.3</td><td>-4.2</td><td>1.0</td><td>-4.0</td><td>6.1E-05</td><td>8.2E-05</td><td>MB231/MB468</td><td>MITO</td></tr>
<tr><td>>hg38_21_+_8433172_8446622_RNA45SN1_45S_50ntFlanks@6786.6806.21|TACGCCTGTCTGAGCGTCGCT</td><td>>hg38_21_+_8433172_8446622_RNA45SN1_45S_50ntFlanks</td><td>6786</td><td>6806</td><td>21</td><td>TACGCCTGTCTGAGCGTCGCT</td><td>442.8</td><td>2.4</td><td>0.6</td><td>4.0</td><td>6.3E-05</td><td>8.2E-05</td><td>BT20/MB468</td><td>NUC</td></tr>
<tr><td>>hg38_21_+_8433172_8446622_RNA45SN1_45S_50ntFlanks@8117.8145.29|CGGGACATGTGGCGTACGGAAGACCCGCT</td><td>>hg38_21_+_8433172_8446622_RNA45SN1_45S_50ntFlanks</td><td>8117</td><td>8145</td><td>29</td><td>CGGGACATGTGGCGTACGGAAGACCCGCT</td><td>298.3</td><td>3.4</td><td>0.8</td><td>4.5</td><td>8.3E-06</td><td>8.2E-05</td><td>BT20/MB231</td><td>MITO</td></tr>
<tr><td>>am_MI0003639|hsa-mir-625&WithFlank&14|+|65471096|65471192@58.80.23&[MIMAT0004808&hsa-miR-625-3p&offsets|0|+1;m-137&14|+|65471153|65471174&offsets|0|+1]|GACTATAGAACTTTCCCCCTCAT</td><td>>am_MI0003639|hsa-mir-625&WithFlank&14|+|65471096|65471192</td><td>58</td><td>80</td><td>23</td><td>GACTATAGAACTTTCCCCCTCAT</td><td>415.0</td><td>-4.5</td><td>1.0</td><td>-4.5</td><td>8.3E-06</td><td>8.2E-05</td><td>BT20/MB231</td><td>MITO</td></tr>
<tr><td>>hg38_21_+_8433172_8446622_RNA45SN1_45S_50ntFlanks@5078.5110.33|CGATAACGAACGAGACTCTGGCATGCTAACTAG</td><td>>hg38_21_+_8433172_8446622_RNA45SN1_45S_50ntFlanks</td><td>5078</td><td>5110</td><td>33</td><td>CGATAACGAACGAGACTCTGGCATGCTAACTAG</td><td>331.8</td><td>-3.4</td><td>0.9</td><td>-4.0</td><td>6.1E-05</td><td>8.2E-05</td><td>MB231/MB468</td><td>NUC</td></tr>
<tr><td>>hg38_21_+_8433172_8446622_RNA45SN1_45S_50ntFlanks@8177.8211.35|TTCTGATCGAGGCCCAGCCCGTGGACGGTGTGAGG</td><td>>hg38_21_+_8433172_8446622_RNA45SN1_45S_50ntFlanks</td><td>8177</td><td>8211</td><td>35</td><td>TTCTGATCGAGGCCCAGCCCGTGGACGGTGTGAGG</td><td>293.6</td><td>-2.7</td><td>0.7</td><td>-4.0</td><td>6.1E-05</td><td>8.2E-05</td><td>MB231/MB468</td><td>NUC</td></tr>
<tr><td>>hg38_21_+_8433172_8446622_RNA45SN1_45S_50ntFlanks@9540.9570.31|CTCTGGTGGAGGTCCGTAGCGGTCCTGACGT</td><td>>hg38_21_+_8433172_8446622_RNA45SN1_45S_50ntFlanks</td><td>9540</td><td>9570</td><td>31</td><td>CTCTGGTGGAGGTCCGTAGCGGTCCTGACGT</td><td>406.3</td><td>-5.2</td><td>1.3</td><td>-4.0</td><td>6.3E-05</td><td>8.2E-05</td><td>BT20/MB468</td><td>NUC</td></tr>
<tr><td>>hg38_21_+_8433172_8446622_RNA45SN1_45S_50ntFlanks@12202.12240.39|CTAAGGCGAGCTCAGGGAGGACAGAAACCTCCCGTGGAG</td><td>>hg38_21_+_8433172_8446622_RNA45SN1_45S_50ntFlanks</td><td>12202</td><td>12240</td><td>39</td><td>CTAAGGCGAGCTCAGGGAGGACAGAAACCTCCCGTGGAG</td><td>207.3</td><td>-5.9</td><td>1.4</td><td>-4.1</td><td>4.5E-05</td><td>8.2E-05</td><td>BT20/MB468</td><td>MITO</td></tr>
<tr><td>>hg38_21_+_8433172_8446622_RNA45SN1_45S_50ntFlanks@11580.11612.33|GAACTGGTGCGGACCAGGGGAATCCGACTGTTT</td><td>>hg38_21_+_8433172_8446622_RNA45SN1_45S_50ntFlanks</td><td>11580</td><td>11612</td><td>33</td><td>GAACTGGTGCGGACCAGGGGAATCCGACTGTTT</td><td>96.7</td><td>-6.1</td><td>1.5</td><td>-4.1</td><td>5.1E-05</td><td>8.2E-05</td><td>MB231/MB468</td><td>MP</td></tr>
<tr><td>>hg38_21_+_8433172_8446622_RNA45SN1_45S_50ntFlanks@3854.3880.27|ACTGTGGTAATTCTAGAGCTAATACAT</td><td>>hg38_21_+_8433172_8446622_RNA45SN1_45S_50ntFlanks</td><td>3854</td><td>3880</td><td>27</td><td>ACTGTGGTAATTCTAGAGCTAATACAT</td><td>302.6</td><td>-5.6</td><td>1.4</td><td>-4.0</td><td>5.1E-05</td><td>8.2E-05</td><td>MB231/MB468</td><td>MP</td></tr>
<tr><td>>hg38_21_+_8433172_8446622_RNA45SN1_45S_50ntFlanks@10821.10839.19|GGAAGTCGGCAAGCCGGAT</td><td>>hg38_21_+_8433172_8446622_RNA45SN1_45S_50ntFlanks</td><td>10821</td><td>10839</td><td>19</td><td>GGAAGTCGGCAAGCCGGAT</td><td>201.8</td><td>-2.5</td><td>0.6</td><td>-4.0</td><td>5.7E-05</td><td>8.2E-05</td><td>MB231/MB468</td><td>CYTO</td></tr>
<tr><td>>hg38_21_+_8433172_8446622_RNA45SN1_45S_50ntFlanks@4550.4572.23|GATACCGCAGCTAGGAATAATGG</td><td>>hg38_21_+_8433172_8446622_RNA45SN1_45S_50ntFlanks</td><td>4550</td><td>4572</td><td>23</td><td>GATACCGCAGCTAGGAATAATGG</td><td>228.5</td><td>-5.7</td><td>1.4</td><td>-4.0</td><td>6.1E-05</td><td>8.2E-05</td><td>MB231/MB468</td><td>NUC</td></tr>
<tr><td>>hg38_21_+_8433172_8446622_RNA45SN1_45S_50ntFlanks@12948.12988.41|GACGACCTGCTTCTGGGTCGGGGTTTCGTACGTAGCAGAGC</td><td>>hg38_21_+_8433172_8446622_RNA45SN1_45S_50ntFlanks</td><td>12948</td><td>12988</td><td>41</td><td>GACGACCTGCTTCTGGGTCGGGGTTTCGTACGTAGCAGAGC</td><td>598.2</td><td>-3.8</td><td>0.8</td><td>-4.5</td><td>6.7E-06</td><td>8.3E-05</td><td>BT20/MB231</td><td>CYTO</td></tr>
<tr><td>>hg38_21_+_8433172_8446622_RNA45SN1_45S_50ntFlanks@12007.12039.33|GCCGCCGGTGAAATACCACTACTCTGATCGTTT</td><td>>hg38_21_+_8433172_8446622_RNA45SN1_45S_50ntFlanks</td><td>12007</td><td>12039</td><td>33</td><td>GCCGCCGGTGAAATACCACTACTCTGATCGTTT</td><td>317.6</td><td>-3.4</td><td>0.8</td><td>-4.0</td><td>6.1E-05</td><td>8.3E-05</td><td>MB231/MB468</td><td>MITO</td></tr>
<tr><td>>hg38_21_+_8433172_8446622_RNA45SN1_45S_50ntFlanks@5515.5533.19|CTAGAGGAAGTAAAAGTCG</td><td>>hg38_21_+_8433172_8446622_RNA45SN1_45S_50ntFlanks</td><td>5515</td><td>5533</td><td>19</td><td>CTAGAGGAAGTAAAAGTCG</td><td>188.5</td><td>-5.6</td><td>1.4</td><td>-4.0</td><td>5.3E-05</td><td>8.3E-05</td><td>BT20/MB468</td><td>CYTO</td></tr>
<tr><td>>hg38_21_+_8433172_8446622_RNA45SN1_45S_50ntFlanks@9938.9956.19|AGCAGGACGGTGGCCATGG</td><td>>hg38_21_+_8433172_8446622_RNA45SN1_45S_50ntFlanks</td><td>9938</td><td>9956</td><td>19</td><td>AGCAGGACGGTGGCCATGG</td><td>779.1</td><td>7.3</td><td>1.7</td><td>4.3</td><td>2.1E-05</td><td>8.3E-05</td><td>BT20/MB468</td><td>MP</td></tr>
<tr><td>>hg38_21_+_8433172_8446622_RNA45SN1_45S_50ntFlanks@11562.11602.41|CGCCTAGCAGCCGACTTAGAACTGGTGCGGACCAGGGGAAT</td><td>>hg38_21_+_8433172_8446622_RNA45SN1_45S_50ntFlanks</td><td>11562</td><td>11602</td><td>41</td><td>CGCCTAGCAGCCGACTTAGAACTGGTGCGGACCAGGGGAAT</td><td>88.8</td><td>9.8</td><td>2.2</td><td>4.5</td><td>7.2E-06</td><td>8.3E-05</td><td>BT20/MB231</td><td>MP</td></tr>
<tr><td>>hg38_21_+_8433172_8446622_RNA45SN1_45S_50ntFlanks@3986.4010.25|GGCTTTGGTGACTCTAGATAACCTC</td><td>>hg38_21_+_8433172_8446622_RNA45SN1_45S_50ntFlanks</td><td>3986</td><td>4010</td><td>25</td><td>GGCTTTGGTGACTCTAGATAACCTC</td><td>416.4</td><td>6.4</td><td>1.6</td><td>4.0</td><td>5.3E-05</td><td>8.3E-05</td><td>BT20/MB468</td><td>CYTO</td></tr>
<tr><td>>hg38_21_+_8433172_8446622_RNA45SN1_45S_50ntFlanks@9033.9059.27|CTCGTGGGGGGGCCGGGCCACCCCTCC</td><td>>hg38_21_+_8433172_8446622_RNA45SN1_45S_50ntFlanks</td><td>9033</td><td>9059</td><td>27</td><td>CTCGTGGGGGGGCCGGGCCACCCCTCC</td><td>1748.1</td><td>-7.1</td><td>1.7</td><td>-4.3</td><td>2.1E-05</td><td>8.3E-05</td><td>BT20/MB468</td><td>MP</td></tr>
<tr><td>>hg38_21_+_8433172_8446622_RNA45SN1_45S_50ntFlanks@3854.3873.20|ACTGTGGTAATTCTAGAGCT</td><td>>hg38_21_+_8433172_8446622_RNA45SN1_45S_50ntFlanks</td><td>3854</td><td>3873</td><td>20</td><td>ACTGTGGTAATTCTAGAGCT</td><td>672.0</td><td>-2.9</td><td>0.7</td><td>-4.0</td><td>5.8E-05</td><td>8.3E-05</td><td>MB231/MB468</td><td>CYTO</td></tr>
<tr><td>>hg38_21_+_8433172_8446622_RNA45SN1_45S_50ntFlanks@9913.9949.37|AGAAAAGGTGTTGGTTGATATAGACAGCAGGACGGTG</td><td>>hg38_21_+_8433172_8446622_RNA45SN1_45S_50ntFlanks</td><td>9913</td><td>9949</td><td>37</td><td>AGAAAAGGTGTTGGTTGATATAGACAGCAGGACGGTG</td><td>82.4</td><td>9.7</td><td>2.2</td><td>4.5</td><td>7.2E-06</td><td>8.3E-05</td><td>BT20/MB231</td><td>MP</td></tr>
<tr><td>>MI0000446|hsa-mir-125b-1&WithFlank&11|-|122099751|122099850@61.80.20(+1A)&[MIMAT0004592&hsa-miR-125b-1-3p&offsets|0|-2(+1A);m-485&11|-|122099769|122099790&offsets|0|-2(+1A)]|ACGGGTTAGGCTCTTGGGAGA</td><td>>MI0000446|hsa-mir-125b-1&WithFlank&11|-|122099751|122099850</td><td>61</td><td>80</td><td>20(+1A)</td><td>ACGGGTTAGGCTCTTGGGAGA</td><td>68.9</td><td>-9.5</td><td>2.1</td><td>-4.5</td><td>7.2E-06</td><td>8.3E-05</td><td>BT20/MB231</td><td>MP</td></tr>
<tr><td>>hg38_21_+_8433172_8446622_RNA45SN1_45S_50ntFlanks@12075.12095.21|GAGGGGCTCTCGCTTCTGGCG</td><td>>hg38_21_+_8433172_8446622_RNA45SN1_45S_50ntFlanks</td><td>12075</td><td>12095</td><td>21</td><td>GAGGGGCTCTCGCTTCTGGCG</td><td>633.3</td><td>-7.0</td><td>1.7</td><td>-4.0</td><td>6.3E-05</td><td>8.3E-05</td><td>BT20/MB468</td><td>NUC</td></tr>
<tr><td>>hg38_21_+_8433172_8446622_RNA45SN1_45S_50ntFlanks@11571.11602.32|GCCGACTTAGAACTGGTGCGGACCAGGGGAAT</td><td>>hg38_21_+_8433172_8446622_RNA45SN1_45S_50ntFlanks</td><td>11571</td><td>11602</td><td>32</td><td>GCCGACTTAGAACTGGTGCGGACCAGGGGAAT</td><td>677.0</td><td>-2.0</td><td>0.5</td><td>-4.0</td><td>6.1E-05</td><td>8.3E-05</td><td>MB231/MB468</td><td>NUC</td></tr>
</table>

<table>
<tr><td>>hg38_21_+_8433172_8446622_RNA45SN1_45S_50ntFlanks@11212.11230.19|CGACTCTGGACGCGAGCCG</td><td>>hg38_21_+_8433172_8446622_RNA45SN1_45S_50ntFlanks</td><td>11212</td><td>11230</td><td>19</td><td>CGACTCTGGACGCGAGCCG</td><td>243.5</td><td>-5.5</td><td>1.4</td><td>-4.0</td><td>5.4E-05</td><td>8.3E-05</td><td>BT20/MB468</td><td>CYTO</td></tr>
<tr><td>>MI0033423|hsa-mir-10399&WithFlank&7|-|139044094|139044163@43.63.21(+1U)&[MIMAT0041630&hsa-miR-10399-3p&offsets|-1|-1(+1U);TJU_CMC_MD2.ID03061.3p-miR&offsets|+3|+2(+1U);m-2657&7|-|139044103|139044124&offsets|+3|+2(+1U)]|TCTCTCGGACAAGCTGTAGGTT</td><td>>MI0033423|hsa-mir-10399&WithFlank&7|-|139044094|139044163</td><td>43</td><td>63</td><td>21(+1U)</td><td>TCTCTCGGACAAGCTGTAGGTT</td><td>247.8</td><td>5.9</td><td>1.5</td><td>4.0</td><td>5.2E-05</td><td>8.3E-05</td><td>MB231/MB468</td><td>MP</td></tr>
<tr><td>>MI0000301|hsa-mir-224&WithFlank&X|-|151958572|151958664@14.34.21(+1A)&[MIMAT0000281&hsa-miR-224-5p&offsets|+1|-3(+1A);m-238&X|-|151958630|151958651&offsets|0|-1(+1A)]|CAAGTCACTAGTGGTTCCGTTA</td><td>>MI0000301|hsa-mir-224&WithFlank&X|-|151958572|151958664</td><td>14</td><td>34</td><td>21(+1A)</td><td>CAAGTCACTAGTGGTTCCGTTA</td><td>465.1</td><td>3.8</td><td>0.9</td><td>4.5</td><td>6.8E-06</td><td>8.3E-05</td><td>BT20/MB231</td><td>CYTO</td></tr>
<tr><td>>hg38_MT_+_1621_3279_RNR2_16S_50ntFlanks@1390.1427.38|GATGGTGCAGCCGCTATTAAAGGTTCGTTTGTTCAACG</td><td>>hg38_MT_+_1621_3279_RNR2_16S_50ntFlanks</td><td>1390</td><td>1427</td><td>38</td><td>GATGGTGCAGCCGCTATTAAAGGTTCGTTTGTTCAACG</td><td>81.9</td><td>6.6</td><td>1.6</td><td>4.0</td><td>6.2E-05</td><td>8.3E-05</td><td>MB231/MB468</td><td>MITO</td></tr>
<tr><td>>hg38_21_+_8433172_8446622_RNA45SN1_45S_50ntFlanks@10783.10801.19|GTGAACAGCCTCTGGCATG</td><td>>hg38_21_+_8433172_8446622_RNA45SN1_45S_50ntFlanks</td><td>10783</td><td>10801</td><td>19</td><td>GTGAACAGCCTCTGGCATG</td><td>520.2</td><td>-5.3</td><td>1.2</td><td>-4.3</td><td>2.1E-05</td><td>8.3E-05</td><td>BT20/MB468</td><td>MP</td></tr>
<tr><td>>hg38_21_+_8433172_8446622_RNA45SN1_45S_50ntFlanks@5090.5120.31|AGACTCTGGCATGCTAACTAGTTACGCGACC</td><td>>hg38_21_+_8433172_8446622_RNA45SN1_45S_50ntFlanks</td><td>5090</td><td>5120</td><td>31</td><td>AGACTCTGGCATGCTAACTAGTTACGCGACC</td><td>52.2</td><td>-8.8</td><td>2.1</td><td>-4.3</td><td>2.1E-05</td><td>8.3E-05</td><td>BT20/MB468</td><td>MP</td></tr>
<tr><td>>hg38_21_+_8433172_8446622_RNA45SN1_45S_50ntFlanks@3840.3873.34|CTCCTACTTGGATAACTGTGGTAATTCTAGAGCT</td><td>>hg38_21_+_8433172_8446622_RNA45SN1_45S_50ntFlanks</td><td>3840</td><td>3873</td><td>34</td><td>CTCCTACTTGGATAACTGTGGTAATTCTAGAGCT</td><td>566.5</td><td>4.1</td><td>0.9</td><td>4.5</td><td>8.0E-06</td><td>8.3E-05</td><td>BT20/MB231</td><td>NUC</td></tr>
<tr><td>>hg38_21_+_8433172_8446622_RNA45SN1_45S_50ntFlanks@11580.11606.27|GAACTGGTGCGGACCAGGGGAATCCGA</td><td>>hg38_21_+_8433172_8446622_RNA45SN1_45S_50ntFlanks</td><td>11580</td><td>11606</td><td>27</td><td>GAACTGGTGCGGACCAGGGGAATCCGA</td><td>214.6</td><td>-3.7</td><td>0.9</td><td>-4.0</td><td>5.8E-05</td><td>8.3E-05</td><td>MB231/MB468</td><td>CYTO</td></tr>
<tr><td>>hg38_21_+_8433172_8446622_RNA45SN1_45S_50ntFlanks@10846.10872.27|CTTCGGGATAAGGATTGGCTCTAAGGG</td><td>>hg38_21_+_8433172_8446622_RNA45SN1_45S_50ntFlanks</td><td>10846</td><td>10872</td><td>27</td><td>CTTCGGGATAAGGATTGGCTCTAAGGG</td><td>183.6</td><td>-2.8</td><td>0.7</td><td>-4.0</td><td>5.4E-05</td><td>8.3E-05</td><td>BT20/MB468</td><td>CYTO</td></tr>
<tr><td>>hg38_21_+_8433172_8446622_RNA45SN1_45S_50ntFlanks@11564.11592.29|CCTAGCAGCCGACTTAGAACTGGTGCGGA</td><td>>hg38_21_+_8433172_8446622_RNA45SN1_45S_50ntFlanks</td><td>11564</td><td>11592</td><td>29</td><td>CCTAGCAGCCGACTTAGAACTGGTGCGGA</td><td>339.8</td><td>-2.8</td><td>0.7</td><td>-4.1</td><td>4.5E-05</td><td>8.3E-05</td><td>BT20/MB468</td><td>MITO</td></tr>
<tr><td>>hg38_21_+_8433172_8446622_RNA45SN1_45S_50ntFlanks@8354.8393.40|TTGAAAAGAACTTTGAAGAGAGAGTTCAAGAGGGCGTGAA</td><td>>hg38_21_+_8433172_8446622_RNA45SN1_45S_50ntFlanks</td><td>8354</td><td>8393</td><td>40</td><td>TTGAAAAGAACTTTGAAGAGAGAGTTCAAGAGGGCGTGAA</td><td>32.0</td><td>8.7</td><td>2.1</td><td>4.2</td><td>2.1E-05</td><td>8.4E-05</td><td>BT20/MB468</td><td>MP</td></tr>
<tr><td>>hg38_MT_+_598_1651_RNR1_12S_50ntFlanks@52.93.42|ATAGGTTTGGTCCTAGCCTTTCTATTAGCTCTTAGTAAGATT</td><td>>hg38_MT_+_598_1651_RNR1_12S_50ntFlanks</td><td>52</td><td>93</td><td>42</td><td>ATAGGTTTGGTCCTAGCCTTTCTATTAGCTCTTAGTAAGATT</td><td>829.2</td><td>4.3</td><td>1.0</td><td>4.1</td><td>4.5E-05</td><td>8.4E-05</td><td>BT20/MB468</td><td>MITO</td></tr>
<tr><td>>hg38_1_-_228634819_228635039_RNA5S12_5S_50ntFlanks@133.170.38|ATGGGAGACCGCCTGGGAATACCGGGTGCTGTAGGCTT</td><td>>hg38_1_-_228634819_228635039_RNA5S12_5S_50ntFlanks</td><td>133</td><td>170</td><td>38</td><td>ATGGGAGACCGCCTGGGAATACCGGGTGCTGTAGGCTT</td><td>25906.9</td><td>4.3</td><td>1.1</td><td>4.1</td><td>4.5E-05</td><td>8.4E-05</td><td>BT20/MB468</td><td>MITO</td></tr>
<tr><td>>hg38_21_+_8433172_8446622_RNA45SN1_45S_50ntFlanks@5494.5516.23|AAGACGGTCGAACTTGACTATCT</td><td>>hg38_21_+_8433172_8446622_RNA45SN1_45S_50ntFlanks</td><td>5494</td><td>5516</td><td>23</td><td>AAGACGGTCGAACTTGACTATCT</td><td>2306.7</td><td>3.5</td><td>0.8</td><td>4.5</td><td>8.4E-06</td><td>8.4E-05</td><td>BT20/MB231</td><td>MITO</td></tr>
<tr><td>>hg38_21_+_8433172_8446622_RNA45SN1_45S_50ntFlanks@12289.12317.29|CCGTGAAAGCGGGGCCTCACGATCCTTCT</td><td>>hg38_21_+_8433172_8446622_RNA45SN1_45S_50ntFlanks</td><td>12289</td><td>12317</td><td>29</td><td>CCGTGAAAGCGGGGCCTCACGATCCTTCT</td><td>618.2</td><td>-5.4</td><td>1.3</td><td>-4.1</td><td>4.5E-05</td><td>8.4E-05</td><td>BT20/MB468</td><td>MITO</td></tr>
<tr><td>>hg38_21_+_8433172_8446622_RNA45SN1_45S_50ntFlanks@11569.11594.26|CAGCCGACTTAGAACTGGTGCGGACC</td><td>>hg38_21_+_8433172_8446622_RNA45SN1_45S_50ntFlanks</td><td>11569</td><td>11594</td><td>26</td><td>CAGCCGACTTAGAACTGGTGCGGACC</td><td>208.7</td><td>-2.3</td><td>0.6</td><td>-4.0</td><td>6.2E-05</td><td>8.4E-05</td><td>MB231/MB468</td><td>NUC</td></tr>
<tr><td>>hg38_MT_+_598_1651_RNR1_12S_50ntFlanks@817.840.24|CGAAGGTGGATTTAGCAGTAAACT</td><td>>hg38_MT_+_598_1651_RNR1_12S_50ntFlanks</td><td>817</td><td>840</td><td>24</td><td>CGAAGGTGGATTTAGCAGTAAACT</td><td>105.9</td><td>7.5</td><td>1.8</td><td>4.2</td><td>2.1E-05</td><td>8.4E-05</td><td>BT20/MB468</td><td>MP</td></tr>
<tr><td>>hg38_21_+_8433172_8446622_RNA45SN1_45S_50ntFlanks@11571.11592.22|GCCGACTTAGAACTGGTGCGGA</td><td>>hg38_21_+_8433172_8446622_RNA45SN1_45S_50ntFlanks</td><td>11571</td><td>11592</td><td>22</td><td>GCCGACTTAGAACTGGTGCGGA</td><td>712.4</td><td>-3.5</td><td>0.8</td><td>-4.2</td><td>2.1E-05</td><td>8.4E-05</td><td>BT20/MB468</td><td>MP</td></tr>
<tr><td>>hg38_MT_+_1621_3279_RNR2_16S_50ntFlanks@341.377.37|AAATAGTGGGAAGATTTATAGGTAGAGGCGACAAACC</td><td>>hg38_MT_+_1621_3279_RNR2_16S_50ntFlanks</td><td>341</td><td>377</td><td>37</td><td>AAATAGTGGGAAGATTTATAGGTAGAGGCGACAAACC</td><td>25.8</td><td>7.9</td><td>2.0</td><td>4.0</td><td>5.2E-05</td><td>8.4E-05</td><td>MB231/MB468</td><td>MP</td></tr>
<tr><td>>hg38_21_+_8433172_8446622_RNA45SN1_45S_50ntFlanks@11771.11791.21|ATTAGTGACGCGCATGAATGG</td><td>>hg38_21_+_8433172_8446622_RNA45SN1_45S_50ntFlanks</td><td>11771</td><td>11791</td><td>21</td><td>ATTAGTGACGCGCATGAATGG</td><td>760.9</td><td>-7.1</td><td>1.8</td><td>-4.0</td><td>6.4E-05</td><td>8.4E-05</td><td>BT20/MB468</td><td>NUC</td></tr>
<tr><td>>hg38_21_+_8433172_8446622_RNA45SN1_45S_50ntFlanks@6646.6679.34|TCGTACGACTCTTAGCGGTGGATCACTCGGCTCG</td><td>>hg38_21_+_8433172_8446622_RNA45SN1_45S_50ntFlanks</td><td>6646</td><td>6679</td><td>34</td><td>TCGTACGACTCTTAGCGGTGGATCACTCGGCTCG</td><td>3339.2</td><td>-4.1</td><td>1.0</td><td>-4.0</td><td>5.2E-05</td><td>8.4E-05</td><td>MB231/MB468</td><td>MP</td></tr>
<tr><td>>hg38_21_+_8433172_8446622_RNA45SN1_45S_50ntFlanks@9788.9819.32|CGCTGGCGTGGAGCCGGGCGTGGAATGCGAGT</td><td>>hg38_21_+_8433172_8446622_RNA45SN1_45S_50ntFlanks</td><td>9788</td><td>9819</td><td>32</td><td>CGCTGGCGTGGAGCCGGGCGTGGAATGCGAGT</td><td>75.8</td><td>-5.8</td><td>1.4</td><td>-4.0</td><td>5.2E-05</td><td>8.4E-05</td><td>MB231/MB468</td><td>MP</td></tr>
<tr><td>>MI0000748|hsa-mir-130b&WithFlank&22|+|21653298|21653391@57.75.19(+1U)&[MIMAT0000691&hsa-miR-130b-3p&offsets|0|-3(+1U);m-205&22|+|21653354|21653375&offsets|0|-3(+1U)]|CAGTGCAATGATGAAAGGGT</td><td>>MI0000748|hsa-mir-130b&WithFlank&22|+|21653298|21653391</td><td>57</td><td>75</td><td>19(+1U)</td><td>CAGTGCAATGATGAAAGGGT</td><td>36.8</td><td>8.4</td><td>2.1</td><td>4.0</td><td>5.2E-05</td><td>8.4E-05</td><td>MB231/MB468</td><td>MP</td></tr>
<tr><td>>MI0000736|hsa-mir-30c-1&WithFlank&1|+|40757278|40757378@23.45.23;MI0000254|hsa-mir-30c-2&WithFlank&6|-|71376954|71377037@13.35.23&[MIMAT0000244&hsa-miR-30c-5p&offsets|0|0;m-57&1|+|40757300|40757323&offsets|0|-1];[MIMAT0000244_1&hsa-miR-30c-5p&offsets|0|0;m-56&6|-|71377002|71377025&offsets|0|-1]|TGTAAACATCCTACACTCTCAGC</td><td>>MI0000736|hsa-mir-30c-1&WithFlank&1|+|40757278|40757378</td><td>23</td><td>45</td><td>23</td><td>TGTAAACATCCTACACTCTCAGC</td><td>45847.1</td><td>-3.5</td><td>0.8</td><td>-4.5</td><td>8.5E-06</td><td>8.4E-05</td><td>BT20/MB231</td><td>MITO</td></tr>
<tr><td>>hg38_21_+_8433172_8446622_RNA45SN1_45S_50ntFlanks@12158.12180.23|GTTTGACTGGGGCGGTACACCTG</td><td>>hg38_21_+_8433172_8446622_RNA45SN1_45S_50ntFlanks</td><td>12158</td><td>12180</td><td>23</td><td>GTTTGACTGGGGCGGTACACCTG</td><td>415.7</td><td>-6.4</td><td>1.5</td><td>-4.2</td><td>2.2E-05</td><td>8.4E-05</td><td>BT20/MB468</td><td>MP</td></tr>
<tr><td>>hg38_21_+_8433172_8446622_RNA45SN1_45S_50ntFlanks@5067.5097.31|CTGGTTAATTCCGATAACGAACGAGACTCTG</td><td>>hg38_21_+_8433172_8446622_RNA45SN1_45S_50ntFlanks</td><td>5067</td><td>5097</td><td>31</td><td>CTGGTTAATTCCGATAACGAACGAGACTCTG</td><td>46.3</td><td>-8.6</td><td>2.0</td><td>-4.2</td><td>2.2E-05</td><td>8.4E-05</td><td>BT20/MB468</td><td>MP</td></tr>
<tr><td>>hg38_21_+_8433172_8446622_RNA45SN1_45S_50ntFlanks@10410.10429.20|TCAGTCGGTCCTGAGAGATG</td><td>>hg38_21_+_8433172_8446622_RNA45SN1_45S_50ntFlanks</td><td>10410</td><td>10429</td><td>20</td><td>TCAGTCGGTCCTGAGAGATG</td><td>1828.0</td><td>-2.3</td><td>0.6</td><td>-4.1</td><td>4.6E-05</td><td>8.4E-05</td><td>BT20/MB468</td><td>MITO</td></tr>
<tr><td>>trna27_GlyCCC_2_-_70476123_70476193@1.32.32,trna34_GlyCCC_16_-_686736_686806@1.32.32|GCGCCGCTGGTGTAGTGGTATCATGCAAGATT</td><td>>trna27_GlyCCC_2_-_70476123_70476193</td><td>1</td><td>32</td><td>32</td><td>GCGCCGCTGGTGTAGTGGTATCATGCAAGATT</td><td>98.6</td><td>6.8</td><td>1.7</td><td>4.0</td><td>5.2E-05</td><td>8.4E-05</td><td>MB231/MB468</td><td>MP</td></tr>
<tr><td>>hg38_21_+_8433172_8446622_RNA45SN1_45S_50ntFlanks@6645.6667.23|CTCGTACGACTCTTAGCGGTGGA</td><td>>hg38_21_+_8433172_8446622_RNA45SN1_45S_50ntFlanks</td><td>6645</td><td>6667</td><td>23</td><td>CTCGTACGACTCTTAGCGGTGGA</td><td>2728.2</td><td>2.9</td><td>0.7</td><td>4.0</td><td>6.2E-05</td><td>8.4E-05</td><td>MB231/MB468</td><td>MITO</td></tr>
<tr><td>>hg38_21_+_8433172_8446622_RNA45SN1_45S_50ntFlanks@11575.11613.39|ACTTAGAACTGGTGCGGACCAGGGGAATCCGACTGTTTA</td><td>>hg38_21_+_8433172_8446622_RNA45SN1_45S_50ntFlanks</td><td>11575</td><td>11613</td><td>39</td><td>ACTTAGAACTGGTGCGGACCAGGGGAATCCGACTGTTTA</td><td>1068.4</td><td>-3.2</td><td>0.8</td><td>-4.0</td><td>6.5E-05</td><td>8.4E-05</td><td>BT20/MB468</td><td>NUC</td></tr>
<tr><td>>hg38_21_+_8433172_8446622_RNA45SN1_45S_50ntFlanks@10401.10430.30|GAACATGGGTCAGTCGGTCCTGAGAGATGG</td><td>>hg38_21_+_8433172_8446622_RNA45SN1_45S_50ntFlanks</td><td>10401</td><td>10430</td><td>30</td><td>GAACATGGGTCAGTCGGTCCTGAGAGATGG</td><td>778.0</td><td>-7.2</td><td>1.8</td><td>-4.0</td><td>6.5E-05</td><td>8.4E-05</td><td>BT20/MB468</td><td>NUC</td></tr>
<tr><td>>hg38_21_+_8433172_8446622_RNA45SN1_45S_50ntFlanks@8528.8551.24|CTTCCCCCGCCGCCCCTCCTCCTC</td><td>>hg38_21_+_8433172_8446622_RNA45SN1_45S_50ntFlanks</td><td>8528</td><td>8551</td><td>24</td><td>CTTCCCCCGCCGCCCCTCCTCCTC</td><td>429.0</td><td>-7.0</td><td>1.7</td><td>-4.0</td><td>6.5E-05</td><td>8.4E-05</td><td>BT20/MB468</td><td>NUC</td></tr>
<tr><td>>hg38_21_+_8433172_8446622_RNA45SN1_45S_50ntFlanks@10397.10424.28|AGTTGAACATGGGTCAGTCGGTCCTGAG</td><td>>hg38_21_+_8433172_8446622_RNA45SN1_45S_50ntFlanks</td><td>10397</td><td>10424</td><td>28</td><td>AGTTGAACATGGGTCAGTCGGTCCTGAG</td><td>3234.8</td><td>3.9</td><td>0.9</td><td>4.5</td><td>8.5E-06</td><td>8.4E-05</td><td>BT20/MB231</td><td>MITO</td></tr>
<tr><td>>hg38_21_+_8433172_8446622_RNA45SN1_45S_50ntFlanks@12990.13014.25|GCTCCCTCGCTGCGATCTATTGAAA</td><td>>hg38_21_+_8433172_8446622_RNA45SN1_45S_50ntFlanks</td><td>12990</td><td>13014</td><td>25</td><td>GCTCCCTCGCTGCGATCTATTGAAA</td><td>579.4</td><td>-3.4</td><td>0.8</td><td>-4.0</td><td>6.3E-05</td><td>8.4E-05</td><td>MB231/MB468</td><td>MITO</td></tr>
<tr><td>>MI0000078|hsa-mir-22&WithFlank&17|-|1713897|1713993@59.77.19(+1A)&[MIMAT0000077&hsa-miR-22-3p&offsets|0|-3(+1A);m-2&17|-|1713914|1713935&offsets|0|-3(+1A)]|AAGCTGCCAGTTGAAGAACA</td><td>>MI0000078|hsa-mir-22&WithFlank&17|-|1713897|1713993</td><td>59</td><td>77</td><td>19(+1A)</td><td>AAGCTGCCAGTTGAAGAACA</td><td>31.1</td><td>8.2</td><td>2.0</td><td>4.0</td><td>5.2E-05</td><td>8.4E-05</td><td>MB231/MB468</td><td>MP</td></tr>
<tr><td>>MI0000087|hsa-mir-29a&WithFlank&7|-|130876741|130876816@48.65.18&[MIMAT0000086&hsa-miR-29a-3p&offsets|0|-4;m-26&7|-|130876749|130876769&offsets|0|-3]|TAGCACCATCTGAAATCG</td><td>>MI0000087|hsa-mir-29a&WithFlank&7|-|130876741|130876816</td><td>48</td><td>65</td><td>18</td><td>TAGCACCATCTGAAATCG</td><td>36.4</td><td>8.4</td><td>2.1</td><td>4.0</td><td>5.2E-05</td><td>8.4E-05</td><td>MB231/MB468</td><td>MP</td></tr>
<tr><td>>hg38_21_+_8433172_8446622_RNA45SN1_45S_50ntFlanks@3787.3810.24|AAACTGCGAATGGCTCATTAAATC</td><td>>hg38_21_+_8433172_8446622_RNA45SN1_45S_50ntFlanks</td><td>3787</td><td>3810</td><td>24</td><td>AAACTGCGAATGGCTCATTAAATC</td><td>548.7</td><td>-3.2</td><td>0.8</td><td>-4.1</td><td>4.6E-05</td><td>8.4E-05</td><td>BT20/MB468</td><td>MITO</td></tr>
<tr><td>>hg38_21_+_8433172_8446622_RNA45SN1_45S_50ntFlanks@3773.3792.20|CGGCCGGTACAGTGAAACTG</td><td>>hg38_21_+_8433172_8446622_RNA45SN1_45S_50ntFlanks</td><td>3773</td><td>3792</td><td>20</td><td>CGGCCGGTACAGTGAAACTG</td><td>185.8</td><td>-3.6</td><td>0.9</td><td>-4.0</td><td>5.5E-05</td><td>8.5E-05</td><td>BT20/MB468</td><td>CYTO</td></tr>
<tr><td>>hg38_21_+_8433172_8446622_RNA45SN1_45S_50ntFlanks@3842.3861.20|CCTACTTGGATAACTGTGGT</td><td>>hg38_21_+_8433172_8446622_RNA45SN1_45S_50ntFlanks</td><td>3842</td><td>3861</td><td>20</td><td>CCTACTTGGATAACTGTGGT</td><td>202.2</td><td>-4.4</td><td>1.1</td><td>-4.0</td><td>5.3E-05</td><td>8.5E-05</td><td>MB231/MB468</td><td>MP</td></tr>
<tr><td>>hg38_21_+_8433172_8446622_RNA45SN1_45S_50ntFlanks@11840.11864.25|CCAAGGGAACGGGCTTGGCGGAATC</td><td>>hg38_21_+_8433172_8446622_RNA45SN1_45S_50ntFlanks</td><td>11840</td><td>11864</td><td>25</td><td>CCAAGGGAACGGGCTTGGCGGAATC</td><td>95.1</td><td>9.9</td><td>2.2</td><td>4.5</td><td>7.4E-06</td><td>8.5E-05</td><td>BT20/MB231</td><td>MP</td></tr>
<tr><td>>MI0000542|hsa-mir-320a&WithFlank&8|-|22244960|22245043@50.68.19(+1U)&[MIMAT0000510&hsa-miR-320a-3p&offsets|+2|-1(+1U);m-60&8|-|22244975|22244996&offsets|+2|-1(+1U)]|AAGCTGGGTTGAGAGGGCGT</td><td>>MI0000542|hsa-mir-320a&WithFlank&8|-|22244960|22245043</td><td>50</td><td>68</td><td>19(+1U)</td><td>AAGCTGGGTTGAGAGGGCGT</td><td>216.4</td><td>7.9</td><td>1.8</td><td>4.2</td><td>2.2E-05</td><td>8.5E-05</td><td>BT20/MB468</td><td>MP</td></tr>
<tr><td>>hg38_21_+_8433172_8446622_RNA45SN1_45S_50ntFlanks@10351.10392.42|CGAGAACTTTGAAGGCCGAAGTGGAGAAGGGTTCCATGTGAA</td><td>>hg38_21_+_8433172_8446622_RNA45SN1_45S_50ntFlanks</td><td>10351</td><td>10392</td><td>42</td><td>CGAGAACTTTGAAGGCCGAAGTGGAGAAGGGTTCCATGTGAA</td><td>256.7</td><td>-5.0</td><td>1.2</td><td>-4.0</td><td>6.3E-05</td><td>8.5E-05</td><td>MB231/MB468</td><td>MITO</td></tr>
<tr><td>>hg38_21_+_8433172_8446622_RNA45SN1_45S_50ntFlanks@11205.11234.30|GCGGCGGCGACTCTGGACGCGAGCCGGGCC</td><td>>hg38_21_+_8433172_8446622_RNA45SN1_45S_50ntFlanks</td><td>11205</td><td>11234</td><td>30</td><td>GCGGCGGCGACTCTGGACGCGAGCCGGGCC</td><td>239.4</td><td>-3.2</td><td>0.8</td><td>-4.0</td><td>5.5E-05</td><td>8.5E-05</td><td>BT20/MB468</td><td>CYTO</td></tr>
<tr><td>>hg38_21_+_8433172_8446622_RNA45SN1_45S_50ntFlanks@10392.10418.27|ACAGCAGTTGAACATGGGTCAGTCGGT</td><td>>hg38_21_+_8433172_8446622_RNA45SN1_45S_50ntFlanks</td><td>10392</td><td>10418</td><td>27</td><td>ACAGCAGTTGAACATGGGTCAGTCGGT</td><td>121.4</td><td>-6.1</td><td>1.5</td><td>-4.0</td><td>5.3E-05</td><td>8.5E-05</td><td>MB231/MB468</td><td>MP</td></tr>
<tr><td>>hg38_21_+_8433172_8446622_RNA45SN1_45S_50ntFlanks@12488.12520.33|AACGTGAGCTGGGTTTAGACCGTCGTGAGACAG</td><td>>hg38_21_+_8433172_8446622_RNA45SN1_45S_50ntFlanks</td><td>12488</td><td>12520</td><td>33</td><td>AACGTGAGCTGGGTTTAGACCGTCGTGAGACAG</td><td>197.1</td><td>-6.2</td><td>1.5</td><td>-4.2</td><td>2.2E-05</td><td>8.5E-05</td><td>BT20/MB468</td><td>MP</td></tr>
<tr><td>>MI0000299|hsa-mir-222&WithFlank&X|-|45747009|45747130@75.93.19(+1U)&[MIMAT0000279&hsa-miR-222-3p&offsets|0|-2(+1U);m-140&X|-|45747033|45747056&offsets|0|-5(+1U)]|AGCTACATCTGGCTACTGGT</td><td>>MI0000299|hsa-mir-222&WithFlank&X|-|45747009|45747130</td><td>75</td><td>93</td><td>19(+1U)</td><td>AGCTACATCTGGCTACTGGT</td><td>29.5</td><td>8.1</td><td>2.0</td><td>4.0</td><td>5.3E-05</td><td>8.5E-05</td><td>MB231/MB468</td><td>MP</td></tr>
</table>

| | | | | | | | | | | | | | |
|---|---|---|---|---|---|---|---|---|---|---|---|---|---|
| >am_trna111_HisGTG_1_-_147774845_147774916@-1G.29.30,trna118_HisGTG_1_-_145396881_145396952@-1G.29.30,trna16_HisGTG_1_+_146544773_146544844@-1G.29.30,trna1_HisGTG_15_+_45493349_45493420@-1G.29.30,trna21_HisGTG_1_+_147753471_147753542@-1G.29.30,trna33_HisGTG_6_+_27125906_27125977@-1G.29.30,trna7_HisGTG_9_-_14433938_14434009@-1G.29.30,trna8_HisGTG_15_-_45492611_45492682@-1G.29.30,trna9_HisGTG_15_-_45490804_45490875@-1G.29.30\|GGCCGTGATCGTATAGTGGTTAGTACTCTG | >am_trna111_HisGTG_1_-_147774845_147774916 | -1G | 29 | 30 | GGCCGTGATCGTATAGTGGTTAGTACTCTG | 283.2 | -4.4 | 1.0 | -4.4 | 8.6E-06 | 8.5E-05 | BT20/MB231 | MITO |
| >hg38_21_+_8433172_8446622_RNA45SN1_45S_50ntFlanks@12324.12346.23\|TTGGGTTTTAAGCAGGAGGTGTC | >hg38_21_+_8433172_8446622_RNA45SN1_45S_50ntFlanks | 12324 | 12346 | 23 | TTGGGTTTTAAGCAGGAGGTGTC | 2219.5 | -6.5 | 1.6 | -4.0 | 5.5E-05 | 8.5E-05 | BT20/MB468 | CYTO |
| >hg38_21_+_8433172_8446622_RNA45SN1_45S_50ntFlanks@12539.12567.29\|ATGATGTGTTGTTGCCATGGTAATCCTGC | >hg38_21_+_8433172_8446622_RNA45SN1_45S_50ntFlanks | 12539 | 12567 | 29 | ATGATGTGTTGTTGCCATGGTAATCCTGC | 539.3 | -5.8 | 1.4 | -4.0 | 5.5E-05 | 8.5E-05 | BT20/MB468 | CYTO |
| >hg38_MT_+_1621_3279_RNR2_16S_50ntFlanks@930.958.29\|AGTGACACATGTTTAACGGCCGCGGTACC | >hg38_MT_+_1621_3279_RNR2_16S_50ntFlanks | 930 | 958 | 29 | AGTGACACATGTTTAACGGCCGCGGTACC | 155.7 | 5.9 | 1.5 | 4.0 | 6.3E-05 | 8.5E-05 | MB231/MB468 | MITO |
| >hg38_21_+_8433172_8446622_RNA45SN1_45S_50ntFlanks@6651.6682.32\|CGACTCTTAGCGGTGGATCACTCGGCTCGTGC | >hg38_21_+_8433172_8446622_RNA45SN1_45S_50ntFlanks | 6651 | 6682 | 32 | CGACTCTTAGCGGTGGATCACTCGGCTCGTGC | 6451.7 | 4.4 | 1.1 | 4.0 | 6.3E-05 | 8.5E-05 | MB231/MB468 | NUC |
| >hg38_21_+_8433172_8446622_RNA45SN1_45S_50ntFlanks@11208.11239.32\|GCGGCGACTCTGGACGCGAGCCGGGCCCTTCC | >hg38_21_+_8433172_8446622_RNA45SN1_45S_50ntFlanks | 11208 | 11239 | 32 | GCGGCGACTCTGGACGCGAGCCGGGCCCTTCC | 308.7 | -4.6 | 1.1 | -4.0 | 6.3E-05 | 8.5E-05 | MB231/MB468 | NUC |
| >hg38_MT_+_598_1651_RNR1_12S_50ntFlanks@853.872.20\|CTTAGTTGAACAGGGCCCTG | >hg38_MT_+_598_1651_RNR1_12S_50ntFlanks | 853 | 872 | 20 | CTTAGTTGAACAGGGCCCTG | 130.6 | 7.9 | 1.8 | 4.2 | 2.2E-05 | 8.5E-05 | BT20/MB468 | MP |
| >hg38_21_+_8433172_8446622_RNA45SN1_45S_50ntFlanks@9536.9571.36\|GAAACTCTGGTGGAGGTCCGTAGCGGTCCTGACGTG | >hg38_21_+_8433172_8446622_RNA45SN1_45S_50ntFlanks | 9536 | 9571 | 36 | GAAACTCTGGTGGAGGTCCGTAGCGGTCCTGACGTG | 2592.7 | -7.7 | 1.9 | -4.0 | 6.5E-05 | 8.5E-05 | BT20/MB468 | NUC |
| >hg38_21_+_8433172_8446622_RNA45SN1_45S_50ntFlanks@9567.9603.37\|ACGTGCAAATCGGTCGTCCGACCTGGGTATAGGGGCG | >hg38_21_+_8433172_8446622_RNA45SN1_45S_50ntFlanks | 9567 | 9603 | 37 | ACGTGCAAATCGGTCGTCCGACCTGGGTATAGGGGCG | 280.2 | -2.9 | 0.7 | -4.0 | 5.5E-05 | 8.5E-05 | BT20/MB468 | CYTO |
| >hg38_21_+_8433172_8446622_RNA45SN1_45S_50ntFlanks@12073.12092.20\|CCGAGGGGCTCTCGCTTCTG | >hg38_21_+_8433172_8446622_RNA45SN1_45S_50ntFlanks | 12073 | 12092 | 20 | CCGAGGGGCTCTCGCTTCTG | 156.5 | -5.3 | 1.3 | -4.1 | 4.6E-05 | 8.5E-05 | BT20/MB468 | MITO |
| >MI0000465\|hsa-mir-191&WithFlank&3\|-\|49020612\|49020715@22.43.22(+1A)&[MIMAT0000440&hsa-miR-191-5p&offsets\|0\|-1(+1A);m-48&3\|-\|49020672\|49020694&offsets\|0\|-1(+1A)]\|CAACGGAATCCCAAAAGCAGCTA | >MI0000465\|hsa-mir-191&WithFlank&3\|-\|49020612\|49020715 | 22 | 43 | 22(+1A) | CAACGGAATCCCAAAAGCAGCTA | 1950.3 | 3.6 | 0.9 | 4.0 | 5.3E-05 | 8.5E-05 | MB231/MB468 | MP |
| >hg38_21_+_8433172_8446622_RNA45SN1_45S_50ntFlanks@3993.4022.30\|GTGACTCTAGATAACCTCGGGCCGATCGCA | >hg38_21_+_8433172_8446622_RNA45SN1_45S_50ntFlanks | 3993 | 4022 | 30 | GTGACTCTAGATAACCTCGGGCCGATCGCA | 800.5 | -4.0 | 1.0 | -4.0 | 6.3E-05 | 8.5E-05 | MB231/MB468 | MITO |
| >MI0000077\|hsa-mir-21&WithFlank&17\|+\|59841260\|59841343@14.31.18&[MIMAT0000076&hsa-miR-21-5p&offsets\|0\|-4;m-1&17\|+\|59841273\|59841294&offsets\|0\|-4]\|TAGCTTATCAGACTGATG | >MI0000077\|hsa-mir-21&WithFlank&17\|+\|59841260\|59841343 | 14 | 31 | 18 | TAGCTTATCAGACTGATG | 1086.3 | 3.6 | 0.8 | 4.5 | 7.0E-06 | 8.5E-05 | BT20/MB231 | CYTO |
| >MI0000804\|hsa-mir-328&WithFlank&16\|-\|67202315\|67202401@54.74.21(+1C)&[MIMAT0000752&hsa-miR-328-3p&offsets\|0\|-1(+1C);m-160&16\|-\|67202327\|67202348&offsets\|0\|-1(+1C)]\|CTGGCCCTCTCTGCCCTTCCGC | >MI0000804\|hsa-mir-328&WithFlank&16\|-\|67202315\|67202401 | 54 | 74 | 21(+1C) | CTGGCCCTCTCTGCCCTTCCGC | 212.7 | 3.7 | 0.8 | 4.5 | 7.0E-06 | 8.5E-05 | BT20/MB231 | CYTO |
| >hg38_21_+_8433172_8446622_RNA45SN1_45S_50ntFlanks@8019.8042.24\|TTAGTCAGCGGAGGAGAAGAAACT | >hg38_21_+_8433172_8446622_RNA45SN1_45S_50ntFlanks | 8019 | 8042 | 24 | TTAGTCAGCGGAGGAGAAGAAACT | 227.9 | -4.2 | 0.9 | -4.5 | 7.0E-06 | 8.5E-05 | BT20/MB231 | CYTO |
| >hg38_1_-_228634819_228635039_RNA5S12_5S_50ntFlanks@104.121.18\|AAGCAGGGTCGGGCCTGG | >hg38_1_-_228634819_228635039_RNA5S12_5S_50ntFlanks | 104 | 121 | 18 | AAGCAGGGTCGGGCCTGG | 41.6 | -8.5 | 2.0 | -4.2 | 2.2E-05 | 8.5E-05 | BT20/MB468 | MP |
| >hg38_21_+_8433172_8446622_RNA45SN1_45S_50ntFlanks@10062.10079.18\|AGTCGAGAGTGGACGGGA | >hg38_21_+_8433172_8446622_RNA45SN1_45S_50ntFlanks | 10062 | 10079 | 18 | AGTCGAGAGTGGACGGGA | 60.3 | -9.8 | 2.4 | -4.0 | 5.3E-05 | 8.5E-05 | MB231/MB468 | MP |
| >hg38_21_+_8433172_8446622_RNA45SN1_45S_50ntFlanks@11208.11254.47\|GCGGCGACTCTGGACGCGAGCCGGGCCCTTCCCGTGGATCGCCCCAG | >hg38_21_+_8433172_8446622_RNA45SN1_45S_50ntFlanks | 11208 | 11254 | 47 | GCGGCGACTCTGGACGCGAGCCGGGCCCTTCCCGTGGATCGCCCCAG | 10380.5 | -4.9 | 1.2 | -4.0 | 6.0E-05 | 8.5E-05 | MB231/MB468 | CYTO |
| >hg38_21_+_8433172_8446622_RNA45SN1_45S_50ntFlanks@4726.4748.23\|TAATCAAGAACGAAAGTCGGAGG | >hg38_21_+_8433172_8446622_RNA45SN1_45S_50ntFlanks | 4726 | 4748 | 23 | TAATCAAGAACGAAAGTCGGAGG | 134.2 | -4.1 | 1.0 | -4.0 | 6.3E-05 | 8.5E-05 | MB231/MB468 | NUC |
| >hg38_21_+_8433172_8446622_RNA45SN1_45S_50ntFlanks@6649.6669.21\|TACGACTCTTAGCGGTGGATC | >hg38_21_+_8433172_8446622_RNA45SN1_45S_50ntFlanks | 6649 | 6669 | 21 | TACGACTCTTAGCGGTGGATC | 412.0 | -3.5 | 0.9 | -4.0 | 5.3E-05 | 8.5E-05 | MB231/MB468 | MP |
| >hg38_21_+_8433172_8446622_RNA45SN1_45S_50ntFlanks@10798.10825.28\|CATGTTGGAACAATGTAGGTAAGGGAAG | >hg38_21_+_8433172_8446622_RNA45SN1_45S_50ntFlanks | 10798 | 10825 | 28 | CATGTTGGAACAATGTAGGTAAGGGAAG | 527.5 | -6.9 | 1.7 | -4.0 | 6.6E-05 | 8.5E-05 | BT20/MB468 | NUC |
| >hg38_21_+_8433172_8446622_RNA45SN1_45S_50ntFlanks@8337.8378.42\|AGTACCGTAAGGGAAAGTTGAAAAGAACTTTGAAGAGAGAGT | >hg38_21_+_8433172_8446622_RNA45SN1_45S_50ntFlanks | 8337 | 8378 | 42 | AGTACCGTAAGGGAAAGTTGAAAAGAACTTTGAAGAGAGAGT | 73.4 | 9.5 | 2.1 | 4.5 | 7.4E-06 | 8.5E-05 | BT20/MB231 | MP |
| >am_trnaMT_ValTAC_MT_+_1602_1670@44.67.24\|ATTTCAACTTAACTTGACCGCTCT | >am_trnaMT_ValTAC_MT_+_1602_1670 | 44 | 67 | 24 | ATTTCAACTTAACTTGACCGCTCT | 598.2 | 5.6 | 1.4 | 4.0 | 5.5E-05 | 8.6E-05 | BT20/MB468 | CYTO |
| >hg38_21_+_8433172_8446622_RNA45SN1_45S_50ntFlanks@6733.6756.24\|CATTGATCATCGACACTTCGAACG | >hg38_21_+_8433172_8446622_RNA45SN1_45S_50ntFlanks | 6733 | 6756 | 24 | CATTGATCATCGACACTTCGAACG | 648.4 | -2.6 | 0.6 | -4.0 | 6.3E-05 | 8.6E-05 | MB231/MB468 | NUC |
| >hg38_21_+_8433172_8446622_RNA45SN1_45S_50ntFlanks@12251.12269.19\|AAAGCTCGCTTGATCTTGA | >hg38_21_+_8433172_8446622_RNA45SN1_45S_50ntFlanks | 12251 | 12269 | 19 | AAAGCTCGCTTGATCTTGA | 510.4 | -6.6 | 1.6 | -4.0 | 6.4E-05 | 8.6E-05 | MB231/MB468 | MITO |
| >hg38_21_+_8433172_8446622_RNA45SN1_45S_50ntFlanks@3813.3852.40\|TTATGGTTCCTTTGGTCGCTCGCTCCTCTCCTACTTGGAT | >hg38_21_+_8433172_8446622_RNA45SN1_45S_50ntFlanks | 3813 | 3852 | 40 | TTATGGTTCCTTTGGTCGCTCGCTCCTCTCCTACTTGGAT | 397.5 | -6.3 | 1.6 | -4.0 | 6.3E-05 | 8.6E-05 | MB231/MB468 | NUC |
| >hg38_21_+_8433172_8446622_RNA45SN1_45S_50ntFlanks@12001.12032.32\|CTGCGGGCCGCCGGTGAAATACCACTACTCTG | >hg38_21_+_8433172_8446622_RNA45SN1_45S_50ntFlanks | 12001 | 12032 | 32 | CTGCGGGCCGCCGGTGAAATACCACTACTCTG | 1851.7 | -7.1 | 1.7 | -4.2 | 2.2E-05 | 8.6E-05 | BT20/MB468 | MP |
| >hg38_21_+_8433172_8446622_RNA45SN1_45S_50ntFlanks@8104.8140.37\|CCGCGGCGGGGCGCGGGACATGTGGCGTACGGAAGAC | >hg38_21_+_8433172_8446622_RNA45SN1_45S_50ntFlanks | 8104 | 8140 | 37 | CCGCGGCGGGGCGCGGGACATGTGGCGTACGGAAGAC | 680.5 | -6.2 | 1.5 | -4.0 | 6.3E-05 | 8.6E-05 | MB231/MB468 | NUC |
| >hg38_21_+_8433172_8446622_RNA45SN1_45S_50ntFlanks@3899.3919.21\|CCTTCGCGGGGGGGATGCGTG | >hg38_21_+_8433172_8446622_RNA45SN1_45S_50ntFlanks | 3899 | 3919 | 21 | CCTTCGCGGGGGGGATGCGTG | 1575.6 | -5.1 | 1.3 | -4.0 | 6.0E-05 | 8.6E-05 | MB231/MB468 | CYTO |
| >hg38_21_+_8433172_8446622_RNA45SN1_45S_50ntFlanks@3771.3792.22\|CACGGCCGGTACAGTGAAACTG | >hg38_21_+_8433172_8446622_RNA45SN1_45S_50ntFlanks | 3771 | 3792 | 22 | CACGGCCGGTACAGTGAAACTG | 396.6 | -3.9 | 1.0 | -4.0 | 6.0E-05 | 8.6E-05 | MB231/MB468 | CYTO |
| >MI0003602\|hsa-mir-590&WithFlank&7\|+\|74191192\|74191300@62.83.22&[MIMAT0004801&hsa-miR-590-3p&offsets\|0\|+1;m-332&7\|+\|74191253\|74191273&offsets\|0\|+1]\|TAATTTTATGTATAAGCTAGTC | >MI0003602\|hsa-mir-590&WithFlank&7\|+\|74191192\|74191300 | 62 | 83 | 22 | TAATTTTATGTATAAGCTAGTC | 63.3 | -9.4 | 2.1 | -4.5 | 7.5E-06 | 8.6E-05 | BT20/MB231 | MP |
| >hg38_21_+_8433172_8446622_RNA45SN1_45S_50ntFlanks@12626.12660.35\|AATGGGGCGAAGCTACCATCTGTGGGATTATGACT | >hg38_21_+_8433172_8446622_RNA45SN1_45S_50ntFlanks | 12626 | 12660 | 35 | AATGGGGCGAAGCTACCATCTGTGGGATTATGACT | 169.9 | -4.8 | 1.2 | -4.0 | 5.4E-05 | 8.6E-05 | MB231/MB468 | MP |
| >hg38_21_+_8433172_8446622_RNA45SN1_45S_50ntFlanks@3843.3860.18\|CTACTTGGATAACTGTGG | >hg38_21_+_8433172_8446622_RNA45SN1_45S_50ntFlanks | 3843 | 3860 | 18 | CTACTTGGATAACTGTGG | 342.7 | -2.8 | 0.7 | -4.0 | 5.6E-05 | 8.6E-05 | BT20/MB468 | CYTO |
| >hg38_21_+_8433172_8446622_RNA45SN1_45S_50ntFlanks@11199.11225.27\|CCGGCGGCGGCGGCGACTCTGGACGCG | >hg38_21_+_8433172_8446622_RNA45SN1_45S_50ntFlanks | 11199 | 11225 | 27 | CCGGCGGCGGCGGCGACTCTGGACGCG | 2932.3 | -5.4 | 1.3 | -4.0 | 5.6E-05 | 8.6E-05 | BT20/MB468 | CYTO |
| >MI0000063\|hsa-let-7b&WithFlank&22\|+\|46113680\|46113774@12.32.21(+1C)&[MIMAT0000063&hsa-let-7b-5p&offsets\|0\|-1(+1C);m-14&22\|+\|46113691\|46113712&offsets\|0\|-1(+1C)]\|TGAGGTAGTAGGTTGTGTGGTC | >MI0000063\|hsa-let-7b&WithFlank&22\|+\|46113680\|46113774 | 12 | 32 | 21(+1C) | TGAGGTAGTAGGTTGTGTGGTC | 3739.8 | 3.4 | 0.8 | 4.0 | 5.4E-05 | 8.6E-05 | MB231/MB468 | MP |
| >trna116_GluCTC_1_-_145399233_145399304@3.33.31,trna59_GluCTC_1_+_249168447_249168518@3.33.31,trna71_GluCTC_1_-_161439189_161439260@3.33.31,trna74_GluCTC_1_-_161431809_161431880@3.33.31,trna77_GluCTC_1_-_161424398_161424469@3.33.31,trna77_GluCTC_6_+_28949976_28950047@3.33.31,trna80_GluCTC_1_-_161417018_161417089@3.33.31,trna87_GluCTC_6_-_126101393_126101464@3.33.31\|CCTGGTGGTCTAGTGGTTAGGATTCGGCGCT | >trna116_GluCTC_1_-_145399233_145399304 | 3 | 33 | 31 | CCTGGTGGTCTAGTGGTTAGGATTCGGCGCT | 289.1 | 6.2 | 1.5 | 4.0 | 5.6E-05 | 8.6E-05 | BT20/MB468 | CYTO |
| >MI0000342\|hsa-mir-200b&WithFlank&1\|+\|1167098\|1167204@63.85.23&[MIMAT0000318&hsa-miR-200b-3p&offsets\|0\|+1;m-93&1\|+\|1167160\|1167182&offsets\|0\|0]\|TAATACTGCCTGGTAATGATGAC | >MI0000342\|hsa-mir-200b&WithFlank&1\|+\|1167098\|1167204 | 63 | 85 | 23 | TAATACTGCCTGGTAATGATGAC | 1323.7 | 3.5 | 0.8 | 4.5 | 7.1E-06 | 8.6E-05 | BT20/MB231 | CYTO |
| >hg38_21_+_8433172_8446622_RNA45SN1_45S_50ntFlanks@4657.4682.26\|CTAGAGGTGAAATTCTTGGACCGGCG | >hg38_21_+_8433172_8446622_RNA45SN1_45S_50ntFlanks | 4657 | 4682 | 26 | CTAGAGGTGAAATTCTTGGACCGGCG | 568.9 | -6.1 | 1.5 | -4.0 | 5.6E-05 | 8.7E-05 | BT20/MB468 | CYTO |
| >trna11_GluTTC_15_-_26327381_26327452@1.33.33,trna3_GluTTC_13_-_45492062_45492133@1.33.33\|TCCCACATGGTCTAGCGGTTAGGATTCCTGGTT | >trna11_GluTTC_15_-_26327381_26327452 | 1 | 33 | 33 | TCCCACATGGTCTAGCGGTTAGGATTCCTGGTT | 1245.1 | 6.7 | 1.6 | 4.2 | 2.2E-05 | 8.7E-05 | BT20/MB468 | MP |
| >hg38_21_+_8433172_8446622_RNA45SN1_45S_50ntFlanks@11578.11613.36\|TAGAACTGGTGCGGACCAGGGGAATCCGACTGTTTA | >hg38_21_+_8433172_8446622_RNA45SN1_45S_50ntFlanks | 11578 | 11613 | 36 | TAGAACTGGTGCGGACCAGGGGAATCCGACTGTTTA | 297.0 | 2.8 | 0.7 | 4.0 | 6.6E-05 | 8.7E-05 | BT20/MB468 | NUC |
| >hg38_21_+_8433172_8446622_RNA45SN1_45S_50ntFlanks@11027.11051.25\|CTCTCTCTCTCTCTCCCCCGCTCCC | >hg38_21_+_8433172_8446622_RNA45SN1_45S_50ntFlanks | 11027 | 11051 | 25 | CTCTCTCTCTCTCTCCCCCGCTCCC | 1596.6 | -5.0 | 1.2 | -4.0 | 5.6E-05 | 8.7E-05 | BT20/MB468 | CYTO |
| >MI0000737\|hsa-mir-200a&WithFlank&1\|+\|1167857\|1167958@60.82.23&[MIMAT0000682&hsa-miR-200a-3p&offsets\|0\|+1;m-142&1\|+\|1167916\|1167938&offsets\|0\|0]\|TAACACTGTCTGGTAACGATGTT | >MI0000737\|hsa-mir-200a&WithFlank&1\|+\|1167857\|1167958 | 60 | 82 | 23 | TAACACTGTCTGGTAACGATGTT | 517.3 | -5.4 | 1.3 | -4.0 | 6.7E-05 | 8.7E-05 | BT20/MB468 | NUC |
| >hg38_21_+_8433172_8446622_RNA45SN1_45S_50ntFlanks@6671.6720.50\|CTCGGCTCGTGCGTCGATGAAGAACGCAGCTAGCTGCGAGAATTAATGTG | >hg38_21_+_8433172_8446622_RNA45SN1_45S_50ntFlanks | 6671 | 6720 | 50 | CTCGGCTCGTGCGTCGATGAAGAACGCAGCTAGCTGCGAGAATTAATGTG | 305.4 | -6.1 | 1.5 | -4.0 | 6.4E-05 | 8.7E-05 | MB231/MB468 | MITO |
| >trnaMT_SerGCT_MT_+_12207_12265@1.40.40\|GAGAAAGCTCACAAGAACTGCTAACTCATGCCCCCATGTC | >trnaMT_SerGCT_MT_+_12207_12265 | 1 | 40 | 40 | GAGAAAGCTCACAAGAACTGCTAACTCATGCCCCCATGTC | 359.3 | 3.4 | 0.9 | 4.0 | 6.4E-05 | 8.7E-05 | MB231/MB468 | MITO |
| >hg38_21_+_8433172_8446622_RNA45SN1_45S_50ntFlanks@3851.3881.31\|ATAACTGTGGTAATTCTAGAGCTAATACATG | >hg38_21_+_8433172_8446622_RNA45SN1_45S_50ntFlanks | 3851 | 3881 | 31 | ATAACTGTGGTAATTCTAGAGCTAATACATG | 157.2 | -5.3 | 1.3 | -4.1 | 4.7E-05 | 8.7E-05 | BT20/MB468 | MITO |
| >hg38_21_+_8433172_8446622_RNA45SN1_45S_50ntFlanks@11255.11283.29\|CTGCGGCGGGCGTCGCGGCCGCCCCCGGG | >hg38_21_+_8433172_8446622_RNA45SN1_45S_50ntFlanks | 11255 | 11283 | 29 | CTGCGGCGGGCGTCGCGGCCGCCCCCGGG | 56.5 | -8.9 | 2.1 | -4.2 | 2.2E-05 | 8.7E-05 | BT20/MB468 | MP |
| >hg38_21_+_8433172_8446622_RNA45SN1_45S_50ntFlanks@7979.8009.31\|ACCTCAGATCAGACGTGGCGACCCGCTGAAT | >hg38_21_+_8433172_8446622_RNA45SN1_45S_50ntFlanks | 7979 | 8009 | 31 | ACCTCAGATCAGACGTGGCGACCCGCTGAAT | 1407.9 | 1.8 | 0.4 | 4.1 | 4.7E-05 | 8.7E-05 | BT20/MB468 | MITO |

<table>
<tr><th></th><th></th><th></th><th></th><th></th><th></th><th></th><th></th><th></th><th></th><th></th><th></th><th></th><th></th></tr>
<tr><td>>hg38_21_+_8433172_8446622_RNA45SN1_45S_50ntFlanks@6739.6770.32|TCATCGACACTTCGAACGCACTTGCGGCCCCG</td><td>>hg38_21_+_8433172_8446622_RNA45SN1_45S_50ntFlanks</td><td>6739</td><td>6770</td><td>32</td><td>TCATCGACACTTCGAACGCACTTGCGGCCCCG</td><td>644.6</td><td>-4.8</td><td>1.2</td><td>-4.1</td><td>4.7E-05</td><td>8.7E-05</td><td>BT20/MB468</td><td>MITO</td></tr>
<tr><td>>hg38_21_+_8433172_8446622_RNA45SN1_45S_50ntFlanks@8106.8139.34|GCGGCGGGGCGCGGGACATGTGGCGTACGGAAGA</td><td>>hg38_21_+_8433172_8446622_RNA45SN1_45S_50ntFlanks</td><td>8106</td><td>8139</td><td>34</td><td>GCGGCGGGGCGCGGGACATGTGGCGTACGGAAGA</td><td>80.0</td><td>9.7</td><td>2.2</td><td>4.5</td><td>7.6E-06</td><td>8.7E-05</td><td>BT20/MB231</td><td>MP</td></tr>
<tr><td>>hg38_21_+_8433172_8446622_RNA45SN1_45S_50ntFlanks@10686.10708.23|GCGGCGTCCGGTGAGCTCTCGCT</td><td>>hg38_21_+_8433172_8446622_RNA45SN1_45S_50ntFlanks</td><td>10686</td><td>10708</td><td>23</td><td>GCGGCGTCCGGTGAGCTCTCGCT</td><td>4439.4</td><td>3.8</td><td>0.9</td><td>4.1</td><td>4.7E-05</td><td>8.7E-05</td><td>BT20/MB468</td><td>MITO</td></tr>
<tr><td>>hg38_MT_+_1621_3279_RNR2_16S_50ntFlanks@729.760.32|GATTAAAACACTGAACTGACAATTAACAGCCC</td><td>>hg38_MT_+_1621_3279_RNR2_16S_50ntFlanks</td><td>729</td><td>760</td><td>32</td><td>GATTAAAACACTGAACTGACAATTAACAGCCC</td><td>27.3</td><td>8.0</td><td>2.0</td><td>4.0</td><td>5.4E-05</td><td>8.7E-05</td><td>MB231/MB468</td><td>MP</td></tr>
<tr><td>>hg38_21_+_8433172_8446622_RNA45SN1_45S_50ntFlanks@10409.10433.25|GTCAGTCGGTCCTGAGAGATGGGCG</td><td>>hg38_21_+_8433172_8446622_RNA45SN1_45S_50ntFlanks</td><td>10409</td><td>10433</td><td>25</td><td>GTCAGTCGGTCCTGAGAGATGGGCG</td><td>264.7</td><td>-5.4</td><td>1.3</td><td>-4.0</td><td>5.6E-05</td><td>8.7E-05</td><td>BT20/MB468</td><td>CYTO</td></tr>
<tr><td>>hg38_21_+_8433172_8446622_RNA45SN1_45S_50ntFlanks@10797.10825.29|GCATGTTGGAACAATGTAGGTAAGGGAAG</td><td>>hg38_21_+_8433172_8446622_RNA45SN1_45S_50ntFlanks</td><td>10797</td><td>10825</td><td>29</td><td>GCATGTTGGAACAATGTAGGTAAGGGAAG</td><td>229.6</td><td>-2.4</td><td>0.6</td><td>-4.1</td><td>4.7E-05</td><td>8.7E-05</td><td>BT20/MB468</td><td>MITO</td></tr>
<tr><td>>hg38_21_+_8433172_8446622_RNA45SN1_45S_50ntFlanks@10726.10753.28|GGGAGAGGGTGTAAATCTCGCGCCGGGC</td><td>>hg38_21_+_8433172_8446622_RNA45SN1_45S_50ntFlanks</td><td>10726</td><td>10753</td><td>28</td><td>GGGAGAGGGTGTAAATCTCGCGCCGGGC</td><td>87.9</td><td>-6.0</td><td>1.5</td><td>-4.0</td><td>5.4E-05</td><td>8.7E-05</td><td>MB231/MB468</td><td>MP</td></tr>
<tr><td>>hg38_21_+_8433172_8446622_RNA45SN1_45S_50ntFlanks@12510.12530.21|TCGTGAGACAGGTTAGTTTTA</td><td>>hg38_21_+_8433172_8446622_RNA45SN1_45S_50ntFlanks</td><td>12510</td><td>12530</td><td>21</td><td>TCGTGAGACAGGTTAGTTTTA</td><td>116.4</td><td>-5.0</td><td>1.2</td><td>-4.0</td><td>5.6E-05</td><td>8.7E-05</td><td>BT20/MB468</td><td>CYTO</td></tr>
<tr><td>>hg38_21_+_8433172_8446622_RNA45SN1_45S_50ntFlanks@11727.11751.25|AGTAACTATGACTCTCTTAAGGTAG</td><td>>hg38_21_+_8433172_8446622_RNA45SN1_45S_50ntFlanks</td><td>11727</td><td>11751</td><td>25</td><td>AGTAACTATGACTCTCTTAAGGTAG</td><td>161.3</td><td>-6.0</td><td>1.5</td><td>-4.0</td><td>5.4E-05</td><td>8.7E-05</td><td>MB231/MB468</td><td>MP</td></tr>
<tr><td>>MI0000264|hsa-mir-7-2&WithFlank&15|+|88611819|88611940@38.61.24;MI0000265|hsa-mir-7-3&WithFlank&19|+|4770664|4770785@37.60.24;MI0000263|hsa-mir-7-1&WithFlank&9|-|83969742|83969863@30.53.24&[MIMAT0000252&hsa-miR-7-5p&offsets|0|0;m-400&15|+|88611856|88611879&offsets|0|0];[MIMAT0000252_1&hsa-miR-7-5p&offsets|0|0;m-402&19|+|4770700|4770723&offsets|0|0];[MIMAT0000252_2&hsa-miR-7-5p&offsets|0|0;m-401&9|-|83969811|83969834&offsets|0|0]|TGGAAGACTAGTGATTTTGTTGTT</td><td>>MI0000264|hsa-mir-7-2&WithFlank&15|+|88611819|88611940</td><td>38</td><td>61</td><td>24</td><td>TGGAAGACTAGTGATTTTGTTGTT</td><td>34362.8</td><td>3.2</td><td>0.8</td><td>4.0</td><td>5.4E-05</td><td>8.7E-05</td><td>MB231/MB468</td><td>MP</td></tr>
<tr><td>>hg38_21_+_8433172_8446622_RNA45SN1_45S_50ntFlanks@12243.12273.31|GAAGGGCAAAAGCTCGCTTGATCTTGATTTT</td><td>>hg38_21_+_8433172_8446622_RNA45SN1_45S_50ntFlanks</td><td>12243</td><td>12273</td><td>31</td><td>GAAGGGCAAAAGCTCGCTTGATCTTGATTTT</td><td>85.9</td><td>-6.0</td><td>1.5</td><td>-4.0</td><td>5.4E-05</td><td>8.7E-05</td><td>MB231/MB468</td><td>MP</td></tr>
<tr><td>>hg38_21_+_8433172_8446622_RNA45SN1_45S_50ntFlanks@11212.11252.41|CGACTCTGGACGCGAGCCGGGCCCTTCCCGTGGATCGCCCC</td><td>>hg38_21_+_8433172_8446622_RNA45SN1_45S_50ntFlanks</td><td>11212</td><td>11252</td><td>41</td><td>CGACTCTGGACGCGAGCCGGGCCCTTCCCGTGGATCGCCCC</td><td>1155.4</td><td>-3.8</td><td>1.0</td><td>-4.0</td><td>6.5E-05</td><td>8.7E-05</td><td>MB231/MB468</td><td>NUC</td></tr>
<tr><td>>MI0000067|hsa-let-7f-1&WithFlank&9|+|94176341|94176439@13.34.22(+1A);MI0000068|hsa-let-7f-2&WithFlank&X|-|53557186|53557280@14.35.22(+1A)&[MIMAT0000067&hsa-let-7f-5p&offsets|0|0(+1A);m-20&9|+|94176353|94176374&offsets|0|0(+1A)];[MIMAT0000067_1&hsa-let-7f-5p&offsets|0|0(+1A);m-19&X|-|53557246|53557267&offsets|0|0(+1A)]|TGAGGTAGTAGATTGTATAGTTA</td><td>>MI0000067|hsa-let-7f-1&WithFlank&9|+|94176341|94176439</td><td>13</td><td>34</td><td>22(+1A)</td><td>TGAGGTAGTAGATTGTATAGTTA</td><td>2331.9</td><td>4.4</td><td>1.0</td><td>4.2</td><td>2.2E-05</td><td>8.7E-05</td><td>BT20/MB468</td><td>MP</td></tr>
<tr><td>>MI0000255|hsa-mir-30d&WithFlank&8|-|134804870|134804951@12.35.24(+1U)&[MIMAT0000245&hsa-miR-30d-5p&offsets|0|+2(+1U);m-21&8|-|134804919|134804940&offsets|0|+2(+1U)]|TGTAAACATCCCCGACTGGAAGCTT</td><td>>MI0000255|hsa-mir-30d&WithFlank&8|-|134804870|134804951</td><td>12</td><td>35</td><td>24(+1U)</td><td>TGTAAACATCCCCGACTGGAAGCTT</td><td>822.4</td><td>4.4</td><td>1.0</td><td>4.2</td><td>2.2E-05</td><td>8.7E-05</td><td>BT20/MB468</td><td>MP</td></tr>
<tr><td>>hg38_21_+_8433172_8446622_RNA45SN1_45S_50ntFlanks@9534.9564.31|AGGAAACTCTGGTGGAGGTCCGTAGCGGTCC</td><td>>hg38_21_+_8433172_8446622_RNA45SN1_45S_50ntFlanks</td><td>9534</td><td>9564</td><td>31</td><td>AGGAAACTCTGGTGGAGGTCCGTAGCGGTCC</td><td>445.7</td><td>-2.5</td><td>0.6</td><td>-4.1</td><td>4.8E-05</td><td>8.7E-05</td><td>BT20/MB468</td><td>MITO</td></tr>
<tr><td>>MI0000079|hsa-mir-23a&WithFlank&19|-|13836581|13836665@54.72.19(+2U)&[MIMAT0000078&hsa-miR-23a-3p&offsets|+3|+1(+2U);m-31&19|-|13836595|13836615&offsets|+3|+1(+2U)]|ACATTGCCAGGGATTTCCATT</td><td>>MI0000079|hsa-mir-23a&WithFlank&19|-|13836581|13836665</td><td>54</td><td>72</td><td>19(+2U)</td><td>ACATTGCCAGGGATTTCCATT</td><td>123.1</td><td>10.2</td><td>2.5</td><td>4.0</td><td>5.5E-05</td><td>8.8E-05</td><td>MB231/MB468</td><td>MP</td></tr>
<tr><td>>hg38_21_+_8433172_8446622_RNA45SN1_45S_50ntFlanks@12488.12521.34|AACGTGAGCTGGGTTTAGACCGTCGTGAGACAGG</td><td>>hg38_21_+_8433172_8446622_RNA45SN1_45S_50ntFlanks</td><td>12488</td><td>12521</td><td>34</td><td>AACGTGAGCTGGGTTTAGACCGTCGTGAGACAGG</td><td>374.5</td><td>-5.6</td><td>1.4</td><td>-4.0</td><td>5.7E-05</td><td>8.8E-05</td><td>BT20/MB468</td><td>CYTO</td></tr>
<tr><td>>hg38_21_+_8433172_8446622_RNA45SN1_45S_50ntFlanks@9533.9550.18|GAGGAAACTCTGGTGGAG</td><td>>hg38_21_+_8433172_8446622_RNA45SN1_45S_50ntFlanks</td><td>9533</td><td>9550</td><td>18</td><td>GAGGAAACTCTGGTGGAG</td><td>731.9</td><td>-7.1</td><td>1.8</td><td>-4.0</td><td>6.7E-05</td><td>8.8E-05</td><td>BT20/MB468</td><td>NUC</td></tr>
<tr><td>>hg38_21_+_8433172_8446622_RNA45SN1_45S_50ntFlanks@9555.9584.30|GTAGCGGTCCTGACGTGCAAATCGGTCGTC</td><td>>hg38_21_+_8433172_8446622_RNA45SN1_45S_50ntFlanks</td><td>9555</td><td>9584</td><td>30</td><td>GTAGCGGTCCTGACGTGCAAATCGGTCGTC</td><td>568.7</td><td>-7.1</td><td>1.8</td><td>-4.0</td><td>6.7E-05</td><td>8.8E-05</td><td>BT20/MB468</td><td>NUC</td></tr>
<tr><td>>trnaMT_HisGTG_MT_+_12138_12206@18.53.36|AAAACATCAGATTGTGAATCTGACAACAGAGGCTTA</td><td>>trnaMT_HisGTG_MT_+_12138_12206</td><td>18</td><td>53</td><td>36</td><td>AAAACATCAGATTGTGAATCTGACAACAGAGGCTTA</td><td>143.7</td><td>10.9</td><td>2.6</td><td>4.2</td><td>2.3E-05</td><td>8.8E-05</td><td>BT20/MB468</td><td>MP</td></tr>
<tr><td>>MI0000809|hsa-mir-151a&WithFlank&8|-|140732558|140732659@53.75.23&[MIMAT0000757&hsa-miR-151a-3p&offsets|0|+2;m-30&8|-|140732587|140732607&offsets|0|+2]|CTAGACTGAAGCTCCTTGAGGAC</td><td>>MI0000809|hsa-mir-151a&WithFlank&8|-|140732558|140732659</td><td>53</td><td>75</td><td>23</td><td>CTAGACTGAAGCTCCTTGAGGAC</td><td>190.6</td><td>4.8</td><td>1.2</td><td>4.0</td><td>5.5E-05</td><td>8.8E-05</td><td>MB231/MB468</td><td>MP</td></tr>
<tr><td>>hg38_21_+_8433172_8446622_RNA45SN1_45S_50ntFlanks@9541.9571.31|TCTGGTGGAGGTCCGTAGCGGTCCTGACGTG</td><td>>hg38_21_+_8433172_8446622_RNA45SN1_45S_50ntFlanks</td><td>9541</td><td>9571</td><td>31</td><td>TCTGGTGGAGGTCCGTAGCGGTCCTGACGTG</td><td>543.0</td><td>-6.9</td><td>1.7</td><td>-4.0</td><td>6.7E-05</td><td>8.8E-05</td><td>BT20/MB468</td><td>NUC</td></tr>
<tr><td>>hg38_21_+_8433172_8446622_RNA45SN1_45S_50ntFlanks@12716.12751.36|GCCTCGGTTGGCCTCGGATAGCCGGTCCCCCGCCTG</td><td>>hg38_21_+_8433172_8446622_RNA45SN1_45S_50ntFlanks</td><td>12716</td><td>12751</td><td>36</td><td>GCCTCGGTTGGCCTCGGATAGCCGGTCCCCCGCCTG</td><td>69.0</td><td>-10.0</td><td>2.5</td><td>-4.0</td><td>5.5E-05</td><td>8.8E-05</td><td>MB231/MB468</td><td>MP</td></tr>
<tr><td>>hg38_21_+_8433172_8446622_RNA45SN1_45S_50ntFlanks@9504.9522.19|GTGAACTATGCCTGGGCAG</td><td>>hg38_21_+_8433172_8446622_RNA45SN1_45S_50ntFlanks</td><td>9504</td><td>9522</td><td>19</td><td>GTGAACTATGCCTGGGCAG</td><td>420.2</td><td>-2.6</td><td>0.6</td><td>-4.1</td><td>4.8E-05</td><td>8.8E-05</td><td>BT20/MB468</td><td>MITO</td></tr>
<tr><td>>hg38_MT_+_1621_3279_RNR2_16S_50ntFlanks@826.854.29|AAGGAAAGGTTAAAAAAAGTAAAAGGAAC</td><td>>hg38_MT_+_1621_3279_RNR2_16S_50ntFlanks</td><td>826</td><td>854</td><td>29</td><td>AAGGAAAGGTTAAAAAAAGTAAAAGGAAC</td><td>27.7</td><td>8.0</td><td>2.0</td><td>4.0</td><td>5.5E-05</td><td>8.8E-05</td><td>MB231/MB468</td><td>MP</td></tr>
<tr><td>>hg38_21_+_8433172_8446622_RNA45SN1_45S_50ntFlanks@11872.11893.22|AAAGAAGACCCTGTTGAGCTTG</td><td>>hg38_21_+_8433172_8446622_RNA45SN1_45S_50ntFlanks</td><td>11872</td><td>11893</td><td>22</td><td>AAAGAAGACCCTGTTGAGCTTG</td><td>258.7</td><td>-2.5</td><td>0.6</td><td>-4.0</td><td>5.7E-05</td><td>8.8E-05</td><td>BT20/MB468</td><td>CYTO</td></tr>
<tr><td>>hg38_21_+_8433172_8446622_RNA45SN1_45S_50ntFlanks@8380.8406.27|CAAGAGGGCGTGAAACCGTTAAGAGGT</td><td>>hg38_21_+_8433172_8446622_RNA45SN1_45S_50ntFlanks</td><td>8380</td><td>8406</td><td>27</td><td>CAAGAGGGCGTGAAACCGTTAAGAGGT</td><td>564.6</td><td>-2.3</td><td>0.6</td><td>-4.0</td><td>6.1E-05</td><td>8.8E-05</td><td>MB231/MB468</td><td>CYTO</td></tr>
<tr><td>>hg38_21_+_8433172_8446622_RNA45SN1_45S_50ntFlanks@4283.4314.32|CTTTAACGAGGATCCATTGGAGGGCAAGTCTG</td><td>>hg38_21_+_8433172_8446622_RNA45SN1_45S_50ntFlanks</td><td>4283</td><td>4314</td><td>32</td><td>CTTTAACGAGGATCCATTGGAGGGCAAGTCTG</td><td>279.4</td><td>-4.0</td><td>1.0</td><td>-4.0</td><td>5.7E-05</td><td>8.8E-05</td><td>BT20/MB468</td><td>CYTO</td></tr>
<tr><td>>hg38_21_+_8433172_8446622_RNA45SN1_45S_50ntFlanks@8342.8380.39|CGTAAGGGAAAGTTGAAAAGAACTTTGAAGAGAGAGTTC</td><td>>hg38_21_+_8433172_8446622_RNA45SN1_45S_50ntFlanks</td><td>8342</td><td>8380</td><td>39</td><td>CGTAAGGGAAAGTTGAAAAGAACTTTGAAGAGAGAGTTC</td><td>26.3</td><td>7.9</td><td>2.0</td><td>4.0</td><td>5.5E-05</td><td>8.8E-05</td><td>MB231/MB468</td><td>MP</td></tr>
<tr><td>>hg38_21_+_8433172_8446622_RNA45SN1_45S_50ntFlanks@4727.4758.32|AATCAAGAACGAAAGTCGGAGGTTCGAAGACG</td><td>>hg38_21_+_8433172_8446622_RNA45SN1_45S_50ntFlanks</td><td>4727</td><td>4758</td><td>32</td><td>AATCAAGAACGAAAGTCGGAGGTTCGAAGACG</td><td>1278.1</td><td>-3.0</td><td>0.7</td><td>-4.0</td><td>6.5E-05</td><td>8.8E-05</td><td>MB231/MB468</td><td>MITO</td></tr>
<tr><td>>hg38_21_+_8433172_8446622_RNA45SN1_45S_50ntFlanks@6661.6679.19|CGGTGGATCACTCGGCTCG</td><td>>hg38_21_+_8433172_8446622_RNA45SN1_45S_50ntFlanks</td><td>6661</td><td>6679</td><td>19</td><td>CGGTGGATCACTCGGCTCG</td><td>367.7</td><td>-1.6</td><td>0.4</td><td>-4.0</td><td>6.8E-05</td><td>8.8E-05</td><td>BT20/MB468</td><td>NUC</td></tr>
<tr><td>>hg38_21_+_8433172_8446622_RNA45SN1_45S_50ntFlanks@5482.5499.18|CGGAGCGCTGAGAAGACG</td><td>>hg38_21_+_8433172_8446622_RNA45SN1_45S_50ntFlanks</td><td>5482</td><td>5499</td><td>18</td><td>CGGAGCGCTGAGAAGACG</td><td>2221.2</td><td>-3.8</td><td>0.9</td><td>-4.2</td><td>2.3E-05</td><td>8.8E-05</td><td>BT20/MB468</td><td>MP</td></tr>
<tr><td>>hg38_21_+_8433172_8446622_RNA45SN1_45S_50ntFlanks@4225.4243.19|AAATAACAATACAGGACTC</td><td>>hg38_21_+_8433172_8446622_RNA45SN1_45S_50ntFlanks</td><td>4225</td><td>4243</td><td>19</td><td>AAATAACAATACAGGACTC</td><td>242.8</td><td>-6.3</td><td>1.5</td><td>-4.2</td><td>2.3E-05</td><td>8.8E-05</td><td>BT20/MB468</td><td>MP</td></tr>
<tr><td>>hg38_21_+_8433172_8446622_RNA45SN1_45S_50ntFlanks@11577.11594.18|TTAGAACTGGTGCGGACC</td><td>>hg38_21_+_8433172_8446622_RNA45SN1_45S_50ntFlanks</td><td>11577</td><td>11594</td><td>18</td><td>TTAGAACTGGTGCGGACC</td><td>388.7</td><td>-2.1</td><td>0.5</td><td>-4.0</td><td>6.5E-05</td><td>8.8E-05</td><td>MB231/MB468</td><td>NUC</td></tr>
<tr><td>>hg38_21_+_8433172_8446622_RNA45SN1_45S_50ntFlanks@10389.10433.45|TGAACAGCAGTTGAACATGGGTCAGTCGGTCCTGAGAGATGGGCG</td><td>>hg38_21_+_8433172_8446622_RNA45SN1_45S_50ntFlanks</td><td>10389</td><td>10433</td><td>45</td><td>TGAACAGCAGTTGAACATGGGTCAGTCGGTCCTGAGAGATGGGCG</td><td>56.1</td><td>-9.7</td><td>2.4</td><td>-4.0</td><td>5.5E-05</td><td>8.8E-05</td><td>MB231/MB468</td><td>MP</td></tr>
<tr><td>>hg38_21_+_8433172_8446622_RNA45SN1_45S_50ntFlanks@5487.5520.34|CGCTGAGAAGACGGTCGAACTTGACTATCTAGAG</td><td>>hg38_21_+_8433172_8446622_RNA45SN1_45S_50ntFlanks</td><td>5487</td><td>5520</td><td>34</td><td>CGCTGAGAAGACGGTCGAACTTGACTATCTAGAG</td><td>114.1</td><td>-6.0</td><td>1.5</td><td>-4.0</td><td>5.5E-05</td><td>8.8E-05</td><td>MB231/MB468</td><td>MP</td></tr>
<tr><td>>trnaMT_PheGAA_MT_+_577_647@36.57.22|AAAATGTTTAGACGGGCTCACA</td><td>>trnaMT_PheGAA_MT_+_577_647</td><td>36</td><td>57</td><td>22</td><td>AAAATGTTTAGACGGGCTCACA</td><td>360.5</td><td>3.7</td><td>0.9</td><td>4.0</td><td>6.6E-05</td><td>8.8E-05</td><td>MB231/MB468</td><td>MITO</td></tr>
<tr><td>>hg38_21_+_8433172_8446622_RNA45SN1_45S_50ntFlanks@10689.10711.23|GCGTCCGGTGAGCTCTCGCTGGC</td><td>>hg38_21_+_8433172_8446622_RNA45SN1_45S_50ntFlanks</td><td>10689</td><td>10711</td><td>23</td><td>GCGTCCGGTGAGCTCTCGCTGGC</td><td>296.1</td><td>-6.8</td><td>1.6</td><td>-4.2</td><td>2.3E-05</td><td>8.8E-05</td><td>BT20/MB468</td><td>MP</td></tr>
<tr><td>>hg38_21_+_8433172_8446622_RNA45SN1_45S_50ntFlanks@9554.9575.22|CGTAGCGGTCCTGACGTGCAAA</td><td>>hg38_21_+_8433172_8446622_RNA45SN1_45S_50ntFlanks</td><td>9554</td><td>9575</td><td>22</td><td>CGTAGCGGTCCTGACGTGCAAA</td><td>1208.7</td><td>-5.6</td><td>1.4</td><td>-4.0</td><td>5.5E-05</td><td>8.8E-05</td><td>MB231/MB468</td><td>MP</td></tr>
<tr><td>>hg38_21_+_8433172_8446622_RNA45SN1_45S_50ntFlanks@7979.8006.28|ACCTCAGATCAGACGTGGCGACCCGCTG</td><td>>hg38_21_+_8433172_8446622_RNA45SN1_45S_50ntFlanks</td><td>7979</td><td>8006</td><td>28</td><td>ACCTCAGATCAGACGTGGCGACCCGCTG</td><td>5208.3</td><td>-4.4</td><td>1.0</td><td>-4.2</td><td>2.3E-05</td><td>8.8E-05</td><td>BT20/MB468</td><td>MP</td></tr>
<tr><td>>hg38_21_+_8433172_8446622_RNA45SN1_45S_50ntFlanks@10841.10874.34|CGTAACTTCGGGATAAGGATTGGCTCTAAGGGCT</td><td>>hg38_21_+_8433172_8446622_RNA45SN1_45S_50ntFlanks</td><td>10841</td><td>10874</td><td>34</td><td>CGTAACTTCGGGATAAGGATTGGCTCTAAGGGCT</td><td>838.4</td><td>-2.9</td><td>0.7</td><td>-4.0</td><td>5.7E-05</td><td>8.8E-05</td><td>BT20/MB468</td><td>CYTO</td></tr>
<tr><td>>trna2_GlyGCC_21_-_18827107_18827177@1.31.31,trna35_GlyGCC_1_+_161413094_161413164@1.31.31,trna37_GlyGCC_1_+_161420467_161420537@1.31.31,trna39_GlyGCC_1_+_161427898_161427968@1.31.31,trna41_GlyGCC_1_+_161435258_161435328@1.31.31|GCATGGGTGGTTCAGTGGTAGAATTCTCGCC</td><td>>trna2_GlyGCC_21_-_18827107_18827177</td><td>1</td><td>31</td><td>31</td><td>GCATGGGTGGTTCAGTGGTAGAATTCTCGCC</td><td>44233.2</td><td>5.5</td><td>1.2</td><td>4.5</td><td>7.7E-06</td><td>8.8E-05</td><td>BT20/MB231</td><td>MP</td></tr>
<tr><td>>MI0000289|hsa-mir-181a-1&WithFlank&1|-|198859038|198859159@30.54.25;MI0000269|hsa-mir-181a-2&WithFlank&9|+|124692436|124692557@45.69.25&[MIMAT0000256&hsa-miR-181a-5p&offsets|0|+2;m-32&1|-|198859108|198859130&offsets|0|+2];[MIMAT0000256_1&hsa-miR-181a-5p&offsets|0|+2;m-32&9|+|124692480|124692502&offsets|0|+2]|AACATTCAACGCTGTCGGTGAGTTT</td><td>>MI0000289|hsa-mir-181a-1&WithFlank&1|-|198859038|198859159</td><td>30</td><td>54</td><td>25</td><td>AACATTCAACGCTGTCGGTGAGTTT</td><td>4462.7</td><td>3.9</td><td>1.0</td><td>4.0</td><td>5.5E-05</td><td>8.8E-05</td><td>MB231/MB468</td><td>MP</td></tr>
<tr><td>>trnaMT_LeuTAA_MT_+_3230_3304@1.42.42|GTTAAGATGGCAGAGCCCGGTAATCGCATAAAACTTAAAACT</td><td>>trnaMT_LeuTAA_MT_+_3230_3304</td><td>1</td><td>42</td><td>42</td><td>GTTAAGATGGCAGAGCCCGGTAATCGCATAAAACTTAAAACT</td><td>32.8</td><td>8.3</td><td>2.0</td><td>4.0</td><td>5.5E-05</td><td>8.8E-05</td><td>MB231/MB468</td><td>MP</td></tr>
<tr><td>>hg38_21_+_8433172_8446622_RNA45SN1_45S_50ntFlanks@9462.9503.42|GGGAGGTGGAGCACGAGCGCACGTGTTAGGACCCGAAAGATG</td><td>>hg38_21_+_8433172_8446622_RNA45SN1_45S_50ntFlanks</td><td>9462</td><td>9503</td><td>42</td><td>GGGAGGTGGAGCACGAGCGCACGTGTTAGGACCCGAAAGATG</td><td>155.4</td><td>-5.7</td><td>1.4</td><td>-4.1</td><td>4.8E-05</td><td>8.9E-05</td><td>BT20/MB468</td><td>MITO</td></tr>
<tr><td>>hg38_MT_+_1621_3279_RNR2_16S_50ntFlanks@761.778.18|AATATCTACAATCAACCA</td><td>>hg38_MT_+_1621_3279_RNR2_16S_50ntFlanks</td><td>761</td><td>778</td><td>18</td><td>AATATCTACAATCAACCA</td><td>25.7</td><td>7.9</td><td>2.0</td><td>4.0</td><td>5.5E-05</td><td>8.9E-05</td><td>MB231/MB468</td><td>MP</td></tr>
<tr><td>>hg38_21_+_8433172_8446622_RNA45SN1_45S_50ntFlanks@5078.5104.27|CGATAACGAACGAGACTCTGGCATGCT</td><td>>hg38_21_+_8433172_8446622_RNA45SN1_45S_50ntFlanks</td><td>5078</td><td>5104</td><td>27</td><td>CGATAACGAACGAGACTCTGGCATGCT</td><td>209.5</td><td>-6.4</td><td>1.6</td><td>-4.0</td><td>5.5E-05</td><td>8.9E-05</td><td>MB231/MB468</td><td>MP</td></tr>
<tr><td>>MI0000542|hsa-mir-320a&WithFlank&8|-|22244960|22245043@48.68.21(+4U)&[MIMAT0000510&hsa-miR-320a-3p&offsets|0|-1(+4U);m-60&8|-|22244975|22244996&offsets|0|-1(+4U)]|AAAAGCTGGGTTGAGAGGGCGTTTT</td><td>>MI0000542|hsa-mir-320a&WithFlank&8|-|22244960|22245043</td><td>48</td><td>68</td><td>21(+4U)</td><td>AAAAGCTGGGTTGAGAGGGCGTTTT</td><td>552.2</td><td>4.9</td><td>1.1</td><td>4.5</td><td>7.4E-06</td><td>8.9E-05</td><td>BT20/MB231</td><td>CYTO</td></tr>
<tr><td>>hg38_21_+_8433172_8446622_RNA45SN1_45S_50ntFlanks@5436.5466.31|GTGAGGCCCTCGGATCGGCCCCGCCGGGGTC</td><td>>hg38_21_+_8433172_8446622_RNA45SN1_45S_50ntFlanks</td><td>5436</td><td>5466</td><td>31</td><td>GTGAGGCCCTCGGATCGGCCCCGCCGGGGTC</td><td>1003.8</td><td>-7.2</td><td>1.8</td><td>-4.0</td><td>6.8E-05</td><td>8.9E-05</td><td>BT20/MB468</td><td>NUC</td></tr>
</table>

<table>
<tr><th></th><th></th><th></th><th></th><th></th><th></th><th></th><th></th><th></th><th></th><th></th><th></th><th></th><th></th></tr>
<tr><td>>hg38_21_+_8433172_8446622_RNA45SN1_45S_50ntFlanks@5485.5509.25|AGCGCTGAGAAGACGGTCGAACTTG</td><td>>hg38_21_+_8433172_8446622_RNA45SN1_45S_50ntFlanks</td><td>5485</td><td>5509</td><td>25</td><td>AGCGCTGAGAAGACGGTCGAACTTG</td><td>331.3</td><td>-3.0</td><td>0.7</td><td>-4.0</td><td>6.2E-05</td><td>8.9E-05</td><td>MB231/MB468</td><td>CYTO</td></tr>
<tr><td>>hg38_21_+_8433172_8446622_RNA45SN1_45S_50ntFlanks@10378.10424.47|AGGGTTCCATGTGAACAGCAGTTGAACATGGGTCAGTCGGTCCTGAG</td><td>>hg38_21_+_8433172_8446622_RNA45SN1_45S_50ntFlanks</td><td>10378</td><td>10424</td><td>47</td><td>AGGGTTCCATGTGAACAGCAGTTGAACATGGGTCAGTCGGTCCTGAG</td><td>267.1</td><td>7.7</td><td>1.8</td><td>4.2</td><td>2.3E-05</td><td>8.9E-05</td><td>BT20/MB468</td><td>MP</td></tr>
<tr><td>>am_trnaMT_HisGTG_MT_+_12138_12206@26.43.18|AGATTGTGAATCTGACAA</td><td>>am_trnaMT_HisGTG_MT_+_12138_12206</td><td>26</td><td>43</td><td>18</td><td>AGATTGTGAATCTGACAA</td><td>297.9</td><td>5.7</td><td>1.4</td><td>4.1</td><td>4.8E-05</td><td>8.9E-05</td><td>BT20/MB468</td><td>MITO</td></tr>
<tr><td>>trnaMT_HisGTG_MT_+_12138_12206@25.59.35|CAGATTGTGAATCTGACAACAGAGGCTTACGACCC</td><td>>trnaMT_HisGTG_MT_+_12138_12206</td><td>25</td><td>59</td><td>35</td><td>CAGATTGTGAATCTGACAACAGAGGCTTACGACCC</td><td>140.8</td><td>10.9</td><td>2.6</td><td>4.2</td><td>2.3E-05</td><td>8.9E-05</td><td>BT20/MB468</td><td>MP</td></tr>
<tr><td>>hg38_21_+_8433172_8446622_RNA45SN1_45S_50ntFlanks@8184.8222.39|CGAGGCCCAGCCCGTGGACGGTGTGAGGCCGGTAGCGGC</td><td>>hg38_21_+_8433172_8446622_RNA45SN1_45S_50ntFlanks</td><td>8184</td><td>8222</td><td>39</td><td>CGAGGCCCAGCCCGTGGACGGTGTGAGGCCGGTAGCGGC</td><td>41.7</td><td>-8.5</td><td>2.0</td><td>-4.2</td><td>2.3E-05</td><td>8.9E-05</td><td>BT20/MB468</td><td>MP</td></tr>
<tr><td>>MI0003137|hsa-mir-193b&WithFlank&16|+|14303961|14304055@57.77.21(+1G)&[MIMAT0002819&hsa-miR-193b-3p&offsets|0|-1(+1G);m-166&16|+|14304017|14304038&offsets|0|-1(+1G)]|AACTGGCCCTCAAAGTCCCGCG</td><td>>MI0003137|hsa-mir-193b&WithFlank&16|+|14303961|14304055</td><td>57</td><td>77</td><td>21(+1G)</td><td>AACTGGCCCTCAAAGTCCCGCG</td><td>177.6</td><td>6.6</td><td>1.5</td><td>4.5</td><td>8.6E-06</td><td>8.9E-05</td><td>BT20/MB231</td><td>NUC</td></tr>
<tr><td>>hg38_21_+_8433172_8446622_RNA45SN1_45S_50ntFlanks@3846.3873.28|CTTGGATAACTGTGGTAATTCTAGAGCT</td><td>>hg38_21_+_8433172_8446622_RNA45SN1_45S_50ntFlanks</td><td>3846</td><td>3873</td><td>28</td><td>CTTGGATAACTGTGGTAATTCTAGAGCT</td><td>161.7</td><td>-6.2</td><td>1.5</td><td>-4.2</td><td>2.3E-05</td><td>8.9E-05</td><td>BT20/MB468</td><td>MP</td></tr>
<tr><td>>hg38_21_+_8433172_8446622_RNA45SN1_45S_50ntFlanks@5487.5513.27|CGCTGAGAAGACGGTCGAACTTGACTA</td><td>>hg38_21_+_8433172_8446622_RNA45SN1_45S_50ntFlanks</td><td>5487</td><td>5513</td><td>27</td><td>CGCTGAGAAGACGGTCGAACTTGACTA</td><td>777.3</td><td>-5.4</td><td>1.3</td><td>-4.2</td><td>2.3E-05</td><td>8.9E-05</td><td>BT20/MB468</td><td>MP</td></tr>
<tr><td>>hg38_21_+_8433172_8446622_RNA45SN1_45S_50ntFlanks@12249.12272.24|CAAAAGCTCGCTTGATCTTGATTT</td><td>>hg38_21_+_8433172_8446622_RNA45SN1_45S_50ntFlanks</td><td>12249</td><td>12272</td><td>24</td><td>CAAAAGCTCGCTTGATCTTGATTT</td><td>312.2</td><td>-4.6</td><td>1.2</td><td>-4.0</td><td>6.6E-05</td><td>8.9E-05</td><td>MB231/MB468</td><td>NUC</td></tr>
<tr><td>>hg38_21_+_8433172_8446622_RNA45SN1_45S_50ntFlanks@10409.10432.24|GTCAGTCGGTCCTGAGAGATGGGC</td><td>>hg38_21_+_8433172_8446622_RNA45SN1_45S_50ntFlanks</td><td>10409</td><td>10432</td><td>24</td><td>GTCAGTCGGTCCTGAGAGATGGGC</td><td>118.6</td><td>-4.6</td><td>1.1</td><td>-4.0</td><td>5.6E-05</td><td>8.9E-05</td><td>MB231/MB468</td><td>MP</td></tr>
<tr><td>>hg38_21_+_8433172_8446622_RNA45SN1_45S_50ntFlanks@12008.12035.28|CCGCCGGTGAAATACCACTACTCTGATC</td><td>>hg38_21_+_8433172_8446622_RNA45SN1_45S_50ntFlanks</td><td>12008</td><td>12035</td><td>28</td><td>CCGCCGGTGAAATACCACTACTCTGATC</td><td>654.6</td><td>-3.6</td><td>0.9</td><td>-4.2</td><td>2.3E-05</td><td>8.9E-05</td><td>BT20/MB468</td><td>MP</td></tr>
<tr><td>>hg38_21_+_8433172_8446622_RNA45SN1_45S_50ntFlanks@4566.4587.22|ATAATGGAATAGGACCGCGGTT</td><td>>hg38_21_+_8433172_8446622_RNA45SN1_45S_50ntFlanks</td><td>4566</td><td>4587</td><td>22</td><td>ATAATGGAATAGGACCGCGGTT</td><td>553.5</td><td>-6.1</td><td>1.5</td><td>-4.0</td><td>6.6E-05</td><td>8.9E-05</td><td>MB231/MB468</td><td>NUC</td></tr>
<tr><td>>hg38_21_+_8433172_8446622_RNA45SN1_45S_50ntFlanks@11876.11909.34|AAGACCCTGTTGAGCTTGACTCTAGTCTGGCACG</td><td>>hg38_21_+_8433172_8446622_RNA45SN1_45S_50ntFlanks</td><td>11876</td><td>11909</td><td>34</td><td>AAGACCCTGTTGAGCTTGACTCTAGTCTGGCACG</td><td>255.0</td><td>-2.4</td><td>0.6</td><td>-4.1</td><td>4.9E-05</td><td>8.9E-05</td><td>BT20/MB468</td><td>MITO</td></tr>
<tr><td>>trna10_ValCAC_5_-_180649395_180649467@35.53.19,trna12_ValAAC_5_-_180645270_180645342@35.53.19,trna132_ValAAC_6_-_27721179_27721251@35.53.19,trna13_ValCAC_19_-_4724647_4724719@35.53.19,trna152_ValCAC_6_-_27248049_27248121@35.53.19,trna157_ValCAC_6_-_27173867_27173939@35.53.19,trna15_ValAAC_5_-_180615416_180615488@35.53.19,trna18_ValCAC_5_-_180529253_180529325@35.53.19,trna2_ValAAC_3_+_169490018_169490090@35.53.19,trna2_ValCAC_5_+_180524070_180524142@35.53.19,trna37_ValAAC_6_+_27203288_27203360@35.53.19,trna4_ValAAC_5_+_180591154_180591226@35.53.19,trna5_ValAAC_5_+_180596610_180596682@35.53.19,trna6_ValCAC_5_+_180600650_180600722@35.53.19,trna85_ValCAC_1_-_161369490_161369562@35.53.19,trna98_ValCAC_1_-_149298555_149298627@35.53.19,trna9_ValCAC_6_+_26538282_26538354@35.53.19|ACACGCGAAAGGTCCCCGG</td><td>>trna10_ValCAC_5_-_180649395_180649467</td><td>35</td><td>53</td><td>19</td><td>ACACGCGAAAGGTCCCCGG</td><td>54.1</td><td>-9.6</td><td>2.4</td><td>-4.0</td><td>5.6E-05</td><td>8.9E-05</td><td>MB231/MB468</td><td>MP</td></tr>
<tr><td>>hg38_21_+_8433172_8446622_RNA45SN1_45S_50ntFlanks@5149.5167.19|TAGAGGGACAAGTGGCGTT</td><td>>hg38_21_+_8433172_8446622_RNA45SN1_45S_50ntFlanks</td><td>5149</td><td>5167</td><td>19</td><td>TAGAGGGACAAGTGGCGTT</td><td>102.8</td><td>10.0</td><td>2.2</td><td>4.5</td><td>7.8E-06</td><td>8.9E-05</td><td>BT20/MB231</td><td>MP</td></tr>
<tr><td>>hg38_21_+_8433172_8446622_RNA45SN1_45S_50ntFlanks@8116.8134.19|GCGGGACATGTGGCGTACG</td><td>>hg38_21_+_8433172_8446622_RNA45SN1_45S_50ntFlanks</td><td>8116</td><td>8134</td><td>19</td><td>GCGGGACATGTGGCGTACG</td><td>306.0</td><td>4.4</td><td>1.0</td><td>4.2</td><td>2.3E-05</td><td>8.9E-05</td><td>BT20/MB468</td><td>MP</td></tr>
<tr><td>>hg38_21_+_8433172_8446622_RNA45SN1_45S_50ntFlanks@11569.11594.26|CAGCCGACTTAGAACTGGTGCGGACC</td><td>>hg38_21_+_8433172_8446622_RNA45SN1_45S_50ntFlanks</td><td>11569</td><td>11594</td><td>26</td><td>CAGCCGACTTAGAACTGGTGCGGACC</td><td>524.2</td><td>-3.1</td><td>0.8</td><td>-4.0</td><td>6.2E-05</td><td>8.9E-05</td><td>MB231/MB468</td><td>CYTO</td></tr>
<tr><td>>hg38_21_+_8433172_8446622_RNA45SN1_45S_50ntFlanks@3854.3898.45|ACTGTGGTAATTCTAGAGCTAATACATGCCGACGGGCGCTGACCC</td><td>>hg38_21_+_8433172_8446622_RNA45SN1_45S_50ntFlanks</td><td>3854</td><td>3898</td><td>45</td><td>ACTGTGGTAATTCTAGAGCTAATACATGCCGACGGGCGCTGACCC</td><td>640.2</td><td>-3.3</td><td>0.8</td><td>-4.0</td><td>6.2E-05</td><td>8.9E-05</td><td>MB231/MB468</td><td>CYTO</td></tr>
<tr><td>>hg38_21_+_8433172_8446622_RNA45SN1_45S_50ntFlanks@7990.8007.18|GACGTGGCGACCCGCTGA</td><td>>hg38_21_+_8433172_8446622_RNA45SN1_45S_50ntFlanks</td><td>7990</td><td>8007</td><td>18</td><td>GACGTGGCGACCCGCTGA</td><td>433.1</td><td>-5.1</td><td>1.3</td><td>-4.0</td><td>6.7E-05</td><td>9.0E-05</td><td>MB231/MB468</td><td>MITO</td></tr>
<tr><td>>MI0000272|hsa-mir-182&WithFlank&7|-|129770377|129770498@29.52.24&[MIMAT0000259&hsa-miR-182-5p&offsets|0|0;m-22&7|-|129770449|129770470&offsets|0|+2]|TTTGGCAATGGTAGAACTCACACT</td><td>>MI0000272|hsa-mir-182&WithFlank&7|-|129770377|129770498</td><td>29</td><td>52</td><td>24</td><td>TTTGGCAATGGTAGAACTCACACT</td><td>4670.6</td><td>3.8</td><td>0.9</td><td>4.0</td><td>6.7E-05</td><td>9.0E-05</td><td>MB231/MB468</td><td>MITO</td></tr>
<tr><td>>MI0000272|hsa-mir-182&WithFlank&7|-|129770377|129770498@29.51.23(+1C)&[MIMAT0000259&hsa-miR-182-5p&offsets|0|-1(+1C);m-22&7|-|129770449|129770470&offsets|0|+1(+1C)]|TTTGGCAATGGTAGAACTCACACC</td><td>>MI0000272|hsa-mir-182&WithFlank&7|-|129770377|129770498</td><td>29</td><td>51</td><td>23(+1C)</td><td>TTTGGCAATGGTAGAACTCACACC</td><td>684.2</td><td>4.1</td><td>1.0</td><td>4.0</td><td>6.7E-05</td><td>9.0E-05</td><td>MB231/MB468</td><td>MITO</td></tr>
<tr><td>>hg38_21_+_8433172_8446622_RNA45SN1_45S_50ntFlanks@10927.10945.19|CGAGGCGCCGCCGCCCCCC</td><td>>hg38_21_+_8433172_8446622_RNA45SN1_45S_50ntFlanks</td><td>10927</td><td>10945</td><td>19</td><td>CGAGGCGCCGCCGCCCCCC</td><td>359.9</td><td>-7.5</td><td>1.8</td><td>-4.2</td><td>2.3E-05</td><td>9.0E-05</td><td>BT20/MB468</td><td>MP</td></tr>
<tr><td>>hg38_21_+_8433172_8446622_RNA45SN1_45S_50ntFlanks@4969.4999.31|ACCCGGCCCGGACACGGACAGGATTGACAGA</td><td>>hg38_21_+_8433172_8446622_RNA45SN1_45S_50ntFlanks</td><td>4969</td><td>4999</td><td>31</td><td>ACCCGGCCCGGACACGGACAGGATTGACAGA</td><td>575.0</td><td>-4.9</td><td>1.2</td><td>-4.0</td><td>6.7E-05</td><td>9.0E-05</td><td>MB231/MB468</td><td>MITO</td></tr>
<tr><td>>hg38_21_+_8433172_8446622_RNA45SN1_45S_50ntFlanks@9454.9491.38|CCGCGCCGGGGAGGTGGAGCACGAGCGCACGTGTTAGG</td><td>>hg38_21_+_8433172_8446622_RNA45SN1_45S_50ntFlanks</td><td>9454</td><td>9491</td><td>38</td><td>CCGCGCCGGGGAGGTGGAGCACGAGCGCACGTGTTAGG</td><td>148.1</td><td>-5.2</td><td>1.3</td><td>-4.1</td><td>4.9E-05</td><td>9.0E-05</td><td>BT20/MB468</td><td>MITO</td></tr>
<tr><td>>MI0000289|hsa-mir-181a-1&WithFlank&1|-|198859038|198859159@30.51.22(+1C);MI0000269|hsa-mir-181a-2&WithFlank&9|+|124692436|124692557@45.66.22(+1C)&[MIMAT0000256&hsa-miR-181a-5p&offsets|0|-1(+1C);m-32&1|-|198859108|198859130&offsets|0|-1(+1C)];[MIMAT0000256_1&hsa-miR-181a-5p&offsets|0|-1(+1C);m-32&9|+|124692480|124692502&offsets|0|-1(+1C)]|AACATTCAACGCTGTCGGTGAGC</td><td>>MI0000289|hsa-mir-181a-1&WithFlank&1|-|198859038|198859159</td><td>30</td><td>51</td><td>22(+1C)</td><td>AACATTCAACGCTGTCGGTGAGC</td><td>1967.0</td><td>3.7</td><td>0.9</td><td>4.0</td><td>5.6E-05</td><td>9.0E-05</td><td>MB231/MB468</td><td>MP</td></tr>
<tr><td>>hg38_21_+_8433172_8446622_RNA45SN1_45S_50ntFlanks@5056.5081.26|GAGCGATTTGTCTGGTTAATTCCGAT</td><td>>hg38_21_+_8433172_8446622_RNA45SN1_45S_50ntFlanks</td><td>5056</td><td>5081</td><td>26</td><td>GAGCGATTTGTCTGGTTAATTCCGAT</td><td>1387.8</td><td>-5.2</td><td>1.3</td><td>-4.0</td><td>5.6E-05</td><td>9.0E-05</td><td>MB231/MB468</td><td>MP</td></tr>
<tr><td>>trnaMT_HisGTG_MT_+_12138_12206@45.72.28|AGAGGCTTACGACCCCTTATTTACCCCA</td><td>>trnaMT_HisGTG_MT_+_12138_12206</td><td>45</td><td>72</td><td>28</td><td>AGAGGCTTACGACCCCTTATTTACCCCA</td><td>25.9</td><td>7.9</td><td>2.0</td><td>4.0</td><td>5.6E-05</td><td>9.0E-05</td><td>MB231/MB468</td><td>MP</td></tr>
<tr><td>>hg38_21_+_8433172_8446622_RNA45SN1_45S_50ntFlanks@10786.10824.39|AACAGCCTCTGGCATGTTGGAACAATGTAGGTAAGGGAA</td><td>>hg38_21_+_8433172_8446622_RNA45SN1_45S_50ntFlanks</td><td>10786</td><td>10824</td><td>39</td><td>AACAGCCTCTGGCATGTTGGAACAATGTAGGTAAGGGAA</td><td>44.2</td><td>-8.5</td><td>2.0</td><td>-4.2</td><td>2.3E-05</td><td>9.0E-05</td><td>BT20/MB468</td><td>MP</td></tr>
<tr><td>>hg38_21_+_8433172_8446622_RNA45SN1_45S_50ntFlanks@12488.12520.33|AACGTGAGCTGGGTTTAGACCGTCGTGAGACAG</td><td>>hg38_21_+_8433172_8446622_RNA45SN1_45S_50ntFlanks</td><td>12488</td><td>12520</td><td>33</td><td>AACGTGAGCTGGGTTTAGACCGTCGTGAGACAG</td><td>195.6</td><td>-5.2</td><td>1.3</td><td>-4.0</td><td>5.9E-05</td><td>9.0E-05</td><td>BT20/MB468</td><td>CYTO</td></tr>
<tr><td>>hg38_21_+_8433172_8446622_RNA45SN1_45S_50ntFlanks@6650.6668.19|ACGACTCTTAGCGGTGGAT</td><td>>hg38_21_+_8433172_8446622_RNA45SN1_45S_50ntFlanks</td><td>6650</td><td>6668</td><td>19</td><td>ACGACTCTTAGCGGTGGAT</td><td>947.4</td><td>3.2</td><td>0.8</td><td>4.2</td><td>2.3E-05</td><td>9.0E-05</td><td>BT20/MB468</td><td>MP</td></tr>
<tr><td>>hg38_21_+_8433172_8446622_RNA45SN1_45S_50ntFlanks@13006.13025.20|CTATTGAAAGTCAGCCCTCG</td><td>>hg38_21_+_8433172_8446622_RNA45SN1_45S_50ntFlanks</td><td>13006</td><td>13025</td><td>20</td><td>CTATTGAAAGTCAGCCCTCG</td><td>155.1</td><td>-6.1</td><td>1.5</td><td>-4.2</td><td>2.3E-05</td><td>9.0E-05</td><td>BT20/MB468</td><td>MP</td></tr>
<tr><td>>MI0000742|hsa-mir-34b&WithFlank&11|+|111512932|111513027@57.78.22&[MIMAT0004676&hsa-miR-34b-3p&offsets|+1|+1;m-585&11|+|111512988|111513009&offsets|0|0]|AATCACTAACTCCACTGCCATC</td><td>>MI0000742|hsa-mir-34b&WithFlank&11|+|111512932|111513027</td><td>57</td><td>78</td><td>22</td><td>AATCACTAACTCCACTGCCATC</td><td>63.9</td><td>-9.4</td><td>2.1</td><td>-4.5</td><td>7.9E-06</td><td>9.0E-05</td><td>BT20/MB231</td><td>MP</td></tr>
<tr><td>>hg38_21_+_8433172_8446622_RNA45SN1_45S_50ntFlanks@10734.10755.22|GTGTAAATCTCGCGCCGGGCCG</td><td>>hg38_21_+_8433172_8446622_RNA45SN1_45S_50ntFlanks</td><td>10734</td><td>10755</td><td>22</td><td>GTGTAAATCTCGCGCCGGGCCG</td><td>142.3</td><td>-5.1</td><td>1.3</td><td>-4.0</td><td>5.9E-05</td><td>9.0E-05</td><td>BT20/MB468</td><td>CYTO</td></tr>
<tr><td>>hg38_21_+_8433172_8446622_RNA45SN1_45S_50ntFlanks@9623.9656.34|TAGTAGCTGGTTCCCTCCGAAGTTTCCCTCAGGA</td><td>>hg38_21_+_8433172_8446622_RNA45SN1_45S_50ntFlanks</td><td>9623</td><td>9656</td><td>34</td><td>TAGTAGCTGGTTCCCTCCGAAGTTTCCCTCAGGA</td><td>308.6</td><td>-1.9</td><td>0.5</td><td>-4.1</td><td>4.9E-05</td><td>9.0E-05</td><td>BT20/MB468</td><td>MITO</td></tr>
<tr><td>>hg38_21_+_8433172_8446622_RNA45SN1_45S_50ntFlanks@8176.8200.25|CTTCTGATCGAGGCCCAGCCCGTGG</td><td>>hg38_21_+_8433172_8446622_RNA45SN1_45S_50ntFlanks</td><td>8176</td><td>8200</td><td>25</td><td>CTTCTGATCGAGGCCCAGCCCGTGG</td><td>159.0</td><td>-6.2</td><td>1.5</td><td>-4.0</td><td>5.6E-05</td><td>9.0E-05</td><td>MB231/MB468</td><td>MP</td></tr>
<tr><td>>hg38_1_-_228634819_228635039_RNA5S12_5S_50ntFlanks@51.77.27|GTCTACGGCCATACCACCCTGAACGCG</td><td>>hg38_1_-_228634819_228635039_RNA5S12_5S_50ntFlanks</td><td>51</td><td>77</td><td>27</td><td>GTCTACGGCCATACCACCCTGAACGCG</td><td>953.4</td><td>3.6</td><td>0.9</td><td>4.0</td><td>6.3E-05</td><td>9.1E-05</td><td>MB231/MB468</td><td>CYTO</td></tr>
<tr><td>>hg38_21_+_8433172_8446622_RNA45SN1_45S_50ntFlanks@4367.4408.42|CAGTTAAAAAGCTCGTAGTTGGATCTTGGGAGCGGGCGGGCG</td><td>>hg38_21_+_8433172_8446622_RNA45SN1_45S_50ntFlanks</td><td>4367</td><td>4408</td><td>42</td><td>CAGTTAAAAAGCTCGTAGTTGGATCTTGGGAGCGGGCGGGCG</td><td>136.4</td><td>-4.9</td><td>1.2</td><td>-4.0</td><td>6.3E-05</td><td>9.1E-05</td><td>MB231/MB468</td><td>CYTO</td></tr>
<tr><td>>hg38_21_+_8433172_8446622_RNA45SN1_45S_50ntFlanks@11669.11710.42|GCCCAGTGCTCTGAATGTCAAAGTGAAGAAATTCAATGAAGC</td><td>>hg38_21_+_8433172_8446622_RNA45SN1_45S_50ntFlanks</td><td>11669</td><td>11710</td><td>42</td><td>GCCCAGTGCTCTGAATGTCAAAGTGAAGAAATTCAATGAAGC</td><td>917.7</td><td>-4.3</td><td>1.0</td><td>-4.5</td><td>7.5E-06</td><td>9.1E-05</td><td>BT20/MB231</td><td>CYTO</td></tr>
<tr><td>>hg38_21_+_8433172_8446622_RNA45SN1_45S_50ntFlanks@10411.10432.22|CAGTCGGTCCTGAGAGATGGGC</td><td>>hg38_21_+_8433172_8446622_RNA45SN1_45S_50ntFlanks</td><td>10411</td><td>10432</td><td>22</td><td>CAGTCGGTCCTGAGAGATGGGC</td><td>92.7</td><td>9.9</td><td>2.2</td><td>4.5</td><td>8.0E-06</td><td>9.1E-05</td><td>BT20/MB231</td><td>MP</td></tr>
<tr><td>>hg38_21_+_8433172_8446622_RNA45SN1_45S_50ntFlanks@10237.10272.36|ACGCTACGCCGCGACGAGTAGGAGGGCCGCTGCGGT</td><td>>hg38_21_+_8433172_8446622_RNA45SN1_45S_50ntFlanks</td><td>10237</td><td>10272</td><td>36</td><td>ACGCTACGCCGCGACGAGTAGGAGGGCCGCTGCGGT</td><td>161.8</td><td>6.9</td><td>1.6</td><td>4.2</td><td>2.3E-05</td><td>9.1E-05</td><td>BT20/MB468</td><td>MP</td></tr>
<tr><td>>MI0000342|hsa-mir-200b&WithFlank&1|+|1167098|1167204@63.85.23(+1U)&[MIMAT0000318&hsa-miR-200b-3p&offsets|0|+1(+1U);m-93&1|+|1167160|1167182&offsets|0|0(+1U)]|TAATACTGCCTGGTAATGATGACT</td><td>>MI0000342|hsa-mir-200b&WithFlank&1|+|1167098|1167204</td><td>63</td><td>85</td><td>23(+1U)</td><td>TAATACTGCCTGGTAATGATGACT</td><td>667.4</td><td>-6.6</td><td>1.6</td><td>-4.2</td><td>2.3E-05</td><td>9.1E-05</td><td>BT20/MB468</td><td>MP</td></tr>
<tr><td>>MI0000273|hsa-mir-183&WithFlank&7|-|129774899|129775020@33.54.22&[MIMAT0000261&hsa-miR-183-5p&offsets|0|0;m-41&7|-|129774966|129774987&offsets|-1|-1]|TATGGCACTGGTAGAATTCACT</td><td>>MI0000273|hsa-mir-183&WithFlank&7|-|129774899|129775020</td><td>33</td><td>54</td><td>22</td><td>TATGGCACTGGTAGAATTCACT</td><td>16043.6</td><td>3.3</td><td>0.7</td><td>4.5</td><td>7.6E-06</td><td>9.1E-05</td><td>BT20/MB231</td><td>CYTO</td></tr>
<tr><td>>MI0000737|hsa-mir-200a&WithFlank&1|+|1167857|1167958@60.82.23(+1U)&[MIMAT0000682&hsa-miR-200a-3p&offsets|0|+1(+1U);m-142&1|+|1167916|1167938&offsets|0|0(+1U)]|TAACACTGTCTGGTAACGATGTTT</td><td>>MI0000737|hsa-mir-200a&WithFlank&1|+|1167857|1167958</td><td>60</td><td>82</td><td>23(+1U)</td><td>TAACACTGTCTGGTAACGATGTTT</td><td>1294.3</td><td>-5.4</td><td>1.3</td><td>-4.0</td><td>5.7E-05</td><td>9.1E-05</td><td>MB231/MB468</td><td>MP</td></tr>
<tr><td>>MI0001445|hsa-mir-423&WithFlank&17|+|30117073|30117178@58.80.23&[MIMAT0001340&hsa-miR-423-3p&offsets|-1|-1;m-100&17|+|30117131|30117153&offsets|-1|-1]|AAGCTCGGTCTGAGGCCCCTCAG</td><td>>MI0001445|hsa-mir-423&WithFlank&17|+|30117073|30117178</td><td>58</td><td>80</td><td>23</td><td>AAGCTCGGTCTGAGGCCCCTCAG</td><td>180.1</td><td>5.1</td><td>1.3</td><td>4.0</td><td>6.8E-05</td><td>9.1E-05</td><td>MB231/MB468</td><td>MITO</td></tr>
<tr><td>>hg38_21_+_8433172_8446622_RNA45SN1_45S_50ntFlanks@11923.11941.19|AGAGGTGTAGAATAAGTGG</td><td>>hg38_21_+_8433172_8446622_RNA45SN1_45S_50ntFlanks</td><td>11923</td><td>11941</td><td>19</td><td>AGAGGTGTAGAATAAGTGG</td><td>33.4</td><td>8.8</td><td>2.1</td><td>4.2</td><td>2.4E-05</td><td>9.1E-05</td><td>BT20/MB468</td><td>MP</td></tr>
</table>

| | | | | | | | | | | | | | |
|---|---|---|---|---|---|---|---|---|---|---|---|---|---|
| >hg38_21_+_8433172_8446622_RNA45SN1_45S_50ntFlanks@12476.12496.21\|CCACTAATAGGGAACGTGAGC | >hg38_21_+_8433172_8446622_RNA45SN1_45S_50ntFlanks | 12476 | 12496 | 21 | CCACTAATAGGGAACGTGAGC | 2217.0 | -2.6 | 0.6 | -4.0 | 6.4E-05 | 9.1E-05 | MB231/MB468 | CYTO |
| >hg38_21_+_8433172_8446622_RNA45SN1_45S_50ntFlanks@12414.12434.21\|CCTTCGATGTCGGCTCTTCCT | >hg38_21_+_8433172_8446622_RNA45SN1_45S_50ntFlanks | 12414 | 12434 | 21 | CCTTCGATGTCGGCTCTTCCT | 435.8 | 3.2 | 0.8 | 4.0 | 6.4E-05 | 9.1E-05 | MB231/MB468 | CYTO |
| >hg38_21_+_8433172_8446622_RNA45SN1_45S_50ntFlanks@10384.10419.36\|CCATGTGAACAGCAGTTGAACATGGGTCAGTCGGTC | >hg38_21_+_8433172_8446622_RNA45SN1_45S_50ntFlanks | 10384 | 10419 | 36 | CCATGTGAACAGCAGTTGAACATGGGTCAGTCGGTC | 121.0 | -4.4 | 1.1 | -4.0 | 6.4E-05 | 9.1E-05 | MB231/MB468 | CYTO |
| >hg38_21_+_8433172_8446622_RNA45SN1_45S_50ntFlanks@6698.6719.22\|AGCTAGCTGCGAGAATTAATGT | >hg38_21_+_8433172_8446622_RNA45SN1_45S_50ntFlanks | 6698 | 6719 | 22 | AGCTAGCTGCGAGAATTAATGT | 1504.5 | -5.6 | 1.4 | -4.0 | 5.7E-05 | 9.1E-05 | MB231/MB468 | MP |
| >hg38_1_-_228634819_228635039_RNA5S12_5S_50ntFlanks@129.171.43\|TTGGATGGGAGACCGCCTGGGAATACCGGGTGCTGTAGGCTTT | >hg38_1_-_228634819_228635039_RNA5S12_5S_50ntFlanks | 129 | 171 | 43 | TTGGATGGGAGACCGCCTGGGAATACCGGGTGCTGTAGGCTTT | 575.0 | 7.1 | 1.7 | 4.2 | 2.4E-05 | 9.1E-05 | BT20/MB468 | MP |
| >MI0000081\|hsa-mir-24-2&WithFlank&19\|-\|13836281\|13836365@56.77.22;MI0000080\|hsa-mir-24-1&WithFlank&9\|+\|95086015\|95086094@50.71.22&[MIMAT0000080&hsa-miR-24-3p&offsets\|0\|0;m-36&19\|-\|13836288\|13836310&offsets\|0\|-1];[MIMAT0000080_1&hsa-miR-24-3p&offsets\|0\|0;m-35&9\|+\|95086064\|95086086&offsets\|0\|-1]\|TGGCTCAGTTCAGCAGGAACAG | >MI0000081\|hsa-mir-24-2&WithFlank&19\|-\|13836281\|13836365 | 56 | 77 | 22 | TGGCTCAGTTCAGCAGGAACAG | 41005.1 | 4.1 | 1.0 | 4.2 | 2.4E-05 | 9.1E-05 | BT20/MB468 | MP |
| >hg38_21_+_8433172_8446622_RNA45SN1_45S_50ntFlanks@8168.8214.47\|CCCAAGTCCTTCTGATCGAGGCCCAGCCCGTGGACGGTGTGAGGCCG | >hg38_21_+_8433172_8446622_RNA45SN1_45S_50ntFlanks | 8168 | 8214 | 47 | CCCAAGTCCTTCTGATCGAGGCCCAGCCCGTGGACGGTGTGAGGCCG | 1107.7 | -5.4 | 1.3 | -4.0 | 6.8E-05 | 9.1E-05 | MB231/MB468 | NUC |
| >trnaMT_ValTAC_MT_+_1602_1670@43.71.29\|GATTTCAACTTAACTTGACCGCTCTGACC | >trnaMT_ValTAC_MT_+_1602_1670 | 43 | 71 | 29 | GATTTCAACTTAACTTGACCGCTCTGACC | 1242.7 | 3.7 | 0.9 | 4.0 | 6.8E-05 | 9.1E-05 | MB231/MB468 | MITO |
| >hg38_21_+_8433172_8446622_RNA45SN1_45S_50ntFlanks@476.500.25\|TTCGTGATCGATGTGGTGACGTCGT | >hg38_21_+_8433172_8446622_RNA45SN1_45S_50ntFlanks | 476 | 500 | 25 | TTCGTGATCGATGTGGTGACGTCGT | 493.4 | 3.9 | 1.0 | 4.0 | 6.8E-05 | 9.1E-05 | MB231/MB468 | NUC |
| >hg38_21_+_8433172_8446622_RNA45SN1_45S_50ntFlanks@12285.12319.35\|CAGACCGTGAAAGCGGGGCCTCACGATCCTTCTGA | >hg38_21_+_8433172_8446622_RNA45SN1_45S_50ntFlanks | 12285 | 12319 | 35 | CAGACCGTGAAAGCGGGGCCTCACGATCCTTCTGA | 482.2 | -3.3 | 0.8 | -4.1 | 5.0E-05 | 9.1E-05 | BT20/MB468 | MITO |
| >hg38_21_+_8433172_8446622_RNA45SN1_45S_50ntFlanks@12071.12093.23\|CCCCGAGGGGCTCTCGCTTCTGG | >hg38_21_+_8433172_8446622_RNA45SN1_45S_50ntFlanks | 12071 | 12093 | 23 | CCCCGAGGGGCTCTCGCTTCTGG | 231.9 | -5.6 | 1.4 | -4.0 | 5.9E-05 | 9.2E-05 | BT20/MB468 | CYTO |
| >hg38_21_+_8433172_8446622_RNA45SN1_45S_50ntFlanks@475.517.43\|CTTCGTGATCGATGTGGTGACGTCGTGCTCTCCCGGGCCGGGT | >hg38_21_+_8433172_8446622_RNA45SN1_45S_50ntFlanks | 475 | 517 | 43 | CTTCGTGATCGATGTGGTGACGTCGTGCTCTCCCGGGCCGGGT | 10955.8 | 3.7 | 0.9 | 4.0 | 6.8E-05 | 9.2E-05 | MB231/MB468 | NUC |
| >hg38_21_+_8433172_8446622_RNA45SN1_45S_50ntFlanks@11580.11606.27\|GAACTGGTGCGGACCAGGGGAATCCGA | >hg38_21_+_8433172_8446622_RNA45SN1_45S_50ntFlanks | 11580 | 11606 | 27 | GAACTGGTGCGGACCAGGGGAATCCGA | 179.4 | -4.0 | 1.0 | -4.0 | 5.9E-05 | 9.2E-05 | BT20/MB468 | CYTO |
| >hg38_21_+_8433172_8446622_RNA45SN1_45S_50ntFlanks@11592.11611.20\|ACCAGGGGAATCCGACTGTT | >hg38_21_+_8433172_8446622_RNA45SN1_45S_50ntFlanks | 11592 | 11611 | 20 | ACCAGGGGAATCCGACTGTT | 517.3 | -3.7 | 0.9 | -4.0 | 6.0E-05 | 9.2E-05 | BT20/MB468 | CYTO |
| >hg38_21_+_8433172_8446622_RNA45SN1_45S_50ntFlanks@12638.12669.32\|CTACCATCTGTGGGATTATGACTGAACGCCTC | >hg38_21_+_8433172_8446622_RNA45SN1_45S_50ntFlanks | 12638 | 12669 | 32 | CTACCATCTGTGGGATTATGACTGAACGCCTC | 600.1 | -3.4 | 0.9 | -4.0 | 6.8E-05 | 9.2E-05 | MB231/MB468 | MITO |
| >hg38_21_+_8433172_8446622_RNA45SN1_45S_50ntFlanks@6661.6696.36\|CGGTGGATCACTCGGCTCGTGCGTCGATGAAGAACG | >hg38_21_+_8433172_8446622_RNA45SN1_45S_50ntFlanks | 6661 | 6696 | 36 | CGGTGGATCACTCGGCTCGTGCGTCGATGAAGAACG | 827.4 | -2.7 | 0.7 | -4.0 | 5.7E-05 | 9.2E-05 | MB231/MB468 | MP |
| >MI0000081\|hsa-mir-24-2&WithFlank&19\|-\|13836281\|13836365@56.74.19(+1A);MI0000080\|hsa-mir-24-1&WithFlank&9\|+\|95086015\|95086094@50.68.19(+1A)&[MIMAT0000080&hsa-miR-24-3p&offsets\|0\|-3(+1A);m-36&19\|-\|13836288\|13836310&offsets\|0\|-4(+1A)];[MIMAT0000080_1&hsa-miR-24-3p&offsets\|0\|-3(+1A);m-35&9\|+\|95086064\|95086086&offsets\|0\|-4(+1A)]\|TGGCTCAGTTCAGCAGGAAA | >MI0000081\|hsa-mir-24-2&WithFlank&19\|-\|13836281\|13836365 | 56 | 74 | 19(+1A) | TGGCTCAGTTCAGCAGGAAA | 414.1 | 3.7 | 0.9 | 4.0 | 5.7E-05 | 9.2E-05 | MB231/MB468 | MP |
| >hg38_21_+_8433172_8446622_RNA45SN1_45S_50ntFlanks@9475.9509.35\|CGAGCGCACGTGTTAGGACCCGAAAGATGGTGAAC | >hg38_21_+_8433172_8446622_RNA45SN1_45S_50ntFlanks | 9475 | 9509 | 35 | CGAGCGCACGTGTTAGGACCCGAAAGATGGTGAAC | 31.5 | 8.7 | 2.1 | 4.2 | 2.4E-05 | 9.2E-05 | BT20/MB468 | MP |
| >hg38_21_+_8433172_8446622_RNA45SN1_45S_50ntFlanks@10910.10932.23\|CGCGCGCCGCGGCTGGACGAGGC | >hg38_21_+_8433172_8446622_RNA45SN1_45S_50ntFlanks | 10910 | 10932 | 23 | CGCGCGCCGCGGCTGGACGAGGC | 87.6 | 9.8 | 2.2 | 4.5 | 8.1E-06 | 9.2E-05 | BT20/MB231 | MP |
| >hg38_21_+_8433172_8446622_RNA45SN1_45S_50ntFlanks@12619.12653.35\|AGGAGCCAATGGGGCGAAGCTACCATCTGTGGGAT | >hg38_21_+_8433172_8446622_RNA45SN1_45S_50ntFlanks | 12619 | 12653 | 35 | AGGAGCCAATGGGGCGAAGCTACCATCTGTGGGAT | 732.6 | -6.7 | 1.6 | -4.2 | 2.4E-05 | 9.2E-05 | BT20/MB468 | MP |
| >hg38_21_+_8433172_8446622_RNA45SN1_45S_50ntFlanks@9791.9817.27\|TGGCGTGGAGCCGGGCGTGGAATGCGA | >hg38_21_+_8433172_8446622_RNA45SN1_45S_50ntFlanks | 9791 | 9817 | 27 | TGGCGTGGAGCCGGGCGTGGAATGCGA | 140.6 | 10.9 | 2.6 | 4.2 | 2.4E-05 | 9.2E-05 | BT20/MB468 | MP |
| >hg38_21_+_8433172_8446622_RNA45SN1_45S_50ntFlanks@12617.12634.18\|TGAGGAGCCAATGGGGCG | >hg38_21_+_8433172_8446622_RNA45SN1_45S_50ntFlanks | 12617 | 12634 | 18 | TGAGGAGCCAATGGGGCG | 37.3 | -8.3 | 2.0 | -4.2 | 2.4E-05 | 9.2E-05 | BT20/MB468 | MP |
| >hg38_21_+_8433172_8446622_RNA45SN1_45S_50ntFlanks@4574.4613.40\|ATAGGACCGCGGTTCTATTTTGTTGGTTTTCGGAACTGAG | >hg38_21_+_8433172_8446622_RNA45SN1_45S_50ntFlanks | 4574 | 4613 | 40 | ATAGGACCGCGGTTCTATTTTGTTGGTTTTCGGAACTGAG | 111.3 | -5.9 | 1.5 | -4.0 | 6.0E-05 | 9.2E-05 | BT20/MB468 | CYTO |
| >hg38_21_+_8433172_8446622_RNA45SN1_45S_50ntFlanks@3899.3919.21\|CCTTCGCGGGGGGGATGCGTG | >hg38_21_+_8433172_8446622_RNA45SN1_45S_50ntFlanks | 3899 | 3919 | 21 | CCTTCGCGGGGGGGATGCGTG | 1316.0 | -4.8 | 1.2 | -4.0 | 6.0E-05 | 9.2E-05 | BT20/MB468 | CYTO |
| >hg38_21_+_8433172_8446622_RNA45SN1_45S_50ntFlanks@12433.12454.22\|CTATCATTGTGAAGCAGAATTC | >hg38_21_+_8433172_8446622_RNA45SN1_45S_50ntFlanks | 12433 | 12454 | 22 | CTATCATTGTGAAGCAGAATTC | 279.6 | -4.5 | 1.1 | -4.0 | 6.9E-05 | 9.3E-05 | MB231/MB468 | NUC |
| >hg38_21_+_8433172_8446622_RNA45SN1_45S_50ntFlanks@10679.10706.28\|GGTTCCGGCGGCGTCCGGTGAGCTCTCG | >hg38_21_+_8433172_8446622_RNA45SN1_45S_50ntFlanks | 10679 | 10706 | 28 | GGTTCCGGCGGCGTCCGGTGAGCTCTCG | 7162.6 | -8.2 | 2.1 | -4.0 | 7.1E-05 | 9.3E-05 | BT20/MB468 | NUC |
| >hg38_21_+_8433172_8446622_RNA45SN1_45S_50ntFlanks@475.513.39\|CTTCGTGATCGATGTGGTGACGTCGTGCTCTCCCGGGCC | >hg38_21_+_8433172_8446622_RNA45SN1_45S_50ntFlanks | 475 | 513 | 39 | CTTCGTGATCGATGTGGTGACGTCGTGCTCTCCCGGGCC | 185.8 | 5.7 | 1.4 | 4.0 | 6.9E-05 | 9.3E-05 | MB231/MB468 | MITO |
| >hg38_21_+_8433172_8446622_RNA45SN1_45S_50ntFlanks@4734.4765.32\|AACGAAAGTCGGAGGTTCGAAGACGATCAGAT | >hg38_21_+_8433172_8446622_RNA45SN1_45S_50ntFlanks | 4734 | 4765 | 32 | AACGAAAGTCGGAGGTTCGAAGACGATCAGAT | 848.9 | 6.1 | 1.4 | 4.2 | 2.4E-05 | 9.3E-05 | BT20/MB468 | MP |
| >hg38_21_+_8433172_8446622_RNA45SN1_45S_50ntFlanks@12999.13039.41\|CTGCGATCTATTGAAAGTCAGCCCTCGACACAAGGGTTTGT | >hg38_21_+_8433172_8446622_RNA45SN1_45S_50ntFlanks | 12999 | 13039 | 41 | CTGCGATCTATTGAAAGTCAGCCCTCGACACAAGGGTTTGT | 5087.1 | 4.5 | 1.1 | 4.0 | 6.0E-05 | 9.3E-05 | BT20/MB468 | CYTO |
| >hg38_21_+_8433172_8446622_RNA45SN1_45S_50ntFlanks@10846.10867.22\|CTTCGGGATAAGGATTGGCTCT | >hg38_21_+_8433172_8446622_RNA45SN1_45S_50ntFlanks | 10846 | 10867 | 22 | CTTCGGGATAAGGATTGGCTCT | 793.4 | -7.1 | 1.8 | -4.0 | 7.1E-05 | 9.3E-05 | BT20/MB468 | NUC |
| >hg38_21_+_8433172_8446622_RNA45SN1_45S_50ntFlanks@11115.11132.18\|CGGCAGGGGCCGGCGGCG | >hg38_21_+_8433172_8446622_RNA45SN1_45S_50ntFlanks | 11115 | 11132 | 18 | CGGCAGGGGCCGGCGGCG | 238.1 | -5.3 | 1.3 | -4.0 | 6.0E-05 | 9.3E-05 | BT20/MB468 | CYTO |
| >hg38_21_+_8433172_8446622_RNA45SN1_45S_50ntFlanks@4390.4423.34\|TCTTGGGAGCGGGCGGGCGGTCCGCCGCGAGGCG | >hg38_21_+_8433172_8446622_RNA45SN1_45S_50ntFlanks | 4390 | 4423 | 34 | TCTTGGGAGCGGGCGGGCGGTCCGCCGCGAGGCG | 206.8 | -5.3 | 1.3 | -4.0 | 6.1E-05 | 9.3E-05 | BT20/MB468 | CYTO |
| >MI0005544\|hsa-mir-147b&WithFlank&15\|+\|45433044\|45433135@55.72.18&[MIMAT0004928&hsa-miR-147b-3p&offsets\|0\|-3;m-575&15\|+\|45433098\|45433118&offsets\|0\|-3]\|GTGTGCGGAAATGCTTCT | >MI0005544\|hsa-mir-147b&WithFlank&15\|+\|45433044\|45433135 | 55 | 72 | 18 | GTGTGCGGAAATGCTTCT | 28.5 | 8.1 | 2.0 | 4.0 | 5.8E-05 | 9.3E-05 | MB231/MB468 | MP |
| >hg38_21_+_8433172_8446622_RNA45SN1_45S_50ntFlanks@12417.12446.30\|TCGATGTCGGCTCTTCCTATCATTGTGAAG | >hg38_21_+_8433172_8446622_RNA45SN1_45S_50ntFlanks | 12417 | 12446 | 30 | TCGATGTCGGCTCTTCCTATCATTGTGAAG | 3161.6 | -7.8 | 2.0 | -4.0 | 7.2E-05 | 9.3E-05 | BT20/MB468 | NUC |
| >hg38_21_+_8433172_8446622_RNA45SN1_45S_50ntFlanks@11565.11603.39\|CTAGCAGCCGACTTAGAACTGGTGCGGACCAGGGGAATC | >hg38_21_+_8433172_8446622_RNA45SN1_45S_50ntFlanks | 11565 | 11603 | 39 | CTAGCAGCCGACTTAGAACTGGTGCGGACCAGGGGAATC | 87.2 | 9.8 | 2.2 | 4.5 | 8.2E-06 | 9.3E-05 | BT20/MB231 | MP |
| >hg38_21_+_8433172_8446622_RNA45SN1_45S_50ntFlanks@8357.8394.38\|AAAAGAACTTTGAAGAGAGAGTTCAAGAGGGCGTGAAA | >hg38_21_+_8433172_8446622_RNA45SN1_45S_50ntFlanks | 8357 | 8394 | 38 | AAAAGAACTTTGAAGAGAGAGTTCAAGAGGGCGTGAAA | 89.6 | 9.8 | 2.2 | 4.5 | 8.2E-06 | 9.3E-05 | BT20/MB231 | MP |
| >hg38_21_+_8433172_8446622_RNA45SN1_45S_50ntFlanks@9603.9629.27\|GAAAGACTAATCGAACCATCTAGTAGC | >hg38_21_+_8433172_8446622_RNA45SN1_45S_50ntFlanks | 9603 | 9629 | 27 | GAAAGACTAATCGAACCATCTAGTAGC | 91.7 | 9.9 | 2.2 | 4.5 | 8.2E-06 | 9.3E-05 | BT20/MB231 | MP |
| >MI0000102\|hsa-mir-100&WithFlank&11\|-\|122152223\|122152314@19.40.22(+3A)&[MIMAT0000098&hsa-miR-100-5p&offsets\|0\|0(+3A);m-27&11\|-\|122152276\|122152296&offsets\|0\|+1(+3A)]\|AACCCGTAGATCCGAACTTGTGAAA | >MI0000102\|hsa-mir-100&WithFlank&11\|-\|122152223\|122152314 | 19 | 40 | 22(+3A) | AACCCGTAGATCCGAACTTGTGAAA | 70.1 | -9.5 | 2.1 | -4.5 | 8.2E-06 | 9.3E-05 | BT20/MB231 | MP |
| >hg38_21_+_8433172_8446622_RNA45SN1_45S_50ntFlanks@6647.6672.26\|CGTACGACTCTTAGCGGTGGATCACT | >hg38_21_+_8433172_8446622_RNA45SN1_45S_50ntFlanks | 6647 | 6672 | 26 | CGTACGACTCTTAGCGGTGGATCACT | 1228.0 | 3.3 | 0.8 | 4.0 | 6.9E-05 | 9.3E-05 | MB231/MB468 | MITO |
| >hg38_MT_+_1621_3279_RNR2_16S_50ntFlanks@727.749.23\|CAGATTAAAACACTGAACTGACA | >hg38_MT_+_1621_3279_RNR2_16S_50ntFlanks | 727 | 749 | 23 | CAGATTAAAACACTGAACTGACA | 28.0 | 8.0 | 2.0 | 4.0 | 5.8E-05 | 9.3E-05 | MB231/MB468 | MP |
| >hg38_21_+_8433172_8446622_RNA45SN1_45S_50ntFlanks@13007.13028.22\|TATTGAAAGTCAGCCCTCGACA | >hg38_21_+_8433172_8446622_RNA45SN1_45S_50ntFlanks | 13007 | 13028 | 22 | TATTGAAAGTCAGCCCTCGACA | 251.5 | -5.9 | 1.5 | -4.0 | 6.9E-05 | 9.3E-05 | MB231/MB468 | MITO |
| >hg38_21_+_8433172_8446622_RNA45SN1_45S_50ntFlanks@12631.12659.29\|GGCGAAGCTACCATCTGTGGGATTATGAC | >hg38_21_+_8433172_8446622_RNA45SN1_45S_50ntFlanks | 12631 | 12659 | 29 | GGCGAAGCTACCATCTGTGGGATTATGAC | 175.2 | -2.9 | 0.7 | -4.0 | 6.9E-05 | 9.3E-05 | MB231/MB468 | NUC |
| >MI0001652\|hsa-mir-450a-1&WithFlank&X\|-\|134540335\|134540437@24.43.20;MI0003187\|hsa-mir-450a-2&WithFlank&X\|-\|134540502\|134540613@29.48.20&[MIMAT0001545&hsa-miR-450a-5p&offsets\|0\|-2;m-288&X\|-\|134540393\|134540414&offsets\|0\|-2];[MIMAT0001545_1&hsa-miR-450a-5p&offsets\|0\|-2;m-287&X\|-\|134540564\|134540585&offsets\|0\|-2]\|TTTTGCGATGTGTTCCTAAT | >MI0001652\|hsa-mir-450a-1&WithFlank&X\|-\|134540335\|134540437 | 24 | 43 | 20 | TTTTGCGATGTGTTCCTAAT | 28.3 | 8.0 | 2.0 | 4.0 | 5.8E-05 | 9.3E-05 | MB231/MB468 | MP |
| >hg38_21_+_8433172_8446622_RNA45SN1_45S_50ntFlanks@12005.12032.28\|GGGCCGCCGGTGAAATACCACTACTCTG | >hg38_21_+_8433172_8446622_RNA45SN1_45S_50ntFlanks | 12005 | 12032 | 28 | GGGCCGCCGGTGAAATACCACTACTCTG | 69.8 | -4.4 | 1.1 | -4.0 | 5.8E-05 | 9.3E-05 | MB231/MB468 | MP |
| >hg38_21_+_8433172_8446622_RNA45SN1_45S_50ntFlanks@11580.11601.22\|GAACTGGTGCGGACCAGGGGAA | >hg38_21_+_8433172_8446622_RNA45SN1_45S_50ntFlanks | 11580 | 11601 | 22 | GAACTGGTGCGGACCAGGGGAA | 1092.3 | -7.1 | 1.7 | -4.2 | 2.4E-05 | 9.3E-05 | BT20/MB468 | MP |
| >hg38_21_+_8433172_8446622_RNA45SN1_45S_50ntFlanks@6646.6685.40\|TCGTACGACTCTTAGCGGTGGATCACTCGGCTCGTGCGTC | >hg38_21_+_8433172_8446622_RNA45SN1_45S_50ntFlanks | 6646 | 6685 | 40 | TCGTACGACTCTTAGCGGTGGATCACTCGGCTCGTGCGTC | 427.8 | 2.6 | 0.7 | 4.0 | 6.5E-05 | 9.3E-05 | MB231/MB468 | CYTO |
| >am_trnaMT_ValTAC_MT_+_1602_1670@50.69.20\|ACTTAACTTGACCGCTCTGA | >am_trnaMT_ValTAC_MT_+_1602_1670 | 50 | 69 | 20 | ACTTAACTTGACCGCTCTGA | 529.6 | -5.6 | 1.4 | -4.0 | 7.2E-05 | 9.3E-05 | BT20/MB468 | NUC |

<table>
<tr><th></th><th></th><th></th><th></th><th></th><th></th><th></th><th></th><th></th><th></th><th></th><th></th><th></th><th></th></tr>
<tr><td>>hg38_21_+_8433172_8446622_RNA45SN1_45S_50ntFlanks@10841.10871.31|CGTAACTTCGGGATAAGGATTGGCTCTAAGG</td><td>>hg38_21_+_8433172_8446622_RNA45SN1_45S_50ntFlanks</td><td>10841</td><td>10871</td><td>31</td><td>CGTAACTTCGGGATAAGGATTGGCTCTAAGG</td><td>1214.1</td><td>-5.7</td><td>1.4</td><td>-4.0</td><td>7.2E-05</td><td>9.3E-05</td><td>BT20/MB468</td><td>NUC</td></tr>
<tr><td>>MI0001448|hsa-mir-425&WithFlank&3|-|49020142|49020240@60.80.21(+1U)&[MIMAT0001343&hsa-miR-425-3p&offsets|-1|-2(+1U);m-258&3|-|49020160|49020181&offsets|0|-1(+1U)]|CATCGGGAATGTCGTGTCCGCT</td><td>>MI0001448|hsa-mir-425&WithFlank&3|-|49020142|49020240</td><td>60</td><td>80</td><td>21(+1U)</td><td>CATCGGGAATGTCGTGTCCGCT</td><td>209.7</td><td>5.1</td><td>1.3</td><td>4.0</td><td>5.8E-05</td><td>9.4E-05</td><td>MB231/MB468</td><td>MP</td></tr>
<tr><td>>hg38_21_+_8433172_8446622_RNA45SN1_45S_50ntFlanks@10682.10710.29|TCCGGCGGCGTCCGGTGAGCTCTCGCTGG</td><td>>hg38_21_+_8433172_8446622_RNA45SN1_45S_50ntFlanks</td><td>10682</td><td>10710</td><td>29</td><td>TCCGGCGGCGTCCGGTGAGCTCTCGCTGG</td><td>130.7</td><td>-6.1</td><td>1.5</td><td>-4.2</td><td>2.4E-05</td><td>9.4E-05</td><td>BT20/MB468</td><td>MP</td></tr>
<tr><td>>MI0000093|hsa-mir-92a-1&WithFlank&13|+|91351308|91351397@54.76.23(+1C)&[MIMAT0000092&hsa-miR-92a-3p&offsets|0|+1(+1C);m-9&13|+|91351361|91351382&offsets|0|+1(+1C)]|TATTGCACTTGTCCCGGCCTGTTC</td><td>>MI0000093|hsa-mir-92a-1&WithFlank&13|+|91351308|91351397</td><td>54</td><td>76</td><td>23(+1C)</td><td>TATTGCACTTGTCCCGGCCTGTTC</td><td>31.2</td><td>8.2</td><td>2.0</td><td>4.0</td><td>5.9E-05</td><td>9.4E-05</td><td>MB231/MB468</td><td>MP</td></tr>
<tr><td>>trnaMT_SerGCT_MT_+_12207_12265@1.19.19|GAGAAAGCTCACAAGAACT</td><td>>trnaMT_SerGCT_MT_+_12207_12265</td><td>1</td><td>19</td><td>19</td><td>GAGAAAGCTCACAAGAACT</td><td>13938.3</td><td>4.8</td><td>1.2</td><td>4.1</td><td>5.1E-05</td><td>9.4E-05</td><td>BT20/MB468</td><td>MITO</td></tr>
<tr><td>>hg38_21_+_8433172_8446622_RNA45SN1_45S_50ntFlanks@6652.6676.25|GACTCTTAGCGGTGGATCACTCGGC</td><td>>hg38_21_+_8433172_8446622_RNA45SN1_45S_50ntFlanks</td><td>6652</td><td>6676</td><td>25</td><td>GACTCTTAGCGGTGGATCACTCGGC</td><td>1508.0</td><td>4.1</td><td>1.0</td><td>4.0</td><td>7.0E-05</td><td>9.4E-05</td><td>MB231/MB468</td><td>NUC</td></tr>
<tr><td>>hg38_21_+_8433172_8446622_RNA45SN1_45S_50ntFlanks@6661.6687.27|CGGTGGATCACTCGGCTCGTGCGTCGA</td><td>>hg38_21_+_8433172_8446622_RNA45SN1_45S_50ntFlanks</td><td>6661</td><td>6687</td><td>27</td><td>CGGTGGATCACTCGGCTCGTGCGTCGA</td><td>1317.0</td><td>2.8</td><td>0.7</td><td>4.0</td><td>7.0E-05</td><td>9.4E-05</td><td>MB231/MB468</td><td>NUC</td></tr>
<tr><td>>hg38_21_+_8433172_8446622_RNA45SN1_45S_50ntFlanks@3773.3799.27|CGGCCGGTACAGTGAAACTGCGAATGG</td><td>>hg38_21_+_8433172_8446622_RNA45SN1_45S_50ntFlanks</td><td>3773</td><td>3799</td><td>27</td><td>CGGCCGGTACAGTGAAACTGCGAATGG</td><td>152.0</td><td>-5.4</td><td>1.3</td><td>-4.0</td><td>7.0E-05</td><td>9.4E-05</td><td>MB231/MB468</td><td>NUC</td></tr>
<tr><td>>hg38_21_+_8433172_8446622_RNA45SN1_45S_50ntFlanks@9483.9520.38|CGTGTTAGGACCCGAAAGATGGTGAACTATGCCTGGGC</td><td>>hg38_21_+_8433172_8446622_RNA45SN1_45S_50ntFlanks</td><td>9483</td><td>9520</td><td>38</td><td>CGTGTTAGGACCCGAAAGATGGTGAACTATGCCTGGGC</td><td>81.3</td><td>9.7</td><td>2.2</td><td>4.5</td><td>8.3E-06</td><td>9.4E-05</td><td>BT20/MB231</td><td>MP</td></tr>
<tr><td>>hg38_21_+_8433172_8446622_RNA45SN1_45S_50ntFlanks@10914.10946.33|CGCCGCGGCTGGACGAGGCGCCGCCGCCCCCCC</td><td>>hg38_21_+_8433172_8446622_RNA45SN1_45S_50ntFlanks</td><td>10914</td><td>10946</td><td>33</td><td>CGCCGCGGCTGGACGAGGCGCCGCCGCCCCCCC</td><td>299.1</td><td>-6.4</td><td>1.5</td><td>-4.2</td><td>2.4E-05</td><td>9.4E-05</td><td>BT20/MB468</td><td>MP</td></tr>
<tr><td>>hg38_21_+_8433172_8446622_RNA45SN1_45S_50ntFlanks@11782.11804.23|GCATGAATGGATGAACGAGATTC</td><td>>hg38_21_+_8433172_8446622_RNA45SN1_45S_50ntFlanks</td><td>11782</td><td>11804</td><td>23</td><td>GCATGAATGGATGAACGAGATTC</td><td>466.8</td><td>-4.7</td><td>1.2</td><td>-4.0</td><td>7.0E-05</td><td>9.4E-05</td><td>MB231/MB468</td><td>NUC</td></tr>
<tr><td>>MI0001445|hsa-mir-423&WithFlank&17|+|30117073|30117178@23.42.20&[MIMAT0004748&hsa-miR-423-5p&offsets|0|-3;m-141&17|+|30117095|30117117&offsets|0|-3]|TGAGGGGCAGAGAGCGAGAC</td><td>>MI0001445|hsa-mir-423&WithFlank&17|+|30117073|30117178</td><td>23</td><td>42</td><td>20</td><td>TGAGGGGCAGAGAGCGAGAC</td><td>341.6</td><td>7.3</td><td>1.7</td><td>4.2</td><td>2.4E-05</td><td>9.4E-05</td><td>BT20/MB468</td><td>MP</td></tr>
<tr><td>>hg38_21_+_8433172_8446622_RNA45SN1_45S_50ntFlanks@11589.11613.25|CGGACCAGGGGAATCCGACTGTTTA</td><td>>hg38_21_+_8433172_8446622_RNA45SN1_45S_50ntFlanks</td><td>11589</td><td>11613</td><td>25</td><td>CGGACCAGGGGAATCCGACTGTTTA</td><td>177.9</td><td>-4.8</td><td>1.2</td><td>-4.0</td><td>5.9E-05</td><td>9.4E-05</td><td>MB231/MB468</td><td>MP</td></tr>
<tr><td>>hg38_21_+_8433172_8446622_RNA45SN1_45S_50ntFlanks@5071.5104.34|TTAATTCCGATAACGAACGAGACTCTGGCATGCT</td><td>>hg38_21_+_8433172_8446622_RNA45SN1_45S_50ntFlanks</td><td>5071</td><td>5104</td><td>34</td><td>TTAATTCCGATAACGAACGAGACTCTGGCATGCT</td><td>54.7</td><td>-9.6</td><td>2.4</td><td>-4.0</td><td>5.9E-05</td><td>9.4E-05</td><td>MB231/MB468</td><td>MP</td></tr>
<tr><td>>hg38_21_+_8433172_8446622_RNA45SN1_45S_50ntFlanks@11562.11587.26|CGCCTAGCAGCCGACTTAGAACTGGT</td><td>>hg38_21_+_8433172_8446622_RNA45SN1_45S_50ntFlanks</td><td>11562</td><td>11587</td><td>26</td><td>CGCCTAGCAGCCGACTTAGAACTGGT</td><td>105.0</td><td>-6.0</td><td>1.5</td><td>-4.0</td><td>5.9E-05</td><td>9.4E-05</td><td>MB231/MB468</td><td>MP</td></tr>
<tr><td>>MI0000061|hsa-let-7a-2&WithFlank&11|-|122146516|122146599@11.31.21;MI0000062|hsa-let-7a-3&WithFlank&22|+|46112743|46112828@10.30.21;MI0000060|hsa-let-7a-1&WithFlank&9|+|94175951|94176042@12.32.21&[MIMAT0000062&hsa-let-7a-5p&offsets|0|-1;m-6&11|-|122146568|122146589&offsets|0|-1];[MIMAT0000062_1&hsa-let-7a-5p&offsets|0|-1;m-5&22|+|46112752|46112773&offsets|0|-1];[MIMAT0000062_2&hsa-let-7a-5p&offsets|0|-1;m-4&9|+|94175962|94175983&offsets|0|-1]|TGAGGTAGTAGGTTGTATAGT</td><td>>MI0000061|hsa-let-7a-2&WithFlank&11|-|122146516|122146599</td><td>11</td><td>31</td><td>21</td><td>TGAGGTAGTAGGTTGTATAGT</td><td>1770.3</td><td>3.3</td><td>0.8</td><td>4.2</td><td>2.4E-05</td><td>9.4E-05</td><td>BT20/MB468</td><td>MP</td></tr>
<tr><td>>hg38_21_+_8433172_8446622_RNA45SN1_45S_50ntFlanks@10392.10426.35|ACAGCAGTTGAACATGGGTCAGTCGGTCCTGAGAG</td><td>>hg38_21_+_8433172_8446622_RNA45SN1_45S_50ntFlanks</td><td>10392</td><td>10426</td><td>35</td><td>ACAGCAGTTGAACATGGGTCAGTCGGTCCTGAGAG</td><td>564.2</td><td>-4.6</td><td>1.1</td><td>-4.0</td><td>6.6E-05</td><td>9.4E-05</td><td>MB231/MB468</td><td>CYTO</td></tr>
<tr><td>>hg38_21_+_8433172_8446622_RNA45SN1_45S_50ntFlanks@10684.10706.23|CGGCGGCGTCCGGTGAGCTCTCG</td><td>>hg38_21_+_8433172_8446622_RNA45SN1_45S_50ntFlanks</td><td>10684</td><td>10706</td><td>23</td><td>CGGCGGCGTCCGGTGAGCTCTCG</td><td>5127.1</td><td>-5.3</td><td>1.3</td><td>-4.0</td><td>6.1E-05</td><td>9.4E-05</td><td>BT20/MB468</td><td>CYTO</td></tr>
<tr><td>>MI0000088|hsa-mir-30a&WithFlank&6|-|71403545|71403627@53.70.18(+1U)&[MIMAT0000088&hsa-miR-30a-3p&offsets|0|-4(+1U);m-38&6|-|71403554|71403575&offsets|0|-4(+1U)]|CTTTCAGTCGGATGTTTGT</td><td>>MI0000088|hsa-mir-30a&WithFlank&6|-|71403545|71403627</td><td>53</td><td>70</td><td>18(+1U)</td><td>CTTTCAGTCGGATGTTTGT</td><td>664.2</td><td>5.2</td><td>1.3</td><td>4.0</td><td>5.9E-05</td><td>9.4E-05</td><td>MB231/MB468</td><td>MP</td></tr>
<tr><td>>trna10_AspGTC_12_-_125424193_125424264@37.74.38,trna12_AspGTC_12_-_125411891_125411962@37.74.38,trna144_AspGTC_6_-_27551236_27551307@37.74.38,trna38_AspGTC_17_-_8125556_8125627@37.74.38,trna45_AspGTC_6_+_27447453_27447524@37.74.38,trna48_AspGTC_6_+_27471523_27471594@37.74.38,trna4_AspGTC_12_+_96429799_96429870@37.74.38,trna69_AspGTC_1_-_161440205_161440276@37.74.38,trna72_AspGTC_1_-_161432824_161432895@37.74.38,trna75_AspGTC_1_-_161425414_161425485@37.74.38,trna78_AspGTC_1_-_161418033_161418104@37.74.38,trna81_AspGTC_1_-_161410615_161410686@37.74.38|ACGCGGGAGACCGGGGTTCGATTCCCCGACGGGGAGCC</td><td>>trna10_AspGTC_12_-_125424193_125424264</td><td>37</td><td>74</td><td>38</td><td>ACGCGGGAGACCGGGGTTCGATTCCCCGACGGGGAGCC</td><td>269.3</td><td>5.6</td><td>1.4</td><td>4.0</td><td>5.9E-05</td><td>9.4E-05</td><td>MB231/MB468</td><td>MP</td></tr>
<tr><td>>hg38_21_+_8433172_8446622_RNA45SN1_45S_50ntFlanks@9597.9634.38|AGGGGCGAAAGACTAATCGAACCATCTAGTAGCTGGTT</td><td>>hg38_21_+_8433172_8446622_RNA45SN1_45S_50ntFlanks</td><td>9597</td><td>9634</td><td>38</td><td>AGGGGCGAAAGACTAATCGAACCATCTAGTAGCTGGTT</td><td>82.6</td><td>9.7</td><td>2.2</td><td>4.5</td><td>8.4E-06</td><td>9.4E-05</td><td>BT20/MB231</td><td>MP</td></tr>
<tr><td>>MI0000065|hsa-let-7d&WithFlank&9|+|94178828|94178926@14.34.21(+1C)&[MIMAT0000065&hsa-let-7d-5p&offsets|0|-1(+1C);m-90&9|+|94178841|94178862&offsets|0|-1(+1C)]|AGAGGTAGTAGGTTGCATAGTC</td><td>>MI0000065|hsa-let-7d&WithFlank&9|+|94178828|94178926</td><td>14</td><td>34</td><td>21(+1C)</td><td>AGAGGTAGTAGGTTGCATAGTC</td><td>867.9</td><td>4.1</td><td>1.0</td><td>4.0</td><td>5.9E-05</td><td>9.4E-05</td><td>MB231/MB468</td><td>MP</td></tr>
<tr><td>>hg38_21_+_8433172_8446622_RNA45SN1_45S_50ntFlanks@4892.4912.21|AAACTTAAAGGAATTGACGGA</td><td>>hg38_21_+_8433172_8446622_RNA45SN1_45S_50ntFlanks</td><td>4892</td><td>4912</td><td>21</td><td>AAACTTAAAGGAATTGACGGA</td><td>104.2</td><td>-2.8</td><td>0.7</td><td>-4.0</td><td>5.9E-05</td><td>9.4E-05</td><td>MB231/MB468</td><td>MP</td></tr>
<tr><td>>hg38_21_+_8433172_8446622_RNA45SN1_45S_50ntFlanks@8380.8398.19|CAAGAGGGCGTGAAACCGT</td><td>>hg38_21_+_8433172_8446622_RNA45SN1_45S_50ntFlanks</td><td>8380</td><td>8398</td><td>19</td><td>CAAGAGGGCGTGAAACCGT</td><td>298.4</td><td>-4.9</td><td>1.2</td><td>-4.0</td><td>5.9E-05</td><td>9.4E-05</td><td>MB231/MB468</td><td>MP</td></tr>
<tr><td>>hg38_21_+_8433172_8446622_RNA45SN1_45S_50ntFlanks@3900.3934.35|CTTCGCGGGGGGGATGCGTGCATTTATCAGATCAA</td><td>>hg38_21_+_8433172_8446622_RNA45SN1_45S_50ntFlanks</td><td>3900</td><td>3934</td><td>35</td><td>CTTCGCGGGGGGGATGCGTGCATTTATCAGATCAA</td><td>41.9</td><td>-8.5</td><td>2.0</td><td>-4.2</td><td>2.5E-05</td><td>9.5E-05</td><td>BT20/MB468</td><td>MP</td></tr>
<tr><td>>hg38_21_+_8433172_8446622_RNA45SN1_45S_50ntFlanks@5444.5467.24|CTCGGATCGGCCCCGCCGGGGTCG</td><td>>hg38_21_+_8433172_8446622_RNA45SN1_45S_50ntFlanks</td><td>5444</td><td>5467</td><td>24</td><td>CTCGGATCGGCCCCGCCGGGGTCG</td><td>724.1</td><td>-5.0</td><td>1.3</td><td>-4.0</td><td>7.0E-05</td><td>9.5E-05</td><td>MB231/MB468</td><td>NUC</td></tr>
<tr><td>>hg38_MT_+_1621_3279_RNR2_16S_50ntFlanks@113.145.33|CCAAATAAAGTATAGGCGATAGAAATTGAAACC</td><td>>hg38_MT_+_1621_3279_RNR2_16S_50ntFlanks</td><td>113</td><td>145</td><td>33</td><td>CCAAATAAAGTATAGGCGATAGAAATTGAAACC</td><td>137.0</td><td>10.8</td><td>2.6</td><td>4.2</td><td>2.5E-05</td><td>9.5E-05</td><td>BT20/MB468</td><td>MP</td></tr>
<tr><td>>hg38_21_+_8433172_8446622_RNA45SN1_45S_50ntFlanks@9198.9225.28|GTTCTCTCGGGGCCACGCGCGCGTCCCC</td><td>>hg38_21_+_8433172_8446622_RNA45SN1_45S_50ntFlanks</td><td>9198</td><td>9225</td><td>28</td><td>GTTCTCTCGGGGCCACGCGCGCGTCCCC</td><td>634.1</td><td>-5.8</td><td>1.4</td><td>-4.0</td><td>5.2E-05</td><td>9.5E-05</td><td>BT20/MB468</td><td>MITO</td></tr>
<tr><td>>hg38_21_+_8433172_8446622_RNA45SN1_45S_50ntFlanks@11784.11803.20|ATGAATGGATGAACGAGATT</td><td>>hg38_21_+_8433172_8446622_RNA45SN1_45S_50ntFlanks</td><td>11784</td><td>11803</td><td>20</td><td>ATGAATGGATGAACGAGATT</td><td>1692.0</td><td>-4.0</td><td>0.9</td><td>-4.2</td><td>2.5E-05</td><td>9.5E-05</td><td>BT20/MB468</td><td>MP</td></tr>
<tr><td>>hg38_21_+_8433172_8446622_RNA45SN1_45S_50ntFlanks@3811.3834.24|AGTTATGGTTCCTTTGGTCGCTCG</td><td>>hg38_21_+_8433172_8446622_RNA45SN1_45S_50ntFlanks</td><td>3811</td><td>3834</td><td>24</td><td>AGTTATGGTTCCTTTGGTCGCTCG</td><td>170.2</td><td>-2.4</td><td>0.6</td><td>-4.0</td><td>6.6E-05</td><td>9.5E-05</td><td>MB231/MB468</td><td>CYTO</td></tr>
<tr><td>>hg38_21_+_8433172_8446622_RNA45SN1_45S_50ntFlanks@12996.13013.18|TCGCTGCGATCTATTGAA</td><td>>hg38_21_+_8433172_8446622_RNA45SN1_45S_50ntFlanks</td><td>12996</td><td>13013</td><td>18</td><td>TCGCTGCGATCTATTGAA</td><td>3393.4</td><td>3.1</td><td>0.8</td><td>4.0</td><td>6.2E-05</td><td>9.5E-05</td><td>BT20/MB468</td><td>CYTO</td></tr>
<tr><td>>hg38_21_+_8433172_8446622_RNA45SN1_45S_50ntFlanks@10395.10439.45|GCAGTTGAACATGGGTCAGTCGGTCCTGAGAGATGGGCGAGCGCC</td><td>>hg38_21_+_8433172_8446622_RNA45SN1_45S_50ntFlanks</td><td>10395</td><td>10439</td><td>45</td><td>GCAGTTGAACATGGGTCAGTCGGTCCTGAGAGATGGGCGAGCGCC</td><td>31.7</td><td>8.7</td><td>2.1</td><td>4.2</td><td>2.5E-05</td><td>9.5E-05</td><td>BT20/MB468</td><td>MP</td></tr>
<tr><td>>trna10_AspGTC_12_-_125424193_125424264@36.70.35,trna12_AspGTC_12_-_125411891_125411962@36.70.35,trna144_AspGTC_6_-_27551236_27551307@36.70.35,trna38_AspGTC_17_-_8125556_8125627@36.70.35,trna45_AspGTC_6_+_27447453_27447524@36.70.35,trna48_AspGTC_6_+_27471523_27471594@36.70.35,trna4_AspGTC_12_+_96429799_96429870@36.70.35,trna69_AspGTC_1_-_161440205_161440276@36.70.35,trna72_AspGTC_1_-_161432824_161432895@36.70.35,trna75_AspGTC_1_-_161425414_161425485@36.70.35,trna78_AspGTC_1_-_161418033_161418104@36.70.35,trna81_AspGTC_1_-_161410615_161410686@36.70.35|CACGCGGGAGACCGGGGTTCGATTCCCCGACGGGG</td><td>>trna10_AspGTC_12_-_125424193_125424264</td><td>36</td><td>70</td><td>35</td><td>CACGCGGGAGACCGGGGTTCGATTCCCCGACGGGG</td><td>839.9</td><td>4.1</td><td>1.0</td><td>4.0</td><td>6.6E-05</td><td>9.5E-05</td><td>MB231/MB468</td><td>CYTO</td></tr>
<tr><td>>hg38_21_+_8433172_8446622_RNA45SN1_45S_50ntFlanks@4049.4066.18|TCGAACGTCTGCCCTATC</td><td>>hg38_21_+_8433172_8446622_RNA45SN1_45S_50ntFlanks</td><td>4049</td><td>4066</td><td>18</td><td>TCGAACGTCTGCCCTATC</td><td>109.5</td><td>-4.4</td><td>1.1</td><td>-4.0</td><td>5.9E-05</td><td>9.5E-05</td><td>MB231/MB468</td><td>MP</td></tr>
<tr><td>>MI0000542|hsa-mir-320a&WithFlank&8|-|22244960|22245043@48.70.23(+1C)&[MIMAT0000510&hsa-miR-320a-3p&offsets|0|+1(+1C);m-60&8|-|22244975|22244996&offsets|0|+1(+1C)]|AAAAGCTGGGTTGAGAGGGCGAAC</td><td>>MI0000542|hsa-mir-320a&WithFlank&8|-|22244960|22245043</td><td>48</td><td>70</td><td>23(+1C)</td><td>AAAAGCTGGGTTGAGAGGGCGAAC</td><td>3113.9</td><td>3.6</td><td>0.8</td><td>4.5</td><td>8.0E-06</td><td>9.5E-05</td><td>BT20/MB231</td><td>CYTO</td></tr>
<tr><td>>hg38_21_+_8433172_8446622_RNA45SN1_45S_50ntFlanks@7978.8016.39|GACCTCAGATCAGACGTGGCGACCCGCTGAATTTAAGCA</td><td>>hg38_21_+_8433172_8446622_RNA45SN1_45S_50ntFlanks</td><td>7978</td><td>8016</td><td>39</td><td>GACCTCAGATCAGACGTGGCGACCCGCTGAATTTAAGCA</td><td>818.9</td><td>-4.6</td><td>1.0</td><td>-4.5</td><td>8.0E-06</td><td>9.5E-05</td><td>BT20/MB231</td><td>CYTO</td></tr>
<tr><td>>hg38_21_+_8433172_8446622_RNA45SN1_45S_50ntFlanks@11578.11605.28|TAGAACTGGTGCGGACCAGGGGAATCCG</td><td>>hg38_21_+_8433172_8446622_RNA45SN1_45S_50ntFlanks</td><td>11578</td><td>11605</td><td>28</td><td>TAGAACTGGTGCGGACCAGGGGAATCCG</td><td>220.8</td><td>-2.2</td><td>0.6</td><td>-4.0</td><td>5.2E-05</td><td>9.5E-05</td><td>BT20/MB468</td><td>MITO</td></tr>
<tr><td>>hg38_21_+_8433172_8446622_RNA45SN1_45S_50ntFlanks@11570.11591.22|AGCCGACTTAGAACTGGTGCGG</td><td>>hg38_21_+_8433172_8446622_RNA45SN1_45S_50ntFlanks</td><td>11570</td><td>11591</td><td>22</td><td>AGCCGACTTAGAACTGGTGCGG</td><td>308.6</td><td>-4.4</td><td>1.1</td><td>-4.0</td><td>7.1E-05</td><td>9.5E-05</td><td>MB231/MB468</td><td>NUC</td></tr>
<tr><td>>hg38_21_+_8433172_8446622_RNA45SN1_45S_50ntFlanks@5493.5516.24|GAAGACGGTCGAACTTGACTATCT</td><td>>hg38_21_+_8433172_8446622_RNA45SN1_45S_50ntFlanks</td><td>5493</td><td>5516</td><td>24</td><td>GAAGACGGTCGAACTTGACTATCT</td><td>1212.8</td><td>3.4</td><td>0.8</td><td>4.4</td><td>9.7E-06</td><td>9.5E-05</td><td>BT20/MB231</td><td>MITO</td></tr>
<tr><td>>hg38_21_+_8433172_8446622_RNA45SN1_45S_50ntFlanks@12997.13035.39|CGCTGCGATCTATTGAAAGTCAGCCCTCGACACAAGGGT</td><td>>hg38_21_+_8433172_8446622_RNA45SN1_45S_50ntFlanks</td><td>12997</td><td>13035</td><td>39</td><td>CGCTGCGATCTATTGAAAGTCAGCCCTCGACACAAGGGT</td><td>222.2</td><td>5.8</td><td>1.5</td><td>4.0</td><td>7.1E-05</td><td>9.5E-05</td><td>MB231/MB468</td><td>NUC</td></tr>
</table>

<table>
<tr><td>>hg38_21_+_8433172_8446622_RNA45SN1_45S_50ntFlanks@12909.12933.25|CCGCACGTTCGTGGGGAACCTGGCG</td><td>>hg38_21_+_8433172_8446622_RNA45SN1_45S_50ntFlanks</td><td>12909</td><td>12933</td><td>25</td><td>CCGCACGTTCGTGGGGAACCTGGCG</td><td>160.1</td><td>-4.5</td><td>1.1</td><td>-4.0</td><td>7.1E-05</td><td>9.5E-05</td><td>MB231/MB468</td><td>NUC</td></tr>
<tr><td>>MI0000477|hsa-mir-146a&WithFlank&5|+|160485346|160485456@27.48.22(+1U)&[MIMAT0000449&hsa-miR-146a-5p&offsets|0|0(+1U);m-129&5|+|160485372|160485393&offsets|0|0(+1U)]|TGAGAACTGAATTCCATGGGTTT</td><td>>MI0000477|hsa-mir-146a&WithFlank&5|+|160485346|160485456</td><td>27</td><td>48</td><td>22(+1U)</td><td>TGAGAACTGAATTCCATGGGTTT</td><td>1354.0</td><td>-9.5</td><td>2.1</td><td>-4.4</td><td>9.8E-06</td><td>9.5E-05</td><td>BT20/MB231</td><td>MITO</td></tr>
<tr><td>>hg38_21_+_8433172_8446622_RNA45SN1_45S_50ntFlanks@12502.12521.20|TTAGACCGTCGTGAGACAGG</td><td>>hg38_21_+_8433172_8446622_RNA45SN1_45S_50ntFlanks</td><td>12502</td><td>12521</td><td>20</td><td>TTAGACCGTCGTGAGACAGG</td><td>40.5</td><td>-8.4</td><td>2.0</td><td>-4.2</td><td>2.5E-05</td><td>9.6E-05</td><td>BT20/MB468</td><td>MP</td></tr>
<tr><td>>hg38_21_+_8433172_8446622_RNA45SN1_45S_50ntFlanks@8379.8400.22|TCAAGAGGGCGTGAAACCGTTA</td><td>>hg38_21_+_8433172_8446622_RNA45SN1_45S_50ntFlanks</td><td>8379</td><td>8400</td><td>22</td><td>TCAAGAGGGCGTGAAACCGTTA</td><td>112.5</td><td>-5.2</td><td>1.3</td><td>-4.0</td><td>5.2E-05</td><td>9.6E-05</td><td>BT20/MB468</td><td>MITO</td></tr>
<tr><td>>hg38_21_+_8433172_8446622_RNA45SN1_45S_50ntFlanks@5086.5110.25|AACGAGACTCTGGCATGCTAACTAG</td><td>>hg38_21_+_8433172_8446622_RNA45SN1_45S_50ntFlanks</td><td>5086</td><td>5110</td><td>25</td><td>AACGAGACTCTGGCATGCTAACTAG</td><td>459.6</td><td>-4.4</td><td>1.1</td><td>-4.0</td><td>6.0E-05</td><td>9.6E-05</td><td>MB231/MB468</td><td>MP</td></tr>
<tr><td>>am_trnaMT_ValTAC_MT_+_1602_1670@35.67.33|ACTTAGGAGATTTCAACTTAACTTGACCGCTCT</td><td>>am_trnaMT_ValTAC_MT_+_1602_1670</td><td>35</td><td>67</td><td>33</td><td>ACTTAGGAGATTTCAACTTAACTTGACCGCTCT</td><td>145.8</td><td>10.9</td><td>2.6</td><td>4.2</td><td>2.5E-05</td><td>9.6E-05</td><td>BT20/MB468</td><td>MP</td></tr>
<tr><td>>MI0000294|hsa-mir-218-1&WithFlank&4|+|20528269|20528390@31.52.22(+1C);MI0000295|hsa-mir-218-2&WithFlank&5|-|168768140|168768261@31.52.22(+1C)&[MIMAT0000275&hsa-miR-218-5p&offsets|0|+1(+1C);m-185&4|+|20528299|20528319&offsets|0|+1(+1C)];[MIMAT0000275_1&hsa-miR-218-5p&offsets|0|+1(+1C);m-187&5|-|168768211|168768231&offsets|0|+1(+1C)]|TTGTGCTTGATCTAACCATGTGC</td><td>>MI0000294|hsa-mir-218-1&WithFlank&4|+|20528269|20528390</td><td>31</td><td>52</td><td>22(+1C)</td><td>TTGTGCTTGATCTAACCATGTGC</td><td>30.1</td><td>8.1</td><td>2.0</td><td>4.0</td><td>6.0E-05</td><td>9.6E-05</td><td>MB231/MB468</td><td>MP</td></tr>
<tr><td>>hg38_21_+_8433172_8446622_RNA45SN1_45S_50ntFlanks@5078.5108.31|CGATAACGAACGAGACTCTGGCATGCTAACT</td><td>>hg38_21_+_8433172_8446622_RNA45SN1_45S_50ntFlanks</td><td>5078</td><td>5108</td><td>31</td><td>CGATAACGAACGAGACTCTGGCATGCTAACT</td><td>214.3</td><td>-6.9</td><td>1.7</td><td>-4.0</td><td>6.0E-05</td><td>9.6E-05</td><td>MB231/MB468</td><td>MP</td></tr>
<tr><td>>hg38_MT_+_598_1651_RNR1_12S_50ntFlanks@818.840.23|GAAGGTGGATTTAGCAGTAAACT</td><td>>hg38_MT_+_598_1651_RNR1_12S_50ntFlanks</td><td>818</td><td>840</td><td>23</td><td>GAAGGTGGATTTAGCAGTAAACT</td><td>212.7</td><td>7.9</td><td>1.9</td><td>4.2</td><td>2.5E-05</td><td>9.6E-05</td><td>BT20/MB468</td><td>MP</td></tr>
<tr><td>>hg38_21_+_8433172_8446622_RNA45SN1_45S_50ntFlanks@6661.6696.36|CGGTGGATCACTCGGCTCGTGCGTCGATGAAGAACG</td><td>>hg38_21_+_8433172_8446622_RNA45SN1_45S_50ntFlanks</td><td>6661</td><td>6696</td><td>36</td><td>CGGTGGATCACTCGGCTCGTGCGTCGATGAAGAACG</td><td>846.7</td><td>2.5</td><td>0.6</td><td>4.0</td><td>7.1E-05</td><td>9.6E-05</td><td>MB231/MB468</td><td>NUC</td></tr>
<tr><td>>hg38_21_+_8433172_8446622_RNA45SN1_45S_50ntFlanks@11600.11621.22|AATCCGACTGTTTAATTAAAAC</td><td>>hg38_21_+_8433172_8446622_RNA45SN1_45S_50ntFlanks</td><td>11600</td><td>11621</td><td>22</td><td>AATCCGACTGTTTAATTAAAAC</td><td>96.0</td><td>7.4</td><td>1.8</td><td>4.2</td><td>2.5E-05</td><td>9.6E-05</td><td>BT20/MB468</td><td>MP</td></tr>
<tr><td>>hg38_21_+_8433172_8446622_RNA45SN1_45S_50ntFlanks@5074.5097.24|ATTCCGATAACGAACGAGACTCTG</td><td>>hg38_21_+_8433172_8446622_RNA45SN1_45S_50ntFlanks</td><td>5074</td><td>5097</td><td>24</td><td>ATTCCGATAACGAACGAGACTCTG</td><td>670.0</td><td>-5.8</td><td>1.5</td><td>-4.0</td><td>7.4E-05</td><td>9.6E-05</td><td>BT20/MB468</td><td>NUC</td></tr>
<tr><td>>hg38_21_+_8433172_8446622_RNA45SN1_45S_50ntFlanks@5188.5205.18|ATAACAGGTCTGTGATGC</td><td>>hg38_21_+_8433172_8446622_RNA45SN1_45S_50ntFlanks</td><td>5188</td><td>5205</td><td>18</td><td>ATAACAGGTCTGTGATGC</td><td>435.9</td><td>-4.8</td><td>1.2</td><td>-4.0</td><td>7.2E-05</td><td>9.6E-05</td><td>MB231/MB468</td><td>MITO</td></tr>
<tr><td>>trnaMT_MetCAT_MT_+_4402_4469@34.71.38|ACCCCGAAAATGTTGGTTATACCCTTCCCGTACTACCA</td><td>>trnaMT_MetCAT_MT_+_4402_4469</td><td>34</td><td>71</td><td>38</td><td>ACCCCGAAAATGTTGGTTATACCCTTCCCGTACTACCA</td><td>637.8</td><td>-4.3</td><td>1.0</td><td>-4.5</td><td>8.1E-06</td><td>9.6E-05</td><td>BT20/MB231</td><td>CYTO</td></tr>
<tr><td>>hg38_21_+_8433172_8446622_RNA45SN1_45S_50ntFlanks@4453.4473.21|TCGGCGCCCCCTCGATGCTCT</td><td>>hg38_21_+_8433172_8446622_RNA45SN1_45S_50ntFlanks</td><td>4453</td><td>4473</td><td>21</td><td>TCGGCGCCCCCTCGATGCTCT</td><td>743.6</td><td>-3.7</td><td>0.8</td><td>-4.5</td><td>8.1E-06</td><td>9.6E-05</td><td>BT20/MB231</td><td>CYTO</td></tr>
<tr><td>>trnaMT_SerGCT_MT_+_12207_12265@1.26.26|GAGAAAGCTCACAAGAACTGCTAACT</td><td>>trnaMT_SerGCT_MT_+_12207_12265</td><td>1</td><td>26</td><td>26</td><td>GAGAAAGCTCACAAGAACTGCTAACT</td><td>1213.2</td><td>4.1</td><td>1.0</td><td>4.0</td><td>6.3E-05</td><td>9.6E-05</td><td>BT20/MB468</td><td>CYTO</td></tr>
<tr><td>>hg38_21_+_8433172_8446622_RNA45SN1_45S_50ntFlanks@11564.11586.23|CCTAGCAGCCGACTTAGAACTGG</td><td>>hg38_21_+_8433172_8446622_RNA45SN1_45S_50ntFlanks</td><td>11564</td><td>11586</td><td>23</td><td>CCTAGCAGCCGACTTAGAACTGG</td><td>429.4</td><td>-4.1</td><td>1.0</td><td>-4.0</td><td>6.3E-05</td><td>9.6E-05</td><td>BT20/MB468</td><td>CYTO</td></tr>
<tr><td>>hg38_21_+_8433172_8446622_RNA45SN1_45S_50ntFlanks@12635.12659.25|AAGCTACCATCTGTGGGATTATGAC</td><td>>hg38_21_+_8433172_8446622_RNA45SN1_45S_50ntFlanks</td><td>12635</td><td>12659</td><td>25</td><td>AAGCTACCATCTGTGGGATTATGAC</td><td>401.4</td><td>-5.7</td><td>1.4</td><td>-4.0</td><td>6.3E-05</td><td>9.6E-05</td><td>BT20/MB468</td><td>CYTO</td></tr>
<tr><td>>MI0000734|hsa-mir-106b&WithFlank&7|-|100093987|100094080@58.75.18(+1U)&[MIMAT0004672&hsa-miR-106b-3p&offsets|0|-4(+1U);m-87&7|-|100094003|100094023&offsets|0|-3(+1U)]|CCGCACTGTGGGTACTTGT</td><td>>MI0000734|hsa-mir-106b&WithFlank&7|-|100093987|100094080</td><td>58</td><td>75</td><td>18(+1U)</td><td>CCGCACTGTGGGTACTTGT</td><td>37.0</td><td>8.4</td><td>2.1</td><td>4.0</td><td>6.0E-05</td><td>9.6E-05</td><td>MB231/MB468</td><td>MP</td></tr>
<tr><td>>am_trnaMT_ValTAC_MT_+_1602_1670@38.62.25|TAGGAGATTTCAACTTAACTTGACC</td><td>>am_trnaMT_ValTAC_MT_+_1602_1670</td><td>38</td><td>62</td><td>25</td><td>TAGGAGATTTCAACTTAACTTGACC</td><td>952.5</td><td>6.0</td><td>1.4</td><td>4.2</td><td>2.5E-05</td><td>9.6E-05</td><td>BT20/MB468</td><td>MP</td></tr>
<tr><td>>hg38_21_+_8433172_8446622_RNA45SN1_45S_50ntFlanks@9198.9222.25|GTTCTCTCGGGGCCACGCGCGCGTC</td><td>>hg38_21_+_8433172_8446622_RNA45SN1_45S_50ntFlanks</td><td>9198</td><td>9222</td><td>25</td><td>GTTCTCTCGGGGCCACGCGCGCGTC</td><td>1441.1</td><td>-7.0</td><td>1.7</td><td>-4.2</td><td>2.5E-05</td><td>9.6E-05</td><td>BT20/MB468</td><td>MP</td></tr>
<tr><td>>hg38_21_+_8433172_8446622_RNA45SN1_45S_50ntFlanks@12406.12435.30|TTTTTGATCCTTCGATGTCGGCTCTTCCTA</td><td>>hg38_21_+_8433172_8446622_RNA45SN1_45S_50ntFlanks</td><td>12406</td><td>12435</td><td>30</td><td>TTTTTGATCCTTCGATGTCGGCTCTTCCTA</td><td>150.7</td><td>-5.2</td><td>1.3</td><td>-4.0</td><td>6.3E-05</td><td>9.7E-05</td><td>BT20/MB468</td><td>CYTO</td></tr>
<tr><td>>hg38_21_+_8433172_8446622_RNA45SN1_45S_50ntFlanks@10845.10871.27|ACTTCGGGATAAGGATTGGCTCTAAGG</td><td>>hg38_21_+_8433172_8446622_RNA45SN1_45S_50ntFlanks</td><td>10845</td><td>10871</td><td>27</td><td>ACTTCGGGATAAGGATTGGCTCTAAGG</td><td>172.5</td><td>-1.8</td><td>0.4</td><td>-4.0</td><td>7.2E-05</td><td>9.7E-05</td><td>MB231/MB468</td><td>NUC</td></tr>
<tr><td>>am_MI0000786|hsa-mir-378a&WithFlank&5|+|149732819|149732896@49.69.21(+1A)&[MIMAT0000732&hsa-miR-378a-3p&offsets|0|-1(+1A);m-44&5|+|149732867|149732888&offsets|0|-1(+1A)]|ACTGGACTTGGAGTCAGAAGGA</td><td>>am_MI0000786|hsa-mir-378a&WithFlank&5|+|149732819|149732896</td><td>49</td><td>69</td><td>21(+1A)</td><td>ACTGGACTTGGAGTCAGAAGGA</td><td>675.4</td><td>3.7</td><td>0.9</td><td>4.0</td><td>6.8E-05</td><td>9.7E-05</td><td>MB231/MB468</td><td>CYTO</td></tr>
<tr><td>>hg38_21_+_8433172_8446622_RNA45SN1_45S_50ntFlanks@10362.10407.46|AAGGCCGAAGTGGAGAAGGGTTCCATGTGAACAGCAGTTGAACATG</td><td>>hg38_21_+_8433172_8446622_RNA45SN1_45S_50ntFlanks</td><td>10362</td><td>10407</td><td>46</td><td>AAGGCCGAAGTGGAGAAGGGTTCCATGTGAACAGCAGTTGAACATG</td><td>225.8</td><td>-4.0</td><td>1.0</td><td>-4.0</td><td>6.8E-05</td><td>9.7E-05</td><td>MB231/MB468</td><td>CYTO</td></tr>
<tr><td>>hg38_21_+_8433172_8446622_RNA45SN1_45S_50ntFlanks@8359.8394.36|AAGAACTTTGAAGAGAGAGTTCAAGAGGGCGTGAAA</td><td>>hg38_21_+_8433172_8446622_RNA45SN1_45S_50ntFlanks</td><td>8359</td><td>8394</td><td>36</td><td>AAGAACTTTGAAGAGAGAGTTCAAGAGGGCGTGAAA</td><td>83.0</td><td>9.7</td><td>2.2</td><td>4.4</td><td>8.6E-06</td><td>9.7E-05</td><td>BT20/MB231</td><td>MP</td></tr>
<tr><td>>MI0000737|hsa-mir-200a&WithFlank&1|+|1167857|1167958@60.82.23&[MIMAT0000682&hsa-miR-200a-3p&offsets|0|+1;m-142&1|+|1167916|1167938&offsets|0|0]|TAACACTGTCTGGTAACGATGTT</td><td>>MI0000737|hsa-mir-200a&WithFlank&1|+|1167857|1167958</td><td>60</td><td>82</td><td>23</td><td>TAACACTGTCTGGTAACGATGTT</td><td>1033.9</td><td>-4.2</td><td>1.1</td><td>-4.0</td><td>7.2E-05</td><td>9.7E-05</td><td>MB231/MB468</td><td>MITO</td></tr>
<tr><td>>hg38_21_+_8433172_8446622_RNA45SN1_45S_50ntFlanks@11321.11344.24|CACGTCTCGTCGCGCGCGCGTCCG</td><td>>hg38_21_+_8433172_8446622_RNA45SN1_45S_50ntFlanks</td><td>11321</td><td>11344</td><td>24</td><td>CACGTCTCGTCGCGCGCGCGTCCG</td><td>24053.7</td><td>-7.3</td><td>1.8</td><td>-4.0</td><td>7.5E-05</td><td>9.7E-05</td><td>BT20/MB468</td><td>NUC</td></tr>
<tr><td>>hg38_21_+_8433172_8446622_RNA45SN1_45S_50ntFlanks@9555.9581.27|GTAGCGGTCCTGACGTGCAAATCGGTC</td><td>>hg38_21_+_8433172_8446622_RNA45SN1_45S_50ntFlanks</td><td>9555</td><td>9581</td><td>27</td><td>GTAGCGGTCCTGACGTGCAAATCGGTC</td><td>317.5</td><td>3.1</td><td>0.8</td><td>4.0</td><td>6.8E-05</td><td>9.7E-05</td><td>MB231/MB468</td><td>CYTO</td></tr>
<tr><td>>MI0003685|hsa-mir-421&WithFlank&X|-|74218371|74218467@57.76.20&[MIMAT0003339&hsa-miR-421&offsets|+3|0;m-309&X|-|74218392|74218414&offsets|+3|0]|AACAGACATTAATTGGGCGC</td><td>>MI0003685|hsa-mir-421&WithFlank&X|-|74218371|74218467</td><td>57</td><td>76</td><td>20</td><td>AACAGACATTAATTGGGCGC</td><td>115.2</td><td>10.1</td><td>2.5</td><td>4.0</td><td>6.1E-05</td><td>9.7E-05</td><td>MB231/MB468</td><td>MP</td></tr>
<tr><td>>hg38_21_+_8433172_8446622_RNA45SN1_45S_50ntFlanks@9545.9569.25|GTGGAGGTCCGTAGCGGTCCTGACG</td><td>>hg38_21_+_8433172_8446622_RNA45SN1_45S_50ntFlanks</td><td>9545</td><td>9569</td><td>25</td><td>GTGGAGGTCCGTAGCGGTCCTGACG</td><td>326.7</td><td>-2.7</td><td>0.7</td><td>-4.0</td><td>5.3E-05</td><td>9.7E-05</td><td>BT20/MB468</td><td>MITO</td></tr>
<tr><td>>hg38_21_+_8433172_8446622_RNA45SN1_45S_50ntFlanks@10786.10825.40|AACAGCCTCTGGCATGTTGGAACAATGTAGGTAAGGGAAG</td><td>>hg38_21_+_8433172_8446622_RNA45SN1_45S_50ntFlanks</td><td>10786</td><td>10825</td><td>40</td><td>AACAGCCTCTGGCATGTTGGAACAATGTAGGTAAGGGAAG</td><td>408.3</td><td>-7.0</td><td>1.7</td><td>-4.2</td><td>2.5E-05</td><td>9.7E-05</td><td>BT20/MB468</td><td>MP</td></tr>
<tr><td>>hg38_21_+_8433172_8446622_RNA45SN1_45S_50ntFlanks@12646.12669.24|TGTGGGATTATGACTGAACGCCTC</td><td>>hg38_21_+_8433172_8446622_RNA45SN1_45S_50ntFlanks</td><td>12646</td><td>12669</td><td>24</td><td>TGTGGGATTATGACTGAACGCCTC</td><td>262.1</td><td>7.0</td><td>1.7</td><td>4.2</td><td>2.5E-05</td><td>9.7E-05</td><td>BT20/MB468</td><td>MP</td></tr>
<tr><td>>hg38_21_+_8433172_8446622_RNA45SN1_45S_50ntFlanks@9566.9595.30|GACGTGCAAATCGGTCGTCCGACCTGGGTA</td><td>>hg38_21_+_8433172_8446622_RNA45SN1_45S_50ntFlanks</td><td>9566</td><td>9595</td><td>30</td><td>GACGTGCAAATCGGTCGTCCGACCTGGGTA</td><td>378.4</td><td>-2.4</td><td>0.6</td><td>-4.0</td><td>6.3E-05</td><td>9.7E-05</td><td>BT20/MB468</td><td>CYTO</td></tr>
<tr><td>>trnaMT_SerGCT_MT_+_12207_12265@1.23.23|GAGAAAGCTCACAAGAACTGCTA</td><td>>trnaMT_SerGCT_MT_+_12207_12265</td><td>1</td><td>23</td><td>23</td><td>GAGAAAGCTCACAAGAACTGCTA</td><td>3881.9</td><td>5.2</td><td>1.3</td><td>4.0</td><td>5.3E-05</td><td>9.7E-05</td><td>BT20/MB468</td><td>MITO</td></tr>
<tr><td>>hg38_21_+_8433172_8446622_RNA45SN1_45S_50ntFlanks@8317.8356.40|CACGAGACCGATAGTCAACAAGTACCGTAAGGGAAAGTTG</td><td>>hg38_21_+_8433172_8446622_RNA45SN1_45S_50ntFlanks</td><td>8317</td><td>8356</td><td>40</td><td>CACGAGACCGATAGTCAACAAGTACCGTAAGGGAAAGTTG</td><td>89.9</td><td>9.8</td><td>2.2</td><td>4.4</td><td>8.6E-06</td><td>9.7E-05</td><td>BT20/MB231</td><td>MP</td></tr>
<tr><td>>hg38_21_+_8433172_8446622_RNA45SN1_45S_50ntFlanks@5484.5509.26|GAGCGCTGAGAAGACGGTCGAACTTG</td><td>>hg38_21_+_8433172_8446622_RNA45SN1_45S_50ntFlanks</td><td>5484</td><td>5509</td><td>26</td><td>GAGCGCTGAGAAGACGGTCGAACTTG</td><td>178.2</td><td>-2.3</td><td>0.6</td><td>-4.0</td><td>5.3E-05</td><td>9.7E-05</td><td>BT20/MB468</td><td>MITO</td></tr>
<tr><td>>hg38_21_+_8433172_8446622_RNA45SN1_45S_50ntFlanks@11211.11252.42|GCGACTCTGGACGCGAGCCGGGCCCTTCCCGTGGATCGCCCC</td><td>>hg38_21_+_8433172_8446622_RNA45SN1_45S_50ntFlanks</td><td>11211</td><td>11252</td><td>42</td><td>GCGACTCTGGACGCGAGCCGGGCCCTTCCCGTGGATCGCCCC</td><td>469.2</td><td>4.0</td><td>0.9</td><td>4.4</td><td>1.0E-05</td><td>9.7E-05</td><td>BT20/MB231</td><td>MITO</td></tr>
<tr><td>>am_trnaMT_PheGAA_MT_+_577_647@40.57.18|TGTTTAGACGGGCTCACA</td><td>>am_trnaMT_PheGAA_MT_+_577_647</td><td>40</td><td>57</td><td>18</td><td>TGTTTAGACGGGCTCACA</td><td>82.5</td><td>3.6</td><td>0.9</td><td>4.0</td><td>7.5E-05</td><td>9.7E-05</td><td>BT20/MB468</td><td>NUC</td></tr>
<tr><td>>hg38_21_+_8433172_8446622_RNA45SN1_45S_50ntFlanks@8115.8143.29|CGCGGGACATGTGGCGTACGGAAGACCCG</td><td>>hg38_21_+_8433172_8446622_RNA45SN1_45S_50ntFlanks</td><td>8115</td><td>8143</td><td>29</td><td>CGCGGGACATGTGGCGTACGGAAGACCCG</td><td>1272.1</td><td>-7.3</td><td>1.8</td><td>-4.0</td><td>7.5E-05</td><td>9.7E-05</td><td>BT20/MB468</td><td>NUC</td></tr>
<tr><td>>hg38_21_+_8433172_8446622_RNA45SN1_45S_50ntFlanks@9625.9650.26|GTAGCTGGTTCCCTCCGAAGTTTCCC</td><td>>hg38_21_+_8433172_8446622_RNA45SN1_45S_50ntFlanks</td><td>9625</td><td>9650</td><td>26</td><td>GTAGCTGGTTCCCTCCGAAGTTTCCC</td><td>125.9</td><td>-5.2</td><td>1.3</td><td>-4.0</td><td>5.3E-05</td><td>9.7E-05</td><td>BT20/MB468</td><td>MITO</td></tr>
<tr><td>>hg38_21_+_8433172_8446622_RNA45SN1_45S_50ntFlanks@12413.12434.22|TCCTTCGATGTCGGCTCTTCCT</td><td>>hg38_21_+_8433172_8446622_RNA45SN1_45S_50ntFlanks</td><td>12413</td><td>12434</td><td>22</td><td>TCCTTCGATGTCGGCTCTTCCT</td><td>829.1</td><td>-2.6</td><td>0.7</td><td>-4.0</td><td>7.3E-05</td><td>9.8E-05</td><td>MB231/MB468</td><td>NUC</td></tr>
<tr><td>>hg38_21_+_8433172_8446622_RNA45SN1_45S_50ntFlanks@10420.10443.24|CTGAGAGATGGGCGAGCGCCGTTC</td><td>>hg38_21_+_8433172_8446622_RNA45SN1_45S_50ntFlanks</td><td>10420</td><td>10443</td><td>24</td><td>CTGAGAGATGGGCGAGCGCCGTTC</td><td>111.4</td><td>-5.2</td><td>1.3</td><td>-4.0</td><td>5.3E-05</td><td>9.8E-05</td><td>BT20/MB468</td><td>MITO</td></tr>
<tr><td>>hg38_21_+_8433172_8446622_RNA45SN1_45S_50ntFlanks@12994.13039.46|CCTCGCTGCGATCTATTGAAAGTCAGCCCTCGACACAAGGGTTTGT</td><td>>hg38_21_+_8433172_8446622_RNA45SN1_45S_50ntFlanks</td><td>12994</td><td>13039</td><td>46</td><td>CCTCGCTGCGATCTATTGAAAGTCAGCCCTCGACACAAGGGTTTGT</td><td>4359.4</td><td>-4.8</td><td>1.2</td><td>-4.0</td><td>7.5E-05</td><td>9.8E-05</td><td>BT20/MB468</td><td>NUC</td></tr>
<tr><td>>MI0000433|hsa-let-7g&WithFlank&3|-|52268272|52268367@11.28.18&[MIMAT0000414&hsa-let-7g-5p&offsets|0|-4;m-58&3|-|52268336|52268357&offsets|0|-4]|TGAGGTAGTAGTTTGTAC</td><td>>MI0000433|hsa-let-7g&WithFlank&3|-|52268272|52268367</td><td>11</td><td>28</td><td>18</td><td>TGAGGTAGTAGTTTGTAC</td><td>917.8</td><td>4.1</td><td>1.0</td><td>4.2</td><td>2.5E-05</td><td>9.8E-05</td><td>BT20/MB468</td><td>MP</td></tr>
<tr><td>>hg38_21_+_8433172_8446622_RNA45SN1_45S_50ntFlanks@11593.11621.29|CCAGGGGAATCCGACTGTTTAATTAAAAC</td><td>>hg38_21_+_8433172_8446622_RNA45SN1_45S_50ntFlanks</td><td>11593</td><td>11621</td><td>29</td><td>CCAGGGGAATCCGACTGTTTAATTAAAAC</td><td>365.1</td><td>-6.6</td><td>1.7</td><td>-4.0</td><td>7.5E-05</td><td>9.8E-05</td><td>BT20/MB468</td><td>NUC</td></tr>
<tr><td>>hg38_21_+_8433172_8446622_RNA45SN1_45S_50ntFlanks@10409.10432.24|GTCAGTCGGTCCTGAGAGATGGGC</td><td>>hg38_21_+_8433172_8446622_RNA45SN1_45S_50ntFlanks</td><td>10409</td><td>10432</td><td>24</td><td>GTCAGTCGGTCCTGAGAGATGGGC</td><td>504.2</td><td>-5.1</td><td>1.3</td><td>-4.0</td><td>7.5E-05</td><td>9.8E-05</td><td>BT20/MB468</td><td>NUC</td></tr>
<tr><td>>hg38_21_+_8433172_8446622_RNA45SN1_45S_50ntFlanks@10394.10430.37|AGCAGTTGAACATGGGTCAGTCGGTCCTGAGAGATGG</td><td>>hg38_21_+_8433172_8446622_RNA45SN1_45S_50ntFlanks</td><td>10394</td><td>10430</td><td>37</td><td>AGCAGTTGAACATGGGTCAGTCGGTCCTGAGAGATGG</td><td>97.2</td><td>-5.0</td><td>1.2</td><td>-4.0</td><td>6.1E-05</td><td>9.8E-05</td><td>MB231/MB468</td><td>MP</td></tr>
<tr><td>>hg38_21_+_8433172_8446622_RNA45SN1_45S_50ntFlanks@4145.4175.31|CCTGAGAAACGGCTACCACATCCAAGGAAGG</td><td>>hg38_21_+_8433172_8446622_RNA45SN1_45S_50ntFlanks</td><td>4145</td><td>4175</td><td>31</td><td>CCTGAGAAACGGCTACCACATCCAAGGAAGG</td><td>60.6</td><td>-9.8</td><td>2.4</td><td>-4.0</td><td>6.1E-05</td><td>9.8E-05</td><td>MB231/MB468</td><td>MP</td></tr>
<tr><td>>hg38_21_+_8433172_8446622_RNA45SN1_45S_50ntFlanks@6678.6696.19|CGTGCGTCGATGAAGAACG</td><td>>hg38_21_+_8433172_8446622_RNA45SN1_45S_50ntFlanks</td><td>6678</td><td>6696</td><td>19</td><td>CGTGCGTCGATGAAGAACG</td><td>225.6</td><td>-4.6</td><td>1.1</td><td>-4.0</td><td>6.1E-05</td><td>9.8E-05</td><td>MB231/MB468</td><td>MP</td></tr>
<tr><td>>hg38_21_+_8433172_8446622_RNA45SN1_45S_50ntFlanks@12406.12429.24|TTTTTGATCCTTCGATGTCGGCTC</td><td>>hg38_21_+_8433172_8446622_RNA45SN1_45S_50ntFlanks</td><td>12406</td><td>12429</td><td>24</td><td>TTTTTGATCCTTCGATGTCGGCTC</td><td>166.3</td><td>-3.6</td><td>0.9</td><td>-4.0</td><td>6.4E-05</td><td>9.8E-05</td><td>BT20/MB468</td><td>CYTO</td></tr>
</table>

<table>
<tr><td>>hg38_21_+_8433172_8446622_RNA45SN1_45S_50ntFlanks@6661.6697.37|CGGTGGATCACTCGGCTCGTGCGTCGATGAAGAACGC</td><td>>hg38_21_+_8433172_8446622_RNA45SN1_45S_50ntFlanks</td><td>6661</td><td>6697</td><td>37</td><td>CGGTGGATCACTCGGCTCGTGCGTCGATGAAGAACGC</td><td>770.1</td><td>-3.1</td><td>0.8</td><td>-4.0</td><td>6.1E-05</td><td>9.8E-05</td><td>MB231/MB468</td><td>MP</td></tr>
<tr><td>>MI0000446|hsa-mir-125b-1&WithFlank&11|-|122099751|122099850@61.82.22(+1A)&[MIMAT0004592&hsa-miR-125b-1-3p&offsets|0|0(+1A);m-485&11|-|122099769|122099790&offsets|0|0(+1A)]|ACGGGTTAGGCTCTTGGGAGCTA</td><td>>MI0000446|hsa-mir-125b-1&WithFlank&11|-|122099751|122099850</td><td>61</td><td>82</td><td>22(+1A)</td><td>ACGGGTTAGGCTCTTGGGAGCTA</td><td>31.4</td><td>8.2</td><td>2.0</td><td>4.0</td><td>6.2E-05</td><td>9.8E-05</td><td>MB231/MB468</td><td>MP</td></tr>
<tr><td>>trna10_LysCTT_16_+_3241501_3241573@4.36.33,trna13_LysCTT_14_-_58706613_58706685@4.36.33,trna2_LysCTT_15_+_79152904_79152976@4.36.33|CGGCTAGCTCAGTCGGTAGAGCATGGGACTCTT</td><td>>trna10_LysCTT_16_+_3241501_3241573</td><td>4</td><td>36</td><td>33</td><td>CGGCTAGCTCAGTCGGTAGAGCATGGGACTCTT</td><td>224.1</td><td>5.1</td><td>1.3</td><td>4.0</td><td>6.2E-05</td><td>9.8E-05</td><td>MB231/MB468</td><td>MP</td></tr>
<tr><td>>hg38_21_+_8433172_8446622_RNA45SN1_45S_50ntFlanks@12928.12956.29|CTGGCGCTAAACCATTCGTAGACGACCTG</td><td>>hg38_21_+_8433172_8446622_RNA45SN1_45S_50ntFlanks</td><td>12928</td><td>12956</td><td>29</td><td>CTGGCGCTAAACCATTCGTAGACGACCTG</td><td>193.5</td><td>-6.2</td><td>1.5</td><td>-4.2</td><td>2.6E-05</td><td>9.8E-05</td><td>BT20/MB468</td><td>MP</td></tr>
<tr><td>>hg38_21_+_8433172_8446622_RNA45SN1_45S_50ntFlanks@10830.10848.19|CAAGCCGGATCCGTAACTT</td><td>>hg38_21_+_8433172_8446622_RNA45SN1_45S_50ntFlanks</td><td>10830</td><td>10848</td><td>19</td><td>CAAGCCGGATCCGTAACTT</td><td>212.1</td><td>-3.8</td><td>0.9</td><td>-4.0</td><td>6.9E-05</td><td>9.8E-05</td><td>MB231/MB468</td><td>CYTO</td></tr>
<tr><td>>hg38_21_+_8433172_8446622_RNA45SN1_45S_50ntFlanks@11565.11602.38|CTAGCAGCCGACTTAGAACTGGTGCGGACCAGGGGAAT</td><td>>hg38_21_+_8433172_8446622_RNA45SN1_45S_50ntFlanks</td><td>11565</td><td>11602</td><td>38</td><td>CTAGCAGCCGACTTAGAACTGGTGCGGACCAGGGGAAT</td><td>441.2</td><td>-6.8</td><td>1.7</td><td>-4.0</td><td>7.6E-05</td><td>9.8E-05</td><td>BT20/MB468</td><td>NUC</td></tr>
<tr><td>>hg38_21_+_8433172_8446622_RNA45SN1_45S_50ntFlanks@4067.4089.23|AACTTTCGATGGTAGTCGCCGTG</td><td>>hg38_21_+_8433172_8446622_RNA45SN1_45S_50ntFlanks</td><td>4067</td><td>4089</td><td>23</td><td>AACTTTCGATGGTAGTCGCCGTG</td><td>315.8</td><td>-2.9</td><td>0.7</td><td>-4.0</td><td>6.2E-05</td><td>9.8E-05</td><td>MB231/MB468</td><td>MP</td></tr>
<tr><td>>hg38_21_+_8433172_8446622_RNA45SN1_45S_50ntFlanks@6664.6699.36|TGGATCACTCGGCTCGTGCGTCGATGAAGAACGCAG</td><td>>hg38_21_+_8433172_8446622_RNA45SN1_45S_50ntFlanks</td><td>6664</td><td>6699</td><td>36</td><td>TGGATCACTCGGCTCGTGCGTCGATGAAGAACGCAG</td><td>358.7</td><td>-4.5</td><td>1.1</td><td>-4.0</td><td>7.3E-05</td><td>9.9E-05</td><td>MB231/MB468</td><td>MITO</td></tr>
<tr><td>>hg38_21_+_8433172_8446622_RNA45SN1_45S_50ntFlanks@12480.12498.19|TAATAGGGAACGTGAGCTG</td><td>>hg38_21_+_8433172_8446622_RNA45SN1_45S_50ntFlanks</td><td>12480</td><td>12498</td><td>19</td><td>TAATAGGGAACGTGAGCTG</td><td>96.4</td><td>-4.5</td><td>1.1</td><td>-4.0</td><td>6.2E-05</td><td>9.9E-05</td><td>MB231/MB468</td><td>MP</td></tr>
<tr><td>>trnaMT_GluTTC_MT_-_14674_14742@38.61.24,trnalookalike8_GluTTC_5_-_93905172_93905240@38.61.24|CATTGGTCGTGGTTGTAGTCCGTG</td><td>>trnaMT_GluTTC_MT_-_14674_14742</td><td>38</td><td>61</td><td>24</td><td>CATTGGTCGTGGTTGTAGTCCGTG</td><td>479.4</td><td>4.7</td><td>1.1</td><td>4.2</td><td>2.6E-05</td><td>9.9E-05</td><td>BT20/MB468</td><td>MP</td></tr>
<tr><td>>hg38_21_+_8433172_8446622_RNA45SN1_45S_50ntFlanks@11641.11667.27|CGGCGGGTGTTGACGCGATGTGATTTC</td><td>>hg38_21_+_8433172_8446622_RNA45SN1_45S_50ntFlanks</td><td>11641</td><td>11667</td><td>27</td><td>CGGCGGGTGTTGACGCGATGTGATTTC</td><td>234.0</td><td>-4.0</td><td>1.0</td><td>-4.0</td><td>6.9E-05</td><td>9.9E-05</td><td>MB231/MB468</td><td>CYTO</td></tr>
<tr><td>>hg38_21_+_8433172_8446622_RNA45SN1_45S_50ntFlanks@12407.12446.40|TTTTGATCCTTCGATGTCGGCTCTTCCTATCATTGTGAAG</td><td>>hg38_21_+_8433172_8446622_RNA45SN1_45S_50ntFlanks</td><td>12407</td><td>12446</td><td>40</td><td>TTTTGATCCTTCGATGTCGGCTCTTCCTATCATTGTGAAG</td><td>1288.3</td><td>-3.0</td><td>0.8</td><td>-4.0</td><td>7.4E-05</td><td>9.9E-05</td><td>MB231/MB468</td><td>NUC</td></tr>
<tr><td>>trna119_LysCTT_1_-_145395522_145395594@3.35.33,trna11_LysCTT_5_-_180648979_180649051@3.35.33,trna13_LysCTT_6_+_26556774_26556846@3.35.33,trna7_LysCTT_16_+_3225692_3225764@3.35.33,trna9_LysCTT_5_+_180634755_180634827@3.35.33|CCGGCTAGCTCAGTCGGTAGAGCATGAGACTCT</td><td>>trna119_LysCTT_1_-_145395522_145395594</td><td>3</td><td>35</td><td>33</td><td>CCGGCTAGCTCAGTCGGTAGAGCATGAGACTCT</td><td>890.8</td><td>-5.0</td><td>1.1</td><td>-4.4</td><td>1.0E-05</td><td>9.9E-05</td><td>BT20/MB231</td><td>MITO</td></tr>
<tr><td>>hg38_21_+_8433172_8446622_RNA45SN1_45S_50ntFlanks@8196.8223.28|CGTGGACGGTGTGAGGCCGGTAGCGGCC</td><td>>hg38_21_+_8433172_8446622_RNA45SN1_45S_50ntFlanks</td><td>8196</td><td>8223</td><td>28</td><td>CGTGGACGGTGTGAGGCCGGTAGCGGCC</td><td>775.2</td><td>-3.7</td><td>0.9</td><td>-4.2</td><td>2.6E-05</td><td>1.0E-04</td><td>BT20/MB468</td><td>MP</td></tr>
<tr><td>>hg38_21_+_8433172_8446622_RNA45SN1_45S_50ntFlanks@10463.10483.21|CTCCGTTGCCCTCGGCCGATC</td><td>>hg38_21_+_8433172_8446622_RNA45SN1_45S_50ntFlanks</td><td>10463</td><td>10483</td><td>21</td><td>CTCCGTTGCCCTCGGCCGATC</td><td>221.9</td><td>-5.5</td><td>1.4</td><td>-4.0</td><td>6.5E-05</td><td>1.0E-04</td><td>BT20/MB468</td><td>CYTO</td></tr>
<tr><td>>hg38_21_+_8433172_8446622_RNA45SN1_45S_50ntFlanks@11670.11699.30|CCCAGTGCTCTGAATGTCAAAGTGAAGAAA</td><td>>hg38_21_+_8433172_8446622_RNA45SN1_45S_50ntFlanks</td><td>11670</td><td>11699</td><td>30</td><td>CCCAGTGCTCTGAATGTCAAAGTGAAGAAA</td><td>214.0</td><td>-3.9</td><td>1.0</td><td>-4.0</td><td>6.5E-05</td><td>1.0E-04</td><td>BT20/MB468</td><td>CYTO</td></tr>
<tr><td>>hg38_21_+_8433172_8446622_RNA45SN1_45S_50ntFlanks@9567.9593.27|ACGTGCAAATCGGTCGTCCGACCTGGG</td><td>>hg38_21_+_8433172_8446622_RNA45SN1_45S_50ntFlanks</td><td>9567</td><td>9593</td><td>27</td><td>ACGTGCAAATCGGTCGTCCGACCTGGG</td><td>162.5</td><td>-5.2</td><td>1.3</td><td>-4.0</td><td>6.5E-05</td><td>1.0E-04</td><td>BT20/MB468</td><td>CYTO</td></tr>
<tr><td>>hg38_21_+_8433172_8446622_RNA45SN1_45S_50ntFlanks@12990.13014.25|GCTCCCTCGCTGCGATCTATTGAAA</td><td>>hg38_21_+_8433172_8446622_RNA45SN1_45S_50ntFlanks</td><td>12990</td><td>13014</td><td>25</td><td>GCTCCCTCGCTGCGATCTATTGAAA</td><td>2137.2</td><td>-3.4</td><td>0.9</td><td>-4.0</td><td>7.4E-05</td><td>1.0E-04</td><td>MB231/MB468</td><td>NUC</td></tr>
<tr><td>>am_trnaMT_ValTAC_MT_+_1602_1670@51.69.19|CTTAACTTGACCGCTCTGA</td><td>>am_trnaMT_ValTAC_MT_+_1602_1670</td><td>51</td><td>69</td><td>19</td><td>CTTAACTTGACCGCTCTGA</td><td>449.7</td><td>-5.3</td><td>1.3</td><td>-4.0</td><td>7.4E-05</td><td>1.0E-04</td><td>MB231/MB468</td><td>NUC</td></tr>
<tr><td>>hg38_21_+_8433172_8446622_RNA45SN1_45S_50ntFlanks@9534.9568.35|AGGAAACTCTGGTGGAGGTCCGTAGCGGTCCTGAC</td><td>>hg38_21_+_8433172_8446622_RNA45SN1_45S_50ntFlanks</td><td>9534</td><td>9568</td><td>35</td><td>AGGAAACTCTGGTGGAGGTCCGTAGCGGTCCTGAC</td><td>187.2</td><td>-6.1</td><td>1.5</td><td>-4.0</td><td>6.2E-05</td><td>1.0E-04</td><td>MB231/MB468</td><td>MP</td></tr>
<tr><td>>hg38_21_+_8433172_8446622_RNA45SN1_45S_50ntFlanks@9930.9948.19|ATATAGACAGCAGGACGGT</td><td>>hg38_21_+_8433172_8446622_RNA45SN1_45S_50ntFlanks</td><td>9930</td><td>9948</td><td>19</td><td>ATATAGACAGCAGGACGGT</td><td>478.5</td><td>4.6</td><td>1.0</td><td>4.4</td><td>8.9E-06</td><td>1.0E-04</td><td>BT20/MB231</td><td>MP</td></tr>
<tr><td>>trna116_GluCTC_1_-_145399233_145399304@1.34.34,trna59_GluCTC_1_+_249168447_249168518@1.34.34,trna71_GluCTC_1_-_161439189_161439260@1.34.34,trna74_GluCTC_1_-_161431809_161431880@1.34.34,trna77_GluCTC_1_-_161424398_161424469@1.34.34,trna77_GluCTC_6_+_28949976_28950047@1.34.34,trna80_GluCTC_1_-_161417018_161417089@1.34.34,trna87_GluCTC_6_-_126101393_126101464@1.34.34|TCCCTGGTGGTCTAGTGGTTAGGATTCGGCGCTC</td><td>>trna116_GluCTC_1_-_145399233_145399304</td><td>1</td><td>34</td><td>34</td><td>TCCCTGGTGGTCTAGTGGTTAGGATTCGGCGCTC</td><td>2905.7</td><td>5.1</td><td>1.2</td><td>4.2</td><td>2.6E-05</td><td>1.0E-04</td><td>BT20/MB468</td><td>MP</td></tr>
<tr><td>>hg38_21_+_8433172_8446622_RNA45SN1_45S_50ntFlanks@11561.11587.27|GCGCCTAGCAGCCGACTTAGAACTGGT</td><td>>hg38_21_+_8433172_8446622_RNA45SN1_45S_50ntFlanks</td><td>11561</td><td>11587</td><td>27</td><td>GCGCCTAGCAGCCGACTTAGAACTGGT</td><td>162.9</td><td>-2.4</td><td>0.6</td><td>-4.0</td><td>7.4E-05</td><td>1.0E-04</td><td>MB231/MB468</td><td>NUC</td></tr>
<tr><td>>MI0003776|hsa-mir-320b-1&WithFlank&1|+|116671740|116671823@48.66.19;MI0003839|hsa-mir-320b-2&WithFlank&1|-|224257034|224257116@47.65.19;MI0003778|hsa-mir-320c-1&WithFlank&18|+|21683512|21683595@48.66.19;MI0008191|hsa-mir-320c-2&WithFlank&18|+|24321669|24321752@48.66.19;MI0000542|hsa-mir-320a&WithFlank&8|-|22244960|22245043@48.66.19&[MIMAT0005792&hsa-miR-320b&offsets|0|-3;m-183&1|+|116671787|116671806&offsets|0|-1];[MIMAT0005792_1&hsa-miR-320b&offsets|0|-3;m-183&1|-|224257051|224257070&offsets|0|-1];[MIMAT0005793&hsa-miR-320c&offsets|0|-1;m-222&18|+|21683559|21683578&offsets|0|-1];[MIMAT0005793_1&hsa-miR-320c&offsets|0|-1;m-219&18|+|24321716|24321735&offsets|0|-1];[MIMAT0000510&hsa-miR-320a-3p&offsets|0|-3;m-60&8|-|22244975|22244996&offsets|0|-3]|AAAAGCTGGGTTGAGAGGG</td><td>>MI0003776|hsa-mir-320b-1&WithFlank&1|+|116671740|116671823</td><td>48</td><td>66</td><td>19</td><td>AAAAGCTGGGTTGAGAGGG</td><td>210.3</td><td>2.9</td><td>0.7</td><td>4.4</td><td>1.0E-05</td><td>1.0E-04</td><td>BT20/MB231</td><td>MITO</td></tr>
<tr><td>>hg38_21_+_8433172_8446622_RNA45SN1_45S_50ntFlanks@10248.10286.39|CGACGAGTAGGAGGGCCGCTGCGGTGAGCCTTGAAGCCT</td><td>>hg38_21_+_8433172_8446622_RNA45SN1_45S_50ntFlanks</td><td>10248</td><td>10286</td><td>39</td><td>CGACGAGTAGGAGGGCCGCTGCGGTGAGCCTTGAAGCCT</td><td>304.5</td><td>3.7</td><td>0.8</td><td>4.4</td><td>1.0E-05</td><td>1.0E-04</td><td>BT20/MB231</td><td>MITO</td></tr>
<tr><td>>hg38_21_+_8433172_8446622_RNA45SN1_45S_50ntFlanks@6655.6689.35|TCTTAGCGGTGGATCACTCGGCTCGTGCGTCGATG</td><td>>hg38_21_+_8433172_8446622_RNA45SN1_45S_50ntFlanks</td><td>6655</td><td>6689</td><td>35</td><td>TCTTAGCGGTGGATCACTCGGCTCGTGCGTCGATG</td><td>629.5</td><td>-5.4</td><td>1.4</td><td>-4.0</td><td>6.3E-05</td><td>1.0E-04</td><td>MB231/MB468</td><td>MP</td></tr>
<tr><td>>hg38_21_+_8433172_8446622_RNA45SN1_45S_50ntFlanks@10822.10865.44|GAAGTCGGCAAGCCGGATCCGTAACTTCGGGATAAGGATTGGCT</td><td>>hg38_21_+_8433172_8446622_RNA45SN1_45S_50ntFlanks</td><td>10822</td><td>10865</td><td>44</td><td>GAAGTCGGCAAGCCGGATCCGTAACTTCGGGATAAGGATTGGCT</td><td>767.7</td><td>-5.8</td><td>1.5</td><td>-4.0</td><td>7.4E-05</td><td>1.0E-04</td><td>MB231/MB468</td><td>NUC</td></tr>
<tr><td>>hg38_21_+_8433172_8446622_RNA45SN1_45S_50ntFlanks@12366.12404.39|TAACTGGCTTGTGGCGGCCAAGCGTTCATAGCGACGTCG</td><td>>hg38_21_+_8433172_8446622_RNA45SN1_45S_50ntFlanks</td><td>12366</td><td>12404</td><td>39</td><td>TAACTGGCTTGTGGCGGCCAAGCGTTCATAGCGACGTCG</td><td>40.1</td><td>-8.4</td><td>2.0</td><td>-4.2</td><td>2.6E-05</td><td>1.0E-04</td><td>BT20/MB468</td><td>MP</td></tr>
<tr><td>>hg38_21_+_8433172_8446622_RNA45SN1_45S_50ntFlanks@5236.5267.32|CACTGACTGGCTCAGCGTGTGCCTACCCTACG</td><td>>hg38_21_+_8433172_8446622_RNA45SN1_45S_50ntFlanks</td><td>5236</td><td>5267</td><td>32</td><td>CACTGACTGGCTCAGCGTGTGCCTACCCTACG</td><td>699.9</td><td>-5.2</td><td>1.3</td><td>-4.0</td><td>5.5E-05</td><td>1.0E-04</td><td>BT20/MB468</td><td>MITO</td></tr>
<tr><td>>MI0003600|hsa-mir-550a-1&WithFlank&7|+|30289788|30289896@28.48.21;MI0003601|hsa-mir-550a-2&WithFlank&7|+|32732975|32733083@28.48.21&[MIMAT0004800&hsa-miR-550a-5p&offsets|0|-2;m-340&7|+|30289815|30289835&offsets|0|0];[MIMAT0004800_1&hsa-miR-550a-5p&offsets|0|-2;m-340&7|+|32733002|32733022&offsets|0|0]|AGTGCCTGAGGGAGTAAGAGC</td><td>>MI0003600|hsa-mir-550a-1&WithFlank&7|+|30289788|30289896</td><td>28</td><td>48</td><td>21</td><td>AGTGCCTGAGGGAGTAAGAGC</td><td>215.5</td><td>6.8</td><td>1.6</td><td>4.2</td><td>2.6E-05</td><td>1.0E-04</td><td>BT20/MB468</td><td>MP</td></tr>
<tr><td>>hg38_21_+_8433172_8446622_RNA45SN1_45S_50ntFlanks@9602.9620.19|CGAAAGACTAATCGAACCA</td><td>>hg38_21_+_8433172_8446622_RNA45SN1_45S_50ntFlanks</td><td>9602</td><td>9620</td><td>19</td><td>CGAAAGACTAATCGAACCA</td><td>53.8</td><td>-9.6</td><td>2.4</td><td>-4.0</td><td>6.3E-05</td><td>1.0E-04</td><td>MB231/MB468</td><td>MP</td></tr>
<tr><td>>hg38_21_+_8433172_8446622_RNA45SN1_45S_50ntFlanks@12406.12436.31|TTTTTGATCCTTCGATGTCGGCTCTTCCTAT</td><td>>hg38_21_+_8433172_8446622_RNA45SN1_45S_50ntFlanks</td><td>12406</td><td>12436</td><td>31</td><td>TTTTTGATCCTTCGATGTCGGCTCTTCCTAT</td><td>331.8</td><td>-4.0</td><td>1.0</td><td>-4.0</td><td>7.0E-05</td><td>1.0E-04</td><td>MB231/MB468</td><td>CYTO</td></tr>
<tr><td>>hg38_21_+_8433172_8446622_RNA45SN1_45S_50ntFlanks@9482.9516.35|ACGTGTTAGGACCCGAAAGATGGTGAACTATGCCT</td><td>>hg38_21_+_8433172_8446622_RNA45SN1_45S_50ntFlanks</td><td>9482</td><td>9516</td><td>35</td><td>ACGTGTTAGGACCCGAAAGATGGTGAACTATGCCT</td><td>77.2</td><td>9.6</td><td>2.2</td><td>4.4</td><td>8.9E-06</td><td>1.0E-04</td><td>BT20/MB231</td><td>MP</td></tr>
<tr><td>>hg38_21_+_8433172_8446622_RNA45SN1_45S_50ntFlanks@11581.11607.27|AACTGGTGCGGACCAGGGGAATCCGAC</td><td>>hg38_21_+_8433172_8446622_RNA45SN1_45S_50ntFlanks</td><td>11581</td><td>11607</td><td>27</td><td>AACTGGTGCGGACCAGGGGAATCCGAC</td><td>425.5</td><td>-2.7</td><td>0.7</td><td>-4.0</td><td>7.0E-05</td><td>1.0E-04</td><td>MB231/MB468</td><td>CYTO</td></tr>
<tr><td>>hg38_21_+_8433172_8446622_RNA45SN1_45S_50ntFlanks@8738.8758.21|CCGTCCGTCCGTCCGTCCGTC</td><td>>hg38_21_+_8433172_8446622_RNA45SN1_45S_50ntFlanks</td><td>8738</td><td>8758</td><td>21</td><td>CCGTCCGTCCGTCCGTCCGTC</td><td>123.2</td><td>-4.2</td><td>1.1</td><td>-4.0</td><td>6.6E-05</td><td>1.0E-04</td><td>BT20/MB468</td><td>CYTO</td></tr>
<tr><td>>hg38_21_+_8433172_8446622_RNA45SN1_45S_50ntFlanks@10843.10867.25|TAACTTCGGGATAAGGATTGGCTCT</td><td>>hg38_21_+_8433172_8446622_RNA45SN1_45S_50ntFlanks</td><td>10843</td><td>10867</td><td>25</td><td>TAACTTCGGGATAAGGATTGGCTCT</td><td>94.1</td><td>-6.1</td><td>1.5</td><td>-4.0</td><td>6.3E-05</td><td>1.0E-04</td><td>MB231/MB468</td><td>MP</td></tr>
<tr><td>>hg38_21_+_8433172_8446622_RNA45SN1_45S_50ntFlanks@11565.11586.22|CTAGCAGCCGACTTAGAACTGG</td><td>>hg38_21_+_8433172_8446622_RNA45SN1_45S_50ntFlanks</td><td>11565</td><td>11586</td><td>22</td><td>CTAGCAGCCGACTTAGAACTGG</td><td>147.1</td><td>-4.6</td><td>1.1</td><td>-4.0</td><td>6.3E-05</td><td>1.0E-04</td><td>MB231/MB468</td><td>MP</td></tr>
<tr><td>>hg38_1_-_228634819_228635039_RNA5S12_5S_50ntFlanks@154.171.18|CCGGGTGCTGTAGGCTTT</td><td>>hg38_1_-_228634819_228635039_RNA5S12_5S_50ntFlanks</td><td>154</td><td>171</td><td>18</td><td>CCGGGTGCTGTAGGCTTT</td><td>3635.6</td><td>-2.9</td><td>0.7</td><td>-4.0</td><td>7.5E-05</td><td>1.0E-04</td><td>MB231/MB468</td><td>MITO</td></tr>
<tr><td>>MI0003591|hsa-mir-584&WithFlank&5|-|149062307|149062415@22.42.21(+2U)&[MIMAT0003249&hsa-miR-584-5p&offsets|0|-1(+2U);m-167&5|-|149062374|149062394&offsets|0|0(+2U)]|TTATGGTTTGCCTGGGACTGATT</td><td>>MI0003591|hsa-mir-584&WithFlank&5|-|149062307|149062415</td><td>22</td><td>42</td><td>21(+2U)</td><td>TTATGGTTTGCCTGGGACTGATT</td><td>130.9</td><td>6.7</td><td>1.7</td><td>4.0</td><td>6.3E-05</td><td>1.0E-04</td><td>MB231/MB468</td><td>MP</td></tr>
<tr><td>>MI0000750|hsa-mir-26a-2&WithFlank&12|-|57824603|57824698@20.41.22(+1C);MI0000083|hsa-mir-26a-1&WithFlank&3|+|37969398|37969486@16.37.22(+1C)&[MIMAT0000082&hsa-miR-26a-5p&offsets|0|0(+1C);m-33&12|-|57824658|57824679&offsets|0|0(+1C)];[MIMAT0000082_1&hsa-miR-26a-5p&offsets|0|0(+1C);m-34&3|+|37969413|37969434&offsets|0|0(+1C)]|TTCAAGTAATCCAGGATAGGCTC</td><td>>MI0000750|hsa-mir-26a-2&WithFlank&12|-|57824603|57824698</td><td>20</td><td>41</td><td>22(+1C)</td><td>TTCAAGTAATCCAGGATAGGCTC</td><td>776.2</td><td>5.4</td><td>1.3</td><td>4.0</td><td>6.3E-05</td><td>1.0E-04</td><td>MB231/MB468</td><td>MP</td></tr>
<tr><td>>hg38_21_+_8433172_8446622_RNA45SN1_45S_50ntFlanks@10355.10383.29|AACTTTGAAGGCCGAAGTGGAGAAGGGTT</td><td>>hg38_21_+_8433172_8446622_RNA45SN1_45S_50ntFlanks</td><td>10355</td><td>10383</td><td>29</td><td>AACTTTGAAGGCCGAAGTGGAGAAGGGTT</td><td>217.2</td><td>-3.8</td><td>1.0</td><td>-4.0</td><td>7.5E-05</td><td>1.0E-04</td><td>MB231/MB468</td><td>MITO</td></tr>
<tr><td>>hg38_21_+_8433172_8446622_RNA45SN1_45S_50ntFlanks@12686.12703.18|CCAGGCGGAACGATACGG</td><td>>hg38_21_+_8433172_8446622_RNA45SN1_45S_50ntFlanks</td><td>12686</td><td>12703</td><td>18</td><td>CCAGGCGGAACGATACGG</td><td>103.3</td><td>-4.9</td><td>1.2</td><td>-4.0</td><td>6.6E-05</td><td>1.0E-04</td><td>BT20/MB468</td><td>CYTO</td></tr>
<tr><td>>trnaMT_GluTTC_MT_-_14674_14742@39.70.32,trnalookalike8_GluTTC_5_-_93905172_93905240@39.70.32|ATTGGTCGTGGTTGTAGTCCGTGCGAGAATAC</td><td>>trnaMT_GluTTC_MT_-_14674_14742</td><td>39</td><td>70</td><td>32</td><td>ATTGGTCGTGGTTGTAGTCCGTGCGAGAATAC</td><td>633.4</td><td>8.4</td><td>2.0</td><td>4.2</td><td>2.6E-05</td><td>1.0E-04</td><td>BT20/MB468</td><td>MP</td></tr>
</table>

| | | | | | | | | | | | | | |
|---|---|---|---|---|---|---|---|---|---|---|---|---|---|
| >MI0003137\|hsa-mir-193b&WithFlank&16\|+\|14303961\|14304055@57.78.22(+1C)&[MIMAT0002819&hsa-miR-193b-3p&offsets\|0\|0(+1C);m-166&16\|+\|14304017\|14304038&offsets\|0\|0(+1C)]\|AACTGGCCCTCAAAGTCCCGCTC | >MI0003137\|hsa-mir-193b&WithFlank&16\|+\|14303961\|14304055 | 57 | 78 | 22(+1C) | AACTGGCCCTCAAAGTCCCGCTC | 216.8 | 7.2 | 1.7 | 4.2 | 2.6E-05 | 1.0E-04 | BT20/MB468 | MP |
| >am_trna2_GlyGCC_21_-_18827107_18827177@3.26.24,trna35_GlyGCC_1_+_161413094_161413164@3.26.24,trna37_GlyGCC_1_+_161420467_161420537@3.26.24,trna39_GlyGCC_1_+_161427898_161427968@3.26.24,trna41_GlyGCC_1_+_161435258_161435328@3.26.24\|ATGGGTGGTTCAGTGGTAGAATTC | >am_trna2_GlyGCC_21_-_18827107_18827177 | 3 | 26 | 24 | ATGGGTGGTTCAGTGGTAGAATTC | 84.0 | 9.7 | 2.2 | 4.4 | 9.0E-06 | 1.0E-04 | BT20/MB231 | MP |
| >hg38_MT_+_1621_3279_RNR2_16S_50ntFlanks@284.305.22\|CCGAAACCAGACGAGCTACCTA | >hg38_MT_+_1621_3279_RNR2_16S_50ntFlanks | 284 | 305 | 22 | CCGAAACCAGACGAGCTACCTA | 133.2 | 7.7 | 1.8 | 4.2 | 2.6E-05 | 1.0E-04 | BT20/MB468 | MP |
| >hg38_21_+_8433172_8446622_RNA45SN1_45S_50ntFlanks@6732.6773.42\|ACATTGATCATCGACACTTCGAACGCACTTGCGGCCCCGGGT | >hg38_21_+_8433172_8446622_RNA45SN1_45S_50ntFlanks | 6732 | 6773 | 42 | ACATTGATCATCGACACTTCGAACGCACTTGCGGCCCCGGGT | 111.4 | 7.4 | 1.8 | 4.2 | 2.6E-05 | 1.0E-04 | BT20/MB468 | MP |
| >MI0000102\|hsa-mir-100&WithFlank&11\|-\|122152223\|122152314@19.39.21(+1A)&[MIMAT0000098&hsa-miR-100-5p&offsets\|0\|-1(+1A);m-27&11\|-\|122152276\|122152296&offsets\|0\|0(+1A)]\|AACCCGTAGATCCGAACTTGTA | >MI0000102\|hsa-mir-100&WithFlank&11\|-\|122152223\|122152314 | 19 | 39 | 21(+1A) | AACCCGTAGATCCGAACTTGTA | 903.0 | 8.3 | 2.0 | 4.0 | 5.5E-05 | 1.0E-04 | BT20/MB468 | MITO |
| >hg38_21_+_8433172_8446622_RNA45SN1_45S_50ntFlanks@11795.11819.25\|AACGAGATTCCCACTGTCCCTACCT | >hg38_21_+_8433172_8446622_RNA45SN1_45S_50ntFlanks | 11795 | 11819 | 25 | AACGAGATTCCCACTGTCCCTACCT | 234.3 | -4.5 | 1.1 | -4.0 | 7.5E-05 | 1.0E-04 | MB231/MB468 | NUC |
| >hg38_21_+_8433172_8446622_RNA45SN1_45S_50ntFlanks@8182.8206.25\|ATCGAGGCCCAGCCCGTGGACGGTG | >hg38_21_+_8433172_8446622_RNA45SN1_45S_50ntFlanks | 8182 | 8206 | 25 | ATCGAGGCCCAGCCCGTGGACGGTG | 524.7 | -5.9 | 1.5 | -4.0 | 7.5E-05 | 1.0E-04 | MB231/MB468 | NUC |
| >hg38_21_+_8433172_8446622_RNA45SN1_45S_50ntFlanks@4297.4314.18\|CATTGGAGGGCAAGTCTG | >hg38_21_+_8433172_8446622_RNA45SN1_45S_50ntFlanks | 4297 | 4314 | 18 | CATTGGAGGGCAAGTCTG | 535.5 | -4.8 | 1.2 | -4.0 | 7.5E-05 | 1.0E-04 | MB231/MB468 | NUC |
| >hg38_21_+_8433172_8446622_RNA45SN1_45S_50ntFlanks@5485.5513.29\|AGCGCTGAGAAGACGGTCGAACTTGACTA | >hg38_21_+_8433172_8446622_RNA45SN1_45S_50ntFlanks | 5485 | 5513 | 29 | AGCGCTGAGAAGACGGTCGAACTTGACTA | 235.6 | -2.5 | 0.6 | -4.0 | 7.1E-05 | 1.0E-04 | MB231/MB468 | CYTO |
| >hg38_21_+_8433172_8446622_RNA45SN1_45S_50ntFlanks@10350.10385.36\|ACGAGAACTTTGAAGGCCGAAGTGGAGAAGGGTTCC | >hg38_21_+_8433172_8446622_RNA45SN1_45S_50ntFlanks | 10350 | 10385 | 36 | ACGAGAACTTTGAAGGCCGAAGTGGAGAAGGGTTCC | 194.5 | -2.7 | 0.7 | -4.0 | 7.5E-05 | 1.0E-04 | MB231/MB468 | NUC |
| >hg38_21_+_8433172_8446622_RNA45SN1_45S_50ntFlanks@4738.4755.18\|AAAGTCGGAGGTTCGAAG | >hg38_21_+_8433172_8446622_RNA45SN1_45S_50ntFlanks | 4738 | 4755 | 18 | AAAGTCGGAGGTTCGAAG | 392.6 | -6.6 | 1.7 | -4.0 | 7.8E-05 | 1.0E-04 | BT20/MB468 | NUC |
| >trnaMT_HisGTG_MT_+_12138_12206@44.72.29\|CAGAGGCTTACGACCCCTTATTTACCCCA | >trnaMT_HisGTG_MT_+_12138_12206 | 44 | 72 | 29 | CAGAGGCTTACGACCCCTTATTTACCCCA | 26.6 | 8.0 | 2.0 | 4.0 | 6.3E-05 | 1.0E-04 | MB231/MB468 | MP |
| >hg38_21_+_8433172_8446622_RNA45SN1_45S_50ntFlanks@5083.5104.22\|ACGAACGAGACTCTGGCATGCT | >hg38_21_+_8433172_8446622_RNA45SN1_45S_50ntFlanks | 5083 | 5104 | 22 | ACGAACGAGACTCTGGCATGCT | 455.3 | -4.5 | 1.1 | -4.0 | 7.6E-05 | 1.0E-04 | MB231/MB468 | MITO |
| >hg38_21_+_8433172_8446622_RNA45SN1_45S_50ntFlanks@6650.6689.40\|ACGACTCTTAGCGGTGGATCACTCGGCTCGTGCGTCGATG | >hg38_21_+_8433172_8446622_RNA45SN1_45S_50ntFlanks | 6650 | 6689 | 40 | ACGACTCTTAGCGGTGGATCACTCGGCTCGTGCGTCGATG | 2086.5 | -3.7 | 0.9 | -4.0 | 6.4E-05 | 1.0E-04 | MB231/MB468 | MP |
| >hg38_21_+_8433172_8446622_RNA45SN1_45S_50ntFlanks@11210.11228.19\|GGCGACTCTGGACGCGAGC | >hg38_21_+_8433172_8446622_RNA45SN1_45S_50ntFlanks | 11210 | 11228 | 19 | GGCGACTCTGGACGCGAGC | 85.4 | -5.9 | 1.5 | -4.0 | 6.4E-05 | 1.0E-04 | MB231/MB468 | MP |
| >hg38_21_+_8433172_8446622_RNA45SN1_45S_50ntFlanks@10393.10425.33\|CAGCAGTTGAACATGGGTCAGTCGGTCCTGAGA | >hg38_21_+_8433172_8446622_RNA45SN1_45S_50ntFlanks | 10393 | 10425 | 33 | CAGCAGTTGAACATGGGTCAGTCGGTCCTGAGA | 91.8 | -6.0 | 1.5 | -4.0 | 6.4E-05 | 1.0E-04 | MB231/MB468 | MP |
| >trnaMT_GluTTC_MT_-_14674_14742@39.68.30,trnalookalike8_GluTTC_5_-_93905172_93905240@39.68.30\|ATTGGTCGTGGTTGTAGTCCGTGCGAGAAT | >trnaMT_GluTTC_MT_-_14674_14742 | 39 | 68 | 30 | ATTGGTCGTGGTTGTAGTCCGTGCGAGAAT | 564.1 | 4.6 | 1.1 | 4.2 | 2.6E-05 | 1.0E-04 | BT20/MB468 | MP |
| >hg38_21_+_8433172_8446622_RNA45SN1_45S_50ntFlanks@9461.9488.28\|GGGGAGGTGGAGCACGAGCGCACGTGTT | >hg38_21_+_8433172_8446622_RNA45SN1_45S_50ntFlanks | 9461 | 9488 | 28 | GGGGAGGTGGAGCACGAGCGCACGTGTT | 71.7 | 9.5 | 2.1 | 4.4 | 9.1E-06 | 1.0E-04 | BT20/MB231 | MP |
| >hg38_21_+_8433172_8446622_RNA45SN1_45S_50ntFlanks@7974.8010.37\|ACGCGACCTCAGATCAGACGTGGCGACCCGCTGAATT | >hg38_21_+_8433172_8446622_RNA45SN1_45S_50ntFlanks | 7974 | 8010 | 37 | ACGCGACCTCAGATCAGACGTGGCGACCCGCTGAATT | 4630.8 | 4.9 | 1.2 | 4.2 | 2.7E-05 | 1.0E-04 | BT20/MB468 | MP |
| >hg38_21_+_8433172_8446622_RNA45SN1_45S_50ntFlanks@11914.11940.27\|AGAGACATGAGAGGTGTAGAATAAGTG | >hg38_21_+_8433172_8446622_RNA45SN1_45S_50ntFlanks | 11914 | 11940 | 27 | AGAGACATGAGAGGTGTAGAATAAGTG | 143.0 | 6.6 | 1.6 | 4.2 | 2.7E-05 | 1.0E-04 | BT20/MB468 | MP |
| >hg38_21_+_8433172_8446622_RNA45SN1_45S_50ntFlanks@12415.12448.34\|CTTCGATGTCGGCTCTTCCTATCATTGTGAAGCA | >hg38_21_+_8433172_8446622_RNA45SN1_45S_50ntFlanks | 12415 | 12448 | 34 | CTTCGATGTCGGCTCTTCCTATCATTGTGAAGCA | 628.3 | 3.3 | 0.8 | 4.0 | 7.6E-05 | 1.0E-04 | MB231/MB468 | NUC |
| >hg38_21_+_8433172_8446622_RNA45SN1_45S_50ntFlanks@10572.10589.18\|AGAAGCCGGCGGGAGCCC | >hg38_21_+_8433172_8446622_RNA45SN1_45S_50ntFlanks | 10572 | 10589 | 18 | AGAAGCCGGCGGGAGCCC | 189.2 | -5.5 | 1.4 | -4.0 | 7.6E-05 | 1.0E-04 | MB231/MB468 | NUC |
| >trna2_GlyGCC_21_-_18827107_18827177@1.33.33,trna35_GlyGCC_1_+_161413094_161413164@1.33.33,trna37_GlyGCC_1_+_161420467_161420537@1.33.33,trna39_GlyGCC_1_+_161427898_161427968@1.33.33,trna41_GlyGCC_1_+_161435258_161435328@1.33.33\|GCATGGGTGGTTCAGTGGTAGAATTCTCGCCTG | >trna2_GlyGCC_21_-_18827107_18827177 | 1 | 33 | 33 | GCATGGGTGGTTCAGTGGTAGAATTCTCGCCTG | 1379.7 | 4.1 | 1.0 | 4.0 | 5.6E-05 | 1.0E-04 | BT20/MB468 | MITO |
| >hg38_21_+_8433172_8446622_RNA45SN1_45S_50ntFlanks@11579.11608.30\|AGAACTGGTGCGGACCAGGGGAATCCGACT | >hg38_21_+_8433172_8446622_RNA45SN1_45S_50ntFlanks | 11579 | 11608 | 30 | AGAACTGGTGCGGACCAGGGGAATCCGACT | 108.3 | -4.6 | 1.1 | -4.0 | 6.4E-05 | 1.0E-04 | MB231/MB468 | MP |
| >hg38_21_+_8433172_8446622_RNA45SN1_45S_50ntFlanks@12004.12035.32\|CGGGCCGCCGGTGAAATACCACTACTCTGATC | >hg38_21_+_8433172_8446622_RNA45SN1_45S_50ntFlanks | 12004 | 12035 | 32 | CGGGCCGCCGGTGAAATACCACTACTCTGATC | 704.3 | 6.0 | 1.4 | 4.4 | 9.9E-06 | 1.0E-04 | BT20/MB231 | NUC |
| >hg38_21_+_8433172_8446622_RNA45SN1_45S_50ntFlanks@4575.4611.37\|TAGGACCGCGGTTCTATTTTGTTGGTTTTCGGAACTG | >hg38_21_+_8433172_8446622_RNA45SN1_45S_50ntFlanks | 4575 | 4611 | 37 | TAGGACCGCGGTTCTATTTTGTTGGTTTTCGGAACTG | 190.6 | -4.5 | 1.1 | -4.0 | 7.6E-05 | 1.0E-04 | MB231/MB468 | NUC |
| >hg38_21_+_8433172_8446622_RNA45SN1_45S_50ntFlanks@9568.9603.36\|CGTGCAAATCGGTCGTCCGACCTGGGTATAGGGGCG | >hg38_21_+_8433172_8446622_RNA45SN1_45S_50ntFlanks | 9568 | 9603 | 36 | CGTGCAAATCGGTCGTCCGACCTGGGTATAGGGGCG | 1405.8 | -2.9 | 0.7 | -4.0 | 5.6E-05 | 1.0E-04 | BT20/MB468 | MITO |
| >hg38_21_+_8433172_8446622_RNA45SN1_45S_50ntFlanks@6651.6686.36\|CGACTCTTAGCGGTGGATCACTCGGCTCGTGCGTCG | >hg38_21_+_8433172_8446622_RNA45SN1_45S_50ntFlanks | 6651 | 6686 | 36 | CGACTCTTAGCGGTGGATCACTCGGCTCGTGCGTCG | 852.6 | 2.9 | 0.7 | 4.0 | 7.6E-05 | 1.0E-04 | MB231/MB468 | NUC |
| >am_trna162_MetCAT_6_-_26758550_26758622@34.54.21,trna164_MetCAT_6_-_26735574_26735646@34.54.21,trna20_MetCAT_16_+_71460396_71460468@34.54.21,trna21_MetCAT_6_+_26701712_26701784@34.54.21,trna27_MetCAT_6_+_26766444_26766516@34.54.21,trna75_MetCAT_6_+_28912352_28912424@34.54.21,trna97_MetCAT_6_-_28921042_28921114@34.54.21\|CATAATCTGAAGGTCCTGAGT | >am_trna162_MetCAT_6_-_26758550_26758622 | 34 | 54 | 21 | CATAATCTGAAGGTCCTGAGT | 437.0 | 4.6 | 1.2 | 4.0 | 7.6E-05 | 1.0E-04 | MB231/MB468 | MITO |
| >hg38_21_+_8433172_8446622_RNA45SN1_45S_50ntFlanks@4730.4752.23\|CAAGAACGAAAGTCGGAGGTTCG | >hg38_21_+_8433172_8446622_RNA45SN1_45S_50ntFlanks | 4730 | 4752 | 23 | CAAGAACGAAAGTCGGAGGTTCG | 297.8 | -3.7 | 0.9 | -4.0 | 7.2E-05 | 1.0E-04 | MB231/MB468 | CYTO |
| >hg38_21_+_8433172_8446622_RNA45SN1_45S_50ntFlanks@11592.11612.21\|ACCAGGGGAATCCGACTGTTT | >hg38_21_+_8433172_8446622_RNA45SN1_45S_50ntFlanks | 11592 | 11612 | 21 | ACCAGGGGAATCCGACTGTTT | 567.4 | -6.8 | 1.6 | -4.2 | 2.7E-05 | 1.0E-04 | BT20/MB468 | MP |
| >hg38_21_+_8433172_8446622_RNA45SN1_45S_50ntFlanks@11875.11898.24\|GAAGACCCTGTTGAGCTTGACTCT | >hg38_21_+_8433172_8446622_RNA45SN1_45S_50ntFlanks | 11875 | 11898 | 24 | GAAGACCCTGTTGAGCTTGACTCT | 343.1 | -3.4 | 0.8 | -4.0 | 7.7E-05 | 1.0E-04 | MB231/MB468 | MITO |
| >MI0000088\|hsa-mir-30a&WithFlank&6\|-\|71403545\|71403627@12.34.23(+1G)&[MIMAT0000087&hsa-miR-30a-5p&offsets\|0\|+1(+1G);m-15&6\|-\|71403595\|71403616&offsets\|0\|+1(+1G)]\|TGTAAACATCCTCGACTGGAAGCG | >MI0000088\|hsa-mir-30a&WithFlank&6\|-\|71403545\|71403627 | 12 | 34 | 23(+1G) | TGTAAACATCCTCGACTGGAAGCG | 673.4 | 4.2 | 1.0 | 4.0 | 7.2E-05 | 1.0E-04 | MB231/MB468 | CYTO |
| >MI0000088\|hsa-mir-30a&WithFlank&6\|-\|71403545\|71403627@53.73.21(+2U)&[MIMAT0000088&hsa-miR-30a-3p&offsets\|0\|-1(+2U);m-38&6\|-\|71403554\|71403575&offsets\|0\|-1(+2U)]\|CTTTCAGTCGGATGTTTGCAGTT | >MI0000088\|hsa-mir-30a&WithFlank&6\|-\|71403545\|71403627 | 53 | 73 | 21(+2U) | CTTTCAGTCGGATGTTTGCAGTT | 876.2 | -4.5 | 1.0 | -4.4 | 9.2E-06 | 1.0E-04 | BT20/MB231 | MP |
| >MI0000073\|hsa-mir-19a&WithFlank&13\|+\|91350885\|91350978@55.78.24&[MIMAT0000073&hsa-miR-19a-3p&offsets\|0\|+1;m-201&13\|+\|91350939\|91350961&offsets\|0\|+1]\|TGTGCAAATCTATGCAAAACTGAT | >MI0000073\|hsa-mir-19a&WithFlank&13\|+\|91350885\|91350978 | 55 | 78 | 24 | TGTGCAAATCTATGCAAAACTGAT | 415.2 | -4.9 | 1.1 | -4.4 | 1.1E-05 | 1.0E-04 | BT20/MB231 | MITO |
| >hg38_21_+_8433172_8446622_RNA45SN1_45S_50ntFlanks@7984.8017.34\|AGATCAGACGTGGCGACCCGCTGAATTTAAGCAT | >hg38_21_+_8433172_8446622_RNA45SN1_45S_50ntFlanks | 7984 | 8017 | 34 | AGATCAGACGTGGCGACCCGCTGAATTTAAGCAT | 350.5 | -3.8 | 1.0 | -4.0 | 7.7E-05 | 1.0E-04 | MB231/MB468 | MITO |
| >hg38_21_+_8433172_8446622_RNA45SN1_45S_50ntFlanks@476.510.35\|TTCGTGATCGATGTGGTGACGTCGTGCTCTCCCGG | >hg38_21_+_8433172_8446622_RNA45SN1_45S_50ntFlanks | 476 | 510 | 35 | TTCGTGATCGATGTGGTGACGTCGTGCTCTCCCGG | 465.8 | 4.8 | 1.2 | 4.0 | 7.7E-05 | 1.0E-04 | MB231/MB468 | MITO |
| >hg38_MT_+_598_1651_RNR1_12S_50ntFlanks@926.946.21\|AACTAAAACCCCTACGCATTT | >hg38_MT_+_598_1651_RNR1_12S_50ntFlanks | 926 | 946 | 21 | AACTAAAACCCCTACGCATTT | 27.8 | 8.0 | 2.0 | 4.0 | 6.5E-05 | 1.0E-04 | MB231/MB468 | MP |
| >trnaMT_GluTTC_MT_-_14674_14742@24.57.34,trnalookalike8_GluTTC_5_-_93905172_93905240@24.57.34\|ATGGTTTTTCATATCATTGGTCGTGGTTGTAGTC | >trnaMT_GluTTC_MT_-_14674_14742 | 24 | 57 | 34 | ATGGTTTTTCATATCATTGGTCGTGGTTGTAGTC | 349.6 | 8.1 | 1.9 | 4.2 | 2.7E-05 | 1.0E-04 | BT20/MB468 | MP |
| >hg38_21_+_8433172_8446622_RNA45SN1_45S_50ntFlanks@9551.9569.19\|GTCCGTAGCGGTCCTGACG | >hg38_21_+_8433172_8446622_RNA45SN1_45S_50ntFlanks | 9551 | 9569 | 19 | GTCCGTAGCGGTCCTGACG | 362.2 | 2.4 | 0.6 | 4.0 | 6.7E-05 | 1.0E-04 | BT20/MB468 | CYTO |
| >hg38_21_+_8433172_8446622_RNA45SN1_45S_50ntFlanks@12409.12437.29\|TTGATCCTTCGATGTCGGCTCTTCCTATC | >hg38_21_+_8433172_8446622_RNA45SN1_45S_50ntFlanks | 12409 | 12437 | 29 | TTGATCCTTCGATGTCGGCTCTTCCTATC | 292.6 | 3.4 | 0.9 | 4.0 | 7.2E-05 | 1.0E-04 | MB231/MB468 | CYTO |
| >hg38_21_+_8433172_8446622_RNA45SN1_45S_50ntFlanks@10107.10127.21\|CGTGTGGTGTGCGTCGGAGGG | >hg38_21_+_8433172_8446622_RNA45SN1_45S_50ntFlanks | 10107 | 10127 | 21 | CGTGTGGTGTGCGTCGGAGGG | 421.8 | -4.2 | 1.1 | -4.0 | 6.7E-05 | 1.0E-04 | BT20/MB468 | CYTO |
| >trnaMT_LysTTT_MT_+_8295_8364@1.38.38,trnalookalike7_LysTTT_1_+_568844_568913@1.38.38\|CACTGTAAAGCTAACTTAGCATTAACCTTTTAAGTTAA | >trnaMT_LysTTT_MT_+_8295_8364 | 1 | 38 | 38 | CACTGTAAAGCTAACTTAGCATTAACCTTTTAAGTTAA | 252.9 | 3.3 | 0.8 | 4.0 | 7.7E-05 | 1.0E-04 | MB231/MB468 | MITO |
| >hg38_21_+_8433172_8446622_RNA45SN1_45S_50ntFlanks@7973.8004.32\|GACGCGACCTCAGATCAGACGTGGCGACCCGC | >hg38_21_+_8433172_8446622_RNA45SN1_45S_50ntFlanks | 7973 | 8004 | 32 | GACGCGACCTCAGATCAGACGTGGCGACCCGC | 259.8 | 4.1 | 1.0 | 4.2 | 2.7E-05 | 1.0E-04 | BT20/MB468 | MP |
| >hg38_21_+_8433172_8446622_RNA45SN1_45S_50ntFlanks@3899.3927.29\|CCTTCGCGGGGGGGATGCGTGCATTTATC | >hg38_21_+_8433172_8446622_RNA45SN1_45S_50ntFlanks | 3899 | 3927 | 29 | CCTTCGCGGGGGGGATGCGTGCATTTATC | 499.3 | -3.1 | 0.8 | -4.0 | 7.2E-05 | 1.0E-04 | MB231/MB468 | CYTO |
| >hg38_21_+_8433172_8446622_RNA45SN1_45S_50ntFlanks@11568.11585.18\|GCAGCCGACTTAGAACTG | >hg38_21_+_8433172_8446622_RNA45SN1_45S_50ntFlanks | 11568 | 11585 | 18 | GCAGCCGACTTAGAACTG | 1278.0 | -1.7 | 0.4 | -4.0 | 6.7E-05 | 1.0E-04 | BT20/MB468 | CYTO |
| >hg38_21_+_8433172_8446622_RNA45SN1_45S_50ntFlanks@12396.12416.21\|GCGACGTCGCTTTTTGATCCT | >hg38_21_+_8433172_8446622_RNA45SN1_45S_50ntFlanks | 12396 | 12416 | 21 | GCGACGTCGCTTTTTGATCCT | 577.0 | -2.6 | 0.7 | -4.0 | 7.7E-05 | 1.0E-04 | MB231/MB468 | NUC |
| >hg38_21_+_8433172_8446622_RNA45SN1_45S_50ntFlanks@10677.10709.33\|GCGGTTCCGGCGGCGTCCGGTGAGCTCTCGCTG | >hg38_21_+_8433172_8446622_RNA45SN1_45S_50ntFlanks | 10677 | 10709 | 33 | GCGGTTCCGGCGGCGTCCGGTGAGCTCTCGCTG | 809.6 | -2.5 | 0.6 | -4.0 | 7.7E-05 | 1.0E-04 | MB231/MB468 | NUC |

| | | | | | | | | | | | | | |
|---|---|---|---|---|---|---|---|---|---|---|---|---|---|
| >hg38_21_+_8433172_8446622_RNA45SN1_45S_50ntFlanks@11594.11613.20\|CAGGGGAATCCGACTGTTTA | >hg38_21_+_8433172_8446622_RNA45SN1_45S_50ntFlanks | 11594 | 11613 | 20 | CAGGGGAATCCGACTGTTTA | 344.2 | -6.2 | 1.5 | -4.0 | 6.5E-05 | 1.0E-04 | MB231/MB468 | MP |
| >hg38_21_+_8433172_8446622_RNA45SN1_45S_50ntFlanks@11209.11252.44\|CGGCGACTCTGGACGCGAGCCGGGCCCTTCCCGTGGATCGCCCC | >hg38_21_+_8433172_8446622_RNA45SN1_45S_50ntFlanks | 11209 | 11252 | 44 | CGGCGACTCTGGACGCGAGCCGGGCCCTTCCCGTGGATCGCCCC | 1793.2 | -5.5 | 1.4 | -4.0 | 7.7E-05 | 1.0E-04 | MB231/MB468 | NUC |
| >hg38_21_+_8433172_8446622_RNA45SN1_45S_50ntFlanks@10768.10792.25\|CAGCAGGTCTCCAAGGTGAACAGCC | >hg38_21_+_8433172_8446622_RNA45SN1_45S_50ntFlanks | 10768 | 10792 | 25 | CAGCAGGTCTCCAAGGTGAACAGCC | 41.0 | -8.4 | 2.0 | -4.2 | 2.7E-05 | 1.0E-04 | BT20/MB468 | MP |
| >hg38_21_+_8433172_8446622_RNA45SN1_45S_50ntFlanks@7975.7997.23\|CGCGACCTCAGATCAGACGTGGC | >hg38_21_+_8433172_8446622_RNA45SN1_45S_50ntFlanks | 7975 | 7997 | 23 | CGCGACCTCAGATCAGACGTGGC | 21211.5 | 4.0 | 1.0 | 4.2 | 2.7E-05 | 1.0E-04 | BT20/MB468 | MP |
| >hg38_21_+_8433172_8446622_RNA45SN1_45S_50ntFlanks@6661.6703.43\|CGGTGGATCACTCGGCTCGTGCGTCGATGAAGAACGCAGCTAG | >hg38_21_+_8433172_8446622_RNA45SN1_45S_50ntFlanks | 6661 | 6703 | 43 | CGGTGGATCACTCGGCTCGTGCGTCGATGAAGAACGCAGCTAG | 709.5 | 2.5 | 0.6 | 4.0 | 5.6E-05 | 1.0E-04 | BT20/MB468 | MITO |
| >hg38_21_+_8433172_8446622_RNA45SN1_45S_50ntFlanks@11575.11593.19\|ACTTAGAACTGGTGCGGAC | >hg38_21_+_8433172_8446622_RNA45SN1_45S_50ntFlanks | 11575 | 11593 | 19 | ACTTAGAACTGGTGCGGAC | 998.3 | -5.4 | 1.3 | -4.2 | 2.7E-05 | 1.0E-04 | BT20/MB468 | MP |
| >hg38_1_-_228634819_228635039_RNA5S12_5S_50ntFlanks@124.170.47\|AGTACTTGGATGGGAGACCGCCTGGGAATACCGGGTGCTGTAGGCTT | >hg38_1_-_228634819_228635039_RNA5S12_5S_50ntFlanks | 124 | 170 | 47 | AGTACTTGGATGGGAGACCGCCTGGGAATACCGGGTGCTGTAGGCTT | 1203.8 | 3.9 | 1.0 | 4.0 | 7.3E-05 | 1.0E-04 | MB231/MB468 | CYTO |
| >hg38_21_+_8433172_8446622_RNA45SN1_45S_50ntFlanks@11235.11259.25\|CTTCCCGTGGATCGCCCCAGCTGCG | >hg38_21_+_8433172_8446622_RNA45SN1_45S_50ntFlanks | 11235 | 11259 | 25 | CTTCCCGTGGATCGCCCCAGCTGCG | 144.8 | -3.0 | 0.8 | -4.0 | 7.3E-05 | 1.0E-04 | MB231/MB468 | CYTO |
| >MI0000078\|hsa-mir-22&WithFlank&17\|-\|1713897\|1713993@59.79.21(+1G)&[MIMAT0000077&hsa-miR-22-3p&offsets\|0\|-1(+1G);m-2&17\|-\|1713914\|1713935&offsets\|0\|-1(+1G)]\|AAGCTGCCAGTTGAAGAACTGG | >MI0000078\|hsa-mir-22&WithFlank&17\|-\|1713897\|1713993 | 59 | 79 | 21(+1G) | AAGCTGCCAGTTGAAGAACTGG | 538.2 | 4.3 | 1.1 | 4.0 | 6.5E-05 | 1.0E-04 | MB231/MB468 | MP |
| >hg38_21_+_8433172_8446622_RNA45SN1_45S_50ntFlanks@3705.3739.35\|TACCTGGTTGATCCTGCCAGTAGCATATGCTTGTC | >hg38_21_+_8433172_8446622_RNA45SN1_45S_50ntFlanks | 3705 | 3739 | 35 | TACCTGGTTGATCCTGCCAGTAGCATATGCTTGTC | 1390.6 | 3.9 | 1.0 | 4.0 | 5.7E-05 | 1.0E-04 | BT20/MB468 | MITO |
| >hg38_21_+_8433172_8446622_RNA45SN1_45S_50ntFlanks@3839.3860.22\|TCTCCTACTTGGATAACTGTGG | >hg38_21_+_8433172_8446622_RNA45SN1_45S_50ntFlanks | 3839 | 3860 | 22 | TCTCCTACTTGGATAACTGTGG | 1041.6 | 3.0 | 0.8 | 4.0 | 6.8E-05 | 1.0E-04 | BT20/MB468 | CYTO |
| >MI0000272\|hsa-mir-182&WithFlank&7\|-\|129770377\|129770498@29.51.23(+1A)&[MIMAT0000259&hsa-miR-182-5p&offsets\|0\|-1(+1A);m-22&7\|-\|129770449\|129770470&offsets\|0\|+1(+1A)]\|TTTGGCAATGGTAGAACTCACACA | >MI0000272\|hsa-mir-182&WithFlank&7\|-\|129770377\|129770498 | 29 | 51 | 23(+1A) | TTTGGCAATGGTAGAACTCACACA | 336.0 | 3.3 | 0.8 | 4.0 | 7.3E-05 | 1.0E-04 | MB231/MB468 | CYTO |
| >hg38_21_+_8433172_8446622_RNA45SN1_45S_50ntFlanks@11656.11685.30\|CGATGTGATTTCTGCCCAGTGCTCTGAATG | >hg38_21_+_8433172_8446622_RNA45SN1_45S_50ntFlanks | 11656 | 11685 | 30 | CGATGTGATTTCTGCCCAGTGCTCTGAATG | 275.3 | -4.9 | 1.2 | -3.9 | 8.0E-05 | 1.0E-04 | BT20/MB468 | NUC |
| >hg38_21_+_8433172_8446622_RNA45SN1_45S_50ntFlanks@6661.6693.33\|CGGTGGATCACTCGGCTCGTGCGTCGATGAAGA | >hg38_21_+_8433172_8446622_RNA45SN1_45S_50ntFlanks | 6661 | 6693 | 33 | CGGTGGATCACTCGGCTCGTGCGTCGATGAAGA | 208.7 | 1.1 | 0.3 | 3.9 | 8.0E-05 | 1.0E-04 | BT20/MB468 | NUC |
| >hg38_21_+_8433172_8446622_RNA45SN1_45S_50ntFlanks@9937.9980.44\|CAGCAGGACGGTGGCCATGGAAGTCGGAATCCGCTAAGGAGTGT | >hg38_21_+_8433172_8446622_RNA45SN1_45S_50ntFlanks | 9937 | 9980 | 44 | CAGCAGGACGGTGGCCATGGAAGTCGGAATCCGCTAAGGAGTGT | 75.0 | 9.6 | 2.2 | 4.4 | 9.3E-06 | 1.0E-04 | BT20/MB231 | MP |
| >MI0000088\|hsa-mir-30a&WithFlank&6\|-\|71403545\|71403627@53.74.22(+1A)&[MIMAT0000088&hsa-miR-30a-3p&offsets\|0\|0(+1A);m-38&6\|-\|71403554\|71403575&offsets\|0\|0(+1A)]\|CTTTCAGTCGGATGTTTGCAGCA | >MI0000088\|hsa-mir-30a&WithFlank&6\|-\|71403545\|71403627 | 53 | 74 | 22(+1A) | CTTTCAGTCGGATGTTTGCAGCA | 159.0 | 6.8 | 1.7 | 4.0 | 7.7E-05 | 1.0E-04 | MB231/MB468 | MITO |
| >hg38_21_+_8433172_8446622_RNA45SN1_45S_50ntFlanks@8380.8406.27\|CAAGAGGGCGTGAAACCGTTAAGAGGT | >hg38_21_+_8433172_8446622_RNA45SN1_45S_50ntFlanks | 8380 | 8406 | 27 | CAAGAGGGCGTGAAACCGTTAAGAGGT | 2002.9 | -3.1 | 0.7 | -4.2 | 2.7E-05 | 1.0E-04 | BT20/MB468 | MP |
| >trna11_SerAGA_8_-_96281885_96281966@1.23.23,trna145_SerAGA_6_-_27521192_27521273@1.23.23,trna147_SerAGA_6_-_27509554_27509635@1.23.23,trna148_SerTGA_6_-_27473607_27473688@1.23.23,trna172_SerTGA_6_-_26312824_26312905@1.23.23,trna35_SerAGA_17_-_8129928_8130009@1.23.23,trna44_SerAGA_6_+_27446591_27446672@1.23.23,trna46_SerAGA_6_+_27463593_27463674@1.23.23,trna47_SerAGA_6_+_27470818_27470899@1.23.23,trna50_SerAGA_6_+_27499987_27500068@1.23.23,trna51_SerTGA_6_+_27513468_27513549@1.23.23,trna5_SerAGA_6_+_26327817_26327898@1.23.23\|GTAGTCGTGGCCGAGTGGTTAAG | >trna11_SerAGA_8_-_96281885_96281966 | 1 | 23 | 23 | GTAGTCGTGGCCGAGTGGTTAAG | 293.1 | 7.1 | 1.7 | 4.2 | 2.7E-05 | 1.0E-04 | BT20/MB468 | MP |
| >hg38_21_+_8433172_8446622_RNA45SN1_45S_50ntFlanks@10797.10816.20\|GCATGTTGGAACAATGTAGG | >hg38_21_+_8433172_8446622_RNA45SN1_45S_50ntFlanks | 10797 | 10816 | 20 | GCATGTTGGAACAATGTAGG | 158.0 | -6.2 | 1.5 | -4.2 | 2.7E-05 | 1.0E-04 | BT20/MB468 | MP |
| >hg38_21_+_8433172_8446622_RNA45SN1_45S_50ntFlanks@12415.12443.29\|CTTCGATGTCGGCTCTTCCTATCATTGTG | >hg38_21_+_8433172_8446622_RNA45SN1_45S_50ntFlanks | 12415 | 12443 | 29 | CTTCGATGTCGGCTCTTCCTATCATTGTG | 853.0 | -6.6 | 1.6 | -4.0 | 6.5E-05 | 1.0E-04 | MB231/MB468 | MP |
| >hg38_21_+_8433172_8446622_RNA45SN1_45S_50ntFlanks@10713.10736.24\|CTTGAAAATCCGGGGGAGAGGGTG | >hg38_21_+_8433172_8446622_RNA45SN1_45S_50ntFlanks | 10713 | 10736 | 24 | CTTGAAAATCCGGGGGAGAGGGTG | 598.6 | -3.9 | 0.9 | -4.2 | 2.7E-05 | 1.0E-04 | BT20/MB468 | MP |
| >hg38_21_+_8433172_8446622_RNA45SN1_45S_50ntFlanks@12423.12455.33\|TCGGCTCTTCCTATCATTGTGAAGCAGAATTCA | >hg38_21_+_8433172_8446622_RNA45SN1_45S_50ntFlanks | 12423 | 12455 | 33 | TCGGCTCTTCCTATCATTGTGAAGCAGAATTCA | 145.2 | -5.3 | 1.3 | -4.0 | 6.8E-05 | 1.0E-04 | BT20/MB468 | CYTO |
| >hg38_21_+_8433172_8446622_RNA45SN1_45S_50ntFlanks@8194.8212.19\|CCCGTGGACGGTGTGAGGC | >hg38_21_+_8433172_8446622_RNA45SN1_45S_50ntFlanks | 8194 | 8212 | 19 | CCCGTGGACGGTGTGAGGC | 519.6 | -2.6 | 0.6 | -4.0 | 6.8E-05 | 1.0E-04 | BT20/MB468 | CYTO |
| >hg38_21_+_8433172_8446622_RNA45SN1_45S_50ntFlanks@8340.8378.39\|ACCGTAAGGGAAAGTTGAAAAGAACTTTGAAGAGAGAGT | >hg38_21_+_8433172_8446622_RNA45SN1_45S_50ntFlanks | 8340 | 8378 | 39 | ACCGTAAGGGAAAGTTGAAAAGAACTTTGAAGAGAGAGT | 74.3 | 9.5 | 2.2 | 4.4 | 9.4E-06 | 1.0E-04 | BT20/MB231 | MP |
| >MI0000107\|hsa-mir-29b-2&WithFlank&1\|-\|207802437\|207802529@57.77.21(+1U);MI0000105\|hsa-mir-29b-1&WithFlank&7\|-\|130877453\|130877545@56.76.21(+1U)&[MIMAT0000100&hsa-miR-29b-3p&offsets\|-1\|-3(+1U);m-81&1\|-\|207802450\|207802472&offsets\|-1\|-3(+1U)];[MIMAT0000100_1&hsa-miR-29b-3p&offsets\|-1\|-3(+1U);m-82&7\|-\|130877467\|130877489&offsets\|-1\|-3(+1U)]\|CTAGCACCATTTGAAATCAGTT | >MI0000107\|hsa-mir-29b-2&WithFlank&1\|-\|207802437\|207802529 | 57 | 77 | 21(+1U) | CTAGCACCATTTGAAATCAGTT | 64.3 | -9.4 | 2.1 | -4.4 | 9.4E-06 | 1.0E-04 | BT20/MB231 | MP |
| >hg38_21_+_8433172_8446622_RNA45SN1_45S_50ntFlanks@5125.5145.21\|AGCGGTCGGCGTCCCCCAACT | >hg38_21_+_8433172_8446622_RNA45SN1_45S_50ntFlanks | 5125 | 5145 | 21 | AGCGGTCGGCGTCCCCCAACT | 190.4 | 6.0 | 1.5 | 3.9 | 8.1E-05 | 1.0E-04 | BT20/MB468 | NUC |
| >trnaMT_PheGAA_MT_+_577_647@56.74.19\|CATCACCCCATAAACACCA | >trnaMT_PheGAA_MT_+_577_647 | 56 | 74 | 19 | CATCACCCCATAAACACCA | 378.1 | 4.1 | 1.0 | 4.0 | 7.8E-05 | 1.0E-04 | MB231/MB468 | NUC |
| >MI0003137\|hsa-mir-193b&WithFlank&16\|+\|14303961\|14304055@57.76.20(+1U)&[MIMAT0002819&hsa-miR-193b-3p&offsets\|0\|-2(+1U);m-166&16\|+\|14304017\|14304038&offsets\|0\|-2(+1U)]\|AACTGGCCCTCAAAGTCCCGT | >MI0003137\|hsa-mir-193b&WithFlank&16\|+\|14303961\|14304055 | 57 | 76 | 20(+1U) | AACTGGCCCTCAAAGTCCCGT | 404.4 | 6.2 | 1.5 | 4.2 | 2.7E-05 | 1.0E-04 | BT20/MB468 | MP |
| >hg38_21_+_8433172_8446622_RNA45SN1_45S_50ntFlanks@11670.11691.22\|CCCAGTGCTCTGAATGTCAAAG | >hg38_21_+_8433172_8446622_RNA45SN1_45S_50ntFlanks | 11670 | 11691 | 22 | CCCAGTGCTCTGAATGTCAAAG | 2668.9 | -6.0 | 1.5 | -3.9 | 8.1E-05 | 1.0E-04 | BT20/MB468 | NUC |
| >MI0000082\|hsa-mir-25&WithFlank&7\|-\|100093554\|100093649@58.76.19(+1A)&[MIMAT0000081&hsa-miR-25-3p&offsets\|0\|-3(+1A);m-23&7\|-\|100093571\|100093592&offsets\|0\|-3(+1A)]\|CATTGCACTTGTCTCGGTCA | >MI0000082\|hsa-mir-25&WithFlank&7\|-\|100093554\|100093649 | 58 | 76 | 19(+1A) | CATTGCACTTGTCTCGGTCA | 35.3 | 8.4 | 2.1 | 4.0 | 6.6E-05 | 1.0E-04 | MB231/MB468 | MP |
| >hg38_21_+_8433172_8446622_RNA45SN1_45S_50ntFlanks@3858.3898.41\|TGGTAATTCTAGAGCTAATACATGCCGACGGGCGCTGACCC | >hg38_21_+_8433172_8446622_RNA45SN1_45S_50ntFlanks | 3858 | 3898 | 41 | TGGTAATTCTAGAGCTAATACATGCCGACGGGCGCTGACCC | 123.2 | -5.5 | 1.4 | -4.0 | 6.8E-05 | 1.0E-04 | BT20/MB468 | CYTO |
| >MI0000264\|hsa-mir-7-2&WithFlank&15\|+\|88611819\|88611940@38.60.23(+1G);MI0000265\|hsa-mir-7-3&WithFlank&19\|+\|4770664\|4770785@37.59.23(+1G);MI0000263\|hsa-mir-7-1&WithFlank&9\|-\|83969742\|83969863@30.52.23(+1G)&[MIMAT0000252&hsa-miR-7-5p&offsets\|0\|-1(+1G);m-400&15\|+\|88611856\|88611879&offsets\|0\|-1(+1G)];[MIMAT0000252_1&hsa-miR-7-5p&offsets\|0\|-1(+1G);m-402&19\|+\|4770700\|4770723&offsets\|0\|-1(+1G)];[MIMAT0000252_2&hsa-miR-7-5p&offsets\|0\|-1(+1G);m-401&9\|-\|83969811\|83969834&offsets\|0\|-1(+1G)]\|TGGAAGACTAGTGATTTTGTTGTG | >MI0000264\|hsa-mir-7-2&WithFlank&15\|+\|88611819\|88611940 | 38 | 60 | 23(+1G) | TGGAAGACTAGTGATTTTGTTGTG | 348.8 | 5.1 | 1.3 | 4.0 | 7.8E-05 | 1.0E-04 | MB231/MB468 | MITO |
| >hg38_21_+_8433172_8446622_RNA45SN1_45S_50ntFlanks@3815.3837.23\|ATGGTTCCTTTGGTCGCTCGCTC | >hg38_21_+_8433172_8446622_RNA45SN1_45S_50ntFlanks | 3815 | 3837 | 23 | ATGGTTCCTTTGGTCGCTCGCTC | 457.2 | -2.7 | 0.7 | -4.0 | 7.8E-05 | 1.0E-04 | MB231/MB468 | NUC |
| >hg38_21_+_8433172_8446622_RNA45SN1_45S_50ntFlanks@5445.5467.23\|TCGGATCGGCCCCGCCGGGGTCG | >hg38_21_+_8433172_8446622_RNA45SN1_45S_50ntFlanks | 5445 | 5467 | 23 | TCGGATCGGCCCCGCCGGGGTCG | 153.7 | -5.3 | 1.3 | -4.0 | 5.7E-05 | 1.0E-04 | BT20/MB468 | MITO |
| >hg38_21_+_8433172_8446622_RNA45SN1_45S_50ntFlanks@6671.6696.26\|CTCGGCTCGTGCGTCGATGAAGAACG | >hg38_21_+_8433172_8446622_RNA45SN1_45S_50ntFlanks | 6671 | 6696 | 26 | CTCGGCTCGTGCGTCGATGAAGAACG | 459.6 | -4.0 | 1.0 | -4.0 | 6.8E-05 | 1.0E-04 | BT20/MB468 | CYTO |
| >MI0000736\|hsa-mir-30c-1&WithFlank&1\|+\|40757278\|40757378@23.44.22(+1U);MI0000254\|hsa-mir-30c-2&WithFlank&6\|-\|71376954\|71377037@13.34.22(+1U)&[MIMAT0000244&hsa-miR-30c-5p&offsets\|0\|-1(+1U);m-57&1\|+\|40757300\|40757323&offsets\|0\|-2(+1U)];[MIMAT0000244_1&hsa-miR-30c-5p&offsets\|0\|-1(+1U);m-56&6\|-\|71377002\|71377025&offsets\|0\|-2(+1U)]\|TGTAAACATCCTACACTCTCAGT | >MI0000736\|hsa-mir-30c-1&WithFlank&1\|+\|40757278\|40757378 | 23 | 44 | 22(+1U) | TGTAAACATCCTACACTCTCAGT | 13502.5 | -3.9 | 0.9 | -4.4 | 1.1E-05 | 1.0E-04 | BT20/MB231 | MITO |
| >hg38_21_+_8433172_8446622_RNA45SN1_45S_50ntFlanks@6698.6720.23\|AGCTAGCTGCGAGAATTAATGTG | >hg38_21_+_8433172_8446622_RNA45SN1_45S_50ntFlanks | 6698 | 6720 | 23 | AGCTAGCTGCGAGAATTAATGTG | 18576.2 | -2.8 | 0.7 | -4.0 | 7.3E-05 | 1.0E-04 | MB231/MB468 | CYTO |
| >hg38_21_+_8433172_8446622_RNA45SN1_45S_50ntFlanks@4006.4039.34\|ACCTCGGGCCGATCGCACGCCCCCCGTGGCGGCG | >hg38_21_+_8433172_8446622_RNA45SN1_45S_50ntFlanks | 4006 | 4039 | 34 | ACCTCGGGCCGATCGCACGCCCCCCGTGGCGGCG | 240.7 | -3.0 | 0.8 | -4.0 | 7.3E-05 | 1.0E-04 | MB231/MB468 | CYTO |
| >hg38_21_+_8433172_8446622_RNA45SN1_45S_50ntFlanks@11561.11585.25\|GCGCCTAGCAGCCGACTTAGAACTG | >hg38_21_+_8433172_8446622_RNA45SN1_45S_50ntFlanks | 11561 | 11585 | 25 | GCGCCTAGCAGCCGACTTAGAACTG | 1634.8 | -4.5 | 1.1 | -4.2 | 2.7E-05 | 1.0E-04 | BT20/MB468 | MP |
| >MI0003685\|hsa-mir-421&WithFlank&X\|-\|74218371\|74218467@57.75.19(+1U)&[MIMAT0003339&hsa-miR-421&offsets\|+3\|-1(+1U);m-309&X\|-\|74218392\|74218414&offsets\|+3\|-1(+1U)]\|AACAGACATTAATTGGGCGT | >MI0003685\|hsa-mir-421&WithFlank&X\|-\|74218371\|74218467 | 57 | 75 | 19(+1U) | AACAGACATTAATTGGGCGT | 111.2 | 10.0 | 2.5 | 4.0 | 6.6E-05 | 1.0E-04 | MB231/MB468 | MP |
| >hg38_MT_+_1621_3279_RNR2_16S_50ntFlanks@51.82.32\|GCTAAACCTAGCCCCAAACCCACTCCACCTTA | >hg38_MT_+_1621_3279_RNR2_16S_50ntFlanks | 51 | 82 | 32 | GCTAAACCTAGCCCCAAACCCACTCCACCTTA | 11978.3 | 5.4 | 1.3 | 4.2 | 2.7E-05 | 1.0E-04 | BT20/MB468 | MP |

<table>
<tr><th></th><th></th><th></th><th></th><th></th><th></th><th></th><th></th><th></th><th></th><th></th><th></th><th></th><th></th><th></th></tr>
<tr><td>>am_MI0005763|hsa-mir-941-1&WithFlank&20|+|63919443|63919526@53.74.22(+1A);MI0005764|hsa-mir-941-2&WithFlank&20|+|63919499|63919582@53.74.22(+1A);MI0005765|hsa-mir-941-3&WithFlank&20|+|63919555|63919638@53.74.22(+1A);MI0005766|hsa-mir-941-4&WithFlank&20|+|63919750|63919833@53.74.22(+1A);MI0031520|hsa-mir-941-5&WithFlank&20|+|63919862|63919945@53.74.22(+1A)&[MIMAT0004984&hsa-miR-941&offsets|0|-1(+1A);m-263&20|+|63919495|63919517&offsets|0|-1(+1A)];[MIMAT0004984_1&hsa-miR-941&offsets|0|-1(+1A);m-263&20|+|63919551|63919573&offsets|0|-1(+1A)];[MIMAT0004984_2&hsa-miR-941&offsets|0|-1(+1A);m-263&20|+|63919607|63919629&offsets|0|-1(+1A)];[MIMAT0004984_3&hsa-miR-941&offsets|0|-1(+1A);m-263&20|+|63919802|63919824&offsets|0|-1(+1A)];[MIMAT0004984_4&hsa-miR-941&offsets|0|-1(+1A);m-263&20|+|63919914|63919936&offsets|0|-1(+1A)]|CACCCGGCTGTGTGCACATGTGA</td><td>>am_MI0005763|hsa-mir-941-1&WithFlank&20|+|63919443|63919526</td><td>53</td><td>74</td><td>22(+1A)</td><td>CACCCGGCTGTGTGCACATGTGA</td><td>369.0</td><td>5.4</td><td>1.3</td><td>4.2</td><td>2.7E-05</td><td>1.0E-04</td><td>BT20/MB468</td><td>MP</td></tr>
<tr><td>>hg38_21_+_8433172_8446622_RNA45SN1_45S_50ntFlanks@10683.10711.29|CCGGCGGCGTCCGGTGAGCTCTCGCTGGC</td><td>>hg38_21_+_8433172_8446622_RNA45SN1_45S_50ntFlanks</td><td>10683</td><td>10711</td><td>29</td><td>CCGGCGGCGTCCGGTGAGCTCTCGCTGGC</td><td>908.0</td><td>-6.7</td><td>1.6</td><td>-4.2</td><td>2.7E-05</td><td>1.0E-04</td><td>BT20/MB468</td><td>MP</td></tr>
<tr><td>>hg38_21_+_8433172_8446622_RNA45SN1_45S_50ntFlanks@8384.8402.19|AGGGCGTGAAACCGTTAAG</td><td>>hg38_21_+_8433172_8446622_RNA45SN1_45S_50ntFlanks</td><td>8384</td><td>8402</td><td>19</td><td>AGGGCGTGAAACCGTTAAG</td><td>401.6</td><td>-3.9</td><td>0.9</td><td>-4.2</td><td>2.7E-05</td><td>1.0E-04</td><td>BT20/MB468</td><td>MP</td></tr>
<tr><td>>hg38_21_+_8433172_8446622_RNA45SN1_45S_50ntFlanks@10348.10369.22|AAACGAGAACTTTGAAGGCCGA</td><td>>hg38_21_+_8433172_8446622_RNA45SN1_45S_50ntFlanks</td><td>10348</td><td>10369</td><td>22</td><td>AAACGAGAACTTTGAAGGCCGA</td><td>560.3</td><td>2.5</td><td>0.6</td><td>4.0</td><td>7.8E-05</td><td>1.0E-04</td><td>MB231/MB468</td><td>NUC</td></tr>
<tr><td>>hg38_21_+_8433172_8446622_RNA45SN1_45S_50ntFlanks@11729.11764.36|TAACTATGACTCTCTTAAGGTAGCCAAATGCCTCGT</td><td>>hg38_21_+_8433172_8446622_RNA45SN1_45S_50ntFlanks</td><td>11729</td><td>11764</td><td>36</td><td>TAACTATGACTCTCTTAAGGTAGCCAAATGCCTCGT</td><td>527.0</td><td>-3.5</td><td>0.9</td><td>-4.0</td><td>7.8E-05</td><td>1.0E-04</td><td>MB231/MB468</td><td>NUC</td></tr>
<tr><td>>hg38_21_+_8433172_8446622_RNA45SN1_45S_50ntFlanks@11580.11611.32|GAACTGGTGCGGACCAGGGGAATCCGACTGTT</td><td>>hg38_21_+_8433172_8446622_RNA45SN1_45S_50ntFlanks</td><td>11580</td><td>11611</td><td>32</td><td>GAACTGGTGCGGACCAGGGGAATCCGACTGTT</td><td>2567.0</td><td>-2.1</td><td>0.5</td><td>-4.0</td><td>5.7E-05</td><td>1.0E-04</td><td>BT20/MB468</td><td>MITO</td></tr>
<tr><td>>trnaMT_SerGCT_MT_+_12207_12265@1.36.36|GAGAAAGCTCACAAGAACTGCTAACTCATGCCCCCA</td><td>>trnaMT_SerGCT_MT_+_12207_12265</td><td>1</td><td>36</td><td>36</td><td>GAGAAAGCTCACAAGAACTGCTAACTCATGCCCCCA</td><td>301.4</td><td>4.2</td><td>1.1</td><td>4.0</td><td>7.8E-05</td><td>1.0E-04</td><td>MB231/MB468</td><td>MITO</td></tr>
<tr><td>>hg38_21_+_8433172_8446622_RNA45SN1_45S_50ntFlanks@11572.11611.40|CCGACTTAGAACTGGTGCGGACCAGGGGAATCCGACTGTT</td><td>>hg38_21_+_8433172_8446622_RNA45SN1_45S_50ntFlanks</td><td>11572</td><td>11611</td><td>40</td><td>CCGACTTAGAACTGGTGCGGACCAGGGGAATCCGACTGTT</td><td>655.7</td><td>-3.0</td><td>0.8</td><td>-4.0</td><td>7.3E-05</td><td>1.0E-04</td><td>MB231/MB468</td><td>CYTO</td></tr>
<tr><td>>hg38_21_+_8433172_8446622_RNA45SN1_45S_50ntFlanks@11664.11685.22|TTTCTGCCCAGTGCTCTGAATG</td><td>>hg38_21_+_8433172_8446622_RNA45SN1_45S_50ntFlanks</td><td>11664</td><td>11685</td><td>22</td><td>TTTCTGCCCAGTGCTCTGAATG</td><td>83.1</td><td>9.7</td><td>2.2</td><td>4.4</td><td>9.4E-06</td><td>1.0E-04</td><td>BT20/MB231</td><td>MP</td></tr>
<tr><td>>MI0000749|hsa-mir-30e&WithFlank&1|+|40754349|40754452@65.85.21(+2U)&[MIMAT0000693&hsa-miR-30e-3p&offsets|0|-1(+2U);m-25&1|+|40754413|40754434&offsets|0|-1(+2U)]|CTTTCAGTCGGATGTTTACAGTT</td><td>>MI0000749|hsa-mir-30e&WithFlank&1|+|40754349|40754452</td><td>65</td><td>85</td><td>21(+2U)</td><td>CTTTCAGTCGGATGTTTACAGTT</td><td>443.5</td><td>3.6</td><td>0.9</td><td>4.0</td><td>7.8E-05</td><td>1.0E-04</td><td>MB231/MB468</td><td>MITO</td></tr>
<tr><td>>hg38_21_+_8433172_8446622_RNA45SN1_45S_50ntFlanks@12993.13012.20|CCCTCGCTGCGATCTATTGA</td><td>>hg38_21_+_8433172_8446622_RNA45SN1_45S_50ntFlanks</td><td>12993</td><td>13012</td><td>20</td><td>CCCTCGCTGCGATCTATTGA</td><td>563.6</td><td>-3.0</td><td>0.8</td><td>-3.9</td><td>7.8E-05</td><td>1.0E-04</td><td>MB231/MB468</td><td>NUC</td></tr>
<tr><td>>hg38_1_-_228634819_228635039_RNA5S12_5S_50ntFlanks@104.131.28|AAGCAGGGTCGGGCCTGGTTAGTACTTG</td><td>>hg38_1_-_228634819_228635039_RNA5S12_5S_50ntFlanks</td><td>104</td><td>131</td><td>28</td><td>AAGCAGGGTCGGGCCTGGTTAGTACTTG</td><td>262.9</td><td>-5.7</td><td>1.4</td><td>-3.9</td><td>7.8E-05</td><td>1.0E-04</td><td>MB231/MB468</td><td>NUC</td></tr>
<tr><td>>hg38_21_+_8433172_8446622_RNA45SN1_45S_50ntFlanks@4641.4663.23|CATTCGTATTGCGCCGCTAGAGG</td><td>>hg38_21_+_8433172_8446622_RNA45SN1_45S_50ntFlanks</td><td>4641</td><td>4663</td><td>23</td><td>CATTCGTATTGCGCCGCTAGAGG</td><td>501.6</td><td>-6.7</td><td>1.6</td><td>-4.2</td><td>2.8E-05</td><td>1.0E-04</td><td>BT20/MB468</td><td>MP</td></tr>
<tr><td>>hg38_21_+_8433172_8446622_RNA45SN1_45S_50ntFlanks@4724.4761.38|ATTAATCAAGAACGAAAGTCGGAGGTTCGAAGACGATC</td><td>>hg38_21_+_8433172_8446622_RNA45SN1_45S_50ntFlanks</td><td>4724</td><td>4761</td><td>38</td><td>ATTAATCAAGAACGAAAGTCGGAGGTTCGAAGACGATC</td><td>383.9</td><td>-5.1</td><td>1.3</td><td>-3.9</td><td>8.1E-05</td><td>1.0E-04</td><td>BT20/MB468</td><td>NUC</td></tr>
<tr><td>>trnaMT_TyrGTA_MT_-_5826_5891@32.69.38|AAATCTAAAGACAGGGGTTAGGCCTCTTTTTACCACCA</td><td>>trnaMT_TyrGTA_MT_-_5826_5891</td><td>32</td><td>69</td><td>38</td><td>AAATCTAAAGACAGGGGTTAGGCCTCTTTTTACCACCA</td><td>317.7</td><td>7.5</td><td>1.9</td><td>3.9</td><td>7.8E-05</td><td>1.0E-04</td><td>MB231/MB468</td><td>MITO</td></tr>
<tr><td>>hg38_21_+_8433172_8446622_RNA45SN1_45S_50ntFlanks@10393.10428.36|CAGCAGTTGAACATGGGTCAGTCGGTCCTGAGAGAT</td><td>>hg38_21_+_8433172_8446622_RNA45SN1_45S_50ntFlanks</td><td>10393</td><td>10428</td><td>36</td><td>CAGCAGTTGAACATGGGTCAGTCGGTCCTGAGAGAT</td><td>137.7</td><td>-6.3</td><td>1.6</td><td>-4.0</td><td>6.6E-05</td><td>1.0E-04</td><td>MB231/MB468</td><td>MP</td></tr>
<tr><td>>hg38_21_+_8433172_8446622_RNA45SN1_45S_50ntFlanks@10922.10941.20|CTGGACGAGGCGCCGCCGCC</td><td>>hg38_21_+_8433172_8446622_RNA45SN1_45S_50ntFlanks</td><td>10922</td><td>10941</td><td>20</td><td>CTGGACGAGGCGCCGCCGCC</td><td>950.7</td><td>-6.5</td><td>1.6</td><td>-3.9</td><td>7.8E-05</td><td>1.1E-04</td><td>MB231/MB468</td><td>NUC</td></tr>
<tr><td>>hg38_21_+_8433172_8446622_RNA45SN1_45S_50ntFlanks@10737.10767.31|TAAATCTCGCGCCGGGCCGTACCCATATCCG</td><td>>hg38_21_+_8433172_8446622_RNA45SN1_45S_50ntFlanks</td><td>10737</td><td>10767</td><td>31</td><td>TAAATCTCGCGCCGGGCCGTACCCATATCCG</td><td>202.7</td><td>-3.9</td><td>1.0</td><td>-4.0</td><td>7.4E-05</td><td>1.1E-04</td><td>MB231/MB468</td><td>CYTO</td></tr>
<tr><td>>MI0000471|hsa-mir-126&WithFlank&9|+|136670596|136670692@57.79.23&[MIMAT0000445&hsa-miR-126-3p&offsets|-1|0;m-40&9|+|136670653|136670674&offsets|-1|0]|CTCGTACCGTGAGTAATAATGCG</td><td>>MI0000471|hsa-mir-126&WithFlank&9|+|136670596|136670692</td><td>57</td><td>79</td><td>23</td><td>CTCGTACCGTGAGTAATAATGCG</td><td>279.2</td><td>6.9</td><td>1.7</td><td>4.0</td><td>6.6E-05</td><td>1.1E-04</td><td>MB231/MB468</td><td>MP</td></tr>
<tr><td>>hg38_21_+_8433172_8446622_RNA45SN1_45S_50ntFlanks@10683.10711.29|CCGGCGGCGTCCGGTGAGCTCTCGCTGGC</td><td>>hg38_21_+_8433172_8446622_RNA45SN1_45S_50ntFlanks</td><td>10683</td><td>10711</td><td>29</td><td>CCGGCGGCGTCCGGTGAGCTCTCGCTGGC</td><td>1591.9</td><td>-5.1</td><td>1.3</td><td>-4.0</td><td>6.9E-05</td><td>1.1E-04</td><td>BT20/MB468</td><td>CYTO</td></tr>
<tr><td>>hg38_21_+_8433172_8446622_RNA45SN1_45S_50ntFlanks@11570.11613.44|AGCCGACTTAGAACTGGTGCGGACCAGGGGAATCCGACTGTTTA</td><td>>hg38_21_+_8433172_8446622_RNA45SN1_45S_50ntFlanks</td><td>11570</td><td>11613</td><td>44</td><td>AGCCGACTTAGAACTGGTGCGGACCAGGGGAATCCGACTGTTTA</td><td>1102.7</td><td>4.0</td><td>1.0</td><td>4.0</td><td>5.8E-05</td><td>1.1E-04</td><td>BT20/MB468</td><td>MITO</td></tr>
<tr><td>>MI0000084|hsa-mir-26b&WithFlank&2|+|218402640|218402728@18.39.22(+2U)&[MIMAT0000083&hsa-miR-26b-5p&offsets|0|+1(+2U);m-53&2|+|218402657|218402678&offsets|0|0(+2U)]|TTCAAGTAATTCAGGATAGGTTTT</td><td>>MI0000084|hsa-mir-26b&WithFlank&2|+|218402640|218402728</td><td>18</td><td>39</td><td>22(+2U)</td><td>TTCAAGTAATTCAGGATAGGTTTT</td><td>67.1</td><td>-9.5</td><td>2.1</td><td>-4.4</td><td>9.5E-06</td><td>1.1E-04</td><td>BT20/MB231</td><td>MP</td></tr>
<tr><td>>hg38_21_+_8433172_8446622_RNA45SN1_45S_50ntFlanks@12940.12965.26|CATTCGTAGACGACCTGCTTCTGGGT</td><td>>hg38_21_+_8433172_8446622_RNA45SN1_45S_50ntFlanks</td><td>12940</td><td>12965</td><td>26</td><td>CATTCGTAGACGACCTGCTTCTGGGT</td><td>187.2</td><td>-2.6</td><td>0.7</td><td>-3.9</td><td>7.9E-05</td><td>1.1E-04</td><td>MB231/MB468</td><td>NUC</td></tr>
<tr><td>>hg38_21_+_8433172_8446622_RNA45SN1_45S_50ntFlanks@6650.6674.25|ACGACTCTTAGCGGTGGATCACTCG</td><td>>hg38_21_+_8433172_8446622_RNA45SN1_45S_50ntFlanks</td><td>6650</td><td>6674</td><td>25</td><td>ACGACTCTTAGCGGTGGATCACTCG</td><td>650.2</td><td>4.6</td><td>1.1</td><td>4.2</td><td>2.8E-05</td><td>1.1E-04</td><td>BT20/MB468</td><td>MP</td></tr>
<tr><td>>MI0000289|hsa-mir-181a-1&WithFlank&1|-|198859038|198859159@30.52.23;MI0000269|hsa-mir-181a-2&WithFlank&9|+|124692436|124692557@45.67.23&[MIMAT0000256&hsa-miR-181a-5p&offsets|0|0;m-32&1|-|198859108|198859130&offsets|0|0];[MIMAT0000256_1&hsa-miR-181a-5p&offsets|0|0;m-32&9|+|124692480|124692502&offsets|0|0]|AACATTCAACGCTGTCGGTGAGT</td><td>>MI0000289|hsa-mir-181a-1&WithFlank&1|-|198859038|198859159</td><td>30</td><td>52</td><td>23</td><td>AACATTCAACGCTGTCGGTGAGT</td><td>12329.7</td><td>3.4</td><td>0.9</td><td>4.0</td><td>6.6E-05</td><td>1.1E-04</td><td>MB231/MB468</td><td>MP</td></tr>
<tr><td>>hg38_21_+_8433172_8446622_RNA45SN1_45S_50ntFlanks@11664.11702.39|TTTCTGCCCAGTGCTCTGAATGTCAAAGTGAAGAAATTC</td><td>>hg38_21_+_8433172_8446622_RNA45SN1_45S_50ntFlanks</td><td>11664</td><td>11702</td><td>39</td><td>TTTCTGCCCAGTGCTCTGAATGTCAAAGTGAAGAAATTC</td><td>238.7</td><td>4.3</td><td>1.1</td><td>4.0</td><td>7.4E-05</td><td>1.1E-04</td><td>MB231/MB468</td><td>CYTO</td></tr>
<tr><td>>hg38_21_+_8433172_8446622_RNA45SN1_45S_50ntFlanks@12638.12659.22|CTACCATCTGTGGGATTATGAC</td><td>>hg38_21_+_8433172_8446622_RNA45SN1_45S_50ntFlanks</td><td>12638</td><td>12659</td><td>22</td><td>CTACCATCTGTGGGATTATGAC</td><td>1849.5</td><td>-5.7</td><td>1.4</td><td>-4.2</td><td>2.8E-05</td><td>1.1E-04</td><td>BT20/MB468</td><td>MP</td></tr>
<tr><td>>MI0000811|hsa-mir-148b&WithFlank&12|+|54337210|54337320@69.90.22&[MIMAT0000759&hsa-miR-148b-3p&offsets|0|0;m-77&12|+|54337278|54337299&offsets|0|0]|TCAGTGCATCACAGAACTTTGT</td><td>>MI0000811|hsa-mir-148b&WithFlank&12|+|54337210|54337320</td><td>69</td><td>90</td><td>22</td><td>TCAGTGCATCACAGAACTTTGT</td><td>14856.9</td><td>3.0</td><td>0.7</td><td>4.0</td><td>6.6E-05</td><td>1.1E-04</td><td>MB231/MB468</td><td>MP</td></tr>
<tr><td>>hg38_21_+_8433172_8446622_RNA45SN1_45S_50ntFlanks@3705.3736.32|TACCTGGTTGATCCTGCCAGTAGCATATGCTT</td><td>>hg38_21_+_8433172_8446622_RNA45SN1_45S_50ntFlanks</td><td>3705</td><td>3736</td><td>32</td><td>TACCTGGTTGATCCTGCCAGTAGCATATGCTT</td><td>991.4</td><td>3.2</td><td>0.7</td><td>4.4</td><td>1.0E-05</td><td>1.1E-04</td><td>BT20/MB231</td><td>NUC</td></tr>
<tr><td>>am_trnaMT_GluTTC_MT_-_14674_14742@27.55.29,trnalookalike8_GluTTC_5_-_93905172_93905240@27.55.29|GTTTTTCATATCATTGGTCGTGGTTGTAG</td><td>>am_trnaMT_GluTTC_MT_-_14674_14742</td><td>27</td><td>55</td><td>29</td><td>GTTTTTCATATCATTGGTCGTGGTTGTAG</td><td>76.0</td><td>-6.3</td><td>1.6</td><td>-4.0</td><td>6.9E-05</td><td>1.1E-04</td><td>BT20/MB468</td><td>CYTO</td></tr>
<tr><td>>trnaMT_GluTTC_MT_-_14674_14742@41.71.31,trnalookalike8_GluTTC_5_-_93905172_93905240@41.71.31|TGGTCGTGGTTGTAGTCCGTGCGAGAATACC</td><td>>trnaMT_GluTTC_MT_-_14674_14742</td><td>41</td><td>71</td><td>31</td><td>TGGTCGTGGTTGTAGTCCGTGCGAGAATACC</td><td>653.5</td><td>-4.0</td><td>0.9</td><td>-4.4</td><td>9.0E-06</td><td>1.1E-04</td><td>BT20/MB231</td><td>CYTO</td></tr>
<tr><td>>hg38_21_+_8433172_8446622_RNA45SN1_45S_50ntFlanks@12003.12029.27|GCGGGCCGCCGGTGAAATACCACTACT</td><td>>hg38_21_+_8433172_8446622_RNA45SN1_45S_50ntFlanks</td><td>12003</td><td>12029</td><td>27</td><td>GCGGGCCGCCGGTGAAATACCACTACT</td><td>1275.2</td><td>-3.8</td><td>0.9</td><td>-4.4</td><td>9.0E-06</td><td>1.1E-04</td><td>BT20/MB231</td><td>CYTO</td></tr>
<tr><td>>hg38_21_+_8433172_8446622_RNA45SN1_45S_50ntFlanks@5470.5514.45|CCACGGCCCTGGCGGAGCGCTGAGAAGACGGTCGAACTTGACTAT</td><td>>hg38_21_+_8433172_8446622_RNA45SN1_45S_50ntFlanks</td><td>5470</td><td>5514</td><td>45</td><td>CCACGGCCCTGGCGGAGCGCTGAGAAGACGGTCGAACTTGACTAT</td><td>556.4</td><td>-6.5</td><td>1.7</td><td>-3.9</td><td>7.9E-05</td><td>1.1E-04</td><td>MB231/MB468</td><td>MITO</td></tr>
<tr><td>>hg38_21_+_8433172_8446622_RNA45SN1_45S_50ntFlanks@9567.9597.31|ACGTGCAAATCGGTCGTCCGACCTGGGTATA</td><td>>hg38_21_+_8433172_8446622_RNA45SN1_45S_50ntFlanks</td><td>9567</td><td>9597</td><td>31</td><td>ACGTGCAAATCGGTCGTCCGACCTGGGTATA</td><td>3036.3</td><td>-7.3</td><td>1.7</td><td>-4.2</td><td>2.8E-05</td><td>1.1E-04</td><td>BT20/MB468</td><td>MP</td></tr>
<tr><td>>hg38_21_+_8433172_8446622_RNA45SN1_45S_50ntFlanks@11769.11803.35|TAATTAGTGACGCGCATGAATGGATGAACGAGATT</td><td>>hg38_21_+_8433172_8446622_RNA45SN1_45S_50ntFlanks</td><td>11769</td><td>11803</td><td>35</td><td>TAATTAGTGACGCGCATGAATGGATGAACGAGATT</td><td>214.2</td><td>-3.8</td><td>0.9</td><td>-4.0</td><td>6.7E-05</td><td>1.1E-04</td><td>MB231/MB468</td><td>MP</td></tr>
<tr><td>>hg38_21_+_8433172_8446622_RNA45SN1_45S_50ntFlanks@4006.4039.34|ACCTCGGGCCGATCGCACGCCCCCCGTGGCGGCG</td><td>>hg38_21_+_8433172_8446622_RNA45SN1_45S_50ntFlanks</td><td>4006</td><td>4039</td><td>34</td><td>ACCTCGGGCCGATCGCACGCCCCCCGTGGCGGCG</td><td>192.7</td><td>-3.9</td><td>1.0</td><td>-4.0</td><td>6.9E-05</td><td>1.1E-04</td><td>BT20/MB468</td><td>CYTO</td></tr>
<tr><td>>hg38_21_+_8433172_8446622_RNA45SN1_45S_50ntFlanks@12432.12455.24|CCTATCATTGTGAAGCAGAATTCA</td><td>>hg38_21_+_8433172_8446622_RNA45SN1_45S_50ntFlanks</td><td>12432</td><td>12455</td><td>24</td><td>CCTATCATTGTGAAGCAGAATTCA</td><td>247.3</td><td>-4.4</td><td>1.1</td><td>-4.0</td><td>5.8E-05</td><td>1.1E-04</td><td>BT20/MB468</td><td>MITO</td></tr>
<tr><td>>hg38_21_+_8433172_8446622_RNA45SN1_45S_50ntFlanks@3846.3865.20|CTTGGATAACTGTGGTAATT</td><td>>hg38_21_+_8433172_8446622_RNA45SN1_45S_50ntFlanks</td><td>3846</td><td>3865</td><td>20</td><td>CTTGGATAACTGTGGTAATT</td><td>156.2</td><td>-2.4</td><td>0.6</td><td>-4.0</td><td>7.5E-05</td><td>1.1E-04</td><td>MB231/MB468</td><td>CYTO</td></tr>
<tr><td>>hg38_21_+_8433172_8446622_RNA45SN1_45S_50ntFlanks@12476.12496.21|CCACTAATAGGGAACGTGAGC</td><td>>hg38_21_+_8433172_8446622_RNA45SN1_45S_50ntFlanks</td><td>12476</td><td>12496</td><td>21</td><td>CCACTAATAGGGAACGTGAGC</td><td>1968.6</td><td>-3.8</td><td>0.9</td><td>-4.2</td><td>2.8E-05</td><td>1.1E-04</td><td>BT20/MB468</td><td>MP</td></tr>
<tr><td>>am_trnaMT_GluTTC_MT_-_14674_14742@27.53.27,trnalookalike8_GluTTC_5_-_93905172_93905240@27.53.27|GTTTTTCATATCATTGGTCGTGGTTGT</td><td>>am_trnaMT_GluTTC_MT_-_14674_14742</td><td>27</td><td>53</td><td>27</td><td>GTTTTTCATATCATTGGTCGTGGTTGT</td><td>202.1</td><td>-4.5</td><td>1.1</td><td>-4.0</td><td>6.7E-05</td><td>1.1E-04</td><td>MB231/MB468</td><td>MP</td></tr>
<tr><td>>hg38_21_+_8433172_8446622_RNA45SN1_45S_50ntFlanks@11672.11693.22|CAGTGCTCTGAATGTCAAAGTG</td><td>>hg38_21_+_8433172_8446622_RNA45SN1_45S_50ntFlanks</td><td>11672</td><td>11693</td><td>22</td><td>CAGTGCTCTGAATGTCAAAGTG</td><td>452.1</td><td>-6.7</td><td>1.7</td><td>-3.9</td><td>8.2E-05</td><td>1.1E-04</td><td>BT20/MB468</td><td>NUC</td></tr>
<tr><td>>hg38_21_+_8433172_8446622_RNA45SN1_45S_50ntFlanks@9306.9323.18|CAAGGAGTCTAACACGTG</td><td>>hg38_21_+_8433172_8446622_RNA45SN1_45S_50ntFlanks</td><td>9306</td><td>9323</td><td>18</td><td>CAAGGAGTCTAACACGTG</td><td>292.2</td><td>-2.5</td><td>0.6</td><td>-4.0</td><td>5.8E-05</td><td>1.1E-04</td><td>BT20/MB468</td><td>MITO</td></tr>
<tr><td>>hg38_21_+_8433172_8446622_RNA45SN1_45S_50ntFlanks@11766.11803.38|ATCTAATTAGTGACGCGCATGAATGGATGAACGAGATT</td><td>>hg38_21_+_8433172_8446622_RNA45SN1_45S_50ntFlanks</td><td>11766</td><td>11803</td><td>38</td><td>ATCTAATTAGTGACGCGCATGAATGGATGAACGAGATT</td><td>177.8</td><td>-6.2</td><td>1.6</td><td>-4.0</td><td>6.7E-05</td><td>1.1E-04</td><td>MB231/MB468</td><td>MP</td></tr>
<tr><td>>hg38_21_+_8433172_8446622_RNA45SN1_45S_50ntFlanks@10349.10390.42|AACGAGAACTTTGAAGGCCGAAGTGGAGAAGGGTTCCATGTG</td><td>>hg38_21_+_8433172_8446622_RNA45SN1_45S_50ntFlanks</td><td>10349</td><td>10390</td><td>42</td><td>AACGAGAACTTTGAAGGCCGAAGTGGAGAAGGGTTCCATGTG</td><td>2312.4</td><td>-3.7</td><td>0.9</td><td>-3.9</td><td>8.0E-05</td><td>1.1E-04</td><td>MB231/MB468</td><td>MITO</td></tr>
<tr><td>>hg38_21_+_8433172_8446622_RNA45SN1_45S_50ntFlanks@8107.8140.34|CGGCGGGGCGCGGGACATGTGGCGTACGGAAGAC</td><td>>hg38_21_+_8433172_8446622_RNA45SN1_45S_50ntFlanks</td><td>8107</td><td>8140</td><td>34</td><td>CGGCGGGGCGCGGGACATGTGGCGTACGGAAGAC</td><td>862.4</td><td>-3.8</td><td>0.9</td><td>-4.2</td><td>2.8E-05</td><td>1.1E-04</td><td>BT20/MB468</td><td>MP</td></tr>
<tr><td>>hg38_21_+_8433172_8446622_RNA45SN1_45S_50ntFlanks@10819.10865.47|AGGGAAGTCGGCAAGCCGGATCCGTAACTTCGGGATAAGGATTGGCT</td><td>>hg38_21_+_8433172_8446622_RNA45SN1_45S_50ntFlanks</td><td>10819</td><td>10865</td><td>47</td><td>AGGGAAGTCGGCAAGCCGGATCCGTAACTTCGGGATAAGGATTGGCT</td><td>315.5</td><td>-4.5</td><td>1.1</td><td>-4.0</td><td>7.5E-05</td><td>1.1E-04</td><td>MB231/MB468</td><td>CYTO</td></tr>
</table>

<table>
<tr><th></th><th></th><th></th><th></th><th></th><th></th><th></th><th></th><th></th><th></th><th></th><th></th><th></th><th></th></tr>
<tr><td>>hg38_21_+_8433172_8446622_RNA45SN1_45S_50ntFlanks@11727.11745.19|AGTAACTATGACTCTCTTA</td><td>>hg38_21_+_8433172_8446622_RNA45SN1_45S_50ntFlanks</td><td>11727</td><td>11745</td><td>19</td><td>AGTAACTATGACTCTCTTA</td><td>351.5</td><td>-5.0</td><td>1.3</td><td>-3.9</td><td>8.2E-05</td><td>1.1E-04</td><td>BT20/MB468</td><td>NUC</td></tr>
<tr><td>>am_trna128_GlyGCC_6_-_27870686_27870756@1.35.35,trna18_GlyGCC_16_+_70822597_70822667@1.35.35,trna19_GlyGCC_16_+_70823410_70823480@1.35.35,trna19_GlyGCC_2_-_157257659_157257729@1.35.35,trna24_GlyGCC_16_-_70812942_70813012@1.35.35,trna25_GlyGCC_16_-_70812114_70812184@1.35.35,trna5_GlyGCC_17_+_8029064_8029134@1.35.35,trna68_GlyGCC_1_-_161493637_161493707@1.35.35|GCATTGGTGGTTCAGTGGTAGAATTCTCGCCTGCC</td><td>>am_trna128_GlyGCC_6_-_27870686_27870756</td><td>1</td><td>35</td><td>35</td><td>GCATTGGTGGTTCAGTGGTAGAATTCTCGCCTGCC</td><td>2443.9</td><td>5.9</td><td>1.4</td><td>4.2</td><td>2.8E-05</td><td>1.1E-04</td><td>BT20/MB468</td><td>MP</td></tr>
<tr><td>>am_trnaMT_ValTAC_MT_+_1602_1670@36.62.27|CTTAGGAGATTTCAACTTAACTTGACC</td><td>>am_trnaMT_ValTAC_MT_+_1602_1670</td><td>36</td><td>62</td><td>27</td><td>CTTAGGAGATTTCAACTTAACTTGACC</td><td>653.7</td><td>6.9</td><td>1.7</td><td>4.2</td><td>2.8E-05</td><td>1.1E-04</td><td>BT20/MB468</td><td>MP</td></tr>
<tr><td>>hg38_21_+_8433172_8446622_RNA45SN1_45S_50ntFlanks@11571.11591.21|GCCGACTTAGAACTGGTGCGG</td><td>>hg38_21_+_8433172_8446622_RNA45SN1_45S_50ntFlanks</td><td>11571</td><td>11591</td><td>21</td><td>GCCGACTTAGAACTGGTGCGG</td><td>196.3</td><td>-4.1</td><td>1.0</td><td>-4.0</td><td>7.0E-05</td><td>1.1E-04</td><td>BT20/MB468</td><td>CYTO</td></tr>
<tr><td>>hg38_21_+_8433172_8446622_RNA45SN1_45S_50ntFlanks@10377.10401.25|AAGGGTTCCATGTGAACAGCAGTTG</td><td>>hg38_21_+_8433172_8446622_RNA45SN1_45S_50ntFlanks</td><td>10377</td><td>10401</td><td>25</td><td>AAGGGTTCCATGTGAACAGCAGTTG</td><td>53.6</td><td>-9.6</td><td>2.4</td><td>-4.0</td><td>6.7E-05</td><td>1.1E-04</td><td>MB231/MB468</td><td>MP</td></tr>
<tr><td>>hg38_MT_+_1621_3279_RNR2_16S_50ntFlanks@1427.1451.25|GATTAAAGTCCTACGTGATCTGAGT</td><td>>hg38_MT_+_1621_3279_RNR2_16S_50ntFlanks</td><td>1427</td><td>1451</td><td>25</td><td>GATTAAAGTCCTACGTGATCTGAGT</td><td>178.1</td><td>4.7</td><td>1.2</td><td>3.9</td><td>8.0E-05</td><td>1.1E-04</td><td>MB231/MB468</td><td>MITO</td></tr>
<tr><td>>MI0003205|hsa-mir-532&WithFlank&X|+|50003142|50003244@26.47.22&[MIMAT0002888&hsa-miR-532-5p&offsets|0|0;m-65&X|+|50003167|50003188&offsets|0|0]|CATGCCTTGAGTGTAGGACCGT</td><td>>MI0003205|hsa-mir-532&WithFlank&X|+|50003142|50003244</td><td>26</td><td>47</td><td>22</td><td>CATGCCTTGAGTGTAGGACCGT</td><td>1592.8</td><td>3.4</td><td>0.9</td><td>3.9</td><td>8.0E-05</td><td>1.1E-04</td><td>MB231/MB468</td><td>MITO</td></tr>
<tr><td>>hg38_21_+_8433172_8446622_RNA45SN1_45S_50ntFlanks@11771.11791.21|ATTAGTGACGCGCATGAATGG</td><td>>hg38_21_+_8433172_8446622_RNA45SN1_45S_50ntFlanks</td><td>11771</td><td>11791</td><td>21</td><td>ATTAGTGACGCGCATGAATGG</td><td>152.4</td><td>-5.8</td><td>1.5</td><td>-4.0</td><td>6.7E-05</td><td>1.1E-04</td><td>MB231/MB468</td><td>MP</td></tr>
<tr><td>>hg38_21_+_8433172_8446622_RNA45SN1_45S_50ntFlanks@4731.4758.28|AAGAACGAAAGTCGGAGGTTCGAAGACG</td><td>>hg38_21_+_8433172_8446622_RNA45SN1_45S_50ntFlanks</td><td>4731</td><td>4758</td><td>28</td><td>AAGAACGAAAGTCGGAGGTTCGAAGACG</td><td>4450.4</td><td>-3.0</td><td>0.7</td><td>-4.0</td><td>5.9E-05</td><td>1.1E-04</td><td>BT20/MB468</td><td>MITO</td></tr>
<tr><td>>hg38_21_+_8433172_8446622_RNA45SN1_45S_50ntFlanks@4002.4042.41|GATAACCTCGGGCCGATCGCACGCCCCCCGTGGCGGCGACG</td><td>>hg38_21_+_8433172_8446622_RNA45SN1_45S_50ntFlanks</td><td>4002</td><td>4042</td><td>41</td><td>GATAACCTCGGGCCGATCGCACGCCCCCCGTGGCGGCGACG</td><td>223.4</td><td>-5.3</td><td>1.3</td><td>-4.0</td><td>5.9E-05</td><td>1.1E-04</td><td>BT20/MB468</td><td>MITO</td></tr>
<tr><td>>MI0000542|hsa-mir-320a&WithFlank&8|-|22244960|22245043@48.70.23(+1G)&[MIMAT0000510&hsa-miR-320a-3p&offsets|0|+1(+1G);m-60&8|-|22244975|22244996&offsets|0|+1(+1G)]|AAAAGCTGGGTTGAGAGGGCGAAG</td><td>>MI0000542|hsa-mir-320a&WithFlank&8|-|22244960|22245043</td><td>48</td><td>70</td><td>23(+1G)</td><td>AAAAGCTGGGTTGAGAGGGCGAAG</td><td>463.4</td><td>2.8</td><td>0.6</td><td>4.4</td><td>1.0E-05</td><td>1.1E-04</td><td>BT20/MB231</td><td>NUC</td></tr>
<tr><td>>hg38_21_+_8433172_8446622_RNA45SN1_45S_50ntFlanks@10843.10866.24|TAACTTCGGGATAAGGATTGGCTC</td><td>>hg38_21_+_8433172_8446622_RNA45SN1_45S_50ntFlanks</td><td>10843</td><td>10866</td><td>24</td><td>TAACTTCGGGATAAGGATTGGCTC</td><td>211.5</td><td>-5.7</td><td>1.4</td><td>-3.9</td><td>8.0E-05</td><td>1.1E-04</td><td>MB231/MB468</td><td>NUC</td></tr>
<tr><td>>trna10_LysCTT_16_+_3241501_3241573@1.28.28,trna119_LysCTT_1_-_145395522_145395594@1.28.28,trna11_LysCTT_5_-_180648979_180649051@1.28.28,trna13_LysCTT_14_-_58706613_58706685@1.28.28,trna13_LysCTT_6_+_26556774_26556846@1.28.28,trna2_LysCTT_15_+_79152904_79152976@1.28.28,trna32_LysCTT_16_-_3207406_3207478@1.28.28,trna7_LysCTT_16_+_3225692_3225764@1.28.28,trna9_LysCTT_5_+_180634755_180634827@1.28.28|GCCCGGCTAGCTCAGTCGGTAGAGCATG</td><td>>trna10_LysCTT_16_+_3241501_3241573</td><td>1</td><td>28</td><td>28</td><td>GCCCGGCTAGCTCAGTCGGTAGAGCATG</td><td>2121.7</td><td>-3.8</td><td>1.0</td><td>-3.9</td><td>8.0E-05</td><td>1.1E-04</td><td>MB231/MB468</td><td>NUC</td></tr>
<tr><td>>MI0000087|hsa-mir-29a&WithFlank&7|-|130876741|130876816@47.67.21(+1A)&[MIMAT0000086&hsa-miR-29a-3p&offsets|-1|-2(+1A);m-26&7|-|130876749|130876769&offsets|-1|-1(+1A)]|CTAGCACCATCTGAAATCGGTA</td><td>>MI0000087|hsa-mir-29a&WithFlank&7|-|130876741|130876816</td><td>47</td><td>67</td><td>21(+1A)</td><td>CTAGCACCATCTGAAATCGGTA</td><td>465.2</td><td>6.3</td><td>1.5</td><td>4.2</td><td>2.8E-05</td><td>1.1E-04</td><td>BT20/MB468</td><td>MP</td></tr>
<tr><td>>hg38_21_+_8433172_8446622_RNA45SN1_45S_50ntFlanks@12479.12509.31|CTAATAGGGAACGTGAGCTGGGTTTAGACCG</td><td>>hg38_21_+_8433172_8446622_RNA45SN1_45S_50ntFlanks</td><td>12479</td><td>12509</td><td>31</td><td>CTAATAGGGAACGTGAGCTGGGTTTAGACCG</td><td>208.1</td><td>-4.7</td><td>1.2</td><td>-4.0</td><td>7.6E-05</td><td>1.1E-04</td><td>MB231/MB468</td><td>CYTO</td></tr>
<tr><td>>hg38_21_+_8433172_8446622_RNA45SN1_45S_50ntFlanks@8121.8140.20|ACATGTGGCGTACGGAAGAC</td><td>>hg38_21_+_8433172_8446622_RNA45SN1_45S_50ntFlanks</td><td>8121</td><td>8140</td><td>20</td><td>ACATGTGGCGTACGGAAGAC</td><td>66.5</td><td>9.4</td><td>2.1</td><td>4.4</td><td>9.7E-06</td><td>1.1E-04</td><td>BT20/MB231</td><td>MP</td></tr>
<tr><td>>hg38_21_+_8433172_8446622_RNA45SN1_45S_50ntFlanks@7981.8009.29|CTCAGATCAGACGTGGCGACCCGCTGAAT</td><td>>hg38_21_+_8433172_8446622_RNA45SN1_45S_50ntFlanks</td><td>7981</td><td>8009</td><td>29</td><td>CTCAGATCAGACGTGGCGACCCGCTGAAT</td><td>394.4</td><td>-3.1</td><td>0.8</td><td>-4.0</td><td>6.8E-05</td><td>1.1E-04</td><td>MB231/MB468</td><td>MP</td></tr>
<tr><td>>trnaMT_PheGAA_MT_+_577_647@35.66.32|GAAAATGTTTAGACGGGCTCACATCACCCCAT</td><td>>trnaMT_PheGAA_MT_+_577_647</td><td>35</td><td>66</td><td>32</td><td>GAAAATGTTTAGACGGGCTCACATCACCCCAT</td><td>141.1</td><td>10.9</td><td>2.6</td><td>4.2</td><td>2.8E-05</td><td>1.1E-04</td><td>BT20/MB468</td><td>MP</td></tr>
<tr><td>>trna108_AlaAGC_6_-_28763741_28763812@58.75.18|TCCCCAGTACCTCCACCA</td><td>>trna108_AlaAGC_6_-_28763741_28763812</td><td>58</td><td>75</td><td>18</td><td>TCCCCAGTACCTCCACCA</td><td>191.1</td><td>3.9</td><td>1.0</td><td>3.9</td><td>8.1E-05</td><td>1.1E-04</td><td>MB231/MB468</td><td>MITO</td></tr>
<tr><td>>hg38_21_+_8433172_8446622_RNA45SN1_45S_50ntFlanks@8196.8222.27|CGTGGACGGTGTGAGGCCGGTAGCGGC</td><td>>hg38_21_+_8433172_8446622_RNA45SN1_45S_50ntFlanks</td><td>8196</td><td>8222</td><td>27</td><td>CGTGGACGGTGTGAGGCCGGTAGCGGC</td><td>429.9</td><td>-3.2</td><td>0.8</td><td>-4.0</td><td>7.1E-05</td><td>1.1E-04</td><td>BT20/MB468</td><td>CYTO</td></tr>
<tr><td>>hg38_21_+_8433172_8446622_RNA45SN1_45S_50ntFlanks@11161.11178.18|CACGGTCCCCCGCGAGGG</td><td>>hg38_21_+_8433172_8446622_RNA45SN1_45S_50ntFlanks</td><td>11161</td><td>11178</td><td>18</td><td>CACGGTCCCCCGCGAGGG</td><td>944.4</td><td>-6.0</td><td>1.5</td><td>-4.0</td><td>5.9E-05</td><td>1.1E-04</td><td>BT20/MB468</td><td>MITO</td></tr>
<tr><td>>hg38_1_-_228634819_228635039_RNA5S12_5S_50ntFlanks@84.101.18|CTCGTCTGATCTCGGAAG</td><td>>hg38_1_-_228634819_228635039_RNA5S12_5S_50ntFlanks</td><td>84</td><td>101</td><td>18</td><td>CTCGTCTGATCTCGGAAG</td><td>2356.6</td><td>-5.2</td><td>1.3</td><td>-4.0</td><td>7.6E-05</td><td>1.1E-04</td><td>MB231/MB468</td><td>CYTO</td></tr>
<tr><td>>MI0000238|hsa-mir-196a-1&WithFlank&17|-|48632484|48632565@13.35.23&[MIMAT0000226_1&hsa-miR-196a-5p&offsets|0|+1;m-268&17|-|48632532|48632553&offsets|0|+1]|TAGGTAGTTTCATGTTGTTGGGC</td><td>>MI0000238|hsa-mir-196a-1&WithFlank&17|-|48632484|48632565</td><td>13</td><td>35</td><td>23</td><td>TAGGTAGTTTCATGTTGTTGGGC</td><td>65.1</td><td>9.4</td><td>2.1</td><td>4.4</td><td>9.8E-06</td><td>1.1E-04</td><td>BT20/MB231</td><td>MP</td></tr>
<tr><td>>hg38_21_+_8433172_8446622_RNA45SN1_45S_50ntFlanks@10686.10706.21|GCGGCGTCCGGTGAGCTCTCG</td><td>>hg38_21_+_8433172_8446622_RNA45SN1_45S_50ntFlanks</td><td>10686</td><td>10706</td><td>21</td><td>GCGGCGTCCGGTGAGCTCTCG</td><td>78.4</td><td>9.6</td><td>2.2</td><td>4.4</td><td>9.8E-06</td><td>1.1E-04</td><td>BT20/MB231</td><td>MP</td></tr>
<tr><td>>am_trnaMT_ValTAC_MT_+_1602_1670@27.62.36|CAACTTACACTTAGGAGATTTCAACTTAACTTGACC</td><td>>am_trnaMT_ValTAC_MT_+_1602_1670</td><td>27</td><td>62</td><td>36</td><td>CAACTTACACTTAGGAGATTTCAACTTAACTTGACC</td><td>279.6</td><td>8.7</td><td>2.1</td><td>4.2</td><td>2.8E-05</td><td>1.1E-04</td><td>BT20/MB468</td><td>MP</td></tr>
<tr><td>>MI0000063|hsa-let-7b&WithFlank&22|+|46113680|46113774@12.33.22(+1C)&[MIMAT0000063&hsa-let-7b-5p&offsets|0|0(+1C);m-14&22|+|46113691|46113712&offsets|0|0(+1C)]|TGAGGTAGTAGGTTGTGTGGTTC</td><td>>MI0000063|hsa-let-7b&WithFlank&22|+|46113680|46113774</td><td>12</td><td>33</td><td>22(+1C)</td><td>TGAGGTAGTAGGTTGTGTGGTTC</td><td>851.3</td><td>2.5</td><td>0.6</td><td>4.4</td><td>1.1E-05</td><td>1.1E-04</td><td>BT20/MB231</td><td>NUC</td></tr>
<tr><td>>hg38_MT_+_1621_3279_RNR2_16S_50ntFlanks@1170.1209.40|AAACCTGCATTAAAAATTTCGGTTGGGGCGACCTCGGAGC</td><td>>hg38_MT_+_1621_3279_RNR2_16S_50ntFlanks</td><td>1170</td><td>1209</td><td>40</td><td>AAACCTGCATTAAAAATTTCGGTTGGGGCGACCTCGGAGC</td><td>25.2</td><td>7.9</td><td>2.0</td><td>4.0</td><td>6.8E-05</td><td>1.1E-04</td><td>MB231/MB468</td><td>MP</td></tr>
<tr><td>>hg38_21_+_8433172_8446622_RNA45SN1_45S_50ntFlanks@10869.10894.26|AGGGCTGGGTCGGTCGGGCTGGGGCG</td><td>>hg38_21_+_8433172_8446622_RNA45SN1_45S_50ntFlanks</td><td>10869</td><td>10894</td><td>26</td><td>AGGGCTGGGTCGGTCGGGCTGGGGCG</td><td>316.8</td><td>-3.5</td><td>0.9</td><td>-4.0</td><td>7.6E-05</td><td>1.1E-04</td><td>MB231/MB468</td><td>CYTO</td></tr>
<tr><td>>MI0006353|hsa-mir-1291&WithFlank&12|-|48654438|48654536@20.40.21&[MIMAT0005881&hsa-miR-1291&offsets|0|-3;m-616&12|-|48654494|48654517&offsets|0|-3]|TGGCCCTGACTGAAGACCAGC</td><td>>MI0006353|hsa-mir-1291&WithFlank&12|-|48654438|48654536</td><td>20</td><td>40</td><td>21</td><td>TGGCCCTGACTGAAGACCAGC</td><td>202.7</td><td>3.8</td><td>0.9</td><td>4.4</td><td>1.1E-05</td><td>1.1E-04</td><td>BT20/MB231</td><td>NUC</td></tr>
<tr><td>>hg38_21_+_8433172_8446622_RNA45SN1_45S_50ntFlanks@7969.8005.37|CCGAGACGCGACCTCAGATCAGACGTGGCGACCCGCT</td><td>>hg38_21_+_8433172_8446622_RNA45SN1_45S_50ntFlanks</td><td>7969</td><td>8005</td><td>37</td><td>CCGAGACGCGACCTCAGATCAGACGTGGCGACCCGCT</td><td>810.6</td><td>4.4</td><td>1.0</td><td>4.4</td><td>1.1E-05</td><td>1.1E-04</td><td>BT20/MB231</td><td>NUC</td></tr>
<tr><td>>hg38_21_+_8433172_8446622_RNA45SN1_45S_50ntFlanks@11585.11623.39|GGTGCGGACCAGGGGAATCCGACTGTTTAATTAAAACAA</td><td>>hg38_21_+_8433172_8446622_RNA45SN1_45S_50ntFlanks</td><td>11585</td><td>11623</td><td>39</td><td>GGTGCGGACCAGGGGAATCCGACTGTTTAATTAAAACAA</td><td>110.2</td><td>10.0</td><td>2.5</td><td>4.0</td><td>6.8E-05</td><td>1.1E-04</td><td>MB231/MB468</td><td>MP</td></tr>
<tr><td>>hg38_21_+_8433172_8446622_RNA45SN1_45S_50ntFlanks@6653.6686.34|ACTCTTAGCGGTGGATCACTCGGCTCGTGCGTCG</td><td>>hg38_21_+_8433172_8446622_RNA45SN1_45S_50ntFlanks</td><td>6653</td><td>6686</td><td>34</td><td>ACTCTTAGCGGTGGATCACTCGGCTCGTGCGTCG</td><td>152.6</td><td>-6.4</td><td>1.6</td><td>-4.0</td><td>6.8E-05</td><td>1.1E-04</td><td>MB231/MB468</td><td>MP</td></tr>
<tr><td>>hg38_21_+_8433172_8446622_RNA45SN1_45S_50ntFlanks@6651.6676.26|CGACTCTTAGCGGTGGATCACTCGGC</td><td>>hg38_21_+_8433172_8446622_RNA45SN1_45S_50ntFlanks</td><td>6651</td><td>6676</td><td>26</td><td>CGACTCTTAGCGGTGGATCACTCGGC</td><td>8365.3</td><td>4.1</td><td>1.0</td><td>3.9</td><td>8.1E-05</td><td>1.1E-04</td><td>MB231/MB468</td><td>NUC</td></tr>
<tr><td>>hg38_21_+_8433172_8446622_RNA45SN1_45S_50ntFlanks@12243.12268.26|GAAGGGCAAAAGCTCGCTTGATCTTG</td><td>>hg38_21_+_8433172_8446622_RNA45SN1_45S_50ntFlanks</td><td>12243</td><td>12268</td><td>26</td><td>GAAGGGCAAAAGCTCGCTTGATCTTG</td><td>401.9</td><td>-5.9</td><td>1.5</td><td>-3.9</td><td>8.1E-05</td><td>1.1E-04</td><td>MB231/MB468</td><td>MITO</td></tr>
<tr><td>>hg38_21_+_8433172_8446622_RNA45SN1_45S_50ntFlanks@8191.8212.22|CAGCCCGTGGACGGTGTGAGGC</td><td>>hg38_21_+_8433172_8446622_RNA45SN1_45S_50ntFlanks</td><td>8191</td><td>8212</td><td>22</td><td>CAGCCCGTGGACGGTGTGAGGC</td><td>409.3</td><td>-4.6</td><td>1.1</td><td>-4.0</td><td>6.0E-05</td><td>1.1E-04</td><td>BT20/MB468</td><td>MITO</td></tr>
<tr><td>>hg38_21_+_8433172_8446622_RNA45SN1_45S_50ntFlanks@3822.3852.31|CTTTGGTCGCTCGCTCCTCTCCTACTTGGAT</td><td>>hg38_21_+_8433172_8446622_RNA45SN1_45S_50ntFlanks</td><td>3822</td><td>3852</td><td>31</td><td>CTTTGGTCGCTCGCTCCTCTCCTACTTGGAT</td><td>460.2</td><td>-6.1</td><td>1.6</td><td>-3.9</td><td>8.1E-05</td><td>1.1E-04</td><td>MB231/MB468</td><td>NUC</td></tr>
<tr><td>>trnaMT_HisGTG_MT_+_12138_12206@18.59.42|AAAACATCAGATTGTGAATCTGACAACAGAGGCTTACGACCC</td><td>>trnaMT_HisGTG_MT_+_12138_12206</td><td>18</td><td>59</td><td>42</td><td>AAAACATCAGATTGTGAATCTGACAACAGAGGCTTACGACCC</td><td>131.2</td><td>10.8</td><td>2.6</td><td>4.2</td><td>2.9E-05</td><td>1.1E-04</td><td>BT20/MB468</td><td>MP</td></tr>
<tr><td>>hg38_21_+_8433172_8446622_RNA45SN1_45S_50ntFlanks@10869.10904.36|AGGGCTGGGTCGGTCGGGCTGGGGCGCGAAGCGGGG</td><td>>hg38_21_+_8433172_8446622_RNA45SN1_45S_50ntFlanks</td><td>10869</td><td>10904</td><td>36</td><td>AGGGCTGGGTCGGTCGGGCTGGGGCGCGAAGCGGGG</td><td>850.0</td><td>-6.3</td><td>1.6</td><td>-3.9</td><td>8.1E-05</td><td>1.1E-04</td><td>MB231/MB468</td><td>NUC</td></tr>
<tr><td>>MI0000076|hsa-mir-20a&WithFlank&13|+|91351059|91351141@14.36.23(+1A)&[MIMAT0000075&hsa-miR-20a-5p&offsets|0|0(+1A);m-96&13|+|91351072|91351094&offsets|0|0(+1A)]|TAAAGTGCTTATAGTGCAGGTAGA</td><td>>MI0000076|hsa-mir-20a&WithFlank&13|+|91351059|91351141</td><td>14</td><td>36</td><td>23(+1A)</td><td>TAAAGTGCTTATAGTGCAGGTAGA</td><td>851.8</td><td>4.0</td><td>1.0</td><td>4.0</td><td>7.7E-05</td><td>1.1E-04</td><td>MB231/MB468</td><td>CYTO</td></tr>
<tr><td>>hg38_21_+_8433172_8446622_RNA45SN1_45S_50ntFlanks@11770.11791.22|AATTAGTGACGCGCATGAATGG</td><td>>hg38_21_+_8433172_8446622_RNA45SN1_45S_50ntFlanks</td><td>11770</td><td>11791</td><td>22</td><td>AATTAGTGACGCGCATGAATGG</td><td>229.1</td><td>-2.7</td><td>0.7</td><td>-3.9</td><td>8.2E-05</td><td>1.1E-04</td><td>MB231/MB468</td><td>NUC</td></tr>
<tr><td>>trnaMT_SerGCT_MT_+_12207_12265@8.30.23|CTCACAAGAACTGCTAACTCATG</td><td>>trnaMT_SerGCT_MT_+_12207_12265</td><td>8</td><td>30</td><td>23</td><td>CTCACAAGAACTGCTAACTCATG</td><td>181.9</td><td>-4.3</td><td>1.1</td><td>-4.0</td><td>7.7E-05</td><td>1.1E-04</td><td>MB231/MB468</td><td>CYTO</td></tr>
<tr><td>>trna13_ValCAC_19_-_4724647_4724719@1.33.33|GTTTCCGTAGTGTAGCGGTTATCACATTCGCCT</td><td>>trna13_ValCAC_19_-_4724647_4724719</td><td>1</td><td>33</td><td>33</td><td>GTTTCCGTAGTGTAGCGGTTATCACATTCGCCT</td><td>122.6</td><td>6.9</td><td>1.7</td><td>4.0</td><td>6.9E-05</td><td>1.1E-04</td><td>MB231/MB468</td><td>MP</td></tr>
<tr><td>>trna10_ValCAC_5_-_180649395_180649467@31.54.24,trna13_ValCAC_19_-_4724647_4724719@31.54.24,trna152_ValCAC_6_-_27248049_27248121@31.54.24,trna157_ValCAC_6_-_27173867_27173939@31.54.24,trna18_ValCAC_5_-_180529253_180529325@31.54.24,trna2_ValCAC_5_+_180524070_180524142@31.54.24,trna6_ValCAC_5_+_180600650_180600722@31.54.24,trna85_ValCAC_1_-_161369490_161369562@31.54.24,trna98_ValCAC_1_-_149298555_149298627@31.54.24,trna9_ValCAC_6_+_26538282_26538354@31.54.24|CCTCACACGCGAAAGGTCCCCGGT</td><td>>trna10_ValCAC_5_-_180649395_180649467</td><td>31</td><td>54</td><td>24</td><td>CCTCACACGCGAAAGGTCCCCGGT</td><td>54.7</td><td>-9.6</td><td>2.4</td><td>-4.0</td><td>6.9E-05</td><td>1.1E-04</td><td>MB231/MB468</td><td>MP</td></tr>
<tr><td>>hg38_21_+_8433172_8446622_RNA45SN1_45S_50ntFlanks@12418.12449.32|CGATGTCGGCTCTTCCTATCATTGTGAAGCAG</td><td>>hg38_21_+_8433172_8446622_RNA45SN1_45S_50ntFlanks</td><td>12418</td><td>12449</td><td>32</td><td>CGATGTCGGCTCTTCCTATCATTGTGAAGCAG</td><td>244.0</td><td>-3.1</td><td>0.8</td><td>-4.0</td><td>6.9E-05</td><td>1.1E-04</td><td>MB231/MB468</td><td>MP</td></tr>
<tr><td>>hg38_21_+_8433172_8446622_RNA45SN1_45S_50ntFlanks@10795.10823.29|TGGCATGTTGGAACAATGTAGGTAAGGGA</td><td>>hg38_21_+_8433172_8446622_RNA45SN1_45S_50ntFlanks</td><td>10795</td><td>10823</td><td>29</td><td>TGGCATGTTGGAACAATGTAGGTAAGGGA</td><td>82.8</td><td>9.7</td><td>2.2</td><td>4.4</td><td>9.9E-06</td><td>1.1E-04</td><td>BT20/MB231</td><td>MP</td></tr>
<tr><td>>hg38_21_+_8433172_8446622_RNA45SN1_45S_50ntFlanks@9793.9819.27|GCGTGGAGCCGGGCGTGGAATGCGAGT</td><td>>hg38_21_+_8433172_8446622_RNA45SN1_45S_50ntFlanks</td><td>9793</td><td>9819</td><td>27</td><td>GCGTGGAGCCGGGCGTGGAATGCGAGT</td><td>68.4</td><td>9.4</td><td>2.1</td><td>4.4</td><td>9.9E-06</td><td>1.1E-04</td><td>BT20/MB231</td><td>MP</td></tr>
</table>

<table>
<tr><th></th><th></th><th></th><th></th><th></th><th></th><th></th><th></th><th></th><th></th><th></th><th></th><th></th><th></th></tr>
<tr><td>>hg38_21_+_8433172_8446622_RNA45SN1_45S_50ntFlanks@11848.11887.40|ACGGGCTTGGCGGAATCAGCGGGGAAAGAAGACCCTGTTG</td><td>>hg38_21_+_8433172_8446622_RNA45SN1_45S_50ntFlanks</td><td>11848</td><td>11887</td><td>40</td><td>ACGGGCTTGGCGGAATCAGCGGGGAAAGAAGACCCTGTTG</td><td>66.7</td><td>9.4</td><td>2.1</td><td>4.4</td><td>9.9E-06</td><td>1.1E-04</td><td>BT20/MB231</td><td>MP</td></tr>
<tr><td>>hg38_21_+_8433172_8446622_RNA45SN1_45S_50ntFlanks@5442.5466.25|CCCTCGGATCGGCCCCGCCGGGGTC</td><td>>hg38_21_+_8433172_8446622_RNA45SN1_45S_50ntFlanks</td><td>5442</td><td>5466</td><td>25</td><td>CCCTCGGATCGGCCCCGCCGGGGTC</td><td>431.6</td><td>-3.0</td><td>0.8</td><td>-4.0</td><td>7.7E-05</td><td>1.1E-04</td><td>MB231/MB468</td><td>CYTO</td></tr>
<tr><td>>hg38_21_+_8433172_8446622_RNA45SN1_45S_50ntFlanks@6722.6747.26|ATTGCAGGACACATTGATCATCGACA</td><td>>hg38_21_+_8433172_8446622_RNA45SN1_45S_50ntFlanks</td><td>6722</td><td>6747</td><td>26</td><td>ATTGCAGGACACATTGATCATCGACA</td><td>178.4</td><td>-2.4</td><td>0.6</td><td>-3.9</td><td>8.2E-05</td><td>1.1E-04</td><td>MB231/MB468</td><td>NUC</td></tr>
<tr><td>>am_trna128_GlyGCC_6_-_27870686_27870756@3.35.33,trna18_GlyGCC_16_+_70822597_70822667@3.35.33,trna19_GlyGCC_16_+_70823410_70823480@3.35.33,trna19_GlyGCC_2_-_157257659_157257729@3.35.33,trna24_GlyGCC_16_-_70812942_70813012@3.35.33,trna25_GlyGCC_16_-_70812114_70812184@3.35.33,trna5_GlyGCC_17_+_8029064_8029134@3.35.33,trna68_GlyGCC_1_-_161493637_161493707@3.35.33|ATTGGTGGTTCAGTGGTAGAATTCTCGCCTGCC</td><td>>am_trna128_GlyGCC_6_-_27870686_27870756</td><td>3</td><td>35</td><td>33</td><td>ATTGGTGGTTCAGTGGTAGAATTCTCGCCTGCC</td><td>276.3</td><td>4.7</td><td>1.2</td><td>4.0</td><td>6.9E-05</td><td>1.1E-04</td><td>MB231/MB468</td><td>MP</td></tr>
<tr><td>>MI0000816|hsa-mir-335&WithFlank&7|+|130496105|130496210@22.42.21(+1A)&[MIMAT0000765&hsa-miR-335-5p&offsets|0|-2(+1A);m-249&7|+|130496126|130496147&offsets|0|-1(+1A)]|TCAAGAGCAATAACGAAAAATA</td><td>>MI0000816|hsa-mir-335&WithFlank&7|+|130496105|130496210</td><td>22</td><td>42</td><td>21(+1A)</td><td>TCAAGAGCAATAACGAAAAATA</td><td>30.3</td><td>8.1</td><td>2.0</td><td>4.0</td><td>6.9E-05</td><td>1.1E-04</td><td>MB231/MB468</td><td>MP</td></tr>
<tr><td>>MI0001733|hsa-mir-452&WithFlank&X|-|151959622|151959718@20.43.24&[MIMAT0001635&hsa-miR-452-5p&offsets|0|+2;m-188&X|-|151959677|151959699&offsets|0|+1]|AACTGTTTGCAGAGGAAACTGAGA</td><td>>MI0001733|hsa-mir-452&WithFlank&X|-|151959622|151959718</td><td>20</td><td>43</td><td>24</td><td>AACTGTTTGCAGAGGAAACTGAGA</td><td>141.9</td><td>7.6</td><td>1.8</td><td>4.2</td><td>2.9E-05</td><td>1.1E-04</td><td>BT20/MB468</td><td>MP</td></tr>
<tr><td>>hg38_21_+_8433172_8446622_RNA45SN1_45S_50ntFlanks@8168.8185.18|CCCAAGTCCTTCTGATCG</td><td>>hg38_21_+_8433172_8446622_RNA45SN1_45S_50ntFlanks</td><td>8168</td><td>8185</td><td>18</td><td>CCCAAGTCCTTCTGATCG</td><td>566.6</td><td>-6.6</td><td>1.6</td><td>-4.2</td><td>2.9E-05</td><td>1.1E-04</td><td>BT20/MB468</td><td>MP</td></tr>
<tr><td>>hg38_21_+_8433172_8446622_RNA45SN1_45S_50ntFlanks@11199.11229.31|CCGGCGGCGGCGGCGACTCTGGACGCGAGCC</td><td>>hg38_21_+_8433172_8446622_RNA45SN1_45S_50ntFlanks</td><td>11199</td><td>11229</td><td>31</td><td>CCGGCGGCGGCGGCGACTCTGGACGCGAGCC</td><td>274.6</td><td>-5.6</td><td>1.4</td><td>-3.9</td><td>8.2E-05</td><td>1.1E-04</td><td>MB231/MB468</td><td>NUC</td></tr>
<tr><td>>MI0000273|hsa-mir-183&WithFlank&7|-|129774899|129775020@33.54.22&[MIMAT0000261&hsa-miR-183-5p&offsets|0|0;m-41&7|-|129774966|129774987&offsets|-1|-1]|TATGGCACTGGTAGAATTCACT</td><td>>MI0000273|hsa-mir-183&WithFlank&7|-|129774899|129775020</td><td>33</td><td>54</td><td>22</td><td>TATGGCACTGGTAGAATTCACT</td><td>4460.0</td><td>2.5</td><td>0.6</td><td>3.9</td><td>8.2E-05</td><td>1.1E-04</td><td>MB231/MB468</td><td>NUC</td></tr>
<tr><td>>hg38_21_+_8433172_8446622_RNA45SN1_45S_50ntFlanks@475.510.36|CTTCGTGATCGATGTGGTGACGTCGTGCTCTCCCGG</td><td>>hg38_21_+_8433172_8446622_RNA45SN1_45S_50ntFlanks</td><td>475</td><td>510</td><td>36</td><td>CTTCGTGATCGATGTGGTGACGTCGTGCTCTCCCGG</td><td>13500.1</td><td>3.0</td><td>0.8</td><td>3.9</td><td>8.2E-05</td><td>1.1E-04</td><td>MB231/MB468</td><td>NUC</td></tr>
<tr><td>>trnaMT_LysTTT_MT_+_8295_8364@37.72.36,trnalookalike7_LysTTT_1_+_568844_568913@37.72.36|AAAGATTAAGAGAACCAACACCTCTTTACAGTGACC</td><td>>trnaMT_LysTTT_MT_+_8295_8364</td><td>37</td><td>72</td><td>36</td><td>AAAGATTAAGAGAACCAACACCTCTTTACAGTGACC</td><td>612.0</td><td>5.1</td><td>1.3</td><td>4.0</td><td>6.9E-05</td><td>1.1E-04</td><td>MB231/MB468</td><td>MP</td></tr>
<tr><td>>hg38_21_+_8433172_8446622_RNA45SN1_45S_50ntFlanks@9345.9368.24|AAAGCCGCCGTGGCGCAATGAAGG</td><td>>hg38_21_+_8433172_8446622_RNA45SN1_45S_50ntFlanks</td><td>9345</td><td>9368</td><td>24</td><td>AAAGCCGCCGTGGCGCAATGAAGG</td><td>37.9</td><td>-8.3</td><td>2.0</td><td>-4.2</td><td>2.9E-05</td><td>1.1E-04</td><td>BT20/MB468</td><td>MP</td></tr>
<tr><td>>MI0000067|hsa-let-7f-1&WithFlank&9|+|94176341|94176439@13.34.22(+1A);MI0000068|hsa-let-7f-2&WithFlank&X|-|53557186|53557280@14.35.22(+1A)&[MIMAT0000067&hsa-let-7f-5p&offsets|0|0(+1A);m-20&9|+|94176353|94176374&offsets|0|0(+1A)];[MIMAT0000067_1&hsa-let-7f-5p&offsets|0|0(+1A);m-19&X|-|53557246|53557267&offsets|0|0(+1A)]|TGAGGTAGTAGATTGTATAGTTA</td><td>>MI0000067|hsa-let-7f-1&WithFlank&9|+|94176341|94176439</td><td>13</td><td>34</td><td>22(+1A)</td><td>TGAGGTAGTAGATTGTATAGTTA</td><td>1831.3</td><td>3.4</td><td>0.9</td><td>4.0</td><td>6.9E-05</td><td>1.1E-04</td><td>MB231/MB468</td><td>MP</td></tr>
<tr><td>>trnaMT_HisGTG_MT_+_12138_12206@19.59.41|AAACATCAGATTGTGAATCTGACAACAGAGGCTTACGACCC</td><td>>trnaMT_HisGTG_MT_+_12138_12206</td><td>19</td><td>59</td><td>41</td><td>AAACATCAGATTGTGAATCTGACAACAGAGGCTTACGACCC</td><td>131.4</td><td>10.8</td><td>2.6</td><td>4.2</td><td>2.9E-05</td><td>1.1E-04</td><td>BT20/MB468</td><td>MP</td></tr>
<tr><td>>hg38_21_+_8433172_8446622_RNA45SN1_45S_50ntFlanks@11564.11603.40|CCTAGCAGCCGACTTAGAACTGGTGCGGACCAGGGGAATC</td><td>>hg38_21_+_8433172_8446622_RNA45SN1_45S_50ntFlanks</td><td>11564</td><td>11603</td><td>40</td><td>CCTAGCAGCCGACTTAGAACTGGTGCGGACCAGGGGAATC</td><td>1692.9</td><td>-4.5</td><td>1.1</td><td>-4.2</td><td>2.9E-05</td><td>1.1E-04</td><td>BT20/MB468</td><td>MP</td></tr>
<tr><td>>hg38_21_+_8433172_8446622_RNA45SN1_45S_50ntFlanks@11208.11250.43|GCGGCGACTCTGGACGCGAGCCGGGCCCTTCCCGTGGATCGCC</td><td>>hg38_21_+_8433172_8446622_RNA45SN1_45S_50ntFlanks</td><td>11208</td><td>11250</td><td>43</td><td>GCGGCGACTCTGGACGCGAGCCGGGCCCTTCCCGTGGATCGCC</td><td>1460.0</td><td>-7.4</td><td>1.9</td><td>-3.9</td><td>8.5E-05</td><td>1.1E-04</td><td>BT20/MB468</td><td>NUC</td></tr>
<tr><td>>hg38_21_+_8433172_8446622_RNA45SN1_45S_50ntFlanks@11578.11600.23|TAGAACTGGTGCGGACCAGGGGA</td><td>>hg38_21_+_8433172_8446622_RNA45SN1_45S_50ntFlanks</td><td>11578</td><td>11600</td><td>23</td><td>TAGAACTGGTGCGGACCAGGGGA</td><td>64.6</td><td>9.3</td><td>2.1</td><td>4.4</td><td>1.0E-05</td><td>1.1E-04</td><td>BT20/MB231</td><td>MP</td></tr>
<tr><td>>MI0000746|hsa-mir-99b&WithFlank&19|+|51692606|51692687@13.32.20(+1A)&[MIMAT0000689&hsa-miR-99b-5p&offsets|0|-2(+1A);m-10&19|+|51692618|51692639&offsets|0|-2(+1A)]|CACCCGTAGAACCGACCTTGA</td><td>>MI0000746|hsa-mir-99b&WithFlank&19|+|51692606|51692687</td><td>13</td><td>32</td><td>20(+1A)</td><td>CACCCGTAGAACCGACCTTGA</td><td>210.4</td><td>2.0</td><td>0.5</td><td>3.9</td><td>8.5E-05</td><td>1.1E-04</td><td>BT20/MB468</td><td>NUC</td></tr>
<tr><td>>hg38_21_+_8433172_8446622_RNA45SN1_45S_50ntFlanks@7975.8017.43|CGCGACCTCAGATCAGACGTGGCGACCCGCTGAATTTAAGCAT</td><td>>hg38_21_+_8433172_8446622_RNA45SN1_45S_50ntFlanks</td><td>7975</td><td>8017</td><td>43</td><td>CGCGACCTCAGATCAGACGTGGCGACCCGCTGAATTTAAGCAT</td><td>1623.4</td><td>5.1</td><td>1.3</td><td>4.0</td><td>7.2E-05</td><td>1.1E-04</td><td>BT20/MB468</td><td>CYTO</td></tr>
<tr><td>>MI0000434|hsa-let-7i&WithFlank&12|+|62603680|62603775@12.32.21(+1G)&[MIMAT0000415&hsa-let-7i-5p&offsets|0|-1(+1G);m-49&12|+|62603691|62603712&offsets|0|-1(+1G)]|TGAGGTAGTAGTTTGTGCTGTG</td><td>>MI0000434|hsa-let-7i&WithFlank&12|+|62603680|62603775</td><td>12</td><td>32</td><td>21(+1G)</td><td>TGAGGTAGTAGTTTGTGCTGTG</td><td>409.4</td><td>4.3</td><td>1.1</td><td>3.9</td><td>8.3E-05</td><td>1.1E-04</td><td>MB231/MB468</td><td>MITO</td></tr>
<tr><td>>MI0000087|hsa-mir-29a&WithFlank&7|-|130876741|130876816@47.67.21&[MIMAT0000086&hsa-miR-29a-3p&offsets|-1|-2;m-26&7|-|130876749|130876769&offsets|-1|-1]|CTAGCACCATCTGAAATCGGT</td><td>>MI0000087|hsa-mir-29a&WithFlank&7|-|130876741|130876816</td><td>47</td><td>67</td><td>21</td><td>CTAGCACCATCTGAAATCGGT</td><td>328.2</td><td>7.1</td><td>1.8</td><td>4.0</td><td>7.0E-05</td><td>1.1E-04</td><td>MB231/MB468</td><td>MP</td></tr>
<tr><td>>hg38_21_+_8433172_8446622_RNA45SN1_45S_50ntFlanks@12586.12604.19|CAGGTTCAGACATTTGGTG</td><td>>hg38_21_+_8433172_8446622_RNA45SN1_45S_50ntFlanks</td><td>12586</td><td>12604</td><td>19</td><td>CAGGTTCAGACATTTGGTG</td><td>120.5</td><td>-6.0</td><td>1.4</td><td>-4.2</td><td>2.9E-05</td><td>1.1E-04</td><td>BT20/MB468</td><td>MP</td></tr>
<tr><td>>MI0000071|hsa-mir-17&WithFlank&13|+|91350599|91350694@20.43.24&[MIMAT0000070&hsa-miR-17-5p&offsets|0|+1;m-88&13|+|91350618|91350640&offsets|0|+1]|CAAAGTGCTTACAGTGCAGGTAGT</td><td>>MI0000071|hsa-mir-17&WithFlank&13|+|91350599|91350694</td><td>20</td><td>43</td><td>24</td><td>CAAAGTGCTTACAGTGCAGGTAGT</td><td>1625.7</td><td>-6.3</td><td>1.4</td><td>-4.4</td><td>1.0E-05</td><td>1.1E-04</td><td>BT20/MB231</td><td>MP</td></tr>
<tr><td>>am_trna112_GlnCTG_1_-_147737382_147737453@24.41.18,trna131_GlnCTG_6_-_27759135_27759206@24.41.18,trna146_GlnCTG_6_-_27515531_27515602@24.41.18,trna15_GlnCTG_1_+_145963304_145963375@24.41.18,trna19_GlnCTG_1_+_147505038_147505109@24.41.18,trna1_GlnCTG_6_+_18836402_18836473@24.41.18,trna22_GlnCTG_1_+_147800937_147801008@24.41.18,trna28_GlnCTG_1_+_149186125_149186196@24.41.18,trna3_GlnCTG_17_+_8023070_8023141@24.41.18,trna42_GlnCTG_6_+_27263212_27263283@24.41.18,trna49_GlnCTG_6_+_27487308_27487379@24.41.18,trna7_GlnCTG_15_-_66161400_66161471@24.41.18,trna99_GlnCTG_6_-_28909378_28909449@24.41.18|ACTCTGGACTCTGAATCC</td><td>>am_trna112_GlnCTG_1_-_147737382_147737453</td><td>24</td><td>41</td><td>18</td><td>ACTCTGGACTCTGAATCC</td><td>232.6</td><td>6.9</td><td>1.7</td><td>4.2</td><td>2.9E-05</td><td>1.1E-04</td><td>BT20/MB468</td><td>MP</td></tr>
<tr><td>>hg38_21_+_8433172_8446622_RNA45SN1_45S_50ntFlanks@4731.4767.37|AAGAACGAAAGTCGGAGGTTCGAAGACGATCAGATAC</td><td>>hg38_21_+_8433172_8446622_RNA45SN1_45S_50ntFlanks</td><td>4731</td><td>4767</td><td>37</td><td>AAGAACGAAAGTCGGAGGTTCGAAGACGATCAGATAC</td><td>311.2</td><td>6.6</td><td>1.6</td><td>4.2</td><td>2.9E-05</td><td>1.1E-04</td><td>BT20/MB468</td><td>MP</td></tr>
<tr><td>>am_trna10_LysCTT_16_+_3241501_3241573@34.51.18,trna119_LysCTT_1_-_145395522_145395594@34.51.18,trna11_LysCTT_5_-_180648979_180649051@34.51.18,trna13_LysCTT_16_+_3246153_3246225@34.51.18,trna13_LysCTT_6_+_26556774_26556846@34.51.18,trna30_LysCTT_16_-_3230555_3230627@34.51.18,trna32_LysCTT_16_-_3207406_3207478@34.51.18,trna5_LysCTT_16_+_3214939_3215011@34.51.18,trna7_LysCTT_16_+_3225692_3225764@34.51.18,trna8_LysCTT_1_+_39970195_39970267@34.51.18,trna9_LysCTT_5_+_180634755_180634827@34.51.18|CTTAATCTCAGGGTCGTG</td><td>>am_trna10_LysCTT_16_+_3241501_3241573</td><td>34</td><td>51</td><td>18</td><td>CTTAATCTCAGGGTCGTG</td><td>220.9</td><td>-6.8</td><td>1.6</td><td>-4.2</td><td>2.9E-05</td><td>1.1E-04</td><td>BT20/MB468</td><td>MP</td></tr>
<tr><td>>hg38_21_+_8433172_8446622_RNA45SN1_45S_50ntFlanks@12365.12384.20|ATAACTGGCTTGTGGCGGCC</td><td>>hg38_21_+_8433172_8446622_RNA45SN1_45S_50ntFlanks</td><td>12365</td><td>12384</td><td>20</td><td>ATAACTGGCTTGTGGCGGCC</td><td>162.6</td><td>-6.5</td><td>1.5</td><td>-4.2</td><td>2.9E-05</td><td>1.1E-04</td><td>BT20/MB468</td><td>MP</td></tr>
<tr><td>>MI0000264|hsa-mir-7-2&WithFlank&15|+|88611819|88611940@38.56.19(+1C);MI0000265|hsa-mir-7-3&WithFlank&19|+|4770664|4770785@37.55.19(+1C);MI0000263|hsa-mir-7-1&WithFlank&9|-|83969742|83969863@30.48.19(+1C)&[MIMAT0000252&hsa-miR-7-5p&offsets|0|-5(+1C);m-400&15|+|88611856|88611879&offsets|0|-5(+1C)];[MIMAT0000252_1&hsa-miR-7-5p&offsets|0|-5(+1C);m-402&19|+|4770700|4770723&offsets|0|-5(+1C)];[MIMAT0000252_2&hsa-miR-7-5p&offsets|0|-5(+1C);m-401&9|-|83969811|83969834&offsets|0|-5(+1C)]|TGGAAGACTAGTGATTTTGC</td><td>>MI0000264|hsa-mir-7-2&WithFlank&15|+|88611819|88611940</td><td>38</td><td>56</td><td>19(+1C)</td><td>TGGAAGACTAGTGATTTTGC</td><td>27.6</td><td>8.0</td><td>2.0</td><td>4.0</td><td>7.0E-05</td><td>1.1E-04</td><td>MB231/MB468</td><td>MP</td></tr>
<tr><td>>trna10_AspGTC_12_-_125424193_125424264@36.74.39,trna12_AspGTC_12_-_125411891_125411962@36.74.39,trna144_AspGTC_6_-_27551236_27551307@36.74.39,trna38_AspGTC_17_-_8125556_8125627@36.74.39,trna45_AspGTC_6_+_27447453_27447524@36.74.39,trna48_AspGTC_6_+_27471523_27471594@36.74.39,trna4_AspGTC_12_+_96429799_96429870@36.74.39,trna69_AspGTC_1_-_161440205_161440276@36.74.39,trna72_AspGTC_1_-_161432824_161432895@36.74.39,trna75_AspGTC_1_-_161425414_161425485@36.74.39,trna78_AspGTC_1_-_161418033_161418104@36.74.39,trna81_AspGTC_1_-_161410615_161410686@36.74.39|CACGCGGGAGACCGGGGTTCGATTCCCCGACGGGGAGCC</td><td>>trna10_AspGTC_12_-_125424193_125424264</td><td>36</td><td>74</td><td>39</td><td>CACGCGGGAGACCGGGGTTCGATTCCCCGACGGGGAGCC</td><td>10716.7</td><td>-4.6</td><td>1.0</td><td>-4.4</td><td>9.5E-06</td><td>1.1E-04</td><td>BT20/MB231</td><td>CYTO</td></tr>
<tr><td>>MI0003591|hsa-mir-584&WithFlank&5|-|149062307|149062415@22.42.21(+1C)&[MIMAT0003249&hsa-miR-584-5p&offsets|0|-1(+1C);m-167&5|-|149062374|149062394&offsets|0|0(+1C)]|TTATGGTTTGCCTGGGACTGAC</td><td>>MI0003591|hsa-mir-584&WithFlank&5|-|149062307|149062415</td><td>22</td><td>42</td><td>21(+1C)</td><td>TTATGGTTTGCCTGGGACTGAC</td><td>29.2</td><td>8.1</td><td>2.0</td><td>4.0</td><td>7.0E-05</td><td>1.1E-04</td><td>MB231/MB468</td><td>MP</td></tr>
<tr><td>>am_trna162_MetCAT_6_-_26758550_26758622@34.54.21,trna164_MetCAT_6_-_26735574_26735646@34.54.21,trna20_MetCAT_16_+_71460396_71460468@34.54.21,trna21_MetCAT_6_+_26701712_26701784@34.54.21,trna27_MetCAT_6_+_26766444_26766516@34.54.21,trna75_MetCAT_6_+_28912352_28912424@34.54.21,trna97_MetCAT_6_-_28921042_28921114@34.54.21|CATAATCTGAAGGTCCTGAGT</td><td>>am_trna162_MetCAT_6_-_26758550_26758622</td><td>34</td><td>54</td><td>21</td><td>CATAATCTGAAGGTCCTGAGT</td><td>567.3</td><td>3.1</td><td>0.8</td><td>3.9</td><td>8.3E-05</td><td>1.1E-04</td><td>MB231/MB468</td><td>NUC</td></tr>
</table>

| | | | | | | | | | | | | | |
|---|---|---|---|---|---|---|---|---|---|---|---|---|---|
| >hg38_21_+_8433172_8446622_RNA45SN1_45S_50ntFlanks@9533.9553.21\|GAGGAAACTCTGGTGGAGGTC | >hg38_21_+_8433172_8446622_RNA45SN1_45S_50ntFlanks | 9533 | 9553 | 21 | GAGGAAACTCTGGTGGAGGTC | 817.4 | -2.1 | 0.5 | -4.0 | 6.1E-05 | 1.1E-04 | BT20/MB468 | MITO |
| >hg38_21_+_8433172_8446622_RNA45SN1_45S_50ntFlanks@10362.10407.46\|AAGGCCGAAGTGGAGAAGGGTTCCATGTGAACAGCAGTTGAACATG | >hg38_21_+_8433172_8446622_RNA45SN1_45S_50ntFlanks | 10362 | 10407 | 46 | AAGGCCGAAGTGGAGAAGGGTTCCATGTGAACAGCAGTTGAACATG | 52.9 | -9.6 | 2.4 | -4.0 | 7.0E-05 | 1.1E-04 | MB231/MB468 | MP |
| >MI0000085\|hsa-mir-27a&WithFlank&19\|-\|13836434\|13836523@57.76.20(+4U)&[MIMAT0000084&hsa-miR-27a-3p&offsets\|0\|-1(+4U);m-43&19\|-\|13836447\|13836467&offsets\|0\|-1(+4U)]\|TTCACAGTGGCTAAGTTCCGTTTT | >MI0000085\|hsa-mir-27a&WithFlank&19\|-\|13836434\|13836523 | 57 | 76 | 20(+4U) | TTCACAGTGGCTAAGTTCCGTTTT | 232.2 | 6.2 | 1.6 | 3.9 | 8.3E-05 | 1.1E-04 | MB231/MB468 | MITO |
| >hg38_21_+_8433172_8446622_RNA45SN1_45S_50ntFlanks@11667.11692.26\|CTGCCCAGTGCTCTGAATGTCAAAGT | >hg38_21_+_8433172_8446622_RNA45SN1_45S_50ntFlanks | 11667 | 11692 | 26 | CTGCCCAGTGCTCTGAATGTCAAAGT | 159.1 | 7.7 | 1.8 | 4.2 | 2.9E-05 | 1.1E-04 | BT20/MB468 | MP |
| >am_trnaMT_PheGAA_MT_+_577_647@40.58.19\|TGTTTAGACGGGCTCACAT | >am_trnaMT_PheGAA_MT_+_577_647 | 40 | 58 | 19 | TGTTTAGACGGGCTCACAT | 120.0 | 3.5 | 0.9 | 4.0 | 7.0E-05 | 1.1E-04 | MB231/MB468 | MP |
| >hg38_1_-_228634819_228635039_RNA5S12_5S_50ntFlanks@134.170.37\|TGGGAGACCGCCTGGGAATACCGGGTGCTGTAGGCTT | >hg38_1_-_228634819_228635039_RNA5S12_5S_50ntFlanks | 134 | 170 | 37 | TGGGAGACCGCCTGGGAATACCGGGTGCTGTAGGCTT | 19162.9 | 3.6 | 0.9 | 4.0 | 7.0E-05 | 1.1E-04 | MB231/MB468 | MP |
| >am_trna119_LysCTT_1_-_145395522_145395594@12.33.22,trna11_LysCTT_5_-_180648979_180649051@12.33.22,trna128_LysCTT_1_-_55423542_55423614@12.33.22,trna13_LysCTT_6_+_26556774_26556846@12.33.22,trna6_LysCTT_19_-_52425396_52425468@12.33.22,trna7_LysCTT_16_+_3225692_3225764@12.33.22,trna9_LysCTT_5_+_180634755_180634827@12.33.22\|TCAGTCGGTAGAGCATGAGACT | >am_trna119_LysCTT_1_-_145395522_145395594 | 12 | 33 | 22 | TCAGTCGGTAGAGCATGAGACT | 28.2 | 8.0 | 2.0 | 4.0 | 7.0E-05 | 1.1E-04 | MB231/MB468 | MP |
| >hg38_21_+_8433172_8446622_RNA45SN1_45S_50ntFlanks@6661.6684.24\|CGGTGGATCACTCGGCTCGTGCGT | >hg38_21_+_8433172_8446622_RNA45SN1_45S_50ntFlanks | 6661 | 6684 | 24 | CGGTGGATCACTCGGCTCGTGCGT | 479.3 | 3.5 | 0.8 | 4.2 | 2.9E-05 | 1.1E-04 | BT20/MB468 | MP |
| >hg38_21_+_8433172_8446622_RNA45SN1_45S_50ntFlanks@13008.13039.32\|ATTGAAAGTCAGCCCTCGACACAAGGGTTTGT | >hg38_21_+_8433172_8446622_RNA45SN1_45S_50ntFlanks | 13008 | 13039 | 32 | ATTGAAAGTCAGCCCTCGACACAAGGGTTTGT | 966.8 | 3.3 | 0.8 | 3.9 | 7.9E-05 | 1.1E-04 | MB231/MB468 | CYTO |
| >hg38_21_+_8433172_8446622_RNA45SN1_45S_50ntFlanks@9534.9574.41\|AGGAAACTCTGGTGGAGGTCCGTAGCGGTCCTGACGTGCAA | >hg38_21_+_8433172_8446622_RNA45SN1_45S_50ntFlanks | 9534 | 9574 | 41 | AGGAAACTCTGGTGGAGGTCCGTAGCGGTCCTGACGTGCAA | 455.2 | -2.8 | 0.7 | -4.0 | 7.3E-05 | 1.1E-04 | BT20/MB468 | CYTO |
| >MI0000446\|hsa-mir-125b-1&WithFlank&11\|-\|122099751\|122099850@21.40.20;MI0000470\|hsa-mir-125b-2&WithFlank&21\|+\|16590231\|16590331@23.42.20&[MIMAT0000423&hsa-miR-125b-5p&offsets\|0\|-2;m-67&11\|-\|122099809\|122099830&offsets\|0\|-2];[MIMAT0000423_1&hsa-miR-125b-5p&offsets\|0\|-2;m-68&21\|+\|16590253\|16590274&offsets\|0\|-2]\|TCCCTGAGACCCTAACTTGT | >MI0000446\|hsa-mir-125b-1&WithFlank&11\|-\|122099751\|122099850 | 21 | 40 | 20 | TCCCTGAGACCCTAACTTGT | 610.0 | 5.2 | 1.3 | 4.0 | 7.0E-05 | 1.1E-04 | MB231/MB468 | MP |
| >hg38_21_+_8433172_8446622_RNA45SN1_45S_50ntFlanks@11561.11600.40\|GCGCCTAGCAGCCGACTTAGAACTGGTGCGGACCAGGGGA | >hg38_21_+_8433172_8446622_RNA45SN1_45S_50ntFlanks | 11561 | 11600 | 40 | GCGCCTAGCAGCCGACTTAGAACTGGTGCGGACCAGGGGA | 384.1 | -4.4 | 1.1 | -3.9 | 7.9E-05 | 1.1E-04 | MB231/MB468 | CYTO |
| >hg38_21_+_8433172_8446622_RNA45SN1_45S_50ntFlanks@9316.9338.23\|AACACGTGCGCGAGTCGGGGGCT | >hg38_21_+_8433172_8446622_RNA45SN1_45S_50ntFlanks | 9316 | 9338 | 23 | AACACGTGCGCGAGTCGGGGGCT | 312.4 | -4.1 | 0.9 | -4.4 | 9.6E-06 | 1.1E-04 | BT20/MB231 | CYTO |
| >hg38_21_+_8433172_8446622_RNA45SN1_45S_50ntFlanks@11660.11685.26\|GTGATTTCTGCCCAGTGCTCTGAATG | >hg38_21_+_8433172_8446622_RNA45SN1_45S_50ntFlanks | 11660 | 11685 | 26 | GTGATTTCTGCCCAGTGCTCTGAATG | 1724.3 | -5.6 | 1.4 | -3.9 | 8.7E-05 | 1.1E-04 | BT20/MB468 | NUC |
| >hg38_21_+_8433172_8446622_RNA45SN1_45S_50ntFlanks@11562.11601.40\|CGCCTAGCAGCCGACTTAGAACTGGTGCGGACCAGGGGAA | >hg38_21_+_8433172_8446622_RNA45SN1_45S_50ntFlanks | 11562 | 11601 | 40 | CGCCTAGCAGCCGACTTAGAACTGGTGCGGACCAGGGGAA | 440.1 | -4.8 | 1.2 | -3.9 | 8.4E-05 | 1.1E-04 | MB231/MB468 | NUC |
| >MI0000542\|hsa-mir-320a&WithFlank&8\|-\|22244960\|22245043@48.69.22(+2U)&[MIMAT0000510&hsa-miR-320a-3p&offsets\|0\|0(+2U);m-60&8\|-\|22244975\|22244996&offsets\|0\|0(+2U)]\|AAAAGCTGGGTTGAGAGGGCGATT | >MI0000542\|hsa-mir-320a&WithFlank&8\|-\|22244960\|22245043 | 48 | 69 | 22(+2U) | AAAAGCTGGGTTGAGAGGGCGATT | 2312.4 | 6.3 | 1.5 | 4.2 | 3.0E-05 | 1.1E-04 | BT20/MB468 | MP |
| >hg38_1_-_228634819_228635039_RNA5S12_5S_50ntFlanks@144.171.28\|CCTGGGAATACCGGGTGCTGTAGGCTTT | >hg38_1_-_228634819_228635039_RNA5S12_5S_50ntFlanks | 144 | 171 | 28 | CCTGGGAATACCGGGTGCTGTAGGCTTT | 4684.2 | -2.7 | 0.7 | -4.0 | 7.1E-05 | 1.1E-04 | MB231/MB468 | MP |
| >hg38_21_+_8433172_8446622_RNA45SN1_45S_50ntFlanks@10869.10892.24\|AGGGCTGGGTCGGTCGGGCTGGGG | >hg38_21_+_8433172_8446622_RNA45SN1_45S_50ntFlanks | 10869 | 10892 | 24 | AGGGCTGGGTCGGTCGGGCTGGGG | 1489.2 | -5.7 | 1.4 | -4.2 | 3.0E-05 | 1.1E-04 | BT20/MB468 | MP |
| >hg38_21_+_8433172_8446622_RNA45SN1_45S_50ntFlanks@476.518.43\|TTCGTGATCGATGTGGTGACGTCGTGCTCTCCCGGGCCGGGTC | >hg38_21_+_8433172_8446622_RNA45SN1_45S_50ntFlanks | 476 | 518 | 43 | TTCGTGATCGATGTGGTGACGTCGTGCTCTCCCGGGCCGGGTC | 7095.6 | 4.1 | 1.0 | 3.9 | 8.4E-05 | 1.1E-04 | MB231/MB468 | NUC |
| >trnaMT_PheGAA_MT_+_577_647@46.74.29\|GACGGGCTCACATCACCCCATAAACACCA | >trnaMT_PheGAA_MT_+_577_647 | 46 | 74 | 29 | GACGGGCTCACATCACCCCATAAACACCA | 283.3 | 5.3 | 1.4 | 3.9 | 8.4E-05 | 1.1E-04 | MB231/MB468 | NUC |
| >MI0005559\|hsa-mir-744&WithFlank&17\|+\|12081893\|12082002@17.37.21&[MIMAT0004945&hsa-miR-744-5p&offsets\|0\|-1;m-172&17\|+\|12081909\|12081929&offsets\|0\|0]\|TGCGGGGCTAGGGCTAACAGC | >MI0005559\|hsa-mir-744&WithFlank&17\|+\|12081893\|12082002 | 17 | 37 | 21 | TGCGGGGCTAGGGCTAACAGC | 3918.8 | 2.9 | 0.7 | 4.4 | 1.2E-05 | 1.1E-04 | BT20/MB231 | MITO |
| >MI0000815\|hsa-mir-339&WithFlank&7\|-\|1022927\|1023032@56.75.20(+1U)&[MIMAT0004702&hsa-miR-339-3p&offsets\|0\|-3(+1U);m-235&7\|-\|1022957\|1022977&offsets\|0\|-1(+1U)]\|TGAGCGCCTCGACGACAGAGT | >MI0000815\|hsa-mir-339&WithFlank&7\|-\|1022927\|1023032 | 56 | 75 | 20(+1U) | TGAGCGCCTCGACGACAGAGT | 323.4 | 8.0 | 1.9 | 4.2 | 3.0E-05 | 1.1E-04 | BT20/MB468 | MP |
| >hg38_21_+_8433172_8446622_RNA45SN1_45S_50ntFlanks@11568.11594.27\|GCAGCCGACTTAGAACTGGTGCGGACC | >hg38_21_+_8433172_8446622_RNA45SN1_45S_50ntFlanks | 11568 | 11594 | 27 | GCAGCCGACTTAGAACTGGTGCGGACC | 274.2 | -2.9 | 0.7 | -4.0 | 7.1E-05 | 1.1E-04 | MB231/MB468 | MP |
| >hg38_21_+_8433172_8446622_RNA45SN1_45S_50ntFlanks@8117.8143.27\|CGGGACATGTGGCGTACGGAAGACCCG | >hg38_21_+_8433172_8446622_RNA45SN1_45S_50ntFlanks | 8117 | 8143 | 27 | CGGGACATGTGGCGTACGGAAGACCCG | 450.4 | -4.4 | 1.1 | -4.2 | 3.0E-05 | 1.1E-04 | BT20/MB468 | MP |
| >hg38_21_+_8433172_8446622_RNA45SN1_45S_50ntFlanks@6661.6698.38\|CGGTGGATCACTCGGCTCGTGCGTCGATGAAGAACGCA | >hg38_21_+_8433172_8446622_RNA45SN1_45S_50ntFlanks | 6661 | 6698 | 38 | CGGTGGATCACTCGGCTCGTGCGTCGATGAAGAACGCA | 488.7 | 2.4 | 0.6 | 4.4 | 1.1E-05 | 1.1E-04 | BT20/MB231 | NUC |
| >hg38_21_+_8433172_8446622_RNA45SN1_45S_50ntFlanks@9561.9597.37\|GTCCTGACGTGCAAATCGGTCGTCCGACCTGGGTATA | >hg38_21_+_8433172_8446622_RNA45SN1_45S_50ntFlanks | 9561 | 9597 | 37 | GTCCTGACGTGCAAATCGGTCGTCCGACCTGGGTATA | 1063.7 | -4.8 | 1.2 | -3.9 | 8.4E-05 | 1.1E-04 | MB231/MB468 | MITO |
| >hg38_21_+_8433172_8446622_RNA45SN1_45S_50ntFlanks@4451.4472.22\|TCTCGGCGCCCCCTCGATGCTC | >hg38_21_+_8433172_8446622_RNA45SN1_45S_50ntFlanks | 4451 | 4472 | 22 | TCTCGGCGCCCCCTCGATGCTC | 87.9 | -5.8 | 1.4 | -4.0 | 7.1E-05 | 1.1E-04 | MB231/MB468 | MP |
| >MI0000285\|hsa-mir-205&WithFlank&1\|+\|209432127\|209432248@40.59.20(+2G)&[MIMAT0000266&hsa-miR-205-5p&offsets\|0\|-2(+2G);m-351&1\|+\|209432166\|209432187&offsets\|0\|-2(+2G)]\|TCCTTCATTCCACCGGAGTCGG | >MI0000285\|hsa-mir-205&WithFlank&1\|+\|209432127\|209432248 | 40 | 59 | 20(+2G) | TCCTTCATTCCACCGGAGTCGG | 60.4 | -9.8 | 2.5 | -4.0 | 7.1E-05 | 1.1E-04 | MB231/MB468 | MP |
| >hg38_21_+_8433172_8446622_RNA45SN1_45S_50ntFlanks@9033.9055.23\|CTCGTGGGGGGGCCGGGCCACCC | >hg38_21_+_8433172_8446622_RNA45SN1_45S_50ntFlanks | 9033 | 9055 | 23 | CTCGTGGGGGGGCCGGGCCACCC | 539.3 | -5.4 | 1.3 | -4.2 | 3.0E-05 | 1.1E-04 | BT20/MB468 | MP |
| >MI0000767\|hsa-mir-365a&WithFlank&16\|+\|14309279\|14309377@22.45.24&[MIMAT0009199&hsa-miR-365a-5p&offsets\|0\|+1;m-595&16\|+\|14309300\|14309322&offsets\|0\|+1]\|AGGGACTTTTGGGGGCAGATGTGT | >MI0000767\|hsa-mir-365a&WithFlank&16\|+\|14309279\|14309377 | 22 | 45 | 24 | AGGGACTTTTGGGGGCAGATGTGT | 168.0 | 3.6 | 0.8 | 4.4 | 9.7E-06 | 1.1E-04 | BT20/MB231 | CYTO |
| >hg38_21_+_8433172_8446622_RNA45SN1_45S_50ntFlanks@5471.5501.31\|CACGGCCCTGGCGGAGCGCTGAGAAGACGGT | >hg38_21_+_8433172_8446622_RNA45SN1_45S_50ntFlanks | 5471 | 5501 | 31 | CACGGCCCTGGCGGAGCGCTGAGAAGACGGT | 803.3 | 3.0 | 0.8 | 3.9 | 7.9E-05 | 1.1E-04 | MB231/MB468 | CYTO |
| >hg38_21_+_8433172_8446622_RNA45SN1_45S_50ntFlanks@10317.10348.32\|GTGCAGATCTTGGTGGTAGTAGCAAATATTCA | >hg38_21_+_8433172_8446622_RNA45SN1_45S_50ntFlanks | 10317 | 10348 | 32 | GTGCAGATCTTGGTGGTAGTAGCAAATATTCA | 83.1 | 9.7 | 2.2 | 4.4 | 1.0E-05 | 1.1E-04 | BT20/MB231 | MP |
| >hg38_21_+_8433172_8446622_RNA45SN1_45S_50ntFlanks@12415.12449.35\|CTTCGATGTCGGCTCTTCCTATCATTGTGAAGCAG | >hg38_21_+_8433172_8446622_RNA45SN1_45S_50ntFlanks | 12415 | 12449 | 35 | CTTCGATGTCGGCTCTTCCTATCATTGTGAAGCAG | 504.1 | 1.7 | 0.4 | 4.0 | 7.4E-05 | 1.1E-04 | BT20/MB468 | CYTO |
| >hg38_21_+_8433172_8446622_RNA45SN1_45S_50ntFlanks@10914.10932.19\|CGCCGCGGCTGGACGAGGC | >hg38_21_+_8433172_8446622_RNA45SN1_45S_50ntFlanks | 10914 | 10932 | 19 | CGCCGCGGCTGGACGAGGC | 569.5 | -6.2 | 1.6 | -4.0 | 7.4E-05 | 1.1E-04 | BT20/MB468 | CYTO |
| >hg38_21_+_8433172_8446622_RNA45SN1_45S_50ntFlanks@6663.6681.19\|GTGGATCACTCGGCTCGTG | >hg38_21_+_8433172_8446622_RNA45SN1_45S_50ntFlanks | 6663 | 6681 | 19 | GTGGATCACTCGGCTCGTG | 768.9 | -4.3 | 1.1 | -4.0 | 7.4E-05 | 1.1E-04 | BT20/MB468 | CYTO |
| >trnaMT_LeuTAA_MT_+_3230_3304@1.25.25\|GTTAAGATGGCAGAGCCCGGTAATC | >trnaMT_LeuTAA_MT_+_3230_3304 | 1 | 25 | 25 | GTTAAGATGGCAGAGCCCGGTAATC | 1138.7 | 6.0 | 1.4 | 4.2 | 3.0E-05 | 1.1E-04 | BT20/MB468 | MP |
| >hg38_21_+_8433172_8446622_RNA45SN1_45S_50ntFlanks@12991.13028.38\|CTCCCTCGCTGCGATCTATTGAAAGTCAGCCCTCGACA | >hg38_21_+_8433172_8446622_RNA45SN1_45S_50ntFlanks | 12991 | 13028 | 38 | CTCCCTCGCTGCGATCTATTGAAAGTCAGCCCTCGACA | 271.5 | -5.5 | 1.4 | -4.0 | 7.4E-05 | 1.1E-04 | BT20/MB468 | CYTO |
| >trnaMT_ValTAC_MT_+_1602_1670@43.72.30\|GATTTCAACTTAACTTGACCGCTCTGACCA | >trnaMT_ValTAC_MT_+_1602_1670 | 43 | 72 | 30 | GATTTCAACTTAACTTGACCGCTCTGACCA | 2737.5 | 3.8 | 1.0 | 3.9 | 8.5E-05 | 1.1E-04 | MB231/MB468 | MITO |
| >hg38_21_+_8433172_8446622_RNA45SN1_45S_50ntFlanks@5073.5104.32\|AATTCCGATAACGAACGAGACTCTGGCATGCT | >hg38_21_+_8433172_8446622_RNA45SN1_45S_50ntFlanks | 5073 | 5104 | 32 | AATTCCGATAACGAACGAGACTCTGGCATGCT | 382.9 | -5.6 | 1.4 | -3.9 | 8.5E-05 | 1.1E-04 | MB231/MB468 | MITO |
| >am_trna116_GluCTC_1_-_145399233_145399304@1.27.27,trna59_GluCTC_1_+_249168447_249168518@1.27.27,trna71_GluCTC_1_-_161439189_161439260@1.27.27,trna74_GluCTC_1_-_161431809_161431880@1.27.27,trna77_GluCTC_1_-_161424398_161424469@1.27.27,trna77_GluCTC_6_+_28949976_28950047@1.27.27,trna80_GluCTC_1_-_161417018_161417089@1.27.27,trna87_GluCTC_6_-_126101393_126101464@1.27.27\|TCCCTGGTGGTCTAGTGGTTAGGATTC | >am_trna116_GluCTC_1_-_145399233_145399304 | 1 | 27 | 27 | TCCCTGGTGGTCTAGTGGTTAGGATTC | 1655.6 | -3.0 | 0.8 | -4.0 | 7.1E-05 | 1.1E-04 | MB231/MB468 | MP |
| >hg38_21_+_8433172_8446622_RNA45SN1_45S_50ntFlanks@10678.10711.34\|CGGTTCCGGCGGCGTCCGGTGAGCTCTCGCTGGC | >hg38_21_+_8433172_8446622_RNA45SN1_45S_50ntFlanks | 10678 | 10711 | 34 | CGGTTCCGGCGGCGTCCGGTGAGCTCTCGCTGGC | 32873.3 | -4.3 | 1.1 | -4.0 | 7.4E-05 | 1.1E-04 | BT20/MB468 | CYTO |
| >hg38_21_+_8433172_8446622_RNA45SN1_45S_50ntFlanks@4415.4438.24\|CGCGAGGCGAGCCACCGCCCGTCC | >hg38_21_+_8433172_8446622_RNA45SN1_45S_50ntFlanks | 4415 | 4438 | 24 | CGCGAGGCGAGCCACCGCCCGTCC | 61.7 | -9.8 | 2.5 | -4.0 | 7.1E-05 | 1.1E-04 | MB231/MB468 | MP |
| >hg38_21_+_8433172_8446622_RNA45SN1_45S_50ntFlanks@12407.12440.34\|TTTTGATCCTTCGATGTCGGCTCTTCCTATCATT | >hg38_21_+_8433172_8446622_RNA45SN1_45S_50ntFlanks | 12407 | 12440 | 34 | TTTTGATCCTTCGATGTCGGCTCTTCCTATCATT | 470.3 | 3.1 | 0.8 | 3.9 | 8.0E-05 | 1.1E-04 | MB231/MB468 | CYTO |
| >hg38_21_+_8433172_8446622_RNA45SN1_45S_50ntFlanks@7970.8009.40\|CGAGACGCGACCTCAGATCAGACGTGGCGACCCGCTGAAT | >hg38_21_+_8433172_8446622_RNA45SN1_45S_50ntFlanks | 7970 | 8009 | 40 | CGAGACGCGACCTCAGATCAGACGTGGCGACCCGCTGAAT | 149.2 | 3.0 | 0.8 | 3.9 | 8.8E-05 | 1.1E-04 | BT20/MB468 | NUC |

| | | | | | | | | | | | | | |
|---|---|---|---|---|---|---|---|---|---|---|---|---|---|
| >MI0000270\|hsa-mir-181b-1&WithFlank&1\|-\|198858867\|198858988@42.65.24;MI0000683\|hsa-mir-181b-2&WithFlank&9\|+\|124693704\|124693804@22.45.24&[MIMAT0000257&hsa-miR-181b-5p&offsets\|0\|+1;m-73&1\|-\|198858925\|198858947&offsets\|0\|+1];[MIMAT0000257_1&hsa-miR-181b-5p&offsets\|0\|+1;m-72&9\|+\|124693725\|124693747&offsets\|0\|+1]\|AACATTCATTGCTGTCGGTGGGTT | >MI0000270\|hsa-mir-181b-1&WithFlank&1\|-\|198858867\|198858988 | 42 | 65 | 24 | AACATTCATTGCTGTCGGTGGGTT | 285.6 | 2.0 | 0.5 | 3.9 | 8.8E-05 | 1.1E-04 | BT20/MB468 | NUC |
| >hg38_21_+_8433172_8446622_RNA45SN1_45S_50ntFlanks@3993.4020.28\|GTGACTCTAGATAACCTCGGGCCGATCG | >hg38_21_+_8433172_8446622_RNA45SN1_45S_50ntFlanks | 3993 | 4020 | 28 | GTGACTCTAGATAACCTCGGGCCGATCG | 213.4 | -4.8 | 1.2 | -4.0 | 7.2E-05 | 1.1E-04 | MB231/MB468 | MP |
| >MI0000782\|hsa-mir-374a&WithFlank&X\|-\|74287280\|74287363@18.38.21&[MIMAT0000727&hsa-miR-374a-5p&offsets\|0\|-1;m-184&X\|-\|74287325\|74287346&offsets\|0\|-1]\|TTATAATACAACCTGATAAGT | >MI0000782\|hsa-mir-374a&WithFlank&X\|-\|74287280\|74287363 | 18 | 38 | 21 | TTATAATACAACCTGATAAGT | 399.7 | 4.6 | 1.2 | 3.9 | 8.5E-05 | 1.1E-04 | MB231/MB468 | MITO |
| >hg38_21_+_8433172_8446622_RNA45SN1_45S_50ntFlanks@12191.12209.19\|ACGCAGGTGTCCTAAGGCG | >hg38_21_+_8433172_8446622_RNA45SN1_45S_50ntFlanks | 12191 | 12209 | 19 | ACGCAGGTGTCCTAAGGCG | 107.4 | -6.1 | 1.5 | -4.0 | 7.2E-05 | 1.1E-04 | MB231/MB468 | MP |
| >hg38_21_+_8433172_8446622_RNA45SN1_45S_50ntFlanks@11217.11234.18\|CTGGACGCGAGCCGGGCC | >hg38_21_+_8433172_8446622_RNA45SN1_45S_50ntFlanks | 11217 | 11234 | 18 | CTGGACGCGAGCCGGGCC | 700.2 | -3.0 | 0.8 | -3.9 | 8.5E-05 | 1.1E-04 | MB231/MB468 | NUC |
| >hg38_21_+_8433172_8446622_RNA45SN1_45S_50ntFlanks@10351.10387.37\|CGAGAACTTTGAAGGCCGAAGTGGAGAAGGGTTCCAT | >hg38_21_+_8433172_8446622_RNA45SN1_45S_50ntFlanks | 10351 | 10387 | 37 | CGAGAACTTTGAAGGCCGAAGTGGAGAAGGGTTCCAT | 143.0 | -5.4 | 1.3 | -4.0 | 6.3E-05 | 1.1E-04 | BT20/MB468 | MITO |
| >hg38_21_+_8433172_8446622_RNA45SN1_45S_50ntFlanks@9564.9593.30\|CTGACGTGCAAATCGGTCGTCCGACCTGGG | >hg38_21_+_8433172_8446622_RNA45SN1_45S_50ntFlanks | 9564 | 9593 | 30 | CTGACGTGCAAATCGGTCGTCCGACCTGGG | 314.2 | -6.4 | 1.6 | -3.9 | 8.8E-05 | 1.1E-04 | BT20/MB468 | NUC |
| >hg38_21_+_8433172_8446622_RNA45SN1_45S_50ntFlanks@4555.4587.33\|CGCAGCTAGGAATAATGGAATAGGACCGCGGTT | >hg38_21_+_8433172_8446622_RNA45SN1_45S_50ntFlanks | 4555 | 4587 | 33 | CGCAGCTAGGAATAATGGAATAGGACCGCGGTT | 189.9 | -2.3 | 0.6 | -4.0 | 7.5E-05 | 1.1E-04 | BT20/MB468 | CYTO |
| >hg38_21_+_8433172_8446622_RNA45SN1_45S_50ntFlanks@9923.9947.25\|TTGGTTGATATAGACAGCAGGACGG | >hg38_21_+_8433172_8446622_RNA45SN1_45S_50ntFlanks | 9923 | 9947 | 25 | TTGGTTGATATAGACAGCAGGACGG | 70.5 | 9.5 | 2.1 | 4.4 | 1.0E-05 | 1.1E-04 | BT20/MB231 | MP |
| >am_trnaMT_ValTAC_MT_+_1602_1670@29.65.37\|ACTTACACTTAGGAGATTTCAACTTAACTTGACCGCT | >am_trnaMT_ValTAC_MT_+_1602_1670 | 29 | 65 | 37 | ACTTACACTTAGGAGATTTCAACTTAACTTGACCGCT | 150.2 | 11.0 | 2.6 | 4.2 | 3.0E-05 | 1.1E-04 | BT20/MB468 | MP |
| >hg38_21_+_8433172_8446622_RNA45SN1_45S_50ntFlanks@9533.9558.26\|GAGGAAACTCTGGTGGAGGTCCGTAG | >hg38_21_+_8433172_8446622_RNA45SN1_45S_50ntFlanks | 9533 | 9558 | 26 | GAGGAAACTCTGGTGGAGGTCCGTAG | 149.6 | -5.9 | 1.5 | -4.0 | 7.2E-05 | 1.1E-04 | MB231/MB468 | MP |
| >trnaMT_MetCAT_MT_+_4402_4469@50.71.22\|TTATACCCTTCCCGTACTACCA | >trnaMT_MetCAT_MT_+_4402_4469 | 50 | 71 | 22 | TTATACCCTTCCCGTACTACCA | 365.2 | 4.1 | 1.0 | 3.9 | 8.6E-05 | 1.1E-04 | MB231/MB468 | NUC |
| >hg38_21_+_8433172_8446622_RNA45SN1_45S_50ntFlanks@12624.12653.30\|CCAATGGGGCGAAGCTACCATCTGTGGGAT | >hg38_21_+_8433172_8446622_RNA45SN1_45S_50ntFlanks | 12624 | 12653 | 30 | CCAATGGGGCGAAGCTACCATCTGTGGGAT | 250.9 | -5.5 | 1.4 | -4.0 | 7.5E-05 | 1.1E-04 | BT20/MB468 | CYTO |
| >hg38_21_+_8433172_8446622_RNA45SN1_45S_50ntFlanks@10410.10442.33\|TCAGTCGGTCCTGAGAGATGGGCGAGCGCCGTT | >hg38_21_+_8433172_8446622_RNA45SN1_45S_50ntFlanks | 10410 | 10442 | 33 | TCAGTCGGTCCTGAGAGATGGGCGAGCGCCGTT | 660.0 | -6.9 | 1.8 | -3.9 | 8.9E-05 | 1.1E-04 | BT20/MB468 | NUC |
| >hg38_21_+_8433172_8446622_RNA45SN1_45S_50ntFlanks@11316.11340.25\|CACCCCACGTCTCGTCGCGCGCGCG | >hg38_21_+_8433172_8446622_RNA45SN1_45S_50ntFlanks | 11316 | 11340 | 25 | CACCCCACGTCTCGTCGCGCGCGCG | 145.0 | -6.3 | 1.5 | -4.2 | 3.0E-05 | 1.1E-04 | BT20/MB468 | MP |
| >hg38_21_+_8433172_8446622_RNA45SN1_45S_50ntFlanks@11668.11710.43\|TGCCCAGTGCTCTGAATGTCAAAGTGAAGAAATTCAATGAAGC | >hg38_21_+_8433172_8446622_RNA45SN1_45S_50ntFlanks | 11668 | 11710 | 43 | TGCCCAGTGCTCTGAATGTCAAAGTGAAGAAATTCAATGAAGC | 774.9 | -4.6 | 1.0 | -4.4 | 9.9E-06 | 1.1E-04 | BT20/MB231 | CYTO |
| >trna116_GluCTC_1_-_145399233_145399304@36.74.39,trna134_GluTTC_1_-_16861774_16861845@36.74.39,trna71_GluCTC_1_-_161439189_161439260@36.74.39,trna74_GluCTC_1_-_161431809_161431880@36.74.39,trna77_GluCTC_1_-_161424398_161424469@36.74.39,trna77_GluCTC_6_+_28949976_28950047@36.74.39,trna80_GluCTC_1_-_161417018_161417089@36.74.39,trna84_GluTTC_1_-_161391883_161391954@36.74.39,trna87_GluCTC_6_-_126101393_126101464@36.74.39\|CACCGCCGCGGCCCGGGTTCGATTCCCGGTCAGGGAACC | >trna116_GluCTC_1_-_145399233_145399304 | 36 | 74 | 39 | CACCGCCGCGGCCCGGGTTCGATTCCCGGTCAGGGAACC | 711.4 | 5.5 | 1.4 | 4.0 | 7.5E-05 | 1.1E-04 | BT20/MB468 | CYTO |
| >MI0000483\|hsa-mir-186&WithFlank&1\|-\|71067625\|71067722@21.41.21(+1A)&[MIMAT0000456&hsa-miR-186-5p&offsets\|0\|-1(+1A);m-84&1\|-\|71067681\|71067702&offsets\|0\|-1(+1A)]\|CAAAGAATTCTCCTTTTGGGCA | >MI0000483\|hsa-mir-186&WithFlank&1\|-\|71067625\|71067722 | 21 | 41 | 21(+1A) | CAAAGAATTCTCCTTTTGGGCA | 69.9 | 4.4 | 1.1 | 3.9 | 8.9E-05 | 1.1E-04 | BT20/MB468 | NUC |
| >hg38_21_+_8433172_8446622_RNA45SN1_45S_50ntFlanks@6645.6662.18\|CTCGTACGACTCTTAGCG | >hg38_21_+_8433172_8446622_RNA45SN1_45S_50ntFlanks | 6645 | 6662 | 18 | CTCGTACGACTCTTAGCG | 5749.0 | 2.7 | 0.7 | 3.9 | 8.1E-05 | 1.1E-04 | MB231/MB468 | CYTO |
| >hg38_21_+_8433172_8446622_RNA45SN1_45S_50ntFlanks@10679.10707.29\|GGTTCCGGCGGCGTCCGGTGAGCTCTCGC | >hg38_21_+_8433172_8446622_RNA45SN1_45S_50ntFlanks | 10679 | 10707 | 29 | GGTTCCGGCGGCGTCCGGTGAGCTCTCGC | 369.6 | -4.9 | 1.2 | -3.9 | 8.6E-05 | 1.1E-04 | MB231/MB468 | MITO |
| >am_trnaMT_PheGAA_MT_+_577_647@42.59.18\|TTTAGACGGGCTCACATC | >am_trnaMT_PheGAA_MT_+_577_647 | 42 | 59 | 18 | TTTAGACGGGCTCACATC | 416.9 | 6.2 | 1.5 | 4.2 | 3.0E-05 | 1.1E-04 | BT20/MB468 | MP |
| >hg38_21_+_8433172_8446622_RNA45SN1_45S_50ntFlanks@3823.3841.19\|TTTGGTCGCTCGCTCCTCT | >hg38_21_+_8433172_8446622_RNA45SN1_45S_50ntFlanks | 3823 | 3841 | 19 | TTTGGTCGCTCGCTCCTCT | 272.6 | 1.9 | 0.5 | 3.9 | 8.1E-05 | 1.1E-04 | MB231/MB468 | CYTO |
| >hg38_21_+_8433172_8446622_RNA45SN1_45S_50ntFlanks@10805.10840.36\|GAACAATGTAGGTAAGGGAAGTCGGCAAGCCGGATC | >hg38_21_+_8433172_8446622_RNA45SN1_45S_50ntFlanks | 10805 | 10840 | 36 | GAACAATGTAGGTAAGGGAAGTCGGCAAGCCGGATC | 255.9 | -3.4 | 0.9 | -3.9 | 8.1E-05 | 1.1E-04 | MB231/MB468 | CYTO |
| >hg38_21_+_8433172_8446622_RNA45SN1_45S_50ntFlanks@12633.12657.25\|CGAAGCTACCATCTGTGGGATTATG | >hg38_21_+_8433172_8446622_RNA45SN1_45S_50ntFlanks | 12633 | 12657 | 25 | CGAAGCTACCATCTGTGGGATTATG | 235.1 | -4.4 | 1.1 | -3.9 | 8.6E-05 | 1.1E-04 | MB231/MB468 | NUC |
| >hg38_21_+_8433172_8446622_RNA45SN1_45S_50ntFlanks@10408.10456.49\|GGTCAGTCGGTCCTGAGAGATGGGCGAGCGCCGTTCCGAAGGGACGGGC | >hg38_21_+_8433172_8446622_RNA45SN1_45S_50ntFlanks | 10408 | 10456 | 49 | GGTCAGTCGGTCCTGAGAGATGGGCGAGCGCCGTTCCGAAGGGACGGGC | 58.0 | -9.7 | 2.5 | -4.0 | 7.2E-05 | 1.1E-04 | MB231/MB468 | MP |
| >hg38_21_+_8433172_8446622_RNA45SN1_45S_50ntFlanks@12002.12030.29\|TGCGGGCCGCCGGTGAAATACCACTACTC | >hg38_21_+_8433172_8446622_RNA45SN1_45S_50ntFlanks | 12002 | 12030 | 29 | TGCGGGCCGCCGGTGAAATACCACTACTC | 936.4 | -5.0 | 1.3 | -3.9 | 8.6E-05 | 1.1E-04 | MB231/MB468 | NUC |
| >hg38_21_+_8433172_8446622_RNA45SN1_45S_50ntFlanks@10728.10755.28\|GAGAGGGTGTAAATCTCGCGCCGGGCCG | >hg38_21_+_8433172_8446622_RNA45SN1_45S_50ntFlanks | 10728 | 10755 | 28 | GAGAGGGTGTAAATCTCGCGCCGGGCCG | 154.8 | -5.1 | 1.3 | -4.0 | 7.5E-05 | 1.1E-04 | BT20/MB468 | CYTO |
| >MI0000081\|hsa-mir-24-2&WithFlank&19\|-\|13836281\|13836365@56.75.20;MI0000080\|hsa-mir-24-1&WithFlank&9\|+\|95086015\|95086094@50.69.20&[MIMAT0000080&hsa-miR-24-3p&offsets\|0\|-2;m-36&19\|-\|13836288\|13836310&offsets\|0\|-3];[MIMAT0000080_1&hsa-miR-24-3p&offsets\|0\|-2;m-35&9\|+\|95086064\|95086086&offsets\|0\|-3]\|TGGCTCAGTTCAGCAGGAAC | >MI0000081\|hsa-mir-24-2&WithFlank&19\|-\|13836281\|13836365 | 56 | 75 | 20 | TGGCTCAGTTCAGCAGGAAC | 27156.2 | 3.2 | 0.8 | 4.0 | 7.3E-05 | 1.1E-04 | MB231/MB468 | MP |
| >hg38_21_+_8433172_8446622_RNA45SN1_45S_50ntFlanks@11854.11887.34\|TTGGCGGAATCAGCGGGGAAAGAAGACCCTGTTG | >hg38_21_+_8433172_8446622_RNA45SN1_45S_50ntFlanks | 11854 | 11887 | 34 | TTGGCGGAATCAGCGGGGAAAGAAGACCCTGTTG | 75.4 | 9.6 | 2.2 | 4.4 | 1.1E-05 | 1.2E-04 | BT20/MB231 | MP |
| >MI0000087\|hsa-mir-29a&WithFlank&7\|-\|130876741\|130876816@48.69.22(+1A)&[MIMAT0000086&hsa-miR-29a-3p&offsets\|0\|0(+1A);m-26&7\|-\|130876749\|130876769&offsets\|0\|+1(+1A)]\|TAGCACCATCTGAAATCGGTTAA | >MI0000087\|hsa-mir-29a&WithFlank&7\|-\|130876741\|130876816 | 48 | 69 | 22(+1A) | TAGCACCATCTGAAATCGGTTAA | 404.6 | 5.0 | 1.3 | 4.0 | 7.3E-05 | 1.2E-04 | MB231/MB468 | MP |
| >MI0000087\|hsa-mir-29a&WithFlank&7\|-\|130876741\|130876816@47.69.23&[MIMAT0000086&hsa-miR-29a-3p&offsets\|-1\|0;m-26&7\|-\|130876749\|130876769&offsets\|-1\|+1]\|CTAGCACCATCTGAAATCGGTTA | >MI0000087\|hsa-mir-29a&WithFlank&7\|-\|130876741\|130876816 | 47 | 69 | 23 | CTAGCACCATCTGAAATCGGTTA | 4308.5 | 5.8 | 1.5 | 4.0 | 7.3E-05 | 1.2E-04 | MB231/MB468 | MP |
| >hg38_21_+_8433172_8446622_RNA45SN1_45S_50ntFlanks@12406.12432.27\|TTTTTGATCCTTCGATGTCGGCTCTTC | >hg38_21_+_8433172_8446622_RNA45SN1_45S_50ntFlanks | 12406 | 12432 | 27 | TTTTTGATCCTTCGATGTCGGCTCTTC | 470.7 | -3.0 | 0.7 | -3.9 | 8.1E-05 | 1.2E-04 | MB231/MB468 | CYTO |
| >am_trnaMT_ValTAC_MT_+_1602_1670@34.67.34\|CACTTAGGAGATTTCAACTTAACTTGACCGCTCT | >am_trnaMT_ValTAC_MT_+_1602_1670 | 34 | 67 | 34 | CACTTAGGAGATTTCAACTTAACTTGACCGCTCT | 140.6 | 10.9 | 2.6 | 4.2 | 3.1E-05 | 1.2E-04 | BT20/MB468 | MP |
| >MI0000063\|hsa-let-7b&WithFlank&22\|+\|46113680\|46113774@12.29.18&[MIMAT0000063&hsa-let-7b-5p&offsets\|0\|-4;m-14&22\|+\|46113691\|46113712&offsets\|0\|-4]\|TGAGGTAGTAGGTTGTGT | >MI0000063\|hsa-let-7b&WithFlank&22\|+\|46113680\|46113774 | 12 | 29 | 18 | TGAGGTAGTAGGTTGTGT | 782.8 | 7.2 | 1.7 | 4.2 | 3.1E-05 | 1.2E-04 | BT20/MB468 | MP |
| >hg38_21_+_8433172_8446622_RNA45SN1_45S_50ntFlanks@11575.11605.31\|ACTTAGAACTGGTGCGGACCAGGGGAATCCG | >hg38_21_+_8433172_8446622_RNA45SN1_45S_50ntFlanks | 11575 | 11605 | 31 | ACTTAGAACTGGTGCGGACCAGGGGAATCCG | 1060.5 | -5.5 | 1.3 | -4.2 | 3.1E-05 | 1.2E-04 | BT20/MB468 | MP |
| >hg38_21_+_8433172_8446622_RNA45SN1_45S_50ntFlanks@4727.4761.35\|AATCAAGAACGAAAGTCGGAGGTTCGAAGACGATC | >hg38_21_+_8433172_8446622_RNA45SN1_45S_50ntFlanks | 4727 | 4761 | 35 | AATCAAGAACGAAAGTCGGAGGTTCGAAGACGATC | 365.6 | 4.0 | 1.0 | 4.2 | 3.1E-05 | 1.2E-04 | BT20/MB468 | MP |
| >hg38_21_+_8433172_8446622_RNA45SN1_45S_50ntFlanks@6648.6667.20\|GTACGACTCTTAGCGGTGGA | >hg38_21_+_8433172_8446622_RNA45SN1_45S_50ntFlanks | 6648 | 6667 | 20 | GTACGACTCTTAGCGGTGGA | 484.8 | 4.0 | 1.0 | 4.2 | 3.1E-05 | 1.2E-04 | BT20/MB468 | MP |
| >hg38_21_+_8433172_8446622_RNA45SN1_45S_50ntFlanks@7975.8006.32\|CGCGACCTCAGATCAGACGTGGCGACCCGCTG | >hg38_21_+_8433172_8446622_RNA45SN1_45S_50ntFlanks | 7975 | 8006 | 32 | CGCGACCTCAGATCAGACGTGGCGACCCGCTG | 44435.6 | -2.9 | 0.7 | -3.9 | 8.7E-05 | 1.2E-04 | MB231/MB468 | NUC |
| >hg38_21_+_8433172_8446622_RNA45SN1_45S_50ntFlanks@475.508.34\|CTTCGTGATCGATGTGGTGACGTCGTGCTCTCCC | >hg38_21_+_8433172_8446622_RNA45SN1_45S_50ntFlanks | 475 | 508 | 34 | CTTCGTGATCGATGTGGTGACGTCGTGCTCTCCC | 2266.4 | 2.6 | 0.7 | 3.9 | 8.7E-05 | 1.2E-04 | MB231/MB468 | NUC |
| >hg38_21_+_8433172_8446622_RNA45SN1_45S_50ntFlanks@12942.12979.38\|TTCGTAGACGACCTGCTTCTGGGTCGGGGTTTCGTACG | >hg38_21_+_8433172_8446622_RNA45SN1_45S_50ntFlanks | 12942 | 12979 | 38 | TTCGTAGACGACCTGCTTCTGGGTCGGGGTTTCGTACG | 37.1 | -8.3 | 2.0 | -4.2 | 3.1E-05 | 1.2E-04 | BT20/MB468 | MP |
| >hg38_21_+_8433172_8446622_RNA45SN1_45S_50ntFlanks@12632.12653.22\|GCGAAGCTACCATCTGTGGGAT | >hg38_21_+_8433172_8446622_RNA45SN1_45S_50ntFlanks | 12632 | 12653 | 22 | GCGAAGCTACCATCTGTGGGAT | 1920.4 | -7.0 | 1.7 | -4.2 | 3.1E-05 | 1.2E-04 | BT20/MB468 | MP |
| >hg38_21_+_8433172_8446622_RNA45SN1_45S_50ntFlanks@9938.9957.20\|AGCAGGACGGTGGCCATGGA | >hg38_21_+_8433172_8446622_RNA45SN1_45S_50ntFlanks | 9938 | 9957 | 20 | AGCAGGACGGTGGCCATGGA | 738.0 | 5.9 | 1.4 | 4.2 | 3.1E-05 | 1.2E-04 | BT20/MB468 | MP |
| >trna11_GluTTC_15_-_26327381_26327452@41.75.35,trna3_GluTTC_13_-_45492062_45492133@41.75.35\|AGGCGGCCCGGGTTCGACTCCCGGTGTGGGAACCA | >trna11_GluTTC_15_-_26327381_26327452 | 41 | 75 | 35 | AGGCGGCCCGGGTTCGACTCCCGGTGTGGGAACCA | 284.7 | 5.2 | 1.3 | 3.9 | 8.7E-05 | 1.2E-04 | MB231/MB468 | MITO |
| >hg38_21_+_8433172_8446622_RNA45SN1_45S_50ntFlanks@11730.11751.22\|AACTATGACTCTCTTAAGGTAG | >hg38_21_+_8433172_8446622_RNA45SN1_45S_50ntFlanks | 11730 | 11751 | 22 | AACTATGACTCTCTTAAGGTAG | 175.1 | -3.5 | 0.9 | -3.9 | 8.2E-05 | 1.2E-04 | MB231/MB468 | CYTO |
| >hg38_21_+_8433172_8446622_RNA45SN1_45S_50ntFlanks@6772.6807.36\|GTTCCTCCCGGGGCTACGCCTGTCTGAGCGTCGCTT | >hg38_21_+_8433172_8446622_RNA45SN1_45S_50ntFlanks | 6772 | 6807 | 36 | GTTCCTCCCGGGGCTACGCCTGTCTGAGCGTCGCTT | 670.5 | -2.8 | 0.7 | -3.9 | 8.7E-05 | 1.2E-04 | MB231/MB468 | NUC |
| >MI0000088\|hsa-mir-30a&WithFlank&6\|-\|71403545\|71403627@53.73.21(+1U)&[MIMAT0000088&hsa-miR-30a-3p&offsets\|0\|-1(+1U);m-38&6\|-\|71403554\|71403575&offsets\|0\|-1(+1U)]\|CTTTCAGTCGGATGTTTGCAGT | >MI0000088\|hsa-mir-30a&WithFlank&6\|-\|71403545\|71403627 | 53 | 73 | 21(+1U) | CTTTCAGTCGGATGTTTGCAGT | 9752.0 | -4.0 | 0.9 | -4.4 | 1.1E-05 | 1.2E-04 | BT20/MB231 | MP |
| >MI0000093\|hsa-mir-92a-1&WithFlank&13\|+\|91351308\|91351397@54.75.22(+2A);MI0000094\|hsa-mir-92a-2&WithFlank&X\|-\|134169532\|134169618@54.75.22(+2A)&[MIMAT0000092&hsa-miR-92a-3p&offsets\|0\|0(+2A);m-9&13\|+\|91351361\|91351382&offsets\|0\|0(+2A)];[MIMAT0000092_1&hsa-miR-92a-3p&offsets\|0\|0(+2A);m-11&X\|-\|134169544\|134169565&offsets\|0\|0(+2A)]\|TATTGCACTTGTCCCGGCCTGTAA | >MI0000093\|hsa-mir-92a-1&WithFlank&13\|+\|91351308\|91351397 | 54 | 75 | 22(+2A) | TATTGCACTTGTCCCGGCCTGTAA | 913.0 | 2.8 | 0.7 | 3.9 | 8.8E-05 | 1.2E-04 | MB231/MB468 | NUC |

<table>
<tr><td>>hg38_21_+_8433172_8446622_RNA45SN1_45S_50ntFlanks@5078.5117.40|CGATAACGAACGAGACTCTGGCATGCTAACTAGTTACGCG</td><td>>hg38_21_+_8433172_8446622_RNA45SN1_45S_50ntFlanks</td><td>5078</td><td>5117</td><td>40</td><td>CGATAACGAACGAGACTCTGGCATGCTAACTAGTTACGCG</td><td>338.6</td><td>-5.0</td><td>1.3</td><td>-3.9</td><td>8.7E-05</td><td>1.2E-04</td><td>MB231/MB468</td><td>MITO</td></tr>
<tr><td>>hg38_21_+_8433172_8446622_RNA45SN1_45S_50ntFlanks@11580.11606.27|GAACTGGTGCGGACCAGGGGAATCCGA</td><td>>hg38_21_+_8433172_8446622_RNA45SN1_45S_50ntFlanks</td><td>11580</td><td>11606</td><td>27</td><td>GAACTGGTGCGGACCAGGGGAATCCGA</td><td>368.7</td><td>-6.2</td><td>1.6</td><td>-4.0</td><td>7.4E-05</td><td>1.2E-04</td><td>MB231/MB468</td><td>MP</td></tr>
<tr><td>>hg38_MT_+_1621_3279_RNR2_16S_50ntFlanks@114.146.33|CAAATAAAGTATAGGCGATAGAAATTGAAACCT</td><td>>hg38_MT_+_1621_3279_RNR2_16S_50ntFlanks</td><td>114</td><td>146</td><td>33</td><td>CAAATAAAGTATAGGCGATAGAAATTGAAACCT</td><td>128.7</td><td>10.8</td><td>2.6</td><td>4.2</td><td>3.1E-05</td><td>1.2E-04</td><td>BT20/MB468</td><td>MP</td></tr>
<tr><td>>hg38_21_+_8433172_8446622_RNA45SN1_45S_50ntFlanks@11569.11605.37|CAGCCGACTTAGAACTGGTGCGGACCAGGGGAATCCG</td><td>>hg38_21_+_8433172_8446622_RNA45SN1_45S_50ntFlanks</td><td>11569</td><td>11605</td><td>37</td><td>CAGCCGACTTAGAACTGGTGCGGACCAGGGGAATCCG</td><td>415.6</td><td>-4.3</td><td>1.1</td><td>-3.9</td><td>8.2E-05</td><td>1.2E-04</td><td>MB231/MB468</td><td>CYTO</td></tr>
<tr><td>>hg38_21_+_8433172_8446622_RNA45SN1_45S_50ntFlanks@11999.12034.36|CCCTGCGGGCCGCCGGTGAAATACCACTACTCTGAT</td><td>>hg38_21_+_8433172_8446622_RNA45SN1_45S_50ntFlanks</td><td>11999</td><td>12034</td><td>36</td><td>CCCTGCGGGCCGCCGGTGAAATACCACTACTCTGAT</td><td>37.9</td><td>-8.3</td><td>2.0</td><td>-4.2</td><td>3.1E-05</td><td>1.2E-04</td><td>BT20/MB468</td><td>MP</td></tr>
<tr><td>>MI0003643|hsa-mir-629&WithFlank&15|-|70079366|70079474@28.48.21&[MIMAT0004810&hsa-miR-629-5p&offsets|0|0;m-150&15|-|70079427|70079447&offsets|0|0]|TGGGTTTACGTTGGGAGAACT</td><td>>MI0003643|hsa-mir-629&WithFlank&15|-|70079366|70079474</td><td>28</td><td>48</td><td>21</td><td>TGGGTTTACGTTGGGAGAACT</td><td>603.2</td><td>2.7</td><td>0.6</td><td>4.4</td><td>1.2E-05</td><td>1.2E-04</td><td>BT20/MB231</td><td>NUC</td></tr>
<tr><td>>hg38_21_+_8433172_8446622_RNA45SN1_45S_50ntFlanks@5491.5516.26|GAGAAGACGGTCGAACTTGACTATCT</td><td>>hg38_21_+_8433172_8446622_RNA45SN1_45S_50ntFlanks</td><td>5491</td><td>5516</td><td>26</td><td>GAGAAGACGGTCGAACTTGACTATCT</td><td>16020.0</td><td>6.0</td><td>1.4</td><td>4.4</td><td>1.2E-05</td><td>1.2E-04</td><td>BT20/MB231</td><td>NUC</td></tr>
<tr><td>>MI0000101|hsa-mir-99a&WithFlank&21|+|16539083|16539175@19.40.22&[MIMAT0000097&hsa-miR-99a-5p&offsets|0|0;m-64&21|+|16539101|16539121&offsets|0|+1]|AACCCGTAGATCCGATCTTGTG</td><td>>MI0000101|hsa-mir-99a&WithFlank&21|+|16539083|16539175</td><td>19</td><td>40</td><td>22</td><td>AACCCGTAGATCCGATCTTGTG</td><td>1127.8</td><td>-6.8</td><td>1.5</td><td>-4.4</td><td>1.2E-05</td><td>1.2E-04</td><td>BT20/MB231</td><td>NUC</td></tr>
<tr><td>>MI0002470|hsa-mir-486-1&WithFlank&8|-|41660435|41660514@10.31.22;MI0023622|hsa-mir-486-2&WithFlank&8|+|41660438|41660513@7.28.22&[MIMAT0002177&hsa-miR-486-5p&offsets|0|0;m-107&8|-|41660484|41660505&offsets|0|0];[MIMAT0002177_1&hsa-miR-486-5p&offsets|0|0;TJU_CMC_MD2.ID03138.5p-miR&offsets|0|-2;m-106&8|+|41660444|41660465&offsets|0|0]|TCCTGTACTGAGCTGCCCCGAG</td><td>>MI0002470|hsa-mir-486-1&WithFlank&8|-|41660435|41660514</td><td>10</td><td>31</td><td>22</td><td>TCCTGTACTGAGCTGCCCCGAG</td><td>283.5</td><td>-4.0</td><td>0.9</td><td>-4.4</td><td>1.0E-05</td><td>1.2E-04</td><td>BT20/MB231</td><td>CYTO</td></tr>
<tr><td>>hg38_21_+_8433172_8446622_RNA45SN1_45S_50ntFlanks@8020.8042.23|TAGTCAGCGGAGGAGAAGAAACT</td><td>>hg38_21_+_8433172_8446622_RNA45SN1_45S_50ntFlanks</td><td>8020</td><td>8042</td><td>23</td><td>TAGTCAGCGGAGGAGAAGAAACT</td><td>423.2</td><td>-4.2</td><td>1.0</td><td>-4.4</td><td>1.0E-05</td><td>1.2E-04</td><td>BT20/MB231</td><td>CYTO</td></tr>
<tr><td>>MI0000077|hsa-mir-21&WithFlank&17|+|59841260|59841343@14.33.20(+1A)&[MIMAT0000076&hsa-miR-21-5p&offsets|0|-2(+1A);m-1&17|+|59841273|59841294&offsets|0|-2(+1A)]|TAGCTTATCAGACTGATGTTA</td><td>>MI0000077|hsa-mir-21&WithFlank&17|+|59841260|59841343</td><td>14</td><td>33</td><td>20(+1A)</td><td>TAGCTTATCAGACTGATGTTA</td><td>3762.4</td><td>3.5</td><td>0.9</td><td>4.0</td><td>7.4E-05</td><td>1.2E-04</td><td>MB231/MB468</td><td>MP</td></tr>
<tr><td>>trnaMT_ValTAC_MT_+_1602_1670@51.71.21|CTTAACTTGACCGCTCTGACC</td><td>>trnaMT_ValTAC_MT_+_1602_1670</td><td>51</td><td>71</td><td>21</td><td>CTTAACTTGACCGCTCTGACC</td><td>1855.7</td><td>-6.9</td><td>1.8</td><td>-3.9</td><td>9.1E-05</td><td>1.2E-04</td><td>BT20/MB468</td><td>NUC</td></tr>
<tr><td>>hg38_21_+_8433172_8446622_RNA45SN1_45S_50ntFlanks@8109.8143.35|GCGGGGCGCGGGACATGTGGCGTACGGAAGACCCG</td><td>>hg38_21_+_8433172_8446622_RNA45SN1_45S_50ntFlanks</td><td>8109</td><td>8143</td><td>35</td><td>GCGGGGCGCGGGACATGTGGCGTACGGAAGACCCG</td><td>163.8</td><td>-5.2</td><td>1.3</td><td>-4.0</td><td>6.4E-05</td><td>1.2E-04</td><td>BT20/MB468</td><td>MITO</td></tr>
<tr><td>>hg38_21_+_8433172_8446622_RNA45SN1_45S_50ntFlanks@12641.12669.29|CCATCTGTGGGATTATGACTGAACGCCTC</td><td>>hg38_21_+_8433172_8446622_RNA45SN1_45S_50ntFlanks</td><td>12641</td><td>12669</td><td>29</td><td>CCATCTGTGGGATTATGACTGAACGCCTC</td><td>503.6</td><td>-3.4</td><td>0.9</td><td>-3.9</td><td>8.8E-05</td><td>1.2E-04</td><td>MB231/MB468</td><td>MITO</td></tr>
<tr><td>>hg38_21_+_8433172_8446622_RNA45SN1_45S_50ntFlanks@12300.12323.24|GGGCCTCACGATCCTTCTGACCTT</td><td>>hg38_21_+_8433172_8446622_RNA45SN1_45S_50ntFlanks</td><td>12300</td><td>12323</td><td>24</td><td>GGGCCTCACGATCCTTCTGACCTT</td><td>59.1</td><td>-9.7</td><td>2.5</td><td>-4.0</td><td>7.4E-05</td><td>1.2E-04</td><td>MB231/MB468</td><td>MP</td></tr>
<tr><td>>hg38_21_+_8433172_8446622_RNA45SN1_45S_50ntFlanks@6651.6694.44|CGACTCTTAGCGGTGGATCACTCGGCTCGTGCGTCGATGAAGAA</td><td>>hg38_21_+_8433172_8446622_RNA45SN1_45S_50ntFlanks</td><td>6651</td><td>6694</td><td>44</td><td>CGACTCTTAGCGGTGGATCACTCGGCTCGTGCGTCGATGAAGAA</td><td>2635.6</td><td>4.0</td><td>1.0</td><td>3.9</td><td>8.8E-05</td><td>1.2E-04</td><td>MB231/MB468</td><td>NUC</td></tr>
<tr><td>>hg38_21_+_8433172_8446622_RNA45SN1_45S_50ntFlanks@9575.9601.27|ATCGGTCGTCCGACCTGGGTATAGGGG</td><td>>hg38_21_+_8433172_8446622_RNA45SN1_45S_50ntFlanks</td><td>9575</td><td>9601</td><td>27</td><td>ATCGGTCGTCCGACCTGGGTATAGGGG</td><td>479.2</td><td>-6.2</td><td>1.6</td><td>-4.0</td><td>7.4E-05</td><td>1.2E-04</td><td>MB231/MB468</td><td>MP</td></tr>
<tr><td>>MI0000542|hsa-mir-320a&WithFlank&8|-|22244960|22245043@49.68.20&[MIMAT0000510&hsa-miR-320a-3p&offsets|+1|-1;m-60&8|-|22244975|22244996&offsets|+1|-1]|AAAGCTGGGTTGAGAGGGCG</td><td>>MI0000542|hsa-mir-320a&WithFlank&8|-|22244960|22245043</td><td>49</td><td>68</td><td>20</td><td>AAAGCTGGGTTGAGAGGGCG</td><td>1187.9</td><td>7.3</td><td>1.7</td><td>4.2</td><td>3.1E-05</td><td>1.2E-04</td><td>BT20/MB468</td><td>MP</td></tr>
<tr><td>>hg38_21_+_8433172_8446622_RNA45SN1_45S_50ntFlanks@10116.10134.19|TGCGTCGGAGGGCGGCGGC</td><td>>hg38_21_+_8433172_8446622_RNA45SN1_45S_50ntFlanks</td><td>10116</td><td>10134</td><td>19</td><td>TGCGTCGGAGGGCGGCGGC</td><td>177.9</td><td>-6.6</td><td>1.6</td><td>-4.2</td><td>3.1E-05</td><td>1.2E-04</td><td>BT20/MB468</td><td>MP</td></tr>
<tr><td>>MI0000542|hsa-mir-320a&WithFlank&8|-|22244960|22245043@48.70.23(+1G)&[MIMAT0000510&hsa-miR-320a-3p&offsets|0|+1(+1G);m-60&8|-|22244975|22244996&offsets|0|+1(+1G)]|AAAAGCTGGGTTGAGAGGGCGAAG</td><td>>MI0000542|hsa-mir-320a&WithFlank&8|-|22244960|22245043</td><td>48</td><td>70</td><td>23(+1G)</td><td>AAAAGCTGGGTTGAGAGGGCGAAG</td><td>249.8</td><td>3.0</td><td>0.7</td><td>4.4</td><td>1.2E-05</td><td>1.2E-04</td><td>BT20/MB231</td><td>MITO</td></tr>
<tr><td>>hg38_21_+_8433172_8446622_RNA45SN1_45S_50ntFlanks@12420.12458.39|ATGTCGGCTCTTCCTATCATTGTGAAGCAGAATTCACCA</td><td>>hg38_21_+_8433172_8446622_RNA45SN1_45S_50ntFlanks</td><td>12420</td><td>12458</td><td>39</td><td>ATGTCGGCTCTTCCTATCATTGTGAAGCAGAATTCACCA</td><td>1347.4</td><td>-3.6</td><td>0.8</td><td>-4.4</td><td>1.2E-05</td><td>1.2E-04</td><td>BT20/MB231</td><td>MITO</td></tr>
<tr><td>>am_trnaMT_TyrGTA_MT_-_5826_5891@13.32.20|GAGTGAAGCATTGGACTGTA</td><td>>am_trnaMT_TyrGTA_MT_-_5826_5891</td><td>13</td><td>32</td><td>20</td><td>GAGTGAAGCATTGGACTGTA</td><td>193.9</td><td>3.0</td><td>0.8</td><td>4.0</td><td>7.4E-05</td><td>1.2E-04</td><td>MB231/MB468</td><td>MP</td></tr>
<tr><td>>hg38_21_+_8433172_8446622_RNA45SN1_45S_50ntFlanks@4728.4752.25|ATCAAGAACGAAAGTCGGAGGTTCG</td><td>>hg38_21_+_8433172_8446622_RNA45SN1_45S_50ntFlanks</td><td>4728</td><td>4752</td><td>25</td><td>ATCAAGAACGAAAGTCGGAGGTTCG</td><td>455.8</td><td>-5.8</td><td>1.5</td><td>-3.9</td><td>8.8E-05</td><td>1.2E-04</td><td>MB231/MB468</td><td>NUC</td></tr>
<tr><td>>hg38_21_+_8433172_8446622_RNA45SN1_45S_50ntFlanks@12241.12266.26|CAGAAGGGCAAAAGCTCGCTTGATCT</td><td>>hg38_21_+_8433172_8446622_RNA45SN1_45S_50ntFlanks</td><td>12241</td><td>12266</td><td>26</td><td>CAGAAGGGCAAAAGCTCGCTTGATCT</td><td>239.2</td><td>-6.0</td><td>1.5</td><td>-4.0</td><td>7.4E-05</td><td>1.2E-04</td><td>MB231/MB468</td><td>MP</td></tr>
<tr><td>>hg38_21_+_8433172_8446622_RNA45SN1_45S_50ntFlanks@5106.5136.31|ACTAGTTACGCGACCCCCGAGCGGTCGGCGT</td><td>>hg38_21_+_8433172_8446622_RNA45SN1_45S_50ntFlanks</td><td>5106</td><td>5136</td><td>31</td><td>ACTAGTTACGCGACCCCCGAGCGGTCGGCGT</td><td>53.4</td><td>-9.6</td><td>2.4</td><td>-4.0</td><td>7.4E-05</td><td>1.2E-04</td><td>MB231/MB468</td><td>MP</td></tr>
<tr><td>>hg38_21_+_8433172_8446622_RNA45SN1_45S_50ntFlanks@10408.10453.46|GGTCAGTCGGTCCTGAGAGATGGGCGAGCGCCGTTCCGAAGGGACG</td><td>>hg38_21_+_8433172_8446622_RNA45SN1_45S_50ntFlanks</td><td>10408</td><td>10453</td><td>46</td><td>GGTCAGTCGGTCCTGAGAGATGGGCGAGCGCCGTTCCGAAGGGACG</td><td>292.4</td><td>-4.9</td><td>1.3</td><td>-3.9</td><td>8.8E-05</td><td>1.2E-04</td><td>MB231/MB468</td><td>MITO</td></tr>
<tr><td>>hg38_21_+_8433172_8446622_RNA45SN1_45S_50ntFlanks@6661.6690.30|CGGTGGATCACTCGGCTCGTGCGTCGATGA</td><td>>hg38_21_+_8433172_8446622_RNA45SN1_45S_50ntFlanks</td><td>6661</td><td>6690</td><td>30</td><td>CGGTGGATCACTCGGCTCGTGCGTCGATGA</td><td>768.7</td><td>4.0</td><td>0.9</td><td>4.4</td><td>1.1E-05</td><td>1.2E-04</td><td>BT20/MB231</td><td>MP</td></tr>
<tr><td>>hg38_21_+_8433172_8446622_RNA45SN1_45S_50ntFlanks@8118.8139.22|GGGACATGTGGCGTACGGAAGA</td><td>>hg38_21_+_8433172_8446622_RNA45SN1_45S_50ntFlanks</td><td>8118</td><td>8139</td><td>22</td><td>GGGACATGTGGCGTACGGAAGA</td><td>675.1</td><td>7.5</td><td>1.8</td><td>4.2</td><td>3.1E-05</td><td>1.2E-04</td><td>BT20/MB468</td><td>MP</td></tr>
<tr><td>>hg38_21_+_8433172_8446622_RNA45SN1_45S_50ntFlanks@6785.6806.22|CTACGCCTGTCTGAGCGTCGCT</td><td>>hg38_21_+_8433172_8446622_RNA45SN1_45S_50ntFlanks</td><td>6785</td><td>6806</td><td>22</td><td>CTACGCCTGTCTGAGCGTCGCT</td><td>443.8</td><td>2.0</td><td>0.5</td><td>3.9</td><td>9.1E-05</td><td>1.2E-04</td><td>BT20/MB468</td><td>NUC</td></tr>
<tr><td>>hg38_21_+_8433172_8446622_RNA45SN1_45S_50ntFlanks@6646.6682.37|TCGTACGACTCTTAGCGGTGGATCACTCGGCTCGTGC</td><td>>hg38_21_+_8433172_8446622_RNA45SN1_45S_50ntFlanks</td><td>6646</td><td>6682</td><td>37</td><td>TCGTACGACTCTTAGCGGTGGATCACTCGGCTCGTGC</td><td>357.3</td><td>4.7</td><td>1.2</td><td>3.9</td><td>8.9E-05</td><td>1.2E-04</td><td>MB231/MB468</td><td>NUC</td></tr>
<tr><td>>hg38_21_+_8433172_8446622_RNA45SN1_45S_50ntFlanks@475.520.46|CTTCGTGATCGATGTGGTGACGTCGTGCTCTCCCGGGCCGGGTCCG</td><td>>hg38_21_+_8433172_8446622_RNA45SN1_45S_50ntFlanks</td><td>475</td><td>520</td><td>46</td><td>CTTCGTGATCGATGTGGTGACGTCGTGCTCTCCCGGGCCGGGTCCG</td><td>484.9</td><td>6.3</td><td>1.6</td><td>4.0</td><td>6.5E-05</td><td>1.2E-04</td><td>BT20/MB468</td><td>MITO</td></tr>
<tr><td>>MI0006444|hsa-mir-1307&WithFlank&10|-|103394247|103394407@87.107.21&[MIMAT0005951&hsa-miR-1307-3p&offsets|+1|0;m-66&10|-|103394300|103394322&offsets|+1|-1]|CTCGGCGTGGCGTCGGTCGTG</td><td>>MI0006444|hsa-mir-1307&WithFlank&10|-|103394247|103394407</td><td>87</td><td>107</td><td>21</td><td>CTCGGCGTGGCGTCGGTCGTG</td><td>157.9</td><td>3.7</td><td>0.9</td><td>3.9</td><td>8.9E-05</td><td>1.2E-04</td><td>MB231/MB468</td><td>MITO</td></tr>
<tr><td>>trnaMT_GlnTTG_MT_-_4329_4400@1.24.24,trnalookalike2_GlnTTG_1_-_564879_564950@1.24.24|TAGGATGGGGTGTGATAGGTGGCA</td><td>>trnaMT_GlnTTG_MT_-_4329_4400</td><td>1</td><td>24</td><td>24</td><td>TAGGATGGGGTGTGATAGGTGGCA</td><td>691.6</td><td>5.2</td><td>1.2</td><td>4.2</td><td>3.2E-05</td><td>1.2E-04</td><td>BT20/MB468</td><td>MP</td></tr>
<tr><td>>hg38_21_+_8433172_8446622_RNA45SN1_45S_50ntFlanks@12477.12498.22|CACTAATAGGGAACGTGAGCTG</td><td>>hg38_21_+_8433172_8446622_RNA45SN1_45S_50ntFlanks</td><td>12477</td><td>12498</td><td>22</td><td>CACTAATAGGGAACGTGAGCTG</td><td>1552.0</td><td>-2.2</td><td>0.6</td><td>-4.0</td><td>7.8E-05</td><td>1.2E-04</td><td>BT20/MB468</td><td>CYTO</td></tr>
<tr><td>>hg38_21_+_8433172_8446622_RNA45SN1_45S_50ntFlanks@11671.11692.22|CCAGTGCTCTGAATGTCAAAGT</td><td>>hg38_21_+_8433172_8446622_RNA45SN1_45S_50ntFlanks</td><td>11671</td><td>11692</td><td>22</td><td>CCAGTGCTCTGAATGTCAAAGT</td><td>200.2</td><td>-3.5</td><td>0.9</td><td>-4.0</td><td>7.5E-05</td><td>1.2E-04</td><td>MB231/MB468</td><td>MP</td></tr>
<tr><td>>hg38_21_+_8433172_8446622_RNA45SN1_45S_50ntFlanks@8189.8214.26|CCCAGCCCGTGGACGGTGTGAGGCCG</td><td>>hg38_21_+_8433172_8446622_RNA45SN1_45S_50ntFlanks</td><td>8189</td><td>8214</td><td>26</td><td>CCCAGCCCGTGGACGGTGTGAGGCCG</td><td>102.9</td><td>-6.0</td><td>1.5</td><td>-4.0</td><td>7.5E-05</td><td>1.2E-04</td><td>MB231/MB468</td><td>MP</td></tr>
<tr><td>>hg38_21_+_8433172_8446622_RNA45SN1_45S_50ntFlanks@12081.12108.28|CTCTCGCTTCTGGCGCCAAGCGCCCGGC</td><td>>hg38_21_+_8433172_8446622_RNA45SN1_45S_50ntFlanks</td><td>12081</td><td>12108</td><td>28</td><td>CTCTCGCTTCTGGCGCCAAGCGCCCGGC</td><td>437.4</td><td>-6.7</td><td>1.7</td><td>-3.9</td><td>9.2E-05</td><td>1.2E-04</td><td>BT20/MB468</td><td>NUC</td></tr>
<tr><td>>hg38_21_+_8433172_8446622_RNA45SN1_45S_50ntFlanks@5473.5491.19|CGGCCCTGGCGGAGCGCTG</td><td>>hg38_21_+_8433172_8446622_RNA45SN1_45S_50ntFlanks</td><td>5473</td><td>5491</td><td>19</td><td>CGGCCCTGGCGGAGCGCTG</td><td>232.0</td><td>-6.0</td><td>1.5</td><td>-4.0</td><td>7.5E-05</td><td>1.2E-04</td><td>MB231/MB468</td><td>MP</td></tr>
<tr><td>>hg38_21_+_8433172_8446622_RNA45SN1_45S_50ntFlanks@6751.6770.20|CGAACGCACTTGCGGCCCCG</td><td>>hg38_21_+_8433172_8446622_RNA45SN1_45S_50ntFlanks</td><td>6751</td><td>6770</td><td>20</td><td>CGAACGCACTTGCGGCCCCG</td><td>610.1</td><td>-6.6</td><td>1.6</td><td>-4.2</td><td>3.2E-05</td><td>1.2E-04</td><td>BT20/MB468</td><td>MP</td></tr>
<tr><td>>hg38_21_+_8433172_8446622_RNA45SN1_45S_50ntFlanks@5472.5516.45|ACGGCCCTGGCGGAGCGCTGAGAAGACGGTCGAACTTGACTATCT</td><td>>hg38_21_+_8433172_8446622_RNA45SN1_45S_50ntFlanks</td><td>5472</td><td>5516</td><td>45</td><td>ACGGCCCTGGCGGAGCGCTGAGAAGACGGTCGAACTTGACTATCT</td><td>1418.0</td><td>7.5</td><td>1.8</td><td>4.2</td><td>3.2E-05</td><td>1.2E-04</td><td>BT20/MB468</td><td>MP</td></tr>
<tr><td>>MI0000087|hsa-mir-29a&WithFlank&7|-|130876741|130876816@48.69.22&[MIMAT0000086&hsa-miR-29a-3p&offsets|0|0;m-26&7|-|130876749|130876769&offsets|0|+1]|TAGCACCATCTGAAATCGGTTA</td><td>>MI0000087|hsa-mir-29a&WithFlank&7|-|130876741|130876816</td><td>48</td><td>69</td><td>22</td><td>TAGCACCATCTGAAATCGGTTA</td><td>30990.7</td><td>6.3</td><td>1.5</td><td>4.2</td><td>3.2E-05</td><td>1.2E-04</td><td>BT20/MB468</td><td>MP</td></tr>
<tr><td>>hg38_21_+_8433172_8446622_RNA45SN1_45S_50ntFlanks@11769.11803.35|TAATTAGTGACGCGCATGAATGGATGAACGAGATT</td><td>>hg38_21_+_8433172_8446622_RNA45SN1_45S_50ntFlanks</td><td>11769</td><td>11803</td><td>35</td><td>TAATTAGTGACGCGCATGAATGGATGAACGAGATT</td><td>721.3</td><td>-2.6</td><td>0.6</td><td>-3.9</td><td>7.8E-05</td><td>1.2E-04</td><td>BT20/MB468</td><td>CYTO</td></tr>
<tr><td>>hg38_21_+_8433172_8446622_RNA45SN1_45S_50ntFlanks@9606.9630.25|AGACTAATCGAACCATCTAGTAGCT</td><td>>hg38_21_+_8433172_8446622_RNA45SN1_45S_50ntFlanks</td><td>9606</td><td>9630</td><td>25</td><td>AGACTAATCGAACCATCTAGTAGCT</td><td>74.9</td><td>9.6</td><td>2.2</td><td>4.4</td><td>1.1E-05</td><td>1.2E-04</td><td>BT20/MB231</td><td>MP</td></tr>
<tr><td>>hg38_21_+_8433172_8446622_RNA45SN1_45S_50ntFlanks@10411.10429.19|CAGTCGGTCCTGAGAGATG</td><td>>hg38_21_+_8433172_8446622_RNA45SN1_45S_50ntFlanks</td><td>10411</td><td>10429</td><td>19</td><td>CAGTCGGTCCTGAGAGATG</td><td>5994.7</td><td>-4.1</td><td>1.0</td><td>-4.2</td><td>3.2E-05</td><td>1.2E-04</td><td>BT20/MB468</td><td>MP</td></tr>
<tr><td>>MI0000087|hsa-mir-29a&WithFlank&7|-|130876741|130876816@47.66.20&[MIMAT0000086&hsa-miR-29a-3p&offsets|-1|-3;m-26&7|-|130876749|130876769&offsets|-1|-2]|CTAGCACCATCTGAAATCGG</td><td>>MI0000087|hsa-mir-29a&WithFlank&7|-|130876741|130876816</td><td>47</td><td>66</td><td>20</td><td>CTAGCACCATCTGAAATCGG</td><td>30.4</td><td>8.1</td><td>2.1</td><td>4.0</td><td>7.5E-05</td><td>1.2E-04</td><td>MB231/MB468</td><td>MP</td></tr>
<tr><td>>hg38_21_+_8433172_8446622_RNA45SN1_45S_50ntFlanks@8169.8197.29|CCAAGTCCTTCTGATCGAGGCCCAGCCCG</td><td>>hg38_21_+_8433172_8446622_RNA45SN1_45S_50ntFlanks</td><td>8169</td><td>8197</td><td>29</td><td>CCAAGTCCTTCTGATCGAGGCCCAGCCCG</td><td>407.4</td><td>-3.5</td><td>0.9</td><td>-4.0</td><td>6.5E-05</td><td>1.2E-04</td><td>BT20/MB468</td><td>MITO</td></tr>
<tr><td>>MI0000273|hsa-mir-183&WithFlank&7|-|129774899|129775020@33.55.23&[MIMAT0000261&hsa-miR-183-5p&offsets|0|+1;m-41&7|-|129774966|129774987&offsets|-1|0]|TATGGCACTGGTAGAATTCACTG</td><td>>MI0000273|hsa-mir-183&WithFlank&7|-|129774899|129775020</td><td>33</td><td>55</td><td>23</td><td>TATGGCACTGGTAGAATTCACTG</td><td>939.2</td><td>2.6</td><td>0.7</td><td>3.9</td><td>9.0E-05</td><td>1.2E-04</td><td>MB231/MB468</td><td>NUC</td></tr>
<tr><td>>hg38_21_+_8433172_8446622_RNA45SN1_45S_50ntFlanks@10869.10902.34|AGGGCTGGGTCGGTCGGGCTGGGGCGCGAAGCGG</td><td>>hg38_21_+_8433172_8446622_RNA45SN1_45S_50ntFlanks</td><td>10869</td><td>10902</td><td>34</td><td>AGGGCTGGGTCGGTCGGGCTGGGGCGCGAAGCGG</td><td>1863.3</td><td>-5.6</td><td>1.3</td><td>-4.2</td><td>3.2E-05</td><td>1.2E-04</td><td>BT20/MB468</td><td>MP</td></tr>
<tr><td>>hg38_21_+_8433172_8446622_RNA45SN1_45S_50ntFlanks@10807.10830.24|ACAATGTAGGTAAGGGAAGTCGGC</td><td>>hg38_21_+_8433172_8446622_RNA45SN1_45S_50ntFlanks</td><td>10807</td><td>10830</td><td>24</td><td>ACAATGTAGGTAAGGGAAGTCGGC</td><td>120.7</td><td>-2.2</td><td>0.6</td><td>-3.9</td><td>7.9E-05</td><td>1.2E-04</td><td>BT20/MB468</td><td>CYTO</td></tr>
</table>

| | | | | | | | | | | | | | |
|---|---|---|---|---|---|---|---|---|---|---|---|---|---|
| >hg38_21_+_8433172_8446622_RNA45SN1_45S_50ntFlanks@12997.13036.40\|CGCTGCGATCTATTGAAAGTCAGCCCTCGACACAAGGGTT | >hg38_21_+_8433172_8446622_RNA45SN1_45S_50ntFlanks | 12997 | 13036 | 40 | CGCTGCGATCTATTGAAAGTCAGCCCTCGACACAAGGGTT | 100.8 | 7.5 | 1.8 | 4.2 | 3.2E-05 | 1.2E-04 | BT20/MB468 | MP |
| >hg38_21_+_8433172_8446622_RNA45SN1_45S_50ntFlanks@10787.10817.31\|ACAGCCTCTGGCATGTTGGAACAATGTAGGT | >hg38_21_+_8433172_8446622_RNA45SN1_45S_50ntFlanks | 10787 | 10817 | 31 | ACAGCCTCTGGCATGTTGGAACAATGTAGGT | 67.4 | 9.4 | 2.1 | 4.4 | 1.1E-05 | 1.2E-04 | BT20/MB231 | MP |
| >hg38_21_+_8433172_8446622_RNA45SN1_45S_50ntFlanks@9572.9601.30\|CAAATCGGTCGTCCGACCTGGGTATAGGGG | >hg38_21_+_8433172_8446622_RNA45SN1_45S_50ntFlanks | 9572 | 9601 | 30 | CAAATCGGTCGTCCGACCTGGGTATAGGGG | 745.7 | -3.6 | 0.9 | -4.2 | 3.2E-05 | 1.2E-04 | BT20/MB468 | MP |
| >hg38_21_+_8433172_8446622_RNA45SN1_45S_50ntFlanks@6647.6667.21\|CGTACGACTCTTAGCGGTGGA | >hg38_21_+_8433172_8446622_RNA45SN1_45S_50ntFlanks | 6647 | 6667 | 21 | CGTACGACTCTTAGCGGTGGA | 578.5 | 2.3 | 0.6 | 3.9 | 9.3E-05 | 1.2E-04 | BT20/MB468 | NUC |
| >hg38_21_+_8433172_8446622_RNA45SN1_45S_50ntFlanks@11584.11603.20\|TGGTGCGGACCAGGGGAATC | >hg38_21_+_8433172_8446622_RNA45SN1_45S_50ntFlanks | 11584 | 11603 | 20 | TGGTGCGGACCAGGGGAATC | 153.0 | -4.9 | 1.2 | -4.0 | 7.6E-05 | 1.2E-04 | MB231/MB468 | MP |
| >hg38_21_+_8433172_8446622_RNA45SN1_45S_50ntFlanks@11573.11593.21\|CGACTTAGAACTGGTGCGGAC | >hg38_21_+_8433172_8446622_RNA45SN1_45S_50ntFlanks | 11573 | 11593 | 21 | CGACTTAGAACTGGTGCGGAC | 300.5 | -2.3 | 0.6 | -3.9 | 7.9E-05 | 1.2E-04 | BT20/MB468 | CYTO |
| >hg38_21_+_8433172_8446622_RNA45SN1_45S_50ntFlanks@11576.11602.27\|CTTAGAACTGGTGCGGACCAGGGGAAT | >hg38_21_+_8433172_8446622_RNA45SN1_45S_50ntFlanks | 11576 | 11602 | 27 | CTTAGAACTGGTGCGGACCAGGGGAAT | 429.6 | -6.4 | 1.6 | -4.2 | 3.2E-05 | 1.2E-04 | BT20/MB468 | MP |
| >hg38_21_+_8433172_8446622_RNA45SN1_45S_50ntFlanks@11924.11950.27\|GAGGTGTAGAATAAGTGGGAGGCCCCC | >hg38_21_+_8433172_8446622_RNA45SN1_45S_50ntFlanks | 11924 | 11950 | 27 | GAGGTGTAGAATAAGTGGGAGGCCCCC | 1510.8 | 6.0 | 1.4 | 4.2 | 3.2E-05 | 1.2E-04 | BT20/MB468 | MP |
| >hg38_21_+_8433172_8446622_RNA45SN1_45S_50ntFlanks@10679.10712.34\|GGTTCCGGCGGCGTCCGGTGAGCTCTCGCTGGCC | >hg38_21_+_8433172_8446622_RNA45SN1_45S_50ntFlanks | 10679 | 10712 | 34 | GGTTCCGGCGGCGTCCGGTGAGCTCTCGCTGGCC | 2160.1 | -5.7 | 1.4 | -4.2 | 3.2E-05 | 1.2E-04 | BT20/MB468 | MP |
| >MI0003562\|hsa-mir-556&WithFlank&1\|+\|162342540\|162342646@22.42.21&[MIMAT0003220&hsa-miR-556-5p&offsets\|0\|-1;m-1043&1\|+\|162342561\|162342581&offsets\|0\|0]\|GATGAGCTCATTGTAATATGA | >MI0003562\|hsa-mir-556&WithFlank&1\|+\|162342540\|162342646 | 22 | 42 | 21 | GATGAGCTCATTGTAATATGA | 29.4 | 8.1 | 2.0 | 4.0 | 7.6E-05 | 1.2E-04 | MB231/MB468 | MP |
| >hg38_21_+_8433172_8446622_RNA45SN1_45S_50ntFlanks@4443.4471.29\|CCCTTGCCTCTCGGCGCCCCCTCGATGCT | >hg38_21_+_8433172_8446622_RNA45SN1_45S_50ntFlanks | 4443 | 4471 | 29 | CCCTTGCCTCTCGGCGCCCCCTCGATGCT | 528.1 | -6.6 | 1.6 | -4.2 | 3.2E-05 | 1.2E-04 | BT20/MB468 | MP |
| >MI0000650\|hsa-mir-200c&WithFlank&12\|+\|6963693\|6963772@50.71.22(+1U)&[MIMAT0000617&hsa-miR-200c-3p&offsets\|0\|-1(+1U);m-54&12\|+\|6963742\|6963764&offsets\|0\|-1(+1U)]\|TAATACTGCCGGGTAATGATGGT | >MI0000650\|hsa-mir-200c&WithFlank&12\|+\|6963693\|6963772 | 50 | 71 | 22(+1U) | TAATACTGCCGGGTAATGATGGT | 1703.8 | 9.2 | 2.1 | 4.4 | 1.3E-05 | 1.2E-04 | BT20/MB231 | MITO |
| >hg38_21_+_8433172_8446622_RNA45SN1_45S_50ntFlanks@12417.12444.28\|TCGATGTCGGCTCTTCCTATCATTGTGA | >hg38_21_+_8433172_8446622_RNA45SN1_45S_50ntFlanks | 12417 | 12444 | 28 | TCGATGTCGGCTCTTCCTATCATTGTGA | 1559.2 | -3.0 | 0.8 | -3.9 | 9.0E-05 | 1.2E-04 | MB231/MB468 | MITO |
| >trna10_AspGTC_12_-_125424193_125424264@35.72.38,trna12_AspGTC_12_-_125411891_125411962@35.72.38,trna144_AspGTC_6_-_27551236_27551307@35.72.38,trna38_AspGTC_17_-_8125556_8125627@35.72.38,trna45_AspGTC_6_+_27447453_27447524@35.72.38,trna48_AspGTC_6_+_27471523_27471594@35.72.38,trna4_AspGTC_12_+_96429799_96429870@35.72.38,trna69_AspGTC_1_-_161440205_161440276@35.72.38,trna72_AspGTC_1_-_161432824_161432895@35.72.38,trna75_AspGTC_1_-_161425414_161425485@35.72.38,trna78_AspGTC_1_-_161418033_161418104@35.72.38,trna81_AspGTC_1_-_161410615_161410686@35.72.38\|TCACGCGGGAGACCGGGGTTCGATTCCCCGACGGGGAG | >trna10_AspGTC_12_-_125424193_125424264 | 35 | 72 | 38 | TCACGCGGGAGACCGGGGTTCGATTCCCCGACGGGGAG | 469.4 | -6.6 | 1.7 | -3.9 | 9.3E-05 | 1.2E-04 | BT20/MB468 | NUC |
| >hg38_21_+_8433172_8446622_RNA45SN1_45S_50ntFlanks@12458.12477.20\|AAGCGTTGGATTGTTCACCC | >hg38_21_+_8433172_8446622_RNA45SN1_45S_50ntFlanks | 12458 | 12477 | 20 | AAGCGTTGGATTGTTCACCC | 524.4 | -3.3 | 0.8 | -4.0 | 7.6E-05 | 1.2E-04 | MB231/MB468 | MP |
| >hg38_21_+_8433172_8446622_RNA45SN1_45S_50ntFlanks@10786.10822.37\|AACAGCCTCTGGCATGTTGGAACAATGTAGGTAAGGG | >hg38_21_+_8433172_8446622_RNA45SN1_45S_50ntFlanks | 10786 | 10822 | 37 | AACAGCCTCTGGCATGTTGGAACAATGTAGGTAAGGG | 475.5 | -4.6 | 1.2 | -3.9 | 7.9E-05 | 1.2E-04 | BT20/MB468 | CYTO |
| >hg38_21_+_8433172_8446622_RNA45SN1_45S_50ntFlanks@11323.11342.20\|CGTCTCGTCGCGCGCGCGTC | >hg38_21_+_8433172_8446622_RNA45SN1_45S_50ntFlanks | 11323 | 11342 | 20 | CGTCTCGTCGCGCGCGCGTC | 19830.4 | -5.8 | 1.5 | -3.9 | 9.3E-05 | 1.2E-04 | BT20/MB468 | NUC |
| >am_trnaMT_PheGAA_MT_+_577_647@40.59.20\|TGTTTAGACGGGCTCACATC | >am_trnaMT_PheGAA_MT_+_577_647 | 40 | 59 | 20 | TGTTTAGACGGGCTCACATC | 326.5 | 6.2 | 1.5 | 4.2 | 3.2E-05 | 1.2E-04 | BT20/MB468 | MP |
| >MI0003760\|hsa-mir-671&WithFlank&7\|+\|151238415\|151238544@35.58.24&[MIMAT0003880&hsa-miR-671-5p&offsets\|0\|+1;m-316&7\|+\|151238449\|151238472&offsets\|0\|0]\|AGGAAGCCCTGGAGGGGCTGGAGG | >MI0003760\|hsa-mir-671&WithFlank&7\|+\|151238415\|151238544 | 35 | 58 | 24 | AGGAAGCCCTGGAGGGGCTGGAGG | 242.1 | 6.1 | 1.6 | 3.9 | 9.0E-05 | 1.2E-04 | MB231/MB468 | MITO |
| >hg38_21_+_8433172_8446622_RNA45SN1_45S_50ntFlanks@4560.4587.28\|CTAGGAATAATGGAATAGGACCGCGGTT | >hg38_21_+_8433172_8446622_RNA45SN1_45S_50ntFlanks | 4560 | 4587 | 28 | CTAGGAATAATGGAATAGGACCGCGGTT | 571.4 | -4.5 | 1.1 | -3.9 | 7.9E-05 | 1.2E-04 | BT20/MB468 | CYTO |
| >hg38_21_+_8433172_8446622_RNA45SN1_45S_50ntFlanks@7991.8016.26\|ACGTGGCGACCCGCTGAATTTAAGCA | >hg38_21_+_8433172_8446622_RNA45SN1_45S_50ntFlanks | 7991 | 8016 | 26 | ACGTGGCGACCCGCTGAATTTAAGCA | 536.2 | -3.9 | 1.0 | -3.9 | 9.3E-05 | 1.2E-04 | BT20/MB468 | NUC |
| >hg38_21_+_8433172_8446622_RNA45SN1_45S_50ntFlanks@6651.6694.44\|CGACTCTTAGCGGTGGATCACTCGGCTCGTGCGTCGATGAAGAA | >hg38_21_+_8433172_8446622_RNA45SN1_45S_50ntFlanks | 6651 | 6694 | 44 | CGACTCTTAGCGGTGGATCACTCGGCTCGTGCGTCGATGAAGAA | 5608.6 | 3.4 | 0.8 | 4.0 | 6.6E-05 | 1.2E-04 | BT20/MB468 | MITO |
| >hg38_21_+_8433172_8446622_RNA45SN1_45S_50ntFlanks@10684.10705.22\|CGGCGGCGTCCGGTGAGCTCTC | >hg38_21_+_8433172_8446622_RNA45SN1_45S_50ntFlanks | 10684 | 10705 | 22 | CGGCGGCGTCCGGTGAGCTCTC | 413.2 | -5.8 | 1.5 | -3.9 | 9.0E-05 | 1.2E-04 | MB231/MB468 | MITO |
| >hg38_21_+_8433172_8446622_RNA45SN1_45S_50ntFlanks@5420.5438.19\|CCGATTGGATGGTTTAGTG | >hg38_21_+_8433172_8446622_RNA45SN1_45S_50ntFlanks | 5420 | 5438 | 19 | CCGATTGGATGGTTTAGTG | 395.9 | -6.4 | 1.5 | -4.2 | 3.2E-05 | 1.2E-04 | BT20/MB468 | MP |
| >hg38_21_+_8433172_8446622_RNA45SN1_45S_50ntFlanks@6732.6777.46\|ACATTGATCATCGACACTTCGAACGCACTTGCGGCCCCGGGTTCCT | >hg38_21_+_8433172_8446622_RNA45SN1_45S_50ntFlanks | 6732 | 6777 | 46 | ACATTGATCATCGACACTTCGAACGCACTTGCGGCCCCGGGTTCCT | 168.8 | -5.1 | 1.2 | -4.4 | 1.0E-05 | 1.2E-04 | BT20/MB231 | CYTO |
| >hg38_21_+_8433172_8446622_RNA45SN1_45S_50ntFlanks@6731.6752.22\|CACATTGATCATCGACACTTCG | >hg38_21_+_8433172_8446622_RNA45SN1_45S_50ntFlanks | 6731 | 6752 | 22 | CACATTGATCATCGACACTTCG | 767.3 | -4.6 | 1.2 | -3.9 | 9.1E-05 | 1.2E-04 | MB231/MB468 | MITO |
| >hg38_21_+_8433172_8446622_RNA45SN1_45S_50ntFlanks@8114.8142.29\|GCGCGGGACATGTGGCGTACGGAAGACCC | >hg38_21_+_8433172_8446622_RNA45SN1_45S_50ntFlanks | 8114 | 8142 | 29 | GCGCGGGACATGTGGCGTACGGAAGACCC | 304.5 | -4.2 | 1.1 | -4.0 | 6.6E-05 | 1.2E-04 | BT20/MB468 | MITO |
| >hg38_21_+_8433172_8446622_RNA45SN1_45S_50ntFlanks@12003.12043.41\|GCGGGCCGCCGGTGAAATACCACTACTCTGATCGTTTTTTC | >hg38_21_+_8433172_8446622_RNA45SN1_45S_50ntFlanks | 12003 | 12043 | 41 | GCGGGCCGCCGGTGAAATACCACTACTCTGATCGTTTTTTC | 866.2 | -5.0 | 1.3 | -4.0 | 6.6E-05 | 1.2E-04 | BT20/MB468 | MITO |
| >hg38_21_+_8433172_8446622_RNA45SN1_45S_50ntFlanks@5186.5218.33\|CAATAACAGGTCTGTGATGCCCTTAGATGTCCG | >hg38_21_+_8433172_8446622_RNA45SN1_45S_50ntFlanks | 5186 | 5218 | 33 | CAATAACAGGTCTGTGATGCCCTTAGATGTCCG | 45.1 | -9.4 | 2.4 | -4.0 | 7.6E-05 | 1.2E-04 | MB231/MB468 | MP |
| >hg38_21_+_8433172_8446622_RNA45SN1_45S_50ntFlanks@9835.9875.41\|TTTGGTAAGCAGAACTGGCGCTGCGGGATGAACCGAACGCC | >hg38_21_+_8433172_8446622_RNA45SN1_45S_50ntFlanks | 9835 | 9875 | 41 | TTTGGTAAGCAGAACTGGCGCTGCGGGATGAACCGAACGCC | 322.1 | 3.9 | 0.9 | 4.4 | 1.3E-05 | 1.2E-04 | BT20/MB231 | MITO |
| >hg38_21_+_8433172_8446622_RNA45SN1_45S_50ntFlanks@11566.11601.36\|TAGCAGCCGACTTAGAACTGGTGCGGACCAGGGGAA | >hg38_21_+_8433172_8446622_RNA45SN1_45S_50ntFlanks | 11566 | 11601 | 36 | TAGCAGCCGACTTAGAACTGGTGCGGACCAGGGGAA | 227.4 | -2.7 | 0.7 | -3.9 | 9.1E-05 | 1.2E-04 | MB231/MB468 | NUC |
| >hg38_MT_+_598_1651_RNR1_12S_50ntFlanks@904.946.43\|AAGTATACTTCAAAGGACATTTAACTAAAACCCCTACGCATTT | >hg38_MT_+_598_1651_RNR1_12S_50ntFlanks | 904 | 946 | 43 | AAGTATACTTCAAAGGACATTTAACTAAAACCCCTACGCATTT | 125.8 | 10.7 | 2.6 | 4.2 | 3.2E-05 | 1.2E-04 | BT20/MB468 | MP |
| >MI0000063\|hsa-let-7b&WithFlank&22\|+\|46113680\|46113774@12.30.19&[MIMAT0000063&hsa-let-7b-5p&offsets\|0\|-3;m-14&22\|+\|46113691\|46113712&offsets\|0\|-3]\|TGAGGTAGTAGGTTGTGTG | >MI0000063\|hsa-let-7b&WithFlank&22\|+\|46113680\|46113774 | 12 | 30 | 19 | TGAGGTAGTAGGTTGTGTG | 184.7 | 3.3 | 0.7 | 4.4 | 1.1E-05 | 1.2E-04 | BT20/MB231 | CYTO |
| >hg38_21_+_8433172_8446622_RNA45SN1_45S_50ntFlanks@11918.11967.50\|ACATGAGAGGTGTAGAATAAGTGGGAGGCCCCCGGCGCCCCCCCGGTGTC | >hg38_21_+_8433172_8446622_RNA45SN1_45S_50ntFlanks | 11918 | 11967 | 50 | ACATGAGAGGTGTAGAATAAGTGGGAGGCCCCCGGCGCCCCCCCGGTGTC | 49.3 | -9.5 | 2.4 | -4.0 | 7.7E-05 | 1.2E-04 | MB231/MB468 | MP |
| >hg38_21_+_8433172_8446622_RNA45SN1_45S_50ntFlanks@11857.11884.28\|GCGGAATCAGCGGGGAAAGAAGACCCTG | >hg38_21_+_8433172_8446622_RNA45SN1_45S_50ntFlanks | 11857 | 11884 | 28 | GCGGAATCAGCGGGGAAAGAAGACCCTG | 260.3 | -6.2 | 1.5 | -4.2 | 3.2E-05 | 1.2E-04 | BT20/MB468 | MP |
| >hg38_21_+_8433172_8446622_RNA45SN1_45S_50ntFlanks@6651.6681.31\|CGACTCTTAGCGGTGGATCACTCGGCTCGTG | >hg38_21_+_8433172_8446622_RNA45SN1_45S_50ntFlanks | 6651 | 6681 | 31 | CGACTCTTAGCGGTGGATCACTCGGCTCGTG | 303539.3 | -3.8 | 1.0 | -3.9 | 9.1E-05 | 1.2E-04 | MB231/MB468 | MITO |
| >hg38_21_+_8433172_8446622_RNA45SN1_45S_50ntFlanks@9623.9647.25\|TAGTAGCTGGTTCCCTCCGAAGTTT | >hg38_21_+_8433172_8446622_RNA45SN1_45S_50ntFlanks | 9623 | 9647 | 25 | TAGTAGCTGGTTCCCTCCGAAGTTT | 281.6 | -5.8 | 1.5 | -3.9 | 9.1E-05 | 1.2E-04 | MB231/MB468 | MITO |
| >hg38_21_+_8433172_8446622_RNA45SN1_45S_50ntFlanks@10355.10383.29\|AACTTTGAAGGCCGAAGTGGAGAAGGGTT | >hg38_21_+_8433172_8446622_RNA45SN1_45S_50ntFlanks | 10355 | 10383 | 29 | AACTTTGAAGGCCGAAGTGGAGAAGGGTT | 169.3 | -2.4 | 0.6 | -3.9 | 8.0E-05 | 1.2E-04 | BT20/MB468 | CYTO |
| >hg38_21_+_8433172_8446622_RNA45SN1_45S_50ntFlanks@11215.11233.19\|CTCTGGACGCGAGCCGGGC | >hg38_21_+_8433172_8446622_RNA45SN1_45S_50ntFlanks | 11215 | 11233 | 19 | CTCTGGACGCGAGCCGGGC | 122.3 | -4.9 | 1.3 | -3.9 | 8.0E-05 | 1.2E-04 | BT20/MB468 | CYTO |
| >hg38_21_+_8433172_8446622_RNA45SN1_45S_50ntFlanks@12324.12349.26\|TTGGGTTTTAAGCAGGAGGTGTCAGA | >hg38_21_+_8433172_8446622_RNA45SN1_45S_50ntFlanks | 12324 | 12349 | 26 | TTGGGTTTTAAGCAGGAGGTGTCAGA | 341.4 | -6.3 | 1.5 | -4.2 | 3.2E-05 | 1.2E-04 | BT20/MB468 | MP |
| >hg38_21_+_8433172_8446622_RNA45SN1_45S_50ntFlanks@3771.3799.29\|CACGGCCGGTACAGTGAAACTGCGAATGG | >hg38_21_+_8433172_8446622_RNA45SN1_45S_50ntFlanks | 3771 | 3799 | 29 | CACGGCCGGTACAGTGAAACTGCGAATGG | 120.8 | -1.7 | 0.4 | -3.9 | 8.0E-05 | 1.2E-04 | BT20/MB468 | CYTO |
| >hg38_21_+_8433172_8446622_RNA45SN1_45S_50ntFlanks@12155.12183.29\|GGAGTTTGACTGGGGCGGTACACCTGTCA | >hg38_21_+_8433172_8446622_RNA45SN1_45S_50ntFlanks | 12155 | 12183 | 29 | GGAGTTTGACTGGGGCGGTACACCTGTCA | 140.2 | -4.6 | 1.2 | -4.0 | 7.7E-05 | 1.2E-04 | MB231/MB468 | MP |
| >am_MI0000809\|hsa-mir-151a&WithFlank&8\|-\|140732558\|140732659@53.70.18(+3G)&[MIMAT0000757&hsa-miR-151a-3p&offsets\|0\|-3(+3G);m-30&8\|-\|140732587\|140732607&offsets\|0\|-3(+3G)]\|CTAGACTGAAGCTCCTTGGGG | >am_MI0000809\|hsa-mir-151a&WithFlank&8\|-\|140732558\|140732659 | 53 | 70 | 18(+3G) | CTAGACTGAAGCTCCTTGGGG | 467.3 | 5.6 | 1.4 | 4.2 | 3.2E-05 | 1.2E-04 | BT20/MB468 | MP |
| >hg38_21_+_8433172_8446622_RNA45SN1_45S_50ntFlanks@10432.10453.22\|CGAGCGCCGTTCCGAAGGGACG | >hg38_21_+_8433172_8446622_RNA45SN1_45S_50ntFlanks | 10432 | 10453 | 22 | CGAGCGCCGTTCCGAAGGGACG | 373.3 | -6.4 | 1.6 | -4.2 | 3.3E-05 | 1.2E-04 | BT20/MB468 | MP |

<table>
<tr><th></th><th></th><th></th><th></th><th></th><th></th><th></th><th></th><th></th><th></th><th></th><th></th><th></th><th></th></tr>
<tr><td>>trna10_AspGTC_12_-_125424193_125424264@37.69.33,trna12_AspGTC_12_-_125411891_125411962@37.69.33,trna144_AspGTC_6_-_27551236_27551307@37.69.33,trna38_AspGTC_17_-_8125556_8125627@37.69.33,trna45_AspGTC_6_+_27447453_27447524@37.69.33,trna48_AspGTC_6_+_27471523_27471594@37.69.33,trna4_AspGTC_12_+_96429799_96429870@37.69.33,trna69_AspGTC_1_-_161440205_161440276@37.69.33,trna72_AspGTC_1_-_161432824_161432895@37.69.33,trna75_AspGTC_1_-_161425414_161425485@37.69.33,trna78_AspGTC_1_-_161418033_161418104@37.69.33,trna81_AspGTC_1_-_161410615_161410686@37.69.33|ACGCGGGAGACCGGGGTTCGATTCCCCGACGGG</td><td>>trna10_AspGTC_12_-_125424193_125424264</td><td>37</td><td>69</td><td>33</td><td>ACGCGGGAGACCGGGGTTCGATTCCCCGACGGG</td><td>1250.4</td><td>-3.1</td><td>0.8</td><td>-4.0</td><td>7.7E-05</td><td>1.2E-04</td><td>MB231/MB468</td><td>MP</td></tr>
<tr><td>>hg38_21_+_8433172_8446622_RNA45SN1_45S_50ntFlanks@8106.8140.35|GCGGCGGGGCGCGGGACATGTGGCGTACGGAAGAC</td><td>>hg38_21_+_8433172_8446622_RNA45SN1_45S_50ntFlanks</td><td>8106</td><td>8140</td><td>35</td><td>GCGGCGGGGCGCGGGACATGTGGCGTACGGAAGAC</td><td>79.2</td><td>9.6</td><td>2.2</td><td>4.4</td><td>1.1E-05</td><td>1.2E-04</td><td>BT20/MB231</td><td>MP</td></tr>
<tr><td>>trnaMT_GluTTC_MT_-_14674_14742@16.57.42,trnalookalike8_GluTTC_5_-_93905172_93905240@16.57.42|ATACAACGATGGTTTTTCATATCATTGGTCGTGGTTGTAGTC</td><td>>trnaMT_GluTTC_MT_-_14674_14742</td><td>16</td><td>57</td><td>42</td><td>ATACAACGATGGTTTTTCATATCATTGGTCGTGGTTGTAGTC</td><td>127.1</td><td>10.7</td><td>2.6</td><td>4.2</td><td>3.3E-05</td><td>1.2E-04</td><td>BT20/MB468</td><td>MP</td></tr>
<tr><td>>hg38_21_+_8433172_8446622_RNA45SN1_45S_50ntFlanks@3782.3800.19|CAGTGAAACTGCGAATGGC</td><td>>hg38_21_+_8433172_8446622_RNA45SN1_45S_50ntFlanks</td><td>3782</td><td>3800</td><td>19</td><td>CAGTGAAACTGCGAATGGC</td><td>557.1</td><td>-4.8</td><td>1.2</td><td>-4.0</td><td>6.7E-05</td><td>1.2E-04</td><td>BT20/MB468</td><td>MITO</td></tr>
<tr><td>>hg38_21_+_8433172_8446622_RNA45SN1_45S_50ntFlanks@12934.12958.25|CTAAACCATTCGTAGACGACCTGCT</td><td>>hg38_21_+_8433172_8446622_RNA45SN1_45S_50ntFlanks</td><td>12934</td><td>12958</td><td>25</td><td>CTAAACCATTCGTAGACGACCTGCT</td><td>264.3</td><td>-1.8</td><td>0.5</td><td>-4.0</td><td>6.7E-05</td><td>1.2E-04</td><td>BT20/MB468</td><td>MITO</td></tr>
<tr><td>>hg38_21_+_8433172_8446622_RNA45SN1_45S_50ntFlanks@4224.4243.20|AAAATAACAATACAGGACTC</td><td>>hg38_21_+_8433172_8446622_RNA45SN1_45S_50ntFlanks</td><td>4224</td><td>4243</td><td>20</td><td>AAAATAACAATACAGGACTC</td><td>261.0</td><td>-6.2</td><td>1.5</td><td>-4.2</td><td>3.3E-05</td><td>1.2E-04</td><td>BT20/MB468</td><td>MP</td></tr>
<tr><td>>hg38_21_+_8433172_8446622_RNA45SN1_45S_50ntFlanks@6661.6678.18|CGGTGGATCACTCGGCTC</td><td>>hg38_21_+_8433172_8446622_RNA45SN1_45S_50ntFlanks</td><td>6661</td><td>6678</td><td>18</td><td>CGGTGGATCACTCGGCTC</td><td>294.5</td><td>2.4</td><td>0.6</td><td>3.9</td><td>9.5E-05</td><td>1.2E-04</td><td>BT20/MB468</td><td>NUC</td></tr>
<tr><td>>hg38_21_+_8433172_8446622_RNA45SN1_45S_50ntFlanks@4770.4793.24|TCGTAGTTCCGACCATAAACGATG</td><td>>hg38_21_+_8433172_8446622_RNA45SN1_45S_50ntFlanks</td><td>4770</td><td>4793</td><td>24</td><td>TCGTAGTTCCGACCATAAACGATG</td><td>290.7</td><td>-2.4</td><td>0.6</td><td>-3.9</td><td>9.2E-05</td><td>1.2E-04</td><td>MB231/MB468</td><td>NUC</td></tr>
<tr><td>>am_trnaMT_GluTTC_MT_-_14674_14742@41.58.18,trnalookalike8_GluTTC_5_-_93905172_93905240@41.58.18|TGGTCGTGGTTGTAGTCC</td><td>>am_trnaMT_GluTTC_MT_-_14674_14742</td><td>41</td><td>58</td><td>18</td><td>TGGTCGTGGTTGTAGTCC</td><td>307.7</td><td>5.4</td><td>1.3</td><td>4.2</td><td>3.3E-05</td><td>1.2E-04</td><td>BT20/MB468</td><td>MP</td></tr>
<tr><td>>hg38_21_+_8433172_8446622_RNA45SN1_45S_50ntFlanks@10816.10840.25|GTAAGGGAAGTCGGCAAGCCGGATC</td><td>>hg38_21_+_8433172_8446622_RNA45SN1_45S_50ntFlanks</td><td>10816</td><td>10840</td><td>25</td><td>GTAAGGGAAGTCGGCAAGCCGGATC</td><td>248.4</td><td>-2.1</td><td>0.5</td><td>-3.9</td><td>8.0E-05</td><td>1.2E-04</td><td>BT20/MB468</td><td>CYTO</td></tr>
<tr><td>>hg38_21_+_8433172_8446622_RNA45SN1_45S_50ntFlanks@4472.4493.22|CTTAGCTGAGTGTCCCGCGGGG</td><td>>hg38_21_+_8433172_8446622_RNA45SN1_45S_50ntFlanks</td><td>4472</td><td>4493</td><td>22</td><td>CTTAGCTGAGTGTCCCGCGGGG</td><td>379.5</td><td>-5.5</td><td>1.4</td><td>-3.9</td><td>8.0E-05</td><td>1.2E-04</td><td>BT20/MB468</td><td>CYTO</td></tr>
<tr><td>>hg38_21_+_8433172_8446622_RNA45SN1_45S_50ntFlanks@9783.9819.37|CGGCTCGCTGGCGTGGAGCCGGGCGTGGAATGCGAGT</td><td>>hg38_21_+_8433172_8446622_RNA45SN1_45S_50ntFlanks</td><td>9783</td><td>9819</td><td>37</td><td>CGGCTCGCTGGCGTGGAGCCGGGCGTGGAATGCGAGT</td><td>31.3</td><td>8.7</td><td>2.1</td><td>4.2</td><td>3.3E-05</td><td>1.2E-04</td><td>BT20/MB468</td><td>MP</td></tr>
<tr><td>>hg38_21_+_8433172_8446622_RNA45SN1_45S_50ntFlanks@10843.10861.19|TAACTTCGGGATAAGGATT</td><td>>hg38_21_+_8433172_8446622_RNA45SN1_45S_50ntFlanks</td><td>10843</td><td>10861</td><td>19</td><td>TAACTTCGGGATAAGGATT</td><td>83.7</td><td>9.7</td><td>2.2</td><td>4.4</td><td>1.1E-05</td><td>1.2E-04</td><td>BT20/MB231</td><td>MP</td></tr>
<tr><td>>hg38_21_+_8433172_8446622_RNA45SN1_45S_50ntFlanks@12635.12661.27|AAGCTACCATCTGTGGGATTATGACTG</td><td>>hg38_21_+_8433172_8446622_RNA45SN1_45S_50ntFlanks</td><td>12635</td><td>12661</td><td>27</td><td>AAGCTACCATCTGTGGGATTATGACTG</td><td>13751.2</td><td>-5.0</td><td>1.2</td><td>-4.2</td><td>3.3E-05</td><td>1.2E-04</td><td>BT20/MB468</td><td>MP</td></tr>
<tr><td>>hg38_21_+_8433172_8446622_RNA45SN1_45S_50ntFlanks@12997.13037.41|CGCTGCGATCTATTGAAAGTCAGCCCTCGACACAAGGGTTT</td><td>>hg38_21_+_8433172_8446622_RNA45SN1_45S_50ntFlanks</td><td>12997</td><td>13037</td><td>41</td><td>CGCTGCGATCTATTGAAAGTCAGCCCTCGACACAAGGGTTT</td><td>251.4</td><td>6.8</td><td>1.7</td><td>3.9</td><td>9.2E-05</td><td>1.2E-04</td><td>MB231/MB468</td><td>NUC</td></tr>
<tr><td>>trna10_GluCTC_3_-_103879524_103879595@16.39.24,trna116_GluCTC_1_-_145399233_145399304@16.39.24,trna59_GluCTC_1_+_249168447_249168518@16.39.24,trna71_GluCTC_1_-_161439189_161439260@16.39.24,trna74_GluCTC_1_-_161431809_161431880@16.39.24,trna77_GluCTC_1_-_161424398_161424469@16.39.24,trna77_GluCTC_6_+_28949976_28950047@16.39.24,trna80_GluCTC_1_-_161417018_161417089@16.39.24,trna87_GluCTC_6_-_126101393_126101464@16.39.24|TGGTTAGGATTCGGCGCTCTCACC</td><td>>trna10_GluCTC_3_-_103879524_103879595</td><td>16</td><td>39</td><td>24</td><td>TGGTTAGGATTCGGCGCTCTCACC</td><td>408.4</td><td>4.6</td><td>1.2</td><td>3.9</td><td>8.6E-05</td><td>1.2E-04</td><td>MB231/MB468</td><td>CYTO</td></tr>
<tr><td>>hg38_21_+_8433172_8446622_RNA45SN1_45S_50ntFlanks@9554.9575.22|CGTAGCGGTCCTGACGTGCAAA</td><td>>hg38_21_+_8433172_8446622_RNA45SN1_45S_50ntFlanks</td><td>9554</td><td>9575</td><td>22</td><td>CGTAGCGGTCCTGACGTGCAAA</td><td>643.2</td><td>-2.2</td><td>0.6</td><td>-3.9</td><td>9.2E-05</td><td>1.2E-04</td><td>MB231/MB468</td><td>NUC</td></tr>
<tr><td>>hg38_21_+_8433172_8446622_RNA45SN1_45S_50ntFlanks@9305.9336.32|CCAAGGAGTCTAACACGTGCGCGAGTCGGGGG</td><td>>hg38_21_+_8433172_8446622_RNA45SN1_45S_50ntFlanks</td><td>9305</td><td>9336</td><td>32</td><td>CCAAGGAGTCTAACACGTGCGCGAGTCGGGGG</td><td>419.3</td><td>-4.4</td><td>1.1</td><td>-4.0</td><td>6.7E-05</td><td>1.2E-04</td><td>BT20/MB468</td><td>MITO</td></tr>
<tr><td>>hg38_21_+_8433172_8446622_RNA45SN1_45S_50ntFlanks@12619.12654.36|AGGAGCCAATGGGGCGAAGCTACCATCTGTGGGATT</td><td>>hg38_21_+_8433172_8446622_RNA45SN1_45S_50ntFlanks</td><td>12619</td><td>12654</td><td>36</td><td>AGGAGCCAATGGGGCGAAGCTACCATCTGTGGGATT</td><td>2039.9</td><td>-2.7</td><td>0.7</td><td>-4.0</td><td>6.7E-05</td><td>1.2E-04</td><td>BT20/MB468</td><td>MITO</td></tr>
<tr><td>>trna10_AspGTC_12_-_125424193_125424264@32.72.41,trna12_AspGTC_12_-_125411891_125411962@32.72.41,trna144_AspGTC_6_-_27551236_27551307@32.72.41,trna38_AspGTC_17_-_8125556_8125627@32.72.41,trna45_AspGTC_6_+_27447453_27447524@32.72.41,trna48_AspGTC_6_+_27471523_27471594@32.72.41,trna4_AspGTC_12_+_96429799_96429870@32.72.41,trna69_AspGTC_1_-_161440205_161440276@32.72.41,trna72_AspGTC_1_-_161432824_161432895@32.72.41,trna75_AspGTC_1_-_161425414_161425485@32.72.41,trna78_AspGTC_1_-_161418033_161418104@32.72.41,trna81_AspGTC_1_-_161410615_161410686@32.72.41|CTGTCACGCGGGAGACCGGGGTTCGATTCCCCGACGGGGAG</td><td>>trna10_AspGTC_12_-_125424193_125424264</td><td>32</td><td>72</td><td>41</td><td>CTGTCACGCGGGAGACCGGGGTTCGATTCCCCGACGGGGAG</td><td>1674.7</td><td>-6.3</td><td>1.6</td><td>-3.9</td><td>8.1E-05</td><td>1.2E-04</td><td>BT20/MB468</td><td>CYTO</td></tr>
<tr><td>>hg38_21_+_8433172_8446622_RNA45SN1_45S_50ntFlanks@10835.10858.24|CGGATCCGTAACTTCGGGATAAGG</td><td>>hg38_21_+_8433172_8446622_RNA45SN1_45S_50ntFlanks</td><td>10835</td><td>10858</td><td>24</td><td>CGGATCCGTAACTTCGGGATAAGG</td><td>355.0</td><td>-1.8</td><td>0.5</td><td>-3.9</td><td>8.1E-05</td><td>1.2E-04</td><td>BT20/MB468</td><td>CYTO</td></tr>
<tr><td>>hg38_21_+_8433172_8446622_RNA45SN1_45S_50ntFlanks@6645.6673.29|CTCGTACGACTCTTAGCGGTGGATCACTC</td><td>>hg38_21_+_8433172_8446622_RNA45SN1_45S_50ntFlanks</td><td>6645</td><td>6673</td><td>29</td><td>CTCGTACGACTCTTAGCGGTGGATCACTC</td><td>193.9</td><td>5.0</td><td>1.3</td><td>3.9</td><td>8.7E-05</td><td>1.2E-04</td><td>MB231/MB468</td><td>CYTO</td></tr>
<tr><td>>hg38_21_+_8433172_8446622_RNA45SN1_45S_50ntFlanks@10466.10488.23|CGTTGCCCTCGGCCGATCGAAAG</td><td>>hg38_21_+_8433172_8446622_RNA45SN1_45S_50ntFlanks</td><td>10466</td><td>10488</td><td>23</td><td>CGTTGCCCTCGGCCGATCGAAAG</td><td>263.1</td><td>-4.6</td><td>1.2</td><td>-3.9</td><td>8.1E-05</td><td>1.2E-04</td><td>BT20/MB468</td><td>CYTO</td></tr>
<tr><td>>hg38_21_+_8433172_8446622_RNA45SN1_45S_50ntFlanks@11589.11612.24|CGGACCAGGGGAATCCGACTGTTT</td><td>>hg38_21_+_8433172_8446622_RNA45SN1_45S_50ntFlanks</td><td>11589</td><td>11612</td><td>24</td><td>CGGACCAGGGGAATCCGACTGTTT</td><td>352.0</td><td>-2.3</td><td>0.6</td><td>-3.9</td><td>9.3E-05</td><td>1.2E-04</td><td>MB231/MB468</td><td>NUC</td></tr>
<tr><td>>hg38_21_+_8433172_8446622_RNA45SN1_45S_50ntFlanks@4103.4121.19|CCACGGGTGACGGGGAATC</td><td>>hg38_21_+_8433172_8446622_RNA45SN1_45S_50ntFlanks</td><td>4103</td><td>4121</td><td>19</td><td>CCACGGGTGACGGGGAATC</td><td>1023.1</td><td>-5.7</td><td>1.4</td><td>-4.2</td><td>3.3E-05</td><td>1.2E-04</td><td>BT20/MB468</td><td>MP</td></tr>
<tr><td>>hg38_21_+_8433172_8446622_RNA45SN1_45S_50ntFlanks@11568.11585.18|GCAGCCGACTTAGAACTG</td><td>>hg38_21_+_8433172_8446622_RNA45SN1_45S_50ntFlanks</td><td>11568</td><td>11585</td><td>18</td><td>GCAGCCGACTTAGAACTG</td><td>1506.3</td><td>-2.9</td><td>0.7</td><td>-3.9</td><td>9.3E-05</td><td>1.2E-04</td><td>MB231/MB468</td><td>MITO</td></tr>
<tr><td>>hg38_21_+_8433172_8446622_RNA45SN1_45S_50ntFlanks@10500.10523.24|CAGATCCCCGAATCCGGAGTGGCG</td><td>>hg38_21_+_8433172_8446622_RNA45SN1_45S_50ntFlanks</td><td>10500</td><td>10523</td><td>24</td><td>CAGATCCCCGAATCCGGAGTGGCG</td><td>52.5</td><td>-9.6</td><td>2.4</td><td>-4.0</td><td>7.8E-05</td><td>1.2E-04</td><td>MB231/MB468</td><td>MP</td></tr>
<tr><td>>hg38_21_+_8433172_8446622_RNA45SN1_45S_50ntFlanks@12940.12960.21|CATTCGTAGACGACCTGCTTC</td><td>>hg38_21_+_8433172_8446622_RNA45SN1_45S_50ntFlanks</td><td>12940</td><td>12960</td><td>21</td><td>CATTCGTAGACGACCTGCTTC</td><td>254.2</td><td>2.2</td><td>0.6</td><td>3.9</td><td>8.7E-05</td><td>1.2E-04</td><td>MB231/MB468</td><td>CYTO</td></tr>
<tr><td>>hg38_21_+_8433172_8446622_RNA45SN1_45S_50ntFlanks@4760.4786.27|TCAGATACCGTCGTAGTTCCGACCATA</td><td>>hg38_21_+_8433172_8446622_RNA45SN1_45S_50ntFlanks</td><td>4760</td><td>4786</td><td>27</td><td>TCAGATACCGTCGTAGTTCCGACCATA</td><td>342.2</td><td>-3.7</td><td>0.9</td><td>-3.9</td><td>8.7E-05</td><td>1.2E-04</td><td>MB231/MB468</td><td>CYTO</td></tr>
<tr><td>>hg38_21_+_8433172_8446622_RNA45SN1_45S_50ntFlanks@6645.6685.41|CTCGTACGACTCTTAGCGGTGGATCACTCGGCTCGTGCGTC</td><td>>hg38_21_+_8433172_8446622_RNA45SN1_45S_50ntFlanks</td><td>6645</td><td>6685</td><td>41</td><td>CTCGTACGACTCTTAGCGGTGGATCACTCGGCTCGTGCGTC</td><td>190.5</td><td>4.0</td><td>1.0</td><td>3.9</td><td>9.6E-05</td><td>1.2E-04</td><td>BT20/MB468</td><td>NUC</td></tr>
<tr><td>>hg38_21_+_8433172_8446622_RNA45SN1_45S_50ntFlanks@12785.12805.21|CCCCGCGCGCGCGGGAGGGCG</td><td>>hg38_21_+_8433172_8446622_RNA45SN1_45S_50ntFlanks</td><td>12785</td><td>12805</td><td>21</td><td>CCCCGCGCGCGCGGGAGGGCG</td><td>2348.9</td><td>-6.4</td><td>1.6</td><td>-4.0</td><td>6.8E-05</td><td>1.2E-04</td><td>BT20/MB468</td><td>MITO</td></tr>
<tr><td>>hg38_21_+_8433172_8446622_RNA45SN1_45S_50ntFlanks@11217.11234.18|CTGGACGCGAGCCGGGCC</td><td>>hg38_21_+_8433172_8446622_RNA45SN1_45S_50ntFlanks</td><td>11217</td><td>11234</td><td>18</td><td>CTGGACGCGAGCCGGGCC</td><td>3191.4</td><td>-4.1</td><td>1.0</td><td>-3.9</td><td>8.1E-05</td><td>1.2E-04</td><td>BT20/MB468</td><td>CYTO</td></tr>
<tr><td>>trna111_HisGTG_1_-_147774845_147774916@-1T.29.30,trna118_HisGTG_1_-_145396881_145396952@-1T.29.30,trna16_HisGTG_1_+_146544773_146544844@-1T.29.30,trna1_HisGTG_15_+_45493349_45493420@-1T.29.30,trna21_HisGTG_1_+_147753471_147753542@-1T.29.30,trna33_HisGTG_6_+_27125906_27125977@-1T.29.30,trna7_HisGTG_9_-_14433938_14434009@-1T.29.30,trna8_HisGTG_15_-_45492611_45492682@-1T.29.30,trna9_HisGTG_15_-_45490804_45490875@-1T.29.30|TGCCGTGATCGTATAGTGGTTAGTACTCTG</td><td>>trna111_HisGTG_1_-_147774845_147774916</td><td>-1T</td><td>29</td><td>30</td><td>TGCCGTGATCGTATAGTGGTTAGTACTCTG</td><td>2237.2</td><td>-7.1</td><td>1.7</td><td>-4.2</td><td>3.3E-05</td><td>1.2E-04</td><td>BT20/MB468</td><td>MP</td></tr>
<tr><td>>hg38_21_+_8433172_8446622_RNA45SN1_45S_50ntFlanks@3838.3874.37|CTCTCCTACTTGGATAACTGTGGTAATTCTAGAGCTA</td><td>>hg38_21_+_8433172_8446622_RNA45SN1_45S_50ntFlanks</td><td>3838</td><td>3874</td><td>37</td><td>CTCTCCTACTTGGATAACTGTGGTAATTCTAGAGCTA</td><td>48.6</td><td>-9.5</td><td>2.4</td><td>-4.0</td><td>7.8E-05</td><td>1.2E-04</td><td>MB231/MB468</td><td>MP</td></tr>
<tr><td>>MI0000270|hsa-mir-181b-1&WithFlank&1|-|198858867|198858988@42.63.22(+1A);MI0000683|hsa-mir-181b-2&WithFlank&9|+|124693704|124693804@22.43.22(+1A)&[MIMAT0000257&hsa-miR-181b-5p&offsets|0|-1(+1A);m-73&1|-|198858925|198858947&offsets|0|-1(+1A)];[MIMAT0000257_1&hsa-miR-181b-5p&offsets|0|-1(+1A);m-72&9|+|124693725|124693747&offsets|0|-1(+1A)]|AACATTCATTGCTGTCGGTGGGA</td><td>>MI0000270|hsa-mir-181b-1&WithFlank&1|-|198858867|198858988</td><td>42</td><td>63</td><td>22(+1A)</td><td>AACATTCATTGCTGTCGGTGGGA</td><td>181.5</td><td>5.2</td><td>1.3</td><td>3.9</td><td>9.3E-05</td><td>1.2E-04</td><td>MB231/MB468</td><td>MITO</td></tr>
<tr><td>>am_trnaMT_TyrGTA_MT_-_5826_5891@1.32.32|GGTAAAATGGCTGAGTGAAGCATTGGACTGTA</td><td>>am_trnaMT_TyrGTA_MT_-_5826_5891</td><td>1</td><td>32</td><td>32</td><td>GGTAAAATGGCTGAGTGAAGCATTGGACTGTA</td><td>2501.7</td><td>4.8</td><td>1.2</td><td>3.9</td><td>9.3E-05</td><td>1.2E-04</td><td>MB231/MB468</td><td>MITO</td></tr>
<tr><td>>hg38_21_+_8433172_8446622_RNA45SN1_45S_50ntFlanks@6721.6746.26|AATTGCAGGACACATTGATCATCGAC</td><td>>hg38_21_+_8433172_8446622_RNA45SN1_45S_50ntFlanks</td><td>6721</td><td>6746</td><td>26</td><td>AATTGCAGGACACATTGATCATCGAC</td><td>1740.6</td><td>-5.1</td><td>1.3</td><td>-3.9</td><td>8.1E-05</td><td>1.2E-04</td><td>BT20/MB468</td><td>CYTO</td></tr>
<tr><td>>hg38_MT_+_1621_3279_RNR2_16S_50ntFlanks@51.83.33|GCTAAACCTAGCCCCAAACCCACTCCACCTTAC</td><td>>hg38_MT_+_1621_3279_RNR2_16S_50ntFlanks</td><td>51</td><td>83</td><td>33</td><td>GCTAAACCTAGCCCCAAACCCACTCCACCTTAC</td><td>236.8</td><td>3.7</td><td>1.0</td><td>3.9</td><td>9.3E-05</td><td>1.2E-04</td><td>MB231/MB468</td><td>MITO</td></tr>
<tr><td>>hg38_21_+_8433172_8446622_RNA45SN1_45S_50ntFlanks@12405.12425.21|CTTTTTGATCCTTCGATGTCG</td><td>>hg38_21_+_8433172_8446622_RNA45SN1_45S_50ntFlanks</td><td>12405</td><td>12425</td><td>21</td><td>CTTTTTGATCCTTCGATGTCG</td><td>578.8</td><td>-6.5</td><td>1.6</td><td>-4.2</td><td>3.3E-05</td><td>1.2E-04</td><td>BT20/MB468</td><td>MP</td></tr>
<tr><td>>hg38_21_+_8433172_8446622_RNA45SN1_45S_50ntFlanks@12420.12438.19|ATGTCGGCTCTTCCTATCA</td><td>>hg38_21_+_8433172_8446622_RNA45SN1_45S_50ntFlanks</td><td>12420</td><td>12438</td><td>19</td><td>ATGTCGGCTCTTCCTATCA</td><td>2136.2</td><td>-3.6</td><td>0.9</td><td>-3.9</td><td>9.6E-05</td><td>1.2E-04</td><td>BT20/MB468</td><td>NUC</td></tr>
</table>

| | | | | | | | | | | | | | |
|---|---|---|---|---|---|---|---|---|---|---|---|---|---|
| >hg38_21_+_8433172_8446622_RNA45SN1_45S_50ntFlanks@12405.12444.40\|CTTTTTGATCCTTCGATGTCGGCTCTTCCTATCATTGTGA | >hg38_21_+_8433172_8446622_RNA45SN1_45S_50ntFlanks | 12405 | 12444 | 40 | CTTTTTGATCCTTCGATGTCGGCTCTTCCTATCATTGTGA | 571.4 | -5.8 | 1.5 | -3.9 | 8.2E-05 | 1.2E-04 | BT20/MB468 | CYTO |
| >hg38_21_+_8433172_8446622_RNA45SN1_45S_50ntFlanks@9930.9957.28\|ATATAGACAGCAGGACGGTGGCCATGGA | >hg38_21_+_8433172_8446622_RNA45SN1_45S_50ntFlanks | 9930 | 9957 | 28 | ATATAGACAGCAGGACGGTGGCCATGGA | 196.8 | -6.4 | 1.6 | -3.9 | 7.8E-05 | 1.2E-04 | MB231/MB468 | MP |
| >MI0000285\|hsa-mir-205&WithFlank&1\|+\|209432127\|209432248@40.60.21(+1U)&[MIMAT0000266&hsa-miR-205-5p&offsets\|0\|-1(+1U);m-351&1\|+\|209432166\|209432187&offsets\|0\|-1(+1U)]\|TCCTTCATTCCACCGGAGTCTT | >MI0000285\|hsa-mir-205&WithFlank&1\|+\|209432127\|209432248 | 40 | 60 | 21(+1U) | TCCTTCATTCCACCGGAGTCTT | 309.7 | 3.6 | 0.9 | 3.9 | 9.6E-05 | 1.2E-04 | BT20/MB468 | NUC |
| >hg38_21_+_8433172_8446622_RNA45SN1_45S_50ntFlanks@10802.10840.39\|TTGGAACAATGTAGGTAAGGGAAGTCGGCAAGCCGGATC | >hg38_21_+_8433172_8446622_RNA45SN1_45S_50ntFlanks | 10802 | 10840 | 39 | TTGGAACAATGTAGGTAAGGGAAGTCGGCAAGCCGGATC | 80.5 | -4.7 | 1.2 | -3.9 | 7.8E-05 | 1.2E-04 | MB231/MB468 | MP |
| >trna10_LysCTT_16_+_3241501_3241573@5.36.32,trna13_LysCTT_14_-_58706613_58706685@5.36.32,trna2_LysCTT_15_+_79152904_79152976@5.36.32\|GGCTAGCTCAGTCGGTAGAGCATGGGACTCTT | >trna10_LysCTT_16_+_3241501_3241573 | 5 | 36 | 32 | GGCTAGCTCAGTCGGTAGAGCATGGGACTCTT | 128.9 | 10.8 | 2.6 | 4.2 | 3.3E-05 | 1.2E-04 | BT20/MB468 | MP |
| >hg38_21_+_8433172_8446622_RNA45SN1_45S_50ntFlanks@475.507.33\|CTTCGTGATCGATGTGGTGACGTCGTGCTCTCC | >hg38_21_+_8433172_8446622_RNA45SN1_45S_50ntFlanks | 475 | 507 | 33 | CTTCGTGATCGATGTGGTGACGTCGTGCTCTCC | 6925.2 | 2.7 | 0.7 | 3.9 | 9.3E-05 | 1.2E-04 | MB231/MB468 | NUC |
| >hg38_21_+_8433172_8446622_RNA45SN1_45S_50ntFlanks@7993.8014.22\|GTGGCGACCCGCTGAATTTAAG | >hg38_21_+_8433172_8446622_RNA45SN1_45S_50ntFlanks | 7993 | 8014 | 22 | GTGGCGACCCGCTGAATTTAAG | 126.7 | -6.0 | 1.5 | -3.9 | 7.8E-05 | 1.2E-04 | MB231/MB468 | MP |
| >trna116_GluCTC_1_-_145399233_145399304@39.74.36,trna134_GluTTC_1_-_16861774_16861845@39.74.36,trna71_GluCTC_1_-_161439189_161439260@39.74.36,trna74_GluCTC_1_-_161431809_161431880@39.74.36,trna77_GluCTC_1_-_161424398_161424469@39.74.36,trna77_GluCTC_6_+_28949976_28950047@39.74.36,trna80_GluCTC_1_-_161417018_161417089@39.74.36,trna84_GluTTC_1_-_161391883_161391954@39.74.36,trna87_GluCTC_6_-_126101393_126101464@39.74.36\|CGCCGCGGCCCGGGTTCGATTCCCGGTCAGGGAACC | >trna116_GluCTC_1_-_145399233_145399304 | 39 | 74 | 36 | CGCCGCGGCCCGGGTTCGATTCCCGGTCAGGGAACC | 34.8 | 8.3 | 2.1 | 3.9 | 7.9E-05 | 1.2E-04 | MB231/MB468 | MP |
| >hg38_21_+_8433172_8446622_RNA45SN1_45S_50ntFlanks@8113.8140.28\|GGCGCGGGACATGTGGCGTACGGAAGAC | >hg38_21_+_8433172_8446622_RNA45SN1_45S_50ntFlanks | 8113 | 8140 | 28 | GGCGCGGGACATGTGGCGTACGGAAGAC | 3850.3 | -4.4 | 1.1 | -4.2 | 3.3E-05 | 1.2E-04 | BT20/MB468 | MP |
| >hg38_21_+_8433172_8446622_RNA45SN1_45S_50ntFlanks@10686.10712.27\|GCGGCGTCCGGTGAGCTCTCGCTGGCC | >hg38_21_+_8433172_8446622_RNA45SN1_45S_50ntFlanks | 10686 | 10712 | 27 | GCGGCGTCCGGTGAGCTCTCGCTGGCC | 476.8 | -2.4 | 0.6 | -3.9 | 9.4E-05 | 1.2E-04 | MB231/MB468 | NUC |
| >hg38_21_+_8433172_8446622_RNA45SN1_45S_50ntFlanks@12419.12441.23\|GATGTCGGCTCTTCCTATCATTG | >hg38_21_+_8433172_8446622_RNA45SN1_45S_50ntFlanks | 12419 | 12441 | 23 | GATGTCGGCTCTTCCTATCATTG | 362.0 | 1.8 | 0.4 | 4.0 | 6.8E-05 | 1.2E-04 | BT20/MB468 | MITO |
| >hg38_21_+_8433172_8446622_RNA45SN1_45S_50ntFlanks@10475.10492.18\|CGGCCGATCGAAAGGGAG | >hg38_21_+_8433172_8446622_RNA45SN1_45S_50ntFlanks | 10475 | 10492 | 18 | CGGCCGATCGAAAGGGAG | 117.6 | -6.1 | 1.5 | -4.1 | 3.3E-05 | 1.2E-04 | BT20/MB468 | MP |
| >MI0000446\|hsa-mir-125b-1&WithFlank&11\|-\|122099751\|122099850@21.40.20;MI0000470\|hsa-mir-125b-2&WithFlank&21\|+\|16590231\|16590331@23.42.20&[MIMAT0000423&hsa-miR-125b-5p&offsets\|0\|-2;m-67&11\|-\|122099809\|122099830&offsets\|0\|-2];[MIMAT0000423_1&hsa-miR-125b-5p&offsets\|0\|-2;m-68&21\|+\|16590253\|16590274&offsets\|0\|-2]\|TCCCTGAGACCCTAACTTGT | >MI0000446\|hsa-mir-125b-1&WithFlank&11\|-\|122099751\|122099850 | 21 | 40 | 20 | TCCCTGAGACCCTAACTTGT | 1641.2 | -5.5 | 1.3 | -4.4 | 1.3E-05 | 1.2E-04 | BT20/MB231 | NUC |
| >hg38_21_+_8433172_8446622_RNA45SN1_45S_50ntFlanks@10186.10209.24\|TCCTCCCCTCCTCCCGCCCACGCC | >hg38_21_+_8433172_8446622_RNA45SN1_45S_50ntFlanks | 10186 | 10209 | 24 | TCCTCCCCTCCTCCCGCCCACGCC | 190.3 | -5.2 | 1.3 | -3.9 | 8.2E-05 | 1.2E-04 | BT20/MB468 | CYTO |
| >MI0000071\|hsa-mir-17&WithFlank&13\|+\|91350599\|91350694@20.41.22&[MIMAT0000070&hsa-miR-17-5p&offsets\|0\|-1;m-88&13\|+\|91350618\|91350640&offsets\|0\|-1]\|CAAAGTGCTTACAGTGCAGGTA | >MI0000071\|hsa-mir-17&WithFlank&13\|+\|91350599\|91350694 | 20 | 41 | 22 | CAAAGTGCTTACAGTGCAGGTA | 1475.0 | -4.1 | 0.9 | -4.4 | 1.1E-05 | 1.2E-04 | BT20/MB231 | MP |
| >trna116_GluCTC_1_-_145399233_145399304@37.74.38,trna134_GluTTC_1_-_16861774_16861845@37.74.38,trna71_GluCTC_1_-_161439189_161439260@37.74.38,trna74_GluCTC_1_-_161431809_161431880@37.74.38,trna77_GluCTC_1_-_161424398_161424469@37.74.38,trna77_GluCTC_6_+_28949976_28950047@37.74.38,trna80_GluCTC_1_-_161417018_161417089@37.74.38,trna84_GluTTC_1_-_161391883_161391954@37.74.38,trna87_GluCTC_6_-_126101393_126101464@37.74.38\|ACCGCCGCGGCCCGGGTTCGATTCCCGGTCAGGGAACC | >trna116_GluCTC_1_-_145399233_145399304 | 37 | 74 | 38 | ACCGCCGCGGCCCGGGTTCGATTCCCGGTCAGGGAACC | 662.6 | 4.8 | 1.2 | 3.9 | 8.2E-05 | 1.2E-04 | BT20/MB468 | CYTO |
| >hg38_21_+_8433172_8446622_RNA45SN1_45S_50ntFlanks@3872.3899.28\|CTAATACATGCCGACGGGCGCTGACCCC | >hg38_21_+_8433172_8446622_RNA45SN1_45S_50ntFlanks | 3872 | 3899 | 28 | CTAATACATGCCGACGGGCGCTGACCCC | 263.0 | -3.7 | 0.9 | -3.9 | 8.2E-05 | 1.2E-04 | BT20/MB468 | CYTO |
| >hg38_21_+_8433172_8446622_RNA45SN1_45S_50ntFlanks@4047.4072.26\|ATTCGAACGTCTGCCCTATCAACTTT | >hg38_21_+_8433172_8446622_RNA45SN1_45S_50ntFlanks | 4047 | 4072 | 26 | ATTCGAACGTCTGCCCTATCAACTTT | 186.3 | -4.2 | 1.1 | -3.9 | 9.4E-05 | 1.2E-04 | MB231/MB468 | NUC |
| >hg38_21_+_8433172_8446622_RNA45SN1_45S_50ntFlanks@12417.12438.22\|TCGATGTCGGCTCTTCCTATCA | >hg38_21_+_8433172_8446622_RNA45SN1_45S_50ntFlanks | 12417 | 12438 | 22 | TCGATGTCGGCTCTTCCTATCA | 778.2 | 1.9 | 0.5 | 3.9 | 8.2E-05 | 1.2E-04 | BT20/MB468 | CYTO |
| >hg38_21_+_8433172_8446622_RNA45SN1_45S_50ntFlanks@11661.11684.24\|TGATTTCTGCCCAGTGCTCTGAAT | >hg38_21_+_8433172_8446622_RNA45SN1_45S_50ntFlanks | 11661 | 11684 | 24 | TGATTTCTGCCCAGTGCTCTGAAT | 162.8 | -5.9 | 1.5 | -3.9 | 7.9E-05 | 1.2E-04 | MB231/MB468 | MP |
| >trna112_GlnCTG_1_-_147737382_147737453@30.50.21,trna131_GlnCTG_6_-_27759135_27759206@30.50.21,trna146_GlnCTG_6_-_27515531_27515602@30.50.21,trna15_GlnCTG_1_+_145963304_145963375@30.50.21,trna19_GlnCTG_1_+_147505038_147505109@30.50.21,trna1_GlnCTG_6_+_18836402_18836473@30.50.21,trna28_GlnCTG_1_+_149186125_149186196@30.50.21,trna3_GlnCTG_17_+_8023070_8023141@30.50.21,trna49_GlnCTG_6_+_27487308_27487379@30.50.21,trna7_GlnCTG_15_-_66161400_66161471@30.50.21,trna99_GlnCTG_6_-_28909378_28909449@30.50.21\|GACTCTGAATCCAGCGATCCG | >trna112_GlnCTG_1_-_147737382_147737453 | 30 | 50 | 21 | GACTCTGAATCCAGCGATCCG | 233.8 | -5.3 | 1.3 | -3.9 | 8.2E-05 | 1.2E-04 | BT20/MB468 | CYTO |
| >hg38_21_+_8433172_8446622_RNA45SN1_45S_50ntFlanks@12606.12625.20\|ATGTGCTTGGCTGAGGAGCC | >hg38_21_+_8433172_8446622_RNA45SN1_45S_50ntFlanks | 12606 | 12625 | 20 | ATGTGCTTGGCTGAGGAGCC | 561.4 | -2.4 | 0.6 | -3.9 | 8.2E-05 | 1.2E-04 | BT20/MB468 | CYTO |
| >hg38_21_+_8433172_8446622_RNA45SN1_45S_50ntFlanks@11710.11727.18\|CGCGGGTAAACGGCGGGA | >hg38_21_+_8433172_8446622_RNA45SN1_45S_50ntFlanks | 11710 | 11727 | 18 | CGCGGGTAAACGGCGGGA | 76.1 | -5.7 | 1.5 | -3.9 | 7.9E-05 | 1.2E-04 | MB231/MB468 | MP |
| >hg38_21_+_8433172_8446622_RNA45SN1_45S_50ntFlanks@9653.9677.25\|AGGATAGCTGGCGCTCTCGCAGACC | >hg38_21_+_8433172_8446622_RNA45SN1_45S_50ntFlanks | 9653 | 9677 | 25 | AGGATAGCTGGCGCTCTCGCAGACC | 178.3 | -5.9 | 1.5 | -3.9 | 7.9E-05 | 1.2E-04 | MB231/MB468 | MP |
| >MI0000803\|hsa-mir-330&WithFlank&19\|-\|45638988\|45639093@64.86.23(+1C)&[MIMAT0000751&hsa-miR-330-3p&offsets\|+1\|+1(+1C);m-347&19\|-\|45639009\|45639030&offsets\|0\|+1(+1C)]\|CAAAGCACACGGCCTGCAGAGAGC | >MI0000803\|hsa-mir-330&WithFlank&19\|-\|45638988\|45639093 | 64 | 86 | 23(+1C) | CAAAGCACACGGCCTGCAGAGAGC | 26.9 | 8.0 | 2.0 | 3.9 | 7.9E-05 | 1.2E-04 | MB231/MB468 | MP |
| >hg38_21_+_8433172_8446622_RNA45SN1_45S_50ntFlanks@8122.8140.19\|CATGTGGCGTACGGAAGAC | >hg38_21_+_8433172_8446622_RNA45SN1_45S_50ntFlanks | 8122 | 8140 | 19 | CATGTGGCGTACGGAAGAC | 67.3 | 9.4 | 2.1 | 4.4 | 1.2E-05 | 1.3E-04 | BT20/MB231 | MP |
| >hg38_21_+_8433172_8446622_RNA45SN1_45S_50ntFlanks@12160.12182.23\|TTGACTGGGGCGGTACACCTGTC | >hg38_21_+_8433172_8446622_RNA45SN1_45S_50ntFlanks | 12160 | 12182 | 23 | TTGACTGGGGCGGTACACCTGTC | 196.8 | -6.4 | 1.5 | -4.1 | 3.4E-05 | 1.3E-04 | BT20/MB468 | MP |
| >MI0000273\|hsa-mir-183&WithFlank&7\|-\|129774899\|129775020@33.55.23&[MIMAT0000261&hsa-miR-183-5p&offsets\|0\|+1;m-41&7\|-\|129774966\|129774987&offsets\|-1\|0]\|TATGGCACTGGTAGAATTCACTG | >MI0000273\|hsa-mir-183&WithFlank&7\|-\|129774899\|129775020 | 33 | 55 | 23 | TATGGCACTGGTAGAATTCACTG | 2658.2 | 3.9 | 0.9 | 4.1 | 3.4E-05 | 1.3E-04 | BT20/MB468 | MP |
| >hg38_21_+_8433172_8446622_RNA45SN1_45S_50ntFlanks@6650.6674.25\|ACGACTCTTAGCGGTGGATCACTCG | >hg38_21_+_8433172_8446622_RNA45SN1_45S_50ntFlanks | 6650 | 6674 | 25 | ACGACTCTTAGCGGTGGATCACTCG | 926.7 | 4.0 | 1.0 | 3.9 | 9.4E-05 | 1.3E-04 | MB231/MB468 | NUC |
| >hg38_21_+_8433172_8446622_RNA45SN1_45S_50ntFlanks@3843.3860.18\|CTACTTGGATAACTGTGG | >hg38_21_+_8433172_8446622_RNA45SN1_45S_50ntFlanks | 3843 | 3860 | 18 | CTACTTGGATAACTGTGG | 157.5 | -4.3 | 1.1 | -3.9 | 9.4E-05 | 1.3E-04 | MB231/MB468 | NUC |
| >hg38_21_+_8433172_8446622_RNA45SN1_45S_50ntFlanks@5186.5204.19\|CAATAACAGGTCTGTGATG | >hg38_21_+_8433172_8446622_RNA45SN1_45S_50ntFlanks | 5186 | 5204 | 19 | CAATAACAGGTCTGTGATG | 453.9 | -6.6 | 1.7 | -4.0 | 6.9E-05 | 1.3E-04 | BT20/MB468 | MITO |
| >hg38_21_+_8433172_8446622_RNA45SN1_45S_50ntFlanks@3910.3935.26\|GGGATGCGTGCATTTATCAGATCAAA | >hg38_21_+_8433172_8446622_RNA45SN1_45S_50ntFlanks | 3910 | 3935 | 26 | GGGATGCGTGCATTTATCAGATCAAA | 43.0 | -8.5 | 2.1 | -4.1 | 3.4E-05 | 1.3E-04 | BT20/MB468 | MP |
| >hg38_21_+_8433172_8446622_RNA45SN1_45S_50ntFlanks@4225.4242.18\|AAATAACAATACAGGACT | >hg38_21_+_8433172_8446622_RNA45SN1_45S_50ntFlanks | 4225 | 4242 | 18 | AAATAACAATACAGGACT | 453.9 | -5.3 | 1.4 | -3.9 | 9.8E-05 | 1.3E-04 | BT20/MB468 | NUC |
| >hg38_21_+_8433172_8446622_RNA45SN1_45S_50ntFlanks@10353.10390.38\|AGAACTTTGAAGGCCGAAGTGGAGAAGGGTTCCATGTG | >hg38_21_+_8433172_8446622_RNA45SN1_45S_50ntFlanks | 10353 | 10390 | 38 | AGAACTTTGAAGGCCGAAGTGGAGAAGGGTTCCATGTG | 985.0 | -5.8 | 1.5 | -3.9 | 9.8E-05 | 1.3E-04 | BT20/MB468 | NUC |
| >hg38_21_+_8433172_8446622_RNA45SN1_45S_50ntFlanks@11661.11680.20\|TGATTTCTGCCCAGTGCTCT | >hg38_21_+_8433172_8446622_RNA45SN1_45S_50ntFlanks | 11661 | 11680 | 20 | TGATTTCTGCCCAGTGCTCT | 629.9 | -3.9 | 1.0 | -3.9 | 8.3E-05 | 1.3E-04 | BT20/MB468 | CYTO |
| >hg38_21_+_8433172_8446622_RNA45SN1_45S_50ntFlanks@9307.9334.28\|AAGGAGTCTAACACGTGCGCGAGTCGGG | >hg38_21_+_8433172_8446622_RNA45SN1_45S_50ntFlanks | 9307 | 9334 | 28 | AAGGAGTCTAACACGTGCGCGAGTCGGG | 485.5 | -5.1 | 1.3 | -3.9 | 8.0E-05 | 1.3E-04 | MB231/MB468 | MP |
| >trnaMT_GluTTC_MT_-_14674_14742@33.72.40,trnalookalike8_GluTTC_5_-_93905172_93905240@33.72.40\|CATATCATTGGTCGTGGTTGTAGTCCGTGCGAGAATACCA | >trnaMT_GluTTC_MT_-_14674_14742 | 33 | 72 | 40 | CATATCATTGGTCGTGGTTGTAGTCCGTGCGAGAATACCA | 906.8 | 5.3 | 1.4 | 3.9 | 8.9E-05 | 1.3E-04 | MB231/MB468 | CYTO |
| >hg38_21_+_8433172_8446622_RNA45SN1_45S_50ntFlanks@10405.10424.20\|ATGGGTCAGTCGGTCCTGAG | >hg38_21_+_8433172_8446622_RNA45SN1_45S_50ntFlanks | 10405 | 10424 | 20 | ATGGGTCAGTCGGTCCTGAG | 669.8 | 3.2 | 0.8 | 3.9 | 9.5E-05 | 1.3E-04 | MB231/MB468 | MITO |
| >hg38_MT_+_1621_3279_RNR2_16S_50ntFlanks@623.663.41\|ATAACTGAACTCCTCACACCCAATTGGACCAATCTATCACC | >hg38_MT_+_1621_3279_RNR2_16S_50ntFlanks | 623 | 663 | 41 | ATAACTGAACTCCTCACACCCAATTGGACCAATCTATCACC | 126.9 | 10.7 | 2.6 | 4.1 | 3.4E-05 | 1.3E-04 | BT20/MB468 | MP |
| >hg38_21_+_8433172_8446622_RNA45SN1_45S_50ntFlanks@11768.11792.25\|CTAATTAGTGACGCGCATGAATGGA | >hg38_21_+_8433172_8446622_RNA45SN1_45S_50ntFlanks | 11768 | 11792 | 25 | CTAATTAGTGACGCGCATGAATGGA | 890.9 | -6.8 | 1.6 | -4.1 | 3.4E-05 | 1.3E-04 | BT20/MB468 | MP |
| >MI0000815\|hsa-mir-339&WithFlank&7\|-\|1022927\|1023032@56.76.21(+1U)&[MIMAT0004702&hsa-miR-339-3p&offsets\|0\|-2(+1U);m-235&7\|-\|1022957\|1022977&offsets\|0\|0(+1U)]\|TGAGCGCCTCGACGACAGAGCT | >MI0000815\|hsa-mir-339&WithFlank&7\|-\|1022927\|1023032 | 56 | 76 | 21(+1U) | TGAGCGCCTCGACGACAGAGCT | 216.1 | 7.9 | 1.9 | 4.1 | 3.4E-05 | 1.3E-04 | BT20/MB468 | MP |

<table>
<tr><td>>hg38_21_+_8433172_8446622_RNA45SN1_45S_50ntFlanks@10237.10273.37|ACGCTACGCCGCGACGAGTAGGAGGGCCGCTGCGGTG</td><td>>hg38_21_+_8433172_8446622_RNA45SN1_45S_50ntFlanks</td><td>10237</td><td>10273</td><td>37</td><td>ACGCTACGCCGCGACGAGTAGGAGGGCCGCTGCGGTG</td><td>958.7</td><td>4.6</td><td>1.1</td><td>4.1</td><td>3.4E-05</td><td>1.3E-04</td><td>BT20/MB468</td><td>MP</td></tr>
<tr><td>>hg38_21_+_8433172_8446622_RNA45SN1_45S_50ntFlanks@5487.5512.26|CGCTGAGAAGACGGTCGAACTTGACT</td><td>>hg38_21_+_8433172_8446622_RNA45SN1_45S_50ntFlanks</td><td>5487</td><td>5512</td><td>26</td><td>CGCTGAGAAGACGGTCGAACTTGACT</td><td>808.6</td><td>3.7</td><td>0.9</td><td>3.9</td><td>8.3E-05</td><td>1.3E-04</td><td>BT20/MB468</td><td>CYTO</td></tr>
<tr><td>>trnaMT_PheGAA_MT_+_577_647@44.74.31|TAGACGGGCTCACATCACCCCATAAACACCA</td><td>>trnaMT_PheGAA_MT_+_577_647</td><td>44</td><td>74</td><td>31</td><td>TAGACGGGCTCACATCACCCCATAAACACCA</td><td>592.2</td><td>4.3</td><td>1.1</td><td>3.9</td><td>9.5E-05</td><td>1.3E-04</td><td>MB231/MB468</td><td>NUC</td></tr>
<tr><td>>hg38_21_+_8433172_8446622_RNA45SN1_45S_50ntFlanks@4279.4314.36|AATCCTTTAACGAGGATCCATTGGAGGGCAAGTCTG</td><td>>hg38_21_+_8433172_8446622_RNA45SN1_45S_50ntFlanks</td><td>4279</td><td>4314</td><td>36</td><td>AATCCTTTAACGAGGATCCATTGGAGGGCAAGTCTG</td><td>156.5</td><td>-2.1</td><td>0.5</td><td>-3.9</td><td>8.4E-05</td><td>1.3E-04</td><td>BT20/MB468</td><td>CYTO</td></tr>
<tr><td>>hg38_21_+_8433172_8446622_RNA45SN1_45S_50ntFlanks@10386.10407.22|ATGTGAACAGCAGTTGAACATG</td><td>>hg38_21_+_8433172_8446622_RNA45SN1_45S_50ntFlanks</td><td>10386</td><td>10407</td><td>22</td><td>ATGTGAACAGCAGTTGAACATG</td><td>51.6</td><td>-9.6</td><td>2.4</td><td>-3.9</td><td>8.0E-05</td><td>1.3E-04</td><td>MB231/MB468</td><td>MP</td></tr>
<tr><td>>hg38_21_+_8433172_8446622_RNA45SN1_45S_50ntFlanks@6647.6689.43|CGTACGACTCTTAGCGGTGGATCACTCGGCTCGTGCGTCGATG</td><td>>hg38_21_+_8433172_8446622_RNA45SN1_45S_50ntFlanks</td><td>6647</td><td>6689</td><td>43</td><td>CGTACGACTCTTAGCGGTGGATCACTCGGCTCGTGCGTCGATG</td><td>5662.6</td><td>-3.5</td><td>0.9</td><td>-3.9</td><td>9.5E-05</td><td>1.3E-04</td><td>MB231/MB468</td><td>MITO</td></tr>
<tr><td>>hg38_21_+_8433172_8446622_RNA45SN1_45S_50ntFlanks@4204.4223.20|ACCCGGGGAGGTAGTGACGA</td><td>>hg38_21_+_8433172_8446622_RNA45SN1_45S_50ntFlanks</td><td>4204</td><td>4223</td><td>20</td><td>ACCCGGGGAGGTAGTGACGA</td><td>337.7</td><td>-6.3</td><td>1.6</td><td>-3.9</td><td>8.0E-05</td><td>1.3E-04</td><td>MB231/MB468</td><td>MP</td></tr>
<tr><td>>hg38_21_+_8433172_8446622_RNA45SN1_45S_50ntFlanks@6778.6806.29|CCCGGGGCTACGCCTGTCTGAGCGTCGCT</td><td>>hg38_21_+_8433172_8446622_RNA45SN1_45S_50ntFlanks</td><td>6778</td><td>6806</td><td>29</td><td>CCCGGGGCTACGCCTGTCTGAGCGTCGCT</td><td>357.8</td><td>1.7</td><td>0.4</td><td>3.9</td><td>9.8E-05</td><td>1.3E-04</td><td>BT20/MB468</td><td>NUC</td></tr>
<tr><td>>hg38_21_+_8433172_8446622_RNA45SN1_45S_50ntFlanks@10362.10391.30|AAGGCCGAAGTGGAGAAGGGTTCCATGTGA</td><td>>hg38_21_+_8433172_8446622_RNA45SN1_45S_50ntFlanks</td><td>10362</td><td>10391</td><td>30</td><td>AAGGCCGAAGTGGAGAAGGGTTCCATGTGA</td><td>149.4</td><td>-6.1</td><td>1.5</td><td>-4.1</td><td>3.4E-05</td><td>1.3E-04</td><td>BT20/MB468</td><td>MP</td></tr>
<tr><td>>trna10_LysCTT_16_+_3241501_3241573@1.40.40,trna13_LysCTT_14_-_58706613_58706685@1.40.40,trna2_LysCTT_15_+_79152904_79152976@1.40.40|GCCCGGCTAGCTCAGTCGGTAGAGCATGGGACTCTTAATC</td><td>>trna10_LysCTT_16_+_3241501_3241573</td><td>1</td><td>40</td><td>40</td><td>GCCCGGCTAGCTCAGTCGGTAGAGCATGGGACTCTTAATC</td><td>719.3</td><td>5.5</td><td>1.4</td><td>3.9</td><td>9.0E-05</td><td>1.3E-04</td><td>MB231/MB468</td><td>CYTO</td></tr>
<tr><td>>hg38_21_+_8433172_8446622_RNA45SN1_45S_50ntFlanks@10794.10812.19|CTGGCATGTTGGAACAATG</td><td>>hg38_21_+_8433172_8446622_RNA45SN1_45S_50ntFlanks</td><td>10794</td><td>10812</td><td>19</td><td>CTGGCATGTTGGAACAATG</td><td>180.3</td><td>-3.6</td><td>0.9</td><td>-3.9</td><td>9.0E-05</td><td>1.3E-04</td><td>MB231/MB468</td><td>CYTO</td></tr>
<tr><td>>hg38_21_+_8433172_8446622_RNA45SN1_45S_50ntFlanks@11666.11687.22|TCTGCCCAGTGCTCTGAATGTC</td><td>>hg38_21_+_8433172_8446622_RNA45SN1_45S_50ntFlanks</td><td>11666</td><td>11687</td><td>22</td><td>TCTGCCCAGTGCTCTGAATGTC</td><td>871.7</td><td>2.9</td><td>0.7</td><td>3.9</td><td>8.4E-05</td><td>1.3E-04</td><td>BT20/MB468</td><td>CYTO</td></tr>
<tr><td>>hg38_21_+_8433172_8446622_RNA45SN1_45S_50ntFlanks@8194.8215.22|CCCGTGGACGGTGTGAGGCCGG</td><td>>hg38_21_+_8433172_8446622_RNA45SN1_45S_50ntFlanks</td><td>8194</td><td>8215</td><td>22</td><td>CCCGTGGACGGTGTGAGGCCGG</td><td>2358.2</td><td>-5.4</td><td>1.4</td><td>-3.9</td><td>8.4E-05</td><td>1.3E-04</td><td>BT20/MB468</td><td>CYTO</td></tr>
<tr><td>>hg38_21_+_8433172_8446622_RNA45SN1_45S_50ntFlanks@7978.8017.40|GACCTCAGATCAGACGTGGCGACCCGCTGAATTTAAGCAT</td><td>>hg38_21_+_8433172_8446622_RNA45SN1_45S_50ntFlanks</td><td>7978</td><td>8017</td><td>40</td><td>GACCTCAGATCAGACGTGGCGACCCGCTGAATTTAAGCAT</td><td>350.4</td><td>5.1</td><td>1.3</td><td>4.0</td><td>7.0E-05</td><td>1.3E-04</td><td>BT20/MB468</td><td>MITO</td></tr>
<tr><td>>hg38_21_+_8433172_8446622_RNA45SN1_45S_50ntFlanks@3926.3959.34|TCAGATCAAAACCAACCCGGTCAGCCCCTCTCCG</td><td>>hg38_21_+_8433172_8446622_RNA45SN1_45S_50ntFlanks</td><td>3926</td><td>3959</td><td>34</td><td>TCAGATCAAAACCAACCCGGTCAGCCCCTCTCCG</td><td>176.9</td><td>-5.2</td><td>1.3</td><td>-4.0</td><td>7.0E-05</td><td>1.3E-04</td><td>BT20/MB468</td><td>MITO</td></tr>
<tr><td>>MI0001445|hsa-mir-423&WithFlank&17|+|30117073|30117178@58.80.23&[MIMAT0001340&hsa-miR-423-3p&offsets|-1|-1;m-100&17|+|30117131|30117153&offsets|-1|-1]|AAGCTCGGTCTGAGGCCCCTCAG</td><td>>MI0001445|hsa-mir-423&WithFlank&17|+|30117073|30117178</td><td>58</td><td>80</td><td>23</td><td>AAGCTCGGTCTGAGGCCCCTCAG</td><td>221.6</td><td>4.9</td><td>1.3</td><td>3.9</td><td>9.6E-05</td><td>1.3E-04</td><td>MB231/MB468</td><td>NUC</td></tr>
<tr><td>>MI0000298|hsa-mir-221&WithFlank&X|-|45746151|45746272@71.93.23(+1C)&[MIMAT0000278&hsa-miR-221-3p&offsets|0|0(+1C);m-91&X|-|45746181|45746202&offsets|0|+1(+1C)]|AGCTACATTGTCTGCTGGGTTTCC</td><td>>MI0000298|hsa-mir-221&WithFlank&X|-|45746151|45746272</td><td>71</td><td>93</td><td>23(+1C)</td><td>AGCTACATTGTCTGCTGGGTTTCC</td><td>2062.5</td><td>-4.2</td><td>0.9</td><td>-4.4</td><td>1.2E-05</td><td>1.3E-04</td><td>BT20/MB231</td><td>MP</td></tr>
<tr><td>>trna17_GluTTC_2_-_204229131_204229201@1.30.30|GCAATGGTGGTTCAGTGGTAGAATTCTCGC</td><td>>trna17_GluTTC_2_-_204229131_204229201</td><td>1</td><td>30</td><td>30</td><td>GCAATGGTGGTTCAGTGGTAGAATTCTCGC</td><td>67.2</td><td>-9.5</td><td>2.2</td><td>-4.4</td><td>1.2E-05</td><td>1.3E-04</td><td>BT20/MB231</td><td>MP</td></tr>
<tr><td>>MI0000087|hsa-mir-29a&WithFlank&7|-|130876741|130876816@48.67.20&[MIMAT0000086&hsa-miR-29a-3p&offsets|0|-2;m-26&7|-|130876749|130876769&offsets|0|-1]|TAGCACCATCTGAAATCGGT</td><td>>MI0000087|hsa-mir-29a&WithFlank&7|-|130876741|130876816</td><td>48</td><td>67</td><td>20</td><td>TAGCACCATCTGAAATCGGT</td><td>1244.2</td><td>6.4</td><td>1.6</td><td>3.9</td><td>8.0E-05</td><td>1.3E-04</td><td>MB231/MB468</td><td>MP</td></tr>
<tr><td>>MI0000065|hsa-let-7d&WithFlank&9|+|94178828|94178926@14.35.22&[MIMAT0000065&hsa-let-7d-5p&offsets|0|0;m-90&9|+|94178841|94178862&offsets|0|0]|AGAGGTAGTAGGTTGCATAGTT</td><td>>MI0000065|hsa-let-7d&WithFlank&9|+|94178828|94178926</td><td>14</td><td>35</td><td>22</td><td>AGAGGTAGTAGGTTGCATAGTT</td><td>5286.8</td><td>3.8</td><td>1.0</td><td>3.9</td><td>8.1E-05</td><td>1.3E-04</td><td>MB231/MB468</td><td>MP</td></tr>
<tr><td>>hg38_21_+_8433172_8446622_RNA45SN1_45S_50ntFlanks@11873.11910.38|AAGAAGACCCTGTTGAGCTTGACTCTAGTCTGGCACGG</td><td>>hg38_21_+_8433172_8446622_RNA45SN1_45S_50ntFlanks</td><td>11873</td><td>11910</td><td>38</td><td>AAGAAGACCCTGTTGAGCTTGACTCTAGTCTGGCACGG</td><td>128.6</td><td>-5.0</td><td>1.3</td><td>-3.9</td><td>8.4E-05</td><td>1.3E-04</td><td>BT20/MB468</td><td>CYTO</td></tr>
<tr><td>>am_MI0000786|hsa-mir-378a&WithFlank&5|+|149732819|149732896@49.70.22(+1A)&[MIMAT0000732&hsa-miR-378a-3p&offsets|0|0(+1A);m-44&5|+|149732867|149732888&offsets|0|0(+1A)]|ACTGGACTTGGAGTCAGAAGGCA</td><td>>am_MI0000786|hsa-mir-378a&WithFlank&5|+|149732819|149732896</td><td>49</td><td>70</td><td>22(+1A)</td><td>ACTGGACTTGGAGTCAGAAGGCA</td><td>961.7</td><td>3.1</td><td>0.8</td><td>3.9</td><td>9.0E-05</td><td>1.3E-04</td><td>MB231/MB468</td><td>CYTO</td></tr>
<tr><td>>hg38_21_+_8433172_8446622_RNA45SN1_45S_50ntFlanks@7989.8009.21|AGACGTGGCGACCCGCTGAAT</td><td>>hg38_21_+_8433172_8446622_RNA45SN1_45S_50ntFlanks</td><td>7989</td><td>8009</td><td>21</td><td>AGACGTGGCGACCCGCTGAAT</td><td>6230.9</td><td>-2.4</td><td>0.6</td><td>-3.9</td><td>8.1E-05</td><td>1.3E-04</td><td>MB231/MB468</td><td>MP</td></tr>
<tr><td>>hg38_21_+_8433172_8446622_RNA45SN1_45S_50ntFlanks@11314.11342.29|CCCACCCCACGTCTCGTCGCGCGCGCGTC</td><td>>hg38_21_+_8433172_8446622_RNA45SN1_45S_50ntFlanks</td><td>11314</td><td>11342</td><td>29</td><td>CCCACCCCACGTCTCGTCGCGCGCGCGTC</td><td>921.3</td><td>-7.1</td><td>1.8</td><td>-3.9</td><td>9.9E-05</td><td>1.3E-04</td><td>BT20/MB468</td><td>NUC</td></tr>
<tr><td>>hg38_21_+_8433172_8446622_RNA45SN1_45S_50ntFlanks@3939.3959.21|AACCCGGTCAGCCCCTCTCCG</td><td>>hg38_21_+_8433172_8446622_RNA45SN1_45S_50ntFlanks</td><td>3939</td><td>3959</td><td>21</td><td>AACCCGGTCAGCCCCTCTCCG</td><td>1062.4</td><td>-3.4</td><td>0.9</td><td>-3.9</td><td>8.4E-05</td><td>1.3E-04</td><td>BT20/MB468</td><td>CYTO</td></tr>
<tr><td>>hg38_21_+_8433172_8446622_RNA45SN1_45S_50ntFlanks@10127.10147.21|GCGGCGGCGGCGGCGGCGGCG</td><td>>hg38_21_+_8433172_8446622_RNA45SN1_45S_50ntFlanks</td><td>10127</td><td>10147</td><td>21</td><td>GCGGCGGCGGCGGCGGCGGCG</td><td>194.8</td><td>-6.3</td><td>1.5</td><td>-4.1</td><td>3.4E-05</td><td>1.3E-04</td><td>BT20/MB468</td><td>MP</td></tr>
<tr><td>>hg38_21_+_8433172_8446622_RNA45SN1_45S_50ntFlanks@8844.8864.21|GCGGCGGCGGCGGCGGCGGCG</td><td>>hg38_21_+_8433172_8446622_RNA45SN1_45S_50ntFlanks</td><td>8844</td><td>8864</td><td>21</td><td>GCGGCGGCGGCGGCGGCGGCG</td><td>194.8</td><td>-6.3</td><td>1.5</td><td>-4.1</td><td>3.4E-05</td><td>1.3E-04</td><td>BT20/MB468</td><td>MP</td></tr>
<tr><td>>hg38_21_+_8433172_8446622_RNA45SN1_45S_50ntFlanks@6652.6680.29|GACTCTTAGCGGTGGATCACTCGGCTCGT</td><td>>hg38_21_+_8433172_8446622_RNA45SN1_45S_50ntFlanks</td><td>6652</td><td>6680</td><td>29</td><td>GACTCTTAGCGGTGGATCACTCGGCTCGT</td><td>764.9</td><td>-2.3</td><td>0.6</td><td>-3.9</td><td>9.9E-05</td><td>1.3E-04</td><td>BT20/MB468</td><td>NUC</td></tr>
<tr><td>>MI0000767|hsa-mir-365a&WithFlank&16|+|14309279|14309377@62.84.23;MI0000769|hsa-mir-365b&WithFlank&17|+|31575405|31575527@74.96.23&[MIMAT0000710&hsa-miR-365a-3p&offsets|0|+1;m-176&16|+|14309340|14309361&offsets|0|+1];[MIMAT0022834&hsa-miR-365b-3p&offsets|0|+1;m-175&17|+|31575478|31575499&offsets|0|+1]|TAATGCCCCTAAAAATCCTTATT</td><td>>MI0000767|hsa-mir-365a&WithFlank&16|+|14309279|14309377</td><td>62</td><td>84</td><td>23</td><td>TAATGCCCCTAAAAATCCTTATT</td><td>757.3</td><td>3.3</td><td>0.8</td><td>4.4</td><td>1.1E-05</td><td>1.3E-04</td><td>BT20/MB231</td><td>CYTO</td></tr>
<tr><td>>hg38_21_+_8433172_8446622_RNA45SN1_45S_50ntFlanks@11769.11806.38|TAATTAGTGACGCGCATGAATGGATGAACGAGATTCCC</td><td>>hg38_21_+_8433172_8446622_RNA45SN1_45S_50ntFlanks</td><td>11769</td><td>11806</td><td>38</td><td>TAATTAGTGACGCGCATGAATGGATGAACGAGATTCCC</td><td>379.1</td><td>-4.7</td><td>1.2</td><td>-3.9</td><td>9.7E-05</td><td>1.3E-04</td><td>MB231/MB468</td><td>NUC</td></tr>
<tr><td>>hg38_21_+_8433172_8446622_RNA45SN1_45S_50ntFlanks@7983.8010.28|CAGATCAGACGTGGCGACCCGCTGAATT</td><td>>hg38_21_+_8433172_8446622_RNA45SN1_45S_50ntFlanks</td><td>7983</td><td>8010</td><td>28</td><td>CAGATCAGACGTGGCGACCCGCTGAATT</td><td>319.1</td><td>-3.3</td><td>0.8</td><td>-3.9</td><td>9.6E-05</td><td>1.3E-04</td><td>MB231/MB468</td><td>MITO</td></tr>
<tr><td>>hg38_21_+_8433172_8446622_RNA45SN1_45S_50ntFlanks@12896.12921.26|CGCCCGTCACGCACCGCACGTTCGTG</td><td>>hg38_21_+_8433172_8446622_RNA45SN1_45S_50ntFlanks</td><td>12896</td><td>12921</td><td>26</td><td>CGCCCGTCACGCACCGCACGTTCGTG</td><td>55.9</td><td>-9.7</td><td>2.5</td><td>-3.9</td><td>8.1E-05</td><td>1.3E-04</td><td>MB231/MB468</td><td>MP</td></tr>
<tr><td>>MI0000487|hsa-mir-193a&WithFlank&17|+|31559990|31560089@27.47.21&[MIMAT0004614&hsa-miR-193a-5p&offsets|0|-1;m-123&17|+|31560016|31560037&offsets|0|-1]|TGGGTCTTTGCGGGCGAGATG</td><td>>MI0000487|hsa-mir-193a&WithFlank&17|+|31559990|31560089</td><td>27</td><td>47</td><td>21</td><td>TGGGTCTTTGCGGGCGAGATG</td><td>1338.3</td><td>6.5</td><td>1.6</td><td>4.1</td><td>3.4E-05</td><td>1.3E-04</td><td>BT20/MB468</td><td>MP</td></tr>
<tr><td>>MI0000063|hsa-let-7b&WithFlank&22|+|46113680|46113774@12.31.20(+1A)&[MIMAT0000063&hsa-let-7b-5p&offsets|0|-2(+1A);m-14&22|+|46113691|46113712&offsets|0|-2(+1A)]|TGAGGTAGTAGGTTGTGTGGA</td><td>>MI0000063|hsa-let-7b&WithFlank&22|+|46113680|46113774</td><td>12</td><td>31</td><td>20(+1A)</td><td>TGAGGTAGTAGGTTGTGTGGA</td><td>502.5</td><td>3.8</td><td>0.9</td><td>4.4</td><td>1.1E-05</td><td>1.3E-04</td><td>BT20/MB231</td><td>CYTO</td></tr>
<tr><td>>hg38_1_-_228634819_228635039_RNA5S12_5S_50ntFlanks@61.89.29|ATACCACCCTGAACGCGCCCGATCTCGTC</td><td>>hg38_1_-_228634819_228635039_RNA5S12_5S_50ntFlanks</td><td>61</td><td>89</td><td>29</td><td>ATACCACCCTGAACGCGCCCGATCTCGTC</td><td>1097.6</td><td>-4.2</td><td>0.9</td><td>-4.4</td><td>1.1E-05</td><td>1.3E-04</td><td>BT20/MB231</td><td>CYTO</td></tr>
<tr><td>>hg38_21_+_8433172_8446622_RNA45SN1_45S_50ntFlanks@8176.8199.24|CTTCTGATCGAGGCCCAGCCCGTG</td><td>>hg38_21_+_8433172_8446622_RNA45SN1_45S_50ntFlanks</td><td>8176</td><td>8199</td><td>24</td><td>CTTCTGATCGAGGCCCAGCCCGTG</td><td>197.6</td><td>-2.0</td><td>0.5</td><td>-3.9</td><td>9.7E-05</td><td>1.3E-04</td><td>MB231/MB468</td><td>NUC</td></tr>
<tr><td>>hg38_21_+_8433172_8446622_RNA45SN1_45S_50ntFlanks@11783.11820.38|CATGAATGGATGAACGAGATTCCCACTGTCCCTACCTA</td><td>>hg38_21_+_8433172_8446622_RNA45SN1_45S_50ntFlanks</td><td>11783</td><td>11820</td><td>38</td><td>CATGAATGGATGAACGAGATTCCCACTGTCCCTACCTA</td><td>231.6</td><td>-6.5</td><td>1.6</td><td>-4.1</td><td>3.5E-05</td><td>1.3E-04</td><td>BT20/MB468</td><td>MP</td></tr>
<tr><td>>trna129_MetCAT_6_-_27870271_27870342@16.33.18,trna142_MetCAT_6_-_27560600_27560671@16.33.18,trna150_MetCAT_6_-_27300764_27300835@16.33.18,trna151_ThrCGT_6_-_27271568_27271639@16.33.18,trna169_MetCAT_6_-_26330529_26330600@16.33.18,trna171_MetCAT_6_-_26313352_26313423@16.33.18,trna20_MetCAT_17_-_80452597_80452668@16.33.18,trna2_MetCAT_6_+_26286754_26286825@16.33.18,trna32_MetCAT_1_+_153643726_153643797@16.33.18,trna61_MetCAT_6_+_27745664_27745735@16.33.18|CGGAAGCGTGCTGGGCCC</td><td>>trna129_MetCAT_6_-_27870271_27870342</td><td>16</td><td>33</td><td>18</td><td>CGGAAGCGTGCTGGGCCC</td><td>128.7</td><td>3.5</td><td>0.9</td><td>3.9</td><td>9.7E-05</td><td>1.3E-04</td><td>MB231/MB468</td><td>MITO</td></tr>
<tr><td>>hg38_21_+_8433172_8446622_RNA45SN1_45S_50ntFlanks@6652.6696.45|GACTCTTAGCGGTGGATCACTCGGCTCGTGCGTCGATGAAGAACG</td><td>>hg38_21_+_8433172_8446622_RNA45SN1_45S_50ntFlanks</td><td>6652</td><td>6696</td><td>45</td><td>GACTCTTAGCGGTGGATCACTCGGCTCGTGCGTCGATGAAGAACG</td><td>4997.8</td><td>2.9</td><td>0.7</td><td>4.0</td><td>7.1E-05</td><td>1.3E-04</td><td>BT20/MB468</td><td>MITO</td></tr>
<tr><td>>hg38_21_+_8433172_8446622_RNA45SN1_45S_50ntFlanks@7982.8008.27|TCAGATCAGACGTGGCGACCCGCTGAA</td><td>>hg38_21_+_8433172_8446622_RNA45SN1_45S_50ntFlanks</td><td>7982</td><td>8008</td><td>27</td><td>TCAGATCAGACGTGGCGACCCGCTGAA</td><td>1326.7</td><td>-2.5</td><td>0.6</td><td>-4.0</td><td>7.1E-05</td><td>1.3E-04</td><td>BT20/MB468</td><td>MITO</td></tr>
<tr><td>>hg38_1_-_228634819_228635039_RNA5S12_5S_50ntFlanks@51.87.37|GTCTACGGCCATACCACCCTGAACGCGCCCGATCTCG</td><td>>hg38_1_-_228634819_228635039_RNA5S12_5S_50ntFlanks</td><td>51</td><td>87</td><td>37</td><td>GTCTACGGCCATACCACCCTGAACGCGCCCGATCTCG</td><td>1417.8</td><td>4.4</td><td>1.1</td><td>3.9</td><td>9.7E-05</td><td>1.3E-04</td><td>MB231/MB468</td><td>NUC</td></tr>
<tr><td>>MI0001445|hsa-mir-423&WithFlank&17|+|30117073|30117178@58.78.21&[MIMAT0001340&hsa-miR-423-3p&offsets|-1|-3;m-100&17|+|30117131|30117153&offsets|-1|-3]|AAGCTCGGTCTGAGGCCCCTC</td><td>>MI0001445|hsa-mir-423&WithFlank&17|+|30117073|30117178</td><td>58</td><td>78</td><td>21</td><td>AAGCTCGGTCTGAGGCCCCTC</td><td>788.4</td><td>5.8</td><td>1.4</td><td>4.1</td><td>3.5E-05</td><td>1.3E-04</td><td>BT20/MB468</td><td>MP</td></tr>
<tr><td>>hg38_21_+_8433172_8446622_RNA45SN1_45S_50ntFlanks@12954.12982.29|CTGCTTCTGGGTCGGGGTTTCGTACGTAG</td><td>>hg38_21_+_8433172_8446622_RNA45SN1_45S_50ntFlanks</td><td>12954</td><td>12982</td><td>29</td><td>CTGCTTCTGGGTCGGGGTTTCGTACGTAG</td><td>156.8</td><td>-5.0</td><td>1.3</td><td>-3.9</td><td>8.5E-05</td><td>1.3E-04</td><td>BT20/MB468</td><td>CYTO</td></tr>
<tr><td>>hg38_21_+_8433172_8446622_RNA45SN1_45S_50ntFlanks@10362.10407.46|AAGGCCGAAGTGGAGAAGGGTTCCATGTGAACAGCAGTTGAACATG</td><td>>hg38_21_+_8433172_8446622_RNA45SN1_45S_50ntFlanks</td><td>10362</td><td>10407</td><td>46</td><td>AAGGCCGAAGTGGAGAAGGGTTCCATGTGAACAGCAGTTGAACATG</td><td>187.6</td><td>-5.7</td><td>1.5</td><td>-3.9</td><td>8.5E-05</td><td>1.3E-04</td><td>BT20/MB468</td><td>CYTO</td></tr>
<tr><td>>hg38_21_+_8433172_8446622_RNA45SN1_45S_50ntFlanks@12841.12862.22|CGGAGTGCCCTTCGTCCTGGGA</td><td>>hg38_21_+_8433172_8446622_RNA45SN1_45S_50ntFlanks</td><td>12841</td><td>12862</td><td>22</td><td>CGGAGTGCCCTTCGTCCTGGGA</td><td>791.2</td><td>-6.3</td><td>1.5</td><td>-4.1</td><td>3.5E-05</td><td>1.3E-04</td><td>BT20/MB468</td><td>MP</td></tr>
</table>

<table>
<tr><th></th><th></th><th></th><th></th><th></th><th></th><th></th><th></th><th></th><th></th><th></th><th></th><th></th><th></th></tr>
<tr><td>>MI0000298|hsa-mir-221&WithFlank&X|-|45746151|45746272@71.93.23(+1U)&[MIMAT0000278&hsa-miR-221-3p&offsets|0|0(+1U);m-91&X|-|45746181|45746202&offsets|0|+1(+1U)]|AGCTACATTGTCTGCTGGGTTTCT</td><td>>MI0000298|hsa-mir-221&WithFlank&X|-|45746151|45746272</td><td>71</td><td>93</td><td>23(+1U)</td><td>AGCTACATTGTCTGCTGGGTTTCT</td><td>11577.6</td><td>-3.9</td><td>0.9</td><td>-4.4</td><td>1.2E-05</td><td>1.3E-04</td><td>BT20/MB231</td><td>MP</td></tr>
<tr><td>>hg38_21_+_8433172_8446622_RNA45SN1_45S_50ntFlanks@5468.5499.32|GCCCACGGCCCTGGCGGAGCGCTGAGAAGACG</td><td>>hg38_21_+_8433172_8446622_RNA45SN1_45S_50ntFlanks</td><td>5468</td><td>5499</td><td>32</td><td>GCCCACGGCCCTGGCGGAGCGCTGAGAAGACG</td><td>1442.2</td><td>-7.4</td><td>1.9</td><td>-3.9</td><td>1.0E-04</td><td>1.3E-04</td><td>BT20/MB468</td><td>NUC</td></tr>
<tr><td>>hg38_21_+_8433172_8446622_RNA45SN1_45S_50ntFlanks@6776.6803.28|CTCCCGGGGCTACGCCTGTCTGAGCGTC</td><td>>hg38_21_+_8433172_8446622_RNA45SN1_45S_50ntFlanks</td><td>6776</td><td>6803</td><td>28</td><td>CTCCCGGGGCTACGCCTGTCTGAGCGTC</td><td>534.1</td><td>-2.1</td><td>0.5</td><td>-3.9</td><td>8.6E-05</td><td>1.3E-04</td><td>BT20/MB468</td><td>CYTO</td></tr>
<tr><td>>am_trnaMT_ValTAC_MT_+_1602_1670@51.68.18|CTTAACTTGACCGCTCTG</td><td>>am_trnaMT_ValTAC_MT_+_1602_1670</td><td>51</td><td>68</td><td>18</td><td>CTTAACTTGACCGCTCTG</td><td>19438.6</td><td>-5.1</td><td>1.3</td><td>-3.9</td><td>8.6E-05</td><td>1.3E-04</td><td>BT20/MB468</td><td>CYTO</td></tr>
<tr><td>>trnaMT_SerGCT_MT_+_12207_12265@1.18.18|GAGAAAGCTCACAAGAAC</td><td>>trnaMT_SerGCT_MT_+_12207_12265</td><td>1</td><td>18</td><td>18</td><td>GAGAAAGCTCACAAGAAC</td><td>7048.7</td><td>5.8</td><td>1.5</td><td>3.9</td><td>1.0E-04</td><td>1.3E-04</td><td>BT20/MB468</td><td>NUC</td></tr>
<tr><td>>hg38_21_+_8433172_8446622_RNA45SN1_45S_50ntFlanks@12075.12093.19|GAGGGGCTCTCGCTTCTGG</td><td>>hg38_21_+_8433172_8446622_RNA45SN1_45S_50ntFlanks</td><td>12075</td><td>12093</td><td>19</td><td>GAGGGGCTCTCGCTTCTGG</td><td>58.6</td><td>-9.7</td><td>2.5</td><td>-3.9</td><td>8.2E-05</td><td>1.3E-04</td><td>MB231/MB468</td><td>MP</td></tr>
<tr><td>>hg38_21_+_8433172_8446622_RNA45SN1_45S_50ntFlanks@12539.12579.41|ATGATGTGTTGTTGCCATGGTAATCCTGCTCAGTACGAGAG</td><td>>hg38_21_+_8433172_8446622_RNA45SN1_45S_50ntFlanks</td><td>12539</td><td>12579</td><td>41</td><td>ATGATGTGTTGTTGCCATGGTAATCCTGCTCAGTACGAGAG</td><td>40.0</td><td>-8.4</td><td>2.0</td><td>-4.1</td><td>3.5E-05</td><td>1.3E-04</td><td>BT20/MB468</td><td>MP</td></tr>
<tr><td>>hg38_21_+_8433172_8446622_RNA45SN1_45S_50ntFlanks@12957.12991.35|CTTCTGGGTCGGGGTTTCGTACGTAGCAGAGCAGC</td><td>>hg38_21_+_8433172_8446622_RNA45SN1_45S_50ntFlanks</td><td>12957</td><td>12991</td><td>35</td><td>CTTCTGGGTCGGGGTTTCGTACGTAGCAGAGCAGC</td><td>102.4</td><td>-4.8</td><td>1.2</td><td>-3.9</td><td>8.6E-05</td><td>1.3E-04</td><td>BT20/MB468</td><td>CYTO</td></tr>
<tr><td>>hg38_21_+_8433172_8446622_RNA45SN1_45S_50ntFlanks@11583.11616.34|CTGGTGCGGACCAGGGGAATCCGACTGTTTAATT</td><td>>hg38_21_+_8433172_8446622_RNA45SN1_45S_50ntFlanks</td><td>11583</td><td>11616</td><td>34</td><td>CTGGTGCGGACCAGGGGAATCCGACTGTTTAATT</td><td>242.0</td><td>2.5</td><td>0.6</td><td>3.9</td><td>9.2E-05</td><td>1.3E-04</td><td>MB231/MB468</td><td>CYTO</td></tr>
<tr><td>>trnaMT_GluTTC_MT_-_14674_14742@34.72.39,trnalookalike8_GluTTC_5_-_93905172_93905240@34.72.39|ATATCATTGGTCGTGGTTGTAGTCCGTGCGAGAATACCA</td><td>>trnaMT_GluTTC_MT_-_14674_14742</td><td>34</td><td>72</td><td>39</td><td>ATATCATTGGTCGTGGTTGTAGTCCGTGCGAGAATACCA</td><td>3872.3</td><td>3.0</td><td>0.8</td><td>3.9</td><td>9.7E-05</td><td>1.3E-04</td><td>MB231/MB468</td><td>MITO</td></tr>
<tr><td>>hg38_21_+_8433172_8446622_RNA45SN1_45S_50ntFlanks@4002.4039.38|GATAACCTCGGGCCGATCGCACGCCCCCCGTGGCGGCG</td><td>>hg38_21_+_8433172_8446622_RNA45SN1_45S_50ntFlanks</td><td>4002</td><td>4039</td><td>38</td><td>GATAACCTCGGGCCGATCGCACGCCCCCCGTGGCGGCG</td><td>169.3</td><td>-5.4</td><td>1.4</td><td>-3.9</td><td>8.6E-05</td><td>1.3E-04</td><td>BT20/MB468</td><td>CYTO</td></tr>
<tr><td>>hg38_21_+_8433172_8446622_RNA45SN1_45S_50ntFlanks@6648.6679.32|GTACGACTCTTAGCGGTGGATCACTCGGCTCG</td><td>>hg38_21_+_8433172_8446622_RNA45SN1_45S_50ntFlanks</td><td>6648</td><td>6679</td><td>32</td><td>GTACGACTCTTAGCGGTGGATCACTCGGCTCG</td><td>544.7</td><td>-4.0</td><td>1.0</td><td>-3.9</td><td>8.2E-05</td><td>1.3E-04</td><td>MB231/MB468</td><td>MP</td></tr>
<tr><td>>hg38_21_+_8433172_8446622_RNA45SN1_45S_50ntFlanks@8176.8202.27|CTTCTGATCGAGGCCCAGCCCGTGGAC</td><td>>hg38_21_+_8433172_8446622_RNA45SN1_45S_50ntFlanks</td><td>8176</td><td>8202</td><td>27</td><td>CTTCTGATCGAGGCCCAGCCCGTGGAC</td><td>1174.5</td><td>2.4</td><td>0.6</td><td>4.0</td><td>7.2E-05</td><td>1.3E-04</td><td>BT20/MB468</td><td>MITO</td></tr>
<tr><td>>hg38_21_+_8433172_8446622_RNA45SN1_45S_50ntFlanks@12971.12992.22|TTTCGTACGTAGCAGAGCAGCT</td><td>>hg38_21_+_8433172_8446622_RNA45SN1_45S_50ntFlanks</td><td>12971</td><td>12992</td><td>22</td><td>TTTCGTACGTAGCAGAGCAGCT</td><td>197.4</td><td>-1.7</td><td>0.4</td><td>-3.9</td><td>8.6E-05</td><td>1.3E-04</td><td>BT20/MB468</td><td>CYTO</td></tr>
<tr><td>>hg38_21_+_8433172_8446622_RNA45SN1_45S_50ntFlanks@6730.6747.18|ACACATTGATCATCGACA</td><td>>hg38_21_+_8433172_8446622_RNA45SN1_45S_50ntFlanks</td><td>6730</td><td>6747</td><td>18</td><td>ACACATTGATCATCGACA</td><td>202.6</td><td>-2.6</td><td>0.7</td><td>-3.9</td><td>9.2E-05</td><td>1.3E-04</td><td>MB231/MB468</td><td>CYTO</td></tr>
<tr><td>>hg38_21_+_8433172_8446622_RNA45SN1_45S_50ntFlanks@3840.3862.23|CTCCTACTTGGATAACTGTGGTA</td><td>>hg38_21_+_8433172_8446622_RNA45SN1_45S_50ntFlanks</td><td>3840</td><td>3862</td><td>23</td><td>CTCCTACTTGGATAACTGTGGTA</td><td>458.7</td><td>4.1</td><td>0.9</td><td>4.4</td><td>1.3E-05</td><td>1.3E-04</td><td>BT20/MB231</td><td>NUC</td></tr>
<tr><td>>hg38_21_+_8433172_8446622_RNA45SN1_45S_50ntFlanks@11581.11602.22|AACTGGTGCGGACCAGGGGAAT</td><td>>hg38_21_+_8433172_8446622_RNA45SN1_45S_50ntFlanks</td><td>11581</td><td>11602</td><td>22</td><td>AACTGGTGCGGACCAGGGGAAT</td><td>433.6</td><td>-4.5</td><td>1.2</td><td>-3.9</td><td>9.8E-05</td><td>1.3E-04</td><td>MB231/MB468</td><td>NUC</td></tr>
<tr><td>>hg38_21_+_8433172_8446622_RNA45SN1_45S_50ntFlanks@11718.11737.20|AACGGCGGGAGTAACTATGA</td><td>>hg38_21_+_8433172_8446622_RNA45SN1_45S_50ntFlanks</td><td>11718</td><td>11737</td><td>20</td><td>AACGGCGGGAGTAACTATGA</td><td>135.8</td><td>-6.1</td><td>1.5</td><td>-4.1</td><td>3.5E-05</td><td>1.3E-04</td><td>BT20/MB468</td><td>MP</td></tr>
<tr><td>>hg38_21_+_8433172_8446622_RNA45SN1_45S_50ntFlanks@11710.11736.27|CGCGGGTAAACGGCGGGAGTAACTATG</td><td>>hg38_21_+_8433172_8446622_RNA45SN1_45S_50ntFlanks</td><td>11710</td><td>11736</td><td>27</td><td>CGCGGGTAAACGGCGGGAGTAACTATG</td><td>638.3</td><td>-5.5</td><td>1.4</td><td>-3.9</td><td>1.0E-04</td><td>1.3E-04</td><td>BT20/MB468</td><td>NUC</td></tr>
<tr><td>>hg38_21_+_8433172_8446622_RNA45SN1_45S_50ntFlanks@11345.11362.18|CTGGGGGCGGGGAGCGGT</td><td>>hg38_21_+_8433172_8446622_RNA45SN1_45S_50ntFlanks</td><td>11345</td><td>11362</td><td>18</td><td>CTGGGGGCGGGGAGCGGT</td><td>1033.5</td><td>-7.0</td><td>1.7</td><td>-4.1</td><td>3.5E-05</td><td>1.3E-04</td><td>BT20/MB468</td><td>MP</td></tr>
<tr><td>>am_trnaMT_SerGCT_MT_+_12207_12265@16.38.23|AACTGCTAACTCATGCCCCCATG</td><td>>am_trnaMT_SerGCT_MT_+_12207_12265</td><td>16</td><td>38</td><td>23</td><td>AACTGCTAACTCATGCCCCCATG</td><td>402.7</td><td>-4.7</td><td>1.2</td><td>-3.9</td><td>8.2E-05</td><td>1.3E-04</td><td>MB231/MB468</td><td>MP</td></tr>
<tr><td>>hg38_21_+_8433172_8446622_RNA45SN1_45S_50ntFlanks@12910.12935.26|CGCACGTTCGTGGGGAACCTGGCGCT</td><td>>hg38_21_+_8433172_8446622_RNA45SN1_45S_50ntFlanks</td><td>12910</td><td>12935</td><td>26</td><td>CGCACGTTCGTGGGGAACCTGGCGCT</td><td>74.7</td><td>-5.9</td><td>1.5</td><td>-3.9</td><td>8.2E-05</td><td>1.3E-04</td><td>MB231/MB468</td><td>MP</td></tr>
<tr><td>>hg38_MT_+_1621_3279_RNR2_16S_50ntFlanks@1185.1215.31|ATTTCGGTTGGGGCGACCTCGGAGCAGAACC</td><td>>hg38_MT_+_1621_3279_RNR2_16S_50ntFlanks</td><td>1185</td><td>1215</td><td>31</td><td>ATTTCGGTTGGGGCGACCTCGGAGCAGAACC</td><td>123.6</td><td>5.6</td><td>1.4</td><td>3.9</td><td>9.8E-05</td><td>1.3E-04</td><td>MB231/MB468</td><td>MITO</td></tr>
<tr><td>>MI0000542|hsa-mir-320a&WithFlank&8|-|22244960|22245043@50.70.21(+1G)&[MIMAT0000510&hsa-miR-320a-3p&offsets|+2|+1(+1G);m-60&8|-|22244975|22244996&offsets|+2|+1(+1G)]|AAGCTGGGTTGAGAGGGCGAAG</td><td>>MI0000542|hsa-mir-320a&WithFlank&8|-|22244960|22245043</td><td>50</td><td>70</td><td>21(+1G)</td><td>AAGCTGGGTTGAGAGGGCGAAG</td><td>58.3</td><td>9.2</td><td>2.1</td><td>4.4</td><td>1.2E-05</td><td>1.3E-04</td><td>BT20/MB231</td><td>MP</td></tr>
<tr><td>>MI0000750|hsa-mir-26a-2&WithFlank&12|-|57824603|57824698@20.40.21;MI0000083|hsa-mir-26a-1&WithFlank&3|+|37969398|37969486@16.36.21&[MIMAT0000082&hsa-miR-26a-5p&offsets|0|-1;m-33&12|-|57824658|57824679&offsets|0|-1];[MIMAT0000082_1&hsa-miR-26a-5p&offsets|0|-1;m-34&3|+|37969413|37969434&offsets|0|-1]|TTCAAGTAATCCAGGATAGGC</td><td>>MI0000750|hsa-mir-26a-2&WithFlank&12|-|57824603|57824698</td><td>20</td><td>40</td><td>21</td><td>TTCAAGTAATCCAGGATAGGC</td><td>2306.0</td><td>-4.4</td><td>1.0</td><td>-4.4</td><td>1.2E-05</td><td>1.3E-04</td><td>BT20/MB231</td><td>MP</td></tr>
<tr><td>>MI0000070|hsa-mir-16-1&WithFlank&13|-|50048967|50049067@20.43.24&[MIMAT0000069&hsa-miR-16-5p&offsets|0|+2;m-62&13|-|50049027|50049048&offsets|0|+2]|TAGCAGCACGTAAATATTGGCGTT</td><td>>MI0000070|hsa-mir-16-1&WithFlank&13|-|50048967|50049067</td><td>20</td><td>43</td><td>24</td><td>TAGCAGCACGTAAATATTGGCGTT</td><td>288.9</td><td>-4.2</td><td>1.0</td><td>-4.4</td><td>1.2E-05</td><td>1.3E-04</td><td>BT20/MB231</td><td>MP</td></tr>
<tr><td>>hg38_21_+_8433172_8446622_RNA45SN1_45S_50ntFlanks@12624.12659.36|CCAATGGGGCGAAGCTACCATCTGTGGGATTATGAC</td><td>>hg38_21_+_8433172_8446622_RNA45SN1_45S_50ntFlanks</td><td>12624</td><td>12659</td><td>36</td><td>CCAATGGGGCGAAGCTACCATCTGTGGGATTATGAC</td><td>117.6</td><td>-4.9</td><td>1.2</td><td>-3.9</td><td>8.6E-05</td><td>1.3E-04</td><td>BT20/MB468</td><td>CYTO</td></tr>
<tr><td>>hg38_21_+_8433172_8446622_RNA45SN1_45S_50ntFlanks@10921.10945.25|GCTGGACGAGGCGCCGCCGCCCCCC</td><td>>hg38_21_+_8433172_8446622_RNA45SN1_45S_50ntFlanks</td><td>10921</td><td>10945</td><td>25</td><td>GCTGGACGAGGCGCCGCCGCCCCCC</td><td>173.7</td><td>-4.1</td><td>1.0</td><td>-3.9</td><td>9.8E-05</td><td>1.3E-04</td><td>MB231/MB468</td><td>NUC</td></tr>
<tr><td>>hg38_21_+_8433172_8446622_RNA45SN1_45S_50ntFlanks@10392.10435.44|ACAGCAGTTGAACATGGGTCAGTCGGTCCTGAGAGATGGGCGAG</td><td>>hg38_21_+_8433172_8446622_RNA45SN1_45S_50ntFlanks</td><td>10392</td><td>10435</td><td>44</td><td>ACAGCAGTTGAACATGGGTCAGTCGGTCCTGAGAGATGGGCGAG</td><td>682.2</td><td>-4.9</td><td>1.2</td><td>-4.0</td><td>7.2E-05</td><td>1.3E-04</td><td>BT20/MB468</td><td>MITO</td></tr>
<tr><td>>hg38_21_+_8433172_8446622_RNA45SN1_45S_50ntFlanks@9572.9604.33|CAAATCGGTCGTCCGACCTGGGTATAGGGGCGA</td><td>>hg38_21_+_8433172_8446622_RNA45SN1_45S_50ntFlanks</td><td>9572</td><td>9604</td><td>33</td><td>CAAATCGGTCGTCCGACCTGGGTATAGGGGCGA</td><td>235.8</td><td>-2.8</td><td>0.7</td><td>-4.0</td><td>7.2E-05</td><td>1.3E-04</td><td>BT20/MB468</td><td>MITO</td></tr>
<tr><td>>hg38_21_+_8433172_8446622_RNA45SN1_45S_50ntFlanks@12995.13037.43|CTCGCTGCGATCTATTGAAAGTCAGCCCTCGACACAAGGGTTT</td><td>>hg38_21_+_8433172_8446622_RNA45SN1_45S_50ntFlanks</td><td>12995</td><td>13037</td><td>43</td><td>CTCGCTGCGATCTATTGAAAGTCAGCCCTCGACACAAGGGTTT</td><td>365.3</td><td>6.2</td><td>1.6</td><td>3.9</td><td>9.9E-05</td><td>1.3E-04</td><td>MB231/MB468</td><td>NUC</td></tr>
<tr><td>>hg38_21_+_8433172_8446622_RNA45SN1_45S_50ntFlanks@10362.10382.21|AAGGCCGAAGTGGAGAAGGGT</td><td>>hg38_21_+_8433172_8446622_RNA45SN1_45S_50ntFlanks</td><td>10362</td><td>10382</td><td>21</td><td>AAGGCCGAAGTGGAGAAGGGT</td><td>536.5</td><td>-5.0</td><td>1.3</td><td>-3.9</td><td>8.3E-05</td><td>1.3E-04</td><td>MB231/MB468</td><td>MP</td></tr>
<tr><td>>hg38_21_+_8433172_8446622_RNA45SN1_45S_50ntFlanks@12636.12657.22|AGCTACCATCTGTGGGATTATG</td><td>>hg38_21_+_8433172_8446622_RNA45SN1_45S_50ntFlanks</td><td>12636</td><td>12657</td><td>22</td><td>AGCTACCATCTGTGGGATTATG</td><td>509.9</td><td>-4.8</td><td>1.2</td><td>-4.0</td><td>7.2E-05</td><td>1.3E-04</td><td>BT20/MB468</td><td>MITO</td></tr>
<tr><td>>hg38_21_+_8433172_8446622_RNA45SN1_45S_50ntFlanks@6700.6719.20|CTAGCTGCGAGAATTAATGT</td><td>>hg38_21_+_8433172_8446622_RNA45SN1_45S_50ntFlanks</td><td>6700</td><td>6719</td><td>20</td><td>CTAGCTGCGAGAATTAATGT</td><td>424.3</td><td>-5.9</td><td>1.5</td><td>-3.9</td><td>9.9E-05</td><td>1.3E-04</td><td>MB231/MB468</td><td>MITO</td></tr>
<tr><td>>hg38_21_+_8433172_8446622_RNA45SN1_45S_50ntFlanks@12006.12032.27|GGCCGCCGGTGAAATACCACTACTCTG</td><td>>hg38_21_+_8433172_8446622_RNA45SN1_45S_50ntFlanks</td><td>12006</td><td>12032</td><td>27</td><td>GGCCGCCGGTGAAATACCACTACTCTG</td><td>788.2</td><td>-5.3</td><td>1.4</td><td>-3.9</td><td>9.9E-05</td><td>1.3E-04</td><td>MB231/MB468</td><td>NUC</td></tr>
<tr><td>>MI0000088|hsa-mir-30a&WithFlank&6|-|71403545|71403627@53.75.23&[MIMAT0000088&hsa-miR-30a-3p&offsets|0|+1;m-38&6|-|71403554|71403575&offsets|0|+1]|CTTTCAGTCGGATGTTTGCAGCT</td><td>>MI0000088|hsa-mir-30a&WithFlank&6|-|71403545|71403627</td><td>53</td><td>75</td><td>23</td><td>CTTTCAGTCGGATGTTTGCAGCT</td><td>262.8</td><td>-3.2</td><td>0.7</td><td>-4.3</td><td>1.4E-05</td><td>1.3E-04</td><td>BT20/MB231</td><td>MITO</td></tr>
<tr><td>>hg38_21_+_8433172_8446622_RNA45SN1_45S_50ntFlanks@8184.8212.29|CGAGGCCCAGCCCGTGGACGGTGTGAGGC</td><td>>hg38_21_+_8433172_8446622_RNA45SN1_45S_50ntFlanks</td><td>8184</td><td>8212</td><td>29</td><td>CGAGGCCCAGCCCGTGGACGGTGTGAGGC</td><td>382.7</td><td>-6.9</td><td>1.7</td><td>-4.1</td><td>3.5E-05</td><td>1.3E-04</td><td>BT20/MB468</td><td>MP</td></tr>
<tr><td>>hg38_21_+_8433172_8446622_RNA45SN1_45S_50ntFlanks@9178.9197.20|CGCCGTCGGGCCCGGGGGAG</td><td>>hg38_21_+_8433172_8446622_RNA45SN1_45S_50ntFlanks</td><td>9178</td><td>9197</td><td>20</td><td>CGCCGTCGGGCCCGGGGGAG</td><td>522.7</td><td>-6.6</td><td>1.6</td><td>-4.1</td><td>3.5E-05</td><td>1.3E-04</td><td>BT20/MB468</td><td>MP</td></tr>
<tr><td>>hg38_21_+_8433172_8446622_RNA45SN1_45S_50ntFlanks@11578.11611.34|TAGAACTGGTGCGGACCAGGGGAATCCGACTGTT</td><td>>hg38_21_+_8433172_8446622_RNA45SN1_45S_50ntFlanks</td><td>11578</td><td>11611</td><td>34</td><td>TAGAACTGGTGCGGACCAGGGGAATCCGACTGTT</td><td>357.9</td><td>3.0</td><td>0.7</td><td>4.4</td><td>1.3E-05</td><td>1.3E-04</td><td>BT20/MB231</td><td>NUC</td></tr>
<tr><td>>MI0000469|hsa-mir-125a&WithFlank&19|+|51693248|51693345@21.41.21(+1C)&[MIMAT0000443&hsa-miR-125a-5p&offsets|0|-3(+1C);m-63&19|+|51693268|51693289&offsets|0|-1(+1C)]|TCCCTGAGACCCTTTAACCTGC</td><td>>MI0000469|hsa-mir-125a&WithFlank&19|+|51693248|51693345</td><td>21</td><td>41</td><td>21(+1C)</td><td>TCCCTGAGACCCTTTAACCTGC</td><td>232.4</td><td>2.9</td><td>0.8</td><td>3.9</td><td>9.9E-05</td><td>1.3E-04</td><td>MB231/MB468</td><td>NUC</td></tr>
<tr><td>>hg38_MT_+_598_1651_RNR1_12S_50ntFlanks@848.876.29|GAGTGCTTAGTTGAACAGGGCCCTGAAGC</td><td>>hg38_MT_+_598_1651_RNR1_12S_50ntFlanks</td><td>848</td><td>876</td><td>29</td><td>GAGTGCTTAGTTGAACAGGGCCCTGAAGC</td><td>139.8</td><td>7.7</td><td>1.9</td><td>4.1</td><td>3.5E-05</td><td>1.3E-04</td><td>BT20/MB468</td><td>MP</td></tr>
<tr><td>>hg38_21_+_8433172_8446622_RNA45SN1_45S_50ntFlanks@12934.12969.36|CTAAACCATTCGTAGACGACCTGCTTCTGGGTCGGG</td><td>>hg38_21_+_8433172_8446622_RNA45SN1_45S_50ntFlanks</td><td>12934</td><td>12969</td><td>36</td><td>CTAAACCATTCGTAGACGACCTGCTTCTGGGTCGGG</td><td>191.8</td><td>-2.2</td><td>0.6</td><td>-4.0</td><td>7.3E-05</td><td>1.3E-04</td><td>BT20/MB468</td><td>MITO</td></tr>
<tr><td>>hg38_21_+_8433172_8446622_RNA45SN1_45S_50ntFlanks@8358.8379.22|AAAGAACTTTGAAGAGAGAGTT</td><td>>hg38_21_+_8433172_8446622_RNA45SN1_45S_50ntFlanks</td><td>8358</td><td>8379</td><td>22</td><td>AAAGAACTTTGAAGAGAGAGTT</td><td>1055.9</td><td>-5.9</td><td>1.4</td><td>-4.1</td><td>3.5E-05</td><td>1.3E-04</td><td>BT20/MB468</td><td>MP</td></tr>
<tr><td>>trna119_LysCTT_1_-_145395522_145395594@1.36.36,trna11_LysCTT_5_-_180648979_180649051@1.36.36,trna13_LysCTT_6_+_26556774_26556846@1.36.36,trna7_LysCTT_16_+_3225692_3225764@1.36.36,trna9_LysCTT_5_+_180634755_180634827@1.36.36|GCCCGGCTAGCTCAGTCGGTAGAGCATGAGACTCTT</td><td>>trna119_LysCTT_1_-_145395522_145395594</td><td>1</td><td>36</td><td>36</td><td>GCCCGGCTAGCTCAGTCGGTAGAGCATGAGACTCTT</td><td>1845.9</td><td>4.8</td><td>1.2</td><td>3.9</td><td>9.9E-05</td><td>1.3E-04</td><td>MB231/MB468</td><td>MITO</td></tr>
<tr><td>>MI0000342|hsa-mir-200b&WithFlank&1|+|1167098|1167204@63.84.22&[MIMAT0000318&hsa-miR-200b-3p&offsets|0|0;m-93&1|+|1167160|1167182&offsets|0|-1]|TAATACTGCCTGGTAATGATGA</td><td>>MI0000342|hsa-mir-200b&WithFlank&1|+|1167098|1167204</td><td>63</td><td>84</td><td>22</td><td>TAATACTGCCTGGTAATGATGA</td><td>344.1</td><td>3.7</td><td>0.8</td><td>4.4</td><td>1.2E-05</td><td>1.3E-04</td><td>BT20/MB231</td><td>CYTO</td></tr>
</table>

| | | | | | | | | | | | | | |
|---|---|---|---|---|---|---|---|---|---|---|---|---|---|
| >trna10_AspGTC_12_-_125424193_125424264@37.72.36,trna12_AspGTC_12_-_125411891_125411962@37.72.36,trna144_AspGTC_6_-_27551236_27551307@37.72.36,trna38_AspGTC_17_-_8125556_8125627@37.72.36,trna45_AspGTC_6_+_27447453_27447524@37.72.36,trna48_AspGTC_6_+_27471523_27471594@37.72.36,trna4_AspGTC_12_+_96429799_96429870@37.72.36,trna69_AspGTC_1_-_161440205_161440276@37.72.36,trna72_AspGTC_1_-_161432824_161432895@37.72.36,trna75_AspGTC_1_-_161425414_161425485@37.72.36,trna78_AspGTC_1_-_161418033_161418104@37.72.36,trna81_AspGTC_1_-_161410615_161410686@37.72.36\|ACGCGGGAGACCGGGGTTCGATTCCCCGACGGGGAG | >trna10_AspGTC_12_-_125424193_125424264 | 37 | 72 | 36 | ACGCGGGAGACCGGGGTTCGATTCCCCGACGGGGAG | 2038.3 | -4.4 | 1.0 | -4.4 | 1.2E-05 | 1.3E-04 | BT20/MB231 | CYTO |
| >hg38_21_+_8433172_8446622_RNA45SN1_45S_50ntFlanks@8012.8046.35\|AAGCATATTAGTCAGCGGAGGAGAAGAAACTAACC | >hg38_21_+_8433172_8446622_RNA45SN1_45S_50ntFlanks | 8012 | 8046 | 35 | AAGCATATTAGTCAGCGGAGGAGAAGAAACTAACC | 242.8 | -3.9 | 0.9 | -4.4 | 1.2E-05 | 1.3E-04 | BT20/MB231 | CYTO |
| >hg38_21_+_8433172_8446622_RNA45SN1_45S_50ntFlanks@5236.5264.29\|CACTGACTGGCTCAGCGTGTGCCTACCCT | >hg38_21_+_8433172_8446622_RNA45SN1_45S_50ntFlanks | 5236 | 5264 | 29 | CACTGACTGGCTCAGCGTGTGCCTACCCT | 92.2 | -5.9 | 1.5 | -3.9 | 8.4E-05 | 1.3E-04 | MB231/MB468 | MP |
| >hg38_21_+_8433172_8446622_RNA45SN1_45S_50ntFlanks@9575.9593.19\|ATCGGTCGTCCGACCTGGG | >hg38_21_+_8433172_8446622_RNA45SN1_45S_50ntFlanks | 9575 | 9593 | 19 | ATCGGTCGTCCGACCTGGG | 182.5 | -6.0 | 1.5 | -3.9 | 8.4E-05 | 1.3E-04 | MB231/MB468 | MP |
| >hg38_21_+_8433172_8446622_RNA45SN1_45S_50ntFlanks@10398.10432.35\|GTTGAACATGGGTCAGTCGGTCCTGAGAGATGGGC | >hg38_21_+_8433172_8446622_RNA45SN1_45S_50ntFlanks | 10398 | 10432 | 35 | GTTGAACATGGGTCAGTCGGTCCTGAGAGATGGGC | 84.6 | 9.7 | 2.2 | 4.4 | 1.2E-05 | 1.3E-04 | BT20/MB231 | MP |
| >MI0000812\|hsa-mir-331&WithFlank&12\|+\|95308414\|95308519@32.51.20(+1U)&[MIMAT0004700&hsa-miR-331-5p&offsets\|0\|-2(+1U);m-377&12\|+\|95308445\|95308465&offsets\|0\|-1(+1U)]\|CTAGGTATGGTCCCAGGGATT | >MI0000812\|hsa-mir-331&WithFlank&12\|+\|95308414\|95308519 | 32 | 51 | 20(+1U) | CTAGGTATGGTCCCAGGGATT | 31.5 | 8.2 | 2.1 | 3.9 | 8.4E-05 | 1.3E-04 | MB231/MB468 | MP |
| >hg38_21_+_8433172_8446622_RNA45SN1_45S_50ntFlanks@11200.11232.33\|CGGCGGCGGCGGCGACTCTGGACGCGAGCCGGG | >hg38_21_+_8433172_8446622_RNA45SN1_45S_50ntFlanks | 11200 | 11232 | 33 | CGGCGGCGGCGGCGACTCTGGACGCGAGCCGGG | 4894.4 | -4.2 | 1.1 | -3.9 | 9.4E-05 | 1.3E-04 | MB231/MB468 | CYTO |
| >MI0000065\|hsa-let-7d&WithFlank&9\|+\|94178828\|94178926@68.88.21(+1A)&[MIMAT0004484&hsa-let-7d-3p&offsets\|0\|-1(+1A);m-98&9\|+\|94178895\|94178916&offsets\|0\|-1(+1A)]\|CTATACGACCTGCTGCCTTTCA | >MI0000065\|hsa-let-7d&WithFlank&9\|+\|94178828\|94178926 | 68 | 88 | 21(+1A) | CTATACGACCTGCTGCCTTTCA | 81.4 | 2.3 | 0.6 | 3.9 | 1.0E-04 | 1.3E-04 | BT20/MB468 | NUC |
| >hg38_21_+_8433172_8446622_RNA45SN1_45S_50ntFlanks@475.497.23\|CTTCGTGATCGATGTGGTGACGT | >hg38_21_+_8433172_8446622_RNA45SN1_45S_50ntFlanks | 475 | 497 | 23 | CTTCGTGATCGATGTGGTGACGT | 4157.7 | 4.6 | 1.2 | 3.9 | 8.4E-05 | 1.3E-04 | MB231/MB468 | MP |
| >trnaMT_HisGTG_MT_+_12138_12206@26.59.34\|AGATTGTGAATCTGACAACAGAGGCTTACGACCC | >trnaMT_HisGTG_MT_+_12138_12206 | 26 | 59 | 34 | AGATTGTGAATCTGACAACAGAGGCTTACGACCC | 123.2 | 10.7 | 2.6 | 4.1 | 3.6E-05 | 1.3E-04 | BT20/MB468 | MP |
| >hg38_21_+_8433172_8446622_RNA45SN1_45S_50ntFlanks@9940.9960.21\|CAGGACGGTGGCCATGGAAGT | >hg38_21_+_8433172_8446622_RNA45SN1_45S_50ntFlanks | 9940 | 9960 | 21 | CAGGACGGTGGCCATGGAAGT | 1296.0 | 4.2 | 1.0 | 4.1 | 3.6E-05 | 1.3E-04 | BT20/MB468 | MP |
| >hg38_21_+_8433172_8446622_RNA45SN1_45S_50ntFlanks@9919.9948.30\|GGTGTTGGTTGATATAGACAGCAGGACGGT | >hg38_21_+_8433172_8446622_RNA45SN1_45S_50ntFlanks | 9919 | 9948 | 30 | GGTGTTGGTTGATATAGACAGCAGGACGGT | 61.8 | 9.3 | 2.1 | 4.4 | 1.2E-05 | 1.3E-04 | BT20/MB231 | MP |
| >hg38_21_+_8433172_8446622_RNA45SN1_45S_50ntFlanks@10350.10382.33\|ACGAGAACTTTGAAGGCCGAAGTGGAGAAGGGT | >hg38_21_+_8433172_8446622_RNA45SN1_45S_50ntFlanks | 10350 | 10382 | 33 | ACGAGAACTTTGAAGGCCGAAGTGGAGAAGGGT | 881.5 | -3.0 | 0.8 | -3.9 | 1.0E-04 | 1.3E-04 | MB231/MB468 | MITO |
| >hg38_21_+_8433172_8446622_RNA45SN1_45S_50ntFlanks@11591.11611.21\|GACCAGGGGAATCCGACTGTT | >hg38_21_+_8433172_8446622_RNA45SN1_45S_50ntFlanks | 11591 | 11611 | 21 | GACCAGGGGAATCCGACTGTT | 313.0 | -6.3 | 1.6 | -3.9 | 8.4E-05 | 1.3E-04 | MB231/MB468 | MP |
| >hg38_21_+_8433172_8446622_RNA45SN1_45S_50ntFlanks@5073.5102.30\|AATTCCGATAACGAACGAGACTCTGGCATG | >hg38_21_+_8433172_8446622_RNA45SN1_45S_50ntFlanks | 5073 | 5102 | 30 | AATTCCGATAACGAACGAGACTCTGGCATG | 690.0 | -5.3 | 1.4 | -3.9 | 1.0E-04 | 1.3E-04 | MB231/MB468 | NUC |
| >hg38_21_+_8433172_8446622_RNA45SN1_45S_50ntFlanks@11857.11886.30\|GCGGAATCAGCGGGGAAAGAAGACCCTGTT | >hg38_21_+_8433172_8446622_RNA45SN1_45S_50ntFlanks | 11857 | 11886 | 30 | GCGGAATCAGCGGGGAAAGAAGACCCTGTT | 74.6 | 9.6 | 2.2 | 4.4 | 1.2E-05 | 1.3E-04 | BT20/MB231 | MP |
| >hg38_21_+_8433172_8446622_RNA45SN1_45S_50ntFlanks@5186.5205.20\|CAATAACAGGTCTGTGATGC | >hg38_21_+_8433172_8446622_RNA45SN1_45S_50ntFlanks | 5186 | 5205 | 20 | CAATAACAGGTCTGTGATGC | 447.5 | -3.2 | 0.8 | -3.9 | 1.0E-04 | 1.3E-04 | MB231/MB468 | NUC |
| >hg38_21_+_8433172_8446622_RNA45SN1_45S_50ntFlanks@10471.10488.18\|CCCTCGGCCGATCGAAAG | >hg38_21_+_8433172_8446622_RNA45SN1_45S_50ntFlanks | 10471 | 10488 | 18 | CCCTCGGCCGATCGAAAG | 657.7 | -3.0 | 0.8 | -4.0 | 7.4E-05 | 1.3E-04 | BT20/MB468 | MITO |
| >hg38_21_+_8433172_8446622_RNA45SN1_45S_50ntFlanks@11273.11293.21\|CCGCCCCCGGGGAGCCCGGCG | >hg38_21_+_8433172_8446622_RNA45SN1_45S_50ntFlanks | 11273 | 11293 | 21 | CCGCCCCCGGGGAGCCCGGCG | 122.1 | -5.1 | 1.3 | -4.0 | 7.4E-05 | 1.3E-04 | BT20/MB468 | MITO |
| >trnaMT_GluTTC_MT_-_14674_14742@26.67.42,trnalookalike8_GluTTC_5_-_93905172_93905240@26.67.42\|GGTTTTTCATATCATTGGTCGTGGTTGTAGTCCGTGCGAGAA | >trnaMT_GluTTC_MT_-_14674_14742 | 26 | 67 | 42 | GGTTTTTCATATCATTGGTCGTGGTTGTAGTCCGTGCGAGAA | 237.7 | 3.1 | 0.7 | 4.3 | 1.4E-05 | 1.3E-04 | BT20/MB231 | MITO |
| >trnaMT_GluTTC_MT_-_14674_14742@16.58.43,trnalookalike8_GluTTC_5_-_93905172_93905240@16.58.43\|ATACAACGATGGTTTTTCATATCATTGGTCGTGGTTGTAGTCC | >trnaMT_GluTTC_MT_-_14674_14742 | 16 | 58 | 43 | ATACAACGATGGTTTTTCATATCATTGGTCGTGGTTGTAGTCC | 122.9 | 10.7 | 2.6 | 4.1 | 3.6E-05 | 1.3E-04 | BT20/MB468 | MP |
| >MI0000085\|hsa-mir-27a&WithFlank&19\|-\|13836434\|13836523@60.77.18&[MIMAT0000084&hsa-miR-27a-3p&offsets\|+3\|0;m-43&19\|-\|13836447\|13836467&offsets\|+3\|0]\|ACAGTGGCTAAGTTCCGC | >MI0000085\|hsa-mir-27a&WithFlank&19\|-\|13836434\|13836523 | 60 | 77 | 18 | ACAGTGGCTAAGTTCCGC | 103.0 | 9.9 | 2.5 | 3.9 | 8.5E-05 | 1.3E-04 | MB231/MB468 | MP |
| >hg38_21_+_8433172_8446622_RNA45SN1_45S_50ntFlanks@12244.12266.23\|AAGGGCAAAAGCTCGCTTGATCT | >hg38_21_+_8433172_8446622_RNA45SN1_45S_50ntFlanks | 12244 | 12266 | 23 | AAGGGCAAAAGCTCGCTTGATCT | 209.8 | -6.0 | 1.5 | -3.9 | 8.5E-05 | 1.3E-04 | MB231/MB468 | MP |
| >trna116_GluCTC_1_-_145399233_145399304@32.71.40,trna71_GluCTC_1_-_161439189_161439260@32.71.40,trna74_GluCTC_1_-_161431809_161431880@32.71.40,trna77_GluCTC_1_-_161424398_161424469@32.71.40,trna77_GluCTC_6_+_28949976_28950047@32.71.40,trna80_GluCTC_1_-_161417018_161417089@32.71.40,trna87_GluCTC_6_-_126101393_126101464@32.71.40\|CTCTCACCGCCGCGGCCCGGGTTCGATTCCCGGTCAGGGA | >trna116_GluCTC_1_-_145399233_145399304 | 32 | 71 | 40 | CTCTCACCGCCGCGGCCCGGGTTCGATTCCCGGTCAGGGA | 325.5 | -6.1 | 1.5 | -4.0 | 7.4E-05 | 1.3E-04 | BT20/MB468 | MITO |
| >hg38_21_+_8433172_8446622_RNA45SN1_45S_50ntFlanks@10807.10828.22\|ACAATGTAGGTAAGGGAAGTCG | >hg38_21_+_8433172_8446622_RNA45SN1_45S_50ntFlanks | 10807 | 10828 | 22 | ACAATGTAGGTAAGGGAAGTCG | 263.0 | -3.3 | 0.8 | -3.9 | 9.5E-05 | 1.3E-04 | MB231/MB468 | CYTO |
| >hg38_21_+_8433172_8446622_RNA45SN1_45S_50ntFlanks@6650.6692.43\|ACGACTCTTAGCGGTGGATCACTCGGCTCGTGCGTCGATGAAG | >hg38_21_+_8433172_8446622_RNA45SN1_45S_50ntFlanks | 6650 | 6692 | 43 | ACGACTCTTAGCGGTGGATCACTCGGCTCGTGCGTCGATGAAG | 904.7 | 3.2 | 0.8 | 3.9 | 1.0E-04 | 1.3E-04 | MB231/MB468 | NUC |
| >hg38_21_+_8433172_8446622_RNA45SN1_45S_50ntFlanks@8378.8402.25\|TTCAAGAGGGCGTGAAACCGTTAAG | >hg38_21_+_8433172_8446622_RNA45SN1_45S_50ntFlanks | 8378 | 8402 | 25 | TTCAAGAGGGCGTGAAACCGTTAAG | 467.8 | -3.6 | 0.9 | -4.1 | 3.6E-05 | 1.3E-04 | BT20/MB468 | MP |
| >trnaMT_SerGCT_MT_+_12207_12265@14.62.49\|AGAACTGCTAACTCATGCCCCCATGTCTAACAACATGGCTTTCTCACCA | >trnaMT_SerGCT_MT_+_12207_12265 | 14 | 62 | 49 | AGAACTGCTAACTCATGCCCCCATGTCTAACAACATGGCTTTCTCACCA | 1160.9 | 6.9 | 1.7 | 4.0 | 7.4E-05 | 1.3E-04 | BT20/MB468 | MITO |
| >hg38_21_+_8433172_8446622_RNA45SN1_45S_50ntFlanks@12423.12443.21\|TCGGCTCTTCCTATCATTGTG | >hg38_21_+_8433172_8446622_RNA45SN1_45S_50ntFlanks | 12423 | 12443 | 21 | TCGGCTCTTCCTATCATTGTG | 406.0 | -4.6 | 1.2 | -3.9 | 1.0E-04 | 1.3E-04 | MB231/MB468 | MITO |
| >hg38_21_+_8433172_8446622_RNA45SN1_45S_50ntFlanks@10248.10279.32\|CGACGAGTAGGAGGGCCGCTGCGGTGAGCCTT | >hg38_21_+_8433172_8446622_RNA45SN1_45S_50ntFlanks | 10248 | 10279 | 32 | CGACGAGTAGGAGGGCCGCTGCGGTGAGCCTT | 842.4 | 3.2 | 0.8 | 3.9 | 1.0E-04 | 1.3E-04 | MB231/MB468 | NUC |
| >hg38_21_+_8433172_8446622_RNA45SN1_45S_50ntFlanks@12997.13036.40\|CGCTGCGATCTATTGAAAGTCAGCCCTCGACACAAGGGTT | >hg38_21_+_8433172_8446622_RNA45SN1_45S_50ntFlanks | 12997 | 13036 | 40 | CGCTGCGATCTATTGAAAGTCAGCCCTCGACACAAGGGTT | 1015.9 | 4.3 | 1.1 | 3.9 | 1.0E-04 | 1.3E-04 | MB231/MB468 | MITO |
| >hg38_21_+_8433172_8446622_RNA45SN1_45S_50ntFlanks@10793.10823.31\|TCTGGCATGTTGGAACAATGTAGGTAAGGGA | >hg38_21_+_8433172_8446622_RNA45SN1_45S_50ntFlanks | 10793 | 10823 | 31 | TCTGGCATGTTGGAACAATGTAGGTAAGGGA | 75.4 | 9.6 | 2.2 | 4.4 | 1.3E-05 | 1.3E-04 | BT20/MB231 | MP |
| >hg38_21_+_8433172_8446622_RNA45SN1_45S_50ntFlanks@12476.12500.25\|CCACTAATAGGGAACGTGAGCTGGG | >hg38_21_+_8433172_8446622_RNA45SN1_45S_50ntFlanks | 12476 | 12500 | 25 | CCACTAATAGGGAACGTGAGCTGGG | 413.0 | -5.3 | 1.3 | -4.1 | 3.6E-05 | 1.3E-04 | BT20/MB468 | MP |
| >MI0000542\|hsa-mir-320a&WithFlank&8\|-\|22244960\|22245043@50.70.21&[MIMAT0000510&hsa-miR-320a-3p&offsets\|+2\|+1;m-60&8\|-\|22244975\|22244996&offsets\|+2\|+1]\|AAGCTGGGTTGAGAGGGCGAA | >MI0000542\|hsa-mir-320a&WithFlank&8\|-\|22244960\|22245043 | 50 | 70 | 21 | AAGCTGGGTTGAGAGGGCGAA | 257.2 | 3.4 | 0.8 | 4.4 | 1.2E-05 | 1.3E-04 | BT20/MB231 | CYTO |
| >hg38_21_+_8433172_8446622_RNA45SN1_45S_50ntFlanks@11770.11791.22\|AATTAGTGACGCGCATGAATGG | >hg38_21_+_8433172_8446622_RNA45SN1_45S_50ntFlanks | 11770 | 11791 | 22 | AATTAGTGACGCGCATGAATGG | 418.8 | -6.6 | 1.7 | -3.9 | 1.0E-04 | 1.3E-04 | BT20/MB468 | NUC |
| >hg38_21_+_8433172_8446622_RNA45SN1_45S_50ntFlanks@12007.12038.32\|GCCGCCGGTGAAATACCACTACTCTGATCGTT | >hg38_21_+_8433172_8446622_RNA45SN1_45S_50ntFlanks | 12007 | 12038 | 32 | GCCGCCGGTGAAATACCACTACTCTGATCGTT | 361.2 | -4.6 | 1.2 | -3.9 | 1.0E-04 | 1.3E-04 | MB231/MB468 | MITO |
| >hg38_1_-_228634819_228635039_RNA5S12_5S_50ntFlanks@51.70.20\|GTCTACGGCCATACCACCCT | >hg38_1_-_228634819_228635039_RNA5S12_5S_50ntFlanks | 51 | 70 | 20 | GTCTACGGCCATACCACCCT | 601.4 | -2.7 | 0.7 | -3.9 | 1.0E-04 | 1.3E-04 | BT20/MB468 | NUC |
| >hg38_21_+_8433172_8446622_RNA45SN1_45S_50ntFlanks@9178.9197.20\|CGCCGTCGGGCCCGGGGGAG | >hg38_21_+_8433172_8446622_RNA45SN1_45S_50ntFlanks | 9178 | 9197 | 20 | CGCCGTCGGGCCCGGGGGAG | 562.8 | -7.1 | 1.8 | -3.9 | 1.0E-04 | 1.3E-04 | BT20/MB468 | NUC |
| >hg38_21_+_8433172_8446622_RNA45SN1_45S_50ntFlanks@12714.12753.40\|GAGCCTCGGTTGGCCTCGGATAGCCGGTCCCCCGCCTGTC | >hg38_21_+_8433172_8446622_RNA45SN1_45S_50ntFlanks | 12714 | 12753 | 40 | GAGCCTCGGTTGGCCTCGGATAGCCGGTCCCCCGCCTGTC | 46.1 | -8.6 | 2.1 | -4.1 | 3.6E-05 | 1.3E-04 | BT20/MB468 | MP |
| >hg38_21_+_8433172_8446622_RNA45SN1_45S_50ntFlanks@12477.12499.23\|CACTAATAGGGAACGTGAGCTGG | >hg38_21_+_8433172_8446622_RNA45SN1_45S_50ntFlanks | 12477 | 12499 | 23 | CACTAATAGGGAACGTGAGCTGG | 171.7 | -4.5 | 1.1 | -3.9 | 8.5E-05 | 1.3E-04 | MB231/MB468 | MP |
| >hg38_21_+_8433172_8446622_RNA45SN1_45S_50ntFlanks@6733.6758.26\|CATTGATCATCGACACTTCGAACGCA | >hg38_21_+_8433172_8446622_RNA45SN1_45S_50ntFlanks | 6733 | 6758 | 26 | CATTGATCATCGACACTTCGAACGCA | 317.0 | -2.0 | 0.5 | -3.9 | 9.5E-05 | 1.3E-04 | MB231/MB468 | CYTO |
| >hg38_21_+_8433172_8446622_RNA45SN1_45S_50ntFlanks@4002.4020.19\|GATAACCTCGGGCCGATCG | >hg38_21_+_8433172_8446622_RNA45SN1_45S_50ntFlanks | 4002 | 4020 | 19 | GATAACCTCGGGCCGATCG | 426.8 | -3.8 | 1.0 | -3.9 | 9.5E-05 | 1.3E-04 | MB231/MB468 | CYTO |
| >hg38_21_+_8433172_8446622_RNA45SN1_45S_50ntFlanks@11583.11609.27\|CTGGTGCGGACCAGGGGAATCCGACTG | >hg38_21_+_8433172_8446622_RNA45SN1_45S_50ntFlanks | 11583 | 11609 | 27 | CTGGTGCGGACCAGGGGAATCCGACTG | 490.7 | -2.0 | 0.5 | -3.9 | 1.0E-04 | 1.3E-04 | MB231/MB468 | NUC |
| >hg38_1_-_228634819_228635039_RNA5S12_5S_50ntFlanks@138.171.34\|AGACCGCCTGGGAATACCGGGTGCTGTAGGCTTT | >hg38_1_-_228634819_228635039_RNA5S12_5S_50ntFlanks | 138 | 171 | 34 | AGACCGCCTGGGAATACCGGGTGCTGTAGGCTTT | 1422.1 | 4.2 | 1.1 | 3.9 | 1.0E-04 | 1.3E-04 | MB231/MB468 | MITO |
| >hg38_21_+_8433172_8446622_RNA45SN1_45S_50ntFlanks@12072.12092.21\|CCCGAGGGGCTCTCGCTTCTG | >hg38_21_+_8433172_8446622_RNA45SN1_45S_50ntFlanks | 12072 | 12092 | 21 | CCCGAGGGGCTCTCGCTTCTG | 542.1 | -4.1 | 1.0 | -3.9 | 8.9E-05 | 1.3E-04 | BT20/MB468 | CYTO |
| >hg38_21_+_8433172_8446622_RNA45SN1_45S_50ntFlanks@10411.10431.21\|CAGTCGGTCCTGAGAGATGGG | >hg38_21_+_8433172_8446622_RNA45SN1_45S_50ntFlanks | 10411 | 10431 | 21 | CAGTCGGTCCTGAGAGATGGG | 1197.4 | -4.6 | 1.2 | -4.0 | 7.5E-05 | 1.4E-04 | BT20/MB468 | MITO |

| | | | | | | | | | | | | | |
|---|---|---|---|---|---|---|---|---|---|---|---|---|---|
| >hg38_21_+_8433172_8446622_RNA45SN1_45S_50ntFlanks@9736.9755.20\|CCGAAACGATCTCAACCTAT | >hg38_21_+_8433172_8446622_RNA45SN1_45S_50ntFlanks | 9736 | 9755 | 20 | CCGAAACGATCTCAACCTAT | 282.3 | -3.2 | 0.8 | -4.0 | 7.5E-05 | 1.4E-04 | BT20/MB468 | MITO |
| >hg38_21_+_8433172_8446622_RNA45SN1_45S_50ntFlanks@11583.11607.25\|CTGGTGCGGACCAGGGGAATCCGAC | >hg38_21_+_8433172_8446622_RNA45SN1_45S_50ntFlanks | 11583 | 11607 | 25 | CTGGTGCGGACCAGGGGAATCCGAC | 270.5 | 3.3 | 0.8 | 4.3 | 1.4E-05 | 1.4E-04 | BT20/MB231 | MITO |
| >hg38_MT_+_1621_3279_RNR2_16S_50ntFlanks@1428.1458.31\|ATTAAAGTCCTACGTGATCTGAGTTCAGACC | >hg38_MT_+_1621_3279_RNR2_16S_50ntFlanks | 1428 | 1458 | 31 | ATTAAAGTCCTACGTGATCTGAGTTCAGACC | 338.4 | 3.6 | 0.9 | 3.9 | 1.0E-04 | 1.4E-04 | MB231/MB468 | NUC |
| >hg38_21_+_8433172_8446622_RNA45SN1_45S_50ntFlanks@10393.10420.28\|CAGCAGTTGAACATGGGTCAGTCGGTCC | >hg38_21_+_8433172_8446622_RNA45SN1_45S_50ntFlanks | 10393 | 10420 | 28 | CAGCAGTTGAACATGGGTCAGTCGGTCC | 202.3 | -5.3 | 1.3 | -4.0 | 7.5E-05 | 1.4E-04 | BT20/MB468 | MITO |
| >trnaMT_GluTTC_MT_-_14674_14742@21.57.37,trnalookalike8_GluTTC_5_-_93905172_93905240@21.57.37\|ACGATGGTTTTTCATATCATTGGTCGTGGTTGTAGTC | >trnaMT_GluTTC_MT_-_14674_14742 | 21 | 57 | 37 | ACGATGGTTTTTCATATCATTGGTCGTGGTTGTAGTC | 500.6 | 8.4 | 2.0 | 4.1 | 3.7E-05 | 1.4E-04 | BT20/MB468 | MP |
| >hg38_21_+_8433172_8446622_RNA45SN1_45S_50ntFlanks@10362.10384.23\|AAGGCCGAAGTGGAGAAGGGTTC | >hg38_21_+_8433172_8446622_RNA45SN1_45S_50ntFlanks | 10362 | 10384 | 23 | AAGGCCGAAGTGGAGAAGGGTTC | 1041.9 | -4.0 | 1.0 | -4.1 | 3.7E-05 | 1.4E-04 | BT20/MB468 | MP |
| >hg38_21_+_8433172_8446622_RNA45SN1_45S_50ntFlanks@4734.4761.28\|AACGAAAGTCGGAGGTTCGAAGACGATC | >hg38_21_+_8433172_8446622_RNA45SN1_45S_50ntFlanks | 4734 | 4761 | 28 | AACGAAAGTCGGAGGTTCGAAGACGATC | 262.5 | 2.0 | 0.5 | 3.9 | 1.0E-04 | 1.4E-04 | MB231/MB468 | NUC |
| >am_MI0000809\|hsa-mir-151a&WithFlank&8\|-\|140732558\|140732659@53.73.21(+3U)&[MIMAT0000757&hsa-miR-151a-3p&offsets\|0\|0(+3U);m-30&8\|-\|140732587\|140732607&offsets\|0\|0(+3U)]\|CTAGACTGAAGCTCCTTGAGGTTT | >am_MI0000809\|hsa-mir-151a&WithFlank&8\|-\|140732558\|140732659 | 53 | 73 | 21(+3U) | CTAGACTGAAGCTCCTTGAGGTTT | 697.5 | 6.0 | 1.5 | 4.1 | 3.7E-05 | 1.4E-04 | BT20/MB468 | MP |
| >hg38_21_+_8433172_8446622_RNA45SN1_45S_50ntFlanks@12459.12477.19\|AGCGTTGGATTGTTCACCC | >hg38_21_+_8433172_8446622_RNA45SN1_45S_50ntFlanks | 12459 | 12477 | 19 | AGCGTTGGATTGTTCACCC | 998.2 | -2.9 | 0.8 | -3.9 | 9.6E-05 | 1.4E-04 | MB231/MB468 | CYTO |
| >MI0000433\|hsa-let-7g&WithFlank&3\|-\|52268272\|52268367@11.30.20(+1C)&[MIMAT0000414&hsa-let-7g-5p&offsets\|0\|-2(+1C);m-58&3\|-\|52268336\|52268357&offsets\|0\|-2(+1C)]\|TGAGGTAGTAGTTTGTACAGC | >MI0000433\|hsa-let-7g&WithFlank&3\|-\|52268272\|52268367 | 11 | 30 | 20(+1C) | TGAGGTAGTAGTTTGTACAGC | 168.8 | 5.1 | 1.3 | 3.9 | 1.0E-04 | 1.4E-04 | MB231/MB468 | MITO |
| >hg38_21_+_8433172_8446622_RNA45SN1_45S_50ntFlanks@11575.11604.30\|ACTTAGAACTGGTGCGGACCAGGGGAATCC | >hg38_21_+_8433172_8446622_RNA45SN1_45S_50ntFlanks | 11575 | 11604 | 30 | ACTTAGAACTGGTGCGGACCAGGGGAATCC | 188.0 | -2.0 | 0.5 | -3.9 | 9.6E-05 | 1.4E-04 | MB231/MB468 | CYTO |
| >hg38_21_+_8433172_8446622_RNA45SN1_45S_50ntFlanks@5055.5081.27\|GGAGCGATTTGTCTGGTTAATTCCGAT | >hg38_21_+_8433172_8446622_RNA45SN1_45S_50ntFlanks | 5055 | 5081 | 27 | GGAGCGATTTGTCTGGTTAATTCCGAT | 761.4 | 8.0 | 1.9 | 4.1 | 3.7E-05 | 1.4E-04 | BT20/MB468 | MP |
| >hg38_21_+_8433172_8446622_RNA45SN1_45S_50ntFlanks@6651.6699.49\|CGACTCTTAGCGGTGGATCACTCGGCTCGTGCGTCGATGAAGAACGCAG | >hg38_21_+_8433172_8446622_RNA45SN1_45S_50ntFlanks | 6651 | 6699 | 49 | CGACTCTTAGCGGTGGATCACTCGGCTCGTGCGTCGATGAAGAACGCAG | 7130.7 | -5.7 | 1.5 | -3.9 | 8.6E-05 | 1.4E-04 | MB231/MB468 | MP |
| >hg38_MT_+_598_1651_RNR1_12S_50ntFlanks@428.456.29\|AAAGTGGCTTTAACATATCTGAACACACA | >hg38_MT_+_598_1651_RNR1_12S_50ntFlanks | 428 | 456 | 29 | AAAGTGGCTTTAACATATCTGAACACACA | 242.7 | 4.9 | 1.3 | 3.9 | 1.0E-04 | 1.4E-04 | MB231/MB468 | MITO |
| >hg38_21_+_8433172_8446622_RNA45SN1_45S_50ntFlanks@11729.11764.36\|TAACTATGACTCTCTTAAGGTAGCCAAATGCCTCGT | >hg38_21_+_8433172_8446622_RNA45SN1_45S_50ntFlanks | 11729 | 11764 | 36 | TAACTATGACTCTCTTAAGGTAGCCAAATGCCTCGT | 350.0 | -5.1 | 1.3 | -3.9 | 1.0E-04 | 1.4E-04 | MB231/MB468 | MITO |
| >hg38_21_+_8433172_8446622_RNA45SN1_45S_50ntFlanks@6721.6745.25\|AATTGCAGGACACATTGATCATCGA | >hg38_21_+_8433172_8446622_RNA45SN1_45S_50ntFlanks | 6721 | 6745 | 25 | AATTGCAGGACACATTGATCATCGA | 269.5 | -4.3 | 1.1 | -4.0 | 7.5E-05 | 1.4E-04 | BT20/MB468 | MITO |
| >hg38_21_+_8433172_8446622_RNA45SN1_45S_50ntFlanks@9546.9572.27\|TGGAGGTCCGTAGCGGTCCTGACGTGC | >hg38_21_+_8433172_8446622_RNA45SN1_45S_50ntFlanks | 9546 | 9572 | 27 | TGGAGGTCCGTAGCGGTCCTGACGTGC | 834.3 | 3.9 | 0.9 | 4.3 | 1.5E-05 | 1.4E-04 | BT20/MB231 | MITO |
| >hg38_21_+_8433172_8446622_RNA45SN1_45S_50ntFlanks@9954.9980.27\|TGGAAGTCGGAATCCGCTAAGGAGTGT | >hg38_21_+_8433172_8446622_RNA45SN1_45S_50ntFlanks | 9954 | 9980 | 27 | TGGAAGTCGGAATCCGCTAAGGAGTGT | 87.1 | 7.3 | 1.8 | 4.1 | 3.7E-05 | 1.4E-04 | BT20/MB468 | MP |
| >MI0000102\|hsa-mir-100&WithFlank&11\|-\|122152223\|122152314@19.37.19&[MIMAT0000098&hsa-miR-100-5p&offsets\|0\|-3;m-27&11\|-\|122152276\|122152296&offsets\|0\|-2]\|AACCCGTAGATCCGAACTT | >MI0000102\|hsa-mir-100&WithFlank&11\|-\|122152223\|122152314 | 19 | 37 | 19 | AACCCGTAGATCCGAACTT | 924.3 | -7.5 | 1.7 | -4.4 | 1.3E-05 | 1.4E-04 | BT20/MB231 | MP |
| >hg38_21_+_8433172_8446622_RNA45SN1_45S_50ntFlanks@12616.12654.39\|CTGAGGAGCCAATGGGGCGAAGCTACCATCTGTGGGATT | >hg38_21_+_8433172_8446622_RNA45SN1_45S_50ntFlanks | 12616 | 12654 | 39 | CTGAGGAGCCAATGGGGCGAAGCTACCATCTGTGGGATT | 656.9 | -3.2 | 0.8 | -3.9 | 9.0E-05 | 1.4E-04 | BT20/MB468 | CYTO |
| >hg38_21_+_8433172_8446622_RNA45SN1_45S_50ntFlanks@12415.12440.26\|CTTCGATGTCGGCTCTTCCTATCATT | >hg38_21_+_8433172_8446622_RNA45SN1_45S_50ntFlanks | 12415 | 12440 | 26 | CTTCGATGTCGGCTCTTCCTATCATT | 1122.7 | 2.7 | 0.7 | 3.9 | 9.7E-05 | 1.4E-04 | MB231/MB468 | CYTO |
| >hg38_21_+_8433172_8446622_RNA45SN1_45S_50ntFlanks@10101.10127.27\|CGCGCGCGTGTGGTGTGCGTCGGAGGG | >hg38_21_+_8433172_8446622_RNA45SN1_45S_50ntFlanks | 10101 | 10127 | 27 | CGCGCGCGTGTGGTGTGCGTCGGAGGG | 1008.4 | -6.1 | 1.5 | -4.0 | 7.6E-05 | 1.4E-04 | BT20/MB468 | MITO |
| >hg38_21_+_8433172_8446622_RNA45SN1_45S_50ntFlanks@10421.10455.35\|TGAGAGATGGGCGAGCGCCGTTCCGAAGGGACGGG | >hg38_21_+_8433172_8446622_RNA45SN1_45S_50ntFlanks | 10421 | 10455 | 35 | TGAGAGATGGGCGAGCGCCGTTCCGAAGGGACGGG | 249.3 | -5.7 | 1.5 | -3.9 | 1.0E-04 | 1.4E-04 | MB231/MB468 | NUC |
| >hg38_21_+_8433172_8446622_RNA45SN1_45S_50ntFlanks@12126.12148.23\|CGACCCGCTCCGGGGACAGTGCC | >hg38_21_+_8433172_8446622_RNA45SN1_45S_50ntFlanks | 12126 | 12148 | 23 | CGACCCGCTCCGGGGACAGTGCC | 166.4 | -5.2 | 1.3 | -3.9 | 9.1E-05 | 1.4E-04 | BT20/MB468 | CYTO |
| >MI0000087\|hsa-mir-29a&WithFlank&7\|-\|130876741\|130876816@48.65.18(+3U)&[MIMAT0000086&hsa-miR-29a-3p&offsets\|0\|-4(+3U);m-26&7\|-\|130876749\|130876769&offsets\|0\|-3(+3U)]\|TAGCACCATCTGAAATCGTTT | >MI0000087\|hsa-mir-29a&WithFlank&7\|-\|130876741\|130876816 | 48 | 65 | 18(+3U) | TAGCACCATCTGAAATCGTTT | 65.8 | -9.4 | 2.2 | -4.4 | 1.3E-05 | 1.4E-04 | BT20/MB231 | MP |
| >hg38_21_+_8433172_8446622_RNA45SN1_45S_50ntFlanks@10903.10939.37\|GGCTGGGCGCGCGCCGCGGCTGGACGAGGCGCCGCCG | >hg38_21_+_8433172_8446622_RNA45SN1_45S_50ntFlanks | 10903 | 10939 | 37 | GGCTGGGCGCGCGCCGCGGCTGGACGAGGCGCCGCCG | 112.6 | -6.0 | 1.5 | -4.1 | 3.7E-05 | 1.4E-04 | BT20/MB468 | MP |
| >hg38_21_+_8433172_8446622_RNA45SN1_45S_50ntFlanks@11537.11585.49\|CCGCGCGGCGCCTCGCCTCGGCCGGCGCCTAGCAGCCGACTTAGAACTG | >hg38_21_+_8433172_8446622_RNA45SN1_45S_50ntFlanks | 11537 | 11585 | 49 | CCGCGCGGCGCCTCGCCTCGGCCGGCGCCTAGCAGCCGACTTAGAACTG | 234.8 | -5.7 | 1.5 | -3.9 | 1.0E-04 | 1.4E-04 | MB231/MB468 | MITO |
| >hg38_21_+_8433172_8446622_RNA45SN1_45S_50ntFlanks@12329.12365.37\|TTTTAAGCAGGAGGTGTCAGAAAAGTTACCACAGGGA | >hg38_21_+_8433172_8446622_RNA45SN1_45S_50ntFlanks | 12329 | 12365 | 37 | TTTTAAGCAGGAGGTGTCAGAAAAGTTACCACAGGGA | 36.2 | -8.3 | 2.0 | -4.1 | 3.7E-05 | 1.4E-04 | BT20/MB468 | MP |
| >hg38_21_+_8433172_8446622_RNA45SN1_45S_50ntFlanks@10380.10424.45\|GGTTCCATGTGAACAGCAGTTGAACATGGGTCAGTCGGTCCTGAG | >hg38_21_+_8433172_8446622_RNA45SN1_45S_50ntFlanks | 10380 | 10424 | 45 | GGTTCCATGTGAACAGCAGTTGAACATGGGTCAGTCGGTCCTGAG | 492.3 | 3.7 | 0.9 | 3.9 | 1.0E-04 | 1.4E-04 | MB231/MB468 | NUC |
| >hg38_21_+_8433172_8446622_RNA45SN1_45S_50ntFlanks@5445.5467.23\|TCGGATCGGCCCCGCCGGGGTCG | >hg38_21_+_8433172_8446622_RNA45SN1_45S_50ntFlanks | 5445 | 5467 | 23 | TCGGATCGGCCCCGCCGGGGTCG | 288.6 | -3.5 | 0.9 | -3.9 | 9.7E-05 | 1.4E-04 | MB231/MB468 | CYTO |
| >hg38_21_+_8433172_8446622_RNA45SN1_45S_50ntFlanks@4724.4755.32\|ATTAATCAAGAACGAAAGTCGGAGGTTCGAAG | >hg38_21_+_8433172_8446622_RNA45SN1_45S_50ntFlanks | 4724 | 4755 | 32 | ATTAATCAAGAACGAAAGTCGGAGGTTCGAAG | 166.2 | -5.3 | 1.4 | -3.9 | 1.0E-04 | 1.4E-04 | MB231/MB468 | NUC |
| >hg38_21_+_8433172_8446622_RNA45SN1_45S_50ntFlanks@4203.4222.20\|GACCCGGGGAGGTAGTGACG | >hg38_21_+_8433172_8446622_RNA45SN1_45S_50ntFlanks | 4203 | 4222 | 20 | GACCCGGGGAGGTAGTGACG | 345.9 | -5.2 | 1.3 | -3.9 | 8.7E-05 | 1.4E-04 | MB231/MB468 | MP |
| >am_MI0000809\|hsa-mir-151a&WithFlank&8\|-\|140732558\|140732659@53.73.21(+1G)&[MIMAT0000757&hsa-miR-151a-3p&offsets\|0\|0(+1G);m-30&8\|-\|140732587\|140732607&offsets\|0\|0(+1G)]\|CTAGACTGAAGCTCCTTGAGGG | >am_MI0000809\|hsa-mir-151a&WithFlank&8\|-\|140732558\|140732659 | 53 | 73 | 21(+1G) | CTAGACTGAAGCTCCTTGAGGG | 148.9 | 5.9 | 1.5 | 3.9 | 1.0E-04 | 1.4E-04 | MB231/MB468 | MITO |
| >MI0000750\|hsa-mir-26a-2&WithFlank&12\|-\|57824603\|57824698@20.41.22(+3U);MI0000083\|hsa-mir-26a-1&WithFlank&3\|+\|37969398\|37969486@16.37.22(+3U)&[MIMAT0000082&hsa-miR-26a-5p&offsets\|0\|0(+3U);m-33&12\|-\|57824658\|57824679&offsets\|0\|0(+3U)];[MIMAT0000082_1&hsa-miR-26a-5p&offsets\|0\|0(+3U);m-34&3\|+\|37969413\|37969434&offsets\|0\|0(+3U)]\|TTCAAGTAATCCAGGATAGGCTTTT | >MI0000750\|hsa-mir-26a-2&WithFlank&12\|-\|57824603\|57824698 | 20 | 41 | 22(+3U) | TTCAAGTAATCCAGGATAGGCTTTT | 407.9 | -3.8 | 0.9 | -4.3 | 1.5E-05 | 1.4E-04 | BT20/MB231 | MITO |
| >hg38_21_+_8433172_8446622_RNA45SN1_45S_50ntFlanks@10791.10823.33\|CCTCTGGCATGTTGGAACAATGTAGGTAAGGGA | >hg38_21_+_8433172_8446622_RNA45SN1_45S_50ntFlanks | 10791 | 10823 | 33 | CCTCTGGCATGTTGGAACAATGTAGGTAAGGGA | 786.6 | 2.2 | 0.6 | 4.0 | 7.6E-05 | 1.4E-04 | BT20/MB468 | MITO |
| >hg38_21_+_8433172_8446622_RNA45SN1_45S_50ntFlanks@10571.10608.38\|GAGAAGCCGGCGGGAGCCCCGGGGAGAGTTCTCTTTTC | >hg38_21_+_8433172_8446622_RNA45SN1_45S_50ntFlanks | 10571 | 10608 | 38 | GAGAAGCCGGCGGGAGCCCCGGGGAGAGTTCTCTTTTC | 149.6 | -5.6 | 1.4 | -4.0 | 7.6E-05 | 1.4E-04 | BT20/MB468 | MITO |
| >hg38_21_+_8433172_8446622_RNA45SN1_45S_50ntFlanks@12141.12162.22\|ACAGTGCCAGGTGGGGAGTTTG | >hg38_21_+_8433172_8446622_RNA45SN1_45S_50ntFlanks | 12141 | 12162 | 22 | ACAGTGCCAGGTGGGGAGTTTG | 171.7 | -6.1 | 1.5 | -4.1 | 3.7E-05 | 1.4E-04 | BT20/MB468 | MP |
| >hg38_21_+_8433172_8446622_RNA45SN1_45S_50ntFlanks@4477.4494.18\|CTGAGTGTCCCGCGGGGC | >hg38_21_+_8433172_8446622_RNA45SN1_45S_50ntFlanks | 4477 | 4494 | 18 | CTGAGTGTCCCGCGGGGC | 165.7 | -6.2 | 1.5 | -4.1 | 3.7E-05 | 1.4E-04 | BT20/MB468 | MP |
| >trna129_MetCAT_6_-_27870271_27870342@33.51.19,trna142_MetCAT_6_-_27560600_27560671@33.51.19,trna150_MetCAT_6_-_27300764_27300835@33.51.19,trna169_MetCAT_6_-_26330529_26330600@33.51.19,trna171_MetCAT_6_-_26313352_26313423@33.51.19,trna20_MetCAT_17_-_80452597_80452668@33.51.19,trna2_MetCAT_6_+_26286754_26286825@33.51.19,trna32_MetCAT_1_+_153643726_153643797@33.51.19,trna61_MetCAT_6_+_27745664_27745735@33.51.19\|CATAACCCAGAGGTCGATG | >trna129_MetCAT_6_-_27870271_27870342 | 33 | 51 | 19 | CATAACCCAGAGGTCGATG | 127.8 | -10.1 | 2.4 | -4.1 | 3.7E-05 | 1.4E-04 | BT20/MB468 | MP |
| >hg38_21_+_8433172_8446622_RNA45SN1_45S_50ntFlanks@6651.6671.21\|CGACTCTTAGCGGTGGATCAC | >hg38_21_+_8433172_8446622_RNA45SN1_45S_50ntFlanks | 6651 | 6671 | 21 | CGACTCTTAGCGGTGGATCAC | 677.6 | 3.4 | 0.9 | 4.0 | 7.6E-05 | 1.4E-04 | BT20/MB468 | MITO |
| >am_trnaMT_HisGTG_MT_+_12138_12206@20.51.32\|AACATCAGATTGTGAATCTGACAACAGAGGCT | >am_trnaMT_HisGTG_MT_+_12138_12206 | 20 | 51 | 32 | AACATCAGATTGTGAATCTGACAACAGAGGCT | 163.2 | 3.6 | 0.9 | 3.9 | 1.0E-04 | 1.4E-04 | MB231/MB468 | MITO |
| >hg38_21_+_8433172_8446622_RNA45SN1_45S_50ntFlanks@10787.10816.30\|ACAGCCTCTGGCATGTTGGAACAATGTAGG | >hg38_21_+_8433172_8446622_RNA45SN1_45S_50ntFlanks | 10787 | 10816 | 30 | ACAGCCTCTGGCATGTTGGAACAATGTAGG | 547.1 | -4.7 | 1.2 | -3.9 | 1.0E-04 | 1.4E-04 | MB231/MB468 | NUC |
| >hg38_21_+_8433172_8446622_RNA45SN1_45S_50ntFlanks@11661.11680.20\|TGATTTCTGCCCAGTGCTCT | >hg38_21_+_8433172_8446622_RNA45SN1_45S_50ntFlanks | 11661 | 11680 | 20 | TGATTTCTGCCCAGTGCTCT | 718.9 | -4.7 | 1.2 | -3.9 | 9.8E-05 | 1.4E-04 | MB231/MB468 | CYTO |
| >trnaMT_MetCAT_MT_+_4402_4469@32.71.40\|ATACCCCGAAAATGTTGGTTATACCCTTCCCGTACTACCA | >trnaMT_MetCAT_MT_+_4402_4469 | 32 | 71 | 40 | ATACCCCGAAAATGTTGGTTATACCCTTCCCGTACTACCA | 169.9 | 6.9 | 1.8 | 3.9 | 1.0E-04 | 1.4E-04 | MB231/MB468 | MITO |
| >am_trna2_GlyGCC_21_-_18827107_18827177@4.26.23,trna35_GlyGCC_1_+_161413094_161413164@4.26.23,trna37_GlyGCC_1_+_161420467_161420537@4.26.23,trna39_GlyGCC_1_+_161427898_161427968@4.26.23,trna41_GlyGCC_1_+_161435258_161435328@4.26.23\|TGGGTGGTTCAGTGGTAGAATTC | >am_trna2_GlyGCC_21_-_18827107_18827177 | 4 | 26 | 23 | TGGGTGGTTCAGTGGTAGAATTC | 57.6 | 9.2 | 2.1 | 4.4 | 1.3E-05 | 1.4E-04 | BT20/MB231 | MP |
| >hg38_21_+_8433172_8446622_RNA45SN1_45S_50ntFlanks@7974.8014.41\|ACGCGACCTCAGATCAGACGTGGCGACCCGCTGAATTTAAG | >hg38_21_+_8433172_8446622_RNA45SN1_45S_50ntFlanks | 7974 | 8014 | 41 | ACGCGACCTCAGATCAGACGTGGCGACCCGCTGAATTTAAG | 1149.5 | 3.1 | 0.8 | 3.9 | 1.0E-04 | 1.4E-04 | MB231/MB468 | NUC |

<table>
<tr><td></td><td></td><td></td><td></td><td></td><td></td><td></td><td></td><td></td><td></td><td></td><td></td><td></td><td></td></tr>
<tr><td>>hg38_21_+_8433172_8446622_RNA45SN1_45S_50ntFlanks@10348.10372.25|AAACGAGAACTTTGAAGGCCGAAGT</td><td>>hg38_21_+_8433172_8446622_RNA45SN1_45S_50ntFlanks</td><td>10348</td><td>10372</td><td>25</td><td>AAACGAGAACTTTGAAGGCCGAAGT</td><td>738.0</td><td>2.7</td><td>0.7</td><td>3.9</td><td>9.8E-05</td><td>1.4E-04</td><td>MB231/MB468</td><td>CYTO</td></tr>
<tr><td>>hg38_21_+_8433172_8446622_RNA45SN1_45S_50ntFlanks@6788.6806.19|CGCCTGTCTGAGCGTCGCT</td><td>>hg38_21_+_8433172_8446622_RNA45SN1_45S_50ntFlanks</td><td>6788</td><td>6806</td><td>19</td><td>CGCCTGTCTGAGCGTCGCT</td><td>1602.8</td><td>3.3</td><td>0.8</td><td>4.1</td><td>3.8E-05</td><td>1.4E-04</td><td>BT20/MB468</td><td>MP</td></tr>
<tr><td>>hg38_21_+_8433172_8446622_RNA45SN1_45S_50ntFlanks@11710.11738.29|CGCGGGTAAACGGCGGGAGTAACTATGAC</td><td>>hg38_21_+_8433172_8446622_RNA45SN1_45S_50ntFlanks</td><td>11710</td><td>11738</td><td>29</td><td>CGCGGGTAAACGGCGGGAGTAACTATGAC</td><td>87.9</td><td>-5.8</td><td>1.5</td><td>-3.9</td><td>8.8E-05</td><td>1.4E-04</td><td>MB231/MB468</td><td>MP</td></tr>
<tr><td>>hg38_21_+_8433172_8446622_RNA45SN1_45S_50ntFlanks@12420.12460.41|ATGTCGGCTCTTCCTATCATTGTGAAGCAGAATTCACCAAG</td><td>>hg38_21_+_8433172_8446622_RNA45SN1_45S_50ntFlanks</td><td>12420</td><td>12460</td><td>41</td><td>ATGTCGGCTCTTCCTATCATTGTGAAGCAGAATTCACCAAG</td><td>279.2</td><td>-3.5</td><td>0.8</td><td>-4.4</td><td>1.2E-05</td><td>1.4E-04</td><td>BT20/MB231</td><td>CYTO</td></tr>
<tr><td>>hg38_21_+_8433172_8446622_RNA45SN1_45S_50ntFlanks@6645.6672.28|CTCGTACGACTCTTAGCGGTGGATCACT</td><td>>hg38_21_+_8433172_8446622_RNA45SN1_45S_50ntFlanks</td><td>6645</td><td>6672</td><td>28</td><td>CTCGTACGACTCTTAGCGGTGGATCACT</td><td>362.1</td><td>3.6</td><td>0.9</td><td>4.1</td><td>3.8E-05</td><td>1.4E-04</td><td>BT20/MB468</td><td>MP</td></tr>
<tr><td>>am_MI0003776|hsa-mir-320b-1&WithFlank&1|+|116671740|116671823@48.65.18;MI0003839|hsa-mir-320b-2&WithFlank&1|-|224257034|224257116@47.64.18;MI0008190|hsa-mir-320d-1&WithFlank&13|-|40727810|40727893@48.65.18;MI0003778|hsa-mir-320c-1&WithFlank&18|+|21683512|21683595@48.65.18;MI0008191|hsa-mir-320c-2&WithFlank&18|+|24321669|24321752@48.65.18;MI0000542|hsa-mir-320a&WithFlank&8|-|22244960|22245043@48.65.18;MI0008192|hsa-mir-320d-2&WithFlank&X|-|140926154|140926237@48.65.18&[MIMAT0005792&hsa-miR-320b&offsets|0|-4;m-183&1|+|116671787|116671806&offsets|0|-2];[MIMAT0005792_1&hsa-miR-320b&offsets|0|-4;m-183&1|-|224257051|224257070&offsets|0|-2];[MIMAT0006764&hsa-miR-320d&offsets|0|-1;m-246&13|-|40727829|40727846&offsets|0|0];[MIMAT0005793&hsa-miR-320c&offsets|0|-2;m-222&18|+|21683559|21683578&offsets|0|-2];[MIMAT0005793_1&hsa-miR-320c&offsets|0|-2;m-219&18|+|24321716|24321735&offsets|0|-2];[MIMAT0000510&hsa-miR-320a-3p&offsets|0|-4;m-60&8|-|22244975|22244996&offsets|0|-4];[MIMAT0006764_1&hsa-miR-320d&offsets|0|-1;m-253&X|-|140926173|140926190&offsets|0|0]|AAAAGCTGGGTTGAGAGG</td><td>>am_MI0003776|hsa-mir-320b-1&WithFlank&1|+|116671740|116671823</td><td>48</td><td>65</td><td>18</td><td>AAAAGCTGGGTTGAGAGG</td><td>368.0</td><td>2.4</td><td>0.6</td><td>4.3</td><td>1.4E-05</td><td>1.4E-04</td><td>BT20/MB231</td><td>NUC</td></tr>
<tr><td>>hg38_21_+_8433172_8446622_RNA45SN1_45S_50ntFlanks@11208.11249.42|GCGGCGACTCTGGACGCGAGCCGGGCCCTTCCCGTGGATCGC</td><td>>hg38_21_+_8433172_8446622_RNA45SN1_45S_50ntFlanks</td><td>11208</td><td>11249</td><td>42</td><td>GCGGCGACTCTGGACGCGAGCCGGGCCCTTCCCGTGGATCGC</td><td>271.2</td><td>-5.6</td><td>1.5</td><td>-3.9</td><td>1.1E-04</td><td>1.4E-04</td><td>MB231/MB468</td><td>NUC</td></tr>
<tr><td>>trna111_HisGTG_1_-_147774845_147774916@-1T.33.34,trna118_HisGTG_1_-_145396881_145396952@-1T.33.34,trna16_HisGTG_1_+_146544773_146544844@-1T.33.34,trna1_HisGTG_15_+_45493349_45493420@-1T.33.34,trna21_HisGTG_1_+_147753471_147753542@-1T.33.34,trna33_HisGTG_6_+_27125906_27125977@-1T.33.34,trna7_HisGTG_9_-_14433938_14434009@-1T.33.34,trna8_HisGTG_15_-_45492611_45492682@-1T.33.34,trna9_HisGTG_15_-_45490804_45490875@-1T.33.34|TGCCGTGATCGTATAGTGGTTAGTACTCTGCGTT</td><td>>trna111_HisGTG_1_-_147774845_147774916</td><td>-1T</td><td>33</td><td>34</td><td>TGCCGTGATCGTATAGTGGTTAGTACTCTGCGTT</td><td>2102.0</td><td>7.8</td><td>1.9</td><td>4.1</td><td>3.8E-05</td><td>1.4E-04</td><td>BT20/MB468</td><td>MP</td></tr>
<tr><td>>hg38_21_+_8433172_8446622_RNA45SN1_45S_50ntFlanks@12415.12453.39|CTTCGATGTCGGCTCTTCCTATCATTGTGAAGCAGAATT</td><td>>hg38_21_+_8433172_8446622_RNA45SN1_45S_50ntFlanks</td><td>12415</td><td>12453</td><td>39</td><td>CTTCGATGTCGGCTCTTCCTATCATTGTGAAGCAGAATT</td><td>192.3</td><td>4.1</td><td>1.1</td><td>3.9</td><td>9.9E-05</td><td>1.4E-04</td><td>MB231/MB468</td><td>CYTO</td></tr>
<tr><td>>hg38_21_+_8433172_8446622_RNA45SN1_45S_50ntFlanks@5497.5526.30|ACGGTCGAACTTGACTATCTAGAGGAAGTA</td><td>>hg38_21_+_8433172_8446622_RNA45SN1_45S_50ntFlanks</td><td>5497</td><td>5526</td><td>30</td><td>ACGGTCGAACTTGACTATCTAGAGGAAGTA</td><td>212.6</td><td>-2.5</td><td>0.6</td><td>-4.0</td><td>7.7E-05</td><td>1.4E-04</td><td>BT20/MB468</td><td>MITO</td></tr>
<tr><td>>hg38_21_+_8433172_8446622_RNA45SN1_45S_50ntFlanks@7974.8002.29|ACGCGACCTCAGATCAGACGTGGCGACCC</td><td>>hg38_21_+_8433172_8446622_RNA45SN1_45S_50ntFlanks</td><td>7974</td><td>8002</td><td>29</td><td>ACGCGACCTCAGATCAGACGTGGCGACCC</td><td>1679.7</td><td>3.8</td><td>0.9</td><td>4.1</td><td>3.8E-05</td><td>1.4E-04</td><td>BT20/MB468</td><td>MP</td></tr>
<tr><td>>hg38_21_+_8433172_8446622_RNA45SN1_45S_50ntFlanks@4372.4389.18|AAAAAGCTCGTAGTTGGA</td><td>>hg38_21_+_8433172_8446622_RNA45SN1_45S_50ntFlanks</td><td>4372</td><td>4389</td><td>18</td><td>AAAAAGCTCGTAGTTGGA</td><td>184.0</td><td>-6.1</td><td>1.6</td><td>-3.9</td><td>8.8E-05</td><td>1.4E-04</td><td>MB231/MB468</td><td>MP</td></tr>
<tr><td>>MI0000108|hsa-mir-103a-2&WithFlank&20|+|3917488|3917577@54.75.22(+1U);MI0000109|hsa-mir-103a-1&WithFlank&5|-|168560890|168560979@54.75.22(+1U)&[MIMAT0000101&hsa-miR-103a-3p&offsets|0|-1(+1U);m-12&20|+|3917541|3917563&offsets|0|-1(+1U)];[MIMAT0000101_1&hsa-miR-103a-3p&offsets|0|-1(+1U);m-13&5|-|168560904|168560926&offsets|0|-1(+1U);m-44220&5|-|168560908|168560926&offsets|0|+3(+1U)]|AGCAGCATTGTACAGGGCTATGT</td><td>>MI0000108|hsa-mir-103a-2&WithFlank&20|+|3917488|3917577</td><td>54</td><td>75</td><td>22(+1U)</td><td>AGCAGCATTGTACAGGGCTATGT</td><td>3302.4</td><td>-4.3</td><td>1.0</td><td>-4.4</td><td>1.3E-05</td><td>1.4E-04</td><td>BT20/MB231</td><td>MP</td></tr>
<tr><td>>hg38_21_+_8433172_8446622_RNA45SN1_45S_50ntFlanks@6661.6688.28|CGGTGGATCACTCGGCTCGTGCGTCGAT</td><td>>hg38_21_+_8433172_8446622_RNA45SN1_45S_50ntFlanks</td><td>6661</td><td>6688</td><td>28</td><td>CGGTGGATCACTCGGCTCGTGCGTCGAT</td><td>5376.7</td><td>5.4</td><td>1.2</td><td>4.4</td><td>1.3E-05</td><td>1.4E-04</td><td>BT20/MB231</td><td>MP</td></tr>
<tr><td>>hg38_21_+_8433172_8446622_RNA45SN1_45S_50ntFlanks@4243.4263.21|CTTTCGAGGCCCTGTAATTGG</td><td>>hg38_21_+_8433172_8446622_RNA45SN1_45S_50ntFlanks</td><td>4243</td><td>4263</td><td>21</td><td>CTTTCGAGGCCCTGTAATTGG</td><td>311.4</td><td>-2.9</td><td>0.7</td><td>-4.0</td><td>7.7E-05</td><td>1.4E-04</td><td>BT20/MB468</td><td>MITO</td></tr>
<tr><td>>hg38_21_+_8433172_8446622_RNA45SN1_45S_50ntFlanks@11755.11777.23|AATGCCTCGTCATCTAATTAGTG</td><td>>hg38_21_+_8433172_8446622_RNA45SN1_45S_50ntFlanks</td><td>11755</td><td>11777</td><td>23</td><td>AATGCCTCGTCATCTAATTAGTG</td><td>156.1</td><td>-5.2</td><td>1.3</td><td>-4.0</td><td>7.7E-05</td><td>1.4E-04</td><td>BT20/MB468</td><td>MITO</td></tr>
<tr><td>>hg38_21_+_8433172_8446622_RNA45SN1_45S_50ntFlanks@12636.12661.26|AGCTACCATCTGTGGGATTATGACTG</td><td>>hg38_21_+_8433172_8446622_RNA45SN1_45S_50ntFlanks</td><td>12636</td><td>12661</td><td>26</td><td>AGCTACCATCTGTGGGATTATGACTG</td><td>298.3</td><td>-2.6</td><td>0.7</td><td>-3.9</td><td>9.9E-05</td><td>1.4E-04</td><td>MB231/MB468</td><td>CYTO</td></tr>
<tr><td>>hg38_21_+_8433172_8446622_RNA45SN1_45S_50ntFlanks@4559.4587.29|GCTAGGAATAATGGAATAGGACCGCGGTT</td><td>>hg38_21_+_8433172_8446622_RNA45SN1_45S_50ntFlanks</td><td>4559</td><td>4587</td><td>29</td><td>GCTAGGAATAATGGAATAGGACCGCGGTT</td><td>130.3</td><td>-3.2</td><td>0.8</td><td>-3.9</td><td>9.9E-05</td><td>1.4E-04</td><td>MB231/MB468</td><td>CYTO</td></tr>
<tr><td>>hg38_21_+_8433172_8446622_RNA45SN1_45S_50ntFlanks@9574.9600.27|AATCGGTCGTCCGACCTGGGTATAGGG</td><td>>hg38_21_+_8433172_8446622_RNA45SN1_45S_50ntFlanks</td><td>9574</td><td>9600</td><td>27</td><td>AATCGGTCGTCCGACCTGGGTATAGGG</td><td>134.6</td><td>-2.8</td><td>0.7</td><td>-3.9</td><td>8.8E-05</td><td>1.4E-04</td><td>MB231/MB468</td><td>MP</td></tr>
<tr><td>>hg38_21_+_8433172_8446622_RNA45SN1_45S_50ntFlanks@10677.10709.33|GCGGTTCCGGCGGCGTCCGGTGAGCTCTCGCTG</td><td>>hg38_21_+_8433172_8446622_RNA45SN1_45S_50ntFlanks</td><td>10677</td><td>10709</td><td>33</td><td>GCGGTTCCGGCGGCGTCCGGTGAGCTCTCGCTG</td><td>370.8</td><td>-2.0</td><td>0.5</td><td>-3.9</td><td>9.9E-05</td><td>1.4E-04</td><td>MB231/MB468</td><td>CYTO</td></tr>
<tr><td>>hg38_MT_+_598_1651_RNR1_12S_50ntFlanks@417.456.40|AATAGACTACGAAAGTGGCTTTAACATATCTGAACACACA</td><td>>hg38_MT_+_598_1651_RNR1_12S_50ntFlanks</td><td>417</td><td>456</td><td>40</td><td>AATAGACTACGAAAGTGGCTTTAACATATCTGAACACACA</td><td>24.0</td><td>7.8</td><td>2.0</td><td>3.9</td><td>8.9E-05</td><td>1.4E-04</td><td>MB231/MB468</td><td>MP</td></tr>
<tr><td>>hg38_21_+_8433172_8446622_RNA45SN1_45S_50ntFlanks@10786.10815.30|AACAGCCTCTGGCATGTTGGAACAATGTAG</td><td>>hg38_21_+_8433172_8446622_RNA45SN1_45S_50ntFlanks</td><td>10786</td><td>10815</td><td>30</td><td>AACAGCCTCTGGCATGTTGGAACAATGTAG</td><td>419.3</td><td>-4.7</td><td>1.2</td><td>-3.9</td><td>9.3E-05</td><td>1.4E-04</td><td>BT20/MB468</td><td>CYTO</td></tr>
<tr><td>>hg38_21_+_8433172_8446622_RNA45SN1_45S_50ntFlanks@8155.8181.27|CGCTCGTGGGGGGCCCAAGTCCTTCTG</td><td>>hg38_21_+_8433172_8446622_RNA45SN1_45S_50ntFlanks</td><td>8155</td><td>8181</td><td>27</td><td>CGCTCGTGGGGGGCCCAAGTCCTTCTG</td><td>151.1</td><td>-3.2</td><td>0.8</td><td>-3.9</td><td>9.9E-05</td><td>1.4E-04</td><td>MB231/MB468</td><td>CYTO</td></tr>
<tr><td>>MI0001445|hsa-mir-423&WithFlank&17|+|30117073|30117178@59.77.19(+1U)&[MIMAT0001340&hsa-miR-423-3p&offsets|0|-4(+1U);m-100&17|+|30117131|30117153&offsets|0|-4(+1U)]|AGCTCGGTCTGAGGCCCCTT</td><td>>MI0001445|hsa-mir-423&WithFlank&17|+|30117073|30117178</td><td>59</td><td>77</td><td>19(+1U)</td><td>AGCTCGGTCTGAGGCCCCTT</td><td>398.6</td><td>6.7</td><td>1.6</td><td>4.1</td><td>3.8E-05</td><td>1.4E-04</td><td>BT20/MB468</td><td>MP</td></tr>
<tr><td>>hg38_21_+_8433172_8446622_RNA45SN1_45S_50ntFlanks@12420.12456.37|ATGTCGGCTCTTCCTATCATTGTGAAGCAGAATTCAC</td><td>>hg38_21_+_8433172_8446622_RNA45SN1_45S_50ntFlanks</td><td>12420</td><td>12456</td><td>37</td><td>ATGTCGGCTCTTCCTATCATTGTGAAGCAGAATTCAC</td><td>2207.3</td><td>-2.7</td><td>0.7</td><td>-3.9</td><td>1.1E-04</td><td>1.4E-04</td><td>MB231/MB468</td><td>NUC</td></tr>
<tr><td>>MI0000063|hsa-let-7b&WithFlank&22|+|46113680|46113774@12.32.21&[MIMAT0000063&hsa-let-7b-5p&offsets|0|-1;m-14&22|+|46113691|46113712&offsets|0|-1]|TGAGGTAGTAGGTTGTGTGGT</td><td>>MI0000063|hsa-let-7b&WithFlank&22|+|46113680|46113774</td><td>12</td><td>32</td><td>21</td><td>TGAGGTAGTAGGTTGTGTGGT</td><td>3888.3</td><td>3.0</td><td>0.8</td><td>3.9</td><td>8.9E-05</td><td>1.4E-04</td><td>MB231/MB468</td><td>MP</td></tr>
<tr><td>>hg38_21_+_8433172_8446622_RNA45SN1_45S_50ntFlanks@6731.6771.41|CACATTGATCATCGACACTTCGAACGCACTTGCGGCCCCGG</td><td>>hg38_21_+_8433172_8446622_RNA45SN1_45S_50ntFlanks</td><td>6731</td><td>6771</td><td>41</td><td>CACATTGATCATCGACACTTCGAACGCACTTGCGGCCCCGG</td><td>295.1</td><td>-5.4</td><td>1.4</td><td>-3.9</td><td>9.3E-05</td><td>1.4E-04</td><td>BT20/MB468</td><td>CYTO</td></tr>
<tr><td>>hg38_21_+_8433172_8446622_RNA45SN1_45S_50ntFlanks@8184.8205.22|CGAGGCCCAGCCCGTGGACGGT</td><td>>hg38_21_+_8433172_8446622_RNA45SN1_45S_50ntFlanks</td><td>8184</td><td>8205</td><td>22</td><td>CGAGGCCCAGCCCGTGGACGGT</td><td>127.5</td><td>-4.5</td><td>1.1</td><td>-3.9</td><td>8.9E-05</td><td>1.4E-04</td><td>MB231/MB468</td><td>MP</td></tr>
<tr><td>>hg38_1_-_228634819_228635039_RNA5S12_5S_50ntFlanks@133.171.39|ATGGGAGACCGCCTGGGAATACCGGGTGCTGTAGGCTTT</td><td>>hg38_1_-_228634819_228635039_RNA5S12_5S_50ntFlanks</td><td>133</td><td>171</td><td>39</td><td>ATGGGAGACCGCCTGGGAATACCGGGTGCTGTAGGCTTT</td><td>8735.1</td><td>4.4</td><td>1.1</td><td>3.9</td><td>9.3E-05</td><td>1.4E-04</td><td>BT20/MB468</td><td>CYTO</td></tr>
<tr><td>>hg38_21_+_8433172_8446622_RNA45SN1_45S_50ntFlanks@8342.8365.24|CGTAAGGGAAAGTTGAAAAGAACT</td><td>>hg38_21_+_8433172_8446622_RNA45SN1_45S_50ntFlanks</td><td>8342</td><td>8365</td><td>24</td><td>CGTAAGGGAAAGTTGAAAAGAACT</td><td>1694.7</td><td>5.7</td><td>1.5</td><td>3.9</td><td>9.3E-05</td><td>1.4E-04</td><td>BT20/MB468</td><td>CYTO</td></tr>
<tr><td>>hg38_21_+_8433172_8446622_RNA45SN1_45S_50ntFlanks@3782.3805.24|CAGTGAAACTGCGAATGGCTCATT</td><td>>hg38_21_+_8433172_8446622_RNA45SN1_45S_50ntFlanks</td><td>3782</td><td>3805</td><td>24</td><td>CAGTGAAACTGCGAATGGCTCATT</td><td>346.0</td><td>-2.5</td><td>0.6</td><td>-3.9</td><td>9.9E-05</td><td>1.4E-04</td><td>MB231/MB468</td><td>CYTO</td></tr>
<tr><td>>hg38_21_+_8433172_8446622_RNA45SN1_45S_50ntFlanks@10346.10382.37|TCAAACGAGAACTTTGAAGGCCGAAGTGGAGAAGGGT</td><td>>hg38_21_+_8433172_8446622_RNA45SN1_45S_50ntFlanks</td><td>10346</td><td>10382</td><td>37</td><td>TCAAACGAGAACTTTGAAGGCCGAAGTGGAGAAGGGT</td><td>537.4</td><td>3.3</td><td>0.9</td><td>3.9</td><td>1.0E-04</td><td>1.4E-04</td><td>MB231/MB468</td><td>CYTO</td></tr>
<tr><td>>hg38_21_+_8433172_8446622_RNA45SN1_45S_50ntFlanks@6670.6698.29|ACTCGGCTCGTGCGTCGATGAAGAACGCA</td><td>>hg38_21_+_8433172_8446622_RNA45SN1_45S_50ntFlanks</td><td>6670</td><td>6698</td><td>29</td><td>ACTCGGCTCGTGCGTCGATGAAGAACGCA</td><td>1253.8</td><td>-3.1</td><td>0.8</td><td>-4.0</td><td>7.8E-05</td><td>1.4E-04</td><td>BT20/MB468</td><td>MITO</td></tr>
<tr><td>>hg38_21_+_8433172_8446622_RNA45SN1_45S_50ntFlanks@8912.8936.25|CGCCGAATCCCGGGGCCGAGGGAGC</td><td>>hg38_21_+_8433172_8446622_RNA45SN1_45S_50ntFlanks</td><td>8912</td><td>8936</td><td>25</td><td>CGCCGAATCCCGGGGCCGAGGGAGC</td><td>947.0</td><td>9.7</td><td>2.5</td><td>3.9</td><td>1.0E-04</td><td>1.4E-04</td><td>MB231/MB468</td><td>CYTO</td></tr>
<tr><td>>hg38_21_+_8433172_8446622_RNA45SN1_45S_50ntFlanks@9475.9506.32|CGAGCGCACGTGTTAGGACCCGAAAGATGGTG</td><td>>hg38_21_+_8433172_8446622_RNA45SN1_45S_50ntFlanks</td><td>9475</td><td>9506</td><td>32</td><td>CGAGCGCACGTGTTAGGACCCGAAAGATGGTG</td><td>73.1</td><td>9.5</td><td>2.2</td><td>4.4</td><td>1.3E-05</td><td>1.4E-04</td><td>BT20/MB231</td><td>MP</td></tr>
<tr><td>>hg38_21_+_8433172_8446622_RNA45SN1_45S_50ntFlanks@10389.10409.21|TGAACAGCAGTTGAACATGGG</td><td>>hg38_21_+_8433172_8446622_RNA45SN1_45S_50ntFlanks</td><td>10389</td><td>10409</td><td>21</td><td>TGAACAGCAGTTGAACATGGG</td><td>403.5</td><td>-6.4</td><td>1.5</td><td>-4.1</td><td>3.8E-05</td><td>1.4E-04</td><td>BT20/MB468</td><td>MP</td></tr>
<tr><td>>trnaMT_PheGAA_MT_+_577_647@53.74.22|TCACATCACCCCATAAACACCA</td><td>>trnaMT_PheGAA_MT_+_577_647</td><td>53</td><td>74</td><td>22</td><td>TCACATCACCCCATAAACACCA</td><td>535.4</td><td>4.8</td><td>1.2</td><td>3.9</td><td>1.0E-04</td><td>1.4E-04</td><td>MB231/MB468</td><td>CYTO</td></tr>
<tr><td>>hg38_21_+_8433172_8446622_RNA45SN1_45S_50ntFlanks@10846.10868.23|CTTCGGGATAAGGATTGGCTCTA</td><td>>hg38_21_+_8433172_8446622_RNA45SN1_45S_50ntFlanks</td><td>10846</td><td>10868</td><td>23</td><td>CTTCGGGATAAGGATTGGCTCTA</td><td>182.2</td><td>-2.3</td><td>0.6</td><td>-4.0</td><td>7.8E-05</td><td>1.4E-04</td><td>BT20/MB468</td><td>MITO</td></tr>
<tr><td>>hg38_21_+_8433172_8446622_RNA45SN1_45S_50ntFlanks@10786.10821.36|AACAGCCTCTGGCATGTTGGAACAATGTAGGTAAGG</td><td>>hg38_21_+_8433172_8446622_RNA45SN1_45S_50ntFlanks</td><td>10786</td><td>10821</td><td>36</td><td>AACAGCCTCTGGCATGTTGGAACAATGTAGGTAAGG</td><td>228.2</td><td>-2.0</td><td>0.5</td><td>-3.9</td><td>1.1E-04</td><td>1.4E-04</td><td>MB231/MB468</td><td>NUC</td></tr>
<tr><td>>hg38_21_+_8433172_8446622_RNA45SN1_45S_50ntFlanks@8342.8364.23|CGTAAGGGAAAGTTGAAAAGAAC</td><td>>hg38_21_+_8433172_8446622_RNA45SN1_45S_50ntFlanks</td><td>8342</td><td>8364</td><td>23</td><td>CGTAAGGGAAAGTTGAAAAGAAC</td><td>136.1</td><td>5.7</td><td>1.5</td><td>3.9</td><td>1.1E-04</td><td>1.4E-04</td><td>MB231/MB468</td><td>NUC</td></tr>
<tr><td>>hg38_21_+_8433172_8446622_RNA45SN1_45S_50ntFlanks@3988.4024.37|CTTTGGTGACTCTAGATAACCTCGGGCCGATCGCACG</td><td>>hg38_21_+_8433172_8446622_RNA45SN1_45S_50ntFlanks</td><td>3988</td><td>4024</td><td>37</td><td>CTTTGGTGACTCTAGATAACCTCGGGCCGATCGCACG</td><td>1334.3</td><td>-5.4</td><td>1.4</td><td>-3.9</td><td>1.1E-04</td><td>1.4E-04</td><td>BT20/MB468</td><td>NUC</td></tr>
</table>

<table>
<tr><th></th><th></th><th></th><th></th><th></th><th></th><th></th><th></th><th></th><th></th><th></th><th></th><th></th><th></th></tr>
<tr><td>>hg38_21_+_8433172_8446622_RNA45SN1_45S_50ntFlanks@6659.6689.31|AGCGGTGGATCACTCGGCTCGTGCGTCGATG</td><td>>hg38_21_+_8433172_8446622_RNA45SN1_45S_50ntFlanks</td><td>6659</td><td>6689</td><td>31</td><td>AGCGGTGGATCACTCGGCTCGTGCGTCGATG</td><td>277.9</td><td>-2.3</td><td>0.6</td><td>-3.9</td><td>1.1E-04</td><td>1.4E-04</td><td>MB231/MB468</td><td>NUC</td></tr>
<tr><td>>hg38_21_+_8433172_8446622_RNA45SN1_45S_50ntFlanks@11630.11669.40|GCGAAGGCCCGCGGCGGGTGTTGACGCGATGTGATTTCTG</td><td>>hg38_21_+_8433172_8446622_RNA45SN1_45S_50ntFlanks</td><td>11630</td><td>11669</td><td>40</td><td>GCGAAGGCCCGCGGCGGGTGTTGACGCGATGTGATTTCTG</td><td>46.8</td><td>-9.4</td><td>2.4</td><td>-3.9</td><td>9.0E-05</td><td>1.4E-04</td><td>MB231/MB468</td><td>MP</td></tr>
<tr><td>>am_trna10_ValCAC_5_-_180649395_180649467@2.32.31,trna12_ValAAC_5_-_180645270_180645342@2.32.31,trna132_ValAAC_6_-_27721179_27721251@2.32.31,trna136_ValAAC_6_-_27648885_27648957@2.32.31,trna139_ValAAC_6_-_27618707_27618779@2.32.31,trna18_ValCAC_5_-_180529253_180529325@2.32.31,trna2_ValAAC_3_+_169490018_169490090@2.32.31,trna2_ValCAC_5_+_180524070_180524142@2.32.31,trna4_ValAAC_5_+_180591154_180591226@2.32.31,trna5_ValAAC_5_+_180596610_180596682@2.32.31,trna6_ValCAC_5_+_180600650_180600722@2.32.31,trna85_ValCAC_1_-_161369490_161369562@2.32.31,trna90_ValCAC_1_-_149684088_149684161@2.32.31,trna98_ValCAC_1_-_149298555_149298627@2.32.31,trna9_ValCAC_6_+_26538282_26538354@2.32.31|TTTCCGTAGTGTAGTGGTTATCACGTTCGCC</td><td>>am_trna10_ValCAC_5_-_180649395_180649467</td><td>2</td><td>32</td><td>31</td><td>TTTCCGTAGTGTAGTGGTTATCACGTTCGCC</td><td>313.4</td><td>-4.6</td><td>1.1</td><td>-4.3</td><td>1.5E-05</td><td>1.4E-04</td><td>BT20/MB231</td><td>MITO</td></tr>
<tr><td>>hg38_21_+_8433172_8446622_RNA45SN1_45S_50ntFlanks@12365.12382.18|ATAACTGGCTTGTGGCGG</td><td>>hg38_21_+_8433172_8446622_RNA45SN1_45S_50ntFlanks</td><td>12365</td><td>12382</td><td>18</td><td>ATAACTGGCTTGTGGCGG</td><td>550.2</td><td>-6.5</td><td>1.6</td><td>-4.1</td><td>3.8E-05</td><td>1.4E-04</td><td>BT20/MB468</td><td>MP</td></tr>
<tr><td>>hg38_21_+_8433172_8446622_RNA45SN1_45S_50ntFlanks@5495.5516.22|AGACGGTCGAACTTGACTATCT</td><td>>hg38_21_+_8433172_8446622_RNA45SN1_45S_50ntFlanks</td><td>5495</td><td>5516</td><td>22</td><td>AGACGGTCGAACTTGACTATCT</td><td>343.3</td><td>7.5</td><td>1.8</td><td>4.1</td><td>3.8E-05</td><td>1.4E-04</td><td>BT20/MB468</td><td>MP</td></tr>
<tr><td>>hg38_21_+_8433172_8446622_RNA45SN1_45S_50ntFlanks@11772.11794.23|TTAGTGACGCGCATGAATGGATG</td><td>>hg38_21_+_8433172_8446622_RNA45SN1_45S_50ntFlanks</td><td>11772</td><td>11794</td><td>23</td><td>TTAGTGACGCGCATGAATGGATG</td><td>1489.1</td><td>-7.3</td><td>1.9</td><td>-3.9</td><td>1.1E-04</td><td>1.4E-04</td><td>BT20/MB468</td><td>NUC</td></tr>
<tr><td>>hg38_MT_+_1621_3279_RNR2_16S_50ntFlanks@51.74.24|GCTAAACCTAGCCCCAAACCCACT</td><td>>hg38_MT_+_1621_3279_RNR2_16S_50ntFlanks</td><td>51</td><td>74</td><td>24</td><td>GCTAAACCTAGCCCCAAACCCACT</td><td>9470.3</td><td>3.7</td><td>1.0</td><td>3.9</td><td>1.1E-04</td><td>1.4E-04</td><td>MB231/MB468</td><td>MITO</td></tr>
<tr><td>>hg38_21_+_8433172_8446622_RNA45SN1_45S_50ntFlanks@9567.9585.19|ACGTGCAAATCGGTCGTCC</td><td>>hg38_21_+_8433172_8446622_RNA45SN1_45S_50ntFlanks</td><td>9567</td><td>9585</td><td>19</td><td>ACGTGCAAATCGGTCGTCC</td><td>325.8</td><td>-5.9</td><td>1.5</td><td>-3.9</td><td>9.4E-05</td><td>1.4E-04</td><td>BT20/MB468</td><td>CYTO</td></tr>
<tr><td>>hg38_21_+_8433172_8446622_RNA45SN1_45S_50ntFlanks@10787.10818.32|ACAGCCTCTGGCATGTTGGAACAATGTAGGTA</td><td>>hg38_21_+_8433172_8446622_RNA45SN1_45S_50ntFlanks</td><td>10787</td><td>10818</td><td>32</td><td>ACAGCCTCTGGCATGTTGGAACAATGTAGGTA</td><td>1203.8</td><td>-2.5</td><td>0.6</td><td>-3.9</td><td>7.9E-05</td><td>1.4E-04</td><td>BT20/MB468</td><td>MITO</td></tr>
<tr><td>>MI0000089|hsa-mir-31&WithFlank&9|-|21512109|21512191@14.34.21&[MIMAT0000089&hsa-miR-31-5p&offsets|0|0;m-360&9|-|21512157|21512178&offsets|0|-1]|AGGCAAGATGCTGGCATAGCT</td><td>>MI0000089|hsa-mir-31&WithFlank&9|-|21512109|21512191</td><td>14</td><td>34</td><td>21</td><td>AGGCAAGATGCTGGCATAGCT</td><td>58.5</td><td>9.2</td><td>2.1</td><td>4.4</td><td>1.3E-05</td><td>1.4E-04</td><td>BT20/MB231</td><td>MP</td></tr>
<tr><td>>hg38_21_+_8433172_8446622_RNA45SN1_45S_50ntFlanks@11565.11588.24|CTAGCAGCCGACTTAGAACTGGTG</td><td>>hg38_21_+_8433172_8446622_RNA45SN1_45S_50ntFlanks</td><td>11565</td><td>11588</td><td>24</td><td>CTAGCAGCCGACTTAGAACTGGTG</td><td>84.2</td><td>9.7</td><td>2.2</td><td>4.4</td><td>1.3E-05</td><td>1.4E-04</td><td>BT20/MB231</td><td>MP</td></tr>
<tr><td>>MI0000298|hsa-mir-221&WithFlank&X|-|45746151|45746272@71.91.21&[MIMAT0000278&hsa-miR-221-3p&offsets|0|-2;m-91&X|-|45746181|45746202&offsets|0|-1]|AGCTACATTGTCTGCTGGGTT</td><td>>MI0000298|hsa-mir-221&WithFlank&X|-|45746151|45746272</td><td>71</td><td>91</td><td>21</td><td>AGCTACATTGTCTGCTGGGTT</td><td>199.5</td><td>2.1</td><td>0.5</td><td>3.9</td><td>1.1E-04</td><td>1.4E-04</td><td>BT20/MB468</td><td>NUC</td></tr>
<tr><td>>hg38_21_+_8433172_8446622_RNA45SN1_45S_50ntFlanks@3850.3872.23|GATAACTGTGGTAATTCTAGAGC</td><td>>hg38_21_+_8433172_8446622_RNA45SN1_45S_50ntFlanks</td><td>3850</td><td>3872</td><td>23</td><td>GATAACTGTGGTAATTCTAGAGC</td><td>399.1</td><td>-5.0</td><td>1.3</td><td>-3.9</td><td>1.1E-04</td><td>1.4E-04</td><td>MB231/MB468</td><td>MITO</td></tr>
<tr><td>>hg38_21_+_8433172_8446622_RNA45SN1_45S_50ntFlanks@12476.12493.18|CCACTAATAGGGAACGTG</td><td>>hg38_21_+_8433172_8446622_RNA45SN1_45S_50ntFlanks</td><td>12476</td><td>12493</td><td>18</td><td>CCACTAATAGGGAACGTG</td><td>364.2</td><td>-6.3</td><td>1.5</td><td>-4.1</td><td>3.8E-05</td><td>1.4E-04</td><td>BT20/MB468</td><td>MP</td></tr>
<tr><td>>hg38_21_+_8433172_8446622_RNA45SN1_45S_50ntFlanks@6735.6772.38|TTGATCATCGACACTTCGAACGCACTTGCGGCCCCGGG</td><td>>hg38_21_+_8433172_8446622_RNA45SN1_45S_50ntFlanks</td><td>6735</td><td>6772</td><td>38</td><td>TTGATCATCGACACTTCGAACGCACTTGCGGCCCCGGG</td><td>56.1</td><td>-9.7</td><td>2.5</td><td>-3.9</td><td>9.0E-05</td><td>1.4E-04</td><td>MB231/MB468</td><td>MP</td></tr>
<tr><td>>hg38_21_+_8433172_8446622_RNA45SN1_45S_50ntFlanks@485.518.34|GATGTGGTGACGTCGTGCTCTCCCGGGCCGGGTC</td><td>>hg38_21_+_8433172_8446622_RNA45SN1_45S_50ntFlanks</td><td>485</td><td>518</td><td>34</td><td>GATGTGGTGACGTCGTGCTCTCCCGGGCCGGGTC</td><td>103.4</td><td>9.9</td><td>2.5</td><td>3.9</td><td>9.0E-05</td><td>1.4E-04</td><td>MB231/MB468</td><td>MP</td></tr>
<tr><td>>hg38_21_+_8433172_8446622_RNA45SN1_45S_50ntFlanks@9940.9968.29|CAGGACGGTGGCCATGGAAGTCGGAATCC</td><td>>hg38_21_+_8433172_8446622_RNA45SN1_45S_50ntFlanks</td><td>9940</td><td>9968</td><td>29</td><td>CAGGACGGTGGCCATGGAAGTCGGAATCC</td><td>624.4</td><td>3.7</td><td>0.9</td><td>4.1</td><td>3.8E-05</td><td>1.4E-04</td><td>BT20/MB468</td><td>MP</td></tr>
<tr><td>>MI0000082|hsa-mir-25&WithFlank&7|-|100093554|100093649@58.79.22(+1U)&[MIMAT0000081&hsa-miR-25-3p&offsets|0|0(+1U);m-23&7|-|100093571|100093592&offsets|0|0(+1U)]|CATTGCACTTGTCTCGGTCTGAT</td><td>>MI0000082|hsa-mir-25&WithFlank&7|-|100093554|100093649</td><td>58</td><td>79</td><td>22(+1U)</td><td>CATTGCACTTGTCTCGGTCTGAT</td><td>424.2</td><td>3.4</td><td>0.9</td><td>3.9</td><td>9.0E-05</td><td>1.4E-04</td><td>MB231/MB468</td><td>MP</td></tr>
<tr><td>>MI0000088|hsa-mir-30a&WithFlank&6|-|71403545|71403627@12.31.20&[MIMAT0000087&hsa-miR-30a-5p&offsets|0|-2;m-15&6|-|71403595|71403616&offsets|0|-2]|TGTAAACATCCTCGACTGGA</td><td>>MI0000088|hsa-mir-30a&WithFlank&6|-|71403545|71403627</td><td>12</td><td>31</td><td>20</td><td>TGTAAACATCCTCGACTGGA</td><td>355.3</td><td>3.4</td><td>0.9</td><td>3.9</td><td>1.1E-04</td><td>1.4E-04</td><td>BT20/MB468</td><td>NUC</td></tr>
<tr><td>>hg38_21_+_8433172_8446622_RNA45SN1_45S_50ntFlanks@11583.11606.24|CTGGTGCGGACCAGGGGAATCCGA</td><td>>hg38_21_+_8433172_8446622_RNA45SN1_45S_50ntFlanks</td><td>11583</td><td>11606</td><td>24</td><td>CTGGTGCGGACCAGGGGAATCCGA</td><td>148.3</td><td>-5.8</td><td>1.5</td><td>-3.9</td><td>9.0E-05</td><td>1.4E-04</td><td>MB231/MB468</td><td>MP</td></tr>
<tr><td>>MI0000746|hsa-mir-99b&WithFlank&19|+|51692606|51692687@51.70.20(+1A)&[MIMAT0004678&hsa-miR-99b-3p&offsets|0|-2(+1A);m-151&19|+|51692656|51692678&offsets|0|-3(+1A)]|CAAGCTCGTGTCTGTGGGTCA</td><td>>MI0000746|hsa-mir-99b&WithFlank&19|+|51692606|51692687</td><td>51</td><td>70</td><td>20(+1A)</td><td>CAAGCTCGTGTCTGTGGGTCA</td><td>145.1</td><td>6.6</td><td>1.6</td><td>4.1</td><td>3.9E-05</td><td>1.4E-04</td><td>BT20/MB468</td><td>MP</td></tr>
<tr><td>>hg38_21_+_8433172_8446622_RNA45SN1_45S_50ntFlanks@10401.10420.20|GAACATGGGTCAGTCGGTCC</td><td>>hg38_21_+_8433172_8446622_RNA45SN1_45S_50ntFlanks</td><td>10401</td><td>10420</td><td>20</td><td>GAACATGGGTCAGTCGGTCC</td><td>187.5</td><td>-2.5</td><td>0.6</td><td>-3.9</td><td>1.0E-04</td><td>1.4E-04</td><td>MB231/MB468</td><td>CYTO</td></tr>
<tr><td>>hg38_21_+_8433172_8446622_RNA45SN1_45S_50ntFlanks@10771.10818.48|CAGGTCTCCAAGGTGAACAGCCTCTGGCATGTTGGAACAATGTAGGTA</td><td>>hg38_21_+_8433172_8446622_RNA45SN1_45S_50ntFlanks</td><td>10771</td><td>10818</td><td>48</td><td>CAGGTCTCCAAGGTGAACAGCCTCTGGCATGTTGGAACAATGTAGGTA</td><td>766.5</td><td>-6.6</td><td>1.7</td><td>-3.9</td><td>1.1E-04</td><td>1.4E-04</td><td>MB231/MB468</td><td>NUC</td></tr>
<tr><td>>hg38_21_+_8433172_8446622_RNA45SN1_45S_50ntFlanks@4163.4186.24|CATCCAAGGAAGGCAGCAGGCGCG</td><td>>hg38_21_+_8433172_8446622_RNA45SN1_45S_50ntFlanks</td><td>4163</td><td>4186</td><td>24</td><td>CATCCAAGGAAGGCAGCAGGCGCG</td><td>257.6</td><td>-5.4</td><td>1.4</td><td>-3.9</td><td>7.9E-05</td><td>1.4E-04</td><td>BT20/MB468</td><td>MITO</td></tr>
<tr><td>>hg38_21_+_8433172_8446622_RNA45SN1_45S_50ntFlanks@12081.12108.28|CTCTCGCTTCTGGCGCCAAGCGCCCGGC</td><td>>hg38_21_+_8433172_8446622_RNA45SN1_45S_50ntFlanks</td><td>12081</td><td>12108</td><td>28</td><td>CTCTCGCTTCTGGCGCCAAGCGCCCGGC</td><td>580.2</td><td>-4.3</td><td>1.1</td><td>-3.9</td><td>7.9E-05</td><td>1.4E-04</td><td>BT20/MB468</td><td>MITO</td></tr>
<tr><td>>hg38_21_+_8433172_8446622_RNA45SN1_45S_50ntFlanks@3993.4012.20|GTGACTCTAGATAACCTCGG</td><td>>hg38_21_+_8433172_8446622_RNA45SN1_45S_50ntFlanks</td><td>3993</td><td>4012</td><td>20</td><td>GTGACTCTAGATAACCTCGG</td><td>104.4</td><td>-4.5</td><td>1.1</td><td>-3.9</td><td>9.0E-05</td><td>1.4E-04</td><td>MB231/MB468</td><td>MP</td></tr>
<tr><td>>MI0000446|hsa-mir-125b-1&WithFlank&11|-|122099751|122099850@21.40.20(+1A);MI0000470|hsa-mir-125b-2&WithFlank&21|+|16590231|16590331@23.42.20(+1A)&[MIMAT0000423&hsa-miR-125b-5p&offsets|0|-2(+1A);m-67&11|-|122099809|122099830&offsets|0|-2(+1A)];[MIMAT0000423_1&hsa-miR-125b-5p&offsets|0|-2(+1A);m-68&21|+|16590253|16590274&offsets|0|-2(+1A)]|TCCCTGAGACCCTAACTTGTA</td><td>>MI0000446|hsa-mir-125b-1&WithFlank&11|-|122099751|122099850</td><td>21</td><td>40</td><td>20(+1A)</td><td>TCCCTGAGACCCTAACTTGTA</td><td>854.2</td><td>5.3</td><td>1.3</td><td>3.9</td><td>9.1E-05</td><td>1.4E-04</td><td>MB231/MB468</td><td>MP</td></tr>
<tr><td>>MI0000239|hsa-mir-197&WithFlank&1|+|109598887|109598973@54.74.21(+1U)&[MIMAT0000227&hsa-miR-197-3p&offsets|0|-1(+1U);m-99&1|+|109598940|109598961&offsets|0|-1(+1U)]|TTCACCACCTTCTCCACCCAGT</td><td>>MI0000239|hsa-mir-197&WithFlank&1|+|109598887|109598973</td><td>54</td><td>74</td><td>21(+1U)</td><td>TTCACCACCTTCTCCACCCAGT</td><td>4951.3</td><td>3.6</td><td>0.8</td><td>4.4</td><td>1.3E-05</td><td>1.4E-04</td><td>BT20/MB231</td><td>CYTO</td></tr>
<tr><td>>hg38_21_+_8433172_8446622_RNA45SN1_45S_50ntFlanks@10349.10382.34|AACGAGAACTTTGAAGGCCGAAGTGGAGAAGGGT</td><td>>hg38_21_+_8433172_8446622_RNA45SN1_45S_50ntFlanks</td><td>10349</td><td>10382</td><td>34</td><td>AACGAGAACTTTGAAGGCCGAAGTGGAGAAGGGT</td><td>857.9</td><td>2.9</td><td>0.7</td><td>3.9</td><td>7.9E-05</td><td>1.4E-04</td><td>BT20/MB468</td><td>MITO</td></tr>
<tr><td>>MI0000811|hsa-mir-148b&WithFlank&12|+|54337210|54337320@69.89.21(+1C)&[MIMAT0000759&hsa-miR-148b-3p&offsets|0|-1(+1C);m-77&12|+|54337278|54337299&offsets|0|-1(+1C)]|TCAGTGCATCACAGAACTTTGC</td><td>>MI0000811|hsa-mir-148b&WithFlank&12|+|54337210|54337320</td><td>69</td><td>89</td><td>21(+1C)</td><td>TCAGTGCATCACAGAACTTTGC</td><td>2220.1</td><td>3.2</td><td>0.8</td><td>3.9</td><td>9.1E-05</td><td>1.4E-04</td><td>MB231/MB468</td><td>MP</td></tr>
<tr><td>>hg38_21_+_8433172_8446622_RNA45SN1_45S_50ntFlanks@10473.10494.22|CTCGGCCGATCGAAAGGGAGTC</td><td>>hg38_21_+_8433172_8446622_RNA45SN1_45S_50ntFlanks</td><td>10473</td><td>10494</td><td>22</td><td>CTCGGCCGATCGAAAGGGAGTC</td><td>124.1</td><td>-4.5</td><td>1.2</td><td>-3.9</td><td>9.1E-05</td><td>1.4E-04</td><td>MB231/MB468</td><td>MP</td></tr>
<tr><td>>hg38_21_+_8433172_8446622_RNA45SN1_45S_50ntFlanks@11572.11597.26|CCGACTTAGAACTGGTGCGGACCAGG</td><td>>hg38_21_+_8433172_8446622_RNA45SN1_45S_50ntFlanks</td><td>11572</td><td>11597</td><td>26</td><td>CCGACTTAGAACTGGTGCGGACCAGG</td><td>706.5</td><td>-3.6</td><td>0.9</td><td>-3.9</td><td>1.0E-04</td><td>1.4E-04</td><td>MB231/MB468</td><td>CYTO</td></tr>
<tr><td>>hg38_21_+_8433172_8446622_RNA45SN1_45S_50ntFlanks@4729.4758.30|TCAAGAACGAAAGTCGGAGGTTCGAAGACG</td><td>>hg38_21_+_8433172_8446622_RNA45SN1_45S_50ntFlanks</td><td>4729</td><td>4758</td><td>30</td><td>TCAAGAACGAAAGTCGGAGGTTCGAAGACG</td><td>3168.3</td><td>-3.2</td><td>0.8</td><td>-4.1</td><td>3.9E-05</td><td>1.4E-04</td><td>BT20/MB468</td><td>MP</td></tr>
<tr><td>>hg38_21_+_8433172_8446622_RNA45SN1_45S_50ntFlanks@7992.8018.27|CGTGGCGACCCGCTGAATTTAAGCATA</td><td>>hg38_21_+_8433172_8446622_RNA45SN1_45S_50ntFlanks</td><td>7992</td><td>8018</td><td>27</td><td>CGTGGCGACCCGCTGAATTTAAGCATA</td><td>434.4</td><td>-2.8</td><td>0.7</td><td>-3.9</td><td>1.1E-04</td><td>1.4E-04</td><td>MB231/MB468</td><td>MITO</td></tr>
<tr><td>>hg38_21_+_8433172_8446622_RNA45SN1_45S_50ntFlanks@10405.10424.20|ATGGGTCAGTCGGTCCTGAG</td><td>>hg38_21_+_8433172_8446622_RNA45SN1_45S_50ntFlanks</td><td>10405</td><td>10424</td><td>20</td><td>ATGGGTCAGTCGGTCCTGAG</td><td>11475.2</td><td>3.6</td><td>0.8</td><td>4.3</td><td>1.5E-05</td><td>1.4E-04</td><td>BT20/MB231</td><td>MITO</td></tr>
<tr><td>>hg38_21_+_8433172_8446622_RNA45SN1_45S_50ntFlanks@3843.3873.31|CTACTTGGATAACTGTGGTAATTCTAGAGCT</td><td>>hg38_21_+_8433172_8446622_RNA45SN1_45S_50ntFlanks</td><td>3843</td><td>3873</td><td>31</td><td>CTACTTGGATAACTGTGGTAATTCTAGAGCT</td><td>302.4</td><td>-5.1</td><td>1.3</td><td>-3.9</td><td>1.1E-04</td><td>1.4E-04</td><td>MB231/MB468</td><td>MITO</td></tr>
<tr><td>>hg38_21_+_8433172_8446622_RNA45SN1_45S_50ntFlanks@11534.11553.20|GTTCCGCGCGGCGCCTCGCC</td><td>>hg38_21_+_8433172_8446622_RNA45SN1_45S_50ntFlanks</td><td>11534</td><td>11553</td><td>20</td><td>GTTCCGCGCGGCGCCTCGCC</td><td>219.7</td><td>-6.2</td><td>1.5</td><td>-4.1</td><td>3.9E-05</td><td>1.4E-04</td><td>BT20/MB468</td><td>MP</td></tr>
<tr><td>>hg38_21_+_8433172_8446622_RNA45SN1_45S_50ntFlanks@9570.9603.34|TGCAAATCGGTCGTCCGACCTGGGTATAGGGGCG</td><td>>hg38_21_+_8433172_8446622_RNA45SN1_45S_50ntFlanks</td><td>9570</td><td>9603</td><td>34</td><td>TGCAAATCGGTCGTCCGACCTGGGTATAGGGGCG</td><td>1953.2</td><td>-3.8</td><td>1.0</td><td>-3.9</td><td>1.1E-04</td><td>1.4E-04</td><td>MB231/MB468</td><td>MITO</td></tr>
<tr><td>>MI0000477|hsa-mir-146a&WithFlank&5|+|160485346|160485456@27.45.19(+2U)&[MIMAT0000449&hsa-miR-146a-5p&offsets|0|-3(+2U);m-129&5|+|160485372|160485393&offsets|0|-3(+2U)]|TGAGAACTGAATTCCATGGTT</td><td>>MI0000477|hsa-mir-146a&WithFlank&5|+|160485346|160485456</td><td>27</td><td>45</td><td>19(+2U)</td><td>TGAGAACTGAATTCCATGGTT</td><td>71.8</td><td>-9.6</td><td>2.2</td><td>-4.4</td><td>1.4E-05</td><td>1.4E-04</td><td>BT20/MB231</td><td>MP</td></tr>
<tr><td>>hg38_21_+_8433172_8446622_RNA45SN1_45S_50ntFlanks@7986.8004.19|ATCAGACGTGGCGACCCGC</td><td>>hg38_21_+_8433172_8446622_RNA45SN1_45S_50ntFlanks</td><td>7986</td><td>8004</td><td>19</td><td>ATCAGACGTGGCGACCCGC</td><td>328.6</td><td>-5.6</td><td>1.4</td><td>-3.9</td><td>1.1E-04</td><td>1.4E-04</td><td>MB231/MB468</td><td>NUC</td></tr>
<tr><td>>hg38_21_+_8433172_8446622_RNA45SN1_45S_50ntFlanks@5473.5509.37|CGGCCCTGGCGGAGCGCTGAGAAGACGGTCGAACTTG</td><td>>hg38_21_+_8433172_8446622_RNA45SN1_45S_50ntFlanks</td><td>5473</td><td>5509</td><td>37</td><td>CGGCCCTGGCGGAGCGCTGAGAAGACGGTCGAACTTG</td><td>398.8</td><td>2.9</td><td>0.8</td><td>3.9</td><td>1.0E-04</td><td>1.4E-04</td><td>MB231/MB468</td><td>CYTO</td></tr>
<tr><td>>hg38_21_+_8433172_8446622_RNA45SN1_45S_50ntFlanks@10401.10431.31|GAACATGGGTCAGTCGGTCCTGAGAGATGGG</td><td>>hg38_21_+_8433172_8446622_RNA45SN1_45S_50ntFlanks</td><td>10401</td><td>10431</td><td>31</td><td>GAACATGGGTCAGTCGGTCCTGAGAGATGGG</td><td>611.3</td><td>-7.0</td><td>1.8</td><td>-3.9</td><td>1.1E-04</td><td>1.4E-04</td><td>BT20/MB468</td><td>NUC</td></tr>
<tr><td>>hg38_21_+_8433172_8446622_RNA45SN1_45S_50ntFlanks@11572.11589.18|CCGACTTAGAACTGGTGC</td><td>>hg38_21_+_8433172_8446622_RNA45SN1_45S_50ntFlanks</td><td>11572</td><td>11589</td><td>18</td><td>CCGACTTAGAACTGGTGC</td><td>517.3</td><td>-3.1</td><td>0.7</td><td>-4.1</td><td>3.9E-05</td><td>1.4E-04</td><td>BT20/MB468</td><td>MP</td></tr>
<tr><td>>MI0000098|hsa-mir-96&WithFlank&7|-|129774686|129774775@15.37.23&[MIMAT0000095&hsa-miR-96-5p&offsets|0|0;m-248&7|-|129774739|129774761&offsets|0|0]|TTTGGCACTAGCACATTTTTGCT</td><td>>MI0000098|hsa-mir-96&WithFlank&7|-|129774686|129774775</td><td>15</td><td>37</td><td>23</td><td>TTTGGCACTAGCACATTTTTGCT</td><td>1391.7</td><td>4.0</td><td>1.0</td><td>3.9</td><td>1.1E-04</td><td>1.4E-04</td><td>MB231/MB468</td><td>MITO</td></tr>
</table>

<table>
<tr><td>>hg38_21_+_8433172_8446622_RNA45SN1_45S_50ntFlanks@4220.4243.24|ACGAAAAATAACAATACAGGACTC</td><td>>hg38_21_+_8433172_8446622_RNA45SN1_45S_50ntFlanks</td><td>4220</td><td>4243</td><td>24</td><td>ACGAAAAATAACAATACAGGACTC</td><td>336.2</td><td>-6.2</td><td>1.5</td><td>-4.1</td><td>3.9E-05</td><td>1.4E-04</td><td>BT20/MB468</td><td>MP</td></tr>
<tr><td>>MI0001448|hsa-mir-425&WithFlank&3|-|49020142|49020240@64.81.18&[MIMAT0001343&hsa-miR-425-3p&offsets|+3|-1;m-258&3|-|49020160|49020181&offsets|+4|0]|GGGAATGTCGTGTCCGCC</td><td>>MI0001448|hsa-mir-425&WithFlank&3|-|49020142|49020240</td><td>64</td><td>81</td><td>18</td><td>GGGAATGTCGTGTCCGCC</td><td>103.2</td><td>9.9</td><td>2.5</td><td>3.9</td><td>9.1E-05</td><td>1.4E-04</td><td>MB231/MB468</td><td>MP</td></tr>
<tr><td>>hg38_21_+_8433172_8446622_RNA45SN1_45S_50ntFlanks@12931.12956.26|GCGCTAAACCATTCGTAGACGACCTG</td><td>>hg38_21_+_8433172_8446622_RNA45SN1_45S_50ntFlanks</td><td>12931</td><td>12956</td><td>26</td><td>GCGCTAAACCATTCGTAGACGACCTG</td><td>474.7</td><td>-4.1</td><td>1.0</td><td>-3.9</td><td>9.5E-05</td><td>1.4E-04</td><td>BT20/MB468</td><td>CYTO</td></tr>
<tr><td>>hg38_21_+_8433172_8446622_RNA45SN1_45S_50ntFlanks@12642.12659.18|CATCTGTGGGATTATGAC</td><td>>hg38_21_+_8433172_8446622_RNA45SN1_45S_50ntFlanks</td><td>12642</td><td>12659</td><td>18</td><td>CATCTGTGGGATTATGAC</td><td>206.9</td><td>-4.3</td><td>1.1</td><td>-3.9</td><td>1.1E-04</td><td>1.4E-04</td><td>MB231/MB468</td><td>NUC</td></tr>
<tr><td>>hg38_21_+_8433172_8446622_RNA45SN1_45S_50ntFlanks@11564.11586.23|CCTAGCAGCCGACTTAGAACTGG</td><td>>hg38_21_+_8433172_8446622_RNA45SN1_45S_50ntFlanks</td><td>11564</td><td>11586</td><td>23</td><td>CCTAGCAGCCGACTTAGAACTGG</td><td>1076.5</td><td>-3.7</td><td>0.9</td><td>-4.1</td><td>3.9E-05</td><td>1.4E-04</td><td>BT20/MB468</td><td>MP</td></tr>
<tr><td>>hg38_21_+_8433172_8446622_RNA45SN1_45S_50ntFlanks@5035.5067.33|CATGGCCGTTCTTAGTTGGTGGAGCGATTTGTC</td><td>>hg38_21_+_8433172_8446622_RNA45SN1_45S_50ntFlanks</td><td>5035</td><td>5067</td><td>33</td><td>CATGGCCGTTCTTAGTTGGTGGAGCGATTTGTC</td><td>470.0</td><td>-5.4</td><td>1.4</td><td>-3.9</td><td>8.0E-05</td><td>1.4E-04</td><td>BT20/MB468</td><td>MITO</td></tr>
<tr><td>>hg38_21_+_8433172_8446622_RNA45SN1_45S_50ntFlanks@9567.9594.28|ACGTGCAAATCGGTCGTCCGACCTGGGT</td><td>>hg38_21_+_8433172_8446622_RNA45SN1_45S_50ntFlanks</td><td>9567</td><td>9594</td><td>28</td><td>ACGTGCAAATCGGTCGTCCGACCTGGGT</td><td>353.6</td><td>-4.5</td><td>1.2</td><td>-3.9</td><td>1.1E-04</td><td>1.4E-04</td><td>MB231/MB468</td><td>NUC</td></tr>
<tr><td>>hg38_21_+_8433172_8446622_RNA45SN1_45S_50ntFlanks@12417.12440.24|TCGATGTCGGCTCTTCCTATCATT</td><td>>hg38_21_+_8433172_8446622_RNA45SN1_45S_50ntFlanks</td><td>12417</td><td>12440</td><td>24</td><td>TCGATGTCGGCTCTTCCTATCATT</td><td>423.9</td><td>1.9</td><td>0.5</td><td>3.9</td><td>8.0E-05</td><td>1.4E-04</td><td>BT20/MB468</td><td>MITO</td></tr>
<tr><td>>MI0000298|hsa-mir-221&WithFlank&X|-|45746151|45746272@71.92.22(+1U)&[MIMAT0000278&hsa-miR-221-3p&offsets|0|-1(+1U);m-91&X|-|45746181|45746202&offsets|0|0(+1U)]|AGCTACATTGTCTGCTGGGTTTT</td><td>>MI0000298|hsa-mir-221&WithFlank&X|-|45746151|45746272</td><td>71</td><td>92</td><td>22(+1U)</td><td>AGCTACATTGTCTGCTGGGTTTT</td><td>307.9</td><td>1.9</td><td>0.5</td><td>3.9</td><td>1.1E-04</td><td>1.4E-04</td><td>BT20/MB468</td><td>NUC</td></tr>
<tr><td>>hg38_MT_+_1621_3279_RNR2_16S_50ntFlanks@51.84.34|GCTAAACCTAGCCCCAAACCCACTCCACCTTACT</td><td>>hg38_MT_+_1621_3279_RNR2_16S_50ntFlanks</td><td>51</td><td>84</td><td>34</td><td>GCTAAACCTAGCCCCAAACCCACTCCACCTTACT</td><td>12786.2</td><td>5.1</td><td>1.2</td><td>4.1</td><td>3.9E-05</td><td>1.4E-04</td><td>BT20/MB468</td><td>MP</td></tr>
<tr><td>>hg38_MT_+_1621_3279_RNR2_16S_50ntFlanks@114.145.32|CAAATAAAGTATAGGCGATAGAAATTGAAACC</td><td>>hg38_MT_+_1621_3279_RNR2_16S_50ntFlanks</td><td>114</td><td>145</td><td>32</td><td>CAAATAAAGTATAGGCGATAGAAATTGAAACC</td><td>248.8</td><td>8.1</td><td>2.0</td><td>4.1</td><td>3.9E-05</td><td>1.4E-04</td><td>BT20/MB468</td><td>MP</td></tr>
<tr><td>>hg38_21_+_8433172_8446622_RNA45SN1_45S_50ntFlanks@8382.8403.22|AGAGGGCGTGAAACCGTTAAGA</td><td>>hg38_21_+_8433172_8446622_RNA45SN1_45S_50ntFlanks</td><td>8382</td><td>8403</td><td>22</td><td>AGAGGGCGTGAAACCGTTAAGA</td><td>124.9</td><td>-4.4</td><td>1.1</td><td>-3.9</td><td>9.2E-05</td><td>1.4E-04</td><td>MB231/MB468</td><td>MP</td></tr>
<tr><td>>hg38_21_+_8433172_8446622_RNA45SN1_45S_50ntFlanks@3835.3872.38|CTCCTCTCCTACTTGGATAACTGTGGTAATTCTAGAGC</td><td>>hg38_21_+_8433172_8446622_RNA45SN1_45S_50ntFlanks</td><td>3835</td><td>3872</td><td>38</td><td>CTCCTCTCCTACTTGGATAACTGTGGTAATTCTAGAGC</td><td>210.3</td><td>-2.2</td><td>0.6</td><td>-3.9</td><td>8.0E-05</td><td>1.4E-04</td><td>BT20/MB468</td><td>MITO</td></tr>
<tr><td>>hg38_21_+_8433172_8446622_RNA45SN1_45S_50ntFlanks@11876.11897.22|AAGACCCTGTTGAGCTTGACTC</td><td>>hg38_21_+_8433172_8446622_RNA45SN1_45S_50ntFlanks</td><td>11876</td><td>11897</td><td>22</td><td>AAGACCCTGTTGAGCTTGACTC</td><td>598.0</td><td>-2.7</td><td>0.7</td><td>-3.9</td><td>1.1E-04</td><td>1.4E-04</td><td>MB231/MB468</td><td>NUC</td></tr>
<tr><td>>MI0000085|hsa-mir-27a&WithFlank&19|-|13836434|13836523@57.76.20(+2U)&[MIMAT0000084&hsa-miR-27a-3p&offsets|0|-1(+2U);m-43&19|-|13836447|13836467&offsets|0|-1(+2U)]|TTCACAGTGGCTAAGTTCCGTT</td><td>>MI0000085|hsa-mir-27a&WithFlank&19|-|13836434|13836523</td><td>57</td><td>76</td><td>20(+2U)</td><td>TTCACAGTGGCTAAGTTCCGTT</td><td>3089.6</td><td>-3.4</td><td>0.8</td><td>-4.3</td><td>1.6E-05</td><td>1.4E-04</td><td>BT20/MB231</td><td>MITO</td></tr>
<tr><td>>am_trnaMT_ValTAC_MT_+_1602_1670@28.67.40|AACTTACACTTAGGAGATTTCAACTTAACTTGACCGCTCT</td><td>>am_trnaMT_ValTAC_MT_+_1602_1670</td><td>28</td><td>67</td><td>40</td><td>AACTTACACTTAGGAGATTTCAACTTAACTTGACCGCTCT</td><td>537.1</td><td>9.9</td><td>2.4</td><td>4.1</td><td>3.9E-05</td><td>1.4E-04</td><td>BT20/MB468</td><td>MP</td></tr>
<tr><td>>MI0000469|hsa-mir-125a&WithFlank&19|+|51693248|51693345@21.42.22(+1A)&[MIMAT0000443&hsa-miR-125a-5p&offsets|0|-2(+1A);m-63&19|+|51693268|51693289&offsets|0|0(+1A)]|TCCCTGAGACCCTTTAACCTGTA</td><td>>MI0000469|hsa-mir-125a&WithFlank&19|+|51693248|51693345</td><td>21</td><td>42</td><td>22(+1A)</td><td>TCCCTGAGACCCTTTAACCTGTA</td><td>790.7</td><td>2.2</td><td>0.6</td><td>3.9</td><td>1.1E-04</td><td>1.4E-04</td><td>BT20/MB468</td><td>NUC</td></tr>
<tr><td>>hg38_21_+_8433172_8446622_RNA45SN1_45S_50ntFlanks@7975.7996.22|CGCGACCTCAGATCAGACGTGG</td><td>>hg38_21_+_8433172_8446622_RNA45SN1_45S_50ntFlanks</td><td>7975</td><td>7996</td><td>22</td><td>CGCGACCTCAGATCAGACGTGG</td><td>4564.5</td><td>-2.5</td><td>0.7</td><td>-3.9</td><td>9.2E-05</td><td>1.4E-04</td><td>MB231/MB468</td><td>MP</td></tr>
<tr><td>>hg38_21_+_8433172_8446622_RNA45SN1_45S_50ntFlanks@10767.10785.19|GCAGCAGGTCTCCAAGGTG</td><td>>hg38_21_+_8433172_8446622_RNA45SN1_45S_50ntFlanks</td><td>10767</td><td>10785</td><td>19</td><td>GCAGCAGGTCTCCAAGGTG</td><td>208.0</td><td>-2.8</td><td>0.7</td><td>-3.9</td><td>9.6E-05</td><td>1.4E-04</td><td>BT20/MB468</td><td>CYTO</td></tr>
<tr><td>>hg38_21_+_8433172_8446622_RNA45SN1_45S_50ntFlanks@9554.9580.27|CGTAGCGGTCCTGACGTGCAAATCGGT</td><td>>hg38_21_+_8433172_8446622_RNA45SN1_45S_50ntFlanks</td><td>9554</td><td>9580</td><td>27</td><td>CGTAGCGGTCCTGACGTGCAAATCGGT</td><td>476.6</td><td>-1.8</td><td>0.5</td><td>-3.9</td><td>1.1E-04</td><td>1.4E-04</td><td>MB231/MB468</td><td>NUC</td></tr>
<tr><td>>MI0003513|hsa-mir-455&WithFlank&9|+|114209428|114209535@22.45.24&[MIMAT0003150&hsa-miR-455-5p&offsets|0|+2;m-232&9|+|114209449|114209470&offsets|0|+2]|TATGTGCCTTTGGACTACATCGTG</td><td>>MI0003513|hsa-mir-455&WithFlank&9|+|114209428|114209535</td><td>22</td><td>45</td><td>24</td><td>TATGTGCCTTTGGACTACATCGTG</td><td>165.9</td><td>-3.1</td><td>0.7</td><td>-4.3</td><td>1.5E-05</td><td>1.4E-04</td><td>BT20/MB231</td><td>NUC</td></tr>
<tr><td>>hg38_21_+_8433172_8446622_RNA45SN1_45S_50ntFlanks@9545.9564.20|GTGGAGGTCCGTAGCGGTCC</td><td>>hg38_21_+_8433172_8446622_RNA45SN1_45S_50ntFlanks</td><td>9545</td><td>9564</td><td>20</td><td>GTGGAGGTCCGTAGCGGTCC</td><td>836.0</td><td>-4.3</td><td>1.1</td><td>-3.9</td><td>1.0E-04</td><td>1.4E-04</td><td>MB231/MB468</td><td>CYTO</td></tr>
<tr><td>>hg38_21_+_8433172_8446622_RNA45SN1_45S_50ntFlanks@11766.11803.38|ATCTAATTAGTGACGCGCATGAATGGATGAACGAGATT</td><td>>hg38_21_+_8433172_8446622_RNA45SN1_45S_50ntFlanks</td><td>11766</td><td>11803</td><td>38</td><td>ATCTAATTAGTGACGCGCATGAATGGATGAACGAGATT</td><td>295.9</td><td>-5.0</td><td>1.3</td><td>-3.9</td><td>1.0E-04</td><td>1.4E-04</td><td>MB231/MB468</td><td>CYTO</td></tr>
<tr><td>>hg38_1_-_228634819_228635039_RNA5S12_5S_50ntFlanks@132.168.37|GATGGGAGACCGCCTGGGAATACCGGGTGCTGTAGGC</td><td>>hg38_1_-_228634819_228635039_RNA5S12_5S_50ntFlanks</td><td>132</td><td>168</td><td>37</td><td>GATGGGAGACCGCCTGGGAATACCGGGTGCTGTAGGC</td><td>443.1</td><td>3.2</td><td>0.8</td><td>3.9</td><td>9.6E-05</td><td>1.4E-04</td><td>BT20/MB468</td><td>CYTO</td></tr>
<tr><td>>hg38_21_+_8433172_8446622_RNA45SN1_45S_50ntFlanks@10409.10432.24|GTCAGTCGGTCCTGAGAGATGGGC</td><td>>hg38_21_+_8433172_8446622_RNA45SN1_45S_50ntFlanks</td><td>10409</td><td>10432</td><td>24</td><td>GTCAGTCGGTCCTGAGAGATGGGC</td><td>1970.9</td><td>8.1</td><td>1.9</td><td>4.3</td><td>1.4E-05</td><td>1.4E-04</td><td>BT20/MB231</td><td>MP</td></tr>
<tr><td>>hg38_21_+_8433172_8446622_RNA45SN1_45S_50ntFlanks@10681.10711.31|TTCCGGCGGCGTCCGGTGAGCTCTCGCTGGC</td><td>>hg38_21_+_8433172_8446622_RNA45SN1_45S_50ntFlanks</td><td>10681</td><td>10711</td><td>31</td><td>TTCCGGCGGCGTCCGGTGAGCTCTCGCTGGC</td><td>1375.7</td><td>-5.2</td><td>1.3</td><td>-3.9</td><td>9.6E-05</td><td>1.4E-04</td><td>BT20/MB468</td><td>CYTO</td></tr>
<tr><td>>hg38_21_+_8433172_8446622_RNA45SN1_45S_50ntFlanks@11873.11910.38|AAGAAGACCCTGTTGAGCTTGACTCTAGTCTGGCACGG</td><td>>hg38_21_+_8433172_8446622_RNA45SN1_45S_50ntFlanks</td><td>11873</td><td>11910</td><td>38</td><td>AAGAAGACCCTGTTGAGCTTGACTCTAGTCTGGCACGG</td><td>487.9</td><td>-4.2</td><td>1.1</td><td>-3.9</td><td>8.1E-05</td><td>1.4E-04</td><td>BT20/MB468</td><td>MITO</td></tr>
<tr><td>>hg38_21_+_8433172_8446622_RNA45SN1_45S_50ntFlanks@6735.6770.36|TTGATCATCGACACTTCGAACGCACTTGCGGCCCCG</td><td>>hg38_21_+_8433172_8446622_RNA45SN1_45S_50ntFlanks</td><td>6735</td><td>6770</td><td>36</td><td>TTGATCATCGACACTTCGAACGCACTTGCGGCCCCG</td><td>437.6</td><td>-4.2</td><td>1.1</td><td>-3.9</td><td>8.1E-05</td><td>1.4E-04</td><td>BT20/MB468</td><td>MITO</td></tr>
<tr><td>>hg38_21_+_8433172_8446622_RNA45SN1_45S_50ntFlanks@8168.8187.20|CCCAAGTCCTTCTGATCGAG</td><td>>hg38_21_+_8433172_8446622_RNA45SN1_45S_50ntFlanks</td><td>8168</td><td>8187</td><td>20</td><td>CCCAAGTCCTTCTGATCGAG</td><td>235.5</td><td>-2.6</td><td>0.7</td><td>-3.9</td><td>8.1E-05</td><td>1.4E-04</td><td>BT20/MB468</td><td>MITO</td></tr>
<tr><td>>hg38_21_+_8433172_8446622_RNA45SN1_45S_50ntFlanks@9595.9612.18|ATAGGGGCGAAAGACTAA</td><td>>hg38_21_+_8433172_8446622_RNA45SN1_45S_50ntFlanks</td><td>9595</td><td>9612</td><td>18</td><td>ATAGGGGCGAAAGACTAA</td><td>84.6</td><td>9.7</td><td>2.2</td><td>4.3</td><td>1.4E-05</td><td>1.5E-04</td><td>BT20/MB231</td><td>MP</td></tr>
<tr><td>>hg38_21_+_8433172_8446622_RNA45SN1_45S_50ntFlanks@3874.3891.18|AATACATGCCGACGGGCG</td><td>>hg38_21_+_8433172_8446622_RNA45SN1_45S_50ntFlanks</td><td>3874</td><td>3891</td><td>18</td><td>AATACATGCCGACGGGCG</td><td>48.7</td><td>-9.5</td><td>2.4</td><td>-3.9</td><td>9.2E-05</td><td>1.5E-04</td><td>MB231/MB468</td><td>MP</td></tr>
<tr><td>>hg38_21_+_8433172_8446622_RNA45SN1_45S_50ntFlanks@4126.4151.26|TTCGATTCCGGAGAGGGAGCCTGAGA</td><td>>hg38_21_+_8433172_8446622_RNA45SN1_45S_50ntFlanks</td><td>4126</td><td>4151</td><td>26</td><td>TTCGATTCCGGAGAGGGAGCCTGAGA</td><td>107.7</td><td>4.5</td><td>1.1</td><td>3.9</td><td>9.2E-05</td><td>1.5E-04</td><td>MB231/MB468</td><td>MP</td></tr>
<tr><td>>MI0000102|hsa-mir-100&WithFlank&11|-|122152223|122152314@19.39.21(+2A)&[MIMAT0000098&hsa-miR-100-5p&offsets|0|-1(+2A);m-27&11|-|122152276|122152296&offsets|0|0(+2A)]|AACCCGTAGATCCGAACTTGTAA</td><td>>MI0000102|hsa-mir-100&WithFlank&11|-|122152223|122152314</td><td>19</td><td>39</td><td>21(+2A)</td><td>AACCCGTAGATCCGAACTTGTAA</td><td>56.3</td><td>-9.2</td><td>2.1</td><td>-4.3</td><td>1.4E-05</td><td>1.5E-04</td><td>BT20/MB231</td><td>MP</td></tr>
<tr><td>>hg38_21_+_8433172_8446622_RNA45SN1_45S_50ntFlanks@9916.9947.32|AAAGGTGTTGGTTGATATAGACAGCAGGACGG</td><td>>hg38_21_+_8433172_8446622_RNA45SN1_45S_50ntFlanks</td><td>9916</td><td>9947</td><td>32</td><td>AAAGGTGTTGGTTGATATAGACAGCAGGACGG</td><td>66.1</td><td>9.4</td><td>2.2</td><td>4.3</td><td>1.4E-05</td><td>1.5E-04</td><td>BT20/MB231</td><td>MP</td></tr>
<tr><td>>hg38_21_+_8433172_8446622_RNA45SN1_45S_50ntFlanks@9819.9838.20|TGCCTAGTGGGCCACTTTTG</td><td>>hg38_21_+_8433172_8446622_RNA45SN1_45S_50ntFlanks</td><td>9819</td><td>9838</td><td>20</td><td>TGCCTAGTGGGCCACTTTTG</td><td>50.9</td><td>-9.5</td><td>2.4</td><td>-3.9</td><td>9.3E-05</td><td>1.5E-04</td><td>MB231/MB468</td><td>MP</td></tr>
<tr><td>>hg38_21_+_8433172_8446622_RNA45SN1_45S_50ntFlanks@12997.13037.41|CGCTGCGATCTATTGAAAGTCAGCCCTCGACACAAGGGTTT</td><td>>hg38_21_+_8433172_8446622_RNA45SN1_45S_50ntFlanks</td><td>12997</td><td>13037</td><td>41</td><td>CGCTGCGATCTATTGAAAGTCAGCCCTCGACACAAGGGTTT</td><td>223.8</td><td>-3.1</td><td>0.7</td><td>-4.3</td><td>1.5E-05</td><td>1.5E-04</td><td>BT20/MB231</td><td>NUC</td></tr>
<tr><td>>hg38_21_+_8433172_8446622_RNA45SN1_45S_50ntFlanks@6659.6688.30|AGCGGTGGATCACTCGGCTCGTGCGTCGAT</td><td>>hg38_21_+_8433172_8446622_RNA45SN1_45S_50ntFlanks</td><td>6659</td><td>6688</td><td>30</td><td>AGCGGTGGATCACTCGGCTCGTGCGTCGAT</td><td>101.4</td><td>-4.7</td><td>1.2</td><td>-3.9</td><td>9.3E-05</td><td>1.5E-04</td><td>MB231/MB468</td><td>MP</td></tr>
<tr><td>>hg38_21_+_8433172_8446622_RNA45SN1_45S_50ntFlanks@8168.8186.19|CCCAAGTCCTTCTGATCGA</td><td>>hg38_21_+_8433172_8446622_RNA45SN1_45S_50ntFlanks</td><td>8168</td><td>8186</td><td>19</td><td>CCCAAGTCCTTCTGATCGA</td><td>612.6</td><td>-3.2</td><td>0.8</td><td>-3.9</td><td>8.1E-05</td><td>1.5E-04</td><td>BT20/MB468</td><td>MITO</td></tr>
<tr><td>>hg38_21_+_8433172_8446622_RNA45SN1_45S_50ntFlanks@11570.11601.32|AGCCGACTTAGAACTGGTGCGGACCAGGGGAA</td><td>>hg38_21_+_8433172_8446622_RNA45SN1_45S_50ntFlanks</td><td>11570</td><td>11601</td><td>32</td><td>AGCCGACTTAGAACTGGTGCGGACCAGGGGAA</td><td>686.0</td><td>-2.6</td><td>0.7</td><td>-3.9</td><td>9.7E-05</td><td>1.5E-04</td><td>BT20/MB468</td><td>CYTO</td></tr>
<tr><td>>hg38_21_+_8433172_8446622_RNA45SN1_45S_50ntFlanks@4723.4748.26|CATTAATCAAGAACGAAAGTCGGAGG</td><td>>hg38_21_+_8433172_8446622_RNA45SN1_45S_50ntFlanks</td><td>4723</td><td>4748</td><td>26</td><td>CATTAATCAAGAACGAAAGTCGGAGG</td><td>496.8</td><td>-6.4</td><td>1.6</td><td>-4.1</td><td>4.0E-05</td><td>1.5E-04</td><td>BT20/MB468</td><td>MP</td></tr>
<tr><td>>hg38_21_+_8433172_8446622_RNA45SN1_45S_50ntFlanks@12641.12661.21|CCATCTGTGGGATTATGACTG</td><td>>hg38_21_+_8433172_8446622_RNA45SN1_45S_50ntFlanks</td><td>12641</td><td>12661</td><td>21</td><td>CCATCTGTGGGATTATGACTG</td><td>297.6</td><td>-2.5</td><td>0.6</td><td>-3.9</td><td>9.7E-05</td><td>1.5E-04</td><td>BT20/MB468</td><td>CYTO</td></tr>
<tr><td>>hg38_21_+_8433172_8446622_RNA45SN1_45S_50ntFlanks@12996.13015.20|TCGCTGCGATCTATTGAAAG</td><td>>hg38_21_+_8433172_8446622_RNA45SN1_45S_50ntFlanks</td><td>12996</td><td>13015</td><td>20</td><td>TCGCTGCGATCTATTGAAAG</td><td>678.6</td><td>3.2</td><td>0.8</td><td>3.9</td><td>9.7E-05</td><td>1.5E-04</td><td>BT20/MB468</td><td>CYTO</td></tr>
<tr><td>>trnaMT_PheGAA_MT_+_577_647@37.57.21|AAATGTTTAGACGGGCTCACA</td><td>>trnaMT_PheGAA_MT_+_577_647</td><td>37</td><td>57</td><td>21</td><td>AAATGTTTAGACGGGCTCACA</td><td>134.6</td><td>6.6</td><td>1.6</td><td>4.1</td><td>4.0E-05</td><td>1.5E-04</td><td>BT20/MB468</td><td>MP</td></tr>
<tr><td>>MI0000071|hsa-mir-17&WithFlank&13|+|91350599|91350694@20.42.23(+1C)&[MIMAT0000070&hsa-miR-17-5p&offsets|0|0(+1C);m-88&13|+|91350618|91350640&offsets|0|0(+1C)]|CAAAGTGCTTACAGTGCAGGTAGC</td><td>>MI0000071|hsa-mir-17&WithFlank&13|+|91350599|91350694</td><td>20</td><td>42</td><td>23(+1C)</td><td>CAAAGTGCTTACAGTGCAGGTAGC</td><td>548.4</td><td>-5.1</td><td>1.2</td><td>-4.3</td><td>1.6E-05</td><td>1.5E-04</td><td>BT20/MB231</td><td>MITO</td></tr>
<tr><td>>hg38_21_+_8433172_8446622_RNA45SN1_45S_50ntFlanks@10681.10706.26|TTCCGGCGGCGTCCGGTGAGCTCTCG</td><td>>hg38_21_+_8433172_8446622_RNA45SN1_45S_50ntFlanks</td><td>10681</td><td>10706</td><td>26</td><td>TTCCGGCGGCGTCCGGTGAGCTCTCG</td><td>970.4</td><td>-5.3</td><td>1.3</td><td>-4.1</td><td>4.0E-05</td><td>1.5E-04</td><td>BT20/MB468</td><td>MP</td></tr>
<tr><td>>hg38_21_+_8433172_8446622_RNA45SN1_45S_50ntFlanks@10841.10864.24|CGTAACTTCGGGATAAGGATTGGC</td><td>>hg38_21_+_8433172_8446622_RNA45SN1_45S_50ntFlanks</td><td>10841</td><td>10864</td><td>24</td><td>CGTAACTTCGGGATAAGGATTGGC</td><td>621.1</td><td>-3.1</td><td>0.8</td><td>-3.9</td><td>8.2E-05</td><td>1.5E-04</td><td>BT20/MB468</td><td>MITO</td></tr>
<tr><td>>MI0000782|hsa-mir-374a&WithFlank&X|-|74287280|74287363@18.39.22&[MIMAT0000727&hsa-miR-374a-5p&offsets|0|0;m-184&X|-|74287325|74287346&offsets|0|0]|TTATAATACAACCTGATAAGTG</td><td>>MI0000782|hsa-mir-374a&WithFlank&X|-|74287280|74287363</td><td>18</td><td>39</td><td>22</td><td>TTATAATACAACCTGATAAGTG</td><td>514.5</td><td>4.3</td><td>1.1</td><td>3.9</td><td>1.1E-04</td><td>1.5E-04</td><td>MB231/MB468</td><td>MITO</td></tr>
<tr><td>>hg38_21_+_8433172_8446622_RNA45SN1_45S_50ntFlanks@6644.6665.22|CCTCGTACGACTCTTAGCGGTG</td><td>>hg38_21_+_8433172_8446622_RNA45SN1_45S_50ntFlanks</td><td>6644</td><td>6665</td><td>22</td><td>CCTCGTACGACTCTTAGCGGTG</td><td>975.3</td><td>3.6</td><td>0.9</td><td>3.9</td><td>1.1E-04</td><td>1.5E-04</td><td>MB231/MB468</td><td>MITO</td></tr>
<tr><td>>MI0001445|hsa-mir-423&WithFlank&17|+|30117073|30117178@59.79.21(+1A)&[MIMAT0001340&hsa-miR-423-3p&offsets|0|-2(+1A);m-100&17|+|30117131|30117153&offsets|0|-2(+1A)]|AGCTCGGTCTGAGGCCCCTCAA</td><td>>MI0001445|hsa-mir-423&WithFlank&17|+|30117073|30117178</td><td>59</td><td>79</td><td>21(+1A)</td><td>AGCTCGGTCTGAGGCCCCTCAA</td><td>93.1</td><td>7.4</td><td>1.8</td><td>4.1</td><td>4.0E-05</td><td>1.5E-04</td><td>BT20/MB468</td><td>MP</td></tr>
<tr><td>>hg38_21_+_8433172_8446622_RNA45SN1_45S_50ntFlanks@8168.8195.28|CCCAAGTCCTTCTGATCGAGGCCCAGCC</td><td>>hg38_21_+_8433172_8446622_RNA45SN1_45S_50ntFlanks</td><td>8168</td><td>8195</td><td>28</td><td>CCCAAGTCCTTCTGATCGAGGCCCAGCC</td><td>259.1</td><td>-5.9</td><td>1.5</td><td>-3.9</td><td>9.3E-05</td><td>1.5E-04</td><td>MB231/MB468</td><td>MP</td></tr>
<tr><td>>MI0000085|hsa-mir-27a&WithFlank&19|-|13836434|13836523@57.75.19(+1U)&[MIMAT0000084&hsa-miR-27a-3p&offsets|0|-2(+1U);m-43&19|-|13836447|13836467&offsets|0|-2(+1U)]|TTCACAGTGGCTAAGTTCCT</td><td>>MI0000085|hsa-mir-27a&WithFlank&19|-|13836434|13836523</td><td>57</td><td>75</td><td>19(+1U)</td><td>TTCACAGTGGCTAAGTTCCT</td><td>532.6</td><td>3.8</td><td>1.0</td><td>3.9</td><td>9.3E-05</td><td>1.5E-04</td><td>MB231/MB468</td><td>MP</td></tr>
</table>

| | | | | | | | | | | | | | |
|---|---|---|---|---|---|---|---|---|---|---|---|---|---|
| >hg38_21_+_8433172_8446622_RNA45SN1_45S_50ntFlanks@11586.11612.27\|GTGCGGACCAGGGGAATCCGACTGTTT | >hg38_21_+_8433172_8446622_RNA45SN1_45S_50ntFlanks | 11586 | 11612 | 27 | GTGCGGACCAGGGGAATCCGACTGTTT | 1307.3 | -2.7 | 0.7 | -3.9 | 1.0E-04 | 1.5E-04 | MB231/MB468 | CYTO |
| >hg38_21_+_8433172_8446622_RNA45SN1_45S_50ntFlanks@4000.4042.43\|TAGATAACCTCGGGCCGATCGCACGCCCCCCGTGGCGGCGACG | >hg38_21_+_8433172_8446622_RNA45SN1_45S_50ntFlanks | 4000 | 4042 | 43 | TAGATAACCTCGGGCCGATCGCACGCCCCCCGTGGCGGCGACG | 684.9 | -3.3 | 0.9 | -3.9 | 1.0E-04 | 1.5E-04 | MB231/MB468 | CYTO |
| >hg38_21_+_8433172_8446622_RNA45SN1_45S_50ntFlanks@3992.4013.22\|GGTGACTCTAGATAACCTCGGG | >hg38_21_+_8433172_8446622_RNA45SN1_45S_50ntFlanks | 3992 | 4013 | 22 | GGTGACTCTAGATAACCTCGGG | 331.3 | -6.4 | 1.7 | -3.9 | 1.1E-04 | 1.5E-04 | BT20/MB468 | NUC |
| >hg38_21_+_8433172_8446622_RNA45SN1_45S_50ntFlanks@11844.11873.30\|GGGAACGGGCTTGGCGGAATCAGCGGGGAA | >hg38_21_+_8433172_8446622_RNA45SN1_45S_50ntFlanks | 11844 | 11873 | 30 | GGGAACGGGCTTGGCGGAATCAGCGGGGAA | 235.5 | -6.1 | 1.5 | -4.1 | 4.0E-05 | 1.5E-04 | BT20/MB468 | MP |
| >MI0000273\|hsa-mir-183&WithFlank&7\|-\|129774899\|129775020@34.56.23&[MIMAT0000261&hsa-miR-183-5p&offsets\|+1\|+2;m-41&7\|-\|129774966\|129774987&offsets\|0\|+1]\|ATGGCACTGGTAGAATTCACTGT | >MI0000273\|hsa-mir-183&WithFlank&7\|-\|129774899\|129775020 | 34 | 56 | 23 | ATGGCACTGGTAGAATTCACTGT | 852.6 | 2.5 | 0.6 | 4.3 | 1.5E-05 | 1.5E-04 | BT20/MB231 | NUC |
| >hg38_21_+_8433172_8446622_RNA45SN1_45S_50ntFlanks@6719.6746.28\|TGAATTGCAGGACACATTGATCATCGAC | >hg38_21_+_8433172_8446622_RNA45SN1_45S_50ntFlanks | 6719 | 6746 | 28 | TGAATTGCAGGACACATTGATCATCGAC | 843.4 | -4.6 | 1.2 | -3.9 | 9.8E-05 | 1.5E-04 | BT20/MB468 | CYTO |
| >MI0000825\|hsa-mir-345&WithFlank&14\|+\|100307853\|100307962@24.45.22&[MIMAT0000772&hsa-miR-345-5p&offsets\|0\|0;m-193&14\|+\|100307876\|100307896&offsets\|0\|+1]\|GCTGACTCCTAGTCCAGGGCTC | >MI0000825\|hsa-mir-345&WithFlank&14\|+\|100307853\|100307962 | 24 | 45 | 22 | GCTGACTCCTAGTCCAGGGCTC | 532.5 | 3.2 | 0.8 | 3.9 | 9.4E-05 | 1.5E-04 | MB231/MB468 | MP |
| >hg38_MT_+_1621_3279_RNR2_16S_50ntFlanks@51.76.26\|GCTAAACCTAGCCCCAAACCCACTCC | >hg38_MT_+_1621_3279_RNR2_16S_50ntFlanks | 51 | 76 | 26 | GCTAAACCTAGCCCCAAACCCACTCC | 563.0 | 3.2 | 0.8 | 3.9 | 1.1E-04 | 1.5E-04 | MB231/MB468 | MITO |
| >hg38_21_+_8433172_8446622_RNA45SN1_45S_50ntFlanks@12997.13038.42\|CGCTGCGATCTATTGAAAGTCAGCCCTCGACACAAGGGTTTG | >hg38_21_+_8433172_8446622_RNA45SN1_45S_50ntFlanks | 12997 | 13038 | 42 | CGCTGCGATCTATTGAAAGTCAGCCCTCGACACAAGGGTTTG | 1317.0 | 6.0 | 1.5 | 3.9 | 1.1E-04 | 1.5E-04 | MB231/MB468 | MITO |
| >hg38_21_+_8433172_8446622_RNA45SN1_45S_50ntFlanks@9313.9338.26\|TCTAACACGTGCGCGAGTCGGGGGCT | >hg38_21_+_8433172_8446622_RNA45SN1_45S_50ntFlanks | 9313 | 9338 | 26 | TCTAACACGTGCGCGAGTCGGGGGCT | 339.0 | -6.4 | 1.7 | -3.9 | 1.1E-04 | 1.5E-04 | BT20/MB468 | NUC |
| >MI0000093\|hsa-mir-92a-1&WithFlank&13\|+\|91351308\|91351397@54.72.19(+1C);MI0000094\|hsa-mir-92a-2&WithFlank&X\|-\|134169532\|134169618@54.72.19(+1C)&[MIMAT0000092&hsa-miR-92a-3p&offsets\|0\|-3(+1C);m-9&13\|+\|91351361\|91351382&offsets\|0\|-3(+1C)];[MIMAT0000092_1&hsa-miR-92a-3p&offsets\|0\|-3(+1C);m-11&X\|-\|134169544\|134169565&offsets\|0\|-3(+1C)]\|TATTGCACTTGTCCCGGCCC | >MI0000093\|hsa-mir-92a-1&WithFlank&13\|+\|91351308\|91351397 | 54 | 72 | 19(+1C) | TATTGCACTTGTCCCGGCCC | 30.4 | 8.1 | 2.1 | 3.9 | 9.4E-05 | 1.5E-04 | MB231/MB468 | MP |
| >hg38_21_+_8433172_8446622_RNA45SN1_45S_50ntFlanks@6652.6686.35\|GACTCTTAGCGGTGGATCACTCGGCTCGTGCGTCG | >hg38_21_+_8433172_8446622_RNA45SN1_45S_50ntFlanks | 6652 | 6686 | 35 | GACTCTTAGCGGTGGATCACTCGGCTCGTGCGTCG | 1083.0 | -3.5 | 0.9 | -3.9 | 9.4E-05 | 1.5E-04 | MB231/MB468 | MP |
| >trnaMT_GluTTC_MT_-_14674_14742@20.63.44,trnalookalike8_GluTTC_5_-_93905172_93905240@20.63.44\|AACGATGGTTTTTCATATCATTGGTCGTGGTTGTAGTCCGTGCG | >trnaMT_GluTTC_MT_-_14674_14742 | 20 | 63 | 44 | AACGATGGTTTTTCATATCATTGGTCGTGGTTGTAGTCCGTGCG | 139.1 | 10.9 | 2.6 | 4.1 | 4.0E-05 | 1.5E-04 | BT20/MB468 | MP |
| >hg38_21_+_8433172_8446622_RNA45SN1_45S_50ntFlanks@5487.5509.23\|CGCTGAGAAGACGGTCGAACTTG | >hg38_21_+_8433172_8446622_RNA45SN1_45S_50ntFlanks | 5487 | 5509 | 23 | CGCTGAGAAGACGGTCGAACTTG | 185.2 | -2.2 | 0.6 | -3.9 | 8.2E-05 | 1.5E-04 | BT20/MB468 | MITO |
| >hg38_21_+_8433172_8446622_RNA45SN1_45S_50ntFlanks@11778.11806.29\|ACGCGCATGAATGGATGAACGAGATTCCC | >hg38_21_+_8433172_8446622_RNA45SN1_45S_50ntFlanks | 11778 | 11806 | 29 | ACGCGCATGAATGGATGAACGAGATTCCC | 224.7 | -2.1 | 0.5 | -3.9 | 1.1E-04 | 1.5E-04 | MB231/MB468 | CYTO |
| >hg38_21_+_8433172_8446622_RNA45SN1_45S_50ntFlanks@11921.11951.31\|TGAGAGGTGTAGAATAAGTGGGAGGCCCCCG | >hg38_21_+_8433172_8446622_RNA45SN1_45S_50ntFlanks | 11921 | 11951 | 31 | TGAGAGGTGTAGAATAAGTGGGAGGCCCCCG | 468.0 | -3.6 | 0.9 | -3.9 | 1.1E-04 | 1.5E-04 | MB231/MB468 | CYTO |
| >hg38_21_+_8433172_8446622_RNA45SN1_45S_50ntFlanks@8190.8214.25\|CCAGCCCGTGGACGGTGTGAGGCCG | >hg38_21_+_8433172_8446622_RNA45SN1_45S_50ntFlanks | 8190 | 8214 | 25 | CCAGCCCGTGGACGGTGTGAGGCCG | 115.1 | -5.8 | 1.5 | -3.9 | 9.4E-05 | 1.5E-04 | MB231/MB468 | MP |
| >trna14_LysTTT_11_-_59327808_59327880@1.24.24,trna2_LysTTT_17_+_8022473_8022545@1.24.24,trna54_LysTTT_1_+_204475655_204475727@1.24.24,trna5_LysTTT_11_+_59323902_59323974@1.24.24,trna62_LysTTT_1_-_204476158_204476230@1.24.24,trna76_LysTTT_6_+_28918806_28918878@1.24.24\|GCCCGGATAGCTCAGTCGGTAGAG | >trna14_LysTTT_11_-_59327808_59327880 | 1 | 24 | 24 | GCCCGGATAGCTCAGTCGGTAGAG | 556.5 | -4.9 | 1.3 | -3.9 | 1.1E-04 | 1.5E-04 | MB231/MB468 | MITO |
| >MI0001448\|hsa-mir-425&WithFlank&3\|-\|49020142\|49020240@60.80.21(+1U)&[MIMAT0001343&hsa-miR-425-3p&offsets\|-1\|-2(+1U);m-258&3\|-\|49020160\|49020181&offsets\|0\|-1(+1U)]\|CATCGGGAATGTCGTGTCCGCT | >MI0001448\|hsa-mir-425&WithFlank&3\|-\|49020142\|49020240 | 60 | 80 | 21(+1U) | CATCGGGAATGTCGTGTCCGCT | 256.1 | 6.1 | 1.5 | 4.1 | 4.0E-05 | 1.5E-04 | BT20/MB468 | MP |
| >hg38_21_+_8433172_8446622_RNA45SN1_45S_50ntFlanks@11771.11794.24\|ATTAGTGACGCGCATGAATGGATG | >hg38_21_+_8433172_8446622_RNA45SN1_45S_50ntFlanks | 11771 | 11794 | 24 | ATTAGTGACGCGCATGAATGGATG | 1924.4 | -7.4 | 1.9 | -3.9 | 1.2E-04 | 1.5E-04 | BT20/MB468 | NUC |
| >hg38_21_+_8433172_8446622_RNA45SN1_45S_50ntFlanks@12283.12318.36\|TACAGACCGTGAAAGCGGGGCCTCACGATCCTTCTG | >hg38_21_+_8433172_8446622_RNA45SN1_45S_50ntFlanks | 12283 | 12318 | 36 | TACAGACCGTGAAAGCGGGGCCTCACGATCCTTCTG | 210.5 | -4.0 | 1.0 | -3.9 | 8.3E-05 | 1.5E-04 | BT20/MB468 | MITO |
| >hg38_21_+_8433172_8446622_RNA45SN1_45S_50ntFlanks@7975.8008.34\|CGCGACCTCAGATCAGACGTGGCGACCCGCTGAA | >hg38_21_+_8433172_8446622_RNA45SN1_45S_50ntFlanks | 7975 | 8008 | 34 | CGCGACCTCAGATCAGACGTGGCGACCCGCTGAA | 450686.0 | 3.2 | 0.8 | 3.9 | 8.3E-05 | 1.5E-04 | BT20/MB468 | MITO |
| >hg38_21_+_8433172_8446622_RNA45SN1_45S_50ntFlanks@12940.12979.40\|CATTCGTAGACGACCTGCTTCTGGGTCGGGGTTTCGTACG | >hg38_21_+_8433172_8446622_RNA45SN1_45S_50ntFlanks | 12940 | 12979 | 40 | CATTCGTAGACGACCTGCTTCTGGGTCGGGGTTTCGTACG | 118.1 | -4.5 | 1.2 | -3.9 | 1.1E-04 | 1.5E-04 | MB231/MB468 | CYTO |
| >hg38_21_+_8433172_8446622_RNA45SN1_45S_50ntFlanks@10855.10874.20\|AAGGATTGGCTCTAAGGGCT | >hg38_21_+_8433172_8446622_RNA45SN1_45S_50ntFlanks | 10855 | 10874 | 20 | AAGGATTGGCTCTAAGGGCT | 89.4 | -5.7 | 1.5 | -3.9 | 9.4E-05 | 1.5E-04 | MB231/MB468 | MP |
| >hg38_21_+_8433172_8446622_RNA45SN1_45S_50ntFlanks@11664.11685.22\|TTTCTGCCCAGTGCTCTGAATG | >hg38_21_+_8433172_8446622_RNA45SN1_45S_50ntFlanks | 11664 | 11685 | 22 | TTTCTGCCCAGTGCTCTGAATG | 314.6 | -2.7 | 0.7 | -3.9 | 9.9E-05 | 1.5E-04 | BT20/MB468 | CYTO |
| >hg38_21_+_8433172_8446622_RNA45SN1_45S_50ntFlanks@12935.12962.28\|TAAACCATTCGTAGACGACCTGCTTCTG | >hg38_21_+_8433172_8446622_RNA45SN1_45S_50ntFlanks | 12935 | 12962 | 28 | TAAACCATTCGTAGACGACCTGCTTCTG | 207.2 | -5.4 | 1.4 | -3.9 | 1.1E-04 | 1.5E-04 | MB231/MB468 | NUC |
| >hg38_21_+_8433172_8446622_RNA45SN1_45S_50ntFlanks@12896.12932.37\|CGCCCGTCACGCACCGCACGTTCGTGGGGAACCTGGC | >hg38_21_+_8433172_8446622_RNA45SN1_45S_50ntFlanks | 12896 | 12932 | 37 | CGCCCGTCACGCACCGCACGTTCGTGGGGAACCTGGC | 97.9 | -5.8 | 1.5 | -3.9 | 9.5E-05 | 1.5E-04 | MB231/MB468 | MP |
| >MI0000816\|hsa-mir-335&WithFlank&7\|+\|130496105\|130496210@22.42.21&[MIMAT0000765&hsa-miR-335-5p&offsets\|0\|-2;m-249&7\|+\|130496126\|130496147&offsets\|0\|-1]\|TCAAGAGCAATAACGAAAAAT | >MI0000816\|hsa-mir-335&WithFlank&7\|+\|130496105\|130496210 | 22 | 42 | 21 | TCAAGAGCAATAACGAAAAAT | 60.6 | -9.3 | 2.1 | -4.3 | 1.4E-05 | 1.5E-04 | BT20/MB231 | MP |
| >MI0003177\|hsa-mir-522&WithFlank&19\|+\|53751205\|53751303@60.80.21&[MIMAT0002868&hsa-miR-522-3p&offsets\|0\|-1;m-909&19\|+\|53751264\|53751285&offsets\|0\|-1]\|AAAATGGTTCCCTTTAGAGTG | >MI0003177\|hsa-mir-522&WithFlank&19\|+\|53751205\|53751303 | 60 | 80 | 21 | AAAATGGTTCCCTTTAGAGTG | 26.9 | 8.0 | 2.0 | 3.9 | 9.5E-05 | 1.5E-04 | MB231/MB468 | MP |
| >hg38_21_+_8433172_8446622_RNA45SN1_45S_50ntFlanks@9545.9574.30\|GTGGAGGTCCGTAGCGGTCCTGACGTGCAA | >hg38_21_+_8433172_8446622_RNA45SN1_45S_50ntFlanks | 9545 | 9574 | 30 | GTGGAGGTCCGTAGCGGTCCTGACGTGCAA | 808.6 | -4.3 | 1.1 | -3.9 | 9.9E-05 | 1.5E-04 | BT20/MB468 | CYTO |
| >am_MI0003639\|hsa-mir-625&WithFlank&14\|+\|65471096\|65471192@58.79.22&[MIMAT0004808&hsa-miR-625-3p&offsets\|0\|0;m-137&14\|+\|65471153\|65471174&offsets\|0\|0]\|GACTATAGAACTTTCCCCCTCA | >am_MI0003639\|hsa-mir-625&WithFlank&14\|+\|65471096\|65471192 | 58 | 79 | 22 | GACTATAGAACTTTCCCCCTCA | 187.5 | 5.7 | 1.5 | 3.9 | 9.5E-05 | 1.5E-04 | MB231/MB468 | MP |
| >hg38_21_+_8433172_8446622_RNA45SN1_45S_50ntFlanks@6731.6752.22\|CACATTGATCATCGACACTTCG | >hg38_21_+_8433172_8446622_RNA45SN1_45S_50ntFlanks | 6731 | 6752 | 22 | CACATTGATCATCGACACTTCG | 564.3 | -4.4 | 1.1 | -3.9 | 9.9E-05 | 1.5E-04 | BT20/MB468 | CYTO |
| >hg38_21_+_8433172_8446622_RNA45SN1_45S_50ntFlanks@10678.10705.28\|CGGTTCCGGCGGCGTCCGGTGAGCTCTC | >hg38_21_+_8433172_8446622_RNA45SN1_45S_50ntFlanks | 10678 | 10705 | 28 | CGGTTCCGGCGGCGTCCGGTGAGCTCTC | 1022.0 | -5.1 | 1.2 | -4.1 | 4.1E-05 | 1.5E-04 | BT20/MB468 | MP |
| >hg38_21_+_8433172_8446622_RNA45SN1_45S_50ntFlanks@12154.12180.27\|GGGAGTTTGACTGGGGCGGTACACCTG | >hg38_21_+_8433172_8446622_RNA45SN1_45S_50ntFlanks | 12154 | 12180 | 27 | GGGAGTTTGACTGGGGCGGTACACCTG | 654.9 | -6.5 | 1.6 | -4.1 | 4.1E-05 | 1.5E-04 | BT20/MB468 | MP |
| >hg38_21_+_8433172_8446622_RNA45SN1_45S_50ntFlanks@10678.10695.18\|CGGTTCCGGCGGCGTCCG | >hg38_21_+_8433172_8446622_RNA45SN1_45S_50ntFlanks | 10678 | 10695 | 18 | CGGTTCCGGCGGCGTCCG | 165.2 | -6.0 | 1.5 | -4.1 | 4.1E-05 | 1.5E-04 | BT20/MB468 | MP |
| >hg38_21_+_8433172_8446622_RNA45SN1_45S_50ntFlanks@11728.11765.38\|GTAACTATGACTCTCTTAAGGTAGCCAAATGCCTCGTC | >hg38_21_+_8433172_8446622_RNA45SN1_45S_50ntFlanks | 11728 | 11765 | 38 | GTAACTATGACTCTCTTAAGGTAGCCAAATGCCTCGTC | 238.7 | -3.0 | 0.8 | -3.9 | 1.1E-04 | 1.5E-04 | MB231/MB468 | NUC |
| >hg38_21_+_8433172_8446622_RNA45SN1_45S_50ntFlanks@12633.12665.33\|CGAAGCTACCATCTGTGGGATTATGACTGAACG | >hg38_21_+_8433172_8446622_RNA45SN1_45S_50ntFlanks | 12633 | 12665 | 33 | CGAAGCTACCATCTGTGGGATTATGACTGAACG | 147.9 | -3.3 | 0.8 | -3.9 | 9.5E-05 | 1.5E-04 | MB231/MB468 | MP |
| >hg38_21_+_8433172_8446622_RNA45SN1_45S_50ntFlanks@10392.10429.38\|ACAGCAGTTGAACATGGGTCAGTCGGTCCTGAGAGATG | >hg38_21_+_8433172_8446622_RNA45SN1_45S_50ntFlanks | 10392 | 10429 | 38 | ACAGCAGTTGAACATGGGTCAGTCGGTCCTGAGAGATG | 2453.6 | -3.7 | 0.9 | -3.9 | 1.0E-04 | 1.5E-04 | BT20/MB468 | CYTO |
| >hg38_21_+_8433172_8446622_RNA45SN1_45S_50ntFlanks@12006.12032.27\|GGCCGCCGGTGAAATACCACTACTCTG | >hg38_21_+_8433172_8446622_RNA45SN1_45S_50ntFlanks | 12006 | 12032 | 27 | GGCCGCCGGTGAAATACCACTACTCTG | 446.4 | -3.7 | 0.9 | -3.9 | 1.1E-04 | 1.5E-04 | MB231/MB468 | CYTO |
| >MI0000063\|hsa-let-7b&WithFlank&22\|+\|46113680\|46113774@12.34.23(+1U)&[MIMAT0000063&hsa-let-7b-5p&offsets\|0\|+1(+1U);m-14&22\|+\|46113691\|46113712&offsets\|0\|+1(+1U)]\|TGAGGTAGTAGGTTGTGTGGTTTT | >MI0000063\|hsa-let-7b&WithFlank&22\|+\|46113680\|46113774 | 12 | 34 | 23(+1U) | TGAGGTAGTAGGTTGTGTGGTTTT | 196.0 | 5.1 | 1.3 | 3.9 | 1.1E-04 | 1.5E-04 | MB231/MB468 | MITO |
| >hg38_21_+_8433172_8446622_RNA45SN1_45S_50ntFlanks@12934.12961.28\|CTAAACCATTCGTAGACGACCTGCTTCT | >hg38_21_+_8433172_8446622_RNA45SN1_45S_50ntFlanks | 12934 | 12961 | 28 | CTAAACCATTCGTAGACGACCTGCTTCT | 404.5 | -4.3 | 1.1 | -3.9 | 1.0E-04 | 1.5E-04 | BT20/MB468 | CYTO |
| >hg38_21_+_8433172_8446622_RNA45SN1_45S_50ntFlanks@10776.10816.41\|CTCCAAGGTGAACAGCCTCTGGCATGTTGGAACAATGTAGG | >hg38_21_+_8433172_8446622_RNA45SN1_45S_50ntFlanks | 10776 | 10816 | 41 | CTCCAAGGTGAACAGCCTCTGGCATGTTGGAACAATGTAGG | 63.9 | 9.3 | 2.2 | 4.3 | 1.4E-05 | 1.5E-04 | BT20/MB231 | MP |
| >hg38_21_+_8433172_8446622_RNA45SN1_45S_50ntFlanks@9299.9331.33\|CACGGACCAAGGAGTCTAACACGTGCGCGAGTC | >hg38_21_+_8433172_8446622_RNA45SN1_45S_50ntFlanks | 9299 | 9331 | 33 | CACGGACCAAGGAGTCTAACACGTGCGCGAGTC | 70.3 | 9.5 | 2.2 | 4.3 | 1.4E-05 | 1.5E-04 | BT20/MB231 | MP |
| >hg38_21_+_8433172_8446622_RNA45SN1_45S_50ntFlanks@4283.4307.25\|CTTTAACGAGGATCCATTGGAGGGC | >hg38_21_+_8433172_8446622_RNA45SN1_45S_50ntFlanks | 4283 | 4307 | 25 | CTTTAACGAGGATCCATTGGAGGGC | 142.5 | -5.1 | 1.3 | -3.9 | 1.0E-04 | 1.5E-04 | BT20/MB468 | CYTO |
| >hg38_21_+_8433172_8446622_RNA45SN1_45S_50ntFlanks@12415.12438.24\|CTTCGATGTCGGCTCTTCCTATCA | >hg38_21_+_8433172_8446622_RNA45SN1_45S_50ntFlanks | 12415 | 12438 | 24 | CTTCGATGTCGGCTCTTCCTATCA | 3102.0 | -4.0 | 1.0 | -3.9 | 1.1E-04 | 1.5E-04 | MB231/MB468 | MITO |
| >trnaMT_PheGAA_MT_+_577_647@35.55.21\|GAAAATGTTTAGACGGGCTCA | >trnaMT_PheGAA_MT_+_577_647 | 35 | 55 | 21 | GAAAATGTTTAGACGGGCTCA | 280.8 | 4.7 | 1.2 | 3.9 | 1.1E-04 | 1.5E-04 | MB231/MB468 | MITO |
| >hg38_21_+_8433172_8446622_RNA45SN1_45S_50ntFlanks@10350.10367.18\|ACGAGAACTTTGAAGGCC | >hg38_21_+_8433172_8446622_RNA45SN1_45S_50ntFlanks | 10350 | 10367 | 18 | ACGAGAACTTTGAAGGCC | 1196.7 | -2.3 | 0.6 | -3.9 | 1.0E-04 | 1.5E-04 | BT20/MB468 | CYTO |
| >hg38_21_+_8433172_8446622_RNA45SN1_45S_50ntFlanks@12934.12957.24\|CTAAACCATTCGTAGACGACCTGC | >hg38_21_+_8433172_8446622_RNA45SN1_45S_50ntFlanks | 12934 | 12957 | 24 | CTAAACCATTCGTAGACGACCTGC | 346.5 | -2.6 | 0.7 | -3.9 | 1.0E-04 | 1.5E-04 | BT20/MB468 | CYTO |

| | | | | | | | | | | | | | |
|---|---|---|---|---|---|---|---|---|---|---|---|---|---|
| >trna10_LysCTT_16_+_3241501_3241573@3.33.31,trna13_LysCTT_14_-_58706613_58706685@3.33.31,trna2_LysCTT_15_+_79152904_79152976@3.33.31\|CCGGCTAGCTCAGTCGGTAGAGCATGGGACT | >trna10_LysCTT_16_+_3241501_3241573 | 3 | 33 | 31 | CCGGCTAGCTCAGTCGGTAGAGCATGGGACT | 314.7 | 4.9 | 1.2 | 3.9 | 9.6E-05 | 1.5E-04 | MB231/MB468 | MP |
| >hg38_21_+_8433172_8446622_RNA45SN1_45S_50ntFlanks@9940.9982.43\|CAGGACGGTGGCCATGGAAGTCGGAATCCGCTAAGGAGTGTGT | >hg38_21_+_8433172_8446622_RNA45SN1_45S_50ntFlanks | 9940 | 9982 | 43 | CAGGACGGTGGCCATGGAAGTCGGAATCCGCTAAGGAGTGTGT | 877.9 | 7.4 | 1.8 | 4.1 | 4.1E-05 | 1.5E-04 | BT20/MB468 | MP |
| >MI0000086\|hsa-mir-28&WithFlank&3\|+\|188688775\|188688872@61.80.20&[MIMAT0004502&hsa-miR-28-3p&offsets\|+1\|-1;m-28&3\|+\|188688834\|188688855&offsets\|+1\|-1]\|ACTAGATTGTGAGCTCCTGG | >MI0000086\|hsa-mir-28&WithFlank&3\|+\|188688775\|188688872 | 61 | 80 | 20 | ACTAGATTGTGAGCTCCTGG | 135.9 | 7.3 | 1.8 | 4.1 | 4.1E-05 | 1.5E-04 | BT20/MB468 | MP |
| >hg38_21_+_8433172_8446622_RNA45SN1_45S_50ntFlanks@3845.3868.24\|ACTTGGATAACTGTGGTAATTCTA | >hg38_21_+_8433172_8446622_RNA45SN1_45S_50ntFlanks | 3845 | 3868 | 24 | ACTTGGATAACTGTGGTAATTCTA | 395.6 | -5.3 | 1.4 | -3.9 | 9.6E-05 | 1.5E-04 | MB231/MB468 | MP |
| >hg38_21_+_8433172_8446622_RNA45SN1_45S_50ntFlanks@7976.7994.19\|GCGACCTCAGATCAGACGT | >hg38_21_+_8433172_8446622_RNA45SN1_45S_50ntFlanks | 7976 | 7994 | 19 | GCGACCTCAGATCAGACGT | 273.5 | 3.5 | 0.8 | 4.1 | 4.1E-05 | 1.5E-04 | BT20/MB468 | MP |
| >hg38_21_+_8433172_8446622_RNA45SN1_45S_50ntFlanks@11873.11905.33\|AAGAAGACCCTGTTGAGCTTGACTCTAGTCTGG | >hg38_21_+_8433172_8446622_RNA45SN1_45S_50ntFlanks | 11873 | 11905 | 33 | AAGAAGACCCTGTTGAGCTTGACTCTAGTCTGG | 1120.5 | -5.7 | 1.4 | -4.1 | 4.1E-05 | 1.5E-04 | BT20/MB468 | MP |
| >hg38_21_+_8433172_8446622_RNA45SN1_45S_50ntFlanks@9569.9595.27\|GTGCAAATCGGTCGTCCGACCTGGGTA | >hg38_21_+_8433172_8446622_RNA45SN1_45S_50ntFlanks | 9569 | 9595 | 27 | GTGCAAATCGGTCGTCCGACCTGGGTA | 217.3 | -2.2 | 0.6 | -3.9 | 1.1E-04 | 1.5E-04 | MB231/MB468 | NUC |
| >hg38_21_+_8433172_8446622_RNA45SN1_45S_50ntFlanks@12251.12284.34\|AAAGCTCGCTTGATCTTGATTTTCAGTACGAATA | >hg38_21_+_8433172_8446622_RNA45SN1_45S_50ntFlanks | 12251 | 12284 | 34 | AAAGCTCGCTTGATCTTGATTTTCAGTACGAATA | 348.7 | -6.4 | 1.7 | -3.9 | 1.2E-04 | 1.5E-04 | BT20/MB468 | NUC |
| >hg38_21_+_8433172_8446622_RNA45SN1_45S_50ntFlanks@8118.8138.21\|GGGACATGTGGCGTACGGAAG | >hg38_21_+_8433172_8446622_RNA45SN1_45S_50ntFlanks | 8118 | 8138 | 21 | GGGACATGTGGCGTACGGAAG | 544.5 | 5.3 | 1.3 | 4.1 | 4.1E-05 | 1.5E-04 | BT20/MB468 | MP |
| >am_trna10_ValCAC_5_-_180649395_180649467@1.31.31,trna12_ValAAC_5_-_180645270_180645342@1.31.31,trna132_ValAAC_6_-_27721179_27721251@1.31.31,trna136_ValAAC_6_-_27648885_27648957@1.31.31,trna139_ValAAC_6_-_27618707_27618779@1.31.31,trna18_ValCAC_5_-_180529253_180529325@1.31.31,trna2_ValAAC_3_+_169490018_169490090@1.31.31,trna2_ValCAC_5_+_180524070_180524142@1.31.31,trna4_ValAAC_5_+_180591154_180591226@1.31.31,trna5_ValAAC_5_+_180596610_180596682@1.31.31,trna6_ValCAC_5_+_180600650_180600722@1.31.31,trna85_ValCAC_1_-_161369490_161369562@1.31.31,trna90_ValCAC_1_-_149684088_149684161@1.31.31,trna98_ValCAC_1_-_149298555_149298627@1.31.31,trna9_ValCAC_6_+_26538282_26538354@1.31.31\|GTTTCCGTAGTGTAGTGGTTATCACGTTCGC | >am_trna10_ValCAC_5_-_180649395_180649467 | 1 | 31 | 31 | GTTTCCGTAGTGTAGTGGTTATCACGTTCGC | 556.0 | -4.0 | 0.9 | -4.3 | 1.7E-05 | 1.5E-04 | BT20/MB231 | MITO |
| >hg38_21_+_8433172_8446622_RNA45SN1_45S_50ntFlanks@11212.11254.43\|CGACTCTGGACGCGAGCCGGGCCCTTCCCGTGGATCGCCCCAG | >hg38_21_+_8433172_8446622_RNA45SN1_45S_50ntFlanks | 11212 | 11254 | 43 | CGACTCTGGACGCGAGCCGGGCCCTTCCCGTGGATCGCCCCAG | 637.9 | -5.3 | 1.3 | -3.9 | 8.4E-05 | 1.5E-04 | BT20/MB468 | MITO |
| >hg38_21_+_8433172_8446622_RNA45SN1_45S_50ntFlanks@6651.6686.36\|CGACTCTTAGCGGTGGATCACTCGGCTCGTGCGTCG | >hg38_21_+_8433172_8446622_RNA45SN1_45S_50ntFlanks | 6651 | 6686 | 36 | CGACTCTTAGCGGTGGATCACTCGGCTCGTGCGTCG | 3425.6 | -2.7 | 0.7 | -3.9 | 1.1E-04 | 1.5E-04 | MB231/MB468 | MITO |
| >hg38_21_+_8433172_8446622_RNA45SN1_45S_50ntFlanks@9032.9059.28\|CCTCGTGGGGGGGCCGGGCCACCCCTCC | >hg38_21_+_8433172_8446622_RNA45SN1_45S_50ntFlanks | 9032 | 9059 | 28 | CCTCGTGGGGGGGCCGGGCCACCCCTCC | 540.9 | -6.4 | 1.6 | -4.1 | 4.1E-05 | 1.5E-04 | BT20/MB468 | MP |
| >hg38_21_+_8433172_8446622_RNA45SN1_45S_50ntFlanks@4857.4876.20\|CTTTGGGTTCCGGGGGGAGT | >hg38_21_+_8433172_8446622_RNA45SN1_45S_50ntFlanks | 4857 | 4876 | 20 | CTTTGGGTTCCGGGGGGAGT | 154.0 | 6.6 | 1.6 | 4.1 | 4.1E-05 | 1.5E-04 | BT20/MB468 | MP |
| >hg38_21_+_8433172_8446622_RNA45SN1_45S_50ntFlanks@12912.12933.22\|CACGTTCGTGGGGAACCTGGCG | >hg38_21_+_8433172_8446622_RNA45SN1_45S_50ntFlanks | 12912 | 12933 | 22 | CACGTTCGTGGGGAACCTGGCG | 1043.5 | -4.9 | 1.3 | -3.9 | 1.0E-04 | 1.5E-04 | BT20/MB468 | CYTO |
| >hg38_21_+_8433172_8446622_RNA45SN1_45S_50ntFlanks@10352.10383.32\|GAGAACTTTGAAGGCCGAAGTGGAGAAGGGTT | >hg38_21_+_8433172_8446622_RNA45SN1_45S_50ntFlanks | 10352 | 10383 | 32 | GAGAACTTTGAAGGCCGAAGTGGAGAAGGGTT | 271.7 | -3.1 | 0.8 | -3.9 | 9.7E-05 | 1.5E-04 | MB231/MB468 | MP |
| >hg38_21_+_8433172_8446622_RNA45SN1_45S_50ntFlanks@10809.10840.32\|AATGTAGGTAAGGGAAGTCGGCAAGCCGGATC | >hg38_21_+_8433172_8446622_RNA45SN1_45S_50ntFlanks | 10809 | 10840 | 32 | AATGTAGGTAAGGGAAGTCGGCAAGCCGGATC | 121.5 | -4.6 | 1.2 | -3.9 | 9.7E-05 | 1.5E-04 | MB231/MB468 | MP |
| >hg38_21_+_8433172_8446622_RNA45SN1_45S_50ntFlanks@10186.10208.23\|TCCTCCCCTCCTCCCGCCCACGC | >hg38_21_+_8433172_8446622_RNA45SN1_45S_50ntFlanks | 10186 | 10208 | 23 | TCCTCCCCTCCTCCCGCCCACGC | 202.3 | -5.2 | 1.3 | -3.9 | 1.0E-04 | 1.5E-04 | BT20/MB468 | CYTO |
| >hg38_21_+_8433172_8446622_RNA45SN1_45S_50ntFlanks@7975.7995.21\|CGCGACCTCAGATCAGACGTG | >hg38_21_+_8433172_8446622_RNA45SN1_45S_50ntFlanks | 7975 | 7995 | 21 | CGCGACCTCAGATCAGACGTG | 7882.2 | -2.9 | 0.7 | -3.9 | 9.7E-05 | 1.5E-04 | MB231/MB468 | MP |
| >hg38_21_+_8433172_8446622_RNA45SN1_45S_50ntFlanks@10855.10875.21\|AAGGATTGGCTCTAAGGGCTG | >hg38_21_+_8433172_8446622_RNA45SN1_45S_50ntFlanks | 10855 | 10875 | 21 | AAGGATTGGCTCTAAGGGCTG | 96.7 | -5.7 | 1.5 | -3.9 | 9.7E-05 | 1.5E-04 | MB231/MB468 | MP |
| >hg38_21_+_8433172_8446622_RNA45SN1_45S_50ntFlanks@4804.4822.19\|ATGCGGCGGCGTTATTCCC | >hg38_21_+_8433172_8446622_RNA45SN1_45S_50ntFlanks | 4804 | 4822 | 19 | ATGCGGCGGCGTTATTCCC | 211.5 | 3.1 | 0.8 | 3.9 | 1.1E-04 | 1.5E-04 | MB231/MB468 | CYTO |
| >hg38_21_+_8433172_8446622_RNA45SN1_45S_50ntFlanks@7978.8011.34\|GACCTCAGATCAGACGTGGCGACCCGCTGAATTT | >hg38_21_+_8433172_8446622_RNA45SN1_45S_50ntFlanks | 7978 | 8011 | 34 | GACCTCAGATCAGACGTGGCGACCCGCTGAATTT | 606.7 | 3.4 | 0.9 | 3.9 | 1.2E-04 | 1.5E-04 | MB231/MB468 | MITO |
| >hg38_21_+_8433172_8446622_RNA45SN1_45S_50ntFlanks@10404.10435.32\|CATGGGTCAGTCGGTCCTGAGAGATGGGCGAG | >hg38_21_+_8433172_8446622_RNA45SN1_45S_50ntFlanks | 10404 | 10435 | 32 | CATGGGTCAGTCGGTCCTGAGAGATGGGCGAG | 142.3 | -5.0 | 1.3 | -3.9 | 8.5E-05 | 1.5E-04 | BT20/MB468 | MITO |
| >hg38_21_+_8433172_8446622_RNA45SN1_45S_50ntFlanks@7979.8004.26\|ACCTCAGATCAGACGTGGCGACCCGC | >hg38_21_+_8433172_8446622_RNA45SN1_45S_50ntFlanks | 7979 | 8004 | 26 | ACCTCAGATCAGACGTGGCGACCCGC | 530.6 | -3.1 | 0.8 | -3.9 | 1.2E-04 | 1.5E-04 | MB231/MB468 | MITO |
| >hg38_21_+_8433172_8446622_RNA45SN1_45S_50ntFlanks@12456.12498.43\|CCAAGCGTTGGATTGTTCACCCACTAATAGGGAACGTGAGCTG | >hg38_21_+_8433172_8446622_RNA45SN1_45S_50ntFlanks | 12456 | 12498 | 43 | CCAAGCGTTGGATTGTTCACCCACTAATAGGGAACGTGAGCTG | 39.2 | -8.4 | 2.0 | -4.1 | 4.2E-05 | 1.5E-04 | BT20/MB468 | MP |
| >hg38_MT_+_1621_3279_RNR2_16S_50ntFlanks@1356.1396.41\|GGTTTACGACCTCGATGTTGGATCAGGACATCCCGATGGTG | >hg38_MT_+_1621_3279_RNR2_16S_50ntFlanks | 1356 | 1396 | 41 | GGTTTACGACCTCGATGTTGGATCAGGACATCCCGATGGTG | 24.6 | 7.8 | 2.0 | 3.9 | 9.7E-05 | 1.5E-04 | MB231/MB468 | MP |
| >hg38_1_-_228634819_228635039_RNA5S12_5S_50ntFlanks@149.169.21\|GAATACCGGGTGCTGTAGGCT | >hg38_1_-_228634819_228635039_RNA5S12_5S_50ntFlanks | 149 | 169 | 21 | GAATACCGGGTGCTGTAGGCT | 330.4 | 3.0 | 0.7 | 4.3 | 1.7E-05 | 1.5E-04 | BT20/MB231 | MITO |
| >hg38_21_+_8433172_8446622_RNA45SN1_45S_50ntFlanks@12990.13016.27\|GCTCCCTCGCTGCGATCTATTGAAAGT | >hg38_21_+_8433172_8446622_RNA45SN1_45S_50ntFlanks | 12990 | 13016 | 27 | GCTCCCTCGCTGCGATCTATTGAAAGT | 168.4 | 2.8 | 0.7 | 3.9 | 1.1E-04 | 1.5E-04 | MB231/MB468 | CYTO |
| >hg38_21_+_8433172_8446622_RNA45SN1_45S_50ntFlanks@11209.11233.25\|CGGCGACTCTGGACGCGAGCCGGGC | >hg38_21_+_8433172_8446622_RNA45SN1_45S_50ntFlanks | 11209 | 11233 | 25 | CGGCGACTCTGGACGCGAGCCGGGC | 1075.5 | -3.9 | 1.0 | -4.1 | 4.2E-05 | 1.5E-04 | BT20/MB468 | MP |
| >hg38_21_+_8433172_8446622_RNA45SN1_45S_50ntFlanks@9714.9735.22\|CGAATGATTAGAGGTCTTGGGG | >hg38_21_+_8433172_8446622_RNA45SN1_45S_50ntFlanks | 9714 | 9735 | 22 | CGAATGATTAGAGGTCTTGGGG | 444.3 | -4.8 | 1.2 | -3.9 | 8.5E-05 | 1.5E-04 | BT20/MB468 | MITO |
| >hg38_21_+_8433172_8446622_RNA45SN1_45S_50ntFlanks@12627.12661.35\|ATGGGGCGAAGCTACCATCTGTGGGATTATGACTG | >hg38_21_+_8433172_8446622_RNA45SN1_45S_50ntFlanks | 12627 | 12661 | 35 | ATGGGGCGAAGCTACCATCTGTGGGATTATGACTG | 327.7 | -3.8 | 1.0 | -3.9 | 1.0E-04 | 1.5E-04 | BT20/MB468 | CYTO |
| >hg38_21_+_8433172_8446622_RNA45SN1_45S_50ntFlanks@3992.4024.33\|GGTGACTCTAGATAACCTCGGGCCGATCGCACG | >hg38_21_+_8433172_8446622_RNA45SN1_45S_50ntFlanks | 3992 | 4024 | 33 | GGTGACTCTAGATAACCTCGGGCCGATCGCACG | 466.5 | 2.9 | 0.7 | 3.9 | 1.1E-04 | 1.5E-04 | MB231/MB468 | CYTO |
| >hg38_21_+_8433172_8446622_RNA45SN1_45S_50ntFlanks@10411.10453.43\|CAGTCGGTCCTGAGAGATGGGCGAGCGCCGTTCCGAAGGGACG | >hg38_21_+_8433172_8446622_RNA45SN1_45S_50ntFlanks | 10411 | 10453 | 43 | CAGTCGGTCCTGAGAGATGGGCGAGCGCCGTTCCGAAGGGACG | 643.2 | -4.3 | 1.0 | -4.1 | 4.2E-05 | 1.5E-04 | BT20/MB468 | MP |
| >hg38_21_+_8433172_8446622_RNA45SN1_45S_50ntFlanks@11562.11600.39\|CGCCTAGCAGCCGACTTAGAACTGGTGCGGACCAGGGGA | >hg38_21_+_8433172_8446622_RNA45SN1_45S_50ntFlanks | 11562 | 11600 | 39 | CGCCTAGCAGCCGACTTAGAACTGGTGCGGACCAGGGGA | 124.4 | -6.3 | 1.6 | -3.9 | 9.8E-05 | 1.5E-04 | MB231/MB468 | MP |
| >hg38_21_+_8433172_8446622_RNA45SN1_45S_50ntFlanks@474.517.44\|TCTTCGTGATCGATGTGGTGACGTCGTGCTCTCCCGGGCCGGGT | >hg38_21_+_8433172_8446622_RNA45SN1_45S_50ntFlanks | 474 | 517 | 44 | TCTTCGTGATCGATGTGGTGACGTCGTGCTCTCCCGGGCCGGGT | 392.5 | 4.0 | 1.0 | 3.9 | 1.2E-04 | 1.5E-04 | MB231/MB468 | NUC |
| >hg38_21_+_8433172_8446622_RNA45SN1_45S_50ntFlanks@10793.10818.26\|TCTGGCATGTTGGAACAATGTAGGTA | >hg38_21_+_8433172_8446622_RNA45SN1_45S_50ntFlanks | 10793 | 10818 | 26 | TCTGGCATGTTGGAACAATGTAGGTA | 353.8 | -5.1 | 1.3 | -4.1 | 4.2E-05 | 1.5E-04 | BT20/MB468 | MP |
| >hg38_21_+_8433172_8446622_RNA45SN1_45S_50ntFlanks@8116.8143.28\|GCGGGACATGTGGCGTACGGAAGACCCG | >hg38_21_+_8433172_8446622_RNA45SN1_45S_50ntFlanks | 8116 | 8143 | 28 | GCGGGACATGTGGCGTACGGAAGACCCG | 334.0 | -3.8 | 0.9 | -4.1 | 4.2E-05 | 1.5E-04 | BT20/MB468 | MP |
| >hg38_21_+_8433172_8446622_RNA45SN1_45S_50ntFlanks@9604.9630.27\|AAAGACTAATCGAACCATCTAGTAGCT | >hg38_21_+_8433172_8446622_RNA45SN1_45S_50ntFlanks | 9604 | 9630 | 27 | AAAGACTAATCGAACCATCTAGTAGCT | 518.1 | 2.8 | 0.7 | 3.9 | 8.6E-05 | 1.5E-04 | BT20/MB468 | MITO |
| >hg38_21_+_8433172_8446622_RNA45SN1_45S_50ntFlanks@9736.9758.23\|CCGAAACGATCTCAACCTATTCT | >hg38_21_+_8433172_8446622_RNA45SN1_45S_50ntFlanks | 9736 | 9758 | 23 | CCGAAACGATCTCAACCTATTCT | 168.7 | -6.1 | 1.5 | -4.1 | 4.2E-05 | 1.5E-04 | BT20/MB468 | MP |
| >trna10_LysCTT_16_+_3241501_3241573@3.36.34,trna13_LysCTT_14_-_58706613_58706685@3.36.34,trna2_LysCTT_15_+_79152904_79152976@3.36.34\|CCGGCTAGCTCAGTCGGTAGAGCATGGGACTCTT | >trna10_LysCTT_16_+_3241501_3241573 | 3 | 36 | 34 | CCGGCTAGCTCAGTCGGTAGAGCATGGGACTCTT | 647.9 | 6.1 | 1.6 | 3.9 | 1.2E-04 | 1.5E-04 | MB231/MB468 | MITO |
| >trna17_ValTAC_11_-_59318102_59318174@1.36.36,trna4_ValTAC_X_-_18693029_18693101@1.36.36\|GGTTCCATAGTGTAGTGGTTATCACGTCTGCTTTAC | >trna17_ValTAC_11_-_59318102_59318174 | 1 | 36 | 36 | GGTTCCATAGTGTAGTGGTTATCACGTCTGCTTTAC | 164.2 | 5.8 | 1.5 | 3.9 | 1.2E-04 | 1.5E-04 | MB231/MB468 | MITO |
| >hg38_21_+_8433172_8446622_RNA45SN1_45S_50ntFlanks@10343.10362.20\|TATTCAAACGAGAACTTTGA | >hg38_21_+_8433172_8446622_RNA45SN1_45S_50ntFlanks | 10343 | 10362 | 20 | TATTCAAACGAGAACTTTGA | 198.2 | 3.3 | 0.8 | 4.1 | 4.2E-05 | 1.5E-04 | BT20/MB468 | MP |
| >trnaMT_PheGAA_MT_+_577_647@35.74.40\|GAAAATGTTTAGACGGGCTCACATCACCCCATAAACACCA | >trnaMT_PheGAA_MT_+_577_647 | 35 | 74 | 40 | GAAAATGTTTAGACGGGCTCACATCACCCCATAAACACCA | 116.5 | 10.6 | 2.6 | 4.1 | 4.2E-05 | 1.5E-04 | BT20/MB468 | MP |
| >hg38_21_+_8433172_8446622_RNA45SN1_45S_50ntFlanks@7974.7995.22\|ACGCGACCTCAGATCAGACGTG | >hg38_21_+_8433172_8446622_RNA45SN1_45S_50ntFlanks | 7974 | 7995 | 22 | ACGCGACCTCAGATCAGACGTG | 2114.7 | 1.7 | 0.4 | 3.9 | 1.2E-04 | 1.5E-04 | MB231/MB468 | NUC |
| >hg38_MT_+_1621_3279_RNR2_16S_50ntFlanks@485.531.47\|GAGGAACAGCTCTTTGGACACTAGGAAAAAACCTTGTAGAGAGAGTA | >hg38_MT_+_1621_3279_RNR2_16S_50ntFlanks | 485 | 531 | 47 | GAGGAACAGCTCTTTGGACACTAGGAAAAAACCTTGTAGAGAGAGTA | 120.7 | 10.7 | 2.6 | 4.1 | 4.2E-05 | 1.5E-04 | BT20/MB468 | MP |
| >hg38_21_+_8433172_8446622_RNA45SN1_45S_50ntFlanks@8184.8210.27\|CGAGGCCCAGCCCGTGGACGGTGTGAG | >hg38_21_+_8433172_8446622_RNA45SN1_45S_50ntFlanks | 8184 | 8210 | 27 | CGAGGCCCAGCCCGTGGACGGTGTGAG | 324.4 | -6.1 | 1.6 | -3.9 | 9.8E-05 | 1.5E-04 | MB231/MB468 | MP |

<table>
<tr><td>>MI0000750|hsa-mir-26a-2&WithFlank&12|-|57824603|57824698@23.40.18(+1A);MI0000083|hsa-mir-26a-1&WithFlank&3|+|37969398|37969486@19.36.18(+1A)&[MIMAT0000082&hsa-miR-26a-5p&offsets|+3|-1(+1A);m-33&12|-|57824658|57824679&offsets|+3|-1(+1A)];[MIMAT0000082_1&hsa-miR-26a-5p&offsets|+3|-1(+1A);m-34&3|+|37969413|37969434&offsets|+3|-1(+1A)]|AAGTAATCCAGGATAGGCA</td><td>>MI0000750|hsa-mir-26a-2&WithFlank&12|-|57824603|57824698</td><td>23</td><td>40</td><td>18(+1A)</td><td>AAGTAATCCAGGATAGGCA</td><td>99.0</td><td>9.8</td><td>2.5</td><td>3.9</td><td>9.8E-05</td><td>1.5E-04</td><td>MB231/MB468</td><td>MP</td></tr>
<tr><td>>hg38_21_+_8433172_8446622_RNA45SN1_45S_50ntFlanks@11921.11951.31|TGAGAGGTGTAGAATAAGTGGGAGGCCCCCG</td><td>>hg38_21_+_8433172_8446622_RNA45SN1_45S_50ntFlanks</td><td>11921</td><td>11951</td><td>31</td><td>TGAGAGGTGTAGAATAAGTGGGAGGCCCCCG</td><td>388.4</td><td>-4.3</td><td>1.1</td><td>-3.9</td><td>1.0E-04</td><td>1.5E-04</td><td>BT20/MB468</td><td>CYTO</td></tr>
<tr><td>>MI0000298|hsa-mir-221&WithFlank&X|-|45746151|45746272@71.88.18(+3U)&[MIMAT0000278&hsa-miR-221-3p&offsets|0|-5(+3U);m-91&X|-|45746181|45746202&offsets|0|-4(+3U)]|AGCTACATTGTCTGCTGGTTT</td><td>>MI0000298|hsa-mir-221&WithFlank&X|-|45746151|45746272</td><td>71</td><td>88</td><td>18(+3U)</td><td>AGCTACATTGTCTGCTGGTTT</td><td>57.0</td><td>-9.2</td><td>2.1</td><td>-4.3</td><td>1.5E-05</td><td>1.5E-04</td><td>BT20/MB231</td><td>MP</td></tr>
<tr><td>>hg38_21_+_8433172_8446622_RNA45SN1_45S_50ntFlanks@4981.5002.22|CACGGACAGGATTGACAGATTG</td><td>>hg38_21_+_8433172_8446622_RNA45SN1_45S_50ntFlanks</td><td>4981</td><td>5002</td><td>22</td><td>CACGGACAGGATTGACAGATTG</td><td>128.9</td><td>-3.5</td><td>0.9</td><td>-3.9</td><td>9.8E-05</td><td>1.5E-04</td><td>MB231/MB468</td><td>MP</td></tr>
<tr><td>>hg38_21_+_8433172_8446622_RNA45SN1_45S_50ntFlanks@8114.8141.28|GCGCGGGACATGTGGCGTACGGAAGACC</td><td>>hg38_21_+_8433172_8446622_RNA45SN1_45S_50ntFlanks</td><td>8114</td><td>8141</td><td>28</td><td>GCGCGGGACATGTGGCGTACGGAAGACC</td><td>220.3</td><td>-5.5</td><td>1.4</td><td>-3.9</td><td>1.2E-04</td><td>1.5E-04</td><td>MB231/MB468</td><td>NUC</td></tr>
<tr><td>>hg38_21_+_8433172_8446622_RNA45SN1_45S_50ntFlanks@9554.9578.25|CGTAGCGGTCCTGACGTGCAAATCG</td><td>>hg38_21_+_8433172_8446622_RNA45SN1_45S_50ntFlanks</td><td>9554</td><td>9578</td><td>25</td><td>CGTAGCGGTCCTGACGTGCAAATCG</td><td>352.9</td><td>-6.5</td><td>1.7</td><td>-3.8</td><td>1.2E-04</td><td>1.5E-04</td><td>BT20/MB468</td><td>NUC</td></tr>
<tr><td>>hg38_21_+_8433172_8446622_RNA45SN1_45S_50ntFlanks@9930.9949.20|ATATAGACAGCAGGACGGTG</td><td>>hg38_21_+_8433172_8446622_RNA45SN1_45S_50ntFlanks</td><td>9930</td><td>9949</td><td>20</td><td>ATATAGACAGCAGGACGGTG</td><td>313.3</td><td>-4.6</td><td>1.2</td><td>-3.9</td><td>9.8E-05</td><td>1.5E-04</td><td>MB231/MB468</td><td>MP</td></tr>
<tr><td>>hg38_21_+_8433172_8446622_RNA45SN1_45S_50ntFlanks@9738.9762.25|GAAACGATCTCAACCTATTCTCAAA</td><td>>hg38_21_+_8433172_8446622_RNA45SN1_45S_50ntFlanks</td><td>9738</td><td>9762</td><td>25</td><td>GAAACGATCTCAACCTATTCTCAAA</td><td>236.5</td><td>-4.3</td><td>1.1</td><td>-3.9</td><td>1.2E-04</td><td>1.5E-04</td><td>MB231/MB468</td><td>NUC</td></tr>
<tr><td>>hg38_21_+_8433172_8446622_RNA45SN1_45S_50ntFlanks@9927.9949.23|TTGATATAGACAGCAGGACGGTG</td><td>>hg38_21_+_8433172_8446622_RNA45SN1_45S_50ntFlanks</td><td>9927</td><td>9949</td><td>23</td><td>TTGATATAGACAGCAGGACGGTG</td><td>251.1</td><td>7.5</td><td>1.8</td><td>4.1</td><td>4.2E-05</td><td>1.5E-04</td><td>BT20/MB468</td><td>MP</td></tr>
<tr><td>>hg38_21_+_8433172_8446622_RNA45SN1_45S_50ntFlanks@3855.3881.27|CTGTGGTAATTCTAGAGCTAATACATG</td><td>>hg38_21_+_8433172_8446622_RNA45SN1_45S_50ntFlanks</td><td>3855</td><td>3881</td><td>27</td><td>CTGTGGTAATTCTAGAGCTAATACATG</td><td>162.2</td><td>-6.1</td><td>1.5</td><td>-4.1</td><td>4.2E-05</td><td>1.5E-04</td><td>BT20/MB468</td><td>MP</td></tr>
<tr><td>>hg38_21_+_8433172_8446622_RNA45SN1_45S_50ntFlanks@6722.6743.22|ATTGCAGGACACATTGATCATC</td><td>>hg38_21_+_8433172_8446622_RNA45SN1_45S_50ntFlanks</td><td>6722</td><td>6743</td><td>22</td><td>ATTGCAGGACACATTGATCATC</td><td>765.7</td><td>-3.3</td><td>0.8</td><td>-3.9</td><td>1.0E-04</td><td>1.5E-04</td><td>BT20/MB468</td><td>CYTO</td></tr>
<tr><td>>hg38_21_+_8433172_8446622_RNA45SN1_45S_50ntFlanks@12936.12962.27|AAACCATTCGTAGACGACCTGCTTCTG</td><td>>hg38_21_+_8433172_8446622_RNA45SN1_45S_50ntFlanks</td><td>12936</td><td>12962</td><td>27</td><td>AAACCATTCGTAGACGACCTGCTTCTG</td><td>158.5</td><td>-3.3</td><td>0.9</td><td>-3.9</td><td>1.1E-04</td><td>1.5E-04</td><td>MB231/MB468</td><td>CYTO</td></tr>
<tr><td>>trna119_LysCTT_1_-_145395522_145395594@1.32.32,trna11_LysCTT_5_-_180648979_180649051@1.32.32,trna13_LysCTT_6_+_26556774_26556846@1.32.32,trna32_LysCTT_16_-_3207406_3207478@1.32.32,trna7_LysCTT_16_+_3225692_3225764@1.32.32,trna9_LysCTT_5_+_180634755_180634827@1.32.32|GCCCGGCTAGCTCAGTCGGTAGAGCATGAGAC</td><td>>trna119_LysCTT_1_-_145395522_145395594</td><td>1</td><td>32</td><td>32</td><td>GCCCGGCTAGCTCAGTCGGTAGAGCATGAGAC</td><td>2569.6</td><td>3.6</td><td>0.9</td><td>3.9</td><td>1.2E-04</td><td>1.5E-04</td><td>MB231/MB468</td><td>MITO</td></tr>
<tr><td>>hg38_21_+_8433172_8446622_RNA45SN1_45S_50ntFlanks@11782.11803.22|GCATGAATGGATGAACGAGATT</td><td>>hg38_21_+_8433172_8446622_RNA45SN1_45S_50ntFlanks</td><td>11782</td><td>11803</td><td>22</td><td>GCATGAATGGATGAACGAGATT</td><td>103.5</td><td>-4.5</td><td>1.2</td><td>-3.9</td><td>9.9E-05</td><td>1.5E-04</td><td>MB231/MB468</td><td>MP</td></tr>
<tr><td>>hg38_21_+_8433172_8446622_RNA45SN1_45S_50ntFlanks@12075.12095.21|GAGGGGCTCTCGCTTCTGGCG</td><td>>hg38_21_+_8433172_8446622_RNA45SN1_45S_50ntFlanks</td><td>12075</td><td>12095</td><td>21</td><td>GAGGGGCTCTCGCTTCTGGCG</td><td>268.4</td><td>-5.7</td><td>1.5</td><td>-3.9</td><td>1.2E-04</td><td>1.5E-04</td><td>MB231/MB468</td><td>MITO</td></tr>
<tr><td>>hg38_MT_+_1621_3279_RNR2_16S_50ntFlanks@1349.1388.40|ACAATAGGGTTTACGACCTCGATGTTGGATCAGGACATCC</td><td>>hg38_MT_+_1621_3279_RNR2_16S_50ntFlanks</td><td>1349</td><td>1388</td><td>40</td><td>ACAATAGGGTTTACGACCTCGATGTTGGATCAGGACATCC</td><td>409.4</td><td>4.5</td><td>1.1</td><td>3.9</td><td>9.9E-05</td><td>1.5E-04</td><td>MB231/MB468</td><td>MP</td></tr>
<tr><td>>hg38_21_+_8433172_8446622_RNA45SN1_45S_50ntFlanks@11933.11951.19|AATAAGTGGGAGGCCCCCG</td><td>>hg38_21_+_8433172_8446622_RNA45SN1_45S_50ntFlanks</td><td>11933</td><td>11951</td><td>19</td><td>AATAAGTGGGAGGCCCCCG</td><td>615.7</td><td>-6.9</td><td>1.7</td><td>-4.1</td><td>4.2E-05</td><td>1.5E-04</td><td>BT20/MB468</td><td>MP</td></tr>
<tr><td>>hg38_1_-_228634819_228635039_RNA5S12_5S_50ntFlanks@132.169.38|GATGGGAGACCGCCTGGGAATACCGGGTGCTGTAGGCT</td><td>>hg38_1_-_228634819_228635039_RNA5S12_5S_50ntFlanks</td><td>132</td><td>169</td><td>38</td><td>GATGGGAGACCGCCTGGGAATACCGGGTGCTGTAGGCT</td><td>14681.2</td><td>3.7</td><td>1.0</td><td>3.9</td><td>1.0E-04</td><td>1.6E-04</td><td>BT20/MB468</td><td>CYTO</td></tr>
<tr><td>>am_trna11_ProAGG_16_+_3241989_3242060@1.28.28,trna12_ProAGG_6_+_26555498_26555569@1.28.28,trna12_ProTGG_11_-_75946869_75946940@1.28.28,trna14_ProTGG_5_-_180615854_180615925@1.28.28,trna22_ProAGG_14_-_21081560_21081631@1.28.28,trna23_ProAGG_14_-_21077495_21077566@1.28.28,trna28_ProTGG_16_-_3234133_3234204@1.28.28,trna29_ProAGG_16_-_3232635_3232706@1.28.28,trna2_ProAGG_7_+_128423504_128423575@1.28.28,trna30_ProCGG_6_+_27059521_27059592@1.28.28,trna37_ProCGG_17_-_8126151_8126222@1.28.28,trna3_ProTGG_16_+_3208923_3208994@1.28.28,trna52_ProCGG_1_+_167683962_167684033@1.28.28,trna65_ProAGG_1_-_167684725_167684796@1.28.28,trna6_ProCGG_16_+_3222049_3222120@1.28.28,trna6_ProTGG_14_+_21152175_21152246@1.28.28,trna8_ProTGG_16_+_3238094_3238165@1.28.28,trna9_ProAGG_11_+_75946557_75946628@1.28.28,trna9_ProAGG_16_+_3239634_3239705@1.28.28|GGCTCGTTGGTCTAGGGGTATGATTCTC</td><td>>am_trna11_ProAGG_16_+_3241989_3242060</td><td>1</td><td>28</td><td>28</td><td>GGCTCGTTGGTCTAGGGGTATGATTCTC</td><td>2579.0</td><td>-5.2</td><td>1.3</td><td>-3.9</td><td>1.0E-04</td><td>1.6E-04</td><td>BT20/MB468</td><td>CYTO</td></tr>
<tr><td>>hg38_21_+_8433172_8446622_RNA45SN1_45S_50ntFlanks@11217.11239.23|CTGGACGCGAGCCGGGCCCTTCC</td><td>>hg38_21_+_8433172_8446622_RNA45SN1_45S_50ntFlanks</td><td>11217</td><td>11239</td><td>23</td><td>CTGGACGCGAGCCGGGCCCTTCC</td><td>381.4</td><td>-5.3</td><td>1.4</td><td>-3.9</td><td>9.9E-05</td><td>1.6E-04</td><td>MB231/MB468</td><td>MP</td></tr>
<tr><td>>hg38_21_+_8433172_8446622_RNA45SN1_45S_50ntFlanks@6652.6690.39|GACTCTTAGCGGTGGATCACTCGGCTCGTGCGTCGATGA</td><td>>hg38_21_+_8433172_8446622_RNA45SN1_45S_50ntFlanks</td><td>6652</td><td>6690</td><td>39</td><td>GACTCTTAGCGGTGGATCACTCGGCTCGTGCGTCGATGA</td><td>3029.2</td><td>3.0</td><td>0.8</td><td>3.9</td><td>8.7E-05</td><td>1.6E-04</td><td>BT20/MB468</td><td>MITO</td></tr>
<tr><td>>hg38_21_+_8433172_8446622_RNA45SN1_45S_50ntFlanks@4760.4783.24|TCAGATACCGTCGTAGTTCCGACC</td><td>>hg38_21_+_8433172_8446622_RNA45SN1_45S_50ntFlanks</td><td>4760</td><td>4783</td><td>24</td><td>TCAGATACCGTCGTAGTTCCGACC</td><td>706.3</td><td>-4.0</td><td>1.0</td><td>-3.9</td><td>8.7E-05</td><td>1.6E-04</td><td>BT20/MB468</td><td>MITO</td></tr>
<tr><td>>MI0000734|hsa-mir-106b&WithFlank&7|-|100093987|100094080@56.76.21&[MIMAT0004672&hsa-miR-106b-3p&offsets|-2|-3;m-87&7|-|100094003|100094023&offsets|-2|-2]|TACCGCACTGTGGGTACTTGC</td><td>>MI0000734|hsa-mir-106b&WithFlank&7|-|100093987|100094080</td><td>56</td><td>76</td><td>21</td><td>TACCGCACTGTGGGTACTTGC</td><td>139.9</td><td>8.0</td><td>1.9</td><td>4.1</td><td>4.3E-05</td><td>1.6E-04</td><td>BT20/MB468</td><td>MP</td></tr>
<tr><td>>hg38_21_+_8433172_8446622_RNA45SN1_45S_50ntFlanks@6773.6797.25|TTCCTCCCGGGGCTACGCCTGTCTG</td><td>>hg38_21_+_8433172_8446622_RNA45SN1_45S_50ntFlanks</td><td>6773</td><td>6797</td><td>25</td><td>TTCCTCCCGGGGCTACGCCTGTCTG</td><td>349.3</td><td>-4.4</td><td>1.1</td><td>-3.9</td><td>8.7E-05</td><td>1.6E-04</td><td>BT20/MB468</td><td>MITO</td></tr>
<tr><td>>hg38_21_+_8433172_8446622_RNA45SN1_45S_50ntFlanks@10048.10078.31|CGGCCGTCGCCGGCAGTCGAGAGTGGACGGG</td><td>>hg38_21_+_8433172_8446622_RNA45SN1_45S_50ntFlanks</td><td>10048</td><td>10078</td><td>31</td><td>CGGCCGTCGCCGGCAGTCGAGAGTGGACGGG</td><td>70.7</td><td>9.5</td><td>2.2</td><td>4.3</td><td>1.5E-05</td><td>1.6E-04</td><td>BT20/MB231</td><td>MP</td></tr>
<tr><td>>hg38_21_+_8433172_8446622_RNA45SN1_45S_50ntFlanks@12478.12498.21|ACTAATAGGGAACGTGAGCTG</td><td>>hg38_21_+_8433172_8446622_RNA45SN1_45S_50ntFlanks</td><td>12478</td><td>12498</td><td>21</td><td>ACTAATAGGGAACGTGAGCTG</td><td>744.4</td><td>-2.4</td><td>0.6</td><td>-3.9</td><td>1.1E-04</td><td>1.6E-04</td><td>MB231/MB468</td><td>CYTO</td></tr>
<tr><td>>MI0000067|hsa-let-7f-1&WithFlank&9|+|94176341|94176439@13.33.21(+1C);MI0000068|hsa-let-7f-2&WithFlank&X|-|53557186|53557280@14.34.21(+1C)&[MIMAT0000067&hsa-let-7f-5p&offsets|0|-1(+1C);m-20&9|+|94176353|94176374&offsets|0|-1(+1C)];[MIMAT0000067_1&hsa-let-7f-5p&offsets|0|-1(+1C);m-19&X|-|53557246|53557267&offsets|0|-1(+1C)]|TGAGGTAGTAGATTGTATAGTC</td><td>>MI0000067|hsa-let-7f-1&WithFlank&9|+|94176341|94176439</td><td>13</td><td>33</td><td>21(+1C)</td><td>TGAGGTAGTAGATTGTATAGTC</td><td>9944.8</td><td>3.8</td><td>1.0</td><td>3.9</td><td>9.9E-05</td><td>1.6E-04</td><td>MB231/MB468</td><td>MP</td></tr>
<tr><td>>hg38_21_+_8433172_8446622_RNA45SN1_45S_50ntFlanks@3985.4026.42|CGGCTTTGGTGACTCTAGATAACCTCGGGCCGATCGCACGCC</td><td>>hg38_21_+_8433172_8446622_RNA45SN1_45S_50ntFlanks</td><td>3985</td><td>4026</td><td>42</td><td>CGGCTTTGGTGACTCTAGATAACCTCGGGCCGATCGCACGCC</td><td>1521.9</td><td>-7.2</td><td>1.8</td><td>-4.1</td><td>4.3E-05</td><td>1.6E-04</td><td>BT20/MB468</td><td>MP</td></tr>
<tr><td>>hg38_21_+_8433172_8446622_RNA45SN1_45S_50ntFlanks@4559.4589.31|GCTAGGAATAATGGAATAGGACCGCGGTTCT</td><td>>hg38_21_+_8433172_8446622_RNA45SN1_45S_50ntFlanks</td><td>4559</td><td>4589</td><td>31</td><td>GCTAGGAATAATGGAATAGGACCGCGGTTCT</td><td>76.2</td><td>9.6</td><td>2.2</td><td>4.3</td><td>1.5E-05</td><td>1.6E-04</td><td>BT20/MB231</td><td>MP</td></tr>
<tr><td>>hg38_21_+_8433172_8446622_RNA45SN1_45S_50ntFlanks@12241.12268.28|CAGAAGGGCAAAAGCTCGCTTGATCTTG</td><td>>hg38_21_+_8433172_8446622_RNA45SN1_45S_50ntFlanks</td><td>12241</td><td>12268</td><td>28</td><td>CAGAAGGGCAAAAGCTCGCTTGATCTTG</td><td>470.1</td><td>-6.5</td><td>1.6</td><td>-4.1</td><td>4.3E-05</td><td>1.6E-04</td><td>BT20/MB468</td><td>MP</td></tr>
<tr><td>>hg38_21_+_8433172_8446622_RNA45SN1_45S_50ntFlanks@10784.10818.35|TGAACAGCCTCTGGCATGTTGGAACAATGTAGGTA</td><td>>hg38_21_+_8433172_8446622_RNA45SN1_45S_50ntFlanks</td><td>10784</td><td>10818</td><td>35</td><td>TGAACAGCCTCTGGCATGTTGGAACAATGTAGGTA</td><td>493.9</td><td>-3.0</td><td>0.8</td><td>-3.9</td><td>1.0E-04</td><td>1.6E-04</td><td>BT20/MB468</td><td>CYTO</td></tr>
<tr><td>>hg38_21_+_8433172_8446622_RNA45SN1_45S_50ntFlanks@9534.9558.25|AGGAAACTCTGGTGGAGGTCCGTAG</td><td>>hg38_21_+_8433172_8446622_RNA45SN1_45S_50ntFlanks</td><td>9534</td><td>9558</td><td>25</td><td>AGGAAACTCTGGTGGAGGTCCGTAG</td><td>449.6</td><td>-5.7</td><td>1.5</td><td>-3.9</td><td>1.0E-04</td><td>1.6E-04</td><td>BT20/MB468</td><td>CYTO</td></tr>
<tr><td>>hg38_21_+_8433172_8446622_RNA45SN1_45S_50ntFlanks@10806.10829.24|AACAATGTAGGTAAGGGAAGTCGG</td><td>>hg38_21_+_8433172_8446622_RNA45SN1_45S_50ntFlanks</td><td>10806</td><td>10829</td><td>24</td><td>AACAATGTAGGTAAGGGAAGTCGG</td><td>199.7</td><td>-3.9</td><td>1.0</td><td>-3.9</td><td>8.7E-05</td><td>1.6E-04</td><td>BT20/MB468</td><td>MITO</td></tr>
<tr><td>>hg38_21_+_8433172_8446622_RNA45SN1_45S_50ntFlanks@6652.6679.28|GACTCTTAGCGGTGGATCACTCGGCTCG</td><td>>hg38_21_+_8433172_8446622_RNA45SN1_45S_50ntFlanks</td><td>6652</td><td>6679</td><td>28</td><td>GACTCTTAGCGGTGGATCACTCGGCTCG</td><td>1118.1</td><td>-4.3</td><td>1.1</td><td>-3.9</td><td>1.0E-04</td><td>1.6E-04</td><td>MB231/MB468</td><td>MP</td></tr>
<tr><td>>MI0000650|hsa-mir-200c&WithFlank&12|+|6963693|6963772@50.72.23(+2U)&[MIMAT0000617&hsa-miR-200c-3p&offsets|0|0(+2U);m-54&12|+|6963742|6963764&offsets|0|0(+2U)]|TAATACTGCCGGGTAATGATGGATT</td><td>>MI0000650|hsa-mir-200c&WithFlank&12|+|6963693|6963772</td><td>50</td><td>72</td><td>23(+2U)</td><td>TAATACTGCCGGGTAATGATGGATT</td><td>126.6</td><td>2.0</td><td>0.5</td><td>3.8</td><td>1.2E-04</td><td>1.6E-04</td><td>BT20/MB468</td><td>NUC</td></tr>
<tr><td>>hg38_21_+_8433172_8446622_RNA45SN1_45S_50ntFlanks@6734.6752.19|ATTGATCATCGACACTTCG</td><td>>hg38_21_+_8433172_8446622_RNA45SN1_45S_50ntFlanks</td><td>6734</td><td>6752</td><td>19</td><td>ATTGATCATCGACACTTCG</td><td>371.9</td><td>-4.5</td><td>1.2</td><td>-3.9</td><td>1.0E-04</td><td>1.6E-04</td><td>MB231/MB468</td><td>MP</td></tr>
<tr><td>>hg38_1_-_228634819_228635039_RNA5S12_5S_50ntFlanks@148.167.20|GGAATACCGGGTGCTGTAGG</td><td>>hg38_1_-_228634819_228635039_RNA5S12_5S_50ntFlanks</td><td>148</td><td>167</td><td>20</td><td>GGAATACCGGGTGCTGTAGG</td><td>135.0</td><td>-5.9</td><td>1.5</td><td>-3.9</td><td>1.0E-04</td><td>1.6E-04</td><td>MB231/MB468</td><td>MP</td></tr>
<tr><td>>MI0000063|hsa-let-7b&WithFlank&22|+|46113680|46113774@12.29.18&[MIMAT0000063&hsa-let-7b-5p&offsets|0|-4;m-14&22|+|46113691|46113712&offsets|0|-4]|TGAGGTAGTAGGTTGTGT</td><td>>MI0000063|hsa-let-7b&WithFlank&22|+|46113680|46113774</td><td>12</td><td>29</td><td>18</td><td>TGAGGTAGTAGGTTGTGT</td><td>452.5</td><td>3.5</td><td>0.8</td><td>4.3</td><td>1.4E-05</td><td>1.6E-04</td><td>BT20/MB231</td><td>CYTO</td></tr>
<tr><td>>hg38_21_+_8433172_8446622_RNA45SN1_45S_50ntFlanks@11540.11586.47|CGCGGCGCCTCGCCTCGGCCGGCGCCTAGCAGCCGACTTAGAACTGG</td><td>>hg38_21_+_8433172_8446622_RNA45SN1_45S_50ntFlanks</td><td>11540</td><td>11586</td><td>47</td><td>CGCGGCGCCTCGCCTCGGCCGGCGCCTAGCAGCCGACTTAGAACTGG</td><td>488.3</td><td>-4.9</td><td>1.1</td><td>-4.3</td><td>1.4E-05</td><td>1.6E-04</td><td>BT20/MB231</td><td>CYTO</td></tr>
<tr><td>>MI0000253|hsa-mir-148a&WithFlank&7|-|25949913|25949992@50.71.22&[MIMAT0000243&hsa-miR-148a-3p&offsets|0|0;m-7&7|-|25949922|25949943&offsets|0|0]|TCAGTGCACTACAGAACTTTGT</td><td>>MI0000253|hsa-mir-148a&WithFlank&7|-|25949913|25949992</td><td>50</td><td>71</td><td>22</td><td>TCAGTGCACTACAGAACTTTGT</td><td>453219.6</td><td>4.4</td><td>1.1</td><td>4.1</td><td>4.3E-05</td><td>1.6E-04</td><td>BT20/MB468</td><td>MP</td></tr>
<tr><td>>trna11_GluTTC_15_-_26327381_26327452@1.30.30,trna3_GluTTC_13_-_45492062_45492133@1.30.30|TCCCACATGGTCTAGCGGTTAGGATTCCTG</td><td>>trna11_GluTTC_15_-_26327381_26327452</td><td>1</td><td>30</td><td>30</td><td>TCCCACATGGTCTAGCGGTTAGGATTCCTG</td><td>1357.3</td><td>-3.0</td><td>0.8</td><td>-3.8</td><td>1.2E-04</td><td>1.6E-04</td><td>MB231/MB468</td><td>MITO</td></tr>
</table>

<table>
<tr><td>>hg38_21_+_8433172_8446622_RNA45SN1_45S_50ntFlanks@8197.8225.29|GTGGACGGTGTGAGGCCGGTAGCGGCCCC</td><td>>hg38_21_+_8433172_8446622_RNA45SN1_45S_50ntFlanks</td><td>8197</td><td>8225</td><td>29</td><td>GTGGACGGTGTGAGGCCGGTAGCGGCCCC</td><td>80.3</td><td>-5.8</td><td>1.5</td><td>-3.9</td><td>1.0E-04</td><td>1.6E-04</td><td>MB231/MB468</td><td>MP</td></tr>
<tr><td>>MI0000086|hsa-mir-28&WithFlank&3|+|188688775|188688872@60.80.21(+1U)&[MIMAT0004502&hsa-miR-28-3p&offsets|0|-1(+1U);m-28&3|+|188688834|188688855&offsets|0|-1(+1U)]|CACTAGATTGTGAGCTCCTGGT</td><td>>MI0000086|hsa-mir-28&WithFlank&3|+|188688775|188688872</td><td>60</td><td>80</td><td>21(+1U)</td><td>CACTAGATTGTGAGCTCCTGGT</td><td>166.1</td><td>6.6</td><td>1.7</td><td>3.9</td><td>1.0E-04</td><td>1.6E-04</td><td>MB231/MB468</td><td>MP</td></tr>
<tr><td>>hg38_21_+_8433172_8446622_RNA45SN1_45S_50ntFlanks@9736.9763.28|CCGAAACGATCTCAACCTATTCTCAAAC</td><td>>hg38_21_+_8433172_8446622_RNA45SN1_45S_50ntFlanks</td><td>9736</td><td>9763</td><td>28</td><td>CCGAAACGATCTCAACCTATTCTCAAAC</td><td>202.8</td><td>-4.1</td><td>1.1</td><td>-3.8</td><td>1.2E-04</td><td>1.6E-04</td><td>MB231/MB468</td><td>NUC</td></tr>
<tr><td>>hg38_21_+_8433172_8446622_RNA45SN1_45S_50ntFlanks@11911.11940.30|TGAAGAGACATGAGAGGTGTAGAATAAGTG</td><td>>hg38_21_+_8433172_8446622_RNA45SN1_45S_50ntFlanks</td><td>11911</td><td>11940</td><td>30</td><td>TGAAGAGACATGAGAGGTGTAGAATAAGTG</td><td>76.6</td><td>9.6</td><td>2.2</td><td>4.3</td><td>1.5E-05</td><td>1.6E-04</td><td>BT20/MB231</td><td>MP</td></tr>
<tr><td>>hg38_21_+_8433172_8446622_RNA45SN1_45S_50ntFlanks@4731.4757.27|AAGAACGAAAGTCGGAGGTTCGAAGAC</td><td>>hg38_21_+_8433172_8446622_RNA45SN1_45S_50ntFlanks</td><td>4731</td><td>4757</td><td>27</td><td>AAGAACGAAAGTCGGAGGTTCGAAGAC</td><td>355.0</td><td>-3.0</td><td>0.8</td><td>-3.9</td><td>1.0E-04</td><td>1.6E-04</td><td>MB231/MB468</td><td>MP</td></tr>
<tr><td>>am_trna2_GlyGCC_21_-_18827107_18827177@1.19.19,trna35_GlyGCC_1_+_161413094_161413164@1.19.19,trna37_GlyGCC_1_+_161420467_161420537@1.19.19,trna39_GlyGCC_1_+_161427898_161427968@1.19.19,trna41_GlyGCC_1_+_161435258_161435328@1.19.19|GCATGGGTGGTTCAGTGGT</td><td>>am_trna2_GlyGCC_21_-_18827107_18827177</td><td>1</td><td>19</td><td>19</td><td>GCATGGGTGGTTCAGTGGT</td><td>582.3</td><td>4.0</td><td>1.0</td><td>3.9</td><td>1.1E-04</td><td>1.6E-04</td><td>BT20/MB468</td><td>CYTO</td></tr>
<tr><td>>hg38_21_+_8433172_8446622_RNA45SN1_45S_50ntFlanks@11571.11592.22|GCCGACTTAGAACTGGTGCGGA</td><td>>hg38_21_+_8433172_8446622_RNA45SN1_45S_50ntFlanks</td><td>11571</td><td>11592</td><td>22</td><td>GCCGACTTAGAACTGGTGCGGA</td><td>303.9</td><td>-3.1</td><td>0.8</td><td>-3.8</td><td>1.2E-04</td><td>1.6E-04</td><td>MB231/MB468</td><td>MITO</td></tr>
<tr><td>>hg38_21_+_8433172_8446622_RNA45SN1_45S_50ntFlanks@6719.6736.18|TGAATTGCAGGACACATT</td><td>>hg38_21_+_8433172_8446622_RNA45SN1_45S_50ntFlanks</td><td>6719</td><td>6736</td><td>18</td><td>TGAATTGCAGGACACATT</td><td>1054.6</td><td>-6.6</td><td>1.6</td><td>-4.1</td><td>4.3E-05</td><td>1.6E-04</td><td>BT20/MB468</td><td>MP</td></tr>
<tr><td>>hg38_21_+_8433172_8446622_RNA45SN1_45S_50ntFlanks@4722.4765.44|TCATTAATCAAGAACGAAAGTCGGAGGTTCGAAGACGATCAGAT</td><td>>hg38_21_+_8433172_8446622_RNA45SN1_45S_50ntFlanks</td><td>4722</td><td>4765</td><td>44</td><td>TCATTAATCAAGAACGAAAGTCGGAGGTTCGAAGACGATCAGAT</td><td>246.0</td><td>4.3</td><td>1.1</td><td>3.9</td><td>1.1E-04</td><td>1.6E-04</td><td>MB231/MB468</td><td>CYTO</td></tr>
<tr><td>>hg38_21_+_8433172_8446622_RNA45SN1_45S_50ntFlanks@4353.4371.19|TATTAAAGTTGCTGCAGTT</td><td>>hg38_21_+_8433172_8446622_RNA45SN1_45S_50ntFlanks</td><td>4353</td><td>4371</td><td>19</td><td>TATTAAAGTTGCTGCAGTT</td><td>325.1</td><td>-2.1</td><td>0.5</td><td>-3.9</td><td>1.1E-04</td><td>1.6E-04</td><td>BT20/MB468</td><td>CYTO</td></tr>
<tr><td>>hg38_21_+_8433172_8446622_RNA45SN1_45S_50ntFlanks@5486.5519.34|GCGCTGAGAAGACGGTCGAACTTGACTATCTAGA</td><td>>hg38_21_+_8433172_8446622_RNA45SN1_45S_50ntFlanks</td><td>5486</td><td>5519</td><td>34</td><td>GCGCTGAGAAGACGGTCGAACTTGACTATCTAGA</td><td>340.2</td><td>7.2</td><td>1.7</td><td>4.3</td><td>1.6E-05</td><td>1.6E-04</td><td>BT20/MB231</td><td>NUC</td></tr>
<tr><td>>hg38_21_+_8433172_8446622_RNA45SN1_45S_50ntFlanks@10871.10893.23|GGCTGGGTCGGTCGGGCTGGGGC</td><td>>hg38_21_+_8433172_8446622_RNA45SN1_45S_50ntFlanks</td><td>10871</td><td>10893</td><td>23</td><td>GGCTGGGTCGGTCGGGCTGGGGC</td><td>73.2</td><td>9.5</td><td>2.2</td><td>4.3</td><td>1.5E-05</td><td>1.6E-04</td><td>BT20/MB231</td><td>MP</td></tr>
<tr><td>>hg38_21_+_8433172_8446622_RNA45SN1_45S_50ntFlanks@10348.10374.27|AAACGAGAACTTTGAAGGCCGAAGTGG</td><td>>hg38_21_+_8433172_8446622_RNA45SN1_45S_50ntFlanks</td><td>10348</td><td>10374</td><td>27</td><td>AAACGAGAACTTTGAAGGCCGAAGTGG</td><td>778.7</td><td>-2.2</td><td>0.6</td><td>-3.9</td><td>1.1E-04</td><td>1.6E-04</td><td>MB231/MB468</td><td>CYTO</td></tr>
<tr><td>>hg38_MT_+_1621_3279_RNR2_16S_50ntFlanks@1374.1414.41|TGGATCAGGACATCCCGATGGTGCAGCCGCTATTAAAGGTT</td><td>>hg38_MT_+_1621_3279_RNR2_16S_50ntFlanks</td><td>1374</td><td>1414</td><td>41</td><td>TGGATCAGGACATCCCGATGGTGCAGCCGCTATTAAAGGTT</td><td>84.2</td><td>6.4</td><td>1.7</td><td>3.8</td><td>1.2E-04</td><td>1.6E-04</td><td>MB231/MB468</td><td>MITO</td></tr>
<tr><td>>am_MI0000809|hsa-mir-151a&WithFlank&8|-|140732558|140732659@53.70.18&[MIMAT0000757&hsa-miR-151a-3p&offsets|0|-3;m-30&8|-|140732587|140732607&offsets|0|-3]|CTAGACTGAAGCTCCTTG</td><td>>am_MI0000809|hsa-mir-151a&WithFlank&8|-|140732558|140732659</td><td>53</td><td>70</td><td>18</td><td>CTAGACTGAAGCTCCTTG</td><td>512.6</td><td>6.0</td><td>1.6</td><td>3.8</td><td>1.2E-04</td><td>1.6E-04</td><td>MB231/MB468</td><td>MITO</td></tr>
<tr><td>>hg38_21_+_8433172_8446622_RNA45SN1_45S_50ntFlanks@7969.7994.26|CCGAGACGCGACCTCAGATCAGACGT</td><td>>hg38_21_+_8433172_8446622_RNA45SN1_45S_50ntFlanks</td><td>7969</td><td>7994</td><td>26</td><td>CCGAGACGCGACCTCAGATCAGACGT</td><td>314.0</td><td>3.0</td><td>0.8</td><td>3.8</td><td>1.2E-04</td><td>1.6E-04</td><td>MB231/MB468</td><td>NUC</td></tr>
<tr><td>>hg38_21_+_8433172_8446622_RNA45SN1_45S_50ntFlanks@4067.4094.28|AACTTTCGATGGTAGTCGCCGTGCCTAC</td><td>>hg38_21_+_8433172_8446622_RNA45SN1_45S_50ntFlanks</td><td>4067</td><td>4094</td><td>28</td><td>AACTTTCGATGGTAGTCGCCGTGCCTAC</td><td>77.8</td><td>-4.5</td><td>1.1</td><td>-3.9</td><td>1.0E-04</td><td>1.6E-04</td><td>MB231/MB468</td><td>MP</td></tr>
<tr><td>>MI0006444|hsa-mir-1307&WithFlank&10|-|103394247|103394407@87.109.23&[MIMAT0005951&hsa-miR-1307-3p&offsets|+1|+2;m-66&10|-|103394300|103394322&offsets|+1|+1]|CTCGGCGTGGCGTCGGTCGTGGT</td><td>>MI0006444|hsa-mir-1307&WithFlank&10|-|103394247|103394407</td><td>87</td><td>109</td><td>23</td><td>CTCGGCGTGGCGTCGGTCGTGGT</td><td>129.9</td><td>6.6</td><td>1.7</td><td>3.8</td><td>1.2E-04</td><td>1.6E-04</td><td>MB231/MB468</td><td>MITO</td></tr>
<tr><td>>MI0000101|hsa-mir-99a&WithFlank&21|+|16539083|16539175@19.40.22(+1A)&[MIMAT0000097&hsa-miR-99a-5p&offsets|0|0(+1A);m-64&21|+|16539101|16539121&offsets|0|+1(+1A)]|AACCCGTAGATCCGATCTTGTGA</td><td>>MI0000101|hsa-mir-99a&WithFlank&21|+|16539083|16539175</td><td>19</td><td>40</td><td>22(+1A)</td><td>AACCCGTAGATCCGATCTTGTGA</td><td>1544.2</td><td>-2.7</td><td>0.7</td><td>-3.9</td><td>1.0E-04</td><td>1.6E-04</td><td>MB231/MB468</td><td>MP</td></tr>
<tr><td>>hg38_21_+_8433172_8446622_RNA45SN1_45S_50ntFlanks@6729.6770.42|GACACATTGATCATCGACACTTCGAACGCACTTGCGGCCCCG</td><td>>hg38_21_+_8433172_8446622_RNA45SN1_45S_50ntFlanks</td><td>6729</td><td>6770</td><td>42</td><td>GACACATTGATCATCGACACTTCGAACGCACTTGCGGCCCCG</td><td>487.2</td><td>-2.6</td><td>0.7</td><td>-3.9</td><td>8.9E-05</td><td>1.6E-04</td><td>BT20/MB468</td><td>MITO</td></tr>
<tr><td>>hg38_MT_+_1621_3279_RNR2_16S_50ntFlanks@1380.1414.35|AGGACATCCCGATGGTGCAGCCGCTATTAAAGGTT</td><td>>hg38_MT_+_1621_3279_RNR2_16S_50ntFlanks</td><td>1380</td><td>1414</td><td>35</td><td>AGGACATCCCGATGGTGCAGCCGCTATTAAAGGTT</td><td>144.2</td><td>4.7</td><td>1.2</td><td>3.8</td><td>1.2E-04</td><td>1.6E-04</td><td>MB231/MB468</td><td>MITO</td></tr>
<tr><td>>hg38_21_+_8433172_8446622_RNA45SN1_45S_50ntFlanks@11666.11701.36|TCTGCCCAGTGCTCTGAATGTCAAAGTGAAGAAATT</td><td>>hg38_21_+_8433172_8446622_RNA45SN1_45S_50ntFlanks</td><td>11666</td><td>11701</td><td>36</td><td>TCTGCCCAGTGCTCTGAATGTCAAAGTGAAGAAATT</td><td>268.8</td><td>2.7</td><td>0.7</td><td>3.9</td><td>1.1E-04</td><td>1.6E-04</td><td>MB231/MB468</td><td>CYTO</td></tr>
<tr><td>>trna111_HisGTG_1_-_147774845_147774916@-1G.32.33,trna118_HisGTG_1_-_145396881_145396952@-1G.32.33,trna16_HisGTG_1_+_146544773_146544844@-1G.32.33,trna1_HisGTG_15_+_45493349_45493420@-1G.32.33,trna21_HisGTG_1_+_147753471_147753542@-1G.32.33,trna33_HisGTG_6_+_27125906_27125977@-1G.32.33,trna7_HisGTG_9_-_14433938_14434009@-1G.32.33,trna8_HisGTG_15_-_45492611_45492682@-1G.32.33,trna9_HisGTG_15_-_45490804_45490875@-1G.32.33|GGCCGTGATCGTATAGTGGTTAGTACTCTGCGT</td><td>>trna111_HisGTG_1_-_147774845_147774916</td><td>-1G</td><td>32</td><td>33</td><td>GGCCGTGATCGTATAGTGGTTAGTACTCTGCGT</td><td>575.2</td><td>8.6</td><td>2.1</td><td>4.1</td><td>4.4E-05</td><td>1.6E-04</td><td>BT20/MB468</td><td>MP</td></tr>
<tr><td>>hg38_21_+_8433172_8446622_RNA45SN1_45S_50ntFlanks@4575.4611.37|TAGGACCGCGGTTCTATTTTGTTGGTTTTCGGAACTG</td><td>>hg38_21_+_8433172_8446622_RNA45SN1_45S_50ntFlanks</td><td>4575</td><td>4611</td><td>37</td><td>TAGGACCGCGGTTCTATTTTGTTGGTTTTCGGAACTG</td><td>117.6</td><td>-6.5</td><td>1.7</td><td>-3.9</td><td>1.0E-04</td><td>1.6E-04</td><td>MB231/MB468</td><td>MP</td></tr>
<tr><td>>hg38_MT_+_598_1651_RNR1_12S_50ntFlanks@845.876.32|GTAGAGTGCTTAGTTGAACAGGGCCCTGAAGC</td><td>>hg38_MT_+_598_1651_RNR1_12S_50ntFlanks</td><td>845</td><td>876</td><td>32</td><td>GTAGAGTGCTTAGTTGAACAGGGCCCTGAAGC</td><td>113.9</td><td>7.7</td><td>1.9</td><td>4.1</td><td>4.4E-05</td><td>1.6E-04</td><td>BT20/MB468</td><td>MP</td></tr>
<tr><td>>hg38_21_+_8433172_8446622_RNA45SN1_45S_50ntFlanks@11923.11943.21|AGAGGTGTAGAATAAGTGGGA</td><td>>hg38_21_+_8433172_8446622_RNA45SN1_45S_50ntFlanks</td><td>11923</td><td>11943</td><td>21</td><td>AGAGGTGTAGAATAAGTGGGA</td><td>80.8</td><td>9.7</td><td>2.2</td><td>4.3</td><td>1.5E-05</td><td>1.6E-04</td><td>BT20/MB231</td><td>MP</td></tr>
<tr><td>>MI0000477|hsa-mir-146a&WithFlank&5|+|160485346|160485456@26.48.23(+1U)&[MIMAT0000449&hsa-miR-146a-5p&offsets|-1|0(+1U);m-129&5|+|160485372|160485393&offsets|-1|0(+1U)]|TTGAGAACTGAATTCCATGGGTTT</td><td>>MI0000477|hsa-mir-146a&WithFlank&5|+|160485346|160485456</td><td>26</td><td>48</td><td>23(+1U)</td><td>TTGAGAACTGAATTCCATGGGTTT</td><td>58.4</td><td>-9.3</td><td>2.1</td><td>-4.3</td><td>1.5E-05</td><td>1.6E-04</td><td>BT20/MB231</td><td>MP</td></tr>
<tr><td>>hg38_21_+_8433172_8446622_RNA45SN1_45S_50ntFlanks@11778.11804.27|ACGCGCATGAATGGATGAACGAGATTC</td><td>>hg38_21_+_8433172_8446622_RNA45SN1_45S_50ntFlanks</td><td>11778</td><td>11804</td><td>27</td><td>ACGCGCATGAATGGATGAACGAGATTC</td><td>1023.2</td><td>-5.4</td><td>1.3</td><td>-4.1</td><td>4.4E-05</td><td>1.6E-04</td><td>BT20/MB468</td><td>MP</td></tr>
<tr><td>>hg38_21_+_8433172_8446622_RNA45SN1_45S_50ntFlanks@12709.12751.43|CCGCGGAGCCTCGGTTGGCCTCGGATAGCCGGTCCCCCGCCTG</td><td>>hg38_21_+_8433172_8446622_RNA45SN1_45S_50ntFlanks</td><td>12709</td><td>12751</td><td>43</td><td>CCGCGGAGCCTCGGTTGGCCTCGGATAGCCGGTCCCCCGCCTG</td><td>44.8</td><td>-8.6</td><td>2.1</td><td>-4.1</td><td>4.4E-05</td><td>1.6E-04</td><td>BT20/MB468</td><td>MP</td></tr>
<tr><td>>hg38_21_+_8433172_8446622_RNA45SN1_45S_50ntFlanks@10381.10424.44|GTTCCATGTGAACAGCAGTTGAACATGGGTCAGTCGGTCCTGAG</td><td>>hg38_21_+_8433172_8446622_RNA45SN1_45S_50ntFlanks</td><td>10381</td><td>10424</td><td>44</td><td>GTTCCATGTGAACAGCAGTTGAACATGGGTCAGTCGGTCCTGAG</td><td>6075.7</td><td>3.7</td><td>0.9</td><td>4.3</td><td>1.7E-05</td><td>1.6E-04</td><td>BT20/MB231</td><td>MITO</td></tr>
<tr><td>>hg38_21_+_8433172_8446622_RNA45SN1_45S_50ntFlanks@10383.10424.42|TCCATGTGAACAGCAGTTGAACATGGGTCAGTCGGTCCTGAG</td><td>>hg38_21_+_8433172_8446622_RNA45SN1_45S_50ntFlanks</td><td>10383</td><td>10424</td><td>42</td><td>TCCATGTGAACAGCAGTTGAACATGGGTCAGTCGGTCCTGAG</td><td>933.8</td><td>4.1</td><td>1.1</td><td>3.8</td><td>1.2E-04</td><td>1.6E-04</td><td>MB231/MB468</td><td>MITO</td></tr>
<tr><td>>trna116_GluCTC_1_-_145399233_145399304@1.33.33,trna59_GluCTC_1_+_249168447_249168518@1.33.33,trna71_GluCTC_1_-_161439189_161439260@1.33.33,trna74_GluCTC_1_-_161431809_161431880@1.33.33,trna77_GluCTC_1_-_161424398_161424469@1.33.33,trna77_GluCTC_6_+_28949976_28950047@1.33.33,trna80_GluCTC_1_-_161417018_161417089@1.33.33,trna87_GluCTC_6_-_126101393_126101464@1.33.33|TCCCTGGTGGTCTAGTGGTTAGGATTCGGCGCT</td><td>>trna116_GluCTC_1_-_145399233_145399304</td><td>1</td><td>33</td><td>33</td><td>TCCCTGGTGGTCTAGTGGTTAGGATTCGGCGCT</td><td>9602.6</td><td>5.5</td><td>1.3</td><td>4.1</td><td>4.4E-05</td><td>1.6E-04</td><td>BT20/MB468</td><td>MP</td></tr>
<tr><td>>hg38_21_+_8433172_8446622_RNA45SN1_45S_50ntFlanks@5486.5514.29|GCGCTGAGAAGACGGTCGAACTTGACTAT</td><td>>hg38_21_+_8433172_8446622_RNA45SN1_45S_50ntFlanks</td><td>5486</td><td>5514</td><td>29</td><td>GCGCTGAGAAGACGGTCGAACTTGACTAT</td><td>1098.6</td><td>-5.5</td><td>1.4</td><td>-3.8</td><td>1.2E-04</td><td>1.6E-04</td><td>BT20/MB468</td><td>NUC</td></tr>
<tr><td>>hg38_21_+_8433172_8446622_RNA45SN1_45S_50ntFlanks@4892.4914.23|AAACTTAAAGGAATTGACGGAAG</td><td>>hg38_21_+_8433172_8446622_RNA45SN1_45S_50ntFlanks</td><td>4892</td><td>4914</td><td>23</td><td>AAACTTAAAGGAATTGACGGAAG</td><td>116.6</td><td>-4.6</td><td>1.2</td><td>-3.9</td><td>1.0E-04</td><td>1.6E-04</td><td>MB231/MB468</td><td>MP</td></tr>
<tr><td>>MI0000298|hsa-mir-221&WithFlank&X|-|45746151|45746272@70.92.23&[MIMAT0000278&hsa-miR-221-3p&offsets|-1|-1;m-91&X|-|45746181|45746202&offsets|-1|0]|CAGCTACATTGTCTGCTGGGTTT</td><td>>MI0000298|hsa-mir-221&WithFlank&X|-|45746151|45746272</td><td>70</td><td>92</td><td>23</td><td>CAGCTACATTGTCTGCTGGGTTT</td><td>56.9</td><td>-9.2</td><td>2.1</td><td>-4.3</td><td>1.5E-05</td><td>1.6E-04</td><td>BT20/MB231</td><td>MP</td></tr>
<tr><td>>hg38_21_+_8433172_8446622_RNA45SN1_45S_50ntFlanks@12420.12445.26|ATGTCGGCTCTTCCTATCATTGTGAA</td><td>>hg38_21_+_8433172_8446622_RNA45SN1_45S_50ntFlanks</td><td>12420</td><td>12445</td><td>26</td><td>ATGTCGGCTCTTCCTATCATTGTGAA</td><td>243.0</td><td>-5.9</td><td>1.5</td><td>-3.9</td><td>1.0E-04</td><td>1.6E-04</td><td>MB231/MB468</td><td>MP</td></tr>
<tr><td>>hg38_21_+_8433172_8446622_RNA45SN1_45S_50ntFlanks@12407.12442.36|TTTTGATCCTTCGATGTCGGCTCTTCCTATCATTGT</td><td>>hg38_21_+_8433172_8446622_RNA45SN1_45S_50ntFlanks</td><td>12407</td><td>12442</td><td>36</td><td>TTTTGATCCTTCGATGTCGGCTCTTCCTATCATTGT</td><td>272.2</td><td>3.5</td><td>0.9</td><td>3.9</td><td>1.1E-04</td><td>1.6E-04</td><td>MB231/MB468</td><td>CYTO</td></tr>
<tr><td>>hg38_21_+_8433172_8446622_RNA45SN1_45S_50ntFlanks@11320.11344.25|CCACGTCTCGTCGCGCGCGCGTCCG</td><td>>hg38_21_+_8433172_8446622_RNA45SN1_45S_50ntFlanks</td><td>11320</td><td>11344</td><td>25</td><td>CCACGTCTCGTCGCGCGCGCGTCCG</td><td>14980.5</td><td>-8.3</td><td>2.2</td><td>-3.8</td><td>1.2E-04</td><td>1.6E-04</td><td>BT20/MB468</td><td>NUC</td></tr>
<tr><td>>MI0000088|hsa-mir-30a&WithFlank&6|-|71403545|71403627@12.30.19(+1U)&[MIMAT0000087&hsa-miR-30a-5p&offsets|0|-3(+1U);m-15&6|-|71403595|71403616&offsets|0|-3(+1U)]|TGTAAACATCCTCGACTGGT</td><td>>MI0000088|hsa-mir-30a&WithFlank&6|-|71403545|71403627</td><td>12</td><td>30</td><td>19(+1U)</td><td>TGTAAACATCCTCGACTGGT</td><td>174.0</td><td>5.7</td><td>1.5</td><td>3.9</td><td>1.0E-04</td><td>1.6E-04</td><td>MB231/MB468</td><td>MP</td></tr>
<tr><td>>hg38_21_+_8433172_8446622_RNA45SN1_45S_50ntFlanks@10350.10378.29|ACGAGAACTTTGAAGGCCGAAGTGGAGAA</td><td>>hg38_21_+_8433172_8446622_RNA45SN1_45S_50ntFlanks</td><td>10350</td><td>10378</td><td>29</td><td>ACGAGAACTTTGAAGGCCGAAGTGGAGAA</td><td>830.3</td><td>-4.3</td><td>1.1</td><td>-4.1</td><td>4.4E-05</td><td>1.6E-04</td><td>BT20/MB468</td><td>MP</td></tr>
<tr><td>>hg38_21_+_8433172_8446622_RNA45SN1_45S_50ntFlanks@9941.9975.35|AGGACGGTGGCCATGGAAGTCGGAATCCGCTAAGG</td><td>>hg38_21_+_8433172_8446622_RNA45SN1_45S_50ntFlanks</td><td>9941</td><td>9975</td><td>35</td><td>AGGACGGTGGCCATGGAAGTCGGAATCCGCTAAGG</td><td>135.6</td><td>7.3</td><td>1.8</td><td>4.1</td><td>4.4E-05</td><td>1.6E-04</td><td>BT20/MB468</td><td>MP</td></tr>
<tr><td>>hg38_21_+_8433172_8446622_RNA45SN1_45S_50ntFlanks@4728.4751.24|ATCAAGAACGAAAGTCGGAGGTTC</td><td>>hg38_21_+_8433172_8446622_RNA45SN1_45S_50ntFlanks</td><td>4728</td><td>4751</td><td>24</td><td>ATCAAGAACGAAAGTCGGAGGTTC</td><td>325.9</td><td>-5.8</td><td>1.5</td><td>-3.8</td><td>1.2E-04</td><td>1.6E-04</td><td>MB231/MB468</td><td>MITO</td></tr>
</table>

<table>
<tr><td>>hg38_21_+_8433172_8446622_RNA45SN1_45S_50ntFlanks@11569.11607.39|CAGCCGACTTAGAACTGGTGCGGACCAGGGGAATCCGAC</td><td>>hg38_21_+_8433172_8446622_RNA45SN1_45S_50ntFlanks</td><td>11569</td><td>11607</td><td>39</td><td>CAGCCGACTTAGAACTGGTGCGGACCAGGGGAATCCGAC</td><td>187.4</td><td>-4.5</td><td>1.2</td><td>-3.8</td><td>1.2E-04</td><td>1.6E-04</td><td>MB231/MB468</td><td>NUC</td></tr>
<tr><td>>MI0000085|hsa-mir-27a&WithFlank&19|-|13836434|13836523@57.76.20(+3A)&[MIMAT0000084&hsa-miR-27a-3p&offsets|0|-1(+3A);m-43&19|-|13836447|13836467&offsets|0|-1(+3A)]|TTCACAGTGGCTAAGTTCCGAAA</td><td>>MI0000085|hsa-mir-27a&WithFlank&19|-|13836434|13836523</td><td>57</td><td>76</td><td>20(+3A)</td><td>TTCACAGTGGCTAAGTTCCGAAA</td><td>59.0</td><td>-9.3</td><td>2.1</td><td>-4.3</td><td>1.5E-05</td><td>1.6E-04</td><td>BT20/MB231</td><td>MP</td></tr>
<tr><td>>trnaMT_HisGTG_MT_+_12138_12206@43.70.28|ACAGAGGCTTACGACCCCTTATTTACCC</td><td>>trnaMT_HisGTG_MT_+_12138_12206</td><td>43</td><td>70</td><td>28</td><td>ACAGAGGCTTACGACCCCTTATTTACCC</td><td>124.0</td><td>10.7</td><td>2.6</td><td>4.1</td><td>4.4E-05</td><td>1.6E-04</td><td>BT20/MB468</td><td>MP</td></tr>
<tr><td>>hg38_21_+_8433172_8446622_RNA45SN1_45S_50ntFlanks@7990.8017.28|GACGTGGCGACCCGCTGAATTTAAGCAT</td><td>>hg38_21_+_8433172_8446622_RNA45SN1_45S_50ntFlanks</td><td>7990</td><td>8017</td><td>28</td><td>GACGTGGCGACCCGCTGAATTTAAGCAT</td><td>175.0</td><td>3.2</td><td>0.8</td><td>3.8</td><td>1.2E-04</td><td>1.6E-04</td><td>MB231/MB468</td><td>NUC</td></tr>
<tr><td>>MI0000269|hsa-mir-181a-2&WithFlank&9|+|124692436|124692557@83.102.20(+1U)&[MIMAT0004558&hsa-miR-181a-2-3p&offsets|0|-2(+1U);m-71&9|+|124692518|124692539&offsets|0|-2(+1U)]|ACCACTGACCGTTGACTGTAT</td><td>>MI0000269|hsa-mir-181a-2&WithFlank&9|+|124692436|124692557</td><td>83</td><td>102</td><td>20(+1U)</td><td>ACCACTGACCGTTGACTGTAT</td><td>26.7</td><td>8.0</td><td>2.0</td><td>3.9</td><td>1.0E-04</td><td>1.6E-04</td><td>MB231/MB468</td><td>MP</td></tr>
<tr><td>>hg38_21_+_8433172_8446622_RNA45SN1_45S_50ntFlanks@12234.12265.32|CGTGGAGCAGAAGGGCAAAAGCTCGCTTGATC</td><td>>hg38_21_+_8433172_8446622_RNA45SN1_45S_50ntFlanks</td><td>12234</td><td>12265</td><td>32</td><td>CGTGGAGCAGAAGGGCAAAAGCTCGCTTGATC</td><td>184.4</td><td>-4.7</td><td>1.2</td><td>-3.9</td><td>1.0E-04</td><td>1.6E-04</td><td>MB231/MB468</td><td>MP</td></tr>
<tr><td>>MI0000266|hsa-mir-10a&WithFlank&17|-|48579832|48579953@69.88.20(+1C)&[MIMAT0004555&hsa-miR-10a-3p&offsets|0|-2(+1C);m-468&17|-|48579865|48579885&offsets|0|-1(+1C)]|CAAATTCGTATCTAGGGGAAC</td><td>>MI0000266|hsa-mir-10a&WithFlank&17|-|48579832|48579953</td><td>69</td><td>88</td><td>20(+1C)</td><td>CAAATTCGTATCTAGGGGAAC</td><td>29.4</td><td>8.1</td><td>2.1</td><td>3.9</td><td>1.0E-04</td><td>1.6E-04</td><td>MB231/MB468</td><td>MP</td></tr>
<tr><td>>hg38_21_+_8433172_8446622_RNA45SN1_45S_50ntFlanks@10513.10533.21|CCGGAGTGGCGGAGATGGGCG</td><td>>hg38_21_+_8433172_8446622_RNA45SN1_45S_50ntFlanks</td><td>10513</td><td>10533</td><td>21</td><td>CCGGAGTGGCGGAGATGGGCG</td><td>864.8</td><td>-4.7</td><td>1.2</td><td>-3.9</td><td>1.1E-04</td><td>1.6E-04</td><td>BT20/MB468</td><td>CYTO</td></tr>
<tr><td>>hg38_21_+_8433172_8446622_RNA45SN1_45S_50ntFlanks@11200.11225.26|CGGCGGCGGCGGCGACTCTGGACGCG</td><td>>hg38_21_+_8433172_8446622_RNA45SN1_45S_50ntFlanks</td><td>11200</td><td>11225</td><td>26</td><td>CGGCGGCGGCGGCGACTCTGGACGCG</td><td>882.7</td><td>-5.2</td><td>1.3</td><td>-4.1</td><td>4.4E-05</td><td>1.6E-04</td><td>BT20/MB468</td><td>MP</td></tr>
<tr><td>>hg38_21_+_8433172_8446622_RNA45SN1_45S_50ntFlanks@12488.12519.32|AACGTGAGCTGGGTTTAGACCGTCGTGAGACA</td><td>>hg38_21_+_8433172_8446622_RNA45SN1_45S_50ntFlanks</td><td>12488</td><td>12519</td><td>32</td><td>AACGTGAGCTGGGTTTAGACCGTCGTGAGACA</td><td>138.6</td><td>-3.9</td><td>1.0</td><td>-3.8</td><td>1.2E-04</td><td>1.6E-04</td><td>MB231/MB468</td><td>NUC</td></tr>
<tr><td>>hg38_21_+_8433172_8446622_RNA45SN1_45S_50ntFlanks@10787.10823.37|ACAGCCTCTGGCATGTTGGAACAATGTAGGTAAGGGA</td><td>>hg38_21_+_8433172_8446622_RNA45SN1_45S_50ntFlanks</td><td>10787</td><td>10823</td><td>37</td><td>ACAGCCTCTGGCATGTTGGAACAATGTAGGTAAGGGA</td><td>76.9</td><td>9.6</td><td>2.2</td><td>4.3</td><td>1.6E-05</td><td>1.6E-04</td><td>BT20/MB231</td><td>MP</td></tr>
<tr><td>>hg38_21_+_8433172_8446622_RNA45SN1_45S_50ntFlanks@3831.3852.22|CTCGCTCCTCTCCTACTTGGAT</td><td>>hg38_21_+_8433172_8446622_RNA45SN1_45S_50ntFlanks</td><td>3831</td><td>3852</td><td>22</td><td>CTCGCTCCTCTCCTACTTGGAT</td><td>268.6</td><td>-6.0</td><td>1.5</td><td>-3.9</td><td>9.0E-05</td><td>1.6E-04</td><td>BT20/MB468</td><td>MITO</td></tr>
<tr><td>>hg38_21_+_8433172_8446622_RNA45SN1_45S_50ntFlanks@11846.11887.42|GAACGGGCTTGGCGGAATCAGCGGGGAAAGAAGACCCTGTTG</td><td>>hg38_21_+_8433172_8446622_RNA45SN1_45S_50ntFlanks</td><td>11846</td><td>11887</td><td>42</td><td>GAACGGGCTTGGCGGAATCAGCGGGGAAAGAAGACCCTGTTG</td><td>64.0</td><td>9.3</td><td>2.2</td><td>4.3</td><td>1.6E-05</td><td>1.6E-04</td><td>BT20/MB231</td><td>MP</td></tr>
<tr><td>>hg38_1_-_228634819_228635039_RNA5S12_5S_50ntFlanks@134.169.36|TGGGAGACCGCCTGGGAATACCGGGTGCTGTAGGCT</td><td>>hg38_1_-_228634819_228635039_RNA5S12_5S_50ntFlanks</td><td>134</td><td>169</td><td>36</td><td>TGGGAGACCGCCTGGGAATACCGGGTGCTGTAGGCT</td><td>2605.5</td><td>4.0</td><td>1.0</td><td>4.1</td><td>4.4E-05</td><td>1.6E-04</td><td>BT20/MB468</td><td>MP</td></tr>
<tr><td>>hg38_21_+_8433172_8446622_RNA45SN1_45S_50ntFlanks@12407.12441.35|TTTTGATCCTTCGATGTCGGCTCTTCCTATCATTG</td><td>>hg38_21_+_8433172_8446622_RNA45SN1_45S_50ntFlanks</td><td>12407</td><td>12441</td><td>35</td><td>TTTTGATCCTTCGATGTCGGCTCTTCCTATCATTG</td><td>394.7</td><td>-1.9</td><td>0.5</td><td>-3.9</td><td>9.0E-05</td><td>1.6E-04</td><td>BT20/MB468</td><td>MITO</td></tr>
<tr><td>>hg38_21_+_8433172_8446622_RNA45SN1_45S_50ntFlanks@10061.10079.19|CAGTCGAGAGTGGACGGGA</td><td>>hg38_21_+_8433172_8446622_RNA45SN1_45S_50ntFlanks</td><td>10061</td><td>10079</td><td>19</td><td>CAGTCGAGAGTGGACGGGA</td><td>146.3</td><td>-5.3</td><td>1.4</td><td>-3.9</td><td>1.0E-04</td><td>1.6E-04</td><td>MB231/MB468</td><td>MP</td></tr>
<tr><td>>hg38_21_+_8433172_8446622_RNA45SN1_45S_50ntFlanks@3783.3800.18|AGTGAAACTGCGAATGGC</td><td>>hg38_21_+_8433172_8446622_RNA45SN1_45S_50ntFlanks</td><td>3783</td><td>3800</td><td>18</td><td>AGTGAAACTGCGAATGGC</td><td>522.5</td><td>-4.2</td><td>1.1</td><td>-3.9</td><td>1.0E-04</td><td>1.6E-04</td><td>MB231/MB468</td><td>MP</td></tr>
<tr><td>>MI0000802|hsa-mir-340&WithFlank&5|-|180015297|180015403@22.43.22&[MIMAT0004692&hsa-miR-340-5p&offsets|0|0;m-220&5|-|180015361|180015382&offsets|0|0]|TTATAAAGCAATGAGACTGATT</td><td>>MI0000802|hsa-mir-340&WithFlank&5|-|180015297|180015403</td><td>22</td><td>43</td><td>22</td><td>TTATAAAGCAATGAGACTGATT</td><td>1124.8</td><td>2.8</td><td>0.7</td><td>3.8</td><td>1.3E-04</td><td>1.6E-04</td><td>BT20/MB468</td><td>NUC</td></tr>
<tr><td>>hg38_21_+_8433172_8446622_RNA45SN1_45S_50ntFlanks@6645.6672.28|CTCGTACGACTCTTAGCGGTGGATCACT</td><td>>hg38_21_+_8433172_8446622_RNA45SN1_45S_50ntFlanks</td><td>6645</td><td>6672</td><td>28</td><td>CTCGTACGACTCTTAGCGGTGGATCACT</td><td>1519.9</td><td>3.3</td><td>0.9</td><td>3.8</td><td>1.2E-04</td><td>1.6E-04</td><td>MB231/MB468</td><td>MITO</td></tr>
<tr><td>>hg38_1_-_228634819_228635039_RNA5S12_5S_50ntFlanks@148.170.23|GGAATACCGGGTGCTGTAGGCTT</td><td>>hg38_1_-_228634819_228635039_RNA5S12_5S_50ntFlanks</td><td>148</td><td>170</td><td>23</td><td>GGAATACCGGGTGCTGTAGGCTT</td><td>2210.6</td><td>3.0</td><td>0.7</td><td>4.3</td><td>1.8E-05</td><td>1.6E-04</td><td>BT20/MB231</td><td>MITO</td></tr>
<tr><td>>am_trna128_GlyGCC_6_-_27870686_27870756@4.35.32,trna18_GlyGCC_16_+_70822597_70822667@4.35.32,trna19_GlyGCC_16_+_70823410_70823480@4.35.32,trna19_GlyGCC_2_-_157257659_157257729@4.35.32,trna24_GlyGCC_16_-_70812942_70813012@4.35.32,trna25_GlyGCC_16_-_70812114_70812184@4.35.32,trna5_GlyGCC_17_+_8029064_8029134@4.35.32,trna68_GlyGCC_1_-_161493637_161493707@4.35.32|TTGGTGGTTCAGTGGTAGAATTCTCGCCTGCC</td><td>>am_trna128_GlyGCC_6_-_27870686_27870756</td><td>4</td><td>35</td><td>32</td><td>TTGGTGGTTCAGTGGTAGAATTCTCGCCTGCC</td><td>331.8</td><td>-4.2</td><td>1.0</td><td>-4.3</td><td>1.8E-05</td><td>1.6E-04</td><td>BT20/MB231</td><td>MITO</td></tr>
<tr><td>>hg38_21_+_8433172_8446622_RNA45SN1_45S_50ntFlanks@10410.10429.20|TCAGTCGGTCCTGAGAGATG</td><td>>hg38_21_+_8433172_8446622_RNA45SN1_45S_50ntFlanks</td><td>10410</td><td>10429</td><td>20</td><td>TCAGTCGGTCCTGAGAGATG</td><td>3018.0</td><td>-3.8</td><td>0.9</td><td>-4.1</td><td>4.5E-05</td><td>1.6E-04</td><td>BT20/MB468</td><td>MP</td></tr>
<tr><td>>hg38_21_+_8433172_8446622_RNA45SN1_45S_50ntFlanks@8106.8141.36|GCGGCGGGGCGCGGGACATGTGGCGTACGGAAGACC</td><td>>hg38_21_+_8433172_8446622_RNA45SN1_45S_50ntFlanks</td><td>8106</td><td>8141</td><td>36</td><td>GCGGCGGGGCGCGGGACATGTGGCGTACGGAAGACC</td><td>506.6</td><td>-6.4</td><td>1.6</td><td>-3.9</td><td>1.0E-04</td><td>1.6E-04</td><td>MB231/MB468</td><td>MP</td></tr>
<tr><td>>hg38_21_+_8433172_8446622_RNA45SN1_45S_50ntFlanks@11568.11611.44|GCAGCCGACTTAGAACTGGTGCGGACCAGGGGAATCCGACTGTT</td><td>>hg38_21_+_8433172_8446622_RNA45SN1_45S_50ntFlanks</td><td>11568</td><td>11611</td><td>44</td><td>GCAGCCGACTTAGAACTGGTGCGGACCAGGGGAATCCGACTGTT</td><td>2996.9</td><td>4.8</td><td>1.2</td><td>3.9</td><td>1.1E-04</td><td>1.6E-04</td><td>BT20/MB468</td><td>CYTO</td></tr>
<tr><td>>hg38_21_+_8433172_8446622_RNA45SN1_45S_50ntFlanks@12633.12652.20|CGAAGCTACCATCTGTGGGA</td><td>>hg38_21_+_8433172_8446622_RNA45SN1_45S_50ntFlanks</td><td>12633</td><td>12652</td><td>20</td><td>CGAAGCTACCATCTGTGGGA</td><td>364.3</td><td>-5.4</td><td>1.4</td><td>-3.9</td><td>1.1E-04</td><td>1.6E-04</td><td>BT20/MB468</td><td>CYTO</td></tr>
<tr><td>>hg38_21_+_8433172_8446622_RNA45SN1_45S_50ntFlanks@3875.3897.23|ATACATGCCGACGGGCGCTGACC</td><td>>hg38_21_+_8433172_8446622_RNA45SN1_45S_50ntFlanks</td><td>3875</td><td>3897</td><td>23</td><td>ATACATGCCGACGGGCGCTGACC</td><td>413.7</td><td>-6.6</td><td>1.7</td><td>-3.8</td><td>1.3E-04</td><td>1.6E-04</td><td>BT20/MB468</td><td>NUC</td></tr>
<tr><td>>hg38_MT_+_598_1651_RNR1_12S_50ntFlanks@916.946.31|AAGGACATTTAACTAAAACCCCTACGCATTT</td><td>>hg38_MT_+_598_1651_RNR1_12S_50ntFlanks</td><td>916</td><td>946</td><td>31</td><td>AAGGACATTTAACTAAAACCCCTACGCATTT</td><td>1890.0</td><td>6.3</td><td>1.5</td><td>4.1</td><td>4.5E-05</td><td>1.6E-04</td><td>BT20/MB468</td><td>MP</td></tr>
<tr><td>>hg38_21_+_8433172_8446622_RNA45SN1_45S_50ntFlanks@9913.9951.39|AGAAAAGGTGTTGGTTGATATAGACAGCAGGACGGTGGC</td><td>>hg38_21_+_8433172_8446622_RNA45SN1_45S_50ntFlanks</td><td>9913</td><td>9951</td><td>39</td><td>AGAAAAGGTGTTGGTTGATATAGACAGCAGGACGGTGGC</td><td>63.7</td><td>9.3</td><td>2.2</td><td>4.3</td><td>1.6E-05</td><td>1.6E-04</td><td>BT20/MB231</td><td>MP</td></tr>
<tr><td>>hg38_21_+_8433172_8446622_RNA45SN1_45S_50ntFlanks@3765.3792.28|AGTACGCACGGCCGGTACAGTGAAACTG</td><td>>hg38_21_+_8433172_8446622_RNA45SN1_45S_50ntFlanks</td><td>3765</td><td>3792</td><td>28</td><td>AGTACGCACGGCCGGTACAGTGAAACTG</td><td>265.6</td><td>-5.9</td><td>1.5</td><td>-3.9</td><td>9.1E-05</td><td>1.6E-04</td><td>BT20/MB468</td><td>MITO</td></tr>
<tr><td>>hg38_21_+_8433172_8446622_RNA45SN1_45S_50ntFlanks@12619.12654.36|AGGAGCCAATGGGGCGAAGCTACCATCTGTGGGATT</td><td>>hg38_21_+_8433172_8446622_RNA45SN1_45S_50ntFlanks</td><td>12619</td><td>12654</td><td>36</td><td>AGGAGCCAATGGGGCGAAGCTACCATCTGTGGGATT</td><td>80.9</td><td>9.7</td><td>2.2</td><td>4.3</td><td>1.6E-05</td><td>1.6E-04</td><td>BT20/MB231</td><td>MP</td></tr>
<tr><td>>MI0000086|hsa-mir-28&WithFlank&3|+|188688775|188688872@60.80.21(+1U)&[MIMAT0004502&hsa-miR-28-3p&offsets|0|-1(+1U);m-28&3|+|188688834|188688855&offsets|0|-1(+1U)]|CACTAGATTGTGAGCTCCTGGT</td><td>>MI0000086|hsa-mir-28&WithFlank&3|+|188688775|188688872</td><td>60</td><td>80</td><td>21(+1U)</td><td>CACTAGATTGTGAGCTCCTGGT</td><td>222.0</td><td>7.6</td><td>1.9</td><td>4.1</td><td>4.5E-05</td><td>1.6E-04</td><td>BT20/MB468</td><td>MP</td></tr>
<tr><td>>am_trnaMT_PheGAA_MT_+_577_647@43.62.20|TTAGACGGGCTCACATCACC</td><td>>am_trnaMT_PheGAA_MT_+_577_647</td><td>43</td><td>62</td><td>20</td><td>TTAGACGGGCTCACATCACC</td><td>249.8</td><td>5.0</td><td>1.2</td><td>4.1</td><td>4.5E-05</td><td>1.6E-04</td><td>BT20/MB468</td><td>MP</td></tr>
<tr><td>>hg38_MT_+_1621_3279_RNR2_16S_50ntFlanks@51.87.37|GCTAAACCTAGCCCCAAACCCACTCCACCTTACTACC</td><td>>hg38_MT_+_1621_3279_RNR2_16S_50ntFlanks</td><td>51</td><td>87</td><td>37</td><td>GCTAAACCTAGCCCCAAACCCACTCCACCTTACTACC</td><td>3218.8</td><td>3.3</td><td>0.8</td><td>3.9</td><td>1.0E-04</td><td>1.6E-04</td><td>MB231/MB468</td><td>MP</td></tr>
<tr><td>>hg38_21_+_8433172_8446622_RNA45SN1_45S_50ntFlanks@10349.10366.18|AACGAGAACTTTGAAGGC</td><td>>hg38_21_+_8433172_8446622_RNA45SN1_45S_50ntFlanks</td><td>10349</td><td>10366</td><td>18</td><td>AACGAGAACTTTGAAGGC</td><td>347.1</td><td>-2.8</td><td>0.7</td><td>-3.8</td><td>1.2E-04</td><td>1.6E-04</td><td>MB231/MB468</td><td>MITO</td></tr>
<tr><td>>hg38_21_+_8433172_8446622_RNA45SN1_45S_50ntFlanks@9604.9622.19|AAAGACTAATCGAACCATC</td><td>>hg38_21_+_8433172_8446622_RNA45SN1_45S_50ntFlanks</td><td>9604</td><td>9622</td><td>19</td><td>AAAGACTAATCGAACCATC</td><td>1217.5</td><td>-2.8</td><td>0.7</td><td>-3.9</td><td>1.1E-04</td><td>1.6E-04</td><td>BT20/MB468</td><td>CYTO</td></tr>
<tr><td>>trnaMT_IleGAT_MT_+_4263_4331@43.71.29|ATAGGAGCTTAAACCCCCTTATTTCTACC</td><td>>trnaMT_IleGAT_MT_+_4263_4331</td><td>43</td><td>71</td><td>29</td><td>ATAGGAGCTTAAACCCCCTTATTTCTACC</td><td>115.0</td><td>10.6</td><td>2.6</td><td>4.1</td><td>4.5E-05</td><td>1.6E-04</td><td>BT20/MB468</td><td>MP</td></tr>
<tr><td>>hg38_21_+_8433172_8446622_RNA45SN1_45S_50ntFlanks@4005.4026.22|AACCTCGGGCCGATCGCACGCC</td><td>>hg38_21_+_8433172_8446622_RNA45SN1_45S_50ntFlanks</td><td>4005</td><td>4026</td><td>22</td><td>AACCTCGGGCCGATCGCACGCC</td><td>262.9</td><td>-5.3</td><td>1.4</td><td>-3.9</td><td>1.1E-04</td><td>1.6E-04</td><td>BT20/MB468</td><td>CYTO</td></tr>
<tr><td>>hg38_21_+_8433172_8446622_RNA45SN1_45S_50ntFlanks@10922.10946.25|CTGGACGAGGCGCCGCCGCCCCCCC</td><td>>hg38_21_+_8433172_8446622_RNA45SN1_45S_50ntFlanks</td><td>10922</td><td>10946</td><td>25</td><td>CTGGACGAGGCGCCGCCGCCCCCCC</td><td>1667.7</td><td>-5.1</td><td>1.3</td><td>-3.9</td><td>1.1E-04</td><td>1.6E-04</td><td>BT20/MB468</td><td>CYTO</td></tr>
<tr><td>>hg38_21_+_8433172_8446622_RNA45SN1_45S_50ntFlanks@5418.5445.28|TACCGATTGGATGGTTTAGTGAGGCCCT</td><td>>hg38_21_+_8433172_8446622_RNA45SN1_45S_50ntFlanks</td><td>5418</td><td>5445</td><td>28</td><td>TACCGATTGGATGGTTTAGTGAGGCCCT</td><td>137.4</td><td>-2.3</td><td>0.6</td><td>-3.9</td><td>1.2E-04</td><td>1.6E-04</td><td>MB231/MB468</td><td>CYTO</td></tr>
<tr><td>>trnaMT_MetCAT_MT_+_4402_4469@34.71.38|ACCCCGAAAATGTTGGTTATACCCTTCCCGTACTACCA</td><td>>trnaMT_MetCAT_MT_+_4402_4469</td><td>34</td><td>71</td><td>38</td><td>ACCCCGAAAATGTTGGTTATACCCTTCCCGTACTACCA</td><td>165.0</td><td>6.9</td><td>1.8</td><td>3.8</td><td>1.2E-04</td><td>1.6E-04</td><td>MB231/MB468</td><td>MITO</td></tr>
<tr><td>>hg38_21_+_8433172_8446622_RNA45SN1_45S_50ntFlanks@3782.3800.19|CAGTGAAACTGCGAATGGC</td><td>>hg38_21_+_8433172_8446622_RNA45SN1_45S_50ntFlanks</td><td>3782</td><td>3800</td><td>19</td><td>CAGTGAAACTGCGAATGGC</td><td>460.8</td><td>-1.6</td><td>0.4</td><td>-3.9</td><td>1.1E-04</td><td>1.6E-04</td><td>BT20/MB468</td><td>CYTO</td></tr>
<tr><td>>hg38_21_+_8433172_8446622_RNA45SN1_45S_50ntFlanks@10877.10902.26|GTCGGTCGGGCTGGGGCGCGAAGCGG</td><td>>hg38_21_+_8433172_8446622_RNA45SN1_45S_50ntFlanks</td><td>10877</td><td>10902</td><td>26</td><td>GTCGGTCGGGCTGGGGCGCGAAGCGG</td><td>164.5</td><td>-5.0</td><td>1.3</td><td>-3.9</td><td>1.1E-04</td><td>1.6E-04</td><td>BT20/MB468</td><td>CYTO</td></tr>
<tr><td>>MI0000086|hsa-mir-28&WithFlank&3|+|188688775|188688872@60.81.22(+2U)&[MIMAT0004502&hsa-miR-28-3p&offsets|0|0(+2U);m-28&3|+|188688834|188688855&offsets|0|0(+2U)]|CACTAGATTGTGAGCTCCTGGATT</td><td>>MI0000086|hsa-mir-28&WithFlank&3|+|188688775|188688872</td><td>60</td><td>81</td><td>22(+2U)</td><td>CACTAGATTGTGAGCTCCTGGATT</td><td>92.1</td><td>6.5</td><td>1.7</td><td>3.9</td><td>1.0E-04</td><td>1.6E-04</td><td>MB231/MB468</td><td>MP</td></tr>
<tr><td>>hg38_21_+_8433172_8446622_RNA45SN1_45S_50ntFlanks@11591.11609.19|GACCAGGGGAATCCGACTG</td><td>>hg38_21_+_8433172_8446622_RNA45SN1_45S_50ntFlanks</td><td>11591</td><td>11609</td><td>19</td><td>GACCAGGGGAATCCGACTG</td><td>69.9</td><td>-5.7</td><td>1.5</td><td>-3.9</td><td>1.0E-04</td><td>1.6E-04</td><td>MB231/MB468</td><td>MP</td></tr>
<tr><td>>hg38_21_+_8433172_8446622_RNA45SN1_45S_50ntFlanks@5078.5108.31|CGATAACGAACGAGACTCTGGCATGCTAACT</td><td>>hg38_21_+_8433172_8446622_RNA45SN1_45S_50ntFlanks</td><td>5078</td><td>5108</td><td>31</td><td>CGATAACGAACGAGACTCTGGCATGCTAACT</td><td>463.2</td><td>-5.6</td><td>1.5</td><td>-3.8</td><td>1.2E-04</td><td>1.6E-04</td><td>MB231/MB468</td><td>MITO</td></tr>
<tr><td>>hg38_21_+_8433172_8446622_RNA45SN1_45S_50ntFlanks@11565.11585.21|CTAGCAGCCGACTTAGAACTG</td><td>>hg38_21_+_8433172_8446622_RNA45SN1_45S_50ntFlanks</td><td>11565</td><td>11585</td><td>21</td><td>CTAGCAGCCGACTTAGAACTG</td><td>309.4</td><td>-4.4</td><td>1.2</td><td>-3.8</td><td>1.2E-04</td><td>1.6E-04</td><td>MB231/MB468</td><td>MITO</td></tr>
<tr><td>>MI0000289|hsa-mir-181a-1&WithFlank&1|-|198859038|198859159@30.51.22(+1C);MI0000269|hsa-mir-181a-2&WithFlank&9|+|124692436|124692557@45.66.22(+1C)&[MIMAT0000256&hsa-miR-181a-5p&offsets|0|-1(+1C);m-32&1|-|198859108|198859130&offsets|0|-1(+1C)];[MIMAT0000256_1&hsa-miR-181a-5p&offsets|0|-1(+1C);m-32&9|+|124692480|124692502&offsets|0|-1(+1C)]|AACATTCAACGCTGTCGGTGAGC</td><td>>MI0000289|hsa-mir-181a-1&WithFlank&1|-|198859038|198859159</td><td>30</td><td>51</td><td>22(+1C)</td><td>AACATTCAACGCTGTCGGTGAGC</td><td>3272.2</td><td>-4.5</td><td>1.0</td><td>-4.3</td><td>1.6E-05</td><td>1.6E-04</td><td>BT20/MB231</td><td>MP</td></tr>
<tr><td>>MI0000088|hsa-mir-30a&WithFlank&6|-|71403545|71403627@53.73.21(+3U)&[MIMAT0000088&hsa-miR-30a-3p&offsets|0|-1(+3U);m-38&6|-|71403554|71403575&offsets|0|-1(+3U)]|CTTTCAGTCGGATGTTTGCAGTTT</td><td>>MI0000088|hsa-mir-30a&WithFlank&6|-|71403545|71403627</td><td>53</td><td>73</td><td>21(+3U)</td><td>CTTTCAGTCGGATGTTTGCAGTTT</td><td>54.8</td><td>-9.2</td><td>2.1</td><td>-4.3</td><td>1.6E-05</td><td>1.6E-04</td><td>BT20/MB231</td><td>MP</td></tr>
</table>

| | | | | | | | | | | | | | |
|---|---|---|---|---|---|---|---|---|---|---|---|---|---|
| >hg38_MT_+_1621_3279_RNR2_16S_50ntFlanks@496.518.23\|CTTTGGACACTAGGAAAAAACCT | >hg38_MT_+_1621_3279_RNR2_16S_50ntFlanks | 496 | 518 | 23 | CTTTGGACACTAGGAAAAAACCT | 127.9 | 7.2 | 1.8 | 4.1 | 4.5E-05 | 1.6E-04 | BT20/MB468 | MP |
| >trna116_GluCTC_1_-_145399233_145399304@32.70.39,trna71_GluCTC_1_-_161439189_161439260@32.70.39,trna74_GluCTC_1_-_161431809_161431880@32.70.39,trna77_GluCTC_1_-_161424398_161424469@32.70.39,trna77_GluCTC_6_+_28949976_28950047@32.70.39,trna80_GluCTC_1_-_161417018_161417089@32.70.39,trna87_GluCTC_6_-_126101393_126101464@32.70.39\|CTCTCACCGCCGCGGCCCGGGTTCGATTCCCGGTCAGGG | >trna116_GluCTC_1_-_145399233_145399304 | 32 | 70 | 39 | CTCTCACCGCCGCGGCCCGGGTTCGATTCCCGGTCAGGG | 1042.8 | -5.2 | 1.3 | -3.9 | 9.2E-05 | 1.6E-04 | BT20/MB468 | MITO |
| >hg38_MT_+_1621_3279_RNR2_16S_50ntFlanks@1380.1414.35\|AGGACATCCCGATGGTGCAGCCGCTATTAAAGGTT | >hg38_MT_+_1621_3279_RNR2_16S_50ntFlanks | 1380 | 1414 | 35 | AGGACATCCCGATGGTGCAGCCGCTATTAAAGGTT | 268.4 | 4.3 | 1.1 | 3.9 | 1.1E-04 | 1.6E-04 | MB231/MB468 | MP |
| >hg38_21_+_8433172_8446622_RNA45SN1_45S_50ntFlanks@8183.8210.28\|TCGAGGCCCAGCCCGTGGACGGTGTGAG | >hg38_21_+_8433172_8446622_RNA45SN1_45S_50ntFlanks | 8183 | 8210 | 28 | TCGAGGCCCAGCCCGTGGACGGTGTGAG | 148.2 | -5.9 | 1.5 | -3.9 | 1.1E-04 | 1.6E-04 | MB231/MB468 | MP |
| >hg38_21_+_8433172_8446622_RNA45SN1_45S_50ntFlanks@10766.10796.31\|CGCAGCAGGTCTCCAAGGTGAACAGCCTCTG | >hg38_21_+_8433172_8446622_RNA45SN1_45S_50ntFlanks | 10766 | 10796 | 31 | CGCAGCAGGTCTCCAAGGTGAACAGCCTCTG | 211.9 | -5.2 | 1.3 | -3.9 | 1.1E-04 | 1.6E-04 | MB231/MB468 | MP |
| >trna27_GlyCCC_2_-_70476123_70476193@1.33.33,trna34_GlyCCC_16_-_686736_686806@1.33.33\|GCGCCGCTGGTGTAGTGGTATCATGCAAGATTC | >trna27_GlyCCC_2_-_70476123_70476193 | 1 | 33 | 33 | GCGCCGCTGGTGTAGTGGTATCATGCAAGATTC | 218.2 | 5.7 | 1.5 | 3.9 | 1.1E-04 | 1.6E-04 | MB231/MB468 | MP |
| >hg38_21_+_8433172_8446622_RNA45SN1_45S_50ntFlanks@4283.4321.39\|CTTTAACGAGGATCCATTGGAGGGCAAGTCTGGTGCCAG | >hg38_21_+_8433172_8446622_RNA45SN1_45S_50ntFlanks | 4283 | 4321 | 39 | CTTTAACGAGGATCCATTGGAGGGCAAGTCTGGTGCCAG | 145.4 | -3.4 | 0.9 | -3.9 | 1.2E-04 | 1.6E-04 | MB231/MB468 | CYTO |
| >hg38_21_+_8433172_8446622_RNA45SN1_45S_50ntFlanks@8384.8403.20\|AGGGCGTGAAACCGTTAAGA | >hg38_21_+_8433172_8446622_RNA45SN1_45S_50ntFlanks | 8384 | 8403 | 20 | AGGGCGTGAAACCGTTAAGA | 133.5 | -4.4 | 1.1 | -3.9 | 1.1E-04 | 1.6E-04 | MB231/MB468 | MP |
| >hg38_21_+_8433172_8446622_RNA45SN1_45S_50ntFlanks@8194.8213.20\|CCCGTGGACGGTGTGAGGCC | >hg38_21_+_8433172_8446622_RNA45SN1_45S_50ntFlanks | 8194 | 8213 | 20 | CCCGTGGACGGTGTGAGGCC | 810.4 | -2.2 | 0.6 | -3.9 | 1.1E-04 | 1.6E-04 | BT20/MB468 | CYTO |
| >trnaMT_SerGCT_MT_+_12207_12265@20.62.43\|GCTAACTCATGCCCCCATGTCTAACAACATGGCTTTCTCACCA | >trnaMT_SerGCT_MT_+_12207_12265 | 20 | 62 | 43 | GCTAACTCATGCCCCCATGTCTAACAACATGGCTTTCTCACCA | 451.4 | 6.3 | 1.6 | 3.9 | 9.2E-05 | 1.6E-04 | BT20/MB468 | MITO |
| >hg38_1_-_228634819_228635039_RNA5S12_5S_50ntFlanks@151.171.21\|ATACCGGGTGCTGTAGGCTTT | >hg38_1_-_228634819_228635039_RNA5S12_5S_50ntFlanks | 151 | 171 | 21 | ATACCGGGTGCTGTAGGCTTT | 1623.4 | 3.6 | 0.9 | 3.8 | 1.3E-04 | 1.6E-04 | MB231/MB468 | NUC |
| >am_trna116_GluCTC_1_-_145399233_145399304@12.29.18,trna25_GluCTC_2_-_71273488_71273560@12.29.18,trna59_GluCTC_1_+_249168447_249168518@12.29.18,trna71_GluCTC_1_-_161439189_161439260@12.29.18,trna74_GluCTC_1_-_161431809_161431880@12.29.18,trna77_GluCTC_1_-_161424398_161424469@12.29.18,trna77_GluCTC_6_+_28949976_28950047@12.29.18,trna80_GluCTC_1_-_161417018_161417089@12.29.18,trna87_GluCTC_6_-_126101393_126101464@12.29.18\|CTAGTGGTTAGGATTCGG | >am_trna116_GluCTC_1_-_145399233_145399304 | 12 | 29 | 18 | CTAGTGGTTAGGATTCGG | 165.7 | -2.5 | 0.7 | -3.9 | 1.1E-04 | 1.6E-04 | BT20/MB468 | CYTO |
| >hg38_21_+_8433172_8446622_RNA45SN1_45S_50ntFlanks@12934.12962.29\|CTAAACCATTCGTAGACGACCTGCTTCTG | >hg38_21_+_8433172_8446622_RNA45SN1_45S_50ntFlanks | 12934 | 12962 | 29 | CTAAACCATTCGTAGACGACCTGCTTCTG | 394.6 | -3.9 | 1.0 | -3.9 | 1.2E-04 | 1.6E-04 | MB231/MB468 | CYTO |
| >hg38_21_+_8433172_8446622_RNA45SN1_45S_50ntFlanks@10067.10086.20\|AGAGTGGACGGGAGCGGCGG | >hg38_21_+_8433172_8446622_RNA45SN1_45S_50ntFlanks | 10067 | 10086 | 20 | AGAGTGGACGGGAGCGGCGG | 74.9 | 9.6 | 2.2 | 4.3 | 1.6E-05 | 1.6E-04 | BT20/MB231 | MP |
| >hg38_21_+_8433172_8446622_RNA45SN1_45S_50ntFlanks@5494.5521.28\|AAGACGGTCGAACTTGACTATCTAGAGG | >hg38_21_+_8433172_8446622_RNA45SN1_45S_50ntFlanks | 5494 | 5521 | 28 | AAGACGGTCGAACTTGACTATCTAGAGG | 175.6 | -4.6 | 1.2 | -3.9 | 1.1E-04 | 1.6E-04 | MB231/MB468 | MP |
| >am_MI0003639\|hsa-mir-625&WithFlank&14\|+\|65471096\|65471192@58.80.23&[MIMAT0004808&hsa-miR-625-3p&offsets\|0\|+1;m-137&14\|+\|65471153\|65471174&offsets\|0\|+1]\|GACTATAGAACTTTCCCCCTCAT | >am_MI0003639\|hsa-mir-625&WithFlank&14\|+\|65471096\|65471192 | 58 | 80 | 23 | GACTATAGAACTTTCCCCCTCAT | 613.1 | -4.0 | 0.9 | -4.3 | 1.6E-05 | 1.6E-04 | BT20/MB231 | MP |
| >hg38_21_+_8433172_8446622_RNA45SN1_45S_50ntFlanks@8196.8226.31\|CGTGGACGGTGTGAGGCCGGTAGCGGCCCCC | >hg38_21_+_8433172_8446622_RNA45SN1_45S_50ntFlanks | 8196 | 8226 | 31 | CGTGGACGGTGTGAGGCCGGTAGCGGCCCCC | 353.9 | -2.9 | 0.8 | -3.8 | 1.3E-04 | 1.6E-04 | MB231/MB468 | NUC |
| >hg38_21_+_8433172_8446622_RNA45SN1_45S_50ntFlanks@10547.10571.25\|AGTGCGGTAACGCGACCGATCCCGG | >hg38_21_+_8433172_8446622_RNA45SN1_45S_50ntFlanks | 10547 | 10571 | 25 | AGTGCGGTAACGCGACCGATCCCGG | 267.5 | -5.0 | 1.3 | -3.9 | 1.1E-04 | 1.7E-04 | MB231/MB468 | MP |
| >hg38_21_+_8433172_8446622_RNA45SN1_45S_50ntFlanks@12161.12181.21\|TGACTGGGGCGGTACACCTGT | >hg38_21_+_8433172_8446622_RNA45SN1_45S_50ntFlanks | 12161 | 12181 | 21 | TGACTGGGGCGGTACACCTGT | 629.4 | -6.4 | 1.6 | -4.1 | 4.6E-05 | 1.7E-04 | BT20/MB468 | MP |
| >hg38_MT_+_1621_3279_RNR2_16S_50ntFlanks@622.652.31\|TATAACTGAACTCCTCACACCCAATTGGACC | >hg38_MT_+_1621_3279_RNR2_16S_50ntFlanks | 622 | 652 | 31 | TATAACTGAACTCCTCACACCCAATTGGACC | 531.4 | 6.0 | 1.5 | 4.1 | 4.6E-05 | 1.7E-04 | BT20/MB468 | MP |
| >trnaMT_GlnTTG_MT_-_4329_4400@1.26.26,trnalookalike2_GlnTTG_1_-_564879_564950@1.26.26\|TAGGATGGGGTGTGATAGGTGGCACG | >trnaMT_GlnTTG_MT_-_4329_4400 | 1 | 26 | 26 | TAGGATGGGGTGTGATAGGTGGCACG | 1339.8 | 5.3 | 1.3 | 4.1 | 4.6E-05 | 1.7E-04 | BT20/MB468 | MP |
| >hg38_21_+_8433172_8446622_RNA45SN1_45S_50ntFlanks@6647.6670.24\|CGTACGACTCTTAGCGGTGGATCA | >hg38_21_+_8433172_8446622_RNA45SN1_45S_50ntFlanks | 6647 | 6670 | 24 | CGTACGACTCTTAGCGGTGGATCA | 579.8 | -3.2 | 0.8 | -4.1 | 4.6E-05 | 1.7E-04 | BT20/MB468 | MP |
| >hg38_21_+_8433172_8446622_RNA45SN1_45S_50ntFlanks@8196.8222.27\|CGTGGACGGTGTGAGGCCGGTAGCGGC | >hg38_21_+_8433172_8446622_RNA45SN1_45S_50ntFlanks | 8196 | 8222 | 27 | CGTGGACGGTGTGAGGCCGGTAGCGGC | 542.0 | -4.8 | 1.2 | -3.9 | 1.1E-04 | 1.7E-04 | MB231/MB468 | MP |
| >hg38_21_+_8433172_8446622_RNA45SN1_45S_50ntFlanks@10391.10422.32\|AACAGCAGTTGAACATGGGTCAGTCGGTCCTG | >hg38_21_+_8433172_8446622_RNA45SN1_45S_50ntFlanks | 10391 | 10422 | 32 | AACAGCAGTTGAACATGGGTCAGTCGGTCCTG | 418.6 | -3.0 | 0.8 | -3.9 | 1.1E-04 | 1.7E-04 | BT20/MB468 | CYTO |
| >trna116_GluCTC_1_-_145399233_145399304@1.31.31,trna59_GluCTC_1_+_249168447_249168518@1.31.31,trna71_GluCTC_1_-_161439189_161439260@1.31.31,trna74_GluCTC_1_-_161431809_161431880@1.31.31,trna77_GluCTC_1_-_161424398_161424469@1.31.31,trna77_GluCTC_6_+_28949976_28950047@1.31.31,trna80_GluCTC_1_-_161417018_161417089@1.31.31,trna87_GluCTC_6_-_126101393_126101464@1.31.31\|TCCCTGGTGGTCTAGTGGTTAGGATTCGGCG | >trna116_GluCTC_1_-_145399233_145399304 | 1 | 31 | 31 | TCCCTGGTGGTCTAGTGGTTAGGATTCGGCG | 292060.2 | -3.5 | 0.9 | -3.8 | 1.2E-04 | 1.7E-04 | MB231/MB468 | CYTO |
| >hg38_21_+_8433172_8446622_RNA45SN1_45S_50ntFlanks@6663.6681.19\|GTGGATCACTCGGCTCGTG | >hg38_21_+_8433172_8446622_RNA45SN1_45S_50ntFlanks | 6663 | 6681 | 19 | GTGGATCACTCGGCTCGTG | 4533.9 | -4.8 | 1.2 | -4.1 | 4.6E-05 | 1.7E-04 | BT20/MB468 | MP |
| >hg38_21_+_8433172_8446622_RNA45SN1_45S_50ntFlanks@11562.11603.42\|CGCCTAGCAGCCGACTTAGAACTGGTGCGGACCAGGGGAATC | >hg38_21_+_8433172_8446622_RNA45SN1_45S_50ntFlanks | 11562 | 11603 | 42 | CGCCTAGCAGCCGACTTAGAACTGGTGCGGACCAGGGGAATC | 337.6 | -4.7 | 1.2 | -3.8 | 1.3E-04 | 1.7E-04 | MB231/MB468 | NUC |
| >hg38_21_+_8433172_8446622_RNA45SN1_45S_50ntFlanks@11873.11899.27\|AAGAAGACCCTGTTGAGCTTGACTCTA | >hg38_21_+_8433172_8446622_RNA45SN1_45S_50ntFlanks | 11873 | 11899 | 27 | AAGAAGACCCTGTTGAGCTTGACTCTA | 491.8 | -5.8 | 1.5 | -3.8 | 1.3E-04 | 1.7E-04 | MB231/MB468 | NUC |
| >hg38_21_+_8433172_8446622_RNA45SN1_45S_50ntFlanks@9533.9562.30\|GAGGAAACTCTGGTGGAGGTCCGTAGCGGT | >hg38_21_+_8433172_8446622_RNA45SN1_45S_50ntFlanks | 9533 | 9562 | 30 | GAGGAAACTCTGGTGGAGGTCCGTAGCGGT | 1354.7 | -7.4 | 1.9 | -3.8 | 1.3E-04 | 1.7E-04 | BT20/MB468 | NUC |
| >hg38_21_+_8433172_8446622_RNA45SN1_45S_50ntFlanks@5057.5082.26\|AGCGATTTGTCTGGTTAATTCCGATA | >hg38_21_+_8433172_8446622_RNA45SN1_45S_50ntFlanks | 5057 | 5082 | 26 | AGCGATTTGTCTGGTTAATTCCGATA | 412.5 | -5.7 | 1.5 | -3.9 | 1.1E-04 | 1.7E-04 | MB231/MB468 | MP |
| >hg38_21_+_8433172_8446622_RNA45SN1_45S_50ntFlanks@10248.10286.39\|CGACGAGTAGGAGGGCCGCTGCGGTGAGCCTTGAAGCCT | >hg38_21_+_8433172_8446622_RNA45SN1_45S_50ntFlanks | 10248 | 10286 | 39 | CGACGAGTAGGAGGGCCGCTGCGGTGAGCCTTGAAGCCT | 309.3 | 3.8 | 0.9 | 4.1 | 4.6E-05 | 1.7E-04 | BT20/MB468 | MP |
| >hg38_21_+_8433172_8446622_RNA45SN1_45S_50ntFlanks@10878.10901.24\|TCGGTCGGGCTGGGGCGCGAAGCG | >hg38_21_+_8433172_8446622_RNA45SN1_45S_50ntFlanks | 10878 | 10901 | 24 | TCGGTCGGGCTGGGGCGCGAAGCG | 354.3 | -3.9 | 1.0 | -3.9 | 1.1E-04 | 1.7E-04 | BT20/MB468 | CYTO |
| >hg38_21_+_8433172_8446622_RNA45SN1_45S_50ntFlanks@3854.3880.27\|ACTGTGGTAATTCTAGAGCTAATACAT | >hg38_21_+_8433172_8446622_RNA45SN1_45S_50ntFlanks | 3854 | 3880 | 27 | ACTGTGGTAATTCTAGAGCTAATACAT | 290.8 | -2.9 | 0.8 | -3.8 | 1.3E-04 | 1.7E-04 | MB231/MB468 | NUC |
| >hg38_21_+_8433172_8446622_RNA45SN1_45S_50ntFlanks@10236.10267.32\|GACGCTACGCCGCGACGAGTAGGAGGGCCGCT | >hg38_21_+_8433172_8446622_RNA45SN1_45S_50ntFlanks | 10236 | 10267 | 32 | GACGCTACGCCGCGACGAGTAGGAGGGCCGCT | 76.1 | -6.0 | 1.5 | -3.9 | 1.1E-04 | 1.7E-04 | MB231/MB468 | MP |
| >hg38_21_+_8433172_8446622_RNA45SN1_45S_50ntFlanks@6690.6720.31\|AAGAACGCAGCTAGCTGCGAGAATTAATGTG | >hg38_21_+_8433172_8446622_RNA45SN1_45S_50ntFlanks | 6690 | 6720 | 31 | AAGAACGCAGCTAGCTGCGAGAATTAATGTG | 764.9 | -6.5 | 1.6 | -4.1 | 4.6E-05 | 1.7E-04 | BT20/MB468 | MP |
| >hg38_21_+_8433172_8446622_RNA45SN1_45S_50ntFlanks@12995.13012.18\|CTCGCTGCGATCTATTGA | >hg38_21_+_8433172_8446622_RNA45SN1_45S_50ntFlanks | 12995 | 13012 | 18 | CTCGCTGCGATCTATTGA | 1054.4 | 2.2 | 0.6 | 3.8 | 1.2E-04 | 1.7E-04 | MB231/MB468 | CYTO |
| >hg38_21_+_8433172_8446622_RNA45SN1_45S_50ntFlanks@8179.8203.25\|CTGATCGAGGCCCAGCCCGTGGACG | >hg38_21_+_8433172_8446622_RNA45SN1_45S_50ntFlanks | 8179 | 8203 | 25 | CTGATCGAGGCCCAGCCCGTGGACG | 256.0 | -3.7 | 0.9 | -3.8 | 1.2E-04 | 1.7E-04 | MB231/MB468 | CYTO |
| >MI0001445\|hsa-mir-423&WithFlank&17\|+\|30117073\|30117178@59.80.22&[MIMAT0001340&hsa-miR-423-3p&offsets\|0\|-1;m-100&17\|+\|30117131\|30117153&offsets\|0\|-1]\|AGCTCGGTCTGAGGCCCCTCAG | >MI0001445\|hsa-mir-423&WithFlank&17\|+\|30117073\|30117178 | 59 | 80 | 22 | AGCTCGGTCTGAGGCCCCTCAG | 505.5 | 3.8 | 1.0 | 3.9 | 1.1E-04 | 1.7E-04 | MB231/MB468 | MP |
| >hg38_21_+_8433172_8446622_RNA45SN1_45S_50ntFlanks@12415.12432.18\|CTTCGATGTCGGCTCTTC | >hg38_21_+_8433172_8446622_RNA45SN1_45S_50ntFlanks | 12415 | 12432 | 18 | CTTCGATGTCGGCTCTTC | 1521.3 | -1.6 | 0.4 | -3.9 | 9.3E-05 | 1.7E-04 | BT20/MB468 | MITO |
| >hg38_21_+_8433172_8446622_RNA45SN1_45S_50ntFlanks@12161.12182.22\|TGACTGGGGCGGTACACCTGTC | >hg38_21_+_8433172_8446622_RNA45SN1_45S_50ntFlanks | 12161 | 12182 | 22 | TGACTGGGGCGGTACACCTGTC | 256.6 | -6.3 | 1.6 | -4.1 | 4.6E-05 | 1.7E-04 | BT20/MB468 | MP |
| >hg38_21_+_8433172_8446622_RNA45SN1_45S_50ntFlanks@4369.4391.23\|GTTAAAAAGCTCGTAGTTGGATC | >hg38_21_+_8433172_8446622_RNA45SN1_45S_50ntFlanks | 4369 | 4391 | 23 | GTTAAAAAGCTCGTAGTTGGATC | 331.5 | -5.1 | 1.3 | -3.8 | 1.3E-04 | 1.7E-04 | BT20/MB468 | NUC |
| >trnaMT_SerGCT_MT_+_12207_12265@1.47.47\|GAGAAAGCTCACAAGAACTGCTAACTCATGCCCCCATGTCTAACAAC | >trnaMT_SerGCT_MT_+_12207_12265 | 1 | 47 | 47 | GAGAAAGCTCACAAGAACTGCTAACTCATGCCCCCATGTCTAACAAC | 347.4 | 6.4 | 1.7 | 3.8 | 1.3E-04 | 1.7E-04 | MB231/MB468 | MITO |
| >hg38_21_+_8433172_8446622_RNA45SN1_45S_50ntFlanks@11729.11748.20\|TAACTATGACTCTCTTAAGG | >hg38_21_+_8433172_8446622_RNA45SN1_45S_50ntFlanks | 11729 | 11748 | 20 | TAACTATGACTCTCTTAAGG | 5084.5 | -2.5 | 0.7 | -3.8 | 1.2E-04 | 1.7E-04 | MB231/MB468 | CYTO |
| >trnaMT_HisGTG_MT_+_12138_12206@45.71.27\|AGAGGCTTACGACCCCTTATTTACCCC | >trnaMT_HisGTG_MT_+_12138_12206 | 45 | 71 | 27 | AGAGGCTTACGACCCCTTATTTACCCC | 23.9 | 7.8 | 2.0 | 3.9 | 1.1E-04 | 1.7E-04 | MB231/MB468 | MP |
| >hg38_21_+_8433172_8446622_RNA45SN1_45S_50ntFlanks@6644.6681.38\|CCTCGTACGACTCTTAGCGGTGGATCACTCGGCTCGTG | >hg38_21_+_8433172_8446622_RNA45SN1_45S_50ntFlanks | 6644 | 6681 | 38 | CCTCGTACGACTCTTAGCGGTGGATCACTCGGCTCGTG | 202.5 | -3.8 | 1.0 | -3.9 | 1.1E-04 | 1.7E-04 | MB231/MB468 | MP |
| >MI0000749\|hsa-mir-30e&WithFlank&1\|+\|40754349\|40754452@65.83.19&[MIMAT0000693&hsa-miR-30e-3p&offsets\|0\|-3;m-25&1\|+\|40754413\|40754434&offsets\|0\|-3]\|CTTTCAGTCGGATGTTTAC | >MI0000749\|hsa-mir-30e&WithFlank&1\|+\|40754349\|40754452 | 65 | 83 | 19 | CTTTCAGTCGGATGTTTAC | 576.4 | -1.8 | 0.5 | -3.8 | 1.3E-04 | 1.7E-04 | BT20/MB468 | NUC |

<table>
<tr><td>>MI0000264|hsa-mir-7-2&WithFlank&15|+|88611819|88611940@38.58.21(+1A);MI0000265|hsa-mir-7-3&WithFlank&19|+|4770664|4770785@37.57.21(+1A);MI0000263|hsa-mir-7-1&WithFlank&9|-|83969742|83969863@30.50.21(+1A)&[MIMAT0000252&hsa-miR-7-5p&offsets|0|-3(+1A);m-400&15|+|88611856|88611879&offsets|0|-3(+1A)];[MIMAT0000252_1&hsa-miR-7-5p&offsets|0|-3(+1A);m-402&19|+|4770700|4770723&offsets|0|-3(+1A)];[MIMAT0000252_2&hsa-miR-7-5p&offsets|0|-3(+1A);m-401&9|-|83969811|83969834&offsets|0|-3(+1A)]|TGGAAGACTAGTGATTTTGTTA</td><td>>MI0000264|hsa-mir-7-2&WithFlank&15|+|88611819|88611940</td><td>38</td><td>58</td><td>21(+1A)</td><td>TGGAAGACTAGTGATTTTGTTA</td><td>242.4</td><td>4.7</td><td>1.2</td><td>3.9</td><td>1.1E-04</td><td>1.7E-04</td><td>MB231/MB468</td><td>MP</td></tr>
<tr><td>>hg38_21_+_8433172_8446622_RNA45SN1_45S_50ntFlanks@3777.3801.25|CGGTACAGTGAAACTGCGAATGGCT</td><td>>hg38_21_+_8433172_8446622_RNA45SN1_45S_50ntFlanks</td><td>3777</td><td>3801</td><td>25</td><td>CGGTACAGTGAAACTGCGAATGGCT</td><td>674.0</td><td>-2.5</td><td>0.7</td><td>-3.8</td><td>1.2E-04</td><td>1.7E-04</td><td>MB231/MB468</td><td>CYTO</td></tr>
<tr><td>>MI0000253|hsa-mir-148a&WithFlank&7|-|25949913|25949992@50.70.21(+1C)&[MIMAT0000243&hsa-miR-148a-3p&offsets|0|-1(+1C);m-7&7|-|25949922|25949943&offsets|0|-1(+1C)]|TCAGTGCACTACAGAACTTTGC</td><td>>MI0000253|hsa-mir-148a&WithFlank&7|-|25949913|25949992</td><td>50</td><td>70</td><td>21(+1C)</td><td>TCAGTGCACTACAGAACTTTGC</td><td>65527.9</td><td>4.6</td><td>1.1</td><td>4.1</td><td>4.6E-05</td><td>1.7E-04</td><td>BT20/MB468</td><td>MP</td></tr>
<tr><td>>am_trna111_HisGTG_1_-_147774845_147774916@-1G.23.24,trna118_HisGTG_1_-_145396881_145396952@-1G.23.24,trna16_HisGTG_1_+_146544773_146544844@-1G.23.24,trna1_HisGTG_15_+_45493349_45493420@-1G.23.24,trna21_HisGTG_1_+_147753471_147753542@-1G.23.24,trna33_HisGTG_6_+_27125906_27125977@-1G.23.24,trna7_HisGTG_9_-_14433938_14434009@-1G.23.24,trna8_HisGTG_15_-_45492611_45492682@-1G.23.24,trna9_HisGTG_15_-_45490804_45490875@-1G.23.24|GGCCGTGATCGTATAGTGGTTAGT</td><td>>am_trna111_HisGTG_1_-_147774845_147774916</td><td>-1G</td><td>23</td><td>24</td><td>GGCCGTGATCGTATAGTGGTTAGT</td><td>1213.0</td><td>-5.6</td><td>1.3</td><td>-4.3</td><td>1.7E-05</td><td>1.7E-04</td><td>BT20/MB231</td><td>NUC</td></tr>
<tr><td>>MI0006444|hsa-mir-1307&WithFlank&10|-|103394247|103394407@86.107.22(+1U)&[MIMAT0005951&hsa-miR-1307-3p&offsets|0|0(+1U);m-66&10|-|103394300|103394322&offsets|0|-1(+1U)]|ACTCGGCGTGGCGTCGGTCGTGT</td><td>>MI0006444|hsa-mir-1307&WithFlank&10|-|103394247|103394407</td><td>86</td><td>107</td><td>22(+1U)</td><td>ACTCGGCGTGGCGTCGGTCGTGT</td><td>397.6</td><td>3.2</td><td>0.8</td><td>3.9</td><td>1.1E-04</td><td>1.7E-04</td><td>MB231/MB468</td><td>MP</td></tr>
<tr><td>>MI0000066|hsa-let-7e&WithFlank&19|+|51692780|51692870@14.35.22&[MIMAT0000066&hsa-let-7e-5p&offsets|0|0;m-50&19|+|51692793|51692814&offsets|0|0]|TGAGGTAGGAGGTTGTATAGTT</td><td>>MI0000066|hsa-let-7e&WithFlank&19|+|51692780|51692870</td><td>14</td><td>35</td><td>22</td><td>TGAGGTAGGAGGTTGTATAGTT</td><td>4229.3</td><td>1.3</td><td>0.3</td><td>3.8</td><td>1.3E-04</td><td>1.7E-04</td><td>BT20/MB468</td><td>NUC</td></tr>
<tr><td>>hg38_21_+_8433172_8446622_RNA45SN1_45S_50ntFlanks@5074.5098.25|ATTCCGATAACGAACGAGACTCTGG</td><td>>hg38_21_+_8433172_8446622_RNA45SN1_45S_50ntFlanks</td><td>5074</td><td>5098</td><td>25</td><td>ATTCCGATAACGAACGAGACTCTGG</td><td>610.1</td><td>-5.4</td><td>1.4</td><td>-3.8</td><td>1.3E-04</td><td>1.7E-04</td><td>MB231/MB468</td><td>NUC</td></tr>
<tr><td>>hg38_21_+_8433172_8446622_RNA45SN1_45S_50ntFlanks@3780.3801.22|TACAGTGAAACTGCGAATGGCT</td><td>>hg38_21_+_8433172_8446622_RNA45SN1_45S_50ntFlanks</td><td>3780</td><td>3801</td><td>22</td><td>TACAGTGAAACTGCGAATGGCT</td><td>335.2</td><td>-4.6</td><td>1.2</td><td>-3.9</td><td>9.4E-05</td><td>1.7E-04</td><td>BT20/MB468</td><td>MITO</td></tr>
<tr><td>>MI0000542|hsa-mir-320a&WithFlank&8|-|22244960|22245043@48.69.22(+1U)&[MIMAT0000510&hsa-miR-320a-3p&offsets|0|0(+1U);m-60&8|-|22244975|22244996&offsets|0|0(+1U)]|AAAAGCTGGGTTGAGAGGGCGAT</td><td>>MI0000542|hsa-mir-320a&WithFlank&8|-|22244960|22245043</td><td>48</td><td>69</td><td>22(+1U)</td><td>AAAAGCTGGGTTGAGAGGGCGAT</td><td>2182.8</td><td>3.9</td><td>1.0</td><td>3.9</td><td>1.1E-04</td><td>1.7E-04</td><td>MB231/MB468</td><td>MP</td></tr>
<tr><td>>MI0003599|hsa-mir-589&WithFlank&7|-|5495813|5495923@30.50.21&[MIMAT0004799&hsa-miR-589-5p&offsets|0|-1;m-162&7|-|5495874|5495894&offsets|0|0]|TGAGAACCACGTCTGCTCTGA</td><td>>MI0003599|hsa-mir-589&WithFlank&7|-|5495813|5495923</td><td>30</td><td>50</td><td>21</td><td>TGAGAACCACGTCTGCTCTGA</td><td>456.9</td><td>5.8</td><td>1.4</td><td>4.1</td><td>4.6E-05</td><td>1.7E-04</td><td>BT20/MB468</td><td>MP</td></tr>
<tr><td>>trnaMT_IleGAT_MT_+_4263_4331@44.72.29|TAGGAGCTTAAACCCCCTTATTTCTACCA</td><td>>trnaMT_IleGAT_MT_+_4263_4331</td><td>44</td><td>72</td><td>29</td><td>TAGGAGCTTAAACCCCCTTATTTCTACCA</td><td>118.4</td><td>10.6</td><td>2.6</td><td>4.1</td><td>4.7E-05</td><td>1.7E-04</td><td>BT20/MB468</td><td>MP</td></tr>
<tr><td>>hg38_21_+_8433172_8446622_RNA45SN1_45S_50ntFlanks@6729.6771.43|GACACATTGATCATCGACACTTCGAACGCACTTGCGGCCCCGG</td><td>>hg38_21_+_8433172_8446622_RNA45SN1_45S_50ntFlanks</td><td>6729</td><td>6771</td><td>43</td><td>GACACATTGATCATCGACACTTCGAACGCACTTGCGGCCCCGG</td><td>477.7</td><td>-3.6</td><td>0.9</td><td>-3.8</td><td>1.3E-04</td><td>1.7E-04</td><td>MB231/MB468</td><td>NUC</td></tr>
<tr><td>>hg38_21_+_8433172_8446622_RNA45SN1_45S_50ntFlanks@5492.5509.18|AGAAGACGGTCGAACTTG</td><td>>hg38_21_+_8433172_8446622_RNA45SN1_45S_50ntFlanks</td><td>5492</td><td>5509</td><td>18</td><td>AGAAGACGGTCGAACTTG</td><td>522.7</td><td>-4.0</td><td>1.0</td><td>-3.8</td><td>1.3E-04</td><td>1.7E-04</td><td>BT20/MB468</td><td>NUC</td></tr>
<tr><td>>hg38_21_+_8433172_8446622_RNA45SN1_45S_50ntFlanks@4241.4258.18|CTCTTTCGAGGCCCTGTA</td><td>>hg38_21_+_8433172_8446622_RNA45SN1_45S_50ntFlanks</td><td>4241</td><td>4258</td><td>18</td><td>CTCTTTCGAGGCCCTGTA</td><td>183.2</td><td>-6.0</td><td>1.5</td><td>-4.1</td><td>4.7E-05</td><td>1.7E-04</td><td>BT20/MB468</td><td>MP</td></tr>
<tr><td>>hg38_MT_+_1621_3279_RNR2_16S_50ntFlanks@621.663.43|ATATAACTGAACTCCTCACACCCAATTGGACCAATCTATCACC</td><td>>hg38_MT_+_1621_3279_RNR2_16S_50ntFlanks</td><td>621</td><td>663</td><td>43</td><td>ATATAACTGAACTCCTCACACCCAATTGGACCAATCTATCACC</td><td>116.7</td><td>10.6</td><td>2.6</td><td>4.1</td><td>4.7E-05</td><td>1.7E-04</td><td>BT20/MB468</td><td>MP</td></tr>
<tr><td>>hg38_21_+_8433172_8446622_RNA45SN1_45S_50ntFlanks@4590.4613.24|ATTTTGTTGGTTTTCGGAACTGAG</td><td>>hg38_21_+_8433172_8446622_RNA45SN1_45S_50ntFlanks</td><td>4590</td><td>4613</td><td>24</td><td>ATTTTGTTGGTTTTCGGAACTGAG</td><td>164.7</td><td>-6.0</td><td>1.5</td><td>-4.1</td><td>4.7E-05</td><td>1.7E-04</td><td>BT20/MB468</td><td>MP</td></tr>
<tr><td>>trna119_LysCTT_1_-_145395522_145395594@1.31.31,trna11_LysCTT_5_-_180648979_180649051@1.31.31,trna13_LysCTT_6_+_26556774_26556846@1.31.31,trna32_LysCTT_16_-_3207406_3207478@1.31.31,trna7_LysCTT_16_+_3225692_3225764@1.31.31,trna9_LysCTT_5_+_180634755_180634827@1.31.31|GCCCGGCTAGCTCAGTCGGTAGAGCATGAGA</td><td>>trna119_LysCTT_1_-_145395522_145395594</td><td>1</td><td>31</td><td>31</td><td>GCCCGGCTAGCTCAGTCGGTAGAGCATGAGA</td><td>13504.3</td><td>-3.6</td><td>0.9</td><td>-3.8</td><td>1.2E-04</td><td>1.7E-04</td><td>MB231/MB468</td><td>CYTO</td></tr>
<tr><td>>hg38_21_+_8433172_8446622_RNA45SN1_45S_50ntFlanks@11667.11693.27|CTGCCCAGTGCTCTGAATGTCAAAGTG</td><td>>hg38_21_+_8433172_8446622_RNA45SN1_45S_50ntFlanks</td><td>11667</td><td>11693</td><td>27</td><td>CTGCCCAGTGCTCTGAATGTCAAAGTG</td><td>304.5</td><td>4.7</td><td>1.2</td><td>3.8</td><td>1.2E-04</td><td>1.7E-04</td><td>MB231/MB468</td><td>CYTO</td></tr>
<tr><td>>MI0005559|hsa-mir-744&WithFlank&17|+|12081893|12082002@17.36.20(+1U)&[MIMAT0004945&hsa-miR-744-5p&offsets|0|-2(+1U);m-172&17|+|12081909|12081929&offsets|0|-1(+1U)]|TGCGGGGCTAGGGCTAACAGT</td><td>>MI0005559|hsa-mir-744&WithFlank&17|+|12081893|12082002</td><td>17</td><td>36</td><td>20(+1U)</td><td>TGCGGGGCTAGGGCTAACAGT</td><td>739.1</td><td>3.0</td><td>0.7</td><td>4.3</td><td>1.9E-05</td><td>1.7E-04</td><td>BT20/MB231</td><td>MITO</td></tr>
<tr><td>>hg38_21_+_8433172_8446622_RNA45SN1_45S_50ntFlanks@11027.11049.23|CTCTCTCTCTCTCTCCCCCGCTC</td><td>>hg38_21_+_8433172_8446622_RNA45SN1_45S_50ntFlanks</td><td>11027</td><td>11049</td><td>23</td><td>CTCTCTCTCTCTCTCCCCCGCTC</td><td>2533.1</td><td>-7.7</td><td>2.0</td><td>-3.8</td><td>1.3E-04</td><td>1.7E-04</td><td>BT20/MB468</td><td>NUC</td></tr>
<tr><td>>hg38_21_+_8433172_8446622_RNA45SN1_45S_50ntFlanks@8184.8211.28|CGAGGCCCAGCCCGTGGACGGTGTGAGG</td><td>>hg38_21_+_8433172_8446622_RNA45SN1_45S_50ntFlanks</td><td>8184</td><td>8211</td><td>28</td><td>CGAGGCCCAGCCCGTGGACGGTGTGAGG</td><td>106.7</td><td>-3.2</td><td>0.8</td><td>-3.9</td><td>1.1E-04</td><td>1.7E-04</td><td>BT20/MB468</td><td>CYTO</td></tr>
<tr><td>>trnaMT_TrpTCA_MT_+_5512_5579@18.35.18,trnalookalike3_TrpTCA_1_+_566062_566129@18.35.18|ACAGACCAAGAGCCTTCA</td><td>>trnaMT_TrpTCA_MT_+_5512_5579</td><td>18</td><td>35</td><td>18</td><td>ACAGACCAAGAGCCTTCA</td><td>195.7</td><td>3.1</td><td>0.8</td><td>3.9</td><td>1.1E-04</td><td>1.7E-04</td><td>MB231/MB468</td><td>MP</td></tr>
<tr><td>>MI0000294|hsa-mir-218-1&WithFlank&4|+|20528269|20528390@31.51.21;MI0000295|hsa-mir-218-2&WithFlank&5|-|168768140|168768261@31.51.21&[MIMAT0000275&hsa-miR-218-5p&offsets|0|0;m-185&4|+|20528299|20528319&offsets|0|0];[MIMAT0000275_1&hsa-miR-218-5p&offsets|0|0;m-187&5|-|168768211|168768231&offsets|0|0]|TTGTGCTTGATCTAACCATGT</td><td>>MI0000294|hsa-mir-218-1&WithFlank&4|+|20528269|20528390</td><td>31</td><td>51</td><td>21</td><td>TTGTGCTTGATCTAACCATGT</td><td>579.7</td><td>7.1</td><td>1.8</td><td>3.9</td><td>1.1E-04</td><td>1.7E-04</td><td>MB231/MB468</td><td>MP</td></tr>
<tr><td>>hg38_21_+_8433172_8446622_RNA45SN1_45S_50ntFlanks@10793.10818.26|TCTGGCATGTTGGAACAATGTAGGTA</td><td>>hg38_21_+_8433172_8446622_RNA45SN1_45S_50ntFlanks</td><td>10793</td><td>10818</td><td>26</td><td>TCTGGCATGTTGGAACAATGTAGGTA</td><td>151.6</td><td>-3.1</td><td>0.8</td><td>-3.8</td><td>1.2E-04</td><td>1.7E-04</td><td>MB231/MB468</td><td>CYTO</td></tr>
<tr><td>>hg38_21_+_8433172_8446622_RNA45SN1_45S_50ntFlanks@9533.9565.33|GAGGAAACTCTGGTGGAGGTCCGTAGCGGTCCT</td><td>>hg38_21_+_8433172_8446622_RNA45SN1_45S_50ntFlanks</td><td>9533</td><td>9565</td><td>33</td><td>GAGGAAACTCTGGTGGAGGTCCGTAGCGGTCCT</td><td>379.4</td><td>-6.6</td><td>1.7</td><td>-3.8</td><td>1.3E-04</td><td>1.7E-04</td><td>BT20/MB468</td><td>NUC</td></tr>
<tr><td>>hg38_MT_+_1621_3279_RNR2_16S_50ntFlanks@1170.1215.46|AAACCTGCATTAAAAATTTCGGTTGGGGCGACCTCGGAGCAGAACC</td><td>>hg38_MT_+_1621_3279_RNR2_16S_50ntFlanks</td><td>1170</td><td>1215</td><td>46</td><td>AAACCTGCATTAAAAATTTCGGTTGGGGCGACCTCGGAGCAGAACC</td><td>112.0</td><td>10.6</td><td>2.6</td><td>4.1</td><td>4.7E-05</td><td>1.7E-04</td><td>BT20/MB468</td><td>MP</td></tr>
<tr><td>>MI0000078|hsa-mir-22&WithFlank&17|-|1713897|1713993@59.81.23(+1U)&[MIMAT0000077&hsa-miR-22-3p&offsets|0|+1(+1U);m-2&17|-|1713914|1713935&offsets|0|+1(+1U)]|AAGCTGCCAGTTGAAGAACTGTTT</td><td>>MI0000078|hsa-mir-22&WithFlank&17|-|1713897|1713993</td><td>59</td><td>81</td><td>23(+1U)</td><td>AAGCTGCCAGTTGAAGAACTGTTT</td><td>27.8</td><td>8.0</td><td>2.1</td><td>3.9</td><td>1.1E-04</td><td>1.7E-04</td><td>MB231/MB468</td><td>MP</td></tr>
<tr><td>>hg38_21_+_8433172_8446622_RNA45SN1_45S_50ntFlanks@8359.8392.34|AAGAACTTTGAAGAGAGAGTTCAAGAGGGCGTGA</td><td>>hg38_21_+_8433172_8446622_RNA45SN1_45S_50ntFlanks</td><td>8359</td><td>8392</td><td>34</td><td>AAGAACTTTGAAGAGAGAGTTCAAGAGGGCGTGA</td><td>74.8</td><td>9.6</td><td>2.2</td><td>4.3</td><td>1.7E-05</td><td>1.7E-04</td><td>BT20/MB231</td><td>MP</td></tr>
<tr><td>>hg38_21_+_8433172_8446622_RNA45SN1_45S_50ntFlanks@12477.12498.22|CACTAATAGGGAACGTGAGCTG</td><td>>hg38_21_+_8433172_8446622_RNA45SN1_45S_50ntFlanks</td><td>12477</td><td>12498</td><td>22</td><td>CACTAATAGGGAACGTGAGCTG</td><td>429.0</td><td>-4.4</td><td>1.1</td><td>-3.8</td><td>1.3E-04</td><td>1.7E-04</td><td>MB231/MB468</td><td>MITO</td></tr>
<tr><td>>hg38_21_+_8433172_8446622_RNA45SN1_45S_50ntFlanks@7984.8006.23|AGATCAGACGTGGCGACCCGCTG</td><td>>hg38_21_+_8433172_8446622_RNA45SN1_45S_50ntFlanks</td><td>7984</td><td>8006</td><td>23</td><td>AGATCAGACGTGGCGACCCGCTG</td><td>1102.5</td><td>-4.7</td><td>1.2</td><td>-3.9</td><td>1.1E-04</td><td>1.7E-04</td><td>BT20/MB468</td><td>CYTO</td></tr>
<tr><td>>hg38_21_+_8433172_8446622_RNA45SN1_45S_50ntFlanks@7975.8005.31|CGCGACCTCAGATCAGACGTGGCGACCCGCT</td><td>>hg38_21_+_8433172_8446622_RNA45SN1_45S_50ntFlanks</td><td>7975</td><td>8005</td><td>31</td><td>CGCGACCTCAGATCAGACGTGGCGACCCGCT</td><td>15426.4</td><td>2.6</td><td>0.7</td><td>3.8</td><td>1.3E-04</td><td>1.7E-04</td><td>MB231/MB468</td><td>NUC</td></tr>
<tr><td>>hg38_21_+_8433172_8446622_RNA45SN1_45S_50ntFlanks@10788.10821.34|CAGCCTCTGGCATGTTGGAACAATGTAGGTAAGG</td><td>>hg38_21_+_8433172_8446622_RNA45SN1_45S_50ntFlanks</td><td>10788</td><td>10821</td><td>34</td><td>CAGCCTCTGGCATGTTGGAACAATGTAGGTAAGG</td><td>130.7</td><td>-4.4</td><td>1.1</td><td>-3.9</td><td>1.1E-04</td><td>1.7E-04</td><td>MB231/MB468</td><td>MP</td></tr>
<tr><td>>hg38_21_+_8433172_8446622_RNA45SN1_45S_50ntFlanks@4049.4066.18|TCGAACGTCTGCCCTATC</td><td>>hg38_21_+_8433172_8446622_RNA45SN1_45S_50ntFlanks</td><td>4049</td><td>4066</td><td>18</td><td>TCGAACGTCTGCCCTATC</td><td>876.2</td><td>2.7</td><td>0.7</td><td>3.8</td><td>1.2E-04</td><td>1.7E-04</td><td>MB231/MB468</td><td>CYTO</td></tr>
<tr><td>>hg38_MT_+_598_1651_RNR1_12S_50ntFlanks@424.469.46|TACGAAAGTGGCTTTAACATATCTGAACACACAATAGCTAAGACCC</td><td>>hg38_MT_+_598_1651_RNR1_12S_50ntFlanks</td><td>424</td><td>469</td><td>46</td><td>TACGAAAGTGGCTTTAACATATCTGAACACACAATAGCTAAGACCC</td><td>116.6</td><td>10.6</td><td>2.6</td><td>4.1</td><td>4.7E-05</td><td>1.7E-04</td><td>BT20/MB468</td><td>MP</td></tr>
<tr><td>>MI0003776|hsa-mir-320b-1&WithFlank&1|+|116671740|116671823@48.68.21(+1U);MI0003839|hsa-mir-320b-2&WithFlank&1|-|224257034|224257116@47.67.21(+1U)&[MIMAT0005792&hsa-miR-320b&offsets|0|-1(+1U);m-183&1|+|116671787|116671806&offsets|0|+1(+1U)];[MIMAT0005792_1&hsa-miR-320b&offsets|0|-1(+1U);m-183&1|-|224257051|224257070&offsets|0|+1(+1U)]|AAAAGCTGGGTTGAGAGGGCAT</td><td>>MI0003776|hsa-mir-320b-1&WithFlank&1|+|116671740|116671823</td><td>48</td><td>68</td><td>21(+1U)</td><td>AAAAGCTGGGTTGAGAGGGCAT</td><td>146.0</td><td>7.6</td><td>1.9</td><td>4.1</td><td>4.7E-05</td><td>1.7E-04</td><td>BT20/MB468</td><td>MP</td></tr>
<tr><td>>hg38_21_+_8433172_8446622_RNA45SN1_45S_50ntFlanks@11564.11585.22|CCTAGCAGCCGACTTAGAACTG</td><td>>hg38_21_+_8433172_8446622_RNA45SN1_45S_50ntFlanks</td><td>11564</td><td>11585</td><td>22</td><td>CCTAGCAGCCGACTTAGAACTG</td><td>1791.8</td><td>-3.9</td><td>1.0</td><td>-4.1</td><td>4.7E-05</td><td>1.7E-04</td><td>BT20/MB468</td><td>MP</td></tr>
<tr><td>>MI0000286|hsa-mir-210&WithFlank&11|-|568083|568204@72.93.22(+1A)&[MIMAT0000267&hsa-miR-210-3p&offsets|0|0(+1A);m-102&11|-|568112|568133&offsets|0|0(+1A)]|CTGTGCGTGTGACAGCGGCTGAA</td><td>>MI0000286|hsa-mir-210&WithFlank&11|-|568083|568204</td><td>72</td><td>93</td><td>22(+1A)</td><td>CTGTGCGTGTGACAGCGGCTGAA</td><td>1282.3</td><td>6.2</td><td>1.5</td><td>4.1</td><td>4.7E-05</td><td>1.7E-04</td><td>BT20/MB468</td><td>MP</td></tr>
<tr><td>>trnaMT_HisGTG_MT_+_12138_12206@27.70.44|GATTGTGAATCTGACAACAGAGGCTTACGACCCCTTATTTACCC</td><td>>trnaMT_HisGTG_MT_+_12138_12206</td><td>27</td><td>70</td><td>44</td><td>GATTGTGAATCTGACAACAGAGGCTTACGACCCCTTATTTACCC</td><td>113.1</td><td>10.6</td><td>2.6</td><td>4.1</td><td>4.7E-05</td><td>1.7E-04</td><td>BT20/MB468</td><td>MP</td></tr>
<tr><td>>hg38_1_-_228634819_228635039_RNA5S12_5S_50ntFlanks@132.169.38|GATGGGAGACCGCCTGGGAATACCGGGTGCTGTAGGCT</td><td>>hg38_1_-_228634819_228635039_RNA5S12_5S_50ntFlanks</td><td>132</td><td>169</td><td>38</td><td>GATGGGAGACCGCCTGGGAATACCGGGTGCTGTAGGCT</td><td>21310.7</td><td>5.7</td><td>1.3</td><td>4.3</td><td>1.7E-05</td><td>1.7E-04</td><td>BT20/MB231</td><td>MP</td></tr>
<tr><td>>MI0003820|hsa-mir-454&WithFlank&17|-|59137752|59137878@70.92.23(+1A)&[MIMAT0003885&hsa-miR-454-3p&offsets|0|0(+1A);m-272&17|-|59137787|59137809&offsets|0|0(+1A)]|TAGTGCAATATTGCTTATAGGGTA</td><td>>MI0003820|hsa-mir-454&WithFlank&17|-|59137752|59137878</td><td>70</td><td>92</td><td>23(+1A)</td><td>TAGTGCAATATTGCTTATAGGGTA</td><td>59.3</td><td>-9.3</td><td>2.2</td><td>-4.3</td><td>1.7E-05</td><td>1.7E-04</td><td>BT20/MB231</td><td>MP</td></tr>
</table>

<table>
<tr><td></td><td></td><td></td><td></td><td></td><td></td><td></td><td></td><td></td><td></td><td></td><td></td><td></td><td></td></tr>
<tr><td>>hg38_21_+_8433172_8446622_RNA45SN1_45S_50ntFlanks@11318.11342.25|CCCCACGTCTCGTCGCGCGCGCGTC</td><td>>hg38_21_+_8433172_8446622_RNA45SN1_45S_50ntFlanks</td><td>11318</td><td>11342</td><td>25</td><td>CCCCACGTCTCGTCGCGCGCGCGTC</td><td>129.2</td><td>-6.1</td><td>1.5</td><td>-4.1</td><td>4.8E-05</td><td>1.7E-04</td><td>BT20/MB468</td><td>MP</td></tr>
<tr><td>>hg38_21_+_8433172_8446622_RNA45SN1_45S_50ntFlanks@10346.10364.19|TCAAACGAGAACTTTGAAG</td><td>>hg38_21_+_8433172_8446622_RNA45SN1_45S_50ntFlanks</td><td>10346</td><td>10364</td><td>19</td><td>TCAAACGAGAACTTTGAAG</td><td>188.6</td><td>-5.9</td><td>1.5</td><td>-3.9</td><td>1.1E-04</td><td>1.7E-04</td><td>MB231/MB468</td><td>MP</td></tr>
<tr><td>>MI0006444|hsa-mir-1307&WithFlank&10|-|103394247|103394407@87.107.21(+1A)&[MIMAT0005951&hsa-miR-1307-3p&offsets|+1|0(+1A);m-66&10|-|103394300|103394322&offsets|+1|-1(+1A)]|CTCGGCGTGGCGTCGGTCGTGA</td><td>>MI0006444|hsa-mir-1307&WithFlank&10|-|103394247|103394407</td><td>87</td><td>107</td><td>21(+1A)</td><td>CTCGGCGTGGCGTCGGTCGTGA</td><td>166.3</td><td>3.7</td><td>0.9</td><td>4.1</td><td>4.8E-05</td><td>1.7E-04</td><td>BT20/MB468</td><td>MP</td></tr>
<tr><td>>hg38_21_+_8433172_8446622_RNA45SN1_45S_50ntFlanks@8381.8404.24|AAGAGGGCGTGAAACCGTTAAGAG</td><td>>hg38_21_+_8433172_8446622_RNA45SN1_45S_50ntFlanks</td><td>8381</td><td>8404</td><td>24</td><td>AAGAGGGCGTGAAACCGTTAAGAG</td><td>210.4</td><td>-5.1</td><td>1.3</td><td>-3.9</td><td>1.2E-04</td><td>1.7E-04</td><td>BT20/MB468</td><td>CYTO</td></tr>
<tr><td>>hg38_21_+_8433172_8446622_RNA45SN1_45S_50ntFlanks@3844.3868.25|TACTTGGATAACTGTGGTAATTCTA</td><td>>hg38_21_+_8433172_8446622_RNA45SN1_45S_50ntFlanks</td><td>3844</td><td>3868</td><td>25</td><td>TACTTGGATAACTGTGGTAATTCTA</td><td>751.1</td><td>1.4</td><td>0.4</td><td>3.9</td><td>1.2E-04</td><td>1.7E-04</td><td>BT20/MB468</td><td>CYTO</td></tr>
<tr><td>>hg38_21_+_8433172_8446622_RNA45SN1_45S_50ntFlanks@8176.8208.33|CTTCTGATCGAGGCCCAGCCCGTGGACGGTGTG</td><td>>hg38_21_+_8433172_8446622_RNA45SN1_45S_50ntFlanks</td><td>8176</td><td>8208</td><td>33</td><td>CTTCTGATCGAGGCCCAGCCCGTGGACGGTGTG</td><td>500.9</td><td>-3.0</td><td>0.7</td><td>-4.1</td><td>4.8E-05</td><td>1.7E-04</td><td>BT20/MB468</td><td>MP</td></tr>
<tr><td>>hg38_21_+_8433172_8446622_RNA45SN1_45S_50ntFlanks@9714.9738.25|CGAATGATTAGAGGTCTTGGGGCCG</td><td>>hg38_21_+_8433172_8446622_RNA45SN1_45S_50ntFlanks</td><td>9714</td><td>9738</td><td>25</td><td>CGAATGATTAGAGGTCTTGGGGCCG</td><td>528.7</td><td>-4.8</td><td>1.2</td><td>-3.9</td><td>9.6E-05</td><td>1.7E-04</td><td>BT20/MB468</td><td>MITO</td></tr>
<tr><td>>hg38_21_+_8433172_8446622_RNA45SN1_45S_50ntFlanks@4008.4042.35|CTCGGGCCGATCGCACGCCCCCCGTGGCGGCGACG</td><td>>hg38_21_+_8433172_8446622_RNA45SN1_45S_50ntFlanks</td><td>4008</td><td>4042</td><td>35</td><td>CTCGGGCCGATCGCACGCCCCCCGTGGCGGCGACG</td><td>157.3</td><td>-2.5</td><td>0.6</td><td>-3.9</td><td>9.6E-05</td><td>1.7E-04</td><td>BT20/MB468</td><td>MITO</td></tr>
<tr><td>>hg38_21_+_8433172_8446622_RNA45SN1_45S_50ntFlanks@12407.12441.35|TTTTGATCCTTCGATGTCGGCTCTTCCTATCATTG</td><td>>hg38_21_+_8433172_8446622_RNA45SN1_45S_50ntFlanks</td><td>12407</td><td>12441</td><td>35</td><td>TTTTGATCCTTCGATGTCGGCTCTTCCTATCATTG</td><td>79.7</td><td>-5.9</td><td>1.5</td><td>-3.9</td><td>1.1E-04</td><td>1.7E-04</td><td>MB231/MB468</td><td>MP</td></tr>
<tr><td>>trnaMT_ThrTGT_MT_+_15888_15953@31.68.38|TTGTAAACCGGAGATGAAAACCTTTTTCCAAGGACACC</td><td>>trnaMT_ThrTGT_MT_+_15888_15953</td><td>31</td><td>68</td><td>38</td><td>TTGTAAACCGGAGATGAAAACCTTTTTCCAAGGACACC</td><td>113.1</td><td>10.6</td><td>2.6</td><td>4.1</td><td>4.8E-05</td><td>1.7E-04</td><td>BT20/MB468</td><td>MP</td></tr>
<tr><td>>hg38_21_+_8433172_8446622_RNA45SN1_45S_50ntFlanks@8184.8211.28|CGAGGCCCAGCCCGTGGACGGTGTGAGG</td><td>>hg38_21_+_8433172_8446622_RNA45SN1_45S_50ntFlanks</td><td>8184</td><td>8211</td><td>28</td><td>CGAGGCCCAGCCCGTGGACGGTGTGAGG</td><td>292.0</td><td>-3.6</td><td>0.9</td><td>-4.1</td><td>4.8E-05</td><td>1.7E-04</td><td>BT20/MB468</td><td>MP</td></tr>
<tr><td>>hg38_21_+_8433172_8446622_RNA45SN1_45S_50ntFlanks@10796.10818.23|GGCATGTTGGAACAATGTAGGTA</td><td>>hg38_21_+_8433172_8446622_RNA45SN1_45S_50ntFlanks</td><td>10796</td><td>10818</td><td>23</td><td>GGCATGTTGGAACAATGTAGGTA</td><td>574.3</td><td>-4.9</td><td>1.2</td><td>-4.1</td><td>4.8E-05</td><td>1.7E-04</td><td>BT20/MB468</td><td>MP</td></tr>
<tr><td>>hg38_21_+_8433172_8446622_RNA45SN1_45S_50ntFlanks@11718.11736.19|AACGGCGGGAGTAACTATG</td><td>>hg38_21_+_8433172_8446622_RNA45SN1_45S_50ntFlanks</td><td>11718</td><td>11736</td><td>19</td><td>AACGGCGGGAGTAACTATG</td><td>348.1</td><td>-3.1</td><td>0.8</td><td>-3.9</td><td>9.6E-05</td><td>1.7E-04</td><td>BT20/MB468</td><td>MITO</td></tr>
<tr><td>>hg38_21_+_8433172_8446622_RNA45SN1_45S_50ntFlanks@11552.11591.40|CCTCGGCCGGCGCCTAGCAGCCGACTTAGAACTGGTGCGG</td><td>>hg38_21_+_8433172_8446622_RNA45SN1_45S_50ntFlanks</td><td>11552</td><td>11591</td><td>40</td><td>CCTCGGCCGGCGCCTAGCAGCCGACTTAGAACTGGTGCGG</td><td>360.3</td><td>-5.6</td><td>1.4</td><td>-3.9</td><td>1.2E-04</td><td>1.7E-04</td><td>BT20/MB468</td><td>CYTO</td></tr>
<tr><td>>hg38_21_+_8433172_8446622_RNA45SN1_45S_50ntFlanks@12712.12755.44|CGGAGCCTCGGTTGGCCTCGGATAGCCGGTCCCCCGCCTGTCCC</td><td>>hg38_21_+_8433172_8446622_RNA45SN1_45S_50ntFlanks</td><td>12712</td><td>12755</td><td>44</td><td>CGGAGCCTCGGTTGGCCTCGGATAGCCGGTCCCCCGCCTGTCCC</td><td>48.2</td><td>-8.7</td><td>2.1</td><td>-4.1</td><td>4.8E-05</td><td>1.7E-04</td><td>BT20/MB468</td><td>MP</td></tr>
<tr><td>>hg38_21_+_8433172_8446622_RNA45SN1_45S_50ntFlanks@8176.8196.21|CTTCTGATCGAGGCCCAGCCC</td><td>>hg38_21_+_8433172_8446622_RNA45SN1_45S_50ntFlanks</td><td>8176</td><td>8196</td><td>21</td><td>CTTCTGATCGAGGCCCAGCCC</td><td>648.6</td><td>-2.0</td><td>0.5</td><td>-3.8</td><td>1.3E-04</td><td>1.7E-04</td><td>MB231/MB468</td><td>NUC</td></tr>
<tr><td>>hg38_21_+_8433172_8446622_RNA45SN1_45S_50ntFlanks@8196.8213.18|CGTGGACGGTGTGAGGCC</td><td>>hg38_21_+_8433172_8446622_RNA45SN1_45S_50ntFlanks</td><td>8196</td><td>8213</td><td>18</td><td>CGTGGACGGTGTGAGGCC</td><td>1046.3</td><td>-2.7</td><td>0.7</td><td>-3.8</td><td>1.3E-04</td><td>1.7E-04</td><td>MB231/MB468</td><td>MITO</td></tr>
<tr><td>>hg38_21_+_8433172_8446622_RNA45SN1_45S_50ntFlanks@9955.9980.26|GGAAGTCGGAATCCGCTAAGGAGTGT</td><td>>hg38_21_+_8433172_8446622_RNA45SN1_45S_50ntFlanks</td><td>9955</td><td>9980</td><td>26</td><td>GGAAGTCGGAATCCGCTAAGGAGTGT</td><td>92.5</td><td>7.2</td><td>1.8</td><td>4.1</td><td>4.8E-05</td><td>1.7E-04</td><td>BT20/MB468</td><td>MP</td></tr>
<tr><td>>hg38_21_+_8433172_8446622_RNA45SN1_45S_50ntFlanks@12394.12420.27|TAGCGACGTCGCTTTTTGATCCTTCGA</td><td>>hg38_21_+_8433172_8446622_RNA45SN1_45S_50ntFlanks</td><td>12394</td><td>12420</td><td>27</td><td>TAGCGACGTCGCTTTTTGATCCTTCGA</td><td>508.0</td><td>-5.3</td><td>1.4</td><td>-3.8</td><td>1.3E-04</td><td>1.7E-04</td><td>MB231/MB468</td><td>MITO</td></tr>
<tr><td>>hg38_21_+_8433172_8446622_RNA45SN1_45S_50ntFlanks@4005.4025.21|AACCTCGGGCCGATCGCACGC</td><td>>hg38_21_+_8433172_8446622_RNA45SN1_45S_50ntFlanks</td><td>4005</td><td>4025</td><td>21</td><td>AACCTCGGGCCGATCGCACGC</td><td>665.6</td><td>-3.7</td><td>1.0</td><td>-3.8</td><td>1.3E-04</td><td>1.7E-04</td><td>MB231/MB468</td><td>NUC</td></tr>
<tr><td>>hg38_21_+_8433172_8446622_RNA45SN1_45S_50ntFlanks@12991.13010.20|CTCCCTCGCTGCGATCTATT</td><td>>hg38_21_+_8433172_8446622_RNA45SN1_45S_50ntFlanks</td><td>12991</td><td>13010</td><td>20</td><td>CTCCCTCGCTGCGATCTATT</td><td>326.3</td><td>-5.3</td><td>1.4</td><td>-3.9</td><td>9.7E-05</td><td>1.7E-04</td><td>BT20/MB468</td><td>MITO</td></tr>
<tr><td>>trna10_LysCTT_16_+_3241501_3241573@3.35.33,trna13_LysCTT_14_-_58706613_58706685@3.35.33,trna2_LysCTT_15_+_79152904_79152976@3.35.33|CCGGCTAGCTCAGTCGGTAGAGCATGGGACTCT</td><td>>trna10_LysCTT_16_+_3241501_3241573</td><td>3</td><td>35</td><td>33</td><td>CCGGCTAGCTCAGTCGGTAGAGCATGGGACTCT</td><td>411.6</td><td>4.9</td><td>1.3</td><td>3.9</td><td>1.1E-04</td><td>1.7E-04</td><td>MB231/MB468</td><td>MP</td></tr>
<tr><td>>hg38_21_+_8433172_8446622_RNA45SN1_45S_50ntFlanks@12419.12454.36|GATGTCGGCTCTTCCTATCATTGTGAAGCAGAATTC</td><td>>hg38_21_+_8433172_8446622_RNA45SN1_45S_50ntFlanks</td><td>12419</td><td>12454</td><td>36</td><td>GATGTCGGCTCTTCCTATCATTGTGAAGCAGAATTC</td><td>483.0</td><td>3.2</td><td>0.8</td><td>3.8</td><td>1.3E-04</td><td>1.7E-04</td><td>MB231/MB468</td><td>NUC</td></tr>
<tr><td>>hg38_21_+_8433172_8446622_RNA45SN1_45S_50ntFlanks@12416.12440.25|TTCGATGTCGGCTCTTCCTATCATT</td><td>>hg38_21_+_8433172_8446622_RNA45SN1_45S_50ntFlanks</td><td>12416</td><td>12440</td><td>25</td><td>TTCGATGTCGGCTCTTCCTATCATT</td><td>359.7</td><td>3.3</td><td>0.9</td><td>3.8</td><td>1.3E-04</td><td>1.7E-04</td><td>MB231/MB468</td><td>MITO</td></tr>
<tr><td>>hg38_21_+_8433172_8446622_RNA45SN1_45S_50ntFlanks@11208.11228.21|GCGGCGACTCTGGACGCGAGC</td><td>>hg38_21_+_8433172_8446622_RNA45SN1_45S_50ntFlanks</td><td>11208</td><td>11228</td><td>21</td><td>GCGGCGACTCTGGACGCGAGC</td><td>383.8</td><td>-3.6</td><td>0.9</td><td>-3.8</td><td>1.3E-04</td><td>1.7E-04</td><td>MB231/MB468</td><td>MITO</td></tr>
<tr><td>>hg38_21_+_8433172_8446622_RNA45SN1_45S_50ntFlanks@9545.9563.19|GTGGAGGTCCGTAGCGGTC</td><td>>hg38_21_+_8433172_8446622_RNA45SN1_45S_50ntFlanks</td><td>9545</td><td>9563</td><td>19</td><td>GTGGAGGTCCGTAGCGGTC</td><td>4240.2</td><td>-5.4</td><td>1.4</td><td>-3.9</td><td>1.2E-04</td><td>1.7E-04</td><td>BT20/MB468</td><td>CYTO</td></tr>
<tr><td>>hg38_21_+_8433172_8446622_RNA45SN1_45S_50ntFlanks@12927.12962.36|CCTGGCGCTAAACCATTCGTAGACGACCTGCTTCTG</td><td>>hg38_21_+_8433172_8446622_RNA45SN1_45S_50ntFlanks</td><td>12927</td><td>12962</td><td>36</td><td>CCTGGCGCTAAACCATTCGTAGACGACCTGCTTCTG</td><td>42.8</td><td>-9.3</td><td>2.4</td><td>-3.9</td><td>1.1E-04</td><td>1.7E-04</td><td>MB231/MB468</td><td>MP</td></tr>
<tr><td>>Antisense_hg38_MT_-_1621_3279_RNR2_16S_50ntFlanks@893.914.22|AGATATTGGGCTGTTAATTGTC</td><td>>Antisense_hg38_MT_-_1621_3279_RNR2_16S_50ntFlanks</td><td>893</td><td>914</td><td>22</td><td>AGATATTGGGCTGTTAATTGTC</td><td>265.6</td><td>3.9</td><td>1.0</td><td>3.8</td><td>1.3E-04</td><td>1.7E-04</td><td>MB231/MB468</td><td>MITO</td></tr>
<tr><td>>hg38_21_+_8433172_8446622_RNA45SN1_45S_50ntFlanks@10791.10820.30|CCTCTGGCATGTTGGAACAATGTAGGTAAG</td><td>>hg38_21_+_8433172_8446622_RNA45SN1_45S_50ntFlanks</td><td>10791</td><td>10820</td><td>30</td><td>CCTCTGGCATGTTGGAACAATGTAGGTAAG</td><td>70.0</td><td>9.5</td><td>2.2</td><td>4.3</td><td>1.7E-05</td><td>1.7E-04</td><td>BT20/MB231</td><td>MP</td></tr>
<tr><td>>MI0000273|hsa-mir-183&WithFlank&7|-|129774899|129775020@34.56.23&[MIMAT0000261&hsa-miR-183-5p&offsets|+1|+2;m-41&7|-|129774966|129774987&offsets|0|+1]|ATGGCACTGGTAGAATTCACTGT</td><td>>MI0000273|hsa-mir-183&WithFlank&7|-|129774899|129775020</td><td>34</td><td>56</td><td>23</td><td>ATGGCACTGGTAGAATTCACTGT</td><td>2247.3</td><td>3.4</td><td>0.8</td><td>4.3</td><td>1.6E-05</td><td>1.7E-04</td><td>BT20/MB231</td><td>CYTO</td></tr>
<tr><td>>MI0000482|hsa-mir-185&WithFlank&22|+|20033133|20033226@21.42.22&[MIMAT0000455&hsa-miR-185-5p&offsets|0|0;m-145&22|+|20033153|20033174&offsets|0|0]|TGGAGAGAAAGGCAGTTCCTGA</td><td>>MI0000482|hsa-mir-185&WithFlank&22|+|20033133|20033226</td><td>21</td><td>42</td><td>22</td><td>TGGAGAGAAAGGCAGTTCCTGA</td><td>2267.1</td><td>3.8</td><td>0.9</td><td>4.3</td><td>1.6E-05</td><td>1.7E-04</td><td>BT20/MB231</td><td>CYTO</td></tr>
<tr><td>>MI0000542|hsa-mir-320a&WithFlank&8|-|22244960|22245043@48.69.22(+3U)&[MIMAT0000510&hsa-miR-320a-3p&offsets|0|0(+3U);m-60&8|-|22244975|22244996&offsets|0|0(+3U)]|AAAAGCTGGGTTGAGAGGGCGATTT</td><td>>MI0000542|hsa-mir-320a&WithFlank&8|-|22244960|22245043</td><td>48</td><td>69</td><td>22(+3U)</td><td>AAAAGCTGGGTTGAGAGGGCGATTT</td><td>697.6</td><td>3.7</td><td>0.9</td><td>4.3</td><td>1.6E-05</td><td>1.7E-04</td><td>BT20/MB231</td><td>CYTO</td></tr>
<tr><td>>hg38_21_+_8433172_8446622_RNA45SN1_45S_50ntFlanks@3846.3869.24|CTTGGATAACTGTGGTAATTCTAG</td><td>>hg38_21_+_8433172_8446622_RNA45SN1_45S_50ntFlanks</td><td>3846</td><td>3869</td><td>24</td><td>CTTGGATAACTGTGGTAATTCTAG</td><td>766.7</td><td>-2.6</td><td>0.7</td><td>-3.8</td><td>1.2E-04</td><td>1.7E-04</td><td>MB231/MB468</td><td>CYTO</td></tr>
<tr><td>>MI0000456|hsa-mir-140&WithFlank&16|+|69933075|69933186@69.91.23&[MIMAT0004597&hsa-miR-140-3p&offsets|+1|+3;m-45&16|+|69933143|69933164&offsets|0|+1]|ACCACAGGGTAGAACCACGGACA</td><td>>MI0000456|hsa-mir-140&WithFlank&16|+|69933075|69933186</td><td>69</td><td>91</td><td>23</td><td>ACCACAGGGTAGAACCACGGACA</td><td>867.6</td><td>6.1</td><td>1.5</td><td>4.1</td><td>4.8E-05</td><td>1.7E-04</td><td>BT20/MB468</td><td>MP</td></tr>
<tr><td>>MI0000815|hsa-mir-339&WithFlank&7|-|1022927|1023032@56.73.18&[MIMAT0004702&hsa-miR-339-3p&offsets|0|-5;m-235&7|-|1022957|1022977&offsets|0|-3]|TGAGCGCCTCGACGACAG</td><td>>MI0000815|hsa-mir-339&WithFlank&7|-|1022927|1023032</td><td>56</td><td>73</td><td>18</td><td>TGAGCGCCTCGACGACAG</td><td>264.6</td><td>8.0</td><td>2.0</td><td>4.1</td><td>4.8E-05</td><td>1.7E-04</td><td>BT20/MB468</td><td>MP</td></tr>
<tr><td>>MI0000255|hsa-mir-30d&WithFlank&8|-|134804870|134804951@12.32.21(+1A)&[MIMAT0000245&hsa-miR-30d-5p&offsets|0|-1(+1A);m-21&8|-|134804919|134804940&offsets|0|-1(+1A)]|TGTAAACATCCCCGACTGGAAA</td><td>>MI0000255|hsa-mir-30d&WithFlank&8|-|134804870|134804951</td><td>12</td><td>32</td><td>21(+1A)</td><td>TGTAAACATCCCCGACTGGAAA</td><td>149.2</td><td>5.8</td><td>1.5</td><td>3.8</td><td>1.3E-04</td><td>1.7E-04</td><td>MB231/MB468</td><td>MITO</td></tr>
<tr><td>>hg38_21_+_8433172_8446622_RNA45SN1_45S_50ntFlanks@5472.5515.44|ACGGCCCTGGCGGAGCGCTGAGAAGACGGTCGAACTTGACTATC</td><td>>hg38_21_+_8433172_8446622_RNA45SN1_45S_50ntFlanks</td><td>5472</td><td>5515</td><td>44</td><td>ACGGCCCTGGCGGAGCGCTGAGAAGACGGTCGAACTTGACTATC</td><td>2593.0</td><td>-3.6</td><td>0.8</td><td>-4.3</td><td>1.6E-05</td><td>1.7E-04</td><td>BT20/MB231</td><td>CYTO</td></tr>
<tr><td>>hg38_21_+_8433172_8446622_RNA45SN1_45S_50ntFlanks@12916.12940.25|TTCGTGGGGAACCTGGCGCTAAACC</td><td>>hg38_21_+_8433172_8446622_RNA45SN1_45S_50ntFlanks</td><td>12916</td><td>12940</td><td>25</td><td>TTCGTGGGGAACCTGGCGCTAAACC</td><td>174.1</td><td>4.5</td><td>1.2</td><td>3.8</td><td>1.2E-04</td><td>1.7E-04</td><td>MB231/MB468</td><td>CYTO</td></tr>
<tr><td>>hg38_21_+_8433172_8446622_RNA45SN1_45S_50ntFlanks@4591.4616.26|TTTTGTTGGTTTTCGGAACTGAGGCC</td><td>>hg38_21_+_8433172_8446622_RNA45SN1_45S_50ntFlanks</td><td>4591</td><td>4616</td><td>26</td><td>TTTTGTTGGTTTTCGGAACTGAGGCC</td><td>241.6</td><td>-4.2</td><td>1.1</td><td>-3.8</td><td>1.3E-04</td><td>1.7E-04</td><td>MB231/MB468</td><td>NUC</td></tr>
<tr><td>>MI0000102|hsa-mir-100&WithFlank&11|-|122152223|122152314@19.39.21(+1C)&[MIMAT0000098&hsa-miR-100-5p&offsets|0|-1(+1C);m-27&11|-|122152276|122152296&offsets|0|0(+1C)]|AACCCGTAGATCCGAACTTGTC</td><td>>MI0000102|hsa-mir-100&WithFlank&11|-|122152223|122152314</td><td>19</td><td>39</td><td>21(+1C)</td><td>AACCCGTAGATCCGAACTTGTC</td><td>2686.6</td><td>-9.3</td><td>2.2</td><td>-4.3</td><td>1.9E-05</td><td>1.7E-04</td><td>BT20/MB231</td><td>MITO</td></tr>
<tr><td>>hg38_21_+_8433172_8446622_RNA45SN1_45S_50ntFlanks@9569.9596.28|GTGCAAATCGGTCGTCCGACCTGGGTAT</td><td>>hg38_21_+_8433172_8446622_RNA45SN1_45S_50ntFlanks</td><td>9569</td><td>9596</td><td>28</td><td>GTGCAAATCGGTCGTCCGACCTGGGTAT</td><td>219.6</td><td>-4.2</td><td>1.1</td><td>-3.8</td><td>1.3E-04</td><td>1.7E-04</td><td>MB231/MB468</td><td>NUC</td></tr>
<tr><td>>am_trnaMT_MetCAT_MT_+_4402_4469@1.42.42|AGTAAGGTCAGCTAAATAAGCTATCGGGCCCATACCCCGAAA</td><td>>am_trnaMT_MetCAT_MT_+_4402_4469</td><td>1</td><td>42</td><td>42</td><td>AGTAAGGTCAGCTAAATAAGCTATCGGGCCCATACCCCGAAA</td><td>113.7</td><td>10.6</td><td>2.6</td><td>4.1</td><td>4.8E-05</td><td>1.7E-04</td><td>BT20/MB468</td><td>MP</td></tr>
<tr><td>>hg38_21_+_8433172_8446622_RNA45SN1_45S_50ntFlanks@6746.6772.27|CACTTCGAACGCACTTGCGGCCCCGGG</td><td>>hg38_21_+_8433172_8446622_RNA45SN1_45S_50ntFlanks</td><td>6746</td><td>6772</td><td>27</td><td>CACTTCGAACGCACTTGCGGCCCCGGG</td><td>236.8</td><td>-6.2</td><td>1.5</td><td>-4.1</td><td>4.8E-05</td><td>1.7E-04</td><td>BT20/MB468</td><td>MP</td></tr>
<tr><td>>hg38_1_-_228634819_228635039_RNA5S12_5S_50ntFlanks@51.89.39|GTCTACGGCCATACCACCCTGAACGCGCCCGATCTCGTC</td><td>>hg38_1_-_228634819_228635039_RNA5S12_5S_50ntFlanks</td><td>51</td><td>89</td><td>39</td><td>GTCTACGGCCATACCACCCTGAACGCGCCCGATCTCGTC</td><td>4065.2</td><td>4.4</td><td>1.1</td><td>4.1</td><td>4.8E-05</td><td>1.7E-04</td><td>BT20/MB468</td><td>MP</td></tr>
<tr><td>>hg38_21_+_8433172_8446622_RNA45SN1_45S_50ntFlanks@9624.9650.27|AGTAGCTGGTTCCCTCCGAAGTTTCCC</td><td>>hg38_21_+_8433172_8446622_RNA45SN1_45S_50ntFlanks</td><td>9624</td><td>9650</td><td>27</td><td>AGTAGCTGGTTCCCTCCGAAGTTTCCC</td><td>172.9</td><td>-5.2</td><td>1.3</td><td>-3.9</td><td>9.8E-05</td><td>1.7E-04</td><td>BT20/MB468</td><td>MITO</td></tr>
<tr><td>>MI0000749|hsa-mir-30e&WithFlank&1|+|40754349|40754452@65.83.19&[MIMAT0000693&hsa-miR-30e-3p&offsets|0|-3;m-25&1|+|40754413|40754434&offsets|0|-3]|CTTTCAGTCGGATGTTTAC</td><td>>MI0000749|hsa-mir-30e&WithFlank&1|+|40754349|40754452</td><td>65</td><td>83</td><td>19</td><td>CTTTCAGTCGGATGTTTAC</td><td>1343.8</td><td>2.7</td><td>0.7</td><td>3.8</td><td>1.3E-04</td><td>1.7E-04</td><td>MB231/MB468</td><td>NUC</td></tr>
<tr><td>>hg38_21_+_8433172_8446622_RNA45SN1_45S_50ntFlanks@11562.11602.41|CGCCTAGCAGCCGACTTAGAACTGGTGCGGACCAGGGGAAT</td><td>>hg38_21_+_8433172_8446622_RNA45SN1_45S_50ntFlanks</td><td>11562</td><td>11602</td><td>41</td><td>CGCCTAGCAGCCGACTTAGAACTGGTGCGGACCAGGGGAAT</td><td>244.9</td><td>-3.2</td><td>0.8</td><td>-3.8</td><td>1.3E-04</td><td>1.7E-04</td><td>MB231/MB468</td><td>NUC</td></tr>
<tr><td>>hg38_21_+_8433172_8446622_RNA45SN1_45S_50ntFlanks@5470.5499.30|CCACGGCCCTGGCGGAGCGCTGAGAAGACG</td><td>>hg38_21_+_8433172_8446622_RNA45SN1_45S_50ntFlanks</td><td>5470</td><td>5499</td><td>30</td><td>CCACGGCCCTGGCGGAGCGCTGAGAAGACG</td><td>1036.9</td><td>-2.9</td><td>0.8</td><td>-3.8</td><td>1.3E-04</td><td>1.7E-04</td><td>MB231/MB468</td><td>CYTO</td></tr>
<tr><td>>hg38_21_+_8433172_8446622_RNA45SN1_45S_50ntFlanks@7974.8012.39|ACGCGACCTCAGATCAGACGTGGCGACCCGCTGAATTTA</td><td>>hg38_21_+_8433172_8446622_RNA45SN1_45S_50ntFlanks</td><td>7974</td><td>8012</td><td>39</td><td>ACGCGACCTCAGATCAGACGTGGCGACCCGCTGAATTTA</td><td>997.8</td><td>3.2</td><td>0.8</td><td>3.9</td><td>1.2E-04</td><td>1.7E-04</td><td>BT20/MB468</td><td>CYTO</td></tr>
<tr><td>>hg38_21_+_8433172_8446622_RNA45SN1_45S_50ntFlanks@11568.11600.33|GCAGCCGACTTAGAACTGGTGCGGACCAGGGGA</td><td>>hg38_21_+_8433172_8446622_RNA45SN1_45S_50ntFlanks</td><td>11568</td><td>11600</td><td>33</td><td>GCAGCCGACTTAGAACTGGTGCGGACCAGGGGA</td><td>2006.3</td><td>-5.8</td><td>1.5</td><td>-3.8</td><td>1.4E-04</td><td>1.7E-04</td><td>BT20/MB468</td><td>NUC</td></tr>
</table>

| | | | | | | | | | | | | | |
|---|---|---|---|---|---|---|---|---|---|---|---|---|---|
| >hg38_21_+_8433172_8446622_RNA45SN1_45S_50ntFlanks@11587.11612.26\|TGCGGACCAGGGGAATCCGACTGTTT | >hg38_21_+_8433172_8446622_RNA45SN1_45S_50ntFlanks | 11587 | 11612 | 26 | TGCGGACCAGGGGAATCCGACTGTTT | 542.5 | -5.0 | 1.3 | -3.9 | 1.1E-04 | 1.7E-04 | MB231/MB468 | MP |
| >hg38_21_+_8433172_8446622_RNA45SN1_45S_50ntFlanks@5494.5512.19\|AAGACGGTCGAACTTGACT | >hg38_21_+_8433172_8446622_RNA45SN1_45S_50ntFlanks | 5494 | 5512 | 19 | AAGACGGTCGAACTTGACT | 120.6 | -4.4 | 1.1 | -3.9 | 1.1E-04 | 1.8E-04 | MB231/MB468 | MP |
| >hg38_21_+_8433172_8446622_RNA45SN1_45S_50ntFlanks@11641.11669.29\|CGGCGGGTGTTGACGCGATGTGATTTCTG | >hg38_21_+_8433172_8446622_RNA45SN1_45S_50ntFlanks | 11641 | 11669 | 29 | CGGCGGGTGTTGACGCGATGTGATTTCTG | 789.5 | -4.5 | 1.2 | -3.9 | 1.2E-04 | 1.8E-04 | BT20/MB468 | CYTO |
| >hg38_21_+_8433172_8446622_RNA45SN1_45S_50ntFlanks@6679.6714.36\|GTGCGTCGATGAAGAACGCAGCTAGCTGCGAGAATT | >hg38_21_+_8433172_8446622_RNA45SN1_45S_50ntFlanks | 6679 | 6714 | 36 | GTGCGTCGATGAAGAACGCAGCTAGCTGCGAGAATT | 91.7 | 7.1 | 1.8 | 4.1 | 4.9E-05 | 1.8E-04 | BT20/MB468 | MP |
| >trnaMT_ValTAC_MT_+_1602_1670@35.71.37\|ACTTAGGAGATTTCAACTTAACTTGACCGCTCTGACC | >trnaMT_ValTAC_MT_+_1602_1670 | 35 | 71 | 37 | ACTTAGGAGATTTCAACTTAACTTGACCGCTCTGACC | 278.5 | 4.8 | 1.2 | 3.9 | 1.1E-04 | 1.8E-04 | MB231/MB468 | MP |
| >hg38_MT_+_1621_3279_RNR2_16S_50ntFlanks@482.504.23\|AAAGAGGAACAGCTCTTTGGACA | >hg38_MT_+_1621_3279_RNR2_16S_50ntFlanks | 482 | 504 | 23 | AAAGAGGAACAGCTCTTTGGACA | 413.2 | 4.0 | 1.0 | 3.8 | 1.3E-04 | 1.8E-04 | MB231/MB468 | MITO |
| >hg38_1_-_228634819_228635039_RNA5S12_5S_50ntFlanks@130.170.41\|TGGATGGGAGACCGCCTGGGAATACCGGGTGCTGTAGGCTT | >hg38_1_-_228634819_228635039_RNA5S12_5S_50ntFlanks | 130 | 170 | 41 | TGGATGGGAGACCGCCTGGGAATACCGGGTGCTGTAGGCTT | 2967.7 | 3.6 | 0.8 | 4.3 | 2.0E-05 | 1.8E-04 | BT20/MB231 | MITO |
| >MI0000070\|hsa-mir-16-1&WithFlank&13\|-\|50048967\|50049067@20.39.20;MI0000115\|hsa-mir-16-2&WithFlank&3\|+\|160404739\|160404831@16.35.20&[MIMAT0000069&hsa-miR-16-5p&offsets\|0\|-2;m-62&13\|-\|50049027\|50049048&offsets\|0\|-2];[MIMAT0000069_1&hsa-miR-16-5p&offsets\|0\|-2;m-61&3\|+\|160404754\|160404775&offsets\|0\|-2]\|TAGCAGCACGTAAATATTGG | >MI0000070\|hsa-mir-16-1&WithFlank&13\|-\|50048967\|50049067 | 20 | 39 | 20 | TAGCAGCACGTAAATATTGG | 207.4 | 5.9 | 1.5 | 3.8 | 1.3E-04 | 1.8E-04 | MB231/MB468 | MITO |
| >hg38_21_+_8433172_8446622_RNA45SN1_45S_50ntFlanks@9960.9994.35\|TCGGAATCCGCTAAGGAGTGTGTAACAACTCACCT | >hg38_21_+_8433172_8446622_RNA45SN1_45S_50ntFlanks | 9960 | 9994 | 35 | TCGGAATCCGCTAAGGAGTGTGTAACAACTCACCT | 336.4 | 3.5 | 0.8 | 4.3 | 2.0E-05 | 1.8E-04 | BT20/MB231 | MITO |
| >MI0000263\|hsa-mir-7-1&WithFlank&9\|-\|83969742\|83969863@30.54.25&[MIMAT0000252_2&hsa-miR-7-5p&offsets\|0\|+1;m-401&9\|-\|83969811\|83969834&offsets\|0\|+1]\|TGGAAGACTAGTGATTTTGTTGTTT | >MI0000263\|hsa-mir-7-1&WithFlank&9\|-\|83969742\|83969863 | 30 | 54 | 25 | TGGAAGACTAGTGATTTTGTTGTTT | 3498.1 | -4.3 | 1.0 | -4.3 | 1.7E-05 | 1.8E-04 | BT20/MB231 | MP |
| >hg38_21_+_8433172_8446622_RNA45SN1_45S_50ntFlanks@8673.8703.31\|CGGTGCGCCGCGACCGGCTCCGGGACGGCTG | >hg38_21_+_8433172_8446622_RNA45SN1_45S_50ntFlanks | 8673 | 8703 | 31 | CGGTGCGCCGCGACCGGCTCCGGGACGGCTG | 65.2 | 9.4 | 2.2 | 4.3 | 1.7E-05 | 1.8E-04 | BT20/MB231 | MP |
| >MI0000093\|hsa-mir-92a-1&WithFlank&13\|+\|91351308\|91351397@54.74.21(+1C);MI0000094\|hsa-mir-92a-2&WithFlank&X\|-\|134169532\|134169618@54.74.21(+1C)&[MIMAT0000092&hsa-miR-92a-3p&offsets\|0\|-1(+1C);m-9&13\|+\|91351361\|91351382&offsets\|0\|-1(+1C)];[MIMAT0000092_1&hsa-miR-92a-3p&offsets\|0\|-1(+1C);m-11&X\|-\|134169544\|134169565&offsets\|0\|-1(+1C)]\|TATTGCACTTGTCCCGGCCTGC | >MI0000093\|hsa-mir-92a-1&WithFlank&13\|+\|91351308\|91351397 | 54 | 74 | 21(+1C) | TATTGCACTTGTCCCGGCCTGC | 7201.9 | 3.2 | 0.8 | 3.9 | 1.1E-04 | 1.8E-04 | MB231/MB468 | MP |
| >hg38_21_+_8433172_8446622_RNA45SN1_45S_50ntFlanks@9486.9510.25\|GTTAGGACCCGAAAGATGGTGAACT | >hg38_21_+_8433172_8446622_RNA45SN1_45S_50ntFlanks | 9486 | 9510 | 25 | GTTAGGACCCGAAAGATGGTGAACT | 77.0 | 9.6 | 2.2 | 4.3 | 1.7E-05 | 1.8E-04 | BT20/MB231 | MP |
| >hg38_21_+_8433172_8446622_RNA45SN1_45S_50ntFlanks@4722.4752.31\|TCATTAATCAAGAACGAAAGTCGGAGGTTCG | >hg38_21_+_8433172_8446622_RNA45SN1_45S_50ntFlanks | 4722 | 4752 | 31 | TCATTAATCAAGAACGAAAGTCGGAGGTTCG | 623.8 | -6.4 | 1.6 | -4.1 | 4.9E-05 | 1.8E-04 | BT20/MB468 | MP |
| >hg38_21_+_8433172_8446622_RNA45SN1_45S_50ntFlanks@11873.11897.25\|AAGAAGACCCTGTTGAGCTTGACTC | >hg38_21_+_8433172_8446622_RNA45SN1_45S_50ntFlanks | 11873 | 11897 | 25 | AAGAAGACCCTGTTGAGCTTGACTC | 732.5 | -4.8 | 1.3 | -3.8 | 1.3E-04 | 1.8E-04 | MB231/MB468 | NUC |
| >trnaMT_PheGAA_MT_+_577_647@32.57.26\|ACTGAAAATGTTTAGACGGGCTCACA | >trnaMT_PheGAA_MT_+_577_647 | 32 | 57 | 26 | ACTGAAAATGTTTAGACGGGCTCACA | 159.4 | 7.6 | 1.9 | 4.1 | 4.9E-05 | 1.8E-04 | BT20/MB468 | MP |
| >hg38_21_+_8433172_8446622_RNA45SN1_45S_50ntFlanks@4047.4073.27\|ATTCGAACGTCTGCCCTATCAACTTTC | >hg38_21_+_8433172_8446622_RNA45SN1_45S_50ntFlanks | 4047 | 4073 | 27 | ATTCGAACGTCTGCCCTATCAACTTTC | 270.3 | -1.8 | 0.5 | -3.8 | 1.2E-04 | 1.8E-04 | BT20/MB468 | CYTO |
| >hg38_21_+_8433172_8446622_RNA45SN1_45S_50ntFlanks@10806.10825.20\|AACAATGTAGGTAAGGGAAG | >hg38_21_+_8433172_8446622_RNA45SN1_45S_50ntFlanks | 10806 | 10825 | 20 | AACAATGTAGGTAAGGGAAG | 247.3 | -6.1 | 1.5 | -4.1 | 4.9E-05 | 1.8E-04 | BT20/MB468 | MP |
| >hg38_21_+_8433172_8446622_RNA45SN1_45S_50ntFlanks@6673.6700.28\|CGGCTCGTGCGTCGATGAAGAACGCAGC | >hg38_21_+_8433172_8446622_RNA45SN1_45S_50ntFlanks | 6673 | 6700 | 28 | CGGCTCGTGCGTCGATGAAGAACGCAGC | 95.8 | 7.3 | 1.8 | 4.1 | 4.9E-05 | 1.8E-04 | BT20/MB468 | MP |
| >hg38_21_+_8433172_8446622_RNA45SN1_45S_50ntFlanks@6660.6700.41\|GCGGTGGATCACTCGGCTCGTGCGTCGATGAAGAACGCAGC | >hg38_21_+_8433172_8446622_RNA45SN1_45S_50ntFlanks | 6660 | 6700 | 41 | GCGGTGGATCACTCGGCTCGTGCGTCGATGAAGAACGCAGC | 111.1 | 10.5 | 2.6 | 4.1 | 4.9E-05 | 1.8E-04 | BT20/MB468 | MP |
| >hg38_21_+_8433172_8446622_RNA45SN1_45S_50ntFlanks@10106.10124.19\|GCGTGTGGTGTGCGTCGGA | >hg38_21_+_8433172_8446622_RNA45SN1_45S_50ntFlanks | 10106 | 10124 | 19 | GCGTGTGGTGTGCGTCGGA | 263.9 | 11.4 | 2.6 | 4.3 | 1.7E-05 | 1.8E-04 | BT20/MB231 | MP |
| >hg38_21_+_8433172_8446622_RNA45SN1_45S_50ntFlanks@9940.9968.29\|CAGGACGGTGGCCATGGAAGTCGGAATCC | >hg38_21_+_8433172_8446622_RNA45SN1_45S_50ntFlanks | 9940 | 9968 | 29 | CAGGACGGTGGCCATGGAAGTCGGAATCC | 405.6 | 4.5 | 1.0 | 4.3 | 2.0E-05 | 1.8E-04 | BT20/MB231 | MITO |
| >MI0000289\|hsa-mir-181a-1&WithFlank&1\|-\|198859038\|198859159@30.48.19(+1U);MI0000269\|hsa-mir-181a-2&WithFlank&9\|+\|124692436\|124692557@45.63.19(+1U)&[MIMAT0000256&hsa-miR-181a-5p&offsets\|0\|-4(+1U);m-32&1\|-\|198859108\|198859130&offsets\|0\|-4(+1U)];[MIMAT0000256_1&hsa-miR-181a-5p&offsets\|0\|-4(+1U);m-32&9\|+\|124692480\|124692502&offsets\|0\|-4(+1U)]\|AACATTCAACGCTGTCGGTT | >MI0000289\|hsa-mir-181a-1&WithFlank&1\|-\|198859038\|198859159 | 30 | 48 | 19(+1U) | AACATTCAACGCTGTCGGTT | 62.6 | -9.4 | 2.2 | -4.3 | 1.8E-05 | 1.8E-04 | BT20/MB231 | MP |
| >hg38_21_+_8433172_8446622_RNA45SN1_45S_50ntFlanks@12007.12041.35\|GCCGCCGGTGAAATACCACTACTCTGATCGTTTTT | >hg38_21_+_8433172_8446622_RNA45SN1_45S_50ntFlanks | 12007 | 12041 | 35 | GCCGCCGGTGAAATACCACTACTCTGATCGTTTTT | 373.8 | -5.1 | 1.3 | -3.8 | 1.4E-04 | 1.8E-04 | MB231/MB468 | MITO |
| >hg38_21_+_8433172_8446622_RNA45SN1_45S_50ntFlanks@4047.4081.35\|ATTCGAACGTCTGCCCTATCAACTTTCGATGGTAG | >hg38_21_+_8433172_8446622_RNA45SN1_45S_50ntFlanks | 4047 | 4081 | 35 | ATTCGAACGTCTGCCCTATCAACTTTCGATGGTAG | 259.7 | -5.4 | 1.4 | -3.8 | 1.2E-04 | 1.8E-04 | BT20/MB468 | CYTO |
| >hg38_21_+_8433172_8446622_RNA45SN1_45S_50ntFlanks@10394.10429.36\|AGCAGTTGAACATGGGTCAGTCGGTCCTGAGAGATG | >hg38_21_+_8433172_8446622_RNA45SN1_45S_50ntFlanks | 10394 | 10429 | 36 | AGCAGTTGAACATGGGTCAGTCGGTCCTGAGAGATG | 338.2 | -3.8 | 1.0 | -3.9 | 1.1E-04 | 1.8E-04 | MB231/MB468 | MP |
| >am_MI0000786\|hsa-mir-378a&WithFlank&5\|+\|149732819\|149732896@49.69.21&[MIMAT0000732&hsa-miR-378a-3p&offsets\|0\|-1;m-44&5\|+\|149732867\|149732888&offsets\|0\|-1]\|ACTGGACTTGGAGTCAGAAGG | >am_MI0000786\|hsa-mir-378a&WithFlank&5\|+\|149732819\|149732896 | 49 | 69 | 21 | ACTGGACTTGGAGTCAGAAGG | 1304.3 | 3.4 | 0.9 | 3.8 | 1.3E-04 | 1.8E-04 | MB231/MB468 | CYTO |
| >hg38_21_+_8433172_8446622_RNA45SN1_45S_50ntFlanks@11581.11602.22\|AACTGGTGCGGACCAGGGGAAT | >hg38_21_+_8433172_8446622_RNA45SN1_45S_50ntFlanks | 11581 | 11602 | 22 | AACTGGTGCGGACCAGGGGAAT | 1545.7 | -2.8 | 0.7 | -3.8 | 1.2E-04 | 1.8E-04 | BT20/MB468 | CYTO |
| >hg38_21_+_8433172_8446622_RNA45SN1_45S_50ntFlanks@4724.4759.36\|ATTAATCAAGAACGAAAGTCGGAGGTTCGAAGACGA | >hg38_21_+_8433172_8446622_RNA45SN1_45S_50ntFlanks | 4724 | 4759 | 36 | ATTAATCAAGAACGAAAGTCGGAGGTTCGAAGACGA | 2693.2 | -3.3 | 0.9 | -3.8 | 1.3E-04 | 1.8E-04 | MB231/MB468 | CYTO |
| >hg38_21_+_8433172_8446622_RNA45SN1_45S_50ntFlanks@6670.6699.30\|ACTCGGCTCGTGCGTCGATGAAGAACGCAG | >hg38_21_+_8433172_8446622_RNA45SN1_45S_50ntFlanks | 6670 | 6699 | 30 | ACTCGGCTCGTGCGTCGATGAAGAACGCAG | 320.3 | -2.6 | 0.7 | -3.8 | 1.3E-04 | 1.8E-04 | MB231/MB468 | CYTO |
| >hg38_21_+_8433172_8446622_RNA45SN1_45S_50ntFlanks@12971.13003.33\|TTTCGTACGTAGCAGAGCAGCTCCCTCGCTGCG | >hg38_21_+_8433172_8446622_RNA45SN1_45S_50ntFlanks | 12971 | 13003 | 33 | TTTCGTACGTAGCAGAGCAGCTCCCTCGCTGCG | 179.4 | -3.1 | 0.8 | -3.8 | 1.2E-04 | 1.8E-04 | BT20/MB468 | CYTO |
| >hg38_21_+_8433172_8446622_RNA45SN1_45S_50ntFlanks@12418.12456.39\|CGATGTCGGCTCTTCCTATCATTGTGAAGCAGAATTCAC | >hg38_21_+_8433172_8446622_RNA45SN1_45S_50ntFlanks | 12418 | 12456 | 39 | CGATGTCGGCTCTTCCTATCATTGTGAAGCAGAATTCAC | 96.8 | -6.1 | 1.6 | -3.9 | 1.1E-04 | 1.8E-04 | MB231/MB468 | MP |
| >hg38_21_+_8433172_8446622_RNA45SN1_45S_50ntFlanks@12478.12499.22\|ACTAATAGGGAACGTGAGCTGG | >hg38_21_+_8433172_8446622_RNA45SN1_45S_50ntFlanks | 12478 | 12499 | 22 | ACTAATAGGGAACGTGAGCTGG | 858.1 | -6.9 | 1.8 | -3.8 | 1.4E-04 | 1.8E-04 | BT20/MB468 | NUC |
| >trnaMT_LysTTT_MT_+_8295_8364@1.47.47,trnalookalike7_LysTTT_1_+_568844_568913@1.47.47\|CACTGTAAAGCTAACTTAGCATTAACCTTTTAAGTTAAAGATTAAGA | >trnaMT_LysTTT_MT_+_8295_8364 | 1 | 47 | 47 | CACTGTAAAGCTAACTTAGCATTAACCTTTTAAGTTAAAGATTAAGA | 377.6 | 6.3 | 1.6 | 3.8 | 1.4E-04 | 1.8E-04 | MB231/MB468 | MITO |
| >MI0000108\|hsa-mir-103a-2&WithFlank&20\|+\|3917488\|3917577@57.76.20;MI0000109\|hsa-mir-103a-1&WithFlank&5\|-\|168560890\|168560979@57.76.20&[MIMAT0000101&hsa-miR-103a-3p&offsets\|+3\|0;m-12&20\|+\|3917541\|3917563&offsets\|+3\|0];[MIMAT0000101_1&hsa-miR-103a-3p&offsets\|+3\|0;m-13&5\|-\|168560904\|168560926&offsets\|+3\|0;m-44220&5\|-\|168560908\|168560926&offsets\|+3\|+4]\|AGCATTGTACAGGGCTATGA | >MI0000108\|hsa-mir-103a-2&WithFlank&20\|+\|3917488\|3917577 | 57 | 76 | 20 | AGCATTGTACAGGGCTATGA | 27.3 | 8.0 | 2.1 | 3.9 | 1.1E-04 | 1.8E-04 | MB231/MB468 | MP |
| >hg38_21_+_8433172_8446622_RNA45SN1_45S_50ntFlanks@8123.8141.19\|ATGTGGCGTACGGAAGACC | >hg38_21_+_8433172_8446622_RNA45SN1_45S_50ntFlanks | 8123 | 8141 | 19 | ATGTGGCGTACGGAAGACC | 236.4 | 3.8 | 0.9 | 4.1 | 5.0E-05 | 1.8E-04 | BT20/MB468 | MP |
| >hg38_1_-_228634819_228635039_RNA5S12_5S_50ntFlanks@148.171.24\|GGAATACCGGGTGCTGTAGGCTTT | >hg38_1_-_228634819_228635039_RNA5S12_5S_50ntFlanks | 148 | 171 | 24 | GGAATACCGGGTGCTGTAGGCTTT | 109018.9 | 7.7 | 1.8 | 4.3 | 1.8E-05 | 1.8E-04 | BT20/MB231 | MP |
| >am_trna111_HisGTG_1_-_147774845_147774916@-1G.18.19,trna118_HisGTG_1_-_145396881_145396952@-1G.18.19,trna16_HisGTG_1_+_146544773_146544844@-1G.18.19,trna1_HisGTG_15_+_45493349_45493420@-1G.18.19,trna21_HisGTG_1_+_147753471_147753542@-1G.18.19,trna33_HisGTG_6_+_27125906_27125977@-1G.18.19,trna7_HisGTG_9_-_14433938_14434009@-1G.18.19,trna8_HisGTG_15_-_45492611_45492682@-1G.18.19,trna9_HisGTG_15_-_45490804_45490875@-1G.18.19\|GGCCGTGATCGTATAGTGG | >am_trna111_HisGTG_1_-_147774845_147774916 | -1G | 18 | 19 | GGCCGTGATCGTATAGTGG | 57.2 | -9.2 | 2.2 | -4.3 | 1.8E-05 | 1.8E-04 | BT20/MB231 | MP |
| >hg38_MT_+_598_1651_RNR1_12S_50ntFlanks@848.872.25\|GAGTGCTTAGTTGAACAGGGCCCTG | >hg38_MT_+_598_1651_RNR1_12S_50ntFlanks | 848 | 872 | 25 | GAGTGCTTAGTTGAACAGGGCCCTG | 115.0 | 10.6 | 2.6 | 4.1 | 5.0E-05 | 1.8E-04 | BT20/MB468 | MP |
| >hg38_21_+_8433172_8446622_RNA45SN1_45S_50ntFlanks@12415.12455.41\|CTTCGATGTCGGCTCTTCCTATCATTGTGAAGCAGAATTCA | >hg38_21_+_8433172_8446622_RNA45SN1_45S_50ntFlanks | 12415 | 12455 | 41 | CTTCGATGTCGGCTCTTCCTATCATTGTGAAGCAGAATTCA | 1508.0 | -3.3 | 0.8 | -3.9 | 1.0E-04 | 1.8E-04 | BT20/MB468 | MITO |
| >hg38_21_+_8433172_8446622_RNA45SN1_45S_50ntFlanks@11564.11590.27\|CCTAGCAGCCGACTTAGAACTGGTGCG | >hg38_21_+_8433172_8446622_RNA45SN1_45S_50ntFlanks | 11564 | 11590 | 27 | CCTAGCAGCCGACTTAGAACTGGTGCG | 177.7 | -5.2 | 1.4 | -3.8 | 1.2E-04 | 1.8E-04 | BT20/MB468 | CYTO |
| >hg38_21_+_8433172_8446622_RNA45SN1_45S_50ntFlanks@10926.10947.22\|ACGAGGCGCCGCCGCCCCCCCC | >hg38_21_+_8433172_8446622_RNA45SN1_45S_50ntFlanks | 10926 | 10947 | 22 | ACGAGGCGCCGCCGCCCCCCCC | 149.9 | -10.3 | 2.5 | -4.1 | 5.0E-05 | 1.8E-04 | BT20/MB468 | MP |
| >hg38_21_+_8433172_8446622_RNA45SN1_45S_50ntFlanks@8176.8195.20\|CTTCTGATCGAGGCCCAGCC | >hg38_21_+_8433172_8446622_RNA45SN1_45S_50ntFlanks | 8176 | 8195 | 20 | CTTCTGATCGAGGCCCAGCC | 4087.2 | 2.1 | 0.5 | 3.8 | 1.2E-04 | 1.8E-04 | BT20/MB468 | CYTO |
| >trnaMT_GluTTC_MT_-_14674_14742@33.71.39,trnalookalike8_GluTTC_5_-_93905172_93905240@33.71.39\|CATATCATTGGTCGTGGTTGTAGTCCGTGCGAGAATACC | >trnaMT_GluTTC_MT_-_14674_14742 | 33 | 71 | 39 | CATATCATTGGTCGTGGTTGTAGTCCGTGCGAGAATACC | 97.1 | 3.4 | 0.9 | 3.8 | 1.4E-04 | 1.8E-04 | MB231/MB468 | MITO |
| >hg38_21_+_8433172_8446622_RNA45SN1_45S_50ntFlanks@11317.11334.18\|ACCCCACGTCTCGTCGCG | >hg38_21_+_8433172_8446622_RNA45SN1_45S_50ntFlanks | 11317 | 11334 | 18 | ACCCCACGTCTCGTCGCG | 606.2 | -4.8 | 1.3 | -3.8 | 1.2E-04 | 1.8E-04 | BT20/MB468 | CYTO |
| >hg38_21_+_8433172_8446622_RNA45SN1_45S_50ntFlanks@9531.9571.41\|CAGAGGAAACTCTGGTGGAGGTCCGTAGCGGTCCTGACGTG | >hg38_21_+_8433172_8446622_RNA45SN1_45S_50ntFlanks | 9531 | 9571 | 41 | CAGAGGAAACTCTGGTGGAGGTCCGTAGCGGTCCTGACGTG | 461.8 | -3.2 | 0.7 | -4.3 | 1.6E-05 | 1.8E-04 | BT20/MB231 | CYTO |

| | | | | | | | | | | | | | |
|---|---|---|---|---|---|---|---|---|---|---|---|---|---|
| >am_trna10_ValCAC_5_-_180649395_180649467@1.33.33,trna12_ValAAC_5_-_180645270_180645342@1.33.33,trna132_ValAAC_6_-_27721179_27721251@1.33.33,trna136_ValAAC_6_-_27648885_27648957@1.33.33,trna139_ValAAC_6_-_27618707_27618779@1.33.33,trna18_ValCAC_5_-_180529253_180529325@1.33.33,trna2_ValAAC_3_+_169490018_169490090@1.33.33,trna2_ValCAC_5_+_180524070_180524142@1.33.33,trna4_ValAAC_5_+_180591154_180591226@1.33.33,trna5_ValAAC_5_+_180596610_180596682@1.33.33,trna6_ValCAC_5_+_180600650_180600722@1.33.33,trna85_ValCAC_1_-_161369490_161369562@1.33.33,trna90_ValCAC_1_-_149684088_149684161@1.33.33,trna98_ValCAC_1_-_149298555_149298627@1.33.33,trna9_ValCAC_6_+_26538282_26538354@1.33.33\|GTTTCCGTAGTGTAGTGGTTATCACGTTCGCCT | >am_trna10_ValCAC_5_-_180649395_180649467 | 1 | 33 | 33 | GTTTCCGTAGTGTAGTGGTTATCACGTTCGCCT | 131.4 | 2.6 | 0.7 | 3.8 | 1.4E-04 | 1.8E-04 | BT20/MB468 | NUC |
| >am_MI0014222\|hsa-mir-548w&WithFlank&16\|+\|26025231\|26025316@15.36.22&[MIMAT0015060&hsa-miR-548w&offsets\|-1\|-2;m-801&16\|+\|26025246\|26025267&offsets\|-1\|-1]\|CAAAAGTAACTGCGGTTTTTGC | >am_MI0014222\|hsa-mir-548w&WithFlank&16\|+\|26025231\|26025316 | 15 | 36 | 22 | CAAAAGTAACTGCGGTTTTTGC | 30.5 | 8.2 | 2.1 | 3.9 | 1.2E-04 | 1.8E-04 | MB231/MB468 | MP |
| >trna10_AspGTC_12_-_125424193_125424264@39.75.37,trna12_AspGTC_12_-_125411891_125411962@39.75.37,trna144_AspGTC_6_-_27551236_27551307@39.75.37,trna38_AspGTC_17_-_8125556_8125627@39.75.37,trna45_AspGTC_6_+_27447453_27447524@39.75.37,trna48_AspGTC_6_+_27471523_27471594@39.75.37,trna4_AspGTC_12_+_96429799_96429870@39.75.37,trna69_AspGTC_1_-_161440205_161440276@39.75.37,trna72_AspGTC_1_-_161432824_161432895@39.75.37,trna75_AspGTC_1_-_161425414_161425485@39.75.37,trna78_AspGTC_1_-_161418033_161418104@39.75.37,trna81_AspGTC_1_-_161410615_161410686@39.75.37\|GCGGGAGACCGGGGTTCGATTCCCCGACGGGGAGCCA | >trna10_AspGTC_12_-_125424193_125424264 | 39 | 75 | 37 | GCGGGAGACCGGGGTTCGATTCCCCGACGGGGAGCCA | 3911.1 | 4.2 | 1.1 | 3.8 | 1.2E-04 | 1.8E-04 | BT20/MB468 | CYTO |
| >MI0000272\|hsa-mir-182&WithFlank&7\|-\|129770377\|129770498@29.51.23(+1C)&[MIMAT0000259&hsa-miR-182-5p&offsets\|0\|-1(+1C);m-22&7\|-\|129770449\|129770470&offsets\|0\|+1(+1C)]\|TTTGGCAATGGTAGAACTCACACC | >MI0000272\|hsa-mir-182&WithFlank&7\|-\|129770377\|129770498 | 29 | 51 | 23(+1C) | TTTGGCAATGGTAGAACTCACACC | 664.8 | 2.9 | 0.7 | 3.8 | 1.3E-04 | 1.8E-04 | MB231/MB468 | CYTO |
| >hg38_21_+_8433172_8446622_RNA45SN1_45S_50ntFlanks@6655.6688.34\|TCTTAGCGGTGGATCACTCGGCTCGTGCGTCGAT | >hg38_21_+_8433172_8446622_RNA45SN1_45S_50ntFlanks | 6655 | 6688 | 34 | TCTTAGCGGTGGATCACTCGGCTCGTGCGTCGAT | 442.0 | -2.9 | 0.7 | -3.8 | 1.4E-04 | 1.8E-04 | MB231/MB468 | MITO |
| >hg38_21_+_8433172_8446622_RNA45SN1_45S_50ntFlanks@11581.11606.26\|AACTGGTGCGGACCAGGGGAATCCGA | >hg38_21_+_8433172_8446622_RNA45SN1_45S_50ntFlanks | 11581 | 11606 | 26 | AACTGGTGCGGACCAGGGGAATCCGA | 158.8 | -4.0 | 1.0 | -3.8 | 1.4E-04 | 1.8E-04 | MB231/MB468 | NUC |
| >trna10_AspGTC_12_-_125424193_125424264@31.75.45,trna12_AspGTC_12_-_125411891_125411962@31.75.45,trna38_AspGTC_17_-_8125556_8125627@31.75.45,trna45_AspGTC_6_+_27447453_27447524@31.75.45,trna48_AspGTC_6_+_27471523_27471594@31.75.45,trna4_AspGTC_12_+_96429799_96429870@31.75.45,trna69_AspGTC_1_-_161440205_161440276@31.75.45,trna72_AspGTC_1_-_161432824_161432895@31.75.45,trna75_AspGTC_1_-_161425414_161425485@31.75.45,trna78_AspGTC_1_-_161418033_161418104@31.75.45,trna81_AspGTC_1_-_161410615_161410686@31.75.45\|CCTGTCACGCGGGAGACCGGGGTTCGATTCCCCGACGGGGAGCCA | >trna10_AspGTC_12_-_125424193_125424264 | 31 | 75 | 45 | CCTGTCACGCGGGAGACCGGGGTTCGATTCCCCGACGGGGAGCCA | 444.8 | -3.8 | 1.0 | -3.8 | 1.3E-04 | 1.8E-04 | MB231/MB468 | CYTO |
| >hg38_21_+_8433172_8446622_RNA45SN1_45S_50ntFlanks@6780.6802.23\|CGGGGCTACGCCTGTCTGAGCGT | >hg38_21_+_8433172_8446622_RNA45SN1_45S_50ntFlanks | 6780 | 6802 | 23 | CGGGGCTACGCCTGTCTGAGCGT | 726.4 | 4.2 | 1.0 | 4.1 | 5.0E-05 | 1.8E-04 | BT20/MB468 | MP |
| >hg38_21_+_8433172_8446622_RNA45SN1_45S_50ntFlanks@9740.9765.26\|AACGATCTCAACCTATTCTCAAACTT | >hg38_21_+_8433172_8446622_RNA45SN1_45S_50ntFlanks | 9740 | 9765 | 26 | AACGATCTCAACCTATTCTCAAACTT | 655.4 | -5.1 | 1.3 | -3.8 | 1.4E-04 | 1.8E-04 | BT20/MB468 | NUC |
| >hg38_21_+_8433172_8446622_RNA45SN1_45S_50ntFlanks@4727.4761.35\|AATCAAGAACGAAAGTCGGAGGTTCGAAGACGATC | >hg38_21_+_8433172_8446622_RNA45SN1_45S_50ntFlanks | 4727 | 4761 | 35 | AATCAAGAACGAAAGTCGGAGGTTCGAAGACGATC | 283.3 | 2.6 | 0.7 | 3.8 | 1.4E-04 | 1.8E-04 | MB231/MB468 | NUC |
| >hg38_21_+_8433172_8446622_RNA45SN1_45S_50ntFlanks@9554.9582.29\|CGTAGCGGTCCTGACGTGCAAATCGGTCG | >hg38_21_+_8433172_8446622_RNA45SN1_45S_50ntFlanks | 9554 | 9582 | 29 | CGTAGCGGTCCTGACGTGCAAATCGGTCG | 590.3 | -2.8 | 0.7 | -3.8 | 1.3E-04 | 1.8E-04 | MB231/MB468 | CYTO |
| >hg38_1_-_228634819_228635039_RNA5S12_5S_50ntFlanks@135.170.36\|GGGAGACCGCCTGGGAATACCGGGTGCTGTAGGCTT | >hg38_1_-_228634819_228635039_RNA5S12_5S_50ntFlanks | 135 | 170 | 36 | GGGAGACCGCCTGGGAATACCGGGTGCTGTAGGCTT | 25956.1 | 3.6 | 0.9 | 4.1 | 5.0E-05 | 1.8E-04 | BT20/MB468 | MP |
| >hg38_MT_+_1621_3279_RNR2_16S_50ntFlanks@505.530.26\|CTAGGAAAAAACCTTGTAGAGAGAGT | >hg38_MT_+_1621_3279_RNR2_16S_50ntFlanks | 505 | 530 | 26 | CTAGGAAAAAACCTTGTAGAGAGAGT | 179.9 | 4.1 | 1.0 | 4.1 | 5.0E-05 | 1.8E-04 | BT20/MB468 | MP |
| >MI0000301\|hsa-mir-224&WithFlank&X\|-\|151958572\|151958664@14.34.21(+1C)&[MIMAT0000281&hsa-miR-224-5p&offsets\|+1\|-3(+1C);m-238&X\|-\|151958630\|151958651&offsets\|0\|-1(+1C)]\|CAAGTCACTAGTGGTTCCGTTC | >MI0000301\|hsa-mir-224&WithFlank&X\|-\|151958572\|151958664 | 14 | 34 | 21(+1C) | CAAGTCACTAGTGGTTCCGTTC | 1398.8 | 3.6 | 0.8 | 4.3 | 1.7E-05 | 1.8E-04 | BT20/MB231 | CYTO |
| >hg38_21_+_8433172_8446622_RNA45SN1_45S_50ntFlanks@12413.12437.25\|TCCTTCGATGTCGGCTCTTCCTATC | >hg38_21_+_8433172_8446622_RNA45SN1_45S_50ntFlanks | 12413 | 12437 | 25 | TCCTTCGATGTCGGCTCTTCCTATC | 657.1 | -5.4 | 1.4 | -3.8 | 1.4E-04 | 1.8E-04 | BT20/MB468 | NUC |
| >am_MI0039740\|hsa-mir-12136&WithFlank&1\|-\|632609\|632691@7.32.26&[MIMAT0049032&hsa-miR-12136&offsets\|0\|+8]\|GAAAAAGTCATGGAGGCCATGGGGTT | >am_MI0039740\|hsa-mir-12136&WithFlank&1\|-\|632609\|632691 | 7 | 32 | 26 | GAAAAAGTCATGGAGGCCATGGGGTT | 1039.6 | 2.8 | 0.7 | 4.3 | 1.9E-05 | 1.8E-04 | BT20/MB231 | NUC |
| >trnaMT_SerTGA_MT_-_7446_7514@1.26.26,trnalookalike6_SerTGA_1_-_567997_568065@1.26.26\|GAAAAAGTCATGGAGGCCATGGGGTT | >trnaMT_SerTGA_MT_-_7446_7514 | 1 | 26 | 26 | GAAAAAGTCATGGAGGCCATGGGGTT | 1039.6 | 2.8 | 0.7 | 4.3 | 1.9E-05 | 1.8E-04 | BT20/MB231 | NUC |
| >trna152_ValCAC_6_-_27248049_27248121@1.34.34\|GCTTCTGTAGTGTAGTGGTTATCACGTTCGCCTC | >trna152_ValCAC_6_-_27248049_27248121 | 1 | 34 | 34 | GCTTCTGTAGTGTAGTGGTTATCACGTTCGCCTC | 3413.8 | -5.0 | 1.2 | -4.3 | 1.9E-05 | 1.8E-04 | BT20/MB231 | NUC |
| >hg38_21_+_8433172_8446622_RNA45SN1_45S_50ntFlanks@12420.12463.44\|ATGTCGGCTCTTCCTATCATTGTGAAGCAGAATTCACCAAGCGT | >hg38_21_+_8433172_8446622_RNA45SN1_45S_50ntFlanks | 12420 | 12463 | 44 | ATGTCGGCTCTTCCTATCATTGTGAAGCAGAATTCACCAAGCGT | 336.8 | -4.8 | 1.1 | -4.3 | 1.7E-05 | 1.8E-04 | BT20/MB231 | CYTO |
| >hg38_MT_+_1621_3279_RNR2_16S_50ntFlanks@1099.1117.19\|GAGAAGACCCTATGGAGCT | >hg38_MT_+_1621_3279_RNR2_16S_50ntFlanks | 1099 | 1117 | 19 | GAGAAGACCCTATGGAGCT | 182.2 | 3.7 | 1.0 | 3.9 | 1.2E-04 | 1.8E-04 | MB231/MB468 | MP |
| >hg38_21_+_8433172_8446622_RNA45SN1_45S_50ntFlanks@6648.6694.47\|GTACGACTCTTAGCGGTGGATCACTCGGCTCGTGCGTCGATGAAGAA | >hg38_21_+_8433172_8446622_RNA45SN1_45S_50ntFlanks | 6648 | 6694 | 47 | GTACGACTCTTAGCGGTGGATCACTCGGCTCGTGCGTCGATGAAGAA | 486.8 | 3.7 | 1.0 | 3.9 | 1.0E-04 | 1.8E-04 | BT20/MB468 | MITO |
| >MI0000061\|hsa-let-7a-2&WithFlank&11\|-\|122146516\|122146599@11.30.20(+1C);MI0000062\|hsa-let-7a-3&WithFlank&22\|+\|46112743\|46112828@10.29.20(+1C);MI0000060\|hsa-let-7a-1&WithFlank&9\|+\|94175951\|94176042@12.31.20(+1C)&[MIMAT0000062&hsa-let-7a-5p&offsets\|0\|-2(+1C);m-6&11\|-\|122146568\|122146589&offsets\|0\|-2(+1C)];[MIMAT0000062_1&hsa-let-7a-5p&offsets\|0\|-2(+1C);m-5&22\|+\|46112752\|46112773&offsets\|0\|-2(+1C)];[MIMAT0000062_2&hsa-let-7a-5p&offsets\|0\|-2(+1C);m-4&9\|+\|94175962\|94175983&offsets\|0\|-2(+1C)]\|TGAGGTAGTAGGTTGTATAGC | >MI0000061\|hsa-let-7a-2&WithFlank&11\|-\|122146516\|122146599 | 11 | 30 | 20(+1C) | TGAGGTAGTAGGTTGTATAGC | 818.0 | 5.0 | 1.3 | 3.9 | 1.2E-04 | 1.8E-04 | MB231/MB468 | MP |
| >trna10_LysCTT_16_+_3241501_3241573@1.28.28,trna119_LysCTT_1_-_145395522_145395594@1.28.28,trna11_LysCTT_5_-_180648979_180649051@1.28.28,trna13_LysCTT_14_-_58706613_58706685@1.28.28,trna13_LysCTT_6_+_26556774_26556846@1.28.28,trna2_LysCTT_15_+_79152904_79152976@1.28.28,trna32_LysCTT_16_-_3207406_3207478@1.28.28,trna7_LysCTT_16_+_3225692_3225764@1.28.28,trna9_LysCTT_5_+_180634755_180634827@1.28.28\|GCCCGGCTAGCTCAGTCGGTAGAGCATG | >trna10_LysCTT_16_+_3241501_3241573 | 1 | 28 | 28 | GCCCGGCTAGCTCAGTCGGTAGAGCATG | 6050.1 | -3.5 | 0.9 | -3.8 | 1.3E-04 | 1.8E-04 | MB231/MB468 | CYTO |
| >hg38_21_+_8433172_8446622_RNA45SN1_45S_50ntFlanks@8178.8208.31\|TCTGATCGAGGCCCAGCCCGTGGACGGTGTG | >hg38_21_+_8433172_8446622_RNA45SN1_45S_50ntFlanks | 8178 | 8208 | 31 | TCTGATCGAGGCCCAGCCCGTGGACGGTGTG | 209.1 | 11.0 | 2.6 | 4.3 | 1.8E-05 | 1.8E-04 | BT20/MB231 | MP |
| >hg38_21_+_8433172_8446622_RNA45SN1_45S_50ntFlanks@8463.8497.35\|CCGTGTCGGCGGCCCGGCGGATCTTTCCCGCCCCC | >hg38_21_+_8433172_8446622_RNA45SN1_45S_50ntFlanks | 8463 | 8497 | 35 | CCGTGTCGGCGGCCCGGCGGATCTTTCCCGCCCCC | 453.9 | -3.6 | 0.9 | -3.9 | 1.0E-04 | 1.8E-04 | BT20/MB468 | MITO |
| >hg38_21_+_8433172_8446622_RNA45SN1_45S_50ntFlanks@11769.11803.35\|TAATTAGTGACGCGCATGAATGGATGAACGAGATT | >hg38_21_+_8433172_8446622_RNA45SN1_45S_50ntFlanks | 11769 | 11803 | 35 | TAATTAGTGACGCGCATGAATGGATGAACGAGATT | 834.6 | -2.9 | 0.8 | -3.8 | 1.3E-04 | 1.8E-04 | MB231/MB468 | CYTO |
| >hg38_MT_+_598_1651_RNR1_12S_50ntFlanks@428.469.42\|AAAGTGGCTTTAACATATCTGAACACACAATAGCTAAGACCC | >hg38_MT_+_598_1651_RNR1_12S_50ntFlanks | 428 | 469 | 42 | AAAGTGGCTTTAACATATCTGAACACACAATAGCTAAGACCC | 710.6 | 7.4 | 1.8 | 4.1 | 5.1E-05 | 1.8E-04 | BT20/MB468 | MP |
| >hg38_21_+_8433172_8446622_RNA45SN1_45S_50ntFlanks@6645.6676.32\|CTCGTACGACTCTTAGCGGTGGATCACTCGGC | >hg38_21_+_8433172_8446622_RNA45SN1_45S_50ntFlanks | 6645 | 6676 | 32 | CTCGTACGACTCTTAGCGGTGGATCACTCGGC | 576.5 | 3.9 | 1.0 | 3.8 | 1.4E-04 | 1.8E-04 | MB231/MB468 | NUC |
| >hg38_21_+_8433172_8446622_RNA45SN1_45S_50ntFlanks@4103.4127.25\|CCACGGGTGACGGGGAATCAGGGTT | >hg38_21_+_8433172_8446622_RNA45SN1_45S_50ntFlanks | 4103 | 4127 | 25 | CCACGGGTGACGGGGAATCAGGGTT | 326.5 | -4.7 | 1.2 | -3.9 | 1.2E-04 | 1.8E-04 | MB231/MB468 | MP |
| >hg38_21_+_8433172_8446622_RNA45SN1_45S_50ntFlanks@10397.10424.28\|AGTTGAACATGGGTCAGTCGGTCCTGAG | >hg38_21_+_8433172_8446622_RNA45SN1_45S_50ntFlanks | 10397 | 10424 | 28 | AGTTGAACATGGGTCAGTCGGTCCTGAG | 1358.4 | 8.2 | 2.0 | 4.1 | 5.1E-05 | 1.8E-04 | BT20/MB468 | MP |
| >hg38_21_+_8433172_8446622_RNA45SN1_45S_50ntFlanks@11561.11601.41\|GCGCCTAGCAGCCGACTTAGAACTGGTGCGGACCAGGGGAA | >hg38_21_+_8433172_8446622_RNA45SN1_45S_50ntFlanks | 11561 | 11601 | 41 | GCGCCTAGCAGCCGACTTAGAACTGGTGCGGACCAGGGGAA | 780.7 | -4.7 | 1.2 | -3.8 | 1.3E-04 | 1.8E-04 | MB231/MB468 | CYTO |
| >hg38_21_+_8433172_8446622_RNA45SN1_45S_50ntFlanks@10878.10902.25\|TCGGTCGGGCTGGGGCGCGAAGCGG | >hg38_21_+_8433172_8446622_RNA45SN1_45S_50ntFlanks | 10878 | 10902 | 25 | TCGGTCGGGCTGGGGCGCGAAGCGG | 401.6 | -4.0 | 1.0 | -3.8 | 1.2E-04 | 1.8E-04 | BT20/MB468 | CYTO |
| >hg38_21_+_8433172_8446622_RNA45SN1_45S_50ntFlanks@11729.11750.22\|TAACTATGACTCTCTTAAGGTA | >hg38_21_+_8433172_8446622_RNA45SN1_45S_50ntFlanks | 11729 | 11750 | 22 | TAACTATGACTCTCTTAAGGTA | 281.7 | -2.4 | 0.6 | -3.9 | 1.0E-04 | 1.8E-04 | BT20/MB468 | MITO |

<table>
<tr><th></th><th></th><th></th><th></th><th></th><th></th><th></th><th></th><th></th><th></th><th></th><th></th><th></th><th></th></tr>
<tr><td>>hg38_21_+_8433172_8446622_RNA45SN1_45S_50ntFlanks@8174.8202.29|TCCTTCTGATCGAGGCCCAGCCCGTGGAC</td><td>>hg38_21_+_8433172_8446622_RNA45SN1_45S_50ntFlanks</td><td>8174</td><td>8202</td><td>29</td><td>TCCTTCTGATCGAGGCCCAGCCCGTGGAC</td><td>165.2</td><td>3.2</td><td>0.8</td><td>3.8</td><td>1.4E-04</td><td>1.8E-04</td><td>MB231/MB468</td><td>NUC</td></tr>
<tr><td>>hg38_21_+_8433172_8446622_RNA45SN1_45S_50ntFlanks@3711.3737.27|GTTGATCCTGCCAGTAGCATATGCTTG</td><td>>hg38_21_+_8433172_8446622_RNA45SN1_45S_50ntFlanks</td><td>3711</td><td>3737</td><td>27</td><td>GTTGATCCTGCCAGTAGCATATGCTTG</td><td>1209.3</td><td>-6.1</td><td>1.6</td><td>-3.8</td><td>1.4E-04</td><td>1.8E-04</td><td>MB231/MB468</td><td>MITO</td></tr>
<tr><td>>hg38_21_+_8433172_8446622_RNA45SN1_45S_50ntFlanks@8176.8193.18|CTTCTGATCGAGGCCCAG</td><td>>hg38_21_+_8433172_8446622_RNA45SN1_45S_50ntFlanks</td><td>8176</td><td>8193</td><td>18</td><td>CTTCTGATCGAGGCCCAG</td><td>359.5</td><td>-2.5</td><td>0.6</td><td>-3.8</td><td>1.2E-04</td><td>1.8E-04</td><td>BT20/MB468</td><td>CYTO</td></tr>
<tr><td>>MI0000816|hsa-mir-335&WithFlank&7|+|130496105|130496210@22.44.23&[MIMAT0000765&hsa-miR-335-5p&offsets|0|0;m-249&7|+|130496126|130496147&offsets|0|+1]|TCAAGAGCAATAACGAAAAATGT</td><td>>MI0000816|hsa-mir-335&WithFlank&7|+|130496105|130496210</td><td>22</td><td>44</td><td>23</td><td>TCAAGAGCAATAACGAAAAATGT</td><td>290.8</td><td>3.2</td><td>0.8</td><td>3.9</td><td>1.2E-04</td><td>1.8E-04</td><td>MB231/MB468</td><td>MP</td></tr>
<tr><td>>MI0000063|hsa-let-7b&WithFlank&22|+|46113680|46113774@12.33.22&[MIMAT0000063&hsa-let-7b-5p&offsets|0|0;m-14&22|+|46113691|46113712&offsets|0|0]|TGAGGTAGTAGGTTGTGTGGTT</td><td>>MI0000063|hsa-let-7b&WithFlank&22|+|46113680|46113774</td><td>12</td><td>33</td><td>22</td><td>TGAGGTAGTAGGTTGTGTGGTT</td><td>23008.8</td><td>3.0</td><td>0.8</td><td>3.9</td><td>1.2E-04</td><td>1.8E-04</td><td>MB231/MB468</td><td>MP</td></tr>
<tr><td>>hg38_21_+_8433172_8446622_RNA45SN1_45S_50ntFlanks@10393.10433.41|CAGCAGTTGAACATGGGTCAGTCGGTCCTGAGAGATGGGCG</td><td>>hg38_21_+_8433172_8446622_RNA45SN1_45S_50ntFlanks</td><td>10393</td><td>10433</td><td>41</td><td>CAGCAGTTGAACATGGGTCAGTCGGTCCTGAGAGATGGGCG</td><td>357.0</td><td>-3.0</td><td>0.8</td><td>-3.8</td><td>1.3E-04</td><td>1.8E-04</td><td>MB231/MB468</td><td>CYTO</td></tr>
<tr><td>>hg38_21_+_8433172_8446622_RNA45SN1_45S_50ntFlanks@12476.12499.24|CCACTAATAGGGAACGTGAGCTGG</td><td>>hg38_21_+_8433172_8446622_RNA45SN1_45S_50ntFlanks</td><td>12476</td><td>12499</td><td>24</td><td>CCACTAATAGGGAACGTGAGCTGG</td><td>357.6</td><td>-5.4</td><td>1.4</td><td>-3.8</td><td>1.2E-04</td><td>1.8E-04</td><td>BT20/MB468</td><td>CYTO</td></tr>
<tr><td>>hg38_21_+_8433172_8446622_RNA45SN1_45S_50ntFlanks@11569.11590.22|CAGCCGACTTAGAACTGGTGCG</td><td>>hg38_21_+_8433172_8446622_RNA45SN1_45S_50ntFlanks</td><td>11569</td><td>11590</td><td>22</td><td>CAGCCGACTTAGAACTGGTGCG</td><td>801.8</td><td>-5.9</td><td>1.4</td><td>-4.0</td><td>5.1E-05</td><td>1.8E-04</td><td>BT20/MB468</td><td>MP</td></tr>
<tr><td>>hg38_21_+_8433172_8446622_RNA45SN1_45S_50ntFlanks@12480.12498.19|TAATAGGGAACGTGAGCTG</td><td>>hg38_21_+_8433172_8446622_RNA45SN1_45S_50ntFlanks</td><td>12480</td><td>12498</td><td>19</td><td>TAATAGGGAACGTGAGCTG</td><td>439.4</td><td>-6.5</td><td>1.7</td><td>-3.8</td><td>1.4E-04</td><td>1.8E-04</td><td>BT20/MB468</td><td>NUC</td></tr>
<tr><td>>hg38_21_+_8433172_8446622_RNA45SN1_45S_50ntFlanks@12407.12426.20|TTTTGATCCTTCGATGTCGG</td><td>>hg38_21_+_8433172_8446622_RNA45SN1_45S_50ntFlanks</td><td>12407</td><td>12426</td><td>20</td><td>TTTTGATCCTTCGATGTCGG</td><td>173.2</td><td>-5.3</td><td>1.4</td><td>-3.8</td><td>1.4E-04</td><td>1.8E-04</td><td>MB231/MB468</td><td>NUC</td></tr>
<tr><td>>hg38_21_+_8433172_8446622_RNA45SN1_45S_50ntFlanks@8384.8406.23|AGGGCGTGAAACCGTTAAGAGGT</td><td>>hg38_21_+_8433172_8446622_RNA45SN1_45S_50ntFlanks</td><td>8384</td><td>8406</td><td>23</td><td>AGGGCGTGAAACCGTTAAGAGGT</td><td>318.5</td><td>2.9</td><td>0.8</td><td>3.8</td><td>1.3E-04</td><td>1.8E-04</td><td>MB231/MB468</td><td>CYTO</td></tr>
<tr><td>>hg38_21_+_8433172_8446622_RNA45SN1_45S_50ntFlanks@9604.9623.20|AAAGACTAATCGAACCATCT</td><td>>hg38_21_+_8433172_8446622_RNA45SN1_45S_50ntFlanks</td><td>9604</td><td>9623</td><td>20</td><td>AAAGACTAATCGAACCATCT</td><td>385.0</td><td>-5.0</td><td>1.3</td><td>-3.8</td><td>1.2E-04</td><td>1.8E-04</td><td>MB231/MB468</td><td>MP</td></tr>
<tr><td>>hg38_21_+_8433172_8446622_RNA45SN1_45S_50ntFlanks@9705.9722.18|CCGGTAAAGCGAATGATT</td><td>>hg38_21_+_8433172_8446622_RNA45SN1_45S_50ntFlanks</td><td>9705</td><td>9722</td><td>18</td><td>CCGGTAAAGCGAATGATT</td><td>117.6</td><td>-5.7</td><td>1.5</td><td>-3.8</td><td>1.2E-04</td><td>1.8E-04</td><td>MB231/MB468</td><td>MP</td></tr>
<tr><td>>hg38_21_+_8433172_8446622_RNA45SN1_45S_50ntFlanks@10736.10755.20|GTAAATCTCGCGCCGGGCCG</td><td>>hg38_21_+_8433172_8446622_RNA45SN1_45S_50ntFlanks</td><td>10736</td><td>10755</td><td>20</td><td>GTAAATCTCGCGCCGGGCCG</td><td>168.8</td><td>-3.8</td><td>1.0</td><td>-3.8</td><td>1.3E-04</td><td>1.8E-04</td><td>MB231/MB468</td><td>CYTO</td></tr>
<tr><td>>am_MI0003776|hsa-mir-320b-1&WithFlank&1|+|116671740|116671823@48.65.18;MI0003839|hsa-mir-320b-2&WithFlank&1|-|224257034|224257116@47.64.18;MI0008190|hsa-mir-320d-1&WithFlank&13|-|40727810|40727893@48.65.18;MI0003778|hsa-mir-320c-1&WithFlank&18|+|21683512|21683595@48.65.18;MI0008191|hsa-mir-320c-2&WithFlank&18|+|24321669|24321752@48.65.18;MI0000542|hsa-mir-320a&WithFlank&8|-|22244960|22245043@48.65.18;MI0008192|hsa-mir-320d-2&WithFlank&X|-|140926154|140926237@48.65.18&[MIMAT0005792&hsa-miR-320b&offsets|0|-4;m-183&1|+|116671787|116671806&offsets|0|-2];[MIMAT0005792_1&hsa-miR-320b&offsets|0|-4;m-183&1|-|224257051|224257070&offsets|0|-2];[MIMAT0006764&hsa-miR-320d&offsets|0|-1;m-246&13|-|40727829|40727846&offsets|0|0];[MIMAT0005793&hsa-miR-320c&offsets|0|-2;m-222&18|+|21683559|21683578&offsets|0|-2];[MIMAT0005793_1&hsa-miR-320c&offsets|0|-2;m-219&18|+|24321716|24321735&offsets|0|-2];[MIMAT0000510&hsa-miR-320a-3p&offsets|0|-4;m-60&8|-|22244975|22244996&offsets|0|-4];[MIMAT0006764_1&hsa-miR-320d&offsets|0|-1;m-253&X|-|140926173|140926190&offsets|0|0]|AAAAGCTGGGTTGAGAGG</td><td>>am_MI0003776|hsa-mir-320b-1&WithFlank&1|+|116671740|116671823</td><td>48</td><td>65</td><td>18</td><td>AAAAGCTGGGTTGAGAGG</td><td>995.1</td><td>3.8</td><td>0.9</td><td>4.3</td><td>1.7E-05</td><td>1.8E-04</td><td>BT20/MB231</td><td>CYTO</td></tr>
<tr><td>>hg38_21_+_8433172_8446622_RNA45SN1_45S_50ntFlanks@11668.11709.42|TGCCCAGTGCTCTGAATGTCAAAGTGAAGAAATTCAATGAAG</td><td>>hg38_21_+_8433172_8446622_RNA45SN1_45S_50ntFlanks</td><td>11668</td><td>11709</td><td>42</td><td>TGCCCAGTGCTCTGAATGTCAAAGTGAAGAAATTCAATGAAG</td><td>958.6</td><td>-4.2</td><td>1.0</td><td>-4.3</td><td>1.7E-05</td><td>1.8E-04</td><td>BT20/MB231</td><td>CYTO</td></tr>
<tr><td>>hg38_21_+_8433172_8446622_RNA45SN1_45S_50ntFlanks@4297.4316.20|CATTGGAGGGCAAGTCTGGT</td><td>>hg38_21_+_8433172_8446622_RNA45SN1_45S_50ntFlanks</td><td>4297</td><td>4316</td><td>20</td><td>CATTGGAGGGCAAGTCTGGT</td><td>2720.6</td><td>3.9</td><td>1.0</td><td>4.0</td><td>5.2E-05</td><td>1.8E-04</td><td>BT20/MB468</td><td>MP</td></tr>
<tr><td>>MI0000750|hsa-mir-26a-2&WithFlank&12|-|57824603|57824698@23.41.19;MI0000083|hsa-mir-26a-1&WithFlank&3|+|37969398|37969486@19.37.19&[MIMAT0000082&hsa-miR-26a-5p&offsets|+3|0;m-33&12|-|57824658|57824679&offsets|+3|0];[MIMAT0000082_1&hsa-miR-26a-5p&offsets|+3|0;m-34&3|+|37969413|37969434&offsets|+3|0]|AAGTAATCCAGGATAGGCT</td><td>>MI0000750|hsa-mir-26a-2&WithFlank&12|-|57824603|57824698</td><td>23</td><td>41</td><td>19</td><td>AAGTAATCCAGGATAGGCT</td><td>970.3</td><td>13.8</td><td>3.6</td><td>3.8</td><td>1.4E-04</td><td>1.8E-04</td><td>MB231/MB468</td><td>NUC</td></tr>
<tr><td>>trnaMT_ValTAC_MT_+_1602_1670@50.72.23|ACTTAACTTGACCGCTCTGACCA</td><td>>trnaMT_ValTAC_MT_+_1602_1670</td><td>50</td><td>72</td><td>23</td><td>ACTTAACTTGACCGCTCTGACCA</td><td>5040.0</td><td>-3.4</td><td>0.9</td><td>-3.8</td><td>1.5E-04</td><td>1.9E-04</td><td>BT20/MB468</td><td>NUC</td></tr>
<tr><td>>hg38_21_+_8433172_8446622_RNA45SN1_45S_50ntFlanks@10409.10431.23|GTCAGTCGGTCCTGAGAGATGGG</td><td>>hg38_21_+_8433172_8446622_RNA45SN1_45S_50ntFlanks</td><td>10409</td><td>10431</td><td>23</td><td>GTCAGTCGGTCCTGAGAGATGGG</td><td>354.0</td><td>-3.6</td><td>0.9</td><td>-3.8</td><td>1.3E-04</td><td>1.9E-04</td><td>MB231/MB468</td><td>CYTO</td></tr>
<tr><td>>hg38_1_-_228634819_228635039_RNA5S12_5S_50ntFlanks@84.101.18|CTCGTCTGATCTCGGAAG</td><td>>hg38_1_-_228634819_228635039_RNA5S12_5S_50ntFlanks</td><td>84</td><td>101</td><td>18</td><td>CTCGTCTGATCTCGGAAG</td><td>391.1</td><td>-4.1</td><td>1.1</td><td>-3.9</td><td>1.0E-04</td><td>1.9E-04</td><td>BT20/MB468</td><td>MITO</td></tr>
<tr><td>>hg38_21_+_8433172_8446622_RNA45SN1_45S_50ntFlanks@9571.9596.26|GCAAATCGGTCGTCCGACCTGGGTAT</td><td>>hg38_21_+_8433172_8446622_RNA45SN1_45S_50ntFlanks</td><td>9571</td><td>9596</td><td>26</td><td>GCAAATCGGTCGTCCGACCTGGGTAT</td><td>383.7</td><td>-5.4</td><td>1.3</td><td>-4.0</td><td>5.2E-05</td><td>1.9E-04</td><td>BT20/MB468</td><td>MP</td></tr>
<tr><td>>hg38_21_+_8433172_8446622_RNA45SN1_45S_50ntFlanks@3985.4004.20|CGGCTTTGGTGACTCTAGAT</td><td>>hg38_21_+_8433172_8446622_RNA45SN1_45S_50ntFlanks</td><td>3985</td><td>4004</td><td>20</td><td>CGGCTTTGGTGACTCTAGAT</td><td>118.6</td><td>-3.4</td><td>0.9</td><td>-3.8</td><td>1.3E-04</td><td>1.9E-04</td><td>BT20/MB468</td><td>CYTO</td></tr>
<tr><td>>hg38_21_+_8433172_8446622_RNA45SN1_45S_50ntFlanks@12942.12959.18|TTCGTAGACGACCTGCTT</td><td>>hg38_21_+_8433172_8446622_RNA45SN1_45S_50ntFlanks</td><td>12942</td><td>12959</td><td>18</td><td>TTCGTAGACGACCTGCTT</td><td>275.7</td><td>-5.8</td><td>1.5</td><td>-3.8</td><td>1.2E-04</td><td>1.9E-04</td><td>MB231/MB468</td><td>MP</td></tr>
<tr><td>>hg38_21_+_8433172_8446622_RNA45SN1_45S_50ntFlanks@12444.12476.33|AAGCAGAATTCACCAAGCGTTGGATTGTTCACC</td><td>>hg38_21_+_8433172_8446622_RNA45SN1_45S_50ntFlanks</td><td>12444</td><td>12476</td><td>33</td><td>AAGCAGAATTCACCAAGCGTTGGATTGTTCACC</td><td>386.5</td><td>-3.7</td><td>1.0</td><td>-3.8</td><td>1.5E-04</td><td>1.9E-04</td><td>BT20/MB468</td><td>NUC</td></tr>
<tr><td>>hg38_MT_+_1621_3279_RNR2_16S_50ntFlanks@1348.1385.38|AACAATAGGGTTTACGACCTCGATGTTGGATCAGGACA</td><td>>hg38_MT_+_1621_3279_RNR2_16S_50ntFlanks</td><td>1348</td><td>1385</td><td>38</td><td>AACAATAGGGTTTACGACCTCGATGTTGGATCAGGACA</td><td>264.4</td><td>4.4</td><td>1.1</td><td>3.8</td><td>1.2E-04</td><td>1.9E-04</td><td>MB231/MB468</td><td>MP</td></tr>
<tr><td>>MI0003628|hsa-mir-615&WithFlank&12|+|54033944|54034051@67.88.22&[MIMAT0003283&hsa-miR-615-3p&offsets|0|0;m-495&12|+|54034010|54034030&offsets|0|+1]|TCCGAGCCTGGGTCTCCCTCTT</td><td>>MI0003628|hsa-mir-615&WithFlank&12|+|54033944|54034051</td><td>67</td><td>88</td><td>22</td><td>TCCGAGCCTGGGTCTCCCTCTT</td><td>325.6</td><td>6.5</td><td>1.6</td><td>4.0</td><td>5.2E-05</td><td>1.9E-04</td><td>BT20/MB468</td><td>MP</td></tr>
<tr><td>>hg38_MT_+_1621_3279_RNR2_16S_50ntFlanks@387.416.30|TGGTGATAGCTGGTTGTCCAAGATAGAATC</td><td>>hg38_MT_+_1621_3279_RNR2_16S_50ntFlanks</td><td>387</td><td>416</td><td>30</td><td>TGGTGATAGCTGGTTGTCCAAGATAGAATC</td><td>143.3</td><td>7.6</td><td>1.9</td><td>4.0</td><td>5.2E-05</td><td>1.9E-04</td><td>BT20/MB468</td><td>MP</td></tr>
<tr><td>>MI0000078|hsa-mir-22&WithFlank&17|-|1713897|1713993@59.78.20(+2U)&[MIMAT0000077&hsa-miR-22-3p&offsets|0|-2(+2U);m-2&17|-|1713914|1713935&offsets|0|-2(+2U)]|AAGCTGCCAGTTGAAGAACTTT</td><td>>MI0000078|hsa-mir-22&WithFlank&17|-|1713897|1713993</td><td>59</td><td>78</td><td>20(+2U)</td><td>AAGCTGCCAGTTGAAGAACTTT</td><td>27.5</td><td>8.0</td><td>2.1</td><td>3.8</td><td>1.2E-04</td><td>1.9E-04</td><td>MB231/MB468</td><td>MP</td></tr>
<tr><td>>hg38_21_+_8433172_8446622_RNA45SN1_45S_50ntFlanks@6651.6674.24|CGACTCTTAGCGGTGGATCACTCG</td><td>>hg38_21_+_8433172_8446622_RNA45SN1_45S_50ntFlanks</td><td>6651</td><td>6674</td><td>24</td><td>CGACTCTTAGCGGTGGATCACTCG</td><td>4467.6</td><td>4.0</td><td>1.0</td><td>3.8</td><td>1.4E-04</td><td>1.9E-04</td><td>MB231/MB468</td><td>NUC</td></tr>
<tr><td>>hg38_21_+_8433172_8446622_RNA45SN1_45S_50ntFlanks@11564.11587.24|CCTAGCAGCCGACTTAGAACTGGT</td><td>>hg38_21_+_8433172_8446622_RNA45SN1_45S_50ntFlanks</td><td>11564</td><td>11587</td><td>24</td><td>CCTAGCAGCCGACTTAGAACTGGT</td><td>215.2</td><td>-1.9</td><td>0.5</td><td>-3.8</td><td>1.4E-04</td><td>1.9E-04</td><td>MB231/MB468</td><td>NUC</td></tr>
<tr><td>>hg38_21_+_8433172_8446622_RNA45SN1_45S_50ntFlanks@8184.8210.27|CGAGGCCCAGCCCGTGGACGGTGTGAG</td><td>>hg38_21_+_8433172_8446622_RNA45SN1_45S_50ntFlanks</td><td>8184</td><td>8210</td><td>27</td><td>CGAGGCCCAGCCCGTGGACGGTGTGAG</td><td>846.5</td><td>-6.9</td><td>1.8</td><td>-3.8</td><td>1.5E-04</td><td>1.9E-04</td><td>BT20/MB468</td><td>NUC</td></tr>
<tr><td>>hg38_21_+_8433172_8446622_RNA45SN1_45S_50ntFlanks@4599.4619.21|GTTTTCGGAACTGAGGCCATG</td><td>>hg38_21_+_8433172_8446622_RNA45SN1_45S_50ntFlanks</td><td>4599</td><td>4619</td><td>21</td><td>GTTTTCGGAACTGAGGCCATG</td><td>88.8</td><td>-5.8</td><td>1.5</td><td>-3.8</td><td>1.2E-04</td><td>1.9E-04</td><td>MB231/MB468</td><td>MP</td></tr>
<tr><td>>MI0000456|hsa-mir-140&WithFlank&16|+|69933075|69933186@69.90.22(+1U)&[MIMAT0004597&hsa-miR-140-3p&offsets|+1|+2(+1U);m-45&16|+|69933143|69933164&offsets|0|0(+1U)]|ACCACAGGGTAGAACCACGGACT</td><td>>MI0000456|hsa-mir-140&WithFlank&16|+|69933075|69933186</td><td>69</td><td>90</td><td>22(+1U)</td><td>ACCACAGGGTAGAACCACGGACT</td><td>898.1</td><td>6.1</td><td>1.5</td><td>4.0</td><td>5.2E-05</td><td>1.9E-04</td><td>BT20/MB468</td><td>MP</td></tr>
<tr><td>>hg38_21_+_8433172_8446622_RNA45SN1_45S_50ntFlanks@11569.11613.45|CAGCCGACTTAGAACTGGTGCGGACCAGGGGAATCCGACTGTTTA</td><td>>hg38_21_+_8433172_8446622_RNA45SN1_45S_50ntFlanks</td><td>11569</td><td>11613</td><td>45</td><td>CAGCCGACTTAGAACTGGTGCGGACCAGGGGAATCCGACTGTTTA</td><td>1112.2</td><td>-4.2</td><td>1.1</td><td>-3.8</td><td>1.4E-04</td><td>1.9E-04</td><td>MB231/MB468</td><td>MITO</td></tr>
<tr><td>>hg38_21_+_8433172_8446622_RNA45SN1_45S_50ntFlanks@9951.9983.33|CCATGGAAGTCGGAATCCGCTAAGGAGTGTGTA</td><td>>hg38_21_+_8433172_8446622_RNA45SN1_45S_50ntFlanks</td><td>9951</td><td>9983</td><td>33</td><td>CCATGGAAGTCGGAATCCGCTAAGGAGTGTGTA</td><td>78.4</td><td>9.6</td><td>2.2</td><td>4.3</td><td>1.9E-05</td><td>1.9E-04</td><td>BT20/MB231</td><td>MP</td></tr>
<tr><td>>MI0000088|hsa-mir-30a&WithFlank&6|-|71403545|71403627@12.35.24&[MIMAT0000087&hsa-miR-30a-5p&offsets|0|+2;m-15&6|-|71403595|71403616&offsets|0|+2]|TGTAAACATCCTCGACTGGAAGCT</td><td>>MI0000088|hsa-mir-30a&WithFlank&6|-|71403545|71403627</td><td>12</td><td>35</td><td>24</td><td>TGTAAACATCCTCGACTGGAAGCT</td><td>153853.5</td><td>-3.3</td><td>0.8</td><td>-4.3</td><td>2.1E-05</td><td>1.9E-04</td><td>BT20/MB231</td><td>MITO</td></tr>
<tr><td>>hg38_21_+_8433172_8446622_RNA45SN1_45S_50ntFlanks@4723.4758.36|CATTAATCAAGAACGAAAGTCGGAGGTTCGAAGACG</td><td>>hg38_21_+_8433172_8446622_RNA45SN1_45S_50ntFlanks</td><td>4723</td><td>4758</td><td>36</td><td>CATTAATCAAGAACGAAAGTCGGAGGTTCGAAGACG</td><td>4366.4</td><td>-2.9</td><td>0.8</td><td>-3.8</td><td>1.3E-04</td><td>1.9E-04</td><td>MB231/MB468</td><td>CYTO</td></tr>
<tr><td>>hg38_21_+_8433172_8446622_RNA45SN1_45S_50ntFlanks@4380.4408.29|CGTAGTTGGATCTTGGGAGCGGGCGGGCG</td><td>>hg38_21_+_8433172_8446622_RNA45SN1_45S_50ntFlanks</td><td>4380</td><td>4408</td><td>29</td><td>CGTAGTTGGATCTTGGGAGCGGGCGGGCG</td><td>142.6</td><td>-3.4</td><td>0.9</td><td>-3.8</td><td>1.3E-04</td><td>1.9E-04</td><td>MB231/MB468</td><td>CYTO</td></tr>
<tr><td>>hg38_21_+_8433172_8446622_RNA45SN1_45S_50ntFlanks@11584.11623.40|TGGTGCGGACCAGGGGAATCCGACTGTTTAATTAAAACAA</td><td>>hg38_21_+_8433172_8446622_RNA45SN1_45S_50ntFlanks</td><td>11584</td><td>11623</td><td>40</td><td>TGGTGCGGACCAGGGGAATCCGACTGTTTAATTAAAACAA</td><td>3408.6</td><td>4.4</td><td>1.0</td><td>4.3</td><td>2.1E-05</td><td>1.9E-04</td><td>BT20/MB231</td><td>MITO</td></tr>
<tr><td>>hg38_21_+_8433172_8446622_RNA45SN1_45S_50ntFlanks@9915.9947.33|AAAAGGTGTTGGTTGATATAGACAGCAGGACGG</td><td>>hg38_21_+_8433172_8446622_RNA45SN1_45S_50ntFlanks</td><td>9915</td><td>9947</td><td>33</td><td>AAAAGGTGTTGGTTGATATAGACAGCAGGACGG</td><td>77.4</td><td>7.1</td><td>1.8</td><td>4.0</td><td>5.2E-05</td><td>1.9E-04</td><td>BT20/MB468</td><td>MP</td></tr>
<tr><td>>hg38_21_+_8433172_8446622_RNA45SN1_45S_50ntFlanks@10854.10873.20|TAAGGATTGGCTCTAAGGGC</td><td>>hg38_21_+_8433172_8446622_RNA45SN1_45S_50ntFlanks</td><td>10854</td><td>10873</td><td>20</td><td>TAAGGATTGGCTCTAAGGGC</td><td>97.0</td><td>-6.1</td><td>1.6</td><td>-3.8</td><td>1.2E-04</td><td>1.9E-04</td><td>MB231/MB468</td><td>MP</td></tr>
<tr><td>>hg38_21_+_8433172_8446622_RNA45SN1_45S_50ntFlanks@12415.12441.27|CTTCGATGTCGGCTCTTCCTATCATTG</td><td>>hg38_21_+_8433172_8446622_RNA45SN1_45S_50ntFlanks</td><td>12415</td><td>12441</td><td>27</td><td>CTTCGATGTCGGCTCTTCCTATCATTG</td><td>231.6</td><td>-5.9</td><td>1.5</td><td>-3.8</td><td>1.2E-04</td><td>1.9E-04</td><td>MB231/MB468</td><td>MP</td></tr>
<tr><td>>hg38_21_+_8433172_8446622_RNA45SN1_45S_50ntFlanks@4007.4024.18|CCTCGGGCCGATCGCACG</td><td>>hg38_21_+_8433172_8446622_RNA45SN1_45S_50ntFlanks</td><td>4007</td><td>4024</td><td>18</td><td>CCTCGGGCCGATCGCACG</td><td>962.8</td><td>-2.5</td><td>0.7</td><td>-3.8</td><td>1.4E-04</td><td>1.9E-04</td><td>MB231/MB468</td><td>CYTO</td></tr>
<tr><td>>hg38_21_+_8433172_8446622_RNA45SN1_45S_50ntFlanks@8114.8141.28|GCGCGGGACATGTGGCGTACGGAAGACC</td><td>>hg38_21_+_8433172_8446622_RNA45SN1_45S_50ntFlanks</td><td>8114</td><td>8141</td><td>28</td><td>GCGCGGGACATGTGGCGTACGGAAGACC</td><td>650.8</td><td>8.0</td><td>1.9</td><td>4.3</td><td>1.9E-05</td><td>1.9E-04</td><td>BT20/MB231</td><td>MP</td></tr>
<tr><td>>trnaMT_PheGAA_MT_+_577_647@36.56.21|AAAATGTTTAGACGGGCTCAC</td><td>>trnaMT_PheGAA_MT_+_577_647</td><td>36</td><td>56</td><td>21</td><td>AAAATGTTTAGACGGGCTCAC</td><td>125.7</td><td>3.2</td><td>0.8</td><td>3.8</td><td>1.2E-04</td><td>1.9E-04</td><td>MB231/MB468</td><td>MP</td></tr>
<tr><td>>trnaMT_GluTTC_MT_-_14674_14742@30.72.43,trnalookalike8_GluTTC_5_-_93905172_93905240@30.72.43|TTTCATATCATTGGTCGTGGTTGTAGTCCGTGCGAGAATACCA</td><td>>trnaMT_GluTTC_MT_-_14674_14742</td><td>30</td><td>72</td><td>43</td><td>TTTCATATCATTGGTCGTGGTTGTAGTCCGTGCGAGAATACCA</td><td>204.2</td><td>4.0</td><td>1.1</td><td>3.8</td><td>1.4E-04</td><td>1.9E-04</td><td>MB231/MB468</td><td>MITO</td></tr>
</table>

| | | | | | | | | | | | | | |
|---|---|---|---|---|---|---|---|---|---|---|---|---|---|
| >MI0000734\|hsa-mir-106b&WithFlank&7\|-\|100093987\|100094080@58.78.21(+1U)&[MIMAT0004672&hsa-miR-106b-3p&offsets\|0\|-1(+1U);m-87&7\|-\|100094003\|100094023&offsets\|0\|0(+1U)]\|CCGCACTGTGGGTACTTGCTGT | >MI0000734\|hsa-mir-106b&WithFlank&7\|-\|100093987\|100094080 | 58 | 78 | 21(+1U) | CCGCACTGTGGGTACTTGCTGT | 826.6 | 3.7 | 1.0 | 3.8 | 1.2E-04 | 1.9E-04 | MB231/MB468 | MP |
| >hg38_21_+_8433172_8446622_RNA45SN1_45S_50ntFlanks@12456.12490.35\|CCAAGCGTTGGATTGTTCACCCACTAATAGGGAAC | >hg38_21_+_8433172_8446622_RNA45SN1_45S_50ntFlanks | 12456 | 12490 | 35 | CCAAGCGTTGGATTGTTCACCCACTAATAGGGAAC | 824.1 | -2.4 | 0.6 | -3.9 | 1.1E-04 | 1.9E-04 | BT20/MB468 | MITO |
| >MI0000439\|hsa-mir-23b&WithFlank&9\|+\|95085202\|95085310@67.86.20&[MIMAT0000418&hsa-miR-23b-3p&offsets\|+3\|0;m-42&9\|+\|95085265\|95085285&offsets\|+3\|+2]\|ACATTGCCAGGGATTACCAC | >MI0000439\|hsa-mir-23b&WithFlank&9\|+\|95085202\|95085310 | 67 | 86 | 20 | ACATTGCCAGGGATTACCAC | 958.9 | 13.8 | 3.6 | 3.8 | 1.4E-04 | 1.9E-04 | MB231/MB468 | NUC |
| >hg38_21_+_8433172_8446622_RNA45SN1_45S_50ntFlanks@10341.10364.24\|AATATTCAAACGAGAACTTTGAAG | >hg38_21_+_8433172_8446622_RNA45SN1_45S_50ntFlanks | 10341 | 10364 | 24 | AATATTCAAACGAGAACTTTGAAG | 72.4 | 9.5 | 2.2 | 4.3 | 1.9E-05 | 1.9E-04 | BT20/MB231 | MP |
| >hg38_21_+_8433172_8446622_RNA45SN1_45S_50ntFlanks@12942.12966.25\|TTCGTAGACGACCTGCTTCTGGGTC | >hg38_21_+_8433172_8446622_RNA45SN1_45S_50ntFlanks | 12942 | 12966 | 25 | TTCGTAGACGACCTGCTTCTGGGTC | 379.4 | -6.4 | 1.6 | -4.0 | 5.3E-05 | 1.9E-04 | BT20/MB468 | MP |
| >hg38_21_+_8433172_8446622_RNA45SN1_45S_50ntFlanks@8196.8224.29\|CGTGGACGGTGTGAGGCCGGTAGCGGCCC | >hg38_21_+_8433172_8446622_RNA45SN1_45S_50ntFlanks | 8196 | 8224 | 29 | CGTGGACGGTGTGAGGCCGGTAGCGGCCC | 607.9 | -3.5 | 0.9 | -4.0 | 5.3E-05 | 1.9E-04 | BT20/MB468 | MP |
| >MI0000085\|hsa-mir-27a&WithFlank&19\|-\|13836434\|13836523@57.74.18;MI0000440\|hsa-mir-27b&WithFlank&9\|+\|95085439\|95085547@67.84.18&[MIMAT0000084&hsa-miR-27a-3p&offsets\|0\|-3;m-43&19\|-\|13836447\|13836467&offsets\|0\|-3];[MIMAT0000419&hsa-miR-27b-3p&offsets\|0\|-3;m-39&9\|+\|95085505\|95085525&offsets\|0\|-3]\|TTCACAGTGGCTAAGTTC | >MI0000085\|hsa-mir-27a&WithFlank&19\|-\|13836434\|13836523 | 57 | 74 | 18 | TTCACAGTGGCTAAGTTC | 105.1 | 2.1 | 0.5 | 3.8 | 1.5E-04 | 1.9E-04 | BT20/MB468 | NUC |
| >MI0000077\|hsa-mir-21&WithFlank&17\|+\|59841260\|59841343@14.32.19&[MIMAT0000076&hsa-miR-21-5p&offsets\|0\|-3;m-1&17\|+\|59841273\|59841294&offsets\|0\|-3]\|TAGCTTATCAGACTGATGT | >MI0000077\|hsa-mir-21&WithFlank&17\|+\|59841260\|59841343 | 14 | 32 | 19 | TAGCTTATCAGACTGATGT | 2628.8 | 4.6 | 1.1 | 4.0 | 5.3E-05 | 1.9E-04 | BT20/MB468 | MP |
| >hg38_21_+_8433172_8446622_RNA45SN1_45S_50ntFlanks@12946.12990.45\|TAGACGACCTGCTTCTGGGTCGGGGTTTCGTACGTAGCAGAGCAG | >hg38_21_+_8433172_8446622_RNA45SN1_45S_50ntFlanks | 12946 | 12990 | 45 | TAGACGACCTGCTTCTGGGTCGGGGTTTCGTACGTAGCAGAGCAG | 586.0 | -5.7 | 1.5 | -3.8 | 1.4E-04 | 1.9E-04 | MB231/MB468 | NUC |
| >am_trnaMT_ValTAC_MT_+_1602_1670@20.57.38\|AAGCACCCAACTTACACTTAGGAGATTTCAACTTAACT | >am_trnaMT_ValTAC_MT_+_1602_1670 | 20 | 57 | 38 | AAGCACCCAACTTACACTTAGGAGATTTCAACTTAACT | 118.5 | 10.6 | 2.6 | 4.0 | 5.3E-05 | 1.9E-04 | BT20/MB468 | MP |
| >hg38_21_+_8433172_8446622_RNA45SN1_45S_50ntFlanks@11767.11794.28\|TCTAATTAGTGACGCGCATGAATGGATG | >hg38_21_+_8433172_8446622_RNA45SN1_45S_50ntFlanks | 11767 | 11794 | 28 | TCTAATTAGTGACGCGCATGAATGGATG | 573.8 | -3.1 | 0.8 | -4.0 | 5.3E-05 | 1.9E-04 | BT20/MB468 | MP |
| >hg38_MT_+_1621_3279_RNR2_16S_50ntFlanks@51.90.40\|GCTAAACCTAGCCCCAAACCCACTCCACCTTACTACCAGA | >hg38_MT_+_1621_3279_RNR2_16S_50ntFlanks | 51 | 90 | 40 | GCTAAACCTAGCCCCAAACCCACTCCACCTTACTACCAGA | 1562.0 | 4.1 | 1.1 | 3.8 | 1.2E-04 | 1.9E-04 | MB231/MB468 | MP |
| >hg38_21_+_8433172_8446622_RNA45SN1_45S_50ntFlanks@6778.6802.25\|CCCGGGGCTACGCCTGTCTGAGCGT | >hg38_21_+_8433172_8446622_RNA45SN1_45S_50ntFlanks | 6778 | 6802 | 25 | CCCGGGGCTACGCCTGTCTGAGCGT | 1329.6 | 3.7 | 1.0 | 3.8 | 1.4E-04 | 1.9E-04 | MB231/MB468 | MITO |
| >hg38_1_-_228634819_228635039_RNA5S12_5S_50ntFlanks@51.98.48\|GTCTACGGCCATACCACCCTGAACGCGCCCGATCTCGTCTGATCTCGG | >hg38_1_-_228634819_228635039_RNA5S12_5S_50ntFlanks | 51 | 98 | 48 | GTCTACGGCCATACCACCCTGAACGCGCCCGATCTCGTCTGATCTCGG | 108.6 | 10.5 | 2.6 | 4.0 | 5.3E-05 | 1.9E-04 | BT20/MB468 | MP |
| >trnaMT_PheGAA_MT_+_577_647@36.54.19\|AAAATGTTTAGACGGGCTC | >trnaMT_PheGAA_MT_+_577_647 | 36 | 54 | 19 | AAAATGTTTAGACGGGCTC | 159.9 | 3.1 | 0.8 | 3.8 | 1.2E-04 | 1.9E-04 | MB231/MB468 | MP |
| >hg38_21_+_8433172_8446622_RNA45SN1_45S_50ntFlanks@10348.10372.25\|AAACGAGAACTTTGAAGGCCGAAGT | >hg38_21_+_8433172_8446622_RNA45SN1_45S_50ntFlanks | 10348 | 10372 | 25 | AAACGAGAACTTTGAAGGCCGAAGT | 453.4 | 2.1 | 0.6 | 3.8 | 1.3E-04 | 1.9E-04 | BT20/MB468 | CYTO |
| >hg38_21_+_8433172_8446622_RNA45SN1_45S_50ntFlanks@12288.12323.36\|ACCGTGAAAGCGGGGCCTCACGATCCTTCTGACCTT | >hg38_21_+_8433172_8446622_RNA45SN1_45S_50ntFlanks | 12288 | 12323 | 36 | ACCGTGAAAGCGGGGCCTCACGATCCTTCTGACCTT | 1027.3 | -3.2 | 0.8 | -3.9 | 1.1E-04 | 1.9E-04 | BT20/MB468 | MITO |
| >hg38_21_+_8433172_8446622_RNA45SN1_45S_50ntFlanks@11771.11804.34\|ATTAGTGACGCGCATGAATGGATGAACGAGATTC | >hg38_21_+_8433172_8446622_RNA45SN1_45S_50ntFlanks | 11771 | 11804 | 34 | ATTAGTGACGCGCATGAATGGATGAACGAGATTC | 117.9 | -4.5 | 1.2 | -3.8 | 1.2E-04 | 1.9E-04 | MB231/MB468 | MP |
| >hg38_21_+_8433172_8446622_RNA45SN1_45S_50ntFlanks@12126.12166.41\|CGACCCGCTCCGGGGACAGTGCCAGGTGGGGAGTTTGACTG | >hg38_21_+_8433172_8446622_RNA45SN1_45S_50ntFlanks | 12126 | 12166 | 41 | CGACCCGCTCCGGGGACAGTGCCAGGTGGGGAGTTTGACTG | 45.1 | -9.4 | 2.4 | -3.8 | 1.2E-04 | 1.9E-04 | MB231/MB468 | MP |
| >hg38_21_+_8433172_8446622_RNA45SN1_45S_50ntFlanks@12413.12441.29\|TCCTTCGATGTCGGCTCTTCCTATCATTG | >hg38_21_+_8433172_8446622_RNA45SN1_45S_50ntFlanks | 12413 | 12441 | 29 | TCCTTCGATGTCGGCTCTTCCTATCATTG | 444.0 | -2.6 | 0.7 | -3.8 | 1.5E-04 | 1.9E-04 | MB231/MB468 | NUC |
| >MI0001445\|hsa-mir-423&WithFlank&17\|+\|30117073\|30117178@23.42.20(+1G)&[MIMAT0004748&hsa-miR-423-5p&offsets\|0\|-3(+1G);m-141&17\|+\|30117095\|30117117&offsets\|0\|-3(+1G)]\|TGAGGGGCAGAGAGCGAGACG | >MI0001445\|hsa-mir-423&WithFlank&17\|+\|30117073\|30117178 | 23 | 42 | 20(+1G) | TGAGGGGCAGAGAGCGAGACG | 236.3 | 7.7 | 1.9 | 4.0 | 5.3E-05 | 1.9E-04 | BT20/MB468 | MP |
| >hg38_21_+_8433172_8446622_RNA45SN1_45S_50ntFlanks@11582.11605.24\|ACTGGTGCGGACCAGGGGAATCCG | >hg38_21_+_8433172_8446622_RNA45SN1_45S_50ntFlanks | 11582 | 11605 | 24 | ACTGGTGCGGACCAGGGGAATCCG | 236.6 | -2.5 | 0.7 | -3.8 | 1.5E-04 | 1.9E-04 | MB231/MB468 | NUC |
| >trna12_ArgCCT_16_+_3243918_3243990@35.54.20,trna18_ArgCCT_17_+_73030001_73030073@35.54.20,trna21_ArgCCT_17_-_73030526_73030598@35.54.20,trna2_ArgCCT_16_+_3202901_3202973@35.54.20,trna3_ArgCCT_7_+_139025446_139025518@35.54.20\|CTAAGCCAGGGATTGTGGGT | >trna12_ArgCCT_16_+_3243918_3243990 | 35 | 54 | 20 | CTAAGCCAGGGATTGTGGGT | 1924.7 | 2.7 | 0.7 | 3.8 | 1.3E-04 | 1.9E-04 | BT20/MB468 | CYTO |
| >hg38_21_+_8433172_8446622_RNA45SN1_45S_50ntFlanks@12454.12476.23\|CACCAAGCGTTGGATTGTTCACC | >hg38_21_+_8433172_8446622_RNA45SN1_45S_50ntFlanks | 12454 | 12476 | 23 | CACCAAGCGTTGGATTGTTCACC | 411.2 | -3.3 | 0.9 | -3.8 | 1.2E-04 | 1.9E-04 | MB231/MB468 | MP |
| >hg38_21_+_8433172_8446622_RNA45SN1_45S_50ntFlanks@12444.12486.43\|AAGCAGAATTCACCAAGCGTTGGATTGTTCACCCACTAATAGG | >hg38_21_+_8433172_8446622_RNA45SN1_45S_50ntFlanks | 12444 | 12486 | 43 | AAGCAGAATTCACCAAGCGTTGGATTGTTCACCCACTAATAGG | 369.6 | -4.1 | 1.1 | -3.8 | 1.5E-04 | 1.9E-04 | MB231/MB468 | MITO |
| >hg38_21_+_8433172_8446622_RNA45SN1_45S_50ntFlanks@10351.10379.29\|CGAGAACTTTGAAGGCCGAAGTGGAGAAG | >hg38_21_+_8433172_8446622_RNA45SN1_45S_50ntFlanks | 10351 | 10379 | 29 | CGAGAACTTTGAAGGCCGAAGTGGAGAAG | 617.4 | -2.2 | 0.6 | -3.9 | 1.1E-04 | 1.9E-04 | BT20/MB468 | MITO |
| >trnaMT_HisGTG_MT_+_12138_12206@32.71.40\|TGAATCTGACAACAGAGGCTTACGACCCCTTATTTACCCC | >trnaMT_HisGTG_MT_+_12138_12206 | 32 | 71 | 40 | TGAATCTGACAACAGAGGCTTACGACCCCTTATTTACCCC | 2418.2 | 3.6 | 1.0 | 3.8 | 1.5E-04 | 1.9E-04 | MB231/MB468 | NUC |
| >MI0000067\|hsa-let-7f-1&WithFlank&9\|+\|94176341\|94176439@13.33.21(+1A);MI0000068\|hsa-let-7f-2&WithFlank&X\|-\|53557186\|53557280@14.34.21(+1A)&[MIMAT0000067&hsa-let-7f-5p&offsets\|0\|-1(+1A);m-20&9\|+\|94176353\|94176374&offsets\|0\|-1(+1A)];[MIMAT0000067_1&hsa-let-7f-5p&offsets\|0\|-1(+1A);m-19&X\|-\|53557246\|53557267&offsets\|0\|-1(+1A)]\|TGAGGTAGTAGATTGTATAGTA | >MI0000067\|hsa-let-7f-1&WithFlank&9\|+\|94176341\|94176439 | 13 | 33 | 21(+1A) | TGAGGTAGTAGATTGTATAGTA | 1069.0 | 2.3 | 0.6 | 3.8 | 1.5E-04 | 1.9E-04 | BT20/MB468 | NUC |
| >hg38_21_+_8433172_8446622_RNA45SN1_45S_50ntFlanks@10682.10712.31\|TCCGGCGGCGTCCGGTGAGCTCTCGCTGGCC | >hg38_21_+_8433172_8446622_RNA45SN1_45S_50ntFlanks | 10682 | 10712 | 31 | TCCGGCGGCGTCCGGTGAGCTCTCGCTGGCC | 724.2 | -2.8 | 0.7 | -3.8 | 1.5E-04 | 1.9E-04 | MB231/MB468 | MITO |
| >hg38_21_+_8433172_8446622_RNA45SN1_45S_50ntFlanks@9564.9598.35\|CTGACGTGCAAATCGGTCGTCCGACCTGGGTATAG | >hg38_21_+_8433172_8446622_RNA45SN1_45S_50ntFlanks | 9564 | 9598 | 35 | CTGACGTGCAAATCGGTCGTCCGACCTGGGTATAG | 183.2 | -4.4 | 1.2 | -3.8 | 1.5E-04 | 1.9E-04 | MB231/MB468 | NUC |
| >hg38_21_+_8433172_8446622_RNA45SN1_45S_50ntFlanks@11315.11338.24\|CCACCCCACGTCTCGTCGCGCGCG | >hg38_21_+_8433172_8446622_RNA45SN1_45S_50ntFlanks | 11315 | 11338 | 24 | CCACCCCACGTCTCGTCGCGCGCG | 2746.0 | -7.6 | 2.0 | -3.8 | 1.5E-04 | 1.9E-04 | BT20/MB468 | NUC |
| >MI0001445\|hsa-mir-423&WithFlank&17\|+\|30117073\|30117178@23.43.21(+1C)&[MIMAT0004748&hsa-miR-423-5p&offsets\|0\|-2(+1C);m-141&17\|+\|30117095\|30117117&offsets\|0\|-2(+1C)]\|TGAGGGGCAGAGAGCGAGACTC | >MI0001445\|hsa-mir-423&WithFlank&17\|+\|30117073\|30117178 | 23 | 43 | 21(+1C) | TGAGGGGCAGAGAGCGAGACTC | 332.3 | 6.1 | 1.6 | 3.8 | 1.2E-04 | 1.9E-04 | MB231/MB468 | MP |
| >hg38_21_+_8433172_8446622_RNA45SN1_45S_50ntFlanks@12274.12307.34\|CAGTACGAATACAGACCGTGAAAGCGGGGCCTCA | >hg38_21_+_8433172_8446622_RNA45SN1_45S_50ntFlanks | 12274 | 12307 | 34 | CAGTACGAATACAGACCGTGAAAGCGGGGCCTCA | 142.0 | -6.4 | 1.6 | -4.0 | 5.4E-05 | 1.9E-04 | BT20/MB468 | MP |
| >hg38_21_+_8433172_8446622_RNA45SN1_45S_50ntFlanks@11579.11606.28\|AGAACTGGTGCGGACCAGGGGAATCCGA | >hg38_21_+_8433172_8446622_RNA45SN1_45S_50ntFlanks | 11579 | 11606 | 28 | AGAACTGGTGCGGACCAGGGGAATCCGA | 216.4 | -2.8 | 0.7 | -3.8 | 1.4E-04 | 1.9E-04 | MB231/MB468 | CYTO |
| >hg38_21_+_8433172_8446622_RNA45SN1_45S_50ntFlanks@9308.9338.31\|AGGAGTCTAACACGTGCGCGAGTCGGGGGCT | >hg38_21_+_8433172_8446622_RNA45SN1_45S_50ntFlanks | 9308 | 9338 | 31 | AGGAGTCTAACACGTGCGCGAGTCGGGGGCT | 589.0 | -4.4 | 1.1 | -3.9 | 1.1E-04 | 1.9E-04 | BT20/MB468 | MITO |
| >hg38_21_+_8433172_8446622_RNA45SN1_45S_50ntFlanks@6746.6770.25\|CACTTCGAACGCACTTGCGGCCCCG | >hg38_21_+_8433172_8446622_RNA45SN1_45S_50ntFlanks | 6746 | 6770 | 25 | CACTTCGAACGCACTTGCGGCCCCG | 2111.1 | -6.9 | 1.7 | -4.0 | 5.4E-05 | 1.9E-04 | BT20/MB468 | MP |
| >hg38_21_+_8433172_8446622_RNA45SN1_45S_50ntFlanks@12399.12443.45\|ACGTCGCTTTTTGATCCTTCGATGTCGGCTCTTCCTATCATTGTG | >hg38_21_+_8433172_8446622_RNA45SN1_45S_50ntFlanks | 12399 | 12443 | 45 | ACGTCGCTTTTTGATCCTTCGATGTCGGCTCTTCCTATCATTGTG | 293.5 | -4.6 | 1.2 | -3.8 | 1.5E-04 | 1.9E-04 | MB231/MB468 | MITO |
| >hg38_21_+_8433172_8446622_RNA45SN1_45S_50ntFlanks@10345.10382.38\|TTCAAACGAGAACTTTGAAGGCCGAAGTGGAGAAGGGT | >hg38_21_+_8433172_8446622_RNA45SN1_45S_50ntFlanks | 10345 | 10382 | 38 | TTCAAACGAGAACTTTGAAGGCCGAAGTGGAGAAGGGT | 145.0 | 5.3 | 1.3 | 4.0 | 5.4E-05 | 1.9E-04 | BT20/MB468 | MP |
| >hg38_21_+_8433172_8446622_RNA45SN1_45S_50ntFlanks@9548.9573.26\|GAGGTCCGTAGCGGTCCTGACGTGCA | >hg38_21_+_8433172_8446622_RNA45SN1_45S_50ntFlanks | 9548 | 9573 | 26 | GAGGTCCGTAGCGGTCCTGACGTGCA | 334.6 | -2.7 | 0.7 | -3.8 | 1.5E-04 | 1.9E-04 | MB231/MB468 | MITO |
| >hg38_21_+_8433172_8446622_RNA45SN1_45S_50ntFlanks@6660.6695.36\|GCGGTGGATCACTCGGCTCGTGCGTCGATGAAGAAC | >hg38_21_+_8433172_8446622_RNA45SN1_45S_50ntFlanks | 6660 | 6695 | 36 | GCGGTGGATCACTCGGCTCGTGCGTCGATGAAGAAC | 415.7 | 4.2 | 1.0 | 4.0 | 5.4E-05 | 1.9E-04 | BT20/MB468 | MP |
| >hg38_21_+_8433172_8446622_RNA45SN1_45S_50ntFlanks@8350.8378.29\|AAAGTTGAAAAGAACTTTGAAGAGAGAGT | >hg38_21_+_8433172_8446622_RNA45SN1_45S_50ntFlanks | 8350 | 8378 | 29 | AAAGTTGAAAAGAACTTTGAAGAGAGAGT | 147.8 | 6.2 | 1.5 | 4.0 | 5.4E-05 | 1.9E-04 | BT20/MB468 | MP |
| >hg38_21_+_8433172_8446622_RNA45SN1_45S_50ntFlanks@475.495.21\|CTTCGTGATCGATGTGGTGAC | >hg38_21_+_8433172_8446622_RNA45SN1_45S_50ntFlanks | 475 | 495 | 21 | CTTCGTGATCGATGTGGTGAC | 816.2 | 5.8 | 1.5 | 3.8 | 1.5E-04 | 1.9E-04 | MB231/MB468 | MITO |
| >hg38_21_+_8433172_8446622_RNA45SN1_45S_50ntFlanks@10685.10710.26\|GGCGGCGTCCGGTGAGCTCTCGCTGG | >hg38_21_+_8433172_8446622_RNA45SN1_45S_50ntFlanks | 10685 | 10710 | 26 | GGCGGCGTCCGGTGAGCTCTCGCTGG | 986.9 | -3.4 | 0.9 | -3.8 | 1.5E-04 | 1.9E-04 | MB231/MB468 | MITO |
| >hg38_21_+_8433172_8446622_RNA45SN1_45S_50ntFlanks@7974.8015.42\|ACGCGACCTCAGATCAGACGTGGCGACCCGCTGAATTTAAGC | >hg38_21_+_8433172_8446622_RNA45SN1_45S_50ntFlanks | 7974 | 8015 | 42 | ACGCGACCTCAGATCAGACGTGGCGACCCGCTGAATTTAAGC | 1025.2 | 3.9 | 1.0 | 3.8 | 1.5E-04 | 1.9E-04 | BT20/MB468 | NUC |
| >am_trna111_HisGTG_1_-_147774845_147774916@3.29.27,trna118_HisGTG_1_-_145396881_145396952@3.29.27,trna16_HisGTG_1_+_146544773_146544844@3.29.27,trna1_HisGTG_15_+_45493349_45493420@3.29.27,trna21_HisGTG_1_+_147753471_147753542@3.29.27,trna33_HisGTG_6_+_27125906_27125977@3.29.27,trna7_HisGTG_9_-_14433938_14434009@3.29.27,trna8_HisGTG_15_-_45492611_45492682@3.29.27,trna9_HisGTG_15_-_45490804_45490875@3.29.27\|CGTGATCGTATAGTGGTTAGTACTCTG | >am_trna111_HisGTG_1_-_147774845_147774916 | 3 | 29 | 27 | CGTGATCGTATAGTGGTTAGTACTCTG | 269.4 | -2.2 | 0.6 | -3.8 | 1.3E-04 | 1.9E-04 | BT20/MB468 | CYTO |
| >hg38_21_+_8433172_8446622_RNA45SN1_45S_50ntFlanks@10689.10707.19\|GCGTCCGGTGAGCTCTCGC | >hg38_21_+_8433172_8446622_RNA45SN1_45S_50ntFlanks | 10689 | 10707 | 19 | GCGTCCGGTGAGCTCTCGC | 273.8 | -4.5 | 1.2 | -3.8 | 1.5E-04 | 1.9E-04 | MB231/MB468 | NUC |

| | | | | | | | | | | | | | |
|---|---|---|---|---|---|---|---|---|---|---|---|---|---|
| >hg38_21_+_8433172_8446622_RNA45SN1_45S_50ntFlanks@6645.6680.36\|CTCGTACGACTCTTAGCGGTGGATCACTCGGCTCGT | >hg38_21_+_8433172_8446622_RNA45SN1_45S_50ntFlanks | 6645 | 6680 | 36 | CTCGTACGACTCTTAGCGGTGGATCACTCGGCTCGT | 4088.5 | 2.2 | 0.6 | 3.9 | 1.1E-04 | 1.9E-04 | BT20/MB468 | MITO |
| >hg38_21_+_8433172_8446622_RNA45SN1_45S_50ntFlanks@12071.12094.24\|CCCCGAGGGGCTCTCGCTTCTGGC | >hg38_21_+_8433172_8446622_RNA45SN1_45S_50ntFlanks | 12071 | 12094 | 24 | CCCCGAGGGGCTCTCGCTTCTGGC | 624.3 | -4.0 | 1.1 | -3.8 | 1.4E-04 | 1.9E-04 | MB231/MB468 | CYTO |
| >hg38_21_+_8433172_8446622_RNA45SN1_45S_50ntFlanks@6644.6661.18\|CCTCGTACGACTCTTAGC | >hg38_21_+_8433172_8446622_RNA45SN1_45S_50ntFlanks | 6644 | 6661 | 18 | CCTCGTACGACTCTTAGC | 287.6 | 3.9 | 1.0 | 4.0 | 5.4E-05 | 1.9E-04 | BT20/MB468 | MP |
| >trna116_GluCTC_1_-_145399233_145399304@2.33.32,trna59_GluCTC_1_+_249168447_249168518@2.33.32,trna71_GluCTC_1_-_161439189_161439260@2.33.32,trna74_GluCTC_1_-_161431809_161431880@2.33.32,trna77_GluCTC_1_-_161424398_161424469@2.33.32,trna77_GluCTC_6_+_28949976_28950047@2.33.32,trna80_GluCTC_1_-_161417018_161417089@2.33.32,trna87_GluCTC_6_-_126101393_126101464@2.33.32\|CCCTGGTGGTCTAGTGGTTAGGATTCGGCGCT | >trna116_GluCTC_1_-_145399233_145399304 | 2 | 33 | 32 | CCCTGGTGGTCTAGTGGTTAGGATTCGGCGCT | 386.2 | 5.6 | 1.5 | 3.8 | 1.4E-04 | 1.9E-04 | MB231/MB468 | CYTO |
| >hg38_21_+_8433172_8446622_RNA45SN1_45S_50ntFlanks@9487.9510.24\|TTAGGACCCGAAAGATGGTGAACT | >hg38_21_+_8433172_8446622_RNA45SN1_45S_50ntFlanks | 9487 | 9510 | 24 | TTAGGACCCGAAAGATGGTGAACT | 68.4 | 9.4 | 2.2 | 4.3 | 1.9E-05 | 1.9E-04 | BT20/MB231 | MP |
| >hg38_21_+_8433172_8446622_RNA45SN1_45S_50ntFlanks@3835.3869.35\|CTCCTCTCCTACTTGGATAACTGTGGTAATTCTAG | >hg38_21_+_8433172_8446622_RNA45SN1_45S_50ntFlanks | 3835 | 3869 | 35 | CTCCTCTCCTACTTGGATAACTGTGGTAATTCTAG | 124.9 | -5.2 | 1.3 | -3.9 | 1.1E-04 | 1.9E-04 | BT20/MB468 | MITO |
| >hg38_21_+_8433172_8446622_RNA45SN1_45S_50ntFlanks@9533.9573.41\|GAGGAAACTCTGGTGGAGGTCCGTAGCGGTCCTGACGTGCA | >hg38_21_+_8433172_8446622_RNA45SN1_45S_50ntFlanks | 9533 | 9573 | 41 | GAGGAAACTCTGGTGGAGGTCCGTAGCGGTCCTGACGTGCA | 406.1 | 2.7 | 0.7 | 3.8 | 1.4E-04 | 1.9E-04 | MB231/MB468 | CYTO |
| >hg38_21_+_8433172_8446622_RNA45SN1_45S_50ntFlanks@10343.10361.19\|TATTCAAACGAGAACTTTG | >hg38_21_+_8433172_8446622_RNA45SN1_45S_50ntFlanks | 10343 | 10361 | 19 | TATTCAAACGAGAACTTTG | 300.6 | -4.1 | 1.1 | -3.8 | 1.4E-04 | 1.9E-04 | MB231/MB468 | CYTO |
| >hg38_21_+_8433172_8446622_RNA45SN1_45S_50ntFlanks@12997.13040.44\|CGCTGCGATCTATTGAAAGTCAGCCCTCGACACAAGGGTTTGTC | >hg38_21_+_8433172_8446622_RNA45SN1_45S_50ntFlanks | 12997 | 13040 | 44 | CGCTGCGATCTATTGAAAGTCAGCCCTCGACACAAGGGTTTGTC | 475.2 | -3.4 | 0.8 | -4.2 | 2.2E-05 | 1.9E-04 | BT20/MB231 | MITO |
| >am_trnaMT_ValTAC_MT_+_1602_1670@28.62.35\|AACTTACACTTAGGAGATTTCAACTTAACTTGACC | >am_trnaMT_ValTAC_MT_+_1602_1670 | 28 | 62 | 35 | AACTTACACTTAGGAGATTTCAACTTAACTTGACC | 776.7 | 6.8 | 1.7 | 4.0 | 5.4E-05 | 1.9E-04 | BT20/MB468 | MP |
| >hg38_21_+_8433172_8446622_RNA45SN1_45S_50ntFlanks@4007.4042.36\|CCTCGGGCCGATCGCACGCCCCCCGTGGCGGCGACG | >hg38_21_+_8433172_8446622_RNA45SN1_45S_50ntFlanks | 4007 | 4042 | 36 | CCTCGGGCCGATCGCACGCCCCCCGTGGCGGCGACG | 432.8 | -2.9 | 0.8 | -3.8 | 1.4E-04 | 1.9E-04 | MB231/MB468 | CYTO |
| >hg38_21_+_8433172_8446622_RNA45SN1_45S_50ntFlanks@12934.12951.18\|CTAAACCATTCGTAGACG | >hg38_21_+_8433172_8446622_RNA45SN1_45S_50ntFlanks | 12934 | 12951 | 18 | CTAAACCATTCGTAGACG | 1350.2 | -6.1 | 1.6 | -3.8 | 1.3E-04 | 1.9E-04 | BT20/MB468 | CYTO |
| >am_trna128_GlyGCC_6_-_27870686_27870756@1.34.34,trna18_GlyGCC_16_+_70822597_70822667@1.34.34,trna19_GlyGCC_16_+_70823410_70823480@1.34.34,trna19_GlyGCC_2_-_157257659_157257729@1.34.34,trna24_GlyGCC_16_-_70812942_70813012@1.34.34,trna25_GlyGCC_16_-_70812114_70812184@1.34.34,trna5_GlyGCC_17_+_8029064_8029134@1.34.34,trna68_GlyGCC_1_-_161493637_161493707@1.34.34\|GCATTGGTGGTTCAGTGGTAGAATTCTCGCCTGC | >am_trna128_GlyGCC_6_-_27870686_27870756 | 1 | 34 | 34 | GCATTGGTGGTTCAGTGGTAGAATTCTCGCCTGC | 888.3 | -5.9 | 1.4 | -4.3 | 2.0E-05 | 1.9E-04 | BT20/MB231 | NUC |
| >hg38_21_+_8433172_8446622_RNA45SN1_45S_50ntFlanks@12947.12988.42\|AGACGACCTGCTTCTGGGTCGGGGTTTCGTACGTAGCAGAGC | >hg38_21_+_8433172_8446622_RNA45SN1_45S_50ntFlanks | 12947 | 12988 | 42 | AGACGACCTGCTTCTGGGTCGGGGTTTCGTACGTAGCAGAGC | 309.4 | -3.7 | 0.9 | -4.3 | 1.8E-05 | 1.9E-04 | BT20/MB231 | CYTO |
| >hg38_21_+_8433172_8446622_RNA45SN1_45S_50ntFlanks@6645.6666.22\|CTCGTACGACTCTTAGCGGTGG | >hg38_21_+_8433172_8446622_RNA45SN1_45S_50ntFlanks | 6645 | 6666 | 22 | CTCGTACGACTCTTAGCGGTGG | 3392.5 | 3.3 | 0.8 | 4.0 | 5.4E-05 | 1.9E-04 | BT20/MB468 | MP |
| >MI0000289\|hsa-mir-181a-1&WithFlank&1\|-\|198859038\|198859159@70.89.20(+1U)&[MIMAT0000270&hsa-miR-181a-3p&offsets\|0\|-2(+1U);m-125&1\|-\|198859069\|198859090&offsets\|0\|-2(+1U)]\|ACCATCGACCGTTGATTGTAT | >MI0000289\|hsa-mir-181a-1&WithFlank&1\|-\|198859038\|198859159 | 70 | 89 | 20(+1U) | ACCATCGACCGTTGATTGTAT | 27.0 | 8.0 | 2.1 | 3.8 | 1.3E-04 | 1.9E-04 | MB231/MB468 | MP |
| >hg38_21_+_8433172_8446622_RNA45SN1_45S_50ntFlanks@7981.8006.26\|CTCAGATCAGACGTGGCGACCCGCTG | >hg38_21_+_8433172_8446622_RNA45SN1_45S_50ntFlanks | 7981 | 8006 | 26 | CTCAGATCAGACGTGGCGACCCGCTG | 5526.5 | -4.5 | 1.1 | -4.0 | 5.5E-05 | 1.9E-04 | BT20/MB468 | MP |
| >hg38_21_+_8433172_8446622_RNA45SN1_45S_50ntFlanks@12657.12674.18\|GACTGAACGCCTCTAAGT | >hg38_21_+_8433172_8446622_RNA45SN1_45S_50ntFlanks | 12657 | 12674 | 18 | GACTGAACGCCTCTAAGT | 331.2 | -5.3 | 1.4 | -3.8 | 1.3E-04 | 1.9E-04 | BT20/MB468 | CYTO |
| >hg38_21_+_8433172_8446622_RNA45SN1_45S_50ntFlanks@9623.9655.33\|TAGTAGCTGGTTCCCTCCGAAGTTTCCCTCAGG | >hg38_21_+_8433172_8446622_RNA45SN1_45S_50ntFlanks | 9623 | 9655 | 33 | TAGTAGCTGGTTCCCTCCGAAGTTTCCCTCAGG | 346.2 | -2.1 | 0.5 | -3.9 | 1.1E-04 | 1.9E-04 | BT20/MB468 | MITO |
| >hg38_21_+_8433172_8446622_RNA45SN1_45S_50ntFlanks@10820.10837.18\|GGGAAGTCGGCAAGCCGG | >hg38_21_+_8433172_8446622_RNA45SN1_45S_50ntFlanks | 10820 | 10837 | 18 | GGGAAGTCGGCAAGCCGG | 255.5 | -4.8 | 1.2 | -3.8 | 1.3E-04 | 1.9E-04 | MB231/MB468 | MP |
| >MI0000255\|hsa-mir-30d&WithFlank&8\|-\|134804870\|134804951@12.35.24(+1C)&[MIMAT0000245&hsa-miR-30d-5p&offsets\|0\|+2(+1C);m-21&8\|-\|134804919\|134804940&offsets\|0\|+2(+1C)]\|TGTAAACATCCCCGACTGGAAGCTC | >MI0000255\|hsa-mir-30d&WithFlank&8\|-\|134804870\|134804951 | 12 | 35 | 24(+1C) | TGTAAACATCCCCGACTGGAAGCTC | 204.2 | 4.6 | 1.2 | 3.8 | 1.3E-04 | 1.9E-04 | MB231/MB468 | MP |
| >hg38_21_+_8433172_8446622_RNA45SN1_45S_50ntFlanks@11452.11471.20\|CTCCTCGGCGCGCGGCGGCG | >hg38_21_+_8433172_8446622_RNA45SN1_45S_50ntFlanks | 11452 | 11471 | 20 | CTCCTCGGCGCGCGGCGGCG | 1196.5 | -7.3 | 1.9 | -3.8 | 1.5E-04 | 1.9E-04 | BT20/MB468 | NUC |
| >hg38_21_+_8433172_8446622_RNA45SN1_45S_50ntFlanks@11564.11611.48\|CCTAGCAGCCGACTTAGAACTGGTGCGGACCAGGGGAATCCGACTGTT | >hg38_21_+_8433172_8446622_RNA45SN1_45S_50ntFlanks | 11564 | 11611 | 48 | CCTAGCAGCCGACTTAGAACTGGTGCGGACCAGGGGAATCCGACTGTT | 444.5 | -6.6 | 1.7 | -3.8 | 1.3E-04 | 1.9E-04 | MB231/MB468 | MP |
| >hg38_21_+_8433172_8446622_RNA45SN1_45S_50ntFlanks@12932.12972.41\|CGCTAAACCATTCGTAGACGACCTGCTTCTGGGTCGGGGTT | >hg38_21_+_8433172_8446622_RNA45SN1_45S_50ntFlanks | 12932 | 12972 | 41 | CGCTAAACCATTCGTAGACGACCTGCTTCTGGGTCGGGGTT | 282.8 | -6.8 | 1.7 | -4.0 | 5.5E-05 | 1.9E-04 | BT20/MB468 | MP |
| >hg38_21_+_8433172_8446622_RNA45SN1_45S_50ntFlanks@10843.10872.30\|TAACTTCGGGATAAGGATTGGCTCTAAGGG | >hg38_21_+_8433172_8446622_RNA45SN1_45S_50ntFlanks | 10843 | 10872 | 30 | TAACTTCGGGATAAGGATTGGCTCTAAGGG | 232.4 | -3.4 | 0.9 | -3.8 | 1.4E-04 | 1.9E-04 | MB231/MB468 | CYTO |
| >hg38_21_+_8433172_8446622_RNA45SN1_45S_50ntFlanks@10388.10424.37\|GTGAACAGCAGTTGAACATGGGTCAGTCGGTCCTGAG | >hg38_21_+_8433172_8446622_RNA45SN1_45S_50ntFlanks | 10388 | 10424 | 37 | GTGAACAGCAGTTGAACATGGGTCAGTCGGTCCTGAG | 1588.3 | 2.7 | 0.7 | 3.8 | 1.5E-04 | 1.9E-04 | MB231/MB468 | NUC |
| >hg38_21_+_8433172_8446622_RNA45SN1_45S_50ntFlanks@7973.7994.22\|GACGCGACCTCAGATCAGACGT | >hg38_21_+_8433172_8446622_RNA45SN1_45S_50ntFlanks | 7973 | 7994 | 22 | GACGCGACCTCAGATCAGACGT | 396.6 | 2.2 | 0.6 | 3.8 | 1.5E-04 | 1.9E-04 | MB231/MB468 | NUC |
| >hg38_21_+_8433172_8446622_RNA45SN1_45S_50ntFlanks@6719.6739.21\|TGAATTGCAGGACACATTGAT | >hg38_21_+_8433172_8446622_RNA45SN1_45S_50ntFlanks | 6719 | 6739 | 21 | TGAATTGCAGGACACATTGAT | 280.3 | -2.8 | 0.7 | -3.9 | 1.1E-04 | 1.9E-04 | BT20/MB468 | MITO |
| >MI0000299\|hsa-mir-222&WithFlank&X\|-\|45747009\|45747130@75.97.23(+1A)&[MIMAT0000279&hsa-miR-222-3p&offsets\|0\|+2(+1A);m-140&X\|-\|45747033\|45747056&offsets\|0\|-1(+1A)]\|AGCTACATCTGGCTACTGGGTCTA | >MI0000299\|hsa-mir-222&WithFlank&X\|-\|45747009\|45747130 | 75 | 97 | 23(+1A) | AGCTACATCTGGCTACTGGGTCTA | 618.0 | 5.0 | 1.3 | 3.8 | 1.3E-04 | 2.0E-04 | MB231/MB468 | MP |
| >trna152_ValCAC_6_-_27248049_27248121@1.35.35\|GCTTCTGTAGTGTAGTGGTTATCACGTTCGCCTCA | >trna152_ValCAC_6_-_27248049_27248121 | 1 | 35 | 35 | GCTTCTGTAGTGTAGTGGTTATCACGTTCGCCTCA | 322.5 | 9.1 | 2.4 | 3.8 | 1.4E-04 | 2.0E-04 | MB231/MB468 | CYTO |
| >MI0000446\|hsa-mir-125b-1&WithFlank&11\|-\|122099751\|122099850@21.41.21;MI0000470\|hsa-mir-125b-2&WithFlank&21\|+\|16590231\|16590331@23.43.21&[MIMAT0000423&hsa-miR-125b-5p&offsets\|0\|-1;m-67&11\|-\|122099809\|122099830&offsets\|0\|-1];[MIMAT0000423_1&hsa-miR-125b-5p&offsets\|0\|-1;m-68&21\|+\|16590253\|16590274&offsets\|0\|-1]\|TCCCTGAGACCCTAACTTGTG | >MI0000446\|hsa-mir-125b-1&WithFlank&11\|-\|122099751\|122099850 | 21 | 41 | 21 | TCCCTGAGACCCTAACTTGTG | 7136.7 | -5.9 | 1.4 | -4.3 | 2.1E-05 | 2.0E-04 | BT20/MB231 | NUC |
| >hg38_21_+_8433172_8446622_RNA45SN1_45S_50ntFlanks@9534.9560.27\|AGGAAACTCTGGTGGAGGTCCGTAGCG | >hg38_21_+_8433172_8446622_RNA45SN1_45S_50ntFlanks | 9534 | 9560 | 27 | AGGAAACTCTGGTGGAGGTCCGTAGCG | 815.4 | -3.2 | 0.8 | -3.8 | 1.4E-04 | 2.0E-04 | MB231/MB468 | CYTO |
| >hg38_21_+_8433172_8446622_RNA45SN1_45S_50ntFlanks@12419.12448.30\|GATGTCGGCTCTTCCTATCATTGTGAAGCA | >hg38_21_+_8433172_8446622_RNA45SN1_45S_50ntFlanks | 12419 | 12448 | 30 | GATGTCGGCTCTTCCTATCATTGTGAAGCA | 912.7 | 3.6 | 1.0 | 3.8 | 1.5E-04 | 2.0E-04 | MB231/MB468 | NUC |
| >hg38_21_+_8433172_8446622_RNA45SN1_45S_50ntFlanks@12456.12488.33\|CCAAGCGTTGGATTGTTCACCCACTAATAGGGA | >hg38_21_+_8433172_8446622_RNA45SN1_45S_50ntFlanks | 12456 | 12488 | 33 | CCAAGCGTTGGATTGTTCACCCACTAATAGGGA | 5350.0 | -7.4 | 1.8 | -4.0 | 5.5E-05 | 2.0E-04 | BT20/MB468 | MP |
| >trna111_HisGTG_1_-_147774845_147774916@-1G.30.31,trna118_HisGTG_1_-_145396881_145396952@-1G.30.31,trna16_HisGTG_1_+_146544773_146544844@-1G.30.31,trna1_HisGTG_15_+_45493349_45493420@-1G.30.31,trna21_HisGTG_1_+_147753471_147753542@-1G.30.31,trna33_HisGTG_6_+_27125906_27125977@-1G.30.31,trna7_HisGTG_9_-_14433938_14434009@-1G.30.31,trna8_HisGTG_15_-_45492611_45492682@-1G.30.31,trna9_HisGTG_15_-_45490804_45490875@-1G.30.31\|GGCCGTGATCGTATAGTGGTTAGTACTCTGC | >trna111_HisGTG_1_-_147774845_147774916 | -1G | 30 | 31 | GGCCGTGATCGTATAGTGGTTAGTACTCTGC | 1190.0 | -3.9 | 0.9 | -4.3 | 2.1E-05 | 2.0E-04 | BT20/MB231 | NUC |
| >hg38_21_+_8433172_8446622_RNA45SN1_45S_50ntFlanks@10783.10801.19\|GTGAACAGCCTCTGGCATG | >hg38_21_+_8433172_8446622_RNA45SN1_45S_50ntFlanks | 10783 | 10801 | 19 | GTGAACAGCCTCTGGCATG | 357.4 | -5.7 | 1.5 | -3.8 | 1.3E-04 | 2.0E-04 | BT20/MB468 | CYTO |
| >hg38_21_+_8433172_8446622_RNA45SN1_45S_50ntFlanks@10473.10491.19\|CTCGGCCGATCGAAAGGGA | >hg38_21_+_8433172_8446622_RNA45SN1_45S_50ntFlanks | 10473 | 10491 | 19 | CTCGGCCGATCGAAAGGGA | 417.9 | 3.6 | 0.9 | 4.0 | 5.5E-05 | 2.0E-04 | BT20/MB468 | MP |
| >hg38_21_+_8433172_8446622_RNA45SN1_45S_50ntFlanks@10786.10817.32\|AACAGCCTCTGGCATGTTGGAACAATGTAGGT | >hg38_21_+_8433172_8446622_RNA45SN1_45S_50ntFlanks | 10786 | 10817 | 32 | AACAGCCTCTGGCATGTTGGAACAATGTAGGT | 674.2 | -2.2 | 0.6 | -3.9 | 1.1E-04 | 2.0E-04 | BT20/MB468 | MITO |
| >hg38_21_+_8433172_8446622_RNA45SN1_45S_50ntFlanks@6651.6683.33\|CGACTCTTAGCGGTGGATCACTCGGCTCGTGCG | >hg38_21_+_8433172_8446622_RNA45SN1_45S_50ntFlanks | 6651 | 6683 | 33 | CGACTCTTAGCGGTGGATCACTCGGCTCGTGCG | 1336.7 | 3.8 | 1.0 | 3.8 | 1.3E-04 | 2.0E-04 | BT20/MB468 | CYTO |
| >hg38_21_+_8433172_8446622_RNA45SN1_45S_50ntFlanks@6732.6770.39\|ACATTGATCATCGACACTTCGAACGCACTTGCGGCCCCG | >hg38_21_+_8433172_8446622_RNA45SN1_45S_50ntFlanks | 6732 | 6770 | 39 | ACATTGATCATCGACACTTCGAACGCACTTGCGGCCCCG | 559.8 | -4.2 | 1.1 | -3.8 | 1.3E-04 | 2.0E-04 | BT20/MB468 | CYTO |
| >MI0000273\|hsa-mir-183&WithFlank&7\|-\|129774899\|129775020@34.55.22(+1C)&[MIMAT0000261&hsa-miR-183-5p&offsets\|+1\|+1(+1C);m-41&7\|-\|129774966\|129774987&offsets\|0\|0(+1C)]\|ATGGCACTGGTAGAATTCACTGC | >MI0000273\|hsa-mir-183&WithFlank&7\|-\|129774899\|129775020 | 34 | 55 | 22(+1C) | ATGGCACTGGTAGAATTCACTGC | 298.3 | 3.4 | 0.8 | 4.3 | 1.8E-05 | 2.0E-04 | BT20/MB231 | CYTO |

<table>
<tr><td>>trna10_AspGTC_12_-_125424193_125424264@32.75.44,trna12_AspGTC_12_-_125411891_125411962@32.75.44,trna144_AspGTC_6_-_27551236_27551307@32.75.44,trna38_AspGTC_17_-_8125556_8125627@32.75.44,trna45_AspGTC_6_+_27447453_27447524@32.75.44,trna48_AspGTC_6_+_27471523_27471594@32.75.44,trna4_AspGTC_12_+_96429799_96429870@32.75.44,trna69_AspGTC_1_-_161440205_161440276@32.75.44,trna72_AspGTC_1_-_161432824_161432895@32.75.44,trna75_AspGTC_1_-_161425414_161425485@32.75.44,trna78_AspGTC_1_-_161418033_161418104@32.75.44,trna81_AspGTC_1_-_161410615_161410686@32.75.44|CTGTCACGCGGGAGACCGGGGTTCGATTCCCCGACGGGGAGCCA</td><td>>trna10_AspGTC_12_-_125424193_125424264</td><td>32</td><td>75</td><td>44</td><td>CTGTCACGCGGGAGACCGGGGTTCGATTCCCCGACGGGGAGCCA</td><td>212.8</td><td>-5.5</td><td>1.4</td><td>-3.8</td><td>1.3E-04</td><td>2.0E-04</td><td>BT20/MB468</td><td>CYTO</td></tr>
<tr><td>>hg38_21_+_8433172_8446622_RNA45SN1_45S_50ntFlanks@6647.6677.31|CGTACGACTCTTAGCGGTGGATCACTCGGCT</td><td>>hg38_21_+_8433172_8446622_RNA45SN1_45S_50ntFlanks</td><td>6647</td><td>6677</td><td>31</td><td>CGTACGACTCTTAGCGGTGGATCACTCGGCT</td><td>739.9</td><td>3.1</td><td>0.8</td><td>4.0</td><td>5.5E-05</td><td>2.0E-04</td><td>BT20/MB468</td><td>MP</td></tr>
<tr><td>>MI0001733|hsa-mir-452&WithFlank&X|-|151959622|151959718@20.41.22&[MIMAT0001635&hsa-miR-452-5p&offsets|0|0;m-188&X|-|151959677|151959699&offsets|0|-1]|AACTGTTTGCAGAGGAAACTGA</td><td>>MI0001733|hsa-mir-452&WithFlank&X|-|151959622|151959718</td><td>20</td><td>41</td><td>22</td><td>AACTGTTTGCAGAGGAAACTGA</td><td>417.1</td><td>5.1</td><td>1.2</td><td>4.3</td><td>2.1E-05</td><td>2.0E-04</td><td>BT20/MB231</td><td>NUC</td></tr>
<tr><td>>hg38_21_+_8433172_8446622_RNA45SN1_45S_50ntFlanks@4727.4758.32|AATCAAGAACGAAAGTCGGAGGTTCGAAGACG</td><td>>hg38_21_+_8433172_8446622_RNA45SN1_45S_50ntFlanks</td><td>4727</td><td>4758</td><td>32</td><td>AATCAAGAACGAAAGTCGGAGGTTCGAAGACG</td><td>721.2</td><td>-1.8</td><td>0.5</td><td>-3.8</td><td>1.5E-04</td><td>2.0E-04</td><td>MB231/MB468</td><td>NUC</td></tr>
<tr><td>>MI0000063|hsa-let-7b&WithFlank&22|+|46113680|46113774@66.86.21&[MIMAT0004482&hsa-let-7b-3p&offsets|0|-1;m-171&22|+|46113745|46113766&offsets|0|-1]|CTATACAACCTACTGCCTTCC</td><td>>MI0000063|hsa-let-7b&WithFlank&22|+|46113680|46113774</td><td>66</td><td>86</td><td>21</td><td>CTATACAACCTACTGCCTTCC</td><td>161.6</td><td>8.0</td><td>2.0</td><td>4.0</td><td>5.6E-05</td><td>2.0E-04</td><td>BT20/MB468</td><td>MP</td></tr>
<tr><td>>hg38_21_+_8433172_8446622_RNA45SN1_45S_50ntFlanks@7985.8003.19|GATCAGACGTGGCGACCCG</td><td>>hg38_21_+_8433172_8446622_RNA45SN1_45S_50ntFlanks</td><td>7985</td><td>8003</td><td>19</td><td>GATCAGACGTGGCGACCCG</td><td>301.6</td><td>-5.6</td><td>1.5</td><td>-3.8</td><td>1.5E-04</td><td>2.0E-04</td><td>MB231/MB468</td><td>NUC</td></tr>
<tr><td>>hg38_21_+_8433172_8446622_RNA45SN1_45S_50ntFlanks@12997.13014.18|CGCTGCGATCTATTGAAA</td><td>>hg38_21_+_8433172_8446622_RNA45SN1_45S_50ntFlanks</td><td>12997</td><td>13014</td><td>18</td><td>CGCTGCGATCTATTGAAA</td><td>3770.4</td><td>2.5</td><td>0.7</td><td>3.8</td><td>1.5E-04</td><td>2.0E-04</td><td>MB231/MB468</td><td>NUC</td></tr>
<tr><td>>hg38_21_+_8433172_8446622_RNA45SN1_45S_50ntFlanks@485.520.36|GATGTGGTGACGTCGTGCTCTCCCGGGCCGGGTCCG</td><td>>hg38_21_+_8433172_8446622_RNA45SN1_45S_50ntFlanks</td><td>485</td><td>520</td><td>36</td><td>GATGTGGTGACGTCGTGCTCTCCCGGGCCGGGTCCG</td><td>112.2</td><td>10.6</td><td>2.6</td><td>4.0</td><td>5.6E-05</td><td>2.0E-04</td><td>BT20/MB468</td><td>MP</td></tr>
<tr><td>>hg38_21_+_8433172_8446622_RNA45SN1_45S_50ntFlanks@9923.9951.29|TTGGTTGATATAGACAGCAGGACGGTGGC</td><td>>hg38_21_+_8433172_8446622_RNA45SN1_45S_50ntFlanks</td><td>9923</td><td>9951</td><td>29</td><td>TTGGTTGATATAGACAGCAGGACGGTGGC</td><td>60.2</td><td>9.2</td><td>2.2</td><td>4.3</td><td>2.0E-05</td><td>2.0E-04</td><td>BT20/MB231</td><td>MP</td></tr>
<tr><td>>MI0000070|hsa-mir-16-1&WithFlank&13|-|50048967|50049067@20.39.20;MI0000115|hsa-mir-16-2&WithFlank&3|+|160404739|160404831@16.35.20&[MIMAT0000069&hsa-miR-16-5p&offsets|0|-2;m-62&13|-|50049027|50049048&offsets|0|-2];[MIMAT0000069_1&hsa-miR-16-5p&offsets|0|-2;m-61&3|+|160404754|160404775&offsets|0|-2]|TAGCAGCACGTAAATATTGG</td><td>>MI0000070|hsa-mir-16-1&WithFlank&13|-|50048967|50049067</td><td>20</td><td>39</td><td>20</td><td>TAGCAGCACGTAAATATTGG</td><td>580.8</td><td>3.8</td><td>1.0</td><td>3.8</td><td>1.3E-04</td><td>2.0E-04</td><td>MB231/MB468</td><td>MP</td></tr>
<tr><td>>hg38_1_-_228634819_228635039_RNA5S12_5S_50ntFlanks@51.71.21|GTCTACGGCCATACCACCCTG</td><td>>hg38_1_-_228634819_228635039_RNA5S12_5S_50ntFlanks</td><td>51</td><td>71</td><td>21</td><td>GTCTACGGCCATACCACCCTG</td><td>752.8</td><td>-3.2</td><td>0.8</td><td>-3.8</td><td>1.5E-04</td><td>2.0E-04</td><td>MB231/MB468</td><td>MITO</td></tr>
<tr><td>>trnaMT_ValTAC_MT_+_1602_1670@51.72.22|CTTAACTTGACCGCTCTGACCA</td><td>>trnaMT_ValTAC_MT_+_1602_1670</td><td>51</td><td>72</td><td>22</td><td>CTTAACTTGACCGCTCTGACCA</td><td>8404.8</td><td>-3.8</td><td>1.0</td><td>-3.8</td><td>1.5E-04</td><td>2.0E-04</td><td>MB231/MB468</td><td>MITO</td></tr>
<tr><td>>hg38_21_+_8433172_8446622_RNA45SN1_45S_50ntFlanks@9806.9830.25|CGTGGAATGCGAGTGCCTAGTGGGC</td><td>>hg38_21_+_8433172_8446622_RNA45SN1_45S_50ntFlanks</td><td>9806</td><td>9830</td><td>25</td><td>CGTGGAATGCGAGTGCCTAGTGGGC</td><td>1041.9</td><td>7.5</td><td>1.8</td><td>4.3</td><td>2.0E-05</td><td>2.0E-04</td><td>BT20/MB231</td><td>MP</td></tr>
<tr><td>>hg38_21_+_8433172_8446622_RNA45SN1_45S_50ntFlanks@11913.11951.39|AAGAGACATGAGAGGTGTAGAATAAGTGGGAGGCCCCCG</td><td>>hg38_21_+_8433172_8446622_RNA45SN1_45S_50ntFlanks</td><td>11913</td><td>11951</td><td>39</td><td>AAGAGACATGAGAGGTGTAGAATAAGTGGGAGGCCCCCG</td><td>178.5</td><td>-5.0</td><td>1.3</td><td>-3.8</td><td>1.3E-04</td><td>2.0E-04</td><td>BT20/MB468</td><td>CYTO</td></tr>
<tr><td>>hg38_21_+_8433172_8446622_RNA45SN1_45S_50ntFlanks@4898.4917.20|AAAGGAATTGACGGAAGGGC</td><td>>hg38_21_+_8433172_8446622_RNA45SN1_45S_50ntFlanks</td><td>4898</td><td>4917</td><td>20</td><td>AAAGGAATTGACGGAAGGGC</td><td>237.1</td><td>-6.9</td><td>1.7</td><td>-4.0</td><td>5.6E-05</td><td>2.0E-04</td><td>BT20/MB468</td><td>MP</td></tr>
<tr><td>>hg38_21_+_8433172_8446622_RNA45SN1_45S_50ntFlanks@12994.13015.22|CCTCGCTGCGATCTATTGAAAG</td><td>>hg38_21_+_8433172_8446622_RNA45SN1_45S_50ntFlanks</td><td>12994</td><td>13015</td><td>22</td><td>CCTCGCTGCGATCTATTGAAAG</td><td>530.0</td><td>-3.4</td><td>0.9</td><td>-3.8</td><td>1.5E-04</td><td>2.0E-04</td><td>MB231/MB468</td><td>MITO</td></tr>
<tr><td>>MI0000750|hsa-mir-26a-2&WithFlank&12|-|57824603|57824698@20.39.20;MI0000083|hsa-mir-26a-1&WithFlank&3|+|37969398|37969486@16.35.20&[MIMAT0000082&hsa-miR-26a-5p&offsets|0|-2;m-33&12|-|57824658|57824679&offsets|0|-2];[MIMAT0000082_1&hsa-miR-26a-5p&offsets|0|-2;m-34&3|+|37969413|37969434&offsets|0|-2]|TTCAAGTAATCCAGGATAGG</td><td>>MI0000750|hsa-mir-26a-2&WithFlank&12|-|57824603|57824698</td><td>20</td><td>39</td><td>20</td><td>TTCAAGTAATCCAGGATAGG</td><td>527.7</td><td>-3.3</td><td>0.8</td><td>-4.2</td><td>2.3E-05</td><td>2.0E-04</td><td>BT20/MB231</td><td>MITO</td></tr>
<tr><td>>am_trnaMT_ValTAC_MT_+_1602_1670@29.62.34|ACTTACACTTAGGAGATTTCAACTTAACTTGACC</td><td>>am_trnaMT_ValTAC_MT_+_1602_1670</td><td>29</td><td>62</td><td>34</td><td>ACTTACACTTAGGAGATTTCAACTTAACTTGACC</td><td>535.2</td><td>7.7</td><td>1.9</td><td>4.0</td><td>5.6E-05</td><td>2.0E-04</td><td>BT20/MB468</td><td>MP</td></tr>
<tr><td>>hg38_1_-_228634819_228635039_RNA5S12_5S_50ntFlanks@140.169.30|ACCGCCTGGGAATACCGGGTGCTGTAGGCT</td><td>>hg38_1_-_228634819_228635039_RNA5S12_5S_50ntFlanks</td><td>140</td><td>169</td><td>30</td><td>ACCGCCTGGGAATACCGGGTGCTGTAGGCT</td><td>5110.1</td><td>3.3</td><td>0.9</td><td>3.8</td><td>1.4E-04</td><td>2.0E-04</td><td>MB231/MB468</td><td>CYTO</td></tr>
<tr><td>>hg38_21_+_8433172_8446622_RNA45SN1_45S_50ntFlanks@8341.8378.38|CCGTAAGGGAAAGTTGAAAAGAACTTTGAAGAGAGAGT</td><td>>hg38_21_+_8433172_8446622_RNA45SN1_45S_50ntFlanks</td><td>8341</td><td>8378</td><td>38</td><td>CCGTAAGGGAAAGTTGAAAAGAACTTTGAAGAGAGAGT</td><td>71.0</td><td>9.5</td><td>2.2</td><td>4.3</td><td>2.0E-05</td><td>2.0E-04</td><td>BT20/MB231</td><td>MP</td></tr>
<tr><td>>hg38_21_+_8433172_8446622_RNA45SN1_45S_50ntFlanks@7990.8010.21|GACGTGGCGACCCGCTGAATT</td><td>>hg38_21_+_8433172_8446622_RNA45SN1_45S_50ntFlanks</td><td>7990</td><td>8010</td><td>21</td><td>GACGTGGCGACCCGCTGAATT</td><td>746.3</td><td>3.2</td><td>0.8</td><td>3.9</td><td>1.1E-04</td><td>2.0E-04</td><td>BT20/MB468</td><td>MITO</td></tr>
<tr><td>>hg38_21_+_8433172_8446622_RNA45SN1_45S_50ntFlanks@9934.9952.19|AGACAGCAGGACGGTGGCC</td><td>>hg38_21_+_8433172_8446622_RNA45SN1_45S_50ntFlanks</td><td>9934</td><td>9952</td><td>19</td><td>AGACAGCAGGACGGTGGCC</td><td>102.6</td><td>7.1</td><td>1.8</td><td>4.0</td><td>5.6E-05</td><td>2.0E-04</td><td>BT20/MB468</td><td>MP</td></tr>
<tr><td>>trna20_GluTTC_2_-_131094701_131094772@1.36.36,trna5_GluTTC_13_-_41634874_41634945@1.36.36|TCCCATATGGTCTAGCGGTTAGGATTCCTGGTTTTC</td><td>>trna20_GluTTC_2_-_131094701_131094772</td><td>1</td><td>36</td><td>36</td><td>TCCCATATGGTCTAGCGGTTAGGATTCCTGGTTTTC</td><td>448.1</td><td>-4.7</td><td>1.1</td><td>-4.2</td><td>2.3E-05</td><td>2.0E-04</td><td>BT20/MB231</td><td>MITO</td></tr>
<tr><td>>trnaMT_SerGCT_MT_+_12207_12265@1.35.35|GAGAAAGCTCACAAGAACTGCTAACTCATGCCCCC</td><td>>trnaMT_SerGCT_MT_+_12207_12265</td><td>1</td><td>35</td><td>35</td><td>GAGAAAGCTCACAAGAACTGCTAACTCATGCCCCC</td><td>194.8</td><td>4.0</td><td>1.1</td><td>3.8</td><td>1.5E-04</td><td>2.0E-04</td><td>MB231/MB468</td><td>MITO</td></tr>
<tr><td>>hg38_21_+_8433172_8446622_RNA45SN1_45S_50ntFlanks@11667.11709.43|CTGCCCAGTGCTCTGAATGTCAAAGTGAAGAAATTCAATGAAG</td><td>>hg38_21_+_8433172_8446622_RNA45SN1_45S_50ntFlanks</td><td>11667</td><td>11709</td><td>43</td><td>CTGCCCAGTGCTCTGAATGTCAAAGTGAAGAAATTCAATGAAG</td><td>199.2</td><td>-4.0</td><td>0.9</td><td>-4.3</td><td>1.9E-05</td><td>2.0E-04</td><td>BT20/MB231</td><td>CYTO</td></tr>
<tr><td>>trna108_AlaAGC_6_-_28763741_28763812@58.75.18|TCCCCAGTACCTCCACCA</td><td>>trna108_AlaAGC_6_-_28763741_28763812</td><td>58</td><td>75</td><td>18</td><td>TCCCCAGTACCTCCACCA</td><td>503.0</td><td>4.0</td><td>1.1</td><td>3.8</td><td>1.5E-04</td><td>2.0E-04</td><td>MB231/MB468</td><td>NUC</td></tr>
<tr><td>>hg38_21_+_8433172_8446622_RNA45SN1_45S_50ntFlanks@12408.12443.36|TTTGATCCTTCGATGTCGGCTCTTCCTATCATTGTG</td><td>>hg38_21_+_8433172_8446622_RNA45SN1_45S_50ntFlanks</td><td>12408</td><td>12443</td><td>36</td><td>TTTGATCCTTCGATGTCGGCTCTTCCTATCATTGTG</td><td>866.5</td><td>-2.2</td><td>0.6</td><td>-3.8</td><td>1.5E-04</td><td>2.0E-04</td><td>MB231/MB468</td><td>NUC</td></tr>
<tr><td>>hg38_21_+_8433172_8446622_RNA45SN1_45S_50ntFlanks@10350.10375.26|ACGAGAACTTTGAAGGCCGAAGTGGA</td><td>>hg38_21_+_8433172_8446622_RNA45SN1_45S_50ntFlanks</td><td>10350</td><td>10375</td><td>26</td><td>ACGAGAACTTTGAAGGCCGAAGTGGA</td><td>150.8</td><td>-2.9</td><td>0.8</td><td>-3.8</td><td>1.5E-04</td><td>2.0E-04</td><td>MB231/MB468</td><td>NUC</td></tr>
<tr><td>>hg38_21_+_8433172_8446622_RNA45SN1_45S_50ntFlanks@10805.10828.24|GAACAATGTAGGTAAGGGAAGTCG</td><td>>hg38_21_+_8433172_8446622_RNA45SN1_45S_50ntFlanks</td><td>10805</td><td>10828</td><td>24</td><td>GAACAATGTAGGTAAGGGAAGTCG</td><td>156.2</td><td>-4.9</td><td>1.3</td><td>-3.8</td><td>1.4E-04</td><td>2.0E-04</td><td>BT20/MB468</td><td>CYTO</td></tr>
<tr><td>>hg38_21_+_8433172_8446622_RNA45SN1_45S_50ntFlanks@10473.10491.19|CTCGGCCGATCGAAAGGGA</td><td>>hg38_21_+_8433172_8446622_RNA45SN1_45S_50ntFlanks</td><td>10473</td><td>10491</td><td>19</td><td>CTCGGCCGATCGAAAGGGA</td><td>560.2</td><td>6.4</td><td>1.5</td><td>4.3</td><td>2.0E-05</td><td>2.0E-04</td><td>BT20/MB231</td><td>MP</td></tr>
<tr><td>>hg38_21_+_8433172_8446622_RNA45SN1_45S_50ntFlanks@10771.10792.22|CAGGTCTCCAAGGTGAACAGCC</td><td>>hg38_21_+_8433172_8446622_RNA45SN1_45S_50ntFlanks</td><td>10771</td><td>10792</td><td>22</td><td>CAGGTCTCCAAGGTGAACAGCC</td><td>187.4</td><td>-4.4</td><td>1.2</td><td>-3.8</td><td>1.5E-04</td><td>2.0E-04</td><td>MB231/MB468</td><td>NUC</td></tr>
<tr><td>>hg38_21_+_8433172_8446622_RNA45SN1_45S_50ntFlanks@10716.10735.20|GAAAATCCGGGGGAGAGGGT</td><td>>hg38_21_+_8433172_8446622_RNA45SN1_45S_50ntFlanks</td><td>10716</td><td>10735</td><td>20</td><td>GAAAATCCGGGGGAGAGGGT</td><td>238.5</td><td>-5.0</td><td>1.3</td><td>-3.8</td><td>1.3E-04</td><td>2.0E-04</td><td>MB231/MB468</td><td>MP</td></tr>
<tr><td>>hg38_21_+_8433172_8446622_RNA45SN1_45S_50ntFlanks@9034.9059.26|TCGTGGGGGGGCCGGGCCACCCCTCC</td><td>>hg38_21_+_8433172_8446622_RNA45SN1_45S_50ntFlanks</td><td>9034</td><td>9059</td><td>26</td><td>TCGTGGGGGGGCCGGGCCACCCCTCC</td><td>187.4</td><td>-5.4</td><td>1.4</td><td>-3.9</td><td>1.1E-04</td><td>2.0E-04</td><td>BT20/MB468</td><td>MITO</td></tr>
<tr><td>>trnaMT_HisGTG_MT_+_12138_12206@26.66.41|AGATTGTGAATCTGACAACAGAGGCTTACGACCCCTTATTT</td><td>>trnaMT_HisGTG_MT_+_12138_12206</td><td>26</td><td>66</td><td>41</td><td>AGATTGTGAATCTGACAACAGAGGCTTACGACCCCTTATTT</td><td>107.8</td><td>10.5</td><td>2.6</td><td>4.0</td><td>5.7E-05</td><td>2.0E-04</td><td>BT20/MB468</td><td>MP</td></tr>
<tr><td>>trna112_GlnCTG_1_-_147737382_147737453@30.50.21,trna131_GlnCTG_6_-_27759135_27759206@30.50.21,trna146_GlnCTG_6_-_27515531_27515602@30.50.21,trna15_GlnCTG_1_+_145963304_145963375@30.50.21,trna19_GlnCTG_1_+_147505038_147505109@30.50.21,trna1_GlnCTG_6_+_18836402_18836473@30.50.21,trna28_GlnCTG_1_+_149186125_149186196@30.50.21,trna3_GlnCTG_17_+_8023070_8023141@30.50.21,trna49_GlnCTG_6_+_27487308_27487379@30.50.21,trna7_GlnCTG_15_-_66161400_66161471@30.50.21,trna99_GlnCTG_6_-_28909378_28909449@30.50.21|GACTCTGAATCCAGCGATCCG</td><td>>trna112_GlnCTG_1_-_147737382_147737453</td><td>30</td><td>50</td><td>21</td><td>GACTCTGAATCCAGCGATCCG</td><td>288.2</td><td>-5.5</td><td>1.4</td><td>-3.9</td><td>1.1E-04</td><td>2.0E-04</td><td>BT20/MB468</td><td>MITO</td></tr>
<tr><td>>hg38_21_+_8433172_8446622_RNA45SN1_45S_50ntFlanks@12640.12659.20|ACCATCTGTGGGATTATGAC</td><td>>hg38_21_+_8433172_8446622_RNA45SN1_45S_50ntFlanks</td><td>12640</td><td>12659</td><td>20</td><td>ACCATCTGTGGGATTATGAC</td><td>1053.3</td><td>-3.1</td><td>0.8</td><td>-3.9</td><td>1.1E-04</td><td>2.0E-04</td><td>BT20/MB468</td><td>MITO</td></tr>
<tr><td>>hg38_21_+_8433172_8446622_RNA45SN1_45S_50ntFlanks@12638.12659.22|CTACCATCTGTGGGATTATGAC</td><td>>hg38_21_+_8433172_8446622_RNA45SN1_45S_50ntFlanks</td><td>12638</td><td>12659</td><td>22</td><td>CTACCATCTGTGGGATTATGAC</td><td>219.0</td><td>-3.0</td><td>0.8</td><td>-3.8</td><td>1.5E-04</td><td>2.0E-04</td><td>MB231/MB468</td><td>NUC</td></tr>
<tr><td>>hg38_21_+_8433172_8446622_RNA45SN1_45S_50ntFlanks@6645.6662.18|CTCGTACGACTCTTAGCG</td><td>>hg38_21_+_8433172_8446622_RNA45SN1_45S_50ntFlanks</td><td>6645</td><td>6662</td><td>18</td><td>CTCGTACGACTCTTAGCG</td><td>5963.1</td><td>3.3</td><td>0.9</td><td>3.8</td><td>1.5E-04</td><td>2.0E-04</td><td>MB231/MB468</td><td>NUC</td></tr>
<tr><td>>trna14_LysTTT_11_-_59327808_59327880@3.36.34,trna2_LysTTT_17_+_8022473_8022545@3.36.34,trna54_LysTTT_1_+_204475655_204475727@3.36.34,trna5_LysTTT_11_+_59323902_59323974@3.36.34,trna62_LysTTT_1_-_204476158_204476230@3.36.34,trna76_LysTTT_6_+_28918806_28918878@3.36.34|CCGGATAGCTCAGTCGGTAGAGCATCAGACTTTT</td><td>>trna14_LysTTT_11_-_59327808_59327880</td><td>3</td><td>36</td><td>34</td><td>CCGGATAGCTCAGTCGGTAGAGCATCAGACTTTT</td><td>303.4</td><td>6.0</td><td>1.6</td><td>3.8</td><td>1.5E-04</td><td>2.0E-04</td><td>MB231/MB468</td><td>MITO</td></tr>
<tr><td>>hg38_21_+_8433172_8446622_RNA45SN1_45S_50ntFlanks@11207.11254.48|GGCGGCGACTCTGGACGCGAGCCGGGCCCTTCCCGTGGATCGCCCCAG</td><td>>hg38_21_+_8433172_8446622_RNA45SN1_45S_50ntFlanks</td><td>11207</td><td>11254</td><td>48</td><td>GGCGGCGACTCTGGACGCGAGCCGGGCCCTTCCCGTGGATCGCCCCAG</td><td>305.0</td><td>-4.8</td><td>1.2</td><td>-3.8</td><td>1.4E-04</td><td>2.0E-04</td><td>BT20/MB468</td><td>CYTO</td></tr>
<tr><td>>hg38_21_+_8433172_8446622_RNA45SN1_45S_50ntFlanks@10344.10388.45|ATTCAAACGAGAACTTTGAAGGCCGAAGTGGAGAAGGGTTCCATG</td><td>>hg38_21_+_8433172_8446622_RNA45SN1_45S_50ntFlanks</td><td>10344</td><td>10388</td><td>45</td><td>ATTCAAACGAGAACTTTGAAGGCCGAAGTGGAGAAGGGTTCCATG</td><td>429.2</td><td>4.6</td><td>1.1</td><td>4.2</td><td>2.3E-05</td><td>2.0E-04</td><td>BT20/MB231</td><td>MITO</td></tr>
<tr><td>>MI0000650|hsa-mir-200c&WithFlank&12|+|6963693|6963772@50.70.21&[MIMAT0000617&hsa-miR-200c-3p&offsets|0|-2;m-54&12|+|6963742|6963764&offsets|0|-2]|TAATACTGCCGGGTAATGATG</td><td>>MI0000650|hsa-mir-200c&WithFlank&12|+|6963693|6963772</td><td>50</td><td>70</td><td>21</td><td>TAATACTGCCGGGTAATGATG</td><td>4158.7</td><td>9.5</td><td>2.2</td><td>4.3</td><td>1.9E-05</td><td>2.0E-04</td><td>BT20/MB231</td><td>CYTO</td></tr>
</table>

| | | | | | | | | | | | | | |
|---|---|---|---|---|---|---|---|---|---|---|---|---|---|
| >MI0003776\|hsa-mir-320b-1&WithFlank&1\|+\|116671740\|116671823@48.67.20(+2U);MI0003839\|hsa-mir-320b-2&WithFlank&1\|-\|224257034\|224257116@47.66.20(+2U);MI0000542\|hsa-mir-320a&WithFlank&8\|-\|22244960\|22245043@48.67.20(+2U)&[MIMAT0005792&hsa-miR-320b&offsets\|0\|-2(+2U);m-183&1\|+\|116671787\|116671806&offsets\|0\|0(+2U)];[MIMAT0005792_1&hsa-miR-320b&offsets\|0\|-2(+2U);m-183&1\|-\|224257051\|224257070&offsets\|0\|0(+2U)];[MIMAT0000510&hsa-miR-320a-3p&offsets\|0\|-2(+2U);m-60&8\|-\|22244975\|22244996&offsets\|0\|-2(+2U)]\|AAAAGCTGGGTTGAGAGGGCTT | >MI0003776\|hsa-mir-320b-1&WithFlank&1\|+\|116671740\|116671823 | 48 | 67 | 20(+2U) | AAAAGCTGGGTTGAGAGGGCTT | 227.0 | 3.9 | 0.9 | 4.3 | 1.9E-05 | 2.0E-04 | BT20/MB231 | CYTO |
| >hg38_21_+_8433172_8446622_RNA45SN1_45S_50ntFlanks@12633.12661.29\|CGAAGCTACCATCTGTGGGATTATGACTG | >hg38_21_+_8433172_8446622_RNA45SN1_45S_50ntFlanks | 12633 | 12661 | 29 | CGAAGCTACCATCTGTGGGATTATGACTG | 535.7 | 8.5 | 2.0 | 4.3 | 2.0E-05 | 2.0E-04 | BT20/MB231 | MP |
| >trnaMT_GluTTC_MT_-_14674_14742@39.61.23,trnalookalike8_GluTTC_5_-_93905172_93905240@39.61.23\|ATTGGTCGTGGTTGTAGTCCGTG | >trnaMT_GluTTC_MT_-_14674_14742 | 39 | 61 | 23 | ATTGGTCGTGGTTGTAGTCCGTG | 184.7 | 3.5 | 0.9 | 3.8 | 1.5E-04 | 2.0E-04 | MB231/MB468 | MITO |
| >hg38_21_+_8433172_8446622_RNA45SN1_45S_50ntFlanks@10407.10424.18\|GGGTCAGTCGGTCCTGAG | >hg38_21_+_8433172_8446622_RNA45SN1_45S_50ntFlanks | 10407 | 10424 | 18 | GGGTCAGTCGGTCCTGAG | 605.8 | 3.6 | 0.8 | 4.2 | 2.3E-05 | 2.0E-04 | BT20/MB231 | MITO |
| >hg38_21_+_8433172_8446622_RNA45SN1_45S_50ntFlanks@474.507.34\|TCTTCGTGATCGATGTGGTGACGTCGTGCTCTCC | >hg38_21_+_8433172_8446622_RNA45SN1_45S_50ntFlanks | 474 | 507 | 34 | TCTTCGTGATCGATGTGGTGACGTCGTGCTCTCC | 223.1 | 2.8 | 0.7 | 3.8 | 1.6E-04 | 2.0E-04 | MB231/MB468 | NUC |
| >MI0000115\|hsa-mir-16-2&WithFlank&3\|+\|160404739\|160404831@59.79.21&[MIMAT0004518&hsa-miR-16-2-3p&offsets\|0\|-1;m-252&3\|+\|160404796\|160404816&offsets\|+1\|+1]\|CCAATATTACTGTGCTGCTTT | >MI0000115\|hsa-mir-16-2&WithFlank&3\|+\|160404739\|160404831 | 59 | 79 | 21 | CCAATATTACTGTGCTGCTTT | 133.5 | 5.8 | 1.5 | 3.8 | 1.5E-04 | 2.0E-04 | MB231/MB468 | MITO |
| >MI0000283\|hsa-mir-203a&WithFlank&14\|+\|104117399\|104117520@72.90.19&[MIMAT0000264&hsa-miR-203a-3p&offsets\|+1\|-2;m-79&14\|+\|104117469\|104117490&offsets\|+1\|-2]\|TGAAATGTTTAGGACCACT | >MI0000283\|hsa-mir-203a&WithFlank&14\|+\|104117399\|104117520 | 72 | 90 | 19 | TGAAATGTTTAGGACCACT | 314.3 | 6.6 | 1.6 | 4.0 | 5.7E-05 | 2.0E-04 | BT20/MB468 | MP |
| >hg38_21_+_8433172_8446622_RNA45SN1_45S_50ntFlanks@6646.6668.23\|TCGTACGACTCTTAGCGGTGGAT | >hg38_21_+_8433172_8446622_RNA45SN1_45S_50ntFlanks | 6646 | 6668 | 23 | TCGTACGACTCTTAGCGGTGGAT | 5419.2 | 2.7 | 0.7 | 4.0 | 5.7E-05 | 2.0E-04 | BT20/MB468 | MP |
| >hg38_21_+_8433172_8446622_RNA45SN1_45S_50ntFlanks@12002.12030.29\|TGCGGGCCGCCGGTGAAATACCACTACTC | >hg38_21_+_8433172_8446622_RNA45SN1_45S_50ntFlanks | 12002 | 12030 | 29 | TGCGGGCCGCCGGTGAAATACCACTACTC | 238.9 | -6.0 | 1.5 | -4.0 | 5.7E-05 | 2.0E-04 | BT20/MB468 | MP |
| >hg38_21_+_8433172_8446622_RNA45SN1_45S_50ntFlanks@10679.10710.32\|GGTTCCGGCGGCGTCCGGTGAGCTCTCGCTGG | >hg38_21_+_8433172_8446622_RNA45SN1_45S_50ntFlanks | 10679 | 10710 | 32 | GGTTCCGGCGGCGTCCGGTGAGCTCTCGCTGG | 1960.5 | 8.2 | 1.9 | 4.3 | 2.0E-05 | 2.0E-04 | BT20/MB231 | MP |
| >hg38_21_+_8433172_8446622_RNA45SN1_45S_50ntFlanks@10845.10862.18\|ACTTCGGGATAAGGATTG | >hg38_21_+_8433172_8446622_RNA45SN1_45S_50ntFlanks | 10845 | 10862 | 18 | ACTTCGGGATAAGGATTG | 228.9 | -6.0 | 1.5 | -4.0 | 5.7E-05 | 2.0E-04 | BT20/MB468 | MP |
| >hg38_21_+_8433172_8446622_RNA45SN1_45S_50ntFlanks@5444.5464.21\|CTCGGATCGGCCCCGCCGGGG | >hg38_21_+_8433172_8446622_RNA45SN1_45S_50ntFlanks | 5444 | 5464 | 21 | CTCGGATCGGCCCCGCCGGGG | 76.7 | -5.9 | 1.5 | -3.8 | 1.3E-04 | 2.0E-04 | MB231/MB468 | MP |
| >MI0006444\|hsa-mir-1307&WithFlank&10\|-\|103394247\|103394407@86.107.22(+1A)&[MIMAT0005951&hsa-miR-1307-3p&offsets\|0\|0(+1A);m-66&10\|-\|103394300\|103394322&offsets\|0\|-1(+1A)]\|ACTCGGCGTGGCGTCGGTCGTGA | >MI0006444\|hsa-mir-1307&WithFlank&10\|-\|103394247\|103394407 | 86 | 107 | 22(+1A) | ACTCGGCGTGGCGTCGGTCGTGA | 253.2 | 3.7 | 0.9 | 4.0 | 5.7E-05 | 2.0E-04 | BT20/MB468 | MP |
| >hg38_21_+_8433172_8446622_RNA45SN1_45S_50ntFlanks@6651.6685.35\|CGACTCTTAGCGGTGGATCACTCGGCTCGTGCGTC | >hg38_21_+_8433172_8446622_RNA45SN1_45S_50ntFlanks | 6651 | 6685 | 35 | CGACTCTTAGCGGTGGATCACTCGGCTCGTGCGTC | 8374.5 | 3.1 | 0.8 | 4.0 | 5.7E-05 | 2.0E-04 | BT20/MB468 | MP |
| >hg38_21_+_8433172_8446622_RNA45SN1_45S_50ntFlanks@12405.12441.37\|CTTTTTGATCCTTCGATGTCGGCTCTTCCTATCATTG | >hg38_21_+_8433172_8446622_RNA45SN1_45S_50ntFlanks | 12405 | 12441 | 37 | CTTTTTGATCCTTCGATGTCGGCTCTTCCTATCATTG | 2654.2 | -4.5 | 1.2 | -3.8 | 1.5E-04 | 2.0E-04 | MB231/MB468 | CYTO |
| >hg38_21_+_8433172_8446622_RNA45SN1_45S_50ntFlanks@9567.9594.28\|ACGTGCAAATCGGTCGTCCGACCTGGGT | >hg38_21_+_8433172_8446622_RNA45SN1_45S_50ntFlanks | 9567 | 9594 | 28 | ACGTGCAAATCGGTCGTCCGACCTGGGT | 580.9 | -3.3 | 0.8 | -4.0 | 5.7E-05 | 2.0E-04 | BT20/MB468 | MP |
| >hg38_21_+_8433172_8446622_RNA45SN1_45S_50ntFlanks@12420.12449.30\|ATGTCGGCTCTTCCTATCATTGTGAAGCAG | >hg38_21_+_8433172_8446622_RNA45SN1_45S_50ntFlanks | 12420 | 12449 | 30 | ATGTCGGCTCTTCCTATCATTGTGAAGCAG | 370.7 | -3.0 | 0.8 | -3.8 | 1.3E-04 | 2.0E-04 | MB231/MB468 | MP |
| >trna10_LysCTT_16_+_3241501_3241573@4.33.30,trna13_LysCTT_14_-_58706613_58706685@4.33.30,trna2_LysCTT_15_+_79152904_79152976@4.33.30\|CGGCTAGCTCAGTCGGTAGAGCATGGGACT | >trna10_LysCTT_16_+_3241501_3241573 | 4 | 33 | 30 | CGGCTAGCTCAGTCGGTAGAGCATGGGACT | 2403.1 | -3.3 | 0.8 | -4.2 | 2.3E-05 | 2.0E-04 | BT20/MB231 | MITO |
| >hg38_21_+_8433172_8446622_RNA45SN1_45S_50ntFlanks@11769.11804.36\|TAATTAGTGACGCGCATGAATGGATGAACGAGATTC | >hg38_21_+_8433172_8446622_RNA45SN1_45S_50ntFlanks | 11769 | 11804 | 36 | TAATTAGTGACGCGCATGAATGGATGAACGAGATTC | 266.2 | -6.1 | 1.6 | -3.8 | 1.3E-04 | 2.0E-04 | MB231/MB468 | MP |
| >hg38_21_+_8433172_8446622_RNA45SN1_45S_50ntFlanks@11727.11747.21\|AGTAACTATGACTCTCTTAAG | >hg38_21_+_8433172_8446622_RNA45SN1_45S_50ntFlanks | 11727 | 11747 | 21 | AGTAACTATGACTCTCTTAAG | 275.3 | -6.3 | 1.7 | -3.8 | 1.6E-04 | 2.0E-04 | BT20/MB468 | NUC |
| >hg38_21_+_8433172_8446622_RNA45SN1_45S_50ntFlanks@7975.8015.41\|CGCGACCTCAGATCAGACGTGGCGACCCGCTGAATTTAAGC | >hg38_21_+_8433172_8446622_RNA45SN1_45S_50ntFlanks | 7975 | 8015 | 41 | CGCGACCTCAGATCAGACGTGGCGACCCGCTGAATTTAAGC | 2437.0 | 4.8 | 1.2 | 4.0 | 5.7E-05 | 2.0E-04 | BT20/MB468 | MP |
| >MI0000076\|hsa-mir-20a&WithFlank&13\|+\|91351059\|91351141@14.36.23&[MIMAT0000075&hsa-miR-20a-5p&offsets\|0\|0;m-96&13\|+\|91351072\|91351094&offsets\|0\|0]\|TAAAGTGCTTATAGTGCAGGTAG | >MI0000076\|hsa-mir-20a&WithFlank&13\|+\|91351059\|91351141 | 14 | 36 | 23 | TAAAGTGCTTATAGTGCAGGTAG | 2720.7 | 3.4 | 0.9 | 3.9 | 1.1E-04 | 2.0E-04 | BT20/MB468 | MITO |
| >hg38_21_+_8433172_8446622_RNA45SN1_45S_50ntFlanks@10679.10706.28\|GGTTCCGGCGGCGTCCGGTGAGCTCTCG | >hg38_21_+_8433172_8446622_RNA45SN1_45S_50ntFlanks | 10679 | 10706 | 28 | GGTTCCGGCGGCGTCCGGTGAGCTCTCG | 3435.1 | -5.2 | 1.3 | -3.9 | 1.1E-04 | 2.0E-04 | BT20/MB468 | MITO |
| >trna16_ValTAC_11_-_59318460_59318532@1.30.30\|GGTTCCATAGTGTAGCGGTTATCACGTCTG | >trna16_ValTAC_11_-_59318460_59318532 | 1 | 30 | 30 | GGTTCCATAGTGTAGCGGTTATCACGTCTG | 269.6 | -5.2 | 1.4 | -3.8 | 1.4E-04 | 2.0E-04 | BT20/MB468 | CYTO |
| >hg38_21_+_8433172_8446622_RNA45SN1_45S_50ntFlanks@12288.12318.31\|ACCGTGAAAGCGGGGCCTCACGATCCTTCTG | >hg38_21_+_8433172_8446622_RNA45SN1_45S_50ntFlanks | 12288 | 12318 | 31 | ACCGTGAAAGCGGGGCCTCACGATCCTTCTG | 272.1 | -5.7 | 1.5 | -3.8 | 1.6E-04 | 2.0E-04 | MB231/MB468 | NUC |
| >hg38_21_+_8433172_8446622_RNA45SN1_45S_50ntFlanks@10829.10858.30\|GCAAGCCGGATCCGTAACTTCGGGATAAGG | >hg38_21_+_8433172_8446622_RNA45SN1_45S_50ntFlanks | 10829 | 10858 | 30 | GCAAGCCGGATCCGTAACTTCGGGATAAGG | 400.2 | -5.7 | 1.5 | -3.8 | 1.4E-04 | 2.0E-04 | BT20/MB468 | CYTO |
| >hg38_21_+_8433172_8446622_RNA45SN1_45S_50ntFlanks@11785.11806.22\|TGAATGGATGAACGAGATTCCC | >hg38_21_+_8433172_8446622_RNA45SN1_45S_50ntFlanks | 11785 | 11806 | 22 | TGAATGGATGAACGAGATTCCC | 645.4 | -6.5 | 1.7 | -3.8 | 1.4E-04 | 2.0E-04 | BT20/MB468 | CYTO |
| >trnaMT_PheGAA_MT_+_577_647@32.59.28\|ACTGAAAATGTTTAGACGGGCTCACATC | >trnaMT_PheGAA_MT_+_577_647 | 32 | 59 | 28 | ACTGAAAATGTTTAGACGGGCTCACATC | 117.0 | 10.6 | 2.6 | 4.0 | 5.7E-05 | 2.0E-04 | BT20/MB468 | MP |
| >hg38_21_+_8433172_8446622_RNA45SN1_45S_50ntFlanks@11579.11609.31\|AGAACTGGTGCGGACCAGGGGAATCCGACTG | >hg38_21_+_8433172_8446622_RNA45SN1_45S_50ntFlanks | 11579 | 11609 | 31 | AGAACTGGTGCGGACCAGGGGAATCCGACTG | 304.9 | -1.5 | 0.4 | -3.8 | 1.4E-04 | 2.0E-04 | BT20/MB468 | CYTO |
| >hg38_21_+_8433172_8446622_RNA45SN1_45S_50ntFlanks@3854.3888.35\|ACTGTGGTAATTCTAGAGCTAATACATGCCGACGG | >hg38_21_+_8433172_8446622_RNA45SN1_45S_50ntFlanks | 3854 | 3888 | 35 | ACTGTGGTAATTCTAGAGCTAATACATGCCGACGG | 176.1 | -4.0 | 1.0 | -3.8 | 1.6E-04 | 2.0E-04 | MB231/MB468 | NUC |
| >hg38_21_+_8433172_8446622_RNA45SN1_45S_50ntFlanks@12712.12744.33\|CGGAGCCTCGGTTGGCCTCGGATAGCCGGTCCC | >hg38_21_+_8433172_8446622_RNA45SN1_45S_50ntFlanks | 12712 | 12744 | 33 | CGGAGCCTCGGTTGGCCTCGGATAGCCGGTCCC | 127.3 | -6.2 | 1.5 | -4.0 | 5.8E-05 | 2.0E-04 | BT20/MB468 | MP |
| >hg38_21_+_8433172_8446622_RNA45SN1_45S_50ntFlanks@12999.13039.41\|CTGCGATCTATTGAAAGTCAGCCCTCGACACAAGGGTTTGT | >hg38_21_+_8433172_8446622_RNA45SN1_45S_50ntFlanks | 12999 | 13039 | 41 | CTGCGATCTATTGAAAGTCAGCCCTCGACACAAGGGTTTGT | 7809.7 | 4.2 | 1.1 | 3.8 | 1.6E-04 | 2.0E-04 | MB231/MB468 | NUC |
| >hg38_21_+_8433172_8446622_RNA45SN1_45S_50ntFlanks@7973.8016.44\|GACGCGACCTCAGATCAGACGTGGCGACCCGCTGAATTTAAGCA | >hg38_21_+_8433172_8446622_RNA45SN1_45S_50ntFlanks | 7973 | 8016 | 44 | GACGCGACCTCAGATCAGACGTGGCGACCCGCTGAATTTAAGCA | 654.5 | 3.7 | 1.0 | 3.8 | 1.6E-04 | 2.0E-04 | MB231/MB468 | MITO |
| >hg38_21_+_8433172_8446622_RNA45SN1_45S_50ntFlanks@8352.8379.28\|AGTTGAAAAGAACTTTGAAGAGAGAGTT | >hg38_21_+_8433172_8446622_RNA45SN1_45S_50ntFlanks | 8352 | 8379 | 28 | AGTTGAAAAGAACTTTGAAGAGAGAGTT | 217.8 | -2.9 | 0.8 | -3.8 | 1.6E-04 | 2.0E-04 | MB231/MB468 | NUC |
| >hg38_21_+_8433172_8446622_RNA45SN1_45S_50ntFlanks@3705.3746.42\|TACCTGGTTGATCCTGCCAGTAGCATATGCTTGTCTCAAAGA | >hg38_21_+_8433172_8446622_RNA45SN1_45S_50ntFlanks | 3705 | 3746 | 42 | TACCTGGTTGATCCTGCCAGTAGCATATGCTTGTCTCAAAGA | 6487.5 | 5.2 | 1.2 | 4.2 | 2.2E-05 | 2.0E-04 | BT20/MB231 | NUC |
| >hg38_21_+_8433172_8446622_RNA45SN1_45S_50ntFlanks@12394.12414.21\|TAGCGACGTCGCTTTTTGATC | >hg38_21_+_8433172_8446622_RNA45SN1_45S_50ntFlanks | 12394 | 12414 | 21 | TAGCGACGTCGCTTTTTGATC | 779.2 | -5.2 | 1.4 | -3.8 | 1.3E-04 | 2.0E-04 | MB231/MB468 | MP |
| >MI0000298\|hsa-mir-221&WithFlank&X\|-\|45746151\|45746272@31.56.26&[MIMAT0004568&hsa-miR-221-5p&offsets\|0\|+4;m-405&X\|-\|45746220\|45746242&offsets\|0\|+3]\|ACCTGGCATACAATGTAGATTTCTGT | >MI0000298\|hsa-mir-221&WithFlank&X\|-\|45746151\|45746272 | 31 | 56 | 26 | ACCTGGCATACAATGTAGATTTCTGT | 1599.7 | -2.8 | 0.7 | -4.2 | 2.2E-05 | 2.0E-04 | BT20/MB231 | NUC |
| >hg38_21_+_8433172_8446622_RNA45SN1_45S_50ntFlanks@6661.6681.21\|CGGTGGATCACTCGGCTCGTG | >hg38_21_+_8433172_8446622_RNA45SN1_45S_50ntFlanks | 6661 | 6681 | 21 | CGGTGGATCACTCGGCTCGTG | 1008.4 | -2.0 | 0.5 | -3.8 | 1.5E-04 | 2.0E-04 | MB231/MB468 | CYTO |
| >MI0000809\|hsa-mir-151a&WithFlank&8\|-\|140732558\|140732659@53.75.23&[MIMAT0000757&hsa-miR-151a-3p&offsets\|0\|+2;m-30&8\|-\|140732587\|140732607&offsets\|0\|+2]\|CTAGACTGAAGCTCCTTGAGGAC | >MI0000809\|hsa-mir-151a&WithFlank&8\|-\|140732558\|140732659 | 53 | 75 | 23 | CTAGACTGAAGCTCCTTGAGGAC | 174.1 | 5.2 | 1.3 | 4.0 | 5.8E-05 | 2.0E-04 | BT20/MB468 | MP |
| >trnaMT_IleGAT_MT_+_4263_4331@24.63.40\|TACTTTGATAGAGTAAATAATAGGAGCTTAAACCCCCTTA | >trnaMT_IleGAT_MT_+_4263_4331 | 24 | 63 | 40 | TACTTTGATAGAGTAAATAATAGGAGCTTAAACCCCCTTA | 107.7 | 10.5 | 2.6 | 4.0 | 5.8E-05 | 2.0E-04 | BT20/MB468 | MP |
| >MI0000270\|hsa-mir-181b-1&WithFlank&1\|-\|198858867\|198858988@42.63.22;MI0000683\|hsa-mir-181b-2&WithFlank&9\|+\|124693704\|124693804@22.43.22&[MIMAT0000257&hsa-miR-181b-5p&offsets\|0\|-1;m-73&1\|-\|198858925\|198858947&offsets\|0\|-1];[MIMAT0000257_1&hsa-miR-181b-5p&offsets\|0\|-1;m-72&9\|+\|124693725\|124693747&offsets\|0\|-1]\|AACATTCATTGCTGTCGGTGGG | >MI0000270\|hsa-mir-181b-1&WithFlank&1\|-\|198858867\|198858988 | 42 | 63 | 22 | AACATTCATTGCTGTCGGTGGG | 225.0 | 5.7 | 1.5 | 3.8 | 1.6E-04 | 2.0E-04 | MB231/MB468 | MITO |
| >hg38_21_+_8433172_8446622_RNA45SN1_45S_50ntFlanks@11876.11898.23\|AAGACCCTGTTGAGCTTGACTCT | >hg38_21_+_8433172_8446622_RNA45SN1_45S_50ntFlanks | 11876 | 11898 | 23 | AAGACCCTGTTGAGCTTGACTCT | 1093.0 | -2.8 | 0.7 | -3.8 | 1.5E-04 | 2.1E-04 | MB231/MB468 | CYTO |
| >MI0000434\|hsa-let-7i&WithFlank&12\|+\|62603680\|62603775@12.32.21(+1C)&[MIMAT0000415&hsa-let-7i-5p&offsets\|0\|-1(+1C);m-49&12\|+\|62603691\|62603712&offsets\|0\|-1(+1C)]\|TGAGGTAGTAGTTTGTGCTGTC | >MI0000434\|hsa-let-7i&WithFlank&12\|+\|62603680\|62603775 | 12 | 32 | 21(+1C) | TGAGGTAGTAGTTTGTGCTGTC | 10085.5 | 4.1 | 1.0 | 4.0 | 5.8E-05 | 2.1E-04 | BT20/MB468 | MP |
| >trnaMT_HisGTG_MT_+_12138_12206@28.70.43\|ATTGTGAATCTGACAACAGAGGCTTACGACCCCTTATTTACCC | >trnaMT_HisGTG_MT_+_12138_12206 | 28 | 70 | 43 | ATTGTGAATCTGACAACAGAGGCTTACGACCCCTTATTTACCC | 107.1 | 10.5 | 2.6 | 4.0 | 5.8E-05 | 2.1E-04 | BT20/MB468 | MP |
| >trna119_LysCTT_1_-_145395522_145395594@1.30.30,trna11_LysCTT_5_-_180648979_180649051@1.30.30,trna13_LysCTT_6_+_26556774_26556846@1.30.30,trna32_LysCTT_16_-_3207406_3207478@1.30.30,trna7_LysCTT_16_+_3225692_3225764@1.30.30,trna9_LysCTT_5_+_180634755_180634827@1.30.30\|GCCCGGCTAGCTCAGTCGGTAGAGCATGAG | >trna119_LysCTT_1_-_145395522_145395594 | 1 | 30 | 30 | GCCCGGCTAGCTCAGTCGGTAGAGCATGAG | 2976.6 | -5.7 | 1.4 | -4.0 | 5.8E-05 | 2.1E-04 | BT20/MB468 | MP |

| | | | | | | | | | | | | | |
|---|---|---|---|---|---|---|---|---|---|---|---|---|---|
| >hg38_21_+_8433172_8446622_RNA45SN1_45S_50ntFlanks@4657.4688.32\|CTAGAGGTGAAATTCTTGGACCGGCGCAAGAC | >hg38_21_+_8433172_8446622_RNA45SN1_45S_50ntFlanks | 4657 | 4688 | 32 | CTAGAGGTGAAATTCTTGGACCGGCGCAAGAC | 369.1 | -6.1 | 1.6 | -3.8 | 1.4E-04 | 2.1E-04 | BT20/MB468 | CYTO |
| >hg38_21_+_8433172_8446622_RNA45SN1_45S_50ntFlanks@12488.12512.25\|AACGTGAGCTGGGTTTAGACCGTCG | >hg38_21_+_8433172_8446622_RNA45SN1_45S_50ntFlanks | 12488 | 12512 | 25 | AACGTGAGCTGGGTTTAGACCGTCG | 219.3 | -4.0 | 1.1 | -3.8 | 1.4E-04 | 2.1E-04 | BT20/MB468 | CYTO |
| >hg38_21_+_8433172_8446622_RNA45SN1_45S_50ntFlanks@10772.10796.25\|AGGTCTCCAAGGTGAACAGCCTCTG | >hg38_21_+_8433172_8446622_RNA45SN1_45S_50ntFlanks | 10772 | 10796 | 25 | AGGTCTCCAAGGTGAACAGCCTCTG | 714.5 | -2.4 | 0.6 | -3.8 | 1.6E-04 | 2.1E-04 | MB231/MB468 | NUC |
| >hg38_21_+_8433172_8446622_RNA45SN1_45S_50ntFlanks@12647.12664.18\|GTGGGATTATGACTGAAC | >hg38_21_+_8433172_8446622_RNA45SN1_45S_50ntFlanks | 12647 | 12664 | 18 | GTGGGATTATGACTGAAC | 400.8 | 9.6 | 2.5 | 3.8 | 1.5E-04 | 2.1E-04 | MB231/MB468 | CYTO |
| >hg38_21_+_8433172_8446622_RNA45SN1_45S_50ntFlanks@7985.8006.22\|GATCAGACGTGGCGACCCGCTG | >hg38_21_+_8433172_8446622_RNA45SN1_45S_50ntFlanks | 7985 | 8006 | 22 | GATCAGACGTGGCGACCCGCTG | 1092.2 | -6.0 | 1.6 | -3.8 | 1.6E-04 | 2.1E-04 | MB231/MB468 | NUC |
| >MI0000076\|hsa-mir-20a&WithFlank&13\|+\|91351059\|91351141@14.35.22(+1A)&[MIMAT0000075&hsa-miR-20a-5p&offsets\|0\|-1(+1A);m-96&13\|+\|91351072\|91351094&offsets\|0\|-1(+1A)]\|TAAAGTGCTTATAGTGCAGGTAA | >MI0000076\|hsa-mir-20a&WithFlank&13\|+\|91351059\|91351141 | 14 | 35 | 22(+1A) | TAAAGTGCTTATAGTGCAGGTAA | 2735.4 | 3.1 | 0.8 | 3.8 | 1.3E-04 | 2.1E-04 | MB231/MB468 | MP |
| >hg38_21_+_8433172_8446622_RNA45SN1_45S_50ntFlanks@9532.9571.40\|AGAGGAAACTCTGGTGGAGGTCCGTAGCGGTCCTGACGTG | >hg38_21_+_8433172_8446622_RNA45SN1_45S_50ntFlanks | 9532 | 9571 | 40 | AGAGGAAACTCTGGTGGAGGTCCGTAGCGGTCCTGACGTG | 1492.8 | -3.5 | 0.8 | -4.3 | 1.9E-05 | 2.1E-04 | BT20/MB231 | CYTO |
| >hg38_21_+_8433172_8446622_RNA45SN1_45S_50ntFlanks@10796.10818.23\|GGCATGTTGGAACAATGTAGGTA | >hg38_21_+_8433172_8446622_RNA45SN1_45S_50ntFlanks | 10796 | 10818 | 23 | GGCATGTTGGAACAATGTAGGTA | 651.0 | -3.9 | 1.0 | -3.8 | 1.5E-04 | 2.1E-04 | MB231/MB468 | CYTO |
| >hg38_21_+_8433172_8446622_RNA45SN1_45S_50ntFlanks@12405.12423.19\|CTTTTTGATCCTTCGATGT | >hg38_21_+_8433172_8446622_RNA45SN1_45S_50ntFlanks | 12405 | 12423 | 19 | CTTTTTGATCCTTCGATGT | 152.7 | -5.9 | 1.5 | -4.0 | 5.8E-05 | 2.1E-04 | BT20/MB468 | MP |
| >hg38_21_+_8433172_8446622_RNA45SN1_45S_50ntFlanks@6644.6662.19\|CCTCGTACGACTCTTAGCG | >hg38_21_+_8433172_8446622_RNA45SN1_45S_50ntFlanks | 6644 | 6662 | 19 | CCTCGTACGACTCTTAGCG | 452.6 | 3.5 | 0.9 | 4.0 | 5.8E-05 | 2.1E-04 | BT20/MB468 | MP |
| >hg38_21_+_8433172_8446622_RNA45SN1_45S_50ntFlanks@12406.12437.32\|TTTTTGATCCTTCGATGTCGGCTCTTCCTATC | >hg38_21_+_8433172_8446622_RNA45SN1_45S_50ntFlanks | 12406 | 12437 | 32 | TTTTTGATCCTTCGATGTCGGCTCTTCCTATC | 841.1 | -2.6 | 0.7 | -3.8 | 1.6E-04 | 2.1E-04 | MB231/MB468 | NUC |
| >hg38_21_+_8433172_8446622_RNA45SN1_45S_50ntFlanks@5527.5550.24\|AAAGTCGTAACAAGGTTTCCGTAG | >hg38_21_+_8433172_8446622_RNA45SN1_45S_50ntFlanks | 5527 | 5550 | 24 | AAAGTCGTAACAAGGTTTCCGTAG | 1008.5 | -6.6 | 1.6 | -4.0 | 5.8E-05 | 2.1E-04 | BT20/MB468 | MP |
| >trnaMT_SerGCT_MT_+_12207_12265@14.41.28\|AGAACTGCTAACTCATGCCCCCATGTCT | >trnaMT_SerGCT_MT_+_12207_12265 | 14 | 41 | 28 | AGAACTGCTAACTCATGCCCCCATGTCT | 380.5 | 4.8 | 1.2 | 4.0 | 5.8E-05 | 2.1E-04 | BT20/MB468 | MP |
| >hg38_21_+_8433172_8446622_RNA45SN1_45S_50ntFlanks@11572.11612.41\|CCGACTTAGAACTGGTGCGGACCAGGGGAATCCGACTGTTT | >hg38_21_+_8433172_8446622_RNA45SN1_45S_50ntFlanks | 11572 | 11612 | 41 | CCGACTTAGAACTGGTGCGGACCAGGGGAATCCGACTGTTT | 3666.5 | -3.9 | 1.0 | -3.8 | 1.6E-04 | 2.1E-04 | MB231/MB468 | NUC |
| >hg38_21_+_8433172_8446622_RNA45SN1_45S_50ntFlanks@6652.6699.48\|GACTCTTAGCGGTGGATCACTCGGCTCGTGCGTCGATGAAGAACGCAG | >hg38_21_+_8433172_8446622_RNA45SN1_45S_50ntFlanks | 6652 | 6699 | 48 | GACTCTTAGCGGTGGATCACTCGGCTCGTGCGTCGATGAAGAACGCAG | 5582.6 | -4.1 | 1.1 | -3.8 | 1.6E-04 | 2.1E-04 | BT20/MB468 | NUC |
| >hg38_MT_+_598_1651_RNR1_12S_50ntFlanks@719.743.25\|TAAAGACGTTAGGTCAAGGTGTAGC | >hg38_MT_+_598_1651_RNR1_12S_50ntFlanks | 719 | 743 | 25 | TAAAGACGTTAGGTCAAGGTGTAGC | 133.4 | -5.5 | 1.4 | -3.8 | 1.4E-04 | 2.1E-04 | BT20/MB468 | CYTO |
| >hg38_21_+_8433172_8446622_RNA45SN1_45S_50ntFlanks@6729.6770.42\|GACACATTGATCATCGACACTTCGAACGCACTTGCGGCCCCG | >hg38_21_+_8433172_8446622_RNA45SN1_45S_50ntFlanks | 6729 | 6770 | 42 | GACACATTGATCATCGACACTTCGAACGCACTTGCGGCCCCG | 217.4 | -5.3 | 1.4 | -3.8 | 1.4E-04 | 2.1E-04 | BT20/MB468 | CYTO |
| >trnaMT_GluTTC_MT_-_14674_14742@40.67.28,trnalookalike8_GluTTC_5_-_93905172_93905240@40.67.28\|TTGGTCGTGGTTGTAGTCCGTGCGAGAA | >trnaMT_GluTTC_MT_-_14674_14742 | 40 | 67 | 28 | TTGGTCGTGGTTGTAGTCCGTGCGAGAA | 174.6 | 6.6 | 1.6 | 4.0 | 5.8E-05 | 2.1E-04 | BT20/MB468 | MP |
| >hg38_21_+_8433172_8446622_RNA45SN1_45S_50ntFlanks@10912.10931.20\|CGCGCCGCGGCTGGACGAGG | >hg38_21_+_8433172_8446622_RNA45SN1_45S_50ntFlanks | 10912 | 10931 | 20 | CGCGCCGCGGCTGGACGAGG | 76.5 | 9.6 | 2.3 | 4.3 | 2.1E-05 | 2.1E-04 | BT20/MB231 | MP |
| >hg38_21_+_8433172_8446622_RNA45SN1_45S_50ntFlanks@8360.8378.19\|AGAACTTTGAAGAGAGAGT | >hg38_21_+_8433172_8446622_RNA45SN1_45S_50ntFlanks | 8360 | 8378 | 19 | AGAACTTTGAAGAGAGAGT | 114.9 | -6.1 | 1.6 | -3.8 | 1.3E-04 | 2.1E-04 | MB231/MB468 | MP |
| >hg38_MT_+_598_1651_RNR1_12S_50ntFlanks@428.470.43\|AAAGTGGCTTTAACATATCTGAACACACAATAGCTAAGACCCA | >hg38_MT_+_598_1651_RNR1_12S_50ntFlanks | 428 | 470 | 43 | AAAGTGGCTTTAACATATCTGAACACACAATAGCTAAGACCCA | 106.9 | 10.5 | 2.6 | 4.0 | 5.8E-05 | 2.1E-04 | BT20/MB468 | MP |
| >hg38_21_+_8433172_8446622_RNA45SN1_45S_50ntFlanks@11564.11593.30\|CCTAGCAGCCGACTTAGAACTGGTGCGGAC | >hg38_21_+_8433172_8446622_RNA45SN1_45S_50ntFlanks | 11564 | 11593 | 30 | CCTAGCAGCCGACTTAGAACTGGTGCGGAC | 273.4 | -6.4 | 1.6 | -4.0 | 5.9E-05 | 2.1E-04 | BT20/MB468 | MP |
| >hg38_21_+_8433172_8446622_RNA45SN1_45S_50ntFlanks@12083.12108.26\|CTCGCTTCTGGCGCCAAGCGCCCGGC | >hg38_21_+_8433172_8446622_RNA45SN1_45S_50ntFlanks | 12083 | 12108 | 26 | CTCGCTTCTGGCGCCAAGCGCCCGGC | 299.2 | -6.1 | 1.5 | -4.0 | 5.9E-05 | 2.1E-04 | BT20/MB468 | MP |
| >hg38_21_+_8433172_8446622_RNA45SN1_45S_50ntFlanks@9602.9622.21\|CGAAAGACTAATCGAACCATC | >hg38_21_+_8433172_8446622_RNA45SN1_45S_50ntFlanks | 9602 | 9622 | 21 | CGAAAGACTAATCGAACCATC | 693.8 | -5.6 | 1.4 | -4.0 | 5.9E-05 | 2.1E-04 | BT20/MB468 | MP |
| >hg38_21_+_8433172_8446622_RNA45SN1_45S_50ntFlanks@4762.4780.19\|AGATACCGTCGTAGTTCCG | >hg38_21_+_8433172_8446622_RNA45SN1_45S_50ntFlanks | 4762 | 4780 | 19 | AGATACCGTCGTAGTTCCG | 653.8 | -2.8 | 0.7 | -3.8 | 1.4E-04 | 2.1E-04 | BT20/MB468 | CYTO |
| >hg38_21_+_8433172_8446622_RNA45SN1_45S_50ntFlanks@10682.10713.32\|TCCGGCGGCGTCCGGTGAGCTCTCGCTGGCCC | >hg38_21_+_8433172_8446622_RNA45SN1_45S_50ntFlanks | 10682 | 10713 | 32 | TCCGGCGGCGTCCGGTGAGCTCTCGCTGGCCC | 118.5 | 3.4 | 0.9 | 3.8 | 1.6E-04 | 2.1E-04 | MB231/MB468 | MITO |
| >hg38_21_+_8433172_8446622_RNA45SN1_45S_50ntFlanks@3705.3735.31\|TACCTGGTTGATCCTGCCAGTAGCATATGCT | >hg38_21_+_8433172_8446622_RNA45SN1_45S_50ntFlanks | 3705 | 3735 | 31 | TACCTGGTTGATCCTGCCAGTAGCATATGCT | 383.5 | 3.8 | 1.0 | 3.8 | 1.6E-04 | 2.1E-04 | MB231/MB468 | MITO |
| >MI0000469\|hsa-mir-125a&WithFlank&19\|+\|51693248\|51693345@21.43.23&[MIMAT0000443&hsa-miR-125a-5p&offsets\|0\|-1;m-63&19\|+\|51693268\|51693289&offsets\|0\|+1]\|TCCCTGAGACCCTTTAACCTGTG | >MI0000469\|hsa-mir-125a&WithFlank&19\|+\|51693248\|51693345 | 21 | 43 | 23 | TCCCTGAGACCCTTTAACCTGTG | 7536.0 | 1.5 | 0.4 | 3.8 | 1.6E-04 | 2.1E-04 | BT20/MB468 | NUC |
| >hg38_21_+_8433172_8446622_RNA45SN1_45S_50ntFlanks@12645.12669.25\|CTGTGGGATTATGACTGAACGCCTC | >hg38_21_+_8433172_8446622_RNA45SN1_45S_50ntFlanks | 12645 | 12669 | 25 | CTGTGGGATTATGACTGAACGCCTC | 566.6 | 3.3 | 0.9 | 3.9 | 1.2E-04 | 2.1E-04 | BT20/MB468 | MITO |
| >hg38_21_+_8433172_8446622_RNA45SN1_45S_50ntFlanks@12476.12498.23\|CCACTAATAGGGAACGTGAGCTG | >hg38_21_+_8433172_8446622_RNA45SN1_45S_50ntFlanks | 12476 | 12498 | 23 | CCACTAATAGGGAACGTGAGCTG | 1879.5 | -4.0 | 1.0 | -4.0 | 5.9E-05 | 2.1E-04 | BT20/MB468 | MP |
| >hg38_21_+_8433172_8446622_RNA45SN1_45S_50ntFlanks@12480.12498.19\|TAATAGGGAACGTGAGCTG | >hg38_21_+_8433172_8446622_RNA45SN1_45S_50ntFlanks | 12480 | 12498 | 19 | TAATAGGGAACGTGAGCTG | 218.5 | -4.5 | 1.2 | -3.8 | 1.6E-04 | 2.1E-04 | MB231/MB468 | NUC |
| >hg38_21_+_8433172_8446622_RNA45SN1_45S_50ntFlanks@11573.11603.31\|CGACTTAGAACTGGTGCGGACCAGGGGAATC | >hg38_21_+_8433172_8446622_RNA45SN1_45S_50ntFlanks | 11573 | 11603 | 31 | CGACTTAGAACTGGTGCGGACCAGGGGAATC | 632.3 | -2.6 | 0.7 | -3.8 | 1.4E-04 | 2.1E-04 | BT20/MB468 | CYTO |
| >am_trna128_GlyGCC_6_-_27870686_27870756@1.25.25,trna133_GlyCCC_1_-_16872434_16872504@1.25.25,trna18_GlyGCC_16_+_70822597_70822667@1.25.25,trna19_GlyGCC_16_+_70823410_70823480@1.25.25,trna19_GlyGCC_2_-_157257659_157257729@1.25.25,trna24_GlyGCC_16_-_70812942_70813012@1.25.25,trna25_GlyGCC_16_-_70812114_70812184@1.25.25,trna4_GlyCCC_1_+_17188416_17188486@1.25.25,trna5_GlyGCC_17_+_8029064_8029134@1.25.25,trna68_GlyGCC_1_-_161493637_161493707@1.25.25\|GCATTGGTGGTTCAGTGGTAGAATT | >am_trna128_GlyGCC_6_-_27870686_27870756 | 1 | 25 | 25 | GCATTGGTGGTTCAGTGGTAGAATT | 2187.1 | 2.4 | 0.6 | 3.8 | 1.5E-04 | 2.1E-04 | MB231/MB468 | CYTO |
| >hg38_MT_+_1621_3279_RNR2_16S_50ntFlanks@833.864.32\|GGTTAAAAAAAGTAAAAGGAACTCGGCAAATC | >hg38_MT_+_1621_3279_RNR2_16S_50ntFlanks | 833 | 864 | 32 | GGTTAAAAAAAGTAAAAGGAACTCGGCAAATC | 104.3 | 7.5 | 1.9 | 4.0 | 5.9E-05 | 2.1E-04 | BT20/MB468 | MP |
| >hg38_21_+_8433172_8446622_RNA45SN1_45S_50ntFlanks@12488.12521.34\|AACGTGAGCTGGGTTTAGACCGTCGTGAGACAGG | >hg38_21_+_8433172_8446622_RNA45SN1_45S_50ntFlanks | 12488 | 12521 | 34 | AACGTGAGCTGGGTTTAGACCGTCGTGAGACAGG | 385.7 | -5.1 | 1.3 | -4.0 | 5.9E-05 | 2.1E-04 | BT20/MB468 | MP |
| >hg38_21_+_8433172_8446622_RNA45SN1_45S_50ntFlanks@10807.10825.19\|ACAATGTAGGTAAGGGAAG | >hg38_21_+_8433172_8446622_RNA45SN1_45S_50ntFlanks | 10807 | 10825 | 19 | ACAATGTAGGTAAGGGAAG | 143.4 | -5.9 | 1.5 | -4.0 | 5.9E-05 | 2.1E-04 | BT20/MB468 | MP |
| >hg38_21_+_8433172_8446622_RNA45SN1_45S_50ntFlanks@8196.8223.28\|CGTGGACGGTGTGAGGCCGGTAGCGGCC | >hg38_21_+_8433172_8446622_RNA45SN1_45S_50ntFlanks | 8196 | 8223 | 28 | CGTGGACGGTGTGAGGCCGGTAGCGGCC | 444.4 | -4.8 | 1.3 | -3.8 | 1.6E-04 | 2.1E-04 | MB231/MB468 | MITO |
| >hg38_21_+_8433172_8446622_RNA45SN1_45S_50ntFlanks@4759.4793.35\|ATCAGATACCGTCGTAGTTCCGACCATAAACGATG | >hg38_21_+_8433172_8446622_RNA45SN1_45S_50ntFlanks | 4759 | 4793 | 35 | ATCAGATACCGTCGTAGTTCCGACCATAAACGATG | 954.0 | -5.5 | 1.4 | -3.8 | 1.5E-04 | 2.1E-04 | MB231/MB468 | CYTO |
| >trna128_GlyGCC_6_-_27870686_27870756@19.38.20,trna19_GlyGCC_16_+_70823410_70823480@19.38.20,trna19_GlyGCC_2_-_157257659_157257729@19.38.20,trna24_GlyGCC_16_-_70812942_70813012@19.38.20,trna25_GlyGCC_16_-_70812114_70812184@19.38.20,trna2_GlyGCC_21_-_18827107_18827177@19.38.20,trna35_GlyGCC_1_+_161413094_161413164@19.38.20,trna37_GlyGCC_1_+_161420467_161420537@19.38.20,trna39_GlyGCC_1_+_161427898_161427968@19.38.20,trna41_GlyGCC_1_+_161435258_161435328@19.38.20,trna5_GlyGCC_17_+_8029064_8029134@19.38.20,trna68_GlyGCC_1_-_161493637_161493707@19.38.20\|TAGAATTCTCGCCTGCCACG | >trna128_GlyGCC_6_-_27870686_27870756 | 19 | 38 | 20 | TAGAATTCTCGCCTGCCACG | 2630.0 | 3.7 | 0.9 | 4.3 | 2.0E-05 | 2.1E-04 | BT20/MB231 | CYTO |
| >trna116_GluCTC_1_-_145399233_145399304@30.70.41,trna71_GluCTC_1_-_161439189_161439260@30.70.41,trna74_GluCTC_1_-_161431809_161431880@30.70.41,trna77_GluCTC_1_-_161424398_161424469@30.70.41,trna77_GluCTC_6_+_28949976_28950047@30.70.41,trna80_GluCTC_1_-_161417018_161417089@30.70.41,trna87_GluCTC_6_-_126101393_126101464@30.70.41\|CGCTCTCACCGCCGCGGCCCGGGTTCGATTCCCGGTCAGGG | >trna116_GluCTC_1_-_145399233_145399304 | 30 | 70 | 41 | CGCTCTCACCGCCGCGGCCCGGGTTCGATTCCCGGTCAGGG | 711.4 | -6.8 | 1.8 | -3.8 | 1.6E-04 | 2.1E-04 | BT20/MB468 | NUC |
| >trnaMT_GlnTTG_MT_-_4329_4400@1.34.34,trnalookalike2_GlnTTG_1_-_564879_564950@1.34.34\|TAGGATGGGGTGTGATAGGTGGCACGGAGAATTT | >trnaMT_GlnTTG_MT_-_4329_4400 | 1 | 34 | 34 | TAGGATGGGGTGTGATAGGTGGCACGGAGAATTT | 889.5 | 6.0 | 1.5 | 4.0 | 5.9E-05 | 2.1E-04 | BT20/MB468 | MP |
| >hg38_21_+_8433172_8446622_RNA45SN1_45S_50ntFlanks@4045.4063.19\|CCATTCGAACGTCTGCCCT | >hg38_21_+_8433172_8446622_RNA45SN1_45S_50ntFlanks | 4045 | 4063 | 19 | CCATTCGAACGTCTGCCCT | 2454.0 | -4.4 | 1.2 | -3.8 | 1.5E-04 | 2.1E-04 | MB231/MB468 | CYTO |
| >hg38_21_+_8433172_8446622_RNA45SN1_45S_50ntFlanks@12481.12498.18\|AATAGGGAACGTGAGCTG | >hg38_21_+_8433172_8446622_RNA45SN1_45S_50ntFlanks | 12481 | 12498 | 18 | AATAGGGAACGTGAGCTG | 323.2 | -4.7 | 1.2 | -3.8 | 1.3E-04 | 2.1E-04 | MB231/MB468 | MP |
| >trnaMT_PheGAA_MT_+_577_647@33.62.30\|CTGAAAATGTTTAGACGGGCTCACATCACC | >trnaMT_PheGAA_MT_+_577_647 | 33 | 62 | 30 | CTGAAAATGTTTAGACGGGCTCACATCACC | 115.6 | 10.6 | 2.6 | 4.0 | 5.9E-05 | 2.1E-04 | BT20/MB468 | MP |
| >hg38_21_+_8433172_8446622_RNA45SN1_45S_50ntFlanks@10796.10817.22\|GGCATGTTGGAACAATGTAGGT | >hg38_21_+_8433172_8446622_RNA45SN1_45S_50ntFlanks | 10796 | 10817 | 22 | GGCATGTTGGAACAATGTAGGT | 238.5 | -5.8 | 1.5 | -3.8 | 1.4E-04 | 2.1E-04 | MB231/MB468 | MP |
| >hg38_21_+_8433172_8446622_RNA45SN1_45S_50ntFlanks@12008.12034.27\|CCGCCGGTGAAATACCACTACTCTGAT | >hg38_21_+_8433172_8446622_RNA45SN1_45S_50ntFlanks | 12008 | 12034 | 27 | CCGCCGGTGAAATACCACTACTCTGAT | 261.2 | -5.5 | 1.5 | -3.8 | 1.6E-04 | 2.1E-04 | MB231/MB468 | NUC |
| >hg38_21_+_8433172_8446622_RNA45SN1_45S_50ntFlanks@12418.12456.39\|CGATGTCGGCTCTTCCTATCATTGTGAAGCAGAATTCAC | >hg38_21_+_8433172_8446622_RNA45SN1_45S_50ntFlanks | 12418 | 12456 | 39 | CGATGTCGGCTCTTCCTATCATTGTGAAGCAGAATTCAC | 249.1 | -3.4 | 0.8 | -4.3 | 2.0E-05 | 2.1E-04 | BT20/MB231 | CYTO |

| | | | | | | | | | | | | | |
|---|---|---|---|---|---|---|---|---|---|---|---|---|---|
| >hg38_21_+_8433172_8446622_RNA45SN1_45S_50ntFlanks@6649.6676.28\|TACGACTCTTAGCGGTGGATCACTCGGC | >hg38_21_+_8433172_8446622_RNA45SN1_45S_50ntFlanks | 6649 | 6676 | 28 | TACGACTCTTAGCGGTGGATCACTCGGC | 288.8 | 3.8 | 1.0 | 3.8 | 1.6E-04 | 2.1E-04 | MB231/MB468 | NUC |
| >hg38_21_+_8433172_8446622_RNA45SN1_45S_50ntFlanks@5471.5499.29\|CACGGCCCTGGCGGAGCGCTGAGAAGACG | >hg38_21_+_8433172_8446622_RNA45SN1_45S_50ntFlanks | 5471 | 5499 | 29 | CACGGCCCTGGCGGAGCGCTGAGAAGACG | 5608.9 | -7.8 | 2.1 | -3.8 | 1.6E-04 | 2.1E-04 | BT20/MB468 | NUC |
| >hg38_21_+_8433172_8446622_RNA45SN1_45S_50ntFlanks@9926.9950.25\|GTTGATATAGACAGCAGGACGGTGG | >hg38_21_+_8433172_8446622_RNA45SN1_45S_50ntFlanks | 9926 | 9950 | 25 | GTTGATATAGACAGCAGGACGGTGG | 54.6 | 9.1 | 2.1 | 4.3 | 2.1E-05 | 2.1E-04 | BT20/MB231 | MP |
| >hg38_21_+_8433172_8446622_RNA45SN1_45S_50ntFlanks@3854.3898.45\|ACTGTGGTAATTCTAGAGCTAATACATGCCGACGGGCGCTGACCC | >hg38_21_+_8433172_8446622_RNA45SN1_45S_50ntFlanks | 3854 | 3898 | 45 | ACTGTGGTAATTCTAGAGCTAATACATGCCGACGGGCGCTGACCC | 507.4 | -5.8 | 1.5 | -3.8 | 1.2E-04 | 2.1E-04 | BT20/MB468 | MITO |
| >MI0000440\|hsa-mir-27b&WithFlank&9\|+\|95085439\|95085547@67.87.21(+2U)&[MIMAT0000419&hsa-miR-27b-3p&offsets\|0\|0(+2U);m-39&9\|+\|95085505\|95085525&offsets\|0\|0(+2U)]\|TTCACAGTGGCTAAGTTCTGCTT | >MI0000440\|hsa-mir-27b&WithFlank&9\|+\|95085439\|95085547 | 67 | 87 | 21(+2U) | TTCACAGTGGCTAAGTTCTGCTT | 198.8 | 3.7 | 0.9 | 4.0 | 6.0E-05 | 2.1E-04 | BT20/MB468 | MP |
| >hg38_21_+_8433172_8446622_RNA45SN1_45S_50ntFlanks@10787.10815.29\|ACAGCCTCTGGCATGTTGGAACAATGTAG | >hg38_21_+_8433172_8446622_RNA45SN1_45S_50ntFlanks | 10787 | 10815 | 29 | ACAGCCTCTGGCATGTTGGAACAATGTAG | 333.2 | -4.4 | 1.2 | -3.8 | 1.6E-04 | 2.1E-04 | MB231/MB468 | NUC |
| >hg38_21_+_8433172_8446622_RNA45SN1_45S_50ntFlanks@9533.9554.22\|GAGGAAACTCTGGTGGAGGTCC | >hg38_21_+_8433172_8446622_RNA45SN1_45S_50ntFlanks | 9533 | 9554 | 22 | GAGGAAACTCTGGTGGAGGTCC | 555.0 | -4.5 | 1.2 | -3.8 | 1.5E-04 | 2.1E-04 | MB231/MB468 | CYTO |
| >hg38_21_+_8433172_8446622_RNA45SN1_45S_50ntFlanks@4723.4766.44\|CATTAATCAAGAACGAAAGTCGGAGGTTCGAAGACGATCAGATA | >hg38_21_+_8433172_8446622_RNA45SN1_45S_50ntFlanks | 4723 | 4766 | 44 | CATTAATCAAGAACGAAAGTCGGAGGTTCGAAGACGATCAGATA | 265.0 | 4.3 | 1.1 | 3.8 | 1.5E-04 | 2.1E-04 | MB231/MB468 | CYTO |
| >hg38_21_+_8433172_8446622_RNA45SN1_45S_50ntFlanks@3850.3872.23\|GATAACTGTGGTAATTCTAGAGC | >hg38_21_+_8433172_8446622_RNA45SN1_45S_50ntFlanks | 3850 | 3872 | 23 | GATAACTGTGGTAATTCTAGAGC | 414.7 | -4.6 | 1.2 | -3.8 | 1.4E-04 | 2.1E-04 | MB231/MB468 | MP |
| >MI0000446\|hsa-mir-125b-1&WithFlank&11\|-\|122099751\|122099850@21.43.23&[MIMAT0000423&hsa-miR-125b-5p&offsets\|0\|+1;m-67&11\|-\|122099809\|122099830&offsets\|0\|+1]\|TCCCTGAGACCCTAACTTGTGAT | >MI0000446\|hsa-mir-125b-1&WithFlank&11\|-\|122099751\|122099850 | 21 | 43 | 23 | TCCCTGAGACCCTAACTTGTGAT | 1250.5 | -6.7 | 1.6 | -4.2 | 2.3E-05 | 2.1E-04 | BT20/MB231 | NUC |
| >hg38_21_+_8433172_8446622_RNA45SN1_45S_50ntFlanks@7973.8009.37\|GACGCGACCTCAGATCAGACGTGGCGACCCGCTGAAT | >hg38_21_+_8433172_8446622_RNA45SN1_45S_50ntFlanks | 7973 | 8009 | 37 | GACGCGACCTCAGATCAGACGTGGCGACCCGCTGAAT | 2931.6 | 2.7 | 0.7 | 3.8 | 1.6E-04 | 2.1E-04 | MB231/MB468 | MITO |
| >hg38_21_+_8433172_8446622_RNA45SN1_45S_50ntFlanks@12071.12106.36\|CCCCGAGGGGCTCTCGCTTCTGGCGCCAAGCGCCCG | >hg38_21_+_8433172_8446622_RNA45SN1_45S_50ntFlanks | 12071 | 12106 | 36 | CCCCGAGGGGCTCTCGCTTCTGGCGCCAAGCGCCCG | 324.2 | -4.4 | 1.2 | -3.8 | 1.4E-04 | 2.1E-04 | BT20/MB468 | CYTO |
| >hg38_21_+_8433172_8446622_RNA45SN1_45S_50ntFlanks@4733.4759.27\|GAACGAAAGTCGGAGGTTCGAAGACGA | >hg38_21_+_8433172_8446622_RNA45SN1_45S_50ntFlanks | 4733 | 4759 | 27 | GAACGAAAGTCGGAGGTTCGAAGACGA | 330.2 | -2.2 | 0.6 | -3.8 | 1.6E-04 | 2.1E-04 | MB231/MB468 | NUC |
| >hg38_21_+_8433172_8446622_RNA45SN1_45S_50ntFlanks@10395.10424.30\|GCAGTTGAACATGGGTCAGTCGGTCCTGAG | >hg38_21_+_8433172_8446622_RNA45SN1_45S_50ntFlanks | 10395 | 10424 | 30 | GCAGTTGAACATGGGTCAGTCGGTCCTGAG | 1287.9 | 2.7 | 0.7 | 3.8 | 1.6E-04 | 2.1E-04 | MB231/MB468 | NUC |
| >hg38_21_+_8433172_8446622_RNA45SN1_45S_50ntFlanks@12950.12988.39\|CGACCTGCTTCTGGGTCGGGGTTTCGTACGTAGCAGAGC | >hg38_21_+_8433172_8446622_RNA45SN1_45S_50ntFlanks | 12950 | 12988 | 39 | CGACCTGCTTCTGGGTCGGGGTTTCGTACGTAGCAGAGC | 300.3 | -2.9 | 0.8 | -3.8 | 1.6E-04 | 2.1E-04 | MB231/MB468 | NUC |
| >MI0000065\|hsa-let-7d&WithFlank&9\|+\|94178828\|94178926@14.36.23&[MIMAT0000065&hsa-let-7d-5p&offsets\|0\|+1;m-90&9\|+\|94178841\|94178862&offsets\|0\|+1]\|AGAGGTAGTAGGTTGCATAGTTT | >MI0000065\|hsa-let-7d&WithFlank&9\|+\|94178828\|94178926 | 14 | 36 | 23 | AGAGGTAGTAGGTTGCATAGTTT | 285.0 | 4.6 | 1.2 | 3.8 | 1.6E-04 | 2.1E-04 | MB231/MB468 | NUC |
| >hg38_21_+_8433172_8446622_RNA45SN1_45S_50ntFlanks@10444.10465.22\|CGAAGGGACGGGCGATGGCCTC | >hg38_21_+_8433172_8446622_RNA45SN1_45S_50ntFlanks | 10444 | 10465 | 22 | CGAAGGGACGGGCGATGGCCTC | 105.8 | -5.7 | 1.5 | -3.8 | 1.4E-04 | 2.1E-04 | MB231/MB468 | MP |
| >hg38_21_+_8433172_8446622_RNA45SN1_45S_50ntFlanks@12006.12031.26\|GGCCGCCGGTGAAATACCACTACTCT | >hg38_21_+_8433172_8446622_RNA45SN1_45S_50ntFlanks | 12006 | 12031 | 26 | GGCCGCCGGTGAAATACCACTACTCT | 127.7 | -6.1 | 1.6 | -3.8 | 1.4E-04 | 2.1E-04 | MB231/MB468 | MP |
| >hg38_21_+_8433172_8446622_RNA45SN1_45S_50ntFlanks@8317.8356.40\|CACGAGACCGATAGTCAACAAGTACCGTAAGGGAAAGTTG | >hg38_21_+_8433172_8446622_RNA45SN1_45S_50ntFlanks | 8317 | 8356 | 40 | CACGAGACCGATAGTCAACAAGTACCGTAAGGGAAAGTTG | 627.9 | -2.2 | 0.6 | -3.8 | 1.2E-04 | 2.1E-04 | BT20/MB468 | MITO |
| >hg38_21_+_8433172_8446622_RNA45SN1_45S_50ntFlanks@11877.11904.28\|AGACCCTGTTGAGCTTGACTCTAGTCTG | >hg38_21_+_8433172_8446622_RNA45SN1_45S_50ntFlanks | 11877 | 11904 | 28 | AGACCCTGTTGAGCTTGACTCTAGTCTG | 150.9 | -6.1 | 1.5 | -4.0 | 6.0E-05 | 2.1E-04 | BT20/MB468 | MP |
| >MI0000749\|hsa-mir-30e&WithFlank&1\|+\|40754349\|40754452@24.46.23(+1U)&[MIMAT0000692&hsa-miR-30e-5p&offsets\|+1\|+2(+1U);m-24&1\|+\|40754371\|40754394&offsets\|+1\|0(+1U)]\|GTAAACATCCTTGACTGGAAGCTT | >MI0000749\|hsa-mir-30e&WithFlank&1\|+\|40754349\|40754452 | 24 | 46 | 23(+1U) | GTAAACATCCTTGACTGGAAGCTT | 186.5 | 3.2 | 0.8 | 3.8 | 1.4E-04 | 2.1E-04 | MB231/MB468 | MP |
| >hg38_21_+_8433172_8446622_RNA45SN1_45S_50ntFlanks@4103.4127.25\|CCACGGGTGACGGGGAATCAGGGTT | >hg38_21_+_8433172_8446622_RNA45SN1_45S_50ntFlanks | 4103 | 4127 | 25 | CCACGGGTGACGGGGAATCAGGGTT | 230.8 | -3.0 | 0.8 | -4.0 | 6.0E-05 | 2.1E-04 | BT20/MB468 | MP |
| >hg38_21_+_8433172_8446622_RNA45SN1_45S_50ntFlanks@12418.12435.18\|CGATGTCGGCTCTTCCTA | >hg38_21_+_8433172_8446622_RNA45SN1_45S_50ntFlanks | 12418 | 12435 | 18 | CGATGTCGGCTCTTCCTA | 1999.0 | -5.4 | 1.4 | -4.0 | 6.0E-05 | 2.1E-04 | BT20/MB468 | MP |
| >MI0000749\|hsa-mir-30e&WithFlank&1\|+\|40754349\|40754452@23.45.23(+1C)&[MIMAT0000692&hsa-miR-30e-5p&offsets\|0\|+1(+1C);m-24&1\|+\|40754371\|40754394&offsets\|0\|-1(+1C)]\|TGTAAACATCCTTGACTGGAAGCC | >MI0000749\|hsa-mir-30e&WithFlank&1\|+\|40754349\|40754452 | 23 | 45 | 23(+1C) | TGTAAACATCCTTGACTGGAAGCC | 1343.2 | 4.3 | 1.2 | 3.8 | 1.6E-04 | 2.1E-04 | MB231/MB468 | MITO |
| >hg38_21_+_8433172_8446622_RNA45SN1_45S_50ntFlanks@11873.11912.40\|AAGAAGACCCTGTTGAGCTTGACTCTAGTCTGGCACGGTG | >hg38_21_+_8433172_8446622_RNA45SN1_45S_50ntFlanks | 11873 | 11912 | 40 | AAGAAGACCCTGTTGAGCTTGACTCTAGTCTGGCACGGTG | 354.6 | -5.2 | 1.3 | -4.0 | 6.0E-05 | 2.1E-04 | BT20/MB468 | MP |
| >hg38_21_+_8433172_8446622_RNA45SN1_45S_50ntFlanks@11908.11931.24\|CGGTGAAGAGACATGAGAGGTGTA | >hg38_21_+_8433172_8446622_RNA45SN1_45S_50ntFlanks | 11908 | 11931 | 24 | CGGTGAAGAGACATGAGAGGTGTA | 62.3 | 9.3 | 2.2 | 4.2 | 2.1E-05 | 2.1E-04 | BT20/MB231 | MP |
| >hg38_21_+_8433172_8446622_RNA45SN1_45S_50ntFlanks@11906.11929.24\|CACGGTGAAGAGACATGAGAGGTG | >hg38_21_+_8433172_8446622_RNA45SN1_45S_50ntFlanks | 11906 | 11929 | 24 | CACGGTGAAGAGACATGAGAGGTG | 138.3 | -3.7 | 1.0 | -3.8 | 1.5E-04 | 2.1E-04 | MB231/MB468 | CYTO |
| >hg38_21_+_8433172_8446622_RNA45SN1_45S_50ntFlanks@11873.11908.36\|AAGAAGACCCTGTTGAGCTTGACTCTAGTCTGGCAC | >hg38_21_+_8433172_8446622_RNA45SN1_45S_50ntFlanks | 11873 | 11908 | 36 | AAGAAGACCCTGTTGAGCTTGACTCTAGTCTGGCAC | 275.1 | -5.3 | 1.4 | -3.8 | 1.4E-04 | 2.1E-04 | BT20/MB468 | CYTO |
| >hg38_21_+_8433172_8446622_RNA45SN1_45S_50ntFlanks@6650.6696.47\|ACGACTCTTAGCGGTGGATCACTCGGCTCGTGCGTCGATGAAGAACG | >hg38_21_+_8433172_8446622_RNA45SN1_45S_50ntFlanks | 6650 | 6696 | 47 | ACGACTCTTAGCGGTGGATCACTCGGCTCGTGCGTCGATGAAGAACG | 2196.3 | 4.3 | 1.1 | 3.8 | 1.6E-04 | 2.1E-04 | MB231/MB468 | NUC |
| >hg38_21_+_8433172_8446622_RNA45SN1_45S_50ntFlanks@10798.10830.33\|CATGTTGGAACAATGTAGGTAAGGGAAGTCGGC | >hg38_21_+_8433172_8446622_RNA45SN1_45S_50ntFlanks | 10798 | 10830 | 33 | CATGTTGGAACAATGTAGGTAAGGGAAGTCGGC | 72.3 | 9.5 | 2.2 | 4.2 | 2.1E-05 | 2.1E-04 | BT20/MB231 | MP |
| >hg38_1_-_228634819_228635039_RNA5S12_5S_50ntFlanks@149.170.22\|GAATACCGGGTGCTGTAGGCTT | >hg38_1_-_228634819_228635039_RNA5S12_5S_50ntFlanks | 149 | 170 | 22 | GAATACCGGGTGCTGTAGGCTT | 1379.8 | 3.3 | 0.9 | 3.8 | 1.6E-04 | 2.1E-04 | MB231/MB468 | NUC |
| >hg38_21_+_8433172_8446622_RNA45SN1_45S_50ntFlanks@11915.11943.29\|GAGACATGAGAGGTGTAGAATAAGTGGGA | >hg38_21_+_8433172_8446622_RNA45SN1_45S_50ntFlanks | 11915 | 11943 | 29 | GAGACATGAGAGGTGTAGAATAAGTGGGA | 69.0 | 9.4 | 2.2 | 4.2 | 2.2E-05 | 2.1E-04 | BT20/MB231 | MP |
| >hg38_21_+_8433172_8446622_RNA45SN1_45S_50ntFlanks@5485.5525.41\|AGCGCTGAGAAGACGGTCGAACTTGACTATCTAGAGGAAGT | >hg38_21_+_8433172_8446622_RNA45SN1_45S_50ntFlanks | 5485 | 5525 | 41 | AGCGCTGAGAAGACGGTCGAACTTGACTATCTAGAGGAAGT | 70.1 | 9.5 | 2.2 | 4.2 | 2.2E-05 | 2.1E-04 | BT20/MB231 | MP |
| >hg38_21_+_8433172_8446622_RNA45SN1_45S_50ntFlanks@10781.10817.37\|AGGTGAACAGCCTCTGGCATGTTGGAACAATGTAGGT | >hg38_21_+_8433172_8446622_RNA45SN1_45S_50ntFlanks | 10781 | 10817 | 37 | AGGTGAACAGCCTCTGGCATGTTGGAACAATGTAGGT | 277.7 | -3.0 | 0.8 | -3.8 | 1.5E-04 | 2.1E-04 | BT20/MB468 | CYTO |
| >MI0000088\|hsa-mir-30a&WithFlank&6\|-\|71403545\|71403627@12.34.23(+1G)&[MIMAT0000087&hsa-miR-30a-5p&offsets\|0\|+1(+1G);m-15&6\|-\|71403595\|71403616&offsets\|0\|+1(+1G)]\|TGTAAACATCCTCGACTGGAAGCG | >MI0000088\|hsa-mir-30a&WithFlank&6\|-\|71403545\|71403627 | 12 | 34 | 23(+1G) | TGTAAACATCCTCGACTGGAAGCG | 5505.1 | 3.2 | 0.8 | 3.8 | 1.4E-04 | 2.1E-04 | MB231/MB468 | MP |
| >hg38_21_+_8433172_8446622_RNA45SN1_45S_50ntFlanks@12407.12441.35\|TTTTGATCCTTCGATGTCGGCTCTTCCTATCATTG | >hg38_21_+_8433172_8446622_RNA45SN1_45S_50ntFlanks | 12407 | 12441 | 35 | TTTTGATCCTTCGATGTCGGCTCTTCCTATCATTG | 786.6 | -4.4 | 1.2 | -3.8 | 1.6E-04 | 2.1E-04 | MB231/MB468 | MITO |
| >hg38_21_+_8433172_8446622_RNA45SN1_45S_50ntFlanks@9524.9571.48\|GCGAAGCCAGAGGAAACTCTGGTGGAGGTCCGTAGCGGTCCTGACGTG | >hg38_21_+_8433172_8446622_RNA45SN1_45S_50ntFlanks | 9524 | 9571 | 48 | GCGAAGCCAGAGGAAACTCTGGTGGAGGTCCGTAGCGGTCCTGACGTG | 242.0 | 2.9 | 0.8 | 3.8 | 1.5E-04 | 2.1E-04 | MB231/MB468 | CYTO |
| >hg38_21_+_8433172_8446622_RNA45SN1_45S_50ntFlanks@11641.11661.21\|CGGCGGGTGTTGACGCGATGT | >hg38_21_+_8433172_8446622_RNA45SN1_45S_50ntFlanks | 11641 | 11661 | 21 | CGGCGGGTGTTGACGCGATGT | 712.3 | -6.5 | 1.6 | -4.0 | 6.1E-05 | 2.1E-04 | BT20/MB468 | MP |
| >MI0000086\|hsa-mir-28&WithFlank&3\|+\|188688775\|188688872@60.81.22(+1U)&[MIMAT0004502&hsa-miR-28-3p&offsets\|0\|0(+1U);m-28&3\|+\|188688834\|188688855&offsets\|0\|0(+1U)]\|CACTAGATTGTGAGCTCCTGGAT | >MI0000086\|hsa-mir-28&WithFlank&3\|+\|188688775\|188688872 | 60 | 81 | 22(+1U) | CACTAGATTGTGAGCTCCTGGAT | 241.3 | 5.4 | 1.3 | 4.0 | 6.1E-05 | 2.1E-04 | BT20/MB468 | MP |
| >hg38_21_+_8433172_8446622_RNA45SN1_45S_50ntFlanks@12607.12625.19\|TGTGCTTGGCTGAGGAGCC | >hg38_21_+_8433172_8446622_RNA45SN1_45S_50ntFlanks | 12607 | 12625 | 19 | TGTGCTTGGCTGAGGAGCC | 293.8 | -3.5 | 0.9 | -3.8 | 1.6E-04 | 2.1E-04 | MB231/MB468 | MITO |
| >hg38_21_+_8433172_8446622_RNA45SN1_45S_50ntFlanks@8353.8392.40\|GTTGAAAAGAACTTTGAAGAGAGAGTTCAAGAGGGCGTGA | >hg38_21_+_8433172_8446622_RNA45SN1_45S_50ntFlanks | 8353 | 8392 | 40 | GTTGAAAAGAACTTTGAAGAGAGAGTTCAAGAGGGCGTGA | 59.5 | 9.2 | 2.2 | 4.2 | 2.2E-05 | 2.1E-04 | BT20/MB231 | MP |
| >hg38_21_+_8433172_8446622_RNA45SN1_45S_50ntFlanks@9788.9818.31\|CGCTGGCGTGGAGCCGGGCGTGGAATGCGAG | >hg38_21_+_8433172_8446622_RNA45SN1_45S_50ntFlanks | 9788 | 9818 | 31 | CGCTGGCGTGGAGCCGGGCGTGGAATGCGAG | 245.0 | 11.3 | 2.7 | 4.2 | 2.2E-05 | 2.1E-04 | BT20/MB231 | MP |
| >hg38_21_+_8433172_8446622_RNA45SN1_45S_50ntFlanks@4976.4999.24\|CCGGACACGGACAGGATTGACAGA | >hg38_21_+_8433172_8446622_RNA45SN1_45S_50ntFlanks | 4976 | 4999 | 24 | CCGGACACGGACAGGATTGACAGA | 518.2 | -3.2 | 0.8 | -4.0 | 6.1E-05 | 2.1E-04 | BT20/MB468 | MP |
| >hg38_21_+_8433172_8446622_RNA45SN1_45S_50ntFlanks@6700.6719.20\|CTAGCTGCGAGAATTAATGT | >hg38_21_+_8433172_8446622_RNA45SN1_45S_50ntFlanks | 6700 | 6719 | 20 | CTAGCTGCGAGAATTAATGT | 670.2 | -2.6 | 0.7 | -3.8 | 1.6E-04 | 2.1E-04 | MB231/MB468 | CYTO |
| >hg38_21_+_8433172_8446622_RNA45SN1_45S_50ntFlanks@12644.12675.32\|TCTGTGGGATTATGACTGAACGCCTCTAAGTC | >hg38_21_+_8433172_8446622_RNA45SN1_45S_50ntFlanks | 12644 | 12675 | 32 | TCTGTGGGATTATGACTGAACGCCTCTAAGTC | 191.1 | -4.3 | 1.1 | -3.8 | 1.7E-04 | 2.1E-04 | MB231/MB468 | NUC |
| >hg38_21_+_8433172_8446622_RNA45SN1_45S_50ntFlanks@4731.4748.18\|AAGAACGAAAGTCGGAGG | >hg38_21_+_8433172_8446622_RNA45SN1_45S_50ntFlanks | 4731 | 4748 | 18 | AAGAACGAAAGTCGGAGG | 695.3 | -3.2 | 0.8 | -4.0 | 6.1E-05 | 2.1E-04 | BT20/MB468 | MP |

<table>
<tr><th></th><th></th><th></th><th></th><th></th><th></th><th></th><th></th><th></th><th></th><th></th><th></th><th></th><th></th></tr>
<tr><td>>trna10_ValCAC_5_-_180649395_180649467@59.76.18,trna12_ValAAC_5_-_180645270_180645342@59.76.18,trna132_ValAAC_6_-_27721179_27721251@59.76.18,trna136_ValAAC_6_-_27648885_27648957@59.76.18,trna15_ValAAC_5_-_180615416_180615488@59.76.18,trna18_ValCAC_5_-_180529253_180529325@59.76.18,trna2_ValAAC_3_+_169490018_169490090@59.76.18,trna2_ValCAC_5_+_180524070_180524142@59.76.18,trna4_ValAAC_5_+_180591154_180591226@59.76.18,trna5_ValAAC_5_+_180596610_180596682@59.76.18,trna6_ValCAC_5_+_180600650_180600722@59.76.18,trna85_ValCAC_1_-_161369490_161369562@59.76.18,trna90_ValCAC_1_-_149684088_149684161@60.77.18,trna9_ValCAC_6_+_26538282_26538354@59.76.18|AACCGGGCGGAAACACCA</td><td>>trna10_ValCAC_5_-_180649395_180649467</td><td>59</td><td>76</td><td>18</td><td>AACCGGGCGGAAACACCA</td><td>224.1</td><td>3.7</td><td>1.0</td><td>3.8</td><td>1.6E-04</td><td>2.1E-04</td><td>MB231/MB468</td><td>MITO</td></tr>
<tr><td>>trna111_HisGTG_1_-_147774845_147774916@-1G.31.32,trna118_HisGTG_1_-_145396881_145396952@-1G.31.32,trna16_HisGTG_1_+_146544773_146544844@-1G.31.32,trna1_HisGTG_15_+_45493349_45493420@-1G.31.32,trna21_HisGTG_1_+_147753471_147753542@-1G.31.32,trna33_HisGTG_6_+_27125906_27125977@-1G.31.32,trna7_HisGTG_9_-_14433938_14434009@-1G.31.32,trna8_HisGTG_15_-_45492611_45492682@-1G.31.32,trna9_HisGTG_15_-_45490804_45490875@-1G.31.32|GGCCGTGATCGTATAGTGGTTAGTACTCTGCG</td><td>>trna111_HisGTG_1_-_147774845_147774916</td><td>-1G</td><td>31</td><td>32</td><td>GGCCGTGATCGTATAGTGGTTAGTACTCTGCG</td><td>3346.5</td><td>-4.5</td><td>1.1</td><td>-4.2</td><td>2.3E-05</td><td>2.2E-04</td><td>BT20/MB231</td><td>NUC</td></tr>
<tr><td>>hg38_21_+_8433172_8446622_RNA45SN1_45S_50ntFlanks@11906.11933.28|CACGGTGAAGAGACATGAGAGGTGTAGA</td><td>>hg38_21_+_8433172_8446622_RNA45SN1_45S_50ntFlanks</td><td>11906</td><td>11933</td><td>28</td><td>CACGGTGAAGAGACATGAGAGGTGTAGA</td><td>69.9</td><td>9.5</td><td>2.2</td><td>4.2</td><td>2.2E-05</td><td>2.2E-04</td><td>BT20/MB231</td><td>MP</td></tr>
<tr><td>>hg38_21_+_8433172_8446622_RNA45SN1_45S_50ntFlanks@9567.9603.37|ACGTGCAAATCGGTCGTCCGACCTGGGTATAGGGGCG</td><td>>hg38_21_+_8433172_8446622_RNA45SN1_45S_50ntFlanks</td><td>9567</td><td>9603</td><td>37</td><td>ACGTGCAAATCGGTCGTCCGACCTGGGTATAGGGGCG</td><td>855.3</td><td>-6.9</td><td>1.8</td><td>-3.8</td><td>1.7E-04</td><td>2.2E-04</td><td>BT20/MB468</td><td>NUC</td></tr>
<tr><td>>hg38_21_+_8433172_8446622_RNA45SN1_45S_50ntFlanks@10786.10826.41|AACAGCCTCTGGCATGTTGGAACAATGTAGGTAAGGGAAGT</td><td>>hg38_21_+_8433172_8446622_RNA45SN1_45S_50ntFlanks</td><td>10786</td><td>10826</td><td>41</td><td>AACAGCCTCTGGCATGTTGGAACAATGTAGGTAAGGGAAGT</td><td>545.3</td><td>-3.6</td><td>1.0</td><td>-3.8</td><td>1.5E-04</td><td>2.2E-04</td><td>BT20/MB468</td><td>CYTO</td></tr>
<tr><td>>MI0000077|hsa-mir-21&WithFlank&17|+|59841260|59841343@14.35.22(+1G)&[MIMAT0000076&hsa-miR-21-5p&offsets|0|0(+1G);m-1&17|+|59841273|59841294&offsets|0|0(+1G)]|TAGCTTATCAGACTGATGTTGAG</td><td>>MI0000077|hsa-mir-21&WithFlank&17|+|59841260|59841343</td><td>14</td><td>35</td><td>22(+1G)</td><td>TAGCTTATCAGACTGATGTTGAG</td><td>1022.8</td><td>4.8</td><td>1.3</td><td>3.8</td><td>1.6E-04</td><td>2.2E-04</td><td>MB231/MB468</td><td>CYTO</td></tr>
<tr><td>>trnaMT_HisGTG_MT_+_12138_12206@25.61.37|CAGATTGTGAATCTGACAACAGAGGCTTACGACCCCT</td><td>>trnaMT_HisGTG_MT_+_12138_12206</td><td>25</td><td>61</td><td>37</td><td>CAGATTGTGAATCTGACAACAGAGGCTTACGACCCCT</td><td>103.4</td><td>10.4</td><td>2.6</td><td>4.0</td><td>6.1E-05</td><td>2.2E-04</td><td>BT20/MB468</td><td>MP</td></tr>
<tr><td>>hg38_21_+_8433172_8446622_RNA45SN1_45S_50ntFlanks@11641.11669.29|CGGCGGGTGTTGACGCGATGTGATTTCTG</td><td>>hg38_21_+_8433172_8446622_RNA45SN1_45S_50ntFlanks</td><td>11641</td><td>11669</td><td>29</td><td>CGGCGGGTGTTGACGCGATGTGATTTCTG</td><td>5831.7</td><td>-7.3</td><td>1.8</td><td>-4.0</td><td>6.1E-05</td><td>2.2E-04</td><td>BT20/MB468</td><td>MP</td></tr>
<tr><td>>hg38_21_+_8433172_8446622_RNA45SN1_45S_50ntFlanks@12622.12654.33|AGCCAATGGGGCGAAGCTACCATCTGTGGGATT</td><td>>hg38_21_+_8433172_8446622_RNA45SN1_45S_50ntFlanks</td><td>12622</td><td>12654</td><td>33</td><td>AGCCAATGGGGCGAAGCTACCATCTGTGGGATT</td><td>198.7</td><td>-2.6</td><td>0.7</td><td>-3.8</td><td>1.5E-04</td><td>2.2E-04</td><td>BT20/MB468</td><td>CYTO</td></tr>
<tr><td>>hg38_21_+_8433172_8446622_RNA45SN1_45S_50ntFlanks@3984.4001.18|GCGGCTTTGGTGACTCTA</td><td>>hg38_21_+_8433172_8446622_RNA45SN1_45S_50ntFlanks</td><td>3984</td><td>4001</td><td>18</td><td>GCGGCTTTGGTGACTCTA</td><td>726.4</td><td>-4.3</td><td>1.1</td><td>-3.8</td><td>1.5E-04</td><td>2.2E-04</td><td>BT20/MB468</td><td>CYTO</td></tr>
<tr><td>>hg38_21_+_8433172_8446622_RNA45SN1_45S_50ntFlanks@11235.11254.20|CTTCCCGTGGATCGCCCCAG</td><td>>hg38_21_+_8433172_8446622_RNA45SN1_45S_50ntFlanks</td><td>11235</td><td>11254</td><td>20</td><td>CTTCCCGTGGATCGCCCCAG</td><td>1582.6</td><td>-6.9</td><td>1.7</td><td>-4.0</td><td>6.1E-05</td><td>2.2E-04</td><td>BT20/MB468</td><td>MP</td></tr>
<tr><td>>hg38_21_+_8433172_8446622_RNA45SN1_45S_50ntFlanks@12625.12659.35|CAATGGGGCGAAGCTACCATCTGTGGGATTATGAC</td><td>>hg38_21_+_8433172_8446622_RNA45SN1_45S_50ntFlanks</td><td>12625</td><td>12659</td><td>35</td><td>CAATGGGGCGAAGCTACCATCTGTGGGATTATGAC</td><td>741.2</td><td>-3.5</td><td>0.9</td><td>-3.8</td><td>1.2E-04</td><td>2.2E-04</td><td>BT20/MB468</td><td>MITO</td></tr>
<tr><td>>hg38_21_+_8433172_8446622_RNA45SN1_45S_50ntFlanks@11580.11613.34|GAACTGGTGCGGACCAGGGGAATCCGACTGTTTA</td><td>>hg38_21_+_8433172_8446622_RNA45SN1_45S_50ntFlanks</td><td>11580</td><td>11613</td><td>34</td><td>GAACTGGTGCGGACCAGGGGAATCCGACTGTTTA</td><td>5568.7</td><td>2.8</td><td>0.7</td><td>3.8</td><td>1.2E-04</td><td>2.2E-04</td><td>BT20/MB468</td><td>MITO</td></tr>
<tr><td>>hg38_21_+_8433172_8446622_RNA45SN1_45S_50ntFlanks@12941.12961.21|ATTCGTAGACGACCTGCTTCT</td><td>>hg38_21_+_8433172_8446622_RNA45SN1_45S_50ntFlanks</td><td>12941</td><td>12961</td><td>21</td><td>ATTCGTAGACGACCTGCTTCT</td><td>485.7</td><td>2.4</td><td>0.6</td><td>3.8</td><td>1.5E-04</td><td>2.2E-04</td><td>BT20/MB468</td><td>CYTO</td></tr>
<tr><td>>hg38_21_+_8433172_8446622_RNA45SN1_45S_50ntFlanks@3850.3881.32|GATAACTGTGGTAATTCTAGAGCTAATACATG</td><td>>hg38_21_+_8433172_8446622_RNA45SN1_45S_50ntFlanks</td><td>3850</td><td>3881</td><td>32</td><td>GATAACTGTGGTAATTCTAGAGCTAATACATG</td><td>208.7</td><td>-4.0</td><td>1.0</td><td>-3.8</td><td>1.2E-04</td><td>2.2E-04</td><td>BT20/MB468</td><td>MITO</td></tr>
<tr><td>>am_trnaMT_PheGAA_MT_+_577_647@40.65.26|TGTTTAGACGGGCTCACATCACCCCA</td><td>>am_trnaMT_PheGAA_MT_+_577_647</td><td>40</td><td>65</td><td>26</td><td>TGTTTAGACGGGCTCACATCACCCCA</td><td>205.1</td><td>8.1</td><td>2.0</td><td>4.0</td><td>6.2E-05</td><td>2.2E-04</td><td>BT20/MB468</td><td>MP</td></tr>
<tr><td>>hg38_21_+_8433172_8446622_RNA45SN1_45S_50ntFlanks@10684.10705.22|CGGCGGCGTCCGGTGAGCTCTC</td><td>>hg38_21_+_8433172_8446622_RNA45SN1_45S_50ntFlanks</td><td>10684</td><td>10705</td><td>22</td><td>CGGCGGCGTCCGGTGAGCTCTC</td><td>1054.4</td><td>-7.0</td><td>1.9</td><td>-3.8</td><td>1.7E-04</td><td>2.2E-04</td><td>BT20/MB468</td><td>NUC</td></tr>
<tr><td>>hg38_21_+_8433172_8446622_RNA45SN1_45S_50ntFlanks@11766.11790.25|ATCTAATTAGTGACGCGCATGAATG</td><td>>hg38_21_+_8433172_8446622_RNA45SN1_45S_50ntFlanks</td><td>11766</td><td>11790</td><td>25</td><td>ATCTAATTAGTGACGCGCATGAATG</td><td>418.0</td><td>-5.0</td><td>1.3</td><td>-3.8</td><td>1.6E-04</td><td>2.2E-04</td><td>MB231/MB468</td><td>CYTO</td></tr>
<tr><td>>MI0000077|hsa-mir-21&WithFlank&17|+|59841260|59841343@14.34.21(+1U)&[MIMAT0000076&hsa-miR-21-5p&offsets|0|-1(+1U);m-1&17|+|59841273|59841294&offsets|0|-1(+1U)]|TAGCTTATCAGACTGATGTTGT</td><td>>MI0000077|hsa-mir-21&WithFlank&17|+|59841260|59841343</td><td>14</td><td>34</td><td>21(+1U)</td><td>TAGCTTATCAGACTGATGTTGT</td><td>395.3</td><td>4.4</td><td>1.2</td><td>3.8</td><td>1.6E-04</td><td>2.2E-04</td><td>MB231/MB468</td><td>CYTO</td></tr>
<tr><td>>hg38_21_+_8433172_8446622_RNA45SN1_45S_50ntFlanks@9554.9595.42|CGTAGCGGTCCTGACGTGCAAATCGGTCGTCCGACCTGGGTA</td><td>>hg38_21_+_8433172_8446622_RNA45SN1_45S_50ntFlanks</td><td>9554</td><td>9595</td><td>42</td><td>CGTAGCGGTCCTGACGTGCAAATCGGTCGTCCGACCTGGGTA</td><td>842.2</td><td>-4.9</td><td>1.3</td><td>-3.8</td><td>1.7E-04</td><td>2.2E-04</td><td>MB231/MB468</td><td>NUC</td></tr>
<tr><td>>hg38_21_+_8433172_8446622_RNA45SN1_45S_50ntFlanks@9526.9570.45|GAAGCCAGAGGAAACTCTGGTGGAGGTCCGTAGCGGTCCTGACGT</td><td>>hg38_21_+_8433172_8446622_RNA45SN1_45S_50ntFlanks</td><td>9526</td><td>9570</td><td>45</td><td>GAAGCCAGAGGAAACTCTGGTGGAGGTCCGTAGCGGTCCTGACGT</td><td>771.1</td><td>-4.0</td><td>1.0</td><td>-4.3</td><td>2.1E-05</td><td>2.2E-04</td><td>BT20/MB231</td><td>CYTO</td></tr>
<tr><td>>MI0000736|hsa-mir-30c-1&WithFlank&1|+|40757278|40757378@23.46.24(+2U);MI0000254|hsa-mir-30c-2&WithFlank&6|-|71376954|71377037@13.36.24(+2U)&[MIMAT0000244&hsa-miR-30c-5p&offsets|0|+1(+2U);m-57&1|+|40757300|40757323&offsets|0|0(+2U)];[MIMAT0000244_1&hsa-miR-30c-5p&offsets|0|+1(+2U);m-56&6|-|71377002|71377025&offsets|0|0(+2U)]|TGTAAACATCCTACACTCTCAGCTTT</td><td>>MI0000736|hsa-mir-30c-1&WithFlank&1|+|40757278|40757378</td><td>23</td><td>46</td><td>24(+2U)</td><td>TGTAAACATCCTACACTCTCAGCTTT</td><td>295.9</td><td>-7.3</td><td>1.7</td><td>-4.2</td><td>2.2E-05</td><td>2.2E-04</td><td>BT20/MB231</td><td>MP</td></tr>
<tr><td>>hg38_21_+_8433172_8446622_RNA45SN1_45S_50ntFlanks@6702.6724.23|AGCTGCGAGAATTAATGTGAATT</td><td>>hg38_21_+_8433172_8446622_RNA45SN1_45S_50ntFlanks</td><td>6702</td><td>6724</td><td>23</td><td>AGCTGCGAGAATTAATGTGAATT</td><td>99.6</td><td>10.4</td><td>2.6</td><td>4.0</td><td>6.2E-05</td><td>2.2E-04</td><td>BT20/MB468</td><td>MP</td></tr>
<tr><td>>am_trnaMT_GluTTC_MT_-_14674_14742@20.54.35,trnalookalike8_GluTTC_5_-_93905172_93905240@20.54.35|AACGATGGTTTTTCATATCATTGGTCGTGGTTGTA</td><td>>am_trnaMT_GluTTC_MT_-_14674_14742</td><td>20</td><td>54</td><td>35</td><td>AACGATGGTTTTTCATATCATTGGTCGTGGTTGTA</td><td>253.5</td><td>7.9</td><td>2.0</td><td>4.0</td><td>6.2E-05</td><td>2.2E-04</td><td>BT20/MB468</td><td>MP</td></tr>
<tr><td>>MI0033423|hsa-mir-10399&WithFlank&7|-|139044094|139044163@43.63.21&[MIMAT0041630&hsa-miR-10399-3p&offsets|-1|-1;TJU_CMC_MD2.ID03061.3p-miR&offsets|+3|+2;m-2657&7|-|139044103|139044124&offsets|+3|+2]|TCTCTCGGACAAGCTGTAGGT</td><td>>MI0033423|hsa-mir-10399&WithFlank&7|-|139044094|139044163</td><td>43</td><td>63</td><td>21</td><td>TCTCTCGGACAAGCTGTAGGT</td><td>52.6</td><td>-9.1</td><td>2.1</td><td>-4.2</td><td>2.2E-05</td><td>2.2E-04</td><td>BT20/MB231</td><td>MP</td></tr>
<tr><td>>hg38_21_+_8433172_8446622_RNA45SN1_45S_50ntFlanks@3828.3860.33|TCGCTCGCTCCTCTCCTACTTGGATAACTGTGG</td><td>>hg38_21_+_8433172_8446622_RNA45SN1_45S_50ntFlanks</td><td>3828</td><td>3860</td><td>33</td><td>TCGCTCGCTCCTCTCCTACTTGGATAACTGTGG</td><td>196.4</td><td>-4.2</td><td>1.1</td><td>-3.8</td><td>1.7E-04</td><td>2.2E-04</td><td>MB231/MB468</td><td>NUC</td></tr>
<tr><td>>hg38_21_+_8433172_8446622_RNA45SN1_45S_50ntFlanks@12913.12935.23|ACGTTCGTGGGGAACCTGGCGCT</td><td>>hg38_21_+_8433172_8446622_RNA45SN1_45S_50ntFlanks</td><td>12913</td><td>12935</td><td>23</td><td>ACGTTCGTGGGGAACCTGGCGCT</td><td>184.9</td><td>-4.2</td><td>1.1</td><td>-3.8</td><td>1.4E-04</td><td>2.2E-04</td><td>MB231/MB468</td><td>MP</td></tr>
<tr><td>>hg38_21_+_8433172_8446622_RNA45SN1_45S_50ntFlanks@10679.10707.29|GGTTCCGGCGGCGTCCGGTGAGCTCTCGC</td><td>>hg38_21_+_8433172_8446622_RNA45SN1_45S_50ntFlanks</td><td>10679</td><td>10707</td><td>29</td><td>GGTTCCGGCGGCGTCCGGTGAGCTCTCGC</td><td>140.8</td><td>-2.4</td><td>0.6</td><td>-3.8</td><td>1.6E-04</td><td>2.2E-04</td><td>MB231/MB468</td><td>CYTO</td></tr>
<tr><td>>hg38_21_+_8433172_8446622_RNA45SN1_45S_50ntFlanks@12652.12675.24|ATTATGACTGAACGCCTCTAAGTC</td><td>>hg38_21_+_8433172_8446622_RNA45SN1_45S_50ntFlanks</td><td>12652</td><td>12675</td><td>24</td><td>ATTATGACTGAACGCCTCTAAGTC</td><td>193.7</td><td>-4.0</td><td>1.0</td><td>-3.8</td><td>1.5E-04</td><td>2.2E-04</td><td>BT20/MB468</td><td>CYTO</td></tr>
<tr><td>>hg38_21_+_8433172_8446622_RNA45SN1_45S_50ntFlanks@10738.10756.19|AAATCTCGCGCCGGGCCGT</td><td>>hg38_21_+_8433172_8446622_RNA45SN1_45S_50ntFlanks</td><td>10738</td><td>10756</td><td>19</td><td>AAATCTCGCGCCGGGCCGT</td><td>959.1</td><td>3.4</td><td>0.9</td><td>3.8</td><td>1.7E-04</td><td>2.2E-04</td><td>MB231/MB468</td><td>NUC</td></tr>
<tr><td>>hg38_21_+_8433172_8446622_RNA45SN1_45S_50ntFlanks@8124.8141.18|TGTGGCGTACGGAAGACC</td><td>>hg38_21_+_8433172_8446622_RNA45SN1_45S_50ntFlanks</td><td>8124</td><td>8141</td><td>18</td><td>TGTGGCGTACGGAAGACC</td><td>53.7</td><td>9.1</td><td>2.1</td><td>4.2</td><td>2.2E-05</td><td>2.2E-04</td><td>BT20/MB231</td><td>MP</td></tr>
<tr><td>>trnaMT_LysTTT_MT_+_8295_8364@48.72.25,trnalookalike7_LysTTT_1_+_568844_568913@48.72.25|GAACCAACACCTCTTTACAGTGACC</td><td>>trnaMT_LysTTT_MT_+_8295_8364</td><td>48</td><td>72</td><td>25</td><td>GAACCAACACCTCTTTACAGTGACC</td><td>22.6</td><td>7.7</td><td>2.0</td><td>3.8</td><td>1.4E-04</td><td>2.2E-04</td><td>MB231/MB468</td><td>MP</td></tr>
<tr><td>>hg38_21_+_8433172_8446622_RNA45SN1_45S_50ntFlanks@10362.10382.21|AAGGCCGAAGTGGAGAAGGGT</td><td>>hg38_21_+_8433172_8446622_RNA45SN1_45S_50ntFlanks</td><td>10362</td><td>10382</td><td>21</td><td>AAGGCCGAAGTGGAGAAGGGT</td><td>386.7</td><td>-2.7</td><td>0.7</td><td>-4.0</td><td>6.2E-05</td><td>2.2E-04</td><td>BT20/MB468</td><td>MP</td></tr>
<tr><td>>hg38_21_+_8433172_8446622_RNA45SN1_45S_50ntFlanks@10729.10755.27|AGAGGGTGTAAATCTCGCGCCGGGCCG</td><td>>hg38_21_+_8433172_8446622_RNA45SN1_45S_50ntFlanks</td><td>10729</td><td>10755</td><td>27</td><td>AGAGGGTGTAAATCTCGCGCCGGGCCG</td><td>271.6</td><td>-5.2</td><td>1.4</td><td>-3.8</td><td>1.5E-04</td><td>2.2E-04</td><td>BT20/MB468</td><td>CYTO</td></tr>
<tr><td>>hg38_21_+_8433172_8446622_RNA45SN1_45S_50ntFlanks@11658.11687.30|ATGTGATTTCTGCCCAGTGCTCTGAATGTC</td><td>>hg38_21_+_8433172_8446622_RNA45SN1_45S_50ntFlanks</td><td>11658</td><td>11687</td><td>30</td><td>ATGTGATTTCTGCCCAGTGCTCTGAATGTC</td><td>265.4</td><td>7.1</td><td>1.8</td><td>4.0</td><td>6.2E-05</td><td>2.2E-04</td><td>BT20/MB468</td><td>MP</td></tr>
<tr><td>>hg38_21_+_8433172_8446622_RNA45SN1_45S_50ntFlanks@10677.10708.32|GCGGTTCCGGCGGCGTCCGGTGAGCTCTCGCT</td><td>>hg38_21_+_8433172_8446622_RNA45SN1_45S_50ntFlanks</td><td>10677</td><td>10708</td><td>32</td><td>GCGGTTCCGGCGGCGTCCGGTGAGCTCTCGCT</td><td>1837.6</td><td>4.1</td><td>1.1</td><td>3.8</td><td>1.2E-04</td><td>2.2E-04</td><td>BT20/MB468</td><td>MITO</td></tr>
<tr><td>>hg38_21_+_8433172_8446622_RNA45SN1_45S_50ntFlanks@5400.5419.20|CACACCGCCCGTCGCTACTA</td><td>>hg38_21_+_8433172_8446622_RNA45SN1_45S_50ntFlanks</td><td>5400</td><td>5419</td><td>20</td><td>CACACCGCCCGTCGCTACTA</td><td>147.4</td><td>-5.4</td><td>1.4</td><td>-3.8</td><td>1.2E-04</td><td>2.2E-04</td><td>BT20/MB468</td><td>MITO</td></tr>
<tr><td>>hg38_21_+_8433172_8446622_RNA45SN1_45S_50ntFlanks@4067.4089.23|AACTTTCGATGGTAGTCGCCGTG</td><td>>hg38_21_+_8433172_8446622_RNA45SN1_45S_50ntFlanks</td><td>4067</td><td>4089</td><td>23</td><td>AACTTTCGATGGTAGTCGCCGTG</td><td>155.6</td><td>-2.9</td><td>0.8</td><td>-3.8</td><td>1.6E-04</td><td>2.2E-04</td><td>MB231/MB468</td><td>CYTO</td></tr>
<tr><td>>hg38_1_-_228634819_228635039_RNA5S12_5S_50ntFlanks@127.171.45|ACTTGGATGGGAGACCGCCTGGGAATACCGGGTGCTGTAGGCTTT</td><td>>hg38_1_-_228634819_228635039_RNA5S12_5S_50ntFlanks</td><td>127</td><td>171</td><td>45</td><td>ACTTGGATGGGAGACCGCCTGGGAATACCGGGTGCTGTAGGCTTT</td><td>708.0</td><td>5.9</td><td>1.6</td><td>3.8</td><td>1.7E-04</td><td>2.2E-04</td><td>MB231/MB468</td><td>NUC</td></tr>
<tr><td>>hg38_21_+_8433172_8446622_RNA45SN1_45S_50ntFlanks@12045.12070.26|CTGACCCGGTGAGGCGGGGGGGCGAG</td><td>>hg38_21_+_8433172_8446622_RNA45SN1_45S_50ntFlanks</td><td>12045</td><td>12070</td><td>26</td><td>CTGACCCGGTGAGGCGGGGGGGCGAG</td><td>451.3</td><td>-5.8</td><td>1.5</td><td>-3.8</td><td>1.2E-04</td><td>2.2E-04</td><td>BT20/MB468</td><td>MITO</td></tr>
<tr><td>>hg38_21_+_8433172_8446622_RNA45SN1_45S_50ntFlanks@5078.5115.38|CGATAACGAACGAGACTCTGGCATGCTAACTAGTTACG</td><td>>hg38_21_+_8433172_8446622_RNA45SN1_45S_50ntFlanks</td><td>5078</td><td>5115</td><td>38</td><td>CGATAACGAACGAGACTCTGGCATGCTAACTAGTTACG</td><td>225.0</td><td>-6.1</td><td>1.6</td><td>-3.8</td><td>1.4E-04</td><td>2.2E-04</td><td>MB231/MB468</td><td>MP</td></tr>
<tr><td>>hg38_21_+_8433172_8446622_RNA45SN1_45S_50ntFlanks@12942.12973.32|TTCGTAGACGACCTGCTTCTGGGTCGGGGTTT</td><td>>hg38_21_+_8433172_8446622_RNA45SN1_45S_50ntFlanks</td><td>12942</td><td>12973</td><td>32</td><td>TTCGTAGACGACCTGCTTCTGGGTCGGGGTTT</td><td>103.9</td><td>-4.9</td><td>1.3</td><td>-3.8</td><td>1.2E-04</td><td>2.2E-04</td><td>BT20/MB468</td><td>MITO</td></tr>
<tr><td>>MI0000065|hsa-let-7d&WithFlank&9|+|94178828|94178926@14.34.21&[MIMAT0000065&hsa-let-7d-5p&offsets|0|-1;m-90&9|+|94178841|94178862&offsets|0|-1]|AGAGGTAGTAGGTTGCATAGT</td><td>>MI0000065|hsa-let-7d&WithFlank&9|+|94178828|94178926</td><td>14</td><td>34</td><td>21</td><td>AGAGGTAGTAGGTTGCATAGT</td><td>407.8</td><td>3.5</td><td>0.9</td><td>4.0</td><td>6.2E-05</td><td>2.2E-04</td><td>BT20/MB468</td><td>MP</td></tr>
</table>

| | | | | | | | | | | | | | |
|---|---|---|---|---|---|---|---|---|---|---|---|---|---|
| >hg38_21_+_8433172_8446622_RNA45SN1_45S_50ntFlanks@10819.10837.19\|AGGGAAGTCGGCAAGCCGG | >hg38_21_+_8433172_8446622_RNA45SN1_45S_50ntFlanks | 10819 | 10837 | 19 | AGGGAAGTCGGCAAGCCGG | 323.7 | -4.9 | 1.3 | -3.8 | 1.7E-04 | 2.2E-04 | MB231/MB468 | NUC |
| >hg38_21_+_8433172_8446622_RNA45SN1_45S_50ntFlanks@4367.4388.22\|CAGTTAAAAAGCTCGTAGTTGG | >hg38_21_+_8433172_8446622_RNA45SN1_45S_50ntFlanks | 4367 | 4388 | 22 | CAGTTAAAAAGCTCGTAGTTGG | 233.8 | -5.5 | 1.5 | -3.8 | 1.5E-04 | 2.2E-04 | BT20/MB468 | CYTO |
| >hg38_21_+_8433172_8446622_RNA45SN1_45S_50ntFlanks@6646.6669.24\|TCGTACGACTCTTAGCGGTGGATC | >hg38_21_+_8433172_8446622_RNA45SN1_45S_50ntFlanks | 6646 | 6669 | 24 | TCGTACGACTCTTAGCGGTGGATC | 114517.8 | 3.0 | 0.7 | 4.2 | 2.5E-05 | 2.2E-04 | BT20/MB231 | MITO |
| >hg38_21_+_8433172_8446622_RNA45SN1_45S_50ntFlanks@6655.6687.33\|TCTTAGCGGTGGATCACTCGGCTCGTGCGTCGA | >hg38_21_+_8433172_8446622_RNA45SN1_45S_50ntFlanks | 6655 | 6687 | 33 | TCTTAGCGGTGGATCACTCGGCTCGTGCGTCGA | 331.2 | 2.9 | 0.8 | 3.8 | 1.7E-04 | 2.2E-04 | MB231/MB468 | NUC |
| >hg38_21_+_8433172_8446622_RNA45SN1_45S_50ntFlanks@6779.6803.25\|CCGGGGCTACGCCTGTCTGAGCGTC | >hg38_21_+_8433172_8446622_RNA45SN1_45S_50ntFlanks | 6779 | 6803 | 25 | CCGGGGCTACGCCTGTCTGAGCGTC | 285.8 | -2.4 | 0.6 | -3.8 | 1.7E-04 | 2.2E-04 | MB231/MB468 | NUC |
| >hg38_21_+_8433172_8446622_RNA45SN1_45S_50ntFlanks@6647.6667.21\|CGTACGACTCTTAGCGGTGGA | >hg38_21_+_8433172_8446622_RNA45SN1_45S_50ntFlanks | 6647 | 6667 | 21 | CGTACGACTCTTAGCGGTGGA | 1870.7 | 2.7 | 0.7 | 3.8 | 1.7E-04 | 2.2E-04 | MB231/MB468 | MITO |
| >hg38_21_+_8433172_8446622_RNA45SN1_45S_50ntFlanks@7984.8003.20\|AGATCAGACGTGGCGACCCG | >hg38_21_+_8433172_8446622_RNA45SN1_45S_50ntFlanks | 7984 | 8003 | 20 | AGATCAGACGTGGCGACCCG | 270.7 | -4.2 | 1.1 | -3.8 | 1.5E-04 | 2.2E-04 | BT20/MB468 | CYTO |
| >MI0000071\|hsa-mir-17&WithFlank&13\|+\|91350599\|91350694@20.41.22(+1A)&[MIMAT0000070&hsa-miR-17-5p&offsets\|0\|-1(+1A);m-88&13\|+\|91350618\|91350640&offsets\|0\|-1(+1A)]\|CAAAGTGCTTACAGTGCAGGTAA | >MI0000071\|hsa-mir-17&WithFlank&13\|+\|91350599\|91350694 | 20 | 41 | 22(+1A) | CAAAGTGCTTACAGTGCAGGTAA | 1474.6 | -3.7 | 0.9 | -4.2 | 2.5E-05 | 2.2E-04 | BT20/MB231 | MITO |
| >hg38_21_+_8433172_8446622_RNA45SN1_45S_50ntFlanks@6722.6758.37\|ATTGCAGGACACATTGATCATCGACACTTCGAACGCA | >hg38_21_+_8433172_8446622_RNA45SN1_45S_50ntFlanks | 6722 | 6758 | 37 | ATTGCAGGACACATTGATCATCGACACTTCGAACGCA | 95.9 | -6.0 | 1.6 | -3.8 | 1.4E-04 | 2.2E-04 | MB231/MB468 | MP |
| >MI0000439\|hsa-mir-23b&WithFlank&9\|+\|95085202\|95085310@67.85.19(+2U)&[MIMAT0000418&hsa-miR-23b-3p&offsets\|+3\|-1(+2U);m-42&9\|+\|95085265\|95085285&offsets\|+3\|+1(+2U)]\|ACATTGCCAGGGATTACCATT | >MI0000439\|hsa-mir-23b&WithFlank&9\|+\|95085202\|95085310 | 67 | 85 | 19(+2U) | ACATTGCCAGGGATTACCATT | 92.5 | 9.8 | 2.6 | 3.8 | 1.4E-04 | 2.2E-04 | MB231/MB468 | MP |
| >hg38_1_-_228634819_228635039_RNA5S12_5S_50ntFlanks@137.171.35\|GAGACCGCCTGGGAATACCGGGTGCTGTAGGCTTT | >hg38_1_-_228634819_228635039_RNA5S12_5S_50ntFlanks | 137 | 171 | 35 | GAGACCGCCTGGGAATACCGGGTGCTGTAGGCTTT | 1407.4 | 4.2 | 1.0 | 4.0 | 6.3E-05 | 2.2E-04 | BT20/MB468 | MP |
| >MI0000088\|hsa-mir-30a&WithFlank&6\|-\|71403545\|71403627@53.73.21(+2U)&[MIMAT0000088&hsa-miR-30a-3p&offsets\|0\|-1(+2U);m-38&6\|-\|71403554\|71403575&offsets\|0\|-1(+2U)]\|CTTTCAGTCGGATGTTTGCAGTT | >MI0000088\|hsa-mir-30a&WithFlank&6\|-\|71403545\|71403627 | 53 | 73 | 21(+2U) | CTTTCAGTCGGATGTTTGCAGTT | 480.8 | 4.9 | 1.3 | 3.8 | 1.4E-04 | 2.2E-04 | MB231/MB468 | MP |
| >hg38_21_+_8433172_8446622_RNA45SN1_45S_50ntFlanks@9300.9336.37\|ACGGACCAAGGAGTCTAACACGTGCGCGAGTCGGGGG | >hg38_21_+_8433172_8446622_RNA45SN1_45S_50ntFlanks | 9300 | 9336 | 37 | ACGGACCAAGGAGTCTAACACGTGCGCGAGTCGGGGG | 64.6 | 9.3 | 2.2 | 4.2 | 2.2E-05 | 2.2E-04 | BT20/MB231 | MP |
| >MI0000265\|hsa-mir-7-3&WithFlank&19\|+\|4770664\|4770785@37.61.25&[MIMAT0000252_1&hsa-miR-7-5p&offsets\|0\|+1;m-402&19\|+\|4770700\|4770723&offsets\|0\|+1]\|TGGAAGACTAGTGATTTTGTTGTTC | >MI0000265\|hsa-mir-7-3&WithFlank&19\|+\|4770664\|4770785 | 37 | 61 | 25 | TGGAAGACTAGTGATTTTGTTGTTC | 501.2 | -4.5 | 1.1 | -4.2 | 2.2E-05 | 2.2E-04 | BT20/MB231 | MP |
| >hg38_21_+_8433172_8446622_RNA45SN1_45S_50ntFlanks@10771.10818.48\|CAGGTCTCCAAGGTGAACAGCCTCTGGCATGTTGGAACAATGTAGGTA | >hg38_21_+_8433172_8446622_RNA45SN1_45S_50ntFlanks | 10771 | 10818 | 48 | CAGGTCTCCAAGGTGAACAGCCTCTGGCATGTTGGAACAATGTAGGTA | 124.8 | -10.0 | 2.5 | -4.0 | 6.3E-05 | 2.2E-04 | BT20/MB468 | MP |
| >hg38_21_+_8433172_8446622_RNA45SN1_45S_50ntFlanks@11592.11612.21\|ACCAGGGGAATCCGACTGTTT | >hg38_21_+_8433172_8446622_RNA45SN1_45S_50ntFlanks | 11592 | 11612 | 21 | ACCAGGGGAATCCGACTGTTT | 465.0 | -4.9 | 1.3 | -3.8 | 1.5E-04 | 2.2E-04 | BT20/MB468 | CYTO |
| >hg38_21_+_8433172_8446622_RNA45SN1_45S_50ntFlanks@10849.10867.19\|CGGGATAAGGATTGGCTCT | >hg38_21_+_8433172_8446622_RNA45SN1_45S_50ntFlanks | 10849 | 10867 | 19 | CGGGATAAGGATTGGCTCT | 449.8 | -6.5 | 1.7 | -3.8 | 1.7E-04 | 2.2E-04 | BT20/MB468 | NUC |
| >am_trnaMT_ValTAC_MT_+_1602_1670@33.66.34\|ACACTTAGGAGATTTCAACTTAACTTGACCGCTC | >am_trnaMT_ValTAC_MT_+_1602_1670 | 33 | 66 | 34 | ACACTTAGGAGATTTCAACTTAACTTGACCGCTC | 112.8 | 10.6 | 2.6 | 4.0 | 6.3E-05 | 2.2E-04 | BT20/MB468 | MP |
| >hg38_21_+_8433172_8446622_RNA45SN1_45S_50ntFlanks@12420.12449.30\|ATGTCGGCTCTTCCTATCATTGTGAAGCAG | >hg38_21_+_8433172_8446622_RNA45SN1_45S_50ntFlanks | 12420 | 12449 | 30 | ATGTCGGCTCTTCCTATCATTGTGAAGCAG | 1023.3 | -1.6 | 0.4 | -3.8 | 1.5E-04 | 2.2E-04 | BT20/MB468 | CYTO |
| >hg38_21_+_8433172_8446622_RNA45SN1_45S_50ntFlanks@7974.7997.24\|ACGCGACCTCAGATCAGACGTGGC | >hg38_21_+_8433172_8446622_RNA45SN1_45S_50ntFlanks | 7974 | 7997 | 24 | ACGCGACCTCAGATCAGACGTGGC | 3184.5 | 3.7 | 0.9 | 4.0 | 6.3E-05 | 2.2E-04 | BT20/MB468 | MP |
| >hg38_21_+_8433172_8446622_RNA45SN1_45S_50ntFlanks@475.501.27\|CTTCGTGATCGATGTGGTGACGTCGTG | >hg38_21_+_8433172_8446622_RNA45SN1_45S_50ntFlanks | 475 | 501 | 27 | CTTCGTGATCGATGTGGTGACGTCGTG | 866.2 | 5.2 | 1.4 | 3.8 | 1.7E-04 | 2.2E-04 | MB231/MB468 | MITO |
| >hg38_21_+_8433172_8446622_RNA45SN1_45S_50ntFlanks@10066.10085.20\|GAGAGTGGACGGGAGCGGCG | >hg38_21_+_8433172_8446622_RNA45SN1_45S_50ntFlanks | 10066 | 10085 | 20 | GAGAGTGGACGGGAGCGGCG | 62.3 | 9.3 | 2.2 | 4.2 | 2.3E-05 | 2.2E-04 | BT20/MB231 | MP |
| >hg38_21_+_8433172_8446622_RNA45SN1_45S_50ntFlanks@6729.6752.24\|GACACATTGATCATCGACACTTCG | >hg38_21_+_8433172_8446622_RNA45SN1_45S_50ntFlanks | 6729 | 6752 | 24 | GACACATTGATCATCGACACTTCG | 79.6 | -5.7 | 1.5 | -3.8 | 1.4E-04 | 2.2E-04 | MB231/MB468 | MP |
| >am_trnaMT_PheGAA_MT_+_577_647@41.62.22\|GTTTAGACGGGCTCACATCACC | >am_trnaMT_PheGAA_MT_+_577_647 | 41 | 62 | 22 | GTTTAGACGGGCTCACATCACC | 190.9 | 2.4 | 0.6 | 3.8 | 1.4E-04 | 2.2E-04 | MB231/MB468 | MP |
| >hg38_21_+_8433172_8446622_RNA45SN1_45S_50ntFlanks@11540.11586.47\|CGCGGCGCCTCGCCTCGGCCGGCGCCTAGCAGCCGACTTAGAACTGG | >hg38_21_+_8433172_8446622_RNA45SN1_45S_50ntFlanks | 11540 | 11586 | 47 | CGCGGCGCCTCGCCTCGGCCGGCGCCTAGCAGCCGACTTAGAACTGG | 539.1 | 3.5 | 0.9 | 3.8 | 1.6E-04 | 2.2E-04 | MB231/MB468 | CYTO |
| >hg38_21_+_8433172_8446622_RNA45SN1_45S_50ntFlanks@6653.6701.49\|ACTCTTAGCGGTGGATCACTCGGCTCGTGCGTCGATGAAGAACGCAGCT | >hg38_21_+_8433172_8446622_RNA45SN1_45S_50ntFlanks | 6653 | 6701 | 49 | ACTCTTAGCGGTGGATCACTCGGCTCGTGCGTCGATGAAGAACGCAGCT | 103.0 | 10.4 | 2.6 | 4.0 | 6.3E-05 | 2.2E-04 | BT20/MB468 | MP |
| >hg38_21_+_8433172_8446622_RNA45SN1_45S_50ntFlanks@12539.12573.35\|ATGATGTGTTGTTGCCATGGTAATCCTGCTCAGTA | >hg38_21_+_8433172_8446622_RNA45SN1_45S_50ntFlanks | 12539 | 12573 | 35 | ATGATGTGTTGTTGCCATGGTAATCCTGCTCAGTA | 484.1 | -4.2 | 1.1 | -3.8 | 1.3E-04 | 2.2E-04 | BT20/MB468 | MITO |
| >MI0000749\|hsa-mir-30e&WithFlank&1\|+\|40754349\|40754452@23.45.23(+1A)&[MIMAT0000692&hsa-miR-30e-5p&offsets\|0\|+1(+1A);m-24&1\|+\|40754371\|40754394&offsets\|0\|-1(+1A)]\|TGTAAACATCCTTGACTGGAAGCA | >MI0000749\|hsa-mir-30e&WithFlank&1\|+\|40754349\|40754452 | 23 | 45 | 23(+1A) | TGTAAACATCCTTGACTGGAAGCA | 781.2 | 4.3 | 1.1 | 3.8 | 1.7E-04 | 2.2E-04 | MB231/MB468 | MITO |
| >hg38_21_+_8433172_8446622_RNA45SN1_45S_50ntFlanks@11323.11343.21\|CGTCTCGTCGCGCGCGCGTCC | >hg38_21_+_8433172_8446622_RNA45SN1_45S_50ntFlanks | 11323 | 11343 | 21 | CGTCTCGTCGCGCGCGCGTCC | 29643.4 | -5.6 | 1.5 | -3.8 | 1.7E-04 | 2.2E-04 | BT20/MB468 | NUC |
| >hg38_21_+_8433172_8446622_RNA45SN1_45S_50ntFlanks@9561.9596.36\|GTCCTGACGTGCAAATCGGTCGTCCGACCTGGGTAT | >hg38_21_+_8433172_8446622_RNA45SN1_45S_50ntFlanks | 9561 | 9596 | 36 | GTCCTGACGTGCAAATCGGTCGTCCGACCTGGGTAT | 169.0 | -5.3 | 1.4 | -3.8 | 1.5E-04 | 2.2E-04 | BT20/MB468 | CYTO |
| >hg38_21_+_8433172_8446622_RNA45SN1_45S_50ntFlanks@8380.8400.21\|CAAGAGGGCGTGAAACCGTTA | >hg38_21_+_8433172_8446622_RNA45SN1_45S_50ntFlanks | 8380 | 8400 | 21 | CAAGAGGGCGTGAAACCGTTA | 4138.2 | -7.3 | 1.8 | -4.0 | 6.3E-05 | 2.2E-04 | BT20/MB468 | MP |
| >trnaMT_PheGAA_MT_+_577_647@43.74.32\|TTAGACGGGCTCACATCACCCCATAAACACCA | >trnaMT_PheGAA_MT_+_577_647 | 43 | 74 | 32 | TTAGACGGGCTCACATCACCCCATAAACACCA | 297.1 | 3.7 | 1.0 | 3.8 | 1.7E-04 | 2.2E-04 | MB231/MB468 | MITO |
| >hg38_21_+_8433172_8446622_RNA45SN1_45S_50ntFlanks@12939.12956.18\|CCATTCGTAGACGACCTG | >hg38_21_+_8433172_8446622_RNA45SN1_45S_50ntFlanks | 12939 | 12956 | 18 | CCATTCGTAGACGACCTG | 493.5 | -2.6 | 0.7 | -3.8 | 1.3E-04 | 2.2E-04 | BT20/MB468 | MITO |
| >hg38_21_+_8433172_8446622_RNA45SN1_45S_50ntFlanks@9938.9960.23\|AGCAGGACGGTGGCCATGGAAGT | >hg38_21_+_8433172_8446622_RNA45SN1_45S_50ntFlanks | 9938 | 9960 | 23 | AGCAGGACGGTGGCCATGGAAGT | 1609.9 | 4.1 | 1.0 | 4.0 | 6.3E-05 | 2.2E-04 | BT20/MB468 | MP |
| >trna116_GluCTC_1_-_145399233_145399304@8.33.26,trna59_GluCTC_1_+_249168447_249168518@8.33.26,trna71_GluCTC_1_-_161439189_161439260@8.33.26,trna74_GluCTC_1_-_161431809_161431880@8.33.26,trna77_GluCTC_1_-_161424398_161424469@8.33.26,trna77_GluCTC_6_+_28949976_28950047@8.33.26,trna80_GluCTC_1_-_161417018_161417089@8.33.26,trna87_GluCTC_6_-_126101393_126101464@8.33.26\|TGGTCTAGTGGTTAGGATTCGGCGCT | >trna116_GluCTC_1_-_145399233_145399304 | 8 | 33 | 26 | TGGTCTAGTGGTTAGGATTCGGCGCT | 367.1 | 6.2 | 1.6 | 3.8 | 1.5E-04 | 2.2E-04 | BT20/MB468 | CYTO |
| >hg38_21_+_8433172_8446622_RNA45SN1_45S_50ntFlanks@5059.5082.24\|CGATTTGTCTGGTTAATTCCGATA | >hg38_21_+_8433172_8446622_RNA45SN1_45S_50ntFlanks | 5059 | 5082 | 24 | CGATTTGTCTGGTTAATTCCGATA | 319.4 | -6.6 | 1.7 | -4.0 | 6.4E-05 | 2.2E-04 | BT20/MB468 | MP |
| >hg38_21_+_8433172_8446622_RNA45SN1_45S_50ntFlanks@6644.6662.19\|CCTCGTACGACTCTTAGCG | >hg38_21_+_8433172_8446622_RNA45SN1_45S_50ntFlanks | 6644 | 6662 | 19 | CCTCGTACGACTCTTAGCG | 1030.8 | 3.9 | 1.0 | 3.8 | 1.7E-04 | 2.2E-04 | MB231/MB468 | MITO |
| >trna10_LysCTT_16_+_3241501_3241573@3.36.34,trna13_LysCTT_14_-_58706613_58706685@3.36.34,trna2_LysCTT_15_+_79152904_79152976@3.36.34\|CCGGCTAGCTCAGTCGGTAGAGCATGGGACTCTT | >trna10_LysCTT_16_+_3241501_3241573 | 3 | 36 | 34 | CCGGCTAGCTCAGTCGGTAGAGCATGGGACTCTT | 716.2 | -3.8 | 0.9 | -4.2 | 2.6E-05 | 2.2E-04 | BT20/MB231 | MITO |
| >hg38_21_+_8433172_8446622_RNA45SN1_45S_50ntFlanks@9454.9488.35\|CCGCGCCGGGGAGGTGGAGCACGAGCGCACGTGTT | >hg38_21_+_8433172_8446622_RNA45SN1_45S_50ntFlanks | 9454 | 9488 | 35 | CCGCGCCGGGGAGGTGGAGCACGAGCGCACGTGTT | 417.6 | -4.1 | 1.1 | -3.8 | 1.3E-04 | 2.2E-04 | BT20/MB468 | MITO |
| >hg38_21_+_8433172_8446622_RNA45SN1_45S_50ntFlanks@12717.12747.31\|CCTCGGTTGGCCTCGGATAGCCGGTCCCCCG | >hg38_21_+_8433172_8446622_RNA45SN1_45S_50ntFlanks | 12717 | 12747 | 31 | CCTCGGTTGGCCTCGGATAGCCGGTCCCCCG | 178.5 | -5.2 | 1.4 | -3.8 | 1.3E-04 | 2.2E-04 | BT20/MB468 | MITO |
| >hg38_21_+_8433172_8446622_RNA45SN1_45S_50ntFlanks@10770.10818.49\|GCAGGTCTCCAAGGTGAACAGCCTCTGGCATGTTGGAACAATGTAGGTA | >hg38_21_+_8433172_8446622_RNA45SN1_45S_50ntFlanks | 10770 | 10818 | 49 | GCAGGTCTCCAAGGTGAACAGCCTCTGGCATGTTGGAACAATGTAGGTA | 160.4 | -4.2 | 1.1 | -3.8 | 1.6E-04 | 2.2E-04 | MB231/MB468 | CYTO |
| >hg38_21_+_8433172_8446622_RNA45SN1_45S_50ntFlanks@10345.10373.29\|TTCAAACGAGAACTTTGAAGGCCGAAGTG | >hg38_21_+_8433172_8446622_RNA45SN1_45S_50ntFlanks | 10345 | 10373 | 29 | TTCAAACGAGAACTTTGAAGGCCGAAGTG | 253.0 | 2.5 | 0.7 | 3.8 | 1.7E-04 | 2.2E-04 | MB231/MB468 | NUC |
| >hg38_21_+_8433172_8446622_RNA45SN1_45S_50ntFlanks@10247.10279.33\|GCGACGAGTAGGAGGGCCGCTGCGGTGAGCCTT | >hg38_21_+_8433172_8446622_RNA45SN1_45S_50ntFlanks | 10247 | 10279 | 33 | GCGACGAGTAGGAGGGCCGCTGCGGTGAGCCTT | 242.3 | 7.3 | 1.8 | 4.0 | 6.4E-05 | 2.2E-04 | BT20/MB468 | MP |
| >hg38_21_+_8433172_8446622_RNA45SN1_45S_50ntFlanks@10856.10878.23\|AGGATTGGCTCTAAGGGCTGGGT | >hg38_21_+_8433172_8446622_RNA45SN1_45S_50ntFlanks | 10856 | 10878 | 23 | AGGATTGGCTCTAAGGGCTGGGT | 64.1 | 9.3 | 2.2 | 4.2 | 2.3E-05 | 2.2E-04 | BT20/MB231 | MP |
| >hg38_21_+_8433172_8446622_RNA45SN1_45S_50ntFlanks@11566.11601.36\|TAGCAGCCGACTTAGAACTGGTGCGGACCAGGGGAA | >hg38_21_+_8433172_8446622_RNA45SN1_45S_50ntFlanks | 11566 | 11601 | 36 | TAGCAGCCGACTTAGAACTGGTGCGGACCAGGGGAA | 416.1 | -6.5 | 1.7 | -3.8 | 1.8E-04 | 2.2E-04 | BT20/MB468 | NUC |
| >hg38_21_+_8433172_8446622_RNA45SN1_45S_50ntFlanks@4982.4999.18\|ACGGACAGGATTGACAGA | >hg38_21_+_8433172_8446622_RNA45SN1_45S_50ntFlanks | 4982 | 4999 | 18 | ACGGACAGGATTGACAGA | 977.5 | -6.1 | 1.6 | -3.8 | 1.7E-04 | 2.2E-04 | MB231/MB468 | MITO |
| >hg38_21_+_8433172_8446622_RNA45SN1_45S_50ntFlanks@10766.10797.32\|CGCAGCAGGTCTCCAAGGTGAACAGCCTCTGG | >hg38_21_+_8433172_8446622_RNA45SN1_45S_50ntFlanks | 10766 | 10797 | 32 | CGCAGCAGGTCTCCAAGGTGAACAGCCTCTGG | 155.9 | -5.3 | 1.4 | -3.8 | 1.7E-04 | 2.2E-04 | MB231/MB468 | NUC |
| >hg38_21_+_8433172_8446622_RNA45SN1_45S_50ntFlanks@12397.12417.21\|CGACGTCGCTTTTTGATCCTT | >hg38_21_+_8433172_8446622_RNA45SN1_45S_50ntFlanks | 12397 | 12417 | 21 | CGACGTCGCTTTTTGATCCTT | 1015.0 | -2.2 | 0.6 | -3.8 | 1.6E-04 | 2.2E-04 | MB231/MB468 | CYTO |
| >hg38_21_+_8433172_8446622_RNA45SN1_45S_50ntFlanks@6731.6753.23\|CACATTGATCATCGACACTTCGA | >hg38_21_+_8433172_8446622_RNA45SN1_45S_50ntFlanks | 6731 | 6753 | 23 | CACATTGATCATCGACACTTCGA | 222.6 | -2.0 | 0.5 | -3.8 | 1.7E-04 | 2.2E-04 | MB231/MB468 | NUC |

<table>
<tr><td>>hg38_21_+_8433172_8446622_RNA45SN1_45S_50ntFlanks@10392.10426.35|ACAGCAGTTGAACATGGGTCAGTCGGTCCTGAGAG</td><td>>hg38_21_+_8433172_8446622_RNA45SN1_45S_50ntFlanks</td><td>10392</td><td>10426</td><td>35</td><td>ACAGCAGTTGAACATGGGTCAGTCGGTCCTGAGAG</td><td>481.5</td><td>-4.9</td><td>1.3</td><td>-3.8</td><td>1.5E-04</td><td>2.2E-04</td><td>BT20/MB468</td><td>CYTO</td></tr>
<tr><td>>hg38_21_+_8433172_8446622_RNA45SN1_45S_50ntFlanks@4661.4691.31|AGGTGAAATTCTTGGACCGGCGCAAGACGGA</td><td>>hg38_21_+_8433172_8446622_RNA45SN1_45S_50ntFlanks</td><td>4661</td><td>4691</td><td>31</td><td>AGGTGAAATTCTTGGACCGGCGCAAGACGGA</td><td>220.4</td><td>-3.1</td><td>0.8</td><td>-3.8</td><td>1.5E-04</td><td>2.2E-04</td><td>MB231/MB468</td><td>MP</td></tr>
<tr><td>>hg38_21_+_8433172_8446622_RNA45SN1_45S_50ntFlanks@11583.11613.31|CTGGTGCGGACCAGGGGAATCCGACTGTTTA</td><td>>hg38_21_+_8433172_8446622_RNA45SN1_45S_50ntFlanks</td><td>11583</td><td>11613</td><td>31</td><td>CTGGTGCGGACCAGGGGAATCCGACTGTTTA</td><td>275.7</td><td>-2.4</td><td>0.6</td><td>-3.8</td><td>1.7E-04</td><td>2.2E-04</td><td>MB231/MB468</td><td>NUC</td></tr>
<tr><td>>am_trna128_GlyGCC_6_-_27870686_27870756@1.36.36,trna18_GlyGCC_16_+_70822597_70822667@1.36.36,trna19_GlyGCC_16_+_70823410_70823480@1.36.36,trna19_GlyGCC_2_-_157257659_157257729@1.36.36,trna24_GlyGCC_16_-_70812942_70813012@1.36.36,trna25_GlyGCC_16_-_70812114_70812184@1.36.36,trna5_GlyGCC_17_+_8029064_8029134@1.36.36,trna68_GlyGCC_1_-_161493637_161493707@1.36.36|GCATTGGTGGTTCAGTGGTAGAATTCTCGCCTGCCA</td><td>>am_trna128_GlyGCC_6_-_27870686_27870756</td><td>1</td><td>36</td><td>36</td><td>GCATTGGTGGTTCAGTGGTAGAATTCTCGCCTGCCA</td><td>310.8</td><td>4.4</td><td>1.2</td><td>3.8</td><td>1.5E-04</td><td>2.2E-04</td><td>MB231/MB468</td><td>MP</td></tr>
<tr><td>>hg38_21_+_8433172_8446622_RNA45SN1_45S_50ntFlanks@12936.12955.20|AAACCATTCGTAGACGACCT</td><td>>hg38_21_+_8433172_8446622_RNA45SN1_45S_50ntFlanks</td><td>12936</td><td>12955</td><td>20</td><td>AAACCATTCGTAGACGACCT</td><td>165.3</td><td>-2.6</td><td>0.7</td><td>-3.8</td><td>1.3E-04</td><td>2.2E-04</td><td>BT20/MB468</td><td>MITO</td></tr>
<tr><td>>hg38_21_+_8433172_8446622_RNA45SN1_45S_50ntFlanks@12941.12965.25|ATTCGTAGACGACCTGCTTCTGGGT</td><td>>hg38_21_+_8433172_8446622_RNA45SN1_45S_50ntFlanks</td><td>12941</td><td>12965</td><td>25</td><td>ATTCGTAGACGACCTGCTTCTGGGT</td><td>224.2</td><td>-2.6</td><td>0.7</td><td>-3.8</td><td>1.7E-04</td><td>2.2E-04</td><td>MB231/MB468</td><td>NUC</td></tr>
<tr><td>>hg38_21_+_8433172_8446622_RNA45SN1_45S_50ntFlanks@12259.12284.26|CTTGATCTTGATTTTCAGTACGAATA</td><td>>hg38_21_+_8433172_8446622_RNA45SN1_45S_50ntFlanks</td><td>12259</td><td>12284</td><td>26</td><td>CTTGATCTTGATTTTCAGTACGAATA</td><td>309.7</td><td>4.4</td><td>1.0</td><td>4.2</td><td>2.6E-05</td><td>2.2E-04</td><td>BT20/MB231</td><td>MITO</td></tr>
<tr><td>>MI0000093|hsa-mir-92a-1&WithFlank&13|+|91351308|91351397@54.75.22;MI0000094|hsa-mir-92a-2&WithFlank&X|-|134169532|134169618@54.75.22&[MIMAT0000092&hsa-miR-92a-3p&offsets|0|0;m-9&13|+|91351361|91351382&offsets|0|0];[MIMAT0000092_1&hsa-miR-92a-3p&offsets|0|0;m-11&X|-|134169544|134169565&offsets|0|0]|TATTGCACTTGTCCCGGCCTGT</td><td>>MI0000093|hsa-mir-92a-1&WithFlank&13|+|91351308|91351397</td><td>54</td><td>75</td><td>22</td><td>TATTGCACTTGTCCCGGCCTGT</td><td>42784.6</td><td>2.9</td><td>0.8</td><td>3.8</td><td>1.5E-04</td><td>2.2E-04</td><td>MB231/MB468</td><td>MP</td></tr>
<tr><td>>hg38_21_+_8433172_8446622_RNA45SN1_45S_50ntFlanks@11190.11208.19|CCCGGGGGGCCGGCGGCGG</td><td>>hg38_21_+_8433172_8446622_RNA45SN1_45S_50ntFlanks</td><td>11190</td><td>11208</td><td>19</td><td>CCCGGGGGGCCGGCGGCGG</td><td>1216.5</td><td>-7.2</td><td>1.9</td><td>-3.7</td><td>1.8E-04</td><td>2.3E-04</td><td>BT20/MB468</td><td>NUC</td></tr>
<tr><td>>hg38_21_+_8433172_8446622_RNA45SN1_45S_50ntFlanks@3842.3868.27|CCTACTTGGATAACTGTGGTAATTCTA</td><td>>hg38_21_+_8433172_8446622_RNA45SN1_45S_50ntFlanks</td><td>3842</td><td>3868</td><td>27</td><td>CCTACTTGGATAACTGTGGTAATTCTA</td><td>486.3</td><td>-4.7</td><td>1.2</td><td>-3.8</td><td>1.7E-04</td><td>2.3E-04</td><td>MB231/MB468</td><td>MITO</td></tr>
<tr><td>>MI0025750|hsa-mir-7974&WithFlank&19|-|11495538|11495628@62.85.24&[MIMAT0031177&hsa-miR-7974&offsets|0|0;m-867&19|-|11495544|11495567&offsets|0|0]|AGGCTGTGATGCTCTCCTGAGCCC</td><td>>MI0025750|hsa-mir-7974&WithFlank&19|-|11495538|11495628</td><td>62</td><td>85</td><td>24</td><td>AGGCTGTGATGCTCTCCTGAGCCC</td><td>1020.8</td><td>-3.8</td><td>1.0</td><td>-3.7</td><td>1.8E-04</td><td>2.3E-04</td><td>BT20/MB468</td><td>NUC</td></tr>
<tr><td>>MI0003776|hsa-mir-320b-1&WithFlank&1|+|116671740|116671823@48.67.20(+1U);MI0003839|hsa-mir-320b-2&WithFlank&1|-|224257034|224257116@47.66.20(+1U);MI0000542|hsa-mir-320a&WithFlank&8|-|22244960|22245043@48.67.20(+1U)&[MIMAT0005792&hsa-miR-320b&offsets|0|-2(+1U);m-183&1|+|116671787|116671806&offsets|0|0(+1U)];[MIMAT0005792_1&hsa-miR-320b&offsets|0|-2(+1U);m-183&1|-|224257051|224257070&offsets|0|0(+1U)];[MIMAT0000510&hsa-miR-320a-3p&offsets|0|-2(+1U);m-60&8|-|22244975|22244996&offsets|0|-2(+1U)]|AAAAGCTGGGTTGAGAGGGCT</td><td>>MI0003776|hsa-mir-320b-1&WithFlank&1|+|116671740|116671823</td><td>48</td><td>67</td><td>20(+1U)</td><td>AAAAGCTGGGTTGAGAGGGCT</td><td>290.6</td><td>2.6</td><td>0.6</td><td>4.2</td><td>2.4E-05</td><td>2.3E-04</td><td>BT20/MB231</td><td>NUC</td></tr>
<tr><td>>hg38_21_+_8433172_8446622_RNA45SN1_45S_50ntFlanks@10463.10497.35|CTCCGTTGCCCTCGGCCGATCGAAAGGGAGTCGGG</td><td>>hg38_21_+_8433172_8446622_RNA45SN1_45S_50ntFlanks</td><td>10463</td><td>10497</td><td>35</td><td>CTCCGTTGCCCTCGGCCGATCGAAAGGGAGTCGGG</td><td>409.9</td><td>-6.2</td><td>1.6</td><td>-4.0</td><td>6.5E-05</td><td>2.3E-04</td><td>BT20/MB468</td><td>MP</td></tr>
<tr><td>>hg38_21_+_8433172_8446622_RNA45SN1_45S_50ntFlanks@12624.12657.34|CCAATGGGGCGAAGCTACCATCTGTGGGATTATG</td><td>>hg38_21_+_8433172_8446622_RNA45SN1_45S_50ntFlanks</td><td>12624</td><td>12657</td><td>34</td><td>CCAATGGGGCGAAGCTACCATCTGTGGGATTATG</td><td>1086.6</td><td>-6.6</td><td>1.7</td><td>-4.0</td><td>6.5E-05</td><td>2.3E-04</td><td>BT20/MB468</td><td>MP</td></tr>
<tr><td>>hg38_21_+_8433172_8446622_RNA45SN1_45S_50ntFlanks@4559.4587.29|GCTAGGAATAATGGAATAGGACCGCGGTT</td><td>>hg38_21_+_8433172_8446622_RNA45SN1_45S_50ntFlanks</td><td>4559</td><td>4587</td><td>29</td><td>GCTAGGAATAATGGAATAGGACCGCGGTT</td><td>122.3</td><td>-1.8</td><td>0.5</td><td>-3.8</td><td>1.5E-04</td><td>2.3E-04</td><td>BT20/MB468</td><td>CYTO</td></tr>
<tr><td>>hg38_MT_+_598_1651_RNR1_12S_50ntFlanks@425.455.31|ACGAAAGTGGCTTTAACATATCTGAACACAC</td><td>>hg38_MT_+_598_1651_RNR1_12S_50ntFlanks</td><td>425</td><td>455</td><td>31</td><td>ACGAAAGTGGCTTTAACATATCTGAACACAC</td><td>177.0</td><td>4.6</td><td>1.2</td><td>3.8</td><td>1.7E-04</td><td>2.3E-04</td><td>MB231/MB468</td><td>MITO</td></tr>
<tr><td>>hg38_21_+_8433172_8446622_RNA45SN1_45S_50ntFlanks@12455.12476.22|ACCAAGCGTTGGATTGTTCACC</td><td>>hg38_21_+_8433172_8446622_RNA45SN1_45S_50ntFlanks</td><td>12455</td><td>12476</td><td>22</td><td>ACCAAGCGTTGGATTGTTCACC</td><td>584.2</td><td>-2.7</td><td>0.7</td><td>-3.8</td><td>1.7E-04</td><td>2.3E-04</td><td>MB231/MB468</td><td>MITO</td></tr>
<tr><td>>hg38_21_+_8433172_8446622_RNA45SN1_45S_50ntFlanks@12156.12182.27|GAGTTTGACTGGGGCGGTACACCTGTC</td><td>>hg38_21_+_8433172_8446622_RNA45SN1_45S_50ntFlanks</td><td>12156</td><td>12182</td><td>27</td><td>GAGTTTGACTGGGGCGGTACACCTGTC</td><td>263.5</td><td>-6.4</td><td>1.6</td><td>-4.0</td><td>6.5E-05</td><td>2.3E-04</td><td>BT20/MB468</td><td>MP</td></tr>
<tr><td>>hg38_21_+_8433172_8446622_RNA45SN1_45S_50ntFlanks@4729.4757.29|TCAAGAACGAAAGTCGGAGGTTCGAAGAC</td><td>>hg38_21_+_8433172_8446622_RNA45SN1_45S_50ntFlanks</td><td>4729</td><td>4757</td><td>29</td><td>TCAAGAACGAAAGTCGGAGGTTCGAAGAC</td><td>78.4</td><td>-5.7</td><td>1.5</td><td>-3.8</td><td>1.5E-04</td><td>2.3E-04</td><td>MB231/MB468</td><td>MP</td></tr>
<tr><td>>hg38_21_+_8433172_8446622_RNA45SN1_45S_50ntFlanks@7990.8016.27|GACGTGGCGACCCGCTGAATTTAAGCA</td><td>>hg38_21_+_8433172_8446622_RNA45SN1_45S_50ntFlanks</td><td>7990</td><td>8016</td><td>27</td><td>GACGTGGCGACCCGCTGAATTTAAGCA</td><td>306.9</td><td>3.8</td><td>1.0</td><td>3.8</td><td>1.8E-04</td><td>2.3E-04</td><td>MB231/MB468</td><td>NUC</td></tr>
<tr><td>>hg38_21_+_8433172_8446622_RNA45SN1_45S_50ntFlanks@12162.12181.20|GACTGGGGCGGTACACCTGT</td><td>>hg38_21_+_8433172_8446622_RNA45SN1_45S_50ntFlanks</td><td>12162</td><td>12181</td><td>20</td><td>GACTGGGGCGGTACACCTGT</td><td>606.5</td><td>-6.3</td><td>1.6</td><td>-4.0</td><td>6.5E-05</td><td>2.3E-04</td><td>BT20/MB468</td><td>MP</td></tr>
<tr><td>>hg38_21_+_8433172_8446622_RNA45SN1_45S_50ntFlanks@8342.8364.23|CGTAAGGGAAAGTTGAAAAGAAC</td><td>>hg38_21_+_8433172_8446622_RNA45SN1_45S_50ntFlanks</td><td>8342</td><td>8364</td><td>23</td><td>CGTAAGGGAAAGTTGAAAAGAAC</td><td>379.6</td><td>6.0</td><td>1.6</td><td>3.8</td><td>1.5E-04</td><td>2.3E-04</td><td>BT20/MB468</td><td>CYTO</td></tr>
<tr><td>>MI0016005|hsa-mir-3615&WithFlank&17|+|74748607|74748705@57.78.22(+1U)&[MIMAT0017994&hsa-miR-3615&offsets|0|+1(+1U);m-271&17|+|74748663|74748684&offsets|0|0(+1U)]|TCTCTCGGCTCCTCGCGGCTCGT</td><td>>MI0016005|hsa-mir-3615&WithFlank&17|+|74748607|74748705</td><td>57</td><td>78</td><td>22(+1U)</td><td>TCTCTCGGCTCCTCGCGGCTCGT</td><td>170.6</td><td>5.0</td><td>1.3</td><td>3.8</td><td>1.7E-04</td><td>2.3E-04</td><td>MB231/MB468</td><td>MITO</td></tr>
<tr><td>>MI0000750|hsa-mir-26a-2&WithFlank&12|-|57824603|57824698@20.40.21(+1G);MI0000083|hsa-mir-26a-1&WithFlank&3|+|37969398|37969486@16.36.21(+1G)&[MIMAT0000082&hsa-miR-26a-5p&offsets|0|-1(+1G);m-33&12|-|57824658|57824679&offsets|0|-1(+1G)];[MIMAT0000082_1&hsa-miR-26a-5p&offsets|0|-1(+1G);m-34&3|+|37969413|37969434&offsets|0|-1(+1G)]|TTCAAGTAATCCAGGATAGGCG</td><td>>MI0000750|hsa-mir-26a-2&WithFlank&12|-|57824603|57824698</td><td>20</td><td>40</td><td>21(+1G)</td><td>TTCAAGTAATCCAGGATAGGCG</td><td>662.9</td><td>1.8</td><td>0.5</td><td>3.7</td><td>1.8E-04</td><td>2.3E-04</td><td>BT20/MB468</td><td>NUC</td></tr>
<tr><td>>hg38_21_+_8433172_8446622_RNA45SN1_45S_50ntFlanks@4776.4793.18|TTCCGACCATAAACGATG</td><td>>hg38_21_+_8433172_8446622_RNA45SN1_45S_50ntFlanks</td><td>4776</td><td>4793</td><td>18</td><td>TTCCGACCATAAACGATG</td><td>75.0</td><td>-4.3</td><td>1.1</td><td>-3.8</td><td>1.5E-04</td><td>2.3E-04</td><td>MB231/MB468</td><td>MP</td></tr>
<tr><td>>MI0000063|hsa-let-7b&WithFlank&22|+|46113680|46113774@66.86.21(+1U)&[MIMAT0004482&hsa-let-7b-3p&offsets|0|-1(+1U);m-171&22|+|46113745|46113766&offsets|0|-1(+1U)]|CTATACAACCTACTGCCTTCCT</td><td>>MI0000063|hsa-let-7b&WithFlank&22|+|46113680|46113774</td><td>66</td><td>86</td><td>21(+1U)</td><td>CTATACAACCTACTGCCTTCCT</td><td>227.0</td><td>4.0</td><td>1.1</td><td>3.8</td><td>1.7E-04</td><td>2.3E-04</td><td>MB231/MB468</td><td>MITO</td></tr>
<tr><td>>hg38_21_+_8433172_8446622_RNA45SN1_45S_50ntFlanks@475.512.38|CTTCGTGATCGATGTGGTGACGTCGTGCTCTCCCGGGC</td><td>>hg38_21_+_8433172_8446622_RNA45SN1_45S_50ntFlanks</td><td>475</td><td>512</td><td>38</td><td>CTTCGTGATCGATGTGGTGACGTCGTGCTCTCCCGGGC</td><td>1554.4</td><td>3.2</td><td>0.8</td><td>3.8</td><td>1.8E-04</td><td>2.3E-04</td><td>MB231/MB468</td><td>NUC</td></tr>
<tr><td>>MI0001445|hsa-mir-423&WithFlank&17|+|30117073|30117178@23.45.23(+1C)&[MIMAT0004748&hsa-miR-423-5p&offsets|0|0(+1C);m-141&17|+|30117095|30117117&offsets|0|0(+1C)]|TGAGGGGCAGAGAGCGAGACTTTC</td><td>>MI0001445|hsa-mir-423&WithFlank&17|+|30117073|30117178</td><td>23</td><td>45</td><td>23(+1C)</td><td>TGAGGGGCAGAGAGCGAGACTTTC</td><td>221.1</td><td>4.7</td><td>1.2</td><td>3.8</td><td>1.5E-04</td><td>2.3E-04</td><td>MB231/MB468</td><td>MP</td></tr>
<tr><td>>MI0000272|hsa-mir-182&WithFlank&7|-|129770377|129770498@29.54.26(+1A)&[MIMAT0000259&hsa-miR-182-5p&offsets|0|+2(+1A);m-22&7|-|129770449|129770470&offsets|0|+4(+1A)]|TTTGGCAATGGTAGAACTCACACTGGA</td><td>>MI0000272|hsa-mir-182&WithFlank&7|-|129770377|129770498</td><td>29</td><td>54</td><td>26(+1A)</td><td>TTTGGCAATGGTAGAACTCACACTGGA</td><td>264.8</td><td>3.2</td><td>0.8</td><td>4.2</td><td>2.2E-05</td><td>2.3E-04</td><td>BT20/MB231</td><td>CYTO</td></tr>
<tr><td>>hg38_21_+_8433172_8446622_RNA45SN1_45S_50ntFlanks@3777.3805.29|CGGTACAGTGAAACTGCGAATGGCTCATT</td><td>>hg38_21_+_8433172_8446622_RNA45SN1_45S_50ntFlanks</td><td>3777</td><td>3805</td><td>29</td><td>CGGTACAGTGAAACTGCGAATGGCTCATT</td><td>398.0</td><td>-5.9</td><td>1.6</td><td>-3.8</td><td>1.7E-04</td><td>2.3E-04</td><td>MB231/MB468</td><td>MITO</td></tr>
<tr><td>>am_trna128_GlyGCC_6_-_27870686_27870756@1.35.35,trna18_GlyGCC_16_+_70822597_70822667@1.35.35,trna19_GlyGCC_16_+_70823410_70823480@1.35.35,trna19_GlyGCC_2_-_157257659_157257729@1.35.35,trna24_GlyGCC_16_-_70812942_70813012@1.35.35,trna25_GlyGCC_16_-_70812114_70812184@1.35.35,trna5_GlyGCC_17_+_8029064_8029134@1.35.35,trna68_GlyGCC_1_-_161493637_161493707@1.35.35|GCATTGGTGGTTCAGTGGTAGAATTCTCGCCTGCC</td><td>>am_trna128_GlyGCC_6_-_27870686_27870756</td><td>1</td><td>35</td><td>35</td><td>GCATTGGTGGTTCAGTGGTAGAATTCTCGCCTGCC</td><td>5679.6</td><td>-3.6</td><td>0.9</td><td>-4.2</td><td>2.7E-05</td><td>2.3E-04</td><td>BT20/MB231</td><td>MITO</td></tr>
<tr><td>>MI0000749|hsa-mir-30e&WithFlank&1|+|40754349|40754452@23.46.24(+2U)&[MIMAT0000692&hsa-miR-30e-5p&offsets|0|+2(+2U);m-24&1|+|40754371|40754394&offsets|0|0(+2U)]|TGTAAACATCCTTGACTGGAAGCTTT</td><td>>MI0000749|hsa-mir-30e&WithFlank&1|+|40754349|40754452</td><td>23</td><td>46</td><td>24(+2U)</td><td>TGTAAACATCCTTGACTGGAAGCTTT</td><td>546.6</td><td>-5.7</td><td>1.4</td><td>-4.2</td><td>2.3E-05</td><td>2.3E-04</td><td>BT20/MB231</td><td>MP</td></tr>
<tr><td>>hg38_21_+_8433172_8446622_RNA45SN1_45S_50ntFlanks@12289.12319.31|CCGTGAAAGCGGGGCCTCACGATCCTTCTGA</td><td>>hg38_21_+_8433172_8446622_RNA45SN1_45S_50ntFlanks</td><td>12289</td><td>12319</td><td>31</td><td>CCGTGAAAGCGGGGCCTCACGATCCTTCTGA</td><td>45.3</td><td>-9.4</td><td>2.5</td><td>-3.8</td><td>1.5E-04</td><td>2.3E-04</td><td>MB231/MB468</td><td>MP</td></tr>
<tr><td>>am_trnaMT_ValTAC_MT_+_1602_1670@38.67.30|TAGGAGATTTCAACTTAACTTGACCGCTCT</td><td>>am_trnaMT_ValTAC_MT_+_1602_1670</td><td>38</td><td>67</td><td>30</td><td>TAGGAGATTTCAACTTAACTTGACCGCTCT</td><td>106.1</td><td>10.5</td><td>2.6</td><td>4.0</td><td>6.5E-05</td><td>2.3E-04</td><td>BT20/MB468</td><td>MP</td></tr>
<tr><td>>hg38_21_+_8433172_8446622_RNA45SN1_45S_50ntFlanks@11564.11600.37|CCTAGCAGCCGACTTAGAACTGGTGCGGACCAGGGGA</td><td>>hg38_21_+_8433172_8446622_RNA45SN1_45S_50ntFlanks</td><td>11564</td><td>11600</td><td>37</td><td>CCTAGCAGCCGACTTAGAACTGGTGCGGACCAGGGGA</td><td>259.4</td><td>3.4</td><td>0.8</td><td>4.2</td><td>2.7E-05</td><td>2.3E-04</td><td>BT20/MB231</td><td>MITO</td></tr>
<tr><td>>MI0000782|hsa-mir-374a&WithFlank&X|-|74287280|74287363@18.39.22&[MIMAT0000727&hsa-miR-374a-5p&offsets|0|0;m-184&X|-|74287325|74287346&offsets|0|0]|TTATAATACAACCTGATAAGTG</td><td>>MI0000782|hsa-mir-374a&WithFlank&X|-|74287280|74287363</td><td>18</td><td>39</td><td>22</td><td>TTATAATACAACCTGATAAGTG</td><td>500.6</td><td>-3.9</td><td>0.9</td><td>-4.2</td><td>2.7E-05</td><td>2.3E-04</td><td>BT20/MB231</td><td>MITO</td></tr>
<tr><td>>hg38_21_+_8433172_8446622_RNA45SN1_45S_50ntFlanks@12850.12871.22|CTTCGTCCTGGGAAACGGGGCG</td><td>>hg38_21_+_8433172_8446622_RNA45SN1_45S_50ntFlanks</td><td>12850</td><td>12871</td><td>22</td><td>CTTCGTCCTGGGAAACGGGGCG</td><td>157.3</td><td>-6.5</td><td>1.6</td><td>-4.0</td><td>6.5E-05</td><td>2.3E-04</td><td>BT20/MB468</td><td>MP</td></tr>
<tr><td>>hg38_21_+_8433172_8446622_RNA45SN1_45S_50ntFlanks@12995.13038.44|CTCGCTGCGATCTATTGAAAGTCAGCCCTCGACACAAGGGTTTG</td><td>>hg38_21_+_8433172_8446622_RNA45SN1_45S_50ntFlanks</td><td>12995</td><td>13038</td><td>44</td><td>CTCGCTGCGATCTATTGAAAGTCAGCCCTCGACACAAGGGTTTG</td><td>1170.9</td><td>5.8</td><td>1.6</td><td>3.8</td><td>1.8E-04</td><td>2.3E-04</td><td>MB231/MB468</td><td>MITO</td></tr>
<tr><td>>MI0000115|hsa-mir-16-2&WithFlank&3|+|160404739|160404831@58.80.23&[MIMAT0004518&hsa-miR-16-2-3p&offsets|-1|0;m-252&3|+|160404796|160404816&offsets|0|+2]|ACCAATATTACTGTGCTGCTTTA</td><td>>MI0000115|hsa-mir-16-2&WithFlank&3|+|160404739|160404831</td><td>58</td><td>80</td><td>23</td><td>ACCAATATTACTGTGCTGCTTTA</td><td>250.9</td><td>4.8</td><td>1.3</td><td>3.8</td><td>1.8E-04</td><td>2.3E-04</td><td>MB231/MB468</td><td>MITO</td></tr>
</table>

<table>
<tr><td>>hg38_21_+_8433172_8446622_RNA45SN1_45S_50ntFlanks@6651.6697.47|CGACTCTTAGCGGTGGATCACTCGGCTCGTGCGTCGATGAAGAACGC</td><td>>hg38_21_+_8433172_8446622_RNA45SN1_45S_50ntFlanks</td><td>6651</td><td>6697</td><td>47</td><td>CGACTCTTAGCGGTGGATCACTCGGCTCGTGCGTCGATGAAGAACGC</td><td>3721.7</td><td>3.2</td><td>0.8</td><td>3.8</td><td>1.7E-04</td><td>2.3E-04</td><td>MB231/MB468</td><td>CYTO</td></tr>
<tr><td>>MI0017294|hsa-mir-4664&WithFlank&8|-|143733077|143733159@50.71.22(+1U)&[MIMAT0019738&hsa-miR-4664-3p&offsets|0|0(+1U);m-1025&8|-|143733094|143733110&offsets|0|+5(+1U)]|CTTCCGGTCTGTGAGCCCCGTCT</td><td>>MI0017294|hsa-mir-4664&WithFlank&8|-|143733077|143733159</td><td>50</td><td>71</td><td>22(+1U)</td><td>CTTCCGGTCTGTGAGCCCCGTCT</td><td>56.1</td><td>9.1</td><td>2.2</td><td>4.2</td><td>2.3E-05</td><td>2.3E-04</td><td>BT20/MB231</td><td>MP</td></tr>
<tr><td>>MI0000239|hsa-mir-197&WithFlank&1|+|109598887|109598973@54.75.22&[MIMAT0000227&hsa-miR-197-3p&offsets|0|0;m-99&1|+|109598940|109598961&offsets|0|0]|TTCACCACCTTCTCCACCCAGC</td><td>>MI0000239|hsa-mir-197&WithFlank&1|+|109598887|109598973</td><td>54</td><td>75</td><td>22</td><td>TTCACCACCTTCTCCACCCAGC</td><td>1511.4</td><td>2.8</td><td>0.7</td><td>3.7</td><td>1.8E-04</td><td>2.3E-04</td><td>MB231/MB468</td><td>NUC</td></tr>
<tr><td>>hg38_21_+_8433172_8446622_RNA45SN1_45S_50ntFlanks@8176.8206.31|CTTCTGATCGAGGCCCAGCCCGTGGACGGTG</td><td>>hg38_21_+_8433172_8446622_RNA45SN1_45S_50ntFlanks</td><td>8176</td><td>8206</td><td>31</td><td>CTTCTGATCGAGGCCCAGCCCGTGGACGGTG</td><td>965.0</td><td>-2.4</td><td>0.6</td><td>-3.8</td><td>1.7E-04</td><td>2.3E-04</td><td>MB231/MB468</td><td>CYTO</td></tr>
<tr><td>>hg38_21_+_8433172_8446622_RNA45SN1_45S_50ntFlanks@12023.12055.33|CACTACTCTGATCGTTTTTTCACTGACCCGGTG</td><td>>hg38_21_+_8433172_8446622_RNA45SN1_45S_50ntFlanks</td><td>12023</td><td>12055</td><td>33</td><td>CACTACTCTGATCGTTTTTTCACTGACCCGGTG</td><td>161.4</td><td>7.2</td><td>1.8</td><td>4.0</td><td>6.6E-05</td><td>2.3E-04</td><td>BT20/MB468</td><td>MP</td></tr>
<tr><td>>hg38_21_+_8433172_8446622_RNA45SN1_45S_50ntFlanks@12433.12454.22|CTATCATTGTGAAGCAGAATTC</td><td>>hg38_21_+_8433172_8446622_RNA45SN1_45S_50ntFlanks</td><td>12433</td><td>12454</td><td>22</td><td>CTATCATTGTGAAGCAGAATTC</td><td>469.0</td><td>-6.4</td><td>1.6</td><td>-4.0</td><td>6.6E-05</td><td>2.3E-04</td><td>BT20/MB468</td><td>MP</td></tr>
<tr><td>>MI0000063|hsa-let-7b&WithFlank&22|+|46113680|46113774@12.29.18&[MIMAT0000063&hsa-let-7b-5p&offsets|0|-4;m-14&22|+|46113691|46113712&offsets|0|-4]|TGAGGTAGTAGGTTGTGT</td><td>>MI0000063|hsa-let-7b&WithFlank&22|+|46113680|46113774</td><td>12</td><td>29</td><td>18</td><td>TGAGGTAGTAGGTTGTGT</td><td>281.7</td><td>2.7</td><td>0.6</td><td>4.2</td><td>2.5E-05</td><td>2.3E-04</td><td>BT20/MB231</td><td>NUC</td></tr>
<tr><td>>hg38_21_+_8433172_8446622_RNA45SN1_45S_50ntFlanks@6661.6690.30|CGGTGGATCACTCGGCTCGTGCGTCGATGA</td><td>>hg38_21_+_8433172_8446622_RNA45SN1_45S_50ntFlanks</td><td>6661</td><td>6690</td><td>30</td><td>CGGTGGATCACTCGGCTCGTGCGTCGATGA</td><td>850.3</td><td>3.6</td><td>0.9</td><td>4.0</td><td>6.6E-05</td><td>2.3E-04</td><td>BT20/MB468</td><td>MP</td></tr>
<tr><td>>hg38_21_+_8433172_8446622_RNA45SN1_45S_50ntFlanks@10802.10823.22|TTGGAACAATGTAGGTAAGGGA</td><td>>hg38_21_+_8433172_8446622_RNA45SN1_45S_50ntFlanks</td><td>10802</td><td>10823</td><td>22</td><td>TTGGAACAATGTAGGTAAGGGA</td><td>295.7</td><td>-3.2</td><td>0.8</td><td>-4.0</td><td>6.6E-05</td><td>2.3E-04</td><td>BT20/MB468</td><td>MP</td></tr>
<tr><td>>hg38_21_+_8433172_8446622_RNA45SN1_45S_50ntFlanks@9482.9505.24|ACGTGTTAGGACCCGAAAGATGGT</td><td>>hg38_21_+_8433172_8446622_RNA45SN1_45S_50ntFlanks</td><td>9482</td><td>9505</td><td>24</td><td>ACGTGTTAGGACCCGAAAGATGGT</td><td>95.2</td><td>10.3</td><td>2.6</td><td>4.0</td><td>6.6E-05</td><td>2.3E-04</td><td>BT20/MB468</td><td>MP</td></tr>
<tr><td>>MI0000077|hsa-mir-21&WithFlank&17|+|59841260|59841343@14.34.21&[MIMAT0000076&hsa-miR-21-5p&offsets|0|-1;m-1&17|+|59841273|59841294&offsets|0|-1]|TAGCTTATCAGACTGATGTTG</td><td>>MI0000077|hsa-mir-21&WithFlank&17|+|59841260|59841343</td><td>14</td><td>34</td><td>21</td><td>TAGCTTATCAGACTGATGTTG</td><td>24462.7</td><td>2.9</td><td>0.8</td><td>3.8</td><td>1.5E-04</td><td>2.3E-04</td><td>MB231/MB468</td><td>MP</td></tr>
<tr><td>>hg38_21_+_8433172_8446622_RNA45SN1_45S_50ntFlanks@9556.9582.27|TAGCGGTCCTGACGTGCAAATCGGTCG</td><td>>hg38_21_+_8433172_8446622_RNA45SN1_45S_50ntFlanks</td><td>9556</td><td>9582</td><td>27</td><td>TAGCGGTCCTGACGTGCAAATCGGTCG</td><td>498.0</td><td>-5.8</td><td>1.5</td><td>-3.8</td><td>1.6E-04</td><td>2.3E-04</td><td>BT20/MB468</td><td>CYTO</td></tr>
<tr><td>>hg38_21_+_8433172_8446622_RNA45SN1_45S_50ntFlanks@10873.10904.32|CTGGGTCGGTCGGGCTGGGGCGCGAAGCGGGG</td><td>>hg38_21_+_8433172_8446622_RNA45SN1_45S_50ntFlanks</td><td>10873</td><td>10904</td><td>32</td><td>CTGGGTCGGTCGGGCTGGGGCGCGAAGCGGGG</td><td>291.1</td><td>-5.2</td><td>1.4</td><td>-3.8</td><td>1.6E-04</td><td>2.3E-04</td><td>BT20/MB468</td><td>CYTO</td></tr>
<tr><td>>hg38_21_+_8433172_8446622_RNA45SN1_45S_50ntFlanks@4243.4271.29|CTTTCGAGGCCCTGTAATTGGAATGAGTC</td><td>>hg38_21_+_8433172_8446622_RNA45SN1_45S_50ntFlanks</td><td>4243</td><td>4271</td><td>29</td><td>CTTTCGAGGCCCTGTAATTGGAATGAGTC</td><td>189.7</td><td>-2.3</td><td>0.6</td><td>-3.7</td><td>1.8E-04</td><td>2.3E-04</td><td>MB231/MB468</td><td>NUC</td></tr>
<tr><td>>hg38_21_+_8433172_8446622_RNA45SN1_45S_50ntFlanks@10347.10376.30|CAAACGAGAACTTTGAAGGCCGAAGTGGAG</td><td>>hg38_21_+_8433172_8446622_RNA45SN1_45S_50ntFlanks</td><td>10347</td><td>10376</td><td>30</td><td>CAAACGAGAACTTTGAAGGCCGAAGTGGAG</td><td>131.2</td><td>-6.3</td><td>1.6</td><td>-3.8</td><td>1.5E-04</td><td>2.3E-04</td><td>MB231/MB468</td><td>MP</td></tr>
<tr><td>>hg38_21_+_8433172_8446622_RNA45SN1_45S_50ntFlanks@10684.10709.26|CGGCGGCGTCCGGTGAGCTCTCGCTG</td><td>>hg38_21_+_8433172_8446622_RNA45SN1_45S_50ntFlanks</td><td>10684</td><td>10709</td><td>26</td><td>CGGCGGCGTCCGGTGAGCTCTCGCTG</td><td>2319.1</td><td>-3.4</td><td>0.9</td><td>-3.8</td><td>1.3E-04</td><td>2.3E-04</td><td>BT20/MB468</td><td>MITO</td></tr>
<tr><td>>hg38_21_+_8433172_8446622_RNA45SN1_45S_50ntFlanks@11569.11611.43|CAGCCGACTTAGAACTGGTGCGGACCAGGGGAATCCGACTGTT</td><td>>hg38_21_+_8433172_8446622_RNA45SN1_45S_50ntFlanks</td><td>11569</td><td>11611</td><td>43</td><td>CAGCCGACTTAGAACTGGTGCGGACCAGGGGAATCCGACTGTT</td><td>600.0</td><td>-4.3</td><td>1.1</td><td>-3.7</td><td>1.8E-04</td><td>2.3E-04</td><td>MB231/MB468</td><td>NUC</td></tr>
<tr><td>>hg38_21_+_8433172_8446622_RNA45SN1_45S_50ntFlanks@9343.9370.28|CGAAAGCCGCCGTGGCGCAATGAAGGTG</td><td>>hg38_21_+_8433172_8446622_RNA45SN1_45S_50ntFlanks</td><td>9343</td><td>9370</td><td>28</td><td>CGAAAGCCGCCGTGGCGCAATGAAGGTG</td><td>1621.9</td><td>-6.8</td><td>1.7</td><td>-4.0</td><td>6.6E-05</td><td>2.3E-04</td><td>BT20/MB468</td><td>MP</td></tr>
<tr><td>>hg38_21_+_8433172_8446622_RNA45SN1_45S_50ntFlanks@8182.8199.18|ATCGAGGCCCAGCCCGTG</td><td>>hg38_21_+_8433172_8446622_RNA45SN1_45S_50ntFlanks</td><td>8182</td><td>8199</td><td>18</td><td>ATCGAGGCCCAGCCCGTG</td><td>116.2</td><td>-9.9</td><td>2.5</td><td>-4.0</td><td>6.6E-05</td><td>2.3E-04</td><td>BT20/MB468</td><td>MP</td></tr>
<tr><td>>hg38_21_+_8433172_8446622_RNA45SN1_45S_50ntFlanks@12141.12158.18|ACAGTGCCAGGTGGGGAG</td><td>>hg38_21_+_8433172_8446622_RNA45SN1_45S_50ntFlanks</td><td>12141</td><td>12158</td><td>18</td><td>ACAGTGCCAGGTGGGGAG</td><td>121.3</td><td>-5.9</td><td>1.5</td><td>-4.0</td><td>6.6E-05</td><td>2.3E-04</td><td>BT20/MB468</td><td>MP</td></tr>
<tr><td>>hg38_21_+_8433172_8446622_RNA45SN1_45S_50ntFlanks@9572.9591.20|CAAATCGGTCGTCCGACCTG</td><td>>hg38_21_+_8433172_8446622_RNA45SN1_45S_50ntFlanks</td><td>9572</td><td>9591</td><td>20</td><td>CAAATCGGTCGTCCGACCTG</td><td>125.9</td><td>-6.2</td><td>1.6</td><td>-4.0</td><td>6.6E-05</td><td>2.3E-04</td><td>BT20/MB468</td><td>MP</td></tr>
<tr><td>>trna11_GluTTC_15_-_26327381_26327452@1.30.30,trna3_GluTTC_13_-_45492062_45492133@1.30.30|TCCCACATGGTCTAGCGGTTAGGATTCCTG</td><td>>trna11_GluTTC_15_-_26327381_26327452</td><td>1</td><td>30</td><td>30</td><td>TCCCACATGGTCTAGCGGTTAGGATTCCTG</td><td>2148.0</td><td>-2.8</td><td>0.7</td><td>-3.8</td><td>1.6E-04</td><td>2.3E-04</td><td>BT20/MB468</td><td>CYTO</td></tr>
<tr><td>>trna10_AlaCGC_6_+_26553731_26553802@1.24.24,trna119_AlaCGC_6_-_28641613_28641684@1.24.24,trna120_AlaAGC_6_-_28626014_28626085@1.24.24,trna13_AlaCGC_2_+_157257281_157257352@1.24.24,trna13_AlaTGC_12_-_125406301_125406372@1.24.24,trna66_AlaTGC_6_+_28611222_28611293@1.24.24,trna8_AlaTGC_12_+_125424512_125424583@1.24.24,trna8_AlaTGC_5_+_180633868_180633939@1.24.24|GGGGATGTAGCTCAGTGGTAGAGC</td><td>>trna10_AlaCGC_6_+_26553731_26553802</td><td>1</td><td>24</td><td>24</td><td>GGGGATGTAGCTCAGTGGTAGAGC</td><td>937.8</td><td>5.2</td><td>1.3</td><td>4.0</td><td>6.6E-05</td><td>2.3E-04</td><td>BT20/MB468</td><td>MP</td></tr>
<tr><td>>hg38_21_+_8433172_8446622_RNA45SN1_45S_50ntFlanks@10329.10362.34|GTGGTAGTAGCAAATATTCAAACGAGAACTTTGA</td><td>>hg38_21_+_8433172_8446622_RNA45SN1_45S_50ntFlanks</td><td>10329</td><td>10362</td><td>34</td><td>GTGGTAGTAGCAAATATTCAAACGAGAACTTTGA</td><td>71.1</td><td>9.5</td><td>2.2</td><td>4.2</td><td>2.4E-05</td><td>2.3E-04</td><td>BT20/MB231</td><td>MP</td></tr>
<tr><td>>hg38_21_+_8433172_8446622_RNA45SN1_45S_50ntFlanks@11886.11904.19|TGAGCTTGACTCTAGTCTG</td><td>>hg38_21_+_8433172_8446622_RNA45SN1_45S_50ntFlanks</td><td>11886</td><td>11904</td><td>19</td><td>TGAGCTTGACTCTAGTCTG</td><td>432.5</td><td>-4.4</td><td>1.2</td><td>-3.8</td><td>1.3E-04</td><td>2.3E-04</td><td>BT20/MB468</td><td>MITO</td></tr>
<tr><td>>hg38_21_+_8433172_8446622_RNA45SN1_45S_50ntFlanks@10516.10533.18|GAGTGGCGGAGATGGGCG</td><td>>hg38_21_+_8433172_8446622_RNA45SN1_45S_50ntFlanks</td><td>10516</td><td>10533</td><td>18</td><td>GAGTGGCGGAGATGGGCG</td><td>259.9</td><td>-3.9</td><td>1.0</td><td>-3.8</td><td>1.6E-04</td><td>2.3E-04</td><td>BT20/MB468</td><td>CYTO</td></tr>
<tr><td>>hg38_21_+_8433172_8446622_RNA45SN1_45S_50ntFlanks@10845.10867.23|ACTTCGGGATAAGGATTGGCTCT</td><td>>hg38_21_+_8433172_8446622_RNA45SN1_45S_50ntFlanks</td><td>10845</td><td>10867</td><td>23</td><td>ACTTCGGGATAAGGATTGGCTCT</td><td>607.5</td><td>-2.5</td><td>0.7</td><td>-3.8</td><td>1.7E-04</td><td>2.3E-04</td><td>MB231/MB468</td><td>CYTO</td></tr>
<tr><td>>hg38_21_+_8433172_8446622_RNA45SN1_45S_50ntFlanks@11580.11606.27|GAACTGGTGCGGACCAGGGGAATCCGA</td><td>>hg38_21_+_8433172_8446622_RNA45SN1_45S_50ntFlanks</td><td>11580</td><td>11606</td><td>27</td><td>GAACTGGTGCGGACCAGGGGAATCCGA</td><td>255.7</td><td>-5.6</td><td>1.5</td><td>-3.7</td><td>1.8E-04</td><td>2.3E-04</td><td>MB231/MB468</td><td>MITO</td></tr>
<tr><td>>MI0000434|hsa-let-7i&WithFlank&12|+|62603680|62603775@12.31.20(+1C)&[MIMAT0000415&hsa-let-7i-5p&offsets|0|-2(+1C);m-49&12|+|62603691|62603712&offsets|0|-2(+1C)]|TGAGGTAGTAGTTTGTGCTGC</td><td>>MI0000434|hsa-let-7i&WithFlank&12|+|62603680|62603775</td><td>12</td><td>31</td><td>20(+1C)</td><td>TGAGGTAGTAGTTTGTGCTGC</td><td>1065.2</td><td>5.3</td><td>1.4</td><td>3.8</td><td>1.5E-04</td><td>2.3E-04</td><td>MB231/MB468</td><td>MP</td></tr>
<tr><td>>MI0015866|hsa-mir-4326&WithFlank&20|+|63286802|63286872@17.40.24(+1U)&[MIMAT0016888&hsa-miR-4326&offsets|0|+4(+1U);m-339&20|+|63286818|63286841&offsets|0|0(+1U)]|TGTTCCTCTGTCTCCCAGACTCTGT</td><td>>MI0015866|hsa-mir-4326&WithFlank&20|+|63286802|63286872</td><td>17</td><td>40</td><td>24(+1U)</td><td>TGTTCCTCTGTCTCCCAGACTCTGT</td><td>88.8</td><td>5.5</td><td>1.4</td><td>3.8</td><td>1.5E-04</td><td>2.3E-04</td><td>MB231/MB468</td><td>MP</td></tr>
<tr><td>>hg38_21_+_8433172_8446622_RNA45SN1_45S_50ntFlanks@5078.5098.21|CGATAACGAACGAGACTCTGG</td><td>>hg38_21_+_8433172_8446622_RNA45SN1_45S_50ntFlanks</td><td>5078</td><td>5098</td><td>21</td><td>CGATAACGAACGAGACTCTGG</td><td>504.5</td><td>-4.0</td><td>1.1</td><td>-3.7</td><td>1.8E-04</td><td>2.3E-04</td><td>MB231/MB468</td><td>NUC</td></tr>
<tr><td>>hg38_21_+_8433172_8446622_RNA45SN1_45S_50ntFlanks@475.511.37|CTTCGTGATCGATGTGGTGACGTCGTGCTCTCCCGGG</td><td>>hg38_21_+_8433172_8446622_RNA45SN1_45S_50ntFlanks</td><td>475</td><td>511</td><td>37</td><td>CTTCGTGATCGATGTGGTGACGTCGTGCTCTCCCGGG</td><td>678.6</td><td>5.0</td><td>1.3</td><td>3.7</td><td>1.8E-04</td><td>2.3E-04</td><td>MB231/MB468</td><td>MITO</td></tr>
<tr><td>>hg38_21_+_8433172_8446622_RNA45SN1_45S_50ntFlanks@10246.10279.34|CGCGACGAGTAGGAGGGCCGCTGCGGTGAGCCTT</td><td>>hg38_21_+_8433172_8446622_RNA45SN1_45S_50ntFlanks</td><td>10246</td><td>10279</td><td>34</td><td>CGCGACGAGTAGGAGGGCCGCTGCGGTGAGCCTT</td><td>85.6</td><td>7.2</td><td>1.8</td><td>4.0</td><td>6.6E-05</td><td>2.3E-04</td><td>BT20/MB468</td><td>MP</td></tr>
<tr><td>>hg38_21_+_8433172_8446622_RNA45SN1_45S_50ntFlanks@3876.3919.44|TACATGCCGACGGGCGCTGACCCCCTTCGCGGGGGGGATGCGTG</td><td>>hg38_21_+_8433172_8446622_RNA45SN1_45S_50ntFlanks</td><td>3876</td><td>3919</td><td>44</td><td>TACATGCCGACGGGCGCTGACCCCCTTCGCGGGGGGGATGCGTG</td><td>102.2</td><td>10.4</td><td>2.6</td><td>4.0</td><td>6.6E-05</td><td>2.3E-04</td><td>BT20/MB468</td><td>MP</td></tr>
<tr><td>>hg38_21_+_8433172_8446622_RNA45SN1_45S_50ntFlanks@9928.9949.22|TGATATAGACAGCAGGACGGTG</td><td>>hg38_21_+_8433172_8446622_RNA45SN1_45S_50ntFlanks</td><td>9928</td><td>9949</td><td>22</td><td>TGATATAGACAGCAGGACGGTG</td><td>245.1</td><td>7.4</td><td>1.8</td><td>4.0</td><td>6.6E-05</td><td>2.3E-04</td><td>BT20/MB468</td><td>MP</td></tr>
<tr><td>>hg38_21_+_8433172_8446622_RNA45SN1_45S_50ntFlanks@11873.11908.36|AAGAAGACCCTGTTGAGCTTGACTCTAGTCTGGCAC</td><td>>hg38_21_+_8433172_8446622_RNA45SN1_45S_50ntFlanks</td><td>11873</td><td>11908</td><td>36</td><td>AAGAAGACCCTGTTGAGCTTGACTCTAGTCTGGCAC</td><td>536.9</td><td>-2.5</td><td>0.7</td><td>-3.8</td><td>1.3E-04</td><td>2.3E-04</td><td>BT20/MB468</td><td>MITO</td></tr>
<tr><td>>hg38_21_+_8433172_8446622_RNA45SN1_45S_50ntFlanks@12476.12496.21|CCACTAATAGGGAACGTGAGC</td><td>>hg38_21_+_8433172_8446622_RNA45SN1_45S_50ntFlanks</td><td>12476</td><td>12496</td><td>21</td><td>CCACTAATAGGGAACGTGAGC</td><td>1114.7</td><td>-2.0</td><td>0.5</td><td>-3.8</td><td>1.3E-04</td><td>2.3E-04</td><td>BT20/MB468</td><td>MITO</td></tr>
<tr><td>>MI0000650|hsa-mir-200c&WithFlank&12|+|6963693|6963772@50.72.23(+1U)&[MIMAT0000617&hsa-miR-200c-3p&offsets|0|0(+1U);m-54&12|+|6963742|6963764&offsets|0|0(+1U)]|TAATACTGCCGGGTAATGATGGAT</td><td>>MI0000650|hsa-mir-200c&WithFlank&12|+|6963693|6963772</td><td>50</td><td>72</td><td>23(+1U)</td><td>TAATACTGCCGGGTAATGATGGAT</td><td>187.6</td><td>-4.6</td><td>1.2</td><td>-3.8</td><td>1.5E-04</td><td>2.3E-04</td><td>MB231/MB468</td><td>MP</td></tr>
<tr><td>>hg38_21_+_8433172_8446622_RNA45SN1_45S_50ntFlanks@8113.8138.26|GGCGCGGGACATGTGGCGTACGGAAG</td><td>>hg38_21_+_8433172_8446622_RNA45SN1_45S_50ntFlanks</td><td>8113</td><td>8138</td><td>26</td><td>GGCGCGGGACATGTGGCGTACGGAAG</td><td>168.4</td><td>-5.5</td><td>1.5</td><td>-3.7</td><td>1.8E-04</td><td>2.3E-04</td><td>MB231/MB468</td><td>NUC</td></tr>
<tr><td>>hg38_21_+_8433172_8446622_RNA45SN1_45S_50ntFlanks@7991.8020.30|ACGTGGCGACCCGCTGAATTTAAGCATATT</td><td>>hg38_21_+_8433172_8446622_RNA45SN1_45S_50ntFlanks</td><td>7991</td><td>8020</td><td>30</td><td>ACGTGGCGACCCGCTGAATTTAAGCATATT</td><td>253.4</td><td>-2.7</td><td>0.7</td><td>-3.7</td><td>1.8E-04</td><td>2.3E-04</td><td>MB231/MB468</td><td>MITO</td></tr>
<tr><td>>hg38_21_+_8433172_8446622_RNA45SN1_45S_50ntFlanks@12997.13038.42|CGCTGCGATCTATTGAAAGTCAGCCCTCGACACAAGGGTTTG</td><td>>hg38_21_+_8433172_8446622_RNA45SN1_45S_50ntFlanks</td><td>12997</td><td>13038</td><td>42</td><td>CGCTGCGATCTATTGAAAGTCAGCCCTCGACACAAGGGTTTG</td><td>758.7</td><td>6.3</td><td>1.7</td><td>3.8</td><td>1.3E-04</td><td>2.3E-04</td><td>BT20/MB468</td><td>MITO</td></tr>
<tr><td>>hg38_21_+_8433172_8446622_RNA45SN1_45S_50ntFlanks@11712.11736.25|CGGGTAAACGGCGGGAGTAACTATG</td><td>>hg38_21_+_8433172_8446622_RNA45SN1_45S_50ntFlanks</td><td>11712</td><td>11736</td><td>25</td><td>CGGGTAAACGGCGGGAGTAACTATG</td><td>173.5</td><td>-4.6</td><td>1.2</td><td>-3.8</td><td>1.5E-04</td><td>2.3E-04</td><td>MB231/MB468</td><td>MP</td></tr>
<tr><td>>hg38_21_+_8433172_8446622_RNA45SN1_45S_50ntFlanks@11620.11660.41|ACAAAGCATCGCGAAGGCCCGCGGCGGGTGTTGACGCGATG</td><td>>hg38_21_+_8433172_8446622_RNA45SN1_45S_50ntFlanks</td><td>11620</td><td>11660</td><td>41</td><td>ACAAAGCATCGCGAAGGCCCGCGGCGGGTGTTGACGCGATG</td><td>188.7</td><td>-5.8</td><td>1.5</td><td>-3.8</td><td>1.3E-04</td><td>2.3E-04</td><td>BT20/MB468</td><td>MITO</td></tr>
<tr><td>>hg38_21_+_8433172_8446622_RNA45SN1_45S_50ntFlanks@7979.8008.30|ACCTCAGATCAGACGTGGCGACCCGCTGAA</td><td>>hg38_21_+_8433172_8446622_RNA45SN1_45S_50ntFlanks</td><td>7979</td><td>8008</td><td>30</td><td>ACCTCAGATCAGACGTGGCGACCCGCTGAA</td><td>2366.5</td><td>-1.9</td><td>0.5</td><td>-3.8</td><td>1.3E-04</td><td>2.3E-04</td><td>BT20/MB468</td><td>MITO</td></tr>
<tr><td>>hg38_MT_+_1621_3279_RNR2_16S_50ntFlanks@51.71.21|GCTAAACCTAGCCCCAAACCC</td><td>>hg38_MT_+_1621_3279_RNR2_16S_50ntFlanks</td><td>51</td><td>71</td><td>21</td><td>GCTAAACCTAGCCCCAAACCC</td><td>168.1</td><td>2.6</td><td>0.7</td><td>3.7</td><td>1.8E-04</td><td>2.3E-04</td><td>BT20/MB468</td><td>NUC</td></tr>
<tr><td>>hg38_21_+_8433172_8446622_RNA45SN1_45S_50ntFlanks@9574.9604.31|AATCGGTCGTCCGACCTGGGTATAGGGGCGA</td><td>>hg38_21_+_8433172_8446622_RNA45SN1_45S_50ntFlanks</td><td>9574</td><td>9604</td><td>31</td><td>AATCGGTCGTCCGACCTGGGTATAGGGGCGA</td><td>411.7</td><td>-3.2</td><td>0.8</td><td>-3.8</td><td>1.5E-04</td><td>2.3E-04</td><td>MB231/MB468</td><td>MP</td></tr>
<tr><td>>hg38_21_+_8433172_8446622_RNA45SN1_45S_50ntFlanks@11317.11338.22|ACCCCACGTCTCGTCGCGCGCG</td><td>>hg38_21_+_8433172_8446622_RNA45SN1_45S_50ntFlanks</td><td>11317</td><td>11338</td><td>22</td><td>ACCCCACGTCTCGTCGCGCGCG</td><td>4700.8</td><td>-7.7</td><td>2.0</td><td>-3.7</td><td>1.8E-04</td><td>2.3E-04</td><td>BT20/MB468</td><td>NUC</td></tr>
<tr><td>>am_trnaMT_ValTAC_MT_+_1602_1670@28.51.24|AACTTACACTTAGGAGATTTCAAC</td><td>>am_trnaMT_ValTAC_MT_+_1602_1670</td><td>28</td><td>51</td><td>24</td><td>AACTTACACTTAGGAGATTTCAAC</td><td>113.3</td><td>10.6</td><td>2.7</td><td>4.0</td><td>6.7E-05</td><td>2.3E-04</td><td>BT20/MB468</td><td>MP</td></tr>
</table>

<table>
<tr><td></td><td></td><td></td><td></td><td></td><td></td><td></td><td></td><td></td><td></td><td></td><td></td><td></td><td></td></tr>
<tr><td>>trna119_LysCTT_1_-_145395522_145395594@3.35.33,trna11_LysCTT_5_-_180648979_180649051@3.35.33,trna13_LysCTT_6_+_26556774_26556846@3.35.33,trna7_LysCTT_16_+_3225692_3225764@3.35.33,trna9_LysCTT_5_+_180634755_180634827@3.35.33|CCGGCTAGCTCAGTCGGTAGAGCATGAGACTCT</td><td>>trna119_LysCTT_1_-_145395522_145395594</td><td>3</td><td>35</td><td>33</td><td>CCGGCTAGCTCAGTCGGTAGAGCATGAGACTCT</td><td>169.0</td><td>5.6</td><td>1.5</td><td>3.8</td><td>1.5E-04</td><td>2.3E-04</td><td>MB231/MB468</td><td>MP</td></tr>
<tr><td>>MI0000065|hsa-let-7d&WithFlank&9|+|94178828|94178926@14.36.23&[MIMAT0000065&hsa-let-7d-5p&offsets|0|+1;m-90&9|+|94178841|94178862&offsets|0|+1]|AGAGGTAGTAGGTTGCATAGTTT</td><td>>MI0000065|hsa-let-7d&WithFlank&9|+|94178828|94178926</td><td>14</td><td>36</td><td>23</td><td>AGAGGTAGTAGGTTGCATAGTTT</td><td>698.4</td><td>5.0</td><td>1.3</td><td>3.8</td><td>1.5E-04</td><td>2.3E-04</td><td>MB231/MB468</td><td>MP</td></tr>
<tr><td>>hg38_21_+_8433172_8446622_RNA45SN1_45S_50ntFlanks@485.521.37|GATGTGGTGACGTCGTGCTCTCCCGGGCCGGGTCCGA</td><td>>hg38_21_+_8433172_8446622_RNA45SN1_45S_50ntFlanks</td><td>485</td><td>521</td><td>37</td><td>GATGTGGTGACGTCGTGCTCTCCCGGGCCGGGTCCGA</td><td>87.3</td><td>9.7</td><td>2.6</td><td>3.8</td><td>1.5E-04</td><td>2.3E-04</td><td>MB231/MB468</td><td>MP</td></tr>
<tr><td>>MI0000093|hsa-mir-92a-1&WithFlank&13|+|91351308|91351397@54.73.20;MI0000094|hsa-mir-92a-2&WithFlank&X|-|134169532|134169618@54.73.20&[MIMAT0000092&hsa-miR-92a-3p&offsets|0|-2;m-9&13|+|91351361|91351382&offsets|0|-2];[MIMAT0000092_1&hsa-miR-92a-3p&offsets|0|-2;m-11&X|-|134169544|134169565&offsets|0|-2]|TATTGCACTTGTCCCGGCCT</td><td>>MI0000093|hsa-mir-92a-1&WithFlank&13|+|91351308|91351397</td><td>54</td><td>73</td><td>20</td><td>TATTGCACTTGTCCCGGCCT</td><td>218.5</td><td>4.8</td><td>1.3</td><td>3.8</td><td>1.5E-04</td><td>2.3E-04</td><td>MB231/MB468</td><td>MP</td></tr>
<tr><td>>hg38_21_+_8433172_8446622_RNA45SN1_45S_50ntFlanks@12602.12626.25|GTGTATGTGCTTGGCTGAGGAGCCA</td><td>>hg38_21_+_8433172_8446622_RNA45SN1_45S_50ntFlanks</td><td>12602</td><td>12626</td><td>25</td><td>GTGTATGTGCTTGGCTGAGGAGCCA</td><td>341.9</td><td>-4.1</td><td>1.1</td><td>-3.8</td><td>1.7E-04</td><td>2.3E-04</td><td>MB231/MB468</td><td>CYTO</td></tr>
<tr><td>>trna10_AspGTC_12_-_125424193_125424264@35.73.39,trna12_AspGTC_12_-_125411891_125411962@35.73.39,trna144_AspGTC_6_-_27551236_27551307@35.73.39,trna38_AspGTC_17_-_8125556_8125627@35.73.39,trna45_AspGTC_6_+_27447453_27447524@35.73.39,trna48_AspGTC_6_+_27471523_27471594@35.73.39,trna4_AspGTC_12_+_96429799_96429870@35.73.39,trna69_AspGTC_1_-_161440205_161440276@35.73.39,trna72_AspGTC_1_-_161432824_161432895@35.73.39,trna75_AspGTC_1_-_161425414_161425485@35.73.39,trna78_AspGTC_1_-_161418033_161418104@35.73.39,trna81_AspGTC_1_-_161410615_161410686@35.73.39|TCACGCGGGAGACCGGGGTTCGATTCCCCGACGGGGAGC</td><td>>trna10_AspGTC_12_-_125424193_125424264</td><td>35</td><td>73</td><td>39</td><td>TCACGCGGGAGACCGGGGTTCGATTCCCCGACGGGGAGC</td><td>4465.9</td><td>-4.7</td><td>1.1</td><td>-4.2</td><td>2.2E-05</td><td>2.3E-04</td><td>BT20/MB231</td><td>CYTO</td></tr>
<tr><td>>MI0000253|hsa-mir-148a&WithFlank&7|-|25949913|25949992@50.71.22&[MIMAT0000243&hsa-miR-148a-3p&offsets|0|0;m-7&7|-|25949922|25949943&offsets|0|0]|TCAGTGCACTACAGAACTTTGT</td><td>>MI0000253|hsa-mir-148a&WithFlank&7|-|25949913|25949992</td><td>50</td><td>71</td><td>22</td><td>TCAGTGCACTACAGAACTTTGT</td><td>97245.9</td><td>3.2</td><td>0.8</td><td>4.2</td><td>2.7E-05</td><td>2.3E-04</td><td>BT20/MB231</td><td>MITO</td></tr>
<tr><td>>MI0000085|hsa-mir-27a&WithFlank&19|-|13836434|13836523@57.76.20(+5U)&[MIMAT0000084&hsa-miR-27a-3p&offsets|0|-1(+5U);m-43&19|-|13836447|13836467&offsets|0|-1(+5U)]|TTCACAGTGGCTAAGTTCCGTTTTT</td><td>>MI0000085|hsa-mir-27a&WithFlank&19|-|13836434|13836523</td><td>57</td><td>76</td><td>20(+5U)</td><td>TTCACAGTGGCTAAGTTCCGTTTTT</td><td>27.6</td><td>8.0</td><td>2.1</td><td>3.8</td><td>1.5E-04</td><td>2.3E-04</td><td>MB231/MB468</td><td>MP</td></tr>
<tr><td>>hg38_21_+_8433172_8446622_RNA45SN1_45S_50ntFlanks@10346.10388.43|TCAAACGAGAACTTTGAAGGCCGAAGTGGAGAAGGGTTCCATG</td><td>>hg38_21_+_8433172_8446622_RNA45SN1_45S_50ntFlanks</td><td>10346</td><td>10388</td><td>43</td><td>TCAAACGAGAACTTTGAAGGCCGAAGTGGAGAAGGGTTCCATG</td><td>606.7</td><td>-3.3</td><td>0.9</td><td>-3.7</td><td>1.8E-04</td><td>2.3E-04</td><td>MB231/MB468</td><td>NUC</td></tr>
<tr><td>>hg38_21_+_8433172_8446622_RNA45SN1_45S_50ntFlanks@5084.5107.24|CGAACGAGACTCTGGCATGCTAAC</td><td>>hg38_21_+_8433172_8446622_RNA45SN1_45S_50ntFlanks</td><td>5084</td><td>5107</td><td>24</td><td>CGAACGAGACTCTGGCATGCTAAC</td><td>192.2</td><td>-5.4</td><td>1.4</td><td>-3.8</td><td>1.5E-04</td><td>2.3E-04</td><td>MB231/MB468</td><td>MP</td></tr>
<tr><td>>trna2_GlyGCC_21_-_18827107_18827177@1.28.28,trna35_GlyGCC_1_+_161413094_161413164@1.28.28,trna37_GlyGCC_1_+_161420467_161420537@1.28.28,trna39_GlyGCC_1_+_161427898_161427968@1.28.28,trna41_GlyGCC_1_+_161435258_161435328@1.28.28|GCATGGGTGGTTCAGTGGTAGAATTCTC</td><td>>trna2_GlyGCC_21_-_18827107_18827177</td><td>1</td><td>28</td><td>28</td><td>GCATGGGTGGTTCAGTGGTAGAATTCTC</td><td>5582.7</td><td>4.3</td><td>1.0</td><td>4.2</td><td>2.2E-05</td><td>2.3E-04</td><td>BT20/MB231</td><td>CYTO</td></tr>
<tr><td>>hg38_21_+_8433172_8446622_RNA45SN1_45S_50ntFlanks@12939.12956.18|CCATTCGTAGACGACCTG</td><td>>hg38_21_+_8433172_8446622_RNA45SN1_45S_50ntFlanks</td><td>12939</td><td>12956</td><td>18</td><td>CCATTCGTAGACGACCTG</td><td>790.3</td><td>-3.6</td><td>0.9</td><td>-4.0</td><td>6.7E-05</td><td>2.3E-04</td><td>BT20/MB468</td><td>MP</td></tr>
<tr><td>>hg38_21_+_8433172_8446622_RNA45SN1_45S_50ntFlanks@8570.8593.24|CTCCGGCGGGTGCGGGGGTGGGCG</td><td>>hg38_21_+_8433172_8446622_RNA45SN1_45S_50ntFlanks</td><td>8570</td><td>8593</td><td>24</td><td>CTCCGGCGGGTGCGGGGGTGGGCG</td><td>156.0</td><td>-4.8</td><td>1.3</td><td>-3.8</td><td>1.6E-04</td><td>2.3E-04</td><td>BT20/MB468</td><td>CYTO</td></tr>
<tr><td>>MI0003591|hsa-mir-584&WithFlank&5|-|149062307|149062415@22.43.22(+1U)&[MIMAT0003249&hsa-miR-584-5p&offsets|0|0(+1U);m-167&5|-|149062374|149062394&offsets|0|+1(+1U)]|TTATGGTTTGCCTGGGACTGAGT</td><td>>MI0003591|hsa-mir-584&WithFlank&5|-|149062307|149062415</td><td>22</td><td>43</td><td>22(+1U)</td><td>TTATGGTTTGCCTGGGACTGAGT</td><td>154.9</td><td>5.3</td><td>1.3</td><td>4.0</td><td>6.7E-05</td><td>2.3E-04</td><td>BT20/MB468</td><td>MP</td></tr>
<tr><td>>hg38_21_+_8433172_8446622_RNA45SN1_45S_50ntFlanks@3842.3874.33|CCTACTTGGATAACTGTGGTAATTCTAGAGCTA</td><td>>hg38_21_+_8433172_8446622_RNA45SN1_45S_50ntFlanks</td><td>3842</td><td>3874</td><td>33</td><td>CCTACTTGGATAACTGTGGTAATTCTAGAGCTA</td><td>94.5</td><td>-4.7</td><td>1.2</td><td>-3.8</td><td>1.5E-04</td><td>2.3E-04</td><td>MB231/MB468</td><td>MP</td></tr>
<tr><td>>hg38_21_+_8433172_8446622_RNA45SN1_45S_50ntFlanks@6652.6695.44|GACTCTTAGCGGTGGATCACTCGGCTCGTGCGTCGATGAAGAAC</td><td>>hg38_21_+_8433172_8446622_RNA45SN1_45S_50ntFlanks</td><td>6652</td><td>6695</td><td>44</td><td>GACTCTTAGCGGTGGATCACTCGGCTCGTGCGTCGATGAAGAAC</td><td>1932.7</td><td>3.2</td><td>0.9</td><td>3.7</td><td>1.8E-04</td><td>2.3E-04</td><td>MB231/MB468</td><td>MITO</td></tr>
<tr><td>>hg38_21_+_8433172_8446622_RNA45SN1_45S_50ntFlanks@11570.11588.19|AGCCGACTTAGAACTGGTG</td><td>>hg38_21_+_8433172_8446622_RNA45SN1_45S_50ntFlanks</td><td>11570</td><td>11588</td><td>19</td><td>AGCCGACTTAGAACTGGTG</td><td>163.3</td><td>-2.5</td><td>0.7</td><td>-3.8</td><td>1.6E-04</td><td>2.3E-04</td><td>BT20/MB468</td><td>CYTO</td></tr>
<tr><td>>MI0000261|hsa-mir-139&WithFlank&11|-|72615057|72615136@13.35.23&[MIMAT0000250&hsa-miR-139-5p&offsets|0|0;m-186&11|-|72615102|72615124&offsets|0|0]|TCTACAGTGCACGTGTCTCCAGT</td><td>>MI0000261|hsa-mir-139&WithFlank&11|-|72615057|72615136</td><td>13</td><td>35</td><td>23</td><td>TCTACAGTGCACGTGTCTCCAGT</td><td>352.2</td><td>7.6</td><td>1.9</td><td>4.0</td><td>6.7E-05</td><td>2.3E-04</td><td>BT20/MB468</td><td>MP</td></tr>
<tr><td>>trna111_HisGTG_1_-_147774845_147774916@1.36.36,trna118_HisGTG_1_-_145396881_145396952@1.36.36,trna16_HisGTG_1_+_146544773_146544844@1.36.36,trna1_HisGTG_15_+_45493349_45493420@1.36.36,trna21_HisGTG_1_+_147753471_147753542@1.36.36,trna33_HisGTG_6_+_27125906_27125977@1.36.36,trna7_HisGTG_9_-_14433938_14434009@1.36.36,trna8_HisGTG_15_-_45492611_45492682@1.36.36,trna9_HisGTG_15_-_45490804_45490875@1.36.36|GCCGTGATCGTATAGTGGTTAGTACTCTGCGTTGTG</td><td>>trna111_HisGTG_1_-_147774845_147774916</td><td>1</td><td>36</td><td>36</td><td>GCCGTGATCGTATAGTGGTTAGTACTCTGCGTTGTG</td><td>767.5</td><td>4.1</td><td>1.0</td><td>4.2</td><td>2.3E-05</td><td>2.3E-04</td><td>BT20/MB231</td><td>CYTO</td></tr>
<tr><td>>hg38_21_+_8433172_8446622_RNA45SN1_45S_50ntFlanks@10868.10902.35|AAGGGCTGGGTCGGTCGGGCTGGGGCGCGAAGCGG</td><td>>hg38_21_+_8433172_8446622_RNA45SN1_45S_50ntFlanks</td><td>10868</td><td>10902</td><td>35</td><td>AAGGGCTGGGTCGGTCGGGCTGGGGCGCGAAGCGG</td><td>490.2</td><td>-6.5</td><td>1.8</td><td>-3.7</td><td>1.8E-04</td><td>2.3E-04</td><td>BT20/MB468</td><td>NUC</td></tr>
<tr><td>>hg38_21_+_8433172_8446622_RNA45SN1_45S_50ntFlanks@8157.8175.19|CTCGTGGGGGGCCCAAGTC</td><td>>hg38_21_+_8433172_8446622_RNA45SN1_45S_50ntFlanks</td><td>8157</td><td>8175</td><td>19</td><td>CTCGTGGGGGGCCCAAGTC</td><td>231.8</td><td>-4.2</td><td>1.1</td><td>-3.8</td><td>1.3E-04</td><td>2.3E-04</td><td>BT20/MB468</td><td>MITO</td></tr>
<tr><td>>hg38_21_+_8433172_8446622_RNA45SN1_45S_50ntFlanks@8114.8141.28|GCGCGGGACATGTGGCGTACGGAAGACC</td><td>>hg38_21_+_8433172_8446622_RNA45SN1_45S_50ntFlanks</td><td>8114</td><td>8141</td><td>28</td><td>GCGCGGGACATGTGGCGTACGGAAGACC</td><td>5014.6</td><td>-3.6</td><td>0.9</td><td>-4.0</td><td>6.8E-05</td><td>2.3E-04</td><td>BT20/MB468</td><td>MP</td></tr>
<tr><td>>hg38_21_+_8433172_8446622_RNA45SN1_45S_50ntFlanks@12454.12473.20|CACCAAGCGTTGGATTGTTC</td><td>>hg38_21_+_8433172_8446622_RNA45SN1_45S_50ntFlanks</td><td>12454</td><td>12473</td><td>20</td><td>CACCAAGCGTTGGATTGTTC</td><td>252.4</td><td>-3.6</td><td>0.9</td><td>-4.0</td><td>6.8E-05</td><td>2.3E-04</td><td>BT20/MB468</td><td>MP</td></tr>
<tr><td>>MI0016887|hsa-mir-4521&WithFlank&17|+|8186939|8187010@10.35.26&[MIMAT0019058&hsa-miR-4521&offsets|0|+4;m-765&17|+|8186948|8186969&offsets|0|+4]|GCTAAGGAAGTCCTGTGCTCAGTTTT</td><td>>MI0016887|hsa-mir-4521&WithFlank&17|+|8186939|8187010</td><td>10</td><td>35</td><td>26</td><td>GCTAAGGAAGTCCTGTGCTCAGTTTT</td><td>842.2</td><td>3.7</td><td>1.0</td><td>3.8</td><td>1.7E-04</td><td>2.4E-04</td><td>MB231/MB468</td><td>CYTO</td></tr>
<tr><td>>MI0000469|hsa-mir-125a&WithFlank&19|+|51693248|51693345@21.42.22&[MIMAT0000443&hsa-miR-125a-5p&offsets|0|-2;m-63&19|+|51693268|51693289&offsets|0|0]|TCCCTGAGACCCTTTAACCTGT</td><td>>MI0000469|hsa-mir-125a&WithFlank&19|+|51693248|51693345</td><td>21</td><td>42</td><td>22</td><td>TCCCTGAGACCCTTTAACCTGT</td><td>2278.4</td><td>2.8</td><td>0.8</td><td>3.7</td><td>1.8E-04</td><td>2.4E-04</td><td>MB231/MB468</td><td>NUC</td></tr>
<tr><td>>hg38_21_+_8433172_8446622_RNA45SN1_45S_50ntFlanks@9572.9598.27|CAAATCGGTCGTCCGACCTGGGTATAG</td><td>>hg38_21_+_8433172_8446622_RNA45SN1_45S_50ntFlanks</td><td>9572</td><td>9598</td><td>27</td><td>CAAATCGGTCGTCCGACCTGGGTATAG</td><td>232.9</td><td>-6.0</td><td>1.5</td><td>-4.0</td><td>6.8E-05</td><td>2.4E-04</td><td>BT20/MB468</td><td>MP</td></tr>
<tr><td>>trnaMT_LysTTT_MT_+_8295_8364@52.72.21,trnalookalike7_LysTTT_1_+_568844_568913@52.72.21|CAACACCTCTTTACAGTGACC</td><td>>trnaMT_LysTTT_MT_+_8295_8364</td><td>52</td><td>72</td><td>21</td><td>CAACACCTCTTTACAGTGACC</td><td>51.9</td><td>-9.1</td><td>2.2</td><td>-4.2</td><td>2.4E-05</td><td>2.4E-04</td><td>BT20/MB231</td><td>MP</td></tr>
<tr><td>>hg38_21_+_8433172_8446622_RNA45SN1_45S_50ntFlanks@9457.9503.47|CGCCGGGGAGGTGGAGCACGAGCGCACGTGTTAGGACCCGAAAGATG</td><td>>hg38_21_+_8433172_8446622_RNA45SN1_45S_50ntFlanks</td><td>9457</td><td>9503</td><td>47</td><td>CGCCGGGGAGGTGGAGCACGAGCGCACGTGTTAGGACCCGAAAGATG</td><td>443.4</td><td>-7.6</td><td>1.9</td><td>-4.0</td><td>6.8E-05</td><td>2.4E-04</td><td>BT20/MB468</td><td>MP</td></tr>
<tr><td>>hg38_21_+_8433172_8446622_RNA45SN1_45S_50ntFlanks@6647.6679.33|CGTACGACTCTTAGCGGTGGATCACTCGGCTCG</td><td>>hg38_21_+_8433172_8446622_RNA45SN1_45S_50ntFlanks</td><td>6647</td><td>6679</td><td>33</td><td>CGTACGACTCTTAGCGGTGGATCACTCGGCTCG</td><td>660.5</td><td>-3.6</td><td>0.9</td><td>-3.8</td><td>1.5E-04</td><td>2.4E-04</td><td>MB231/MB468</td><td>MP</td></tr>
<tr><td>>hg38_21_+_8433172_8446622_RNA45SN1_45S_50ntFlanks@10482.10500.19|TCGAAAGGGAGTCGGGTTC</td><td>>hg38_21_+_8433172_8446622_RNA45SN1_45S_50ntFlanks</td><td>10482</td><td>10500</td><td>19</td><td>TCGAAAGGGAGTCGGGTTC</td><td>194.1</td><td>-2.6</td><td>0.7</td><td>-3.8</td><td>1.7E-04</td><td>2.4E-04</td><td>MB231/MB468</td><td>CYTO</td></tr>
<tr><td>>hg38_21_+_8433172_8446622_RNA45SN1_45S_50ntFlanks@6651.6672.22|CGACTCTTAGCGGTGGATCACT</td><td>>hg38_21_+_8433172_8446622_RNA45SN1_45S_50ntFlanks</td><td>6651</td><td>6672</td><td>22</td><td>CGACTCTTAGCGGTGGATCACT</td><td>1469.4</td><td>3.5</td><td>0.9</td><td>4.0</td><td>6.8E-05</td><td>2.4E-04</td><td>BT20/MB468</td><td>MP</td></tr>
<tr><td>>hg38_21_+_8433172_8446622_RNA45SN1_45S_50ntFlanks@10349.10376.28|AACGAGAACTTTGAAGGCCGAAGTGGAG</td><td>>hg38_21_+_8433172_8446622_RNA45SN1_45S_50ntFlanks</td><td>10349</td><td>10376</td><td>28</td><td>AACGAGAACTTTGAAGGCCGAAGTGGAG</td><td>614.3</td><td>-1.7</td><td>0.5</td><td>-3.8</td><td>1.6E-04</td><td>2.4E-04</td><td>BT20/MB468</td><td>CYTO</td></tr>
<tr><td>>MI0000081|hsa-mir-24-2&WithFlank&19|-|13836281|13836365@56.74.19(+1U);MI0000080|hsa-mir-24-1&WithFlank&9|+|95086015|95086094@50.68.19(+1U)&[MIMAT0000080&hsa-miR-24-3p&offsets|0|-3(+1U);m-36&19|-|13836288|13836310&offsets|0|-4(+1U)];[MIMAT0000080_1&hsa-miR-24-3p&offsets|0|-3(+1U);m-35&9|+|95086064|95086086&offsets|0|-4(+1U)]|TGGCTCAGTTCAGCAGGAAT</td><td>>MI0000081|hsa-mir-24-2&WithFlank&19|-|13836281|13836365</td><td>56</td><td>74</td><td>19(+1U)</td><td>TGGCTCAGTTCAGCAGGAAT</td><td>2721.5</td><td>4.3</td><td>1.1</td><td>4.0</td><td>6.8E-05</td><td>2.4E-04</td><td>BT20/MB468</td><td>MP</td></tr>
<tr><td>>hg38_21_+_8433172_8446622_RNA45SN1_45S_50ntFlanks@11767.11804.38|TCTAATTAGTGACGCGCATGAATGGATGAACGAGATTC</td><td>>hg38_21_+_8433172_8446622_RNA45SN1_45S_50ntFlanks</td><td>11767</td><td>11804</td><td>38</td><td>TCTAATTAGTGACGCGCATGAATGGATGAACGAGATTC</td><td>222.5</td><td>-4.1</td><td>1.1</td><td>-3.7</td><td>1.8E-04</td><td>2.4E-04</td><td>MB231/MB468</td><td>NUC</td></tr>
<tr><td>>hg38_21_+_8433172_8446622_RNA45SN1_45S_50ntFlanks@12001.12030.30|CTGCGGGCCGCCGGTGAAATACCACTACTC</td><td>>hg38_21_+_8433172_8446622_RNA45SN1_45S_50ntFlanks</td><td>12001</td><td>12030</td><td>30</td><td>CTGCGGGCCGCCGGTGAAATACCACTACTC</td><td>646.0</td><td>-5.9</td><td>1.6</td><td>-3.7</td><td>1.8E-04</td><td>2.4E-04</td><td>MB231/MB468</td><td>NUC</td></tr>
<tr><td>>MI0000273|hsa-mir-183&WithFlank&7|-|129774899|129775020@33.53.21(+1C)&[MIMAT0000261&hsa-miR-183-5p&offsets|0|-1(+1C);m-41&7|-|129774966|129774987&offsets|-1|-2(+1C)]|TATGGCACTGGTAGAATTCACC</td><td>>MI0000273|hsa-mir-183&WithFlank&7|-|129774899|129775020</td><td>33</td><td>53</td><td>21(+1C)</td><td>TATGGCACTGGTAGAATTCACC</td><td>353.7</td><td>3.6</td><td>1.0</td><td>3.7</td><td>1.8E-04</td><td>2.4E-04</td><td>MB231/MB468</td><td>MITO</td></tr>
<tr><td>>hg38_21_+_8433172_8446622_RNA45SN1_45S_50ntFlanks@10369.10406.38|AAGTGGAGAAGGGTTCCATGTGAACAGCAGTTGAACAT</td><td>>hg38_21_+_8433172_8446622_RNA45SN1_45S_50ntFlanks</td><td>10369</td><td>10406</td><td>38</td><td>AAGTGGAGAAGGGTTCCATGTGAACAGCAGTTGAACAT</td><td>166.1</td><td>-5.0</td><td>1.3</td><td>-3.8</td><td>1.6E-04</td><td>2.4E-04</td><td>BT20/MB468</td><td>CYTO</td></tr>
<tr><td>>hg38_21_+_8433172_8446622_RNA45SN1_45S_50ntFlanks@12635.12657.23|AAGCTACCATCTGTGGGATTATG</td><td>>hg38_21_+_8433172_8446622_RNA45SN1_45S_50ntFlanks</td><td>12635</td><td>12657</td><td>23</td><td>AAGCTACCATCTGTGGGATTATG</td><td>231.5</td><td>-4.4</td><td>1.2</td><td>-3.7</td><td>1.8E-04</td><td>2.4E-04</td><td>MB231/MB468</td><td>NUC</td></tr>
<tr><td>>am_MI0000786|hsa-mir-378a&WithFlank&5|+|149732819|149732896@49.70.22(+1U)&[MIMAT0000732&hsa-miR-378a-3p&offsets|0|0(+1U);m-44&5|+|149732867|149732888&offsets|0|0(+1U)]|ACTGGACTTGGAGTCAGAAGGCT</td><td>>am_MI0000786|hsa-mir-378a&WithFlank&5|+|149732819|149732896</td><td>49</td><td>70</td><td>22(+1U)</td><td>ACTGGACTTGGAGTCAGAAGGCT</td><td>1495.5</td><td>2.4</td><td>0.7</td><td>3.7</td><td>1.8E-04</td><td>2.4E-04</td><td>MB231/MB468</td><td>NUC</td></tr>
</table>

| | | | | | | | | | | | | | |
|---|---|---|---|---|---|---|---|---|---|---|---|---|---|
| >hg38_21_+_8433172_8446622_RNA45SN1_45S_50ntFlanks@10423.10453.31\|AGAGATGGGCGAGCGCCGTTCCGAAGGGACG | >hg38_21_+_8433172_8446622_RNA45SN1_45S_50ntFlanks | 10423 | 10453 | 31 | AGAGATGGGCGAGCGCCGTTCCGAAGGGACG | 102.2 | -5.7 | 1.5 | -3.8 | 1.5E-04 | 2.4E-04 | MB231/MB468 | MP |
| >hg38_MT_+_1621_3279_RNR2_16S_50ntFlanks@1425.1448.24\|ACGATTAAAGTCCTACGTGATCTG | >hg38_MT_+_1621_3279_RNR2_16S_50ntFlanks | 1425 | 1448 | 24 | ACGATTAAAGTCCTACGTGATCTG | 255.8 | 4.0 | 1.1 | 3.8 | 1.5E-04 | 2.4E-04 | MB231/MB468 | MP |
| >am_trna111_HisGTG_1_-_147774845_147774916@-1G.26.27,trna118_HisGTG_1_-_145396881_145396952@-1G.26.27,trna16_HisGTG_1_+_146544773_146544844@-1G.26.27,trna1_HisGTG_15_+_45493349_45493420@-1G.26.27,trna21_HisGTG_1_+_147753471_147753542@-1G.26.27,trna33_HisGTG_6_+_27125906_27125977@-1G.26.27,trna7_HisGTG_9_-_14433938_14434009@-1G.26.27,trna8_HisGTG_15_-_45492611_45492682@-1G.26.27,trna9_HisGTG_15_-_45490804_45490875@-1G.26.27\|GGCCGTGATCGTATAGTGGTTAGTACT | >am_trna111_HisGTG_1_-_147774845_147774916 | -1G | 26 | 27 | GGCCGTGATCGTATAGTGGTTAGTACT | 926.7 | 5.0 | 1.2 | 4.0 | 6.8E-05 | 2.4E-04 | BT20/MB468 | MP |
| >hg38_21_+_8433172_8446622_RNA45SN1_45S_50ntFlanks@6788.6807.20\|CGCCTGTCTGAGCGTCGCTT | >hg38_21_+_8433172_8446622_RNA45SN1_45S_50ntFlanks | 6788 | 6807 | 20 | CGCCTGTCTGAGCGTCGCTT | 688.8 | 4.6 | 1.2 | 4.0 | 6.8E-05 | 2.4E-04 | BT20/MB468 | MP |
| >hg38_21_+_8433172_8446622_RNA45SN1_45S_50ntFlanks@12479.12500.22\|CTAATAGGGAACGTGAGCTGGG | >hg38_21_+_8433172_8446622_RNA45SN1_45S_50ntFlanks | 12479 | 12500 | 22 | CTAATAGGGAACGTGAGCTGGG | 1166.6 | -7.0 | 1.9 | -3.7 | 1.9E-04 | 2.4E-04 | BT20/MB468 | NUC |
| >hg38_21_+_8433172_8446622_RNA45SN1_45S_50ntFlanks@10684.10712.29\|CGGCGGCGTCCGGTGAGCTCTCGCTGGCC | >hg38_21_+_8433172_8446622_RNA45SN1_45S_50ntFlanks | 10684 | 10712 | 29 | CGGCGGCGTCCGGTGAGCTCTCGCTGGCC | 542.9 | -4.1 | 1.1 | -3.8 | 1.6E-04 | 2.4E-04 | BT20/MB468 | CYTO |
| >hg38_21_+_8433172_8446622_RNA45SN1_45S_50ntFlanks@475.506.32\|CTTCGTGATCGATGTGGTGACGTCGTGCTCTC | >hg38_21_+_8433172_8446622_RNA45SN1_45S_50ntFlanks | 475 | 506 | 32 | CTTCGTGATCGATGTGGTGACGTCGTGCTCTC | 266.9 | 3.0 | 0.8 | 3.8 | 1.5E-04 | 2.4E-04 | MB231/MB468 | MP |
| >hg38_21_+_8433172_8446622_RNA45SN1_45S_50ntFlanks@9861.9885.25\|GATGAACCGAACGCCGGGTTAAGGC | >hg38_21_+_8433172_8446622_RNA45SN1_45S_50ntFlanks | 9861 | 9885 | 25 | GATGAACCGAACGCCGGGTTAAGGC | 183.7 | 7.3 | 1.8 | 4.0 | 6.8E-05 | 2.4E-04 | BT20/MB468 | MP |
| >trna10_AspGTC_12_-_125424193_125424264@35.75.41,trna12_AspGTC_12_-_125411891_125411962@35.75.41,trna144_AspGTC_6_-_27551236_27551307@35.75.41,trna38_AspGTC_17_-_8125556_8125627@35.75.41,trna45_AspGTC_6_+_27447453_27447524@35.75.41,trna48_AspGTC_6_+_27471523_27471594@35.75.41,trna4_AspGTC_12_+_96429799_96429870@35.75.41,trna69_AspGTC_1_-_161440205_161440276@35.75.41,trna72_AspGTC_1_-_161432824_161432895@35.75.41,trna75_AspGTC_1_-_161425414_161425485@35.75.41,trna78_AspGTC_1_-_161418033_161418104@35.75.41,trna81_AspGTC_1_-_161410615_161410686@35.75.41\|TCACGCGGGAGACCGGGGTTCGATTCCCCGACGGGGAGCCA | >trna10_AspGTC_12_-_125424193_125424264 | 35 | 75 | 41 | TCACGCGGGAGACCGGGGTTCGATTCCCCGACGGGGAGCCA | 342.3 | 4.1 | 1.1 | 3.7 | 1.8E-04 | 2.4E-04 | MB231/MB468 | MITO |
| >hg38_21_+_8433172_8446622_RNA45SN1_45S_50ntFlanks@10234.10262.29\|CGGACGCTACGCCGCGACGAGTAGGAGGG | >hg38_21_+_8433172_8446622_RNA45SN1_45S_50ntFlanks | 10234 | 10262 | 29 | CGGACGCTACGCCGCGACGAGTAGGAGGG | 391.4 | -3.3 | 0.8 | -4.0 | 6.9E-05 | 2.4E-04 | BT20/MB468 | MP |
| >hg38_21_+_8433172_8446622_RNA45SN1_45S_50ntFlanks@11564.11594.31\|CCTAGCAGCCGACTTAGAACTGGTGCGGACC | >hg38_21_+_8433172_8446622_RNA45SN1_45S_50ntFlanks | 11564 | 11594 | 31 | CCTAGCAGCCGACTTAGAACTGGTGCGGACC | 59.8 | 9.2 | 2.2 | 4.2 | 2.4E-05 | 2.4E-04 | BT20/MB231 | MP |
| >hg38_21_+_8433172_8446622_RNA45SN1_45S_50ntFlanks@9299.9322.24\|CACGGACCAAGGAGTCTAACACGT | >hg38_21_+_8433172_8446622_RNA45SN1_45S_50ntFlanks | 9299 | 9322 | 24 | CACGGACCAAGGAGTCTAACACGT | 64.1 | 9.3 | 2.2 | 4.2 | 2.5E-05 | 2.4E-04 | BT20/MB231 | MP |
| >trnaMT_SerGCT_MT_+_12207_12265@14.40.27\|AGAACTGCTAACTCATGCCCCCATGTC | >trnaMT_SerGCT_MT_+_12207_12265 | 14 | 40 | 27 | AGAACTGCTAACTCATGCCCCCATGTC | 415.4 | 4.4 | 1.1 | 4.0 | 6.9E-05 | 2.4E-04 | BT20/MB468 | MP |
| >hg38_21_+_8433172_8446622_RNA45SN1_45S_50ntFlanks@5471.5500.30\|CACGGCCCTGGCGGAGCGCTGAGAAGACGG | >hg38_21_+_8433172_8446622_RNA45SN1_45S_50ntFlanks | 5471 | 5500 | 30 | CACGGCCCTGGCGGAGCGCTGAGAAGACGG | 367.0 | 2.8 | 0.7 | 3.8 | 1.7E-04 | 2.4E-04 | MB231/MB468 | CYTO |
| >MI0000085\|hsa-mir-27a&WithFlank&19\|-\|13836434\|13836523@57.76.20(+2U)&[MIMAT0000084&hsa-miR-27a-3p&offsets\|0\|-1(+2U);m-43&19\|-\|13836447\|13836467&offsets\|0\|-1(+2U)]\|TTCACAGTGGCTAAGTTCCGTT | >MI0000085\|hsa-mir-27a&WithFlank&19\|-\|13836434\|13836523 | 57 | 76 | 20(+2U) | TTCACAGTGGCTAAGTTCCGTT | 638.6 | 3.0 | 0.8 | 3.8 | 1.4E-04 | 2.4E-04 | BT20/MB468 | MITO |
| >hg38_21_+_8433172_8446622_RNA45SN1_45S_50ntFlanks@12415.12441.27\|CTTCGATGTCGGCTCTTCCTATCATTG | >hg38_21_+_8433172_8446622_RNA45SN1_45S_50ntFlanks | 12415 | 12441 | 27 | CTTCGATGTCGGCTCTTCCTATCATTG | 2368.6 | -2.3 | 0.6 | -3.8 | 1.4E-04 | 2.4E-04 | BT20/MB468 | MITO |
| >hg38_21_+_8433172_8446622_RNA45SN1_45S_50ntFlanks@4666.4692.27\|AAATTCTTGGACCGGCGCAAGACGGAC | >hg38_21_+_8433172_8446622_RNA45SN1_45S_50ntFlanks | 4666 | 4692 | 27 | AAATTCTTGGACCGGCGCAAGACGGAC | 87.1 | -4.3 | 1.1 | -3.8 | 1.5E-04 | 2.4E-04 | MB231/MB468 | MP |
| >hg38_21_+_8433172_8446622_RNA45SN1_45S_50ntFlanks@5444.5466.23\|CTCGGATCGGCCCCGCCGGGGTC | >hg38_21_+_8433172_8446622_RNA45SN1_45S_50ntFlanks | 5444 | 5466 | 23 | CTCGGATCGGCCCCGCCGGGGTC | 40104.0 | -7.1 | 1.9 | -3.7 | 1.9E-04 | 2.4E-04 | BT20/MB468 | NUC |
| >hg38_MT_+_598_1651_RNR1_12S_50ntFlanks@979.1003.25\|GTACTGGAAAGTGCACTTGGACGAA | >hg38_MT_+_598_1651_RNR1_12S_50ntFlanks | 979 | 1003 | 25 | GTACTGGAAAGTGCACTTGGACGAA | 23.2 | 7.8 | 2.1 | 3.8 | 1.5E-04 | 2.4E-04 | MB231/MB468 | MP |
| >MI0000255\|hsa-mir-30d&WithFlank&8\|-\|134804870\|134804951@12.33.22(+1U)&[MIMAT0000245&hsa-miR-30d-5p&offsets\|0\|0(+1U);m-21&8\|-\|134804919\|134804940&offsets\|0\|0(+1U)]\|TGTAAACATCCCCGACTGGAAGT | >MI0000255\|hsa-mir-30d&WithFlank&8\|-\|134804870\|134804951 | 12 | 33 | 22(+1U) | TGTAAACATCCCCGACTGGAAGT | 1194.6 | 3.1 | 0.8 | 3.8 | 1.5E-04 | 2.4E-04 | MB231/MB468 | MP |
| >MI0003776\|hsa-mir-320b-1&WithFlank&1\|+\|116671740\|116671823@47.67.21(+1U);MI0003839\|hsa-mir-320b-2&WithFlank&1\|-\|224257034\|224257116@46.66.21(+1U);MI0000542\|hsa-mir-320a&WithFlank&8\|-\|22244960\|22245043@47.67.21(+1U)&[MIMAT0005792&hsa-miR-320b&offsets\|-1\|-2(+1U);m-183&1\|+\|116671787\|116671806&offsets\|-1\|0(+1U)];[MIMAT0005792_1&hsa-miR-320b&offsets\|-1\|-2(+1U);m-183&1\|-\|224257051\|224257070&offsets\|-1\|0(+1U)];[MIMAT0000510&hsa-miR-320a-3p&offsets\|-1\|-2(+1U);m-60&8\|-\|22244975\|22244996&offsets\|-1\|-2(+1U)]\|GAAAAGCTGGGTTGAGAGGGCT | >MI0003776\|hsa-mir-320b-1&WithFlank&1\|+\|116671740\|116671823 | 47 | 67 | 21(+1U) | GAAAAGCTGGGTTGAGAGGGCT | 153.4 | 7.4 | 1.8 | 4.0 | 6.9E-05 | 2.4E-04 | BT20/MB468 | MP |
| >hg38_21_+_8433172_8446622_RNA45SN1_45S_50ntFlanks@6652.6688.37\|GACTCTTAGCGGTGGATCACTCGGCTCGTGCGTCGAT | >hg38_21_+_8433172_8446622_RNA45SN1_45S_50ntFlanks | 6652 | 6688 | 37 | GACTCTTAGCGGTGGATCACTCGGCTCGTGCGTCGAT | 388.9 | 2.0 | 0.5 | 3.8 | 1.7E-04 | 2.4E-04 | MB231/MB468 | CYTO |
| >MI0000816\|hsa-mir-335&WithFlank&7\|+\|130496105\|130496210@58.78.21&[MIMAT0004703&hsa-miR-335-3p&offsets\|0\|-1;m-159&7\|+\|130496162\|130496183&offsets\|0\|-1]\|TTTTTCATTATTGCTCCTGAC | >MI0000816\|hsa-mir-335&WithFlank&7\|+\|130496105\|130496210 | 58 | 78 | 21 | TTTTTCATTATTGCTCCTGAC | 468.0 | -3.6 | 0.9 | -4.2 | 2.8E-05 | 2.4E-04 | BT20/MB231 | MITO |
| >hg38_21_+_8433172_8446622_RNA45SN1_45S_50ntFlanks@6651.6668.18\|CGACTCTTAGCGGTGGAT | >hg38_21_+_8433172_8446622_RNA45SN1_45S_50ntFlanks | 6651 | 6668 | 18 | CGACTCTTAGCGGTGGAT | 5556.8 | 3.0 | 0.8 | 4.0 | 6.9E-05 | 2.4E-04 | BT20/MB468 | MP |
| >MI0000457\|hsa-mir-141&WithFlank&12\|+\|6964091\|6964197@65.86.22&[MIMAT0000432&hsa-miR-141-3p&offsets\|0\|0;m-134&12\|+\|6964155\|6964175&offsets\|0\|+1]\|TAACACTGTCTGGTAAAGATGG | >MI0000457\|hsa-mir-141&WithFlank&12\|+\|6964091\|6964197 | 65 | 86 | 22 | TAACACTGTCTGGTAAAGATGG | 47.3 | -9.4 | 2.5 | -3.8 | 1.6E-04 | 2.4E-04 | MB231/MB468 | MP |
| >hg38_21_+_8433172_8446622_RNA45SN1_45S_50ntFlanks@6645.6679.35\|CTCGTACGACTCTTAGCGGTGGATCACTCGGCTCG | >hg38_21_+_8433172_8446622_RNA45SN1_45S_50ntFlanks | 6645 | 6679 | 35 | CTCGTACGACTCTTAGCGGTGGATCACTCGGCTCG | 744.6 | -3.8 | 1.0 | -3.8 | 1.6E-04 | 2.4E-04 | MB231/MB468 | MP |
| >hg38_21_+_8433172_8446622_RNA45SN1_45S_50ntFlanks@3815.3841.27\|ATGGTTCCTTTGGTCGCTCGCTCCTCT | >hg38_21_+_8433172_8446622_RNA45SN1_45S_50ntFlanks | 3815 | 3841 | 27 | ATGGTTCCTTTGGTCGCTCGCTCCTCT | 289.3 | -2.2 | 0.6 | -3.7 | 1.9E-04 | 2.4E-04 | MB231/MB468 | NUC |
| >trnaMT_GluTTC_MT_-_14674_14742@39.66.28,trnalookalike8_GluTTC_5_-_93905172_93905240@39.66.28\|ATTGGTCGTGGTTGTAGTCCGTGCGAGA | >trnaMT_GluTTC_MT_-_14674_14742 | 39 | 66 | 28 | ATTGGTCGTGGTTGTAGTCCGTGCGAGA | 177.6 | 4.6 | 1.2 | 3.7 | 1.8E-04 | 2.4E-04 | MB231/MB468 | MITO |
| >MI0000750\|hsa-mir-26a-2&WithFlank&12\|-\|57824603\|57824698@23.41.19;MI0000083\|hsa-mir-26a-1&WithFlank&3\|+\|37969398\|37969486@19.37.19&[MIMAT0000082&hsa-miR-26a-5p&offsets\|+3\|0;m-33&12\|-\|57824658\|57824679&offsets\|+3\|0];[MIMAT0000082_1&hsa-miR-26a-5p&offsets\|+3\|0;m-34&3\|+\|37969413\|37969434&offsets\|+3\|0]\|AAGTAATCCAGGATAGGCT | >MI0000750\|hsa-mir-26a-2&WithFlank&12\|-\|57824603\|57824698 | 23 | 41 | 19 | AAGTAATCCAGGATAGGCT | 790.7 | 13.6 | 3.6 | 3.8 | 1.7E-04 | 2.4E-04 | MB231/MB468 | CYTO |
| >hg38_21_+_8433172_8446622_RNA45SN1_45S_50ntFlanks@3874.3899.26\|AATACATGCCGACGGGCGCTGACCCC | >hg38_21_+_8433172_8446622_RNA45SN1_45S_50ntFlanks | 3874 | 3899 | 26 | AATACATGCCGACGGGCGCTGACCCC | 698.5 | -2.6 | 0.7 | -3.8 | 1.6E-04 | 2.4E-04 | BT20/MB468 | CYTO |
| >hg38_21_+_8433172_8446622_RNA45SN1_45S_50ntFlanks@5479.5512.34\|TGGCGGAGCGCTGAGAAGACGGTCGAACTTGACT | >hg38_21_+_8433172_8446622_RNA45SN1_45S_50ntFlanks | 5479 | 5512 | 34 | TGGCGGAGCGCTGAGAAGACGGTCGAACTTGACT | 72.5 | 7.0 | 1.8 | 4.0 | 6.9E-05 | 2.4E-04 | BT20/MB468 | MP |
| >MI0000433\|hsa-let-7g&WithFlank&3\|-\|52268272\|52268367@11.32.22(+1G)&[MIMAT0000414&hsa-let-7g-5p&offsets\|0\|0(+1G);m-58&3\|-\|52268336\|52268357&offsets\|0\|0(+1G)]\|TGAGGTAGTAGTTTGTACAGTTG | >MI0000433\|hsa-let-7g&WithFlank&3\|-\|52268272\|52268367 | 11 | 32 | 22(+1G) | TGAGGTAGTAGTTTGTACAGTTG | 165.1 | 6.5 | 1.6 | 4.0 | 6.9E-05 | 2.4E-04 | BT20/MB468 | MP |
| >hg38_21_+_8433172_8446622_RNA45SN1_45S_50ntFlanks@12416.12441.26\|TTCGATGTCGGCTCTTCCTATCATTG | >hg38_21_+_8433172_8446622_RNA45SN1_45S_50ntFlanks | 12416 | 12441 | 26 | TTCGATGTCGGCTCTTCCTATCATTG | 2813.3 | 2.8 | 0.7 | 3.8 | 1.7E-04 | 2.4E-04 | MB231/MB468 | CYTO |
| >hg38_21_+_8433172_8446622_RNA45SN1_45S_50ntFlanks@6772.6803.32\|GTTCCTCCCGGGGCTACGCCTGTCTGAGCGTC | >hg38_21_+_8433172_8446622_RNA45SN1_45S_50ntFlanks | 6772 | 6803 | 32 | GTTCCTCCCGGGGCTACGCCTGTCTGAGCGTC | 1041.3 | -4.5 | 1.2 | -3.8 | 1.4E-04 | 2.4E-04 | BT20/MB468 | MITO |
| >hg38_21_+_8433172_8446622_RNA45SN1_45S_50ntFlanks@11572.11604.33\|CCGACTTAGAACTGGTGCGGACCAGGGGAATCC | >hg38_21_+_8433172_8446622_RNA45SN1_45S_50ntFlanks | 11572 | 11604 | 33 | CCGACTTAGAACTGGTGCGGACCAGGGGAATCC | 436.4 | -1.9 | 0.5 | -3.7 | 1.9E-04 | 2.4E-04 | MB231/MB468 | NUC |
| >hg38_21_+_8433172_8446622_RNA45SN1_45S_50ntFlanks@10822.10844.23\|GAAGTCGGCAAGCCGGATCCGTA | >hg38_21_+_8433172_8446622_RNA45SN1_45S_50ntFlanks | 10822 | 10844 | 23 | GAAGTCGGCAAGCCGGATCCGTA | 168.5 | -4.6 | 1.2 | -3.8 | 1.6E-04 | 2.4E-04 | MB231/MB468 | MP |
| >MI0000063\|hsa-let-7b&WithFlank&22\|+\|46113680\|46113774@12.30.19&[MIMAT0000063&hsa-let-7b-5p&offsets\|0\|-3;m-14&22\|+\|46113691\|46113712&offsets\|0\|-3]\|TGAGGTAGTAGGTTGTGTG | >MI0000063\|hsa-let-7b&WithFlank&22\|+\|46113680\|46113774 | 12 | 30 | 19 | TGAGGTAGTAGGTTGTGTG | 322.3 | 5.4 | 1.4 | 4.0 | 6.9E-05 | 2.4E-04 | BT20/MB468 | MP |
| >hg38_21_+_8433172_8446622_RNA45SN1_45S_50ntFlanks@8380.8415.36\|CAAGAGGGCGTGAAACCGTTAAGAGGTAAACGGGTG | >hg38_21_+_8433172_8446622_RNA45SN1_45S_50ntFlanks | 8380 | 8415 | 36 | CAAGAGGGCGTGAAACCGTTAAGAGGTAAACGGGTG | 213.2 | -6.2 | 1.5 | -4.0 | 6.9E-05 | 2.4E-04 | BT20/MB468 | MP |
| >hg38_21_+_8433172_8446622_RNA45SN1_45S_50ntFlanks@4130.4161.32\|ATTCCGGAGAGGGAGCCTGAGAAACGGCTACC | >hg38_21_+_8433172_8446622_RNA45SN1_45S_50ntFlanks | 4130 | 4161 | 32 | ATTCCGGAGAGGGAGCCTGAGAAACGGCTACC | 80.4 | -4.4 | 1.2 | -3.8 | 1.6E-04 | 2.4E-04 | MB231/MB468 | MP |
| >hg38_21_+_8433172_8446622_RNA45SN1_45S_50ntFlanks@9300.9332.33\|ACGGACCAAGGAGTCTAACACGTGCGCGAGTCG | >hg38_21_+_8433172_8446622_RNA45SN1_45S_50ntFlanks | 9300 | 9332 | 33 | ACGGACCAAGGAGTCTAACACGTGCGCGAGTCG | 66.6 | 9.4 | 2.2 | 4.2 | 2.5E-05 | 2.4E-04 | BT20/MB231 | MP |
| >MI0000063\|hsa-let-7b&WithFlank&22\|+\|46113680\|46113774@12.32.21(+1A)&[MIMAT0000063&hsa-let-7b-5p&offsets\|0\|-1(+1A);m-14&22\|+\|46113691\|46113712&offsets\|0\|-1(+1A)]\|TGAGGTAGTAGGTTGTGTGGTA | >MI0000063\|hsa-let-7b&WithFlank&22\|+\|46113680\|46113774 | 12 | 32 | 21(+1A) | TGAGGTAGTAGGTTGTGTGGTA | 4850.4 | 3.7 | 0.9 | 4.2 | 2.3E-05 | 2.4E-04 | BT20/MB231 | CYTO |

| | | | | | | | | | | | | | |
|---|---|---|---|---|---|---|---|---|---|---|---|---|---|
| >hg38_21_+_8433172_8446622_RNA45SN1_45S_50ntFlanks@8172.8214.43\|AGTCCTTCTGATCGAGGCCCAGCCCGTGGACGGTGTGAGGCCG | >hg38_21_+_8433172_8446622_RNA45SN1_45S_50ntFlanks | 8172 | 8214 | 43 | AGTCCTTCTGATCGAGGCCCAGCCCGTGGACGGTGTGAGGCCG | 232.5 | -3.6 | 1.0 | -3.7 | 1.9E-04 | 2.4E-04 | MB231/MB468 | MITO |
| >hg38_21_+_8433172_8446622_RNA45SN1_45S_50ntFlanks@9533.9569.37\|GAGGAAACTCTGGTGGAGGTCCGTAGCGGTCCTGACG | >hg38_21_+_8433172_8446622_RNA45SN1_45S_50ntFlanks | 9533 | 9569 | 37 | GAGGAAACTCTGGTGGAGGTCCGTAGCGGTCCTGACG | 1746.7 | -7.3 | 2.0 | -3.7 | 1.9E-04 | 2.4E-04 | BT20/MB468 | NUC |
| >MI0000434\|hsa-let-7i&WithFlank&12\|+\|62603680\|62603775@12.33.22&[MIMAT0000415&hsa-let-7i-5p&offsets\|0\|0;m-49&12\|+\|62603691\|62603712&offsets\|0\|0]\|TGAGGTAGTAGTTTGTGCTGTT | >MI0000434\|hsa-let-7i&WithFlank&12\|+\|62603680\|62603775 | 12 | 33 | 22 | TGAGGTAGTAGTTTGTGCTGTT | 70831.5 | 3.8 | 1.0 | 4.0 | 7.0E-05 | 2.4E-04 | BT20/MB468 | MP |
| >hg38_21_+_8433172_8446622_RNA45SN1_45S_50ntFlanks@6648.6680.33\|GTACGACTCTTAGCGGTGGATCACTCGGCTCGT | >hg38_21_+_8433172_8446622_RNA45SN1_45S_50ntFlanks | 6648 | 6680 | 33 | GTACGACTCTTAGCGGTGGATCACTCGGCTCGT | 525.9 | -3.1 | 0.8 | -3.8 | 1.6E-04 | 2.4E-04 | MB231/MB468 | MP |
| >hg38_21_+_8433172_8446622_RNA45SN1_45S_50ntFlanks@10441.10460.20\|TTCCGAAGGGACGGGCGATG | >hg38_21_+_8433172_8446622_RNA45SN1_45S_50ntFlanks | 10441 | 10460 | 20 | TTCCGAAGGGACGGGCGATG | 1152.9 | -7.1 | 1.9 | -3.7 | 1.9E-04 | 2.4E-04 | BT20/MB468 | NUC |
| >hg38_21_+_8433172_8446622_RNA45SN1_45S_50ntFlanks@11578.11611.34\|TAGAACTGGTGCGGACCAGGGGAATCCGACTGTT | >hg38_21_+_8433172_8446622_RNA45SN1_45S_50ntFlanks | 11578 | 11611 | 34 | TAGAACTGGTGCGGACCAGGGGAATCCGACTGTT | 391.9 | 2.8 | 0.7 | 4.2 | 2.9E-05 | 2.4E-04 | BT20/MB231 | MITO |
| >hg38_21_+_8433172_8446622_RNA45SN1_45S_50ntFlanks@9940.9971.32\|CAGGACGGTGGCCATGGAAGTCGGAATCCGCT | >hg38_21_+_8433172_8446622_RNA45SN1_45S_50ntFlanks | 9940 | 9971 | 32 | CAGGACGGTGGCCATGGAAGTCGGAATCCGCT | 140.6 | 7.1 | 1.8 | 4.0 | 7.0E-05 | 2.4E-04 | BT20/MB468 | MP |
| >hg38_21_+_8433172_8446622_RNA45SN1_45S_50ntFlanks@12412.12443.32\|ATCCTTCGATGTCGGCTCTTCCTATCATTGTG | >hg38_21_+_8433172_8446622_RNA45SN1_45S_50ntFlanks | 12412 | 12443 | 32 | ATCCTTCGATGTCGGCTCTTCCTATCATTGTG | 1154.7 | -6.0 | 1.6 | -3.7 | 1.9E-04 | 2.4E-04 | MB231/MB468 | MITO |
| >hg38_21_+_8433172_8446622_RNA45SN1_45S_50ntFlanks@12247.12272.26\|GGCAAAAGCTCGCTTGATCTTGATTT | >hg38_21_+_8433172_8446622_RNA45SN1_45S_50ntFlanks | 12247 | 12272 | 26 | GGCAAAAGCTCGCTTGATCTTGATTT | 81.9 | -5.7 | 1.5 | -3.8 | 1.6E-04 | 2.4E-04 | MB231/MB468 | MP |
| >hg38_MT_+_598_1651_RNR1_12S_50ntFlanks@570.611.42\|AAGGACCTGGCGGTGCTTCATATCCCTCTAGAGGAGCCTGTT | >hg38_MT_+_598_1651_RNR1_12S_50ntFlanks | 570 | 611 | 42 | AAGGACCTGGCGGTGCTTCATATCCCTCTAGAGGAGCCTGTT | 219.1 | 5.7 | 1.5 | 3.7 | 1.9E-04 | 2.4E-04 | MB231/MB468 | MITO |
| >hg38_21_+_8433172_8446622_RNA45SN1_45S_50ntFlanks@4000.4023.24\|TAGATAACCTCGGGCCGATCGCAC | >hg38_21_+_8433172_8446622_RNA45SN1_45S_50ntFlanks | 4000 | 4023 | 24 | TAGATAACCTCGGGCCGATCGCAC | 741.2 | -3.9 | 1.0 | -3.7 | 1.9E-04 | 2.4E-04 | BT20/MB468 | NUC |
| >hg38_21_+_8433172_8446622_RNA45SN1_45S_50ntFlanks@6777.6806.30\|TCCCGGGGCTACGCCTGTCTGAGCGTCGCT | >hg38_21_+_8433172_8446622_RNA45SN1_45S_50ntFlanks | 6777 | 6806 | 30 | TCCCGGGGCTACGCCTGTCTGAGCGTCGCT | 144.9 | 1.2 | 0.3 | 3.7 | 1.9E-04 | 2.4E-04 | BT20/MB468 | NUC |
| >hg38_21_+_8433172_8446622_RNA45SN1_45S_50ntFlanks@6649.6676.28\|TACGACTCTTAGCGGTGGATCACTCGGC | >hg38_21_+_8433172_8446622_RNA45SN1_45S_50ntFlanks | 6649 | 6676 | 28 | TACGACTCTTAGCGGTGGATCACTCGGC | 255.2 | 4.1 | 1.0 | 4.0 | 7.0E-05 | 2.4E-04 | BT20/MB468 | MP |
| >MI0000063\|hsa-let-7b&WithFlank&22\|+\|46113680\|46113774@12.31.20(+1C)&[MIMAT0000063&hsa-let-7b-5p&offsets\|0\|-2(+1C);m-14&22\|+\|46113691\|46113712&offsets\|0\|-2(+1C)]\|TGAGGTAGTAGGTTGTGTGGC | >MI0000063\|hsa-let-7b&WithFlank&22\|+\|46113680\|46113774 | 12 | 31 | 20(+1C) | TGAGGTAGTAGGTTGTGTGGC | 602.0 | 3.3 | 0.9 | 3.8 | 1.6E-04 | 2.4E-04 | MB231/MB468 | MP |
| >hg38_21_+_8433172_8446622_RNA45SN1_45S_50ntFlanks@11674.11692.19\|GTGCTCTGAATGTCAAAGT | >hg38_21_+_8433172_8446622_RNA45SN1_45S_50ntFlanks | 11674 | 11692 | 19 | GTGCTCTGAATGTCAAAGT | 322.7 | 3.2 | 0.8 | 4.2 | 2.9E-05 | 2.4E-04 | BT20/MB231 | MITO |
| >hg38_21_+_8433172_8446622_RNA45SN1_45S_50ntFlanks@9736.9767.32\|CCGAAACGATCTCAACCTATTCTCAAACTTTA | >hg38_21_+_8433172_8446622_RNA45SN1_45S_50ntFlanks | 9736 | 9767 | 32 | CCGAAACGATCTCAACCTATTCTCAAACTTTA | 579.7 | -2.6 | 0.7 | -3.8 | 1.4E-04 | 2.4E-04 | BT20/MB468 | MITO |
| >MI0003667\|hsa-mir-652&WithFlank&X\|+\|110055323\|110055432@67.87.21(+1U)&[MIMAT0003322&hsa-miR-652-3p&offsets\|0\|0(+1U);m-148&X\|+\|110055389\|110055410&offsets\|0\|-1(+1U)]\|AATGGCGCCACTAGGGTTGTGT | >MI0003667\|hsa-mir-652&WithFlank&X\|+\|110055323\|110055432 | 67 | 87 | 21(+1U) | AATGGCGCCACTAGGGTTGTGT | 1822.8 | 5.0 | 1.3 | 4.0 | 7.0E-05 | 2.4E-04 | BT20/MB468 | MP |
| >MI0000077\|hsa-mir-21&WithFlank&17\|+\|59841260\|59841343@15.36.22(+1A)&[MIMAT0000076&hsa-miR-21-5p&offsets\|+1\|+1(+1A);m-1&17\|+\|59841273\|59841294&offsets\|+1\|+1(+1A)]\|AGCTTATCAGACTGATGTTGACA | >MI0000077\|hsa-mir-21&WithFlank&17\|+\|59841260\|59841343 | 15 | 36 | 22(+1A) | AGCTTATCAGACTGATGTTGACA | 323.3 | 3.3 | 0.8 | 4.2 | 2.4E-05 | 2.4E-04 | BT20/MB231 | CYTO |
| >hg38_21_+_8433172_8446622_RNA45SN1_45S_50ntFlanks@3835.3869.35\|CTCCTCTCCTACTTGGATAACTGTGGTAATTCTAG | >hg38_21_+_8433172_8446622_RNA45SN1_45S_50ntFlanks | 3835 | 3869 | 35 | CTCCTCTCCTACTTGGATAACTGTGGTAATTCTAG | 204.8 | -2.7 | 0.7 | -3.8 | 1.7E-04 | 2.4E-04 | BT20/MB468 | CYTO |
| >hg38_21_+_8433172_8446622_RNA45SN1_45S_50ntFlanks@11233.11251.19\|CCCTTCCCGTGGATCGCCC | >hg38_21_+_8433172_8446622_RNA45SN1_45S_50ntFlanks | 11233 | 11251 | 19 | CCCTTCCCGTGGATCGCCC | 204.2 | -5.6 | 1.5 | -3.8 | 1.7E-04 | 2.4E-04 | BT20/MB468 | CYTO |
| >MI0000542\|hsa-mir-320a&WithFlank&8\|-\|22244960\|22245043@48.68.21(+2U)&[MIMAT0000510&hsa-miR-320a-3p&offsets\|0\|-1(+2U);m-60&8\|-\|22244975\|22244996&offsets\|0\|-1(+2U)]\|AAAAGCTGGGTTGAGAGGGCGTT | >MI0000542\|hsa-mir-320a&WithFlank&8\|-\|22244960\|22245043 | 48 | 68 | 21(+2U) | AAAAGCTGGGTTGAGAGGGCGTT | 194.8 | 6.6 | 1.8 | 3.7 | 1.9E-04 | 2.4E-04 | MB231/MB468 | MITO |
| >hg38_21_+_8433172_8446622_RNA45SN1_45S_50ntFlanks@6721.6751.31\|AATTGCAGGACACATTGATCATCGACACTTC | >hg38_21_+_8433172_8446622_RNA45SN1_45S_50ntFlanks | 6721 | 6751 | 31 | AATTGCAGGACACATTGATCATCGACACTTC | 244.9 | -4.2 | 1.1 | -3.7 | 1.9E-04 | 2.4E-04 | MB231/MB468 | MITO |
| >hg38_21_+_8433172_8446622_RNA45SN1_45S_50ntFlanks@7976.8006.31\|GCGACCTCAGATCAGACGTGGCGACCCGCTG | >hg38_21_+_8433172_8446622_RNA45SN1_45S_50ntFlanks | 7976 | 8006 | 31 | GCGACCTCAGATCAGACGTGGCGACCCGCTG | 653.7 | -4.5 | 1.2 | -3.7 | 1.9E-04 | 2.4E-04 | MB231/MB468 | MITO |
| >MI0000440\|hsa-mir-27b&WithFlank&9\|+\|95085439\|95085547@67.86.20(+1U)&[MIMAT0000419&hsa-miR-27b-3p&offsets\|0\|-1(+1U);m-39&9\|+\|95085505\|95085525&offsets\|0\|-1(+1U)]\|TTCACAGTGGCTAAGTTCTGT | >MI0000440\|hsa-mir-27b&WithFlank&9\|+\|95085439\|95085547 | 67 | 86 | 20(+1U) | TTCACAGTGGCTAAGTTCTGT | 4403.8 | 4.1 | 1.0 | 4.0 | 7.0E-05 | 2.4E-04 | BT20/MB468 | MP |
| >hg38_21_+_8433172_8446622_RNA45SN1_45S_50ntFlanks@12286.12318.33\|AGACCGTGAAAGCGGGGCCTCACGATCCTTCTG | >hg38_21_+_8433172_8446622_RNA45SN1_45S_50ntFlanks | 12286 | 12318 | 33 | AGACCGTGAAAGCGGGGCCTCACGATCCTTCTG | 191.8 | -5.6 | 1.5 | -3.8 | 1.4E-04 | 2.4E-04 | BT20/MB468 | MITO |
| >am_trna111_HisGTG_1_-_147774845_147774916@1.26.26,trna118_HisGTG_1_-_145396881_145396952@1.26.26,trna16_HisGTG_1_+_146544773_146544844@1.26.26,trna1_HisGTG_15_+_45493349_45493420@1.26.26,trna21_HisGTG_1_+_147753471_147753542@1.26.26,trna33_HisGTG_6_+_27125906_27125977@1.26.26,trna7_HisGTG_9_-_14433938_14434009@1.26.26,trna8_HisGTG_15_-_45492611_45492682@1.26.26,trna9_HisGTG_15_-_45490804_45490875@1.26.26\|GCCGTGATCGTATAGTGGTTAGTACT | >am_trna111_HisGTG_1_-_147774845_147774916 | 1 | 26 | 26 | GCCGTGATCGTATAGTGGTTAGTACT | 314.5 | -2.7 | 0.7 | -3.7 | 1.9E-04 | 2.4E-04 | BT20/MB468 | NUC |
| >hg38_21_+_8433172_8446622_RNA45SN1_45S_50ntFlanks@11873.11903.31\|AAGAAGACCCTGTTGAGCTTGACTCTAGTCT | >hg38_21_+_8433172_8446622_RNA45SN1_45S_50ntFlanks | 11873 | 11903 | 31 | AAGAAGACCCTGTTGAGCTTGACTCTAGTCT | 475.9 | -3.4 | 0.9 | -4.0 | 7.0E-05 | 2.4E-04 | BT20/MB468 | MP |
| >hg38_21_+_8433172_8446622_RNA45SN1_45S_50ntFlanks@3853.3872.20\|AACTGTGGTAATTCTAGAGC | >hg38_21_+_8433172_8446622_RNA45SN1_45S_50ntFlanks | 3853 | 3872 | 20 | AACTGTGGTAATTCTAGAGC | 340.5 | -6.1 | 1.5 | -4.0 | 7.0E-05 | 2.4E-04 | BT20/MB468 | MP |
| >hg38_21_+_8433172_8446622_RNA45SN1_45S_50ntFlanks@9621.9648.28\|TCTAGTAGCTGGTTCCCTCCGAAGTTTC | >hg38_21_+_8433172_8446622_RNA45SN1_45S_50ntFlanks | 9621 | 9648 | 28 | TCTAGTAGCTGGTTCCCTCCGAAGTTTC | 239.2 | -6.2 | 1.6 | -4.0 | 7.0E-05 | 2.4E-04 | BT20/MB468 | MP |
| >hg38_MT_+_1621_3279_RNR2_16S_50ntFlanks@488.517.30\|GAACAGCTCTTTGGACACTAGGAAAAAACC | >hg38_MT_+_1621_3279_RNR2_16S_50ntFlanks | 488 | 517 | 30 | GAACAGCTCTTTGGACACTAGGAAAAAACC | 137.5 | 4.5 | 1.2 | 3.7 | 1.9E-04 | 2.4E-04 | MB231/MB468 | MITO |
| >MI0000264\|hsa-mir-7-2&WithFlank&15\|+\|88611819\|88611940@38.60.23(+1G);MI0000265\|hsa-mir-7-3&WithFlank&19\|+\|4770664\|4770785@37.59.23(+1G);MI0000263\|hsa-mir-7-1&WithFlank&9\|-\|83969742\|83969863@30.52.23(+1G)&[MIMAT0000252&hsa-miR-7-5p&offsets\|0\|-1(+1G);m-400&15\|+\|88611856\|88611879&offsets\|0\|-1(+1G)];[MIMAT0000252_1&hsa-miR-7-5p&offsets\|0\|-1(+1G);m-402&19\|+\|4770700\|4770723&offsets\|0\|-1(+1G)];[MIMAT0000252_2&hsa-miR-7-5p&offsets\|0\|-1(+1G);m-401&9\|-\|83969811\|83969834&offsets\|0\|-1(+1G)]\|TGGAAGACTAGTGATTTTGTTGTG | >MI0000264\|hsa-mir-7-2&WithFlank&15\|+\|88611819\|88611940 | 38 | 60 | 23(+1G) | TGGAAGACTAGTGATTTTGTTGTG | 2156.7 | 4.1 | 1.0 | 4.2 | 2.4E-05 | 2.4E-04 | BT20/MB231 | CYTO |
| >hg38_21_+_8433172_8446622_RNA45SN1_45S_50ntFlanks@12290.12323.34\|CGTGAAAGCGGGGCCTCACGATCCTTCTGACCTT | >hg38_21_+_8433172_8446622_RNA45SN1_45S_50ntFlanks | 12290 | 12323 | 34 | CGTGAAAGCGGGGCCTCACGATCCTTCTGACCTT | 295.9 | -4.5 | 1.2 | -3.8 | 1.4E-04 | 2.4E-04 | BT20/MB468 | MITO |
| >hg38_21_+_8433172_8446622_RNA45SN1_45S_50ntFlanks@10797.10817.21\|GCATGTTGGAACAATGTAGGT | >hg38_21_+_8433172_8446622_RNA45SN1_45S_50ntFlanks | 10797 | 10817 | 21 | GCATGTTGGAACAATGTAGGT | 290.0 | -5.0 | 1.3 | -3.7 | 1.9E-04 | 2.4E-04 | BT20/MB468 | NUC |
| >hg38_21_+_8433172_8446622_RNA45SN1_45S_50ntFlanks@7970.8010.41\|CGAGACGCGACCTCAGATCAGACGTGGCGACCCGCTGAATT | >hg38_21_+_8433172_8446622_RNA45SN1_45S_50ntFlanks | 7970 | 8010 | 41 | CGAGACGCGACCTCAGATCAGACGTGGCGACCCGCTGAATT | 328.0 | 3.5 | 0.9 | 3.7 | 1.9E-04 | 2.4E-04 | BT20/MB468 | NUC |
| >hg38_21_+_8433172_8446622_RNA45SN1_45S_50ntFlanks@6671.6694.24\|CTCGGCTCGTGCGTCGATGAAGAA | >hg38_21_+_8433172_8446622_RNA45SN1_45S_50ntFlanks | 6671 | 6694 | 24 | CTCGGCTCGTGCGTCGATGAAGAA | 110.5 | -5.6 | 1.5 | -3.8 | 1.6E-04 | 2.4E-04 | MB231/MB468 | MP |
| >hg38_21_+_8433172_8446622_RNA45SN1_45S_50ntFlanks@10841.10872.32\|CGTAACTTCGGGATAAGGATTGGCTCTAAGGG | >hg38_21_+_8433172_8446622_RNA45SN1_45S_50ntFlanks | 10841 | 10872 | 32 | CGTAACTTCGGGATAAGGATTGGCTCTAAGGG | 1142.2 | -2.7 | 0.7 | -3.7 | 1.8E-04 | 2.4E-04 | MB231/MB468 | CYTO |
| >MI0000115\|hsa-mir-16-2&WithFlank&3\|+\|160404739\|160404831@58.79.22&[MIMAT0004518&hsa-miR-16-2-3p&offsets\|-1\|-1;m-252&3\|+\|160404796\|160404816&offsets\|0\|+1]\|ACCAATATTACTGTGCTGCTTT | >MI0000115\|hsa-mir-16-2&WithFlank&3\|+\|160404739\|160404831 | 58 | 79 | 22 | ACCAATATTACTGTGCTGCTTT | 994.3 | 3.5 | 0.9 | 3.8 | 1.6E-04 | 2.5E-04 | MB231/MB468 | MP |
| >hg38_21_+_8433172_8446622_RNA45SN1_45S_50ntFlanks@5083.5108.26\|ACGAACGAGACTCTGGCATGCTAACT | >hg38_21_+_8433172_8446622_RNA45SN1_45S_50ntFlanks | 5083 | 5108 | 26 | ACGAACGAGACTCTGGCATGCTAACT | 190.8 | -4.1 | 1.1 | -3.8 | 1.6E-04 | 2.5E-04 | MB231/MB468 | MP |
| >hg38_21_+_8433172_8446622_RNA45SN1_45S_50ntFlanks@11209.11250.42\|CGGCGACTCTGGACGCGAGCCGGGCCCTTCCCGTGGATCGCC | >hg38_21_+_8433172_8446622_RNA45SN1_45S_50ntFlanks | 11209 | 11250 | 42 | CGGCGACTCTGGACGCGAGCCGGGCCCTTCCCGTGGATCGCC | 483.9 | -5.9 | 1.6 | -3.7 | 1.9E-04 | 2.5E-04 | MB231/MB468 | NUC |
| >hg38_21_+_8433172_8446622_RNA45SN1_45S_50ntFlanks@11783.11802.20\|CATGAATGGATGAACGAGAT | >hg38_21_+_8433172_8446622_RNA45SN1_45S_50ntFlanks | 11783 | 11802 | 20 | CATGAATGGATGAACGAGAT | 228.6 | -2.8 | 0.7 | -3.7 | 1.8E-04 | 2.5E-04 | MB231/MB468 | CYTO |
| >hg38_21_+_8433172_8446622_RNA45SN1_45S_50ntFlanks@10349.10372.24\|AACGAGAACTTTGAAGGCCGAAGT | >hg38_21_+_8433172_8446622_RNA45SN1_45S_50ntFlanks | 10349 | 10372 | 24 | AACGAGAACTTTGAAGGCCGAAGT | 512.7 | 1.9 | 0.5 | 3.8 | 1.7E-04 | 2.5E-04 | BT20/MB468 | CYTO |

| | | | | | | | | | | | | | |
|---|---|---|---|---|---|---|---|---|---|---|---|---|---|
| >am_trna10_ValCAC_5_-_180649395_180649467@1.19.19,trna12_ValAAC_5_-_180645270_180645342@1.19.19,trna132_ValAAC_6_-_27721179_27721251@1.19.19,trna133_ValCAC_6_-_27696327_27696399@1.19.19,trna136_ValAAC_6_-_27648885_27648957@1.19.19,trna139_ValAAC_6_-_27618707_27618779@1.19.19,trna15_ValAAC_5_-_180615416_180615488@1.19.19,trna18_ValCAC_5_-_180529253_180529325@1.19.19,trna2_ValAAC_3_+_169490018_169490090@1.19.19,trna2_ValCAC_5_+_180524070_180524142@1.19.19,trna37_ValAAC_6_+_27203288_27203360@1.19.19,trna4_ValAAC_5_+_180591154_180591226@1.19.19,trna5_ValAAC_5_+_180596610_180596682@1.19.19,trna6_ValCAC_5_+_180600650_180600722@1.19.19,trna85_ValCAC_1_-_161369490_161369562@1.19.19,trna90_ValCAC_1_-_149684088_149684161@1.19.19,trna98_ValCAC_1_-_149298555_149298627@1.19.19,trna9_ValCAC_6_+_26538282_26538354@1.19.19\|GTTTCCGTAGTGTAGTGGT | >am_trna10_ValCAC_5_-_180649395_180649467 | 1 | 19 | 19 | GTTTCCGTAGTGTAGTGGT | 227.5 | 4.0 | 1.1 | 3.7 | 1.9E-04 | 2.5E-04 | MB231/MB468 | MITO |
| >MI0001445\|hsa-mir-423&WithFlank&17\|+\|30117073\|30117178@59.77.19(+1A)&[MIMAT0001340&hsa-miR-423-3p&offsets\|0\|-4(+1A);m-100&17\|+\|30117131\|30117153&offsets\|0\|-4(+1A)]\|AGCTCGGTCTGAGGCCCCTA | >MI0001445\|hsa-mir-423&WithFlank&17\|+\|30117073\|30117178 | 59 | 77 | 19(+1A) | AGCTCGGTCTGAGGCCCCTA | 123.4 | 6.3 | 1.6 | 4.0 | 7.1E-05 | 2.5E-04 | BT20/MB468 | MP |
| >hg38_21_+_8433172_8446622_RNA45SN1_45S_50ntFlanks@11892.11922.31\|TGACTCTAGTCTGGCACGGTGAAGAGACATG | >hg38_21_+_8433172_8446622_RNA45SN1_45S_50ntFlanks | 11892 | 11922 | 31 | TGACTCTAGTCTGGCACGGTGAAGAGACATG | 128.4 | 7.2 | 1.8 | 4.0 | 7.1E-05 | 2.5E-04 | BT20/MB468 | MP |
| >trna15_ValAAC_5_-_180615416_180615488@1.33.33\|GTTTCCGTAGTGTAGTGGTCATCACGTTCGCCT | >trna15_ValAAC_5_-_180615416_180615488 | 1 | 33 | 33 | GTTTCCGTAGTGTAGTGGTCATCACGTTCGCCT | 821.2 | 10.1 | 2.7 | 3.7 | 1.8E-04 | 2.5E-04 | MB231/MB468 | CYTO |
| >MI0000447\|hsa-mir-128-1&WithFlank&2\|+\|135665391\|135665484@56.78.23&[MIMAT0000424&hsa-miR-128-3p&offsets\|0\|+2;m-108&2\|+\|135665446\|135665466&offsets\|0\|+2]\|TCACAGTGAACCGGTCTCTTTTT | >MI0000447\|hsa-mir-128-1&WithFlank&2\|+\|135665391\|135665484 | 56 | 78 | 23 | TCACAGTGAACCGGTCTCTTTTT | 329.5 | 3.0 | 0.8 | 3.7 | 1.9E-04 | 2.5E-04 | MB231/MB468 | NUC |
| >MI0005566\|hsa-mir-374b&WithFlank&X\|-\|74218541\|74218624@17.37.21(+1A)&[MIMAT0004955&hsa-miR-374b-5p&offsets\|0\|-1(+1A);m-153&X\|-\|74218587\|74218608&offsets\|0\|-1(+1A)]\|ATATAATACAACCTGCTAAGTA | >MI0005566\|hsa-mir-374b&WithFlank&X\|-\|74218541\|74218624 | 17 | 37 | 21(+1A) | ATATAATACAACCTGCTAAGTA | 342.2 | 6.0 | 1.6 | 3.7 | 1.9E-04 | 2.5E-04 | MB231/MB468 | MITO |
| >hg38_21_+_8433172_8446622_RNA45SN1_45S_50ntFlanks@12638.12657.20\|CTACCATCTGTGGGATTATG | >hg38_21_+_8433172_8446622_RNA45SN1_45S_50ntFlanks | 12638 | 12657 | 20 | CTACCATCTGTGGGATTATG | 670.1 | -4.0 | 1.1 | -3.8 | 1.4E-04 | 2.5E-04 | BT20/MB468 | MITO |
| >MI0000456\|hsa-mir-140&WithFlank&16\|+\|69933075\|69933186@69.91.23&[MIMAT0004597&hsa-miR-140-3p&offsets\|+1\|+3;m-45&16\|+\|69933143\|69933164&offsets\|0\|+1]\|ACCACAGGGTAGAACCACGGACA | >MI0000456\|hsa-mir-140&WithFlank&16\|+\|69933075\|69933186 | 69 | 91 | 23 | ACCACAGGGTAGAACCACGGACA | 473.7 | 4.6 | 1.2 | 3.8 | 1.6E-04 | 2.5E-04 | MB231/MB468 | MP |
| >hg38_21_+_8433172_8446622_RNA45SN1_45S_50ntFlanks@9852.9899.48\|GCGCTGCGGGATGAACCGAACGCCGGGTTAAGGCGCCCGATGCCGACG | >hg38_21_+_8433172_8446622_RNA45SN1_45S_50ntFlanks | 9852 | 9899 | 48 | GCGCTGCGGGATGAACCGAACGCCGGGTTAAGGCGCCCGATGCCGACG | 429.8 | -12.5 | 3.4 | -3.7 | 1.9E-04 | 2.5E-04 | MB231/MB468 | MITO |
| >hg38_21_+_8433172_8446622_RNA45SN1_45S_50ntFlanks@11571.11592.22\|GCCGACTTAGAACTGGTGCGGA | >hg38_21_+_8433172_8446622_RNA45SN1_45S_50ntFlanks | 11571 | 11592 | 22 | GCCGACTTAGAACTGGTGCGGA | 222.2 | -3.0 | 0.8 | -3.8 | 1.7E-04 | 2.5E-04 | BT20/MB468 | CYTO |
| >MI0000101\|hsa-mir-99a&WithFlank&21\|+\|16539083\|16539175@19.38.20(+1A)&[MIMAT0000097&hsa-miR-99a-5p&offsets\|0\|-2(+1A);m-64&21\|+\|16539101\|16539121&offsets\|0\|-1(+1A)]\|AACCCGTAGATCCGATCTTGA | >MI0000101\|hsa-mir-99a&WithFlank&21\|+\|16539083\|16539175 | 19 | 38 | 20(+1A) | AACCCGTAGATCCGATCTTGA | 56.5 | -9.2 | 2.2 | -4.2 | 2.5E-05 | 2.5E-04 | BT20/MB231 | MP |
| >hg38_21_+_8433172_8446622_RNA45SN1_45S_50ntFlanks@6731.6770.40\|CACATTGATCATCGACACTTCGAACGCACTTGCGGCCCCG | >hg38_21_+_8433172_8446622_RNA45SN1_45S_50ntFlanks | 6731 | 6770 | 40 | CACATTGATCATCGACACTTCGAACGCACTTGCGGCCCCG | 812.8 | -5.8 | 1.5 | -3.8 | 1.7E-04 | 2.5E-04 | BT20/MB468 | CYTO |
| >hg38_21_+_8433172_8446622_RNA45SN1_45S_50ntFlanks@9835.9854.20\|TTTGGTAAGCAGAACTGGCG | >hg38_21_+_8433172_8446622_RNA45SN1_45S_50ntFlanks | 9835 | 9854 | 20 | TTTGGTAAGCAGAACTGGCG | 221.3 | 11.1 | 2.6 | 4.2 | 2.6E-05 | 2.5E-04 | BT20/MB231 | MP |
| >hg38_21_+_8433172_8446622_RNA45SN1_45S_50ntFlanks@11235.11259.25\|CTTCCCGTGGATCGCCCCAGCTGCG | >hg38_21_+_8433172_8446622_RNA45SN1_45S_50ntFlanks | 11235 | 11259 | 25 | CTTCCCGTGGATCGCCCCAGCTGCG | 190.3 | -2.4 | 0.7 | -3.7 | 1.9E-04 | 2.5E-04 | MB231/MB468 | NUC |
| >hg38_21_+_8433172_8446622_RNA45SN1_45S_50ntFlanks@11574.11603.30\|GACTTAGAACTGGTGCGGACCAGGGGAATC | >hg38_21_+_8433172_8446622_RNA45SN1_45S_50ntFlanks | 11574 | 11603 | 30 | GACTTAGAACTGGTGCGGACCAGGGGAATC | 263.0 | -5.5 | 1.5 | -3.7 | 1.9E-04 | 2.5E-04 | MB231/MB468 | NUC |
| >MI0000542\|hsa-mir-320a&WithFlank&8\|-\|22244960\|22245043@49.68.20&[MIMAT0000510&hsa-miR-320a-3p&offsets\|+1\|-1;m-60&8\|-\|22244975\|22244996&offsets\|+1\|-1]\|AAAGCTGGGTTGAGAGGGCG | >MI0000542\|hsa-mir-320a&WithFlank&8\|-\|22244960\|22245043 | 49 | 68 | 20 | AAAGCTGGGTTGAGAGGGCG | 752.3 | 3.5 | 0.8 | 4.2 | 2.4E-05 | 2.5E-04 | BT20/MB231 | CYTO |
| >hg38_21_+_8433172_8446622_RNA45SN1_45S_50ntFlanks@5485.5506.22\|AGCGCTGAGAAGACGGTCGAAC | >hg38_21_+_8433172_8446622_RNA45SN1_45S_50ntFlanks | 5485 | 5506 | 22 | AGCGCTGAGAAGACGGTCGAAC | 454.8 | 4.3 | 1.0 | 4.2 | 2.4E-05 | 2.5E-04 | BT20/MB231 | CYTO |
| >hg38_21_+_8433172_8446622_RNA45SN1_45S_50ntFlanks@6719.6752.34\|TGAATTGCAGGACACATTGATCATCGACACTTCG | >hg38_21_+_8433172_8446622_RNA45SN1_45S_50ntFlanks | 6719 | 6752 | 34 | TGAATTGCAGGACACATTGATCATCGACACTTCG | 226.0 | -3.7 | 1.0 | -3.7 | 1.8E-04 | 2.5E-04 | MB231/MB468 | CYTO |
| >hg38_21_+_8433172_8446622_RNA45SN1_45S_50ntFlanks@11027.11055.29\|CTCTCTCTCTCTCTCCCCCGCTCCCCGTC | >hg38_21_+_8433172_8446622_RNA45SN1_45S_50ntFlanks | 11027 | 11055 | 29 | CTCTCTCTCTCTCTCCCCCGCTCCCCGTC | 339.7 | -5.7 | 1.5 | -3.7 | 1.9E-04 | 2.5E-04 | MB231/MB468 | NUC |
| >hg38_21_+_8433172_8446622_RNA45SN1_45S_50ntFlanks@7984.8008.25\|AGATCAGACGTGGCGACCCGCTGAA | >hg38_21_+_8433172_8446622_RNA45SN1_45S_50ntFlanks | 7984 | 8008 | 25 | AGATCAGACGTGGCGACCCGCTGAA | 1149.4 | -5.5 | 1.5 | -3.7 | 2.0E-04 | 2.5E-04 | BT20/MB468 | NUC |
| >hg38_21_+_8433172_8446622_RNA45SN1_45S_50ntFlanks@10239.10264.26\|GCTACGCCGCGACGAGTAGGAGGGCC | >hg38_21_+_8433172_8446622_RNA45SN1_45S_50ntFlanks | 10239 | 10264 | 26 | GCTACGCCGCGACGAGTAGGAGGGCC | 106.7 | 6.4 | 1.6 | 4.0 | 7.2E-05 | 2.5E-04 | BT20/MB468 | MP |
| >hg38_21_+_8433172_8446622_RNA45SN1_45S_50ntFlanks@10784.10818.35\|TGAACAGCCTCTGGCATGTTGGAACAATGTAGGTA | >hg38_21_+_8433172_8446622_RNA45SN1_45S_50ntFlanks | 10784 | 10818 | 35 | TGAACAGCCTCTGGCATGTTGGAACAATGTAGGTA | 386.9 | -1.8 | 0.5 | -3.8 | 1.4E-04 | 2.5E-04 | BT20/MB468 | MITO |
| >hg38_21_+_8433172_8446622_RNA45SN1_45S_50ntFlanks@7975.7996.22\|CGCGACCTCAGATCAGACGTGG | >hg38_21_+_8433172_8446622_RNA45SN1_45S_50ntFlanks | 7975 | 7996 | 22 | CGCGACCTCAGATCAGACGTGG | 27636.7 | 3.4 | 0.9 | 3.8 | 1.4E-04 | 2.5E-04 | BT20/MB468 | MITO |
| >hg38_21_+_8433172_8446622_RNA45SN1_45S_50ntFlanks@12405.12438.34\|CTTTTTGATCCTTCGATGTCGGCTCTTCCTATCA | >hg38_21_+_8433172_8446622_RNA45SN1_45S_50ntFlanks | 12405 | 12438 | 34 | CTTTTTGATCCTTCGATGTCGGCTCTTCCTATCA | 904.7 | -2.9 | 0.8 | -3.8 | 1.4E-04 | 2.5E-04 | BT20/MB468 | MITO |
| >hg38_21_+_8433172_8446622_RNA45SN1_45S_50ntFlanks@7975.7997.23\|CGCGACCTCAGATCAGACGTGGC | >hg38_21_+_8433172_8446622_RNA45SN1_45S_50ntFlanks | 7975 | 7997 | 23 | CGCGACCTCAGATCAGACGTGGC | 85910.2 | 3.5 | 0.8 | 4.2 | 2.7E-05 | 2.5E-04 | BT20/MB231 | NUC |
| >hg38_21_+_8433172_8446622_RNA45SN1_45S_50ntFlanks@10352.10369.18\|GAGAACTTTGAAGGCCGA | >hg38_21_+_8433172_8446622_RNA45SN1_45S_50ntFlanks | 10352 | 10369 | 18 | GAGAACTTTGAAGGCCGA | 153.5 | 3.2 | 0.9 | 3.7 | 1.9E-04 | 2.5E-04 | MB231/MB468 | MITO |
| >hg38_21_+_8433172_8446622_RNA45SN1_45S_50ntFlanks@12444.12476.33\|AAGCAGAATTCACCAAGCGTTGGATTGTTCACC | >hg38_21_+_8433172_8446622_RNA45SN1_45S_50ntFlanks | 12444 | 12476 | 33 | AAGCAGAATTCACCAAGCGTTGGATTGTTCACC | 916.8 | -3.1 | 0.8 | -3.8 | 1.6E-04 | 2.5E-04 | MB231/MB468 | MP |
| >hg38_21_+_8433172_8446622_RNA45SN1_45S_50ntFlanks@12007.12031.25\|GCCGCCGGTGAAATACCACTACTCT | >hg38_21_+_8433172_8446622_RNA45SN1_45S_50ntFlanks | 12007 | 12031 | 25 | GCCGCCGGTGAAATACCACTACTCT | 1008.4 | -5.3 | 1.3 | -4.0 | 7.2E-05 | 2.5E-04 | BT20/MB468 | MP |
| >hg38_21_+_8433172_8446622_RNA45SN1_45S_50ntFlanks@10349.10384.36\|AACGAGAACTTTGAAGGCCGAAGTGGAGAAGGGTTC | >hg38_21_+_8433172_8446622_RNA45SN1_45S_50ntFlanks | 10349 | 10384 | 36 | AACGAGAACTTTGAAGGCCGAAGTGGAGAAGGGTTC | 847.1 | -3.5 | 1.0 | -3.7 | 1.9E-04 | 2.5E-04 | MB231/MB468 | MITO |
| >am_trnaMT_ValTAC_MT_+_1602_1670@30.54.25\|CTTACACTTAGGAGATTTCAACTTA | >am_trnaMT_ValTAC_MT_+_1602_1670 | 30 | 54 | 25 | CTTACACTTAGGAGATTTCAACTTA | 339.4 | -4.7 | 1.3 | -3.7 | 1.9E-04 | 2.5E-04 | MB231/MB468 | MITO |
| >hg38_21_+_8433172_8446622_RNA45SN1_45S_50ntFlanks@11918.11967.50\|ACATGAGAGGTGTAGAATAAGTGGGAGGCCCCCGGCGCCCCCCCGGTGTC | >hg38_21_+_8433172_8446622_RNA45SN1_45S_50ntFlanks | 11918 | 11967 | 50 | ACATGAGAGGTGTAGAATAAGTGGGAGGCCCCCGGCGCCCCCCCGGTGTC | 605.6 | -5.4 | 1.5 | -3.7 | 1.8E-04 | 2.5E-04 | MB231/MB468 | CYTO |
| >hg38_21_+_8433172_8446622_RNA45SN1_45S_50ntFlanks@6652.6678.27\|GACTCTTAGCGGTGGATCACTCGGCTC | >hg38_21_+_8433172_8446622_RNA45SN1_45S_50ntFlanks | 6652 | 6678 | 27 | GACTCTTAGCGGTGGATCACTCGGCTC | 8813.5 | 5.8 | 1.4 | 4.2 | 2.6E-05 | 2.5E-04 | BT20/MB231 | MP |
| >hg38_21_+_8433172_8446622_RNA45SN1_45S_50ntFlanks@4679.4707.29\|GGCGCAAGACGGACCAGAGCGAAAGCATT | >hg38_21_+_8433172_8446622_RNA45SN1_45S_50ntFlanks | 4679 | 4707 | 29 | GGCGCAAGACGGACCAGAGCGAAAGCATT | 210.4 | 6.7 | 1.7 | 4.0 | 7.2E-05 | 2.5E-04 | BT20/MB468 | MP |
| >hg38_21_+_8433172_8446622_RNA45SN1_45S_50ntFlanks@9496.9525.30\|GAAAGATGGTGAACTATGCCTGGGCAGGGC | >hg38_21_+_8433172_8446622_RNA45SN1_45S_50ntFlanks | 9496 | 9525 | 30 | GAAAGATGGTGAACTATGCCTGGGCAGGGC | 248.7 | 6.6 | 1.7 | 4.0 | 7.2E-05 | 2.5E-04 | BT20/MB468 | MP |
| >am_MI0000809\|hsa-mir-151a&WithFlank&8\|-\|140732558\|140732659@54.73.20&[MIMAT0000757&hsa-miR-151a-3p&offsets\|+1\|0;m-30&8\|-\|140732587\|140732607&offsets\|+1\|0]\|TAGACTGAAGCTCCTTGAGG | >am_MI0000809\|hsa-mir-151a&WithFlank&8\|-\|140732558\|140732659 | 54 | 73 | 20 | TAGACTGAAGCTCCTTGAGG | 124.5 | 6.3 | 1.6 | 4.0 | 7.2E-05 | 2.5E-04 | BT20/MB468 | MP |
| >hg38_21_+_8433172_8446622_RNA45SN1_45S_50ntFlanks@11200.11227.28\|CGGCGGCGGCGGCGACTCTGGACGCGAG | >hg38_21_+_8433172_8446622_RNA45SN1_45S_50ntFlanks | 11200 | 11227 | 28 | CGGCGGCGGCGGCGACTCTGGACGCGAG | 381.4 | -6.3 | 1.7 | -3.7 | 2.0E-04 | 2.5E-04 | BT20/MB468 | NUC |
| >MI0000446\|hsa-mir-125b-1&WithFlank&11\|-\|122099751\|122099850@61.82.22(+2U)&[MIMAT0004592&hsa-miR-125b-1-3p&offsets\|0\|0(+2U);m-485&11\|-\|122099769\|122099790&offsets\|0\|0(+2U)]\|ACGGGTTAGGCTCTTGGGAGCTTT | >MI0000446\|hsa-mir-125b-1&WithFlank&11\|-\|122099751\|122099850 | 61 | 82 | 22(+2U) | ACGGGTTAGGCTCTTGGGAGCTTT | 24.9 | 7.9 | 2.1 | 3.8 | 1.6E-04 | 2.5E-04 | MB231/MB468 | MP |
| >hg38_21_+_8433172_8446622_RNA45SN1_45S_50ntFlanks@9554.9595.42\|CGTAGCGGTCCTGACGTGCAAATCGGTCGTCCGACCTGGGTA | >hg38_21_+_8433172_8446622_RNA45SN1_45S_50ntFlanks | 9554 | 9595 | 42 | CGTAGCGGTCCTGACGTGCAAATCGGTCGTCCGACCTGGGTA | 328.4 | -5.8 | 1.5 | -3.8 | 1.7E-04 | 2.5E-04 | BT20/MB468 | CYTO |
| >MI0000086\|hsa-mir-28&WithFlank&3\|+\|188688775\|188688872@60.82.23(+1U)&[MIMAT0004502&hsa-miR-28-3p&offsets\|0\|+1(+1U);m-28&3\|+\|188688834\|188688855&offsets\|0\|+1(+1U)]\|CACTAGATTGTGAGCTCCTGGAGT | >MI0000086\|hsa-mir-28&WithFlank&3\|+\|188688775\|188688872 | 60 | 82 | 23(+1U) | CACTAGATTGTGAGCTCCTGGAGT | 510.0 | 7.1 | 1.8 | 4.0 | 7.2E-05 | 2.5E-04 | BT20/MB468 | MP |
| >hg38_21_+_8433172_8446622_RNA45SN1_45S_50ntFlanks@8348.8378.31\|GGAAAGTTGAAAAGAACTTTGAAGAGAGAGT | >hg38_21_+_8433172_8446622_RNA45SN1_45S_50ntFlanks | 8348 | 8378 | 31 | GGAAAGTTGAAAAGAACTTTGAAGAGAGAGT | 153.7 | 6.2 | 1.6 | 4.0 | 7.2E-05 | 2.5E-04 | BT20/MB468 | MP |
| >hg38_21_+_8433172_8446622_RNA45SN1_45S_50ntFlanks@8170.8214.45\|CAAGTCCTTCTGATCGAGGCCCAGCCCGTGGACGGTGTGAGGCCG | >hg38_21_+_8433172_8446622_RNA45SN1_45S_50ntFlanks | 8170 | 8214 | 45 | CAAGTCCTTCTGATCGAGGCCCAGCCCGTGGACGGTGTGAGGCCG | 716.4 | -5.6 | 1.4 | -4.0 | 7.2E-05 | 2.5E-04 | BT20/MB468 | MP |
| >trnaMT_GluTTC_MT_-_14674_14742@39.68.30,trnalookalike8_GluTTC_5_-_93905172_93905240@39.68.30\|ATTGGTCGTGGTTGTAGTCCGTGCGAGAAT | >trnaMT_GluTTC_MT_-_14674_14742 | 39 | 68 | 30 | ATTGGTCGTGGTTGTAGTCCGTGCGAGAAT | 155.0 | 3.6 | 1.0 | 3.7 | 1.9E-04 | 2.5E-04 | MB231/MB468 | MITO |
| >hg38_21_+_8433172_8446622_RNA45SN1_45S_50ntFlanks@5484.5516.33\|GAGCGCTGAGAAGACGGTCGAACTTGACTATCT | >hg38_21_+_8433172_8446622_RNA45SN1_45S_50ntFlanks | 5484 | 5516 | 33 | GAGCGCTGAGAAGACGGTCGAACTTGACTATCT | 268.6 | -5.0 | 1.3 | -3.8 | 1.6E-04 | 2.5E-04 | MB231/MB468 | MP |

<table>
<tr><th></th><th></th><th></th><th></th><th></th><th></th><th></th><th></th><th></th><th></th><th></th><th></th><th></th><th></th></tr>
<tr><td>>trna2_GlyGCC_21_-_18827107_18827177@1.35.35,trna35_GlyGCC_1_+_161413094_161413164@1.35.35,trna37_GlyGCC_1_+_161420467_161420537@1.35.35,trna39_GlyGCC_1_+_161427898_161427968@1.35.35,trna41_GlyGCC_1_+_161435258_161435328@1.35.35|GCATGGGTGGTTCAGTGGTAGAATTCTCGCCTGCC</td><td>>trna2_GlyGCC_21_-_18827107_18827177</td><td>1</td><td>35</td><td>35</td><td>GCATGGGTGGTTCAGTGGTAGAATTCTCGCCTGCC</td><td>605.1</td><td>5.6</td><td>1.4</td><td>4.0</td><td>7.3E-05</td><td>2.5E-04</td><td>BT20/MB468</td><td>MP</td></tr>
<tr><td>>hg38_MT_+_1621_3279_RNR2_16S_50ntFlanks@1352.1396.45|ATAGGGTTTACGACCTCGATGTTGGATCAGGACATCCCGATGGTG</td><td>>hg38_MT_+_1621_3279_RNR2_16S_50ntFlanks</td><td>1352</td><td>1396</td><td>45</td><td>ATAGGGTTTACGACCTCGATGTTGGATCAGGACATCCCGATGGTG</td><td>98.7</td><td>10.4</td><td>2.6</td><td>4.0</td><td>7.3E-05</td><td>2.5E-04</td><td>BT20/MB468</td><td>MP</td></tr>
<tr><td>>MI0000085|hsa-mir-27a&WithFlank&19|-|13836434|13836523@57.77.21&[MIMAT0000084&hsa-miR-27a-3p&offsets|0|0;m-43&19|-|13836447|13836467&offsets|0|0]|TTCACAGTGGCTAAGTTCCGC</td><td>>MI0000085|hsa-mir-27a&WithFlank&19|-|13836434|13836523</td><td>57</td><td>77</td><td>21</td><td>TTCACAGTGGCTAAGTTCCGC</td><td>374.2</td><td>2.5</td><td>0.7</td><td>3.7</td><td>2.0E-04</td><td>2.5E-04</td><td>BT20/MB468</td><td>NUC</td></tr>
<tr><td>>am_MI0000809|hsa-mir-151a&WithFlank&8|-|140732558|140732659@52.73.22&[MIMAT0000757&hsa-miR-151a-3p&offsets|-1|0;m-30&8|-|140732587|140732607&offsets|-1|0]|ACTAGACTGAAGCTCCTTGAGG</td><td>>am_MI0000809|hsa-mir-151a&WithFlank&8|-|140732558|140732659</td><td>52</td><td>73</td><td>22</td><td>ACTAGACTGAAGCTCCTTGAGG</td><td>703.9</td><td>3.1</td><td>0.8</td><td>3.8</td><td>1.6E-04</td><td>2.5E-04</td><td>MB231/MB468</td><td>MP</td></tr>
<tr><td>>hg38_1_-_228634819_228635039_RNA5S12_5S_50ntFlanks@137.170.34|GAGACCGCCTGGGAATACCGGGTGCTGTAGGCTT</td><td>>hg38_1_-_228634819_228635039_RNA5S12_5S_50ntFlanks</td><td>137</td><td>170</td><td>34</td><td>GAGACCGCCTGGGAATACCGGGTGCTGTAGGCTT</td><td>641.5</td><td>2.7</td><td>0.7</td><td>4.0</td><td>7.3E-05</td><td>2.5E-04</td><td>BT20/MB468</td><td>MP</td></tr>
<tr><td>>hg38_21_+_8433172_8446622_RNA45SN1_45S_50ntFlanks@11795.11821.27|AACGAGATTCCCACTGTCCCTACCTAC</td><td>>hg38_21_+_8433172_8446622_RNA45SN1_45S_50ntFlanks</td><td>11795</td><td>11821</td><td>27</td><td>AACGAGATTCCCACTGTCCCTACCTAC</td><td>386.4</td><td>-6.0</td><td>1.6</td><td>-3.7</td><td>2.0E-04</td><td>2.5E-04</td><td>MB231/MB468</td><td>NUC</td></tr>
<tr><td>>hg38_MT_+_1621_3279_RNR2_16S_50ntFlanks@1438.1477.40|TACGTGATCTGAGTTCAGACCGGAGTAATCCAGGTCGGTT</td><td>>hg38_MT_+_1621_3279_RNR2_16S_50ntFlanks</td><td>1438</td><td>1477</td><td>40</td><td>TACGTGATCTGAGTTCAGACCGGAGTAATCCAGGTCGGTT</td><td>169.3</td><td>4.4</td><td>1.2</td><td>3.7</td><td>2.0E-04</td><td>2.5E-04</td><td>MB231/MB468</td><td>NUC</td></tr>
<tr><td>>hg38_21_+_8433172_8446622_RNA45SN1_45S_50ntFlanks@12412.12441.30|ATCCTTCGATGTCGGCTCTTCCTATCATTG</td><td>>hg38_21_+_8433172_8446622_RNA45SN1_45S_50ntFlanks</td><td>12412</td><td>12441</td><td>30</td><td>ATCCTTCGATGTCGGCTCTTCCTATCATTG</td><td>480.5</td><td>-5.8</td><td>1.6</td><td>-3.7</td><td>1.9E-04</td><td>2.5E-04</td><td>MB231/MB468</td><td>MITO</td></tr>
<tr><td>>hg38_21_+_8433172_8446622_RNA45SN1_45S_50ntFlanks@5472.5493.22|ACGGCCCTGGCGGAGCGCTGAG</td><td>>hg38_21_+_8433172_8446622_RNA45SN1_45S_50ntFlanks</td><td>5472</td><td>5493</td><td>22</td><td>ACGGCCCTGGCGGAGCGCTGAG</td><td>376.2</td><td>2.9</td><td>0.8</td><td>3.7</td><td>1.8E-04</td><td>2.5E-04</td><td>MB231/MB468</td><td>CYTO</td></tr>
<tr><td>>hg38_21_+_8433172_8446622_RNA45SN1_45S_50ntFlanks@12644.12661.18|TCTGTGGGATTATGACTG</td><td>>hg38_21_+_8433172_8446622_RNA45SN1_45S_50ntFlanks</td><td>12644</td><td>12661</td><td>18</td><td>TCTGTGGGATTATGACTG</td><td>328.0</td><td>-2.6</td><td>0.7</td><td>-3.8</td><td>1.7E-04</td><td>2.5E-04</td><td>BT20/MB468</td><td>CYTO</td></tr>
<tr><td>>hg38_21_+_8433172_8446622_RNA45SN1_45S_50ntFlanks@12458.12475.18|AAGCGTTGGATTGTTCAC</td><td>>hg38_21_+_8433172_8446622_RNA45SN1_45S_50ntFlanks</td><td>12458</td><td>12475</td><td>18</td><td>AAGCGTTGGATTGTTCAC</td><td>654.5</td><td>-4.8</td><td>1.3</td><td>-3.8</td><td>1.7E-04</td><td>2.5E-04</td><td>BT20/MB468</td><td>CYTO</td></tr>
<tr><td>>trnaMT_HisGTG_MT_+_12138_12206@25.70.46|CAGATTGTGAATCTGACAACAGAGGCTTACGACCCCTTATTTACCC</td><td>>trnaMT_HisGTG_MT_+_12138_12206</td><td>25</td><td>70</td><td>46</td><td>CAGATTGTGAATCTGACAACAGAGGCTTACGACCCCTTATTTACCC</td><td>97.6</td><td>10.4</td><td>2.6</td><td>4.0</td><td>7.3E-05</td><td>2.5E-04</td><td>BT20/MB468</td><td>MP</td></tr>
<tr><td>>hg38_21_+_8433172_8446622_RNA45SN1_45S_50ntFlanks@11879.11904.26|ACCCTGTTGAGCTTGACTCTAGTCTG</td><td>>hg38_21_+_8433172_8446622_RNA45SN1_45S_50ntFlanks</td><td>11879</td><td>11904</td><td>26</td><td>ACCCTGTTGAGCTTGACTCTAGTCTG</td><td>128.6</td><td>-3.3</td><td>0.9</td><td>-3.8</td><td>1.7E-04</td><td>2.5E-04</td><td>BT20/MB468</td><td>CYTO</td></tr>
<tr><td>>hg38_1_-_228634819_228635039_RNA5S12_5S_50ntFlanks@51.82.32|GTCTACGGCCATACCACCCTGAACGCGCCCGA</td><td>>hg38_1_-_228634819_228635039_RNA5S12_5S_50ntFlanks</td><td>51</td><td>82</td><td>32</td><td>GTCTACGGCCATACCACCCTGAACGCGCCCGA</td><td>474.6</td><td>3.2</td><td>0.9</td><td>3.8</td><td>1.4E-04</td><td>2.5E-04</td><td>BT20/MB468</td><td>MITO</td></tr>
<tr><td>>hg38_21_+_8433172_8446622_RNA45SN1_45S_50ntFlanks@476.508.33|TTCGTGATCGATGTGGTGACGTCGTGCTCTCCC</td><td>>hg38_21_+_8433172_8446622_RNA45SN1_45S_50ntFlanks</td><td>476</td><td>508</td><td>33</td><td>TTCGTGATCGATGTGGTGACGTCGTGCTCTCCC</td><td>170.5</td><td>4.5</td><td>1.2</td><td>3.7</td><td>1.9E-04</td><td>2.5E-04</td><td>MB231/MB468</td><td>MITO</td></tr>
<tr><td>>MI0000081|hsa-mir-24-2&WithFlank&19|-|13836281|13836365@56.76.21;MI0000080|hsa-mir-24-1&WithFlank&9|+|95086015|95086094@50.70.21&[MIMAT0000080&hsa-miR-24-3p&offsets|0|-1;m-36&19|-|13836288|13836310&offsets|0|-2];[MIMAT0000080_1&hsa-miR-24-3p&offsets|0|-1;m-35&9|+|95086064|95086086&offsets|0|-2]|TGGCTCAGTTCAGCAGGAACA</td><td>>MI0000081|hsa-mir-24-2&WithFlank&19|-|13836281|13836365</td><td>56</td><td>76</td><td>21</td><td>TGGCTCAGTTCAGCAGGAACA</td><td>6282.5</td><td>3.4</td><td>0.9</td><td>4.0</td><td>7.3E-05</td><td>2.5E-04</td><td>BT20/MB468</td><td>MP</td></tr>
<tr><td>>hg38_21_+_8433172_8446622_RNA45SN1_45S_50ntFlanks@8179.8203.25|CTGATCGAGGCCCAGCCCGTGGACG</td><td>>hg38_21_+_8433172_8446622_RNA45SN1_45S_50ntFlanks</td><td>8179</td><td>8203</td><td>25</td><td>CTGATCGAGGCCCAGCCCGTGGACG</td><td>181.8</td><td>-2.2</td><td>0.6</td><td>-3.8</td><td>1.4E-04</td><td>2.5E-04</td><td>BT20/MB468</td><td>MITO</td></tr>
<tr><td>>hg38_21_+_8433172_8446622_RNA45SN1_45S_50ntFlanks@6661.6710.50|CGGTGGATCACTCGGCTCGTGCGTCGATGAAGAACGCAGCTAGCTGCGAG</td><td>>hg38_21_+_8433172_8446622_RNA45SN1_45S_50ntFlanks</td><td>6661</td><td>6710</td><td>50</td><td>CGGTGGATCACTCGGCTCGTGCGTCGATGAAGAACGCAGCTAGCTGCGAG</td><td>3162.2</td><td>4.4</td><td>1.0</td><td>4.2</td><td>3.0E-05</td><td>2.5E-04</td><td>BT20/MB231</td><td>MITO</td></tr>
<tr><td>>hg38_21_+_8433172_8446622_RNA45SN1_45S_50ntFlanks@4283.4314.32|CTTTAACGAGGATCCATTGGAGGGCAAGTCTG</td><td>>hg38_21_+_8433172_8446622_RNA45SN1_45S_50ntFlanks</td><td>4283</td><td>4314</td><td>32</td><td>CTTTAACGAGGATCCATTGGAGGGCAAGTCTG</td><td>350.1</td><td>-3.1</td><td>0.8</td><td>-3.7</td><td>1.8E-04</td><td>2.5E-04</td><td>MB231/MB468</td><td>CYTO</td></tr>
<tr><td>>am_MI0000786|hsa-mir-378a&WithFlank&5|+|149732819|149732896@49.70.22&[MIMAT0000732&hsa-miR-378a-3p&offsets|0|0;m-44&5|+|149732867|149732888&offsets|0|0]|ACTGGACTTGGAGTCAGAAGGC</td><td>>am_MI0000786|hsa-mir-378a&WithFlank&5|+|149732819|149732896</td><td>49</td><td>70</td><td>22</td><td>ACTGGACTTGGAGTCAGAAGGC</td><td>6842.7</td><td>3.0</td><td>0.8</td><td>3.7</td><td>2.0E-04</td><td>2.5E-04</td><td>MB231/MB468</td><td>MITO</td></tr>
<tr><td>>hg38_21_+_8433172_8446622_RNA45SN1_45S_50ntFlanks@7971.8009.39|GAGACGCGACCTCAGATCAGACGTGGCGACCCGCTGAAT</td><td>>hg38_21_+_8433172_8446622_RNA45SN1_45S_50ntFlanks</td><td>7971</td><td>8009</td><td>39</td><td>GAGACGCGACCTCAGATCAGACGTGGCGACCCGCTGAAT</td><td>102.7</td><td>7.1</td><td>1.8</td><td>4.0</td><td>7.3E-05</td><td>2.5E-04</td><td>BT20/MB468</td><td>MP</td></tr>
<tr><td>>trnaMT_GluTTC_MT_-_14674_14742@19.57.39,trnalookalike8_GluTTC_5_-_93905172_93905240@19.57.39|CAACGATGGTTTTTCATATCATTGGTCGTGGTTGTAGTC</td><td>>trnaMT_GluTTC_MT_-_14674_14742</td><td>19</td><td>57</td><td>39</td><td>CAACGATGGTTTTTCATATCATTGGTCGTGGTTGTAGTC</td><td>104.7</td><td>10.5</td><td>2.6</td><td>4.0</td><td>7.3E-05</td><td>2.5E-04</td><td>BT20/MB468</td><td>MP</td></tr>
<tr><td>>hg38_21_+_8433172_8446622_RNA45SN1_45S_50ntFlanks@11664.11685.22|TTTCTGCCCAGTGCTCTGAATG</td><td>>hg38_21_+_8433172_8446622_RNA45SN1_45S_50ntFlanks</td><td>11664</td><td>11685</td><td>22</td><td>TTTCTGCCCAGTGCTCTGAATG</td><td>357.0</td><td>-3.7</td><td>1.0</td><td>-3.7</td><td>1.8E-04</td><td>2.5E-04</td><td>MB231/MB468</td><td>CYTO</td></tr>
<tr><td>>hg38_21_+_8433172_8446622_RNA45SN1_45S_50ntFlanks@4122.4145.24|AGGGTTCGATTCCGGAGAGGGAGC</td><td>>hg38_21_+_8433172_8446622_RNA45SN1_45S_50ntFlanks</td><td>4122</td><td>4145</td><td>24</td><td>AGGGTTCGATTCCGGAGAGGGAGC</td><td>125.7</td><td>-2.5</td><td>0.7</td><td>-3.7</td><td>1.8E-04</td><td>2.5E-04</td><td>MB231/MB468</td><td>CYTO</td></tr>
<tr><td>>hg38_MT_+_1621_3279_RNR2_16S_50ntFlanks@760.778.19|CAATATCTACAATCAACCA</td><td>>hg38_MT_+_1621_3279_RNR2_16S_50ntFlanks</td><td>760</td><td>778</td><td>19</td><td>CAATATCTACAATCAACCA</td><td>21.8</td><td>7.7</td><td>2.0</td><td>3.8</td><td>1.6E-04</td><td>2.5E-04</td><td>MB231/MB468</td><td>MP</td></tr>
<tr><td>>trnaMT_GluTTC_MT_-_14674_14742@40.59.20,trnalookalike8_GluTTC_5_-_93905172_93905240@40.59.20|TTGGTCGTGGTTGTAGTCCG</td><td>>trnaMT_GluTTC_MT_-_14674_14742</td><td>40</td><td>59</td><td>20</td><td>TTGGTCGTGGTTGTAGTCCG</td><td>175.2</td><td>5.2</td><td>1.3</td><td>4.0</td><td>7.4E-05</td><td>2.5E-04</td><td>BT20/MB468</td><td>MP</td></tr>
<tr><td>>hg38_21_+_8433172_8446622_RNA45SN1_45S_50ntFlanks@4414.4438.25|CCGCGAGGCGAGCCACCGCCCGTCC</td><td>>hg38_21_+_8433172_8446622_RNA45SN1_45S_50ntFlanks</td><td>4414</td><td>4438</td><td>25</td><td>CCGCGAGGCGAGCCACCGCCCGTCC</td><td>109.8</td><td>-9.9</td><td>2.5</td><td>-4.0</td><td>7.4E-05</td><td>2.5E-04</td><td>BT20/MB468</td><td>MP</td></tr>
<tr><td>>hg38_21_+_8433172_8446622_RNA45SN1_45S_50ntFlanks@6697.6718.22|CAGCTAGCTGCGAGAATTAATG</td><td>>hg38_21_+_8433172_8446622_RNA45SN1_45S_50ntFlanks</td><td>6697</td><td>6718</td><td>22</td><td>CAGCTAGCTGCGAGAATTAATG</td><td>441.3</td><td>-2.0</td><td>0.5</td><td>-3.8</td><td>1.7E-04</td><td>2.5E-04</td><td>BT20/MB468</td><td>CYTO</td></tr>
<tr><td>>hg38_21_+_8433172_8446622_RNA45SN1_45S_50ntFlanks@12418.12452.35|CGATGTCGGCTCTTCCTATCATTGTGAAGCAGAAT</td><td>>hg38_21_+_8433172_8446622_RNA45SN1_45S_50ntFlanks</td><td>12418</td><td>12452</td><td>35</td><td>CGATGTCGGCTCTTCCTATCATTGTGAAGCAGAAT</td><td>385.6</td><td>2.8</td><td>0.7</td><td>3.7</td><td>1.8E-04</td><td>2.5E-04</td><td>MB231/MB468</td><td>CYTO</td></tr>
<tr><td>>hg38_21_+_8433172_8446622_RNA45SN1_45S_50ntFlanks@5491.5515.25|GAGAAGACGGTCGAACTTGACTATC</td><td>>hg38_21_+_8433172_8446622_RNA45SN1_45S_50ntFlanks</td><td>5491</td><td>5515</td><td>25</td><td>GAGAAGACGGTCGAACTTGACTATC</td><td>234.6</td><td>3.6</td><td>0.9</td><td>4.0</td><td>7.4E-05</td><td>2.5E-04</td><td>BT20/MB468</td><td>MP</td></tr>
<tr><td>>hg38_21_+_8433172_8446622_RNA45SN1_45S_50ntFlanks@11668.11708.41|TGCCCAGTGCTCTGAATGTCAAAGTGAAGAAATTCAATGAA</td><td>>hg38_21_+_8433172_8446622_RNA45SN1_45S_50ntFlanks</td><td>11668</td><td>11708</td><td>41</td><td>TGCCCAGTGCTCTGAATGTCAAAGTGAAGAAATTCAATGAA</td><td>365.3</td><td>-3.7</td><td>0.9</td><td>-4.2</td><td>2.5E-05</td><td>2.5E-04</td><td>BT20/MB231</td><td>CYTO</td></tr>
<tr><td>>hg38_21_+_8433172_8446622_RNA45SN1_45S_50ntFlanks@11208.11225.18|GCGGCGACTCTGGACGCG</td><td>>hg38_21_+_8433172_8446622_RNA45SN1_45S_50ntFlanks</td><td>11208</td><td>11225</td><td>18</td><td>GCGGCGACTCTGGACGCG</td><td>1264.0</td><td>-3.3</td><td>0.9</td><td>-3.8</td><td>1.4E-04</td><td>2.5E-04</td><td>BT20/MB468</td><td>MITO</td></tr>
<tr><td>>hg38_21_+_8433172_8446622_RNA45SN1_45S_50ntFlanks@12002.12032.31|TGCGGGCCGCCGGTGAAATACCACTACTCTG</td><td>>hg38_21_+_8433172_8446622_RNA45SN1_45S_50ntFlanks</td><td>12002</td><td>12032</td><td>31</td><td>TGCGGGCCGCCGGTGAAATACCACTACTCTG</td><td>4999.4</td><td>-6.6</td><td>1.8</td><td>-3.8</td><td>1.7E-04</td><td>2.5E-04</td><td>BT20/MB468</td><td>CYTO</td></tr>
<tr><td>>trnaMT_SerGCT_MT_+_12207_12265@13.53.41|AAGAACTGCTAACTCATGCCCCCATGTCTAACAACATGGCT</td><td>>trnaMT_SerGCT_MT_+_12207_12265</td><td>13</td><td>53</td><td>41</td><td>AAGAACTGCTAACTCATGCCCCCATGTCTAACAACATGGCT</td><td>98.9</td><td>10.4</td><td>2.6</td><td>4.0</td><td>7.4E-05</td><td>2.5E-04</td><td>BT20/MB468</td><td>MP</td></tr>
<tr><td>>MI0000114|hsa-mir-107&WithFlank&10|-|89592741|89592833@56.75.20(+1G);MI0000108|hsa-mir-103a-2&WithFlank&20|+|3917488|3917577@54.73.20(+1G);MI0000109|hsa-mir-103a-1&WithFlank&5|-|168560890|168560979@54.73.20(+1G)&[MIMAT0000104&hsa-miR-107&offsets|0|-3(+1G);m-78&10|-|89592758|89592778&offsets|0|-1(+1G)];[MIMAT0000101&hsa-miR-103a-3p&offsets|0|-3(+1G);m-12&20|+|3917541|3917563&offsets|0|-3(+1G)];[MIMAT0000101_1&hsa-miR-103a-3p&offsets|0|-3(+1G);m-13&5|-|168560904|168560926&offsets|0|-3(+1G);m-44220&5|-|168560908|168560926&offsets|0|+1(+1G)]|AGCAGCATTGTACAGGGCTAG</td><td>>MI0000114|hsa-mir-107&WithFlank&10|-|89592741|89592833</td><td>56</td><td>75</td><td>20(+1G)</td><td>AGCAGCATTGTACAGGGCTAG</td><td>26.2</td><td>7.9</td><td>2.1</td><td>3.8</td><td>1.7E-04</td><td>2.5E-04</td><td>MB231/MB468</td><td>MP</td></tr>
<tr><td>>hg38_21_+_8433172_8446622_RNA45SN1_45S_50ntFlanks@8176.8201.26|CTTCTGATCGAGGCCCAGCCCGTGGA</td><td>>hg38_21_+_8433172_8446622_RNA45SN1_45S_50ntFlanks</td><td>8176</td><td>8201</td><td>26</td><td>CTTCTGATCGAGGCCCAGCCCGTGGA</td><td>457.7</td><td>-2.3</td><td>0.6</td><td>-3.8</td><td>1.7E-04</td><td>2.5E-04</td><td>BT20/MB468</td><td>CYTO</td></tr>
<tr><td>>hg38_21_+_8433172_8446622_RNA45SN1_45S_50ntFlanks@7968.7991.24|TCCGAGACGCGACCTCAGATCAGA</td><td>>hg38_21_+_8433172_8446622_RNA45SN1_45S_50ntFlanks</td><td>7968</td><td>7991</td><td>24</td><td>TCCGAGACGCGACCTCAGATCAGA</td><td>544.9</td><td>3.8</td><td>1.0</td><td>3.7</td><td>2.0E-04</td><td>2.5E-04</td><td>MB231/MB468</td><td>NUC</td></tr>
<tr><td>>hg38_21_+_8433172_8446622_RNA45SN1_45S_50ntFlanks@10869.10894.26|AGGGCTGGGTCGGTCGGGCTGGGGCG</td><td>>hg38_21_+_8433172_8446622_RNA45SN1_45S_50ntFlanks</td><td>10869</td><td>10894</td><td>26</td><td>AGGGCTGGGTCGGTCGGGCTGGGGCG</td><td>1098.7</td><td>-3.9</td><td>1.0</td><td>-4.0</td><td>7.4E-05</td><td>2.5E-04</td><td>BT20/MB468</td><td>MP</td></tr>
<tr><td>>hg38_21_+_8433172_8446622_RNA45SN1_45S_50ntFlanks@6747.6772.26|ACTTCGAACGCACTTGCGGCCCCGGG</td><td>>hg38_21_+_8433172_8446622_RNA45SN1_45S_50ntFlanks</td><td>6747</td><td>6772</td><td>26</td><td>ACTTCGAACGCACTTGCGGCCCCGGG</td><td>142.4</td><td>-6.0</td><td>1.5</td><td>-4.0</td><td>7.4E-05</td><td>2.5E-04</td><td>BT20/MB468</td><td>MP</td></tr>
<tr><td>>MI0000465|hsa-mir-191&WithFlank&3|-|49020612|49020715@22.44.23&[MIMAT0000440&hsa-miR-191-5p&offsets|0|0;m-48&3|-|49020672|49020694&offsets|0|0]|CAACGGAATCCCAAAAGCAGCTG</td><td>>MI0000465|hsa-mir-191&WithFlank&3|-|49020612|49020715</td><td>22</td><td>44</td><td>23</td><td>CAACGGAATCCCAAAAGCAGCTG</td><td>13022.8</td><td>3.0</td><td>0.8</td><td>3.8</td><td>1.7E-04</td><td>2.5E-04</td><td>MB231/MB468</td><td>MP</td></tr>
<tr><td>>hg38_21_+_8433172_8446622_RNA45SN1_45S_50ntFlanks@4005.4023.19|AACCTCGGGCCGATCGCAC</td><td>>hg38_21_+_8433172_8446622_RNA45SN1_45S_50ntFlanks</td><td>4005</td><td>4023</td><td>19</td><td>AACCTCGGGCCGATCGCAC</td><td>1143.4</td><td>3.3</td><td>0.9</td><td>3.8</td><td>1.7E-04</td><td>2.5E-04</td><td>BT20/MB468</td><td>CYTO</td></tr>
<tr><td>>MI0006444|hsa-mir-1307&WithFlank&10|-|103394247|103394407@86.108.23&[MIMAT0005951&hsa-miR-1307-3p&offsets|0|+1;m-66&10|-|103394300|103394322&offsets|0|0]|ACTCGGCGTGGCGTCGGTCGTGG</td><td>>MI0006444|hsa-mir-1307&WithFlank&10|-|103394247|103394407</td><td>86</td><td>108</td><td>23</td><td>ACTCGGCGTGGCGTCGGTCGTGG</td><td>1477.7</td><td>3.7</td><td>0.9</td><td>4.0</td><td>7.4E-05</td><td>2.5E-04</td><td>BT20/MB468</td><td>MP</td></tr>
<tr><td>>hg38_21_+_8433172_8446622_RNA45SN1_45S_50ntFlanks@9305.9337.33|CCAAGGAGTCTAACACGTGCGCGAGTCGGGGGC</td><td>>hg38_21_+_8433172_8446622_RNA45SN1_45S_50ntFlanks</td><td>9305</td><td>9337</td><td>33</td><td>CCAAGGAGTCTAACACGTGCGCGAGTCGGGGGC</td><td>301.5</td><td>-5.3</td><td>1.4</td><td>-3.8</td><td>1.5E-04</td><td>2.5E-04</td><td>BT20/MB468</td><td>MITO</td></tr>
<tr><td>>hg38_21_+_8433172_8446622_RNA45SN1_45S_50ntFlanks@10420.10453.34|CTGAGAGATGGGCGAGCGCCGTTCCGAAGGGACG</td><td>>hg38_21_+_8433172_8446622_RNA45SN1_45S_50ntFlanks</td><td>10420</td><td>10453</td><td>34</td><td>CTGAGAGATGGGCGAGCGCCGTTCCGAAGGGACG</td><td>1827.8</td><td>-2.7</td><td>0.7</td><td>-3.7</td><td>1.9E-04</td><td>2.5E-04</td><td>MB231/MB468</td><td>CYTO</td></tr>
<tr><td>>hg38_MT_+_1621_3279_RNR2_16S_50ntFlanks@1095.1137.43|AGACGAGAAGACCCTATGGAGCTTTAATTTATTAATGCAAACA</td><td>>hg38_MT_+_1621_3279_RNR2_16S_50ntFlanks</td><td>1095</td><td>1137</td><td>43</td><td>AGACGAGAAGACCCTATGGAGCTTTAATTTATTAATGCAAACA</td><td>98.3</td><td>10.4</td><td>2.6</td><td>4.0</td><td>7.4E-05</td><td>2.6E-04</td><td>BT20/MB468</td><td>MP</td></tr>
<tr><td>>hg38_21_+_8433172_8446622_RNA45SN1_45S_50ntFlanks@7992.8019.28|CGTGGCGACCCGCTGAATTTAAGCATAT</td><td>>hg38_21_+_8433172_8446622_RNA45SN1_45S_50ntFlanks</td><td>7992</td><td>8019</td><td>28</td><td>CGTGGCGACCCGCTGAATTTAAGCATAT</td><td>140.5</td><td>4.5</td><td>1.2</td><td>3.7</td><td>2.0E-04</td><td>2.6E-04</td><td>MB231/MB468</td><td>NUC</td></tr>
<tr><td>>hg38_21_+_8433172_8446622_RNA45SN1_45S_50ntFlanks@10853.10878.26|ATAAGGATTGGCTCTAAGGGCTGGGT</td><td>>hg38_21_+_8433172_8446622_RNA45SN1_45S_50ntFlanks</td><td>10853</td><td>10878</td><td>26</td><td>ATAAGGATTGGCTCTAAGGGCTGGGT</td><td>64.9</td><td>9.4</td><td>2.2</td><td>4.2</td><td>2.6E-05</td><td>2.6E-04</td><td>BT20/MB231</td><td>MP</td></tr>
</table>

| | | | | | | | | | | | | | |
|---|---|---|---|---|---|---|---|---|---|---|---|---|---|
| >hg38_21_+_8433172_8446622_RNA45SN1_45S_50ntFlanks@11200.11225.26\|CGGCGGCGGCGGCGACTCTGGACGCG | >hg38_21_+_8433172_8446622_RNA45SN1_45S_50ntFlanks | 11200 | 11225 | 26 | CGGCGGCGGCGGCGACTCTGGACGCG | 540.1 | -2.5 | 0.7 | -3.8 | 1.5E-04 | 2.6E-04 | BT20/MB468 | MITO |
| >hg38_MT_+_598_1651_RNR1_12S_50ntFlanks@446.480.35\|CTGAACACACAATAGCTAAGACCCAAACTGGGATT | >hg38_MT_+_598_1651_RNR1_12S_50ntFlanks | 446 | 480 | 35 | CTGAACACACAATAGCTAAGACCCAAACTGGGATT | 155.4 | 4.7 | 1.3 | 3.7 | 2.0E-04 | 2.6E-04 | MB231/MB468 | MITO |
| >hg38_21_+_8433172_8446622_RNA45SN1_45S_50ntFlanks@11321.11344.24\|CACGTCTCGTCGCGCGCGCGTCCG | >hg38_21_+_8433172_8446622_RNA45SN1_45S_50ntFlanks | 11321 | 11344 | 24 | CACGTCTCGTCGCGCGCGCGTCCG | 6493.9 | -5.1 | 1.4 | -3.8 | 1.7E-04 | 2.6E-04 | BT20/MB468 | CYTO |
| >hg38_21_+_8433172_8446622_RNA45SN1_45S_50ntFlanks@10792.10818.27\|CTCTGGCATGTTGGAACAATGTAGGTA | >hg38_21_+_8433172_8446622_RNA45SN1_45S_50ntFlanks | 10792 | 10818 | 27 | CTCTGGCATGTTGGAACAATGTAGGTA | 359.0 | -5.2 | 1.3 | -4.0 | 7.4E-05 | 2.6E-04 | BT20/MB468 | MP |
| >hg38_21_+_8433172_8446622_RNA45SN1_45S_50ntFlanks@9544.9570.27\|GGTGGAGGTCCGTAGCGGTCCTGACGT | >hg38_21_+_8433172_8446622_RNA45SN1_45S_50ntFlanks | 9544 | 9570 | 27 | GGTGGAGGTCCGTAGCGGTCCTGACGT | 1035.1 | -5.4 | 1.5 | -3.7 | 2.0E-04 | 2.6E-04 | BT20/MB468 | NUC |
| >hg38_21_+_8433172_8446622_RNA45SN1_45S_50ntFlanks@6647.6681.35\|CGTACGACTCTTAGCGGTGGATCACTCGGCTCGTG | >hg38_21_+_8433172_8446622_RNA45SN1_45S_50ntFlanks | 6647 | 6681 | 35 | CGTACGACTCTTAGCGGTGGATCACTCGGCTCGTG | 21894.1 | -3.4 | 0.9 | -3.7 | 2.0E-04 | 2.6E-04 | MB231/MB468 | MITO |
| >hg38_21_+_8433172_8446622_RNA45SN1_45S_50ntFlanks@6648.6681.34\|GTACGACTCTTAGCGGTGGATCACTCGGCTCGTG | >hg38_21_+_8433172_8446622_RNA45SN1_45S_50ntFlanks | 6648 | 6681 | 34 | GTACGACTCTTAGCGGTGGATCACTCGGCTCGTG | 22543.5 | -3.5 | 0.9 | -3.7 | 2.0E-04 | 2.6E-04 | MB231/MB468 | MITO |
| >am_trna162_MetCAT_6_-_26758550_26758622@34.54.21,trna164_MetCAT_6_-_26735574_26735646@34.54.21,trna20_MetCAT_16_+_71460396_71460468@34.54.21,trna21_MetCAT_6_+_26701712_26701784@34.54.21,trna27_MetCAT_6_+_26766444_26766516@34.54.21,trna75_MetCAT_6_+_28912352_28912424@34.54.21,trna97_MetCAT_6_-_28921042_28921114@34.54.21\|CATAATCTGAAGGTCCTGAGT | >am_trna162_MetCAT_6_-_26758550_26758622 | 34 | 54 | 21 | CATAATCTGAAGGTCCTGAGT | 381.8 | 3.0 | 0.8 | 3.7 | 1.9E-04 | 2.6E-04 | MB231/MB468 | CYTO |
| >hg38_21_+_8433172_8446622_RNA45SN1_45S_50ntFlanks@12937.12954.18\|AACCATTCGTAGACGACC | >hg38_21_+_8433172_8446622_RNA45SN1_45S_50ntFlanks | 12937 | 12954 | 18 | AACCATTCGTAGACGACC | 76.7 | -5.7 | 1.5 | -3.8 | 1.7E-04 | 2.6E-04 | MB231/MB468 | MP |
| >MI0003137\|hsa-mir-193b&WithFlank&16\|+\|14303961\|14304055@19.40.22(+1U)&[MIMAT0004767&hsa-miR-193b-5p&offsets\|-1\|-1(+1U);m-343&16\|+\|14303980\|14304001&offsets\|-1\|-1(+1U)]\|TCGGGGTTTTGAGGGCGAGATGT | >MI0003137\|hsa-mir-193b&WithFlank&16\|+\|14303961\|14304055 | 19 | 40 | 22(+1U) | TCGGGGTTTTGAGGGCGAGATGT | 52.7 | 9.1 | 2.2 | 4.2 | 2.7E-05 | 2.6E-04 | BT20/MB231 | MP |
| >hg38_21_+_8433172_8446622_RNA45SN1_45S_50ntFlanks@3813.3843.31\|TTATGGTTCCTTTGGTCGCTCGCTCCTCTCC | >hg38_21_+_8433172_8446622_RNA45SN1_45S_50ntFlanks | 3813 | 3843 | 31 | TTATGGTTCCTTTGGTCGCTCGCTCCTCTCC | 243.7 | -5.4 | 1.5 | -3.7 | 2.0E-04 | 2.6E-04 | MB231/MB468 | NUC |
| >trnaMT_GluTTC_MT_-_14674_14742@26.57.32,trnalookalike8_GluTTC_5_-_93905172_93905240@26.57.32\|GGTTTTTCATATCATTGGTCGTGGTTGTAGTC | >trnaMT_GluTTC_MT_-_14674_14742 | 26 | 57 | 32 | GGTTTTTCATATCATTGGTCGTGGTTGTAGTC | 596.1 | 7.0 | 1.8 | 4.0 | 7.5E-05 | 2.6E-04 | BT20/MB468 | MP |
| >MI0000088\|hsa-mir-30a&WithFlank&6\|-\|71403545\|71403627@53.73.21(+1A)&[MIMAT0000088&hsa-miR-30a-3p&offsets\|0\|-1(+1A);m-38&6\|-\|71403554\|71403575&offsets\|0\|-1(+1A)]\|CTTTCAGTCGGATGTTTGCAGA | >MI0000088\|hsa-mir-30a&WithFlank&6\|-\|71403545\|71403627 | 53 | 73 | 21(+1A) | CTTTCAGTCGGATGTTTGCAGA | 964.2 | -4.2 | 1.0 | -4.2 | 2.7E-05 | 2.6E-04 | BT20/MB231 | MP |
| >hg38_21_+_8433172_8446622_RNA45SN1_45S_50ntFlanks@6772.6797.26\|GTTCCTCCCGGGGCTACGCCTGTCTG | >hg38_21_+_8433172_8446622_RNA45SN1_45S_50ntFlanks | 6772 | 6797 | 26 | GTTCCTCCCGGGGCTACGCCTGTCTG | 181.0 | -4.9 | 1.3 | -3.8 | 1.8E-04 | 2.6E-04 | BT20/MB468 | CYTO |
| >hg38_21_+_8433172_8446622_RNA45SN1_45S_50ntFlanks@12658.12675.18\|ACTGAACGCCTCTAAGTC | >hg38_21_+_8433172_8446622_RNA45SN1_45S_50ntFlanks | 12658 | 12675 | 18 | ACTGAACGCCTCTAAGTC | 627.0 | -2.5 | 0.7 | -3.8 | 1.5E-04 | 2.6E-04 | BT20/MB468 | MITO |
| >MI0000090\|hsa-mir-32&WithFlank&9\|-\|109046223\|109046304@12.32.21&[MIMAT0000090&hsa-miR-32-5p&offsets\|0\|-1;m-208&9\|-\|109046273\|109046293&offsets\|0\|0]\|TATTGCACATTACTAAGTTGC | >MI0000090\|hsa-mir-32&WithFlank&9\|-\|109046223\|109046304 | 12 | 32 | 21 | TATTGCACATTACTAAGTTGC | 207.9 | 3.1 | 0.8 | 3.8 | 1.7E-04 | 2.6E-04 | MB231/MB468 | MP |
| >hg38_21_+_8433172_8446622_RNA45SN1_45S_50ntFlanks@11206.11227.22\|CGGCGGCGACTCTGGACGCGAG | >hg38_21_+_8433172_8446622_RNA45SN1_45S_50ntFlanks | 11206 | 11227 | 22 | CGGCGGCGACTCTGGACGCGAG | 159.9 | -5.7 | 1.5 | -3.8 | 1.7E-04 | 2.6E-04 | MB231/MB468 | MP |
| >MI0000067\|hsa-let-7f-1&WithFlank&9\|+\|94176341\|94176439@13.34.22(+1C);MI0000068\|hsa-let-7f-2&WithFlank&X\|-\|53557186\|53557280@14.35.22(+1C)&[MIMAT0000067&hsa-let-7f-5p&offsets\|0\|0(+1C);m-20&9\|+\|94176353\|94176374&offsets\|0\|0(+1C)];[MIMAT0000067_1&hsa-let-7f-5p&offsets\|0\|0(+1C);m-19&X\|-\|53557246\|53557267&offsets\|0\|0(+1C)]\|TGAGGTAGTAGATTGTATAGTTC | >MI0000067\|hsa-let-7f-1&WithFlank&9\|+\|94176341\|94176439 | 13 | 34 | 22(+1C) | TGAGGTAGTAGATTGTATAGTTC | 165.0 | 5.7 | 1.5 | 3.7 | 2.0E-04 | 2.6E-04 | MB231/MB468 | MITO |
| >hg38_1_-_228634819_228635039_RNA5S12_5S_50ntFlanks@129.169.41\|TTGGATGGGAGACCGCCTGGGAATACCGGGTGCTGTAGGCT | >hg38_1_-_228634819_228635039_RNA5S12_5S_50ntFlanks | 129 | 169 | 41 | TTGGATGGGAGACCGCCTGGGAATACCGGGTGCTGTAGGCT | 387.6 | 3.3 | 0.8 | 4.2 | 3.1E-05 | 2.6E-04 | BT20/MB231 | MITO |
| >hg38_MT_+_598_1651_RNR1_12S_50ntFlanks@972.1004.33\|GGTAAGTGTACTGGAAAGTGCACTTGGACGAAC | >hg38_MT_+_598_1651_RNR1_12S_50ntFlanks | 972 | 1004 | 33 | GGTAAGTGTACTGGAAAGTGCACTTGGACGAAC | 747.1 | -3.7 | 0.9 | -4.2 | 3.1E-05 | 2.6E-04 | BT20/MB231 | MITO |
| >hg38_21_+_8433172_8446622_RNA45SN1_45S_50ntFlanks@11873.11903.31\|AAGAAGACCCTGTTGAGCTTGACTCTAGTCT | >hg38_21_+_8433172_8446622_RNA45SN1_45S_50ntFlanks | 11873 | 11903 | 31 | AAGAAGACCCTGTTGAGCTTGACTCTAGTCT | 178.4 | -5.3 | 1.4 | -3.7 | 2.0E-04 | 2.6E-04 | MB231/MB468 | NUC |
| >hg38_21_+_8433172_8446622_RNA45SN1_45S_50ntFlanks@11729.11752.24\|TAACTATGACTCTCTTAAGGTAGC | >hg38_21_+_8433172_8446622_RNA45SN1_45S_50ntFlanks | 11729 | 11752 | 24 | TAACTATGACTCTCTTAAGGTAGC | 295.4 | -3.0 | 0.8 | -3.8 | 1.5E-04 | 2.6E-04 | BT20/MB468 | MITO |
| >hg38_21_+_8433172_8446622_RNA45SN1_45S_50ntFlanks@10411.10440.30\|CAGTCGGTCCTGAGAGATGGGCGAGCGCCG | >hg38_21_+_8433172_8446622_RNA45SN1_45S_50ntFlanks | 10411 | 10440 | 30 | CAGTCGGTCCTGAGAGATGGGCGAGCGCCG | 214.0 | -4.0 | 1.1 | -3.8 | 1.5E-04 | 2.6E-04 | BT20/MB468 | MITO |
| >hg38_21_+_8433172_8446622_RNA45SN1_45S_50ntFlanks@10802.10825.24\|TTGGAACAATGTAGGTAAGGGAAG | >hg38_21_+_8433172_8446622_RNA45SN1_45S_50ntFlanks | 10802 | 10825 | 24 | TTGGAACAATGTAGGTAAGGGAAG | 226.8 | -6.0 | 1.5 | -4.0 | 7.5E-05 | 2.6E-04 | BT20/MB468 | MP |
| >hg38_21_+_8433172_8446622_RNA45SN1_45S_50ntFlanks@6670.6698.29\|ACTCGGCTCGTGCGTCGATGAAGAACGCA | >hg38_21_+_8433172_8446622_RNA45SN1_45S_50ntFlanks | 6670 | 6698 | 29 | ACTCGGCTCGTGCGTCGATGAAGAACGCA | 172.5 | -3.0 | 0.8 | -3.7 | 1.9E-04 | 2.6E-04 | MB231/MB468 | CYTO |
| >hg38_21_+_8433172_8446622_RNA45SN1_45S_50ntFlanks@9736.9759.24\|CCGAAACGATCTCAACCTATTCTC | >hg38_21_+_8433172_8446622_RNA45SN1_45S_50ntFlanks | 9736 | 9759 | 24 | CCGAAACGATCTCAACCTATTCTC | 481.8 | -5.2 | 1.4 | -3.8 | 1.5E-04 | 2.6E-04 | BT20/MB468 | MITO |
| >hg38_21_+_8433172_8446622_RNA45SN1_45S_50ntFlanks@11562.11588.27\|CGCCTAGCAGCCGACTTAGAACTGGTG | >hg38_21_+_8433172_8446622_RNA45SN1_45S_50ntFlanks | 11562 | 11588 | 27 | CGCCTAGCAGCCGACTTAGAACTGGTG | 373.4 | -3.9 | 1.0 | -3.7 | 1.9E-04 | 2.6E-04 | MB231/MB468 | CYTO |
| >hg38_21_+_8433172_8446622_RNA45SN1_45S_50ntFlanks@4738.4758.21\|AAAGTCGGAGGTTCGAAGACG | >hg38_21_+_8433172_8446622_RNA45SN1_45S_50ntFlanks | 4738 | 4758 | 21 | AAAGTCGGAGGTTCGAAGACG | 13795.5 | -2.2 | 0.6 | -3.8 | 1.7E-04 | 2.6E-04 | MB231/MB468 | MP |
| >am_trna2_GlyGCC_21_-_18827107_18827177@1.27.27,trna35_GlyGCC_1_+_161413094_161413164@1.27.27,trna37_GlyGCC_1_+_161420467_161420537@1.27.27,trna39_GlyGCC_1_+_161427898_161427968@1.27.27,trna41_GlyGCC_1_+_161435258_161435328@1.27.27\|GCATGGGTGGTTCAGTGGTAGAATTCT | >am_trna2_GlyGCC_21_-_18827107_18827177 | 1 | 27 | 27 | GCATGGGTGGTTCAGTGGTAGAATTCT | 4341.9 | 2.6 | 0.7 | 3.7 | 1.8E-04 | 2.6E-04 | BT20/MB468 | CYTO |
| >MI0000067\|hsa-let-7f-1&WithFlank&9\|+\|94176341\|94176439@13.30.18;MI0000068\|hsa-let-7f-2&WithFlank&X\|-\|53557186\|53557280@14.31.18&[MIMAT0000067&hsa-let-7f-5p&offsets\|0\|-4;m-20&9\|+\|94176353\|94176374&offsets\|0\|-4];[MIMAT0000067_1&hsa-let-7f-5p&offsets\|0\|-4;m-19&X\|-\|53557246\|53557267&offsets\|0\|-4]\|TGAGGTAGTAGATTGTAT | >MI0000067\|hsa-let-7f-1&WithFlank&9\|+\|94176341\|94176439 | 13 | 30 | 18 | TGAGGTAGTAGATTGTAT | 1197.9 | 3.6 | 0.9 | 4.0 | 7.5E-05 | 2.6E-04 | BT20/MB468 | MP |
| >MI0000299\|hsa-mir-222&WithFlank&X\|-\|45747009\|45747130@75.99.25(+4U)&[MIMAT0000279&hsa-miR-222-3p&offsets\|0\|+4(+4U);m-140&X\|-\|45747033\|45747056&offsets\|0\|+1(+4U)]\|AGCTACATCTGGCTACTGGGTCTCTTTTT | >MI0000299\|hsa-mir-222&WithFlank&X\|-\|45747009\|45747130 | 75 | 99 | 25(+4U) | AGCTACATCTGGCTACTGGGTCTCTTTTT | 562.0 | -3.3 | 0.8 | -4.2 | 2.5E-05 | 2.6E-04 | BT20/MB231 | CYTO |
| >hg38_21_+_8433172_8446622_RNA45SN1_45S_50ntFlanks@12433.12452.20\|CTATCATTGTGAAGCAGAAT | >hg38_21_+_8433172_8446622_RNA45SN1_45S_50ntFlanks | 12433 | 12452 | 20 | CTATCATTGTGAAGCAGAAT | 99.9 | -4.3 | 1.1 | -3.8 | 1.7E-04 | 2.6E-04 | MB231/MB468 | MP |
| >hg38_21_+_8433172_8446622_RNA45SN1_45S_50ntFlanks@7981.7998.18\|CTCAGATCAGACGTGGCG | >hg38_21_+_8433172_8446622_RNA45SN1_45S_50ntFlanks | 7981 | 7998 | 18 | CTCAGATCAGACGTGGCG | 94.4 | -4.8 | 1.3 | -3.7 | 1.8E-04 | 2.6E-04 | BT20/MB468 | CYTO |
| >hg38_21_+_8433172_8446622_RNA45SN1_45S_50ntFlanks@4762.4780.19\|AGATACCGTCGTAGTTCCG | >hg38_21_+_8433172_8446622_RNA45SN1_45S_50ntFlanks | 4762 | 4780 | 19 | AGATACCGTCGTAGTTCCG | 726.1 | -4.8 | 1.3 | -3.7 | 2.0E-04 | 2.6E-04 | MB231/MB468 | MITO |
| >am_trnaMT_MetCAT_MT_+_4402_4469@1.47.47\|AGTAAGGTCAGCTAAATAAGCTATCGGGCCCATACCCCGAAAATGTT | >am_trnaMT_MetCAT_MT_+_4402_4469 | 1 | 47 | 47 | AGTAAGGTCAGCTAAATAAGCTATCGGGCCCATACCCCGAAAATGTT | 618.5 | 6.1 | 1.6 | 3.7 | 2.0E-04 | 2.6E-04 | MB231/MB468 | MITO |
| >hg38_21_+_8433172_8446622_RNA45SN1_45S_50ntFlanks@5184.5204.21\|AGCAATAACAGGTCTGTGATG | >hg38_21_+_8433172_8446622_RNA45SN1_45S_50ntFlanks | 5184 | 5204 | 21 | AGCAATAACAGGTCTGTGATG | 142.5 | -4.0 | 1.1 | -3.8 | 1.7E-04 | 2.6E-04 | MB231/MB468 | MP |
| >hg38_21_+_8433172_8446622_RNA45SN1_45S_50ntFlanks@4449.4470.22\|CCTCTCGGCGCCCCCTCGATGC | >hg38_21_+_8433172_8446622_RNA45SN1_45S_50ntFlanks | 4449 | 4470 | 22 | CCTCTCGGCGCCCCCTCGATGC | 142.2 | -5.0 | 1.3 | -3.8 | 1.5E-04 | 2.6E-04 | BT20/MB468 | MITO |
| >MI0000264\|hsa-mir-7-2&WithFlank&15\|+\|88611819\|88611940@38.60.23(+1A);MI0000265\|hsa-mir-7-3&WithFlank&19\|+\|4770664\|4770785@37.59.23(+1A);MI0000263\|hsa-mir-7-1&WithFlank&9\|-\|83969742\|83969863@30.52.23(+1A)&[MIMAT0000252&hsa-miR-7-5p&offsets\|0\|-1(+1A);m-400&15\|+\|88611856\|88611879&offsets\|0\|-1(+1A)];[MIMAT0000252_1&hsa-miR-7-5p&offsets\|0\|-1(+1A);m-402&19\|+\|4770700\|4770723&offsets\|0\|-1(+1A)];[MIMAT0000252_2&hsa-miR-7-5p&offsets\|0\|-1(+1A);m-401&9\|-\|83969811\|83969834&offsets\|0\|-1(+1A)]\|TGGAAGACTAGTGATTTTGTTGTA | >MI0000264\|hsa-mir-7-2&WithFlank&15\|+\|88611819\|88611940 | 38 | 60 | 23(+1A) | TGGAAGACTAGTGATTTTGTTGTA | 5810.5 | 4.1 | 1.0 | 4.2 | 2.6E-05 | 2.6E-04 | BT20/MB231 | CYTO |
| >hg38_21_+_8433172_8446622_RNA45SN1_45S_50ntFlanks@12008.12034.27\|CCGCCGGTGAAATACCACTACTCTGAT | >hg38_21_+_8433172_8446622_RNA45SN1_45S_50ntFlanks | 12008 | 12034 | 27 | CCGCCGGTGAAATACCACTACTCTGAT | 316.9 | -6.1 | 1.5 | -4.0 | 7.6E-05 | 2.6E-04 | BT20/MB468 | MP |
| >hg38_21_+_8433172_8446622_RNA45SN1_45S_50ntFlanks@4044.4063.20\|CCCATTCGAACGTCTGCCCT | >hg38_21_+_8433172_8446622_RNA45SN1_45S_50ntFlanks | 4044 | 4063 | 20 | CCCATTCGAACGTCTGCCCT | 195.5 | -5.0 | 1.3 | -3.7 | 1.8E-04 | 2.6E-04 | BT20/MB468 | CYTO |
| >hg38_21_+_8433172_8446622_RNA45SN1_45S_50ntFlanks@6745.6762.18\|ACACTTCGAACGCACTTG | >hg38_21_+_8433172_8446622_RNA45SN1_45S_50ntFlanks | 6745 | 6762 | 18 | ACACTTCGAACGCACTTG | 219.7 | -5.1 | 1.4 | -3.7 | 1.8E-04 | 2.6E-04 | BT20/MB468 | CYTO |
| >hg38_MT_+_1621_3279_RNR2_16S_50ntFlanks@825.859.35\|TAAGGAAAGGTTAAAAAAAGTAAAAGGAACTCGGC | >hg38_MT_+_1621_3279_RNR2_16S_50ntFlanks | 825 | 859 | 35 | TAAGGAAAGGTTAAAAAAAGTAAAAGGAACTCGGC | 22.0 | 7.7 | 2.0 | 3.8 | 1.7E-04 | 2.6E-04 | MB231/MB468 | MP |
| >hg38_21_+_8433172_8446622_RNA45SN1_45S_50ntFlanks@9778.9819.42\|AAGCCCGGCTCGCTGGCGTGGAGCCGGGCGTGGAATGCGAGT | >hg38_21_+_8433172_8446622_RNA45SN1_45S_50ntFlanks | 9778 | 9819 | 42 | AAGCCCGGCTCGCTGGCGTGGAGCCGGGCGTGGAATGCGAGT | 51.4 | 9.0 | 2.1 | 4.2 | 2.7E-05 | 2.6E-04 | BT20/MB231 | MP |
| >hg38_21_+_8433172_8446622_RNA45SN1_45S_50ntFlanks@475.519.45\|CTTCGTGATCGATGTGGTGACGTCGTGCTCTCCCGGGCCGGGTCC | >hg38_21_+_8433172_8446622_RNA45SN1_45S_50ntFlanks | 475 | 519 | 45 | CTTCGTGATCGATGTGGTGACGTCGTGCTCTCCCGGGCCGGGTCC | 1407.7 | 4.2 | 1.1 | 3.7 | 2.0E-04 | 2.6E-04 | MB231/MB468 | NUC |

<table>
<tr><td>>hg38_21_+_8433172_8446622_RNA45SN1_45S_50ntFlanks@10391.10428.38|AACAGCAGTTGAACATGGGTCAGTCGGTCCTGAGAGAT</td><td>>hg38_21_+_8433172_8446622_RNA45SN1_45S_50ntFlanks</td><td>10391</td><td>10428</td><td>38</td><td>AACAGCAGTTGAACATGGGTCAGTCGGTCCTGAGAGAT</td><td>108.4</td><td>-6.0</td><td>1.6</td><td>-3.8</td><td>1.7E-04</td><td>2.6E-04</td><td>MB231/MB468</td><td>MP</td></tr>
<tr><td>>hg38_21_+_8433172_8446622_RNA45SN1_45S_50ntFlanks@10336.10373.38|TAGCAAATATTCAAACGAGAACTTTGAAGGCCGAAGTG</td><td>>hg38_21_+_8433172_8446622_RNA45SN1_45S_50ntFlanks</td><td>10336</td><td>10373</td><td>38</td><td>TAGCAAATATTCAAACGAGAACTTTGAAGGCCGAAGTG</td><td>72.5</td><td>7.0</td><td>1.8</td><td>4.0</td><td>7.6E-05</td><td>2.6E-04</td><td>BT20/MB468</td><td>MP</td></tr>
<tr><td>>hg38_21_+_8433172_8446622_RNA45SN1_45S_50ntFlanks@9493.9520.28|CCCGAAAGATGGTGAACTATGCCTGGGC</td><td>>hg38_21_+_8433172_8446622_RNA45SN1_45S_50ntFlanks</td><td>9493</td><td>9520</td><td>28</td><td>CCCGAAAGATGGTGAACTATGCCTGGGC</td><td>63.3</td><td>9.3</td><td>2.2</td><td>4.2</td><td>2.7E-05</td><td>2.6E-04</td><td>BT20/MB231</td><td>MP</td></tr>
<tr><td>>MI0000737|hsa-mir-200a&WithFlank&1|+|1167857|1167958@60.80.21&[MIMAT0000682&hsa-miR-200a-3p&offsets|0|-1;m-142&1|+|1167916|1167938&offsets|0|-2]|TAACACTGTCTGGTAACGATG</td><td>>MI0000737|hsa-mir-200a&WithFlank&1|+|1167857|1167958</td><td>60</td><td>80</td><td>21</td><td>TAACACTGTCTGGTAACGATG</td><td>356.0</td><td>-6.1</td><td>1.5</td><td>-4.0</td><td>7.6E-05</td><td>2.6E-04</td><td>BT20/MB468</td><td>MP</td></tr>
<tr><td>>MI0000081|hsa-mir-24-2&WithFlank&19|-|13836281|13836365@56.77.22(+1U);MI0000080|hsa-mir-24-1&WithFlank&9|+|95086015|95086094@50.71.22(+1U)&[MIMAT0000080&hsa-miR-24-3p&offsets|0|0(+1U);m-36&19|-|13836288|13836310&offsets|0|-1(+1U)];[MIMAT0000080_1&hsa-miR-24-3p&offsets|0|0(+1U);m-35&9|+|95086064|95086086&offsets|0|-1(+1U)]|TGGCTCAGTTCAGCAGGAACAGT</td><td>>MI0000081|hsa-mir-24-2&WithFlank&19|-|13836281|13836365</td><td>56</td><td>77</td><td>22(+1U)</td><td>TGGCTCAGTTCAGCAGGAACAGT</td><td>17344.7</td><td>-4.0</td><td>0.9</td><td>-4.2</td><td>2.7E-05</td><td>2.6E-04</td><td>BT20/MB231</td><td>MP</td></tr>
<tr><td>>hg38_21_+_8433172_8446622_RNA45SN1_45S_50ntFlanks@6723.6743.21|TTGCAGGACACATTGATCATC</td><td>>hg38_21_+_8433172_8446622_RNA45SN1_45S_50ntFlanks</td><td>6723</td><td>6743</td><td>21</td><td>TTGCAGGACACATTGATCATC</td><td>236.9</td><td>-4.5</td><td>1.2</td><td>-3.8</td><td>1.7E-04</td><td>2.6E-04</td><td>MB231/MB468</td><td>MP</td></tr>
<tr><td>>hg38_21_+_8433172_8446622_RNA45SN1_45S_50ntFlanks@8114.8138.25|GCGCGGGACATGTGGCGTACGGAAG</td><td>>hg38_21_+_8433172_8446622_RNA45SN1_45S_50ntFlanks</td><td>8114</td><td>8138</td><td>25</td><td>GCGCGGGACATGTGGCGTACGGAAG</td><td>1081.2</td><td>-7.1</td><td>1.9</td><td>-3.7</td><td>2.1E-04</td><td>2.6E-04</td><td>BT20/MB468</td><td>NUC</td></tr>
<tr><td>>MI0005531|hsa-mir-450b&WithFlank&X|-|134540179|134540268@17.37.21&[MIMAT0004909&hsa-miR-450b-5p&offsets|0|-1;m-266&X|-|134540232|134540252&offsets|0|0]|TTTTGCAATATGTTCCTGAAT</td><td>>MI0005531|hsa-mir-450b&WithFlank&X|-|134540179|134540268</td><td>17</td><td>37</td><td>21</td><td>TTTTGCAATATGTTCCTGAAT</td><td>109.3</td><td>5.4</td><td>1.4</td><td>3.8</td><td>1.7E-04</td><td>2.6E-04</td><td>MB231/MB468</td><td>MP</td></tr>
<tr><td>>hg38_21_+_8433172_8446622_RNA45SN1_45S_50ntFlanks@6721.6745.25|AATTGCAGGACACATTGATCATCGA</td><td>>hg38_21_+_8433172_8446622_RNA45SN1_45S_50ntFlanks</td><td>6721</td><td>6745</td><td>25</td><td>AATTGCAGGACACATTGATCATCGA</td><td>2243.5</td><td>-6.3</td><td>1.7</td><td>-3.7</td><td>1.8E-04</td><td>2.6E-04</td><td>BT20/MB468</td><td>CYTO</td></tr>
<tr><td>>hg38_21_+_8433172_8446622_RNA45SN1_45S_50ntFlanks@10357.10378.22|CTTTGAAGGCCGAAGTGGAGAA</td><td>>hg38_21_+_8433172_8446622_RNA45SN1_45S_50ntFlanks</td><td>10357</td><td>10378</td><td>22</td><td>CTTTGAAGGCCGAAGTGGAGAA</td><td>164.3</td><td>-4.2</td><td>1.1</td><td>-3.7</td><td>2.0E-04</td><td>2.6E-04</td><td>MB231/MB468</td><td>NUC</td></tr>
<tr><td>>MI0000088|hsa-mir-30a&WithFlank&6|-|71403545|71403627@12.33.22&[MIMAT0000087&hsa-miR-30a-5p&offsets|0|0;m-15&6|-|71403595|71403616&offsets|0|0]|TGTAAACATCCTCGACTGGAAG</td><td>>MI0000088|hsa-mir-30a&WithFlank&6|-|71403545|71403627</td><td>12</td><td>33</td><td>22</td><td>TGTAAACATCCTCGACTGGAAG</td><td>9516.5</td><td>2.6</td><td>0.7</td><td>3.8</td><td>1.7E-04</td><td>2.6E-04</td><td>MB231/MB468</td><td>MP</td></tr>
<tr><td>>hg38_21_+_8433172_8446622_RNA45SN1_45S_50ntFlanks@8362.8391.30|AACTTTGAAGAGAGAGTTCAAGAGGGCGTG</td><td>>hg38_21_+_8433172_8446622_RNA45SN1_45S_50ntFlanks</td><td>8362</td><td>8391</td><td>30</td><td>AACTTTGAAGAGAGAGTTCAAGAGGGCGTG</td><td>168.2</td><td>7.6</td><td>1.9</td><td>4.0</td><td>7.6E-05</td><td>2.6E-04</td><td>BT20/MB468</td><td>MP</td></tr>
<tr><td>>hg38_21_+_8433172_8446622_RNA45SN1_45S_50ntFlanks@12418.12446.29|CGATGTCGGCTCTTCCTATCATTGTGAAG</td><td>>hg38_21_+_8433172_8446622_RNA45SN1_45S_50ntFlanks</td><td>12418</td><td>12446</td><td>29</td><td>CGATGTCGGCTCTTCCTATCATTGTGAAG</td><td>4091.1</td><td>-2.4</td><td>0.6</td><td>-3.7</td><td>1.9E-04</td><td>2.6E-04</td><td>MB231/MB468</td><td>CYTO</td></tr>
<tr><td>>hg38_21_+_8433172_8446622_RNA45SN1_45S_50ntFlanks@7980.8006.27|CCTCAGATCAGACGTGGCGACCCGCTG</td><td>>hg38_21_+_8433172_8446622_RNA45SN1_45S_50ntFlanks</td><td>7980</td><td>8006</td><td>27</td><td>CCTCAGATCAGACGTGGCGACCCGCTG</td><td>2110.4</td><td>-3.7</td><td>1.0</td><td>-3.7</td><td>1.9E-04</td><td>2.6E-04</td><td>MB231/MB468</td><td>CYTO</td></tr>
<tr><td>>hg38_21_+_8433172_8446622_RNA45SN1_45S_50ntFlanks@12202.12220.19|CTAAGGCGAGCTCAGGGAG</td><td>>hg38_21_+_8433172_8446622_RNA45SN1_45S_50ntFlanks</td><td>12202</td><td>12220</td><td>19</td><td>CTAAGGCGAGCTCAGGGAG</td><td>636.5</td><td>-5.0</td><td>1.3</td><td>-3.8</td><td>1.5E-04</td><td>2.6E-04</td><td>BT20/MB468</td><td>MITO</td></tr>
<tr><td>>hg38_21_+_8433172_8446622_RNA45SN1_45S_50ntFlanks@12926.12956.31|ACCTGGCGCTAAACCATTCGTAGACGACCTG</td><td>>hg38_21_+_8433172_8446622_RNA45SN1_45S_50ntFlanks</td><td>12926</td><td>12956</td><td>31</td><td>ACCTGGCGCTAAACCATTCGTAGACGACCTG</td><td>443.3</td><td>-4.4</td><td>1.2</td><td>-3.8</td><td>1.5E-04</td><td>2.6E-04</td><td>BT20/MB468</td><td>MITO</td></tr>
<tr><td>>hg38_21_+_8433172_8446622_RNA45SN1_45S_50ntFlanks@10910.10928.19|CGCGCGCCGCGGCTGGACG</td><td>>hg38_21_+_8433172_8446622_RNA45SN1_45S_50ntFlanks</td><td>10910</td><td>10928</td><td>19</td><td>CGCGCGCCGCGGCTGGACG</td><td>306.5</td><td>-3.4</td><td>0.9</td><td>-3.7</td><td>2.0E-04</td><td>2.6E-04</td><td>MB231/MB468</td><td>NUC</td></tr>
<tr><td>>hg38_21_+_8433172_8446622_RNA45SN1_45S_50ntFlanks@9929.9948.20|GATATAGACAGCAGGACGGT</td><td>>hg38_21_+_8433172_8446622_RNA45SN1_45S_50ntFlanks</td><td>9929</td><td>9948</td><td>20</td><td>GATATAGACAGCAGGACGGT</td><td>1020.4</td><td>4.6</td><td>1.1</td><td>4.2</td><td>2.7E-05</td><td>2.6E-04</td><td>BT20/MB231</td><td>MP</td></tr>
<tr><td>>am_trnaMT_HisGTG_MT_+_12138_12206@19.50.32|AAACATCAGATTGTGAATCTGACAACAGAGGC</td><td>>am_trnaMT_HisGTG_MT_+_12138_12206</td><td>19</td><td>50</td><td>32</td><td>AAACATCAGATTGTGAATCTGACAACAGAGGC</td><td>23.4</td><td>7.8</td><td>2.1</td><td>3.8</td><td>1.7E-04</td><td>2.6E-04</td><td>MB231/MB468</td><td>MP</td></tr>
<tr><td>>hg38_21_+_8433172_8446622_RNA45SN1_45S_50ntFlanks@12633.12661.29|CGAAGCTACCATCTGTGGGATTATGACTG</td><td>>hg38_21_+_8433172_8446622_RNA45SN1_45S_50ntFlanks</td><td>12633</td><td>12661</td><td>29</td><td>CGAAGCTACCATCTGTGGGATTATGACTG</td><td>5946.7</td><td>-4.5</td><td>1.1</td><td>-4.0</td><td>7.7E-05</td><td>2.6E-04</td><td>BT20/MB468</td><td>MP</td></tr>
<tr><td>>hg38_21_+_8433172_8446622_RNA45SN1_45S_50ntFlanks@11029.11050.22|CTCTCTCTCTCTCCCCCGCTCC</td><td>>hg38_21_+_8433172_8446622_RNA45SN1_45S_50ntFlanks</td><td>11029</td><td>11050</td><td>22</td><td>CTCTCTCTCTCTCCCCCGCTCC</td><td>318.6</td><td>-5.9</td><td>1.6</td><td>-3.7</td><td>2.0E-04</td><td>2.6E-04</td><td>MB231/MB468</td><td>NUC</td></tr>
<tr><td>>MI0000088|hsa-mir-30a&WithFlank&6|-|71403545|71403627@12.35.24(+1C)&[MIMAT0000087&hsa-miR-30a-5p&offsets|0|+2(+1C);m-15&6|-|71403595|71403616&offsets|0|+2(+1C)]|TGTAAACATCCTCGACTGGAAGCTC</td><td>>MI0000088|hsa-mir-30a&WithFlank&6|-|71403545|71403627</td><td>12</td><td>35</td><td>24(+1C)</td><td>TGTAAACATCCTCGACTGGAAGCTC</td><td>5123.0</td><td>-4.4</td><td>1.1</td><td>-4.2</td><td>2.7E-05</td><td>2.6E-04</td><td>BT20/MB231</td><td>MP</td></tr>
<tr><td>>hg38_21_+_8433172_8446622_RNA45SN1_45S_50ntFlanks@6648.6685.38|GTACGACTCTTAGCGGTGGATCACTCGGCTCGTGCGTC</td><td>>hg38_21_+_8433172_8446622_RNA45SN1_45S_50ntFlanks</td><td>6648</td><td>6685</td><td>38</td><td>GTACGACTCTTAGCGGTGGATCACTCGGCTCGTGCGTC</td><td>358.6</td><td>4.2</td><td>1.1</td><td>4.0</td><td>7.7E-05</td><td>2.6E-04</td><td>BT20/MB468</td><td>MP</td></tr>
<tr><td>>hg38_21_+_8433172_8446622_RNA45SN1_45S_50ntFlanks@12406.12432.27|TTTTTGATCCTTCGATGTCGGCTCTTC</td><td>>hg38_21_+_8433172_8446622_RNA45SN1_45S_50ntFlanks</td><td>12406</td><td>12432</td><td>27</td><td>TTTTTGATCCTTCGATGTCGGCTCTTC</td><td>379.6</td><td>-4.3</td><td>1.1</td><td>-3.7</td><td>1.8E-04</td><td>2.6E-04</td><td>BT20/MB468</td><td>CYTO</td></tr>
<tr><td>>hg38_21_+_8433172_8446622_RNA45SN1_45S_50ntFlanks@8191.8214.24|CAGCCCGTGGACGGTGTGAGGCCG</td><td>>hg38_21_+_8433172_8446622_RNA45SN1_45S_50ntFlanks</td><td>8191</td><td>8214</td><td>24</td><td>CAGCCCGTGGACGGTGTGAGGCCG</td><td>656.4</td><td>-6.4</td><td>1.6</td><td>-4.0</td><td>7.7E-05</td><td>2.6E-04</td><td>BT20/MB468</td><td>MP</td></tr>
<tr><td>>hg38_21_+_8433172_8446622_RNA45SN1_45S_50ntFlanks@8115.8138.24|CGCGGGACATGTGGCGTACGGAAG</td><td>>hg38_21_+_8433172_8446622_RNA45SN1_45S_50ntFlanks</td><td>8115</td><td>8138</td><td>24</td><td>CGCGGGACATGTGGCGTACGGAAG</td><td>6541.4</td><td>-4.5</td><td>1.1</td><td>-4.0</td><td>7.7E-05</td><td>2.6E-04</td><td>BT20/MB468</td><td>MP</td></tr>
<tr><td>>hg38_MT_+_1621_3279_RNR2_16S_50ntFlanks@1508.1551.44|AAAGGACAAGAGAAATAAGGCCTACTTCACAAAGCGCCTTCCCC</td><td>>hg38_MT_+_1621_3279_RNR2_16S_50ntFlanks</td><td>1508</td><td>1551</td><td>44</td><td>AAAGGACAAGAGAAATAAGGCCTACTTCACAAAGCGCCTTCCCC</td><td>95.9</td><td>10.3</td><td>2.6</td><td>4.0</td><td>7.7E-05</td><td>2.6E-04</td><td>BT20/MB468</td><td>MP</td></tr>
<tr><td>>hg38_MT_+_1621_3279_RNR2_16S_50ntFlanks@252.282.31|TAACTTTGCAAGGAGAGCCAAAGCTAAGACC</td><td>>hg38_MT_+_1621_3279_RNR2_16S_50ntFlanks</td><td>252</td><td>282</td><td>31</td><td>TAACTTTGCAAGGAGAGCCAAAGCTAAGACC</td><td>101.7</td><td>10.4</td><td>2.6</td><td>4.0</td><td>7.7E-05</td><td>2.6E-04</td><td>BT20/MB468</td><td>MP</td></tr>
<tr><td>>hg38_21_+_8433172_8446622_RNA45SN1_45S_50ntFlanks@12416.12455.40|TTCGATGTCGGCTCTTCCTATCATTGTGAAGCAGAATTCA</td><td>>hg38_21_+_8433172_8446622_RNA45SN1_45S_50ntFlanks</td><td>12416</td><td>12455</td><td>40</td><td>TTCGATGTCGGCTCTTCCTATCATTGTGAAGCAGAATTCA</td><td>693.8</td><td>-7.2</td><td>1.9</td><td>-3.7</td><td>2.1E-04</td><td>2.6E-04</td><td>BT20/MB468</td><td>NUC</td></tr>
<tr><td>>hg38_21_+_8433172_8446622_RNA45SN1_45S_50ntFlanks@7984.8016.33|AGATCAGACGTGGCGACCCGCTGAATTTAAGCA</td><td>>hg38_21_+_8433172_8446622_RNA45SN1_45S_50ntFlanks</td><td>7984</td><td>8016</td><td>33</td><td>AGATCAGACGTGGCGACCCGCTGAATTTAAGCA</td><td>323.2</td><td>-3.3</td><td>0.9</td><td>-3.7</td><td>2.0E-04</td><td>2.6E-04</td><td>MB231/MB468</td><td>MITO</td></tr>
<tr><td>>hg38_21_+_8433172_8446622_RNA45SN1_45S_50ntFlanks@11314.11338.25|CCCACCCCACGTCTCGTCGCGCGCG</td><td>>hg38_21_+_8433172_8446622_RNA45SN1_45S_50ntFlanks</td><td>11314</td><td>11338</td><td>25</td><td>CCCACCCCACGTCTCGTCGCGCGCG</td><td>593.7</td><td>-6.7</td><td>1.8</td><td>-3.7</td><td>2.1E-04</td><td>2.6E-04</td><td>BT20/MB468</td><td>NUC</td></tr>
<tr><td>>hg38_21_+_8433172_8446622_RNA45SN1_45S_50ntFlanks@12602.12626.25|GTGTATGTGCTTGGCTGAGGAGCCA</td><td>>hg38_21_+_8433172_8446622_RNA45SN1_45S_50ntFlanks</td><td>12602</td><td>12626</td><td>25</td><td>GTGTATGTGCTTGGCTGAGGAGCCA</td><td>280.4</td><td>-4.1</td><td>1.1</td><td>-3.7</td><td>1.8E-04</td><td>2.6E-04</td><td>BT20/MB468</td><td>CYTO</td></tr>
<tr><td>>MI0000750|hsa-mir-26a-2&WithFlank&12|-|57824603|57824698@20.40.21(+1G);MI0000083|hsa-mir-26a-1&WithFlank&3|+|37969398|37969486@16.36.21(+1G)&[MIMAT0000082&hsa-miR-26a-5p&offsets|0|-1(+1G);m-33&12|-|57824658|57824679&offsets|0|-1(+1G)];[MIMAT0000082_1&hsa-miR-26a-5p&offsets|0|-1(+1G);m-34&3|+|37969413|37969434&offsets|0|-1(+1G)]|TTCAAGTAATCCAGGATAGGCG</td><td>>MI0000750|hsa-mir-26a-2&WithFlank&12|-|57824603|57824698</td><td>20</td><td>40</td><td>21(+1G)</td><td>TTCAAGTAATCCAGGATAGGCG</td><td>1763.7</td><td>3.7</td><td>1.0</td><td>3.7</td><td>1.9E-04</td><td>2.6E-04</td><td>MB231/MB468</td><td>CYTO</td></tr>
<tr><td>>hg38_21_+_8433172_8446622_RNA45SN1_45S_50ntFlanks@7973.8011.39|GACGCGACCTCAGATCAGACGTGGCGACCCGCTGAATTT</td><td>>hg38_21_+_8433172_8446622_RNA45SN1_45S_50ntFlanks</td><td>7973</td><td>8011</td><td>39</td><td>GACGCGACCTCAGATCAGACGTGGCGACCCGCTGAATTT</td><td>2257.4</td><td>2.6</td><td>0.7</td><td>3.7</td><td>2.0E-04</td><td>2.6E-04</td><td>MB231/MB468</td><td>MITO</td></tr>
<tr><td>>hg38_21_+_8433172_8446622_RNA45SN1_45S_50ntFlanks@8176.8212.37|CTTCTGATCGAGGCCCAGCCCGTGGACGGTGTGAGGC</td><td>>hg38_21_+_8433172_8446622_RNA45SN1_45S_50ntFlanks</td><td>8176</td><td>8212</td><td>37</td><td>CTTCTGATCGAGGCCCAGCCCGTGGACGGTGTGAGGC</td><td>958.0</td><td>-3.9</td><td>1.0</td><td>-4.0</td><td>7.7E-05</td><td>2.6E-04</td><td>BT20/MB468</td><td>MP</td></tr>
<tr><td>>hg38_21_+_8433172_8446622_RNA45SN1_45S_50ntFlanks@4207.4242.36|CGGGGAGGTAGTGACGAAAAATAACAATACAGGACT</td><td>>hg38_21_+_8433172_8446622_RNA45SN1_45S_50ntFlanks</td><td>4207</td><td>4242</td><td>36</td><td>CGGGGAGGTAGTGACGAAAAATAACAATACAGGACT</td><td>146.8</td><td>7.3</td><td>1.8</td><td>4.0</td><td>7.7E-05</td><td>2.6E-04</td><td>BT20/MB468</td><td>MP</td></tr>
<tr><td>>hg38_21_+_8433172_8446622_RNA45SN1_45S_50ntFlanks@11209.11233.25|CGGCGACTCTGGACGCGAGCCGGGC</td><td>>hg38_21_+_8433172_8446622_RNA45SN1_45S_50ntFlanks</td><td>11209</td><td>11233</td><td>25</td><td>CGGCGACTCTGGACGCGAGCCGGGC</td><td>2820.0</td><td>-5.9</td><td>1.6</td><td>-3.7</td><td>2.1E-04</td><td>2.6E-04</td><td>BT20/MB468</td><td>NUC</td></tr>
<tr><td>>hg38_21_+_8433172_8446622_RNA45SN1_45S_50ntFlanks@4724.4752.29|ATTAATCAAGAACGAAAGTCGGAGGTTCG</td><td>>hg38_21_+_8433172_8446622_RNA45SN1_45S_50ntFlanks</td><td>4724</td><td>4752</td><td>29</td><td>ATTAATCAAGAACGAAAGTCGGAGGTTCG</td><td>928.2</td><td>-6.4</td><td>1.6</td><td>-4.0</td><td>7.7E-05</td><td>2.6E-04</td><td>BT20/MB468</td><td>MP</td></tr>
<tr><td>>hg38_21_+_8433172_8446622_RNA45SN1_45S_50ntFlanks@11718.11738.21|AACGGCGGGAGTAACTATGAC</td><td>>hg38_21_+_8433172_8446622_RNA45SN1_45S_50ntFlanks</td><td>11718</td><td>11738</td><td>21</td><td>AACGGCGGGAGTAACTATGAC</td><td>263.4</td><td>-4.1</td><td>1.1</td><td>-3.8</td><td>1.5E-04</td><td>2.7E-04</td><td>BT20/MB468</td><td>MITO</td></tr>
<tr><td>>hg38_21_+_8433172_8446622_RNA45SN1_45S_50ntFlanks@10866.10883.18|CTAAGGGCTGGGTCGGTC</td><td>>hg38_21_+_8433172_8446622_RNA45SN1_45S_50ntFlanks</td><td>10866</td><td>10883</td><td>18</td><td>CTAAGGGCTGGGTCGGTC</td><td>254.1</td><td>-3.6</td><td>1.0</td><td>-3.7</td><td>1.8E-04</td><td>2.7E-04</td><td>BT20/MB468</td><td>CYTO</td></tr>
<tr><td>>hg38_MT_+_1621_3279_RNR2_16S_50ntFlanks@1094.1117.24|AAGACGAGAAGACCCTATGGAGCT</td><td>>hg38_MT_+_1621_3279_RNR2_16S_50ntFlanks</td><td>1094</td><td>1117</td><td>24</td><td>AAGACGAGAAGACCCTATGGAGCT</td><td>155.3</td><td>3.5</td><td>0.9</td><td>4.0</td><td>7.8E-05</td><td>2.7E-04</td><td>BT20/MB468</td><td>MP</td></tr>
<tr><td>>trnaMT_HisGTG_MT_+_12138_12206@26.65.40|AGATTGTGAATCTGACAACAGAGGCTTACGACCCCTTATT</td><td>>trnaMT_HisGTG_MT_+_12138_12206</td><td>26</td><td>65</td><td>40</td><td>AGATTGTGAATCTGACAACAGAGGCTTACGACCCCTTATT</td><td>100.7</td><td>10.4</td><td>2.6</td><td>4.0</td><td>7.8E-05</td><td>2.7E-04</td><td>BT20/MB468</td><td>MP</td></tr>
<tr><td>>hg38_1_-_228634819_228635039_RNA5S12_5S_50ntFlanks@63.89.27|ACCACCCTGAACGCGCCCGATCTCGTC</td><td>>hg38_1_-_228634819_228635039_RNA5S12_5S_50ntFlanks</td><td>63</td><td>89</td><td>27</td><td>ACCACCCTGAACGCGCCCGATCTCGTC</td><td>361.2</td><td>-3.4</td><td>0.8</td><td>-4.2</td><td>2.6E-05</td><td>2.7E-04</td><td>BT20/MB231</td><td>CYTO</td></tr>
<tr><td>>hg38_21_+_8433172_8446622_RNA45SN1_45S_50ntFlanks@12996.13040.45|TCGCTGCGATCTATTGAAAGTCAGCCCTCGACACAAGGGTTTGTC</td><td>>hg38_21_+_8433172_8446622_RNA45SN1_45S_50ntFlanks</td><td>12996</td><td>13040</td><td>45</td><td>TCGCTGCGATCTATTGAAAGTCAGCCCTCGACACAAGGGTTTGTC</td><td>476.2</td><td>6.8</td><td>1.8</td><td>3.7</td><td>2.1E-04</td><td>2.7E-04</td><td>MB231/MB468</td><td>MITO</td></tr>
<tr><td>>hg38_1_-_228634819_228635039_RNA5S12_5S_50ntFlanks@146.169.24|TGGGAATACCGGGTGCTGTAGGCT</td><td>>hg38_1_-_228634819_228635039_RNA5S12_5S_50ntFlanks</td><td>146</td><td>169</td><td>24</td><td>TGGGAATACCGGGTGCTGTAGGCT</td><td>356.6</td><td>2.9</td><td>0.7</td><td>4.2</td><td>3.2E-05</td><td>2.7E-04</td><td>BT20/MB231</td><td>MITO</td></tr>
<tr><td>>hg38_21_+_8433172_8446622_RNA45SN1_45S_50ntFlanks@6660.6689.30|GCGGTGGATCACTCGGCTCGTGCGTCGATG</td><td>>hg38_21_+_8433172_8446622_RNA45SN1_45S_50ntFlanks</td><td>6660</td><td>6689</td><td>30</td><td>GCGGTGGATCACTCGGCTCGTGCGTCGATG</td><td>542.8</td><td>3.2</td><td>0.8</td><td>4.2</td><td>3.2E-05</td><td>2.7E-04</td><td>BT20/MB231</td><td>MITO</td></tr>
<tr><td>>MI0000487|hsa-mir-193a&WithFlank&17|+|31559990|31560089@27.46.20(+1A)&[MIMAT0004614&hsa-miR-193a-5p&offsets|0|-2(+1A);m-123&17|+|31560016|31560037&offsets|0|-2(+1A)]|TGGGTCTTTGCGGGCGAGATA</td><td>>MI0000487|hsa-mir-193a&WithFlank&17|+|31559990|31560089</td><td>27</td><td>46</td><td>20(+1A)</td><td>TGGGTCTTTGCGGGCGAGATA</td><td>191.2</td><td>7.7</td><td>1.9</td><td>4.0</td><td>7.8E-05</td><td>2.7E-04</td><td>BT20/MB468</td><td>MP</td></tr>
<tr><td>>MI0000285|hsa-mir-205&WithFlank&1|+|209432127|209432248@40.62.23&[MIMAT0000266&hsa-miR-205-5p&offsets|0|+1;m-351&1|+|209432166|209432187&offsets|0|+1]|TCCTTCATTCCACCGGAGTCTGT</td><td>>MI0000285|hsa-mir-205&WithFlank&1|+|209432127|209432248</td><td>40</td><td>62</td><td>23</td><td>TCCTTCATTCCACCGGAGTCTGT</td><td>1057.2</td><td>-2.9</td><td>0.8</td><td>-3.7</td><td>1.9E-04</td><td>2.7E-04</td><td>MB231/MB468</td><td>CYTO</td></tr>
</table>

| | | | | | | | | | | | | | |
|---|---|---|---|---|---|---|---|---|---|---|---|---|---|
| >hg38_21_+_8433172_8446622_RNA45SN1_45S_50ntFlanks@4725.4751.27\|TTAATCAAGAACGAAAGTCGGAGGTTC | >hg38_21_+_8433172_8446622_RNA45SN1_45S_50ntFlanks | 4725 | 4751 | 27 | TTAATCAAGAACGAAAGTCGGAGGTTC | 153.9 | -5.8 | 1.5 | -4.0 | 7.8E-05 | 2.7E-04 | BT20/MB468 | MP |
| >hg38_21_+_8433172_8446622_RNA45SN1_45S_50ntFlanks@12934.12959.26\|CTAAACCATTCGTAGACGACCTGCTT | >hg38_21_+_8433172_8446622_RNA45SN1_45S_50ntFlanks | 12934 | 12959 | 26 | CTAAACCATTCGTAGACGACCTGCTT | 606.2 | -2.5 | 0.7 | -3.7 | 1.9E-04 | 2.7E-04 | MB231/MB468 | CYTO |
| >hg38_21_+_8433172_8446622_RNA45SN1_45S_50ntFlanks@6647.6674.28\|CGTACGACTCTTAGCGGTGGATCACTCG | >hg38_21_+_8433172_8446622_RNA45SN1_45S_50ntFlanks | 6647 | 6674 | 28 | CGTACGACTCTTAGCGGTGGATCACTCG | 316.5 | 3.2 | 0.8 | 4.0 | 7.8E-05 | 2.7E-04 | BT20/MB468 | MP |
| >hg38_MT_+_1621_3279_RNR2_16S_50ntFlanks@931.961.31\|GTGACACATGTTTAACGGCCGCGGTACCCTA | >hg38_MT_+_1621_3279_RNR2_16S_50ntFlanks | 931 | 961 | 31 | GTGACACATGTTTAACGGCCGCGGTACCCTA | 23.4 | 7.8 | 2.1 | 3.8 | 1.7E-04 | 2.7E-04 | MB231/MB468 | MP |
| >hg38_21_+_8433172_8446622_RNA45SN1_45S_50ntFlanks@3882.3900.19\|CCGACGGGCGCTGACCCCC | >hg38_21_+_8433172_8446622_RNA45SN1_45S_50ntFlanks | 3882 | 3900 | 19 | CCGACGGGCGCTGACCCCC | 74.3 | -5.5 | 1.5 | -3.8 | 1.7E-04 | 2.7E-04 | MB231/MB468 | MP |
| >am_trnaMT_ValTAC_MT_+_1602_1670@49.66.18\|AACTTAACTTGACCGCTC | >am_trnaMT_ValTAC_MT_+_1602_1670 | 49 | 66 | 18 | AACTTAACTTGACCGCTC | 376.6 | 3.8 | 0.9 | 4.2 | 2.6E-05 | 2.7E-04 | BT20/MB231 | CYTO |
| >hg38_21_+_8433172_8446622_RNA45SN1_45S_50ntFlanks@12435.12454.20\|ATCATTGTGAAGCAGAATTC | >hg38_21_+_8433172_8446622_RNA45SN1_45S_50ntFlanks | 12435 | 12454 | 20 | ATCATTGTGAAGCAGAATTC | 1066.4 | -6.3 | 1.7 | -3.8 | 1.7E-04 | 2.7E-04 | MB231/MB468 | MP |
| >trna10_AspGTC_12_-_125424193_125424264@31.71.41,trna12_AspGTC_12_-_125411891_125411962@31.71.41,trna38_AspGTC_17_-_8125556_8125627@31.71.41,trna45_AspGTC_6_+_27447453_27447524@31.71.41,trna48_AspGTC_6_+_27471523_27471594@31.71.41,trna4_AspGTC_12_+_96429799_96429870@31.71.41,trna69_AspGTC_1_-_161440205_161440276@31.71.41,trna72_AspGTC_1_-_161432824_161432895@31.71.41,trna75_AspGTC_1_-_161425414_161425485@31.71.41,trna78_AspGTC_1_-_161418033_161418104@31.71.41,trna81_AspGTC_1_-_161410615_161410686@31.71.41\|CCTGTCACGCGGGAGACCGGGGTTCGATTCCCCGACGGGGA | >trna10_AspGTC_12_-_125424193_125424264 | 31 | 71 | 41 | CCTGTCACGCGGGAGACCGGGGTTCGATTCCCCGACGGGGA | 202.7 | -5.2 | 1.4 | -3.7 | 1.8E-04 | 2.7E-04 | BT20/MB468 | CYTO |
| >MI0001445\|hsa-mir-423&WithFlank&17\|+\|30117073\|30117178@23.42.20(+1C)&[MIMAT0004748&hsa-miR-423-5p&offsets\|0\|-3(+1C);m-141&17\|+\|30117095\|30117117&offsets\|0\|-3(+1C)]\|TGAGGGGCAGAGAGCGAGACC | >MI0001445\|hsa-mir-423&WithFlank&17\|+\|30117073\|30117178 | 23 | 42 | 20(+1C) | TGAGGGGCAGAGAGCGAGACC | 392.0 | 4.8 | 1.3 | 3.8 | 1.7E-04 | 2.7E-04 | MB231/MB468 | MP |
| >hg38_21_+_8433172_8446622_RNA45SN1_45S_50ntFlanks@10574.10608.35\|AAGCCGGCGGGAGCCCCGGGGAGAGTTCTCTTTTC | >hg38_21_+_8433172_8446622_RNA45SN1_45S_50ntFlanks | 10574 | 10608 | 35 | AAGCCGGCGGGAGCCCCGGGGAGAGTTCTCTTTTC | 310.4 | -4.0 | 1.1 | -3.8 | 1.5E-04 | 2.7E-04 | BT20/MB468 | MITO |
| >trna10_LysCTT_16_+_3241501_3241573@3.34.32,trna13_LysCTT_14_-_58706613_58706685@3.34.32,trna2_LysCTT_15_+_79152904_79152976@3.34.32\|CCGGCTAGCTCAGTCGGTAGAGCATGGGACTC | >trna10_LysCTT_16_+_3241501_3241573 | 3 | 34 | 32 | CCGGCTAGCTCAGTCGGTAGAGCATGGGACTC | 114.8 | 6.5 | 1.7 | 3.8 | 1.7E-04 | 2.7E-04 | MB231/MB468 | MP |
| >hg38_21_+_8433172_8446622_RNA45SN1_45S_50ntFlanks@8019.8046.28\|TTAGTCAGCGGAGGAGAAGAAACTAACC | >hg38_21_+_8433172_8446622_RNA45SN1_45S_50ntFlanks | 8019 | 8046 | 28 | TTAGTCAGCGGAGGAGAAGAAACTAACC | 316.3 | -4.7 | 1.1 | -4.2 | 2.7E-05 | 2.7E-04 | BT20/MB231 | CYTO |
| >hg38_21_+_8433172_8446622_RNA45SN1_45S_50ntFlanks@9531.9573.43\|CAGAGGAAACTCTGGTGGAGGTCCGTAGCGGTCCTGACGTGCA | >hg38_21_+_8433172_8446622_RNA45SN1_45S_50ntFlanks | 9531 | 9573 | 43 | CAGAGGAAACTCTGGTGGAGGTCCGTAGCGGTCCTGACGTGCA | 178.9 | -3.3 | 0.8 | -4.2 | 2.7E-05 | 2.7E-04 | BT20/MB231 | CYTO |
| >hg38_21_+_8433172_8446622_RNA45SN1_45S_50ntFlanks@9572.9603.32\|CAAATCGGTCGTCCGACCTGGGTATAGGGGCG | >hg38_21_+_8433172_8446622_RNA45SN1_45S_50ntFlanks | 9572 | 9603 | 32 | CAAATCGGTCGTCCGACCTGGGTATAGGGGCG | 4859.2 | -4.4 | 1.1 | -4.0 | 7.8E-05 | 2.7E-04 | BT20/MB468 | MP |
| >MI0000736\|hsa-mir-30c-1&WithFlank&1\|+\|40757278\|40757378@23.43.21;MI0000254\|hsa-mir-30c-2&WithFlank&6\|-\|71376954\|71377037@13.33.21&[MIMAT0000244&hsa-miR-30c-5p&offsets\|0\|-2;m-57&1\|+\|40757300\|40757323&offsets\|0\|-3];[MIMAT0000244_1&hsa-miR-30c-5p&offsets\|0\|-2;m-56&6\|-\|71377002\|71377025&offsets\|0\|-3]\|TGTAAACATCCTACACTCTCA | >MI0000736\|hsa-mir-30c-1&WithFlank&1\|+\|40757278\|40757378 | 23 | 43 | 21 | TGTAAACATCCTACACTCTCA | 270.0 | 4.9 | 1.3 | 3.8 | 1.7E-04 | 2.7E-04 | MB231/MB468 | MP |
| >hg38_21_+_8433172_8446622_RNA45SN1_45S_50ntFlanks@485.522.38\|GATGTGGTGACGTCGTGCTCTCCCGGGCCGGGTCCGAG | >hg38_21_+_8433172_8446622_RNA45SN1_45S_50ntFlanks | 485 | 522 | 38 | GATGTGGTGACGTCGTGCTCTCCCGGGCCGGGTCCGAG | 86.2 | 9.7 | 2.6 | 3.8 | 1.7E-04 | 2.7E-04 | MB231/MB468 | MP |
| >hg38_21_+_8433172_8446622_RNA45SN1_45S_50ntFlanks@11768.11804.37\|CTAATTAGTGACGCGCATGAATGGATGAACGAGATTC | >hg38_21_+_8433172_8446622_RNA45SN1_45S_50ntFlanks | 11768 | 11804 | 37 | CTAATTAGTGACGCGCATGAATGGATGAACGAGATTC | 326.4 | -2.5 | 0.6 | -3.8 | 1.5E-04 | 2.7E-04 | BT20/MB468 | MITO |
| >hg38_21_+_8433172_8446622_RNA45SN1_45S_50ntFlanks@11569.11604.36\|CAGCCGACTTAGAACTGGTGCGGACCAGGGGAATCC | >hg38_21_+_8433172_8446622_RNA45SN1_45S_50ntFlanks | 11569 | 11604 | 36 | CAGCCGACTTAGAACTGGTGCGGACCAGGGGAATCC | 316.8 | -4.4 | 1.2 | -3.7 | 2.1E-04 | 2.7E-04 | MB231/MB468 | MITO |
| >hg38_21_+_8433172_8446622_RNA45SN1_45S_50ntFlanks@12455.12474.20\|ACCAAGCGTTGGATTGTTCA | >hg38_21_+_8433172_8446622_RNA45SN1_45S_50ntFlanks | 12455 | 12474 | 20 | ACCAAGCGTTGGATTGTTCA | 138.6 | -6.1 | 1.5 | -3.9 | 7.8E-05 | 2.7E-04 | BT20/MB468 | MP |
| >hg38_21_+_8433172_8446622_RNA45SN1_45S_50ntFlanks@9122.9152.31\|CCGCGCGCGGGTCGGGGGGCGGGGCGGACTG | >hg38_21_+_8433172_8446622_RNA45SN1_45S_50ntFlanks | 9122 | 9152 | 31 | CCGCGCGCGGGTCGGGGGGCGGGGCGGACTG | 178.8 | -5.5 | 1.5 | -3.8 | 1.5E-04 | 2.7E-04 | BT20/MB468 | MITO |
| >hg38_MT_+_1621_3279_RNR2_16S_50ntFlanks@1363.1401.39\|GACCTCGATGTTGGATCAGGACATCCCGATGGTGCAGCC | >hg38_MT_+_1621_3279_RNR2_16S_50ntFlanks | 1363 | 1401 | 39 | GACCTCGATGTTGGATCAGGACATCCCGATGGTGCAGCC | 22.8 | 7.7 | 2.1 | 3.8 | 1.7E-04 | 2.7E-04 | MB231/MB468 | MP |
| >hg38_21_+_8433172_8446622_RNA45SN1_45S_50ntFlanks@11918.11940.23\|ACATGAGAGGTGTAGAATAAGTG | >hg38_21_+_8433172_8446622_RNA45SN1_45S_50ntFlanks | 11918 | 11940 | 23 | ACATGAGAGGTGTAGAATAAGTG | 329.9 | 3.6 | 0.9 | 3.9 | 7.8E-05 | 2.7E-04 | BT20/MB468 | MP |
| >hg38_21_+_8433172_8446622_RNA45SN1_45S_50ntFlanks@10792.10815.24\|CTCTGGCATGTTGGAACAATGTAG | >hg38_21_+_8433172_8446622_RNA45SN1_45S_50ntFlanks | 10792 | 10815 | 24 | CTCTGGCATGTTGGAACAATGTAG | 237.8 | -2.7 | 0.7 | -3.7 | 2.1E-04 | 2.7E-04 | MB231/MB468 | NUC |
| >hg38_1_-_228634819_228635039_RNA5S12_5S_50ntFlanks@153.170.18\|ACCGGGTGCTGTAGGCTT | >hg38_1_-_228634819_228635039_RNA5S12_5S_50ntFlanks | 153 | 170 | 18 | ACCGGGTGCTGTAGGCTT | 64908.4 | 4.7 | 1.2 | 3.9 | 7.9E-05 | 2.7E-04 | BT20/MB468 | MP |
| >hg38_21_+_8433172_8446622_RNA45SN1_45S_50ntFlanks@11669.11702.34\|GCCCAGTGCTCTGAATGTCAAAGTGAAGAAATTC | >hg38_21_+_8433172_8446622_RNA45SN1_45S_50ntFlanks | 11669 | 11702 | 34 | GCCCAGTGCTCTGAATGTCAAAGTGAAGAAATTC | 908.6 | -3.4 | 0.9 | -3.7 | 2.1E-04 | 2.7E-04 | BT20/MB468 | NUC |
| >am_MI0000086\|hsa-mir-28&WithFlank&3\|+\|188688775\|188688872@60.77.18(+1A)&[MIMAT0004502&hsa-miR-28-3p&offsets\|0\|-4(+1A);m-28&3\|+\|188688834\|188688855&offsets\|0\|-4(+1A)]\|CACTAGATTGTGAGCTCCA | >am_MI0000086\|hsa-mir-28&WithFlank&3\|+\|188688775\|188688872 | 60 | 77 | 18(+1A) | CACTAGATTGTGAGCTCCA | 25.8 | 7.9 | 2.1 | 3.8 | 1.7E-04 | 2.7E-04 | MB231/MB468 | MP |
| >hg38_21_+_8433172_8446622_RNA45SN1_45S_50ntFlanks@5492.5510.19\|AGAAGACGGTCGAACTTGA | >hg38_21_+_8433172_8446622_RNA45SN1_45S_50ntFlanks | 5492 | 5510 | 19 | AGAAGACGGTCGAACTTGA | 250.4 | -2.7 | 0.7 | -3.7 | 2.0E-04 | 2.7E-04 | MB231/MB468 | CYTO |
| >hg38_21_+_8433172_8446622_RNA45SN1_45S_50ntFlanks@11602.11629.28\|TCCGACTGTTTAATTAAAACAAAGCATC | >hg38_21_+_8433172_8446622_RNA45SN1_45S_50ntFlanks | 11602 | 11629 | 28 | TCCGACTGTTTAATTAAAACAAAGCATC | 75.9 | 9.6 | 2.3 | 4.2 | 2.8E-05 | 2.7E-04 | BT20/MB231 | MP |
| >trna146_GlnCTG_6_-_27515531_27515602@14.41.28,trna1_GlnCTG_6_+_18836402_18836473@14.41.28,trna3_GlnCTG_17_+_8023070_8023141@14.41.28,trna42_GlnCTG_6_+_27263212_27263283@14.41.28,trna49_GlnCTG_6_+_27487308_27487379@14.41.28,trna7_GlnCTG_15_-_66161400_66161471@14.41.28,trna99_GlnCTG_6_-_28909378_28909449@14.41.28\|AATGGTTAGCACTCTGGACTCTGAATCC | >trna146_GlnCTG_6_-_27515531_27515602 | 14 | 41 | 28 | AATGGTTAGCACTCTGGACTCTGAATCC | 100.7 | 10.4 | 2.6 | 3.9 | 7.9E-05 | 2.7E-04 | BT20/MB468 | MP |
| >hg38_21_+_8433172_8446622_RNA45SN1_45S_50ntFlanks@9535.9571.37\|GGAAACTCTGGTGGAGGTCCGTAGCGGTCCTGACGTG | >hg38_21_+_8433172_8446622_RNA45SN1_45S_50ntFlanks | 9535 | 9571 | 37 | GGAAACTCTGGTGGAGGTCCGTAGCGGTCCTGACGTG | 857.9 | -3.5 | 0.8 | -4.2 | 2.7E-05 | 2.7E-04 | BT20/MB231 | CYTO |
| >hg38_21_+_8433172_8446622_RNA45SN1_45S_50ntFlanks@8012.8042.31\|AAGCATATTAGTCAGCGGAGGAGAAGAAACT | >hg38_21_+_8433172_8446622_RNA45SN1_45S_50ntFlanks | 8012 | 8042 | 31 | AAGCATATTAGTCAGCGGAGGAGAAGAAACT | 283.9 | -3.9 | 0.9 | -4.2 | 2.7E-05 | 2.7E-04 | BT20/MB231 | CYTO |
| >MI0000264\|hsa-mir-7-2&WithFlank&15\|+\|88611819\|88611940@38.57.20(+1C);MI0000265\|hsa-mir-7-3&WithFlank&19\|+\|4770664\|4770785@37.56.20(+1C);MI0000263\|hsa-mir-7-1&WithFlank&9\|-\|83969742\|83969863@30.49.20(+1C)&[MIMAT0000252&hsa-miR-7-5p&offsets\|0\|-4(+1C);m-400&15\|+\|88611856\|88611879&offsets\|0\|-4(+1C)];[MIMAT0000252_1&hsa-miR-7-5p&offsets\|0\|-4(+1C);m-402&19\|+\|4770700\|4770723&offsets\|0\|-4(+1C)];[MIMAT0000252_2&hsa-miR-7-5p&offsets\|0\|-4(+1C);m-401&9\|-\|83969811\|83969834&offsets\|0\|-4(+1C)]\|TGGAAGACTAGTGATTTTGTC | >MI0000264\|hsa-mir-7-2&WithFlank&15\|+\|88611819\|88611940 | 38 | 57 | 20(+1C) | TGGAAGACTAGTGATTTTGTC | 272.1 | 4.5 | 1.2 | 3.8 | 1.8E-04 | 2.7E-04 | MB231/MB468 | MP |
| >hg38_21_+_8433172_8446622_RNA45SN1_45S_50ntFlanks@10349.10376.28\|AACGAGAACTTTGAAGGCCGAAGTGGAG | >hg38_21_+_8433172_8446622_RNA45SN1_45S_50ntFlanks | 10349 | 10376 | 28 | AACGAGAACTTTGAAGGCCGAAGTGGAG | 1308.8 | -4.5 | 1.1 | -3.9 | 7.9E-05 | 2.7E-04 | BT20/MB468 | MP |
| >hg38_21_+_8433172_8446622_RNA45SN1_45S_50ntFlanks@9252.9273.22\|CACGGGGTCGGCGGCGACGTCG | >hg38_21_+_8433172_8446622_RNA45SN1_45S_50ntFlanks | 9252 | 9273 | 22 | CACGGGGTCGGCGGCGACGTCG | 267.7 | -3.0 | 0.8 | -3.8 | 1.5E-04 | 2.7E-04 | BT20/MB468 | MITO |
| >hg38_21_+_8433172_8446622_RNA45SN1_45S_50ntFlanks@9564.9594.31\|CTGACGTGCAAATCGGTCGTCCGACCTGGGT | >hg38_21_+_8433172_8446622_RNA45SN1_45S_50ntFlanks | 9564 | 9594 | 31 | CTGACGTGCAAATCGGTCGTCCGACCTGGGT | 911.3 | -2.0 | 0.5 | -3.8 | 1.5E-04 | 2.7E-04 | BT20/MB468 | MITO |
| >hg38_21_+_8433172_8446622_RNA45SN1_45S_50ntFlanks@8528.8549.22\|CTTCCCCCGCCGCCCCTCCTCC | >hg38_21_+_8433172_8446622_RNA45SN1_45S_50ntFlanks | 8528 | 8549 | 22 | CTTCCCCCGCCGCCCCTCCTCC | 340.9 | -6.6 | 1.8 | -3.7 | 2.1E-04 | 2.7E-04 | BT20/MB468 | NUC |
| >hg38_21_+_8433172_8446622_RNA45SN1_45S_50ntFlanks@11586.11609.24\|GTGCGGACCAGGGGAATCCGACTG | >hg38_21_+_8433172_8446622_RNA45SN1_45S_50ntFlanks | 11586 | 11609 | 24 | GTGCGGACCAGGGGAATCCGACTG | 590.7 | -2.7 | 0.7 | -3.7 | 1.9E-04 | 2.7E-04 | BT20/MB468 | CYTO |
| >hg38_21_+_8433172_8446622_RNA45SN1_45S_50ntFlanks@6649.6674.26\|TACGACTCTTAGCGGTGGATCACTCG | >hg38_21_+_8433172_8446622_RNA45SN1_45S_50ntFlanks | 6649 | 6674 | 26 | TACGACTCTTAGCGGTGGATCACTCG | 322.7 | 3.7 | 1.0 | 3.7 | 2.1E-04 | 2.7E-04 | MB231/MB468 | NUC |
| >MI0000101\|hsa-mir-99a&WithFlank&21\|+\|16539083\|16539175@19.40.22(+1A)&[MIMAT0000097&hsa-miR-99a-5p&offsets\|0\|0(+1A);m-64&21\|+\|16539101\|16539121&offsets\|0\|+1(+1A)]\|AACCCGTAGATCCGATCTTGTGA | >MI0000101\|hsa-mir-99a&WithFlank&21\|+\|16539083\|16539175 | 19 | 40 | 22(+1A) | AACCCGTAGATCCGATCTTGTGA | 959.8 | -6.7 | 1.7 | -3.9 | 7.9E-05 | 2.7E-04 | BT20/MB468 | MP |
| >MI0003188\|hsa-mir-503&WithFlank&X\|-\|134546322\|134546404@12.29.18(+1C)&[MIMAT0002874&hsa-miR-503-5p&offsets\|0\|-5(+1C);m-297&X\|-\|134546371\|134546393&offsets\|0\|-5(+1C)]\|TAGCAGCGGGAACAGTTCC | >MI0003188\|hsa-mir-503&WithFlank&X\|-\|134546322\|134546404 | 12 | 29 | 18(+1C) | TAGCAGCGGGAACAGTTCC | 25.6 | 7.9 | 2.1 | 3.8 | 1.8E-04 | 2.7E-04 | MB231/MB468 | MP |
| >hg38_1_-_228634819_228635039_RNA5S12_5S_50ntFlanks@136.171.36\|GGAGACCGCCTGGGAATACCGGGTGCTGTAGGCTTT | >hg38_1_-_228634819_228635039_RNA5S12_5S_50ntFlanks | 136 | 171 | 36 | GGAGACCGCCTGGGAATACCGGGTGCTGTAGGCTTT | 2193.7 | 4.1 | 1.1 | 3.7 | 1.9E-04 | 2.7E-04 | BT20/MB468 | CYTO |
| >hg38_21_+_8433172_8446622_RNA45SN1_45S_50ntFlanks@11709.11728.20\|GCGCGGGTAAACGGCGGGAG | >hg38_21_+_8433172_8446622_RNA45SN1_45S_50ntFlanks | 11709 | 11728 | 20 | GCGCGGGTAAACGGCGGGAG | 165.6 | -5.2 | 1.4 | -3.7 | 2.1E-04 | 2.7E-04 | MB231/MB468 | NUC |

| | | | | | | | | | | | | | |
|---|---|---|---|---|---|---|---|---|---|---|---|---|---|
| >hg38_21_+_8433172_8446622_RNA45SN1_45S_50ntFlanks@11315.11342.28\|CCACCCCACGTCTCGTCGCGCGCGCGTC | >hg38_21_+_8433172_8446622_RNA45SN1_45S_50ntFlanks | 11315 | 11342 | 28 | CCACCCCACGTCTCGTCGCGCGCGCGTC | 3277.2 | -7.5 | 2.0 | -3.7 | 2.1E-04 | 2.7E-04 | BT20/MB468 | NUC |
| >hg38_MT_+_1621_3279_RNR2_16S_50ntFlanks@113.146.34\|CCAAATAAAGTATAGGCGATAGAAATTGAAACCT | >hg38_MT_+_1621_3279_RNR2_16S_50ntFlanks | 113 | 146 | 34 | CCAAATAAAGTATAGGCGATAGAAATTGAAACCT | 95.4 | 10.3 | 2.6 | 3.9 | 7.9E-05 | 2.7E-04 | BT20/MB468 | MP |
| >hg38_21_+_8433172_8446622_RNA45SN1_45S_50ntFlanks@3841.3868.28\|TCCTACTTGGATAACTGTGGTAATTCTA | >hg38_21_+_8433172_8446622_RNA45SN1_45S_50ntFlanks | 3841 | 3868 | 28 | TCCTACTTGGATAACTGTGGTAATTCTA | 128.2 | -6.5 | 1.7 | -3.8 | 1.8E-04 | 2.7E-04 | MB231/MB468 | MP |
| >hg38_21_+_8433172_8446622_RNA45SN1_45S_50ntFlanks@3831.3860.30\|CTCGCTCCTCTCCTACTTGGATAACTGTGG | >hg38_21_+_8433172_8446622_RNA45SN1_45S_50ntFlanks | 3831 | 3860 | 30 | CTCGCTCCTCTCCTACTTGGATAACTGTGG | 334.6 | -2.6 | 0.7 | -3.8 | 1.6E-04 | 2.7E-04 | BT20/MB468 | MITO |
| >hg38_21_+_8433172_8446622_RNA45SN1_45S_50ntFlanks@10806.10826.21\|AACAATGTAGGTAAGGGAAGT | >hg38_21_+_8433172_8446622_RNA45SN1_45S_50ntFlanks | 10806 | 10826 | 21 | AACAATGTAGGTAAGGGAAGT | 318.3 | -5.3 | 1.4 | -3.7 | 1.9E-04 | 2.7E-04 | BT20/MB468 | CYTO |
| >MI0000088\|hsa-mir-30a&WithFlank&6\|-\|71403545\|71403627@13.34.22(+1C)&[MIMAT0000087&hsa-miR-30a-5p&offsets\|+1\|+1(+1C);m-15&6\|-\|71403595\|71403616&offsets\|+1\|+1(+1C)]\|GTAAACATCCTCGACTGGAAGCC | >MI0000088\|hsa-mir-30a&WithFlank&6\|-\|71403545\|71403627 | 13 | 34 | 22(+1C) | GTAAACATCCTCGACTGGAAGCC | 559.4 | 4.6 | 1.2 | 3.7 | 1.8E-04 | 2.7E-04 | MB231/MB468 | MP |
| >trnaMT_GluTTC_MT_-_14674_14742@39.66.28,trnalookalike8_GluTTC_5_-_93905172_93905240@39.66.28\|ATTGGTCGTGGTTGTAGTCCGTGCGAGA | >trnaMT_GluTTC_MT_-_14674_14742 | 39 | 66 | 28 | ATTGGTCGTGGTTGTAGTCCGTGCGAGA | 207.0 | 3.1 | 0.8 | 3.7 | 1.8E-04 | 2.7E-04 | MB231/MB468 | MP |
| >trnaMT_IleGAT_MT_+_4263_4331@42.68.27\|AATAGGAGCTTAAACCCCCTTATTTCT | >trnaMT_IleGAT_MT_+_4263_4331 | 42 | 68 | 27 | AATAGGAGCTTAAACCCCCTTATTTCT | 95.1 | 10.3 | 2.6 | 3.9 | 8.0E-05 | 2.7E-04 | BT20/MB468 | MP |
| >trnaMT_PheGAA_MT_+_577_647@54.74.21\|CACATCACCCCATAAACACCA | >trnaMT_PheGAA_MT_+_577_647 | 54 | 74 | 21 | CACATCACCCCATAAACACCA | 1090.5 | -4.4 | 1.2 | -3.7 | 2.2E-04 | 2.7E-04 | BT20/MB468 | NUC |
| >hg38_21_+_8433172_8446622_RNA45SN1_45S_50ntFlanks@11039.11060.22\|CTCCCCCGCTCCCCGTCCTCCC | >hg38_21_+_8433172_8446622_RNA45SN1_45S_50ntFlanks | 11039 | 11060 | 22 | CTCCCCCGCTCCCCGTCCTCCC | 445.3 | -6.1 | 1.6 | -3.7 | 2.1E-04 | 2.7E-04 | MB231/MB468 | NUC |
| >trnaMT_IleGAT_MT_+_4263_4331@43.72.30\|ATAGGAGCTTAAACCCCCTTATTTCTACCA | >trnaMT_IleGAT_MT_+_4263_4331 | 43 | 72 | 30 | ATAGGAGCTTAAACCCCCTTATTTCTACCA | 98.9 | 10.4 | 2.6 | 3.9 | 8.0E-05 | 2.7E-04 | BT20/MB468 | MP |
| >hg38_21_+_8433172_8446622_RNA45SN1_45S_50ntFlanks@4160.4177.18\|CCACATCCAAGGAAGGCA | >hg38_21_+_8433172_8446622_RNA45SN1_45S_50ntFlanks | 4160 | 4177 | 18 | CCACATCCAAGGAAGGCA | 90.6 | -9.6 | 2.4 | -3.9 | 8.0E-05 | 2.7E-04 | BT20/MB468 | MP |
| >hg38_21_+_8433172_8446622_RNA45SN1_45S_50ntFlanks@7985.8009.25\|GATCAGACGTGGCGACCCGCTGAAT | >hg38_21_+_8433172_8446622_RNA45SN1_45S_50ntFlanks | 7985 | 8009 | 25 | GATCAGACGTGGCGACCCGCTGAAT | 681.0 | -6.1 | 1.6 | -3.7 | 1.8E-04 | 2.7E-04 | MB231/MB468 | MP |
| >hg38_21_+_8433172_8446622_RNA45SN1_45S_50ntFlanks@7971.7988.18\|GAGACGCGACCTCAGATC | >hg38_21_+_8433172_8446622_RNA45SN1_45S_50ntFlanks | 7971 | 7988 | 18 | GAGACGCGACCTCAGATC | 914.6 | 4.2 | 1.1 | 3.9 | 8.0E-05 | 2.7E-04 | BT20/MB468 | MP |
| >hg38_MT_+_598_1651_RNR1_12S_50ntFlanks@843.876.34\|GAGTAGAGTGCTTAGTTGAACAGGGCCCTGAAGC | >hg38_MT_+_598_1651_RNR1_12S_50ntFlanks | 843 | 876 | 34 | GAGTAGAGTGCTTAGTTGAACAGGGCCCTGAAGC | 181.1 | 7.7 | 2.0 | 3.9 | 8.0E-05 | 2.7E-04 | BT20/MB468 | MP |
| >hg38_MT_+_598_1651_RNR1_12S_50ntFlanks@428.468.41\|AAAGTGGCTTTAACATATCTGAACACACAATAGCTAAGACC | >hg38_MT_+_598_1651_RNR1_12S_50ntFlanks | 428 | 468 | 41 | AAAGTGGCTTTAACATATCTGAACACACAATAGCTAAGACC | 959.6 | 6.2 | 1.6 | 3.9 | 8.0E-05 | 2.7E-04 | BT20/MB468 | MP |
| >hg38_21_+_8433172_8446622_RNA45SN1_45S_50ntFlanks@4367.4400.34\|CAGTTAAAAAGCTCGTAGTTGGATCTTGGGAGCG | >hg38_21_+_8433172_8446622_RNA45SN1_45S_50ntFlanks | 4367 | 4400 | 34 | CAGTTAAAAAGCTCGTAGTTGGATCTTGGGAGCG | 137.6 | -2.8 | 0.8 | -3.7 | 1.9E-04 | 2.7E-04 | BT20/MB468 | CYTO |
| >hg38_21_+_8433172_8446622_RNA45SN1_45S_50ntFlanks@7991.8014.24\|ACGTGGCGACCCGCTGAATTTAAG | >hg38_21_+_8433172_8446622_RNA45SN1_45S_50ntFlanks | 7991 | 8014 | 24 | ACGTGGCGACCCGCTGAATTTAAG | 1849.2 | -2.7 | 0.7 | -3.8 | 1.6E-04 | 2.7E-04 | BT20/MB468 | MITO |
| >trnaMT_ValTAC_MT_+_1602_1670@55.72.18\|ACTTGACCGCTCTGACCA | >trnaMT_ValTAC_MT_+_1602_1670 | 55 | 72 | 18 | ACTTGACCGCTCTGACCA | 3326.6 | 3.2 | 0.9 | 3.7 | 2.1E-04 | 2.7E-04 | MB231/MB468 | NUC |
| >hg38_21_+_8433172_8446622_RNA45SN1_45S_50ntFlanks@11669.11708.40\|GCCCAGTGCTCTGAATGTCAAAGTGAAGAAATTCAATGAA | >hg38_21_+_8433172_8446622_RNA45SN1_45S_50ntFlanks | 11669 | 11708 | 40 | GCCCAGTGCTCTGAATGTCAAAGTGAAGAAATTCAATGAA | 448.0 | -3.5 | 0.8 | -4.2 | 2.7E-05 | 2.7E-04 | BT20/MB231 | CYTO |
| >hg38_21_+_8433172_8446622_RNA45SN1_45S_50ntFlanks@9027.9063.37\|CGTCTCCTCGTGGGGGGGCCGGGCCACCCCTCCCACG | >hg38_21_+_8433172_8446622_RNA45SN1_45S_50ntFlanks | 9027 | 9063 | 37 | CGTCTCCTCGTGGGGGGGCCGGGCCACCCCTCCCACG | 278.5 | -5.5 | 1.5 | -3.8 | 1.6E-04 | 2.7E-04 | BT20/MB468 | MITO |
| >MI0000071\|hsa-mir-17&WithFlank&13\|+\|91350599\|91350694@20.42.23&[MIMAT0000070&hsa-miR-17-5p&offsets\|0\|0;m-88&13\|+\|91350618\|91350640&offsets\|0\|0]\|CAAAGTGCTTACAGTGCAGGTAG | >MI0000071\|hsa-mir-17&WithFlank&13\|+\|91350599\|91350694 | 20 | 42 | 23 | CAAAGTGCTTACAGTGCAGGTAG | 3285.6 | 3.8 | 1.0 | 3.7 | 2.0E-04 | 2.7E-04 | MB231/MB468 | CYTO |
| >hg38_21_+_8433172_8446622_RNA45SN1_45S_50ntFlanks@11586.11623.38\|GTGCGGACCAGGGGAATCCGACTGTTTAATTAAAACAA | >hg38_21_+_8433172_8446622_RNA45SN1_45S_50ntFlanks | 11586 | 11623 | 38 | GTGCGGACCAGGGGAATCCGACTGTTTAATTAAAACAA | 186.4 | 7.4 | 1.9 | 3.9 | 8.0E-05 | 2.7E-04 | BT20/MB468 | MP |
| >hg38_21_+_8433172_8446622_RNA45SN1_45S_50ntFlanks@6671.6696.26\|CTCGGCTCGTGCGTCGATGAAGAACG | >hg38_21_+_8433172_8446622_RNA45SN1_45S_50ntFlanks | 6671 | 6696 | 26 | CTCGGCTCGTGCGTCGATGAAGAACG | 690.3 | -2.1 | 0.6 | -3.7 | 2.1E-04 | 2.7E-04 | MB231/MB468 | NUC |
| >trna10_AspGTC_12_-_125424193_125424264@40.72.33,trna12_AspGTC_12_-_125411891_125411962@40.72.33,trna144_AspGTC_6_-_27551236_27551307@40.72.33,trna38_AspGTC_17_-_8125556_8125627@40.72.33,trna45_AspGTC_6_+_27447453_27447524@40.72.33,trna48_AspGTC_6_+_27471523_27471594@40.72.33,trna4_AspGTC_12_+_96429799_96429870@40.72.33,trna69_AspGTC_1_-_161440205_161440276@40.72.33,trna72_AspGTC_1_-_161432824_161432895@40.72.33,trna75_AspGTC_1_-_161425414_161425485@40.72.33,trna78_AspGTC_1_-_161418033_161418104@40.72.33,trna81_AspGTC_1_-_161410615_161410686@40.72.33\|CGGGAGACCGGGGTTCGATTCCCCGACGGGGAG | >trna10_AspGTC_12_-_125424193_125424264 | 40 | 72 | 33 | CGGGAGACCGGGGTTCGATTCCCCGACGGGGAG | 195.4 | -3.3 | 0.9 | -3.7 | 1.8E-04 | 2.7E-04 | MB231/MB468 | MP |
| >hg38_21_+_8433172_8446622_RNA45SN1_45S_50ntFlanks@12455.12487.33\|ACCAAGCGTTGGATTGTTCACCCACTAATAGGG | >hg38_21_+_8433172_8446622_RNA45SN1_45S_50ntFlanks | 12455 | 12487 | 33 | ACCAAGCGTTGGATTGTTCACCCACTAATAGGG | 245.9 | -3.7 | 1.0 | -3.7 | 2.0E-04 | 2.7E-04 | MB231/MB468 | CYTO |
| >hg38_21_+_8433172_8446622_RNA45SN1_45S_50ntFlanks@6646.6666.21\|TCGTACGACTCTTAGCGGTGG | >hg38_21_+_8433172_8446622_RNA45SN1_45S_50ntFlanks | 6646 | 6666 | 21 | TCGTACGACTCTTAGCGGTGG | 19000.5 | 3.1 | 0.8 | 3.7 | 1.9E-04 | 2.7E-04 | BT20/MB468 | CYTO |
| >hg38_21_+_8433172_8446622_RNA45SN1_45S_50ntFlanks@8185.8207.23\|GAGGCCCAGCCCGTGGACGGTGT | >hg38_21_+_8433172_8446622_RNA45SN1_45S_50ntFlanks | 8185 | 8207 | 23 | GAGGCCCAGCCCGTGGACGGTGT | 314.0 | 8.1 | 2.1 | 3.9 | 8.1E-05 | 2.7E-04 | BT20/MB468 | MP |
| >hg38_MT_+_598_1651_RNR1_12S_50ntFlanks@848.873.26\|GAGTGCTTAGTTGAACAGGGCCCTGA | >hg38_MT_+_598_1651_RNR1_12S_50ntFlanks | 848 | 873 | 26 | GAGTGCTTAGTTGAACAGGGCCCTGA | 101.8 | 10.4 | 2.6 | 3.9 | 8.1E-05 | 2.7E-04 | BT20/MB468 | MP |
| >hg38_MT_+_1621_3279_RNR2_16S_50ntFlanks@1369.1396.28\|GATGTTGGATCAGGACATCCCGATGGTG | >hg38_MT_+_1621_3279_RNR2_16S_50ntFlanks | 1369 | 1396 | 28 | GATGTTGGATCAGGACATCCCGATGGTG | 315.4 | 3.0 | 0.8 | 3.7 | 1.8E-04 | 2.7E-04 | MB231/MB468 | MP |
| >MI0000542\|hsa-mir-320a&WithFlank&8\|-\|22244960\|22245043@48.71.24(+1U)&[MIMAT0000510&hsa-miR-320a-3p&offsets\|0\|+2(+1U);m-60&8\|-\|22244975\|22244996&offsets\|0\|+2(+1U)]\|AAAAGCTGGGTTGAGAGGGCGAAAT | >MI0000542\|hsa-mir-320a&WithFlank&8\|-\|22244960\|22245043 | 48 | 71 | 24(+1U) | AAAAGCTGGGTTGAGAGGGCGAAAT | 1706.2 | 3.4 | 0.8 | 4.2 | 2.8E-05 | 2.7E-04 | BT20/MB231 | CYTO |
| >trnaMT_LeuTAA_MT_+_3230_3304@18.42.25\|CGGTAATCGCATAAAACTTAAAACT | >trnaMT_LeuTAA_MT_+_3230_3304 | 18 | 42 | 25 | CGGTAATCGCATAAAACTTAAAACT | 428.2 | 3.5 | 0.8 | 4.2 | 2.8E-05 | 2.7E-04 | BT20/MB231 | CYTO |
| >hg38_21_+_8433172_8446622_RNA45SN1_45S_50ntFlanks@10786.10821.36\|AACAGCCTCTGGCATGTTGGAACAATGTAGGTAAGG | >hg38_21_+_8433172_8446622_RNA45SN1_45S_50ntFlanks | 10786 | 10821 | 36 | AACAGCCTCTGGCATGTTGGAACAATGTAGGTAAGG | 655.6 | -2.1 | 0.6 | -3.8 | 1.6E-04 | 2.7E-04 | BT20/MB468 | MITO |
| >hg38_21_+_8433172_8446622_RNA45SN1_45S_50ntFlanks@9504.9531.28\|GTGAACTATGCCTGGGCAGGGCGAAGCC | >hg38_21_+_8433172_8446622_RNA45SN1_45S_50ntFlanks | 9504 | 9531 | 28 | GTGAACTATGCCTGGGCAGGGCGAAGCC | 231.7 | -2.8 | 0.7 | -3.8 | 1.6E-04 | 2.7E-04 | BT20/MB468 | MITO |
| >hg38_21_+_8433172_8446622_RNA45SN1_45S_50ntFlanks@11572.11589.18\|CCGACTTAGAACTGGTGC | >hg38_21_+_8433172_8446622_RNA45SN1_45S_50ntFlanks | 11572 | 11589 | 18 | CCGACTTAGAACTGGTGC | 372.9 | -2.0 | 0.5 | -3.7 | 2.0E-04 | 2.7E-04 | MB231/MB468 | CYTO |
| >hg38_21_+_8433172_8446622_RNA45SN1_45S_50ntFlanks@11778.11805.28\|ACGCGCATGAATGGATGAACGAGATTCC | >hg38_21_+_8433172_8446622_RNA45SN1_45S_50ntFlanks | 11778 | 11805 | 28 | ACGCGCATGAATGGATGAACGAGATTCC | 119.6 | -5.5 | 1.5 | -3.7 | 1.8E-04 | 2.7E-04 | MB231/MB468 | MP |
| >trnaMT_PheGAA_MT_+_577_647@36.54.19\|AAAATGTTTAGACGGGCTC | >trnaMT_PheGAA_MT_+_577_647 | 36 | 54 | 19 | AAAATGTTTAGACGGGCTC | 247.7 | 5.7 | 1.5 | 3.7 | 2.1E-04 | 2.7E-04 | MB231/MB468 | MITO |
| >hg38_21_+_8433172_8446622_RNA45SN1_45S_50ntFlanks@8111.8132.22\|GGGGCGCGGGACATGTGGCGTA | >hg38_21_+_8433172_8446622_RNA45SN1_45S_50ntFlanks | 8111 | 8132 | 22 | GGGGCGCGGGACATGTGGCGTA | 120.5 | 7.3 | 1.9 | 3.9 | 8.1E-05 | 2.7E-04 | BT20/MB468 | MP |
| >hg38_21_+_8433172_8446622_RNA45SN1_45S_50ntFlanks@11667.11701.35\|CTGCCCAGTGCTCTGAATGTCAAAGTGAAGAAATT | >hg38_21_+_8433172_8446622_RNA45SN1_45S_50ntFlanks | 11667 | 11701 | 35 | CTGCCCAGTGCTCTGAATGTCAAAGTGAAGAAATT | 97.6 | 7.2 | 1.8 | 3.9 | 8.1E-05 | 2.8E-04 | BT20/MB468 | MP |
| >MI0003667\|hsa-mir-652&WithFlank&X\|+\|110055323\|110055432@67.86.20(+1A)&[MIMAT0003322&hsa-miR-652-3p&offsets\|0\|-1(+1A);m-148&X\|+\|110055389\|110055410&offsets\|0\|-2(+1A)]\|AATGGCGCCACTAGGGTTGTA | >MI0003667\|hsa-mir-652&WithFlank&X\|+\|110055323\|110055432 | 67 | 86 | 20(+1A) | AATGGCGCCACTAGGGTTGTA | 200.9 | 5.7 | 1.4 | 3.9 | 8.1E-05 | 2.8E-04 | BT20/MB468 | MP |
| >hg38_21_+_8433172_8446622_RNA45SN1_45S_50ntFlanks@10485.10505.21\|AAAGGGAGTCGGGTTCAGATC | >hg38_21_+_8433172_8446622_RNA45SN1_45S_50ntFlanks | 10485 | 10505 | 21 | AAAGGGAGTCGGGTTCAGATC | 176.0 | 2.3 | 0.6 | 3.7 | 2.0E-04 | 2.8E-04 | MB231/MB468 | CYTO |
| >hg38_21_+_8433172_8446622_RNA45SN1_45S_50ntFlanks@5486.5514.29\|GCGCTGAGAAGACGGTCGAACTTGACTAT | >hg38_21_+_8433172_8446622_RNA45SN1_45S_50ntFlanks | 5486 | 5514 | 29 | GCGCTGAGAAGACGGTCGAACTTGACTAT | 540.5 | -5.8 | 1.6 | -3.7 | 2.2E-04 | 2.8E-04 | MB231/MB468 | NUC |
| >trnaMT_HisGTG_MT_+_12138_12206@20.61.42\|AACATCAGATTGTGAATCTGACAACAGAGGCTTACGACCCCT | >trnaMT_HisGTG_MT_+_12138_12206 | 20 | 61 | 42 | AACATCAGATTGTGAATCTGACAACAGAGGCTTACGACCCCT | 96.8 | 10.3 | 2.6 | 3.9 | 8.1E-05 | 2.8E-04 | BT20/MB468 | MP |
| >hg38_21_+_8433172_8446622_RNA45SN1_45S_50ntFlanks@10866.10904.39\|CTAAGGGCTGGGTCGGTCGGGCTGGGGCGCGAAGCGGGG | >hg38_21_+_8433172_8446622_RNA45SN1_45S_50ntFlanks | 10866 | 10904 | 39 | CTAAGGGCTGGGTCGGTCGGGCTGGGGCGCGAAGCGGGG | 147.5 | -6.0 | 1.5 | -3.9 | 8.1E-05 | 2.8E-04 | BT20/MB468 | MP |
| >hg38_21_+_8433172_8446622_RNA45SN1_45S_50ntFlanks@9564.9598.35\|CTGACGTGCAAATCGGTCGTCCGACCTGGGTATAG | >hg38_21_+_8433172_8446622_RNA45SN1_45S_50ntFlanks | 9564 | 9598 | 35 | CTGACGTGCAAATCGGTCGTCCGACCTGGGTATAG | 365.0 | -2.6 | 0.7 | -3.8 | 1.6E-04 | 2.8E-04 | BT20/MB468 | MITO |
| >hg38_21_+_8433172_8446622_RNA45SN1_45S_50ntFlanks@6644.6681.38\|CCTCGTACGACTCTTAGCGGTGGATCACTCGGCTCGTG | >hg38_21_+_8433172_8446622_RNA45SN1_45S_50ntFlanks | 6644 | 6681 | 38 | CCTCGTACGACTCTTAGCGGTGGATCACTCGGCTCGTG | 641.8 | -2.8 | 0.7 | -4.2 | 3.0E-05 | 2.8E-04 | BT20/MB231 | NUC |
| >MI0000294\|hsa-mir-218-1&WithFlank&4\|+\|20528269\|20528390@31.51.21(+1A);MI0000295\|hsa-mir-218-2&WithFlank&5\|-\|168768140\|168768261@31.51.21(+1A)&[MIMAT0000275&hsa-miR-218-5p&offsets\|0\|0(+1A);m-185&4\|+\|20528299\|20528319&offsets\|0\|0(+1A)];[MIMAT0000275_1&hsa-miR-218-5p&offsets\|0\|0(+1A);m-187&5\|-\|168768211\|168768231&offsets\|0\|0(+1A)]\|TTGTGCTTGATCTAACCATGTA | >MI0000294\|hsa-mir-218-1&WithFlank&4\|+\|20528269\|20528390 | 31 | 51 | 21(+1A) | TTGTGCTTGATCTAACCATGTA | 57.0 | -9.2 | 2.2 | -4.2 | 2.9E-05 | 2.8E-04 | BT20/MB231 | MP |
| >hg38_21_+_8433172_8446622_RNA45SN1_45S_50ntFlanks@12008.12041.34\|CCGCCGGTGAAATACCACTACTCTGATCGTTTTT | >hg38_21_+_8433172_8446622_RNA45SN1_45S_50ntFlanks | 12008 | 12041 | 34 | CCGCCGGTGAAATACCACTACTCTGATCGTTTTT | 555.3 | -4.6 | 1.2 | -3.8 | 1.6E-04 | 2.8E-04 | BT20/MB468 | MITO |

<table>
<tr><td>>MI0000542|hsa-mir-320a&WithFlank&8|-|22244960|22245043@49.70.22(+1C)&[MIMAT0000510&hsa-miR-320a-3p&offsets|+1|+1(+1C);m-60&8|-|22244975|22244996&offsets|+1|+1(+1C)]|AAAGCTGGGTTGAGAGGGCGAAC</td><td>>MI0000542|hsa-mir-320a&WithFlank&8|-|22244960|22245043</td><td>49</td><td>70</td><td>22(+1C)</td><td>AAAGCTGGGTTGAGAGGGCGAAC</td><td>149.0</td><td>7.2</td><td>1.8</td><td>3.9</td><td>8.1E-05</td><td>2.8E-04</td><td>BT20/MB468</td><td>MP</td></tr>
<tr><td>>hg38_21_+_8433172_8446622_RNA45SN1_45S_50ntFlanks@10347.10368.22|CAAACGAGAACTTTGAAGGCCG</td><td>>hg38_21_+_8433172_8446622_RNA45SN1_45S_50ntFlanks</td><td>10347</td><td>10368</td><td>22</td><td>CAAACGAGAACTTTGAAGGCCG</td><td>454.8</td><td>-2.5</td><td>0.7</td><td>-3.7</td><td>2.2E-04</td><td>2.8E-04</td><td>MB231/MB468</td><td>NUC</td></tr>
<tr><td>>hg38_21_+_8433172_8446622_RNA45SN1_45S_50ntFlanks@7977.8007.31|CGACCTCAGATCAGACGTGGCGACCCGCTGA</td><td>>hg38_21_+_8433172_8446622_RNA45SN1_45S_50ntFlanks</td><td>7977</td><td>8007</td><td>31</td><td>CGACCTCAGATCAGACGTGGCGACCCGCTGA</td><td>152.4</td><td>-5.6</td><td>1.5</td><td>-3.7</td><td>1.8E-04</td><td>2.8E-04</td><td>MB231/MB468</td><td>MP</td></tr>
<tr><td>>hg38_21_+_8433172_8446622_RNA45SN1_45S_50ntFlanks@10771.10816.46|CAGGTCTCCAAGGTGAACAGCCTCTGGCATGTTGGAACAATGTAGG</td><td>>hg38_21_+_8433172_8446622_RNA45SN1_45S_50ntFlanks</td><td>10771</td><td>10816</td><td>46</td><td>CAGGTCTCCAAGGTGAACAGCCTCTGGCATGTTGGAACAATGTAGG</td><td>441.9</td><td>-3.2</td><td>0.9</td><td>-3.7</td><td>2.0E-04</td><td>2.8E-04</td><td>MB231/MB468</td><td>CYTO</td></tr>
<tr><td>>hg38_21_+_8433172_8446622_RNA45SN1_45S_50ntFlanks@4589.4616.28|TATTTTGTTGGTTTTCGGAACTGAGGCC</td><td>>hg38_21_+_8433172_8446622_RNA45SN1_45S_50ntFlanks</td><td>4589</td><td>4616</td><td>28</td><td>TATTTTGTTGGTTTTCGGAACTGAGGCC</td><td>151.8</td><td>-4.0</td><td>1.1</td><td>-3.7</td><td>2.2E-04</td><td>2.8E-04</td><td>MB231/MB468</td><td>NUC</td></tr>
<tr><td>>hg38_MT_+_598_1651_RNR1_12S_50ntFlanks@638.665.28|TCACCACCTCTTGCTCAGCCTATATACC</td><td>>hg38_MT_+_598_1651_RNR1_12S_50ntFlanks</td><td>638</td><td>665</td><td>28</td><td>TCACCACCTCTTGCTCAGCCTATATACC</td><td>127.6</td><td>5.5</td><td>1.5</td><td>3.7</td><td>2.1E-04</td><td>2.8E-04</td><td>MB231/MB468</td><td>MITO</td></tr>
<tr><td>>MI0000471|hsa-mir-126&WithFlank&9|+|136670596|136670692@58.79.22(+1U)&[MIMAT0000445&hsa-miR-126-3p&offsets|0|0(+1U);m-40&9|+|136670653|136670674&offsets|0|0(+1U)]|TCGTACCGTGAGTAATAATGCGT</td><td>>MI0000471|hsa-mir-126&WithFlank&9|+|136670596|136670692</td><td>58</td><td>79</td><td>22(+1U)</td><td>TCGTACCGTGAGTAATAATGCGT</td><td>94.7</td><td>10.3</td><td>2.6</td><td>3.9</td><td>8.2E-05</td><td>2.8E-04</td><td>BT20/MB468</td><td>MP</td></tr>
<tr><td>>MI0000082|hsa-mir-25&WithFlank&7|-|100093554|100093649@58.79.22(+2A)&[MIMAT0000081&hsa-miR-25-3p&offsets|0|0(+2A);m-23&7|-|100093571|100093592&offsets|0|0(+2A)]|CATTGCACTTGTCTCGGTCTGAAA</td><td>>MI0000082|hsa-mir-25&WithFlank&7|-|100093554|100093649</td><td>58</td><td>79</td><td>22(+2A)</td><td>CATTGCACTTGTCTCGGTCTGAAA</td><td>174.5</td><td>3.4</td><td>0.9</td><td>3.7</td><td>2.1E-04</td><td>2.8E-04</td><td>MB231/MB468</td><td>MITO</td></tr>
<tr><td>>hg38_21_+_8433172_8446622_RNA45SN1_45S_50ntFlanks@10725.10755.31|GGGGAGAGGGTGTAAATCTCGCGCCGGGCCG</td><td>>hg38_21_+_8433172_8446622_RNA45SN1_45S_50ntFlanks</td><td>10725</td><td>10755</td><td>31</td><td>GGGGAGAGGGTGTAAATCTCGCGCCGGGCCG</td><td>186.0</td><td>-5.0</td><td>1.3</td><td>-3.7</td><td>1.9E-04</td><td>2.8E-04</td><td>BT20/MB468</td><td>CYTO</td></tr>
<tr><td>>hg38_21_+_8433172_8446622_RNA45SN1_45S_50ntFlanks@7991.8017.27|ACGTGGCGACCCGCTGAATTTAAGCAT</td><td>>hg38_21_+_8433172_8446622_RNA45SN1_45S_50ntFlanks</td><td>7991</td><td>8017</td><td>27</td><td>ACGTGGCGACCCGCTGAATTTAAGCAT</td><td>307.7</td><td>-3.3</td><td>0.8</td><td>-3.9</td><td>8.2E-05</td><td>2.8E-04</td><td>BT20/MB468</td><td>MP</td></tr>
<tr><td>>MI0000286|hsa-mir-210&WithFlank&11|-|568083|568204@72.93.22(+2A)&[MIMAT0000267&hsa-miR-210-3p&offsets|0|0(+2A);m-102&11|-|568112|568133&offsets|0|0(+2A)]|CTGTGCGTGTGACAGCGGCTGAAA</td><td>>MI0000286|hsa-mir-210&WithFlank&11|-|568083|568204</td><td>72</td><td>93</td><td>22(+2A)</td><td>CTGTGCGTGTGACAGCGGCTGAAA</td><td>2125.5</td><td>4.1</td><td>1.0</td><td>4.2</td><td>2.8E-05</td><td>2.8E-04</td><td>BT20/MB231</td><td>CYTO</td></tr>
<tr><td>>trnaMT_GluTTC_MT_-_14674_14742@20.57.38,trnalookalike8_GluTTC_5_-_93905172_93905240@20.57.38|AACGATGGTTTTTCATATCATTGGTCGTGGTTGTAGTC</td><td>>trnaMT_GluTTC_MT_-_14674_14742</td><td>20</td><td>57</td><td>38</td><td>AACGATGGTTTTTCATATCATTGGTCGTGGTTGTAGTC</td><td>199.4</td><td>7.9</td><td>2.0</td><td>3.9</td><td>8.2E-05</td><td>2.8E-04</td><td>BT20/MB468</td><td>MP</td></tr>
<tr><td>>hg38_21_+_8433172_8446622_RNA45SN1_45S_50ntFlanks@3920.3959.40|CATTTATCAGATCAAAACCAACCCGGTCAGCCCCTCTCCG</td><td>>hg38_21_+_8433172_8446622_RNA45SN1_45S_50ntFlanks</td><td>3920</td><td>3959</td><td>40</td><td>CATTTATCAGATCAAAACCAACCCGGTCAGCCCCTCTCCG</td><td>383.9</td><td>-5.3</td><td>1.4</td><td>-3.7</td><td>2.0E-04</td><td>2.8E-04</td><td>MB231/MB468</td><td>CYTO</td></tr>
<tr><td>>hg38_21_+_8433172_8446622_RNA45SN1_45S_50ntFlanks@3903.3935.33|CGCGGGGGGGATGCGTGCATTTATCAGATCAAA</td><td>>hg38_21_+_8433172_8446622_RNA45SN1_45S_50ntFlanks</td><td>3903</td><td>3935</td><td>33</td><td>CGCGGGGGGGATGCGTGCATTTATCAGATCAAA</td><td>37.0</td><td>-8.3</td><td>2.1</td><td>-3.9</td><td>8.2E-05</td><td>2.8E-04</td><td>BT20/MB468</td><td>MP</td></tr>
<tr><td>>MI0000683|hsa-mir-181b-2&WithFlank&9|+|124693704|124693804@22.46.25&[MIMAT0000257_1&hsa-miR-181b-5p&offsets|0|+2;m-72&9|+|124693725|124693747&offsets|0|+2]|AACATTCATTGCTGTCGGTGGGTTT</td><td>>MI0000683|hsa-mir-181b-2&WithFlank&9|+|124693704|124693804</td><td>22</td><td>46</td><td>25</td><td>AACATTCATTGCTGTCGGTGGGTTT</td><td>1706.7</td><td>-4.3</td><td>1.0</td><td>-4.2</td><td>2.9E-05</td><td>2.8E-04</td><td>BT20/MB231</td><td>MP</td></tr>
<tr><td>>hg38_21_+_8433172_8446622_RNA45SN1_45S_50ntFlanks@9716.9737.22|AATGATTAGAGGTCTTGGGGCC</td><td>>hg38_21_+_8433172_8446622_RNA45SN1_45S_50ntFlanks</td><td>9716</td><td>9737</td><td>22</td><td>AATGATTAGAGGTCTTGGGGCC</td><td>381.0</td><td>-2.8</td><td>0.7</td><td>-3.7</td><td>2.0E-04</td><td>2.8E-04</td><td>MB231/MB468</td><td>CYTO</td></tr>
<tr><td>>hg38_21_+_8433172_8446622_RNA45SN1_45S_50ntFlanks@11212.11231.20|CGACTCTGGACGCGAGCCGG</td><td>>hg38_21_+_8433172_8446622_RNA45SN1_45S_50ntFlanks</td><td>11212</td><td>11231</td><td>20</td><td>CGACTCTGGACGCGAGCCGG</td><td>291.2</td><td>-3.0</td><td>0.8</td><td>-3.9</td><td>8.2E-05</td><td>2.8E-04</td><td>BT20/MB468</td><td>MP</td></tr>
<tr><td>>hg38_21_+_8433172_8446622_RNA45SN1_45S_50ntFlanks@10788.10812.25|CAGCCTCTGGCATGTTGGAACAATG</td><td>>hg38_21_+_8433172_8446622_RNA45SN1_45S_50ntFlanks</td><td>10788</td><td>10812</td><td>25</td><td>CAGCCTCTGGCATGTTGGAACAATG</td><td>62.6</td><td>9.3</td><td>2.2</td><td>4.2</td><td>2.9E-05</td><td>2.8E-04</td><td>BT20/MB231</td><td>MP</td></tr>
<tr><td>>MI0001446|hsa-mir-424&WithFlank&X|-|134546608|134546717@54.74.21&[MIMAT0004749&hsa-miR-424-3p&offsets|0|0;m-305&X|-|134546644|134546664&offsets|0|0]|CAAAACGTGAGGCGCTGCTAT</td><td>>MI0001446|hsa-mir-424&WithFlank&X|-|134546608|134546717</td><td>54</td><td>74</td><td>21</td><td>CAAAACGTGAGGCGCTGCTAT</td><td>134.8</td><td>5.8</td><td>1.6</td><td>3.7</td><td>1.8E-04</td><td>2.8E-04</td><td>MB231/MB468</td><td>MP</td></tr>
<tr><td>>MI0000462|hsa-mir-152&WithFlank&17|-|48037155|48037253@60.80.21(+1U)&[MIMAT0000438&hsa-miR-152-3p&offsets|0|0(+1U);m-97&17|-|48037174|48037194&offsets|0|0(+1U)]|TCAGTGCATGACAGAACTTGGT</td><td>>MI0000462|hsa-mir-152&WithFlank&17|-|48037155|48037253</td><td>60</td><td>80</td><td>21(+1U)</td><td>TCAGTGCATGACAGAACTTGGT</td><td>190.0</td><td>6.7</td><td>1.7</td><td>3.9</td><td>8.2E-05</td><td>2.8E-04</td><td>BT20/MB468</td><td>MP</td></tr>
<tr><td>>hg38_21_+_8433172_8446622_RNA45SN1_45S_50ntFlanks@11604.11627.24|CGACTGTTTAATTAAAACAAAGCA</td><td>>hg38_21_+_8433172_8446622_RNA45SN1_45S_50ntFlanks</td><td>11604</td><td>11627</td><td>24</td><td>CGACTGTTTAATTAAAACAAAGCA</td><td>155.1</td><td>3.6</td><td>0.9</td><td>3.9</td><td>8.2E-05</td><td>2.8E-04</td><td>BT20/MB468</td><td>MP</td></tr>
<tr><td>>hg38_MT_+_1621_3279_RNR2_16S_50ntFlanks@495.526.32|TCTTTGGACACTAGGAAAAAACCTTGTAGAGA</td><td>>hg38_MT_+_1621_3279_RNR2_16S_50ntFlanks</td><td>495</td><td>526</td><td>32</td><td>TCTTTGGACACTAGGAAAAAACCTTGTAGAGA</td><td>89.6</td><td>7.3</td><td>1.9</td><td>3.9</td><td>8.2E-05</td><td>2.8E-04</td><td>BT20/MB468</td><td>MP</td></tr>
<tr><td>>hg38_21_+_8433172_8446622_RNA45SN1_45S_50ntFlanks@10927.10947.21|CGAGGCGCCGCCGCCCCCCCC</td><td>>hg38_21_+_8433172_8446622_RNA45SN1_45S_50ntFlanks</td><td>10927</td><td>10947</td><td>21</td><td>CGAGGCGCCGCCGCCCCCCCC</td><td>120.9</td><td>-10.0</td><td>2.5</td><td>-3.9</td><td>8.2E-05</td><td>2.8E-04</td><td>BT20/MB468</td><td>MP</td></tr>
<tr><td>>hg38_21_+_8433172_8446622_RNA45SN1_45S_50ntFlanks@10794.10817.24|CTGGCATGTTGGAACAATGTAGGT</td><td>>hg38_21_+_8433172_8446622_RNA45SN1_45S_50ntFlanks</td><td>10794</td><td>10817</td><td>24</td><td>CTGGCATGTTGGAACAATGTAGGT</td><td>368.5</td><td>-2.6</td><td>0.7</td><td>-3.8</td><td>1.6E-04</td><td>2.8E-04</td><td>BT20/MB468</td><td>MITO</td></tr>
<tr><td>>MI0000447|hsa-mir-128-1&WithFlank&2|+|135665391|135665484@56.77.22&[MIMAT0000424&hsa-miR-128-3p&offsets|0|+1;m-108&2|+|135665446|135665466&offsets|0|+1]|TCACAGTGAACCGGTCTCTTTT</td><td>>MI0000447|hsa-mir-128-1&WithFlank&2|+|135665391|135665484</td><td>56</td><td>77</td><td>22</td><td>TCACAGTGAACCGGTCTCTTTT</td><td>2832.8</td><td>3.0</td><td>0.8</td><td>3.7</td><td>2.2E-04</td><td>2.8E-04</td><td>MB231/MB468</td><td>NUC</td></tr>
<tr><td>>hg38_21_+_8433172_8446622_RNA45SN1_45S_50ntFlanks@4203.4223.21|GACCCGGGGAGGTAGTGACGA</td><td>>hg38_21_+_8433172_8446622_RNA45SN1_45S_50ntFlanks</td><td>4203</td><td>4223</td><td>21</td><td>GACCCGGGGAGGTAGTGACGA</td><td>490.8</td><td>-6.6</td><td>1.8</td><td>-3.7</td><td>2.2E-04</td><td>2.8E-04</td><td>BT20/MB468</td><td>NUC</td></tr>
<tr><td>>hg38_21_+_8433172_8446622_RNA45SN1_45S_50ntFlanks@11792.11810.19|ATGAACGAGATTCCCACTG</td><td>>hg38_21_+_8433172_8446622_RNA45SN1_45S_50ntFlanks</td><td>11792</td><td>11810</td><td>19</td><td>ATGAACGAGATTCCCACTG</td><td>278.1</td><td>-5.7</td><td>1.5</td><td>-3.7</td><td>1.9E-04</td><td>2.8E-04</td><td>BT20/MB468</td><td>CYTO</td></tr>
<tr><td>>hg38_21_+_8433172_8446622_RNA45SN1_45S_50ntFlanks@7987.8011.25|TCAGACGTGGCGACCCGCTGAATTT</td><td>>hg38_21_+_8433172_8446622_RNA45SN1_45S_50ntFlanks</td><td>7987</td><td>8011</td><td>25</td><td>TCAGACGTGGCGACCCGCTGAATTT</td><td>66.4</td><td>-5.7</td><td>1.5</td><td>-3.7</td><td>1.8E-04</td><td>2.8E-04</td><td>MB231/MB468</td><td>MP</td></tr>
<tr><td>>hg38_21_+_8433172_8446622_RNA45SN1_45S_50ntFlanks@10689.10706.18|GCGTCCGGTGAGCTCTCG</td><td>>hg38_21_+_8433172_8446622_RNA45SN1_45S_50ntFlanks</td><td>10689</td><td>10706</td><td>18</td><td>GCGTCCGGTGAGCTCTCG</td><td>1413.0</td><td>-6.7</td><td>1.7</td><td>-3.9</td><td>8.3E-05</td><td>2.8E-04</td><td>BT20/MB468</td><td>MP</td></tr>
<tr><td>>am_trnaMT_ValTAC_MT_+_1602_1670@28.67.40|AACTTACACTTAGGAGATTTCAACTTAACTTGACCGCTCT</td><td>>am_trnaMT_ValTAC_MT_+_1602_1670</td><td>28</td><td>67</td><td>40</td><td>AACTTACACTTAGGAGATTTCAACTTAACTTGACCGCTCT</td><td>75.2</td><td>6.2</td><td>1.7</td><td>3.7</td><td>1.8E-04</td><td>2.8E-04</td><td>MB231/MB468</td><td>MP</td></tr>
<tr><td>>hg38_21_+_8433172_8446622_RNA45SN1_45S_50ntFlanks@8659.8677.19|CATTTCCACCGCGGCGGTG</td><td>>hg38_21_+_8433172_8446622_RNA45SN1_45S_50ntFlanks</td><td>8659</td><td>8677</td><td>19</td><td>CATTTCCACCGCGGCGGTG</td><td>45.8</td><td>-9.4</td><td>2.5</td><td>-3.7</td><td>1.8E-04</td><td>2.8E-04</td><td>MB231/MB468</td><td>MP</td></tr>
<tr><td>>MI0000115|hsa-mir-16-2&WithFlank&3|+|160404739|160404831@59.80.22&[MIMAT0004518&hsa-miR-16-2-3p&offsets|0|0;m-252&3|+|160404796|160404816&offsets|+1|+2]|CCAATATTACTGTGCTGCTTTA</td><td>>MI0000115|hsa-mir-16-2&WithFlank&3|+|160404739|160404831</td><td>59</td><td>80</td><td>22</td><td>CCAATATTACTGTGCTGCTTTA</td><td>166.4</td><td>4.6</td><td>1.2</td><td>3.7</td><td>2.2E-04</td><td>2.8E-04</td><td>MB231/MB468</td><td>MITO</td></tr>
<tr><td>>hg38_21_+_8433172_8446622_RNA45SN1_45S_50ntFlanks@11576.11600.25|CTTAGAACTGGTGCGGACCAGGGGA</td><td>>hg38_21_+_8433172_8446622_RNA45SN1_45S_50ntFlanks</td><td>11576</td><td>11600</td><td>25</td><td>CTTAGAACTGGTGCGGACCAGGGGA</td><td>424.4</td><td>-5.3</td><td>1.4</td><td>-3.9</td><td>8.3E-05</td><td>2.8E-04</td><td>BT20/MB468</td><td>MP</td></tr>
<tr><td>>MI0000760|hsa-mir-361&WithFlank&X|-|85903630|85903713@51.73.23&[MIMAT0004682&hsa-miR-361-3p&offsets|0|0;m-124&X|-|85903642|85903663&offsets|0|+1]|TCCCCCAGGTGTGATTCTGATTT</td><td>>MI0000760|hsa-mir-361&WithFlank&X|-|85903630|85903713</td><td>51</td><td>73</td><td>23</td><td>TCCCCCAGGTGTGATTCTGATTT</td><td>195.7</td><td>5.6</td><td>1.5</td><td>3.7</td><td>2.2E-04</td><td>2.8E-04</td><td>MB231/MB468</td><td>MITO</td></tr>
<tr><td>>hg38_21_+_8433172_8446622_RNA45SN1_45S_50ntFlanks@6653.6681.29|ACTCTTAGCGGTGGATCACTCGGCTCGTG</td><td>>hg38_21_+_8433172_8446622_RNA45SN1_45S_50ntFlanks</td><td>6653</td><td>6681</td><td>29</td><td>ACTCTTAGCGGTGGATCACTCGGCTCGTG</td><td>6204.9</td><td>-3.5</td><td>0.9</td><td>-3.7</td><td>2.2E-04</td><td>2.8E-04</td><td>MB231/MB468</td><td>MITO</td></tr>
<tr><td>>hg38_21_+_8433172_8446622_RNA45SN1_45S_50ntFlanks@10677.10709.33|GCGGTTCCGGCGGCGTCCGGTGAGCTCTCGCTG</td><td>>hg38_21_+_8433172_8446622_RNA45SN1_45S_50ntFlanks</td><td>10677</td><td>10709</td><td>33</td><td>GCGGTTCCGGCGGCGTCCGGTGAGCTCTCGCTG</td><td>146.1</td><td>-3.4</td><td>0.9</td><td>-3.7</td><td>1.8E-04</td><td>2.8E-04</td><td>MB231/MB468</td><td>MP</td></tr>
<tr><td>>hg38_21_+_8433172_8446622_RNA45SN1_45S_50ntFlanks@11573.11602.30|CGACTTAGAACTGGTGCGGACCAGGGGAAT</td><td>>hg38_21_+_8433172_8446622_RNA45SN1_45S_50ntFlanks</td><td>11573</td><td>11602</td><td>30</td><td>CGACTTAGAACTGGTGCGGACCAGGGGAAT</td><td>537.8</td><td>-2.5</td><td>0.7</td><td>-3.7</td><td>1.9E-04</td><td>2.8E-04</td><td>BT20/MB468</td><td>CYTO</td></tr>
<tr><td>>hg38_1_-_228634819_228635039_RNA5S12_5S_50ntFlanks@85.126.42|TCGTCTGATCTCGGAAGCTAAGCAGGGTCGGGCCTGGTTAGT</td><td>>hg38_1_-_228634819_228635039_RNA5S12_5S_50ntFlanks</td><td>85</td><td>126</td><td>42</td><td>TCGTCTGATCTCGGAAGCTAAGCAGGGTCGGGCCTGGTTAGT</td><td>51.5</td><td>9.0</td><td>2.2</td><td>4.2</td><td>3.0E-05</td><td>2.8E-04</td><td>BT20/MB231</td><td>MP</td></tr>
<tr><td>>hg38_21_+_8433172_8446622_RNA45SN1_45S_50ntFlanks@10914.10931.18|CGCCGCGGCTGGACGAGG</td><td>>hg38_21_+_8433172_8446622_RNA45SN1_45S_50ntFlanks</td><td>10914</td><td>10931</td><td>18</td><td>CGCCGCGGCTGGACGAGG</td><td>64.9</td><td>9.4</td><td>2.2</td><td>4.2</td><td>2.9E-05</td><td>2.8E-04</td><td>BT20/MB231</td><td>MP</td></tr>
<tr><td>>hg38_21_+_8433172_8446622_RNA45SN1_45S_50ntFlanks@9592.9610.19|GGTATAGGGGCGAAAGACT</td><td>>hg38_21_+_8433172_8446622_RNA45SN1_45S_50ntFlanks</td><td>9592</td><td>9610</td><td>19</td><td>GGTATAGGGGCGAAAGACT</td><td>85.6</td><td>7.2</td><td>1.8</td><td>3.9</td><td>8.3E-05</td><td>2.8E-04</td><td>BT20/MB468</td><td>MP</td></tr>
<tr><td>>hg38_21_+_8433172_8446622_RNA45SN1_45S_50ntFlanks@6651.6685.35|CGACTCTTAGCGGTGGATCACTCGGCTCGTGCGTC</td><td>>hg38_21_+_8433172_8446622_RNA45SN1_45S_50ntFlanks</td><td>6651</td><td>6685</td><td>35</td><td>CGACTCTTAGCGGTGGATCACTCGGCTCGTGCGTC</td><td>1249.5</td><td>1.4</td><td>0.4</td><td>3.7</td><td>2.2E-04</td><td>2.8E-04</td><td>BT20/MB468</td><td>NUC</td></tr>
<tr><td>>MI0000273|hsa-mir-183&WithFlank&7|-|129774899|129775020@34.54.21&[MIMAT0000261&hsa-miR-183-5p&offsets|+1|0;m-41&7|-|129774966|129774987&offsets|0|-1]|ATGGCACTGGTAGAATTCACT</td><td>>MI0000273|hsa-mir-183&WithFlank&7|-|129774899|129775020</td><td>34</td><td>54</td><td>21</td><td>ATGGCACTGGTAGAATTCACT</td><td>709.9</td><td>2.5</td><td>0.7</td><td>3.7</td><td>2.2E-04</td><td>2.8E-04</td><td>MB231/MB468</td><td>NUC</td></tr>
<tr><td>>hg38_21_+_8433172_8446622_RNA45SN1_45S_50ntFlanks@11766.11803.38|ATCTAATTAGTGACGCGCATGAATGGATGAACGAGATT</td><td>>hg38_21_+_8433172_8446622_RNA45SN1_45S_50ntFlanks</td><td>11766</td><td>11803</td><td>38</td><td>ATCTAATTAGTGACGCGCATGAATGGATGAACGAGATT</td><td>453.8</td><td>-5.5</td><td>1.5</td><td>-3.7</td><td>2.2E-04</td><td>2.8E-04</td><td>MB231/MB468</td><td>NUC</td></tr>
<tr><td>>hg38_21_+_8433172_8446622_RNA45SN1_45S_50ntFlanks@9736.9753.18|CCGAAACGATCTCAACCT</td><td>>hg38_21_+_8433172_8446622_RNA45SN1_45S_50ntFlanks</td><td>9736</td><td>9753</td><td>18</td><td>CCGAAACGATCTCAACCT</td><td>157.5</td><td>-4.3</td><td>1.2</td><td>-3.7</td><td>2.2E-04</td><td>2.8E-04</td><td>MB231/MB468</td><td>NUC</td></tr>
<tr><td>>hg38_21_+_8433172_8446622_RNA45SN1_45S_50ntFlanks@10682.10712.31|TCCGGCGGCGTCCGGTGAGCTCTCGCTGGCC</td><td>>hg38_21_+_8433172_8446622_RNA45SN1_45S_50ntFlanks</td><td>10682</td><td>10712</td><td>31</td><td>TCCGGCGGCGTCCGGTGAGCTCTCGCTGGCC</td><td>431.3</td><td>-2.5</td><td>0.7</td><td>-3.7</td><td>2.2E-04</td><td>2.8E-04</td><td>MB231/MB468</td><td>NUC</td></tr>
<tr><td>>hg38_21_+_8433172_8446622_RNA45SN1_45S_50ntFlanks@11896.11935.40|TCTAGTCTGGCACGGTGAAGAGACATGAGAGGTGTAGAAT</td><td>>hg38_21_+_8433172_8446622_RNA45SN1_45S_50ntFlanks</td><td>11896</td><td>11935</td><td>40</td><td>TCTAGTCTGGCACGGTGAAGAGACATGAGAGGTGTAGAAT</td><td>71.5</td><td>9.5</td><td>2.3</td><td>4.2</td><td>3.0E-05</td><td>2.8E-04</td><td>BT20/MB231</td><td>MP</td></tr>
<tr><td>>hg38_21_+_8433172_8446622_RNA45SN1_45S_50ntFlanks@6646.6676.31|TCGTACGACTCTTAGCGGTGGATCACTCGGC</td><td>>hg38_21_+_8433172_8446622_RNA45SN1_45S_50ntFlanks</td><td>6646</td><td>6676</td><td>31</td><td>TCGTACGACTCTTAGCGGTGGATCACTCGGC</td><td>2394.0</td><td>3.7</td><td>1.0</td><td>3.7</td><td>2.2E-04</td><td>2.8E-04</td><td>MB231/MB468</td><td>NUC</td></tr>
<tr><td>>hg38_21_+_8433172_8446622_RNA45SN1_45S_50ntFlanks@4733.4771.39|GAACGAAAGTCGGAGGTTCGAAGACGATCAGATACCGTC</td><td>>hg38_21_+_8433172_8446622_RNA45SN1_45S_50ntFlanks</td><td>4733</td><td>4771</td><td>39</td><td>GAACGAAAGTCGGAGGTTCGAAGACGATCAGATACCGTC</td><td>76.0</td><td>7.1</td><td>1.8</td><td>3.9</td><td>8.3E-05</td><td>2.8E-04</td><td>BT20/MB468</td><td>MP</td></tr>
<tr><td>>trnaMT_HisGTG_MT_+_12138_12206@26.70.45|AGATTGTGAATCTGACAACAGAGGCTTACGACCCCTTATTTACCC</td><td>>trnaMT_HisGTG_MT_+_12138_12206</td><td>26</td><td>70</td><td>45</td><td>AGATTGTGAATCTGACAACAGAGGCTTACGACCCCTTATTTACCC</td><td>93.8</td><td>10.3</td><td>2.6</td><td>3.9</td><td>8.3E-05</td><td>2.8E-04</td><td>BT20/MB468</td><td>MP</td></tr>
<tr><td>>hg38_21_+_8433172_8446622_RNA45SN1_45S_50ntFlanks@9566.9595.30|GACGTGCAAATCGGTCGTCCGACCTGGGTA</td><td>>hg38_21_+_8433172_8446622_RNA45SN1_45S_50ntFlanks</td><td>9566</td><td>9595</td><td>30</td><td>GACGTGCAAATCGGTCGTCCGACCTGGGTA</td><td>201.6</td><td>-5.6</td><td>1.5</td><td>-3.7</td><td>1.8E-04</td><td>2.8E-04</td><td>MB231/MB468</td><td>MP</td></tr>
</table>

| | | | | | | | | | | | | | |
|---|---|---|---|---|---|---|---|---|---|---|---|---|---|
| >hg38_21_+_8433172_8446622_RNA45SN1_45S_50ntFlanks@4685.4722.38\|AGACGGACCAGAGCGAAAGCATTTGCCAAGAATGTTTT | >hg38_21_+_8433172_8446622_RNA45SN1_45S_50ntFlanks | 4685 | 4722 | 38 | AGACGGACCAGAGCGAAAGCATTTGCCAAGAATGTTTT | 167.7 | -5.1 | 1.4 | -3.7 | 1.8E-04 | 2.8E-04 | MB231/MB468 | MP |
| >hg38_21_+_8433172_8446622_RNA45SN1_45S_50ntFlanks@11209.11234.26\|CGGCGACTCTGGACGCGAGCCGGGCC | >hg38_21_+_8433172_8446622_RNA45SN1_45S_50ntFlanks | 11209 | 11234 | 26 | CGGCGACTCTGGACGCGAGCCGGGCC | 3907.7 | -5.3 | 1.4 | -3.7 | 1.9E-04 | 2.8E-04 | BT20/MB468 | CYTO |
| >MI0000086\|hsa-mir-28&WithFlank&3\|+\|188688775\|188688872@60.82.23(+1U)&[MIMAT0004502&hsa-miR-28-3p&offsets\|0\|+1(+1U);m-28&3\|+\|188688834\|188688855&offsets\|0\|+1(+1U)]\|CACTAGATTGTGAGCTCCTGGAGT | >MI0000086\|hsa-mir-28&WithFlank&3\|+\|188688775\|188688872 | 60 | 82 | 23(+1U) | CACTAGATTGTGAGCTCCTGGAGT | 328.1 | 5.8 | 1.5 | 3.7 | 1.8E-04 | 2.8E-04 | MB231/MB468 | MP |
| >hg38_MT_+_598_1651_RNR1_12S_50ntFlanks@853.877.25\|CTTAGTTGAACAGGGCCCTGAAGCG | >hg38_MT_+_598_1651_RNR1_12S_50ntFlanks | 853 | 877 | 25 | CTTAGTTGAACAGGGCCCTGAAGCG | 128.4 | 7.6 | 1.9 | 3.9 | 8.3E-05 | 2.8E-04 | BT20/MB468 | MP |
| >am_trnaMT_ValTAC_MT_+_1602_1670@20.48.29\|AAGCACCCAACTTACACTTAGGAGATTTC | >am_trnaMT_ValTAC_MT_+_1602_1670 | 20 | 48 | 29 | AAGCACCCAACTTACACTTAGGAGATTTC | 101.9 | 10.4 | 2.6 | 3.9 | 8.3E-05 | 2.8E-04 | BT20/MB468 | MP |
| >hg38_21_+_8433172_8446622_RNA45SN1_45S_50ntFlanks@12476.12493.18\|CCACTAATAGGGAACGTG | >hg38_21_+_8433172_8446622_RNA45SN1_45S_50ntFlanks | 12476 | 12493 | 18 | CCACTAATAGGGAACGTG | 169.2 | -2.8 | 0.7 | -3.8 | 1.6E-04 | 2.8E-04 | BT20/MB468 | MITO |
| >hg38_21_+_8433172_8446622_RNA45SN1_45S_50ntFlanks@10535.10567.33\|CGCGAGGCGTCCAGTGCGGTAACGCGACCGATC | >hg38_21_+_8433172_8446622_RNA45SN1_45S_50ntFlanks | 10535 | 10567 | 33 | CGCGAGGCGTCCAGTGCGGTAACGCGACCGATC | 227.2 | -5.9 | 1.5 | -3.9 | 8.4E-05 | 2.8E-04 | BT20/MB468 | MP |
| >hg38_21_+_8433172_8446622_RNA45SN1_45S_50ntFlanks@12934.12952.19\|CTAAACCATTCGTAGACGA | >hg38_21_+_8433172_8446622_RNA45SN1_45S_50ntFlanks | 12934 | 12952 | 19 | CTAAACCATTCGTAGACGA | 206.8 | -2.7 | 0.7 | -3.8 | 1.6E-04 | 2.8E-04 | BT20/MB468 | MITO |
| >hg38_21_+_8433172_8446622_RNA45SN1_45S_50ntFlanks@12420.12439.20\|ATGTCGGCTCTTCCTATCAT | >hg38_21_+_8433172_8446622_RNA45SN1_45S_50ntFlanks | 12420 | 12439 | 20 | ATGTCGGCTCTTCCTATCAT | 2582.2 | -3.6 | 1.0 | -3.7 | 2.2E-04 | 2.8E-04 | BT20/MB468 | NUC |
| >hg38_21_+_8433172_8446622_RNA45SN1_45S_50ntFlanks@5497.5514.18\|ACGGTCGAACTTGACTAT | >hg38_21_+_8433172_8446622_RNA45SN1_45S_50ntFlanks | 5497 | 5514 | 18 | ACGGTCGAACTTGACTAT | 95.3 | -9.7 | 2.5 | -3.9 | 8.4E-05 | 2.8E-04 | BT20/MB468 | MP |
| >hg38_21_+_8433172_8446622_RNA45SN1_45S_50ntFlanks@11561.11592.32\|GCGCCTAGCAGCCGACTTAGAACTGGTGCGGA | >hg38_21_+_8433172_8446622_RNA45SN1_45S_50ntFlanks | 11561 | 11592 | 32 | GCGCCTAGCAGCCGACTTAGAACTGGTGCGGA | 1825.4 | -6.9 | 1.8 | -3.9 | 8.4E-05 | 2.8E-04 | BT20/MB468 | MP |
| >hg38_21_+_8433172_8446622_RNA45SN1_45S_50ntFlanks@475.515.41\|CTTCGTGATCGATGTGGTGACGTCGTGCTCTCCCGGGCCGG | >hg38_21_+_8433172_8446622_RNA45SN1_45S_50ntFlanks | 475 | 515 | 41 | CTTCGTGATCGATGTGGTGACGTCGTGCTCTCCCGGGCCGG | 488.8 | 5.8 | 1.6 | 3.7 | 2.2E-04 | 2.8E-04 | MB231/MB468 | MITO |
| >trnaMT_ProTGG_MT_-_15956_16023@47.70.24\|GAGTTAAAGACTTTTTCTCTGACC | >trnaMT_ProTGG_MT_-_15956_16023 | 47 | 70 | 24 | GAGTTAAAGACTTTTTCTCTGACC | 182.7 | 3.0 | 0.8 | 3.7 | 1.9E-04 | 2.8E-04 | MB231/MB468 | MP |
| >trnaMT_GluTTC_MT_-_14674_14742@39.59.21,trnalookalike8_GluTTC_5_-_93905172_93905240@39.59.21\|ATTGGTCGTGGTTGTAGTCCG | >trnaMT_GluTTC_MT_-_14674_14742 | 39 | 59 | 21 | ATTGGTCGTGGTTGTAGTCCG | 226.0 | 5.3 | 1.3 | 3.9 | 8.4E-05 | 2.8E-04 | BT20/MB468 | MP |
| >hg38_21_+_8433172_8446622_RNA45SN1_45S_50ntFlanks@12324.12347.24\|TTGGGTTTTAAGCAGGAGGTGTCA | >hg38_21_+_8433172_8446622_RNA45SN1_45S_50ntFlanks | 12324 | 12347 | 24 | TTGGGTTTTAAGCAGGAGGTGTCA | 151.6 | -5.8 | 1.5 | -3.9 | 8.4E-05 | 2.8E-04 | BT20/MB468 | MP |
| >hg38_21_+_8433172_8446622_RNA45SN1_45S_50ntFlanks@9925.9947.23\|GGTTGATATAGACAGCAGGACGG | >hg38_21_+_8433172_8446622_RNA45SN1_45S_50ntFlanks | 9925 | 9947 | 23 | GGTTGATATAGACAGCAGGACGG | 57.3 | 9.2 | 2.2 | 4.2 | 3.0E-05 | 2.8E-04 | BT20/MB231 | MP |
| >hg38_21_+_8433172_8446622_RNA45SN1_45S_50ntFlanks@7974.8001.28\|ACGCGACCTCAGATCAGACGTGGCGACC | >hg38_21_+_8433172_8446622_RNA45SN1_45S_50ntFlanks | 7974 | 8001 | 28 | ACGCGACCTCAGATCAGACGTGGCGACC | 528.1 | 3.4 | 0.8 | 4.2 | 3.1E-05 | 2.8E-04 | BT20/MB231 | NUC |
| >MI0000342\|hsa-mir-200b&WithFlank&1\|+\|1167098\|1167204@63.83.21&[MIMAT0000318&hsa-miR-200b-3p&offsets\|0\|-1;m-93&1\|+\|1167160\|1167182&offsets\|0\|-2]\|TAATACTGCCTGGTAATGATG | >MI0000342\|hsa-mir-200b&WithFlank&1\|+\|1167098\|1167204 | 63 | 83 | 21 | TAATACTGCCTGGTAATGATG | 1091.3 | -3.9 | 1.0 | -3.9 | 8.4E-05 | 2.9E-04 | BT20/MB468 | MP |
| >hg38_21_+_8433172_8446622_RNA45SN1_45S_50ntFlanks@10807.10829.23\|ACAATGTAGGTAAGGGAAGTCGG | >hg38_21_+_8433172_8446622_RNA45SN1_45S_50ntFlanks | 10807 | 10829 | 23 | ACAATGTAGGTAAGGGAAGTCGG | 185.4 | -4.6 | 1.2 | -3.7 | 1.9E-04 | 2.9E-04 | MB231/MB468 | MP |
| >MI0003560\|hsa-mir-92b&WithFlank&1\|+\|155195171\|155195278@67.86.20(+1A)&[MIMAT0003218&hsa-miR-92b-3p&offsets\|0\|-2(+1A);m-115&1\|+\|155195237\|155195258&offsets\|0\|-2(+1A)]\|TATTGCACTCGTCCCGGCCTA | >MI0003560\|hsa-mir-92b&WithFlank&1\|+\|155195171\|155195278 | 67 | 86 | 20(+1A) | TATTGCACTCGTCCCGGCCTA | 152.9 | 7.9 | 2.0 | 3.9 | 8.4E-05 | 2.9E-04 | BT20/MB468 | MP |
| >hg38_21_+_8433172_8446622_RNA45SN1_45S_50ntFlanks@10678.10708.31\|CGGTTCCGGCGGCGTCCGGTGAGCTCTCGCT | >hg38_21_+_8433172_8446622_RNA45SN1_45S_50ntFlanks | 10678 | 10708 | 31 | CGGTTCCGGCGGCGTCCGGTGAGCTCTCGCT | 22488.8 | -3.7 | 1.0 | -3.8 | 1.6E-04 | 2.9E-04 | BT20/MB468 | MITO |
| >MI0000085\|hsa-mir-27a&WithFlank&19\|-\|13836434\|13836523@57.76.20(+1U)&[MIMAT0000084&hsa-miR-27a-3p&offsets\|0\|-1(+1U);m-43&19\|-\|13836447\|13836467&offsets\|0\|-1(+1U)]\|TTCACAGTGGCTAAGTTCCGT | >MI0000085\|hsa-mir-27a&WithFlank&19\|-\|13836434\|13836523 | 57 | 76 | 20(+1U) | TTCACAGTGGCTAAGTTCCGT | 12576.2 | 3.0 | 0.8 | 3.7 | 1.9E-04 | 2.9E-04 | MB231/MB468 | MP |
| >hg38_21_+_8433172_8446622_RNA45SN1_45S_50ntFlanks@5485.5509.25\|AGCGCTGAGAAGACGGTCGAACTTG | >hg38_21_+_8433172_8446622_RNA45SN1_45S_50ntFlanks | 5485 | 5509 | 25 | AGCGCTGAGAAGACGGTCGAACTTG | 479.3 | -3.6 | 1.0 | -3.7 | 2.3E-04 | 2.9E-04 | BT20/MB468 | NUC |
| >trnaMT_PheGAA_MT_+_577_647@42.74.33\|TTTAGACGGGCTCACATCACCCCATAAACACCA | >trnaMT_PheGAA_MT_+_577_647 | 42 | 74 | 33 | TTTAGACGGGCTCACATCACCCCATAAACACCA | 632.0 | 4.1 | 1.1 | 3.7 | 2.2E-04 | 2.9E-04 | MB231/MB468 | MITO |
| >hg38_21_+_8433172_8446622_RNA45SN1_45S_50ntFlanks@4728.4751.24\|ATCAAGAACGAAAGTCGGAGGTTC | >hg38_21_+_8433172_8446622_RNA45SN1_45S_50ntFlanks | 4728 | 4751 | 24 | ATCAAGAACGAAAGTCGGAGGTTC | 1866.9 | -6.4 | 1.7 | -3.7 | 1.9E-04 | 2.9E-04 | MB231/MB468 | MP |
| >hg38_21_+_8433172_8446622_RNA45SN1_45S_50ntFlanks@12360.12384.25\|CAGGGATAACTGGCTTGTGGCGGCC | >hg38_21_+_8433172_8446622_RNA45SN1_45S_50ntFlanks | 12360 | 12384 | 25 | CAGGGATAACTGGCTTGTGGCGGCC | 245.5 | -2.4 | 0.6 | -3.8 | 1.6E-04 | 2.9E-04 | BT20/MB468 | MITO |
| >hg38_21_+_8433172_8446622_RNA45SN1_45S_50ntFlanks@10316.10352.37\|GGTGCAGATCTTGGTGGTAGTAGCAAATATTCAAACG | >hg38_21_+_8433172_8446622_RNA45SN1_45S_50ntFlanks | 10316 | 10352 | 37 | GGTGCAGATCTTGGTGGTAGTAGCAAATATTCAAACG | 65.2 | 9.4 | 2.2 | 4.2 | 3.0E-05 | 2.9E-04 | BT20/MB231 | MP |
| >trnaMT_PheGAA_MT_+_577_647@22.65.44\|AAAGCAATACACTGAAAATGTTTAGACGGGCTCACATCACCCCA | >trnaMT_PheGAA_MT_+_577_647 | 22 | 65 | 44 | AAAGCAATACACTGAAAATGTTTAGACGGGCTCACATCACCCCA | 93.0 | 10.3 | 2.6 | 3.9 | 8.5E-05 | 2.9E-04 | BT20/MB468 | MP |
| >hg38_21_+_8433172_8446622_RNA45SN1_45S_50ntFlanks@5467.5498.32\|GGCCCACGGCCCTGGCGGAGCGCTGAGAAGAC | >hg38_21_+_8433172_8446622_RNA45SN1_45S_50ntFlanks | 5467 | 5498 | 32 | GGCCCACGGCCCTGGCGGAGCGCTGAGAAGAC | 174.1 | -3.6 | 1.0 | -3.7 | 2.1E-04 | 2.9E-04 | MB231/MB468 | CYTO |
| >trnaMT_PheGAA_MT_+_577_647@44.74.31\|TAGACGGGCTCACATCACCCCATAAACACCA | >trnaMT_PheGAA_MT_+_577_647 | 44 | 74 | 31 | TAGACGGGCTCACATCACCCCATAAACACCA | 322.0 | 5.0 | 1.3 | 3.7 | 2.0E-04 | 2.9E-04 | BT20/MB468 | CYTO |
| >hg38_21_+_8433172_8446622_RNA45SN1_45S_50ntFlanks@10409.10429.21\|GTCAGTCGGTCCTGAGAGATG | >hg38_21_+_8433172_8446622_RNA45SN1_45S_50ntFlanks | 10409 | 10429 | 21 | GTCAGTCGGTCCTGAGAGATG | 1515.5 | 7.5 | 1.8 | 4.2 | 3.0E-05 | 2.9E-04 | BT20/MB231 | MP |
| >hg38_21_+_8433172_8446622_RNA45SN1_45S_50ntFlanks@12941.12960.20\|ATTCGTAGACGACCTGCTTC | >hg38_21_+_8433172_8446622_RNA45SN1_45S_50ntFlanks | 12941 | 12960 | 20 | ATTCGTAGACGACCTGCTTC | 335.5 | 1.9 | 0.5 | 3.7 | 2.2E-04 | 2.9E-04 | MB231/MB468 | NUC |
| >hg38_21_+_8433172_8446622_RNA45SN1_45S_50ntFlanks@3985.4008.24\|CGGCTTTGGTGACTCTAGATAACC | >hg38_21_+_8433172_8446622_RNA45SN1_45S_50ntFlanks | 3985 | 4008 | 24 | CGGCTTTGGTGACTCTAGATAACC | 193.6 | -4.1 | 1.1 | -3.7 | 2.2E-04 | 2.9E-04 | MB231/MB468 | NUC |
| >hg38_MT_+_598_1651_RNR1_12S_50ntFlanks@424.462.39\|TACGAAAGTGGCTTTAACATATCTGAACACACAATAGCT | >hg38_MT_+_598_1651_RNR1_12S_50ntFlanks | 424 | 462 | 39 | TACGAAAGTGGCTTTAACATATCTGAACACACAATAGCT | 21.8 | 7.7 | 2.1 | 3.7 | 1.9E-04 | 2.9E-04 | MB231/MB468 | MP |
| >hg38_21_+_8433172_8446622_RNA45SN1_45S_50ntFlanks@11901.11920.20\|TCTGGCACGGTGAAGAGACA | >hg38_21_+_8433172_8446622_RNA45SN1_45S_50ntFlanks | 11901 | 11920 | 20 | TCTGGCACGGTGAAGAGACA | 231.4 | -2.5 | 0.7 | -3.7 | 2.3E-04 | 2.9E-04 | MB231/MB468 | NUC |
| >hg38_21_+_8433172_8446622_RNA45SN1_45S_50ntFlanks@7985.8015.31\|GATCAGACGTGGCGACCCGCTGAATTTAAGC | >hg38_21_+_8433172_8446622_RNA45SN1_45S_50ntFlanks | 7985 | 8015 | 31 | GATCAGACGTGGCGACCCGCTGAATTTAAGC | 93.5 | -5.6 | 1.5 | -3.7 | 1.9E-04 | 2.9E-04 | MB231/MB468 | MP |
| >hg38_21_+_8433172_8446622_RNA45SN1_45S_50ntFlanks@8168.8197.30\|CCCAAGTCCTTCTGATCGAGGCCCAGCCCG | >hg38_21_+_8433172_8446622_RNA45SN1_45S_50ntFlanks | 8168 | 8197 | 30 | CCCAAGTCCTTCTGATCGAGGCCCAGCCCG | 600.7 | -5.9 | 1.6 | -3.8 | 1.7E-04 | 2.9E-04 | BT20/MB468 | MITO |
| >hg38_21_+_8433172_8446622_RNA45SN1_45S_50ntFlanks@6734.6753.20\|ATTGATCATCGACACTTCGA | >hg38_21_+_8433172_8446622_RNA45SN1_45S_50ntFlanks | 6734 | 6753 | 20 | ATTGATCATCGACACTTCGA | 234.5 | -4.3 | 1.2 | -3.7 | 1.9E-04 | 2.9E-04 | MB231/MB468 | MP |
| >hg38_21_+_8433172_8446622_RNA45SN1_45S_50ntFlanks@11569.11594.26\|CAGCCGACTTAGAACTGGTGCGGACC | >hg38_21_+_8433172_8446622_RNA45SN1_45S_50ntFlanks | 11569 | 11594 | 26 | CAGCCGACTTAGAACTGGTGCGGACC | 722.7 | -3.3 | 0.9 | -3.9 | 8.5E-05 | 2.9E-04 | BT20/MB468 | MP |
| >hg38_21_+_8433172_8446622_RNA45SN1_45S_50ntFlanks@11656.11687.32\|CGATGTGATTTCTGCCCAGTGCTCTGAATGTC | >hg38_21_+_8433172_8446622_RNA45SN1_45S_50ntFlanks | 11656 | 11687 | 32 | CGATGTGATTTCTGCCCAGTGCTCTGAATGTC | 243.1 | 5.7 | 1.5 | 3.9 | 8.5E-05 | 2.9E-04 | BT20/MB468 | MP |
| >hg38_21_+_8433172_8446622_RNA45SN1_45S_50ntFlanks@3705.3746.42\|TACCTGGTTGATCCTGCCAGTAGCATATGCTTGTCTCAAAGA | >hg38_21_+_8433172_8446622_RNA45SN1_45S_50ntFlanks | 3705 | 3746 | 42 | TACCTGGTTGATCCTGCCAGTAGCATATGCTTGTCTCAAAGA | 1825.2 | 4.4 | 1.2 | 3.7 | 2.3E-04 | 2.9E-04 | BT20/MB468 | NUC |
| >trnaMT_PheGAA_MT_+_577_647@22.55.34\|AAAGCAATACACTGAAAATGTTTAGACGGGCTCA | >trnaMT_PheGAA_MT_+_577_647 | 22 | 55 | 34 | AAAGCAATACACTGAAAATGTTTAGACGGGCTCA | 94.7 | 10.3 | 2.6 | 3.9 | 8.6E-05 | 2.9E-04 | BT20/MB468 | MP |
| >hg38_21_+_8433172_8446622_RNA45SN1_45S_50ntFlanks@6731.6771.41\|CACATTGATCATCGACACTTCGAACGCACTTGCGGCCCCGG | >hg38_21_+_8433172_8446622_RNA45SN1_45S_50ntFlanks | 6731 | 6771 | 41 | CACATTGATCATCGACACTTCGAACGCACTTGCGGCCCCGG | 362.5 | -3.9 | 1.0 | -3.7 | 2.1E-04 | 2.9E-04 | MB231/MB468 | CYTO |
| >MI0000825\|hsa-mir-345&WithFlank&14\|+\|100307853\|100307962@24.45.22(+1A)&[MIMAT0000772&hsa-miR-345-5p&offsets\|0\|0(+1A);m-193&14\|+\|100307876\|100307896&offsets\|0\|+1(+1A)]\|GCTGACTCCTAGTCCAGGGCTCA | >MI0000825\|hsa-mir-345&WithFlank&14\|+\|100307853\|100307962 | 24 | 45 | 22(+1A) | GCTGACTCCTAGTCCAGGGCTCA | 28.0 | 8.0 | 2.2 | 3.7 | 1.9E-04 | 2.9E-04 | MB231/MB468 | MP |
| >hg38_21_+_8433172_8446622_RNA45SN1_45S_50ntFlanks@10391.10430.40\|AACAGCAGTTGAACATGGGTCAGTCGGTCCTGAGAGATGG | >hg38_21_+_8433172_8446622_RNA45SN1_45S_50ntFlanks | 10391 | 10430 | 40 | AACAGCAGTTGAACATGGGTCAGTCGGTCCTGAGAGATGG | 625.7 | -4.6 | 1.2 | -3.7 | 2.0E-04 | 2.9E-04 | BT20/MB468 | CYTO |
| >hg38_21_+_8433172_8446622_RNA45SN1_45S_50ntFlanks@11212.11254.43\|CGACTCTGGACGCGAGCCGGGCCCTTCCCGTGGATCGCCCCAG | >hg38_21_+_8433172_8446622_RNA45SN1_45S_50ntFlanks | 11212 | 11254 | 43 | CGACTCTGGACGCGAGCCGGGCCCTTCCCGTGGATCGCCCCAG | 361.7 | -5.4 | 1.4 | -3.7 | 2.0E-04 | 2.9E-04 | BT20/MB468 | CYTO |
| >MI0000487\|hsa-mir-193a&WithFlank&17\|+\|31559990\|31560089@27.48.22&[MIMAT0004614&hsa-miR-193a-5p&offsets\|0\|0;m-123&17\|+\|31560016\|31560037&offsets\|0\|0]\|TGGGTCTTTGCGGGCGAGATGA | >MI0000487\|hsa-mir-193a&WithFlank&17\|+\|31559990\|31560089 | 27 | 48 | 22 | TGGGTCTTTGCGGGCGAGATGA | 840.5 | 4.5 | 1.1 | 4.2 | 3.1E-05 | 2.9E-04 | BT20/MB231 | MP |
| >hg38_21_+_8433172_8446622_RNA45SN1_45S_50ntFlanks@12437.12472.36\|CATTGTGAAGCAGAATTCACCAAGCGTTGGATTGTT | >hg38_21_+_8433172_8446622_RNA45SN1_45S_50ntFlanks | 12437 | 12472 | 36 | CATTGTGAAGCAGAATTCACCAAGCGTTGGATTGTT | 130.4 | -6.1 | 1.6 | -3.7 | 1.9E-04 | 2.9E-04 | MB231/MB468 | MP |
| >hg38_21_+_8433172_8446622_RNA45SN1_45S_50ntFlanks@10846.10870.25\|CTTCGGGATAAGGATTGGCTCTAAG | >hg38_21_+_8433172_8446622_RNA45SN1_45S_50ntFlanks | 10846 | 10870 | 25 | CTTCGGGATAAGGATTGGCTCTAAG | 198.9 | -2.3 | 0.6 | -3.7 | 2.3E-04 | 2.9E-04 | MB231/MB468 | NUC |
| >hg38_21_+_8433172_8446622_RNA45SN1_45S_50ntFlanks@4283.4321.39\|CTTTAACGAGGATCCATTGGAGGGCAAGTCTGGTGCCAG | >hg38_21_+_8433172_8446622_RNA45SN1_45S_50ntFlanks | 4283 | 4321 | 39 | CTTTAACGAGGATCCATTGGAGGGCAAGTCTGGTGCCAG | 116.0 | -5.0 | 1.3 | -3.7 | 2.0E-04 | 2.9E-04 | BT20/MB468 | CYTO |
| >MI0000542\|hsa-mir-320a&WithFlank&8\|-\|22244960\|22245043@49.70.22(+2U)&[MIMAT0000510&hsa-miR-320a-3p&offsets\|+1\|+1(+2U);m-60&8\|-\|22244975\|22244996&offsets\|+1\|+1(+2U)]\|AAAGCTGGGTTGAGAGGGCGAATT | >MI0000542\|hsa-mir-320a&WithFlank&8\|-\|22244960\|22245043 | 49 | 70 | 22(+2U) | AAAGCTGGGTTGAGAGGGCGAATT | 94.8 | 6.2 | 1.6 | 3.9 | 8.6E-05 | 2.9E-04 | BT20/MB468 | MP |
| >trna10_LysCTT_16_+_3241501_3241573@5.35.31,trna13_LysCTT_14_-_58706613_58706685@5.35.31,trna2_LysCTT_15_+_79152904_79152976@5.35.31\|GGCTAGCTCAGTCGGTAGAGCATGGGACTCT | >trna10_LysCTT_16_+_3241501_3241573 | 5 | 35 | 31 | GGCTAGCTCAGTCGGTAGAGCATGGGACTCT | 855.5 | -3.5 | 0.8 | -4.1 | 3.5E-05 | 2.9E-04 | BT20/MB231 | MITO |

<table>
<tr><td>>hg38_1_-_228634819_228635039_RNA5S12_5S_50ntFlanks@124.147.24|AGTACTTGGATGGGAGACCGCCTG</td><td>>hg38_1_-_228634819_228635039_RNA5S12_5S_50ntFlanks</td><td>124</td><td>147</td><td>24</td><td>AGTACTTGGATGGGAGACCGCCTG</td><td>165.8</td><td>-4.5</td><td>1.2</td><td>-3.7</td><td>2.1E-04</td><td>2.9E-04</td><td>MB231/MB468</td><td>CYTO</td></tr>
<tr><td>>hg38_21_+_8433172_8446622_RNA45SN1_45S_50ntFlanks@12995.13014.20|CTCGCTGCGATCTATTGAAA</td><td>>hg38_21_+_8433172_8446622_RNA45SN1_45S_50ntFlanks</td><td>12995</td><td>13014</td><td>20</td><td>CTCGCTGCGATCTATTGAAA</td><td>859.6</td><td>2.8</td><td>0.8</td><td>3.7</td><td>2.1E-04</td><td>2.9E-04</td><td>MB231/MB468</td><td>CYTO</td></tr>
<tr><td>>hg38_21_+_8433172_8446622_RNA45SN1_45S_50ntFlanks@10912.10932.21|CGCGCCGCGGCTGGACGAGGC</td><td>>hg38_21_+_8433172_8446622_RNA45SN1_45S_50ntFlanks</td><td>10912</td><td>10932</td><td>21</td><td>CGCGCCGCGGCTGGACGAGGC</td><td>556.9</td><td>-4.8</td><td>1.2</td><td>-3.9</td><td>8.6E-05</td><td>2.9E-04</td><td>BT20/MB468</td><td>MP</td></tr>
<tr><td>>hg38_21_+_8433172_8446622_RNA45SN1_45S_50ntFlanks@6652.6673.22|GACTCTTAGCGGTGGATCACTC</td><td>>hg38_21_+_8433172_8446622_RNA45SN1_45S_50ntFlanks</td><td>6652</td><td>6673</td><td>22</td><td>GACTCTTAGCGGTGGATCACTC</td><td>541.5</td><td>4.2</td><td>1.1</td><td>3.9</td><td>8.6E-05</td><td>2.9E-04</td><td>BT20/MB468</td><td>MP</td></tr>
<tr><td>>hg38_21_+_8433172_8446622_RNA45SN1_45S_50ntFlanks@9533.9553.21|GAGGAAACTCTGGTGGAGGTC</td><td>>hg38_21_+_8433172_8446622_RNA45SN1_45S_50ntFlanks</td><td>9533</td><td>9553</td><td>21</td><td>GAGGAAACTCTGGTGGAGGTC</td><td>3879.6</td><td>-3.9</td><td>1.0</td><td>-3.9</td><td>8.6E-05</td><td>2.9E-04</td><td>BT20/MB468</td><td>MP</td></tr>
<tr><td>>hg38_21_+_8433172_8446622_RNA45SN1_45S_50ntFlanks@11867.11893.27|CGGGGAAAGAAGACCCTGTTGAGCTTG</td><td>>hg38_21_+_8433172_8446622_RNA45SN1_45S_50ntFlanks</td><td>11867</td><td>11893</td><td>27</td><td>CGGGGAAAGAAGACCCTGTTGAGCTTG</td><td>69.8</td><td>9.5</td><td>2.3</td><td>4.2</td><td>3.1E-05</td><td>2.9E-04</td><td>BT20/MB231</td><td>MP</td></tr>
<tr><td>>hg38_21_+_8433172_8446622_RNA45SN1_45S_50ntFlanks@11190.11207.18|CCCGGGGGGCCGGCGGCG</td><td>>hg38_21_+_8433172_8446622_RNA45SN1_45S_50ntFlanks</td><td>11190</td><td>11207</td><td>18</td><td>CCCGGGGGGCCGGCGGCG</td><td>2083.5</td><td>-7.3</td><td>2.0</td><td>-3.7</td><td>2.3E-04</td><td>2.9E-04</td><td>BT20/MB468</td><td>NUC</td></tr>
<tr><td>>hg38_21_+_8433172_8446622_RNA45SN1_45S_50ntFlanks@10133.10159.27|GCGGCGGCGGCGGCGGGGGTGTGGGGT</td><td>>hg38_21_+_8433172_8446622_RNA45SN1_45S_50ntFlanks</td><td>10133</td><td>10159</td><td>27</td><td>GCGGCGGCGGCGGCGGGGGTGTGGGGT</td><td>764.4</td><td>7.6</td><td>1.9</td><td>3.9</td><td>8.6E-05</td><td>2.9E-04</td><td>BT20/MB468</td><td>MP</td></tr>
<tr><td>>hg38_21_+_8433172_8446622_RNA45SN1_45S_50ntFlanks@4734.4761.28|AACGAAAGTCGGAGGTTCGAAGACGATC</td><td>>hg38_21_+_8433172_8446622_RNA45SN1_45S_50ntFlanks</td><td>4734</td><td>4761</td><td>28</td><td>AACGAAAGTCGGAGGTTCGAAGACGATC</td><td>1154.6</td><td>3.6</td><td>0.9</td><td>3.9</td><td>8.6E-05</td><td>2.9E-04</td><td>BT20/MB468</td><td>MP</td></tr>
<tr><td>>am_trnaMT_ValTAC_MT_+_1602_1670@41.59.19|GAGATTTCAACTTAACTTG</td><td>>am_trnaMT_ValTAC_MT_+_1602_1670</td><td>41</td><td>59</td><td>19</td><td>GAGATTTCAACTTAACTTG</td><td>69.1</td><td>-5.6</td><td>1.5</td><td>-3.7</td><td>1.9E-04</td><td>2.9E-04</td><td>MB231/MB468</td><td>MP</td></tr>
<tr><td>>MI0000279|hsa-mir-196a-2&WithFlank&12|+|53991732|53991853@31.52.22(+1U);MI0000238|hsa-mir-196a-1&WithFlank&17|-|48632484|48632565@13.34.22(+1U)&[MIMAT0000226&hsa-miR-196a-5p&offsets|0|0(+1U);m-261&12|+|53991762|53991783&offsets|0|0(+1U)];[MIMAT0000226_1&hsa-miR-196a-5p&offsets|0|0(+1U);m-268&17|-|48632532|48632553&offsets|0|0(+1U)]|TAGGTAGTTTCATGTTGTTGGGT</td><td>>MI0000279|hsa-mir-196a-2&WithFlank&12|+|53991732|53991853</td><td>31</td><td>52</td><td>22(+1U)</td><td>TAGGTAGTTTCATGTTGTTGGGT</td><td>562.5</td><td>4.1</td><td>1.0</td><td>3.9</td><td>8.6E-05</td><td>2.9E-04</td><td>BT20/MB468</td><td>MP</td></tr>
<tr><td>>hg38_21_+_8433172_8446622_RNA45SN1_45S_50ntFlanks@12996.13038.43|TCGCTGCGATCTATTGAAAGTCAGCCCTCGACACAAGGGTTTG</td><td>>hg38_21_+_8433172_8446622_RNA45SN1_45S_50ntFlanks</td><td>12996</td><td>13038</td><td>43</td><td>TCGCTGCGATCTATTGAAAGTCAGCCCTCGACACAAGGGTTTG</td><td>93.7</td><td>10.3</td><td>2.6</td><td>3.9</td><td>8.6E-05</td><td>2.9E-04</td><td>BT20/MB468</td><td>MP</td></tr>
<tr><td>>hg38_21_+_8433172_8446622_RNA45SN1_45S_50ntFlanks@10388.10409.22|GTGAACAGCAGTTGAACATGGG</td><td>>hg38_21_+_8433172_8446622_RNA45SN1_45S_50ntFlanks</td><td>10388</td><td>10409</td><td>22</td><td>GTGAACAGCAGTTGAACATGGG</td><td>113.2</td><td>-6.1</td><td>1.5</td><td>-3.9</td><td>8.6E-05</td><td>2.9E-04</td><td>BT20/MB468</td><td>MP</td></tr>
<tr><td>>trnaMT_ValTAC_MT_+_1602_1670@33.72.40|ACACTTAGGAGATTTCAACTTAACTTGACCGCTCTGACCA</td><td>>trnaMT_ValTAC_MT_+_1602_1670</td><td>33</td><td>72</td><td>40</td><td>ACACTTAGGAGATTTCAACTTAACTTGACCGCTCTGACCA</td><td>398.8</td><td>4.4</td><td>1.2</td><td>3.7</td><td>2.3E-04</td><td>2.9E-04</td><td>MB231/MB468</td><td>MITO</td></tr>
<tr><td>>hg38_MT_+_598_1651_RNR1_12S_50ntFlanks@428.469.42|AAAGTGGCTTTAACATATCTGAACACACAATAGCTAAGACCC</td><td>>hg38_MT_+_598_1651_RNR1_12S_50ntFlanks</td><td>428</td><td>469</td><td>42</td><td>AAAGTGGCTTTAACATATCTGAACACACAATAGCTAAGACCC</td><td>203.8</td><td>-4.8</td><td>1.1</td><td>-4.2</td><td>3.0E-05</td><td>2.9E-04</td><td>BT20/MB231</td><td>CYTO</td></tr>
<tr><td>>MI0000434|hsa-let-7i&WithFlank&12|+|62603680|62603775@12.31.20(+1A)&[MIMAT0000415&hsa-let-7i-5p&offsets|0|-2(+1A);m-49&12|+|62603691|62603712&offsets|0|-2(+1A)]|TGAGGTAGTAGTTTGTGCTGA</td><td>>MI0000434|hsa-let-7i&WithFlank&12|+|62603680|62603775</td><td>12</td><td>31</td><td>20(+1A)</td><td>TGAGGTAGTAGTTTGTGCTGA</td><td>260.2</td><td>6.0</td><td>1.6</td><td>3.7</td><td>1.9E-04</td><td>2.9E-04</td><td>MB231/MB468</td><td>MP</td></tr>
<tr><td>>hg38_21_+_8433172_8446622_RNA45SN1_45S_50ntFlanks@10822.10840.19|GAAGTCGGCAAGCCGGATC</td><td>>hg38_21_+_8433172_8446622_RNA45SN1_45S_50ntFlanks</td><td>10822</td><td>10840</td><td>19</td><td>GAAGTCGGCAAGCCGGATC</td><td>412.2</td><td>-5.4</td><td>1.4</td><td>-3.9</td><td>8.7E-05</td><td>2.9E-04</td><td>BT20/MB468</td><td>MP</td></tr>
<tr><td>>hg38_21_+_8433172_8446622_RNA45SN1_45S_50ntFlanks@12028.12055.28|CTCTGATCGTTTTTTCACTGACCCGGTG</td><td>>hg38_21_+_8433172_8446622_RNA45SN1_45S_50ntFlanks</td><td>12028</td><td>12055</td><td>28</td><td>CTCTGATCGTTTTTTCACTGACCCGGTG</td><td>152.0</td><td>7.1</td><td>1.8</td><td>3.9</td><td>8.7E-05</td><td>2.9E-04</td><td>BT20/MB468</td><td>MP</td></tr>
<tr><td>>hg38_21_+_8433172_8446622_RNA45SN1_45S_50ntFlanks@11564.11600.37|CCTAGCAGCCGACTTAGAACTGGTGCGGACCAGGGGA</td><td>>hg38_21_+_8433172_8446622_RNA45SN1_45S_50ntFlanks</td><td>11564</td><td>11600</td><td>37</td><td>CCTAGCAGCCGACTTAGAACTGGTGCGGACCAGGGGA</td><td>660.0</td><td>-4.1</td><td>1.1</td><td>-3.7</td><td>2.1E-04</td><td>2.9E-04</td><td>MB231/MB468</td><td>CYTO</td></tr>
<tr><td>>hg38_1_-_228634819_228635039_RNA5S12_5S_50ntFlanks@133.170.38|ATGGGAGACCGCCTGGGAATACCGGGTGCTGTAGGCTT</td><td>>hg38_1_-_228634819_228635039_RNA5S12_5S_50ntFlanks</td><td>133</td><td>170</td><td>38</td><td>ATGGGAGACCGCCTGGGAATACCGGGTGCTGTAGGCTT</td><td>33306.3</td><td>3.9</td><td>1.0</td><td>3.9</td><td>8.7E-05</td><td>2.9E-04</td><td>BT20/MB468</td><td>MP</td></tr>
<tr><td>>MI0003581|hsa-mir-574&WithFlank&4|+|38868026|38868133@67.87.21&[MIMAT0003239&hsa-miR-574-3p&offsets|0|-1;m-149&4|+|38868092|38868113&offsets|0|-1]|CACGCTCATGCACACACCCAC</td><td>>MI0003581|hsa-mir-574&WithFlank&4|+|38868026|38868133</td><td>67</td><td>87</td><td>21</td><td>CACGCTCATGCACACACCCAC</td><td>310.7</td><td>3.1</td><td>0.8</td><td>3.7</td><td>1.9E-04</td><td>2.9E-04</td><td>MB231/MB468</td><td>MP</td></tr>
<tr><td>>hg38_21_+_8433172_8446622_RNA45SN1_45S_50ntFlanks@11572.11604.33|CCGACTTAGAACTGGTGCGGACCAGGGGAATCC</td><td>>hg38_21_+_8433172_8446622_RNA45SN1_45S_50ntFlanks</td><td>11572</td><td>11604</td><td>33</td><td>CCGACTTAGAACTGGTGCGGACCAGGGGAATCC</td><td>295.7</td><td>-4.5</td><td>1.2</td><td>-3.7</td><td>1.9E-04</td><td>2.9E-04</td><td>MB231/MB468</td><td>MP</td></tr>
<tr><td>>hg38_21_+_8433172_8446622_RNA45SN1_45S_50ntFlanks@10344.10382.39|ATTCAAACGAGAACTTTGAAGGCCGAAGTGGAGAAGGGT</td><td>>hg38_21_+_8433172_8446622_RNA45SN1_45S_50ntFlanks</td><td>10344</td><td>10382</td><td>39</td><td>ATTCAAACGAGAACTTTGAAGGCCGAAGTGGAGAAGGGT</td><td>109.1</td><td>7.0</td><td>1.8</td><td>3.9</td><td>8.7E-05</td><td>2.9E-04</td><td>BT20/MB468</td><td>MP</td></tr>
<tr><td>>MI0000433|hsa-let-7g&WithFlank&3|-|52268272|52268367@10.32.23&[MIMAT0000414&hsa-let-7g-5p&offsets|-1|0;m-58&3|-|52268336|52268357&offsets|-1|0]|CTGAGGTAGTAGTTTGTACAGTT</td><td>>MI0000433|hsa-let-7g&WithFlank&3|-|52268272|52268367</td><td>10</td><td>32</td><td>23</td><td>CTGAGGTAGTAGTTTGTACAGTT</td><td>177.1</td><td>4.7</td><td>1.3</td><td>3.7</td><td>2.3E-04</td><td>2.9E-04</td><td>MB231/MB468</td><td>MITO</td></tr>
<tr><td>>hg38_21_+_8433172_8446622_RNA45SN1_45S_50ntFlanks@10845.10874.30|ACTTCGGGATAAGGATTGGCTCTAAGGGCT</td><td>>hg38_21_+_8433172_8446622_RNA45SN1_45S_50ntFlanks</td><td>10845</td><td>10874</td><td>30</td><td>ACTTCGGGATAAGGATTGGCTCTAAGGGCT</td><td>188.5</td><td>-5.3</td><td>1.4</td><td>-3.7</td><td>2.3E-04</td><td>2.9E-04</td><td>MB231/MB468</td><td>NUC</td></tr>
<tr><td>>MI0000465|hsa-mir-191&WithFlank&3|-|49020612|49020715@22.43.22(+1U)&[MIMAT0000440&hsa-miR-191-5p&offsets|0|-1(+1U);m-48&3|-|49020672|49020694&offsets|0|-1(+1U)]|CAACGGAATCCCAAAAGCAGCTT</td><td>>MI0000465|hsa-mir-191&WithFlank&3|-|49020612|49020715</td><td>22</td><td>43</td><td>22(+1U)</td><td>CAACGGAATCCCAAAAGCAGCTT</td><td>463.3</td><td>3.5</td><td>0.9</td><td>3.7</td><td>1.9E-04</td><td>2.9E-04</td><td>MB231/MB468</td><td>MP</td></tr>
<tr><td>>hg38_21_+_8433172_8446622_RNA45SN1_45S_50ntFlanks@10798.10826.29|CATGTTGGAACAATGTAGGTAAGGGAAGT</td><td>>hg38_21_+_8433172_8446622_RNA45SN1_45S_50ntFlanks</td><td>10798</td><td>10826</td><td>29</td><td>CATGTTGGAACAATGTAGGTAAGGGAAGT</td><td>243.5</td><td>-4.1</td><td>1.1</td><td>-3.7</td><td>2.3E-04</td><td>2.9E-04</td><td>MB231/MB468</td><td>NUC</td></tr>
<tr><td>>hg38_21_+_8433172_8446622_RNA45SN1_45S_50ntFlanks@6719.6740.22|TGAATTGCAGGACACATTGATC</td><td>>hg38_21_+_8433172_8446622_RNA45SN1_45S_50ntFlanks</td><td>6719</td><td>6740</td><td>22</td><td>TGAATTGCAGGACACATTGATC</td><td>1027.8</td><td>-2.3</td><td>0.6</td><td>-3.8</td><td>1.7E-04</td><td>2.9E-04</td><td>BT20/MB468</td><td>MITO</td></tr>
<tr><td>>hg38_21_+_8433172_8446622_RNA45SN1_45S_50ntFlanks@4102.4121.20|ACCACGGGTGACGGGGAATC</td><td>>hg38_21_+_8433172_8446622_RNA45SN1_45S_50ntFlanks</td><td>4102</td><td>4121</td><td>20</td><td>ACCACGGGTGACGGGGAATC</td><td>643.4</td><td>-5.1</td><td>1.4</td><td>-3.8</td><td>1.7E-04</td><td>2.9E-04</td><td>BT20/MB468</td><td>MITO</td></tr>
<tr><td>>hg38_21_+_8433172_8446622_RNA45SN1_45S_50ntFlanks@12457.12474.18|CAAGCGTTGGATTGTTCA</td><td>>hg38_21_+_8433172_8446622_RNA45SN1_45S_50ntFlanks</td><td>12457</td><td>12474</td><td>18</td><td>CAAGCGTTGGATTGTTCA</td><td>220.6</td><td>-4.0</td><td>1.1</td><td>-3.7</td><td>2.3E-04</td><td>2.9E-04</td><td>MB231/MB468</td><td>NUC</td></tr>
<tr><td>>hg38_21_+_8433172_8446622_RNA45SN1_45S_50ntFlanks@10410.10440.31|TCAGTCGGTCCTGAGAGATGGGCGAGCGCCG</td><td>>hg38_21_+_8433172_8446622_RNA45SN1_45S_50ntFlanks</td><td>10410</td><td>10440</td><td>31</td><td>TCAGTCGGTCCTGAGAGATGGGCGAGCGCCG</td><td>131.4</td><td>-2.0</td><td>0.5</td><td>-3.8</td><td>1.7E-04</td><td>2.9E-04</td><td>BT20/MB468</td><td>MITO</td></tr>
<tr><td>>hg38_21_+_8433172_8446622_RNA45SN1_45S_50ntFlanks@6733.6756.24|CATTGATCATCGACACTTCGAACG</td><td>>hg38_21_+_8433172_8446622_RNA45SN1_45S_50ntFlanks</td><td>6733</td><td>6756</td><td>24</td><td>CATTGATCATCGACACTTCGAACG</td><td>754.1</td><td>-2.3</td><td>0.6</td><td>-3.7</td><td>2.1E-04</td><td>2.9E-04</td><td>MB231/MB468</td><td>CYTO</td></tr>
<tr><td>>MI0001733|hsa-mir-452&WithFlank&X|-|151959622|151959718@23.44.22&[MIMAT0001635&hsa-miR-452-5p&offsets|+3|+3;m-188&X|-|151959677|151959699&offsets|+3|+2]|TGTTTGCAGAGGAAACTGAGAC</td><td>>MI0001733|hsa-mir-452&WithFlank&X|-|151959622|151959718</td><td>23</td><td>44</td><td>22</td><td>TGTTTGCAGAGGAAACTGAGAC</td><td>432.0</td><td>2.6</td><td>0.6</td><td>4.2</td><td>3.2E-05</td><td>2.9E-04</td><td>BT20/MB231</td><td>NUC</td></tr>
<tr><td>>hg38_21_+_8433172_8446622_RNA45SN1_45S_50ntFlanks@4297.4314.18|CATTGGAGGGCAAGTCTG</td><td>>hg38_21_+_8433172_8446622_RNA45SN1_45S_50ntFlanks</td><td>4297</td><td>4314</td><td>18</td><td>CATTGGAGGGCAAGTCTG</td><td>1624.1</td><td>8.9</td><td>2.1</td><td>4.2</td><td>3.1E-05</td><td>2.9E-04</td><td>BT20/MB231</td><td>MP</td></tr>
<tr><td>>hg38_21_+_8433172_8446622_RNA45SN1_45S_50ntFlanks@8117.8134.18|CGGGACATGTGGCGTACG</td><td>>hg38_21_+_8433172_8446622_RNA45SN1_45S_50ntFlanks</td><td>8117</td><td>8134</td><td>18</td><td>CGGGACATGTGGCGTACG</td><td>519.0</td><td>7.5</td><td>1.8</td><td>4.2</td><td>3.1E-05</td><td>2.9E-04</td><td>BT20/MB231</td><td>MP</td></tr>
<tr><td>>hg38_21_+_8433172_8446622_RNA45SN1_45S_50ntFlanks@9941.9980.40|AGGACGGTGGCCATGGAAGTCGGAATCCGCTAAGGAGTGT</td><td>>hg38_21_+_8433172_8446622_RNA45SN1_45S_50ntFlanks</td><td>9941</td><td>9980</td><td>40</td><td>AGGACGGTGGCCATGGAAGTCGGAATCCGCTAAGGAGTGT</td><td>1201.1</td><td>6.8</td><td>1.6</td><td>4.2</td><td>3.1E-05</td><td>2.9E-04</td><td>BT20/MB231</td><td>MP</td></tr>
<tr><td>>trna2_GlyGCC_21_-_18827107_18827177@1.33.33,trna35_GlyGCC_1_+_161413094_161413164@1.33.33,trna37_GlyGCC_1_+_161420467_161420537@1.33.33,trna39_GlyGCC_1_+_161427898_161427968@1.33.33,trna41_GlyGCC_1_+_161435258_161435328@1.33.33|GCATGGGTGGTTCAGTGGTAGAATTCTCGCCTG</td><td>>trna2_GlyGCC_21_-_18827107_18827177</td><td>1</td><td>33</td><td>33</td><td>GCATGGGTGGTTCAGTGGTAGAATTCTCGCCTG</td><td>597.1</td><td>3.6</td><td>0.9</td><td>4.1</td><td>3.5E-05</td><td>2.9E-04</td><td>BT20/MB231</td><td>MITO</td></tr>
<tr><td>>hg38_21_+_8433172_8446622_RNA45SN1_45S_50ntFlanks@3987.4023.37|GCTTTGGTGACTCTAGATAACCTCGGGCCGATCGCAC</td><td>>hg38_21_+_8433172_8446622_RNA45SN1_45S_50ntFlanks</td><td>3987</td><td>4023</td><td>37</td><td>GCTTTGGTGACTCTAGATAACCTCGGGCCGATCGCAC</td><td>120.2</td><td>3.3</td><td>0.9</td><td>3.7</td><td>2.3E-04</td><td>2.9E-04</td><td>BT20/MB468</td><td>NUC</td></tr>
<tr><td>>hg38_21_+_8433172_8446622_RNA45SN1_45S_50ntFlanks@9533.9571.39|GAGGAAACTCTGGTGGAGGTCCGTAGCGGTCCTGACGTG</td><td>>hg38_21_+_8433172_8446622_RNA45SN1_45S_50ntFlanks</td><td>9533</td><td>9571</td><td>39</td><td>GAGGAAACTCTGGTGGAGGTCCGTAGCGGTCCTGACGTG</td><td>2305.0</td><td>-2.4</td><td>0.6</td><td>-3.7</td><td>2.0E-04</td><td>2.9E-04</td><td>BT20/MB468</td><td>CYTO</td></tr>
<tr><td>>hg38_21_+_8433172_8446622_RNA45SN1_45S_50ntFlanks@6652.6690.39|GACTCTTAGCGGTGGATCACTCGGCTCGTGCGTCGATGA</td><td>>hg38_21_+_8433172_8446622_RNA45SN1_45S_50ntFlanks</td><td>6652</td><td>6690</td><td>39</td><td>GACTCTTAGCGGTGGATCACTCGGCTCGTGCGTCGATGA</td><td>1064.7</td><td>2.7</td><td>0.7</td><td>3.7</td><td>2.3E-04</td><td>2.9E-04</td><td>MB231/MB468</td><td>NUC</td></tr>
<tr><td>>hg38_21_+_8433172_8446622_RNA45SN1_45S_50ntFlanks@11571.11591.21|GCCGACTTAGAACTGGTGCGG</td><td>>hg38_21_+_8433172_8446622_RNA45SN1_45S_50ntFlanks</td><td>11571</td><td>11591</td><td>21</td><td>GCCGACTTAGAACTGGTGCGG</td><td>235.5</td><td>-3.5</td><td>1.0</td><td>-3.7</td><td>2.1E-04</td><td>2.9E-04</td><td>MB231/MB468</td><td>CYTO</td></tr>
<tr><td>>MI0000103|hsa-mir-101-1&WithFlank&1|-|65058428|65058514@53.73.21(+2U);MI0000739|hsa-mir-101-2&WithFlank&9|+|4850291|4850381@55.75.21(+2U)&[MIMAT0000099&hsa-miR-101-3p&offsets|0|0(+2U);m-17&1|-|65058443|65058462&offsets|0|+1(+2U)];[MIMAT0000099_1&hsa-miR-101-3p&offsets|0|0(+2U);m-16&9|+|4850345|4850364&offsets|0|+1(+2U)]|TACAGTACTGTGATAACTGAATT</td><td>>MI0000103|hsa-mir-101-1&WithFlank&1|-|65058428|65058514</td><td>53</td><td>73</td><td>21(+2U)</td><td>TACAGTACTGTGATAACTGAATT</td><td>75.7</td><td>6.4</td><td>1.7</td><td>3.7</td><td>1.9E-04</td><td>2.9E-04</td><td>MB231/MB468</td><td>MP</td></tr>
<tr><td>>hg38_21_+_8433172_8446622_RNA45SN1_45S_50ntFlanks@9539.9570.32|ACTCTGGTGGAGGTCCGTAGCGGTCCTGACGT</td><td>>hg38_21_+_8433172_8446622_RNA45SN1_45S_50ntFlanks</td><td>9539</td><td>9570</td><td>32</td><td>ACTCTGGTGGAGGTCCGTAGCGGTCCTGACGT</td><td>490.9</td><td>-5.1</td><td>1.4</td><td>-3.7</td><td>2.3E-04</td><td>2.9E-04</td><td>BT20/MB468</td><td>NUC</td></tr>
<tr><td>>hg38_21_+_8433172_8446622_RNA45SN1_45S_50ntFlanks@12710.12747.38|CGCGGAGCCTCGGTTGGCCTCGGATAGCCGGTCCCCCG</td><td>>hg38_21_+_8433172_8446622_RNA45SN1_45S_50ntFlanks</td><td>12710</td><td>12747</td><td>38</td><td>CGCGGAGCCTCGGTTGGCCTCGGATAGCCGGTCCCCCG</td><td>188.8</td><td>-6.1</td><td>1.6</td><td>-3.9</td><td>8.8E-05</td><td>2.9E-04</td><td>BT20/MB468</td><td>MP</td></tr>
<tr><td>>hg38_21_+_8433172_8446622_RNA45SN1_45S_50ntFlanks@12991.13027.37|CTCCCTCGCTGCGATCTATTGAAAGTCAGCCCTCGAC</td><td>>hg38_21_+_8433172_8446622_RNA45SN1_45S_50ntFlanks</td><td>12991</td><td>13027</td><td>37</td><td>CTCCCTCGCTGCGATCTATTGAAAGTCAGCCCTCGAC</td><td>356.0</td><td>-5.5</td><td>1.5</td><td>-3.7</td><td>2.0E-04</td><td>2.9E-04</td><td>BT20/MB468</td><td>CYTO</td></tr>
<tr><td>>hg38_21_+_8433172_8446622_RNA45SN1_45S_50ntFlanks@10349.10369.21|AACGAGAACTTTGAAGGCCGA</td><td>>hg38_21_+_8433172_8446622_RNA45SN1_45S_50ntFlanks</td><td>10349</td><td>10369</td><td>21</td><td>AACGAGAACTTTGAAGGCCGA</td><td>754.4</td><td>2.4</td><td>0.6</td><td>3.7</td><td>2.2E-04</td><td>2.9E-04</td><td>MB231/MB468</td><td>CYTO</td></tr>
<tr><td>>hg38_21_+_8433172_8446622_RNA45SN1_45S_50ntFlanks@3853.3872.20|AACTGTGGTAATTCTAGAGC</td><td>>hg38_21_+_8433172_8446622_RNA45SN1_45S_50ntFlanks</td><td>3853</td><td>3872</td><td>20</td><td>AACTGTGGTAATTCTAGAGC</td><td>202.8</td><td>-3.7</td><td>1.0</td><td>-3.7</td><td>2.2E-04</td><td>2.9E-04</td><td>MB231/MB468</td><td>CYTO</td></tr>
<tr><td>>hg38_21_+_8433172_8446622_RNA45SN1_45S_50ntFlanks@9571.9594.24|GCAAATCGGTCGTCCGACCTGGGT</td><td>>hg38_21_+_8433172_8446622_RNA45SN1_45S_50ntFlanks</td><td>9571</td><td>9594</td><td>24</td><td>GCAAATCGGTCGTCCGACCTGGGT</td><td>131.8</td><td>-4.4</td><td>1.2</td><td>-3.7</td><td>1.9E-04</td><td>2.9E-04</td><td>MB231/MB468</td><td>MP</td></tr>
</table>

<table>
<tr><td></td><td></td><td></td><td></td><td></td><td></td><td></td><td></td><td></td><td></td><td></td><td></td><td></td><td></td></tr>
<tr><td>>hg38_MT_+_1621_3279_RNR2_16S_50ntFlanks@618.663.46|AACATATAACTGAACTCCTCACACCCAATTGGACCAATCTATCACC</td><td>>hg38_MT_+_1621_3279_RNR2_16S_50ntFlanks</td><td>618</td><td>663</td><td>46</td><td>AACATATAACTGAACTCCTCACACCCAATTGGACCAATCTATCACC</td><td>93.4</td><td>10.3</td><td>2.6</td><td>3.9</td><td>8.8E-05</td><td>3.0E-04</td><td>BT20/MB468</td><td>MP</td></tr>
<tr><td>>trna111_HisGTG_1_-_147774845_147774916@1.31.31,trna118_HisGTG_1_-_145396881_145396952@1.31.31,trna16_HisGTG_1_+_146544773_146544844@1.31.31,trna1_HisGTG_15_+_45493349_45493420@1.31.31,trna21_HisGTG_1_+_147753471_147753542@1.31.31,trna33_HisGTG_6_+_27125906_27125977@1.31.31,trna7_HisGTG_9_-_14433938_14434009@1.31.31,trna8_HisGTG_15_-_45492611_45492682@1.31.31,trna9_HisGTG_15_-_45490804_45490875@1.31.31|GCCGTGATCGTATAGTGGTTAGTACTCTGCG</td><td>>trna111_HisGTG_1_-_147774845_147774916</td><td>1</td><td>31</td><td>31</td><td>GCCGTGATCGTATAGTGGTTAGTACTCTGCG</td><td>6568.5</td><td>-3.8</td><td>1.0</td><td>-3.9</td><td>8.8E-05</td><td>3.0E-04</td><td>BT20/MB468</td><td>MP</td></tr>
<tr><td>>hg38_21_+_8433172_8446622_RNA45SN1_45S_50ntFlanks@7978.8016.39|GACCTCAGATCAGACGTGGCGACCCGCTGAATTTAAGCA</td><td>>hg38_21_+_8433172_8446622_RNA45SN1_45S_50ntFlanks</td><td>7978</td><td>8016</td><td>39</td><td>GACCTCAGATCAGACGTGGCGACCCGCTGAATTTAAGCA</td><td>788.3</td><td>9.1</td><td>2.5</td><td>3.7</td><td>2.2E-04</td><td>3.0E-04</td><td>MB231/MB468</td><td>CYTO</td></tr>
<tr><td>>MI0000434|hsa-let-7i&WithFlank&12|+|62603680|62603775@12.33.22(+1A)&[MIMAT0000415&hsa-let-7i-5p&offsets|0|0(+1A);m-49&12|+|62603691|62603712&offsets|0|0(+1A)]|TGAGGTAGTAGTTTGTGCTGTTA</td><td>>MI0000434|hsa-let-7i&WithFlank&12|+|62603680|62603775</td><td>12</td><td>33</td><td>22(+1A)</td><td>TGAGGTAGTAGTTTGTGCTGTTA</td><td>10104.8</td><td>4.2</td><td>1.1</td><td>3.9</td><td>8.8E-05</td><td>3.0E-04</td><td>BT20/MB468</td><td>MP</td></tr>
<tr><td>>MI0003667|hsa-mir-652&WithFlank&X|+|110055323|110055432@67.87.21(+1U)&[MIMAT0003322&hsa-miR-652-3p&offsets|0|0(+1U);m-148&X|+|110055389|110055410&offsets|0|-1(+1U)]|AATGGCGCCACTAGGGTTGTGT</td><td>>MI0003667|hsa-mir-652&WithFlank&X|+|110055323|110055432</td><td>67</td><td>87</td><td>21(+1U)</td><td>AATGGCGCCACTAGGGTTGTGT</td><td>577.0</td><td>3.0</td><td>0.7</td><td>4.2</td><td>3.3E-05</td><td>3.0E-04</td><td>BT20/MB231</td><td>NUC</td></tr>
<tr><td>>hg38_21_+_8433172_8446622_RNA45SN1_45S_50ntFlanks@11772.11794.23|TTAGTGACGCGCATGAATGGATG</td><td>>hg38_21_+_8433172_8446622_RNA45SN1_45S_50ntFlanks</td><td>11772</td><td>11794</td><td>23</td><td>TTAGTGACGCGCATGAATGGATG</td><td>811.7</td><td>-2.8</td><td>0.7</td><td>-3.7</td><td>2.3E-04</td><td>3.0E-04</td><td>MB231/MB468</td><td>NUC</td></tr>
<tr><td>>hg38_21_+_8433172_8446622_RNA45SN1_45S_50ntFlanks@12347.12375.29|AGAAAAGTTACCACAGGGATAACTGGCTT</td><td>>hg38_21_+_8433172_8446622_RNA45SN1_45S_50ntFlanks</td><td>12347</td><td>12375</td><td>29</td><td>AGAAAAGTTACCACAGGGATAACTGGCTT</td><td>123.7</td><td>-10.0</td><td>2.6</td><td>-3.9</td><td>8.8E-05</td><td>3.0E-04</td><td>BT20/MB468</td><td>MP</td></tr>
<tr><td>>hg38_21_+_8433172_8446622_RNA45SN1_45S_50ntFlanks@11668.11685.18|TGCCCAGTGCTCTGAATG</td><td>>hg38_21_+_8433172_8446622_RNA45SN1_45S_50ntFlanks</td><td>11668</td><td>11685</td><td>18</td><td>TGCCCAGTGCTCTGAATG</td><td>656.4</td><td>-2.7</td><td>0.7</td><td>-3.7</td><td>2.3E-04</td><td>3.0E-04</td><td>MB231/MB468</td><td>MITO</td></tr>
<tr><td>>hg38_21_+_8433172_8446622_RNA45SN1_45S_50ntFlanks@9574.9604.31|AATCGGTCGTCCGACCTGGGTATAGGGGCGA</td><td>>hg38_21_+_8433172_8446622_RNA45SN1_45S_50ntFlanks</td><td>9574</td><td>9604</td><td>31</td><td>AATCGGTCGTCCGACCTGGGTATAGGGGCGA</td><td>459.6</td><td>-2.8</td><td>0.8</td><td>-3.7</td><td>2.3E-04</td><td>3.0E-04</td><td>MB231/MB468</td><td>MITO</td></tr>
<tr><td>>MI0000440|hsa-mir-27b&WithFlank&9|+|95085439|95085547@67.88.22&[MIMAT0000419&hsa-miR-27b-3p&offsets|0|+1;m-39&9|+|95085505|95085525&offsets|0|+1]|TTCACAGTGGCTAAGTTCTGCA</td><td>>MI0000440|hsa-mir-27b&WithFlank&9|+|95085439|95085547</td><td>67</td><td>88</td><td>22</td><td>TTCACAGTGGCTAAGTTCTGCA</td><td>1041.3</td><td>2.8</td><td>0.8</td><td>3.7</td><td>1.9E-04</td><td>3.0E-04</td><td>MB231/MB468</td><td>MP</td></tr>
<tr><td>>hg38_21_+_8433172_8446622_RNA45SN1_45S_50ntFlanks@4002.4022.21|GATAACCTCGGGCCGATCGCA</td><td>>hg38_21_+_8433172_8446622_RNA45SN1_45S_50ntFlanks</td><td>4002</td><td>4022</td><td>21</td><td>GATAACCTCGGGCCGATCGCA</td><td>450.7</td><td>-2.4</td><td>0.6</td><td>-3.7</td><td>2.0E-04</td><td>3.0E-04</td><td>BT20/MB468</td><td>CYTO</td></tr>
<tr><td>>hg38_MT_+_1621_3279_RNR2_16S_50ntFlanks@1221.1261.41|TCCGAGCAGTACATGCTAAGACTTCACCAGTCAAAGCGAAC</td><td>>hg38_MT_+_1621_3279_RNR2_16S_50ntFlanks</td><td>1221</td><td>1261</td><td>41</td><td>TCCGAGCAGTACATGCTAAGACTTCACCAGTCAAAGCGAAC</td><td>21.3</td><td>7.6</td><td>2.0</td><td>3.7</td><td>1.9E-04</td><td>3.0E-04</td><td>MB231/MB468</td><td>MP</td></tr>
<tr><td>>hg38_21_+_8433172_8446622_RNA45SN1_45S_50ntFlanks@4102.4120.19|ACCACGGGTGACGGGGAAT</td><td>>hg38_21_+_8433172_8446622_RNA45SN1_45S_50ntFlanks</td><td>4102</td><td>4120</td><td>19</td><td>ACCACGGGTGACGGGGAAT</td><td>346.4</td><td>-2.5</td><td>0.7</td><td>-3.7</td><td>2.2E-04</td><td>3.0E-04</td><td>MB231/MB468</td><td>CYTO</td></tr>
<tr><td>>MI0000746|hsa-mir-99b&WithFlank&19|+|51692606|51692687@51.71.21(+1A)&[MIMAT0004678&hsa-miR-99b-3p&offsets|0|-1(+1A);m-151&19|+|51692656|51692678&offsets|0|-2(+1A)]|CAAGCTCGTGTCTGTGGGTCCA</td><td>>MI0000746|hsa-mir-99b&WithFlank&19|+|51692606|51692687</td><td>51</td><td>71</td><td>21(+1A)</td><td>CAAGCTCGTGTCTGTGGGTCCA</td><td>786.6</td><td>3.3</td><td>0.8</td><td>4.2</td><td>3.0E-05</td><td>3.0E-04</td><td>BT20/MB231</td><td>CYTO</td></tr>
<tr><td>>hg38_21_+_8433172_8446622_RNA45SN1_45S_50ntFlanks@11567.11611.45|AGCAGCCGACTTAGAACTGGTGCGGACCAGGGGAATCCGACTGTT</td><td>>hg38_21_+_8433172_8446622_RNA45SN1_45S_50ntFlanks</td><td>11567</td><td>11611</td><td>45</td><td>AGCAGCCGACTTAGAACTGGTGCGGACCAGGGGAATCCGACTGTT</td><td>5191.6</td><td>4.1</td><td>1.1</td><td>3.8</td><td>1.7E-04</td><td>3.0E-04</td><td>BT20/MB468</td><td>MITO</td></tr>
<tr><td>>hg38_21_+_8433172_8446622_RNA45SN1_45S_50ntFlanks@9554.9580.27|CGTAGCGGTCCTGACGTGCAAATCGGT</td><td>>hg38_21_+_8433172_8446622_RNA45SN1_45S_50ntFlanks</td><td>9554</td><td>9580</td><td>27</td><td>CGTAGCGGTCCTGACGTGCAAATCGGT</td><td>811.6</td><td>-6.7</td><td>1.8</td><td>-3.7</td><td>2.4E-04</td><td>3.0E-04</td><td>BT20/MB468</td><td>NUC</td></tr>
<tr><td>>hg38_21_+_8433172_8446622_RNA45SN1_45S_50ntFlanks@9469.9488.20|GGAGCACGAGCGCACGTGTT</td><td>>hg38_21_+_8433172_8446622_RNA45SN1_45S_50ntFlanks</td><td>9469</td><td>9488</td><td>20</td><td>GGAGCACGAGCGCACGTGTT</td><td>52.5</td><td>9.0</td><td>2.2</td><td>4.2</td><td>3.2E-05</td><td>3.0E-04</td><td>BT20/MB231</td><td>MP</td></tr>
<tr><td>>hg38_21_+_8433172_8446622_RNA45SN1_45S_50ntFlanks@3853.3898.46|AACTGTGGTAATTCTAGAGCTAATACATGCCGACGGGCGCTGACCC</td><td>>hg38_21_+_8433172_8446622_RNA45SN1_45S_50ntFlanks</td><td>3853</td><td>3898</td><td>46</td><td>AACTGTGGTAATTCTAGAGCTAATACATGCCGACGGGCGCTGACCC</td><td>334.6</td><td>-5.6</td><td>1.5</td><td>-3.8</td><td>1.7E-04</td><td>3.0E-04</td><td>BT20/MB468</td><td>MITO</td></tr>
<tr><td>>MI0001733|hsa-mir-452&WithFlank&X|-|151959622|151959718@20.40.21&[MIMAT0001635&hsa-miR-452-5p&offsets|0|-1;m-188&X|-|151959677|151959699&offsets|0|-2]|AACTGTTTGCAGAGGAAACTG</td><td>>MI0001733|hsa-mir-452&WithFlank&X|-|151959622|151959718</td><td>20</td><td>40</td><td>21</td><td>AACTGTTTGCAGAGGAAACTG</td><td>91.8</td><td>10.3</td><td>2.6</td><td>3.9</td><td>8.9E-05</td><td>3.0E-04</td><td>BT20/MB468</td><td>MP</td></tr>
<tr><td>>hg38_21_+_8433172_8446622_RNA45SN1_45S_50ntFlanks@11579.11605.27|AGAACTGGTGCGGACCAGGGGAATCCG</td><td>>hg38_21_+_8433172_8446622_RNA45SN1_45S_50ntFlanks</td><td>11579</td><td>11605</td><td>27</td><td>AGAACTGGTGCGGACCAGGGGAATCCG</td><td>988.5</td><td>-4.3</td><td>1.1</td><td>-3.8</td><td>1.7E-04</td><td>3.0E-04</td><td>BT20/MB468</td><td>MITO</td></tr>
<tr><td>>hg38_21_+_8433172_8446622_RNA45SN1_45S_50ntFlanks@11205.11225.21|GCGGCGGCGACTCTGGACGCG</td><td>>hg38_21_+_8433172_8446622_RNA45SN1_45S_50ntFlanks</td><td>11205</td><td>11225</td><td>21</td><td>GCGGCGGCGACTCTGGACGCG</td><td>192.0</td><td>-5.3</td><td>1.4</td><td>-3.7</td><td>2.3E-04</td><td>3.0E-04</td><td>MB231/MB468</td><td>NUC</td></tr>
<tr><td>>MI0000088|hsa-mir-30a&WithFlank&6|-|71403545|71403627@13.35.23&[MIMAT0000087&hsa-miR-30a-5p&offsets|+1|+2;m-15&6|-|71403595|71403616&offsets|+1|+2]|GTAAACATCCTCGACTGGAAGCT</td><td>>MI0000088|hsa-mir-30a&WithFlank&6|-|71403545|71403627</td><td>13</td><td>35</td><td>23</td><td>GTAAACATCCTCGACTGGAAGCT</td><td>2496.4</td><td>4.4</td><td>1.1</td><td>3.9</td><td>8.9E-05</td><td>3.0E-04</td><td>BT20/MB468</td><td>MP</td></tr>
<tr><td>>hg38_21_+_8433172_8446622_RNA45SN1_45S_50ntFlanks@8737.8758.22|CCCGTCCGTCCGTCCGTCCGTC</td><td>>hg38_21_+_8433172_8446622_RNA45SN1_45S_50ntFlanks</td><td>8737</td><td>8758</td><td>22</td><td>CCCGTCCGTCCGTCCGTCCGTC</td><td>198.9</td><td>-3.9</td><td>1.0</td><td>-3.8</td><td>1.7E-04</td><td>3.0E-04</td><td>BT20/MB468</td><td>MITO</td></tr>
<tr><td>>hg38_21_+_8433172_8446622_RNA45SN1_45S_50ntFlanks@11212.11252.41|CGACTCTGGACGCGAGCCGGGCCCTTCCCGTGGATCGCCCC</td><td>>hg38_21_+_8433172_8446622_RNA45SN1_45S_50ntFlanks</td><td>11212</td><td>11252</td><td>41</td><td>CGACTCTGGACGCGAGCCGGGCCCTTCCCGTGGATCGCCCC</td><td>243.1</td><td>3.6</td><td>0.9</td><td>4.1</td><td>3.6E-05</td><td>3.0E-04</td><td>BT20/MB231</td><td>MITO</td></tr>
<tr><td>>hg38_21_+_8433172_8446622_RNA45SN1_45S_50ntFlanks@10866.10884.19|CTAAGGGCTGGGTCGGTCG</td><td>>hg38_21_+_8433172_8446622_RNA45SN1_45S_50ntFlanks</td><td>10866</td><td>10884</td><td>19</td><td>CTAAGGGCTGGGTCGGTCG</td><td>746.4</td><td>-5.6</td><td>1.5</td><td>-3.7</td><td>2.1E-04</td><td>3.0E-04</td><td>BT20/MB468</td><td>CYTO</td></tr>
<tr><td>>hg38_21_+_8433172_8446622_RNA45SN1_45S_50ntFlanks@10678.10713.36|CGGTTCCGGCGGCGTCCGGTGAGCTCTCGCTGGCCC</td><td>>hg38_21_+_8433172_8446622_RNA45SN1_45S_50ntFlanks</td><td>10678</td><td>10713</td><td>36</td><td>CGGTTCCGGCGGCGTCCGGTGAGCTCTCGCTGGCCC</td><td>241.8</td><td>-4.7</td><td>1.3</td><td>-3.7</td><td>2.3E-04</td><td>3.0E-04</td><td>MB231/MB468</td><td>MITO</td></tr>
<tr><td>>hg38_21_+_8433172_8446622_RNA45SN1_45S_50ntFlanks@12243.12272.30|GAAGGGCAAAAGCTCGCTTGATCTTGATTT</td><td>>hg38_21_+_8433172_8446622_RNA45SN1_45S_50ntFlanks</td><td>12243</td><td>12272</td><td>30</td><td>GAAGGGCAAAAGCTCGCTTGATCTTGATTT</td><td>295.6</td><td>3.4</td><td>0.9</td><td>3.7</td><td>2.2E-04</td><td>3.0E-04</td><td>MB231/MB468</td><td>CYTO</td></tr>
<tr><td>>hg38_21_+_8433172_8446622_RNA45SN1_45S_50ntFlanks@6730.6756.27|ACACATTGATCATCGACACTTCGAACG</td><td>>hg38_21_+_8433172_8446622_RNA45SN1_45S_50ntFlanks</td><td>6730</td><td>6756</td><td>27</td><td>ACACATTGATCATCGACACTTCGAACG</td><td>225.4</td><td>-1.6</td><td>0.4</td><td>-3.7</td><td>2.2E-04</td><td>3.0E-04</td><td>MB231/MB468</td><td>CYTO</td></tr>
<tr><td>>hg38_21_+_8433172_8446622_RNA45SN1_45S_50ntFlanks@4727.4752.26|AATCAAGAACGAAAGTCGGAGGTTCG</td><td>>hg38_21_+_8433172_8446622_RNA45SN1_45S_50ntFlanks</td><td>4727</td><td>4752</td><td>26</td><td>AATCAAGAACGAAAGTCGGAGGTTCG</td><td>314.5</td><td>-3.3</td><td>0.9</td><td>-3.9</td><td>8.9E-05</td><td>3.0E-04</td><td>BT20/MB468</td><td>MP</td></tr>
<tr><td>>hg38_21_+_8433172_8446622_RNA45SN1_45S_50ntFlanks@6650.6681.32|ACGACTCTTAGCGGTGGATCACTCGGCTCGTG</td><td>>hg38_21_+_8433172_8446622_RNA45SN1_45S_50ntFlanks</td><td>6650</td><td>6681</td><td>32</td><td>ACGACTCTTAGCGGTGGATCACTCGGCTCGTG</td><td>20480.8</td><td>-3.4</td><td>0.9</td><td>-3.7</td><td>2.3E-04</td><td>3.0E-04</td><td>MB231/MB468</td><td>MITO</td></tr>
<tr><td>>hg38_21_+_8433172_8446622_RNA45SN1_45S_50ntFlanks@10666.10706.41|TGGAAAGCGTCGCGGTTCCGGCGGCGTCCGGTGAGCTCTCG</td><td>>hg38_21_+_8433172_8446622_RNA45SN1_45S_50ntFlanks</td><td>10666</td><td>10706</td><td>41</td><td>TGGAAAGCGTCGCGGTTCCGGCGGCGTCCGGTGAGCTCTCG</td><td>286.5</td><td>-5.8</td><td>1.6</td><td>-3.7</td><td>2.4E-04</td><td>3.0E-04</td><td>MB231/MB468</td><td>NUC</td></tr>
<tr><td>>hg38_MT_+_598_1651_RNR1_12S_50ntFlanks@628.665.38|CCGATCAACCTCACCACCTCTTGCTCAGCCTATATACC</td><td>>hg38_MT_+_598_1651_RNR1_12S_50ntFlanks</td><td>628</td><td>665</td><td>38</td><td>CCGATCAACCTCACCACCTCTTGCTCAGCCTATATACC</td><td>114.7</td><td>6.2</td><td>1.7</td><td>3.7</td><td>2.3E-04</td><td>3.0E-04</td><td>MB231/MB468</td><td>MITO</td></tr>
<tr><td>>hg38_21_+_8433172_8446622_RNA45SN1_45S_50ntFlanks@6660.6681.22|GCGGTGGATCACTCGGCTCGTG</td><td>>hg38_21_+_8433172_8446622_RNA45SN1_45S_50ntFlanks</td><td>6660</td><td>6681</td><td>22</td><td>GCGGTGGATCACTCGGCTCGTG</td><td>2096.5</td><td>-4.3</td><td>1.2</td><td>-3.7</td><td>2.3E-04</td><td>3.0E-04</td><td>MB231/MB468</td><td>MITO</td></tr>
<tr><td>>hg38_21_+_8433172_8446622_RNA45SN1_45S_50ntFlanks@12447.12476.30|CAGAATTCACCAAGCGTTGGATTGTTCACC</td><td>>hg38_21_+_8433172_8446622_RNA45SN1_45S_50ntFlanks</td><td>12447</td><td>12476</td><td>30</td><td>CAGAATTCACCAAGCGTTGGATTGTTCACC</td><td>470.7</td><td>-5.4</td><td>1.4</td><td>-3.9</td><td>9.0E-05</td><td>3.0E-04</td><td>BT20/MB468</td><td>MP</td></tr>
<tr><td>>hg38_MT_+_598_1651_RNR1_12S_50ntFlanks@52.90.39|ATAGGTTTGGTCCTAGCCTTTCTATTAGCTCTTAGTAAG</td><td>>hg38_MT_+_598_1651_RNR1_12S_50ntFlanks</td><td>52</td><td>90</td><td>39</td><td>ATAGGTTTGGTCCTAGCCTTTCTATTAGCTCTTAGTAAG</td><td>858.4</td><td>3.0</td><td>0.8</td><td>3.7</td><td>2.3E-04</td><td>3.0E-04</td><td>MB231/MB468</td><td>MITO</td></tr>
<tr><td>>hg38_21_+_8433172_8446622_RNA45SN1_45S_50ntFlanks@3861.3898.38|TAATTCTAGAGCTAATACATGCCGACGGGCGCTGACCC</td><td>>hg38_21_+_8433172_8446622_RNA45SN1_45S_50ntFlanks</td><td>3861</td><td>3898</td><td>38</td><td>TAATTCTAGAGCTAATACATGCCGACGGGCGCTGACCC</td><td>86.8</td><td>-9.5</td><td>2.4</td><td>-3.9</td><td>9.0E-05</td><td>3.0E-04</td><td>BT20/MB468</td><td>MP</td></tr>
<tr><td>>hg38_21_+_8433172_8446622_RNA45SN1_45S_50ntFlanks@12274.12306.33|CAGTACGAATACAGACCGTGAAAGCGGGGCCTC</td><td>>hg38_21_+_8433172_8446622_RNA45SN1_45S_50ntFlanks</td><td>12274</td><td>12306</td><td>33</td><td>CAGTACGAATACAGACCGTGAAAGCGGGGCCTC</td><td>270.5</td><td>-6.1</td><td>1.6</td><td>-3.9</td><td>9.0E-05</td><td>3.0E-04</td><td>BT20/MB468</td><td>MP</td></tr>
<tr><td>>hg38_21_+_8433172_8446622_RNA45SN1_45S_50ntFlanks@12940.12979.40|CATTCGTAGACGACCTGCTTCTGGGTCGGGGTTTCGTACG</td><td>>hg38_21_+_8433172_8446622_RNA45SN1_45S_50ntFlanks</td><td>12940</td><td>12979</td><td>40</td><td>CATTCGTAGACGACCTGCTTCTGGGTCGGGGTTTCGTACG</td><td>99.1</td><td>-5.2</td><td>1.4</td><td>-3.7</td><td>2.1E-04</td><td>3.0E-04</td><td>BT20/MB468</td><td>CYTO</td></tr>
<tr><td>>hg38_21_+_8433172_8446622_RNA45SN1_45S_50ntFlanks@9552.9572.21|TCCGTAGCGGTCCTGACGTGC</td><td>>hg38_21_+_8433172_8446622_RNA45SN1_45S_50ntFlanks</td><td>9552</td><td>9572</td><td>21</td><td>TCCGTAGCGGTCCTGACGTGC</td><td>254.0</td><td>-2.0</td><td>0.6</td><td>-3.7</td><td>2.1E-04</td><td>3.0E-04</td><td>BT20/MB468</td><td>CYTO</td></tr>
<tr><td>>hg38_21_+_8433172_8446622_RNA45SN1_45S_50ntFlanks@12072.12097.26|CCCGAGGGGCTCTCGCTTCTGGCGCC</td><td>>hg38_21_+_8433172_8446622_RNA45SN1_45S_50ntFlanks</td><td>12072</td><td>12097</td><td>26</td><td>CCCGAGGGGCTCTCGCTTCTGGCGCC</td><td>693.0</td><td>-2.8</td><td>0.8</td><td>-3.7</td><td>2.1E-04</td><td>3.0E-04</td><td>BT20/MB468</td><td>CYTO</td></tr>
<tr><td>>am_trnaMT_HisGTG_MT_+_12138_12206@20.51.32|AACATCAGATTGTGAATCTGACAACAGAGGCT</td><td>>am_trnaMT_HisGTG_MT_+_12138_12206</td><td>20</td><td>51</td><td>32</td><td>AACATCAGATTGTGAATCTGACAACAGAGGCT</td><td>942.0</td><td>5.9</td><td>1.5</td><td>3.9</td><td>9.0E-05</td><td>3.0E-04</td><td>BT20/MB468</td><td>MP</td></tr>
<tr><td>>MI0000101|hsa-mir-99a&WithFlank&21|+|16539083|16539175@19.39.21&[MIMAT0000097&hsa-miR-99a-5p&offsets|0|-1;m-64&21|+|16539101|16539121&offsets|0|0]|AACCCGTAGATCCGATCTTGT</td><td>>MI0000101|hsa-mir-99a&WithFlank&21|+|16539083|16539175</td><td>19</td><td>39</td><td>21</td><td>AACCCGTAGATCCGATCTTGT</td><td>1229.5</td><td>-6.6</td><td>1.6</td><td>-4.1</td><td>3.3E-05</td><td>3.0E-04</td><td>BT20/MB231</td><td>NUC</td></tr>
<tr><td>>hg38_21_+_8433172_8446622_RNA45SN1_45S_50ntFlanks@9567.9601.35|ACGTGCAAATCGGTCGTCCGACCTGGGTATAGGGG</td><td>>hg38_21_+_8433172_8446622_RNA45SN1_45S_50ntFlanks</td><td>9567</td><td>9601</td><td>35</td><td>ACGTGCAAATCGGTCGTCCGACCTGGGTATAGGGG</td><td>228.9</td><td>-6.1</td><td>1.6</td><td>-3.7</td><td>2.0E-04</td><td>3.0E-04</td><td>MB231/MB468</td><td>MP</td></tr>
<tr><td>>hg38_21_+_8433172_8446622_RNA45SN1_45S_50ntFlanks@11885.11905.21|TTGAGCTTGACTCTAGTCTGG</td><td>>hg38_21_+_8433172_8446622_RNA45SN1_45S_50ntFlanks</td><td>11885</td><td>11905</td><td>21</td><td>TTGAGCTTGACTCTAGTCTGG</td><td>595.6</td><td>-6.3</td><td>1.6</td><td>-3.9</td><td>9.0E-05</td><td>3.0E-04</td><td>BT20/MB468</td><td>MP</td></tr>
<tr><td>>MI0000342|hsa-mir-200b&WithFlank&1|+|1167098|1167204@63.84.22&[MIMAT0000318&hsa-miR-200b-3p&offsets|0|0;m-93&1|+|1167160|1167182&offsets|0|-1]|TAATACTGCCTGGTAATGATGA</td><td>>MI0000342|hsa-mir-200b&WithFlank&1|+|1167098|1167204</td><td>63</td><td>84</td><td>22</td><td>TAATACTGCCTGGTAATGATGA</td><td>2747.3</td><td>-4.1</td><td>1.0</td><td>-3.9</td><td>9.0E-05</td><td>3.0E-04</td><td>BT20/MB468</td><td>MP</td></tr>
<tr><td>>hg38_21_+_8433172_8446622_RNA45SN1_45S_50ntFlanks@11576.11607.32|CTTAGAACTGGTGCGGACCAGGGGAATCCGAC</td><td>>hg38_21_+_8433172_8446622_RNA45SN1_45S_50ntFlanks</td><td>11576</td><td>11607</td><td>32</td><td>CTTAGAACTGGTGCGGACCAGGGGAATCCGAC</td><td>475.9</td><td>-2.7</td><td>0.7</td><td>-3.7</td><td>2.1E-04</td><td>3.0E-04</td><td>BT20/MB468</td><td>CYTO</td></tr>
<tr><td>>hg38_21_+_8433172_8446622_RNA45SN1_45S_50ntFlanks@6672.6699.28|TCGGCTCGTGCGTCGATGAAGAACGCAG</td><td>>hg38_21_+_8433172_8446622_RNA45SN1_45S_50ntFlanks</td><td>6672</td><td>6699</td><td>28</td><td>TCGGCTCGTGCGTCGATGAAGAACGCAG</td><td>474.0</td><td>-3.9</td><td>1.0</td><td>-3.7</td><td>2.3E-04</td><td>3.0E-04</td><td>MB231/MB468</td><td>MITO</td></tr>
<tr><td>>hg38_21_+_8433172_8446622_RNA45SN1_45S_50ntFlanks@12636.12657.22|AGCTACCATCTGTGGGATTATG</td><td>>hg38_21_+_8433172_8446622_RNA45SN1_45S_50ntFlanks</td><td>12636</td><td>12657</td><td>22</td><td>AGCTACCATCTGTGGGATTATG</td><td>740.6</td><td>-6.3</td><td>1.6</td><td>-3.9</td><td>9.0E-05</td><td>3.0E-04</td><td>BT20/MB468</td><td>MP</td></tr>
</table>

| | | | | | | | | | | | | | |
|---|---|---|---|---|---|---|---|---|---|---|---|---|---|
| >hg38_21_+_8433172_8446622_RNA45SN1_45S_50ntFlanks@11730.11751.22\|AACTATGACTCTCTTAAGGTAG | >hg38_21_+_8433172_8446622_RNA45SN1_45S_50ntFlanks | 11730 | 11751 | 22 | AACTATGACTCTCTTAAGGTAG | 147.7 | -3.6 | 1.0 | -3.7 | 2.1E-04 | 3.0E-04 | BT20/MB468 | CYTO |
| >hg38_21_+_8433172_8446622_RNA45SN1_45S_50ntFlanks@6660.6686.27\|GCGGTGGATCACTCGGCTCGTGCGTCG | >hg38_21_+_8433172_8446622_RNA45SN1_45S_50ntFlanks | 6660 | 6686 | 27 | GCGGTGGATCACTCGGCTCGTGCGTCG | 655.4 | -4.6 | 1.3 | -3.7 | 2.4E-04 | 3.0E-04 | MB231/MB468 | MITO |
| >MI0001445\|hsa-mir-423&WithFlank&17\|+\|30117073\|30117178@59.81.23(+2U)&[MIMAT0001340&hsa-miR-423-3p&offsets\|0\|0(+2U);m-100&17\|+\|30117131\|30117153&offsets\|0\|0(+2U)]\|AGCTCGGTCTGAGGCCCCTCAGTTT | >MI0001445\|hsa-mir-423&WithFlank&17\|+\|30117073\|30117178 | 59 | 81 | 23(+2U) | AGCTCGGTCTGAGGCCCCTCAGTTT | 183.7 | 5.8 | 1.6 | 3.7 | 2.4E-04 | 3.0E-04 | MB231/MB468 | MITO |
| >hg38_21_+_8433172_8446622_RNA45SN1_45S_50ntFlanks@11208.11250.43\|GCGGCGACTCTGGACGCGAGCCGGGCCCTTCCCGTGGATCGCC | >hg38_21_+_8433172_8446622_RNA45SN1_45S_50ntFlanks | 11208 | 11250 | 43 | GCGGCGACTCTGGACGCGAGCCGGGCCCTTCCCGTGGATCGCC | 528.6 | -4.8 | 1.3 | -3.7 | 2.1E-04 | 3.0E-04 | BT20/MB468 | CYTO |
| >hg38_21_+_8433172_8446622_RNA45SN1_45S_50ntFlanks@9496.9515.20\|GAAAGATGGTGAACTATGCC | >hg38_21_+_8433172_8446622_RNA45SN1_45S_50ntFlanks | 9496 | 9515 | 20 | GAAAGATGGTGAACTATGCC | 95.5 | 7.0 | 1.8 | 3.9 | 9.1E-05 | 3.0E-04 | BT20/MB468 | MP |
| >hg38_MT_+_1621_3279_RNR2_16S_50ntFlanks@1092.1120.29\|GCAAGACGAGAAGACCCTATGGAGCTTTA | >hg38_MT_+_1621_3279_RNR2_16S_50ntFlanks | 1092 | 1120 | 29 | GCAAGACGAGAAGACCCTATGGAGCTTTA | 369.0 | 3.9 | 1.0 | 3.9 | 9.1E-05 | 3.0E-04 | BT20/MB468 | MP |
| >am_trna128_GlyGCC_6_-_27870686_27870756@1.27.27,trna133_GlyCCC_1_-_16872434_16872504@1.27.27,trna18_GlyGCC_16_+_70822597_70822667@1.27.27,trna19_GlyGCC_16_+_70823410_70823480@1.27.27,trna19_GlyGCC_2_-_157257659_157257729@1.27.27,trna24_GlyGCC_16_-_70812942_70813012@1.27.27,trna25_GlyGCC_16_-_70812114_70812184@1.27.27,trna4_GlyCCC_1_+_17188416_17188486@1.27.27,trna5_GlyGCC_17_+_8029064_8029134@1.27.27,trna68_GlyGCC_1_-_161493637_161493707@1.27.27\|GCATTGGTGGTTCAGTGGTAGAATTCT | >am_trna128_GlyGCC_6_-_27870686_27870756 | 1 | 27 | 27 | GCATTGGTGGTTCAGTGGTAGAATTCT | 2302.4 | -2.5 | 0.7 | -3.7 | 2.4E-04 | 3.0E-04 | MB231/MB468 | NUC |
| >hg38_21_+_8433172_8446622_RNA45SN1_45S_50ntFlanks@11879.11904.26\|ACCCTGTTGAGCTTGACTCTAGTCTG | >hg38_21_+_8433172_8446622_RNA45SN1_45S_50ntFlanks | 11879 | 11904 | 26 | ACCCTGTTGAGCTTGACTCTAGTCTG | 199.9 | -4.0 | 1.1 | -3.7 | 2.4E-04 | 3.0E-04 | MB231/MB468 | NUC |
| >am_trnaMT_LeuTAG_MT_+_12266_12336@42.61.20\|AAAATTTTGGTGCAACTCCA | >am_trnaMT_LeuTAG_MT_+_12266_12336 | 42 | 61 | 20 | AAAATTTTGGTGCAACTCCA | 126.0 | -4.3 | 1.2 | -3.7 | 2.0E-04 | 3.0E-04 | MB231/MB468 | MP |
| >hg38_21_+_8433172_8446622_RNA45SN1_45S_50ntFlanks@9146.9165.20\|CGGACTGTCCCCAGTGCGCC | >hg38_21_+_8433172_8446622_RNA45SN1_45S_50ntFlanks | 9146 | 9165 | 20 | CGGACTGTCCCCAGTGCGCC | 355.2 | -3.8 | 1.0 | -3.9 | 9.1E-05 | 3.0E-04 | BT20/MB468 | MP |
| >hg38_21_+_8433172_8446622_RNA45SN1_45S_50ntFlanks@10350.10385.36\|ACGAGAACTTTGAAGGCCGAAGTGGAGAAGGGTTCC | >hg38_21_+_8433172_8446622_RNA45SN1_45S_50ntFlanks | 10350 | 10385 | 36 | ACGAGAACTTTGAAGGCCGAAGTGGAGAAGGGTTCC | 364.1 | -6.4 | 1.7 | -3.7 | 2.4E-04 | 3.0E-04 | BT20/MB468 | NUC |
| >hg38_21_+_8433172_8446622_RNA45SN1_45S_50ntFlanks@12420.12458.39\|ATGTCGGCTCTTCCTATCATTGTGAAGCAGAATTCACCA | >hg38_21_+_8433172_8446622_RNA45SN1_45S_50ntFlanks | 12420 | 12458 | 39 | ATGTCGGCTCTTCCTATCATTGTGAAGCAGAATTCACCA | 600.6 | -5.3 | 1.5 | -3.7 | 2.4E-04 | 3.0E-04 | BT20/MB468 | NUC |
| >MI0000446\|hsa-mir-125b-1&WithFlank&11\|-\|122099751\|122099850@21.42.22;MI0000470\|hsa-mir-125b-2&WithFlank&21\|+\|16590231\|16590331@23.44.22&[MIMAT0000423&hsa-miR-125b-5p&offsets\|0\|0;m-67&11\|-\|122099809\|122099830&offsets\|0\|0];[MIMAT0000423_1&hsa-miR-125b-5p&offsets\|0\|0;m-68&21\|+\|16590253\|16590274&offsets\|0\|0]\|TCCCTGAGACCCTAACTTGTGA | >MI0000446\|hsa-mir-125b-1&WithFlank&11\|-\|122099751\|122099850 | 21 | 42 | 22 | TCCCTGAGACCCTAACTTGTGA | 26774.2 | -7.2 | 1.7 | -4.2 | 3.2E-05 | 3.0E-04 | BT20/MB231 | MP |
| >am_MI0000809\|hsa-mir-151a&WithFlank&8\|-\|140732558\|140732659@53.72.20(+3U)&[MIMAT0000757&hsa-miR-151a-3p&offsets\|0\|-1(+3U);m-30&8\|-\|140732587\|140732607&offsets\|0\|-1(+3U)]\|CTAGACTGAAGCTCCTTGAGTTT | >am_MI0000809\|hsa-mir-151a&WithFlank&8\|-\|140732558\|140732659 | 53 | 72 | 20(+3U) | CTAGACTGAAGCTCCTTGAGTTT | 68.5 | 6.2 | 1.7 | 3.7 | 2.0E-04 | 3.0E-04 | MB231/MB468 | MP |
| >hg38_21_+_8433172_8446622_RNA45SN1_45S_50ntFlanks@485.523.39\|GATGTGGTGACGTCGTGCTCTCCCGGGCCGGGTCCGAGC | >hg38_21_+_8433172_8446622_RNA45SN1_45S_50ntFlanks | 485 | 523 | 39 | GATGTGGTGACGTCGTGCTCTCCCGGGCCGGGTCCGAGC | 88.0 | 9.7 | 2.6 | 3.7 | 2.0E-04 | 3.0E-04 | MB231/MB468 | MP |
| >hg38_21_+_8433172_8446622_RNA45SN1_45S_50ntFlanks@11235.11255.21\|CTTCCCGTGGATCGCCCCAGC | >hg38_21_+_8433172_8446622_RNA45SN1_45S_50ntFlanks | 11235 | 11255 | 21 | CTTCCCGTGGATCGCCCCAGC | 312.4 | -3.9 | 1.0 | -3.7 | 2.1E-04 | 3.0E-04 | BT20/MB468 | CYTO |
| >trnaMT_GluTTC_MT_-_14674_14742@39.57.19,trnalookalike8_GluTTC_5_-_93905172_93905240@39.57.19\|ATTGGTCGTGGTTGTAGTC | >trnaMT_GluTTC_MT_-_14674_14742 | 39 | 57 | 19 | ATTGGTCGTGGTTGTAGTC | 189.6 | 4.6 | 1.3 | 3.7 | 2.4E-04 | 3.0E-04 | MB231/MB468 | MITO |
| >MI0000767\|hsa-mir-365a&WithFlank&16\|+\|14309279\|14309377@62.82.21(+1C);MI0000769\|hsa-mir-365b&WithFlank&17\|+\|31575405\|31575527@74.94.21(+1C)&[MIMAT0000710&hsa-miR-365a-3p&offsets\|0\|-1(+1C);m-176&16\|+\|14309340\|14309361&offsets\|0\|-1(+1C)];[MIMAT0022834&hsa-miR-365b-3p&offsets\|0\|-1(+1C);m-175&17\|+\|31575478\|31575499&offsets\|0\|-1(+1C)]\|TAATGCCCCTAAAAATCCTTAC | >MI0000767\|hsa-mir-365a&WithFlank&16\|+\|14309279\|14309377 | 62 | 82 | 21(+1C) | TAATGCCCCTAAAAATCCTTAC | 240.5 | 4.3 | 1.0 | 4.2 | 3.1E-05 | 3.0E-04 | BT20/MB231 | CYTO |
| >hg38_21_+_8433172_8446622_RNA45SN1_45S_50ntFlanks@3940.3959.20\|ACCCGGTCAGCCCCTCTCCG | >hg38_21_+_8433172_8446622_RNA45SN1_45S_50ntFlanks | 3940 | 3959 | 20 | ACCCGGTCAGCCCCTCTCCG | 302.4 | -3.9 | 1.0 | -3.7 | 2.1E-04 | 3.0E-04 | BT20/MB468 | CYTO |
| >hg38_21_+_8433172_8446622_RNA45SN1_45S_50ntFlanks@7974.8009.36\|ACGCGACCTCAGATCAGACGTGGCGACCCGCTGAAT | >hg38_21_+_8433172_8446622_RNA45SN1_45S_50ntFlanks | 7974 | 8009 | 36 | ACGCGACCTCAGATCAGACGTGGCGACCCGCTGAAT | 12426.8 | 4.1 | 1.1 | 3.9 | 9.1E-05 | 3.1E-04 | BT20/MB468 | MP |
| >hg38_21_+_8433172_8446622_RNA45SN1_45S_50ntFlanks@12991.13010.20\|CTCCCTCGCTGCGATCTATT | >hg38_21_+_8433172_8446622_RNA45SN1_45S_50ntFlanks | 12991 | 13010 | 20 | CTCCCTCGCTGCGATCTATT | 627.4 | -4.0 | 1.1 | -3.7 | 2.2E-04 | 3.1E-04 | MB231/MB468 | CYTO |
| >am_trna128_GlyGCC_6_-_27870686_27870756@1.36.36,trna18_GlyGCC_16_+_70822597_70822667@1.36.36,trna19_GlyGCC_16_+_70823410_70823480@1.36.36,trna19_GlyGCC_2_-_157257659_157257729@1.36.36,trna24_GlyGCC_16_-_70812942_70813012@1.36.36,trna25_GlyGCC_16_-_70812114_70812184@1.36.36,trna5_GlyGCC_17_+_8029064_8029134@1.36.36,trna68_GlyGCC_1_-_161493637_161493707@1.36.36\|GCATTGGTGGTTCAGTGGTAGAATTCTCGCCTGCCA | >am_trna128_GlyGCC_6_-_27870686_27870756 | 1 | 36 | 36 | GCATTGGTGGTTCAGTGGTAGAATTCTCGCCTGCCA | 698.2 | 6.2 | 1.6 | 3.9 | 9.1E-05 | 3.1E-04 | BT20/MB468 | MP |
| >trnaMT_HisGTG_MT_+_12138_12206@26.71.46\|AGATTGTGAATCTGACAACAGAGGCTTACGACCCCTTATTTACCCC | >trnaMT_HisGTG_MT_+_12138_12206 | 26 | 71 | 46 | AGATTGTGAATCTGACAACAGAGGCTTACGACCCCTTATTTACCCC | 93.2 | 10.3 | 2.6 | 3.9 | 9.1E-05 | 3.1E-04 | BT20/MB468 | MP |
| >hg38_21_+_8433172_8446622_RNA45SN1_45S_50ntFlanks@10343.10369.27\|TATTCAAACGAGAACTTTGAAGGCCGA | >hg38_21_+_8433172_8446622_RNA45SN1_45S_50ntFlanks | 10343 | 10369 | 27 | TATTCAAACGAGAACTTTGAAGGCCGA | 101.8 | 7.1 | 1.8 | 3.9 | 9.1E-05 | 3.1E-04 | BT20/MB468 | MP |
| >trna27_GlyCCC_2_-_70476123_70476193@1.34.34,trna34_GlyCCC_16_-_686736_686806@1.34.34\|GCGCCGCTGGTGTAGTGGTATCATGCAAGATTCC | >trna27_GlyCCC_2_-_70476123_70476193 | 1 | 34 | 34 | GCGCCGCTGGTGTAGTGGTATCATGCAAGATTCC | 96.7 | 10.3 | 2.6 | 3.9 | 9.1E-05 | 3.1E-04 | BT20/MB468 | MP |
| >hg38_21_+_8433172_8446622_RNA45SN1_45S_50ntFlanks@4212.4242.31\|AGGTAGTGACGAAAAATAACAATACAGGACT | >hg38_21_+_8433172_8446622_RNA45SN1_45S_50ntFlanks | 4212 | 4242 | 31 | AGGTAGTGACGAAAAATAACAATACAGGACT | 146.9 | -3.6 | 1.0 | -3.7 | 2.0E-04 | 3.1E-04 | MB231/MB468 | MP |
| >hg38_21_+_8433172_8446622_RNA45SN1_45S_50ntFlanks@4738.4755.18\|AAAGTCGGAGGTTCGAAG | >hg38_21_+_8433172_8446622_RNA45SN1_45S_50ntFlanks | 4738 | 4755 | 18 | AAAGTCGGAGGTTCGAAG | 226.3 | 2.1 | 0.6 | 3.7 | 2.2E-04 | 3.1E-04 | MB231/MB468 | CYTO |
| >hg38_21_+_8433172_8446622_RNA45SN1_45S_50ntFlanks@12416.12460.45\|TTCGATGTCGGCTCTTCCTATCATTGTGAAGCAGAATTCACCAAG | >hg38_21_+_8433172_8446622_RNA45SN1_45S_50ntFlanks | 12416 | 12460 | 45 | TTCGATGTCGGCTCTTCCTATCATTGTGAAGCAGAATTCACCAAG | 160.3 | -4.8 | 1.1 | -4.2 | 3.1E-05 | 3.1E-04 | BT20/MB231 | CYTO |
| >hg38_21_+_8433172_8446622_RNA45SN1_45S_50ntFlanks@10510.10534.25\|AATCCGGAGTGGCGGAGATGGGCGC | >hg38_21_+_8433172_8446622_RNA45SN1_45S_50ntFlanks | 10510 | 10534 | 25 | AATCCGGAGTGGCGGAGATGGGCGC | 169.2 | -5.0 | 1.3 | -3.8 | 1.8E-04 | 3.1E-04 | BT20/MB468 | MITO |
| >MI0000077\|hsa-mir-21&WithFlank&17\|+\|59841260\|59841343@52.72.21&[MIMAT0004494&hsa-miR-21-3p&offsets\|0\|0;m-52&17\|+\|59841311\|59841332&offsets\|0\|-1]\|CAACACCAGTCGATGGGCTGT | >MI0000077\|hsa-mir-21&WithFlank&17\|+\|59841260\|59841343 | 52 | 72 | 21 | CAACACCAGTCGATGGGCTGT | 2303.0 | 5.3 | 1.4 | 3.9 | 9.1E-05 | 3.1E-04 | BT20/MB468 | MP |
| >MI0000542\|hsa-mir-320a&WithFlank&8\|-\|22244960\|22245043@49.69.21(+1U)&[MIMAT0000510&hsa-miR-320a-3p&offsets\|+1\|0(+1U);m-60&8\|-\|22244975\|22244996&offsets\|+1\|0(+1U)]\|AAAGCTGGGTTGAGAGGGCGAT | >MI0000542\|hsa-mir-320a&WithFlank&8\|-\|22244960\|22245043 | 49 | 69 | 21(+1U) | AAAGCTGGGTTGAGAGGGCGAT | 984.3 | 3.5 | 0.8 | 4.2 | 3.1E-05 | 3.1E-04 | BT20/MB231 | CYTO |
| >MI0003137\|hsa-mir-193b&WithFlank&16\|+\|14303961\|14304055@57.76.20&[MIMAT0002819&hsa-miR-193b-3p&offsets\|0\|-2;m-166&16\|+\|14304017\|14304038&offsets\|0\|-2]\|AACTGGCCCTCAAAGTCCCG | >MI0003137\|hsa-mir-193b&WithFlank&16\|+\|14303961\|14304055 | 57 | 76 | 20 | AACTGGCCCTCAAAGTCCCG | 337.5 | 7.0 | 1.8 | 3.9 | 9.2E-05 | 3.1E-04 | BT20/MB468 | MP |
| >hg38_21_+_8433172_8446622_RNA45SN1_45S_50ntFlanks@12128.12146.19\|ACCCGCTCCGGGGACAGTG | >hg38_21_+_8433172_8446622_RNA45SN1_45S_50ntFlanks | 12128 | 12146 | 19 | ACCCGCTCCGGGGACAGTG | 96.4 | -5.8 | 1.5 | -3.9 | 9.2E-05 | 3.1E-04 | BT20/MB468 | MP |
| >hg38_21_+_8433172_8446622_RNA45SN1_45S_50ntFlanks@11583.11600.18\|CTGGTGCGGACCAGGGGA | >hg38_21_+_8433172_8446622_RNA45SN1_45S_50ntFlanks | 11583 | 11600 | 18 | CTGGTGCGGACCAGGGGA | 655.6 | -6.6 | 1.8 | -3.7 | 2.4E-04 | 3.1E-04 | BT20/MB468 | NUC |
| >hg38_21_+_8433172_8446622_RNA45SN1_45S_50ntFlanks@8189.8212.24\|CCCAGCCCGTGGACGGTGTGAGGC | >hg38_21_+_8433172_8446622_RNA45SN1_45S_50ntFlanks | 8189 | 8212 | 24 | CCCAGCCCGTGGACGGTGTGAGGC | 441.6 | -6.2 | 1.7 | -3.7 | 2.1E-04 | 3.1E-04 | BT20/MB468 | CYTO |
| >hg38_21_+_8433172_8446622_RNA45SN1_45S_50ntFlanks@10473.10490.18\|CTCGGCCGATCGAAAGGG | >hg38_21_+_8433172_8446622_RNA45SN1_45S_50ntFlanks | 10473 | 10490 | 18 | CTCGGCCGATCGAAAGGG | 194.4 | -4.2 | 1.1 | -3.7 | 2.4E-04 | 3.1E-04 | MB231/MB468 | NUC |
| >hg38_21_+_8433172_8446622_RNA45SN1_45S_50ntFlanks@5492.5516.25\|AGAAGACGGTCGAACTTGACTATCT | >hg38_21_+_8433172_8446622_RNA45SN1_45S_50ntFlanks | 5492 | 5516 | 25 | AGAAGACGGTCGAACTTGACTATCT | 20007.0 | 5.5 | 1.3 | 4.1 | 3.4E-05 | 3.1E-04 | BT20/MB231 | NUC |
| >am_trnaMT_ValTAC_MT_+_1602_1670@50.69.20\|ACTTAACTTGACCGCTCTGA | >am_trnaMT_ValTAC_MT_+_1602_1670 | 50 | 69 | 20 | ACTTAACTTGACCGCTCTGA | 252.0 | -3.1 | 0.9 | -3.7 | 2.4E-04 | 3.1E-04 | MB231/MB468 | NUC |
| >am_trnaMT_ValTAC_MT_+_1602_1670@29.57.29\|ACTTACACTTAGGAGATTTCAACTTAACT | >am_trnaMT_ValTAC_MT_+_1602_1670 | 29 | 57 | 29 | ACTTACACTTAGGAGATTTCAACTTAACT | 374.2 | 6.1 | 1.6 | 3.9 | 9.2E-05 | 3.1E-04 | BT20/MB468 | MP |
| >hg38_MT_+_1621_3279_RNR2_16S_50ntFlanks@1175.1215.41\|TGCATTAAAAATTTCGGTTGGGGCGACCTCGGAGCAGAACC | >hg38_MT_+_1621_3279_RNR2_16S_50ntFlanks | 1175 | 1215 | 41 | TGCATTAAAAATTTCGGTTGGGGCGACCTCGGAGCAGAACC | 230.4 | 7.9 | 2.0 | 3.9 | 9.2E-05 | 3.1E-04 | BT20/MB468 | MP |
| >hg38_21_+_8433172_8446622_RNA45SN1_45S_50ntFlanks@6786.6806.21\|TACGCCTGTCTGAGCGTCGCT | >hg38_21_+_8433172_8446622_RNA45SN1_45S_50ntFlanks | 6786 | 6806 | 21 | TACGCCTGTCTGAGCGTCGCT | 816.0 | 3.7 | 0.9 | 3.9 | 9.2E-05 | 3.1E-04 | BT20/MB468 | MP |
| >hg38_21_+_8433172_8446622_RNA45SN1_45S_50ntFlanks@12007.12042.36\|GCCGCCGGTGAAATACCACTACTCTGATCGTTTTTT | >hg38_21_+_8433172_8446622_RNA45SN1_45S_50ntFlanks | 12007 | 12042 | 36 | GCCGCCGGTGAAATACCACTACTCTGATCGTTTTTT | 752.7 | -7.2 | 2.0 | -3.7 | 2.4E-04 | 3.1E-04 | BT20/MB468 | NUC |
| >hg38_21_+_8433172_8446622_RNA45SN1_45S_50ntFlanks@4976.5000.25\|CCGGACACGGACAGGATTGACAGAT | >hg38_21_+_8433172_8446622_RNA45SN1_45S_50ntFlanks | 4976 | 5000 | 25 | CCGGACACGGACAGGATTGACAGAT | 96.2 | -5.8 | 1.5 | -3.9 | 9.2E-05 | 3.1E-04 | BT20/MB468 | MP |
| >hg38_21_+_8433172_8446622_RNA45SN1_45S_50ntFlanks@11565.11591.27\|CTAGCAGCCGACTTAGAACTGGTGCGG | >hg38_21_+_8433172_8446622_RNA45SN1_45S_50ntFlanks | 11565 | 11591 | 27 | CTAGCAGCCGACTTAGAACTGGTGCGG | 120.3 | -6.1 | 1.6 | -3.9 | 9.2E-05 | 3.1E-04 | BT20/MB468 | MP |
| >hg38_21_+_8433172_8446622_RNA45SN1_45S_50ntFlanks@7988.8014.27\|CAGACGTGGCGACCCGCTGAATTTAAG | >hg38_21_+_8433172_8446622_RNA45SN1_45S_50ntFlanks | 7988 | 8014 | 27 | CAGACGTGGCGACCCGCTGAATTTAAG | 1154.0 | -5.3 | 1.4 | -3.9 | 9.2E-05 | 3.1E-04 | BT20/MB468 | MP |
| >hg38_21_+_8433172_8446622_RNA45SN1_45S_50ntFlanks@4731.4768.38\|AAGAACGAAAGTCGGAGGTTCGAAGACGATCAGATACC | >hg38_21_+_8433172_8446622_RNA45SN1_45S_50ntFlanks | 4731 | 4768 | 38 | AAGAACGAAAGTCGGAGGTTCGAAGACGATCAGATACC | 224.3 | 6.2 | 1.6 | 3.9 | 9.2E-05 | 3.1E-04 | BT20/MB468 | MP |
| >hg38_21_+_8433172_8446622_RNA45SN1_45S_50ntFlanks@12717.12747.31\|CCTCGGTTGGCCTCGGATAGCCGGTCCCCCG | >hg38_21_+_8433172_8446622_RNA45SN1_45S_50ntFlanks | 12717 | 12747 | 31 | CCTCGGTTGGCCTCGGATAGCCGGTCCCCCG | 844.7 | -6.3 | 1.6 | -3.9 | 9.2E-05 | 3.1E-04 | BT20/MB468 | MP |

<table>
<tr><td></td><td></td><td></td><td></td><td></td><td></td><td></td><td></td><td></td><td></td><td></td><td></td><td></td><td></td></tr>
<tr><td>>hg38_21_+_8433172_8446622_RNA45SN1_45S_50ntFlanks@12163.12182.20|ACTGGGGCGGTACACCTGTC</td><td>>hg38_21_+_8433172_8446622_RNA45SN1_45S_50ntFlanks</td><td>12163</td><td>12182</td><td>20</td><td>ACTGGGGCGGTACACCTGTC</td><td>498.8</td><td>-4.2</td><td>1.1</td><td>-3.7</td><td>2.1E-04</td><td>3.1E-04</td><td>BT20/MB468</td><td>CYTO</td></tr>
<tr><td>>hg38_21_+_8433172_8446622_RNA45SN1_45S_50ntFlanks@5423.5443.21|ATTGGATGGTTTAGTGAGGCC</td><td>>hg38_21_+_8433172_8446622_RNA45SN1_45S_50ntFlanks</td><td>5423</td><td>5443</td><td>21</td><td>ATTGGATGGTTTAGTGAGGCC</td><td>179.7</td><td>-5.0</td><td>1.3</td><td>-3.7</td><td>2.1E-04</td><td>3.1E-04</td><td>BT20/MB468</td><td>CYTO</td></tr>
<tr><td>>hg38_21_+_8433172_8446622_RNA45SN1_45S_50ntFlanks@9563.9595.33|CCTGACGTGCAAATCGGTCGTCCGACCTGGGTA</td><td>>hg38_21_+_8433172_8446622_RNA45SN1_45S_50ntFlanks</td><td>9563</td><td>9595</td><td>33</td><td>CCTGACGTGCAAATCGGTCGTCCGACCTGGGTA</td><td>160.6</td><td>-3.0</td><td>0.8</td><td>-3.7</td><td>2.3E-04</td><td>3.1E-04</td><td>MB231/MB468</td><td>CYTO</td></tr>
<tr><td>>hg38_MT_+_1621_3279_RNR2_16S_50ntFlanks@617.656.40|AAACATATAACTGAACTCCTCACACCCAATTGGACCAATC</td><td>>hg38_MT_+_1621_3279_RNR2_16S_50ntFlanks</td><td>617</td><td>656</td><td>40</td><td>AAACATATAACTGAACTCCTCACACCCAATTGGACCAATC</td><td>93.7</td><td>10.3</td><td>2.6</td><td>3.9</td><td>9.2E-05</td><td>3.1E-04</td><td>BT20/MB468</td><td>MP</td></tr>
<tr><td>>hg38_MT_+_1621_3279_RNR2_16S_50ntFlanks@486.518.33|AGGAACAGCTCTTTGGACACTAGGAAAAAACCT</td><td>>hg38_MT_+_1621_3279_RNR2_16S_50ntFlanks</td><td>486</td><td>518</td><td>33</td><td>AGGAACAGCTCTTTGGACACTAGGAAAAAACCT</td><td>188.4</td><td>2.9</td><td>0.8</td><td>3.7</td><td>2.4E-04</td><td>3.1E-04</td><td>MB231/MB468</td><td>MITO</td></tr>
<tr><td>>hg38_21_+_8433172_8446622_RNA45SN1_45S_50ntFlanks@12420.12457.38|ATGTCGGCTCTTCCTATCATTGTGAAGCAGAATTCACC</td><td>>hg38_21_+_8433172_8446622_RNA45SN1_45S_50ntFlanks</td><td>12420</td><td>12457</td><td>38</td><td>ATGTCGGCTCTTCCTATCATTGTGAAGCAGAATTCACC</td><td>1253.6</td><td>-3.2</td><td>0.8</td><td>-4.1</td><td>3.7E-05</td><td>3.1E-04</td><td>BT20/MB231</td><td>MITO</td></tr>
<tr><td>>hg38_21_+_8433172_8446622_RNA45SN1_45S_50ntFlanks@12457.12496.40|CAAGCGTTGGATTGTTCACCCACTAATAGGGAACGTGAGC</td><td>>hg38_21_+_8433172_8446622_RNA45SN1_45S_50ntFlanks</td><td>12457</td><td>12496</td><td>40</td><td>CAAGCGTTGGATTGTTCACCCACTAATAGGGAACGTGAGC</td><td>347.6</td><td>2.1</td><td>0.6</td><td>3.7</td><td>2.3E-04</td><td>3.1E-04</td><td>MB231/MB468</td><td>CYTO</td></tr>
<tr><td>>MI0000061|hsa-let-7a-2&WithFlank&11|-|122146516|122146599@11.31.21(+1G);MI0000062|hsa-let-7a-3&WithFlank&22|+|46112743|46112828@10.30.21(+1G);MI0000060|hsa-let-7a-1&WithFlank&9|+|94175951|94176042@12.32.21(+1G)&[MIMAT0000062&hsa-let-7a-5p&offsets|0|-1(+1G);m-6&11|-|122146568|122146589&offsets|0|-1(+1G)];[MIMAT0000062_1&hsa-let-7a-5p&offsets|0|-1(+1G);m-5&22|+|46112752|46112773&offsets|0|-1(+1G)];[MIMAT0000062_2&hsa-let-7a-5p&offsets|0|-1(+1G);m-4&9|+|94175962|94175983&offsets|0|-1(+1G)]|TGAGGTAGTAGGTTGTATAGTG</td><td>>MI0000061|hsa-let-7a-2&WithFlank&11|-|122146516|122146599</td><td>11</td><td>31</td><td>21(+1G)</td><td>TGAGGTAGTAGGTTGTATAGTG</td><td>407.0</td><td>2.4</td><td>0.7</td><td>3.7</td><td>2.5E-04</td><td>3.1E-04</td><td>BT20/MB468</td><td>NUC</td></tr>
<tr><td>>hg38_21_+_8433172_8446622_RNA45SN1_45S_50ntFlanks@11233.11253.21|CCCTTCCCGTGGATCGCCCCA</td><td>>hg38_21_+_8433172_8446622_RNA45SN1_45S_50ntFlanks</td><td>11233</td><td>11253</td><td>21</td><td>CCCTTCCCGTGGATCGCCCCA</td><td>128.6</td><td>-6.0</td><td>1.5</td><td>-3.9</td><td>9.3E-05</td><td>3.1E-04</td><td>BT20/MB468</td><td>MP</td></tr>
<tr><td>>hg38_21_+_8433172_8446622_RNA45SN1_45S_50ntFlanks@10389.10407.19|TGAACAGCAGTTGAACATG</td><td>>hg38_21_+_8433172_8446622_RNA45SN1_45S_50ntFlanks</td><td>10389</td><td>10407</td><td>19</td><td>TGAACAGCAGTTGAACATG</td><td>379.7</td><td>-2.3</td><td>0.6</td><td>-3.7</td><td>1.8E-04</td><td>3.1E-04</td><td>BT20/MB468</td><td>MITO</td></tr>
<tr><td>>MI0000767|hsa-mir-365a&WithFlank&16|+|14309279|14309377@22.45.24&[MIMAT0009199&hsa-miR-365a-5p&offsets|0|+1;m-595&16|+|14309300|14309322&offsets|0|+1]|AGGGACTTTTGGGGGCAGATGTGT</td><td>>MI0000767|hsa-mir-365a&WithFlank&16|+|14309279|14309377</td><td>22</td><td>45</td><td>24</td><td>AGGGACTTTTGGGGGCAGATGTGT</td><td>220.2</td><td>6.3</td><td>1.6</td><td>3.9</td><td>9.3E-05</td><td>3.1E-04</td><td>BT20/MB468</td><td>MP</td></tr>
<tr><td>>MI0000077|hsa-mir-21&WithFlank&17|+|59841260|59841343@13.36.24&[MIMAT0000076&hsa-miR-21-5p&offsets|-1|+1;m-1&17|+|59841273|59841294&offsets|-1|+1]|GTAGCTTATCAGACTGATGTTGAC</td><td>>MI0000077|hsa-mir-21&WithFlank&17|+|59841260|59841343</td><td>13</td><td>36</td><td>24</td><td>GTAGCTTATCAGACTGATGTTGAC</td><td>410.9</td><td>-4.0</td><td>1.0</td><td>-4.2</td><td>3.3E-05</td><td>3.1E-04</td><td>BT20/MB231</td><td>MP</td></tr>
<tr><td>>hg38_MT_+_1621_3279_RNR2_16S_50ntFlanks@644.677.34|AATTGGACCAATCTATCACCCTATAGAAGAACTA</td><td>>hg38_MT_+_1621_3279_RNR2_16S_50ntFlanks</td><td>644</td><td>677</td><td>34</td><td>AATTGGACCAATCTATCACCCTATAGAAGAACTA</td><td>118.0</td><td>7.1</td><td>1.8</td><td>3.9</td><td>9.3E-05</td><td>3.1E-04</td><td>BT20/MB468</td><td>MP</td></tr>
<tr><td>>hg38_21_+_8433172_8446622_RNA45SN1_45S_50ntFlanks@11718.11736.19|AACGGCGGGAGTAACTATG</td><td>>hg38_21_+_8433172_8446622_RNA45SN1_45S_50ntFlanks</td><td>11718</td><td>11736</td><td>19</td><td>AACGGCGGGAGTAACTATG</td><td>946.7</td><td>-3.5</td><td>1.0</td><td>-3.7</td><td>2.3E-04</td><td>3.1E-04</td><td>MB231/MB468</td><td>CYTO</td></tr>
<tr><td>>hg38_21_+_8433172_8446622_RNA45SN1_45S_50ntFlanks@7974.8016.43|ACGCGACCTCAGATCAGACGTGGCGACCCGCTGAATTTAAGCA</td><td>>hg38_21_+_8433172_8446622_RNA45SN1_45S_50ntFlanks</td><td>7974</td><td>8016</td><td>43</td><td>ACGCGACCTCAGATCAGACGTGGCGACCCGCTGAATTTAAGCA</td><td>6241.8</td><td>3.3</td><td>0.9</td><td>3.7</td><td>2.4E-04</td><td>3.1E-04</td><td>MB231/MB468</td><td>MITO</td></tr>
<tr><td>>hg38_21_+_8433172_8446622_RNA45SN1_45S_50ntFlanks@10845.10866.22|ACTTCGGGATAAGGATTGGCTC</td><td>>hg38_21_+_8433172_8446622_RNA45SN1_45S_50ntFlanks</td><td>10845</td><td>10866</td><td>22</td><td>ACTTCGGGATAAGGATTGGCTC</td><td>349.1</td><td>-4.2</td><td>1.1</td><td>-3.7</td><td>2.1E-04</td><td>3.1E-04</td><td>BT20/MB468</td><td>CYTO</td></tr>
<tr><td>>hg38_21_+_8433172_8446622_RNA45SN1_45S_50ntFlanks@11199.11226.28|CCGGCGGCGGCGGCGACTCTGGACGCGA</td><td>>hg38_21_+_8433172_8446622_RNA45SN1_45S_50ntFlanks</td><td>11199</td><td>11226</td><td>28</td><td>CCGGCGGCGGCGGCGACTCTGGACGCGA</td><td>95.3</td><td>-4.6</td><td>1.2</td><td>-3.7</td><td>2.1E-04</td><td>3.1E-04</td><td>BT20/MB468</td><td>CYTO</td></tr>
<tr><td>>hg38_21_+_8433172_8446622_RNA45SN1_45S_50ntFlanks@10678.10713.36|CGGTTCCGGCGGCGTCCGGTGAGCTCTCGCTGGCCC</td><td>>hg38_21_+_8433172_8446622_RNA45SN1_45S_50ntFlanks</td><td>10678</td><td>10713</td><td>36</td><td>CGGTTCCGGCGGCGTCCGGTGAGCTCTCGCTGGCCC</td><td>75.1</td><td>-5.5</td><td>1.5</td><td>-3.7</td><td>2.0E-04</td><td>3.1E-04</td><td>MB231/MB468</td><td>MP</td></tr>
<tr><td>>hg38_21_+_8433172_8446622_RNA45SN1_45S_50ntFlanks@12444.12486.43|AAGCAGAATTCACCAAGCGTTGGATTGTTCACCCACTAATAGG</td><td>>hg38_21_+_8433172_8446622_RNA45SN1_45S_50ntFlanks</td><td>12444</td><td>12486</td><td>43</td><td>AAGCAGAATTCACCAAGCGTTGGATTGTTCACCCACTAATAGG</td><td>921.6</td><td>-3.8</td><td>1.0</td><td>-3.7</td><td>2.4E-04</td><td>3.1E-04</td><td>MB231/MB468</td><td>NUC</td></tr>
<tr><td>>hg38_21_+_8433172_8446622_RNA45SN1_45S_50ntFlanks@9535.9553.19|GGAAACTCTGGTGGAGGTC</td><td>>hg38_21_+_8433172_8446622_RNA45SN1_45S_50ntFlanks</td><td>9535</td><td>9553</td><td>19</td><td>GGAAACTCTGGTGGAGGTC</td><td>362.5</td><td>-5.5</td><td>1.5</td><td>-3.7</td><td>2.1E-04</td><td>3.1E-04</td><td>BT20/MB468</td><td>CYTO</td></tr>
<tr><td>>hg38_21_+_8433172_8446622_RNA45SN1_45S_50ntFlanks@10408.10443.36|GGTCAGTCGGTCCTGAGAGATGGGCGAGCGCCGTTC</td><td>>hg38_21_+_8433172_8446622_RNA45SN1_45S_50ntFlanks</td><td>10408</td><td>10443</td><td>36</td><td>GGTCAGTCGGTCCTGAGAGATGGGCGAGCGCCGTTC</td><td>939.1</td><td>-6.8</td><td>1.7</td><td>-3.9</td><td>9.3E-05</td><td>3.1E-04</td><td>BT20/MB468</td><td>MP</td></tr>
<tr><td>>hg38_21_+_8433172_8446622_RNA45SN1_45S_50ntFlanks@11572.11593.22|CCGACTTAGAACTGGTGCGGAC</td><td>>hg38_21_+_8433172_8446622_RNA45SN1_45S_50ntFlanks</td><td>11572</td><td>11593</td><td>22</td><td>CCGACTTAGAACTGGTGCGGAC</td><td>1051.0</td><td>-5.0</td><td>1.3</td><td>-3.9</td><td>9.3E-05</td><td>3.1E-04</td><td>BT20/MB468</td><td>MP</td></tr>
<tr><td>>hg38_21_+_8433172_8446622_RNA45SN1_45S_50ntFlanks@12946.12964.19|TAGACGACCTGCTTCTGGG</td><td>>hg38_21_+_8433172_8446622_RNA45SN1_45S_50ntFlanks</td><td>12946</td><td>12964</td><td>19</td><td>TAGACGACCTGCTTCTGGG</td><td>162.5</td><td>-4.0</td><td>1.1</td><td>-3.7</td><td>2.4E-04</td><td>3.1E-04</td><td>MB231/MB468</td><td>NUC</td></tr>
<tr><td>>trnaMT_PheGAA_MT_+_577_647@39.59.21|ATGTTTAGACGGGCTCACATC</td><td>>trnaMT_PheGAA_MT_+_577_647</td><td>39</td><td>59</td><td>21</td><td>ATGTTTAGACGGGCTCACATC</td><td>550.3</td><td>4.6</td><td>1.2</td><td>3.9</td><td>9.3E-05</td><td>3.1E-04</td><td>BT20/MB468</td><td>MP</td></tr>
<tr><td>>hg38_21_+_8433172_8446622_RNA45SN1_45S_50ntFlanks@9549.9572.24|AGGTCCGTAGCGGTCCTGACGTGC</td><td>>hg38_21_+_8433172_8446622_RNA45SN1_45S_50ntFlanks</td><td>9549</td><td>9572</td><td>24</td><td>AGGTCCGTAGCGGTCCTGACGTGC</td><td>192.6</td><td>4.9</td><td>1.3</td><td>3.9</td><td>9.3E-05</td><td>3.1E-04</td><td>BT20/MB468</td><td>MP</td></tr>
<tr><td>>hg38_21_+_8433172_8446622_RNA45SN1_45S_50ntFlanks@10801.10818.18|GTTGGAACAATGTAGGTA</td><td>>hg38_21_+_8433172_8446622_RNA45SN1_45S_50ntFlanks</td><td>10801</td><td>10818</td><td>18</td><td>GTTGGAACAATGTAGGTA</td><td>662.9</td><td>-4.9</td><td>1.3</td><td>-3.7</td><td>2.2E-04</td><td>3.1E-04</td><td>BT20/MB468</td><td>CYTO</td></tr>
<tr><td>>hg38_21_+_8433172_8446622_RNA45SN1_45S_50ntFlanks@10905.10935.31|CTGGGCGCGCGCCGCGGCTGGACGAGGCGCC</td><td>>hg38_21_+_8433172_8446622_RNA45SN1_45S_50ntFlanks</td><td>10905</td><td>10935</td><td>31</td><td>CTGGGCGCGCGCCGCGGCTGGACGAGGCGCC</td><td>200.4</td><td>-5.3</td><td>1.4</td><td>-3.7</td><td>2.2E-04</td><td>3.1E-04</td><td>BT20/MB468</td><td>CYTO</td></tr>
<tr><td>>hg38_21_+_8433172_8446622_RNA45SN1_45S_50ntFlanks@11578.11611.34|TAGAACTGGTGCGGACCAGGGGAATCCGACTGTT</td><td>>hg38_21_+_8433172_8446622_RNA45SN1_45S_50ntFlanks</td><td>11578</td><td>11611</td><td>34</td><td>TAGAACTGGTGCGGACCAGGGGAATCCGACTGTT</td><td>277.1</td><td>-4.7</td><td>1.3</td><td>-3.7</td><td>2.0E-04</td><td>3.1E-04</td><td>MB231/MB468</td><td>MP</td></tr>
<tr><td>>MI0003599|hsa-mir-589&WithFlank&7|-|5495813|5495923@30.52.23&[MIMAT0004799&hsa-miR-589-5p&offsets|0|+1;m-162&7|-|5495874|5495894&offsets|0|+2]|TGAGAACCACGTCTGCTCTGAGC</td><td>>MI0003599|hsa-mir-589&WithFlank&7|-|5495813|5495923</td><td>30</td><td>52</td><td>23</td><td>TGAGAACCACGTCTGCTCTGAGC</td><td>381.6</td><td>6.5</td><td>1.7</td><td>3.9</td><td>9.4E-05</td><td>3.1E-04</td><td>BT20/MB468</td><td>MP</td></tr>
<tr><td>>hg38_21_+_8433172_8446622_RNA45SN1_45S_50ntFlanks@10238.10263.26|CGCTACGCCGCGACGAGTAGGAGGGC</td><td>>hg38_21_+_8433172_8446622_RNA45SN1_45S_50ntFlanks</td><td>10238</td><td>10263</td><td>26</td><td>CGCTACGCCGCGACGAGTAGGAGGGC</td><td>735.7</td><td>3.9</td><td>1.0</td><td>3.9</td><td>9.4E-05</td><td>3.1E-04</td><td>BT20/MB468</td><td>MP</td></tr>
<tr><td>>hg38_21_+_8433172_8446622_RNA45SN1_45S_50ntFlanks@10679.10707.29|GGTTCCGGCGGCGTCCGGTGAGCTCTCGC</td><td>>hg38_21_+_8433172_8446622_RNA45SN1_45S_50ntFlanks</td><td>10679</td><td>10707</td><td>29</td><td>GGTTCCGGCGGCGTCCGGTGAGCTCTCGC</td><td>144.9</td><td>-3.9</td><td>1.1</td><td>-3.7</td><td>2.4E-04</td><td>3.1E-04</td><td>MB231/MB468</td><td>NUC</td></tr>
<tr><td>>hg38_21_+_8433172_8446622_RNA45SN1_45S_50ntFlanks@10843.10868.26|TAACTTCGGGATAAGGATTGGCTCTA</td><td>>hg38_21_+_8433172_8446622_RNA45SN1_45S_50ntFlanks</td><td>10843</td><td>10868</td><td>26</td><td>TAACTTCGGGATAAGGATTGGCTCTA</td><td>281.4</td><td>-4.8</td><td>1.2</td><td>-3.9</td><td>9.4E-05</td><td>3.1E-04</td><td>BT20/MB468</td><td>MP</td></tr>
<tr><td>>hg38_21_+_8433172_8446622_RNA45SN1_45S_50ntFlanks@13007.13024.18|TATTGAAAGTCAGCCCTC</td><td>>hg38_21_+_8433172_8446622_RNA45SN1_45S_50ntFlanks</td><td>13007</td><td>13024</td><td>18</td><td>TATTGAAAGTCAGCCCTC</td><td>148.4</td><td>-4.9</td><td>1.3</td><td>-3.7</td><td>2.2E-04</td><td>3.1E-04</td><td>BT20/MB468</td><td>CYTO</td></tr>
<tr><td>>hg38_21_+_8433172_8446622_RNA45SN1_45S_50ntFlanks@10807.10828.22|ACAATGTAGGTAAGGGAAGTCG</td><td>>hg38_21_+_8433172_8446622_RNA45SN1_45S_50ntFlanks</td><td>10807</td><td>10828</td><td>22</td><td>ACAATGTAGGTAAGGGAAGTCG</td><td>235.9</td><td>-5.5</td><td>1.5</td><td>-3.7</td><td>2.5E-04</td><td>3.1E-04</td><td>MB231/MB468</td><td>NUC</td></tr>
<tr><td>>hg38_21_+_8433172_8446622_RNA45SN1_45S_50ntFlanks@4050.4082.33|CGAACGTCTGCCCTATCAACTTTCGATGGTAGT</td><td>>hg38_21_+_8433172_8446622_RNA45SN1_45S_50ntFlanks</td><td>4050</td><td>4082</td><td>33</td><td>CGAACGTCTGCCCTATCAACTTTCGATGGTAGT</td><td>130.8</td><td>7.3</td><td>1.9</td><td>3.9</td><td>9.4E-05</td><td>3.1E-04</td><td>BT20/MB468</td><td>MP</td></tr>
<tr><td>>hg38_MT_+_598_1651_RNR1_12S_50ntFlanks@916.950.35|AAGGACATTTAACTAAAACCCCTACGCATTTATAT</td><td>>hg38_MT_+_598_1651_RNR1_12S_50ntFlanks</td><td>916</td><td>950</td><td>35</td><td>AAGGACATTTAACTAAAACCCCTACGCATTTATAT</td><td>758.0</td><td>7.2</td><td>1.8</td><td>3.9</td><td>9.4E-05</td><td>3.1E-04</td><td>BT20/MB468</td><td>MP</td></tr>
<tr><td>>hg38_21_+_8433172_8446622_RNA45SN1_45S_50ntFlanks@10357.10383.27|CTTTGAAGGCCGAAGTGGAGAAGGGTT</td><td>>hg38_21_+_8433172_8446622_RNA45SN1_45S_50ntFlanks</td><td>10357</td><td>10383</td><td>27</td><td>CTTTGAAGGCCGAAGTGGAGAAGGGTT</td><td>585.3</td><td>-3.3</td><td>0.8</td><td>-3.9</td><td>9.4E-05</td><td>3.1E-04</td><td>BT20/MB468</td><td>MP</td></tr>
<tr><td>>hg38_21_+_8433172_8446622_RNA45SN1_45S_50ntFlanks@10402.10433.32|AACATGGGTCAGTCGGTCCTGAGAGATGGGCG</td><td>>hg38_21_+_8433172_8446622_RNA45SN1_45S_50ntFlanks</td><td>10402</td><td>10433</td><td>32</td><td>AACATGGGTCAGTCGGTCCTGAGAGATGGGCG</td><td>581.6</td><td>-3.3</td><td>0.9</td><td>-3.9</td><td>9.4E-05</td><td>3.1E-04</td><td>BT20/MB468</td><td>MP</td></tr>
<tr><td>>trna111_HisGTG_1_-_147774845_147774916@-1T.34.35,trna118_HisGTG_1_-_145396881_145396952@-1T.34.35,trna16_HisGTG_1_+_146544773_146544844@-1T.34.35,trna1_HisGTG_15_+_45493349_45493420@-1T.34.35,trna21_HisGTG_1_+_147753471_147753542@-1T.34.35,trna33_HisGTG_6_+_27125906_27125977@-1T.34.35,trna7_HisGTG_9_-_14433938_14434009@-1T.34.35,trna8_HisGTG_15_-_45492611_45492682@-1T.34.35,trna9_HisGTG_15_-_45490804_45490875@-1T.34.35|TGCCGTGATCGTATAGTGGTTAGTACTCTGCGTTG</td><td>>trna111_HisGTG_1_-_147774845_147774916</td><td>-1T</td><td>34</td><td>35</td><td>TGCCGTGATCGTATAGTGGTTAGTACTCTGCGTTG</td><td>1141.0</td><td>4.4</td><td>1.2</td><td>3.7</td><td>2.5E-04</td><td>3.1E-04</td><td>MB231/MB468</td><td>NUC</td></tr>
<tr><td>>MI0000088|hsa-mir-30a&WithFlank&6|-|71403545|71403627@54.75.22&[MIMAT0000088&hsa-miR-30a-3p&offsets|+1|+1;m-38&6|-|71403554|71403575&offsets|+1|+1]|TTTCAGTCGGATGTTTGCAGCT</td><td>>MI0000088|hsa-mir-30a&WithFlank&6|-|71403545|71403627</td><td>54</td><td>75</td><td>22</td><td>TTTCAGTCGGATGTTTGCAGCT</td><td>623.0</td><td>5.1</td><td>1.4</td><td>3.7</td><td>2.1E-04</td><td>3.1E-04</td><td>MB231/MB468</td><td>MP</td></tr>
<tr><td>>hg38_21_+_8433172_8446622_RNA45SN1_45S_50ntFlanks@10409.10426.18|GTCAGTCGGTCCTGAGAG</td><td>>hg38_21_+_8433172_8446622_RNA45SN1_45S_50ntFlanks</td><td>10409</td><td>10426</td><td>18</td><td>GTCAGTCGGTCCTGAGAG</td><td>155.9</td><td>-3.4</td><td>0.9</td><td>-3.7</td><td>2.2E-04</td><td>3.1E-04</td><td>BT20/MB468</td><td>CYTO</td></tr>
<tr><td>>hg38_21_+_8433172_8446622_RNA45SN1_45S_50ntFlanks@5490.5526.37|TGAGAAGACGGTCGAACTTGACTATCTAGAGGAAGTA</td><td>>hg38_21_+_8433172_8446622_RNA45SN1_45S_50ntFlanks</td><td>5490</td><td>5526</td><td>37</td><td>TGAGAAGACGGTCGAACTTGACTATCTAGAGGAAGTA</td><td>359.2</td><td>6.9</td><td>1.8</td><td>3.9</td><td>9.4E-05</td><td>3.1E-04</td><td>BT20/MB468</td><td>MP</td></tr>
<tr><td>>hg38_1_-_228634819_228635039_RNA5S12_5S_50ntFlanks@85.109.25|TCGTCTGATCTCGGAAGCTAAGCAG</td><td>>hg38_1_-_228634819_228635039_RNA5S12_5S_50ntFlanks</td><td>85</td><td>109</td><td>25</td><td>TCGTCTGATCTCGGAAGCTAAGCAG</td><td>307.1</td><td>5.8</td><td>1.5</td><td>3.9</td><td>9.4E-05</td><td>3.1E-04</td><td>BT20/MB468</td><td>MP</td></tr>
<tr><td>>hg38_21_+_8433172_8446622_RNA45SN1_45S_50ntFlanks@12605.12623.19|TATGTGCTTGGCTGAGGAG</td><td>>hg38_21_+_8433172_8446622_RNA45SN1_45S_50ntFlanks</td><td>12605</td><td>12623</td><td>19</td><td>TATGTGCTTGGCTGAGGAG</td><td>402.4</td><td>-4.0</td><td>1.1</td><td>-3.7</td><td>2.2E-04</td><td>3.1E-04</td><td>BT20/MB468</td><td>CYTO</td></tr>
<tr><td>>hg38_21_+_8433172_8446622_RNA45SN1_45S_50ntFlanks@4003.4039.37|ATAACCTCGGGCCGATCGCACGCCCCCCGTGGCGGCG</td><td>>hg38_21_+_8433172_8446622_RNA45SN1_45S_50ntFlanks</td><td>4003</td><td>4039</td><td>37</td><td>ATAACCTCGGGCCGATCGCACGCCCCCCGTGGCGGCG</td><td>217.1</td><td>-4.0</td><td>1.1</td><td>-3.7</td><td>2.2E-04</td><td>3.1E-04</td><td>BT20/MB468</td><td>CYTO</td></tr>
<tr><td>>hg38_21_+_8433172_8446622_RNA45SN1_45S_50ntFlanks@12437.12476.40|CATTGTGAAGCAGAATTCACCAAGCGTTGGATTGTTCACC</td><td>>hg38_21_+_8433172_8446622_RNA45SN1_45S_50ntFlanks</td><td>12437</td><td>12476</td><td>40</td><td>CATTGTGAAGCAGAATTCACCAAGCGTTGGATTGTTCACC</td><td>105.0</td><td>-6.2</td><td>1.7</td><td>-3.7</td><td>2.1E-04</td><td>3.1E-04</td><td>MB231/MB468</td><td>MP</td></tr>
<tr><td>>trnaMT_IleGAT_MT_+_4263_4331@43.68.26|ATAGGAGCTTAAACCCCCTTATTTCT</td><td>>trnaMT_IleGAT_MT_+_4263_4331</td><td>43</td><td>68</td><td>26</td><td>ATAGGAGCTTAAACCCCCTTATTTCT</td><td>90.6</td><td>10.2</td><td>2.6</td><td>3.9</td><td>9.5E-05</td><td>3.1E-04</td><td>BT20/MB468</td><td>MP</td></tr>
<tr><td>>hg38_21_+_8433172_8446622_RNA45SN1_45S_50ntFlanks@10668.10706.39|GAAAGCGTCGCGGTTCCGGCGGCGTCCGGTGAGCTCTCG</td><td>>hg38_21_+_8433172_8446622_RNA45SN1_45S_50ntFlanks</td><td>10668</td><td>10706</td><td>39</td><td>GAAAGCGTCGCGGTTCCGGCGGCGTCCGGTGAGCTCTCG</td><td>564.0</td><td>-5.1</td><td>1.4</td><td>-3.7</td><td>2.2E-04</td><td>3.1E-04</td><td>BT20/MB468</td><td>CYTO</td></tr>
</table>

| | | | | | | | | | | | | | |
|---|---|---|---|---|---|---|---|---|---|---|---|---|---|
| >hg38_21_+_8433172_8446622_RNA45SN1_45S_50ntFlanks@12957.12982.26\|CTTCTGGGTCGGGGTTTCGTACGTAG | >hg38_21_+_8433172_8446622_RNA45SN1_45S_50ntFlanks | 12957 | 12982 | 26 | CTTCTGGGTCGGGGTTTCGTACGTAG | 1556.3 | -6.0 | 1.6 | -3.7 | 2.2E-04 | 3.1E-04 | BT20/MB468 | CYTO |
| >MI0000085\|hsa-mir-27a&WithFlank&19\|-\|13836434\|13836523@57.76.20(+4U)&[MIMAT0000084&hsa-miR-27a-3p&offsets\|0\|-1(+4U);m-43&19\|-\|13836447\|13836467&offsets\|0\|-1(+4U)]\|TTCACAGTGGCTAAGTTCCGTTTT | >MI0000085\|hsa-mir-27a&WithFlank&19\|-\|13836434\|13836523 | 57 | 76 | 20(+4U) | TTCACAGTGGCTAAGTTCCGTTTT | 161.5 | 5.7 | 1.5 | 3.7 | 2.1E-04 | 3.2E-04 | MB231/MB468 | MP |
| >trna14_LysTTT_11_-_59327808_59327880@1.31.31,trna2_LysTTT_17_+_8022473_8022545@1.31.31,trna54_LysTTT_1_+_204475655_204475727@1.31.31,trna5_LysTTT_11_+_59323902_59323974@1.31.31,trna62_LysTTT_1_-_204476158_204476230@1.31.31,trna76_LysTTT_6_+_28918806_28918878@1.31.31\|GCCCGGATAGCTCAGTCGGTAGAGCATCAGA | >trna14_LysTTT_11_-_59327808_59327880 | 1 | 31 | 31 | GCCCGGATAGCTCAGTCGGTAGAGCATCAGA | 409.4 | -3.5 | 1.0 | -3.7 | 2.1E-04 | 3.2E-04 | MB231/MB468 | MP |
| >MI0000266\|hsa-mir-10a&WithFlank&17\|-\|48579832\|48579953@28.46.19&[MIMAT0000253&hsa-miR-10a-5p&offsets\|0\|-4;m-18&17\|-\|48579905\|48579926&offsets\|0\|-3]\|TACCCTGTAGATCCGAATT | >MI0000266\|hsa-mir-10a&WithFlank&17\|-\|48579832\|48579953 | 28 | 46 | 19 | TACCCTGTAGATCCGAATT | 95.0 | 10.3 | 2.6 | 3.9 | 9.5E-05 | 3.2E-04 | BT20/MB468 | MP |
| >hg38_21_+_8433172_8446622_RNA45SN1_45S_50ntFlanks@4046.4069.24\|CATTCGAACGTCTGCCCTATCAAC | >hg38_21_+_8433172_8446622_RNA45SN1_45S_50ntFlanks | 4046 | 4069 | 24 | CATTCGAACGTCTGCCCTATCAAC | 185.6 | -2.5 | 0.7 | -3.7 | 2.5E-04 | 3.2E-04 | MB231/MB468 | NUC |
| >hg38_MT_+_598_1651_RNR1_12S_50ntFlanks@425.462.38\|ACGAAAGTGGCTTTAACATATCTGAACACACAATAGCT | >hg38_MT_+_598_1651_RNR1_12S_50ntFlanks | 425 | 462 | 38 | ACGAAAGTGGCTTTAACATATCTGAACACACAATAGCT | 22.1 | 7.7 | 2.1 | 3.7 | 2.1E-04 | 3.2E-04 | MB231/MB468 | MP |
| >hg38_21_+_8433172_8446622_RNA45SN1_45S_50ntFlanks@6650.6699.50\|ACGACTCTTAGCGGTGGATCACTCGGCTCGTGCGTCGATGAAGAACGCAG | >hg38_21_+_8433172_8446622_RNA45SN1_45S_50ntFlanks | 6650 | 6699 | 50 | ACGACTCTTAGCGGTGGATCACTCGGCTCGTGCGTCGATGAAGAACGCAG | 2227.3 | -4.4 | 1.2 | -3.7 | 2.5E-04 | 3.2E-04 | BT20/MB468 | NUC |
| >hg38_21_+_8433172_8446622_RNA45SN1_45S_50ntFlanks@10186.10209.24\|TCCTCCCCTCCTCCCGCCCACGCC | >hg38_21_+_8433172_8446622_RNA45SN1_45S_50ntFlanks | 10186 | 10209 | 24 | TCCTCCCCTCCTCCCGCCCACGCC | 119.4 | -10.0 | 2.6 | -3.9 | 9.5E-05 | 3.2E-04 | BT20/MB468 | MP |
| >hg38_21_+_8433172_8446622_RNA45SN1_45S_50ntFlanks@3843.3871.29\|CTACTTGGATAACTGTGGTAATTCTAGAG | >hg38_21_+_8433172_8446622_RNA45SN1_45S_50ntFlanks | 3843 | 3871 | 29 | CTACTTGGATAACTGTGGTAATTCTAGAG | 634.5 | -3.1 | 0.8 | -3.7 | 2.3E-04 | 3.2E-04 | MB231/MB468 | CYTO |
| >trna111_HisGTG_1_-_147774845_147774916@-1T.22.23,trna118_HisGTG_1_-_145396881_145396952@-1T.22.23,trna16_HisGTG_1_+_146544773_146544844@-1T.22.23,trna1_HisGTG_15_+_45493349_45493420@-1T.22.23,trna21_HisGTG_1_+_147753471_147753542@-1T.22.23,trna33_HisGTG_6_+_27125906_27125977@-1T.22.23,trna7_HisGTG_9_-_14433938_14434009@-1T.22.23,trna8_HisGTG_15_-_45492611_45492682@-1T.22.23,trna9_HisGTG_15_-_45490804_45490875@-1T.22.23\|TGCCGTGATCGTATAGTGGTTAG | >trna111_HisGTG_1_-_147774845_147774916 | -1T | 22 | 23 | TGCCGTGATCGTATAGTGGTTAG | 277.7 | -3.1 | 0.7 | -4.1 | 3.9E-05 | 3.2E-04 | BT20/MB231 | MITO |
| >MI0001445\|hsa-mir-423&WithFlank&17\|+\|30117073\|30117178@25.46.22&[MIMAT0004748&hsa-miR-423-5p&offsets\|+2\|+1;m-141&17\|+\|30117095\|30117117&offsets\|+2\|+1]\|AGGGGCAGAGAGCGAGACTTTT | >MI0001445\|hsa-mir-423&WithFlank&17\|+\|30117073\|30117178 | 25 | 46 | 22 | AGGGGCAGAGAGCGAGACTTTT | 178.2 | 6.5 | 1.7 | 3.9 | 9.5E-05 | 3.2E-04 | BT20/MB468 | MP |
| >hg38_21_+_8433172_8446622_RNA45SN1_45S_50ntFlanks@10685.10707.23\|GGCGGCGTCCGGTGAGCTCTCGC | >hg38_21_+_8433172_8446622_RNA45SN1_45S_50ntFlanks | 10685 | 10707 | 23 | GGCGGCGTCCGGTGAGCTCTCGC | 255.4 | 6.8 | 1.7 | 3.9 | 9.5E-05 | 3.2E-04 | BT20/MB468 | MP |
| >MI0000085\|hsa-mir-27a&WithFlank&19\|-\|13836434\|13836523@57.75.19(+1C)&[MIMAT0000084&hsa-miR-27a-3p&offsets\|0\|-2(+1C);m-43&19\|-\|13836447\|13836467&offsets\|0\|-2(+1C)]\|TTCACAGTGGCTAAGTTCCC | >MI0000085\|hsa-mir-27a&WithFlank&19\|-\|13836434\|13836523 | 57 | 75 | 19(+1C) | TTCACAGTGGCTAAGTTCCC | 154.9 | 5.9 | 1.6 | 3.7 | 2.5E-04 | 3.2E-04 | MB231/MB468 | MITO |
| >hg38_21_+_8433172_8446622_RNA45SN1_45S_50ntFlanks@12076.12097.22\|AGGGGCTCTCGCTTCTGGCGCC | >hg38_21_+_8433172_8446622_RNA45SN1_45S_50ntFlanks | 12076 | 12097 | 22 | AGGGGCTCTCGCTTCTGGCGCC | 583.7 | -6.5 | 1.8 | -3.7 | 2.5E-04 | 3.2E-04 | BT20/MB468 | NUC |
| >hg38_21_+_8433172_8446622_RNA45SN1_45S_50ntFlanks@4005.4022.18\|AACCTCGGGCCGATCGCA | >hg38_21_+_8433172_8446622_RNA45SN1_45S_50ntFlanks | 4005 | 4022 | 18 | AACCTCGGGCCGATCGCA | 1638.9 | -7.2 | 2.0 | -3.7 | 2.5E-04 | 3.2E-04 | BT20/MB468 | NUC |
| >hg38_21_+_8433172_8446622_RNA45SN1_45S_50ntFlanks@3842.3860.19\|CCTACTTGGATAACTGTGG | >hg38_21_+_8433172_8446622_RNA45SN1_45S_50ntFlanks | 3842 | 3860 | 19 | CCTACTTGGATAACTGTGG | 166.3 | -6.1 | 1.6 | -3.9 | 9.6E-05 | 3.2E-04 | BT20/MB468 | MP |
| >hg38_21_+_8433172_8446622_RNA45SN1_45S_50ntFlanks@9548.9574.27\|GAGGTCCGTAGCGGTCCTGACGTGCAA | >hg38_21_+_8433172_8446622_RNA45SN1_45S_50ntFlanks | 9548 | 9574 | 27 | GAGGTCCGTAGCGGTCCTGACGTGCAA | 645.2 | -6.6 | 1.8 | -3.7 | 2.5E-04 | 3.2E-04 | BT20/MB468 | NUC |
| >hg38_21_+_8433172_8446622_RNA45SN1_45S_50ntFlanks@5486.5526.41\|GCGCTGAGAAGACGGTCGAACTTGACTATCTAGAGGAAGTA | >hg38_21_+_8433172_8446622_RNA45SN1_45S_50ntFlanks | 5486 | 5526 | 41 | GCGCTGAGAAGACGGTCGAACTTGACTATCTAGAGGAAGTA | 69.6 | 9.5 | 2.3 | 4.1 | 3.4E-05 | 3.2E-04 | BT20/MB231 | MP |
| >hg38_21_+_8433172_8446622_RNA45SN1_45S_50ntFlanks@6661.6699.39\|CGGTGGATCACTCGGCTCGTGCGTCGATGAAGAACGCAG | >hg38_21_+_8433172_8446622_RNA45SN1_45S_50ntFlanks | 6661 | 6699 | 39 | CGGTGGATCACTCGGCTCGTGCGTCGATGAAGAACGCAG | 2910.6 | 4.1 | 1.0 | 4.1 | 3.4E-05 | 3.2E-04 | BT20/MB231 | MP |
| >hg38_21_+_8433172_8446622_RNA45SN1_45S_50ntFlanks@5071.5097.27\|TTAATTCCGATAACGAACGAGACTCTG | >hg38_21_+_8433172_8446622_RNA45SN1_45S_50ntFlanks | 5071 | 5097 | 27 | TTAATTCCGATAACGAACGAGACTCTG | 500.3 | -3.4 | 0.9 | -3.7 | 2.5E-04 | 3.2E-04 | MB231/MB468 | NUC |
| >MI0003591\|hsa-mir-584&WithFlank&5\|-\|149062307\|149062415@22.39.18(+1C)&[MIMAT0003249&hsa-miR-584-5p&offsets\|0\|-4(+1C);m-167&5\|-\|149062374\|149062394&offsets\|0\|-3(+1C)]\|TTATGGTTTGCCTGGGACC | >MI0003591\|hsa-mir-584&WithFlank&5\|-\|149062307\|149062415 | 22 | 39 | 18(+1C) | TTATGGTTTGCCTGGGACC | 207.8 | 7.5 | 1.9 | 3.9 | 9.6E-05 | 3.2E-04 | BT20/MB468 | MP |
| >am_MI0016846\|hsa-mir-4485&WithFlank&11\|-\|10508264\|10508332@34.52.19&[MIMAT0019019&hsa-miR-4485-3p&offsets\|-3\|-4;m-604&11\|-\|10508277\|10508296&offsets\|-3\|-4]\|GTTTAACGGCCGCGGTACC | >am_MI0016846\|hsa-mir-4485&WithFlank&11\|-\|10508264\|10508332 | 34 | 52 | 19 | GTTTAACGGCCGCGGTACC | 170.0 | 5.4 | 1.5 | 3.7 | 2.5E-04 | 3.2E-04 | MB231/MB468 | MITO |
| >hg38_MT_+_1621_3279_RNR2_16S_50ntFlanks@940.958.19\|GTTTAACGGCCGCGGTACC | >hg38_MT_+_1621_3279_RNR2_16S_50ntFlanks | 940 | 958 | 19 | GTTTAACGGCCGCGGTACC | 170.0 | 5.4 | 1.5 | 3.7 | 2.5E-04 | 3.2E-04 | MB231/MB468 | MITO |
| >hg38_21_+_8433172_8446622_RNA45SN1_45S_50ntFlanks@12407.12446.40\|TTTTGATCCTTCGATGTCGGCTCTTCCTATCATTGTGAAG | >hg38_21_+_8433172_8446622_RNA45SN1_45S_50ntFlanks | 12407 | 12446 | 40 | TTTTGATCCTTCGATGTCGGCTCTTCCTATCATTGTGAAG | 313.8 | -2.2 | 0.6 | -3.7 | 2.2E-04 | 3.2E-04 | BT20/MB468 | CYTO |
| >hg38_21_+_8433172_8446622_RNA45SN1_45S_50ntFlanks@11661.11681.21\|TGATTTCTGCCCAGTGCTCTG | >hg38_21_+_8433172_8446622_RNA45SN1_45S_50ntFlanks | 11661 | 11681 | 21 | TGATTTCTGCCCAGTGCTCTG | 1724.8 | -2.4 | 0.6 | -3.7 | 1.8E-04 | 3.2E-04 | BT20/MB468 | MITO |
| >hg38_21_+_8433172_8446622_RNA45SN1_45S_50ntFlanks@9304.9338.35\|ACCAAGGAGTCTAACACGTGCGCGAGTCGGGGGCT | >hg38_21_+_8433172_8446622_RNA45SN1_45S_50ntFlanks | 9304 | 9338 | 35 | ACCAAGGAGTCTAACACGTGCGCGAGTCGGGGGCT | 637.3 | -5.5 | 1.5 | -3.7 | 1.8E-04 | 3.2E-04 | BT20/MB468 | MITO |
| >hg38_21_+_8433172_8446622_RNA45SN1_45S_50ntFlanks@10794.10815.22\|CTGGCATGTTGGAACAATGTAG | >hg38_21_+_8433172_8446622_RNA45SN1_45S_50ntFlanks | 10794 | 10815 | 22 | CTGGCATGTTGGAACAATGTAG | 225.3 | -5.9 | 1.5 | -3.9 | 9.6E-05 | 3.2E-04 | BT20/MB468 | MP |
| >hg38_21_+_8433172_8446622_RNA45SN1_45S_50ntFlanks@11027.11054.28\|CTCTCTCTCTCTCTCCCCCGCTCCCCGT | >hg38_21_+_8433172_8446622_RNA45SN1_45S_50ntFlanks | 11027 | 11054 | 28 | CTCTCTCTCTCTCTCCCCCGCTCCCCGT | 217.4 | -5.5 | 1.5 | -3.7 | 2.5E-04 | 3.2E-04 | MB231/MB468 | NUC |
| >hg38_21_+_8433172_8446622_RNA45SN1_45S_50ntFlanks@10404.10422.19\|CATGGGTCAGTCGGTCCTG | >hg38_21_+_8433172_8446622_RNA45SN1_45S_50ntFlanks | 10404 | 10422 | 19 | CATGGGTCAGTCGGTCCTG | 272.0 | -5.3 | 1.4 | -3.7 | 2.2E-04 | 3.2E-04 | BT20/MB468 | CYTO |
| >hg38_21_+_8433172_8446622_RNA45SN1_45S_50ntFlanks@11580.11604.25\|GAACTGGTGCGGACCAGGGGAATCC | >hg38_21_+_8433172_8446622_RNA45SN1_45S_50ntFlanks | 11580 | 11604 | 25 | GAACTGGTGCGGACCAGGGGAATCC | 186.1 | -5.8 | 1.6 | -3.7 | 2.1E-04 | 3.2E-04 | MB231/MB468 | MP |
| >hg38_21_+_8433172_8446622_RNA45SN1_45S_50ntFlanks@7983.8009.27\|CAGATCAGACGTGGCGACCCGCTGAAT | >hg38_21_+_8433172_8446622_RNA45SN1_45S_50ntFlanks | 7983 | 8009 | 27 | CAGATCAGACGTGGCGACCCGCTGAAT | 197.5 | -2.2 | 0.6 | -3.7 | 2.5E-04 | 3.2E-04 | MB231/MB468 | NUC |
| >hg38_21_+_8433172_8446622_RNA45SN1_45S_50ntFlanks@9298.9338.41\|ACACGGACCAAGGAGTCTAACACGTGCGCGAGTCGGGGGCT | >hg38_21_+_8433172_8446622_RNA45SN1_45S_50ntFlanks | 9298 | 9338 | 41 | ACACGGACCAAGGAGTCTAACACGTGCGCGAGTCGGGGGCT | 167.9 | -4.8 | 1.3 | -3.7 | 2.1E-04 | 3.2E-04 | MB231/MB468 | MP |
| >trnaMT_ValTAC_MT_+_1602_1670@41.71.31\|GAGATTTCAACTTAACTTGACCGCTCTGACC | >trnaMT_ValTAC_MT_+_1602_1670 | 41 | 71 | 31 | GAGATTTCAACTTAACTTGACCGCTCTGACC | 237.1 | 4.2 | 1.1 | 3.7 | 2.5E-04 | 3.2E-04 | MB231/MB468 | NUC |
| >hg38_21_+_8433172_8446622_RNA45SN1_45S_50ntFlanks@10344.10388.45\|ATTCAAACGAGAACTTTGAAGGCCGAAGTGGAGAAGGGTTCCATG | >hg38_21_+_8433172_8446622_RNA45SN1_45S_50ntFlanks | 10344 | 10388 | 45 | ATTCAAACGAGAACTTTGAAGGCCGAAGTGGAGAAGGGTTCCATG | 140.1 | 7.2 | 1.9 | 3.9 | 9.6E-05 | 3.2E-04 | BT20/MB468 | MP |
| >hg38_21_+_8433172_8446622_RNA45SN1_45S_50ntFlanks@4657.4690.34\|CTAGAGGTGAAATTCTTGGACCGGCGCAAGACGG | >hg38_21_+_8433172_8446622_RNA45SN1_45S_50ntFlanks | 4657 | 4690 | 34 | CTAGAGGTGAAATTCTTGGACCGGCGCAAGACGG | 144.9 | -6.2 | 1.6 | -3.9 | 9.6E-05 | 3.2E-04 | BT20/MB468 | MP |
| >hg38_21_+_8433172_8446622_RNA45SN1_45S_50ntFlanks@12439.12480.42\|TTGTGAAGCAGAATTCACCAAGCGTTGGATTGTTCACCCACT | >hg38_21_+_8433172_8446622_RNA45SN1_45S_50ntFlanks | 12439 | 12480 | 42 | TTGTGAAGCAGAATTCACCAAGCGTTGGATTGTTCACCCACT | 89.9 | 10.2 | 2.6 | 3.9 | 9.6E-05 | 3.2E-04 | BT20/MB468 | MP |
| >hg38_21_+_8433172_8446622_RNA45SN1_45S_50ntFlanks@5238.5255.18\|CTGACTGGCTCAGCGTGT | >hg38_21_+_8433172_8446622_RNA45SN1_45S_50ntFlanks | 5238 | 5255 | 18 | CTGACTGGCTCAGCGTGT | 208.8 | 3.7 | 1.0 | 3.9 | 9.6E-05 | 3.2E-04 | BT20/MB468 | MP |
| >trnaMT_HisGTG_MT_+_12138_12206@26.69.44\|AGATTGTGAATCTGACAACAGAGGCTTACGACCCCTTATTTACC | >trnaMT_HisGTG_MT_+_12138_12206 | 26 | 69 | 44 | AGATTGTGAATCTGACAACAGAGGCTTACGACCCCTTATTTACC | 91.1 | 10.3 | 2.6 | 3.9 | 9.6E-05 | 3.2E-04 | BT20/MB468 | MP |
| >hg38_21_+_8433172_8446622_RNA45SN1_45S_50ntFlanks@9915.9937.23\|AAAAGGTGTTGGTTGATATAGAC | >hg38_21_+_8433172_8446622_RNA45SN1_45S_50ntFlanks | 9915 | 9937 | 23 | AAAAGGTGTTGGTTGATATAGAC | 61.1 | 9.3 | 2.2 | 4.1 | 3.4E-05 | 3.2E-04 | BT20/MB231 | MP |
| >hg38_21_+_8433172_8446622_RNA45SN1_45S_50ntFlanks@3872.3899.28\|CTAATACATGCCGACGGGCGCTGACCCC | >hg38_21_+_8433172_8446622_RNA45SN1_45S_50ntFlanks | 3872 | 3899 | 28 | CTAATACATGCCGACGGGCGCTGACCCC | 222.4 | -5.3 | 1.5 | -3.7 | 2.5E-04 | 3.2E-04 | MB231/MB468 | NUC |
| >hg38_21_+_8433172_8446622_RNA45SN1_45S_50ntFlanks@11564.11590.27\|CCTAGCAGCCGACTTAGAACTGGTGCG | >hg38_21_+_8433172_8446622_RNA45SN1_45S_50ntFlanks | 11564 | 11590 | 27 | CCTAGCAGCCGACTTAGAACTGGTGCG | 140.5 | -3.8 | 1.0 | -3.7 | 2.5E-04 | 3.2E-04 | MB231/MB468 | NUC |
| >MI0000434\|hsa-let-7i&WithFlank&12\|+\|62603680\|62603775@12.31.20(+1C)&[MIMAT0000415&hsa-let-7i-5p&offsets\|0\|-2(+1C);m-49&12\|+\|62603691\|62603712&offsets\|0\|-2(+1C)]\|TGAGGTAGTAGTTTGTGCTGC | >MI0000434\|hsa-let-7i&WithFlank&12\|+\|62603680\|62603775 | 12 | 31 | 20(+1C) | TGAGGTAGTAGTTTGTGCTGC | 237.6 | 6.5 | 1.8 | 3.7 | 2.5E-04 | 3.2E-04 | MB231/MB468 | MITO |
| >hg38_MT_+_598_1651_RNR1_12S_50ntFlanks@441.480.40\|CATATCTGAACACACAATAGCTAAGACCCAAACTGGGATT | >hg38_MT_+_598_1651_RNR1_12S_50ntFlanks | 441 | 480 | 40 | CATATCTGAACACACAATAGCTAAGACCCAAACTGGGATT | 217.8 | 5.5 | 1.5 | 3.7 | 2.5E-04 | 3.2E-04 | MB231/MB468 | MITO |
| >hg38_21_+_8433172_8446622_RNA45SN1_45S_50ntFlanks@12488.12509.22\|AACGTGAGCTGGGTTTAGACCG | >hg38_21_+_8433172_8446622_RNA45SN1_45S_50ntFlanks | 12488 | 12509 | 22 | AACGTGAGCTGGGTTTAGACCG | 2109.4 | -6.8 | 1.8 | -3.9 | 9.7E-05 | 3.2E-04 | BT20/MB468 | MP |
| >hg38_21_+_8433172_8446622_RNA45SN1_45S_50ntFlanks@8105.8135.31\|CGCGGCGGGGCGCGGGACATGTGGCGTACGG | >hg38_21_+_8433172_8446622_RNA45SN1_45S_50ntFlanks | 8105 | 8135 | 31 | CGCGGCGGGGCGCGGGACATGTGGCGTACGG | 240.5 | 7.1 | 1.7 | 4.1 | 3.4E-05 | 3.2E-04 | BT20/MB231 | MP |

<table>
<tr><td>>MI0000093|hsa-mir-92a-1&WithFlank&13|+|91351308|91351397@54.76.23(+1C)&[MIMAT0000092&hsa-miR-92a-3p&offsets|0|+1(+1C);m-9&13|+|91351361|91351382&offsets|0|+1(+1C)]|TATTGCACTTGTCCCGGCCTGTTC</td><td>>MI0000093|hsa-mir-92a-1&WithFlank&13|+|91351308|91351397</td><td>54</td><td>76</td><td>23(+1C)</td><td>TATTGCACTTGTCCCGGCCTGTTC</td><td>54.9</td><td>-9.2</td><td>2.2</td><td>-4.1</td><td>3.4E-05</td><td>3.2E-04</td><td>BT20/MB231</td><td>MP</td></tr>
<tr><td>>hg38_21_+_8433172_8446622_RNA45SN1_45S_50ntFlanks@10482.10506.25|TCGAAAGGGAGTCGGGTTCAGATCC</td><td>>hg38_21_+_8433172_8446622_RNA45SN1_45S_50ntFlanks</td><td>10482</td><td>10506</td><td>25</td><td>TCGAAAGGGAGTCGGGTTCAGATCC</td><td>133.4</td><td>-5.8</td><td>1.6</td><td>-3.7</td><td>2.1E-04</td><td>3.2E-04</td><td>MB231/MB468</td><td>MP</td></tr>
<tr><td>>hg38_21_+_8433172_8446622_RNA45SN1_45S_50ntFlanks@10246.10272.27|CGCGACGAGTAGGAGGGCCGCTGCGGT</td><td>>hg38_21_+_8433172_8446622_RNA45SN1_45S_50ntFlanks</td><td>10246</td><td>10272</td><td>27</td><td>CGCGACGAGTAGGAGGGCCGCTGCGGT</td><td>63.3</td><td>9.3</td><td>2.2</td><td>4.1</td><td>3.4E-05</td><td>3.2E-04</td><td>BT20/MB231</td><td>MP</td></tr>
<tr><td>>hg38_21_+_8433172_8446622_RNA45SN1_45S_50ntFlanks@12420.12455.36|ATGTCGGCTCTTCCTATCATTGTGAAGCAGAATTCA</td><td>>hg38_21_+_8433172_8446622_RNA45SN1_45S_50ntFlanks</td><td>12420</td><td>12455</td><td>36</td><td>ATGTCGGCTCTTCCTATCATTGTGAAGCAGAATTCA</td><td>474.2</td><td>-3.4</td><td>0.8</td><td>-4.1</td><td>3.9E-05</td><td>3.2E-04</td><td>BT20/MB231</td><td>MITO</td></tr>
<tr><td>>hg38_21_+_8433172_8446622_RNA45SN1_45S_50ntFlanks@4047.4069.23|ATTCGAACGTCTGCCCTATCAAC</td><td>>hg38_21_+_8433172_8446622_RNA45SN1_45S_50ntFlanks</td><td>4047</td><td>4069</td><td>23</td><td>ATTCGAACGTCTGCCCTATCAAC</td><td>512.0</td><td>-3.7</td><td>1.0</td><td>-3.7</td><td>1.9E-04</td><td>3.2E-04</td><td>BT20/MB468</td><td>MITO</td></tr>
<tr><td>>hg38_21_+_8433172_8446622_RNA45SN1_45S_50ntFlanks@9738.9762.25|GAAACGATCTCAACCTATTCTCAAA</td><td>>hg38_21_+_8433172_8446622_RNA45SN1_45S_50ntFlanks</td><td>9738</td><td>9762</td><td>25</td><td>GAAACGATCTCAACCTATTCTCAAA</td><td>482.7</td><td>-3.7</td><td>1.0</td><td>-3.7</td><td>1.9E-04</td><td>3.2E-04</td><td>BT20/MB468</td><td>MITO</td></tr>
<tr><td>>hg38_21_+_8433172_8446622_RNA45SN1_45S_50ntFlanks@10807.10837.31|ACAATGTAGGTAAGGGAAGTCGGCAAGCCGG</td><td>>hg38_21_+_8433172_8446622_RNA45SN1_45S_50ntFlanks</td><td>10807</td><td>10837</td><td>31</td><td>ACAATGTAGGTAAGGGAAGTCGGCAAGCCGG</td><td>89.7</td><td>-5.6</td><td>1.5</td><td>-3.7</td><td>2.1E-04</td><td>3.2E-04</td><td>MB231/MB468</td><td>MP</td></tr>
<tr><td>>hg38_21_+_8433172_8446622_RNA45SN1_45S_50ntFlanks@6645.6670.26|CTCGTACGACTCTTAGCGGTGGATCA</td><td>>hg38_21_+_8433172_8446622_RNA45SN1_45S_50ntFlanks</td><td>6645</td><td>6670</td><td>26</td><td>CTCGTACGACTCTTAGCGGTGGATCA</td><td>549.4</td><td>-3.1</td><td>0.8</td><td>-3.9</td><td>9.7E-05</td><td>3.2E-04</td><td>BT20/MB468</td><td>MP</td></tr>
<tr><td>>MI0005566|hsa-mir-374b&WithFlank&X|-|74218541|74218624@17.34.18&[MIMAT0004955&hsa-miR-374b-5p&offsets|0|-4;m-153&X|-|74218587|74218608&offsets|0|-4]|ATATAATACAACCTGCTA</td><td>>MI0005566|hsa-mir-374b&WithFlank&X|-|74218541|74218624</td><td>17</td><td>34</td><td>18</td><td>ATATAATACAACCTGCTA</td><td>179.8</td><td>5.9</td><td>1.6</td><td>3.7</td><td>2.1E-04</td><td>3.2E-04</td><td>MB231/MB468</td><td>MP</td></tr>
<tr><td>>hg38_21_+_8433172_8446622_RNA45SN1_45S_50ntFlanks@7989.8009.21|AGACGTGGCGACCCGCTGAAT</td><td>>hg38_21_+_8433172_8446622_RNA45SN1_45S_50ntFlanks</td><td>7989</td><td>8009</td><td>21</td><td>AGACGTGGCGACCCGCTGAAT</td><td>2535.9</td><td>3.8</td><td>1.0</td><td>3.7</td><td>2.5E-04</td><td>3.2E-04</td><td>MB231/MB468</td><td>NUC</td></tr>
<tr><td>>hg38_21_+_8433172_8446622_RNA45SN1_45S_50ntFlanks@6778.6803.26|CCCGGGGCTACGCCTGTCTGAGCGTC</td><td>>hg38_21_+_8433172_8446622_RNA45SN1_45S_50ntFlanks</td><td>6778</td><td>6803</td><td>26</td><td>CCCGGGGCTACGCCTGTCTGAGCGTC</td><td>450.0</td><td>-3.2</td><td>0.9</td><td>-3.7</td><td>2.6E-04</td><td>3.2E-04</td><td>BT20/MB468</td><td>NUC</td></tr>
<tr><td>>hg38_MT_+_598_1651_RNR1_12S_50ntFlanks@429.468.40|AAGTGGCTTTAACATATCTGAACACACAATAGCTAAGACC</td><td>>hg38_MT_+_598_1651_RNR1_12S_50ntFlanks</td><td>429</td><td>468</td><td>40</td><td>AAGTGGCTTTAACATATCTGAACACACAATAGCTAAGACC</td><td>151.1</td><td>7.5</td><td>1.9</td><td>3.9</td><td>9.7E-05</td><td>3.2E-04</td><td>BT20/MB468</td><td>MP</td></tr>
<tr><td>>hg38_21_+_8433172_8446622_RNA45SN1_45S_50ntFlanks@485.520.36|GATGTGGTGACGTCGTGCTCTCCCGGGCCGGGTCCG</td><td>>hg38_21_+_8433172_8446622_RNA45SN1_45S_50ntFlanks</td><td>485</td><td>520</td><td>36</td><td>GATGTGGTGACGTCGTGCTCTCCCGGGCCGGGTCCG</td><td>80.0</td><td>9.5</td><td>2.6</td><td>3.7</td><td>2.1E-04</td><td>3.2E-04</td><td>MB231/MB468</td><td>MP</td></tr>
<tr><td>>hg38_21_+_8433172_8446622_RNA45SN1_45S_50ntFlanks@10362.10384.23|AAGGCCGAAGTGGAGAAGGGTTC</td><td>>hg38_21_+_8433172_8446622_RNA45SN1_45S_50ntFlanks</td><td>10362</td><td>10384</td><td>23</td><td>AAGGCCGAAGTGGAGAAGGGTTC</td><td>583.9</td><td>-4.3</td><td>1.2</td><td>-3.7</td><td>2.2E-04</td><td>3.2E-04</td><td>BT20/MB468</td><td>CYTO</td></tr>
<tr><td>>hg38_21_+_8433172_8446622_RNA45SN1_45S_50ntFlanks@5485.5509.25|AGCGCTGAGAAGACGGTCGAACTTG</td><td>>hg38_21_+_8433172_8446622_RNA45SN1_45S_50ntFlanks</td><td>5485</td><td>5509</td><td>25</td><td>AGCGCTGAGAAGACGGTCGAACTTG</td><td>477.3</td><td>6.8</td><td>1.6</td><td>4.1</td><td>3.5E-05</td><td>3.2E-04</td><td>BT20/MB231</td><td>MP</td></tr>
<tr><td>>MI0000071|hsa-mir-17&WithFlank&13|+|91350599|91350694@20.42.23(+1G)&[MIMAT0000070&hsa-miR-17-5p&offsets|0|0(+1G);m-88&13|+|91350618|91350640&offsets|0|0(+1G)]|CAAAGTGCTTACAGTGCAGGTAGG</td><td>>MI0000071|hsa-mir-17&WithFlank&13|+|91350599|91350694</td><td>20</td><td>42</td><td>23(+1G)</td><td>CAAAGTGCTTACAGTGCAGGTAGG</td><td>53.8</td><td>-9.1</td><td>2.2</td><td>-4.1</td><td>3.5E-05</td><td>3.2E-04</td><td>BT20/MB231</td><td>MP</td></tr>
<tr><td>>MI0000683|hsa-mir-181b-2&WithFlank&9|+|124693704|124693804@22.46.25(+1U)&[MIMAT0000257_1&hsa-miR-181b-5p&offsets|0|+2(+1U);m-72&9|+|124693725|124693747&offsets|0|+2(+1U)]|AACATTCATTGCTGTCGGTGGGTTTT</td><td>>MI0000683|hsa-mir-181b-2&WithFlank&9|+|124693704|124693804</td><td>22</td><td>46</td><td>25(+1U)</td><td>AACATTCATTGCTGTCGGTGGGTTTT</td><td>811.9</td><td>-5.7</td><td>1.4</td><td>-4.1</td><td>3.5E-05</td><td>3.2E-04</td><td>BT20/MB231</td><td>MP</td></tr>
<tr><td>>hg38_21_+_8433172_8446622_RNA45SN1_45S_50ntFlanks@11583.11611.29|CTGGTGCGGACCAGGGGAATCCGACTGTT</td><td>>hg38_21_+_8433172_8446622_RNA45SN1_45S_50ntFlanks</td><td>11583</td><td>11611</td><td>29</td><td>CTGGTGCGGACCAGGGGAATCCGACTGTT</td><td>4301.0</td><td>-3.0</td><td>0.8</td><td>-3.7</td><td>2.6E-04</td><td>3.2E-04</td><td>BT20/MB468</td><td>NUC</td></tr>
<tr><td>>hg38_21_+_8433172_8446622_RNA45SN1_45S_50ntFlanks@10351.10381.31|CGAGAACTTTGAAGGCCGAAGTGGAGAAGGG</td><td>>hg38_21_+_8433172_8446622_RNA45SN1_45S_50ntFlanks</td><td>10351</td><td>10381</td><td>31</td><td>CGAGAACTTTGAAGGCCGAAGTGGAGAAGGG</td><td>461.3</td><td>-2.2</td><td>0.6</td><td>-3.7</td><td>1.9E-04</td><td>3.2E-04</td><td>BT20/MB468</td><td>MITO</td></tr>
<tr><td>>hg38_21_+_8433172_8446622_RNA45SN1_45S_50ntFlanks@6653.6676.24|ACTCTTAGCGGTGGATCACTCGGC</td><td>>hg38_21_+_8433172_8446622_RNA45SN1_45S_50ntFlanks</td><td>6653</td><td>6676</td><td>24</td><td>ACTCTTAGCGGTGGATCACTCGGC</td><td>305.1</td><td>3.4</td><td>0.9</td><td>3.7</td><td>2.5E-04</td><td>3.2E-04</td><td>MB231/MB468</td><td>MITO</td></tr>
<tr><td>>MI0000079|hsa-mir-23a&WithFlank&19|-|13836581|13836665@54.72.19(+1U)&[MIMAT0000078&hsa-miR-23a-3p&offsets|+3|+1(+1U);m-31&19|-|13836595|13836615&offsets|+3|+1(+1U)]|ACATTGCCAGGGATTTCCAT</td><td>>MI0000079|hsa-mir-23a&WithFlank&19|-|13836581|13836665</td><td>54</td><td>72</td><td>19(+1U)</td><td>ACATTGCCAGGGATTTCCAT</td><td>669.9</td><td>13.3</td><td>3.6</td><td>3.7</td><td>2.5E-04</td><td>3.2E-04</td><td>MB231/MB468</td><td>NUC</td></tr>
<tr><td>>MI0000446|hsa-mir-125b-1&WithFlank&11|-|122099751|122099850@61.82.22(+1A)&[MIMAT0004592&hsa-miR-125b-1-3p&offsets|0|0(+1A);m-485&11|-|122099769|122099790&offsets|0|0(+1A)]|ACGGGTTAGGCTCTTGGGAGCTA</td><td>>MI0000446|hsa-mir-125b-1&WithFlank&11|-|122099751|122099850</td><td>61</td><td>82</td><td>22(+1A)</td><td>ACGGGTTAGGCTCTTGGGAGCTA</td><td>55.5</td><td>-9.2</td><td>2.2</td><td>-4.1</td><td>3.5E-05</td><td>3.2E-04</td><td>BT20/MB231</td><td>MP</td></tr>
<tr><td>>hg38_21_+_8433172_8446622_RNA45SN1_45S_50ntFlanks@6789.6806.18|GCCTGTCTGAGCGTCGCT</td><td>>hg38_21_+_8433172_8446622_RNA45SN1_45S_50ntFlanks</td><td>6789</td><td>6806</td><td>18</td><td>GCCTGTCTGAGCGTCGCT</td><td>490.1</td><td>3.6</td><td>0.9</td><td>3.9</td><td>9.8E-05</td><td>3.2E-04</td><td>BT20/MB468</td><td>MP</td></tr>
<tr><td>>hg38_21_+_8433172_8446622_RNA45SN1_45S_50ntFlanks@10361.10388.28|GAAGGCCGAAGTGGAGAAGGGTTCCATG</td><td>>hg38_21_+_8433172_8446622_RNA45SN1_45S_50ntFlanks</td><td>10361</td><td>10388</td><td>28</td><td>GAAGGCCGAAGTGGAGAAGGGTTCCATG</td><td>571.4</td><td>-5.4</td><td>1.4</td><td>-3.9</td><td>9.8E-05</td><td>3.2E-04</td><td>BT20/MB468</td><td>MP</td></tr>
<tr><td>>trnaMT_ValTAC_MT_+_1602_1670@30.72.43|CTTACACTTAGGAGATTTCAACTTAACTTGACCGCTCTGACCA</td><td>>trnaMT_ValTAC_MT_+_1602_1670</td><td>30</td><td>72</td><td>43</td><td>CTTACACTTAGGAGATTTCAACTTAACTTGACCGCTCTGACCA</td><td>776.5</td><td>3.2</td><td>0.9</td><td>3.7</td><td>1.9E-04</td><td>3.2E-04</td><td>BT20/MB468</td><td>MITO</td></tr>
<tr><td>>hg38_21_+_8433172_8446622_RNA45SN1_45S_50ntFlanks@7981.8010.30|CTCAGATCAGACGTGGCGACCCGCTGAATT</td><td>>hg38_21_+_8433172_8446622_RNA45SN1_45S_50ntFlanks</td><td>7981</td><td>8010</td><td>30</td><td>CTCAGATCAGACGTGGCGACCCGCTGAATT</td><td>95.2</td><td>-5.4</td><td>1.5</td><td>-3.7</td><td>2.1E-04</td><td>3.2E-04</td><td>MB231/MB468</td><td>MP</td></tr>
<tr><td>>hg38_21_+_8433172_8446622_RNA45SN1_45S_50ntFlanks@9465.9503.39|AGGTGGAGCACGAGCGCACGTGTTAGGACCCGAAAGATG</td><td>>hg38_21_+_8433172_8446622_RNA45SN1_45S_50ntFlanks</td><td>9465</td><td>9503</td><td>39</td><td>AGGTGGAGCACGAGCGCACGTGTTAGGACCCGAAAGATG</td><td>759.3</td><td>-4.5</td><td>1.2</td><td>-3.7</td><td>1.9E-04</td><td>3.2E-04</td><td>BT20/MB468</td><td>MITO</td></tr>
<tr><td>>hg38_21_+_8433172_8446622_RNA45SN1_45S_50ntFlanks@3854.3875.22|ACTGTGGTAATTCTAGAGCTAA</td><td>>hg38_21_+_8433172_8446622_RNA45SN1_45S_50ntFlanks</td><td>3854</td><td>3875</td><td>22</td><td>ACTGTGGTAATTCTAGAGCTAA</td><td>68.1</td><td>-5.5</td><td>1.5</td><td>-3.7</td><td>2.1E-04</td><td>3.2E-04</td><td>MB231/MB468</td><td>MP</td></tr>
<tr><td>>am_trnaMT_ValTAC_MT_+_1602_1670@28.52.25|AACTTACACTTAGGAGATTTCAACT</td><td>>am_trnaMT_ValTAC_MT_+_1602_1670</td><td>28</td><td>52</td><td>25</td><td>AACTTACACTTAGGAGATTTCAACT</td><td>99.4</td><td>10.4</td><td>2.7</td><td>3.9</td><td>9.8E-05</td><td>3.2E-04</td><td>BT20/MB468</td><td>MP</td></tr>
<tr><td>>hg38_21_+_8433172_8446622_RNA45SN1_45S_50ntFlanks@10349.10372.24|AACGAGAACTTTGAAGGCCGAAGT</td><td>>hg38_21_+_8433172_8446622_RNA45SN1_45S_50ntFlanks</td><td>10349</td><td>10372</td><td>24</td><td>AACGAGAACTTTGAAGGCCGAAGT</td><td>878.1</td><td>2.6</td><td>0.7</td><td>3.7</td><td>2.4E-04</td><td>3.3E-04</td><td>MB231/MB468</td><td>CYTO</td></tr>
<tr><td>>hg38_21_+_8433172_8446622_RNA45SN1_45S_50ntFlanks@10819.10837.19|AGGGAAGTCGGCAAGCCGG</td><td>>hg38_21_+_8433172_8446622_RNA45SN1_45S_50ntFlanks</td><td>10819</td><td>10837</td><td>19</td><td>AGGGAAGTCGGCAAGCCGG</td><td>568.4</td><td>-3.9</td><td>1.1</td><td>-3.7</td><td>2.4E-04</td><td>3.3E-04</td><td>MB231/MB468</td><td>CYTO</td></tr>
<tr><td>>hg38_21_+_8433172_8446622_RNA45SN1_45S_50ntFlanks@6734.6773.40|ATTGATCATCGACACTTCGAACGCACTTGCGGCCCCGGGT</td><td>>hg38_21_+_8433172_8446622_RNA45SN1_45S_50ntFlanks</td><td>6734</td><td>6773</td><td>40</td><td>ATTGATCATCGACACTTCGAACGCACTTGCGGCCCCGGGT</td><td>303.6</td><td>2.5</td><td>0.7</td><td>3.7</td><td>2.6E-04</td><td>3.3E-04</td><td>BT20/MB468</td><td>NUC</td></tr>
<tr><td>>hg38_21_+_8433172_8446622_RNA45SN1_45S_50ntFlanks@4325.4342.18|CCGCGGTAATTCCAGCTC</td><td>>hg38_21_+_8433172_8446622_RNA45SN1_45S_50ntFlanks</td><td>4325</td><td>4342</td><td>18</td><td>CCGCGGTAATTCCAGCTC</td><td>90.9</td><td>-5.7</td><td>1.5</td><td>-3.9</td><td>9.8E-05</td><td>3.3E-04</td><td>BT20/MB468</td><td>MP</td></tr>
<tr><td>>hg38_21_+_8433172_8446622_RNA45SN1_45S_50ntFlanks@6722.6760.39|ATTGCAGGACACATTGATCATCGACACTTCGAACGCACT</td><td>>hg38_21_+_8433172_8446622_RNA45SN1_45S_50ntFlanks</td><td>6722</td><td>6760</td><td>39</td><td>ATTGCAGGACACATTGATCATCGACACTTCGAACGCACT</td><td>182.1</td><td>2.5</td><td>0.7</td><td>3.7</td><td>2.4E-04</td><td>3.3E-04</td><td>MB231/MB468</td><td>CYTO</td></tr>
<tr><td>>hg38_21_+_8433172_8446622_RNA45SN1_45S_50ntFlanks@10391.10409.19|AACAGCAGTTGAACATGGG</td><td>>hg38_21_+_8433172_8446622_RNA45SN1_45S_50ntFlanks</td><td>10391</td><td>10409</td><td>19</td><td>AACAGCAGTTGAACATGGG</td><td>2907.2</td><td>-6.3</td><td>1.7</td><td>-3.7</td><td>2.3E-04</td><td>3.3E-04</td><td>BT20/MB468</td><td>CYTO</td></tr>
<tr><td>>hg38_21_+_8433172_8446622_RNA45SN1_45S_50ntFlanks@6723.6744.22|TTGCAGGACACATTGATCATCG</td><td>>hg38_21_+_8433172_8446622_RNA45SN1_45S_50ntFlanks</td><td>6723</td><td>6744</td><td>22</td><td>TTGCAGGACACATTGATCATCG</td><td>199.6</td><td>-5.9</td><td>1.5</td><td>-3.9</td><td>9.9E-05</td><td>3.3E-04</td><td>BT20/MB468</td><td>MP</td></tr>
<tr><td>>hg38_21_+_8433172_8446622_RNA45SN1_45S_50ntFlanks@10677.10709.33|GCGGTTCCGGCGGCGTCCGGTGAGCTCTCGCTG</td><td>>hg38_21_+_8433172_8446622_RNA45SN1_45S_50ntFlanks</td><td>10677</td><td>10709</td><td>33</td><td>GCGGTTCCGGCGGCGTCCGGTGAGCTCTCGCTG</td><td>778.3</td><td>-2.7</td><td>0.7</td><td>-3.7</td><td>2.5E-04</td><td>3.3E-04</td><td>MB231/MB468</td><td>MITO</td></tr>
<tr><td>>hg38_21_+_8433172_8446622_RNA45SN1_45S_50ntFlanks@10471.10488.18|CCCTCGGCCGATCGAAAG</td><td>>hg38_21_+_8433172_8446622_RNA45SN1_45S_50ntFlanks</td><td>10471</td><td>10488</td><td>18</td><td>CCCTCGGCCGATCGAAAG</td><td>718.8</td><td>-2.6</td><td>0.7</td><td>-3.7</td><td>2.4E-04</td><td>3.3E-04</td><td>MB231/MB468</td><td>CYTO</td></tr>
<tr><td>>hg38_21_+_8433172_8446622_RNA45SN1_45S_50ntFlanks@11570.11592.23|AGCCGACTTAGAACTGGTGCGGA</td><td>>hg38_21_+_8433172_8446622_RNA45SN1_45S_50ntFlanks</td><td>11570</td><td>11592</td><td>23</td><td>AGCCGACTTAGAACTGGTGCGGA</td><td>1041.2</td><td>-3.8</td><td>1.0</td><td>-3.9</td><td>9.9E-05</td><td>3.3E-04</td><td>BT20/MB468</td><td>MP</td></tr>
<tr><td>>hg38_MT_+_1621_3279_RNR2_16S_50ntFlanks@714.749.36|CATAAGCCTGCGTCAGATTAAAACACTGAACTGACA</td><td>>hg38_MT_+_1621_3279_RNR2_16S_50ntFlanks</td><td>714</td><td>749</td><td>36</td><td>CATAAGCCTGCGTCAGATTAAAACACTGAACTGACA</td><td>21.6</td><td>7.7</td><td>2.1</td><td>3.7</td><td>2.1E-04</td><td>3.3E-04</td><td>MB231/MB468</td><td>MP</td></tr>
<tr><td>>MI0016887|hsa-mir-4521&WithFlank&17|+|8186939|8187010@10.35.26&[MIMAT0019058&hsa-miR-4521&offsets|0|+4;m-765&17|+|8186948|8186969&offsets|0|+4]|GCTAAGGAAGTCCTGTGCTCAGTTTT</td><td>>MI0016887|hsa-mir-4521&WithFlank&17|+|8186939|8187010</td><td>10</td><td>35</td><td>26</td><td>GCTAAGGAAGTCCTGTGCTCAGTTTT</td><td>140.5</td><td>3.1</td><td>0.9</td><td>3.7</td><td>2.6E-04</td><td>3.3E-04</td><td>BT20/MB468</td><td>NUC</td></tr>
<tr><td>>hg38_21_+_8433172_8446622_RNA45SN1_45S_50ntFlanks@8079.8105.27|AAGAGCCCAGCGCCGAATCCCCGCCCC</td><td>>hg38_21_+_8433172_8446622_RNA45SN1_45S_50ntFlanks</td><td>8079</td><td>8105</td><td>27</td><td>AAGAGCCCAGCGCCGAATCCCCGCCCC</td><td>243.0</td><td>-5.2</td><td>1.4</td><td>-3.7</td><td>1.9E-04</td><td>3.3E-04</td><td>BT20/MB468</td><td>MITO</td></tr>
<tr><td>>hg38_21_+_8433172_8446622_RNA45SN1_45S_50ntFlanks@4970.4999.30|CCCGGCCCGGACACGGACAGGATTGACAGA</td><td>>hg38_21_+_8433172_8446622_RNA45SN1_45S_50ntFlanks</td><td>4970</td><td>4999</td><td>30</td><td>CCCGGCCCGGACACGGACAGGATTGACAGA</td><td>455.1</td><td>-4.1</td><td>1.1</td><td>-3.7</td><td>2.5E-04</td><td>3.3E-04</td><td>MB231/MB468</td><td>MITO</td></tr>
<tr><td>>hg38_21_+_8433172_8446622_RNA45SN1_45S_50ntFlanks@10793.10812.20|TCTGGCATGTTGGAACAATG</td><td>>hg38_21_+_8433172_8446622_RNA45SN1_45S_50ntFlanks</td><td>10793</td><td>10812</td><td>20</td><td>TCTGGCATGTTGGAACAATG</td><td>68.1</td><td>9.4</td><td>2.3</td><td>4.1</td><td>3.5E-05</td><td>3.3E-04</td><td>BT20/MB231</td><td>MP</td></tr>
<tr><td>>MI0000294|hsa-mir-218-1&WithFlank&4|+|20528269|20528390@31.52.22(+1C);MI0000295|hsa-mir-218-2&WithFlank&5|-|168768140|168768261@31.52.22(+1C)&[MIMAT0000275&hsa-miR-218-5p&offsets|0|+1(+1C);m-185&4|+|20528299|20528319&offsets|0|+1(+1C)];[MIMAT0000275_1&hsa-miR-218-5p&offsets|0|+1(+1C);m-187&5|-|168768211|168768231&offsets|0|+1(+1C)]|TTGTGCTTGATCTAACCATGTGC</td><td>>MI0000294|hsa-mir-218-1&WithFlank&4|+|20528269|20528390</td><td>31</td><td>52</td><td>22(+1C)</td><td>TTGTGCTTGATCTAACCATGTGC</td><td>53.6</td><td>-9.1</td><td>2.2</td><td>-4.1</td><td>3.5E-05</td><td>3.3E-04</td><td>BT20/MB231</td><td>MP</td></tr>
<tr><td>>hg38_21_+_8433172_8446622_RNA45SN1_45S_50ntFlanks@3985.4004.20|CGGCTTTGGTGACTCTAGAT</td><td>>hg38_21_+_8433172_8446622_RNA45SN1_45S_50ntFlanks</td><td>3985</td><td>4004</td><td>20</td><td>CGGCTTTGGTGACTCTAGAT</td><td>266.0</td><td>-6.1</td><td>1.6</td><td>-3.9</td><td>9.9E-05</td><td>3.3E-04</td><td>BT20/MB468</td><td>MP</td></tr>
<tr><td>>hg38_21_+_8433172_8446622_RNA45SN1_45S_50ntFlanks@7989.8015.27|AGACGTGGCGACCCGCTGAATTTAAGC</td><td>>hg38_21_+_8433172_8446622_RNA45SN1_45S_50ntFlanks</td><td>7989</td><td>8015</td><td>27</td><td>AGACGTGGCGACCCGCTGAATTTAAGC</td><td>298.8</td><td>4.7</td><td>1.3</td><td>3.7</td><td>2.6E-04</td><td>3.3E-04</td><td>MB231/MB468</td><td>NUC</td></tr>
</table>

<table>
<tr><td></td><td></td><td></td><td></td><td></td><td></td><td></td><td></td><td></td><td></td><td></td><td></td><td></td><td></td></tr>
<tr><td>>am_trna119_LysCTT_1_-_145395522_145395594@15.33.19,trna11_LysCTT_5_-_180648979_180649051@15.33.19,trna128_LysCTT_1_-_55423542_55423614@15.33.19,trna13_LysCTT_6_+_26556774_26556846@15.33.19,trna6_LysCTT_19_-_52425396_52425468@15.33.19,trna7_LysCTT_16_+_3225692_3225764@15.33.19,trna9_LysCTT_5_+_180634755_180634827@15.33.19|GTCGGTAGAGCATGAGACT</td><td>>am_trna119_LysCTT_1_-_145395522_145395594</td><td>15</td><td>33</td><td>19</td><td>GTCGGTAGAGCATGAGACT</td><td>50.9</td><td>-9.1</td><td>2.2</td><td>-4.1</td><td>3.5E-05</td><td>3.3E-04</td><td>BT20/MB231</td><td>MP</td></tr>
<tr><td>>am_MI0000786|hsa-mir-378a&WithFlank&5|+|149732819|149732896@49.69.21(+1A)&[MIMAT0000732&hsa-miR-378a-3p&offsets|0|-1(+1A);m-44&5|+|149732867|149732888&offsets|0|-1(+1A)]|ACTGGACTTGGAGTCAGAAGGA</td><td>>am_MI0000786|hsa-mir-378a&WithFlank&5|+|149732819|149732896</td><td>49</td><td>69</td><td>21(+1A)</td><td>ACTGGACTTGGAGTCAGAAGGA</td><td>586.5</td><td>3.3</td><td>0.9</td><td>3.7</td><td>2.6E-04</td><td>3.3E-04</td><td>MB231/MB468</td><td>MITO</td></tr>
<tr><td>>hg38_21_+_8433172_8446622_RNA45SN1_45S_50ntFlanks@10835.10858.24|CGGATCCGTAACTTCGGGATAAGG</td><td>>hg38_21_+_8433172_8446622_RNA45SN1_45S_50ntFlanks</td><td>10835</td><td>10858</td><td>24</td><td>CGGATCCGTAACTTCGGGATAAGG</td><td>498.4</td><td>-3.5</td><td>0.9</td><td>-3.7</td><td>2.2E-04</td><td>3.3E-04</td><td>MB231/MB468</td><td>MP</td></tr>
<tr><td>>hg38_21_+_8433172_8446622_RNA45SN1_45S_50ntFlanks@8178.8208.31|TCTGATCGAGGCCCAGCCCGTGGACGGTGTG</td><td>>hg38_21_+_8433172_8446622_RNA45SN1_45S_50ntFlanks</td><td>8178</td><td>8208</td><td>31</td><td>TCTGATCGAGGCCCAGCCCGTGGACGGTGTG</td><td>243.6</td><td>-2.9</td><td>0.8</td><td>-3.7</td><td>2.6E-04</td><td>3.3E-04</td><td>MB231/MB468</td><td>NUC</td></tr>
<tr><td>>hg38_21_+_8433172_8446622_RNA45SN1_45S_50ntFlanks@4003.4020.18|ATAACCTCGGGCCGATCG</td><td>>hg38_21_+_8433172_8446622_RNA45SN1_45S_50ntFlanks</td><td>4003</td><td>4020</td><td>18</td><td>ATAACCTCGGGCCGATCG</td><td>692.9</td><td>-4.2</td><td>1.1</td><td>-3.7</td><td>2.4E-04</td><td>3.3E-04</td><td>MB231/MB468</td><td>CYTO</td></tr>
<tr><td>>hg38_21_+_8433172_8446622_RNA45SN1_45S_50ntFlanks@7967.7988.22|CTCCGAGACGCGACCTCAGATC</td><td>>hg38_21_+_8433172_8446622_RNA45SN1_45S_50ntFlanks</td><td>7967</td><td>7988</td><td>22</td><td>CTCCGAGACGCGACCTCAGATC</td><td>73.9</td><td>2.4</td><td>0.7</td><td>3.7</td><td>2.6E-04</td><td>3.3E-04</td><td>BT20/MB468</td><td>NUC</td></tr>
<tr><td>>MI0000782|hsa-mir-374a&WithFlank&X|-|74287280|74287363@18.39.22&[MIMAT0000727&hsa-miR-374a-5p&offsets|0|0;m-184&X|-|74287325|74287346&offsets|0|0]|TTATAATACAACCTGATAAGTG</td><td>>MI0000782|hsa-mir-374a&WithFlank&X|-|74287280|74287363</td><td>18</td><td>39</td><td>22</td><td>TTATAATACAACCTGATAAGTG</td><td>694.1</td><td>-4.3</td><td>1.0</td><td>-4.1</td><td>3.6E-05</td><td>3.3E-04</td><td>BT20/MB231</td><td>MP</td></tr>
<tr><td>>hg38_21_+_8433172_8446622_RNA45SN1_45S_50ntFlanks@7975.8015.41|CGCGACCTCAGATCAGACGTGGCGACCCGCTGAATTTAAGC</td><td>>hg38_21_+_8433172_8446622_RNA45SN1_45S_50ntFlanks</td><td>7975</td><td>8015</td><td>41</td><td>CGCGACCTCAGATCAGACGTGGCGACCCGCTGAATTTAAGC</td><td>18819.8</td><td>3.5</td><td>0.9</td><td>3.7</td><td>2.6E-04</td><td>3.3E-04</td><td>MB231/MB468</td><td>MITO</td></tr>
<tr><td>>hg38_21_+_8433172_8446622_RNA45SN1_45S_50ntFlanks@11919.11948.30|CATGAGAGGTGTAGAATAAGTGGGAGGCCC</td><td>>hg38_21_+_8433172_8446622_RNA45SN1_45S_50ntFlanks</td><td>11919</td><td>11948</td><td>30</td><td>CATGAGAGGTGTAGAATAAGTGGGAGGCCC</td><td>214.2</td><td>-4.2</td><td>1.1</td><td>-3.7</td><td>2.3E-04</td><td>3.3E-04</td><td>BT20/MB468</td><td>CYTO</td></tr>
<tr><td>>hg38_21_+_8433172_8446622_RNA45SN1_45S_50ntFlanks@9570.9601.32|TGCAAATCGGTCGTCCGACCTGGGTATAGGGG</td><td>>hg38_21_+_8433172_8446622_RNA45SN1_45S_50ntFlanks</td><td>9570</td><td>9601</td><td>32</td><td>TGCAAATCGGTCGTCCGACCTGGGTATAGGGG</td><td>324.5</td><td>-4.4</td><td>1.2</td><td>-3.7</td><td>2.6E-04</td><td>3.3E-04</td><td>MB231/MB468</td><td>MITO</td></tr>
<tr><td>>hg38_21_+_8433172_8446622_RNA45SN1_45S_50ntFlanks@11561.11582.22|GCGCCTAGCAGCCGACTTAGAA</td><td>>hg38_21_+_8433172_8446622_RNA45SN1_45S_50ntFlanks</td><td>11561</td><td>11582</td><td>22</td><td>GCGCCTAGCAGCCGACTTAGAA</td><td>105.4</td><td>-5.8</td><td>1.5</td><td>-3.9</td><td>9.9E-05</td><td>3.3E-04</td><td>BT20/MB468</td><td>MP</td></tr>
<tr><td>>hg38_MT_+_1621_3279_RNR2_16S_50ntFlanks@1389.1414.26|CGATGGTGCAGCCGCTATTAAAGGTT</td><td>>hg38_MT_+_1621_3279_RNR2_16S_50ntFlanks</td><td>1389</td><td>1414</td><td>26</td><td>CGATGGTGCAGCCGCTATTAAAGGTT</td><td>423.6</td><td>4.0</td><td>1.1</td><td>3.7</td><td>2.2E-04</td><td>3.3E-04</td><td>MB231/MB468</td><td>MP</td></tr>
<tr><td>>am_trnaMT_HisGTG_MT_+_12138_12206@26.51.26|AGATTGTGAATCTGACAACAGAGGCT</td><td>>am_trnaMT_HisGTG_MT_+_12138_12206</td><td>26</td><td>51</td><td>26</td><td>AGATTGTGAATCTGACAACAGAGGCT</td><td>221.6</td><td>6.7</td><td>1.7</td><td>3.9</td><td>1.0E-04</td><td>3.3E-04</td><td>BT20/MB468</td><td>MP</td></tr>
<tr><td>>hg38_21_+_8433172_8446622_RNA45SN1_45S_50ntFlanks@12415.12441.27|CTTCGATGTCGGCTCTTCCTATCATTG</td><td>>hg38_21_+_8433172_8446622_RNA45SN1_45S_50ntFlanks</td><td>12415</td><td>12441</td><td>27</td><td>CTTCGATGTCGGCTCTTCCTATCATTG</td><td>3044.4</td><td>-2.9</td><td>0.8</td><td>-3.7</td><td>2.6E-04</td><td>3.3E-04</td><td>MB231/MB468</td><td>NUC</td></tr>
<tr><td>>hg38_21_+_8433172_8446622_RNA45SN1_45S_50ntFlanks@6701.6718.18|TAGCTGCGAGAATTAATG</td><td>>hg38_21_+_8433172_8446622_RNA45SN1_45S_50ntFlanks</td><td>6701</td><td>6718</td><td>18</td><td>TAGCTGCGAGAATTAATG</td><td>134.7</td><td>-2.3</td><td>0.6</td><td>-3.7</td><td>2.3E-04</td><td>3.3E-04</td><td>BT20/MB468</td><td>CYTO</td></tr>
<tr><td>>hg38_21_+_8433172_8446622_RNA45SN1_45S_50ntFlanks@11570.11592.23|AGCCGACTTAGAACTGGTGCGGA</td><td>>hg38_21_+_8433172_8446622_RNA45SN1_45S_50ntFlanks</td><td>11570</td><td>11592</td><td>23</td><td>AGCCGACTTAGAACTGGTGCGGA</td><td>221.2</td><td>-2.8</td><td>0.8</td><td>-3.7</td><td>2.3E-04</td><td>3.3E-04</td><td>BT20/MB468</td><td>CYTO</td></tr>
<tr><td>>trna10_AlaCGC_6_+_26553731_26553802@1.18.18,trna119_AlaCGC_6_-_28641613_28641684@1.18.18,trna120_AlaAGC_6_-_28626014_28626085@1.18.18,trna13_AlaCGC_2_+_157257281_157257352@1.18.18,trna13_AlaTGC_12_-_125406301_125406372@1.18.18,trna66_AlaTGC_6_+_28611222_28611293@1.18.18,trna8_AlaTGC_12_+_125424512_125424583@1.18.18,trna8_AlaTGC_5_+_180633868_180633939@1.18.18|GGGGATGTAGCTCAGTGG</td><td>>trna10_AlaCGC_6_+_26553731_26553802</td><td>1</td><td>18</td><td>18</td><td>GGGGATGTAGCTCAGTGG</td><td>320.7</td><td>3.2</td><td>0.9</td><td>3.7</td><td>2.2E-04</td><td>3.3E-04</td><td>MB231/MB468</td><td>MP</td></tr>
<tr><td>>trnaMT_LysTTT_MT_+_8295_8364@44.72.29,trnalookalike7_LysTTT_1_+_568844_568913@44.72.29|AAGAGAACCAACACCTCTTTACAGTGACC</td><td>>trnaMT_LysTTT_MT_+_8295_8364</td><td>44</td><td>72</td><td>29</td><td>AAGAGAACCAACACCTCTTTACAGTGACC</td><td>375.7</td><td>3.0</td><td>0.8</td><td>3.7</td><td>2.2E-04</td><td>3.3E-04</td><td>MB231/MB468</td><td>MP</td></tr>
<tr><td>>hg38_21_+_8433172_8446622_RNA45SN1_45S_50ntFlanks@6733.6753.21|CATTGATCATCGACACTTCGA</td><td>>hg38_21_+_8433172_8446622_RNA45SN1_45S_50ntFlanks</td><td>6733</td><td>6753</td><td>21</td><td>CATTGATCATCGACACTTCGA</td><td>361.9</td><td>-2.4</td><td>0.7</td><td>-3.7</td><td>2.4E-04</td><td>3.3E-04</td><td>MB231/MB468</td><td>CYTO</td></tr>
<tr><td>>hg38_21_+_8433172_8446622_RNA45SN1_45S_50ntFlanks@10571.10608.38|GAGAAGCCGGCGGGAGCCCCGGGGAGAGTTCTCTTTTC</td><td>>hg38_21_+_8433172_8446622_RNA45SN1_45S_50ntFlanks</td><td>10571</td><td>10608</td><td>38</td><td>GAGAAGCCGGCGGGAGCCCCGGGGAGAGTTCTCTTTTC</td><td>332.9</td><td>-5.3</td><td>1.4</td><td>-3.7</td><td>2.3E-04</td><td>3.3E-04</td><td>BT20/MB468</td><td>CYTO</td></tr>
<tr><td>>hg38_21_+_8433172_8446622_RNA45SN1_45S_50ntFlanks@3853.3898.46|AACTGTGGTAATTCTAGAGCTAATACATGCCGACGGGCGCTGACCC</td><td>>hg38_21_+_8433172_8446622_RNA45SN1_45S_50ntFlanks</td><td>3853</td><td>3898</td><td>46</td><td>AACTGTGGTAATTCTAGAGCTAATACATGCCGACGGGCGCTGACCC</td><td>104.6</td><td>-9.8</td><td>2.5</td><td>-3.9</td><td>1.0E-04</td><td>3.3E-04</td><td>BT20/MB468</td><td>MP</td></tr>
<tr><td>>hg38_21_+_8433172_8446622_RNA45SN1_45S_50ntFlanks@6733.6752.20|CATTGATCATCGACACTTCG</td><td>>hg38_21_+_8433172_8446622_RNA45SN1_45S_50ntFlanks</td><td>6733</td><td>6752</td><td>20</td><td>CATTGATCATCGACACTTCG</td><td>978.1</td><td>-4.0</td><td>1.0</td><td>-3.9</td><td>1.0E-04</td><td>3.3E-04</td><td>BT20/MB468</td><td>MP</td></tr>
<tr><td>>hg38_21_+_8433172_8446622_RNA45SN1_45S_50ntFlanks@10907.10930.24|GGGCGCGCGCCGCGGCTGGACGAG</td><td>>hg38_21_+_8433172_8446622_RNA45SN1_45S_50ntFlanks</td><td>10907</td><td>10930</td><td>24</td><td>GGGCGCGCGCCGCGGCTGGACGAG</td><td>210.8</td><td>6.6</td><td>1.7</td><td>3.9</td><td>1.0E-04</td><td>3.3E-04</td><td>BT20/MB468</td><td>MP</td></tr>
<tr><td>>hg38_21_+_8433172_8446622_RNA45SN1_45S_50ntFlanks@6652.6677.26|GACTCTTAGCGGTGGATCACTCGGCT</td><td>>hg38_21_+_8433172_8446622_RNA45SN1_45S_50ntFlanks</td><td>6652</td><td>6677</td><td>26</td><td>GACTCTTAGCGGTGGATCACTCGGCT</td><td>1012.8</td><td>3.1</td><td>0.8</td><td>3.7</td><td>2.4E-04</td><td>3.3E-04</td><td>MB231/MB468</td><td>CYTO</td></tr>
<tr><td>>hg38_21_+_8433172_8446622_RNA45SN1_45S_50ntFlanks@6651.6688.38|CGACTCTTAGCGGTGGATCACTCGGCTCGTGCGTCGAT</td><td>>hg38_21_+_8433172_8446622_RNA45SN1_45S_50ntFlanks</td><td>6651</td><td>6688</td><td>38</td><td>CGACTCTTAGCGGTGGATCACTCGGCTCGTGCGTCGAT</td><td>2721.8</td><td>3.0</td><td>0.8</td><td>3.7</td><td>2.6E-04</td><td>3.3E-04</td><td>MB231/MB468</td><td>NUC</td></tr>
<tr><td>>hg38_21_+_8433172_8446622_RNA45SN1_45S_50ntFlanks@12405.12430.26|CTTTTTGATCCTTCGATGTCGGCTCT</td><td>>hg38_21_+_8433172_8446622_RNA45SN1_45S_50ntFlanks</td><td>12405</td><td>12430</td><td>26</td><td>CTTTTTGATCCTTCGATGTCGGCTCT</td><td>239.7</td><td>-2.3</td><td>0.6</td><td>-3.7</td><td>1.9E-04</td><td>3.3E-04</td><td>BT20/MB468</td><td>MITO</td></tr>
<tr><td>>hg38_21_+_8433172_8446622_RNA45SN1_45S_50ntFlanks@9300.9338.39|ACGGACCAAGGAGTCTAACACGTGCGCGAGTCGGGGGCT</td><td>>hg38_21_+_8433172_8446622_RNA45SN1_45S_50ntFlanks</td><td>9300</td><td>9338</td><td>39</td><td>ACGGACCAAGGAGTCTAACACGTGCGCGAGTCGGGGGCT</td><td>142.4</td><td>-3.4</td><td>0.9</td><td>-3.7</td><td>2.2E-04</td><td>3.3E-04</td><td>MB231/MB468</td><td>MP</td></tr>
<tr><td>>MI0000253|hsa-mir-148a&WithFlank&7|-|25949913|25949992@50.73.24&[MIMAT0000243&hsa-miR-148a-3p&offsets|0|+2;m-7&7|-|25949922|25949943&offsets|0|+2]|TCAGTGCACTACAGAACTTTGTCT</td><td>>MI0000253|hsa-mir-148a&WithFlank&7|-|25949913|25949992</td><td>50</td><td>73</td><td>24</td><td>TCAGTGCACTACAGAACTTTGTCT</td><td>366.8</td><td>5.1</td><td>1.2</td><td>4.1</td><td>3.7E-05</td><td>3.3E-04</td><td>BT20/MB231</td><td>NUC</td></tr>
<tr><td>>hg38_21_+_8433172_8446622_RNA45SN1_45S_50ntFlanks@11581.11604.24|AACTGGTGCGGACCAGGGGAATCC</td><td>>hg38_21_+_8433172_8446622_RNA45SN1_45S_50ntFlanks</td><td>11581</td><td>11604</td><td>24</td><td>AACTGGTGCGGACCAGGGGAATCC</td><td>241.6</td><td>-2.6</td><td>0.7</td><td>-3.7</td><td>2.6E-04</td><td>3.3E-04</td><td>MB231/MB468</td><td>NUC</td></tr>
<tr><td>>hg38_MT_+_1621_3279_RNR2_16S_50ntFlanks@1092.1117.26|GCAAGACGAGAAGACCCTATGGAGCT</td><td>>hg38_MT_+_1621_3279_RNR2_16S_50ntFlanks</td><td>1092</td><td>1117</td><td>26</td><td>GCAAGACGAGAAGACCCTATGGAGCT</td><td>194.7</td><td>3.9</td><td>1.1</td><td>3.7</td><td>2.2E-04</td><td>3.3E-04</td><td>MB231/MB468</td><td>MP</td></tr>
<tr><td>>hg38_21_+_8433172_8446622_RNA45SN1_45S_50ntFlanks@6648.6693.46|GTACGACTCTTAGCGGTGGATCACTCGGCTCGTGCGTCGATGAAGA</td><td>>hg38_21_+_8433172_8446622_RNA45SN1_45S_50ntFlanks</td><td>6648</td><td>6693</td><td>46</td><td>GTACGACTCTTAGCGGTGGATCACTCGGCTCGTGCGTCGATGAAGA</td><td>411.0</td><td>4.1</td><td>1.1</td><td>3.7</td><td>2.6E-04</td><td>3.3E-04</td><td>MB231/MB468</td><td>NUC</td></tr>
<tr><td>>hg38_21_+_8433172_8446622_RNA45SN1_45S_50ntFlanks@3986.4010.25|GGCTTTGGTGACTCTAGATAACCTC</td><td>>hg38_21_+_8433172_8446622_RNA45SN1_45S_50ntFlanks</td><td>3986</td><td>4010</td><td>25</td><td>GGCTTTGGTGACTCTAGATAACCTC</td><td>448.4</td><td>3.1</td><td>0.8</td><td>4.1</td><td>4.1E-05</td><td>3.3E-04</td><td>BT20/MB231</td><td>MITO</td></tr>
<tr><td>>MI0000440|hsa-mir-27b&WithFlank&9|+|95085439|95085547@66.86.21(+1U)&[MIMAT0000419&hsa-miR-27b-3p&offsets|-1|-1(+1U);m-39&9|+|95085505|95085525&offsets|-1|-1(+1U)]|GTTCACAGTGGCTAAGTTCTGT</td><td>>MI0000440|hsa-mir-27b&WithFlank&9|+|95085439|95085547</td><td>66</td><td>86</td><td>21(+1U)</td><td>GTTCACAGTGGCTAAGTTCTGT</td><td>25.0</td><td>7.9</td><td>2.1</td><td>3.7</td><td>2.2E-04</td><td>3.3E-04</td><td>MB231/MB468</td><td>MP</td></tr>
<tr><td>>hg38_21_+_8433172_8446622_RNA45SN1_45S_50ntFlanks@485.531.47|GATGTGGTGACGTCGTGCTCTCCCGGGCCGGGTCCGAGCCGCGACGG</td><td>>hg38_21_+_8433172_8446622_RNA45SN1_45S_50ntFlanks</td><td>485</td><td>531</td><td>47</td><td>GATGTGGTGACGTCGTGCTCTCCCGGGCCGGGTCCGAGCCGCGACGG</td><td>92.2</td><td>10.3</td><td>2.6</td><td>3.9</td><td>1.0E-04</td><td>3.3E-04</td><td>BT20/MB468</td><td>MP</td></tr>
<tr><td>>hg38_21_+_8433172_8446622_RNA45SN1_45S_50ntFlanks@6650.6682.33|ACGACTCTTAGCGGTGGATCACTCGGCTCGTGC</td><td>>hg38_21_+_8433172_8446622_RNA45SN1_45S_50ntFlanks</td><td>6650</td><td>6682</td><td>33</td><td>ACGACTCTTAGCGGTGGATCACTCGGCTCGTGC</td><td>146.1</td><td>3.7</td><td>0.9</td><td>3.9</td><td>1.0E-04</td><td>3.3E-04</td><td>BT20/MB468</td><td>MP</td></tr>
<tr><td>>hg38_MT_+_1621_3279_RNR2_16S_50ntFlanks@489.518.30|AACAGCTCTTTGGACACTAGGAAAAAACCT</td><td>>hg38_MT_+_1621_3279_RNR2_16S_50ntFlanks</td><td>489</td><td>518</td><td>30</td><td>AACAGCTCTTTGGACACTAGGAAAAAACCT</td><td>85.0</td><td>7.0</td><td>1.8</td><td>3.9</td><td>1.0E-04</td><td>3.3E-04</td><td>BT20/MB468</td><td>MP</td></tr>
<tr><td>>hg38_21_+_8433172_8446622_RNA45SN1_45S_50ntFlanks@10805.10822.18|GAACAATGTAGGTAAGGG</td><td>>hg38_21_+_8433172_8446622_RNA45SN1_45S_50ntFlanks</td><td>10805</td><td>10822</td><td>18</td><td>GAACAATGTAGGTAAGGG</td><td>169.2</td><td>-5.8</td><td>1.5</td><td>-3.9</td><td>1.0E-04</td><td>3.3E-04</td><td>BT20/MB468</td><td>MP</td></tr>
<tr><td>>hg38_21_+_8433172_8446622_RNA45SN1_45S_50ntFlanks@11576.11608.33|CTTAGAACTGGTGCGGACCAGGGGAATCCGACT</td><td>>hg38_21_+_8433172_8446622_RNA45SN1_45S_50ntFlanks</td><td>11576</td><td>11608</td><td>33</td><td>CTTAGAACTGGTGCGGACCAGGGGAATCCGACT</td><td>80.8</td><td>-5.8</td><td>1.6</td><td>-3.7</td><td>2.2E-04</td><td>3.3E-04</td><td>MB231/MB468</td><td>MP</td></tr>
<tr><td>>hg38_21_+_8433172_8446622_RNA45SN1_45S_50ntFlanks@10796.10823.28|GGCATGTTGGAACAATGTAGGTAAGGGA</td><td>>hg38_21_+_8433172_8446622_RNA45SN1_45S_50ntFlanks</td><td>10796</td><td>10823</td><td>28</td><td>GGCATGTTGGAACAATGTAGGTAAGGGA</td><td>63.1</td><td>9.3</td><td>2.3</td><td>4.1</td><td>3.6E-05</td><td>3.3E-04</td><td>BT20/MB231</td><td>MP</td></tr>
<tr><td>>hg38_21_+_8433172_8446622_RNA45SN1_45S_50ntFlanks@6649.6681.33|TACGACTCTTAGCGGTGGATCACTCGGCTCGTG</td><td>>hg38_21_+_8433172_8446622_RNA45SN1_45S_50ntFlanks</td><td>6649</td><td>6681</td><td>33</td><td>TACGACTCTTAGCGGTGGATCACTCGGCTCGTG</td><td>312.2</td><td>-2.2</td><td>0.6</td><td>-3.7</td><td>2.4E-04</td><td>3.3E-04</td><td>MB231/MB468</td><td>CYTO</td></tr>
<tr><td>>trnaMT_ThrTGT_MT_+_15888_15953@31.69.39|TTGTAAACCGGAGATGAAAACCTTTTTCCAAGGACACCA</td><td>>trnaMT_ThrTGT_MT_+_15888_15953</td><td>31</td><td>69</td><td>39</td><td>TTGTAAACCGGAGATGAAAACCTTTTTCCAAGGACACCA</td><td>1101.8</td><td>6.5</td><td>1.7</td><td>3.9</td><td>1.0E-04</td><td>3.3E-04</td><td>BT20/MB468</td><td>MP</td></tr>
<tr><td>>MI0000746|hsa-mir-99b&WithFlank&19|+|51692606|51692687@51.70.20(+1A)&[MIMAT0004678&hsa-miR-99b-3p&offsets|0|-2(+1A);m-151&19|+|51692656|51692678&offsets|0|-3(+1A)]|CAAGCTCGTGTCTGTGGGTCA</td><td>>MI0000746|hsa-mir-99b&WithFlank&19|+|51692606|51692687</td><td>51</td><td>70</td><td>20(+1A)</td><td>CAAGCTCGTGTCTGTGGGTCA</td><td>256.5</td><td>3.6</td><td>0.9</td><td>4.1</td><td>3.4E-05</td><td>3.3E-04</td><td>BT20/MB231</td><td>CYTO</td></tr>
<tr><td>>MI0000440|hsa-mir-27b&WithFlank&9|+|95085439|95085547@66.87.22&[MIMAT0000419&hsa-miR-27b-3p&offsets|-1|0;m-39&9|+|95085505|95085525&offsets|-1|0]|GTTCACAGTGGCTAAGTTCTGC</td><td>>MI0000440|hsa-mir-27b&WithFlank&9|+|95085439|95085547</td><td>66</td><td>87</td><td>22</td><td>GTTCACAGTGGCTAAGTTCTGC</td><td>162.5</td><td>5.6</td><td>1.5</td><td>3.7</td><td>2.6E-04</td><td>3.3E-04</td><td>MB231/MB468</td><td>MITO</td></tr>
<tr><td>>MI0000737|hsa-mir-200a&WithFlank&1|+|1167857|1167958@22.43.22&[MIMAT0001620&hsa-miR-200a-5p&offsets|0|0;m-154&1|+|1167878|1167899&offsets|0|0]|CATCTTACCGGACAGTGCTGGA</td><td>>MI0000737|hsa-mir-200a&WithFlank&1|+|1167857|1167958</td><td>22</td><td>43</td><td>22</td><td>CATCTTACCGGACAGTGCTGGA</td><td>228.7</td><td>-4.1</td><td>1.1</td><td>-3.6</td><td>2.6E-04</td><td>3.3E-04</td><td>MB231/MB468</td><td>NUC</td></tr>
<tr><td>>MI0000289|hsa-mir-181a-1&WithFlank&1|-|198859038|198859159@30.50.21;MI0000269|hsa-mir-181a-2&WithFlank&9|+|124692436|124692557@45.65.21&[MIMAT0000256&hsa-miR-181a-5p&offsets|0|-2;m-32&1|-|198859108|198859130&offsets|0|-2];[MIMAT0000256_1&hsa-miR-181a-5p&offsets|0|-2;m-32&9|+|124692480|124692502&offsets|0|-2]|AACATTCAACGCTGTCGGTGA</td><td>>MI0000289|hsa-mir-181a-1&WithFlank&1|-|198859038|198859159</td><td>30</td><td>50</td><td>21</td><td>AACATTCAACGCTGTCGGTGA</td><td>404.0</td><td>3.6</td><td>1.0</td><td>3.7</td><td>2.6E-04</td><td>3.3E-04</td><td>MB231/MB468</td><td>MITO</td></tr>
<tr><td>>hg38_21_+_8433172_8446622_RNA45SN1_45S_50ntFlanks@12046.12070.25|TGACCCGGTGAGGCGGGGGGGCGAG</td><td>>hg38_21_+_8433172_8446622_RNA45SN1_45S_50ntFlanks</td><td>12046</td><td>12070</td><td>25</td><td>TGACCCGGTGAGGCGGGGGGGCGAG</td><td>226.3</td><td>-5.9</td><td>1.6</td><td>-3.7</td><td>1.9E-04</td><td>3.3E-04</td><td>BT20/MB468</td><td>MITO</td></tr>
<tr><td>>trnaMT_PheGAA_MT_+_577_647@35.68.34|GAAAATGTTTAGACGGGCTCACATCACCCCATAA</td><td>>trnaMT_PheGAA_MT_+_577_647</td><td>35</td><td>68</td><td>34</td><td>GAAAATGTTTAGACGGGCTCACATCACCCCATAA</td><td>96.6</td><td>10.3</td><td>2.7</td><td>3.9</td><td>1.0E-04</td><td>3.3E-04</td><td>BT20/MB468</td><td>MP</td></tr>
<tr><td>>hg38_21_+_8433172_8446622_RNA45SN1_45S_50ntFlanks@6745.6772.28|ACACTTCGAACGCACTTGCGGCCCCGGG</td><td>>hg38_21_+_8433172_8446622_RNA45SN1_45S_50ntFlanks</td><td>6745</td><td>6772</td><td>28</td><td>ACACTTCGAACGCACTTGCGGCCCCGGG</td><td>903.1</td><td>-6.5</td><td>1.7</td><td>-3.9</td><td>1.0E-04</td><td>3.3E-04</td><td>BT20/MB468</td><td>MP</td></tr>
<tr><td>>hg38_21_+_8433172_8446622_RNA45SN1_45S_50ntFlanks@3837.3860.24|CCTCTCCTACTTGGATAACTGTGG</td><td>>hg38_21_+_8433172_8446622_RNA45SN1_45S_50ntFlanks</td><td>3837</td><td>3860</td><td>24</td><td>CCTCTCCTACTTGGATAACTGTGG</td><td>351.1</td><td>-4.8</td><td>1.3</td><td>-3.7</td><td>1.9E-04</td><td>3.3E-04</td><td>BT20/MB468</td><td>MITO</td></tr>
</table>

<table>
<tr><td>>hg38_21_+_8433172_8446622_RNA45SN1_45S_50ntFlanks@4975.4994.20|CCCGGACACGGACAGGATTG</td><td>>hg38_21_+_8433172_8446622_RNA45SN1_45S_50ntFlanks</td><td>4975</td><td>4994</td><td>20</td><td>CCCGGACACGGACAGGATTG</td><td>745.6</td><td>-6.4</td><td>1.6</td><td>-3.9</td><td>1.0E-04</td><td>3.3E-04</td><td>BT20/MB468</td><td>MP</td></tr>
<tr><td>>hg38_21_+_8433172_8446622_RNA45SN1_45S_50ntFlanks@9938.9964.27|AGCAGGACGGTGGCCATGGAAGTCGGA</td><td>>hg38_21_+_8433172_8446622_RNA45SN1_45S_50ntFlanks</td><td>9938</td><td>9964</td><td>27</td><td>AGCAGGACGGTGGCCATGGAAGTCGGA</td><td>77.9</td><td>7.0</td><td>1.8</td><td>3.9</td><td>1.0E-04</td><td>3.3E-04</td><td>BT20/MB468</td><td>MP</td></tr>
<tr><td>>MI0000737|hsa-mir-200a&WithFlank&1|+|1167857|1167958@60.81.22&[MIMAT0000682&hsa-miR-200a-3p&offsets|0|0;m-142&1|+|1167916|1167938&offsets|0|-1]|TAACACTGTCTGGTAACGATGT</td><td>>MI0000737|hsa-mir-200a&WithFlank&1|+|1167857|1167958</td><td>60</td><td>81</td><td>22</td><td>TAACACTGTCTGGTAACGATGT</td><td>1219.4</td><td>-5.3</td><td>1.4</td><td>-3.9</td><td>1.0E-04</td><td>3.3E-04</td><td>BT20/MB468</td><td>MP</td></tr>
<tr><td>>MI0000433|hsa-let-7g&WithFlank&3|-|52268272|52268367@11.32.22(+1C)&[MIMAT0000414&hsa-let-7g-5p&offsets|0|0(+1C);m-58&3|-|52268336|52268357&offsets|0|0(+1C)]|TGAGGTAGTAGTTTGTACAGTTC</td><td>>MI0000433|hsa-let-7g&WithFlank&3|-|52268272|52268367</td><td>11</td><td>32</td><td>22(+1C)</td><td>TGAGGTAGTAGTTTGTACAGTTC</td><td>363.4</td><td>6.1</td><td>1.6</td><td>3.7</td><td>2.2E-04</td><td>3.3E-04</td><td>MB231/MB468</td><td>MP</td></tr>
<tr><td>>MI0000070|hsa-mir-16-1&WithFlank&13|-|50048967|50049067@20.39.20(+1U);MI0000115|hsa-mir-16-2&WithFlank&3|+|160404739|160404831@16.35.20(+1U)&[MIMAT0000069&hsa-miR-16-5p&offsets|0|-2(+1U);m-62&13|-|50049027|50049048&offsets|0|-2(+1U)];[MIMAT0000069_1&hsa-miR-16-5p&offsets|0|-2(+1U);m-61&3|+|160404754|160404775&offsets|0|-2(+1U)]|TAGCAGCACGTAAATATTGGT</td><td>>MI0000070|hsa-mir-16-1&WithFlank&13|-|50048967|50049067</td><td>20</td><td>39</td><td>20(+1U)</td><td>TAGCAGCACGTAAATATTGGT</td><td>863.9</td><td>3.6</td><td>1.0</td><td>3.7</td><td>2.2E-04</td><td>3.3E-04</td><td>MB231/MB468</td><td>MP</td></tr>
<tr><td>>hg38_21_+_8433172_8446622_RNA45SN1_45S_50ntFlanks@12539.12560.22|ATGATGTGTTGTTGCCATGGTA</td><td>>hg38_21_+_8433172_8446622_RNA45SN1_45S_50ntFlanks</td><td>12539</td><td>12560</td><td>22</td><td>ATGATGTGTTGTTGCCATGGTA</td><td>207.5</td><td>-4.0</td><td>1.1</td><td>-3.7</td><td>1.9E-04</td><td>3.3E-04</td><td>BT20/MB468</td><td>MITO</td></tr>
<tr><td>>hg38_21_+_8433172_8446622_RNA45SN1_45S_50ntFlanks@10681.10710.30|TTCCGGCGGCGTCCGGTGAGCTCTCGCTGG</td><td>>hg38_21_+_8433172_8446622_RNA45SN1_45S_50ntFlanks</td><td>10681</td><td>10710</td><td>30</td><td>TTCCGGCGGCGTCCGGTGAGCTCTCGCTGG</td><td>2022.1</td><td>-5.2</td><td>1.4</td><td>-3.7</td><td>2.3E-04</td><td>3.3E-04</td><td>BT20/MB468</td><td>CYTO</td></tr>
<tr><td>>hg38_21_+_8433172_8446622_RNA45SN1_45S_50ntFlanks@12458.12478.21|AAGCGTTGGATTGTTCACCCA</td><td>>hg38_21_+_8433172_8446622_RNA45SN1_45S_50ntFlanks</td><td>12458</td><td>12478</td><td>21</td><td>AAGCGTTGGATTGTTCACCCA</td><td>1184.8</td><td>-2.6</td><td>0.7</td><td>-3.7</td><td>2.3E-04</td><td>3.3E-04</td><td>BT20/MB468</td><td>CYTO</td></tr>
<tr><td>>hg38_21_+_8433172_8446622_RNA45SN1_45S_50ntFlanks@6742.6771.30|TCGACACTTCGAACGCACTTGCGGCCCCGG</td><td>>hg38_21_+_8433172_8446622_RNA45SN1_45S_50ntFlanks</td><td>6742</td><td>6771</td><td>30</td><td>TCGACACTTCGAACGCACTTGCGGCCCCGG</td><td>170.0</td><td>-5.0</td><td>1.4</td><td>-3.7</td><td>2.3E-04</td><td>3.3E-04</td><td>BT20/MB468</td><td>CYTO</td></tr>
<tr><td>>hg38_21_+_8433172_8446622_RNA45SN1_45S_50ntFlanks@4353.4391.39|TATTAAAGTTGCTGCAGTTAAAAAGCTCGTAGTTGGATC</td><td>>hg38_21_+_8433172_8446622_RNA45SN1_45S_50ntFlanks</td><td>4353</td><td>4391</td><td>39</td><td>TATTAAAGTTGCTGCAGTTAAAAAGCTCGTAGTTGGATC</td><td>124.9</td><td>-6.2</td><td>1.7</td><td>-3.7</td><td>2.2E-04</td><td>3.3E-04</td><td>MB231/MB468</td><td>MP</td></tr>
<tr><td>>hg38_21_+_8433172_8446622_RNA45SN1_45S_50ntFlanks@8178.8204.27|TCTGATCGAGGCCCAGCCCGTGGACGG</td><td>>hg38_21_+_8433172_8446622_RNA45SN1_45S_50ntFlanks</td><td>8178</td><td>8204</td><td>27</td><td>TCTGATCGAGGCCCAGCCCGTGGACGG</td><td>109.6</td><td>-4.4</td><td>1.2</td><td>-3.7</td><td>2.2E-04</td><td>3.4E-04</td><td>MB231/MB468</td><td>MP</td></tr>
<tr><td>>hg38_21_+_8433172_8446622_RNA45SN1_45S_50ntFlanks@11879.11898.20|ACCCTGTTGAGCTTGACTCT</td><td>>hg38_21_+_8433172_8446622_RNA45SN1_45S_50ntFlanks</td><td>11879</td><td>11898</td><td>20</td><td>ACCCTGTTGAGCTTGACTCT</td><td>236.6</td><td>3.4</td><td>0.8</td><td>4.1</td><td>4.1E-05</td><td>3.4E-04</td><td>BT20/MB231</td><td>MITO</td></tr>
<tr><td>>hg38_21_+_8433172_8446622_RNA45SN1_45S_50ntFlanks@3987.4010.24|GCTTTGGTGACTCTAGATAACCTC</td><td>>hg38_21_+_8433172_8446622_RNA45SN1_45S_50ntFlanks</td><td>3987</td><td>4010</td><td>24</td><td>GCTTTGGTGACTCTAGATAACCTC</td><td>207.9</td><td>3.1</td><td>0.8</td><td>4.1</td><td>4.1E-05</td><td>3.4E-04</td><td>BT20/MB231</td><td>MITO</td></tr>
<tr><td>>MI0005544|hsa-mir-147b&WithFlank&15|+|45433044|45433135@55.72.18&[MIMAT0004928&hsa-miR-147b-3p&offsets|0|-3;m-575&15|+|45433098|45433118&offsets|0|-3]|GTGTGCGGAAATGCTTCT</td><td>>MI0005544|hsa-mir-147b&WithFlank&15|+|45433044|45433135</td><td>55</td><td>72</td><td>18</td><td>GTGTGCGGAAATGCTTCT</td><td>51.2</td><td>-9.1</td><td>2.2</td><td>-4.1</td><td>3.6E-05</td><td>3.4E-04</td><td>BT20/MB231</td><td>MP</td></tr>
<tr><td>>hg38_21_+_8433172_8446622_RNA45SN1_45S_50ntFlanks@4511.4528.18|GAAAAAATTAGAGTGTTC</td><td>>hg38_21_+_8433172_8446622_RNA45SN1_45S_50ntFlanks</td><td>4511</td><td>4528</td><td>18</td><td>GAAAAAATTAGAGTGTTC</td><td>210.4</td><td>-6.1</td><td>1.7</td><td>-3.7</td><td>2.2E-04</td><td>3.4E-04</td><td>MB231/MB468</td><td>MP</td></tr>
<tr><td>>am_trna130_GlnTTG_6_-_27763640_27763711@13.30.18,trna146_GlnCTG_6_-_27515531_27515602@13.30.18,trna16_GlnTTG_17_+_47269890_47269961@13.30.18,trna173_GlnTTG_6_-_26311975_26312046@13.30.18,trna174_GlnTTG_6_-_26311424_26311495@13.30.18,trna1_GlnCTG_6_+_18836402_18836473@13.30.18,trna3_GlnCTG_17_+_8023070_8023141@13.30.18,trna42_GlnCTG_6_+_27263212_27263283@13.30.18,trna49_GlnCTG_6_+_27487308_27487379@13.30.18,trna64_GlnTTG_6_+_28557156_28557227@13.30.18,trna7_GlnCTG_15_-_66161400_66161471@13.30.18,trna84_GlnTTG_6_+_145503859_145503930@13.30.18,trna99_GlnCTG_6_-_28909378_28909449@13.30.18|TAATGGTTAGCACTCTGG</td><td>>am_trna130_GlnTTG_6_-_27763640_27763711</td><td>13</td><td>30</td><td>18</td><td>TAATGGTTAGCACTCTGG</td><td>183.0</td><td>-3.5</td><td>1.0</td><td>-3.7</td><td>2.5E-04</td><td>3.4E-04</td><td>MB231/MB468</td><td>CYTO</td></tr>
<tr><td>>MI0000066|hsa-let-7e&WithFlank&19|+|51692780|51692870@14.34.21(+1A)&[MIMAT0000066&hsa-let-7e-5p&offsets|0|-1(+1A);m-50&19|+|51692793|51692814&offsets|0|-1(+1A)]|TGAGGTAGGAGGTTGTATAGTA</td><td>>MI0000066|hsa-let-7e&WithFlank&19|+|51692780|51692870</td><td>14</td><td>34</td><td>21(+1A)</td><td>TGAGGTAGGAGGTTGTATAGTA</td><td>96.8</td><td>2.1</td><td>0.6</td><td>3.6</td><td>2.7E-04</td><td>3.4E-04</td><td>BT20/MB468</td><td>NUC</td></tr>
<tr><td>>hg38_21_+_8433172_8446622_RNA45SN1_45S_50ntFlanks@6740.6770.31|CATCGACACTTCGAACGCACTTGCGGCCCCG</td><td>>hg38_21_+_8433172_8446622_RNA45SN1_45S_50ntFlanks</td><td>6740</td><td>6770</td><td>31</td><td>CATCGACACTTCGAACGCACTTGCGGCCCCG</td><td>2001.2</td><td>-6.7</td><td>1.7</td><td>-3.9</td><td>1.0E-04</td><td>3.4E-04</td><td>BT20/MB468</td><td>MP</td></tr>
<tr><td>>hg38_21_+_8433172_8446622_RNA45SN1_45S_50ntFlanks@6680.6718.39|TGCGTCGATGAAGAACGCAGCTAGCTGCGAGAATTAATG</td><td>>hg38_21_+_8433172_8446622_RNA45SN1_45S_50ntFlanks</td><td>6680</td><td>6718</td><td>39</td><td>TGCGTCGATGAAGAACGCAGCTAGCTGCGAGAATTAATG</td><td>197.3</td><td>-2.9</td><td>0.8</td><td>-3.7</td><td>1.9E-04</td><td>3.4E-04</td><td>BT20/MB468</td><td>MITO</td></tr>
<tr><td>>hg38_21_+_8433172_8446622_RNA45SN1_45S_50ntFlanks@10346.10367.22|TCAAACGAGAACTTTGAAGGCC</td><td>>hg38_21_+_8433172_8446622_RNA45SN1_45S_50ntFlanks</td><td>10346</td><td>10367</td><td>22</td><td>TCAAACGAGAACTTTGAAGGCC</td><td>385.7</td><td>-4.6</td><td>1.2</td><td>-3.7</td><td>2.2E-04</td><td>3.4E-04</td><td>MB231/MB468</td><td>MP</td></tr>
<tr><td>>hg38_21_+_8433172_8446622_RNA45SN1_45S_50ntFlanks@7973.8004.32|GACGCGACCTCAGATCAGACGTGGCGACCCGC</td><td>>hg38_21_+_8433172_8446622_RNA45SN1_45S_50ntFlanks</td><td>7973</td><td>8004</td><td>32</td><td>GACGCGACCTCAGATCAGACGTGGCGACCCGC</td><td>1296.8</td><td>4.0</td><td>1.0</td><td>4.1</td><td>3.8E-05</td><td>3.4E-04</td><td>BT20/MB231</td><td>NUC</td></tr>
<tr><td>>hg38_21_+_8433172_8446622_RNA45SN1_45S_50ntFlanks@11583.11612.30|CTGGTGCGGACCAGGGGAATCCGACTGTTT</td><td>>hg38_21_+_8433172_8446622_RNA45SN1_45S_50ntFlanks</td><td>11583</td><td>11612</td><td>30</td><td>CTGGTGCGGACCAGGGGAATCCGACTGTTT</td><td>1107.3</td><td>-3.1</td><td>0.8</td><td>-3.9</td><td>1.0E-04</td><td>3.4E-04</td><td>BT20/MB468</td><td>MP</td></tr>
<tr><td>>MI0003137|hsa-mir-193b&WithFlank&16|+|14303961|14304055@57.77.21(+1A)&[MIMAT0002819&hsa-miR-193b-3p&offsets|0|-1(+1A);m-166&16|+|14304017|14304038&offsets|0|-1(+1A)]|AACTGGCCCTCAAAGTCCCGCA</td><td>>MI0003137|hsa-mir-193b&WithFlank&16|+|14303961|14304055</td><td>57</td><td>77</td><td>21(+1A)</td><td>AACTGGCCCTCAAAGTCCCGCA</td><td>602.9</td><td>5.7</td><td>1.5</td><td>3.9</td><td>1.0E-04</td><td>3.4E-04</td><td>BT20/MB468</td><td>MP</td></tr>
<tr><td>>hg38_21_+_8433172_8446622_RNA45SN1_45S_50ntFlanks@10547.10576.30|AGTGCGGTAACGCGACCGATCCCGGAGAAG</td><td>>hg38_21_+_8433172_8446622_RNA45SN1_45S_50ntFlanks</td><td>10547</td><td>10576</td><td>30</td><td>AGTGCGGTAACGCGACCGATCCCGGAGAAG</td><td>227.2</td><td>3.4</td><td>0.8</td><td>4.1</td><td>4.1E-05</td><td>3.4E-04</td><td>BT20/MB231</td><td>MITO</td></tr>
<tr><td>>hg38_21_+_8433172_8446622_RNA45SN1_45S_50ntFlanks@12417.12458.42|TCGATGTCGGCTCTTCCTATCATTGTGAAGCAGAATTCACCA</td><td>>hg38_21_+_8433172_8446622_RNA45SN1_45S_50ntFlanks</td><td>12417</td><td>12458</td><td>42</td><td>TCGATGTCGGCTCTTCCTATCATTGTGAAGCAGAATTCACCA</td><td>244.2</td><td>4.2</td><td>1.2</td><td>3.7</td><td>2.5E-04</td><td>3.4E-04</td><td>MB231/MB468</td><td>CYTO</td></tr>
<tr><td>>hg38_21_+_8433172_8446622_RNA45SN1_45S_50ntFlanks@6741.6770.30|ATCGACACTTCGAACGCACTTGCGGCCCCG</td><td>>hg38_21_+_8433172_8446622_RNA45SN1_45S_50ntFlanks</td><td>6741</td><td>6770</td><td>30</td><td>ATCGACACTTCGAACGCACTTGCGGCCCCG</td><td>2054.9</td><td>-6.7</td><td>1.7</td><td>-3.9</td><td>1.0E-04</td><td>3.4E-04</td><td>BT20/MB468</td><td>MP</td></tr>
<tr><td>>hg38_21_+_8433172_8446622_RNA45SN1_45S_50ntFlanks@6721.6756.36|AATTGCAGGACACATTGATCATCGACACTTCGAACG</td><td>>hg38_21_+_8433172_8446622_RNA45SN1_45S_50ntFlanks</td><td>6721</td><td>6756</td><td>36</td><td>AATTGCAGGACACATTGATCATCGACACTTCGAACG</td><td>667.3</td><td>-2.3</td><td>0.6</td><td>-3.7</td><td>1.9E-04</td><td>3.4E-04</td><td>BT20/MB468</td><td>MITO</td></tr>
<tr><td>>hg38_21_+_8433172_8446622_RNA45SN1_45S_50ntFlanks@8176.8220.45|CTTCTGATCGAGGCCCAGCCCGTGGACGGTGTGAGGCCGGTAGCG</td><td>>hg38_21_+_8433172_8446622_RNA45SN1_45S_50ntFlanks</td><td>8176</td><td>8220</td><td>45</td><td>CTTCTGATCGAGGCCCAGCCCGTGGACGGTGTGAGGCCGGTAGCG</td><td>196.0</td><td>-4.9</td><td>1.3</td><td>-3.7</td><td>2.2E-04</td><td>3.4E-04</td><td>MB231/MB468</td><td>MP</td></tr>
<tr><td>>hg38_21_+_8433172_8446622_RNA45SN1_45S_50ntFlanks@11562.11591.30|CGCCTAGCAGCCGACTTAGAACTGGTGCGG</td><td>>hg38_21_+_8433172_8446622_RNA45SN1_45S_50ntFlanks</td><td>11562</td><td>11591</td><td>30</td><td>CGCCTAGCAGCCGACTTAGAACTGGTGCGG</td><td>419.6</td><td>-5.3</td><td>1.4</td><td>-3.9</td><td>1.0E-04</td><td>3.4E-04</td><td>BT20/MB468</td><td>MP</td></tr>
<tr><td>>am_trnaMT_ArgTCG_MT_+_10405_10469@28.48.21|ATTTCGACTCATTAAATTATG</td><td>>am_trnaMT_ArgTCG_MT_+_10405_10469</td><td>28</td><td>48</td><td>21</td><td>ATTTCGACTCATTAAATTATG</td><td>580.6</td><td>-5.2</td><td>1.4</td><td>-3.7</td><td>2.3E-04</td><td>3.4E-04</td><td>BT20/MB468</td><td>CYTO</td></tr>
<tr><td>>hg38_21_+_8433172_8446622_RNA45SN1_45S_50ntFlanks@9567.9592.26|ACGTGCAAATCGGTCGTCCGACCTGG</td><td>>hg38_21_+_8433172_8446622_RNA45SN1_45S_50ntFlanks</td><td>9567</td><td>9592</td><td>26</td><td>ACGTGCAAATCGGTCGTCCGACCTGG</td><td>713.0</td><td>-5.1</td><td>1.3</td><td>-3.9</td><td>1.0E-04</td><td>3.4E-04</td><td>BT20/MB468</td><td>MP</td></tr>
<tr><td>>hg38_21_+_8433172_8446622_RNA45SN1_45S_50ntFlanks@10687.10706.20|CGGCGTCCGGTGAGCTCTCG</td><td>>hg38_21_+_8433172_8446622_RNA45SN1_45S_50ntFlanks</td><td>10687</td><td>10706</td><td>20</td><td>CGGCGTCCGGTGAGCTCTCG</td><td>2746.0</td><td>-5.7</td><td>1.5</td><td>-3.9</td><td>1.0E-04</td><td>3.4E-04</td><td>BT20/MB468</td><td>MP</td></tr>
<tr><td>>hg38_21_+_8433172_8446622_RNA45SN1_45S_50ntFlanks@10686.10707.22|GCGGCGTCCGGTGAGCTCTCGC</td><td>>hg38_21_+_8433172_8446622_RNA45SN1_45S_50ntFlanks</td><td>10686</td><td>10707</td><td>22</td><td>GCGGCGTCCGGTGAGCTCTCGC</td><td>368.8</td><td>4.3</td><td>1.1</td><td>3.9</td><td>1.0E-04</td><td>3.4E-04</td><td>BT20/MB468</td><td>MP</td></tr>
<tr><td>>hg38_21_+_8433172_8446622_RNA45SN1_45S_50ntFlanks@8346.8378.33|AGGGAAAGTTGAAAAGAACTTTGAAGAGAGAGT</td><td>>hg38_21_+_8433172_8446622_RNA45SN1_45S_50ntFlanks</td><td>8346</td><td>8378</td><td>33</td><td>AGGGAAAGTTGAAAAGAACTTTGAAGAGAGAGT</td><td>173.6</td><td>6.0</td><td>1.6</td><td>3.9</td><td>1.0E-04</td><td>3.4E-04</td><td>BT20/MB468</td><td>MP</td></tr>
<tr><td>>hg38_21_+_8433172_8446622_RNA45SN1_45S_50ntFlanks@9714.9735.22|CGAATGATTAGAGGTCTTGGGG</td><td>>hg38_21_+_8433172_8446622_RNA45SN1_45S_50ntFlanks</td><td>9714</td><td>9735</td><td>22</td><td>CGAATGATTAGAGGTCTTGGGG</td><td>208.8</td><td>-5.3</td><td>1.4</td><td>-3.7</td><td>2.4E-04</td><td>3.4E-04</td><td>BT20/MB468</td><td>CYTO</td></tr>
<tr><td>>MI0000255|hsa-mir-30d&WithFlank&8|-|134804870|134804951@12.35.24(+3U)&[MIMAT0000245&hsa-miR-30d-5p&offsets|0|+2(+3U);m-21&8|-|134804919|134804940&offsets|0|+2(+3U)]|TGTAAACATCCCCGACTGGAAGCTTTT</td><td>>MI0000255|hsa-mir-30d&WithFlank&8|-|134804870|134804951</td><td>12</td><td>35</td><td>24(+3U)</td><td>TGTAAACATCCCCGACTGGAAGCTTTT</td><td>499.3</td><td>-3.2</td><td>0.8</td><td>-4.1</td><td>4.2E-05</td><td>3.4E-04</td><td>BT20/MB231</td><td>MITO</td></tr>
<tr><td>>hg38_21_+_8433172_8446622_RNA45SN1_45S_50ntFlanks@9937.9979.43|CAGCAGGACGGTGGCCATGGAAGTCGGAATCCGCTAAGGAGTG</td><td>>hg38_21_+_8433172_8446622_RNA45SN1_45S_50ntFlanks</td><td>9937</td><td>9979</td><td>43</td><td>CAGCAGGACGGTGGCCATGGAAGTCGGAATCCGCTAAGGAGTG</td><td>60.3</td><td>9.2</td><td>2.2</td><td>4.1</td><td>3.7E-05</td><td>3.4E-04</td><td>BT20/MB231</td><td>MP</td></tr>
<tr><td>>MI0000289|hsa-mir-181a-1&WithFlank&1|-|198859038|198859159@30.49.20;MI0000269|hsa-mir-181a-2&WithFlank&9|+|124692436|124692557@45.64.20&[MIMAT0000256&hsa-miR-181a-5p&offsets|0|-3;m-32&1|-|198859108|198859130&offsets|0|-3];[MIMAT0000256_1&hsa-miR-181a-5p&offsets|0|-3;m-32&9|+|124692480|124692502&offsets|0|-3]|AACATTCAACGCTGTCGGTG</td><td>>MI0000289|hsa-mir-181a-1&WithFlank&1|-|198859038|198859159</td><td>30</td><td>49</td><td>20</td><td>AACATTCAACGCTGTCGGTG</td><td>905.7</td><td>3.2</td><td>0.9</td><td>3.7</td><td>2.2E-04</td><td>3.4E-04</td><td>MB231/MB468</td><td>MP</td></tr>
<tr><td>>hg38_21_+_8433172_8446622_RNA45SN1_45S_50ntFlanks@6661.6679.19|CGGTGGATCACTCGGCTCG</td><td>>hg38_21_+_8433172_8446622_RNA45SN1_45S_50ntFlanks</td><td>6661</td><td>6679</td><td>19</td><td>CGGTGGATCACTCGGCTCG</td><td>1372.8</td><td>3.5</td><td>1.0</td><td>3.7</td><td>2.4E-04</td><td>3.4E-04</td><td>BT20/MB468</td><td>CYTO</td></tr>
<tr><td>>hg38_MT_+_1621_3279_RNR2_16S_50ntFlanks@644.663.20|AATTGGACCAATCTATCACC</td><td>>hg38_MT_+_1621_3279_RNR2_16S_50ntFlanks</td><td>644</td><td>663</td><td>20</td><td>AATTGGACCAATCTATCACC</td><td>123.0</td><td>3.0</td><td>0.8</td><td>3.7</td><td>2.2E-04</td><td>3.4E-04</td><td>MB231/MB468</td><td>MP</td></tr>
<tr><td>>hg38_21_+_8433172_8446622_RNA45SN1_45S_50ntFlanks@8105.8143.39|CGCGGCGGGGCGCGGGACATGTGGCGTACGGAAGACCCG</td><td>>hg38_21_+_8433172_8446622_RNA45SN1_45S_50ntFlanks</td><td>8105</td><td>8143</td><td>39</td><td>CGCGGCGGGGCGCGGGACATGTGGCGTACGGAAGACCCG</td><td>18848.6</td><td>-5.8</td><td>1.6</td><td>-3.7</td><td>2.0E-04</td><td>3.4E-04</td><td>BT20/MB468</td><td>MITO</td></tr>
<tr><td>>hg38_21_+_8433172_8446622_RNA45SN1_45S_50ntFlanks@10808.10840.33|CAATGTAGGTAAGGGAAGTCGGCAAGCCGGATC</td><td>>hg38_21_+_8433172_8446622_RNA45SN1_45S_50ntFlanks</td><td>10808</td><td>10840</td><td>33</td><td>CAATGTAGGTAAGGGAAGTCGGCAAGCCGGATC</td><td>1468.9</td><td>-4.2</td><td>1.1</td><td>-3.9</td><td>1.0E-04</td><td>3.4E-04</td><td>BT20/MB468</td><td>MP</td></tr>
<tr><td>>trnaMT_PheGAA_MT_+_577_647@39.56.18|ATGTTTAGACGGGCTCAC</td><td>>trnaMT_PheGAA_MT_+_577_647</td><td>39</td><td>56</td><td>18</td><td>ATGTTTAGACGGGCTCAC</td><td>1337.8</td><td>9.2</td><td>2.5</td><td>3.7</td><td>2.5E-04</td><td>3.4E-04</td><td>MB231/MB468</td><td>CYTO</td></tr>
<tr><td>>hg38_21_+_8433172_8446622_RNA45SN1_45S_50ntFlanks@10510.10533.24|AATCCGGAGTGGCGGAGATGGGCG</td><td>>hg38_21_+_8433172_8446622_RNA45SN1_45S_50ntFlanks</td><td>10510</td><td>10533</td><td>24</td><td>AATCCGGAGTGGCGGAGATGGGCG</td><td>8270.8</td><td>-4.8</td><td>1.2</td><td>-3.9</td><td>1.0E-04</td><td>3.4E-04</td><td>BT20/MB468</td><td>MP</td></tr>
<tr><td>>hg38_21_+_8433172_8446622_RNA45SN1_45S_50ntFlanks@9528.9570.43|AGCCAGAGGAAACTCTGGTGGAGGTCCGTAGCGGTCCTGACGT</td><td>>hg38_21_+_8433172_8446622_RNA45SN1_45S_50ntFlanks</td><td>9528</td><td>9570</td><td>43</td><td>AGCCAGAGGAAACTCTGGTGGAGGTCCGTAGCGGTCCTGACGT</td><td>60.3</td><td>9.2</td><td>2.2</td><td>4.1</td><td>3.7E-05</td><td>3.4E-04</td><td>BT20/MB231</td><td>MP</td></tr>
<tr><td>>hg38_21_+_8433172_8446622_RNA45SN1_45S_50ntFlanks@8171.8212.42|AAGTCCTTCTGATCGAGGCCCAGCCCGTGGACGGTGTGAGGC</td><td>>hg38_21_+_8433172_8446622_RNA45SN1_45S_50ntFlanks</td><td>8171</td><td>8212</td><td>42</td><td>AAGTCCTTCTGATCGAGGCCCAGCCCGTGGACGGTGTGAGGC</td><td>302.5</td><td>-2.2</td><td>0.6</td><td>-3.7</td><td>2.4E-04</td><td>3.4E-04</td><td>BT20/MB468</td><td>CYTO</td></tr>
</table>

<table>
<tr><th></th><th></th><th></th><th></th><th></th><th></th><th></th><th></th><th></th><th></th><th></th><th></th><th></th><th></th></tr>
<tr><td>>hg38_21_+_8433172_8446622_RNA45SN1_45S_50ntFlanks@12463.12487.25|TTGGATTGTTCACCCACTAATAGGG</td><td>>hg38_21_+_8433172_8446622_RNA45SN1_45S_50ntFlanks</td><td>12463</td><td>12487</td><td>25</td><td>TTGGATTGTTCACCCACTAATAGGG</td><td>223.6</td><td>-2.8</td><td>0.8</td><td>-3.7</td><td>2.5E-04</td><td>3.4E-04</td><td>MB231/MB468</td><td>CYTO</td></tr>
<tr><td>>MI0000542|hsa-mir-320a&WithFlank&8|-|22244960|22245043@49.68.20(+1U)&[MIMAT0000510&hsa-miR-320a-3p&offsets|+1|-1(+1U);m-60&8|-|22244975|22244996&offsets|+1|-1(+1U)]|AAAGCTGGGTTGAGAGGGCGT</td><td>>MI0000542|hsa-mir-320a&WithFlank&8|-|22244960|22245043</td><td>49</td><td>68</td><td>20(+1U)</td><td>AAAGCTGGGTTGAGAGGGCGT</td><td>751.7</td><td>3.4</td><td>0.8</td><td>4.1</td><td>3.5E-05</td><td>3.4E-04</td><td>BT20/MB231</td><td>CYTO</td></tr>
<tr><td>>trna15_ValAAC_5_-_180615416_180615488@1.29.29|GTTTCCGTAGTGTAGTGGTCATCACGTTC</td><td>>trna15_ValAAC_5_-_180615416_180615488</td><td>1</td><td>29</td><td>29</td><td>GTTTCCGTAGTGTAGTGGTCATCACGTTC</td><td>1861.1</td><td>-6.2</td><td>1.7</td><td>-3.7</td><td>2.4E-04</td><td>3.4E-04</td><td>BT20/MB468</td><td>CYTO</td></tr>
<tr><td>>hg38_21_+_8433172_8446622_RNA45SN1_45S_50ntFlanks@9536.9571.36|GAAACTCTGGTGGAGGTCCGTAGCGGTCCTGACGTG</td><td>>hg38_21_+_8433172_8446622_RNA45SN1_45S_50ntFlanks</td><td>9536</td><td>9571</td><td>36</td><td>GAAACTCTGGTGGAGGTCCGTAGCGGTCCTGACGTG</td><td>587.3</td><td>-3.4</td><td>0.8</td><td>-4.1</td><td>3.5E-05</td><td>3.4E-04</td><td>BT20/MB231</td><td>CYTO</td></tr>
<tr><td>>hg38_21_+_8433172_8446622_RNA45SN1_45S_50ntFlanks@7977.8008.32|CGACCTCAGATCAGACGTGGCGACCCGCTGAA</td><td>>hg38_21_+_8433172_8446622_RNA45SN1_45S_50ntFlanks</td><td>7977</td><td>8008</td><td>32</td><td>CGACCTCAGATCAGACGTGGCGACCCGCTGAA</td><td>1483.9</td><td>-2.8</td><td>0.8</td><td>-3.6</td><td>2.7E-04</td><td>3.4E-04</td><td>MB231/MB468</td><td>MITO</td></tr>
<tr><td>>hg38_21_+_8433172_8446622_RNA45SN1_45S_50ntFlanks@7983.8003.21|CAGATCAGACGTGGCGACCCG</td><td>>hg38_21_+_8433172_8446622_RNA45SN1_45S_50ntFlanks</td><td>7983</td><td>8003</td><td>21</td><td>CAGATCAGACGTGGCGACCCG</td><td>343.5</td><td>7.8</td><td>1.9</td><td>4.1</td><td>3.7E-05</td><td>3.4E-04</td><td>BT20/MB231</td><td>MP</td></tr>
<tr><td>>hg38_21_+_8433172_8446622_RNA45SN1_45S_50ntFlanks@3846.3869.24|CTTGGATAACTGTGGTAATTCTAG</td><td>>hg38_21_+_8433172_8446622_RNA45SN1_45S_50ntFlanks</td><td>3846</td><td>3869</td><td>24</td><td>CTTGGATAACTGTGGTAATTCTAG</td><td>519.5</td><td>-4.8</td><td>1.2</td><td>-3.9</td><td>1.0E-04</td><td>3.4E-04</td><td>BT20/MB468</td><td>MP</td></tr>
<tr><td>>hg38_MT_+_598_1651_RNR1_12S_50ntFlanks@848.868.21|GAGTGCTTAGTTGAACAGGGC</td><td>>hg38_MT_+_598_1651_RNR1_12S_50ntFlanks</td><td>848</td><td>868</td><td>21</td><td>GAGTGCTTAGTTGAACAGGGC</td><td>99.3</td><td>7.0</td><td>1.8</td><td>3.9</td><td>1.0E-04</td><td>3.4E-04</td><td>BT20/MB468</td><td>MP</td></tr>
<tr><td>>hg38_21_+_8433172_8446622_RNA45SN1_45S_50ntFlanks@11602.11620.19|TCCGACTGTTTAATTAAAA</td><td>>hg38_21_+_8433172_8446622_RNA45SN1_45S_50ntFlanks</td><td>11602</td><td>11620</td><td>19</td><td>TCCGACTGTTTAATTAAAA</td><td>375.9</td><td>-3.2</td><td>0.8</td><td>-3.9</td><td>1.0E-04</td><td>3.4E-04</td><td>BT20/MB468</td><td>MP</td></tr>
<tr><td>>am_trnaMT_ValTAC_MT_+_1602_1670@31.67.37|TTACACTTAGGAGATTTCAACTTAACTTGACCGCTCT</td><td>>am_trnaMT_ValTAC_MT_+_1602_1670</td><td>31</td><td>67</td><td>37</td><td>TTACACTTAGGAGATTTCAACTTAACTTGACCGCTCT</td><td>95.9</td><td>10.3</td><td>2.7</td><td>3.9</td><td>1.0E-04</td><td>3.4E-04</td><td>BT20/MB468</td><td>MP</td></tr>
<tr><td>>trnaMT_IleGAT_MT_+_4263_4331@25.63.39|ACTTTGATAGAGTAAATAATAGGAGCTTAAACCCCCTTA</td><td>>trnaMT_IleGAT_MT_+_4263_4331</td><td>25</td><td>63</td><td>39</td><td>ACTTTGATAGAGTAAATAATAGGAGCTTAAACCCCCTTA</td><td>78.0</td><td>9.5</td><td>2.6</td><td>3.7</td><td>2.2E-04</td><td>3.4E-04</td><td>MB231/MB468</td><td>MP</td></tr>
<tr><td>>hg38_21_+_8433172_8446622_RNA45SN1_45S_50ntFlanks@8175.8216.42|CCTTCTGATCGAGGCCCAGCCCGTGGACGGTGTGAGGCCGGT</td><td>>hg38_21_+_8433172_8446622_RNA45SN1_45S_50ntFlanks</td><td>8175</td><td>8216</td><td>42</td><td>CCTTCTGATCGAGGCCCAGCCCGTGGACGGTGTGAGGCCGGT</td><td>262.5</td><td>5.7</td><td>1.6</td><td>3.7</td><td>2.2E-04</td><td>3.4E-04</td><td>MB231/MB468</td><td>MP</td></tr>
<tr><td>>hg38_21_+_8433172_8446622_RNA45SN1_45S_50ntFlanks@6648.6668.21|GTACGACTCTTAGCGGTGGAT</td><td>>hg38_21_+_8433172_8446622_RNA45SN1_45S_50ntFlanks</td><td>6648</td><td>6668</td><td>21</td><td>GTACGACTCTTAGCGGTGGAT</td><td>443.5</td><td>1.8</td><td>0.5</td><td>3.6</td><td>2.7E-04</td><td>3.4E-04</td><td>BT20/MB468</td><td>NUC</td></tr>
<tr><td>>trna116_GluCTC_1_-_145399233_145399304@1.35.35,trna59_GluCTC_1_+_249168447_249168518@1.35.35,trna71_GluCTC_1_-_161439189_161439260@1.35.35,trna74_GluCTC_1_-_161431809_161431880@1.35.35,trna77_GluCTC_1_-_161424398_161424469@1.35.35,trna77_GluCTC_6_+_28949976_28950047@1.35.35,trna80_GluCTC_1_-_161417018_161417089@1.35.35,trna87_GluCTC_6_-_126101393_126101464@1.35.35|TCCCTGGTGGTCTAGTGGTTAGGATTCGGCGCTCT</td><td>>trna116_GluCTC_1_-_145399233_145399304</td><td>1</td><td>35</td><td>35</td><td>TCCCTGGTGGTCTAGTGGTTAGGATTCGGCGCTCT</td><td>2055.4</td><td>5.3</td><td>1.4</td><td>3.9</td><td>1.0E-04</td><td>3.4E-04</td><td>BT20/MB468</td><td>MP</td></tr>
<tr><td>>trnaMT_GluTTC_MT_-_14674_14742@31.71.41,trnalookalike8_GluTTC_5_-_93905172_93905240@31.71.41|TTCATATCATTGGTCGTGGTTGTAGTCCGTGCGAGAATACC</td><td>>trnaMT_GluTTC_MT_-_14674_14742</td><td>31</td><td>71</td><td>41</td><td>TTCATATCATTGGTCGTGGTTGTAGTCCGTGCGAGAATACC</td><td>169.8</td><td>5.4</td><td>1.5</td><td>3.6</td><td>2.7E-04</td><td>3.4E-04</td><td>MB231/MB468</td><td>MITO</td></tr>
<tr><td>>MI0000473|hsa-mir-129-2&WithFlank&11|+|43581388|43581489@21.41.21;MI0000252|hsa-mir-129-1&WithFlank&7|+|128207866|128207949@11.31.21&[MIMAT0000242&hsa-miR-129-5p&offsets|0|0;m-434&11|+|43581408|43581428&offsets|0|0];[MIMAT0000242_1&hsa-miR-129-5p&offsets|0|0;m-434&7|+|128207876|128207896&offsets|0|0]|CTTTTTGCGGTCTGGGCTTGC</td><td>>MI0000473|hsa-mir-129-2&WithFlank&11|+|43581388|43581489</td><td>21</td><td>41</td><td>21</td><td>CTTTTTGCGGTCTGGGCTTGC</td><td>184.9</td><td>3.3</td><td>0.8</td><td>4.1</td><td>3.6E-05</td><td>3.4E-04</td><td>BT20/MB231</td><td>CYTO</td></tr>
<tr><td>>hg38_21_+_8433172_8446622_RNA45SN1_45S_50ntFlanks@5086.5110.25|AACGAGACTCTGGCATGCTAACTAG</td><td>>hg38_21_+_8433172_8446622_RNA45SN1_45S_50ntFlanks</td><td>5086</td><td>5110</td><td>25</td><td>AACGAGACTCTGGCATGCTAACTAG</td><td>481.6</td><td>-3.9</td><td>1.1</td><td>-3.6</td><td>2.7E-04</td><td>3.4E-04</td><td>MB231/MB468</td><td>MITO</td></tr>
<tr><td>>hg38_21_+_8433172_8446622_RNA45SN1_45S_50ntFlanks@11211.11252.42|GCGACTCTGGACGCGAGCCGGGCCCTTCCCGTGGATCGCCCC</td><td>>hg38_21_+_8433172_8446622_RNA45SN1_45S_50ntFlanks</td><td>11211</td><td>11252</td><td>42</td><td>GCGACTCTGGACGCGAGCCGGGCCCTTCCCGTGGATCGCCCC</td><td>658.7</td><td>3.5</td><td>0.9</td><td>3.7</td><td>2.0E-04</td><td>3.4E-04</td><td>BT20/MB468</td><td>MITO</td></tr>
<tr><td>>hg38_21_+_8433172_8446622_RNA45SN1_45S_50ntFlanks@4557.4589.33|CAGCTAGGAATAATGGAATAGGACCGCGGTTCT</td><td>>hg38_21_+_8433172_8446622_RNA45SN1_45S_50ntFlanks</td><td>4557</td><td>4589</td><td>33</td><td>CAGCTAGGAATAATGGAATAGGACCGCGGTTCT</td><td>72.4</td><td>-5.7</td><td>1.5</td><td>-3.7</td><td>2.3E-04</td><td>3.4E-04</td><td>MB231/MB468</td><td>MP</td></tr>
<tr><td>>hg38_21_+_8433172_8446622_RNA45SN1_45S_50ntFlanks@4759.4783.25|ATCAGATACCGTCGTAGTTCCGACC</td><td>>hg38_21_+_8433172_8446622_RNA45SN1_45S_50ntFlanks</td><td>4759</td><td>4783</td><td>25</td><td>ATCAGATACCGTCGTAGTTCCGACC</td><td>1264.0</td><td>-5.4</td><td>1.5</td><td>-3.7</td><td>2.5E-04</td><td>3.4E-04</td><td>MB231/MB468</td><td>CYTO</td></tr>
<tr><td>>hg38_21_+_8433172_8446622_RNA45SN1_45S_50ntFlanks@9477.9497.21|AGCGCACGTGTTAGGACCCGA</td><td>>hg38_21_+_8433172_8446622_RNA45SN1_45S_50ntFlanks</td><td>9477</td><td>9497</td><td>21</td><td>AGCGCACGTGTTAGGACCCGA</td><td>1672.6</td><td>-3.6</td><td>0.9</td><td>-4.1</td><td>3.6E-05</td><td>3.4E-04</td><td>BT20/MB231</td><td>CYTO</td></tr>
<tr><td>>MI0000750|hsa-mir-26a-2&WithFlank&12|-|57824603|57824698@20.41.22(+2A);MI0000083|hsa-mir-26a-1&WithFlank&3|+|37969398|37969486@16.37.22(+2A)&[MIMAT0000082&hsa-miR-26a-5p&offsets|0|0(+2A);m-33&12|-|57824658|57824679&offsets|0|0(+2A)];[MIMAT0000082_1&hsa-miR-26a-5p&offsets|0|0(+2A);m-34&3|+|37969413|37969434&offsets|0|0(+2A)]|TTCAAGTAATCCAGGATAGGCTAA</td><td>>MI0000750|hsa-mir-26a-2&WithFlank&12|-|57824603|57824698</td><td>20</td><td>41</td><td>22(+2A)</td><td>TTCAAGTAATCCAGGATAGGCTAA</td><td>507.7</td><td>3.4</td><td>0.9</td><td>3.7</td><td>2.3E-04</td><td>3.4E-04</td><td>MB231/MB468</td><td>MP</td></tr>
<tr><td>>MI0000750|hsa-mir-26a-2&WithFlank&12|-|57824603|57824698@20.42.23;MI0000083|hsa-mir-26a-1&WithFlank&3|+|37969398|37969486@16.38.23&[MIMAT0000082&hsa-miR-26a-5p&offsets|0|+1;m-33&12|-|57824658|57824679&offsets|0|+1];[MIMAT0000082_1&hsa-miR-26a-5p&offsets|0|+1;m-34&3|+|37969413|37969434&offsets|0|+1]|TTCAAGTAATCCAGGATAGGCTG</td><td>>MI0000750|hsa-mir-26a-2&WithFlank&12|-|57824603|57824698</td><td>20</td><td>42</td><td>23</td><td>TTCAAGTAATCCAGGATAGGCTG</td><td>408.2</td><td>3.2</td><td>0.9</td><td>3.7</td><td>2.3E-04</td><td>3.4E-04</td><td>MB231/MB468</td><td>MP</td></tr>
<tr><td>>hg38_21_+_8433172_8446622_RNA45SN1_45S_50ntFlanks@11569.11591.23|CAGCCGACTTAGAACTGGTGCGG</td><td>>hg38_21_+_8433172_8446622_RNA45SN1_45S_50ntFlanks</td><td>11569</td><td>11591</td><td>23</td><td>CAGCCGACTTAGAACTGGTGCGG</td><td>126.0</td><td>-2.2</td><td>0.6</td><td>-3.7</td><td>2.0E-04</td><td>3.4E-04</td><td>BT20/MB468</td><td>MITO</td></tr>
<tr><td>>hg38_1_-_228634819_228635039_RNA5S12_5S_50ntFlanks@134.171.38|TGGGAGACCGCCTGGGAATACCGGGTGCTGTAGGCTTT</td><td>>hg38_1_-_228634819_228635039_RNA5S12_5S_50ntFlanks</td><td>134</td><td>171</td><td>38</td><td>TGGGAGACCGCCTGGGAATACCGGGTGCTGTAGGCTTT</td><td>4441.8</td><td>3.9</td><td>1.0</td><td>3.9</td><td>1.0E-04</td><td>3.4E-04</td><td>BT20/MB468</td><td>MP</td></tr>
<tr><td>>MI0000081|hsa-mir-24-2&WithFlank&19|-|13836281|13836365@56.75.20;MI0000080|hsa-mir-24-1&WithFlank&9|+|95086015|95086094@50.69.20&[MIMAT0000080&hsa-miR-24-3p&offsets|0|-2;m-36&19|-|13836288|13836310&offsets|0|-3];[MIMAT0000080_1&hsa-miR-24-3p&offsets|0|-2;m-35&9|+|95086064|95086086&offsets|0|-3]|TGGCTCAGTTCAGCAGGAAC</td><td>>MI0000081|hsa-mir-24-2&WithFlank&19|-|13836281|13836365</td><td>56</td><td>75</td><td>20</td><td>TGGCTCAGTTCAGCAGGAAC</td><td>28239.9</td><td>3.9</td><td>1.0</td><td>3.9</td><td>1.0E-04</td><td>3.4E-04</td><td>BT20/MB468</td><td>MP</td></tr>
<tr><td>>hg38_21_+_8433172_8446622_RNA45SN1_45S_50ntFlanks@485.513.29|GATGTGGTGACGTCGTGCTCTCCCGGGCC</td><td>>hg38_21_+_8433172_8446622_RNA45SN1_45S_50ntFlanks</td><td>485</td><td>513</td><td>29</td><td>GATGTGGTGACGTCGTGCTCTCCCGGGCC</td><td>641.9</td><td>13.2</td><td>3.6</td><td>3.6</td><td>2.7E-04</td><td>3.4E-04</td><td>MB231/MB468</td><td>NUC</td></tr>
<tr><td>>hg38_21_+_8433172_8446622_RNA45SN1_45S_50ntFlanks@8357.8376.20|AAAAGAACTTTGAAGAGAGA</td><td>>hg38_21_+_8433172_8446622_RNA45SN1_45S_50ntFlanks</td><td>8357</td><td>8376</td><td>20</td><td>AAAAGAACTTTGAAGAGAGA</td><td>281.9</td><td>-3.6</td><td>1.0</td><td>-3.7</td><td>2.5E-04</td><td>3.4E-04</td><td>MB231/MB468</td><td>CYTO</td></tr>
<tr><td>>hg38_21_+_8433172_8446622_RNA45SN1_45S_50ntFlanks@4223.4242.20|AAAAATAACAATACAGGACT</td><td>>hg38_21_+_8433172_8446622_RNA45SN1_45S_50ntFlanks</td><td>4223</td><td>4242</td><td>20</td><td>AAAAATAACAATACAGGACT</td><td>166.7</td><td>-4.0</td><td>1.1</td><td>-3.6</td><td>2.7E-04</td><td>3.4E-04</td><td>MB231/MB468</td><td>NUC</td></tr>
<tr><td>>hg38_21_+_8433172_8446622_RNA45SN1_45S_50ntFlanks@11586.11609.24|GTGCGGACCAGGGGAATCCGACTG</td><td>>hg38_21_+_8433172_8446622_RNA45SN1_45S_50ntFlanks</td><td>11586</td><td>11609</td><td>24</td><td>GTGCGGACCAGGGGAATCCGACTG</td><td>671.2</td><td>-3.6</td><td>0.9</td><td>-3.9</td><td>1.1E-04</td><td>3.4E-04</td><td>BT20/MB468</td><td>MP</td></tr>
<tr><td>>hg38_21_+_8433172_8446622_RNA45SN1_45S_50ntFlanks@12397.12418.22|CGACGTCGCTTTTTGATCCTTC</td><td>>hg38_21_+_8433172_8446622_RNA45SN1_45S_50ntFlanks</td><td>12397</td><td>12418</td><td>22</td><td>CGACGTCGCTTTTTGATCCTTC</td><td>409.0</td><td>1.9</td><td>0.5</td><td>3.7</td><td>2.0E-04</td><td>3.4E-04</td><td>BT20/MB468</td><td>MITO</td></tr>
<tr><td>>hg38_21_+_8433172_8446622_RNA45SN1_45S_50ntFlanks@10534.10570.37|CCGCGAGGCGTCCAGTGCGGTAACGCGACCGATCCCG</td><td>>hg38_21_+_8433172_8446622_RNA45SN1_45S_50ntFlanks</td><td>10534</td><td>10570</td><td>37</td><td>CCGCGAGGCGTCCAGTGCGGTAACGCGACCGATCCCG</td><td>210.7</td><td>-5.4</td><td>1.5</td><td>-3.7</td><td>2.4E-04</td><td>3.4E-04</td><td>BT20/MB468</td><td>CYTO</td></tr>
<tr><td>>hg38_MT_+_598_1651_RNR1_12S_50ntFlanks@52.83.32|ATAGGTTTGGTCCTAGCCTTTCTATTAGCTCT</td><td>>hg38_MT_+_598_1651_RNR1_12S_50ntFlanks</td><td>52</td><td>83</td><td>32</td><td>ATAGGTTTGGTCCTAGCCTTTCTATTAGCTCT</td><td>92.2</td><td>2.2</td><td>0.6</td><td>3.6</td><td>2.8E-04</td><td>3.5E-04</td><td>BT20/MB468</td><td>NUC</td></tr>
<tr><td>>MI0000456|hsa-mir-140&WithFlank&16|+|69933075|69933186@68.91.24&[MIMAT0004597&hsa-miR-140-3p&offsets|0|+3;m-45&16|+|69933143|69933164&offsets|-1|+1]|TACCACAGGGTAGAACCACGGACA</td><td>>MI0000456|hsa-mir-140&WithFlank&16|+|69933075|69933186</td><td>68</td><td>91</td><td>24</td><td>TACCACAGGGTAGAACCACGGACA</td><td>186.8</td><td>4.3</td><td>1.2</td><td>3.7</td><td>2.3E-04</td><td>3.5E-04</td><td>MB231/MB468</td><td>MP</td></tr>
<tr><td>>am_MI0015825|hsa-mir-378c&WithFlank&10|-|130962582|130962674@17.37.21&[MIMAT0016847&hsa-miR-378c&offsets|0|-4;m-130&10|-|130962639|130962658&offsets|0|+1]|ACTGGACTTGGAGTCAGAAGA</td><td>>am_MI0015825|hsa-mir-378c&WithFlank&10|-|130962582|130962674</td><td>17</td><td>37</td><td>21</td><td>ACTGGACTTGGAGTCAGAAGA</td><td>118.3</td><td>1.6</td><td>0.4</td><td>3.6</td><td>2.8E-04</td><td>3.5E-04</td><td>BT20/MB468</td><td>NUC</td></tr>
<tr><td>>hg38_21_+_8433172_8446622_RNA45SN1_45S_50ntFlanks@10514.10531.18|CGGAGTGGCGGAGATGGG</td><td>>hg38_21_+_8433172_8446622_RNA45SN1_45S_50ntFlanks</td><td>10514</td><td>10531</td><td>18</td><td>CGGAGTGGCGGAGATGGG</td><td>197.4</td><td>11.0</td><td>2.7</td><td>4.1</td><td>3.8E-05</td><td>3.5E-04</td><td>BT20/MB231</td><td>MP</td></tr>
<tr><td>>MI0000114|hsa-mir-107&WithFlank&10|-|89592741|89592833@56.78.23&[MIMAT0000104&hsa-miR-107&offsets|0|0;m-78&10|-|89592758|89592778&offsets|0|+2]|AGCAGCATTGTACAGGGCTATCA</td><td>>MI0000114|hsa-mir-107&WithFlank&10|-|89592741|89592833</td><td>56</td><td>78</td><td>23</td><td>AGCAGCATTGTACAGGGCTATCA</td><td>391.9</td><td>3.7</td><td>1.0</td><td>3.7</td><td>2.5E-04</td><td>3.5E-04</td><td>MB231/MB468</td><td>CYTO</td></tr>
<tr><td>>MI0000264|hsa-mir-7-2&WithFlank&15|+|88611819|88611940@38.58.21(+1U);MI0000265|hsa-mir-7-3&WithFlank&19|+|4770664|4770785@37.57.21(+1U);MI0000263|hsa-mir-7-1&WithFlank&9|-|83969742|83969863@30.50.21(+1U)&[MIMAT0000252&hsa-miR-7-5p&offsets|0|-3(+1U);m-400&15|+|88611856|88611879&offsets|0|-3(+1U)];[MIMAT0000252_1&hsa-miR-7-5p&offsets|0|-3(+1U);m-402&19|+|4770700|4770723&offsets|0|-3(+1U)];[MIMAT0000252_2&hsa-miR-7-5p&offsets|0|-3(+1U);m-401&9|-|83969811|83969834&offsets|0|-3(+1U)]|TGGAAGACTAGTGATTTTGTTT</td><td>>MI0000264|hsa-mir-7-2&WithFlank&15|+|88611819|88611940</td><td>38</td><td>58</td><td>21(+1U)</td><td>TGGAAGACTAGTGATTTTGTTT</td><td>95.0</td><td>6.3</td><td>1.7</td><td>3.7</td><td>2.3E-04</td><td>3.5E-04</td><td>MB231/MB468</td><td>MP</td></tr>
<tr><td>>hg38_21_+_8433172_8446622_RNA45SN1_45S_50ntFlanks@10848.10865.18|TCGGGATAAGGATTGGCT</td><td>>hg38_21_+_8433172_8446622_RNA45SN1_45S_50ntFlanks</td><td>10848</td><td>10865</td><td>18</td><td>TCGGGATAAGGATTGGCT</td><td>87.0</td><td>-5.6</td><td>1.5</td><td>-3.7</td><td>2.3E-04</td><td>3.5E-04</td><td>MB231/MB468</td><td>MP</td></tr>
<tr><td>>hg38_21_+_8433172_8446622_RNA45SN1_45S_50ntFlanks@5489.5522.34|CTGAGAAGACGGTCGAACTTGACTATCTAGAGGA</td><td>>hg38_21_+_8433172_8446622_RNA45SN1_45S_50ntFlanks</td><td>5489</td><td>5522</td><td>34</td><td>CTGAGAAGACGGTCGAACTTGACTATCTAGAGGA</td><td>74.1</td><td>-5.6</td><td>1.5</td><td>-3.7</td><td>2.3E-04</td><td>3.5E-04</td><td>MB231/MB468</td><td>MP</td></tr>
<tr><td>>hg38_21_+_8433172_8446622_RNA45SN1_45S_50ntFlanks@12455.12487.33|ACCAAGCGTTGGATTGTTCACCCACTAATAGGG</td><td>>hg38_21_+_8433172_8446622_RNA45SN1_45S_50ntFlanks</td><td>12455</td><td>12487</td><td>33</td><td>ACCAAGCGTTGGATTGTTCACCCACTAATAGGG</td><td>325.8</td><td>-4.2</td><td>1.1</td><td>-3.6</td><td>2.7E-04</td><td>3.5E-04</td><td>MB231/MB468</td><td>NUC</td></tr>
<tr><td>>hg38_21_+_8433172_8446622_RNA45SN1_45S_50ntFlanks@12456.12497.42|CCAAGCGTTGGATTGTTCACCCACTAATAGGGAACGTGAGCT</td><td>>hg38_21_+_8433172_8446622_RNA45SN1_45S_50ntFlanks</td><td>12456</td><td>12497</td><td>42</td><td>CCAAGCGTTGGATTGTTCACCCACTAATAGGGAACGTGAGCT</td><td>225.0</td><td>-4.5</td><td>1.2</td><td>-3.6</td><td>2.7E-04</td><td>3.5E-04</td><td>MB231/MB468</td><td>MITO</td></tr>
<tr><td>>MI0000301|hsa-mir-224&WithFlank&X|-|151958572|151958664@14.35.22&[MIMAT0000281&hsa-miR-224-5p&offsets|+1|-2;m-238&X|-|151958630|151958651&offsets|0|0]|CAAGTCACTAGTGGTTCCGTTT</td><td>>MI0000301|hsa-mir-224&WithFlank&X|-|151958572|151958664</td><td>14</td><td>35</td><td>22</td><td>CAAGTCACTAGTGGTTCCGTTT</td><td>645.0</td><td>6.0</td><td>1.6</td><td>3.7</td><td>2.3E-04</td><td>3.5E-04</td><td>MB231/MB468</td><td>MP</td></tr>
</table>

<table>
<tr><td>>am_MI0000809|hsa-mir-151a&WithFlank&8|-|140732558|140732659@53.72.20(+1C)&[MIMAT0000757&hsa-miR-151a-3p&offsets|0|-1(+1C);m-30&8|-|140732587|140732607&offsets|0|-1(+1C)]|CTAGACTGAAGCTCCTTGAGC</td><td>>am_MI0000809|hsa-mir-151a&WithFlank&8|-|140732558|140732659</td><td>53</td><td>72</td><td>20(+1C)</td><td>CTAGACTGAAGCTCCTTGAGC</td><td>525.6</td><td>4.6</td><td>1.3</td><td>3.7</td><td>2.3E-04</td><td>3.5E-04</td><td>MB231/MB468</td><td>MP</td></tr>
<tr><td>>hg38_21_+_8433172_8446622_RNA45SN1_45S_50ntFlanks@485.523.39|GATGTGGTGACGTCGTGCTCTCCCGGGCCGGGTCCGAGC</td><td>>hg38_21_+_8433172_8446622_RNA45SN1_45S_50ntFlanks</td><td>485</td><td>523</td><td>39</td><td>GATGTGGTGACGTCGTGCTCTCCCGGGCCGGGTCCGAGC</td><td>637.6</td><td>13.2</td><td>3.6</td><td>3.6</td><td>2.7E-04</td><td>3.5E-04</td><td>MB231/MB468</td><td>NUC</td></tr>
<tr><td>>trnaMT_PheGAA_MT_+_577_647@42.74.33|TTTAGACGGGCTCACATCACCCCATAAACACCA</td><td>>trnaMT_PheGAA_MT_+_577_647</td><td>42</td><td>74</td><td>33</td><td>TTTAGACGGGCTCACATCACCCCATAAACACCA</td><td>1493.3</td><td>5.2</td><td>1.3</td><td>3.9</td><td>1.1E-04</td><td>3.5E-04</td><td>BT20/MB468</td><td>MP</td></tr>
<tr><td>>hg38_21_+_8433172_8446622_RNA45SN1_45S_50ntFlanks@12991.13016.26|CTCCCTCGCTGCGATCTATTGAAAGT</td><td>>hg38_21_+_8433172_8446622_RNA45SN1_45S_50ntFlanks</td><td>12991</td><td>13016</td><td>26</td><td>CTCCCTCGCTGCGATCTATTGAAAGT</td><td>266.6</td><td>-2.0</td><td>0.5</td><td>-3.7</td><td>2.0E-04</td><td>3.5E-04</td><td>BT20/MB468</td><td>MITO</td></tr>
<tr><td>>MI0000447|hsa-mir-128-1&WithFlank&2|+|135665391|135665484@56.74.19;MI0000727|hsa-mir-128-2&WithFlank&3|+|35744470|35744565@58.76.19&[MIMAT0000424&hsa-miR-128-3p&offsets|0|-2;m-108&2|+|135665446|135665466&offsets|0|-2];[MIMAT0000424_1&hsa-miR-128-3p&offsets|0|-2;m-117&3|+|35744527|35744547&offsets|0|-2]|TCACAGTGAACCGGTCTCT</td><td>>MI0000447|hsa-mir-128-1&WithFlank&2|+|135665391|135665484</td><td>56</td><td>74</td><td>19</td><td>TCACAGTGAACCGGTCTCT</td><td>150.1</td><td>3.8</td><td>1.0</td><td>3.6</td><td>2.7E-04</td><td>3.5E-04</td><td>MB231/MB468</td><td>MITO</td></tr>
<tr><td>>hg38_21_+_8433172_8446622_RNA45SN1_45S_50ntFlanks@6651.6696.46|CGACTCTTAGCGGTGGATCACTCGGCTCGTGCGTCGATGAAGAACG</td><td>>hg38_21_+_8433172_8446622_RNA45SN1_45S_50ntFlanks</td><td>6651</td><td>6696</td><td>46</td><td>CGACTCTTAGCGGTGGATCACTCGGCTCGTGCGTCGATGAAGAACG</td><td>28223.6</td><td>3.8</td><td>1.0</td><td>3.6</td><td>2.7E-04</td><td>3.5E-04</td><td>MB231/MB468</td><td>NUC</td></tr>
<tr><td>>hg38_21_+_8433172_8446622_RNA45SN1_45S_50ntFlanks@3993.4023.31|GTGACTCTAGATAACCTCGGGCCGATCGCAC</td><td>>hg38_21_+_8433172_8446622_RNA45SN1_45S_50ntFlanks</td><td>3993</td><td>4023</td><td>31</td><td>GTGACTCTAGATAACCTCGGGCCGATCGCAC</td><td>770.0</td><td>2.7</td><td>0.7</td><td>3.6</td><td>2.7E-04</td><td>3.5E-04</td><td>MB231/MB468</td><td>NUC</td></tr>
<tr><td>>hg38_21_+_8433172_8446622_RNA45SN1_45S_50ntFlanks@11778.11804.27|ACGCGCATGAATGGATGAACGAGATTC</td><td>>hg38_21_+_8433172_8446622_RNA45SN1_45S_50ntFlanks</td><td>11778</td><td>11804</td><td>27</td><td>ACGCGCATGAATGGATGAACGAGATTC</td><td>301.0</td><td>-4.3</td><td>1.2</td><td>-3.6</td><td>2.8E-04</td><td>3.5E-04</td><td>MB231/MB468</td><td>NUC</td></tr>
<tr><td>>hg38_21_+_8433172_8446622_RNA45SN1_45S_50ntFlanks@6661.6692.32|CGGTGGATCACTCGGCTCGTGCGTCGATGAAG</td><td>>hg38_21_+_8433172_8446622_RNA45SN1_45S_50ntFlanks</td><td>6661</td><td>6692</td><td>32</td><td>CGGTGGATCACTCGGCTCGTGCGTCGATGAAG</td><td>1310.0</td><td>2.1</td><td>0.6</td><td>3.7</td><td>2.0E-04</td><td>3.5E-04</td><td>BT20/MB468</td><td>MITO</td></tr>
<tr><td>>hg38_21_+_8433172_8446622_RNA45SN1_45S_50ntFlanks@5379.5413.35|ATTAAGTCCCTGCCCTTTGTACACACCGCCCGTCG</td><td>>hg38_21_+_8433172_8446622_RNA45SN1_45S_50ntFlanks</td><td>5379</td><td>5413</td><td>35</td><td>ATTAAGTCCCTGCCCTTTGTACACACCGCCCGTCG</td><td>188.1</td><td>-5.6</td><td>1.5</td><td>-3.7</td><td>2.0E-04</td><td>3.5E-04</td><td>BT20/MB468</td><td>MITO</td></tr>
<tr><td>>trna111_HisGTG_1_-_147774845_147774916@-1T.25.26,trna118_HisGTG_1_-_145396881_145396952@-1T.25.26,trna16_HisGTG_1_+_146544773_146544844@-1T.25.26,trna1_HisGTG_15_+_45493349_45493420@-1T.25.26,trna21_HisGTG_1_+_147753471_147753542@-1T.25.26,trna33_HisGTG_6_+_27125906_27125977@-1T.25.26,trna7_HisGTG_9_-_14433938_14434009@-1T.25.26,trna8_HisGTG_15_-_45492611_45492682@-1T.25.26,trna9_HisGTG_15_-_45490804_45490875@-1T.25.26|TGCCGTGATCGTATAGTGGTTAGTAC</td><td>>trna111_HisGTG_1_-_147774845_147774916</td><td>-1T</td><td>25</td><td>26</td><td>TGCCGTGATCGTATAGTGGTTAGTAC</td><td>393.9</td><td>3.2</td><td>0.9</td><td>3.6</td><td>2.7E-04</td><td>3.5E-04</td><td>MB231/MB468</td><td>MITO</td></tr>
<tr><td>>MI0003820|hsa-mir-454&WithFlank&17|-|59137752|59137878@70.92.23&[MIMAT0003885&hsa-miR-454-3p&offsets|0|0;m-272&17|-|59137787|59137809&offsets|0|0]|TAGTGCAATATTGCTTATAGGGT</td><td>>MI0003820|hsa-mir-454&WithFlank&17|-|59137752|59137878</td><td>70</td><td>92</td><td>23</td><td>TAGTGCAATATTGCTTATAGGGT</td><td>479.0</td><td>3.2</td><td>0.9</td><td>3.7</td><td>2.3E-04</td><td>3.5E-04</td><td>MB231/MB468</td><td>MP</td></tr>
<tr><td>>hg38_21_+_8433172_8446622_RNA45SN1_45S_50ntFlanks@4971.5002.32|CCGGCCCGGACACGGACAGGATTGACAGATTG</td><td>>hg38_21_+_8433172_8446622_RNA45SN1_45S_50ntFlanks</td><td>4971</td><td>5002</td><td>32</td><td>CCGGCCCGGACACGGACAGGATTGACAGATTG</td><td>84.7</td><td>-5.9</td><td>1.6</td><td>-3.7</td><td>2.3E-04</td><td>3.5E-04</td><td>MB231/MB468</td><td>MP</td></tr>
<tr><td>>hg38_21_+_8433172_8446622_RNA45SN1_45S_50ntFlanks@3992.4026.35|GGTGACTCTAGATAACCTCGGGCCGATCGCACGCC</td><td>>hg38_21_+_8433172_8446622_RNA45SN1_45S_50ntFlanks</td><td>3992</td><td>4026</td><td>35</td><td>GGTGACTCTAGATAACCTCGGGCCGATCGCACGCC</td><td>201.2</td><td>-2.2</td><td>0.6</td><td>-3.7</td><td>2.0E-04</td><td>3.5E-04</td><td>BT20/MB468</td><td>MITO</td></tr>
<tr><td>>hg38_21_+_8433172_8446622_RNA45SN1_45S_50ntFlanks@9564.9603.40|CTGACGTGCAAATCGGTCGTCCGACCTGGGTATAGGGGCG</td><td>>hg38_21_+_8433172_8446622_RNA45SN1_45S_50ntFlanks</td><td>9564</td><td>9603</td><td>40</td><td>CTGACGTGCAAATCGGTCGTCCGACCTGGGTATAGGGGCG</td><td>2362.9</td><td>-3.8</td><td>1.1</td><td>-3.6</td><td>2.7E-04</td><td>3.5E-04</td><td>MB231/MB468</td><td>MITO</td></tr>
<tr><td>>am_trnaMT_MetCAT_MT_+_4402_4469@1.22.22|AGTAAGGTCAGCTAAATAAGCT</td><td>>am_trnaMT_MetCAT_MT_+_4402_4469</td><td>1</td><td>22</td><td>22</td><td>AGTAAGGTCAGCTAAATAAGCT</td><td>810.9</td><td>4.5</td><td>1.2</td><td>3.7</td><td>2.0E-04</td><td>3.5E-04</td><td>BT20/MB468</td><td>MITO</td></tr>
<tr><td>>hg38_21_+_8433172_8446622_RNA45SN1_45S_50ntFlanks@6664.6696.33|TGGATCACTCGGCTCGTGCGTCGATGAAGAACG</td><td>>hg38_21_+_8433172_8446622_RNA45SN1_45S_50ntFlanks</td><td>6664</td><td>6696</td><td>33</td><td>TGGATCACTCGGCTCGTGCGTCGATGAAGAACG</td><td>140.6</td><td>-5.5</td><td>1.5</td><td>-3.7</td><td>2.3E-04</td><td>3.5E-04</td><td>MB231/MB468</td><td>MP</td></tr>
<tr><td>>hg38_21_+_8433172_8446622_RNA45SN1_45S_50ntFlanks@6654.6681.28|CTCTTAGCGGTGGATCACTCGGCTCGTG</td><td>>hg38_21_+_8433172_8446622_RNA45SN1_45S_50ntFlanks</td><td>6654</td><td>6681</td><td>28</td><td>CTCTTAGCGGTGGATCACTCGGCTCGTG</td><td>1534.2</td><td>-3.4</td><td>0.9</td><td>-3.9</td><td>1.1E-04</td><td>3.5E-04</td><td>BT20/MB468</td><td>MP</td></tr>
<tr><td>>MI0000299|hsa-mir-222&WithFlank&X|-|45747009|45747130@75.96.22(+1A)&[MIMAT0000279&hsa-miR-222-3p&offsets|0|+1(+1A);m-140&X|-|45747033|45747056&offsets|0|-2(+1A)]|AGCTACATCTGGCTACTGGGTCA</td><td>>MI0000299|hsa-mir-222&WithFlank&X|-|45747009|45747130</td><td>75</td><td>96</td><td>22(+1A)</td><td>AGCTACATCTGGCTACTGGGTCA</td><td>1043.2</td><td>5.1</td><td>1.4</td><td>3.7</td><td>2.3E-04</td><td>3.5E-04</td><td>MB231/MB468</td><td>MP</td></tr>
<tr><td>>trnaMT_GluTTC_MT_-_14674_14742@18.57.40,trnalookalike8_GluTTC_5_-_93905172_93905240@18.57.40|ACAACGATGGTTTTTCATATCATTGGTCGTGGTTGTAGTC</td><td>>trnaMT_GluTTC_MT_-_14674_14742</td><td>18</td><td>57</td><td>40</td><td>ACAACGATGGTTTTTCATATCATTGGTCGTGGTTGTAGTC</td><td>92.7</td><td>10.3</td><td>2.7</td><td>3.9</td><td>1.1E-04</td><td>3.5E-04</td><td>BT20/MB468</td><td>MP</td></tr>
<tr><td>>hg38_21_+_8433172_8446622_RNA45SN1_45S_50ntFlanks@6645.6675.31|CTCGTACGACTCTTAGCGGTGGATCACTCGG</td><td>>hg38_21_+_8433172_8446622_RNA45SN1_45S_50ntFlanks</td><td>6645</td><td>6675</td><td>31</td><td>CTCGTACGACTCTTAGCGGTGGATCACTCGG</td><td>382.7</td><td>2.6</td><td>0.7</td><td>3.7</td><td>2.6E-04</td><td>3.5E-04</td><td>MB231/MB468</td><td>CYTO</td></tr>
<tr><td>>hg38_21_+_8433172_8446622_RNA45SN1_45S_50ntFlanks@11316.11336.21|CACCCCACGTCTCGTCGCGCG</td><td>>hg38_21_+_8433172_8446622_RNA45SN1_45S_50ntFlanks</td><td>11316</td><td>11336</td><td>21</td><td>CACCCCACGTCTCGTCGCGCG</td><td>6440.5</td><td>-7.7</td><td>2.1</td><td>-3.6</td><td>2.8E-04</td><td>3.5E-04</td><td>BT20/MB468</td><td>NUC</td></tr>
<tr><td>>trna10_LysCTT_16_+_3241501_3241573@1.33.33,trna13_LysCTT_14_-_58706613_58706685@1.33.33,trna2_LysCTT_15_+_79152904_79152976@1.33.33|GCCCGGCTAGCTCAGTCGGTAGAGCATGGGACT</td><td>>trna10_LysCTT_16_+_3241501_3241573</td><td>1</td><td>33</td><td>33</td><td>GCCCGGCTAGCTCAGTCGGTAGAGCATGGGACT</td><td>327.1</td><td>4.6</td><td>1.3</td><td>3.7</td><td>2.3E-04</td><td>3.5E-04</td><td>MB231/MB468</td><td>MP</td></tr>
<tr><td>>hg38_21_+_8433172_8446622_RNA45SN1_45S_50ntFlanks@10686.10710.25|GCGGCGTCCGGTGAGCTCTCGCTGG</td><td>>hg38_21_+_8433172_8446622_RNA45SN1_45S_50ntFlanks</td><td>10686</td><td>10710</td><td>25</td><td>GCGGCGTCCGGTGAGCTCTCGCTGG</td><td>7228.7</td><td>-6.3</td><td>1.7</td><td>-3.6</td><td>2.8E-04</td><td>3.5E-04</td><td>BT20/MB468</td><td>NUC</td></tr>
<tr><td>>MI0000471|hsa-mir-126&WithFlank&9|+|136670596|136670692@21.41.21&[MIMAT0000444&hsa-miR-126-5p&offsets|0|0;m-114&9|+|136670616|136670636&offsets|0|0]|CATTATTACTTTTGGTACGCG</td><td>>MI0000471|hsa-mir-126&WithFlank&9|+|136670596|136670692</td><td>21</td><td>41</td><td>21</td><td>CATTATTACTTTTGGTACGCG</td><td>93.1</td><td>7.4</td><td>1.9</td><td>3.9</td><td>1.1E-04</td><td>3.5E-04</td><td>BT20/MB468</td><td>MP</td></tr>
<tr><td>>am_MI0003583|hsa-mir-576&WithFlank&4|+|109488692|109488801@61.82.22(+1U)&[MIMAT0004796&hsa-miR-576-3p&offsets|0|0(+1U);m-524&4|+|109488752|109488772&offsets|0|+1(+1U)]|AAGATGTGGAAAAATTGGAATCT</td><td>>am_MI0003583|hsa-mir-576&WithFlank&4|+|109488692|109488801</td><td>61</td><td>82</td><td>22(+1U)</td><td>AAGATGTGGAAAAATTGGAATCT</td><td>381.3</td><td>5.8</td><td>1.5</td><td>3.9</td><td>1.1E-04</td><td>3.5E-04</td><td>BT20/MB468</td><td>MP</td></tr>
<tr><td>>MI0000456|hsa-mir-140&WithFlank&16|+|69933075|69933186@69.90.22&[MIMAT0004597&hsa-miR-140-3p&offsets|+1|+2;m-45&16|+|69933143|69933164&offsets|0|0]|ACCACAGGGTAGAACCACGGAC</td><td>>MI0000456|hsa-mir-140&WithFlank&16|+|69933075|69933186</td><td>69</td><td>90</td><td>22</td><td>ACCACAGGGTAGAACCACGGAC</td><td>1153.9</td><td>6.0</td><td>1.6</td><td>3.9</td><td>1.1E-04</td><td>3.5E-04</td><td>BT20/MB468</td><td>MP</td></tr>
<tr><td>>hg38_21_+_8433172_8446622_RNA45SN1_45S_50ntFlanks@10731.10752.22|AGGGTGTAAATCTCGCGCCGGG</td><td>>hg38_21_+_8433172_8446622_RNA45SN1_45S_50ntFlanks</td><td>10731</td><td>10752</td><td>22</td><td>AGGGTGTAAATCTCGCGCCGGG</td><td>492.6</td><td>-5.0</td><td>1.3</td><td>-3.9</td><td>1.1E-04</td><td>3.5E-04</td><td>BT20/MB468</td><td>MP</td></tr>
<tr><td>>hg38_1_-_228634819_228635039_RNA5S12_5S_50ntFlanks@142.171.30|CGCCTGGGAATACCGGGTGCTGTAGGCTTT</td><td>>hg38_1_-_228634819_228635039_RNA5S12_5S_50ntFlanks</td><td>142</td><td>171</td><td>30</td><td>CGCCTGGGAATACCGGGTGCTGTAGGCTTT</td><td>1856.7</td><td>-2.5</td><td>0.7</td><td>-3.7</td><td>2.3E-04</td><td>3.5E-04</td><td>MB231/MB468</td><td>MP</td></tr>
<tr><td>>hg38_21_+_8433172_8446622_RNA45SN1_45S_50ntFlanks@12423.12455.33|TCGGCTCTTCCTATCATTGTGAAGCAGAATTCA</td><td>>hg38_21_+_8433172_8446622_RNA45SN1_45S_50ntFlanks</td><td>12423</td><td>12455</td><td>33</td><td>TCGGCTCTTCCTATCATTGTGAAGCAGAATTCA</td><td>183.8</td><td>-3.4</td><td>0.9</td><td>-3.7</td><td>2.6E-04</td><td>3.5E-04</td><td>MB231/MB468</td><td>CYTO</td></tr>
<tr><td>>hg38_21_+_8433172_8446622_RNA45SN1_45S_50ntFlanks@12159.12178.20|TTTGACTGGGGCGGTACACC</td><td>>hg38_21_+_8433172_8446622_RNA45SN1_45S_50ntFlanks</td><td>12159</td><td>12178</td><td>20</td><td>TTTGACTGGGGCGGTACACC</td><td>331.1</td><td>-4.5</td><td>1.2</td><td>-3.7</td><td>2.5E-04</td><td>3.5E-04</td><td>BT20/MB468</td><td>CYTO</td></tr>
<tr><td>>hg38_21_+_8433172_8446622_RNA45SN1_45S_50ntFlanks@4899.4918.20|AAGGAATTGACGGAAGGGCA</td><td>>hg38_21_+_8433172_8446622_RNA45SN1_45S_50ntFlanks</td><td>4899</td><td>4918</td><td>20</td><td>AAGGAATTGACGGAAGGGCA</td><td>154.0</td><td>-6.3</td><td>1.6</td><td>-3.9</td><td>1.1E-04</td><td>3.5E-04</td><td>BT20/MB468</td><td>MP</td></tr>
<tr><td>>hg38_21_+_8433172_8446622_RNA45SN1_45S_50ntFlanks@10868.10901.34|AAGGGCTGGGTCGGTCGGGCTGGGGCGCGAAGCG</td><td>>hg38_21_+_8433172_8446622_RNA45SN1_45S_50ntFlanks</td><td>10868</td><td>10901</td><td>34</td><td>AAGGGCTGGGTCGGTCGGGCTGGGGCGCGAAGCG</td><td>92.8</td><td>-5.6</td><td>1.5</td><td>-3.7</td><td>2.3E-04</td><td>3.5E-04</td><td>MB231/MB468</td><td>MP</td></tr>
<tr><td>>hg38_21_+_8433172_8446622_RNA45SN1_45S_50ntFlanks@11669.11703.35|GCCCAGTGCTCTGAATGTCAAAGTGAAGAAATTCA</td><td>>hg38_21_+_8433172_8446622_RNA45SN1_45S_50ntFlanks</td><td>11669</td><td>11703</td><td>35</td><td>GCCCAGTGCTCTGAATGTCAAAGTGAAGAAATTCA</td><td>338.2</td><td>2.5</td><td>0.7</td><td>3.6</td><td>2.8E-04</td><td>3.5E-04</td><td>MB231/MB468</td><td>NUC</td></tr>
<tr><td>>hg38_21_+_8433172_8446622_RNA45SN1_45S_50ntFlanks@10794.10823.30|CTGGCATGTTGGAACAATGTAGGTAAGGGA</td><td>>hg38_21_+_8433172_8446622_RNA45SN1_45S_50ntFlanks</td><td>10794</td><td>10823</td><td>30</td><td>CTGGCATGTTGGAACAATGTAGGTAAGGGA</td><td>168.5</td><td>-3.6</td><td>1.0</td><td>-3.7</td><td>2.5E-04</td><td>3.5E-04</td><td>BT20/MB468</td><td>CYTO</td></tr>
<tr><td>>MI0000077|hsa-mir-21&WithFlank&17|+|59841260|59841343@14.39.26(+1U)&[MIMAT0000076&hsa-miR-21-5p&offsets|0|+4(+1U);m-1&17|+|59841273|59841294&offsets|0|+4(+1U)]|TAGCTTATCAGACTGATGTTGACTGTT</td><td>>MI0000077|hsa-mir-21&WithFlank&17|+|59841260|59841343</td><td>14</td><td>39</td><td>26(+1U)</td><td>TAGCTTATCAGACTGATGTTGACTGTT</td><td>174.5</td><td>2.2</td><td>0.6</td><td>3.6</td><td>2.8E-04</td><td>3.5E-04</td><td>BT20/MB468</td><td>NUC</td></tr>
<tr><td>>hg38_21_+_8433172_8446622_RNA45SN1_45S_50ntFlanks@10680.10709.30|GTTCCGGCGGCGTCCGGTGAGCTCTCGCTG</td><td>>hg38_21_+_8433172_8446622_RNA45SN1_45S_50ntFlanks</td><td>10680</td><td>10709</td><td>30</td><td>GTTCCGGCGGCGTCCGGTGAGCTCTCGCTG</td><td>6364.2</td><td>-7.7</td><td>2.1</td><td>-3.6</td><td>2.8E-04</td><td>3.5E-04</td><td>BT20/MB468</td><td>NUC</td></tr>
<tr><td>>hg38_21_+_8433172_8446622_RNA45SN1_45S_50ntFlanks@12072.12096.25|CCCGAGGGGCTCTCGCTTCTGGCGC</td><td>>hg38_21_+_8433172_8446622_RNA45SN1_45S_50ntFlanks</td><td>12072</td><td>12096</td><td>25</td><td>CCCGAGGGGCTCTCGCTTCTGGCGC</td><td>706.4</td><td>-4.5</td><td>1.2</td><td>-3.7</td><td>2.5E-04</td><td>3.6E-04</td><td>BT20/MB468</td><td>CYTO</td></tr>
<tr><td>>hg38_21_+_8433172_8446622_RNA45SN1_45S_50ntFlanks@6741.6770.30|ATCGACACTTCGAACGCACTTGCGGCCCCG</td><td>>hg38_21_+_8433172_8446622_RNA45SN1_45S_50ntFlanks</td><td>6741</td><td>6770</td><td>30</td><td>ATCGACACTTCGAACGCACTTGCGGCCCCG</td><td>1474.4</td><td>-6.0</td><td>1.6</td><td>-3.7</td><td>2.5E-04</td><td>3.6E-04</td><td>BT20/MB468</td><td>CYTO</td></tr>
<tr><td>>hg38_21_+_8433172_8446622_RNA45SN1_45S_50ntFlanks@8170.8202.33|CAAGTCCTTCTGATCGAGGCCCAGCCCGTGGAC</td><td>>hg38_21_+_8433172_8446622_RNA45SN1_45S_50ntFlanks</td><td>8170</td><td>8202</td><td>33</td><td>CAAGTCCTTCTGATCGAGGCCCAGCCCGTGGAC</td><td>101.1</td><td>-5.7</td><td>1.5</td><td>-3.7</td><td>2.3E-04</td><td>3.6E-04</td><td>MB231/MB468</td><td>MP</td></tr>
<tr><td>>am_MI0000786|hsa-mir-378a&WithFlank&5|+|149732819|149732896@50.70.21&[MIMAT0000732&hsa-miR-378a-3p&offsets|+1|0;m-44&5|+|149732867|149732888&offsets|+1|0]|CTGGACTTGGAGTCAGAAGGC</td><td>>am_MI0000786|hsa-mir-378a&WithFlank&5|+|149732819|149732896</td><td>50</td><td>70</td><td>21</td><td>CTGGACTTGGAGTCAGAAGGC</td><td>915.0</td><td>2.1</td><td>0.6</td><td>3.6</td><td>2.8E-04</td><td>3.6E-04</td><td>MB231/MB468</td><td>NUC</td></tr>
<tr><td>>hg38_21_+_8433172_8446622_RNA45SN1_45S_50ntFlanks@12412.12436.25|ATCCTTCGATGTCGGCTCTTCCTAT</td><td>>hg38_21_+_8433172_8446622_RNA45SN1_45S_50ntFlanks</td><td>12412</td><td>12436</td><td>25</td><td>ATCCTTCGATGTCGGCTCTTCCTAT</td><td>478.5</td><td>-5.2</td><td>1.4</td><td>-3.7</td><td>2.6E-04</td><td>3.6E-04</td><td>MB231/MB468</td><td>CYTO</td></tr>
<tr><td>>hg38_21_+_8433172_8446622_RNA45SN1_45S_50ntFlanks@10869.10890.22|AGGGCTGGGTCGGTCGGGCTGG</td><td>>hg38_21_+_8433172_8446622_RNA45SN1_45S_50ntFlanks</td><td>10869</td><td>10890</td><td>22</td><td>AGGGCTGGGTCGGTCGGGCTGG</td><td>2236.5</td><td>-7.3</td><td>2.0</td><td>-3.6</td><td>2.8E-04</td><td>3.6E-04</td><td>BT20/MB468</td><td>NUC</td></tr>
<tr><td>>hg38_21_+_8433172_8446622_RNA45SN1_45S_50ntFlanks@11589.11612.24|CGGACCAGGGGAATCCGACTGTTT</td><td>>hg38_21_+_8433172_8446622_RNA45SN1_45S_50ntFlanks</td><td>11589</td><td>11612</td><td>24</td><td>CGGACCAGGGGAATCCGACTGTTT</td><td>527.5</td><td>3.8</td><td>0.9</td><td>4.1</td><td>4.0E-05</td><td>3.6E-04</td><td>BT20/MB231</td><td>NUC</td></tr>
</table>

| | | | | | | | | | | | | | |
|---|---|---|---|---|---|---|---|---|---|---|---|---|---|
| >hg38_21_+_8433172_8446622_RNA45SN1_45S_50ntFlanks@6651.6700.50\|CGACTCTTAGCGGTGGATCACTCGGCTCGTGCGTCGATGAAGAACGCAGC | >hg38_21_+_8433172_8446622_RNA45SN1_45S_50ntFlanks | 6651 | 6700 | 50 | CGACTCTTAGCGGTGGATCACTCGGCTCGTGCGTCGATGAAGAACGCAGC | 1208.7 | 5.4 | 1.5 | 3.7 | 2.5E-04 | 3.6E-04 | BT20/MB468 | CYTO |
| >hg38_21_+_8433172_8446622_RNA45SN1_45S_50ntFlanks@9534.9559.26\|AGGAAACTCTGGTGGAGGTCCGTAGC | >hg38_21_+_8433172_8446622_RNA45SN1_45S_50ntFlanks | 9534 | 9559 | 26 | AGGAAACTCTGGTGGAGGTCCGTAGC | 244.7 | -3.5 | 0.9 | -3.6 | 2.6E-04 | 3.6E-04 | MB231/MB468 | CYTO |
| >hg38_1_-_228634819_228635039_RNA5S12_5S_50ntFlanks@147.170.24\|GGGAATACCGGGTGCTGTAGGCTT | >hg38_1_-_228634819_228635039_RNA5S12_5S_50ntFlanks | 147 | 170 | 24 | GGGAATACCGGGTGCTGTAGGCTT | 1818.5 | 2.5 | 0.7 | 3.6 | 2.8E-04 | 3.6E-04 | MB231/MB468 | NUC |
| >hg38_21_+_8433172_8446622_RNA45SN1_45S_50ntFlanks@10393.10432.40\|CAGCAGTTGAACATGGGTCAGTCGGTCCTGAGAGATGGGC | >hg38_21_+_8433172_8446622_RNA45SN1_45S_50ntFlanks | 10393 | 10432 | 40 | CAGCAGTTGAACATGGGTCAGTCGGTCCTGAGAGATGGGC | 200.4 | -4.4 | 1.2 | -3.6 | 2.8E-04 | 3.6E-04 | MB231/MB468 | NUC |
| >hg38_21_+_8433172_8446622_RNA45SN1_45S_50ntFlanks@12539.12563.25\|ATGATGTGTTGTTGCCATGGTAATC | >hg38_21_+_8433172_8446622_RNA45SN1_45S_50ntFlanks | 12539 | 12563 | 25 | ATGATGTGTTGTTGCCATGGTAATC | 354.3 | -2.4 | 0.6 | -3.7 | 2.1E-04 | 3.6E-04 | BT20/MB468 | MITO |
| >hg38_21_+_8433172_8446622_RNA45SN1_45S_50ntFlanks@6650.6669.20\|ACGACTCTTAGCGGTGGATC | >hg38_21_+_8433172_8446622_RNA45SN1_45S_50ntFlanks | 6650 | 6669 | 20 | ACGACTCTTAGCGGTGGATC | 370.0 | 1.8 | 0.5 | 3.6 | 2.8E-04 | 3.6E-04 | MB231/MB468 | NUC |
| >hg38_21_+_8433172_8446622_RNA45SN1_45S_50ntFlanks@6648.6689.42\|GTACGACTCTTAGCGGTGGATCACTCGGCTCGTGCGTCGATG | >hg38_21_+_8433172_8446622_RNA45SN1_45S_50ntFlanks | 6648 | 6689 | 42 | GTACGACTCTTAGCGGTGGATCACTCGGCTCGTGCGTCGATG | 2262.0 | -3.8 | 1.0 | -3.7 | 2.4E-04 | 3.6E-04 | MB231/MB468 | MP |
| >hg38_21_+_8433172_8446622_RNA45SN1_45S_50ntFlanks@10687.10706.20\|CGGCGTCCGGTGAGCTCTCG | >hg38_21_+_8433172_8446622_RNA45SN1_45S_50ntFlanks | 10687 | 10706 | 20 | CGGCGTCCGGTGAGCTCTCG | 4081.5 | -4.8 | 1.3 | -3.7 | 2.5E-04 | 3.6E-04 | BT20/MB468 | CYTO |
| >hg38_21_+_8433172_8446622_RNA45SN1_45S_50ntFlanks@10683.10712.30\|CCGGCGGCGTCCGGTGAGCTCTCGCTGGCC | >hg38_21_+_8433172_8446622_RNA45SN1_45S_50ntFlanks | 10683 | 10712 | 30 | CCGGCGGCGTCCGGTGAGCTCTCGCTGGCC | 834.7 | -5.2 | 1.3 | -3.9 | 1.1E-04 | 3.6E-04 | BT20/MB468 | MP |
| >hg38_21_+_8433172_8446622_RNA45SN1_45S_50ntFlanks@12081.12110.30\|CTCTCGCTTCTGGCGCCAAGCGCCCGGCCG | >hg38_21_+_8433172_8446622_RNA45SN1_45S_50ntFlanks | 12081 | 12110 | 30 | CTCTCGCTTCTGGCGCCAAGCGCCCGGCCG | 590.3 | -6.3 | 1.6 | -3.9 | 1.1E-04 | 3.6E-04 | BT20/MB468 | MP |
| >hg38_21_+_8433172_8446622_RNA45SN1_45S_50ntFlanks@8382.8402.21\|AGAGGGCGTGAAACCGTTAAG | >hg38_21_+_8433172_8446622_RNA45SN1_45S_50ntFlanks | 8382 | 8402 | 21 | AGAGGGCGTGAAACCGTTAAG | 526.8 | -3.4 | 0.9 | -3.9 | 1.1E-04 | 3.6E-04 | BT20/MB468 | MP |
| >hg38_21_+_8433172_8446622_RNA45SN1_45S_50ntFlanks@11604.11626.23\|CGACTGTTTAATTAAAACAAAGC | >hg38_21_+_8433172_8446622_RNA45SN1_45S_50ntFlanks | 11604 | 11626 | 23 | CGACTGTTTAATTAAAACAAAGC | 170.0 | 3.6 | 0.9 | 3.9 | 1.1E-04 | 3.6E-04 | BT20/MB468 | MP |
| >MI0003820\|hsa-mir-454&WithFlank&17\|-\|59137752\|59137878@70.93.24&[MIMAT0003885&hsa-miR-454-3p&offsets\|0\|+1;m-272&17\|-\|59137787\|59137809&offsets\|0\|+1]\|TAGTGCAATATTGCTTATAGGGTT | >MI0003820\|hsa-mir-454&WithFlank&17\|-\|59137752\|59137878 | 70 | 93 | 24 | TAGTGCAATATTGCTTATAGGGTT | 884.5 | -3.4 | 0.8 | -4.1 | 4.5E-05 | 3.6E-04 | BT20/MB231 | MITO |
| >hg38_21_+_8433172_8446622_RNA45SN1_45S_50ntFlanks@12933.12956.24\|GCTAAACCATTCGTAGACGACCTG | >hg38_21_+_8433172_8446622_RNA45SN1_45S_50ntFlanks | 12933 | 12956 | 24 | GCTAAACCATTCGTAGACGACCTG | 428.6 | -2.0 | 0.5 | -3.7 | 2.1E-04 | 3.6E-04 | BT20/MB468 | MITO |
| >hg38_21_+_8433172_8446622_RNA45SN1_45S_50ntFlanks@11217.11250.34\|CTGGACGCGAGCCGGGCCCTTCCCGTGGATCGCC | >hg38_21_+_8433172_8446622_RNA45SN1_45S_50ntFlanks | 11217 | 11250 | 34 | CTGGACGCGAGCCGGGCCCTTCCCGTGGATCGCC | 366.3 | -4.6 | 1.3 | -3.7 | 2.5E-04 | 3.6E-04 | BT20/MB468 | CYTO |
| >hg38_21_+_8433172_8446622_RNA45SN1_45S_50ntFlanks@3874.3897.24\|AATACATGCCGACGGGCGCTGACC | >hg38_21_+_8433172_8446622_RNA45SN1_45S_50ntFlanks | 3874 | 3897 | 24 | AATACATGCCGACGGGCGCTGACC | 491.2 | -5.0 | 1.4 | -3.6 | 2.9E-04 | 3.6E-04 | BT20/MB468 | NUC |
| >hg38_21_+_8433172_8446622_RNA45SN1_45S_50ntFlanks@12289.12319.31\|CCGTGAAAGCGGGGCCTCACGATCCTTCTGA | >hg38_21_+_8433172_8446622_RNA45SN1_45S_50ntFlanks | 12289 | 12319 | 31 | CCGTGAAAGCGGGGCCTCACGATCCTTCTGA | 837.9 | -3.8 | 1.0 | -3.7 | 2.1E-04 | 3.6E-04 | BT20/MB468 | MITO |
| >hg38_21_+_8433172_8446622_RNA45SN1_45S_50ntFlanks@4567.4606.40\|TAATGGAATAGGACCGCGGTTCTATTTTGTTGGTTTTCGG | >hg38_21_+_8433172_8446622_RNA45SN1_45S_50ntFlanks | 4567 | 4606 | 40 | TAATGGAATAGGACCGCGGTTCTATTTTGTTGGTTTTCGG | 87.2 | 10.2 | 2.6 | 3.9 | 1.1E-04 | 3.6E-04 | BT20/MB468 | MP |
| >hg38_21_+_8433172_8446622_RNA45SN1_45S_50ntFlanks@10798.10826.29\|CATGTTGGAACAATGTAGGTAAGGGAAGT | >hg38_21_+_8433172_8446622_RNA45SN1_45S_50ntFlanks | 10798 | 10826 | 29 | CATGTTGGAACAATGTAGGTAAGGGAAGT | 194.7 | -2.2 | 0.6 | -3.6 | 2.7E-04 | 3.6E-04 | MB231/MB468 | CYTO |
| >hg38_21_+_8433172_8446622_RNA45SN1_45S_50ntFlanks@9947.9968.22\|GTGGCCATGGAAGTCGGAATCC | >hg38_21_+_8433172_8446622_RNA45SN1_45S_50ntFlanks | 9947 | 9968 | 22 | GTGGCCATGGAAGTCGGAATCC | 177.7 | 7.4 | 1.9 | 3.9 | 1.1E-04 | 3.6E-04 | BT20/MB468 | MP |
| >hg38_21_+_8433172_8446622_RNA45SN1_45S_50ntFlanks@10350.10391.42\|ACGAGAACTTTGAAGGCCGAAGTGGAGAAGGGTTCCATGTGA | >hg38_21_+_8433172_8446622_RNA45SN1_45S_50ntFlanks | 10350 | 10391 | 42 | ACGAGAACTTTGAAGGCCGAAGTGGAGAAGGGTTCCATGTGA | 652.3 | -4.6 | 1.3 | -3.6 | 2.8E-04 | 3.6E-04 | MB231/MB468 | MITO |
| >MI0000253\|hsa-mir-148a&WithFlank&7\|-\|25949913\|25949992@12.31.20&[MIMAT0004549&hsa-miR-148a-5p&offsets\|0\|-2;m-189&7\|-\|25949960\|25949981&offsets\|0\|-2]\|AAAGTTCTGAGACACTCCGA | >MI0000253\|hsa-mir-148a&WithFlank&7\|-\|25949913\|25949992 | 12 | 31 | 20 | AAAGTTCTGAGACACTCCGA | 114.5 | 7.2 | 1.9 | 3.9 | 1.1E-04 | 3.6E-04 | BT20/MB468 | MP |
| >hg38_21_+_8433172_8446622_RNA45SN1_45S_50ntFlanks@12406.12436.31\|TTTTTGATCCTTCGATGTCGGCTCTTCCTAT | >hg38_21_+_8433172_8446622_RNA45SN1_45S_50ntFlanks | 12406 | 12436 | 31 | TTTTTGATCCTTCGATGTCGGCTCTTCCTAT | 282.4 | -4.1 | 1.1 | -3.7 | 2.5E-04 | 3.6E-04 | BT20/MB468 | CYTO |
| >hg38_21_+_8433172_8446622_RNA45SN1_45S_50ntFlanks@6646.6663.18\|TCGTACGACTCTTAGCGG | >hg38_21_+_8433172_8446622_RNA45SN1_45S_50ntFlanks | 6646 | 6663 | 18 | TCGTACGACTCTTAGCGG | 15397.5 | 2.6 | 0.7 | 3.6 | 2.7E-04 | 3.6E-04 | MB231/MB468 | CYTO |
| >trnaMT_GluTTC_MT_-_14674_14742@38.61.24,trnalookalike8_GluTTC_5_-_93905172_93905240@38.61.24\|CATTGGTCGTGGTTGTAGTCCGTG | >trnaMT_GluTTC_MT_-_14674_14742 | 38 | 61 | 24 | CATTGGTCGTGGTTGTAGTCCGTG | 131.7 | 3.4 | 0.9 | 3.6 | 2.8E-04 | 3.6E-04 | MB231/MB468 | MITO |
| >hg38_21_+_8433172_8446622_RNA45SN1_45S_50ntFlanks@11593.11613.21\|CCAGGGGAATCCGACTGTTTA | >hg38_21_+_8433172_8446622_RNA45SN1_45S_50ntFlanks | 11593 | 11613 | 21 | CCAGGGGAATCCGACTGTTTA | 577.3 | -2.3 | 0.6 | -3.7 | 2.1E-04 | 3.6E-04 | BT20/MB468 | MITO |
| >MI0000446\|hsa-mir-125b-1&WithFlank&11\|-\|122099751\|122099850@21.42.22(+2A);MI0000470\|hsa-mir-125b-2&WithFlank&21\|+\|16590231\|16590331@23.44.22(+2A)&[MIMAT0000423&hsa-miR-125b-5p&offsets\|0\|0(+2A);m-67&11\|-\|122099809\|122099830&offsets\|0\|0(+2A)];[MIMAT0000423_1&hsa-miR-125b-5p&offsets\|0\|0(+2A);m-68&21\|+\|16590253\|16590274&offsets\|0\|0(+2A)]\|TCCCTGAGACCCTAACTTGTGAAA | >MI0000446\|hsa-mir-125b-1&WithFlank&11\|-\|122099751\|122099850 | 21 | 42 | 22(+2A) | TCCCTGAGACCCTAACTTGTGAAA | 549.7 | -4.7 | 1.2 | -4.1 | 4.1E-05 | 3.6E-04 | BT20/MB231 | NUC |
| >trnaMT_PheGAA_MT_+_577_647@28.59.32\|ATACACTGAAAATGTTTAGACGGGCTCACATC | >trnaMT_PheGAA_MT_+_577_647 | 28 | 59 | 32 | ATACACTGAAAATGTTTAGACGGGCTCACATC | 92.1 | 10.3 | 2.7 | 3.9 | 1.1E-04 | 3.6E-04 | BT20/MB468 | MP |
| >am_trnaMT_ValTAC_MT_+_1602_1670@49.67.19\|AACTTAACTTGACCGCTCT | >am_trnaMT_ValTAC_MT_+_1602_1670 | 49 | 67 | 19 | AACTTAACTTGACCGCTCT | 5321.7 | 3.6 | 0.9 | 4.1 | 3.8E-05 | 3.6E-04 | BT20/MB231 | CYTO |
| >hg38_MT_+_598_1651_RNR1_12S_50ntFlanks@904.929.26\|AAGTATACTTCAAAGGACATTTAACT | >hg38_MT_+_598_1651_RNR1_12S_50ntFlanks | 904 | 929 | 26 | AAGTATACTTCAAAGGACATTTAACT | 363.8 | 4.2 | 1.2 | 3.7 | 2.4E-04 | 3.6E-04 | MB231/MB468 | MP |
| >hg38_21_+_8433172_8446622_RNA45SN1_45S_50ntFlanks@6646.6672.27\|TCGTACGACTCTTAGCGGTGGATCACT | >hg38_21_+_8433172_8446622_RNA45SN1_45S_50ntFlanks | 6646 | 6672 | 27 | TCGTACGACTCTTAGCGGTGGATCACT | 5639.1 | 3.0 | 0.8 | 3.6 | 2.8E-04 | 3.6E-04 | MB231/MB468 | MITO |
| >hg38_21_+_8433172_8446622_RNA45SN1_45S_50ntFlanks@11315.11343.29\|CCACCCCACGTCTCGTCGCGCGCGCGTCC | >hg38_21_+_8433172_8446622_RNA45SN1_45S_50ntFlanks | 11315 | 11343 | 29 | CCACCCCACGTCTCGTCGCGCGCGCGTCC | 1985.9 | -7.3 | 2.0 | -3.6 | 2.9E-04 | 3.6E-04 | BT20/MB468 | NUC |
| >hg38_21_+_8433172_8446622_RNA45SN1_45S_50ntFlanks@10909.10930.22\|GCGCGCGCCGCGGCTGGACGAG | >hg38_21_+_8433172_8446622_RNA45SN1_45S_50ntFlanks | 10909 | 10930 | 22 | GCGCGCGCCGCGGCTGGACGAG | 59.2 | 9.2 | 2.2 | 4.1 | 4.0E-05 | 3.6E-04 | BT20/MB231 | MP |
| >MI0000079\|hsa-mir-23a&WithFlank&19\|-\|13836581\|13836665@54.73.20(+1A)&[MIMAT0000078&hsa-miR-23a-3p&offsets\|+3\|+2(+1A);m-31&19\|-\|13836595\|13836615&offsets\|+3\|+2(+1A)]\|ACATTGCCAGGGATTTCCAAA | >MI0000079\|hsa-mir-23a&WithFlank&19\|-\|13836581\|13836665 | 54 | 73 | 20(+1A) | ACATTGCCAGGGATTTCCAAA | 599.7 | 13.2 | 3.6 | 3.6 | 2.7E-04 | 3.6E-04 | MB231/MB468 | CYTO |
| >hg38_21_+_8433172_8446622_RNA45SN1_45S_50ntFlanks@10350.10391.42\|ACGAGAACTTTGAAGGCCGAAGTGGAGAAGGGTTCCATGTGA | >hg38_21_+_8433172_8446622_RNA45SN1_45S_50ntFlanks | 10350 | 10391 | 42 | ACGAGAACTTTGAAGGCCGAAGTGGAGAAGGGTTCCATGTGA | 570.1 | -3.4 | 0.9 | -3.6 | 2.9E-04 | 3.6E-04 | MB231/MB468 | NUC |
| >am_trna119_LysCTT_1_-_145395522_145395594@14.31.18,trna11_LysCTT_5_-_180648979_180649051@14.31.18,trna128_LysCTT_1_-_55423542_55423614@14.31.18,trna13_LysCTT_6_+_26556774_26556846@14.31.18,trna32_LysCTT_16_-_3207406_3207478@14.31.18,trna6_LysCTT_19_-_52425396_52425468@14.31.18,trna7_LysCTT_16_+_3225692_3225764@14.31.18,trna9_LysCTT_5_+_180634755_180634827@14.31.18\|AGTCGGTAGAGCATGAGA | >am_trna119_LysCTT_1_-_145395522_145395594 | 14 | 31 | 18 | AGTCGGTAGAGCATGAGA | 608.0 | -5.8 | 1.6 | -3.6 | 2.9E-04 | 3.6E-04 | MB231/MB468 | NUC |
| >hg38_21_+_8433172_8446622_RNA45SN1_45S_50ntFlanks@11664.11685.22\|TTTCTGCCCAGTGCTCTGAATG | >hg38_21_+_8433172_8446622_RNA45SN1_45S_50ntFlanks | 11664 | 11685 | 22 | TTTCTGCCCAGTGCTCTGAATG | 199.4 | -2.4 | 0.7 | -3.6 | 2.9E-04 | 3.6E-04 | MB231/MB468 | NUC |
| >hg38_21_+_8433172_8446622_RNA45SN1_45S_50ntFlanks@10348.10388.41\|AAACGAGAACTTTGAAGGCCGAAGTGGAGAAGGGTTCCATG | >hg38_21_+_8433172_8446622_RNA45SN1_45S_50ntFlanks | 10348 | 10388 | 41 | AAACGAGAACTTTGAAGGCCGAAGTGGAGAAGGGTTCCATG | 205.0 | -3.5 | 1.0 | -3.7 | 2.4E-04 | 3.6E-04 | MB231/MB468 | MP |
| >hg38_21_+_8433172_8446622_RNA45SN1_45S_50ntFlanks@10349.10388.40\|AACGAGAACTTTGAAGGCCGAAGTGGAGAAGGGTTCCATG | >hg38_21_+_8433172_8446622_RNA45SN1_45S_50ntFlanks | 10349 | 10388 | 40 | AACGAGAACTTTGAAGGCCGAAGTGGAGAAGGGTTCCATG | 1365.3 | -2.9 | 0.8 | -3.6 | 2.7E-04 | 3.7E-04 | MB231/MB468 | CYTO |
| >hg38_21_+_8433172_8446622_RNA45SN1_45S_50ntFlanks@12970.12991.22\|GTTTCGTACGTAGCAGAGCAGC | >hg38_21_+_8433172_8446622_RNA45SN1_45S_50ntFlanks | 12970 | 12991 | 22 | GTTTCGTACGTAGCAGAGCAGC | 145.6 | -3.8 | 1.0 | -3.7 | 2.5E-04 | 3.7E-04 | BT20/MB468 | CYTO |
| >hg38_21_+_8433172_8446622_RNA45SN1_45S_50ntFlanks@11843.11861.19\|AGGGAACGGGCTTGGCGGA | >hg38_21_+_8433172_8446622_RNA45SN1_45S_50ntFlanks | 11843 | 11861 | 19 | AGGGAACGGGCTTGGCGGA | 64.0 | 9.3 | 2.3 | 4.1 | 4.0E-05 | 3.7E-04 | BT20/MB231 | MP |
| >hg38_21_+_8433172_8446622_RNA45SN1_45S_50ntFlanks@12935.12960.26\|TAAACCATTCGTAGACGACCTGCTTC | >hg38_21_+_8433172_8446622_RNA45SN1_45S_50ntFlanks | 12935 | 12960 | 26 | TAAACCATTCGTAGACGACCTGCTTC | 322.4 | -2.2 | 0.6 | -3.6 | 2.9E-04 | 3.7E-04 | MB231/MB468 | NUC |
| >hg38_21_+_8433172_8446622_RNA45SN1_45S_50ntFlanks@6653.6700.48\|ACTCTTAGCGGTGGATCACTCGGCTCGTGCGTCGATGAAGAACGCAGC | >hg38_21_+_8433172_8446622_RNA45SN1_45S_50ntFlanks | 6653 | 6700 | 48 | ACTCTTAGCGGTGGATCACTCGGCTCGTGCGTCGATGAAGAACGCAGC | 221.2 | 5.5 | 1.5 | 3.6 | 2.9E-04 | 3.7E-04 | MB231/MB468 | MITO |
| >hg38_21_+_8433172_8446622_RNA45SN1_45S_50ntFlanks@10392.10433.42\|ACAGCAGTTGAACATGGGTCAGTCGGTCCTGAGAGATGGGCG | >hg38_21_+_8433172_8446622_RNA45SN1_45S_50ntFlanks | 10392 | 10433 | 42 | ACAGCAGTTGAACATGGGTCAGTCGGTCCTGAGAGATGGGCG | 424.6 | -3.0 | 0.8 | -3.7 | 2.1E-04 | 3.7E-04 | BT20/MB468 | MITO |
| >MI0003776\|hsa-mir-320b-1&WithFlank&1\|+\|116671740\|116671823@49.67.19(+1U);MI0003839\|hsa-mir-320b-2&WithFlank&1\|-\|224257034\|224257116@48.66.19(+1U);MI0000542\|hsa-mir-320a&WithFlank&8\|-\|22244960\|22245043@49.67.19(+1U)&[MIMAT0005792&hsa-miR-320b&offsets\|+1\|-2(+1U);m-183&1\|+\|116671787\|116671806&offsets\|+1\|0(+1U)];[MIMAT0005792_1&hsa-miR-320b&offsets\|+1\|-2(+1U);m-183&1\|-\|224257051\|224257070&offsets\|+1\|0(+1U)];[MIMAT0000510&hsa-miR-320a-3p&offsets\|+1\|-2(+1U);m-60&8\|-\|22244975\|22244996&offsets\|+1\|-2(+1U)]\|AAAGCTGGGTTGAGAGGGCT | >MI0003776\|hsa-mir-320b-1&WithFlank&1\|+\|116671740\|116671823 | 49 | 67 | 19(+1U) | AAAGCTGGGTTGAGAGGGCT | 108.5 | 7.4 | 1.9 | 3.9 | 1.1E-04 | 3.7E-04 | BT20/MB468 | MP |
| >MI0000285\|hsa-mir-205&WithFlank&1\|+\|209432127\|209432248@40.61.22&[MIMAT0000266&hsa-miR-205-5p&offsets\|0\|0;m-351&1\|+\|209432166\|209432187&offsets\|0\|0]\|TCCTTCATTCCACCGGAGTCTG | >MI0000285\|hsa-mir-205&WithFlank&1\|+\|209432127\|209432248 | 40 | 61 | 22 | TCCTTCATTCCACCGGAGTCTG | 10438.6 | 2.5 | 0.7 | 3.6 | 2.9E-04 | 3.7E-04 | BT20/MB468 | NUC |

<table>
<tr><td>>hg38_21_+_8433172_8446622_RNA45SN1_45S_50ntFlanks@12405.12445.41|CTTTTTGATCCTTCGATGTCGGCTCTTCCTATCATTGTGAA</td><td>>hg38_21_+_8433172_8446622_RNA45SN1_45S_50ntFlanks</td><td>12405</td><td>12445</td><td>41</td><td>CTTTTTGATCCTTCGATGTCGGCTCTTCCTATCATTGTGAA</td><td>172.9</td><td>-3.7</td><td>1.0</td><td>-3.6</td><td>2.7E-04</td><td>3.7E-04</td><td>MB231/MB468</td><td>CYTO</td></tr>
<tr><td>>hg38_21_+_8433172_8446622_RNA45SN1_45S_50ntFlanks@7977.8003.27|CGACCTCAGATCAGACGTGGCGACCCG</td><td>>hg38_21_+_8433172_8446622_RNA45SN1_45S_50ntFlanks</td><td>7977</td><td>8003</td><td>27</td><td>CGACCTCAGATCAGACGTGGCGACCCG</td><td>359.4</td><td>-3.3</td><td>0.9</td><td>-3.6</td><td>2.9E-04</td><td>3.7E-04</td><td>MB231/MB468</td><td>MITO</td></tr>
<tr><td>>hg38_21_+_8433172_8446622_RNA45SN1_45S_50ntFlanks@4102.4120.19|ACCACGGGTGACGGGGAAT</td><td>>hg38_21_+_8433172_8446622_RNA45SN1_45S_50ntFlanks</td><td>4102</td><td>4120</td><td>19</td><td>ACCACGGGTGACGGGGAAT</td><td>305.8</td><td>-4.5</td><td>1.2</td><td>-3.6</td><td>2.9E-04</td><td>3.7E-04</td><td>MB231/MB468</td><td>NUC</td></tr>
<tr><td>>hg38_21_+_8433172_8446622_RNA45SN1_45S_50ntFlanks@8357.8376.20|AAAAGAACTTTGAAGAGAGA</td><td>>hg38_21_+_8433172_8446622_RNA45SN1_45S_50ntFlanks</td><td>8357</td><td>8376</td><td>20</td><td>AAAAGAACTTTGAAGAGAGA</td><td>550.3</td><td>-3.2</td><td>0.9</td><td>-3.6</td><td>2.9E-04</td><td>3.7E-04</td><td>MB231/MB468</td><td>NUC</td></tr>
<tr><td>>MI0000063|hsa-let-7b&WithFlank&22|+|46113680|46113774@12.30.19(+1U)&[MIMAT0000063&hsa-let-7b-5p&offsets|0|-3(+1U);m-14&22|+|46113691|46113712&offsets|0|-3(+1U)]|TGAGGTAGTAGGTTGTGTGT</td><td>>MI0000063|hsa-let-7b&WithFlank&22|+|46113680|46113774</td><td>12</td><td>30</td><td>19(+1U)</td><td>TGAGGTAGTAGGTTGTGTGT</td><td>144.3</td><td>6.3</td><td>1.6</td><td>3.9</td><td>1.1E-04</td><td>3.7E-04</td><td>BT20/MB468</td><td>MP</td></tr>
<tr><td>>hg38_21_+_8433172_8446622_RNA45SN1_45S_50ntFlanks@11575.11613.39|ACTTAGAACTGGTGCGGACCAGGGGAATCCGACTGTTTA</td><td>>hg38_21_+_8433172_8446622_RNA45SN1_45S_50ntFlanks</td><td>11575</td><td>11613</td><td>39</td><td>ACTTAGAACTGGTGCGGACCAGGGGAATCCGACTGTTTA</td><td>1984.1</td><td>-3.7</td><td>1.0</td><td>-3.6</td><td>2.9E-04</td><td>3.7E-04</td><td>MB231/MB468</td><td>MITO</td></tr>
<tr><td>>hg38_21_+_8433172_8446622_RNA45SN1_45S_50ntFlanks@5084.5108.25|CGAACGAGACTCTGGCATGCTAACT</td><td>>hg38_21_+_8433172_8446622_RNA45SN1_45S_50ntFlanks</td><td>5084</td><td>5108</td><td>25</td><td>CGAACGAGACTCTGGCATGCTAACT</td><td>335.2</td><td>-3.4</td><td>0.9</td><td>-3.6</td><td>2.9E-04</td><td>3.7E-04</td><td>MB231/MB468</td><td>NUC</td></tr>
<tr><td>>hg38_21_+_8433172_8446622_RNA45SN1_45S_50ntFlanks@10821.10838.18|GGAAGTCGGCAAGCCGGA</td><td>>hg38_21_+_8433172_8446622_RNA45SN1_45S_50ntFlanks</td><td>10821</td><td>10838</td><td>18</td><td>GGAAGTCGGCAAGCCGGA</td><td>1006.5</td><td>-3.2</td><td>0.8</td><td>-3.9</td><td>1.1E-04</td><td>3.7E-04</td><td>BT20/MB468</td><td>MP</td></tr>
<tr><td>>hg38_21_+_8433172_8446622_RNA45SN1_45S_50ntFlanks@4006.4025.20|ACCTCGGGCCGATCGCACGC</td><td>>hg38_21_+_8433172_8446622_RNA45SN1_45S_50ntFlanks</td><td>4006</td><td>4025</td><td>20</td><td>ACCTCGGGCCGATCGCACGC</td><td>339.9</td><td>-5.4</td><td>1.4</td><td>-3.7</td><td>2.1E-04</td><td>3.7E-04</td><td>BT20/MB468</td><td>MITO</td></tr>
<tr><td>>am_trna11_ProAGG_16_+_3241989_3242060@1.31.31,trna12_ProAGG_6_+_26555498_26555569@1.31.31,trna14_ProTGG_5_-_180615854_180615925@1.31.31,trna22_ProAGG_14_-_21081560_21081631@1.31.31,trna23_ProAGG_14_-_21077495_21077566@1.31.31,trna28_ProTGG_16_-_3234133_3234204@1.31.31,trna29_ProAGG_16_-_3232635_3232706@1.31.31,trna2_ProAGG_7_+_128423504_128423575@1.31.31,trna30_ProCGG_6_+_27059521_27059592@1.31.31,trna37_ProCGG_17_-_8126151_8126222@1.31.31,trna3_ProTGG_16_+_3208923_3208994@1.31.31,trna52_ProCGG_1_+_167683962_167684033@1.31.31,trna65_ProAGG_1_-_167684725_167684796@1.31.31,trna6_ProCGG_16_+_3222049_3222120@1.31.31,trna6_ProTGG_14_+_21152175_21152246@1.31.31,trna8_ProTGG_16_+_3238094_3238165@1.31.31,trna9_ProAGG_11_+_75946557_75946628@1.31.31,trna9_ProAGG_16_+_3239634_3239705@1.31.31|GGCTCGTTGGTCTAGGGGTATGATTCTCGCT</td><td>>am_trna11_ProAGG_16_+_3241989_3242060</td><td>1</td><td>31</td><td>31</td><td>GGCTCGTTGGTCTAGGGGTATGATTCTCGCT</td><td>1248.7</td><td>5.7</td><td>1.6</td><td>3.7</td><td>2.6E-04</td><td>3.7E-04</td><td>BT20/MB468</td><td>CYTO</td></tr>
<tr><td>>hg38_21_+_8433172_8446622_RNA45SN1_45S_50ntFlanks@12996.13038.43|TCGCTGCGATCTATTGAAAGTCAGCCCTCGACACAAGGGTTTG</td><td>>hg38_21_+_8433172_8446622_RNA45SN1_45S_50ntFlanks</td><td>12996</td><td>13038</td><td>43</td><td>TCGCTGCGATCTATTGAAAGTCAGCCCTCGACACAAGGGTTTG</td><td>775.7</td><td>6.2</td><td>1.7</td><td>3.7</td><td>2.1E-04</td><td>3.7E-04</td><td>BT20/MB468</td><td>MITO</td></tr>
<tr><td>>hg38_21_+_8433172_8446622_RNA45SN1_45S_50ntFlanks@11668.11702.35|TGCCCAGTGCTCTGAATGTCAAAGTGAAGAAATTC</td><td>>hg38_21_+_8433172_8446622_RNA45SN1_45S_50ntFlanks</td><td>11668</td><td>11702</td><td>35</td><td>TGCCCAGTGCTCTGAATGTCAAAGTGAAGAAATTC</td><td>1872.2</td><td>2.2</td><td>0.6</td><td>3.7</td><td>2.6E-04</td><td>3.7E-04</td><td>BT20/MB468</td><td>CYTO</td></tr>
<tr><td>>hg38_21_+_8433172_8446622_RNA45SN1_45S_50ntFlanks@9483.9505.23|CGTGTTAGGACCCGAAAGATGGT</td><td>>hg38_21_+_8433172_8446622_RNA45SN1_45S_50ntFlanks</td><td>9483</td><td>9505</td><td>23</td><td>CGTGTTAGGACCCGAAAGATGGT</td><td>128.0</td><td>7.1</td><td>1.8</td><td>3.9</td><td>1.1E-04</td><td>3.7E-04</td><td>BT20/MB468</td><td>MP</td></tr>
<tr><td>>hg38_21_+_8433172_8446622_RNA45SN1_45S_50ntFlanks@10346.10366.21|TCAAACGAGAACTTTGAAGGC</td><td>>hg38_21_+_8433172_8446622_RNA45SN1_45S_50ntFlanks</td><td>10346</td><td>10366</td><td>21</td><td>TCAAACGAGAACTTTGAAGGC</td><td>207.0</td><td>-4.4</td><td>1.2</td><td>-3.7</td><td>2.4E-04</td><td>3.7E-04</td><td>MB231/MB468</td><td>MP</td></tr>
<tr><td>>hg38_21_+_8433172_8446622_RNA45SN1_45S_50ntFlanks@4202.4224.23|CGACCCGGGGAGGTAGTGACGAA</td><td>>hg38_21_+_8433172_8446622_RNA45SN1_45S_50ntFlanks</td><td>4202</td><td>4224</td><td>23</td><td>CGACCCGGGGAGGTAGTGACGAA</td><td>89.0</td><td>-5.5</td><td>1.5</td><td>-3.7</td><td>2.4E-04</td><td>3.7E-04</td><td>MB231/MB468</td><td>MP</td></tr>
<tr><td>>MI0000086|hsa-mir-28&WithFlank&3|+|188688775|188688872@61.82.22&[MIMAT0004502&hsa-miR-28-3p&offsets|+1|+1;m-28&3|+|188688834|188688855&offsets|+1|+1]|ACTAGATTGTGAGCTCCTGGAG</td><td>>MI0000086|hsa-mir-28&WithFlank&3|+|188688775|188688872</td><td>61</td><td>82</td><td>22</td><td>ACTAGATTGTGAGCTCCTGGAG</td><td>486.9</td><td>6.5</td><td>1.7</td><td>3.9</td><td>1.1E-04</td><td>3.7E-04</td><td>BT20/MB468</td><td>MP</td></tr>
<tr><td>>hg38_21_+_8433172_8446622_RNA45SN1_45S_50ntFlanks@8171.8211.41|AAGTCCTTCTGATCGAGGCCCAGCCCGTGGACGGTGTGAGG</td><td>>hg38_21_+_8433172_8446622_RNA45SN1_45S_50ntFlanks</td><td>8171</td><td>8211</td><td>41</td><td>AAGTCCTTCTGATCGAGGCCCAGCCCGTGGACGGTGTGAGG</td><td>71.5</td><td>-9.2</td><td>2.4</td><td>-3.9</td><td>1.1E-04</td><td>3.7E-04</td><td>BT20/MB468</td><td>MP</td></tr>
<tr><td>>MI0000809|hsa-mir-151a&WithFlank&8|-|140732558|140732659@53.74.22(+2U)&[MIMAT0000757&hsa-miR-151a-3p&offsets|0|+1(+2U);m-30&8|-|140732587|140732607&offsets|0|+1(+2U)]|CTAGACTGAAGCTCCTTGAGGATT</td><td>>MI0000809|hsa-mir-151a&WithFlank&8|-|140732558|140732659</td><td>53</td><td>74</td><td>22(+2U)</td><td>CTAGACTGAAGCTCCTTGAGGATT</td><td>662.1</td><td>4.6</td><td>1.3</td><td>3.7</td><td>2.4E-04</td><td>3.7E-04</td><td>MB231/MB468</td><td>MP</td></tr>
<tr><td>>hg38_21_+_8433172_8446622_RNA45SN1_45S_50ntFlanks@10842.10865.24|GTAACTTCGGGATAAGGATTGGCT</td><td>>hg38_21_+_8433172_8446622_RNA45SN1_45S_50ntFlanks</td><td>10842</td><td>10865</td><td>24</td><td>GTAACTTCGGGATAAGGATTGGCT</td><td>575.7</td><td>-4.2</td><td>1.2</td><td>-3.7</td><td>2.6E-04</td><td>3.7E-04</td><td>BT20/MB468</td><td>CYTO</td></tr>
<tr><td>>hg38_21_+_8433172_8446622_RNA45SN1_45S_50ntFlanks@12635.12659.25|AAGCTACCATCTGTGGGATTATGAC</td><td>>hg38_21_+_8433172_8446622_RNA45SN1_45S_50ntFlanks</td><td>12635</td><td>12659</td><td>25</td><td>AAGCTACCATCTGTGGGATTATGAC</td><td>501.8</td><td>-4.0</td><td>1.1</td><td>-3.6</td><td>2.7E-04</td><td>3.7E-04</td><td>MB231/MB468</td><td>CYTO</td></tr>
<tr><td>>MI0000471|hsa-mir-126&WithFlank&9|+|136670596|136670692@57.79.23&[MIMAT0000445&hsa-miR-126-3p&offsets|-1|0;m-40&9|+|136670653|136670674&offsets|-1|0]|CTCGTACCGTGAGTAATAATGCG</td><td>>MI0000471|hsa-mir-126&WithFlank&9|+|136670596|136670692</td><td>57</td><td>79</td><td>23</td><td>CTCGTACCGTGAGTAATAATGCG</td><td>166.3</td><td>7.0</td><td>1.8</td><td>3.9</td><td>1.1E-04</td><td>3.7E-04</td><td>BT20/MB468</td><td>MP</td></tr>
<tr><td>>hg38_21_+_8433172_8446622_RNA45SN1_45S_50ntFlanks@11785.11806.22|TGAATGGATGAACGAGATTCCC</td><td>>hg38_21_+_8433172_8446622_RNA45SN1_45S_50ntFlanks</td><td>11785</td><td>11806</td><td>22</td><td>TGAATGGATGAACGAGATTCCC</td><td>295.7</td><td>-4.3</td><td>1.2</td><td>-3.6</td><td>2.9E-04</td><td>3.7E-04</td><td>MB231/MB468</td><td>NUC</td></tr>
<tr><td>>hg38_21_+_8433172_8446622_RNA45SN1_45S_50ntFlanks@11029.11051.23|CTCTCTCTCTCTCCCCCGCTCCC</td><td>>hg38_21_+_8433172_8446622_RNA45SN1_45S_50ntFlanks</td><td>11029</td><td>11051</td><td>23</td><td>CTCTCTCTCTCTCCCCCGCTCCC</td><td>222.1</td><td>-3.7</td><td>1.0</td><td>-3.7</td><td>2.6E-04</td><td>3.7E-04</td><td>BT20/MB468</td><td>CYTO</td></tr>
<tr><td>>hg38_21_+_8433172_8446622_RNA45SN1_45S_50ntFlanks@7983.8016.34|CAGATCAGACGTGGCGACCCGCTGAATTTAAGCA</td><td>>hg38_21_+_8433172_8446622_RNA45SN1_45S_50ntFlanks</td><td>7983</td><td>8016</td><td>34</td><td>CAGATCAGACGTGGCGACCCGCTGAATTTAAGCA</td><td>147.9</td><td>-5.6</td><td>1.5</td><td>-3.7</td><td>2.4E-04</td><td>3.7E-04</td><td>MB231/MB468</td><td>MP</td></tr>
<tr><td>>hg38_21_+_8433172_8446622_RNA45SN1_45S_50ntFlanks@6662.6689.28|GGTGGATCACTCGGCTCGTGCGTCGATG</td><td>>hg38_21_+_8433172_8446622_RNA45SN1_45S_50ntFlanks</td><td>6662</td><td>6689</td><td>28</td><td>GGTGGATCACTCGGCTCGTGCGTCGATG</td><td>650.0</td><td>2.0</td><td>0.5</td><td>3.7</td><td>2.1E-04</td><td>3.7E-04</td><td>BT20/MB468</td><td>MITO</td></tr>
<tr><td>>hg38_21_+_8433172_8446622_RNA45SN1_45S_50ntFlanks@3835.3862.28|CTCCTCTCCTACTTGGATAACTGTGGTA</td><td>>hg38_21_+_8433172_8446622_RNA45SN1_45S_50ntFlanks</td><td>3835</td><td>3862</td><td>28</td><td>CTCCTCTCCTACTTGGATAACTGTGGTA</td><td>663.1</td><td>-4.1</td><td>1.1</td><td>-3.7</td><td>2.1E-04</td><td>3.7E-04</td><td>BT20/MB468</td><td>MITO</td></tr>
<tr><td>>hg38_21_+_8433172_8446622_RNA45SN1_45S_50ntFlanks@9837.9868.32|TGGTAAGCAGAACTGGCGCTGCGGGATGAACC</td><td>>hg38_21_+_8433172_8446622_RNA45SN1_45S_50ntFlanks</td><td>9837</td><td>9868</td><td>32</td><td>TGGTAAGCAGAACTGGCGCTGCGGGATGAACC</td><td>57.8</td><td>9.2</td><td>2.2</td><td>4.1</td><td>4.0E-05</td><td>3.7E-04</td><td>BT20/MB231</td><td>MP</td></tr>
<tr><td>>MI0000061|hsa-let-7a-2&WithFlank&11|-|122146516|122146599@11.32.22(+1A);MI0000062|hsa-let-7a-3&WithFlank&22|+|46112743|46112828@10.31.22(+1A);MI0000060|hsa-let-7a-1&WithFlank&9|+|94175951|94176042@12.33.22(+1A)&[MIMAT0000062&hsa-let-7a-5p&offsets|0|0(+1A);m-6&11|-|122146568|122146589&offsets|0|0(+1A)];[MIMAT0000062_1&hsa-let-7a-5p&offsets|0|0(+1A);m-5&22|+|46112752|46112773&offsets|0|0(+1A)];[MIMAT0000062_2&hsa-let-7a-5p&offsets|0|0(+1A);m-4&9|+|94175962|94175983&offsets|0|0(+1A)]|TGAGGTAGTAGGTTGTATAGTTA</td><td>>MI0000061|hsa-let-7a-2&WithFlank&11|-|122146516|122146599</td><td>11</td><td>32</td><td>22(+1A)</td><td>TGAGGTAGTAGGTTGTATAGTTA</td><td>540.8</td><td>1.7</td><td>0.5</td><td>3.6</td><td>3.0E-04</td><td>3.7E-04</td><td>BT20/MB468</td><td>NUC</td></tr>
<tr><td>>hg38_21_+_8433172_8446622_RNA45SN1_45S_50ntFlanks@9532.9571.40|AGAGGAAACTCTGGTGGAGGTCCGTAGCGGTCCTGACGTG</td><td>>hg38_21_+_8433172_8446622_RNA45SN1_45S_50ntFlanks</td><td>9532</td><td>9571</td><td>40</td><td>AGAGGAAACTCTGGTGGAGGTCCGTAGCGGTCCTGACGTG</td><td>193.2</td><td>3.8</td><td>1.1</td><td>3.6</td><td>2.9E-04</td><td>3.7E-04</td><td>MB231/MB468</td><td>NUC</td></tr>
<tr><td>>hg38_21_+_8433172_8446622_RNA45SN1_45S_50ntFlanks@12163.12180.18|ACTGGGGCGGTACACCTG</td><td>>hg38_21_+_8433172_8446622_RNA45SN1_45S_50ntFlanks</td><td>12163</td><td>12180</td><td>18</td><td>ACTGGGGCGGTACACCTG</td><td>574.5</td><td>-5.1</td><td>1.4</td><td>-3.6</td><td>2.7E-04</td><td>3.7E-04</td><td>MB231/MB468</td><td>CYTO</td></tr>
<tr><td>>hg38_21_+_8433172_8446622_RNA45SN1_45S_50ntFlanks@5467.5484.18|GGCCCACGGCCCTGGCGG</td><td>>hg38_21_+_8433172_8446622_RNA45SN1_45S_50ntFlanks</td><td>5467</td><td>5484</td><td>18</td><td>GGCCCACGGCCCTGGCGG</td><td>314.0</td><td>-5.2</td><td>1.4</td><td>-3.7</td><td>2.2E-04</td><td>3.7E-04</td><td>BT20/MB468</td><td>MITO</td></tr>
<tr><td>>hg38_21_+_8433172_8446622_RNA45SN1_45S_50ntFlanks@6661.6684.24|CGGTGGATCACTCGGCTCGTGCGT</td><td>>hg38_21_+_8433172_8446622_RNA45SN1_45S_50ntFlanks</td><td>6661</td><td>6684</td><td>24</td><td>CGGTGGATCACTCGGCTCGTGCGT</td><td>214.2</td><td>1.3</td><td>0.4</td><td>3.6</td><td>3.0E-04</td><td>3.7E-04</td><td>BT20/MB468</td><td>NUC</td></tr>
<tr><td>>hg38_21_+_8433172_8446622_RNA45SN1_45S_50ntFlanks@7981.8006.26|CTCAGATCAGACGTGGCGACCCGCTG</td><td>>hg38_21_+_8433172_8446622_RNA45SN1_45S_50ntFlanks</td><td>7981</td><td>8006</td><td>26</td><td>CTCAGATCAGACGTGGCGACCCGCTG</td><td>1851.4</td><td>-4.5</td><td>1.2</td><td>-3.6</td><td>2.7E-04</td><td>3.7E-04</td><td>MB231/MB468</td><td>CYTO</td></tr>
<tr><td>>hg38_21_+_8433172_8446622_RNA45SN1_45S_50ntFlanks@11567.11584.18|AGCAGCCGACTTAGAACT</td><td>>hg38_21_+_8433172_8446622_RNA45SN1_45S_50ntFlanks</td><td>11567</td><td>11584</td><td>18</td><td>AGCAGCCGACTTAGAACT</td><td>504.5</td><td>3.2</td><td>0.8</td><td>3.9</td><td>1.1E-04</td><td>3.7E-04</td><td>BT20/MB468</td><td>MP</td></tr>
<tr><td>>hg38_21_+_8433172_8446622_RNA45SN1_45S_50ntFlanks@11920.11950.31|ATGAGAGGTGTAGAATAAGTGGGAGGCCCCC</td><td>>hg38_21_+_8433172_8446622_RNA45SN1_45S_50ntFlanks</td><td>11920</td><td>11950</td><td>31</td><td>ATGAGAGGTGTAGAATAAGTGGGAGGCCCCC</td><td>567.1</td><td>-3.8</td><td>0.9</td><td>-4.1</td><td>3.9E-05</td><td>3.7E-04</td><td>BT20/MB231</td><td>CYTO</td></tr>
<tr><td>>hg38_21_+_8433172_8446622_RNA45SN1_45S_50ntFlanks@11849.11873.25|CGGGCTTGGCGGAATCAGCGGGGAA</td><td>>hg38_21_+_8433172_8446622_RNA45SN1_45S_50ntFlanks</td><td>11849</td><td>11873</td><td>25</td><td>CGGGCTTGGCGGAATCAGCGGGGAA</td><td>429.6</td><td>-5.1</td><td>1.3</td><td>-3.9</td><td>1.1E-04</td><td>3.7E-04</td><td>BT20/MB468</td><td>MP</td></tr>
<tr><td>>hg38_21_+_8433172_8446622_RNA45SN1_45S_50ntFlanks@3881.3898.18|GCCGACGGGCGCTGACCC</td><td>>hg38_21_+_8433172_8446622_RNA45SN1_45S_50ntFlanks</td><td>3881</td><td>3898</td><td>18</td><td>GCCGACGGGCGCTGACCC</td><td>450.0</td><td>-3.4</td><td>0.9</td><td>-3.6</td><td>2.7E-04</td><td>3.7E-04</td><td>MB231/MB468</td><td>CYTO</td></tr>
<tr><td>>hg38_21_+_8433172_8446622_RNA45SN1_45S_50ntFlanks@12406.12434.29|TTTTTGATCCTTCGATGTCGGCTCTTCCT</td><td>>hg38_21_+_8433172_8446622_RNA45SN1_45S_50ntFlanks</td><td>12406</td><td>12434</td><td>29</td><td>TTTTTGATCCTTCGATGTCGGCTCTTCCT</td><td>87.4</td><td>-5.6</td><td>1.5</td><td>-3.7</td><td>2.5E-04</td><td>3.7E-04</td><td>MB231/MB468</td><td>MP</td></tr>
<tr><td>>trna2_GlyGCC_21_-_18827107_18827177@4.31.28,trna35_GlyGCC_1_+_161413094_161413164@4.31.28,trna37_GlyGCC_1_+_161420467_161420537@4.31.28,trna39_GlyGCC_1_+_161427898_161427968@4.31.28,trna41_GlyGCC_1_+_161435258_161435328@4.31.28|TGGGTGGTTCAGTGGTAGAATTCTCGCC</td><td>>trna2_GlyGCC_21_-_18827107_18827177</td><td>4</td><td>31</td><td>28</td><td>TGGGTGGTTCAGTGGTAGAATTCTCGCC</td><td>51.8</td><td>9.0</td><td>2.2</td><td>4.1</td><td>4.1E-05</td><td>3.7E-04</td><td>BT20/MB231</td><td>MP</td></tr>
<tr><td>>hg38_21_+_8433172_8446622_RNA45SN1_45S_50ntFlanks@12008.12039.32|CCGCCGGTGAAATACCACTACTCTGATCGTTT</td><td>>hg38_21_+_8433172_8446622_RNA45SN1_45S_50ntFlanks</td><td>12008</td><td>12039</td><td>32</td><td>CCGCCGGTGAAATACCACTACTCTGATCGTTT</td><td>398.9</td><td>-3.1</td><td>0.9</td><td>-3.7</td><td>2.6E-04</td><td>3.7E-04</td><td>BT20/MB468</td><td>CYTO</td></tr>
<tr><td>>hg38_21_+_8433172_8446622_RNA45SN1_45S_50ntFlanks@10239.10261.23|GCTACGCCGCGACGAGTAGGAGG</td><td>>hg38_21_+_8433172_8446622_RNA45SN1_45S_50ntFlanks</td><td>10239</td><td>10261</td><td>23</td><td>GCTACGCCGCGACGAGTAGGAGG</td><td>175.9</td><td>3.6</td><td>0.9</td><td>3.9</td><td>1.1E-04</td><td>3.7E-04</td><td>BT20/MB468</td><td>MP</td></tr>
<tr><td>>MI0000067|hsa-let-7f-1&WithFlank&9|+|94176341|94176439@69.89.21(+1U)&[MIMAT0004486&hsa-let-7f-1-3p&offsets|0|-1(+1U);m-275&9|+|94176409|94176430&offsets|0|-1(+1U)]|CTATACAATCTATTGCCTTCCT</td><td>>MI0000067|hsa-let-7f-1&WithFlank&9|+|94176341|94176439</td><td>69</td><td>89</td><td>21(+1U)</td><td>CTATACAATCTATTGCCTTCCT</td><td>446.1</td><td>4.4</td><td>1.2</td><td>3.9</td><td>1.1E-04</td><td>3.7E-04</td><td>BT20/MB468</td><td>MP</td></tr>
<tr><td>>hg38_21_+_8433172_8446622_RNA45SN1_45S_50ntFlanks@12841.12869.29|CGGAGTGCCCTTCGTCCTGGGAAACGGGG</td><td>>hg38_21_+_8433172_8446622_RNA45SN1_45S_50ntFlanks</td><td>12841</td><td>12869</td><td>29</td><td>CGGAGTGCCCTTCGTCCTGGGAAACGGGG</td><td>98.8</td><td>-9.7</td><td>2.5</td><td>-3.9</td><td>1.1E-04</td><td>3.7E-04</td><td>BT20/MB468</td><td>MP</td></tr>
<tr><td>>hg38_21_+_8433172_8446622_RNA45SN1_45S_50ntFlanks@3705.3742.38|TACCTGGTTGATCCTGCCAGTAGCATATGCTTGTCTCA</td><td>>hg38_21_+_8433172_8446622_RNA45SN1_45S_50ntFlanks</td><td>3705</td><td>3742</td><td>38</td><td>TACCTGGTTGATCCTGCCAGTAGCATATGCTTGTCTCA</td><td>1361.8</td><td>3.2</td><td>0.9</td><td>3.7</td><td>2.2E-04</td><td>3.7E-04</td><td>BT20/MB468</td><td>MITO</td></tr>
</table>

| | | | | | | | | | | | | | |
|---|---|---|---|---|---|---|---|---|---|---|---|---|---|
| >hg38_21_+_8433172_8446622_RNA45SN1_45S_50ntFlanks@8176.8199.24\|CTTCTGATCGAGGCCCAGCCCGTG | >hg38_21_+_8433172_8446622_RNA45SN1_45S_50ntFlanks | 8176 | 8199 | 24 | CTTCTGATCGAGGCCCAGCCCGTG | 350.2 | -6.1 | 1.7 | -3.6 | 3.0E-04 | 3.7E-04 | BT20/MB468 | NUC |
| >MI0000272\|hsa-mir-182&WithFlank&7\|-\|129770377\|129770498@29.50.22&[MIMAT0000259&hsa-miR-182-5p&offsets\|0\|-2;m-22&7\|-\|129770449\|129770470&offsets\|0\|0]\|TTTGGCAATGGTAGAACTCACA | >MI0000272\|hsa-mir-182&WithFlank&7\|-\|129770377\|129770498 | 29 | 50 | 22 | TTTGGCAATGGTAGAACTCACA | 295.3 | 3.3 | 0.9 | 3.6 | 2.9E-04 | 3.7E-04 | MB231/MB468 | MITO |
| >hg38_21_+_8433172_8446622_RNA45SN1_45S_50ntFlanks@11199.11224.26\|CCGGCGGCGGCGGCGACTCTGGACGC | >hg38_21_+_8433172_8446622_RNA45SN1_45S_50ntFlanks | 11199 | 11224 | 26 | CCGGCGGCGGCGGCGACTCTGGACGC | 223.0 | -5.0 | 1.4 | -3.7 | 2.6E-04 | 3.7E-04 | BT20/MB468 | CYTO |
| >MI0000108\|hsa-mir-103a-2&WithFlank&20\|+\|3917488\|3917577@54.76.23;MI0000109\|hsa-mir-103a-1&WithFlank&5\|-\|168560890\|168560979@54.76.23&[MIMAT0000101&hsa-miR-103a-3p&offsets\|0\|0;m-12&20\|+\|3917541\|3917563&offsets\|0\|0];[MIMAT0000101_1&hsa-miR-103a-3p&offsets\|0\|0;m-13&5\|-\|168560904\|168560926&offsets\|0\|0;m-44220&5\|-\|168560908\|168560926&offsets\|0\|+4]\|AGCAGCATTGTACAGGGCTATGA | >MI0000108\|hsa-mir-103a-2&WithFlank&20\|+\|3917488\|3917577 | 54 | 76 | 23 | AGCAGCATTGTACAGGGCTATGA | 49452.6 | -3.5 | 0.9 | -4.1 | 4.1E-05 | 3.7E-04 | BT20/MB231 | MP |
| >hg38_21_+_8433172_8446622_RNA45SN1_45S_50ntFlanks@11896.11922.27\|TCTAGTCTGGCACGGTGAAGAGACATG | >hg38_21_+_8433172_8446622_RNA45SN1_45S_50ntFlanks | 11896 | 11922 | 27 | TCTAGTCTGGCACGGTGAAGAGACATG | 103.2 | 6.3 | 1.6 | 3.9 | 1.1E-04 | 3.7E-04 | BT20/MB468 | MP |
| >hg38_21_+_8433172_8446622_RNA45SN1_45S_50ntFlanks@10186.10207.22\|TCCTCCCCTCCTCCCGCCCACG | >hg38_21_+_8433172_8446622_RNA45SN1_45S_50ntFlanks | 10186 | 10207 | 22 | TCCTCCCCTCCTCCCGCCCACG | 112.6 | -9.9 | 2.6 | -3.9 | 1.1E-04 | 3.7E-04 | BT20/MB468 | MP |
| >hg38_21_+_8433172_8446622_RNA45SN1_45S_50ntFlanks@11913.11939.27\|AAGAGACATGAGAGGTGTAGAATAAGT | >hg38_21_+_8433172_8446622_RNA45SN1_45S_50ntFlanks | 11913 | 11939 | 27 | AAGAGACATGAGAGGTGTAGAATAAGT | 54.7 | 9.1 | 2.2 | 4.1 | 4.1E-05 | 3.7E-04 | BT20/MB231 | MP |
| >MI0000272\|hsa-mir-182&WithFlank&7\|-\|129770377\|129770498@29.53.25(+1C)&[MIMAT0000259&hsa-miR-182-5p&offsets\|0\|+1(+1C);m-22&7\|-\|129770449\|129770470&offsets\|0\|+3(+1C)]\|TTTGGCAATGGTAGAACTCACACTGC | >MI0000272\|hsa-mir-182&WithFlank&7\|-\|129770377\|129770498 | 29 | 53 | 25(+1C) | TTTGGCAATGGTAGAACTCACACTGC | 432.0 | 3.3 | 0.8 | 4.1 | 3.9E-05 | 3.7E-04 | BT20/MB231 | CYTO |
| >hg38_21_+_8433172_8446622_RNA45SN1_45S_50ntFlanks@12655.12673.19\|ATGACTGAACGCCTCTAAG | >hg38_21_+_8433172_8446622_RNA45SN1_45S_50ntFlanks | 12655 | 12673 | 19 | ATGACTGAACGCCTCTAAG | 229.3 | -5.0 | 1.4 | -3.7 | 2.6E-04 | 3.7E-04 | BT20/MB468 | CYTO |
| >hg38_21_+_8433172_8446622_RNA45SN1_45S_50ntFlanks@10797.10830.34\|GCATGTTGGAACAATGTAGGTAAGGGAAGTCGGC | >hg38_21_+_8433172_8446622_RNA45SN1_45S_50ntFlanks | 10797 | 10830 | 34 | GCATGTTGGAACAATGTAGGTAAGGGAAGTCGGC | 144.9 | -4.3 | 1.2 | -3.7 | 2.5E-04 | 3.7E-04 | MB231/MB468 | MP |
| >hg38_21_+_8433172_8446622_RNA45SN1_45S_50ntFlanks@6652.6677.26\|GACTCTTAGCGGTGGATCACTCGGCT | >hg38_21_+_8433172_8446622_RNA45SN1_45S_50ntFlanks | 6652 | 6677 | 26 | GACTCTTAGCGGTGGATCACTCGGCT | 3703.3 | 3.8 | 1.0 | 3.9 | 1.2E-04 | 3.7E-04 | BT20/MB468 | MP |
| >MI0000767\|hsa-mir-365a&WithFlank&16\|+\|14309279\|14309377@22.43.22&[MIMAT0009199&hsa-miR-365a-5p&offsets\|0\|-1;m-595&16\|+\|14309300\|14309322&offsets\|0\|-1]\|AGGGACTTTTGGGGGCAGATGT | >MI0000767\|hsa-mir-365a&WithFlank&16\|+\|14309279\|14309377 | 22 | 43 | 22 | AGGGACTTTTGGGGGCAGATGT | 367.4 | 4.0 | 1.0 | 4.1 | 4.1E-05 | 3.7E-04 | BT20/MB231 | MP |
| >hg38_21_+_8433172_8446622_RNA45SN1_45S_50ntFlanks@12934.12965.32\|CTAAACCATTCGTAGACGACCTGCTTCTGGGT | >hg38_21_+_8433172_8446622_RNA45SN1_45S_50ntFlanks | 12934 | 12965 | 32 | CTAAACCATTCGTAGACGACCTGCTTCTGGGT | 691.0 | -4.7 | 1.3 | -3.6 | 3.0E-04 | 3.8E-04 | MB231/MB468 | NUC |
| >hg38_21_+_8433172_8446622_RNA45SN1_45S_50ntFlanks@3769.3791.23\|CGCACGGCCGGTACAGTGAAACT | >hg38_21_+_8433172_8446622_RNA45SN1_45S_50ntFlanks | 3769 | 3791 | 23 | CGCACGGCCGGTACAGTGAAACT | 299.8 | 2.7 | 0.7 | 3.6 | 2.8E-04 | 3.8E-04 | MB231/MB468 | CYTO |
| >hg38_21_+_8433172_8446622_RNA45SN1_45S_50ntFlanks@12934.12954.21\|CTAAACCATTCGTAGACGACC | >hg38_21_+_8433172_8446622_RNA45SN1_45S_50ntFlanks | 12934 | 12954 | 21 | CTAAACCATTCGTAGACGACC | 569.5 | -2.5 | 0.7 | -3.6 | 2.8E-04 | 3.8E-04 | MB231/MB468 | CYTO |
| >MI0000434\|hsa-let-7i&WithFlank&12\|+\|62603680\|62603775@12.33.22(+1A)&[MIMAT0000415&hsa-let-7i-5p&offsets\|0\|0(+1A);m-49&12\|+\|62603691\|62603712&offsets\|0\|0(+1A)]\|TGAGGTAGTAGTTTGTGCTGTTA | >MI0000434\|hsa-let-7i&WithFlank&12\|+\|62603680\|62603775 | 12 | 33 | 22(+1A) | TGAGGTAGTAGTTTGTGCTGTTA | 7969.0 | 3.1 | 0.8 | 3.7 | 2.5E-04 | 3.8E-04 | MB231/MB468 | MP |
| >trnaMT_PheGAA_MT_+_577_647@35.65.31\|GAAAATGTTTAGACGGGCTCACATCACCCCA | >trnaMT_PheGAA_MT_+_577_647 | 35 | 65 | 31 | GAAAATGTTTAGACGGGCTCACATCACCCCA | 1122.7 | 7.3 | 1.9 | 3.9 | 1.2E-04 | 3.8E-04 | BT20/MB468 | MP |
| >hg38_21_+_8433172_8446622_RNA45SN1_45S_50ntFlanks@11778.11804.27\|ACGCGCATGAATGGATGAACGAGATTC | >hg38_21_+_8433172_8446622_RNA45SN1_45S_50ntFlanks | 11778 | 11804 | 27 | ACGCGCATGAATGGATGAACGAGATTC | 200.9 | -5.1 | 1.4 | -3.6 | 2.6E-04 | 3.8E-04 | BT20/MB468 | CYTO |
| >hg38_1_-_228634819_228635039_RNA5S12_5S_50ntFlanks@132.169.38\|GATGGGAGACCGCCTGGGAATACCGGGTGCTGTAGGCT | >hg38_1_-_228634819_228635039_RNA5S12_5S_50ntFlanks | 132 | 169 | 38 | GATGGGAGACCGCCTGGGAATACCGGGTGCTGTAGGCT | 1923.5 | -2.6 | 0.7 | -3.7 | 2.5E-04 | 3.8E-04 | MB231/MB468 | MP |
| >hg38_21_+_8433172_8446622_RNA45SN1_45S_50ntFlanks@9604.9623.20\|AAAGACTAATCGAACCATCT | >hg38_21_+_8433172_8446622_RNA45SN1_45S_50ntFlanks | 9604 | 9623 | 20 | AAAGACTAATCGAACCATCT | 226.2 | -3.2 | 0.9 | -3.6 | 3.0E-04 | 3.8E-04 | MB231/MB468 | MITO |
| >hg38_21_+_8433172_8446622_RNA45SN1_45S_50ntFlanks@12251.12288.38\|AAAGCTCGCTTGATCTTGATTTTCAGTACGAATACAGA | >hg38_21_+_8433172_8446622_RNA45SN1_45S_50ntFlanks | 12251 | 12288 | 38 | AAAGCTCGCTTGATCTTGATTTTCAGTACGAATACAGA | 345.3 | -5.6 | 1.5 | -3.6 | 3.0E-04 | 3.8E-04 | MB231/MB468 | MITO |
| >hg38_21_+_8433172_8446622_RNA45SN1_45S_50ntFlanks@7982.8009.28\|TCAGATCAGACGTGGCGACCCGCTGAAT | >hg38_21_+_8433172_8446622_RNA45SN1_45S_50ntFlanks | 7982 | 8009 | 28 | TCAGATCAGACGTGGCGACCCGCTGAAT | 481.5 | -2.7 | 0.8 | -3.6 | 3.0E-04 | 3.8E-04 | MB231/MB468 | MITO |
| >MI0000803\|hsa-mir-330&WithFlank&19\|-\|45638988\|45639093@64.86.23(+1U)&[MIMAT0000751&hsa-miR-330-3p&offsets\|+1\|+1(+1U);m-347&19\|-\|45639009\|45639030&offsets\|0\|+1(+1U)]\|CAAAGCACACGGCCTGCAGAGAGT | >MI0000803\|hsa-mir-330&WithFlank&19\|-\|45638988\|45639093 | 64 | 86 | 23(+1U) | CAAAGCACACGGCCTGCAGAGAGT | 233.6 | 5.3 | 1.4 | 3.9 | 1.2E-04 | 3.8E-04 | BT20/MB468 | MP |
| >MI0006273\|hsa-mir-1180&WithFlank&17\|-\|19344500\|19344580@47.68.22&[MIMAT0005825&hsa-miR-1180-3p&offsets\|0\|0;m-214&17\|-\|19344513\|19344534&offsets\|0\|0]\|TTTCCGGCTCGCGTGGGTGTGT | >MI0006273\|hsa-mir-1180&WithFlank&17\|-\|19344500\|19344580 | 47 | 68 | 22 | TTTCCGGCTCGCGTGGGTGTGT | 497.3 | 3.1 | 0.9 | 3.6 | 3.0E-04 | 3.8E-04 | MB231/MB468 | NUC |
| >trna111_HisGTG_1_-_147774845_147774916@1.31.31,trna118_HisGTG_1_-_145396881_145396952@1.31.31,trna16_HisGTG_1_+_146544773_146544844@1.31.31,trna1_HisGTG_15_+_45493349_45493420@1.31.31,trna21_HisGTG_1_+_147753471_147753542@1.31.31,trna33_HisGTG_6_+_27125906_27125977@1.31.31,trna7_HisGTG_9_-_14433938_14434009@1.31.31,trna8_HisGTG_15_-_45492611_45492682@1.31.31,trna9_HisGTG_15_-_45490804_45490875@1.31.31\|GCCGTGATCGTATAGTGGTTAGTACTCTGCG | >trna111_HisGTG_1_-_147774845_147774916 | 1 | 31 | 31 | GCCGTGATCGTATAGTGGTTAGTACTCTGCG | 36660.9 | -3.0 | 0.8 | -3.6 | 2.6E-04 | 3.8E-04 | BT20/MB468 | CYTO |
| >hg38_21_+_8433172_8446622_RNA45SN1_45S_50ntFlanks@12618.12654.37\|GAGGAGCCAATGGGGCGAAGCTACCATCTGTGGGATT | >hg38_21_+_8433172_8446622_RNA45SN1_45S_50ntFlanks | 12618 | 12654 | 37 | GAGGAGCCAATGGGGCGAAGCTACCATCTGTGGGATT | 485.2 | -6.4 | 1.8 | -3.6 | 3.0E-04 | 3.8E-04 | BT20/MB468 | NUC |
| >hg38_21_+_8433172_8446622_RNA45SN1_45S_50ntFlanks@11783.11802.20\|CATGAATGGATGAACGAGAT | >hg38_21_+_8433172_8446622_RNA45SN1_45S_50ntFlanks | 11783 | 11802 | 20 | CATGAATGGATGAACGAGAT | 189.2 | -2.8 | 0.8 | -3.6 | 2.7E-04 | 3.8E-04 | BT20/MB468 | CYTO |
| >hg38_21_+_8433172_8446622_RNA45SN1_45S_50ntFlanks@10872.10902.31\|GCTGGGTCGGTCGGGCTGGGGCGCGAAGCGG | >hg38_21_+_8433172_8446622_RNA45SN1_45S_50ntFlanks | 10872 | 10902 | 31 | GCTGGGTCGGTCGGGCTGGGGCGCGAAGCGG | 163.4 | -5.2 | 1.4 | -3.6 | 3.0E-04 | 3.8E-04 | MB231/MB468 | NUC |
| >hg38_21_+_8433172_8446622_RNA45SN1_45S_50ntFlanks@9544.9574.31\|GGTGGAGGTCCGTAGCGGTCCTGACGTGCAA | >hg38_21_+_8433172_8446622_RNA45SN1_45S_50ntFlanks | 9544 | 9574 | 31 | GGTGGAGGTCCGTAGCGGTCCTGACGTGCAA | 329.9 | -2.9 | 0.8 | -3.6 | 3.0E-04 | 3.8E-04 | MB231/MB468 | NUC |
| >MI0000825\|hsa-mir-345&WithFlank&14\|+\|100307853\|100307962@24.45.22(+1U)&[MIMAT0000772&hsa-miR-345-5p&offsets\|0\|0(+1U);m-193&14\|+\|100307876\|100307896&offsets\|0\|+1(+1U)]\|GCTGACTCCTAGTCCAGGGCTCT | >MI0000825\|hsa-mir-345&WithFlank&14\|+\|100307853\|100307962 | 24 | 45 | 22(+1U) | GCTGACTCCTAGTCCAGGGCTCT | 216.5 | 3.3 | 0.9 | 3.7 | 2.5E-04 | 3.8E-04 | MB231/MB468 | MP |
| >hg38_21_+_8433172_8446622_RNA45SN1_45S_50ntFlanks@4588.4614.27\|CTATTTTGTTGGTTTTCGGAACTGAGG | >hg38_21_+_8433172_8446622_RNA45SN1_45S_50ntFlanks | 4588 | 4614 | 27 | CTATTTTGTTGGTTTTCGGAACTGAGG | 210.7 | -5.0 | 1.4 | -3.7 | 2.2E-04 | 3.8E-04 | BT20/MB468 | MITO |
| >hg38_21_+_8433172_8446622_RNA45SN1_45S_50ntFlanks@7975.8005.31\|CGCGACCTCAGATCAGACGTGGCGACCCGCT | >hg38_21_+_8433172_8446622_RNA45SN1_45S_50ntFlanks | 7975 | 8005 | 31 | CGCGACCTCAGATCAGACGTGGCGACCCGCT | 467935.4 | 3.0 | 0.7 | 4.1 | 4.7E-05 | 3.8E-04 | BT20/MB231 | MITO |
| >hg38_21_+_8433172_8446622_RNA45SN1_45S_50ntFlanks@10403.10420.18\|ACATGGGTCAGTCGGTCC | >hg38_21_+_8433172_8446622_RNA45SN1_45S_50ntFlanks | 10403 | 10420 | 18 | ACATGGGTCAGTCGGTCC | 173.6 | -4.9 | 1.3 | -3.6 | 2.7E-04 | 3.8E-04 | BT20/MB468 | CYTO |
| >hg38_21_+_8433172_8446622_RNA45SN1_45S_50ntFlanks@11882.11904.23\|CTGTTGAGCTTGACTCTAGTCTG | >hg38_21_+_8433172_8446622_RNA45SN1_45S_50ntFlanks | 11882 | 11904 | 23 | CTGTTGAGCTTGACTCTAGTCTG | 1004.0 | -4.9 | 1.3 | -3.9 | 1.2E-04 | 3.8E-04 | BT20/MB468 | MP |
| >hg38_21_+_8433172_8446622_RNA45SN1_45S_50ntFlanks@5485.5509.25\|AGCGCTGAGAAGACGGTCGAACTTG | >hg38_21_+_8433172_8446622_RNA45SN1_45S_50ntFlanks | 5485 | 5509 | 25 | AGCGCTGAGAAGACGGTCGAACTTG | 226.4 | -4.1 | 1.1 | -3.6 | 3.0E-04 | 3.8E-04 | MB231/MB468 | NUC |
| >hg38_21_+_8433172_8446622_RNA45SN1_45S_50ntFlanks@10784.10823.40\|TGAACAGCCTCTGGCATGTTGGAACAATGTAGGTAAGGGA | >hg38_21_+_8433172_8446622_RNA45SN1_45S_50ntFlanks | 10784 | 10823 | 40 | TGAACAGCCTCTGGCATGTTGGAACAATGTAGGTAAGGGA | 58.8 | 9.2 | 2.2 | 4.1 | 4.2E-05 | 3.8E-04 | BT20/MB231 | MP |
| >hg38_21_+_8433172_8446622_RNA45SN1_45S_50ntFlanks@11671.11691.21\|CCAGTGCTCTGAATGTCAAAG | >hg38_21_+_8433172_8446622_RNA45SN1_45S_50ntFlanks | 11671 | 11691 | 21 | CCAGTGCTCTGAATGTCAAAG | 784.9 | -2.7 | 0.7 | -3.6 | 3.0E-04 | 3.8E-04 | MB231/MB468 | NUC |
| >MI0000299\|hsa-mir-222&WithFlank&X\|-\|45747009\|45747130@75.99.25(+1U)&[MIMAT0000279&hsa-miR-222-3p&offsets\|0\|+4(+1U);m-140&X\|-\|45747033\|45747056&offsets\|0\|+1(+1U)]\|AGCTACATCTGGCTACTGGGTCTCTT | >MI0000299\|hsa-mir-222&WithFlank&X\|-\|45747009\|45747130 | 75 | 99 | 25(+1U) | AGCTACATCTGGCTACTGGGTCTCTT | 584.9 | 4.7 | 1.3 | 3.7 | 2.5E-04 | 3.8E-04 | MB231/MB468 | MP |
| >hg38_21_+_8433172_8446622_RNA45SN1_45S_50ntFlanks@10903.10935.33\|GGCTGGGCGCGCGCCGCGGCTGGACGAGGCGCC | >hg38_21_+_8433172_8446622_RNA45SN1_45S_50ntFlanks | 10903 | 10935 | 33 | GGCTGGGCGCGCGCCGCGGCTGGACGAGGCGCC | 136.6 | -5.4 | 1.5 | -3.6 | 2.7E-04 | 3.8E-04 | BT20/MB468 | CYTO |
| >MI0003776\|hsa-mir-320b-1&WithFlank&1\|+\|116671740\|116671823@48.67.20(+2U);MI0003839\|hsa-mir-320b-2&WithFlank&1\|-\|224257034\|224257116@47.66.20(+2U);MI0000542\|hsa-mir-320a&WithFlank&8\|-\|22244960\|22245043@48.67.20(+2U)&[MIMAT0005792&hsa-miR-320b&offsets\|0\|-2(+2U);m-183&1\|+\|116671787\|116671806&offsets\|0\|0(+2U)];[MIMAT0005792_1&hsa-miR-320b&offsets\|0\|-2(+2U);m-183&1\|-\|224257051\|224257070&offsets\|0\|0(+2U)];[MIMAT0000510&hsa-miR-320a-3p&offsets\|0\|-2(+2U);m-60&8\|-\|22244975\|22244996&offsets\|0\|-2(+2U)]\|AAAAGCTGGGTTGAGAGGGCTT | >MI0003776\|hsa-mir-320b-1&WithFlank&1\|+\|116671740\|116671823 | 48 | 67 | 20(+2U) | AAAAGCTGGGTTGAGAGGGCTT | 120.8 | 7.3 | 1.9 | 3.9 | 1.2E-04 | 3.8E-04 | BT20/MB468 | MP |
| >am_MI0016746\|hsa-mir-548o-2&WithFlank&20\|+\|38516557\|38516638@51.71.21(+1U);MI0006402\|hsa-mir-548o&WithFlank&7\|-\|102405736\|102405861@93.113.21(+1U)&[MIMAT0005919&hsa-miR-548o-3p&offsets\|0\|-1(+1U);m-431&20\|+\|38516607\|38516628&offsets\|0\|-1(+1U)];[MIMAT0005919_1&hsa-miR-548o-3p&offsets\|0\|-1(+1U);m-430&7\|-\|102405748\|102405769&offsets\|0\|-1(+1U)]\|CCAAAACTGCAGTTACTTTTGT | >am_MI0016746\|hsa-mir-548o-2&WithFlank&20\|+\|38516557\|38516638 | 51 | 71 | 21(+1U) | CCAAAACTGCAGTTACTTTTGT | 134.0 | 4.3 | 1.2 | 3.7 | 2.5E-04 | 3.8E-04 | MB231/MB468 | MP |
| >trnaMT_ProTGG_MT_-_15956_16023@40.58.19\|AATGGTGGAGTTAAAGACT | >trnaMT_ProTGG_MT_-_15956_16023 | 40 | 58 | 19 | AATGGTGGAGTTAAAGACT | 382.7 | 2.6 | 0.7 | 3.6 | 3.0E-04 | 3.8E-04 | MB231/MB468 | MITO |

| | | | | | | | | | | | | | |
|---|---|---|---|---|---|---|---|---|---|---|---|---|---|
| >hg38_21_+_8433172_8446622_RNA45SN1_45S_50ntFlanks@10681.10709.29\|TTCCGGCGGCGTCCGGTGAGCTCTCGCTG | >hg38_21_+_8433172_8446622_RNA45SN1_45S_50ntFlanks | 10681 | 10709 | 29 | TTCCGGCGGCGTCCGGTGAGCTCTCGCTG | 212.0 | -3.8 | 1.0 | -3.6 | 2.7E-04 | 3.8E-04 | BT20/MB468 | CYTO |
| >hg38_21_+_8433172_8446622_RNA45SN1_45S_50ntFlanks@12999.13028.30\|CTGCGATCTATTGAAAGTCAGCCCTCGACA | >hg38_21_+_8433172_8446622_RNA45SN1_45S_50ntFlanks | 12999 | 13028 | 30 | CTGCGATCTATTGAAAGTCAGCCCTCGACA | 89.2 | -5.4 | 1.5 | -3.7 | 2.5E-04 | 3.8E-04 | MB231/MB468 | MP |
| >hg38_21_+_8433172_8446622_RNA45SN1_45S_50ntFlanks@9623.9648.26\|TAGTAGCTGGTTCCCTCCGAAGTTTC | >hg38_21_+_8433172_8446622_RNA45SN1_45S_50ntFlanks | 9623 | 9648 | 26 | TAGTAGCTGGTTCCCTCCGAAGTTTC | 148.3 | -3.9 | 1.1 | -3.6 | 3.0E-04 | 3.8E-04 | MB231/MB468 | NUC |
| >hg38_21_+_8433172_8446622_RNA45SN1_45S_50ntFlanks@10907.10928.22\|GGGCGCGCGCCGCGGCTGGACG | >hg38_21_+_8433172_8446622_RNA45SN1_45S_50ntFlanks | 10907 | 10928 | 22 | GGGCGCGCGCCGCGGCTGGACG | 99.3 | 7.2 | 1.9 | 3.9 | 1.2E-04 | 3.8E-04 | BT20/MB468 | MP |
| >hg38_21_+_8433172_8446622_RNA45SN1_45S_50ntFlanks@11209.11234.26\|CGGCGACTCTGGACGCGAGCCGGGCC | >hg38_21_+_8433172_8446622_RNA45SN1_45S_50ntFlanks | 11209 | 11234 | 26 | CGGCGACTCTGGACGCGAGCCGGGCC | 1022.7 | -2.6 | 0.7 | -3.7 | 2.2E-04 | 3.8E-04 | BT20/MB468 | MITO |
| >hg38_21_+_8433172_8446622_RNA45SN1_45S_50ntFlanks@5084.5112.29\|CGAACGAGACTCTGGCATGCTAACTAGTT | >hg38_21_+_8433172_8446622_RNA45SN1_45S_50ntFlanks | 5084 | 5112 | 29 | CGAACGAGACTCTGGCATGCTAACTAGTT | 610.3 | -4.1 | 1.1 | -3.7 | 2.2E-04 | 3.8E-04 | BT20/MB468 | MITO |
| >hg38_21_+_8433172_8446622_RNA45SN1_45S_50ntFlanks@6719.6758.40\|TGAATTGCAGGACACATTGATCATCGACACTTCGAACGCA | >hg38_21_+_8433172_8446622_RNA45SN1_45S_50ntFlanks | 6719 | 6758 | 40 | TGAATTGCAGGACACATTGATCATCGACACTTCGAACGCA | 223.1 | -2.7 | 0.7 | -3.6 | 2.7E-04 | 3.8E-04 | BT20/MB468 | CYTO |
| >hg38_21_+_8433172_8446622_RNA45SN1_45S_50ntFlanks@9534.9569.36\|AGGAAACTCTGGTGGAGGTCCGTAGCGGTCCTGACG | >hg38_21_+_8433172_8446622_RNA45SN1_45S_50ntFlanks | 9534 | 9569 | 36 | AGGAAACTCTGGTGGAGGTCCGTAGCGGTCCTGACG | 1225.3 | -2.5 | 0.7 | -3.7 | 2.2E-04 | 3.8E-04 | BT20/MB468 | MITO |
| >trnaMT_HisGTG_MT_+_12138_12206@18.52.35\|AAAACATCAGATTGTGAATCTGACAACAGAGGCTT | >trnaMT_HisGTG_MT_+_12138_12206 | 18 | 52 | 35 | AAAACATCAGATTGTGAATCTGACAACAGAGGCTT | 86.2 | 10.2 | 2.6 | 3.9 | 1.2E-04 | 3.8E-04 | BT20/MB468 | MP |
| >hg38_21_+_8433172_8446622_RNA45SN1_45S_50ntFlanks@10914.10938.25\|CGCCGCGGCTGGACGAGGCGCCGCC | >hg38_21_+_8433172_8446622_RNA45SN1_45S_50ntFlanks | 10914 | 10938 | 25 | CGCCGCGGCTGGACGAGGCGCCGCC | 161.8 | -5.7 | 1.5 | -3.9 | 1.2E-04 | 3.8E-04 | BT20/MB468 | MP |
| >hg38_21_+_8433172_8446622_RNA45SN1_45S_50ntFlanks@12417.12437.21\|TCGATGTCGGCTCTTCCTATC | >hg38_21_+_8433172_8446622_RNA45SN1_45S_50ntFlanks | 12417 | 12437 | 21 | TCGATGTCGGCTCTTCCTATC | 1752.5 | -1.7 | 0.5 | -3.7 | 2.2E-04 | 3.8E-04 | BT20/MB468 | MITO |
| >hg38_21_+_8433172_8446622_RNA45SN1_45S_50ntFlanks@11785.11805.21\|TGAATGGATGAACGAGATTCC | >hg38_21_+_8433172_8446622_RNA45SN1_45S_50ntFlanks | 11785 | 11805 | 21 | TGAATGGATGAACGAGATTCC | 640.2 | -4.8 | 1.2 | -3.8 | 1.2E-04 | 3.8E-04 | BT20/MB468 | MP |
| >hg38_21_+_8433172_8446622_RNA45SN1_45S_50ntFlanks@10186.10208.23\|TCCTCCCCTCCTCCCGCCCACGC | >hg38_21_+_8433172_8446622_RNA45SN1_45S_50ntFlanks | 10186 | 10208 | 23 | TCCTCCCCTCCTCCCGCCCACGC | 115.3 | -9.9 | 2.6 | -3.8 | 1.2E-04 | 3.8E-04 | BT20/MB468 | MP |
| >hg38_21_+_8433172_8446622_RNA45SN1_45S_50ntFlanks@3829.3852.24\|CGCTCGCTCCTCTCCTACTTGGAT | >hg38_21_+_8433172_8446622_RNA45SN1_45S_50ntFlanks | 3829 | 3852 | 24 | CGCTCGCTCCTCTCCTACTTGGAT | 220.8 | -4.4 | 1.2 | -3.6 | 3.0E-04 | 3.8E-04 | MB231/MB468 | NUC |
| >hg38_21_+_8433172_8446622_RNA45SN1_45S_50ntFlanks@9837.9864.28\|TGGTAAGCAGAACTGGCGCTGCGGGATG | >hg38_21_+_8433172_8446622_RNA45SN1_45S_50ntFlanks | 9837 | 9864 | 28 | TGGTAAGCAGAACTGGCGCTGCGGGATG | 76.1 | 7.1 | 1.8 | 3.8 | 1.2E-04 | 3.8E-04 | BT20/MB468 | MP |
| >hg38_21_+_8433172_8446622_RNA45SN1_45S_50ntFlanks@11718.11742.25\|AACGGCGGGAGTAACTATGACTCTC | >hg38_21_+_8433172_8446622_RNA45SN1_45S_50ntFlanks | 11718 | 11742 | 25 | AACGGCGGGAGTAACTATGACTCTC | 937.3 | -5.2 | 1.3 | -3.8 | 1.2E-04 | 3.8E-04 | BT20/MB468 | MP |
| >hg38_1_-_228634819_228635039_RNA5S12_5S_50ntFlanks@51.82.32\|GTCTACGGCCATACCACCCTGAACGCGCCCGA | >hg38_1_-_228634819_228635039_RNA5S12_5S_50ntFlanks | 51 | 82 | 32 | GTCTACGGCCATACCACCCTGAACGCGCCCGA | 1101.8 | -2.2 | 0.6 | -3.6 | 3.1E-04 | 3.8E-04 | BT20/MB468 | NUC |
| >hg38_MT_+_1621_3279_RNR2_16S_50ntFlanks@287.310.24\|AAACCAGACGAGCTACCTAAGAAC | >hg38_MT_+_1621_3279_RNR2_16S_50ntFlanks | 287 | 310 | 24 | AAACCAGACGAGCTACCTAAGAAC | 92.9 | 7.2 | 1.9 | 3.8 | 1.2E-04 | 3.8E-04 | BT20/MB468 | MP |
| >hg38_21_+_8433172_8446622_RNA45SN1_45S_50ntFlanks@8170.8214.45\|CAAGTCCTTCTGATCGAGGCCCAGCCCGTGGACGGTGTGAGGCCG | >hg38_21_+_8433172_8446622_RNA45SN1_45S_50ntFlanks | 8170 | 8214 | 45 | CAAGTCCTTCTGATCGAGGCCCAGCCCGTGGACGGTGTGAGGCCG | 1723.4 | -5.2 | 1.4 | -3.6 | 2.7E-04 | 3.8E-04 | BT20/MB468 | CYTO |
| >hg38_21_+_8433172_8446622_RNA45SN1_45S_50ntFlanks@11767.11790.24\|TCTAATTAGTGACGCGCATGAATG | >hg38_21_+_8433172_8446622_RNA45SN1_45S_50ntFlanks | 11767 | 11790 | 24 | TCTAATTAGTGACGCGCATGAATG | 348.6 | -4.1 | 1.1 | -3.6 | 3.1E-04 | 3.8E-04 | MB231/MB468 | NUC |
| >hg38_21_+_8433172_8446622_RNA45SN1_45S_50ntFlanks@12408.12434.27\|TTTGATCCTTCGATGTCGGCTCTTCCT | >hg38_21_+_8433172_8446622_RNA45SN1_45S_50ntFlanks | 12408 | 12434 | 27 | TTTGATCCTTCGATGTCGGCTCTTCCT | 398.4 | 1.6 | 0.4 | 3.7 | 2.3E-04 | 3.9E-04 | BT20/MB468 | MITO |
| >hg38_21_+_8433172_8446622_RNA45SN1_45S_50ntFlanks@12999.13027.29\|CTGCGATCTATTGAAAGTCAGCCCTCGAC | >hg38_21_+_8433172_8446622_RNA45SN1_45S_50ntFlanks | 12999 | 13027 | 29 | CTGCGATCTATTGAAAGTCAGCCCTCGAC | 348.3 | -4.1 | 1.0 | -4.1 | 4.4E-05 | 3.9E-04 | BT20/MB231 | NUC |
| >MI0000803\|hsa-mir-330&WithFlank&19\|-\|45638988\|45639093@65.87.23&[MIMAT0000751&hsa-miR-330-3p&offsets\|+2\|+2;m-347&19\|-\|45639009\|45639030&offsets\|+1\|+2]\|AAAGCACACGGCCTGCAGAGAGG | >MI0000803\|hsa-mir-330&WithFlank&19\|-\|45638988\|45639093 | 65 | 87 | 23 | AAAGCACACGGCCTGCAGAGAGG | 86.5 | 7.3 | 1.9 | 3.8 | 1.2E-04 | 3.9E-04 | BT20/MB468 | MP |
| >hg38_21_+_8433172_8446622_RNA45SN1_45S_50ntFlanks@11629.11662.34\|CGCGAAGGCCCGCGGCGGGTGTTGACGCGATGTG | >hg38_21_+_8433172_8446622_RNA45SN1_45S_50ntFlanks | 11629 | 11662 | 34 | CGCGAAGGCCCGCGGCGGGTGTTGACGCGATGTG | 559.1 | -3.4 | 0.9 | -3.8 | 1.2E-04 | 3.9E-04 | BT20/MB468 | MP |
| >hg38_21_+_8433172_8446622_RNA45SN1_45S_50ntFlanks@6656.6681.26\|CTTAGCGGTGGATCACTCGGCTCGTG | >hg38_21_+_8433172_8446622_RNA45SN1_45S_50ntFlanks | 6656 | 6681 | 26 | CTTAGCGGTGGATCACTCGGCTCGTG | 3836.3 | -4.2 | 1.1 | -3.8 | 1.2E-04 | 3.9E-04 | BT20/MB468 | MP |
| >hg38_21_+_8433172_8446622_RNA45SN1_45S_50ntFlanks@12445.12484.40\|AGCAGAATTCACCAAGCGTTGGATTGTTCACCCACTAATA | >hg38_21_+_8433172_8446622_RNA45SN1_45S_50ntFlanks | 12445 | 12484 | 40 | AGCAGAATTCACCAAGCGTTGGATTGTTCACCCACTAATA | 214.2 | -2.8 | 0.8 | -3.6 | 3.1E-04 | 3.9E-04 | MB231/MB468 | NUC |
| >hg38_21_+_8433172_8446622_RNA45SN1_45S_50ntFlanks@4570.4607.38\|TGGAATAGGACCGCGGTTCTATTTTGTTGGTTTTCGGA | >hg38_21_+_8433172_8446622_RNA45SN1_45S_50ntFlanks | 4570 | 4607 | 38 | TGGAATAGGACCGCGGTTCTATTTTGTTGGTTTTCGGA | 85.4 | 10.2 | 2.6 | 3.8 | 1.2E-04 | 3.9E-04 | BT20/MB468 | MP |
| >hg38_21_+_8433172_8446622_RNA45SN1_45S_50ntFlanks@10788.10807.20\|CAGCCTCTGGCATGTTGGAA | >hg38_21_+_8433172_8446622_RNA45SN1_45S_50ntFlanks | 10788 | 10807 | 20 | CAGCCTCTGGCATGTTGGAA | 89.3 | -5.7 | 1.5 | -3.8 | 1.2E-04 | 3.9E-04 | BT20/MB468 | MP |
| >trnaMT_GlyTCC_MT_+_9991_10058@15.33.19\|AATAGTACCGTTAACTTCC | >trnaMT_GlyTCC_MT_+_9991_10058 | 15 | 33 | 19 | AATAGTACCGTTAACTTCC | 90.8 | 10.3 | 2.7 | 3.8 | 1.2E-04 | 3.9E-04 | BT20/MB468 | MP |
| >hg38_21_+_8433172_8446622_RNA45SN1_45S_50ntFlanks@12444.12485.42\|AAGCAGAATTCACCAAGCGTTGGATTGTTCACCCACTAATAG | >hg38_21_+_8433172_8446622_RNA45SN1_45S_50ntFlanks | 12444 | 12485 | 42 | AAGCAGAATTCACCAAGCGTTGGATTGTTCACCCACTAATAG | 299.3 | -4.6 | 1.3 | -3.6 | 3.0E-04 | 3.9E-04 | MB231/MB468 | MITO |
| >am_trnaMT_SerGCT_MT_+_12207_12265@16.50.35\|AACTGCTAACTCATGCCCCCATGTCTAACAACATG | >am_trnaMT_SerGCT_MT_+_12207_12265 | 16 | 50 | 35 | AACTGCTAACTCATGCCCCCATGTCTAACAACATG | 194.4 | -4.5 | 1.2 | -3.6 | 3.1E-04 | 3.9E-04 | MB231/MB468 | NUC |
| >hg38_21_+_8433172_8446622_RNA45SN1_45S_50ntFlanks@3922.3959.38\|TTTATCAGATCAAAACCAACCCGGTCAGCCCCTCTCCG | >hg38_21_+_8433172_8446622_RNA45SN1_45S_50ntFlanks | 3922 | 3959 | 38 | TTTATCAGATCAAAACCAACCCGGTCAGCCCCTCTCCG | 173.5 | -5.0 | 1.4 | -3.7 | 2.3E-04 | 3.9E-04 | BT20/MB468 | MITO |
| >hg38_MT_+_598_1651_RNR1_12S_50ntFlanks@428.468.41\|AAAGTGGCTTTAACATATCTGAACACACAATAGCTAAGACC | >hg38_MT_+_598_1651_RNR1_12S_50ntFlanks | 428 | 468 | 41 | AAAGTGGCTTTAACATATCTGAACACACAATAGCTAAGACC | 410.7 | 4.5 | 1.2 | 3.7 | 2.6E-04 | 3.9E-04 | MB231/MB468 | MP |
| >hg38_21_+_8433172_8446622_RNA45SN1_45S_50ntFlanks@10780.10818.39\|AAGGTGAACAGCCTCTGGCATGTTGGAACAATGTAGGTA | >hg38_21_+_8433172_8446622_RNA45SN1_45S_50ntFlanks | 10780 | 10818 | 39 | AAGGTGAACAGCCTCTGGCATGTTGGAACAATGTAGGTA | 170.4 | -2.2 | 0.6 | -3.6 | 2.9E-04 | 3.9E-04 | MB231/MB468 | CYTO |
| >hg38_21_+_8433172_8446622_RNA45SN1_45S_50ntFlanks@10404.10424.21\|CATGGGTCAGTCGGTCCTGAG | >hg38_21_+_8433172_8446622_RNA45SN1_45S_50ntFlanks | 10404 | 10424 | 21 | CATGGGTCAGTCGGTCCTGAG | 1613.6 | 2.6 | 0.7 | 3.6 | 2.9E-04 | 3.9E-04 | MB231/MB468 | CYTO |
| >hg38_21_+_8433172_8446622_RNA45SN1_45S_50ntFlanks@10797.10820.24\|GCATGTTGGAACAATGTAGGTAAG | >hg38_21_+_8433172_8446622_RNA45SN1_45S_50ntFlanks | 10797 | 10820 | 24 | GCATGTTGGAACAATGTAGGTAAG | 263.2 | -4.4 | 1.2 | -3.7 | 2.6E-04 | 3.9E-04 | MB231/MB468 | MP |
| >trnaMT_PheGAA_MT_+_577_647@39.57.19\|ATGTTTAGACGGGCTCACA | >trnaMT_PheGAA_MT_+_577_647 | 39 | 57 | 19 | ATGTTTAGACGGGCTCACA | 830.3 | 2.7 | 0.8 | 3.6 | 3.0E-04 | 3.9E-04 | MB231/MB468 | MITO |
| >trna111_HisGTG_1_-_147774845_147774916@-1T.24.25,trna118_HisGTG_1_-_145396881_145396952@-1T.24.25,trna16_HisGTG_1_+_146544773_146544844@-1T.24.25,trna1_HisGTG_15_+_45493349_45493420@-1T.24.25,trna21_HisGTG_1_+_147753471_147753542@-1T.24.25,trna33_HisGTG_6_+_27125906_27125977@-1T.24.25,trna7_HisGTG_9_-_14433938_14434009@-1T.24.25,trna8_HisGTG_15_-_45492611_45492682@-1T.24.25,trna9_HisGTG_15_-_45490804_45490875@-1T.24.25\|TGCCGTGATCGTATAGTGGTTAGTA | >trna111_HisGTG_1_-_147774845_147774916 | -1T | 24 | 25 | TGCCGTGATCGTATAGTGGTTAGTA | 177.7 | 3.2 | 0.9 | 3.6 | 3.0E-04 | 3.9E-04 | MB231/MB468 | MITO |
| >MI0000079\|hsa-mir-23a&WithFlank&19\|-\|13836581\|13836665@54.72.19(+2U)&[MIMAT0000078&hsa-miR-23a-3p&offsets\|+3\|+1(+2U);m-31&19\|-\|13836595\|13836615&offsets\|+3\|+1(+2U)]\|ACATTGCCAGGGATTTCCATT | >MI0000079\|hsa-mir-23a&WithFlank&19\|-\|13836581\|13836665 | 54 | 72 | 19(+2U) | ACATTGCCAGGGATTTCCATT | 590.4 | 13.1 | 3.6 | 3.6 | 3.1E-04 | 3.9E-04 | MB231/MB468 | NUC |
| >hg38_21_+_8433172_8446622_RNA45SN1_45S_50ntFlanks@10229.10263.35\|CCCCGCGGACGCTACGCCGCGACGAGTAGGAGGGC | >hg38_21_+_8433172_8446622_RNA45SN1_45S_50ntFlanks | 10229 | 10263 | 35 | CCCCGCGGACGCTACGCCGCGACGAGTAGGAGGGC | 150.9 | 4.3 | 1.1 | 3.8 | 1.2E-04 | 3.9E-04 | BT20/MB468 | MP |
| >hg38_21_+_8433172_8446622_RNA45SN1_45S_50ntFlanks@12410.12437.28\|TGATCCTTCGATGTCGGCTCTTCCTATC | >hg38_21_+_8433172_8446622_RNA45SN1_45S_50ntFlanks | 12410 | 12437 | 28 | TGATCCTTCGATGTCGGCTCTTCCTATC | 196.5 | 4.1 | 1.1 | 3.6 | 2.9E-04 | 3.9E-04 | MB231/MB468 | CYTO |
| >hg38_21_+_8433172_8446622_RNA45SN1_45S_50ntFlanks@5078.5097.20\|CGATAACGAACGAGACTCTG | >hg38_21_+_8433172_8446622_RNA45SN1_45S_50ntFlanks | 5078 | 5097 | 20 | CGATAACGAACGAGACTCTG | 640.6 | -5.3 | 1.5 | -3.6 | 3.1E-04 | 3.9E-04 | BT20/MB468 | NUC |
| >hg38_21_+_8433172_8446622_RNA45SN1_45S_50ntFlanks@9352.9370.19\|CCGTGGCGCAATGAAGGTG | >hg38_21_+_8433172_8446622_RNA45SN1_45S_50ntFlanks | 9352 | 9370 | 19 | CCGTGGCGCAATGAAGGTG | 293.1 | -4.0 | 1.1 | -3.6 | 2.9E-04 | 3.9E-04 | MB231/MB468 | CYTO |
| >trnaMT_GluTTC_MT_-_14674_14742@24.72.49,trnalookalike8_GluTTC_5_-_93905172_93905240@24.72.49\|ATGGTTTTTCATATCATTGGTCGTGGTTGTAGTCCGTGCGAGAATACCA | >trnaMT_GluTTC_MT_-_14674_14742 | 24 | 72 | 49 | ATGGTTTTTCATATCATTGGTCGTGGTTGTAGTCCGTGCGAGAATACCA | 146.5 | 6.7 | 1.9 | 3.6 | 3.0E-04 | 3.9E-04 | MB231/MB468 | MITO |
| >hg38_21_+_8433172_8446622_RNA45SN1_45S_50ntFlanks@7984.8009.26\|AGATCAGACGTGGCGACCCGCTGAAT | >hg38_21_+_8433172_8446622_RNA45SN1_45S_50ntFlanks | 7984 | 8009 | 26 | AGATCAGACGTGGCGACCCGCTGAAT | 901.3 | -3.5 | 1.0 | -3.6 | 3.0E-04 | 3.9E-04 | MB231/MB468 | MITO |
| >hg38_21_+_8433172_8446622_RNA45SN1_45S_50ntFlanks@9740.9758.19\|AACGATCTCAACCTATTCT | >hg38_21_+_8433172_8446622_RNA45SN1_45S_50ntFlanks | 9740 | 9758 | 19 | AACGATCTCAACCTATTCT | 497.2 | -4.8 | 1.3 | -3.6 | 3.1E-04 | 3.9E-04 | BT20/MB468 | NUC |
| >hg38_21_+_8433172_8446622_RNA45SN1_45S_50ntFlanks@4002.4020.19\|GATAACCTCGGGCCGATCG | >hg38_21_+_8433172_8446622_RNA45SN1_45S_50ntFlanks | 4002 | 4020 | 19 | GATAACCTCGGGCCGATCG | 323.0 | -5.3 | 1.4 | -3.7 | 2.3E-04 | 3.9E-04 | BT20/MB468 | MITO |
| >hg38_21_+_8433172_8446622_RNA45SN1_45S_50ntFlanks@12419.12436.18\|GATGTCGGCTCTTCCTAT | >hg38_21_+_8433172_8446622_RNA45SN1_45S_50ntFlanks | 12419 | 12436 | 18 | GATGTCGGCTCTTCCTAT | 2212.5 | -2.3 | 0.6 | -3.6 | 2.7E-04 | 3.9E-04 | BT20/MB468 | CYTO |
| >hg38_21_+_8433172_8446622_RNA45SN1_45S_50ntFlanks@6646.6667.22\|TCGTACGACTCTTAGCGGTGGA | >hg38_21_+_8433172_8446622_RNA45SN1_45S_50ntFlanks | 6646 | 6667 | 22 | TCGTACGACTCTTAGCGGTGGA | 12472.3 | 2.8 | 0.7 | 4.1 | 4.1E-05 | 3.9E-04 | BT20/MB231 | CYTO |
| >trna128_GlyGCC_6_-_27870686_27870756@1.40.40,trna19_GlyGCC_16_+_70823410_70823480@1.40.40,trna19_GlyGCC_2_-_157257659_157257729@1.40.40,trna24_GlyGCC_16_-_70812942_70813012@1.40.40,trna25_GlyGCC_16_-_70812114_70812184@1.40.40,trna5_GlyGCC_17_+_8029064_8029134@1.40.40,trna68_GlyGCC_1_-_161493637_161493707@1.40.40\|GCATTGGTGGTTCAGTGGTAGAATTCTCGCCTGCCACGCG | >trna128_GlyGCC_6_-_27870686_27870756 | 1 | 40 | 40 | GCATTGGTGGTTCAGTGGTAGAATTCTCGCCTGCCACGCG | 930.1 | -4.0 | 1.0 | -4.1 | 4.1E-05 | 3.9E-04 | BT20/MB231 | CYTO |

| | | | | | | | | | | | | | |
|---|---|---|---|---|---|---|---|---|---|---|---|---|---|
| >hg38_21_+_8433172_8446622_RNA45SN1_45S_50ntFlanks@12417.12456.40\|TCGATGTCGGCTCTTCCTATCATTGTGAAGCAGAATTCAC | >hg38_21_+_8433172_8446622_RNA45SN1_45S_50ntFlanks | 12417 | 12456 | 40 | TCGATGTCGGCTCTTCCTATCATTGTGAAGCAGAATTCAC | 163.4 | -3.6 | 0.9 | -4.1 | 4.1E-05 | 3.9E-04 | BT20/MB231 | CYTO |
| >MI0000065\|hsa-let-7d&WithFlank&9\|+\|94178828\|94178926@14.31.18&[MIMAT0000065&hsa-let-7d-5p&offsets\|0\|-4;m-90&9\|+\|94178841\|94178862&offsets\|0\|-4]\|AGAGGTAGTAGGTTGCAT | >MI0000065\|hsa-let-7d&WithFlank&9\|+\|94178828\|94178926 | 14 | 31 | 18 | AGAGGTAGTAGGTTGCAT | 130.6 | 5.6 | 1.6 | 3.6 | 3.1E-04 | 3.9E-04 | MB231/MB468 | MITO |
| >hg38_21_+_8433172_8446622_RNA45SN1_45S_50ntFlanks@10694.10711.18\|CGGTGAGCTCTCGCTGGC | >hg38_21_+_8433172_8446622_RNA45SN1_45S_50ntFlanks | 10694 | 10711 | 18 | CGGTGAGCTCTCGCTGGC | 1141.9 | -5.1 | 1.4 | -3.6 | 3.1E-04 | 3.9E-04 | MB231/MB468 | MITO |
| >MI0000746\|hsa-mir-99b&WithFlank&19\|+\|51692606\|51692687@51.71.21(+1U)&[MIMAT0004678&hsa-miR-99b-3p&offsets\|0\|-1(+1U);m-151&19\|+\|51692656\|51692678&offsets\|0\|-2(+1U)]\|CAAGCTCGTGTCTGTGGGTCCT | >MI0000746\|hsa-mir-99b&WithFlank&19\|+\|51692606\|51692687 | 51 | 71 | 21(+1U) | CAAGCTCGTGTCTGTGGGTCCT | 110.0 | 6.9 | 1.8 | 3.8 | 1.2E-04 | 3.9E-04 | BT20/MB468 | MP |
| >MI0003137\|hsa-mir-193b&WithFlank&16\|+\|14303961\|14304055@57.80.24&[MIMAT0002819&hsa-miR-193b-3p&offsets\|0\|+2;m-166&16\|+\|14304017\|14304038&offsets\|0\|+2]\|AACTGGCCCTCAAAGTCCCGCTTT | >MI0003137\|hsa-mir-193b&WithFlank&16\|+\|14303961\|14304055 | 57 | 80 | 24 | AACTGGCCCTCAAAGTCCCGCTTT | 168.3 | 7.6 | 2.0 | 3.8 | 1.2E-04 | 3.9E-04 | BT20/MB468 | MP |
| >MI0000087\|hsa-mir-29a&WithFlank&7\|-\|130876741\|130876816@48.69.22(+1C)&[MIMAT0000086&hsa-miR-29a-3p&offsets\|0\|0(+1C);m-26&7\|-\|130876749\|130876769&offsets\|0\|+1(+1C)]\|TAGCACCATCTGAAATCGGTTAC | >MI0000087\|hsa-mir-29a&WithFlank&7\|-\|130876741\|130876816 | 48 | 69 | 22(+1C) | TAGCACCATCTGAAATCGGTTAC | 86.6 | 10.2 | 2.6 | 3.8 | 1.2E-04 | 3.9E-04 | BT20/MB468 | MP |
| >hg38_1_-_228634819_228635039_RNA5S12_5S_50ntFlanks@133.169.37\|ATGGGAGACCGCCTGGGAATACCGGGTGCTGTAGGCT | >hg38_1_-_228634819_228635039_RNA5S12_5S_50ntFlanks | 133 | 169 | 37 | ATGGGAGACCGCCTGGGAATACCGGGTGCTGTAGGCT | 2973.6 | 4.0 | 1.1 | 3.7 | 2.3E-04 | 3.9E-04 | BT20/MB468 | MITO |
| >hg38_21_+_8433172_8446622_RNA45SN1_45S_50ntFlanks@4734.4752.19\|AACGAAAGTCGGAGGTTCG | >hg38_21_+_8433172_8446622_RNA45SN1_45S_50ntFlanks | 4734 | 4752 | 19 | AACGAAAGTCGGAGGTTCG | 528.6 | -3.4 | 0.9 | -3.8 | 1.2E-04 | 3.9E-04 | BT20/MB468 | MP |
| >hg38_21_+_8433172_8446622_RNA45SN1_45S_50ntFlanks@12029.12056.28\|TCTGATCGTTTTTTCACTGACCCGGTGA | >hg38_21_+_8433172_8446622_RNA45SN1_45S_50ntFlanks | 12029 | 12056 | 28 | TCTGATCGTTTTTTCACTGACCCGGTGA | 75.8 | 10.0 | 2.6 | 3.8 | 1.2E-04 | 3.9E-04 | BT20/MB468 | MP |
| >hg38_21_+_8433172_8446622_RNA45SN1_45S_50ntFlanks@12638.12669.32\|CTACCATCTGTGGGATTATGACTGAACGCCTC | >hg38_21_+_8433172_8446622_RNA45SN1_45S_50ntFlanks | 12638 | 12669 | 32 | CTACCATCTGTGGGATTATGACTGAACGCCTC | 117.8 | -4.4 | 1.2 | -3.7 | 2.6E-04 | 3.9E-04 | MB231/MB468 | MP |
| >hg38_21_+_8433172_8446622_RNA45SN1_45S_50ntFlanks@6650.6699.50\|ACGACTCTTAGCGGTGGATCACTCGGCTCGTGCGTCGATGAAGAACGCAG | >hg38_21_+_8433172_8446622_RNA45SN1_45S_50ntFlanks | 6650 | 6699 | 50 | ACGACTCTTAGCGGTGGATCACTCGGCTCGTGCGTCGATGAAGAACGCAG | 4448.4 | -3.4 | 0.9 | -3.6 | 3.1E-04 | 3.9E-04 | MB231/MB468 | MITO |
| >hg38_21_+_8433172_8446622_RNA45SN1_45S_50ntFlanks@12635.12669.35\|AAGCTACCATCTGTGGGATTATGACTGAACGCCTC | >hg38_21_+_8433172_8446622_RNA45SN1_45S_50ntFlanks | 12635 | 12669 | 35 | AAGCTACCATCTGTGGGATTATGACTGAACGCCTC | 160.5 | -3.2 | 0.9 | -3.7 | 2.6E-04 | 3.9E-04 | MB231/MB468 | MP |
| >hg38_21_+_8433172_8446622_RNA45SN1_45S_50ntFlanks@12241.12266.26\|CAGAAGGGCAAAAGCTCGCTTGATCT | >hg38_21_+_8433172_8446622_RNA45SN1_45S_50ntFlanks | 12241 | 12266 | 26 | CAGAAGGGCAAAAGCTCGCTTGATCT | 163.5 | -5.8 | 1.5 | -3.8 | 1.2E-04 | 3.9E-04 | BT20/MB468 | MP |
| >hg38_21_+_8433172_8446622_RNA45SN1_45S_50ntFlanks@3841.3865.25\|TCCTACTTGGATAACTGTGGTAATT | >hg38_21_+_8433172_8446622_RNA45SN1_45S_50ntFlanks | 3841 | 3865 | 25 | TCCTACTTGGATAACTGTGGTAATT | 289.0 | 3.0 | 0.7 | 4.1 | 4.2E-05 | 3.9E-04 | BT20/MB231 | CYTO |
| >hg38_21_+_8433172_8446622_RNA45SN1_45S_50ntFlanks@3775.3794.20\|GCCGGTACAGTGAAACTGCG | >hg38_21_+_8433172_8446622_RNA45SN1_45S_50ntFlanks | 3775 | 3794 | 20 | GCCGGTACAGTGAAACTGCG | 227.7 | -3.5 | 1.0 | -3.6 | 2.8E-04 | 3.9E-04 | BT20/MB468 | CYTO |
| >MI0000079\|hsa-mir-23a&WithFlank&19\|-\|13836581\|13836665@54.73.20(+1U)&[MIMAT0000078&hsa-miR-23a-3p&offsets\|+3\|+2(+1U);m-31&19\|-\|13836595\|13836615&offsets\|+3\|+2(+1U)]\|ACATTGCCAGGGATTTCCAAT | >MI0000079\|hsa-mir-23a&WithFlank&19\|-\|13836581\|13836665 | 54 | 73 | 20(+1U) | ACATTGCCAGGGATTTCCAAT | 83.5 | 10.1 | 2.6 | 3.8 | 1.2E-04 | 3.9E-04 | BT20/MB468 | MP |
| >hg38_21_+_8433172_8446622_RNA45SN1_45S_50ntFlanks@3811.3836.26\|AGTTATGGTTCCTTTGGTCGCTCGCT | >hg38_21_+_8433172_8446622_RNA45SN1_45S_50ntFlanks | 3811 | 3836 | 26 | AGTTATGGTTCCTTTGGTCGCTCGCT | 285.9 | -4.2 | 1.2 | -3.6 | 3.1E-04 | 3.9E-04 | MB231/MB468 | NUC |
| >MI0000067\|hsa-let-7f-1&WithFlank&9\|+\|94176341\|94176439@13.34.22;MI0000068\|hsa-let-7f-2&WithFlank&X\|-\|53557186\|53557280@14.35.22&[MIMAT0000067&hsa-let-7f-5p&offsets\|0\|0;m-20&9\|+\|94176353\|94176374&offsets\|0\|0];[MIMAT0000067_1&hsa-let-7f-5p&offsets\|0\|0;m-19&X\|-\|53557246\|53557267&offsets\|0\|0]\|TGAGGTAGTAGATTGTATAGTT | >MI0000067\|hsa-let-7f-1&WithFlank&9\|+\|94176341\|94176439 | 13 | 34 | 22 | TGAGGTAGTAGATTGTATAGTT | 60967.5 | 3.4 | 0.9 | 3.7 | 2.6E-04 | 3.9E-04 | MB231/MB468 | MP |
| >hg38_21_+_8433172_8446622_RNA45SN1_45S_50ntFlanks@8104.8142.39\|CCGCGGCGGGGCGCGGGACATGTGGCGTACGGAAGACCC | >hg38_21_+_8433172_8446622_RNA45SN1_45S_50ntFlanks | 8104 | 8142 | 39 | CCGCGGCGGGGCGCGGGACATGTGGCGTACGGAAGACCC | 450.4 | -6.4 | 1.8 | -3.6 | 3.1E-04 | 3.9E-04 | BT20/MB468 | NUC |
| >hg38_21_+_8433172_8446622_RNA45SN1_45S_50ntFlanks@9454.9491.38\|CCGCGCCGGGGAGGTGGAGCACGAGCGCACGTGTTAGG | >hg38_21_+_8433172_8446622_RNA45SN1_45S_50ntFlanks | 9454 | 9491 | 38 | CCGCGCCGGGGAGGTGGAGCACGAGCGCACGTGTTAGG | 201.9 | -5.9 | 1.5 | -3.8 | 1.2E-04 | 3.9E-04 | BT20/MB468 | MP |
| >hg38_21_+_8433172_8446622_RNA45SN1_45S_50ntFlanks@11783.11818.36\|CATGAATGGATGAACGAGATTCCCACTGTCCCTACC | >hg38_21_+_8433172_8446622_RNA45SN1_45S_50ntFlanks | 11783 | 11818 | 36 | CATGAATGGATGAACGAGATTCCCACTGTCCCTACC | 118.9 | -4.5 | 1.2 | -3.7 | 2.6E-04 | 3.9E-04 | MB231/MB468 | MP |
| >hg38_21_+_8433172_8446622_RNA45SN1_45S_50ntFlanks@11208.11249.42\|GCGGCGACTCTGGACGCGAGCCGGGCCCTTCCCGTGGATCGC | >hg38_21_+_8433172_8446622_RNA45SN1_45S_50ntFlanks | 11208 | 11249 | 42 | GCGGCGACTCTGGACGCGAGCCGGGCCCTTCCCGTGGATCGC | 550.6 | -6.4 | 1.8 | -3.6 | 3.2E-04 | 3.9E-04 | BT20/MB468 | NUC |
| >hg38_21_+_8433172_8446622_RNA45SN1_45S_50ntFlanks@11326.11345.20\|CTCGTCGCGCGCGCGTCCGC | >hg38_21_+_8433172_8446622_RNA45SN1_45S_50ntFlanks | 11326 | 11345 | 20 | CTCGTCGCGCGCGCGTCCGC | 1014.0 | -4.2 | 1.0 | -4.1 | 4.9E-05 | 3.9E-04 | BT20/MB231 | MITO |
| >MI0000465\|hsa-mir-191&WithFlank&3\|-\|49020612\|49020715@23.44.22&[MIMAT0000440&hsa-miR-191-5p&offsets\|+1\|0;m-48&3\|-\|49020672\|49020694&offsets\|+1\|0]\|AACGGAATCCCAAAAGCAGCTG | >MI0000465\|hsa-mir-191&WithFlank&3\|-\|49020612\|49020715 | 23 | 44 | 22 | AACGGAATCCCAAAAGCAGCTG | 389.2 | 3.9 | 1.0 | 3.8 | 1.2E-04 | 3.9E-04 | BT20/MB468 | MP |
| >hg38_21_+_8433172_8446622_RNA45SN1_45S_50ntFlanks@9792.9816.25\|GGCGTGGAGCCGGGCGTGGAATGCG | >hg38_21_+_8433172_8446622_RNA45SN1_45S_50ntFlanks | 9792 | 9816 | 25 | GGCGTGGAGCCGGGCGTGGAATGCG | 183.0 | 7.1 | 1.9 | 3.8 | 1.2E-04 | 3.9E-04 | BT20/MB468 | MP |
| >hg38_21_+_8433172_8446622_RNA45SN1_45S_50ntFlanks@4370.4391.22\|TTAAAAAGCTCGTAGTTGGATC | >hg38_21_+_8433172_8446622_RNA45SN1_45S_50ntFlanks | 4370 | 4391 | 22 | TTAAAAAGCTCGTAGTTGGATC | 164.1 | -4.4 | 1.2 | -3.6 | 3.1E-04 | 3.9E-04 | MB231/MB468 | NUC |
| >hg38_21_+_8433172_8446622_RNA45SN1_45S_50ntFlanks@3711.3729.19\|GTTGATCCTGCCAGTAGCA | >hg38_21_+_8433172_8446622_RNA45SN1_45S_50ntFlanks | 3711 | 3729 | 19 | GTTGATCCTGCCAGTAGCA | 226.6 | -4.5 | 1.2 | -3.7 | 2.6E-04 | 3.9E-04 | MB231/MB468 | MP |
| >MI0000273\|hsa-mir-183&WithFlank&7\|-\|129774899\|129775020@34.53.20&[MIMAT0000261&hsa-miR-183-5p&offsets\|+1\|-1;m-41&7\|-\|129774966\|129774987&offsets\|0\|-2]\|ATGGCACTGGTAGAATTCAC | >MI0000273\|hsa-mir-183&WithFlank&7\|-\|129774899\|129775020 | 34 | 53 | 20 | ATGGCACTGGTAGAATTCAC | 243.5 | 6.5 | 1.7 | 3.8 | 1.2E-04 | 4.0E-04 | BT20/MB468 | MP |
| >hg38_21_+_8433172_8446622_RNA45SN1_45S_50ntFlanks@8692.8715.24\|CCGGGACGGCTGGGAAGGCCCGGC | >hg38_21_+_8433172_8446622_RNA45SN1_45S_50ntFlanks | 8692 | 8715 | 24 | CCGGGACGGCTGGGAAGGCCCGGC | 86.3 | 10.2 | 2.6 | 3.8 | 1.2E-04 | 4.0E-04 | BT20/MB468 | MP |
| >hg38_21_+_8433172_8446622_RNA45SN1_45S_50ntFlanks@11027.11050.24\|CTCTCTCTCTCTCTCCCCCGCTCC | >hg38_21_+_8433172_8446622_RNA45SN1_45S_50ntFlanks | 11027 | 11050 | 24 | CTCTCTCTCTCTCTCCCCCGCTCC | 1396.5 | -4.5 | 1.3 | -3.6 | 2.8E-04 | 4.0E-04 | BT20/MB468 | CYTO |
| >MI0003834\|hsa-mir-769&WithFlank&19\|+\|46018926\|46019055@36.57.22(+1U)&[MIMAT0003886&hsa-miR-769-5p&offsets\|0\|0(+1U);m-216&19\|+\|46018961\|46018982&offsets\|0\|0(+1U)]\|TGAGACCTCTGGGTTCTGAGCTT | >MI0003834\|hsa-mir-769&WithFlank&19\|+\|46018926\|46019055 | 36 | 57 | 22(+1U) | TGAGACCTCTGGGTTCTGAGCTT | 345.7 | 2.9 | 0.8 | 3.7 | 2.6E-04 | 4.0E-04 | MB231/MB468 | MP |
| >hg38_21_+_8433172_8446622_RNA45SN1_45S_50ntFlanks@6660.6696.37\|GCGGTGGATCACTCGGCTCGTGCGTCGATGAAGAACG | >hg38_21_+_8433172_8446622_RNA45SN1_45S_50ntFlanks | 6660 | 6696 | 37 | GCGGTGGATCACTCGGCTCGTGCGTCGATGAAGAACG | 479.6 | 2.3 | 0.6 | 3.7 | 2.3E-04 | 4.0E-04 | BT20/MB468 | MITO |
| >hg38_21_+_8433172_8446622_RNA45SN1_45S_50ntFlanks@6774.6806.33\|TCCTCCCGGGGCTACGCCTGTCTGAGCGTCGCT | >hg38_21_+_8433172_8446622_RNA45SN1_45S_50ntFlanks | 6774 | 6806 | 33 | TCCTCCCGGGGCTACGCCTGTCTGAGCGTCGCT | 2670.4 | 3.9 | 1.1 | 3.6 | 3.1E-04 | 4.0E-04 | MB231/MB468 | MITO |
| >hg38_21_+_8433172_8446622_RNA45SN1_45S_50ntFlanks@4927.4944.18\|AGTGGAGCCTGCGGCTTA | >hg38_21_+_8433172_8446622_RNA45SN1_45S_50ntFlanks | 4927 | 4944 | 18 | AGTGGAGCCTGCGGCTTA | 410.3 | 3.5 | 1.0 | 3.7 | 2.3E-04 | 4.0E-04 | BT20/MB468 | MITO |
| >hg38_21_+_8433172_8446622_RNA45SN1_45S_50ntFlanks@476.523.48\|TTCGTGATCGATGTGGTGACGTCGTGCTCTCCCGGGCCGGGTCCGAGC | >hg38_21_+_8433172_8446622_RNA45SN1_45S_50ntFlanks | 476 | 523 | 48 | TTCGTGATCGATGTGGTGACGTCGTGCTCTCCCGGGCCGGGTCCGAGC | 10555.8 | 4.7 | 1.3 | 3.6 | 3.1E-04 | 4.0E-04 | MB231/MB468 | NUC |
| >MI0003562\|hsa-mir-556&WithFlank&1\|+\|162342540\|162342646@22.42.21&[MIMAT0003220&hsa-miR-556-5p&offsets\|0\|-1;m-1043&1\|+\|162342561\|162342581&offsets\|0\|0]\|GATGAGCTCATTGTAATATGA | >MI0003562\|hsa-mir-556&WithFlank&1\|+\|162342540\|162342646 | 22 | 42 | 21 | GATGAGCTCATTGTAATATGA | 52.3 | -9.1 | 2.2 | -4.1 | 4.4E-05 | 4.0E-04 | BT20/MB231 | MP |
| >MI0000061\|hsa-let-7a-2&WithFlank&11\|-\|122146516\|122146599@11.33.23;MI0000062\|hsa-let-7a-3&WithFlank&22\|+\|46112743\|46112828@10.32.23;MI0000060\|hsa-let-7a-1&WithFlank&9\|+\|94175951\|94176042@12.34.23&[MIMAT0000062&hsa-let-7a-5p&offsets\|0\|+1;m-6&11\|-\|122146568\|122146589&offsets\|0\|+1];[MIMAT0000062_1&hsa-let-7a-5p&offsets\|0\|+1;m-5&22\|+\|46112752\|46112773&offsets\|0\|+1];[MIMAT0000062_2&hsa-let-7a-5p&offsets\|0\|+1;m-4&9\|+\|94175962\|94175983&offsets\|0\|+1]\|TGAGGTAGTAGGTTGTATAGTTT | >MI0000061\|hsa-let-7a-2&WithFlank&11\|-\|122146516\|122146599 | 11 | 33 | 23 | TGAGGTAGTAGGTTGTATAGTTT | 1320.8 | 3.7 | 1.0 | 3.8 | 1.2E-04 | 4.0E-04 | BT20/MB468 | MP |
| >hg38_21_+_8433172_8446622_RNA45SN1_45S_50ntFlanks@10866.10884.19\|CTAAGGGCTGGGTCGGTCG | >hg38_21_+_8433172_8446622_RNA45SN1_45S_50ntFlanks | 10866 | 10884 | 19 | CTAAGGGCTGGGTCGGTCG | 266.9 | -3.9 | 1.1 | -3.7 | 2.3E-04 | 4.0E-04 | BT20/MB468 | MITO |
| >hg38_21_+_8433172_8446622_RNA45SN1_45S_50ntFlanks@12604.12625.22\|GTATGTGCTTGGCTGAGGAGCC | >hg38_21_+_8433172_8446622_RNA45SN1_45S_50ntFlanks | 12604 | 12625 | 22 | GTATGTGCTTGGCTGAGGAGCC | 231.9 | 2.7 | 0.7 | 3.6 | 2.9E-04 | 4.0E-04 | MB231/MB468 | CYTO |
| >MI0000542\|hsa-mir-320a&WithFlank&8\|-\|22244960\|22245043@50.69.20(+1U)&[MIMAT0000510&hsa-miR-320a-3p&offsets\|+2\|0(+1U);m-60&8\|-\|22244975\|22244996&offsets\|+2\|0(+1U)]\|AAGCTGGGTTGAGAGGGCGAT | >MI0000542\|hsa-mir-320a&WithFlank&8\|-\|22244960\|22245043 | 50 | 69 | 20(+1U) | AAGCTGGGTTGAGAGGGCGAT | 227.2 | 7.4 | 1.9 | 3.8 | 1.2E-04 | 4.0E-04 | BT20/MB468 | MP |
| >hg38_21_+_8433172_8446622_RNA45SN1_45S_50ntFlanks@476.509.34\|TTCGTGATCGATGTGGTGACGTCGTGCTCTCCCG | >hg38_21_+_8433172_8446622_RNA45SN1_45S_50ntFlanks | 476 | 509 | 34 | TTCGTGATCGATGTGGTGACGTCGTGCTCTCCCG | 465.2 | 4.6 | 1.3 | 3.6 | 3.1E-04 | 4.0E-04 | MB231/MB468 | MITO |
| >MI0000085\|hsa-mir-27a&WithFlank&19\|-\|13836434\|13836523@57.77.21&[MIMAT0000084&hsa-miR-27a-3p&offsets\|0\|0;m-43&19\|-\|13836447\|13836467&offsets\|0\|0]\|TTCACAGTGGCTAAGTTCCGC | >MI0000085\|hsa-mir-27a&WithFlank&19\|-\|13836434\|13836523 | 57 | 77 | 21 | TTCACAGTGGCTAAGTTCCGC | 3742.9 | -3.4 | 0.8 | -4.1 | 5.0E-05 | 4.0E-04 | BT20/MB231 | MITO |
| >MI0000261\|hsa-mir-139&WithFlank&11\|-\|72615057\|72615136@13.34.22(+1C)&[MIMAT0000250&hsa-miR-139-5p&offsets\|0\|-1(+1C);m-186&11\|-\|72615102\|72615124&offsets\|0\|-1(+1C)]\|TCTACAGTGCACGTGTCTCCAGC | >MI0000261\|hsa-mir-139&WithFlank&11\|-\|72615057\|72615136 | 13 | 34 | 22(+1C) | TCTACAGTGCACGTGTCTCCAGC | 281.6 | -3.4 | 0.8 | -4.1 | 5.0E-05 | 4.0E-04 | BT20/MB231 | MITO |
| >am_trnaMT_HisGTG_MT_+_12138_12206@19.51.33\|AAACATCAGATTGTGAATCTGACAACAGAGGCT | >am_trnaMT_HisGTG_MT_+_12138_12206 | 19 | 51 | 33 | AAACATCAGATTGTGAATCTGACAACAGAGGCT | 127.6 | 3.4 | 0.9 | 3.6 | 3.2E-04 | 4.0E-04 | BT20/MB468 | NUC |
| >hg38_21_+_8433172_8446622_RNA45SN1_45S_50ntFlanks@9535.9569.35\|GGAAACTCTGGTGGAGGTCCGTAGCGGTCCTGACG | >hg38_21_+_8433172_8446622_RNA45SN1_45S_50ntFlanks | 9535 | 9569 | 35 | GGAAACTCTGGTGGAGGTCCGTAGCGGTCCTGACG | 245.8 | -3.1 | 0.8 | -3.6 | 3.2E-04 | 4.0E-04 | MB231/MB468 | NUC |
| >hg38_21_+_8433172_8446622_RNA45SN1_45S_50ntFlanks@10683.10712.30\|CCGGCGGCGTCCGGTGAGCTCTCGCTGGCC | >hg38_21_+_8433172_8446622_RNA45SN1_45S_50ntFlanks | 10683 | 10712 | 30 | CCGGCGGCGTCCGGTGAGCTCTCGCTGGCC | 2050.4 | -4.7 | 1.3 | -3.6 | 2.8E-04 | 4.0E-04 | BT20/MB468 | CYTO |
| >trnaMT_SerGCT_MT_+_12207_12265@8.31.24\|CTCACAAGAACTGCTAACTCATGC | >trnaMT_SerGCT_MT_+_12207_12265 | 8 | 31 | 24 | CTCACAAGAACTGCTAACTCATGC | 125.3 | -4.6 | 1.3 | -3.6 | 2.6E-04 | 4.0E-04 | MB231/MB468 | MP |

| | | | | | | | | | | | | | |
|---|---|---|---|---|---|---|---|---|---|---|---|---|---|
| >trna17_ValTAC_11_-_59318102_59318174@1.34.34,trna4_ValTAC_X_-_18693029_18693101@1.34.34\|GGTTCCATAGTGTAGTGGTTATCACGTCTGCTTT | >trna17_ValTAC_11_-_59318102_59318174 | 1 | 34 | 34 | GGTTCCATAGTGTAGTGGTTATCACGTCTGCTTT | 580.6 | 9.5 | 2.6 | 3.6 | 2.9E-04 | 4.0E-04 | MB231/MB468 | CYTO |
| >hg38_21_+_8433172_8446622_RNA45SN1_45S_50ntFlanks@9527.9570.44\|AAGCCAGAGGAAACTCTGGTGGAGGTCCGTAGCGGTCCTGACGT | >hg38_21_+_8433172_8446622_RNA45SN1_45S_50ntFlanks | 9527 | 9570 | 44 | AAGCCAGAGGAAACTCTGGTGGAGGTCCGTAGCGGTCCTGACGT | 58.8 | 9.2 | 2.3 | 4.1 | 4.4E-05 | 4.0E-04 | BT20/MB231 | MP |
| >hg38_21_+_8433172_8446622_RNA45SN1_45S_50ntFlanks@3987.4026.40\|GCTTTGGTGACTCTAGATAACCTCGGGCCGATCGCACGCC | >hg38_21_+_8433172_8446622_RNA45SN1_45S_50ntFlanks | 3987 | 4026 | 40 | GCTTTGGTGACTCTAGATAACCTCGGGCCGATCGCACGCC | 458.7 | -2.6 | 0.7 | -3.7 | 2.3E-04 | 4.0E-04 | BT20/MB468 | MITO |
| >hg38_21_+_8433172_8446622_RNA45SN1_45S_50ntFlanks@9939.9982.44\|GCAGGACGGTGGCCATGGAAGTCGGAATCCGCTAAGGAGTGTGT | >hg38_21_+_8433172_8446622_RNA45SN1_45S_50ntFlanks | 9939 | 9982 | 44 | GCAGGACGGTGGCCATGGAAGTCGGAATCCGCTAAGGAGTGTGT | 101.8 | 7.3 | 1.9 | 3.8 | 1.2E-04 | 4.0E-04 | BT20/MB468 | MP |
| >hg38_21_+_8433172_8446622_RNA45SN1_45S_50ntFlanks@6721.6742.22\|AATTGCAGGACACATTGATCAT | >hg38_21_+_8433172_8446622_RNA45SN1_45S_50ntFlanks | 6721 | 6742 | 22 | AATTGCAGGACACATTGATCAT | 518.0 | -5.7 | 1.6 | -3.6 | 2.8E-04 | 4.0E-04 | BT20/MB468 | CYTO |
| >trnaMT_SerGCT_MT_+_12207_12265@20.62.43\|GCTAACTCATGCCCCCATGTCTAACAACATGGCTTTCTCACCA | >trnaMT_SerGCT_MT_+_12207_12265 | 20 | 62 | 43 | GCTAACTCATGCCCCCATGTCTAACAACATGGCTTTCTCACCA | 87.6 | 10.2 | 2.7 | 3.8 | 1.2E-04 | 4.0E-04 | BT20/MB468 | MP |
| >hg38_21_+_8433172_8446622_RNA45SN1_45S_50ntFlanks@11704.11740.37\|ATGAAGCGCGGGTAAACGGCGGGAGTAACTATGACTC | >hg38_21_+_8433172_8446622_RNA45SN1_45S_50ntFlanks | 11704 | 11740 | 37 | ATGAAGCGCGGGTAAACGGCGGGAGTAACTATGACTC | 85.9 | -6.0 | 1.7 | -3.6 | 2.6E-04 | 4.0E-04 | MB231/MB468 | MP |
| >hg38_21_+_8433172_8446622_RNA45SN1_45S_50ntFlanks@4264.4282.19\|AATGAGTCCACTTTAAATC | >hg38_21_+_8433172_8446622_RNA45SN1_45S_50ntFlanks | 4264 | 4282 | 19 | AATGAGTCCACTTTAAATC | 120.3 | -5.7 | 1.5 | -3.8 | 1.2E-04 | 4.0E-04 | BT20/MB468 | MP |
| >MI0000072\|hsa-mir-18a&WithFlank&13\|+\|91350745\|91350827@53.73.21&[MIMAT0002891&hsa-miR-18a-3p&offsets\|0\|-2;m-389&13\|+\|91350797\|91350819&offsets\|0\|-2]\|ACTGCCCTAAGTGCTCCTTCT | >MI0000072\|hsa-mir-18a&WithFlank&13\|+\|91350745\|91350827 | 53 | 73 | 21 | ACTGCCCTAAGTGCTCCTTCT | 24.6 | 7.8 | 2.1 | 3.6 | 2.6E-04 | 4.0E-04 | MB231/MB468 | MP |
| >hg38_21_+_8433172_8446622_RNA45SN1_45S_50ntFlanks@9736.9753.18\|CCGAAACGATCTCAACCT | >hg38_21_+_8433172_8446622_RNA45SN1_45S_50ntFlanks | 9736 | 9753 | 18 | CCGAAACGATCTCAACCT | 289.1 | -5.5 | 1.5 | -3.6 | 2.8E-04 | 4.0E-04 | BT20/MB468 | CYTO |
| >hg38_21_+_8433172_8446622_RNA45SN1_45S_50ntFlanks@10360.10383.24\|TGAAGGCCGAAGTGGAGAAGGGTT | >hg38_21_+_8433172_8446622_RNA45SN1_45S_50ntFlanks | 10360 | 10383 | 24 | TGAAGGCCGAAGTGGAGAAGGGTT | 155.6 | -5.5 | 1.5 | -3.6 | 2.6E-04 | 4.0E-04 | MB231/MB468 | MP |
| >hg38_21_+_8433172_8446622_RNA45SN1_45S_50ntFlanks@9565.9595.31\|TGACGTGCAAATCGGTCGTCCGACCTGGGTA | >hg38_21_+_8433172_8446622_RNA45SN1_45S_50ntFlanks | 9565 | 9595 | 31 | TGACGTGCAAATCGGTCGTCCGACCTGGGTA | 1174.2 | -2.8 | 0.8 | -3.6 | 3.0E-04 | 4.0E-04 | MB231/MB468 | CYTO |
| >hg38_21_+_8433172_8446622_RNA45SN1_45S_50ntFlanks@3899.3927.29\|CCTTCGCGGGGGGGATGCGTGCATTTATC | >hg38_21_+_8433172_8446622_RNA45SN1_45S_50ntFlanks | 3899 | 3927 | 29 | CCTTCGCGGGGGGGATGCGTGCATTTATC | 714.2 | -2.9 | 0.8 | -3.7 | 2.3E-04 | 4.0E-04 | BT20/MB468 | MITO |
| >hg38_21_+_8433172_8446622_RNA45SN1_45S_50ntFlanks@10737.10755.19\|TAAATCTCGCGCCGGGCCG | >hg38_21_+_8433172_8446622_RNA45SN1_45S_50ntFlanks | 10737 | 10755 | 19 | TAAATCTCGCGCCGGGCCG | 746.1 | -5.6 | 1.6 | -3.6 | 2.8E-04 | 4.0E-04 | BT20/MB468 | CYTO |
| >MI0006444\|hsa-mir-1307&WithFlank&10\|-\|103394247\|103394407@86.107.22(+1U)&[MIMAT0005951&hsa-miR-1307-3p&offsets\|0\|0(+1U);m-66&10\|-\|103394300\|103394322&offsets\|0\|-1(+1U)]\|ACTCGGCGTGGCGTCGGTCGTGT | >MI0006444\|hsa-mir-1307&WithFlank&10\|-\|103394247\|103394407 | 86 | 107 | 22(+1U) | ACTCGGCGTGGCGTCGGTCGTGT | 322.1 | 3.6 | 0.9 | 3.8 | 1.2E-04 | 4.0E-04 | BT20/MB468 | MP |
| >hg38_21_+_8433172_8446622_RNA45SN1_45S_50ntFlanks@10869.10889.21\|AGGGCTGGGTCGGTCGGGCTG | >hg38_21_+_8433172_8446622_RNA45SN1_45S_50ntFlanks | 10869 | 10889 | 21 | AGGGCTGGGTCGGTCGGGCTG | 747.5 | -3.3 | 0.9 | -3.6 | 2.8E-04 | 4.0E-04 | BT20/MB468 | CYTO |
| >hg38_MT_+_598_1651_RNR1_12S_50ntFlanks@417.462.46\|AATAGACTACGAAAGTGGCTTTAACATATCTGAACACACAATAGCT | >hg38_MT_+_598_1651_RNR1_12S_50ntFlanks | 417 | 462 | 46 | AATAGACTACGAAAGTGGCTTTAACATATCTGAACACACAATAGCT | 84.7 | 10.2 | 2.6 | 3.8 | 1.2E-04 | 4.0E-04 | BT20/MB468 | MP |
| >trnaMT_GluTTC_MT_-_14674_14742@40.62.23,trnalookalike8_GluTTC_5_-_93905172_93905240@40.62.23\|TTGGTCGTGGTTGTAGTCCGTGC | >trnaMT_GluTTC_MT_-_14674_14742 | 40 | 62 | 23 | TTGGTCGTGGTTGTAGTCCGTGC | 239.9 | 5.4 | 1.4 | 3.8 | 1.2E-04 | 4.0E-04 | BT20/MB468 | MP |
| >MI0000095\|hsa-mir-93&WithFlank&7\|-\|100093762\|100093853@17.39.23&[MIMAT0000093&hsa-miR-93-5p&offsets\|0\|0;m-29&7\|-\|100093815\|100093837&offsets\|0\|0]\|CAAAGTGCTGTTCGTGCAGGTAG | >MI0000095\|hsa-mir-93&WithFlank&7\|-\|100093762\|100093853 | 17 | 39 | 23 | CAAAGTGCTGTTCGTGCAGGTAG | 3306.8 | 1.9 | 0.5 | 3.6 | 3.2E-04 | 4.0E-04 | BT20/MB468 | NUC |
| >hg38_21_+_8433172_8446622_RNA45SN1_45S_50ntFlanks@10781.10802.22\|AGGTGAACAGCCTCTGGCATGT | >hg38_21_+_8433172_8446622_RNA45SN1_45S_50ntFlanks | 10781 | 10802 | 22 | AGGTGAACAGCCTCTGGCATGT | 391.2 | -4.8 | 1.3 | -3.6 | 3.0E-04 | 4.0E-04 | MB231/MB468 | CYTO |
| >MI0000466\|hsa-mir-9-1&WithFlank&1\|-\|156420335\|156420435@22.43.22;MI0000468\|hsa-mir-9-3&WithFlank&15\|+\|89368011\|89368112@22.43.22;MI0000467\|hsa-mir-9-2&WithFlank&5\|-\|88666847\|88666945@22.43.22&[MIMAT0000441&hsa-miR-9-5p&offsets\|0\|-1;m-104&1\|-\|156420392\|156420414&offsets\|0\|-1];[MIMAT0000441_1&hsa-miR-9-5p&offsets\|0\|-1;m-105&15\|+\|89368032\|89368054&offsets\|0\|-1];[MIMAT0000441_2&hsa-miR-9-5p&offsets\|0\|-1;m-104&5\|-\|88666902\|88666924&offsets\|0\|-1]\|TCTTTGGTTATCTAGCTGTATG | >MI0000466\|hsa-mir-9-1&WithFlank&1\|-\|156420335\|156420435 | 22 | 43 | 22 | TCTTTGGTTATCTAGCTGTATG | 245.2 | 5.8 | 1.6 | 3.6 | 3.1E-04 | 4.0E-04 | MB231/MB468 | MITO |
| >hg38_21_+_8433172_8446622_RNA45SN1_45S_50ntFlanks@12420.12447.28\|ATGTCGGCTCTTCCTATCATTGTGAAGC | >hg38_21_+_8433172_8446622_RNA45SN1_45S_50ntFlanks | 12420 | 12447 | 28 | ATGTCGGCTCTTCCTATCATTGTGAAGC | 993.3 | -3.3 | 0.8 | -4.1 | 5.0E-05 | 4.0E-04 | BT20/MB231 | MITO |
| >hg38_21_+_8433172_8446622_RNA45SN1_45S_50ntFlanks@10344.10373.30\|ATTCAAACGAGAACTTTGAAGGCCGAAGTG | >hg38_21_+_8433172_8446622_RNA45SN1_45S_50ntFlanks | 10344 | 10373 | 30 | ATTCAAACGAGAACTTTGAAGGCCGAAGTG | 181.5 | 4.9 | 1.3 | 3.8 | 1.2E-04 | 4.0E-04 | BT20/MB468 | MP |
| >hg38_1_-_228634819_228635039_RNA5S12_5S_50ntFlanks@147.170.24\|GGGAATACCGGGTGCTGTAGGCTT | >hg38_1_-_228634819_228635039_RNA5S12_5S_50ntFlanks | 147 | 170 | 24 | GGGAATACCGGGTGCTGTAGGCTT | 2957.6 | 2.9 | 0.7 | 4.1 | 5.1E-05 | 4.0E-04 | BT20/MB231 | MITO |
| >hg38_MT_+_1621_3279_RNR2_16S_50ntFlanks@617.663.47\|AAACATATAACTGAACTCCTCACACCCAATTGGACCAATCTATCACC | >hg38_MT_+_1621_3279_RNR2_16S_50ntFlanks | 617 | 663 | 47 | AAACATATAACTGAACTCCTCACACCCAATTGGACCAATCTATCACC | 82.9 | 10.1 | 2.6 | 3.8 | 1.2E-04 | 4.0E-04 | BT20/MB468 | MP |
| >hg38_21_+_8433172_8446622_RNA45SN1_45S_50ntFlanks@6650.6682.33\|ACGACTCTTAGCGGTGGATCACTCGGCTCGTGC | >hg38_21_+_8433172_8446622_RNA45SN1_45S_50ntFlanks | 6650 | 6682 | 33 | ACGACTCTTAGCGGTGGATCACTCGGCTCGTGC | 731.0 | 3.8 | 1.0 | 3.6 | 3.2E-04 | 4.0E-04 | MB231/MB468 | MITO |
| >hg38_21_+_8433172_8446622_RNA45SN1_45S_50ntFlanks@3845.3868.24\|ACTTGGATAACTGTGGTAATTCTA | >hg38_21_+_8433172_8446622_RNA45SN1_45S_50ntFlanks | 3845 | 3868 | 24 | ACTTGGATAACTGTGGTAATTCTA | 461.4 | -4.4 | 1.2 | -3.6 | 3.2E-04 | 4.0E-04 | MB231/MB468 | MITO |
| >hg38_21_+_8433172_8446622_RNA45SN1_45S_50ntFlanks@10473.10493.21\|CTCGGCCGATCGAAAGGGAGT | >hg38_21_+_8433172_8446622_RNA45SN1_45S_50ntFlanks | 10473 | 10493 | 21 | CTCGGCCGATCGAAAGGGAGT | 118.2 | -2.8 | 0.8 | -3.6 | 2.7E-04 | 4.0E-04 | MB231/MB468 | MP |
| >hg38_21_+_8433172_8446622_RNA45SN1_45S_50ntFlanks@6781.6803.23\|GGGGCTACGCCTGTCTGAGCGTC | >hg38_21_+_8433172_8446622_RNA45SN1_45S_50ntFlanks | 6781 | 6803 | 23 | GGGGCTACGCCTGTCTGAGCGTC | 170.7 | -4.4 | 1.2 | -3.6 | 2.7E-04 | 4.0E-04 | MB231/MB468 | MP |
| >hg38_21_+_8433172_8446622_RNA45SN1_45S_50ntFlanks@11783.11813.31\|CATGAATGGATGAACGAGATTCCCACTGTCC | >hg38_21_+_8433172_8446622_RNA45SN1_45S_50ntFlanks | 11783 | 11813 | 31 | CATGAATGGATGAACGAGATTCCCACTGTCC | 59.4 | 9.2 | 2.3 | 4.1 | 4.5E-05 | 4.0E-04 | BT20/MB231 | MP |
| >hg38_21_+_8433172_8446622_RNA45SN1_45S_50ntFlanks@10392.10433.42\|ACAGCAGTTGAACATGGGTCAGTCGGTCCTGAGAGATGGGCG | >hg38_21_+_8433172_8446622_RNA45SN1_45S_50ntFlanks | 10392 | 10433 | 42 | ACAGCAGTTGAACATGGGTCAGTCGGTCCTGAGAGATGGGCG | 570.5 | -4.8 | 1.3 | -3.6 | 2.8E-04 | 4.0E-04 | BT20/MB468 | CYTO |
| >MI0000816\|hsa-mir-335&WithFlank&7\|+\|130496105\|130496210@58.78.21&[MIMAT0004703&hsa-miR-335-3p&offsets\|0\|-1;m-159&7\|+\|130496162\|130496183&offsets\|0\|-1]\|TTTTTCATTATTGCTCCTGAC | >MI0000816\|hsa-mir-335&WithFlank&7\|+\|130496105\|130496210 | 58 | 78 | 21 | TTTTTCATTATTGCTCCTGAC | 176.4 | 3.4 | 0.9 | 3.6 | 2.7E-04 | 4.0E-04 | MB231/MB468 | MP |
| >MI0000749\|hsa-mir-30e&WithFlank&1\|+\|40754349\|40754452@23.46.24&[MIMAT0000692&hsa-miR-30e-5p&offsets\|0\|+2;m-24&1\|+\|40754371\|40754394&offsets\|0\|0]\|TGTAAACATCCTTGACTGGAAGCT | >MI0000749\|hsa-mir-30e&WithFlank&1\|+\|40754349\|40754452 | 23 | 46 | 24 | TGTAAACATCCTTGACTGGAAGCT | 7708.7 | 3.9 | 1.1 | 3.6 | 3.2E-04 | 4.0E-04 | MB231/MB468 | MITO |
| >hg38_21_+_8433172_8446622_RNA45SN1_45S_50ntFlanks@11876.11907.32\|AAGACCCTGTTGAGCTTGACTCTAGTCTGGCA | >hg38_21_+_8433172_8446622_RNA45SN1_45S_50ntFlanks | 11876 | 11907 | 32 | AAGACCCTGTTGAGCTTGACTCTAGTCTGGCA | 152.8 | -3.8 | 1.0 | -3.7 | 2.4E-04 | 4.0E-04 | BT20/MB468 | MITO |
| >hg38_21_+_8433172_8446622_RNA45SN1_45S_50ntFlanks@8176.8194.19\|CTTCTGATCGAGGCCCAGC | >hg38_21_+_8433172_8446622_RNA45SN1_45S_50ntFlanks | 8176 | 8194 | 19 | CTTCTGATCGAGGCCCAGC | 759.4 | 2.5 | 0.7 | 3.6 | 3.0E-04 | 4.0E-04 | MB231/MB468 | CYTO |
| >hg38_21_+_8433172_8446622_RNA45SN1_45S_50ntFlanks@12998.13039.42\|GCTGCGATCTATTGAAAGTCAGCCCTCGACACAAGGGTTTGT | >hg38_21_+_8433172_8446622_RNA45SN1_45S_50ntFlanks | 12998 | 13039 | 42 | GCTGCGATCTATTGAAAGTCAGCCCTCGACACAAGGGTTTGT | 5718.2 | 4.1 | 1.1 | 3.6 | 3.2E-04 | 4.0E-04 | MB231/MB468 | MITO |
| >hg38_21_+_8433172_8446622_RNA45SN1_45S_50ntFlanks@5193.5225.33\|AGGTCTGTGATGCCCTTAGATGTCCGGGGCTGC | >hg38_21_+_8433172_8446622_RNA45SN1_45S_50ntFlanks | 5193 | 5225 | 33 | AGGTCTGTGATGCCCTTAGATGTCCGGGGCTGC | 71.2 | 7.0 | 1.8 | 3.8 | 1.3E-04 | 4.0E-04 | BT20/MB468 | MP |
| >trnaMT_GluTTC_MT_-_14674_14742@47.71.25,trnalookalike8_GluTTC_5_-_93905172_93905240@47.71.25\|TGGTTGTAGTCCGTGCGAGAATACC | >trnaMT_GluTTC_MT_-_14674_14742 | 47 | 71 | 25 | TGGTTGTAGTCCGTGCGAGAATACC | 142.4 | 2.8 | 0.8 | 3.6 | 2.7E-04 | 4.0E-04 | MB231/MB468 | MP |
| >hg38_21_+_8433172_8446622_RNA45SN1_45S_50ntFlanks@8113.8138.26\|GGCGCGGGACATGTGGCGTACGGAAG | >hg38_21_+_8433172_8446622_RNA45SN1_45S_50ntFlanks | 8113 | 8138 | 26 | GGCGCGGGACATGTGGCGTACGGAAG | 394.6 | 8.0 | 2.0 | 4.1 | 4.5E-05 | 4.0E-04 | BT20/MB231 | MP |
| >hg38_21_+_8433172_8446622_RNA45SN1_45S_50ntFlanks@4160.4184.25\|CCACATCCAAGGAAGGCAGCAGGCG | >hg38_21_+_8433172_8446622_RNA45SN1_45S_50ntFlanks | 4160 | 4184 | 25 | CCACATCCAAGGAAGGCAGCAGGCG | 396.2 | -6.2 | 1.6 | -3.8 | 1.3E-04 | 4.1E-04 | BT20/MB468 | MP |
| >MI0000466\|hsa-mir-9-1&WithFlank&1\|-\|156420335\|156420435@22.44.23;MI0000468\|hsa-mir-9-3&WithFlank&15\|+\|89368011\|89368112@22.44.23;MI0000467\|hsa-mir-9-2&WithFlank&5\|-\|88666847\|88666945@22.44.23&[MIMAT0000441&hsa-miR-9-5p&offsets\|0\|0;m-104&1\|-\|156420392\|156420414&offsets\|0\|0];[MIMAT0000441_1&hsa-miR-9-5p&offsets\|0\|0;m-105&15\|+\|89368032\|89368054&offsets\|0\|0];[MIMAT0000441_2&hsa-miR-9-5p&offsets\|0\|0;m-104&5\|-\|88666902\|88666924&offsets\|0\|0]\|TCTTTGGTTATCTAGCTGTATGA | >MI0000466\|hsa-mir-9-1&WithFlank&1\|-\|156420335\|156420435 | 22 | 44 | 23 | TCTTTGGTTATCTAGCTGTATGA | 332.7 | 3.6 | 0.9 | 3.8 | 1.3E-04 | 4.1E-04 | BT20/MB468 | MP |
| >MI0000082\|hsa-mir-25&WithFlank&7\|-\|100093554\|100093649@58.77.20(+1U)&[MIMAT0000081&hsa-miR-25-3p&offsets\|0\|-2(+1U);m-23&7\|-\|100093571\|100093592&offsets\|0\|-2(+1U)]\|CATTGCACTTGTCTCGGTCTT | >MI0000082\|hsa-mir-25&WithFlank&7\|-\|100093554\|100093649 | 58 | 77 | 20(+1U) | CATTGCACTTGTCTCGGTCTT | 149.0 | 4.6 | 1.3 | 3.6 | 2.7E-04 | 4.1E-04 | MB231/MB468 | MP |
| >hg38_1_-_228634819_228635039_RNA5S12_5S_50ntFlanks@144.170.27\|CCTGGGAATACCGGGTGCTGTAGGCTT | >hg38_1_-_228634819_228635039_RNA5S12_5S_50ntFlanks | 144 | 170 | 27 | CCTGGGAATACCGGGTGCTGTAGGCTT | 4155.1 | -2.5 | 0.7 | -3.6 | 2.7E-04 | 4.1E-04 | MB231/MB468 | MP |
| >MI0000087\|hsa-mir-29a&WithFlank&7\|-\|130876741\|130876816@47.70.24&[MIMAT0000086&hsa-miR-29a-3p&offsets\|-1\|+1;m-26&7\|-\|130876749\|130876769&offsets\|-1\|+2]\|CTAGCACCATCTGAAATCGGTTAT | >MI0000087\|hsa-mir-29a&WithFlank&7\|-\|130876741\|130876816 | 47 | 70 | 24 | CTAGCACCATCTGAAATCGGTTAT | 177.8 | 7.6 | 2.0 | 3.8 | 1.3E-04 | 4.1E-04 | BT20/MB468 | MP |
| >hg38_MT_+_1621_3279_RNR2_16S_50ntFlanks@1370.1415.46\|ATGTTGGATCAGGACATCCCGATGGTGCAGCCGCTATTAAAGGTTC | >hg38_MT_+_1621_3279_RNR2_16S_50ntFlanks | 1370 | 1415 | 46 | ATGTTGGATCAGGACATCCCGATGGTGCAGCCGCTATTAAAGGTTC | 85.1 | 10.2 | 2.6 | 3.8 | 1.3E-04 | 4.1E-04 | BT20/MB468 | MP |
| >hg38_21_+_8433172_8446622_RNA45SN1_45S_50ntFlanks@12182.12211.30\|CAAACGGTAACGCAGGTGTCCTAAGGCGAG | >hg38_21_+_8433172_8446622_RNA45SN1_45S_50ntFlanks | 12182 | 12211 | 30 | CAAACGGTAACGCAGGTGTCCTAAGGCGAG | 105.5 | -5.8 | 1.5 | -3.8 | 1.3E-04 | 4.1E-04 | BT20/MB468 | MP |
| >hg38_21_+_8433172_8446622_RNA45SN1_45S_50ntFlanks@12007.12043.37\|GCCGCCGGTGAAATACCACTACTCTGATCGTTTTTTC | >hg38_21_+_8433172_8446622_RNA45SN1_45S_50ntFlanks | 12007 | 12043 | 37 | GCCGCCGGTGAAATACCACTACTCTGATCGTTTTTTC | 1055.2 | -6.1 | 1.6 | -3.8 | 1.3E-04 | 4.1E-04 | BT20/MB468 | MP |

| | | | | | | | | | | | | | |
|---|---|---|---|---|---|---|---|---|---|---|---|---|---|
| >hg38_21_+_8433172_8446622_RNA45SN1_45S_50ntFlanks@12285.12318.34\|CAGACCGTGAAAGCGGGGCCTCACGATCCTTCTG | >hg38_21_+_8433172_8446622_RNA45SN1_45S_50ntFlanks | 12285 | 12318 | 34 | CAGACCGTGAAAGCGGGGCCTCACGATCCTTCTG | 91.8 | -5.7 | 1.5 | -3.8 | 1.3E-04 | 4.1E-04 | BT20/MB468 | MP |
| >hg38_MT_+_598_1651_RNR1_12S_50ntFlanks@442.469.28\|ATATCTGAACACACAATAGCTAAGACCC | >hg38_MT_+_598_1651_RNR1_12S_50ntFlanks | 442 | 469 | 28 | ATATCTGAACACACAATAGCTAAGACCC | 143.0 | 4.8 | 1.3 | 3.6 | 2.7E-04 | 4.1E-04 | MB231/MB468 | MP |
| >hg38_21_+_8433172_8446622_RNA45SN1_45S_50ntFlanks@476.517.42\|TTCGTGATCGATGTGGTGACGTCGTGCTCTCCCGGGCCGGGT | >hg38_21_+_8433172_8446622_RNA45SN1_45S_50ntFlanks | 476 | 517 | 42 | TTCGTGATCGATGTGGTGACGTCGTGCTCTCCCGGGCCGGGT | 550.9 | 5.4 | 1.5 | 3.6 | 3.2E-04 | 4.1E-04 | MB231/MB468 | MITO |
| >hg38_21_+_8433172_8446622_RNA45SN1_45S_50ntFlanks@11561.11602.42\|GCGCCTAGCAGCCGACTTAGAACTGGTGCGGACCAGGGGAAT | >hg38_21_+_8433172_8446622_RNA45SN1_45S_50ntFlanks | 11561 | 11602 | 42 | GCGCCTAGCAGCCGACTTAGAACTGGTGCGGACCAGGGGAAT | 414.4 | -5.5 | 1.5 | -3.6 | 2.9E-04 | 4.1E-04 | BT20/MB468 | CYTO |
| >hg38_21_+_8433172_8446622_RNA45SN1_45S_50ntFlanks@11849.11871.23\|CGGGCTTGGCGGAATCAGCGGGG | >hg38_21_+_8433172_8446622_RNA45SN1_45S_50ntFlanks | 11849 | 11871 | 23 | CGGGCTTGGCGGAATCAGCGGGG | 424.4 | -3.3 | 0.9 | -3.8 | 1.3E-04 | 4.1E-04 | BT20/MB468 | MP |
| >hg38_21_+_8433172_8446622_RNA45SN1_45S_50ntFlanks@12076.12097.22\|AGGGGCTCTCGCTTCTGGCGCC | >hg38_21_+_8433172_8446622_RNA45SN1_45S_50ntFlanks | 12076 | 12097 | 22 | AGGGGCTCTCGCTTCTGGCGCC | 399.5 | -2.3 | 0.6 | -3.6 | 2.9E-04 | 4.1E-04 | BT20/MB468 | CYTO |
| >am_trnaMT_ValTAC_MT_+_1602_1670@33.65.33\|ACACTTAGGAGATTTCAACTTAACTTGACCGCT | >am_trnaMT_ValTAC_MT_+_1602_1670 | 33 | 65 | 33 | ACACTTAGGAGATTTCAACTTAACTTGACCGCT | 89.9 | 10.2 | 2.7 | 3.8 | 1.3E-04 | 4.1E-04 | BT20/MB468 | MP |
| >MI0000255\|hsa-mir-30d&WithFlank&8\|-\|134804870\|134804951@12.33.22&[MIMAT0000245&hsa-miR-30d-5p&offsets\|0\|0;m-21&8\|-\|134804919\|134804940&offsets\|0\|0]\|TGTAAACATCCCCGACTGGAAG | >MI0000255\|hsa-mir-30d&WithFlank&8\|-\|134804870\|134804951 | 12 | 33 | 22 | TGTAAACATCCCCGACTGGAAG | 1464.0 | 3.8 | 1.0 | 3.8 | 1.3E-04 | 4.1E-04 | BT20/MB468 | MP |
| >hg38_21_+_8433172_8446622_RNA45SN1_45S_50ntFlanks@11576.11612.37\|CTTAGAACTGGTGCGGACCAGGGGAATCCGACTGTTT | >hg38_21_+_8433172_8446622_RNA45SN1_45S_50ntFlanks | 11576 | 11612 | 37 | CTTAGAACTGGTGCGGACCAGGGGAATCCGACTGTTT | 303.4 | -3.9 | 1.1 | -3.6 | 3.0E-04 | 4.1E-04 | MB231/MB468 | CYTO |
| >hg38_21_+_8433172_8446622_RNA45SN1_45S_50ntFlanks@6646.6682.37\|TCGTACGACTCTTAGCGGTGGATCACTCGGCTCGTGC | >hg38_21_+_8433172_8446622_RNA45SN1_45S_50ntFlanks | 6646 | 6682 | 37 | TCGTACGACTCTTAGCGGTGGATCACTCGGCTCGTGC | 581.2 | 4.0 | 1.0 | 3.8 | 1.3E-04 | 4.1E-04 | BT20/MB468 | MP |
| >hg38_21_+_8433172_8446622_RNA45SN1_45S_50ntFlanks@12934.12954.21\|CTAAACCATTCGTAGACGACC | >hg38_21_+_8433172_8446622_RNA45SN1_45S_50ntFlanks | 12934 | 12954 | 21 | CTAAACCATTCGTAGACGACC | 444.1 | -4.0 | 1.1 | -3.6 | 2.9E-04 | 4.1E-04 | BT20/MB468 | CYTO |
| >MI0001445\|hsa-mir-423&WithFlank&17\|+\|30117073\|30117178@59.80.22(+1A)&[MIMAT0001340&hsa-miR-423-3p&offsets\|0\|-1(+1A);m-100&17\|+\|30117131\|30117153&offsets\|0\|-1(+1A)]\|AGCTCGGTCTGAGGCCCCTCAGA | >MI0001445\|hsa-mir-423&WithFlank&17\|+\|30117073\|30117178 | 59 | 80 | 22(+1A) | AGCTCGGTCTGAGGCCCCTCAGA | 235.1 | 2.1 | 0.6 | 3.6 | 3.3E-04 | 4.1E-04 | BT20/MB468 | NUC |
| >trna10_AspGTC_12_-_125424193_125424264@37.70.34,trna12_AspGTC_12_-_125411891_125411962@37.70.34,trna144_AspGTC_6_-_27551236_27551307@37.70.34,trna38_AspGTC_17_-_8125556_8125627@37.70.34,trna45_AspGTC_6_+_27447453_27447524@37.70.34,trna48_AspGTC_6_+_27471523_27471594@37.70.34,trna4_AspGTC_12_+_96429799_96429870@37.70.34,trna69_AspGTC_1_-_161440205_161440276@37.70.34,trna72_AspGTC_1_-_161432824_161432895@37.70.34,trna75_AspGTC_1_-_161425414_161425485@37.70.34,trna78_AspGTC_1_-_161418033_161418104@37.70.34,trna81_AspGTC_1_-_161410615_161410686@37.70.34\|ACGCGGGAGACCGGGGTTCGATTCCCCGACGGGG | >trna10_AspGTC_12_-_125424193_125424264 | 37 | 70 | 34 | ACGCGGGAGACCGGGGTTCGATTCCCCGACGGGG | 1168.8 | -4.1 | 1.0 | -4.1 | 4.3E-05 | 4.1E-04 | BT20/MB231 | CYTO |
| >trna10_LysCTT_16_+_3241501_3241573@6.35.30,trna13_LysCTT_14_-_58706613_58706685@6.35.30,trna2_LysCTT_15_+_79152904_79152976@6.35.30,trna4_LysCTT_18_-_43669271_43669343@6.35.30\|GCTAGCTCAGTCGGTAGAGCATGGGACTCT | >trna10_LysCTT_16_+_3241501_3241573 | 6 | 35 | 30 | GCTAGCTCAGTCGGTAGAGCATGGGACTCT | 276.4 | -4.3 | 1.1 | -4.0 | 5.2E-05 | 4.1E-04 | BT20/MB231 | MITO |
| >hg38_21_+_8433172_8446622_RNA45SN1_45S_50ntFlanks@9300.9331.32\|ACGGACCAAGGAGTCTAACACGTGCGCGAGTC | >hg38_21_+_8433172_8446622_RNA45SN1_45S_50ntFlanks | 9300 | 9331 | 32 | ACGGACCAAGGAGTCTAACACGTGCGCGAGTC | 89.8 | 6.9 | 1.8 | 3.8 | 1.3E-04 | 4.1E-04 | BT20/MB468 | MP |
| >MI0000253\|hsa-mir-148a&WithFlank&7\|-\|25949913\|25949992@50.70.21(+1A)&[MIMAT0000243&hsa-miR-148a-3p&offsets\|0\|-1(+1A);m-7&7\|-\|25949922\|25949943&offsets\|0\|-1(+1A)]\|TCAGTGCACTACAGAACTTTGA | >MI0000253\|hsa-mir-148a&WithFlank&7\|-\|25949913\|25949992 | 50 | 70 | 21(+1A) | TCAGTGCACTACAGAACTTTGA | 14380.5 | 4.7 | 1.2 | 3.8 | 1.3E-04 | 4.1E-04 | BT20/MB468 | MP |
| >MI0003776\|hsa-mir-320b-1&WithFlank&1\|+\|116671740\|116671823@48.66.19(+3A);MI0003839\|hsa-mir-320b-2&WithFlank&1\|-\|224257034\|224257116@47.65.19(+3A);MI0003778\|hsa-mir-320c-1&WithFlank&18\|+\|21683512\|21683595@48.66.19(+3A);MI0008191\|hsa-mir-320c-2&WithFlank&18\|+\|24321669\|24321752@48.66.19(+3A);MI0000542\|hsa-mir-320a&WithFlank&8\|-\|22244960\|22245043@48.66.19(+3A)&[MIMAT0005792&hsa-miR-320b&offsets\|0\|-3(+3A);m-183&1\|+\|116671787\|116671806&offsets\|0\|-1(+3A)];[MIMAT0005792_1&hsa-miR-320b&offsets\|0\|-3(+3A);m-183&1\|-\|224257051\|224257070&offsets\|0\|-1(+3A)];[MIMAT0005793&hsa-miR-320c&offsets\|0\|-1(+3A);m-222&18\|+\|21683559\|21683578&offsets\|0\|-1(+3A)];[MIMAT0005793_1&hsa-miR-320c&offsets\|0\|-1(+3A);m-219&18\|+\|24321716\|24321735&offsets\|0\|-1(+3A)];[MIMAT0000510&hsa-miR-320a-3p&offsets\|0\|-3(+3A);m-60&8\|-\|22244975\|22244996&offsets\|0\|-3(+3A)]\|AAAAGCTGGGTTGAGAGGGAAA | >MI0003776\|hsa-mir-320b-1&WithFlank&1\|+\|116671740\|116671823 | 48 | 66 | 19(+3A) | AAAAGCTGGGTTGAGAGGGAAA | 51.8 | 9.0 | 2.2 | 4.1 | 4.5E-05 | 4.1E-04 | BT20/MB231 | MP |
| >am_trnaMT_GluTTC_MT_-_14674_14742@24.53.30,trnalookalike8_GluTTC_5_-_93905172_93905240@24.53.30\|ATGGTTTTTCATATCATTGGTCGTGGTTGT | >am_trnaMT_GluTTC_MT_-_14674_14742 | 24 | 53 | 30 | ATGGTTTTTCATATCATTGGTCGTGGTTGT | 80.2 | 7.1 | 1.9 | 3.8 | 1.3E-04 | 4.1E-04 | BT20/MB468 | MP |
| >hg38_21_+_8433172_8446622_RNA45SN1_45S_50ntFlanks@12289.12322.34\|CCGTGAAAGCGGGGCCTCACGATCCTTCTGACCT | >hg38_21_+_8433172_8446622_RNA45SN1_45S_50ntFlanks | 12289 | 12322 | 34 | CCGTGAAAGCGGGGCCTCACGATCCTTCTGACCT | 358.3 | -4.0 | 1.1 | -3.7 | 2.4E-04 | 4.1E-04 | BT20/MB468 | MITO |
| >hg38_21_+_8433172_8446622_RNA45SN1_45S_50ntFlanks@11323.11344.22\|CGTCTCGTCGCGCGCGCGTCCG | >hg38_21_+_8433172_8446622_RNA45SN1_45S_50ntFlanks | 11323 | 11344 | 22 | CGTCTCGTCGCGCGCGCGTCCG | 41578.3 | -8.3 | 2.3 | -3.6 | 3.3E-04 | 4.1E-04 | BT20/MB468 | NUC |
| >trna152_ValCAC_6_-_27248049_27248121@3.36.34\|TTCTGTAGTGTAGTGGTTATCACGTTCGCCTCAC | >trna152_ValCAC_6_-_27248049_27248121 | 3 | 36 | 34 | TTCTGTAGTGTAGTGGTTATCACGTTCGCCTCAC | 233.1 | -4.0 | 1.0 | -4.0 | 5.2E-05 | 4.1E-04 | BT20/MB231 | MITO |
| >hg38_21_+_8433172_8446622_RNA45SN1_45S_50ntFlanks@12405.12440.36\|CTTTTTGATCCTTCGATGTCGGCTCTTCCTATCATT | >hg38_21_+_8433172_8446622_RNA45SN1_45S_50ntFlanks | 12405 | 12440 | 36 | CTTTTTGATCCTTCGATGTCGGCTCTTCCTATCATT | 386.5 | -3.7 | 1.0 | -3.6 | 2.9E-04 | 4.1E-04 | BT20/MB468 | CYTO |
| >hg38_21_+_8433172_8446622_RNA45SN1_45S_50ntFlanks@12007.12037.31\|GCCGCCGGTGAAATACCACTACTCTGATCGT | >hg38_21_+_8433172_8446622_RNA45SN1_45S_50ntFlanks | 12007 | 12037 | 31 | GCCGCCGGTGAAATACCACTACTCTGATCGT | 753.9 | 3.2 | 0.9 | 3.7 | 2.4E-04 | 4.1E-04 | BT20/MB468 | MITO |
| >hg38_21_+_8433172_8446622_RNA45SN1_45S_50ntFlanks@11320.11343.24\|CCACGTCTCGTCGCGCGCGCGTCC | >hg38_21_+_8433172_8446622_RNA45SN1_45S_50ntFlanks | 11320 | 11343 | 24 | CCACGTCTCGTCGCGCGCGCGTCC | 385.0 | -4.4 | 1.2 | -3.6 | 3.0E-04 | 4.1E-04 | MB231/MB468 | CYTO |
| >hg38_21_+_8433172_8446622_RNA45SN1_45S_50ntFlanks@11208.11251.44\|GCGGCGACTCTGGACGCGAGCCGGGCCCTTCCCGTGGATCGCCC | >hg38_21_+_8433172_8446622_RNA45SN1_45S_50ntFlanks | 11208 | 11251 | 44 | GCGGCGACTCTGGACGCGAGCCGGGCCCTTCCCGTGGATCGCCC | 264.8 | -4.4 | 1.2 | -3.6 | 2.9E-04 | 4.1E-04 | BT20/MB468 | CYTO |
| >hg38_21_+_8433172_8446622_RNA45SN1_45S_50ntFlanks@11920.11950.31\|ATGAGAGGTGTAGAATAAGTGGGAGGCCCCC | >hg38_21_+_8433172_8446622_RNA45SN1_45S_50ntFlanks | 11920 | 11950 | 31 | ATGAGAGGTGTAGAATAAGTGGGAGGCCCCC | 478.4 | -2.2 | 0.6 | -3.6 | 2.9E-04 | 4.1E-04 | BT20/MB468 | CYTO |
| >hg38_21_+_8433172_8446622_RNA45SN1_45S_50ntFlanks@4498.4517.20\|AAGCGTTTACTTTGAAAAAA | >hg38_21_+_8433172_8446622_RNA45SN1_45S_50ntFlanks | 4498 | 4517 | 20 | AAGCGTTTACTTTGAAAAAA | 293.0 | 3.3 | 0.8 | 4.1 | 4.4E-05 | 4.1E-04 | BT20/MB231 | CYTO |
| >hg38_21_+_8433172_8446622_RNA45SN1_45S_50ntFlanks@12456.12499.44\|CCAAGCGTTGGATTGTTCACCCACTAATAGGGAACGTGAGCTGG | >hg38_21_+_8433172_8446622_RNA45SN1_45S_50ntFlanks | 12456 | 12499 | 44 | CCAAGCGTTGGATTGTTCACCCACTAATAGGGAACGTGAGCTGG | 624.3 | -5.0 | 1.4 | -3.6 | 2.9E-04 | 4.1E-04 | BT20/MB468 | CYTO |
| >hg38_21_+_8433172_8446622_RNA45SN1_45S_50ntFlanks@10357.10388.32\|CTTTGAAGGCCGAAGTGGAGAAGGGTTCCATG | >hg38_21_+_8433172_8446622_RNA45SN1_45S_50ntFlanks | 10357 | 10388 | 32 | CTTTGAAGGCCGAAGTGGAGAAGGGTTCCATG | 416.9 | -5.0 | 1.3 | -3.8 | 1.3E-04 | 4.1E-04 | BT20/MB468 | MP |
| >hg38_21_+_8433172_8446622_RNA45SN1_45S_50ntFlanks@3835.3859.25\|CTCCTCTCCTACTTGGATAACTGTG | >hg38_21_+_8433172_8446622_RNA45SN1_45S_50ntFlanks | 3835 | 3859 | 25 | CTCCTCTCCTACTTGGATAACTGTG | 229.7 | -3.3 | 0.9 | -3.6 | 3.0E-04 | 4.1E-04 | MB231/MB468 | CYTO |
| >hg38_21_+_8433172_8446622_RNA45SN1_45S_50ntFlanks@9855.9899.45\|CTGCGGGATGAACCGAACGCCGGGTTAAGGCGCCCGATGCCGACG | >hg38_21_+_8433172_8446622_RNA45SN1_45S_50ntFlanks | 9855 | 9899 | 45 | CTGCGGGATGAACCGAACGCCGGGTTAAGGCGCCCGATGCCGACG | 317.4 | -6.3 | 1.7 | -3.6 | 3.2E-04 | 4.1E-04 | MB231/MB468 | MITO |
| >MI0000456\|hsa-mir-140&WithFlank&16\|+\|69933075\|69933186@69.89.21(+1U)&[MIMAT0004597&hsa-miR-140-3p&offsets\|+1\|+1(+1U);m-45&16\|+\|69933143\|69933164&offsets\|0\|-1(+1U)]\|ACCACAGGGTAGAACCACGGAT | >MI0000456\|hsa-mir-140&WithFlank&16\|+\|69933075\|69933186 | 69 | 89 | 21(+1U) | ACCACAGGGTAGAACCACGGAT | 183.3 | 7.7 | 2.0 | 3.8 | 1.3E-04 | 4.1E-04 | BT20/MB468 | MP |
| >hg38_MT_+_1621_3279_RNR2_16S_50ntFlanks@1367.1396.30\|TCGATGTTGGATCAGGACATCCCGATGGTG | >hg38_MT_+_1621_3279_RNR2_16S_50ntFlanks | 1367 | 1396 | 30 | TCGATGTTGGATCAGGACATCCCGATGGTG | 490.3 | 4.2 | 1.1 | 3.8 | 1.3E-04 | 4.1E-04 | BT20/MB468 | MP |
| >hg38_21_+_8433172_8446622_RNA45SN1_45S_50ntFlanks@7984.8014.31\|AGATCAGACGTGGCGACCCGCTGAATTTAAG | >hg38_21_+_8433172_8446622_RNA45SN1_45S_50ntFlanks | 7984 | 8014 | 31 | AGATCAGACGTGGCGACCCGCTGAATTTAAG | 96.7 | -5.7 | 1.5 | -3.8 | 1.3E-04 | 4.1E-04 | BT20/MB468 | MP |
| >trnaMT_SerGCT_MT_+_12207_12265@1.18.18\|GAGAAAGCTCACAAGAAC | >trnaMT_SerGCT_MT_+_12207_12265 | 1 | 18 | 18 | GAGAAAGCTCACAAGAAC | 86370.5 | -3.8 | 0.9 | -4.0 | 5.3E-05 | 4.1E-04 | BT20/MB231 | MITO |
| >hg38_21_+_8433172_8446622_RNA45SN1_45S_50ntFlanks@10396.10424.29\|CAGTTGAACATGGGTCAGTCGGTCCTGAG | >hg38_21_+_8433172_8446622_RNA45SN1_45S_50ntFlanks | 10396 | 10424 | 29 | CAGTTGAACATGGGTCAGTCGGTCCTGAG | 20940.0 | 3.5 | 0.9 | 4.0 | 5.3E-05 | 4.1E-04 | BT20/MB231 | MITO |
| >MI0000085\|hsa-mir-27a&WithFlank&19\|-\|13836434\|13836523@57.75.19(+1U)&[MIMAT0000084&hsa-miR-27a-3p&offsets\|0\|-2(+1U);m-43&19\|-\|13836447\|13836467&offsets\|0\|-2(+1U)]\|TTCACAGTGGCTAAGTTCCT | >MI0000085\|hsa-mir-27a&WithFlank&19\|-\|13836434\|13836523 | 57 | 75 | 19(+1U) | TTCACAGTGGCTAAGTTCCT | 533.1 | -3.3 | 0.8 | -4.0 | 5.2E-05 | 4.1E-04 | BT20/MB231 | MITO |
| >hg38_21_+_8433172_8446622_RNA45SN1_45S_50ntFlanks@11541.11584.44\|GCGGCGCCTCGCCTCGGCCGGCGCCTAGCAGCCGACTTAGAACT | >hg38_21_+_8433172_8446622_RNA45SN1_45S_50ntFlanks | 11541 | 11584 | 44 | GCGGCGCCTCGCCTCGGCCGGCGCCTAGCAGCCGACTTAGAACT | 421.7 | -3.2 | 0.8 | -4.1 | 4.4E-05 | 4.1E-04 | BT20/MB231 | CYTO |
| >hg38_21_+_8433172_8446622_RNA45SN1_45S_50ntFlanks@8171.8195.25\|AAGTCCTTCTGATCGAGGCCCAGCC | >hg38_21_+_8433172_8446622_RNA45SN1_45S_50ntFlanks | 8171 | 8195 | 25 | AAGTCCTTCTGATCGAGGCCCAGCC | 192.6 | -2.4 | 0.7 | -3.6 | 2.9E-04 | 4.1E-04 | BT20/MB468 | CYTO |
| >hg38_21_+_8433172_8446622_RNA45SN1_45S_50ntFlanks@3993.4010.18\|GTGACTCTAGATAACCTC | >hg38_21_+_8433172_8446622_RNA45SN1_45S_50ntFlanks | 3993 | 4010 | 18 | GTGACTCTAGATAACCTC | 423.6 | 2.6 | 0.7 | 3.6 | 3.1E-04 | 4.1E-04 | MB231/MB468 | CYTO |
| >hg38_MT_+_1621_3279_RNR2_16S_50ntFlanks@1349.1385.37\|ACAATAGGGTTTACGACCTCGATGTTGGATCAGGACA | >hg38_MT_+_1621_3279_RNR2_16S_50ntFlanks | 1349 | 1385 | 37 | ACAATAGGGTTTACGACCTCGATGTTGGATCAGGACA | 159.4 | 5.2 | 1.4 | 3.6 | 3.2E-04 | 4.1E-04 | MB231/MB468 | MITO |
| >hg38_21_+_8433172_8446622_RNA45SN1_45S_50ntFlanks@11572.11611.40\|CCGACTTAGAACTGGTGCGGACCAGGGGAATCCGACTGTT | >hg38_21_+_8433172_8446622_RNA45SN1_45S_50ntFlanks | 11572 | 11611 | 40 | CCGACTTAGAACTGGTGCGGACCAGGGGAATCCGACTGTT | 2899.9 | -2.9 | 0.8 | -3.7 | 2.4E-04 | 4.1E-04 | BT20/MB468 | MITO |
| >hg38_21_+_8433172_8446622_RNA45SN1_45S_50ntFlanks@11571.11592.22\|GCCGACTTAGAACTGGTGCGGA | >hg38_21_+_8433172_8446622_RNA45SN1_45S_50ntFlanks | 11571 | 11592 | 22 | GCCGACTTAGAACTGGTGCGGA | 264.1 | -2.8 | 0.8 | -3.6 | 3.1E-04 | 4.1E-04 | MB231/MB468 | CYTO |

| | | | | | | | | | | | | | |
|---|---|---|---|---|---|---|---|---|---|---|---|---|---|
| >hg38_21_+_8433172_8446622_RNA45SN1_45S_50ntFlanks@12479.12497.19\|CTAATAGGGAACGTGAGCT | >hg38_21_+_8433172_8446622_RNA45SN1_45S_50ntFlanks | 12479 | 12497 | 19 | CTAATAGGGAACGTGAGCT | 3949.2 | 2.6 | 0.7 | 3.6 | 2.9E-04 | 4.1E-04 | BT20/MB468 | CYTO |
| >trnaMT_PheGAA_MT_+_577_647@22.59.38\|AAAGCAATACACTGAAAATGTTTAGACGGGCTCACATC | >trnaMT_PheGAA_MT_+_577_647 | 22 | 59 | 38 | AAAGCAATACACTGAAAATGTTTAGACGGGCTCACATC | 90.0 | 10.2 | 2.7 | 3.8 | 1.3E-04 | 4.1E-04 | BT20/MB468 | MP |
| >hg38_21_+_8433172_8446622_RNA45SN1_45S_50ntFlanks@11480.11502.23\|CGGCGGAGGGGCCGCGGGCCGGT | >hg38_21_+_8433172_8446622_RNA45SN1_45S_50ntFlanks | 11480 | 11502 | 23 | CGGCGGAGGGGCCGCGGGCCGGT | 161.8 | -3.7 | 1.0 | -3.6 | 3.1E-04 | 4.1E-04 | MB231/MB468 | CYTO |
| >hg38_21_+_8433172_8446622_RNA45SN1_45S_50ntFlanks@10390.10424.35\|GAACAGCAGTTGAACATGGGTCAGTCGGTCCTGAG | >hg38_21_+_8433172_8446622_RNA45SN1_45S_50ntFlanks | 10390 | 10424 | 35 | GAACAGCAGTTGAACATGGGTCAGTCGGTCCTGAG | 5672.4 | 3.5 | 0.9 | 4.0 | 5.3E-05 | 4.1E-04 | BT20/MB231 | MITO |
| >hg38_21_+_8433172_8446622_RNA45SN1_45S_50ntFlanks@12420.12457.38\|ATGTCGGCTCTTCCTATCATTGTGAAGCAGAATTCACC | >hg38_21_+_8433172_8446622_RNA45SN1_45S_50ntFlanks | 12420 | 12457 | 38 | ATGTCGGCTCTTCCTATCATTGTGAAGCAGAATTCACC | 2812.2 | -3.2 | 0.8 | -4.1 | 4.4E-05 | 4.1E-04 | BT20/MB231 | CYTO |
| >hg38_21_+_8433172_8446622_RNA45SN1_45S_50ntFlanks@12934.12966.33\|CTAAACCATTCGTAGACGACCTGCTTCTGGGTC | >hg38_21_+_8433172_8446622_RNA45SN1_45S_50ntFlanks | 12934 | 12966 | 33 | CTAAACCATTCGTAGACGACCTGCTTCTGGGTC | 353.0 | -5.6 | 1.6 | -3.6 | 3.3E-04 | 4.1E-04 | MB231/MB468 | NUC |
| >am_trnaMT_GluTTC_MT_-_14674_14742@26.54.29,trnalookalike8_GluTTC_5_-_93905172_93905240@26.54.29\|GGTTTTTCATATCATTGGTCGTGGTTGTA | >am_trnaMT_GluTTC_MT_-_14674_14742 | 26 | 54 | 29 | GGTTTTTCATATCATTGGTCGTGGTTGTA | 296.1 | 7.2 | 1.9 | 3.8 | 1.3E-04 | 4.1E-04 | BT20/MB468 | MP |
| >hg38_21_+_8433172_8446622_RNA45SN1_45S_50ntFlanks@6734.6756.23\|ATTGATCATCGACACTTCGAACG | >hg38_21_+_8433172_8446622_RNA45SN1_45S_50ntFlanks | 6734 | 6756 | 23 | ATTGATCATCGACACTTCGAACG | 433.3 | -2.8 | 0.8 | -3.6 | 3.3E-04 | 4.1E-04 | BT20/MB468 | NUC |
| >hg38_21_+_8433172_8446622_RNA45SN1_45S_50ntFlanks@10687.10709.23\|CGGCGTCCGGTGAGCTCTCGCTG | >hg38_21_+_8433172_8446622_RNA45SN1_45S_50ntFlanks | 10687 | 10709 | 23 | CGGCGTCCGGTGAGCTCTCGCTG | 2373.5 | -2.8 | 0.8 | -3.7 | 2.4E-04 | 4.2E-04 | BT20/MB468 | MITO |
| >trnaMT_GluTTC_MT_-_14674_14742@39.58.20,trnalookalike8_GluTTC_5_-_93905172_93905240@39.58.20\|ATTGGTCGTGGTTGTAGTCC | >trnaMT_GluTTC_MT_-_14674_14742 | 39 | 58 | 20 | ATTGGTCGTGGTTGTAGTCC | 253.5 | 4.5 | 1.3 | 3.6 | 3.3E-04 | 4.2E-04 | MB231/MB468 | MITO |
| >hg38_21_+_8433172_8446622_RNA45SN1_45S_50ntFlanks@10727.10755.29\|GGAGAGGGTGTAAATCTCGCGCCGGGCCG | >hg38_21_+_8433172_8446622_RNA45SN1_45S_50ntFlanks | 10727 | 10755 | 29 | GGAGAGGGTGTAAATCTCGCGCCGGGCCG | 595.4 | -5.5 | 1.5 | -3.6 | 2.9E-04 | 4.2E-04 | BT20/MB468 | CYTO |
| >hg38_21_+_8433172_8446622_RNA45SN1_45S_50ntFlanks@9603.9622.20\|GAAAGACTAATCGAACCATC | >hg38_21_+_8433172_8446622_RNA45SN1_45S_50ntFlanks | 9603 | 9622 | 20 | GAAAGACTAATCGAACCATC | 186.2 | -4.5 | 1.2 | -3.6 | 2.7E-04 | 4.2E-04 | MB231/MB468 | MP |
| >hg38_21_+_8433172_8446622_RNA45SN1_45S_50ntFlanks@8177.8203.27\|TTCTGATCGAGGCCCAGCCCGTGGACG | >hg38_21_+_8433172_8446622_RNA45SN1_45S_50ntFlanks | 8177 | 8203 | 27 | TTCTGATCGAGGCCCAGCCCGTGGACG | 406.5 | 2.0 | 0.5 | 3.7 | 2.4E-04 | 4.2E-04 | BT20/MB468 | MITO |
| >trnaMT_ValTAC_MT_+_1602_1670@47.72.26\|TCAACTTAACTTGACCGCTCTGACCA | >trnaMT_ValTAC_MT_+_1602_1670 | 47 | 72 | 26 | TCAACTTAACTTGACCGCTCTGACCA | 3205.9 | 3.5 | 1.0 | 3.7 | 2.4E-04 | 4.2E-04 | BT20/MB468 | MITO |
| >MI0000299\|hsa-mir-222&WithFlank&X\|-\|45747009\|45747130@75.99.25(+2U)&[MIMAT0000279&hsa-miR-222-3p&offsets\|0\|+4(+2U);m-140&X\|-\|45747033\|45747056&offsets\|0\|+1(+2U)]\|AGCTACATCTGGCTACTGGGTCTCTTT | >MI0000299\|hsa-mir-222&WithFlank&X\|-\|45747009\|45747130 | 75 | 99 | 25(+2U) | AGCTACATCTGGCTACTGGGTCTCTTT | 3064.4 | -3.5 | 0.9 | -4.0 | 5.3E-05 | 4.2E-04 | BT20/MB231 | MITO |
| >hg38_21_+_8433172_8446622_RNA45SN1_45S_50ntFlanks@10384.10424.41\|CCATGTGAACAGCAGTTGAACATGGGTCAGTCGGTCCTGAG | >hg38_21_+_8433172_8446622_RNA45SN1_45S_50ntFlanks | 10384 | 10424 | 41 | CCATGTGAACAGCAGTTGAACATGGGTCAGTCGGTCCTGAG | 1283.0 | 2.4 | 0.7 | 3.6 | 3.1E-04 | 4.2E-04 | MB231/MB468 | CYTO |
| >hg38_21_+_8433172_8446622_RNA45SN1_45S_50ntFlanks@4047.4069.23\|ATTCGAACGTCTGCCCTATCAAC | >hg38_21_+_8433172_8446622_RNA45SN1_45S_50ntFlanks | 4047 | 4069 | 23 | ATTCGAACGTCTGCCCTATCAAC | 274.9 | -2.4 | 0.7 | -3.6 | 3.3E-04 | 4.2E-04 | MB231/MB468 | NUC |
| >trnaMT_PheGAA_MT_+_577_647@45.73.29\|AGACGGGCTCACATCACCCCATAAACACC | >trnaMT_PheGAA_MT_+_577_647 | 45 | 73 | 29 | AGACGGGCTCACATCACCCCATAAACACC | 664.4 | -4.2 | 1.2 | -3.6 | 3.3E-04 | 4.2E-04 | MB231/MB468 | MITO |
| >hg38_21_+_8433172_8446622_RNA45SN1_45S_50ntFlanks@12456.12500.45\|CCAAGCGTTGGATTGTTCACCCACTAATAGGGAACGTGAGCTGGG | >hg38_21_+_8433172_8446622_RNA45SN1_45S_50ntFlanks | 12456 | 12500 | 45 | CCAAGCGTTGGATTGTTCACCCACTAATAGGGAACGTGAGCTGGG | 264.1 | -3.1 | 0.9 | -3.6 | 3.1E-04 | 4.2E-04 | MB231/MB468 | CYTO |
| >hg38_21_+_8433172_8446622_RNA45SN1_45S_50ntFlanks@11670.11701.32\|CCCAGTGCTCTGAATGTCAAAGTGAAGAAATT | >hg38_21_+_8433172_8446622_RNA45SN1_45S_50ntFlanks | 11670 | 11701 | 32 | CCCAGTGCTCTGAATGTCAAAGTGAAGAAATT | 540.6 | -2.1 | 0.6 | -3.6 | 3.1E-04 | 4.2E-04 | MB231/MB468 | CYTO |
| >hg38_21_+_8433172_8446622_RNA45SN1_45S_50ntFlanks@6661.6696.36\|CGGTGGATCACTCGGCTCGTGCGTCGATGAAGAACG | >hg38_21_+_8433172_8446622_RNA45SN1_45S_50ntFlanks | 6661 | 6696 | 36 | CGGTGGATCACTCGGCTCGTGCGTCGATGAAGAACG | 2149.9 | 2.4 | 0.6 | 3.7 | 2.4E-04 | 4.2E-04 | BT20/MB468 | MITO |
| >hg38_21_+_8433172_8446622_RNA45SN1_45S_50ntFlanks@4620.4639.20\|ATTAAGAGGGACGGCCGGGG | >hg38_21_+_8433172_8446622_RNA45SN1_45S_50ntFlanks | 4620 | 4639 | 20 | ATTAAGAGGGACGGCCGGGG | 190.2 | -3.8 | 1.0 | -3.6 | 2.8E-04 | 4.2E-04 | MB231/MB468 | MP |
| >MI0003129\|hsa-mir-146b&WithFlank&10\|+\|102436506\|102436590@15.32.18&[MIMAT0002809&hsa-miR-146b-5p&offsets\|0\|-5;m-70&10\|+\|102436520\|102436543&offsets\|0\|-6]\|TGAGAACTGAATTCCATA | >MI0003129\|hsa-mir-146b&WithFlank&10\|+\|102436506\|102436590 | 15 | 32 | 18 | TGAGAACTGAATTCCATA | 72.5 | 9.4 | 2.6 | 3.6 | 2.8E-04 | 4.2E-04 | MB231/MB468 | MP |
| >hg38_21_+_8433172_8446622_RNA45SN1_45S_50ntFlanks@12083.12106.24\|CTCGCTTCTGGCGCCAAGCGCCCG | >hg38_21_+_8433172_8446622_RNA45SN1_45S_50ntFlanks | 12083 | 12106 | 24 | CTCGCTTCTGGCGCCAAGCGCCCG | 237.3 | -5.3 | 1.5 | -3.6 | 2.9E-04 | 4.2E-04 | BT20/MB468 | CYTO |
| >hg38_21_+_8433172_8446622_RNA45SN1_45S_50ntFlanks@11233.11254.22\|CCCTTCCCGTGGATCGCCCCAG | >hg38_21_+_8433172_8446622_RNA45SN1_45S_50ntFlanks | 11233 | 11254 | 22 | CCCTTCCCGTGGATCGCCCCAG | 1393.6 | -6.0 | 1.7 | -3.6 | 3.0E-04 | 4.2E-04 | BT20/MB468 | CYTO |
| >hg38_21_+_8433172_8446622_RNA45SN1_45S_50ntFlanks@11604.11622.19\|CGACTGTTTAATTAAAACA | >hg38_21_+_8433172_8446622_RNA45SN1_45S_50ntFlanks | 11604 | 11622 | 19 | CGACTGTTTAATTAAAACA | 385.0 | 3.7 | 0.9 | 4.0 | 5.4E-05 | 4.2E-04 | BT20/MB231 | MITO |
| >hg38_21_+_8433172_8446622_RNA45SN1_45S_50ntFlanks@8195.8215.21\|CCGTGGACGGTGTGAGGCCGG | >hg38_21_+_8433172_8446622_RNA45SN1_45S_50ntFlanks | 8195 | 8215 | 21 | CCGTGGACGGTGTGAGGCCGG | 224.8 | -2.4 | 0.7 | -3.6 | 3.1E-04 | 4.2E-04 | MB231/MB468 | CYTO |
| >trna3_ProTGG_14_+_21101165_21101236@1.29.29\|GGCTCGTTGGTCTAGTGGTATGATTCTCG | >trna3_ProTGG_14_+_21101165_21101236 | 1 | 29 | 29 | GGCTCGTTGGTCTAGTGGTATGATTCTCG | 150.0 | -4.8 | 1.3 | -3.6 | 3.0E-04 | 4.2E-04 | BT20/MB468 | CYTO |
| >MI0000115\|hsa-mir-16-2&WithFlank&3\|+\|160404739\|160404831@58.79.22(+1U)&[MIMAT0004518&hsa-miR-16-2-3p&offsets\|-1\|-1(+1U);m-252&3\|+\|160404796\|160404816&offsets\|0\|+1(+1U)]\|ACCAATATTACTGTGCTGCTTTT | >MI0000115\|hsa-mir-16-2&WithFlank&3\|+\|160404739\|160404831 | 58 | 79 | 22(+1U) | ACCAATATTACTGTGCTGCTTTT | 152.3 | 5.7 | 1.6 | 3.6 | 3.3E-04 | 4.2E-04 | MB231/MB468 | MITO |
| >hg38_MT_+_1621_3279_RNR2_16S_50ntFlanks@51.90.40\|GCTAAACCTAGCCCCAAACCCACTCCACCTTACTACCAGA | >hg38_MT_+_1621_3279_RNR2_16S_50ntFlanks | 51 | 90 | 40 | GCTAAACCTAGCCCCAAACCCACTCCACCTTACTACCAGA | 3359.8 | 5.6 | 1.5 | 3.8 | 1.3E-04 | 4.2E-04 | BT20/MB468 | MP |
| >hg38_21_+_8433172_8446622_RNA45SN1_45S_50ntFlanks@12627.12669.43\|ATGGGGCGAAGCTACCATCTGTGGGATTATGACTGAACGCCTC | >hg38_21_+_8433172_8446622_RNA45SN1_45S_50ntFlanks | 12627 | 12669 | 43 | ATGGGGCGAAGCTACCATCTGTGGGATTATGACTGAACGCCTC | 584.7 | -3.3 | 0.9 | -3.6 | 3.3E-04 | 4.2E-04 | MB231/MB468 | MITO |
| >MI0000266\|hsa-mir-10a&WithFlank&17\|-\|48579832\|48579953@29.49.21(+1U)&[MIMAT0000253&hsa-miR-10a-5p&offsets\|+1\|-1(+1U);m-18&17\|-\|48579905\|48579926&offsets\|+1\|0(+1U)]\|ACCCTGTAGATCCGAATTTGTT | >MI0000266\|hsa-mir-10a&WithFlank&17\|-\|48579832\|48579953 | 29 | 49 | 21(+1U) | ACCCTGTAGATCCGAATTTGTT | 207.9 | 3.0 | 0.7 | 4.1 | 4.5E-05 | 4.2E-04 | BT20/MB231 | CYTO |
| >hg38_21_+_8433172_8446622_RNA45SN1_45S_50ntFlanks@6723.6740.18\|TTGCAGGACACATTGATC | >hg38_21_+_8433172_8446622_RNA45SN1_45S_50ntFlanks | 6723 | 6740 | 18 | TTGCAGGACACATTGATC | 283.8 | -5.7 | 1.6 | -3.6 | 2.8E-04 | 4.2E-04 | MB231/MB468 | MP |
| >hg38_21_+_8433172_8446622_RNA45SN1_45S_50ntFlanks@12785.12817.33\|CCCCGCGCGCGCGGGAGGGCGCGTGCCCCGCCG | >hg38_21_+_8433172_8446622_RNA45SN1_45S_50ntFlanks | 12785 | 12817 | 33 | CCCCGCGCGCGCGGGAGGGCGCGTGCCCCGCCG | 203.7 | -5.1 | 1.4 | -3.7 | 2.5E-04 | 4.2E-04 | BT20/MB468 | MITO |
| >hg38_21_+_8433172_8446622_RNA45SN1_45S_50ntFlanks@10845.10872.28\|ACTTCGGGATAAGGATTGGCTCTAAGGG | >hg38_21_+_8433172_8446622_RNA45SN1_45S_50ntFlanks | 10845 | 10872 | 28 | ACTTCGGGATAAGGATTGGCTCTAAGGG | 209.5 | -3.8 | 1.0 | -3.6 | 3.0E-04 | 4.2E-04 | BT20/MB468 | CYTO |
| >hg38_21_+_8433172_8446622_RNA45SN1_45S_50ntFlanks@6753.6770.18\|AACGCACTTGCGGCCCCG | >hg38_21_+_8433172_8446622_RNA45SN1_45S_50ntFlanks | 6753 | 6770 | 18 | AACGCACTTGCGGCCCCG | 91.3 | -4.9 | 1.3 | -3.7 | 2.5E-04 | 4.2E-04 | BT20/MB468 | MITO |
| >MI0000085\|hsa-mir-27a&WithFlank&19\|-\|13836434\|13836523@57.75.19(+4U)&[MIMAT0000084&hsa-miR-27a-3p&offsets\|0\|-2(+4U);m-43&19\|-\|13836447\|13836467&offsets\|0\|-2(+4U)]\|TTCACAGTGGCTAAGTTCCTTTT | >MI0000085\|hsa-mir-27a&WithFlank&19\|-\|13836434\|13836523 | 57 | 75 | 19(+4U) | TTCACAGTGGCTAAGTTCCTTTT | 50.9 | -9.1 | 2.2 | -4.1 | 4.7E-05 | 4.2E-04 | BT20/MB231 | MP |
| >MI0006444\|hsa-mir-1307&WithFlank&10\|-\|103394247\|103394407@87.108.22&[MIMAT0005951&hsa-miR-1307-3p&offsets\|+1\|+1;m-66&10\|-\|103394300\|103394322&offsets\|+1\|0]\|CTCGGCGTGGCGTCGGTCGTGG | >MI0006444\|hsa-mir-1307&WithFlank&10\|-\|103394247\|103394407 | 87 | 108 | 22 | CTCGGCGTGGCGTCGGTCGTGG | 1053.8 | 3.5 | 0.9 | 3.8 | 1.3E-04 | 4.2E-04 | BT20/MB468 | MP |
| >trnaMT_GluTTC_MT_-_14674_14742@38.66.29,trnalookalike8_GluTTC_5_-_93905172_93905240@38.66.29\|CATTGGTCGTGGTTGTAGTCCGTGCGAGA | >trnaMT_GluTTC_MT_-_14674_14742 | 38 | 66 | 29 | CATTGGTCGTGGTTGTAGTCCGTGCGAGA | 375.9 | 4.4 | 1.2 | 3.8 | 1.3E-04 | 4.2E-04 | BT20/MB468 | MP |
| >hg38_21_+_8433172_8446622_RNA45SN1_45S_50ntFlanks@9561.9596.36\|GTCCTGACGTGCAAATCGGTCGTCCGACCTGGGTAT | >hg38_21_+_8433172_8446622_RNA45SN1_45S_50ntFlanks | 9561 | 9596 | 36 | GTCCTGACGTGCAAATCGGTCGTCCGACCTGGGTAT | 205.7 | -3.7 | 1.0 | -3.6 | 3.1E-04 | 4.2E-04 | MB231/MB468 | CYTO |
| >trna10_AspGTC_12_-_125424193_125424264@30.72.43,trna12_AspGTC_12_-_125411891_125411962@30.72.43,trna38_AspGTC_17_-_8125556_8125627@30.72.43,trna45_AspGTC_6_+_27447453_27447524@30.72.43,trna48_AspGTC_6_+_27471523_27471594@30.72.43,trna4_AspGTC_12_+_96429799_96429870@30.72.43,trna69_AspGTC_1_-_161440205_161440276@30.72.43,trna72_AspGTC_1_-_161432824_161432895@30.72.43,trna75_AspGTC_1_-_161425414_161425485@30.72.43,trna78_AspGTC_1_-_161418033_161418104@30.72.43,trna81_AspGTC_1_-_161410615_161410686@30.72.43\|GCCTGTCACGCGGGAGACCGGGGTTCGATTCCCCGACGGGGAG | >trna10_AspGTC_12_-_125424193_125424264 | 30 | 72 | 43 | GCCTGTCACGCGGGAGACCGGGGTTCGATTCCCCGACGGGGAG | 356.5 | -3.9 | 1.1 | -3.6 | 3.0E-04 | 4.2E-04 | BT20/MB468 | CYTO |
| >hg38_21_+_8433172_8446622_RNA45SN1_45S_50ntFlanks@10411.10445.35\|CAGTCGGTCCTGAGAGATGGGCGAGCGCCGTTCCG | >hg38_21_+_8433172_8446622_RNA45SN1_45S_50ntFlanks | 10411 | 10445 | 35 | CAGTCGGTCCTGAGAGATGGGCGAGCGCCGTTCCG | 168.2 | -3.3 | 0.9 | -3.6 | 3.0E-04 | 4.2E-04 | BT20/MB468 | CYTO |
| >hg38_21_+_8433172_8446622_RNA45SN1_45S_50ntFlanks@6779.6803.25\|CCGGGGCTACGCCTGTCTGAGCGTC | >hg38_21_+_8433172_8446622_RNA45SN1_45S_50ntFlanks | 6779 | 6803 | 25 | CCGGGGCTACGCCTGTCTGAGCGTC | 309.0 | -1.4 | 0.4 | -3.6 | 3.0E-04 | 4.2E-04 | BT20/MB468 | CYTO |
| >hg38_21_+_8433172_8446622_RNA45SN1_45S_50ntFlanks@11933.11954.22\|AATAAGTGGGAGGCCCCCGGCG | >hg38_21_+_8433172_8446622_RNA45SN1_45S_50ntFlanks | 11933 | 11954 | 22 | AATAAGTGGGAGGCCCCCGGCG | 613.6 | -5.1 | 1.3 | -3.8 | 1.3E-04 | 4.2E-04 | BT20/MB468 | MP |
| >hg38_21_+_8433172_8446622_RNA45SN1_45S_50ntFlanks@476.522.47\|TTCGTGATCGATGTGGTGACGTCGTGCTCTCCCGGGCCGGGTCCGAG | >hg38_21_+_8433172_8446622_RNA45SN1_45S_50ntFlanks | 476 | 522 | 47 | TTCGTGATCGATGTGGTGACGTCGTGCTCTCCCGGGCCGGGTCCGAG | 2888.8 | 4.2 | 1.2 | 3.6 | 3.4E-04 | 4.2E-04 | MB231/MB468 | NUC |
| >MI0000108\|hsa-mir-103a-2&WithFlank&20\|+\|3917488\|3917577@54.76.23(+2U);MI0000109\|hsa-mir-103a-1&WithFlank&5\|-\|168560890\|168560979@54.76.23(+2U)&[MIMAT0000101&hsa-miR-103a-3p&offsets\|0\|0(+2U);m-12&20\|+\|3917541\|3917563&offsets\|0\|0(+2U)];[MIMAT0000101_1&hsa-miR-103a-3p&offsets\|0\|0(+2U);m-13&5\|-\|168560904\|168560926&offsets\|0\|0(+2U);m-44220&5\|-\|168560908\|168560926&offsets\|0\|+4(+2U)]\|AGCAGCATTGTACAGGGCTATGATT | >MI0000108\|hsa-mir-103a-2&WithFlank&20\|+\|3917488\|3917577 | 54 | 76 | 23(+2U) | AGCAGCATTGTACAGGGCTATGATT | 1194.2 | -3.2 | 0.8 | -4.0 | 5.5E-05 | 4.2E-04 | BT20/MB231 | MITO |
| >hg38_21_+_8433172_8446622_RNA45SN1_45S_50ntFlanks@6687.6720.34\|ATGAAGAACGCAGCTAGCTGCGAGAATTAATGTG | >hg38_21_+_8433172_8446622_RNA45SN1_45S_50ntFlanks | 6687 | 6720 | 34 | ATGAAGAACGCAGCTAGCTGCGAGAATTAATGTG | 349.7 | -3.6 | 1.0 | -3.6 | 3.1E-04 | 4.2E-04 | MB231/MB468 | CYTO |

<table>
<tr><th></th><th></th><th></th><th></th><th></th><th></th><th></th><th></th><th></th><th></th><th></th><th></th><th></th><th></th></tr>
<tr><td>>hg38_21_+_8433172_8446622_RNA45SN1_45S_50ntFlanks@11583.11611.29|CTGGTGCGGACCAGGGGAATCCGACTGTT</td><td>>hg38_21_+_8433172_8446622_RNA45SN1_45S_50ntFlanks</td><td>11583</td><td>11611</td><td>29</td><td>CTGGTGCGGACCAGGGGAATCCGACTGTT</td><td>3072.4</td><td>-3.1</td><td>0.8</td><td>-3.8</td><td>1.3E-04</td><td>4.2E-04</td><td>BT20/MB468</td><td>MP</td></tr>
<tr><td>>trna111_HisGTG_1_-_147774845_147774916@-1G.33.34,trna118_HisGTG_1_-_145396881_145396952@-1G.33.34,trna16_HisGTG_1_+_146544773_146544844@-1G.33.34,trna1_HisGTG_15_+_45493349_45493420@-1G.33.34,trna21_HisGTG_1_+_147753471_147753542@-1G.33.34,trna33_HisGTG_6_+_27125906_27125977@-1G.33.34,trna7_HisGTG_9_-_14433938_14434009@-1G.33.34,trna8_HisGTG_15_-_45492611_45492682@-1G.33.34,trna9_HisGTG_15_-_45490804_45490875@-1G.33.34|GGCCGTGATCGTATAGTGGTTAGTACTCTGCGTT</td><td>>trna111_HisGTG_1_-_147774845_147774916</td><td>-1G</td><td>33</td><td>34</td><td>GGCCGTGATCGTATAGTGGTTAGTACTCTGCGTT</td><td>25258.8</td><td>-4.5</td><td>1.1</td><td>-4.1</td><td>4.8E-05</td><td>4.2E-04</td><td>BT20/MB231</td><td>NUC</td></tr>
<tr><td>>hg38_MT_+_598_1651_RNR1_12S_50ntFlanks@428.469.42|AAAGTGGCTTTAACATATCTGAACACACAATAGCTAAGACCC</td><td>>hg38_MT_+_598_1651_RNR1_12S_50ntFlanks</td><td>428</td><td>469</td><td>42</td><td>AAAGTGGCTTTAACATATCTGAACACACAATAGCTAAGACCC</td><td>235.9</td><td>5.4</td><td>1.5</td><td>3.6</td><td>2.8E-04</td><td>4.2E-04</td><td>MB231/MB468</td><td>MP</td></tr>
<tr><td>>hg38_21_+_8433172_8446622_RNA45SN1_45S_50ntFlanks@10391.10420.30|AACAGCAGTTGAACATGGGTCAGTCGGTCC</td><td>>hg38_21_+_8433172_8446622_RNA45SN1_45S_50ntFlanks</td><td>10391</td><td>10420</td><td>30</td><td>AACAGCAGTTGAACATGGGTCAGTCGGTCC</td><td>362.8</td><td>-3.9</td><td>1.1</td><td>-3.7</td><td>2.5E-04</td><td>4.2E-04</td><td>BT20/MB468</td><td>MITO</td></tr>
<tr><td>>hg38_21_+_8433172_8446622_RNA45SN1_45S_50ntFlanks@10357.10377.21|CTTTGAAGGCCGAAGTGGAGA</td><td>>hg38_21_+_8433172_8446622_RNA45SN1_45S_50ntFlanks</td><td>10357</td><td>10377</td><td>21</td><td>CTTTGAAGGCCGAAGTGGAGA</td><td>176.9</td><td>-3.2</td><td>0.9</td><td>-3.6</td><td>3.2E-04</td><td>4.2E-04</td><td>MB231/MB468</td><td>CYTO</td></tr>
<tr><td>>hg38_21_+_8433172_8446622_RNA45SN1_45S_50ntFlanks@4884.4905.22|CAAAGCTGAAACTTAAAGGAAT</td><td>>hg38_21_+_8433172_8446622_RNA45SN1_45S_50ntFlanks</td><td>4884</td><td>4905</td><td>22</td><td>CAAAGCTGAAACTTAAAGGAAT</td><td>87.1</td><td>-5.4</td><td>1.5</td><td>-3.6</td><td>2.8E-04</td><td>4.2E-04</td><td>MB231/MB468</td><td>MP</td></tr>
<tr><td>>hg38_21_+_8433172_8446622_RNA45SN1_45S_50ntFlanks@11781.11802.22|CGCATGAATGGATGAACGAGAT</td><td>>hg38_21_+_8433172_8446622_RNA45SN1_45S_50ntFlanks</td><td>11781</td><td>11802</td><td>22</td><td>CGCATGAATGGATGAACGAGAT</td><td>121.0</td><td>-5.6</td><td>1.5</td><td>-3.6</td><td>2.8E-04</td><td>4.2E-04</td><td>MB231/MB468</td><td>MP</td></tr>
<tr><td>>hg38_21_+_8433172_8446622_RNA45SN1_45S_50ntFlanks@7979.8012.34|ACCTCAGATCAGACGTGGCGACCCGCTGAATTTA</td><td>>hg38_21_+_8433172_8446622_RNA45SN1_45S_50ntFlanks</td><td>7979</td><td>8012</td><td>34</td><td>ACCTCAGATCAGACGTGGCGACCCGCTGAATTTA</td><td>438.0</td><td>-2.8</td><td>0.8</td><td>-3.6</td><td>3.4E-04</td><td>4.2E-04</td><td>MB231/MB468</td><td>NUC</td></tr>
<tr><td>>hg38_21_+_8433172_8446622_RNA45SN1_45S_50ntFlanks@10869.10889.21|AGGGCTGGGTCGGTCGGGCTG</td><td>>hg38_21_+_8433172_8446622_RNA45SN1_45S_50ntFlanks</td><td>10869</td><td>10889</td><td>21</td><td>AGGGCTGGGTCGGTCGGGCTG</td><td>603.6</td><td>-3.5</td><td>0.9</td><td>-3.8</td><td>1.3E-04</td><td>4.3E-04</td><td>BT20/MB468</td><td>MP</td></tr>
<tr><td>>hg38_21_+_8433172_8446622_RNA45SN1_45S_50ntFlanks@11781.11804.24|CGCATGAATGGATGAACGAGATTC</td><td>>hg38_21_+_8433172_8446622_RNA45SN1_45S_50ntFlanks</td><td>11781</td><td>11804</td><td>24</td><td>CGCATGAATGGATGAACGAGATTC</td><td>1074.3</td><td>-2.6</td><td>0.7</td><td>-3.7</td><td>2.5E-04</td><td>4.3E-04</td><td>BT20/MB468</td><td>MITO</td></tr>
<tr><td>>MI0000063|hsa-let-7b&WithFlank&22|+|46113680|46113774@12.32.21(+1A)&[MIMAT0000063&hsa-let-7b-5p&offsets|0|-1(+1A);m-14&22|+|46113691|46113712&offsets|0|-1(+1A)]|TGAGGTAGTAGGTTGTGTGGTA</td><td>>MI0000063|hsa-let-7b&WithFlank&22|+|46113680|46113774</td><td>12</td><td>32</td><td>21(+1A)</td><td>TGAGGTAGTAGGTTGTGTGGTA</td><td>1160.1</td><td>3.3</td><td>0.9</td><td>3.6</td><td>2.8E-04</td><td>4.3E-04</td><td>MB231/MB468</td><td>MP</td></tr>
<tr><td>>hg38_21_+_8433172_8446622_RNA45SN1_45S_50ntFlanks@3854.3888.35|ACTGTGGTAATTCTAGAGCTAATACATGCCGACGG</td><td>>hg38_21_+_8433172_8446622_RNA45SN1_45S_50ntFlanks</td><td>3854</td><td>3888</td><td>35</td><td>ACTGTGGTAATTCTAGAGCTAATACATGCCGACGG</td><td>350.6</td><td>-6.1</td><td>1.7</td><td>-3.6</td><td>3.4E-04</td><td>4.3E-04</td><td>BT20/MB468</td><td>NUC</td></tr>
<tr><td>>hg38_21_+_8433172_8446622_RNA45SN1_45S_50ntFlanks@4378.4398.21|CTCGTAGTTGGATCTTGGGAG</td><td>>hg38_21_+_8433172_8446622_RNA45SN1_45S_50ntFlanks</td><td>4378</td><td>4398</td><td>21</td><td>CTCGTAGTTGGATCTTGGGAG</td><td>154.1</td><td>-5.8</td><td>1.5</td><td>-3.8</td><td>1.3E-04</td><td>4.3E-04</td><td>BT20/MB468</td><td>MP</td></tr>
<tr><td>>am_trnaMT_PheGAA_MT_+_577_647@44.62.19|TAGACGGGCTCACATCACC</td><td>>am_trnaMT_PheGAA_MT_+_577_647</td><td>44</td><td>62</td><td>19</td><td>TAGACGGGCTCACATCACC</td><td>937.9</td><td>4.4</td><td>1.2</td><td>3.8</td><td>1.3E-04</td><td>4.3E-04</td><td>BT20/MB468</td><td>MP</td></tr>
<tr><td>>MI0003137|hsa-mir-193b&WithFlank&16|+|14303961|14304055@57.77.21&[MIMAT0002819&hsa-miR-193b-3p&offsets|0|-1;m-166&16|+|14304017|14304038&offsets|0|-1]|AACTGGCCCTCAAAGTCCCGC</td><td>>MI0003137|hsa-mir-193b&WithFlank&16|+|14303961|14304055</td><td>57</td><td>77</td><td>21</td><td>AACTGGCCCTCAAAGTCCCGC</td><td>1307.9</td><td>6.0</td><td>1.6</td><td>3.8</td><td>1.3E-04</td><td>4.3E-04</td><td>BT20/MB468</td><td>MP</td></tr>
<tr><td>>MI0000446|hsa-mir-125b-1&WithFlank&11|-|122099751|122099850@21.38.18(+1A);MI0000470|hsa-mir-125b-2&WithFlank&21|+|16590231|16590331@23.40.18(+1A)&[MIMAT0000423&hsa-miR-125b-5p&offsets|0|-4(+1A);m-67&11|-|122099809|122099830&offsets|0|-4(+1A)];[MIMAT0000423_1&hsa-miR-125b-5p&offsets|0|-4(+1A);m-68&21|+|16590253|16590274&offsets|0|-4(+1A)]|TCCCTGAGACCCTAACTTA</td><td>>MI0000446|hsa-mir-125b-1&WithFlank&11|-|122099751|122099850</td><td>21</td><td>38</td><td>18(+1A)</td><td>TCCCTGAGACCCTAACTTA</td><td>72.1</td><td>9.4</td><td>2.6</td><td>3.6</td><td>2.8E-04</td><td>4.3E-04</td><td>MB231/MB468</td><td>MP</td></tr>
<tr><td>>trnaMT_PheGAA_MT_+_577_647@39.60.22|ATGTTTAGACGGGCTCACATCA</td><td>>trnaMT_PheGAA_MT_+_577_647</td><td>39</td><td>60</td><td>22</td><td>ATGTTTAGACGGGCTCACATCA</td><td>197.0</td><td>4.6</td><td>1.3</td><td>3.6</td><td>3.3E-04</td><td>4.3E-04</td><td>MB231/MB468</td><td>MITO</td></tr>
<tr><td>>MI0000301|hsa-mir-224&WithFlank&X|-|151958572|151958664@14.35.22&[MIMAT0000281&hsa-miR-224-5p&offsets|+1|-2;m-238&X|-|151958630|151958651&offsets|0|0]|CAAGTCACTAGTGGTTCCGTTT</td><td>>MI0000301|hsa-mir-224&WithFlank&X|-|151958572|151958664</td><td>14</td><td>35</td><td>22</td><td>CAAGTCACTAGTGGTTCCGTTT</td><td>9144.0</td><td>2.8</td><td>0.7</td><td>4.1</td><td>4.9E-05</td><td>4.3E-04</td><td>BT20/MB231</td><td>NUC</td></tr>
<tr><td>>hg38_21_+_8433172_8446622_RNA45SN1_45S_50ntFlanks@12457.12490.34|CAAGCGTTGGATTGTTCACCCACTAATAGGGAAC</td><td>>hg38_21_+_8433172_8446622_RNA45SN1_45S_50ntFlanks</td><td>12457</td><td>12490</td><td>34</td><td>CAAGCGTTGGATTGTTCACCCACTAATAGGGAAC</td><td>130.2</td><td>-5.6</td><td>1.5</td><td>-3.6</td><td>2.8E-04</td><td>4.3E-04</td><td>MB231/MB468</td><td>MP</td></tr>
<tr><td>>MI0003667|hsa-mir-652&WithFlank&X|+|110055323|110055432@67.86.20&[MIMAT0003322&hsa-miR-652-3p&offsets|0|-1;m-148&X|+|110055389|110055410&offsets|0|-2]|AATGGCGCCACTAGGGTTGT</td><td>>MI0003667|hsa-mir-652&WithFlank&X|+|110055323|110055432</td><td>67</td><td>86</td><td>20</td><td>AATGGCGCCACTAGGGTTGT</td><td>747.0</td><td>6.1</td><td>1.6</td><td>3.8</td><td>1.3E-04</td><td>4.3E-04</td><td>BT20/MB468</td><td>MP</td></tr>
<tr><td>>hg38_21_+_8433172_8446622_RNA45SN1_45S_50ntFlanks@7978.8008.31|GACCTCAGATCAGACGTGGCGACCCGCTGAA</td><td>>hg38_21_+_8433172_8446622_RNA45SN1_45S_50ntFlanks</td><td>7978</td><td>8008</td><td>31</td><td>GACCTCAGATCAGACGTGGCGACCCGCTGAA</td><td>190.7</td><td>-2.8</td><td>0.8</td><td>-3.6</td><td>2.8E-04</td><td>4.3E-04</td><td>MB231/MB468</td><td>MP</td></tr>
<tr><td>>hg38_21_+_8433172_8446622_RNA45SN1_45S_50ntFlanks@12290.12318.29|CGTGAAAGCGGGGCCTCACGATCCTTCTG</td><td>>hg38_21_+_8433172_8446622_RNA45SN1_45S_50ntFlanks</td><td>12290</td><td>12318</td><td>29</td><td>CGTGAAAGCGGGGCCTCACGATCCTTCTG</td><td>334.7</td><td>-5.8</td><td>1.6</td><td>-3.7</td><td>2.5E-04</td><td>4.3E-04</td><td>BT20/MB468</td><td>MITO</td></tr>
<tr><td>>hg38_21_+_8433172_8446622_RNA45SN1_45S_50ntFlanks@10711.10736.26|CCCTTGAAAATCCGGGGGAGAGGGTG</td><td>>hg38_21_+_8433172_8446622_RNA45SN1_45S_50ntFlanks</td><td>10711</td><td>10736</td><td>26</td><td>CCCTTGAAAATCCGGGGGAGAGGGTG</td><td>177.8</td><td>-3.7</td><td>1.0</td><td>-3.6</td><td>3.2E-04</td><td>4.3E-04</td><td>MB231/MB468</td><td>CYTO</td></tr>
<tr><td>>MI0001733|hsa-mir-452&WithFlank&X|-|151959622|151959718@23.43.21&[MIMAT0001635&hsa-miR-452-5p&offsets|+3|+2;m-188&X|-|151959677|151959699&offsets|+3|+1]|TGTTTGCAGAGGAAACTGAGA</td><td>>MI0001733|hsa-mir-452&WithFlank&X|-|151959622|151959718</td><td>23</td><td>43</td><td>21</td><td>TGTTTGCAGAGGAAACTGAGA</td><td>109.3</td><td>6.2</td><td>1.7</td><td>3.6</td><td>2.8E-04</td><td>4.3E-04</td><td>MB231/MB468</td><td>MP</td></tr>
<tr><td>>hg38_21_+_8433172_8446622_RNA45SN1_45S_50ntFlanks@476.524.49|TTCGTGATCGATGTGGTGACGTCGTGCTCTCCCGGGCCGGGTCCGAGCC</td><td>>hg38_21_+_8433172_8446622_RNA45SN1_45S_50ntFlanks</td><td>476</td><td>524</td><td>49</td><td>TTCGTGATCGATGTGGTGACGTCGTGCTCTCCCGGGCCGGGTCCGAGCC</td><td>7994.3</td><td>5.1</td><td>1.4</td><td>3.6</td><td>3.4E-04</td><td>4.3E-04</td><td>MB231/MB468</td><td>NUC</td></tr>
<tr><td>>hg38_21_+_8433172_8446622_RNA45SN1_45S_50ntFlanks@10384.10429.46|CCATGTGAACAGCAGTTGAACATGGGTCAGTCGGTCCTGAGAGATG</td><td>>hg38_21_+_8433172_8446622_RNA45SN1_45S_50ntFlanks</td><td>10384</td><td>10429</td><td>46</td><td>CCATGTGAACAGCAGTTGAACATGGGTCAGTCGGTCCTGAGAGATG</td><td>537.0</td><td>-4.6</td><td>1.3</td><td>-3.6</td><td>3.4E-04</td><td>4.3E-04</td><td>MB231/MB468</td><td>NUC</td></tr>
<tr><td>>MI0003776|hsa-mir-320b-1&WithFlank&1|+|116671740|116671823@47.68.22;MI0003839|hsa-mir-320b-2&WithFlank&1|-|224257034|224257116@46.67.22&[MIMAT0005792&hsa-miR-320b&offsets|-1|-1;m-183&1|+|116671787|116671806&offsets|-1|+1];[MIMAT0005792_1&hsa-miR-320b&offsets|-1|-1;m-183&1|-|224257051|224257070&offsets|-1|+1]|GAAAAGCTGGGTTGAGAGGGCA</td><td>>MI0003776|hsa-mir-320b-1&WithFlank&1|+|116671740|116671823</td><td>47</td><td>68</td><td>22</td><td>GAAAAGCTGGGTTGAGAGGGCA</td><td>83.3</td><td>7.2</td><td>1.9</td><td>3.8</td><td>1.3E-04</td><td>4.3E-04</td><td>BT20/MB468</td><td>MP</td></tr>
<tr><td>>hg38_21_+_8433172_8446622_RNA45SN1_45S_50ntFlanks@8176.8215.40|CTTCTGATCGAGGCCCAGCCCGTGGACGGTGTGAGGCCGG</td><td>>hg38_21_+_8433172_8446622_RNA45SN1_45S_50ntFlanks</td><td>8176</td><td>8215</td><td>40</td><td>CTTCTGATCGAGGCCCAGCCCGTGGACGGTGTGAGGCCGG</td><td>375.4</td><td>-4.6</td><td>1.3</td><td>-3.6</td><td>3.0E-04</td><td>4.3E-04</td><td>BT20/MB468</td><td>CYTO</td></tr>
<tr><td>>hg38_21_+_8433172_8446622_RNA45SN1_45S_50ntFlanks@10401.10419.19|GAACATGGGTCAGTCGGTC</td><td>>hg38_21_+_8433172_8446622_RNA45SN1_45S_50ntFlanks</td><td>10401</td><td>10419</td><td>19</td><td>GAACATGGGTCAGTCGGTC</td><td>121.9</td><td>-5.7</td><td>1.6</td><td>-3.6</td><td>2.8E-04</td><td>4.3E-04</td><td>MB231/MB468</td><td>MP</td></tr>
<tr><td>>hg38_21_+_8433172_8446622_RNA45SN1_45S_50ntFlanks@4726.4759.34|TAATCAAGAACGAAAGTCGGAGGTTCGAAGACGA</td><td>>hg38_21_+_8433172_8446622_RNA45SN1_45S_50ntFlanks</td><td>4726</td><td>4759</td><td>34</td><td>TAATCAAGAACGAAAGTCGGAGGTTCGAAGACGA</td><td>1005.0</td><td>-2.5</td><td>0.7</td><td>-3.6</td><td>3.0E-04</td><td>4.3E-04</td><td>BT20/MB468</td><td>CYTO</td></tr>
<tr><td>>hg38_21_+_8433172_8446622_RNA45SN1_45S_50ntFlanks@8381.8402.22|AAGAGGGCGTGAAACCGTTAAG</td><td>>hg38_21_+_8433172_8446622_RNA45SN1_45S_50ntFlanks</td><td>8381</td><td>8402</td><td>22</td><td>AAGAGGGCGTGAAACCGTTAAG</td><td>897.2</td><td>-5.7</td><td>1.6</td><td>-3.6</td><td>3.0E-04</td><td>4.3E-04</td><td>BT20/MB468</td><td>CYTO</td></tr>
<tr><td>>hg38_1_-_228634819_228635039_RNA5S12_5S_50ntFlanks@89.130.42|CTGATCTCGGAAGCTAAGCAGGGTCGGGCCTGGTTAGTACTT</td><td>>hg38_1_-_228634819_228635039_RNA5S12_5S_50ntFlanks</td><td>89</td><td>130</td><td>42</td><td>CTGATCTCGGAAGCTAAGCAGGGTCGGGCCTGGTTAGTACTT</td><td>77.4</td><td>10.0</td><td>2.6</td><td>3.8</td><td>1.3E-04</td><td>4.3E-04</td><td>BT20/MB468</td><td>MP</td></tr>
<tr><td>>hg38_21_+_8433172_8446622_RNA45SN1_45S_50ntFlanks@11694.11717.24|AAGAAATTCAATGAAGCGCGGGTA</td><td>>hg38_21_+_8433172_8446622_RNA45SN1_45S_50ntFlanks</td><td>11694</td><td>11717</td><td>24</td><td>AAGAAATTCAATGAAGCGCGGGTA</td><td>383.4</td><td>-6.3</td><td>1.6</td><td>-3.8</td><td>1.3E-04</td><td>4.3E-04</td><td>BT20/MB468</td><td>MP</td></tr>
<tr><td>>hg38_21_+_8433172_8446622_RNA45SN1_45S_50ntFlanks@10059.10076.18|GGCAGTCGAGAGTGGACG</td><td>>hg38_21_+_8433172_8446622_RNA45SN1_45S_50ntFlanks</td><td>10059</td><td>10076</td><td>18</td><td>GGCAGTCGAGAGTGGACG</td><td>140.0</td><td>4.9</td><td>1.3</td><td>3.8</td><td>1.3E-04</td><td>4.3E-04</td><td>BT20/MB468</td><td>MP</td></tr>
<tr><td>>hg38_21_+_8433172_8446622_RNA45SN1_45S_50ntFlanks@10821.10865.45|GGAAGTCGGCAAGCCGGATCCGTAACTTCGGGATAAGGATTGGCT</td><td>>hg38_21_+_8433172_8446622_RNA45SN1_45S_50ntFlanks</td><td>10821</td><td>10865</td><td>45</td><td>GGAAGTCGGCAAGCCGGATCCGTAACTTCGGGATAAGGATTGGCT</td><td>530.5</td><td>-6.0</td><td>1.6</td><td>-3.6</td><td>3.0E-04</td><td>4.3E-04</td><td>BT20/MB468</td><td>CYTO</td></tr>
<tr><td>>hg38_21_+_8433172_8446622_RNA45SN1_45S_50ntFlanks@10794.10817.24|CTGGCATGTTGGAACAATGTAGGT</td><td>>hg38_21_+_8433172_8446622_RNA45SN1_45S_50ntFlanks</td><td>10794</td><td>10817</td><td>24</td><td>CTGGCATGTTGGAACAATGTAGGT</td><td>267.2</td><td>-2.4</td><td>0.7</td><td>-3.6</td><td>3.0E-04</td><td>4.3E-04</td><td>BT20/MB468</td><td>CYTO</td></tr>
<tr><td>>hg38_21_+_8433172_8446622_RNA45SN1_45S_50ntFlanks@10351.10373.23|CGAGAACTTTGAAGGCCGAAGTG</td><td>>hg38_21_+_8433172_8446622_RNA45SN1_45S_50ntFlanks</td><td>10351</td><td>10373</td><td>23</td><td>CGAGAACTTTGAAGGCCGAAGTG</td><td>904.5</td><td>-3.8</td><td>1.0</td><td>-3.6</td><td>3.2E-04</td><td>4.3E-04</td><td>MB231/MB468</td><td>CYTO</td></tr>
<tr><td>>hg38_21_+_8433172_8446622_RNA45SN1_45S_50ntFlanks@12971.12989.19|TTTCGTACGTAGCAGAGCA</td><td>>hg38_21_+_8433172_8446622_RNA45SN1_45S_50ntFlanks</td><td>12971</td><td>12989</td><td>19</td><td>TTTCGTACGTAGCAGAGCA</td><td>381.6</td><td>-3.5</td><td>1.0</td><td>-3.6</td><td>3.2E-04</td><td>4.3E-04</td><td>MB231/MB468</td><td>CYTO</td></tr>
<tr><td>>MI0000273|hsa-mir-183&WithFlank&7|-|129774899|129775020@34.55.22&[MIMAT0000261&hsa-miR-183-5p&offsets|+1|+1;m-41&7|-|129774966|129774987&offsets|0|0]|ATGGCACTGGTAGAATTCACTG</td><td>>MI0000273|hsa-mir-183&WithFlank&7|-|129774899|129775020</td><td>34</td><td>55</td><td>22</td><td>ATGGCACTGGTAGAATTCACTG</td><td>578.2</td><td>2.5</td><td>0.7</td><td>3.6</td><td>3.4E-04</td><td>4.3E-04</td><td>MB231/MB468</td><td>NUC</td></tr>
<tr><td>>hg38_21_+_8433172_8446622_RNA45SN1_45S_50ntFlanks@6650.6698.49|ACGACTCTTAGCGGTGGATCACTCGGCTCGTGCGTCGATGAAGAACGCA</td><td>>hg38_21_+_8433172_8446622_RNA45SN1_45S_50ntFlanks</td><td>6650</td><td>6698</td><td>49</td><td>ACGACTCTTAGCGGTGGATCACTCGGCTCGTGCGTCGATGAAGAACGCA</td><td>3211.5</td><td>-3.9</td><td>1.1</td><td>-3.6</td><td>3.5E-04</td><td>4.3E-04</td><td>BT20/MB468</td><td>NUC</td></tr>
<tr><td>>hg38_21_+_8433172_8446622_RNA45SN1_45S_50ntFlanks@8388.8405.18|CGTGAAACCGTTAAGAGG</td><td>>hg38_21_+_8433172_8446622_RNA45SN1_45S_50ntFlanks</td><td>8388</td><td>8405</td><td>18</td><td>CGTGAAACCGTTAAGAGG</td><td>400.2</td><td>-5.5</td><td>1.5</td><td>-3.6</td><td>3.0E-04</td><td>4.3E-04</td><td>BT20/MB468</td><td>CYTO</td></tr>
<tr><td>>hg38_21_+_8433172_8446622_RNA45SN1_45S_50ntFlanks@7985.8017.33|GATCAGACGTGGCGACCCGCTGAATTTAAGCAT</td><td>>hg38_21_+_8433172_8446622_RNA45SN1_45S_50ntFlanks</td><td>7985</td><td>8017</td><td>33</td><td>GATCAGACGTGGCGACCCGCTGAATTTAAGCAT</td><td>241.4</td><td>-5.8</td><td>1.6</td><td>-3.6</td><td>2.9E-04</td><td>4.3E-04</td><td>MB231/MB468</td><td>MP</td></tr>
<tr><td>>hg38_21_+_8433172_8446622_RNA45SN1_45S_50ntFlanks@9203.9224.22|CTCGGGGCCACGCGCGCGTCCC</td><td>>hg38_21_+_8433172_8446622_RNA45SN1_45S_50ntFlanks</td><td>9203</td><td>9224</td><td>22</td><td>CTCGGGGCCACGCGCGCGTCCC</td><td>405.2</td><td>-6.1</td><td>1.6</td><td>-3.8</td><td>1.4E-04</td><td>4.3E-04</td><td>BT20/MB468</td><td>MP</td></tr>
<tr><td>>hg38_21_+_8433172_8446622_RNA45SN1_45S_50ntFlanks@9792.9831.40|GGCGTGGAGCCGGGCGTGGAATGCGAGTGCCTAGTGGGCC</td><td>>hg38_21_+_8433172_8446622_RNA45SN1_45S_50ntFlanks</td><td>9792</td><td>9831</td><td>40</td><td>GGCGTGGAGCCGGGCGTGGAATGCGAGTGCCTAGTGGGCC</td><td>182.1</td><td>10.8</td><td>2.7</td><td>4.1</td><td>4.8E-05</td><td>4.3E-04</td><td>BT20/MB231</td><td>MP</td></tr>
</table>

| | | | | | | | | | | | | | |
|---|---|---|---|---|---|---|---|---|---|---|---|---|---|
| >trna116_GluCTC_1_-_145399233_145399304@36.70.35,trna134_GluTTC_1_-_16861774_16861845@36.70.35,trna71_GluCTC_1_-_161439189_161439260@36.70.35,trna74_GluCTC_1_-_161431809_161431880@36.70.35,trna77_GluCTC_1_-_161424398_161424469@36.70.35,trna77_GluCTC_6_+_28949976_28950047@36.70.35,trna80_GluCTC_1_-_161417018_161417089@36.70.35,trna84_GluTTC_1_-_161391883_161391954@36.70.35,trna87_GluCTC_6_-_126101393_126101464@36.70.35\|CACCGCCGCGGCCCGGGTTCGATTCCCGGTCAGGG | >trna116_GluCTC_1_-_145399233_145399304 | 36 | 70 | 35 | CACCGCCGCGGCCCGGGTTCGATTCCCGGTCAGGG | 346.5 | -3.8 | 1.1 | -3.6 | 3.0E-04 | 4.3E-04 | BT20/MB468 | CYTO |
| >hg38_21_+_8433172_8446622_RNA45SN1_45S_50ntFlanks@10391.10410.20\|AACAGCAGTTGAACATGGGT | >hg38_21_+_8433172_8446622_RNA45SN1_45S_50ntFlanks | 10391 | 10410 | 20 | AACAGCAGTTGAACATGGGT | 1754.6 | -5.1 | 1.3 | -3.8 | 1.4E-04 | 4.3E-04 | BT20/MB468 | MP |
| >trnaMT_HisGTG_MT_+_12138_12206@27.71.45\|GATTGTGAATCTGACAACAGAGGCTTACGACCCCTTATTTACCCC | >trnaMT_HisGTG_MT_+_12138_12206 | 27 | 71 | 45 | GATTGTGAATCTGACAACAGAGGCTTACGACCCCTTATTTACCCC | 81.0 | 10.1 | 2.6 | 3.8 | 1.4E-04 | 4.3E-04 | BT20/MB468 | MP |
| >hg38_21_+_8433172_8446622_RNA45SN1_45S_50ntFlanks@12405.12444.40\|CTTTTTGATCCTTCGATGTCGGCTCTTCCTATCATTGTGA | >hg38_21_+_8433172_8446622_RNA45SN1_45S_50ntFlanks | 12405 | 12444 | 40 | CTTTTTGATCCTTCGATGTCGGCTCTTCCTATCATTGTGA | 732.1 | -2.6 | 0.7 | -3.7 | 2.5E-04 | 4.3E-04 | BT20/MB468 | MITO |
| >hg38_21_+_8433172_8446622_RNA45SN1_45S_50ntFlanks@10411.10442.32\|CAGTCGGTCCTGAGAGATGGGCGAGCGCCGTT | >hg38_21_+_8433172_8446622_RNA45SN1_45S_50ntFlanks | 10411 | 10442 | 32 | CAGTCGGTCCTGAGAGATGGGCGAGCGCCGTT | 601.4 | -2.8 | 0.8 | -3.7 | 2.5E-04 | 4.3E-04 | BT20/MB468 | MITO |
| >hg38_21_+_8433172_8446622_RNA45SN1_45S_50ntFlanks@6723.6770.48\|TTGCAGGACACATTGATCATCGACACTTCGAACGCACTTGCGGCCCCG | >hg38_21_+_8433172_8446622_RNA45SN1_45S_50ntFlanks | 6723 | 6770 | 48 | TTGCAGGACACATTGATCATCGACACTTCGAACGCACTTGCGGCCCCG | 255.5 | -5.8 | 1.6 | -3.6 | 3.4E-04 | 4.3E-04 | MB231/MB468 | MITO |
| >hg38_21_+_8433172_8446622_RNA45SN1_45S_50ntFlanks@12407.12436.30\|TTTTGATCCTTCGATGTCGGCTCTTCCTAT | >hg38_21_+_8433172_8446622_RNA45SN1_45S_50ntFlanks | 12407 | 12436 | 30 | TTTTGATCCTTCGATGTCGGCTCTTCCTAT | 1030.5 | -4.3 | 1.2 | -3.6 | 3.1E-04 | 4.3E-04 | BT20/MB468 | CYTO |
| >hg38_21_+_8433172_8446622_RNA45SN1_45S_50ntFlanks@10423.10453.31\|AGAGATGGGCGAGCGCCGTTCCGAAGGGACG | >hg38_21_+_8433172_8446622_RNA45SN1_45S_50ntFlanks | 10423 | 10453 | 31 | AGAGATGGGCGAGCGCCGTTCCGAAGGGACG | 326.0 | -3.7 | 0.9 | -4.1 | 4.7E-05 | 4.3E-04 | BT20/MB231 | CYTO |
| >MI0003129\|hsa-mir-146b&WithFlank&10\|+\|102436506\|102436590@15.38.24&[MIMAT0002809&hsa-miR-146b-5p&offsets\|0\|+1;m-70&10\|+\|102436520\|102436543&offsets\|0\|0]\|TGAGAACTGAATTCCATAGGCTGT | >MI0003129\|hsa-mir-146b&WithFlank&10\|+\|102436506\|102436590 | 15 | 38 | 24 | TGAGAACTGAATTCCATAGGCTGT | 284.9 | 3.8 | 1.1 | 3.6 | 3.4E-04 | 4.3E-04 | MB231/MB468 | MITO |
| >hg38_1_-_228634819_228635039_RNA5S12_5S_50ntFlanks@51.93.43\|GTCTACGGCCATACCACCCTGAACGCGCCCGATCTCGTCTGAT | >hg38_1_-_228634819_228635039_RNA5S12_5S_50ntFlanks | 51 | 93 | 43 | GTCTACGGCCATACCACCCTGAACGCGCCCGATCTCGTCTGAT | 361.1 | 3.4 | 0.9 | 3.6 | 3.4E-04 | 4.3E-04 | MB231/MB468 | NUC |
| >hg38_21_+_8433172_8446622_RNA45SN1_45S_50ntFlanks@8912.8936.25\|CGCCGAATCCCGGGGCCGAGGGAGC | >hg38_21_+_8433172_8446622_RNA45SN1_45S_50ntFlanks | 8912 | 8936 | 25 | CGCCGAATCCCGGGGCCGAGGGAGC | 138.8 | 6.3 | 1.6 | 3.8 | 1.4E-04 | 4.3E-04 | BT20/MB468 | MP |
| >MI0000086\|hsa-mir-28&WithFlank&3\|+\|188688775\|188688872@60.81.22(+1A)&[MIMAT0004502&hsa-miR-28-3p&offsets\|0\|0(+1A);m-28&3\|+\|188688834\|188688855&offsets\|0\|0(+1A)]\|CACTAGATTGTGAGCTCCTGGAA | >MI0000086\|hsa-mir-28&WithFlank&3\|+\|188688775\|188688872 | 60 | 81 | 22(+1A) | CACTAGATTGTGAGCTCCTGGAA | 378.0 | 4.5 | 1.2 | 3.6 | 2.9E-04 | 4.3E-04 | MB231/MB468 | MP |
| >trnaMT_PheGAA_MT_+_577_647@36.60.25\|AAAATGTTTAGACGGGCTCACATCA | >trnaMT_PheGAA_MT_+_577_647 | 36 | 60 | 25 | AAAATGTTTAGACGGGCTCACATCA | 143.1 | 6.3 | 1.8 | 3.6 | 3.4E-04 | 4.3E-04 | MB231/MB468 | MITO |
| >hg38_21_+_8433172_8446622_RNA45SN1_45S_50ntFlanks@11209.11250.42\|CGGCGACTCTGGACGCGAGCCGGGCCCTTCCCGTGGATCGCC | >hg38_21_+_8433172_8446622_RNA45SN1_45S_50ntFlanks | 11209 | 11250 | 42 | CGGCGACTCTGGACGCGAGCCGGGCCCTTCCCGTGGATCGCC | 977.4 | -6.8 | 1.9 | -3.6 | 3.5E-04 | 4.3E-04 | BT20/MB468 | NUC |
| >hg38_21_+_8433172_8446622_RNA45SN1_45S_50ntFlanks@12243.12272.30\|GAAGGGCAAAAGCTCGCTTGATCTTGATTT | >hg38_21_+_8433172_8446622_RNA45SN1_45S_50ntFlanks | 12243 | 12272 | 30 | GAAGGGCAAAAGCTCGCTTGATCTTGATTT | 647.9 | -3.7 | 1.0 | -3.6 | 3.4E-04 | 4.4E-04 | MB231/MB468 | MITO |
| >MI0000270\|hsa-mir-181b-1&WithFlank&1\|-\|198858867\|198858988@42.61.20;MI0000683\|hsa-mir-181b-2&WithFlank&9\|+\|124693704\|124693804@22.41.20&[MIMAT0000257&hsa-miR-181b-5p&offsets\|0\|-3;m-73&1\|-\|198858925\|198858947&offsets\|0\|-3];[MIMAT0000257_1&hsa-miR-181b-5p&offsets\|0\|-3;m-72&9\|+\|124693725\|124693747&offsets\|0\|-3]\|AACATTCATTGCTGTCGGTG | >MI0000270\|hsa-mir-181b-1&WithFlank&1\|-\|198858867\|198858988 | 42 | 61 | 20 | AACATTCATTGCTGTCGGTG | 162.6 | 3.7 | 1.0 | 3.6 | 3.4E-04 | 4.4E-04 | MB231/MB468 | MITO |
| >hg38_21_+_8433172_8446622_RNA45SN1_45S_50ntFlanks@12481.12500.20\|AATAGGGAACGTGAGCTGGG | >hg38_21_+_8433172_8446622_RNA45SN1_45S_50ntFlanks | 12481 | 12500 | 20 | AATAGGGAACGTGAGCTGGG | 180.2 | -3.1 | 0.8 | -3.6 | 2.9E-04 | 4.4E-04 | MB231/MB468 | MP |
| >hg38_21_+_8433172_8446622_RNA45SN1_45S_50ntFlanks@10393.10427.35\|CAGCAGTTGAACATGGGTCAGTCGGTCCTGAGAGA | >hg38_21_+_8433172_8446622_RNA45SN1_45S_50ntFlanks | 10393 | 10427 | 35 | CAGCAGTTGAACATGGGTCAGTCGGTCCTGAGAGA | 92.6 | -5.5 | 1.5 | -3.6 | 2.9E-04 | 4.4E-04 | MB231/MB468 | MP |
| >hg38_21_+_8433172_8446622_RNA45SN1_45S_50ntFlanks@12941.12958.18\|ATTCGTAGACGACCTGCT | >hg38_21_+_8433172_8446622_RNA45SN1_45S_50ntFlanks | 12941 | 12958 | 18 | ATTCGTAGACGACCTGCT | 580.8 | 2.2 | 0.6 | 3.6 | 3.2E-04 | 4.4E-04 | MB231/MB468 | CYTO |
| >hg38_21_+_8433172_8446622_RNA45SN1_45S_50ntFlanks@12243.12268.26\|GAAGGGCAAAAGCTCGCTTGATCTTG | >hg38_21_+_8433172_8446622_RNA45SN1_45S_50ntFlanks | 12243 | 12268 | 26 | GAAGGGCAAAAGCTCGCTTGATCTTG | 371.3 | -6.3 | 1.8 | -3.6 | 3.5E-04 | 4.4E-04 | BT20/MB468 | NUC |
| >hg38_21_+_8433172_8446622_RNA45SN1_45S_50ntFlanks@8341.8373.33\|CCGTAAGGGAAAGTTGAAAAGAACTTTGAAGAG | >hg38_21_+_8433172_8446622_RNA45SN1_45S_50ntFlanks | 8341 | 8373 | 33 | CCGTAAGGGAAAGTTGAAAAGAACTTTGAAGAG | 323.8 | 7.3 | 1.9 | 3.8 | 1.4E-04 | 4.4E-04 | BT20/MB468 | MP |
| >hg38_21_+_8433172_8446622_RNA45SN1_45S_50ntFlanks@10905.10926.22\|CTGGGCGCGCGCCGCGGCTGGA | >hg38_21_+_8433172_8446622_RNA45SN1_45S_50ntFlanks | 10905 | 10926 | 22 | CTGGGCGCGCGCCGCGGCTGGA | 104.8 | -4.9 | 1.3 | -3.7 | 2.6E-04 | 4.4E-04 | BT20/MB468 | MITO |
| >trnaMT_ValTAC_MT_+_1602_1670@42.70.29\|AGATTTCAACTTAACTTGACCGCTCTGAC | >trnaMT_ValTAC_MT_+_1602_1670 | 42 | 70 | 29 | AGATTTCAACTTAACTTGACCGCTCTGAC | 584.3 | 3.8 | 1.1 | 3.6 | 3.4E-04 | 4.4E-04 | MB231/MB468 | MITO |
| >hg38_21_+_8433172_8446622_RNA45SN1_45S_50ntFlanks@10687.10707.21\|CGGCGTCCGGTGAGCTCTCGC | >hg38_21_+_8433172_8446622_RNA45SN1_45S_50ntFlanks | 10687 | 10707 | 21 | CGGCGTCCGGTGAGCTCTCGC | 336.8 | -4.6 | 1.3 | -3.6 | 3.5E-04 | 4.4E-04 | BT20/MB468 | NUC |
| >MI0000088\|hsa-mir-30a&WithFlank&6\|-\|71403545\|71403627@54.73.20(+1A)&[MIMAT0000088&hsa-miR-30a-3p&offsets\|+1\|-1(+1A);m-38&6\|-\|71403554\|71403575&offsets\|+1\|-1(+1A)]\|TTTCAGTCGGATGTTTGCAGA | >MI0000088\|hsa-mir-30a&WithFlank&6\|-\|71403545\|71403627 | 54 | 73 | 20(+1A) | TTTCAGTCGGATGTTTGCAGA | 206.5 | 5.5 | 1.5 | 3.6 | 2.9E-04 | 4.4E-04 | MB231/MB468 | MP |
| >hg38_21_+_8433172_8446622_RNA45SN1_45S_50ntFlanks@6651.6678.28\|CGACTCTTAGCGGTGGATCACTCGGCTC | >hg38_21_+_8433172_8446622_RNA45SN1_45S_50ntFlanks | 6651 | 6678 | 28 | CGACTCTTAGCGGTGGATCACTCGGCTC | 27388.9 | 5.6 | 1.4 | 4.1 | 4.9E-05 | 4.4E-04 | BT20/MB231 | MP |
| >trna119_LysCTT_1_-_145395522_145395594@1.35.35,trna11_LysCTT_5_-_180648979_180649051@1.35.35,trna13_LysCTT_6_+_26556774_26556846@1.35.35,trna7_LysCTT_16_+_3225692_3225764@1.35.35,trna9_LysCTT_5_+_180634755_180634827@1.35.35\|GCCCGGCTAGCTCAGTCGGTAGAGCATGAGACTCT | >trna119_LysCTT_1_-_145395522_145395594 | 1 | 35 | 35 | GCCCGGCTAGCTCAGTCGGTAGAGCATGAGACTCT | 1372.3 | 5.1 | 1.3 | 3.8 | 1.4E-04 | 4.4E-04 | BT20/MB468 | MP |
| >am_MI0003581\|hsa-mir-574&WithFlank&4\|+\|38868026\|38868133@31.53.23&[MIMAT0004795&hsa-miR-574-5p&offsets\|0\|0;m-213&4\|+\|38868056\|38868078&offsets\|0\|0]\|TGAGTGTGTGTGTGTGAGTGTGT | >am_MI0003581\|hsa-mir-574&WithFlank&4\|+\|38868026\|38868133 | 31 | 53 | 23 | TGAGTGTGTGTGTGTGAGTGTGT | 995.5 | 3.8 | 1.0 | 3.8 | 1.4E-04 | 4.4E-04 | BT20/MB468 | MP |
| >hg38_21_+_8433172_8446622_RNA45SN1_45S_50ntFlanks@6652.6674.23\|GACTCTTAGCGGTGGATCACTCG | >hg38_21_+_8433172_8446622_RNA45SN1_45S_50ntFlanks | 6652 | 6674 | 23 | GACTCTTAGCGGTGGATCACTCG | 365.2 | 4.0 | 1.1 | 3.6 | 3.5E-04 | 4.4E-04 | MB231/MB468 | NUC |
| >hg38_21_+_8433172_8446622_RNA45SN1_45S_50ntFlanks@3931.3959.29\|TCAAAACCAACCCGGTCAGCCCCTCTCCG | >hg38_21_+_8433172_8446622_RNA45SN1_45S_50ntFlanks | 3931 | 3959 | 29 | TCAAAACCAACCCGGTCAGCCCCTCTCCG | 370.1 | -5.2 | 1.4 | -3.7 | 2.6E-04 | 4.4E-04 | BT20/MB468 | MITO |
| >hg38_21_+_8433172_8446622_RNA45SN1_45S_50ntFlanks@10685.10710.26\|GGCGGCGTCCGGTGAGCTCTCGCTGG | >hg38_21_+_8433172_8446622_RNA45SN1_45S_50ntFlanks | 10685 | 10710 | 26 | GGCGGCGTCCGGTGAGCTCTCGCTGG | 945.3 | -2.5 | 0.7 | -3.6 | 3.5E-04 | 4.4E-04 | MB231/MB468 | NUC |
| >hg38_21_+_8433172_8446622_RNA45SN1_45S_50ntFlanks@8355.8395.41\|TGAAAAGAACTTTGAAGAGAGAGTTCAAGAGGGCGTGAAAC | >hg38_21_+_8433172_8446622_RNA45SN1_45S_50ntFlanks | 8355 | 8395 | 41 | TGAAAAGAACTTTGAAGAGAGAGTTCAAGAGGGCGTGAAAC | 60.6 | 9.3 | 2.3 | 4.1 | 4.9E-05 | 4.4E-04 | BT20/MB231 | MP |
| >hg38_21_+_8433172_8446622_RNA45SN1_45S_50ntFlanks@10404.10424.21\|CATGGGTCAGTCGGTCCTGAG | >hg38_21_+_8433172_8446622_RNA45SN1_45S_50ntFlanks | 10404 | 10424 | 21 | CATGGGTCAGTCGGTCCTGAG | 4894.4 | 3.5 | 1.0 | 3.6 | 2.9E-04 | 4.4E-04 | MB231/MB468 | MP |
| >hg38_21_+_8433172_8446622_RNA45SN1_45S_50ntFlanks@8176.8212.37\|CTTCTGATCGAGGCCCAGCCCGTGGACGGTGTGAGGC | >hg38_21_+_8433172_8446622_RNA45SN1_45S_50ntFlanks | 8176 | 8212 | 37 | CTTCTGATCGAGGCCCAGCCCGTGGACGGTGTGAGGC | 1383.3 | -3.9 | 1.1 | -3.6 | 2.9E-04 | 4.4E-04 | MB231/MB468 | MP |
| >hg38_21_+_8433172_8446622_RNA45SN1_45S_50ntFlanks@9569.9595.27\|GTGCAAATCGGTCGTCCGACCTGGGTA | >hg38_21_+_8433172_8446622_RNA45SN1_45S_50ntFlanks | 9569 | 9595 | 27 | GTGCAAATCGGTCGTCCGACCTGGGTA | 485.1 | -4.3 | 1.2 | -3.6 | 3.5E-04 | 4.4E-04 | MB231/MB468 | MITO |
| >MI0000542\|hsa-mir-320a&WithFlank&8\|-\|22244960\|22245043@48.70.23(+2U)&[MIMAT0000510&hsa-miR-320a-3p&offsets\|0\|+1(+2U);m-60&8\|-\|22244975\|22244996&offsets\|0\|+1(+2U)]\|AAAAGCTGGGTTGAGAGGGCGAATT | >MI0000542\|hsa-mir-320a&WithFlank&8\|-\|22244960\|22245043 | 48 | 70 | 23(+2U) | AAAAGCTGGGTTGAGAGGGCGAATT | 1505.2 | 3.1 | 0.8 | 4.1 | 4.8E-05 | 4.4E-04 | BT20/MB231 | CYTO |
| >hg38_21_+_8433172_8446622_RNA45SN1_45S_50ntFlanks@3813.3834.22\|TTATGGTTCCTTTGGTCGCTCG | >hg38_21_+_8433172_8446622_RNA45SN1_45S_50ntFlanks | 3813 | 3834 | 22 | TTATGGTTCCTTTGGTCGCTCG | 313.3 | -4.5 | 1.2 | -3.7 | 2.6E-04 | 4.4E-04 | BT20/MB468 | MITO |
| >MI0003780\|hsa-mir-1296&WithFlank&10\|-\|63372951\|63373054@22.42.21(+1U)&[MIMAT0005794&hsa-miR-1296-5p&offsets\|0\|-1(+1U);m-304&10\|-\|63373012\|63373033&offsets\|0\|-1(+1U)]\|TTAGGGCCCTGGCTCCATCTCT | >MI0003780\|hsa-mir-1296&WithFlank&10\|-\|63372951\|63373054 | 22 | 42 | 21(+1U) | TTAGGGCCCTGGCTCCATCTCT | 159.5 | 4.7 | 1.3 | 3.6 | 2.9E-04 | 4.4E-04 | MB231/MB468 | MP |
| >hg38_21_+_8433172_8446622_RNA45SN1_45S_50ntFlanks@6641.6683.43\|CGACCTCGTACGACTCTTAGCGGTGGATCACTCGGCTCGTGCG | >hg38_21_+_8433172_8446622_RNA45SN1_45S_50ntFlanks | 6641 | 6683 | 43 | CGACCTCGTACGACTCTTAGCGGTGGATCACTCGGCTCGTGCG | 431.1 | -3.6 | 0.9 | -4.1 | 5.0E-05 | 4.4E-04 | BT20/MB231 | NUC |
| >hg38_21_+_8433172_8446622_RNA45SN1_45S_50ntFlanks@6719.6744.26\|TGAATTGCAGGACACATTGATCATCG | >hg38_21_+_8433172_8446622_RNA45SN1_45S_50ntFlanks | 6719 | 6744 | 26 | TGAATTGCAGGACACATTGATCATCG | 3270.5 | -6.8 | 1.8 | -3.8 | 1.4E-04 | 4.4E-04 | BT20/MB468 | MP |
| >hg38_21_+_8433172_8446622_RNA45SN1_45S_50ntFlanks@5492.5526.35\|AGAAGACGGTCGAACTTGACTATCTAGAGGAAGTA | >hg38_21_+_8433172_8446622_RNA45SN1_45S_50ntFlanks | 5492 | 5526 | 35 | AGAAGACGGTCGAACTTGACTATCTAGAGGAAGTA | 893.9 | -3.4 | 1.0 | -3.6 | 3.3E-04 | 4.4E-04 | MB231/MB468 | CYTO |
| >hg38_1_-_228634819_228635039_RNA5S12_5S_50ntFlanks@132.170.39\|GATGGGAGACCGCCTGGGAATACCGGGTGCTGTAGGCTT | >hg38_1_-_228634819_228635039_RNA5S12_5S_50ntFlanks | 132 | 170 | 39 | GATGGGAGACCGCCTGGGAATACCGGGTGCTGTAGGCTT | 47790.8 | 3.8 | 0.9 | 4.0 | 5.7E-05 | 4.4E-04 | BT20/MB231 | MITO |
| >hg38_21_+_8433172_8446622_RNA45SN1_45S_50ntFlanks@6784.6803.20\|GCTACGCCTGTCTGAGCGTC | >hg38_21_+_8433172_8446622_RNA45SN1_45S_50ntFlanks | 6784 | 6803 | 20 | GCTACGCCTGTCTGAGCGTC | 379.4 | -5.7 | 1.6 | -3.7 | 2.6E-04 | 4.4E-04 | BT20/MB468 | MITO |
| >hg38_21_+_8433172_8446622_RNA45SN1_45S_50ntFlanks@12257.12284.28\|CGCTTGATCTTGATTTTCAGTACGAATA | >hg38_21_+_8433172_8446622_RNA45SN1_45S_50ntFlanks | 12257 | 12284 | 28 | CGCTTGATCTTGATTTTCAGTACGAATA | 111.0 | -6.1 | 1.7 | -3.6 | 2.9E-04 | 4.4E-04 | MB231/MB468 | MP |

| | | | | | | | | | | | | | |
|---|---|---|---|---|---|---|---|---|---|---|---|---|---|
| >hg38_21_+_8433172_8446622_RNA45SN1_45S_50ntFlanks@10059.10076.18\|GGCAGTCGAGAGTGGACG | >hg38_21_+_8433172_8446622_RNA45SN1_45S_50ntFlanks | 10059 | 10076 | 18 | GGCAGTCGAGAGTGGACG | 203.6 | 7.0 | 1.7 | 4.1 | 4.9E-05 | 4.4E-04 | BT20/MB231 | MP |
| >hg38_21_+_8433172_8446622_RNA45SN1_45S_50ntFlanks@4722.4748.27\|TCATTAATCAAGAACGAAAGTCGGAGG | >hg38_21_+_8433172_8446622_RNA45SN1_45S_50ntFlanks | 4722 | 4748 | 27 | TCATTAATCAAGAACGAAAGTCGGAGG | 666.2 | -3.1 | 0.9 | -3.6 | 3.3E-04 | 4.4E-04 | MB231/MB468 | CYTO |
| >MI0000283\|hsa-mir-203a&WithFlank&14\|+\|104117399\|104117520@71.91.21(+1A)&[MIMAT0000264&hsa-miR-203a-3p&offsets\|0\|-1(+1A);m-79&14\|+\|104117469\|104117490&offsets\|0\|-1(+1A)]\|GTGAAATGTTTAGGACCACTAA | >MI0000283\|hsa-mir-203a&WithFlank&14\|+\|104117399\|104117520 | 71 | 91 | 21(+1A) | GTGAAATGTTTAGGACCACTAA | 95.1 | 7.2 | 1.9 | 3.8 | 1.4E-04 | 4.4E-04 | BT20/MB468 | MP |
| >hg38_21_+_8433172_8446622_RNA45SN1_45S_50ntFlanks@11317.11344.28\|ACCCCACGTCTCGTCGCGCGCGCGTCCG | >hg38_21_+_8433172_8446622_RNA45SN1_45S_50ntFlanks | 11317 | 11344 | 28 | ACCCCACGTCTCGTCGCGCGCGCGTCCG | 10108.9 | -5.6 | 1.5 | -3.7 | 2.6E-04 | 4.4E-04 | BT20/MB468 | MITO |
| >hg38_21_+_8433172_8446622_RNA45SN1_45S_50ntFlanks@12416.12459.44\|TTCGATGTCGGCTCTTCCTATCATTGTGAAGCAGAATTCACCAA | >hg38_21_+_8433172_8446622_RNA45SN1_45S_50ntFlanks | 12416 | 12459 | 44 | TTCGATGTCGGCTCTTCCTATCATTGTGAAGCAGAATTCACCAA | 169.9 | -4.6 | 1.1 | -4.1 | 4.8E-05 | 4.4E-04 | BT20/MB231 | CYTO |
| >hg38_21_+_8433172_8446622_RNA45SN1_45S_50ntFlanks@6644.6662.19\|CCTCGTACGACTCTTAGCG | >hg38_21_+_8433172_8446622_RNA45SN1_45S_50ntFlanks | 6644 | 6662 | 19 | CCTCGTACGACTCTTAGCG | 189.5 | 2.8 | 0.8 | 3.6 | 3.3E-04 | 4.4E-04 | MB231/MB468 | CYTO |
| >hg38_21_+_8433172_8446622_RNA45SN1_45S_50ntFlanks@3989.4023.35\|TTTGGTGACTCTAGATAACCTCGGGCCGATCGCAC | >hg38_21_+_8433172_8446622_RNA45SN1_45S_50ntFlanks | 3989 | 4023 | 35 | TTTGGTGACTCTAGATAACCTCGGGCCGATCGCAC | 289.9 | 2.6 | 0.7 | 3.6 | 3.5E-04 | 4.4E-04 | MB231/MB468 | NUC |
| >hg38_21_+_8433172_8446622_RNA45SN1_45S_50ntFlanks@9792.9817.26\|GGCGTGGAGCCGGGCGTGGAATGCGA | >hg38_21_+_8433172_8446622_RNA45SN1_45S_50ntFlanks | 9792 | 9817 | 26 | GGCGTGGAGCCGGGCGTGGAATGCGA | 176.9 | 10.8 | 2.7 | 4.1 | 4.9E-05 | 4.4E-04 | BT20/MB231 | MP |
| >hg38_21_+_8433172_8446622_RNA45SN1_45S_50ntFlanks@12083.12107.25\|CTCGCTTCTGGCGCCAAGCGCCCGG | >hg38_21_+_8433172_8446622_RNA45SN1_45S_50ntFlanks | 12083 | 12107 | 25 | CTCGCTTCTGGCGCCAAGCGCCCGG | 369.1 | -5.2 | 1.4 | -3.7 | 2.6E-04 | 4.4E-04 | BT20/MB468 | MITO |
| >hg38_21_+_8433172_8446622_RNA45SN1_45S_50ntFlanks@10349.10376.28\|AACGAGAACTTTGAAGGCCGAAGTGGAG | >hg38_21_+_8433172_8446622_RNA45SN1_45S_50ntFlanks | 10349 | 10376 | 28 | AACGAGAACTTTGAAGGCCGAAGTGGAG | 491.5 | -4.3 | 1.2 | -3.6 | 3.5E-04 | 4.4E-04 | MB231/MB468 | MITO |
| >hg38_21_+_8433172_8446622_RNA45SN1_45S_50ntFlanks@11703.11744.42\|AATGAAGCGCGGGTAAACGGCGGGAGTAACTATGACTCTCTT | >hg38_21_+_8433172_8446622_RNA45SN1_45S_50ntFlanks | 11703 | 11744 | 42 | AATGAAGCGCGGGTAAACGGCGGGAGTAACTATGACTCTCTT | 190.2 | -4.7 | 1.3 | -3.6 | 3.5E-04 | 4.4E-04 | MB231/MB468 | MITO |
| >hg38_21_+_8433172_8446622_RNA45SN1_45S_50ntFlanks@12004.12035.32\|CGGGCCGCCGGTGAAATACCACTACTCTGATC | >hg38_21_+_8433172_8446622_RNA45SN1_45S_50ntFlanks | 12004 | 12035 | 32 | CGGGCCGCCGGTGAAATACCACTACTCTGATC | 214.4 | 3.5 | 1.0 | 3.6 | 3.6E-04 | 4.5E-04 | BT20/MB468 | NUC |
| >hg38_21_+_8433172_8446622_RNA45SN1_45S_50ntFlanks@10357.10390.34\|CTTTGAAGGCCGAAGTGGAGAAGGGTTCCATGTG | >hg38_21_+_8433172_8446622_RNA45SN1_45S_50ntFlanks | 10357 | 10390 | 34 | CTTTGAAGGCCGAAGTGGAGAAGGGTTCCATGTG | 140.2 | -1.9 | 0.5 | -3.7 | 2.6E-04 | 4.5E-04 | BT20/MB468 | MITO |
| >hg38_21_+_8433172_8446622_RNA45SN1_45S_50ntFlanks@12319.12346.28\|ACCTTTTGGGTTTTAAGCAGGAGGTGTC | >hg38_21_+_8433172_8446622_RNA45SN1_45S_50ntFlanks | 12319 | 12346 | 28 | ACCTTTTGGGTTTTAAGCAGGAGGTGTC | 78.4 | -9.4 | 2.5 | -3.8 | 1.4E-04 | 4.5E-04 | BT20/MB468 | MP |
| >hg38_21_+_8433172_8446622_RNA45SN1_45S_50ntFlanks@7973.8010.38\|GACGCGACCTCAGATCAGACGTGGCGACCCGCTGAATT | >hg38_21_+_8433172_8446622_RNA45SN1_45S_50ntFlanks | 7973 | 8010 | 38 | GACGCGACCTCAGATCAGACGTGGCGACCCGCTGAATT | 520.2 | 4.6 | 1.2 | 3.8 | 1.4E-04 | 4.5E-04 | BT20/MB468 | MP |
| >hg38_21_+_8433172_8446622_RNA45SN1_45S_50ntFlanks@7984.8002.19\|AGATCAGACGTGGCGACCC | >hg38_21_+_8433172_8446622_RNA45SN1_45S_50ntFlanks | 7984 | 8002 | 19 | AGATCAGACGTGGCGACCC | 97.8 | -5.9 | 1.6 | -3.6 | 3.0E-04 | 4.5E-04 | MB231/MB468 | MP |
| >trnaMT_HisGTG_MT_+_12138_12206@27.65.39\|GATTGTGAATCTGACAACAGAGGCTTACGACCCCTTATT | >trnaMT_HisGTG_MT_+_12138_12206 | 27 | 65 | 39 | GATTGTGAATCTGACAACAGAGGCTTACGACCCCTTATT | 87.1 | 10.2 | 2.7 | 3.8 | 1.4E-04 | 4.5E-04 | BT20/MB468 | MP |
| >MI0000473\|hsa-mir-129-2&WithFlank&11\|+\|43581388\|43581489@21.41.21;MI0000252\|hsa-mir-129-1&WithFlank&7\|+\|128207866\|128207949@11.31.21&[MIMAT0000242&hsa-miR-129-5p&offsets\|0\|0;m-434&11\|+\|43581408\|43581428&offsets\|0\|0];[MIMAT0000242_1&hsa-miR-129-5p&offsets\|0\|0;m-434&7\|+\|128207876\|128207896&offsets\|0\|0]\|CTTTTTGCGGTCTGGGCTTGC | >MI0000473\|hsa-mir-129-2&WithFlank&11\|+\|43581388\|43581489 | 21 | 41 | 21 | CTTTTTGCGGTCTGGGCTTGC | 199.8 | 6.2 | 1.6 | 3.8 | 1.4E-04 | 4.5E-04 | BT20/MB468 | MP |
| >hg38_MT_+_598_1651_RNR1_12S_50ntFlanks@369.404.36\|AATAAAGCTAAAACTCACCTGAGTTGTAAAAAACTC | >hg38_MT_+_598_1651_RNR1_12S_50ntFlanks | 369 | 404 | 36 | AATAAAGCTAAAACTCACCTGAGTTGTAAAAAACTC | 840.7 | 2.9 | 0.8 | 3.6 | 3.5E-04 | 4.5E-04 | MB231/MB468 | MITO |
| >hg38_21_+_8433172_8446622_RNA45SN1_45S_50ntFlanks@7968.7994.27\|TCCGAGACGCGACCTCAGATCAGACGT | >hg38_21_+_8433172_8446622_RNA45SN1_45S_50ntFlanks | 7968 | 7994 | 27 | TCCGAGACGCGACCTCAGATCAGACGT | 1022.0 | 2.9 | 0.8 | 3.6 | 3.6E-04 | 4.5E-04 | MB231/MB468 | NUC |
| >MI0001445\|hsa-mir-423&WithFlank&17\|+\|30117073\|30117178@59.81.23(+1U)&[MIMAT0001340&hsa-miR-423-3p&offsets\|0\|0(+1U);m-100&17\|+\|30117131\|30117153&offsets\|0\|0(+1U)]\|AGCTCGGTCTGAGGCCCCTCAGTT | >MI0001445\|hsa-mir-423&WithFlank&17\|+\|30117073\|30117178 | 59 | 81 | 23(+1U) | AGCTCGGTCTGAGGCCCCTCAGTT | 298.3 | 5.0 | 1.3 | 3.8 | 1.4E-04 | 4.5E-04 | BT20/MB468 | MP |
| >hg38_21_+_8433172_8446622_RNA45SN1_45S_50ntFlanks@11784.11805.22\|ATGAATGGATGAACGAGATTCC | >hg38_21_+_8433172_8446622_RNA45SN1_45S_50ntFlanks | 11784 | 11805 | 22 | ATGAATGGATGAACGAGATTCC | 628.7 | -3.3 | 0.9 | -3.8 | 1.4E-04 | 4.5E-04 | BT20/MB468 | MP |
| >hg38_21_+_8433172_8446622_RNA45SN1_45S_50ntFlanks@12500.12518.19\|GTTTAGACCGTCGTGAGAC | >hg38_21_+_8433172_8446622_RNA45SN1_45S_50ntFlanks | 12500 | 12518 | 19 | GTTTAGACCGTCGTGAGAC | 103.0 | -5.8 | 1.5 | -3.8 | 1.4E-04 | 4.5E-04 | BT20/MB468 | MP |
| >hg38_21_+_8433172_8446622_RNA45SN1_45S_50ntFlanks@11564.11611.48\|CCTAGCAGCCGACTTAGAACTGGTGCGGACCAGGGGAATCCGACTGTT | >hg38_21_+_8433172_8446622_RNA45SN1_45S_50ntFlanks | 11564 | 11611 | 48 | CCTAGCAGCCGACTTAGAACTGGTGCGGACCAGGGGAATCCGACTGTT | 1950.9 | 4.9 | 1.2 | 4.0 | 5.8E-05 | 4.5E-04 | BT20/MB231 | MITO |
| >hg38_21_+_8433172_8446622_RNA45SN1_45S_50ntFlanks@9575.9597.23\|ATCGGTCGTCCGACCTGGGTATA | >hg38_21_+_8433172_8446622_RNA45SN1_45S_50ntFlanks | 9575 | 9597 | 23 | ATCGGTCGTCCGACCTGGGTATA | 276.7 | -5.2 | 1.5 | -3.6 | 3.6E-04 | 4.5E-04 | MB231/MB468 | NUC |
| >hg38_21_+_8433172_8446622_RNA45SN1_45S_50ntFlanks@10410.10430.21\|TCAGTCGGTCCTGAGAGATGG | >hg38_21_+_8433172_8446622_RNA45SN1_45S_50ntFlanks | 10410 | 10430 | 21 | TCAGTCGGTCCTGAGAGATGG | 1870.8 | -5.2 | 1.4 | -3.8 | 1.4E-04 | 4.5E-04 | BT20/MB468 | MP |
| >MI0000108\|hsa-mir-103a-2&WithFlank&20\|+\|3917488\|3917577@54.77.24(+1U);MI0000109\|hsa-mir-103a-1&WithFlank&5\|-\|168560890\|168560979@54.77.24(+1U)&[MIMAT0000101&hsa-miR-103a-3p&offsets\|0\|+1(+1U);m-12&20\|+\|3917541\|3917563&offsets\|0\|+1(+1U)];[MIMAT0000101_1&hsa-miR-103a-3p&offsets\|0\|+1(+1U);m-13&5\|-\|168560904\|168560926&offsets\|0\|+1(+1U);m-44220&5\|-\|168560908\|168560926&offsets\|0\|+5(+1U)]\|AGCAGCATTGTACAGGGCTATGAAT | >MI0000108\|hsa-mir-103a-2&WithFlank&20\|+\|3917488\|3917577 | 54 | 77 | 24(+1U) | AGCAGCATTGTACAGGGCTATGAAT | 730.3 | -3.4 | 0.9 | -4.0 | 5.8E-05 | 4.5E-04 | BT20/MB231 | MITO |
| >MI0000299\|hsa-mir-222&WithFlank&X\|-\|45747009\|45747130@75.94.20(+1C)&[MIMAT0000279&hsa-miR-222-3p&offsets\|0\|-1(+1C);m-140&X\|-\|45747033\|45747056&offsets\|0\|-4(+1C)]\|AGCTACATCTGGCTACTGGGC | >MI0000299\|hsa-mir-222&WithFlank&X\|-\|45747009\|45747130 | 75 | 94 | 20(+1C) | AGCTACATCTGGCTACTGGGC | 574.2 | 4.2 | 1.1 | 3.8 | 1.4E-04 | 4.5E-04 | BT20/MB468 | MP |
| >hg38_21_+_8433172_8446622_RNA45SN1_45S_50ntFlanks@8380.8404.25\|CAAGAGGGCGTGAAACCGTTAAGAG | >hg38_21_+_8433172_8446622_RNA45SN1_45S_50ntFlanks | 8380 | 8404 | 25 | CAAGAGGGCGTGAAACCGTTAAGAG | 496.3 | -5.4 | 1.5 | -3.6 | 3.2E-04 | 4.5E-04 | BT20/MB468 | CYTO |
| >hg38_21_+_8433172_8446622_RNA45SN1_45S_50ntFlanks@10882.10902.21\|TCGGGCTGGGGCGCGAAGCGG | >hg38_21_+_8433172_8446622_RNA45SN1_45S_50ntFlanks | 10882 | 10902 | 21 | TCGGGCTGGGGCGCGAAGCGG | 714.7 | -6.6 | 1.8 | -3.6 | 3.6E-04 | 4.5E-04 | BT20/MB468 | NUC |
| >hg38_21_+_8433172_8446622_RNA45SN1_45S_50ntFlanks@10393.10428.36\|CAGCAGTTGAACATGGGTCAGTCGGTCCTGAGAGAT | >hg38_21_+_8433172_8446622_RNA45SN1_45S_50ntFlanks | 10393 | 10428 | 36 | CAGCAGTTGAACATGGGTCAGTCGGTCCTGAGAGAT | 245.3 | -2.5 | 0.7 | -3.6 | 3.6E-04 | 4.5E-04 | MB231/MB468 | NUC |
| >hg38_MT_+_598_1651_RNR1_12S_50ntFlanks@425.452.28\|ACGAAAGTGGCTTTAACATATCTGAACA | >hg38_MT_+_598_1651_RNR1_12S_50ntFlanks | 425 | 452 | 28 | ACGAAAGTGGCTTTAACATATCTGAACA | 393.1 | 4.7 | 1.2 | 3.8 | 1.4E-04 | 4.5E-04 | BT20/MB468 | MP |
| >hg38_21_+_8433172_8446622_RNA45SN1_45S_50ntFlanks@10908.10930.23\|GGCGCGCGCCGCGGCTGGACGAG | >hg38_21_+_8433172_8446622_RNA45SN1_45S_50ntFlanks | 10908 | 10930 | 23 | GGCGCGCGCCGCGGCTGGACGAG | 65.9 | 9.4 | 2.3 | 4.1 | 5.0E-05 | 4.5E-04 | BT20/MB231 | MP |
| >hg38_21_+_8433172_8446622_RNA45SN1_45S_50ntFlanks@9543.9571.29\|TGGTGGAGGTCCGTAGCGGTCCTGACGTG | >hg38_21_+_8433172_8446622_RNA45SN1_45S_50ntFlanks | 9543 | 9571 | 29 | TGGTGGAGGTCCGTAGCGGTCCTGACGTG | 157.4 | -4.3 | 1.2 | -3.6 | 3.0E-04 | 4.5E-04 | MB231/MB468 | MP |
| >am_trna128_GlyGCC_6_-_27870686_27870756@7.35.29,trna18_GlyGCC_16_+_70822597_70822667@7.35.29,trna19_GlyGCC_16_+_70823410_70823480@7.35.29,trna19_GlyGCC_2_-_157257659_157257729@7.35.29,trna24_GlyGCC_16_-_70812942_70813012@7.35.29,trna25_GlyGCC_16_-_70812114_70812184@7.35.29,trna2_GlyGCC_21_-_18827107_18827177@7.35.29,trna35_GlyGCC_1_+_161413094_161413164@7.35.29,trna37_GlyGCC_1_+_161420467_161420537@7.35.29,trna39_GlyGCC_1_+_161427898_161427968@7.35.29,trna41_GlyGCC_1_+_161435258_161435328@7.35.29,trna5_GlyGCC_17_+_8029064_8029134@7.35.29,trna68_GlyGCC_1_-_161493637_161493707@7.35.29\|GTGGTTCAGTGGTAGAATTCTCGCCTGCC | >am_trna128_GlyGCC_6_-_27870686_27870756 | 7 | 35 | 29 | GTGGTTCAGTGGTAGAATTCTCGCCTGCC | 577.3 | -3.0 | 0.7 | -4.0 | 5.8E-05 | 4.5E-04 | BT20/MB231 | MITO |
| >hg38_21_+_8433172_8446622_RNA45SN1_45S_50ntFlanks@7989.8008.20\|AGACGTGGCGACCCGCTGAA | >hg38_21_+_8433172_8446622_RNA45SN1_45S_50ntFlanks | 7989 | 8008 | 20 | AGACGTGGCGACCCGCTGAA | 3173.7 | -3.6 | 1.0 | -3.6 | 3.6E-04 | 4.5E-04 | BT20/MB468 | NUC |
| >trna116_GluCTC_1_-_145399233_145399304@13.31.19,trna25_GluCTC_2_-_71273488_71273560@13.31.19,trna59_GluCTC_1_+_249168447_249168518@13.31.19,trna71_GluCTC_1_-_161439189_161439260@13.31.19,trna74_GluCTC_1_-_161431809_161431880@13.31.19,trna77_GluCTC_1_-_161424398_161424469@13.31.19,trna77_GluCTC_6_+_28949976_28950047@13.31.19,trna80_GluCTC_1_-_161417018_161417089@13.31.19,trna87_GluCTC_6_-_126101393_126101464@13.31.19\|TAGTGGTTAGGATTCGGCG | >trna116_GluCTC_1_-_145399233_145399304 | 13 | 31 | 19 | TAGTGGTTAGGATTCGGCG | 291.5 | -6.1 | 1.7 | -3.6 | 3.6E-04 | 4.5E-04 | BT20/MB468 | NUC |
| >hg38_21_+_8433172_8446622_RNA45SN1_45S_50ntFlanks@6649.6672.24\|TACGACTCTTAGCGGTGGATCACT | >hg38_21_+_8433172_8446622_RNA45SN1_45S_50ntFlanks | 6649 | 6672 | 24 | TACGACTCTTAGCGGTGGATCACT | 532.5 | 3.1 | 0.9 | 3.6 | 3.5E-04 | 4.5E-04 | MB231/MB468 | MITO |
| >hg38_21_+_8433172_8446622_RNA45SN1_45S_50ntFlanks@10389.10409.21\|TGAACAGCAGTTGAACATGGG | >hg38_21_+_8433172_8446622_RNA45SN1_45S_50ntFlanks | 10389 | 10409 | 21 | TGAACAGCAGTTGAACATGGG | 857.6 | -5.7 | 1.6 | -3.6 | 3.2E-04 | 4.5E-04 | BT20/MB468 | CYTO |
| >hg38_21_+_8433172_8446622_RNA45SN1_45S_50ntFlanks@6652.6682.31\|GACTCTTAGCGGTGGATCACTCGGCTCGTGC | >hg38_21_+_8433172_8446622_RNA45SN1_45S_50ntFlanks | 6652 | 6682 | 31 | GACTCTTAGCGGTGGATCACTCGGCTCGTGC | 507.0 | 4.7 | 1.3 | 3.6 | 3.3E-04 | 4.5E-04 | MB231/MB468 | CYTO |
| >hg38_21_+_8433172_8446622_RNA45SN1_45S_50ntFlanks@12420.12446.27\|ATGTCGGCTCTTCCTATCATTGTGAAG | >hg38_21_+_8433172_8446622_RNA45SN1_45S_50ntFlanks | 12420 | 12446 | 27 | ATGTCGGCTCTTCCTATCATTGTGAAG | 476.8 | -3.4 | 0.9 | -4.0 | 5.9E-05 | 4.5E-04 | BT20/MB231 | MITO |
| >hg38_21_+_8433172_8446622_RNA45SN1_45S_50ntFlanks@6719.6750.32\|TGAATTGCAGGACACATTGATCATCGACACTT | >hg38_21_+_8433172_8446622_RNA45SN1_45S_50ntFlanks | 6719 | 6750 | 32 | TGAATTGCAGGACACATTGATCATCGACACTT | 92.6 | -4.7 | 1.3 | -3.6 | 3.2E-04 | 4.5E-04 | BT20/MB468 | CYTO |
| >hg38_21_+_8433172_8446622_RNA45SN1_45S_50ntFlanks@11784.11806.23\|ATGAATGGATGAACGAGATTCCC | >hg38_21_+_8433172_8446622_RNA45SN1_45S_50ntFlanks | 11784 | 11806 | 23 | ATGAATGGATGAACGAGATTCCC | 425.8 | -4.2 | 1.2 | -3.6 | 3.6E-04 | 4.5E-04 | MB231/MB468 | NUC |

<table>
<tr><td>>hg38_21_+_8433172_8446622_RNA45SN1_45S_50ntFlanks@3863.3898.36|ATTCTAGAGCTAATACATGCCGACGGGCGCTGACCC</td><td>>hg38_21_+_8433172_8446622_RNA45SN1_45S_50ntFlanks</td><td>3863</td><td>3898</td><td>36</td><td>ATTCTAGAGCTAATACATGCCGACGGGCGCTGACCC</td><td>148.2</td><td>-6.3</td><td>1.7</td><td>-3.8</td><td>1.4E-04</td><td>4.5E-04</td><td>BT20/MB468</td><td>MP</td></tr>
<tr><td>>hg38_MT_+_1621_3279_RNR2_16S_50ntFlanks@1000.1038.39|GGGACCTGTATGAATGGCTCCACGAGGGTTCAGCTGTCT</td><td>>hg38_MT_+_1621_3279_RNR2_16S_50ntFlanks</td><td>1000</td><td>1038</td><td>39</td><td>GGGACCTGTATGAATGGCTCCACGAGGGTTCAGCTGTCT</td><td>84.7</td><td>10.2</td><td>2.7</td><td>3.8</td><td>1.4E-04</td><td>4.5E-04</td><td>BT20/MB468</td><td>MP</td></tr>
<tr><td>>hg38_21_+_8433172_8446622_RNA45SN1_45S_50ntFlanks@11913.11942.30|AAGAGACATGAGAGGTGTAGAATAAGTGGG</td><td>>hg38_21_+_8433172_8446622_RNA45SN1_45S_50ntFlanks</td><td>11913</td><td>11942</td><td>30</td><td>AAGAGACATGAGAGGTGTAGAATAAGTGGG</td><td>66.3</td><td>9.4</td><td>2.3</td><td>4.1</td><td>5.0E-05</td><td>4.5E-04</td><td>BT20/MB231</td><td>MP</td></tr>
<tr><td>>hg38_21_+_8433172_8446622_RNA45SN1_45S_50ntFlanks@10391.10433.43|AACAGCAGTTGAACATGGGTCAGTCGGTCCTGAGAGATGGGCG</td><td>>hg38_21_+_8433172_8446622_RNA45SN1_45S_50ntFlanks</td><td>10391</td><td>10433</td><td>43</td><td>AACAGCAGTTGAACATGGGTCAGTCGGTCCTGAGAGATGGGCG</td><td>406.2</td><td>-4.6</td><td>1.3</td><td>-3.6</td><td>3.2E-04</td><td>4.5E-04</td><td>BT20/MB468</td><td>CYTO</td></tr>
<tr><td>>hg38_21_+_8433172_8446622_RNA45SN1_45S_50ntFlanks@7980.8010.31|CCTCAGATCAGACGTGGCGACCCGCTGAATT</td><td>>hg38_21_+_8433172_8446622_RNA45SN1_45S_50ntFlanks</td><td>7980</td><td>8010</td><td>31</td><td>CCTCAGATCAGACGTGGCGACCCGCTGAATT</td><td>271.9</td><td>-2.8</td><td>0.8</td><td>-3.6</td><td>3.6E-04</td><td>4.5E-04</td><td>MB231/MB468</td><td>MITO</td></tr>
<tr><td>>hg38_21_+_8433172_8446622_RNA45SN1_45S_50ntFlanks@10805.10822.18|GAACAATGTAGGTAAGGG</td><td>>hg38_21_+_8433172_8446622_RNA45SN1_45S_50ntFlanks</td><td>10805</td><td>10822</td><td>18</td><td>GAACAATGTAGGTAAGGG</td><td>167.5</td><td>-4.9</td><td>1.3</td><td>-3.6</td><td>2.7E-04</td><td>4.5E-04</td><td>BT20/MB468</td><td>MITO</td></tr>
<tr><td>>hg38_21_+_8433172_8446622_RNA45SN1_45S_50ntFlanks@9300.9330.31|ACGGACCAAGGAGTCTAACACGTGCGCGAGT</td><td>>hg38_21_+_8433172_8446622_RNA45SN1_45S_50ntFlanks</td><td>9300</td><td>9330</td><td>31</td><td>ACGGACCAAGGAGTCTAACACGTGCGCGAGT</td><td>145.3</td><td>7.0</td><td>1.8</td><td>3.8</td><td>1.4E-04</td><td>4.5E-04</td><td>BT20/MB468</td><td>MP</td></tr>
<tr><td>>hg38_21_+_8433172_8446622_RNA45SN1_45S_50ntFlanks@10513.10545.33|CCGGAGTGGCGGAGATGGGCGCCGCGAGGCGTC</td><td>>hg38_21_+_8433172_8446622_RNA45SN1_45S_50ntFlanks</td><td>10513</td><td>10545</td><td>33</td><td>CCGGAGTGGCGGAGATGGGCGCCGCGAGGCGTC</td><td>81.9</td><td>-5.5</td><td>1.5</td><td>-3.6</td><td>3.0E-04</td><td>4.5E-04</td><td>MB231/MB468</td><td>MP</td></tr>
<tr><td>>hg38_MT_+_1621_3279_RNR2_16S_50ntFlanks@1181.1215.35|AAAAATTTCGGTTGGGGCGACCTCGGAGCAGAACC</td><td>>hg38_MT_+_1621_3279_RNR2_16S_50ntFlanks</td><td>1181</td><td>1215</td><td>35</td><td>AAAAATTTCGGTTGGGGCGACCTCGGAGCAGAACC</td><td>147.9</td><td>7.3</td><td>1.9</td><td>3.8</td><td>1.4E-04</td><td>4.5E-04</td><td>BT20/MB468</td><td>MP</td></tr>
<tr><td>>hg38_21_+_8433172_8446622_RNA45SN1_45S_50ntFlanks@12073.12091.19|CCGAGGGGCTCTCGCTTCT</td><td>>hg38_21_+_8433172_8446622_RNA45SN1_45S_50ntFlanks</td><td>12073</td><td>12091</td><td>19</td><td>CCGAGGGGCTCTCGCTTCT</td><td>101.8</td><td>-5.6</td><td>1.5</td><td>-3.8</td><td>1.4E-04</td><td>4.5E-04</td><td>BT20/MB468</td><td>MP</td></tr>
<tr><td>>hg38_21_+_8433172_8446622_RNA45SN1_45S_50ntFlanks@12909.12931.23|CCGCACGTTCGTGGGGAACCTGG</td><td>>hg38_21_+_8433172_8446622_RNA45SN1_45S_50ntFlanks</td><td>12909</td><td>12931</td><td>23</td><td>CCGCACGTTCGTGGGGAACCTGG</td><td>100.2</td><td>-5.4</td><td>1.5</td><td>-3.6</td><td>3.0E-04</td><td>4.5E-04</td><td>MB231/MB468</td><td>MP</td></tr>
<tr><td>>MI0000273|hsa-mir-183&WithFlank&7|-|129774899|129775020@33.55.23(+1C)&[MIMAT0000261&hsa-miR-183-5p&offsets|0|+1(+1C);m-41&7|-|129774966|129774987&offsets|-1|0(+1C)]|TATGGCACTGGTAGAATTCACTGC</td><td>>MI0000273|hsa-mir-183&WithFlank&7|-|129774899|129775020</td><td>33</td><td>55</td><td>23(+1C)</td><td>TATGGCACTGGTAGAATTCACTGC</td><td>206.8</td><td>3.1</td><td>0.8</td><td>4.1</td><td>4.9E-05</td><td>4.5E-04</td><td>BT20/MB231</td><td>CYTO</td></tr>
<tr><td>>hg38_21_+_8433172_8446622_RNA45SN1_45S_50ntFlanks@10877.10902.26|GTCGGTCGGGCTGGGGCGCGAAGCGG</td><td>>hg38_21_+_8433172_8446622_RNA45SN1_45S_50ntFlanks</td><td>10877</td><td>10902</td><td>26</td><td>GTCGGTCGGGCTGGGGCGCGAAGCGG</td><td>243.0</td><td>-5.3</td><td>1.5</td><td>-3.6</td><td>3.6E-04</td><td>4.5E-04</td><td>MB231/MB468</td><td>NUC</td></tr>
<tr><td>>hg38_21_+_8433172_8446622_RNA45SN1_45S_50ntFlanks@6739.6772.34|TCATCGACACTTCGAACGCACTTGCGGCCCCGGG</td><td>>hg38_21_+_8433172_8446622_RNA45SN1_45S_50ntFlanks</td><td>6739</td><td>6772</td><td>34</td><td>TCATCGACACTTCGAACGCACTTGCGGCCCCGGG</td><td>339.6</td><td>-2.7</td><td>0.8</td><td>-3.6</td><td>2.7E-04</td><td>4.5E-04</td><td>BT20/MB468</td><td>MITO</td></tr>
<tr><td>>MI0001445|hsa-mir-423&WithFlank&17|+|30117073|30117178@59.78.20&[MIMAT0001340&hsa-miR-423-3p&offsets|0|-3;m-100&17|+|30117131|30117153&offsets|0|-3]|AGCTCGGTCTGAGGCCCCTC</td><td>>MI0001445|hsa-mir-423&WithFlank&17|+|30117073|30117178</td><td>59</td><td>78</td><td>20</td><td>AGCTCGGTCTGAGGCCCCTC</td><td>974.7</td><td>3.6</td><td>1.0</td><td>3.6</td><td>3.0E-04</td><td>4.5E-04</td><td>MB231/MB468</td><td>MP</td></tr>
<tr><td>>hg38_21_+_8433172_8446622_RNA45SN1_45S_50ntFlanks@3876.3899.24|TACATGCCGACGGGCGCTGACCCC</td><td>>hg38_21_+_8433172_8446622_RNA45SN1_45S_50ntFlanks</td><td>3876</td><td>3899</td><td>24</td><td>TACATGCCGACGGGCGCTGACCCC</td><td>202.5</td><td>-2.6</td><td>0.7</td><td>-3.6</td><td>3.6E-04</td><td>4.5E-04</td><td>MB231/MB468</td><td>NUC</td></tr>
<tr><td>>MI0000086|hsa-mir-28&WithFlank&3|+|188688775|188688872@61.82.22(+1C)&[MIMAT0004502&hsa-miR-28-3p&offsets|+1|+1(+1C);m-28&3|+|188688834|188688855&offsets|+1|+1(+1C)]|ACTAGATTGTGAGCTCCTGGAGC</td><td>>MI0000086|hsa-mir-28&WithFlank&3|+|188688775|188688872</td><td>61</td><td>82</td><td>22(+1C)</td><td>ACTAGATTGTGAGCTCCTGGAGC</td><td>85.7</td><td>7.1</td><td>1.9</td><td>3.8</td><td>1.4E-04</td><td>4.5E-04</td><td>BT20/MB468</td><td>MP</td></tr>
<tr><td>>trna116_GluCTC_1_-_145399233_145399304@1.36.36,trna59_GluCTC_1_+_249168447_249168518@1.36.36,trna71_GluCTC_1_-_161439189_161439260@1.36.36,trna74_GluCTC_1_-_161431809_161431880@1.36.36,trna77_GluCTC_1_-_161424398_161424469@1.36.36,trna77_GluCTC_6_+_28949976_28950047@1.36.36,trna80_GluCTC_1_-_161417018_161417089@1.36.36,trna87_GluCTC_6_-_126101393_126101464@1.36.36|TCCCTGGTGGTCTAGTGGTTAGGATTCGGCGCTCTC</td><td>>trna116_GluCTC_1_-_145399233_145399304</td><td>1</td><td>36</td><td>36</td><td>TCCCTGGTGGTCTAGTGGTTAGGATTCGGCGCTCTC</td><td>1345.4</td><td>5.9</td><td>1.6</td><td>3.8</td><td>1.4E-04</td><td>4.5E-04</td><td>BT20/MB468</td><td>MP</td></tr>
<tr><td>>trna10_LysCTT_16_+_3241501_3241573@4.33.30,trna13_LysCTT_14_-_58706613_58706685@4.33.30,trna2_LysCTT_15_+_79152904_79152976@4.33.30|CGGCTAGCTCAGTCGGTAGAGCATGGGACT</td><td>>trna10_LysCTT_16_+_3241501_3241573</td><td>4</td><td>33</td><td>30</td><td>CGGCTAGCTCAGTCGGTAGAGCATGGGACT</td><td>345.8</td><td>4.2</td><td>1.1</td><td>3.8</td><td>1.4E-04</td><td>4.5E-04</td><td>BT20/MB468</td><td>MP</td></tr>
<tr><td>>hg38_21_+_8433172_8446622_RNA45SN1_45S_50ntFlanks@10350.10372.23|ACGAGAACTTTGAAGGCCGAAGT</td><td>>hg38_21_+_8433172_8446622_RNA45SN1_45S_50ntFlanks</td><td>10350</td><td>10372</td><td>23</td><td>ACGAGAACTTTGAAGGCCGAAGT</td><td>1229.5</td><td>-3.0</td><td>0.8</td><td>-3.6</td><td>3.0E-04</td><td>4.5E-04</td><td>MB231/MB468</td><td>MP</td></tr>
<tr><td>>hg38_21_+_8433172_8446622_RNA45SN1_45S_50ntFlanks@10391.10420.30|AACAGCAGTTGAACATGGGTCAGTCGGTCC</td><td>>hg38_21_+_8433172_8446622_RNA45SN1_45S_50ntFlanks</td><td>10391</td><td>10420</td><td>30</td><td>AACAGCAGTTGAACATGGGTCAGTCGGTCC</td><td>496.2</td><td>-5.3</td><td>1.5</td><td>-3.6</td><td>3.2E-04</td><td>4.5E-04</td><td>BT20/MB468</td><td>CYTO</td></tr>
<tr><td>>hg38_21_+_8433172_8446622_RNA45SN1_45S_50ntFlanks@12456.12496.41|CCAAGCGTTGGATTGTTCACCCACTAATAGGGAACGTGAGC</td><td>>hg38_21_+_8433172_8446622_RNA45SN1_45S_50ntFlanks</td><td>12456</td><td>12496</td><td>41</td><td>CCAAGCGTTGGATTGTTCACCCACTAATAGGGAACGTGAGC</td><td>311.0</td><td>-4.6</td><td>1.3</td><td>-3.6</td><td>3.0E-04</td><td>4.5E-04</td><td>MB231/MB468</td><td>MP</td></tr>
<tr><td>>trna116_GluCTC_1_-_145399233_145399304@36.71.36,trna134_GluTTC_1_-_16861774_16861845@36.71.36,trna71_GluCTC_1_-_161439189_161439260@36.71.36,trna74_GluCTC_1_-_161431809_161431880@36.71.36,trna77_GluCTC_1_-_161424398_161424469@36.71.36,trna77_GluCTC_6_+_28949976_28950047@36.71.36,trna80_GluCTC_1_-_161417018_161417089@36.71.36,trna84_GluTTC_1_-_161391883_161391954@36.71.36,trna87_GluCTC_6_-_126101393_126101464@36.71.36|CACCGCCGCGGCCCGGGTTCGATTCCCGGTCAGGGA</td><td>>trna116_GluCTC_1_-_145399233_145399304</td><td>36</td><td>71</td><td>36</td><td>CACCGCCGCGGCCCGGGTTCGATTCCCGGTCAGGGA</td><td>125.5</td><td>-3.9</td><td>1.1</td><td>-3.6</td><td>3.2E-04</td><td>4.5E-04</td><td>BT20/MB468</td><td>CYTO</td></tr>
<tr><td>>hg38_MT_+_598_1651_RNR1_12S_50ntFlanks@367.404.38|CCAATAAAGCTAAAACTCACCTGAGTTGTAAAAAACTC</td><td>>hg38_MT_+_598_1651_RNR1_12S_50ntFlanks</td><td>367</td><td>404</td><td>38</td><td>CCAATAAAGCTAAAACTCACCTGAGTTGTAAAAAACTC</td><td>848.4</td><td>3.5</td><td>1.0</td><td>3.6</td><td>3.6E-04</td><td>4.5E-04</td><td>MB231/MB468</td><td>MITO</td></tr>
<tr><td>>MI0000746|hsa-mir-99b&WithFlank&19|+|51692606|51692687@13.32.20(+1A)&[MIMAT0000689&hsa-miR-99b-5p&offsets|0|-2(+1A);m-10&19|+|51692618|51692639&offsets|0|-2(+1A)]|CACCCGTAGAACCGACCTTGA</td><td>>MI0000746|hsa-mir-99b&WithFlank&19|+|51692606|51692687</td><td>13</td><td>32</td><td>20(+1A)</td><td>CACCCGTAGAACCGACCTTGA</td><td>511.7</td><td>5.3</td><td>1.4</td><td>3.8</td><td>1.4E-04</td><td>4.5E-04</td><td>BT20/MB468</td><td>MP</td></tr>
<tr><td>>hg38_21_+_8433172_8446622_RNA45SN1_45S_50ntFlanks@12397.12414.18|CGACGTCGCTTTTTGATC</td><td>>hg38_21_+_8433172_8446622_RNA45SN1_45S_50ntFlanks</td><td>12397</td><td>12414</td><td>18</td><td>CGACGTCGCTTTTTGATC</td><td>951.4</td><td>-1.5</td><td>0.4</td><td>-3.6</td><td>3.6E-04</td><td>4.5E-04</td><td>MB231/MB468</td><td>NUC</td></tr>
<tr><td>>MI0000439|hsa-mir-23b&WithFlank&9|+|95085202|95085310@67.84.18&[MIMAT0000418&hsa-miR-23b-3p&offsets|+3|-2;m-42&9|+|95085265|95085285&offsets|+3|0]|ACATTGCCAGGGATTACC</td><td>>MI0000439|hsa-mir-23b&WithFlank&9|+|95085202|95085310</td><td>67</td><td>84</td><td>18</td><td>ACATTGCCAGGGATTACC</td><td>530.6</td><td>12.9</td><td>3.6</td><td>3.6</td><td>3.6E-04</td><td>4.6E-04</td><td>MB231/MB468</td><td>NUC</td></tr>
<tr><td>>hg38_21_+_8433172_8446622_RNA45SN1_45S_50ntFlanks@3835.3861.27|CTCCTCTCCTACTTGGATAACTGTGGT</td><td>>hg38_21_+_8433172_8446622_RNA45SN1_45S_50ntFlanks</td><td>3835</td><td>3861</td><td>27</td><td>CTCCTCTCCTACTTGGATAACTGTGGT</td><td>680.4</td><td>-3.9</td><td>1.1</td><td>-3.6</td><td>2.7E-04</td><td>4.6E-04</td><td>BT20/MB468</td><td>MITO</td></tr>
<tr><td>>hg38_21_+_8433172_8446622_RNA45SN1_45S_50ntFlanks@6776.6802.27|CTCCCGGGGCTACGCCTGTCTGAGCGT</td><td>>hg38_21_+_8433172_8446622_RNA45SN1_45S_50ntFlanks</td><td>6776</td><td>6802</td><td>27</td><td>CTCCCGGGGCTACGCCTGTCTGAGCGT</td><td>593.8</td><td>2.6</td><td>0.7</td><td>3.6</td><td>3.4E-04</td><td>4.6E-04</td><td>MB231/MB468</td><td>CYTO</td></tr>
<tr><td>>hg38_21_+_8433172_8446622_RNA45SN1_45S_50ntFlanks@10829.10858.30|GCAAGCCGGATCCGTAACTTCGGGATAAGG</td><td>>hg38_21_+_8433172_8446622_RNA45SN1_45S_50ntFlanks</td><td>10829</td><td>10858</td><td>30</td><td>GCAAGCCGGATCCGTAACTTCGGGATAAGG</td><td>485.9</td><td>-4.0</td><td>1.1</td><td>-3.6</td><td>3.4E-04</td><td>4.6E-04</td><td>MB231/MB468</td><td>CYTO</td></tr>
<tr><td>>MI0000478|hsa-mir-149&WithFlank&2|+|240455995|240456095@21.42.22&[MIMAT0000450&hsa-miR-149-5p&offsets|0|-1;m-195&2|+|240456015|240456037&offsets|0|-1]|TCTGGCTCCGTGTCTTCACTCC</td><td>>MI0000478|hsa-mir-149&WithFlank&2|+|240455995|240456095</td><td>21</td><td>42</td><td>22</td><td>TCTGGCTCCGTGTCTTCACTCC</td><td>1269.1</td><td>2.9</td><td>0.7</td><td>4.0</td><td>6.0E-05</td><td>4.6E-04</td><td>BT20/MB231</td><td>MITO</td></tr>
<tr><td>>trnaMT_ValTAC_MT_+_1602_1670@36.72.37|CTTAGGAGATTTCAACTTAACTTGACCGCTCTGACCA</td><td>>trnaMT_ValTAC_MT_+_1602_1670</td><td>36</td><td>72</td><td>37</td><td>CTTAGGAGATTTCAACTTAACTTGACCGCTCTGACCA</td><td>589.5</td><td>6.3</td><td>1.6</td><td>3.8</td><td>1.4E-04</td><td>4.6E-04</td><td>BT20/MB468</td><td>MP</td></tr>
<tr><td>>hg38_21_+_8433172_8446622_RNA45SN1_45S_50ntFlanks@12151.12180.30|GTGGGGAGTTTGACTGGGGCGGTACACCTG</td><td>>hg38_21_+_8433172_8446622_RNA45SN1_45S_50ntFlanks</td><td>12151</td><td>12180</td><td>30</td><td>GTGGGGAGTTTGACTGGGGCGGTACACCTG</td><td>772.5</td><td>-6.2</td><td>1.6</td><td>-3.8</td><td>1.4E-04</td><td>4.6E-04</td><td>BT20/MB468</td><td>MP</td></tr>
<tr><td>>am_MI0000809|hsa-mir-151a&WithFlank&8|-|140732558|140732659@53.71.19(+1A)&[MIMAT0000757&hsa-miR-151a-3p&offsets|0|-2(+1A);m-30&8|-|140732587|140732607&offsets|0|-2(+1A)]|CTAGACTGAAGCTCCTTGAA</td><td>>am_MI0000809|hsa-mir-151a&WithFlank&8|-|140732558|140732659</td><td>53</td><td>71</td><td>19(+1A)</td><td>CTAGACTGAAGCTCCTTGAA</td><td>554.4</td><td>3.0</td><td>0.7</td><td>4.1</td><td>5.0E-05</td><td>4.6E-04</td><td>BT20/MB231</td><td>CYTO</td></tr>
<tr><td>>trnaMT_PheGAA_MT_+_577_647@38.55.18|AATGTTTAGACGGGCTCA</td><td>>trnaMT_PheGAA_MT_+_577_647</td><td>38</td><td>55</td><td>18</td><td>AATGTTTAGACGGGCTCA</td><td>247.1</td><td>7.3</td><td>1.9</td><td>3.8</td><td>1.4E-04</td><td>4.6E-04</td><td>BT20/MB468</td><td>MP</td></tr>
<tr><td>>MI0000071|hsa-mir-17&WithFlank&13|+|91350599|91350694@20.42.23&[MIMAT0000070&hsa-miR-17-5p&offsets|0|0;m-88&13|+|91350618|91350640&offsets|0|0]|CAAAGTGCTTACAGTGCAGGTAG</td><td>>MI0000071|hsa-mir-17&WithFlank&13|+|91350599|91350694</td><td>20</td><td>42</td><td>23</td><td>CAAAGTGCTTACAGTGCAGGTAG</td><td>12401.7</td><td>-3.5</td><td>0.9</td><td>-4.0</td><td>6.0E-05</td><td>4.6E-04</td><td>BT20/MB231</td><td>MITO</td></tr>
<tr><td>>hg38_21_+_8433172_8446622_RNA45SN1_45S_50ntFlanks@6648.6691.44|GTACGACTCTTAGCGGTGGATCACTCGGCTCGTGCGTCGATGAA</td><td>>hg38_21_+_8433172_8446622_RNA45SN1_45S_50ntFlanks</td><td>6648</td><td>6691</td><td>44</td><td>GTACGACTCTTAGCGGTGGATCACTCGGCTCGTGCGTCGATGAA</td><td>247.4</td><td>4.8</td><td>1.3</td><td>3.6</td><td>3.6E-04</td><td>4.6E-04</td><td>MB231/MB468</td><td>NUC</td></tr>
<tr><td>>MI0000081|hsa-mir-24-2&WithFlank&19|-|13836281|13836365@56.77.22(+1C);MI0000080|hsa-mir-24-1&WithFlank&9|+|95086015|95086094@50.71.22(+1C)&[MIMAT0000080&hsa-miR-24-3p&offsets|0|0(+1C);m-36&19|-|13836288|13836310&offsets|0|-1(+1C)];[MIMAT0000080_1&hsa-miR-24-3p&offsets|0|0(+1C);m-35&9|+|95086064|95086086&offsets|0|-1(+1C)]|TGGCTCAGTTCAGCAGGAACAGC</td><td>>MI0000081|hsa-mir-24-2&WithFlank&19|-|13836281|13836365</td><td>56</td><td>77</td><td>22(+1C)</td><td>TGGCTCAGTTCAGCAGGAACAGC</td><td>2661.0</td><td>-4.3</td><td>1.1</td><td>-4.1</td><td>5.1E-05</td><td>4.6E-04</td><td>BT20/MB231</td><td>MP</td></tr>
<tr><td>>hg38_21_+_8433172_8446622_RNA45SN1_45S_50ntFlanks@9545.9563.19|GTGGAGGTCCGTAGCGGTC</td><td>>hg38_21_+_8433172_8446622_RNA45SN1_45S_50ntFlanks</td><td>9545</td><td>9563</td><td>19</td><td>GTGGAGGTCCGTAGCGGTC</td><td>626.1</td><td>-4.1</td><td>1.1</td><td>-3.6</td><td>2.7E-04</td><td>4.6E-04</td><td>BT20/MB468</td><td>MITO</td></tr>
<tr><td>>trnaMT_SerGCT_MT_+_12207_12265@1.25.25|GAGAAAGCTCACAAGAACTGCTAAC</td><td>>trnaMT_SerGCT_MT_+_12207_12265</td><td>1</td><td>25</td><td>25</td><td>GAGAAAGCTCACAAGAACTGCTAAC</td><td>37412.0</td><td>2.7</td><td>0.8</td><td>3.6</td><td>3.0E-04</td><td>4.6E-04</td><td>MB231/MB468</td><td>MP</td></tr>
<tr><td>>hg38_MT_+_598_1651_RNR1_12S_50ntFlanks@908.950.43|ATACTTCAAAGGACATTTAACTAAAACCCCTACGCATTTATAT</td><td>>hg38_MT_+_598_1651_RNR1_12S_50ntFlanks</td><td>908</td><td>950</td><td>43</td><td>ATACTTCAAAGGACATTTAACTAAAACCCCTACGCATTTATAT</td><td>83.0</td><td>10.1</td><td>2.7</td><td>3.8</td><td>1.4E-04</td><td>4.6E-04</td><td>BT20/MB468</td><td>MP</td></tr>
</table>

| | | | | | | | | | | | | | |
|---|---|---|---|---|---|---|---|---|---|---|---|---|---|
| >trna10_AspGTC_12_-_125424193_125424264@35.69.35,trna12_AspGTC_12_-_125411891_125411962@35.69.35,trna144_AspGTC_6_-_27551236_27551307@35.69.35,trna38_AspGTC_17_-_8125556_8125627@35.69.35,trna45_AspGTC_6_+_27447453_27447524@35.69.35,trna48_AspGTC_6_+_27471523_27471594@35.69.35,trna4_AspGTC_12_+_96429799_96429870@35.69.35,trna69_AspGTC_1_-_161440205_161440276@35.69.35,trna72_AspGTC_1_-_161432824_161432895@35.69.35,trna75_AspGTC_1_-_161425414_161425485@35.69.35,trna78_AspGTC_1_-_161418033_161418104@35.69.35,trna81_AspGTC_1_-_161410615_161410686@35.69.35\|TCACGCGGGAGACCGGGGTTCGATTCCCCGACGGG | >trna10_AspGTC_12_-_125424193_125424264 | 35 | 69 | 35 | TCACGCGGGAGACCGGGGTTCGATTCCCCGACGGG | 395.2 | -3.1 | 0.9 | -3.6 | 3.0E-04 | 4.6E-04 | MB231/MB468 | MP |
| >hg38_21_+_8433172_8446622_RNA45SN1_45S_50ntFlanks@6721.6752.32\|AATTGCAGGACACATTGATCATCGACACTTCG | >hg38_21_+_8433172_8446622_RNA45SN1_45S_50ntFlanks | 6721 | 6752 | 32 | AATTGCAGGACACATTGATCATCGACACTTCG | 473.1 | -1.8 | 0.5 | -3.6 | 2.7E-04 | 4.6E-04 | BT20/MB468 | MITO |
| >hg38_21_+_8433172_8446622_RNA45SN1_45S_50ntFlanks@10356.10383.28\|ACTTTGAAGGCCGAAGTGGAGAAGGGTT | >hg38_21_+_8433172_8446622_RNA45SN1_45S_50ntFlanks | 10356 | 10383 | 28 | ACTTTGAAGGCCGAAGTGGAGAAGGGTT | 109.8 | -2.9 | 0.8 | -3.6 | 3.0E-04 | 4.6E-04 | MB231/MB468 | MP |
| >hg38_21_+_8433172_8446622_RNA45SN1_45S_50ntFlanks@12283.12323.41\|TACAGACCGTGAAAGCGGGGCCTCACGATCCTTCTGACCTT | >hg38_21_+_8433172_8446622_RNA45SN1_45S_50ntFlanks | 12283 | 12323 | 41 | TACAGACCGTGAAAGCGGGGCCTCACGATCCTTCTGACCTT | 165.0 | -4.8 | 1.3 | -3.6 | 2.7E-04 | 4.6E-04 | BT20/MB468 | MITO |
| >hg38_21_+_8433172_8446622_RNA45SN1_45S_50ntFlanks@4006.4024.19\|ACCTCGGGCCGATCGCACG | >hg38_21_+_8433172_8446622_RNA45SN1_45S_50ntFlanks | 4006 | 4024 | 19 | ACCTCGGGCCGATCGCACG | 466.0 | -5.3 | 1.5 | -3.6 | 2.7E-04 | 4.6E-04 | BT20/MB468 | MITO |
| >hg38_21_+_8433172_8446622_RNA45SN1_45S_50ntFlanks@9739.9769.31\|AAACGATCTCAACCTATTCTCAAACTTTAAA | >hg38_21_+_8433172_8446622_RNA45SN1_45S_50ntFlanks | 9739 | 9769 | 31 | AAACGATCTCAACCTATTCTCAAACTTTAAA | 353.9 | -4.9 | 1.4 | -3.6 | 3.7E-04 | 4.6E-04 | BT20/MB468 | NUC |
| >hg38_21_+_8433172_8446622_RNA45SN1_45S_50ntFlanks@10248.10279.32\|CGACGAGTAGGAGGGCCGCTGCGGTGAGCCTT | >hg38_21_+_8433172_8446622_RNA45SN1_45S_50ntFlanks | 10248 | 10279 | 32 | CGACGAGTAGGAGGGCCGCTGCGGTGAGCCTT | 289.3 | 4.8 | 1.3 | 3.8 | 1.5E-04 | 4.6E-04 | BT20/MB468 | MP |
| >MI0000736\|hsa-mir-30c-1&WithFlank&1\|+\|40757278\|40757378@23.48.26;MI0000254\|hsa-mir-30c-2&WithFlank&6\|-\|71376954\|71377037@13.38.26&[MIMAT0000244&hsa-miR-30c-5p&offsets\|0\|+3;m-57&1\|+\|40757300\|40757323&offsets\|0\|+2];[MIMAT0000244_1&hsa-miR-30c-5p&offsets\|0\|+3;m-56&6\|-\|71377002\|71377025&offsets\|0\|+2]\|TGTAAACATCCTACACTCTCAGCTGT | >MI0000736\|hsa-mir-30c-1&WithFlank&1\|+\|40757278\|40757378 | 23 | 48 | 26 | TGTAAACATCCTACACTCTCAGCTGT | 96.6 | 6.5 | 1.8 | 3.6 | 3.1E-04 | 4.6E-04 | MB231/MB468 | MP |
| >hg38_21_+_8433172_8446622_RNA45SN1_45S_50ntFlanks@10771.10815.45\|CAGGTCTCCAAGGTGAACAGCCTCTGGCATGTTGGAACAATGTAG | >hg38_21_+_8433172_8446622_RNA45SN1_45S_50ntFlanks | 10771 | 10815 | 45 | CAGGTCTCCAAGGTGAACAGCCTCTGGCATGTTGGAACAATGTAG | 354.0 | -5.7 | 1.6 | -3.6 | 3.7E-04 | 4.6E-04 | MB231/MB468 | NUC |
| >hg38_21_+_8433172_8446622_RNA45SN1_45S_50ntFlanks@12927.12951.25\|CCTGGCGCTAAACCATTCGTAGACG | >hg38_21_+_8433172_8446622_RNA45SN1_45S_50ntFlanks | 12927 | 12951 | 25 | CCTGGCGCTAAACCATTCGTAGACG | 235.6 | -3.0 | 0.8 | -3.6 | 2.7E-04 | 4.6E-04 | BT20/MB468 | MITO |
| >MI0000088\|hsa-mir-30a&WithFlank&6\|-\|71403545\|71403627@12.33.22(+1A)&[MIMAT0000087&hsa-miR-30a-5p&offsets\|0\|0(+1A);m-15&6\|-\|71403595\|71403616&offsets\|0\|0(+1A)]\|TGTAAACATCCTCGACTGGAAGA | >MI0000088\|hsa-mir-30a&WithFlank&6\|-\|71403545\|71403627 | 12 | 33 | 22(+1A) | TGTAAACATCCTCGACTGGAAGA | 3174.2 | 3.3 | 0.9 | 3.6 | 3.1E-04 | 4.6E-04 | MB231/MB468 | MP |
| >hg38_21_+_8433172_8446622_RNA45SN1_45S_50ntFlanks@11874.11903.30\|AGAAGACCCTGTTGAGCTTGACTCTAGTCT | >hg38_21_+_8433172_8446622_RNA45SN1_45S_50ntFlanks | 11874 | 11903 | 30 | AGAAGACCCTGTTGAGCTTGACTCTAGTCT | 176.2 | -4.9 | 1.4 | -3.6 | 3.3E-04 | 4.6E-04 | BT20/MB468 | CYTO |
| >hg38_21_+_8433172_8446622_RNA45SN1_45S_50ntFlanks@10409.10443.35\|GTCAGTCGGTCCTGAGAGATGGGCGAGCGCCGTTC | >hg38_21_+_8433172_8446622_RNA45SN1_45S_50ntFlanks | 10409 | 10443 | 35 | GTCAGTCGGTCCTGAGAGATGGGCGAGCGCCGTTC | 134.0 | -1.9 | 0.5 | -3.6 | 3.3E-04 | 4.6E-04 | BT20/MB468 | CYTO |
| >MI0000803\|hsa-mir-330&WithFlank&19\|-\|45638988\|45639093@63.86.24(+1U)&[MIMAT0000751&hsa-miR-330-3p&offsets\|0\|+1(+1U);m-347&19\|-\|45639009\|45639030&offsets\|-1\|+1(+1U)]\|GCAAAGCACACGGCCTGCAGAGAGT | >MI0000803\|hsa-mir-330&WithFlank&19\|-\|45638988\|45639093 | 63 | 86 | 24(+1U) | GCAAAGCACACGGCCTGCAGAGAGT | 218.8 | 3.8 | 1.0 | 3.8 | 1.5E-04 | 4.6E-04 | BT20/MB468 | MP |
| >hg38_21_+_8433172_8446622_RNA45SN1_45S_50ntFlanks@12044.12061.18\|ACTGACCCGGTGAGGCGG | >hg38_21_+_8433172_8446622_RNA45SN1_45S_50ntFlanks | 12044 | 12061 | 18 | ACTGACCCGGTGAGGCGG | 109.2 | -9.9 | 2.6 | -3.8 | 1.5E-04 | 4.6E-04 | BT20/MB468 | MP |
| >hg38_21_+_8433172_8446622_RNA45SN1_45S_50ntFlanks@3999.4024.26\|CTAGATAACCTCGGGCCGATCGCACG | >hg38_21_+_8433172_8446622_RNA45SN1_45S_50ntFlanks | 3999 | 4024 | 26 | CTAGATAACCTCGGGCCGATCGCACG | 273.0 | -2.2 | 0.6 | -3.6 | 3.4E-04 | 4.6E-04 | MB231/MB468 | CYTO |
| >hg38_21_+_8433172_8446622_RNA45SN1_45S_50ntFlanks@11708.11736.29\|AGCGCGGGTAAACGGCGGGAGTAACTATG | >hg38_21_+_8433172_8446622_RNA45SN1_45S_50ntFlanks | 11708 | 11736 | 29 | AGCGCGGGTAAACGGCGGGAGTAACTATG | 218.5 | 3.4 | 1.0 | 3.6 | 3.4E-04 | 4.6E-04 | MB231/MB468 | CYTO |
| >hg38_21_+_8433172_8446622_RNA45SN1_45S_50ntFlanks@7973.8013.41\|GACGCGACCTCAGATCAGACGTGGCGACCCGCTGAATTTAA | >hg38_21_+_8433172_8446622_RNA45SN1_45S_50ntFlanks | 7973 | 8013 | 41 | GACGCGACCTCAGATCAGACGTGGCGACCCGCTGAATTTAA | 260.5 | 3.4 | 0.9 | 3.6 | 3.6E-04 | 4.6E-04 | MB231/MB468 | MITO |
| >hg38_21_+_8433172_8446622_RNA45SN1_45S_50ntFlanks@11209.11249.41\|CGGCGACTCTGGACGCGAGCCGGGCCCTTCCCGTGGATCGC | >hg38_21_+_8433172_8446622_RNA45SN1_45S_50ntFlanks | 11209 | 11249 | 41 | CGGCGACTCTGGACGCGAGCCGGGCCCTTCCCGTGGATCGC | 463.7 | -6.3 | 1.8 | -3.6 | 3.7E-04 | 4.6E-04 | BT20/MB468 | NUC |
| >MI0001733\|hsa-mir-452&WithFlank&X\|-\|151959622\|151959718@20.43.24&[MIMAT0001635&hsa-miR-452-5p&offsets\|0\|+2;m-188&X\|-\|151959677\|151959699&offsets\|0\|+1]\|AACTGTTTGCAGAGGAAACTGAGA | >MI0001733\|hsa-mir-452&WithFlank&X\|-\|151959622\|151959718 | 20 | 43 | 24 | AACTGTTTGCAGAGGAAACTGAGA | 1080.8 | 3.9 | 1.0 | 4.1 | 5.0E-05 | 4.6E-04 | BT20/MB231 | CYTO |
| >hg38_21_+_8433172_8446622_RNA45SN1_45S_50ntFlanks@8358.8391.34\|AAAGAACTTTGAAGAGAGAGTTCAAGAGGGCGTG | >hg38_21_+_8433172_8446622_RNA45SN1_45S_50ntFlanks | 8358 | 8391 | 34 | AAAGAACTTTGAAGAGAGAGTTCAAGAGGGCGTG | 144.6 | 3.2 | 0.8 | 3.8 | 1.5E-04 | 4.6E-04 | BT20/MB468 | MP |
| >hg38_21_+_8433172_8446622_RNA45SN1_45S_50ntFlanks@10678.10710.33\|CGGTTCCGGCGGCGTCCGGTGAGCTCTCGCTGG | >hg38_21_+_8433172_8446622_RNA45SN1_45S_50ntFlanks | 10678 | 10710 | 33 | CGGTTCCGGCGGCGTCCGGTGAGCTCTCGCTGG | 3986.2 | 6.4 | 1.6 | 4.0 | 5.2E-05 | 4.6E-04 | BT20/MB231 | MP |
| >hg38_21_+_8433172_8446622_RNA45SN1_45S_50ntFlanks@9563.9595.33\|CCTGACGTGCAAATCGGTCGTCCGACCTGGGTA | >hg38_21_+_8433172_8446622_RNA45SN1_45S_50ntFlanks | 9563 | 9595 | 33 | CCTGACGTGCAAATCGGTCGTCCGACCTGGGTA | 132.8 | -3.3 | 0.9 | -3.6 | 3.3E-04 | 4.6E-04 | BT20/MB468 | CYTO |
| >hg38_21_+_8433172_8446622_RNA45SN1_45S_50ntFlanks@10571.10589.19\|GAGAAGCCGGCGGGAGCCC | >hg38_21_+_8433172_8446622_RNA45SN1_45S_50ntFlanks | 10571 | 10589 | 19 | GAGAAGCCGGCGGGAGCCC | 171.4 | -4.9 | 1.4 | -3.6 | 3.3E-04 | 4.6E-04 | BT20/MB468 | CYTO |
| >hg38_21_+_8433172_8446622_RNA45SN1_45S_50ntFlanks@4000.4023.24\|TAGATAACCTCGGGCCGATCGCAC | >hg38_21_+_8433172_8446622_RNA45SN1_45S_50ntFlanks | 4000 | 4023 | 24 | TAGATAACCTCGGGCCGATCGCAC | 407.6 | -2.1 | 0.6 | -3.6 | 3.7E-04 | 4.6E-04 | MB231/MB468 | NUC |
| >hg38_21_+_8433172_8446622_RNA45SN1_45S_50ntFlanks@10389.10422.34\|TGAACAGCAGTTGAACATGGGTCAGTCGGTCCTG | >hg38_21_+_8433172_8446622_RNA45SN1_45S_50ntFlanks | 10389 | 10422 | 34 | TGAACAGCAGTTGAACATGGGTCAGTCGGTCCTG | 261.6 | -4.3 | 1.2 | -3.6 | 3.7E-04 | 4.6E-04 | MB231/MB468 | NUC |
| >MI0000433\|hsa-let-7g&WithFlank&3\|-\|52268272\|52268367@11.31.21(+1G)&[MIMAT0000414&hsa-let-7g-5p&offsets\|0\|-1(+1G);m-58&3\|-\|52268336\|52268357&offsets\|0\|-1(+1G)]\|TGAGGTAGTAGTTTGTACAGTG | >MI0000433\|hsa-let-7g&WithFlank&3\|-\|52268272\|52268367 | 11 | 31 | 21(+1G) | TGAGGTAGTAGTTTGTACAGTG | 232.7 | 3.5 | 1.0 | 3.6 | 3.7E-04 | 4.6E-04 | MB231/MB468 | MITO |
| >hg38_21_+_8433172_8446622_RNA45SN1_45S_50ntFlanks@6651.6688.38\|CGACTCTTAGCGGTGGATCACTCGGCTCGTGCGTCGAT | >hg38_21_+_8433172_8446622_RNA45SN1_45S_50ntFlanks | 6651 | 6688 | 38 | CGACTCTTAGCGGTGGATCACTCGGCTCGTGCGTCGAT | 17887.2 | 2.3 | 0.6 | 3.6 | 2.7E-04 | 4.6E-04 | BT20/MB468 | MITO |
| >hg38_21_+_8433172_8446622_RNA45SN1_45S_50ntFlanks@8172.8214.43\|AGTCCTTCTGATCGAGGCCCAGCCCGTGGACGGTGTGAGGCCG | >hg38_21_+_8433172_8446622_RNA45SN1_45S_50ntFlanks | 8172 | 8214 | 43 | AGTCCTTCTGATCGAGGCCCAGCCCGTGGACGGTGTGAGGCCG | 361.6 | -6.2 | 1.7 | -3.6 | 3.7E-04 | 4.6E-04 | BT20/MB468 | NUC |
| >hg38_21_+_8433172_8446622_RNA45SN1_45S_50ntFlanks@6719.6749.31\|TGAATTGCAGGACACATTGATCATCGACACT | >hg38_21_+_8433172_8446622_RNA45SN1_45S_50ntFlanks | 6719 | 6749 | 31 | TGAATTGCAGGACACATTGATCATCGACACT | 258.3 | -4.0 | 1.1 | -3.6 | 3.3E-04 | 4.6E-04 | BT20/MB468 | CYTO |
| >trnaMT_GluTTC_MT_-_14674_14742@35.72.38,trnalookalike8_GluTTC_5_-_93905172_93905240@35.72.38\|TATCATTGGTCGTGGTTGTAGTCCGTGCGAGAATACCA | >trnaMT_GluTTC_MT_-_14674_14742 | 35 | 72 | 38 | TATCATTGGTCGTGGTTGTAGTCCGTGCGAGAATACCA | 10088.7 | -3.8 | 0.9 | -4.1 | 5.1E-05 | 4.7E-04 | BT20/MB231 | CYTO |
| >hg38_21_+_8433172_8446622_RNA45SN1_45S_50ntFlanks@475.508.34\|CTTCGTGATCGATGTGGTGACGTCGTGCTCTCCC | >hg38_21_+_8433172_8446622_RNA45SN1_45S_50ntFlanks | 475 | 508 | 34 | CTTCGTGATCGATGTGGTGACGTCGTGCTCTCCC | 77.9 | 10.0 | 2.6 | 3.8 | 1.5E-04 | 4.7E-04 | BT20/MB468 | MP |
| >hg38_MT_+_1621_3279_RNR2_16S_50ntFlanks@999.1038.40\|AGGGACCTGTATGAATGGCTCCACGAGGGTTCAGCTGTCT | >hg38_MT_+_1621_3279_RNR2_16S_50ntFlanks | 999 | 1038 | 40 | AGGGACCTGTATGAATGGCTCCACGAGGGTTCAGCTGTCT | 81.1 | 10.1 | 2.7 | 3.8 | 1.5E-04 | 4.7E-04 | BT20/MB468 | MP |
| >MI0000746\|hsa-mir-99b&WithFlank&19\|+\|51692606\|51692687@51.71.21(+1A)&[MIMAT0004678&hsa-miR-99b-3p&offsets\|0\|-1(+1A);m-151&19\|+\|51692656\|51692678&offsets\|0\|-2(+1A)]\|CAAGCTCGTGTCTGTGGGTCCA | >MI0000746\|hsa-mir-99b&WithFlank&19\|+\|51692606\|51692687 | 51 | 71 | 21(+1A) | CAAGCTCGTGTCTGTGGGTCCA | 121.2 | 2.5 | 0.7 | 3.6 | 3.7E-04 | 4.7E-04 | BT20/MB468 | NUC |
| >am_trna128_GlyGCC_6_-_27870686_27870756@1.25.25,trna133_GlyCCC_1_-_16872434_16872504@1.25.25,trna18_GlyGCC_16_+_70822597_70822667@1.25.25,trna19_GlyGCC_16_+_70823410_70823480@1.25.25,trna19_GlyGCC_2_-_157257659_157257729@1.25.25,trna24_GlyGCC_16_-_70812942_70813012@1.25.25,trna25_GlyGCC_16_-_70812114_70812184@1.25.25,trna4_GlyCCC_1_+_17188416_17188486@1.25.25,trna5_GlyGCC_17_+_8029064_8029134@1.25.25,trna68_GlyGCC_1_-_161493637_161493707@1.25.25\|GCATTGGTGGTTCAGTGGTAGAATT | >am_trna128_GlyGCC_6_-_27870686_27870756 | 1 | 25 | 25 | GCATTGGTGGTTCAGTGGTAGAATT | 1666.5 | 2.2 | 0.6 | 3.6 | 3.3E-04 | 4.7E-04 | BT20/MB468 | CYTO |
| >trna111_HisGTG_1_-_147774845_147774916@-1G.34.35,trna118_HisGTG_1_-_145396881_145396952@-1G.34.35,trna16_HisGTG_1_+_146544773_146544844@-1G.34.35,trna1_HisGTG_15_+_45493349_45493420@-1G.34.35,trna21_HisGTG_1_+_147753471_147753542@-1G.34.35,trna33_HisGTG_6_+_27125906_27125977@-1G.34.35,trna7_HisGTG_9_-_14433938_14434009@-1G.34.35,trna8_HisGTG_15_-_45492611_45492682@-1G.34.35,trna9_HisGTG_15_-_45490804_45490875@-1G.34.35\|GGCCGTGATCGTATAGTGGTTAGTACTCTGCGTTG | >trna111_HisGTG_1_-_147774845_147774916 | -1G | 34 | 35 | GGCCGTGATCGTATAGTGGTTAGTACTCTGCGTTG | 49321.0 | -4.8 | 1.2 | -4.1 | 5.1E-05 | 4.7E-04 | BT20/MB231 | CYTO |
| >MI0000465\|hsa-mir-191&WithFlank&3\|-\|49020612\|49020715@22.44.23&[MIMAT0000440&hsa-miR-191-5p&offsets\|0\|0;m-48&3\|-\|49020672\|49020694&offsets\|0\|0]\|CAACGGAATCCCAAAAGCAGCTG | >MI0000465\|hsa-mir-191&WithFlank&3\|-\|49020612\|49020715 | 22 | 44 | 23 | CAACGGAATCCCAAAAGCAGCTG | 15524.4 | 3.9 | 1.0 | 3.8 | 1.5E-04 | 4.7E-04 | BT20/MB468 | MP |
| >hg38_21_+_8433172_8446622_RNA45SN1_45S_50ntFlanks@12413.12443.31\|TCCTTCGATGTCGGCTCTTCCTATCATTGTG | >hg38_21_+_8433172_8446622_RNA45SN1_45S_50ntFlanks | 12413 | 12443 | 31 | TCCTTCGATGTCGGCTCTTCCTATCATTGTG | 425.6 | -2.1 | 0.6 | -3.6 | 3.3E-04 | 4.7E-04 | BT20/MB468 | CYTO |

<table>
<tr><th></th><th></th><th></th><th></th><th></th><th></th><th></th><th></th><th></th><th></th><th></th><th></th><th></th><th></th></tr>
<tr><td>>MI0000433|hsa-let-7g&WithFlank&3|-|52268272|52268367@10.32.23&[MIMAT0000414&hsa-let-7g-5p&offsets|-1|0;m-58&3|-|52268336|52268357&offsets|-1|0]|CTGAGGTAGTAGTTTGTACAGTT</td><td>>MI0000433|hsa-let-7g&WithFlank&3|-|52268272|52268367</td><td>10</td><td>32</td><td>23</td><td>CTGAGGTAGTAGTTTGTACAGTT</td><td>152.0</td><td>1.5</td><td>0.4</td><td>3.6</td><td>3.8E-04</td><td>4.7E-04</td><td>BT20/MB468</td><td>NUC</td></tr>
<tr><td>>hg38_21_+_8433172_8446622_RNA45SN1_45S_50ntFlanks@10349.10384.36|AACGAGAACTTTGAAGGCCGAAGTGGAGAAGGGTTC</td><td>>hg38_21_+_8433172_8446622_RNA45SN1_45S_50ntFlanks</td><td>10349</td><td>10384</td><td>36</td><td>AACGAGAACTTTGAAGGCCGAAGTGGAGAAGGGTTC</td><td>1036.0</td><td>-3.9</td><td>1.0</td><td>-3.8</td><td>1.5E-04</td><td>4.7E-04</td><td>BT20/MB468</td><td>MP</td></tr>
<tr><td>>hg38_21_+_8433172_8446622_RNA45SN1_45S_50ntFlanks@10866.10894.29|CTAAGGGCTGGGTCGGTCGGGCTGGGGCG</td><td>>hg38_21_+_8433172_8446622_RNA45SN1_45S_50ntFlanks</td><td>10866</td><td>10894</td><td>29</td><td>CTAAGGGCTGGGTCGGTCGGGCTGGGGCG</td><td>463.0</td><td>-4.0</td><td>1.1</td><td>-3.6</td><td>2.8E-04</td><td>4.7E-04</td><td>BT20/MB468</td><td>MITO</td></tr>
<tr><td>>hg38_21_+_8433172_8446622_RNA45SN1_45S_50ntFlanks@11569.11599.31|CAGCCGACTTAGAACTGGTGCGGACCAGGGG</td><td>>hg38_21_+_8433172_8446622_RNA45SN1_45S_50ntFlanks</td><td>11569</td><td>11599</td><td>31</td><td>CAGCCGACTTAGAACTGGTGCGGACCAGGGG</td><td>492.1</td><td>-2.7</td><td>0.8</td><td>-3.6</td><td>3.5E-04</td><td>4.7E-04</td><td>MB231/MB468</td><td>CYTO</td></tr>
<tr><td>>hg38_21_+_8433172_8446622_RNA45SN1_45S_50ntFlanks@11577.11609.33|TTAGAACTGGTGCGGACCAGGGGAATCCGACTG</td><td>>hg38_21_+_8433172_8446622_RNA45SN1_45S_50ntFlanks</td><td>11577</td><td>11609</td><td>33</td><td>TTAGAACTGGTGCGGACCAGGGGAATCCGACTG</td><td>176.1</td><td>-3.1</td><td>0.9</td><td>-3.6</td><td>3.5E-04</td><td>4.7E-04</td><td>MB231/MB468</td><td>CYTO</td></tr>
<tr><td>>hg38_21_+_8433172_8446622_RNA45SN1_45S_50ntFlanks@9533.9566.34|GAGGAAACTCTGGTGGAGGTCCGTAGCGGTCCTG</td><td>>hg38_21_+_8433172_8446622_RNA45SN1_45S_50ntFlanks</td><td>9533</td><td>9566</td><td>34</td><td>GAGGAAACTCTGGTGGAGGTCCGTAGCGGTCCTG</td><td>305.6</td><td>-5.4</td><td>1.5</td><td>-3.6</td><td>3.3E-04</td><td>4.7E-04</td><td>BT20/MB468</td><td>CYTO</td></tr>
<tr><td>>hg38_21_+_8433172_8446622_RNA45SN1_45S_50ntFlanks@5489.5512.24|CTGAGAAGACGGTCGAACTTGACT</td><td>>hg38_21_+_8433172_8446622_RNA45SN1_45S_50ntFlanks</td><td>5489</td><td>5512</td><td>24</td><td>CTGAGAAGACGGTCGAACTTGACT</td><td>442.2</td><td>-2.5</td><td>0.7</td><td>-3.6</td><td>3.1E-04</td><td>4.7E-04</td><td>MB231/MB468</td><td>MP</td></tr>
<tr><td>>hg38_21_+_8433172_8446622_RNA45SN1_45S_50ntFlanks@12934.12962.29|CTAAACCATTCGTAGACGACCTGCTTCTG</td><td>>hg38_21_+_8433172_8446622_RNA45SN1_45S_50ntFlanks</td><td>12934</td><td>12962</td><td>29</td><td>CTAAACCATTCGTAGACGACCTGCTTCTG</td><td>153.8</td><td>-2.1</td><td>0.6</td><td>-3.6</td><td>2.8E-04</td><td>4.7E-04</td><td>BT20/MB468</td><td>MITO</td></tr>
<tr><td>>hg38_21_+_8433172_8446622_RNA45SN1_45S_50ntFlanks@12458.12476.19|AAGCGTTGGATTGTTCACC</td><td>>hg38_21_+_8433172_8446622_RNA45SN1_45S_50ntFlanks</td><td>12458</td><td>12476</td><td>19</td><td>AAGCGTTGGATTGTTCACC</td><td>1446.9</td><td>-2.6</td><td>0.7</td><td>-3.6</td><td>3.7E-04</td><td>4.7E-04</td><td>MB231/MB468</td><td>MITO</td></tr>
<tr><td>>MI0000469|hsa-mir-125a&WithFlank&19|+|51693248|51693345@21.41.21(+1C)&[MIMAT0000443&hsa-miR-125a-5p&offsets|0|-3(+1C);m-63&19|+|51693268|51693289&offsets|0|-1(+1C)]|TCCCTGAGACCCTTTAACCTGC</td><td>>MI0000469|hsa-mir-125a&WithFlank&19|+|51693248|51693345</td><td>21</td><td>41</td><td>21(+1C)</td><td>TCCCTGAGACCCTTTAACCTGC</td><td>220.3</td><td>3.6</td><td>1.0</td><td>3.6</td><td>3.7E-04</td><td>4.7E-04</td><td>MB231/MB468</td><td>MITO</td></tr>
<tr><td>>hg38_21_+_8433172_8446622_RNA45SN1_45S_50ntFlanks@11873.11903.31|AAGAAGACCCTGTTGAGCTTGACTCTAGTCT</td><td>>hg38_21_+_8433172_8446622_RNA45SN1_45S_50ntFlanks</td><td>11873</td><td>11903</td><td>31</td><td>AAGAAGACCCTGTTGAGCTTGACTCTAGTCT</td><td>215.4</td><td>-4.0</td><td>1.1</td><td>-3.6</td><td>3.3E-04</td><td>4.7E-04</td><td>BT20/MB468</td><td>CYTO</td></tr>
<tr><td>>MI0000087|hsa-mir-29a&WithFlank&7|-|130876741|130876816@47.65.19(+3U)&[MIMAT0000086&hsa-miR-29a-3p&offsets|-1|-4(+3U);m-26&7|-|130876749|130876769&offsets|-1|-3(+3U)]|CTAGCACCATCTGAAATCGTTT</td><td>>MI0000087|hsa-mir-29a&WithFlank&7|-|130876741|130876816</td><td>47</td><td>65</td><td>19(+3U)</td><td>CTAGCACCATCTGAAATCGTTT</td><td>24.6</td><td>7.8</td><td>2.2</td><td>3.6</td><td>3.1E-04</td><td>4.7E-04</td><td>MB231/MB468</td><td>MP</td></tr>
<tr><td>>MI0000299|hsa-mir-222&WithFlank&X|-|45747009|45747130@75.98.24&[MIMAT0000279&hsa-miR-222-3p&offsets|0|+3;m-140&X|-|45747033|45747056&offsets|0|0]|AGCTACATCTGGCTACTGGGTCTC</td><td>>MI0000299|hsa-mir-222&WithFlank&X|-|45747009|45747130</td><td>75</td><td>98</td><td>24</td><td>AGCTACATCTGGCTACTGGGTCTC</td><td>3429.6</td><td>3.8</td><td>1.0</td><td>3.8</td><td>1.5E-04</td><td>4.7E-04</td><td>BT20/MB468</td><td>MP</td></tr>
<tr><td>>trnaMT_HisGTG_MT_+_12138_12206@20.59.40|AACATCAGATTGTGAATCTGACAACAGAGGCTTACGACCC</td><td>>trnaMT_HisGTG_MT_+_12138_12206</td><td>20</td><td>59</td><td>40</td><td>AACATCAGATTGTGAATCTGACAACAGAGGCTTACGACCC</td><td>79.4</td><td>10.1</td><td>2.7</td><td>3.8</td><td>1.5E-04</td><td>4.7E-04</td><td>BT20/MB468</td><td>MP</td></tr>
<tr><td>>hg38_21_+_8433172_8446622_RNA45SN1_45S_50ntFlanks@11836.11872.37|ACAGCCAAGGGAACGGGCTTGGCGGAATCAGCGGGGA</td><td>>hg38_21_+_8433172_8446622_RNA45SN1_45S_50ntFlanks</td><td>11836</td><td>11872</td><td>37</td><td>ACAGCCAAGGGAACGGGCTTGGCGGAATCAGCGGGGA</td><td>158.0</td><td>-5.9</td><td>1.6</td><td>-3.8</td><td>1.5E-04</td><td>4.7E-04</td><td>BT20/MB468</td><td>MP</td></tr>
<tr><td>>MI0000085|hsa-mir-27a&WithFlank&19|-|13836434|13836523@57.75.19(+1C)&[MIMAT0000084&hsa-miR-27a-3p&offsets|0|-2(+1C);m-43&19|-|13836447|13836467&offsets|0|-2(+1C)]|TTCACAGTGGCTAAGTTCCC</td><td>>MI0000085|hsa-mir-27a&WithFlank&19|-|13836434|13836523</td><td>57</td><td>75</td><td>19(+1C)</td><td>TTCACAGTGGCTAAGTTCCC</td><td>223.4</td><td>3.5</td><td>1.0</td><td>3.6</td><td>3.1E-04</td><td>4.7E-04</td><td>MB231/MB468</td><td>MP</td></tr>
<tr><td>>hg38_21_+_8433172_8446622_RNA45SN1_45S_50ntFlanks@9533.9563.31|GAGGAAACTCTGGTGGAGGTCCGTAGCGGTC</td><td>>hg38_21_+_8433172_8446622_RNA45SN1_45S_50ntFlanks</td><td>9533</td><td>9563</td><td>31</td><td>GAGGAAACTCTGGTGGAGGTCCGTAGCGGTC</td><td>325.9</td><td>-1.9</td><td>0.5</td><td>-3.6</td><td>2.8E-04</td><td>4.7E-04</td><td>BT20/MB468</td><td>MITO</td></tr>
<tr><td>>hg38_21_+_8433172_8446622_RNA45SN1_45S_50ntFlanks@11564.11601.38|CCTAGCAGCCGACTTAGAACTGGTGCGGACCAGGGGAA</td><td>>hg38_21_+_8433172_8446622_RNA45SN1_45S_50ntFlanks</td><td>11564</td><td>11601</td><td>38</td><td>CCTAGCAGCCGACTTAGAACTGGTGCGGACCAGGGGAA</td><td>3238.1</td><td>-4.1</td><td>1.1</td><td>-3.8</td><td>1.5E-04</td><td>4.7E-04</td><td>BT20/MB468</td><td>MP</td></tr>
<tr><td>>hg38_21_+_8433172_8446622_RNA45SN1_45S_50ntFlanks@9167.9197.31|CGGGCGGGTCGCGCCGTCGGGCCCGGGGGAG</td><td>>hg38_21_+_8433172_8446622_RNA45SN1_45S_50ntFlanks</td><td>9167</td><td>9197</td><td>31</td><td>CGGGCGGGTCGCGCCGTCGGGCCCGGGGGAG</td><td>430.9</td><td>-6.4</td><td>1.8</td><td>-3.6</td><td>3.8E-04</td><td>4.7E-04</td><td>BT20/MB468</td><td>NUC</td></tr>
<tr><td>>hg38_21_+_8433172_8446622_RNA45SN1_45S_50ntFlanks@7974.8015.42|ACGCGACCTCAGATCAGACGTGGCGACCCGCTGAATTTAAGC</td><td>>hg38_21_+_8433172_8446622_RNA45SN1_45S_50ntFlanks</td><td>7974</td><td>8015</td><td>42</td><td>ACGCGACCTCAGATCAGACGTGGCGACCCGCTGAATTTAAGC</td><td>2881.6</td><td>3.6</td><td>1.0</td><td>3.6</td><td>3.7E-04</td><td>4.7E-04</td><td>MB231/MB468</td><td>MITO</td></tr>
<tr><td>>MI0000234|hsa-mir-192&WithFlank&11|-|64891131|64891252@31.51.21(+1U)&[MIMAT0000222&hsa-miR-192-5p&offsets|+1|+1(+1U);m-59&11|-|64891203|64891223&offsets|+1|+1(+1U)]|TGACCTATGAATTGACAGCCAT</td><td>>MI0000234|hsa-mir-192&WithFlank&11|-|64891131|64891252</td><td>31</td><td>51</td><td>21(+1U)</td><td>TGACCTATGAATTGACAGCCAT</td><td>196.4</td><td>4.8</td><td>1.3</td><td>3.6</td><td>3.7E-04</td><td>4.7E-04</td><td>MB231/MB468</td><td>MITO</td></tr>
<tr><td>>hg38_MT_+_598_1651_RNR1_12S_50ntFlanks@424.468.45|TACGAAAGTGGCTTTAACATATCTGAACACACAATAGCTAAGACC</td><td>>hg38_MT_+_598_1651_RNR1_12S_50ntFlanks</td><td>424</td><td>468</td><td>45</td><td>TACGAAAGTGGCTTTAACATATCTGAACACACAATAGCTAAGACC</td><td>83.8</td><td>10.1</td><td>2.7</td><td>3.8</td><td>1.5E-04</td><td>4.7E-04</td><td>BT20/MB468</td><td>MP</td></tr>
<tr><td>>hg38_21_+_8433172_8446622_RNA45SN1_45S_50ntFlanks@9932.9951.20|ATAGACAGCAGGACGGTGGC</td><td>>hg38_21_+_8433172_8446622_RNA45SN1_45S_50ntFlanks</td><td>9932</td><td>9951</td><td>20</td><td>ATAGACAGCAGGACGGTGGC</td><td>966.6</td><td>8.0</td><td>2.0</td><td>4.0</td><td>5.3E-05</td><td>4.7E-04</td><td>BT20/MB231</td><td>MP</td></tr>
<tr><td>>hg38_21_+_8433172_8446622_RNA45SN1_45S_50ntFlanks@8176.8214.39|CTTCTGATCGAGGCCCAGCCCGTGGACGGTGTGAGGCCG</td><td>>hg38_21_+_8433172_8446622_RNA45SN1_45S_50ntFlanks</td><td>8176</td><td>8214</td><td>39</td><td>CTTCTGATCGAGGCCCAGCCCGTGGACGGTGTGAGGCCG</td><td>744.9</td><td>-3.5</td><td>1.0</td><td>-3.6</td><td>3.3E-04</td><td>4.7E-04</td><td>BT20/MB468</td><td>CYTO</td></tr>
<tr><td>>hg38_21_+_8433172_8446622_RNA45SN1_45S_50ntFlanks@12251.12273.23|AAAGCTCGCTTGATCTTGATTTT</td><td>>hg38_21_+_8433172_8446622_RNA45SN1_45S_50ntFlanks</td><td>12251</td><td>12273</td><td>23</td><td>AAAGCTCGCTTGATCTTGATTTT</td><td>361.9</td><td>-3.9</td><td>1.1</td><td>-3.6</td><td>2.8E-04</td><td>4.7E-04</td><td>BT20/MB468</td><td>MITO</td></tr>
<tr><td>>hg38_21_+_8433172_8446622_RNA45SN1_45S_50ntFlanks@11580.11613.34|GAACTGGTGCGGACCAGGGGAATCCGACTGTTTA</td><td>>hg38_21_+_8433172_8446622_RNA45SN1_45S_50ntFlanks</td><td>11580</td><td>11613</td><td>34</td><td>GAACTGGTGCGGACCAGGGGAATCCGACTGTTTA</td><td>546.1</td><td>-2.8</td><td>0.8</td><td>-3.6</td><td>3.1E-04</td><td>4.7E-04</td><td>MB231/MB468</td><td>MP</td></tr>
<tr><td>>hg38_21_+_8433172_8446622_RNA45SN1_45S_50ntFlanks@12407.12429.23|TTTTGATCCTTCGATGTCGGCTC</td><td>>hg38_21_+_8433172_8446622_RNA45SN1_45S_50ntFlanks</td><td>12407</td><td>12429</td><td>23</td><td>TTTTGATCCTTCGATGTCGGCTC</td><td>432.2</td><td>-6.2</td><td>1.7</td><td>-3.6</td><td>3.8E-04</td><td>4.7E-04</td><td>BT20/MB468</td><td>NUC</td></tr>
<tr><td>>hg38_21_+_8433172_8446622_RNA45SN1_45S_50ntFlanks@12407.12440.34|TTTTGATCCTTCGATGTCGGCTCTTCCTATCATT</td><td>>hg38_21_+_8433172_8446622_RNA45SN1_45S_50ntFlanks</td><td>12407</td><td>12440</td><td>34</td><td>TTTTGATCCTTCGATGTCGGCTCTTCCTATCATT</td><td>310.6</td><td>2.7</td><td>0.8</td><td>3.6</td><td>3.3E-04</td><td>4.7E-04</td><td>BT20/MB468</td><td>CYTO</td></tr>
<tr><td>>hg38_21_+_8433172_8446622_RNA45SN1_45S_50ntFlanks@10808.10839.32|CAATGTAGGTAAGGGAAGTCGGCAAGCCGGAT</td><td>>hg38_21_+_8433172_8446622_RNA45SN1_45S_50ntFlanks</td><td>10808</td><td>10839</td><td>32</td><td>CAATGTAGGTAAGGGAAGTCGGCAAGCCGGAT</td><td>115.2</td><td>-4.7</td><td>1.3</td><td>-3.6</td><td>2.8E-04</td><td>4.7E-04</td><td>BT20/MB468</td><td>MITO</td></tr>
<tr><td>>trnaMT_HisGTG_MT_+_12138_12206@27.69.43|GATTGTGAATCTGACAACAGAGGCTTACGACCCCTTATTTACC</td><td>>trnaMT_HisGTG_MT_+_12138_12206</td><td>27</td><td>69</td><td>43</td><td>GATTGTGAATCTGACAACAGAGGCTTACGACCCCTTATTTACC</td><td>80.1</td><td>10.1</td><td>2.7</td><td>3.8</td><td>1.5E-04</td><td>4.7E-04</td><td>BT20/MB468</td><td>MP</td></tr>
<tr><td>>hg38_21_+_8433172_8446622_RNA45SN1_45S_50ntFlanks@9198.9225.28|GTTCTCTCGGGGCCACGCGCGCGTCCCC</td><td>>hg38_21_+_8433172_8446622_RNA45SN1_45S_50ntFlanks</td><td>9198</td><td>9225</td><td>28</td><td>GTTCTCTCGGGGCCACGCGCGCGTCCCC</td><td>344.9</td><td>-6.0</td><td>1.6</td><td>-3.8</td><td>1.5E-04</td><td>4.7E-04</td><td>BT20/MB468</td><td>MP</td></tr>
<tr><td>>hg38_21_+_8433172_8446622_RNA45SN1_45S_50ntFlanks@8180.8211.32|TGATCGAGGCCCAGCCCGTGGACGGTGTGAGG</td><td>>hg38_21_+_8433172_8446622_RNA45SN1_45S_50ntFlanks</td><td>8180</td><td>8211</td><td>32</td><td>TGATCGAGGCCCAGCCCGTGGACGGTGTGAGG</td><td>169.7</td><td>-2.9</td><td>0.8</td><td>-3.8</td><td>1.5E-04</td><td>4.7E-04</td><td>BT20/MB468</td><td>MP</td></tr>
<tr><td>>MI0000471|hsa-mir-126&WithFlank&9|+|136670596|136670692@58.77.20(+1U)&[MIMAT0000445&hsa-miR-126-3p&offsets|0|-2(+1U);m-40&9|+|136670653|136670674&offsets|0|-2(+1U)]|TCGTACCGTGAGTAATAATGT</td><td>>MI0000471|hsa-mir-126&WithFlank&9|+|136670596|136670692</td><td>58</td><td>77</td><td>20(+1U)</td><td>TCGTACCGTGAGTAATAATGT</td><td>443.9</td><td>7.5</td><td>2.0</td><td>3.8</td><td>1.5E-04</td><td>4.7E-04</td><td>BT20/MB468</td><td>MP</td></tr>
<tr><td>>hg38_21_+_8433172_8446622_RNA45SN1_45S_50ntFlanks@12606.12624.19|ATGTGCTTGGCTGAGGAGC</td><td>>hg38_21_+_8433172_8446622_RNA45SN1_45S_50ntFlanks</td><td>12606</td><td>12624</td><td>19</td><td>ATGTGCTTGGCTGAGGAGC</td><td>798.9</td><td>-1.8</td><td>0.5</td><td>-3.6</td><td>3.8E-04</td><td>4.7E-04</td><td>MB231/MB468</td><td>NUC</td></tr>
<tr><td>>MI0000084|hsa-mir-26b&WithFlank&2|+|218402640|218402728@18.39.22(+1U)&[MIMAT0000083&hsa-miR-26b-5p&offsets|0|+1(+1U);m-53&2|+|218402657|218402678&offsets|0|0(+1U)]|TTCAAGTAATTCAGGATAGGTTT</td><td>>MI0000084|hsa-mir-26b&WithFlank&2|+|218402640|218402728</td><td>18</td><td>39</td><td>22(+1U)</td><td>TTCAAGTAATTCAGGATAGGTTT</td><td>804.9</td><td>-3.3</td><td>0.8</td><td>-4.0</td><td>6.2E-05</td><td>4.7E-04</td><td>BT20/MB231</td><td>MITO</td></tr>
<tr><td>>hg38_21_+_8433172_8446622_RNA45SN1_45S_50ntFlanks@10404.10422.19|CATGGGTCAGTCGGTCCTG</td><td>>hg38_21_+_8433172_8446622_RNA45SN1_45S_50ntFlanks</td><td>10404</td><td>10422</td><td>19</td><td>CATGGGTCAGTCGGTCCTG</td><td>476.2</td><td>-3.2</td><td>0.8</td><td>-3.8</td><td>1.5E-04</td><td>4.7E-04</td><td>BT20/MB468</td><td>MP</td></tr>
<tr><td>>MI0000082|hsa-mir-25&WithFlank&7|-|100093554|100093649@58.78.21(+1G)&[MIMAT0000081&hsa-miR-25-3p&offsets|0|-1(+1G);m-23&7|-|100093571|100093592&offsets|0|-1(+1G)]|CATTGCACTTGTCTCGGTCTGG</td><td>>MI0000082|hsa-mir-25&WithFlank&7|-|100093554|100093649</td><td>58</td><td>78</td><td>21(+1G)</td><td>CATTGCACTTGTCTCGGTCTGG</td><td>673.4</td><td>4.1</td><td>1.1</td><td>3.8</td><td>1.5E-04</td><td>4.8E-04</td><td>BT20/MB468</td><td>MP</td></tr>
<tr><td>>hg38_21_+_8433172_8446622_RNA45SN1_45S_50ntFlanks@7976.8009.34|GCGACCTCAGATCAGACGTGGCGACCCGCTGAAT</td><td>>hg38_21_+_8433172_8446622_RNA45SN1_45S_50ntFlanks</td><td>7976</td><td>8009</td><td>34</td><td>GCGACCTCAGATCAGACGTGGCGACCCGCTGAAT</td><td>347.4</td><td>2.6</td><td>0.7</td><td>3.6</td><td>3.8E-04</td><td>4.8E-04</td><td>BT20/MB468</td><td>NUC</td></tr>
<tr><td>>hg38_21_+_8433172_8446622_RNA45SN1_45S_50ntFlanks@3705.3723.19|TACCTGGTTGATCCTGCCA</td><td>>hg38_21_+_8433172_8446622_RNA45SN1_45S_50ntFlanks</td><td>3705</td><td>3723</td><td>19</td><td>TACCTGGTTGATCCTGCCA</td><td>711.1</td><td>3.3</td><td>0.9</td><td>3.6</td><td>2.8E-04</td><td>4.8E-04</td><td>BT20/MB468</td><td>MITO</td></tr>
<tr><td>>hg38_21_+_8433172_8446622_RNA45SN1_45S_50ntFlanks@8674.8702.29|GGTGCGCCGCGACCGGCTCCGGGACGGCT</td><td>>hg38_21_+_8433172_8446622_RNA45SN1_45S_50ntFlanks</td><td>8674</td><td>8702</td><td>29</td><td>GGTGCGCCGCGACCGGCTCCGGGACGGCT</td><td>73.6</td><td>5.9</td><td>1.6</td><td>3.8</td><td>1.5E-04</td><td>4.8E-04</td><td>BT20/MB468</td><td>MP</td></tr>
<tr><td>>hg38_1_-_228634819_228635039_RNA5S12_5S_50ntFlanks@145.171.27|CTGGGAATACCGGGTGCTGTAGGCTTT</td><td>>hg38_1_-_228634819_228635039_RNA5S12_5S_50ntFlanks</td><td>145</td><td>171</td><td>27</td><td>CTGGGAATACCGGGTGCTGTAGGCTTT</td><td>1054.7</td><td>-2.8</td><td>0.8</td><td>-3.6</td><td>3.2E-04</td><td>4.8E-04</td><td>MB231/MB468</td><td>MP</td></tr>
<tr><td>>MI0006273|hsa-mir-1180&WithFlank&17|-|19344500|19344580@47.66.20&[MIMAT0005825&hsa-miR-1180-3p&offsets|0|-2;m-214&17|-|19344513|19344534&offsets|0|-2]|TTTCCGGCTCGCGTGGGTGT</td><td>>MI0006273|hsa-mir-1180&WithFlank&17|-|19344500|19344580</td><td>47</td><td>66</td><td>20</td><td>TTTCCGGCTCGCGTGGGTGT</td><td>931.7</td><td>3.4</td><td>0.8</td><td>4.0</td><td>5.2E-05</td><td>4.8E-04</td><td>BT20/MB231</td><td>CYTO</td></tr>
<tr><td>>hg38_21_+_8433172_8446622_RNA45SN1_45S_50ntFlanks@11571.11605.35|GCCGACTTAGAACTGGTGCGGACCAGGGGAATCCG</td><td>>hg38_21_+_8433172_8446622_RNA45SN1_45S_50ntFlanks</td><td>11571</td><td>11605</td><td>35</td><td>GCCGACTTAGAACTGGTGCGGACCAGGGGAATCCG</td><td>505.5</td><td>-6.3</td><td>1.8</td><td>-3.6</td><td>3.8E-04</td><td>4.8E-04</td><td>BT20/MB468</td><td>NUC</td></tr>
<tr><td>>hg38_21_+_8433172_8446622_RNA45SN1_45S_50ntFlanks@9533.9552.20|GAGGAAACTCTGGTGGAGGT</td><td>>hg38_21_+_8433172_8446622_RNA45SN1_45S_50ntFlanks</td><td>9533</td><td>9552</td><td>20</td><td>GAGGAAACTCTGGTGGAGGT</td><td>770.5</td><td>-6.6</td><td>1.8</td><td>-3.6</td><td>3.8E-04</td><td>4.8E-04</td><td>BT20/MB468</td><td>NUC</td></tr>
<tr><td>>MI0000342|hsa-mir-200b&WithFlank&1|+|1167098|1167204@63.84.22(+1U)&[MIMAT0000318&hsa-miR-200b-3p&offsets|0|0(+1U);m-93&1|+|1167160|1167182&offsets|0|-1(+1U)]|TAATACTGCCTGGTAATGATGAT</td><td>>MI0000342|hsa-mir-200b&WithFlank&1|+|1167098|1167204</td><td>63</td><td>84</td><td>22(+1U)</td><td>TAATACTGCCTGGTAATGATGAT</td><td>177.9</td><td>-2.7</td><td>0.8</td><td>-3.6</td><td>3.5E-04</td><td>4.8E-04</td><td>MB231/MB468</td><td>CYTO</td></tr>
<tr><td>>hg38_21_+_8433172_8446622_RNA45SN1_45S_50ntFlanks@11564.11602.39|CCTAGCAGCCGACTTAGAACTGGTGCGGACCAGGGGAAT</td><td>>hg38_21_+_8433172_8446622_RNA45SN1_45S_50ntFlanks</td><td>11564</td><td>11602</td><td>39</td><td>CCTAGCAGCCGACTTAGAACTGGTGCGGACCAGGGGAAT</td><td>3415.3</td><td>-2.8</td><td>0.8</td><td>-3.6</td><td>2.8E-04</td><td>4.8E-04</td><td>BT20/MB468</td><td>MITO</td></tr>
<tr><td>>hg38_21_+_8433172_8446622_RNA45SN1_45S_50ntFlanks@11235.11268.34|CTTCCCGTGGATCGCCCCAGCTGCGGCGGGCGTC</td><td>>hg38_21_+_8433172_8446622_RNA45SN1_45S_50ntFlanks</td><td>11235</td><td>11268</td><td>34</td><td>CTTCCCGTGGATCGCCCCAGCTGCGGCGGGCGTC</td><td>972.5</td><td>-6.8</td><td>1.9</td><td>-3.6</td><td>3.8E-04</td><td>4.8E-04</td><td>BT20/MB468</td><td>NUC</td></tr>
</table>

<table>
<tr><td>>hg38_21_+_8433172_8446622_RNA45SN1_45S_50ntFlanks@9021.9039.19|CGCCGGCGTCTCCTCGTGG</td><td>>hg38_21_+_8433172_8446622_RNA45SN1_45S_50ntFlanks</td><td>9021</td><td>9039</td><td>19</td><td>CGCCGGCGTCTCCTCGTGG</td><td>371.5</td><td>-6.2</td><td>1.7</td><td>-3.6</td><td>3.8E-04</td><td>4.8E-04</td><td>BT20/MB468</td><td>NUC</td></tr>
<tr><td>>hg38_21_+_8433172_8446622_RNA45SN1_45S_50ntFlanks@11671.11705.35|CCAGTGCTCTGAATGTCAAAGTGAAGAAATTCAAT</td><td>>hg38_21_+_8433172_8446622_RNA45SN1_45S_50ntFlanks</td><td>11671</td><td>11705</td><td>35</td><td>CCAGTGCTCTGAATGTCAAAGTGAAGAAATTCAAT</td><td>62.0</td><td>9.3</td><td>2.3</td><td>4.0</td><td>5.4E-05</td><td>4.8E-04</td><td>BT20/MB231</td><td>MP</td></tr>
<tr><td>>hg38_21_+_8433172_8446622_RNA45SN1_45S_50ntFlanks@3705.3741.37|TACCTGGTTGATCCTGCCAGTAGCATATGCTTGTCTC</td><td>>hg38_21_+_8433172_8446622_RNA45SN1_45S_50ntFlanks</td><td>3705</td><td>3741</td><td>37</td><td>TACCTGGTTGATCCTGCCAGTAGCATATGCTTGTCTC</td><td>1767.3</td><td>3.0</td><td>0.8</td><td>3.6</td><td>2.8E-04</td><td>4.8E-04</td><td>BT20/MB468</td><td>MITO</td></tr>
<tr><td>>hg38_21_+_8433172_8446622_RNA45SN1_45S_50ntFlanks@9941.9959.19|AGGACGGTGGCCATGGAAG</td><td>>hg38_21_+_8433172_8446622_RNA45SN1_45S_50ntFlanks</td><td>9941</td><td>9959</td><td>19</td><td>AGGACGGTGGCCATGGAAG</td><td>71.9</td><td>6.8</td><td>1.8</td><td>3.8</td><td>1.5E-04</td><td>4.8E-04</td><td>BT20/MB468</td><td>MP</td></tr>
<tr><td>>hg38_21_+_8433172_8446622_RNA45SN1_45S_50ntFlanks@10794.10828.35|CTGGCATGTTGGAACAATGTAGGTAAGGGAAGTCG</td><td>>hg38_21_+_8433172_8446622_RNA45SN1_45S_50ntFlanks</td><td>10794</td><td>10828</td><td>35</td><td>CTGGCATGTTGGAACAATGTAGGTAAGGGAAGTCG</td><td>159.2</td><td>-5.7</td><td>1.5</td><td>-3.8</td><td>1.5E-04</td><td>4.8E-04</td><td>BT20/MB468</td><td>MP</td></tr>
<tr><td>>hg38_21_+_8433172_8446622_RNA45SN1_45S_50ntFlanks@12934.12952.19|CTAAACCATTCGTAGACGA</td><td>>hg38_21_+_8433172_8446622_RNA45SN1_45S_50ntFlanks</td><td>12934</td><td>12952</td><td>19</td><td>CTAAACCATTCGTAGACGA</td><td>478.9</td><td>-6.0</td><td>1.6</td><td>-3.8</td><td>1.5E-04</td><td>4.8E-04</td><td>BT20/MB468</td><td>MP</td></tr>
<tr><td>>hg38_21_+_8433172_8446622_RNA45SN1_45S_50ntFlanks@9569.9596.28|GTGCAAATCGGTCGTCCGACCTGGGTAT</td><td>>hg38_21_+_8433172_8446622_RNA45SN1_45S_50ntFlanks</td><td>9569</td><td>9596</td><td>28</td><td>GTGCAAATCGGTCGTCCGACCTGGGTAT</td><td>536.8</td><td>-6.2</td><td>1.6</td><td>-3.8</td><td>1.5E-04</td><td>4.8E-04</td><td>BT20/MB468</td><td>MP</td></tr>
<tr><td>>hg38_21_+_8433172_8446622_RNA45SN1_45S_50ntFlanks@6671.6690.20|CTCGGCTCGTGCGTCGATGA</td><td>>hg38_21_+_8433172_8446622_RNA45SN1_45S_50ntFlanks</td><td>6671</td><td>6690</td><td>20</td><td>CTCGGCTCGTGCGTCGATGA</td><td>364.1</td><td>-6.3</td><td>1.7</td><td>-3.6</td><td>2.8E-04</td><td>4.8E-04</td><td>BT20/MB468</td><td>MITO</td></tr>
<tr><td>>hg38_21_+_8433172_8446622_RNA45SN1_45S_50ntFlanks@12607.12626.20|TGTGCTTGGCTGAGGAGCCA</td><td>>hg38_21_+_8433172_8446622_RNA45SN1_45S_50ntFlanks</td><td>12607</td><td>12626</td><td>20</td><td>TGTGCTTGGCTGAGGAGCCA</td><td>236.8</td><td>-3.3</td><td>0.9</td><td>-3.6</td><td>3.6E-04</td><td>4.8E-04</td><td>MB231/MB468</td><td>CYTO</td></tr>
<tr><td>>hg38_21_+_8433172_8446622_RNA45SN1_45S_50ntFlanks@5526.5550.25|AAAAGTCGTAACAAGGTTTCCGTAG</td><td>>hg38_21_+_8433172_8446622_RNA45SN1_45S_50ntFlanks</td><td>5526</td><td>5550</td><td>25</td><td>AAAAGTCGTAACAAGGTTTCCGTAG</td><td>142.6</td><td>-3.6</td><td>1.0</td><td>-3.6</td><td>3.4E-04</td><td>4.8E-04</td><td>BT20/MB468</td><td>CYTO</td></tr>
<tr><td>>hg38_21_+_8433172_8446622_RNA45SN1_45S_50ntFlanks@6702.6720.19|AGCTGCGAGAATTAATGTG</td><td>>hg38_21_+_8433172_8446622_RNA45SN1_45S_50ntFlanks</td><td>6702</td><td>6720</td><td>19</td><td>AGCTGCGAGAATTAATGTG</td><td>356.7</td><td>-2.6</td><td>0.7</td><td>-3.6</td><td>3.8E-04</td><td>4.8E-04</td><td>MB231/MB468</td><td>MITO</td></tr>
<tr><td>>trnaMT_ProTGG_MT_-_15956_16023@39.58.20|TAATGGTGGAGTTAAAGACT</td><td>>trnaMT_ProTGG_MT_-_15956_16023</td><td>39</td><td>58</td><td>20</td><td>TAATGGTGGAGTTAAAGACT</td><td>155.9</td><td>4.2</td><td>1.2</td><td>3.6</td><td>3.2E-04</td><td>4.8E-04</td><td>MB231/MB468</td><td>MP</td></tr>
<tr><td>>trnaMT_GluTTC_MT_-_14674_14742@39.67.29,trnalookalike8_GluTTC_5_-_93905172_93905240@39.67.29|ATTGGTCGTGGTTGTAGTCCGTGCGAGAA</td><td>>trnaMT_GluTTC_MT_-_14674_14742</td><td>39</td><td>67</td><td>29</td><td>ATTGGTCGTGGTTGTAGTCCGTGCGAGAA</td><td>191.7</td><td>6.4</td><td>1.7</td><td>3.8</td><td>1.5E-04</td><td>4.8E-04</td><td>BT20/MB468</td><td>MP</td></tr>
<tr><td>>MI0003776|hsa-mir-320b-1&WithFlank&1|+|116671740|116671823@47.67.21;MI0003839|hsa-mir-320b-2&WithFlank&1|-|224257034|224257116@46.66.21;MI0000542|hsa-mir-320a&WithFlank&8|-|22244960|22245043@47.67.21&[MIMAT0005792&hsa-miR-320b&offsets|-1|-2;m-183&1|+|116671787|116671806&offsets|-1|0];[MIMAT0005792_1&hsa-miR-320b&offsets|-1|-2;m-183&1|-|224257051|224257070&offsets|-1|0];[MIMAT0000510&hsa-miR-320a-3p&offsets|-1|-2;m-60&8|-|22244975|22244996&offsets|-1|-2]|GAAAAGCTGGGTTGAGAGGGC</td><td>>MI0003776|hsa-mir-320b-1&WithFlank&1|+|116671740|116671823</td><td>47</td><td>67</td><td>21</td><td>GAAAAGCTGGGTTGAGAGGGC</td><td>113.7</td><td>7.1</td><td>1.9</td><td>3.8</td><td>1.5E-04</td><td>4.8E-04</td><td>BT20/MB468</td><td>MP</td></tr>
<tr><td>>hg38_21_+_8433172_8446622_RNA45SN1_45S_50ntFlanks@475.524.50|CTTCGTGATCGATGTGGTGACGTCGTGCTCTCCCGGGCCGGGTCCGAGCC</td><td>>hg38_21_+_8433172_8446622_RNA45SN1_45S_50ntFlanks</td><td>475</td><td>524</td><td>50</td><td>CTTCGTGATCGATGTGGTGACGTCGTGCTCTCCCGGGCCGGGTCCGAGCC</td><td>323.6</td><td>5.7</td><td>1.6</td><td>3.6</td><td>3.8E-04</td><td>4.8E-04</td><td>MB231/MB468</td><td>MITO</td></tr>
<tr><td>>hg38_21_+_8433172_8446622_RNA45SN1_45S_50ntFlanks@10317.10340.24|GTGCAGATCTTGGTGGTAGTAGCA</td><td>>hg38_21_+_8433172_8446622_RNA45SN1_45S_50ntFlanks</td><td>10317</td><td>10340</td><td>24</td><td>GTGCAGATCTTGGTGGTAGTAGCA</td><td>839.4</td><td>5.8</td><td>1.4</td><td>4.0</td><td>5.4E-05</td><td>4.8E-04</td><td>BT20/MB231</td><td>MP</td></tr>
<tr><td>>hg38_21_+_8433172_8446622_RNA45SN1_45S_50ntFlanks@8371.8391.21|GAGAGAGTTCAAGAGGGCGTG</td><td>>hg38_21_+_8433172_8446622_RNA45SN1_45S_50ntFlanks</td><td>8371</td><td>8391</td><td>21</td><td>GAGAGAGTTCAAGAGGGCGTG</td><td>54.0</td><td>9.1</td><td>2.2</td><td>4.0</td><td>5.4E-05</td><td>4.8E-04</td><td>BT20/MB231</td><td>MP</td></tr>
<tr><td>>hg38_21_+_8433172_8446622_RNA45SN1_45S_50ntFlanks@12408.12441.34|TTTGATCCTTCGATGTCGGCTCTTCCTATCATTG</td><td>>hg38_21_+_8433172_8446622_RNA45SN1_45S_50ntFlanks</td><td>12408</td><td>12441</td><td>34</td><td>TTTGATCCTTCGATGTCGGCTCTTCCTATCATTG</td><td>494.3</td><td>-6.3</td><td>1.8</td><td>-3.5</td><td>3.9E-04</td><td>4.8E-04</td><td>BT20/MB468</td><td>NUC</td></tr>
<tr><td>>am_trnaMT_GluTTC_MT_-_14674_14742@21.54.34,trnalookalike8_GluTTC_5_-_93905172_93905240@21.54.34|ACGATGGTTTTTCATATCATTGGTCGTGGTTGTA</td><td>>am_trnaMT_GluTTC_MT_-_14674_14742</td><td>21</td><td>54</td><td>34</td><td>ACGATGGTTTTTCATATCATTGGTCGTGGTTGTA</td><td>640.3</td><td>7.9</td><td>2.1</td><td>3.8</td><td>1.5E-04</td><td>4.8E-04</td><td>BT20/MB468</td><td>MP</td></tr>
<tr><td>>hg38_21_+_8433172_8446622_RNA45SN1_45S_50ntFlanks@12324.12342.19|TTGGGTTTTAAGCAGGAGG</td><td>>hg38_21_+_8433172_8446622_RNA45SN1_45S_50ntFlanks</td><td>12324</td><td>12342</td><td>19</td><td>TTGGGTTTTAAGCAGGAGG</td><td>397.2</td><td>-5.3</td><td>1.5</td><td>-3.6</td><td>3.4E-04</td><td>4.8E-04</td><td>BT20/MB468</td><td>CYTO</td></tr>
<tr><td>>trnaMT_GluTTC_MT_-_14674_14742@26.58.33,trnalookalike8_GluTTC_5_-_93905172_93905240@26.58.33|GGTTTTTCATATCATTGGTCGTGGTTGTAGTCC</td><td>>trnaMT_GluTTC_MT_-_14674_14742</td><td>26</td><td>58</td><td>33</td><td>GGTTTTTCATATCATTGGTCGTGGTTGTAGTCC</td><td>359.6</td><td>3.0</td><td>0.7</td><td>4.0</td><td>6.3E-05</td><td>4.8E-04</td><td>BT20/MB231</td><td>MITO</td></tr>
<tr><td>>hg38_21_+_8433172_8446622_RNA45SN1_45S_50ntFlanks@10392.10435.44|ACAGCAGTTGAACATGGGTCAGTCGGTCCTGAGAGATGGGCGAG</td><td>>hg38_21_+_8433172_8446622_RNA45SN1_45S_50ntFlanks</td><td>10392</td><td>10435</td><td>44</td><td>ACAGCAGTTGAACATGGGTCAGTCGGTCCTGAGAGATGGGCGAG</td><td>239.4</td><td>-6.6</td><td>1.7</td><td>-3.8</td><td>1.5E-04</td><td>4.8E-04</td><td>BT20/MB468</td><td>MP</td></tr>
<tr><td>>hg38_21_+_8433172_8446622_RNA45SN1_45S_50ntFlanks@12407.12442.36|TTTTGATCCTTCGATGTCGGCTCTTCCTATCATTGT</td><td>>hg38_21_+_8433172_8446622_RNA45SN1_45S_50ntFlanks</td><td>12407</td><td>12442</td><td>36</td><td>TTTTGATCCTTCGATGTCGGCTCTTCCTATCATTGT</td><td>582.9</td><td>-5.3</td><td>1.5</td><td>-3.5</td><td>3.9E-04</td><td>4.8E-04</td><td>BT20/MB468</td><td>NUC</td></tr>
<tr><td>>hg38_21_+_8433172_8446622_RNA45SN1_45S_50ntFlanks@6646.6681.36|TCGTACGACTCTTAGCGGTGGATCACTCGGCTCGTG</td><td>>hg38_21_+_8433172_8446622_RNA45SN1_45S_50ntFlanks</td><td>6646</td><td>6681</td><td>36</td><td>TCGTACGACTCTTAGCGGTGGATCACTCGGCTCGTG</td><td>109098.1</td><td>-3.5</td><td>1.0</td><td>-3.6</td><td>3.8E-04</td><td>4.8E-04</td><td>MB231/MB468</td><td>MITO</td></tr>
<tr><td>>MI0000289|hsa-mir-181a-1&WithFlank&1|-|198859038|198859159@30.51.22;MI0000269|hsa-mir-181a-2&WithFlank&9|+|124692436|124692557@45.66.22&[MIMAT0000256&hsa-miR-181a-5p&offsets|0|-1;m-32&1|-|198859108|198859130&offsets|0|-1];[MIMAT0000256_1&hsa-miR-181a-5p&offsets|0|-1;m-32&9|+|124692480|124692502&offsets|0|-1]|AACATTCAACGCTGTCGGTGAG</td><td>>MI0000289|hsa-mir-181a-1&WithFlank&1|-|198859038|198859159</td><td>30</td><td>51</td><td>22</td><td>AACATTCAACGCTGTCGGTGAG</td><td>694.0</td><td>3.1</td><td>0.9</td><td>3.6</td><td>3.2E-04</td><td>4.8E-04</td><td>MB231/MB468</td><td>MP</td></tr>
<tr><td>>hg38_21_+_8433172_8446622_RNA45SN1_45S_50ntFlanks@9531.9572.42|CAGAGGAAACTCTGGTGGAGGTCCGTAGCGGTCCTGACGTGC</td><td>>hg38_21_+_8433172_8446622_RNA45SN1_45S_50ntFlanks</td><td>9531</td><td>9572</td><td>42</td><td>CAGAGGAAACTCTGGTGGAGGTCCGTAGCGGTCCTGACGTGC</td><td>559.3</td><td>3.8</td><td>0.9</td><td>4.0</td><td>6.3E-05</td><td>4.8E-04</td><td>BT20/MB231</td><td>MITO</td></tr>
<tr><td>>hg38_21_+_8433172_8446622_RNA45SN1_45S_50ntFlanks@10587.10607.21|CCCCGGGGAGAGTTCTCTTTT</td><td>>hg38_21_+_8433172_8446622_RNA45SN1_45S_50ntFlanks</td><td>10587</td><td>10607</td><td>21</td><td>CCCCGGGGAGAGTTCTCTTTT</td><td>262.2</td><td>-5.2</td><td>1.4</td><td>-3.6</td><td>3.4E-04</td><td>4.8E-04</td><td>BT20/MB468</td><td>CYTO</td></tr>
<tr><td>>hg38_21_+_8433172_8446622_RNA45SN1_45S_50ntFlanks@9955.9981.27|GGAAGTCGGAATCCGCTAAGGAGTGTG</td><td>>hg38_21_+_8433172_8446622_RNA45SN1_45S_50ntFlanks</td><td>9955</td><td>9981</td><td>27</td><td>GGAAGTCGGAATCCGCTAAGGAGTGTG</td><td>98.1</td><td>7.0</td><td>1.9</td><td>3.8</td><td>1.5E-04</td><td>4.8E-04</td><td>BT20/MB468</td><td>MP</td></tr>
<tr><td>>hg38_21_+_8433172_8446622_RNA45SN1_45S_50ntFlanks@9484.9513.30|GTGTTAGGACCCGAAAGATGGTGAACTATG</td><td>>hg38_21_+_8433172_8446622_RNA45SN1_45S_50ntFlanks</td><td>9484</td><td>9513</td><td>30</td><td>GTGTTAGGACCCGAAAGATGGTGAACTATG</td><td>59.2</td><td>9.2</td><td>2.3</td><td>4.0</td><td>5.4E-05</td><td>4.8E-04</td><td>BT20/MB231</td><td>MP</td></tr>
<tr><td>>hg38_21_+_8433172_8446622_RNA45SN1_45S_50ntFlanks@10350.10367.18|ACGAGAACTTTGAAGGCC</td><td>>hg38_21_+_8433172_8446622_RNA45SN1_45S_50ntFlanks</td><td>10350</td><td>10367</td><td>18</td><td>ACGAGAACTTTGAAGGCC</td><td>1282.5</td><td>-3.9</td><td>1.0</td><td>-3.8</td><td>1.5E-04</td><td>4.8E-04</td><td>BT20/MB468</td><td>MP</td></tr>
<tr><td>>hg38_21_+_8433172_8446622_RNA45SN1_45S_50ntFlanks@12405.12438.34|CTTTTTGATCCTTCGATGTCGGCTCTTCCTATCA</td><td>>hg38_21_+_8433172_8446622_RNA45SN1_45S_50ntFlanks</td><td>12405</td><td>12438</td><td>34</td><td>CTTTTTGATCCTTCGATGTCGGCTCTTCCTATCA</td><td>180.0</td><td>-2.3</td><td>0.6</td><td>-3.6</td><td>3.4E-04</td><td>4.8E-04</td><td>BT20/MB468</td><td>CYTO</td></tr>
<tr><td>>hg38_21_+_8433172_8446622_RNA45SN1_45S_50ntFlanks@6719.6745.27|TGAATTGCAGGACACATTGATCATCGA</td><td>>hg38_21_+_8433172_8446622_RNA45SN1_45S_50ntFlanks</td><td>6719</td><td>6745</td><td>27</td><td>TGAATTGCAGGACACATTGATCATCGA</td><td>262.5</td><td>-6.0</td><td>1.6</td><td>-3.8</td><td>1.5E-04</td><td>4.8E-04</td><td>BT20/MB468</td><td>MP</td></tr>
<tr><td>>hg38_21_+_8433172_8446622_RNA45SN1_45S_50ntFlanks@7988.8014.27|CAGACGTGGCGACCCGCTGAATTTAAG</td><td>>hg38_21_+_8433172_8446622_RNA45SN1_45S_50ntFlanks</td><td>7988</td><td>8014</td><td>27</td><td>CAGACGTGGCGACCCGCTGAATTTAAG</td><td>531.9</td><td>-5.2</td><td>1.5</td><td>-3.5</td><td>3.9E-04</td><td>4.8E-04</td><td>BT20/MB468</td><td>NUC</td></tr>
<tr><td>>hg38_21_+_8433172_8446622_RNA45SN1_45S_50ntFlanks@4000.4024.25|TAGATAACCTCGGGCCGATCGCACG</td><td>>hg38_21_+_8433172_8446622_RNA45SN1_45S_50ntFlanks</td><td>4000</td><td>4024</td><td>25</td><td>TAGATAACCTCGGGCCGATCGCACG</td><td>300.8</td><td>-5.1</td><td>1.4</td><td>-3.6</td><td>2.9E-04</td><td>4.8E-04</td><td>BT20/MB468</td><td>MITO</td></tr>
<tr><td>>hg38_21_+_8433172_8446622_RNA45SN1_45S_50ntFlanks@11576.11613.38|CTTAGAACTGGTGCGGACCAGGGGAATCCGACTGTTTA</td><td>>hg38_21_+_8433172_8446622_RNA45SN1_45S_50ntFlanks</td><td>11576</td><td>11613</td><td>38</td><td>CTTAGAACTGGTGCGGACCAGGGGAATCCGACTGTTTA</td><td>1037.1</td><td>3.4</td><td>0.8</td><td>4.0</td><td>6.4E-05</td><td>4.8E-04</td><td>BT20/MB231</td><td>MITO</td></tr>
<tr><td>>hg38_21_+_8433172_8446622_RNA45SN1_45S_50ntFlanks@7973.7991.19|GACGCGACCTCAGATCAGA</td><td>>hg38_21_+_8433172_8446622_RNA45SN1_45S_50ntFlanks</td><td>7973</td><td>7991</td><td>19</td><td>GACGCGACCTCAGATCAGA</td><td>1006.8</td><td>2.6</td><td>0.7</td><td>3.6</td><td>3.8E-04</td><td>4.8E-04</td><td>MB231/MB468</td><td>NUC</td></tr>
<tr><td>>hg38_21_+_8433172_8446622_RNA45SN1_45S_50ntFlanks@4241.4264.24|CTCTTTCGAGGCCCTGTAATTGGA</td><td>>hg38_21_+_8433172_8446622_RNA45SN1_45S_50ntFlanks</td><td>4241</td><td>4264</td><td>24</td><td>CTCTTTCGAGGCCCTGTAATTGGA</td><td>102.1</td><td>-4.6</td><td>1.3</td><td>-3.6</td><td>3.4E-04</td><td>4.8E-04</td><td>BT20/MB468</td><td>CYTO</td></tr>
<tr><td>>hg38_21_+_8433172_8446622_RNA45SN1_45S_50ntFlanks@12029.12059.31|TCTGATCGTTTTTTCACTGACCCGGTGAGGC</td><td>>hg38_21_+_8433172_8446622_RNA45SN1_45S_50ntFlanks</td><td>12029</td><td>12059</td><td>31</td><td>TCTGATCGTTTTTTCACTGACCCGGTGAGGC</td><td>172.2</td><td>10.8</td><td>2.7</td><td>4.0</td><td>5.4E-05</td><td>4.8E-04</td><td>BT20/MB231</td><td>MP</td></tr>
<tr><td>>MI0001445|hsa-mir-423&WithFlank&17|+|30117073|30117178@59.80.22(+1A)&[MIMAT0001340&hsa-miR-423-3p&offsets|0|-1(+1A);m-100&17|+|30117131|30117153&offsets|0|-1(+1A)]|AGCTCGGTCTGAGGCCCCTCAGA</td><td>>MI0001445|hsa-mir-423&WithFlank&17|+|30117073|30117178</td><td>59</td><td>80</td><td>22(+1A)</td><td>AGCTCGGTCTGAGGCCCCTCAGA</td><td>326.1</td><td>3.1</td><td>0.9</td><td>3.6</td><td>3.2E-04</td><td>4.8E-04</td><td>MB231/MB468</td><td>MP</td></tr>
<tr><td>>am_trna10_ValCAC_5_-_180649395_180649467@1.35.35,trna18_ValCAC_5_-_180529253_180529325@1.35.35,trna2_ValCAC_5_+_180524070_180524142@1.35.35,trna6_ValCAC_5_+_180600650_180600722@1.35.35,trna85_ValCAC_1_-_161369490_161369562@1.35.35,trna90_ValCAC_1_-_149684088_149684161@1.35.35,trna98_ValCAC_1_-_149298555_149298627@1.35.35,trna9_ValCAC_6_+_26538282_26538354@1.35.35|GTTTCCGTAGTGTAGTGGTTATCACGTTCGCCTCA</td><td>>am_trna10_ValCAC_5_-_180649395_180649467</td><td>1</td><td>35</td><td>35</td><td>GTTTCCGTAGTGTAGTGGTTATCACGTTCGCCTCA</td><td>355.7</td><td>8.9</td><td>2.5</td><td>3.6</td><td>3.6E-04</td><td>4.8E-04</td><td>MB231/MB468</td><td>CYTO</td></tr>
<tr><td>>am_MI0003581|hsa-mir-574&WithFlank&4|+|38868026|38868133@31.52.22(+1C)&[MIMAT0004795&hsa-miR-574-5p&offsets|0|-1(+1C);m-213&4|+|38868056|38868078&offsets|0|-1(+1C)]|TGAGTGTGTGTGTGTGAGTGTGC</td><td>>am_MI0003581|hsa-mir-574&WithFlank&4|+|38868026|38868133</td><td>31</td><td>52</td><td>22(+1C)</td><td>TGAGTGTGTGTGTGTGAGTGTGC</td><td>257.8</td><td>4.4</td><td>1.2</td><td>3.6</td><td>3.2E-04</td><td>4.8E-04</td><td>MB231/MB468</td><td>MP</td></tr>
<tr><td>>hg38_21_+_8433172_8446622_RNA45SN1_45S_50ntFlanks@6652.6692.41|GACTCTTAGCGGTGGATCACTCGGCTCGTGCGTCGATGAAG</td><td>>hg38_21_+_8433172_8446622_RNA45SN1_45S_50ntFlanks</td><td>6652</td><td>6692</td><td>41</td><td>GACTCTTAGCGGTGGATCACTCGGCTCGTGCGTCGATGAAG</td><td>1443.6</td><td>-1.7</td><td>0.5</td><td>-3.5</td><td>3.9E-04</td><td>4.8E-04</td><td>BT20/MB468</td><td>NUC</td></tr>
<tr><td>>hg38_21_+_8433172_8446622_RNA45SN1_45S_50ntFlanks@11872.11893.22|AAAGAAGACCCTGTTGAGCTTG</td><td>>hg38_21_+_8433172_8446622_RNA45SN1_45S_50ntFlanks</td><td>11872</td><td>11893</td><td>22</td><td>AAAGAAGACCCTGTTGAGCTTG</td><td>248.6</td><td>-4.1</td><td>1.2</td><td>-3.5</td><td>3.9E-04</td><td>4.8E-04</td><td>MB231/MB468</td><td>NUC</td></tr>
<tr><td>>hg38_21_+_8433172_8446622_RNA45SN1_45S_50ntFlanks@10685.10710.26|GGCGGCGTCCGGTGAGCTCTCGCTGG</td><td>>hg38_21_+_8433172_8446622_RNA45SN1_45S_50ntFlanks</td><td>10685</td><td>10710</td><td>26</td><td>GGCGGCGTCCGGTGAGCTCTCGCTGG</td><td>289.0</td><td>-3.4</td><td>0.9</td><td>-3.6</td><td>3.2E-04</td><td>4.8E-04</td><td>MB231/MB468</td><td>MP</td></tr>
<tr><td>>hg38_21_+_8433172_8446622_RNA45SN1_45S_50ntFlanks@10391.10427.37|AACAGCAGTTGAACATGGGTCAGTCGGTCCTGAGAGA</td><td>>hg38_21_+_8433172_8446622_RNA45SN1_45S_50ntFlanks</td><td>10391</td><td>10427</td><td>37</td><td>AACAGCAGTTGAACATGGGTCAGTCGGTCCTGAGAGA</td><td>70.9</td><td>-5.8</td><td>1.6</td><td>-3.6</td><td>3.2E-04</td><td>4.8E-04</td><td>MB231/MB468</td><td>MP</td></tr>
<tr><td>>hg38_21_+_8433172_8446622_RNA45SN1_45S_50ntFlanks@10409.10433.25|GTCAGTCGGTCCTGAGAGATGGGCG</td><td>>hg38_21_+_8433172_8446622_RNA45SN1_45S_50ntFlanks</td><td>10409</td><td>10433</td><td>25</td><td>GTCAGTCGGTCCTGAGAGATGGGCG</td><td>171.9</td><td>-3.8</td><td>1.1</td><td>-3.5</td><td>3.9E-04</td><td>4.8E-04</td><td>MB231/MB468</td><td>NUC</td></tr>
<tr><td>>hg38_MT_+_1621_3279_RNR2_16S_50ntFlanks@115.155.41|AAATAAAGTATAGGCGATAGAAATTGAAACCTGGCGCAATA</td><td>>hg38_MT_+_1621_3279_RNR2_16S_50ntFlanks</td><td>115</td><td>155</td><td>41</td><td>AAATAAAGTATAGGCGATAGAAATTGAAACCTGGCGCAATA</td><td>78.8</td><td>10.0</td><td>2.7</td><td>3.8</td><td>1.5E-04</td><td>4.9E-04</td><td>BT20/MB468</td><td>MP</td></tr>
</table>

<table>
<tr><td>>hg38_21_+_8433172_8446622_RNA45SN1_45S_50ntFlanks@12933.12960.28|GCTAAACCATTCGTAGACGACCTGCTTC</td><td>>hg38_21_+_8433172_8446622_RNA45SN1_45S_50ntFlanks</td><td>12933</td><td>12960</td><td>28</td><td>GCTAAACCATTCGTAGACGACCTGCTTC</td><td>261.0</td><td>2.5</td><td>0.7</td><td>3.6</td><td>3.6E-04</td><td>4.9E-04</td><td>MB231/MB468</td><td>CYTO</td></tr>
<tr><td>>MI0000086|hsa-mir-28&WithFlank&3|+|188688775|188688872@60.81.22(+2A)&[MIMAT0004502&hsa-miR-28-3p&offsets|0|0(+2A);m-28&3|+|188688834|188688855&offsets|0|0(+2A)]|CACTAGATTGTGAGCTCCTGGAAA</td><td>>MI0000086|hsa-mir-28&WithFlank&3|+|188688775|188688872</td><td>60</td><td>81</td><td>22(+2A)</td><td>CACTAGATTGTGAGCTCCTGGAAA</td><td>65.8</td><td>6.2</td><td>1.7</td><td>3.6</td><td>3.2E-04</td><td>4.9E-04</td><td>MB231/MB468</td><td>MP</td></tr>
<tr><td>>am_trnaMT_ProTGG_MT_-_15956_16023@53.70.18|AAGACTTTTTCTCTGACC</td><td>>am_trnaMT_ProTGG_MT_-_15956_16023</td><td>53</td><td>70</td><td>18</td><td>AAGACTTTTTCTCTGACC</td><td>472.3</td><td>3.8</td><td>1.1</td><td>3.5</td><td>3.9E-04</td><td>4.9E-04</td><td>MB231/MB468</td><td>NUC</td></tr>
<tr><td>>hg38_21_+_8433172_8446622_RNA45SN1_45S_50ntFlanks@6663.6686.24|GTGGATCACTCGGCTCGTGCGTCG</td><td>>hg38_21_+_8433172_8446622_RNA45SN1_45S_50ntFlanks</td><td>6663</td><td>6686</td><td>24</td><td>GTGGATCACTCGGCTCGTGCGTCG</td><td>190.0</td><td>-3.1</td><td>0.9</td><td>-3.6</td><td>3.6E-04</td><td>4.9E-04</td><td>MB231/MB468</td><td>CYTO</td></tr>
<tr><td>>hg38_21_+_8433172_8446622_RNA45SN1_45S_50ntFlanks@5490.5516.27|TGAGAAGACGGTCGAACTTGACTATCT</td><td>>hg38_21_+_8433172_8446622_RNA45SN1_45S_50ntFlanks</td><td>5490</td><td>5516</td><td>27</td><td>TGAGAAGACGGTCGAACTTGACTATCT</td><td>171.1</td><td>4.6</td><td>1.3</td><td>3.6</td><td>3.8E-04</td><td>4.9E-04</td><td>MB231/MB468</td><td>MITO</td></tr>
<tr><td>>hg38_21_+_8433172_8446622_RNA45SN1_45S_50ntFlanks@10921.10941.21|GCTGGACGAGGCGCCGCCGCC</td><td>>hg38_21_+_8433172_8446622_RNA45SN1_45S_50ntFlanks</td><td>10921</td><td>10941</td><td>21</td><td>GCTGGACGAGGCGCCGCCGCC</td><td>210.9</td><td>-5.5</td><td>1.5</td><td>-3.5</td><td>3.9E-04</td><td>4.9E-04</td><td>MB231/MB468</td><td>NUC</td></tr>
<tr><td>>hg38_21_+_8433172_8446622_RNA45SN1_45S_50ntFlanks@6660.6683.24|GCGGTGGATCACTCGGCTCGTGCG</td><td>>hg38_21_+_8433172_8446622_RNA45SN1_45S_50ntFlanks</td><td>6660</td><td>6683</td><td>24</td><td>GCGGTGGATCACTCGGCTCGTGCG</td><td>884.0</td><td>-2.7</td><td>0.7</td><td>-4.0</td><td>5.6E-05</td><td>4.9E-04</td><td>BT20/MB231</td><td>NUC</td></tr>
<tr><td>>hg38_1_-_228634819_228635039_RNA5S12_5S_50ntFlanks@151.171.21|ATACCGGGTGCTGTAGGCTTT</td><td>>hg38_1_-_228634819_228635039_RNA5S12_5S_50ntFlanks</td><td>151</td><td>171</td><td>21</td><td>ATACCGGGTGCTGTAGGCTTT</td><td>2298.6</td><td>-2.5</td><td>0.7</td><td>-3.6</td><td>3.8E-04</td><td>4.9E-04</td><td>MB231/MB468</td><td>MITO</td></tr>
<tr><td>>hg38_21_+_8433172_8446622_RNA45SN1_45S_50ntFlanks@10681.10711.31|TTCCGGCGGCGTCCGGTGAGCTCTCGCTGGC</td><td>>hg38_21_+_8433172_8446622_RNA45SN1_45S_50ntFlanks</td><td>10681</td><td>10711</td><td>31</td><td>TTCCGGCGGCGTCCGGTGAGCTCTCGCTGGC</td><td>219.7</td><td>-5.7</td><td>1.5</td><td>-3.8</td><td>1.5E-04</td><td>4.9E-04</td><td>BT20/MB468</td><td>MP</td></tr>
<tr><td>>hg38_21_+_8433172_8446622_RNA45SN1_45S_50ntFlanks@11579.11612.34|AGAACTGGTGCGGACCAGGGGAATCCGACTGTTT</td><td>>hg38_21_+_8433172_8446622_RNA45SN1_45S_50ntFlanks</td><td>11579</td><td>11612</td><td>34</td><td>AGAACTGGTGCGGACCAGGGGAATCCGACTGTTT</td><td>114.0</td><td>-5.7</td><td>1.6</td><td>-3.6</td><td>3.2E-04</td><td>4.9E-04</td><td>MB231/MB468</td><td>MP</td></tr>
<tr><td>>hg38_21_+_8433172_8446622_RNA45SN1_45S_50ntFlanks@3878.3898.21|CATGCCGACGGGCGCTGACCC</td><td>>hg38_21_+_8433172_8446622_RNA45SN1_45S_50ntFlanks</td><td>3878</td><td>3898</td><td>21</td><td>CATGCCGACGGGCGCTGACCC</td><td>699.0</td><td>-6.6</td><td>1.9</td><td>-3.5</td><td>3.9E-04</td><td>4.9E-04</td><td>BT20/MB468</td><td>NUC</td></tr>
<tr><td>>hg38_21_+_8433172_8446622_RNA45SN1_45S_50ntFlanks@10843.10861.19|TAACTTCGGGATAAGGATT</td><td>>hg38_21_+_8433172_8446622_RNA45SN1_45S_50ntFlanks</td><td>10843</td><td>10861</td><td>19</td><td>TAACTTCGGGATAAGGATT</td><td>403.9</td><td>-3.5</td><td>1.0</td><td>-3.6</td><td>3.6E-04</td><td>4.9E-04</td><td>MB231/MB468</td><td>CYTO</td></tr>
<tr><td>>hg38_21_+_8433172_8446622_RNA45SN1_45S_50ntFlanks@10849.10867.19|CGGGATAAGGATTGGCTCT</td><td>>hg38_21_+_8433172_8446622_RNA45SN1_45S_50ntFlanks</td><td>10849</td><td>10867</td><td>19</td><td>CGGGATAAGGATTGGCTCT</td><td>86.1</td><td>-5.5</td><td>1.5</td><td>-3.6</td><td>3.2E-04</td><td>4.9E-04</td><td>MB231/MB468</td><td>MP</td></tr>
<tr><td>>MI0000542|hsa-mir-320a&WithFlank&8|-|22244960|22245043@48.68.21(+1C)&[MIMAT0000510&hsa-miR-320a-3p&offsets|0|-1(+1C);m-60&8|-|22244975|22244996&offsets|0|-1(+1C)]|AAAAGCTGGGTTGAGAGGGCGC</td><td>>MI0000542|hsa-mir-320a&WithFlank&8|-|22244960|22245043</td><td>48</td><td>68</td><td>21(+1C)</td><td>AAAAGCTGGGTTGAGAGGGCGC</td><td>634.9</td><td>4.9</td><td>1.3</td><td>3.6</td><td>3.2E-04</td><td>4.9E-04</td><td>MB231/MB468</td><td>MP</td></tr>
<tr><td>>hg38_21_+_8433172_8446622_RNA45SN1_45S_50ntFlanks@6645.6673.29|CTCGTACGACTCTTAGCGGTGGATCACTC</td><td>>hg38_21_+_8433172_8446622_RNA45SN1_45S_50ntFlanks</td><td>6645</td><td>6673</td><td>29</td><td>CTCGTACGACTCTTAGCGGTGGATCACTC</td><td>250.2</td><td>5.7</td><td>1.6</td><td>3.6</td><td>3.5E-04</td><td>4.9E-04</td><td>BT20/MB468</td><td>CYTO</td></tr>
<tr><td>>hg38_21_+_8433172_8446622_RNA45SN1_45S_50ntFlanks@6661.6701.41|CGGTGGATCACTCGGCTCGTGCGTCGATGAAGAACGCAGCT</td><td>>hg38_21_+_8433172_8446622_RNA45SN1_45S_50ntFlanks</td><td>6661</td><td>6701</td><td>41</td><td>CGGTGGATCACTCGGCTCGTGCGTCGATGAAGAACGCAGCT</td><td>820.5</td><td>2.9</td><td>0.8</td><td>3.5</td><td>3.9E-04</td><td>4.9E-04</td><td>MB231/MB468</td><td>MITO</td></tr>
<tr><td>>hg38_MT_+_1621_3279_RNR2_16S_50ntFlanks@1347.1388.42|CAACAATAGGGTTTACGACCTCGATGTTGGATCAGGACATCC</td><td>>hg38_MT_+_1621_3279_RNR2_16S_50ntFlanks</td><td>1347</td><td>1388</td><td>42</td><td>CAACAATAGGGTTTACGACCTCGATGTTGGATCAGGACATCC</td><td>78.8</td><td>10.0</td><td>2.7</td><td>3.8</td><td>1.6E-04</td><td>4.9E-04</td><td>BT20/MB468</td><td>MP</td></tr>
<tr><td>>hg38_MT_+_1621_3279_RNR2_16S_50ntFlanks@1426.1451.26|CGATTAAAGTCCTACGTGATCTGAGT</td><td>>hg38_MT_+_1621_3279_RNR2_16S_50ntFlanks</td><td>1426</td><td>1451</td><td>26</td><td>CGATTAAAGTCCTACGTGATCTGAGT</td><td>75.9</td><td>3.0</td><td>0.8</td><td>3.6</td><td>3.3E-04</td><td>4.9E-04</td><td>MB231/MB468</td><td>MP</td></tr>
<tr><td>>trna10_LysCTT_16_+_3241501_3241573@3.36.34,trna13_LysCTT_14_-_58706613_58706685@3.36.34,trna2_LysCTT_15_+_79152904_79152976@3.36.34|CCGGCTAGCTCAGTCGGTAGAGCATGGGACTCTT</td><td>>trna10_LysCTT_16_+_3241501_3241573</td><td>3</td><td>36</td><td>34</td><td>CCGGCTAGCTCAGTCGGTAGAGCATGGGACTCTT</td><td>281.5</td><td>4.7</td><td>1.3</td><td>3.6</td><td>3.3E-04</td><td>4.9E-04</td><td>MB231/MB468</td><td>MP</td></tr>
<tr><td>>hg38_21_+_8433172_8446622_RNA45SN1_45S_50ntFlanks@6733.6770.38|CATTGATCATCGACACTTCGAACGCACTTGCGGCCCCG</td><td>>hg38_21_+_8433172_8446622_RNA45SN1_45S_50ntFlanks</td><td>6733</td><td>6770</td><td>38</td><td>CATTGATCATCGACACTTCGAACGCACTTGCGGCCCCG</td><td>1103.9</td><td>-3.9</td><td>1.1</td><td>-3.6</td><td>3.7E-04</td><td>4.9E-04</td><td>MB231/MB468</td><td>CYTO</td></tr>
<tr><td>>MI0000440|hsa-mir-27b&WithFlank&9|+|95085439|95085547@67.86.20(+1A)&[MIMAT0000419&hsa-miR-27b-3p&offsets|0|-1(+1A);m-39&9|+|95085505|95085525&offsets|0|-1(+1A)]|TTCACAGTGGCTAAGTTCTGA</td><td>>MI0000440|hsa-mir-27b&WithFlank&9|+|95085439|95085547</td><td>67</td><td>86</td><td>20(+1A)</td><td>TTCACAGTGGCTAAGTTCTGA</td><td>1315.1</td><td>3.8</td><td>1.0</td><td>3.8</td><td>1.6E-04</td><td>4.9E-04</td><td>BT20/MB468</td><td>MP</td></tr>
<tr><td>>hg38_21_+_8433172_8446622_RNA45SN1_45S_50ntFlanks@12995.13013.19|CTCGCTGCGATCTATTGAA</td><td>>hg38_21_+_8433172_8446622_RNA45SN1_45S_50ntFlanks</td><td>12995</td><td>13013</td><td>19</td><td>CTCGCTGCGATCTATTGAA</td><td>492.6</td><td>-2.7</td><td>0.7</td><td>-4.0</td><td>5.7E-05</td><td>4.9E-04</td><td>BT20/MB231</td><td>NUC</td></tr>
<tr><td>>hg38_1_-_228634819_228635039_RNA5S12_5S_50ntFlanks@51.89.39|GTCTACGGCCATACCACCCTGAACGCGCCCGATCTCGTC</td><td>>hg38_1_-_228634819_228635039_RNA5S12_5S_50ntFlanks</td><td>51</td><td>89</td><td>39</td><td>GTCTACGGCCATACCACCCTGAACGCGCCCGATCTCGTC</td><td>3284.9</td><td>3.7</td><td>1.0</td><td>3.6</td><td>3.3E-04</td><td>4.9E-04</td><td>MB231/MB468</td><td>MP</td></tr>
<tr><td>>MI0000283|hsa-mir-203a&WithFlank&14|+|104117399|104117520@72.93.22&[MIMAT0000264&hsa-miR-203a-3p&offsets|+1|+1;m-79&14|+|104117469|104117490&offsets|+1|+1]|TGAAATGTTTAGGACCACTAGA</td><td>>MI0000283|hsa-mir-203a&WithFlank&14|+|104117399|104117520</td><td>72</td><td>93</td><td>22</td><td>TGAAATGTTTAGGACCACTAGA</td><td>171.6</td><td>6.2</td><td>1.6</td><td>3.8</td><td>1.6E-04</td><td>4.9E-04</td><td>BT20/MB468</td><td>MP</td></tr>
<tr><td>>hg38_21_+_8433172_8446622_RNA45SN1_45S_50ntFlanks@10468.10497.30|TTGCCCTCGGCCGATCGAAAGGGAGTCGGG</td><td>>hg38_21_+_8433172_8446622_RNA45SN1_45S_50ntFlanks</td><td>10468</td><td>10497</td><td>30</td><td>TTGCCCTCGGCCGATCGAAAGGGAGTCGGG</td><td>250.1</td><td>-5.8</td><td>1.5</td><td>-3.8</td><td>1.6E-04</td><td>4.9E-04</td><td>BT20/MB468</td><td>MP</td></tr>
<tr><td>>hg38_21_+_8433172_8446622_RNA45SN1_45S_50ntFlanks@4287.4316.30|AACGAGGATCCATTGGAGGGCAAGTCTGGT</td><td>>hg38_21_+_8433172_8446622_RNA45SN1_45S_50ntFlanks</td><td>4287</td><td>4316</td><td>30</td><td>AACGAGGATCCATTGGAGGGCAAGTCTGGT</td><td>542.8</td><td>7.8</td><td>1.9</td><td>4.0</td><td>5.5E-05</td><td>4.9E-04</td><td>BT20/MB231</td><td>MP</td></tr>
<tr><td>>hg38_21_+_8433172_8446622_RNA45SN1_45S_50ntFlanks@11027.11054.28|CTCTCTCTCTCTCTCCCCCGCTCCCCGT</td><td>>hg38_21_+_8433172_8446622_RNA45SN1_45S_50ntFlanks</td><td>11027</td><td>11054</td><td>28</td><td>CTCTCTCTCTCTCTCCCCCGCTCCCCGT</td><td>225.4</td><td>-3.2</td><td>0.9</td><td>-3.6</td><td>3.7E-04</td><td>4.9E-04</td><td>MB231/MB468</td><td>CYTO</td></tr>
<tr><td>>hg38_21_+_8433172_8446622_RNA45SN1_45S_50ntFlanks@9939.9968.30|GCAGGACGGTGGCCATGGAAGTCGGAATCC</td><td>>hg38_21_+_8433172_8446622_RNA45SN1_45S_50ntFlanks</td><td>9939</td><td>9968</td><td>30</td><td>GCAGGACGGTGGCCATGGAAGTCGGAATCC</td><td>1401.4</td><td>8.2</td><td>2.0</td><td>4.0</td><td>5.6E-05</td><td>4.9E-04</td><td>BT20/MB231</td><td>MP</td></tr>
<tr><td>>trnaMT_GluTTC_MT_-_14674_14742@40.65.26,trnalookalike8_GluTTC_5_-_93905172_93905240@40.65.26|TTGGTCGTGGTTGTAGTCCGTGCGAG</td><td>>trnaMT_GluTTC_MT_-_14674_14742</td><td>40</td><td>65</td><td>26</td><td>TTGGTCGTGGTTGTAGTCCGTGCGAG</td><td>411.0</td><td>4.3</td><td>1.1</td><td>3.8</td><td>1.6E-04</td><td>4.9E-04</td><td>BT20/MB468</td><td>MP</td></tr>
<tr><td>>hg38_21_+_8433172_8446622_RNA45SN1_45S_50ntFlanks@10348.10384.37|AAACGAGAACTTTGAAGGCCGAAGTGGAGAAGGGTTC</td><td>>hg38_21_+_8433172_8446622_RNA45SN1_45S_50ntFlanks</td><td>10348</td><td>10384</td><td>37</td><td>AAACGAGAACTTTGAAGGCCGAAGTGGAGAAGGGTTC</td><td>397.4</td><td>-2.8</td><td>0.8</td><td>-3.5</td><td>3.9E-04</td><td>4.9E-04</td><td>MB231/MB468</td><td>NUC</td></tr>
<tr><td>>hg38_21_+_8433172_8446622_RNA45SN1_45S_50ntFlanks@6650.6688.39|ACGACTCTTAGCGGTGGATCACTCGGCTCGTGCGTCGAT</td><td>>hg38_21_+_8433172_8446622_RNA45SN1_45S_50ntFlanks</td><td>6650</td><td>6688</td><td>39</td><td>ACGACTCTTAGCGGTGGATCACTCGGCTCGTGCGTCGAT</td><td>532.7</td><td>2.8</td><td>0.8</td><td>3.6</td><td>2.9E-04</td><td>4.9E-04</td><td>BT20/MB468</td><td>MITO</td></tr>
<tr><td>>hg38_21_+_8433172_8446622_RNA45SN1_45S_50ntFlanks@7985.8018.34|GATCAGACGTGGCGACCCGCTGAATTTAAGCATA</td><td>>hg38_21_+_8433172_8446622_RNA45SN1_45S_50ntFlanks</td><td>7985</td><td>8018</td><td>34</td><td>GATCAGACGTGGCGACCCGCTGAATTTAAGCATA</td><td>142.9</td><td>-5.8</td><td>1.6</td><td>-3.6</td><td>3.3E-04</td><td>4.9E-04</td><td>MB231/MB468</td><td>MP</td></tr>
<tr><td>>hg38_21_+_8433172_8446622_RNA45SN1_45S_50ntFlanks@6762.6806.45|GCGGCCCCGGGTTCCTCCCGGGGCTACGCCTGTCTGAGCGTCGCT</td><td>>hg38_21_+_8433172_8446622_RNA45SN1_45S_50ntFlanks</td><td>6762</td><td>6806</td><td>45</td><td>GCGGCCCCGGGTTCCTCCCGGGGCTACGCCTGTCTGAGCGTCGCT</td><td>212.4</td><td>5.2</td><td>1.5</td><td>3.5</td><td>3.9E-04</td><td>4.9E-04</td><td>MB231/MB468</td><td>MITO</td></tr>
<tr><td>>hg38_MT_+_1621_3279_RNR2_16S_50ntFlanks@1427.1458.32|GATTAAAGTCCTACGTGATCTGAGTTCAGACC</td><td>>hg38_MT_+_1621_3279_RNR2_16S_50ntFlanks</td><td>1427</td><td>1458</td><td>32</td><td>GATTAAAGTCCTACGTGATCTGAGTTCAGACC</td><td>327.2</td><td>4.3</td><td>1.2</td><td>3.5</td><td>3.9E-04</td><td>4.9E-04</td><td>MB231/MB468</td><td>MITO</td></tr>
<tr><td>>trnaMT_SerGCT_MT_+_12207_12265@14.47.34|AGAACTGCTAACTCATGCCCCCATGTCTAACAAC</td><td>>trnaMT_SerGCT_MT_+_12207_12265</td><td>14</td><td>47</td><td>34</td><td>AGAACTGCTAACTCATGCCCCCATGTCTAACAAC</td><td>95.1</td><td>6.1</td><td>1.7</td><td>3.5</td><td>3.9E-04</td><td>4.9E-04</td><td>MB231/MB468</td><td>MITO</td></tr>
<tr><td>>hg38_21_+_8433172_8446622_RNA45SN1_45S_50ntFlanks@10770.10796.27|GCAGGTCTCCAAGGTGAACAGCCTCTG</td><td>>hg38_21_+_8433172_8446622_RNA45SN1_45S_50ntFlanks</td><td>10770</td><td>10796</td><td>27</td><td>GCAGGTCTCCAAGGTGAACAGCCTCTG</td><td>740.6</td><td>-3.0</td><td>0.8</td><td>-3.5</td><td>3.9E-04</td><td>4.9E-04</td><td>MB231/MB468</td><td>MITO</td></tr>
<tr><td>>hg38_21_+_8433172_8446622_RNA45SN1_45S_50ntFlanks@8351.8378.28|AAGTTGAAAAGAACTTTGAAGAGAGAGT</td><td>>hg38_21_+_8433172_8446622_RNA45SN1_45S_50ntFlanks</td><td>8351</td><td>8378</td><td>28</td><td>AAGTTGAAAAGAACTTTGAAGAGAGAGT</td><td>172.2</td><td>4.8</td><td>1.3</td><td>3.8</td><td>1.6E-04</td><td>4.9E-04</td><td>BT20/MB468</td><td>MP</td></tr>
<tr><td>>hg38_21_+_8433172_8446622_RNA45SN1_45S_50ntFlanks@11210.11252.43|GGCGACTCTGGACGCGAGCCGGGCCCTTCCCGTGGATCGCCCC</td><td>>hg38_21_+_8433172_8446622_RNA45SN1_45S_50ntFlanks</td><td>11210</td><td>11252</td><td>43</td><td>GGCGACTCTGGACGCGAGCCGGGCCCTTCCCGTGGATCGCCCC</td><td>487.7</td><td>-4.9</td><td>1.4</td><td>-3.5</td><td>4.0E-04</td><td>4.9E-04</td><td>BT20/MB468</td><td>NUC</td></tr>
<tr><td>>hg38_21_+_8433172_8446622_RNA45SN1_45S_50ntFlanks@12417.12442.26|TCGATGTCGGCTCTTCCTATCATTGT</td><td>>hg38_21_+_8433172_8446622_RNA45SN1_45S_50ntFlanks</td><td>12417</td><td>12442</td><td>26</td><td>TCGATGTCGGCTCTTCCTATCATTGT</td><td>994.5</td><td>2.4</td><td>0.7</td><td>3.6</td><td>3.7E-04</td><td>4.9E-04</td><td>MB231/MB468</td><td>CYTO</td></tr>
<tr><td>>hg38_MT_+_1621_3279_RNR2_16S_50ntFlanks@1348.1389.42|AACAATAGGGTTTACGACCTCGATGTTGGATCAGGACATCCC</td><td>>hg38_MT_+_1621_3279_RNR2_16S_50ntFlanks</td><td>1348</td><td>1389</td><td>42</td><td>AACAATAGGGTTTACGACCTCGATGTTGGATCAGGACATCCC</td><td>78.2</td><td>10.0</td><td>2.7</td><td>3.8</td><td>1.6E-04</td><td>5.0E-04</td><td>BT20/MB468</td><td>MP</td></tr>
<tr><td>>hg38_21_+_8433172_8446622_RNA45SN1_45S_50ntFlanks@10384.10405.22|CCATGTGAACAGCAGTTGAACA</td><td>>hg38_21_+_8433172_8446622_RNA45SN1_45S_50ntFlanks</td><td>10384</td><td>10405</td><td>22</td><td>CCATGTGAACAGCAGTTGAACA</td><td>193.3</td><td>-5.1</td><td>1.4</td><td>-3.6</td><td>3.5E-04</td><td>5.0E-04</td><td>BT20/MB468</td><td>CYTO</td></tr>
<tr><td>>MI0000447|hsa-mir-128-1&WithFlank&2|+|135665391|135665484@56.75.20;MI0000727|hsa-mir-128-2&WithFlank&3|+|35744470|35744565@58.77.20&[MIMAT0000424&hsa-miR-128-3p&offsets|0|-1;m-108&2|+|135665446|135665466&offsets|0|-1];[MIMAT0000424_1&hsa-miR-128-3p&offsets|0|-1;m-117&3|+|35744527|35744547&offsets|0|-1]|TCACAGTGAACCGGTCTCTT</td><td>>MI0000447|hsa-mir-128-1&WithFlank&2|+|135665391|135665484</td><td>56</td><td>75</td><td>20</td><td>TCACAGTGAACCGGTCTCTT</td><td>697.1</td><td>3.3</td><td>0.9</td><td>3.5</td><td>3.9E-04</td><td>5.0E-04</td><td>MB231/MB468</td><td>MITO</td></tr>
<tr><td>>hg38_21_+_8433172_8446622_RNA45SN1_45S_50ntFlanks@4759.4786.28|ATCAGATACCGTCGTAGTTCCGACCATA</td><td>>hg38_21_+_8433172_8446622_RNA45SN1_45S_50ntFlanks</td><td>4759</td><td>4786</td><td>28</td><td>ATCAGATACCGTCGTAGTTCCGACCATA</td><td>230.2</td><td>-5.0</td><td>1.4</td><td>-3.5</td><td>4.0E-04</td><td>5.0E-04</td><td>MB231/MB468</td><td>NUC</td></tr>
<tr><td>>hg38_MT_+_1621_3279_RNR2_16S_50ntFlanks@739.777.39|CTGAACTGACAATTAACAGCCCAATATCTACAATCAACC</td><td>>hg38_MT_+_1621_3279_RNR2_16S_50ntFlanks</td><td>739</td><td>777</td><td>39</td><td>CTGAACTGACAATTAACAGCCCAATATCTACAATCAACC</td><td>84.5</td><td>10.1</td><td>2.7</td><td>3.8</td><td>1.6E-04</td><td>5.0E-04</td><td>BT20/MB468</td><td>MP</td></tr>
<tr><td>>hg38_21_+_8433172_8446622_RNA45SN1_45S_50ntFlanks@6761.6806.46|TGCGGCCCCGGGTTCCTCCCGGGGCTACGCCTGTCTGAGCGTCGCT</td><td>>hg38_21_+_8433172_8446622_RNA45SN1_45S_50ntFlanks</td><td>6761</td><td>6806</td><td>46</td><td>TGCGGCCCCGGGTTCCTCCCGGGGCTACGCCTGTCTGAGCGTCGCT</td><td>336.8</td><td>4.3</td><td>1.2</td><td>3.5</td><td>3.9E-04</td><td>5.0E-04</td><td>MB231/MB468</td><td>MITO</td></tr>
<tr><td>>hg38_21_+_8433172_8446622_RNA45SN1_45S_50ntFlanks@12444.12486.43|AAGCAGAATTCACCAAGCGTTGGATTGTTCACCCACTAATAGG</td><td>>hg38_21_+_8433172_8446622_RNA45SN1_45S_50ntFlanks</td><td>12444</td><td>12486</td><td>43</td><td>AAGCAGAATTCACCAAGCGTTGGATTGTTCACCCACTAATAGG</td><td>159.7</td><td>-5.7</td><td>1.6</td><td>-3.6</td><td>2.9E-04</td><td>5.0E-04</td><td>BT20/MB468</td><td>MITO</td></tr>
<tr><td>>trna5_AspGTC_12_+_98897281_98897352@35.74.40|TCACGCGGGAGACCGGGGTTCAATTCCCCGACGGGGAGCC</td><td>>trna5_AspGTC_12_+_98897281_98897352</td><td>35</td><td>74</td><td>40</td><td>TCACGCGGGAGACCGGGGTTCAATTCCCCGACGGGGAGCC</td><td>2423.9</td><td>-7.4</td><td>1.8</td><td>-4.0</td><td>5.5E-05</td><td>5.0E-04</td><td>BT20/MB231</td><td>CYTO</td></tr>
<tr><td>>MI0000093|hsa-mir-92a-1&WithFlank&13|+|91351308|91351397@54.74.21(+1A);MI0000094|hsa-mir-92a-2&WithFlank&X|-|134169532|134169618@54.74.21(+1A)&[MIMAT0000092&hsa-miR-92a-3p&offsets|0|-1(+1A);m-9&13|+|91351361|91351382&offsets|0|-1(+1A)];[MIMAT0000092_1&hsa-miR-92a-3p&offsets|0|-1(+1A);m-11&X|-|134169544|134169565&offsets|0|-1(+1A)]|TATTGCACTTGTCCCGGCCTGA</td><td>>MI0000093|hsa-mir-92a-1&WithFlank&13|+|91351308|91351397</td><td>54</td><td>74</td><td>21(+1A)</td><td>TATTGCACTTGTCCCGGCCTGA</td><td>1903.8</td><td>3.3</td><td>0.9</td><td>3.6</td><td>3.3E-04</td><td>5.0E-04</td><td>MB231/MB468</td><td>MP</td></tr>
</table>

<table>
<tr><th></th><th></th><th></th><th></th><th></th><th></th><th></th><th></th><th></th><th></th><th></th><th></th><th></th><th></th></tr>
<tr><td>>trna119_LysCTT_1_-_145395522_145395594@4.34.31,trna11_LysCTT_5_-_180648979_180649051@4.34.31,trna13_LysCTT_6_+_26556774_26556846@4.34.31,trna7_LysCTT_16_+_3225692_3225764@4.34.31,trna9_LysCTT_5_+_180634755_180634827@4.34.31|CGGCTAGCTCAGTCGGTAGAGCATGAGACTC</td><td>>trna119_LysCTT_1_-_145395522_145395594</td><td>4</td><td>34</td><td>31</td><td>CGGCTAGCTCAGTCGGTAGAGCATGAGACTC</td><td>136.7</td><td>5.4</td><td>1.5</td><td>3.6</td><td>3.3E-04</td><td>5.0E-04</td><td>MB231/MB468</td><td>MP</td></tr>
<tr><td>>MI0000070|hsa-mir-16-1&WithFlank&13|-|50048967|50049067@20.39.20;MI0000115|hsa-mir-16-2&WithFlank&3|+|160404739|160404831@16.35.20&[MIMAT0000069&hsa-miR-16-5p&offsets|0|-2;m-62&13|-|50049027|50049048&offsets|0|-2];[MIMAT0000069_1&hsa-miR-16-5p&offsets|0|-2;m-61&3|+|160404754|160404775&offsets|0|-2]|TAGCAGCACGTAAATATTGG</td><td>>MI0000070|hsa-mir-16-1&WithFlank&13|-|50048967|50049067</td><td>20</td><td>39</td><td>20</td><td>TAGCAGCACGTAAATATTGG</td><td>636.1</td><td>4.6</td><td>1.2</td><td>3.8</td><td>1.6E-04</td><td>5.0E-04</td><td>BT20/MB468</td><td>MP</td></tr>
<tr><td>>hg38_21_+_8433172_8446622_RNA45SN1_45S_50ntFlanks@5482.5516.35|CGGAGCGCTGAGAAGACGGTCGAACTTGACTATCT</td><td>>hg38_21_+_8433172_8446622_RNA45SN1_45S_50ntFlanks</td><td>5482</td><td>5516</td><td>35</td><td>CGGAGCGCTGAGAAGACGGTCGAACTTGACTATCT</td><td>173.6</td><td>3.9</td><td>1.0</td><td>3.8</td><td>1.6E-04</td><td>5.0E-04</td><td>BT20/MB468</td><td>MP</td></tr>
<tr><td>>hg38_21_+_8433172_8446622_RNA45SN1_45S_50ntFlanks@4739.4758.20|AAGTCGGAGGTTCGAAGACG</td><td>>hg38_21_+_8433172_8446622_RNA45SN1_45S_50ntFlanks</td><td>4739</td><td>4758</td><td>20</td><td>AAGTCGGAGGTTCGAAGACG</td><td>185.0</td><td>-4.1</td><td>1.1</td><td>-3.6</td><td>2.9E-04</td><td>5.0E-04</td><td>BT20/MB468</td><td>MITO</td></tr>
<tr><td>>hg38_21_+_8433172_8446622_RNA45SN1_45S_50ntFlanks@7973.8015.43|GACGCGACCTCAGATCAGACGTGGCGACCCGCTGAATTTAAGC</td><td>>hg38_21_+_8433172_8446622_RNA45SN1_45S_50ntFlanks</td><td>7973</td><td>8015</td><td>43</td><td>GACGCGACCTCAGATCAGACGTGGCGACCCGCTGAATTTAAGC</td><td>521.4</td><td>5.9</td><td>1.6</td><td>3.6</td><td>3.0E-04</td><td>5.0E-04</td><td>BT20/MB468</td><td>MITO</td></tr>
<tr><td>>hg38_21_+_8433172_8446622_RNA45SN1_45S_50ntFlanks@10922.10945.24|CTGGACGAGGCGCCGCCGCCCCCC</td><td>>hg38_21_+_8433172_8446622_RNA45SN1_45S_50ntFlanks</td><td>10922</td><td>10945</td><td>24</td><td>CTGGACGAGGCGCCGCCGCCCCCC</td><td>1290.6</td><td>-6.8</td><td>1.9</td><td>-3.5</td><td>4.0E-04</td><td>5.0E-04</td><td>BT20/MB468</td><td>NUC</td></tr>
<tr><td>>hg38_21_+_8433172_8446622_RNA45SN1_45S_50ntFlanks@7974.7994.21|ACGCGACCTCAGATCAGACGT</td><td>>hg38_21_+_8433172_8446622_RNA45SN1_45S_50ntFlanks</td><td>7974</td><td>7994</td><td>21</td><td>ACGCGACCTCAGATCAGACGT</td><td>852.1</td><td>1.9</td><td>0.5</td><td>3.5</td><td>4.0E-04</td><td>5.0E-04</td><td>MB231/MB468</td><td>NUC</td></tr>
<tr><td>>hg38_21_+_8433172_8446622_RNA45SN1_45S_50ntFlanks@11212.11250.39|CGACTCTGGACGCGAGCCGGGCCCTTCCCGTGGATCGCC</td><td>>hg38_21_+_8433172_8446622_RNA45SN1_45S_50ntFlanks</td><td>11212</td><td>11250</td><td>39</td><td>CGACTCTGGACGCGAGCCGGGCCCTTCCCGTGGATCGCC</td><td>454.7</td><td>-2.9</td><td>0.8</td><td>-3.6</td><td>3.7E-04</td><td>5.0E-04</td><td>MB231/MB468</td><td>CYTO</td></tr>
<tr><td>>hg38_21_+_8433172_8446622_RNA45SN1_45S_50ntFlanks@10347.10368.22|CAAACGAGAACTTTGAAGGCCG</td><td>>hg38_21_+_8433172_8446622_RNA45SN1_45S_50ntFlanks</td><td>10347</td><td>10368</td><td>22</td><td>CAAACGAGAACTTTGAAGGCCG</td><td>556.3</td><td>-4.0</td><td>1.1</td><td>-3.8</td><td>1.6E-04</td><td>5.0E-04</td><td>BT20/MB468</td><td>MP</td></tr>
<tr><td>>trnaMT_MetCAT_MT_+_4402_4469@36.71.36|CCCGAAAATGTTGGTTATACCCTTCCCGTACTACCA</td><td>>trnaMT_MetCAT_MT_+_4402_4469</td><td>36</td><td>71</td><td>36</td><td>CCCGAAAATGTTGGTTATACCCTTCCCGTACTACCA</td><td>141.6</td><td>3.7</td><td>1.0</td><td>3.5</td><td>4.0E-04</td><td>5.0E-04</td><td>MB231/MB468</td><td>MITO</td></tr>
<tr><td>>trnaMT_GluTTC_MT_-_14674_14742@38.57.20,trnalookalike8_GluTTC_5_-_93905172_93905240@38.57.20|CATTGGTCGTGGTTGTAGTC</td><td>>trnaMT_GluTTC_MT_-_14674_14742</td><td>38</td><td>57</td><td>20</td><td>CATTGGTCGTGGTTGTAGTC</td><td>190.1</td><td>5.3</td><td>1.4</td><td>3.8</td><td>1.6E-04</td><td>5.0E-04</td><td>BT20/MB468</td><td>MP</td></tr>
<tr><td>>hg38_21_+_8433172_8446622_RNA45SN1_45S_50ntFlanks@11873.11902.30|AAGAAGACCCTGTTGAGCTTGACTCTAGTC</td><td>>hg38_21_+_8433172_8446622_RNA45SN1_45S_50ntFlanks</td><td>11873</td><td>11902</td><td>30</td><td>AAGAAGACCCTGTTGAGCTTGACTCTAGTC</td><td>286.4</td><td>-4.2</td><td>1.2</td><td>-3.5</td><td>4.0E-04</td><td>5.0E-04</td><td>MB231/MB468</td><td>NUC</td></tr>
<tr><td>>hg38_21_+_8433172_8446622_RNA45SN1_45S_50ntFlanks@3875.3897.23|ATACATGCCGACGGGCGCTGACC</td><td>>hg38_21_+_8433172_8446622_RNA45SN1_45S_50ntFlanks</td><td>3875</td><td>3897</td><td>23</td><td>ATACATGCCGACGGGCGCTGACC</td><td>228.7</td><td>-2.5</td><td>0.7</td><td>-3.5</td><td>4.0E-04</td><td>5.0E-04</td><td>MB231/MB468</td><td>NUC</td></tr>
<tr><td>>hg38_21_+_8433172_8446622_RNA45SN1_45S_50ntFlanks@10391.10418.28|AACAGCAGTTGAACATGGGTCAGTCGGT</td><td>>hg38_21_+_8433172_8446622_RNA45SN1_45S_50ntFlanks</td><td>10391</td><td>10418</td><td>28</td><td>AACAGCAGTTGAACATGGGTCAGTCGGT</td><td>185.2</td><td>-5.2</td><td>1.5</td><td>-3.5</td><td>4.0E-04</td><td>5.0E-04</td><td>MB231/MB468</td><td>NUC</td></tr>
<tr><td>>hg38_MT_+_598_1651_RNR1_12S_50ntFlanks@416.462.47|AAATAGACTACGAAAGTGGCTTTAACATATCTGAACACACAATAGCT</td><td>>hg38_MT_+_598_1651_RNR1_12S_50ntFlanks</td><td>416</td><td>462</td><td>47</td><td>AAATAGACTACGAAAGTGGCTTTAACATATCTGAACACACAATAGCT</td><td>77.5</td><td>10.0</td><td>2.7</td><td>3.8</td><td>1.6E-04</td><td>5.0E-04</td><td>BT20/MB468</td><td>MP</td></tr>
<tr><td>>hg38_21_+_8433172_8446622_RNA45SN1_45S_50ntFlanks@11657.11685.29|GATGTGATTTCTGCCCAGTGCTCTGAATG</td><td>>hg38_21_+_8433172_8446622_RNA45SN1_45S_50ntFlanks</td><td>11657</td><td>11685</td><td>29</td><td>GATGTGATTTCTGCCCAGTGCTCTGAATG</td><td>575.8</td><td>5.3</td><td>1.4</td><td>3.8</td><td>1.6E-04</td><td>5.0E-04</td><td>BT20/MB468</td><td>MP</td></tr>
<tr><td>>MI0001445|hsa-mir-423&WithFlank&17|+|30117073|30117178@23.44.22&[MIMAT0004748&hsa-miR-423-5p&offsets|0|-1;m-141&17|+|30117095|30117117&offsets|0|-1]|TGAGGGGCAGAGAGCGAGACTT</td><td>>MI0001445|hsa-mir-423&WithFlank&17|+|30117073|30117178</td><td>23</td><td>44</td><td>22</td><td>TGAGGGGCAGAGAGCGAGACTT</td><td>1701.9</td><td>5.0</td><td>1.4</td><td>3.6</td><td>3.3E-04</td><td>5.0E-04</td><td>MB231/MB468</td><td>MP</td></tr>
<tr><td>>trnaMT_ThrTGT_MT_+_15888_15953@37.69.33|ACCGGAGATGAAAACCTTTTTCCAAGGACACCA</td><td>>trnaMT_ThrTGT_MT_+_15888_15953</td><td>37</td><td>69</td><td>33</td><td>ACCGGAGATGAAAACCTTTTTCCAAGGACACCA</td><td>257.4</td><td>4.8</td><td>1.4</td><td>3.5</td><td>4.0E-04</td><td>5.0E-04</td><td>MB231/MB468</td><td>MITO</td></tr>
<tr><td>>hg38_21_+_8433172_8446622_RNA45SN1_45S_50ntFlanks@11718.11736.19|AACGGCGGGAGTAACTATG</td><td>>hg38_21_+_8433172_8446622_RNA45SN1_45S_50ntFlanks</td><td>11718</td><td>11736</td><td>19</td><td>AACGGCGGGAGTAACTATG</td><td>3567.7</td><td>-7.3</td><td>2.1</td><td>-3.5</td><td>4.1E-04</td><td>5.0E-04</td><td>BT20/MB468</td><td>NUC</td></tr>
<tr><td>>am_trnaMT_ValTAC_MT_+_1602_1670@51.68.18|CTTAACTTGACCGCTCTG</td><td>>am_trnaMT_ValTAC_MT_+_1602_1670</td><td>51</td><td>68</td><td>18</td><td>CTTAACTTGACCGCTCTG</td><td>10633.5</td><td>-6.1</td><td>1.6</td><td>-3.8</td><td>1.6E-04</td><td>5.0E-04</td><td>BT20/MB468</td><td>MP</td></tr>
<tr><td>>MI0000299|hsa-mir-222&WithFlank&X|-|45747009|45747130@75.97.23(+1U)&[MIMAT0000279&hsa-miR-222-3p&offsets|0|+2(+1U);m-140&X|-|45747033|45747056&offsets|0|-1(+1U)]|AGCTACATCTGGCTACTGGGTCTT</td><td>>MI0000299|hsa-mir-222&WithFlank&X|-|45747009|45747130</td><td>75</td><td>97</td><td>23(+1U)</td><td>AGCTACATCTGGCTACTGGGTCTT</td><td>706.4</td><td>3.9</td><td>1.0</td><td>3.8</td><td>1.6E-04</td><td>5.0E-04</td><td>BT20/MB468</td><td>MP</td></tr>
<tr><td>>hg38_21_+_8433172_8446622_RNA45SN1_45S_50ntFlanks@10403.10429.27|ACATGGGTCAGTCGGTCCTGAGAGATG</td><td>>hg38_21_+_8433172_8446622_RNA45SN1_45S_50ntFlanks</td><td>10403</td><td>10429</td><td>27</td><td>ACATGGGTCAGTCGGTCCTGAGAGATG</td><td>549.5</td><td>-2.0</td><td>0.6</td><td>-3.6</td><td>3.8E-04</td><td>5.0E-04</td><td>MB231/MB468</td><td>CYTO</td></tr>
<tr><td>>hg38_21_+_8433172_8446622_RNA45SN1_45S_50ntFlanks@12974.12992.19|CGTACGTAGCAGAGCAGCT</td><td>>hg38_21_+_8433172_8446622_RNA45SN1_45S_50ntFlanks</td><td>12974</td><td>12992</td><td>19</td><td>CGTACGTAGCAGAGCAGCT</td><td>294.1</td><td>-3.6</td><td>1.0</td><td>-3.6</td><td>3.6E-04</td><td>5.0E-04</td><td>BT20/MB468</td><td>CYTO</td></tr>
<tr><td>>hg38_21_+_8433172_8446622_RNA45SN1_45S_50ntFlanks@6655.6681.27|TCTTAGCGGTGGATCACTCGGCTCGTG</td><td>>hg38_21_+_8433172_8446622_RNA45SN1_45S_50ntFlanks</td><td>6655</td><td>6681</td><td>27</td><td>TCTTAGCGGTGGATCACTCGGCTCGTG</td><td>4486.5</td><td>-3.5</td><td>1.0</td><td>-3.5</td><td>4.0E-04</td><td>5.0E-04</td><td>MB231/MB468</td><td>MITO</td></tr>
<tr><td>>hg38_21_+_8433172_8446622_RNA45SN1_45S_50ntFlanks@8176.8214.39|CTTCTGATCGAGGCCCAGCCCGTGGACGGTGTGAGGCCG</td><td>>hg38_21_+_8433172_8446622_RNA45SN1_45S_50ntFlanks</td><td>8176</td><td>8214</td><td>39</td><td>CTTCTGATCGAGGCCCAGCCCGTGGACGGTGTGAGGCCG</td><td>1890.2</td><td>-5.6</td><td>1.5</td><td>-3.8</td><td>1.6E-04</td><td>5.0E-04</td><td>BT20/MB468</td><td>MP</td></tr>
<tr><td>>trna119_LysCTT_1_-_145395522_145395594@3.34.32,trna11_LysCTT_5_-_180648979_180649051@3.34.32,trna13_LysCTT_6_+_26556774_26556846@3.34.32,trna7_LysCTT_16_+_3225692_3225764@3.34.32,trna9_LysCTT_5_+_180634755_180634827@3.34.32|CCGGCTAGCTCAGTCGGTAGAGCATGAGACTC</td><td>>trna119_LysCTT_1_-_145395522_145395594</td><td>3</td><td>34</td><td>32</td><td>CCGGCTAGCTCAGTCGGTAGAGCATGAGACTC</td><td>235.0</td><td>-6.2</td><td>1.5</td><td>-4.0</td><td>5.8E-05</td><td>5.0E-04</td><td>BT20/MB231</td><td>NUC</td></tr>
<tr><td>>hg38_21_+_8433172_8446622_RNA45SN1_45S_50ntFlanks@8105.8138.34|CGCGGCGGGGCGCGGGACATGTGGCGTACGGAAG</td><td>>hg38_21_+_8433172_8446622_RNA45SN1_45S_50ntFlanks</td><td>8105</td><td>8138</td><td>34</td><td>CGCGGCGGGGCGCGGGACATGTGGCGTACGGAAG</td><td>1169.1</td><td>-5.4</td><td>1.5</td><td>-3.5</td><td>4.1E-04</td><td>5.0E-04</td><td>BT20/MB468</td><td>NUC</td></tr>
<tr><td>>trnaMT_PheGAA_MT_+_577_647@28.54.27|ATACACTGAAAATGTTTAGACGGGCTC</td><td>>trnaMT_PheGAA_MT_+_577_647</td><td>28</td><td>54</td><td>27</td><td>ATACACTGAAAATGTTTAGACGGGCTC</td><td>77.0</td><td>10.0</td><td>2.7</td><td>3.8</td><td>1.6E-04</td><td>5.0E-04</td><td>BT20/MB468</td><td>MP</td></tr>
<tr><td>>hg38_21_+_8433172_8446622_RNA45SN1_45S_50ntFlanks@5078.5102.25|CGATAACGAACGAGACTCTGGCATG</td><td>>hg38_21_+_8433172_8446622_RNA45SN1_45S_50ntFlanks</td><td>5078</td><td>5102</td><td>25</td><td>CGATAACGAACGAGACTCTGGCATG</td><td>458.9</td><td>-5.8</td><td>1.6</td><td>-3.6</td><td>3.0E-04</td><td>5.0E-04</td><td>BT20/MB468</td><td>MITO</td></tr>
<tr><td>>hg38_21_+_8433172_8446622_RNA45SN1_45S_50ntFlanks@9576.9603.28|TCGGTCGTCCGACCTGGGTATAGGGGCG</td><td>>hg38_21_+_8433172_8446622_RNA45SN1_45S_50ntFlanks</td><td>9576</td><td>9603</td><td>28</td><td>TCGGTCGTCCGACCTGGGTATAGGGGCG</td><td>148.4</td><td>-5.6</td><td>1.6</td><td>-3.6</td><td>3.4E-04</td><td>5.0E-04</td><td>MB231/MB468</td><td>MP</td></tr>
<tr><td>>hg38_21_+_8433172_8446622_RNA45SN1_45S_50ntFlanks@11671.11688.18|CCAGTGCTCTGAATGTCA</td><td>>hg38_21_+_8433172_8446622_RNA45SN1_45S_50ntFlanks</td><td>11671</td><td>11688</td><td>18</td><td>CCAGTGCTCTGAATGTCA</td><td>789.5</td><td>-4.6</td><td>1.3</td><td>-3.6</td><td>3.6E-04</td><td>5.0E-04</td><td>BT20/MB468</td><td>CYTO</td></tr>
<tr><td>>hg38_21_+_8433172_8446622_RNA45SN1_45S_50ntFlanks@9928.9951.24|TGATATAGACAGCAGGACGGTGGC</td><td>>hg38_21_+_8433172_8446622_RNA45SN1_45S_50ntFlanks</td><td>9928</td><td>9951</td><td>24</td><td>TGATATAGACAGCAGGACGGTGGC</td><td>186.3</td><td>7.0</td><td>1.8</td><td>3.8</td><td>1.6E-04</td><td>5.1E-04</td><td>BT20/MB468</td><td>MP</td></tr>
<tr><td>>hg38_21_+_8433172_8446622_RNA45SN1_45S_50ntFlanks@12419.12459.41|GATGTCGGCTCTTCCTATCATTGTGAAGCAGAATTCACCAA</td><td>>hg38_21_+_8433172_8446622_RNA45SN1_45S_50ntFlanks</td><td>12419</td><td>12459</td><td>41</td><td>GATGTCGGCTCTTCCTATCATTGTGAAGCAGAATTCACCAA</td><td>211.9</td><td>5.4</td><td>1.5</td><td>3.5</td><td>4.0E-04</td><td>5.1E-04</td><td>MB231/MB468</td><td>MITO</td></tr>
<tr><td>>hg38_21_+_8433172_8446622_RNA45SN1_45S_50ntFlanks@4226.4243.18|AATAACAATACAGGACTC</td><td>>hg38_21_+_8433172_8446622_RNA45SN1_45S_50ntFlanks</td><td>4226</td><td>4243</td><td>18</td><td>AATAACAATACAGGACTC</td><td>135.2</td><td>-4.4</td><td>1.2</td><td>-3.6</td><td>3.4E-04</td><td>5.1E-04</td><td>MB231/MB468</td><td>MP</td></tr>
<tr><td>>hg38_21_+_8433172_8446622_RNA45SN1_45S_50ntFlanks@12990.13012.23|GCTCCCTCGCTGCGATCTATTGA</td><td>>hg38_21_+_8433172_8446622_RNA45SN1_45S_50ntFlanks</td><td>12990</td><td>13012</td><td>23</td><td>GCTCCCTCGCTGCGATCTATTGA</td><td>1631.6</td><td>-3.3</td><td>0.9</td><td>-3.5</td><td>4.1E-04</td><td>5.1E-04</td><td>MB231/MB468</td><td>NUC</td></tr>
<tr><td>>MI0001445|hsa-mir-423&WithFlank&17|+|30117073|30117178@23.46.24(+1U)&[MIMAT0004748&hsa-miR-423-5p&offsets|0|+1(+1U);m-141&17|+|30117095|30117117&offsets|0|+1(+1U)]|TGAGGGGCAGAGAGCGAGACTTTTT</td><td>>MI0001445|hsa-mir-423&WithFlank&17|+|30117073|30117178</td><td>23</td><td>46</td><td>24(+1U)</td><td>TGAGGGGCAGAGAGCGAGACTTTTT</td><td>159.2</td><td>6.1</td><td>1.6</td><td>3.8</td><td>1.6E-04</td><td>5.1E-04</td><td>BT20/MB468</td><td>MP</td></tr>
<tr><td>>hg38_21_+_8433172_8446622_RNA45SN1_45S_50ntFlanks@10238.10260.23|CGCTACGCCGCGACGAGTAGGAG</td><td>>hg38_21_+_8433172_8446622_RNA45SN1_45S_50ntFlanks</td><td>10238</td><td>10260</td><td>23</td><td>CGCTACGCCGCGACGAGTAGGAG</td><td>155.4</td><td>-5.5</td><td>1.5</td><td>-3.6</td><td>3.4E-04</td><td>5.1E-04</td><td>MB231/MB468</td><td>MP</td></tr>
<tr><td>>hg38_21_+_8433172_8446622_RNA45SN1_45S_50ntFlanks@11663.11692.30|ATTTCTGCCCAGTGCTCTGAATGTCAAAGT</td><td>>hg38_21_+_8433172_8446622_RNA45SN1_45S_50ntFlanks</td><td>11663</td><td>11692</td><td>30</td><td>ATTTCTGCCCAGTGCTCTGAATGTCAAAGT</td><td>93.9</td><td>7.0</td><td>1.8</td><td>3.8</td><td>1.6E-04</td><td>5.1E-04</td><td>BT20/MB468</td><td>MP</td></tr>
<tr><td>>hg38_21_+_8433172_8446622_RNA45SN1_45S_50ntFlanks@10843.10862.20|TAACTTCGGGATAAGGATTG</td><td>>hg38_21_+_8433172_8446622_RNA45SN1_45S_50ntFlanks</td><td>10843</td><td>10862</td><td>20</td><td>TAACTTCGGGATAAGGATTG</td><td>169.6</td><td>-5.8</td><td>1.5</td><td>-3.8</td><td>1.6E-04</td><td>5.1E-04</td><td>BT20/MB468</td><td>MP</td></tr>
<tr><td>>trnaMT_LeuTAA_MT_+_3230_3304@1.35.35|GTTAAGATGGCAGAGCCCGGTAATCGCATAAAACT</td><td>>trnaMT_LeuTAA_MT_+_3230_3304</td><td>1</td><td>35</td><td>35</td><td>GTTAAGATGGCAGAGCCCGGTAATCGCATAAAACT</td><td>154.0</td><td>4.7</td><td>1.3</td><td>3.5</td><td>4.0E-04</td><td>5.1E-04</td><td>MB231/MB468</td><td>MITO</td></tr>
<tr><td>>MI0000542|hsa-mir-320a&WithFlank&8|-|22244960|22245043@49.69.21(+2U)&[MIMAT0000510&hsa-miR-320a-3p&offsets|+1|0(+2U);m-60&8|-|22244975|22244996&offsets|+1|0(+2U)]|AAAGCTGGGTTGAGAGGGCGATT</td><td>>MI0000542|hsa-mir-320a&WithFlank&8|-|22244960|22245043</td><td>49</td><td>69</td><td>21(+2U)</td><td>AAAGCTGGGTTGAGAGGGCGATT</td><td>466.3</td><td>3.2</td><td>0.8</td><td>4.0</td><td>5.6E-05</td><td>5.1E-04</td><td>BT20/MB231</td><td>CYTO</td></tr>
<tr><td>>hg38_MT_+_598_1651_RNR1_12S_50ntFlanks@427.469.43|GAAAGTGGCTTTAACATATCTGAACACACAATAGCTAAGACCC</td><td>>hg38_MT_+_598_1651_RNR1_12S_50ntFlanks</td><td>427</td><td>469</td><td>43</td><td>GAAAGTGGCTTTAACATATCTGAACACACAATAGCTAAGACCC</td><td>103.4</td><td>6.1</td><td>1.7</td><td>3.5</td><td>4.0E-04</td><td>5.1E-04</td><td>MB231/MB468</td><td>MITO</td></tr>
<tr><td>>hg38_21_+_8433172_8446622_RNA45SN1_45S_50ntFlanks@3899.3937.39|CCTTCGCGGGGGGGATGCGTGCATTTATCAGATCAAAAC</td><td>>hg38_21_+_8433172_8446622_RNA45SN1_45S_50ntFlanks</td><td>3899</td><td>3937</td><td>39</td><td>CCTTCGCGGGGGGGATGCGTGCATTTATCAGATCAAAAC</td><td>500.0</td><td>-4.7</td><td>1.3</td><td>-3.6</td><td>3.8E-04</td><td>5.1E-04</td><td>MB231/MB468</td><td>CYTO</td></tr>
<tr><td>>hg38_21_+_8433172_8446622_RNA45SN1_45S_50ntFlanks@4046.4069.24|CATTCGAACGTCTGCCCTATCAAC</td><td>>hg38_21_+_8433172_8446622_RNA45SN1_45S_50ntFlanks</td><td>4046</td><td>4069</td><td>24</td><td>CATTCGAACGTCTGCCCTATCAAC</td><td>342.4</td><td>-4.7</td><td>1.3</td><td>-3.5</td><td>4.1E-04</td><td>5.1E-04</td><td>BT20/MB468</td><td>NUC</td></tr>
<tr><td>>hg38_21_+_8433172_8446622_RNA45SN1_45S_50ntFlanks@11882.11900.19|CTGTTGAGCTTGACTCTAG</td><td>>hg38_21_+_8433172_8446622_RNA45SN1_45S_50ntFlanks</td><td>11882</td><td>11900</td><td>19</td><td>CTGTTGAGCTTGACTCTAG</td><td>179.8</td><td>-5.9</td><td>1.6</td><td>-3.8</td><td>1.6E-04</td><td>5.1E-04</td><td>BT20/MB468</td><td>MP</td></tr>
<tr><td>>MI0000101|hsa-mir-99a&WithFlank&21|+|16539083|16539175@19.40.22&[MIMAT0000097&hsa-miR-99a-5p&offsets|0|0;m-64&21|+|16539101|16539121&offsets|0|+1]|AACCCGTAGATCCGATCTTGTG</td><td>>MI0000101|hsa-mir-99a&WithFlank&21|+|16539083|16539175</td><td>19</td><td>40</td><td>22</td><td>AACCCGTAGATCCGATCTTGTG</td><td>1362.4</td><td>-8.3</td><td>2.1</td><td>-4.0</td><td>6.7E-05</td><td>5.1E-04</td><td>BT20/MB231</td><td>MITO</td></tr>
<tr><td>>hg38_21_+_8433172_8446622_RNA45SN1_45S_50ntFlanks@7990.8019.30|GACGTGGCGACCCGCTGAATTTAAGCATAT</td><td>>hg38_21_+_8433172_8446622_RNA45SN1_45S_50ntFlanks</td><td>7990</td><td>8019</td><td>30</td><td>GACGTGGCGACCCGCTGAATTTAAGCATAT</td><td>255.8</td><td>5.0</td><td>1.4</td><td>3.5</td><td>4.1E-04</td><td>5.1E-04</td><td>MB231/MB468</td><td>NUC</td></tr>
<tr><td>>hg38_21_+_8433172_8446622_RNA45SN1_45S_50ntFlanks@10866.10884.19|CTAAGGGCTGGGTCGGTCG</td><td>>hg38_21_+_8433172_8446622_RNA45SN1_45S_50ntFlanks</td><td>10866</td><td>10884</td><td>19</td><td>CTAAGGGCTGGGTCGGTCG</td><td>637.8</td><td>-4.9</td><td>1.3</td><td>-3.8</td><td>1.6E-04</td><td>5.1E-04</td><td>BT20/MB468</td><td>MP</td></tr>
<tr><td>>MI0016003|hsa-mir-3613&WithFlank&13|-|49996409|49996507@22.43.22&[MIMAT0017990&hsa-miR-3613-5p&offsets|0|0;m-259&13|-|49996465|49996486&offsets|0|0]|TGTTGTACTTTTTTTTTTGTTC</td><td>>MI0016003|hsa-mir-3613&WithFlank&13|-|49996409|49996507</td><td>22</td><td>43</td><td>22</td><td>TGTTGTACTTTTTTTTTTGTTC</td><td>122.3</td><td>3.5</td><td>1.0</td><td>3.5</td><td>4.1E-04</td><td>5.1E-04</td><td>BT20/MB468</td><td>NUC</td></tr>
</table>

<table>
<tr><td>>MI0000108|hsa-mir-103a-2&WithFlank&20|+|3917488|3917577@54.76.23(+1U);MI0000109|hsa-mir-103a-1&WithFlank&5|-|168560890|168560979@54.76.23(+1U)&[MIMAT0000101&hsa-miR-103a-3p&offsets|0|0(+1U);m-12&20|+|3917541|3917563&offsets|0|0(+1U)];[MIMAT0000101_1&hsa-miR-103a-3p&offsets|0|0(+1U);m-13&5|-|168560904|168560926&offsets|0|0(+1U);m-44220&5|-|168560908|168560926&offsets|0|+4(+1U)]|AGCAGCATTGTACAGGGCTATGAT</td><td>>MI0000108|hsa-mir-103a-2&WithFlank&20|+|3917488|3917577</td><td>54</td><td>76</td><td>23(+1U)</td><td>AGCAGCATTGTACAGGGCTATGAT</td><td>1240.9</td><td>-3.7</td><td>0.9</td><td>-4.0</td><td>5.7E-05</td><td>5.1E-04</td><td>BT20/MB231</td><td>MP</td></tr>
<tr><td>>hg38_21_+_8433172_8446622_RNA45SN1_45S_50ntFlanks@8196.8218.23|CGTGGACGGTGTGAGGCCGGTAG</td><td>>hg38_21_+_8433172_8446622_RNA45SN1_45S_50ntFlanks</td><td>8196</td><td>8218</td><td>23</td><td>CGTGGACGGTGTGAGGCCGGTAG</td><td>164.5</td><td>-4.3</td><td>1.2</td><td>-3.6</td><td>3.4E-04</td><td>5.1E-04</td><td>MB231/MB468</td><td>MP</td></tr>
<tr><td>>hg38_21_+_8433172_8446622_RNA45SN1_45S_50ntFlanks@12416.12445.30|TTCGATGTCGGCTCTTCCTATCATTGTGAA</td><td>>hg38_21_+_8433172_8446622_RNA45SN1_45S_50ntFlanks</td><td>12416</td><td>12445</td><td>30</td><td>TTCGATGTCGGCTCTTCCTATCATTGTGAA</td><td>389.9</td><td>2.3</td><td>0.7</td><td>3.5</td><td>4.1E-04</td><td>5.1E-04</td><td>MB231/MB468</td><td>NUC</td></tr>
<tr><td>>hg38_21_+_8433172_8446622_RNA45SN1_45S_50ntFlanks@3838.3872.35|CTCTCCTACTTGGATAACTGTGGTAATTCTAGAGC</td><td>>hg38_21_+_8433172_8446622_RNA45SN1_45S_50ntFlanks</td><td>3838</td><td>3872</td><td>35</td><td>CTCTCCTACTTGGATAACTGTGGTAATTCTAGAGC</td><td>325.9</td><td>-4.0</td><td>1.1</td><td>-3.5</td><td>4.1E-04</td><td>5.1E-04</td><td>MB231/MB468</td><td>NUC</td></tr>
<tr><td>>trnaMT_SerGCT_MT_+_12207_12265@23.62.40|AACTCATGCCCCCATGTCTAACAACATGGCTTTCTCACCA</td><td>>trnaMT_SerGCT_MT_+_12207_12265</td><td>23</td><td>62</td><td>40</td><td>AACTCATGCCCCCATGTCTAACAACATGGCTTTCTCACCA</td><td>226.9</td><td>3.5</td><td>1.0</td><td>3.5</td><td>4.0E-04</td><td>5.1E-04</td><td>MB231/MB468</td><td>MITO</td></tr>
<tr><td>>trna116_GluCTC_1_-_145399233_145399304@39.73.35,trna134_GluTTC_1_-_16861774_16861845@39.73.35,trna71_GluCTC_1_-_161439189_161439260@39.73.35,trna74_GluCTC_1_-_161431809_161431880@39.73.35,trna77_GluCTC_1_-_161424398_161424469@39.73.35,trna77_GluCTC_6_+_28949976_28950047@39.73.35,trna80_GluCTC_1_-_161417018_161417089@39.73.35,trna84_GluTTC_1_-_161391883_161391954@39.73.35,trna87_GluCTC_6_-_126101393_126101464@39.73.35|CGCCGCGGCCCGGGTTCGATTCCCGGTCAGGGAAC</td><td>>trna116_GluCTC_1_-_145399233_145399304</td><td>39</td><td>73</td><td>35</td><td>CGCCGCGGCCCGGGTTCGATTCCCGGTCAGGGAAC</td><td>71.5</td><td>9.4</td><td>2.6</td><td>3.6</td><td>3.4E-04</td><td>5.1E-04</td><td>MB231/MB468</td><td>MP</td></tr>
<tr><td>>hg38_21_+_8433172_8446622_RNA45SN1_45S_50ntFlanks@11844.11872.29|GGGAACGGGCTTGGCGGAATCAGCGGGGA</td><td>>hg38_21_+_8433172_8446622_RNA45SN1_45S_50ntFlanks</td><td>11844</td><td>11872</td><td>29</td><td>GGGAACGGGCTTGGCGGAATCAGCGGGGA</td><td>865.4</td><td>-6.2</td><td>1.6</td><td>-3.8</td><td>1.6E-04</td><td>5.1E-04</td><td>BT20/MB468</td><td>MP</td></tr>
<tr><td>>am_MI0003639|hsa-mir-625&WithFlank&14|+|65471096|65471192@58.78.21(+1U)&[MIMAT0004808&hsa-miR-625-3p&offsets|0|-1(+1U);m-137&14|+|65471153|65471174&offsets|0|-1(+1U)]|GACTATAGAACTTTCCCCCTCT</td><td>>am_MI0003639|hsa-mir-625&WithFlank&14|+|65471096|65471192</td><td>58</td><td>78</td><td>21(+1U)</td><td>GACTATAGAACTTTCCCCCTCT</td><td>216.8</td><td>-3.0</td><td>0.7</td><td>-4.0</td><td>5.9E-05</td><td>5.1E-04</td><td>BT20/MB231</td><td>NUC</td></tr>
<tr><td>>hg38_21_+_8433172_8446622_RNA45SN1_45S_50ntFlanks@12995.13012.18|CTCGCTGCGATCTATTGA</td><td>>hg38_21_+_8433172_8446622_RNA45SN1_45S_50ntFlanks</td><td>12995</td><td>13012</td><td>18</td><td>CTCGCTGCGATCTATTGA</td><td>439.3</td><td>-3.1</td><td>0.8</td><td>-4.0</td><td>5.9E-05</td><td>5.1E-04</td><td>BT20/MB231</td><td>NUC</td></tr>
<tr><td>>hg38_MT_+_598_1651_RNR1_12S_50ntFlanks@908.946.39|ATACTTCAAAGGACATTTAACTAAAACCCCTACGCATTT</td><td>>hg38_MT_+_598_1651_RNR1_12S_50ntFlanks</td><td>908</td><td>946</td><td>39</td><td>ATACTTCAAAGGACATTTAACTAAAACCCCTACGCATTT</td><td>485.6</td><td>6.9</td><td>1.8</td><td>3.8</td><td>1.6E-04</td><td>5.1E-04</td><td>BT20/MB468</td><td>MP</td></tr>
<tr><td>>MI0000063|hsa-let-7b&WithFlank&22|+|46113680|46113774@12.31.20(+1A)&[MIMAT0000063&hsa-let-7b-5p&offsets|0|-2(+1A);m-14&22|+|46113691|46113712&offsets|0|-2(+1A)]|TGAGGTAGTAGGTTGTGTGGA</td><td>>MI0000063|hsa-let-7b&WithFlank&22|+|46113680|46113774</td><td>12</td><td>31</td><td>20(+1A)</td><td>TGAGGTAGTAGGTTGTGTGGA</td><td>410.1</td><td>5.4</td><td>1.4</td><td>3.8</td><td>1.6E-04</td><td>5.1E-04</td><td>BT20/MB468</td><td>MP</td></tr>
<tr><td>>hg38_21_+_8433172_8446622_RNA45SN1_45S_50ntFlanks@12476.12500.25|CCACTAATAGGGAACGTGAGCTGGG</td><td>>hg38_21_+_8433172_8446622_RNA45SN1_45S_50ntFlanks</td><td>12476</td><td>12500</td><td>25</td><td>CCACTAATAGGGAACGTGAGCTGGG</td><td>652.9</td><td>-4.3</td><td>1.2</td><td>-3.6</td><td>3.6E-04</td><td>5.1E-04</td><td>BT20/MB468</td><td>CYTO</td></tr>
<tr><td>>hg38_21_+_8433172_8446622_RNA45SN1_45S_50ntFlanks@8201.8223.23|ACGGTGTGAGGCCGGTAGCGGCC</td><td>>hg38_21_+_8433172_8446622_RNA45SN1_45S_50ntFlanks</td><td>8201</td><td>8223</td><td>23</td><td>ACGGTGTGAGGCCGGTAGCGGCC</td><td>89.4</td><td>-5.6</td><td>1.5</td><td>-3.8</td><td>1.6E-04</td><td>5.1E-04</td><td>BT20/MB468</td><td>MP</td></tr>
<tr><td>>hg38_MT_+_598_1651_RNR1_12S_50ntFlanks@853.870.18|CTTAGTTGAACAGGGCCC</td><td>>hg38_MT_+_598_1651_RNR1_12S_50ntFlanks</td><td>853</td><td>870</td><td>18</td><td>CTTAGTTGAACAGGGCCC</td><td>76.3</td><td>10.0</td><td>2.7</td><td>3.8</td><td>1.6E-04</td><td>5.1E-04</td><td>BT20/MB468</td><td>MP</td></tr>
<tr><td>>trnaMT_TrpTCA_MT_+_5512_5579@53.71.19,trnalookalike3_TrpTCA_1_+_566062_566129@53.71.19|CAATACTTAATTTCTGCCA</td><td>>trnaMT_TrpTCA_MT_+_5512_5579</td><td>53</td><td>71</td><td>19</td><td>CAATACTTAATTTCTGCCA</td><td>680.5</td><td>-3.7</td><td>1.0</td><td>-3.8</td><td>1.6E-04</td><td>5.1E-04</td><td>BT20/MB468</td><td>MP</td></tr>
<tr><td>>hg38_21_+_8433172_8446622_RNA45SN1_45S_50ntFlanks@11200.11234.35|CGGCGGCGGCGGCGACTCTGGACGCGAGCCGGGCC</td><td>>hg38_21_+_8433172_8446622_RNA45SN1_45S_50ntFlanks</td><td>11200</td><td>11234</td><td>35</td><td>CGGCGGCGGCGGCGACTCTGGACGCGAGCCGGGCC</td><td>107.0</td><td>-5.8</td><td>1.6</td><td>-3.8</td><td>1.6E-04</td><td>5.1E-04</td><td>BT20/MB468</td><td>MP</td></tr>
<tr><td>>hg38_21_+_8433172_8446622_RNA45SN1_45S_50ntFlanks@12934.12953.20|CTAAACCATTCGTAGACGAC</td><td>>hg38_21_+_8433172_8446622_RNA45SN1_45S_50ntFlanks</td><td>12934</td><td>12953</td><td>20</td><td>CTAAACCATTCGTAGACGAC</td><td>654.8</td><td>-6.1</td><td>1.6</td><td>-3.8</td><td>1.6E-04</td><td>5.1E-04</td><td>BT20/MB468</td><td>MP</td></tr>
<tr><td>>hg38_21_+_8433172_8446622_RNA45SN1_45S_50ntFlanks@11848.11886.39|ACGGGCTTGGCGGAATCAGCGGGGAAAGAAGACCCTGTT</td><td>>hg38_21_+_8433172_8446622_RNA45SN1_45S_50ntFlanks</td><td>11848</td><td>11886</td><td>39</td><td>ACGGGCTTGGCGGAATCAGCGGGGAAAGAAGACCCTGTT</td><td>61.6</td><td>6.8</td><td>1.8</td><td>3.8</td><td>1.6E-04</td><td>5.1E-04</td><td>BT20/MB468</td><td>MP</td></tr>
<tr><td>>hg38_21_+_8433172_8446622_RNA45SN1_45S_50ntFlanks@7974.7994.21|ACGCGACCTCAGATCAGACGT</td><td>>hg38_21_+_8433172_8446622_RNA45SN1_45S_50ntFlanks</td><td>7974</td><td>7994</td><td>21</td><td>ACGCGACCTCAGATCAGACGT</td><td>6148.4</td><td>3.5</td><td>0.9</td><td>4.0</td><td>6.0E-05</td><td>5.1E-04</td><td>BT20/MB231</td><td>NUC</td></tr>
<tr><td>>hg38_21_+_8433172_8446622_RNA45SN1_45S_50ntFlanks@11674.11692.19|GTGCTCTGAATGTCAAAGT</td><td>>hg38_21_+_8433172_8446622_RNA45SN1_45S_50ntFlanks</td><td>11674</td><td>11692</td><td>19</td><td>GTGCTCTGAATGTCAAAGT</td><td>86.6</td><td>6.8</td><td>1.8</td><td>3.8</td><td>1.6E-04</td><td>5.1E-04</td><td>BT20/MB468</td><td>MP</td></tr>
<tr><td>>hg38_21_+_8433172_8446622_RNA45SN1_45S_50ntFlanks@10839.10865.27|TCCGTAACTTCGGGATAAGGATTGGCT</td><td>>hg38_21_+_8433172_8446622_RNA45SN1_45S_50ntFlanks</td><td>10839</td><td>10865</td><td>27</td><td>TCCGTAACTTCGGGATAAGGATTGGCT</td><td>344.3</td><td>-5.9</td><td>1.6</td><td>-3.8</td><td>1.6E-04</td><td>5.1E-04</td><td>BT20/MB468</td><td>MP</td></tr>
<tr><td>>hg38_21_+_8433172_8446622_RNA45SN1_45S_50ntFlanks@12000.12031.32|CCTGCGGGCCGCCGGTGAAATACCACTACTCT</td><td>>hg38_21_+_8433172_8446622_RNA45SN1_45S_50ntFlanks</td><td>12000</td><td>12031</td><td>32</td><td>CCTGCGGGCCGCCGGTGAAATACCACTACTCT</td><td>149.6</td><td>-5.0</td><td>1.4</td><td>-3.5</td><td>4.1E-04</td><td>5.1E-04</td><td>MB231/MB468</td><td>NUC</td></tr>
<tr><td>>hg38_21_+_8433172_8446622_RNA45SN1_45S_50ntFlanks@12941.12964.24|ATTCGTAGACGACCTGCTTCTGGG</td><td>>hg38_21_+_8433172_8446622_RNA45SN1_45S_50ntFlanks</td><td>12941</td><td>12964</td><td>24</td><td>ATTCGTAGACGACCTGCTTCTGGG</td><td>192.3</td><td>-4.0</td><td>1.1</td><td>-3.5</td><td>4.1E-04</td><td>5.1E-04</td><td>MB231/MB468</td><td>NUC</td></tr>
<tr><td>>am_MI0000239|hsa-mir-197&WithFlank&1|+|109598887|109598973@54.71.18(+1U)&[MIMAT0000227&hsa-miR-197-3p&offsets|0|-4(+1U);m-99&1|+|109598940|109598961&offsets|0|-4(+1U)]|TTCACCACCTTCTCCACCT</td><td>>am_MI0000239|hsa-mir-197&WithFlank&1|+|109598887|109598973</td><td>54</td><td>71</td><td>18(+1U)</td><td>TTCACCACCTTCTCCACCT</td><td>344.8</td><td>3.2</td><td>0.8</td><td>4.0</td><td>5.7E-05</td><td>5.1E-04</td><td>BT20/MB231</td><td>CYTO</td></tr>
<tr><td>>trna111_HisGTG_1_-_147774845_147774916@-1T.26.27,trna118_HisGTG_1_-_145396881_145396952@-1T.26.27,trna16_HisGTG_1_+_146544773_146544844@-1T.26.27,trna1_HisGTG_15_+_45493349_45493420@-1T.26.27,trna21_HisGTG_1_+_147753471_147753542@-1T.26.27,trna33_HisGTG_6_+_27125906_27125977@-1T.26.27,trna7_HisGTG_9_-_14433938_14434009@-1T.26.27,trna8_HisGTG_15_-_45492611_45492682@-1T.26.27,trna9_HisGTG_15_-_45490804_45490875@-1T.26.27|TGCCGTGATCGTATAGTGGTTAGTACT</td><td>>trna111_HisGTG_1_-_147774845_147774916</td><td>-1T</td><td>26</td><td>27</td><td>TGCCGTGATCGTATAGTGGTTAGTACT</td><td>352.4</td><td>3.2</td><td>0.9</td><td>3.6</td><td>3.4E-04</td><td>5.1E-04</td><td>MB231/MB468</td><td>MP</td></tr>
<tr><td>>hg38_21_+_8433172_8446622_RNA45SN1_45S_50ntFlanks@6650.6672.23|ACGACTCTTAGCGGTGGATCACT</td><td>>hg38_21_+_8433172_8446622_RNA45SN1_45S_50ntFlanks</td><td>6650</td><td>6672</td><td>23</td><td>ACGACTCTTAGCGGTGGATCACT</td><td>876.9</td><td>3.2</td><td>0.9</td><td>3.5</td><td>4.1E-04</td><td>5.1E-04</td><td>MB231/MB468</td><td>MITO</td></tr>
<tr><td>>hg38_21_+_8433172_8446622_RNA45SN1_45S_50ntFlanks@11785.11803.19|TGAATGGATGAACGAGATT</td><td>>hg38_21_+_8433172_8446622_RNA45SN1_45S_50ntFlanks</td><td>11785</td><td>11803</td><td>19</td><td>TGAATGGATGAACGAGATT</td><td>1702.5</td><td>-3.6</td><td>1.0</td><td>-3.6</td><td>3.7E-04</td><td>5.1E-04</td><td>BT20/MB468</td><td>CYTO</td></tr>
<tr><td>>hg38_21_+_8433172_8446622_RNA45SN1_45S_50ntFlanks@12633.12669.37|CGAAGCTACCATCTGTGGGATTATGACTGAACGCCTC</td><td>>hg38_21_+_8433172_8446622_RNA45SN1_45S_50ntFlanks</td><td>12633</td><td>12669</td><td>37</td><td>CGAAGCTACCATCTGTGGGATTATGACTGAACGCCTC</td><td>792.6</td><td>-3.1</td><td>0.9</td><td>-3.5</td><td>4.1E-04</td><td>5.1E-04</td><td>MB231/MB468</td><td>MITO</td></tr>
<tr><td>>hg38_21_+_8433172_8446622_RNA45SN1_45S_50ntFlanks@12410.12433.24|TGATCCTTCGATGTCGGCTCTTCC</td><td>>hg38_21_+_8433172_8446622_RNA45SN1_45S_50ntFlanks</td><td>12410</td><td>12433</td><td>24</td><td>TGATCCTTCGATGTCGGCTCTTCC</td><td>260.1</td><td>3.4</td><td>0.9</td><td>3.6</td><td>3.7E-04</td><td>5.1E-04</td><td>BT20/MB468</td><td>CYTO</td></tr>
<tr><td>>hg38_21_+_8433172_8446622_RNA45SN1_45S_50ntFlanks@12423.12441.19|TCGGCTCTTCCTATCATTG</td><td>>hg38_21_+_8433172_8446622_RNA45SN1_45S_50ntFlanks</td><td>12423</td><td>12441</td><td>19</td><td>TCGGCTCTTCCTATCATTG</td><td>422.9</td><td>-2.6</td><td>0.7</td><td>-3.5</td><td>4.1E-04</td><td>5.1E-04</td><td>MB231/MB468</td><td>NUC</td></tr>
<tr><td>>hg38_21_+_8433172_8446622_RNA45SN1_45S_50ntFlanks@11561.11593.33|GCGCCTAGCAGCCGACTTAGAACTGGTGCGGAC</td><td>>hg38_21_+_8433172_8446622_RNA45SN1_45S_50ntFlanks</td><td>11561</td><td>11593</td><td>33</td><td>GCGCCTAGCAGCCGACTTAGAACTGGTGCGGAC</td><td>398.0</td><td>-6.1</td><td>1.6</td><td>-3.8</td><td>1.7E-04</td><td>5.2E-04</td><td>BT20/MB468</td><td>MP</td></tr>
<tr><td>>am_trnaMT_ProTGG_MT_-_15956_16023@52.70.19|AAAGACTTTTTCTCTGACC</td><td>>am_trnaMT_ProTGG_MT_-_15956_16023</td><td>52</td><td>70</td><td>19</td><td>AAAGACTTTTTCTCTGACC</td><td>1391.2</td><td>3.4</td><td>0.9</td><td>3.6</td><td>3.4E-04</td><td>5.2E-04</td><td>MB231/MB468</td><td>MP</td></tr>
<tr><td>>trnaMT_ValTAC_MT_+_1602_1670@49.70.22|AACTTAACTTGACCGCTCTGAC</td><td>>trnaMT_ValTAC_MT_+_1602_1670</td><td>49</td><td>70</td><td>22</td><td>AACTTAACTTGACCGCTCTGAC</td><td>216.6</td><td>5.2</td><td>1.4</td><td>3.8</td><td>1.7E-04</td><td>5.2E-04</td><td>BT20/MB468</td><td>MP</td></tr>
<tr><td>>hg38_21_+_8433172_8446622_RNA45SN1_45S_50ntFlanks@12907.12932.26|CACCGCACGTTCGTGGGGAACCTGGC</td><td>>hg38_21_+_8433172_8446622_RNA45SN1_45S_50ntFlanks</td><td>12907</td><td>12932</td><td>26</td><td>CACCGCACGTTCGTGGGGAACCTGGC</td><td>320.1</td><td>-5.9</td><td>1.6</td><td>-3.6</td><td>3.4E-04</td><td>5.2E-04</td><td>MB231/MB468</td><td>MP</td></tr>
<tr><td>>hg38_21_+_8433172_8446622_RNA45SN1_45S_50ntFlanks@9792.9817.26|GGCGTGGAGCCGGGCGTGGAATGCGA</td><td>>hg38_21_+_8433172_8446622_RNA45SN1_45S_50ntFlanks</td><td>9792</td><td>9817</td><td>26</td><td>GGCGTGGAGCCGGGCGTGGAATGCGA</td><td>70.4</td><td>9.9</td><td>2.6</td><td>3.8</td><td>1.7E-04</td><td>5.2E-04</td><td>BT20/MB468</td><td>MP</td></tr>
<tr><td>>hg38_21_+_8433172_8446622_RNA45SN1_45S_50ntFlanks@3705.3744.40|TACCTGGTTGATCCTGCCAGTAGCATATGCTTGTCTCAAA</td><td>>hg38_21_+_8433172_8446622_RNA45SN1_45S_50ntFlanks</td><td>3705</td><td>3744</td><td>40</td><td>TACCTGGTTGATCCTGCCAGTAGCATATGCTTGTCTCAAA</td><td>1353.9</td><td>3.2</td><td>0.8</td><td>4.0</td><td>6.0E-05</td><td>5.2E-04</td><td>BT20/MB231</td><td>NUC</td></tr>
<tr><td>>MI0000542|hsa-mir-320a&WithFlank&8|-|22244960|22245043@48.68.21(+2U)&[MIMAT0000510&hsa-miR-320a-3p&offsets|0|-1(+2U);m-60&8|-|22244975|22244996&offsets|0|-1(+2U)]|AAAAGCTGGGTTGAGAGGGCGTT</td><td>>MI0000542|hsa-mir-320a&WithFlank&8|-|22244960|22245043</td><td>48</td><td>68</td><td>21(+2U)</td><td>AAAAGCTGGGTTGAGAGGGCGTT</td><td>2283.0</td><td>2.6</td><td>0.7</td><td>4.0</td><td>6.1E-05</td><td>5.2E-04</td><td>BT20/MB231</td><td>NUC</td></tr>
<tr><td>>MI0003591|hsa-mir-584&WithFlank&5|-|149062307|149062415@22.40.19(+1A)&[MIMAT0003249&hsa-miR-584-5p&offsets|0|-3(+1A);m-167&5|-|149062374|149062394&offsets|0|-2(+1A)]|TTATGGTTTGCCTGGGACTA</td><td>>MI0003591|hsa-mir-584&WithFlank&5|-|149062307|149062415</td><td>22</td><td>40</td><td>19(+1A)</td><td>TTATGGTTTGCCTGGGACTA</td><td>84.4</td><td>6.3</td><td>1.7</td><td>3.6</td><td>3.4E-04</td><td>5.2E-04</td><td>MB231/MB468</td><td>MP</td></tr>
<tr><td>>hg38_21_+_8433172_8446622_RNA45SN1_45S_50ntFlanks@9544.9572.29|GGTGGAGGTCCGTAGCGGTCCTGACGTGC</td><td>>hg38_21_+_8433172_8446622_RNA45SN1_45S_50ntFlanks</td><td>9544</td><td>9572</td><td>29</td><td>GGTGGAGGTCCGTAGCGGTCCTGACGTGC</td><td>1759.2</td><td>-6.9</td><td>2.0</td><td>-3.5</td><td>4.2E-04</td><td>5.2E-04</td><td>BT20/MB468</td><td>NUC</td></tr>
<tr><td>>MI0000114|hsa-mir-107&WithFlank&10|-|89592741|89592833@56.77.22&[MIMAT0000104&hsa-miR-107&offsets|0|-1;m-78&10|-|89592758|89592778&offsets|0|+1]|AGCAGCATTGTACAGGGCTATC</td><td>>MI0000114|hsa-mir-107&WithFlank&10|-|89592741|89592833</td><td>56</td><td>77</td><td>22</td><td>AGCAGCATTGTACAGGGCTATC</td><td>151.2</td><td>2.3</td><td>0.7</td><td>3.5</td><td>4.2E-04</td><td>5.2E-04</td><td>BT20/MB468</td><td>NUC</td></tr>
<tr><td>>MI0000440|hsa-mir-27b&WithFlank&9|+|95085439|95085547@67.87.21&[MIMAT0000419&hsa-miR-27b-3p&offsets|0|0;m-39&9|+|95085505|95085525&offsets|0|0]|TTCACAGTGGCTAAGTTCTGC</td><td>>MI0000440|hsa-mir-27b&WithFlank&9|+|95085439|95085547</td><td>67</td><td>87</td><td>21</td><td>TTCACAGTGGCTAAGTTCTGC</td><td>18135.7</td><td>3.5</td><td>0.9</td><td>3.8</td><td>1.7E-04</td><td>5.2E-04</td><td>BT20/MB468</td><td>MP</td></tr>
<tr><td>>MI0000063|hsa-let-7b&WithFlank&22|+|46113680|46113774@12.31.20&[MIMAT0000063&hsa-let-7b-5p&offsets|0|-2;m-14&22|+|46113691|46113712&offsets|0|-2]|TGAGGTAGTAGGTTGTGTGG</td><td>>MI0000063|hsa-let-7b&WithFlank&22|+|46113680|46113774</td><td>12</td><td>31</td><td>20</td><td>TGAGGTAGTAGGTTGTGTGG</td><td>1130.6</td><td>3.1</td><td>0.8</td><td>4.0</td><td>5.7E-05</td><td>5.2E-04</td><td>BT20/MB231</td><td>CYTO</td></tr>
<tr><td>>hg38_21_+_8433172_8446622_RNA45SN1_45S_50ntFlanks@11894.11912.19|ACTCTAGTCTGGCACGGTG</td><td>>hg38_21_+_8433172_8446622_RNA45SN1_45S_50ntFlanks</td><td>11894</td><td>11912</td><td>19</td><td>ACTCTAGTCTGGCACGGTG</td><td>434.1</td><td>-5.9</td><td>1.6</td><td>-3.8</td><td>1.7E-04</td><td>5.2E-04</td><td>BT20/MB468</td><td>MP</td></tr>
<tr><td>>hg38_21_+_8433172_8446622_RNA45SN1_45S_50ntFlanks@10907.10932.26|GGGCGCGCGCCGCGGCTGGACGAGGC</td><td>>hg38_21_+_8433172_8446622_RNA45SN1_45S_50ntFlanks</td><td>10907</td><td>10932</td><td>26</td><td>GGGCGCGCGCCGCGGCTGGACGAGGC</td><td>114.5</td><td>6.2</td><td>1.6</td><td>3.8</td><td>1.7E-04</td><td>5.2E-04</td><td>BT20/MB468</td><td>MP</td></tr>
<tr><td>>hg38_21_+_8433172_8446622_RNA45SN1_45S_50ntFlanks@11667.11706.40|CTGCCCAGTGCTCTGAATGTCAAAGTGAAGAAATTCAATG</td><td>>hg38_21_+_8433172_8446622_RNA45SN1_45S_50ntFlanks</td><td>11667</td><td>11706</td><td>40</td><td>CTGCCCAGTGCTCTGAATGTCAAAGTGAAGAAATTCAATG</td><td>60.8</td><td>6.7</td><td>1.8</td><td>3.8</td><td>1.7E-04</td><td>5.2E-04</td><td>BT20/MB468</td><td>MP</td></tr>
</table>

| | | | | | | | | | | | | | |
|---|---|---|---|---|---|---|---|---|---|---|---|---|---|
| >hg38_21_+_8433172_8446622_RNA45SN1_45S_50ntFlanks@6655.6688.34\|TCTTAGCGGTGGATCACTCGGCTCGTGCGTCGAT | >hg38_21_+_8433172_8446622_RNA45SN1_45S_50ntFlanks | 6655 | 6688 | 34 | TCTTAGCGGTGGATCACTCGGCTCGTGCGTCGAT | 711.4 | 2.2 | 0.6 | 3.6 | 3.1E-04 | 5.2E-04 | BT20/MB468 | MITO |
| >hg38_21_+_8433172_8446622_RNA45SN1_45S_50ntFlanks@10786.10827.42\|AACAGCCTCTGGCATGTTGGAACAATGTAGGTAAGGGAAGTC | >hg38_21_+_8433172_8446622_RNA45SN1_45S_50ntFlanks | 10786 | 10827 | 42 | AACAGCCTCTGGCATGTTGGAACAATGTAGGTAAGGGAAGTC | 156.8 | -5.3 | 1.5 | -3.5 | 4.2E-04 | 5.2E-04 | MB231/MB468 | NUC |
| >MI0000542\|hsa-mir-320a&WithFlank&8\|-\|22244960\|22245043@49.70.22(+1U)&[MIMAT0000510&hsa-miR-320a-3p&offsets\|+1\|+1(+1U);m-60&8\|-\|22244975\|22244996&offsets\|+1\|+1(+1U)]\|AAAGCTGGGTTGAGAGGGCGAAT | >MI0000542\|hsa-mir-320a&WithFlank&8\|-\|22244960\|22245043 | 49 | 70 | 22(+1U) | AAAGCTGGGTTGAGAGGGCGAAT | 1112.8 | 3.1 | 0.8 | 4.0 | 5.8E-05 | 5.2E-04 | BT20/MB231 | CYTO |
| >hg38_21_+_8433172_8446622_RNA45SN1_45S_50ntFlanks@12416.12454.39\|TTCGATGTCGGCTCTTCCTATCATTGTGAAGCAGAATTC | >hg38_21_+_8433172_8446622_RNA45SN1_45S_50ntFlanks | 12416 | 12454 | 39 | TTCGATGTCGGCTCTTCCTATCATTGTGAAGCAGAATTC | 506.5 | -3.3 | 0.8 | -4.0 | 5.8E-05 | 5.2E-04 | BT20/MB231 | CYTO |
| >hg38_21_+_8433172_8446622_RNA45SN1_45S_50ntFlanks@11583.11602.20\|CTGGTGCGGACCAGGGGAAT | >hg38_21_+_8433172_8446622_RNA45SN1_45S_50ntFlanks | 11583 | 11602 | 20 | CTGGTGCGGACCAGGGGAAT | 1176.9 | -2.7 | 0.8 | -3.5 | 3.9E-04 | 5.2E-04 | MB231/MB468 | CYTO |
| >trnaMT_GluTTC_MT_-_14674_14742@39.72.34,trnalookalike8_GluTTC_5_-_93905172_93905240@39.72.34\|ATTGGTCGTGGTTGTAGTCCGTGCGAGAATACCA | >trnaMT_GluTTC_MT_-_14674_14742 | 39 | 72 | 34 | ATTGGTCGTGGTTGTAGTCCGTGCGAGAATACCA | 33161.1 | 4.4 | 1.2 | 3.8 | 1.7E-04 | 5.2E-04 | BT20/MB468 | MP |
| >trnaMT_HisGTG_MT_+_12138_12206@44.71.28\|CAGAGGCTTACGACCCCTTATTTACCCC | >trnaMT_HisGTG_MT_+_12138_12206 | 44 | 71 | 28 | CAGAGGCTTACGACCCCTTATTTACCCC | 78.9 | 10.0 | 2.7 | 3.8 | 1.7E-04 | 5.2E-04 | BT20/MB468 | MP |
| >hg38_21_+_8433172_8446622_RNA45SN1_45S_50ntFlanks@10349.10377.29\|AACGAGAACTTTGAAGGCCGAAGTGGAGA | >hg38_21_+_8433172_8446622_RNA45SN1_45S_50ntFlanks | 10349 | 10377 | 29 | AACGAGAACTTTGAAGGCCGAAGTGGAGA | 2443.4 | -3.9 | 1.0 | -3.8 | 1.7E-04 | 5.2E-04 | BT20/MB468 | MP |
| >MI0000088\|hsa-mir-30a&WithFlank&6\|-\|71403545\|71403627@12.34.23(+1G)&[MIMAT0000087&hsa-miR-30a-5p&offsets\|0\|+1(+1G);m-15&6\|-\|71403595\|71403616&offsets\|0\|+1(+1G)]\|TGTAAACATCCTCGACTGGAAGCG | >MI0000088\|hsa-mir-30a&WithFlank&6\|-\|71403545\|71403627 | 12 | 34 | 23(+1G) | TGTAAACATCCTCGACTGGAAGCG | 3156.8 | -3.5 | 0.9 | -4.0 | 6.9E-05 | 5.2E-04 | BT20/MB231 | MITO |
| >hg38_21_+_8433172_8446622_RNA45SN1_45S_50ntFlanks@12147.12173.27\|CCAGGTGGGGAGTTTGACTGGGGCGGT | >hg38_21_+_8433172_8446622_RNA45SN1_45S_50ntFlanks | 12147 | 12173 | 27 | CCAGGTGGGGAGTTTGACTGGGGCGGT | 56.0 | 9.1 | 2.3 | 4.0 | 5.9E-05 | 5.2E-04 | BT20/MB231 | MP |
| >hg38_21_+_8433172_8446622_RNA45SN1_45S_50ntFlanks@5486.5521.36\|GCGCTGAGAAGACGGTCGAACTTGACTATCTAGAGG | >hg38_21_+_8433172_8446622_RNA45SN1_45S_50ntFlanks | 5486 | 5521 | 36 | GCGCTGAGAAGACGGTCGAACTTGACTATCTAGAGG | 143.3 | -5.5 | 1.5 | -3.6 | 3.5E-04 | 5.2E-04 | MB231/MB468 | MP |
| >hg38_21_+_8433172_8446622_RNA45SN1_45S_50ntFlanks@10830.10855.26\|CAAGCCGGATCCGTAACTTCGGGATA | >hg38_21_+_8433172_8446622_RNA45SN1_45S_50ntFlanks | 10830 | 10855 | 26 | CAAGCCGGATCCGTAACTTCGGGATA | 188.4 | -6.1 | 1.6 | -3.8 | 1.7E-04 | 5.2E-04 | BT20/MB468 | MP |
| >hg38_MT_+_1621_3279_RNR2_16S_50ntFlanks@621.656.36\|ATATAACTGAACTCCTCACACCCAATTGGACCAATC | >hg38_MT_+_1621_3279_RNR2_16S_50ntFlanks | 621 | 656 | 36 | ATATAACTGAACTCCTCACACCCAATTGGACCAATC | 78.5 | 10.0 | 2.7 | 3.8 | 1.7E-04 | 5.2E-04 | BT20/MB468 | MP |
| >MI0000088\|hsa-mir-30a&WithFlank&6\|-\|71403545\|71403627@12.34.23(+1C)&[MIMAT0000087&hsa-miR-30a-5p&offsets\|0\|+1(+1C);m-15&6\|-\|71403595\|71403616&offsets\|0\|+1(+1C)]\|TGTAAACATCCTCGACTGGAAGCC | >MI0000088\|hsa-mir-30a&WithFlank&6\|-\|71403545\|71403627 | 12 | 34 | 23(+1C) | TGTAAACATCCTCGACTGGAAGCC | 2057.4 | 3.7 | 1.1 | 3.5 | 4.2E-04 | 5.2E-04 | BT20/MB468 | NUC |
| >hg38_21_+_8433172_8446622_RNA45SN1_45S_50ntFlanks@12637.12657.21\|GCTACCATCTGTGGGATTATG | >hg38_21_+_8433172_8446622_RNA45SN1_45S_50ntFlanks | 12637 | 12657 | 21 | GCTACCATCTGTGGGATTATG | 532.4 | -4.6 | 1.2 | -3.8 | 1.7E-04 | 5.2E-04 | BT20/MB468 | MP |
| >hg38_21_+_8433172_8446622_RNA45SN1_45S_50ntFlanks@11561.11591.31\|GCGCCTAGCAGCCGACTTAGAACTGGTGCGG | >hg38_21_+_8433172_8446622_RNA45SN1_45S_50ntFlanks | 11561 | 11591 | 31 | GCGCCTAGCAGCCGACTTAGAACTGGTGCGG | 986.3 | -6.3 | 1.7 | -3.8 | 1.7E-04 | 5.2E-04 | BT20/MB468 | MP |
| >hg38_21_+_8433172_8446622_RNA45SN1_45S_50ntFlanks@9575.9605.31\|ATCGGTCGTCCGACCTGGGTATAGGGGCGAA | >hg38_21_+_8433172_8446622_RNA45SN1_45S_50ntFlanks | 9575 | 9605 | 31 | ATCGGTCGTCCGACCTGGGTATAGGGGCGAA | 620.6 | -6.0 | 1.7 | -3.6 | 3.5E-04 | 5.2E-04 | MB231/MB468 | MP |
| >hg38_21_+_8433172_8446622_RNA45SN1_45S_50ntFlanks@12630.12654.25\|GGGCGAAGCTACCATCTGTGGGATT | >hg38_21_+_8433172_8446622_RNA45SN1_45S_50ntFlanks | 12630 | 12654 | 25 | GGGCGAAGCTACCATCTGTGGGATT | 308.3 | -4.1 | 1.1 | -3.6 | 3.1E-04 | 5.2E-04 | BT20/MB468 | MITO |
| >hg38_21_+_8433172_8446622_RNA45SN1_45S_50ntFlanks@11569.11600.32\|CAGCCGACTTAGAACTGGTGCGGACCAGGGGA | >hg38_21_+_8433172_8446622_RNA45SN1_45S_50ntFlanks | 11569 | 11600 | 32 | CAGCCGACTTAGAACTGGTGCGGACCAGGGGA | 1973.1 | -3.0 | 0.8 | -3.5 | 3.9E-04 | 5.2E-04 | MB231/MB468 | CYTO |
| >am_MI0000809\|hsa-mir-151a&WithFlank&8\|-\|140732558\|140732659@53.72.20(+2U)&[MIMAT0000757&hsa-miR-151a-3p&offsets\|0\|-1(+2U);m-30&8\|-\|140732587\|140732607&offsets\|0\|-1(+2U)]\|CTAGACTGAAGCTCCTTGAGTT | >am_MI0000809\|hsa-mir-151a&WithFlank&8\|-\|140732558\|140732659 | 53 | 72 | 20(+2U) | CTAGACTGAAGCTCCTTGAGTT | 136.5 | 6.2 | 1.7 | 3.6 | 3.5E-04 | 5.2E-04 | MB231/MB468 | MP |
| >hg38_MT_+_598_1651_RNR1_12S_50ntFlanks@561.605.45\|TAAAACTCAAAGGACCTGGCGGTGCTTCATATCCCTCTAGAGGAG | >hg38_MT_+_598_1651_RNR1_12S_50ntFlanks | 561 | 605 | 45 | TAAAACTCAAAGGACCTGGCGGTGCTTCATATCCCTCTAGAGGAG | 124.8 | 5.2 | 1.5 | 3.5 | 4.2E-04 | 5.2E-04 | MB231/MB468 | MITO |
| >MI0000264\|hsa-mir-7-2&WithFlank&15\|+\|88611819\|88611940@38.59.22(+1A);MI0000265\|hsa-mir-7-3&WithFlank&19\|+\|4770664\|4770785@37.58.22(+1A);MI0000263\|hsa-mir-7-1&WithFlank&9\|-\|83969742\|83969863@30.51.22(+1A)&[MIMAT0000252&hsa-miR-7-5p&offsets\|0\|-2(+1A);m-400&15\|+\|88611856\|88611879&offsets\|0\|-2(+1A)];[MIMAT0000252_1&hsa-miR-7-5p&offsets\|0\|-2(+1A);m-402&19\|+\|4770700\|4770723&offsets\|0\|-2(+1A)];[MIMAT0000252_2&hsa-miR-7-5p&offsets\|0\|-2(+1A);m-401&9\|-\|83969811\|83969834&offsets\|0\|-2(+1A)]\|TGGAAGACTAGTGATTTTGTTGA | >MI0000264\|hsa-mir-7-2&WithFlank&15\|+\|88611819\|88611940 | 38 | 59 | 22(+1A) | TGGAAGACTAGTGATTTTGTTGA | 679.8 | 4.3 | 1.1 | 4.0 | 5.9E-05 | 5.3E-04 | BT20/MB231 | CYTO |
| >hg38_21_+_8433172_8446622_RNA45SN1_45S_50ntFlanks@7969.8011.43\|CCGAGACGCGACCTCAGATCAGACGTGGCGACCCGCTGAATTT | >hg38_21_+_8433172_8446622_RNA45SN1_45S_50ntFlanks | 7969 | 8011 | 43 | CCGAGACGCGACCTCAGATCAGACGTGGCGACCCGCTGAATTT | 297.2 | 5.9 | 1.7 | 3.5 | 4.2E-04 | 5.3E-04 | MB231/MB468 | NUC |
| >trnaMT_HisGTG_MT_+_12138_12206@28.66.39\|ATTGTGAATCTGACAACAGAGGCTTACGACCCCTTATTT | >trnaMT_HisGTG_MT_+_12138_12206 | 28 | 66 | 39 | ATTGTGAATCTGACAACAGAGGCTTACGACCCCTTATTT | 149.1 | -4.7 | 1.3 | -3.5 | 4.2E-04 | 5.3E-04 | MB231/MB468 | NUC |
| >hg38_21_+_8433172_8446622_RNA45SN1_45S_50ntFlanks@12473.12496.24\|CACCCACTAATAGGGAACGTGAGC | >hg38_21_+_8433172_8446622_RNA45SN1_45S_50ntFlanks | 12473 | 12496 | 24 | CACCCACTAATAGGGAACGTGAGC | 552.8 | 2.6 | 0.7 | 3.5 | 3.9E-04 | 5.3E-04 | MB231/MB468 | CYTO |
| >hg38_21_+_8433172_8446622_RNA45SN1_45S_50ntFlanks@13001.13027.27\|GCGATCTATTGAAAGTCAGCCCTCGAC | >hg38_21_+_8433172_8446622_RNA45SN1_45S_50ntFlanks | 13001 | 13027 | 27 | GCGATCTATTGAAAGTCAGCCCTCGAC | 366.7 | -3.8 | 1.1 | -3.5 | 3.9E-04 | 5.3E-04 | MB231/MB468 | CYTO |
| >MI0000746\|hsa-mir-99b&WithFlank&19\|+\|51692606\|51692687@51.70.20(+1U)&[MIMAT0004678&hsa-miR-99b-3p&offsets\|0\|-2(+1U);m-151&19\|+\|51692656\|51692678&offsets\|0\|-3(+1U)]\|CAAGCTCGTGTCTGTGGGTCT | >MI0000746\|hsa-mir-99b&WithFlank&19\|+\|51692606\|51692687 | 51 | 70 | 20(+1U) | CAAGCTCGTGTCTGTGGGTCT | 515.4 | 3.2 | 0.8 | 4.0 | 5.9E-05 | 5.3E-04 | BT20/MB231 | CYTO |
| >MI0000746\|hsa-mir-99b&WithFlank&19\|+\|51692606\|51692687@51.72.22(+1A)&[MIMAT0004678&hsa-miR-99b-3p&offsets\|0\|0(+1A);m-151&19\|+\|51692656\|51692678&offsets\|0\|-1(+1A)]\|CAAGCTCGTGTCTGTGGGTCCGA | >MI0000746\|hsa-mir-99b&WithFlank&19\|+\|51692606\|51692687 | 51 | 72 | 22(+1A) | CAAGCTCGTGTCTGTGGGTCCGA | 1648.9 | 3.6 | 0.9 | 4.0 | 5.9E-05 | 5.3E-04 | BT20/MB231 | CYTO |
| >hg38_21_+_8433172_8446622_RNA45SN1_45S_50ntFlanks@10841.10868.28\|CGTAACTTCGGGATAAGGATTGGCTCTA | >hg38_21_+_8433172_8446622_RNA45SN1_45S_50ntFlanks | 10841 | 10868 | 28 | CGTAACTTCGGGATAAGGATTGGCTCTA | 8898.3 | -3.3 | 0.9 | -3.5 | 3.9E-04 | 5.3E-04 | MB231/MB468 | CYTO |
| >hg38_21_+_8433172_8446622_RNA45SN1_45S_50ntFlanks@3840.3862.23\|CTCCTACTTGGATAACTGTGGTA | >hg38_21_+_8433172_8446622_RNA45SN1_45S_50ntFlanks | 3840 | 3862 | 23 | CTCCTACTTGGATAACTGTGGTA | 357.2 | 3.0 | 0.8 | 3.6 | 3.1E-04 | 5.3E-04 | BT20/MB468 | MITO |
| >hg38_21_+_8433172_8446622_RNA45SN1_45S_50ntFlanks@6648.6677.30\|GTACGACTCTTAGCGGTGGATCACTCGGCT | >hg38_21_+_8433172_8446622_RNA45SN1_45S_50ntFlanks | 6648 | 6677 | 30 | GTACGACTCTTAGCGGTGGATCACTCGGCT | 785.1 | 3.0 | 0.9 | 3.5 | 4.2E-04 | 5.3E-04 | MB231/MB468 | NUC |
| >hg38_MT_+_1621_3279_RNR2_16S_50ntFlanks@1431.1453.23\|AAAGTCCTACGTGATCTGAGTTC | >hg38_MT_+_1621_3279_RNR2_16S_50ntFlanks | 1431 | 1453 | 23 | AAAGTCCTACGTGATCTGAGTTC | 337.4 | 3.2 | 0.9 | 3.6 | 3.5E-04 | 5.3E-04 | MB231/MB468 | MP |
| >trna111_HisGTG_1_-_147774845_147774916@1.32.32,trna118_HisGTG_1_-_145396881_145396952@1.32.32,trna16_HisGTG_1_+_146544773_146544844@1.32.32,trna1_HisGTG_15_+_45493349_45493420@1.32.32,trna21_HisGTG_1_+_147753471_147753542@1.32.32,trna33_HisGTG_6_+_27125906_27125977@1.32.32,trna7_HisGTG_9_-_14433938_14434009@1.32.32,trna8_HisGTG_15_-_45492611_45492682@1.32.32,trna9_HisGTG_15_-_45490804_45490875@1.32.32\|GCCGTGATCGTATAGTGGTTAGTACTCTGCGT | >trna111_HisGTG_1_-_147774845_147774916 | 1 | 32 | 32 | GCCGTGATCGTATAGTGGTTAGTACTCTGCGT | 309.9 | 6.6 | 1.9 | 3.6 | 3.5E-04 | 5.3E-04 | MB231/MB468 | MP |
| >hg38_21_+_8433172_8446622_RNA45SN1_45S_50ntFlanks@11209.11254.46\|CGGCGACTCTGGACGCGAGCCGGGCCCTTCCCGTGGATCGCCCCAG | >hg38_21_+_8433172_8446622_RNA45SN1_45S_50ntFlanks | 11209 | 11254 | 46 | CGGCGACTCTGGACGCGAGCCGGGCCCTTCCCGTGGATCGCCCCAG | 393.6 | 4.0 | 1.0 | 4.0 | 7.0E-05 | 5.3E-04 | BT20/MB231 | MITO |
| >hg38_21_+_8433172_8446622_RNA45SN1_45S_50ntFlanks@7975.8006.32\|CGCGACCTCAGATCAGACGTGGCGACCCGCTG | >hg38_21_+_8433172_8446622_RNA45SN1_45S_50ntFlanks | 7975 | 8006 | 32 | CGCGACCTCAGATCAGACGTGGCGACCCGCTG | 149509.9 | 2.9 | 0.7 | 4.0 | 7.1E-05 | 5.3E-04 | BT20/MB231 | MITO |
| >hg38_21_+_8433172_8446622_RNA45SN1_45S_50ntFlanks@6670.6700.31\|ACTCGGCTCGTGCGTCGATGAAGAACGCAGC | >hg38_21_+_8433172_8446622_RNA45SN1_45S_50ntFlanks | 6670 | 6700 | 31 | ACTCGGCTCGTGCGTCGATGAAGAACGCAGC | 191.5 | 3.5 | 0.9 | 3.8 | 1.7E-04 | 5.3E-04 | BT20/MB468 | MP |
| >hg38_21_+_8433172_8446622_RNA45SN1_45S_50ntFlanks@10798.10825.28\|CATGTTGGAACAATGTAGGTAAGGGAAG | >hg38_21_+_8433172_8446622_RNA45SN1_45S_50ntFlanks | 10798 | 10825 | 28 | CATGTTGGAACAATGTAGGTAAGGGAAG | 338.1 | -5.9 | 1.6 | -3.8 | 1.7E-04 | 5.3E-04 | BT20/MB468 | MP |
| >hg38_21_+_8433172_8446622_RNA45SN1_45S_50ntFlanks@11923.11941.19\|AGAGGTGTAGAATAAGTGG | >hg38_21_+_8433172_8446622_RNA45SN1_45S_50ntFlanks | 11923 | 11941 | 19 | AGAGGTGTAGAATAAGTGG | 62.5 | 9.3 | 2.3 | 4.0 | 6.0E-05 | 5.3E-04 | BT20/MB231 | MP |
| >hg38_21_+_8433172_8446622_RNA45SN1_45S_50ntFlanks@12413.12443.31\|TCCTTCGATGTCGGCTCTTCCTATCATTGTG | >hg38_21_+_8433172_8446622_RNA45SN1_45S_50ntFlanks | 12413 | 12443 | 31 | TCCTTCGATGTCGGCTCTTCCTATCATTGTG | 886.8 | -2.9 | 0.8 | -3.5 | 4.2E-04 | 5.3E-04 | MB231/MB468 | NUC |
| >MI0000273\|hsa-mir-183&WithFlank&7\|-\|129774899\|129775020@33.53.21&[MIMAT0000261&hsa-miR-183-5p&offsets\|0\|-1;m-41&7\|-\|129774966\|129774987&offsets\|-1\|-2]\|TATGGCACTGGTAGAATTCAC | >MI0000273\|hsa-mir-183&WithFlank&7\|-\|129774899\|129775020 | 33 | 53 | 21 | TATGGCACTGGTAGAATTCAC | 386.2 | 3.3 | 0.8 | 4.0 | 5.9E-05 | 5.3E-04 | BT20/MB231 | CYTO |
| >MI0000273\|hsa-mir-183&WithFlank&7\|-\|129774899\|129775020@33.55.23&[MIMAT0000261&hsa-miR-183-5p&offsets\|0\|+1;m-41&7\|-\|129774966\|129774987&offsets\|-1\|0]\|TATGGCACTGGTAGAATTCACTG | >MI0000273\|hsa-mir-183&WithFlank&7\|-\|129774899\|129775020 | 33 | 55 | 23 | TATGGCACTGGTAGAATTCACTG | 494.6 | 3.1 | 0.9 | 3.5 | 4.2E-04 | 5.3E-04 | MB231/MB468 | MITO |
| >hg38_21_+_8433172_8446622_RNA45SN1_45S_50ntFlanks@9533.9570.38\|GAGGAAACTCTGGTGGAGGTCCGTAGCGGTCCTGACGT | >hg38_21_+_8433172_8446622_RNA45SN1_45S_50ntFlanks | 9533 | 9570 | 38 | GAGGAAACTCTGGTGGAGGTCCGTAGCGGTCCTGACGT | 888.8 | -2.5 | 0.7 | -3.5 | 4.2E-04 | 5.3E-04 | MB231/MB468 | NUC |
| >hg38_21_+_8433172_8446622_RNA45SN1_45S_50ntFlanks@8196.8217.22\|CGTGGACGGTGTGAGGCCGGTA | >hg38_21_+_8433172_8446622_RNA45SN1_45S_50ntFlanks | 8196 | 8217 | 22 | CGTGGACGGTGTGAGGCCGGTA | 330.8 | -3.3 | 0.9 | -3.8 | 1.7E-04 | 5.3E-04 | BT20/MB468 | MP |
| >hg38_21_+_8433172_8446622_RNA45SN1_45S_50ntFlanks@12994.13036.43\|CCTCGCTGCGATCTATTGAAAGTCAGCCCTCGACACAAGGGTT | >hg38_21_+_8433172_8446622_RNA45SN1_45S_50ntFlanks | 12994 | 13036 | 43 | CCTCGCTGCGATCTATTGAAAGTCAGCCCTCGACACAAGGGTT | 431.2 | 4.7 | 1.3 | 3.6 | 3.1E-04 | 5.3E-04 | BT20/MB468 | MITO |
| >hg38_21_+_8433172_8446622_RNA45SN1_45S_50ntFlanks@11921.11948.28\|TGAGAGGTGTAGAATAAGTGGGAGGCCC | >hg38_21_+_8433172_8446622_RNA45SN1_45S_50ntFlanks | 11921 | 11948 | 28 | TGAGAGGTGTAGAATAAGTGGGAGGCCC | 387.0 | -3.7 | 1.1 | -3.5 | 4.0E-04 | 5.3E-04 | MB231/MB468 | CYTO |

<table>
<tr><td></td><td></td><td></td><td></td><td></td><td></td><td></td><td></td><td></td><td></td><td></td><td></td><td></td><td></td></tr>
<tr><td>>trnaMT_ValTAC_MT_+_1602_1670@27.71.45|CAACTTACACTTAGGAGATTTCAACTTAACTTGACCGCTCTGACC</td><td>>trnaMT_ValTAC_MT_+_1602_1670</td><td>27</td><td>71</td><td>45</td><td>CAACTTACACTTAGGAGATTTCAACTTAACTTGACCGCTCTGACC</td><td>75.6</td><td>10.0</td><td>2.7</td><td>3.8</td><td>1.7E-04</td><td>5.3E-04</td><td>BT20/MB468</td><td>MP</td></tr>
<tr><td>>hg38_21_+_8433172_8446622_RNA45SN1_45S_50ntFlanks@6719.6756.38|TGAATTGCAGGACACATTGATCATCGACACTTCGAACG</td><td>>hg38_21_+_8433172_8446622_RNA45SN1_45S_50ntFlanks</td><td>6719</td><td>6756</td><td>38</td><td>TGAATTGCAGGACACATTGATCATCGACACTTCGAACG</td><td>167.6</td><td>-2.3</td><td>0.6</td><td>-3.6</td><td>3.8E-04</td><td>5.3E-04</td><td>BT20/MB468</td><td>CYTO</td></tr>
<tr><td>>MI0003513|hsa-mir-455&WithFlank&9|+|114209428|114209535@22.43.22&[MIMAT0003150&hsa-miR-455-5p&offsets|0|0;m-232&9|+|114209449|114209470&offsets|0|0]|TATGTGCCTTTGGACTACATCG</td><td>>MI0003513|hsa-mir-455&WithFlank&9|+|114209428|114209535</td><td>22</td><td>43</td><td>22</td><td>TATGTGCCTTTGGACTACATCG</td><td>572.8</td><td>-5.5</td><td>1.4</td><td>-4.0</td><td>6.0E-05</td><td>5.3E-04</td><td>BT20/MB231</td><td>MP</td></tr>
<tr><td>>hg38_21_+_8433172_8446622_RNA45SN1_45S_50ntFlanks@11212.11250.39|CGACTCTGGACGCGAGCCGGGCCCTTCCCGTGGATCGCC</td><td>>hg38_21_+_8433172_8446622_RNA45SN1_45S_50ntFlanks</td><td>11212</td><td>11250</td><td>39</td><td>CGACTCTGGACGCGAGCCGGGCCCTTCCCGTGGATCGCC</td><td>382.6</td><td>-2.9</td><td>0.8</td><td>-3.6</td><td>3.8E-04</td><td>5.3E-04</td><td>BT20/MB468</td><td>CYTO</td></tr>
<tr><td>>MI0000342|hsa-mir-200b&WithFlank&1|+|1167098|1167204@63.83.21&[MIMAT0000318&hsa-miR-200b-3p&offsets|0|-1;m-93&1|+|1167160|1167182&offsets|0|-2]|TAATACTGCCTGGTAATGATG</td><td>>MI0000342|hsa-mir-200b&WithFlank&1|+|1167098|1167204</td><td>63</td><td>83</td><td>21</td><td>TAATACTGCCTGGTAATGATG</td><td>365.8</td><td>-4.5</td><td>1.3</td><td>-3.5</td><td>4.2E-04</td><td>5.3E-04</td><td>MB231/MB468</td><td>MITO</td></tr>
<tr><td>>hg38_21_+_8433172_8446622_RNA45SN1_45S_50ntFlanks@7982.8012.31|TCAGATCAGACGTGGCGACCCGCTGAATTTA</td><td>>hg38_21_+_8433172_8446622_RNA45SN1_45S_50ntFlanks</td><td>7982</td><td>8012</td><td>31</td><td>TCAGATCAGACGTGGCGACCCGCTGAATTTA</td><td>168.5</td><td>-5.8</td><td>1.5</td><td>-3.8</td><td>1.7E-04</td><td>5.3E-04</td><td>BT20/MB468</td><td>MP</td></tr>
<tr><td>>am_trna117_GlyTCC_1_-_145397864_145397935@1.30.30,trna45_GlyTCC_1_+_161500903_161500974@1.30.30,trna70_GlyTCC_1_-_161439547_161439618@1.30.30,trna73_GlyTCC_1_-_161432166_161432237@1.30.30,trna76_GlyTCC_1_-_161424756_161424827@1.30.30,trna79_GlyTCC_1_-_161417375_161417446@1.30.30|GCGTTGGTGGTATAGTGGTGAGCATAGCTG</td><td>>am_trna117_GlyTCC_1_-_145397864_145397935</td><td>1</td><td>30</td><td>30</td><td>GCGTTGGTGGTATAGTGGTGAGCATAGCTG</td><td>720.3</td><td>-4.5</td><td>1.2</td><td>-3.6</td><td>3.1E-04</td><td>5.3E-04</td><td>BT20/MB468</td><td>MITO</td></tr>
<tr><td>>hg38_21_+_8433172_8446622_RNA45SN1_45S_50ntFlanks@12409.12446.38|TTGATCCTTCGATGTCGGCTCTTCCTATCATTGTGAAG</td><td>>hg38_21_+_8433172_8446622_RNA45SN1_45S_50ntFlanks</td><td>12409</td><td>12446</td><td>38</td><td>TTGATCCTTCGATGTCGGCTCTTCCTATCATTGTGAAG</td><td>213.6</td><td>2.8</td><td>0.8</td><td>3.5</td><td>4.0E-04</td><td>5.3E-04</td><td>MB231/MB468</td><td>CYTO</td></tr>
<tr><td>>hg38_21_+_8433172_8446622_RNA45SN1_45S_50ntFlanks@11644.11691.48|CGGGTGTTGACGCGATGTGATTTCTGCCCAGTGCTCTGAATGTCAAAG</td><td>>hg38_21_+_8433172_8446622_RNA45SN1_45S_50ntFlanks</td><td>11644</td><td>11691</td><td>48</td><td>CGGGTGTTGACGCGATGTGATTTCTGCCCAGTGCTCTGAATGTCAAAG</td><td>392.8</td><td>13.1</td><td>3.6</td><td>3.6</td><td>3.2E-04</td><td>5.3E-04</td><td>BT20/MB468</td><td>MITO</td></tr>
<tr><td>>hg38_21_+_8433172_8446622_RNA45SN1_45S_50ntFlanks@10782.10801.20|GGTGAACAGCCTCTGGCATG</td><td>>hg38_21_+_8433172_8446622_RNA45SN1_45S_50ntFlanks</td><td>10782</td><td>10801</td><td>20</td><td>GGTGAACAGCCTCTGGCATG</td><td>289.5</td><td>-2.3</td><td>0.7</td><td>-3.6</td><td>3.2E-04</td><td>5.3E-04</td><td>BT20/MB468</td><td>MITO</td></tr>
<tr><td>>hg38_21_+_8433172_8446622_RNA45SN1_45S_50ntFlanks@6645.6682.38|CTCGTACGACTCTTAGCGGTGGATCACTCGGCTCGTGC</td><td>>hg38_21_+_8433172_8446622_RNA45SN1_45S_50ntFlanks</td><td>6645</td><td>6682</td><td>38</td><td>CTCGTACGACTCTTAGCGGTGGATCACTCGGCTCGTGC</td><td>341.8</td><td>3.5</td><td>1.0</td><td>3.5</td><td>4.2E-04</td><td>5.3E-04</td><td>MB231/MB468</td><td>MITO</td></tr>
<tr><td>>hg38_21_+_8433172_8446622_RNA45SN1_45S_50ntFlanks@9535.9563.29|GGAAACTCTGGTGGAGGTCCGTAGCGGTC</td><td>>hg38_21_+_8433172_8446622_RNA45SN1_45S_50ntFlanks</td><td>9535</td><td>9563</td><td>29</td><td>GGAAACTCTGGTGGAGGTCCGTAGCGGTC</td><td>163.7</td><td>-3.0</td><td>0.9</td><td>-3.5</td><td>4.0E-04</td><td>5.3E-04</td><td>MB231/MB468</td><td>CYTO</td></tr>
<tr><td>>hg38_21_+_8433172_8446622_RNA45SN1_45S_50ntFlanks@11562.11600.39|CGCCTAGCAGCCGACTTAGAACTGGTGCGGACCAGGGGA</td><td>>hg38_21_+_8433172_8446622_RNA45SN1_45S_50ntFlanks</td><td>11562</td><td>11600</td><td>39</td><td>CGCCTAGCAGCCGACTTAGAACTGGTGCGGACCAGGGGA</td><td>209.1</td><td>-2.8</td><td>0.8</td><td>-3.5</td><td>4.3E-04</td><td>5.3E-04</td><td>MB231/MB468</td><td>NUC</td></tr>
<tr><td>>hg38_21_+_8433172_8446622_RNA45SN1_45S_50ntFlanks@9545.9564.20|GTGGAGGTCCGTAGCGGTCC</td><td>>hg38_21_+_8433172_8446622_RNA45SN1_45S_50ntFlanks</td><td>9545</td><td>9564</td><td>20</td><td>GTGGAGGTCCGTAGCGGTCC</td><td>688.9</td><td>-5.5</td><td>1.6</td><td>-3.6</td><td>3.8E-04</td><td>5.3E-04</td><td>BT20/MB468</td><td>CYTO</td></tr>
<tr><td>>hg38_21_+_8433172_8446622_RNA45SN1_45S_50ntFlanks@11670.11687.18|CCCAGTGCTCTGAATGTC</td><td>>hg38_21_+_8433172_8446622_RNA45SN1_45S_50ntFlanks</td><td>11670</td><td>11687</td><td>18</td><td>CCCAGTGCTCTGAATGTC</td><td>1207.0</td><td>-2.6</td><td>0.7</td><td>-3.6</td><td>3.8E-04</td><td>5.3E-04</td><td>BT20/MB468</td><td>CYTO</td></tr>
<tr><td>>MI0000093|hsa-mir-92a-1&WithFlank&13|+|91351308|91351397@54.73.20(+1A);MI0000094|hsa-mir-92a-2&WithFlank&X|-|134169532|134169618@54.73.20(+1A)&[MIMAT0000092&hsa-miR-92a-3p&offsets|0|-2(+1A);m-9&13|+|91351361|91351382&offsets|0|-2(+1A)];[MIMAT0000092_1&hsa-miR-92a-3p&offsets|0|-2(+1A);m-11&X|-|134169544|134169565&offsets|0|-2(+1A)]|TATTGCACTTGTCCCGGCCTA</td><td>>MI0000093|hsa-mir-92a-1&WithFlank&13|+|91351308|91351397</td><td>54</td><td>73</td><td>20(+1A)</td><td>TATTGCACTTGTCCCGGCCTA</td><td>198.5</td><td>5.7</td><td>1.6</td><td>3.6</td><td>3.6E-04</td><td>5.3E-04</td><td>MB231/MB468</td><td>MP</td></tr>
<tr><td>>hg38_21_+_8433172_8446622_RNA45SN1_45S_50ntFlanks@7968.8005.38|TCCGAGACGCGACCTCAGATCAGACGTGGCGACCCGCT</td><td>>hg38_21_+_8433172_8446622_RNA45SN1_45S_50ntFlanks</td><td>7968</td><td>8005</td><td>38</td><td>TCCGAGACGCGACCTCAGATCAGACGTGGCGACCCGCT</td><td>1704.7</td><td>3.9</td><td>1.0</td><td>4.0</td><td>6.3E-05</td><td>5.3E-04</td><td>BT20/MB231</td><td>NUC</td></tr>
<tr><td>>hg38_21_+_8433172_8446622_RNA45SN1_45S_50ntFlanks@7975.8007.33|CGCGACCTCAGATCAGACGTGGCGACCCGCTGA</td><td>>hg38_21_+_8433172_8446622_RNA45SN1_45S_50ntFlanks</td><td>7975</td><td>8007</td><td>33</td><td>CGCGACCTCAGATCAGACGTGGCGACCCGCTGA</td><td>37826.4</td><td>2.9</td><td>0.7</td><td>4.0</td><td>6.3E-05</td><td>5.3E-04</td><td>BT20/MB231</td><td>NUC</td></tr>
<tr><td>>trna116_GluCTC_1_-_145399233_145399304@35.75.41,trna134_GluTTC_1_-_16861774_16861845@35.75.41,trna71_GluCTC_1_-_161439189_161439260@35.75.41,trna74_GluCTC_1_-_161431809_161431880@35.75.41,trna77_GluCTC_1_-_161424398_161424469@35.75.41,trna77_GluCTC_6_+_28949976_28950047@35.75.41,trna80_GluCTC_1_-_161417018_161417089@35.75.41,trna84_GluTTC_1_-_161391883_161391954@35.75.41,trna87_GluCTC_6_-_126101393_126101464@35.75.41|TCACCGCCGCGGCCCGGGTTCGATTCCCGGTCAGGGAACCA</td><td>>trna116_GluCTC_1_-_145399233_145399304</td><td>35</td><td>75</td><td>41</td><td>TCACCGCCGCGGCCCGGGTTCGATTCCCGGTCAGGGAACCA</td><td>3695.0</td><td>-4.0</td><td>1.0</td><td>-4.0</td><td>6.0E-05</td><td>5.3E-04</td><td>BT20/MB231</td><td>CYTO</td></tr>
<tr><td>>hg38_21_+_8433172_8446622_RNA45SN1_45S_50ntFlanks@4732.4767.36|AGAACGAAAGTCGGAGGTTCGAAGACGATCAGATAC</td><td>>hg38_21_+_8433172_8446622_RNA45SN1_45S_50ntFlanks</td><td>4732</td><td>4767</td><td>36</td><td>AGAACGAAAGTCGGAGGTTCGAAGACGATCAGATAC</td><td>184.0</td><td>6.1</td><td>1.6</td><td>3.8</td><td>1.7E-04</td><td>5.3E-04</td><td>BT20/MB468</td><td>MP</td></tr>
<tr><td>>hg38_21_+_8433172_8446622_RNA45SN1_45S_50ntFlanks@8380.8407.28|CAAGAGGGCGTGAAACCGTTAAGAGGTA</td><td>>hg38_21_+_8433172_8446622_RNA45SN1_45S_50ntFlanks</td><td>8380</td><td>8407</td><td>28</td><td>CAAGAGGGCGTGAAACCGTTAAGAGGTA</td><td>394.3</td><td>-3.2</td><td>0.8</td><td>-3.8</td><td>1.7E-04</td><td>5.3E-04</td><td>BT20/MB468</td><td>MP</td></tr>
<tr><td>>trnaMT_GluTTC_MT_-_14674_14742@13.57.45,trnalookalike8_GluTTC_5_-_93905172_93905240@13.57.45|GAAATACAACGATGGTTTTTCATATCATTGGTCGTGGTTGTAGTC</td><td>>trnaMT_GluTTC_MT_-_14674_14742</td><td>13</td><td>57</td><td>45</td><td>GAAATACAACGATGGTTTTTCATATCATTGGTCGTGGTTGTAGTC</td><td>78.1</td><td>10.0</td><td>2.7</td><td>3.8</td><td>1.7E-04</td><td>5.3E-04</td><td>BT20/MB468</td><td>MP</td></tr>
<tr><td>>hg38_21_+_8433172_8446622_RNA45SN1_45S_50ntFlanks@5467.5484.18|GGCCCACGGCCCTGGCGG</td><td>>hg38_21_+_8433172_8446622_RNA45SN1_45S_50ntFlanks</td><td>5467</td><td>5484</td><td>18</td><td>GGCCCACGGCCCTGGCGG</td><td>226.3</td><td>-3.8</td><td>1.1</td><td>-3.6</td><td>3.8E-04</td><td>5.3E-04</td><td>BT20/MB468</td><td>CYTO</td></tr>
<tr><td>>hg38_21_+_8433172_8446622_RNA45SN1_45S_50ntFlanks@3841.3868.28|TCCTACTTGGATAACTGTGGTAATTCTA</td><td>>hg38_21_+_8433172_8446622_RNA45SN1_45S_50ntFlanks</td><td>3841</td><td>3868</td><td>28</td><td>TCCTACTTGGATAACTGTGGTAATTCTA</td><td>383.1</td><td>2.3</td><td>0.7</td><td>3.6</td><td>3.2E-04</td><td>5.3E-04</td><td>BT20/MB468</td><td>MITO</td></tr>
<tr><td>>hg38_21_+_8433172_8446622_RNA45SN1_45S_50ntFlanks@6721.6749.29|AATTGCAGGACACATTGATCATCGACACT</td><td>>hg38_21_+_8433172_8446622_RNA45SN1_45S_50ntFlanks</td><td>6721</td><td>6749</td><td>29</td><td>AATTGCAGGACACATTGATCATCGACACT</td><td>157.0</td><td>-5.5</td><td>1.5</td><td>-3.6</td><td>3.6E-04</td><td>5.4E-04</td><td>MB231/MB468</td><td>MP</td></tr>
<tr><td>>hg38_21_+_8433172_8446622_RNA45SN1_45S_50ntFlanks@4370.4391.22|TTAAAAAGCTCGTAGTTGGATC</td><td>>hg38_21_+_8433172_8446622_RNA45SN1_45S_50ntFlanks</td><td>4370</td><td>4391</td><td>22</td><td>TTAAAAAGCTCGTAGTTGGATC</td><td>246.1</td><td>-2.1</td><td>0.6</td><td>-3.5</td><td>4.0E-04</td><td>5.4E-04</td><td>MB231/MB468</td><td>CYTO</td></tr>
<tr><td>>hg38_21_+_8433172_8446622_RNA45SN1_45S_50ntFlanks@8425.8469.45|CAGTCCGCCCGGAGGATTCAACCCGGCGGCGGGTCCGGCCGTGTC</td><td>>hg38_21_+_8433172_8446622_RNA45SN1_45S_50ntFlanks</td><td>8425</td><td>8469</td><td>45</td><td>CAGTCCGCCCGGAGGATTCAACCCGGCGGCGGGTCCGGCCGTGTC</td><td>222.5</td><td>-5.2</td><td>1.4</td><td>-3.6</td><td>3.2E-04</td><td>5.4E-04</td><td>BT20/MB468</td><td>MITO</td></tr>
<tr><td>>trna119_LysCTT_1_-_145395522_145395594@1.30.30,trna11_LysCTT_5_-_180648979_180649051@1.30.30,trna13_LysCTT_6_+_26556774_26556846@1.30.30,trna32_LysCTT_16_-_3207406_3207478@1.30.30,trna7_LysCTT_16_+_3225692_3225764@1.30.30,trna9_LysCTT_5_+_180634755_180634827@1.30.30|GCCCGGCTAGCTCAGTCGGTAGAGCATGAG</td><td>>trna119_LysCTT_1_-_145395522_145395594</td><td>1</td><td>30</td><td>30</td><td>GCCCGGCTAGCTCAGTCGGTAGAGCATGAG</td><td>4897.7</td><td>-5.2</td><td>1.4</td><td>-3.6</td><td>3.2E-04</td><td>5.4E-04</td><td>BT20/MB468</td><td>MITO</td></tr>
<tr><td>>MI0000446|hsa-mir-125b-1&WithFlank&11|-|122099751|122099850@61.82.22(+1U)&[MIMAT0004592&hsa-miR-125b-1-3p&offsets|0|0(+1U);m-485&11|-|122099769|122099790&offsets|0|0(+1U)]|ACGGGTTAGGCTCTTGGGAGCTT</td><td>>MI0000446|hsa-mir-125b-1&WithFlank&11|-|122099751|122099850</td><td>61</td><td>82</td><td>22(+1U)</td><td>ACGGGTTAGGCTCTTGGGAGCTT</td><td>427.6</td><td>-6.8</td><td>1.7</td><td>-4.0</td><td>6.1E-05</td><td>5.4E-04</td><td>BT20/MB231</td><td>MP</td></tr>
<tr><td>>hg38_21_+_8433172_8446622_RNA45SN1_45S_50ntFlanks@6661.6706.46|CGGTGGATCACTCGGCTCGTGCGTCGATGAAGAACGCAGCTAGCTG</td><td>>hg38_21_+_8433172_8446622_RNA45SN1_45S_50ntFlanks</td><td>6661</td><td>6706</td><td>46</td><td>CGGTGGATCACTCGGCTCGTGCGTCGATGAAGAACGCAGCTAGCTG</td><td>9830.4</td><td>3.5</td><td>0.9</td><td>4.0</td><td>7.2E-05</td><td>5.4E-04</td><td>BT20/MB231</td><td>MITO</td></tr>
<tr><td>>MI0005559|hsa-mir-744&WithFlank&17|+|12081893|12082002@17.37.21(+1U)&[MIMAT0004945&hsa-miR-744-5p&offsets|0|-1(+1U);m-172&17|+|12081909|12081929&offsets|0|0(+1U)]|TGCGGGGCTAGGGCTAACAGCT</td><td>>MI0005559|hsa-mir-744&WithFlank&17|+|12081893|12082002</td><td>17</td><td>37</td><td>21(+1U)</td><td>TGCGGGGCTAGGGCTAACAGCT</td><td>61.2</td><td>6.7</td><td>1.8</td><td>3.8</td><td>1.7E-04</td><td>5.4E-04</td><td>BT20/MB468</td><td>MP</td></tr>
<tr><td>>hg38_MT_+_598_1651_RNR1_12S_50ntFlanks@849.877.29|AGTGCTTAGTTGAACAGGGCCCTGAAGCG</td><td>>hg38_MT_+_598_1651_RNR1_12S_50ntFlanks</td><td>849</td><td>877</td><td>29</td><td>AGTGCTTAGTTGAACAGGGCCCTGAAGCG</td><td>103.7</td><td>7.3</td><td>1.9</td><td>3.8</td><td>1.7E-04</td><td>5.4E-04</td><td>BT20/MB468</td><td>MP</td></tr>
<tr><td>>hg38_21_+_8433172_8446622_RNA45SN1_45S_50ntFlanks@8102.8143.42|CCCCGCGGCGGGGCGCGGGACATGTGGCGTACGGAAGACCCG</td><td>>hg38_21_+_8433172_8446622_RNA45SN1_45S_50ntFlanks</td><td>8102</td><td>8143</td><td>42</td><td>CCCCGCGGCGGGGCGCGGGACATGTGGCGTACGGAAGACCCG</td><td>66.9</td><td>-9.1</td><td>2.4</td><td>-3.8</td><td>1.7E-04</td><td>5.4E-04</td><td>BT20/MB468</td><td>MP</td></tr>
<tr><td>>trnaMT_PheGAA_MT_+_577_647@35.60.26|GAAAATGTTTAGACGGGCTCACATCA</td><td>>trnaMT_PheGAA_MT_+_577_647</td><td>35</td><td>60</td><td>26</td><td>GAAAATGTTTAGACGGGCTCACATCA</td><td>291.2</td><td>5.4</td><td>1.5</td><td>3.5</td><td>4.3E-04</td><td>5.4E-04</td><td>MB231/MB468</td><td>MITO</td></tr>
<tr><td>>am_MI0000086|hsa-mir-28&WithFlank&3|+|188688775|188688872@20.39.20&[MIMAT0000085&hsa-miR-28-5p&offsets|0|-2;m-133&3|+|188688794|188688815&offsets|0|-2]|AAGGAGCTCACAGTCTATTG</td><td>>am_MI0000086|hsa-mir-28&WithFlank&3|+|188688775|188688872</td><td>20</td><td>39</td><td>20</td><td>AAGGAGCTCACAGTCTATTG</td><td>551.4</td><td>5.3</td><td>1.4</td><td>3.8</td><td>1.7E-04</td><td>5.4E-04</td><td>BT20/MB468</td><td>MP</td></tr>
<tr><td>>MI0005566|hsa-mir-374b&WithFlank&X|-|74218541|74218624@17.36.20&[MIMAT0004955&hsa-miR-374b-5p&offsets|0|-2;m-153&X|-|74218587|74218608&offsets|0|-2]|ATATAATACAACCTGCTAAG</td><td>>MI0005566|hsa-mir-374b&WithFlank&X|-|74218541|74218624</td><td>17</td><td>36</td><td>20</td><td>ATATAATACAACCTGCTAAG</td><td>52.0</td><td>-9.1</td><td>2.3</td><td>-4.0</td><td>6.1E-05</td><td>5.4E-04</td><td>BT20/MB231</td><td>MP</td></tr>
<tr><td>>hg38_MT_+_598_1651_RNR1_12S_50ntFlanks@416.454.39|AAATAGACTACGAAAGTGGCTTTAACATATCTGAACACA</td><td>>hg38_MT_+_598_1651_RNR1_12S_50ntFlanks</td><td>416</td><td>454</td><td>39</td><td>AAATAGACTACGAAAGTGGCTTTAACATATCTGAACACA</td><td>79.5</td><td>10.1</td><td>2.7</td><td>3.8</td><td>1.7E-04</td><td>5.4E-04</td><td>BT20/MB468</td><td>MP</td></tr>
<tr><td>>hg38_21_+_8433172_8446622_RNA45SN1_45S_50ntFlanks@11768.11806.39|CTAATTAGTGACGCGCATGAATGGATGAACGAGATTCCC</td><td>>hg38_21_+_8433172_8446622_RNA45SN1_45S_50ntFlanks</td><td>11768</td><td>11806</td><td>39</td><td>CTAATTAGTGACGCGCATGAATGGATGAACGAGATTCCC</td><td>191.5</td><td>-4.9</td><td>1.4</td><td>-3.6</td><td>3.8E-04</td><td>5.4E-04</td><td>BT20/MB468</td><td>CYTO</td></tr>
<tr><td>>hg38_21_+_8433172_8446622_RNA45SN1_45S_50ntFlanks@12243.12272.30|GAAGGGCAAAAGCTCGCTTGATCTTGATTT</td><td>>hg38_21_+_8433172_8446622_RNA45SN1_45S_50ntFlanks</td><td>12243</td><td>12272</td><td>30</td><td>GAAGGGCAAAAGCTCGCTTGATCTTGATTT</td><td>144.9</td><td>-4.5</td><td>1.2</td><td>-3.6</td><td>3.6E-04</td><td>5.4E-04</td><td>MB231/MB468</td><td>MP</td></tr>
<tr><td>>hg38_1_-_228634819_228635039_RNA5S12_5S_50ntFlanks@128.170.43|CTTGGATGGGAGACCGCCTGGGAATACCGGGTGCTGTAGGCTT</td><td>>hg38_1_-_228634819_228635039_RNA5S12_5S_50ntFlanks</td><td>128</td><td>170</td><td>43</td><td>CTTGGATGGGAGACCGCCTGGGAATACCGGGTGCTGTAGGCTT</td><td>1424.4</td><td>3.6</td><td>0.9</td><td>4.0</td><td>7.2E-05</td><td>5.4E-04</td><td>BT20/MB231</td><td>MITO</td></tr>
<tr><td>>hg38_21_+_8433172_8446622_RNA45SN1_45S_50ntFlanks@7978.8016.39|GACCTCAGATCAGACGTGGCGACCCGCTGAATTTAAGCA</td><td>>hg38_21_+_8433172_8446622_RNA45SN1_45S_50ntFlanks</td><td>7978</td><td>8016</td><td>39</td><td>GACCTCAGATCAGACGTGGCGACCCGCTGAATTTAAGCA</td><td>376.6</td><td>4.2</td><td>1.2</td><td>3.5</td><td>4.3E-04</td><td>5.4E-04</td><td>MB231/MB468</td><td>MITO</td></tr>
<tr><td>>MI0000299|hsa-mir-222&WithFlank&X|-|45747009|45747130@75.99.25(+3U)&[MIMAT0000279&hsa-miR-222-3p&offsets|0|+4(+3U);m-140&X|-|45747033|45747056&offsets|0|+1(+3U)]|AGCTACATCTGGCTACTGGGTCTCTTTT</td><td>>MI0000299|hsa-mir-222&WithFlank&X|-|45747009|45747130</td><td>75</td><td>99</td><td>25(+3U)</td><td>AGCTACATCTGGCTACTGGGTCTCTTTT</td><td>171.5</td><td>6.4</td><td>1.8</td><td>3.6</td><td>3.6E-04</td><td>5.4E-04</td><td>MB231/MB468</td><td>MP</td></tr>
<tr><td>>MI0000433|hsa-let-7g&WithFlank&3|-|52268272|52268367@11.31.21(+1C)&[MIMAT0000414&hsa-let-7g-5p&offsets|0|-1(+1C);m-58&3|-|52268336|52268357&offsets|0|-1(+1C)]|TGAGGTAGTAGTTTGTACAGTC</td><td>>MI0000433|hsa-let-7g&WithFlank&3|-|52268272|52268367</td><td>11</td><td>31</td><td>21(+1C)</td><td>TGAGGTAGTAGTTTGTACAGTC</td><td>2165.7</td><td>3.5</td><td>0.9</td><td>3.8</td><td>1.7E-04</td><td>5.4E-04</td><td>BT20/MB468</td><td>MP</td></tr>
</table>

<table>
<tr><td>>trna2_GlyGCC_21_-_18827107_18827177@1.29.29,trna35_GlyGCC_1_+_161413094_161413164@1.29.29,trna37_GlyGCC_1_+_161420467_161420537@1.29.29,trna39_GlyGCC_1_+_161427898_161427968@1.29.29,trna41_GlyGCC_1_+_161435258_161435328@1.29.29|GCATGGGTGGTTCAGTGGTAGAATTCTCG</td><td>>trna2_GlyGCC_21_-_18827107_18827177</td><td>1</td><td>29</td><td>29</td><td>GCATGGGTGGTTCAGTGGTAGAATTCTCG</td><td>12550.3</td><td>-3.6</td><td>1.0</td><td>-3.8</td><td>1.7E-04</td><td>5.4E-04</td><td>BT20/MB468</td><td>MP</td></tr>
<tr><td>>MI0000101|hsa-mir-99a&WithFlank&21|+|16539083|16539175@19.40.22&[MIMAT0000097&hsa-miR-99a-5p&offsets|0|0;m-64&21|+|16539101|16539121&offsets|0|+1]|AACCCGTAGATCCGATCTTGTG</td><td>>MI0000101|hsa-mir-99a&WithFlank&21|+|16539083|16539175</td><td>19</td><td>40</td><td>22</td><td>AACCCGTAGATCCGATCTTGTG</td><td>1390.2</td><td>-7.5</td><td>1.9</td><td>-4.0</td><td>6.1E-05</td><td>5.4E-04</td><td>BT20/MB231</td><td>MP</td></tr>
<tr><td>>hg38_21_+_8433172_8446622_RNA45SN1_45S_50ntFlanks@11583.11609.27|CTGGTGCGGACCAGGGGAATCCGACTG</td><td>>hg38_21_+_8433172_8446622_RNA45SN1_45S_50ntFlanks</td><td>11583</td><td>11609</td><td>27</td><td>CTGGTGCGGACCAGGGGAATCCGACTG</td><td>812.3</td><td>-2.9</td><td>0.8</td><td>-3.8</td><td>1.7E-04</td><td>5.4E-04</td><td>BT20/MB468</td><td>MP</td></tr>
<tr><td>>hg38_21_+_8433172_8446622_RNA45SN1_45S_50ntFlanks@4590.4615.26|ATTTTGTTGGTTTTCGGAACTGAGGC</td><td>>hg38_21_+_8433172_8446622_RNA45SN1_45S_50ntFlanks</td><td>4590</td><td>4615</td><td>26</td><td>ATTTTGTTGGTTTTCGGAACTGAGGC</td><td>129.3</td><td>-3.9</td><td>1.1</td><td>-3.5</td><td>4.3E-04</td><td>5.4E-04</td><td>MB231/MB468</td><td>NUC</td></tr>
<tr><td>>hg38_21_+_8433172_8446622_RNA45SN1_45S_50ntFlanks@9345.9370.26|AAAGCCGCCGTGGCGCAATGAAGGTG</td><td>>hg38_21_+_8433172_8446622_RNA45SN1_45S_50ntFlanks</td><td>9345</td><td>9370</td><td>26</td><td>AAAGCCGCCGTGGCGCAATGAAGGTG</td><td>749.4</td><td>-6.3</td><td>1.7</td><td>-3.8</td><td>1.7E-04</td><td>5.4E-04</td><td>BT20/MB468</td><td>MP</td></tr>
<tr><td>>hg38_21_+_8433172_8446622_RNA45SN1_45S_50ntFlanks@8197.8215.19|GTGGACGGTGTGAGGCCGG</td><td>>hg38_21_+_8433172_8446622_RNA45SN1_45S_50ntFlanks</td><td>8197</td><td>8215</td><td>19</td><td>GTGGACGGTGTGAGGCCGG</td><td>444.1</td><td>-3.9</td><td>1.1</td><td>-3.5</td><td>4.0E-04</td><td>5.4E-04</td><td>MB231/MB468</td><td>CYTO</td></tr>
<tr><td>>hg38_21_+_8433172_8446622_RNA45SN1_45S_50ntFlanks@12711.12742.32|GCGGAGCCTCGGTTGGCCTCGGATAGCCGGTC</td><td>>hg38_21_+_8433172_8446622_RNA45SN1_45S_50ntFlanks</td><td>12711</td><td>12742</td><td>32</td><td>GCGGAGCCTCGGTTGGCCTCGGATAGCCGGTC</td><td>375.2</td><td>-6.0</td><td>1.6</td><td>-3.8</td><td>1.7E-04</td><td>5.4E-04</td><td>BT20/MB468</td><td>MP</td></tr>
<tr><td>>hg38_21_+_8433172_8446622_RNA45SN1_45S_50ntFlanks@5417.5443.27|CTACCGATTGGATGGTTTAGTGAGGCC</td><td>>hg38_21_+_8433172_8446622_RNA45SN1_45S_50ntFlanks</td><td>5417</td><td>5443</td><td>27</td><td>CTACCGATTGGATGGTTTAGTGAGGCC</td><td>281.1</td><td>-3.3</td><td>0.9</td><td>-3.5</td><td>4.0E-04</td><td>5.4E-04</td><td>MB231/MB468</td><td>CYTO</td></tr>
<tr><td>>am_MI0014222|hsa-mir-548w&WithFlank&16|+|26025231|26025316@15.36.22&[MIMAT0015060&hsa-miR-548w&offsets|-1|-2;m-801&16|+|26025246|26025267&offsets|-1|-1]|CAAAAGTAACTGCGGTTTTTGC</td><td>>am_MI0014222|hsa-mir-548w&WithFlank&16|+|26025231|26025316</td><td>15</td><td>36</td><td>22</td><td>CAAAAGTAACTGCGGTTTTTGC</td><td>53.5</td><td>-9.1</td><td>2.3</td><td>-4.0</td><td>6.1E-05</td><td>5.4E-04</td><td>BT20/MB231</td><td>MP</td></tr>
<tr><td>>hg38_21_+_8433172_8446622_RNA45SN1_45S_50ntFlanks@12007.12034.28|GCCGCCGGTGAAATACCACTACTCTGAT</td><td>>hg38_21_+_8433172_8446622_RNA45SN1_45S_50ntFlanks</td><td>12007</td><td>12034</td><td>28</td><td>GCCGCCGGTGAAATACCACTACTCTGAT</td><td>459.4</td><td>2.7</td><td>0.7</td><td>3.6</td><td>3.2E-04</td><td>5.4E-04</td><td>BT20/MB468</td><td>MITO</td></tr>
<tr><td>>MI0000253|hsa-mir-148a&WithFlank&7|-|25949913|25949992@50.70.21(+1C)&[MIMAT0000243&hsa-miR-148a-3p&offsets|0|-1(+1C);m-7&7|-|25949922|25949943&offsets|0|-1(+1C)]|TCAGTGCACTACAGAACTTTGC</td><td>>MI0000253|hsa-mir-148a&WithFlank&7|-|25949913|25949992</td><td>50</td><td>70</td><td>21(+1C)</td><td>TCAGTGCACTACAGAACTTTGC</td><td>7402.4</td><td>2.7</td><td>0.8</td><td>3.5</td><td>4.4E-04</td><td>5.4E-04</td><td>BT20/MB468</td><td>NUC</td></tr>
<tr><td>>hg38_21_+_8433172_8446622_RNA45SN1_45S_50ntFlanks@11578.11602.25|TAGAACTGGTGCGGACCAGGGGAAT</td><td>>hg38_21_+_8433172_8446622_RNA45SN1_45S_50ntFlanks</td><td>11578</td><td>11602</td><td>25</td><td>TAGAACTGGTGCGGACCAGGGGAAT</td><td>149.7</td><td>-3.8</td><td>1.1</td><td>-3.5</td><td>4.3E-04</td><td>5.4E-04</td><td>MB231/MB468</td><td>NUC</td></tr>
<tr><td>>hg38_21_+_8433172_8446622_RNA45SN1_45S_50ntFlanks@9532.9573.42|AGAGGAAACTCTGGTGGAGGTCCGTAGCGGTCCTGACGTGCA</td><td>>hg38_21_+_8433172_8446622_RNA45SN1_45S_50ntFlanks</td><td>9532</td><td>9573</td><td>42</td><td>AGAGGAAACTCTGGTGGAGGTCCGTAGCGGTCCTGACGTGCA</td><td>355.6</td><td>-3.1</td><td>0.8</td><td>-4.0</td><td>6.1E-05</td><td>5.4E-04</td><td>BT20/MB231</td><td>CYTO</td></tr>
<tr><td>>hg38_21_+_8433172_8446622_RNA45SN1_45S_50ntFlanks@10819.10839.21|AGGGAAGTCGGCAAGCCGGAT</td><td>>hg38_21_+_8433172_8446622_RNA45SN1_45S_50ntFlanks</td><td>10819</td><td>10839</td><td>21</td><td>AGGGAAGTCGGCAAGCCGGAT</td><td>834.0</td><td>-3.6</td><td>1.0</td><td>-3.8</td><td>1.7E-04</td><td>5.4E-04</td><td>BT20/MB468</td><td>MP</td></tr>
<tr><td>>hg38_21_+_8433172_8446622_RNA45SN1_45S_50ntFlanks@11661.11687.27|TGATTTCTGCCCAGTGCTCTGAATGTC</td><td>>hg38_21_+_8433172_8446622_RNA45SN1_45S_50ntFlanks</td><td>11661</td><td>11687</td><td>27</td><td>TGATTTCTGCCCAGTGCTCTGAATGTC</td><td>171.4</td><td>-4.2</td><td>1.2</td><td>-3.6</td><td>3.6E-04</td><td>5.4E-04</td><td>MB231/MB468</td><td>MP</td></tr>
<tr><td>>hg38_21_+_8433172_8446622_RNA45SN1_45S_50ntFlanks@11209.11249.41|CGGCGACTCTGGACGCGAGCCGGGCCCTTCCCGTGGATCGC</td><td>>hg38_21_+_8433172_8446622_RNA45SN1_45S_50ntFlanks</td><td>11209</td><td>11249</td><td>41</td><td>CGGCGACTCTGGACGCGAGCCGGGCCCTTCCCGTGGATCGC</td><td>230.2</td><td>-5.2</td><td>1.5</td><td>-3.5</td><td>4.4E-04</td><td>5.4E-04</td><td>MB231/MB468</td><td>NUC</td></tr>
<tr><td>>hg38_21_+_8433172_8446622_RNA45SN1_45S_50ntFlanks@7978.8016.39|GACCTCAGATCAGACGTGGCGACCCGCTGAATTTAAGCA</td><td>>hg38_21_+_8433172_8446622_RNA45SN1_45S_50ntFlanks</td><td>7978</td><td>8016</td><td>39</td><td>GACCTCAGATCAGACGTGGCGACCCGCTGAATTTAAGCA</td><td>330.9</td><td>5.2</td><td>1.5</td><td>3.5</td><td>4.4E-04</td><td>5.4E-04</td><td>MB231/MB468</td><td>NUC</td></tr>
<tr><td>>MI0000079|hsa-mir-23a&WithFlank&19|-|13836581|13836665@54.71.18(+1G)&[MIMAT0000078&hsa-miR-23a-3p&offsets|+3|0(+1G);m-31&19|-|13836595|13836615&offsets|+3|0(+1G)]|ACATTGCCAGGGATTTCCG</td><td>>MI0000079|hsa-mir-23a&WithFlank&19|-|13836581|13836665</td><td>54</td><td>71</td><td>18(+1G)</td><td>ACATTGCCAGGGATTTCCG</td><td>457.3</td><td>12.8</td><td>3.6</td><td>3.5</td><td>4.1E-04</td><td>5.4E-04</td><td>MB231/MB468</td><td>CYTO</td></tr>
<tr><td>>hg38_21_+_8433172_8446622_RNA45SN1_45S_50ntFlanks@476.514.39|TTCGTGATCGATGTGGTGACGTCGTGCTCTCCCGGGCCG</td><td>>hg38_21_+_8433172_8446622_RNA45SN1_45S_50ntFlanks</td><td>476</td><td>514</td><td>39</td><td>TTCGTGATCGATGTGGTGACGTCGTGCTCTCCCGGGCCG</td><td>1067.4</td><td>3.5</td><td>1.0</td><td>3.5</td><td>4.4E-04</td><td>5.4E-04</td><td>MB231/MB468</td><td>NUC</td></tr>
<tr><td>>trnaMT_PheGAA_MT_+_577_647@36.64.29|AAAATGTTTAGACGGGCTCACATCACCCC</td><td>>trnaMT_PheGAA_MT_+_577_647</td><td>36</td><td>64</td><td>29</td><td>AAAATGTTTAGACGGGCTCACATCACCCC</td><td>345.7</td><td>9.1</td><td>2.4</td><td>3.8</td><td>1.8E-04</td><td>5.4E-04</td><td>BT20/MB468</td><td>MP</td></tr>
<tr><td>>MI0000077|hsa-mir-21&WithFlank&17|+|59841260|59841343@14.39.26&[MIMAT0000076&hsa-miR-21-5p&offsets|0|+4;m-1&17|+|59841273|59841294&offsets|0|+4]|TAGCTTATCAGACTGATGTTGACTGT</td><td>>MI0000077|hsa-mir-21&WithFlank&17|+|59841260|59841343</td><td>14</td><td>39</td><td>26</td><td>TAGCTTATCAGACTGATGTTGACTGT</td><td>312.7</td><td>2.8</td><td>0.7</td><td>4.0</td><td>6.5E-05</td><td>5.4E-04</td><td>BT20/MB231</td><td>NUC</td></tr>
<tr><td>>hg38_21_+_8433172_8446622_RNA45SN1_45S_50ntFlanks@12995.13014.20|CTCGCTGCGATCTATTGAAA</td><td>>hg38_21_+_8433172_8446622_RNA45SN1_45S_50ntFlanks</td><td>12995</td><td>13014</td><td>20</td><td>CTCGCTGCGATCTATTGAAA</td><td>351.8</td><td>-2.7</td><td>0.7</td><td>-4.0</td><td>6.5E-05</td><td>5.4E-04</td><td>BT20/MB231</td><td>NUC</td></tr>
<tr><td>>hg38_21_+_8433172_8446622_RNA45SN1_45S_50ntFlanks@9736.9765.30|CCGAAACGATCTCAACCTATTCTCAAACTT</td><td>>hg38_21_+_8433172_8446622_RNA45SN1_45S_50ntFlanks</td><td>9736</td><td>9765</td><td>30</td><td>CCGAAACGATCTCAACCTATTCTCAAACTT</td><td>109.3</td><td>-5.0</td><td>1.4</td><td>-3.5</td><td>3.9E-04</td><td>5.4E-04</td><td>BT20/MB468</td><td>CYTO</td></tr>
<tr><td>>hg38_21_+_8433172_8446622_RNA45SN1_45S_50ntFlanks@11326.11344.19|CTCGTCGCGCGCGCGTCCG</td><td>>hg38_21_+_8433172_8446622_RNA45SN1_45S_50ntFlanks</td><td>11326</td><td>11344</td><td>19</td><td>CTCGTCGCGCGCGCGTCCG</td><td>85.8</td><td>-5.7</td><td>1.5</td><td>-3.8</td><td>1.8E-04</td><td>5.4E-04</td><td>BT20/MB468</td><td>MP</td></tr>
<tr><td>>MI0000086|hsa-mir-28&WithFlank&3|+|188688775|188688872@60.79.20&[MIMAT0004502&hsa-miR-28-3p&offsets|0|-2;m-28&3|+|188688834|188688855&offsets|0|-2]|CACTAGATTGTGAGCTCCTG</td><td>>MI0000086|hsa-mir-28&WithFlank&3|+|188688775|188688872</td><td>60</td><td>79</td><td>20</td><td>CACTAGATTGTGAGCTCCTG</td><td>139.1</td><td>6.2</td><td>1.6</td><td>3.8</td><td>1.8E-04</td><td>5.4E-04</td><td>BT20/MB468</td><td>MP</td></tr>
<tr><td>>hg38_21_+_8433172_8446622_RNA45SN1_45S_50ntFlanks@9791.9813.23|TGGCGTGGAGCCGGGCGTGGAAT</td><td>>hg38_21_+_8433172_8446622_RNA45SN1_45S_50ntFlanks</td><td>9791</td><td>9813</td><td>23</td><td>TGGCGTGGAGCCGGGCGTGGAAT</td><td>87.7</td><td>6.9</td><td>1.8</td><td>3.8</td><td>1.8E-04</td><td>5.4E-04</td><td>BT20/MB468</td><td>MP</td></tr>
<tr><td>>hg38_1_-_228634819_228635039_RNA5S12_5S_50ntFlanks@132.170.39|GATGGGAGACCGCCTGGGAATACCGGGTGCTGTAGGCTT</td><td>>hg38_1_-_228634819_228635039_RNA5S12_5S_50ntFlanks</td><td>132</td><td>170</td><td>39</td><td>GATGGGAGACCGCCTGGGAATACCGGGTGCTGTAGGCTT</td><td>110557.0</td><td>3.8</td><td>1.0</td><td>3.8</td><td>1.8E-04</td><td>5.4E-04</td><td>BT20/MB468</td><td>MP</td></tr>
<tr><td>>hg38_21_+_8433172_8446622_RNA45SN1_45S_50ntFlanks@12935.12953.19|TAAACCATTCGTAGACGAC</td><td>>hg38_21_+_8433172_8446622_RNA45SN1_45S_50ntFlanks</td><td>12935</td><td>12953</td><td>19</td><td>TAAACCATTCGTAGACGAC</td><td>421.1</td><td>-5.4</td><td>1.5</td><td>-3.5</td><td>3.9E-04</td><td>5.5E-04</td><td>BT20/MB468</td><td>CYTO</td></tr>
<tr><td>>hg38_21_+_8433172_8446622_RNA45SN1_45S_50ntFlanks@10411.10429.19|CAGTCGGTCCTGAGAGATG</td><td>>hg38_21_+_8433172_8446622_RNA45SN1_45S_50ntFlanks</td><td>10411</td><td>10429</td><td>19</td><td>CAGTCGGTCCTGAGAGATG</td><td>611.5</td><td>7.7</td><td>1.9</td><td>4.0</td><td>6.2E-05</td><td>5.5E-04</td><td>BT20/MB231</td><td>MP</td></tr>
<tr><td>>hg38_21_+_8433172_8446622_RNA45SN1_45S_50ntFlanks@9533.9567.35|GAGGAAACTCTGGTGGAGGTCCGTAGCGGTCCTGA</td><td>>hg38_21_+_8433172_8446622_RNA45SN1_45S_50ntFlanks</td><td>9533</td><td>9567</td><td>35</td><td>GAGGAAACTCTGGTGGAGGTCCGTAGCGGTCCTGA</td><td>90.5</td><td>-5.6</td><td>1.6</td><td>-3.6</td><td>3.6E-04</td><td>5.5E-04</td><td>MB231/MB468</td><td>MP</td></tr>
<tr><td>>am_trnaMT_HisGTG_MT_+_12138_12206@18.51.34|AAAACATCAGATTGTGAATCTGACAACAGAGGCT</td><td>>am_trnaMT_HisGTG_MT_+_12138_12206</td><td>18</td><td>51</td><td>34</td><td>AAAACATCAGATTGTGAATCTGACAACAGAGGCT</td><td>245.0</td><td>5.5</td><td>1.5</td><td>3.6</td><td>3.6E-04</td><td>5.5E-04</td><td>MB231/MB468</td><td>MP</td></tr>
<tr><td>>hg38_21_+_8433172_8446622_RNA45SN1_45S_50ntFlanks@10802.10826.25|TTGGAACAATGTAGGTAAGGGAAGT</td><td>>hg38_21_+_8433172_8446622_RNA45SN1_45S_50ntFlanks</td><td>10802</td><td>10826</td><td>25</td><td>TTGGAACAATGTAGGTAAGGGAAGT</td><td>368.3</td><td>-5.9</td><td>1.6</td><td>-3.8</td><td>1.8E-04</td><td>5.5E-04</td><td>BT20/MB468</td><td>MP</td></tr>
<tr><td>>am_trnaMT_SerGCT_MT_+_12207_12265@13.32.20|AAGAACTGCTAACTCATGCC</td><td>>am_trnaMT_SerGCT_MT_+_12207_12265</td><td>13</td><td>32</td><td>20</td><td>AAGAACTGCTAACTCATGCC</td><td>277.9</td><td>3.2</td><td>0.8</td><td>4.0</td><td>6.2E-05</td><td>5.5E-04</td><td>BT20/MB231</td><td>CYTO</td></tr>
<tr><td>>hg38_21_+_8433172_8446622_RNA45SN1_45S_50ntFlanks@10786.10821.36|AACAGCCTCTGGCATGTTGGAACAATGTAGGTAAGG</td><td>>hg38_21_+_8433172_8446622_RNA45SN1_45S_50ntFlanks</td><td>10786</td><td>10821</td><td>36</td><td>AACAGCCTCTGGCATGTTGGAACAATGTAGGTAAGG</td><td>302.4</td><td>-2.4</td><td>0.7</td><td>-3.5</td><td>3.9E-04</td><td>5.5E-04</td><td>BT20/MB468</td><td>CYTO</td></tr>
<tr><td>>hg38_21_+_8433172_8446622_RNA45SN1_45S_50ntFlanks@12419.12458.40|GATGTCGGCTCTTCCTATCATTGTGAAGCAGAATTCACCA</td><td>>hg38_21_+_8433172_8446622_RNA45SN1_45S_50ntFlanks</td><td>12419</td><td>12458</td><td>40</td><td>GATGTCGGCTCTTCCTATCATTGTGAAGCAGAATTCACCA</td><td>314.0</td><td>4.6</td><td>1.3</td><td>3.5</td><td>4.4E-04</td><td>5.5E-04</td><td>MB231/MB468</td><td>NUC</td></tr>
<tr><td>>hg38_21_+_8433172_8446622_RNA45SN1_45S_50ntFlanks@12417.12454.38|TCGATGTCGGCTCTTCCTATCATTGTGAAGCAGAATTC</td><td>>hg38_21_+_8433172_8446622_RNA45SN1_45S_50ntFlanks</td><td>12417</td><td>12454</td><td>38</td><td>TCGATGTCGGCTCTTCCTATCATTGTGAAGCAGAATTC</td><td>314.0</td><td>-3.1</td><td>0.8</td><td>-4.0</td><td>6.2E-05</td><td>5.5E-04</td><td>BT20/MB231</td><td>CYTO</td></tr>
<tr><td>>hg38_21_+_8433172_8446622_RNA45SN1_45S_50ntFlanks@12418.12455.38|CGATGTCGGCTCTTCCTATCATTGTGAAGCAGAATTCA</td><td>>hg38_21_+_8433172_8446622_RNA45SN1_45S_50ntFlanks</td><td>12418</td><td>12455</td><td>38</td><td>CGATGTCGGCTCTTCCTATCATTGTGAAGCAGAATTCA</td><td>304.7</td><td>-3.0</td><td>0.7</td><td>-4.0</td><td>6.2E-05</td><td>5.5E-04</td><td>BT20/MB231</td><td>CYTO</td></tr>
<tr><td>>hg38_21_+_8433172_8446622_RNA45SN1_45S_50ntFlanks@10352.10371.20|GAGAACTTTGAAGGCCGAAG</td><td>>hg38_21_+_8433172_8446622_RNA45SN1_45S_50ntFlanks</td><td>10352</td><td>10371</td><td>20</td><td>GAGAACTTTGAAGGCCGAAG</td><td>220.9</td><td>-2.6</td><td>0.7</td><td>-4.0</td><td>6.5E-05</td><td>5.5E-04</td><td>BT20/MB231</td><td>NUC</td></tr>
<tr><td>>hg38_21_+_8433172_8446622_RNA45SN1_45S_50ntFlanks@7974.8006.33|ACGCGACCTCAGATCAGACGTGGCGACCCGCTG</td><td>>hg38_21_+_8433172_8446622_RNA45SN1_45S_50ntFlanks</td><td>7974</td><td>8006</td><td>33</td><td>ACGCGACCTCAGATCAGACGTGGCGACCCGCTG</td><td>6075.9</td><td>-2.6</td><td>0.7</td><td>-3.5</td><td>4.4E-04</td><td>5.5E-04</td><td>MB231/MB468</td><td>NUC</td></tr>
<tr><td>>MI0000433|hsa-let-7g&WithFlank&3|-|52268272|52268367@11.32.22&[MIMAT0000414&hsa-let-7g-5p&offsets|0|0;m-58&3|-|52268336|52268357&offsets|0|0]|TGAGGTAGTAGTTTGTACAGTT</td><td>>MI0000433|hsa-let-7g&WithFlank&3|-|52268272|52268367</td><td>11</td><td>32</td><td>22</td><td>TGAGGTAGTAGTTTGTACAGTT</td><td>14592.8</td><td>3.3</td><td>0.9</td><td>3.7</td><td>1.8E-04</td><td>5.5E-04</td><td>BT20/MB468</td><td>MP</td></tr>
<tr><td>>hg38_21_+_8433172_8446622_RNA45SN1_45S_50ntFlanks@11694.11728.35|AAGAAATTCAATGAAGCGCGGGTAAACGGCGGGAG</td><td>>hg38_21_+_8433172_8446622_RNA45SN1_45S_50ntFlanks</td><td>11694</td><td>11728</td><td>35</td><td>AAGAAATTCAATGAAGCGCGGGTAAACGGCGGGAG</td><td>153.9</td><td>-4.8</td><td>1.3</td><td>-3.5</td><td>3.9E-04</td><td>5.5E-04</td><td>BT20/MB468</td><td>CYTO</td></tr>
<tr><td>>hg38_21_+_8433172_8446622_RNA45SN1_45S_50ntFlanks@12420.12453.34|ATGTCGGCTCTTCCTATCATTGTGAAGCAGAATT</td><td>>hg38_21_+_8433172_8446622_RNA45SN1_45S_50ntFlanks</td><td>12420</td><td>12453</td><td>34</td><td>ATGTCGGCTCTTCCTATCATTGTGAAGCAGAATT</td><td>297.9</td><td>-5.0</td><td>1.4</td><td>-3.5</td><td>4.4E-04</td><td>5.5E-04</td><td>BT20/MB468</td><td>NUC</td></tr>
<tr><td>>hg38_21_+_8433172_8446622_RNA45SN1_45S_50ntFlanks@7990.8014.25|GACGTGGCGACCCGCTGAATTTAAG</td><td>>hg38_21_+_8433172_8446622_RNA45SN1_45S_50ntFlanks</td><td>7990</td><td>8014</td><td>25</td><td>GACGTGGCGACCCGCTGAATTTAAG</td><td>311.5</td><td>-2.5</td><td>0.7</td><td>-3.5</td><td>4.3E-04</td><td>5.5E-04</td><td>MB231/MB468</td><td>MITO</td></tr>
<tr><td>>hg38_21_+_8433172_8446622_RNA45SN1_45S_50ntFlanks@6721.6760.40|AATTGCAGGACACATTGATCATCGACACTTCGAACGCACT</td><td>>hg38_21_+_8433172_8446622_RNA45SN1_45S_50ntFlanks</td><td>6721</td><td>6760</td><td>40</td><td>AATTGCAGGACACATTGATCATCGACACTTCGAACGCACT</td><td>691.7</td><td>-4.4</td><td>1.2</td><td>-3.5</td><td>4.4E-04</td><td>5.5E-04</td><td>MB231/MB468</td><td>MITO</td></tr>
<tr><td>>hg38_21_+_8433172_8446622_RNA45SN1_45S_50ntFlanks@12685.12711.27|CCCAGGCGGAACGATACGGCAGCGCCG</td><td>>hg38_21_+_8433172_8446622_RNA45SN1_45S_50ntFlanks</td><td>12685</td><td>12711</td><td>27</td><td>CCCAGGCGGAACGATACGGCAGCGCCG</td><td>459.9</td><td>-4.3</td><td>1.2</td><td>-3.6</td><td>3.3E-04</td><td>5.5E-04</td><td>BT20/MB468</td><td>MITO</td></tr>
<tr><td>>hg38_21_+_8433172_8446622_RNA45SN1_45S_50ntFlanks@13006.13039.34|CTATTGAAAGTCAGCCCTCGACACAAGGGTTTGT</td><td>>hg38_21_+_8433172_8446622_RNA45SN1_45S_50ntFlanks</td><td>13006</td><td>13039</td><td>34</td><td>CTATTGAAAGTCAGCCCTCGACACAAGGGTTTGT</td><td>577.6</td><td>3.1</td><td>0.9</td><td>3.5</td><td>4.1E-04</td><td>5.5E-04</td><td>MB231/MB468</td><td>CYTO</td></tr>
<tr><td>>hg38_21_+_8433172_8446622_RNA45SN1_45S_50ntFlanks@12728.12747.20|CTCGGATAGCCGGTCCCCCG</td><td>>hg38_21_+_8433172_8446622_RNA45SN1_45S_50ntFlanks</td><td>12728</td><td>12747</td><td>20</td><td>CTCGGATAGCCGGTCCCCCG</td><td>119.1</td><td>-4.4</td><td>1.2</td><td>-3.5</td><td>4.1E-04</td><td>5.5E-04</td><td>MB231/MB468</td><td>CYTO</td></tr>
<tr><td>>hg38_21_+_8433172_8446622_RNA45SN1_45S_50ntFlanks@12244.12268.25|AAGGGCAAAAGCTCGCTTGATCTTG</td><td>>hg38_21_+_8433172_8446622_RNA45SN1_45S_50ntFlanks</td><td>12244</td><td>12268</td><td>25</td><td>AAGGGCAAAAGCTCGCTTGATCTTG</td><td>636.0</td><td>-3.6</td><td>1.0</td><td>-3.7</td><td>1.8E-04</td><td>5.5E-04</td><td>BT20/MB468</td><td>MP</td></tr>
</table>

| | | | | | | | | | | | | | |
|---|---|---|---|---|---|---|---|---|---|---|---|---|---|
| >trna10_AspGTC_12_-_125424193_125424264@37.75.39,trna12_AspGTC_12_-_125411891_125411962@37.75.39,trna144_AspGTC_6_-_27551236_27551307@37.75.39,trna38_AspGTC_17_-_8125556_8125627@37.75.39,trna45_AspGTC_6_+_27447453_27447524@37.75.39,trna48_AspGTC_6_+_27471523_27471594@37.75.39,trna4_AspGTC_12_+_96429799_96429870@37.75.39,trna69_AspGTC_1_-_161440205_161440276@37.75.39,trna72_AspGTC_1_-_161432824_161432895@37.75.39,trna75_AspGTC_1_-_161425414_161425485@37.75.39,trna78_AspGTC_1_-_161418033_161418104@37.75.39,trna81_AspGTC_1_-_161410615_161410686@37.75.39\|ACGCGGGAGACCGGGGTTCGATTCCCCGACGGGGAGCCA | >trna10_AspGTC_12_-_125424193_125424264 | 37 | 75 | 39 | ACGCGGGAGACCGGGGTTCGATTCCCCGACGGGGAGCCA | 21701.0 | -4.9 | 1.2 | -4.0 | 6.3E-05 | 5.5E-04 | BT20/MB231 | CYTO |
| >hg38_21_+_8433172_8446622_RNA45SN1_45S_50ntFlanks@12251.12289.39\|AAAGCTCGCTTGATCTTGATTTTCAGTACGAATACAGAC | >hg38_21_+_8433172_8446622_RNA45SN1_45S_50ntFlanks | 12251 | 12289 | 39 | AAAGCTCGCTTGATCTTGATTTTCAGTACGAATACAGAC | 502.1 | -3.5 | 0.9 | -4.0 | 6.3E-05 | 5.5E-04 | BT20/MB231 | CYTO |
| >MI0000077\|hsa-mir-21&WithFlank&17\|+\|59841260\|59841343@14.35.22(+1G)&[MIMAT0000076&hsa-miR-21-5p&offsets\|0\|0(+1G);m-1&17\|+\|59841273\|59841294&offsets\|0\|0(+1G)]\|TAGCTTATCAGACTGATGTTGAG | >MI0000077\|hsa-mir-21&WithFlank&17\|+\|59841260\|59841343 | 14 | 35 | 22(+1G) | TAGCTTATCAGACTGATGTTGAG | 2507.2 | -3.4 | 0.9 | -4.0 | 7.4E-05 | 5.5E-04 | BT20/MB231 | MITO |
| >hg38_21_+_8433172_8446622_RNA45SN1_45S_50ntFlanks@6652.6686.35\|GACTCTTAGCGGTGGATCACTCGGCTCGTGCGTCG | >hg38_21_+_8433172_8446622_RNA45SN1_45S_50ntFlanks | 6652 | 6686 | 35 | GACTCTTAGCGGTGGATCACTCGGCTCGTGCGTCG | 3435.0 | 2.5 | 0.7 | 3.5 | 4.4E-04 | 5.5E-04 | MB231/MB468 | NUC |
| >MI0000438\|hsa-mir-15b&WithFlank&3\|+\|160404582\|160404691@26.46.21&[MIMAT0000417&hsa-miR-15b-5p&offsets\|0\|-1;m-110&3\|+\|160404607\|160404628&offsets\|0\|-1]\|TAGCAGCACATCATGGTTTAC | >MI0000438\|hsa-mir-15b&WithFlank&3\|+\|160404582\|160404691 | 26 | 46 | 21 | TAGCAGCACATCATGGTTTAC | 254.1 | 2.0 | 0.6 | 3.5 | 4.5E-04 | 5.5E-04 | BT20/MB468 | NUC |
| >hg38_21_+_8433172_8446622_RNA45SN1_45S_50ntFlanks@12938.12956.19\|ACCATTCGTAGACGACCTG | >hg38_21_+_8433172_8446622_RNA45SN1_45S_50ntFlanks | 12938 | 12956 | 19 | ACCATTCGTAGACGACCTG | 694.5 | -1.9 | 0.5 | -3.6 | 3.3E-04 | 5.5E-04 | BT20/MB468 | MITO |
| >MI0000085\|hsa-mir-27a&WithFlank&19\|-\|13836434\|13836523@57.76.20(+1U)&[MIMAT0000084&hsa-miR-27a-3p&offsets\|0\|-1(+1U);m-43&19\|-\|13836447\|13836467&offsets\|0\|-1(+1U)]\|TTCACAGTGGCTAAGTTCCGT | >MI0000085\|hsa-mir-27a&WithFlank&19\|-\|13836434\|13836523 | 57 | 76 | 20(+1U) | TTCACAGTGGCTAAGTTCCGT | 8782.4 | -3.1 | 0.8 | -4.0 | 7.4E-05 | 5.5E-04 | BT20/MB231 | MITO |
| >MI0000078\|hsa-mir-22&WithFlank&17\|-\|1713897\|1713993@59.81.23(+1U)&[MIMAT0000077&hsa-miR-22-3p&offsets\|0\|+1(+1U);m-2&17\|-\|1713914\|1713935&offsets\|0\|+1(+1U)]\|AAGCTGCCAGTTGAAGAACTGTTT | >MI0000078\|hsa-mir-22&WithFlank&17\|-\|1713897\|1713993 | 59 | 81 | 23(+1U) | AAGCTGCCAGTTGAAGAACTGTTT | 49.3 | -9.0 | 2.3 | -4.0 | 6.3E-05 | 5.5E-04 | BT20/MB231 | MP |
| >hg38_21_+_8433172_8446622_RNA45SN1_45S_50ntFlanks@8381.8405.25\|AAGAGGGCGTGAAACCGTTAAGAGG | >hg38_21_+_8433172_8446622_RNA45SN1_45S_50ntFlanks | 8381 | 8405 | 25 | AAGAGGGCGTGAAACCGTTAAGAGG | 461.8 | -3.1 | 0.9 | -3.5 | 4.4E-04 | 5.5E-04 | MB231/MB468 | NUC |
| >hg38_MT_+_1621_3279_RNR2_16S_50ntFlanks@51.74.24\|GCTAAACCTAGCCCCAAACCCACT | >hg38_MT_+_1621_3279_RNR2_16S_50ntFlanks | 51 | 74 | 24 | GCTAAACCTAGCCCCAAACCCACT | 396.3 | 1.8 | 0.5 | 3.5 | 4.5E-04 | 5.5E-04 | BT20/MB468 | NUC |
| >MI0001445\|hsa-mir-423&WithFlank&17\|+\|30117073\|30117178@23.41.19&[MIMAT0004748&hsa-miR-423-5p&offsets\|0\|-4;m-141&17\|+\|30117095\|30117117&offsets\|0\|-4]\|TGAGGGGCAGAGAGCGAGA | >MI0001445\|hsa-mir-423&WithFlank&17\|+\|30117073\|30117178 | 23 | 41 | 19 | TGAGGGGCAGAGAGCGAGA | 313.3 | 7.4 | 2.0 | 3.7 | 1.8E-04 | 5.5E-04 | BT20/MB468 | MP |
| >hg38_21_+_8433172_8446622_RNA45SN1_45S_50ntFlanks@6741.6772.32\|ATCGACACTTCGAACGCACTTGCGGCCCCGGG | >hg38_21_+_8433172_8446622_RNA45SN1_45S_50ntFlanks | 6741 | 6772 | 32 | ATCGACACTTCGAACGCACTTGCGGCCCCGGG | 360.2 | -6.2 | 1.7 | -3.7 | 1.8E-04 | 5.5E-04 | BT20/MB468 | MP |
| >MI0000264\|hsa-mir-7-2&WithFlank&15\|+\|88611819\|88611940@38.59.22(+1G);MI0000265\|hsa-mir-7-3&WithFlank&19\|+\|4770664\|4770785@37.58.22(+1G);MI0000263\|hsa-mir-7-1&WithFlank&9\|-\|83969742\|83969863@30.51.22(+1G)&[MIMAT0000252&hsa-miR-7-5p&offsets\|0\|-2(+1G);m-400&15\|+\|88611856\|88611879&offsets\|0\|-2(+1G)];[MIMAT0000252_1&hsa-miR-7-5p&offsets\|0\|-2(+1G);m-402&19\|+\|4770700\|4770723&offsets\|0\|-2(+1G)];[MIMAT0000252_2&hsa-miR-7-5p&offsets\|0\|-2(+1G);m-401&9\|-\|83969811\|83969834&offsets\|0\|-2(+1G)]\|TGGAAGACTAGTGATTTTGTTGG | >MI0000264\|hsa-mir-7-2&WithFlank&15\|+\|88611819\|88611940 | 38 | 59 | 22(+1G) | TGGAAGACTAGTGATTTTGTTGG | 126.1 | 2.6 | 0.7 | 3.6 | 3.7E-04 | 5.5E-04 | MB231/MB468 | MP |
| >hg38_21_+_8433172_8446622_RNA45SN1_45S_50ntFlanks@3993.4025.33\|GTGACTCTAGATAACCTCGGGCCGATCGCACGC | >hg38_21_+_8433172_8446622_RNA45SN1_45S_50ntFlanks | 3993 | 4025 | 33 | GTGACTCTAGATAACCTCGGGCCGATCGCACGC | 279.7 | -3.0 | 0.8 | -3.6 | 3.3E-04 | 5.5E-04 | BT20/MB468 | MITO |
| >hg38_1_-_228634819_228635039_RNA5S12_5S_50ntFlanks@141.170.30\|CCGCCTGGGAATACCGGGTGCTGTAGGCTT | >hg38_1_-_228634819_228635039_RNA5S12_5S_50ntFlanks | 141 | 170 | 30 | CCGCCTGGGAATACCGGGTGCTGTAGGCTT | 3295.4 | -2.4 | 0.7 | -3.6 | 3.7E-04 | 5.5E-04 | MB231/MB468 | MP |
| >hg38_21_+_8433172_8446622_RNA45SN1_45S_50ntFlanks@4685.4720.36\|AGACGGACCAGAGCGAAAGCATTTGCCAAGAATGTT | >hg38_21_+_8433172_8446622_RNA45SN1_45S_50ntFlanks | 4685 | 4720 | 36 | AGACGGACCAGAGCGAAAGCATTTGCCAAGAATGTT | 574.7 | 4.3 | 1.2 | 3.5 | 4.1E-04 | 5.5E-04 | MB231/MB468 | CYTO |
| >hg38_21_+_8433172_8446622_RNA45SN1_45S_50ntFlanks@10681.10708.28\|TTCCGGCGGCGTCCGGTGAGCTCTCGCT | >hg38_21_+_8433172_8446622_RNA45SN1_45S_50ntFlanks | 10681 | 10708 | 28 | TTCCGGCGGCGTCCGGTGAGCTCTCGCT | 121.4 | -5.6 | 1.5 | -3.7 | 1.8E-04 | 5.5E-04 | BT20/MB468 | MP |
| >hg38_21_+_8433172_8446622_RNA45SN1_45S_50ntFlanks@4067.4089.23\|AACTTTCGATGGTAGTCGCCGTG | >hg38_21_+_8433172_8446622_RNA45SN1_45S_50ntFlanks | 4067 | 4089 | 23 | AACTTTCGATGGTAGTCGCCGTG | 207.8 | -3.1 | 0.8 | -3.7 | 1.8E-04 | 5.6E-04 | BT20/MB468 | MP |
| >am_trnaMT_ValTAC_MT_+_1602_1670@50.68.19\|ACTTAACTTGACCGCTCTG | >am_trnaMT_ValTAC_MT_+_1602_1670 | 50 | 68 | 19 | ACTTAACTTGACCGCTCTG | 4274.6 | -4.0 | 1.1 | -3.5 | 4.2E-04 | 5.6E-04 | MB231/MB468 | CYTO |
| >hg38_21_+_8433172_8446622_RNA45SN1_45S_50ntFlanks@5492.5516.25\|AGAAGACGGTCGAACTTGACTATCT | >hg38_21_+_8433172_8446622_RNA45SN1_45S_50ntFlanks | 5492 | 5516 | 25 | AGAAGACGGTCGAACTTGACTATCT | 812.2 | 2.0 | 0.6 | 3.5 | 4.5E-04 | 5.6E-04 | MB231/MB468 | NUC |
| >hg38_21_+_8433172_8446622_RNA45SN1_45S_50ntFlanks@6645.6677.33\|CTCGTACGACTCTTAGCGGTGGATCACTCGGCT | >hg38_21_+_8433172_8446622_RNA45SN1_45S_50ntFlanks | 6645 | 6677 | 33 | CTCGTACGACTCTTAGCGGTGGATCACTCGGCT | 682.4 | 2.6 | 0.7 | 3.5 | 4.5E-04 | 5.6E-04 | MB231/MB468 | NUC |
| >MI0000760\|hsa-mir-361&WithFlank&X\|-\|85903630\|85903713@12.33.22&[MIMAT0000703&hsa-miR-361-5p&offsets\|0\|0;m-89&X\|-\|85903681\|85903702&offsets\|0\|0]\|TTATCAGAATCTCCAGGGGTAC | >MI0000760\|hsa-mir-361&WithFlank&X\|-\|85903630\|85903713 | 12 | 33 | 22 | TTATCAGAATCTCCAGGGGTAC | 273.5 | 3.0 | 0.8 | 3.6 | 3.7E-04 | 5.6E-04 | MB231/MB468 | MP |
| >MI0000734\|hsa-mir-106b&WithFlank&7\|-\|100093987\|100094080@58.78.21&[MIMAT0004672&hsa-miR-106b-3p&offsets\|0\|-1;m-87&7\|-\|100094003\|100094023&offsets\|0\|0]\|CCGCACTGTGGGTACTTGCTG | >MI0000734\|hsa-mir-106b&WithFlank&7\|-\|100093987\|100094080 | 58 | 78 | 21 | CCGCACTGTGGGTACTTGCTG | 1078.6 | 3.5 | 1.0 | 3.6 | 3.7E-04 | 5.6E-04 | MB231/MB468 | MP |
| >am_MI0000809\|hsa-mir-151a&WithFlank&8\|-\|140732558\|140732659@53.73.21(+1G)&[MIMAT0000757&hsa-miR-151a-3p&offsets\|0\|0(+1G);m-30&8\|-\|140732587\|140732607&offsets\|0\|0(+1G)]\|CTAGACTGAAGCTCCTTGAGGG | >am_MI0000809\|hsa-mir-151a&WithFlank&8\|-\|140732558\|140732659 | 53 | 73 | 21(+1G) | CTAGACTGAAGCTCCTTGAGGG | 302.0 | 5.1 | 1.4 | 3.7 | 1.8E-04 | 5.6E-04 | BT20/MB468 | MP |
| >hg38_21_+_8433172_8446622_RNA45SN1_45S_50ntFlanks@11776.11794.19\|TGACGCGCATGAATGGATG | >hg38_21_+_8433172_8446622_RNA45SN1_45S_50ntFlanks | 11776 | 11794 | 19 | TGACGCGCATGAATGGATG | 443.4 | 2.5 | 0.7 | 3.5 | 4.2E-04 | 5.6E-04 | MB231/MB468 | CYTO |
| >hg38_21_+_8433172_8446622_RNA45SN1_45S_50ntFlanks@12994.13013.20\|CCTCGCTGCGATCTATTGAA | >hg38_21_+_8433172_8446622_RNA45SN1_45S_50ntFlanks | 12994 | 13013 | 20 | CCTCGCTGCGATCTATTGAA | 1336.2 | -2.7 | 0.8 | -3.5 | 4.2E-04 | 5.6E-04 | MB231/MB468 | CYTO |
| >hg38_1_-_228634819_228635039_RNA5S12_5S_50ntFlanks@84.109.26\|CTCGTCTGATCTCGGAAGCTAAGCAG | >hg38_1_-_228634819_228635039_RNA5S12_5S_50ntFlanks | 84 | 109 | 26 | CTCGTCTGATCTCGGAAGCTAAGCAG | 510.4 | -6.3 | 1.8 | -3.5 | 4.5E-04 | 5.6E-04 | BT20/MB468 | NUC |
| >hg38_21_+_8433172_8446622_RNA45SN1_45S_50ntFlanks@9761.9786.26\|AACTTTAAATGGGTAAGAAGCCCGGC | >hg38_21_+_8433172_8446622_RNA45SN1_45S_50ntFlanks | 9761 | 9786 | 26 | AACTTTAAATGGGTAAGAAGCCCGGC | 98.6 | -9.7 | 2.6 | -3.7 | 1.8E-04 | 5.6E-04 | BT20/MB468 | MP |
| >hg38_21_+_8433172_8446622_RNA45SN1_45S_50ntFlanks@11575.11604.30\|ACTTAGAACTGGTGCGGACCAGGGGAATCC | >hg38_21_+_8433172_8446622_RNA45SN1_45S_50ntFlanks | 11575 | 11604 | 30 | ACTTAGAACTGGTGCGGACCAGGGGAATCC | 93.9 | -5.6 | 1.6 | -3.6 | 3.7E-04 | 5.6E-04 | MB231/MB468 | MP |
| >hg38_21_+_8433172_8446622_RNA45SN1_45S_50ntFlanks@12935.12953.19\|TAAACCATTCGTAGACGAC | >hg38_21_+_8433172_8446622_RNA45SN1_45S_50ntFlanks | 12935 | 12953 | 19 | TAAACCATTCGTAGACGAC | 184.3 | -4.0 | 1.1 | -3.5 | 4.5E-04 | 5.6E-04 | MB231/MB468 | NUC |
| >hg38_21_+_8433172_8446622_RNA45SN1_45S_50ntFlanks@12934.12966.33\|CTAAACCATTCGTAGACGACCTGCTTCTGGGTC | >hg38_21_+_8433172_8446622_RNA45SN1_45S_50ntFlanks | 12934 | 12966 | 33 | CTAAACCATTCGTAGACGACCTGCTTCTGGGTC | 215.6 | -6.3 | 1.7 | -3.7 | 1.8E-04 | 5.6E-04 | BT20/MB468 | MP |
| >hg38_21_+_8433172_8446622_RNA45SN1_45S_50ntFlanks@9568.9585.18\|CGTGCAAATCGGTCGTCC | >hg38_21_+_8433172_8446622_RNA45SN1_45S_50ntFlanks | 9568 | 9585 | 18 | CGTGCAAATCGGTCGTCC | 278.7 | -3.7 | 1.1 | -3.5 | 4.2E-04 | 5.6E-04 | MB231/MB468 | CYTO |
| >hg38_21_+_8433172_8446622_RNA45SN1_45S_50ntFlanks@10798.10823.26\|CATGTTGGAACAATGTAGGTAAGGGA | >hg38_21_+_8433172_8446622_RNA45SN1_45S_50ntFlanks | 10798 | 10823 | 26 | CATGTTGGAACAATGTAGGTAAGGGA | 206.2 | -1.8 | 0.5 | -3.6 | 3.3E-04 | 5.6E-04 | BT20/MB468 | MITO |
| >hg38_21_+_8433172_8446622_RNA45SN1_45S_50ntFlanks@10805.10840.36\|GAACAATGTAGGTAAGGGAAGTCGGCAAGCCGGATC | >hg38_21_+_8433172_8446622_RNA45SN1_45S_50ntFlanks | 10805 | 10840 | 36 | GAACAATGTAGGTAAGGGAAGTCGGCAAGCCGGATC | 305.7 | -3.8 | 1.1 | -3.6 | 3.3E-04 | 5.6E-04 | BT20/MB468 | MITO |
| >hg38_21_+_8433172_8446622_RNA45SN1_45S_50ntFlanks@12988.13013.26\|CAGCTCCCTCGCTGCGATCTATTGAA | >hg38_21_+_8433172_8446622_RNA45SN1_45S_50ntFlanks | 12988 | 13013 | 26 | CAGCTCCCTCGCTGCGATCTATTGAA | 1124.7 | -3.2 | 0.9 | -3.6 | 3.3E-04 | 5.6E-04 | BT20/MB468 | MITO |
| >hg38_21_+_8433172_8446622_RNA45SN1_45S_50ntFlanks@11999.12033.35\|CCCTGCGGGCCGCCGGTGAAATACCACTACTCTGA | >hg38_21_+_8433172_8446622_RNA45SN1_45S_50ntFlanks | 11999 | 12033 | 35 | CCCTGCGGGCCGCCGGTGAAATACCACTACTCTGA | 109.7 | -5.8 | 1.5 | -3.7 | 1.8E-04 | 5.6E-04 | BT20/MB468 | MP |
| >hg38_21_+_8433172_8446622_RNA45SN1_45S_50ntFlanks@9653.9677.25\|AGGATAGCTGGCGCTCTCGCAGACC | >hg38_21_+_8433172_8446622_RNA45SN1_45S_50ntFlanks | 9653 | 9677 | 25 | AGGATAGCTGGCGCTCTCGCAGACC | 944.9 | -5.7 | 1.6 | -3.5 | 4.5E-04 | 5.6E-04 | BT20/MB468 | NUC |
| >hg38_21_+_8433172_8446622_RNA45SN1_45S_50ntFlanks@12652.12675.24\|ATTATGACTGAACGCCTCTAAGTC | >hg38_21_+_8433172_8446622_RNA45SN1_45S_50ntFlanks | 12652 | 12675 | 24 | ATTATGACTGAACGCCTCTAAGTC | 140.6 | -5.0 | 1.4 | -3.5 | 4.5E-04 | 5.6E-04 | MB231/MB468 | NUC |
| >hg38_21_+_8433172_8446622_RNA45SN1_45S_50ntFlanks@5073.5093.21\|AATTCCGATAACGAACGAGAC | >hg38_21_+_8433172_8446622_RNA45SN1_45S_50ntFlanks | 5073 | 5093 | 21 | AATTCCGATAACGAACGAGAC | 428.0 | -6.5 | 1.8 | -3.6 | 3.3E-04 | 5.6E-04 | BT20/MB468 | MITO |
| >hg38_21_+_8433172_8446622_RNA45SN1_45S_50ntFlanks@11586.11613.28\|GTGCGGACCAGGGGAATCCGACTGTTTA | >hg38_21_+_8433172_8446622_RNA45SN1_45S_50ntFlanks | 11586 | 11613 | 28 | GTGCGGACCAGGGGAATCCGACTGTTTA | 225.2 | -2.9 | 0.8 | -3.5 | 4.5E-04 | 5.6E-04 | MB231/MB468 | NUC |
| >MI0000067\|hsa-let-7f-1&WithFlank&9\|+\|94176341\|94176439@13.33.21(+1G);MI0000068\|hsa-let-7f-2&WithFlank&X\|-\|53557186\|53557280@14.34.21(+1G)&[MIMAT0000067&hsa-let-7f-5p&offsets\|0\|-1(+1G);m-20&9\|+\|94176353\|94176374&offsets\|0\|-1(+1G)];[MIMAT0000067_1&hsa-let-7f-5p&offsets\|0\|-1(+1G);m-19&X\|-\|53557246\|53557267&offsets\|0\|-1(+1G)]\|TGAGGTAGTAGATTGTATAGTG | >MI0000067\|hsa-let-7f-1&WithFlank&9\|+\|94176341\|94176439 | 13 | 33 | 21(+1G) | TGAGGTAGTAGATTGTATAGTG | 343.6 | 3.8 | 1.1 | 3.5 | 4.5E-04 | 5.6E-04 | MB231/MB468 | MITO |
| >hg38_21_+_8433172_8446622_RNA45SN1_45S_50ntFlanks@9556.9573.18\|TAGCGGTCCTGACGTGCA | >hg38_21_+_8433172_8446622_RNA45SN1_45S_50ntFlanks | 9556 | 9573 | 18 | TAGCGGTCCTGACGTGCA | 445.0 | -2.5 | 0.7 | -3.5 | 4.2E-04 | 5.6E-04 | MB231/MB468 | CYTO |
| >hg38_21_+_8433172_8446622_RNA45SN1_45S_50ntFlanks@11892.11922.31\|TGACTCTAGTCTGGCACGGTGAAGAGACATG | >hg38_21_+_8433172_8446622_RNA45SN1_45S_50ntFlanks | 11892 | 11922 | 31 | TGACTCTAGTCTGGCACGGTGAAGAGACATG | 334.5 | 3.2 | 0.8 | 4.0 | 7.6E-05 | 5.6E-04 | BT20/MB231 | MITO |

| | | | | | | | | | | | | | |
|---|---|---|---|---|---|---|---|---|---|---|---|---|---|
| >hg38_21_+_8433172_8446622_RNA45SN1_45S_50ntFlanks@8196.8213.18\|CGTGGACGGTGTGAGGCC | >hg38_21_+_8433172_8446622_RNA45SN1_45S_50ntFlanks | 8196 | 8213 | 18 | CGTGGACGGTGTGAGGCC | 8690.5 | 3.2 | 0.9 | 3.5 | 4.2E-04 | 5.6E-04 | MB231/MB468 | CYTO |
| >trnaMT_HisGTG_MT_+_12138_12206@27.66.40\|GATTGTGAATCTGACAACAGAGGCTTACGACCCCTTATTT | >trnaMT_HisGTG_MT_+_12138_12206 | 27 | 66 | 40 | GATTGTGAATCTGACAACAGAGGCTTACGACCCCTTATTT | 75.6 | 10.0 | 2.7 | 3.7 | 1.8E-04 | 5.6E-04 | BT20/MB468 | MP |
| >hg38_1_-_228634819_228635039_RNA5S12_5S_50ntFlanks@133.169.37\|ATGGGAGACCGCCTGGGAATACCGGGTGCTGTAGGCT | >hg38_1_-_228634819_228635039_RNA5S12_5S_50ntFlanks | 133 | 169 | 37 | ATGGGAGACCGCCTGGGAATACCGGGTGCTGTAGGCT | 2433.5 | 2.8 | 0.8 | 3.5 | 4.2E-04 | 5.6E-04 | MB231/MB468 | CYTO |
| >hg38_21_+_8433172_8446622_RNA45SN1_45S_50ntFlanks@9535.9571.37\|GGAAACTCTGGTGGAGGTCCGTAGCGGTCCTGACGTG | >hg38_21_+_8433172_8446622_RNA45SN1_45S_50ntFlanks | 9535 | 9571 | 37 | GGAAACTCTGGTGGAGGTCCGTAGCGGTCCTGACGTG | 307.8 | -3.1 | 0.9 | -3.5 | 4.5E-04 | 5.6E-04 | MB231/MB468 | MITO |
| >MI0000439\|hsa-mir-23b&WithFlank&9\|+\|95085202\|95085310@67.85.19(+1U)&[MIMAT0000418&hsa-miR-23b-3p&offsets\|+3\|-1(+1U);m-42&9\|+\|95085265\|95085285&offsets\|+3\|+1(+1U)]\|ACATTGCCAGGGATTACCAT | >MI0000439\|hsa-mir-23b&WithFlank&9\|+\|95085202\|95085310 | 67 | 85 | 19(+1U) | ACATTGCCAGGGATTACCAT | 68.1 | 9.3 | 2.6 | 3.6 | 3.8E-04 | 5.6E-04 | MB231/MB468 | MP |
| >hg38_21_+_8433172_8446622_RNA45SN1_45S_50ntFlanks@10901.10930.30\|GGGGCTGGGCGCGCGCCGCGGCTGGACGAG | >hg38_21_+_8433172_8446622_RNA45SN1_45S_50ntFlanks | 10901 | 10930 | 30 | GGGGCTGGGCGCGCGCCGCGGCTGGACGAG | 60.6 | 9.3 | 2.3 | 4.0 | 6.4E-05 | 5.6E-04 | BT20/MB231 | MP |
| >MI0000239\|hsa-mir-197&WithFlank&1\|+\|109598887\|109598973@54.74.21&[MIMAT0000227&hsa-miR-197-3p&offsets\|0\|-1;m-99&1\|+\|109598940\|109598961&offsets\|0\|-1]\|TTCACCACCTTCTCCACCCAG | >MI0000239\|hsa-mir-197&WithFlank&1\|+\|109598887\|109598973 | 54 | 74 | 21 | TTCACCACCTTCTCCACCCAG | 713.0 | 3.1 | 0.8 | 4.0 | 6.5E-05 | 5.6E-04 | BT20/MB231 | CYTO |
| >trna5_AspGTC_12_+_98897281_98897352@36.75.40\|CACGCGGGAGACCGGGGTTCAATTCCCCGACGGGGAGCCA | >trna5_AspGTC_12_+_98897281_98897352 | 36 | 75 | 40 | CACGCGGGAGACCGGGGTTCAATTCCCCGACGGGGAGCCA | 3316.2 | -4.5 | 1.1 | -4.0 | 6.5E-05 | 5.6E-04 | BT20/MB231 | CYTO |
| >hg38_21_+_8433172_8446622_RNA45SN1_45S_50ntFlanks@11589.11611.23\|CGGACCAGGGGAATCCGACTGTT | >hg38_21_+_8433172_8446622_RNA45SN1_45S_50ntFlanks | 11589 | 11611 | 23 | CGGACCAGGGGAATCCGACTGTT | 638.8 | -2.0 | 0.6 | -3.5 | 4.5E-04 | 5.6E-04 | MB231/MB468 | NUC |
| >hg38_21_+_8433172_8446622_RNA45SN1_45S_50ntFlanks@475.514.40\|CTTCGTGATCGATGTGGTGACGTCGTGCTCTCCCGGGCCG | >hg38_21_+_8433172_8446622_RNA45SN1_45S_50ntFlanks | 475 | 514 | 40 | CTTCGTGATCGATGTGGTGACGTCGTGCTCTCCCGGGCCG | 7016.2 | 3.4 | 1.0 | 3.5 | 4.5E-04 | 5.6E-04 | MB231/MB468 | NUC |
| >hg38_21_+_8433172_8446622_RNA45SN1_45S_50ntFlanks@11212.11251.40\|CGACTCTGGACGCGAGCCGGGCCCTTCCCGTGGATCGCCC | >hg38_21_+_8433172_8446622_RNA45SN1_45S_50ntFlanks | 11212 | 11251 | 40 | CGACTCTGGACGCGAGCCGGGCCCTTCCCGTGGATCGCCC | 387.9 | -3.0 | 0.9 | -3.5 | 4.5E-04 | 5.6E-04 | MB231/MB468 | NUC |
| >hg38_21_+_8433172_8446622_RNA45SN1_45S_50ntFlanks@10393.10430.38\|CAGCAGTTGAACATGGGTCAGTCGGTCCTGAGAGATGG | >hg38_21_+_8433172_8446622_RNA45SN1_45S_50ntFlanks | 10393 | 10430 | 38 | CAGCAGTTGAACATGGGTCAGTCGGTCCTGAGAGATGG | 354.8 | -3.7 | 1.0 | -3.7 | 1.8E-04 | 5.6E-04 | BT20/MB468 | MP |
| >hg38_21_+_8433172_8446622_RNA45SN1_45S_50ntFlanks@4297.4318.22\|CATTGGAGGGCAAGTCTGGTGC | >hg38_21_+_8433172_8446622_RNA45SN1_45S_50ntFlanks | 4297 | 4318 | 22 | CATTGGAGGGCAAGTCTGGTGC | 198.6 | -2.2 | 0.6 | -3.5 | 4.2E-04 | 5.6E-04 | MB231/MB468 | CYTO |
| >hg38_21_+_8433172_8446622_RNA45SN1_45S_50ntFlanks@5491.5525.35\|GAGAAGACGGTCGAACTTGACTATCTAGAGGAAGT | >hg38_21_+_8433172_8446622_RNA45SN1_45S_50ntFlanks | 5491 | 5525 | 35 | GAGAAGACGGTCGAACTTGACTATCTAGAGGAAGT | 258.3 | 5.9 | 1.6 | 3.7 | 1.8E-04 | 5.6E-04 | BT20/MB468 | MP |
| >hg38_21_+_8433172_8446622_RNA45SN1_45S_50ntFlanks@6661.6697.37\|CGGTGGATCACTCGGCTCGTGCGTCGATGAAGAACGC | >hg38_21_+_8433172_8446622_RNA45SN1_45S_50ntFlanks | 6661 | 6697 | 37 | CGGTGGATCACTCGGCTCGTGCGTCGATGAAGAACGC | 2386.9 | -2.5 | 0.7 | -3.5 | 4.5E-04 | 5.6E-04 | MB231/MB468 | MITO |
| >hg38_21_+_8433172_8446622_RNA45SN1_45S_50ntFlanks@13001.13027.27\|GCGATCTATTGAAAGTCAGCCCTCGAC | >hg38_21_+_8433172_8446622_RNA45SN1_45S_50ntFlanks | 13001 | 13027 | 27 | GCGATCTATTGAAAGTCAGCCCTCGAC | 308.4 | -4.2 | 1.2 | -3.5 | 4.0E-04 | 5.7E-04 | BT20/MB468 | CYTO |
| >MI0000063\|hsa-let-7b&WithFlank&22\|+\|46113680\|46113774@12.32.21(+1C)&[MIMAT0000063&hsa-let-7b-5p&offsets\|0\|-1(+1C);m-14&22\|+\|46113691\|46113712&offsets\|0\|-1(+1C)]\|TGAGGTAGTAGGTTGTGTGGTC | >MI0000063\|hsa-let-7b&WithFlank&22\|+\|46113680\|46113774 | 12 | 32 | 21(+1C) | TGAGGTAGTAGGTTGTGTGGTC | 9666.0 | 3.3 | 0.8 | 4.0 | 6.5E-05 | 5.7E-04 | BT20/MB231 | CYTO |
| >MI0000746\|hsa-mir-99b&WithFlank&19\|+\|51692606\|51692687@13.34.22(+1U)&[MIMAT0000689&hsa-miR-99b-5p&offsets\|0\|0(+1U);m-10&19\|+\|51692618\|51692639&offsets\|0\|0(+1U)]\|CACCCGTAGAACCGACCTTGCGT | >MI0000746\|hsa-mir-99b&WithFlank&19\|+\|51692606\|51692687 | 13 | 34 | 22(+1U) | CACCCGTAGAACCGACCTTGCGT | 810.5 | 2.8 | 0.8 | 3.6 | 3.8E-04 | 5.7E-04 | MB231/MB468 | MP |
| >hg38_21_+_8433172_8446622_RNA45SN1_45S_50ntFlanks@11704.11736.33\|ATGAAGCGCGGGTAAACGGCGGGAGTAACTATG | >hg38_21_+_8433172_8446622_RNA45SN1_45S_50ntFlanks | 11704 | 11736 | 33 | ATGAAGCGCGGGTAAACGGCGGGAGTAACTATG | 202.5 | -4.5 | 1.3 | -3.6 | 3.8E-04 | 5.7E-04 | MB231/MB468 | MP |
| >MI0003591\|hsa-mir-584&WithFlank&5\|-\|149062307\|149062415@22.44.23&[MIMAT0003249&hsa-miR-584-5p&offsets\|0\|+1;m-167&5\|-\|149062374\|149062394&offsets\|0\|+2]\|TTATGGTTTGCCTGGGACTGAGG | >MI0003591\|hsa-mir-584&WithFlank&5\|-\|149062307\|149062415 | 22 | 44 | 23 | TTATGGTTTGCCTGGGACTGAGG | 235.3 | 6.1 | 1.6 | 3.7 | 1.8E-04 | 5.7E-04 | BT20/MB468 | MP |
| >trnaMT_GluTTC_MT_-_14674_14742@39.71.33,trnalookalike8_GluTTC_5_-_93905172_93905240@39.71.33\|ATTGGTCGTGGTTGTAGTCCGTGCGAGAATACC | >trnaMT_GluTTC_MT_-_14674_14742 | 39 | 71 | 33 | ATTGGTCGTGGTTGTAGTCCGTGCGAGAATACC | 2454.8 | 4.4 | 1.2 | 3.7 | 1.8E-04 | 5.7E-04 | BT20/MB468 | MP |
| >hg38_21_+_8433172_8446622_RNA45SN1_45S_50ntFlanks@11322.11344.23\|ACGTCTCGTCGCGCGCGCGTCCG | >hg38_21_+_8433172_8446622_RNA45SN1_45S_50ntFlanks | 11322 | 11344 | 23 | ACGTCTCGTCGCGCGCGCGTCCG | 2343.1 | -5.5 | 1.5 | -3.7 | 1.8E-04 | 5.7E-04 | BT20/MB468 | MP |
| >hg38_21_+_8433172_8446622_RNA45SN1_45S_50ntFlanks@5487.5512.26\|CGCTGAGAAGACGGTCGAACTTGACT | >hg38_21_+_8433172_8446622_RNA45SN1_45S_50ntFlanks | 5487 | 5512 | 26 | CGCTGAGAAGACGGTCGAACTTGACT | 402.2 | -3.5 | 0.9 | -3.7 | 1.8E-04 | 5.7E-04 | BT20/MB468 | MP |
| >hg38_21_+_8433172_8446622_RNA45SN1_45S_50ntFlanks@10386.10424.39\|ATGTGAACAGCAGTTGAACATGGGTCAGTCGGTCCTGAG | >hg38_21_+_8433172_8446622_RNA45SN1_45S_50ntFlanks | 10386 | 10424 | 39 | ATGTGAACAGCAGTTGAACATGGGTCAGTCGGTCCTGAG | 550.5 | 3.2 | 0.9 | 3.5 | 4.5E-04 | 5.7E-04 | MB231/MB468 | MITO |
| >hg38_21_+_8433172_8446622_RNA45SN1_45S_50ntFlanks@9941.9968.28\|AGGACGGTGGCCATGGAAGTCGGAATCC | >hg38_21_+_8433172_8446622_RNA45SN1_45S_50ntFlanks | 9941 | 9968 | 28 | AGGACGGTGGCCATGGAAGTCGGAATCC | 2253.4 | 5.7 | 1.4 | 4.0 | 6.5E-05 | 5.7E-04 | BT20/MB231 | MP |
| >hg38_21_+_8433172_8446622_RNA45SN1_45S_50ntFlanks@10786.10821.36\|AACAGCCTCTGGCATGTTGGAACAATGTAGGTAAGG | >hg38_21_+_8433172_8446622_RNA45SN1_45S_50ntFlanks | 10786 | 10821 | 36 | AACAGCCTCTGGCATGTTGGAACAATGTAGGTAAGG | 363.1 | -2.3 | 0.7 | -3.5 | 4.3E-04 | 5.7E-04 | MB231/MB468 | CYTO |
| >MI0000079\|hsa-mir-23a&WithFlank&19\|-\|13836581\|13836665@54.74.21&[MIMAT0000078&hsa-miR-23a-3p&offsets\|+3\|+3;m-31&19\|-\|13836595\|13836615&offsets\|+3\|+3]\|ACATTGCCAGGGATTTCCAAC | >MI0000079\|hsa-mir-23a&WithFlank&19\|-\|13836581\|13836665 | 54 | 74 | 21 | ACATTGCCAGGGATTTCCAAC | 69.0 | 9.3 | 2.6 | 3.6 | 3.8E-04 | 5.7E-04 | MB231/MB468 | MP |
| >MI0000746\|hsa-mir-99b&WithFlank&19\|+\|51692606\|51692687@13.34.22(+1U)&[MIMAT0000689&hsa-miR-99b-5p&offsets\|0\|0(+1U);m-10&19\|+\|51692618\|51692639&offsets\|0\|0(+1U)]\|CACCCGTAGAACCGACCTTGCGT | >MI0000746\|hsa-mir-99b&WithFlank&19\|+\|51692606\|51692687 | 13 | 34 | 22(+1U) | CACCCGTAGAACCGACCTTGCGT | 621.2 | 3.1 | 0.8 | 3.7 | 1.8E-04 | 5.7E-04 | BT20/MB468 | MP |
| >hg38_21_+_8433172_8446622_RNA45SN1_45S_50ntFlanks@10683.10706.24\|CCGGCGGCGTCCGGTGAGCTCTCG | >hg38_21_+_8433172_8446622_RNA45SN1_45S_50ntFlanks | 10683 | 10706 | 24 | CCGGCGGCGTCCGGTGAGCTCTCG | 5738.1 | -6.9 | 1.9 | -3.7 | 1.8E-04 | 5.7E-04 | BT20/MB468 | MP |
| >hg38_21_+_8433172_8446622_RNA45SN1_45S_50ntFlanks@476.506.31\|TTCGTGATCGATGTGGTGACGTCGTGCTCTC | >hg38_21_+_8433172_8446622_RNA45SN1_45S_50ntFlanks | 476 | 506 | 31 | TTCGTGATCGATGTGGTGACGTCGTGCTCTC | 173.6 | 5.2 | 1.4 | 3.7 | 1.8E-04 | 5.7E-04 | BT20/MB468 | MP |
| >hg38_21_+_8433172_8446622_RNA45SN1_45S_50ntFlanks@12029.12059.31\|TCTGATCGTTTTTTCACTGACCCGGTGAGGC | >hg38_21_+_8433172_8446622_RNA45SN1_45S_50ntFlanks | 12029 | 12059 | 31 | TCTGATCGTTTTTTCACTGACCCGGTGAGGC | 68.3 | 9.8 | 2.6 | 3.7 | 1.8E-04 | 5.7E-04 | BT20/MB468 | MP |
| >hg38_21_+_8433172_8446622_RNA45SN1_45S_50ntFlanks@6760.6806.47\|TTGCGGCCCCGGGTTCCTCCCGGGGCTACGCCTGTCTGAGCGTCGCT | >hg38_21_+_8433172_8446622_RNA45SN1_45S_50ntFlanks | 6760 | 6806 | 47 | TTGCGGCCCCGGGTTCCTCCCGGGGCTACGCCTGTCTGAGCGTCGCT | 775.5 | 9.0 | 2.5 | 3.5 | 4.3E-04 | 5.7E-04 | MB231/MB468 | CYTO |
| >MI0000088\|hsa-mir-30a&WithFlank&6\|-\|71403545\|71403627@13.34.22(+1A)&[MIMAT0000087&hsa-miR-30a-5p&offsets\|+1\|+1(+1A);m-15&6\|-\|71403595\|71403616&offsets\|+1\|+1(+1A)]\|GTAAACATCCTCGACTGGAAGCA | >MI0000088\|hsa-mir-30a&WithFlank&6\|-\|71403545\|71403627 | 13 | 34 | 22(+1A) | GTAAACATCCTCGACTGGAAGCA | 367.4 | 4.5 | 1.3 | 3.6 | 3.8E-04 | 5.7E-04 | MB231/MB468 | MP |
| >hg38_21_+_8433172_8446622_RNA45SN1_45S_50ntFlanks@6653.6698.46\|ACTCTTAGCGGTGGATCACTCGGCTCGTGCGTCGATGAAGAACGCA | >hg38_21_+_8433172_8446622_RNA45SN1_45S_50ntFlanks | 6653 | 6698 | 46 | ACTCTTAGCGGTGGATCACTCGGCTCGTGCGTCGATGAAGAACGCA | 549.3 | -3.2 | 0.9 | -3.5 | 4.6E-04 | 5.7E-04 | BT20/MB468 | NUC |
| >MI0000264\|hsa-mir-7-2&WithFlank&15\|+\|88611819\|88611940@38.60.23(+1C);MI0000265\|hsa-mir-7-3&WithFlank&19\|+\|4770664\|4770785@37.59.23(+1C);MI0000263\|hsa-mir-7-1&WithFlank&9\|-\|83969742\|83969863@30.52.23(+1C)&[MIMAT0000252&hsa-miR-7-5p&offsets\|0\|-1(+1C);m-400&15\|+\|88611856\|88611879&offsets\|0\|-1(+1C)];[MIMAT0000252_1&hsa-miR-7-5p&offsets\|0\|-1(+1C);m-402&19\|+\|4770700\|4770723&offsets\|0\|-1(+1C)];[MIMAT0000252_2&hsa-miR-7-5p&offsets\|0\|-1(+1C);m-401&9\|-\|83969811\|83969834&offsets\|0\|-1(+1C)]\|TGGAAGACTAGTGATTTTGTTGTC | >MI0000264\|hsa-mir-7-2&WithFlank&15\|+\|88611819\|88611940 | 38 | 60 | 23(+1C) | TGGAAGACTAGTGATTTTGTTGTC | 13798.2 | 3.5 | 0.9 | 4.0 | 6.6E-05 | 5.7E-04 | BT20/MB231 | CYTO |
| >hg38_21_+_8433172_8446622_RNA45SN1_45S_50ntFlanks@12444.12483.40\|AAGCAGAATTCACCAAGCGTTGGATTGTTCACCCACTAAT | >hg38_21_+_8433172_8446622_RNA45SN1_45S_50ntFlanks | 12444 | 12483 | 40 | AAGCAGAATTCACCAAGCGTTGGATTGTTCACCCACTAAT | 384.9 | -2.9 | 0.8 | -3.5 | 4.6E-04 | 5.7E-04 | MB231/MB468 | NUC |
| >hg38_21_+_8433172_8446622_RNA45SN1_45S_50ntFlanks@12412.12455.44\|ATCCTTCGATGTCGGCTCTTCCTATCATTGTGAAGCAGAATTCA | >hg38_21_+_8433172_8446622_RNA45SN1_45S_50ntFlanks | 12412 | 12455 | 44 | ATCCTTCGATGTCGGCTCTTCCTATCATTGTGAAGCAGAATTCA | 353.4 | -5.8 | 1.6 | -3.5 | 4.5E-04 | 5.7E-04 | MB231/MB468 | MITO |
| >hg38_21_+_8433172_8446622_RNA45SN1_45S_50ntFlanks@9739.9757.19\|AAACGATCTCAACCTATTC | >hg38_21_+_8433172_8446622_RNA45SN1_45S_50ntFlanks | 9739 | 9757 | 19 | AAACGATCTCAACCTATTC | 949.8 | -6.2 | 1.7 | -3.7 | 1.9E-04 | 5.7E-04 | BT20/MB468 | MP |
| >hg38_21_+_8433172_8446622_RNA45SN1_45S_50ntFlanks@3984.4004.21\|GCGGCTTTGGTGACTCTAGAT | >hg38_21_+_8433172_8446622_RNA45SN1_45S_50ntFlanks | 3984 | 4004 | 21 | GCGGCTTTGGTGACTCTAGAT | 130.3 | -5.7 | 1.5 | -3.7 | 1.9E-04 | 5.7E-04 | BT20/MB468 | MP |
| >hg38_21_+_8433172_8446622_RNA45SN1_45S_50ntFlanks@12914.12933.20\|CGTTCGTGGGGAACCTGGCG | >hg38_21_+_8433172_8446622_RNA45SN1_45S_50ntFlanks | 12914 | 12933 | 20 | CGTTCGTGGGGAACCTGGCG | 426.0 | -3.8 | 1.1 | -3.5 | 4.3E-04 | 5.7E-04 | MB231/MB468 | CYTO |
| >trnaMT_GluTTC_MT_-_14674_14742@34.63.30,trnalookalike8_GluTTC_5_-_93905172_93905240@34.63.30\|ATATCATTGGTCGTGGTTGTAGTCCGTGCG | >trnaMT_GluTTC_MT_-_14674_14742 | 34 | 63 | 30 | ATATCATTGGTCGTGGTTGTAGTCCGTGCG | 255.4 | 4.4 | 1.2 | 3.7 | 1.9E-04 | 5.7E-04 | BT20/MB468 | MP |
| >hg38_21_+_8433172_8446622_RNA45SN1_45S_50ntFlanks@10572.10607.36\|AGAAGCCGGCGGGAGCCCCGGGGAGAGTTCTCTTTT | >hg38_21_+_8433172_8446622_RNA45SN1_45S_50ntFlanks | 10572 | 10607 | 36 | AGAAGCCGGCGGGAGCCCCGGGGAGAGTTCTCTTTT | 191.8 | -2.2 | 0.6 | -3.6 | 3.4E-04 | 5.7E-04 | BT20/MB468 | MITO |
| >hg38_21_+_8433172_8446622_RNA45SN1_45S_50ntFlanks@12418.12456.39\|CGATGTCGGCTCTTCCTATCATTGTGAAGCAGAATTCAC | >hg38_21_+_8433172_8446622_RNA45SN1_45S_50ntFlanks | 12418 | 12456 | 39 | CGATGTCGGCTCTTCCTATCATTGTGAAGCAGAATTCAC | 1894.3 | -3.2 | 0.9 | -3.5 | 4.5E-04 | 5.7E-04 | MB231/MB468 | MITO |
| >hg38_21_+_8433172_8446622_RNA45SN1_45S_50ntFlanks@475.521.47\|CTTCGTGATCGATGTGGTGACGTCGTGCTCTCCCGGGCCGGGTCCGA | >hg38_21_+_8433172_8446622_RNA45SN1_45S_50ntFlanks | 475 | 521 | 47 | CTTCGTGATCGATGTGGTGACGTCGTGCTCTCCCGGGCCGGGTCCGA | 1078.2 | 6.2 | 1.7 | 3.6 | 3.4E-04 | 5.7E-04 | BT20/MB468 | MITO |
| >hg38_21_+_8433172_8446622_RNA45SN1_45S_50ntFlanks@12415.12440.26\|CTTCGATGTCGGCTCTTCCTATCATT | >hg38_21_+_8433172_8446622_RNA45SN1_45S_50ntFlanks | 12415 | 12440 | 26 | CTTCGATGTCGGCTCTTCCTATCATT | 693.3 | 2.1 | 0.6 | 3.5 | 4.1E-04 | 5.7E-04 | BT20/MB468 | CYTO |
| >hg38_21_+_8433172_8446622_RNA45SN1_45S_50ntFlanks@11568.11592.25\|GCAGCCGACTTAGAACTGGTGCGGA | >hg38_21_+_8433172_8446622_RNA45SN1_45S_50ntFlanks | 11568 | 11592 | 25 | GCAGCCGACTTAGAACTGGTGCGGA | 1588.9 | -5.7 | 1.5 | -3.7 | 1.9E-04 | 5.7E-04 | BT20/MB468 | MP |

<table>
<tr><td>>hg38_21_+_8433172_8446622_RNA45SN1_45S_50ntFlanks@11318.11338.21|CCCCACGTCTCGTCGCGCGCG</td><td>>hg38_21_+_8433172_8446622_RNA45SN1_45S_50ntFlanks</td><td>11318</td><td>11338</td><td>21</td><td>CCCCACGTCTCGTCGCGCGCG</td><td>1823.5</td><td>-7.0</td><td>2.0</td><td>-3.5</td><td>4.6E-04</td><td>5.7E-04</td><td>BT20/MB468</td><td>NUC</td></tr>
<tr><td>>hg38_21_+_8433172_8446622_RNA45SN1_45S_50ntFlanks@7981.7999.19|CTCAGATCAGACGTGGCGA</td><td>>hg38_21_+_8433172_8446622_RNA45SN1_45S_50ntFlanks</td><td>7981</td><td>7999</td><td>19</td><td>CTCAGATCAGACGTGGCGA</td><td>169.5</td><td>-5.0</td><td>1.4</td><td>-3.5</td><td>4.6E-04</td><td>5.7E-04</td><td>MB231/MB468</td><td>NUC</td></tr>
<tr><td>>hg38_21_+_8433172_8446622_RNA45SN1_45S_50ntFlanks@5494.5525.32|AAGACGGTCGAACTTGACTATCTAGAGGAAGT</td><td>>hg38_21_+_8433172_8446622_RNA45SN1_45S_50ntFlanks</td><td>5494</td><td>5525</td><td>32</td><td>AAGACGGTCGAACTTGACTATCTAGAGGAAGT</td><td>277.5</td><td>-4.3</td><td>1.2</td><td>-3.6</td><td>3.8E-04</td><td>5.7E-04</td><td>MB231/MB468</td><td>MP</td></tr>
<tr><td>>hg38_21_+_8433172_8446622_RNA45SN1_45S_50ntFlanks@10101.10126.26|CGCGCGCGTGTGGTGTGCGTCGGAGG</td><td>>hg38_21_+_8433172_8446622_RNA45SN1_45S_50ntFlanks</td><td>10101</td><td>10126</td><td>26</td><td>CGCGCGCGTGTGGTGTGCGTCGGAGG</td><td>547.3</td><td>-6.5</td><td>1.8</td><td>-3.5</td><td>4.7E-04</td><td>5.7E-04</td><td>BT20/MB468</td><td>NUC</td></tr>
<tr><td>>hg38_21_+_8433172_8446622_RNA45SN1_45S_50ntFlanks@5237.5281.45|ACTGACTGGCTCAGCGTGTGCCTACCCTACGCCGGCAGGCGCGGG</td><td>>hg38_21_+_8433172_8446622_RNA45SN1_45S_50ntFlanks</td><td>5237</td><td>5281</td><td>45</td><td>ACTGACTGGCTCAGCGTGTGCCTACCCTACGCCGGCAGGCGCGGG</td><td>75.1</td><td>10.0</td><td>2.7</td><td>3.7</td><td>1.9E-04</td><td>5.7E-04</td><td>BT20/MB468</td><td>MP</td></tr>
<tr><td>>hg38_21_+_8433172_8446622_RNA45SN1_45S_50ntFlanks@11661.11678.18|TGATTTCTGCCCAGTGCT</td><td>>hg38_21_+_8433172_8446622_RNA45SN1_45S_50ntFlanks</td><td>11661</td><td>11678</td><td>18</td><td>TGATTTCTGCCCAGTGCT</td><td>90.5</td><td>-5.7</td><td>1.5</td><td>-3.7</td><td>1.9E-04</td><td>5.8E-04</td><td>BT20/MB468</td><td>MP</td></tr>
<tr><td>>hg38_21_+_8433172_8446622_RNA45SN1_45S_50ntFlanks@12001.12043.43|CTGCGGGCCGCCGGTGAAATACCACTACTCTGATCGTTTTTTC</td><td>>hg38_21_+_8433172_8446622_RNA45SN1_45S_50ntFlanks</td><td>12001</td><td>12043</td><td>43</td><td>CTGCGGGCCGCCGGTGAAATACCACTACTCTGATCGTTTTTTC</td><td>84.8</td><td>-9.5</td><td>2.5</td><td>-3.7</td><td>1.9E-04</td><td>5.8E-04</td><td>BT20/MB468</td><td>MP</td></tr>
<tr><td>>trna13_ValCAC_19_-_4724647_4724719@1.33.33|GTTTCCGTAGTGTAGCGGTTATCACATTCGCCT</td><td>>trna13_ValCAC_19_-_4724647_4724719</td><td>1</td><td>33</td><td>33</td><td>GTTTCCGTAGTGTAGCGGTTATCACATTCGCCT</td><td>419.0</td><td>-4.2</td><td>1.1</td><td>-4.0</td><td>6.7E-05</td><td>5.8E-04</td><td>BT20/MB231</td><td>CYTO</td></tr>
<tr><td>>hg38_21_+_8433172_8446622_RNA45SN1_45S_50ntFlanks@6656.6689.34|CTTAGCGGTGGATCACTCGGCTCGTGCGTCGATG</td><td>>hg38_21_+_8433172_8446622_RNA45SN1_45S_50ntFlanks</td><td>6656</td><td>6689</td><td>34</td><td>CTTAGCGGTGGATCACTCGGCTCGTGCGTCGATG</td><td>211.8</td><td>-3.2</td><td>0.9</td><td>-3.5</td><td>4.3E-04</td><td>5.8E-04</td><td>MB231/MB468</td><td>CYTO</td></tr>
<tr><td>>MI0000108|hsa-mir-103a-2&WithFlank&20|+|3917488|3917577@57.76.20;MI0000109|hsa-mir-103a-1&WithFlank&5|-|168560890|168560979@57.76.20&[MIMAT0000101&hsa-miR-103a-3p&offsets|+3|0;m-12&20|+|3917541|3917563&offsets|+3|0];[MIMAT0000101_1&hsa-miR-103a-3p&offsets|+3|0;m-13&5|-|168560904|168560926&offsets|+3|0;m-44220&5|-|168560908|168560926&offsets|+3|+4]|AGCATTGTACAGGGCTATGA</td><td>>MI0000108|hsa-mir-103a-2&WithFlank&20|+|3917488|3917577</td><td>57</td><td>76</td><td>20</td><td>AGCATTGTACAGGGCTATGA</td><td>48.6</td><td>-9.0</td><td>2.3</td><td>-4.0</td><td>6.6E-05</td><td>5.8E-04</td><td>BT20/MB231</td><td>MP</td></tr>
<tr><td>>hg38_21_+_8433172_8446622_RNA45SN1_45S_50ntFlanks@6671.6696.26|CTCGGCTCGTGCGTCGATGAAGAACG</td><td>>hg38_21_+_8433172_8446622_RNA45SN1_45S_50ntFlanks</td><td>6671</td><td>6696</td><td>26</td><td>CTCGGCTCGTGCGTCGATGAAGAACG</td><td>557.1</td><td>-3.5</td><td>1.0</td><td>-3.5</td><td>4.3E-04</td><td>5.8E-04</td><td>MB231/MB468</td><td>CYTO</td></tr>
<tr><td>>hg38_21_+_8433172_8446622_RNA45SN1_45S_50ntFlanks@10403.10431.29|ACATGGGTCAGTCGGTCCTGAGAGATGGG</td><td>>hg38_21_+_8433172_8446622_RNA45SN1_45S_50ntFlanks</td><td>10403</td><td>10431</td><td>29</td><td>ACATGGGTCAGTCGGTCCTGAGAGATGGG</td><td>197.2</td><td>-5.3</td><td>1.5</td><td>-3.5</td><td>4.7E-04</td><td>5.8E-04</td><td>MB231/MB468</td><td>NUC</td></tr>
<tr><td>>hg38_21_+_8433172_8446622_RNA45SN1_45S_50ntFlanks@6652.6697.46|GACTCTTAGCGGTGGATCACTCGGCTCGTGCGTCGATGAAGAACGC</td><td>>hg38_21_+_8433172_8446622_RNA45SN1_45S_50ntFlanks</td><td>6652</td><td>6697</td><td>46</td><td>GACTCTTAGCGGTGGATCACTCGGCTCGTGCGTCGATGAAGAACGC</td><td>782.4</td><td>2.8</td><td>0.8</td><td>3.5</td><td>4.3E-04</td><td>5.8E-04</td><td>MB231/MB468</td><td>CYTO</td></tr>
<tr><td>>hg38_21_+_8433172_8446622_RNA45SN1_45S_50ntFlanks@8357.8376.20|AAAAGAACTTTGAAGAGAGA</td><td>>hg38_21_+_8433172_8446622_RNA45SN1_45S_50ntFlanks</td><td>8357</td><td>8376</td><td>20</td><td>AAAAGAACTTTGAAGAGAGA</td><td>253.0</td><td>-2.6</td><td>0.7</td><td>-3.5</td><td>4.1E-04</td><td>5.8E-04</td><td>BT20/MB468</td><td>CYTO</td></tr>
<tr><td>>hg38_21_+_8433172_8446622_RNA45SN1_45S_50ntFlanks@3789.3814.26|ACTGCGAATGGCTCATTAAATCAGTT</td><td>>hg38_21_+_8433172_8446622_RNA45SN1_45S_50ntFlanks</td><td>3789</td><td>3814</td><td>26</td><td>ACTGCGAATGGCTCATTAAATCAGTT</td><td>59.3</td><td>-5.5</td><td>1.5</td><td>-3.5</td><td>3.9E-04</td><td>5.8E-04</td><td>MB231/MB468</td><td>MP</td></tr>
<tr><td>>hg38_21_+_8433172_8446622_RNA45SN1_45S_50ntFlanks@6663.6686.24|GTGGATCACTCGGCTCGTGCGTCG</td><td>>hg38_21_+_8433172_8446622_RNA45SN1_45S_50ntFlanks</td><td>6663</td><td>6686</td><td>24</td><td>GTGGATCACTCGGCTCGTGCGTCG</td><td>213.4</td><td>3.9</td><td>1.0</td><td>3.9</td><td>7.9E-05</td><td>5.8E-04</td><td>BT20/MB231</td><td>MITO</td></tr>
<tr><td>>MI0000811|hsa-mir-148b&WithFlank&12|+|54337210|54337320@69.90.22(+3U)&[MIMAT0000759&hsa-miR-148b-3p&offsets|0|0(+3U);m-77&12|+|54337278|54337299&offsets|0|0(+3U)]|TCAGTGCATCACAGAACTTTGTTTT</td><td>>MI0000811|hsa-mir-148b&WithFlank&12|+|54337210|54337320</td><td>69</td><td>90</td><td>22(+3U)</td><td>TCAGTGCATCACAGAACTTTGTTTT</td><td>334.4</td><td>-3.0</td><td>0.8</td><td>-3.9</td><td>7.9E-05</td><td>5.8E-04</td><td>BT20/MB231</td><td>MITO</td></tr>
<tr><td>>hg38_21_+_8433172_8446622_RNA45SN1_45S_50ntFlanks@6649.6672.24|TACGACTCTTAGCGGTGGATCACT</td><td>>hg38_21_+_8433172_8446622_RNA45SN1_45S_50ntFlanks</td><td>6649</td><td>6672</td><td>24</td><td>TACGACTCTTAGCGGTGGATCACT</td><td>134.0</td><td>2.1</td><td>0.6</td><td>3.5</td><td>4.7E-04</td><td>5.8E-04</td><td>BT20/MB468</td><td>NUC</td></tr>
<tr><td>>hg38_21_+_8433172_8446622_RNA45SN1_45S_50ntFlanks@8168.8213.46|CCCAAGTCCTTCTGATCGAGGCCCAGCCCGTGGACGGTGTGAGGCC</td><td>>hg38_21_+_8433172_8446622_RNA45SN1_45S_50ntFlanks</td><td>8168</td><td>8213</td><td>46</td><td>CCCAAGTCCTTCTGATCGAGGCCCAGCCCGTGGACGGTGTGAGGCC</td><td>511.9</td><td>-4.5</td><td>1.3</td><td>-3.5</td><td>4.7E-04</td><td>5.8E-04</td><td>MB231/MB468</td><td>NUC</td></tr>
<tr><td>>hg38_21_+_8433172_8446622_RNA45SN1_45S_50ntFlanks@6730.6771.42|ACACATTGATCATCGACACTTCGAACGCACTTGCGGCCCCGG</td><td>>hg38_21_+_8433172_8446622_RNA45SN1_45S_50ntFlanks</td><td>6730</td><td>6771</td><td>42</td><td>ACACATTGATCATCGACACTTCGAACGCACTTGCGGCCCCGG</td><td>136.8</td><td>-3.3</td><td>0.9</td><td>-3.5</td><td>4.4E-04</td><td>5.8E-04</td><td>MB231/MB468</td><td>CYTO</td></tr>
<tr><td>>hg38_21_+_8433172_8446622_RNA45SN1_45S_50ntFlanks@12991.13036.46|CTCCCTCGCTGCGATCTATTGAAAGTCAGCCCTCGACACAAGGGTT</td><td>>hg38_21_+_8433172_8446622_RNA45SN1_45S_50ntFlanks</td><td>12991</td><td>13036</td><td>46</td><td>CTCCCTCGCTGCGATCTATTGAAAGTCAGCCCTCGACACAAGGGTT</td><td>398.1</td><td>-6.1</td><td>1.7</td><td>-3.5</td><td>4.6E-04</td><td>5.8E-04</td><td>MB231/MB468</td><td>MITO</td></tr>
<tr><td>>hg38_21_+_8433172_8446622_RNA45SN1_45S_50ntFlanks@8176.8214.39|CTTCTGATCGAGGCCCAGCCCGTGGACGGTGTGAGGCCG</td><td>>hg38_21_+_8433172_8446622_RNA45SN1_45S_50ntFlanks</td><td>8176</td><td>8214</td><td>39</td><td>CTTCTGATCGAGGCCCAGCCCGTGGACGGTGTGAGGCCG</td><td>677.3</td><td>-6.8</td><td>1.9</td><td>-3.5</td><td>4.7E-04</td><td>5.8E-04</td><td>BT20/MB468</td><td>NUC</td></tr>
<tr><td>>MI0000298|hsa-mir-221&WithFlank&X|-|45746151|45746272@71.93.23&[MIMAT0000278&hsa-miR-221-3p&offsets|0|0;m-91&X|-|45746181|45746202&offsets|0|+1]|AGCTACATTGTCTGCTGGGTTTC</td><td>>MI0000298|hsa-mir-221&WithFlank&X|-|45746151|45746272</td><td>71</td><td>93</td><td>23</td><td>AGCTACATTGTCTGCTGGGTTTC</td><td>30508.0</td><td>-2.0</td><td>0.5</td><td>-4.0</td><td>7.0E-05</td><td>5.8E-04</td><td>BT20/MB231</td><td>NUC</td></tr>
<tr><td>>hg38_21_+_8433172_8446622_RNA45SN1_45S_50ntFlanks@9573.9598.26|AAATCGGTCGTCCGACCTGGGTATAG</td><td>>hg38_21_+_8433172_8446622_RNA45SN1_45S_50ntFlanks</td><td>9573</td><td>9598</td><td>26</td><td>AAATCGGTCGTCCGACCTGGGTATAG</td><td>98.6</td><td>-5.6</td><td>1.5</td><td>-3.7</td><td>1.9E-04</td><td>5.8E-04</td><td>BT20/MB468</td><td>MP</td></tr>
<tr><td>>MI0003820|hsa-mir-454&WithFlank&17|-|59137752|59137878@70.93.24(+1C)&[MIMAT0003885&hsa-miR-454-3p&offsets|0|+1(+1C);m-272&17|-|59137787|59137809&offsets|0|+1(+1C)]|TAGTGCAATATTGCTTATAGGGTTC</td><td>>MI0003820|hsa-mir-454&WithFlank&17|-|59137752|59137878</td><td>70</td><td>93</td><td>24(+1C)</td><td>TAGTGCAATATTGCTTATAGGGTTC</td><td>100.3</td><td>6.3</td><td>1.8</td><td>3.5</td><td>3.9E-04</td><td>5.8E-04</td><td>MB231/MB468</td><td>MP</td></tr>
<tr><td>>trnaMT_SerGCT_MT_+_12207_12265@1.19.19|GAGAAAGCTCACAAGAACT</td><td>>trnaMT_SerGCT_MT_+_12207_12265</td><td>1</td><td>19</td><td>19</td><td>GAGAAAGCTCACAAGAACT</td><td>369.0</td><td>5.1</td><td>1.4</td><td>3.5</td><td>4.4E-04</td><td>5.8E-04</td><td>MB231/MB468</td><td>CYTO</td></tr>
<tr><td>>MI0000063|hsa-let-7b&WithFlank&22|+|46113680|46113774@12.33.22(+2A)&[MIMAT0000063&hsa-let-7b-5p&offsets|0|0(+2A);m-14&22|+|46113691|46113712&offsets|0|0(+2A)]|TGAGGTAGTAGGTTGTGTGGTTAA</td><td>>MI0000063|hsa-let-7b&WithFlank&22|+|46113680|46113774</td><td>12</td><td>33</td><td>22(+2A)</td><td>TGAGGTAGTAGGTTGTGTGGTTAA</td><td>440.2</td><td>5.3</td><td>1.4</td><td>3.7</td><td>1.9E-04</td><td>5.8E-04</td><td>BT20/MB468</td><td>MP</td></tr>
<tr><td>>hg38_21_+_8433172_8446622_RNA45SN1_45S_50ntFlanks@12454.12493.40|CACCAAGCGTTGGATTGTTCACCCACTAATAGGGAACGTG</td><td>>hg38_21_+_8433172_8446622_RNA45SN1_45S_50ntFlanks</td><td>12454</td><td>12493</td><td>40</td><td>CACCAAGCGTTGGATTGTTCACCCACTAATAGGGAACGTG</td><td>83.7</td><td>7.0</td><td>1.9</td><td>3.7</td><td>1.9E-04</td><td>5.8E-04</td><td>BT20/MB468</td><td>MP</td></tr>
<tr><td>>hg38_21_+_8433172_8446622_RNA45SN1_45S_50ntFlanks@5048.5081.34|AGTTGGTGGAGCGATTTGTCTGGTTAATTCCGAT</td><td>>hg38_21_+_8433172_8446622_RNA45SN1_45S_50ntFlanks</td><td>5048</td><td>5081</td><td>34</td><td>AGTTGGTGGAGCGATTTGTCTGGTTAATTCCGAT</td><td>107.7</td><td>7.0</td><td>1.9</td><td>3.7</td><td>1.9E-04</td><td>5.8E-04</td><td>BT20/MB468</td><td>MP</td></tr>
<tr><td>>MI0000746|hsa-mir-99b&WithFlank&19|+|51692606|51692687@51.71.21(+1U)&[MIMAT0004678&hsa-miR-99b-3p&offsets|0|-1(+1U);m-151&19|+|51692656|51692678&offsets|0|-2(+1U)]|CAAGCTCGTGTCTGTGGGTCCT</td><td>>MI0000746|hsa-mir-99b&WithFlank&19|+|51692606|51692687</td><td>51</td><td>71</td><td>21(+1U)</td><td>CAAGCTCGTGTCTGTGGGTCCT</td><td>192.0</td><td>3.0</td><td>0.7</td><td>4.0</td><td>6.8E-05</td><td>5.8E-04</td><td>BT20/MB231</td><td>CYTO</td></tr>
<tr><td>>hg38_21_+_8433172_8446622_RNA45SN1_45S_50ntFlanks@10841.10870.30|CGTAACTTCGGGATAAGGATTGGCTCTAAG</td><td>>hg38_21_+_8433172_8446622_RNA45SN1_45S_50ntFlanks</td><td>10841</td><td>10870</td><td>30</td><td>CGTAACTTCGGGATAAGGATTGGCTCTAAG</td><td>607.8</td><td>-6.3</td><td>1.8</td><td>-3.5</td><td>4.7E-04</td><td>5.9E-04</td><td>BT20/MB468</td><td>NUC</td></tr>
<tr><td>>hg38_21_+_8433172_8446622_RNA45SN1_45S_50ntFlanks@12133.12150.18|CTCCGGGGACAGTGCCAG</td><td>>hg38_21_+_8433172_8446622_RNA45SN1_45S_50ntFlanks</td><td>12133</td><td>12150</td><td>18</td><td>CTCCGGGGACAGTGCCAG</td><td>811.5</td><td>-6.3</td><td>1.7</td><td>-3.7</td><td>1.9E-04</td><td>5.9E-04</td><td>BT20/MB468</td><td>MP</td></tr>
<tr><td>>hg38_21_+_8433172_8446622_RNA45SN1_45S_50ntFlanks@10349.10384.36|AACGAGAACTTTGAAGGCCGAAGTGGAGAAGGGTTC</td><td>>hg38_21_+_8433172_8446622_RNA45SN1_45S_50ntFlanks</td><td>10349</td><td>10384</td><td>36</td><td>AACGAGAACTTTGAAGGCCGAAGTGGAGAAGGGTTC</td><td>1417.1</td><td>-5.0</td><td>1.4</td><td>-3.5</td><td>3.9E-04</td><td>5.9E-04</td><td>MB231/MB468</td><td>MP</td></tr>
<tr><td>>hg38_21_+_8433172_8446622_RNA45SN1_45S_50ntFlanks@10766.10796.31|CGCAGCAGGTCTCCAAGGTGAACAGCCTCTG</td><td>>hg38_21_+_8433172_8446622_RNA45SN1_45S_50ntFlanks</td><td>10766</td><td>10796</td><td>31</td><td>CGCAGCAGGTCTCCAAGGTGAACAGCCTCTG</td><td>150.4</td><td>-6.2</td><td>1.7</td><td>-3.7</td><td>1.9E-04</td><td>5.9E-04</td><td>BT20/MB468</td><td>MP</td></tr>
<tr><td>>hg38_21_+_8433172_8446622_RNA45SN1_45S_50ntFlanks@3853.3883.31|AACTGTGGTAATTCTAGAGCTAATACATGCC</td><td>>hg38_21_+_8433172_8446622_RNA45SN1_45S_50ntFlanks</td><td>3853</td><td>3883</td><td>31</td><td>AACTGTGGTAATTCTAGAGCTAATACATGCC</td><td>163.8</td><td>-3.0</td><td>0.9</td><td>-3.5</td><td>3.9E-04</td><td>5.9E-04</td><td>MB231/MB468</td><td>MP</td></tr>
<tr><td>>MI0000440|hsa-mir-27b&WithFlank&9|+|95085439|95085547@67.85.19&[MIMAT0000419&hsa-miR-27b-3p&offsets|0|-2;m-39&9|+|95085505|95085525&offsets|0|-2]|TTCACAGTGGCTAAGTTCT</td><td>>MI0000440|hsa-mir-27b&WithFlank&9|+|95085439|95085547</td><td>67</td><td>85</td><td>19</td><td>TTCACAGTGGCTAAGTTCT</td><td>486.3</td><td>1.9</td><td>0.5</td><td>3.5</td><td>4.8E-04</td><td>5.9E-04</td><td>BT20/MB468</td><td>NUC</td></tr>
<tr><td>>trna10_AspGTC_12_-_125424193_125424264@1.29.29,trna12_AspGTC_12_-_125411891_125411962@1.29.29,trna38_AspGTC_17_-_8125556_8125627@1.29.29,trna45_AspGTC_6_+_27447453_27447524@1.29.29,trna48_AspGTC_6_+_27471523_27471594@1.29.29,trna4_AspGTC_12_+_96429799_96429870@1.29.29,trna69_AspGTC_1_-_161440205_161440276@1.29.29,trna72_AspGTC_1_-_161432824_161432895@1.29.29,trna75_AspGTC_1_-_161425414_161425485@1.29.29,trna78_AspGTC_1_-_161418033_161418104@1.29.29,trna81_AspGTC_1_-_161410615_161410686@1.29.29|TCCTCGTTAGTATAGTGGTGAGTATCCCC</td><td>>trna10_AspGTC_12_-_125424193_125424264</td><td>1</td><td>29</td><td>29</td><td>TCCTCGTTAGTATAGTGGTGAGTATCCCC</td><td>504.0</td><td>-5.3</td><td>1.5</td><td>-3.5</td><td>4.2E-04</td><td>5.9E-04</td><td>BT20/MB468</td><td>CYTO</td></tr>
<tr><td>>hg38_21_+_8433172_8446622_RNA45SN1_45S_50ntFlanks@4924.4944.21|AGGAGTGGAGCCTGCGGCTTA</td><td>>hg38_21_+_8433172_8446622_RNA45SN1_45S_50ntFlanks</td><td>4924</td><td>4944</td><td>21</td><td>AGGAGTGGAGCCTGCGGCTTA</td><td>1176.8</td><td>7.2</td><td>1.8</td><td>4.0</td><td>6.7E-05</td><td>5.9E-04</td><td>BT20/MB231</td><td>MP</td></tr>
<tr><td>>MI0000750|hsa-mir-26a-2&WithFlank&12|-|57824603|57824698@23.41.19;MI0000083|hsa-mir-26a-1&WithFlank&3|+|37969398|37969486@19.37.19&[MIMAT0000082&hsa-miR-26a-5p&offsets|+3|0;m-33&12|-|57824658|57824679&offsets|+3|0];[MIMAT0000082_1&hsa-miR-26a-5p&offsets|+3|0;m-34&3|+|37969413|37969434&offsets|+3|0]|AAGTAATCCAGGATAGGCT</td><td>>MI0000750|hsa-mir-26a-2&WithFlank&12|-|57824603|57824698</td><td>23</td><td>41</td><td>19</td><td>AAGTAATCCAGGATAGGCT</td><td>803.2</td><td>12.9</td><td>3.6</td><td>3.5</td><td>3.9E-04</td><td>5.9E-04</td><td>MB231/MB468</td><td>MP</td></tr>
<tr><td>>trnaMT_HisGTG_MT_+_12138_12206@44.71.28|CAGAGGCTTACGACCCCTTATTTACCCC</td><td>>trnaMT_HisGTG_MT_+_12138_12206</td><td>44</td><td>71</td><td>28</td><td>CAGAGGCTTACGACCCCTTATTTACCCC</td><td>20.1</td><td>7.5</td><td>2.1</td><td>3.5</td><td>3.9E-04</td><td>5.9E-04</td><td>MB231/MB468</td><td>MP</td></tr>
<tr><td>>hg38_21_+_8433172_8446622_RNA45SN1_45S_50ntFlanks@12605.12634.30|TATGTGCTTGGCTGAGGAGCCAATGGGGCG</td><td>>hg38_21_+_8433172_8446622_RNA45SN1_45S_50ntFlanks</td><td>12605</td><td>12634</td><td>30</td><td>TATGTGCTTGGCTGAGGAGCCAATGGGGCG</td><td>177.0</td><td>-3.8</td><td>1.1</td><td>-3.5</td><td>4.2E-04</td><td>5.9E-04</td><td>BT20/MB468</td><td>CYTO</td></tr>
<tr><td>>MI0000439|hsa-mir-23b&WithFlank&9|+|95085202|95085310@67.86.20&[MIMAT0000418&hsa-miR-23b-3p&offsets|+3|0;m-42&9|+|95085265|95085285&offsets|+3|+2]|ACATTGCCAGGGATTACCAC</td><td>>MI0000439|hsa-mir-23b&WithFlank&9|+|95085202|95085310</td><td>67</td><td>86</td><td>20</td><td>ACATTGCCAGGGATTACCAC</td><td>715.2</td><td>12.7</td><td>3.6</td><td>3.5</td><td>4.7E-04</td><td>5.9E-04</td><td>MB231/MB468</td><td>MITO</td></tr>
<tr><td>>MI0000065|hsa-let-7d&WithFlank&9|+|94178828|94178926@14.31.18&[MIMAT0000065&hsa-let-7d-5p&offsets|0|-4;m-90&9|+|94178841|94178862&offsets|0|-4]|AGAGGTAGTAGGTTGCAT</td><td>>MI0000065|hsa-let-7d&WithFlank&9|+|94178828|94178926</td><td>14</td><td>31</td><td>18</td><td>AGAGGTAGTAGGTTGCAT</td><td>398.0</td><td>6.0</td><td>1.7</td><td>3.5</td><td>3.9E-04</td><td>5.9E-04</td><td>MB231/MB468</td><td>MP</td></tr>
<tr><td>>hg38_21_+_8433172_8446622_RNA45SN1_45S_50ntFlanks@4759.4786.28|ATCAGATACCGTCGTAGTTCCGACCATA</td><td>>hg38_21_+_8433172_8446622_RNA45SN1_45S_50ntFlanks</td><td>4759</td><td>4786</td><td>28</td><td>ATCAGATACCGTCGTAGTTCCGACCATA</td><td>1344.6</td><td>-6.1</td><td>1.7</td><td>-3.5</td><td>3.9E-04</td><td>5.9E-04</td><td>MB231/MB468</td><td>MP</td></tr>
<tr><td>>hg38_1_-_228634819_228635039_RNA5S12_5S_50ntFlanks@140.169.30|ACCGCCTGGGAATACCGGGTGCTGTAGGCT</td><td>>hg38_1_-_228634819_228635039_RNA5S12_5S_50ntFlanks</td><td>140</td><td>169</td><td>30</td><td>ACCGCCTGGGAATACCGGGTGCTGTAGGCT</td><td>2459.7</td><td>3.2</td><td>0.9</td><td>3.5</td><td>4.7E-04</td><td>5.9E-04</td><td>MB231/MB468</td><td>NUC</td></tr>
<tr><td>>hg38_21_+_8433172_8446622_RNA45SN1_45S_50ntFlanks@4103.4121.19|CCACGGGTGACGGGGAATC</td><td>>hg38_21_+_8433172_8446622_RNA45SN1_45S_50ntFlanks</td><td>4103</td><td>4121</td><td>19</td><td>CCACGGGTGACGGGGAATC</td><td>615.0</td><td>-4.1</td><td>1.2</td><td>-3.5</td><td>4.2E-04</td><td>5.9E-04</td><td>BT20/MB468</td><td>CYTO</td></tr>
</table>

<table>
<tr><th></th><th></th><th></th><th></th><th></th><th></th><th></th><th></th><th></th><th></th><th></th><th></th><th></th><th></th></tr>
<tr><td>>trnaMT_HisGTG_MT_+_12138_12206@26.63.38|AGATTGTGAATCTGACAACAGAGGCTTACGACCCCTTA</td><td>>trnaMT_HisGTG_MT_+_12138_12206</td><td>26</td><td>63</td><td>38</td><td>AGATTGTGAATCTGACAACAGAGGCTTACGACCCCTTA</td><td>147.9</td><td>5.4</td><td>1.6</td><td>3.5</td><td>4.7E-04</td><td>5.9E-04</td><td>MB231/MB468</td><td>MITO</td></tr>
<tr><td>>hg38_21_+_8433172_8446622_RNA45SN1_45S_50ntFlanks@4223.4243.21|AAAAATAACAATACAGGACTC</td><td>>hg38_21_+_8433172_8446622_RNA45SN1_45S_50ntFlanks</td><td>4223</td><td>4243</td><td>21</td><td>AAAAATAACAATACAGGACTC</td><td>91.7</td><td>-5.6</td><td>1.5</td><td>-3.7</td><td>1.9E-04</td><td>5.9E-04</td><td>BT20/MB468</td><td>MP</td></tr>
<tr><td>>trnaMT_SerGCT_MT_+_12207_12265@24.62.39|ACTCATGCCCCCATGTCTAACAACATGGCTTTCTCACCA</td><td>>trnaMT_SerGCT_MT_+_12207_12265</td><td>24</td><td>62</td><td>39</td><td>ACTCATGCCCCCATGTCTAACAACATGGCTTTCTCACCA</td><td>159.2</td><td>3.4</td><td>1.0</td><td>3.5</td><td>4.7E-04</td><td>5.9E-04</td><td>MB231/MB468</td><td>MITO</td></tr>
<tr><td>>MI0000447|hsa-mir-128-1&WithFlank&2|+|135665391|135665484@56.75.20(+1C);MI0000727|hsa-mir-128-2&WithFlank&3|+|35744470|35744565@58.77.20(+1C)&[MIMAT0000424&hsa-miR-128-3p&offsets|0|-1(+1C);m-108&2|+|135665446|135665466&offsets|0|-1(+1C)];[MIMAT0000424_1&hsa-miR-128-3p&offsets|0|-1(+1C);m-117&3|+|35744527|35744547&offsets|0|-1(+1C)]|TCACAGTGAACCGGTCTCTTC</td><td>>MI0000447|hsa-mir-128-1&WithFlank&2|+|135665391|135665484</td><td>56</td><td>75</td><td>20(+1C)</td><td>TCACAGTGAACCGGTCTCTTC</td><td>654.0</td><td>2.8</td><td>0.8</td><td>3.5</td><td>4.8E-04</td><td>5.9E-04</td><td>MB231/MB468</td><td>NUC</td></tr>
<tr><td>>hg38_21_+_8433172_8446622_RNA45SN1_45S_50ntFlanks@11233.11255.23|CCCTTCCCGTGGATCGCCCCAGC</td><td>>hg38_21_+_8433172_8446622_RNA45SN1_45S_50ntFlanks</td><td>11233</td><td>11255</td><td>23</td><td>CCCTTCCCGTGGATCGCCCCAGC</td><td>143.4</td><td>-6.3</td><td>1.7</td><td>-3.7</td><td>1.9E-04</td><td>5.9E-04</td><td>BT20/MB468</td><td>MP</td></tr>
<tr><td>>hg38_21_+_8433172_8446622_RNA45SN1_45S_50ntFlanks@12479.12497.19|CTAATAGGGAACGTGAGCT</td><td>>hg38_21_+_8433172_8446622_RNA45SN1_45S_50ntFlanks</td><td>12479</td><td>12497</td><td>19</td><td>CTAATAGGGAACGTGAGCT</td><td>3539.1</td><td>2.1</td><td>0.6</td><td>3.5</td><td>4.4E-04</td><td>5.9E-04</td><td>MB231/MB468</td><td>CYTO</td></tr>
<tr><td>>hg38_21_+_8433172_8446622_RNA45SN1_45S_50ntFlanks@10351.10373.23|CGAGAACTTTGAAGGCCGAAGTG</td><td>>hg38_21_+_8433172_8446622_RNA45SN1_45S_50ntFlanks</td><td>10351</td><td>10373</td><td>23</td><td>CGAGAACTTTGAAGGCCGAAGTG</td><td>2189.3</td><td>-4.3</td><td>1.2</td><td>-3.7</td><td>1.9E-04</td><td>5.9E-04</td><td>BT20/MB468</td><td>MP</td></tr>
<tr><td>>hg38_21_+_8433172_8446622_RNA45SN1_45S_50ntFlanks@9534.9561.28|AGGAAACTCTGGTGGAGGTCCGTAGCGG</td><td>>hg38_21_+_8433172_8446622_RNA45SN1_45S_50ntFlanks</td><td>9534</td><td>9561</td><td>28</td><td>AGGAAACTCTGGTGGAGGTCCGTAGCGG</td><td>2467.2</td><td>-5.3</td><td>1.4</td><td>-3.7</td><td>1.9E-04</td><td>5.9E-04</td><td>BT20/MB468</td><td>MP</td></tr>
<tr><td>>hg38_21_+_8433172_8446622_RNA45SN1_45S_50ntFlanks@12416.12444.29|TTCGATGTCGGCTCTTCCTATCATTGTGA</td><td>>hg38_21_+_8433172_8446622_RNA45SN1_45S_50ntFlanks</td><td>12416</td><td>12444</td><td>29</td><td>TTCGATGTCGGCTCTTCCTATCATTGTGA</td><td>509.6</td><td>-2.5</td><td>0.7</td><td>-3.5</td><td>4.8E-04</td><td>5.9E-04</td><td>BT20/MB468</td><td>NUC</td></tr>
<tr><td>>hg38_21_+_8433172_8446622_RNA45SN1_45S_50ntFlanks@13008.13028.21|ATTGAAAGTCAGCCCTCGACA</td><td>>hg38_21_+_8433172_8446622_RNA45SN1_45S_50ntFlanks</td><td>13008</td><td>13028</td><td>21</td><td>ATTGAAAGTCAGCCCTCGACA</td><td>770.8</td><td>-4.8</td><td>1.3</td><td>-3.5</td><td>4.2E-04</td><td>5.9E-04</td><td>BT20/MB468</td><td>CYTO</td></tr>
<tr><td>>hg38_MT_+_1621_3279_RNR2_16S_50ntFlanks@486.517.32|AGGAACAGCTCTTTGGACACTAGGAAAAAACC</td><td>>hg38_MT_+_1621_3279_RNR2_16S_50ntFlanks</td><td>486</td><td>517</td><td>32</td><td>AGGAACAGCTCTTTGGACACTAGGAAAAAACC</td><td>236.1</td><td>4.1</td><td>1.1</td><td>3.7</td><td>1.9E-04</td><td>5.9E-04</td><td>BT20/MB468</td><td>MP</td></tr>
<tr><td>>hg38_21_+_8433172_8446622_RNA45SN1_45S_50ntFlanks@7984.8012.29|AGATCAGACGTGGCGACCCGCTGAATTTA</td><td>>hg38_21_+_8433172_8446622_RNA45SN1_45S_50ntFlanks</td><td>7984</td><td>8012</td><td>29</td><td>AGATCAGACGTGGCGACCCGCTGAATTTA</td><td>364.1</td><td>-5.9</td><td>1.6</td><td>-3.7</td><td>1.9E-04</td><td>5.9E-04</td><td>BT20/MB468</td><td>MP</td></tr>
<tr><td>>hg38_21_+_8433172_8446622_RNA45SN1_45S_50ntFlanks@11572.11605.34|CCGACTTAGAACTGGTGCGGACCAGGGGAATCCG</td><td>>hg38_21_+_8433172_8446622_RNA45SN1_45S_50ntFlanks</td><td>11572</td><td>11605</td><td>34</td><td>CCGACTTAGAACTGGTGCGGACCAGGGGAATCCG</td><td>2461.8</td><td>-3.9</td><td>1.0</td><td>-3.7</td><td>1.9E-04</td><td>5.9E-04</td><td>BT20/MB468</td><td>MP</td></tr>
<tr><td>>MI0000063|hsa-let-7b&WithFlank&22|+|46113680|46113774@12.33.22(+1A)&[MIMAT0000063&hsa-let-7b-5p&offsets|0|0(+1A);m-14&22|+|46113691|46113712&offsets|0|0(+1A)]|TGAGGTAGTAGGTTGTGTGGTTA</td><td>>MI0000063|hsa-let-7b&WithFlank&22|+|46113680|46113774</td><td>12</td><td>33</td><td>22(+1A)</td><td>TGAGGTAGTAGGTTGTGTGGTTA</td><td>3449.9</td><td>3.2</td><td>0.8</td><td>4.0</td><td>6.9E-05</td><td>5.9E-04</td><td>BT20/MB231</td><td>CYTO</td></tr>
<tr><td>>hg38_21_+_8433172_8446622_RNA45SN1_45S_50ntFlanks@11604.11626.23|CGACTGTTTAATTAAAACAAAGC</td><td>>hg38_21_+_8433172_8446622_RNA45SN1_45S_50ntFlanks</td><td>11604</td><td>11626</td><td>23</td><td>CGACTGTTTAATTAAAACAAAGC</td><td>384.1</td><td>3.4</td><td>0.9</td><td>3.9</td><td>8.1E-05</td><td>5.9E-04</td><td>BT20/MB231</td><td>MITO</td></tr>
<tr><td>>hg38_21_+_8433172_8446622_RNA45SN1_45S_50ntFlanks@12599.12626.28|TTGGTGTATGTGCTTGGCTGAGGAGCCA</td><td>>hg38_21_+_8433172_8446622_RNA45SN1_45S_50ntFlanks</td><td>12599</td><td>12626</td><td>28</td><td>TTGGTGTATGTGCTTGGCTGAGGAGCCA</td><td>138.2</td><td>-3.8</td><td>1.1</td><td>-3.5</td><td>4.2E-04</td><td>5.9E-04</td><td>BT20/MB468</td><td>CYTO</td></tr>
<tr><td>>trnaMT_LeuTAA_MT_+_3230_3304@17.44.28|CCGGTAATCGCATAAAACTTAAAACTTT</td><td>>trnaMT_LeuTAA_MT_+_3230_3304</td><td>17</td><td>44</td><td>28</td><td>CCGGTAATCGCATAAAACTTAAAACTTT</td><td>127.6</td><td>-6.0</td><td>1.7</td><td>-3.5</td><td>4.0E-04</td><td>5.9E-04</td><td>MB231/MB468</td><td>MP</td></tr>
<tr><td>>hg38_21_+_8433172_8446622_RNA45SN1_45S_50ntFlanks@12995.13036.42|CTCGCTGCGATCTATTGAAAGTCAGCCCTCGACACAAGGGTT</td><td>>hg38_21_+_8433172_8446622_RNA45SN1_45S_50ntFlanks</td><td>12995</td><td>13036</td><td>42</td><td>CTCGCTGCGATCTATTGAAAGTCAGCCCTCGACACAAGGGTT</td><td>829.3</td><td>4.4</td><td>1.2</td><td>3.5</td><td>4.7E-04</td><td>5.9E-04</td><td>MB231/MB468</td><td>MITO</td></tr>
<tr><td>>hg38_21_+_8433172_8446622_RNA45SN1_45S_50ntFlanks@11933.11950.18|AATAAGTGGGAGGCCCCC</td><td>>hg38_21_+_8433172_8446622_RNA45SN1_45S_50ntFlanks</td><td>11933</td><td>11950</td><td>18</td><td>AATAAGTGGGAGGCCCCC</td><td>678.2</td><td>-6.3</td><td>1.7</td><td>-3.7</td><td>1.9E-04</td><td>5.9E-04</td><td>BT20/MB468</td><td>MP</td></tr>
<tr><td>>hg38_21_+_8433172_8446622_RNA45SN1_45S_50ntFlanks@9575.9601.27|ATCGGTCGTCCGACCTGGGTATAGGGG</td><td>>hg38_21_+_8433172_8446622_RNA45SN1_45S_50ntFlanks</td><td>9575</td><td>9601</td><td>27</td><td>ATCGGTCGTCCGACCTGGGTATAGGGG</td><td>419.9</td><td>-5.4</td><td>1.6</td><td>-3.5</td><td>4.7E-04</td><td>5.9E-04</td><td>MB231/MB468</td><td>MITO</td></tr>
<tr><td>>hg38_21_+_8433172_8446622_RNA45SN1_45S_50ntFlanks@6655.6695.41|TCTTAGCGGTGGATCACTCGGCTCGTGCGTCGATGAAGAAC</td><td>>hg38_21_+_8433172_8446622_RNA45SN1_45S_50ntFlanks</td><td>6655</td><td>6695</td><td>41</td><td>TCTTAGCGGTGGATCACTCGGCTCGTGCGTCGATGAAGAAC</td><td>81.5</td><td>6.8</td><td>1.8</td><td>3.7</td><td>1.9E-04</td><td>6.0E-04</td><td>BT20/MB468</td><td>MP</td></tr>
<tr><td>>hg38_21_+_8433172_8446622_RNA45SN1_45S_50ntFlanks@12418.12458.41|CGATGTCGGCTCTTCCTATCATTGTGAAGCAGAATTCACCA</td><td>>hg38_21_+_8433172_8446622_RNA45SN1_45S_50ntFlanks</td><td>12418</td><td>12458</td><td>41</td><td>CGATGTCGGCTCTTCCTATCATTGTGAAGCAGAATTCACCA</td><td>429.4</td><td>-3.2</td><td>0.8</td><td>-4.0</td><td>6.9E-05</td><td>6.0E-04</td><td>BT20/MB231</td><td>CYTO</td></tr>
<tr><td>>MI0005561|hsa-mir-877&WithFlank&6|+|30584326|30584423@7.29.23&[MIMAT0004949&hsa-miR-877-5p&offsets|0|+3;m-382&6|+|30584332|30584354&offsets|0|0]|GTAGAGGAGATGGCGCAGGGGAC</td><td>>MI0005561|hsa-mir-877&WithFlank&6|+|30584326|30584423</td><td>7</td><td>29</td><td>23</td><td>GTAGAGGAGATGGCGCAGGGGAC</td><td>262.7</td><td>5.0</td><td>1.3</td><td>3.7</td><td>1.9E-04</td><td>6.0E-04</td><td>BT20/MB468</td><td>MP</td></tr>
<tr><td>>hg38_21_+_8433172_8446622_RNA45SN1_45S_50ntFlanks@11209.11230.22|CGGCGACTCTGGACGCGAGCCG</td><td>>hg38_21_+_8433172_8446622_RNA45SN1_45S_50ntFlanks</td><td>11209</td><td>11230</td><td>22</td><td>CGGCGACTCTGGACGCGAGCCG</td><td>1072.7</td><td>-5.0</td><td>1.3</td><td>-3.7</td><td>1.9E-04</td><td>6.0E-04</td><td>BT20/MB468</td><td>MP</td></tr>
<tr><td>>hg38_21_+_8433172_8446622_RNA45SN1_45S_50ntFlanks@11719.11742.24|ACGGCGGGAGTAACTATGACTCTC</td><td>>hg38_21_+_8433172_8446622_RNA45SN1_45S_50ntFlanks</td><td>11719</td><td>11742</td><td>24</td><td>ACGGCGGGAGTAACTATGACTCTC</td><td>143.3</td><td>-5.4</td><td>1.5</td><td>-3.5</td><td>4.0E-04</td><td>6.0E-04</td><td>MB231/MB468</td><td>MP</td></tr>
<tr><td>>MI0000273|hsa-mir-183&WithFlank&7|-|129774899|129775020@34.54.21&[MIMAT0000261&hsa-miR-183-5p&offsets|+1|0;m-41&7|-|129774966|129774987&offsets|0|-1]|ATGGCACTGGTAGAATTCACT</td><td>>MI0000273|hsa-mir-183&WithFlank&7|-|129774899|129775020</td><td>34</td><td>54</td><td>21</td><td>ATGGCACTGGTAGAATTCACT</td><td>3815.2</td><td>2.7</td><td>0.7</td><td>3.9</td><td>8.1E-05</td><td>6.0E-04</td><td>BT20/MB231</td><td>MITO</td></tr>
<tr><td>>hg38_21_+_8433172_8446622_RNA45SN1_45S_50ntFlanks@6661.6709.49|CGGTGGATCACTCGGCTCGTGCGTCGATGAAGAACGCAGCTAGCTGCGA</td><td>>hg38_21_+_8433172_8446622_RNA45SN1_45S_50ntFlanks</td><td>6661</td><td>6709</td><td>49</td><td>CGGTGGATCACTCGGCTCGTGCGTCGATGAAGAACGCAGCTAGCTGCGA</td><td>67.6</td><td>6.7</td><td>1.8</td><td>3.7</td><td>2.0E-04</td><td>6.0E-04</td><td>BT20/MB468</td><td>MP</td></tr>
<tr><td>>hg38_21_+_8433172_8446622_RNA45SN1_45S_50ntFlanks@10347.10383.37|CAAACGAGAACTTTGAAGGCCGAAGTGGAGAAGGGTT</td><td>>hg38_21_+_8433172_8446622_RNA45SN1_45S_50ntFlanks</td><td>10347</td><td>10383</td><td>37</td><td>CAAACGAGAACTTTGAAGGCCGAAGTGGAGAAGGGTT</td><td>431.4</td><td>2.2</td><td>0.6</td><td>3.5</td><td>4.5E-04</td><td>6.0E-04</td><td>MB231/MB468</td><td>CYTO</td></tr>
<tr><td>>MI0003205|hsa-mir-532&WithFlank&X|+|50003142|50003244@26.46.21(+1C)&[MIMAT0002888&hsa-miR-532-5p&offsets|0|-1(+1C);m-65&X|+|50003167|50003188&offsets|0|-1(+1C)]|CATGCCTTGAGTGTAGGACCGC</td><td>>MI0003205|hsa-mir-532&WithFlank&X|+|50003142|50003244</td><td>26</td><td>46</td><td>21(+1C)</td><td>CATGCCTTGAGTGTAGGACCGC</td><td>216.1</td><td>3.5</td><td>1.0</td><td>3.5</td><td>4.7E-04</td><td>6.0E-04</td><td>MB231/MB468</td><td>MITO</td></tr>
<tr><td>>hg38_21_+_8433172_8446622_RNA45SN1_45S_50ntFlanks@10124.10149.26|AGGGCGGCGGCGGCGGCGGCGGCGGG</td><td>>hg38_21_+_8433172_8446622_RNA45SN1_45S_50ntFlanks</td><td>10124</td><td>10149</td><td>26</td><td>AGGGCGGCGGCGGCGGCGGCGGCGGG</td><td>24.5</td><td>7.8</td><td>2.2</td><td>3.5</td><td>4.0E-04</td><td>6.0E-04</td><td>MB231/MB468</td><td>MP</td></tr>
<tr><td>>hg38_MT_+_1621_3279_RNR2_16S_50ntFlanks@695.727.33|ATGAAAACATTCTCCTCCGCATAAGCCTGCGTC</td><td>>hg38_MT_+_1621_3279_RNR2_16S_50ntFlanks</td><td>695</td><td>727</td><td>33</td><td>ATGAAAACATTCTCCTCCGCATAAGCCTGCGTC</td><td>77.1</td><td>10.0</td><td>2.7</td><td>3.7</td><td>2.0E-04</td><td>6.0E-04</td><td>BT20/MB468</td><td>MP</td></tr>
<tr><td>>hg38_21_+_8433172_8446622_RNA45SN1_45S_50ntFlanks@5485.5518.34|AGCGCTGAGAAGACGGTCGAACTTGACTATCTAG</td><td>>hg38_21_+_8433172_8446622_RNA45SN1_45S_50ntFlanks</td><td>5485</td><td>5518</td><td>34</td><td>AGCGCTGAGAAGACGGTCGAACTTGACTATCTAG</td><td>1253.9</td><td>-6.8</td><td>1.9</td><td>-3.5</td><td>4.8E-04</td><td>6.0E-04</td><td>BT20/MB468</td><td>NUC</td></tr>
<tr><td>>hg38_21_+_8433172_8446622_RNA45SN1_45S_50ntFlanks@6645.6682.38|CTCGTACGACTCTTAGCGGTGGATCACTCGGCTCGTGC</td><td>>hg38_21_+_8433172_8446622_RNA45SN1_45S_50ntFlanks</td><td>6645</td><td>6682</td><td>38</td><td>CTCGTACGACTCTTAGCGGTGGATCACTCGGCTCGTGC</td><td>203.9</td><td>4.2</td><td>1.1</td><td>3.7</td><td>2.0E-04</td><td>6.0E-04</td><td>BT20/MB468</td><td>MP</td></tr>
<tr><td>>hg38_21_+_8433172_8446622_RNA45SN1_45S_50ntFlanks@6646.6675.30|TCGTACGACTCTTAGCGGTGGATCACTCGG</td><td>>hg38_21_+_8433172_8446622_RNA45SN1_45S_50ntFlanks</td><td>6646</td><td>6675</td><td>30</td><td>TCGTACGACTCTTAGCGGTGGATCACTCGG</td><td>3071.6</td><td>-2.0</td><td>0.6</td><td>-3.5</td><td>4.9E-04</td><td>6.0E-04</td><td>BT20/MB468</td><td>NUC</td></tr>
<tr><td>>MI0000067|hsa-let-7f-1&WithFlank&9|+|94176341|94176439@13.33.21(+1A);MI0000068|hsa-let-7f-2&WithFlank&X|-|53557186|53557280@14.34.21(+1A)&[MIMAT0000067&hsa-let-7f-5p&offsets|0|-1(+1A);m-20&9|+|94176353|94176374&offsets|0|-1(+1A)];[MIMAT0000067_1&hsa-let-7f-5p&offsets|0|-1(+1A);m-19&X|-|53557246|53557267&offsets|0|-1(+1A)]|TGAGGTAGTAGATTGTATAGTA</td><td>>MI0000067|hsa-let-7f-1&WithFlank&9|+|94176341|94176439</td><td>13</td><td>33</td><td>21(+1A)</td><td>TGAGGTAGTAGATTGTATAGTA</td><td>1847.4</td><td>3.8</td><td>1.1</td><td>3.5</td><td>4.0E-04</td><td>6.0E-04</td><td>MB231/MB468</td><td>MP</td></tr>
<tr><td>>trnaMT_HisGTG_MT_+_12138_12206@32.61.30|TGAATCTGACAACAGAGGCTTACGACCCCT</td><td>>trnaMT_HisGTG_MT_+_12138_12206</td><td>32</td><td>61</td><td>30</td><td>TGAATCTGACAACAGAGGCTTACGACCCCT</td><td>1269.0</td><td>5.6</td><td>1.6</td><td>3.6</td><td>3.6E-04</td><td>6.0E-04</td><td>BT20/MB468</td><td>MITO</td></tr>
<tr><td>>hg38_21_+_8433172_8446622_RNA45SN1_45S_50ntFlanks@6663.6689.27|GTGGATCACTCGGCTCGTGCGTCGATG</td><td>>hg38_21_+_8433172_8446622_RNA45SN1_45S_50ntFlanks</td><td>6663</td><td>6689</td><td>27</td><td>GTGGATCACTCGGCTCGTGCGTCGATG</td><td>701.9</td><td>-2.7</td><td>0.8</td><td>-3.5</td><td>4.3E-04</td><td>6.0E-04</td><td>BT20/MB468</td><td>CYTO</td></tr>
<tr><td>>hg38_21_+_8433172_8446622_RNA45SN1_45S_50ntFlanks@11679.11702.24|CTGAATGTCAAAGTGAAGAAATTC</td><td>>hg38_21_+_8433172_8446622_RNA45SN1_45S_50ntFlanks</td><td>11679</td><td>11702</td><td>24</td><td>CTGAATGTCAAAGTGAAGAAATTC</td><td>185.2</td><td>-5.7</td><td>1.5</td><td>-3.7</td><td>2.0E-04</td><td>6.0E-04</td><td>BT20/MB468</td><td>MP</td></tr>
<tr><td>>hg38_MT_+_598_1651_RNR1_12S_50ntFlanks@915.950.36|AAAGGACATTTAACTAAAACCCCTACGCATTTATAT</td><td>>hg38_MT_+_598_1651_RNR1_12S_50ntFlanks</td><td>915</td><td>950</td><td>36</td><td>AAAGGACATTTAACTAAAACCCCTACGCATTTATAT</td><td>483.6</td><td>6.8</td><td>1.8</td><td>3.7</td><td>2.0E-04</td><td>6.0E-04</td><td>BT20/MB468</td><td>MP</td></tr>
<tr><td>>hg38_21_+_8433172_8446622_RNA45SN1_45S_50ntFlanks@8168.8195.28|CCCAAGTCCTTCTGATCGAGGCCCAGCC</td><td>>hg38_21_+_8433172_8446622_RNA45SN1_45S_50ntFlanks</td><td>8168</td><td>8195</td><td>28</td><td>CCCAAGTCCTTCTGATCGAGGCCCAGCC</td><td>173.6</td><td>-5.6</td><td>1.5</td><td>-3.7</td><td>2.0E-04</td><td>6.0E-04</td><td>BT20/MB468</td><td>MP</td></tr>
<tr><td>>hg38_21_+_8433172_8446622_RNA45SN1_45S_50ntFlanks@12417.12455.39|TCGATGTCGGCTCTTCCTATCATTGTGAAGCAGAATTCA</td><td>>hg38_21_+_8433172_8446622_RNA45SN1_45S_50ntFlanks</td><td>12417</td><td>12455</td><td>39</td><td>TCGATGTCGGCTCTTCCTATCATTGTGAAGCAGAATTCA</td><td>208.5</td><td>-3.3</td><td>0.8</td><td>-4.0</td><td>7.0E-05</td><td>6.0E-04</td><td>BT20/MB231</td><td>CYTO</td></tr>
<tr><td>>hg38_21_+_8433172_8446622_RNA45SN1_45S_50ntFlanks@3853.3874.22|AACTGTGGTAATTCTAGAGCTA</td><td>>hg38_21_+_8433172_8446622_RNA45SN1_45S_50ntFlanks</td><td>3853</td><td>3874</td><td>22</td><td>AACTGTGGTAATTCTAGAGCTA</td><td>331.6</td><td>-5.0</td><td>1.4</td><td>-3.5</td><td>4.8E-04</td><td>6.0E-04</td><td>MB231/MB468</td><td>MITO</td></tr>
<tr><td>>hg38_21_+_8433172_8446622_RNA45SN1_45S_50ntFlanks@10482.10499.18|TCGAAAGGGAGTCGGGTT</td><td>>hg38_21_+_8433172_8446622_RNA45SN1_45S_50ntFlanks</td><td>10482</td><td>10499</td><td>18</td><td>TCGAAAGGGAGTCGGGTT</td><td>174.5</td><td>-2.3</td><td>0.6</td><td>-3.5</td><td>4.3E-04</td><td>6.0E-04</td><td>BT20/MB468</td><td>CYTO</td></tr>
<tr><td>>hg38_21_+_8433172_8446622_RNA45SN1_45S_50ntFlanks@4722.4758.37|TCATTAATCAAGAACGAAAGTCGGAGGTTCGAAGACG</td><td>>hg38_21_+_8433172_8446622_RNA45SN1_45S_50ntFlanks</td><td>4722</td><td>4758</td><td>37</td><td>TCATTAATCAAGAACGAAAGTCGGAGGTTCGAAGACG</td><td>5921.5</td><td>-4.8</td><td>1.3</td><td>-3.7</td><td>2.0E-04</td><td>6.0E-04</td><td>BT20/MB468</td><td>MP</td></tr>
<tr><td>>hg38_21_+_8433172_8446622_RNA45SN1_45S_50ntFlanks@8176.8206.31|CTTCTGATCGAGGCCCAGCCCGTGGACGGTG</td><td>>hg38_21_+_8433172_8446622_RNA45SN1_45S_50ntFlanks</td><td>8176</td><td>8206</td><td>31</td><td>CTTCTGATCGAGGCCCAGCCCGTGGACGGTG</td><td>399.8</td><td>-1.4</td><td>0.4</td><td>-3.6</td><td>3.6E-04</td><td>6.0E-04</td><td>BT20/MB468</td><td>MITO</td></tr>
<tr><td>>hg38_21_+_8433172_8446622_RNA45SN1_45S_50ntFlanks@12015.12035.21|TGAAATACCACTACTCTGATC</td><td>>hg38_21_+_8433172_8446622_RNA45SN1_45S_50ntFlanks</td><td>12015</td><td>12035</td><td>21</td><td>TGAAATACCACTACTCTGATC</td><td>57.7</td><td>9.2</td><td>2.3</td><td>4.0</td><td>6.9E-05</td><td>6.0E-04</td><td>BT20/MB231</td><td>MP</td></tr>
<tr><td>>hg38_21_+_8433172_8446622_RNA45SN1_45S_50ntFlanks@12420.12458.39|ATGTCGGCTCTTCCTATCATTGTGAAGCAGAATTCACCA</td><td>>hg38_21_+_8433172_8446622_RNA45SN1_45S_50ntFlanks</td><td>12420</td><td>12458</td><td>39</td><td>ATGTCGGCTCTTCCTATCATTGTGAAGCAGAATTCACCA</td><td>1202.9</td><td>-3.2</td><td>0.8</td><td>-4.0</td><td>7.0E-05</td><td>6.0E-04</td><td>BT20/MB231</td><td>CYTO</td></tr>
<tr><td>>hg38_21_+_8433172_8446622_RNA45SN1_45S_50ntFlanks@8176.8212.37|CTTCTGATCGAGGCCCAGCCCGTGGACGGTGTGAGGC</td><td>>hg38_21_+_8433172_8446622_RNA45SN1_45S_50ntFlanks</td><td>8176</td><td>8212</td><td>37</td><td>CTTCTGATCGAGGCCCAGCCCGTGGACGGTGTGAGGC</td><td>660.5</td><td>-3.2</td><td>0.9</td><td>-3.5</td><td>4.8E-04</td><td>6.0E-04</td><td>MB231/MB468</td><td>MITO</td></tr>
<tr><td>>hg38_21_+_8433172_8446622_RNA45SN1_45S_50ntFlanks@9554.9586.33|CGTAGCGGTCCTGACGTGCAAATCGGTCGTCCG</td><td>>hg38_21_+_8433172_8446622_RNA45SN1_45S_50ntFlanks</td><td>9554</td><td>9586</td><td>33</td><td>CGTAGCGGTCCTGACGTGCAAATCGGTCGTCCG</td><td>61.0</td><td>-5.4</td><td>1.5</td><td>-3.5</td><td>4.0E-04</td><td>6.0E-04</td><td>MB231/MB468</td><td>MP</td></tr>
</table>

| | | | | | | | | | | | | | |
|---|---|---|---|---|---|---|---|---|---|---|---|---|---|
| >hg38_21_+_8433172_8446622_RNA45SN1_45S_50ntFlanks@9795.9819.25\|GTGGAGCCGGGCGTGGAATGCGAGT | >hg38_21_+_8433172_8446622_RNA45SN1_45S_50ntFlanks | 9795 | 9819 | 25 | GTGGAGCCGGGCGTGGAATGCGAGT | 92.7 | 6.9 | 1.9 | 3.7 | 2.0E-04 | 6.0E-04 | BT20/MB468 | MP |
| >hg38_21_+_8433172_8446622_RNA45SN1_45S_50ntFlanks@11209.11251.43\|CGGCGACTCTGGACGCGAGCCGGGCCCTTCCCGTGGATCGCCC | >hg38_21_+_8433172_8446622_RNA45SN1_45S_50ntFlanks | 11209 | 11251 | 43 | CGGCGACTCTGGACGCGAGCCGGGCCCTTCCCGTGGATCGCCC | 200.7 | -4.0 | 1.1 | -3.5 | 4.5E-04 | 6.0E-04 | MB231/MB468 | CYTO |
| >hg38_21_+_8433172_8446622_RNA45SN1_45S_50ntFlanks@9478.9497.20\|GCGCACGTGTTAGGACCCGA | >hg38_21_+_8433172_8446622_RNA45SN1_45S_50ntFlanks | 9478 | 9497 | 20 | GCGCACGTGTTAGGACCCGA | 1044.1 | -3.4 | 0.9 | -4.0 | 7.1E-05 | 6.0E-04 | BT20/MB231 | CYTO |
| >hg38_21_+_8433172_8446622_RNA45SN1_45S_50ntFlanks@9621.9648.28\|TCTAGTAGCTGGTTCCCTCCGAAGTTTC | >hg38_21_+_8433172_8446622_RNA45SN1_45S_50ntFlanks | 9621 | 9648 | 28 | TCTAGTAGCTGGTTCCCTCCGAAGTTTC | 166.9 | -5.0 | 1.4 | -3.5 | 4.9E-04 | 6.0E-04 | MB231/MB468 | NUC |
| >hg38_21_+_8433172_8446622_RNA45SN1_45S_50ntFlanks@10684.10709.26\|CGGCGGCGTCCGGTGAGCTCTCGCTG | >hg38_21_+_8433172_8446622_RNA45SN1_45S_50ntFlanks | 10684 | 10709 | 26 | CGGCGGCGTCCGGTGAGCTCTCGCTG | 1451.9 | -4.2 | 1.2 | -3.5 | 4.3E-04 | 6.0E-04 | BT20/MB468 | CYTO |
| >hg38_21_+_8433172_8446622_RNA45SN1_45S_50ntFlanks@9802.9819.18\|CGGGCGTGGAATGCGAGT | >hg38_21_+_8433172_8446622_RNA45SN1_45S_50ntFlanks | 9802 | 9819 | 18 | CGGGCGTGGAATGCGAGT | 83.9 | 5.8 | 1.5 | 3.7 | 2.0E-04 | 6.0E-04 | BT20/MB468 | MP |
| >MI0000086\|hsa-mir-28&WithFlank&3\|+\|188688775\|188688872@61.82.22(+1A)&[MIMAT0004502&hsa-miR-28-3p&offsets\|+1\|+1(+1A);m-28&3\|+\|188688834\|188688855&offsets\|+1\|+1(+1A)]\|ACTAGATTGTGAGCTCCTGGAGA | >MI0000086\|hsa-mir-28&WithFlank&3\|+\|188688775\|188688872 | 61 | 82 | 22(+1A) | ACTAGATTGTGAGCTCCTGGAGA | 294.4 | 3.2 | 0.8 | 4.0 | 7.1E-05 | 6.0E-04 | BT20/MB231 | CYTO |
| >MI0000542\|hsa-mir-320a&WithFlank&8\|-\|22244960\|22245043@48.68.21&[MIMAT0000510&hsa-miR-320a-3p&offsets\|0\|-1;m-60&8\|-\|22244975\|22244996&offsets\|0\|-1]\|AAAAGCTGGGTTGAGAGGGCG | >MI0000542\|hsa-mir-320a&WithFlank&8\|-\|22244960\|22245043 | 48 | 68 | 21 | AAAAGCTGGGTTGAGAGGGCG | 3859.3 | 2.3 | 0.6 | 4.0 | 7.3E-05 | 6.0E-04 | BT20/MB231 | NUC |
| >hg38_21_+_8433172_8446622_RNA45SN1_45S_50ntFlanks@11663.11706.44\|ATTTCTGCCCAGTGCTCTGAATGTCAAAGTGAAGAAATTCAATG | >hg38_21_+_8433172_8446622_RNA45SN1_45S_50ntFlanks | 11663 | 11706 | 44 | ATTTCTGCCCAGTGCTCTGAATGTCAAAGTGAAGAAATTCAATG | 195.4 | -3.7 | 0.9 | -4.0 | 7.1E-05 | 6.0E-04 | BT20/MB231 | CYTO |
| >hg38_21_+_8433172_8446622_RNA45SN1_45S_50ntFlanks@6649.6666.18\|TACGACTCTTAGCGGTGG | >hg38_21_+_8433172_8446622_RNA45SN1_45S_50ntFlanks | 6649 | 6666 | 18 | TACGACTCTTAGCGGTGG | 140.3 | -3.0 | 0.8 | -3.5 | 4.0E-04 | 6.0E-04 | MB231/MB468 | MP |
| >hg38_21_+_8433172_8446622_RNA45SN1_45S_50ntFlanks@12324.12344.21\|TTGGGTTTTAAGCAGGAGGTG | >hg38_21_+_8433172_8446622_RNA45SN1_45S_50ntFlanks | 12324 | 12344 | 21 | TTGGGTTTTAAGCAGGAGGTG | 1765.1 | -6.4 | 1.7 | -3.7 | 2.0E-04 | 6.0E-04 | BT20/MB468 | MP |
| >MI0000063\|hsa-let-7b&WithFlank&22\|+\|46113680\|46113774@12.34.23&[MIMAT0000063&hsa-let-7b-5p&offsets\|0\|+1;m-14&22\|+\|46113691\|46113712&offsets\|0\|+1]\|TGAGGTAGTAGGTTGTGTGGTTT | >MI0000063\|hsa-let-7b&WithFlank&22\|+\|46113680\|46113774 | 12 | 34 | 23 | TGAGGTAGTAGGTTGTGTGGTTT | 7052.0 | 2.1 | 0.5 | 4.0 | 7.3E-05 | 6.1E-04 | BT20/MB231 | NUC |
| >hg38_21_+_8433172_8446622_RNA45SN1_45S_50ntFlanks@6656.6680.25\|CTTAGCGGTGGATCACTCGGCTCGT | >hg38_21_+_8433172_8446622_RNA45SN1_45S_50ntFlanks | 6656 | 6680 | 25 | CTTAGCGGTGGATCACTCGGCTCGT | 297.6 | -5.3 | 1.5 | -3.5 | 4.8E-04 | 6.1E-04 | MB231/MB468 | MITO |
| >hg38_21_+_8433172_8446622_RNA45SN1_45S_50ntFlanks@11586.11605.20\|GTGCGGACCAGGGGAATCCG | >hg38_21_+_8433172_8446622_RNA45SN1_45S_50ntFlanks | 11586 | 11605 | 20 | GTGCGGACCAGGGGAATCCG | 607.2 | -6.3 | 1.7 | -3.7 | 2.0E-04 | 6.1E-04 | BT20/MB468 | MP |
| >hg38_21_+_8433172_8446622_RNA45SN1_45S_50ntFlanks@8198.8223.26\|TGGACGGTGTGAGGCCGGTAGCGGCC | >hg38_21_+_8433172_8446622_RNA45SN1_45S_50ntFlanks | 8198 | 8223 | 26 | TGGACGGTGTGAGGCCGGTAGCGGCC | 587.2 | -4.0 | 1.1 | -3.5 | 4.3E-04 | 6.1E-04 | BT20/MB468 | CYTO |
| >hg38_21_+_8433172_8446622_RNA45SN1_45S_50ntFlanks@12407.12435.29\|TTTTGATCCTTCGATGTCGGCTCTTCCTA | >hg38_21_+_8433172_8446622_RNA45SN1_45S_50ntFlanks | 12407 | 12435 | 29 | TTTTGATCCTTCGATGTCGGCTCTTCCTA | 605.2 | -3.8 | 1.1 | -3.5 | 4.6E-04 | 6.1E-04 | MB231/MB468 | CYTO |
| >hg38_21_+_8433172_8446622_RNA45SN1_45S_50ntFlanks@4976.5001.26\|CCGGACACGGACAGGATTGACAGATT | >hg38_21_+_8433172_8446622_RNA45SN1_45S_50ntFlanks | 4976 | 5001 | 26 | CCGGACACGGACAGGATTGACAGATT | 350.6 | 2.6 | 0.7 | 4.0 | 7.3E-05 | 6.1E-04 | BT20/MB231 | NUC |
| >trnaMT_GluTTC_MT_-_14674_14742@46.70.25,trnalookalike8_GluTTC_5_-_93905172_93905240@46.70.25\|GTGGTTGTAGTCCGTGCGAGAATAC | >trnaMT_GluTTC_MT_-_14674_14742 | 46 | 70 | 25 | GTGGTTGTAGTCCGTGCGAGAATAC | 252.9 | 7.2 | 1.9 | 3.7 | 2.0E-04 | 6.1E-04 | BT20/MB468 | MP |
| >hg38_1_-_228634819_228635039_RNA5S12_5S_50ntFlanks@127.169.43\|ACTTGGATGGGAGACCGCCTGGGAATACCGGGTGCTGTAGGCT | >hg38_1_-_228634819_228635039_RNA5S12_5S_50ntFlanks | 127 | 169 | 43 | ACTTGGATGGGAGACCGCCTGGGAATACCGGGTGCTGTAGGCT | 349.7 | 3.5 | 0.9 | 3.9 | 8.3E-05 | 6.1E-04 | BT20/MB231 | MITO |
| >hg38_21_+_8433172_8446622_RNA45SN1_45S_50ntFlanks@475.517.43\|CTTCGTGATCGATGTGGTGACGTCGTGCTCTCCCGGGCCGGGT | >hg38_21_+_8433172_8446622_RNA45SN1_45S_50ntFlanks | 475 | 517 | 43 | CTTCGTGATCGATGTGGTGACGTCGTGCTCTCCCGGGCCGGGT | 1294.3 | 5.6 | 1.6 | 3.5 | 4.8E-04 | 6.1E-04 | MB231/MB468 | MITO |
| >hg38_21_+_8433172_8446622_RNA45SN1_45S_50ntFlanks@476.495.20\|TTCGTGATCGATGTGGTGAC | >hg38_21_+_8433172_8446622_RNA45SN1_45S_50ntFlanks | 476 | 495 | 20 | TTCGTGATCGATGTGGTGAC | 484.3 | 5.4 | 1.5 | 3.5 | 4.8E-04 | 6.1E-04 | MB231/MB468 | MITO |
| >hg38_21_+_8433172_8446622_RNA45SN1_45S_50ntFlanks@12393.12419.27\|ATAGCGACGTCGCTTTTTGATCCTTCG | >hg38_21_+_8433172_8446622_RNA45SN1_45S_50ntFlanks | 12393 | 12419 | 27 | ATAGCGACGTCGCTTTTTGATCCTTCG | 449.8 | -5.5 | 1.6 | -3.5 | 4.9E-04 | 6.1E-04 | MB231/MB468 | MITO |
| >hg38_21_+_8433172_8446622_RNA45SN1_45S_50ntFlanks@10848.10865.18\|TCGGGATAAGGATTGGCT | >hg38_21_+_8433172_8446622_RNA45SN1_45S_50ntFlanks | 10848 | 10865 | 18 | TCGGGATAAGGATTGGCT | 445.7 | -2.5 | 0.7 | -3.5 | 4.6E-04 | 6.1E-04 | MB231/MB468 | CYTO |
| >hg38_21_+_8433172_8446622_RNA45SN1_45S_50ntFlanks@8175.8203.29\|CCTTCTGATCGAGGCCCAGCCCGTGGACG | >hg38_21_+_8433172_8446622_RNA45SN1_45S_50ntFlanks | 8175 | 8203 | 29 | CCTTCTGATCGAGGCCCAGCCCGTGGACG | 215.8 | 2.5 | 0.7 | 3.5 | 4.6E-04 | 6.1E-04 | MB231/MB468 | CYTO |
| >hg38_21_+_8433172_8446622_RNA45SN1_45S_50ntFlanks@3775.3794.20\|GCCGGTACAGTGAAACTGCG | >hg38_21_+_8433172_8446622_RNA45SN1_45S_50ntFlanks | 3775 | 3794 | 20 | GCCGGTACAGTGAAACTGCG | 275.5 | -3.2 | 0.9 | -3.5 | 4.6E-04 | 6.1E-04 | MB231/MB468 | CYTO |
| >hg38_21_+_8433172_8446622_RNA45SN1_45S_50ntFlanks@11452.11475.24\|CTCCTCGGCGCGCGGCGGCGGCGG | >hg38_21_+_8433172_8446622_RNA45SN1_45S_50ntFlanks | 11452 | 11475 | 24 | CTCCTCGGCGCGCGGCGGCGGCGG | 473.6 | -6.3 | 1.8 | -3.5 | 5.0E-04 | 6.1E-04 | BT20/MB468 | NUC |
| >hg38_21_+_8433172_8446622_RNA45SN1_45S_50ntFlanks@5472.5499.28\|ACGGCCCTGGCGGAGCGCTGAGAAGACG | >hg38_21_+_8433172_8446622_RNA45SN1_45S_50ntFlanks | 5472 | 5499 | 28 | ACGGCCCTGGCGGAGCGCTGAGAAGACG | 7521.4 | 2.3 | 0.7 | 3.5 | 4.6E-04 | 6.1E-04 | MB231/MB468 | CYTO |
| >MI0000542\|hsa-mir-320a&WithFlank&8\|-\|22244960\|22245043@48.70.23(+1C)&[MIMAT0000510&hsa-miR-320a-3p&offsets\|0\|+1(+1C);m-60&8\|-\|22244975\|22244996&offsets\|0\|+1(+1C)]\|AAAAGCTGGGTTGAGAGGGCGAAC | >MI0000542\|hsa-mir-320a&WithFlank&8\|-\|22244960\|22245043 | 48 | 70 | 23(+1C) | AAAAGCTGGGTTGAGAGGGCGAAC | 315.1 | 4.5 | 1.3 | 3.5 | 4.1E-04 | 6.1E-04 | MB231/MB468 | MP |
| >hg38_21_+_8433172_8446622_RNA45SN1_45S_50ntFlanks@4764.4781.18\|ATACCGTCGTAGTTCCGA | >hg38_21_+_8433172_8446622_RNA45SN1_45S_50ntFlanks | 4764 | 4781 | 18 | ATACCGTCGTAGTTCCGA | 194.7 | -3.7 | 1.0 | -3.5 | 4.4E-04 | 6.1E-04 | BT20/MB468 | CYTO |
| >hg38_21_+_8433172_8446622_RNA45SN1_45S_50ntFlanks@10587.10608.22\|CCCCGGGGAGAGTTCTCTTTTC | >hg38_21_+_8433172_8446622_RNA45SN1_45S_50ntFlanks | 10587 | 10608 | 22 | CCCCGGGGAGAGTTCTCTTTTC | 248.1 | -5.1 | 1.4 | -3.5 | 4.4E-04 | 6.1E-04 | BT20/MB468 | CYTO |
| >MI0000298\|hsa-mir-221&WithFlank&X\|-\|45746151\|45746272@31.54.24&[MIMAT0004568&hsa-miR-221-5p&offsets\|0\|+2;m-405&X\|-\|45746220\|45746242&offsets\|0\|+1]\|ACCTGGCATACAATGTAGATTTCT | >MI0000298\|hsa-mir-221&WithFlank&X\|-\|45746151\|45746272 | 31 | 54 | 24 | ACCTGGCATACAATGTAGATTTCT | 2247.1 | -3.2 | 0.8 | -3.9 | 8.4E-05 | 6.1E-04 | BT20/MB231 | MITO |
| >hg38_MT_+_598_1651_RNR1_12S_50ntFlanks@915.946.32\|AAAGGACATTTAACTAAAACCCCTACGCATTT | >hg38_MT_+_598_1651_RNR1_12S_50ntFlanks | 915 | 946 | 32 | AAAGGACATTTAACTAAAACCCCTACGCATTT | 1226.9 | 5.7 | 1.5 | 3.7 | 2.0E-04 | 6.1E-04 | BT20/MB468 | MP |
| >hg38_21_+_8433172_8446622_RNA45SN1_45S_50ntFlanks@5486.5518.33\|GCGCTGAGAAGACGGTCGAACTTGACTATCTAG | >hg38_21_+_8433172_8446622_RNA45SN1_45S_50ntFlanks | 5486 | 5518 | 33 | GCGCTGAGAAGACGGTCGAACTTGACTATCTAG | 379.9 | -5.9 | 1.6 | -3.7 | 2.0E-04 | 6.1E-04 | BT20/MB468 | MP |
| >hg38_21_+_8433172_8446622_RNA45SN1_45S_50ntFlanks@9777.9819.43\|GAAGCCCGGCTCGCTGGCGTGGAGCCGGGCGTGGAATGCGAGT | >hg38_21_+_8433172_8446622_RNA45SN1_45S_50ntFlanks | 9777 | 9819 | 43 | GAAGCCCGGCTCGCTGGCGTGGAGCCGGGCGTGGAATGCGAGT | 45.2 | 8.8 | 2.2 | 4.0 | 7.0E-05 | 6.1E-04 | BT20/MB231 | MP |
| >am_trnaMT_PheGAA_MT_+_577_647@41.60.20\|GTTTAGACGGGCTCACATCA | >am_trnaMT_PheGAA_MT_+_577_647 | 41 | 60 | 20 | GTTTAGACGGGCTCACATCA | 268.9 | 5.4 | 1.5 | 3.7 | 2.0E-04 | 6.1E-04 | BT20/MB468 | MP |
| >am_trnaMT_TyrGTA_MT_-_5826_5891@1.33.33\|GGTAAAATGGCTGAGTGAAGCATTGGACTGTAA | >am_trnaMT_TyrGTA_MT_-_5826_5891 | 1 | 33 | 33 | GGTAAAATGGCTGAGTGAAGCATTGGACTGTAA | 491.9 | 5.1 | 1.5 | 3.5 | 4.9E-04 | 6.1E-04 | MB231/MB468 | MITO |
| >hg38_21_+_8433172_8446622_RNA45SN1_45S_50ntFlanks@11913.11940.28\|AAGAGACATGAGAGGTGTAGAATAAGTG | >hg38_21_+_8433172_8446622_RNA45SN1_45S_50ntFlanks | 11913 | 11940 | 28 | AAGAGACATGAGAGGTGTAGAATAAGTG | 104.9 | 5.9 | 1.6 | 3.7 | 2.0E-04 | 6.1E-04 | BT20/MB468 | MP |
| >hg38_21_+_8433172_8446622_RNA45SN1_45S_50ntFlanks@9533.9561.29\|GAGGAAACTCTGGTGGAGGTCCGTAGCGG | >hg38_21_+_8433172_8446622_RNA45SN1_45S_50ntFlanks | 9533 | 9561 | 29 | GAGGAAACTCTGGTGGAGGTCCGTAGCGG | 442.1 | -4.3 | 1.2 | -3.5 | 4.4E-04 | 6.1E-04 | BT20/MB468 | CYTO |
| >hg38_21_+_8433172_8446622_RNA45SN1_45S_50ntFlanks@11808.11830.23\|CTGTCCCTACCTACTATCCAGCG | >hg38_21_+_8433172_8446622_RNA45SN1_45S_50ntFlanks | 11808 | 11830 | 23 | CTGTCCCTACCTACTATCCAGCG | 167.9 | -3.8 | 1.1 | -3.5 | 5.0E-04 | 6.1E-04 | MB231/MB468 | NUC |
| >MI0000746\|hsa-mir-99b&WithFlank&19\|+\|51692606\|51692687@13.33.21(+1A)&[MIMAT0000689&hsa-miR-99b-5p&offsets\|0\|-1(+1A);m-10&19\|+\|51692618\|51692639&offsets\|0\|-1(+1A)]\|CACCCGTAGAACCGACCTTGCA | >MI0000746\|hsa-mir-99b&WithFlank&19\|+\|51692606\|51692687 | 13 | 33 | 21(+1A) | CACCCGTAGAACCGACCTTGCA | 1582.4 | 2.8 | 0.8 | 3.7 | 2.0E-04 | 6.1E-04 | BT20/MB468 | MP |
| >hg38_21_+_8433172_8446622_RNA45SN1_45S_50ntFlanks@10513.10530.18\|CCGGAGTGGCGGAGATGG | >hg38_21_+_8433172_8446622_RNA45SN1_45S_50ntFlanks | 10513 | 10530 | 18 | CCGGAGTGGCGGAGATGG | 491.9 | 7.6 | 1.9 | 4.0 | 7.1E-05 | 6.1E-04 | BT20/MB231 | MP |
| >hg38_21_+_8433172_8446622_RNA45SN1_45S_50ntFlanks@11671.11693.23\|CCAGTGCTCTGAATGTCAAAGTG | >hg38_21_+_8433172_8446622_RNA45SN1_45S_50ntFlanks | 11671 | 11693 | 23 | CCAGTGCTCTGAATGTCAAAGTG | 1161.6 | -4.2 | 1.2 | -3.5 | 4.4E-04 | 6.1E-04 | BT20/MB468 | CYTO |
| >hg38_21_+_8433172_8446622_RNA45SN1_45S_50ntFlanks@10233.10273.41\|GCGGACGCTACGCCGCGACGAGTAGGAGGGCCGCTGCGGTG | >hg38_21_+_8433172_8446622_RNA45SN1_45S_50ntFlanks | 10233 | 10273 | 41 | GCGGACGCTACGCCGCGACGAGTAGGAGGGCCGCTGCGGTG | 123.0 | -5.7 | 1.6 | -3.5 | 4.1E-04 | 6.1E-04 | MB231/MB468 | MP |
| >hg38_21_+_8433172_8446622_RNA45SN1_45S_50ntFlanks@4494.4511.18\|CCCGAAGCGTTTACTTTG | >hg38_21_+_8433172_8446622_RNA45SN1_45S_50ntFlanks | 4494 | 4511 | 18 | CCCGAAGCGTTTACTTTG | 158.0 | -6.6 | 1.8 | -3.7 | 2.0E-04 | 6.1E-04 | BT20/MB468 | MP |
| >hg38_21_+_8433172_8446622_RNA45SN1_45S_50ntFlanks@10407.10424.18\|GGGTCAGTCGGTCCTGAG | >hg38_21_+_8433172_8446622_RNA45SN1_45S_50ntFlanks | 10407 | 10424 | 18 | GGGTCAGTCGGTCCTGAG | 412.7 | -3.3 | 0.9 | -3.5 | 4.9E-04 | 6.2E-04 | MB231/MB468 | MITO |
| >hg38_21_+_8433172_8446622_RNA45SN1_45S_50ntFlanks@4738.4765.28\|AAAGTCGGAGGTTCGAAGACGATCAGAT | >hg38_21_+_8433172_8446622_RNA45SN1_45S_50ntFlanks | 4738 | 4765 | 28 | AAAGTCGGAGGTTCGAAGACGATCAGAT | 412.0 | 2.8 | 0.8 | 3.7 | 2.0E-04 | 6.2E-04 | BT20/MB468 | MP |
| >hg38_21_+_8433172_8446622_RNA45SN1_45S_50ntFlanks@9481.9505.25\|CACGTGTTAGGACCCGAAAGATGGT | >hg38_21_+_8433172_8446622_RNA45SN1_45S_50ntFlanks | 9481 | 9505 | 25 | CACGTGTTAGGACCCGAAAGATGGT | 157.8 | 10.6 | 2.7 | 4.0 | 7.1E-05 | 6.2E-04 | BT20/MB231 | MP |
| >hg38_21_+_8433172_8446622_RNA45SN1_45S_50ntFlanks@11206.11229.24\|CGGCGGCGACTCTGGACGCGAGCC | >hg38_21_+_8433172_8446622_RNA45SN1_45S_50ntFlanks | 11206 | 11229 | 24 | CGGCGGCGACTCTGGACGCGAGCC | 55.2 | 9.1 | 2.3 | 4.0 | 7.1E-05 | 6.2E-04 | BT20/MB231 | MP |
| >MI0000434\|hsa-let-7i&WithFlank&12\|+\|62603680\|62603775@12.30.19(+1U)&[MIMAT0000415&hsa-let-7i-5p&offsets\|0\|-3(+1U);m-49&12\|+\|62603691\|62603712&offsets\|0\|-3(+1U)]\|TGAGGTAGTAGTTTGTGCTT | >MI0000434\|hsa-let-7i&WithFlank&12\|+\|62603680\|62603775 | 12 | 30 | 19(+1U) | TGAGGTAGTAGTTTGTGCTT | 23.3 | 7.8 | 2.2 | 3.5 | 4.1E-04 | 6.2E-04 | MB231/MB468 | MP |
| >trnaMT_HisGTG_MT_+_12138_12206@25.69.45\|CAGATTGTGAATCTGACAACAGAGGCTTACGACCCCTTATTTACC | >trnaMT_HisGTG_MT_+_12138_12206 | 25 | 69 | 45 | CAGATTGTGAATCTGACAACAGAGGCTTACGACCCCTTATTTACC | 71.5 | 9.9 | 2.7 | 3.7 | 2.0E-04 | 6.2E-04 | BT20/MB468 | MP |
| >hg38_21_+_8433172_8446622_RNA45SN1_45S_50ntFlanks@12622.12654.33\|AGCCAATGGGGCGAAGCTACCATCTGTGGGATT | >hg38_21_+_8433172_8446622_RNA45SN1_45S_50ntFlanks | 12622 | 12654 | 33 | AGCCAATGGGGCGAAGCTACCATCTGTGGGATT | 313.9 | -4.6 | 1.2 | -3.7 | 2.0E-04 | 6.2E-04 | BT20/MB468 | MP |
| >MI0000301\|hsa-mir-224&WithFlank&X\|-\|151958572\|151958664@14.35.22&[MIMAT0000281&hsa-miR-224-5p&offsets\|+1\|-2;m-238&X\|-\|151958630\|151958651&offsets\|0\|0]\|CAAGTCACTAGTGGTTCCGTTT | >MI0000301\|hsa-mir-224&WithFlank&X\|-\|151958572\|151958664 | 14 | 35 | 22 | CAAGTCACTAGTGGTTCCGTTT | 13481.5 | 3.4 | 0.9 | 4.0 | 7.3E-05 | 6.2E-04 | BT20/MB231 | CYTO |
| >trnaMT_PheGAA_MT_+_577_647@36.55.20\|AAAATGTTTAGACGGGCTCA | >trnaMT_PheGAA_MT_+_577_647 | 36 | 55 | 20 | AAAATGTTTAGACGGGCTCA | 132.2 | 4.3 | 1.2 | 3.5 | 4.1E-04 | 6.2E-04 | MB231/MB468 | MP |
| >MI0000441\|hsa-mir-30b&WithFlank&8\|-\|134800514\|134800613@23.43.21(+1A)&[MIMAT0000420&hsa-miR-30b-5p&offsets\|0\|-1(+1A);m-76&8\|-\|134800570\|134800591&offsets\|0\|-1(+1A)]\|TGTAAACATCCTACACTCAGCA | >MI0000441\|hsa-mir-30b&WithFlank&8\|-\|134800514\|134800613 | 23 | 43 | 21(+1A) | TGTAAACATCCTACACTCAGCA | 1074.6 | -4.1 | 1.1 | -3.9 | 8.5E-05 | 6.2E-04 | BT20/MB231 | MITO |

| | | | | | | | | | | | | | |
|---|---|---|---|---|---|---|---|---|---|---|---|---|---|
| >hg38_21_+_8433172_8446622_RNA45SN1_45S_50ntFlanks@6650.6695.46\|ACGACTCTTAGCGGTGGATCACTCGGCTCGTGCGTCGATGAAGAAC | >hg38_21_+_8433172_8446622_RNA45SN1_45S_50ntFlanks | 6650 | 6695 | 46 | ACGACTCTTAGCGGTGGATCACTCGGCTCGTGCGTCGATGAAGAAC | 921.9 | 3.6 | 1.0 | 3.5 | 4.9E-04 | 6.2E-04 | MB231/MB468 | MITO |
| >hg38_21_+_8433172_8446622_RNA45SN1_45S_50ntFlanks@4728.4752.25\|ATCAAGAACGAAAGTCGGAGGTTCG | >hg38_21_+_8433172_8446622_RNA45SN1_45S_50ntFlanks | 4728 | 4752 | 25 | ATCAAGAACGAAAGTCGGAGGTTCG | 494.4 | -4.9 | 1.4 | -3.5 | 4.6E-04 | 6.2E-04 | MB231/MB468 | CYTO |
| >trna134_GluTTC_1_-_16861774_16861845@1.33.33,trna5_GluTTC_1_+_17199078_17199149@1.33.33,trna84_GluTTC_1_-_161391883_161391954@1.33.33,trna94_GluTTC_1_-_149664355_149664427@1.33.33\|TCCCTGGTGGTCTAGTGGCTAGGATTCGGCGCT | >trna134_GluTTC_1_-_16861774_16861845 | 1 | 33 | 33 | TCCCTGGTGGTCTAGTGGCTAGGATTCGGCGCT | 388.6 | 7.5 | 2.0 | 3.7 | 2.0E-04 | 6.2E-04 | BT20/MB468 | MP |
| >trnaMT_HisGTG_MT_+_12138_12206@32.62.31\|TGAATCTGACAACAGAGGCTTACGACCCCTT | >trnaMT_HisGTG_MT_+_12138_12206 | 32 | 62 | 31 | TGAATCTGACAACAGAGGCTTACGACCCCTT | 972.0 | 5.6 | 1.6 | 3.6 | 3.7E-04 | 6.2E-04 | BT20/MB468 | MITO |
| >hg38_21_+_8433172_8446622_RNA45SN1_45S_50ntFlanks@8172.8214.43\|AGTCCTTCTGATCGAGGCCCAGCCCGTGGACGGTGTGAGGCCG | >hg38_21_+_8433172_8446622_RNA45SN1_45S_50ntFlanks | 8172 | 8214 | 43 | AGTCCTTCTGATCGAGGCCCAGCCCGTGGACGGTGTGAGGCCG | 89.2 | -5.8 | 1.6 | -3.5 | 4.1E-04 | 6.2E-04 | MB231/MB468 | MP |
| >MI0006444\|hsa-mir-1307&WithFlank&10\|-\|103394247\|103394407@86.108.23(+1A)&[MIMAT0005951&hsa-miR-1307-3p&offsets\|0\|+1(+1A);m-66&10\|-\|103394300\|103394322&offsets\|0\|0(+1A)]\|ACTCGGCGTGGCGTCGGTCGTGGA | >MI0006444\|hsa-mir-1307&WithFlank&10\|-\|103394247\|103394407 | 86 | 108 | 23(+1A) | ACTCGGCGTGGCGTCGGTCGTGGA | 121.5 | 4.4 | 1.3 | 3.5 | 4.9E-04 | 6.2E-04 | MB231/MB468 | MITO |
| >hg38_21_+_8433172_8446622_RNA45SN1_45S_50ntFlanks@13007.13039.33\|TATTGAAAGTCAGCCCTCGACACAAGGGTTTGT | >hg38_21_+_8433172_8446622_RNA45SN1_45S_50ntFlanks | 13007 | 13039 | 33 | TATTGAAAGTCAGCCCTCGACACAAGGGTTTGT | 1306.7 | 3.0 | 0.9 | 3.5 | 4.9E-04 | 6.2E-04 | MB231/MB468 | MITO |
| >am_trnaMT_HisGTG_MT_+_12138_12206@27.51.25\|GATTGTGAATCTGACAACAGAGGCT | >am_trnaMT_HisGTG_MT_+_12138_12206 | 27 | 51 | 25 | GATTGTGAATCTGACAACAGAGGCT | 225.1 | 6.6 | 1.8 | 3.7 | 2.0E-04 | 6.2E-04 | BT20/MB468 | MP |
| >MI0000465\|hsa-mir-191&WithFlank&3\|-\|49020612\|49020715@22.39.18&[MIMAT0000440&hsa-miR-191-5p&offsets\|0\|-5;m-48&3\|-\|49020672\|49020694&offsets\|0\|-5]\|CAACGGAATCCCAAAAGC | >MI0000465\|hsa-mir-191&WithFlank&3\|-\|49020612\|49020715 | 22 | 39 | 18 | CAACGGAATCCCAAAAGC | 244.4 | 3.4 | 1.0 | 3.5 | 5.0E-04 | 6.2E-04 | MB231/MB468 | NUC |
| >MI0000439\|hsa-mir-23b&WithFlank&9\|+\|95085202\|95085310@67.86.20(+1C)&[MIMAT0000418&hsa-miR-23b-3p&offsets\|+3\|0(+1C);m-42&9\|+\|95085265\|95085285&offsets\|+3\|+2(+1C)]\|ACATTGCCAGGGATTACCACC | >MI0000439\|hsa-mir-23b&WithFlank&9\|+\|95085202\|95085310 | 67 | 86 | 20(+1C) | ACATTGCCAGGGATTACCACC | 428.9 | 12.6 | 3.6 | 3.5 | 5.0E-04 | 6.2E-04 | MB231/MB468 | NUC |
| >hg38_21_+_8433172_8446622_RNA45SN1_45S_50ntFlanks@12510.12527.18\|TCGTGAGACAGGTTAGTT | >hg38_21_+_8433172_8446622_RNA45SN1_45S_50ntFlanks | 12510 | 12527 | 18 | TCGTGAGACAGGTTAGTT | 145.6 | -3.7 | 1.0 | -3.6 | 3.7E-04 | 6.2E-04 | BT20/MB468 | MITO |
| >trnaMT_PheGAA_MT_+_577_647@33.55.23\|CTGAAAATGTTTAGACGGGCTCA | >trnaMT_PheGAA_MT_+_577_647 | 33 | 55 | 23 | CTGAAAATGTTTAGACGGGCTCA | 113.0 | 7.1 | 1.9 | 3.7 | 2.0E-04 | 6.2E-04 | BT20/MB468 | MP |
| >hg38_21_+_8433172_8446622_RNA45SN1_45S_50ntFlanks@5083.5100.18\|ACGAACGAGACTCTGGCA | >hg38_21_+_8433172_8446622_RNA45SN1_45S_50ntFlanks | 5083 | 5100 | 18 | ACGAACGAGACTCTGGCA | 251.0 | -4.0 | 1.1 | -3.5 | 4.1E-04 | 6.2E-04 | MB231/MB468 | MP |
| >trnaMT_SerGCT_MT_+_12207_12265@19.62.44\|TGCTAACTCATGCCCCCATGTCTAACAACATGGCTTTCTCACCA | >trnaMT_SerGCT_MT_+_12207_12265 | 19 | 62 | 44 | TGCTAACTCATGCCCCCATGTCTAACAACATGGCTTTCTCACCA | 76.8 | 10.0 | 2.7 | 3.7 | 2.0E-04 | 6.2E-04 | BT20/MB468 | MP |
| >MI0000088\|hsa-mir-30a&WithFlank&6\|-\|71403545\|71403627@12.34.23(+1C)&[MIMAT0000087&hsa-miR-30a-5p&offsets\|0\|+1(+1C);m-15&6\|-\|71403595\|71403616&offsets\|0\|+1(+1C)]\|TGTAAACATCCTCGACTGGAAGCC | >MI0000088\|hsa-mir-30a&WithFlank&6\|-\|71403545\|71403627 | 12 | 34 | 23(+1C) | TGTAAACATCCTCGACTGGAAGCC | 26290.8 | -3.5 | 0.9 | -3.9 | 8.5E-05 | 6.2E-04 | BT20/MB231 | MITO |
| >hg38_21_+_8433172_8446622_RNA45SN1_45S_50ntFlanks@10247.10286.40\|GCGACGAGTAGGAGGGCCGCTGCGGTGAGCCTTGAAGCCT | >hg38_21_+_8433172_8446622_RNA45SN1_45S_50ntFlanks | 10247 | 10286 | 40 | GCGACGAGTAGGAGGGCCGCTGCGGTGAGCCTTGAAGCCT | 169.6 | 6.8 | 1.8 | 3.7 | 2.0E-04 | 6.2E-04 | BT20/MB468 | MP |
| >MI0000465\|hsa-mir-191&WithFlank&3\|-\|49020612\|49020715@22.43.22(+1A)&[MIMAT0000440&hsa-miR-191-5p&offsets\|0\|-1(+1A);m-48&3\|-\|49020672\|49020694&offsets\|0\|-1(+1A)]\|CAACGGAATCCCAAAAGCAGCTA | >MI0000465\|hsa-mir-191&WithFlank&3\|-\|49020612\|49020715 | 22 | 43 | 22(+1A) | CAACGGAATCCCAAAAGCAGCTA | 1643.2 | 3.9 | 1.1 | 3.7 | 2.0E-04 | 6.2E-04 | BT20/MB468 | MP |
| >hg38_21_+_8433172_8446622_RNA45SN1_45S_50ntFlanks@10401.10439.39\|GAACATGGGTCAGTCGGTCCTGAGAGATGGGCGAGCGCC | >hg38_21_+_8433172_8446622_RNA45SN1_45S_50ntFlanks | 10401 | 10439 | 39 | GAACATGGGTCAGTCGGTCCTGAGAGATGGGCGAGCGCC | 386.3 | -3.6 | 0.9 | -4.0 | 7.4E-05 | 6.2E-04 | BT20/MB231 | CYTO |
| >hg38_21_+_8433172_8446622_RNA45SN1_45S_50ntFlanks@8192.8213.22\|AGCCCGTGGACGGTGTGAGGCC | >hg38_21_+_8433172_8446622_RNA45SN1_45S_50ntFlanks | 8192 | 8213 | 22 | AGCCCGTGGACGGTGTGAGGCC | 275.9 | -4.3 | 1.2 | -3.5 | 4.2E-04 | 6.2E-04 | MB231/MB468 | MP |
| >hg38_21_+_8433172_8446622_RNA45SN1_45S_50ntFlanks@3988.4013.26\|CTTTGGTGACTCTAGATAACCTCGGG | >hg38_21_+_8433172_8446622_RNA45SN1_45S_50ntFlanks | 3988 | 4013 | 26 | CTTTGGTGACTCTAGATAACCTCGGG | 177.9 | -5.0 | 1.4 | -3.5 | 5.0E-04 | 6.2E-04 | MB231/MB468 | NUC |
| >hg38_21_+_8433172_8446622_RNA45SN1_45S_50ntFlanks@6644.6665.22\|CCTCGTACGACTCTTAGCGGTG | >hg38_21_+_8433172_8446622_RNA45SN1_45S_50ntFlanks | 6644 | 6665 | 22 | CCTCGTACGACTCTTAGCGGTG | 266.6 | 3.8 | 1.1 | 3.5 | 4.7E-04 | 6.2E-04 | MB231/MB468 | CYTO |
| >hg38_21_+_8433172_8446622_RNA45SN1_45S_50ntFlanks@12444.12483.40\|AAGCAGAATTCACCAAGCGTTGGATTGTTCACCCACTAAT | >hg38_21_+_8433172_8446622_RNA45SN1_45S_50ntFlanks | 12444 | 12483 | 40 | AAGCAGAATTCACCAAGCGTTGGATTGTTCACCCACTAAT | 242.6 | -3.4 | 1.0 | -3.5 | 5.0E-04 | 6.2E-04 | MB231/MB468 | MITO |
| >hg38_21_+_8433172_8446622_RNA45SN1_45S_50ntFlanks@6700.6719.20\|CTAGCTGCGAGAATTAATGT | >hg38_21_+_8433172_8446622_RNA45SN1_45S_50ntFlanks | 6700 | 6719 | 20 | CTAGCTGCGAGAATTAATGT | 440.3 | -2.9 | 0.8 | -3.5 | 5.0E-04 | 6.2E-04 | MB231/MB468 | NUC |
| >hg38_MT_+_1621_3279_RNR2_16S_50ntFlanks@1370.1397.28\|ATGTTGGATCAGGACATCCCGATGGTGC | >hg38_MT_+_1621_3279_RNR2_16S_50ntFlanks | 1370 | 1397 | 28 | ATGTTGGATCAGGACATCCCGATGGTGC | 163.3 | 5.2 | 1.4 | 3.7 | 2.0E-04 | 6.2E-04 | BT20/MB468 | MP |
| >hg38_21_+_8433172_8446622_RNA45SN1_45S_50ntFlanks@11208.11226.19\|GCGGCGACTCTGGACGCGA | >hg38_21_+_8433172_8446622_RNA45SN1_45S_50ntFlanks | 11208 | 11226 | 19 | GCGGCGACTCTGGACGCGA | 197.3 | -4.5 | 1.2 | -3.7 | 2.0E-04 | 6.2E-04 | BT20/MB468 | MP |
| >MI0000750\|hsa-mir-26a-2&WithFlank&12\|-\|57824603\|57824698@20.41.22(+2U);MI0000083\|hsa-mir-26a-1&WithFlank&3\|+\|37969398\|37969486@16.37.22(+2U)&[MIMAT0000082&hsa-miR-26a-5p&offsets\|0\|0(+2U);m-33&12\|-\|57824658\|57824679&offsets\|0\|0(+2U)];[MIMAT0000082_1&hsa-miR-26a-5p&offsets\|0\|0(+2U);m-34&3\|+\|37969413\|37969434&offsets\|0\|0(+2U)]\|TTCAAGTAATCCAGGATAGGCTTT | >MI0000750\|hsa-mir-26a-2&WithFlank&12\|-\|57824603\|57824698 | 20 | 41 | 22(+2U) | TTCAAGTAATCCAGGATAGGCTTT | 5403.9 | -3.4 | 0.9 | -3.9 | 8.6E-05 | 6.2E-04 | BT20/MB231 | MITO |
| >trna10_LysCTT_16_+_3241501_3241573@5.34.30,trna13_LysCTT_14_-_58706613_58706685@5.34.30,trna2_LysCTT_15_+_79152904_79152976@5.34.30\|GGCTAGCTCAGTCGGTAGAGCATGGGACTC | >trna10_LysCTT_16_+_3241501_3241573 | 5 | 34 | 30 | GGCTAGCTCAGTCGGTAGAGCATGGGACTC | 192.1 | -4.0 | 1.0 | -3.9 | 8.6E-05 | 6.2E-04 | BT20/MB231 | MITO |
| >trnaMT_ValTAC_MT_+_1602_1670@34.71.38\|CACTTAGGAGATTTCAACTTAACTTGACCGCTCTGACC | >trnaMT_ValTAC_MT_+_1602_1670 | 34 | 71 | 38 | CACTTAGGAGATTTCAACTTAACTTGACCGCTCTGACC | 107.2 | 7.3 | 2.0 | 3.7 | 2.0E-04 | 6.2E-04 | BT20/MB468 | MP |
| >MI0000650\|hsa-mir-200c&WithFlank&12\|+\|6963693\|6963772@50.72.23&[MIMAT0000617&hsa-miR-200c-3p&offsets\|0\|0;m-54&12\|+\|6963742\|6963764&offsets\|0\|0]\|TAATACTGCCGGGTAATGATGGA | >MI0000650\|hsa-mir-200c&WithFlank&12\|+\|6963693\|6963772 | 50 | 72 | 23 | TAATACTGCCGGGTAATGATGGA | 1562.2 | -3.2 | 0.9 | -3.5 | 5.0E-04 | 6.2E-04 | MB231/MB468 | MITO |
| >hg38_21_+_8433172_8446622_RNA45SN1_45S_50ntFlanks@10402.10443.42\|AACATGGGTCAGTCGGTCCTGAGAGATGGGCGAGCGCCGTTC | >hg38_21_+_8433172_8446622_RNA45SN1_45S_50ntFlanks | 10402 | 10443 | 42 | AACATGGGTCAGTCGGTCCTGAGAGATGGGCGAGCGCCGTTC | 272.4 | -5.3 | 1.5 | -3.5 | 5.0E-04 | 6.2E-04 | MB231/MB468 | MITO |
| >hg38_21_+_8433172_8446622_RNA45SN1_45S_50ntFlanks@10794.10814.21\|CTGGCATGTTGGAACAATGTA | >hg38_21_+_8433172_8446622_RNA45SN1_45S_50ntFlanks | 10794 | 10814 | 21 | CTGGCATGTTGGAACAATGTA | 55.2 | 9.1 | 2.3 | 4.0 | 7.2E-05 | 6.2E-04 | BT20/MB231 | MP |
| >hg38_21_+_8433172_8446622_RNA45SN1_45S_50ntFlanks@4003.4038.36\|ATAACCTCGGGCCGATCGCACGCCCCCCGTGGCGGC | >hg38_21_+_8433172_8446622_RNA45SN1_45S_50ntFlanks | 4003 | 4038 | 36 | ATAACCTCGGGCCGATCGCACGCCCCCCGTGGCGGC | 110.6 | -2.4 | 0.7 | -3.5 | 4.5E-04 | 6.2E-04 | BT20/MB468 | CYTO |
| >hg38_21_+_8433172_8446622_RNA45SN1_45S_50ntFlanks@10393.10435.43\|CAGCAGTTGAACATGGGTCAGTCGGTCCTGAGAGATGGGCGAG | >hg38_21_+_8433172_8446622_RNA45SN1_45S_50ntFlanks | 10393 | 10435 | 43 | CAGCAGTTGAACATGGGTCAGTCGGTCCTGAGAGATGGGCGAG | 457.4 | -4.3 | 1.2 | -3.6 | 3.7E-04 | 6.2E-04 | BT20/MB468 | MITO |
| >hg38_21_+_8433172_8446622_RNA45SN1_45S_50ntFlanks@10680.10707.28\|GTTCCGGCGGCGTCCGGTGAGCTCTCGC | >hg38_21_+_8433172_8446622_RNA45SN1_45S_50ntFlanks | 10680 | 10707 | 28 | GTTCCGGCGGCGTCCGGTGAGCTCTCGC | 144.8 | -5.5 | 1.5 | -3.5 | 4.2E-04 | 6.2E-04 | MB231/MB468 | MP |
| >hg38_21_+_8433172_8446622_RNA45SN1_45S_50ntFlanks@11847.11886.40\|AACGGGCTTGGCGGAATCAGCGGGGAAAGAAGACCCTGTT | >hg38_21_+_8433172_8446622_RNA45SN1_45S_50ntFlanks | 11847 | 11886 | 40 | AACGGGCTTGGCGGAATCAGCGGGGAAAGAAGACCCTGTT | 82.0 | 6.8 | 1.8 | 3.7 | 2.1E-04 | 6.2E-04 | BT20/MB468 | MP |
| >trnaMT_GluTTC_MT_-_14674_14742@35.72.38,trnalookalike8_GluTTC_5_-_93905172_93905240@35.72.38\|TATCATTGGTCGTGGTTGTAGTCCGTGCGAGAATACCA | >trnaMT_GluTTC_MT_-_14674_14742 | 35 | 72 | 38 | TATCATTGGTCGTGGTTGTAGTCCGTGCGAGAATACCA | 5425.8 | -3.6 | 0.9 | -4.0 | 7.6E-05 | 6.2E-04 | BT20/MB231 | NUC |
| >hg38_1_-_228634819_228635039_RNA5S12_5S_50ntFlanks@136.169.34\|GGAGACCGCCTGGGAATACCGGGTGCTGTAGGCT | >hg38_1_-_228634819_228635039_RNA5S12_5S_50ntFlanks | 136 | 169 | 34 | GGAGACCGCCTGGGAATACCGGGTGCTGTAGGCT | 2448.4 | 5.6 | 1.4 | 4.0 | 7.2E-05 | 6.2E-04 | BT20/MB231 | MP |
| >hg38_21_+_8433172_8446622_RNA45SN1_45S_50ntFlanks@9501.9523.23\|ATGGTGAACTATGCCTGGGCAGG | >hg38_21_+_8433172_8446622_RNA45SN1_45S_50ntFlanks | 9501 | 9523 | 23 | ATGGTGAACTATGCCTGGGCAGG | 160.1 | 10.7 | 2.7 | 4.0 | 7.2E-05 | 6.2E-04 | BT20/MB231 | MP |
| >hg38_21_+_8433172_8446622_RNA45SN1_45S_50ntFlanks@10868.10902.35\|AAGGGCTGGGTCGGTCGGGCTGGGGCGCGAAGCGG | >hg38_21_+_8433172_8446622_RNA45SN1_45S_50ntFlanks | 10868 | 10902 | 35 | AAGGGCTGGGTCGGTCGGGCTGGGGCGCGAAGCGG | 240.4 | -5.2 | 1.5 | -3.5 | 5.1E-04 | 6.3E-04 | MB231/MB468 | NUC |
| >MI0000077\|hsa-mir-21&WithFlank&17\|+\|59841260\|59841343@14.36.23(+1G)&[MIMAT0000076&hsa-miR-21-5p&offsets\|0\|+1(+1G);m-1&17\|+\|59841273\|59841294&offsets\|0\|+1(+1G)]\|TAGCTTATCAGACTGATGTTGACG | >MI0000077\|hsa-mir-21&WithFlank&17\|+\|59841260\|59841343 | 14 | 36 | 23(+1G) | TAGCTTATCAGACTGATGTTGACG | 4086.3 | 3.6 | 1.0 | 3.5 | 4.7E-04 | 6.3E-04 | MB231/MB468 | CYTO |
| >hg38_21_+_8433172_8446622_RNA45SN1_45S_50ntFlanks@10349.10390.42\|AACGAGAACTTTGAAGGCCGAAGTGGAGAAGGGTTCCATGTG | >hg38_21_+_8433172_8446622_RNA45SN1_45S_50ntFlanks | 10349 | 10390 | 42 | AACGAGAACTTTGAAGGCCGAAGTGGAGAAGGGTTCCATGTG | 1844.0 | -3.1 | 0.9 | -3.5 | 5.1E-04 | 6.3E-04 | MB231/MB468 | NUC |
| >hg38_21_+_8433172_8446622_RNA45SN1_45S_50ntFlanks@12996.13013.18\|TCGCTGCGATCTATTGAA | >hg38_21_+_8433172_8446622_RNA45SN1_45S_50ntFlanks | 12996 | 13013 | 18 | TCGCTGCGATCTATTGAA | 3225.9 | 2.2 | 0.6 | 3.5 | 5.1E-04 | 6.3E-04 | MB231/MB468 | NUC |
| >MI0000289\|hsa-mir-181a-1&WithFlank&1\|-\|198859038\|198859159@70.89.20(+1U)&[MIMAT0000270&hsa-miR-181a-3p&offsets\|0\|-2(+1U);m-125&1\|-\|198859069\|198859090&offsets\|0\|-2(+1U)]\|ACCATCGACCGTTGATTGTAT | >MI0000289\|hsa-mir-181a-1&WithFlank&1\|-\|198859038\|198859159 | 70 | 89 | 20(+1U) | ACCATCGACCGTTGATTGTAT | 47.9 | -9.0 | 2.3 | -4.0 | 7.3E-05 | 6.3E-04 | BT20/MB231 | MP |
| >hg38_21_+_8433172_8446622_RNA45SN1_45S_50ntFlanks@5055.5081.27\|GGAGCGATTTGTCTGGTTAATTCCGAT | >hg38_21_+_8433172_8446622_RNA45SN1_45S_50ntFlanks | 5055 | 5081 | 27 | GGAGCGATTTGTCTGGTTAATTCCGAT | 898.1 | 3.8 | 1.0 | 4.0 | 7.3E-05 | 6.3E-04 | BT20/MB231 | MP |
| >hg38_21_+_8433172_8446622_RNA45SN1_45S_50ntFlanks@8176.8214.39\|CTTCTGATCGAGGCCCAGCCCGTGGACGGTGTGAGGCCG | >hg38_21_+_8433172_8446622_RNA45SN1_45S_50ntFlanks | 8176 | 8214 | 39 | CTTCTGATCGAGGCCCAGCCCGTGGACGGTGTGAGGCCG | 334.1 | -3.0 | 0.9 | -3.5 | 5.1E-04 | 6.3E-04 | MB231/MB468 | NUC |
| >hg38_21_+_8433172_8446622_RNA45SN1_45S_50ntFlanks@12459.12478.20\|AGCGTTGGATTGTTCACCCA | >hg38_21_+_8433172_8446622_RNA45SN1_45S_50ntFlanks | 12459 | 12478 | 20 | AGCGTTGGATTGTTCACCCA | 1011.6 | -6.6 | 1.9 | -3.5 | 5.1E-04 | 6.3E-04 | BT20/MB468 | NUC |
| >trnaMT_LeuTAA_MT_+_3230_3304@18.44.27\|CGGTAATCGCATAAAACTTAAAACTTT | >trnaMT_LeuTAA_MT_+_3230_3304 | 18 | 44 | 27 | CGGTAATCGCATAAAACTTAAAACTTT | 611.0 | -3.2 | 0.9 | -3.5 | 5.1E-04 | 6.3E-04 | MB231/MB468 | NUC |
| >hg38_21_+_8433172_8446622_RNA45SN1_45S_50ntFlanks@13000.13039.40\|TGCGATCTATTGAAAGTCAGCCCTCGACACAAGGGTTTGT | >hg38_21_+_8433172_8446622_RNA45SN1_45S_50ntFlanks | 13000 | 13039 | 40 | TGCGATCTATTGAAAGTCAGCCCTCGACACAAGGGTTTGT | 421.3 | 4.4 | 1.2 | 3.5 | 4.5E-04 | 6.3E-04 | BT20/MB468 | CYTO |
| >MI0000085\|hsa-mir-27a&WithFlank&19\|-\|13836434\|13836523@57.76.20&[MIMAT0000084&hsa-miR-27a-3p&offsets\|0\|-1;m-43&19\|-\|13836447\|13836467&offsets\|0\|-1]\|TTCACAGTGGCTAAGTTCCG | >MI0000085\|hsa-mir-27a&WithFlank&19\|-\|13836434\|13836523 | 57 | 76 | 20 | TTCACAGTGGCTAAGTTCCG | 16166.2 | 3.0 | 0.9 | 3.5 | 4.2E-04 | 6.3E-04 | MB231/MB468 | MP |

<table>
<tr><td>>hg38_21_+_8433172_8446622_RNA45SN1_45S_50ntFlanks@11583.11602.20|CTGGTGCGGACCAGGGGAAT</td><td>>hg38_21_+_8433172_8446622_RNA45SN1_45S_50ntFlanks</td><td>11583</td><td>11602</td><td>20</td><td>CTGGTGCGGACCAGGGGAAT</td><td>1011.3</td><td>-2.7</td><td>0.8</td><td>-3.5</td><td>4.5E-04</td><td>6.3E-04</td><td>BT20/MB468</td><td>CYTO</td></tr>
<tr><td>>hg38_21_+_8433172_8446622_RNA45SN1_45S_50ntFlanks@9156.9187.32|CCAGTGCGCCCCGGGCGGGTCGCGCCGTCGGG</td><td>>hg38_21_+_8433172_8446622_RNA45SN1_45S_50ntFlanks</td><td>9156</td><td>9187</td><td>32</td><td>CCAGTGCGCCCCGGGCGGGTCGCGCCGTCGGG</td><td>217.8</td><td>-4.9</td><td>1.4</td><td>-3.6</td><td>3.8E-04</td><td>6.3E-04</td><td>BT20/MB468</td><td>MITO</td></tr>
<tr><td>>hg38_21_+_8433172_8446622_RNA45SN1_45S_50ntFlanks@10391.10435.45|AACAGCAGTTGAACATGGGTCAGTCGGTCCTGAGAGATGGGCGAG</td><td>>hg38_21_+_8433172_8446622_RNA45SN1_45S_50ntFlanks</td><td>10391</td><td>10435</td><td>45</td><td>AACAGCAGTTGAACATGGGTCAGTCGGTCCTGAGAGATGGGCGAG</td><td>538.1</td><td>-4.7</td><td>1.3</td><td>-3.6</td><td>3.8E-04</td><td>6.3E-04</td><td>BT20/MB468</td><td>MITO</td></tr>
<tr><td>>MI0000456|hsa-mir-140&WithFlank&16|+|69933075|69933186@69.90.22(+1C)&[MIMAT0004597&hsa-miR-140-3p&offsets|+1|+2(+1C);m-45&16|+|69933143|69933164&offsets|0|0(+1C)]|ACCACAGGGTAGAACCACGGACC</td><td>>MI0000456|hsa-mir-140&WithFlank&16|+|69933075|69933186</td><td>69</td><td>90</td><td>22(+1C)</td><td>ACCACAGGGTAGAACCACGGACC</td><td>123.1</td><td>7.3</td><td>2.0</td><td>3.7</td><td>2.1E-04</td><td>6.3E-04</td><td>BT20/MB468</td><td>MP</td></tr>
<tr><td>>hg38_21_+_8433172_8446622_RNA45SN1_45S_50ntFlanks@8342.8378.37|CGTAAGGGAAAGTTGAAAAGAACTTTGAAGAGAGAGT</td><td>>hg38_21_+_8433172_8446622_RNA45SN1_45S_50ntFlanks</td><td>8342</td><td>8378</td><td>37</td><td>CGTAAGGGAAAGTTGAAAAGAACTTTGAAGAGAGAGT</td><td>299.7</td><td>6.1</td><td>1.6</td><td>3.7</td><td>2.1E-04</td><td>6.3E-04</td><td>BT20/MB468</td><td>MP</td></tr>
<tr><td>>hg38_21_+_8433172_8446622_RNA45SN1_45S_50ntFlanks@6739.6771.33|TCATCGACACTTCGAACGCACTTGCGGCCCCGG</td><td>>hg38_21_+_8433172_8446622_RNA45SN1_45S_50ntFlanks</td><td>6739</td><td>6771</td><td>33</td><td>TCATCGACACTTCGAACGCACTTGCGGCCCCGG</td><td>134.9</td><td>-4.6</td><td>1.3</td><td>-3.5</td><td>4.5E-04</td><td>6.3E-04</td><td>BT20/MB468</td><td>CYTO</td></tr>
<tr><td>>hg38_21_+_8433172_8446622_RNA45SN1_45S_50ntFlanks@6721.6757.37|AATTGCAGGACACATTGATCATCGACACTTCGAACGC</td><td>>hg38_21_+_8433172_8446622_RNA45SN1_45S_50ntFlanks</td><td>6721</td><td>6757</td><td>37</td><td>AATTGCAGGACACATTGATCATCGACACTTCGAACGC</td><td>220.5</td><td>-4.8</td><td>1.4</td><td>-3.5</td><td>4.5E-04</td><td>6.3E-04</td><td>BT20/MB468</td><td>CYTO</td></tr>
<tr><td>>hg38_21_+_8433172_8446622_RNA45SN1_45S_50ntFlanks@6651.6687.37|CGACTCTTAGCGGTGGATCACTCGGCTCGTGCGTCGA</td><td>>hg38_21_+_8433172_8446622_RNA45SN1_45S_50ntFlanks</td><td>6651</td><td>6687</td><td>37</td><td>CGACTCTTAGCGGTGGATCACTCGGCTCGTGCGTCGA</td><td>2876.2</td><td>2.7</td><td>0.8</td><td>3.6</td><td>3.8E-04</td><td>6.3E-04</td><td>BT20/MB468</td><td>MITO</td></tr>
<tr><td>>hg38_MT_+_1621_3279_RNR2_16S_50ntFlanks@51.70.20|GCTAAACCTAGCCCCAAACC</td><td>>hg38_MT_+_1621_3279_RNR2_16S_50ntFlanks</td><td>51</td><td>70</td><td>20</td><td>GCTAAACCTAGCCCCAAACC</td><td>2372.0</td><td>2.7</td><td>0.8</td><td>3.5</td><td>4.2E-04</td><td>6.3E-04</td><td>MB231/MB468</td><td>MP</td></tr>
<tr><td>>hg38_21_+_8433172_8446622_RNA45SN1_45S_50ntFlanks@12418.12437.20|CGATGTCGGCTCTTCCTATC</td><td>>hg38_21_+_8433172_8446622_RNA45SN1_45S_50ntFlanks</td><td>12418</td><td>12437</td><td>20</td><td>CGATGTCGGCTCTTCCTATC</td><td>8550.2</td><td>-1.9</td><td>0.5</td><td>-3.5</td><td>4.8E-04</td><td>6.3E-04</td><td>MB231/MB468</td><td>CYTO</td></tr>
<tr><td>>hg38_21_+_8433172_8446622_RNA45SN1_45S_50ntFlanks@9544.9569.26|GGTGGAGGTCCGTAGCGGTCCTGACG</td><td>>hg38_21_+_8433172_8446622_RNA45SN1_45S_50ntFlanks</td><td>9544</td><td>9569</td><td>26</td><td>GGTGGAGGTCCGTAGCGGTCCTGACG</td><td>588.7</td><td>-6.3</td><td>1.8</td><td>-3.5</td><td>5.2E-04</td><td>6.4E-04</td><td>BT20/MB468</td><td>NUC</td></tr>
<tr><td>>am_trna111_HisGTG_1_-_147774845_147774916@1.26.26,trna118_HisGTG_1_-_145396881_145396952@1.26.26,trna16_HisGTG_1_+_146544773_146544844@1.26.26,trna1_HisGTG_15_+_45493349_45493420@1.26.26,trna21_HisGTG_1_+_147753471_147753542@1.26.26,trna33_HisGTG_6_+_27125906_27125977@1.26.26,trna7_HisGTG_9_-_14433938_14434009@1.26.26,trna8_HisGTG_15_-_45492611_45492682@1.26.26,trna9_HisGTG_15_-_45490804_45490875@1.26.26|GCCGTGATCGTATAGTGGTTAGTACT</td><td>>am_trna111_HisGTG_1_-_147774845_147774916</td><td>1</td><td>26</td><td>26</td><td>GCCGTGATCGTATAGTGGTTAGTACT</td><td>424.0</td><td>4.3</td><td>1.1</td><td>3.7</td><td>2.1E-04</td><td>6.4E-04</td><td>BT20/MB468</td><td>MP</td></tr>
<tr><td>>hg38_21_+_8433172_8446622_RNA45SN1_45S_50ntFlanks@11584.11612.29|TGGTGCGGACCAGGGGAATCCGACTGTTT</td><td>>hg38_21_+_8433172_8446622_RNA45SN1_45S_50ntFlanks</td><td>11584</td><td>11612</td><td>29</td><td>TGGTGCGGACCAGGGGAATCCGACTGTTT</td><td>261.1</td><td>3.0</td><td>0.9</td><td>3.5</td><td>5.2E-04</td><td>6.4E-04</td><td>BT20/MB468</td><td>NUC</td></tr>
<tr><td>>hg38_21_+_8433172_8446622_RNA45SN1_45S_50ntFlanks@9533.9551.19|GAGGAAACTCTGGTGGAGG</td><td>>hg38_21_+_8433172_8446622_RNA45SN1_45S_50ntFlanks</td><td>9533</td><td>9551</td><td>19</td><td>GAGGAAACTCTGGTGGAGG</td><td>872.6</td><td>-6.6</td><td>1.9</td><td>-3.5</td><td>5.2E-04</td><td>6.4E-04</td><td>BT20/MB468</td><td>NUC</td></tr>
<tr><td>>hg38_21_+_8433172_8446622_RNA45SN1_45S_50ntFlanks@3997.4022.26|CTCTAGATAACCTCGGGCCGATCGCA</td><td>>hg38_21_+_8433172_8446622_RNA45SN1_45S_50ntFlanks</td><td>3997</td><td>4022</td><td>26</td><td>CTCTAGATAACCTCGGGCCGATCGCA</td><td>502.2</td><td>-6.4</td><td>1.8</td><td>-3.5</td><td>5.2E-04</td><td>6.4E-04</td><td>BT20/MB468</td><td>NUC</td></tr>
<tr><td>>hg38_21_+_8433172_8446622_RNA45SN1_45S_50ntFlanks@6700.6719.20|CTAGCTGCGAGAATTAATGT</td><td>>hg38_21_+_8433172_8446622_RNA45SN1_45S_50ntFlanks</td><td>6700</td><td>6719</td><td>20</td><td>CTAGCTGCGAGAATTAATGT</td><td>182.9</td><td>-3.8</td><td>1.1</td><td>-3.6</td><td>3.8E-04</td><td>6.4E-04</td><td>BT20/MB468</td><td>MITO</td></tr>
<tr><td>>hg38_21_+_8433172_8446622_RNA45SN1_45S_50ntFlanks@3985.4004.20|CGGCTTTGGTGACTCTAGAT</td><td>>hg38_21_+_8433172_8446622_RNA45SN1_45S_50ntFlanks</td><td>3985</td><td>4004</td><td>20</td><td>CGGCTTTGGTGACTCTAGAT</td><td>138.5</td><td>-4.0</td><td>1.1</td><td>-3.5</td><td>4.8E-04</td><td>6.4E-04</td><td>MB231/MB468</td><td>CYTO</td></tr>
<tr><td>>hg38_21_+_8433172_8446622_RNA45SN1_45S_50ntFlanks@4380.4410.31|CGTAGTTGGATCTTGGGAGCGGGCGGGCGGT</td><td>>hg38_21_+_8433172_8446622_RNA45SN1_45S_50ntFlanks</td><td>4380</td><td>4410</td><td>31</td><td>CGTAGTTGGATCTTGGGAGCGGGCGGGCGGT</td><td>195.1</td><td>-5.0</td><td>1.5</td><td>-3.5</td><td>5.1E-04</td><td>6.4E-04</td><td>MB231/MB468</td><td>NUC</td></tr>
<tr><td>>am_trna119_LysCTT_1_-_145395522_145395594@13.33.21,trna11_LysCTT_5_-_180648979_180649051@13.33.21,trna128_LysCTT_1_-_55423542_55423614@13.33.21,trna13_LysCTT_6_+_26556774_26556846@13.33.21,trna6_LysCTT_19_-_52425396_52425468@13.33.21,trna7_LysCTT_16_+_3225692_3225764@13.33.21,trna9_LysCTT_5_+_180634755_180634827@13.33.21|CAGTCGGTAGAGCATGAGACT</td><td>>am_trna119_LysCTT_1_-_145395522_145395594</td><td>13</td><td>33</td><td>21</td><td>CAGTCGGTAGAGCATGAGACT</td><td>207.2</td><td>4.6</td><td>1.3</td><td>3.5</td><td>4.3E-04</td><td>6.4E-04</td><td>MB231/MB468</td><td>MP</td></tr>
<tr><td>>hg38_21_+_8433172_8446622_RNA45SN1_45S_50ntFlanks@12407.12429.23|TTTTGATCCTTCGATGTCGGCTC</td><td>>hg38_21_+_8433172_8446622_RNA45SN1_45S_50ntFlanks</td><td>12407</td><td>12429</td><td>23</td><td>TTTTGATCCTTCGATGTCGGCTC</td><td>172.6</td><td>-4.3</td><td>1.2</td><td>-3.5</td><td>4.3E-04</td><td>6.4E-04</td><td>MB231/MB468</td><td>MP</td></tr>
<tr><td>>hg38_21_+_8433172_8446622_RNA45SN1_45S_50ntFlanks@10369.10407.39|AAGTGGAGAAGGGTTCCATGTGAACAGCAGTTGAACATG</td><td>>hg38_21_+_8433172_8446622_RNA45SN1_45S_50ntFlanks</td><td>10369</td><td>10407</td><td>39</td><td>AAGTGGAGAAGGGTTCCATGTGAACAGCAGTTGAACATG</td><td>130.4</td><td>-4.7</td><td>1.3</td><td>-3.6</td><td>3.8E-04</td><td>6.4E-04</td><td>BT20/MB468</td><td>MITO</td></tr>
<tr><td>>hg38_21_+_8433172_8446622_RNA45SN1_45S_50ntFlanks@10425.10443.19|AGATGGGCGAGCGCCGTTC</td><td>>hg38_21_+_8433172_8446622_RNA45SN1_45S_50ntFlanks</td><td>10425</td><td>10443</td><td>19</td><td>AGATGGGCGAGCGCCGTTC</td><td>86.1</td><td>-9.5</td><td>2.6</td><td>-3.7</td><td>2.1E-04</td><td>6.4E-04</td><td>BT20/MB468</td><td>MP</td></tr>
<tr><td>>hg38_21_+_8433172_8446622_RNA45SN1_45S_50ntFlanks@6645.6674.30|CTCGTACGACTCTTAGCGGTGGATCACTCG</td><td>>hg38_21_+_8433172_8446622_RNA45SN1_45S_50ntFlanks</td><td>6645</td><td>6674</td><td>30</td><td>CTCGTACGACTCTTAGCGGTGGATCACTCG</td><td>1700.8</td><td>3.4</td><td>1.0</td><td>3.5</td><td>5.2E-04</td><td>6.4E-04</td><td>MB231/MB468</td><td>NUC</td></tr>
<tr><td>>MI0000298|hsa-mir-221&WithFlank&X|-|45746151|45746272@71.94.24&[MIMAT0000278&hsa-miR-221-3p&offsets|0|+1;m-91&X|-|45746181|45746202&offsets|0|+2]|AGCTACATTGTCTGCTGGGTTTCA</td><td>>MI0000298|hsa-mir-221&WithFlank&X|-|45746151|45746272</td><td>71</td><td>94</td><td>24</td><td>AGCTACATTGTCTGCTGGGTTTCA</td><td>1736.1</td><td>-2.2</td><td>0.5</td><td>-4.0</td><td>7.8E-05</td><td>6.4E-04</td><td>BT20/MB231</td><td>NUC</td></tr>
<tr><td>>hg38_21_+_8433172_8446622_RNA45SN1_45S_50ntFlanks@3840.3867.28|CTCCTACTTGGATAACTGTGGTAATTCT</td><td>>hg38_21_+_8433172_8446622_RNA45SN1_45S_50ntFlanks</td><td>3840</td><td>3867</td><td>28</td><td>CTCCTACTTGGATAACTGTGGTAATTCT</td><td>321.3</td><td>2.9</td><td>0.7</td><td>4.0</td><td>7.8E-05</td><td>6.4E-04</td><td>BT20/MB231</td><td>NUC</td></tr>
<tr><td>>MI0000446|hsa-mir-125b-1&WithFlank&11|-|122099751|122099850@21.39.19;MI0000470|hsa-mir-125b-2&WithFlank&21|+|16590231|16590331@23.41.19&[MIMAT0000423&hsa-miR-125b-5p&offsets|0|-3;m-67&11|-|122099809|122099830&offsets|0|-3];[MIMAT0000423_1&hsa-miR-125b-5p&offsets|0|-3;m-68&21|+|16590253|16590274&offsets|0|-3]|TCCCTGAGACCCTAACTTG</td><td>>MI0000446|hsa-mir-125b-1&WithFlank&11|-|122099751|122099850</td><td>21</td><td>39</td><td>19</td><td>TCCCTGAGACCCTAACTTG</td><td>1007.3</td><td>-6.4</td><td>1.6</td><td>-4.0</td><td>7.8E-05</td><td>6.4E-04</td><td>BT20/MB231</td><td>NUC</td></tr>
<tr><td>>trna119_LysCTT_1_-_145395522_145395594@4.36.33,trna11_LysCTT_5_-_180648979_180649051@4.36.33,trna13_LysCTT_6_+_26556774_26556846@4.36.33,trna7_LysCTT_16_+_3225692_3225764@4.36.33,trna9_LysCTT_5_+_180634755_180634827@4.36.33|CGGCTAGCTCAGTCGGTAGAGCATGAGACTCTT</td><td>>trna119_LysCTT_1_-_145395522_145395594</td><td>4</td><td>36</td><td>33</td><td>CGGCTAGCTCAGTCGGTAGAGCATGAGACTCTT</td><td>560.7</td><td>4.7</td><td>1.3</td><td>3.5</td><td>4.3E-04</td><td>6.4E-04</td><td>MB231/MB468</td><td>MP</td></tr>
<tr><td>>trna17_ValTAC_11_-_59318102_59318174@3.36.34,trna4_ValTAC_X_-_18693029_18693101@3.36.34|TTCCATAGTGTAGTGGTTATCACGTCTGCTTTAC</td><td>>trna17_ValTAC_11_-_59318102_59318174</td><td>3</td><td>36</td><td>34</td><td>TTCCATAGTGTAGTGGTTATCACGTCTGCTTTAC</td><td>151.8</td><td>5.1</td><td>1.5</td><td>3.5</td><td>5.1E-04</td><td>6.4E-04</td><td>MB231/MB468</td><td>MITO</td></tr>
<tr><td>>hg38_21_+_8433172_8446622_RNA45SN1_45S_50ntFlanks@11273.11293.21|CCGCCCCCGGGGAGCCCGGCG</td><td>>hg38_21_+_8433172_8446622_RNA45SN1_45S_50ntFlanks</td><td>11273</td><td>11293</td><td>21</td><td>CCGCCCCCGGGGAGCCCGGCG</td><td>273.5</td><td>-5.0</td><td>1.4</td><td>-3.5</td><td>4.6E-04</td><td>6.4E-04</td><td>BT20/MB468</td><td>CYTO</td></tr>
<tr><td>>hg38_21_+_8433172_8446622_RNA45SN1_45S_50ntFlanks@11708.11728.21|AGCGCGGGTAAACGGCGGGAG</td><td>>hg38_21_+_8433172_8446622_RNA45SN1_45S_50ntFlanks</td><td>11708</td><td>11728</td><td>21</td><td>AGCGCGGGTAAACGGCGGGAG</td><td>79.6</td><td>-5.5</td><td>1.5</td><td>-3.7</td><td>2.1E-04</td><td>6.4E-04</td><td>BT20/MB468</td><td>MP</td></tr>
<tr><td>>hg38_21_+_8433172_8446622_RNA45SN1_45S_50ntFlanks@9544.9571.28|GGTGGAGGTCCGTAGCGGTCCTGACGTG</td><td>>hg38_21_+_8433172_8446622_RNA45SN1_45S_50ntFlanks</td><td>9544</td><td>9571</td><td>28</td><td>GGTGGAGGTCCGTAGCGGTCCTGACGTG</td><td>1352.7</td><td>-4.7</td><td>1.3</td><td>-3.5</td><td>4.3E-04</td><td>6.4E-04</td><td>MB231/MB468</td><td>MP</td></tr>
<tr><td>>hg38_21_+_8433172_8446622_RNA45SN1_45S_50ntFlanks@10821.10840.20|GGAAGTCGGCAAGCCGGATC</td><td>>hg38_21_+_8433172_8446622_RNA45SN1_45S_50ntFlanks</td><td>10821</td><td>10840</td><td>20</td><td>GGAAGTCGGCAAGCCGGATC</td><td>4122.3</td><td>-4.2</td><td>1.1</td><td>-3.7</td><td>2.1E-04</td><td>6.4E-04</td><td>BT20/MB468</td><td>MP</td></tr>
<tr><td>>MI0003129|hsa-mir-146b&WithFlank&10|+|102436506|102436590@15.32.18&[MIMAT0002809&hsa-miR-146b-5p&offsets|0|-5;m-70&10|+|102436520|102436543&offsets|0|-6]|TGAGAACTGAATTCCATA</td><td>>MI0003129|hsa-mir-146b&WithFlank&10|+|102436506|102436590</td><td>15</td><td>32</td><td>18</td><td>TGAGAACTGAATTCCATA</td><td>123.6</td><td>-10.3</td><td>2.6</td><td>-4.0</td><td>7.4E-05</td><td>6.4E-04</td><td>BT20/MB231</td><td>MP</td></tr>
<tr><td>>hg38_21_+_8433172_8446622_RNA45SN1_45S_50ntFlanks@10843.10863.21|TAACTTCGGGATAAGGATTGG</td><td>>hg38_21_+_8433172_8446622_RNA45SN1_45S_50ntFlanks</td><td>10843</td><td>10863</td><td>21</td><td>TAACTTCGGGATAAGGATTGG</td><td>125.4</td><td>-5.7</td><td>1.5</td><td>-3.7</td><td>2.1E-04</td><td>6.4E-04</td><td>BT20/MB468</td><td>MP</td></tr>
<tr><td>>MI0000457|hsa-mir-141&WithFlank&12|+|6964091|6964197@65.86.22&[MIMAT0000432&hsa-miR-141-3p&offsets|0|0;m-134&12|+|6964155|6964175&offsets|0|+1]|TAACACTGTCTGGTAAAGATGG</td><td>>MI0000457|hsa-mir-141&WithFlank&12|+|6964091|6964197</td><td>65</td><td>86</td><td>22</td><td>TAACACTGTCTGGTAAAGATGG</td><td>115.9</td><td>10.2</td><td>2.6</td><td>4.0</td><td>7.4E-05</td><td>6.4E-04</td><td>BT20/MB231</td><td>MP</td></tr>
<tr><td>>hg38_21_+_8433172_8446622_RNA45SN1_45S_50ntFlanks@10680.10706.27|GTTCCGGCGGCGTCCGGTGAGCTCTCG</td><td>>hg38_21_+_8433172_8446622_RNA45SN1_45S_50ntFlanks</td><td>10680</td><td>10706</td><td>27</td><td>GTTCCGGCGGCGTCCGGTGAGCTCTCG</td><td>6901.6</td><td>-5.5</td><td>1.5</td><td>-3.7</td><td>2.1E-04</td><td>6.4E-04</td><td>BT20/MB468</td><td>MP</td></tr>
<tr><td>>MI0000071|hsa-mir-17&WithFlank&13|+|91350599|91350694@20.41.22(+1A)&[MIMAT0000070&hsa-miR-17-5p&offsets|0|-1(+1A);m-88&13|+|91350618|91350640&offsets|0|-1(+1A)]|CAAAGTGCTTACAGTGCAGGTAA</td><td>>MI0000071|hsa-mir-17&WithFlank&13|+|91350599|91350694</td><td>20</td><td>41</td><td>22(+1A)</td><td>CAAAGTGCTTACAGTGCAGGTAA</td><td>351.1</td><td>4.0</td><td>1.2</td><td>3.5</td><td>4.8E-04</td><td>6.4E-04</td><td>MB231/MB468</td><td>CYTO</td></tr>
<tr><td>>hg38_21_+_8433172_8446622_RNA45SN1_45S_50ntFlanks@12420.12450.31|ATGTCGGCTCTTCCTATCATTGTGAAGCAGA</td><td>>hg38_21_+_8433172_8446622_RNA45SN1_45S_50ntFlanks</td><td>12420</td><td>12450</td><td>31</td><td>ATGTCGGCTCTTCCTATCATTGTGAAGCAGA</td><td>608.2</td><td>-3.1</td><td>0.8</td><td>-3.9</td><td>8.9E-05</td><td>6.4E-04</td><td>BT20/MB231</td><td>MITO</td></tr>
<tr><td>>hg38_21_+_8433172_8446622_RNA45SN1_45S_50ntFlanks@12420.12452.33|ATGTCGGCTCTTCCTATCATTGTGAAGCAGAAT</td><td>>hg38_21_+_8433172_8446622_RNA45SN1_45S_50ntFlanks</td><td>12420</td><td>12452</td><td>33</td><td>ATGTCGGCTCTTCCTATCATTGTGAAGCAGAAT</td><td>802.6</td><td>-3.0</td><td>0.8</td><td>-3.9</td><td>8.9E-05</td><td>6.4E-04</td><td>BT20/MB231</td><td>MITO</td></tr>
<tr><td>>hg38_21_+_8433172_8446622_RNA45SN1_45S_50ntFlanks@12997.13037.41|CGCTGCGATCTATTGAAAGTCAGCCCTCGACACAAGGGTTT</td><td>>hg38_21_+_8433172_8446622_RNA45SN1_45S_50ntFlanks</td><td>12997</td><td>13037</td><td>41</td><td>CGCTGCGATCTATTGAAAGTCAGCCCTCGACACAAGGGTTT</td><td>472.9</td><td>-2.9</td><td>0.8</td><td>-3.9</td><td>9.0E-05</td><td>6.4E-04</td><td>BT20/MB231</td><td>MITO</td></tr>
<tr><td>>hg38_21_+_8433172_8446622_RNA45SN1_45S_50ntFlanks@12444.12481.38|AAGCAGAATTCACCAAGCGTTGGATTGTTCACCCACTA</td><td>>hg38_21_+_8433172_8446622_RNA45SN1_45S_50ntFlanks</td><td>12444</td><td>12481</td><td>38</td><td>AAGCAGAATTCACCAAGCGTTGGATTGTTCACCCACTA</td><td>191.8</td><td>-4.4</td><td>1.3</td><td>-3.5</td><td>5.1E-04</td><td>6.4E-04</td><td>MB231/MB468</td><td>MITO</td></tr>
<tr><td>>MI0005767|hsa-mir-942&WithFlank&1|+|117094637|117094734@21.41.21&[MIMAT0004985&hsa-miR-942-5p&offsets|+2|+1;m-292&1|+|117094655|117094677&offsets|+2|0]|TTCTCTGTTTTGGCCATGTGT</td><td>>MI0005767|hsa-mir-942&WithFlank&1|+|117094637|117094734</td><td>21</td><td>41</td><td>21</td><td>TTCTCTGTTTTGGCCATGTGT</td><td>105.8</td><td>7.1</td><td>1.9</td><td>3.7</td><td>2.1E-04</td><td>6.4E-04</td><td>BT20/MB468</td><td>MP</td></tr>
<tr><td>>hg38_21_+_8433172_8446622_RNA45SN1_45S_50ntFlanks@12437.12474.38|CATTGTGAAGCAGAATTCACCAAGCGTTGGATTGTTCA</td><td>>hg38_21_+_8433172_8446622_RNA45SN1_45S_50ntFlanks</td><td>12437</td><td>12474</td><td>38</td><td>CATTGTGAAGCAGAATTCACCAAGCGTTGGATTGTTCA</td><td>229.6</td><td>-6.4</td><td>1.7</td><td>-3.7</td><td>2.1E-04</td><td>6.4E-04</td><td>BT20/MB468</td><td>MP</td></tr>
<tr><td>>hg38_21_+_8433172_8446622_RNA45SN1_45S_50ntFlanks@11564.11581.18|CCTAGCAGCCGACTTAGA</td><td>>hg38_21_+_8433172_8446622_RNA45SN1_45S_50ntFlanks</td><td>11564</td><td>11581</td><td>18</td><td>CCTAGCAGCCGACTTAGA</td><td>323.4</td><td>-4.7</td><td>1.3</td><td>-3.7</td><td>2.1E-04</td><td>6.4E-04</td><td>BT20/MB468</td><td>MP</td></tr>
<tr><td>>hg38_21_+_8433172_8446622_RNA45SN1_45S_50ntFlanks@3874.3899.26|AATACATGCCGACGGGCGCTGACCCC</td><td>>hg38_21_+_8433172_8446622_RNA45SN1_45S_50ntFlanks</td><td>3874</td><td>3899</td><td>26</td><td>AATACATGCCGACGGGCGCTGACCCC</td><td>1350.2</td><td>-5.6</td><td>1.6</td><td>-3.5</td><td>5.2E-04</td><td>6.4E-04</td><td>BT20/MB468</td><td>NUC</td></tr>
</table>

| | | | | | | | | | | | | | |
|---|---|---|---|---|---|---|---|---|---|---|---|---|---|
| >MI0000446\|hsa-mir-125b-1&WithFlank&11\|-\|122099751\|122099850@21.38.18(+1A);MI0000470\|hsa-mir-125b-2&WithFlank&21\|+\|16590231\|16590331@23.40.18(+1A)&[MIMAT0000423&hsa-miR-125b-5p&offsets\|0\|-4(+1A);m-67&11\|-\|122099809\|122099830&offsets\|0\|-4(+1A)];[MIMAT0000423_1&hsa-miR-125b-5p&offsets\|0\|-4(+1A);m-68&21\|+\|16590253\|16590274&offsets\|0\|-4(+1A)]\|TCCCTGAGACCCTAACTTA | >MI0000446\|hsa-mir-125b-1&WithFlank&11\|-\|122099751\|122099850 | 21 | 38 | 18(+1A) | TCCCTGAGACCCTAACTTA | 123.4 | -10.3 | 2.6 | -4.0 | 7.5E-05 | 6.4E-04 | BT20/MB231 | MP |
| >trna14_LysTTT_11_-_59327808_59327880@1.24.24,trna2_LysTTT_17_+_8022473_8022545@1.24.24,trna54_LysTTT_1_+_204475655_204475727@1.24.24,trna5_LysTTT_11_+_59323902_59323974@1.24.24,trna62_LysTTT_1_-_204476158_204476230@1.24.24,trna76_LysTTT_6_+_28918806_28918878@1.24.24\|GCCCGGATAGCTCAGTCGGTAGAG | >trna14_LysTTT_11_-_59327808_59327880 | 1 | 24 | 24 | GCCCGGATAGCTCAGTCGGTAGAG | 302.9 | -4.1 | 1.2 | -3.5 | 5.2E-04 | 6.4E-04 | MB231/MB468 | NUC |
| >MI0000289\|hsa-mir-181a-1&WithFlank&1\|-\|198859038\|198859159@30.53.24(+1C);MI0000269\|hsa-mir-181a-2&WithFlank&9\|+\|124692436\|124692557@45.68.24(+1C)&[MIMAT0000256&hsa-miR-181a-5p&offsets\|0\|+1(+1C);m-32&1\|-\|198859108\|198859130&offsets\|0\|+1(+1C)];[MIMAT0000256_1&hsa-miR-181a-5p&offsets\|0\|+1(+1C);m-32&9\|+\|124692480\|124692502&offsets\|0\|+1(+1C)]\|AACATTCAACGCTGTCGGTGAGTTC | >MI0000289\|hsa-mir-181a-1&WithFlank&1\|-\|198859038\|198859159 | 30 | 53 | 24(+1C) | AACATTCAACGCTGTCGGTGAGTTC | 1252.9 | -5.6 | 1.4 | -4.0 | 7.5E-05 | 6.4E-04 | BT20/MB231 | MP |
| >am_trnaMT_PheGAA_MT_+_577_647@40.65.26\|TGTTTAGACGGGCTCACATCACCCCA | >am_trnaMT_PheGAA_MT_+_577_647 | 40 | 65 | 26 | TGTTTAGACGGGCTCACATCACCCCA | 76.1 | 6.1 | 1.7 | 3.5 | 4.3E-04 | 6.4E-04 | MB231/MB468 | MP |
| >hg38_21_+_8433172_8446622_RNA45SN1_45S_50ntFlanks@10731.10757.27\|AGGGTGTAAATCTCGCGCCGGGCCGTA | >hg38_21_+_8433172_8446622_RNA45SN1_45S_50ntFlanks | 10731 | 10757 | 27 | AGGGTGTAAATCTCGCGCCGGGCCGTA | 144.1 | -2.9 | 0.8 | -3.7 | 2.1E-04 | 6.5E-04 | BT20/MB468 | MP |
| >hg38_21_+_8433172_8446622_RNA45SN1_45S_50ntFlanks@12243.12275.33\|GAAGGGCAAAAGCTCGCTTGATCTTGATTTTCA | >hg38_21_+_8433172_8446622_RNA45SN1_45S_50ntFlanks | 12243 | 12275 | 33 | GAAGGGCAAAAGCTCGCTTGATCTTGATTTTCA | 144.1 | 7.3 | 2.1 | 3.5 | 4.9E-04 | 6.5E-04 | MB231/MB468 | CYTO |
| >MI0000439\|hsa-mir-23b&WithFlank&9\|+\|95085202\|95085310@67.85.19(+2U)&[MIMAT0000418&hsa-miR-23b-3p&offsets\|+3\|-1(+2U);m-42&9\|+\|95085265\|95085285&offsets\|+3\|+1(+2U)]\|ACATTGCCAGGGATTACCATT | >MI0000439\|hsa-mir-23b&WithFlank&9\|+\|95085202\|95085310 | 67 | 85 | 19(+2U) | ACATTGCCAGGGATTACCATT | 416.2 | 12.6 | 3.6 | 3.5 | 5.2E-04 | 6.5E-04 | MB231/MB468 | NUC |
| >am_trna106_PheGAA_6_-_28775610_28775682@31.52.22,trna109_PheGAA_6_-_28758499_28758571@31.52.22,trna11_PheGAA_12_-_125412389_125412461@31.52.22,trna13_PheGAA_11_-_59333853_59333925@31.52.22,trna14_PheGAA_19_-_1383361_1383433@31.52.22,trna15_PheGAA_11_-_59324970_59325042@31.52.22,trna1_PheGAA_13_-_95201904_95201976@31.52.22,trna72_PheGAA_6_+_28732164_28732236@31.52.22,trna96_PheGAA_6_-_28949449_28949521@31.52.22\|ACTGAAGATCTAAAGGTCCCTG | >am_trna106_PheGAA_6_-_28775610_28775682 | 31 | 52 | 22 | ACTGAAGATCTAAAGGTCCCTG | 148.2 | -4.7 | 1.3 | -3.5 | 4.6E-04 | 6.5E-04 | BT20/MB468 | CYTO |
| >hg38_21_+_8433172_8446622_RNA45SN1_45S_50ntFlanks@9504.9527.24\|GTGAACTATGCCTGGGCAGGGCGA | >hg38_21_+_8433172_8446622_RNA45SN1_45S_50ntFlanks | 9504 | 9527 | 24 | GTGAACTATGCCTGGGCAGGGCGA | 276.6 | -2.8 | 0.8 | -3.5 | 3.9E-04 | 6.5E-04 | BT20/MB468 | MITO |
| >MI0003628\|hsa-mir-615&WithFlank&12\|+\|54033944\|54034051@67.86.20&[MIMAT0003283&hsa-miR-615-3p&offsets\|0\|-2;m-495&12\|+\|54034010\|54034030&offsets\|0\|-1]\|TCCGAGCCTGGGTCTCCCTC | >MI0003628\|hsa-mir-615&WithFlank&12\|+\|54033944\|54034051 | 67 | 86 | 20 | TCCGAGCCTGGGTCTCCCTC | 89.3 | 6.7 | 1.8 | 3.7 | 2.1E-04 | 6.5E-04 | BT20/MB468 | MP |
| >MI0000079\|hsa-mir-23a&WithFlank&19\|-\|13836581\|13836665@54.71.18(+1U)&[MIMAT0000078&hsa-miR-23a-3p&offsets\|+3\|0(+1U);m-31&19\|-\|13836595\|13836615&offsets\|+3\|0(+1U)]\|ACATTGCCAGGGATTTCCT | >MI0000079\|hsa-mir-23a&WithFlank&19\|-\|13836581\|13836665 | 54 | 71 | 18(+1U) | ACATTGCCAGGGATTTCCT | 65.3 | 9.3 | 2.6 | 3.5 | 4.3E-04 | 6.5E-04 | MB231/MB468 | MP |
| >hg38_21_+_8433172_8446622_RNA45SN1_45S_50ntFlanks@10341.10369.29\|AATATTCAAACGAGAACTTTGAAGGCCGA | >hg38_21_+_8433172_8446622_RNA45SN1_45S_50ntFlanks | 10341 | 10369 | 29 | AATATTCAAACGAGAACTTTGAAGGCCGA | 100.6 | 5.8 | 1.6 | 3.7 | 2.1E-04 | 6.5E-04 | BT20/MB468 | MP |
| >hg38_21_+_8433172_8446622_RNA45SN1_45S_50ntFlanks@9250.9271.22\|CGCACGGGGTCGGCGGCGACGT | >hg38_21_+_8433172_8446622_RNA45SN1_45S_50ntFlanks | 9250 | 9271 | 22 | CGCACGGGGTCGGCGGCGACGT | 94.4 | -2.9 | 0.8 | -3.5 | 4.3E-04 | 6.5E-04 | MB231/MB468 | MP |
| >MI0000471\|hsa-mir-126&WithFlank&9\|+\|136670596\|136670692@58.77.20&[MIMAT0000445&hsa-miR-126-3p&offsets\|0\|-2;m-40&9\|+\|136670653\|136670674&offsets\|0\|-2]\|TCGTACCGTGAGTAATAATG | >MI0000471\|hsa-mir-126&WithFlank&9\|+\|136670596\|136670692 | 58 | 77 | 20 | TCGTACCGTGAGTAATAATG | 305.0 | 7.2 | 1.9 | 3.7 | 2.1E-04 | 6.5E-04 | BT20/MB468 | MP |
| >am_trnaMT_ProTGG_MT_-_15956_16023@52.71.20\|AAAGACTTTTTCTCTGACCA | >am_trnaMT_ProTGG_MT_-_15956_16023 | 52 | 71 | 20 | AAAGACTTTTTCTCTGACCA | 2401.2 | 3.2 | 0.9 | 3.5 | 5.2E-04 | 6.5E-04 | MB231/MB468 | MITO |
| >trna10_AspGTC_12_-_125424193_125424264@32.75.44,trna12_AspGTC_12_-_125411891_125411962@32.75.44,trna144_AspGTC_6_-_27551236_27551307@32.75.44,trna38_AspGTC_17_-_8125556_8125627@32.75.44,trna45_AspGTC_6_+_27447453_27447524@32.75.44,trna48_AspGTC_6_+_27471523_27471594@32.75.44,trna4_AspGTC_12_+_96429799_96429870@32.75.44,trna69_AspGTC_1_-_161440205_161440276@32.75.44,trna72_AspGTC_1_-_161432824_161432895@32.75.44,trna75_AspGTC_1_-_161425414_161425485@32.75.44,trna78_AspGTC_1_-_161418033_161418104@32.75.44,trna81_AspGTC_1_-_161410615_161410686@32.75.44\|CTGTCACGCGGGAGACCGGGGTTCGATTCCCCGACGGGGAGCCA | >trna10_AspGTC_12_-_125424193_125424264 | 32 | 75 | 44 | CTGTCACGCGGGAGACCGGGGTTCGATTCCCCGACGGGGAGCCA | 268.5 | -3.5 | 1.0 | -3.5 | 4.9E-04 | 6.5E-04 | MB231/MB468 | CYTO |
| >hg38_21_+_8433172_8446622_RNA45SN1_45S_50ntFlanks@11602.11640.39\|TCCGACTGTTTAATTAAAACAAAGCATCGCGAAGGCCCG | >hg38_21_+_8433172_8446622_RNA45SN1_45S_50ntFlanks | 11602 | 11640 | 39 | TCCGACTGTTTAATTAAAACAAAGCATCGCGAAGGCCCG | 96.9 | -5.0 | 1.4 | -3.5 | 3.9E-04 | 6.5E-04 | BT20/MB468 | MITO |
| >hg38_21_+_8433172_8446622_RNA45SN1_45S_50ntFlanks@12459.12477.19\|AGCGTTGGATTGTTCACCC | >hg38_21_+_8433172_8446622_RNA45SN1_45S_50ntFlanks | 12459 | 12477 | 19 | AGCGTTGGATTGTTCACCC | 190.3 | -3.8 | 1.1 | -3.5 | 3.9E-04 | 6.5E-04 | BT20/MB468 | MITO |
| >hg38_21_+_8433172_8446622_RNA45SN1_45S_50ntFlanks@12957.12989.33\|CTTCTGGGTCGGGGTTTCGTACGTAGCAGAGCA | >hg38_21_+_8433172_8446622_RNA45SN1_45S_50ntFlanks | 12957 | 12989 | 33 | CTTCTGGGTCGGGGTTTCGTACGTAGCAGAGCA | 191.5 | -3.0 | 0.9 | -3.5 | 4.9E-04 | 6.5E-04 | MB231/MB468 | CYTO |
| >hg38_21_+_8433172_8446622_RNA45SN1_45S_50ntFlanks@6646.6677.32\|TCGTACGACTCTTAGCGGTGGATCACTCGGCT | >hg38_21_+_8433172_8446622_RNA45SN1_45S_50ntFlanks | 6646 | 6677 | 32 | TCGTACGACTCTTAGCGGTGGATCACTCGGCT | 3186.9 | 2.7 | 0.8 | 3.5 | 5.2E-04 | 6.5E-04 | MB231/MB468 | NUC |
| >MI0000466\|hsa-mir-9-1&WithFlank&1\|-\|156420335\|156420435@22.42.21(+1A);MI0000468\|hsa-mir-9-3&WithFlank&15\|+\|89368011\|89368112@22.42.21(+1A);MI0000467\|hsa-mir-9-2&WithFlank&5\|-\|88666847\|88666945@22.42.21(+1A)&[MIMAT0000441&hsa-miR-9-5p&offsets\|0\|-2(+1A);m-104&1\|-\|156420392\|156420414&offsets\|0\|-2(+1A)];[MIMAT0000441_1&hsa-miR-9-5p&offsets\|0\|-2(+1A);m-105&15\|+\|89368032\|89368054&offsets\|0\|-2(+1A)];[MIMAT0000441_2&hsa-miR-9-5p&offsets\|0\|-2(+1A);m-104&5\|-\|88666902\|88666924&offsets\|0\|-2(+1A)]\|TCTTTGGTTATCTAGCTGTATA | >MI0000466\|hsa-mir-9-1&WithFlank&1\|-\|156420335\|156420435 | 22 | 42 | 21(+1A) | TCTTTGGTTATCTAGCTGTATA | 158.0 | 4.2 | 1.1 | 4.0 | 7.7E-05 | 6.5E-04 | BT20/MB231 | CYTO |
| >hg38_21_+_8433172_8446622_RNA45SN1_45S_50ntFlanks@11701.11728.28\|TCAATGAAGCGCGGGTAAACGGCGGGAG | >hg38_21_+_8433172_8446622_RNA45SN1_45S_50ntFlanks | 11701 | 11728 | 28 | TCAATGAAGCGCGGGTAAACGGCGGGAG | 127.5 | -5.8 | 1.6 | -3.7 | 2.1E-04 | 6.5E-04 | BT20/MB468 | MP |
| >hg38_21_+_8433172_8446622_RNA45SN1_45S_50ntFlanks@9545.9569.25\|GTGGAGGTCCGTAGCGGTCCTGACG | >hg38_21_+_8433172_8446622_RNA45SN1_45S_50ntFlanks | 9545 | 9569 | 25 | GTGGAGGTCCGTAGCGGTCCTGACG | 523.2 | -4.9 | 1.3 | -3.7 | 2.1E-04 | 6.5E-04 | BT20/MB468 | MP |
| >hg38_21_+_8433172_8446622_RNA45SN1_45S_50ntFlanks@12418.12460.43\|CGATGTCGGCTCTTCCTATCATTGTGAAGCAGAATTCACCAAG | >hg38_21_+_8433172_8446622_RNA45SN1_45S_50ntFlanks | 12418 | 12460 | 43 | CGATGTCGGCTCTTCCTATCATTGTGAAGCAGAATTCACCAAG | 162.3 | -4.1 | 1.2 | -3.5 | 4.6E-04 | 6.5E-04 | BT20/MB468 | CYTO |
| >trna2_GlyGCC_21_-_18827107_18827177@1.33.33,trna35_GlyGCC_1_+_161413094_161413164@1.33.33,trna37_GlyGCC_1_+_161420467_161420537@1.33.33,trna39_GlyGCC_1_+_161427898_161427968@1.33.33,trna41_GlyGCC_1_+_161435258_161435328@1.33.33\|GCATGGGTGGTTCAGTGGTAGAATTCTCGCCTG | >trna2_GlyGCC_21_-_18827107_18827177 | 1 | 33 | 33 | GCATGGGTGGTTCAGTGGTAGAATTCTCGCCTG | 1703.7 | 5.0 | 1.3 | 3.7 | 2.1E-04 | 6.5E-04 | BT20/MB468 | MP |
| >hg38_21_+_8433172_8446622_RNA45SN1_45S_50ntFlanks@11603.11620.18\|CCGACTGTTTAATTAAAA | >hg38_21_+_8433172_8446622_RNA45SN1_45S_50ntFlanks | 11603 | 11620 | 18 | CCGACTGTTTAATTAAAA | 623.8 | -2.9 | 0.8 | -3.5 | 4.7E-04 | 6.5E-04 | BT20/MB468 | CYTO |
| >MI0003140\|hsa-mir-512-1&WithFlank&19\|+\|53666673\|53666768@57.78.22;MI0003141\|hsa-mir-512-2&WithFlank&19\|+\|53669151\|53669260@63.84.22&[MIMAT0002823&hsa-miR-512-3p&offsets\|0\|0;m-849&19\|+\|53666729\|53666751&offsets\|0\|-1];[MIMAT0002823_1&hsa-miR-512-3p&offsets\|0\|0;m-849&19\|+\|53669213\|53669235&offsets\|0\|-1]\|AAGTGCTGTCATAGCTGAGGTC | >MI0003140\|hsa-mir-512-1&WithFlank&19\|+\|53666673\|53666768 | 57 | 78 | 22 | AAGTGCTGTCATAGCTGAGGTC | 46.4 | 8.9 | 2.2 | 4.0 | 7.6E-05 | 6.5E-04 | BT20/MB231 | MP |
| >hg38_21_+_8433172_8446622_RNA45SN1_45S_50ntFlanks@12394.12422.29\|TAGCGACGTCGCTTTTTGATCCTTCGATG | >hg38_21_+_8433172_8446622_RNA45SN1_45S_50ntFlanks | 12394 | 12422 | 29 | TAGCGACGTCGCTTTTTGATCCTTCGATG | 282.0 | -4.4 | 1.3 | -3.5 | 5.2E-04 | 6.5E-04 | MB231/MB468 | MITO |
| >MI0000286\|hsa-mir-210&WithFlank&11\|-\|568083\|568204@72.93.22&[MIMAT0000267&hsa-miR-210-3p&offsets\|0\|0;m-102&11\|-\|568112\|568133&offsets\|0\|0]\|CTGTGCGTGTGACAGCGGCTGA | >MI0000286\|hsa-mir-210&WithFlank&11\|-\|568083\|568204 | 72 | 93 | 22 | CTGTGCGTGTGACAGCGGCTGA | 758.2 | 4.0 | 1.0 | 4.0 | 7.8E-05 | 6.5E-04 | BT20/MB231 | CYTO |
| >trnaMT_LeuTAA_MT_+_3230_3304@19.44.26\|GGTAATCGCATAAAACTTAAAACTTT | >trnaMT_LeuTAA_MT_+_3230_3304 | 19 | 44 | 26 | GGTAATCGCATAAAACTTAAAACTTT | 414.9 | 3.1 | 0.8 | 4.0 | 7.8E-05 | 6.5E-04 | BT20/MB231 | CYTO |
| >hg38_21_+_8433172_8446622_RNA45SN1_45S_50ntFlanks@7976.8011.36\|GCGACCTCAGATCAGACGTGGCGACCCGCTGAATTT | >hg38_21_+_8433172_8446622_RNA45SN1_45S_50ntFlanks | 7976 | 8011 | 36 | GCGACCTCAGATCAGACGTGGCGACCCGCTGAATTT | 1765.3 | -3.1 | 0.8 | -4.0 | 7.8E-05 | 6.5E-04 | BT20/MB231 | CYTO |
| >hg38_21_+_8433172_8446622_RNA45SN1_45S_50ntFlanks@9306.9323.18\|CAAGGAGTCTAACACGTG | >hg38_21_+_8433172_8446622_RNA45SN1_45S_50ntFlanks | 9306 | 9323 | 18 | CAAGGAGTCTAACACGTG | 980.9 | -3.8 | 1.0 | -3.7 | 2.2E-04 | 6.5E-04 | BT20/MB468 | MP |
| >hg38_MT_+_1621_3279_RNR2_16S_50ntFlanks@1367.1385.19\|TCGATGTTGGATCAGGACA | >hg38_MT_+_1621_3279_RNR2_16S_50ntFlanks | 1367 | 1385 | 19 | TCGATGTTGGATCAGGACA | 266.0 | 3.6 | 1.0 | 3.5 | 4.4E-04 | 6.5E-04 | MB231/MB468 | MP |
| >MI0000434\|hsa-let-7i&WithFlank&12\|+\|62603680\|62603775@12.33.22(+2U)&[MIMAT0000415&hsa-let-7i-5p&offsets\|0\|0(+2U);m-49&12\|+\|62603691\|62603712&offsets\|0\|0(+2U)]\|TGAGGTAGTAGTTTGTGCTGTTTT | >MI0000434\|hsa-let-7i&WithFlank&12\|+\|62603680\|62603775 | 12 | 33 | 22(+2U) | TGAGGTAGTAGTTTGTGCTGTTTT | 118.9 | 6.2 | 1.8 | 3.5 | 5.2E-04 | 6.5E-04 | MB231/MB468 | MITO |
| >hg38_21_+_8433172_8446622_RNA45SN1_45S_50ntFlanks@7975.8018.44\|CGCGACCTCAGATCAGACGTGGCGACCCGCTGAATTTAAGCATA | >hg38_21_+_8433172_8446622_RNA45SN1_45S_50ntFlanks | 7975 | 8018 | 44 | CGCGACCTCAGATCAGACGTGGCGACCCGCTGAATTTAAGCATA | 1093.8 | 6.0 | 1.7 | 3.5 | 4.7E-04 | 6.5E-04 | BT20/MB468 | CYTO |
| >MI0000272\|hsa-mir-182&WithFlank&7\|-\|129770377\|129770498@29.50.22(+1A)&[MIMAT0000259&hsa-miR-182-5p&offsets\|0\|-2(+1A);m-22&7\|-\|129770449\|129770470&offsets\|0\|0(+1A)]\|TTTGGCAATGGTAGAACTCACAA | >MI0000272\|hsa-mir-182&WithFlank&7\|-\|129770377\|129770498 | 29 | 50 | 22(+1A) | TTTGGCAATGGTAGAACTCACAA | 210.5 | 7.0 | 1.8 | 3.9 | 7.8E-05 | 6.5E-04 | BT20/MB231 | CYTO |
| >MI0000542\|hsa-mir-320a&WithFlank&8\|-\|22244960\|22245043@49.69.21(+2U)&[MIMAT0000510&hsa-miR-320a-3p&offsets\|+1\|0(+2U);m-60&8\|-\|22244975\|22244996&offsets\|+1\|0(+2U)]\|AAAGCTGGGTTGAGAGGGCGATT | >MI0000542\|hsa-mir-320a&WithFlank&8\|-\|22244960\|22245043 | 49 | 69 | 21(+2U) | AAAGCTGGGTTGAGAGGGCGATT | 270.6 | 7.3 | 2.0 | 3.7 | 2.2E-04 | 6.5E-04 | BT20/MB468 | MP |

<table>
<tr><td>>hg38_21_+_8433172_8446622_RNA45SN1_45S_50ntFlanks@11924.11947.24|GAGGTGTAGAATAAGTGGGAGGCC</td><td>>hg38_21_+_8433172_8446622_RNA45SN1_45S_50ntFlanks</td><td>11924</td><td>11947</td><td>24</td><td>GAGGTGTAGAATAAGTGGGAGGCC</td><td>157.2</td><td>10.6</td><td>2.7</td><td>4.0</td><td>7.6E-05</td><td>6.5E-04</td><td>BT20/MB231</td><td>MP</td></tr>
<tr><td>>hg38_21_+_8433172_8446622_RNA45SN1_45S_50ntFlanks@10726.10755.30|GGGAGAGGGTGTAAATCTCGCGCCGGGCCG</td><td>>hg38_21_+_8433172_8446622_RNA45SN1_45S_50ntFlanks</td><td>10726</td><td>10755</td><td>30</td><td>GGGAGAGGGTGTAAATCTCGCGCCGGGCCG</td><td>139.7</td><td>-4.7</td><td>1.3</td><td>-3.5</td><td>4.7E-04</td><td>6.5E-04</td><td>BT20/MB468</td><td>CYTO</td></tr>
<tr><td>>hg38_21_+_8433172_8446622_RNA45SN1_45S_50ntFlanks@6733.6770.38|CATTGATCATCGACACTTCGAACGCACTTGCGGCCCCG</td><td>>hg38_21_+_8433172_8446622_RNA45SN1_45S_50ntFlanks</td><td>6733</td><td>6770</td><td>38</td><td>CATTGATCATCGACACTTCGAACGCACTTGCGGCCCCG</td><td>899.7</td><td>-5.5</td><td>1.6</td><td>-3.5</td><td>4.7E-04</td><td>6.5E-04</td><td>BT20/MB468</td><td>CYTO</td></tr>
<tr><td>>hg38_21_+_8433172_8446622_RNA45SN1_45S_50ntFlanks@6652.6689.38|GACTCTTAGCGGTGGATCACTCGGCTCGTGCGTCGATG</td><td>>hg38_21_+_8433172_8446622_RNA45SN1_45S_50ntFlanks</td><td>6652</td><td>6689</td><td>38</td><td>GACTCTTAGCGGTGGATCACTCGGCTCGTGCGTCGATG</td><td>7441.9</td><td>-3.8</td><td>1.1</td><td>-3.5</td><td>4.4E-04</td><td>6.5E-04</td><td>MB231/MB468</td><td>MP</td></tr>
<tr><td>>hg38_21_+_8433172_8446622_RNA45SN1_45S_50ntFlanks@11671.11707.37|CCAGTGCTCTGAATGTCAAAGTGAAGAAATTCAATGA</td><td>>hg38_21_+_8433172_8446622_RNA45SN1_45S_50ntFlanks</td><td>11671</td><td>11707</td><td>37</td><td>CCAGTGCTCTGAATGTCAAAGTGAAGAAATTCAATGA</td><td>628.1</td><td>-2.2</td><td>0.6</td><td>-3.5</td><td>5.3E-04</td><td>6.5E-04</td><td>MB231/MB468</td><td>NUC</td></tr>
<tr><td>>hg38_21_+_8433172_8446622_RNA45SN1_45S_50ntFlanks@6719.6747.29|TGAATTGCAGGACACATTGATCATCGACA</td><td>>hg38_21_+_8433172_8446622_RNA45SN1_45S_50ntFlanks</td><td>6719</td><td>6747</td><td>29</td><td>TGAATTGCAGGACACATTGATCATCGACA</td><td>1319.6</td><td>-3.0</td><td>0.8</td><td>-3.5</td><td>4.9E-04</td><td>6.5E-04</td><td>MB231/MB468</td><td>CYTO</td></tr>
<tr><td>>hg38_1_-_228634819_228635039_RNA5S12_5S_50ntFlanks@154.172.19|CCGGGTGCTGTAGGCTTTT</td><td>>hg38_1_-_228634819_228635039_RNA5S12_5S_50ntFlanks</td><td>154</td><td>172</td><td>19</td><td>CCGGGTGCTGTAGGCTTTT</td><td>184.3</td><td>-5.2</td><td>1.5</td><td>-3.5</td><td>4.4E-04</td><td>6.6E-04</td><td>MB231/MB468</td><td>MP</td></tr>
<tr><td>>hg38_21_+_8433172_8446622_RNA45SN1_45S_50ntFlanks@9914.9938.25|GAAAAGGTGTTGGTTGATATAGACA</td><td>>hg38_21_+_8433172_8446622_RNA45SN1_45S_50ntFlanks</td><td>9914</td><td>9938</td><td>25</td><td>GAAAAGGTGTTGGTTGATATAGACA</td><td>109.3</td><td>5.8</td><td>1.6</td><td>3.7</td><td>2.2E-04</td><td>6.6E-04</td><td>BT20/MB468</td><td>MP</td></tr>
<tr><td>>hg38_21_+_8433172_8446622_RNA45SN1_45S_50ntFlanks@10349.10379.31|AACGAGAACTTTGAAGGCCGAAGTGGAGAAG</td><td>>hg38_21_+_8433172_8446622_RNA45SN1_45S_50ntFlanks</td><td>10349</td><td>10379</td><td>31</td><td>AACGAGAACTTTGAAGGCCGAAGTGGAGAAG</td><td>464.5</td><td>1.9</td><td>0.5</td><td>3.5</td><td>4.7E-04</td><td>6.6E-04</td><td>BT20/MB468</td><td>CYTO</td></tr>
<tr><td>>MI0000804|hsa-mir-328&WithFlank&16|-|67202315|67202401@54.75.22&[MIMAT0000752&hsa-miR-328-3p&offsets|0|0;m-160&16|-|67202327|67202348&offsets|0|0]|CTGGCCCTCTCTGCCCTTCCGT</td><td>>MI0000804|hsa-mir-328&WithFlank&16|-|67202315|67202401</td><td>54</td><td>75</td><td>22</td><td>CTGGCCCTCTCTGCCCTTCCGT</td><td>1550.4</td><td>3.3</td><td>0.8</td><td>3.9</td><td>7.9E-05</td><td>6.6E-04</td><td>BT20/MB231</td><td>CYTO</td></tr>
<tr><td>>trna134_GluTTC_1_-_16861774_16861845@1.33.33,trna5_GluTTC_1_+_17199078_17199149@1.33.33,trna84_GluTTC_1_-_161391883_161391954@1.33.33,trna94_GluTTC_1_-_149664355_149664427@1.33.33|TCCCTGGTGGTCTAGTGGCTAGGATTCGGCGCT</td><td>>trna134_GluTTC_1_-_16861774_16861845</td><td>1</td><td>33</td><td>33</td><td>TCCCTGGTGGTCTAGTGGCTAGGATTCGGCGCT</td><td>6747.7</td><td>3.1</td><td>0.8</td><td>3.9</td><td>7.9E-05</td><td>6.6E-04</td><td>BT20/MB231</td><td>CYTO</td></tr>
<tr><td>>hg38_21_+_8433172_8446622_RNA45SN1_45S_50ntFlanks@8182.8223.42|ATCGAGGCCCAGCCCGTGGACGGTGTGAGGCCGGTAGCGGCC</td><td>>hg38_21_+_8433172_8446622_RNA45SN1_45S_50ntFlanks</td><td>8182</td><td>8223</td><td>42</td><td>ATCGAGGCCCAGCCCGTGGACGGTGTGAGGCCGGTAGCGGCC</td><td>376.9</td><td>-4.8</td><td>1.4</td><td>-3.5</td><td>5.0E-04</td><td>6.6E-04</td><td>MB231/MB468</td><td>CYTO</td></tr>
<tr><td>>hg38_21_+_8433172_8446622_RNA45SN1_45S_50ntFlanks@8356.8377.22|GAAAAGAACTTTGAAGAGAGAG</td><td>>hg38_21_+_8433172_8446622_RNA45SN1_45S_50ntFlanks</td><td>8356</td><td>8377</td><td>22</td><td>GAAAAGAACTTTGAAGAGAGAG</td><td>1771.3</td><td>-6.7</td><td>1.8</td><td>-3.7</td><td>2.2E-04</td><td>6.6E-04</td><td>BT20/MB468</td><td>MP</td></tr>
<tr><td>>trna152_ValCAC_6_-_27248049_27248121@1.36.36|GCTTCTGTAGTGTAGTGGTTATCACGTTCGCCTCAC</td><td>>trna152_ValCAC_6_-_27248049_27248121</td><td>1</td><td>36</td><td>36</td><td>GCTTCTGTAGTGTAGTGGTTATCACGTTCGCCTCAC</td><td>1973.5</td><td>-3.9</td><td>1.0</td><td>-3.9</td><td>7.9E-05</td><td>6.6E-04</td><td>BT20/MB231</td><td>CYTO</td></tr>
<tr><td>>hg38_21_+_8433172_8446622_RNA45SN1_45S_50ntFlanks@12418.12456.39|CGATGTCGGCTCTTCCTATCATTGTGAAGCAGAATTCAC</td><td>>hg38_21_+_8433172_8446622_RNA45SN1_45S_50ntFlanks</td><td>12418</td><td>12456</td><td>39</td><td>CGATGTCGGCTCTTCCTATCATTGTGAAGCAGAATTCAC</td><td>854.8</td><td>-2.8</td><td>0.8</td><td>-3.5</td><td>3.9E-04</td><td>6.6E-04</td><td>BT20/MB468</td><td>MITO</td></tr>
<tr><td>>trna111_HisGTG_1_-_147774845_147774916@1.34.34,trna118_HisGTG_1_-_145396881_145396952@1.34.34,trna16_HisGTG_1_+_146544773_146544844@1.34.34,trna1_HisGTG_15_+_45493349_45493420@1.34.34,trna21_HisGTG_1_+_147753471_147753542@1.34.34,trna33_HisGTG_6_+_27125906_27125977@1.34.34,trna7_HisGTG_9_-_14433938_14434009@1.34.34,trna8_HisGTG_15_-_45492611_45492682@1.34.34,trna9_HisGTG_15_-_45490804_45490875@1.34.34|GCCGTGATCGTATAGTGGTTAGTACTCTGCGTTG</td><td>>trna111_HisGTG_1_-_147774845_147774916</td><td>1</td><td>34</td><td>34</td><td>GCCGTGATCGTATAGTGGTTAGTACTCTGCGTTG</td><td>588.6</td><td>3.7</td><td>1.0</td><td>3.5</td><td>4.4E-04</td><td>6.6E-04</td><td>MB231/MB468</td><td>MP</td></tr>
<tr><td>>hg38_21_+_8433172_8446622_RNA45SN1_45S_50ntFlanks@10686.10711.26|GCGGCGTCCGGTGAGCTCTCGCTGGC</td><td>>hg38_21_+_8433172_8446622_RNA45SN1_45S_50ntFlanks</td><td>10686</td><td>10711</td><td>26</td><td>GCGGCGTCCGGTGAGCTCTCGCTGGC</td><td>498.3</td><td>-5.6</td><td>1.6</td><td>-3.5</td><td>4.4E-04</td><td>6.6E-04</td><td>MB231/MB468</td><td>MP</td></tr>
<tr><td>>hg38_21_+_8433172_8446622_RNA45SN1_45S_50ntFlanks@8113.8140.28|GGCGCGGGACATGTGGCGTACGGAAGAC</td><td>>hg38_21_+_8433172_8446622_RNA45SN1_45S_50ntFlanks</td><td>8113</td><td>8140</td><td>28</td><td>GGCGCGGGACATGTGGCGTACGGAAGAC</td><td>384.4</td><td>-4.0</td><td>1.1</td><td>-3.5</td><td>3.9E-04</td><td>6.6E-04</td><td>BT20/MB468</td><td>MITO</td></tr>
<tr><td>>hg38_21_+_8433172_8446622_RNA45SN1_45S_50ntFlanks@9307.9330.24|AAGGAGTCTAACACGTGCGCGAGT</td><td>>hg38_21_+_8433172_8446622_RNA45SN1_45S_50ntFlanks</td><td>9307</td><td>9330</td><td>24</td><td>AAGGAGTCTAACACGTGCGCGAGT</td><td>382.8</td><td>3.8</td><td>1.0</td><td>3.7</td><td>2.2E-04</td><td>6.6E-04</td><td>BT20/MB468</td><td>MP</td></tr>
<tr><td>>hg38_21_+_8433172_8446622_RNA45SN1_45S_50ntFlanks@4683.4722.40|CAAGACGGACCAGAGCGAAAGCATTTGCCAAGAATGTTTT</td><td>>hg38_21_+_8433172_8446622_RNA45SN1_45S_50ntFlanks</td><td>4683</td><td>4722</td><td>40</td><td>CAAGACGGACCAGAGCGAAAGCATTTGCCAAGAATGTTTT</td><td>187.9</td><td>-2.8</td><td>0.8</td><td>-3.5</td><td>5.3E-04</td><td>6.6E-04</td><td>MB231/MB468</td><td>NUC</td></tr>
<tr><td>>MI0000065|hsa-let-7d&WithFlank&9|+|94178828|94178926@14.34.21(+1A)&[MIMAT0000065&hsa-let-7d-5p&offsets|0|-1(+1A);m-90&9|+|94178841|94178862&offsets|0|-1(+1A)]|AGAGGTAGTAGGTTGCATAGTA</td><td>>MI0000065|hsa-let-7d&WithFlank&9|+|94178828|94178926</td><td>14</td><td>34</td><td>21(+1A)</td><td>AGAGGTAGTAGGTTGCATAGTA</td><td>161.7</td><td>4.6</td><td>1.3</td><td>3.5</td><td>4.4E-04</td><td>6.6E-04</td><td>MB231/MB468</td><td>MP</td></tr>
<tr><td>>hg38_21_+_8433172_8446622_RNA45SN1_45S_50ntFlanks@11579.11602.24|AGAACTGGTGCGGACCAGGGGAAT</td><td>>hg38_21_+_8433172_8446622_RNA45SN1_45S_50ntFlanks</td><td>11579</td><td>11602</td><td>24</td><td>AGAACTGGTGCGGACCAGGGGAAT</td><td>777.5</td><td>-2.9</td><td>0.8</td><td>-3.5</td><td>4.7E-04</td><td>6.6E-04</td><td>BT20/MB468</td><td>CYTO</td></tr>
<tr><td>>hg38_MT_+_1621_3279_RNR2_16S_50ntFlanks@643.676.34|CAATTGGACCAATCTATCACCCTATAGAAGAACT</td><td>>hg38_MT_+_1621_3279_RNR2_16S_50ntFlanks</td><td>643</td><td>676</td><td>34</td><td>CAATTGGACCAATCTATCACCCTATAGAAGAACT</td><td>127.8</td><td>6.0</td><td>1.7</td><td>3.5</td><td>5.3E-04</td><td>6.6E-04</td><td>MB231/MB468</td><td>MITO</td></tr>
<tr><td>>trnaMT_SerGCT_MT_+_12207_12265@14.62.49|AGAACTGCTAACTCATGCCCCCATGTCTAACAACATGGCTTTCTCACCA</td><td>>trnaMT_SerGCT_MT_+_12207_12265</td><td>14</td><td>62</td><td>49</td><td>AGAACTGCTAACTCATGCCCCCATGTCTAACAACATGGCTTTCTCACCA</td><td>720.8</td><td>7.2</td><td>1.9</td><td>3.7</td><td>2.2E-04</td><td>6.6E-04</td><td>BT20/MB468</td><td>MP</td></tr>
<tr><td>>hg38_21_+_8433172_8446622_RNA45SN1_45S_50ntFlanks@12204.12223.20|AAGGCGAGCTCAGGGAGGAC</td><td>>hg38_21_+_8433172_8446622_RNA45SN1_45S_50ntFlanks</td><td>12204</td><td>12223</td><td>20</td><td>AAGGCGAGCTCAGGGAGGAC</td><td>496.1</td><td>-3.5</td><td>0.9</td><td>-3.9</td><td>8.0E-05</td><td>6.6E-04</td><td>BT20/MB231</td><td>CYTO</td></tr>
<tr><td>>trnaMT_PheGAA_MT_+_577_647@1.19.19|GTTTATGTAGCTTACCTCC</td><td>>trnaMT_PheGAA_MT_+_577_647</td><td>1</td><td>19</td><td>19</td><td>GTTTATGTAGCTTACCTCC</td><td>996.0</td><td>2.8</td><td>0.8</td><td>3.5</td><td>5.3E-04</td><td>6.6E-04</td><td>MB231/MB468</td><td>MITO</td></tr>
<tr><td>>hg38_21_+_8433172_8446622_RNA45SN1_45S_50ntFlanks@11572.11606.35|CCGACTTAGAACTGGTGCGGACCAGGGGAATCCGA</td><td>>hg38_21_+_8433172_8446622_RNA45SN1_45S_50ntFlanks</td><td>11572</td><td>11606</td><td>35</td><td>CCGACTTAGAACTGGTGCGGACCAGGGGAATCCGA</td><td>474.0</td><td>-4.3</td><td>1.2</td><td>-3.5</td><td>5.3E-04</td><td>6.6E-04</td><td>MB231/MB468</td><td>NUC</td></tr>
<tr><td>>MI0000273|hsa-mir-183&WithFlank&7|-|129774899|129775020@33.54.22&[MIMAT0000261&hsa-miR-183-5p&offsets|0|0;m-41&7|-|129774966|129774987&offsets|-1|-1]|TATGGCACTGGTAGAATTCACT</td><td>>MI0000273|hsa-mir-183&WithFlank&7|-|129774899|129775020</td><td>33</td><td>54</td><td>22</td><td>TATGGCACTGGTAGAATTCACT</td><td>2411.4</td><td>3.2</td><td>0.9</td><td>3.5</td><td>5.3E-04</td><td>6.6E-04</td><td>MB231/MB468</td><td>MITO</td></tr>
<tr><td>>hg38_MT_+_1621_3279_RNR2_16S_50ntFlanks@1508.1550.43|AAAGGACAAGAGAAATAAGGCCTACTTCACAAAGCGCCTTCCC</td><td>>hg38_MT_+_1621_3279_RNR2_16S_50ntFlanks</td><td>1508</td><td>1550</td><td>43</td><td>AAAGGACAAGAGAAATAAGGCCTACTTCACAAAGCGCCTTCCC</td><td>69.9</td><td>9.9</td><td>2.7</td><td>3.7</td><td>2.2E-04</td><td>6.6E-04</td><td>BT20/MB468</td><td>MP</td></tr>
<tr><td>>MI0000298|hsa-mir-221&WithFlank&X|-|45746151|45746272@71.92.22(+1U)&[MIMAT0000278&hsa-miR-221-3p&offsets|0|-1(+1U);m-91&X|-|45746181|45746202&offsets|0|0(+1U)]|AGCTACATTGTCTGCTGGGTTTT</td><td>>MI0000298|hsa-mir-221&WithFlank&X|-|45746151|45746272</td><td>71</td><td>92</td><td>22(+1U)</td><td>AGCTACATTGTCTGCTGGGTTTT</td><td>7236.2</td><td>-3.0</td><td>0.8</td><td>-3.9</td><td>9.3E-05</td><td>6.6E-04</td><td>BT20/MB231</td><td>MITO</td></tr>
<tr><td>>hg38_21_+_8433172_8446622_RNA45SN1_45S_50ntFlanks@9534.9572.39|AGGAAACTCTGGTGGAGGTCCGTAGCGGTCCTGACGTGC</td><td>>hg38_21_+_8433172_8446622_RNA45SN1_45S_50ntFlanks</td><td>9534</td><td>9572</td><td>39</td><td>AGGAAACTCTGGTGGAGGTCCGTAGCGGTCCTGACGTGC</td><td>653.1</td><td>4.6</td><td>1.2</td><td>3.9</td><td>9.3E-05</td><td>6.6E-04</td><td>BT20/MB231</td><td>MITO</td></tr>
<tr><td>>trna119_LysCTT_1_-_145395522_145395594@5.35.31,trna11_LysCTT_5_-_180648979_180649051@5.35.31,trna13_LysCTT_6_+_26556774_26556846@5.35.31,trna7_LysCTT_16_+_3225692_3225764@5.35.31,trna9_LysCTT_5_+_180634755_180634827@5.35.31|GGCTAGCTCAGTCGGTAGAGCATGAGACTCT</td><td>>trna119_LysCTT_1_-_145395522_145395594</td><td>5</td><td>35</td><td>31</td><td>GGCTAGCTCAGTCGGTAGAGCATGAGACTCT</td><td>216.1</td><td>4.5</td><td>1.3</td><td>3.5</td><td>4.4E-04</td><td>6.6E-04</td><td>MB231/MB468</td><td>MP</td></tr>
<tr><td>>hg38_21_+_8433172_8446622_RNA45SN1_45S_50ntFlanks@12419.12436.18|GATGTCGGCTCTTCCTAT</td><td>>hg38_21_+_8433172_8446622_RNA45SN1_45S_50ntFlanks</td><td>12419</td><td>12436</td><td>18</td><td>GATGTCGGCTCTTCCTAT</td><td>349.4</td><td>-5.6</td><td>1.6</td><td>-3.5</td><td>4.4E-04</td><td>6.6E-04</td><td>MB231/MB468</td><td>MP</td></tr>
<tr><td>>hg38_1_-_228634819_228635039_RNA5S12_5S_50ntFlanks@134.169.36|TGGGAGACCGCCTGGGAATACCGGGTGCTGTAGGCT</td><td>>hg38_1_-_228634819_228635039_RNA5S12_5S_50ntFlanks</td><td>134</td><td>169</td><td>36</td><td>TGGGAGACCGCCTGGGAATACCGGGTGCTGTAGGCT</td><td>1417.7</td><td>3.2</td><td>0.9</td><td>3.5</td><td>5.3E-04</td><td>6.6E-04</td><td>MB231/MB468</td><td>MITO</td></tr>
<tr><td>>MI0000746|hsa-mir-99b&WithFlank&19|+|51692606|51692687@13.34.22&[MIMAT0000689&hsa-miR-99b-5p&offsets|0|0;m-10&19|+|51692618|51692639&offsets|0|0]|CACCCGTAGAACCGACCTTGCG</td><td>>MI0000746|hsa-mir-99b&WithFlank&19|+|51692606|51692687</td><td>13</td><td>34</td><td>22</td><td>CACCCGTAGAACCGACCTTGCG</td><td>9396.2</td><td>2.8</td><td>0.8</td><td>3.7</td><td>2.2E-04</td><td>6.6E-04</td><td>BT20/MB468</td><td>MP</td></tr>
<tr><td>>hg38_21_+_8433172_8446622_RNA45SN1_45S_50ntFlanks@4066.4095.30|CAACTTTCGATGGTAGTCGCCGTGCCTACC</td><td>>hg38_21_+_8433172_8446622_RNA45SN1_45S_50ntFlanks</td><td>4066</td><td>4095</td><td>30</td><td>CAACTTTCGATGGTAGTCGCCGTGCCTACC</td><td>89.3</td><td>-5.6</td><td>1.5</td><td>-3.7</td><td>2.2E-04</td><td>6.6E-04</td><td>BT20/MB468</td><td>MP</td></tr>
<tr><td>>MI0000434|hsa-let-7i&WithFlank&12|+|62603680|62603775@12.31.20&[MIMAT0000415&hsa-let-7i-5p&offsets|0|-2;m-49&12|+|62603691|62603712&offsets|0|-2]|TGAGGTAGTAGTTTGTGCTG</td><td>>MI0000434|hsa-let-7i&WithFlank&12|+|62603680|62603775</td><td>12</td><td>31</td><td>20</td><td>TGAGGTAGTAGTTTGTGCTG</td><td>411.7</td><td>4.8</td><td>1.4</td><td>3.5</td><td>4.5E-04</td><td>6.6E-04</td><td>MB231/MB468</td><td>MP</td></tr>
<tr><td>>hg38_21_+_8433172_8446622_RNA45SN1_45S_50ntFlanks@11843.11872.30|AGGGAACGGGCTTGGCGGAATCAGCGGGGA</td><td>>hg38_21_+_8433172_8446622_RNA45SN1_45S_50ntFlanks</td><td>11843</td><td>11872</td><td>30</td><td>AGGGAACGGGCTTGGCGGAATCAGCGGGGA</td><td>193.5</td><td>-4.8</td><td>1.4</td><td>-3.5</td><td>4.8E-04</td><td>6.6E-04</td><td>BT20/MB468</td><td>CYTO</td></tr>
<tr><td>>hg38_21_+_8433172_8446622_RNA45SN1_45S_50ntFlanks@3853.3873.21|AACTGTGGTAATTCTAGAGCT</td><td>>hg38_21_+_8433172_8446622_RNA45SN1_45S_50ntFlanks</td><td>3853</td><td>3873</td><td>21</td><td>AACTGTGGTAATTCTAGAGCT</td><td>550.8</td><td>-4.9</td><td>1.3</td><td>-3.7</td><td>2.2E-04</td><td>6.6E-04</td><td>BT20/MB468</td><td>MP</td></tr>
<tr><td>>trnaMT_PheGAA_MT_+_577_647@28.65.38|ATACACTGAAAATGTTTAGACGGGCTCACATCACCCCA</td><td>>trnaMT_PheGAA_MT_+_577_647</td><td>28</td><td>65</td><td>38</td><td>ATACACTGAAAATGTTTAGACGGGCTCACATCACCCCA</td><td>69.8</td><td>9.9</td><td>2.7</td><td>3.7</td><td>2.2E-04</td><td>6.6E-04</td><td>BT20/MB468</td><td>MP</td></tr>
<tr><td>>hg38_21_+_8433172_8446622_RNA45SN1_45S_50ntFlanks@8105.8140.36|CGCGGCGGGGCGCGGGACATGTGGCGTACGGAAGAC</td><td>>hg38_21_+_8433172_8446622_RNA45SN1_45S_50ntFlanks</td><td>8105</td><td>8140</td><td>36</td><td>CGCGGCGGGGCGCGGGACATGTGGCGTACGGAAGAC</td><td>10556.0</td><td>-3.7</td><td>1.0</td><td>-3.7</td><td>2.2E-04</td><td>6.6E-04</td><td>BT20/MB468</td><td>MP</td></tr>
<tr><td>>MI0000746|hsa-mir-99b&WithFlank&19|+|51692606|51692687@13.33.21&[MIMAT0000689&hsa-miR-99b-5p&offsets|0|-1;m-10&19|+|51692618|51692639&offsets|0|-1]|CACCCGTAGAACCGACCTTGC</td><td>>MI0000746|hsa-mir-99b&WithFlank&19|+|51692606|51692687</td><td>13</td><td>33</td><td>21</td><td>CACCCGTAGAACCGACCTTGC</td><td>3135.3</td><td>2.3</td><td>0.7</td><td>3.5</td><td>4.5E-04</td><td>6.6E-04</td><td>MB231/MB468</td><td>MP</td></tr>
<tr><td>>hg38_21_+_8433172_8446622_RNA45SN1_45S_50ntFlanks@11641.11668.28|CGGCGGGTGTTGACGCGATGTGATTTCT</td><td>>hg38_21_+_8433172_8446622_RNA45SN1_45S_50ntFlanks</td><td>11641</td><td>11668</td><td>28</td><td>CGGCGGGTGTTGACGCGATGTGATTTCT</td><td>1290.7</td><td>-6.4</td><td>1.7</td><td>-3.7</td><td>2.2E-04</td><td>6.7E-04</td><td>BT20/MB468</td><td>MP</td></tr>
<tr><td>>hg38_21_+_8433172_8446622_RNA45SN1_45S_50ntFlanks@3843.3881.39|CTACTTGGATAACTGTGGTAATTCTAGAGCTAATACATG</td><td>>hg38_21_+_8433172_8446622_RNA45SN1_45S_50ntFlanks</td><td>3843</td><td>3881</td><td>39</td><td>CTACTTGGATAACTGTGGTAATTCTAGAGCTAATACATG</td><td>62.3</td><td>-9.0</td><td>2.4</td><td>-3.7</td><td>2.2E-04</td><td>6.7E-04</td><td>BT20/MB468</td><td>MP</td></tr>
<tr><td>>hg38_21_+_8433172_8446622_RNA45SN1_45S_50ntFlanks@7988.8013.26|CAGACGTGGCGACCCGCTGAATTTAA</td><td>>hg38_21_+_8433172_8446622_RNA45SN1_45S_50ntFlanks</td><td>7988</td><td>8013</td><td>26</td><td>CAGACGTGGCGACCCGCTGAATTTAA</td><td>332.5</td><td>-3.1</td><td>0.8</td><td>-3.7</td><td>2.2E-04</td><td>6.7E-04</td><td>BT20/MB468</td><td>MP</td></tr>
<tr><td>>hg38_21_+_8433172_8446622_RNA45SN1_45S_50ntFlanks@9928.9957.30|TGATATAGACAGCAGGACGGTGGCCATGGA</td><td>>hg38_21_+_8433172_8446622_RNA45SN1_45S_50ntFlanks</td><td>9928</td><td>9957</td><td>30</td><td>TGATATAGACAGCAGGACGGTGGCCATGGA</td><td>61.8</td><td>6.8</td><td>1.8</td><td>3.7</td><td>2.2E-04</td><td>6.7E-04</td><td>BT20/MB468</td><td>MP</td></tr>
<tr><td>>hg38_21_+_8433172_8446622_RNA45SN1_45S_50ntFlanks@10694.10711.18|CGGTGAGCTCTCGCTGGC</td><td>>hg38_21_+_8433172_8446622_RNA45SN1_45S_50ntFlanks</td><td>10694</td><td>10711</td><td>18</td><td>CGGTGAGCTCTCGCTGGC</td><td>910.8</td><td>-6.2</td><td>1.7</td><td>-3.7</td><td>2.2E-04</td><td>6.7E-04</td><td>BT20/MB468</td><td>MP</td></tr>
<tr><td>>hg38_21_+_8433172_8446622_RNA45SN1_45S_50ntFlanks@10427.10456.30|ATGGGCGAGCGCCGTTCCGAAGGGACGGGC</td><td>>hg38_21_+_8433172_8446622_RNA45SN1_45S_50ntFlanks</td><td>10427</td><td>10456</td><td>30</td><td>ATGGGCGAGCGCCGTTCCGAAGGGACGGGC</td><td>1942.6</td><td>-2.1</td><td>0.6</td><td>-3.5</td><td>5.0E-04</td><td>6.7E-04</td><td>MB231/MB468</td><td>CYTO</td></tr>
</table>

<table>
<tr><td>>hg38_21_+_8433172_8446622_RNA45SN1_45S_50ntFlanks@10834.10865.32|CCGGATCCGTAACTTCGGGATAAGGATTGGCT</td><td>>hg38_21_+_8433172_8446622_RNA45SN1_45S_50ntFlanks</td><td>10834</td><td>10865</td><td>32</td><td>CCGGATCCGTAACTTCGGGATAAGGATTGGCT</td><td>472.4</td><td>-6.4</td><td>1.8</td><td>-3.5</td><td>4.5E-04</td><td>6.7E-04</td><td>MB231/MB468</td><td>MP</td></tr>
<tr><td>>hg38_21_+_8433172_8446622_RNA45SN1_45S_50ntFlanks@10349.10379.31|AACGAGAACTTTGAAGGCCGAAGTGGAGAAG</td><td>>hg38_21_+_8433172_8446622_RNA45SN1_45S_50ntFlanks</td><td>10349</td><td>10379</td><td>31</td><td>AACGAGAACTTTGAAGGCCGAAGTGGAGAAG</td><td>480.1</td><td>-3.0</td><td>0.9</td><td>-3.5</td><td>4.5E-04</td><td>6.7E-04</td><td>MB231/MB468</td><td>MP</td></tr>
<tr><td>>hg38_21_+_8433172_8446622_RNA45SN1_45S_50ntFlanks@4975.4999.25|CCCGGACACGGACAGGATTGACAGA</td><td>>hg38_21_+_8433172_8446622_RNA45SN1_45S_50ntFlanks</td><td>4975</td><td>4999</td><td>25</td><td>CCCGGACACGGACAGGATTGACAGA</td><td>454.5</td><td>-3.4</td><td>1.0</td><td>-3.5</td><td>4.0E-04</td><td>6.7E-04</td><td>BT20/MB468</td><td>MITO</td></tr>
<tr><td>>hg38_21_+_8433172_8446622_RNA45SN1_45S_50ntFlanks@6742.6770.29|TCGACACTTCGAACGCACTTGCGGCCCCG</td><td>>hg38_21_+_8433172_8446622_RNA45SN1_45S_50ntFlanks</td><td>6742</td><td>6770</td><td>29</td><td>TCGACACTTCGAACGCACTTGCGGCCCCG</td><td>307.7</td><td>-3.9</td><td>1.1</td><td>-3.5</td><td>4.0E-04</td><td>6.7E-04</td><td>BT20/MB468</td><td>MITO</td></tr>
<tr><td>>hg38_21_+_8433172_8446622_RNA45SN1_45S_50ntFlanks@12626.12659.34|AATGGGGCGAAGCTACCATCTGTGGGATTATGAC</td><td>>hg38_21_+_8433172_8446622_RNA45SN1_45S_50ntFlanks</td><td>12626</td><td>12659</td><td>34</td><td>AATGGGGCGAAGCTACCATCTGTGGGATTATGAC</td><td>327.1</td><td>-2.3</td><td>0.7</td><td>-3.5</td><td>4.8E-04</td><td>6.7E-04</td><td>BT20/MB468</td><td>CYTO</td></tr>
<tr><td>>hg38_21_+_8433172_8446622_RNA45SN1_45S_50ntFlanks@6678.6714.37|CGTGCGTCGATGAAGAACGCAGCTAGCTGCGAGAATT</td><td>>hg38_21_+_8433172_8446622_RNA45SN1_45S_50ntFlanks</td><td>6678</td><td>6714</td><td>37</td><td>CGTGCGTCGATGAAGAACGCAGCTAGCTGCGAGAATT</td><td>79.1</td><td>6.7</td><td>1.8</td><td>3.7</td><td>2.2E-04</td><td>6.7E-04</td><td>BT20/MB468</td><td>MP</td></tr>
<tr><td>>hg38_21_+_8433172_8446622_RNA45SN1_45S_50ntFlanks@12676.12700.25|AGAATCCCGCCCAGGCGGAACGATA</td><td>>hg38_21_+_8433172_8446622_RNA45SN1_45S_50ntFlanks</td><td>12676</td><td>12700</td><td>25</td><td>AGAATCCCGCCCAGGCGGAACGATA</td><td>79.2</td><td>-9.4</td><td>2.5</td><td>-3.7</td><td>2.2E-04</td><td>6.7E-04</td><td>BT20/MB468</td><td>MP</td></tr>
<tr><td>>hg38_1_-_228634819_228635039_RNA5S12_5S_50ntFlanks@147.171.25|GGGAATACCGGGTGCTGTAGGCTTT</td><td>>hg38_1_-_228634819_228635039_RNA5S12_5S_50ntFlanks</td><td>147</td><td>171</td><td>25</td><td>GGGAATACCGGGTGCTGTAGGCTTT</td><td>2866.9</td><td>-2.5</td><td>0.7</td><td>-3.5</td><td>4.5E-04</td><td>6.7E-04</td><td>MB231/MB468</td><td>MP</td></tr>
<tr><td>>hg38_21_+_8433172_8446622_RNA45SN1_45S_50ntFlanks@4573.4616.44|AATAGGACCGCGGTTCTATTTTGTTGGTTTTCGGAACTGAGGCC</td><td>>hg38_21_+_8433172_8446622_RNA45SN1_45S_50ntFlanks</td><td>4573</td><td>4616</td><td>44</td><td>AATAGGACCGCGGTTCTATTTTGTTGGTTTTCGGAACTGAGGCC</td><td>157.7</td><td>-3.8</td><td>1.1</td><td>-3.5</td><td>5.0E-04</td><td>6.7E-04</td><td>MB231/MB468</td><td>CYTO</td></tr>
<tr><td>>hg38_21_+_8433172_8446622_RNA45SN1_45S_50ntFlanks@6739.6770.32|TCATCGACACTTCGAACGCACTTGCGGCCCCG</td><td>>hg38_21_+_8433172_8446622_RNA45SN1_45S_50ntFlanks</td><td>6739</td><td>6770</td><td>32</td><td>TCATCGACACTTCGAACGCACTTGCGGCCCCG</td><td>242.5</td><td>-5.0</td><td>1.4</td><td>-3.5</td><td>4.8E-04</td><td>6.7E-04</td><td>BT20/MB468</td><td>CYTO</td></tr>
<tr><td>>hg38_MT_+_1621_3279_RNR2_16S_50ntFlanks@1367.1396.30|TCGATGTTGGATCAGGACATCCCGATGGTG</td><td>>hg38_MT_+_1621_3279_RNR2_16S_50ntFlanks</td><td>1367</td><td>1396</td><td>30</td><td>TCGATGTTGGATCAGGACATCCCGATGGTG</td><td>430.5</td><td>3.5</td><td>1.0</td><td>3.5</td><td>4.5E-04</td><td>6.7E-04</td><td>MB231/MB468</td><td>MP</td></tr>
<tr><td>>hg38_21_+_8433172_8446622_RNA45SN1_45S_50ntFlanks@6748.6772.25|CTTCGAACGCACTTGCGGCCCCGGG</td><td>>hg38_21_+_8433172_8446622_RNA45SN1_45S_50ntFlanks</td><td>6748</td><td>6772</td><td>25</td><td>CTTCGAACGCACTTGCGGCCCCGGG</td><td>120.1</td><td>-5.7</td><td>1.5</td><td>-3.7</td><td>2.2E-04</td><td>6.7E-04</td><td>BT20/MB468</td><td>MP</td></tr>
<tr><td>>hg38_21_+_8433172_8446622_RNA45SN1_45S_50ntFlanks@9544.9563.20|GGTGGAGGTCCGTAGCGGTC</td><td>>hg38_21_+_8433172_8446622_RNA45SN1_45S_50ntFlanks</td><td>9544</td><td>9563</td><td>20</td><td>GGTGGAGGTCCGTAGCGGTC</td><td>320.1</td><td>-3.8</td><td>1.1</td><td>-3.5</td><td>4.8E-04</td><td>6.7E-04</td><td>BT20/MB468</td><td>CYTO</td></tr>
<tr><td>>hg38_21_+_8433172_8446622_RNA45SN1_45S_50ntFlanks@12626.12652.27|AATGGGGCGAAGCTACCATCTGTGGGA</td><td>>hg38_21_+_8433172_8446622_RNA45SN1_45S_50ntFlanks</td><td>12626</td><td>12652</td><td>27</td><td>AATGGGGCGAAGCTACCATCTGTGGGA</td><td>282.8</td><td>-5.0</td><td>1.4</td><td>-3.5</td><td>4.0E-04</td><td>6.7E-04</td><td>BT20/MB468</td><td>MITO</td></tr>
<tr><td>>hg38_21_+_8433172_8446622_RNA45SN1_45S_50ntFlanks@10841.10873.33|CGTAACTTCGGGATAAGGATTGGCTCTAAGGGC</td><td>>hg38_21_+_8433172_8446622_RNA45SN1_45S_50ntFlanks</td><td>10841</td><td>10873</td><td>33</td><td>CGTAACTTCGGGATAAGGATTGGCTCTAAGGGC</td><td>186.4</td><td>-5.7</td><td>1.6</td><td>-3.5</td><td>4.5E-04</td><td>6.7E-04</td><td>MB231/MB468</td><td>MP</td></tr>
<tr><td>>hg38_21_+_8433172_8446622_RNA45SN1_45S_50ntFlanks@11853.11871.19|CTTGGCGGAATCAGCGGGG</td><td>>hg38_21_+_8433172_8446622_RNA45SN1_45S_50ntFlanks</td><td>11853</td><td>11871</td><td>19</td><td>CTTGGCGGAATCAGCGGGG</td><td>84.9</td><td>-5.6</td><td>1.5</td><td>-3.7</td><td>2.2E-04</td><td>6.7E-04</td><td>BT20/MB468</td><td>MP</td></tr>
<tr><td>>MI0016005|hsa-mir-3615&WithFlank&17|+|74748607|74748705@57.78.22(+2U)&[MIMAT0017994&hsa-miR-3615&offsets|0|+1(+2U);m-271&17|+|74748663|74748684&offsets|0|0(+2U)]|TCTCTCGGCTCCTCGCGGCTCGTT</td><td>>MI0016005|hsa-mir-3615&WithFlank&17|+|74748607|74748705</td><td>57</td><td>78</td><td>22(+2U)</td><td>TCTCTCGGCTCCTCGCGGCTCGTT</td><td>135.6</td><td>5.3</td><td>1.5</td><td>3.5</td><td>5.4E-04</td><td>6.7E-04</td><td>MB231/MB468</td><td>MITO</td></tr>
<tr><td>>am_trnaMT_SerGCT_MT_+_12207_12265@13.38.26|AAGAACTGCTAACTCATGCCCCCATG</td><td>>am_trnaMT_SerGCT_MT_+_12207_12265</td><td>13</td><td>38</td><td>26</td><td>AAGAACTGCTAACTCATGCCCCCATG</td><td>480.0</td><td>-2.8</td><td>0.8</td><td>-3.5</td><td>5.4E-04</td><td>6.7E-04</td><td>MB231/MB468</td><td>MITO</td></tr>
<tr><td>>hg38_MT_+_598_1651_RNR1_12S_50ntFlanks@910.950.41|ACTTCAAAGGACATTTAACTAAAACCCCTACGCATTTATAT</td><td>>hg38_MT_+_598_1651_RNR1_12S_50ntFlanks</td><td>910</td><td>950</td><td>41</td><td>ACTTCAAAGGACATTTAACTAAAACCCCTACGCATTTATAT</td><td>71.7</td><td>9.9</td><td>2.7</td><td>3.7</td><td>2.2E-04</td><td>6.7E-04</td><td>BT20/MB468</td><td>MP</td></tr>
<tr><td>>hg38_21_+_8433172_8446622_RNA45SN1_45S_50ntFlanks@12442.12473.32|TGAAGCAGAATTCACCAAGCGTTGGATTGTTC</td><td>>hg38_21_+_8433172_8446622_RNA45SN1_45S_50ntFlanks</td><td>12442</td><td>12473</td><td>32</td><td>TGAAGCAGAATTCACCAAGCGTTGGATTGTTC</td><td>243.3</td><td>-4.3</td><td>1.2</td><td>-3.5</td><td>4.5E-04</td><td>6.7E-04</td><td>MB231/MB468</td><td>MP</td></tr>
<tr><td>>MI0005561|hsa-mir-877&WithFlank&6|+|30584326|30584423@7.28.22(+1U)&[MIMAT0004949&hsa-miR-877-5p&offsets|0|+2(+1U);m-382&6|+|30584332|30584354&offsets|0|-1(+1U)]|GTAGAGGAGATGGCGCAGGGGAT</td><td>>MI0005561|hsa-mir-877&WithFlank&6|+|30584326|30584423</td><td>7</td><td>28</td><td>22(+1U)</td><td>GTAGAGGAGATGGCGCAGGGGAT</td><td>87.1</td><td>6.2</td><td>1.7</td><td>3.7</td><td>2.2E-04</td><td>6.7E-04</td><td>BT20/MB468</td><td>MP</td></tr>
<tr><td>>hg38_21_+_8433172_8446622_RNA45SN1_45S_50ntFlanks@8184.8211.28|CGAGGCCCAGCCCGTGGACGGTGTGAGG</td><td>>hg38_21_+_8433172_8446622_RNA45SN1_45S_50ntFlanks</td><td>8184</td><td>8211</td><td>28</td><td>CGAGGCCCAGCCCGTGGACGGTGTGAGG</td><td>130.6</td><td>-2.7</td><td>0.8</td><td>-3.5</td><td>5.1E-04</td><td>6.7E-04</td><td>MB231/MB468</td><td>CYTO</td></tr>
<tr><td>>hg38_21_+_8433172_8446622_RNA45SN1_45S_50ntFlanks@7974.7996.23|ACGCGACCTCAGATCAGACGTGG</td><td>>hg38_21_+_8433172_8446622_RNA45SN1_45S_50ntFlanks</td><td>7974</td><td>7996</td><td>23</td><td>ACGCGACCTCAGATCAGACGTGG</td><td>414.2</td><td>2.4</td><td>0.7</td><td>3.5</td><td>5.4E-04</td><td>6.7E-04</td><td>MB231/MB468</td><td>NUC</td></tr>
<tr><td>>hg38_21_+_8433172_8446622_RNA45SN1_45S_50ntFlanks@11703.11736.34|AATGAAGCGCGGGTAAACGGCGGGAGTAACTATG</td><td>>hg38_21_+_8433172_8446622_RNA45SN1_45S_50ntFlanks</td><td>11703</td><td>11736</td><td>34</td><td>AATGAAGCGCGGGTAAACGGCGGGAGTAACTATG</td><td>402.7</td><td>-2.4</td><td>0.7</td><td>-3.5</td><td>5.4E-04</td><td>6.7E-04</td><td>MB231/MB468</td><td>NUC</td></tr>
<tr><td>>hg38_21_+_8433172_8446622_RNA45SN1_45S_50ntFlanks@10059.10079.21|GGCAGTCGAGAGTGGACGGGA</td><td>>hg38_21_+_8433172_8446622_RNA45SN1_45S_50ntFlanks</td><td>10059</td><td>10079</td><td>21</td><td>GGCAGTCGAGAGTGGACGGGA</td><td>127.5</td><td>6.8</td><td>1.8</td><td>3.7</td><td>2.2E-04</td><td>6.7E-04</td><td>BT20/MB468</td><td>MP</td></tr>
<tr><td>>hg38_21_+_8433172_8446622_RNA45SN1_45S_50ntFlanks@13002.13030.29|CGATCTATTGAAAGTCAGCCCTCGACACA</td><td>>hg38_21_+_8433172_8446622_RNA45SN1_45S_50ntFlanks</td><td>13002</td><td>13030</td><td>29</td><td>CGATCTATTGAAAGTCAGCCCTCGACACA</td><td>257.3</td><td>-4.5</td><td>1.3</td><td>-3.5</td><td>5.1E-04</td><td>6.7E-04</td><td>MB231/MB468</td><td>CYTO</td></tr>
<tr><td>>hg38_21_+_8433172_8446622_RNA45SN1_45S_50ntFlanks@3853.3884.32|AACTGTGGTAATTCTAGAGCTAATACATGCCG</td><td>>hg38_21_+_8433172_8446622_RNA45SN1_45S_50ntFlanks</td><td>3853</td><td>3884</td><td>32</td><td>AACTGTGGTAATTCTAGAGCTAATACATGCCG</td><td>671.3</td><td>2.3</td><td>0.7</td><td>3.5</td><td>4.8E-04</td><td>6.7E-04</td><td>BT20/MB468</td><td>CYTO</td></tr>
<tr><td>>hg38_1_-_228634819_228635039_RNA5S12_5S_50ntFlanks@149.171.23|GAATACCGGGTGCTGTAGGCTTT</td><td>>hg38_1_-_228634819_228635039_RNA5S12_5S_50ntFlanks</td><td>149</td><td>171</td><td>23</td><td>GAATACCGGGTGCTGTAGGCTTT</td><td>1606.3</td><td>2.0</td><td>0.6</td><td>3.5</td><td>4.0E-04</td><td>6.7E-04</td><td>BT20/MB468</td><td>MITO</td></tr>
<tr><td>>hg38_21_+_8433172_8446622_RNA45SN1_45S_50ntFlanks@10481.10500.20|ATCGAAAGGGAGTCGGGTTC</td><td>>hg38_21_+_8433172_8446622_RNA45SN1_45S_50ntFlanks</td><td>10481</td><td>10500</td><td>20</td><td>ATCGAAAGGGAGTCGGGTTC</td><td>64.9</td><td>-9.1</td><td>2.5</td><td>-3.7</td><td>2.2E-04</td><td>6.7E-04</td><td>BT20/MB468</td><td>MP</td></tr>
<tr><td>>hg38_21_+_8433172_8446622_RNA45SN1_45S_50ntFlanks@9935.9955.21|GACAGCAGGACGGTGGCCATG</td><td>>hg38_21_+_8433172_8446622_RNA45SN1_45S_50ntFlanks</td><td>9935</td><td>9955</td><td>21</td><td>GACAGCAGGACGGTGGCCATG</td><td>120.3</td><td>5.8</td><td>1.6</td><td>3.7</td><td>2.2E-04</td><td>6.7E-04</td><td>BT20/MB468</td><td>MP</td></tr>
<tr><td>>trna10_AspGTC_12_-_125424193_125424264@1.32.32,trna12_AspGTC_12_-_125411891_125411962@1.32.32,trna38_AspGTC_17_-_8125556_8125627@1.32.32,trna45_AspGTC_6_+_27447453_27447524@1.32.32,trna48_AspGTC_6_+_27471523_27471594@1.32.32,trna4_AspGTC_12_+_96429799_96429870@1.32.32,trna69_AspGTC_1_-_161440205_161440276@1.32.32,trna72_AspGTC_1_-_161432824_161432895@1.32.32,trna75_AspGTC_1_-_161425414_161425485@1.32.32,trna78_AspGTC_1_-_161418033_161418104@1.32.32,trna81_AspGTC_1_-_161410615_161410686@1.32.32|TCCTCGTTAGTATAGTGGTGAGTATCCCCGCC</td><td>>trna10_AspGTC_12_-_125424193_125424264</td><td>1</td><td>32</td><td>32</td><td>TCCTCGTTAGTATAGTGGTGAGTATCCCCGCC</td><td>136.5</td><td>5.3</td><td>1.5</td><td>3.5</td><td>5.4E-04</td><td>6.7E-04</td><td>MB231/MB468</td><td>MITO</td></tr>
<tr><td>>MI0000088|hsa-mir-30a&WithFlank&6|-|71403545|71403627@12.32.21&[MIMAT0000087&hsa-miR-30a-5p&offsets|0|-1;m-15&6|-|71403595|71403616&offsets|0|-1]|TGTAAACATCCTCGACTGGAA</td><td>>MI0000088|hsa-mir-30a&WithFlank&6|-|71403545|71403627</td><td>12</td><td>32</td><td>21</td><td>TGTAAACATCCTCGACTGGAA</td><td>172.2</td><td>3.2</td><td>0.9</td><td>3.5</td><td>5.5E-04</td><td>6.7E-04</td><td>BT20/MB468</td><td>NUC</td></tr>
<tr><td>>hg38_21_+_8433172_8446622_RNA45SN1_45S_50ntFlanks@12997.13015.19|CGCTGCGATCTATTGAAAG</td><td>>hg38_21_+_8433172_8446622_RNA45SN1_45S_50ntFlanks</td><td>12997</td><td>13015</td><td>19</td><td>CGCTGCGATCTATTGAAAG</td><td>767.8</td><td>2.9</td><td>0.8</td><td>3.5</td><td>4.8E-04</td><td>6.8E-04</td><td>BT20/MB468</td><td>CYTO</td></tr>
<tr><td>>hg38_21_+_8433172_8446622_RNA45SN1_45S_50ntFlanks@3845.3868.24|ACTTGGATAACTGTGGTAATTCTA</td><td>>hg38_21_+_8433172_8446622_RNA45SN1_45S_50ntFlanks</td><td>3845</td><td>3868</td><td>24</td><td>ACTTGGATAACTGTGGTAATTCTA</td><td>953.9</td><td>1.6</td><td>0.5</td><td>3.5</td><td>4.8E-04</td><td>6.8E-04</td><td>BT20/MB468</td><td>CYTO</td></tr>
<tr><td>>hg38_21_+_8433172_8446622_RNA45SN1_45S_50ntFlanks@10427.10455.29|ATGGGCGAGCGCCGTTCCGAAGGGACGGG</td><td>>hg38_21_+_8433172_8446622_RNA45SN1_45S_50ntFlanks</td><td>10427</td><td>10455</td><td>29</td><td>ATGGGCGAGCGCCGTTCCGAAGGGACGGG</td><td>182.5</td><td>-4.7</td><td>1.4</td><td>-3.5</td><td>4.8E-04</td><td>6.8E-04</td><td>BT20/MB468</td><td>CYTO</td></tr>
<tr><td>>MI0000286|hsa-mir-210&WithFlank&11|-|568083|568204@72.93.22(+1A)&[MIMAT0000267&hsa-miR-210-3p&offsets|0|0(+1A);m-102&11|-|568112|568133&offsets|0|0(+1A)]|CTGTGCGTGTGACAGCGGCTGAA</td><td>>MI0000286|hsa-mir-210&WithFlank&11|-|568083|568204</td><td>72</td><td>93</td><td>22(+1A)</td><td>CTGTGCGTGTGACAGCGGCTGAA</td><td>713.8</td><td>4.0</td><td>1.0</td><td>3.9</td><td>8.2E-05</td><td>6.8E-04</td><td>BT20/MB231</td><td>CYTO</td></tr>
<tr><td>>hg38_21_+_8433172_8446622_RNA45SN1_45S_50ntFlanks@5084.5115.32|CGAACGAGACTCTGGCATGCTAACTAGTTACG</td><td>>hg38_21_+_8433172_8446622_RNA45SN1_45S_50ntFlanks</td><td>5084</td><td>5115</td><td>32</td><td>CGAACGAGACTCTGGCATGCTAACTAGTTACG</td><td>435.2</td><td>-3.1</td><td>0.9</td><td>-3.5</td><td>5.5E-04</td><td>6.8E-04</td><td>MB231/MB468</td><td>NUC</td></tr>
<tr><td>>hg38_21_+_8433172_8446622_RNA45SN1_45S_50ntFlanks@11921.11950.30|TGAGAGGTGTAGAATAAGTGGGAGGCCCCC</td><td>>hg38_21_+_8433172_8446622_RNA45SN1_45S_50ntFlanks</td><td>11921</td><td>11950</td><td>30</td><td>TGAGAGGTGTAGAATAAGTGGGAGGCCCCC</td><td>435.1</td><td>-3.2</td><td>0.9</td><td>-3.5</td><td>4.9E-04</td><td>6.8E-04</td><td>BT20/MB468</td><td>CYTO</td></tr>
<tr><td>>trnaMT_GluTTC_MT_-_14674_14742@21.66.46,trnalookalike8_GluTTC_5_-_93905172_93905240@21.66.46|ACGATGGTTTTTCATATCATTGGTCGTGGTTGTAGTCCGTGCGAGA</td><td>>trnaMT_GluTTC_MT_-_14674_14742</td><td>21</td><td>66</td><td>46</td><td>ACGATGGTTTTTCATATCATTGGTCGTGGTTGTAGTCCGTGCGAGA</td><td>70.1</td><td>9.9</td><td>2.7</td><td>3.7</td><td>2.2E-04</td><td>6.8E-04</td><td>BT20/MB468</td><td>MP</td></tr>
<tr><td>>hg38_21_+_8433172_8446622_RNA45SN1_45S_50ntFlanks@4728.4758.31|ATCAAGAACGAAAGTCGGAGGTTCGAAGACG</td><td>>hg38_21_+_8433172_8446622_RNA45SN1_45S_50ntFlanks</td><td>4728</td><td>4758</td><td>31</td><td>ATCAAGAACGAAAGTCGGAGGTTCGAAGACG</td><td>3723.9</td><td>-6.1</td><td>1.8</td><td>-3.5</td><td>5.5E-04</td><td>6.8E-04</td><td>MB231/MB468</td><td>NUC</td></tr>
<tr><td>>MI0005561|hsa-mir-877&WithFlank&6|+|30584326|30584423@7.26.20&[MIMAT0004949&hsa-miR-877-5p&offsets|0|0;m-382&6|+|30584332|30584354&offsets|0|-3]|GTAGAGGAGATGGCGCAGGG</td><td>>MI0005561|hsa-mir-877&WithFlank&6|+|30584326|30584423</td><td>7</td><td>26</td><td>20</td><td>GTAGAGGAGATGGCGCAGGG</td><td>383.3</td><td>5.4</td><td>1.5</td><td>3.7</td><td>2.2E-04</td><td>6.8E-04</td><td>BT20/MB468</td><td>MP</td></tr>
<tr><td>>hg38_21_+_8433172_8446622_RNA45SN1_45S_50ntFlanks@10571.10591.21|GAGAAGCCGGCGGGAGCCCCG</td><td>>hg38_21_+_8433172_8446622_RNA45SN1_45S_50ntFlanks</td><td>10571</td><td>10591</td><td>21</td><td>GAGAAGCCGGCGGGAGCCCCG</td><td>259.5</td><td>-3.9</td><td>1.1</td><td>-3.5</td><td>4.1E-04</td><td>6.8E-04</td><td>BT20/MB468</td><td>MITO</td></tr>
<tr><td>>hg38_21_+_8433172_8446622_RNA45SN1_45S_50ntFlanks@8107.8143.37|CGGCGGGGCGCGGGACATGTGGCGTACGGAAGACCCG</td><td>>hg38_21_+_8433172_8446622_RNA45SN1_45S_50ntFlanks</td><td>8107</td><td>8143</td><td>37</td><td>CGGCGGGGCGCGGGACATGTGGCGTACGGAAGACCCG</td><td>556.4</td><td>-4.3</td><td>1.2</td><td>-3.5</td><td>4.1E-04</td><td>6.8E-04</td><td>BT20/MB468</td><td>MITO</td></tr>
<tr><td>>hg38_21_+_8433172_8446622_RNA45SN1_45S_50ntFlanks@7975.8010.36|CGCGACCTCAGATCAGACGTGGCGACCCGCTGAATT</td><td>>hg38_21_+_8433172_8446622_RNA45SN1_45S_50ntFlanks</td><td>7975</td><td>8010</td><td>36</td><td>CGCGACCTCAGATCAGACGTGGCGACCCGCTGAATT</td><td>121716.3</td><td>2.5</td><td>0.7</td><td>3.5</td><td>5.4E-04</td><td>6.8E-04</td><td>MB231/MB468</td><td>MITO</td></tr>
<tr><td>>hg38_21_+_8433172_8446622_RNA45SN1_45S_50ntFlanks@10351.10387.37|CGAGAACTTTGAAGGCCGAAGTGGAGAAGGGTTCCAT</td><td>>hg38_21_+_8433172_8446622_RNA45SN1_45S_50ntFlanks</td><td>10351</td><td>10387</td><td>37</td><td>CGAGAACTTTGAAGGCCGAAGTGGAGAAGGGTTCCAT</td><td>210.5</td><td>-5.2</td><td>1.5</td><td>-3.5</td><td>5.5E-04</td><td>6.8E-04</td><td>MB231/MB468</td><td>NUC</td></tr>
</table>

<table>
<tr><td>>MI0000081|hsa-mir-24-2&WithFlank&19|-|13836281|13836365@56.76.21(+1U);MI0000080|hsa-mir-24-1&WithFlank&9|+|95086015|95086094@50.70.21(+1U)&[MIMAT0000080&hsa-miR-24-3p&offsets|0|-1(+1U);m-36&19|-|13836288|13836310&offsets|0|-2(+1U)];[MIMAT0000080_1&hsa-miR-24-3p&offsets|0|-1(+1U);m-35&9|+|95086064|95086086&offsets|0|-2(+1U)]|TGGCTCAGTTCAGCAGGAACAT</td><td>>MI0000081|hsa-mir-24-2&WithFlank&19|-|13836281|13836365</td><td>56</td><td>76</td><td>21(+1U)</td><td>TGGCTCAGTTCAGCAGGAACAT</td><td>144.6</td><td>4.6</td><td>1.3</td><td>3.5</td><td>5.4E-04</td><td>6.8E-04</td><td>MB231/MB468</td><td>MITO</td></tr>
<tr><td>>MI0000542|hsa-mir-320a&WithFlank&8|-|22244960|22245043@48.72.25(+1U)&[MIMAT0000510&hsa-miR-320a-3p&offsets|0|+3(+1U);m-60&8|-|22244975|22244996&offsets|0|+3(+1U)]|AAAAGCTGGGTTGAGAGGGCGAAAAT</td><td>>MI0000542|hsa-mir-320a&WithFlank&8|-|22244960|22245043</td><td>48</td><td>72</td><td>25(+1U)</td><td>AAAAGCTGGGTTGAGAGGGCGAAAAT</td><td>71.2</td><td>6.7</td><td>1.8</td><td>3.7</td><td>2.3E-04</td><td>6.8E-04</td><td>BT20/MB468</td><td>MP</td></tr>
<tr><td>>hg38_21_+_8433172_8446622_RNA45SN1_45S_50ntFlanks@6644.6685.42|CCTCGTACGACTCTTAGCGGTGGATCACTCGGCTCGTGCGTC</td><td>>hg38_21_+_8433172_8446622_RNA45SN1_45S_50ntFlanks</td><td>6644</td><td>6685</td><td>42</td><td>CCTCGTACGACTCTTAGCGGTGGATCACTCGGCTCGTGCGTC</td><td>102.9</td><td>6.0</td><td>1.6</td><td>3.7</td><td>2.3E-04</td><td>6.8E-04</td><td>BT20/MB468</td><td>MP</td></tr>
<tr><td>>hg38_21_+_8433172_8446622_RNA45SN1_45S_50ntFlanks@5485.5514.30|AGCGCTGAGAAGACGGTCGAACTTGACTAT</td><td>>hg38_21_+_8433172_8446622_RNA45SN1_45S_50ntFlanks</td><td>5485</td><td>5514</td><td>30</td><td>AGCGCTGAGAAGACGGTCGAACTTGACTAT</td><td>12180.4</td><td>-4.7</td><td>1.3</td><td>-3.7</td><td>2.3E-04</td><td>6.8E-04</td><td>BT20/MB468</td><td>MP</td></tr>
<tr><td>>hg38_21_+_8433172_8446622_RNA45SN1_45S_50ntFlanks@6646.6685.40|TCGTACGACTCTTAGCGGTGGATCACTCGGCTCGTGCGTC</td><td>>hg38_21_+_8433172_8446622_RNA45SN1_45S_50ntFlanks</td><td>6646</td><td>6685</td><td>40</td><td>TCGTACGACTCTTAGCGGTGGATCACTCGGCTCGTGCGTC</td><td>725.6</td><td>2.7</td><td>0.8</td><td>3.5</td><td>5.5E-04</td><td>6.8E-04</td><td>MB231/MB468</td><td>NUC</td></tr>
<tr><td>>MI0000067|hsa-let-7f-1&WithFlank&9|+|94176341|94176439@13.34.22(+1A);MI0000068|hsa-let-7f-2&WithFlank&X|-|53557186|53557280@14.35.22(+1A)&[MIMAT0000067&hsa-let-7f-5p&offsets|0|0(+1A);m-20&9|+|94176353|94176374&offsets|0|0(+1A)];[MIMAT0000067_1&hsa-let-7f-5p&offsets|0|0(+1A);m-19&X|-|53557246|53557267&offsets|0|0(+1A)]|TGAGGTAGTAGATTGTATAGTTA</td><td>>MI0000067|hsa-let-7f-1&WithFlank&9|+|94176341|94176439</td><td>13</td><td>34</td><td>22(+1A)</td><td>TGAGGTAGTAGATTGTATAGTTA</td><td>876.1</td><td>1.5</td><td>0.4</td><td>3.5</td><td>5.5E-04</td><td>6.8E-04</td><td>BT20/MB468</td><td>NUC</td></tr>
<tr><td>>hg38_21_+_8433172_8446622_RNA45SN1_45S_50ntFlanks@10345.10388.44|TTCAAACGAGAACTTTGAAGGCCGAAGTGGAGAAGGGTTCCATG</td><td>>hg38_21_+_8433172_8446622_RNA45SN1_45S_50ntFlanks</td><td>10345</td><td>10388</td><td>44</td><td>TTCAAACGAGAACTTTGAAGGCCGAAGTGGAGAAGGGTTCCATG</td><td>396.6</td><td>3.9</td><td>1.0</td><td>3.9</td><td>9.5E-05</td><td>6.8E-04</td><td>BT20/MB231</td><td>MITO</td></tr>
<tr><td>>MI0000101|hsa-mir-99a&WithFlank&21|+|16539083|16539175@19.40.22(+1A)&[MIMAT0000097&hsa-miR-99a-5p&offsets|0|0(+1A);m-64&21|+|16539101|16539121&offsets|0|+1(+1A)]|AACCCGTAGATCCGATCTTGTGA</td><td>>MI0000101|hsa-mir-99a&WithFlank&21|+|16539083|16539175</td><td>19</td><td>40</td><td>22(+1A)</td><td>AACCCGTAGATCCGATCTTGTGA</td><td>291.4</td><td>-3.8</td><td>1.1</td><td>-3.5</td><td>4.1E-04</td><td>6.8E-04</td><td>BT20/MB468</td><td>MITO</td></tr>
<tr><td>>hg38_21_+_8433172_8446622_RNA45SN1_45S_50ntFlanks@8190.8213.24|CCAGCCCGTGGACGGTGTGAGGCC</td><td>>hg38_21_+_8433172_8446622_RNA45SN1_45S_50ntFlanks</td><td>8190</td><td>8213</td><td>24</td><td>CCAGCCCGTGGACGGTGTGAGGCC</td><td>305.3</td><td>-4.4</td><td>1.3</td><td>-3.5</td><td>4.6E-04</td><td>6.8E-04</td><td>MB231/MB468</td><td>MP</td></tr>
<tr><td>>trnaMT_TyrGTA_MT_-_5826_5891@23.69.47|TTGGACTGTAAATCTAAAGACAGGGGTTAGGCCTCTTTTTACCACCA</td><td>>trnaMT_TyrGTA_MT_-_5826_5891</td><td>23</td><td>69</td><td>47</td><td>TTGGACTGTAAATCTAAAGACAGGGGTTAGGCCTCTTTTTACCACCA</td><td>69.2</td><td>9.9</td><td>2.7</td><td>3.7</td><td>2.3E-04</td><td>6.8E-04</td><td>BT20/MB468</td><td>MP</td></tr>
<tr><td>>hg38_21_+_8433172_8446622_RNA45SN1_45S_50ntFlanks@11849.11873.25|CGGGCTTGGCGGAATCAGCGGGGAA</td><td>>hg38_21_+_8433172_8446622_RNA45SN1_45S_50ntFlanks</td><td>11849</td><td>11873</td><td>25</td><td>CGGGCTTGGCGGAATCAGCGGGGAA</td><td>159.8</td><td>-4.9</td><td>1.4</td><td>-3.5</td><td>5.5E-04</td><td>6.8E-04</td><td>MB231/MB468</td><td>NUC</td></tr>
<tr><td>>hg38_21_+_8433172_8446622_RNA45SN1_45S_50ntFlanks@10421.10456.36|TGAGAGATGGGCGAGCGCCGTTCCGAAGGGACGGGC</td><td>>hg38_21_+_8433172_8446622_RNA45SN1_45S_50ntFlanks</td><td>10421</td><td>10456</td><td>36</td><td>TGAGAGATGGGCGAGCGCCGTTCCGAAGGGACGGGC</td><td>537.0</td><td>-3.6</td><td>0.9</td><td>-3.9</td><td>8.3E-05</td><td>6.8E-04</td><td>BT20/MB231</td><td>CYTO</td></tr>
<tr><td>>hg38_21_+_8433172_8446622_RNA45SN1_45S_50ntFlanks@11919.11967.49|CATGAGAGGTGTAGAATAAGTGGGAGGCCCCCGGCGCCCCCCCGGTGTC</td><td>>hg38_21_+_8433172_8446622_RNA45SN1_45S_50ntFlanks</td><td>11919</td><td>11967</td><td>49</td><td>CATGAGAGGTGTAGAATAAGTGGGAGGCCCCCGGCGCCCCCCCGGTGTC</td><td>148.9</td><td>-4.3</td><td>1.2</td><td>-3.5</td><td>5.1E-04</td><td>6.8E-04</td><td>MB231/MB468</td><td>CYTO</td></tr>
<tr><td>>hg38_21_+_8433172_8446622_RNA45SN1_45S_50ntFlanks@10737.10755.19|TAAATCTCGCGCCGGGCCG</td><td>>hg38_21_+_8433172_8446622_RNA45SN1_45S_50ntFlanks</td><td>10737</td><td>10755</td><td>19</td><td>TAAATCTCGCGCCGGGCCG</td><td>915.1</td><td>-4.3</td><td>1.3</td><td>-3.5</td><td>5.1E-04</td><td>6.8E-04</td><td>MB231/MB468</td><td>CYTO</td></tr>
<tr><td>>hg38_21_+_8433172_8446622_RNA45SN1_45S_50ntFlanks@8317.8356.40|CACGAGACCGATAGTCAACAAGTACCGTAAGGGAAAGTTG</td><td>>hg38_21_+_8433172_8446622_RNA45SN1_45S_50ntFlanks</td><td>8317</td><td>8356</td><td>40</td><td>CACGAGACCGATAGTCAACAAGTACCGTAAGGGAAAGTTG</td><td>436.8</td><td>-3.0</td><td>0.9</td><td>-3.5</td><td>4.9E-04</td><td>6.8E-04</td><td>BT20/MB468</td><td>CYTO</td></tr>
<tr><td>>hg38_21_+_8433172_8446622_RNA45SN1_45S_50ntFlanks@10349.10381.33|AACGAGAACTTTGAAGGCCGAAGTGGAGAAGGG</td><td>>hg38_21_+_8433172_8446622_RNA45SN1_45S_50ntFlanks</td><td>10349</td><td>10381</td><td>33</td><td>AACGAGAACTTTGAAGGCCGAAGTGGAGAAGGG</td><td>599.6</td><td>-6.3</td><td>1.8</td><td>-3.5</td><td>5.5E-04</td><td>6.8E-04</td><td>BT20/MB468</td><td>NUC</td></tr>
<tr><td>>hg38_21_+_8433172_8446622_RNA45SN1_45S_50ntFlanks@6741.6771.31|ATCGACACTTCGAACGCACTTGCGGCCCCGG</td><td>>hg38_21_+_8433172_8446622_RNA45SN1_45S_50ntFlanks</td><td>6741</td><td>6771</td><td>31</td><td>ATCGACACTTCGAACGCACTTGCGGCCCCGG</td><td>562.8</td><td>-4.0</td><td>1.1</td><td>-3.5</td><td>4.9E-04</td><td>6.8E-04</td><td>BT20/MB468</td><td>CYTO</td></tr>
<tr><td>>hg38_21_+_8433172_8446622_RNA45SN1_45S_50ntFlanks@10350.10383.34|ACGAGAACTTTGAAGGCCGAAGTGGAGAAGGGTT</td><td>>hg38_21_+_8433172_8446622_RNA45SN1_45S_50ntFlanks</td><td>10350</td><td>10383</td><td>34</td><td>ACGAGAACTTTGAAGGCCGAAGTGGAGAAGGGTT</td><td>12385.9</td><td>-4.4</td><td>1.2</td><td>-3.7</td><td>2.3E-04</td><td>6.8E-04</td><td>BT20/MB468</td><td>MP</td></tr>
<tr><td>>hg38_21_+_8433172_8446622_RNA45SN1_45S_50ntFlanks@12990.13015.26|GCTCCCTCGCTGCGATCTATTGAAAG</td><td>>hg38_21_+_8433172_8446622_RNA45SN1_45S_50ntFlanks</td><td>12990</td><td>13015</td><td>26</td><td>GCTCCCTCGCTGCGATCTATTGAAAG</td><td>311.7</td><td>-2.8</td><td>0.8</td><td>-3.5</td><td>4.1E-04</td><td>6.8E-04</td><td>BT20/MB468</td><td>MITO</td></tr>
<tr><td>>hg38_21_+_8433172_8446622_RNA45SN1_45S_50ntFlanks@11731.11751.21|ACTATGACTCTCTTAAGGTAG</td><td>>hg38_21_+_8433172_8446622_RNA45SN1_45S_50ntFlanks</td><td>11731</td><td>11751</td><td>21</td><td>ACTATGACTCTCTTAAGGTAG</td><td>216.1</td><td>-4.9</td><td>1.4</td><td>-3.5</td><td>4.9E-04</td><td>6.8E-04</td><td>BT20/MB468</td><td>CYTO</td></tr>
<tr><td>>hg38_21_+_8433172_8446622_RNA45SN1_45S_50ntFlanks@3843.3868.26|CTACTTGGATAACTGTGGTAATTCTA</td><td>>hg38_21_+_8433172_8446622_RNA45SN1_45S_50ntFlanks</td><td>3843</td><td>3868</td><td>26</td><td>CTACTTGGATAACTGTGGTAATTCTA</td><td>327.4</td><td>-2.2</td><td>0.6</td><td>-3.5</td><td>5.5E-04</td><td>6.8E-04</td><td>MB231/MB468</td><td>NUC</td></tr>
<tr><td>>MI0000107|hsa-mir-29b-2&WithFlank&1|-|207802437|207802529@58.77.20;MI0000105|hsa-mir-29b-1&WithFlank&7|-|130877453|130877545@57.76.20&[MIMAT0000100&hsa-miR-29b-3p&offsets|0|-3;m-81&1|-|207802450|207802472&offsets|0|-3];[MIMAT0000100_1&hsa-miR-29b-3p&offsets|0|-3;m-82&7|-|130877467|130877489&offsets|0|-3]|TAGCACCATTTGAAATCAGT</td><td>>MI0000107|hsa-mir-29b-2&WithFlank&1|-|207802437|207802529</td><td>58</td><td>77</td><td>20</td><td>TAGCACCATTTGAAATCAGT</td><td>113.2</td><td>5.4</td><td>1.5</td><td>3.5</td><td>4.6E-04</td><td>6.8E-04</td><td>MB231/MB468</td><td>MP</td></tr>
<tr><td>>hg38_21_+_8433172_8446622_RNA45SN1_45S_50ntFlanks@10392.10422.31|ACAGCAGTTGAACATGGGTCAGTCGGTCCTG</td><td>>hg38_21_+_8433172_8446622_RNA45SN1_45S_50ntFlanks</td><td>10392</td><td>10422</td><td>31</td><td>ACAGCAGTTGAACATGGGTCAGTCGGTCCTG</td><td>610.1</td><td>-4.6</td><td>1.3</td><td>-3.5</td><td>4.9E-04</td><td>6.8E-04</td><td>BT20/MB468</td><td>CYTO</td></tr>
<tr><td>>MI0000289|hsa-mir-181a-1&WithFlank&1|-|198859038|198859159@70.90.21(+1U)&[MIMAT0000270&hsa-miR-181a-3p&offsets|0|-1(+1U);m-125&1|-|198859069|198859090&offsets|0|-1(+1U)]|ACCATCGACCGTTGATTGTACT</td><td>>MI0000289|hsa-mir-181a-1&WithFlank&1|-|198859038|198859159</td><td>70</td><td>90</td><td>21(+1U)</td><td>ACCATCGACCGTTGATTGTACT</td><td>290.9</td><td>4.4</td><td>1.3</td><td>3.5</td><td>4.6E-04</td><td>6.8E-04</td><td>MB231/MB468</td><td>MP</td></tr>
<tr><td>>hg38_21_+_8433172_8446622_RNA45SN1_45S_50ntFlanks@3840.3865.26|CTCCTACTTGGATAACTGTGGTAATT</td><td>>hg38_21_+_8433172_8446622_RNA45SN1_45S_50ntFlanks</td><td>3840</td><td>3865</td><td>26</td><td>CTCCTACTTGGATAACTGTGGTAATT</td><td>112.5</td><td>2.2</td><td>0.6</td><td>3.5</td><td>5.5E-04</td><td>6.8E-04</td><td>BT20/MB468</td><td>NUC</td></tr>
<tr><td>>hg38_21_+_8433172_8446622_RNA45SN1_45S_50ntFlanks@12416.12455.40|TTCGATGTCGGCTCTTCCTATCATTGTGAAGCAGAATTCA</td><td>>hg38_21_+_8433172_8446622_RNA45SN1_45S_50ntFlanks</td><td>12416</td><td>12455</td><td>40</td><td>TTCGATGTCGGCTCTTCCTATCATTGTGAAGCAGAATTCA</td><td>275.0</td><td>-3.3</td><td>0.8</td><td>-3.9</td><td>8.3E-05</td><td>6.8E-04</td><td>BT20/MB231</td><td>CYTO</td></tr>
<tr><td>>MI0003820|hsa-mir-454&WithFlank&17|-|59137752|59137878@70.94.25&[MIMAT0003885&hsa-miR-454-3p&offsets|0|+2;m-272&17|-|59137787|59137809&offsets|0|+2]|TAGTGCAATATTGCTTATAGGGTTT</td><td>>MI0003820|hsa-mir-454&WithFlank&17|-|59137752|59137878</td><td>70</td><td>94</td><td>25</td><td>TAGTGCAATATTGCTTATAGGGTTT</td><td>670.2</td><td>4.7</td><td>1.3</td><td>3.5</td><td>4.6E-04</td><td>6.8E-04</td><td>MB231/MB468</td><td>MP</td></tr>
<tr><td>>MI0000095|hsa-mir-93&WithFlank&7|-|100093762|100093853@17.36.20(+1C)&[MIMAT0000093&hsa-miR-93-5p&offsets|0|-3(+1C);m-29&7|-|100093815|100093837&offsets|0|-3(+1C)]|CAAAGTGCTGTTCGTGCAGGC</td><td>>MI0000095|hsa-mir-93&WithFlank&7|-|100093762|100093853</td><td>17</td><td>36</td><td>20(+1C)</td><td>CAAAGTGCTGTTCGTGCAGGC</td><td>714.3</td><td>3.3</td><td>1.0</td><td>3.5</td><td>4.6E-04</td><td>6.8E-04</td><td>MB231/MB468</td><td>MP</td></tr>
<tr><td>>MI0000077|hsa-mir-21&WithFlank&17|+|59841260|59841343@14.31.18(+1C)&[MIMAT0000076&hsa-miR-21-5p&offsets|0|-4(+1C);m-1&17|+|59841273|59841294&offsets|0|-4(+1C)]|TAGCTTATCAGACTGATGC</td><td>>MI0000077|hsa-mir-21&WithFlank&17|+|59841260|59841343</td><td>14</td><td>31</td><td>18(+1C)</td><td>TAGCTTATCAGACTGATGC</td><td>369.2</td><td>4.6</td><td>1.3</td><td>3.7</td><td>2.3E-04</td><td>6.8E-04</td><td>BT20/MB468</td><td>MP</td></tr>
<tr><td>>MI0000272|hsa-mir-182&WithFlank&7|-|129770377|129770498@29.47.19&[MIMAT0000259&hsa-miR-182-5p&offsets|0|-5;m-22&7|-|129770449|129770470&offsets|0|-3]|TTTGGCAATGGTAGAACTC</td><td>>MI0000272|hsa-mir-182&WithFlank&7|-|129770377|129770498</td><td>29</td><td>47</td><td>19</td><td>TTTGGCAATGGTAGAACTC</td><td>177.8</td><td>4.3</td><td>1.1</td><td>3.9</td><td>8.3E-05</td><td>6.8E-04</td><td>BT20/MB231</td><td>CYTO</td></tr>
<tr><td>>MI0017308|hsa-mir-4677&WithFlank&1|+|243346170|243346261@56.76.21&[MIMAT0019761&hsa-miR-4677-3p&offsets|0|-1;m-295&1|+|243346225|243346246&offsets|0|-1]|TCTGTGAGACCAAAGAACTAC</td><td>>MI0017308|hsa-mir-4677&WithFlank&1|+|243346170|243346261</td><td>56</td><td>76</td><td>21</td><td>TCTGTGAGACCAAAGAACTAC</td><td>78.3</td><td>7.0</td><td>1.9</td><td>3.7</td><td>2.3E-04</td><td>6.8E-04</td><td>BT20/MB468</td><td>MP</td></tr>
<tr><td>>hg38_21_+_8433172_8446622_RNA45SN1_45S_50ntFlanks@9574.9605.32|AATCGGTCGTCCGACCTGGGTATAGGGGCGAA</td><td>>hg38_21_+_8433172_8446622_RNA45SN1_45S_50ntFlanks</td><td>9574</td><td>9605</td><td>32</td><td>AATCGGTCGTCCGACCTGGGTATAGGGGCGAA</td><td>1284.9</td><td>-2.9</td><td>0.8</td><td>-3.5</td><td>4.6E-04</td><td>6.8E-04</td><td>MB231/MB468</td><td>MP</td></tr>
<tr><td>>hg38_21_+_8433172_8446622_RNA45SN1_45S_50ntFlanks@10319.10354.36|GCAGATCTTGGTGGTAGTAGCAAATATTCAAACGAG</td><td>>hg38_21_+_8433172_8446622_RNA45SN1_45S_50ntFlanks</td><td>10319</td><td>10354</td><td>36</td><td>GCAGATCTTGGTGGTAGTAGCAAATATTCAAACGAG</td><td>56.6</td><td>6.6</td><td>1.8</td><td>3.7</td><td>2.3E-04</td><td>6.8E-04</td><td>BT20/MB468</td><td>MP</td></tr>
<tr><td>>hg38_21_+_8433172_8446622_RNA45SN1_45S_50ntFlanks@10239.10259.21|GCTACGCCGCGACGAGTAGGA</td><td>>hg38_21_+_8433172_8446622_RNA45SN1_45S_50ntFlanks</td><td>10239</td><td>10259</td><td>21</td><td>GCTACGCCGCGACGAGTAGGA</td><td>855.0</td><td>5.2</td><td>1.3</td><td>3.9</td><td>8.0E-05</td><td>6.9E-04</td><td>BT20/MB231</td><td>MP</td></tr>
<tr><td>>hg38_21_+_8433172_8446622_RNA45SN1_45S_50ntFlanks@10905.10941.37|CTGGGCGCGCGCCGCGGCTGGACGAGGCGCCGCCGCC</td><td>>hg38_21_+_8433172_8446622_RNA45SN1_45S_50ntFlanks</td><td>10905</td><td>10941</td><td>37</td><td>CTGGGCGCGCGCCGCGGCTGGACGAGGCGCCGCCGCC</td><td>339.4</td><td>-5.4</td><td>1.5</td><td>-3.5</td><td>4.9E-04</td><td>6.9E-04</td><td>BT20/MB468</td><td>CYTO</td></tr>
<tr><td>>hg38_21_+_8433172_8446622_RNA45SN1_45S_50ntFlanks@7979.8003.25|ACCTCAGATCAGACGTGGCGACCCG</td><td>>hg38_21_+_8433172_8446622_RNA45SN1_45S_50ntFlanks</td><td>7979</td><td>8003</td><td>25</td><td>ACCTCAGATCAGACGTGGCGACCCG</td><td>516.4</td><td>-4.1</td><td>1.2</td><td>-3.5</td><td>4.9E-04</td><td>6.9E-04</td><td>BT20/MB468</td><td>CYTO</td></tr>
<tr><td>>MI0000063|hsa-let-7b&WithFlank&22|+|46113680|46113774@66.86.21(+1U)&[MIMAT0004482&hsa-let-7b-3p&offsets|0|-1(+1U);m-171&22|+|46113745|46113766&offsets|0|-1(+1U)]|CTATACAACCTACTGCCTTCCT</td><td>>MI0000063|hsa-let-7b&WithFlank&22|+|46113680|46113774</td><td>66</td><td>86</td><td>21(+1U)</td><td>CTATACAACCTACTGCCTTCCT</td><td>183.5</td><td>3.2</td><td>0.9</td><td>3.5</td><td>4.6E-04</td><td>6.9E-04</td><td>MB231/MB468</td><td>MP</td></tr>
<tr><td>>hg38_21_+_8433172_8446622_RNA45SN1_45S_50ntFlanks@9005.9026.22|CTCCCCCGCGGGGGCGCGCCGG</td><td>>hg38_21_+_8433172_8446622_RNA45SN1_45S_50ntFlanks</td><td>9005</td><td>9026</td><td>22</td><td>CTCCCCCGCGGGGGCGCGCCGG</td><td>667.4</td><td>-5.5</td><td>1.6</td><td>-3.5</td><td>4.1E-04</td><td>6.9E-04</td><td>BT20/MB468</td><td>MITO</td></tr>
<tr><td>>hg38_21_+_8433172_8446622_RNA45SN1_45S_50ntFlanks@10411.10456.46|CAGTCGGTCCTGAGAGATGGGCGAGCGCCGTTCCGAAGGGACGGGC</td><td>>hg38_21_+_8433172_8446622_RNA45SN1_45S_50ntFlanks</td><td>10411</td><td>10456</td><td>46</td><td>CAGTCGGTCCTGAGAGATGGGCGAGCGCCGTTCCGAAGGGACGGGC</td><td>686.5</td><td>-4.1</td><td>1.2</td><td>-3.5</td><td>4.1E-04</td><td>6.9E-04</td><td>BT20/MB468</td><td>MITO</td></tr>
<tr><td>>MI0006273|hsa-mir-1180&WithFlank&17|-|19344500|19344580@47.67.21(+1C)&[MIMAT0005825&hsa-miR-1180-3p&offsets|0|-1(+1C);m-214&17|-|19344513|19344534&offsets|0|-1(+1C)]|TTTCCGGCTCGCGTGGGTGTGC</td><td>>MI0006273|hsa-mir-1180&WithFlank&17|-|19344500|19344580</td><td>47</td><td>67</td><td>21(+1C)</td><td>TTTCCGGCTCGCGTGGGTGTGC</td><td>68.7</td><td>2.7</td><td>0.8</td><td>3.5</td><td>5.6E-04</td><td>6.9E-04</td><td>BT20/MB468</td><td>NUC</td></tr>
<tr><td>>trnaMT_AsnGTT_MT_-_5657_5729@1.39.39,trnalookalike5_AsnGTT_1_-_566207_566279@1.39.39|TAGATTGAAGCCAGTTGATTAGGGTGCTTAGCTGTTAAC</td><td>>trnaMT_AsnGTT_MT_-_5657_5729</td><td>1</td><td>39</td><td>39</td><td>TAGATTGAAGCCAGTTGATTAGGGTGCTTAGCTGTTAAC</td><td>158.5</td><td>5.3</td><td>1.5</td><td>3.5</td><td>5.5E-04</td><td>6.9E-04</td><td>MB231/MB468</td><td>MITO</td></tr>
<tr><td>>hg38_21_+_8433172_8446622_RNA45SN1_45S_50ntFlanks@9554.9573.20|CGTAGCGGTCCTGACGTGCA</td><td>>hg38_21_+_8433172_8446622_RNA45SN1_45S_50ntFlanks</td><td>9554</td><td>9573</td><td>20</td><td>CGTAGCGGTCCTGACGTGCA</td><td>4241.0</td><td>-3.6</td><td>1.0</td><td>-3.7</td><td>2.3E-04</td><td>6.9E-04</td><td>BT20/MB468</td><td>MP</td></tr>
<tr><td>>MI0000079|hsa-mir-23a&WithFlank&19|-|13836581|13836665@51.70.20(+1U)&[MIMAT0000078&hsa-miR-23a-3p&offsets|0|-1(+1U);m-31&19|-|13836595|13836615&offsets|0|-1(+1U)]|ATCACATTGCCAGGGATTTCT</td><td>>MI0000079|hsa-mir-23a&WithFlank&19|-|13836581|13836665</td><td>51</td><td>70</td><td>20(+1U)</td><td>ATCACATTGCCAGGGATTTCT</td><td>63.6</td><td>-9.1</td><td>2.5</td><td>-3.7</td><td>2.3E-04</td><td>6.9E-04</td><td>BT20/MB468</td><td>MP</td></tr>
</table>

| | | | | | | | | | | | | | |
|---|---|---|---|---|---|---|---|---|---|---|---|---|---|
| >hg38_21_+_8433172_8446622_RNA45SN1_45S_50ntFlanks@12417.12434.18\|TCGATGTCGGCTCTTCCT | >hg38_21_+_8433172_8446622_RNA45SN1_45S_50ntFlanks | 12417 | 12434 | 18 | TCGATGTCGGCTCTTCCT | 1563.7 | -1.4 | 0.4 | -3.5 | 4.1E-04 | 6.9E-04 | BT20/MB468 | MITO |
| >hg38_21_+_8433172_8446622_RNA45SN1_45S_50ntFlanks@6721.6760.40\|AATTGCAGGACACATTGATCATCGACACTTCGAACGCACT | >hg38_21_+_8433172_8446622_RNA45SN1_45S_50ntFlanks | 6721 | 6760 | 40 | AATTGCAGGACACATTGATCATCGACACTTCGAACGCACT | 164.5 | -4.6 | 1.3 | -3.5 | 4.6E-04 | 6.9E-04 | MB231/MB468 | MP |
| >MI0000090\|hsa-mir-32&WithFlank&9\|-\|109046223\|109046304@53.73.21&[MIMAT0004505&hsa-miR-32-3p&offsets\|0\|-1;m-518&9\|-\|109046232\|109046252&offsets\|0\|0]\|CAATTTAGTGTGTGTGATATT | >MI0000090\|hsa-mir-32&WithFlank&9\|-\|109046223\|109046304 | 53 | 73 | 21 | CAATTTAGTGTGTGTGATATT | 163.7 | -3.1 | 0.8 | -3.9 | 9.7E-05 | 6.9E-04 | BT20/MB231 | MITO |
| >trnaMT_ValTAC_MT_+_1602_1670@50.72.23\|ACTTAACTTGACCGCTCTGACCA | >trnaMT_ValTAC_MT_+_1602_1670 | 50 | 72 | 23 | ACTTAACTTGACCGCTCTGACCA | 15131.4 | 4.4 | 1.3 | 3.5 | 4.1E-04 | 6.9E-04 | BT20/MB468 | MITO |
| >hg38_MT_+_1621_3279_RNR2_16S_50ntFlanks@1368.1405.38\|CGATGTTGGATCAGGACATCCCGATGGTGCAGCCGCTA | >hg38_MT_+_1621_3279_RNR2_16S_50ntFlanks | 1368 | 1405 | 38 | CGATGTTGGATCAGGACATCCCGATGGTGCAGCCGCTA | 19.3 | 7.5 | 2.1 | 3.5 | 4.6E-04 | 6.9E-04 | MB231/MB468 | MP |
| >trna116_GluCTC_1_-_145399233_145399304@1.31.31,trna59_GluCTC_1_+_249168447_249168518@1.31.31,trna71_GluCTC_1_-_161439189_161439260@1.31.31,trna74_GluCTC_1_-_161431809_161431880@1.31.31,trna77_GluCTC_1_-_161424398_161424469@1.31.31,trna77_GluCTC_6_+_28949976_28950047@1.31.31,trna80_GluCTC_1_-_161417018_161417089@1.31.31,trna87_GluCTC_6_-_126101393_126101464@1.31.31\|TCCCTGGTGGTCTAGTGGTTAGGATTCGGCG | >trna116_GluCTC_1_-_145399233_145399304 | 1 | 31 | 31 | TCCCTGGTGGTCTAGTGGTTAGGATTCGGCG | 21120.4 | -2.8 | 0.8 | -3.5 | 5.5E-04 | 6.9E-04 | MB231/MB468 | MITO |
| >hg38_21_+_8433172_8446622_RNA45SN1_45S_50ntFlanks@12184.12211.28\|AACGGTAACGCAGGTGTCCTAAGGCGAG | >hg38_21_+_8433172_8446622_RNA45SN1_45S_50ntFlanks | 12184 | 12211 | 28 | AACGGTAACGCAGGTGTCCTAAGGCGAG | 120.5 | -5.6 | 1.5 | -3.7 | 2.3E-04 | 6.9E-04 | BT20/MB468 | MP |
| >hg38_1_-_228634819_228635039_RNA5S12_5S_50ntFlanks@84.102.19\|CTCGTCTGATCTCGGAAGC | >hg38_1_-_228634819_228635039_RNA5S12_5S_50ntFlanks | 84 | 102 | 19 | CTCGTCTGATCTCGGAAGC | 582.8 | -4.3 | 1.2 | -3.5 | 5.2E-04 | 6.9E-04 | MB231/MB468 | CYTO |
| >MI0003591\|hsa-mir-584&WithFlank&5\|-\|149062307\|149062415@22.42.21(+2A)&[MIMAT0003249&hsa-miR-584-5p&offsets\|0\|-1(+2A);m-167&5\|-\|149062374\|149062394&offsets\|0\|0(+2A)]\|TTATGGTTTGCCTGGGACTGAAA | >MI0003591\|hsa-mir-584&WithFlank&5\|-\|149062307\|149062415 | 22 | 42 | 21(+2A) | TTATGGTTTGCCTGGGACTGAAA | 77.6 | 5.8 | 1.6 | 3.7 | 2.3E-04 | 6.9E-04 | BT20/MB468 | MP |
| >MI0000095\|hsa-mir-93&WithFlank&7\|-\|100093762\|100093853@17.37.21&[MIMAT0000093&hsa-miR-93-5p&offsets\|0\|-2;m-29&7\|-\|100093815\|100093837&offsets\|0\|-2]\|CAAAGTGCTGTTCGTGCAGGT | >MI0000095\|hsa-mir-93&WithFlank&7\|-\|100093762\|100093853 | 17 | 37 | 21 | CAAAGTGCTGTTCGTGCAGGT | 4114.2 | 3.0 | 0.9 | 3.5 | 4.6E-04 | 6.9E-04 | MB231/MB468 | MP |
| >hg38_21_+_8433172_8446622_RNA45SN1_45S_50ntFlanks@7986.8016.31\|ATCAGACGTGGCGACCCGCTGAATTTAAGCA | >hg38_21_+_8433172_8446622_RNA45SN1_45S_50ntFlanks | 7986 | 8016 | 31 | ATCAGACGTGGCGACCCGCTGAATTTAAGCA | 1601.5 | -5.8 | 1.7 | -3.5 | 5.5E-04 | 6.9E-04 | MB231/MB468 | MITO |
| >hg38_21_+_8433172_8446622_RNA45SN1_45S_50ntFlanks@10678.10708.31\|CGGTTCCGGCGGCGTCCGGTGAGCTCTCGCT | >hg38_21_+_8433172_8446622_RNA45SN1_45S_50ntFlanks | 10678 | 10708 | 31 | CGGTTCCGGCGGCGTCCGGTGAGCTCTCGCT | 9772.6 | -3.3 | 0.9 | -3.5 | 5.0E-04 | 6.9E-04 | BT20/MB468 | CYTO |
| >hg38_21_+_8433172_8446622_RNA45SN1_45S_50ntFlanks@11784.11803.20\|ATGAATGGATGAACGAGATT | >hg38_21_+_8433172_8446622_RNA45SN1_45S_50ntFlanks | 11784 | 11803 | 20 | ATGAATGGATGAACGAGATT | 1244.0 | -3.1 | 0.9 | -3.5 | 5.0E-04 | 6.9E-04 | BT20/MB468 | CYTO |
| >hg38_21_+_8433172_8446622_RNA45SN1_45S_50ntFlanks@11874.11903.30\|AGAAGACCCTGTTGAGCTTGACTCTAGTCT | >hg38_21_+_8433172_8446622_RNA45SN1_45S_50ntFlanks | 11874 | 11903 | 30 | AGAAGACCCTGTTGAGCTTGACTCTAGTCT | 484.9 | -3.3 | 0.9 | -3.7 | 2.3E-04 | 6.9E-04 | BT20/MB468 | MP |
| >hg38_1_-_228634819_228635039_RNA5S12_5S_50ntFlanks@133.170.38\|ATGGGAGACCGCCTGGGAATACCGGGTGCTGTAGGCTT | >hg38_1_-_228634819_228635039_RNA5S12_5S_50ntFlanks | 133 | 170 | 38 | ATGGGAGACCGCCTGGGAATACCGGGTGCTGTAGGCTT | 12281.2 | 3.0 | 0.9 | 3.5 | 5.2E-04 | 6.9E-04 | MB231/MB468 | CYTO |
| >hg38_21_+_8433172_8446622_RNA45SN1_45S_50ntFlanks@12418.12444.27\|CGATGTCGGCTCTTCCTATCATTGTGA | >hg38_21_+_8433172_8446622_RNA45SN1_45S_50ntFlanks | 12418 | 12444 | 27 | CGATGTCGGCTCTTCCTATCATTGTGA | 1834.4 | -2.2 | 0.6 | -3.5 | 5.0E-04 | 6.9E-04 | BT20/MB468 | CYTO |
| >trna27_GlyCCC_2_-_70476123_70476193@1.28.28,trna34_GlyCCC_16_-_686736_686806@1.28.28\|GCGCCGCTGGTGTAGTGGTATCATGCAA | >trna27_GlyCCC_2_-_70476123_70476193 | 1 | 28 | 28 | GCGCCGCTGGTGTAGTGGTATCATGCAA | 86.2 | -9.5 | 2.6 | -3.7 | 2.3E-04 | 6.9E-04 | BT20/MB468 | MP |
| >hg38_21_+_8433172_8446622_RNA45SN1_45S_50ntFlanks@7974.7996.23\|ACGCGACCTCAGATCAGACGTGG | >hg38_21_+_8433172_8446622_RNA45SN1_45S_50ntFlanks | 7974 | 7996 | 23 | ACGCGACCTCAGATCAGACGTGG | 504.6 | -2.2 | 0.6 | -3.5 | 4.7E-04 | 6.9E-04 | MB231/MB468 | MP |
| >hg38_21_+_8433172_8446622_RNA45SN1_45S_50ntFlanks@11703.11740.38\|AATGAAGCGCGGGTAAACGGCGGGAGTAACTATGACTC | >hg38_21_+_8433172_8446622_RNA45SN1_45S_50ntFlanks | 11703 | 11740 | 38 | AATGAAGCGCGGGTAAACGGCGGGAGTAACTATGACTC | 95.0 | -5.7 | 1.6 | -3.5 | 4.7E-04 | 6.9E-04 | MB231/MB468 | MP |
| >MI0000089\|hsa-mir-31&WithFlank&9\|-\|21512109\|21512191@14.33.20&[MIMAT0000089&hsa-miR-31-5p&offsets\|0\|-1;m-360&9\|-\|21512157\|21512178&offsets\|0\|-2]\|AGGCAAGATGCTGGCATAGC | >MI0000089\|hsa-mir-31&WithFlank&9\|-\|21512109\|21512191 | 14 | 33 | 20 | AGGCAAGATGCTGGCATAGC | 45.4 | 8.8 | 2.2 | 3.9 | 8.1E-05 | 6.9E-04 | BT20/MB231 | MP |
| >hg38_21_+_8433172_8446622_RNA45SN1_45S_50ntFlanks@12416.12457.42\|TTCGATGTCGGCTCTTCCTATCATTGTGAAGCAGAATTCACC | >hg38_21_+_8433172_8446622_RNA45SN1_45S_50ntFlanks | 12416 | 12457 | 42 | TTCGATGTCGGCTCTTCCTATCATTGTGAAGCAGAATTCACC | 491.7 | -3.8 | 1.0 | -3.9 | 8.5E-05 | 7.0E-04 | BT20/MB231 | NUC |
| >MI0000063\|hsa-let-7b&WithFlank&22\|+\|46113680\|46113774@12.32.21(+1G)&[MIMAT0000063&hsa-let-7b-5p&offsets\|0\|-1(+1G);m-14&22\|+\|46113691\|46113712&offsets\|0\|-1(+1G)]\|TGAGGTAGTAGGTTGTGTGGTG | >MI0000063\|hsa-let-7b&WithFlank&22\|+\|46113680\|46113774 | 12 | 32 | 21(+1G) | TGAGGTAGTAGGTTGTGTGGTG | 802.1 | 4.4 | 1.2 | 3.7 | 2.3E-04 | 7.0E-04 | BT20/MB468 | MP |
| >hg38_21_+_8433172_8446622_RNA45SN1_45S_50ntFlanks@12083.12108.26\|CTCGCTTCTGGCGCCAAGCGCCCGGC | >hg38_21_+_8433172_8446622_RNA45SN1_45S_50ntFlanks | 12083 | 12108 | 26 | CTCGCTTCTGGCGCCAAGCGCCCGGC | 329.2 | -4.0 | 1.1 | -3.5 | 4.2E-04 | 7.0E-04 | BT20/MB468 | MITO |
| >trna116_GluCTC_1_-_145399233_145399304@30.71.42,trna71_GluCTC_1_-_161439189_161439260@30.71.42,trna74_GluCTC_1_-_161431809_161431880@30.71.42,trna77_GluCTC_1_-_161424398_161424469@30.71.42,trna77_GluCTC_6_+_28949976_28950047@30.71.42,trna80_GluCTC_1_-_161417018_161417089@30.71.42,trna87_GluCTC_6_-_126101393_126101464@30.71.42\|CGCTCTCACCGCCGCGGCCCGGGTTCGATTCCCGGTCAGGGA | >trna116_GluCTC_1_-_145399233_145399304 | 30 | 71 | 42 | CGCTCTCACCGCCGCGGCCCGGGTTCGATTCCCGGTCAGGGA | 354.2 | -6.2 | 1.8 | -3.4 | 5.7E-04 | 7.0E-04 | BT20/MB468 | NUC |
| >hg38_21_+_8433172_8446622_RNA45SN1_45S_50ntFlanks@4727.4751.25\|AATCAAGAACGAAAGTCGGAGGTTC | >hg38_21_+_8433172_8446622_RNA45SN1_45S_50ntFlanks | 4727 | 4751 | 25 | AATCAAGAACGAAAGTCGGAGGTTC | 455.8 | -3.1 | 0.8 | -3.7 | 2.3E-04 | 7.0E-04 | BT20/MB468 | MP |
| >hg38_1_-_228634819_228635039_RNA5S12_5S_50ntFlanks@51.90.40\|GTCTACGGCCATACCACCCTGAACGCGCCCGATCTCGTCT | >hg38_1_-_228634819_228635039_RNA5S12_5S_50ntFlanks | 51 | 90 | 40 | GTCTACGGCCATACCACCCTGAACGCGCCCGATCTCGTCT | 145.2 | 2.9 | 0.8 | 3.5 | 5.6E-04 | 7.0E-04 | MB231/MB468 | MITO |
| >hg38_21_+_8433172_8446622_RNA45SN1_45S_50ntFlanks@12957.12981.25\|CTTCTGGGTCGGGGTTTCGTACGTA | >hg38_21_+_8433172_8446622_RNA45SN1_45S_50ntFlanks | 12957 | 12981 | 25 | CTTCTGGGTCGGGGTTTCGTACGTA | 181.4 | -3.4 | 1.0 | -3.5 | 5.0E-04 | 7.0E-04 | BT20/MB468 | CYTO |
| >trna4_ArgTCG_15_+_89878304_89878376@31.52.22\|ACTTCGGATCAGAAGATTGCAG | >trna4_ArgTCG_15_+_89878304_89878376 | 31 | 52 | 22 | ACTTCGGATCAGAAGATTGCAG | 89.0 | -9.6 | 2.6 | -3.7 | 2.3E-04 | 7.0E-04 | BT20/MB468 | MP |
| >hg38_21_+_8433172_8446622_RNA45SN1_45S_50ntFlanks@6649.6668.20\|TACGACTCTTAGCGGTGGAT | >hg38_21_+_8433172_8446622_RNA45SN1_45S_50ntFlanks | 6649 | 6668 | 20 | TACGACTCTTAGCGGTGGAT | 244.0 | 1.8 | 0.5 | 3.4 | 5.7E-04 | 7.0E-04 | BT20/MB468 | NUC |
| >hg38_21_+_8433172_8446622_RNA45SN1_45S_50ntFlanks@11206.11225.20\|CGGCGGCGACTCTGGACGCG | >hg38_21_+_8433172_8446622_RNA45SN1_45S_50ntFlanks | 11206 | 11225 | 20 | CGGCGGCGACTCTGGACGCG | 257.2 | -3.6 | 1.0 | -3.5 | 5.0E-04 | 7.0E-04 | BT20/MB468 | CYTO |
| >hg38_21_+_8433172_8446622_RNA45SN1_45S_50ntFlanks@5186.5226.41\|CAATAACAGGTCTGTGATGCCCTTAGATGTCCGGGGCTGCA | >hg38_21_+_8433172_8446622_RNA45SN1_45S_50ntFlanks | 5186 | 5226 | 41 | CAATAACAGGTCTGTGATGCCCTTAGATGTCCGGGGCTGCA | 75.5 | -9.3 | 2.5 | -3.7 | 2.3E-04 | 7.0E-04 | BT20/MB468 | MP |
| >hg38_MT_+_1621_3279_RNR2_16S_50ntFlanks@1348.1385.38\|AACAATAGGGTTTACGACCTCGATGTTGGATCAGGACA | >hg38_MT_+_1621_3279_RNR2_16S_50ntFlanks | 1348 | 1385 | 38 | AACAATAGGGTTTACGACCTCGATGTTGGATCAGGACA | 395.1 | 5.4 | 1.5 | 3.7 | 2.3E-04 | 7.0E-04 | BT20/MB468 | MP |
| >hg38_21_+_8433172_8446622_RNA45SN1_45S_50ntFlanks@8191.8215.25\|CAGCCCGTGGACGGTGTGAGGCCGG | >hg38_21_+_8433172_8446622_RNA45SN1_45S_50ntFlanks | 8191 | 8215 | 25 | CAGCCCGTGGACGGTGTGAGGCCGG | 798.4 | -2.2 | 0.6 | -3.5 | 4.2E-04 | 7.0E-04 | BT20/MB468 | MITO |
| >hg38_21_+_8433172_8446622_RNA45SN1_45S_50ntFlanks@12626.12660.35\|AATGGGGCGAAGCTACCATCTGTGGGATTATGACT | >hg38_21_+_8433172_8446622_RNA45SN1_45S_50ntFlanks | 12626 | 12660 | 35 | AATGGGGCGAAGCTACCATCTGTGGGATTATGACT | 406.9 | -4.3 | 1.3 | -3.5 | 5.6E-04 | 7.0E-04 | MB231/MB468 | MITO |
| >am_trnaMT_PheGAA_MT_+_577_647@42.60.19\|TTTAGACGGGCTCACATCA | >am_trnaMT_PheGAA_MT_+_577_647 | 42 | 60 | 19 | TTTAGACGGGCTCACATCA | 143.7 | 7.4 | 2.0 | 3.7 | 2.3E-04 | 7.0E-04 | BT20/MB468 | MP |
| >am_MI0003583\|hsa-mir-576&WithFlank&4\|+\|109488692\|109488801@61.82.22(+1U)&[MIMAT0004796&hsa-miR-576-3p&offsets\|0\|0(+1U);m-524&4\|+\|109488752\|109488772&offsets\|0\|+1(+1U)]\|AAGATGTGGAAAAATTGGAATCT | >am_MI0003583\|hsa-mir-576&WithFlank&4\|+\|109488692\|109488801 | 61 | 82 | 22(+1U) | AAGATGTGGAAAAATTGGAATCT | 207.1 | 4.3 | 1.2 | 3.5 | 4.7E-04 | 7.0E-04 | MB231/MB468 | MP |
| >hg38_21_+_8433172_8446622_RNA45SN1_45S_50ntFlanks@7979.8007.29\|ACCTCAGATCAGACGTGGCGACCCGCTGA | >hg38_21_+_8433172_8446622_RNA45SN1_45S_50ntFlanks | 7979 | 8007 | 29 | ACCTCAGATCAGACGTGGCGACCCGCTGA | 314.9 | -2.8 | 0.8 | -3.7 | 2.3E-04 | 7.0E-04 | BT20/MB468 | MP |
| >hg38_21_+_8433172_8446622_RNA45SN1_45S_50ntFlanks@11704.11736.33\|ATGAAGCGCGGGTAAACGGCGGGAGTAACTATG | >hg38_21_+_8433172_8446622_RNA45SN1_45S_50ntFlanks | 11704 | 11736 | 33 | ATGAAGCGCGGGTAAACGGCGGGAGTAACTATG | 283.6 | -2.6 | 0.8 | -3.4 | 5.7E-04 | 7.0E-04 | MB231/MB468 | NUC |
| >hg38_21_+_8433172_8446622_RNA45SN1_45S_50ntFlanks@485.527.43\|GATGTGGTGACGTCGTGCTCTCCCGGGCCGGGTCCGAGCCGCG | >hg38_21_+_8433172_8446622_RNA45SN1_45S_50ntFlanks | 485 | 527 | 43 | GATGTGGTGACGTCGTGCTCTCCCGGGCCGGGTCCGAGCCGCG | 67.1 | 9.3 | 2.7 | 3.5 | 4.7E-04 | 7.0E-04 | MB231/MB468 | MP |
| >hg38_21_+_8433172_8446622_RNA45SN1_45S_50ntFlanks@12973.12997.25\|TCGTACGTAGCAGAGCAGCTCCCTC | >hg38_21_+_8433172_8446622_RNA45SN1_45S_50ntFlanks | 12973 | 12997 | 25 | TCGTACGTAGCAGAGCAGCTCCCTC | 236.2 | -3.0 | 0.8 | -3.9 | 8.6E-05 | 7.0E-04 | BT20/MB231 | CYTO |
| >hg38_21_+_8433172_8446622_RNA45SN1_45S_50ntFlanks@6723.6743.21\|TTGCAGGACACATTGATCATC | >hg38_21_+_8433172_8446622_RNA45SN1_45S_50ntFlanks | 6723 | 6743 | 21 | TTGCAGGACACATTGATCATC | 670.8 | -2.8 | 0.8 | -3.4 | 5.7E-04 | 7.0E-04 | MB231/MB468 | NUC |
| >hg38_21_+_8433172_8446622_RNA45SN1_45S_50ntFlanks@12008.12041.34\|CCGCCGGTGAAATACCACTACTCTGATCGTTTTT | >hg38_21_+_8433172_8446622_RNA45SN1_45S_50ntFlanks | 12008 | 12041 | 34 | CCGCCGGTGAAATACCACTACTCTGATCGTTTTT | 277.6 | -5.2 | 1.5 | -3.5 | 5.1E-04 | 7.0E-04 | BT20/MB468 | CYTO |
| >hg38_MT_+_1621_3279_RNR2_16S_50ntFlanks@1184.1209.26\|AATTTCGGTTGGGGCGACCTCGGAGC | >hg38_MT_+_1621_3279_RNR2_16S_50ntFlanks | 1184 | 1209 | 26 | AATTTCGGTTGGGGCGACCTCGGAGC | 82.2 | 7.0 | 1.9 | 3.7 | 2.3E-04 | 7.0E-04 | BT20/MB468 | MP |
| >MI0000650\|hsa-mir-200c&WithFlank&12\|+\|6963693\|6963772@50.69.20(+1U)&[MIMAT0000617&hsa-miR-200c-3p&offsets\|0\|-3(+1U);m-54&12\|+\|6963742\|6963764&offsets\|0\|-3(+1U)]\|TAATACTGCCGGGTAATGATT | >MI0000650\|hsa-mir-200c&WithFlank&12\|+\|6963693\|6963772 | 50 | 69 | 20(+1U) | TAATACTGCCGGGTAATGATT | 43.9 | 8.8 | 2.2 | 3.9 | 8.2E-05 | 7.0E-04 | BT20/MB231 | MP |
| >MI0005566\|hsa-mir-374b&WithFlank&X\|-\|74218541\|74218624@17.38.22&[MIMAT0004955&hsa-miR-374b-5p&offsets\|0\|0;m-153&X\|-\|74218587\|74218608&offsets\|0\|0]\|ATATAATACAACCTGCTAAGTG | >MI0005566\|hsa-mir-374b&WithFlank&X\|-\|74218541\|74218624 | 17 | 38 | 22 | ATATAATACAACCTGCTAAGTG | 874.9 | 3.5 | 1.0 | 3.5 | 4.7E-04 | 7.0E-04 | MB231/MB468 | MP |
| >hg38_21_+_8433172_8446622_RNA45SN1_45S_50ntFlanks@12710.12745.36\|CGCGGAGCCTCGGTTGGCCTCGGATAGCCGGTCCCC | >hg38_21_+_8433172_8446622_RNA45SN1_45S_50ntFlanks | 12710 | 12745 | 36 | CGCGGAGCCTCGGTTGGCCTCGGATAGCCGGTCCCC | 322.5 | -5.9 | 1.6 | -3.7 | 2.3E-04 | 7.0E-04 | BT20/MB468 | MP |
| >hg38_21_+_8433172_8446622_RNA45SN1_45S_50ntFlanks@9569.9586.18\|GTGCAAATCGGTCGTCCG | >hg38_21_+_8433172_8446622_RNA45SN1_45S_50ntFlanks | 9569 | 9586 | 18 | GTGCAAATCGGTCGTCCG | 93.7 | -4.0 | 1.2 | -3.5 | 4.7E-04 | 7.0E-04 | MB231/MB468 | MP |

<table>
<tr><th></th><th></th><th></th><th></th><th></th><th></th><th></th><th></th><th></th><th></th><th></th><th></th><th></th><th></th></tr>
<tr><td>>hg38_21_+_8433172_8446622_RNA45SN1_45S_50ntFlanks@11718.11742.25|AACGGCGGGAGTAACTATGACTCTC</td><td>>hg38_21_+_8433172_8446622_RNA45SN1_45S_50ntFlanks</td><td>11718</td><td>11742</td><td>25</td><td>AACGGCGGGAGTAACTATGACTCTC</td><td>341.2</td><td>-2.3</td><td>0.7</td><td>-3.5</td><td>4.2E-04</td><td>7.0E-04</td><td>BT20/MB468</td><td>MITO</td></tr>
<tr><td>>trna10_AlaCGC_6_+_26553731_26553802@1.18.18,trna119_AlaCGC_6_-_28641613_28641684@1.18.18,trna120_AlaAGC_6_-_28626014_28626085@1.18.18,trna13_AlaCGC_2_+_157257281_157257352@1.18.18,trna13_AlaTGC_12_-_125406301_125406372@1.18.18,trna66_AlaTGC_6_+_28611222_28611293@1.18.18,trna8_AlaTGC_12_+_125424512_125424583@1.18.18,trna8_AlaTGC_5_+_180633868_180633939@1.18.18|GGGGATGTAGCTCAGTGG</td><td>>trna10_AlaCGC_6_+_26553731_26553802</td><td>1</td><td>18</td><td>18</td><td>GGGGATGTAGCTCAGTGG</td><td>341.7</td><td>4.0</td><td>1.1</td><td>3.7</td><td>2.4E-04</td><td>7.0E-04</td><td>BT20/MB468</td><td>MP</td></tr>
<tr><td>>trnaMT_PheGAA_MT_+_577_647@37.55.19|AAATGTTTAGACGGGCTCA</td><td>>trnaMT_PheGAA_MT_+_577_647</td><td>37</td><td>55</td><td>19</td><td>AAATGTTTAGACGGGCTCA</td><td>177.5</td><td>6.1</td><td>1.7</td><td>3.7</td><td>2.4E-04</td><td>7.0E-04</td><td>BT20/MB468</td><td>MP</td></tr>
<tr><td>>hg38_21_+_8433172_8446622_RNA45SN1_45S_50ntFlanks@6748.6770.23|CTTCGAACGCACTTGCGGCCCCG</td><td>>hg38_21_+_8433172_8446622_RNA45SN1_45S_50ntFlanks</td><td>6748</td><td>6770</td><td>23</td><td>CTTCGAACGCACTTGCGGCCCCG</td><td>960.8</td><td>-6.2</td><td>1.7</td><td>-3.7</td><td>2.4E-04</td><td>7.0E-04</td><td>BT20/MB468</td><td>MP</td></tr>
<tr><td>>hg38_21_+_8433172_8446622_RNA45SN1_45S_50ntFlanks@10684.10706.23|CGGCGGCGTCCGGTGAGCTCTCG</td><td>>hg38_21_+_8433172_8446622_RNA45SN1_45S_50ntFlanks</td><td>10684</td><td>10706</td><td>23</td><td>CGGCGGCGTCCGGTGAGCTCTCG</td><td>3581.8</td><td>-5.5</td><td>1.5</td><td>-3.7</td><td>2.4E-04</td><td>7.0E-04</td><td>BT20/MB468</td><td>MP</td></tr>
<tr><td>>MI0000736|hsa-mir-30c-1&WithFlank&1|+|40757278|40757378@23.46.24(+2U);MI0000254|hsa-mir-30c-2&WithFlank&6|-|71376954|71377037@13.36.24(+2U)&[MIMAT0000244&hsa-miR-30c-5p&offsets|0|+1(+2U);m-57&1|+|40757300|40757323&offsets|0|0(+2U)];[MIMAT0000244_1&hsa-miR-30c-5p&offsets|0|+1(+2U);m-56&6|-|71377002|71377025&offsets|0|0(+2U)]|TGTAAACATCCTACACTCTCAGCTTT</td><td>>MI0000736|hsa-mir-30c-1&WithFlank&1|+|40757278|40757378</td><td>23</td><td>46</td><td>24(+2U)</td><td>TGTAAACATCCTACACTCTCAGCTTT</td><td>169.6</td><td>5.6</td><td>1.6</td><td>3.5</td><td>4.7E-04</td><td>7.1E-04</td><td>MB231/MB468</td><td>MP</td></tr>
<tr><td>>hg38_21_+_8433172_8446622_RNA45SN1_45S_50ntFlanks@11821.11844.24|CTATCCAGCGAAACCACAGCCAAG</td><td>>hg38_21_+_8433172_8446622_RNA45SN1_45S_50ntFlanks</td><td>11821</td><td>11844</td><td>24</td><td>CTATCCAGCGAAACCACAGCCAAG</td><td>156.7</td><td>-4.6</td><td>1.3</td><td>-3.5</td><td>5.1E-04</td><td>7.1E-04</td><td>BT20/MB468</td><td>CYTO</td></tr>
<tr><td>>hg38_21_+_8433172_8446622_RNA45SN1_45S_50ntFlanks@5188.5226.39|ATAACAGGTCTGTGATGCCCTTAGATGTCCGGGGCTGCA</td><td>>hg38_21_+_8433172_8446622_RNA45SN1_45S_50ntFlanks</td><td>5188</td><td>5226</td><td>39</td><td>ATAACAGGTCTGTGATGCCCTTAGATGTCCGGGGCTGCA</td><td>327.8</td><td>-4.7</td><td>1.3</td><td>-3.5</td><td>4.3E-04</td><td>7.1E-04</td><td>BT20/MB468</td><td>MITO</td></tr>
<tr><td>>hg38_1_-_228634819_228635039_RNA5S12_5S_50ntFlanks@132.170.39|GATGGGAGACCGCCTGGGAATACCGGGTGCTGTAGGCTT</td><td>>hg38_1_-_228634819_228635039_RNA5S12_5S_50ntFlanks</td><td>132</td><td>170</td><td>39</td><td>GATGGGAGACCGCCTGGGAATACCGGGTGCTGTAGGCTT</td><td>245844.1</td><td>5.3</td><td>1.3</td><td>3.9</td><td>8.3E-05</td><td>7.1E-04</td><td>BT20/MB231</td><td>MP</td></tr>
<tr><td>>hg38_21_+_8433172_8446622_RNA45SN1_45S_50ntFlanks@11923.11950.28|AGAGGTGTAGAATAAGTGGGAGGCCCCC</td><td>>hg38_21_+_8433172_8446622_RNA45SN1_45S_50ntFlanks</td><td>11923</td><td>11950</td><td>28</td><td>AGAGGTGTAGAATAAGTGGGAGGCCCCC</td><td>375.3</td><td>7.5</td><td>1.9</td><td>3.9</td><td>8.3E-05</td><td>7.1E-04</td><td>BT20/MB231</td><td>MP</td></tr>
<tr><td>>trna10_LysCTT_16_+_3241501_3241573@3.33.31,trna13_LysCTT_14_-_58706613_58706685@3.33.31,trna2_LysCTT_15_+_79152904_79152976@3.33.31|CCGGCTAGCTCAGTCGGTAGAGCATGGGACT</td><td>>trna10_LysCTT_16_+_3241501_3241573</td><td>3</td><td>33</td><td>31</td><td>CCGGCTAGCTCAGTCGGTAGAGCATGGGACT</td><td>292.3</td><td>5.5</td><td>1.5</td><td>3.7</td><td>2.4E-04</td><td>7.1E-04</td><td>BT20/MB468</td><td>MP</td></tr>
<tr><td>>MI0000255|hsa-mir-30d&WithFlank&8|-|134804870|134804951@12.34.23&[MIMAT0000245&hsa-miR-30d-5p&offsets|0|+1;m-21&8|-|134804919|134804940&offsets|0|+1]|TGTAAACATCCCCGACTGGAAGC</td><td>>MI0000255|hsa-mir-30d&WithFlank&8|-|134804870|134804951</td><td>12</td><td>34</td><td>23</td><td>TGTAAACATCCCCGACTGGAAGC</td><td>3495.0</td><td>2.4</td><td>0.7</td><td>3.5</td><td>4.8E-04</td><td>7.1E-04</td><td>MB231/MB468</td><td>MP</td></tr>
<tr><td>>MI0005561|hsa-mir-877&WithFlank&6|+|30584326|30584423@8.29.22(+1U)&[MIMAT0004949&hsa-miR-877-5p&offsets|+1|+3(+1U);m-382&6|+|30584332|30584354&offsets|+1|0(+1U)]|TAGAGGAGATGGCGCAGGGGACT</td><td>>MI0005561|hsa-mir-877&WithFlank&6|+|30584326|30584423</td><td>8</td><td>29</td><td>22(+1U)</td><td>TAGAGGAGATGGCGCAGGGGACT</td><td>70.8</td><td>9.9</td><td>2.7</td><td>3.7</td><td>2.4E-04</td><td>7.1E-04</td><td>BT20/MB468</td><td>MP</td></tr>
<tr><td>>hg38_21_+_8433172_8446622_RNA45SN1_45S_50ntFlanks@6647.6685.39|CGTACGACTCTTAGCGGTGGATCACTCGGCTCGTGCGTC</td><td>>hg38_21_+_8433172_8446622_RNA45SN1_45S_50ntFlanks</td><td>6647</td><td>6685</td><td>39</td><td>CGTACGACTCTTAGCGGTGGATCACTCGGCTCGTGCGTC</td><td>722.7</td><td>4.6</td><td>1.3</td><td>3.7</td><td>2.4E-04</td><td>7.1E-04</td><td>BT20/MB468</td><td>MP</td></tr>
<tr><td>>hg38_21_+_8433172_8446622_RNA45SN1_45S_50ntFlanks@4882.4902.21|TGCAAAGCTGAAACTTAAAGG</td><td>>hg38_21_+_8433172_8446622_RNA45SN1_45S_50ntFlanks</td><td>4882</td><td>4902</td><td>21</td><td>TGCAAAGCTGAAACTTAAAGG</td><td>82.6</td><td>6.8</td><td>1.8</td><td>3.7</td><td>2.4E-04</td><td>7.1E-04</td><td>BT20/MB468</td><td>MP</td></tr>
<tr><td>>hg38_21_+_8433172_8446622_RNA45SN1_45S_50ntFlanks@6723.6740.18|TTGCAGGACACATTGATC</td><td>>hg38_21_+_8433172_8446622_RNA45SN1_45S_50ntFlanks</td><td>6723</td><td>6740</td><td>18</td><td>TTGCAGGACACATTGATC</td><td>210.9</td><td>-2.9</td><td>0.8</td><td>-3.7</td><td>2.4E-04</td><td>7.1E-04</td><td>BT20/MB468</td><td>MP</td></tr>
<tr><td>>MI0000469|hsa-mir-125a&WithFlank&19|+|51693248|51693345@21.42.22&[MIMAT0000443&hsa-miR-125a-5p&offsets|0|-2;m-63&19|+|51693268|51693289&offsets|0|0]|TCCCTGAGACCCTTTAACCTGT</td><td>>MI0000469|hsa-mir-125a&WithFlank&19|+|51693248|51693345</td><td>21</td><td>42</td><td>22</td><td>TCCCTGAGACCCTTTAACCTGT</td><td>494.6</td><td>3.3</td><td>0.9</td><td>3.7</td><td>2.4E-04</td><td>7.1E-04</td><td>BT20/MB468</td><td>MP</td></tr>
<tr><td>>hg38_21_+_8433172_8446622_RNA45SN1_45S_50ntFlanks@10107.10127.21|CGTGTGGTGTGCGTCGGAGGG</td><td>>hg38_21_+_8433172_8446622_RNA45SN1_45S_50ntFlanks</td><td>10107</td><td>10127</td><td>21</td><td>CGTGTGGTGTGCGTCGGAGGG</td><td>697.8</td><td>-4.6</td><td>1.3</td><td>-3.7</td><td>2.4E-04</td><td>7.1E-04</td><td>BT20/MB468</td><td>MP</td></tr>
<tr><td>>hg38_21_+_8433172_8446622_RNA45SN1_45S_50ntFlanks@3835.3861.27|CTCCTCTCCTACTTGGATAACTGTGGT</td><td>>hg38_21_+_8433172_8446622_RNA45SN1_45S_50ntFlanks</td><td>3835</td><td>3861</td><td>27</td><td>CTCCTCTCCTACTTGGATAACTGTGGT</td><td>367.7</td><td>-1.8</td><td>0.5</td><td>-3.5</td><td>5.4E-04</td><td>7.1E-04</td><td>MB231/MB468</td><td>CYTO</td></tr>
<tr><td>>hg38_1_-_228634819_228635039_RNA5S12_5S_50ntFlanks@127.170.44|ACTTGGATGGGAGACCGCCTGGGAATACCGGGTGCTGTAGGCTT</td><td>>hg38_1_-_228634819_228635039_RNA5S12_5S_50ntFlanks</td><td>127</td><td>170</td><td>44</td><td>ACTTGGATGGGAGACCGCCTGGGAATACCGGGTGCTGTAGGCTT</td><td>592.9</td><td>6.3</td><td>1.7</td><td>3.7</td><td>2.4E-04</td><td>7.1E-04</td><td>BT20/MB468</td><td>MP</td></tr>
<tr><td>>hg38_MT_+_598_1651_RNR1_12S_50ntFlanks@365.404.40|CCCCAATAAAGCTAAAACTCACCTGAGTTGTAAAAAACTC</td><td>>hg38_MT_+_598_1651_RNR1_12S_50ntFlanks</td><td>365</td><td>404</td><td>40</td><td>CCCCAATAAAGCTAAAACTCACCTGAGTTGTAAAAAACTC</td><td>245.0</td><td>3.9</td><td>1.1</td><td>3.4</td><td>5.7E-04</td><td>7.1E-04</td><td>MB231/MB468</td><td>MITO</td></tr>
<tr><td>>hg38_21_+_8433172_8446622_RNA45SN1_45S_50ntFlanks@5073.5112.40|AATTCCGATAACGAACGAGACTCTGGCATGCTAACTAGTT</td><td>>hg38_21_+_8433172_8446622_RNA45SN1_45S_50ntFlanks</td><td>5073</td><td>5112</td><td>40</td><td>AATTCCGATAACGAACGAGACTCTGGCATGCTAACTAGTT</td><td>174.9</td><td>-2.3</td><td>0.7</td><td>-3.5</td><td>4.3E-04</td><td>7.1E-04</td><td>BT20/MB468</td><td>MITO</td></tr>
<tr><td>>hg38_21_+_8433172_8446622_RNA45SN1_45S_50ntFlanks@6650.6667.18|ACGACTCTTAGCGGTGGA</td><td>>hg38_21_+_8433172_8446622_RNA45SN1_45S_50ntFlanks</td><td>6650</td><td>6667</td><td>18</td><td>ACGACTCTTAGCGGTGGA</td><td>282.2</td><td>3.5</td><td>0.9</td><td>3.7</td><td>2.4E-04</td><td>7.1E-04</td><td>BT20/MB468</td><td>MP</td></tr>
<tr><td>>hg38_21_+_8433172_8446622_RNA45SN1_45S_50ntFlanks@4126.4151.26|TTCGATTCCGGAGAGGGAGCCTGAGA</td><td>>hg38_21_+_8433172_8446622_RNA45SN1_45S_50ntFlanks</td><td>4126</td><td>4151</td><td>26</td><td>TTCGATTCCGGAGAGGGAGCCTGAGA</td><td>119.2</td><td>5.2</td><td>1.4</td><td>3.7</td><td>2.4E-04</td><td>7.1E-04</td><td>BT20/MB468</td><td>MP</td></tr>
<tr><td>>hg38_21_+_8433172_8446622_RNA45SN1_45S_50ntFlanks@10425.10455.31|AGATGGGCGAGCGCCGTTCCGAAGGGACGGG</td><td>>hg38_21_+_8433172_8446622_RNA45SN1_45S_50ntFlanks</td><td>10425</td><td>10455</td><td>31</td><td>AGATGGGCGAGCGCCGTTCCGAAGGGACGGG</td><td>128.6</td><td>-4.6</td><td>1.3</td><td>-3.5</td><td>5.1E-04</td><td>7.1E-04</td><td>BT20/MB468</td><td>CYTO</td></tr>
<tr><td>>MI0005559|hsa-mir-744&WithFlank&17|+|12081893|12082002@17.34.18&[MIMAT0004945&hsa-miR-744-5p&offsets|0|-4;m-172&17|+|12081909|12081929&offsets|0|-3]|TGCGGGGCTAGGGCTAAC</td><td>>MI0005559|hsa-mir-744&WithFlank&17|+|12081893|12082002</td><td>17</td><td>34</td><td>18</td><td>TGCGGGGCTAGGGCTAAC</td><td>303.3</td><td>7.0</td><td>1.9</td><td>3.7</td><td>2.4E-04</td><td>7.1E-04</td><td>BT20/MB468</td><td>MP</td></tr>
<tr><td>>hg38_21_+_8433172_8446622_RNA45SN1_45S_50ntFlanks@12406.12443.38|TTTTTGATCCTTCGATGTCGGCTCTTCCTATCATTGTG</td><td>>hg38_21_+_8433172_8446622_RNA45SN1_45S_50ntFlanks</td><td>12406</td><td>12443</td><td>38</td><td>TTTTTGATCCTTCGATGTCGGCTCTTCCTATCATTGTG</td><td>343.0</td><td>-4.2</td><td>1.2</td><td>-3.4</td><td>5.7E-04</td><td>7.1E-04</td><td>MB231/MB468</td><td>MITO</td></tr>
<tr><td>>hg38_21_+_8433172_8446622_RNA45SN1_45S_50ntFlanks@7973.7997.25|GACGCGACCTCAGATCAGACGTGGC</td><td>>hg38_21_+_8433172_8446622_RNA45SN1_45S_50ntFlanks</td><td>7973</td><td>7997</td><td>25</td><td>GACGCGACCTCAGATCAGACGTGGC</td><td>280.1</td><td>3.2</td><td>0.9</td><td>3.4</td><td>5.7E-04</td><td>7.1E-04</td><td>MB231/MB468</td><td>MITO</td></tr>
<tr><td>>hg38_21_+_8433172_8446622_RNA45SN1_45S_50ntFlanks@9574.9602.29|AATCGGTCGTCCGACCTGGGTATAGGGGC</td><td>>hg38_21_+_8433172_8446622_RNA45SN1_45S_50ntFlanks</td><td>9574</td><td>9602</td><td>29</td><td>AATCGGTCGTCCGACCTGGGTATAGGGGC</td><td>832.0</td><td>2.1</td><td>0.6</td><td>3.5</td><td>4.3E-04</td><td>7.1E-04</td><td>BT20/MB468</td><td>MITO</td></tr>
<tr><td>>hg38_21_+_8433172_8446622_RNA45SN1_45S_50ntFlanks@12083.12106.24|CTCGCTTCTGGCGCCAAGCGCCCG</td><td>>hg38_21_+_8433172_8446622_RNA45SN1_45S_50ntFlanks</td><td>12083</td><td>12106</td><td>24</td><td>CTCGCTTCTGGCGCCAAGCGCCCG</td><td>473.1</td><td>-5.3</td><td>1.5</td><td>-3.5</td><td>4.3E-04</td><td>7.1E-04</td><td>BT20/MB468</td><td>MITO</td></tr>
<tr><td>>hg38_21_+_8433172_8446622_RNA45SN1_45S_50ntFlanks@4002.4025.24|GATAACCTCGGGCCGATCGCACGC</td><td>>hg38_21_+_8433172_8446622_RNA45SN1_45S_50ntFlanks</td><td>4002</td><td>4025</td><td>24</td><td>GATAACCTCGGGCCGATCGCACGC</td><td>320.8</td><td>-5.1</td><td>1.4</td><td>-3.5</td><td>4.3E-04</td><td>7.1E-04</td><td>BT20/MB468</td><td>MITO</td></tr>
<tr><td>>hg38_21_+_8433172_8446622_RNA45SN1_45S_50ntFlanks@5124.5158.35|GAGCGGTCGGCGTCCCCCAACTTCTTAGAGGGACA</td><td>>hg38_21_+_8433172_8446622_RNA45SN1_45S_50ntFlanks</td><td>5124</td><td>5158</td><td>35</td><td>GAGCGGTCGGCGTCCCCCAACTTCTTAGAGGGACA</td><td>267.6</td><td>7.9</td><td>2.3</td><td>3.5</td><td>5.4E-04</td><td>7.1E-04</td><td>MB231/MB468</td><td>CYTO</td></tr>
<tr><td>>hg38_21_+_8433172_8446622_RNA45SN1_45S_50ntFlanks@5050.5081.32|TTGGTGGAGCGATTTGTCTGGTTAATTCCGAT</td><td>>hg38_21_+_8433172_8446622_RNA45SN1_45S_50ntFlanks</td><td>5050</td><td>5081</td><td>32</td><td>TTGGTGGAGCGATTTGTCTGGTTAATTCCGAT</td><td>44.0</td><td>8.8</td><td>2.2</td><td>3.9</td><td>8.4E-05</td><td>7.1E-04</td><td>BT20/MB231</td><td>MP</td></tr>
<tr><td>>hg38_21_+_8433172_8446622_RNA45SN1_45S_50ntFlanks@12942.12962.21|TTCGTAGACGACCTGCTTCTG</td><td>>hg38_21_+_8433172_8446622_RNA45SN1_45S_50ntFlanks</td><td>12942</td><td>12962</td><td>21</td><td>TTCGTAGACGACCTGCTTCTG</td><td>349.4</td><td>-2.2</td><td>0.6</td><td>-3.5</td><td>5.4E-04</td><td>7.2E-04</td><td>MB231/MB468</td><td>CYTO</td></tr>
<tr><td>>hg38_21_+_8433172_8446622_RNA45SN1_45S_50ntFlanks@3855.3881.27|CTGTGGTAATTCTAGAGCTAATACATG</td><td>>hg38_21_+_8433172_8446622_RNA45SN1_45S_50ntFlanks</td><td>3855</td><td>3881</td><td>27</td><td>CTGTGGTAATTCTAGAGCTAATACATG</td><td>217.1</td><td>-3.6</td><td>1.0</td><td>-3.5</td><td>5.4E-04</td><td>7.2E-04</td><td>MB231/MB468</td><td>CYTO</td></tr>
<tr><td>>trnaMT_GluTTC_MT_-_14674_14742@35.71.37,trnalookalike8_GluTTC_5_-_93905172_93905240@35.71.37|TATCATTGGTCGTGGTTGTAGTCCGTGCGAGAATACC</td><td>>trnaMT_GluTTC_MT_-_14674_14742</td><td>35</td><td>71</td><td>37</td><td>TATCATTGGTCGTGGTTGTAGTCCGTGCGAGAATACC</td><td>438.6</td><td>-3.5</td><td>0.9</td><td>-3.9</td><td>8.8E-05</td><td>7.2E-04</td><td>BT20/MB231</td><td>NUC</td></tr>
<tr><td>>trna152_ValCAC_6_-_27248049_27248121@1.31.31|GCTTCTGTAGTGTAGTGGTTATCACGTTCGC</td><td>>trna152_ValCAC_6_-_27248049_27248121</td><td>1</td><td>31</td><td>31</td><td>GCTTCTGTAGTGTAGTGGTTATCACGTTCGC</td><td>228.0</td><td>-2.6</td><td>0.8</td><td>-3.5</td><td>5.2E-04</td><td>7.2E-04</td><td>BT20/MB468</td><td>CYTO</td></tr>
<tr><td>>hg38_21_+_8433172_8446622_RNA45SN1_45S_50ntFlanks@3854.3887.34|ACTGTGGTAATTCTAGAGCTAATACATGCCGACG</td><td>>hg38_21_+_8433172_8446622_RNA45SN1_45S_50ntFlanks</td><td>3854</td><td>3887</td><td>34</td><td>ACTGTGGTAATTCTAGAGCTAATACATGCCGACG</td><td>365.9</td><td>-6.0</td><td>1.7</td><td>-3.4</td><td>5.8E-04</td><td>7.2E-04</td><td>BT20/MB468</td><td>NUC</td></tr>
<tr><td>>hg38_21_+_8433172_8446622_RNA45SN1_45S_50ntFlanks@11207.11227.21|GGCGGCGACTCTGGACGCGAG</td><td>>hg38_21_+_8433172_8446622_RNA45SN1_45S_50ntFlanks</td><td>11207</td><td>11227</td><td>21</td><td>GGCGGCGACTCTGGACGCGAG</td><td>174.4</td><td>-3.5</td><td>1.0</td><td>-3.5</td><td>5.2E-04</td><td>7.2E-04</td><td>BT20/MB468</td><td>CYTO</td></tr>
<tr><td>>hg38_21_+_8433172_8446622_RNA45SN1_45S_50ntFlanks@4983.5018.36|CGGACAGGATTGACAGATTGATAGCTCTTTCTCGAT</td><td>>hg38_21_+_8433172_8446622_RNA45SN1_45S_50ntFlanks</td><td>4983</td><td>5018</td><td>36</td><td>CGGACAGGATTGACAGATTGATAGCTCTTTCTCGAT</td><td>89.3</td><td>6.9</td><td>1.9</td><td>3.7</td><td>2.4E-04</td><td>7.2E-04</td><td>BT20/MB468</td><td>MP</td></tr>
<tr><td>>hg38_21_+_8433172_8446622_RNA45SN1_45S_50ntFlanks@11919.11948.30|CATGAGAGGTGTAGAATAAGTGGGAGGCCC</td><td>>hg38_21_+_8433172_8446622_RNA45SN1_45S_50ntFlanks</td><td>11919</td><td>11948</td><td>30</td><td>CATGAGAGGTGTAGAATAAGTGGGAGGCCC</td><td>730.9</td><td>-3.6</td><td>1.0</td><td>-3.7</td><td>2.4E-04</td><td>7.2E-04</td><td>BT20/MB468</td><td>MP</td></tr>
<tr><td>>hg38_21_+_8433172_8446622_RNA45SN1_45S_50ntFlanks@8379.8405.27|TCAAGAGGGCGTGAAACCGTTAAGAGG</td><td>>hg38_21_+_8433172_8446622_RNA45SN1_45S_50ntFlanks</td><td>8379</td><td>8405</td><td>27</td><td>TCAAGAGGGCGTGAAACCGTTAAGAGG</td><td>241.6</td><td>-2.7</td><td>0.7</td><td>-3.7</td><td>2.4E-04</td><td>7.2E-04</td><td>BT20/MB468</td><td>MP</td></tr>
<tr><td>>hg38_21_+_8433172_8446622_RNA45SN1_45S_50ntFlanks@10402.10433.32|AACATGGGTCAGTCGGTCCTGAGAGATGGGCG</td><td>>hg38_21_+_8433172_8446622_RNA45SN1_45S_50ntFlanks</td><td>10402</td><td>10433</td><td>32</td><td>AACATGGGTCAGTCGGTCCTGAGAGATGGGCG</td><td>268.5</td><td>-1.9</td><td>0.5</td><td>-3.5</td><td>4.3E-04</td><td>7.2E-04</td><td>BT20/MB468</td><td>MITO</td></tr>
<tr><td>>hg38_21_+_8433172_8446622_RNA45SN1_45S_50ntFlanks@10508.10533.26|CGAATCCGGAGTGGCGGAGATGGGCG</td><td>>hg38_21_+_8433172_8446622_RNA45SN1_45S_50ntFlanks</td><td>10508</td><td>10533</td><td>26</td><td>CGAATCCGGAGTGGCGGAGATGGGCG</td><td>1205.1</td><td>-3.5</td><td>1.0</td><td>-3.7</td><td>2.4E-04</td><td>7.2E-04</td><td>BT20/MB468</td><td>MP</td></tr>
<tr><td>>hg38_21_+_8433172_8446622_RNA45SN1_45S_50ntFlanks@11873.11902.30|AAGAAGACCCTGTTGAGCTTGACTCTAGTC</td><td>>hg38_21_+_8433172_8446622_RNA45SN1_45S_50ntFlanks</td><td>11873</td><td>11902</td><td>30</td><td>AAGAAGACCCTGTTGAGCTTGACTCTAGTC</td><td>324.1</td><td>-2.4</td><td>0.7</td><td>-3.5</td><td>5.2E-04</td><td>7.2E-04</td><td>BT20/MB468</td><td>CYTO</td></tr>
<tr><td>>hg38_21_+_8433172_8446622_RNA45SN1_45S_50ntFlanks@11552.11588.37|CCTCGGCCGGCGCCTAGCAGCCGACTTAGAACTGGTG</td><td>>hg38_21_+_8433172_8446622_RNA45SN1_45S_50ntFlanks</td><td>11552</td><td>11588</td><td>37</td><td>CCTCGGCCGGCGCCTAGCAGCCGACTTAGAACTGGTG</td><td>201.7</td><td>-3.1</td><td>0.9</td><td>-3.5</td><td>5.4E-04</td><td>7.2E-04</td><td>MB231/MB468</td><td>CYTO</td></tr>
</table>

| | | | | | | | | | | | | | |
|---|---|---|---|---|---|---|---|---|---|---|---|---|---|
| >MI0003581\|hsa-mir-574&WithFlank&4\|+\|38868026\|38868133@67.88.22&[MIMAT0003239&hsa-miR-574-3p&offsets\|0\|0;m-149&4\|+\|38868092\|38868113&offsets\|0\|0]\|CACGCTCATGCACACACCCACA | >MI0003581\|hsa-mir-574&WithFlank&4\|+\|38868026\|38868133 | 67 | 88 | 22 | CACGCTCATGCACACACCCACA | 774.5 | 3.0 | 0.9 | 3.5 | 4.8E-04 | 7.2E-04 | MB231/MB468 | MP |
| >hg38_21_+_8433172_8446622_RNA45SN1_45S_50ntFlanks@485.506.22\|GATGTGGTGACGTCGTGCTCTC | >hg38_21_+_8433172_8446622_RNA45SN1_45S_50ntFlanks | 485 | 506 | 22 | GATGTGGTGACGTCGTGCTCTC | 699.1 | 12.7 | 3.6 | 3.5 | 4.8E-04 | 7.2E-04 | MB231/MB468 | MP |
| >trnaMT_HisGTG_MT_+_12138_12206@27.59.33\|GATTGTGAATCTGACAACAGAGGCTTACGACCC | >trnaMT_HisGTG_MT_+_12138_12206 | 27 | 59 | 33 | GATTGTGAATCTGACAACAGAGGCTTACGACCC | 68.1 | 9.8 | 2.7 | 3.7 | 2.4E-04 | 7.2E-04 | BT20/MB468 | MP |
| >hg38_21_+_8433172_8446622_RNA45SN1_45S_50ntFlanks@7982.8008.27\|TCAGATCAGACGTGGCGACCCGCTGAA | >hg38_21_+_8433172_8446622_RNA45SN1_45S_50ntFlanks | 7982 | 8008 | 27 | TCAGATCAGACGTGGCGACCCGCTGAA | 1072.3 | -5.0 | 1.4 | -3.7 | 2.4E-04 | 7.2E-04 | BT20/MB468 | MP |
| >MI0000063\|hsa-let-7b&WithFlank&22\|+\|46113680\|46113774@12.33.22(+1A)&[MIMAT0000063&hsa-let-7b-5p&offsets\|0\|0(+1A);m-14&22\|+\|46113691\|46113712&offsets\|0\|0(+1A)]\|TGAGGTAGTAGGTTGTGTGGTTA | >MI0000063\|hsa-let-7b&WithFlank&22\|+\|46113680\|46113774 | 12 | 33 | 22(+1A) | TGAGGTAGTAGGTTGTGTGGTTA | 1624.2 | 2.6 | 0.8 | 3.5 | 4.8E-04 | 7.2E-04 | MB231/MB468 | MP |
| >hg38_21_+_8433172_8446622_RNA45SN1_45S_50ntFlanks@7976.8005.30\|GCGACCTCAGATCAGACGTGGCGACCCGCT | >hg38_21_+_8433172_8446622_RNA45SN1_45S_50ntFlanks | 7976 | 8005 | 30 | GCGACCTCAGATCAGACGTGGCGACCCGCT | 170.2 | 2.7 | 0.8 | 3.4 | 5.8E-04 | 7.2E-04 | BT20/MB468 | NUC |
| >hg38_21_+_8433172_8446622_RNA45SN1_45S_50ntFlanks@3843.3860.18\|CTACTTGGATAACTGTGG | >hg38_21_+_8433172_8446622_RNA45SN1_45S_50ntFlanks | 3843 | 3860 | 18 | CTACTTGGATAACTGTGG | 324.2 | -6.0 | 1.6 | -3.7 | 2.4E-04 | 7.2E-04 | BT20/MB468 | MP |
| >hg38_21_+_8433172_8446622_RNA45SN1_45S_50ntFlanks@12159.12178.20\|TTTGACTGGGGCGGTACACC | >hg38_21_+_8433172_8446622_RNA45SN1_45S_50ntFlanks | 12159 | 12178 | 20 | TTTGACTGGGGCGGTACACC | 410.4 | -4.1 | 1.2 | -3.5 | 5.4E-04 | 7.2E-04 | MB231/MB468 | CYTO |
| >hg38_21_+_8433172_8446622_RNA45SN1_45S_50ntFlanks@5073.5113.41\|AATTCCGATAACGAACGAGACTCTGGCATGCTAACTAGTTA | >hg38_21_+_8433172_8446622_RNA45SN1_45S_50ntFlanks | 5073 | 5113 | 41 | AATTCCGATAACGAACGAGACTCTGGCATGCTAACTAGTTA | 205.3 | -5.0 | 1.4 | -3.5 | 4.3E-04 | 7.2E-04 | BT20/MB468 | MITO |
| >trnaMT_LeuTAA_MT_+_3230_3304@16.44.29\|CCCGGTAATCGCATAAAACTTAAAACTTT | >trnaMT_LeuTAA_MT_+_3230_3304 | 16 | 44 | 29 | CCCGGTAATCGCATAAAACTTAAAACTTT | 1073.2 | -3.0 | 0.9 | -3.5 | 5.2E-04 | 7.2E-04 | BT20/MB468 | CYTO |
| >hg38_21_+_8433172_8446622_RNA45SN1_45S_50ntFlanks@11670.11697.28\|CCCAGTGCTCTGAATGTCAAAGTGAAGA | >hg38_21_+_8433172_8446622_RNA45SN1_45S_50ntFlanks | 11670 | 11697 | 28 | CCCAGTGCTCTGAATGTCAAAGTGAAGA | 613.3 | -6.3 | 1.8 | -3.4 | 5.9E-04 | 7.2E-04 | BT20/MB468 | NUC |
| >hg38_21_+_8433172_8446622_RNA45SN1_45S_50ntFlanks@8176.8202.27\|CTTCTGATCGAGGCCCAGCCCGTGGAC | >hg38_21_+_8433172_8446622_RNA45SN1_45S_50ntFlanks | 8176 | 8202 | 27 | CTTCTGATCGAGGCCCAGCCCGTGGAC | 2692.0 | 2.6 | 0.7 | 3.5 | 5.5E-04 | 7.2E-04 | MB231/MB468 | CYTO |
| >trna14_LysTTT_11_-_59327808_59327880@1.26.26,trna2_LysTTT_17_+_8022473_8022545@1.26.26,trna54_LysTTT_1_+_204475655_204475727@1.26.26,trna5_LysTTT_11_+_59323902_59323974@1.26.26,trna62_LysTTT_1_-_204476158_204476230@1.26.26,trna76_LysTTT_6_+_28918806_28918878@1.26.26\|GCCCGGATAGCTCAGTCGGTAGAGCA | >trna14_LysTTT_11_-_59327808_59327880 | 1 | 26 | 26 | GCCCGGATAGCTCAGTCGGTAGAGCA | 73.7 | -5.4 | 1.6 | -3.5 | 4.9E-04 | 7.2E-04 | MB231/MB468 | MP |
| >hg38_21_+_8433172_8446622_RNA45SN1_45S_50ntFlanks@12615.12634.20\|GCTGAGGAGCCAATGGGGCG | >hg38_21_+_8433172_8446622_RNA45SN1_45S_50ntFlanks | 12615 | 12634 | 20 | GCTGAGGAGCCAATGGGGCG | 115.0 | -4.5 | 1.3 | -3.5 | 5.2E-04 | 7.2E-04 | BT20/MB468 | CYTO |
| >hg38_21_+_8433172_8446622_RNA45SN1_45S_50ntFlanks@10786.10825.40\|AACAGCCTCTGGCATGTTGGAACAATGTAGGTAAGGGAAG | >hg38_21_+_8433172_8446622_RNA45SN1_45S_50ntFlanks | 10786 | 10825 | 40 | AACAGCCTCTGGCATGTTGGAACAATGTAGGTAAGGGAAG | 390.9 | -5.3 | 1.5 | -3.5 | 5.2E-04 | 7.2E-04 | BT20/MB468 | CYTO |
| >hg38_21_+_8433172_8446622_RNA45SN1_45S_50ntFlanks@12071.12096.26\|CCCCGAGGGGCTCTCGCTTCTGGCGC | >hg38_21_+_8433172_8446622_RNA45SN1_45S_50ntFlanks | 12071 | 12096 | 26 | CCCCGAGGGGCTCTCGCTTCTGGCGC | 724.4 | -4.6 | 1.3 | -3.5 | 5.2E-04 | 7.2E-04 | BT20/MB468 | CYTO |
| >hg38_21_+_8433172_8446622_RNA45SN1_45S_50ntFlanks@9306.9335.30\|CAAGGAGTCTAACACGTGCGCGAGTCGGGG | >hg38_21_+_8433172_8446622_RNA45SN1_45S_50ntFlanks | 9306 | 9335 | 30 | CAAGGAGTCTAACACGTGCGCGAGTCGGGG | 884.7 | -3.4 | 0.9 | -3.7 | 2.4E-04 | 7.2E-04 | BT20/MB468 | MP |
| >hg38_21_+_8433172_8446622_RNA45SN1_45S_50ntFlanks@6648.6670.23\|GTACGACTCTTAGCGGTGGATCA | >hg38_21_+_8433172_8446622_RNA45SN1_45S_50ntFlanks | 6648 | 6670 | 23 | GTACGACTCTTAGCGGTGGATCA | 144.2 | -5.8 | 1.6 | -3.7 | 2.4E-04 | 7.2E-04 | BT20/MB468 | MP |
| >hg38_21_+_8433172_8446622_RNA45SN1_45S_50ntFlanks@11876.11893.18\|AAGACCCTGTTGAGCTTG | >hg38_21_+_8433172_8446622_RNA45SN1_45S_50ntFlanks | 11876 | 11893 | 18 | AAGACCCTGTTGAGCTTG | 804.9 | -2.1 | 0.6 | -3.5 | 4.4E-04 | 7.2E-04 | BT20/MB468 | MITO |
| >MI0000085\|hsa-mir-27a&WithFlank&19\|-\|13836434\|13836523@57.76.20(+5U)&[MIMAT0000084&hsa-miR-27a-3p&offsets\|0\|-1(+5U);m-43&19\|-\|13836447\|13836467&offsets\|0\|-1(+5U)]\|TTCACAGTGGCTAAGTTCCGTTTTT | >MI0000085\|hsa-mir-27a&WithFlank&19\|-\|13836434\|13836523 | 57 | 76 | 20(+5U) | TTCACAGTGGCTAAGTTCCGTTTTT | 48.4 | -9.0 | 2.3 | -3.9 | 8.5E-05 | 7.2E-04 | BT20/MB231 | MP |
| >trna14_LysTTT_11_-_59327808_59327880@1.30.30,trna2_LysTTT_17_+_8022473_8022545@1.30.30,trna54_LysTTT_1_+_204475655_204475727@1.30.30,trna5_LysTTT_11_+_59323902_59323974@1.30.30,trna62_LysTTT_1_-_204476158_204476230@1.30.30,trna76_LysTTT_6_+_28918806_28918878@1.30.30\|GCCCGGATAGCTCAGTCGGTAGAGCATCAG | >trna14_LysTTT_11_-_59327808_59327880 | 1 | 30 | 30 | GCCCGGATAGCTCAGTCGGTAGAGCATCAG | 415.8 | -6.2 | 1.7 | -3.7 | 2.4E-04 | 7.2E-04 | BT20/MB468 | MP |
| >hg38_MT_+_598_1651_RNR1_12S_50ntFlanks@442.468.27\|ATATCTGAACACACAATAGCTAAGACC | >hg38_MT_+_598_1651_RNR1_12S_50ntFlanks | 442 | 468 | 27 | ATATCTGAACACACAATAGCTAAGACC | 152.9 | 4.6 | 1.3 | 3.5 | 4.9E-04 | 7.2E-04 | MB231/MB468 | MP |
| >trnaMT_IleGAT_MT_+_4263_4331@42.71.30\|AATAGGAGCTTAAACCCCCTTATTTCTACC | >trnaMT_IleGAT_MT_+_4263_4331 | 42 | 71 | 30 | AATAGGAGCTTAAACCCCCTTATTTCTACC | 67.8 | 9.8 | 2.7 | 3.7 | 2.4E-04 | 7.2E-04 | BT20/MB468 | MP |
| >hg38_21_+_8433172_8446622_RNA45SN1_45S_50ntFlanks@10806.10830.25\|AACAATGTAGGTAAGGGAAGTCGGC | >hg38_21_+_8433172_8446622_RNA45SN1_45S_50ntFlanks | 10806 | 10830 | 25 | AACAATGTAGGTAAGGGAAGTCGGC | 150.0 | -2.0 | 0.6 | -3.5 | 4.4E-04 | 7.2E-04 | BT20/MB468 | MITO |
| >am_trna128_GlyGCC_6_-_27870686_27870756@1.30.30,trna133_GlyCCC_1_-_16872434_16872504@1.30.30,trna18_GlyGCC_16_+_70822597_70822667@1.30.30,trna19_GlyGCC_16_+_70823410_70823480@1.30.30,trna19_GlyGCC_2_-_157257659_157257729@1.30.30,trna24_GlyGCC_16_-_70812942_70813012@1.30.30,trna25_GlyGCC_16_-_70812114_70812184@1.30.30,trna4_GlyCCC_1_+_17188416_17188486@1.30.30,trna5_GlyGCC_17_+_8029064_8029134@1.30.30,trna68_GlyGCC_1_-_161493637_161493707@1.30.30\|GCATTGGTGGTTCAGTGGTAGAATTCTCGC | >am_trna128_GlyGCC_6_-_27870686_27870756 | 1 | 30 | 30 | GCATTGGTGGTTCAGTGGTAGAATTCTCGC | 14094.2 | -3.4 | 0.9 | -3.9 | 8.9E-05 | 7.2E-04 | BT20/MB231 | CYTO |
| >hg38_21_+_8433172_8446622_RNA45SN1_45S_50ntFlanks@10391.10419.29\|AACAGCAGTTGAACATGGGTCAGTCGGTC | >hg38_21_+_8433172_8446622_RNA45SN1_45S_50ntFlanks | 10391 | 10419 | 29 | AACAGCAGTTGAACATGGGTCAGTCGGTC | 286.0 | -5.8 | 1.6 | -3.7 | 2.4E-04 | 7.2E-04 | BT20/MB468 | MP |
| >MI0000086\|hsa-mir-28&WithFlank&3\|+\|188688775\|188688872@20.40.21(+1A)&[MIMAT0000085&hsa-miR-28-5p&offsets\|0\|-1(+1A);m-133&3\|+\|188688794\|188688815&offsets\|0\|-1(+1A)]\|AAGGAGCTCACAGTCTATTGAA | >MI0000086\|hsa-mir-28&WithFlank&3\|+\|188688775\|188688872 | 20 | 40 | 21(+1A) | AAGGAGCTCACAGTCTATTGAA | 139.0 | 5.4 | 1.6 | 3.5 | 4.9E-04 | 7.3E-04 | MB231/MB468 | MP |
| >hg38_21_+_8433172_8446622_RNA45SN1_45S_50ntFlanks@10772.10796.25\|AGGTCTCCAAGGTGAACAGCCTCTG | >hg38_21_+_8433172_8446622_RNA45SN1_45S_50ntFlanks | 10772 | 10796 | 25 | AGGTCTCCAAGGTGAACAGCCTCTG | 520.9 | -4.1 | 1.2 | -3.4 | 5.8E-04 | 7.3E-04 | MB231/MB468 | MITO |
| >trnaMT_HisGTG_MT_+_12138_12206@42.70.29\|AACAGAGGCTTACGACCCCTTATTTACCC | >trnaMT_HisGTG_MT_+_12138_12206 | 42 | 70 | 29 | AACAGAGGCTTACGACCCCTTATTTACCC | 71.2 | 9.9 | 2.7 | 3.7 | 2.4E-04 | 7.3E-04 | BT20/MB468 | MP |
| >trna11_GluTTC_15_-_26327381_26327452@3.36.34,trna3_GluTTC_13_-_45492062_45492133@3.36.34\|CCACATGGTCTAGCGGTTAGGATTCCTGGTTTTC | >trna11_GluTTC_15_-_26327381_26327452 | 3 | 36 | 34 | CCACATGGTCTAGCGGTTAGGATTCCTGGTTTTC | 268.8 | -4.0 | 1.0 | -3.9 | 1.0E-04 | 7.3E-04 | BT20/MB231 | MITO |
| >hg38_21_+_8433172_8446622_RNA45SN1_45S_50ntFlanks@9033.9076.44\|CTCGTGGGGGGGCCGGGCCACCCCTCCCACGGCGCGACCGCTCT | >hg38_21_+_8433172_8446622_RNA45SN1_45S_50ntFlanks | 9033 | 9076 | 44 | CTCGTGGGGGGGCCGGGCCACCCCTCCCACGGCGCGACCGCTCT | 187.7 | -5.3 | 1.5 | -3.4 | 5.9E-04 | 7.3E-04 | MB231/MB468 | NUC |
| >hg38_21_+_8433172_8446622_RNA45SN1_45S_50ntFlanks@10131.10149.19\|CGGCGGCGGCGGCGGCGGG | >hg38_21_+_8433172_8446622_RNA45SN1_45S_50ntFlanks | 10131 | 10149 | 19 | CGGCGGCGGCGGCGGCGGG | 528.9 | -5.4 | 1.5 | -3.5 | 4.4E-04 | 7.3E-04 | BT20/MB468 | MITO |
| >hg38_21_+_8433172_8446622_RNA45SN1_45S_50ntFlanks@8848.8866.19\|CGGCGGCGGCGGCGGCGGG | >hg38_21_+_8433172_8446622_RNA45SN1_45S_50ntFlanks | 8848 | 8866 | 19 | CGGCGGCGGCGGCGGCGGG | 528.9 | -5.4 | 1.5 | -3.5 | 4.4E-04 | 7.3E-04 | BT20/MB468 | MITO |
| >hg38_21_+_8433172_8446622_RNA45SN1_45S_50ntFlanks@11570.11613.44\|AGCCGACTTAGAACTGGTGCGGACCAGGGGAATCCGACTGTTTA | >hg38_21_+_8433172_8446622_RNA45SN1_45S_50ntFlanks | 11570 | 11613 | 44 | AGCCGACTTAGAACTGGTGCGGACCAGGGGAATCCGACTGTTTA | 837.2 | 4.0 | 1.0 | 3.9 | 1.0E-04 | 7.3E-04 | BT20/MB231 | MITO |
| >hg38_21_+_8433172_8446622_RNA45SN1_45S_50ntFlanks@9794.9811.18\|CGTGGAGCCGGGCGTGGA | >hg38_21_+_8433172_8446622_RNA45SN1_45S_50ntFlanks | 9794 | 9811 | 18 | CGTGGAGCCGGGCGTGGA | 110.5 | 5.7 | 1.6 | 3.7 | 2.4E-04 | 7.3E-04 | BT20/MB468 | MP |
| >hg38_21_+_8433172_8446622_RNA45SN1_45S_50ntFlanks@5190.5234.45\|AACAGGTCTGTGATGCCCTTAGATGTCCGGGGCTGCACGCGCGCT | >hg38_21_+_8433172_8446622_RNA45SN1_45S_50ntFlanks | 5190 | 5234 | 45 | AACAGGTCTGTGATGCCCTTAGATGTCCGGGGCTGCACGCGCGCT | 67.9 | 9.8 | 2.7 | 3.7 | 2.4E-04 | 7.3E-04 | BT20/MB468 | MP |
| >hg38_21_+_8433172_8446622_RNA45SN1_45S_50ntFlanks@8176.8202.27\|CTTCTGATCGAGGCCCAGCCCGTGGAC | >hg38_21_+_8433172_8446622_RNA45SN1_45S_50ntFlanks | 8176 | 8202 | 27 | CTTCTGATCGAGGCCCAGCCCGTGGAC | 673.8 | -3.1 | 0.9 | -3.5 | 4.9E-04 | 7.3E-04 | MB231/MB468 | MP |
| >hg38_21_+_8433172_8446622_RNA45SN1_45S_50ntFlanks@6660.6688.29\|GCGGTGGATCACTCGGCTCGTGCGTCGAT | >hg38_21_+_8433172_8446622_RNA45SN1_45S_50ntFlanks | 6660 | 6688 | 29 | GCGGTGGATCACTCGGCTCGTGCGTCGAT | 186.8 | -4.3 | 1.2 | -3.5 | 4.9E-04 | 7.3E-04 | MB231/MB468 | MP |
| >hg38_21_+_8433172_8446622_RNA45SN1_45S_50ntFlanks@12626.12661.36\|AATGGGGCGAAGCTACCATCTGTGGGATTATGACTG | >hg38_21_+_8433172_8446622_RNA45SN1_45S_50ntFlanks | 12626 | 12661 | 36 | AATGGGGCGAAGCTACCATCTGTGGGATTATGACTG | 2654.8 | -4.4 | 1.2 | -3.7 | 2.4E-04 | 7.3E-04 | BT20/MB468 | MP |
| >hg38_21_+_8433172_8446622_RNA45SN1_45S_50ntFlanks@10572.10591.20\|AGAAGCCGGCGGGAGCCCCG | >hg38_21_+_8433172_8446622_RNA45SN1_45S_50ntFlanks | 10572 | 10591 | 20 | AGAAGCCGGCGGGAGCCCCG | 800.6 | -5.5 | 1.6 | -3.5 | 5.3E-04 | 7.3E-04 | BT20/MB468 | CYTO |
| >hg38_21_+_8433172_8446622_RNA45SN1_45S_50ntFlanks@9572.9605.34\|CAAATCGGTCGTCCGACCTGGGTATAGGGGCGAA | >hg38_21_+_8433172_8446622_RNA45SN1_45S_50ntFlanks | 9572 | 9605 | 34 | CAAATCGGTCGTCCGACCTGGGTATAGGGGCGAA | 226.2 | -3.3 | 1.0 | -3.5 | 5.5E-04 | 7.3E-04 | MB231/MB468 | CYTO |
| >hg38_21_+_8433172_8446622_RNA45SN1_45S_50ntFlanks@6651.6692.42\|CGACTCTTAGCGGTGGATCACTCGGCTCGTGCGTCGATGAAG | >hg38_21_+_8433172_8446622_RNA45SN1_45S_50ntFlanks | 6651 | 6692 | 42 | CGACTCTTAGCGGTGGATCACTCGGCTCGTGCGTCGATGAAG | 7987.4 | 2.6 | 0.7 | 3.4 | 5.9E-04 | 7.3E-04 | MB231/MB468 | NUC |
| >trnaMT_LysTTT_MT_+_8295_8364@54.72.19,trnalookalike7_LysTTT_1_+_568844_568913@54.72.19\|ACACCTCTTTACAGTGACC | >trnaMT_LysTTT_MT_+_8295_8364 | 54 | 72 | 19 | ACACCTCTTTACAGTGACC | 205.6 | 4.3 | 1.2 | 3.5 | 4.9E-04 | 7.3E-04 | MB231/MB468 | MP |
| >hg38_21_+_8433172_8446622_RNA45SN1_45S_50ntFlanks@12914.12933.20\|CGTTCGTGGGGAACCTGGCG | >hg38_21_+_8433172_8446622_RNA45SN1_45S_50ntFlanks | 12914 | 12933 | 20 | CGTTCGTGGGGAACCTGGCG | 354.0 | -2.0 | 0.6 | -3.5 | 4.4E-04 | 7.3E-04 | BT20/MB468 | MITO |
| >hg38_21_+_8433172_8446622_RNA45SN1_45S_50ntFlanks@11669.11687.19\|GCCCAGTGCTCTGAATGTC | >hg38_21_+_8433172_8446622_RNA45SN1_45S_50ntFlanks | 11669 | 11687 | 19 | GCCCAGTGCTCTGAATGTC | 210.1 | 2.6 | 0.8 | 3.4 | 5.8E-04 | 7.3E-04 | MB231/MB468 | MITO |
| >hg38_21_+_8433172_8446622_RNA45SN1_45S_50ntFlanks@11581.11613.33\|AACTGGTGCGGACCAGGGGAATCCGACTGTTTA | >hg38_21_+_8433172_8446622_RNA45SN1_45S_50ntFlanks | 11581 | 11613 | 33 | AACTGGTGCGGACCAGGGGAATCCGACTGTTTA | 269.9 | -2.1 | 0.6 | -3.4 | 5.9E-04 | 7.3E-04 | MB231/MB468 | NUC |
| >MI0000286\|hsa-mir-210&WithFlank&11\|-\|568083\|568204@72.93.22(+1C)&[MIMAT0000267&hsa-miR-210-3p&offsets\|0\|0(+1C);m-102&11\|-\|568112\|568133&offsets\|0\|0(+1C)]\|CTGTGCGTGTGACAGCGGCTGAC | >MI0000286\|hsa-mir-210&WithFlank&11\|-\|568083\|568204 | 72 | 93 | 22(+1C) | CTGTGCGTGTGACAGCGGCTGAC | 196.1 | 7.2 | 2.0 | 3.7 | 2.5E-04 | 7.3E-04 | BT20/MB468 | MP |
| >hg38_21_+_8433172_8446622_RNA45SN1_45S_50ntFlanks@9162.9187.26\|CGCCCCGGGCGGGTCGCGCCGTCGGG | >hg38_21_+_8433172_8446622_RNA45SN1_45S_50ntFlanks | 9162 | 9187 | 26 | CGCCCCGGGCGGGTCGCGCCGTCGGG | 128.5 | -5.7 | 1.5 | -3.7 | 2.5E-04 | 7.3E-04 | BT20/MB468 | MP |

<table>
<tr><td>>hg38_21_+_8433172_8446622_RNA45SN1_45S_50ntFlanks@5059.5098.40|CGATTTGTCTGGTTAATTCCGATAACGAACGAGACTCTGG</td><td>>hg38_21_+_8433172_8446622_RNA45SN1_45S_50ntFlanks</td><td>5059</td><td>5098</td><td>40</td><td>CGATTTGTCTGGTTAATTCCGATAACGAACGAGACTCTGG</td><td>999.3</td><td>-4.5</td><td>1.3</td><td>-3.4</td><td>5.9E-04</td><td>7.3E-04</td><td>MB231/MB468</td><td>NUC</td></tr>
<tr><td>>hg38_21_+_8433172_8446622_RNA45SN1_45S_50ntFlanks@5074.5110.37|ATTCCGATAACGAACGAGACTCTGGCATGCTAACTAG</td><td>>hg38_21_+_8433172_8446622_RNA45SN1_45S_50ntFlanks</td><td>5074</td><td>5110</td><td>37</td><td>ATTCCGATAACGAACGAGACTCTGGCATGCTAACTAG</td><td>208.2</td><td>-4.3</td><td>1.2</td><td>-3.5</td><td>4.4E-04</td><td>7.3E-04</td><td>BT20/MB468</td><td>MITO</td></tr>
<tr><td>>hg38_21_+_8433172_8446622_RNA45SN1_45S_50ntFlanks@11710.11744.35|CGCGGGTAAACGGCGGGAGTAACTATGACTCTCTT</td><td>>hg38_21_+_8433172_8446622_RNA45SN1_45S_50ntFlanks</td><td>11710</td><td>11744</td><td>35</td><td>CGCGGGTAAACGGCGGGAGTAACTATGACTCTCTT</td><td>81.2</td><td>-5.3</td><td>1.5</td><td>-3.5</td><td>4.9E-04</td><td>7.3E-04</td><td>MB231/MB468</td><td>MP</td></tr>
<tr><td>>hg38_21_+_8433172_8446622_RNA45SN1_45S_50ntFlanks@10236.10263.28|GACGCTACGCCGCGACGAGTAGGAGGGC</td><td>>hg38_21_+_8433172_8446622_RNA45SN1_45S_50ntFlanks</td><td>10236</td><td>10263</td><td>28</td><td>GACGCTACGCCGCGACGAGTAGGAGGGC</td><td>80.2</td><td>-2.7</td><td>0.8</td><td>-3.5</td><td>4.9E-04</td><td>7.3E-04</td><td>MB231/MB468</td><td>MP</td></tr>
<tr><td>>hg38_21_+_8433172_8446622_RNA45SN1_45S_50ntFlanks@9940.9968.29|CAGGACGGTGGCCATGGAAGTCGGAATCC</td><td>>hg38_21_+_8433172_8446622_RNA45SN1_45S_50ntFlanks</td><td>9940</td><td>9968</td><td>29</td><td>CAGGACGGTGGCCATGGAAGTCGGAATCC</td><td>1201.0</td><td>5.3</td><td>1.3</td><td>3.9</td><td>8.6E-05</td><td>7.3E-04</td><td>BT20/MB231</td><td>MP</td></tr>
<tr><td>>hg38_21_+_8433172_8446622_RNA45SN1_45S_50ntFlanks@10914.10931.18|CGCCGCGGCTGGACGAGG</td><td>>hg38_21_+_8433172_8446622_RNA45SN1_45S_50ntFlanks</td><td>10914</td><td>10931</td><td>18</td><td>CGCCGCGGCTGGACGAGG</td><td>663.6</td><td>-4.5</td><td>1.3</td><td>-3.5</td><td>5.3E-04</td><td>7.3E-04</td><td>BT20/MB468</td><td>CYTO</td></tr>
<tr><td>>hg38_MT_+_598_1651_RNR1_12S_50ntFlanks@52.83.32|ATAGGTTTGGTCCTAGCCTTTCTATTAGCTCT</td><td>>hg38_MT_+_598_1651_RNR1_12S_50ntFlanks</td><td>52</td><td>83</td><td>32</td><td>ATAGGTTTGGTCCTAGCCTTTCTATTAGCTCT</td><td>3976.5</td><td>4.0</td><td>1.1</td><td>3.5</td><td>4.4E-04</td><td>7.3E-04</td><td>BT20/MB468</td><td>MITO</td></tr>
<tr><td>>hg38_MT_+_1621_3279_RNR2_16S_50ntFlanks@496.521.26|CTTTGGACACTAGGAAAAAACCTTGT</td><td>>hg38_MT_+_1621_3279_RNR2_16S_50ntFlanks</td><td>496</td><td>521</td><td>26</td><td>CTTTGGACACTAGGAAAAAACCTTGT</td><td>147.6</td><td>7.0</td><td>1.9</td><td>3.7</td><td>2.5E-04</td><td>7.3E-04</td><td>BT20/MB468</td><td>MP</td></tr>
<tr><td>>am_trna2_GlyTCC_19_+_4724082_4724153@1.30.30|GCGTTGGTGGTATAGTGGTTAGCATAGCTG</td><td>>am_trna2_GlyTCC_19_+_4724082_4724153</td><td>1</td><td>30</td><td>30</td><td>GCGTTGGTGGTATAGTGGTTAGCATAGCTG</td><td>363.1</td><td>-5.8</td><td>1.6</td><td>-3.7</td><td>2.5E-04</td><td>7.3E-04</td><td>BT20/MB468</td><td>MP</td></tr>
<tr><td>>trnaMT_GlnTTG_MT_-_4329_4400@1.23.23,trnalookalike2_GlnTTG_1_-_564879_564950@1.23.23|TAGGATGGGGTGTGATAGGTGGC</td><td>>trnaMT_GlnTTG_MT_-_4329_4400</td><td>1</td><td>23</td><td>23</td><td>TAGGATGGGGTGTGATAGGTGGC</td><td>682.8</td><td>4.8</td><td>1.3</td><td>3.7</td><td>2.5E-04</td><td>7.3E-04</td><td>BT20/MB468</td><td>MP</td></tr>
<tr><td>>hg38_21_+_8433172_8446622_RNA45SN1_45S_50ntFlanks@4732.4765.34|AGAACGAAAGTCGGAGGTTCGAAGACGATCAGAT</td><td>>hg38_21_+_8433172_8446622_RNA45SN1_45S_50ntFlanks</td><td>4732</td><td>4765</td><td>34</td><td>AGAACGAAAGTCGGAGGTTCGAAGACGATCAGAT</td><td>360.6</td><td>3.4</td><td>0.9</td><td>3.7</td><td>2.5E-04</td><td>7.3E-04</td><td>BT20/MB468</td><td>MP</td></tr>
<tr><td>>hg38_21_+_8433172_8446622_RNA45SN1_45S_50ntFlanks@6773.6802.30|TTCCTCCCGGGGCTACGCCTGTCTGAGCGT</td><td>>hg38_21_+_8433172_8446622_RNA45SN1_45S_50ntFlanks</td><td>6773</td><td>6802</td><td>30</td><td>TTCCTCCCGGGGCTACGCCTGTCTGAGCGT</td><td>93.3</td><td>-5.2</td><td>1.5</td><td>-3.5</td><td>4.9E-04</td><td>7.3E-04</td><td>MB231/MB468</td><td>MP</td></tr>
<tr><td>>hg38_21_+_8433172_8446622_RNA45SN1_45S_50ntFlanks@12008.12043.36|CCGCCGGTGAAATACCACTACTCTGATCGTTTTTTC</td><td>>hg38_21_+_8433172_8446622_RNA45SN1_45S_50ntFlanks</td><td>12008</td><td>12043</td><td>36</td><td>CCGCCGGTGAAATACCACTACTCTGATCGTTTTTTC</td><td>3881.4</td><td>-6.6</td><td>1.9</td><td>-3.5</td><td>5.3E-04</td><td>7.3E-04</td><td>BT20/MB468</td><td>CYTO</td></tr>
<tr><td>>hg38_21_+_8433172_8446622_RNA45SN1_45S_50ntFlanks@6721.6753.33|AATTGCAGGACACATTGATCATCGACACTTCGA</td><td>>hg38_21_+_8433172_8446622_RNA45SN1_45S_50ntFlanks</td><td>6721</td><td>6753</td><td>33</td><td>AATTGCAGGACACATTGATCATCGACACTTCGA</td><td>134.1</td><td>-2.6</td><td>0.8</td><td>-3.5</td><td>5.6E-04</td><td>7.3E-04</td><td>MB231/MB468</td><td>CYTO</td></tr>
<tr><td>>hg38_21_+_8433172_8446622_RNA45SN1_45S_50ntFlanks@6702.6720.19|AGCTGCGAGAATTAATGTG</td><td>>hg38_21_+_8433172_8446622_RNA45SN1_45S_50ntFlanks</td><td>6702</td><td>6720</td><td>19</td><td>AGCTGCGAGAATTAATGTG</td><td>349.9</td><td>-2.1</td><td>0.6</td><td>-3.5</td><td>5.3E-04</td><td>7.3E-04</td><td>BT20/MB468</td><td>CYTO</td></tr>
<tr><td>>MI0000650|hsa-mir-200c&WithFlank&12|+|6963693|6963772@50.71.22(+2U)&[MIMAT0000617&hsa-miR-200c-3p&offsets|0|-1(+2U);m-54&12|+|6963742|6963764&offsets|0|-1(+2U)]|TAATACTGCCGGGTAATGATGGTT</td><td>>MI0000650|hsa-mir-200c&WithFlank&12|+|6963693|6963772</td><td>50</td><td>71</td><td>22(+2U)</td><td>TAATACTGCCGGGTAATGATGGTT</td><td>367.3</td><td>5.8</td><td>1.6</td><td>3.7</td><td>2.5E-04</td><td>7.3E-04</td><td>BT20/MB468</td><td>MP</td></tr>
<tr><td>>hg38_21_+_8433172_8446622_RNA45SN1_45S_50ntFlanks@8177.8206.30|TTCTGATCGAGGCCCAGCCCGTGGACGGTG</td><td>>hg38_21_+_8433172_8446622_RNA45SN1_45S_50ntFlanks</td><td>8177</td><td>8206</td><td>30</td><td>TTCTGATCGAGGCCCAGCCCGTGGACGGTG</td><td>842.2</td><td>-6.5</td><td>1.9</td><td>-3.4</td><td>6.0E-04</td><td>7.3E-04</td><td>BT20/MB468</td><td>NUC</td></tr>
<tr><td>>hg38_21_+_8433172_8446622_RNA45SN1_45S_50ntFlanks@6644.6668.25|CCTCGTACGACTCTTAGCGGTGGAT</td><td>>hg38_21_+_8433172_8446622_RNA45SN1_45S_50ntFlanks</td><td>6644</td><td>6668</td><td>25</td><td>CCTCGTACGACTCTTAGCGGTGGAT</td><td>292.6</td><td>3.0</td><td>0.9</td><td>3.4</td><td>5.9E-04</td><td>7.4E-04</td><td>MB231/MB468</td><td>MITO</td></tr>
<tr><td>>hg38_21_+_8433172_8446622_RNA45SN1_45S_50ntFlanks@12642.12659.18|CATCTGTGGGATTATGAC</td><td>>hg38_21_+_8433172_8446622_RNA45SN1_45S_50ntFlanks</td><td>12642</td><td>12659</td><td>18</td><td>CATCTGTGGGATTATGAC</td><td>310.0</td><td>-2.9</td><td>0.8</td><td>-3.5</td><td>5.3E-04</td><td>7.4E-04</td><td>BT20/MB468</td><td>CYTO</td></tr>
<tr><td>>hg38_21_+_8433172_8446622_RNA45SN1_45S_50ntFlanks@6661.6689.29|CGGTGGATCACTCGGCTCGTGCGTCGATG</td><td>>hg38_21_+_8433172_8446622_RNA45SN1_45S_50ntFlanks</td><td>6661</td><td>6689</td><td>29</td><td>CGGTGGATCACTCGGCTCGTGCGTCGATG</td><td>4665.6</td><td>-1.8</td><td>0.5</td><td>-3.4</td><td>6.0E-04</td><td>7.4E-04</td><td>MB231/MB468</td><td>NUC</td></tr>
<tr><td>>MI0000299|hsa-mir-222&WithFlank&X|-|45747009|45747130@75.97.23(+2U)&[MIMAT0000279&hsa-miR-222-3p&offsets|0|+2(+2U);m-140&X|-|45747033|45747056&offsets|0|-1(+2U)]|AGCTACATCTGGCTACTGGGTCTTT</td><td>>MI0000299|hsa-mir-222&WithFlank&X|-|45747009|45747130</td><td>75</td><td>97</td><td>23(+2U)</td><td>AGCTACATCTGGCTACTGGGTCTTT</td><td>554.9</td><td>-4.0</td><td>1.0</td><td>-3.9</td><td>8.7E-05</td><td>7.4E-04</td><td>BT20/MB231</td><td>MP</td></tr>
<tr><td>>hg38_21_+_8433172_8446622_RNA45SN1_45S_50ntFlanks@8355.8375.21|TGAAAAGAACTTTGAAGAGAG</td><td>>hg38_21_+_8433172_8446622_RNA45SN1_45S_50ntFlanks</td><td>8355</td><td>8375</td><td>21</td><td>TGAAAAGAACTTTGAAGAGAG</td><td>9415.2</td><td>5.1</td><td>1.3</td><td>3.9</td><td>8.7E-05</td><td>7.4E-04</td><td>BT20/MB231</td><td>MP</td></tr>
<tr><td>>hg38_21_+_8433172_8446622_RNA45SN1_45S_50ntFlanks@12419.12443.25|GATGTCGGCTCTTCCTATCATTGTG</td><td>>hg38_21_+_8433172_8446622_RNA45SN1_45S_50ntFlanks</td><td>12419</td><td>12443</td><td>25</td><td>GATGTCGGCTCTTCCTATCATTGTG</td><td>3609.8</td><td>2.3</td><td>0.7</td><td>3.5</td><td>5.6E-04</td><td>7.4E-04</td><td>MB231/MB468</td><td>CYTO</td></tr>
<tr><td>>trna130_GlnTTG_6_-_27763640_27763711@31.52.22,trna16_GlnTTG_17_+_47269890_47269961@31.52.22,trna173_GlnTTG_6_-_26311975_26312046@31.52.22,trna174_GlnTTG_6_-_26311424_26311495@31.52.22|ACTTTGAATCCAGCGATCCGAG</td><td>>trna130_GlnTTG_6_-_27763640_27763711</td><td>31</td><td>52</td><td>22</td><td>ACTTTGAATCCAGCGATCCGAG</td><td>72.3</td><td>-9.3</td><td>2.5</td><td>-3.7</td><td>2.5E-04</td><td>7.4E-04</td><td>BT20/MB468</td><td>MP</td></tr>
<tr><td>>MI0000085|hsa-mir-27a&WithFlank&19|-|13836434|13836523@57.75.19(+1A)&[MIMAT0000084&hsa-miR-27a-3p&offsets|0|-2(+1A);m-43&19|-|13836447|13836467&offsets|0|-2(+1A)]|TTCACAGTGGCTAAGTTCCA</td><td>>MI0000085|hsa-mir-27a&WithFlank&19|-|13836434|13836523</td><td>57</td><td>75</td><td>19(+1A)</td><td>TTCACAGTGGCTAAGTTCCA</td><td>3550.8</td><td>-3.1</td><td>0.8</td><td>-3.9</td><td>1.0E-04</td><td>7.4E-04</td><td>BT20/MB231</td><td>MITO</td></tr>
<tr><td>>hg38_21_+_8433172_8446622_RNA45SN1_45S_50ntFlanks@11585.11623.39|GGTGCGGACCAGGGGAATCCGACTGTTTAATTAAAACAA</td><td>>hg38_21_+_8433172_8446622_RNA45SN1_45S_50ntFlanks</td><td>11585</td><td>11623</td><td>39</td><td>GGTGCGGACCAGGGGAATCCGACTGTTTAATTAAAACAA</td><td>1090.2</td><td>3.8</td><td>1.0</td><td>3.9</td><td>1.0E-04</td><td>7.4E-04</td><td>BT20/MB231</td><td>MITO</td></tr>
<tr><td>>hg38_21_+_8433172_8446622_RNA45SN1_45S_50ntFlanks@10819.10840.22|AGGGAAGTCGGCAAGCCGGATC</td><td>>hg38_21_+_8433172_8446622_RNA45SN1_45S_50ntFlanks</td><td>10819</td><td>10840</td><td>22</td><td>AGGGAAGTCGGCAAGCCGGATC</td><td>7333.6</td><td>-4.3</td><td>1.2</td><td>-3.7</td><td>2.5E-04</td><td>7.4E-04</td><td>BT20/MB468</td><td>MP</td></tr>
<tr><td>>hg38_21_+_8433172_8446622_RNA45SN1_45S_50ntFlanks@7978.8009.32|GACCTCAGATCAGACGTGGCGACCCGCTGAAT</td><td>>hg38_21_+_8433172_8446622_RNA45SN1_45S_50ntFlanks</td><td>7978</td><td>8009</td><td>32</td><td>GACCTCAGATCAGACGTGGCGACCCGCTGAAT</td><td>273.5</td><td>3.2</td><td>0.9</td><td>3.7</td><td>2.5E-04</td><td>7.4E-04</td><td>BT20/MB468</td><td>MP</td></tr>
<tr><td>>MI0000439|hsa-mir-23b&WithFlank&9|+|95085202|95085310@67.85.19&[MIMAT0000418&hsa-miR-23b-3p&offsets|+3|-1;m-42&9|+|95085265|95085285&offsets|+3|+1]|ACATTGCCAGGGATTACCA</td><td>>MI0000439|hsa-mir-23b&WithFlank&9|+|95085202|95085310</td><td>67</td><td>85</td><td>19</td><td>ACATTGCCAGGGATTACCA</td><td>63.9</td><td>9.2</td><td>2.6</td><td>3.5</td><td>5.0E-04</td><td>7.4E-04</td><td>MB231/MB468</td><td>MP</td></tr>
<tr><td>>hg38_21_+_8433172_8446622_RNA45SN1_45S_50ntFlanks@3843.3871.29|CTACTTGGATAACTGTGGTAATTCTAGAG</td><td>>hg38_21_+_8433172_8446622_RNA45SN1_45S_50ntFlanks</td><td>3843</td><td>3871</td><td>29</td><td>CTACTTGGATAACTGTGGTAATTCTAGAG</td><td>370.9</td><td>-3.2</td><td>0.9</td><td>-3.5</td><td>4.5E-04</td><td>7.4E-04</td><td>BT20/MB468</td><td>MITO</td></tr>
<tr><td>>trnaMT_TyrGTA_MT_-_5826_5891@33.69.37|AATCTAAAGACAGGGGTTAGGCCTCTTTTTACCACCA</td><td>>trnaMT_TyrGTA_MT_-_5826_5891</td><td>33</td><td>69</td><td>37</td><td>AATCTAAAGACAGGGGTTAGGCCTCTTTTTACCACCA</td><td>210.7</td><td>5.9</td><td>1.7</td><td>3.4</td><td>5.9E-04</td><td>7.4E-04</td><td>MB231/MB468</td><td>MITO</td></tr>
<tr><td>>hg38_MT_+_1621_3279_RNR2_16S_50ntFlanks@999.1029.31|AGGGACCTGTATGAATGGCTCCACGAGGGTT</td><td>>hg38_MT_+_1621_3279_RNR2_16S_50ntFlanks</td><td>999</td><td>1029</td><td>31</td><td>AGGGACCTGTATGAATGGCTCCACGAGGGTT</td><td>68.3</td><td>6.9</td><td>1.9</td><td>3.7</td><td>2.5E-04</td><td>7.4E-04</td><td>BT20/MB468</td><td>MP</td></tr>
<tr><td>>hg38_21_+_8433172_8446622_RNA45SN1_45S_50ntFlanks@12126.12150.25|CGACCCGCTCCGGGGACAGTGCCAG</td><td>>hg38_21_+_8433172_8446622_RNA45SN1_45S_50ntFlanks</td><td>12126</td><td>12150</td><td>25</td><td>CGACCCGCTCCGGGGACAGTGCCAG</td><td>555.4</td><td>-5.3</td><td>1.5</td><td>-3.5</td><td>5.4E-04</td><td>7.4E-04</td><td>BT20/MB468</td><td>CYTO</td></tr>
<tr><td>>hg38_21_+_8433172_8446622_RNA45SN1_45S_50ntFlanks@9567.9603.37|ACGTGCAAATCGGTCGTCCGACCTGGGTATAGGGGCG</td><td>>hg38_21_+_8433172_8446622_RNA45SN1_45S_50ntFlanks</td><td>9567</td><td>9603</td><td>37</td><td>ACGTGCAAATCGGTCGTCCGACCTGGGTATAGGGGCG</td><td>460.1</td><td>-2.4</td><td>0.7</td><td>-3.4</td><td>6.0E-04</td><td>7.4E-04</td><td>MB231/MB468</td><td>NUC</td></tr>
<tr><td>>hg38_21_+_8433172_8446622_RNA45SN1_45S_50ntFlanks@6648.6676.29|GTACGACTCTTAGCGGTGGATCACTCGGC</td><td>>hg38_21_+_8433172_8446622_RNA45SN1_45S_50ntFlanks</td><td>6648</td><td>6676</td><td>29</td><td>GTACGACTCTTAGCGGTGGATCACTCGGC</td><td>2933.2</td><td>3.1</td><td>0.9</td><td>3.4</td><td>6.0E-04</td><td>7.4E-04</td><td>MB231/MB468</td><td>MITO</td></tr>
<tr><td>>hg38_21_+_8433172_8446622_RNA45SN1_45S_50ntFlanks@3987.4024.38|GCTTTGGTGACTCTAGATAACCTCGGGCCGATCGCACG</td><td>>hg38_21_+_8433172_8446622_RNA45SN1_45S_50ntFlanks</td><td>3987</td><td>4024</td><td>38</td><td>GCTTTGGTGACTCTAGATAACCTCGGGCCGATCGCACG</td><td>657.2</td><td>-2.6</td><td>0.7</td><td>-3.5</td><td>4.5E-04</td><td>7.4E-04</td><td>BT20/MB468</td><td>MITO</td></tr>
<tr><td>>hg38_21_+_8433172_8446622_RNA45SN1_45S_50ntFlanks@10808.10829.22|CAATGTAGGTAAGGGAAGTCGG</td><td>>hg38_21_+_8433172_8446622_RNA45SN1_45S_50ntFlanks</td><td>10808</td><td>10829</td><td>22</td><td>CAATGTAGGTAAGGGAAGTCGG</td><td>289.3</td><td>-5.7</td><td>1.6</td><td>-3.7</td><td>2.5E-04</td><td>7.4E-04</td><td>BT20/MB468</td><td>MP</td></tr>
<tr><td>>MI0000456|hsa-mir-140&WithFlank&16|+|69933075|69933186@68.88.21&[MIMAT0004597&hsa-miR-140-3p&offsets|0|0;m-45&16|+|69933143|69933164&offsets|-1|-2]|TACCACAGGGTAGAACCACGG</td><td>>MI0000456|hsa-mir-140&WithFlank&16|+|69933075|69933186</td><td>68</td><td>88</td><td>21</td><td>TACCACAGGGTAGAACCACGG</td><td>271.8</td><td>6.3</td><td>1.8</td><td>3.5</td><td>5.0E-04</td><td>7.4E-04</td><td>MB231/MB468</td><td>MP</td></tr>
<tr><td>>hg38_21_+_8433172_8446622_RNA45SN1_45S_50ntFlanks@12418.12439.22|CGATGTCGGCTCTTCCTATCAT</td><td>>hg38_21_+_8433172_8446622_RNA45SN1_45S_50ntFlanks</td><td>12418</td><td>12439</td><td>22</td><td>CGATGTCGGCTCTTCCTATCAT</td><td>869.0</td><td>-2.9</td><td>0.8</td><td>-3.4</td><td>6.0E-04</td><td>7.4E-04</td><td>MB231/MB468</td><td>MITO</td></tr>
<tr><td>>MI0000815|hsa-mir-339&WithFlank&7|-|1022927|1023032@56.78.23&[MIMAT0004702&hsa-miR-339-3p&offsets|0|0;m-235&7|-|1022957|1022977&offsets|0|+2]|TGAGCGCCTCGACGACAGAGCCG</td><td>>MI0000815|hsa-mir-339&WithFlank&7|-|1022927|1023032</td><td>56</td><td>78</td><td>23</td><td>TGAGCGCCTCGACGACAGAGCCG</td><td>634.3</td><td>5.7</td><td>1.6</td><td>3.7</td><td>2.5E-04</td><td>7.4E-04</td><td>BT20/MB468</td><td>MP</td></tr>
<tr><td>>MI0000066|hsa-let-7e&WithFlank&19|+|51692780|51692870@14.34.21&[MIMAT0000066&hsa-let-7e-5p&offsets|0|-1;m-50&19|+|51692793|51692814&offsets|0|-1]|TGAGGTAGGAGGTTGTATAGT</td><td>>MI0000066|hsa-let-7e&WithFlank&19|+|51692780|51692870</td><td>14</td><td>34</td><td>21</td><td>TGAGGTAGGAGGTTGTATAGT</td><td>446.3</td><td>3.1</td><td>0.8</td><td>3.7</td><td>2.5E-04</td><td>7.4E-04</td><td>BT20/MB468</td><td>MP</td></tr>
<tr><td>>hg38_MT_+_598_1651_RNR1_12S_50ntFlanks@917.950.34|AGGACATTTAACTAAAACCCCTACGCATTTATAT</td><td>>hg38_MT_+_598_1651_RNR1_12S_50ntFlanks</td><td>917</td><td>950</td><td>34</td><td>AGGACATTTAACTAAAACCCCTACGCATTTATAT</td><td>409.6</td><td>6.9</td><td>1.9</td><td>3.7</td><td>2.5E-04</td><td>7.5E-04</td><td>BT20/MB468</td><td>MP</td></tr>
<tr><td>>hg38_1_-_228634819_228635039_RNA5S12_5S_50ntFlanks@133.169.37|ATGGGAGACCGCCTGGGAATACCGGGTGCTGTAGGCT</td><td>>hg38_1_-_228634819_228635039_RNA5S12_5S_50ntFlanks</td><td>133</td><td>169</td><td>37</td><td>ATGGGAGACCGCCTGGGAATACCGGGTGCTGTAGGCT</td><td>7904.3</td><td>5.2</td><td>1.3</td><td>3.9</td><td>8.8E-05</td><td>7.5E-04</td><td>BT20/MB231</td><td>MP</td></tr>
<tr><td>>trnaMT_MetCAT_MT_+_4402_4469@38.70.33|CGAAAATGTTGGTTATACCCTTCCCGTACTACC</td><td>>trnaMT_MetCAT_MT_+_4402_4469</td><td>38</td><td>70</td><td>33</td><td>CGAAAATGTTGGTTATACCCTTCCCGTACTACC</td><td>67.2</td><td>9.8</td><td>2.7</td><td>3.7</td><td>2.5E-04</td><td>7.5E-04</td><td>BT20/MB468</td><td>MP</td></tr>
<tr><td>>hg38_21_+_8433172_8446622_RNA45SN1_45S_50ntFlanks@8176.8216.41|CTTCTGATCGAGGCCCAGCCCGTGGACGGTGTGAGGCCGGT</td><td>>hg38_21_+_8433172_8446622_RNA45SN1_45S_50ntFlanks</td><td>8176</td><td>8216</td><td>41</td><td>CTTCTGATCGAGGCCCAGCCCGTGGACGGTGTGAGGCCGGT</td><td>280.8</td><td>-3.5</td><td>1.0</td><td>-3.5</td><td>5.0E-04</td><td>7.5E-04</td><td>MB231/MB468</td><td>MP</td></tr>
<tr><td>>hg38_21_+_8433172_8446622_RNA45SN1_45S_50ntFlanks@11920.11948.29|ATGAGAGGTGTAGAATAAGTGGGAGGCCC</td><td>>hg38_21_+_8433172_8446622_RNA45SN1_45S_50ntFlanks</td><td>11920</td><td>11948</td><td>29</td><td>ATGAGAGGTGTAGAATAAGTGGGAGGCCC</td><td>144.4</td><td>10.5</td><td>2.7</td><td>3.9</td><td>8.8E-05</td><td>7.5E-04</td><td>BT20/MB231</td><td>MP</td></tr>
<tr><td>>MI0000088|hsa-mir-30a&WithFlank&6|-|71403545|71403627@12.35.24(+3U)&[MIMAT0000087&hsa-miR-30a-5p&offsets|0|+2(+3U);m-15&6|-|71403595|71403616&offsets|0|+2(+3U)]|TGTAAACATCCTCGACTGGAAGCTTTT</td><td>>MI0000088|hsa-mir-30a&WithFlank&6|-|71403545|71403627</td><td>12</td><td>35</td><td>24(+3U)</td><td>TGTAAACATCCTCGACTGGAAGCTTTT</td><td>449.7</td><td>-7.9</td><td>2.0</td><td>-3.9</td><td>1.1E-04</td><td>7.5E-04</td><td>BT20/MB231</td><td>MITO</td></tr>
<tr><td>>hg38_21_+_8433172_8446622_RNA45SN1_45S_50ntFlanks@10407.10424.18|GGGTCAGTCGGTCCTGAG</td><td>>hg38_21_+_8433172_8446622_RNA45SN1_45S_50ntFlanks</td><td>10407</td><td>10424</td><td>18</td><td>GGGTCAGTCGGTCCTGAG</td><td>938.9</td><td>2.7</td><td>0.8</td><td>3.5</td><td>4.5E-04</td><td>7.5E-04</td><td>BT20/MB468</td><td>MITO</td></tr>
<tr><td>>hg38_21_+_8433172_8446622_RNA45SN1_45S_50ntFlanks@12300.12323.24|GGGCCTCACGATCCTTCTGACCTT</td><td>>hg38_21_+_8433172_8446622_RNA45SN1_45S_50ntFlanks</td><td>12300</td><td>12323</td><td>24</td><td>GGGCCTCACGATCCTTCTGACCTT</td><td>907.6</td><td>-4.2</td><td>1.2</td><td>-3.5</td><td>4.5E-04</td><td>7.5E-04</td><td>BT20/MB468</td><td>MITO</td></tr>
<tr><td>>hg38_21_+_8433172_8446622_RNA45SN1_45S_50ntFlanks@7968.8008.41|TCCGAGACGCGACCTCAGATCAGACGTGGCGACCCGCTGAA</td><td>>hg38_21_+_8433172_8446622_RNA45SN1_45S_50ntFlanks</td><td>7968</td><td>8008</td><td>41</td><td>TCCGAGACGCGACCTCAGATCAGACGTGGCGACCCGCTGAA</td><td>81.2</td><td>2.8</td><td>0.8</td><td>3.4</td><td>6.1E-04</td><td>7.5E-04</td><td>BT20/MB468</td><td>NUC</td></tr>
<tr><td>>am_trnaMT_SerGCT_MT_+_12207_12265@16.38.23|AACTGCTAACTCATGCCCCCATG</td><td>>am_trnaMT_SerGCT_MT_+_12207_12265</td><td>16</td><td>38</td><td>23</td><td>AACTGCTAACTCATGCCCCCATG</td><td>161.4</td><td>-4.1</td><td>1.2</td><td>-3.4</td><td>5.7E-04</td><td>7.5E-04</td><td>MB231/MB468</td><td>CYTO</td></tr>
<tr><td>>MI0000088|hsa-mir-30a&WithFlank&6|-|71403545|71403627@53.74.22&[MIMAT0000088&hsa-miR-30a-3p&offsets|0|0;m-38&6|-|71403554|71403575&offsets|0|0]|CTTTCAGTCGGATGTTTGCAGC</td><td>>MI0000088|hsa-mir-30a&WithFlank&6|-|71403545|71403627</td><td>53</td><td>74</td><td>22</td><td>CTTTCAGTCGGATGTTTGCAGC</td><td>7222.5</td><td>3.1</td><td>0.9</td><td>3.5</td><td>5.0E-04</td><td>7.5E-04</td><td>MB231/MB468</td><td>MP</td></tr>
</table>

<table>
<tr><th></th><th></th><th></th><th></th><th></th><th></th><th></th><th></th><th></th><th></th><th></th><th></th><th></th><th></th></tr>
<tr><td>>hg38_21_+_8433172_8446622_RNA45SN1_45S_50ntFlanks@5059.5081.23|CGATTTGTCTGGTTAATTCCGAT</td><td>>hg38_21_+_8433172_8446622_RNA45SN1_45S_50ntFlanks</td><td>5059</td><td>5081</td><td>23</td><td>CGATTTGTCTGGTTAATTCCGAT</td><td>311.8</td><td>-3.7</td><td>1.1</td><td>-3.5</td><td>5.0E-04</td><td>7.5E-04</td><td>MB231/MB468</td><td>MP</td></tr>
<tr><td>>hg38_21_+_8433172_8446622_RNA45SN1_45S_50ntFlanks@8168.8214.47|CCCAAGTCCTTCTGATCGAGGCCCAGCCCGTGGACGGTGTGAGGCCG</td><td>>hg38_21_+_8433172_8446622_RNA45SN1_45S_50ntFlanks</td><td>8168</td><td>8214</td><td>47</td><td>CCCAAGTCCTTCTGATCGAGGCCCAGCCCGTGGACGGTGTGAGGCCG</td><td>188.5</td><td>-4.3</td><td>1.2</td><td>-3.5</td><td>5.4E-04</td><td>7.5E-04</td><td>BT20/MB468</td><td>CYTO</td></tr>
<tr><td>>hg38_21_+_8433172_8446622_RNA45SN1_45S_50ntFlanks@12916.12933.18|TTCGTGGGGAACCTGGCG</td><td>>hg38_21_+_8433172_8446622_RNA45SN1_45S_50ntFlanks</td><td>12916</td><td>12933</td><td>18</td><td>TTCGTGGGGAACCTGGCG</td><td>393.8</td><td>-6.2</td><td>1.8</td><td>-3.4</td><td>6.1E-04</td><td>7.5E-04</td><td>BT20/MB468</td><td>NUC</td></tr>
<tr><td>>am_trnaMT_GluTTC_MT_-_14674_14742@27.48.22,trnalookalike8_GluTTC_5_-_93905172_93905240@27.48.22|GTTTTTCATATCATTGGTCGTG</td><td>>am_trnaMT_GluTTC_MT_-_14674_14742</td><td>27</td><td>48</td><td>22</td><td>GTTTTTCATATCATTGGTCGTG</td><td>186.5</td><td>-5.4</td><td>1.5</td><td>-3.5</td><td>5.1E-04</td><td>7.5E-04</td><td>MB231/MB468</td><td>MP</td></tr>
<tr><td>>hg38_21_+_8433172_8446622_RNA45SN1_45S_50ntFlanks@10361.10383.23|GAAGGCCGAAGTGGAGAAGGGTT</td><td>>hg38_21_+_8433172_8446622_RNA45SN1_45S_50ntFlanks</td><td>10361</td><td>10383</td><td>23</td><td>GAAGGCCGAAGTGGAGAAGGGTT</td><td>188.7</td><td>-3.8</td><td>1.1</td><td>-3.5</td><td>4.5E-04</td><td>7.5E-04</td><td>BT20/MB468</td><td>MITO</td></tr>
<tr><td>>hg38_21_+_8433172_8446622_RNA45SN1_45S_50ntFlanks@12399.12437.39|ACGTCGCTTTTTGATCCTTCGATGTCGGCTCTTCCTATC</td><td>>hg38_21_+_8433172_8446622_RNA45SN1_45S_50ntFlanks</td><td>12399</td><td>12437</td><td>39</td><td>ACGTCGCTTTTTGATCCTTCGATGTCGGCTCTTCCTATC</td><td>388.4</td><td>-2.9</td><td>0.8</td><td>-3.4</td><td>6.0E-04</td><td>7.5E-04</td><td>MB231/MB468</td><td>MITO</td></tr>
<tr><td>>hg38_21_+_8433172_8446622_RNA45SN1_45S_50ntFlanks@12419.12446.28|GATGTCGGCTCTTCCTATCATTGTGAAG</td><td>>hg38_21_+_8433172_8446622_RNA45SN1_45S_50ntFlanks</td><td>12419</td><td>12446</td><td>28</td><td>GATGTCGGCTCTTCCTATCATTGTGAAG</td><td>2401.7</td><td>2.2</td><td>0.6</td><td>3.4</td><td>5.7E-04</td><td>7.5E-04</td><td>MB231/MB468</td><td>CYTO</td></tr>
<tr><td>>hg38_21_+_8433172_8446622_RNA45SN1_45S_50ntFlanks@4547.4566.20|CTGGATACCGCAGCTAGGAA</td><td>>hg38_21_+_8433172_8446622_RNA45SN1_45S_50ntFlanks</td><td>4547</td><td>4566</td><td>20</td><td>CTGGATACCGCAGCTAGGAA</td><td>180.1</td><td>-6.2</td><td>1.7</td><td>-3.7</td><td>2.5E-04</td><td>7.5E-04</td><td>BT20/MB468</td><td>MP</td></tr>
<tr><td>>hg38_1_-_228634819_228635039_RNA5S12_5S_50ntFlanks@139.170.32|GACCGCCTGGGAATACCGGGTGCTGTAGGCTT</td><td>>hg38_1_-_228634819_228635039_RNA5S12_5S_50ntFlanks</td><td>139</td><td>170</td><td>32</td><td>GACCGCCTGGGAATACCGGGTGCTGTAGGCTT</td><td>1816.5</td><td>2.8</td><td>0.8</td><td>3.7</td><td>2.5E-04</td><td>7.5E-04</td><td>BT20/MB468</td><td>MP</td></tr>
<tr><td>>MI0000809|hsa-mir-151a&WithFlank&8|-|140732558|140732659@51.70.20&[MIMAT0000757&hsa-miR-151a-3p&offsets|-2|-3;m-30&8|-|140732587|140732607&offsets|-2|-3]|TACTAGACTGAAGCTCCTTG</td><td>>MI0000809|hsa-mir-151a&WithFlank&8|-|140732558|140732659</td><td>51</td><td>70</td><td>20</td><td>TACTAGACTGAAGCTCCTTG</td><td>197.8</td><td>6.1</td><td>1.7</td><td>3.7</td><td>2.5E-04</td><td>7.5E-04</td><td>BT20/MB468</td><td>MP</td></tr>
<tr><td>>hg38_21_+_8433172_8446622_RNA45SN1_45S_50ntFlanks@10727.10754.28|GGAGAGGGTGTAAATCTCGCGCCGGGCC</td><td>>hg38_21_+_8433172_8446622_RNA45SN1_45S_50ntFlanks</td><td>10727</td><td>10754</td><td>28</td><td>GGAGAGGGTGTAAATCTCGCGCCGGGCC</td><td>136.0</td><td>-5.6</td><td>1.5</td><td>-3.7</td><td>2.5E-04</td><td>7.5E-04</td><td>BT20/MB468</td><td>MP</td></tr>
<tr><td>>hg38_21_+_8433172_8446622_RNA45SN1_45S_50ntFlanks@10798.10815.18|CATGTTGGAACAATGTAG</td><td>>hg38_21_+_8433172_8446622_RNA45SN1_45S_50ntFlanks</td><td>10798</td><td>10815</td><td>18</td><td>CATGTTGGAACAATGTAG</td><td>101.1</td><td>-5.5</td><td>1.5</td><td>-3.7</td><td>2.5E-04</td><td>7.5E-04</td><td>BT20/MB468</td><td>MP</td></tr>
<tr><td>>hg38_21_+_8433172_8446622_RNA45SN1_45S_50ntFlanks@13007.13025.19|TATTGAAAGTCAGCCCTCG</td><td>>hg38_21_+_8433172_8446622_RNA45SN1_45S_50ntFlanks</td><td>13007</td><td>13025</td><td>19</td><td>TATTGAAAGTCAGCCCTCG</td><td>102.9</td><td>-5.5</td><td>1.5</td><td>-3.7</td><td>2.5E-04</td><td>7.5E-04</td><td>BT20/MB468</td><td>MP</td></tr>
<tr><td>>hg38_21_+_8433172_8446622_RNA45SN1_45S_50ntFlanks@9787.9819.33|TCGCTGGCGTGGAGCCGGGCGTGGAATGCGAGT</td><td>>hg38_21_+_8433172_8446622_RNA45SN1_45S_50ntFlanks</td><td>9787</td><td>9819</td><td>33</td><td>TCGCTGGCGTGGAGCCGGGCGTGGAATGCGAGT</td><td>517.9</td><td>7.5</td><td>1.9</td><td>3.9</td><td>8.9E-05</td><td>7.5E-04</td><td>BT20/MB231</td><td>MP</td></tr>
<tr><td>>hg38_21_+_8433172_8446622_RNA45SN1_45S_50ntFlanks@12417.12456.40|TCGATGTCGGCTCTTCCTATCATTGTGAAGCAGAATTCAC</td><td>>hg38_21_+_8433172_8446622_RNA45SN1_45S_50ntFlanks</td><td>12417</td><td>12456</td><td>40</td><td>TCGATGTCGGCTCTTCCTATCATTGTGAAGCAGAATTCAC</td><td>206.7</td><td>-2.6</td><td>0.8</td><td>-3.5</td><td>5.4E-04</td><td>7.5E-04</td><td>BT20/MB468</td><td>CYTO</td></tr>
<tr><td>>hg38_21_+_8433172_8446622_RNA45SN1_45S_50ntFlanks@5467.5497.31|GGCCCACGGCCCTGGCGGAGCGCTGAGAAGA</td><td>>hg38_21_+_8433172_8446622_RNA45SN1_45S_50ntFlanks</td><td>5467</td><td>5497</td><td>31</td><td>GGCCCACGGCCCTGGCGGAGCGCTGAGAAGA</td><td>184.8</td><td>-3.8</td><td>1.1</td><td>-3.4</td><td>5.7E-04</td><td>7.5E-04</td><td>MB231/MB468</td><td>CYTO</td></tr>
<tr><td>>hg38_21_+_8433172_8446622_RNA45SN1_45S_50ntFlanks@12416.12460.45|TTCGATGTCGGCTCTTCCTATCATTGTGAAGCAGAATTCACCAAG</td><td>>hg38_21_+_8433172_8446622_RNA45SN1_45S_50ntFlanks</td><td>12416</td><td>12460</td><td>45</td><td>TTCGATGTCGGCTCTTCCTATCATTGTGAAGCAGAATTCACCAAG</td><td>264.2</td><td>5.4</td><td>1.6</td><td>3.4</td><td>6.0E-04</td><td>7.5E-04</td><td>MB231/MB468</td><td>MITO</td></tr>
<tr><td>>am_trna111_HisGTG_1_-_147774845_147774916@2.29.28,trna118_HisGTG_1_-_145396881_145396952@2.29.28,trna16_HisGTG_1_+_146544773_146544844@2.29.28,trna1_HisGTG_15_+_45493349_45493420@2.29.28,trna21_HisGTG_1_+_147753471_147753542@2.29.28,trna33_HisGTG_6_+_27125906_27125977@2.29.28,trna7_HisGTG_9_-_14433938_14434009@2.29.28,trna8_HisGTG_15_-_45492611_45492682@2.29.28,trna9_HisGTG_15_-_45490804_45490875@2.29.28|CCGTGATCGTATAGTGGTTAGTACTCTG</td><td>>am_trna111_HisGTG_1_-_147774845_147774916</td><td>2</td><td>29</td><td>28</td><td>CCGTGATCGTATAGTGGTTAGTACTCTG</td><td>309.0</td><td>-5.9</td><td>1.7</td><td>-3.4</td><td>6.1E-04</td><td>7.5E-04</td><td>BT20/MB468</td><td>NUC</td></tr>
<tr><td>>hg38_21_+_8433172_8446622_RNA45SN1_45S_50ntFlanks@10905.10935.31|CTGGGCGCGCGCCGCGGCTGGACGAGGCGCC</td><td>>hg38_21_+_8433172_8446622_RNA45SN1_45S_50ntFlanks</td><td>10905</td><td>10935</td><td>31</td><td>CTGGGCGCGCGCCGCGGCTGGACGAGGCGCC</td><td>371.3</td><td>-6.0</td><td>1.8</td><td>-3.4</td><td>6.1E-04</td><td>7.5E-04</td><td>BT20/MB468</td><td>NUC</td></tr>
<tr><td>>MI0000542|hsa-mir-320a&WithFlank&8|-|22244960|22245043@48.70.23(+1U)&[MIMAT0000510&hsa-miR-320a-3p&offsets|0|+1(+1U);m-60&8|-|22244975|22244996&offsets|0|+1(+1U)]|AAAAGCTGGGTTGAGAGGGCGAAT</td><td>>MI0000542|hsa-mir-320a&WithFlank&8|-|22244960|22245043</td><td>48</td><td>70</td><td>23(+1U)</td><td>AAAAGCTGGGTTGAGAGGGCGAAT</td><td>22662.6</td><td>3.1</td><td>0.8</td><td>3.9</td><td>9.3E-05</td><td>7.5E-04</td><td>BT20/MB231</td><td>CYTO</td></tr>
<tr><td>>MI0000077|hsa-mir-21&WithFlank&17|+|59841260|59841343@52.73.22&[MIMAT0004494&hsa-miR-21-3p&offsets|0|+1;m-52&17|+|59841311|59841332&offsets|0|0]|CAACACCAGTCGATGGGCTGTC</td><td>>MI0000077|hsa-mir-21&WithFlank&17|+|59841260|59841343</td><td>52</td><td>73</td><td>22</td><td>CAACACCAGTCGATGGGCTGTC</td><td>1592.2</td><td>4.9</td><td>1.4</td><td>3.7</td><td>2.6E-04</td><td>7.5E-04</td><td>BT20/MB468</td><td>MP</td></tr>
<tr><td>>am_trna2_GlyGCC_21_-_18827107_18827177@1.26.26,trna35_GlyGCC_1_+_161413094_161413164@1.26.26,trna37_GlyGCC_1_+_161420467_161420537@1.26.26,trna39_GlyGCC_1_+_161427898_161427968@1.26.26,trna41_GlyGCC_1_+_161435258_161435328@1.26.26|GCATGGGTGGTTCAGTGGTAGAATTC</td><td>>am_trna2_GlyGCC_21_-_18827107_18827177</td><td>1</td><td>26</td><td>26</td><td>GCATGGGTGGTTCAGTGGTAGAATTC</td><td>2459.4</td><td>6.5</td><td>1.7</td><td>3.9</td><td>8.9E-05</td><td>7.5E-04</td><td>BT20/MB231</td><td>MP</td></tr>
<tr><td>>hg38_21_+_8433172_8446622_RNA45SN1_45S_50ntFlanks@10849.10866.18|CGGGATAAGGATTGGCTC</td><td>>hg38_21_+_8433172_8446622_RNA45SN1_45S_50ntFlanks</td><td>10849</td><td>10866</td><td>18</td><td>CGGGATAAGGATTGGCTC</td><td>362.2</td><td>-2.9</td><td>0.8</td><td>-3.4</td><td>5.7E-04</td><td>7.6E-04</td><td>MB231/MB468</td><td>CYTO</td></tr>
<tr><td>>hg38_21_+_8433172_8446622_RNA45SN1_45S_50ntFlanks@11580.11609.30|GAACTGGTGCGGACCAGGGGAATCCGACTG</td><td>>hg38_21_+_8433172_8446622_RNA45SN1_45S_50ntFlanks</td><td>11580</td><td>11609</td><td>30</td><td>GAACTGGTGCGGACCAGGGGAATCCGACTG</td><td>785.4</td><td>-1.8</td><td>0.5</td><td>-3.4</td><td>5.7E-04</td><td>7.6E-04</td><td>MB231/MB468</td><td>CYTO</td></tr>
<tr><td>>hg38_21_+_8433172_8446622_RNA45SN1_45S_50ntFlanks@9621.9655.35|TCTAGTAGCTGGTTCCCTCCGAAGTTTCCCTCAGG</td><td>>hg38_21_+_8433172_8446622_RNA45SN1_45S_50ntFlanks</td><td>9621</td><td>9655</td><td>35</td><td>TCTAGTAGCTGGTTCCCTCCGAAGTTTCCCTCAGG</td><td>132.3</td><td>-2.4</td><td>0.7</td><td>-3.5</td><td>4.6E-04</td><td>7.6E-04</td><td>BT20/MB468</td><td>MITO</td></tr>
<tr><td>>hg38_21_+_8433172_8446622_RNA45SN1_45S_50ntFlanks@11587.11605.19|TGCGGACCAGGGGAATCCG</td><td>>hg38_21_+_8433172_8446622_RNA45SN1_45S_50ntFlanks</td><td>11587</td><td>11605</td><td>19</td><td>TGCGGACCAGGGGAATCCG</td><td>244.7</td><td>-6.1</td><td>1.7</td><td>-3.7</td><td>2.6E-04</td><td>7.6E-04</td><td>BT20/MB468</td><td>MP</td></tr>
<tr><td>>trnaMT_HisGTG_MT_+_12138_12206@26.72.47|AGATTGTGAATCTGACAACAGAGGCTTACGACCCCTTATTTACCCCA</td><td>>trnaMT_HisGTG_MT_+_12138_12206</td><td>26</td><td>72</td><td>47</td><td>AGATTGTGAATCTGACAACAGAGGCTTACGACCCCTTATTTACCCCA</td><td>121.4</td><td>6.4</td><td>1.9</td><td>3.4</td><td>6.1E-04</td><td>7.6E-04</td><td>MB231/MB468</td><td>MITO</td></tr>
<tr><td>>hg38_21_+_8433172_8446622_RNA45SN1_45S_50ntFlanks@11552.11588.37|CCTCGGCCGGCGCCTAGCAGCCGACTTAGAACTGGTG</td><td>>hg38_21_+_8433172_8446622_RNA45SN1_45S_50ntFlanks</td><td>11552</td><td>11588</td><td>37</td><td>CCTCGGCCGGCGCCTAGCAGCCGACTTAGAACTGGTG</td><td>690.3</td><td>-2.7</td><td>0.8</td><td>-3.4</td><td>6.1E-04</td><td>7.6E-04</td><td>MB231/MB468</td><td>MITO</td></tr>
<tr><td>>MI0000746|hsa-mir-99b&WithFlank&19|+|51692606|51692687@13.31.19&[MIMAT0000689&hsa-miR-99b-5p&offsets|0|-3;m-10&19|+|51692618|51692639&offsets|0|-3]|CACCCGTAGAACCGACCTT</td><td>>MI0000746|hsa-mir-99b&WithFlank&19|+|51692606|51692687</td><td>13</td><td>31</td><td>19</td><td>CACCCGTAGAACCGACCTT</td><td>253.5</td><td>6.3</td><td>1.7</td><td>3.7</td><td>2.6E-04</td><td>7.6E-04</td><td>BT20/MB468</td><td>MP</td></tr>
<tr><td>>trna14_LysTTT_11_-_59327808_59327880@1.30.30,trna2_LysTTT_17_+_8022473_8022545@1.30.30,trna54_LysTTT_1_+_204475655_204475727@1.30.30,trna5_LysTTT_11_+_59323902_59323974@1.30.30,trna62_LysTTT_1_-_204476158_204476230@1.30.30,trna76_LysTTT_6_+_28918806_28918878@1.30.30|GCCCGGATAGCTCAGTCGGTAGAGCATCAG</td><td>>trna14_LysTTT_11_-_59327808_59327880</td><td>1</td><td>30</td><td>30</td><td>GCCCGGATAGCTCAGTCGGTAGAGCATCAG</td><td>920.9</td><td>-5.7</td><td>1.6</td><td>-3.5</td><td>5.5E-04</td><td>7.6E-04</td><td>BT20/MB468</td><td>CYTO</td></tr>
<tr><td>>hg38_21_+_8433172_8446622_RNA45SN1_45S_50ntFlanks@11027.11049.23|CTCTCTCTCTCTCTCCCCCGCTC</td><td>>hg38_21_+_8433172_8446622_RNA45SN1_45S_50ntFlanks</td><td>11027</td><td>11049</td><td>23</td><td>CTCTCTCTCTCTCTCCCCCGCTC</td><td>810.5</td><td>-4.3</td><td>1.2</td><td>-3.5</td><td>5.5E-04</td><td>7.6E-04</td><td>BT20/MB468</td><td>CYTO</td></tr>
<tr><td>>hg38_21_+_8433172_8446622_RNA45SN1_45S_50ntFlanks@12999.13027.29|CTGCGATCTATTGAAAGTCAGCCCTCGAC</td><td>>hg38_21_+_8433172_8446622_RNA45SN1_45S_50ntFlanks</td><td>12999</td><td>13027</td><td>29</td><td>CTGCGATCTATTGAAAGTCAGCCCTCGAC</td><td>410.7</td><td>-2.2</td><td>0.6</td><td>-3.5</td><td>4.6E-04</td><td>7.6E-04</td><td>BT20/MB468</td><td>MITO</td></tr>
<tr><td>>hg38_21_+_8433172_8446622_RNA45SN1_45S_50ntFlanks@11577.11601.25|TTAGAACTGGTGCGGACCAGGGGAA</td><td>>hg38_21_+_8433172_8446622_RNA45SN1_45S_50ntFlanks</td><td>11577</td><td>11601</td><td>25</td><td>TTAGAACTGGTGCGGACCAGGGGAA</td><td>366.0</td><td>-4.2</td><td>1.2</td><td>-3.5</td><td>5.5E-04</td><td>7.6E-04</td><td>BT20/MB468</td><td>CYTO</td></tr>
<tr><td>>hg38_21_+_8433172_8446622_RNA45SN1_45S_50ntFlanks@4726.4748.23|TAATCAAGAACGAAAGTCGGAGG</td><td>>hg38_21_+_8433172_8446622_RNA45SN1_45S_50ntFlanks</td><td>4726</td><td>4748</td><td>23</td><td>TAATCAAGAACGAAAGTCGGAGG</td><td>169.0</td><td>-5.7</td><td>1.6</td><td>-3.7</td><td>2.6E-04</td><td>7.6E-04</td><td>BT20/MB468</td><td>MP</td></tr>
<tr><td>>hg38_21_+_8433172_8446622_RNA45SN1_45S_50ntFlanks@6652.6688.37|GACTCTTAGCGGTGGATCACTCGGCTCGTGCGTCGAT</td><td>>hg38_21_+_8433172_8446622_RNA45SN1_45S_50ntFlanks</td><td>6652</td><td>6688</td><td>37</td><td>GACTCTTAGCGGTGGATCACTCGGCTCGTGCGTCGAT</td><td>3205.9</td><td>2.8</td><td>0.8</td><td>3.4</td><td>6.2E-04</td><td>7.6E-04</td><td>MB231/MB468</td><td>NUC</td></tr>
<tr><td>>trnaMT_ValTAC_MT_+_1602_1670@53.72.20|TAACTTGACCGCTCTGACCA</td><td>>trnaMT_ValTAC_MT_+_1602_1670</td><td>53</td><td>72</td><td>20</td><td>TAACTTGACCGCTCTGACCA</td><td>784.6</td><td>3.7</td><td>1.0</td><td>3.7</td><td>2.6E-04</td><td>7.6E-04</td><td>BT20/MB468</td><td>MP</td></tr>
<tr><td>>hg38_21_+_8433172_8446622_RNA45SN1_45S_50ntFlanks@10409.10426.18|GTCAGTCGGTCCTGAGAG</td><td>>hg38_21_+_8433172_8446622_RNA45SN1_45S_50ntFlanks</td><td>10409</td><td>10426</td><td>18</td><td>GTCAGTCGGTCCTGAGAG</td><td>189.2</td><td>-2.9</td><td>0.9</td><td>-3.4</td><td>5.8E-04</td><td>7.6E-04</td><td>MB231/MB468</td><td>CYTO</td></tr>
<tr><td>>hg38_21_+_8433172_8446622_RNA45SN1_45S_50ntFlanks@5056.5083.28|GAGCGATTTGTCTGGTTAATTCCGATAA</td><td>>hg38_21_+_8433172_8446622_RNA45SN1_45S_50ntFlanks</td><td>5056</td><td>5083</td><td>28</td><td>GAGCGATTTGTCTGGTTAATTCCGATAA</td><td>77.9</td><td>-10.1</td><td>2.9</td><td>-3.5</td><td>5.1E-04</td><td>7.6E-04</td><td>MB231/MB468</td><td>MP</td></tr>
<tr><td>>hg38_21_+_8433172_8446622_RNA45SN1_45S_50ntFlanks@9788.9820.33|CGCTGGCGTGGAGCCGGGCGTGGAATGCGAGTG</td><td>>hg38_21_+_8433172_8446622_RNA45SN1_45S_50ntFlanks</td><td>9788</td><td>9820</td><td>33</td><td>CGCTGGCGTGGAGCCGGGCGTGGAATGCGAGTG</td><td>699.3</td><td>8.0</td><td>2.0</td><td>3.9</td><td>9.0E-05</td><td>7.6E-04</td><td>BT20/MB231</td><td>MP</td></tr>
<tr><td>>hg38_21_+_8433172_8446622_RNA45SN1_45S_50ntFlanks@10410.10443.34|TCAGTCGGTCCTGAGAGATGGGCGAGCGCCGTTC</td><td>>hg38_21_+_8433172_8446622_RNA45SN1_45S_50ntFlanks</td><td>10410</td><td>10443</td><td>34</td><td>TCAGTCGGTCCTGAGAGATGGGCGAGCGCCGTTC</td><td>865.2</td><td>-4.9</td><td>1.3</td><td>-3.7</td><td>2.6E-04</td><td>7.6E-04</td><td>BT20/MB468</td><td>MP</td></tr>
<tr><td>>hg38_21_+_8433172_8446622_RNA45SN1_45S_50ntFlanks@5071.5110.40|TTAATTCCGATAACGAACGAGACTCTGGCATGCTAACTAG</td><td>>hg38_21_+_8433172_8446622_RNA45SN1_45S_50ntFlanks</td><td>5071</td><td>5110</td><td>40</td><td>TTAATTCCGATAACGAACGAGACTCTGGCATGCTAACTAG</td><td>78.2</td><td>-10.2</td><td>2.9</td><td>-3.5</td><td>5.1E-04</td><td>7.6E-04</td><td>MB231/MB468</td><td>MP</td></tr>
<tr><td>>MI0000265|hsa-mir-7-3&WithFlank&19|+|4770664|4770785@37.61.25&[MIMAT0000252_1&hsa-miR-7-5p&offsets|0|+1;m-402&19|+|4770700|4770723&offsets|0|+1]|TGGAAGACTAGTGATTTTGTTGTTC</td><td>>MI0000265|hsa-mir-7-3&WithFlank&19|+|4770664|4770785</td><td>37</td><td>61</td><td>25</td><td>TGGAAGACTAGTGATTTTGTTGTTC</td><td>278.6</td><td>4.4</td><td>1.3</td><td>3.5</td><td>5.1E-04</td><td>7.6E-04</td><td>MB231/MB468</td><td>MP</td></tr>
<tr><td>>hg38_21_+_8433172_8446622_RNA45SN1_45S_50ntFlanks@10238.10261.24|CGCTACGCCGCGACGAGTAGGAGG</td><td>>hg38_21_+_8433172_8446622_RNA45SN1_45S_50ntFlanks</td><td>10238</td><td>10261</td><td>24</td><td>CGCTACGCCGCGACGAGTAGGAGG</td><td>350.6</td><td>5.6</td><td>1.4</td><td>3.9</td><td>9.0E-05</td><td>7.6E-04</td><td>BT20/MB231</td><td>MP</td></tr>
<tr><td>>hg38_21_+_8433172_8446622_RNA45SN1_45S_50ntFlanks@8171.8195.25|AAGTCCTTCTGATCGAGGCCCAGCC</td><td>>hg38_21_+_8433172_8446622_RNA45SN1_45S_50ntFlanks</td><td>8171</td><td>8195</td><td>25</td><td>AAGTCCTTCTGATCGAGGCCCAGCC</td><td>338.7</td><td>-6.0</td><td>1.6</td><td>-3.7</td><td>2.6E-04</td><td>7.6E-04</td><td>BT20/MB468</td><td>MP</td></tr>
<tr><td>>hg38_21_+_8433172_8446622_RNA45SN1_45S_50ntFlanks@11670.11692.23|CCCAGTGCTCTGAATGTCAAAGT</td><td>>hg38_21_+_8433172_8446622_RNA45SN1_45S_50ntFlanks</td><td>11670</td><td>11692</td><td>23</td><td>CCCAGTGCTCTGAATGTCAAAGT</td><td>332.3</td><td>3.0</td><td>0.8</td><td>3.9</td><td>1.1E-04</td><td>7.6E-04</td><td>BT20/MB231</td><td>MITO</td></tr>
<tr><td>>hg38_21_+_8433172_8446622_RNA45SN1_45S_50ntFlanks@5472.5489.18|ACGGCCCTGGCGGAGCGC</td><td>>hg38_21_+_8433172_8446622_RNA45SN1_45S_50ntFlanks</td><td>5472</td><td>5489</td><td>18</td><td>ACGGCCCTGGCGGAGCGC</td><td>264.0</td><td>5.6</td><td>1.6</td><td>3.5</td><td>5.1E-04</td><td>7.6E-04</td><td>MB231/MB468</td><td>MP</td></tr>
<tr><td>>hg38_21_+_8433172_8446622_RNA45SN1_45S_50ntFlanks@4588.4607.20|CTATTTTGTTGGTTTTCGGA</td><td>>hg38_21_+_8433172_8446622_RNA45SN1_45S_50ntFlanks</td><td>4588</td><td>4607</td><td>20</td><td>CTATTTTGTTGGTTTTCGGA</td><td>196.5</td><td>-2.8</td><td>0.8</td><td>-3.4</td><td>5.8E-04</td><td>7.6E-04</td><td>MB231/MB468</td><td>CYTO</td></tr>
<tr><td>>hg38_21_+_8433172_8446622_RNA45SN1_45S_50ntFlanks@9167.9197.31|CGGGCGGGTCGCGCCGTCGGGCCCGGGGGAG</td><td>>hg38_21_+_8433172_8446622_RNA45SN1_45S_50ntFlanks</td><td>9167</td><td>9197</td><td>31</td><td>CGGGCGGGTCGCGCCGTCGGGCCCGGGGGAG</td><td>943.4</td><td>-6.3</td><td>1.7</td><td>-3.7</td><td>2.6E-04</td><td>7.6E-04</td><td>BT20/MB468</td><td>MP</td></tr>
<tr><td>>MI0000087|hsa-mir-29a&WithFlank&7|-|130876741|130876816@48.69.22(+1A)&[MIMAT0000086&hsa-miR-29a-3p&offsets|0|0(+1A);m-26&7|-|130876749|130876769&offsets|0|+1(+1A)]|TAGCACCATCTGAAATCGGTTAA</td><td>>MI0000087|hsa-mir-29a&WithFlank&7|-|130876741|130876816</td><td>48</td><td>69</td><td>22(+1A)</td><td>TAGCACCATCTGAAATCGGTTAA</td><td>554.6</td><td>6.0</td><td>1.7</td><td>3.7</td><td>2.6E-04</td><td>7.6E-04</td><td>BT20/MB468</td><td>MP</td></tr>
</table>

<table>
<tr><td>>hg38_21_+_8433172_8446622_RNA45SN1_45S_50ntFlanks@7973.8009.37|GACGCGACCTCAGATCAGACGTGGCGACCCGCTGAAT</td><td>>hg38_21_+_8433172_8446622_RNA45SN1_45S_50ntFlanks</td><td>7973</td><td>8009</td><td>37</td><td>GACGCGACCTCAGATCAGACGTGGCGACCCGCTGAAT</td><td>1759.3</td><td>4.1</td><td>1.1</td><td>3.7</td><td>2.6E-04</td><td>7.6E-04</td><td>BT20/MB468</td><td>MP</td></tr>
<tr><td>>hg38_21_+_8433172_8446622_RNA45SN1_45S_50ntFlanks@10841.10868.28|CGTAACTTCGGGATAAGGATTGGCTCTA</td><td>>hg38_21_+_8433172_8446622_RNA45SN1_45S_50ntFlanks</td><td>10841</td><td>10868</td><td>28</td><td>CGTAACTTCGGGATAAGGATTGGCTCTA</td><td>7894.6</td><td>-2.6</td><td>0.8</td><td>-3.5</td><td>5.5E-04</td><td>7.6E-04</td><td>BT20/MB468</td><td>CYTO</td></tr>
<tr><td>>hg38_21_+_8433172_8446622_RNA45SN1_45S_50ntFlanks@9623.9658.36|TAGTAGCTGGTTCCCTCCGAAGTTTCCCTCAGGATA</td><td>>hg38_21_+_8433172_8446622_RNA45SN1_45S_50ntFlanks</td><td>9623</td><td>9658</td><td>36</td><td>TAGTAGCTGGTTCCCTCCGAAGTTTCCCTCAGGATA</td><td>302.9</td><td>-2.4</td><td>0.7</td><td>-3.4</td><td>6.2E-04</td><td>7.6E-04</td><td>MB231/MB468</td><td>NUC</td></tr>
<tr><td>>hg38_21_+_8433172_8446622_RNA45SN1_45S_50ntFlanks@5073.5098.26|AATTCCGATAACGAACGAGACTCTGG</td><td>>hg38_21_+_8433172_8446622_RNA45SN1_45S_50ntFlanks</td><td>5073</td><td>5098</td><td>26</td><td>AATTCCGATAACGAACGAGACTCTGG</td><td>1989.3</td><td>-5.3</td><td>1.5</td><td>-3.5</td><td>4.6E-04</td><td>7.6E-04</td><td>BT20/MB468</td><td>MITO</td></tr>
<tr><td>>hg38_21_+_8433172_8446622_RNA45SN1_45S_50ntFlanks@10391.10424.34|AACAGCAGTTGAACATGGGTCAGTCGGTCCTGAG</td><td>>hg38_21_+_8433172_8446622_RNA45SN1_45S_50ntFlanks</td><td>10391</td><td>10424</td><td>34</td><td>AACAGCAGTTGAACATGGGTCAGTCGGTCCTGAG</td><td>7644.3</td><td>3.8</td><td>1.0</td><td>3.7</td><td>2.6E-04</td><td>7.6E-04</td><td>BT20/MB468</td><td>MP</td></tr>
<tr><td>>hg38_21_+_8433172_8446622_RNA45SN1_45S_50ntFlanks@9478.9497.20|GCGCACGTGTTAGGACCCGA</td><td>>hg38_21_+_8433172_8446622_RNA45SN1_45S_50ntFlanks</td><td>9478</td><td>9497</td><td>20</td><td>GCGCACGTGTTAGGACCCGA</td><td>1250.8</td><td>8.9</td><td>2.6</td><td>3.4</td><td>5.8E-04</td><td>7.7E-04</td><td>MB231/MB468</td><td>CYTO</td></tr>
<tr><td>>hg38_21_+_8433172_8446622_RNA45SN1_45S_50ntFlanks@12636.12659.24|AGCTACCATCTGTGGGATTATGAC</td><td>>hg38_21_+_8433172_8446622_RNA45SN1_45S_50ntFlanks</td><td>12636</td><td>12659</td><td>24</td><td>AGCTACCATCTGTGGGATTATGAC</td><td>240.9</td><td>-3.7</td><td>1.1</td><td>-3.4</td><td>5.8E-04</td><td>7.7E-04</td><td>MB231/MB468</td><td>CYTO</td></tr>
<tr><td>>hg38_21_+_8433172_8446622_RNA45SN1_45S_50ntFlanks@6689.6720.32|GAAGAACGCAGCTAGCTGCGAGAATTAATGTG</td><td>>hg38_21_+_8433172_8446622_RNA45SN1_45S_50ntFlanks</td><td>6689</td><td>6720</td><td>32</td><td>GAAGAACGCAGCTAGCTGCGAGAATTAATGTG</td><td>76.2</td><td>-5.5</td><td>1.5</td><td>-3.7</td><td>2.6E-04</td><td>7.7E-04</td><td>BT20/MB468</td><td>MP</td></tr>
<tr><td>>hg38_21_+_8433172_8446622_RNA45SN1_45S_50ntFlanks@11325.11344.20|TCTCGTCGCGCGCGCGTCCG</td><td>>hg38_21_+_8433172_8446622_RNA45SN1_45S_50ntFlanks</td><td>11325</td><td>11344</td><td>20</td><td>TCTCGTCGCGCGCGCGTCCG</td><td>1836.8</td><td>-4.3</td><td>1.2</td><td>-3.4</td><td>6.2E-04</td><td>7.7E-04</td><td>BT20/MB468</td><td>NUC</td></tr>
<tr><td>>hg38_21_+_8433172_8446622_RNA45SN1_45S_50ntFlanks@10394.10429.36|AGCAGTTGAACATGGGTCAGTCGGTCCTGAGAGATG</td><td>>hg38_21_+_8433172_8446622_RNA45SN1_45S_50ntFlanks</td><td>10394</td><td>10429</td><td>36</td><td>AGCAGTTGAACATGGGTCAGTCGGTCCTGAGAGATG</td><td>425.4</td><td>-1.9</td><td>0.6</td><td>-3.4</td><td>6.2E-04</td><td>7.7E-04</td><td>MB231/MB468</td><td>NUC</td></tr>
<tr><td>>hg38_21_+_8433172_8446622_RNA45SN1_45S_50ntFlanks@10791.10818.28|CCTCTGGCATGTTGGAACAATGTAGGTA</td><td>>hg38_21_+_8433172_8446622_RNA45SN1_45S_50ntFlanks</td><td>10791</td><td>10818</td><td>28</td><td>CCTCTGGCATGTTGGAACAATGTAGGTA</td><td>228.5</td><td>-2.3</td><td>0.7</td><td>-3.4</td><td>5.8E-04</td><td>7.7E-04</td><td>MB231/MB468</td><td>CYTO</td></tr>
<tr><td>>hg38_21_+_8433172_8446622_RNA45SN1_45S_50ntFlanks@8192.8210.19|AGCCCGTGGACGGTGTGAG</td><td>>hg38_21_+_8433172_8446622_RNA45SN1_45S_50ntFlanks</td><td>8192</td><td>8210</td><td>19</td><td>AGCCCGTGGACGGTGTGAG</td><td>127.6</td><td>-5.5</td><td>1.6</td><td>-3.5</td><td>5.2E-04</td><td>7.7E-04</td><td>MB231/MB468</td><td>MP</td></tr>
<tr><td>>MI0003560|hsa-mir-92b&WithFlank&1|+|155195171|155195278@67.88.22(+2U)&[MIMAT0003218&hsa-miR-92b-3p&offsets|0|0(+2U);m-115&1|+|155195237|155195258&offsets|0|0(+2U)]|TATTGCACTCGTCCCGGCCTCCTT</td><td>>MI0003560|hsa-mir-92b&WithFlank&1|+|155195171|155195278</td><td>67</td><td>88</td><td>22(+2U)</td><td>TATTGCACTCGTCCCGGCCTCCTT</td><td>142.6</td><td>7.4</td><td>2.0</td><td>3.7</td><td>2.6E-04</td><td>7.7E-04</td><td>BT20/MB468</td><td>MP</td></tr>
<tr><td>>hg38_21_+_8433172_8446622_RNA45SN1_45S_50ntFlanks@9533.9571.39|GAGGAAACTCTGGTGGAGGTCCGTAGCGGTCCTGACGTG</td><td>>hg38_21_+_8433172_8446622_RNA45SN1_45S_50ntFlanks</td><td>9533</td><td>9571</td><td>39</td><td>GAGGAAACTCTGGTGGAGGTCCGTAGCGGTCCTGACGTG</td><td>2496.8</td><td>-3.8</td><td>1.0</td><td>-3.9</td><td>9.4E-05</td><td>7.7E-04</td><td>BT20/MB231</td><td>CYTO</td></tr>
<tr><td>>hg38_21_+_8433172_8446622_RNA45SN1_45S_50ntFlanks@12957.12980.24|CTTCTGGGTCGGGGTTTCGTACGT</td><td>>hg38_21_+_8433172_8446622_RNA45SN1_45S_50ntFlanks</td><td>12957</td><td>12980</td><td>24</td><td>CTTCTGGGTCGGGGTTTCGTACGT</td><td>144.9</td><td>-4.6</td><td>1.3</td><td>-3.5</td><td>5.6E-04</td><td>7.7E-04</td><td>BT20/MB468</td><td>CYTO</td></tr>
<tr><td>>hg38_21_+_8433172_8446622_RNA45SN1_45S_50ntFlanks@12474.12497.24|ACCCACTAATAGGGAACGTGAGCT</td><td>>hg38_21_+_8433172_8446622_RNA45SN1_45S_50ntFlanks</td><td>12474</td><td>12497</td><td>24</td><td>ACCCACTAATAGGGAACGTGAGCT</td><td>548.9</td><td>2.5</td><td>0.7</td><td>3.4</td><td>5.8E-04</td><td>7.7E-04</td><td>MB231/MB468</td><td>CYTO</td></tr>
<tr><td>>trna10_AspGTC_12_-_125424193_125424264@38.71.34,trna12_AspGTC_12_-_125411891_125411962@38.71.34,trna144_AspGTC_6_-_27551236_27551307@38.71.34,trna38_AspGTC_17_-_8125556_8125627@38.71.34,trna45_AspGTC_6_+_27447453_27447524@38.71.34,trna48_AspGTC_6_+_27471523_27471594@38.71.34,trna4_AspGTC_12_+_96429799_96429870@38.71.34,trna69_AspGTC_1_-_161440205_161440276@38.71.34,trna72_AspGTC_1_-_161432824_161432895@38.71.34,trna75_AspGTC_1_-_161425414_161425485@38.71.34,trna78_AspGTC_1_-_161418033_161418104@38.71.34,trna81_AspGTC_1_-_161410615_161410686@38.71.34|CGCGGGAGACCGGGGTTCGATTCCCCGACGGGGA</td><td>>trna10_AspGTC_12_-_125424193_125424264</td><td>38</td><td>71</td><td>34</td><td>CGCGGGAGACCGGGGTTCGATTCCCCGACGGGGA</td><td>461.1</td><td>2.8</td><td>0.8</td><td>3.4</td><td>5.8E-04</td><td>7.7E-04</td><td>MB231/MB468</td><td>CYTO</td></tr>
<tr><td>>hg38_21_+_8433172_8446622_RNA45SN1_45S_50ntFlanks@10829.10852.24|GCAAGCCGGATCCGTAACTTCGGG</td><td>>hg38_21_+_8433172_8446622_RNA45SN1_45S_50ntFlanks</td><td>10829</td><td>10852</td><td>24</td><td>GCAAGCCGGATCCGTAACTTCGGG</td><td>331.1</td><td>-4.3</td><td>1.2</td><td>-3.5</td><td>5.6E-04</td><td>7.7E-04</td><td>BT20/MB468</td><td>CYTO</td></tr>
<tr><td>>hg38_21_+_8433172_8446622_RNA45SN1_45S_50ntFlanks@10440.10462.23|GTTCCGAAGGGACGGGCGATGGC</td><td>>hg38_21_+_8433172_8446622_RNA45SN1_45S_50ntFlanks</td><td>10440</td><td>10462</td><td>23</td><td>GTTCCGAAGGGACGGGCGATGGC</td><td>367.4</td><td>2.7</td><td>0.8</td><td>3.4</td><td>5.8E-04</td><td>7.7E-04</td><td>MB231/MB468</td><td>CYTO</td></tr>
<tr><td>>hg38_21_+_8433172_8446622_RNA45SN1_45S_50ntFlanks@8916.8936.21|GAATCCCGGGGCCGAGGGAGC</td><td>>hg38_21_+_8433172_8446622_RNA45SN1_45S_50ntFlanks</td><td>8916</td><td>8936</td><td>21</td><td>GAATCCCGGGGCCGAGGGAGC</td><td>84.9</td><td>6.8</td><td>1.9</td><td>3.7</td><td>2.6E-04</td><td>7.7E-04</td><td>BT20/MB468</td><td>MP</td></tr>
<tr><td>>hg38_21_+_8433172_8446622_RNA45SN1_45S_50ntFlanks@4509.4528.20|TTGAAAAAATTAGAGTGTTC</td><td>>hg38_21_+_8433172_8446622_RNA45SN1_45S_50ntFlanks</td><td>4509</td><td>4528</td><td>20</td><td>TTGAAAAAATTAGAGTGTTC</td><td>149.6</td><td>-4.1</td><td>1.2</td><td>-3.5</td><td>5.2E-04</td><td>7.7E-04</td><td>MB231/MB468</td><td>MP</td></tr>
<tr><td>>hg38_MT_+_1621_3279_RNR2_16S_50ntFlanks@617.642.26|AAACATATAACTGAACTCCTCACACC</td><td>>hg38_MT_+_1621_3279_RNR2_16S_50ntFlanks</td><td>617</td><td>642</td><td>26</td><td>AAACATATAACTGAACTCCTCACACC</td><td>212.6</td><td>4.4</td><td>1.3</td><td>3.4</td><td>6.2E-04</td><td>7.7E-04</td><td>MB231/MB468</td><td>MITO</td></tr>
<tr><td>>MI0000085|hsa-mir-27a&WithFlank&19|-|13836434|13836523@57.76.20(+1A)&[MIMAT0000084&hsa-miR-27a-3p&offsets|0|-1(+1A);m-43&19|-|13836447|13836467&offsets|0|-1(+1A)]|TTCACAGTGGCTAAGTTCCGA</td><td>>MI0000085|hsa-mir-27a&WithFlank&19|-|13836434|13836523</td><td>57</td><td>76</td><td>20(+1A)</td><td>TTCACAGTGGCTAAGTTCCGA</td><td>5040.1</td><td>2.8</td><td>0.8</td><td>3.5</td><td>5.2E-04</td><td>7.7E-04</td><td>MB231/MB468</td><td>MP</td></tr>
<tr><td>>hg38_21_+_8433172_8446622_RNA45SN1_45S_50ntFlanks@5085.5112.28|GAACGAGACTCTGGCATGCTAACTAGTT</td><td>>hg38_21_+_8433172_8446622_RNA45SN1_45S_50ntFlanks</td><td>5085</td><td>5112</td><td>28</td><td>GAACGAGACTCTGGCATGCTAACTAGTT</td><td>752.1</td><td>5.7</td><td>1.5</td><td>3.9</td><td>9.1E-05</td><td>7.7E-04</td><td>BT20/MB231</td><td>MP</td></tr>
<tr><td>>hg38_21_+_8433172_8446622_RNA45SN1_45S_50ntFlanks@3705.3746.42|TACCTGGTTGATCCTGCCAGTAGCATATGCTTGTCTCAAAGA</td><td>>hg38_21_+_8433172_8446622_RNA45SN1_45S_50ntFlanks</td><td>3705</td><td>3746</td><td>42</td><td>TACCTGGTTGATCCTGCCAGTAGCATATGCTTGTCTCAAAGA</td><td>1437.1</td><td>2.7</td><td>0.8</td><td>3.5</td><td>4.7E-04</td><td>7.7E-04</td><td>BT20/MB468</td><td>MITO</td></tr>
<tr><td>>MI0000281|hsa-mir-199a-2&WithFlank&1|-|172144529|172144650@37.59.23;MI0000242|hsa-mir-199a-1&WithFlank&19|-|10817420|10817502@12.34.23&[MIMAT0000231&hsa-miR-199a-5p&offsets|0|0;m-86&1|-|172144592|172144614&offsets|0|0];[MIMAT0000231_1&hsa-miR-199a-5p&offsets|0|0;m-85&19|-|10817469|10817491&offsets|0|0]|CCCAGTGTTCAGACTACCTGTTC</td><td>>MI0000281|hsa-mir-199a-2&WithFlank&1|-|172144529|172144650</td><td>37</td><td>59</td><td>23</td><td>CCCAGTGTTCAGACTACCTGTTC</td><td>179.1</td><td>7.3</td><td>2.0</td><td>3.7</td><td>2.6E-04</td><td>7.7E-04</td><td>BT20/MB468</td><td>MP</td></tr>
<tr><td>>hg38_21_+_8433172_8446622_RNA45SN1_45S_50ntFlanks@11569.11606.38|CAGCCGACTTAGAACTGGTGCGGACCAGGGGAATCCGA</td><td>>hg38_21_+_8433172_8446622_RNA45SN1_45S_50ntFlanks</td><td>11569</td><td>11606</td><td>38</td><td>CAGCCGACTTAGAACTGGTGCGGACCAGGGGAATCCGA</td><td>257.3</td><td>-5.3</td><td>1.5</td><td>-3.4</td><td>6.3E-04</td><td>7.7E-04</td><td>MB231/MB468</td><td>NUC</td></tr>
<tr><td>>MI0000063|hsa-let-7b&WithFlank&22|+|46113680|46113774@12.31.20(+1G)&[MIMAT0000063&hsa-let-7b-5p&offsets|0|-2(+1G);m-14&22|+|46113691|46113712&offsets|0|-2(+1G)]|TGAGGTAGTAGGTTGTGTGGG</td><td>>MI0000063|hsa-let-7b&WithFlank&22|+|46113680|46113774</td><td>12</td><td>31</td><td>20(+1G)</td><td>TGAGGTAGTAGGTTGTGTGGG</td><td>238.5</td><td>3.3</td><td>0.9</td><td>3.9</td><td>9.6E-05</td><td>7.7E-04</td><td>BT20/MB231</td><td>CYTO</td></tr>
<tr><td>>trna58_LeuCAA_1_+_249168054_249168159@48.65.18|CCGGATGGAGGCGTGGGT</td><td>>trna58_LeuCAA_1_+_249168054_249168159</td><td>48</td><td>65</td><td>18</td><td>CCGGATGGAGGCGTGGGT</td><td>411.1</td><td>3.5</td><td>0.9</td><td>3.9</td><td>9.5E-05</td><td>7.7E-04</td><td>BT20/MB231</td><td>CYTO</td></tr>
<tr><td>>hg38_21_+_8433172_8446622_RNA45SN1_45S_50ntFlanks@10786.10823.38|AACAGCCTCTGGCATGTTGGAACAATGTAGGTAAGGGA</td><td>>hg38_21_+_8433172_8446622_RNA45SN1_45S_50ntFlanks</td><td>10786</td><td>10823</td><td>38</td><td>AACAGCCTCTGGCATGTTGGAACAATGTAGGTAAGGGA</td><td>676.8</td><td>-3.6</td><td>1.0</td><td>-3.7</td><td>2.6E-04</td><td>7.7E-04</td><td>BT20/MB468</td><td>MP</td></tr>
<tr><td>>MI0000088|hsa-mir-30a&WithFlank&6|-|71403545|71403627@12.34.23(+1A)&[MIMAT0000087&hsa-miR-30a-5p&offsets|0|+1(+1A);m-15&6|-|71403595|71403616&offsets|0|+1(+1A)]|TGTAAACATCCTCGACTGGAAGCA</td><td>>MI0000088|hsa-mir-30a&WithFlank&6|-|71403545|71403627</td><td>12</td><td>34</td><td>23(+1A)</td><td>TGTAAACATCCTCGACTGGAAGCA</td><td>15539.3</td><td>-3.4</td><td>0.9</td><td>-3.9</td><td>1.1E-04</td><td>7.7E-04</td><td>BT20/MB231</td><td>MITO</td></tr>
<tr><td>>hg38_21_+_8433172_8446622_RNA45SN1_45S_50ntFlanks@12007.12036.30|GCCGCCGGTGAAATACCACTACTCTGATCG</td><td>>hg38_21_+_8433172_8446622_RNA45SN1_45S_50ntFlanks</td><td>12007</td><td>12036</td><td>30</td><td>GCCGCCGGTGAAATACCACTACTCTGATCG</td><td>1234.5</td><td>3.2</td><td>0.9</td><td>3.5</td><td>4.7E-04</td><td>7.7E-04</td><td>BT20/MB468</td><td>MITO</td></tr>
<tr><td>>hg38_21_+_8433172_8446622_RNA45SN1_45S_50ntFlanks@5082.5110.29|AACGAACGAGACTCTGGCATGCTAACTAG</td><td>>hg38_21_+_8433172_8446622_RNA45SN1_45S_50ntFlanks</td><td>5082</td><td>5110</td><td>29</td><td>AACGAACGAGACTCTGGCATGCTAACTAG</td><td>504.4</td><td>-5.5</td><td>1.6</td><td>-3.5</td><td>5.2E-04</td><td>7.8E-04</td><td>MB231/MB468</td><td>MP</td></tr>
<tr><td>>hg38_21_+_8433172_8446622_RNA45SN1_45S_50ntFlanks@10393.10429.37|CAGCAGTTGAACATGGGTCAGTCGGTCCTGAGAGATG</td><td>>hg38_21_+_8433172_8446622_RNA45SN1_45S_50ntFlanks</td><td>10393</td><td>10429</td><td>37</td><td>CAGCAGTTGAACATGGGTCAGTCGGTCCTGAGAGATG</td><td>1352.0</td><td>-2.8</td><td>0.8</td><td>-3.4</td><td>5.6E-04</td><td>7.8E-04</td><td>BT20/MB468</td><td>CYTO</td></tr>
<tr><td>>MI0000085|hsa-mir-27a&WithFlank&19|-|13836434|13836523@57.77.21(+1G)&[MIMAT0000084&hsa-miR-27a-3p&offsets|0|0(+1G);m-43&19|-|13836447|13836467&offsets|0|0(+1G)]|TTCACAGTGGCTAAGTTCCGCG</td><td>>MI0000085|hsa-mir-27a&WithFlank&19|-|13836434|13836523</td><td>57</td><td>77</td><td>21(+1G)</td><td>TTCACAGTGGCTAAGTTCCGCG</td><td>46.3</td><td>-8.9</td><td>2.3</td><td>-3.9</td><td>9.2E-05</td><td>7.8E-04</td><td>BT20/MB231</td><td>MP</td></tr>
<tr><td>>hg38_21_+_8433172_8446622_RNA45SN1_45S_50ntFlanks@11890.11907.18|CTTGACTCTAGTCTGGCA</td><td>>hg38_21_+_8433172_8446622_RNA45SN1_45S_50ntFlanks</td><td>11890</td><td>11907</td><td>18</td><td>CTTGACTCTAGTCTGGCA</td><td>478.0</td><td>-2.8</td><td>0.8</td><td>-3.5</td><td>4.7E-04</td><td>7.8E-04</td><td>BT20/MB468</td><td>MITO</td></tr>
<tr><td>>hg38_21_+_8433172_8446622_RNA45SN1_45S_50ntFlanks@6736.6770.35|TGATCATCGACACTTCGAACGCACTTGCGGCCCCG</td><td>>hg38_21_+_8433172_8446622_RNA45SN1_45S_50ntFlanks</td><td>6736</td><td>6770</td><td>35</td><td>TGATCATCGACACTTCGAACGCACTTGCGGCCCCG</td><td>145.4</td><td>-4.7</td><td>1.3</td><td>-3.5</td><td>4.7E-04</td><td>7.8E-04</td><td>BT20/MB468</td><td>MITO</td></tr>
<tr><td>>hg38_21_+_8433172_8446622_RNA45SN1_45S_50ntFlanks@8168.8187.20|CCCAAGTCCTTCTGATCGAG</td><td>>hg38_21_+_8433172_8446622_RNA45SN1_45S_50ntFlanks</td><td>8168</td><td>8187</td><td>20</td><td>CCCAAGTCCTTCTGATCGAG</td><td>177.4</td><td>-5.5</td><td>1.5</td><td>-3.6</td><td>2.6E-04</td><td>7.8E-04</td><td>BT20/MB468</td><td>MP</td></tr>
<tr><td>>hg38_21_+_8433172_8446622_RNA45SN1_45S_50ntFlanks@11209.11232.24|CGGCGACTCTGGACGCGAGCCGGG</td><td>>hg38_21_+_8433172_8446622_RNA45SN1_45S_50ntFlanks</td><td>11209</td><td>11232</td><td>24</td><td>CGGCGACTCTGGACGCGAGCCGGG</td><td>7199.3</td><td>-2.8</td><td>0.8</td><td>-3.5</td><td>4.7E-04</td><td>7.8E-04</td><td>BT20/MB468</td><td>MITO</td></tr>
<tr><td>>hg38_21_+_8433172_8446622_RNA45SN1_45S_50ntFlanks@6653.6701.49|ACTCTTAGCGGTGGATCACTCGGCTCGTGCGTCGATGAAGAACGCAGCT</td><td>>hg38_21_+_8433172_8446622_RNA45SN1_45S_50ntFlanks</td><td>6653</td><td>6701</td><td>49</td><td>ACTCTTAGCGGTGGATCACTCGGCTCGTGCGTCGATGAAGAACGCAGCT</td><td>176.9</td><td>5.5</td><td>1.6</td><td>3.4</td><td>6.2E-04</td><td>7.8E-04</td><td>MB231/MB468</td><td>MITO</td></tr>
<tr><td>>hg38_MT_+_1621_3279_RNR2_16S_50ntFlanks@1217.1248.32|AACCTCCGAGCAGTACATGCTAAGACTTCACC</td><td>>hg38_MT_+_1621_3279_RNR2_16S_50ntFlanks</td><td>1217</td><td>1248</td><td>32</td><td>AACCTCCGAGCAGTACATGCTAAGACTTCACC</td><td>187.7</td><td>5.1</td><td>1.5</td><td>3.4</td><td>6.2E-04</td><td>7.8E-04</td><td>MB231/MB468</td><td>MITO</td></tr>
<tr><td>>hg38_21_+_8433172_8446622_RNA45SN1_45S_50ntFlanks@6698.6719.22|AGCTAGCTGCGAGAATTAATGT</td><td>>hg38_21_+_8433172_8446622_RNA45SN1_45S_50ntFlanks</td><td>6698</td><td>6719</td><td>22</td><td>AGCTAGCTGCGAGAATTAATGT</td><td>917.9</td><td>-2.3</td><td>0.7</td><td>-3.4</td><td>5.9E-04</td><td>7.8E-04</td><td>MB231/MB468</td><td>CYTO</td></tr>
<tr><td>>hg38_21_+_8433172_8446622_RNA45SN1_45S_50ntFlanks@11767.11803.37|TCTAATTAGTGACGCGCATGAATGGATGAACGAGATT</td><td>>hg38_21_+_8433172_8446622_RNA45SN1_45S_50ntFlanks</td><td>11767</td><td>11803</td><td>37</td><td>TCTAATTAGTGACGCGCATGAATGGATGAACGAGATT</td><td>469.1</td><td>-3.5</td><td>1.0</td><td>-3.4</td><td>5.9E-04</td><td>7.8E-04</td><td>MB231/MB468</td><td>CYTO</td></tr>
<tr><td>>hg38_21_+_8433172_8446622_RNA45SN1_45S_50ntFlanks@476.505.30|TTCGTGATCGATGTGGTGACGTCGTGCTCT</td><td>>hg38_21_+_8433172_8446622_RNA45SN1_45S_50ntFlanks</td><td>476</td><td>505</td><td>30</td><td>TTCGTGATCGATGTGGTGACGTCGTGCTCT</td><td>329.4</td><td>5.2</td><td>1.4</td><td>3.6</td><td>2.6E-04</td><td>7.8E-04</td><td>BT20/MB468</td><td>MP</td></tr>
<tr><td>>hg38_21_+_8433172_8446622_RNA45SN1_45S_50ntFlanks@6647.6676.30|CGTACGACTCTTAGCGGTGGATCACTCGGC</td><td>>hg38_21_+_8433172_8446622_RNA45SN1_45S_50ntFlanks</td><td>6647</td><td>6676</td><td>30</td><td>CGTACGACTCTTAGCGGTGGATCACTCGGC</td><td>1408.8</td><td>3.1</td><td>0.9</td><td>3.4</td><td>6.2E-04</td><td>7.8E-04</td><td>MB231/MB468</td><td>MITO</td></tr>
<tr><td>>trnaMT_GlnTTG_MT_-_4329_4400@1.33.33,trnalookalike2_GlnTTG_1_-_564879_564950@1.33.33|TAGGATGGGGTGTGATAGGTGGCACGGAGAATT</td><td>>trnaMT_GlnTTG_MT_-_4329_4400</td><td>1</td><td>33</td><td>33</td><td>TAGGATGGGGTGTGATAGGTGGCACGGAGAATT</td><td>949.5</td><td>4.8</td><td>1.3</td><td>3.6</td><td>2.6E-04</td><td>7.8E-04</td><td>BT20/MB468</td><td>MP</td></tr>
</table>

| | | | | | | | | | | | | | |
|---|---|---|---|---|---|---|---|---|---|---|---|---|---|
| >hg38_21_+_8433172_8446622_RNA45SN1_45S_50ntFlanks@11561.11599.39|GCGCCTAGCAGCCGACTTAGAACTGGTGCGGACCAGGGG | >hg38_21_+_8433172_8446622_RNA45SN1_45S_50ntFlanks | 11561 | 11599 | 39 | GCGCCTAGCAGCCGACTTAGAACTGGTGCGGACCAGGGG | 166.6 | -4.9 | 1.4 | -3.4 | 6.3E-04 | 7.8E-04 | MB231/MB468 | NUC |
| >MI0000102|hsa-mir-100&WithFlank&11|-|122152223|122152314@19.37.19&[MIMAT0000098&hsa-miR-100-5p&offsets|0|-3;m-27&11|-|122152276|122152296&offsets|0|-2]|AACCCGTAGATCCGAACTT | >MI0000102|hsa-mir-100&WithFlank&11|-|122152223|122152314 | 19 | 37 | 19 | AACCCGTAGATCCGAACTT | 249.7 | -5.7 | 1.5 | -3.9 | 9.6E-05 | 7.8E-04 | BT20/MB231 | NUC |
| >MI0003591|hsa-mir-584&WithFlank&5|-|149062307|149062415@22.39.18(+1A)&[MIMAT0003249&hsa-miR-584-5p&offsets|0|-4(+1A);m-167&5|-|149062374|149062394&offsets|0|-3(+1A)]|TTATGGTTTGCCTGGGACA | >MI0003591|hsa-mir-584&WithFlank&5|-|149062307|149062415 | 22 | 39 | 18(+1A) | TTATGGTTTGCCTGGGACA | 87.3 | 6.2 | 1.8 | 3.5 | 5.3E-04 | 7.8E-04 | MB231/MB468 | MP |
| >MI0000077|hsa-mir-21&WithFlank&17|+|59841260|59841343@14.33.20(+1U)&[MIMAT0000076&hsa-miR-21-5p&offsets|0|-2(+1U);m-1&17|+|59841273|59841294&offsets|0|-2(+1U)]|TAGCTTATCAGACTGATGTTT | >MI0000077|hsa-mir-21&WithFlank&17|+|59841260|59841343 | 14 | 33 | 20(+1U) | TAGCTTATCAGACTGATGTTT | 556.5 | 3.6 | 1.0 | 3.5 | 5.3E-04 | 7.8E-04 | MB231/MB468 | MP |
| >hg38_21_+_8433172_8446622_RNA45SN1_45S_50ntFlanks@10468.10495.28|TTGCCCTCGGCCGATCGAAAGGGAGTCG | >hg38_21_+_8433172_8446622_RNA45SN1_45S_50ntFlanks | 10468 | 10495 | 28 | TTGCCCTCGGCCGATCGAAAGGGAGTCG | 220.8 | -2.9 | 0.8 | -3.4 | 5.7E-04 | 7.8E-04 | BT20/MB468 | CYTO |
| >hg38_MT_+_598_1651_RNR1_12S_50ntFlanks@416.455.40|AAATAGACTACGAAAGTGGCTTTAACATATCTGAACACAC | >hg38_MT_+_598_1651_RNR1_12S_50ntFlanks | 416 | 455 | 40 | AAATAGACTACGAAAGTGGCTTTAACATATCTGAACACAC | 136.4 | 5.9 | 1.7 | 3.4 | 6.3E-04 | 7.8E-04 | MB231/MB468 | MITO |
| >hg38_21_+_8433172_8446622_RNA45SN1_45S_50ntFlanks@9835.9856.22|TTTGGTAAGCAGAACTGGCGCT | >hg38_21_+_8433172_8446622_RNA45SN1_45S_50ntFlanks | 9835 | 9856 | 22 | TTTGGTAAGCAGAACTGGCGCT | 60.1 | 9.7 | 2.6 | 3.6 | 2.6E-04 | 7.8E-04 | BT20/MB468 | MP |
| >MI0000542|hsa-mir-320a&WithFlank&8|-|22244960|22245043@49.70.22&[MIMAT0000510&hsa-miR-320a-3p&offsets|+1|+1;m-60&8|-|22244975|22244996&offsets|+1|+1]|AAAGCTGGGTTGAGAGGGCGAA | >MI0000542|hsa-mir-320a&WithFlank&8|-|22244960|22245043 | 49 | 70 | 22 | AAAGCTGGGTTGAGAGGGCGAA | 263.3 | 2.7 | 0.7 | 3.9 | 1.1E-04 | 7.8E-04 | BT20/MB231 | MITO |
| >MI0000088|hsa-mir-30a&WithFlank&6|-|71403545|71403627@53.73.21(+2U)&[MIMAT0000088&hsa-miR-30a-3p&offsets|0|-1(+2U);m-38&6|-|71403554|71403575&offsets|0|-1(+2U)]|CTTTCAGTCGGATGTTTGCAGTT | >MI0000088|hsa-mir-30a&WithFlank&6|-|71403545|71403627 | 53 | 73 | 21(+2U) | CTTTCAGTCGGATGTTTGCAGTT | 468.1 | -2.8 | 0.7 | -3.9 | 1.1E-04 | 7.8E-04 | BT20/MB231 | MITO |
| >hg38_21_+_8433172_8446622_RNA45SN1_45S_50ntFlanks@8192.8212.21|AGCCCGTGGACGGTGTGAGGC | >hg38_21_+_8433172_8446622_RNA45SN1_45S_50ntFlanks | 8192 | 8212 | 21 | AGCCCGTGGACGGTGTGAGGC | 288.6 | -2.7 | 0.8 | -3.4 | 6.4E-04 | 7.8E-04 | MB231/MB468 | NUC |
| >hg38_21_+_8433172_8446622_RNA45SN1_45S_50ntFlanks@12227.12249.23|AACCTCCCGTGGAGCAGAAGGGC | >hg38_21_+_8433172_8446622_RNA45SN1_45S_50ntFlanks | 12227 | 12249 | 23 | AACCTCCCGTGGAGCAGAAGGGC | 65.0 | 6.8 | 1.9 | 3.6 | 2.7E-04 | 7.8E-04 | BT20/MB468 | MP |
| >hg38_21_+_8433172_8446622_RNA45SN1_45S_50ntFlanks@11217.11248.32|CTGGACGCGAGCCGGGCCCTTCCCGTGGATCG | >hg38_21_+_8433172_8446622_RNA45SN1_45S_50ntFlanks | 11217 | 11248 | 32 | CTGGACGCGAGCCGGGCCCTTCCCGTGGATCG | 517.3 | -3.2 | 0.9 | -3.4 | 6.4E-04 | 7.8E-04 | MB231/MB468 | NUC |
| >trnaMT_GluTTC_MT_-_14674_14742@45.72.28,trnalookalike8_GluTTC_5_-_93905172_93905240@45.72.28|CGTGGTTGTAGTCCGTGCGAGAATACCA | >trnaMT_GluTTC_MT_-_14674_14742 | 45 | 72 | 28 | CGTGGTTGTAGTCCGTGCGAGAATACCA | 561.1 | 2.6 | 0.8 | 3.4 | 6.3E-04 | 7.8E-04 | MB231/MB468 | MITO |
| >trnaMT_PheGAA_MT_+_577_647@36.74.39|AAAATGTTTAGACGGGCTCACATCACCCCATAAACACCA | >trnaMT_PheGAA_MT_+_577_647 | 36 | 74 | 39 | AAAATGTTTAGACGGGCTCACATCACCCCATAAACACCA | 66.0 | 9.8 | 2.7 | 3.6 | 2.7E-04 | 7.8E-04 | BT20/MB468 | MP |
| >hg38_21_+_8433172_8446622_RNA45SN1_45S_50ntFlanks@9484.9510.27|GTGTTAGGACCCGAAAGATGGTGAACT | >hg38_21_+_8433172_8446622_RNA45SN1_45S_50ntFlanks | 9484 | 9510 | 27 | GTGTTAGGACCCGAAAGATGGTGAACT | 294.3 | 7.3 | 1.9 | 3.9 | 9.3E-05 | 7.8E-04 | BT20/MB231 | MP |
| >hg38_21_+_8433172_8446622_RNA45SN1_45S_50ntFlanks@10680.10708.29|GTTCCGGCGGCGTCCGGTGAGCTCTCGCT | >hg38_21_+_8433172_8446622_RNA45SN1_45S_50ntFlanks | 10680 | 10708 | 29 | GTTCCGGCGGCGTCCGGTGAGCTCTCGCT | 6685.0 | -4.1 | 1.1 | -3.6 | 2.7E-04 | 7.8E-04 | BT20/MB468 | MP |
| >hg38_21_+_8433172_8446622_RNA45SN1_45S_50ntFlanks@10137.10158.22|CGGCGGCGGCGGGGGTGTGGGG | >hg38_21_+_8433172_8446622_RNA45SN1_45S_50ntFlanks | 10137 | 10158 | 22 | CGGCGGCGGCGGGGGTGTGGGG | 95.5 | -5.7 | 1.6 | -3.6 | 2.7E-04 | 7.8E-04 | BT20/MB468 | MP |
| >hg38_21_+_8433172_8446622_RNA45SN1_45S_50ntFlanks@11575.11611.37|ACTTAGAACTGGTGCGGACCAGGGGAATCCGACTGTT | >hg38_21_+_8433172_8446622_RNA45SN1_45S_50ntFlanks | 11575 | 11611 | 37 | ACTTAGAACTGGTGCGGACCAGGGGAATCCGACTGTT | 336.6 | -3.5 | 1.0 | -3.4 | 5.9E-04 | 7.8E-04 | MB231/MB468 | CYTO |
| >hg38_21_+_8433172_8446622_RNA45SN1_45S_50ntFlanks@10128.10145.18|CGGCGGCGGCGGCGGCGG | >hg38_21_+_8433172_8446622_RNA45SN1_45S_50ntFlanks | 10128 | 10145 | 18 | CGGCGGCGGCGGCGGCGG | 311.5 | -5.8 | 1.6 | -3.6 | 2.7E-04 | 7.8E-04 | BT20/MB468 | MP |
| >hg38_21_+_8433172_8446622_RNA45SN1_45S_50ntFlanks@10131.10148.18|CGGCGGCGGCGGCGGCGG | >hg38_21_+_8433172_8446622_RNA45SN1_45S_50ntFlanks | 10131 | 10148 | 18 | CGGCGGCGGCGGCGGCGG | 311.5 | -5.8 | 1.6 | -3.6 | 2.7E-04 | 7.8E-04 | BT20/MB468 | MP |
| >hg38_21_+_8433172_8446622_RNA45SN1_45S_50ntFlanks@10392.10419.28|ACAGCAGTTGAACATGGGTCAGTCGGTC | >hg38_21_+_8433172_8446622_RNA45SN1_45S_50ntFlanks | 10392 | 10419 | 28 | ACAGCAGTTGAACATGGGTCAGTCGGTC | 376.7 | -4.6 | 1.3 | -3.6 | 2.7E-04 | 7.8E-04 | BT20/MB468 | MP |
| >hg38_21_+_8433172_8446622_RNA45SN1_45S_50ntFlanks@8824.8841.18|CGGCGGCGGCGGCGGCGG | >hg38_21_+_8433172_8446622_RNA45SN1_45S_50ntFlanks | 8824 | 8841 | 18 | CGGCGGCGGCGGCGGCGG | 311.5 | -5.8 | 1.6 | -3.6 | 2.7E-04 | 7.8E-04 | BT20/MB468 | MP |
| >hg38_21_+_8433172_8446622_RNA45SN1_45S_50ntFlanks@8845.8862.18|CGGCGGCGGCGGCGGCGG | >hg38_21_+_8433172_8446622_RNA45SN1_45S_50ntFlanks | 8845 | 8862 | 18 | CGGCGGCGGCGGCGGCGG | 311.5 | -5.8 | 1.6 | -3.6 | 2.7E-04 | 7.8E-04 | BT20/MB468 | MP |
| >hg38_21_+_8433172_8446622_RNA45SN1_45S_50ntFlanks@8848.8865.18|CGGCGGCGGCGGCGGCGG | >hg38_21_+_8433172_8446622_RNA45SN1_45S_50ntFlanks | 8848 | 8865 | 18 | CGGCGGCGGCGGCGGCGG | 311.5 | -5.8 | 1.6 | -3.6 | 2.7E-04 | 7.8E-04 | BT20/MB468 | MP |
| >hg38_21_+_8433172_8446622_RNA45SN1_45S_50ntFlanks@8354.8393.40|TTGAAAAGAACTTTGAAGAGAGAGTTCAAGAGGGCGTGAA | >hg38_21_+_8433172_8446622_RNA45SN1_45S_50ntFlanks | 8354 | 8393 | 40 | TTGAAAAGAACTTTGAAGAGAGAGTTCAAGAGGGCGTGAA | 52.6 | 9.0 | 2.3 | 3.9 | 9.3E-05 | 7.8E-04 | BT20/MB231 | MP |
| >hg38_21_+_8433172_8446622_RNA45SN1_45S_50ntFlanks@9835.9856.22|TTTGGTAAGCAGAACTGGCGCT | >hg38_21_+_8433172_8446622_RNA45SN1_45S_50ntFlanks | 9835 | 9856 | 22 | TTTGGTAAGCAGAACTGGCGCT | 140.2 | 10.5 | 2.7 | 3.9 | 9.3E-05 | 7.8E-04 | BT20/MB231 | MP |
| >MI0000446|hsa-mir-125b-1&WithFlank&11|-|122099751|122099850@61.81.21&[MIMAT0004592&hsa-miR-125b-1-3p&offsets|0|-1;m-485&11|-|122099769|122099790&offsets|0|-1]|ACGGGTTAGGCTCTTGGGAGC | >MI0000446|hsa-mir-125b-1&WithFlank&11|-|122099751|122099850 | 61 | 81 | 21 | ACGGGTTAGGCTCTTGGGAGC | 550.7 | -6.8 | 1.7 | -3.9 | 9.3E-05 | 7.8E-04 | BT20/MB231 | MP |
| >hg38_21_+_8433172_8446622_RNA45SN1_45S_50ntFlanks@6645.6687.43|CTCGTACGACTCTTAGCGGTGGATCACTCGGCTCGTGCGTCGA | >hg38_21_+_8433172_8446622_RNA45SN1_45S_50ntFlanks | 6645 | 6687 | 43 | CTCGTACGACTCTTAGCGGTGGATCACTCGGCTCGTGCGTCGA | 199.8 | 3.3 | 1.0 | 3.4 | 6.3E-04 | 7.8E-04 | MB231/MB468 | MITO |
| >hg38_21_+_8433172_8446622_RNA45SN1_45S_50ntFlanks@10852.10873.22|GATAAGGATTGGCTCTAAGGGC | >hg38_21_+_8433172_8446622_RNA45SN1_45S_50ntFlanks | 10852 | 10873 | 22 | GATAAGGATTGGCTCTAAGGGC | 120.3 | 6.7 | 1.8 | 3.6 | 2.7E-04 | 7.8E-04 | BT20/MB468 | MP |
| >hg38_21_+_8433172_8446622_RNA45SN1_45S_50ntFlanks@12479.12499.21|CTAATAGGGAACGTGAGCTGG | >hg38_21_+_8433172_8446622_RNA45SN1_45S_50ntFlanks | 12479 | 12499 | 21 | CTAATAGGGAACGTGAGCTGG | 1117.1 | -6.5 | 1.9 | -3.4 | 6.4E-04 | 7.8E-04 | BT20/MB468 | NUC |
| >MI0000088|hsa-mir-30a&WithFlank&6|-|71403545|71403627@12.35.24(+1U)&[MIMAT0000087&hsa-miR-30a-5p&offsets|0|+2(+1U);m-15&6|-|71403595|71403616&offsets|0|+2(+1U)]|TGTAAACATCCTCGACTGGAAGCTT | >MI0000088|hsa-mir-30a&WithFlank&6|-|71403545|71403627 | 12 | 35 | 24(+1U) | TGTAAACATCCTCGACTGGAAGCTT | 35622.7 | -4.3 | 1.1 | -3.9 | 9.4E-05 | 7.8E-04 | BT20/MB231 | MP |
| >hg38_21_+_8433172_8446622_RNA45SN1_45S_50ntFlanks@7987.8012.26|TCAGACGTGGCGACCCGCTGAATTTA | >hg38_21_+_8433172_8446622_RNA45SN1_45S_50ntFlanks | 7987 | 8012 | 26 | TCAGACGTGGCGACCCGCTGAATTTA | 122.5 | -5.6 | 1.5 | -3.6 | 2.7E-04 | 7.8E-04 | BT20/MB468 | MP |
| >hg38_21_+_8433172_8446622_RNA45SN1_45S_50ntFlanks@12407.12432.26|TTTTGATCCTTCGATGTCGGCTCTTC | >hg38_21_+_8433172_8446622_RNA45SN1_45S_50ntFlanks | 12407 | 12432 | 26 | TTTTGATCCTTCGATGTCGGCTCTTC | 935.1 | -2.8 | 0.8 | -3.4 | 5.7E-04 | 7.8E-04 | BT20/MB468 | CYTO |
| >hg38_21_+_8433172_8446622_RNA45SN1_45S_50ntFlanks@9564.9591.28|CTGACGTGCAAATCGGTCGTCCGACCTG | >hg38_21_+_8433172_8446622_RNA45SN1_45S_50ntFlanks | 9564 | 9591 | 28 | CTGACGTGCAAATCGGTCGTCCGACCTG | 216.2 | -2.3 | 0.7 | -3.4 | 6.0E-04 | 7.8E-04 | MB231/MB468 | CYTO |
| >hg38_21_+_8433172_8446622_RNA45SN1_45S_50ntFlanks@10614.10639.26|GAAGGGCAGGGCGCCCTGGAATGGGT | >hg38_21_+_8433172_8446622_RNA45SN1_45S_50ntFlanks | 10614 | 10639 | 26 | GAAGGGCAGGGCGCCCTGGAATGGGT | 61.6 | 9.2 | 2.6 | 3.5 | 5.3E-04 | 7.8E-04 | MB231/MB468 | MP |
| >MI0000298|hsa-mir-221&WithFlank&X|-|45746151|45746272@71.92.22(+2U)&[MIMAT0000278&hsa-miR-221-3p&offsets|0|-1(+2U);m-91&X|-|45746181|45746202&offsets|0|0(+2U)]|AGCTACATTGTCTGCTGGGTTTTT | >MI0000298|hsa-mir-221&WithFlank&X|-|45746151|45746272 | 71 | 92 | 22(+2U) | AGCTACATTGTCTGCTGGGTTTTT | 315.3 | -2.8 | 0.7 | -3.9 | 9.7E-05 | 7.9E-04 | BT20/MB231 | NUC |
| >hg38_21_+_8433172_8446622_RNA45SN1_45S_50ntFlanks@5059.5099.41|CGATTTGTCTGGTTAATTCCGATAACGAACGAGACTCTGGC | >hg38_21_+_8433172_8446622_RNA45SN1_45S_50ntFlanks | 5059 | 5099 | 41 | CGATTTGTCTGGTTAATTCCGATAACGAACGAGACTCTGGC | 77.6 | -10.1 | 2.9 | -3.5 | 5.3E-04 | 7.9E-04 | MB231/MB468 | MP |
| >trna17_ValTAC_11_-_59318102_59318174@1.30.30,trna4_ValTAC_X_-_18693029_18693101@1.30.30|GGTTCCATAGTGTAGTGGTTATCACGTCTG | >trna17_ValTAC_11_-_59318102_59318174 | 1 | 30 | 30 | GGTTCCATAGTGTAGTGGTTATCACGTCTG | 708.4 | -5.5 | 1.6 | -3.4 | 5.7E-04 | 7.9E-04 | BT20/MB468 | CYTO |
| >hg38_21_+_8433172_8446622_RNA45SN1_45S_50ntFlanks@10351.10381.31|CGAGAACTTTGAAGGCCGAAGTGGAGAAGGG | >hg38_21_+_8433172_8446622_RNA45SN1_45S_50ntFlanks | 10351 | 10381 | 31 | CGAGAACTTTGAAGGCCGAAGTGGAGAAGGG | 1632.3 | -3.5 | 1.0 | -3.6 | 2.7E-04 | 7.9E-04 | BT20/MB468 | MP |
| >hg38_21_+_8433172_8446622_RNA45SN1_45S_50ntFlanks@4047.4069.23|ATTCGAACGTCTGCCCTATCAAC | >hg38_21_+_8433172_8446622_RNA45SN1_45S_50ntFlanks | 4047 | 4069 | 23 | ATTCGAACGTCTGCCCTATCAAC | 270.3 | 2.4 | 0.7 | 3.4 | 6.0E-04 | 7.9E-04 | MB231/MB468 | CYTO |
| >hg38_21_+_8433172_8446622_RNA45SN1_45S_50ntFlanks@12243.12280.38|GAAGGGCAAAAGCTCGCTTGATCTTGATTTTCAGTACG | >hg38_21_+_8433172_8446622_RNA45SN1_45S_50ntFlanks | 12243 | 12280 | 38 | GAAGGGCAAAAGCTCGCTTGATCTTGATTTTCAGTACG | 190.3 | -3.9 | 1.0 | -3.9 | 9.7E-05 | 7.9E-04 | BT20/MB231 | CYTO |
| >hg38_21_+_8433172_8446622_RNA45SN1_45S_50ntFlanks@7973.7992.20|GACGCGACCTCAGATCAGAC | >hg38_21_+_8433172_8446622_RNA45SN1_45S_50ntFlanks | 7973 | 7992 | 20 | GACGCGACCTCAGATCAGAC | 1307.9 | 2.2 | 0.6 | 3.4 | 6.4E-04 | 7.9E-04 | MB231/MB468 | NUC |
| >hg38_21_+_8433172_8446622_RNA45SN1_45S_50ntFlanks@10841.10859.19|CGTAACTTCGGGATAAGGA | >hg38_21_+_8433172_8446622_RNA45SN1_45S_50ntFlanks | 10841 | 10859 | 19 | CGTAACTTCGGGATAAGGA | 434.3 | -6.0 | 1.8 | -3.4 | 6.4E-04 | 7.9E-04 | BT20/MB468 | NUC |
| >am_MI0003581|hsa-mir-574&WithFlank&4|+|38868026|38868133@31.55.25&[MIMAT0004795&hsa-miR-574-5p&offsets|0|+2;m-213&4|+|38868056|38868078&offsets|0|+2]|TGAGTGTGTGTGTGTGAGTGTGTGT | >am_MI0003581|hsa-mir-574&WithFlank&4|+|38868026|38868133 | 31 | 55 | 25 | TGAGTGTGTGTGTGTGAGTGTGTGT | 1221.0 | -4.5 | 1.1 | -3.9 | 9.4E-05 | 7.9E-04 | BT20/MB231 | MP |
| >hg38_21_+_8433172_8446622_RNA45SN1_45S_50ntFlanks@10351.10375.25|CGAGAACTTTGAAGGCCGAAGTGGA | >hg38_21_+_8433172_8446622_RNA45SN1_45S_50ntFlanks | 10351 | 10375 | 25 | CGAGAACTTTGAAGGCCGAAGTGGA | 683.0 | -6.5 | 1.8 | -3.6 | 2.7E-04 | 7.9E-04 | BT20/MB468 | MP |
| >hg38_21_+_8433172_8446622_RNA45SN1_45S_50ntFlanks@8381.8400.20|AAGAGGGCGTGAAACCGTTA | >hg38_21_+_8433172_8446622_RNA45SN1_45S_50ntFlanks | 8381 | 8400 | 20 | AAGAGGGCGTGAAACCGTTA | 2262.5 | -6.7 | 1.8 | -3.6 | 2.7E-04 | 7.9E-04 | BT20/MB468 | MP |
| >hg38_21_+_8433172_8446622_RNA45SN1_45S_50ntFlanks@7979.8009.31|ACCTCAGATCAGACGTGGCGACCCGCTGAAT | >hg38_21_+_8433172_8446622_RNA45SN1_45S_50ntFlanks | 7979 | 8009 | 31 | ACCTCAGATCAGACGTGGCGACCCGCTGAAT | 1080.9 | -2.0 | 0.6 | -3.4 | 6.3E-04 | 7.9E-04 | MB231/MB468 | MITO |
| >hg38_21_+_8433172_8446622_RNA45SN1_45S_50ntFlanks@11703.11726.24|AATGAAGCGCGGGTAAACGGCGGG | >hg38_21_+_8433172_8446622_RNA45SN1_45S_50ntFlanks | 11703 | 11726 | 24 | AATGAAGCGCGGGTAAACGGCGGG | 260.5 | -2.2 | 0.6 | -3.4 | 6.0E-04 | 7.9E-04 | MB231/MB468 | CYTO |
| >MI0000749|hsa-mir-30e&WithFlank&1|+|40754349|40754452@23.46.24(+1U)&[MIMAT0000692&hsa-miR-30e-5p&offsets|0|+2(+1U);m-24&1|+|40754371|40754394&offsets|0|0(+1U)]|TGTAAACATCCTTGACTGGAAGCTT | >MI0000749|hsa-mir-30e&WithFlank&1|+|40754349|40754452 | 23 | 46 | 24(+1U) | TGTAAACATCCTTGACTGGAAGCTT | 656.8 | -3.5 | 0.9 | -3.9 | 1.1E-04 | 7.9E-04 | BT20/MB231 | MITO |

<table>
<tr><td>>hg38_21_+_8433172_8446622_RNA45SN1_45S_50ntFlanks@12418.12458.41|CGATGTCGGCTCTTCCTATCATTGTGAAGCAGAATTCACCA</td><td>>hg38_21_+_8433172_8446622_RNA45SN1_45S_50ntFlanks</td><td>12418</td><td>12458</td><td>41</td><td>CGATGTCGGCTCTTCCTATCATTGTGAAGCAGAATTCACCA</td><td>625.0</td><td>-6.8</td><td>2.0</td><td>-3.4</td><td>6.4E-04</td><td>7.9E-04</td><td>BT20/MB468</td><td>NUC</td></tr>
<tr><td>>hg38_21_+_8433172_8446622_RNA45SN1_45S_50ntFlanks@12605.12624.20|TATGTGCTTGGCTGAGGAGC</td><td>>hg38_21_+_8433172_8446622_RNA45SN1_45S_50ntFlanks</td><td>12605</td><td>12624</td><td>20</td><td>TATGTGCTTGGCTGAGGAGC</td><td>1296.3</td><td>-1.9</td><td>0.6</td><td>-3.4</td><td>5.7E-04</td><td>7.9E-04</td><td>BT20/MB468</td><td>CYTO</td></tr>
<tr><td>>hg38_21_+_8433172_8446622_RNA45SN1_45S_50ntFlanks@4919.4941.23|CCACCAGGAGTGGAGCCTGCGGC</td><td>>hg38_21_+_8433172_8446622_RNA45SN1_45S_50ntFlanks</td><td>4919</td><td>4941</td><td>23</td><td>CCACCAGGAGTGGAGCCTGCGGC</td><td>1038.7</td><td>-4.1</td><td>1.2</td><td>-3.5</td><td>4.8E-04</td><td>7.9E-04</td><td>BT20/MB468</td><td>MITO</td></tr>
<tr><td>>MI0000087|hsa-mir-29a&WithFlank&7|-|130876741|130876816@48.69.22(+1G)&[MIMAT0000086&hsa-miR-29a-3p&offsets|0|0(+1G);m-26&7|-|130876749|130876769&offsets|0|+1(+1G)]|TAGCACCATCTGAAATCGGTTAG</td><td>>MI0000087|hsa-mir-29a&WithFlank&7|-|130876741|130876816</td><td>48</td><td>69</td><td>22(+1G)</td><td>TAGCACCATCTGAAATCGGTTAG</td><td>67.1</td><td>9.8</td><td>2.7</td><td>3.6</td><td>2.7E-04</td><td>7.9E-04</td><td>BT20/MB468</td><td>MP</td></tr>
<tr><td>>hg38_21_+_8433172_8446622_RNA45SN1_45S_50ntFlanks@11316.11334.19|CACCCCACGTCTCGTCGCG</td><td>>hg38_21_+_8433172_8446622_RNA45SN1_45S_50ntFlanks</td><td>11316</td><td>11334</td><td>19</td><td>CACCCCACGTCTCGTCGCG</td><td>1955.5</td><td>-7.0</td><td>2.1</td><td>-3.4</td><td>6.5E-04</td><td>7.9E-04</td><td>BT20/MB468</td><td>NUC</td></tr>
<tr><td>>MI0003137|hsa-mir-193b&WithFlank&16|+|14303961|14304055@57.77.21(+1C)&[MIMAT0002819&hsa-miR-193b-3p&offsets|0|-1(+1C);m-166&16|+|14304017|14304038&offsets|0|-1(+1C)]|AACTGGCCCTCAAAGTCCCGCC</td><td>>MI0003137|hsa-mir-193b&WithFlank&16|+|14303961|14304055</td><td>57</td><td>77</td><td>21(+1C)</td><td>AACTGGCCCTCAAAGTCCCGCC</td><td>1007.5</td><td>5.5</td><td>1.5</td><td>3.6</td><td>2.7E-04</td><td>7.9E-04</td><td>BT20/MB468</td><td>MP</td></tr>
<tr><td>>am_trnaMT_PheGAA_MT_+_577_647@42.63.22|TTTAGACGGGCTCACATCACCC</td><td>>am_trnaMT_PheGAA_MT_+_577_647</td><td>42</td><td>63</td><td>22</td><td>TTTAGACGGGCTCACATCACCC</td><td>117.0</td><td>5.5</td><td>1.5</td><td>3.6</td><td>2.7E-04</td><td>7.9E-04</td><td>BT20/MB468</td><td>MP</td></tr>
<tr><td>>hg38_21_+_8433172_8446622_RNA45SN1_45S_50ntFlanks@9493.9525.33|CCCGAAAGATGGTGAACTATGCCTGGGCAGGGC</td><td>>hg38_21_+_8433172_8446622_RNA45SN1_45S_50ntFlanks</td><td>9493</td><td>9525</td><td>33</td><td>CCCGAAAGATGGTGAACTATGCCTGGGCAGGGC</td><td>115.2</td><td>5.2</td><td>1.4</td><td>3.6</td><td>2.7E-04</td><td>7.9E-04</td><td>BT20/MB468</td><td>MP</td></tr>
<tr><td>>hg38_21_+_8433172_8446622_RNA45SN1_45S_50ntFlanks@12912.12935.24|CACGTTCGTGGGGAACCTGGCGCT</td><td>>hg38_21_+_8433172_8446622_RNA45SN1_45S_50ntFlanks</td><td>12912</td><td>12935</td><td>24</td><td>CACGTTCGTGGGGAACCTGGCGCT</td><td>236.1</td><td>-4.6</td><td>1.4</td><td>-3.4</td><td>6.0E-04</td><td>7.9E-04</td><td>MB231/MB468</td><td>CYTO</td></tr>
<tr><td>>hg38_21_+_8433172_8446622_RNA45SN1_45S_50ntFlanks@10687.10705.19|CGGCGTCCGGTGAGCTCTC</td><td>>hg38_21_+_8433172_8446622_RNA45SN1_45S_50ntFlanks</td><td>10687</td><td>10705</td><td>19</td><td>CGGCGTCCGGTGAGCTCTC</td><td>242.8</td><td>-3.4</td><td>1.0</td><td>-3.4</td><td>5.7E-04</td><td>7.9E-04</td><td>BT20/MB468</td><td>CYTO</td></tr>
<tr><td>>hg38_21_+_8433172_8446622_RNA45SN1_45S_50ntFlanks@11668.11687.20|TGCCCAGTGCTCTGAATGTC</td><td>>hg38_21_+_8433172_8446622_RNA45SN1_45S_50ntFlanks</td><td>11668</td><td>11687</td><td>20</td><td>TGCCCAGTGCTCTGAATGTC</td><td>213.7</td><td>2.8</td><td>0.8</td><td>3.4</td><td>6.4E-04</td><td>7.9E-04</td><td>MB231/MB468</td><td>MITO</td></tr>
<tr><td>>hg38_21_+_8433172_8446622_RNA45SN1_45S_50ntFlanks@7979.8010.32|ACCTCAGATCAGACGTGGCGACCCGCTGAATT</td><td>>hg38_21_+_8433172_8446622_RNA45SN1_45S_50ntFlanks</td><td>7979</td><td>8010</td><td>32</td><td>ACCTCAGATCAGACGTGGCGACCCGCTGAATT</td><td>161.5</td><td>-2.7</td><td>0.8</td><td>-3.5</td><td>5.3E-04</td><td>7.9E-04</td><td>MB231/MB468</td><td>MP</td></tr>
<tr><td>>MI0001445|hsa-mir-423&WithFlank&17|+|30117073|30117178@23.45.23(+1G)&[MIMAT0004748&hsa-miR-423-5p&offsets|0|0(+1G);m-141&17|+|30117095|30117117&offsets|0|0(+1G)]|TGAGGGGCAGAGAGCGAGACTTTG</td><td>>MI0001445|hsa-mir-423&WithFlank&17|+|30117073|30117178</td><td>23</td><td>45</td><td>23(+1G)</td><td>TGAGGGGCAGAGAGCGAGACTTTG</td><td>100.7</td><td>7.0</td><td>1.9</td><td>3.6</td><td>2.7E-04</td><td>7.9E-04</td><td>BT20/MB468</td><td>MP</td></tr>
<tr><td>>hg38_21_+_8433172_8446622_RNA45SN1_45S_50ntFlanks@12001.12030.30|CTGCGGGCCGCCGGTGAAATACCACTACTC</td><td>>hg38_21_+_8433172_8446622_RNA45SN1_45S_50ntFlanks</td><td>12001</td><td>12030</td><td>30</td><td>CTGCGGGCCGCCGGTGAAATACCACTACTC</td><td>240.0</td><td>-2.9</td><td>0.8</td><td>-3.4</td><td>6.0E-04</td><td>7.9E-04</td><td>MB231/MB468</td><td>CYTO</td></tr>
<tr><td>>hg38_MT_+_1621_3279_RNR2_16S_50ntFlanks@460.492.33|TTGTAAATTTAACTGTTAGTCCAAAGAGGAACA</td><td>>hg38_MT_+_1621_3279_RNR2_16S_50ntFlanks</td><td>460</td><td>492</td><td>33</td><td>TTGTAAATTTAACTGTTAGTCCAAAGAGGAACA</td><td>293.3</td><td>4.5</td><td>1.3</td><td>3.4</td><td>6.4E-04</td><td>7.9E-04</td><td>MB231/MB468</td><td>MITO</td></tr>
<tr><td>>MI0000289|hsa-mir-181a-1&WithFlank&1|-|198859038|198859159@30.52.23;MI0000269|hsa-mir-181a-2&WithFlank&9|+|124692436|124692557@45.67.23&[MIMAT0000256&hsa-miR-181a-5p&offsets|0|0;m-32&1|-|198859108|198859130&offsets|0|0];[MIMAT0000256_1&hsa-miR-181a-5p&offsets|0|0;m-32&9|+|124692480|124692502&offsets|0|0]|AACATTCAACGCTGTCGGTGAGT</td><td>>MI0000289|hsa-mir-181a-1&WithFlank&1|-|198859038|198859159</td><td>30</td><td>52</td><td>23</td><td>AACATTCAACGCTGTCGGTGAGT</td><td>7593.1</td><td>-3.1</td><td>0.8</td><td>-3.9</td><td>1.1E-04</td><td>7.9E-04</td><td>BT20/MB231</td><td>MITO</td></tr>
<tr><td>>hg38_21_+_8433172_8446622_RNA45SN1_45S_50ntFlanks@12933.12953.21|GCTAAACCATTCGTAGACGAC</td><td>>hg38_21_+_8433172_8446622_RNA45SN1_45S_50ntFlanks</td><td>12933</td><td>12953</td><td>21</td><td>GCTAAACCATTCGTAGACGAC</td><td>256.1</td><td>-5.1</td><td>1.5</td><td>-3.4</td><td>5.8E-04</td><td>7.9E-04</td><td>BT20/MB468</td><td>CYTO</td></tr>
<tr><td>>hg38_MT_+_1621_3279_RNR2_16S_50ntFlanks@1338.1374.37|AGTCCATATCAACAATAGGGTTTACGACCTCGATGTT</td><td>>hg38_MT_+_1621_3279_RNR2_16S_50ntFlanks</td><td>1338</td><td>1374</td><td>37</td><td>AGTCCATATCAACAATAGGGTTTACGACCTCGATGTT</td><td>69.1</td><td>9.9</td><td>2.7</td><td>3.6</td><td>2.7E-04</td><td>7.9E-04</td><td>BT20/MB468</td><td>MP</td></tr>
<tr><td>>hg38_21_+_8433172_8446622_RNA45SN1_45S_50ntFlanks@11872.11893.22|AAAGAAGACCCTGTTGAGCTTG</td><td>>hg38_21_+_8433172_8446622_RNA45SN1_45S_50ntFlanks</td><td>11872</td><td>11893</td><td>22</td><td>AAAGAAGACCCTGTTGAGCTTG</td><td>298.8</td><td>-3.0</td><td>0.9</td><td>-3.4</td><td>6.1E-04</td><td>8.0E-04</td><td>MB231/MB468</td><td>CYTO</td></tr>
<tr><td>>MI0006273|hsa-mir-1180&WithFlank&17|-|19344500|19344580@47.67.21(+1C)&[MIMAT0005825&hsa-miR-1180-3p&offsets|0|-1(+1C);m-214&17|-|19344513|19344534&offsets|0|-1(+1C)]|TTTCCGGCTCGCGTGGGTGTGC</td><td>>MI0006273|hsa-mir-1180&WithFlank&17|-|19344500|19344580</td><td>47</td><td>67</td><td>21(+1C)</td><td>TTTCCGGCTCGCGTGGGTGTGC</td><td>519.0</td><td>3.2</td><td>0.8</td><td>3.9</td><td>9.9E-05</td><td>8.0E-04</td><td>BT20/MB231</td><td>CYTO</td></tr>
<tr><td>>MI0000446|hsa-mir-125b-1&WithFlank&11|-|122099751|122099850@61.82.22(+2U)&[MIMAT0004592&hsa-miR-125b-1-3p&offsets|0|0(+2U);m-485&11|-|122099769|122099790&offsets|0|0(+2U)]|ACGGGTTAGGCTCTTGGGAGCTTT</td><td>>MI0000446|hsa-mir-125b-1&WithFlank&11|-|122099751|122099850</td><td>61</td><td>82</td><td>22(+2U)</td><td>ACGGGTTAGGCTCTTGGGAGCTTT</td><td>44.6</td><td>-8.9</td><td>2.3</td><td>-3.9</td><td>9.5E-05</td><td>8.0E-04</td><td>BT20/MB231</td><td>MP</td></tr>
<tr><td>>MI0000433|hsa-let-7g&WithFlank&3|-|52268272|52268367@11.31.21(+1G)&[MIMAT0000414&hsa-let-7g-5p&offsets|0|-1(+1G);m-58&3|-|52268336|52268357&offsets|0|-1(+1G)]|TGAGGTAGTAGTTTGTACAGTG</td><td>>MI0000433|hsa-let-7g&WithFlank&3|-|52268272|52268367</td><td>11</td><td>31</td><td>21(+1G)</td><td>TGAGGTAGTAGTTTGTACAGTG</td><td>442.7</td><td>3.5</td><td>1.0</td><td>3.5</td><td>5.4E-04</td><td>8.0E-04</td><td>MB231/MB468</td><td>MP</td></tr>
<tr><td>>hg38_MT_+_1621_3279_RNR2_16S_50ntFlanks@1338.1373.36|AGTCCATATCAACAATAGGGTTTACGACCTCGATGT</td><td>>hg38_MT_+_1621_3279_RNR2_16S_50ntFlanks</td><td>1338</td><td>1373</td><td>36</td><td>AGTCCATATCAACAATAGGGTTTACGACCTCGATGT</td><td>137.8</td><td>7.4</td><td>2.0</td><td>3.6</td><td>2.7E-04</td><td>8.0E-04</td><td>BT20/MB468</td><td>MP</td></tr>
<tr><td>>trnaMT_ValTAC_MT_+_1602_1670@51.71.21|CTTAACTTGACCGCTCTGACC</td><td>>trnaMT_ValTAC_MT_+_1602_1670</td><td>51</td><td>71</td><td>21</td><td>CTTAACTTGACCGCTCTGACC</td><td>2048.6</td><td>-3.4</td><td>1.0</td><td>-3.4</td><td>6.4E-04</td><td>8.0E-04</td><td>MB231/MB468</td><td>MITO</td></tr>
<tr><td>>hg38_21_+_8433172_8446622_RNA45SN1_45S_50ntFlanks@6653.6686.34|ACTCTTAGCGGTGGATCACTCGGCTCGTGCGTCG</td><td>>hg38_21_+_8433172_8446622_RNA45SN1_45S_50ntFlanks</td><td>6653</td><td>6686</td><td>34</td><td>ACTCTTAGCGGTGGATCACTCGGCTCGTGCGTCG</td><td>1209.5</td><td>-3.0</td><td>0.9</td><td>-3.4</td><td>6.4E-04</td><td>8.0E-04</td><td>MB231/MB468</td><td>MITO</td></tr>
<tr><td>>MI0000446|hsa-mir-125b-1&WithFlank&11|-|122099751|122099850@21.40.20(+1U);MI0000470|hsa-mir-125b-2&WithFlank&21|+|16590231|16590331@23.42.20(+1U)&[MIMAT0000423&hsa-miR-125b-5p&offsets|0|-2(+1U);m-67&11|-|122099809|122099830&offsets|0|-2(+1U)];[MIMAT0000423_1&hsa-miR-125b-5p&offsets|0|-2(+1U);m-68&21|+|16590253|16590274&offsets|0|-2(+1U)]|TCCCTGAGACCCTAACTTGTT</td><td>>MI0000446|hsa-mir-125b-1&WithFlank&11|-|122099751|122099850</td><td>21</td><td>40</td><td>20(+1U)</td><td>TCCCTGAGACCCTAACTTGTT</td><td>86.0</td><td>6.3</td><td>1.8</td><td>3.5</td><td>5.4E-04</td><td>8.0E-04</td><td>MB231/MB468</td><td>MP</td></tr>
<tr><td>>MI0000439|hsa-mir-23b&WithFlank&9|+|95085202|95085310@67.84.18&[MIMAT0000418&hsa-miR-23b-3p&offsets|+3|-2;m-42&9|+|95085265|95085285&offsets|+3|0]|ACATTGCCAGGGATTACC</td><td>>MI0000439|hsa-mir-23b&WithFlank&9|+|95085202|95085310</td><td>67</td><td>84</td><td>18</td><td>ACATTGCCAGGGATTACC</td><td>576.1</td><td>12.4</td><td>3.6</td><td>3.4</td><td>6.4E-04</td><td>8.0E-04</td><td>MB231/MB468</td><td>MITO</td></tr>
<tr><td>>trnaMT_GluTTC_MT_-_14674_14742@45.70.26,trnalookalike8_GluTTC_5_-_93905172_93905240@45.70.26|CGTGGTTGTAGTCCGTGCGAGAATAC</td><td>>trnaMT_GluTTC_MT_-_14674_14742</td><td>45</td><td>70</td><td>26</td><td>CGTGGTTGTAGTCCGTGCGAGAATAC</td><td>201.1</td><td>4.1</td><td>1.2</td><td>3.4</td><td>6.4E-04</td><td>8.0E-04</td><td>MB231/MB468</td><td>MITO</td></tr>
<tr><td>>trnaMT_SerGCT_MT_+_12207_12265@1.18.18|GAGAAAGCTCACAAGAAC</td><td>>trnaMT_SerGCT_MT_+_12207_12265</td><td>1</td><td>18</td><td>18</td><td>GAGAAAGCTCACAAGAAC</td><td>168672.6</td><td>-3.8</td><td>1.0</td><td>-3.9</td><td>9.6E-05</td><td>8.0E-04</td><td>BT20/MB231</td><td>MP</td></tr>
<tr><td>>hg38_21_+_8433172_8446622_RNA45SN1_45S_50ntFlanks@9475.9509.35|CGAGCGCACGTGTTAGGACCCGAAAGATGGTGAAC</td><td>>hg38_21_+_8433172_8446622_RNA45SN1_45S_50ntFlanks</td><td>9475</td><td>9509</td><td>35</td><td>CGAGCGCACGTGTTAGGACCCGAAAGATGGTGAAC</td><td>52.6</td><td>9.0</td><td>2.3</td><td>3.9</td><td>9.6E-05</td><td>8.0E-04</td><td>BT20/MB231</td><td>MP</td></tr>
<tr><td>>trna119_LysCTT_1_-_145395522_145395594@1.36.36,trna11_LysCTT_5_-_180648979_180649051@1.36.36,trna13_LysCTT_6_+_26556774_26556846@1.36.36,trna7_LysCTT_16_+_3225692_3225764@1.36.36,trna9_LysCTT_5_+_180634755_180634827@1.36.36|GCCCGGCTAGCTCAGTCGGTAGAGCATGAGACTCTT</td><td>>trna119_LysCTT_1_-_145395522_145395594</td><td>1</td><td>36</td><td>36</td><td>GCCCGGCTAGCTCAGTCGGTAGAGCATGAGACTCTT</td><td>774.9</td><td>4.9</td><td>1.4</td><td>3.6</td><td>2.7E-04</td><td>8.0E-04</td><td>BT20/MB468</td><td>MP</td></tr>
<tr><td>>hg38_21_+_8433172_8446622_RNA45SN1_45S_50ntFlanks@8170.8215.46|CAAGTCCTTCTGATCGAGGCCCAGCCCGTGGACGGTGTGAGGCCGG</td><td>>hg38_21_+_8433172_8446622_RNA45SN1_45S_50ntFlanks</td><td>8170</td><td>8215</td><td>46</td><td>CAAGTCCTTCTGATCGAGGCCCAGCCCGTGGACGGTGTGAGGCCGG</td><td>361.4</td><td>-6.6</td><td>1.8</td><td>-3.6</td><td>2.7E-04</td><td>8.0E-04</td><td>BT20/MB468</td><td>MP</td></tr>
<tr><td>>hg38_21_+_8433172_8446622_RNA45SN1_45S_50ntFlanks@12324.12350.27|TTGGGTTTTAAGCAGGAGGTGTCAGAA</td><td>>hg38_21_+_8433172_8446622_RNA45SN1_45S_50ntFlanks</td><td>12324</td><td>12350</td><td>27</td><td>TTGGGTTTTAAGCAGGAGGTGTCAGAA</td><td>196.6</td><td>-5.6</td><td>1.5</td><td>-3.6</td><td>2.7E-04</td><td>8.0E-04</td><td>BT20/MB468</td><td>MP</td></tr>
<tr><td>>hg38_21_+_8433172_8446622_RNA45SN1_45S_50ntFlanks@4498.4520.23|AAGCGTTTACTTTGAAAAAATTA</td><td>>hg38_21_+_8433172_8446622_RNA45SN1_45S_50ntFlanks</td><td>4498</td><td>4520</td><td>23</td><td>AAGCGTTTACTTTGAAAAAATTA</td><td>753.8</td><td>3.5</td><td>0.9</td><td>3.9</td><td>1.0E-04</td><td>8.0E-04</td><td>BT20/MB231</td><td>CYTO</td></tr>
<tr><td>>hg38_21_+_8433172_8446622_RNA45SN1_45S_50ntFlanks@12419.12456.38|GATGTCGGCTCTTCCTATCATTGTGAAGCAGAATTCAC</td><td>>hg38_21_+_8433172_8446622_RNA45SN1_45S_50ntFlanks</td><td>12419</td><td>12456</td><td>38</td><td>GATGTCGGCTCTTCCTATCATTGTGAAGCAGAATTCAC</td><td>1198.5</td><td>-3.2</td><td>0.8</td><td>-3.9</td><td>1.0E-04</td><td>8.0E-04</td><td>BT20/MB231</td><td>CYTO</td></tr>
<tr><td>>hg38_21_+_8433172_8446622_RNA45SN1_45S_50ntFlanks@10349.10388.40|AACGAGAACTTTGAAGGCCGAAGTGGAGAAGGGTTCCATG</td><td>>hg38_21_+_8433172_8446622_RNA45SN1_45S_50ntFlanks</td><td>10349</td><td>10388</td><td>40</td><td>AACGAGAACTTTGAAGGCCGAAGTGGAGAAGGGTTCCATG</td><td>1151.8</td><td>-3.2</td><td>0.9</td><td>-3.4</td><td>5.8E-04</td><td>8.0E-04</td><td>BT20/MB468</td><td>CYTO</td></tr>
<tr><td>>hg38_21_+_8433172_8446622_RNA45SN1_45S_50ntFlanks@9955.9979.25|GGAAGTCGGAATCCGCTAAGGAGTG</td><td>>hg38_21_+_8433172_8446622_RNA45SN1_45S_50ntFlanks</td><td>9955</td><td>9979</td><td>25</td><td>GGAAGTCGGAATCCGCTAAGGAGTG</td><td>326.6</td><td>7.4</td><td>1.9</td><td>3.9</td><td>9.6E-05</td><td>8.0E-04</td><td>BT20/MB231</td><td>MP</td></tr>
<tr><td>>hg38_21_+_8433172_8446622_RNA45SN1_45S_50ntFlanks@11769.11794.26|TAATTAGTGACGCGCATGAATGGATG</td><td>>hg38_21_+_8433172_8446622_RNA45SN1_45S_50ntFlanks</td><td>11769</td><td>11794</td><td>26</td><td>TAATTAGTGACGCGCATGAATGGATG</td><td>462.2</td><td>-2.8</td><td>0.8</td><td>-3.6</td><td>2.7E-04</td><td>8.0E-04</td><td>BT20/MB468</td><td>MP</td></tr>
<tr><td>>hg38_MT_+_1621_3279_RNR2_16S_50ntFlanks@617.643.27|AAACATATAACTGAACTCCTCACACCC</td><td>>hg38_MT_+_1621_3279_RNR2_16S_50ntFlanks</td><td>617</td><td>643</td><td>27</td><td>AAACATATAACTGAACTCCTCACACCC</td><td>124.0</td><td>5.1</td><td>1.5</td><td>3.4</td><td>6.4E-04</td><td>8.0E-04</td><td>MB231/MB468</td><td>MITO</td></tr>
<tr><td>>MI0000456|hsa-mir-140&WithFlank&16|+|69933075|69933186@70.90.21(+1U)&[MIMAT0004597&hsa-miR-140-3p&offsets|+2|+2(+1U);m-45&16|+|69933143|69933164&offsets|+1|0(+1U)]|CCACAGGGTAGAACCACGGACT</td><td>>MI0000456|hsa-mir-140&WithFlank&16|+|69933075|69933186</td><td>70</td><td>90</td><td>21(+1U)</td><td>CCACAGGGTAGAACCACGGACT</td><td>65.2</td><td>9.8</td><td>2.7</td><td>3.6</td><td>2.7E-04</td><td>8.0E-04</td><td>BT20/MB468</td><td>MP</td></tr>
<tr><td>>hg38_21_+_8433172_8446622_RNA45SN1_45S_50ntFlanks@11322.11343.22|ACGTCTCGTCGCGCGCGCGTCC</td><td>>hg38_21_+_8433172_8446622_RNA45SN1_45S_50ntFlanks</td><td>11322</td><td>11343</td><td>22</td><td>ACGTCTCGTCGCGCGCGCGTCC</td><td>6823.9</td><td>-4.3</td><td>1.3</td><td>-3.4</td><td>6.5E-04</td><td>8.0E-04</td><td>BT20/MB468</td><td>NUC</td></tr>
<tr><td>>MI0000301|hsa-mir-224&WithFlank&X|-|151958572|151958664@14.34.21&[MIMAT0000281&hsa-miR-224-5p&offsets|+1|-3;m-238&X|-|151958630|151958651&offsets|0|-1]|CAAGTCACTAGTGGTTCCGTT</td><td>>MI0000301|hsa-mir-224&WithFlank&X|-|151958572|151958664</td><td>14</td><td>34</td><td>21</td><td>CAAGTCACTAGTGGTTCCGTT</td><td>1482.9</td><td>3.0</td><td>0.8</td><td>3.9</td><td>1.0E-04</td><td>8.0E-04</td><td>BT20/MB231</td><td>CYTO</td></tr>
<tr><td>>hg38_21_+_8433172_8446622_RNA45SN1_45S_50ntFlanks@12158.12183.26|GTTTGACTGGGGCGGTACACCTGTCA</td><td>>hg38_21_+_8433172_8446622_RNA45SN1_45S_50ntFlanks</td><td>12158</td><td>12183</td><td>26</td><td>GTTTGACTGGGGCGGTACACCTGTCA</td><td>82.7</td><td>-5.6</td><td>1.5</td><td>-3.6</td><td>2.7E-04</td><td>8.0E-04</td><td>BT20/MB468</td><td>MP</td></tr>
<tr><td>>trna16_TyrGTA_14_-_21131351_21131444@59.76.18,trna17_TyrGTA_14_-_21128117_21128210@59.76.18,trna19_TyrGTA_14_-_21121258_21121351@59.76.18,trna2_TyrGTA_2_+_27273650_27273738@59.76.18,trna4_TyrGTA_8_+_67025602_67025694@59.76.18,trna5_TyrGTA_14_+_21151432_21151520@59.76.18,trna5_TyrGTA_8_+_67026223_67026311@59.76.18|TTCCGGCTCGAAGGACCA</td><td>>trna16_TyrGTA_14_-_21131351_21131444</td><td>59</td><td>76</td><td>18</td><td>TTCCGGCTCGAAGGACCA</td><td>320.0</td><td>2.9</td><td>0.8</td><td>3.4</td><td>6.6E-04</td><td>8.0E-04</td><td>BT20/MB468</td><td>NUC</td></tr>
<tr><td>>hg38_21_+_8433172_8446622_RNA45SN1_45S_50ntFlanks@11924.11947.24|GAGGTGTAGAATAAGTGGGAGGCC</td><td>>hg38_21_+_8433172_8446622_RNA45SN1_45S_50ntFlanks</td><td>11924</td><td>11947</td><td>24</td><td>GAGGTGTAGAATAAGTGGGAGGCC</td><td>61.3</td><td>9.7</td><td>2.7</td><td>3.6</td><td>2.7E-04</td><td>8.0E-04</td><td>BT20/MB468</td><td>MP</td></tr>
</table>

<table>
<tr><td>>MI0003137|hsa-mir-193b&WithFlank&16|+|14303961|14304055@20.41.22&[MIMAT0004767&hsa-miR-193b-5p&offsets|0|0;m-343&16|+|14303980|14304001&offsets|0|0]|CGGGGTTTTGAGGGCGAGATGA</td><td>>MI0003137|hsa-mir-193b&WithFlank&16|+|14303961|14304055</td><td>20</td><td>41</td><td>22</td><td>CGGGGTTTTGAGGGCGAGATGA</td><td>304.5</td><td>6.5</td><td>1.7</td><td>3.9</td><td>9.7E-05</td><td>8.1E-04</td><td>BT20/MB231</td><td>MP</td></tr>
<tr><td>>hg38_21_+_8433172_8446622_RNA45SN1_45S_50ntFlanks@10782.10801.20|GGTGAACAGCCTCTGGCATG</td><td>>hg38_21_+_8433172_8446622_RNA45SN1_45S_50ntFlanks</td><td>10782</td><td>10801</td><td>20</td><td>GGTGAACAGCCTCTGGCATG</td><td>413.6</td><td>-3.3</td><td>0.9</td><td>-3.6</td><td>2.8E-04</td><td>8.1E-04</td><td>BT20/MB468</td><td>MP</td></tr>
<tr><td>>hg38_21_+_8433172_8446622_RNA45SN1_45S_50ntFlanks@11569.11586.18|CAGCCGACTTAGAACTGG</td><td>>hg38_21_+_8433172_8446622_RNA45SN1_45S_50ntFlanks</td><td>11569</td><td>11586</td><td>18</td><td>CAGCCGACTTAGAACTGG</td><td>1309.2</td><td>-2.3</td><td>0.7</td><td>-3.4</td><td>6.1E-04</td><td>8.1E-04</td><td>MB231/MB468</td><td>CYTO</td></tr>
<tr><td>>hg38_1_-_228634819_228635039_RNA5S12_5S_50ntFlanks@130.171.42|TGGATGGGAGACCGCCTGGGAATACCGGGTGCTGTAGGCTTT</td><td>>hg38_1_-_228634819_228635039_RNA5S12_5S_50ntFlanks</td><td>130</td><td>171</td><td>42</td><td>TGGATGGGAGACCGCCTGGGAATACCGGGTGCTGTAGGCTTT</td><td>189.7</td><td>4.6</td><td>1.3</td><td>3.5</td><td>5.5E-04</td><td>8.1E-04</td><td>MB231/MB468</td><td>MP</td></tr>
<tr><td>>MI0000456|hsa-mir-140&WithFlank&16|+|69933075|69933186@68.91.24&[MIMAT0004597&hsa-miR-140-3p&offsets|0|+3;m-45&16|+|69933143|69933164&offsets|-1|+1]|TACCACAGGGTAGAACCACGGACA</td><td>>MI0000456|hsa-mir-140&WithFlank&16|+|69933075|69933186</td><td>68</td><td>91</td><td>24</td><td>TACCACAGGGTAGAACCACGGACA</td><td>199.7</td><td>5.1</td><td>1.4</td><td>3.6</td><td>2.8E-04</td><td>8.1E-04</td><td>BT20/MB468</td><td>MP</td></tr>
<tr><td>>trna2_GlyGCC_21_-_18827107_18827177@1.30.30,trna35_GlyGCC_1_+_161413094_161413164@1.30.30,trna37_GlyGCC_1_+_161420467_161420537@1.30.30,trna39_GlyGCC_1_+_161427898_161427968@1.30.30,trna41_GlyGCC_1_+_161435258_161435328@1.30.30|GCATGGGTGGTTCAGTGGTAGAATTCTCGC</td><td>>trna2_GlyGCC_21_-_18827107_18827177</td><td>1</td><td>30</td><td>30</td><td>GCATGGGTGGTTCAGTGGTAGAATTCTCGC</td><td>2832.9</td><td>3.4</td><td>0.9</td><td>3.9</td><td>1.0E-04</td><td>8.1E-04</td><td>BT20/MB231</td><td>CYTO</td></tr>
<tr><td>>hg38_21_+_8433172_8446622_RNA45SN1_45S_50ntFlanks@9027.9049.23|CGTCTCCTCGTGGGGGGGCCGGG</td><td>>hg38_21_+_8433172_8446622_RNA45SN1_45S_50ntFlanks</td><td>9027</td><td>9049</td><td>23</td><td>CGTCTCCTCGTGGGGGGGCCGGG</td><td>66.2</td><td>-9.1</td><td>2.5</td><td>-3.6</td><td>2.8E-04</td><td>8.1E-04</td><td>BT20/MB468</td><td>MP</td></tr>
<tr><td>>hg38_21_+_8433172_8446622_RNA45SN1_45S_50ntFlanks@8170.8201.32|CAAGTCCTTCTGATCGAGGCCCAGCCCGTGGA</td><td>>hg38_21_+_8433172_8446622_RNA45SN1_45S_50ntFlanks</td><td>8170</td><td>8201</td><td>32</td><td>CAAGTCCTTCTGATCGAGGCCCAGCCCGTGGA</td><td>270.1</td><td>-3.9</td><td>1.1</td><td>-3.5</td><td>4.9E-04</td><td>8.1E-04</td><td>BT20/MB468</td><td>MITO</td></tr>
<tr><td>>hg38_21_+_8433172_8446622_RNA45SN1_45S_50ntFlanks@11103.11121.19|GAAGGGTCGGGGCGGCAGG</td><td>>hg38_21_+_8433172_8446622_RNA45SN1_45S_50ntFlanks</td><td>11103</td><td>11121</td><td>19</td><td>GAAGGGTCGGGGCGGCAGG</td><td>143.5</td><td>10.5</td><td>2.7</td><td>3.9</td><td>9.7E-05</td><td>8.1E-04</td><td>BT20/MB231</td><td>MP</td></tr>
<tr><td>>hg38_21_+_8433172_8446622_RNA45SN1_45S_50ntFlanks@5419.5443.25|ACCGATTGGATGGTTTAGTGAGGCC</td><td>>hg38_21_+_8433172_8446622_RNA45SN1_45S_50ntFlanks</td><td>5419</td><td>5443</td><td>25</td><td>ACCGATTGGATGGTTTAGTGAGGCC</td><td>114.5</td><td>-3.2</td><td>0.9</td><td>-3.4</td><td>5.9E-04</td><td>8.1E-04</td><td>BT20/MB468</td><td>CYTO</td></tr>
<tr><td>>MI0000433|hsa-let-7g&WithFlank&3|-|52268272|52268367@11.30.20(+1C)&[MIMAT0000414&hsa-let-7g-5p&offsets|0|-2(+1C);m-58&3|-|52268336|52268357&offsets|0|-2(+1C)]|TGAGGTAGTAGTTTGTACAGC</td><td>>MI0000433|hsa-let-7g&WithFlank&3|-|52268272|52268367</td><td>11</td><td>30</td><td>20(+1C)</td><td>TGAGGTAGTAGTTTGTACAGC</td><td>456.5</td><td>5.0</td><td>1.4</td><td>3.5</td><td>5.5E-04</td><td>8.1E-04</td><td>MB231/MB468</td><td>MP</td></tr>
<tr><td>>hg38_21_+_8433172_8446622_RNA45SN1_45S_50ntFlanks@10687.10712.26|CGGCGTCCGGTGAGCTCTCGCTGGCC</td><td>>hg38_21_+_8433172_8446622_RNA45SN1_45S_50ntFlanks</td><td>10687</td><td>10712</td><td>26</td><td>CGGCGTCCGGTGAGCTCTCGCTGGCC</td><td>605.0</td><td>-4.0</td><td>1.2</td><td>-3.5</td><td>4.9E-04</td><td>8.1E-04</td><td>BT20/MB468</td><td>MITO</td></tr>
<tr><td>>hg38_21_+_8433172_8446622_RNA45SN1_45S_50ntFlanks@3899.3937.39|CCTTCGCGGGGGGGATGCGTGCATTTATCAGATCAAAAC</td><td>>hg38_21_+_8433172_8446622_RNA45SN1_45S_50ntFlanks</td><td>3899</td><td>3937</td><td>39</td><td>CCTTCGCGGGGGGGATGCGTGCATTTATCAGATCAAAAC</td><td>588.0</td><td>-5.3</td><td>1.5</td><td>-3.5</td><td>4.9E-04</td><td>8.1E-04</td><td>BT20/MB468</td><td>MITO</td></tr>
<tr><td>>am_MI0003639|hsa-mir-625&WithFlank&14|+|65471096|65471192@58.78.21(+1U)&[MIMAT0004808&hsa-miR-625-3p&offsets|0|-1(+1U);m-137&14|+|65471153|65471174&offsets|0|-1(+1U)]|GACTATAGAACTTTCCCCCTCT</td><td>>am_MI0003639|hsa-mir-625&WithFlank&14|+|65471096|65471192</td><td>58</td><td>78</td><td>21(+1U)</td><td>GACTATAGAACTTTCCCCCTCT</td><td>183.0</td><td>5.3</td><td>1.5</td><td>3.5</td><td>5.5E-04</td><td>8.1E-04</td><td>MB231/MB468</td><td>MP</td></tr>
<tr><td>>hg38_21_+_8433172_8446622_RNA45SN1_45S_50ntFlanks@7983.8009.27|CAGATCAGACGTGGCGACCCGCTGAAT</td><td>>hg38_21_+_8433172_8446622_RNA45SN1_45S_50ntFlanks</td><td>7983</td><td>8009</td><td>27</td><td>CAGATCAGACGTGGCGACCCGCTGAAT</td><td>814.2</td><td>-2.7</td><td>0.7</td><td>-3.6</td><td>2.8E-04</td><td>8.1E-04</td><td>BT20/MB468</td><td>MP</td></tr>
<tr><td>>MI0000273|hsa-mir-183&WithFlank&7|-|129774899|129775020@33.54.22&[MIMAT0000261&hsa-miR-183-5p&offsets|0|0;m-41&7|-|129774966|129774987&offsets|-1|-1]|TATGGCACTGGTAGAATTCACT</td><td>>MI0000273|hsa-mir-183&WithFlank&7|-|129774899|129775020</td><td>33</td><td>54</td><td>22</td><td>TATGGCACTGGTAGAATTCACT</td><td>12675.4</td><td>2.0</td><td>0.5</td><td>3.9</td><td>1.0E-04</td><td>8.1E-04</td><td>BT20/MB231</td><td>NUC</td></tr>
<tr><td>>trnaMT_ValTAC_MT_+_1602_1670@51.72.22|CTTAACTTGACCGCTCTGACCA</td><td>>trnaMT_ValTAC_MT_+_1602_1670</td><td>51</td><td>72</td><td>22</td><td>CTTAACTTGACCGCTCTGACCA</td><td>5647.0</td><td>-3.2</td><td>0.9</td><td>-3.4</td><td>6.6E-04</td><td>8.1E-04</td><td>MB231/MB468</td><td>NUC</td></tr>
<tr><td>>hg38_21_+_8433172_8446622_RNA45SN1_45S_50ntFlanks@10349.10384.36|AACGAGAACTTTGAAGGCCGAAGTGGAGAAGGGTTC</td><td>>hg38_21_+_8433172_8446622_RNA45SN1_45S_50ntFlanks</td><td>10349</td><td>10384</td><td>36</td><td>AACGAGAACTTTGAAGGCCGAAGTGGAGAAGGGTTC</td><td>759.3</td><td>-1.3</td><td>0.4</td><td>-3.4</td><td>5.9E-04</td><td>8.1E-04</td><td>BT20/MB468</td><td>CYTO</td></tr>
<tr><td>>hg38_21_+_8433172_8446622_RNA45SN1_45S_50ntFlanks@10683.10709.27|CCGGCGGCGTCCGGTGAGCTCTCGCTG</td><td>>hg38_21_+_8433172_8446622_RNA45SN1_45S_50ntFlanks</td><td>10683</td><td>10709</td><td>27</td><td>CCGGCGGCGTCCGGTGAGCTCTCGCTG</td><td>558.5</td><td>-4.0</td><td>1.2</td><td>-3.4</td><td>5.9E-04</td><td>8.1E-04</td><td>BT20/MB468</td><td>CYTO</td></tr>
<tr><td>>hg38_21_+_8433172_8446622_RNA45SN1_45S_50ntFlanks@7977.7995.19|CGACCTCAGATCAGACGTG</td><td>>hg38_21_+_8433172_8446622_RNA45SN1_45S_50ntFlanks</td><td>7977</td><td>7995</td><td>19</td><td>CGACCTCAGATCAGACGTG</td><td>366.1</td><td>-2.3</td><td>0.7</td><td>-3.4</td><td>6.2E-04</td><td>8.1E-04</td><td>MB231/MB468</td><td>CYTO</td></tr>
<tr><td>>hg38_21_+_8433172_8446622_RNA45SN1_45S_50ntFlanks@9305.9338.34|CCAAGGAGTCTAACACGTGCGCGAGTCGGGGGCT</td><td>>hg38_21_+_8433172_8446622_RNA45SN1_45S_50ntFlanks</td><td>9305</td><td>9338</td><td>34</td><td>CCAAGGAGTCTAACACGTGCGCGAGTCGGGGGCT</td><td>2457.5</td><td>-6.5</td><td>1.8</td><td>-3.6</td><td>2.8E-04</td><td>8.1E-04</td><td>BT20/MB468</td><td>MP</td></tr>
<tr><td>>hg38_21_+_8433172_8446622_RNA45SN1_45S_50ntFlanks@8341.8375.35|CCGTAAGGGAAAGTTGAAAAGAACTTTGAAGAGAG</td><td>>hg38_21_+_8433172_8446622_RNA45SN1_45S_50ntFlanks</td><td>8341</td><td>8375</td><td>35</td><td>CCGTAAGGGAAAGTTGAAAAGAACTTTGAAGAGAG</td><td>1097.7</td><td>7.6</td><td>2.1</td><td>3.6</td><td>2.8E-04</td><td>8.1E-04</td><td>BT20/MB468</td><td>MP</td></tr>
<tr><td>>hg38_21_+_8433172_8446622_RNA45SN1_45S_50ntFlanks@9738.9760.23|GAAACGATCTCAACCTATTCTCA</td><td>>hg38_21_+_8433172_8446622_RNA45SN1_45S_50ntFlanks</td><td>9738</td><td>9760</td><td>23</td><td>GAAACGATCTCAACCTATTCTCA</td><td>106.0</td><td>-3.2</td><td>0.9</td><td>-3.4</td><td>5.9E-04</td><td>8.1E-04</td><td>BT20/MB468</td><td>CYTO</td></tr>
<tr><td>>hg38_21_+_8433172_8446622_RNA45SN1_45S_50ntFlanks@10872.10890.19|GCTGGGTCGGTCGGGCTGG</td><td>>hg38_21_+_8433172_8446622_RNA45SN1_45S_50ntFlanks</td><td>10872</td><td>10890</td><td>19</td><td>GCTGGGTCGGTCGGGCTGG</td><td>95.5</td><td>-4.3</td><td>1.3</td><td>-3.5</td><td>5.5E-04</td><td>8.1E-04</td><td>MB231/MB468</td><td>MP</td></tr>
<tr><td>>trnaMT_GlnTTG_MT_-_4329_4400@1.36.36,trnalookalike2_GlnTTG_1_-_564879_564950@1.36.36|TAGGATGGGGTGTGATAGGTGGCACGGAGAATTTTG</td><td>>trnaMT_GlnTTG_MT_-_4329_4400</td><td>1</td><td>36</td><td>36</td><td>TAGGATGGGGTGTGATAGGTGGCACGGAGAATTTTG</td><td>978.4</td><td>6.0</td><td>1.7</td><td>3.6</td><td>2.8E-04</td><td>8.1E-04</td><td>BT20/MB468</td><td>MP</td></tr>
<tr><td>>hg38_21_+_8433172_8446622_RNA45SN1_45S_50ntFlanks@8118.8142.25|GGGACATGTGGCGTACGGAAGACCC</td><td>>hg38_21_+_8433172_8446622_RNA45SN1_45S_50ntFlanks</td><td>8118</td><td>8142</td><td>25</td><td>GGGACATGTGGCGTACGGAAGACCC</td><td>121.6</td><td>6.9</td><td>1.9</td><td>3.6</td><td>2.8E-04</td><td>8.1E-04</td><td>BT20/MB468</td><td>MP</td></tr>
<tr><td>>hg38_21_+_8433172_8446622_RNA45SN1_45S_50ntFlanks@12488.12505.18|AACGTGAGCTGGGTTTAG</td><td>>hg38_21_+_8433172_8446622_RNA45SN1_45S_50ntFlanks</td><td>12488</td><td>12505</td><td>18</td><td>AACGTGAGCTGGGTTTAG</td><td>104.4</td><td>-5.8</td><td>1.6</td><td>-3.6</td><td>2.8E-04</td><td>8.1E-04</td><td>BT20/MB468</td><td>MP</td></tr>
<tr><td>>MI0006273|hsa-mir-1180&WithFlank&17|-|19344500|19344580@47.66.20&[MIMAT0005825&hsa-miR-1180-3p&offsets|0|-2;m-214&17|-|19344513|19344534&offsets|0|-2]|TTTCCGGCTCGCGTGGGTGT</td><td>>MI0006273|hsa-mir-1180&WithFlank&17|-|19344500|19344580</td><td>47</td><td>66</td><td>20</td><td>TTTCCGGCTCGCGTGGGTGT</td><td>488.5</td><td>5.0</td><td>1.4</td><td>3.6</td><td>2.8E-04</td><td>8.1E-04</td><td>BT20/MB468</td><td>MP</td></tr>
<tr><td>>hg38_21_+_8433172_8446622_RNA45SN1_45S_50ntFlanks@5237.5285.49|ACTGACTGGCTCAGCGTGTGCCTACCCTACGCCGGCAGGCGCGGGTAAC</td><td>>hg38_21_+_8433172_8446622_RNA45SN1_45S_50ntFlanks</td><td>5237</td><td>5285</td><td>49</td><td>ACTGACTGGCTCAGCGTGTGCCTACCCTACGCCGGCAGGCGCGGGTAAC</td><td>65.0</td><td>9.8</td><td>2.7</td><td>3.6</td><td>2.8E-04</td><td>8.1E-04</td><td>BT20/MB468</td><td>MP</td></tr>
<tr><td>>hg38_21_+_8433172_8446622_RNA45SN1_45S_50ntFlanks@10802.10820.19|TTGGAACAATGTAGGTAAG</td><td>>hg38_21_+_8433172_8446622_RNA45SN1_45S_50ntFlanks</td><td>10802</td><td>10820</td><td>19</td><td>TTGGAACAATGTAGGTAAG</td><td>119.2</td><td>-5.6</td><td>1.5</td><td>-3.6</td><td>2.8E-04</td><td>8.1E-04</td><td>BT20/MB468</td><td>MP</td></tr>
<tr><td>>hg38_21_+_8433172_8446622_RNA45SN1_45S_50ntFlanks@11666.11688.23|TCTGCCCAGTGCTCTGAATGTCA</td><td>>hg38_21_+_8433172_8446622_RNA45SN1_45S_50ntFlanks</td><td>11666</td><td>11688</td><td>23</td><td>TCTGCCCAGTGCTCTGAATGTCA</td><td>408.3</td><td>3.1</td><td>0.9</td><td>3.4</td><td>6.2E-04</td><td>8.1E-04</td><td>MB231/MB468</td><td>CYTO</td></tr>
<tr><td>>am_trnaMT_HisGTG_MT_+_12138_12206@18.37.20|AAAACATCAGATTGTGAATC</td><td>>am_trnaMT_HisGTG_MT_+_12138_12206</td><td>18</td><td>37</td><td>20</td><td>AAAACATCAGATTGTGAATC</td><td>68.0</td><td>9.8</td><td>2.7</td><td>3.6</td><td>2.8E-04</td><td>8.2E-04</td><td>BT20/MB468</td><td>MP</td></tr>
<tr><td>>hg38_21_+_8433172_8446622_RNA45SN1_45S_50ntFlanks@11666.11693.28|TCTGCCCAGTGCTCTGAATGTCAAAGTG</td><td>>hg38_21_+_8433172_8446622_RNA45SN1_45S_50ntFlanks</td><td>11666</td><td>11693</td><td>28</td><td>TCTGCCCAGTGCTCTGAATGTCAAAGTG</td><td>267.2</td><td>3.2</td><td>0.9</td><td>3.4</td><td>5.9E-04</td><td>8.2E-04</td><td>BT20/MB468</td><td>CYTO</td></tr>
<tr><td>>hg38_MT_+_1621_3279_RNR2_16S_50ntFlanks@1425.1451.27|ACGATTAAAGTCCTACGTGATCTGAGT</td><td>>hg38_MT_+_1621_3279_RNR2_16S_50ntFlanks</td><td>1425</td><td>1451</td><td>27</td><td>ACGATTAAAGTCCTACGTGATCTGAGT</td><td>202.0</td><td>4.7</td><td>1.4</td><td>3.5</td><td>5.5E-04</td><td>8.2E-04</td><td>MB231/MB468</td><td>MP</td></tr>
<tr><td>>hg38_21_+_8433172_8446622_RNA45SN1_45S_50ntFlanks@6773.6802.30|TTCCTCCCGGGGCTACGCCTGTCTGAGCGT</td><td>>hg38_21_+_8433172_8446622_RNA45SN1_45S_50ntFlanks</td><td>6773</td><td>6802</td><td>30</td><td>TTCCTCCCGGGGCTACGCCTGTCTGAGCGT</td><td>684.7</td><td>-2.7</td><td>0.8</td><td>-3.4</td><td>6.6E-04</td><td>8.2E-04</td><td>MB231/MB468</td><td>MITO</td></tr>
<tr><td>>hg38_21_+_8433172_8446622_RNA45SN1_45S_50ntFlanks@13005.13039.35|TCTATTGAAAGTCAGCCCTCGACACAAGGGTTTGT</td><td>>hg38_21_+_8433172_8446622_RNA45SN1_45S_50ntFlanks</td><td>13005</td><td>13039</td><td>35</td><td>TCTATTGAAAGTCAGCCCTCGACACAAGGGTTTGT</td><td>230.0</td><td>3.1</td><td>0.9</td><td>3.4</td><td>6.6E-04</td><td>8.2E-04</td><td>MB231/MB468</td><td>NUC</td></tr>
<tr><td>>hg38_21_+_8433172_8446622_RNA45SN1_45S_50ntFlanks@11577.11609.33|TTAGAACTGGTGCGGACCAGGGGAATCCGACTG</td><td>>hg38_21_+_8433172_8446622_RNA45SN1_45S_50ntFlanks</td><td>11577</td><td>11609</td><td>33</td><td>TTAGAACTGGTGCGGACCAGGGGAATCCGACTG</td><td>164.4</td><td>-1.8</td><td>0.5</td><td>-3.4</td><td>5.9E-04</td><td>8.2E-04</td><td>BT20/MB468</td><td>CYTO</td></tr>
<tr><td>>hg38_21_+_8433172_8446622_RNA45SN1_45S_50ntFlanks@10822.10847.26|GAAGTCGGCAAGCCGGATCCGTAACT</td><td>>hg38_21_+_8433172_8446622_RNA45SN1_45S_50ntFlanks</td><td>10822</td><td>10847</td><td>26</td><td>GAAGTCGGCAAGCCGGATCCGTAACT</td><td>262.0</td><td>1.7</td><td>0.5</td><td>3.4</td><td>6.2E-04</td><td>8.2E-04</td><td>MB231/MB468</td><td>CYTO</td></tr>
<tr><td>>hg38_21_+_8433172_8446622_RNA45SN1_45S_50ntFlanks@11534.11551.18|GTTCCGCGCGGCGCCTCG</td><td>>hg38_21_+_8433172_8446622_RNA45SN1_45S_50ntFlanks</td><td>11534</td><td>11551</td><td>18</td><td>GTTCCGCGCGGCGCCTCG</td><td>1338.1</td><td>-5.8</td><td>1.7</td><td>-3.4</td><td>5.9E-04</td><td>8.2E-04</td><td>BT20/MB468</td><td>CYTO</td></tr>
<tr><td>>hg38_21_+_8433172_8446622_RNA45SN1_45S_50ntFlanks@6661.6700.40|CGGTGGATCACTCGGCTCGTGCGTCGATGAAGAACGCAGC</td><td>>hg38_21_+_8433172_8446622_RNA45SN1_45S_50ntFlanks</td><td>6661</td><td>6700</td><td>40</td><td>CGGTGGATCACTCGGCTCGTGCGTCGATGAAGAACGCAGC</td><td>290.1</td><td>5.0</td><td>1.5</td><td>3.4</td><td>6.2E-04</td><td>8.2E-04</td><td>MB231/MB468</td><td>CYTO</td></tr>
<tr><td>>hg38_21_+_8433172_8446622_RNA45SN1_45S_50ntFlanks@10829.10855.27|GCAAGCCGGATCCGTAACTTCGGGATA</td><td>>hg38_21_+_8433172_8446622_RNA45SN1_45S_50ntFlanks</td><td>10829</td><td>10855</td><td>27</td><td>GCAAGCCGGATCCGTAACTTCGGGATA</td><td>218.8</td><td>-3.9</td><td>1.1</td><td>-3.4</td><td>5.9E-04</td><td>8.2E-04</td><td>BT20/MB468</td><td>CYTO</td></tr>
<tr><td>>hg38_21_+_8433172_8446622_RNA45SN1_45S_50ntFlanks@9549.9570.22|AGGTCCGTAGCGGTCCTGACGT</td><td>>hg38_21_+_8433172_8446622_RNA45SN1_45S_50ntFlanks</td><td>9549</td><td>9570</td><td>22</td><td>AGGTCCGTAGCGGTCCTGACGT</td><td>324.2</td><td>3.8</td><td>1.0</td><td>3.9</td><td>9.8E-05</td><td>8.2E-04</td><td>BT20/MB231</td><td>MP</td></tr>
<tr><td>>hg38_21_+_8433172_8446622_RNA45SN1_45S_50ntFlanks@3705.3736.32|TACCTGGTTGATCCTGCCAGTAGCATATGCTT</td><td>>hg38_21_+_8433172_8446622_RNA45SN1_45S_50ntFlanks</td><td>3705</td><td>3736</td><td>32</td><td>TACCTGGTTGATCCTGCCAGTAGCATATGCTT</td><td>1130.6</td><td>3.3</td><td>1.0</td><td>3.4</td><td>6.6E-04</td><td>8.2E-04</td><td>MB231/MB468</td><td>MITO</td></tr>
<tr><td>>MI0000469|hsa-mir-125a&WithFlank&19|+|51693248|51693345@21.43.23&[MIMAT0000443&hsa-miR-125a-5p&offsets|0|-1;m-63&19|+|51693268|51693289&offsets|0|+1]|TCCCTGAGACCCTTTAACCTGTG</td><td>>MI0000469|hsa-mir-125a&WithFlank&19|+|51693248|51693345</td><td>21</td><td>43</td><td>23</td><td>TCCCTGAGACCCTTTAACCTGTG</td><td>3219.2</td><td>3.2</td><td>0.9</td><td>3.6</td><td>2.8E-04</td><td>8.2E-04</td><td>BT20/MB468</td><td>MP</td></tr>
<tr><td>>hg38_21_+_8433172_8446622_RNA45SN1_45S_50ntFlanks@7979.7996.18|ACCTCAGATCAGACGTGG</td><td>>hg38_21_+_8433172_8446622_RNA45SN1_45S_50ntFlanks</td><td>7979</td><td>7996</td><td>18</td><td>ACCTCAGATCAGACGTGG</td><td>178.4</td><td>-5.7</td><td>1.6</td><td>-3.6</td><td>2.8E-04</td><td>8.2E-04</td><td>BT20/MB468</td><td>MP</td></tr>
<tr><td>>hg38_21_+_8433172_8446622_RNA45SN1_45S_50ntFlanks@8915.8936.22|CGAATCCCGGGGCCGAGGGAGC</td><td>>hg38_21_+_8433172_8446622_RNA45SN1_45S_50ntFlanks</td><td>8915</td><td>8936</td><td>22</td><td>CGAATCCCGGGGCCGAGGGAGC</td><td>81.4</td><td>6.0</td><td>1.6</td><td>3.6</td><td>2.8E-04</td><td>8.2E-04</td><td>BT20/MB468</td><td>MP</td></tr>
<tr><td>>hg38_1_-_228634819_228635039_RNA5S12_5S_50ntFlanks@149.169.21|GAATACCGGGTGCTGTAGGCT</td><td>>hg38_1_-_228634819_228635039_RNA5S12_5S_50ntFlanks</td><td>149</td><td>169</td><td>21</td><td>GAATACCGGGTGCTGTAGGCT</td><td>249.9</td><td>-2.7</td><td>0.8</td><td>-3.4</td><td>6.6E-04</td><td>8.2E-04</td><td>MB231/MB468</td><td>MITO</td></tr>
<tr><td>>hg38_21_+_8433172_8446622_RNA45SN1_45S_50ntFlanks@11580.11609.30|GAACTGGTGCGGACCAGGGGAATCCGACTG</td><td>>hg38_21_+_8433172_8446622_RNA45SN1_45S_50ntFlanks</td><td>11580</td><td>11609</td><td>30</td><td>GAACTGGTGCGGACCAGGGGAATCCGACTG</td><td>579.5</td><td>-2.1</td><td>0.6</td><td>-3.4</td><td>6.7E-04</td><td>8.2E-04</td><td>MB231/MB468</td><td>NUC</td></tr>
<tr><td>>hg38_21_+_8433172_8446622_RNA45SN1_45S_50ntFlanks@12605.12623.19|TATGTGCTTGGCTGAGGAG</td><td>>hg38_21_+_8433172_8446622_RNA45SN1_45S_50ntFlanks</td><td>12605</td><td>12623</td><td>19</td><td>TATGTGCTTGGCTGAGGAG</td><td>491.8</td><td>-3.5</td><td>1.0</td><td>-3.4</td><td>6.3E-04</td><td>8.2E-04</td><td>MB231/MB468</td><td>CYTO</td></tr>
<tr><td>>hg38_21_+_8433172_8446622_RNA45SN1_45S_50ntFlanks@11572.11605.34|CCGACTTAGAACTGGTGCGGACCAGGGGAATCCG</td><td>>hg38_21_+_8433172_8446622_RNA45SN1_45S_50ntFlanks</td><td>11572</td><td>11605</td><td>34</td><td>CCGACTTAGAACTGGTGCGGACCAGGGGAATCCG</td><td>301.0</td><td>7.5</td><td>1.9</td><td>3.9</td><td>9.9E-05</td><td>8.2E-04</td><td>BT20/MB231</td><td>MP</td></tr>
<tr><td>>hg38_21_+_8433172_8446622_RNA45SN1_45S_50ntFlanks@9837.9854.18|TGGTAAGCAGAACTGGCG</td><td>>hg38_21_+_8433172_8446622_RNA45SN1_45S_50ntFlanks</td><td>9837</td><td>9854</td><td>18</td><td>TGGTAAGCAGAACTGGCG</td><td>141.4</td><td>10.5</td><td>2.7</td><td>3.9</td><td>9.9E-05</td><td>8.2E-04</td><td>BT20/MB231</td><td>MP</td></tr>
<tr><td>>hg38_21_+_8433172_8446622_RNA45SN1_45S_50ntFlanks@5485.5514.30|AGCGCTGAGAAGACGGTCGAACTTGACTAT</td><td>>hg38_21_+_8433172_8446622_RNA45SN1_45S_50ntFlanks</td><td>5485</td><td>5514</td><td>30</td><td>AGCGCTGAGAAGACGGTCGAACTTGACTAT</td><td>5077.3</td><td>-2.8</td><td>0.8</td><td>-3.4</td><td>6.0E-04</td><td>8.3E-04</td><td>BT20/MB468</td><td>CYTO</td></tr>
</table>

<table>
<tr><td></td><td></td><td></td><td></td><td></td><td></td><td></td><td></td><td></td><td></td><td></td><td></td><td></td><td></td></tr>
<tr><td>>hg38_21_+_8433172_8446622_RNA45SN1_45S_50ntFlanks@11666.11701.36|TCTGCCCAGTGCTCTGAATGTCAAAGTGAAGAAATT</td><td>>hg38_21_+_8433172_8446622_RNA45SN1_45S_50ntFlanks</td><td>11666</td><td>11701</td><td>36</td><td>TCTGCCCAGTGCTCTGAATGTCAAAGTGAAGAAATT</td><td>477.4</td><td>-4.7</td><td>1.4</td><td>-3.4</td><td>6.7E-04</td><td>8.3E-04</td><td>BT20/MB468</td><td>NUC</td></tr>
<tr><td>>hg38_21_+_8433172_8446622_RNA45SN1_45S_50ntFlanks@4567.4589.23|TAATGGAATAGGACCGCGGTTCT</td><td>>hg38_21_+_8433172_8446622_RNA45SN1_45S_50ntFlanks</td><td>4567</td><td>4589</td><td>23</td><td>TAATGGAATAGGACCGCGGTTCT</td><td>272.2</td><td>-3.5</td><td>1.0</td><td>-3.4</td><td>6.0E-04</td><td>8.3E-04</td><td>BT20/MB468</td><td>CYTO</td></tr>
<tr><td>>trna10_AspGTC_12_-_125424193_125424264@37.71.35,trna12_AspGTC_12_-_125411891_125411962@37.71.35,trna144_AspGTC_6_-_27551236_27551307@37.71.35,trna38_AspGTC_17_-_8125556_8125627@37.71.35,trna45_AspGTC_6_+_27447453_27447524@37.71.35,trna48_AspGTC_6_+_27471523_27471594@37.71.35,trna4_AspGTC_12_+_96429799_96429870@37.71.35,trna69_AspGTC_1_-_161440205_161440276@37.71.35,trna72_AspGTC_1_-_161432824_161432895@37.71.35,trna75_AspGTC_1_-_161425414_161425485@37.71.35,trna78_AspGTC_1_-_161418033_161418104@37.71.35,trna81_AspGTC_1_-_161410615_161410686@37.71.35|ACGCGGGAGACCGGGGTTCGATTCCCCGACGGGGA</td><td>>trna10_AspGTC_12_-_125424193_125424264</td><td>37</td><td>71</td><td>35</td><td>ACGCGGGAGACCGGGGTTCGATTCCCCGACGGGGA</td><td>504.6</td><td>-3.0</td><td>0.9</td><td>-3.5</td><td>5.6E-04</td><td>8.3E-04</td><td>MB231/MB468</td><td>MP</td></tr>
<tr><td>>MI0000103|hsa-mir-101-1&WithFlank&1|-|65058428|65058514@53.72.20;MI0000739|hsa-mir-101-2&WithFlank&9|+|4850291|4850381@55.74.20&[MIMAT0000099&hsa-miR-101-3p&offsets|0|-1;m-17&1|-|65058443|65058462&offsets|0|0];[MIMAT0000099_1&hsa-miR-101-3p&offsets|0|-1;m-16&9|+|4850345|4850364&offsets|0|0]|TACAGTACTGTGATAACTGA</td><td>>MI0000103|hsa-mir-101-1&WithFlank&1|-|65058428|65058514</td><td>53</td><td>72</td><td>20</td><td>TACAGTACTGTGATAACTGA</td><td>1764.9</td><td>2.4</td><td>0.7</td><td>3.5</td><td>5.6E-04</td><td>8.3E-04</td><td>MB231/MB468</td><td>MP</td></tr>
<tr><td>>hg38_21_+_8433172_8446622_RNA45SN1_45S_50ntFlanks@10392.10420.29|ACAGCAGTTGAACATGGGTCAGTCGGTCC</td><td>>hg38_21_+_8433172_8446622_RNA45SN1_45S_50ntFlanks</td><td>10392</td><td>10420</td><td>29</td><td>ACAGCAGTTGAACATGGGTCAGTCGGTCC</td><td>369.5</td><td>-5.1</td><td>1.5</td><td>-3.4</td><td>6.0E-04</td><td>8.3E-04</td><td>BT20/MB468</td><td>CYTO</td></tr>
<tr><td>>hg38_21_+_8433172_8446622_RNA45SN1_45S_50ntFlanks@3782.3812.31|CAGTGAAACTGCGAATGGCTCATTAAATCAG</td><td>>hg38_21_+_8433172_8446622_RNA45SN1_45S_50ntFlanks</td><td>3782</td><td>3812</td><td>31</td><td>CAGTGAAACTGCGAATGGCTCATTAAATCAG</td><td>240.2</td><td>-6.3</td><td>1.7</td><td>-3.6</td><td>2.8E-04</td><td>8.3E-04</td><td>BT20/MB468</td><td>MP</td></tr>
<tr><td>>hg38_21_+_8433172_8446622_RNA45SN1_45S_50ntFlanks@12641.12659.19|CCATCTGTGGGATTATGAC</td><td>>hg38_21_+_8433172_8446622_RNA45SN1_45S_50ntFlanks</td><td>12641</td><td>12659</td><td>19</td><td>CCATCTGTGGGATTATGAC</td><td>1313.2</td><td>-4.1</td><td>1.1</td><td>-3.6</td><td>2.8E-04</td><td>8.3E-04</td><td>BT20/MB468</td><td>MP</td></tr>
<tr><td>>hg38_21_+_8433172_8446622_RNA45SN1_45S_50ntFlanks@12418.12445.28|CGATGTCGGCTCTTCCTATCATTGTGAA</td><td>>hg38_21_+_8433172_8446622_RNA45SN1_45S_50ntFlanks</td><td>12418</td><td>12445</td><td>28</td><td>CGATGTCGGCTCTTCCTATCATTGTGAA</td><td>191.4</td><td>-5.3</td><td>1.5</td><td>-3.5</td><td>5.6E-04</td><td>8.3E-04</td><td>MB231/MB468</td><td>MP</td></tr>
<tr><td>>MI0000299|hsa-mir-222&WithFlank&X|-|45747009|45747130@75.95.21(+1U)&[MIMAT0000279&hsa-miR-222-3p&offsets|0|0(+1U);m-140&X|-|45747033|45747056&offsets|0|-3(+1U)]|AGCTACATCTGGCTACTGGGTT</td><td>>MI0000299|hsa-mir-222&WithFlank&X|-|45747009|45747130</td><td>75</td><td>95</td><td>21(+1U)</td><td>AGCTACATCTGGCTACTGGGTT</td><td>452.7</td><td>4.5</td><td>1.3</td><td>3.5</td><td>5.6E-04</td><td>8.3E-04</td><td>MB231/MB468</td><td>MP</td></tr>
<tr><td>>hg38_21_+_8433172_8446622_RNA45SN1_45S_50ntFlanks@10346.10370.25|TCAAACGAGAACTTTGAAGGCCGAA</td><td>>hg38_21_+_8433172_8446622_RNA45SN1_45S_50ntFlanks</td><td>10346</td><td>10370</td><td>25</td><td>TCAAACGAGAACTTTGAAGGCCGAA</td><td>197.6</td><td>-5.5</td><td>1.6</td><td>-3.5</td><td>5.6E-04</td><td>8.3E-04</td><td>MB231/MB468</td><td>MP</td></tr>
<tr><td>>MI0000736|hsa-mir-30c-1&WithFlank&1|+|40757278|40757378@23.42.20;MI0000254|hsa-mir-30c-2&WithFlank&6|-|71376954|71377037@13.32.20&[MIMAT0000244&hsa-miR-30c-5p&offsets|0|-3;m-57&1|+|40757300|40757323&offsets|0|-4];[MIMAT0000244_1&hsa-miR-30c-5p&offsets|0|-3;m-56&6|-|71377002|71377025&offsets|0|-4]|TGTAAACATCCTACACTCTC</td><td>>MI0000736|hsa-mir-30c-1&WithFlank&1|+|40757278|40757378</td><td>23</td><td>42</td><td>20</td><td>TGTAAACATCCTACACTCTC</td><td>282.9</td><td>3.7</td><td>1.0</td><td>3.6</td><td>2.8E-04</td><td>8.3E-04</td><td>BT20/MB468</td><td>MP</td></tr>
<tr><td>>hg38_21_+_8433172_8446622_RNA45SN1_45S_50ntFlanks@10351.10372.22|CGAGAACTTTGAAGGCCGAAGT</td><td>>hg38_21_+_8433172_8446622_RNA45SN1_45S_50ntFlanks</td><td>10351</td><td>10372</td><td>22</td><td>CGAGAACTTTGAAGGCCGAAGT</td><td>426.9</td><td>-1.9</td><td>0.6</td><td>-3.4</td><td>6.3E-04</td><td>8.3E-04</td><td>MB231/MB468</td><td>CYTO</td></tr>
<tr><td>>hg38_21_+_8433172_8446622_RNA45SN1_45S_50ntFlanks@6650.6677.28|ACGACTCTTAGCGGTGGATCACTCGGCT</td><td>>hg38_21_+_8433172_8446622_RNA45SN1_45S_50ntFlanks</td><td>6650</td><td>6677</td><td>28</td><td>ACGACTCTTAGCGGTGGATCACTCGGCT</td><td>896.8</td><td>2.9</td><td>0.8</td><td>3.4</td><td>6.8E-04</td><td>8.3E-04</td><td>MB231/MB468</td><td>NUC</td></tr>
<tr><td>>hg38_21_+_8433172_8446622_RNA45SN1_45S_50ntFlanks@6660.6689.30|GCGGTGGATCACTCGGCTCGTGCGTCGATG</td><td>>hg38_21_+_8433172_8446622_RNA45SN1_45S_50ntFlanks</td><td>6660</td><td>6689</td><td>30</td><td>GCGGTGGATCACTCGGCTCGTGCGTCGATG</td><td>558.2</td><td>-2.0</td><td>0.6</td><td>-3.4</td><td>6.8E-04</td><td>8.3E-04</td><td>MB231/MB468</td><td>NUC</td></tr>
<tr><td>>hg38_21_+_8433172_8446622_RNA45SN1_45S_50ntFlanks@9740.9765.26|AACGATCTCAACCTATTCTCAAACTT</td><td>>hg38_21_+_8433172_8446622_RNA45SN1_45S_50ntFlanks</td><td>9740</td><td>9765</td><td>26</td><td>AACGATCTCAACCTATTCTCAAACTT</td><td>1558.7</td><td>-2.7</td><td>0.8</td><td>-3.5</td><td>5.0E-04</td><td>8.3E-04</td><td>BT20/MB468</td><td>MITO</td></tr>
<tr><td>>hg38_21_+_8433172_8446622_RNA45SN1_45S_50ntFlanks@9604.9621.18|AAAGACTAATCGAACCAT</td><td>>hg38_21_+_8433172_8446622_RNA45SN1_45S_50ntFlanks</td><td>9604</td><td>9621</td><td>18</td><td>AAAGACTAATCGAACCAT</td><td>188.6</td><td>-3.6</td><td>1.0</td><td>-3.5</td><td>5.0E-04</td><td>8.3E-04</td><td>BT20/MB468</td><td>MITO</td></tr>
<tr><td>>MI0000076|hsa-mir-20a&WithFlank&13|+|91351059|91351141@14.36.23&[MIMAT0000075&hsa-miR-20a-5p&offsets|0|0;m-96&13|+|91351072|91351094&offsets|0|0]|TAAAGTGCTTATAGTGCAGGTAG</td><td>>MI0000076|hsa-mir-20a&WithFlank&13|+|91351059|91351141</td><td>14</td><td>36</td><td>23</td><td>TAAAGTGCTTATAGTGCAGGTAG</td><td>2904.7</td><td>3.6</td><td>1.0</td><td>3.4</td><td>6.3E-04</td><td>8.3E-04</td><td>MB231/MB468</td><td>CYTO</td></tr>
<tr><td>>hg38_21_+_8433172_8446622_RNA45SN1_45S_50ntFlanks@12822.12848.27|CCGGGACCGGGGTCCGGTGCGGAGTGC</td><td>>hg38_21_+_8433172_8446622_RNA45SN1_45S_50ntFlanks</td><td>12822</td><td>12848</td><td>27</td><td>CCGGGACCGGGGTCCGGTGCGGAGTGC</td><td>72.2</td><td>-9.3</td><td>2.6</td><td>-3.6</td><td>2.9E-04</td><td>8.3E-04</td><td>BT20/MB468</td><td>MP</td></tr>
<tr><td>>hg38_21_+_8433172_8446622_RNA45SN1_45S_50ntFlanks@3840.3873.34|CTCCTACTTGGATAACTGTGGTAATTCTAGAGCT</td><td>>hg38_21_+_8433172_8446622_RNA45SN1_45S_50ntFlanks</td><td>3840</td><td>3873</td><td>34</td><td>CTCCTACTTGGATAACTGTGGTAATTCTAGAGCT</td><td>99.7</td><td>-4.8</td><td>1.4</td><td>-3.4</td><td>5.6E-04</td><td>8.3E-04</td><td>MB231/MB468</td><td>MP</td></tr>
<tr><td>>hg38_21_+_8433172_8446622_RNA45SN1_45S_50ntFlanks@12003.12029.27|GCGGGCCGCCGGTGAAATACCACTACT</td><td>>hg38_21_+_8433172_8446622_RNA45SN1_45S_50ntFlanks</td><td>12003</td><td>12029</td><td>27</td><td>GCGGGCCGCCGGTGAAATACCACTACT</td><td>98.3</td><td>-5.5</td><td>1.5</td><td>-3.6</td><td>2.9E-04</td><td>8.3E-04</td><td>BT20/MB468</td><td>MP</td></tr>
<tr><td>>MI0000102|hsa-mir-100&WithFlank&11|-|122152223|122152314@19.39.21&[MIMAT0000098&hsa-miR-100-5p&offsets|0|-1;m-27&11|-|122152276|122152296&offsets|0|0]|AACCCGTAGATCCGAACTTGT</td><td>>MI0000102|hsa-mir-100&WithFlank&11|-|122152223|122152314</td><td>19</td><td>39</td><td>21</td><td>AACCCGTAGATCCGAACTTGT</td><td>1452.8</td><td>7.7</td><td>2.2</td><td>3.5</td><td>5.0E-04</td><td>8.3E-04</td><td>BT20/MB468</td><td>MITO</td></tr>
<tr><td>>trnaMT_GluTTC_MT_-_14674_14742@40.57.18,trnalookalike8_GluTTC_5_-_93905172_93905240@40.57.18|TTGGTCGTGGTTGTAGTC</td><td>>trnaMT_GluTTC_MT_-_14674_14742</td><td>40</td><td>57</td><td>18</td><td>TTGGTCGTGGTTGTAGTC</td><td>159.2</td><td>4.2</td><td>1.2</td><td>3.4</td><td>6.7E-04</td><td>8.3E-04</td><td>MB231/MB468</td><td>MITO</td></tr>
<tr><td>>hg38_21_+_8433172_8446622_RNA45SN1_45S_50ntFlanks@6648.6674.27|GTACGACTCTTAGCGGTGGATCACTCG</td><td>>hg38_21_+_8433172_8446622_RNA45SN1_45S_50ntFlanks</td><td>6648</td><td>6674</td><td>27</td><td>GTACGACTCTTAGCGGTGGATCACTCG</td><td>229.8</td><td>3.6</td><td>1.0</td><td>3.6</td><td>2.9E-04</td><td>8.3E-04</td><td>BT20/MB468</td><td>MP</td></tr>
<tr><td>>hg38_21_+_8433172_8446622_RNA45SN1_45S_50ntFlanks@12045.12071.27|CTGACCCGGTGAGGCGGGGGGGCGAGC</td><td>>hg38_21_+_8433172_8446622_RNA45SN1_45S_50ntFlanks</td><td>12045</td><td>12071</td><td>27</td><td>CTGACCCGGTGAGGCGGGGGGGCGAGC</td><td>273.0</td><td>-4.1</td><td>1.2</td><td>-3.5</td><td>5.1E-04</td><td>8.3E-04</td><td>BT20/MB468</td><td>MITO</td></tr>
<tr><td>>hg38_21_+_8433172_8446622_RNA45SN1_45S_50ntFlanks@6668.6686.19|TCACTCGGCTCGTGCGTCG</td><td>>hg38_21_+_8433172_8446622_RNA45SN1_45S_50ntFlanks</td><td>6668</td><td>6686</td><td>19</td><td>TCACTCGGCTCGTGCGTCG</td><td>99.7</td><td>-4.6</td><td>1.3</td><td>-3.5</td><td>5.1E-04</td><td>8.3E-04</td><td>BT20/MB468</td><td>MITO</td></tr>
<tr><td>>hg38_21_+_8433172_8446622_RNA45SN1_45S_50ntFlanks@10782.10818.37|GGTGAACAGCCTCTGGCATGTTGGAACAATGTAGGTA</td><td>>hg38_21_+_8433172_8446622_RNA45SN1_45S_50ntFlanks</td><td>10782</td><td>10818</td><td>37</td><td>GGTGAACAGCCTCTGGCATGTTGGAACAATGTAGGTA</td><td>369.4</td><td>-4.1</td><td>1.2</td><td>-3.4</td><td>6.1E-04</td><td>8.3E-04</td><td>BT20/MB468</td><td>CYTO</td></tr>
<tr><td>>hg38_21_+_8433172_8446622_RNA45SN1_45S_50ntFlanks@9313.9338.26|TCTAACACGTGCGCGAGTCGGGGGCT</td><td>>hg38_21_+_8433172_8446622_RNA45SN1_45S_50ntFlanks</td><td>9313</td><td>9338</td><td>26</td><td>TCTAACACGTGCGCGAGTCGGGGGCT</td><td>421.8</td><td>-2.5</td><td>0.7</td><td>-3.5</td><td>5.1E-04</td><td>8.3E-04</td><td>BT20/MB468</td><td>MITO</td></tr>
<tr><td>>hg38_21_+_8433172_8446622_RNA45SN1_45S_50ntFlanks@6650.6667.18|ACGACTCTTAGCGGTGGA</td><td>>hg38_21_+_8433172_8446622_RNA45SN1_45S_50ntFlanks</td><td>6650</td><td>6667</td><td>18</td><td>ACGACTCTTAGCGGTGGA</td><td>338.1</td><td>3.1</td><td>0.8</td><td>3.9</td><td>1.0E-04</td><td>8.3E-04</td><td>BT20/MB231</td><td>CYTO</td></tr>
<tr><td>>hg38_21_+_8433172_8446622_RNA45SN1_45S_50ntFlanks@6652.6692.41|GACTCTTAGCGGTGGATCACTCGGCTCGTGCGTCGATGAAG</td><td>>hg38_21_+_8433172_8446622_RNA45SN1_45S_50ntFlanks</td><td>6652</td><td>6692</td><td>41</td><td>GACTCTTAGCGGTGGATCACTCGGCTCGTGCGTCGATGAAG</td><td>6122.1</td><td>2.1</td><td>0.6</td><td>3.5</td><td>5.1E-04</td><td>8.3E-04</td><td>BT20/MB468</td><td>MITO</td></tr>
<tr><td>>hg38_21_+_8433172_8446622_RNA45SN1_45S_50ntFlanks@11579.11602.24|AGAACTGGTGCGGACCAGGGGAAT</td><td>>hg38_21_+_8433172_8446622_RNA45SN1_45S_50ntFlanks</td><td>11579</td><td>11602</td><td>24</td><td>AGAACTGGTGCGGACCAGGGGAAT</td><td>324.4</td><td>-4.0</td><td>1.2</td><td>-3.4</td><td>6.8E-04</td><td>8.3E-04</td><td>MB231/MB468</td><td>NUC</td></tr>
<tr><td>>hg38_21_+_8433172_8446622_RNA45SN1_45S_50ntFlanks@11728.11745.18|GTAACTATGACTCTCTTA</td><td>>hg38_21_+_8433172_8446622_RNA45SN1_45S_50ntFlanks</td><td>11728</td><td>11745</td><td>18</td><td>GTAACTATGACTCTCTTA</td><td>770.3</td><td>-6.4</td><td>1.9</td><td>-3.4</td><td>6.8E-04</td><td>8.3E-04</td><td>BT20/MB468</td><td>NUC</td></tr>
<tr><td>>hg38_21_+_8433172_8446622_RNA45SN1_45S_50ntFlanks@6698.6721.24|AGCTAGCTGCGAGAATTAATGTGA</td><td>>hg38_21_+_8433172_8446622_RNA45SN1_45S_50ntFlanks</td><td>6698</td><td>6721</td><td>24</td><td>AGCTAGCTGCGAGAATTAATGTGA</td><td>293.2</td><td>-4.2</td><td>1.2</td><td>-3.4</td><td>5.7E-04</td><td>8.4E-04</td><td>MB231/MB468</td><td>MP</td></tr>
<tr><td>>hg38_21_+_8433172_8446622_RNA45SN1_45S_50ntFlanks@10808.10831.24|CAATGTAGGTAAGGGAAGTCGGCA</td><td>>hg38_21_+_8433172_8446622_RNA45SN1_45S_50ntFlanks</td><td>10808</td><td>10831</td><td>24</td><td>CAATGTAGGTAAGGGAAGTCGGCA</td><td>196.3</td><td>-2.9</td><td>0.8</td><td>-3.6</td><td>2.9E-04</td><td>8.4E-04</td><td>BT20/MB468</td><td>MP</td></tr>
<tr><td>>hg38_21_+_8433172_8446622_RNA45SN1_45S_50ntFlanks@6653.6695.43|ACTCTTAGCGGTGGATCACTCGGCTCGTGCGTCGATGAAGAAC</td><td>>hg38_21_+_8433172_8446622_RNA45SN1_45S_50ntFlanks</td><td>6653</td><td>6695</td><td>43</td><td>ACTCTTAGCGGTGGATCACTCGGCTCGTGCGTCGATGAAGAAC</td><td>84.7</td><td>6.8</td><td>1.9</td><td>3.6</td><td>2.9E-04</td><td>8.4E-04</td><td>BT20/MB468</td><td>MP</td></tr>
<tr><td>>hg38_21_+_8433172_8446622_RNA45SN1_45S_50ntFlanks@12436.12473.38|TCATTGTGAAGCAGAATTCACCAAGCGTTGGATTGTTC</td><td>>hg38_21_+_8433172_8446622_RNA45SN1_45S_50ntFlanks</td><td>12436</td><td>12473</td><td>38</td><td>TCATTGTGAAGCAGAATTCACCAAGCGTTGGATTGTTC</td><td>239.5</td><td>-5.0</td><td>1.4</td><td>-3.6</td><td>2.9E-04</td><td>8.4E-04</td><td>BT20/MB468</td><td>MP</td></tr>
<tr><td>>MI0000289|hsa-mir-181a-1&WithFlank&1|-|198859038|198859159@70.91.22&[MIMAT0000270&hsa-miR-181a-3p&offsets|0|0;m-125&1|-|198859069|198859090&offsets|0|0]|ACCATCGACCGTTGATTGTACC</td><td>>MI0000289|hsa-mir-181a-1&WithFlank&1|-|198859038|198859159</td><td>70</td><td>91</td><td>22</td><td>ACCATCGACCGTTGATTGTACC</td><td>1921.0</td><td>-4.1</td><td>1.0</td><td>-3.9</td><td>1.1E-04</td><td>8.4E-04</td><td>BT20/MB231</td><td>CYTO</td></tr>
<tr><td>>trnaMT_GluTTC_MT_-_14674_14742@33.72.40,trnalookalike8_GluTTC_5_-_93905172_93905240@33.72.40|CATATCATTGGTCGTGGTTGTAGTCCGTGCGAGAATACCA</td><td>>trnaMT_GluTTC_MT_-_14674_14742</td><td>33</td><td>72</td><td>40</td><td>CATATCATTGGTCGTGGTTGTAGTCCGTGCGAGAATACCA</td><td>585.9</td><td>2.5</td><td>0.7</td><td>3.4</td><td>6.7E-04</td><td>8.4E-04</td><td>MB231/MB468</td><td>MITO</td></tr>
<tr><td>>MI0000076|hsa-mir-20a&WithFlank&13|+|91351059|91351141@14.33.20(+1C)&[MIMAT0000075&hsa-miR-20a-5p&offsets|0|-3(+1C);m-96&13|+|91351072|91351094&offsets|0|-3(+1C)]|TAAAGTGCTTATAGTGCAGGC</td><td>>MI0000076|hsa-mir-20a&WithFlank&13|+|91351059|91351141</td><td>14</td><td>33</td><td>20(+1C)</td><td>TAAAGTGCTTATAGTGCAGGC</td><td>554.3</td><td>-5.4</td><td>1.4</td><td>-3.9</td><td>1.0E-04</td><td>8.4E-04</td><td>BT20/MB231</td><td>MP</td></tr>
<tr><td>>hg38_21_+_8433172_8446622_RNA45SN1_45S_50ntFlanks@11783.11802.20|CATGAATGGATGAACGAGAT</td><td>>hg38_21_+_8433172_8446622_RNA45SN1_45S_50ntFlanks</td><td>11783</td><td>11802</td><td>20</td><td>CATGAATGGATGAACGAGAT</td><td>514.7</td><td>-1.7</td><td>0.5</td><td>-3.4</td><td>6.8E-04</td><td>8.4E-04</td><td>MB231/MB468</td><td>NUC</td></tr>
<tr><td>>am_trna10_ValCAC_5_-_180649395_180649467@5.29.25,trna12_ValAAC_5_-_180645270_180645342@5.29.25,trna132_ValAAC_6_-_27721179_27721251@5.29.25,trna136_ValAAC_6_-_27648885_27648957@5.29.25,trna139_ValAAC_6_-_27618707_27618779@5.29.25,trna18_ValCAC_5_-_180529253_180529325@5.29.25,trna2_ValAAC_3_+_169490018_169490090@5.29.25,trna2_ValCAC_5_+_180524070_180524142@5.29.25,trna4_ValAAC_5_+_180591154_180591226@5.29.25,trna5_ValAAC_5_+_180596610_180596682@5.29.25,trna6_ValCAC_5_+_180600650_180600722@5.29.25,trna85_ValCAC_1_-_161369490_161369562@5.29.25,trna90_ValCAC_1_-_149684088_149684161@5.29.25,trna98_ValCAC_1_-_149298555_149298627@5.29.25,trna9_ValCAC_6_+_26538282_26538354@5.29.25|CCGTAGTGTAGTGGTTATCACGTTC</td><td>>am_trna10_ValCAC_5_-_180649395_180649467</td><td>5</td><td>29</td><td>25</td><td>CCGTAGTGTAGTGGTTATCACGTTC</td><td>143.3</td><td>-5.5</td><td>1.6</td><td>-3.4</td><td>5.7E-04</td><td>8.4E-04</td><td>MB231/MB468</td><td>MP</td></tr>
<tr><td>>hg38_21_+_8433172_8446622_RNA45SN1_45S_50ntFlanks@11668.11701.34|TGCCCAGTGCTCTGAATGTCAAAGTGAAGAAATT</td><td>>hg38_21_+_8433172_8446622_RNA45SN1_45S_50ntFlanks</td><td>11668</td><td>11701</td><td>34</td><td>TGCCCAGTGCTCTGAATGTCAAAGTGAAGAAATT</td><td>594.0</td><td>2.4</td><td>0.7</td><td>3.4</td><td>6.4E-04</td><td>8.4E-04</td><td>MB231/MB468</td><td>CYTO</td></tr>
</table>

| | | | | | | | | | | | | | |
|---|---|---|---|---|---|---|---|---|---|---|---|---|---|
| >MI0000342\|hsa-mir-200b&WithFlank&1\|+\|1167098\|1167204@63.84.22(+1U)&[MIMAT0000318&hsa-miR-200b-3p&offsets\|0\|0(+1U);m-93&1\|+\|1167160\|1167182&offsets\|0\|-1(+1U)]\|TAATACTGCCTGGTAATGATGAT | >MI0000342\|hsa-mir-200b&WithFlank&1\|+\|1167098\|1167204 | 63 | 84 | 22(+1U) | TAATACTGCCTGGTAATGATGAT | 364.0 | -4.4 | 1.3 | -3.4 | 5.7E-04 | 8.4E-04 | MB231/MB468 | MP |
| >hg38_21_+_8433172_8446622_RNA45SN1_45S_50ntFlanks@10686.10711.26\|GCGGCGTCCGGTGAGCTCTCGCTGGC | >hg38_21_+_8433172_8446622_RNA45SN1_45S_50ntFlanks | 10686 | 10711 | 26 | GCGGCGTCCGGTGAGCTCTCGCTGGC | 382.0 | -2.7 | 0.8 | -3.6 | 2.9E-04 | 8.4E-04 | BT20/MB468 | MP |
| >hg38_21_+_8433172_8446622_RNA45SN1_45S_50ntFlanks@9791.9816.26\|TGGCGTGGAGCCGGGCGTGGAATGCG | >hg38_21_+_8433172_8446622_RNA45SN1_45S_50ntFlanks | 9791 | 9816 | 26 | TGGCGTGGAGCCGGGCGTGGAATGCG | 197.6 | 7.3 | 2.0 | 3.6 | 2.9E-04 | 8.4E-04 | BT20/MB468 | MP |
| >hg38_21_+_8433172_8446622_RNA45SN1_45S_50ntFlanks@6761.6806.46\|TGCGGCCCCGGGTTCCTCCCGGGGCTACGCCTGTCTGAGCGTCGCT | >hg38_21_+_8433172_8446622_RNA45SN1_45S_50ntFlanks | 6761 | 6806 | 46 | TGCGGCCCCGGGTTCCTCCCGGGGCTACGCCTGTCTGAGCGTCGCT | 339.2 | 8.7 | 2.5 | 3.4 | 6.1E-04 | 8.4E-04 | BT20/MB468 | CYTO |
| >hg38_21_+_8433172_8446622_RNA45SN1_45S_50ntFlanks@10059.10077.19\|GGCAGTCGAGAGTGGACGG | >hg38_21_+_8433172_8446622_RNA45SN1_45S_50ntFlanks | 10059 | 10077 | 19 | GGCAGTCGAGAGTGGACGG | 353.8 | 5.9 | 1.5 | 3.9 | 1.0E-04 | 8.4E-04 | BT20/MB231 | MP |
| >hg38_MT_+_598_1651_RNR1_12S_50ntFlanks@447.468.22\|TGAACACACAATAGCTAAGACC | >hg38_MT_+_598_1651_RNR1_12S_50ntFlanks | 447 | 468 | 22 | TGAACACACAATAGCTAAGACC | 64.9 | 9.8 | 2.7 | 3.6 | 2.9E-04 | 8.4E-04 | BT20/MB468 | MP |
| >hg38_21_+_8433172_8446622_RNA45SN1_45S_50ntFlanks@4685.4722.38\|AGACGGACCAGAGCGAAAGCATTTGCCAAGAATGTTTT | >hg38_21_+_8433172_8446622_RNA45SN1_45S_50ntFlanks | 4685 | 4722 | 38 | AGACGGACCAGAGCGAAAGCATTTGCCAAGAATGTTTT | 146.9 | -2.5 | 0.7 | -3.4 | 6.8E-04 | 8.4E-04 | MB231/MB468 | NUC |
| >hg38_21_+_8433172_8446622_RNA45SN1_45S_50ntFlanks@5077.5108.32\|CCGATAACGAACGAGACTCTGGCATGCTAACT | >hg38_21_+_8433172_8446622_RNA45SN1_45S_50ntFlanks | 5077 | 5108 | 32 | CCGATAACGAACGAGACTCTGGCATGCTAACT | 109.8 | -6.1 | 1.8 | -3.4 | 5.7E-04 | 8.4E-04 | MB231/MB468 | MP |
| >hg38_21_+_8433172_8446622_RNA45SN1_45S_50ntFlanks@11846.11872.27\|GAACGGGCTTGGCGGAATCAGCGGGGA | >hg38_21_+_8433172_8446622_RNA45SN1_45S_50ntFlanks | 11846 | 11872 | 27 | GAACGGGCTTGGCGGAATCAGCGGGGA | 142.5 | -2.0 | 0.6 | -3.5 | 5.1E-04 | 8.4E-04 | BT20/MB468 | MITO |
| >hg38_21_+_8433172_8446622_RNA45SN1_45S_50ntFlanks@9836.9864.29\|TTGGTAAGCAGAACTGGCGCTGCGGGATG | >hg38_21_+_8433172_8446622_RNA45SN1_45S_50ntFlanks | 9836 | 9864 | 29 | TTGGTAAGCAGAACTGGCGCTGCGGGATG | 139.1 | 10.5 | 2.7 | 3.9 | 1.0E-04 | 8.4E-04 | BT20/MB231 | MP |
| >hg38_21_+_8433172_8446622_RNA45SN1_45S_50ntFlanks@10391.10418.28\|AACAGCAGTTGAACATGGGTCAGTCGGT | >hg38_21_+_8433172_8446622_RNA45SN1_45S_50ntFlanks | 10391 | 10418 | 28 | AACAGCAGTTGAACATGGGTCAGTCGGT | 128.2 | -3.2 | 0.9 | -3.4 | 6.1E-04 | 8.4E-04 | BT20/MB468 | CYTO |
| >hg38_21_+_8433172_8446622_RNA45SN1_45S_50ntFlanks@11568.11586.19\|GCAGCCGACTTAGAACTGG | >hg38_21_+_8433172_8446622_RNA45SN1_45S_50ntFlanks | 11568 | 11586 | 19 | GCAGCCGACTTAGAACTGG | 1518.4 | -2.8 | 0.8 | -3.4 | 5.7E-04 | 8.4E-04 | MB231/MB468 | MP |
| >hg38_21_+_8433172_8446622_RNA45SN1_45S_50ntFlanks@9569.9597.29\|GTGCAAATCGGTCGTCCGACCTGGGTATA | >hg38_21_+_8433172_8446622_RNA45SN1_45S_50ntFlanks | 9569 | 9597 | 29 | GTGCAAATCGGTCGTCCGACCTGGGTATA | 825.8 | -3.7 | 1.0 | -3.6 | 2.9E-04 | 8.4E-04 | BT20/MB468 | MP |
| >hg38_21_+_8433172_8446622_RNA45SN1_45S_50ntFlanks@9565.9596.32\|TGACGTGCAAATCGGTCGTCCGACCTGGGTAT | >hg38_21_+_8433172_8446622_RNA45SN1_45S_50ntFlanks | 9565 | 9596 | 32 | TGACGTGCAAATCGGTCGTCCGACCTGGGTAT | 2336.3 | -4.0 | 1.1 | -3.6 | 2.9E-04 | 8.4E-04 | BT20/MB468 | MP |
| >hg38_21_+_8433172_8446622_RNA45SN1_45S_50ntFlanks@3815.3834.20\|ATGGTTCCTTTGGTCGCTCG | >hg38_21_+_8433172_8446622_RNA45SN1_45S_50ntFlanks | 3815 | 3834 | 20 | ATGGTTCCTTTGGTCGCTCG | 199.1 | -2.5 | 0.7 | -3.4 | 6.9E-04 | 8.4E-04 | MB231/MB468 | NUC |
| >MI0000065\|hsa-let-7d&WithFlank&9\|+\|94178828\|94178926@14.35.22&[MIMAT0000065&hsa-let-7d-5p&offsets\|0\|0;m-90&9\|+\|94178841\|94178862&offsets\|0\|0]\|AGAGGTAGTAGGTTGCATAGTT | >MI0000065\|hsa-let-7d&WithFlank&9\|+\|94178828\|94178926 | 14 | 35 | 22 | AGAGGTAGTAGGTTGCATAGTT | 3106.7 | 3.6 | 1.0 | 3.6 | 2.9E-04 | 8.4E-04 | BT20/MB468 | MP |
| >hg38_21_+_8433172_8446622_RNA45SN1_45S_50ntFlanks@6721.6770.50\|AATTGCAGGACACATTGATCATCGACACTTCGAACGCACTTGCGGCCCCG | >hg38_21_+_8433172_8446622_RNA45SN1_45S_50ntFlanks | 6721 | 6770 | 50 | AATTGCAGGACACATTGATCATCGACACTTCGAACGCACTTGCGGCCCCG | 949.6 | -6.1 | 1.8 | -3.4 | 6.9E-04 | 8.4E-04 | MB231/MB468 | NUC |
| >trnaMT_LeuTAA_MT_+_3230_3304@19.44.26\|GGTAATCGCATAAAACTTAAAACTTT | >trnaMT_LeuTAA_MT_+_3230_3304 | 19 | 44 | 26 | GGTAATCGCATAAAACTTAAAACTTT | 685.9 | 4.7 | 1.4 | 3.4 | 6.1E-04 | 8.4E-04 | BT20/MB468 | CYTO |
| >hg38_21_+_8433172_8446622_RNA45SN1_45S_50ntFlanks@485.510.26\|GATGTGGTGACGTCGTGCTCTCCCGG | >hg38_21_+_8433172_8446622_RNA45SN1_45S_50ntFlanks | 485 | 510 | 26 | GATGTGGTGACGTCGTGCTCTCCCGG | 65.9 | 9.8 | 2.7 | 3.6 | 2.9E-04 | 8.4E-04 | BT20/MB468 | MP |
| >trna2_GlyGCC_21_-_18827107_18827177@3.31.29,trna35_GlyGCC_1_+_161413094_161413164@3.31.29,trna37_GlyGCC_1_+_161420467_161420537@3.31.29,trna39_GlyGCC_1_+_161427898_161427968@3.31.29,trna41_GlyGCC_1_+_161435258_161435328@3.31.29\|ATGGGTGGTTCAGTGGTAGAATTCTCGCC | >trna2_GlyGCC_21_-_18827107_18827177 | 3 | 31 | 29 | ATGGGTGGTTCAGTGGTAGAATTCTCGCC | 146.4 | 5.9 | 1.6 | 3.6 | 2.9E-04 | 8.5E-04 | BT20/MB468 | MP |
| >hg38_MT_+_1621_3279_RNR2_16S_50ntFlanks@119.145.27\|AAAGTATAGGCGATAGAAATTGAAACC | >hg38_MT_+_1621_3279_RNR2_16S_50ntFlanks | 119 | 145 | 27 | AAAGTATAGGCGATAGAAATTGAAACC | 66.4 | 9.8 | 2.7 | 3.6 | 2.9E-04 | 8.5E-04 | BT20/MB468 | MP |
| >MI0000077\|hsa-mir-21&WithFlank&17\|+\|59841260\|59841343@14.35.22(+1A)&[MIMAT0000076&hsa-miR-21-5p&offsets\|0\|0(+1A);m-1&17\|+\|59841273\|59841294&offsets\|0\|0(+1A)]\|TAGCTTATCAGACTGATGTTGAA | >MI0000077\|hsa-mir-21&WithFlank&17\|+\|59841260\|59841343 | 14 | 35 | 22(+1A) | TAGCTTATCAGACTGATGTTGAA | 12942.7 | 3.3 | 0.9 | 3.4 | 5.7E-04 | 8.5E-04 | MB231/MB468 | MP |
| >hg38_MT_+_598_1651_RNR1_12S_50ntFlanks@634.657.24\|AACCTCACCACCTCTTGCTCAGCC | >hg38_MT_+_598_1651_RNR1_12S_50ntFlanks | 634 | 657 | 24 | AACCTCACCACCTCTTGCTCAGCC | 186.7 | 7.0 | 1.9 | 3.6 | 2.9E-04 | 8.5E-04 | BT20/MB468 | MP |
| >hg38_21_+_8433172_8446622_RNA45SN1_45S_50ntFlanks@10140.10157.18\|CGGCGGCGGGGGTGTGGG | >hg38_21_+_8433172_8446622_RNA45SN1_45S_50ntFlanks | 10140 | 10157 | 18 | CGGCGGCGGGGGTGTGGG | 137.4 | -5.6 | 1.5 | -3.6 | 2.9E-04 | 8.5E-04 | BT20/MB468 | MP |
| >trna111_HisGTG_1_-_147774845_147774916@1.34.34,trna118_HisGTG_1_-_145396881_145396952@1.34.34,trna16_HisGTG_1_+_146544773_146544844@1.34.34,trna1_HisGTG_15_+_45493349_45493420@1.34.34,trna21_HisGTG_1_+_147753471_147753542@1.34.34,trna33_HisGTG_6_+_27125906_27125977@1.34.34,trna7_HisGTG_9_-_14433938_14434009@1.34.34,trna8_HisGTG_15_-_45492611_45492682@1.34.34,trna9_HisGTG_15_-_45490804_45490875@1.34.34\|GCCGTGATCGTATAGTGGTTAGTACTCTGCGTTG | >trna111_HisGTG_1_-_147774845_147774916 | 1 | 34 | 34 | GCCGTGATCGTATAGTGGTTAGTACTCTGCGTTG | 1620.8 | 4.1 | 1.2 | 3.4 | 6.9E-04 | 8.5E-04 | MB231/MB468 | NUC |
| >hg38_21_+_8433172_8446622_RNA45SN1_45S_50ntFlanks@12419.12457.39\|GATGTCGGCTCTTCCTATCATTGTGAAGCAGAATTCACC | >hg38_21_+_8433172_8446622_RNA45SN1_45S_50ntFlanks | 12419 | 12457 | 39 | GATGTCGGCTCTTCCTATCATTGTGAAGCAGAATTCACC | 2688.0 | -3.0 | 0.8 | -3.9 | 1.1E-04 | 8.5E-04 | BT20/MB231 | CYTO |
| >hg38_21_+_8433172_8446622_RNA45SN1_45S_50ntFlanks@10786.10812.27\|AACAGCCTCTGGCATGTTGGAACAATG | >hg38_21_+_8433172_8446622_RNA45SN1_45S_50ntFlanks | 10786 | 10812 | 27 | AACAGCCTCTGGCATGTTGGAACAATG | 359.5 | -2.7 | 0.8 | -3.4 | 6.5E-04 | 8.5E-04 | MB231/MB468 | CYTO |
| >hg38_21_+_8433172_8446622_RNA45SN1_45S_50ntFlanks@11578.11603.26\|TAGAACTGGTGCGGACCAGGGGAATC | >hg38_21_+_8433172_8446622_RNA45SN1_45S_50ntFlanks | 11578 | 11603 | 26 | TAGAACTGGTGCGGACCAGGGGAATC | 348.3 | -2.1 | 0.6 | -3.5 | 5.2E-04 | 8.5E-04 | BT20/MB468 | MITO |
| >hg38_MT_+_598_1651_RNR1_12S_50ntFlanks@366.404.39\|CCCAATAAAGCTAAAACTCACCTGAGTTGTAAAAAACTC | >hg38_MT_+_598_1651_RNR1_12S_50ntFlanks | 366 | 404 | 39 | CCCAATAAAGCTAAAACTCACCTGAGTTGTAAAAAACTC | 489.6 | 3.6 | 1.1 | 3.4 | 6.8E-04 | 8.5E-04 | MB231/MB468 | MITO |
| >MI0000273\|hsa-mir-183&WithFlank&7\|-\|129774899\|129775020@33.53.21&[MIMAT0000261&hsa-miR-183-5p&offsets\|0\|-1;m-41&7\|-\|129774966\|129774987&offsets\|-1\|-2]\|TATGGCACTGGTAGAATTCAC | >MI0000273\|hsa-mir-183&WithFlank&7\|-\|129774899\|129775020 | 33 | 53 | 21 | TATGGCACTGGTAGAATTCAC | 395.4 | 5.2 | 1.4 | 3.6 | 2.9E-04 | 8.5E-04 | BT20/MB468 | MP |
| >MI0000542\|hsa-mir-320a&WithFlank&8\|-\|22244960\|22245043@49.69.21(+1G)&[MIMAT0000510&hsa-miR-320a-3p&offsets\|+1\|0(+1G);m-60&8\|-\|22244975\|22244996&offsets\|+1\|0(+1G)]\|AAAGCTGGGTTGAGAGGGCGAG | >MI0000542\|hsa-mir-320a&WithFlank&8\|-\|22244960\|22245043 | 49 | 69 | 21(+1G) | AAAGCTGGGTTGAGAGGGCGAG | 76.7 | 6.9 | 1.9 | 3.6 | 2.9E-04 | 8.5E-04 | BT20/MB468 | MP |
| >hg38_21_+_8433172_8446622_RNA45SN1_45S_50ntFlanks@6721.6752.32\|AATTGCAGGACACATTGATCATCGACACTTCG | >hg38_21_+_8433172_8446622_RNA45SN1_45S_50ntFlanks | 6721 | 6752 | 32 | AATTGCAGGACACATTGATCATCGACACTTCG | 278.3 | -3.7 | 1.1 | -3.4 | 6.2E-04 | 8.5E-04 | BT20/MB468 | CYTO |
| >hg38_21_+_8433172_8446622_RNA45SN1_45S_50ntFlanks@12645.12669.25\|CTGTGGGATTATGACTGAACGCCTC | >hg38_21_+_8433172_8446622_RNA45SN1_45S_50ntFlanks | 12645 | 12669 | 25 | CTGTGGGATTATGACTGAACGCCTC | 586.8 | 7.0 | 1.8 | 3.9 | 1.0E-04 | 8.5E-04 | BT20/MB231 | MP |
| >MI0000439\|hsa-mir-23b&WithFlank&9\|+\|95085202\|95085310@67.86.20(+1A)&[MIMAT0000418&hsa-miR-23b-3p&offsets\|+3\|0(+1A);m-42&9\|+\|95085265\|95085285&offsets\|+3\|+2(+1A)]\|ACATTGCCAGGGATTACCACA | >MI0000439\|hsa-mir-23b&WithFlank&9\|+\|95085202\|95085310 | 67 | 86 | 20(+1A) | ACATTGCCAGGGATTACCACA | 550.7 | 12.3 | 3.6 | 3.4 | 6.9E-04 | 8.5E-04 | MB231/MB468 | MITO |
| >hg38_21_+_8433172_8446622_RNA45SN1_45S_50ntFlanks@3881.3898.18\|GCCGACGGGCGCTGACCC | >hg38_21_+_8433172_8446622_RNA45SN1_45S_50ntFlanks | 3881 | 3898 | 18 | GCCGACGGGCGCTGACCC | 282.5 | -5.1 | 1.5 | -3.5 | 5.2E-04 | 8.5E-04 | BT20/MB468 | MITO |
| >hg38_MT_+_598_1651_RNR1_12S_50ntFlanks@52.87.36\|ATAGGTTTGGTCCTAGCCTTTCTATTAGCTCTTAGT | >hg38_MT_+_598_1651_RNR1_12S_50ntFlanks | 52 | 87 | 36 | ATAGGTTTGGTCCTAGCCTTTCTATTAGCTCTTAGT | 174.0 | 2.0 | 0.6 | 3.4 | 7.0E-04 | 8.5E-04 | BT20/MB468 | NUC |
| >hg38_21_+_8433172_8446622_RNA45SN1_45S_50ntFlanks@476.511.36\|TTCGTGATCGATGTGGTGACGTCGTGCTCTCCCGGG | >hg38_21_+_8433172_8446622_RNA45SN1_45S_50ntFlanks | 476 | 511 | 36 | TTCGTGATCGATGTGGTGACGTCGTGCTCTCCCGGG | 276.3 | 5.1 | 1.5 | 3.4 | 6.9E-04 | 8.5E-04 | MB231/MB468 | MITO |
| >hg38_MT_+_1621_3279_RNR2_16S_50ntFlanks@622.664.43\|TATAACTGAACTCCTCACACCCAATTGGACCAATCTATCACCC | >hg38_MT_+_1621_3279_RNR2_16S_50ntFlanks | 622 | 664 | 43 | TATAACTGAACTCCTCACACCCAATTGGACCAATCTATCACCC | 66.1 | 9.8 | 2.7 | 3.6 | 2.9E-04 | 8.5E-04 | BT20/MB468 | MP |
| >MI0000447\|hsa-mir-128-1&WithFlank&2\|+\|135665391\|135665484@56.77.22&[MIMAT0000424&hsa-miR-128-3p&offsets\|0\|+1;m-108&2\|+\|135665446\|135665466&offsets\|0\|+1]\|TCACAGTGAACCGGTCTCTTTT | >MI0000447\|hsa-mir-128-1&WithFlank&2\|+\|135665391\|135665484 | 56 | 77 | 22 | TCACAGTGAACCGGTCTCTTTT | 2048.3 | 3.0 | 0.9 | 3.4 | 6.9E-04 | 8.5E-04 | MB231/MB468 | MITO |
| >hg38_21_+_8433172_8446622_RNA45SN1_45S_50ntFlanks@6653.6680.28\|ACTCTTAGCGGTGGATCACTCGGCTCGT | >hg38_21_+_8433172_8446622_RNA45SN1_45S_50ntFlanks | 6653 | 6680 | 28 | ACTCTTAGCGGTGGATCACTCGGCTCGT | 189.8 | -4.6 | 1.3 | -3.4 | 5.8E-04 | 8.5E-04 | MB231/MB468 | MP |
| >hg38_21_+_8433172_8446622_RNA45SN1_45S_50ntFlanks@12420.12443.24\|ATGTCGGCTCTTCCTATCATTGTG | >hg38_21_+_8433172_8446622_RNA45SN1_45S_50ntFlanks | 12420 | 12443 | 24 | ATGTCGGCTCTTCCTATCATTGTG | 1281.0 | -3.7 | 1.0 | -3.6 | 2.9E-04 | 8.5E-04 | BT20/MB468 | MP |
| >hg38_21_+_8433172_8446622_RNA45SN1_45S_50ntFlanks@12691.12708.18\|CGGAACGATACGGCAGCG | >hg38_21_+_8433172_8446622_RNA45SN1_45S_50ntFlanks | 12691 | 12708 | 18 | CGGAACGATACGGCAGCG | 241.5 | -5.9 | 1.6 | -3.6 | 2.9E-04 | 8.5E-04 | BT20/MB468 | MP |
| >hg38_21_+_8433172_8446622_RNA45SN1_45S_50ntFlanks@11561.11603.43\|GCGCCTAGCAGCCGACTTAGAACTGGTGCGGACCAGGGGAATC | >hg38_21_+_8433172_8446622_RNA45SN1_45S_50ntFlanks | 11561 | 11603 | 43 | GCGCCTAGCAGCCGACTTAGAACTGGTGCGGACCAGGGGAATC | 535.5 | -4.1 | 1.2 | -3.4 | 6.5E-04 | 8.6E-04 | MB231/MB468 | CYTO |
| >hg38_21_+_8433172_8446622_RNA45SN1_45S_50ntFlanks@12599.12626.28\|TTGGTGTATGTGCTTGGCTGAGGAGCCA | >hg38_21_+_8433172_8446622_RNA45SN1_45S_50ntFlanks | 12599 | 12626 | 28 | TTGGTGTATGTGCTTGGCTGAGGAGCCA | 169.3 | -3.6 | 1.1 | -3.4 | 6.5E-04 | 8.6E-04 | MB231/MB468 | CYTO |
| >hg38_21_+_8433172_8446622_RNA45SN1_45S_50ntFlanks@12957.12975.19\|CTTCTGGGTCGGGGTTTCG | >hg38_21_+_8433172_8446622_RNA45SN1_45S_50ntFlanks | 12957 | 12975 | 19 | CTTCTGGGTCGGGGTTTCG | 165.9 | -3.3 | 1.0 | -3.4 | 6.2E-04 | 8.6E-04 | BT20/MB468 | CYTO |
| >hg38_21_+_8433172_8446622_RNA45SN1_45S_50ntFlanks@12841.12863.23\|CGGAGTGCCCTTCGTCCTGGGAA | >hg38_21_+_8433172_8446622_RNA45SN1_45S_50ntFlanks | 12841 | 12863 | 23 | CGGAGTGCCCTTCGTCCTGGGAA | 63.0 | -9.1 | 2.5 | -3.6 | 2.9E-04 | 8.6E-04 | BT20/MB468 | MP |
| >am_MI0003581\|hsa-mir-574&WithFlank&4\|+\|38868026\|38868133@31.55.25&[MIMAT0004795&hsa-miR-574-5p&offsets\|0\|+2;m-213&4\|+\|38868056\|38868078&offsets\|0\|+2]\|TGAGTGTGTGTGTGTGAGTGTGTGT | >am_MI0003581\|hsa-mir-574&WithFlank&4\|+\|38868026\|38868133 | 31 | 55 | 25 | TGAGTGTGTGTGTGTGAGTGTGTGT | 945.4 | -3.2 | 0.8 | -3.8 | 1.2E-04 | 8.6E-04 | BT20/MB231 | MITO |
| >hg38_21_+_8433172_8446622_RNA45SN1_45S_50ntFlanks@6653.6696.44\|ACTCTTAGCGGTGGATCACTCGGCTCGTGCGTCGATGAAGAACG | >hg38_21_+_8433172_8446622_RNA45SN1_45S_50ntFlanks | 6653 | 6696 | 44 | ACTCTTAGCGGTGGATCACTCGGCTCGTGCGTCGATGAAGAACG | 243.4 | -3.2 | 0.8 | -3.9 | 1.1E-04 | 8.6E-04 | BT20/MB231 | CYTO |

| | | | | | | | | | | | | | |
|---|---|---|---|---|---|---|---|---|---|---|---|---|---|
| >hg38_MT_+_1621_3279_RNR2_16S_50ntFlanks@1091.1120.30\|AGCAAGACGAGAAGACCCTATGGAGCTTTA | >hg38_MT_+_1621_3279_RNR2_16S_50ntFlanks | 1091 | 1120 | 30 | AGCAAGACGAGAAGACCCTATGGAGCTTTA | 193.5 | 3.9 | 1.1 | 3.6 | 3.0E-04 | 8.6E-04 | BT20/MB468 | MP |
| >MI0000301\|hsa-mir-224&WithFlank&X\|-\|151958572\|151958664@14.35.22(+1U)&[MIMAT0000281&hsa-miR-224-5p&offsets\|+1\|-2(+1U);m-238&X\|-\|151958630\|151958651&offsets\|0\|0(+1U)]\|CAAGTCACTAGTGGTTCCGTTTT | >MI0000301\|hsa-mir-224&WithFlank&X\|-\|151958572\|151958664 | 14 | 35 | 22(+1U) | CAAGTCACTAGTGGTTCCGTTTT | 259.6 | 3.3 | 0.9 | 3.9 | 1.1E-04 | 8.6E-04 | BT20/MB231 | CYTO |
| >hg38_21_+_8433172_8446622_RNA45SN1_45S_50ntFlanks@5473.5509.37\|CGGCCCTGGCGGAGCGCTGAGAAGACGGTCGAACTTG | >hg38_21_+_8433172_8446622_RNA45SN1_45S_50ntFlanks | 5473 | 5509 | 37 | CGGCCCTGGCGGAGCGCTGAGAAGACGGTCGAACTTG | 126.5 | -2.8 | 0.8 | -3.4 | 5.8E-04 | 8.6E-04 | MB231/MB468 | MP |
| >hg38_21_+_8433172_8446622_RNA45SN1_45S_50ntFlanks@10240.10262.23\|CTACGCCGCGACGAGTAGGAGGG | >hg38_21_+_8433172_8446622_RNA45SN1_45S_50ntFlanks | 10240 | 10262 | 23 | CTACGCCGCGACGAGTAGGAGGG | 259.6 | -4.3 | 1.3 | -3.4 | 6.9E-04 | 8.6E-04 | MB231/MB468 | MITO |
| >hg38_21_+_8433172_8446622_RNA45SN1_45S_50ntFlanks@5472.5521.50\|ACGGCCCTGGCGGAGCGCTGAGAAGACGGTCGAACTTGACTATCTAGAGG | >hg38_21_+_8433172_8446622_RNA45SN1_45S_50ntFlanks | 5472 | 5521 | 50 | ACGGCCCTGGCGGAGCGCTGAGAAGACGGTCGAACTTGACTATCTAGAGG | 340.7 | -3.9 | 1.0 | -3.9 | 1.1E-04 | 8.6E-04 | BT20/MB231 | CYTO |
| >hg38_21_+_8433172_8446622_RNA45SN1_45S_50ntFlanks@3854.3882.29\|ACTGTGGTAATTCTAGAGCTAATACATGC | >hg38_21_+_8433172_8446622_RNA45SN1_45S_50ntFlanks | 3854 | 3882 | 29 | ACTGTGGTAATTCTAGAGCTAATACATGC | 233.1 | -3.9 | 1.1 | -3.4 | 7.0E-04 | 8.6E-04 | MB231/MB468 | NUC |
| >am_trnaMT_ValTAC_MT_+_1602_1670@35.66.32\|ACTTAGGAGATTTCAACTTAACTTGACCGCTC | >am_trnaMT_ValTAC_MT_+_1602_1670 | 35 | 66 | 32 | ACTTAGGAGATTTCAACTTAACTTGACCGCTC | 66.2 | 9.8 | 2.7 | 3.6 | 3.0E-04 | 8.6E-04 | BT20/MB468 | MP |
| >MI0000082\|hsa-mir-25&WithFlank&7\|-\|100093554\|100093649@58.77.20(+1A)&[MIMAT0000081&hsa-miR-25-3p&offsets\|0\|-2(+1A);m-23&7\|-\|100093571\|100093592&offsets\|0\|-2(+1A)]\|CATTGCACTTGTCTCGGTCTA | >MI0000082\|hsa-mir-25&WithFlank&7\|-\|100093554\|100093649 | 58 | 77 | 20(+1A) | CATTGCACTTGTCTCGGTCTA | 862.3 | 4.6 | 1.3 | 3.4 | 5.8E-04 | 8.6E-04 | MB231/MB468 | MP |
| >hg38_21_+_8433172_8446622_RNA45SN1_45S_50ntFlanks@11584.11616.33\|TGGTGCGGACCAGGGGAATCCGACTGTTTAATT | >hg38_21_+_8433172_8446622_RNA45SN1_45S_50ntFlanks | 11584 | 11616 | 33 | TGGTGCGGACCAGGGGAATCCGACTGTTTAATT | 355.4 | 4.0 | 1.2 | 3.4 | 7.0E-04 | 8.6E-04 | BT20/MB468 | NUC |
| >am_trnaMT_ValTAC_MT_+_1602_1670@1.39.39\|CAGAGTGTAGCTTAACACAAAGCACCCAACTTACACTTA | >am_trnaMT_ValTAC_MT_+_1602_1670 | 1 | 39 | 39 | CAGAGTGTAGCTTAACACAAAGCACCCAACTTACACTTA | 330.4 | 6.9 | 1.9 | 3.6 | 3.0E-04 | 8.6E-04 | BT20/MB468 | MP |
| >hg38_21_+_8433172_8446622_RNA45SN1_45S_50ntFlanks@3853.3873.21\|AACTGTGGTAATTCTAGAGCT | >hg38_21_+_8433172_8446622_RNA45SN1_45S_50ntFlanks | 3853 | 3873 | 21 | AACTGTGGTAATTCTAGAGCT | 450.4 | -3.9 | 1.1 | -3.4 | 6.9E-04 | 8.6E-04 | MB231/MB468 | MITO |
| >trnaMT_SerGCT_MT_+_12207_12265@1.38.38\|GAGAAAGCTCACAAGAACTGCTAACTCATGCCCCCATG | >trnaMT_SerGCT_MT_+_12207_12265 | 1 | 38 | 38 | GAGAAAGCTCACAAGAACTGCTAACTCATGCCCCCATG | 1179.1 | -3.4 | 1.0 | -3.4 | 7.0E-04 | 8.6E-04 | BT20/MB468 | NUC |
| >hg38_21_+_8433172_8446622_RNA45SN1_45S_50ntFlanks@12457.12486.30\|CAAGCGTTGGATTGTTCACCCACTAATAGG | >hg38_21_+_8433172_8446622_RNA45SN1_45S_50ntFlanks | 12457 | 12486 | 30 | CAAGCGTTGGATTGTTCACCCACTAATAGG | 512.5 | -4.2 | 1.2 | -3.4 | 7.0E-04 | 8.6E-04 | MB231/MB468 | NUC |
| >hg38_21_+_8433172_8446622_RNA45SN1_45S_50ntFlanks@4683.4720.38\|CAAGACGGACCAGAGCGAAAGCATTTGCCAAGAATGTT | >hg38_21_+_8433172_8446622_RNA45SN1_45S_50ntFlanks | 4683 | 4720 | 38 | CAAGACGGACCAGAGCGAAAGCATTTGCCAAGAATGTT | 478.0 | 4.3 | 1.3 | 3.4 | 6.6E-04 | 8.6E-04 | MB231/MB468 | CYTO |
| >hg38_21_+_8433172_8446622_RNA45SN1_45S_50ntFlanks@11658.11687.30\|ATGTGATTTCTGCCCAGTGCTCTGAATGTC | >hg38_21_+_8433172_8446622_RNA45SN1_45S_50ntFlanks | 11658 | 11687 | 30 | ATGTGATTTCTGCCCAGTGCTCTGAATGTC | 755.9 | 8.1 | 2.1 | 3.9 | 1.0E-04 | 8.6E-04 | BT20/MB231 | MP |
| >hg38_21_+_8433172_8446622_RNA45SN1_45S_50ntFlanks@6650.6678.29\|ACGACTCTTAGCGGTGGATCACTCGGCTC | >hg38_21_+_8433172_8446622_RNA45SN1_45S_50ntFlanks | 6650 | 6678 | 29 | ACGACTCTTAGCGGTGGATCACTCGGCTC | 945.0 | 2.5 | 0.8 | 3.4 | 7.0E-04 | 8.6E-04 | MB231/MB468 | NUC |
| >hg38_21_+_8433172_8446622_RNA45SN1_45S_50ntFlanks@10717.10734.18\|AAAATCCGGGGGAGAGGG | >hg38_21_+_8433172_8446622_RNA45SN1_45S_50ntFlanks | 10717 | 10734 | 18 | AAAATCCGGGGGAGAGGG | 150.0 | -5.4 | 1.6 | -3.4 | 5.8E-04 | 8.6E-04 | MB231/MB468 | MP |
| >hg38_21_+_8433172_8446622_RNA45SN1_45S_50ntFlanks@11583.11602.20\|CTGGTGCGGACCAGGGGAAT | >hg38_21_+_8433172_8446622_RNA45SN1_45S_50ntFlanks | 11583 | 11602 | 20 | CTGGTGCGGACCAGGGGAAT | 1570.8 | -5.4 | 1.6 | -3.4 | 7.0E-04 | 8.6E-04 | BT20/MB468 | NUC |
| >MI0000063\|hsa-let-7b&WithFlank&22\|+\|46113680\|46113774@12.32.21(+2A)&[MIMAT0000063&hsa-let-7b-5p&offsets\|0\|-1(+2A);m-14&22\|+\|46113691\|46113712&offsets\|0\|-1(+2A)]\|TGAGGTAGTAGGTTGTGTGGTAA | >MI0000063\|hsa-let-7b&WithFlank&22\|+\|46113680\|46113774 | 12 | 32 | 21(+2A) | TGAGGTAGTAGGTTGTGTGGTAA | 165.0 | 5.9 | 1.6 | 3.6 | 3.0E-04 | 8.6E-04 | BT20/MB468 | MP |
| >hg38_21_+_8433172_8446622_RNA45SN1_45S_50ntFlanks@11581.11612.32\|AACTGGTGCGGACCAGGGGAATCCGACTGTTT | >hg38_21_+_8433172_8446622_RNA45SN1_45S_50ntFlanks | 11581 | 11612 | 32 | AACTGGTGCGGACCAGGGGAATCCGACTGTTT | 1672.0 | -2.1 | 0.6 | -3.4 | 6.3E-04 | 8.6E-04 | BT20/MB468 | CYTO |
| >MI0005566\|hsa-mir-374b&WithFlank&X\|-\|74218541\|74218624@17.38.22&[MIMAT0004955&hsa-miR-374b-5p&offsets\|0\|0;m-153&X\|-\|74218587\|74218608&offsets\|0\|0]\|ATATAATACAACCTGCTAAGTG | >MI0005566\|hsa-mir-374b&WithFlank&X\|-\|74218541\|74218624 | 17 | 38 | 22 | ATATAATACAACCTGCTAAGTG | 280.7 | 4.7 | 1.4 | 3.4 | 6.6E-04 | 8.6E-04 | MB231/MB468 | CYTO |
| >hg38_21_+_8433172_8446622_RNA45SN1_45S_50ntFlanks@9507.9527.21\|AACTATGCCTGGGCAGGGCGA | >hg38_21_+_8433172_8446622_RNA45SN1_45S_50ntFlanks | 9507 | 9527 | 21 | AACTATGCCTGGGCAGGGCGA | 72.0 | -9.3 | 2.6 | -3.6 | 3.0E-04 | 8.6E-04 | BT20/MB468 | MP |
| >MI0000270\|hsa-mir-181b-1&WithFlank&1\|-\|198858867\|198858988@42.64.23;MI0000683\|hsa-mir-181b-2&WithFlank&9\|+\|124693704\|124693804@22.44.23&[MIMAT0000257&hsa-miR-181b-5p&offsets\|0\|0;m-73&1\|-\|198858925\|198858947&offsets\|0\|0];[MIMAT0000257_1&hsa-miR-181b-5p&offsets\|0\|0;m-72&9\|+\|124693725\|124693747&offsets\|0\|0]\|AACATTCATTGCTGTCGGTGGGT | >MI0000270\|hsa-mir-181b-1&WithFlank&1\|-\|198858867\|198858988 | 42 | 64 | 23 | AACATTCATTGCTGTCGGTGGGT | 2129.6 | 2.9 | 0.9 | 3.4 | 5.9E-04 | 8.7E-04 | MB231/MB468 | MP |
| >hg38_21_+_8433172_8446622_RNA45SN1_45S_50ntFlanks@11641.11664.24\|CGGCGGGTGTTGACGCGATGTGAT | >hg38_21_+_8433172_8446622_RNA45SN1_45S_50ntFlanks | 11641 | 11664 | 24 | CGGCGGGTGTTGACGCGATGTGAT | 280.4 | -3.8 | 1.1 | -3.4 | 6.3E-04 | 8.7E-04 | BT20/MB468 | CYTO |
| >hg38_21_+_8433172_8446622_RNA45SN1_45S_50ntFlanks@6652.6692.41\|GACTCTTAGCGGTGGATCACTCGGCTCGTGCGTCGATGAAG | >hg38_21_+_8433172_8446622_RNA45SN1_45S_50ntFlanks | 6652 | 6692 | 41 | GACTCTTAGCGGTGGATCACTCGGCTCGTGCGTCGATGAAG | 2684.1 | 2.3 | 0.7 | 3.4 | 7.1E-04 | 8.7E-04 | MB231/MB468 | NUC |
| >hg38_21_+_8433172_8446622_RNA45SN1_45S_50ntFlanks@6658.6681.24\|TAGCGGTGGATCACTCGGCTCGTG | >hg38_21_+_8433172_8446622_RNA45SN1_45S_50ntFlanks | 6658 | 6681 | 24 | TAGCGGTGGATCACTCGGCTCGTG | 121.0 | -3.0 | 0.9 | -3.4 | 6.3E-04 | 8.7E-04 | BT20/MB468 | CYTO |
| >hg38_21_+_8433172_8446622_RNA45SN1_45S_50ntFlanks@6650.6677.28\|ACGACTCTTAGCGGTGGATCACTCGGCT | >hg38_21_+_8433172_8446622_RNA45SN1_45S_50ntFlanks | 6650 | 6677 | 28 | ACGACTCTTAGCGGTGGATCACTCGGCT | 999.2 | 3.5 | 1.0 | 3.6 | 3.0E-04 | 8.7E-04 | BT20/MB468 | MP |
| >hg38_21_+_8433172_8446622_RNA45SN1_45S_50ntFlanks@12412.12434.23\|ATCCTTCGATGTCGGCTCTTCCT | >hg38_21_+_8433172_8446622_RNA45SN1_45S_50ntFlanks | 12412 | 12434 | 23 | ATCCTTCGATGTCGGCTCTTCCT | 670.4 | -5.4 | 1.6 | -3.4 | 7.0E-04 | 8.7E-04 | MB231/MB468 | MITO |
| >hg38_21_+_8433172_8446622_RNA45SN1_45S_50ntFlanks@10362.10384.23\|AAGGCCGAAGTGGAGAAGGGTTC | >hg38_21_+_8433172_8446622_RNA45SN1_45S_50ntFlanks | 10362 | 10384 | 23 | AAGGCCGAAGTGGAGAAGGGTTC | 276.0 | -4.9 | 1.4 | -3.5 | 5.3E-04 | 8.7E-04 | BT20/MB468 | MITO |
| >hg38_21_+_8433172_8446622_RNA45SN1_45S_50ntFlanks@11567.11592.26\|AGCAGCCGACTTAGAACTGGTGCGGA | >hg38_21_+_8433172_8446622_RNA45SN1_45S_50ntFlanks | 11567 | 11592 | 26 | AGCAGCCGACTTAGAACTGGTGCGGA | 173.9 | -4.0 | 1.2 | -3.4 | 6.3E-04 | 8.7E-04 | BT20/MB468 | CYTO |
| >hg38_21_+_8433172_8446622_RNA45SN1_45S_50ntFlanks@6651.6668.18\|CGACTCTTAGCGGTGGAT | >hg38_21_+_8433172_8446622_RNA45SN1_45S_50ntFlanks | 6651 | 6668 | 18 | CGACTCTTAGCGGTGGAT | 2837.2 | 1.7 | 0.5 | 3.4 | 6.6E-04 | 8.7E-04 | MB231/MB468 | CYTO |
| >hg38_21_+_8433172_8446622_RNA45SN1_45S_50ntFlanks@8168.8205.38\|CCCAAGTCCTTCTGATCGAGGCCCAGCCCGTGGACGGT | >hg38_21_+_8433172_8446622_RNA45SN1_45S_50ntFlanks | 8168 | 8205 | 38 | CCCAAGTCCTTCTGATCGAGGCCCAGCCCGTGGACGGT | 62.4 | -5.6 | 1.6 | -3.4 | 5.9E-04 | 8.7E-04 | MB231/MB468 | MP |
| >hg38_21_+_8433172_8446622_RNA45SN1_45S_50ntFlanks@7992.8013.22\|CGTGGCGACCCGCTGAATTTAA | >hg38_21_+_8433172_8446622_RNA45SN1_45S_50ntFlanks | 7992 | 8013 | 22 | CGTGGCGACCCGCTGAATTTAA | 303.8 | -2.6 | 0.8 | -3.4 | 7.0E-04 | 8.7E-04 | MB231/MB468 | MITO |
| >hg38_21_+_8433172_8446622_RNA45SN1_45S_50ntFlanks@9500.9520.21\|GATGGTGAACTATGCCTGGGC | >hg38_21_+_8433172_8446622_RNA45SN1_45S_50ntFlanks | 9500 | 9520 | 21 | GATGGTGAACTATGCCTGGGC | 85.4 | 5.8 | 1.6 | 3.6 | 3.0E-04 | 8.7E-04 | BT20/MB468 | MP |
| >MI0000812\|hsa-mir-331&WithFlank&12\|+\|95308414\|95308519@30.52.23&[MIMAT0004700&hsa-miR-331-5p&offsets\|-2\|-1;TJU_CMC_MD2.ID00734.5p-miR&offsets\|+18\|+21;m-2747&12\|+\|95308425\|95308444&offsets\|+18\|+21;m-377&12\|+\|95308445\|95308465&offsets\|-2\|0]\|TTCTAGGTATGGTCCCAGGGATC | >MI0000812\|hsa-mir-331&WithFlank&12\|+\|95308414\|95308519 | 30 | 52 | 23 | TTCTAGGTATGGTCCCAGGGATC | 65.0 | 9.8 | 2.7 | 3.6 | 3.0E-04 | 8.7E-04 | BT20/MB468 | MP |
| >hg38_21_+_8433172_8446622_RNA45SN1_45S_50ntFlanks@10350.10391.42\|ACGAGAACTTTGAAGGCCGAAGTGGAGAAGGGTTCCATGTGA | >hg38_21_+_8433172_8446622_RNA45SN1_45S_50ntFlanks | 10350 | 10391 | 42 | ACGAGAACTTTGAAGGCCGAAGTGGAGAAGGGTTCCATGTGA | 306.3 | -2.7 | 0.8 | -3.5 | 5.3E-04 | 8.7E-04 | BT20/MB468 | MITO |
| >hg38_21_+_8433172_8446622_RNA45SN1_45S_50ntFlanks@9783.9819.37\|CGGCTCGCTGGCGTGGAGCCGGGCGTGGAATGCGAGT | >hg38_21_+_8433172_8446622_RNA45SN1_45S_50ntFlanks | 9783 | 9819 | 37 | CGGCTCGCTGGCGTGGAGCCGGGCGTGGAATGCGAGT | 53.6 | 9.1 | 2.3 | 3.9 | 1.1E-04 | 8.7E-04 | BT20/MB231 | MP |
| >hg38_21_+_8433172_8446622_RNA45SN1_45S_50ntFlanks@10349.10371.23\|AACGAGAACTTTGAAGGCCGAAG | >hg38_21_+_8433172_8446622_RNA45SN1_45S_50ntFlanks | 10349 | 10371 | 23 | AACGAGAACTTTGAAGGCCGAAG | 1019.9 | 1.7 | 0.5 | 3.5 | 5.3E-04 | 8.7E-04 | BT20/MB468 | MITO |
| >MI0000299\|hsa-mir-222&WithFlank&X\|-\|45747009\|45747130@75.97.23(+3U)&[MIMAT0000279&hsa-miR-222-3p&offsets\|0\|+2(+3U);m-140&X\|-\|45747033\|45747056&offsets\|0\|-1(+3U)]\|AGCTACATCTGGCTACTGGGTCTTTT | >MI0000299\|hsa-mir-222&WithFlank&X\|-\|45747009\|45747130 | 75 | 97 | 23(+3U) | AGCTACATCTGGCTACTGGGTCTTTT | 412.6 | -4.2 | 1.1 | -3.8 | 1.3E-04 | 8.7E-04 | BT20/MB231 | MITO |
| >hg38_21_+_8433172_8446622_RNA45SN1_45S_50ntFlanks@11922.11940.19\|GAGAGGTGTAGAATAAGTG | >hg38_21_+_8433172_8446622_RNA45SN1_45S_50ntFlanks | 11922 | 11940 | 19 | GAGAGGTGTAGAATAAGTG | 59.0 | 6.7 | 1.9 | 3.6 | 3.0E-04 | 8.7E-04 | BT20/MB468 | MP |
| >hg38_21_+_8433172_8446622_RNA45SN1_45S_50ntFlanks@4003.4020.18\|ATAACCTCGGGCCGATCG | >hg38_21_+_8433172_8446622_RNA45SN1_45S_50ntFlanks | 4003 | 4020 | 18 | ATAACCTCGGGCCGATCG | 742.7 | -5.4 | 1.6 | -3.5 | 5.3E-04 | 8.7E-04 | BT20/MB468 | MITO |
| >hg38_21_+_8433172_8446622_RNA45SN1_45S_50ntFlanks@6648.6672.25\|GTACGACTCTTAGCGGTGGATCACT | >hg38_21_+_8433172_8446622_RNA45SN1_45S_50ntFlanks | 6648 | 6672 | 25 | GTACGACTCTTAGCGGTGGATCACT | 771.0 | 2.9 | 0.9 | 3.4 | 7.0E-04 | 8.7E-04 | MB231/MB468 | MITO |
| >hg38_21_+_8433172_8446622_RNA45SN1_45S_50ntFlanks@11781.11803.23\|CGCATGAATGGATGAACGAGATT | >hg38_21_+_8433172_8446622_RNA45SN1_45S_50ntFlanks | 11781 | 11803 | 23 | CGCATGAATGGATGAACGAGATT | 1854.8 | -3.8 | 1.1 | -3.6 | 3.0E-04 | 8.7E-04 | BT20/MB468 | MP |
| >hg38_21_+_8433172_8446622_RNA45SN1_45S_50ntFlanks@7983.8012.30\|CAGATCAGACGTGGCGACCCGCTGAATTTA | >hg38_21_+_8433172_8446622_RNA45SN1_45S_50ntFlanks | 7983 | 8012 | 30 | CAGATCAGACGTGGCGACCCGCTGAATTTA | 500.2 | -5.9 | 1.6 | -3.6 | 3.0E-04 | 8.7E-04 | BT20/MB468 | MP |
| >hg38_21_+_8433172_8446622_RNA45SN1_45S_50ntFlanks@9203.9222.20\|CTCGGGGCCACGCGCGCGTC | >hg38_21_+_8433172_8446622_RNA45SN1_45S_50ntFlanks | 9203 | 9222 | 20 | CTCGGGGCCACGCGCGCGTC | 1302.2 | -5.2 | 1.4 | -3.6 | 3.0E-04 | 8.7E-04 | BT20/MB468 | MP |
| >hg38_21_+_8433172_8446622_RNA45SN1_45S_50ntFlanks@4099.4121.23\|GTGACCACGGGTGACGGGGAATC | >hg38_21_+_8433172_8446622_RNA45SN1_45S_50ntFlanks | 4099 | 4121 | 23 | GTGACCACGGGTGACGGGGAATC | 173.9 | -4.7 | 1.4 | -3.4 | 6.4E-04 | 8.7E-04 | BT20/MB468 | CYTO |
| >hg38_21_+_8433172_8446622_RNA45SN1_45S_50ntFlanks@3862.3898.37\|AATTCTAGAGCTAATACATGCCGACGGGCGCTGACCC | >hg38_21_+_8433172_8446622_RNA45SN1_45S_50ntFlanks | 3862 | 3898 | 37 | AATTCTAGAGCTAATACATGCCGACGGGCGCTGACCC | 130.8 | -4.6 | 1.3 | -3.5 | 5.3E-04 | 8.7E-04 | BT20/MB468 | MITO |
| >hg38_21_+_8433172_8446622_RNA45SN1_45S_50ntFlanks@9561.9597.37\|GTCCTGACGTGCAAATCGGTCGTCCGACCTGGGTATA | >hg38_21_+_8433172_8446622_RNA45SN1_45S_50ntFlanks | 9561 | 9597 | 37 | GTCCTGACGTGCAAATCGGTCGTCCGACCTGGGTATA | 506.7 | -2.4 | 0.7 | -3.5 | 5.3E-04 | 8.7E-04 | BT20/MB468 | MITO |
| >hg38_21_+_8433172_8446622_RNA45SN1_45S_50ntFlanks@9926.9948.23\|GTTGATATAGACAGCAGGACGGT | >hg38_21_+_8433172_8446622_RNA45SN1_45S_50ntFlanks | 9926 | 9948 | 23 | GTTGATATAGACAGCAGGACGGT | 90.6 | 6.6 | 1.8 | 3.6 | 3.0E-04 | 8.7E-04 | BT20/MB468 | MP |

| | | | | | | | | | | | | | |
|---|---|---|---|---|---|---|---|---|---|---|---|---|---|
| >trnaMT_GluTTC_MT_-_14674_14742@24.65.42,trnalookalike8_GluTTC_5_-_93905172_93905240@24.65.42\|ATGGTTTTTCATATCATTGGTCGTGGTTGTAGTCCGTGCGAG | >trnaMT_GluTTC_MT_-_14674_14742 | 24 | 65 | 42 | ATGGTTTTTCATATCATTGGTCGTGGTTGTAGTCCGTGCGAG | 65.3 | 9.8 | 2.7 | 3.6 | 3.0E-04 | 8.7E-04 | BT20/MB468 | MP |
| >hg38_21_+_8433172_8446622_RNA45SN1_45S_50ntFlanks@6653.6677.25\|ACTCTTAGCGGTGGATCACTCGGCT | >hg38_21_+_8433172_8446622_RNA45SN1_45S_50ntFlanks | 6653 | 6677 | 25 | ACTCTTAGCGGTGGATCACTCGGCT | 372.0 | 2.7 | 0.8 | 3.4 | 7.1E-04 | 8.7E-04 | MB231/MB468 | NUC |
| >hg38_21_+_8433172_8446622_RNA45SN1_45S_50ntFlanks@4104.4121.18\|CACGGGTGACGGGGAATC | >hg38_21_+_8433172_8446622_RNA45SN1_45S_50ntFlanks | 4104 | 4121 | 18 | CACGGGTGACGGGGAATC | 157.5 | -3.3 | 1.0 | -3.4 | 6.4E-04 | 8.7E-04 | BT20/MB468 | CYTO |
| >MI0000077\|hsa-mir-21&WithFlank&17\|+\|59841260\|59841343@14.35.22(+1G)&[MIMAT0000076&hsa-miR-21-5p&offsets\|0\|0(+1G);m-1&17\|+\|59841273\|59841294&offsets\|0\|0(+1G)]\|TAGCTTATCAGACTGATGTTGAG | >MI0000077\|hsa-mir-21&WithFlank&17\|+\|59841260\|59841343 | 14 | 35 | 22(+1G) | TAGCTTATCAGACTGATGTTGAG | 1281.4 | 3.7 | 1.1 | 3.4 | 5.9E-04 | 8.7E-04 | MB231/MB468 | MP |
| >hg38_21_+_8433172_8446622_RNA45SN1_45S_50ntFlanks@13007.13028.22\|TATTGAAAGTCAGCCCTCGACA | >hg38_21_+_8433172_8446622_RNA45SN1_45S_50ntFlanks | 13007 | 13028 | 22 | TATTGAAAGTCAGCCCTCGACA | 337.9 | -4.0 | 1.2 | -3.4 | 7.1E-04 | 8.7E-04 | MB231/MB468 | NUC |
| >hg38_21_+_8433172_8446622_RNA45SN1_45S_50ntFlanks@6719.6749.31\|TGAATTGCAGGACACATTGATCATCGACACT | >hg38_21_+_8433172_8446622_RNA45SN1_45S_50ntFlanks | 6719 | 6749 | 31 | TGAATTGCAGGACACATTGATCATCGACACT | 318.9 | -3.3 | 1.0 | -3.4 | 6.7E-04 | 8.7E-04 | MB231/MB468 | CYTO |
| >MI0000103\|hsa-mir-101-1&WithFlank&1\|-\|65058428\|65058514@53.74.22;MI0000739\|hsa-mir-101-2&WithFlank&9\|+\|4850291\|4850381@55.76.22&[MIMAT0000099&hsa-miR-101-3p&offsets\|0\|+1;m-17&1\|-\|65058443\|65058462&offsets\|0\|+2];[MIMAT0000099_1&hsa-miR-101-3p&offsets\|0\|+1;m-16&9\|+\|4850345\|4850364&offsets\|0\|+2]\|TACAGTACTGTGATAACTGAAG | >MI0000103\|hsa-mir-101-1&WithFlank&1\|-\|65058428\|65058514 | 53 | 74 | 22 | TACAGTACTGTGATAACTGAAG | 695.5 | 3.2 | 0.9 | 3.4 | 7.2E-04 | 8.8E-04 | BT20/MB468 | NUC |
| >hg38_MT_+_1621_3279_RNR2_16S_50ntFlanks@51.81.31\|GCTAAACCTAGCCCCAAACCCACTCCACCTT | >hg38_MT_+_1621_3279_RNR2_16S_50ntFlanks | 51 | 81 | 31 | GCTAAACCTAGCCCCAAACCCACTCCACCTT | 3604.4 | 4.5 | 1.2 | 3.6 | 3.0E-04 | 8.8E-04 | BT20/MB468 | MP |
| >trnaMT_HisGTG_MT_+_12138_12206@26.63.38\|AGATTGTGAATCTGACAACAGAGGCTTACGACCCCTTA | >trnaMT_HisGTG_MT_+_12138_12206 | 26 | 63 | 38 | AGATTGTGAATCTGACAACAGAGGCTTACGACCCCTTA | 380.9 | 5.4 | 1.5 | 3.6 | 3.0E-04 | 8.8E-04 | BT20/MB468 | MP |
| >hg38_21_+_8433172_8446622_RNA45SN1_45S_50ntFlanks@4615.4633.19\|CCATGATTAAGAGGGACGG | >hg38_21_+_8433172_8446622_RNA45SN1_45S_50ntFlanks | 4615 | 4633 | 19 | CCATGATTAAGAGGGACGG | 155.8 | -5.5 | 1.5 | -3.6 | 3.0E-04 | 8.8E-04 | BT20/MB468 | MP |
| >hg38_21_+_8433172_8446622_RNA45SN1_45S_50ntFlanks@11566.11584.19\|TAGCAGCCGACTTAGAACT | >hg38_21_+_8433172_8446622_RNA45SN1_45S_50ntFlanks | 11566 | 11584 | 19 | TAGCAGCCGACTTAGAACT | 134.4 | 4.6 | 1.3 | 3.6 | 3.0E-04 | 8.8E-04 | BT20/MB468 | MP |
| >hg38_21_+_8433172_8446622_RNA45SN1_45S_50ntFlanks@12394.12419.26\|TAGCGACGTCGCTTTTTGATCCTTCG | >hg38_21_+_8433172_8446622_RNA45SN1_45S_50ntFlanks | 12394 | 12419 | 26 | TAGCGACGTCGCTTTTTGATCCTTCG | 962.3 | -2.7 | 0.8 | -3.4 | 7.2E-04 | 8.8E-04 | MB231/MB468 | NUC |
| >trna10_LysCTT_16_+_3241501_3241573@5.35.31,trna13_LysCTT_14_-_58706613_58706685@5.35.31,trna2_LysCTT_15_+_79152904_79152976@5.35.31\|GGCTAGCTCAGTCGGTAGAGCATGGGACTCT | >trna10_LysCTT_16_+_3241501_3241573 | 5 | 35 | 31 | GGCTAGCTCAGTCGGTAGAGCATGGGACTCT | 64.2 | 9.8 | 2.7 | 3.6 | 3.0E-04 | 8.8E-04 | BT20/MB468 | MP |
| >am_MI0003581\|hsa-mir-574&WithFlank&4\|+\|38868026\|38868133@31.54.24&[MIMAT0004795&hsa-miR-574-5p&offsets\|0\|+1;m-213&4\|+\|38868056\|38868078&offsets\|0\|+1]\|TGAGTGTGTGTGTGTGAGTGTGTG | >am_MI0003581\|hsa-mir-574&WithFlank&4\|+\|38868026\|38868133 | 31 | 54 | 24 | TGAGTGTGTGTGTGTGAGTGTGTG | 505.0 | 2.8 | 0.8 | 3.4 | 5.9E-04 | 8.8E-04 | MB231/MB468 | MP |
| >MI0003591\|hsa-mir-584&WithFlank&5\|-\|149062307\|149062415@22.43.22&[MIMAT0003249&hsa-miR-584-5p&offsets\|0\|0;m-167&5\|-\|149062374\|149062394&offsets\|0\|+1]\|TTATGGTTTGCCTGGGACTGAG | >MI0003591\|hsa-mir-584&WithFlank&5\|-\|149062307\|149062415 | 22 | 43 | 22 | TTATGGTTTGCCTGGGACTGAG | 261.7 | 5.0 | 1.4 | 3.6 | 3.0E-04 | 8.8E-04 | BT20/MB468 | MP |
| >hg38_21_+_8433172_8446622_RNA45SN1_45S_50ntFlanks@12939.12972.34\|CCATTCGTAGACGACCTGCTTCTGGGTCGGGGTT | >hg38_21_+_8433172_8446622_RNA45SN1_45S_50ntFlanks | 12939 | 12972 | 34 | CCATTCGTAGACGACCTGCTTCTGGGTCGGGGTT | 134.8 | -5.7 | 1.6 | -3.6 | 3.0E-04 | 8.8E-04 | BT20/MB468 | MP |
| >hg38_21_+_8433172_8446622_RNA45SN1_45S_50ntFlanks@6773.6806.34\|TTCCTCCCGGGGCTACGCCTGTCTGAGCGTCGCT | >hg38_21_+_8433172_8446622_RNA45SN1_45S_50ntFlanks | 6773 | 6806 | 34 | TTCCTCCCGGGGCTACGCCTGTCTGAGCGTCGCT | 1122.6 | 2.6 | 0.8 | 3.4 | 6.4E-04 | 8.8E-04 | BT20/MB468 | CYTO |
| >hg38_21_+_8433172_8446622_RNA45SN1_45S_50ntFlanks@9604.9630.27\|AAAGACTAATCGAACCATCTAGTAGCT | >hg38_21_+_8433172_8446622_RNA45SN1_45S_50ntFlanks | 9604 | 9630 | 27 | AAAGACTAATCGAACCATCTAGTAGCT | 200.2 | -4.0 | 1.2 | -3.4 | 6.0E-04 | 8.8E-04 | MB231/MB468 | MP |
| >hg38_21_+_8433172_8446622_RNA45SN1_45S_50ntFlanks@11579.11606.28\|AGAACTGGTGCGGACCAGGGGAATCCGA | >hg38_21_+_8433172_8446622_RNA45SN1_45S_50ntFlanks | 11579 | 11606 | 28 | AGAACTGGTGCGGACCAGGGGAATCCGA | 170.2 | -3.8 | 1.1 | -3.4 | 7.2E-04 | 8.8E-04 | MB231/MB468 | NUC |
| >hg38_21_+_8433172_8446622_RNA45SN1_45S_50ntFlanks@6646.6663.18\|TCGTACGACTCTTAGCGG | >hg38_21_+_8433172_8446622_RNA45SN1_45S_50ntFlanks | 6646 | 6663 | 18 | TCGTACGACTCTTAGCGG | 8353.6 | -2.0 | 0.6 | -3.4 | 6.0E-04 | 8.8E-04 | MB231/MB468 | MP |
| >hg38_21_+_8433172_8446622_RNA45SN1_45S_50ntFlanks@11682.11702.21\|AATGTCAAAGTGAAGAAATTC | >hg38_21_+_8433172_8446622_RNA45SN1_45S_50ntFlanks | 11682 | 11702 | 21 | AATGTCAAAGTGAAGAAATTC | 170.4 | -2.4 | 0.7 | -3.4 | 7.2E-04 | 8.8E-04 | MB231/MB468 | NUC |
| >hg38_21_+_8433172_8446622_RNA45SN1_45S_50ntFlanks@12409.12432.24\|TTGATCCTTCGATGTCGGCTCTTC | >hg38_21_+_8433172_8446622_RNA45SN1_45S_50ntFlanks | 12409 | 12432 | 24 | TTGATCCTTCGATGTCGGCTCTTC | 136.4 | -5.4 | 1.6 | -3.4 | 6.0E-04 | 8.8E-04 | MB231/MB468 | MP |
| >hg38_21_+_8433172_8446622_RNA45SN1_45S_50ntFlanks@10687.10707.21\|CGGCGTCCGGTGAGCTCTCGC | >hg38_21_+_8433172_8446622_RNA45SN1_45S_50ntFlanks | 10687 | 10707 | 21 | CGGCGTCCGGTGAGCTCTCGC | 123.0 | -2.8 | 0.8 | -3.4 | 6.7E-04 | 8.8E-04 | MB231/MB468 | CYTO |
| >hg38_1_-_228634819_228635039_RNA5S12_5S_50ntFlanks@129.169.41\|TTGGATGGGAGACCGCCTGGGAATACCGGGTGCTGTAGGCT | >hg38_1_-_228634819_228635039_RNA5S12_5S_50ntFlanks | 129 | 169 | 41 | TTGGATGGGAGACCGCCTGGGAATACCGGGTGCTGTAGGCT | 558.2 | 6.3 | 1.8 | 3.6 | 3.0E-04 | 8.8E-04 | BT20/MB468 | MP |
| >hg38_21_+_8433172_8446622_RNA45SN1_45S_50ntFlanks@6648.6665.18\|GTACGACTCTTAGCGGTG | >hg38_21_+_8433172_8446622_RNA45SN1_45S_50ntFlanks | 6648 | 6665 | 18 | GTACGACTCTTAGCGGTG | 337.6 | 2.3 | 0.7 | 3.4 | 7.2E-04 | 8.8E-04 | BT20/MB468 | NUC |
| >MI0000093\|hsa-mir-92a-1&WithFlank&13\|+\|91351308\|91351397@54.76.23(+1U)&[MIMAT0000092&hsa-miR-92a-3p&offsets\|0\|+1(+1U);m-9&13\|+\|91351361\|91351382&offsets\|0\|+1(+1U)]\|TATTGCACTTGTCCCGGCCTGTTT | >MI0000093\|hsa-mir-92a-1&WithFlank&13\|+\|91351308\|91351397 | 54 | 76 | 23(+1U) | TATTGCACTTGTCCCGGCCTGTTT | 333.9 | -4.2 | 1.1 | -3.9 | 1.1E-04 | 8.8E-04 | BT20/MB231 | MP |
| >hg38_21_+_8433172_8446622_RNA45SN1_45S_50ntFlanks@12913.12932.20\|ACGTTCGTGGGGAACCTGGC | >hg38_21_+_8433172_8446622_RNA45SN1_45S_50ntFlanks | 12913 | 12932 | 20 | ACGTTCGTGGGGAACCTGGC | 96.9 | -5.4 | 1.6 | -3.4 | 6.0E-04 | 8.8E-04 | MB231/MB468 | MP |
| >MI0000093\|hsa-mir-92a-1&WithFlank&13\|+\|91351308\|91351397@54.73.20(+1U);MI0000094\|hsa-mir-92a-2&WithFlank&X\|-\|134169532\|134169618@54.73.20(+1U)&[MIMAT0000092&hsa-miR-92a-3p&offsets\|0\|-2(+1U);m-9&13\|+\|91351361\|91351382&offsets\|0\|-2(+1U)];[MIMAT0000092_1&hsa-miR-92a-3p&offsets\|0\|-2(+1U);m-11&X\|-\|134169544\|134169565&offsets\|0\|-2(+1U)]\|TATTGCACTTGTCCCGGCCTT | >MI0000093\|hsa-mir-92a-1&WithFlank&13\|+\|91351308\|91351397 | 54 | 73 | 20(+1U) | TATTGCACTTGTCCCGGCCTT | 130.3 | 5.3 | 1.5 | 3.4 | 6.0E-04 | 8.8E-04 | MB231/MB468 | MP |
| >hg38_21_+_8433172_8446622_RNA45SN1_45S_50ntFlanks@10866.10902.37\|CTAAGGGCTGGGTCGGTCGGGCTGGGGCGCGAAGCGG | >hg38_21_+_8433172_8446622_RNA45SN1_45S_50ntFlanks | 10866 | 10902 | 37 | CTAAGGGCTGGGTCGGTCGGGCTGGGGCGCGAAGCGG | 1223.6 | -5.5 | 1.6 | -3.5 | 5.4E-04 | 8.8E-04 | BT20/MB468 | MITO |
| >am_trnaMT_ValTAC_MT_+_1602_1670@48.67.20\|CAACTTAACTTGACCGCTCT | >am_trnaMT_ValTAC_MT_+_1602_1670 | 48 | 67 | 20 | CAACTTAACTTGACCGCTCT | 216.0 | 4.5 | 1.2 | 3.6 | 3.1E-04 | 8.8E-04 | BT20/MB468 | MP |
| >hg38_21_+_8433172_8446622_RNA45SN1_45S_50ntFlanks@10922.10945.24\|CTGGACGAGGCGCCGCCGCCCCCC | >hg38_21_+_8433172_8446622_RNA45SN1_45S_50ntFlanks | 10922 | 10945 | 24 | CTGGACGAGGCGCCGCCGCCCCCC | 1023.7 | -4.2 | 1.2 | -3.4 | 6.4E-04 | 8.8E-04 | BT20/MB468 | CYTO |
| >hg38_21_+_8433172_8446622_RNA45SN1_45S_50ntFlanks@8082.8100.19\|AGCCCAGCGCCGAATCCCC | >hg38_21_+_8433172_8446622_RNA45SN1_45S_50ntFlanks | 8082 | 8100 | 19 | AGCCCAGCGCCGAATCCCC | 108.0 | 10.1 | 2.6 | 3.9 | 1.1E-04 | 8.8E-04 | BT20/MB231 | MP |
| >hg38_MT_+_598_1651_RNR1_12S_50ntFlanks@849.873.25\|AGTGCTTAGTTGAACAGGGCCCTGA | >hg38_MT_+_598_1651_RNR1_12S_50ntFlanks | 849 | 873 | 25 | AGTGCTTAGTTGAACAGGGCCCTGA | 81.6 | 7.2 | 2.0 | 3.6 | 3.1E-04 | 8.8E-04 | BT20/MB468 | MP |
| >hg38_21_+_8433172_8446622_RNA45SN1_45S_50ntFlanks@6654.6681.28\|CTCTTAGCGGTGGATCACTCGGCTCGTG | >hg38_21_+_8433172_8446622_RNA45SN1_45S_50ntFlanks | 6654 | 6681 | 28 | CTCTTAGCGGTGGATCACTCGGCTCGTG | 923.9 | -4.1 | 1.2 | -3.4 | 6.5E-04 | 8.8E-04 | BT20/MB468 | CYTO |
| >hg38_21_+_8433172_8446622_RNA45SN1_45S_50ntFlanks@5490.5516.27\|TGAGAAGACGGTCGAACTTGACTATCT | >hg38_21_+_8433172_8446622_RNA45SN1_45S_50ntFlanks | 5490 | 5516 | 27 | TGAGAAGACGGTCGAACTTGACTATCT | 3773.6 | 3.6 | 0.9 | 3.8 | 1.3E-04 | 8.8E-04 | BT20/MB231 | MITO |
| >hg38_21_+_8433172_8446622_RNA45SN1_45S_50ntFlanks@11564.11584.21\|CCTAGCAGCCGACTTAGAACT | >hg38_21_+_8433172_8446622_RNA45SN1_45S_50ntFlanks | 11564 | 11584 | 21 | CCTAGCAGCCGACTTAGAACT | 223.4 | -4.0 | 1.2 | -3.4 | 6.0E-04 | 8.8E-04 | MB231/MB468 | MP |
| >hg38_21_+_8433172_8446622_RNA45SN1_45S_50ntFlanks@476.518.43\|TTCGTGATCGATGTGGTGACGTCGTGCTCTCCCGGGCCGGGTC | >hg38_21_+_8433172_8446622_RNA45SN1_45S_50ntFlanks | 476 | 518 | 43 | TTCGTGATCGATGTGGTGACGTCGTGCTCTCCCGGGCCGGGTC | 297.0 | 5.1 | 1.5 | 3.4 | 7.1E-04 | 8.8E-04 | MB231/MB468 | MITO |
| >trnaMT_PheGAA_MT_+_577_647@36.64.29\|AAAATGTTTAGACGGGCTCACATCACCCC | >trnaMT_PheGAA_MT_+_577_647 | 36 | 64 | 29 | AAAATGTTTAGACGGGCTCACATCACCCC | 74.9 | 6.0 | 1.7 | 3.4 | 6.0E-04 | 8.8E-04 | MB231/MB468 | MP |
| >hg38_21_+_8433172_8446622_RNA45SN1_45S_50ntFlanks@10808.10837.30\|CAATGTAGGTAAGGGAAGTCGGCAAGCCGG | >hg38_21_+_8433172_8446622_RNA45SN1_45S_50ntFlanks | 10808 | 10837 | 30 | CAATGTAGGTAAGGGAAGTCGGCAAGCCGG | 246.1 | -4.8 | 1.3 | -3.6 | 3.1E-04 | 8.9E-04 | BT20/MB468 | MP |
| >hg38_21_+_8433172_8446622_RNA45SN1_45S_50ntFlanks@12836.12861.26\|CGGTGCGGAGTGCCCTTCGTCCTGGG | >hg38_21_+_8433172_8446622_RNA45SN1_45S_50ntFlanks | 12836 | 12861 | 26 | CGGTGCGGAGTGCCCTTCGTCCTGGG | 181.9 | -4.5 | 1.3 | -3.6 | 3.1E-04 | 8.9E-04 | BT20/MB468 | MP |
| >trnaMT_IleGAT_MT_+_4263_4331@42.72.31\|AATAGGAGCTTAAACCCCCTTATTTCTACCA | >trnaMT_IleGAT_MT_+_4263_4331 | 42 | 72 | 31 | AATAGGAGCTTAAACCCCCTTATTTCTACCA | 64.1 | 9.7 | 2.7 | 3.6 | 3.1E-04 | 8.9E-04 | BT20/MB468 | MP |
| >hg38_1_-_228634819_228635039_RNA5S12_5S_50ntFlanks@136.170.35\|GGAGACCGCCTGGGAATACCGGGTGCTGTAGGCTT | >hg38_1_-_228634819_228635039_RNA5S12_5S_50ntFlanks | 136 | 170 | 35 | GGAGACCGCCTGGGAATACCGGGTGCTGTAGGCTT | 8898.8 | 3.8 | 1.1 | 3.6 | 3.1E-04 | 8.9E-04 | BT20/MB468 | MP |
| >MI0000062\|hsa-let-7a-3&WithFlank&22\|+\|46112743\|46112828@58.79.22;MI0000060\|hsa-let-7a-1&WithFlank&9\|+\|94175951\|94176042@63.84.22&[MIMAT0004481&hsa-let-7a-3p&offsets\|0\|+1;m-180&22\|+\|46112800\|46112821&offsets\|0\|0];[MIMAT0004481_1&hsa-let-7a-3p&offsets\|0\|+1;m-181&9\|+\|94176013\|94176034&offsets\|0\|0]\|CTATACAATCTACTGTCTTTCC | >MI0000062\|hsa-let-7a-3&WithFlank&22\|+\|46112743\|46112828 | 58 | 79 | 22 | CTATACAATCTACTGTCTTTCC | 220.5 | 2.9 | 0.8 | 3.4 | 6.0E-04 | 8.9E-04 | MB231/MB468 | MP |
| >hg38_21_+_8433172_8446622_RNA45SN1_45S_50ntFlanks@11874.11909.36\|AGAAGACCCTGTTGAGCTTGACTCTAGTCTGGCACG | >hg38_21_+_8433172_8446622_RNA45SN1_45S_50ntFlanks | 11874 | 11909 | 36 | AGAAGACCCTGTTGAGCTTGACTCTAGTCTGGCACG | 252.9 | -5.1 | 1.5 | -3.4 | 7.2E-04 | 8.9E-04 | MB231/MB468 | NUC |
| >hg38_21_+_8433172_8446622_RNA45SN1_45S_50ntFlanks@4733.4767.35\|GAACGAAAGTCGGAGGTTCGAAGACGATCAGATAC | >hg38_21_+_8433172_8446622_RNA45SN1_45S_50ntFlanks | 4733 | 4767 | 35 | GAACGAAAGTCGGAGGTTCGAAGACGATCAGATAC | 190.4 | 5.8 | 1.6 | 3.6 | 3.1E-04 | 8.9E-04 | BT20/MB468 | MP |
| >hg38_21_+_8433172_8446622_RNA45SN1_45S_50ntFlanks@11320.11342.23\|CCACGTCTCGTCGCGCGCGCGTC | >hg38_21_+_8433172_8446622_RNA45SN1_45S_50ntFlanks | 11320 | 11342 | 23 | CCACGTCTCGTCGCGCGCGCGTC | 1535.5 | -4.4 | 1.3 | -3.4 | 6.5E-04 | 8.9E-04 | BT20/MB468 | CYTO |
| >MI0000077\|hsa-mir-21&WithFlank&17\|+\|59841260\|59841343@52.74.23&[MIMAT0004494&hsa-miR-21-3p&offsets\|0\|+2;m-52&17\|+\|59841311\|59841332&offsets\|0\|+1]\|CAACACCAGTCGATGGGCTGTCT | >MI0000077\|hsa-mir-21&WithFlank&17\|+\|59841260\|59841343 | 52 | 74 | 23 | CAACACCAGTCGATGGGCTGTCT | 496.9 | 4.3 | 1.3 | 3.4 | 6.0E-04 | 8.9E-04 | MB231/MB468 | MP |
| >hg38_21_+_8433172_8446622_RNA45SN1_45S_50ntFlanks@485.525.41\|GATGTGGTGACGTCGTGCTCTCCCGGGCCGGGTCCGAGCCG | >hg38_21_+_8433172_8446622_RNA45SN1_45S_50ntFlanks | 485 | 525 | 41 | GATGTGGTGACGTCGTGCTCTCCCGGGCCGGGTCCGAGCCG | 59.3 | 9.1 | 2.7 | 3.4 | 6.0E-04 | 8.9E-04 | MB231/MB468 | MP |
| >hg38_MT_+_1621_3279_RNR2_16S_50ntFlanks@51.98.48\|GCTAAACCTAGCCCCAAACCCACTCCACCTTACTACCAGACAACCTTA | >hg38_MT_+_1621_3279_RNR2_16S_50ntFlanks | 51 | 98 | 48 | GCTAAACCTAGCCCCAAACCCACTCCACCTTACTACCAGACAACCTTA | 775.0 | 4.4 | 1.3 | 3.4 | 7.2E-04 | 8.9E-04 | MB231/MB468 | MITO |
| >hg38_21_+_8433172_8446622_RNA45SN1_45S_50ntFlanks@12417.12453.37\|TCGATGTCGGCTCTTCCTATCATTGTGAAGCAGAATT | >hg38_21_+_8433172_8446622_RNA45SN1_45S_50ntFlanks | 12417 | 12453 | 37 | TCGATGTCGGCTCTTCCTATCATTGTGAAGCAGAATT | 350.5 | 2.7 | 0.8 | 3.4 | 6.8E-04 | 8.9E-04 | MB231/MB468 | CYTO |

| | | | | | | | | | | | | | |
|---|---|---|---|---|---|---|---|---|---|---|---|---|---|
| >hg38_21_+_8433172_8446622_RNA45SN1_45S_50ntFlanks@8104.8138.35\|CCGCGGCGGGGCGCGGGACATGTGGCGTACGGAAG | >hg38_21_+_8433172_8446622_RNA45SN1_45S_50ntFlanks | 8104 | 8138 | 35 | CCGCGGCGGGGCGCGGGACATGTGGCGTACGGAAG | 3153.5 | -3.5 | 1.0 | -3.6 | 3.1E-04 | 8.9E-04 | BT20/MB468 | MP |
| >hg38_21_+_8433172_8446622_RNA45SN1_45S_50ntFlanks@12417.12443.27\|TCGATGTCGGCTCTTCCTATCATTGTG | >hg38_21_+_8433172_8446622_RNA45SN1_45S_50ntFlanks | 12417 | 12443 | 27 | TCGATGTCGGCTCTTCCTATCATTGTG | 9042.0 | -2.3 | 0.7 | -3.4 | 6.8E-04 | 8.9E-04 | MB231/MB468 | CYTO |
| >hg38_MT_+_1621_3279_RNR2_16S_50ntFlanks@761.785.25\|AATATCTACAATCAACCAACAAGTC | >hg38_MT_+_1621_3279_RNR2_16S_50ntFlanks | 761 | 785 | 25 | AATATCTACAATCAACCAACAAGTC | 57.7 | 9.1 | 2.6 | 3.4 | 6.1E-04 | 8.9E-04 | MB231/MB468 | MP |
| >hg38_21_+_8433172_8446622_RNA45SN1_45S_50ntFlanks@5515.5534.20\|CTAGAGGAAGTAAAAGTCGT | >hg38_21_+_8433172_8446622_RNA45SN1_45S_50ntFlanks | 5515 | 5534 | 20 | CTAGAGGAAGTAAAAGTCGT | 183.6 | -2.6 | 0.8 | -3.4 | 6.5E-04 | 8.9E-04 | BT20/MB468 | CYTO |
| >hg38_21_+_8433172_8446622_RNA45SN1_45S_50ntFlanks@11540.11580.41\|CGCGGCGCCTCGCCTCGGCCGGCGCCTAGCAGCCGACTTAG | >hg38_21_+_8433172_8446622_RNA45SN1_45S_50ntFlanks | 11540 | 11580 | 41 | CGCGGCGCCTCGCCTCGGCCGGCGCCTAGCAGCCGACTTAG | 1961.0 | -2.7 | 0.8 | -3.4 | 7.2E-04 | 8.9E-04 | MB231/MB468 | MITO |
| >hg38_21_+_8433172_8446622_RNA45SN1_45S_50ntFlanks@12420.12452.33\|ATGTCGGCTCTTCCTATCATTGTGAAGCAGAAT | >hg38_21_+_8433172_8446622_RNA45SN1_45S_50ntFlanks | 12420 | 12452 | 33 | ATGTCGGCTCTTCCTATCATTGTGAAGCAGAAT | 487.2 | 2.7 | 0.8 | 3.4 | 7.2E-04 | 8.9E-04 | MB231/MB468 | MITO |
| >hg38_21_+_8433172_8446622_RNA45SN1_45S_50ntFlanks@12405.12442.38\|CTTTTTGATCCTTCGATGTCGGCTCTTCCTATCATTGT | >hg38_21_+_8433172_8446622_RNA45SN1_45S_50ntFlanks | 12405 | 12442 | 38 | CTTTTTGATCCTTCGATGTCGGCTCTTCCTATCATTGT | 364.4 | -4.1 | 1.2 | -3.4 | 7.2E-04 | 8.9E-04 | MB231/MB468 | MITO |
| >MI0000084\|hsa-mir-26b&WithFlank&2\|+\|218402640\|218402728@21.39.19&[MIMAT0000083&hsa-miR-26b-5p&offsets\|+3\|+1;m-53&2\|+\|218402657\|218402678&offsets\|+3\|0]\|AAGTAATTCAGGATAGGTT | >MI0000084\|hsa-mir-26b&WithFlank&2\|+\|218402640\|218402728 | 21 | 39 | 19 | AAGTAATTCAGGATAGGTT | 58.0 | 9.1 | 2.6 | 3.4 | 6.1E-04 | 8.9E-04 | MB231/MB468 | MP |
| >hg38_21_+_8433172_8446622_RNA45SN1_45S_50ntFlanks@5467.5514.48\|GGCCCACGGCCCTGGCGGAGCGCTGAGAAGACGGTCGAACTTGACTAT | >hg38_21_+_8433172_8446622_RNA45SN1_45S_50ntFlanks | 5467 | 5514 | 48 | GGCCCACGGCCCTGGCGGAGCGCTGAGAAGACGGTCGAACTTGACTAT | 888.4 | -4.7 | 1.4 | -3.4 | 6.8E-04 | 8.9E-04 | MB231/MB468 | CYTO |
| >hg38_21_+_8433172_8446622_RNA45SN1_45S_50ntFlanks@10266.10284.19\|CTGCGGTGAGCCTTGAAGC | >hg38_21_+_8433172_8446622_RNA45SN1_45S_50ntFlanks | 10266 | 10284 | 19 | CTGCGGTGAGCCTTGAAGC | 77.4 | 6.7 | 1.8 | 3.6 | 3.1E-04 | 8.9E-04 | BT20/MB468 | MP |
| >hg38_MT_+_1621_3279_RNR2_16S_50ntFlanks@1094.1117.24\|AAGACGAGAAGACCCTATGGAGCT | >hg38_MT_+_1621_3279_RNR2_16S_50ntFlanks | 1094 | 1117 | 24 | AAGACGAGAAGACCCTATGGAGCT | 125.6 | 2.5 | 0.7 | 3.4 | 6.1E-04 | 8.9E-04 | MB231/MB468 | MP |
| >hg38_21_+_8433172_8446622_RNA45SN1_45S_50ntFlanks@11641.11662.22\|CGGCGGGTGTTGACGCGATGTG | >hg38_21_+_8433172_8446622_RNA45SN1_45S_50ntFlanks | 11641 | 11662 | 22 | CGGCGGGTGTTGACGCGATGTG | 449.7 | -3.9 | 1.1 | -3.4 | 6.5E-04 | 8.9E-04 | BT20/MB468 | CYTO |
| >hg38_21_+_8433172_8446622_RNA45SN1_45S_50ntFlanks@11575.11601.27\|ACTTAGAACTGGTGCGGACCAGGGGAA | >hg38_21_+_8433172_8446622_RNA45SN1_45S_50ntFlanks | 11575 | 11601 | 27 | ACTTAGAACTGGTGCGGACCAGGGGAA | 1007.1 | -6.2 | 1.7 | -3.6 | 3.1E-04 | 8.9E-04 | BT20/MB468 | MP |
| >MI0000063\|hsa-let-7b&WithFlank&22\|+\|46113680\|46113774@12.35.24&[MIMAT0000063&hsa-let-7b-5p&offsets\|0\|+2;m-14&22\|+\|46113691\|46113712&offsets\|0\|+2]\|TGAGGTAGTAGGTTGTGTGGTTTC | >MI0000063\|hsa-let-7b&WithFlank&22\|+\|46113680\|46113774 | 12 | 35 | 24 | TGAGGTAGTAGGTTGTGTGGTTTC | 128.5 | 7.0 | 1.9 | 3.6 | 3.1E-04 | 9.0E-04 | BT20/MB468 | MP |
| >hg38_21_+_8433172_8446622_RNA45SN1_45S_50ntFlanks@11320.11338.19\|CCACGTCTCGTCGCGCGCG | >hg38_21_+_8433172_8446622_RNA45SN1_45S_50ntFlanks | 11320 | 11338 | 19 | CCACGTCTCGTCGCGCGCG | 240.2 | -4.9 | 1.4 | -3.4 | 6.5E-04 | 9.0E-04 | BT20/MB468 | CYTO |
| >hg38_21_+_8433172_8446622_RNA45SN1_45S_50ntFlanks@9032.9056.25\|CCTCGTGGGGGGGCCGGGCCACCCC | >hg38_21_+_8433172_8446622_RNA45SN1_45S_50ntFlanks | 9032 | 9056 | 25 | CCTCGTGGGGGGGCCGGGCCACCCC | 136.7 | -5.1 | 1.5 | -3.5 | 5.5E-04 | 9.0E-04 | BT20/MB468 | MITO |
| >hg38_1_-_228634819_228635039_RNA5S12_5S_50ntFlanks@150.167.18\|AATACCGGGTGCTGTAGG | >hg38_1_-_228634819_228635039_RNA5S12_5S_50ntFlanks | 150 | 167 | 18 | AATACCGGGTGCTGTAGG | 87.0 | -5.5 | 1.6 | -3.4 | 6.1E-04 | 9.0E-04 | MB231/MB468 | MP |
| >hg38_21_+_8433172_8446622_RNA45SN1_45S_50ntFlanks@9970.9988.19\|CTAAGGAGTGTGTAACAAC | >hg38_21_+_8433172_8446622_RNA45SN1_45S_50ntFlanks | 9970 | 9988 | 19 | CTAAGGAGTGTGTAACAAC | 140.4 | 10.5 | 2.7 | 3.9 | 1.1E-04 | 9.0E-04 | BT20/MB231 | MP |
| >hg38_MT_+_1621_3279_RNR2_16S_50ntFlanks@1095.1117.23\|AGACGAGAAGACCCTATGGAGCT | >hg38_MT_+_1621_3279_RNR2_16S_50ntFlanks | 1095 | 1117 | 23 | AGACGAGAAGACCCTATGGAGCT | 315.3 | 3.2 | 0.9 | 3.4 | 6.1E-04 | 9.0E-04 | MB231/MB468 | MP |
| >am_trna111_HisGTG_1_-_147774845_147774916@-1G.26.27,trna118_HisGTG_1_-_145396881_145396952@-1G.26.27,trna16_HisGTG_1_+_146544773_146544844@-1G.26.27,trna1_HisGTG_15_+_45493349_45493420@-1G.26.27,trna21_HisGTG_1_+_147753471_147753542@-1G.26.27,trna33_HisGTG_6_+_27125906_27125977@-1G.26.27,trna7_HisGTG_9_-_14433938_14434009@-1G.26.27,trna8_HisGTG_15_-_45492611_45492682@-1G.26.27,trna9_HisGTG_15_-_45490804_45490875@-1G.26.27\|GGCCGTGATCGTATAGTGGTTAGTACT | >am_trna111_HisGTG_1_-_147774845_147774916 | -1G | 26 | 27 | GGCCGTGATCGTATAGTGGTTAGTACT | 632.8 | -3.2 | 1.0 | -3.4 | 7.3E-04 | 9.0E-04 | BT20/MB468 | NUC |
| >hg38_21_+_8433172_8446622_RNA45SN1_45S_50ntFlanks@4068.4094.27\|ACTTTCGATGGTAGTCGCCGTGCCTAC | >hg38_21_+_8433172_8446622_RNA45SN1_45S_50ntFlanks | 4068 | 4094 | 27 | ACTTTCGATGGTAGTCGCCGTGCCTAC | 93.1 | -5.3 | 1.5 | -3.4 | 6.1E-04 | 9.0E-04 | MB231/MB468 | MP |
| >MI0000809\|hsa-mir-151a&WithFlank&8\|-\|140732558\|140732659@53.74.22(+1G)&[MIMAT0000757&hsa-miR-151a-3p&offsets\|0\|+1(+1G);m-30&8\|-\|140732587\|140732607&offsets\|0\|+1(+1G)]\|CTAGACTGAAGCTCCTTGAGGAG | >MI0000809\|hsa-mir-151a&WithFlank&8\|-\|140732558\|140732659 | 53 | 74 | 22(+1G) | CTAGACTGAAGCTCCTTGAGGAG | 205.1 | 2.9 | 0.8 | 3.9 | 1.1E-04 | 9.0E-04 | BT20/MB231 | CYTO |
| >hg38_21_+_8433172_8446622_RNA45SN1_45S_50ntFlanks@9419.9453.35\|TCCGCCGAGGGCGCACCACCGGCCCGTCTCGCCCG | >hg38_21_+_8433172_8446622_RNA45SN1_45S_50ntFlanks | 9419 | 9453 | 35 | TCCGCCGAGGGCGCACCACCGGCCCGTCTCGCCCG | 144.4 | -5.5 | 1.5 | -3.6 | 3.1E-04 | 9.0E-04 | BT20/MB468 | MP |
| >hg38_21_+_8433172_8446622_RNA45SN1_45S_50ntFlanks@10797.10828.32\|GCATGTTGGAACAATGTAGGTAAGGGAAGTCG | >hg38_21_+_8433172_8446622_RNA45SN1_45S_50ntFlanks | 10797 | 10828 | 32 | GCATGTTGGAACAATGTAGGTAAGGGAAGTCG | 245.0 | -4.3 | 1.2 | -3.6 | 3.1E-04 | 9.0E-04 | BT20/MB468 | MP |
| >hg38_1_-_228634819_228635039_RNA5S12_5S_50ntFlanks@51.91.41\|GTCTACGGCCATACCACCCTGAACGCGCCCGATCTCGTCTG | >hg38_1_-_228634819_228635039_RNA5S12_5S_50ntFlanks | 51 | 91 | 41 | GTCTACGGCCATACCACCCTGAACGCGCCCGATCTCGTCTG | 4027.0 | 7.0 | 1.9 | 3.6 | 3.1E-04 | 9.0E-04 | BT20/MB468 | MP |
| >MI0000102\|hsa-mir-100&WithFlank&11\|-\|122152223\|122152314@19.39.21&[MIMAT0000098&hsa-miR-100-5p&offsets\|0\|-1;m-27&11\|-\|122152276\|122152296&offsets\|0\|0]\|AACCCGTAGATCCGAACTTGT | >MI0000102\|hsa-mir-100&WithFlank&11\|-\|122152223\|122152314 | 19 | 39 | 21 | AACCCGTAGATCCGAACTTGT | 3308.7 | 6.2 | 1.7 | 3.6 | 3.1E-04 | 9.0E-04 | BT20/MB468 | MP |
| >hg38_21_+_8433172_8446622_RNA45SN1_45S_50ntFlanks@11854.11886.33\|TTGGCGGAATCAGCGGGGAAAGAAGACCCTGTT | >hg38_21_+_8433172_8446622_RNA45SN1_45S_50ntFlanks | 11854 | 11886 | 33 | TTGGCGGAATCAGCGGGGAAAGAAGACCCTGTT | 64.3 | 6.6 | 1.8 | 3.6 | 3.1E-04 | 9.0E-04 | BT20/MB468 | MP |
| >MI0000079\|hsa-mir-23a&WithFlank&19\|-\|13836581\|13836665@54.73.20(+2U)&[MIMAT0000078&hsa-miR-23a-3p&offsets\|+3\|+2(+2U);m-31&19\|-\|13836595\|13836615&offsets\|+3\|+2(+2U)]\|ACATTGCCAGGGATTTCCAATT | >MI0000079\|hsa-mir-23a&WithFlank&19\|-\|13836581\|13836665 | 54 | 73 | 20(+2U) | ACATTGCCAGGGATTTCCAATT | 328.6 | 12.3 | 3.6 | 3.4 | 7.4E-04 | 9.0E-04 | MB231/MB468 | NUC |
| >trnaMT_HisGTG_MT_+_12138_12206@46.70.25\|GAGGCTTACGACCCCTTATTTACCC | >trnaMT_HisGTG_MT_+_12138_12206 | 46 | 70 | 25 | GAGGCTTACGACCCCTTATTTACCC | 63.7 | 9.7 | 2.7 | 3.6 | 3.1E-04 | 9.0E-04 | BT20/MB468 | MP |
| >hg38_21_+_8433172_8446622_RNA45SN1_45S_50ntFlanks@12911.12933.23\|GCACGTTCGTGGGGAACCTGGCG | >hg38_21_+_8433172_8446622_RNA45SN1_45S_50ntFlanks | 12911 | 12933 | 23 | GCACGTTCGTGGGGAACCTGGCG | 76.0 | -5.3 | 1.6 | -3.4 | 6.1E-04 | 9.0E-04 | MB231/MB468 | MP |
| >hg38_21_+_8433172_8446622_RNA45SN1_45S_50ntFlanks@6653.6688.36\|ACTCTTAGCGGTGGATCACTCGGCTCGTGCGTCGAT | >hg38_21_+_8433172_8446622_RNA45SN1_45S_50ntFlanks | 6653 | 6688 | 36 | ACTCTTAGCGGTGGATCACTCGGCTCGTGCGTCGAT | 2327.5 | 2.7 | 0.7 | 3.8 | 1.3E-04 | 9.0E-04 | BT20/MB231 | MITO |
| >hg38_MT_+_1621_3279_RNR2_16S_50ntFlanks@1351.1396.46\|AATAGGGTTTACGACCTCGATGTTGGATCAGGACATCCCGATGGTG | >hg38_MT_+_1621_3279_RNR2_16S_50ntFlanks | 1351 | 1396 | 46 | AATAGGGTTTACGACCTCGATGTTGGATCAGGACATCCCGATGGTG | 62.8 | 9.7 | 2.7 | 3.6 | 3.1E-04 | 9.0E-04 | BT20/MB468 | MP |
| >hg38_21_+_8433172_8446622_RNA45SN1_45S_50ntFlanks@5425.5466.42\|TGGATGGTTTAGTGAGGCCCTCGGATCGGCCCCGCCGGGGTC | >hg38_21_+_8433172_8446622_RNA45SN1_45S_50ntFlanks | 5425 | 5466 | 42 | TGGATGGTTTAGTGAGGCCCTCGGATCGGCCCCGCCGGGGTC | 51.0 | 9.0 | 2.3 | 3.9 | 1.1E-04 | 9.0E-04 | BT20/MB231 | MP |
| >hg38_21_+_8433172_8446622_RNA45SN1_45S_50ntFlanks@7992.8013.22\|CGTGGCGACCCGCTGAATTTAA | >hg38_21_+_8433172_8446622_RNA45SN1_45S_50ntFlanks | 7992 | 8013 | 22 | CGTGGCGACCCGCTGAATTTAA | 63.1 | -5.4 | 1.6 | -3.4 | 6.1E-04 | 9.0E-04 | MB231/MB468 | MP |
| >hg38_21_+_8433172_8446622_RNA45SN1_45S_50ntFlanks@7976.8008.33\|GCGACCTCAGATCAGACGTGGCGACCCGCTGAA | >hg38_21_+_8433172_8446622_RNA45SN1_45S_50ntFlanks | 7976 | 8008 | 33 | GCGACCTCAGATCAGACGTGGCGACCCGCTGAA | 830.3 | 3.0 | 0.9 | 3.4 | 7.4E-04 | 9.1E-04 | MB231/MB468 | NUC |
| >hg38_21_+_8433172_8446622_RNA45SN1_45S_50ntFlanks@12624.12661.38\|CCAATGGGGCGAAGCTACCATCTGTGGGATTATGACTG | >hg38_21_+_8433172_8446622_RNA45SN1_45S_50ntFlanks | 12624 | 12661 | 38 | CCAATGGGGCGAAGCTACCATCTGTGGGATTATGACTG | 1690.0 | -4.1 | 1.1 | -3.6 | 3.1E-04 | 9.1E-04 | BT20/MB468 | MP |
| >hg38_21_+_8433172_8446622_RNA45SN1_45S_50ntFlanks@4759.4785.27\|ATCAGATACCGTCGTAGTTCCGACCAT | >hg38_21_+_8433172_8446622_RNA45SN1_45S_50ntFlanks | 4759 | 4785 | 27 | ATCAGATACCGTCGTAGTTCCGACCAT | 95.6 | -5.3 | 1.6 | -3.4 | 6.1E-04 | 9.1E-04 | MB231/MB468 | MP |
| >am_MI0000086\|hsa-mir-28&WithFlank&3\|+\|188688775\|188688872@20.39.20&[MIMAT0000085&hsa-miR-28-5p&offsets\|0\|-2;m-133&3\|+\|188688794\|188688815&offsets\|0\|-2]\|AAGGAGCTCACAGTCTATTG | >am_MI0000086\|hsa-mir-28&WithFlank&3\|+\|188688775\|188688872 | 20 | 39 | 20 | AAGGAGCTCACAGTCTATTG | 572.1 | 4.7 | 1.4 | 3.4 | 6.2E-04 | 9.1E-04 | MB231/MB468 | MP |
| >hg38_21_+_8433172_8446622_RNA45SN1_45S_50ntFlanks@10686.10712.27\|GCGGCGTCCGGTGAGCTCTCGCTGGCC | >hg38_21_+_8433172_8446622_RNA45SN1_45S_50ntFlanks | 10686 | 10712 | 27 | GCGGCGTCCGGTGAGCTCTCGCTGGCC | 240.7 | -2.6 | 0.7 | -3.4 | 6.2E-04 | 9.1E-04 | MB231/MB468 | MP |
| >hg38_21_+_8433172_8446622_RNA45SN1_45S_50ntFlanks@6647.6664.18\|CGTACGACTCTTAGCGGT | >hg38_21_+_8433172_8446622_RNA45SN1_45S_50ntFlanks | 6647 | 6664 | 18 | CGTACGACTCTTAGCGGT | 4594.7 | 2.3 | 0.7 | 3.4 | 7.3E-04 | 9.1E-04 | MB231/MB468 | MITO |
| >MI0000478\|hsa-mir-149&WithFlank&2\|+\|240455995\|240456095@21.43.23(+1U)&[MIMAT0000450&hsa-miR-149-5p&offsets\|0\|0(+1U);m-195&2\|+\|240456015\|240456037&offsets\|0\|0(+1U)]\|TCTGGCTCCGTGTCTTCACTCCCT | >MI0000478\|hsa-mir-149&WithFlank&2\|+\|240455995\|240456095 | 21 | 43 | 23(+1U) | TCTGGCTCCGTGTCTTCACTCCCT | 242.6 | 3.6 | 0.9 | 3.9 | 1.2E-04 | 9.1E-04 | BT20/MB231 | CYTO |
| >MI0000440\|hsa-mir-27b&WithFlank&9\|+\|95085439\|95085547@70.87.18&[MIMAT0000419&hsa-miR-27b-3p&offsets\|+3\|0;m-39&9\|+\|95085505\|95085525&offsets\|+3\|0]\|ACAGTGGCTAAGTTCTGC | >MI0000440\|hsa-mir-27b&WithFlank&9\|+\|95085439\|95085547 | 70 | 87 | 18 | ACAGTGGCTAAGTTCTGC | 327.0 | 12.2 | 3.6 | 3.4 | 7.4E-04 | 9.1E-04 | MB231/MB468 | NUC |
| >hg38_21_+_8433172_8446622_RNA45SN1_45S_50ntFlanks@9566.9595.30\|GACGTGCAAATCGGTCGTCCGACCTGGGTA | >hg38_21_+_8433172_8446622_RNA45SN1_45S_50ntFlanks | 9566 | 9595 | 30 | GACGTGCAAATCGGTCGTCCGACCTGGGTA | 880.5 | -6.4 | 1.9 | -3.4 | 7.4E-04 | 9.1E-04 | BT20/MB468 | NUC |
| >trna111_HisGTG_1_-_147774845_147774916@-1G.31.32,trna118_HisGTG_1_-_145396881_145396952@-1G.31.32,trna16_HisGTG_1_+_146544773_146544844@-1G.31.32,trna1_HisGTG_15_+_45493349_45493420@-1G.31.32,trna21_HisGTG_1_+_147753471_147753542@-1G.31.32,trna33_HisGTG_6_+_27125906_27125977@-1G.31.32,trna7_HisGTG_9_-_14433938_14434009@-1G.31.32,trna8_HisGTG_15_-_45492611_45492682@-1G.31.32,trna9_HisGTG_15_-_45490804_45490875@-1G.31.32\|GGCCGTGATCGTATAGTGGTTAGTACTCTGCG | >trna111_HisGTG_1_-_147774845_147774916 | -1G | 31 | 32 | GGCCGTGATCGTATAGTGGTTAGTACTCTGCG | 17291.0 | -4.6 | 1.2 | -3.9 | 1.1E-04 | 9.1E-04 | BT20/MB231 | MP |
| >hg38_21_+_8433172_8446622_RNA45SN1_45S_50ntFlanks@10410.10429.20\|TCAGTCGGTCCTGAGAGATG | >hg38_21_+_8433172_8446622_RNA45SN1_45S_50ntFlanks | 10410 | 10429 | 20 | TCAGTCGGTCCTGAGAGATG | 344.8 | 6.3 | 1.6 | 3.9 | 1.1E-04 | 9.1E-04 | BT20/MB231 | MP |
| >hg38_21_+_8433172_8446622_RNA45SN1_45S_50ntFlanks@12894.12927.34\|CTCGCCCGTCACGCACCGCACGTTCGTGGGGAAC | >hg38_21_+_8433172_8446622_RNA45SN1_45S_50ntFlanks | 12894 | 12927 | 34 | CTCGCCCGTCACGCACCGCACGTTCGTGGGGAAC | 110.7 | -5.6 | 1.5 | -3.6 | 3.2E-04 | 9.1E-04 | BT20/MB468 | MP |
| >hg38_21_+_8433172_8446622_RNA45SN1_45S_50ntFlanks@9566.9595.30\|GACGTGCAAATCGGTCGTCCGACCTGGGTA | >hg38_21_+_8433172_8446622_RNA45SN1_45S_50ntFlanks | 9566 | 9595 | 30 | GACGTGCAAATCGGTCGTCCGACCTGGGTA | 710.9 | -4.2 | 1.2 | -3.4 | 7.3E-04 | 9.1E-04 | MB231/MB468 | MITO |

<table>
<tr><th></th><th></th><th></th><th></th><th></th><th></th><th></th><th></th><th></th><th></th><th></th><th></th><th></th><th></th></tr>
<tr><td>>hg38_21_+_8433172_8446622_RNA45SN1_45S_50ntFlanks@7980.8003.24|CCTCAGATCAGACGTGGCGACCCG</td><td>>hg38_21_+_8433172_8446622_RNA45SN1_45S_50ntFlanks</td><td>7980</td><td>8003</td><td>24</td><td>CCTCAGATCAGACGTGGCGACCCG</td><td>303.6</td><td>-3.8</td><td>1.1</td><td>-3.4</td><td>6.6E-04</td><td>9.1E-04</td><td>BT20/MB468</td><td>CYTO</td></tr>
<tr><td>>MI0000434|hsa-let-7i&WithFlank&12|+|62603680|62603775@12.32.21(+1A)&[MIMAT0000415&hsa-let-7i-5p&offsets|0|-1(+1A);m-49&12|+|62603691|62603712&offsets|0|-1(+1A)]|TGAGGTAGTAGTTTGTGCTGTA</td><td>>MI0000434|hsa-let-7i&WithFlank&12|+|62603680|62603775</td><td>12</td><td>32</td><td>21(+1A)</td><td>TGAGGTAGTAGTTTGTGCTGTA</td><td>2792.0</td><td>4.0</td><td>1.1</td><td>3.6</td><td>3.2E-04</td><td>9.1E-04</td><td>BT20/MB468</td><td>MP</td></tr>
<tr><td>>hg38_MT_+_1621_3279_RNR2_16S_50ntFlanks@382.411.30|GAGCCTGGTGATAGCTGGTTGTCCAAGATA</td><td>>hg38_MT_+_1621_3279_RNR2_16S_50ntFlanks</td><td>382</td><td>411</td><td>30</td><td>GAGCCTGGTGATAGCTGGTTGTCCAAGATA</td><td>64.4</td><td>9.8</td><td>2.7</td><td>3.6</td><td>3.2E-04</td><td>9.1E-04</td><td>BT20/MB468</td><td>MP</td></tr>
<tr><td>>trna116_GluCTC_1_-_145399233_145399304@7.31.25,trna59_GluCTC_1_+_249168447_249168518@7.31.25,trna71_GluCTC_1_-_161439189_161439260@7.31.25,trna74_GluCTC_1_-_161431809_161431880@7.31.25,trna77_GluCTC_1_-_161424398_161424469@7.31.25,trna77_GluCTC_6_+_28949976_28950047@7.31.25,trna80_GluCTC_1_-_161417018_161417089@7.31.25,trna87_GluCTC_6_-_126101393_126101464@7.31.25|GTGGTCTAGTGGTTAGGATTCGGCG</td><td>>trna116_GluCTC_1_-_145399233_145399304</td><td>7</td><td>31</td><td>25</td><td>GTGGTCTAGTGGTTAGGATTCGGCG</td><td>315.8</td><td>-5.8</td><td>1.7</td><td>-3.4</td><td>7.4E-04</td><td>9.1E-04</td><td>BT20/MB468</td><td>NUC</td></tr>
<tr><td>>MI0000446|hsa-mir-125b-1&WithFlank&11|-|122099751|122099850@61.82.22&[MIMAT0004592&hsa-miR-125b-1-3p&offsets|0|0;m-485&11|-|122099769|122099790&offsets|0|0]|ACGGGTTAGGCTCTTGGGAGCT</td><td>>MI0000446|hsa-mir-125b-1&WithFlank&11|-|122099751|122099850</td><td>61</td><td>82</td><td>22</td><td>ACGGGTTAGGCTCTTGGGAGCT</td><td>1664.2</td><td>-7.1</td><td>1.8</td><td>-3.9</td><td>1.1E-04</td><td>9.1E-04</td><td>BT20/MB231</td><td>MP</td></tr>
<tr><td>>hg38_21_+_8433172_8446622_RNA45SN1_45S_50ntFlanks@10808.10831.24|CAATGTAGGTAAGGGAAGTCGGCA</td><td>>hg38_21_+_8433172_8446622_RNA45SN1_45S_50ntFlanks</td><td>10808</td><td>10831</td><td>24</td><td>CAATGTAGGTAAGGGAAGTCGGCA</td><td>256.5</td><td>-3.8</td><td>1.1</td><td>-3.5</td><td>5.5E-04</td><td>9.1E-04</td><td>BT20/MB468</td><td>MITO</td></tr>
<tr><td>>hg38_21_+_8433172_8446622_RNA45SN1_45S_50ntFlanks@10345.10388.44|TTCAAACGAGAACTTTGAAGGCCGAAGTGGAGAAGGGTTCCATG</td><td>>hg38_21_+_8433172_8446622_RNA45SN1_45S_50ntFlanks</td><td>10345</td><td>10388</td><td>44</td><td>TTCAAACGAGAACTTTGAAGGCCGAAGTGGAGAAGGGTTCCATG</td><td>473.2</td><td>4.7</td><td>1.4</td><td>3.5</td><td>5.6E-04</td><td>9.1E-04</td><td>BT20/MB468</td><td>MITO</td></tr>
<tr><td>>hg38_21_+_8433172_8446622_RNA45SN1_45S_50ntFlanks@12081.12106.26|CTCTCGCTTCTGGCGCCAAGCGCCCG</td><td>>hg38_21_+_8433172_8446622_RNA45SN1_45S_50ntFlanks</td><td>12081</td><td>12106</td><td>26</td><td>CTCTCGCTTCTGGCGCCAAGCGCCCG</td><td>536.0</td><td>-4.0</td><td>1.2</td><td>-3.5</td><td>5.6E-04</td><td>9.1E-04</td><td>BT20/MB468</td><td>MITO</td></tr>
<tr><td>>MI0000815|hsa-mir-339&WithFlank&7|-|1022927|1023032@56.76.21&[MIMAT0004702&hsa-miR-339-3p&offsets|0|-2;m-235&7|-|1022957|1022977&offsets|0|0]|TGAGCGCCTCGACGACAGAGC</td><td>>MI0000815|hsa-mir-339&WithFlank&7|-|1022927|1023032</td><td>56</td><td>76</td><td>21</td><td>TGAGCGCCTCGACGACAGAGC</td><td>330.1</td><td>2.7</td><td>0.8</td><td>3.4</td><td>6.2E-04</td><td>9.1E-04</td><td>MB231/MB468</td><td>MP</td></tr>
<tr><td>>hg38_1_-_228634819_228635039_RNA5S12_5S_50ntFlanks@139.169.31|GACCGCCTGGGAATACCGGGTGCTGTAGGCT</td><td>>hg38_1_-_228634819_228635039_RNA5S12_5S_50ntFlanks</td><td>139</td><td>169</td><td>31</td><td>GACCGCCTGGGAATACCGGGTGCTGTAGGCT</td><td>1661.5</td><td>3.2</td><td>0.9</td><td>3.6</td><td>3.2E-04</td><td>9.1E-04</td><td>BT20/MB468</td><td>MP</td></tr>
<tr><td>>hg38_21_+_8433172_8446622_RNA45SN1_45S_50ntFlanks@10869.10890.22|AGGGCTGGGTCGGTCGGGCTGG</td><td>>hg38_21_+_8433172_8446622_RNA45SN1_45S_50ntFlanks</td><td>10869</td><td>10890</td><td>22</td><td>AGGGCTGGGTCGGTCGGGCTGG</td><td>801.7</td><td>-3.0</td><td>0.8</td><td>-3.6</td><td>3.2E-04</td><td>9.1E-04</td><td>BT20/MB468</td><td>MP</td></tr>
<tr><td>>hg38_21_+_8433172_8446622_RNA45SN1_45S_50ntFlanks@5472.5512.41|ACGGCCCTGGCGGAGCGCTGAGAAGACGGTCGAACTTGACT</td><td>>hg38_21_+_8433172_8446622_RNA45SN1_45S_50ntFlanks</td><td>5472</td><td>5512</td><td>41</td><td>ACGGCCCTGGCGGAGCGCTGAGAAGACGGTCGAACTTGACT</td><td>503.3</td><td>5.3</td><td>1.5</td><td>3.6</td><td>3.2E-04</td><td>9.1E-04</td><td>BT20/MB468</td><td>MP</td></tr>
<tr><td>>hg38_21_+_8433172_8446622_RNA45SN1_45S_50ntFlanks@10797.10825.29|GCATGTTGGAACAATGTAGGTAAGGGAAG</td><td>>hg38_21_+_8433172_8446622_RNA45SN1_45S_50ntFlanks</td><td>10797</td><td>10825</td><td>29</td><td>GCATGTTGGAACAATGTAGGTAAGGGAAG</td><td>406.8</td><td>-5.8</td><td>1.6</td><td>-3.6</td><td>3.2E-04</td><td>9.1E-04</td><td>BT20/MB468</td><td>MP</td></tr>
<tr><td>>hg38_21_+_8433172_8446622_RNA45SN1_45S_50ntFlanks@9504.9523.20|GTGAACTATGCCTGGGCAGG</td><td>>hg38_21_+_8433172_8446622_RNA45SN1_45S_50ntFlanks</td><td>9504</td><td>9523</td><td>20</td><td>GTGAACTATGCCTGGGCAGG</td><td>2878.7</td><td>-4.3</td><td>1.2</td><td>-3.6</td><td>3.2E-04</td><td>9.1E-04</td><td>BT20/MB468</td><td>MP</td></tr>
<tr><td>>hg38_21_+_8433172_8446622_RNA45SN1_45S_50ntFlanks@4732.4752.21|AGAACGAAAGTCGGAGGTTCG</td><td>>hg38_21_+_8433172_8446622_RNA45SN1_45S_50ntFlanks</td><td>4732</td><td>4752</td><td>21</td><td>AGAACGAAAGTCGGAGGTTCG</td><td>163.8</td><td>-2.1</td><td>0.6</td><td>-3.4</td><td>7.5E-04</td><td>9.1E-04</td><td>MB231/MB468</td><td>NUC</td></tr>
<tr><td>>hg38_21_+_8433172_8446622_RNA45SN1_45S_50ntFlanks@12463.12487.25|TTGGATTGTTCACCCACTAATAGGG</td><td>>hg38_21_+_8433172_8446622_RNA45SN1_45S_50ntFlanks</td><td>12463</td><td>12487</td><td>25</td><td>TTGGATTGTTCACCCACTAATAGGG</td><td>191.4</td><td>-2.8</td><td>0.8</td><td>-3.4</td><td>6.7E-04</td><td>9.1E-04</td><td>BT20/MB468</td><td>CYTO</td></tr>
<tr><td>>hg38_21_+_8433172_8446622_RNA45SN1_45S_50ntFlanks@4102.4127.26|ACCACGGGTGACGGGGAATCAGGGTT</td><td>>hg38_21_+_8433172_8446622_RNA45SN1_45S_50ntFlanks</td><td>4102</td><td>4127</td><td>26</td><td>ACCACGGGTGACGGGGAATCAGGGTT</td><td>409.0</td><td>-3.6</td><td>1.1</td><td>-3.4</td><td>7.4E-04</td><td>9.1E-04</td><td>MB231/MB468</td><td>MITO</td></tr>
<tr><td>>trnaMT_PheGAA_MT_+_577_647@37.65.29|AAATGTTTAGACGGGCTCACATCACCCCA</td><td>>trnaMT_PheGAA_MT_+_577_647</td><td>37</td><td>65</td><td>29</td><td>AAATGTTTAGACGGGCTCACATCACCCCA</td><td>62.4</td><td>9.7</td><td>2.7</td><td>3.6</td><td>3.2E-04</td><td>9.1E-04</td><td>BT20/MB468</td><td>MP</td></tr>
<tr><td>>hg38_21_+_8433172_8446622_RNA45SN1_45S_50ntFlanks@4090.4111.22|CCTACCATGGTGACCACGGGTG</td><td>>hg38_21_+_8433172_8446622_RNA45SN1_45S_50ntFlanks</td><td>4090</td><td>4111</td><td>22</td><td>CCTACCATGGTGACCACGGGTG</td><td>63.9</td><td>-9.1</td><td>2.5</td><td>-3.6</td><td>3.2E-04</td><td>9.1E-04</td><td>BT20/MB468</td><td>MP</td></tr>
<tr><td>>hg38_21_+_8433172_8446622_RNA45SN1_45S_50ntFlanks@9266.9285.20|CGACGTCGGCTACCCACCCG</td><td>>hg38_21_+_8433172_8446622_RNA45SN1_45S_50ntFlanks</td><td>9266</td><td>9285</td><td>20</td><td>CGACGTCGGCTACCCACCCG</td><td>328.2</td><td>-3.3</td><td>1.0</td><td>-3.4</td><td>7.5E-04</td><td>9.1E-04</td><td>MB231/MB468</td><td>NUC</td></tr>
<tr><td>>trnaMT_GluTTC_MT_-_14674_14742@46.71.26,trnalookalike8_GluTTC_5_-_93905172_93905240@46.71.26|GTGGTTGTAGTCCGTGCGAGAATACC</td><td>>trnaMT_GluTTC_MT_-_14674_14742</td><td>46</td><td>71</td><td>26</td><td>GTGGTTGTAGTCCGTGCGAGAATACC</td><td>247.3</td><td>3.1</td><td>0.9</td><td>3.4</td><td>6.2E-04</td><td>9.1E-04</td><td>MB231/MB468</td><td>MP</td></tr>
<tr><td>>hg38_MT_+_598_1651_RNR1_12S_50ntFlanks@917.946.30|AGGACATTTAACTAAAACCCCTACGCATTT</td><td>>hg38_MT_+_598_1651_RNR1_12S_50ntFlanks</td><td>917</td><td>946</td><td>30</td><td>AGGACATTTAACTAAAACCCCTACGCATTT</td><td>797.9</td><td>6.6</td><td>1.8</td><td>3.6</td><td>3.2E-04</td><td>9.2E-04</td><td>BT20/MB468</td><td>MP</td></tr>
<tr><td>>hg38_21_+_8433172_8446622_RNA45SN1_45S_50ntFlanks@7988.8018.31|CAGACGTGGCGACCCGCTGAATTTAAGCATA</td><td>>hg38_21_+_8433172_8446622_RNA45SN1_45S_50ntFlanks</td><td>7988</td><td>8018</td><td>31</td><td>CAGACGTGGCGACCCGCTGAATTTAAGCATA</td><td>325.1</td><td>-4.6</td><td>1.4</td><td>-3.4</td><td>7.5E-04</td><td>9.2E-04</td><td>BT20/MB468</td><td>NUC</td></tr>
<tr><td>>hg38_21_+_8433172_8446622_RNA45SN1_45S_50ntFlanks@8319.8356.38|CGAGACCGATAGTCAACAAGTACCGTAAGGGAAAGTTG</td><td>>hg38_21_+_8433172_8446622_RNA45SN1_45S_50ntFlanks</td><td>8319</td><td>8356</td><td>38</td><td>CGAGACCGATAGTCAACAAGTACCGTAAGGGAAAGTTG</td><td>319.7</td><td>-4.1</td><td>1.2</td><td>-3.4</td><td>7.5E-04</td><td>9.2E-04</td><td>MB231/MB468</td><td>NUC</td></tr>
<tr><td>>hg38_21_+_8433172_8446622_RNA45SN1_45S_50ntFlanks@9534.9562.29|AGGAAACTCTGGTGGAGGTCCGTAGCGGT</td><td>>hg38_21_+_8433172_8446622_RNA45SN1_45S_50ntFlanks</td><td>9534</td><td>9562</td><td>29</td><td>AGGAAACTCTGGTGGAGGTCCGTAGCGGT</td><td>2284.8</td><td>-5.0</td><td>1.4</td><td>-3.6</td><td>3.2E-04</td><td>9.2E-04</td><td>BT20/MB468</td><td>MP</td></tr>
<tr><td>>hg38_21_+_8433172_8446622_RNA45SN1_45S_50ntFlanks@10099.10117.19|CGCGCGCGCGTGTGGTGTG</td><td>>hg38_21_+_8433172_8446622_RNA45SN1_45S_50ntFlanks</td><td>10099</td><td>10117</td><td>19</td><td>CGCGCGCGCGTGTGGTGTG</td><td>2888.0</td><td>-5.2</td><td>1.5</td><td>-3.6</td><td>3.2E-04</td><td>9.2E-04</td><td>BT20/MB468</td><td>MP</td></tr>
<tr><td>>hg38_21_+_8433172_8446622_RNA45SN1_45S_50ntFlanks@6671.6697.27|CTCGGCTCGTGCGTCGATGAAGAACGC</td><td>>hg38_21_+_8433172_8446622_RNA45SN1_45S_50ntFlanks</td><td>6671</td><td>6697</td><td>27</td><td>CTCGGCTCGTGCGTCGATGAAGAACGC</td><td>883.3</td><td>-3.6</td><td>1.0</td><td>-3.6</td><td>3.2E-04</td><td>9.2E-04</td><td>BT20/MB468</td><td>MP</td></tr>
<tr><td>>hg38_21_+_8433172_8446622_RNA45SN1_45S_50ntFlanks@8170.8196.27|CAAGTCCTTCTGATCGAGGCCCAGCCC</td><td>>hg38_21_+_8433172_8446622_RNA45SN1_45S_50ntFlanks</td><td>8170</td><td>8196</td><td>27</td><td>CAAGTCCTTCTGATCGAGGCCCAGCCC</td><td>302.4</td><td>-3.9</td><td>1.1</td><td>-3.5</td><td>5.6E-04</td><td>9.2E-04</td><td>BT20/MB468</td><td>MITO</td></tr>
<tr><td>>hg38_21_+_8433172_8446622_RNA45SN1_45S_50ntFlanks@4726.4752.27|TAATCAAGAACGAAAGTCGGAGGTTCG</td><td>>hg38_21_+_8433172_8446622_RNA45SN1_45S_50ntFlanks</td><td>4726</td><td>4752</td><td>27</td><td>TAATCAAGAACGAAAGTCGGAGGTTCG</td><td>285.6</td><td>-5.6</td><td>1.6</td><td>-3.6</td><td>3.2E-04</td><td>9.2E-04</td><td>BT20/MB468</td><td>MP</td></tr>
<tr><td>>hg38_21_+_8433172_8446622_RNA45SN1_45S_50ntFlanks@12607.12625.19|TGTGCTTGGCTGAGGAGCC</td><td>>hg38_21_+_8433172_8446622_RNA45SN1_45S_50ntFlanks</td><td>12607</td><td>12625</td><td>19</td><td>TGTGCTTGGCTGAGGAGCC</td><td>423.5</td><td>-1.7</td><td>0.5</td><td>-3.4</td><td>7.5E-04</td><td>9.2E-04</td><td>MB231/MB468</td><td>NUC</td></tr>
<tr><td>>MI0000434|hsa-let-7i&WithFlank&12|+|62603680|62603775@12.33.22&[MIMAT0000415&hsa-let-7i-5p&offsets|0|0;m-49&12|+|62603691|62603712&offsets|0|0]|TGAGGTAGTAGTTTGTGCTGTT</td><td>>MI0000434|hsa-let-7i&WithFlank&12|+|62603680|62603775</td><td>12</td><td>33</td><td>22</td><td>TGAGGTAGTAGTTTGTGCTGTT</td><td>30950.3</td><td>1.5</td><td>0.5</td><td>3.4</td><td>7.5E-04</td><td>9.2E-04</td><td>BT20/MB468</td><td>NUC</td></tr>
<tr><td>>hg38_21_+_8433172_8446622_RNA45SN1_45S_50ntFlanks@6661.6682.22|CGGTGGATCACTCGGCTCGTGC</td><td>>hg38_21_+_8433172_8446622_RNA45SN1_45S_50ntFlanks</td><td>6661</td><td>6682</td><td>22</td><td>CGGTGGATCACTCGGCTCGTGC</td><td>381.6</td><td>3.8</td><td>1.0</td><td>3.9</td><td>1.1E-04</td><td>9.2E-04</td><td>BT20/MB231</td><td>MP</td></tr>
<tr><td>>hg38_21_+_8433172_8446622_RNA45SN1_45S_50ntFlanks@12685.12708.24|CCCAGGCGGAACGATACGGCAGCG</td><td>>hg38_21_+_8433172_8446622_RNA45SN1_45S_50ntFlanks</td><td>12685</td><td>12708</td><td>24</td><td>CCCAGGCGGAACGATACGGCAGCG</td><td>140.9</td><td>-4.9</td><td>1.4</td><td>-3.4</td><td>5.6E-04</td><td>9.2E-04</td><td>BT20/MB468</td><td>MITO</td></tr>
<tr><td>>MI0000465|hsa-mir-191&WithFlank&3|-|49020612|49020715@22.39.18&[MIMAT0000440&hsa-miR-191-5p&offsets|0|-5;m-48&3|-|49020672|49020694&offsets|0|-5]|CAACGGAATCCCAAAAGC</td><td>>MI0000465|hsa-mir-191&WithFlank&3|-|49020612|49020715</td><td>22</td><td>39</td><td>18</td><td>CAACGGAATCCCAAAAGC</td><td>305.0</td><td>3.9</td><td>1.1</td><td>3.6</td><td>3.2E-04</td><td>9.2E-04</td><td>BT20/MB468</td><td>MP</td></tr>
<tr><td>>hg38_21_+_8433172_8446622_RNA45SN1_45S_50ntFlanks@11580.11608.29|GAACTGGTGCGGACCAGGGGAATCCGACT</td><td>>hg38_21_+_8433172_8446622_RNA45SN1_45S_50ntFlanks</td><td>11580</td><td>11608</td><td>29</td><td>GAACTGGTGCGGACCAGGGGAATCCGACT</td><td>167.0</td><td>-4.1</td><td>1.2</td><td>-3.4</td><td>6.3E-04</td><td>9.2E-04</td><td>MB231/MB468</td><td>MP</td></tr>
<tr><td>>hg38_21_+_8433172_8446622_RNA45SN1_45S_50ntFlanks@10781.10814.34|AGGTGAACAGCCTCTGGCATGTTGGAACAATGTA</td><td>>hg38_21_+_8433172_8446622_RNA45SN1_45S_50ntFlanks</td><td>10781</td><td>10814</td><td>34</td><td>AGGTGAACAGCCTCTGGCATGTTGGAACAATGTA</td><td>165.9</td><td>-2.6</td><td>0.8</td><td>-3.4</td><td>6.8E-04</td><td>9.2E-04</td><td>BT20/MB468</td><td>CYTO</td></tr>
<tr><td>>hg38_MT_+_598_1651_RNR1_12S_50ntFlanks@416.452.37|AAATAGACTACGAAAGTGGCTTTAACATATCTGAACA</td><td>>hg38_MT_+_598_1651_RNR1_12S_50ntFlanks</td><td>416</td><td>452</td><td>37</td><td>AAATAGACTACGAAAGTGGCTTTAACATATCTGAACA</td><td>64.1</td><td>9.7</td><td>2.7</td><td>3.6</td><td>3.2E-04</td><td>9.2E-04</td><td>BT20/MB468</td><td>MP</td></tr>
<tr><td>>hg38_1_-_228634819_228635039_RNA5S12_5S_50ntFlanks@152.169.18|TACCGGGTGCTGTAGGCT</td><td>>hg38_1_-_228634819_228635039_RNA5S12_5S_50ntFlanks</td><td>152</td><td>169</td><td>18</td><td>TACCGGGTGCTGTAGGCT</td><td>289.5</td><td>3.0</td><td>0.9</td><td>3.4</td><td>7.6E-04</td><td>9.2E-04</td><td>MB231/MB468</td><td>NUC</td></tr>
<tr><td>>hg38_21_+_8433172_8446622_RNA45SN1_45S_50ntFlanks@9736.9758.23|CCGAAACGATCTCAACCTATTCT</td><td>>hg38_21_+_8433172_8446622_RNA45SN1_45S_50ntFlanks</td><td>9736</td><td>9758</td><td>23</td><td>CCGAAACGATCTCAACCTATTCT</td><td>406.7</td><td>-5.5</td><td>1.6</td><td>-3.4</td><td>5.6E-04</td><td>9.3E-04</td><td>BT20/MB468</td><td>MITO</td></tr>
<tr><td>>trnaMT_MetCAT_MT_+_4402_4469@43.71.29|ATGTTGGTTATACCCTTCCCGTACTACCA</td><td>>trnaMT_MetCAT_MT_+_4402_4469</td><td>43</td><td>71</td><td>29</td><td>ATGTTGGTTATACCCTTCCCGTACTACCA</td><td>195.9</td><td>2.6</td><td>0.8</td><td>3.4</td><td>6.3E-04</td><td>9.3E-04</td><td>MB231/MB468</td><td>MP</td></tr>
<tr><td>>hg38_21_+_8433172_8446622_RNA45SN1_45S_50ntFlanks@4759.4790.32|ATCAGATACCGTCGTAGTTCCGACCATAAACG</td><td>>hg38_21_+_8433172_8446622_RNA45SN1_45S_50ntFlanks</td><td>4759</td><td>4790</td><td>32</td><td>ATCAGATACCGTCGTAGTTCCGACCATAAACG</td><td>61.4</td><td>-5.3</td><td>1.6</td><td>-3.4</td><td>6.3E-04</td><td>9.3E-04</td><td>MB231/MB468</td><td>MP</td></tr>
<tr><td>>hg38_MT_+_598_1651_RNR1_12S_50ntFlanks@430.468.39|AGTGGCTTTAACATATCTGAACACACAATAGCTAAGACC</td><td>>hg38_MT_+_598_1651_RNR1_12S_50ntFlanks</td><td>430</td><td>468</td><td>39</td><td>AGTGGCTTTAACATATCTGAACACACAATAGCTAAGACC</td><td>88.1</td><td>6.0</td><td>1.8</td><td>3.4</td><td>6.3E-04</td><td>9.3E-04</td><td>MB231/MB468</td><td>MP</td></tr>
<tr><td>>hg38_MT_+_1621_3279_RNR2_16S_50ntFlanks@51.97.47|GCTAAACCTAGCCCCAAACCCACTCCACCTTACTACCAGACAACCTT</td><td>>hg38_MT_+_1621_3279_RNR2_16S_50ntFlanks</td><td>51</td><td>97</td><td>47</td><td>GCTAAACCTAGCCCCAAACCCACTCCACCTTACTACCAGACAACCTT</td><td>576.9</td><td>4.3</td><td>1.3</td><td>3.4</td><td>7.5E-04</td><td>9.3E-04</td><td>MB231/MB468</td><td>MITO</td></tr>
<tr><td>>hg38_21_+_8433172_8446622_RNA45SN1_45S_50ntFlanks@10682.10705.24|TCCGGCGGCGTCCGGTGAGCTCTC</td><td>>hg38_21_+_8433172_8446622_RNA45SN1_45S_50ntFlanks</td><td>10682</td><td>10705</td><td>24</td><td>TCCGGCGGCGTCCGGTGAGCTCTC</td><td>162.4</td><td>-2.2</td><td>0.7</td><td>-3.4</td><td>7.1E-04</td><td>9.3E-04</td><td>MB231/MB468</td><td>CYTO</td></tr>
<tr><td>>hg38_21_+_8433172_8446622_RNA45SN1_45S_50ntFlanks@4970.4999.30|CCCGGCCCGGACACGGACAGGATTGACAGA</td><td>>hg38_21_+_8433172_8446622_RNA45SN1_45S_50ntFlanks</td><td>4970</td><td>4999</td><td>30</td><td>CCCGGCCCGGACACGGACAGGATTGACAGA</td><td>274.4</td><td>-4.7</td><td>1.4</td><td>-3.4</td><td>6.3E-04</td><td>9.3E-04</td><td>MB231/MB468</td><td>MP</td></tr>
<tr><td>>hg38_21_+_8433172_8446622_RNA45SN1_45S_50ntFlanks@6669.6696.28|CACTCGGCTCGTGCGTCGATGAAGAACG</td><td>>hg38_21_+_8433172_8446622_RNA45SN1_45S_50ntFlanks</td><td>6669</td><td>6696</td><td>28</td><td>CACTCGGCTCGTGCGTCGATGAAGAACG</td><td>195.9</td><td>-2.7</td><td>0.8</td><td>-3.6</td><td>3.2E-04</td><td>9.3E-04</td><td>BT20/MB468</td><td>MP</td></tr>
<tr><td>>hg38_21_+_8433172_8446622_RNA45SN1_45S_50ntFlanks@6646.6674.29|TCGTACGACTCTTAGCGGTGGATCACTCG</td><td>>hg38_21_+_8433172_8446622_RNA45SN1_45S_50ntFlanks</td><td>6646</td><td>6674</td><td>29</td><td>TCGTACGACTCTTAGCGGTGGATCACTCG</td><td>2105.5</td><td>3.3</td><td>0.9</td><td>3.6</td><td>3.2E-04</td><td>9.3E-04</td><td>BT20/MB468</td><td>MP</td></tr>
<tr><td>>hg38_21_+_8433172_8446622_RNA45SN1_45S_50ntFlanks@10461.10488.28|GCCTCCGTTGCCCTCGGCCGATCGAAAG</td><td>>hg38_21_+_8433172_8446622_RNA45SN1_45S_50ntFlanks</td><td>10461</td><td>10488</td><td>28</td><td>GCCTCCGTTGCCCTCGGCCGATCGAAAG</td><td>280.4</td><td>-3.4</td><td>1.0</td><td>-3.4</td><td>7.6E-04</td><td>9.3E-04</td><td>MB231/MB468</td><td>NUC</td></tr>
<tr><td>>hg38_21_+_8433172_8446622_RNA45SN1_45S_50ntFlanks@9498.9525.28|AAGATGGTGAACTATGCCTGGGCAGGGC</td><td>>hg38_21_+_8433172_8446622_RNA45SN1_45S_50ntFlanks</td><td>9498</td><td>9525</td><td>28</td><td>AAGATGGTGAACTATGCCTGGGCAGGGC</td><td>92.5</td><td>6.8</td><td>1.9</td><td>3.6</td><td>3.2E-04</td><td>9.3E-04</td><td>BT20/MB468</td><td>MP</td></tr>
<tr><td>>hg38_21_+_8433172_8446622_RNA45SN1_45S_50ntFlanks@10781.10814.34|AGGTGAACAGCCTCTGGCATGTTGGAACAATGTA</td><td>>hg38_21_+_8433172_8446622_RNA45SN1_45S_50ntFlanks</td><td>10781</td><td>10814</td><td>34</td><td>AGGTGAACAGCCTCTGGCATGTTGGAACAATGTA</td><td>168.1</td><td>-3.7</td><td>1.1</td><td>-3.4</td><td>7.6E-04</td><td>9.3E-04</td><td>MB231/MB468</td><td>NUC</td></tr>
</table>

<table>
<tr><td>>hg38_21_+_8433172_8446622_RNA45SN1_45S_50ntFlanks@6648.6678.31|GTACGACTCTTAGCGGTGGATCACTCGGCTC</td><td>>hg38_21_+_8433172_8446622_RNA45SN1_45S_50ntFlanks</td><td>6648</td><td>6678</td><td>31</td><td>GTACGACTCTTAGCGGTGGATCACTCGGCTC</td><td>982.1</td><td>2.8</td><td>0.8</td><td>3.4</td><td>7.6E-04</td><td>9.3E-04</td><td>MB231/MB468</td><td>NUC</td></tr>
<tr><td>>MI0001733|hsa-mir-452&WithFlank&X|-|151959622|151959718@20.42.23(+1U)&[MIMAT0001635&hsa-miR-452-5p&offsets|0|+1(+1U);m-188&X|-|151959677|151959699&offsets|0|0(+1U)]|AACTGTTTGCAGAGGAAACTGAGT</td><td>>MI0001733|hsa-mir-452&WithFlank&X|-|151959622|151959718</td><td>20</td><td>42</td><td>23(+1U)</td><td>AACTGTTTGCAGAGGAAACTGAGT</td><td>261.3</td><td>2.7</td><td>0.7</td><td>3.9</td><td>1.2E-04</td><td>9.3E-04</td><td>BT20/MB231</td><td>NUC</td></tr>
<tr><td>>MI0003780|hsa-mir-1296&WithFlank&10|-|63372951|63373054@22.42.21(+1U)&[MIMAT0005794&hsa-miR-1296-5p&offsets|0|-1(+1U);m-304&10|-|63373012|63373033&offsets|0|-1(+1U)]|TTAGGGCCCTGGCTCCATCTCT</td><td>>MI0003780|hsa-mir-1296&WithFlank&10|-|63372951|63373054</td><td>22</td><td>42</td><td>21(+1U)</td><td>TTAGGGCCCTGGCTCCATCTCT</td><td>142.8</td><td>5.2</td><td>1.4</td><td>3.6</td><td>3.2E-04</td><td>9.3E-04</td><td>BT20/MB468</td><td>MP</td></tr>
<tr><td>>hg38_21_+_8433172_8446622_RNA45SN1_45S_50ntFlanks@5443.5466.24|CCTCGGATCGGCCCCGCCGGGGTC</td><td>>hg38_21_+_8433172_8446622_RNA45SN1_45S_50ntFlanks</td><td>5443</td><td>5466</td><td>24</td><td>CCTCGGATCGGCCCCGCCGGGGTC</td><td>1003.8</td><td>-2.4</td><td>0.7</td><td>-3.4</td><td>7.1E-04</td><td>9.3E-04</td><td>MB231/MB468</td><td>CYTO</td></tr>
<tr><td>>hg38_21_+_8433172_8446622_RNA45SN1_45S_50ntFlanks@12456.12483.28|CCAAGCGTTGGATTGTTCACCCACTAAT</td><td>>hg38_21_+_8433172_8446622_RNA45SN1_45S_50ntFlanks</td><td>12456</td><td>12483</td><td>28</td><td>CCAAGCGTTGGATTGTTCACCCACTAAT</td><td>901.1</td><td>-6.0</td><td>1.7</td><td>-3.6</td><td>3.3E-04</td><td>9.3E-04</td><td>BT20/MB468</td><td>MP</td></tr>
<tr><td>>MI0000298|hsa-mir-221&WithFlank&X|-|45746151|45746272@71.92.22(+1A)&[MIMAT0000278&hsa-miR-221-3p&offsets|0|-1(+1A);m-91&X|-|45746181|45746202&offsets|0|0(+1A)]|AGCTACATTGTCTGCTGGGTTTA</td><td>>MI0000298|hsa-mir-221&WithFlank&X|-|45746151|45746272</td><td>71</td><td>92</td><td>22(+1A)</td><td>AGCTACATTGTCTGCTGGGTTTA</td><td>97.6</td><td>1.5</td><td>0.5</td><td>3.4</td><td>7.6E-04</td><td>9.3E-04</td><td>BT20/MB468</td><td>NUC</td></tr>
<tr><td>>hg38_21_+_8433172_8446622_RNA45SN1_45S_50ntFlanks@9543.9571.29|TGGTGGAGGTCCGTAGCGGTCCTGACGTG</td><td>>hg38_21_+_8433172_8446622_RNA45SN1_45S_50ntFlanks</td><td>9543</td><td>9571</td><td>29</td><td>TGGTGGAGGTCCGTAGCGGTCCTGACGTG</td><td>1057.3</td><td>-6.5</td><td>1.9</td><td>-3.4</td><td>7.6E-04</td><td>9.3E-04</td><td>BT20/MB468</td><td>NUC</td></tr>
<tr><td>>hg38_21_+_8433172_8446622_RNA45SN1_45S_50ntFlanks@5087.5120.34|ACGAGACTCTGGCATGCTAACTAGTTACGCGACC</td><td>>hg38_21_+_8433172_8446622_RNA45SN1_45S_50ntFlanks</td><td>5087</td><td>5120</td><td>34</td><td>ACGAGACTCTGGCATGCTAACTAGTTACGCGACC</td><td>105.9</td><td>-4.7</td><td>1.4</td><td>-3.4</td><td>6.4E-04</td><td>9.3E-04</td><td>MB231/MB468</td><td>MP</td></tr>
<tr><td>>hg38_21_+_8433172_8446622_RNA45SN1_45S_50ntFlanks@4724.4766.43|ATTAATCAAGAACGAAAGTCGGAGGTTCGAAGACGATCAGATA</td><td>>hg38_21_+_8433172_8446622_RNA45SN1_45S_50ntFlanks</td><td>4724</td><td>4766</td><td>43</td><td>ATTAATCAAGAACGAAAGTCGGAGGTTCGAAGACGATCAGATA</td><td>258.2</td><td>3.9</td><td>1.1</td><td>3.4</td><td>7.2E-04</td><td>9.4E-04</td><td>MB231/MB468</td><td>CYTO</td></tr>
<tr><td>>hg38_21_+_8433172_8446622_RNA45SN1_45S_50ntFlanks@6700.6717.18|CTAGCTGCGAGAATTAAT</td><td>>hg38_21_+_8433172_8446622_RNA45SN1_45S_50ntFlanks</td><td>6700</td><td>6717</td><td>18</td><td>CTAGCTGCGAGAATTAAT</td><td>225.6</td><td>-4.8</td><td>1.4</td><td>-3.4</td><td>6.8E-04</td><td>9.4E-04</td><td>BT20/MB468</td><td>CYTO</td></tr>
<tr><td>>hg38_21_+_8433172_8446622_RNA45SN1_45S_50ntFlanks@5190.5236.47|AACAGGTCTGTGATGCCCTTAGATGTCCGGGGCTGCACGCGCGCTAC</td><td>>hg38_21_+_8433172_8446622_RNA45SN1_45S_50ntFlanks</td><td>5190</td><td>5236</td><td>47</td><td>AACAGGTCTGTGATGCCCTTAGATGTCCGGGGCTGCACGCGCGCTAC</td><td>62.2</td><td>9.7</td><td>2.7</td><td>3.6</td><td>3.3E-04</td><td>9.4E-04</td><td>BT20/MB468</td><td>MP</td></tr>
<tr><td>>hg38_MT_+_598_1651_RNR1_12S_50ntFlanks@424.455.32|TACGAAAGTGGCTTTAACATATCTGAACACAC</td><td>>hg38_MT_+_598_1651_RNR1_12S_50ntFlanks</td><td>424</td><td>455</td><td>32</td><td>TACGAAAGTGGCTTTAACATATCTGAACACAC</td><td>154.4</td><td>4.2</td><td>1.2</td><td>3.4</td><td>7.6E-04</td><td>9.4E-04</td><td>MB231/MB468</td><td>MITO</td></tr>
<tr><td>>hg38_21_+_8433172_8446622_RNA45SN1_45S_50ntFlanks@9568.9594.27|CGTGCAAATCGGTCGTCCGACCTGGGT</td><td>>hg38_21_+_8433172_8446622_RNA45SN1_45S_50ntFlanks</td><td>9568</td><td>9594</td><td>27</td><td>CGTGCAAATCGGTCGTCCGACCTGGGT</td><td>591.9</td><td>-2.2</td><td>0.6</td><td>-3.4</td><td>5.7E-04</td><td>9.4E-04</td><td>BT20/MB468</td><td>MITO</td></tr>
<tr><td>>hg38_21_+_8433172_8446622_RNA45SN1_45S_50ntFlanks@11710.11740.31|CGCGGGTAAACGGCGGGAGTAACTATGACTC</td><td>>hg38_21_+_8433172_8446622_RNA45SN1_45S_50ntFlanks</td><td>11710</td><td>11740</td><td>31</td><td>CGCGGGTAAACGGCGGGAGTAACTATGACTC</td><td>459.9</td><td>-2.7</td><td>0.8</td><td>-3.4</td><td>7.7E-04</td><td>9.4E-04</td><td>MB231/MB468</td><td>NUC</td></tr>
<tr><td>>hg38_21_+_8433172_8446622_RNA45SN1_45S_50ntFlanks@7982.8004.23|TCAGATCAGACGTGGCGACCCGC</td><td>>hg38_21_+_8433172_8446622_RNA45SN1_45S_50ntFlanks</td><td>7982</td><td>8004</td><td>23</td><td>TCAGATCAGACGTGGCGACCCGC</td><td>418.5</td><td>-2.1</td><td>0.6</td><td>-3.4</td><td>5.7E-04</td><td>9.4E-04</td><td>BT20/MB468</td><td>MITO</td></tr>
<tr><td>>MI0000803|hsa-mir-330&WithFlank&19|-|45638988|45639093@64.87.24&[MIMAT0000751&hsa-miR-330-3p&offsets|+1|+2;m-347&19|-|45639009|45639030&offsets|0|+2]|CAAAGCACACGGCCTGCAGAGAGG</td><td>>MI0000803|hsa-mir-330&WithFlank&19|-|45638988|45639093</td><td>64</td><td>87</td><td>24</td><td>CAAAGCACACGGCCTGCAGAGAGG</td><td>258.1</td><td>6.2</td><td>1.7</td><td>3.6</td><td>3.3E-04</td><td>9.4E-04</td><td>BT20/MB468</td><td>MP</td></tr>
<tr><td>>hg38_21_+_8433172_8446622_RNA45SN1_45S_50ntFlanks@8356.8377.22|GAAAAGAACTTTGAAGAGAGAG</td><td>>hg38_21_+_8433172_8446622_RNA45SN1_45S_50ntFlanks</td><td>8356</td><td>8377</td><td>22</td><td>GAAAAGAACTTTGAAGAGAGAG</td><td>209.7</td><td>-4.8</td><td>1.4</td><td>-3.4</td><td>6.9E-04</td><td>9.4E-04</td><td>BT20/MB468</td><td>CYTO</td></tr>
<tr><td>>hg38_21_+_8433172_8446622_RNA45SN1_45S_50ntFlanks@6730.6770.41|ACACATTGATCATCGACACTTCGAACGCACTTGCGGCCCCG</td><td>>hg38_21_+_8433172_8446622_RNA45SN1_45S_50ntFlanks</td><td>6730</td><td>6770</td><td>41</td><td>ACACATTGATCATCGACACTTCGAACGCACTTGCGGCCCCG</td><td>337.0</td><td>-3.5</td><td>1.0</td><td>-3.4</td><td>7.2E-04</td><td>9.4E-04</td><td>MB231/MB468</td><td>CYTO</td></tr>
<tr><td>>hg38_21_+_8433172_8446622_RNA45SN1_45S_50ntFlanks@3994.4013.20|TGACTCTAGATAACCTCGGG</td><td>>hg38_21_+_8433172_8446622_RNA45SN1_45S_50ntFlanks</td><td>3994</td><td>4013</td><td>20</td><td>TGACTCTAGATAACCTCGGG</td><td>203.8</td><td>-1.7</td><td>0.5</td><td>-3.4</td><td>6.9E-04</td><td>9.4E-04</td><td>BT20/MB468</td><td>CYTO</td></tr>
<tr><td>>hg38_21_+_8433172_8446622_RNA45SN1_45S_50ntFlanks@11905.11940.36|GCACGGTGAAGAGACATGAGAGGTGTAGAATAAGTG</td><td>>hg38_21_+_8433172_8446622_RNA45SN1_45S_50ntFlanks</td><td>11905</td><td>11940</td><td>36</td><td>GCACGGTGAAGAGACATGAGAGGTGTAGAATAAGTG</td><td>132.9</td><td>10.4</td><td>2.7</td><td>3.9</td><td>1.1E-04</td><td>9.4E-04</td><td>BT20/MB231</td><td>MP</td></tr>
<tr><td>>hg38_21_+_8433172_8446622_RNA45SN1_45S_50ntFlanks@11670.11688.19|CCCAGTGCTCTGAATGTCA</td><td>>hg38_21_+_8433172_8446622_RNA45SN1_45S_50ntFlanks</td><td>11670</td><td>11688</td><td>19</td><td>CCCAGTGCTCTGAATGTCA</td><td>783.1</td><td>-3.1</td><td>0.9</td><td>-3.4</td><td>7.2E-04</td><td>9.4E-04</td><td>MB231/MB468</td><td>CYTO</td></tr>
<tr><td>>trnaMT_GluTTC_MT_-_14674_14742@41.72.32,trnalookalike8_GluTTC_5_-_93905172_93905240@41.72.32|TGGTCGTGGTTGTAGTCCGTGCGAGAATACCA</td><td>>trnaMT_GluTTC_MT_-_14674_14742</td><td>41</td><td>72</td><td>32</td><td>TGGTCGTGGTTGTAGTCCGTGCGAGAATACCA</td><td>1870.2</td><td>-2.4</td><td>0.7</td><td>-3.4</td><td>7.7E-04</td><td>9.4E-04</td><td>BT20/MB468</td><td>NUC</td></tr>
<tr><td>>MI0000803|hsa-mir-330&WithFlank&19|-|45638988|45639093@64.86.23(+1U)&[MIMAT0000751&hsa-miR-330-3p&offsets|+1|+1(+1U);m-347&19|-|45639009|45639030&offsets|0|+1(+1U)]|CAAAGCACACGGCCTGCAGAGAGT</td><td>>MI0000803|hsa-mir-330&WithFlank&19|-|45638988|45639093</td><td>64</td><td>86</td><td>23(+1U)</td><td>CAAAGCACACGGCCTGCAGAGAGT</td><td>181.2</td><td>4.2</td><td>1.2</td><td>3.4</td><td>6.4E-04</td><td>9.4E-04</td><td>MB231/MB468</td><td>MP</td></tr>
<tr><td>>hg38_21_+_8433172_8446622_RNA45SN1_45S_50ntFlanks@10552.10581.30|GGTAACGCGACCGATCCCGGAGAAGCCGGC</td><td>>hg38_21_+_8433172_8446622_RNA45SN1_45S_50ntFlanks</td><td>10552</td><td>10581</td><td>30</td><td>GGTAACGCGACCGATCCCGGAGAAGCCGGC</td><td>61.1</td><td>9.7</td><td>2.7</td><td>3.6</td><td>3.3E-04</td><td>9.4E-04</td><td>BT20/MB468</td><td>MP</td></tr>
<tr><td>>MI0006273|hsa-mir-1180&WithFlank&17|-|19344500|19344580@47.67.21(+1C)&[MIMAT0005825&hsa-miR-1180-3p&offsets|0|-1(+1C);m-214&17|-|19344513|19344534&offsets|0|-1(+1C)]|TTTCCGGCTCGCGTGGGTGTGC</td><td>>MI0006273|hsa-mir-1180&WithFlank&17|-|19344500|19344580</td><td>47</td><td>67</td><td>21(+1C)</td><td>TTTCCGGCTCGCGTGGGTGTGC</td><td>232.3</td><td>2.3</td><td>0.6</td><td>3.8</td><td>1.2E-04</td><td>9.4E-04</td><td>BT20/MB231</td><td>NUC</td></tr>
<tr><td>>hg38_21_+_8433172_8446622_RNA45SN1_45S_50ntFlanks@7967.8005.39|CTCCGAGACGCGACCTCAGATCAGACGTGGCGACCCGCT</td><td>>hg38_21_+_8433172_8446622_RNA45SN1_45S_50ntFlanks</td><td>7967</td><td>8005</td><td>39</td><td>CTCCGAGACGCGACCTCAGATCAGACGTGGCGACCCGCT</td><td>2202.1</td><td>4.7</td><td>1.2</td><td>3.9</td><td>1.2E-04</td><td>9.4E-04</td><td>BT20/MB231</td><td>NUC</td></tr>
<tr><td>>hg38_21_+_8433172_8446622_RNA45SN1_45S_50ntFlanks@12456.12487.32|CCAAGCGTTGGATTGTTCACCCACTAATAGGG</td><td>>hg38_21_+_8433172_8446622_RNA45SN1_45S_50ntFlanks</td><td>12456</td><td>12487</td><td>32</td><td>CCAAGCGTTGGATTGTTCACCCACTAATAGGG</td><td>16142.1</td><td>-7.0</td><td>2.0</td><td>-3.6</td><td>3.3E-04</td><td>9.4E-04</td><td>BT20/MB468</td><td>MP</td></tr>
<tr><td>>hg38_21_+_8433172_8446622_RNA45SN1_45S_50ntFlanks@11907.11933.27|ACGGTGAAGAGACATGAGAGGTGTAGA</td><td>>hg38_21_+_8433172_8446622_RNA45SN1_45S_50ntFlanks</td><td>11907</td><td>11933</td><td>27</td><td>ACGGTGAAGAGACATGAGAGGTGTAGA</td><td>52.7</td><td>9.0</td><td>2.3</td><td>3.9</td><td>1.2E-04</td><td>9.4E-04</td><td>BT20/MB231</td><td>MP</td></tr>
<tr><td>>hg38_21_+_8433172_8446622_RNA45SN1_45S_50ntFlanks@9568.9595.28|CGTGCAAATCGGTCGTCCGACCTGGGTA</td><td>>hg38_21_+_8433172_8446622_RNA45SN1_45S_50ntFlanks</td><td>9568</td><td>9595</td><td>28</td><td>CGTGCAAATCGGTCGTCCGACCTGGGTA</td><td>1601.4</td><td>-5.1</td><td>1.4</td><td>-3.6</td><td>3.3E-04</td><td>9.4E-04</td><td>BT20/MB468</td><td>MP</td></tr>
<tr><td>>hg38_21_+_8433172_8446622_RNA45SN1_45S_50ntFlanks@9534.9570.37|AGGAAACTCTGGTGGAGGTCCGTAGCGGTCCTGACGT</td><td>>hg38_21_+_8433172_8446622_RNA45SN1_45S_50ntFlanks</td><td>9534</td><td>9570</td><td>37</td><td>AGGAAACTCTGGTGGAGGTCCGTAGCGGTCCTGACGT</td><td>290.6</td><td>3.1</td><td>0.8</td><td>3.8</td><td>1.4E-04</td><td>9.4E-04</td><td>BT20/MB231</td><td>MITO</td></tr>
<tr><td>>hg38_21_+_8433172_8446622_RNA45SN1_45S_50ntFlanks@10014.10038.25|AAAATGGATGGCGCTGGAGCGTCGG</td><td>>hg38_21_+_8433172_8446622_RNA45SN1_45S_50ntFlanks</td><td>10014</td><td>10038</td><td>25</td><td>AAAATGGATGGCGCTGGAGCGTCGG</td><td>74.5</td><td>-10.1</td><td>3.0</td><td>-3.4</td><td>6.4E-04</td><td>9.4E-04</td><td>MB231/MB468</td><td>MP</td></tr>
<tr><td>>MI0000263|hsa-mir-7-1&WithFlank&9|-|83969742|83969863@30.55.26&[MIMAT0000252_2&hsa-miR-7-5p&offsets|0|+2;m-401&9|-|83969811|83969834&offsets|0|+2]|TGGAAGACTAGTGATTTTGTTGTTTT</td><td>>MI0000263|hsa-mir-7-1&WithFlank&9|-|83969742|83969863</td><td>30</td><td>55</td><td>26</td><td>TGGAAGACTAGTGATTTTGTTGTTTT</td><td>321.2</td><td>-2.9</td><td>0.8</td><td>-3.8</td><td>1.4E-04</td><td>9.4E-04</td><td>BT20/MB231</td><td>MITO</td></tr>
<tr><td>>MI0000266|hsa-mir-10a&WithFlank&17|-|48579832|48579953@28.49.22(+1A)&[MIMAT0000253&hsa-miR-10a-5p&offsets|0|-1(+1A);m-18&17|-|48579905|48579926&offsets|0|0(+1A)]|TACCCTGTAGATCCGAATTTGTA</td><td>>MI0000266|hsa-mir-10a&WithFlank&17|-|48579832|48579953</td><td>28</td><td>49</td><td>22(+1A)</td><td>TACCCTGTAGATCCGAATTTGTA</td><td>3111.8</td><td>3.0</td><td>0.8</td><td>3.8</td><td>1.2E-04</td><td>9.5E-04</td><td>BT20/MB231</td><td>CYTO</td></tr>
<tr><td>>hg38_21_+_8433172_8446622_RNA45SN1_45S_50ntFlanks@10921.10946.26|GCTGGACGAGGCGCCGCCGCCCCCCC</td><td>>hg38_21_+_8433172_8446622_RNA45SN1_45S_50ntFlanks</td><td>10921</td><td>10946</td><td>26</td><td>GCTGGACGAGGCGCCGCCGCCCCCCC</td><td>333.9</td><td>-3.7</td><td>1.1</td><td>-3.4</td><td>6.9E-04</td><td>9.5E-04</td><td>BT20/MB468</td><td>CYTO</td></tr>
<tr><td>>hg38_21_+_8433172_8446622_RNA45SN1_45S_50ntFlanks@8196.8223.28|CGTGGACGGTGTGAGGCCGGTAGCGGCC</td><td>>hg38_21_+_8433172_8446622_RNA45SN1_45S_50ntFlanks</td><td>8196</td><td>8223</td><td>28</td><td>CGTGGACGGTGTGAGGCCGGTAGCGGCC</td><td>868.6</td><td>-2.2</td><td>0.6</td><td>-3.4</td><td>6.9E-04</td><td>9.5E-04</td><td>BT20/MB468</td><td>CYTO</td></tr>
<tr><td>>MI0000079|hsa-mir-23a&WithFlank&19|-|13836581|13836665@54.71.18(+1U)&[MIMAT0000078&hsa-miR-23a-3p&offsets|+3|0(+1U);m-31&19|-|13836595|13836615&offsets|+3|0(+1U)]|ACATTGCCAGGGATTTCCT</td><td>>MI0000079|hsa-mir-23a&WithFlank&19|-|13836581|13836665</td><td>54</td><td>71</td><td>18(+1U)</td><td>ACATTGCCAGGGATTTCCT</td><td>308.9</td><td>12.3</td><td>3.6</td><td>3.4</td><td>7.2E-04</td><td>9.5E-04</td><td>MB231/MB468</td><td>CYTO</td></tr>
<tr><td>>hg38_21_+_8433172_8446622_RNA45SN1_45S_50ntFlanks@3985.4027.43|CGGCTTTGGTGACTCTAGATAACCTCGGGCCGATCGCACGCCC</td><td>>hg38_21_+_8433172_8446622_RNA45SN1_45S_50ntFlanks</td><td>3985</td><td>4027</td><td>43</td><td>CGGCTTTGGTGACTCTAGATAACCTCGGGCCGATCGCACGCCC</td><td>101.8</td><td>-5.1</td><td>1.5</td><td>-3.4</td><td>5.8E-04</td><td>9.5E-04</td><td>BT20/MB468</td><td>MITO</td></tr>
<tr><td>>MI0000070|hsa-mir-16-1&WithFlank&13|-|50048967|50049067@20.37.18;MI0000115|hsa-mir-16-2&WithFlank&3|+|160404739|160404831@16.33.18&[MIMAT0000069&hsa-miR-16-5p&offsets|0|-4;m-62&13|-|50049027|50049048&offsets|0|-4];[MIMAT0000069_1&hsa-miR-16-5p&offsets|0|-4;m-61&3|+|160404754|160404775&offsets|0|-4]|TAGCAGCACGTAAATATT</td><td>>MI0000070|hsa-mir-16-1&WithFlank&13|-|50048967|50049067</td><td>20</td><td>37</td><td>18</td><td>TAGCAGCACGTAAATATT</td><td>269.8</td><td>6.5</td><td>1.8</td><td>3.6</td><td>3.3E-04</td><td>9.5E-04</td><td>BT20/MB468</td><td>MP</td></tr>
<tr><td>>hg38_21_+_8433172_8446622_RNA45SN1_45S_50ntFlanks@3769.3786.18|CGCACGGCCGGTACAGTG</td><td>>hg38_21_+_8433172_8446622_RNA45SN1_45S_50ntFlanks</td><td>3769</td><td>3786</td><td>18</td><td>CGCACGGCCGGTACAGTG</td><td>200.0</td><td>-4.0</td><td>1.2</td><td>-3.4</td><td>7.8E-04</td><td>9.5E-04</td><td>MB231/MB468</td><td>NUC</td></tr>
<tr><td>>MI0000683|hsa-mir-181b-2&WithFlank&9|+|124693704|124693804@22.46.25&[MIMAT0000257_1&hsa-miR-181b-5p&offsets|0|+2;m-72&9|+|124693725|124693747&offsets|0|+2]|AACATTCATTGCTGTCGGTGGGTTT</td><td>>MI0000683|hsa-mir-181b-2&WithFlank&9|+|124693704|124693804</td><td>22</td><td>46</td><td>25</td><td>AACATTCATTGCTGTCGGTGGGTTT</td><td>1703.2</td><td>-3.2</td><td>0.8</td><td>-3.8</td><td>1.4E-04</td><td>9.5E-04</td><td>BT20/MB231</td><td>MITO</td></tr>
<tr><td>>hg38_21_+_8433172_8446622_RNA45SN1_45S_50ntFlanks@12409.12433.25|TTGATCCTTCGATGTCGGCTCTTCC</td><td>>hg38_21_+_8433172_8446622_RNA45SN1_45S_50ntFlanks</td><td>12409</td><td>12433</td><td>25</td><td>TTGATCCTTCGATGTCGGCTCTTCC</td><td>831.4</td><td>3.2</td><td>0.9</td><td>3.4</td><td>7.3E-04</td><td>9.5E-04</td><td>MB231/MB468</td><td>CYTO</td></tr>
<tr><td>>trnaMT_PheGAA_MT_+_577_647@55.74.20|ACATCACCCCATAAACACCA</td><td>>trnaMT_PheGAA_MT_+_577_647</td><td>55</td><td>74</td><td>20</td><td>ACATCACCCCATAAACACCA</td><td>391.0</td><td>4.9</td><td>1.5</td><td>3.4</td><td>7.8E-04</td><td>9.5E-04</td><td>MB231/MB468</td><td>NUC</td></tr>
<tr><td>>hg38_21_+_8433172_8446622_RNA45SN1_45S_50ntFlanks@12408.12433.26|TTTGATCCTTCGATGTCGGCTCTTCC</td><td>>hg38_21_+_8433172_8446622_RNA45SN1_45S_50ntFlanks</td><td>12408</td><td>12433</td><td>26</td><td>TTTGATCCTTCGATGTCGGCTCTTCC</td><td>1177.9</td><td>3.3</td><td>1.0</td><td>3.4</td><td>7.3E-04</td><td>9.5E-04</td><td>MB231/MB468</td><td>CYTO</td></tr>
<tr><td>>hg38_21_+_8433172_8446622_RNA45SN1_45S_50ntFlanks@4047.4064.18|ATTCGAACGTCTGCCCTA</td><td>>hg38_21_+_8433172_8446622_RNA45SN1_45S_50ntFlanks</td><td>4047</td><td>4064</td><td>18</td><td>ATTCGAACGTCTGCCCTA</td><td>361.8</td><td>-5.7</td><td>1.6</td><td>-3.6</td><td>3.3E-04</td><td>9.5E-04</td><td>BT20/MB468</td><td>MP</td></tr>
<tr><td>>hg38_21_+_8433172_8446622_RNA45SN1_45S_50ntFlanks@12499.12520.22|GGTTTAGACCGTCGTGAGACAG</td><td>>hg38_21_+_8433172_8446622_RNA45SN1_45S_50ntFlanks</td><td>12499</td><td>12520</td><td>22</td><td>GGTTTAGACCGTCGTGAGACAG</td><td>70.0</td><td>-9.2</td><td>2.6</td><td>-3.6</td><td>3.3E-04</td><td>9.5E-04</td><td>BT20/MB468</td><td>MP</td></tr>
<tr><td>>hg38_21_+_8433172_8446622_RNA45SN1_45S_50ntFlanks@10852.10879.28|GATAAGGATTGGCTCTAAGGGCTGGGTC</td><td>>hg38_21_+_8433172_8446622_RNA45SN1_45S_50ntFlanks</td><td>10852</td><td>10879</td><td>28</td><td>GATAAGGATTGGCTCTAAGGGCTGGGTC</td><td>146.2</td><td>6.7</td><td>1.9</td><td>3.6</td><td>3.3E-04</td><td>9.5E-04</td><td>BT20/MB468</td><td>MP</td></tr>
<tr><td>>hg38_21_+_8433172_8446622_RNA45SN1_45S_50ntFlanks@9953.9982.30|ATGGAAGTCGGAATCCGCTAAGGAGTGTGT</td><td>>hg38_21_+_8433172_8446622_RNA45SN1_45S_50ntFlanks</td><td>9953</td><td>9982</td><td>30</td><td>ATGGAAGTCGGAATCCGCTAAGGAGTGTGT</td><td>55.7</td><td>6.6</td><td>1.8</td><td>3.6</td><td>3.3E-04</td><td>9.5E-04</td><td>BT20/MB468</td><td>MP</td></tr>
<tr><td>>hg38_21_+_8433172_8446622_RNA45SN1_45S_50ntFlanks@10466.10488.23|CGTTGCCCTCGGCCGATCGAAAG</td><td>>hg38_21_+_8433172_8446622_RNA45SN1_45S_50ntFlanks</td><td>10466</td><td>10488</td><td>23</td><td>CGTTGCCCTCGGCCGATCGAAAG</td><td>324.1</td><td>-2.9</td><td>0.8</td><td>-3.4</td><td>5.8E-04</td><td>9.5E-04</td><td>BT20/MB468</td><td>MITO</td></tr>
</table>

<table>
<tr><td>>am_MI0005763|hsa-mir-941-1&WithFlank&20|+|63919443|63919526@53.74.22(+1A);MI0005764|hsa-mir-941-2&WithFlank&20|+|63919499|63919582@53.74.22(+1A);MI0005765|hsa-mir-941-3&WithFlank&20|+|63919555|63919638@53.74.22(+1A);MI0005766|hsa-mir-941-4&WithFlank&20|+|63919750|63919833@53.74.22(+1A);MI0031520|hsa-mir-941-5&WithFlank&20|+|63919862|63919945@53.74.22(+1A)&[MIMAT0004984&hsa-miR-941&offsets|0|-1(+1A);m-263&20|+|63919495|63919517&offsets|0|-1(+1A)];[MIMAT0004984_1&hsa-miR-941&offsets|0|-1(+1A);m-263&20|+|63919551|63919573&offsets|0|-1(+1A)];[MIMAT0004984_2&hsa-miR-941&offsets|0|-1(+1A);m-263&20|+|63919607|63919629&offsets|0|-1(+1A)];[MIMAT0004984_3&hsa-miR-941&offsets|0|-1(+1A);m-263&20|+|63919802|63919824&offsets|0|-1(+1A)];[MIMAT0004984_4&hsa-miR-941&offsets|0|-1(+1A);m-263&20|+|63919914|63919936&offsets|0|-1(+1A)]|CACCCGGCTGTGTGCACATGTGA</td><td>>am_MI0005763|hsa-mir-941-1&WithFlank&20|+|63919443|63919526</td><td>53</td><td>74</td><td>22(+1A)</td><td>CACCCGGCTGTGTGCACATGTGA</td><td>266.9</td><td>4.2</td><td>1.2</td><td>3.4</td><td>6.5E-04</td><td>9.5E-04</td><td>MB231/MB468</td><td>MP</td></tr>
<tr><td>>hg38_21_+_8433172_8446622_RNA45SN1_45S_50ntFlanks@10906.10926.21|TGGGCGCGCGCCGCGGCTGGA</td><td>>hg38_21_+_8433172_8446622_RNA45SN1_45S_50ntFlanks</td><td>10906</td><td>10926</td><td>21</td><td>TGGGCGCGCGCCGCGGCTGGA</td><td>190.0</td><td>-2.4</td><td>0.7</td><td>-3.4</td><td>7.0E-04</td><td>9.5E-04</td><td>BT20/MB468</td><td>CYTO</td></tr>
<tr><td>>hg38_21_+_8433172_8446622_RNA45SN1_45S_50ntFlanks@12714.12747.34|GAGCCTCGGTTGGCCTCGGATAGCCGGTCCCCCG</td><td>>hg38_21_+_8433172_8446622_RNA45SN1_45S_50ntFlanks</td><td>12714</td><td>12747</td><td>34</td><td>GAGCCTCGGTTGGCCTCGGATAGCCGGTCCCCCG</td><td>716.0</td><td>-2.9</td><td>0.9</td><td>-3.4</td><td>7.7E-04</td><td>9.5E-04</td><td>MB231/MB468</td><td>MITO</td></tr>
<tr><td>>MI0003667|hsa-mir-652&WithFlank&X|+|110055323|110055432@67.87.21&[MIMAT0003322&hsa-miR-652-3p&offsets|0|0;m-148&X|+|110055389|110055410&offsets|0|-1]|AATGGCGCCACTAGGGTTGTG</td><td>>MI0003667|hsa-mir-652&WithFlank&X|+|110055323|110055432</td><td>67</td><td>87</td><td>21</td><td>AATGGCGCCACTAGGGTTGTG</td><td>1385.2</td><td>5.5</td><td>1.5</td><td>3.6</td><td>3.3E-04</td><td>9.6E-04</td><td>BT20/MB468</td><td>MP</td></tr>
<tr><td>>hg38_21_+_8433172_8446622_RNA45SN1_45S_50ntFlanks@6662.6689.28|GGTGGATCACTCGGCTCGTGCGTCGATG</td><td>>hg38_21_+_8433172_8446622_RNA45SN1_45S_50ntFlanks</td><td>6662</td><td>6689</td><td>28</td><td>GGTGGATCACTCGGCTCGTGCGTCGATG</td><td>509.4</td><td>7.1</td><td>1.8</td><td>3.9</td><td>1.2E-04</td><td>9.6E-04</td><td>BT20/MB231</td><td>MP</td></tr>
<tr><td>>MI0000746|hsa-mir-99b&WithFlank&19|+|51692606|51692687@13.33.21(+1A)&[MIMAT0000689&hsa-miR-99b-5p&offsets|0|-1(+1A);m-10&19|+|51692618|51692639&offsets|0|-1(+1A)]|CACCCGTAGAACCGACCTTGCA</td><td>>MI0000746|hsa-mir-99b&WithFlank&19|+|51692606|51692687</td><td>13</td><td>33</td><td>21(+1A)</td><td>CACCCGTAGAACCGACCTTGCA</td><td>1870.5</td><td>2.5</td><td>0.7</td><td>3.4</td><td>6.5E-04</td><td>9.6E-04</td><td>MB231/MB468</td><td>MP</td></tr>
<tr><td>>trna16_TyrGTA_14_-_21131351_21131444@59.76.18,trna17_TyrGTA_14_-_21128117_21128210@59.76.18,trna19_TyrGTA_14_-_21121258_21121351@59.76.18,trna2_TyrGTA_2_+_27273650_27273738@59.76.18,trna4_TyrGTA_8_+_67025602_67025694@59.76.18,trna5_TyrGTA_14_+_21151432_21151520@59.76.18,trna5_TyrGTA_8_+_67026223_67026311@59.76.18|TTCCGGCTCGAAGGACCA</td><td>>trna16_TyrGTA_14_-_21131351_21131444</td><td>59</td><td>76</td><td>18</td><td>TTCCGGCTCGAAGGACCA</td><td>387.7</td><td>3.2</td><td>1.0</td><td>3.4</td><td>7.7E-04</td><td>9.6E-04</td><td>MB231/MB468</td><td>MITO</td></tr>
<tr><td>>hg38_21_+_8433172_8446622_RNA45SN1_45S_50ntFlanks@7983.8003.21|CAGATCAGACGTGGCGACCCG</td><td>>hg38_21_+_8433172_8446622_RNA45SN1_45S_50ntFlanks</td><td>7983</td><td>8003</td><td>21</td><td>CAGATCAGACGTGGCGACCCG</td><td>491.8</td><td>-3.9</td><td>1.2</td><td>-3.4</td><td>7.3E-04</td><td>9.6E-04</td><td>MB231/MB468</td><td>CYTO</td></tr>
<tr><td>>MI0000466|hsa-mir-9-1&WithFlank&1|-|156420335|156420435@22.43.22;MI0000468|hsa-mir-9-3&WithFlank&15|+|89368011|89368112@22.43.22;MI0000467|hsa-mir-9-2&WithFlank&5|-|88666847|88666945@22.43.22&[MIMAT0000441&hsa-miR-9-5p&offsets|0|-1;m-104&1|-|156420392|156420414&offsets|0|-1];[MIMAT0000441_1&hsa-miR-9-5p&offsets|0|-1;m-105&15|+|89368032|89368054&offsets|0|-1];[MIMAT0000441_2&hsa-miR-9-5p&offsets|0|-1;m-104&5|-|88666902|88666924&offsets|0|-1]|TCTTTGGTTATCTAGCTGTATG</td><td>>MI0000466|hsa-mir-9-1&WithFlank&1|-|156420335|156420435</td><td>22</td><td>43</td><td>22</td><td>TCTTTGGTTATCTAGCTGTATG</td><td>1277.6</td><td>3.5</td><td>0.9</td><td>3.8</td><td>1.2E-04</td><td>9.6E-04</td><td>BT20/MB231</td><td>CYTO</td></tr>
<tr><td>>trnaMT_PheGAA_MT_+_577_647@27.74.48|AATACACTGAAAATGTTTAGACGGGCTCACATCACCCCATAAACACCA</td><td>>trnaMT_PheGAA_MT_+_577_647</td><td>27</td><td>74</td><td>48</td><td>AATACACTGAAAATGTTTAGACGGGCTCACATCACCCCATAAACACCA</td><td>62.7</td><td>9.7</td><td>2.7</td><td>3.6</td><td>3.4E-04</td><td>9.6E-04</td><td>BT20/MB468</td><td>MP</td></tr>
<tr><td>>hg38_21_+_8433172_8446622_RNA45SN1_45S_50ntFlanks@5486.5506.21|GCGCTGAGAAGACGGTCGAAC</td><td>>hg38_21_+_8433172_8446622_RNA45SN1_45S_50ntFlanks</td><td>5486</td><td>5506</td><td>21</td><td>GCGCTGAGAAGACGGTCGAAC</td><td>195.1</td><td>-4.2</td><td>1.2</td><td>-3.4</td><td>6.5E-04</td><td>9.6E-04</td><td>MB231/MB468</td><td>MP</td></tr>
<tr><td>>hg38_21_+_8433172_8446622_RNA45SN1_45S_50ntFlanks@11205.11232.28|GCGGCGGCGACTCTGGACGCGAGCCGGG</td><td>>hg38_21_+_8433172_8446622_RNA45SN1_45S_50ntFlanks</td><td>11205</td><td>11232</td><td>28</td><td>GCGGCGGCGACTCTGGACGCGAGCCGGG</td><td>492.4</td><td>-5.4</td><td>1.6</td><td>-3.4</td><td>7.9E-04</td><td>9.6E-04</td><td>MB231/MB468</td><td>NUC</td></tr>
<tr><td>>hg38_21_+_8433172_8446622_RNA45SN1_45S_50ntFlanks@3830.3872.43|GCTCGCTCCTCTCCTACTTGGATAACTGTGGTAATTCTAGAGC</td><td>>hg38_21_+_8433172_8446622_RNA45SN1_45S_50ntFlanks</td><td>3830</td><td>3872</td><td>43</td><td>GCTCGCTCCTCTCCTACTTGGATAACTGTGGTAATTCTAGAGC</td><td>307.6</td><td>-3.5</td><td>1.0</td><td>-3.4</td><td>7.9E-04</td><td>9.6E-04</td><td>MB231/MB468</td><td>NUC</td></tr>
<tr><td>>hg38_21_+_8433172_8446622_RNA45SN1_45S_50ntFlanks@4367.4402.36|CAGTTAAAAAGCTCGTAGTTGGATCTTGGGAGCGGG</td><td>>hg38_21_+_8433172_8446622_RNA45SN1_45S_50ntFlanks</td><td>4367</td><td>4402</td><td>36</td><td>CAGTTAAAAAGCTCGTAGTTGGATCTTGGGAGCGGG</td><td>134.9</td><td>-3.3</td><td>1.0</td><td>-3.4</td><td>7.3E-04</td><td>9.6E-04</td><td>MB231/MB468</td><td>CYTO</td></tr>
<tr><td>>MI0000077|hsa-mir-21&WithFlank&17|+|59841260|59841343@14.36.23(+1C)&[MIMAT0000076&hsa-miR-21-5p&offsets|0|+1(+1C);m-1&17|+|59841273|59841294&offsets|0|+1(+1C)]|TAGCTTATCAGACTGATGTTGACC</td><td>>MI0000077|hsa-mir-21&WithFlank&17|+|59841260|59841343</td><td>14</td><td>36</td><td>23(+1C)</td><td>TAGCTTATCAGACTGATGTTGACC</td><td>3312.3</td><td>3.2</td><td>0.9</td><td>3.6</td><td>3.4E-04</td><td>9.6E-04</td><td>BT20/MB468</td><td>MP</td></tr>
<tr><td>>hg38_21_+_8433172_8446622_RNA45SN1_45S_50ntFlanks@9495.9525.31|CGAAAGATGGTGAACTATGCCTGGGCAGGGC</td><td>>hg38_21_+_8433172_8446622_RNA45SN1_45S_50ntFlanks</td><td>9495</td><td>9525</td><td>31</td><td>CGAAAGATGGTGAACTATGCCTGGGCAGGGC</td><td>245.4</td><td>5.1</td><td>1.4</td><td>3.6</td><td>3.4E-04</td><td>9.6E-04</td><td>BT20/MB468</td><td>MP</td></tr>
<tr><td>>trna152_ValCAC_6_-_27248049_27248121@1.30.30|GCTTCTGTAGTGTAGTGGTTATCACGTTCG</td><td>>trna152_ValCAC_6_-_27248049_27248121</td><td>1</td><td>30</td><td>30</td><td>GCTTCTGTAGTGTAGTGGTTATCACGTTCG</td><td>340.6</td><td>-3.8</td><td>1.1</td><td>-3.4</td><td>7.0E-04</td><td>9.6E-04</td><td>BT20/MB468</td><td>CYTO</td></tr>
<tr><td>>hg38_21_+_8433172_8446622_RNA45SN1_45S_50ntFlanks@7983.8016.34|CAGATCAGACGTGGCGACCCGCTGAATTTAAGCA</td><td>>hg38_21_+_8433172_8446622_RNA45SN1_45S_50ntFlanks</td><td>7983</td><td>8016</td><td>34</td><td>CAGATCAGACGTGGCGACCCGCTGAATTTAAGCA</td><td>373.1</td><td>-2.9</td><td>0.9</td><td>-3.4</td><td>7.8E-04</td><td>9.6E-04</td><td>MB231/MB468</td><td>MITO</td></tr>
<tr><td>>MI0003137|hsa-mir-193b&WithFlank&16|+|14303961|14304055@57.75.19(+1A)&[MIMAT0002819&hsa-miR-193b-3p&offsets|0|-3(+1A);m-166&16|+|14304017|14304038&offsets|0|-3(+1A)]|AACTGGCCCTCAAAGTCCCA</td><td>>MI0003137|hsa-mir-193b&WithFlank&16|+|14303961|14304055</td><td>57</td><td>75</td><td>19(+1A)</td><td>AACTGGCCCTCAAAGTCCCA</td><td>62.4</td><td>9.7</td><td>2.7</td><td>3.6</td><td>3.4E-04</td><td>9.6E-04</td><td>BT20/MB468</td><td>MP</td></tr>
<tr><td>>hg38_21_+_8433172_8446622_RNA45SN1_45S_50ntFlanks@8242.8270.29|GGTCTTCCCGGAGTCGGGTTGCTTGGGAA</td><td>>hg38_21_+_8433172_8446622_RNA45SN1_45S_50ntFlanks</td><td>8242</td><td>8270</td><td>29</td><td>GGTCTTCCCGGAGTCGGGTTGCTTGGGAA</td><td>135.4</td><td>10.4</td><td>2.7</td><td>3.9</td><td>1.2E-04</td><td>9.6E-04</td><td>BT20/MB231</td><td>MP</td></tr>
<tr><td>>hg38_21_+_8433172_8446622_RNA45SN1_45S_50ntFlanks@3789.3810.22|ACTGCGAATGGCTCATTAAATC</td><td>>hg38_21_+_8433172_8446622_RNA45SN1_45S_50ntFlanks</td><td>3789</td><td>3810</td><td>22</td><td>ACTGCGAATGGCTCATTAAATC</td><td>315.6</td><td>-1.6</td><td>0.5</td><td>-3.4</td><td>7.0E-04</td><td>9.6E-04</td><td>BT20/MB468</td><td>CYTO</td></tr>
<tr><td>>MI0000456|hsa-mir-140&WithFlank&16|+|69933075|69933186@69.89.21(+1A)&[MIMAT0004597&hsa-miR-140-3p&offsets|+1|+1(+1A);m-45&16|+|69933143|69933164&offsets|0|-1(+1A)]|ACCACAGGGTAGAACCACGGAA</td><td>>MI0000456|hsa-mir-140&WithFlank&16|+|69933075|69933186</td><td>69</td><td>89</td><td>21(+1A)</td><td>ACCACAGGGTAGAACCACGGAA</td><td>119.3</td><td>6.0</td><td>1.8</td><td>3.4</td><td>6.5E-04</td><td>9.6E-04</td><td>MB231/MB468</td><td>MP</td></tr>
<tr><td>>hg38_21_+_8433172_8446622_RNA45SN1_45S_50ntFlanks@7981.8011.31|CTCAGATCAGACGTGGCGACCCGCTGAATTT</td><td>>hg38_21_+_8433172_8446622_RNA45SN1_45S_50ntFlanks</td><td>7981</td><td>8011</td><td>31</td><td>CTCAGATCAGACGTGGCGACCCGCTGAATTT</td><td>318.7</td><td>-2.6</td><td>0.8</td><td>-3.4</td><td>7.8E-04</td><td>9.6E-04</td><td>MB231/MB468</td><td>MITO</td></tr>
<tr><td>>hg38_21_+_8433172_8446622_RNA45SN1_45S_50ntFlanks@7975.8010.36|CGCGACCTCAGATCAGACGTGGCGACCCGCTGAATT</td><td>>hg38_21_+_8433172_8446622_RNA45SN1_45S_50ntFlanks</td><td>7975</td><td>8010</td><td>36</td><td>CGCGACCTCAGATCAGACGTGGCGACCCGCTGAATT</td><td>464526.6</td><td>3.0</td><td>0.8</td><td>3.8</td><td>1.2E-04</td><td>9.6E-04</td><td>BT20/MB231</td><td>NUC</td></tr>
<tr><td>>hg38_21_+_8433172_8446622_RNA45SN1_45S_50ntFlanks@11208.11227.20|GCGGCGACTCTGGACGCGAG</td><td>>hg38_21_+_8433172_8446622_RNA45SN1_45S_50ntFlanks</td><td>11208</td><td>11227</td><td>20</td><td>GCGGCGACTCTGGACGCGAG</td><td>2601.6</td><td>-5.1</td><td>1.4</td><td>-3.6</td><td>3.4E-04</td><td>9.6E-04</td><td>BT20/MB468</td><td>MP</td></tr>
<tr><td>>hg38_21_+_8433172_8446622_RNA45SN1_45S_50ntFlanks@7988.8017.30|CAGACGTGGCGACCCGCTGAATTTAAGCAT</td><td>>hg38_21_+_8433172_8446622_RNA45SN1_45S_50ntFlanks</td><td>7988</td><td>8017</td><td>30</td><td>CAGACGTGGCGACCCGCTGAATTTAAGCAT</td><td>793.1</td><td>-2.3</td><td>0.7</td><td>-3.4</td><td>5.9E-04</td><td>9.6E-04</td><td>BT20/MB468</td><td>MITO</td></tr>
<tr><td>>hg38_21_+_8433172_8446622_RNA45SN1_45S_50ntFlanks@7975.7997.23|CGCGACCTCAGATCAGACGTGGC</td><td>>hg38_21_+_8433172_8446622_RNA45SN1_45S_50ntFlanks</td><td>7975</td><td>7997</td><td>23</td><td>CGCGACCTCAGATCAGACGTGGC</td><td>159463.5</td><td>2.7</td><td>0.7</td><td>3.8</td><td>1.4E-04</td><td>9.6E-04</td><td>BT20/MB231</td><td>MITO</td></tr>
<tr><td>>hg38_1_-_228634819_228635039_RNA5S12_5S_50ntFlanks@140.171.32|ACCGCCTGGGAATACCGGGTGCTGTAGGCTTT</td><td>>hg38_1_-_228634819_228635039_RNA5S12_5S_50ntFlanks</td><td>140</td><td>171</td><td>32</td><td>ACCGCCTGGGAATACCGGGTGCTGTAGGCTTT</td><td>8752.4</td><td>-3.0</td><td>0.9</td><td>-3.4</td><td>6.6E-04</td><td>9.6E-04</td><td>MB231/MB468</td><td>MP</td></tr>
<tr><td>>MI0000439|hsa-mir-23b&WithFlank&9|+|95085202|95085310@67.86.20(+1A)&[MIMAT0000418&hsa-miR-23b-3p&offsets|+3|0(+1A);m-42&9|+|95085265|95085285&offsets|+3|+2(+1A)]|ACATTGCCAGGGATTACCACA</td><td>>MI0000439|hsa-mir-23b&WithFlank&9|+|95085202|95085310</td><td>67</td><td>86</td><td>20(+1A)</td><td>ACATTGCCAGGGATTACCACA</td><td>313.0</td><td>12.2</td><td>3.6</td><td>3.4</td><td>7.9E-04</td><td>9.6E-04</td><td>MB231/MB468</td><td>NUC</td></tr>
<tr><td>>hg38_21_+_8433172_8446622_RNA45SN1_45S_50ntFlanks@4724.4764.41|ATTAATCAAGAACGAAAGTCGGAGGTTCGAAGACGATCAGA</td><td>>hg38_21_+_8433172_8446622_RNA45SN1_45S_50ntFlanks</td><td>4724</td><td>4764</td><td>41</td><td>ATTAATCAAGAACGAAAGTCGGAGGTTCGAAGACGATCAGA</td><td>72.4</td><td>-10.0</td><td>2.9</td><td>-3.4</td><td>6.6E-04</td><td>9.6E-04</td><td>MB231/MB468</td><td>MP</td></tr>
<tr><td>>MI0000750|hsa-mir-26a-2&WithFlank&12|-|57824603|57824698@20.41.22(+3U);MI0000083|hsa-mir-26a-1&WithFlank&3|+|37969398|37969486@16.37.22(+3U)&[MIMAT0000082&hsa-miR-26a-5p&offsets|0|0(+3U);m-33&12|-|57824658|57824679&offsets|0|0(+3U)];[MIMAT0000082_1&hsa-miR-26a-5p&offsets|0|0(+3U);m-34&3|+|37969413|37969434&offsets|0|0(+3U)]|TTCAAGTAATCCAGGATAGGCTTTT</td><td>>MI0000750|hsa-mir-26a-2&WithFlank&12|-|57824603|57824698</td><td>20</td><td>41</td><td>22(+3U)</td><td>TTCAAGTAATCCAGGATAGGCTTTT</td><td>203.0</td><td>6.3</td><td>1.8</td><td>3.4</td><td>6.6E-04</td><td>9.7E-04</td><td>MB231/MB468</td><td>MP</td></tr>
<tr><td>>hg38_21_+_8433172_8446622_RNA45SN1_45S_50ntFlanks@5492.5511.20|AGAAGACGGTCGAACTTGAC</td><td>>hg38_21_+_8433172_8446622_RNA45SN1_45S_50ntFlanks</td><td>5492</td><td>5511</td><td>20</td><td>AGAAGACGGTCGAACTTGAC</td><td>111.3</td><td>-2.9</td><td>0.8</td><td>-3.4</td><td>6.6E-04</td><td>9.7E-04</td><td>MB231/MB468</td><td>MP</td></tr>
<tr><td>>hg38_21_+_8433172_8446622_RNA45SN1_45S_50ntFlanks@6781.6802.22|GGGGCTACGCCTGTCTGAGCGT</td><td>>hg38_21_+_8433172_8446622_RNA45SN1_45S_50ntFlanks</td><td>6781</td><td>6802</td><td>22</td><td>GGGGCTACGCCTGTCTGAGCGT</td><td>384.2</td><td>5.2</td><td>1.4</td><td>3.6</td><td>3.4E-04</td><td>9.7E-04</td><td>BT20/MB468</td><td>MP</td></tr>
<tr><td>>hg38_21_+_8433172_8446622_RNA45SN1_45S_50ntFlanks@3979.4001.23|CGCCGGCGGCTTTGGTGACTCTA</td><td>>hg38_21_+_8433172_8446622_RNA45SN1_45S_50ntFlanks</td><td>3979</td><td>4001</td><td>23</td><td>CGCCGGCGGCTTTGGTGACTCTA</td><td>318.9</td><td>-3.2</td><td>0.9</td><td>-3.4</td><td>5.9E-04</td><td>9.7E-04</td><td>BT20/MB468</td><td>MITO</td></tr>
<tr><td>>hg38_21_+_8433172_8446622_RNA45SN1_45S_50ntFlanks@5472.5506.35|ACGGCCCTGGCGGAGCGCTGAGAAGACGGTCGAAC</td><td>>hg38_21_+_8433172_8446622_RNA45SN1_45S_50ntFlanks</td><td>5472</td><td>5506</td><td>35</td><td>ACGGCCCTGGCGGAGCGCTGAGAAGACGGTCGAAC</td><td>249.5</td><td>3.1</td><td>0.9</td><td>3.4</td><td>6.6E-04</td><td>9.7E-04</td><td>MB231/MB468</td><td>MP</td></tr>
<tr><td>>hg38_21_+_8433172_8446622_RNA45SN1_45S_50ntFlanks@10229.10255.27|CCCCGCGGACGCTACGCCGCGACGAGT</td><td>>hg38_21_+_8433172_8446622_RNA45SN1_45S_50ntFlanks</td><td>10229</td><td>10255</td><td>27</td><td>CCCCGCGGACGCTACGCCGCGACGAGT</td><td>61.2</td><td>9.7</td><td>2.7</td><td>3.6</td><td>3.4E-04</td><td>9.7E-04</td><td>BT20/MB468</td><td>MP</td></tr>
<tr><td>>hg38_21_+_8433172_8446622_RNA45SN1_45S_50ntFlanks@10127.10159.33|GCGGCGGCGGCGGCGGCGGCGGGGGTGTGGGGT</td><td>>hg38_21_+_8433172_8446622_RNA45SN1_45S_50ntFlanks</td><td>10127</td><td>10159</td><td>33</td><td>GCGGCGGCGGCGGCGGCGGCGGGGGTGTGGGGT</td><td>63.1</td><td>9.7</td><td>2.7</td><td>3.6</td><td>3.4E-04</td><td>9.7E-04</td><td>BT20/MB468</td><td>MP</td></tr>
<tr><td>>hg38_21_+_8433172_8446622_RNA45SN1_45S_50ntFlanks@11319.11338.20|CCCACGTCTCGTCGCGCGCG</td><td>>hg38_21_+_8433172_8446622_RNA45SN1_45S_50ntFlanks</td><td>11319</td><td>11338</td><td>20</td><td>CCCACGTCTCGTCGCGCGCG</td><td>310.2</td><td>-5.8</td><td>1.7</td><td>-3.4</td><td>7.9E-04</td><td>9.7E-04</td><td>BT20/MB468</td><td>NUC</td></tr>
<tr><td>>MI0000088|hsa-mir-30a&WithFlank&6|-|71403545|71403627@12.35.24(+3U)&[MIMAT0000087&hsa-miR-30a-5p&offsets|0|+2(+3U);m-15&6|-|71403595|71403616&offsets|0|+2(+3U)]|TGTAAACATCCTCGACTGGAAGCTTTT</td><td>>MI0000088|hsa-mir-30a&WithFlank&6|-|71403545|71403627</td><td>12</td><td>35</td><td>24(+3U)</td><td>TGTAAACATCCTCGACTGGAAGCTTTT</td><td>234.9</td><td>5.2</td><td>1.5</td><td>3.4</td><td>6.6E-04</td><td>9.7E-04</td><td>MB231/MB468</td><td>MP</td></tr>
</table>

<table>
<tr><td>>hg38_MT_+_1621_3279_RNR2_16S_50ntFlanks@1170.1210.41|AAACCTGCATTAAAAATTTCGGTTGGGGCGACCTCGGAGCA</td><td>>hg38_MT_+_1621_3279_RNR2_16S_50ntFlanks</td><td>1170</td><td>1210</td><td>41</td><td>AAACCTGCATTAAAAATTTCGGTTGGGGCGACCTCGGAGCA</td><td>82.4</td><td>5.9</td><td>1.7</td><td>3.4</td><td>7.8E-04</td><td>9.7E-04</td><td>MB231/MB468</td><td>MITO</td></tr>
<tr><td>>hg38_21_+_8433172_8446622_RNA45SN1_45S_50ntFlanks@12932.12966.35|CGCTAAACCATTCGTAGACGACCTGCTTCTGGGTC</td><td>>hg38_21_+_8433172_8446622_RNA45SN1_45S_50ntFlanks</td><td>12932</td><td>12966</td><td>35</td><td>CGCTAAACCATTCGTAGACGACCTGCTTCTGGGTC</td><td>167.1</td><td>-2.9</td><td>0.9</td><td>-3.4</td><td>7.4E-04</td><td>9.7E-04</td><td>MB231/MB468</td><td>CYTO</td></tr>
<tr><td>>hg38_21_+_8433172_8446622_RNA45SN1_45S_50ntFlanks@8170.8218.49|CAAGTCCTTCTGATCGAGGCCCAGCCCGTGGACGGTGTGAGGCCGGTAG</td><td>>hg38_21_+_8433172_8446622_RNA45SN1_45S_50ntFlanks</td><td>8170</td><td>8218</td><td>49</td><td>CAAGTCCTTCTGATCGAGGCCCAGCCCGTGGACGGTGTGAGGCCGGTAG</td><td>261.3</td><td>-4.7</td><td>1.4</td><td>-3.4</td><td>8.0E-04</td><td>9.7E-04</td><td>MB231/MB468</td><td>NUC</td></tr>
<tr><td>>hg38_21_+_8433172_8446622_RNA45SN1_45S_50ntFlanks@11579.11600.22|AGAACTGGTGCGGACCAGGGGA</td><td>>hg38_21_+_8433172_8446622_RNA45SN1_45S_50ntFlanks</td><td>11579</td><td>11600</td><td>22</td><td>AGAACTGGTGCGGACCAGGGGA</td><td>260.7</td><td>-4.4</td><td>1.3</td><td>-3.4</td><td>7.1E-04</td><td>9.7E-04</td><td>BT20/MB468</td><td>CYTO</td></tr>
<tr><td>>trna111_HisGTG_1_-_147774845_147774916@-1G.35.36,trna118_HisGTG_1_-_145396881_145396952@-1G.35.36,trna16_HisGTG_1_+_146544773_146544844@-1G.35.36,trna1_HisGTG_15_+_45493349_45493420@-1G.35.36,trna21_HisGTG_1_+_147753471_147753542@-1G.35.36,trna33_HisGTG_6_+_27125906_27125977@-1G.35.36,trna7_HisGTG_9_-_14433938_14434009@-1G.35.36,trna8_HisGTG_15_-_45492611_45492682@-1G.35.36,trna9_HisGTG_15_-_45490804_45490875@-1G.35.36|GGCCGTGATCGTATAGTGGTTAGTACTCTGCGTTGT</td><td>>trna111_HisGTG_1_-_147774845_147774916</td><td>-1G</td><td>35</td><td>36</td><td>GGCCGTGATCGTATAGTGGTTAGTACTCTGCGTTGT</td><td>1445.7</td><td>-3.1</td><td>0.8</td><td>-3.8</td><td>1.4E-04</td><td>9.7E-04</td><td>BT20/MB231</td><td>MITO</td></tr>
<tr><td>>hg38_1_-_228634819_228635039_RNA5S12_5S_50ntFlanks@51.68.18|GTCTACGGCCATACCACC</td><td>>hg38_1_-_228634819_228635039_RNA5S12_5S_50ntFlanks</td><td>51</td><td>68</td><td>18</td><td>GTCTACGGCCATACCACC</td><td>360.0</td><td>-3.7</td><td>1.1</td><td>-3.4</td><td>6.6E-04</td><td>9.7E-04</td><td>MB231/MB468</td><td>MP</td></tr>
<tr><td>>hg38_21_+_8433172_8446622_RNA45SN1_45S_50ntFlanks@10393.10431.39|CAGCAGTTGAACATGGGTCAGTCGGTCCTGAGAGATGGG</td><td>>hg38_21_+_8433172_8446622_RNA45SN1_45S_50ntFlanks</td><td>10393</td><td>10431</td><td>39</td><td>CAGCAGTTGAACATGGGTCAGTCGGTCCTGAGAGATGGG</td><td>202.2</td><td>-3.2</td><td>1.0</td><td>-3.4</td><td>7.4E-04</td><td>9.7E-04</td><td>MB231/MB468</td><td>CYTO</td></tr>
<tr><td>>MI0000298|hsa-mir-221&WithFlank&X|-|45746151|45746272@31.54.24&[MIMAT0004568&hsa-miR-221-5p&offsets|0|+2;m-405&X|-|45746220|45746242&offsets|0|+1]|ACCTGGCATACAATGTAGATTTCT</td><td>>MI0000298|hsa-mir-221&WithFlank&X|-|45746151|45746272</td><td>31</td><td>54</td><td>24</td><td>ACCTGGCATACAATGTAGATTTCT</td><td>1397.4</td><td>-2.4</td><td>0.6</td><td>-3.8</td><td>1.2E-04</td><td>9.7E-04</td><td>BT20/MB231</td><td>NUC</td></tr>
<tr><td>>hg38_21_+_8433172_8446622_RNA45SN1_45S_50ntFlanks@9156.9188.33|CCAGTGCGCCCCGGGCGGGTCGCGCCGTCGGGC</td><td>>hg38_21_+_8433172_8446622_RNA45SN1_45S_50ntFlanks</td><td>9156</td><td>9188</td><td>33</td><td>CCAGTGCGCCCCGGGCGGGTCGCGCCGTCGGGC</td><td>893.2</td><td>-4.8</td><td>1.3</td><td>-3.8</td><td>1.2E-04</td><td>9.7E-04</td><td>BT20/MB231</td><td>NUC</td></tr>
<tr><td>>hg38_21_+_8433172_8446622_RNA45SN1_45S_50ntFlanks@9761.9787.27|AACTTTAAATGGGTAAGAAGCCCGGCT</td><td>>hg38_21_+_8433172_8446622_RNA45SN1_45S_50ntFlanks</td><td>9761</td><td>9787</td><td>27</td><td>AACTTTAAATGGGTAAGAAGCCCGGCT</td><td>73.1</td><td>-9.3</td><td>2.6</td><td>-3.6</td><td>3.4E-04</td><td>9.7E-04</td><td>BT20/MB468</td><td>MP</td></tr>
<tr><td>>hg38_21_+_8433172_8446622_RNA45SN1_45S_50ntFlanks@6719.6746.28|TGAATTGCAGGACACATTGATCATCGAC</td><td>>hg38_21_+_8433172_8446622_RNA45SN1_45S_50ntFlanks</td><td>6719</td><td>6746</td><td>28</td><td>TGAATTGCAGGACACATTGATCATCGAC</td><td>1039.1</td><td>-4.9</td><td>1.4</td><td>-3.6</td><td>3.4E-04</td><td>9.7E-04</td><td>BT20/MB468</td><td>MP</td></tr>
<tr><td>>MI0000440|hsa-mir-27b&WithFlank&9|+|95085439|95085547@70.87.18&[MIMAT0000419&hsa-miR-27b-3p&offsets|+3|0;m-39&9|+|95085505|95085525&offsets|+3|0]|ACAGTGGCTAAGTTCTGC</td><td>>MI0000440|hsa-mir-27b&WithFlank&9|+|95085439|95085547</td><td>70</td><td>87</td><td>18</td><td>ACAGTGGCTAAGTTCTGC</td><td>500.8</td><td>12.2</td><td>3.6</td><td>3.4</td><td>7.9E-04</td><td>9.7E-04</td><td>MB231/MB468</td><td>MITO</td></tr>
<tr><td>>MI0000289|hsa-mir-181a-1&WithFlank&1|-|198859038|198859159@30.51.22(+1A);MI0000269|hsa-mir-181a-2&WithFlank&9|+|124692436|124692557@45.66.22(+1A)&[MIMAT0000256&hsa-miR-181a-5p&offsets|0|-1(+1A);m-32&1|-|198859108|198859130&offsets|0|-1(+1A)];[MIMAT0000256_1&hsa-miR-181a-5p&offsets|0|-1(+1A);m-32&9|+|124692480|124692502&offsets|0|-1(+1A)]|AACATTCAACGCTGTCGGTGAGA</td><td>>MI0000289|hsa-mir-181a-1&WithFlank&1|-|198859038|198859159</td><td>30</td><td>51</td><td>22(+1A)</td><td>AACATTCAACGCTGTCGGTGAGA</td><td>601.3</td><td>2.8</td><td>0.8</td><td>3.4</td><td>6.6E-04</td><td>9.7E-04</td><td>MB231/MB468</td><td>MP</td></tr>
<tr><td>>hg38_21_+_8433172_8446622_RNA45SN1_45S_50ntFlanks@10401.10426.26|GAACATGGGTCAGTCGGTCCTGAGAG</td><td>>hg38_21_+_8433172_8446622_RNA45SN1_45S_50ntFlanks</td><td>10401</td><td>10426</td><td>26</td><td>GAACATGGGTCAGTCGGTCCTGAGAG</td><td>215.1</td><td>-5.4</td><td>1.6</td><td>-3.4</td><td>6.6E-04</td><td>9.7E-04</td><td>MB231/MB468</td><td>MP</td></tr>
<tr><td>>MI0000456|hsa-mir-140&WithFlank&16|+|69933075|69933186@68.89.22&[MIMAT0004597&hsa-miR-140-3p&offsets|0|+1;m-45&16|+|69933143|69933164&offsets|-1|-1]|TACCACAGGGTAGAACCACGGA</td><td>>MI0000456|hsa-mir-140&WithFlank&16|+|69933075|69933186</td><td>68</td><td>89</td><td>22</td><td>TACCACAGGGTAGAACCACGGA</td><td>388.4</td><td>5.2</td><td>1.4</td><td>3.6</td><td>3.4E-04</td><td>9.7E-04</td><td>BT20/MB468</td><td>MP</td></tr>
<tr><td>>hg38_21_+_8433172_8446622_RNA45SN1_45S_50ntFlanks@8112.8138.27|GGGCGCGGGACATGTGGCGTACGGAAG</td><td>>hg38_21_+_8433172_8446622_RNA45SN1_45S_50ntFlanks</td><td>8112</td><td>8138</td><td>27</td><td>GGGCGCGGGACATGTGGCGTACGGAAG</td><td>1581.0</td><td>-3.5</td><td>1.0</td><td>-3.6</td><td>3.4E-04</td><td>9.7E-04</td><td>BT20/MB468</td><td>MP</td></tr>
<tr><td>>hg38_21_+_8433172_8446622_RNA45SN1_45S_50ntFlanks@11321.11338.18|CACGTCTCGTCGCGCGCG</td><td>>hg38_21_+_8433172_8446622_RNA45SN1_45S_50ntFlanks</td><td>11321</td><td>11338</td><td>18</td><td>CACGTCTCGTCGCGCGCG</td><td>1110.8</td><td>-4.5</td><td>1.3</td><td>-3.4</td><td>7.1E-04</td><td>9.8E-04</td><td>BT20/MB468</td><td>CYTO</td></tr>
<tr><td>>hg38_21_+_8433172_8446622_RNA45SN1_45S_50ntFlanks@9571.9597.27|GCAAATCGGTCGTCCGACCTGGGTATA</td><td>>hg38_21_+_8433172_8446622_RNA45SN1_45S_50ntFlanks</td><td>9571</td><td>9597</td><td>27</td><td>GCAAATCGGTCGTCCGACCTGGGTATA</td><td>288.5</td><td>-5.8</td><td>1.6</td><td>-3.6</td><td>3.4E-04</td><td>9.8E-04</td><td>BT20/MB468</td><td>MP</td></tr>
<tr><td>>MI0003760|hsa-mir-671&WithFlank&7|+|151238415|151238544@74.94.21&[MIMAT0004819&hsa-miR-671-3p&offsets|0|0;m-323&7|+|151238488|151238508&offsets|0|0]|TCCGGTTCTCAGGGCTCCACC</td><td>>MI0003760|hsa-mir-671&WithFlank&7|+|151238415|151238544</td><td>74</td><td>94</td><td>21</td><td>TCCGGTTCTCAGGGCTCCACC</td><td>129.2</td><td>4.3</td><td>1.3</td><td>3.4</td><td>7.9E-04</td><td>9.8E-04</td><td>MB231/MB468</td><td>MITO</td></tr>
<tr><td>>hg38_21_+_8433172_8446622_RNA45SN1_45S_50ntFlanks@11660.11685.26|GTGATTTCTGCCCAGTGCTCTGAATG</td><td>>hg38_21_+_8433172_8446622_RNA45SN1_45S_50ntFlanks</td><td>11660</td><td>11685</td><td>26</td><td>GTGATTTCTGCCCAGTGCTCTGAATG</td><td>2225.4</td><td>-2.9</td><td>0.8</td><td>-3.4</td><td>6.0E-04</td><td>9.8E-04</td><td>BT20/MB468</td><td>MITO</td></tr>
<tr><td>>hg38_21_+_8433172_8446622_RNA45SN1_45S_50ntFlanks@485.511.27|GATGTGGTGACGTCGTGCTCTCCCGGG</td><td>>hg38_21_+_8433172_8446622_RNA45SN1_45S_50ntFlanks</td><td>485</td><td>511</td><td>27</td><td>GATGTGGTGACGTCGTGCTCTCCCGGG</td><td>56.1</td><td>9.0</td><td>2.7</td><td>3.4</td><td>6.6E-04</td><td>9.8E-04</td><td>MB231/MB468</td><td>MP</td></tr>
<tr><td>>hg38_21_+_8433172_8446622_RNA45SN1_45S_50ntFlanks@11215.11234.20|CTCTGGACGCGAGCCGGGCC</td><td>>hg38_21_+_8433172_8446622_RNA45SN1_45S_50ntFlanks</td><td>11215</td><td>11234</td><td>20</td><td>CTCTGGACGCGAGCCGGGCC</td><td>128.0</td><td>-5.8</td><td>1.6</td><td>-3.6</td><td>3.4E-04</td><td>9.8E-04</td><td>BT20/MB468</td><td>MP</td></tr>
<tr><td>>hg38_21_+_8433172_8446622_RNA45SN1_45S_50ntFlanks@11552.11592.41|CCTCGGCCGGCGCCTAGCAGCCGACTTAGAACTGGTGCGGA</td><td>>hg38_21_+_8433172_8446622_RNA45SN1_45S_50ntFlanks</td><td>11552</td><td>11592</td><td>41</td><td>CCTCGGCCGGCGCCTAGCAGCCGACTTAGAACTGGTGCGGA</td><td>284.1</td><td>-4.1</td><td>1.2</td><td>-3.4</td><td>7.2E-04</td><td>9.8E-04</td><td>BT20/MB468</td><td>CYTO</td></tr>
<tr><td>>am_trna111_HisGTG_1_-_147774845_147774916@-1G.23.24,trna118_HisGTG_1_-_145396881_145396952@-1G.23.24,trna16_HisGTG_1_+_146544773_146544844@-1G.23.24,trna1_HisGTG_15_+_45493349_45493420@-1G.23.24,trna21_HisGTG_1_+_147753471_147753542@-1G.23.24,trna33_HisGTG_6_+_27125906_27125977@-1G.23.24,trna7_HisGTG_9_-_14433938_14434009@-1G.23.24,trna8_HisGTG_15_-_45492611_45492682@-1G.23.24,trna9_HisGTG_15_-_45490804_45490875@-1G.23.24|GGCCGTGATCGTATAGTGGTTAGT</td><td>>am_trna111_HisGTG_1_-_147774845_147774916</td><td>-1G</td><td>23</td><td>24</td><td>GGCCGTGATCGTATAGTGGTTAGT</td><td>873.0</td><td>-6.1</td><td>1.6</td><td>-3.8</td><td>1.2E-04</td><td>9.8E-04</td><td>BT20/MB231</td><td>MP</td></tr>
<tr><td>>hg38_21_+_8433172_8446622_RNA45SN1_45S_50ntFlanks@12073.12110.38|CCGAGGGGCTCTCGCTTCTGGCGCCAAGCGCCCGGCCG</td><td>>hg38_21_+_8433172_8446622_RNA45SN1_45S_50ntFlanks</td><td>12073</td><td>12110</td><td>38</td><td>CCGAGGGGCTCTCGCTTCTGGCGCCAAGCGCCCGGCCG</td><td>196.0</td><td>-4.9</td><td>1.5</td><td>-3.4</td><td>7.2E-04</td><td>9.8E-04</td><td>BT20/MB468</td><td>CYTO</td></tr>
<tr><td>>hg38_21_+_8433172_8446622_RNA45SN1_45S_50ntFlanks@475.520.46|CTTCGTGATCGATGTGGTGACGTCGTGCTCTCCCGGGCCGGGTCCG</td><td>>hg38_21_+_8433172_8446622_RNA45SN1_45S_50ntFlanks</td><td>475</td><td>520</td><td>46</td><td>CTTCGTGATCGATGTGGTGACGTCGTGCTCTCCCGGGCCGGGTCCG</td><td>3043.3</td><td>3.8</td><td>1.1</td><td>3.4</td><td>8.0E-04</td><td>9.8E-04</td><td>MB231/MB468</td><td>NUC</td></tr>
<tr><td>>hg38_21_+_8433172_8446622_RNA45SN1_45S_50ntFlanks@11455.11477.23|CTCGGCGCGCGGCGGCGGCGGCG</td><td>>hg38_21_+_8433172_8446622_RNA45SN1_45S_50ntFlanks</td><td>11455</td><td>11477</td><td>23</td><td>CTCGGCGCGCGGCGGCGGCGGCG</td><td>197.1</td><td>-4.8</td><td>1.4</td><td>-3.4</td><td>7.2E-04</td><td>9.8E-04</td><td>BT20/MB468</td><td>CYTO</td></tr>
<tr><td>>trna143_LysTTT_6_-_27559593_27559665@13.30.18,trna149_LysTTT_6_-_27302769_27302841@13.30.18,trna14_LysTTT_11_-_59327808_59327880@13.30.18,trna2_LysTTT_17_+_8022473_8022545@13.30.18,trna54_LysTTT_1_+_204475655_204475727@13.30.18,trna5_LysTTT_11_+_59323902_59323974@13.30.18,trna62_LysTTT_1_-_204476158_204476230@13.30.18,trna76_LysTTT_6_+_28918806_28918878@13.30.18|CAGTCGGTAGAGCATCAG</td><td>>trna143_LysTTT_6_-_27559593_27559665</td><td>13</td><td>30</td><td>18</td><td>CAGTCGGTAGAGCATCAG</td><td>240.5</td><td>-5.7</td><td>1.6</td><td>-3.6</td><td>3.4E-04</td><td>9.8E-04</td><td>BT20/MB468</td><td>MP</td></tr>
<tr><td>>hg38_21_+_8433172_8446622_RNA45SN1_45S_50ntFlanks@12165.12182.18|TGGGGCGGTACACCTGTC</td><td>>hg38_21_+_8433172_8446622_RNA45SN1_45S_50ntFlanks</td><td>12165</td><td>12182</td><td>18</td><td>TGGGGCGGTACACCTGTC</td><td>367.5</td><td>-3.2</td><td>0.9</td><td>-3.6</td><td>3.4E-04</td><td>9.8E-04</td><td>BT20/MB468</td><td>MP</td></tr>
<tr><td>>hg38_21_+_8433172_8446622_RNA45SN1_45S_50ntFlanks@11217.11234.18|CTGGACGCGAGCCGGGCC</td><td>>hg38_21_+_8433172_8446622_RNA45SN1_45S_50ntFlanks</td><td>11217</td><td>11234</td><td>18</td><td>CTGGACGCGAGCCGGGCC</td><td>1295.9</td><td>-5.4</td><td>1.6</td><td>-3.4</td><td>8.0E-04</td><td>9.8E-04</td><td>BT20/MB468</td><td>NUC</td></tr>
<tr><td>>hg38_21_+_8433172_8446622_RNA45SN1_45S_50ntFlanks@5085.5104.20|GAACGAGACTCTGGCATGCT</td><td>>hg38_21_+_8433172_8446622_RNA45SN1_45S_50ntFlanks</td><td>5085</td><td>5104</td><td>20</td><td>GAACGAGACTCTGGCATGCT</td><td>361.9</td><td>5.6</td><td>1.6</td><td>3.6</td><td>3.4E-04</td><td>9.8E-04</td><td>BT20/MB468</td><td>MP</td></tr>
<tr><td>>trnaMT_PheGAA_MT_+_577_647@41.74.34|GTTTAGACGGGCTCACATCACCCCATAAACACCA</td><td>>trnaMT_PheGAA_MT_+_577_647</td><td>41</td><td>74</td><td>34</td><td>GTTTAGACGGGCTCACATCACCCCATAAACACCA</td><td>61.1</td><td>9.7</td><td>2.7</td><td>3.6</td><td>3.4E-04</td><td>9.8E-04</td><td>BT20/MB468</td><td>MP</td></tr>
<tr><td>>hg38_21_+_8433172_8446622_RNA45SN1_45S_50ntFlanks@10240.10259.20|CTACGCCGCGACGAGTAGGA</td><td>>hg38_21_+_8433172_8446622_RNA45SN1_45S_50ntFlanks</td><td>10240</td><td>10259</td><td>20</td><td>CTACGCCGCGACGAGTAGGA</td><td>143.5</td><td>2.3</td><td>0.7</td><td>3.4</td><td>8.0E-04</td><td>9.8E-04</td><td>BT20/MB468</td><td>NUC</td></tr>
<tr><td>>trnaMT_GluTTC_MT_-_14674_14742@40.70.31,trnalookalike8_GluTTC_5_-_93905172_93905240@40.70.31|TTGGTCGTGGTTGTAGTCCGTGCGAGAATAC</td><td>>trnaMT_GluTTC_MT_-_14674_14742</td><td>40</td><td>70</td><td>31</td><td>TTGGTCGTGGTTGTAGTCCGTGCGAGAATAC</td><td>536.0</td><td>7.4</td><td>2.1</td><td>3.6</td><td>3.4E-04</td><td>9.8E-04</td><td>BT20/MB468</td><td>MP</td></tr>
<tr><td>>MI0000299|hsa-mir-222&WithFlank&X|-|45747009|45747130@75.99.25(+2U)&[MIMAT0000279&hsa-miR-222-3p&offsets|0|+4(+2U);m-140&X|-|45747033|45747056&offsets|0|+1(+2U)]|AGCTACATCTGGCTACTGGGTCTCTTT</td><td>>MI0000299|hsa-mir-222&WithFlank&X|-|45747009|45747130</td><td>75</td><td>99</td><td>25(+2U)</td><td>AGCTACATCTGGCTACTGGGTCTCTTT</td><td>1712.8</td><td>-3.1</td><td>0.8</td><td>-3.8</td><td>1.2E-04</td><td>9.8E-04</td><td>BT20/MB231</td><td>NUC</td></tr>
<tr><td>>MI0000071|hsa-mir-17&WithFlank&13|+|91350599|91350694@20.42.23(+1A)&[MIMAT0000070&hsa-miR-17-5p&offsets|0|0(+1A);m-88&13|+|91350618|91350640&offsets|0|0(+1A)]|CAAAGTGCTTACAGTGCAGGTAGA</td><td>>MI0000071|hsa-mir-17&WithFlank&13|+|91350599|91350694</td><td>20</td><td>42</td><td>23(+1A)</td><td>CAAAGTGCTTACAGTGCAGGTAGA</td><td>5344.0</td><td>-3.2</td><td>0.8</td><td>-3.8</td><td>1.5E-04</td><td>9.8E-04</td><td>BT20/MB231</td><td>MITO</td></tr>
<tr><td>>trna10_LysCTT_16_+_3241501_3241573@1.36.36,trna13_LysCTT_14_-_58706613_58706685@1.36.36,trna2_LysCTT_15_+_79152904_79152976@1.36.36|GCCCGGCTAGCTCAGTCGGTAGAGCATGGGACTCTT</td><td>>trna10_LysCTT_16_+_3241501_3241573</td><td>1</td><td>36</td><td>36</td><td>GCCCGGCTAGCTCAGTCGGTAGAGCATGGGACTCTT</td><td>5198.2</td><td>-3.8</td><td>1.0</td><td>-3.8</td><td>1.5E-04</td><td>9.8E-04</td><td>BT20/MB231</td><td>MITO</td></tr>
<tr><td>>hg38_21_+_8433172_8446622_RNA45SN1_45S_50ntFlanks@4569.4587.19|ATGGAATAGGACCGCGGTT</td><td>>hg38_21_+_8433172_8446622_RNA45SN1_45S_50ntFlanks</td><td>4569</td><td>4587</td><td>19</td><td>ATGGAATAGGACCGCGGTT</td><td>319.5</td><td>-2.0</td><td>0.6</td><td>-3.4</td><td>7.5E-04</td><td>9.8E-04</td><td>MB231/MB468</td><td>CYTO</td></tr>
<tr><td>>hg38_21_+_8433172_8446622_RNA45SN1_45S_50ntFlanks@12274.12309.36|CAGTACGAATACAGACCGTGAAAGCGGGGCCTCACG</td><td>>hg38_21_+_8433172_8446622_RNA45SN1_45S_50ntFlanks</td><td>12274</td><td>12309</td><td>36</td><td>CAGTACGAATACAGACCGTGAAAGCGGGGCCTCACG</td><td>91.0</td><td>-5.7</td><td>1.6</td><td>-3.6</td><td>3.5E-04</td><td>9.8E-04</td><td>BT20/MB468</td><td>MP</td></tr>
<tr><td>>trnaMT_PheGAA_MT_+_577_647@39.57.19|ATGTTTAGACGGGCTCACA</td><td>>trnaMT_PheGAA_MT_+_577_647</td><td>39</td><td>57</td><td>19</td><td>ATGTTTAGACGGGCTCACA</td><td>784.8</td><td>3.2</td><td>0.9</td><td>3.6</td><td>3.5E-04</td><td>9.8E-04</td><td>BT20/MB468</td><td>MP</td></tr>
<tr><td>>hg38_21_+_8433172_8446622_RNA45SN1_45S_50ntFlanks@12937.12956.20|AACCATTCGTAGACGACCTG</td><td>>hg38_21_+_8433172_8446622_RNA45SN1_45S_50ntFlanks</td><td>12937</td><td>12956</td><td>20</td><td>AACCATTCGTAGACGACCTG</td><td>2368.4</td><td>-4.1</td><td>1.2</td><td>-3.6</td><td>3.5E-04</td><td>9.8E-04</td><td>BT20/MB468</td><td>MP</td></tr>
<tr><td>>hg38_21_+_8433172_8446622_RNA45SN1_45S_50ntFlanks@9623.9654.32|TAGTAGCTGGTTCCCTCCGAAGTTTCCCTCAG</td><td>>hg38_21_+_8433172_8446622_RNA45SN1_45S_50ntFlanks</td><td>9623</td><td>9654</td><td>32</td><td>TAGTAGCTGGTTCCCTCCGAAGTTTCCCTCAG</td><td>184.6</td><td>-3.5</td><td>1.0</td><td>-3.4</td><td>6.0E-04</td><td>9.9E-04</td><td>BT20/MB468</td><td>MITO</td></tr>
<tr><td>>hg38_21_+_8433172_8446622_RNA45SN1_45S_50ntFlanks@12991.13015.25|CTCCCTCGCTGCGATCTATTGAAAG</td><td>>hg38_21_+_8433172_8446622_RNA45SN1_45S_50ntFlanks</td><td>12991</td><td>13015</td><td>25</td><td>CTCCCTCGCTGCGATCTATTGAAAG</td><td>2351.2</td><td>-4.6</td><td>1.4</td><td>-3.4</td><td>7.2E-04</td><td>9.9E-04</td><td>BT20/MB468</td><td>CYTO</td></tr>
<tr><td>>hg38_21_+_8433172_8446622_RNA45SN1_45S_50ntFlanks@11905.11940.36|GCACGGTGAAGAGACATGAGAGGTGTAGAATAAGTG</td><td>>hg38_21_+_8433172_8446622_RNA45SN1_45S_50ntFlanks</td><td>11905</td><td>11940</td><td>36</td><td>GCACGGTGAAGAGACATGAGAGGTGTAGAATAAGTG</td><td>55.8</td><td>9.6</td><td>2.7</td><td>3.6</td><td>3.5E-04</td><td>9.9E-04</td><td>BT20/MB468</td><td>MP</td></tr>
</table>

<table>
<tr><td>>hg38_21_+_8433172_8446622_RNA45SN1_45S_50ntFlanks@10348.10377.30|AAACGAGAACTTTGAAGGCCGAAGTGGAGA</td><td>>hg38_21_+_8433172_8446622_RNA45SN1_45S_50ntFlanks</td><td>10348</td><td>10377</td><td>30</td><td>AAACGAGAACTTTGAAGGCCGAAGTGGAGA</td><td>564.8</td><td>-4.6</td><td>1.3</td><td>-3.4</td><td>6.7E-04</td><td>9.9E-04</td><td>MB231/MB468</td><td>MP</td></tr>
<tr><td>>hg38_21_+_8433172_8446622_RNA45SN1_45S_50ntFlanks@10791.10818.28|CCTCTGGCATGTTGGAACAATGTAGGTA</td><td>>hg38_21_+_8433172_8446622_RNA45SN1_45S_50ntFlanks</td><td>10791</td><td>10818</td><td>28</td><td>CCTCTGGCATGTTGGAACAATGTAGGTA</td><td>229.7</td><td>-3.0</td><td>0.8</td><td>-3.6</td><td>3.5E-04</td><td>9.9E-04</td><td>BT20/MB468</td><td>MP</td></tr>
<tr><td>>hg38_21_+_8433172_8446622_RNA45SN1_45S_50ntFlanks@12415.12446.32|CTTCGATGTCGGCTCTTCCTATCATTGTGAAG</td><td>>hg38_21_+_8433172_8446622_RNA45SN1_45S_50ntFlanks</td><td>12415</td><td>12446</td><td>32</td><td>CTTCGATGTCGGCTCTTCCTATCATTGTGAAG</td><td>177.8</td><td>-5.5</td><td>1.6</td><td>-3.4</td><td>6.7E-04</td><td>9.9E-04</td><td>MB231/MB468</td><td>MP</td></tr>
<tr><td>>MI0000734|hsa-mir-106b&WithFlank&7|-|100093987|100094080@18.38.21&[MIMAT0000680&hsa-miR-106b-5p&offsets|0|0;m-103&7|-|100094043|100094063&offsets|0|0]|TAAAGTGCTGACAGTGCAGAT</td><td>>MI0000734|hsa-mir-106b&WithFlank&7|-|100093987|100094080</td><td>18</td><td>38</td><td>21</td><td>TAAAGTGCTGACAGTGCAGAT</td><td>1388.1</td><td>-3.6</td><td>0.9</td><td>-3.8</td><td>1.5E-04</td><td>9.9E-04</td><td>BT20/MB231</td><td>MITO</td></tr>
<tr><td>>hg38_MT_+_1621_3279_RNR2_16S_50ntFlanks@1184.1216.33|AATTTCGGTTGGGGCGACCTCGGAGCAGAACCC</td><td>>hg38_MT_+_1621_3279_RNR2_16S_50ntFlanks</td><td>1184</td><td>1216</td><td>33</td><td>AATTTCGGTTGGGGCGACCTCGGAGCAGAACCC</td><td>140.5</td><td>4.2</td><td>1.2</td><td>3.4</td><td>8.0E-04</td><td>9.9E-04</td><td>MB231/MB468</td><td>MITO</td></tr>
<tr><td>>am_trnaMT_ValTAC_MT_+_1602_1670@44.68.25|ATTTCAACTTAACTTGACCGCTCTG</td><td>>am_trnaMT_ValTAC_MT_+_1602_1670</td><td>44</td><td>68</td><td>25</td><td>ATTTCAACTTAACTTGACCGCTCTG</td><td>589.7</td><td>-3.8</td><td>1.1</td><td>-3.4</td><td>8.0E-04</td><td>9.9E-04</td><td>MB231/MB468</td><td>MITO</td></tr>
<tr><td>>trnaMT_GluTTC_MT_-_14674_14742@38.66.29,trnalookalike8_GluTTC_5_-_93905172_93905240@38.66.29|CATTGGTCGTGGTTGTAGTCCGTGCGAGA</td><td>>trnaMT_GluTTC_MT_-_14674_14742</td><td>38</td><td>66</td><td>29</td><td>CATTGGTCGTGGTTGTAGTCCGTGCGAGA</td><td>184.4</td><td>3.2</td><td>1.0</td><td>3.4</td><td>8.0E-04</td><td>9.9E-04</td><td>MB231/MB468</td><td>MITO</td></tr>
<tr><td>>hg38_21_+_8433172_8446622_RNA45SN1_45S_50ntFlanks@6661.6685.25|CGGTGGATCACTCGGCTCGTGCGTC</td><td>>hg38_21_+_8433172_8446622_RNA45SN1_45S_50ntFlanks</td><td>6661</td><td>6685</td><td>25</td><td>CGGTGGATCACTCGGCTCGTGCGTC</td><td>494.5</td><td>1.2</td><td>0.4</td><td>3.3</td><td>8.1E-04</td><td>9.9E-04</td><td>BT20/MB468</td><td>NUC</td></tr>
<tr><td>>MI0000283|hsa-mir-203a&WithFlank&14|+|104117399|104117520@72.92.21(+1U)&[MIMAT0000264&hsa-miR-203a-3p&offsets|+1|0(+1U);m-79&14|+|104117469|104117490&offsets|+1|0(+1U)]|TGAAATGTTTAGGACCACTAGT</td><td>>MI0000283|hsa-mir-203a&WithFlank&14|+|104117399|104117520</td><td>72</td><td>92</td><td>21(+1U)</td><td>TGAAATGTTTAGGACCACTAGT</td><td>202.1</td><td>5.9</td><td>1.6</td><td>3.6</td><td>3.5E-04</td><td>9.9E-04</td><td>BT20/MB468</td><td>MP</td></tr>
<tr><td>>trnaMT_GluTTC_MT_-_14674_14742@38.65.28,trnalookalike8_GluTTC_5_-_93905172_93905240@38.65.28|CATTGGTCGTGGTTGTAGTCCGTGCGAG</td><td>>trnaMT_GluTTC_MT_-_14674_14742</td><td>38</td><td>65</td><td>28</td><td>CATTGGTCGTGGTTGTAGTCCGTGCGAG</td><td>212.8</td><td>4.3</td><td>1.2</td><td>3.6</td><td>3.5E-04</td><td>9.9E-04</td><td>BT20/MB468</td><td>MP</td></tr>
<tr><td>>hg38_21_+_8433172_8446622_RNA45SN1_45S_50ntFlanks@11703.11726.24|AATGAAGCGCGGGTAAACGGCGGG</td><td>>hg38_21_+_8433172_8446622_RNA45SN1_45S_50ntFlanks</td><td>11703</td><td>11726</td><td>24</td><td>AATGAAGCGCGGGTAAACGGCGGG</td><td>219.4</td><td>-2.2</td><td>0.6</td><td>-3.4</td><td>7.3E-04</td><td>9.9E-04</td><td>BT20/MB468</td><td>CYTO</td></tr>
<tr><td>>hg38_21_+_8433172_8446622_RNA45SN1_45S_50ntFlanks@8423.8467.45|CGCAGTCCGCCCGGAGGATTCAACCCGGCGGCGGGTCCGGCCGTG</td><td>>hg38_21_+_8433172_8446622_RNA45SN1_45S_50ntFlanks</td><td>8423</td><td>8467</td><td>45</td><td>CGCAGTCCGCCCGGAGGATTCAACCCGGCGGCGGGTCCGGCCGTG</td><td>299.3</td><td>-5.2</td><td>1.5</td><td>-3.4</td><td>6.1E-04</td><td>9.9E-04</td><td>BT20/MB468</td><td>MITO</td></tr>
<tr><td>>hg38_21_+_8433172_8446622_RNA45SN1_45S_50ntFlanks@11209.11254.46|CGGCGACTCTGGACGCGAGCCGGGCCCTTCCCGTGGATCGCCCCAG</td><td>>hg38_21_+_8433172_8446622_RNA45SN1_45S_50ntFlanks</td><td>11209</td><td>11254</td><td>46</td><td>CGGCGACTCTGGACGCGAGCCGGGCCCTTCCCGTGGATCGCCCCAG</td><td>9172.6</td><td>-3.7</td><td>1.1</td><td>-3.4</td><td>6.1E-04</td><td>9.9E-04</td><td>BT20/MB468</td><td>MITO</td></tr>
<tr><td>>hg38_21_+_8433172_8446622_RNA45SN1_45S_50ntFlanks@11316.11336.21|CACCCCACGTCTCGTCGCGCG</td><td>>hg38_21_+_8433172_8446622_RNA45SN1_45S_50ntFlanks</td><td>11316</td><td>11336</td><td>21</td><td>CACCCCACGTCTCGTCGCGCG</td><td>2659.8</td><td>-3.8</td><td>1.1</td><td>-3.4</td><td>6.1E-04</td><td>9.9E-04</td><td>BT20/MB468</td><td>MITO</td></tr>
<tr><td>>hg38_21_+_8433172_8446622_RNA45SN1_45S_50ntFlanks@7971.7988.18|GAGACGCGACCTCAGATC</td><td>>hg38_21_+_8433172_8446622_RNA45SN1_45S_50ntFlanks</td><td>7971</td><td>7988</td><td>18</td><td>GAGACGCGACCTCAGATC</td><td>498.9</td><td>2.9</td><td>0.9</td><td>3.3</td><td>8.1E-04</td><td>9.9E-04</td><td>BT20/MB468</td><td>NUC</td></tr>
<tr><td>>hg38_21_+_8433172_8446622_RNA45SN1_45S_50ntFlanks@10852.10873.22|GATAAGGATTGGCTCTAAGGGC</td><td>>hg38_21_+_8433172_8446622_RNA45SN1_45S_50ntFlanks</td><td>10852</td><td>10873</td><td>22</td><td>GATAAGGATTGGCTCTAAGGGC</td><td>242.7</td><td>7.1</td><td>1.8</td><td>3.8</td><td>1.2E-04</td><td>9.9E-04</td><td>BT20/MB231</td><td>MP</td></tr>
<tr><td>>hg38_21_+_8433172_8446622_RNA45SN1_45S_50ntFlanks@6734.6770.37|ATTGATCATCGACACTTCGAACGCACTTGCGGCCCCG</td><td>>hg38_21_+_8433172_8446622_RNA45SN1_45S_50ntFlanks</td><td>6734</td><td>6770</td><td>37</td><td>ATTGATCATCGACACTTCGAACGCACTTGCGGCCCCG</td><td>463.1</td><td>-2.3</td><td>0.7</td><td>-3.4</td><td>7.6E-04</td><td>9.9E-04</td><td>MB231/MB468</td><td>CYTO</td></tr>
<tr><td>>hg38_21_+_8433172_8446622_RNA45SN1_45S_50ntFlanks@12406.12431.26|TTTTTGATCCTTCGATGTCGGCTCTT</td><td>>hg38_21_+_8433172_8446622_RNA45SN1_45S_50ntFlanks</td><td>12406</td><td>12431</td><td>26</td><td>TTTTTGATCCTTCGATGTCGGCTCTT</td><td>338.8</td><td>-4.0</td><td>1.2</td><td>-3.3</td><td>8.2E-04</td><td>9.9E-04</td><td>MB231/MB468</td><td>NUC</td></tr>
<tr><td>>MI0033423|hsa-mir-10399&WithFlank&7|-|139044094|139044163@45.63.19(+2U)&[MIMAT0041630&hsa-miR-10399-3p&offsets|+1|-1(+2U);TJU_CMC_MD2.ID03061.3p-miR&offsets|+5|+2(+2U);m-2657&7|-|139044103|139044124&offsets|+5|+2(+2U)]|TCTCGGACAAGCTGTAGGTTT</td><td>>MI0033423|hsa-mir-10399&WithFlank&7|-|139044094|139044163</td><td>45</td><td>63</td><td>19(+2U)</td><td>TCTCGGACAAGCTGTAGGTTT</td><td>428.4</td><td>-5.2</td><td>1.4</td><td>-3.8</td><td>1.2E-04</td><td>9.9E-04</td><td>BT20/MB231</td><td>MP</td></tr>
<tr><td>>MI0000085|hsa-mir-27a&WithFlank&19|-|13836434|13836523@57.76.20(+2U)&[MIMAT0000084&hsa-miR-27a-3p&offsets|0|-1(+2U);m-43&19|-|13836447|13836467&offsets|0|-1(+2U)]|TTCACAGTGGCTAAGTTCCGTT</td><td>>MI0000085|hsa-mir-27a&WithFlank&19|-|13836434|13836523</td><td>57</td><td>76</td><td>20(+2U)</td><td>TTCACAGTGGCTAAGTTCCGTT</td><td>6178.7</td><td>-3.9</td><td>1.0</td><td>-3.8</td><td>1.2E-04</td><td>9.9E-04</td><td>BT20/MB231</td><td>MP</td></tr>
<tr><td>>hg38_21_+_8433172_8446622_RNA45SN1_45S_50ntFlanks@11667.11705.39|CTGCCCAGTGCTCTGAATGTCAAAGTGAAGAAATTCAAT</td><td>>hg38_21_+_8433172_8446622_RNA45SN1_45S_50ntFlanks</td><td>11667</td><td>11705</td><td>39</td><td>CTGCCCAGTGCTCTGAATGTCAAAGTGAAGAAATTCAAT</td><td>236.3</td><td>-2.7</td><td>0.7</td><td>-3.8</td><td>1.3E-04</td><td>9.9E-04</td><td>BT20/MB231</td><td>CYTO</td></tr>
<tr><td>>hg38_21_+_8433172_8446622_RNA45SN1_45S_50ntFlanks@11669.11692.24|GCCCAGTGCTCTGAATGTCAAAGT</td><td>>hg38_21_+_8433172_8446622_RNA45SN1_45S_50ntFlanks</td><td>11669</td><td>11692</td><td>24</td><td>GCCCAGTGCTCTGAATGTCAAAGT</td><td>147.5</td><td>5.9</td><td>1.7</td><td>3.6</td><td>3.5E-04</td><td>9.9E-04</td><td>BT20/MB468</td><td>MP</td></tr>
<tr><td>>hg38_21_+_8433172_8446622_RNA45SN1_45S_50ntFlanks@11572.11606.35|CCGACTTAGAACTGGTGCGGACCAGGGGAATCCGA</td><td>>hg38_21_+_8433172_8446622_RNA45SN1_45S_50ntFlanks</td><td>11572</td><td>11606</td><td>35</td><td>CCGACTTAGAACTGGTGCGGACCAGGGGAATCCGA</td><td>276.3</td><td>-2.2</td><td>0.6</td><td>-3.4</td><td>7.3E-04</td><td>1.0E-03</td><td>BT20/MB468</td><td>CYTO</td></tr>
<tr><td>>hg38_21_+_8433172_8446622_RNA45SN1_45S_50ntFlanks@6645.6679.35|CTCGTACGACTCTTAGCGGTGGATCACTCGGCTCG</td><td>>hg38_21_+_8433172_8446622_RNA45SN1_45S_50ntFlanks</td><td>6645</td><td>6679</td><td>35</td><td>CTCGTACGACTCTTAGCGGTGGATCACTCGGCTCG</td><td>878.4</td><td>2.6</td><td>0.8</td><td>3.4</td><td>7.3E-04</td><td>1.0E-03</td><td>BT20/MB468</td><td>CYTO</td></tr>
<tr><td>>trna119_LysCTT_1_-_145395522_145395594@1.29.29,trna11_LysCTT_5_-_180648979_180649051@1.29.29,trna13_LysCTT_6_+_26556774_26556846@1.29.29,trna32_LysCTT_16_-_3207406_3207478@1.29.29,trna7_LysCTT_16_+_3225692_3225764@1.29.29,trna9_LysCTT_5_+_180634755_180634827@1.29.29|GCCCGGCTAGCTCAGTCGGTAGAGCATGA</td><td>>trna119_LysCTT_1_-_145395522_145395594</td><td>1</td><td>29</td><td>29</td><td>GCCCGGCTAGCTCAGTCGGTAGAGCATGA</td><td>482.4</td><td>-4.7</td><td>1.3</td><td>-3.6</td><td>3.5E-04</td><td>1.0E-03</td><td>BT20/MB468</td><td>MP</td></tr>
<tr><td>>hg38_21_+_8433172_8446622_RNA45SN1_45S_50ntFlanks@4452.4472.21|CTCGGCGCCCCCTCGATGCTC</td><td>>hg38_21_+_8433172_8446622_RNA45SN1_45S_50ntFlanks</td><td>4452</td><td>4472</td><td>21</td><td>CTCGGCGCCCCCTCGATGCTC</td><td>197.5</td><td>-5.5</td><td>1.5</td><td>-3.6</td><td>3.5E-04</td><td>1.0E-03</td><td>BT20/MB468</td><td>MP</td></tr>
<tr><td>>hg38_21_+_8433172_8446622_RNA45SN1_45S_50ntFlanks@11845.11873.29|GGAACGGGCTTGGCGGAATCAGCGGGGAA</td><td>>hg38_21_+_8433172_8446622_RNA45SN1_45S_50ntFlanks</td><td>11845</td><td>11873</td><td>29</td><td>GGAACGGGCTTGGCGGAATCAGCGGGGAA</td><td>151.5</td><td>-4.6</td><td>1.3</td><td>-3.4</td><td>6.1E-04</td><td>1.0E-03</td><td>BT20/MB468</td><td>MITO</td></tr>
<tr><td>>hg38_21_+_8433172_8446622_RNA45SN1_45S_50ntFlanks@10716.10736.21|GAAAATCCGGGGGAGAGGGTG</td><td>>hg38_21_+_8433172_8446622_RNA45SN1_45S_50ntFlanks</td><td>10716</td><td>10736</td><td>21</td><td>GAAAATCCGGGGGAGAGGGTG</td><td>249.6</td><td>-3.2</td><td>0.9</td><td>-3.6</td><td>3.5E-04</td><td>1.0E-03</td><td>BT20/MB468</td><td>MP</td></tr>
<tr><td>>hg38_21_+_8433172_8446622_RNA45SN1_45S_50ntFlanks@11580.11613.34|GAACTGGTGCGGACCAGGGGAATCCGACTGTTTA</td><td>>hg38_21_+_8433172_8446622_RNA45SN1_45S_50ntFlanks</td><td>11580</td><td>11613</td><td>34</td><td>GAACTGGTGCGGACCAGGGGAATCCGACTGTTTA</td><td>2284.9</td><td>-2.8</td><td>0.8</td><td>-3.4</td><td>8.1E-04</td><td>1.0E-03</td><td>MB231/MB468</td><td>MITO</td></tr>
<tr><td>>hg38_21_+_8433172_8446622_RNA45SN1_45S_50ntFlanks@9545.9576.32|GTGGAGGTCCGTAGCGGTCCTGACGTGCAAAT</td><td>>hg38_21_+_8433172_8446622_RNA45SN1_45S_50ntFlanks</td><td>9545</td><td>9576</td><td>32</td><td>GTGGAGGTCCGTAGCGGTCCTGACGTGCAAAT</td><td>204.0</td><td>2.0</td><td>0.6</td><td>3.4</td><td>7.7E-04</td><td>1.0E-03</td><td>MB231/MB468</td><td>CYTO</td></tr>
<tr><td>>hg38_21_+_8433172_8446622_RNA45SN1_45S_50ntFlanks@12932.12970.39|CGCTAAACCATTCGTAGACGACCTGCTTCTGGGTCGGGG</td><td>>hg38_21_+_8433172_8446622_RNA45SN1_45S_50ntFlanks</td><td>12932</td><td>12970</td><td>39</td><td>CGCTAAACCATTCGTAGACGACCTGCTTCTGGGTCGGGG</td><td>197.5</td><td>-4.9</td><td>1.5</td><td>-3.4</td><td>7.3E-04</td><td>1.0E-03</td><td>BT20/MB468</td><td>CYTO</td></tr>
<tr><td>>hg38_21_+_8433172_8446622_RNA45SN1_45S_50ntFlanks@12367.12384.18|AACTGGCTTGTGGCGGCC</td><td>>hg38_21_+_8433172_8446622_RNA45SN1_45S_50ntFlanks</td><td>12367</td><td>12384</td><td>18</td><td>AACTGGCTTGTGGCGGCC</td><td>137.9</td><td>-3.9</td><td>1.1</td><td>-3.4</td><td>6.1E-04</td><td>1.0E-03</td><td>BT20/MB468</td><td>MITO</td></tr>
<tr><td>>trna14_LysTTT_11_-_59327808_59327880@1.24.24,trna2_LysTTT_17_+_8022473_8022545@1.24.24,trna54_LysTTT_1_+_204475655_204475727@1.24.24,trna5_LysTTT_11_+_59323902_59323974@1.24.24,trna62_LysTTT_1_-_204476158_204476230@1.24.24,trna76_LysTTT_6_+_28918806_28918878@1.24.24|GCCCGGATAGCTCAGTCGGTAGAG</td><td>>trna14_LysTTT_11_-_59327808_59327880</td><td>1</td><td>24</td><td>24</td><td>GCCCGGATAGCTCAGTCGGTAGAG</td><td>108.1</td><td>-5.2</td><td>1.5</td><td>-3.4</td><td>6.8E-04</td><td>1.0E-03</td><td>MB231/MB468</td><td>MP</td></tr>
<tr><td>>MI0000081|hsa-mir-24-2&WithFlank&19|-|13836281|13836365@56.74.19(+1A);MI0000080|hsa-mir-24-1&WithFlank&9|+|95086015|95086094@50.68.19(+1A)&[MIMAT0000080&hsa-miR-24-3p&offsets|0|-3(+1A);m-36&19|-|13836288|13836310&offsets|0|-4(+1A)];[MIMAT0000080_1&hsa-miR-24-3p&offsets|0|-3(+1A);m-35&9|+|95086064|95086086&offsets|0|-4(+1A)]|TGGCTCAGTTCAGCAGGAAA</td><td>>MI0000081|hsa-mir-24-2&WithFlank&19|-|13836281|13836365</td><td>56</td><td>74</td><td>19(+1A)</td><td>TGGCTCAGTTCAGCAGGAAA</td><td>59.2</td><td>2.8</td><td>0.8</td><td>3.3</td><td>8.2E-04</td><td>1.0E-03</td><td>BT20/MB468</td><td>NUC</td></tr>
<tr><td>>MI0005561|hsa-mir-877&WithFlank&6|+|30584326|30584423@7.28.22&[MIMAT0004949&hsa-miR-877-5p&offsets|0|+2;m-382&6|+|30584332|30584354&offsets|0|-1]|GTAGAGGAGATGGCGCAGGGGA</td><td>>MI0005561|hsa-mir-877&WithFlank&6|+|30584326|30584423</td><td>7</td><td>28</td><td>22</td><td>GTAGAGGAGATGGCGCAGGGGA</td><td>248.6</td><td>5.0</td><td>1.4</td><td>3.6</td><td>3.5E-04</td><td>1.0E-03</td><td>BT20/MB468</td><td>MP</td></tr>
<tr><td>>trnaMT_ValTAC_MT_+_1602_1670@39.70.32|AGGAGATTTCAACTTAACTTGACCGCTCTGAC</td><td>>trnaMT_ValTAC_MT_+_1602_1670</td><td>39</td><td>70</td><td>32</td><td>AGGAGATTTCAACTTAACTTGACCGCTCTGAC</td><td>62.6</td><td>9.7</td><td>2.7</td><td>3.6</td><td>3.5E-04</td><td>1.0E-03</td><td>BT20/MB468</td><td>MP</td></tr>
<tr><td>>hg38_21_+_8433172_8446622_RNA45SN1_45S_50ntFlanks@12991.13017.27|CTCCCTCGCTGCGATCTATTGAAAGTC</td><td>>hg38_21_+_8433172_8446622_RNA45SN1_45S_50ntFlanks</td><td>12991</td><td>13017</td><td>27</td><td>CTCCCTCGCTGCGATCTATTGAAAGTC</td><td>215.5</td><td>-2.7</td><td>0.8</td><td>-3.4</td><td>6.1E-04</td><td>1.0E-03</td><td>BT20/MB468</td><td>MITO</td></tr>
<tr><td>>trnaMT_GluTTC_MT_-_14674_14742@20.61.42,trnalookalike8_GluTTC_5_-_93905172_93905240@20.61.42|AACGATGGTTTTTCATATCATTGGTCGTGGTTGTAGTCCGTG</td><td>>trnaMT_GluTTC_MT_-_14674_14742</td><td>20</td><td>61</td><td>42</td><td>AACGATGGTTTTTCATATCATTGGTCGTGGTTGTAGTCCGTG</td><td>63.5</td><td>9.7</td><td>2.7</td><td>3.6</td><td>3.5E-04</td><td>1.0E-03</td><td>BT20/MB468</td><td>MP</td></tr>
<tr><td>>hg38_21_+_8433172_8446622_RNA45SN1_45S_50ntFlanks@10458.10476.19|ATGGCCTCCGTTGCCCTCG</td><td>>hg38_21_+_8433172_8446622_RNA45SN1_45S_50ntFlanks</td><td>10458</td><td>10476</td><td>19</td><td>ATGGCCTCCGTTGCCCTCG</td><td>55.6</td><td>-8.9</td><td>2.5</td><td>-3.6</td><td>3.5E-04</td><td>1.0E-03</td><td>BT20/MB468</td><td>MP</td></tr>
<tr><td>>hg38_MT_+_598_1651_RNR1_12S_50ntFlanks@371.404.34|TAAAGCTAAAACTCACCTGAGTTGTAAAAAACTC</td><td>>hg38_MT_+_598_1651_RNR1_12S_50ntFlanks</td><td>371</td><td>404</td><td>34</td><td>TAAAGCTAAAACTCACCTGAGTTGTAAAAAACTC</td><td>413.5</td><td>3.2</td><td>0.9</td><td>3.3</td><td>8.1E-04</td><td>1.0E-03</td><td>MB231/MB468</td><td>MITO</td></tr>
<tr><td>>hg38_21_+_8433172_8446622_RNA45SN1_45S_50ntFlanks@11199.11225.27|CCGGCGGCGGCGGCGACTCTGGACGCG</td><td>>hg38_21_+_8433172_8446622_RNA45SN1_45S_50ntFlanks</td><td>11199</td><td>11225</td><td>27</td><td>CCGGCGGCGGCGGCGACTCTGGACGCG</td><td>1102.9</td><td>-6.2</td><td>1.7</td><td>-3.6</td><td>3.5E-04</td><td>1.0E-03</td><td>BT20/MB468</td><td>MP</td></tr>
<tr><td>>MI0000749|hsa-mir-30e&WithFlank&1|+|40754349|40754452@65.86.22&[MIMAT0000693&hsa-miR-30e-3p&offsets|0|0;m-25&1|+|40754413|40754434&offsets|0|0]|CTTTCAGTCGGATGTTTACAGC</td><td>>MI0000749|hsa-mir-30e&WithFlank&1|+|40754349|40754452</td><td>65</td><td>86</td><td>22</td><td>CTTTCAGTCGGATGTTTACAGC</td><td>1484.4</td><td>2.8</td><td>0.8</td><td>3.3</td><td>8.1E-04</td><td>1.0E-03</td><td>MB231/MB468</td><td>MITO</td></tr>
<tr><td>>hg38_21_+_8433172_8446622_RNA45SN1_45S_50ntFlanks@9572.9601.30|CAAATCGGTCGTCCGACCTGGGTATAGGGG</td><td>>hg38_21_+_8433172_8446622_RNA45SN1_45S_50ntFlanks</td><td>9572</td><td>9601</td><td>30</td><td>CAAATCGGTCGTCCGACCTGGGTATAGGGG</td><td>204.8</td><td>-3.7</td><td>1.1</td><td>-3.3</td><td>8.2E-04</td><td>1.0E-03</td><td>MB231/MB468</td><td>NUC</td></tr>
<tr><td>>MI0000299|hsa-mir-222&WithFlank&X|-|45747009|45747130@75.97.23(+2U)&[MIMAT0000279&hsa-miR-222-3p&offsets|0|+2(+2U);m-140&X|-|45747033|45747056&offsets|0|-1(+2U)]|AGCTACATCTGGCTACTGGGTCTTT</td><td>>MI0000299|hsa-mir-222&WithFlank&X|-|45747009|45747130</td><td>75</td><td>97</td><td>23(+2U)</td><td>AGCTACATCTGGCTACTGGGTCTTT</td><td>300.0</td><td>5.5</td><td>1.6</td><td>3.4</td><td>6.9E-04</td><td>1.0E-03</td><td>MB231/MB468</td><td>MP</td></tr>
<tr><td>>hg38_21_+_8433172_8446622_RNA45SN1_45S_50ntFlanks@3854.3884.31|ACTGTGGTAATTCTAGAGCTAATACATGCCG</td><td>>hg38_21_+_8433172_8446622_RNA45SN1_45S_50ntFlanks</td><td>3854</td><td>3884</td><td>31</td><td>ACTGTGGTAATTCTAGAGCTAATACATGCCG</td><td>114.1</td><td>-5.7</td><td>1.6</td><td>-3.6</td><td>3.5E-04</td><td>1.0E-03</td><td>BT20/MB468</td><td>MP</td></tr>
<tr><td>>hg38_21_+_8433172_8446622_RNA45SN1_45S_50ntFlanks@10392.10430.39|ACAGCAGTTGAACATGGGTCAGTCGGTCCTGAGAGATGG</td><td>>hg38_21_+_8433172_8446622_RNA45SN1_45S_50ntFlanks</td><td>10392</td><td>10430</td><td>39</td><td>ACAGCAGTTGAACATGGGTCAGTCGGTCCTGAGAGATGG</td><td>790.3</td><td>-5.6</td><td>1.7</td><td>-3.4</td><td>7.4E-04</td><td>1.0E-03</td><td>BT20/MB468</td><td>CYTO</td></tr>
</table>

<table>
<tr><td>>hg38_21_+_8433172_8446622_RNA45SN1_45S_50ntFlanks@11907.11939.33|ACGGTGAAGAGACATGAGAGGTGTAGAATAAGT</td><td>>hg38_21_+_8433172_8446622_RNA45SN1_45S_50ntFlanks</td><td>11907</td><td>11939</td><td>33</td><td>ACGGTGAAGAGACATGAGAGGTGTAGAATAAGT</td><td>54.8</td><td>9.5</td><td>2.7</td><td>3.6</td><td>3.6E-04</td><td>1.0E-03</td><td>BT20/MB468</td><td>MP</td></tr>
<tr><td>>hg38_21_+_8433172_8446622_RNA45SN1_45S_50ntFlanks@10349.10390.42|AACGAGAACTTTGAAGGCCGAAGTGGAGAAGGGTTCCATGTG</td><td>>hg38_21_+_8433172_8446622_RNA45SN1_45S_50ntFlanks</td><td>10349</td><td>10390</td><td>42</td><td>AACGAGAACTTTGAAGGCCGAAGTGGAGAAGGGTTCCATGTG</td><td>1130.8</td><td>-2.6</td><td>0.8</td><td>-3.4</td><td>7.4E-04</td><td>1.0E-03</td><td>BT20/MB468</td><td>CYTO</td></tr>
<tr><td>>hg38_21_+_8433172_8446622_RNA45SN1_45S_50ntFlanks@10393.10419.27|CAGCAGTTGAACATGGGTCAGTCGGTC</td><td>>hg38_21_+_8433172_8446622_RNA45SN1_45S_50ntFlanks</td><td>10393</td><td>10419</td><td>27</td><td>CAGCAGTTGAACATGGGTCAGTCGGTC</td><td>310.4</td><td>-3.5</td><td>1.0</td><td>-3.4</td><td>7.7E-04</td><td>1.0E-03</td><td>MB231/MB468</td><td>CYTO</td></tr>
<tr><td>>MI0000082|hsa-mir-25&WithFlank&7|-|100093554|100093649@58.77.20(+1U)&[MIMAT0000081&hsa-miR-25-3p&offsets|0|-2(+1U);m-23&7|-|100093571|100093592&offsets|0|-2(+1U)]|CATTGCACTTGTCTCGGTCTT</td><td>>MI0000082|hsa-mir-25&WithFlank&7|-|100093554|100093649</td><td>58</td><td>77</td><td>20(+1U)</td><td>CATTGCACTTGTCTCGGTCTT</td><td>131.8</td><td>5.1</td><td>1.4</td><td>3.6</td><td>3.6E-04</td><td>1.0E-03</td><td>BT20/MB468</td><td>MP</td></tr>
<tr><td>>hg38_21_+_8433172_8446622_RNA45SN1_45S_50ntFlanks@10402.10424.23|AACATGGGTCAGTCGGTCCTGAG</td><td>>hg38_21_+_8433172_8446622_RNA45SN1_45S_50ntFlanks</td><td>10402</td><td>10424</td><td>23</td><td>AACATGGGTCAGTCGGTCCTGAG</td><td>14192.9</td><td>3.4</td><td>0.9</td><td>3.8</td><td>1.5E-04</td><td>1.0E-03</td><td>BT20/MB231</td><td>MITO</td></tr>
<tr><td>>hg38_21_+_8433172_8446622_RNA45SN1_45S_50ntFlanks@11323.11343.21|CGTCTCGTCGCGCGCGCGTCC</td><td>>hg38_21_+_8433172_8446622_RNA45SN1_45S_50ntFlanks</td><td>11323</td><td>11343</td><td>21</td><td>CGTCTCGTCGCGCGCGCGTCC</td><td>5988.4</td><td>-3.4</td><td>1.0</td><td>-3.4</td><td>7.4E-04</td><td>1.0E-03</td><td>BT20/MB468</td><td>CYTO</td></tr>
<tr><td>>hg38_21_+_8433172_8446622_RNA45SN1_45S_50ntFlanks@4592.4616.25|TTTGTTGGTTTTCGGAACTGAGGCC</td><td>>hg38_21_+_8433172_8446622_RNA45SN1_45S_50ntFlanks</td><td>4592</td><td>4616</td><td>25</td><td>TTTGTTGGTTTTCGGAACTGAGGCC</td><td>180.0</td><td>-3.7</td><td>1.1</td><td>-3.3</td><td>8.3E-04</td><td>1.0E-03</td><td>MB231/MB468</td><td>NUC</td></tr>
<tr><td>>hg38_21_+_8433172_8446622_RNA45SN1_45S_50ntFlanks@11598.11622.25|GGAATCCGACTGTTTAATTAAAACA</td><td>>hg38_21_+_8433172_8446622_RNA45SN1_45S_50ntFlanks</td><td>11598</td><td>11622</td><td>25</td><td>GGAATCCGACTGTTTAATTAAAACA</td><td>71.1</td><td>6.6</td><td>1.8</td><td>3.6</td><td>3.6E-04</td><td>1.0E-03</td><td>BT20/MB468</td><td>MP</td></tr>
<tr><td>>hg38_21_+_8433172_8446622_RNA45SN1_45S_50ntFlanks@11666.11685.20|TCTGCCCAGTGCTCTGAATG</td><td>>hg38_21_+_8433172_8446622_RNA45SN1_45S_50ntFlanks</td><td>11666</td><td>11685</td><td>20</td><td>TCTGCCCAGTGCTCTGAATG</td><td>3192.3</td><td>-3.1</td><td>0.9</td><td>-3.4</td><td>7.4E-04</td><td>1.0E-03</td><td>BT20/MB468</td><td>CYTO</td></tr>
<tr><td>>MI0000734|hsa-mir-106b&WithFlank&7|-|100093987|100094080@56.77.22&[MIMAT0004672&hsa-miR-106b-3p&offsets|-2|-2;m-87&7|-|100094003|100094023&offsets|-2|-1]|TACCGCACTGTGGGTACTTGCT</td><td>>MI0000734|hsa-mir-106b&WithFlank&7|-|100093987|100094080</td><td>56</td><td>77</td><td>22</td><td>TACCGCACTGTGGGTACTTGCT</td><td>601.1</td><td>6.5</td><td>1.8</td><td>3.6</td><td>3.6E-04</td><td>1.0E-03</td><td>BT20/MB468</td><td>MP</td></tr>
<tr><td>>hg38_21_+_8433172_8446622_RNA45SN1_45S_50ntFlanks@9308.9335.28|AGGAGTCTAACACGTGCGCGAGTCGGGG</td><td>>hg38_21_+_8433172_8446622_RNA45SN1_45S_50ntFlanks</td><td>9308</td><td>9335</td><td>28</td><td>AGGAGTCTAACACGTGCGCGAGTCGGGG</td><td>833.1</td><td>-3.5</td><td>1.0</td><td>-3.6</td><td>3.6E-04</td><td>1.0E-03</td><td>BT20/MB468</td><td>MP</td></tr>
<tr><td>>hg38_21_+_8433172_8446622_RNA45SN1_45S_50ntFlanks@12408.12431.24|TTTGATCCTTCGATGTCGGCTCTT</td><td>>hg38_21_+_8433172_8446622_RNA45SN1_45S_50ntFlanks</td><td>12408</td><td>12431</td><td>24</td><td>TTTGATCCTTCGATGTCGGCTCTT</td><td>1172.4</td><td>-2.6</td><td>0.8</td><td>-3.4</td><td>7.8E-04</td><td>1.0E-03</td><td>MB231/MB468</td><td>CYTO</td></tr>
<tr><td>>hg38_21_+_8433172_8446622_RNA45SN1_45S_50ntFlanks@12324.12346.23|TTGGGTTTTAAGCAGGAGGTGTC</td><td>>hg38_21_+_8433172_8446622_RNA45SN1_45S_50ntFlanks</td><td>12324</td><td>12346</td><td>23</td><td>TTGGGTTTTAAGCAGGAGGTGTC</td><td>3261.1</td><td>-6.5</td><td>1.8</td><td>-3.6</td><td>3.6E-04</td><td>1.0E-03</td><td>BT20/MB468</td><td>MP</td></tr>
<tr><td>>hg38_21_+_8433172_8446622_RNA45SN1_45S_50ntFlanks@11583.11601.19|CTGGTGCGGACCAGGGGAA</td><td>>hg38_21_+_8433172_8446622_RNA45SN1_45S_50ntFlanks</td><td>11583</td><td>11601</td><td>19</td><td>CTGGTGCGGACCAGGGGAA</td><td>1357.0</td><td>-3.3</td><td>1.0</td><td>-3.4</td><td>7.4E-04</td><td>1.0E-03</td><td>BT20/MB468</td><td>CYTO</td></tr>
<tr><td>>hg38_21_+_8433172_8446622_RNA45SN1_45S_50ntFlanks@3842.3872.31|CCTACTTGGATAACTGTGGTAATTCTAGAGC</td><td>>hg38_21_+_8433172_8446622_RNA45SN1_45S_50ntFlanks</td><td>3842</td><td>3872</td><td>31</td><td>CCTACTTGGATAACTGTGGTAATTCTAGAGC</td><td>124.0</td><td>-4.1</td><td>1.2</td><td>-3.4</td><td>6.9E-04</td><td>1.0E-03</td><td>MB231/MB468</td><td>MP</td></tr>
<tr><td>>hg38_21_+_8433172_8446622_RNA45SN1_45S_50ntFlanks@3843.3862.20|CTACTTGGATAACTGTGGTA</td><td>>hg38_21_+_8433172_8446622_RNA45SN1_45S_50ntFlanks</td><td>3843</td><td>3862</td><td>20</td><td>CTACTTGGATAACTGTGGTA</td><td>437.6</td><td>2.3</td><td>0.7</td><td>3.4</td><td>7.4E-04</td><td>1.0E-03</td><td>BT20/MB468</td><td>CYTO</td></tr>
<tr><td>>trnaMT_MetCAT_MT_+_4402_4469@37.71.35|CCGAAAATGTTGGTTATACCCTTCCCGTACTACCA</td><td>>trnaMT_MetCAT_MT_+_4402_4469</td><td>37</td><td>71</td><td>35</td><td>CCGAAAATGTTGGTTATACCCTTCCCGTACTACCA</td><td>255.3</td><td>3.1</td><td>0.9</td><td>3.3</td><td>8.2E-04</td><td>1.0E-03</td><td>MB231/MB468</td><td>MITO</td></tr>
<tr><td>>hg38_21_+_8433172_8446622_RNA45SN1_45S_50ntFlanks@6763.6806.44|CGGCCCCGGGTTCCTCCCGGGGCTACGCCTGTCTGAGCGTCGCT</td><td>>hg38_21_+_8433172_8446622_RNA45SN1_45S_50ntFlanks</td><td>6763</td><td>6806</td><td>44</td><td>CGGCCCCGGGTTCCTCCCGGGGCTACGCCTGTCTGAGCGTCGCT</td><td>196.9</td><td>5.0</td><td>1.5</td><td>3.3</td><td>8.2E-04</td><td>1.0E-03</td><td>MB231/MB468</td><td>MITO</td></tr>
<tr><td>>trnaMT_SerGCT_MT_+_12207_12265@1.41.41|GAGAAAGCTCACAAGAACTGCTAACTCATGCCCCCATGTCT</td><td>>trnaMT_SerGCT_MT_+_12207_12265</td><td>1</td><td>41</td><td>41</td><td>GAGAAAGCTCACAAGAACTGCTAACTCATGCCCCCATGTCT</td><td>400.6</td><td>3.2</td><td>1.0</td><td>3.3</td><td>8.2E-04</td><td>1.0E-03</td><td>MB231/MB468</td><td>MITO</td></tr>
<tr><td>>trnaMT_PheGAA_MT_+_577_647@35.62.28|GAAAATGTTTAGACGGGCTCACATCACC</td><td>>trnaMT_PheGAA_MT_+_577_647</td><td>35</td><td>62</td><td>28</td><td>GAAAATGTTTAGACGGGCTCACATCACC</td><td>252.7</td><td>3.5</td><td>1.0</td><td>3.4</td><td>6.9E-04</td><td>1.0E-03</td><td>MB231/MB468</td><td>MP</td></tr>
<tr><td>>trnaMT_PheGAA_MT_+_577_647@55.74.20|ACATCACCCCATAAACACCA</td><td>>trnaMT_PheGAA_MT_+_577_647</td><td>55</td><td>74</td><td>20</td><td>ACATCACCCCATAAACACCA</td><td>233.7</td><td>4.4</td><td>1.3</td><td>3.3</td><td>8.2E-04</td><td>1.0E-03</td><td>MB231/MB468</td><td>MITO</td></tr>
<tr><td>>hg38_21_+_8433172_8446622_RNA45SN1_45S_50ntFlanks@9504.9521.18|GTGAACTATGCCTGGGCA</td><td>>hg38_21_+_8433172_8446622_RNA45SN1_45S_50ntFlanks</td><td>9504</td><td>9521</td><td>18</td><td>GTGAACTATGCCTGGGCA</td><td>1054.8</td><td>-3.8</td><td>1.1</td><td>-3.6</td><td>3.6E-04</td><td>1.0E-03</td><td>BT20/MB468</td><td>MP</td></tr>
<tr><td>>hg38_21_+_8433172_8446622_RNA45SN1_45S_50ntFlanks@4917.4940.24|CACCACCAGGAGTGGAGCCTGCGG</td><td>>hg38_21_+_8433172_8446622_RNA45SN1_45S_50ntFlanks</td><td>4917</td><td>4940</td><td>24</td><td>CACCACCAGGAGTGGAGCCTGCGG</td><td>231.4</td><td>-5.2</td><td>1.5</td><td>-3.4</td><td>6.2E-04</td><td>1.0E-03</td><td>BT20/MB468</td><td>MITO</td></tr>
<tr><td>>hg38_21_+_8433172_8446622_RNA45SN1_45S_50ntFlanks@10872.10891.20|GCTGGGTCGGTCGGGCTGGG</td><td>>hg38_21_+_8433172_8446622_RNA45SN1_45S_50ntFlanks</td><td>10872</td><td>10891</td><td>20</td><td>GCTGGGTCGGTCGGGCTGGG</td><td>521.3</td><td>7.8</td><td>2.0</td><td>3.8</td><td>1.3E-04</td><td>1.0E-03</td><td>BT20/MB231</td><td>MP</td></tr>
<tr><td>>hg38_21_+_8433172_8446622_RNA45SN1_45S_50ntFlanks@4203.4222.20|GACCCGGGGAGGTAGTGACG</td><td>>hg38_21_+_8433172_8446622_RNA45SN1_45S_50ntFlanks</td><td>4203</td><td>4222</td><td>20</td><td>GACCCGGGGAGGTAGTGACG</td><td>2485.6</td><td>7.7</td><td>2.0</td><td>3.8</td><td>1.3E-04</td><td>1.0E-03</td><td>BT20/MB231</td><td>MP</td></tr>
<tr><td>>hg38_21_+_8433172_8446622_RNA45SN1_45S_50ntFlanks@11718.11736.19|AACGGCGGGAGTAACTATG</td><td>>hg38_21_+_8433172_8446622_RNA45SN1_45S_50ntFlanks</td><td>11718</td><td>11736</td><td>19</td><td>AACGGCGGGAGTAACTATG</td><td>815.9</td><td>-3.4</td><td>1.0</td><td>-3.4</td><td>7.5E-04</td><td>1.0E-03</td><td>BT20/MB468</td><td>CYTO</td></tr>
<tr><td>>hg38_21_+_8433172_8446622_RNA45SN1_45S_50ntFlanks@11553.11591.39|CTCGGCCGGCGCCTAGCAGCCGACTTAGAACTGGTGCGG</td><td>>hg38_21_+_8433172_8446622_RNA45SN1_45S_50ntFlanks</td><td>11553</td><td>11591</td><td>39</td><td>CTCGGCCGGCGCCTAGCAGCCGACTTAGAACTGGTGCGG</td><td>258.0</td><td>-4.2</td><td>1.3</td><td>-3.3</td><td>8.4E-04</td><td>1.0E-03</td><td>MB231/MB468</td><td>NUC</td></tr>
<tr><td>>MI0000298|hsa-mir-221&WithFlank&X|-|45746151|45746272@71.94.24&[MIMAT0000278&hsa-miR-221-3p&offsets|0|+1;m-91&X|-|45746181|45746202&offsets|0|+2]|AGCTACATTGTCTGCTGGGTTTCA</td><td>>MI0000298|hsa-mir-221&WithFlank&X|-|45746151|45746272</td><td>71</td><td>94</td><td>24</td><td>AGCTACATTGTCTGCTGGGTTTCA</td><td>1908.7</td><td>2.9</td><td>0.9</td><td>3.4</td><td>6.9E-04</td><td>1.0E-03</td><td>MB231/MB468</td><td>MP</td></tr>
<tr><td>>MI0000456|hsa-mir-140&WithFlank&16|+|69933075|69933186@29.49.21(+1A)&[MIMAT0000431&hsa-miR-140-5p&offsets|0|-1(+1A);m-173&16|+|69933103|69933124&offsets|0|-1(+1A)]|CAGTGGTTTTACCCTATGGTAA</td><td>>MI0000456|hsa-mir-140&WithFlank&16|+|69933075|69933186</td><td>29</td><td>49</td><td>21(+1A)</td><td>CAGTGGTTTTACCCTATGGTAA</td><td>203.7</td><td>6.4</td><td>1.9</td><td>3.4</td><td>6.9E-04</td><td>1.0E-03</td><td>MB231/MB468</td><td>MP</td></tr>
<tr><td>>hg38_21_+_8433172_8446622_RNA45SN1_45S_50ntFlanks@12945.12962.18|GTAGACGACCTGCTTCTG</td><td>>hg38_21_+_8433172_8446622_RNA45SN1_45S_50ntFlanks</td><td>12945</td><td>12962</td><td>18</td><td>GTAGACGACCTGCTTCTG</td><td>110.9</td><td>-5.3</td><td>1.6</td><td>-3.4</td><td>6.9E-04</td><td>1.0E-03</td><td>MB231/MB468</td><td>MP</td></tr>
<tr><td>>hg38_21_+_8433172_8446622_RNA45SN1_45S_50ntFlanks@8344.8378.35|TAAGGGAAAGTTGAAAAGAACTTTGAAGAGAGAGT</td><td>>hg38_21_+_8433172_8446622_RNA45SN1_45S_50ntFlanks</td><td>8344</td><td>8378</td><td>35</td><td>TAAGGGAAAGTTGAAAAGAACTTTGAAGAGAGAGT</td><td>93.2</td><td>6.6</td><td>1.9</td><td>3.6</td><td>3.6E-04</td><td>1.0E-03</td><td>BT20/MB468</td><td>MP</td></tr>
<tr><td>>hg38_21_+_8433172_8446622_RNA45SN1_45S_50ntFlanks@10389.10410.22|TGAACAGCAGTTGAACATGGGT</td><td>>hg38_21_+_8433172_8446622_RNA45SN1_45S_50ntFlanks</td><td>10389</td><td>10410</td><td>22</td><td>TGAACAGCAGTTGAACATGGGT</td><td>1088.4</td><td>-4.7</td><td>1.3</td><td>-3.6</td><td>3.6E-04</td><td>1.0E-03</td><td>BT20/MB468</td><td>MP</td></tr>
<tr><td>>hg38_21_+_8433172_8446622_RNA45SN1_45S_50ntFlanks@6646.6674.29|TCGTACGACTCTTAGCGGTGGATCACTCG</td><td>>hg38_21_+_8433172_8446622_RNA45SN1_45S_50ntFlanks</td><td>6646</td><td>6674</td><td>29</td><td>TCGTACGACTCTTAGCGGTGGATCACTCG</td><td>2891.6</td><td>3.5</td><td>1.1</td><td>3.3</td><td>8.4E-04</td><td>1.0E-03</td><td>MB231/MB468</td><td>NUC</td></tr>
<tr><td>>hg38_21_+_8433172_8446622_RNA45SN1_45S_50ntFlanks@8079.8101.23|AAGAGCCCAGCGCCGAATCCCCG</td><td>>hg38_21_+_8433172_8446622_RNA45SN1_45S_50ntFlanks</td><td>8079</td><td>8101</td><td>23</td><td>AAGAGCCCAGCGCCGAATCCCCG</td><td>534.5</td><td>-5.1</td><td>1.4</td><td>-3.6</td><td>3.6E-04</td><td>1.0E-03</td><td>BT20/MB468</td><td>MP</td></tr>
<tr><td>>hg38_21_+_8433172_8446622_RNA45SN1_45S_50ntFlanks@6722.6752.31|ATTGCAGGACACATTGATCATCGACACTTCG</td><td>>hg38_21_+_8433172_8446622_RNA45SN1_45S_50ntFlanks</td><td>6722</td><td>6752</td><td>31</td><td>ATTGCAGGACACATTGATCATCGACACTTCG</td><td>81.6</td><td>-5.2</td><td>1.5</td><td>-3.4</td><td>6.9E-04</td><td>1.0E-03</td><td>MB231/MB468</td><td>MP</td></tr>
<tr><td>>MI0001448|hsa-mir-425&WithFlank&3|-|49020142|49020240@64.82.19&[MIMAT0001343&hsa-miR-425-3p&offsets|+3|0;m-258&3|-|49020160|49020181&offsets|+4|+1]|GGGAATGTCGTGTCCGCCC</td><td>>MI0001448|hsa-mir-425&WithFlank&3|-|49020142|49020240</td><td>64</td><td>82</td><td>19</td><td>GGGAATGTCGTGTCCGCCC</td><td>56.6</td><td>9.0</td><td>2.7</td><td>3.4</td><td>6.9E-04</td><td>1.0E-03</td><td>MB231/MB468</td><td>MP</td></tr>
<tr><td>>MI0000802|hsa-mir-340&WithFlank&5|-|180015297|180015403@22.43.22&[MIMAT0004692&hsa-miR-340-5p&offsets|0|0;m-220&5|-|180015361|180015382&offsets|0|0]|TTATAAAGCAATGAGACTGATT</td><td>>MI0000802|hsa-mir-340&WithFlank&5|-|180015297|180015403</td><td>22</td><td>43</td><td>22</td><td>TTATAAAGCAATGAGACTGATT</td><td>1140.2</td><td>3.7</td><td>1.0</td><td>3.8</td><td>1.3E-04</td><td>1.0E-03</td><td>BT20/MB231</td><td>CYTO</td></tr>
<tr><td>>trnaMT_PheGAA_MT_+_577_647@28.53.26|ATACACTGAAAATGTTTAGACGGGCT</td><td>>trnaMT_PheGAA_MT_+_577_647</td><td>28</td><td>53</td><td>26</td><td>ATACACTGAAAATGTTTAGACGGGCT</td><td>158.9</td><td>6.2</td><td>1.7</td><td>3.6</td><td>3.6E-04</td><td>1.0E-03</td><td>BT20/MB468</td><td>MP</td></tr>
<tr><td>>hg38_21_+_8433172_8446622_RNA45SN1_45S_50ntFlanks@12539.12564.26|ATGATGTGTTGTTGCCATGGTAATCC</td><td>>hg38_21_+_8433172_8446622_RNA45SN1_45S_50ntFlanks</td><td>12539</td><td>12564</td><td>26</td><td>ATGATGTGTTGTTGCCATGGTAATCC</td><td>81.3</td><td>-5.4</td><td>1.5</td><td>-3.6</td><td>3.6E-04</td><td>1.0E-03</td><td>BT20/MB468</td><td>MP</td></tr>
<tr><td>>hg38_21_+_8433172_8446622_RNA45SN1_45S_50ntFlanks@8197.8215.19|GTGGACGGTGTGAGGCCGG</td><td>>hg38_21_+_8433172_8446622_RNA45SN1_45S_50ntFlanks</td><td>8197</td><td>8215</td><td>19</td><td>GTGGACGGTGTGAGGCCGG</td><td>381.3</td><td>-3.1</td><td>0.9</td><td>-3.6</td><td>3.6E-04</td><td>1.0E-03</td><td>BT20/MB468</td><td>MP</td></tr>
<tr><td>>hg38_21_+_8433172_8446622_RNA45SN1_45S_50ntFlanks@10729.10755.27|AGAGGGTGTAAATCTCGCGCCGGGCCG</td><td>>hg38_21_+_8433172_8446622_RNA45SN1_45S_50ntFlanks</td><td>10729</td><td>10755</td><td>27</td><td>AGAGGGTGTAAATCTCGCGCCGGGCCG</td><td>154.3</td><td>-3.5</td><td>1.0</td><td>-3.4</td><td>6.3E-04</td><td>1.0E-03</td><td>BT20/MB468</td><td>MITO</td></tr>
<tr><td>>hg38_21_+_8433172_8446622_RNA45SN1_45S_50ntFlanks@5473.5501.29|CGGCCCTGGCGGAGCGCTGAGAAGACGGT</td><td>>hg38_21_+_8433172_8446622_RNA45SN1_45S_50ntFlanks</td><td>5473</td><td>5501</td><td>29</td><td>CGGCCCTGGCGGAGCGCTGAGAAGACGGT</td><td>242.0</td><td>3.4</td><td>1.0</td><td>3.6</td><td>3.6E-04</td><td>1.0E-03</td><td>BT20/MB468</td><td>MP</td></tr>
<tr><td>>hg38_21_+_8433172_8446622_RNA45SN1_45S_50ntFlanks@10871.10892.22|GGCTGGGTCGGTCGGGCTGGGG</td><td>>hg38_21_+_8433172_8446622_RNA45SN1_45S_50ntFlanks</td><td>10871</td><td>10892</td><td>22</td><td>GGCTGGGTCGGTCGGGCTGGGG</td><td>154.5</td><td>-3.4</td><td>1.0</td><td>-3.4</td><td>7.5E-04</td><td>1.0E-03</td><td>BT20/MB468</td><td>CYTO</td></tr>
<tr><td>>MI0000101|hsa-mir-99a&WithFlank&21|+|16539083|16539175@19.40.22(+1A)&[MIMAT0000097&hsa-miR-99a-5p&offsets|0|0(+1A);m-64&21|+|16539101|16539121&offsets|0|+1(+1A)]|AACCCGTAGATCCGATCTTGTGA</td><td>>MI0000101|hsa-mir-99a&WithFlank&21|+|16539083|16539175</td><td>19</td><td>40</td><td>22(+1A)</td><td>AACCCGTAGATCCGATCTTGTGA</td><td>317.5</td><td>-6.5</td><td>1.7</td><td>-3.8</td><td>1.3E-04</td><td>1.0E-03</td><td>BT20/MB231</td><td>MP</td></tr>
<tr><td>>MI0000273|hsa-mir-183&WithFlank&7|-|129774899|129775020@34.53.20(+1C)&[MIMAT0000261&hsa-miR-183-5p&offsets|+1|-1(+1C);m-41&7|-|129774966|129774987&offsets|0|-2(+1C)]|ATGGCACTGGTAGAATTCACC</td><td>>MI0000273|hsa-mir-183&WithFlank&7|-|129774899|129775020</td><td>34</td><td>53</td><td>20(+1C)</td><td>ATGGCACTGGTAGAATTCACC</td><td>520.4</td><td>2.7</td><td>0.7</td><td>3.8</td><td>1.5E-04</td><td>1.0E-03</td><td>BT20/MB231</td><td>MITO</td></tr>
<tr><td>>trnaMT_GluTTC_MT_-_14674_14742@28.71.44,trnalookalike8_GluTTC_5_-_93905172_93905240@28.71.44|TTTTTCATATCATTGGTCGTGGTTGTAGTCCGTGCGAGAATACC</td><td>>trnaMT_GluTTC_MT_-_14674_14742</td><td>28</td><td>71</td><td>44</td><td>TTTTTCATATCATTGGTCGTGGTTGTAGTCCGTGCGAGAATACC</td><td>160.7</td><td>-3.6</td><td>0.9</td><td>-3.8</td><td>1.5E-04</td><td>1.0E-03</td><td>BT20/MB231</td><td>MITO</td></tr>
<tr><td>>hg38_21_+_8433172_8446622_RNA45SN1_45S_50ntFlanks@10128.10159.32|CGGCGGCGGCGGCGGCGGCGGGGGTGTGGGGT</td><td>>hg38_21_+_8433172_8446622_RNA45SN1_45S_50ntFlanks</td><td>10128</td><td>10159</td><td>32</td><td>CGGCGGCGGCGGCGGCGGCGGGGGTGTGGGGT</td><td>60.6</td><td>9.7</td><td>2.7</td><td>3.6</td><td>3.6E-04</td><td>1.0E-03</td><td>BT20/MB468</td><td>MP</td></tr>
<tr><td>>hg38_21_+_8433172_8446622_RNA45SN1_45S_50ntFlanks@6719.6758.40|TGAATTGCAGGACACATTGATCATCGACACTTCGAACGCA</td><td>>hg38_21_+_8433172_8446622_RNA45SN1_45S_50ntFlanks</td><td>6719</td><td>6758</td><td>40</td><td>TGAATTGCAGGACACATTGATCATCGACACTTCGAACGCA</td><td>346.0</td><td>-2.8</td><td>0.8</td><td>-3.4</td><td>6.3E-04</td><td>1.0E-03</td><td>BT20/MB468</td><td>MITO</td></tr>
<tr><td>>MI0000446|hsa-mir-125b-1&WithFlank&11|-|122099751|122099850@61.80.20(+1U)&[MIMAT0004592&hsa-miR-125b-1-3p&offsets|0|-2(+1U);m-485&11|-|122099769|122099790&offsets|0|-2(+1U)]|ACGGGTTAGGCTCTTGGGAGT</td><td>>MI0000446|hsa-mir-125b-1&WithFlank&11|-|122099751|122099850</td><td>61</td><td>80</td><td>20(+1U)</td><td>ACGGGTTAGGCTCTTGGGAGT</td><td>1257.8</td><td>-5.8</td><td>1.5</td><td>-3.8</td><td>1.3E-04</td><td>1.0E-03</td><td>BT20/MB231</td><td>MP</td></tr>
<tr><td>>hg38_21_+_8433172_8446622_RNA45SN1_45S_50ntFlanks@11672.11703.32|CAGTGCTCTGAATGTCAAAGTGAAGAAATTCA</td><td>>hg38_21_+_8433172_8446622_RNA45SN1_45S_50ntFlanks</td><td>11672</td><td>11703</td><td>32</td><td>CAGTGCTCTGAATGTCAAAGTGAAGAAATTCA</td><td>277.8</td><td>-2.6</td><td>0.7</td><td>-3.8</td><td>1.3E-04</td><td>1.0E-03</td><td>BT20/MB231</td><td>NUC</td></tr>
<tr><td>>trnaMT_GluTTC_MT_-_14674_14742@40.66.27,trnalookalike8_GluTTC_5_-_93905172_93905240@40.66.27|TTGGTCGTGGTTGTAGTCCGTGCGAGA</td><td>>trnaMT_GluTTC_MT_-_14674_14742</td><td>40</td><td>66</td><td>27</td><td>TTGGTCGTGGTTGTAGTCCGTGCGAGA</td><td>252.4</td><td>2.9</td><td>0.8</td><td>3.4</td><td>7.0E-04</td><td>1.0E-03</td><td>MB231/MB468</td><td>MP</td></tr>
<tr><td>>hg38_21_+_8433172_8446622_RNA45SN1_45S_50ntFlanks@11604.11623.20|CGACTGTTTAATTAAAACAA</td><td>>hg38_21_+_8433172_8446622_RNA45SN1_45S_50ntFlanks</td><td>11604</td><td>11623</td><td>20</td><td>CGACTGTTTAATTAAAACAA</td><td>273.7</td><td>-1.3</td><td>0.4</td><td>-3.3</td><td>8.4E-04</td><td>1.0E-03</td><td>BT20/MB468</td><td>NUC</td></tr>
</table>

<table>
<tr><td>>trnaMT_SerGCT_MT_+_12207_12265@14.53.40|AGAACTGCTAACTCATGCCCCCATGTCTAACAACATGGCT</td><td>>trnaMT_SerGCT_MT_+_12207_12265</td><td>14</td><td>53</td><td>40</td><td>AGAACTGCTAACTCATGCCCCCATGTCTAACAACATGGCT</td><td>60.6</td><td>9.7</td><td>2.7</td><td>3.6</td><td>3.6E-04</td><td>1.0E-03</td><td>BT20/MB468</td><td>MP</td></tr>
<tr><td>>hg38_21_+_8433172_8446622_RNA45SN1_45S_50ntFlanks@10410.10432.23|TCAGTCGGTCCTGAGAGATGGGC</td><td>>hg38_21_+_8433172_8446622_RNA45SN1_45S_50ntFlanks</td><td>10410</td><td>10432</td><td>23</td><td>TCAGTCGGTCCTGAGAGATGGGC</td><td>757.1</td><td>-3.0</td><td>0.8</td><td>-3.6</td><td>3.6E-04</td><td>1.0E-03</td><td>BT20/MB468</td><td>MP</td></tr>
<tr><td>>MI0000289|hsa-mir-181a-1&WithFlank&1|-|198859038|198859159@30.51.22(+1C);MI0000269|hsa-mir-181a-2&WithFlank&9|+|124692436|124692557@45.66.22(+1C)&[MIMAT0000256&hsa-miR-181a-5p&offsets|0|-1(+1C);m-32&1|-|198859108|198859130&offsets|0|-1(+1C)];[MIMAT0000256_1&hsa-miR-181a-5p&offsets|0|-1(+1C);m-32&9|+|124692480|124692502&offsets|0|-1(+1C)]|AACATTCAACGCTGTCGGTGAGC</td><td>>MI0000289|hsa-mir-181a-1&WithFlank&1|-|198859038|198859159</td><td>30</td><td>51</td><td>22(+1C)</td><td>AACATTCAACGCTGTCGGTGAGC</td><td>1059.4</td><td>-3.2</td><td>0.8</td><td>-3.8</td><td>1.5E-04</td><td>1.0E-03</td><td>BT20/MB231</td><td>MITO</td></tr>
<tr><td>>hg38_21_+_8433172_8446622_RNA45SN1_45S_50ntFlanks@9921.9949.29|TGTTGGTTGATATAGACAGCAGGACGGTG</td><td>>hg38_21_+_8433172_8446622_RNA45SN1_45S_50ntFlanks</td><td>9921</td><td>9949</td><td>29</td><td>TGTTGGTTGATATAGACAGCAGGACGGTG</td><td>59.7</td><td>6.5</td><td>1.8</td><td>3.6</td><td>3.6E-04</td><td>1.0E-03</td><td>BT20/MB468</td><td>MP</td></tr>
<tr><td>>hg38_21_+_8433172_8446622_RNA45SN1_45S_50ntFlanks@12476.12499.24|CCACTAATAGGGAACGTGAGCTGG</td><td>>hg38_21_+_8433172_8446622_RNA45SN1_45S_50ntFlanks</td><td>12476</td><td>12499</td><td>24</td><td>CCACTAATAGGGAACGTGAGCTGG</td><td>207.1</td><td>-5.6</td><td>1.6</td><td>-3.6</td><td>3.6E-04</td><td>1.0E-03</td><td>BT20/MB468</td><td>MP</td></tr>
<tr><td>>hg38_21_+_8433172_8446622_RNA45SN1_45S_50ntFlanks@11572.11592.21|CCGACTTAGAACTGGTGCGGA</td><td>>hg38_21_+_8433172_8446622_RNA45SN1_45S_50ntFlanks</td><td>11572</td><td>11592</td><td>21</td><td>CCGACTTAGAACTGGTGCGGA</td><td>307.7</td><td>-5.9</td><td>1.8</td><td>-3.3</td><td>8.4E-04</td><td>1.0E-03</td><td>BT20/MB468</td><td>NUC</td></tr>
<tr><td>>hg38_21_+_8433172_8446622_RNA45SN1_45S_50ntFlanks@7969.8008.40|CCGAGACGCGACCTCAGATCAGACGTGGCGACCCGCTGAA</td><td>>hg38_21_+_8433172_8446622_RNA45SN1_45S_50ntFlanks</td><td>7969</td><td>8008</td><td>40</td><td>CCGAGACGCGACCTCAGATCAGACGTGGCGACCCGCTGAA</td><td>64.2</td><td>2.6</td><td>0.8</td><td>3.3</td><td>8.4E-04</td><td>1.0E-03</td><td>BT20/MB468</td><td>NUC</td></tr>
<tr><td>>MI0000088|hsa-mir-30a&WithFlank&6|-|71403545|71403627@54.74.21(+1A)&[MIMAT0000088&hsa-miR-30a-3p&offsets|+1|0(+1A);m-38&6|-|71403554|71403575&offsets|+1|0(+1A)]|TTTCAGTCGGATGTTTGCAGCA</td><td>>MI0000088|hsa-mir-30a&WithFlank&6|-|71403545|71403627</td><td>54</td><td>74</td><td>21(+1A)</td><td>TTTCAGTCGGATGTTTGCAGCA</td><td>138.0</td><td>6.2</td><td>1.8</td><td>3.4</td><td>7.0E-04</td><td>1.0E-03</td><td>MB231/MB468</td><td>MP</td></tr>
<tr><td>>hg38_21_+_8433172_8446622_RNA45SN1_45S_50ntFlanks@10366.10383.18|CCGAAGTGGAGAAGGGTT</td><td>>hg38_21_+_8433172_8446622_RNA45SN1_45S_50ntFlanks</td><td>10366</td><td>10383</td><td>18</td><td>CCGAAGTGGAGAAGGGTT</td><td>122.3</td><td>-4.5</td><td>1.3</td><td>-3.4</td><td>7.6E-04</td><td>1.0E-03</td><td>BT20/MB468</td><td>CYTO</td></tr>
<tr><td>>trna111_HisGTG_1_-_147774845_147774916@-1T.23.24,trna118_HisGTG_1_-_145396881_145396952@-1T.23.24,trna16_HisGTG_1_+_146544773_146544844@-1T.23.24,trna1_HisGTG_15_+_45493349_45493420@-1T.23.24,trna21_HisGTG_1_+_147753471_147753542@-1T.23.24,trna33_HisGTG_6_+_27125906_27125977@-1T.23.24,trna7_HisGTG_9_-_14433938_14434009@-1T.23.24,trna8_HisGTG_15_-_45492611_45492682@-1T.23.24,trna9_HisGTG_15_-_45490804_45490875@-1T.23.24|TGCCGTGATCGTATAGTGGTTAGT</td><td>>trna111_HisGTG_1_-_147774845_147774916</td><td>-1T</td><td>23</td><td>24</td><td>TGCCGTGATCGTATAGTGGTTAGT</td><td>1651.6</td><td>-3.4</td><td>0.9</td><td>-3.8</td><td>1.3E-04</td><td>1.0E-03</td><td>BT20/MB231</td><td>CYTO</td></tr>
<tr><td>>hg38_21_+_8433172_8446622_RNA45SN1_45S_50ntFlanks@4735.4765.31|ACGAAAGTCGGAGGTTCGAAGACGATCAGAT</td><td>>hg38_21_+_8433172_8446622_RNA45SN1_45S_50ntFlanks</td><td>4735</td><td>4765</td><td>31</td><td>ACGAAAGTCGGAGGTTCGAAGACGATCAGAT</td><td>136.4</td><td>3.8</td><td>1.1</td><td>3.6</td><td>3.7E-04</td><td>1.0E-03</td><td>BT20/MB468</td><td>MP</td></tr>
<tr><td>>hg38_21_+_8433172_8446622_RNA45SN1_45S_50ntFlanks@12007.12042.36|GCCGCCGGTGAAATACCACTACTCTGATCGTTTTTT</td><td>>hg38_21_+_8433172_8446622_RNA45SN1_45S_50ntFlanks</td><td>12007</td><td>12042</td><td>36</td><td>GCCGCCGGTGAAATACCACTACTCTGATCGTTTTTT</td><td>371.3</td><td>-5.7</td><td>1.7</td><td>-3.3</td><td>8.5E-04</td><td>1.0E-03</td><td>MB231/MB468</td><td>NUC</td></tr>
<tr><td>>hg38_21_+_8433172_8446622_RNA45SN1_45S_50ntFlanks@3989.4023.35|TTTGGTGACTCTAGATAACCTCGGGCCGATCGCAC</td><td>>hg38_21_+_8433172_8446622_RNA45SN1_45S_50ntFlanks</td><td>3989</td><td>4023</td><td>35</td><td>TTTGGTGACTCTAGATAACCTCGGGCCGATCGCAC</td><td>151.8</td><td>3.7</td><td>1.0</td><td>3.6</td><td>3.7E-04</td><td>1.0E-03</td><td>BT20/MB468</td><td>MP</td></tr>
<tr><td>>hg38_21_+_8433172_8446622_RNA45SN1_45S_50ntFlanks@7968.8011.44|TCCGAGACGCGACCTCAGATCAGACGTGGCGACCCGCTGAATTT</td><td>>hg38_21_+_8433172_8446622_RNA45SN1_45S_50ntFlanks</td><td>7968</td><td>8011</td><td>44</td><td>TCCGAGACGCGACCTCAGATCAGACGTGGCGACCCGCTGAATTT</td><td>305.6</td><td>5.0</td><td>1.5</td><td>3.3</td><td>8.5E-04</td><td>1.0E-03</td><td>MB231/MB468</td><td>NUC</td></tr>
<tr><td>>hg38_21_+_8433172_8446622_RNA45SN1_45S_50ntFlanks@10420.10453.34|CTGAGAGATGGGCGAGCGCCGTTCCGAAGGGACG</td><td>>hg38_21_+_8433172_8446622_RNA45SN1_45S_50ntFlanks</td><td>10420</td><td>10453</td><td>34</td><td>CTGAGAGATGGGCGAGCGCCGTTCCGAAGGGACG</td><td>227.3</td><td>-4.8</td><td>1.4</td><td>-3.4</td><td>6.3E-04</td><td>1.0E-03</td><td>BT20/MB468</td><td>MITO</td></tr>
<tr><td>>hg38_21_+_8433172_8446622_RNA45SN1_45S_50ntFlanks@8176.8195.20|CTTCTGATCGAGGCCCAGCC</td><td>>hg38_21_+_8433172_8446622_RNA45SN1_45S_50ntFlanks</td><td>8176</td><td>8195</td><td>20</td><td>CTTCTGATCGAGGCCCAGCC</td><td>5409.3</td><td>2.2</td><td>0.7</td><td>3.4</td><td>7.9E-04</td><td>1.0E-03</td><td>MB231/MB468</td><td>CYTO</td></tr>
<tr><td>>hg38_21_+_8433172_8446622_RNA45SN1_45S_50ntFlanks@12487.12509.23|GAACGTGAGCTGGGTTTAGACCG</td><td>>hg38_21_+_8433172_8446622_RNA45SN1_45S_50ntFlanks</td><td>12487</td><td>12509</td><td>23</td><td>GAACGTGAGCTGGGTTTAGACCG</td><td>306.7</td><td>-4.2</td><td>1.3</td><td>-3.3</td><td>8.5E-04</td><td>1.0E-03</td><td>MB231/MB468</td><td>NUC</td></tr>
<tr><td>>trnaMT_PheGAA_MT_+_577_647@39.60.22|ATGTTTAGACGGGCTCACATCA</td><td>>trnaMT_PheGAA_MT_+_577_647</td><td>39</td><td>60</td><td>22</td><td>ATGTTTAGACGGGCTCACATCA</td><td>190.9</td><td>4.4</td><td>1.2</td><td>3.6</td><td>3.7E-04</td><td>1.0E-03</td><td>BT20/MB468</td><td>MP</td></tr>
<tr><td>>hg38_21_+_8433172_8446622_RNA45SN1_45S_50ntFlanks@9506.9526.21|GAACTATGCCTGGGCAGGGCG</td><td>>hg38_21_+_8433172_8446622_RNA45SN1_45S_50ntFlanks</td><td>9506</td><td>9526</td><td>21</td><td>GAACTATGCCTGGGCAGGGCG</td><td>61.0</td><td>-9.0</td><td>2.5</td><td>-3.6</td><td>3.7E-04</td><td>1.0E-03</td><td>BT20/MB468</td><td>MP</td></tr>
<tr><td>>hg38_MT_+_1621_3279_RNR2_16S_50ntFlanks@1370.1395.26|ATGTTGGATCAGGACATCCCGATGGT</td><td>>hg38_MT_+_1621_3279_RNR2_16S_50ntFlanks</td><td>1370</td><td>1395</td><td>26</td><td>ATGTTGGATCAGGACATCCCGATGGT</td><td>294.7</td><td>3.5</td><td>1.0</td><td>3.4</td><td>7.1E-04</td><td>1.0E-03</td><td>MB231/MB468</td><td>MP</td></tr>
<tr><td>>hg38_21_+_8433172_8446622_RNA45SN1_45S_50ntFlanks@9930.9951.22|ATATAGACAGCAGGACGGTGGC</td><td>>hg38_21_+_8433172_8446622_RNA45SN1_45S_50ntFlanks</td><td>9930</td><td>9951</td><td>22</td><td>ATATAGACAGCAGGACGGTGGC</td><td>549.5</td><td>7.2</td><td>1.9</td><td>3.8</td><td>1.3E-04</td><td>1.0E-03</td><td>BT20/MB231</td><td>MP</td></tr>
<tr><td>>hg38_21_+_8433172_8446622_RNA45SN1_45S_50ntFlanks@10823.10865.43|AAGTCGGCAAGCCGGATCCGTAACTTCGGGATAAGGATTGGCT</td><td>>hg38_21_+_8433172_8446622_RNA45SN1_45S_50ntFlanks</td><td>10823</td><td>10865</td><td>43</td><td>AAGTCGGCAAGCCGGATCCGTAACTTCGGGATAAGGATTGGCT</td><td>55.6</td><td>-8.9</td><td>2.5</td><td>-3.6</td><td>3.7E-04</td><td>1.0E-03</td><td>BT20/MB468</td><td>MP</td></tr>
<tr><td>>MI0000816|hsa-mir-335&WithFlank&7|+|130496105|130496210@58.78.21(+1U)&[MIMAT0004703&hsa-miR-335-3p&offsets|0|-1(+1U);m-159&7|+|130496162|130496183&offsets|0|-1(+1U)]|TTTTTCATTATTGCTCCTGACT</td><td>>MI0000816|hsa-mir-335&WithFlank&7|+|130496105|130496210</td><td>58</td><td>78</td><td>21(+1U)</td><td>TTTTTCATTATTGCTCCTGACT</td><td>340.9</td><td>4.4</td><td>1.3</td><td>3.4</td><td>7.1E-04</td><td>1.0E-03</td><td>MB231/MB468</td><td>MP</td></tr>
<tr><td>>MI0000264|hsa-mir-7-2&WithFlank&15|+|88611819|88611940@38.57.20;MI0000265|hsa-mir-7-3&WithFlank&19|+|4770664|4770785@37.56.20;MI0000263|hsa-mir-7-1&WithFlank&9|-|83969742|83969863@30.49.20&[MIMAT0000252&hsa-miR-7-5p&offsets|0|-4;m-400&15|+|88611856|88611879&offsets|0|-4];[MIMAT0000252_1&hsa-miR-7-5p&offsets|0|-4;m-402&19|+|4770700|4770723&offsets|0|-4];[MIMAT0000252_2&hsa-miR-7-5p&offsets|0|-4;m-401&9|-|83969811|83969834&offsets|0|-4]|TGGAAGACTAGTGATTTTGT</td><td>>MI0000264|hsa-mir-7-2&WithFlank&15|+|88611819|88611940</td><td>38</td><td>57</td><td>20</td><td>TGGAAGACTAGTGATTTTGT</td><td>208.9</td><td>4.1</td><td>1.2</td><td>3.4</td><td>7.1E-04</td><td>1.0E-03</td><td>MB231/MB468</td><td>MP</td></tr>
<tr><td>>hg38_21_+_8433172_8446622_RNA45SN1_45S_50ntFlanks@11200.11225.26|CGGCGGCGGCGGCGACTCTGGACGCG</td><td>>hg38_21_+_8433172_8446622_RNA45SN1_45S_50ntFlanks</td><td>11200</td><td>11225</td><td>26</td><td>CGGCGGCGGCGGCGACTCTGGACGCG</td><td>804.6</td><td>-4.5</td><td>1.3</td><td>-3.3</td><td>8.5E-04</td><td>1.0E-03</td><td>MB231/MB468</td><td>NUC</td></tr>
<tr><td>>hg38_21_+_8433172_8446622_RNA45SN1_45S_50ntFlanks@4762.4780.19|AGATACCGTCGTAGTTCCG</td><td>>hg38_21_+_8433172_8446622_RNA45SN1_45S_50ntFlanks</td><td>4762</td><td>4780</td><td>19</td><td>AGATACCGTCGTAGTTCCG</td><td>397.0</td><td>-4.4</td><td>1.3</td><td>-3.4</td><td>7.1E-04</td><td>1.0E-03</td><td>MB231/MB468</td><td>MP</td></tr>
<tr><td>>hg38_21_+_8433172_8446622_RNA45SN1_45S_50ntFlanks@11564.11584.21|CCTAGCAGCCGACTTAGAACT</td><td>>hg38_21_+_8433172_8446622_RNA45SN1_45S_50ntFlanks</td><td>11564</td><td>11584</td><td>21</td><td>CCTAGCAGCCGACTTAGAACT</td><td>585.9</td><td>1.7</td><td>0.5</td><td>3.4</td><td>6.4E-04</td><td>1.0E-03</td><td>BT20/MB468</td><td>MITO</td></tr>
<tr><td>>MI0000102|hsa-mir-100&WithFlank&11|-|122152223|122152314@19.37.19(+1A)&[MIMAT0000098&hsa-miR-100-5p&offsets|0|-3(+1A);m-27&11|-|122152276|122152296&offsets|0|-2(+1A)]|AACCCGTAGATCCGAACTTA</td><td>>MI0000102|hsa-mir-100&WithFlank&11|-|122152223|122152314</td><td>19</td><td>37</td><td>19(+1A)</td><td>AACCCGTAGATCCGAACTTA</td><td>1554.2</td><td>-5.1</td><td>1.3</td><td>-3.8</td><td>1.3E-04</td><td>1.0E-03</td><td>BT20/MB231</td><td>NUC</td></tr>
<tr><td>>hg38_MT_+_598_1651_RNR1_12S_50ntFlanks@905.929.25|AGTATACTTCAAAGGACATTTAACT</td><td>>hg38_MT_+_598_1651_RNR1_12S_50ntFlanks</td><td>905</td><td>929</td><td>25</td><td>AGTATACTTCAAAGGACATTTAACT</td><td>369.1</td><td>4.1</td><td>1.2</td><td>3.3</td><td>8.4E-04</td><td>1.0E-03</td><td>MB231/MB468</td><td>MITO</td></tr>
<tr><td>>hg38_21_+_8433172_8446622_RNA45SN1_45S_50ntFlanks@3858.3898.41|TGGTAATTCTAGAGCTAATACATGCCGACGGGCGCTGACCC</td><td>>hg38_21_+_8433172_8446622_RNA45SN1_45S_50ntFlanks</td><td>3858</td><td>3898</td><td>41</td><td>TGGTAATTCTAGAGCTAATACATGCCGACGGGCGCTGACCC</td><td>68.9</td><td>-10.0</td><td>2.9</td><td>-3.4</td><td>7.1E-04</td><td>1.0E-03</td><td>MB231/MB468</td><td>MP</td></tr>
<tr><td>>hg38_21_+_8433172_8446622_RNA45SN1_45S_50ntFlanks@8179.8203.25|CTGATCGAGGCCCAGCCCGTGGACG</td><td>>hg38_21_+_8433172_8446622_RNA45SN1_45S_50ntFlanks</td><td>8179</td><td>8203</td><td>25</td><td>CTGATCGAGGCCCAGCCCGTGGACG</td><td>234.0</td><td>-2.4</td><td>0.7</td><td>-3.4</td><td>7.7E-04</td><td>1.0E-03</td><td>BT20/MB468</td><td>CYTO</td></tr>
<tr><td>>am_trna116_GluCTC_1_-_145399233_145399304@1.30.30,trna59_GluCTC_1_+_249168447_249168518@1.30.30,trna71_GluCTC_1_-_161439189_161439260@1.30.30,trna74_GluCTC_1_-_161431809_161431880@1.30.30,trna77_GluCTC_1_-_161424398_161424469@1.30.30,trna77_GluCTC_6_+_28949976_28950047@1.30.30,trna80_GluCTC_1_-_161417018_161417089@1.30.30,trna87_GluCTC_6_-_126101393_126101464@1.30.30|TCCCTGGTGGTCTAGTGGTTAGGATTCGGC</td><td>>am_trna116_GluCTC_1_-_145399233_145399304</td><td>1</td><td>30</td><td>30</td><td>TCCCTGGTGGTCTAGTGGTTAGGATTCGGC</td><td>19805.3</td><td>-3.4</td><td>1.0</td><td>-3.3</td><td>8.6E-04</td><td>1.0E-03</td><td>MB231/MB468</td><td>NUC</td></tr>
<tr><td>>hg38_21_+_8433172_8446622_RNA45SN1_45S_50ntFlanks@3990.4023.34|TTGGTGACTCTAGATAACCTCGGGCCGATCGCAC</td><td>>hg38_21_+_8433172_8446622_RNA45SN1_45S_50ntFlanks</td><td>3990</td><td>4023</td><td>34</td><td>TTGGTGACTCTAGATAACCTCGGGCCGATCGCAC</td><td>630.6</td><td>2.1</td><td>0.6</td><td>3.3</td><td>8.6E-04</td><td>1.0E-03</td><td>MB231/MB468</td><td>NUC</td></tr>
<tr><td>>hg38_21_+_8433172_8446622_RNA45SN1_45S_50ntFlanks@8342.8365.24|CGTAAGGGAAAGTTGAAAAGAACT</td><td>>hg38_21_+_8433172_8446622_RNA45SN1_45S_50ntFlanks</td><td>8342</td><td>8365</td><td>24</td><td>CGTAAGGGAAAGTTGAAAAGAACT</td><td>147.8</td><td>5.9</td><td>1.8</td><td>3.3</td><td>8.4E-04</td><td>1.0E-03</td><td>MB231/MB468</td><td>MITO</td></tr>
<tr><td>>hg38_21_+_8433172_8446622_RNA45SN1_45S_50ntFlanks@11321.11342.22|CACGTCTCGTCGCGCGCGCGTC</td><td>>hg38_21_+_8433172_8446622_RNA45SN1_45S_50ntFlanks</td><td>11321</td><td>11342</td><td>22</td><td>CACGTCTCGTCGCGCGCGCGTC</td><td>2108.5</td><td>-3.4</td><td>1.0</td><td>-3.4</td><td>7.7E-04</td><td>1.0E-03</td><td>BT20/MB468</td><td>CYTO</td></tr>
<tr><td>>hg38_21_+_8433172_8446622_RNA45SN1_45S_50ntFlanks@12159.12179.21|TTTGACTGGGGCGGTACACCT</td><td>>hg38_21_+_8433172_8446622_RNA45SN1_45S_50ntFlanks</td><td>12159</td><td>12179</td><td>21</td><td>TTTGACTGGGGCGGTACACCT</td><td>5937.8</td><td>-6.7</td><td>1.9</td><td>-3.6</td><td>3.7E-04</td><td>1.0E-03</td><td>BT20/MB468</td><td>MP</td></tr>
<tr><td>>trna2_GlyGCC_21_-_18827107_18827177@53.74.22,trna35_GlyGCC_1_+_161413094_161413164@53.74.22,trna37_GlyGCC_1_+_161420467_161420537@53.74.22,trna39_GlyGCC_1_+_161427898_161427968@53.74.22,trna41_GlyGCC_1_+_161435258_161435328@53.74.22|TCGATTCCCGGCCCATGCACCA</td><td>>trna2_GlyGCC_21_-_18827107_18827177</td><td>53</td><td>74</td><td>22</td><td>TCGATTCCCGGCCCATGCACCA</td><td>306.1</td><td>7.2</td><td>1.9</td><td>3.8</td><td>1.3E-04</td><td>1.0E-03</td><td>BT20/MB231</td><td>CYTO</td></tr>
<tr><td>>hg38_21_+_8433172_8446622_RNA45SN1_45S_50ntFlanks@6647.6675.29|CGTACGACTCTTAGCGGTGGATCACTCGG</td><td>>hg38_21_+_8433172_8446622_RNA45SN1_45S_50ntFlanks</td><td>6647</td><td>6675</td><td>29</td><td>CGTACGACTCTTAGCGGTGGATCACTCGG</td><td>876.0</td><td>2.6</td><td>0.8</td><td>3.3</td><td>8.6E-04</td><td>1.0E-03</td><td>MB231/MB468</td><td>NUC</td></tr>
<tr><td>>hg38_21_+_8433172_8446622_RNA45SN1_45S_50ntFlanks@9954.9978.25|TGGAAGTCGGAATCCGCTAAGGAGT</td><td>>hg38_21_+_8433172_8446622_RNA45SN1_45S_50ntFlanks</td><td>9954</td><td>9978</td><td>25</td><td>TGGAAGTCGGAATCCGCTAAGGAGT</td><td>219.1</td><td>2.9</td><td>0.8</td><td>3.6</td><td>3.7E-04</td><td>1.0E-03</td><td>BT20/MB468</td><td>MP</td></tr>
<tr><td>>hg38_21_+_8433172_8446622_RNA45SN1_45S_50ntFlanks@12332.12365.34|TAAGCAGGAGGTGTCAGAAAAGTTACCACAGGGA</td><td>>hg38_21_+_8433172_8446622_RNA45SN1_45S_50ntFlanks</td><td>12332</td><td>12365</td><td>34</td><td>TAAGCAGGAGGTGTCAGAAAAGTTACCACAGGGA</td><td>108.8</td><td>-5.6</td><td>1.6</td><td>-3.6</td><td>3.7E-04</td><td>1.0E-03</td><td>BT20/MB468</td><td>MP</td></tr>
<tr><td>>hg38_21_+_8433172_8446622_RNA45SN1_45S_50ntFlanks@9696.9722.27|CAGTTTTATCCGGTAAAGCGAATGATT</td><td>>hg38_21_+_8433172_8446622_RNA45SN1_45S_50ntFlanks</td><td>9696</td><td>9722</td><td>27</td><td>CAGTTTTATCCGGTAAAGCGAATGATT</td><td>147.6</td><td>-5.4</td><td>1.6</td><td>-3.4</td><td>6.4E-04</td><td>1.0E-03</td><td>BT20/MB468</td><td>MITO</td></tr>
<tr><td>>hg38_21_+_8433172_8446622_RNA45SN1_45S_50ntFlanks@10681.10708.28|TTCCGGCGGCGTCCGGTGAGCTCTCGCT</td><td>>hg38_21_+_8433172_8446622_RNA45SN1_45S_50ntFlanks</td><td>10681</td><td>10708</td><td>28</td><td>TTCCGGCGGCGTCCGGTGAGCTCTCGCT</td><td>322.4</td><td>-5.1</td><td>1.5</td><td>-3.4</td><td>7.7E-04</td><td>1.0E-03</td><td>BT20/MB468</td><td>CYTO</td></tr>
<tr><td>>hg38_21_+_8433172_8446622_RNA45SN1_45S_50ntFlanks@11668.11687.20|TGCCCAGTGCTCTGAATGTC</td><td>>hg38_21_+_8433172_8446622_RNA45SN1_45S_50ntFlanks</td><td>11668</td><td>11687</td><td>20</td><td>TGCCCAGTGCTCTGAATGTC</td><td>335.8</td><td>3.8</td><td>1.1</td><td>3.6</td><td>3.7E-04</td><td>1.0E-03</td><td>BT20/MB468</td><td>MP</td></tr>
</table>

| | | | | | | | | | | | | | |
|---|---|---|---|---|---|---|---|---|---|---|---|---|---|
| >hg38_21_+_8433172_8446622_RNA45SN1_45S_50ntFlanks@11999.12034.36\|CCCTGCGGGCCGCCGGTGAAATACCACTACTCTGAT | >hg38_21_+_8433172_8446622_RNA45SN1_45S_50ntFlanks | 11999 | 12034 | 36 | CCCTGCGGGCCGCCGGTGAAATACCACTACTCTGAT | 195.5 | -4.9 | 1.5 | -3.4 | 7.7E-04 | 1.0E-03 | BT20/MB468 | CYTO |
| >MI0000081\|hsa-mir-24-2&WithFlank&19\|-\|13836281\|13836365@56.74.19(+1A);MI0000080\|hsa-mir-24-1&WithFlank&9\|+\|95086015\|95086094@50.68.19(+1A)&[MIMAT0000080&hsa-miR-24-3p&offsets\|0\|-3(+1A);m-36&19\|-\|13836288\|13836310&offsets\|0\|-4(+1A)];[MIMAT0000080_1&hsa-miR-24-3p&offsets\|0\|-3(+1A);m-35&9\|+\|95086064\|95086086&offsets\|0\|-4(+1A)]\|TGGCTCAGTTCAGCAGGAAA | >MI0000081\|hsa-mir-24-2&WithFlank&19\|-\|13836281\|13836365 | 56 | 74 | 19(+1A) | TGGCTCAGTTCAGCAGGAAA | 336.2 | 4.0 | 1.1 | 3.6 | 3.7E-04 | 1.0E-03 | BT20/MB468 | MP |
| >hg38_21_+_8433172_8446622_RNA45SN1_45S_50ntFlanks@10350.10375.26\|ACGAGAACTTTGAAGGCCGAAGTGGA | >hg38_21_+_8433172_8446622_RNA45SN1_45S_50ntFlanks | 10350 | 10375 | 26 | ACGAGAACTTTGAAGGCCGAAGTGGA | 248.0 | -6.1 | 1.7 | -3.6 | 3.7E-04 | 1.0E-03 | BT20/MB468 | MP |
| >MI0000478\|hsa-mir-149&WithFlank&2\|+\|240455995\|240456095@21.42.22(+1A)&[MIMAT0000450&hsa-miR-149-5p&offsets\|0\|-1(+1A);m-195&2\|+\|240456015\|240456037&offsets\|0\|-1(+1A)]\|TCTGGCTCCGTGTCTTCACTCCA | >MI0000478\|hsa-mir-149&WithFlank&2\|+\|240455995\|240456095 | 21 | 42 | 22(+1A) | TCTGGCTCCGTGTCTTCACTCCA | 340.9 | 3.3 | 0.9 | 3.8 | 1.6E-04 | 1.0E-03 | BT20/MB231 | MITO |
| >MI0000079\|hsa-mir-23a&WithFlank&19\|-\|13836581\|13836665@54.71.18(+1C)&[MIMAT0000078&hsa-miR-23a-3p&offsets\|+3\|0(+1C);m-31&19\|-\|13836595\|13836615&offsets\|+3\|0(+1C)]\|ACATTGCCAGGGATTTCCC | >MI0000079\|hsa-mir-23a&WithFlank&19\|-\|13836581\|13836665 | 54 | 71 | 18(+1C) | ACATTGCCAGGGATTTCCC | 57.0 | 9.1 | 2.7 | 3.4 | 7.1E-04 | 1.0E-03 | MB231/MB468 | MP |
| >MI0000542\|hsa-mir-320a&WithFlank&8\|-\|22244960\|22245043@48.68.21(+2U)&[MIMAT0000510&hsa-miR-320a-3p&offsets\|0\|-1(+2U);m-60&8\|-\|22244975\|22244996&offsets\|0\|-1(+2U)]\|AAAAGCTGGGTTGAGAGGGCGTT | >MI0000542\|hsa-mir-320a&WithFlank&8\|-\|22244960\|22245043 | 48 | 68 | 21(+2U) | AAAAGCTGGGTTGAGAGGGCGTT | 749.7 | 4.6 | 1.4 | 3.4 | 7.1E-04 | 1.0E-03 | MB231/MB468 | MP |
| >hg38_21_+_8433172_8446622_RNA45SN1_45S_50ntFlanks@9575.9605.31\|ATCGGTCGTCCGACCTGGGTATAGGGGCGAA | >hg38_21_+_8433172_8446622_RNA45SN1_45S_50ntFlanks | 9575 | 9605 | 31 | ATCGGTCGTCCGACCTGGGTATAGGGGCGAA | 646.9 | -5.4 | 1.6 | -3.3 | 8.5E-04 | 1.0E-03 | MB231/MB468 | MITO |
| >MI0000253\|hsa-mir-148a&WithFlank&7\|-\|25949913\|25949992@50.70.21(+1A)&[MIMAT0000243&hsa-miR-148a-3p&offsets\|0\|-1(+1A);m-7&7\|-\|25949922\|25949943&offsets\|0\|-1(+1A)]\|TCAGTGCACTACAGAACTTTGA | >MI0000253\|hsa-mir-148a&WithFlank&7\|-\|25949913\|25949992 | 50 | 70 | 21(+1A) | TCAGTGCACTACAGAACTTTGA | 2056.6 | 3.1 | 0.8 | 3.8 | 1.6E-04 | 1.0E-03 | BT20/MB231 | MITO |
| >hg38_1_-_228634819_228635039_RNA5S12_5S_50ntFlanks@146.170.25\|TGGGAATACCGGGTGCTGTAGGCTT | >hg38_1_-_228634819_228635039_RNA5S12_5S_50ntFlanks | 146 | 170 | 25 | TGGGAATACCGGGTGCTGTAGGCTT | 1959.5 | 2.6 | 0.7 | 3.8 | 1.6E-04 | 1.0E-03 | BT20/MB231 | MITO |
| >hg38_21_+_8433172_8446622_RNA45SN1_45S_50ntFlanks@6722.6747.26\|ATTGCAGGACACATTGATCATCGACA | >hg38_21_+_8433172_8446622_RNA45SN1_45S_50ntFlanks | 6722 | 6747 | 26 | ATTGCAGGACACATTGATCATCGACA | 253.5 | -3.8 | 1.1 | -3.3 | 8.5E-04 | 1.0E-03 | MB231/MB468 | MITO |
| >MI0000286\|hsa-mir-210&WithFlank&11\|-\|568083\|568204@72.93.22(+1A)&[MIMAT0000267&hsa-miR-210-3p&offsets\|0\|0(+1A);m-102&11\|-\|568112\|568133&offsets\|0\|0(+1A)]\|CTGTGCGTGTGACAGCGGCTGAA | >MI0000286\|hsa-mir-210&WithFlank&11\|-\|568083\|568204 | 72 | 93 | 22(+1A) | CTGTGCGTGTGACAGCGGCTGAA | 810.1 | 3.4 | 0.9 | 3.8 | 1.3E-04 | 1.1E-03 | BT20/MB231 | NUC |
| >trna111_HisGTG_1_-_147774845_147774916@-1A.29.30,trna118_HisGTG_1_-_145396881_145396952@-1A.29.30,trna16_HisGTG_1_+_146544773_146544844@-1A.29.30,trna1_HisGTG_15_+_45493349_45493420@-1A.29.30,trna21_HisGTG_1_+_147753471_147753542@-1A.29.30,trna33_HisGTG_6_+_27125906_27125977@-1A.29.30,trna7_HisGTG_9_-_14433938_14434009@-1A.29.30,trna8_HisGTG_15_-_45492611_45492682@-1A.29.30,trna9_HisGTG_15_-_45490804_45490875@-1A.29.30\|AGCCGTGATCGTATAGTGGTTAGTACTCTG | >trna111_HisGTG_1_-_147774845_147774916 | -1A | 29 | 30 | AGCCGTGATCGTATAGTGGTTAGTACTCTG | 246.7 | -5.6 | 1.6 | -3.6 | 3.7E-04 | 1.1E-03 | BT20/MB468 | MP |
| >MI0005567\|hsa-mir-760&WithFlank&1\|+\|93846826\|93846917@55.75.21(+1U)&[MIMAT0004957&hsa-miR-760&offsets\|0\|+1(+1U);m-448&1\|+\|93846880\|93846901&offsets\|0\|-1(+1U)]\|CGGCTCTGGGTCTGTGGGGAGT | >MI0005567\|hsa-mir-760&WithFlank&1\|+\|93846826\|93846917 | 55 | 75 | 21(+1U) | CGGCTCTGGGTCTGTGGGGAGT | 334.9 | 6.0 | 1.7 | 3.6 | 3.7E-04 | 1.1E-03 | BT20/MB468 | MP |
| >hg38_21_+_8433172_8446622_RNA45SN1_45S_50ntFlanks@5083.5110.28\|ACGAACGAGACTCTGGCATGCTAACTAG | >hg38_21_+_8433172_8446622_RNA45SN1_45S_50ntFlanks | 5083 | 5110 | 28 | ACGAACGAGACTCTGGCATGCTAACTAG | 520.7 | -3.6 | 1.0 | -3.4 | 6.5E-04 | 1.1E-03 | BT20/MB468 | MITO |
| >hg38_21_+_8433172_8446622_RNA45SN1_45S_50ntFlanks@12928.12953.26\|CTGGCGCTAAACCATTCGTAGACGAC | >hg38_21_+_8433172_8446622_RNA45SN1_45S_50ntFlanks | 12928 | 12953 | 26 | CTGGCGCTAAACCATTCGTAGACGAC | 165.8 | -2.0 | 0.6 | -3.4 | 6.5E-04 | 1.1E-03 | BT20/MB468 | MITO |
| >hg38_21_+_8433172_8446622_RNA45SN1_45S_50ntFlanks@9623.9646.24\|TAGTAGCTGGTTCCCTCCGAAGTT | >hg38_21_+_8433172_8446622_RNA45SN1_45S_50ntFlanks | 9623 | 9646 | 24 | TAGTAGCTGGTTCCCTCCGAAGTT | 337.7 | -4.2 | 1.3 | -3.3 | 8.5E-04 | 1.1E-03 | MB231/MB468 | MITO |
| >hg38_21_+_8433172_8446622_RNA45SN1_45S_50ntFlanks@8659.8699.41\|CATTTCCACCGCGGCGGTGCGCCGCGACCGGCTCCGGGACG | >hg38_21_+_8433172_8446622_RNA45SN1_45S_50ntFlanks | 8659 | 8699 | 41 | CATTTCCACCGCGGCGGTGCGCCGCGACCGGCTCCGGGACG | 118.8 | -5.3 | 1.5 | -3.4 | 6.5E-04 | 1.1E-03 | BT20/MB468 | MITO |
| >MI0003137\|hsa-mir-193b&WithFlank&16\|+\|14303961\|14304055@57.76.20(+1A)&[MIMAT0002819&hsa-miR-193b-3p&offsets\|0\|-2(+1A);m-166&16\|+\|14304017\|14304038&offsets\|0\|-2(+1A)]\|AACTGGCCCTCAAAGTCCCGA | >MI0003137\|hsa-mir-193b&WithFlank&16\|+\|14303961\|14304055 | 57 | 76 | 20(+1A) | AACTGGCCCTCAAAGTCCCGA | 106.6 | 7.1 | 2.0 | 3.6 | 3.7E-04 | 1.1E-03 | BT20/MB468 | MP |
| >MI0000095\|hsa-mir-93&WithFlank&7\|-\|100093762\|100093853@17.40.24&[MIMAT0000093&hsa-miR-93-5p&offsets\|0\|+1;m-29&7\|-\|100093815\|100093837&offsets\|0\|+1]\|CAAAGTGCTGTTCGTGCAGGTAGT | >MI0000095\|hsa-mir-93&WithFlank&7\|-\|100093762\|100093853 | 17 | 40 | 24 | CAAAGTGCTGTTCGTGCAGGTAGT | 1938.3 | -3.7 | 1.0 | -3.8 | 1.3E-04 | 1.1E-03 | BT20/MB231 | MP |
| >am_trna117_GlyTCC_1_-_145397864_145397935@1.32.32,trna45_GlyTCC_1_+_161500903_161500974@1.32.32,trna70_GlyTCC_1_-_161439547_161439618@1.32.32,trna73_GlyTCC_1_-_161432166_161432237@1.32.32,trna76_GlyTCC_1_-_161424756_161424827@1.32.32,trna79_GlyTCC_1_-_161417375_161417446@1.32.32\|GCGTTGGTGGTATAGTGGTGAGCATAGCTGCC | >am_trna117_GlyTCC_1_-_145397864_145397935 | 1 | 32 | 32 | GCGTTGGTGGTATAGTGGTGAGCATAGCTGCC | 441.2 | 4.8 | 1.3 | 3.6 | 3.8E-04 | 1.1E-03 | BT20/MB468 | MP |
| >hg38_21_+_8433172_8446622_RNA45SN1_45S_50ntFlanks@6661.6678.18\|CGGTGGATCACTCGGCTC | >hg38_21_+_8433172_8446622_RNA45SN1_45S_50ntFlanks | 6661 | 6678 | 18 | CGGTGGATCACTCGGCTC | 2077.9 | 2.9 | 0.8 | 3.8 | 1.6E-04 | 1.1E-03 | BT20/MB231 | MITO |
| >hg38_21_+_8433172_8446622_RNA45SN1_45S_50ntFlanks@11778.11803.26\|ACGCGCATGAATGGATGAACGAGATT | >hg38_21_+_8433172_8446622_RNA45SN1_45S_50ntFlanks | 11778 | 11803 | 26 | ACGCGCATGAATGGATGAACGAGATT | 286.9 | -4.3 | 1.3 | -3.3 | 8.6E-04 | 1.1E-03 | MB231/MB468 | MITO |
| >hg38_21_+_8433172_8446622_RNA45SN1_45S_50ntFlanks@9546.9571.26\|TGGAGGTCCGTAGCGGTCCTGACGTG | >hg38_21_+_8433172_8446622_RNA45SN1_45S_50ntFlanks | 9546 | 9571 | 26 | TGGAGGTCCGTAGCGGTCCTGACGTG | 484.1 | -2.5 | 0.8 | -3.3 | 8.6E-04 | 1.1E-03 | MB231/MB468 | MITO |
| >hg38_21_+_8433172_8446622_RNA45SN1_45S_50ntFlanks@12156.12180.25\|GAGTTTGACTGGGGCGGTACACCTG | >hg38_21_+_8433172_8446622_RNA45SN1_45S_50ntFlanks | 12156 | 12180 | 25 | GAGTTTGACTGGGGCGGTACACCTG | 110.7 | -4.5 | 1.3 | -3.4 | 6.5E-04 | 1.1E-03 | BT20/MB468 | MITO |
| >trnaMT_GluTTC_MT_-_14674_14742@40.58.19,trnalookalike8_GluTTC_5_-_93905172_93905240@40.58.19\|TTGGTCGTGGTTGTAGTCC | >trnaMT_GluTTC_MT_-_14674_14742 | 40 | 58 | 19 | TTGGTCGTGGTTGTAGTCC | 384.0 | 3.2 | 0.9 | 3.4 | 7.2E-04 | 1.1E-03 | MB231/MB468 | MP |
| >hg38_21_+_8433172_8446622_RNA45SN1_45S_50ntFlanks@5485.5506.22\|AGCGCTGAGAAGACGGTCGAAC | >hg38_21_+_8433172_8446622_RNA45SN1_45S_50ntFlanks | 5485 | 5506 | 22 | AGCGCTGAGAAGACGGTCGAAC | 726.4 | 3.3 | 1.0 | 3.4 | 7.8E-04 | 1.1E-03 | BT20/MB468 | CYTO |
| >hg38_21_+_8433172_8446622_RNA45SN1_45S_50ntFlanks@9562.9595.34\|TCCTGACGTGCAAATCGGTCGTCCGACCTGGGTA | >hg38_21_+_8433172_8446622_RNA45SN1_45S_50ntFlanks | 9562 | 9595 | 34 | TCCTGACGTGCAAATCGGTCGTCCGACCTGGGTA | 489.1 | -2.9 | 0.8 | -3.4 | 6.5E-04 | 1.1E-03 | BT20/MB468 | MITO |
| >hg38_1_-_228634819_228635039_RNA5S12_5S_50ntFlanks@151.170.20\|ATACCGGGTGCTGTAGGCTT | >hg38_1_-_228634819_228635039_RNA5S12_5S_50ntFlanks | 151 | 170 | 20 | ATACCGGGTGCTGTAGGCTT | 1435.7 | 3.0 | 0.9 | 3.3 | 8.7E-04 | 1.1E-03 | MB231/MB468 | NUC |
| >hg38_21_+_8433172_8446622_RNA45SN1_45S_50ntFlanks@8176.8215.40\|CTTCTGATCGAGGCCCAGCCCGTGGACGGTGTGAGGCCGG | >hg38_21_+_8433172_8446622_RNA45SN1_45S_50ntFlanks | 8176 | 8215 | 40 | CTTCTGATCGAGGCCCAGCCCGTGGACGGTGTGAGGCCGG | 856.9 | -5.3 | 1.5 | -3.6 | 3.8E-04 | 1.1E-03 | BT20/MB468 | MP |
| >hg38_21_+_8433172_8446622_RNA45SN1_45S_50ntFlanks@12404.12434.31\|GCTTTTTGATCCTTCGATGTCGGCTCTTCCT | >hg38_21_+_8433172_8446622_RNA45SN1_45S_50ntFlanks | 12404 | 12434 | 31 | GCTTTTTGATCCTTCGATGTCGGCTCTTCCT | 1208.6 | -2.6 | 0.8 | -3.3 | 8.7E-04 | 1.1E-03 | MB231/MB468 | NUC |
| >MI0000272\|hsa-mir-182&WithFlank&7\|-\|129770377\|129770498@29.51.23&[MIMAT0000259&hsa-miR-182-5p&offsets\|0\|-1;m-22&7\|-\|129770449\|129770470&offsets\|0\|+1]\|TTTGGCAATGGTAGAACTCACAC | >MI0000272\|hsa-mir-182&WithFlank&7\|-\|129770377\|129770498 | 29 | 51 | 23 | TTTGGCAATGGTAGAACTCACAC | 200.1 | 3.4 | 1.0 | 3.3 | 8.6E-04 | 1.1E-03 | MB231/MB468 | MITO |
| >hg38_21_+_8433172_8446622_RNA45SN1_45S_50ntFlanks@10679.10705.27\|GGTTCCGGCGGCGTCCGGTGAGCTCTC | >hg38_21_+_8433172_8446622_RNA45SN1_45S_50ntFlanks | 10679 | 10705 | 27 | GGTTCCGGCGGCGTCCGGTGAGCTCTC | 537.1 | -6.1 | 1.8 | -3.3 | 8.7E-04 | 1.1E-03 | BT20/MB468 | NUC |
| >hg38_21_+_8433172_8446622_RNA45SN1_45S_50ntFlanks@9554.9582.29\|CGTAGCGGTCCTGACGTGCAAATCGGTCG | >hg38_21_+_8433172_8446622_RNA45SN1_45S_50ntFlanks | 9554 | 9582 | 29 | CGTAGCGGTCCTGACGTGCAAATCGGTCG | 274.7 | -1.8 | 0.5 | -3.4 | 6.5E-04 | 1.1E-03 | BT20/MB468 | MITO |
| >hg38_21_+_8433172_8446622_RNA45SN1_45S_50ntFlanks@11102.11121.20\|AGAAGGGTCGGGGCGGCAGG | >hg38_21_+_8433172_8446622_RNA45SN1_45S_50ntFlanks | 11102 | 11121 | 20 | AGAAGGGTCGGGGCGGCAGG | 131.2 | 10.4 | 2.7 | 3.8 | 1.3E-04 | 1.1E-03 | BT20/MB231 | MP |
| >hg38_21_+_8433172_8446622_RNA45SN1_45S_50ntFlanks@12603.12624.22\|TGTATGTGCTTGGCTGAGGAGC | >hg38_21_+_8433172_8446622_RNA45SN1_45S_50ntFlanks | 12603 | 12624 | 22 | TGTATGTGCTTGGCTGAGGAGC | 399.4 | 4.0 | 1.2 | 3.3 | 8.1E-04 | 1.1E-03 | MB231/MB468 | CYTO |
| >hg38_21_+_8433172_8446622_RNA45SN1_45S_50ntFlanks@3771.3798.28\|CACGGCCGGTACAGTGAAACTGCGAATG | >hg38_21_+_8433172_8446622_RNA45SN1_45S_50ntFlanks | 3771 | 3798 | 28 | CACGGCCGGTACAGTGAAACTGCGAATG | 131.4 | -5.3 | 1.6 | -3.4 | 7.3E-04 | 1.1E-03 | MB231/MB468 | MP |
| >hg38_21_+_8433172_8446622_RNA45SN1_45S_50ntFlanks@11564.11601.38\|CCTAGCAGCCGACTTAGAACTGGTGCGGACCAGGGGAA | >hg38_21_+_8433172_8446622_RNA45SN1_45S_50ntFlanks | 11564 | 11601 | 38 | CCTAGCAGCCGACTTAGAACTGGTGCGGACCAGGGGAA | 1401.9 | -4.4 | 1.3 | -3.4 | 7.8E-04 | 1.1E-03 | BT20/MB468 | CYTO |
| >hg38_21_+_8433172_8446622_RNA45SN1_45S_50ntFlanks@4557.4576.20\|CAGCTAGGAATAATGGAATA | >hg38_21_+_8433172_8446622_RNA45SN1_45S_50ntFlanks | 4557 | 4576 | 20 | CAGCTAGGAATAATGGAATA | 159.3 | -2.5 | 0.7 | -3.3 | 8.7E-04 | 1.1E-03 | MB231/MB468 | NUC |
| >hg38_21_+_8433172_8446622_RNA45SN1_45S_50ntFlanks@11924.11949.26\|GAGGTGTAGAATAAGTGGGAGGCCCC | >hg38_21_+_8433172_8446622_RNA45SN1_45S_50ntFlanks | 11924 | 11949 | 26 | GAGGTGTAGAATAAGTGGGAGGCCCC | 575.8 | 4.8 | 1.3 | 3.6 | 3.8E-04 | 1.1E-03 | BT20/MB468 | MP |
| >hg38_21_+_8433172_8446622_RNA45SN1_45S_50ntFlanks@10780.10801.22\|AAGGTGAACAGCCTCTGGCATG | >hg38_21_+_8433172_8446622_RNA45SN1_45S_50ntFlanks | 10780 | 10801 | 22 | AAGGTGAACAGCCTCTGGCATG | 245.2 | -1.6 | 0.5 | -3.3 | 8.2E-04 | 1.1E-03 | MB231/MB468 | CYTO |
| >hg38_21_+_8433172_8446622_RNA45SN1_45S_50ntFlanks@7974.7996.23\|ACGCGACCTCAGATCAGACGTGG | >hg38_21_+_8433172_8446622_RNA45SN1_45S_50ntFlanks | 7974 | 7996 | 23 | ACGCGACCTCAGATCAGACGTGG | 1471.6 | 2.8 | 0.8 | 3.4 | 6.5E-04 | 1.1E-03 | BT20/MB468 | MITO |
| >hg38_21_+_8433172_8446622_RNA45SN1_45S_50ntFlanks@6648.6695.48\|GTACGACTCTTAGCGGTGGATCACTCGGCTCGTGCGTCGATGAAGAAC | >hg38_21_+_8433172_8446622_RNA45SN1_45S_50ntFlanks | 6648 | 6695 | 48 | GTACGACTCTTAGCGGTGGATCACTCGGCTCGTGCGTCGATGAAGAAC | 1206.2 | 3.8 | 1.1 | 3.3 | 8.6E-04 | 1.1E-03 | MB231/MB468 | MITO |

| | | | | | | | | | | | | | |
|---|---|---|---|---|---|---|---|---|---|---|---|---|---|
| >hg38_MT_+_1621_3279_RNR2_16S_50ntFlanks@1182.1215.34\|AAAATTTCGGTTGGGGCGACCTCGGAGCAGAACC | >hg38_MT_+_1621_3279_RNR2_16S_50ntFlanks | 1182 | 1215 | 34 | AAAATTTCGGTTGGGGCGACCTCGGAGCAGAACC | 280.6 | 5.0 | 1.4 | 3.6 | 3.8E-04 | 1.1E-03 | BT20/MB468 | MP |
| >hg38_21_+_8433172_8446622_RNA45SN1_45S_50ntFlanks@10350.10379.30\|ACGAGAACTTTGAAGGCCGAAGTGGAGAAG | >hg38_21_+_8433172_8446622_RNA45SN1_45S_50ntFlanks | 10350 | 10379 | 30 | ACGAGAACTTTGAAGGCCGAAGTGGAGAAG | 514.2 | -3.9 | 1.2 | -3.3 | 8.7E-04 | 1.1E-03 | MB231/MB468 | MITO |
| >hg38_21_+_8433172_8446622_RNA45SN1_45S_50ntFlanks@11641.11665.25\|CGGCGGGTGTTGACGCGATGTGATT | >hg38_21_+_8433172_8446622_RNA45SN1_45S_50ntFlanks | 11641 | 11665 | 25 | CGGCGGGTGTTGACGCGATGTGATT | 1170.1 | -4.4 | 1.3 | -3.4 | 7.8E-04 | 1.1E-03 | BT20/MB468 | CYTO |
| >MI0000809\|hsa-mir-151a&WithFlank&8\|-\|140732558\|140732659@53.75.23(+1U)&[MIMAT0000757&hsa-miR-151a-3p&offsets\|0\|+2(+1U);m-30&8\|-\|140732587\|140732607&offsets\|0\|+2(+1U)]\|CTAGACTGAAGCTCCTTGAGGACT | >MI0000809\|hsa-mir-151a&WithFlank&8\|-\|140732558\|140732659 | 53 | 75 | 23(+1U) | CTAGACTGAAGCTCCTTGAGGACT | 190.3 | 5.0 | 1.4 | 3.6 | 3.8E-04 | 1.1E-03 | BT20/MB468 | MP |
| >MI0015866\|hsa-mir-4326&WithFlank&20\|+\|63286802\|63286872@17.40.24&[MIMAT0016888&hsa-miR-4326&offsets\|0\|+4;m-339&20\|+\|63286818\|63286841&offsets\|0\|0]\|TGTTCCTCTGTCTCCCAGACTCTG | >MI0015866\|hsa-mir-4326&WithFlank&20\|+\|63286802\|63286872 | 17 | 40 | 24 | TGTTCCTCTGTCTCCCAGACTCTG | 439.6 | 4.9 | 1.4 | 3.6 | 3.8E-04 | 1.1E-03 | BT20/MB468 | MP |
| >hg38_21_+_8433172_8446622_RNA45SN1_45S_50ntFlanks@11203.11229.27\|CGGCGGCGGCGACTCTGGACGCGAGCC | >hg38_21_+_8433172_8446622_RNA45SN1_45S_50ntFlanks | 11203 | 11229 | 27 | CGGCGGCGGCGACTCTGGACGCGAGCC | 120.0 | 10.2 | 2.7 | 3.8 | 1.3E-04 | 1.1E-03 | BT20/MB231 | MP |
| >hg38_21_+_8433172_8446622_RNA45SN1_45S_50ntFlanks@8105.8141.37\|CGCGGCGGGGCGCGGGACATGTGGCGTACGGAAGACC | >hg38_21_+_8433172_8446622_RNA45SN1_45S_50ntFlanks | 8105 | 8141 | 37 | CGCGGCGGGGCGCGGGACATGTGGCGTACGGAAGACC | 964.5 | 4.7 | 1.2 | 3.8 | 1.3E-04 | 1.1E-03 | BT20/MB231 | MP |
| >hg38_21_+_8433172_8446622_RNA45SN1_45S_50ntFlanks@11577.11611.35\|TTAGAACTGGTGCGGACCAGGGGAATCCGACTGTT | >hg38_21_+_8433172_8446622_RNA45SN1_45S_50ntFlanks | 11577 | 11611 | 35 | TTAGAACTGGTGCGGACCAGGGGAATCCGACTGTT | 258.4 | -3.2 | 0.9 | -3.3 | 8.2E-04 | 1.1E-03 | MB231/MB468 | CYTO |
| >MI0000746\|hsa-mir-99b&WithFlank&19\|+\|51692606\|51692687@13.32.20(+2G)&[MIMAT0000689&hsa-miR-99b-5p&offsets\|0\|-2(+2G);m-10&19\|+\|51692618\|51692639&offsets\|0\|-2(+2G)]\|CACCCGTAGAACCGACCTTGGG | >MI0000746\|hsa-mir-99b&WithFlank&19\|+\|51692606\|51692687 | 13 | 32 | 20(+2G) | CACCCGTAGAACCGACCTTGGG | 343.9 | 3.5 | 0.9 | 3.8 | 1.4E-04 | 1.1E-03 | BT20/MB231 | CYTO |
| >trna152_ValCAC_6_-_27248049_27248121@1.34.34\|GCTTCTGTAGTGTAGTGGTTATCACGTTCGCCTC | >trna152_ValCAC_6_-_27248049_27248121 | 1 | 34 | 34 | GCTTCTGTAGTGTAGTGGTTATCACGTTCGCCTC | 13469.6 | -4.1 | 1.1 | -3.8 | 1.4E-04 | 1.1E-03 | BT20/MB231 | CYTO |
| >hg38_21_+_8433172_8446622_RNA45SN1_45S_50ntFlanks@5492.5512.21\|AGAAGACGGTCGAACTTGACT | >hg38_21_+_8433172_8446622_RNA45SN1_45S_50ntFlanks | 5492 | 5512 | 21 | AGAAGACGGTCGAACTTGACT | 2168.0 | 4.3 | 1.1 | 3.8 | 1.4E-04 | 1.1E-03 | BT20/MB231 | CYTO |
| >hg38_21_+_8433172_8446622_RNA45SN1_45S_50ntFlanks@11567.11586.20\|AGCAGCCGACTTAGAACTGG | >hg38_21_+_8433172_8446622_RNA45SN1_45S_50ntFlanks | 11567 | 11586 | 20 | AGCAGCCGACTTAGAACTGG | 1396.3 | 2.5 | 0.8 | 3.3 | 8.8E-04 | 1.1E-03 | MB231/MB468 | NUC |
| >trnaMT_LeuTAA_MT_+_3230_3304@22.44.23\|AATCGCATAAAACTTAAAACTTT | >trnaMT_LeuTAA_MT_+_3230_3304 | 22 | 44 | 23 | AATCGCATAAAACTTAAAACTTT | 79.3 | -5.4 | 1.6 | -3.4 | 7.3E-04 | 1.1E-03 | MB231/MB468 | MP |
| >hg38_21_+_8433172_8446622_RNA45SN1_45S_50ntFlanks@4007.4042.36\|CCTCGGGCCGATCGCACGCCCCCCGTGGCGGCGACG | >hg38_21_+_8433172_8446622_RNA45SN1_45S_50ntFlanks | 4007 | 4042 | 36 | CCTCGGGCCGATCGCACGCCCCCCGTGGCGGCGACG | 64.4 | -5.2 | 1.5 | -3.4 | 7.3E-04 | 1.1E-03 | MB231/MB468 | MP |
| >MI0000063\|hsa-let-7b&WithFlank&22\|+\|46113680\|46113774@12.32.21(+1G)&[MIMAT0000063&hsa-let-7b-5p&offsets\|0\|-1(+1G);m-14&22\|+\|46113691\|46113712&offsets\|0\|-1(+1G)]\|TGAGGTAGTAGGTTGTGTGGTG | >MI0000063\|hsa-let-7b&WithFlank&22\|+\|46113680\|46113774 | 12 | 32 | 21(+1G) | TGAGGTAGTAGGTTGTGTGGTG | 1515.3 | 3.7 | 1.0 | 3.8 | 1.4E-04 | 1.1E-03 | BT20/MB231 | CYTO |
| >hg38_21_+_8433172_8446622_RNA45SN1_45S_50ntFlanks@10351.10370.20\|CGAGAACTTTGAAGGCCGAA | >hg38_21_+_8433172_8446622_RNA45SN1_45S_50ntFlanks | 10351 | 10370 | 20 | CGAGAACTTTGAAGGCCGAA | 222.2 | -2.4 | 0.7 | -3.3 | 8.8E-04 | 1.1E-03 | MB231/MB468 | NUC |
| >hg38_21_+_8433172_8446622_RNA45SN1_45S_50ntFlanks@5486.5512.27\|GCGCTGAGAAGACGGTCGAACTTGACT | >hg38_21_+_8433172_8446622_RNA45SN1_45S_50ntFlanks | 5486 | 5512 | 27 | GCGCTGAGAAGACGGTCGAACTTGACT | 1208.1 | 4.0 | 1.1 | 3.8 | 1.4E-04 | 1.1E-03 | BT20/MB231 | CYTO |
| >hg38_21_+_8433172_8446622_RNA45SN1_45S_50ntFlanks@11575.11604.30\|ACTTAGAACTGGTGCGGACCAGGGGAATCC | >hg38_21_+_8433172_8446622_RNA45SN1_45S_50ntFlanks | 11575 | 11604 | 30 | ACTTAGAACTGGTGCGGACCAGGGGAATCC | 158.0 | -2.0 | 0.6 | -3.4 | 7.9E-04 | 1.1E-03 | BT20/MB468 | CYTO |
| >hg38_21_+_8433172_8446622_RNA45SN1_45S_50ntFlanks@4738.4767.30\|AAAGTCGGAGGTTCGAAGACGATCAGATAC | >hg38_21_+_8433172_8446622_RNA45SN1_45S_50ntFlanks | 4738 | 4767 | 30 | AAAGTCGGAGGTTCGAAGACGATCAGATAC | 185.7 | 3.3 | 0.9 | 3.6 | 3.8E-04 | 1.1E-03 | BT20/MB468 | MP |
| >hg38_21_+_8433172_8446622_RNA45SN1_45S_50ntFlanks@7989.8008.20\|AGACGTGGCGACCCGCTGAA | >hg38_21_+_8433172_8446622_RNA45SN1_45S_50ntFlanks | 7989 | 8008 | 20 | AGACGTGGCGACCCGCTGAA | 6633.1 | -2.5 | 0.7 | -3.4 | 6.6E-04 | 1.1E-03 | BT20/MB468 | MITO |
| >hg38_21_+_8433172_8446622_RNA45SN1_45S_50ntFlanks@9485.9505.21\|TGTTAGGACCCGAAAGATGGT | >hg38_21_+_8433172_8446622_RNA45SN1_45S_50ntFlanks | 9485 | 9505 | 21 | TGTTAGGACCCGAAAGATGGT | 69.4 | 6.5 | 1.8 | 3.6 | 3.8E-04 | 1.1E-03 | BT20/MB468 | MP |
| >MI0000469\|hsa-mir-125a&WithFlank&19\|+\|51693248\|51693345@21.41.21(+1C)&[MIMAT0000443&hsa-miR-125a-5p&offsets\|0\|-3(+1C);m-63&19\|+\|51693268\|51693289&offsets\|0\|-1(+1C)]\|TCCCTGAGACCCTTTAACCTGC | >MI0000469\|hsa-mir-125a&WithFlank&19\|+\|51693248\|51693345 | 21 | 41 | 21(+1C) | TCCCTGAGACCCTTTAACCTGC | 243.8 | 1.7 | 0.5 | 3.3 | 8.8E-04 | 1.1E-03 | BT20/MB468 | NUC |
| >MI0000239\|hsa-mir-197&WithFlank&1\|+\|109598887\|109598973@57.75.19&[MIMAT0000227&hsa-miR-197-3p&offsets\|+3\|0;m-99&1\|+\|109598940\|109598961&offsets\|+3\|0]\|ACCACCTTCTCCACCCAGC | >MI0000239\|hsa-mir-197&WithFlank&1\|+\|109598887\|109598973 | 57 | 75 | 19 | ACCACCTTCTCCACCCAGC | 250.8 | 4.1 | 1.1 | 3.8 | 1.4E-04 | 1.1E-03 | BT20/MB231 | CYTO |
| >hg38_21_+_8433172_8446622_RNA45SN1_45S_50ntFlanks@11575.11600.26\|ACTTAGAACTGGTGCGGACCAGGGGA | >hg38_21_+_8433172_8446622_RNA45SN1_45S_50ntFlanks | 11575 | 11600 | 26 | ACTTAGAACTGGTGCGGACCAGGGGA | 640.2 | -4.7 | 1.3 | -3.6 | 3.8E-04 | 1.1E-03 | BT20/MB468 | MP |
| >hg38_21_+_8433172_8446622_RNA45SN1_45S_50ntFlanks@10392.10418.27\|ACAGCAGTTGAACATGGGTCAGTCGGT | >hg38_21_+_8433172_8446622_RNA45SN1_45S_50ntFlanks | 10392 | 10418 | 27 | ACAGCAGTTGAACATGGGTCAGTCGGT | 192.6 | -2.5 | 0.7 | -3.4 | 7.9E-04 | 1.1E-03 | BT20/MB468 | CYTO |
| >hg38_21_+_8433172_8446622_RNA45SN1_45S_50ntFlanks@11583.11600.18\|CTGGTGCGGACCAGGGGA | >hg38_21_+_8433172_8446622_RNA45SN1_45S_50ntFlanks | 11583 | 11600 | 18 | CTGGTGCGGACCAGGGGA | 396.3 | -4.6 | 1.3 | -3.6 | 3.8E-04 | 1.1E-03 | BT20/MB468 | MP |
| >hg38_21_+_8433172_8446622_RNA45SN1_45S_50ntFlanks@12641.12661.21\|CCATCTGTGGGATTATGACTG | >hg38_21_+_8433172_8446622_RNA45SN1_45S_50ntFlanks | 12641 | 12661 | 21 | CCATCTGTGGGATTATGACTG | 364.1 | -2.3 | 0.7 | -3.3 | 8.3E-04 | 1.1E-03 | MB231/MB468 | CYTO |
| >hg38_21_+_8433172_8446622_RNA45SN1_45S_50ntFlanks@8169.8214.46\|CCAAGTCCTTCTGATCGAGGCCCAGCCCGTGGACGGTGTGAGGCCG | >hg38_21_+_8433172_8446622_RNA45SN1_45S_50ntFlanks | 8169 | 8214 | 46 | CCAAGTCCTTCTGATCGAGGCCCAGCCCGTGGACGGTGTGAGGCCG | 513.2 | -4.3 | 1.3 | -3.3 | 8.9E-04 | 1.1E-03 | MB231/MB468 | NUC |
| >am_trnaMT_GluTTC_MT_-_14674_14742@20.53.34,trnalookalike8_GluTTC_5_-_93905172_93905240@20.53.34\|AACGATGGTTTTTCATATCATTGGTCGTGGTTGT | >am_trnaMT_GluTTC_MT_-_14674_14742 | 20 | 53 | 34 | AACGATGGTTTTTCATATCATTGGTCGTGGTTGT | 116.3 | 7.1 | 2.0 | 3.6 | 3.8E-04 | 1.1E-03 | BT20/MB468 | MP |
| >trnaMT_ValTAC_MT_+_1602_1670@33.70.38\|ACACTTAGGAGATTTCAACTTAACTTGACCGCTCTGAC | >trnaMT_ValTAC_MT_+_1602_1670 | 33 | 70 | 38 | ACACTTAGGAGATTTCAACTTAACTTGACCGCTCTGAC | 61.9 | 9.7 | 2.7 | 3.6 | 3.8E-04 | 1.1E-03 | BT20/MB468 | MP |
| >hg38_21_+_8433172_8446622_RNA45SN1_45S_50ntFlanks@10247.10276.30\|GCGACGAGTAGGAGGGCCGCTGCGGTGAGC | >hg38_21_+_8433172_8446622_RNA45SN1_45S_50ntFlanks | 10247 | 10276 | 30 | GCGACGAGTAGGAGGGCCGCTGCGGTGAGC | 100.4 | 6.6 | 1.9 | 3.6 | 3.8E-04 | 1.1E-03 | BT20/MB468 | MP |
| >hg38_MT_+_1621_3279_RNR2_16S_50ntFlanks@617.652.36\|AAACATATAACTGAACTCCTCACACCCAATTGGACC | >hg38_MT_+_1621_3279_RNR2_16S_50ntFlanks | 617 | 652 | 36 | AAACATATAACTGAACTCCTCACACCCAATTGGACC | 356.4 | 5.6 | 1.6 | 3.6 | 3.8E-04 | 1.1E-03 | BT20/MB468 | MP |
| >hg38_21_+_8433172_8446622_RNA45SN1_45S_50ntFlanks@11853.11872.20\|CTTGGCGGAATCAGCGGGGA | >hg38_21_+_8433172_8446622_RNA45SN1_45S_50ntFlanks | 11853 | 11872 | 20 | CTTGGCGGAATCAGCGGGGA | 312.4 | -5.6 | 1.6 | -3.6 | 3.8E-04 | 1.1E-03 | BT20/MB468 | MP |
| >hg38_21_+_8433172_8446622_RNA45SN1_45S_50ntFlanks@10786.10821.36\|AACAGCCTCTGGCATGTTGGAACAATGTAGGTAAGG | >hg38_21_+_8433172_8446622_RNA45SN1_45S_50ntFlanks | 10786 | 10821 | 36 | AACAGCCTCTGGCATGTTGGAACAATGTAGGTAAGG | 403.0 | -5.9 | 1.8 | -3.3 | 8.8E-04 | 1.1E-03 | BT20/MB468 | NUC |
| >hg38_21_+_8433172_8446622_RNA45SN1_45S_50ntFlanks@5431.5466.36\|GTTTAGTGAGGCCCTCGGATCGGCCCCGCCGGGGTC | >hg38_21_+_8433172_8446622_RNA45SN1_45S_50ntFlanks | 5431 | 5466 | 36 | GTTTAGTGAGGCCCTCGGATCGGCCCCGCCGGGGTC | 429.1 | -5.9 | 1.8 | -3.3 | 8.8E-04 | 1.1E-03 | BT20/MB468 | NUC |
| >hg38_21_+_8433172_8446622_RNA45SN1_45S_50ntFlanks@8117.8135.19\|CGGGACATGTGGCGTACGG | >hg38_21_+_8433172_8446622_RNA45SN1_45S_50ntFlanks | 8117 | 8135 | 19 | CGGGACATGTGGCGTACGG | 210.2 | 4.7 | 1.3 | 3.6 | 3.8E-04 | 1.1E-03 | BT20/MB468 | MP |
| >hg38_21_+_8433172_8446622_RNA45SN1_45S_50ntFlanks@7976.8010.35\|GCGACCTCAGATCAGACGTGGCGACCCGCTGAATT | >hg38_21_+_8433172_8446622_RNA45SN1_45S_50ntFlanks | 7976 | 8010 | 35 | GCGACCTCAGATCAGACGTGGCGACCCGCTGAATT | 283.0 | 2.6 | 0.8 | 3.3 | 8.8E-04 | 1.1E-03 | MB231/MB468 | MITO |
| >hg38_21_+_8433172_8446622_RNA45SN1_45S_50ntFlanks@6651.6698.48\|CGACTCTTAGCGGTGGATCACTCGGCTCGTGCGTCGATGAAGAACGCA | >hg38_21_+_8433172_8446622_RNA45SN1_45S_50ntFlanks | 6651 | 6698 | 48 | CGACTCTTAGCGGTGGATCACTCGGCTCGTGCGTCGATGAAGAACGCA | 11450.4 | -3.5 | 1.0 | -3.3 | 8.9E-04 | 1.1E-03 | BT20/MB468 | NUC |
| >MI0000342\|hsa-mir-200b&WithFlank&1\|+\|1167098\|1167204@63.85.23(+1U)&[MIMAT0000318&hsa-miR-200b-3p&offsets\|0\|+1(+1U);m-93&1\|+\|1167160\|1167182&offsets\|0\|0(+1U)]\|TAATACTGCCTGGTAATGATGACT | >MI0000342\|hsa-mir-200b&WithFlank&1\|+\|1167098\|1167204 | 63 | 85 | 23(+1U) | TAATACTGCCTGGTAATGATGACT | 1529.1 | -3.3 | 1.0 | -3.3 | 8.8E-04 | 1.1E-03 | MB231/MB468 | MITO |
| >hg38_21_+_8433172_8446622_RNA45SN1_45S_50ntFlanks@7983.8018.36\|CAGATCAGACGTGGCGACCCGCTGAATTTAAGCATA | >hg38_21_+_8433172_8446622_RNA45SN1_45S_50ntFlanks | 7983 | 8018 | 36 | CAGATCAGACGTGGCGACCCGCTGAATTTAAGCATA | 315.9 | -2.9 | 0.9 | -3.3 | 8.8E-04 | 1.1E-03 | MB231/MB468 | MITO |
| >hg38_21_+_8433172_8446622_RNA45SN1_45S_50ntFlanks@9933.9951.19\|TAGACAGCAGGACGGTGGC | >hg38_21_+_8433172_8446622_RNA45SN1_45S_50ntFlanks | 9933 | 9951 | 19 | TAGACAGCAGGACGGTGGC | 183.8 | 5.8 | 1.6 | 3.6 | 3.8E-04 | 1.1E-03 | BT20/MB468 | MP |
| >hg38_21_+_8433172_8446622_RNA45SN1_45S_50ntFlanks@8123.8141.19\|ATGTGGCGTACGGAAGACC | >hg38_21_+_8433172_8446622_RNA45SN1_45S_50ntFlanks | 8123 | 8141 | 19 | ATGTGGCGTACGGAAGACC | 213.3 | 5.4 | 1.4 | 3.8 | 1.3E-04 | 1.1E-03 | BT20/MB231 | MP |
| >hg38_21_+_8433172_8446622_RNA45SN1_45S_50ntFlanks@11784.11805.22\|ATGAATGGATGAACGAGATTCC | >hg38_21_+_8433172_8446622_RNA45SN1_45S_50ntFlanks | 11784 | 11805 | 22 | ATGAATGGATGAACGAGATTCC | 599.2 | -4.6 | 1.4 | -3.4 | 7.9E-04 | 1.1E-03 | BT20/MB468 | CYTO |
| >hg38_21_+_8433172_8446622_RNA45SN1_45S_50ntFlanks@11279.11296.18\|CCGGGGAGCCCGGCGGGC | >hg38_21_+_8433172_8446622_RNA45SN1_45S_50ntFlanks | 11279 | 11296 | 18 | CCGGGGAGCCCGGCGGGC | 56.7 | 9.0 | 2.7 | 3.4 | 7.4E-04 | 1.1E-03 | MB231/MB468 | MP |
| >hg38_21_+_8433172_8446622_RNA45SN1_45S_50ntFlanks@10059.10078.20\|GGCAGTCGAGAGTGGACGGG | >hg38_21_+_8433172_8446622_RNA45SN1_45S_50ntFlanks | 10059 | 10078 | 20 | GGCAGTCGAGAGTGGACGGG | 205.6 | 4.6 | 1.3 | 3.6 | 3.8E-04 | 1.1E-03 | BT20/MB468 | MP |
| >hg38_21_+_8433172_8446622_RNA45SN1_45S_50ntFlanks@7983.8005.23\|CAGATCAGACGTGGCGACCCGCT | >hg38_21_+_8433172_8446622_RNA45SN1_45S_50ntFlanks | 7983 | 8005 | 23 | CAGATCAGACGTGGCGACCCGCT | 573.4 | 7.5 | 2.0 | 3.8 | 1.3E-04 | 1.1E-03 | BT20/MB231 | MP |
| >hg38_21_+_8433172_8446622_RNA45SN1_45S_50ntFlanks@12415.12437.23\|CTTCGATGTCGGCTCTTCCTATC | >hg38_21_+_8433172_8446622_RNA45SN1_45S_50ntFlanks | 12415 | 12437 | 23 | CTTCGATGTCGGCTCTTCCTATC | 3748.2 | -2.1 | 0.6 | -3.3 | 8.9E-04 | 1.1E-03 | MB231/MB468 | NUC |
| >MI0001733\|hsa-mir-452&WithFlank&X\|-\|151959622\|151959718@58.79.22&[MIMAT0001636&hsa-miR-452-3p&offsets\|-6\|-6;m-457&X\|-\|151959639\|151959660&offsets\|-1\|-1]\|CTCAGTCTCATCTGCAAAGAAG | >MI0001733\|hsa-mir-452&WithFlank&X\|-\|151959622\|151959718 | 58 | 79 | 22 | CTCAGTCTCATCTGCAAAGAAG | 58.8 | 9.6 | 2.7 | 3.6 | 3.9E-04 | 1.1E-03 | BT20/MB468 | MP |
| >hg38_21_+_8433172_8446622_RNA45SN1_45S_50ntFlanks@9739.9769.31\|AAACGATCTCAACCTATTCTCAAACTTTAAA | >hg38_21_+_8433172_8446622_RNA45SN1_45S_50ntFlanks | 9739 | 9769 | 31 | AAACGATCTCAACCTATTCTCAAACTTTAAA | 183.4 | -2.6 | 0.8 | -3.3 | 8.9E-04 | 1.1E-03 | MB231/MB468 | NUC |
| >hg38_21_+_8433172_8446622_RNA45SN1_45S_50ntFlanks@6652.6695.44\|GACTCTTAGCGGTGGATCACTCGGCTCGTGCGTCGATGAAGAAC | >hg38_21_+_8433172_8446622_RNA45SN1_45S_50ntFlanks | 6652 | 6695 | 44 | GACTCTTAGCGGTGGATCACTCGGCTCGTGCGTCGATGAAGAAC | 1858.3 | 5.3 | 1.5 | 3.5 | 3.9E-04 | 1.1E-03 | BT20/MB468 | MP |
| >hg38_21_+_8433172_8446622_RNA45SN1_45S_50ntFlanks@3840.3859.20\|CTCCTACTTGGATAACTGTG | >hg38_21_+_8433172_8446622_RNA45SN1_45S_50ntFlanks | 3840 | 3859 | 20 | CTCCTACTTGGATAACTGTG | 361.3 | -3.2 | 1.0 | -3.3 | 8.8E-04 | 1.1E-03 | MB231/MB468 | MITO |
| >hg38_21_+_8433172_8446622_RNA45SN1_45S_50ntFlanks@5472.5513.42\|ACGGCCCTGGCGGAGCGCTGAGAAGACGGTCGAACTTGACTA | >hg38_21_+_8433172_8446622_RNA45SN1_45S_50ntFlanks | 5472 | 5513 | 42 | ACGGCCCTGGCGGAGCGCTGAGAAGACGGTCGAACTTGACTA | 289.0 | 3.2 | 1.0 | 3.3 | 8.9E-04 | 1.1E-03 | MB231/MB468 | NUC |
| >hg38_MT_+_1621_3279_RNR2_16S_50ntFlanks@285.305.21\|CGAAACCAGACGAGCTACCTA | >hg38_MT_+_1621_3279_RNR2_16S_50ntFlanks | 285 | 305 | 21 | CGAAACCAGACGAGCTACCTA | 214.8 | 5.0 | 1.4 | 3.5 | 3.9E-04 | 1.1E-03 | BT20/MB468 | MP |

<table>
<tr><td>>hg38_21_+_8433172_8446622_RNA45SN1_45S_50ntFlanks@4370.4389.20|TTAAAAAGCTCGTAGTTGGA</td><td>>hg38_21_+_8433172_8446622_RNA45SN1_45S_50ntFlanks</td><td>4370</td><td>4389</td><td>20</td><td>TTAAAAAGCTCGTAGTTGGA</td><td>260.3</td><td>-4.0</td><td>1.2</td><td>-3.4</td><td>8.0E-04</td><td>1.1E-03</td><td>BT20/MB468</td><td>CYTO</td></tr>
<tr><td>>hg38_21_+_8433172_8446622_RNA45SN1_45S_50ntFlanks@6648.6680.33|GTACGACTCTTAGCGGTGGATCACTCGGCTCGT</td><td>>hg38_21_+_8433172_8446622_RNA45SN1_45S_50ntFlanks</td><td>6648</td><td>6680</td><td>33</td><td>GTACGACTCTTAGCGGTGGATCACTCGGCTCGT</td><td>1179.7</td><td>2.5</td><td>0.8</td><td>3.3</td><td>8.9E-04</td><td>1.1E-03</td><td>MB231/MB468</td><td>NUC</td></tr>
<tr><td>>hg38_21_+_8433172_8446622_RNA45SN1_45S_50ntFlanks@12487.12509.23|GAACGTGAGCTGGGTTTAGACCG</td><td>>hg38_21_+_8433172_8446622_RNA45SN1_45S_50ntFlanks</td><td>12487</td><td>12509</td><td>23</td><td>GAACGTGAGCTGGGTTTAGACCG</td><td>606.6</td><td>-6.1</td><td>1.8</td><td>-3.3</td><td>8.9E-04</td><td>1.1E-03</td><td>BT20/MB468</td><td>NUC</td></tr>
<tr><td>>hg38_21_+_8433172_8446622_RNA45SN1_45S_50ntFlanks@5472.5489.18|ACGGCCCTGGCGGAGCGC</td><td>>hg38_21_+_8433172_8446622_RNA45SN1_45S_50ntFlanks</td><td>5472</td><td>5489</td><td>18</td><td>ACGGCCCTGGCGGAGCGC</td><td>185.5</td><td>5.9</td><td>1.7</td><td>3.5</td><td>3.9E-04</td><td>1.1E-03</td><td>BT20/MB468</td><td>MP</td></tr>
<tr><td>>hg38_21_+_8433172_8446622_RNA45SN1_45S_50ntFlanks@5080.5104.25|ATAACGAACGAGACTCTGGCATGCT</td><td>>hg38_21_+_8433172_8446622_RNA45SN1_45S_50ntFlanks</td><td>5080</td><td>5104</td><td>25</td><td>ATAACGAACGAGACTCTGGCATGCT</td><td>178.7</td><td>-5.6</td><td>1.7</td><td>-3.4</td><td>7.4E-04</td><td>1.1E-03</td><td>MB231/MB468</td><td>MP</td></tr>
<tr><td>>trna10_LysCTT_16_+_3241501_3241573@5.36.32,trna13_LysCTT_14_-_58706613_58706685@5.36.32,trna2_LysCTT_15_+_79152904_79152976@5.36.32|GGCTAGCTCAGTCGGTAGAGCATGGGACTCTT</td><td>>trna10_LysCTT_16_+_3241501_3241573</td><td>5</td><td>36</td><td>32</td><td>GGCTAGCTCAGTCGGTAGAGCATGGGACTCTT</td><td>620.0</td><td>-3.3</td><td>0.9</td><td>-3.8</td><td>1.6E-04</td><td>1.1E-03</td><td>BT20/MB231</td><td>MITO</td></tr>
<tr><td>>hg38_21_+_8433172_8446622_RNA45SN1_45S_50ntFlanks@9551.9569.19|GTCCGTAGCGGTCCTGACG</td><td>>hg38_21_+_8433172_8446622_RNA45SN1_45S_50ntFlanks</td><td>9551</td><td>9569</td><td>19</td><td>GTCCGTAGCGGTCCTGACG</td><td>256.6</td><td>2.2</td><td>0.7</td><td>3.3</td><td>9.0E-04</td><td>1.1E-03</td><td>MB231/MB468</td><td>NUC</td></tr>
<tr><td>>hg38_MT_+_598_1651_RNR1_12S_50ntFlanks@843.873.31|GAGTAGAGTGCTTAGTTGAACAGGGCCCTGA</td><td>>hg38_MT_+_598_1651_RNR1_12S_50ntFlanks</td><td>843</td><td>873</td><td>31</td><td>GAGTAGAGTGCTTAGTTGAACAGGGCCCTGA</td><td>115.2</td><td>6.9</td><td>2.0</td><td>3.5</td><td>3.9E-04</td><td>1.1E-03</td><td>BT20/MB468</td><td>MP</td></tr>
<tr><td>>hg38_21_+_8433172_8446622_RNA45SN1_45S_50ntFlanks@11783.11821.39|CATGAATGGATGAACGAGATTCCCACTGTCCCTACCTAC</td><td>>hg38_21_+_8433172_8446622_RNA45SN1_45S_50ntFlanks</td><td>11783</td><td>11821</td><td>39</td><td>CATGAATGGATGAACGAGATTCCCACTGTCCCTACCTAC</td><td>157.0</td><td>-2.6</td><td>0.8</td><td>-3.4</td><td>6.7E-04</td><td>1.1E-03</td><td>BT20/MB468</td><td>MITO</td></tr>
<tr><td>>MI0000063|hsa-let-7b&WithFlank&22|+|46113680|46113774@12.31.20(+1C)&[MIMAT0000063&hsa-let-7b-5p&offsets|0|-2(+1C);m-14&22|+|46113691|46113712&offsets|0|-2(+1C)]|TGAGGTAGTAGGTTGTGTGGC</td><td>>MI0000063|hsa-let-7b&WithFlank&22|+|46113680|46113774</td><td>12</td><td>31</td><td>20(+1C)</td><td>TGAGGTAGTAGGTTGTGTGGC</td><td>1135.9</td><td>3.2</td><td>0.8</td><td>3.8</td><td>1.4E-04</td><td>1.1E-03</td><td>BT20/MB231</td><td>CYTO</td></tr>
<tr><td>>hg38_21_+_8433172_8446622_RNA45SN1_45S_50ntFlanks@8229.8251.23|CGCGCCGGGCCCGGGTCTTCCCG</td><td>>hg38_21_+_8433172_8446622_RNA45SN1_45S_50ntFlanks</td><td>8229</td><td>8251</td><td>23</td><td>CGCGCCGGGCCCGGGTCTTCCCG</td><td>738.0</td><td>-3.6</td><td>0.9</td><td>-3.8</td><td>1.4E-04</td><td>1.1E-03</td><td>BT20/MB231</td><td>CYTO</td></tr>
<tr><td>>hg38_21_+_8433172_8446622_RNA45SN1_45S_50ntFlanks@6661.6697.37|CGGTGGATCACTCGGCTCGTGCGTCGATGAAGAACGC</td><td>>hg38_21_+_8433172_8446622_RNA45SN1_45S_50ntFlanks</td><td>6661</td><td>6697</td><td>37</td><td>CGGTGGATCACTCGGCTCGTGCGTCGATGAAGAACGC</td><td>1775.1</td><td>3.7</td><td>1.0</td><td>3.8</td><td>1.4E-04</td><td>1.1E-03</td><td>BT20/MB231</td><td>MP</td></tr>
<tr><td>>hg38_21_+_8433172_8446622_RNA45SN1_45S_50ntFlanks@9459.9506.48|CCGGGGAGGTGGAGCACGAGCGCACGTGTTAGGACCCGAAAGATGGTG</td><td>>hg38_21_+_8433172_8446622_RNA45SN1_45S_50ntFlanks</td><td>9459</td><td>9506</td><td>48</td><td>CCGGGGAGGTGGAGCACGAGCGCACGTGTTAGGACCCGAAAGATGGTG</td><td>67.5</td><td>-9.9</td><td>2.9</td><td>-3.4</td><td>7.4E-04</td><td>1.1E-03</td><td>MB231/MB468</td><td>MP</td></tr>
<tr><td>>hg38_21_+_8433172_8446622_RNA45SN1_45S_50ntFlanks@12444.12481.38|AAGCAGAATTCACCAAGCGTTGGATTGTTCACCCACTA</td><td>>hg38_21_+_8433172_8446622_RNA45SN1_45S_50ntFlanks</td><td>12444</td><td>12481</td><td>38</td><td>AAGCAGAATTCACCAAGCGTTGGATTGTTCACCCACTA</td><td>169.2</td><td>-2.1</td><td>0.6</td><td>-3.3</td><td>9.0E-04</td><td>1.1E-03</td><td>MB231/MB468</td><td>NUC</td></tr>
<tr><td>>hg38_21_+_8433172_8446622_RNA45SN1_45S_50ntFlanks@10841.10871.31|CGTAACTTCGGGATAAGGATTGGCTCTAAGG</td><td>>hg38_21_+_8433172_8446622_RNA45SN1_45S_50ntFlanks</td><td>10841</td><td>10871</td><td>31</td><td>CGTAACTTCGGGATAAGGATTGGCTCTAAGG</td><td>659.7</td><td>-2.4</td><td>0.7</td><td>-3.3</td><td>9.0E-04</td><td>1.1E-03</td><td>MB231/MB468</td><td>NUC</td></tr>
<tr><td>>MI0000063|hsa-let-7b&WithFlank&22|+|46113680|46113774@66.85.20(+1U)&[MIMAT0004482&hsa-let-7b-3p&offsets|0|-2(+1U);m-171&22|+|46113745|46113766&offsets|0|-2(+1U)]|CTATACAACCTACTGCCTTCT</td><td>>MI0000063|hsa-let-7b&WithFlank&22|+|46113680|46113774</td><td>66</td><td>85</td><td>20(+1U)</td><td>CTATACAACCTACTGCCTTCT</td><td>59.4</td><td>9.6</td><td>2.7</td><td>3.5</td><td>3.9E-04</td><td>1.1E-03</td><td>BT20/MB468</td><td>MP</td></tr>
<tr><td>>MI0000457|hsa-mir-141&WithFlank&12|+|6964091|6964197@65.84.20(+1A)&[MIMAT0000432&hsa-miR-141-3p&offsets|0|-2(+1A);m-134&12|+|6964155|6964175&offsets|0|-1(+1A)]|TAACACTGTCTGGTAAAGATA</td><td>>MI0000457|hsa-mir-141&WithFlank&12|+|6964091|6964197</td><td>65</td><td>84</td><td>20(+1A)</td><td>TAACACTGTCTGGTAAAGATA</td><td>99.9</td><td>10.0</td><td>2.6</td><td>3.8</td><td>1.4E-04</td><td>1.1E-03</td><td>BT20/MB231</td><td>MP</td></tr>
<tr><td>>hg38_21_+_8433172_8446622_RNA45SN1_45S_50ntFlanks@9508.9526.19|ACTATGCCTGGGCAGGGCG</td><td>>hg38_21_+_8433172_8446622_RNA45SN1_45S_50ntFlanks</td><td>9508</td><td>9526</td><td>19</td><td>ACTATGCCTGGGCAGGGCG</td><td>72.4</td><td>-9.3</td><td>2.6</td><td>-3.5</td><td>3.9E-04</td><td>1.1E-03</td><td>BT20/MB468</td><td>MP</td></tr>
<tr><td>>hg38_21_+_8433172_8446622_RNA45SN1_45S_50ntFlanks@4760.4793.34|TCAGATACCGTCGTAGTTCCGACCATAAACGATG</td><td>>hg38_21_+_8433172_8446622_RNA45SN1_45S_50ntFlanks</td><td>4760</td><td>4793</td><td>34</td><td>TCAGATACCGTCGTAGTTCCGACCATAAACGATG</td><td>507.7</td><td>-2.5</td><td>0.7</td><td>-3.3</td><td>8.4E-04</td><td>1.1E-03</td><td>MB231/MB468</td><td>CYTO</td></tr>
<tr><td>>hg38_21_+_8433172_8446622_RNA45SN1_45S_50ntFlanks@10808.10837.30|CAATGTAGGTAAGGGAAGTCGGCAAGCCGG</td><td>>hg38_21_+_8433172_8446622_RNA45SN1_45S_50ntFlanks</td><td>10808</td><td>10837</td><td>30</td><td>CAATGTAGGTAAGGGAAGTCGGCAAGCCGG</td><td>255.8</td><td>-3.4</td><td>1.0</td><td>-3.3</td><td>8.4E-04</td><td>1.1E-03</td><td>MB231/MB468</td><td>CYTO</td></tr>
<tr><td>>hg38_21_+_8433172_8446622_RNA45SN1_45S_50ntFlanks@10341.10362.22|AATATTCAAACGAGAACTTTGA</td><td>>hg38_21_+_8433172_8446622_RNA45SN1_45S_50ntFlanks</td><td>10341</td><td>10362</td><td>22</td><td>AATATTCAAACGAGAACTTTGA</td><td>127.2</td><td>4.6</td><td>1.3</td><td>3.5</td><td>3.9E-04</td><td>1.1E-03</td><td>BT20/MB468</td><td>MP</td></tr>
<tr><td>>hg38_21_+_8433172_8446622_RNA45SN1_45S_50ntFlanks@9573.9604.32|AAATCGGTCGTCCGACCTGGGTATAGGGGCGA</td><td>>hg38_21_+_8433172_8446622_RNA45SN1_45S_50ntFlanks</td><td>9573</td><td>9604</td><td>32</td><td>AAATCGGTCGTCCGACCTGGGTATAGGGGCGA</td><td>266.0</td><td>-4.0</td><td>1.2</td><td>-3.3</td><td>8.9E-04</td><td>1.1E-03</td><td>MB231/MB468</td><td>MITO</td></tr>
<tr><td>>trna111_HisGTG_1_-_147774845_147774916@-1T.26.27,trna118_HisGTG_1_-_145396881_145396952@-1T.26.27,trna16_HisGTG_1_+_146544773_146544844@-1T.26.27,trna1_HisGTG_15_+_45493349_45493420@-1T.26.27,trna21_HisGTG_1_+_147753471_147753542@-1T.26.27,trna33_HisGTG_6_+_27125906_27125977@-1T.26.27,trna7_HisGTG_9_-_14433938_14434009@-1T.26.27,trna8_HisGTG_15_-_45492611_45492682@-1T.26.27,trna9_HisGTG_15_-_45490804_45490875@-1T.26.27|TGCCGTGATCGTATAGTGGTTAGTACT</td><td>>trna111_HisGTG_1_-_147774845_147774916</td><td>-1T</td><td>26</td><td>27</td><td>TGCCGTGATCGTATAGTGGTTAGTACT</td><td>325.0</td><td>3.7</td><td>1.1</td><td>3.5</td><td>3.9E-04</td><td>1.1E-03</td><td>BT20/MB468</td><td>MP</td></tr>
<tr><td>>hg38_21_+_8433172_8446622_RNA45SN1_45S_50ntFlanks@12417.12440.24|TCGATGTCGGCTCTTCCTATCATT</td><td>>hg38_21_+_8433172_8446622_RNA45SN1_45S_50ntFlanks</td><td>12417</td><td>12440</td><td>24</td><td>TCGATGTCGGCTCTTCCTATCATT</td><td>1513.0</td><td>2.0</td><td>0.6</td><td>3.3</td><td>8.4E-04</td><td>1.1E-03</td><td>MB231/MB468</td><td>CYTO</td></tr>
<tr><td>>hg38_21_+_8433172_8446622_RNA45SN1_45S_50ntFlanks@10362.10379.18|AAGGCCGAAGTGGAGAAG</td><td>>hg38_21_+_8433172_8446622_RNA45SN1_45S_50ntFlanks</td><td>10362</td><td>10379</td><td>18</td><td>AAGGCCGAAGTGGAGAAG</td><td>194.0</td><td>-3.2</td><td>1.0</td><td>-3.3</td><td>8.4E-04</td><td>1.1E-03</td><td>MB231/MB468</td><td>CYTO</td></tr>
<tr><td>>hg38_21_+_8433172_8446622_RNA45SN1_45S_50ntFlanks@11315.11343.29|CCACCCCACGTCTCGTCGCGCGCGCGTCC</td><td>>hg38_21_+_8433172_8446622_RNA45SN1_45S_50ntFlanks</td><td>11315</td><td>11343</td><td>29</td><td>CCACCCCACGTCTCGTCGCGCGCGCGTCC</td><td>170.1</td><td>-3.4</td><td>1.0</td><td>-3.4</td><td>8.0E-04</td><td>1.1E-03</td><td>BT20/MB468</td><td>CYTO</td></tr>
<tr><td>>trna111_HisGTG_1_-_147774845_147774916@-1T.31.32,trna118_HisGTG_1_-_145396881_145396952@-1T.31.32,trna16_HisGTG_1_+_146544773_146544844@-1T.31.32,trna1_HisGTG_15_+_45493349_45493420@-1T.31.32,trna21_HisGTG_1_+_147753471_147753542@-1T.31.32,trna33_HisGTG_6_+_27125906_27125977@-1T.31.32,trna7_HisGTG_9_-_14433938_14434009@-1T.31.32,trna8_HisGTG_15_-_45492611_45492682@-1T.31.32,trna9_HisGTG_15_-_45490804_45490875@-1T.31.32|TGCCGTGATCGTATAGTGGTTAGTACTCTGCG</td><td>>trna111_HisGTG_1_-_147774845_147774916</td><td>-1T</td><td>31</td><td>32</td><td>TGCCGTGATCGTATAGTGGTTAGTACTCTGCG</td><td>21040.7</td><td>-3.3</td><td>1.0</td><td>-3.4</td><td>8.1E-04</td><td>1.1E-03</td><td>BT20/MB468</td><td>CYTO</td></tr>
<tr><td>>hg38_21_+_8433172_8446622_RNA45SN1_45S_50ntFlanks@12654.12675.22|TATGACTGAACGCCTCTAAGTC</td><td>>hg38_21_+_8433172_8446622_RNA45SN1_45S_50ntFlanks</td><td>12654</td><td>12675</td><td>22</td><td>TATGACTGAACGCCTCTAAGTC</td><td>583.9</td><td>-5.8</td><td>1.6</td><td>-3.5</td><td>3.9E-04</td><td>1.1E-03</td><td>BT20/MB468</td><td>MP</td></tr>
<tr><td>>MI0003560|hsa-mir-92b&WithFlank&1|+|155195171|155195278@26.49.24&[MIMAT0004792&hsa-miR-92b-5p&offsets|0|+2;m-581&1|+|155195196|155195219&offsets|0|0]|AGGGACGGGACGCGGTGCAGTGTT</td><td>>MI0003560|hsa-mir-92b&WithFlank&1|+|155195171|155195278</td><td>26</td><td>49</td><td>24</td><td>AGGGACGGGACGCGGTGCAGTGTT</td><td>120.2</td><td>6.7</td><td>1.9</td><td>3.5</td><td>3.9E-04</td><td>1.1E-03</td><td>BT20/MB468</td><td>MP</td></tr>
<tr><td>>hg38_21_+_8433172_8446622_RNA45SN1_45S_50ntFlanks@12640.12667.28|ACCATCTGTGGGATTATGACTGAACGCC</td><td>>hg38_21_+_8433172_8446622_RNA45SN1_45S_50ntFlanks</td><td>12640</td><td>12667</td><td>28</td><td>ACCATCTGTGGGATTATGACTGAACGCC</td><td>148.1</td><td>3.5</td><td>1.0</td><td>3.3</td><td>8.9E-04</td><td>1.1E-03</td><td>MB231/MB468</td><td>MITO</td></tr>
<tr><td>>hg38_21_+_8433172_8446622_RNA45SN1_45S_50ntFlanks@11668.11700.33|TGCCCAGTGCTCTGAATGTCAAAGTGAAGAAAT</td><td>>hg38_21_+_8433172_8446622_RNA45SN1_45S_50ntFlanks</td><td>11668</td><td>11700</td><td>33</td><td>TGCCCAGTGCTCTGAATGTCAAAGTGAAGAAAT</td><td>244.3</td><td>-2.0</td><td>0.6</td><td>-3.3</td><td>9.0E-04</td><td>1.1E-03</td><td>MB231/MB468</td><td>NUC</td></tr>
<tr><td>>hg38_21_+_8433172_8446622_RNA45SN1_45S_50ntFlanks@10781.10802.22|AGGTGAACAGCCTCTGGCATGT</td><td>>hg38_21_+_8433172_8446622_RNA45SN1_45S_50ntFlanks</td><td>10781</td><td>10802</td><td>22</td><td>AGGTGAACAGCCTCTGGCATGT</td><td>337.8</td><td>-4.1</td><td>1.2</td><td>-3.3</td><td>8.1E-04</td><td>1.1E-03</td><td>BT20/MB468</td><td>CYTO</td></tr>
<tr><td>>trna128_GlyGCC_6_-_27870686_27870756@20.38.19,trna19_GlyGCC_16_+_70823410_70823480@20.38.19,trna19_GlyGCC_2_-_157257659_157257729@20.38.19,trna24_GlyGCC_16_-_70812942_70813012@20.38.19,trna25_GlyGCC_16_-_70812114_70812184@20.38.19,trna2_GlyGCC_21_-_18827107_18827177@20.38.19,trna35_GlyGCC_1_+_161413094_161413164@20.38.19,trna37_GlyGCC_1_+_161420467_161420537@20.38.19,trna39_GlyGCC_1_+_161427898_161427968@20.38.19,trna41_GlyGCC_1_+_161435258_161435328@20.38.19,trna5_GlyGCC_17_+_8029064_8029134@20.38.19,trna68_GlyGCC_1_-_161493637_161493707@20.38.19|AGAATTCTCGCCTGCCACG</td><td>>trna128_GlyGCC_6_-_27870686_27870756</td><td>20</td><td>38</td><td>19</td><td>AGAATTCTCGCCTGCCACG</td><td>298.8</td><td>3.2</td><td>0.8</td><td>3.8</td><td>1.4E-04</td><td>1.1E-03</td><td>BT20/MB231</td><td>CYTO</td></tr>
<tr><td>>hg38_21_+_8433172_8446622_RNA45SN1_45S_50ntFlanks@11918.11939.22|ACATGAGAGGTGTAGAATAAGT</td><td>>hg38_21_+_8433172_8446622_RNA45SN1_45S_50ntFlanks</td><td>11918</td><td>11939</td><td>22</td><td>ACATGAGAGGTGTAGAATAAGT</td><td>160.8</td><td>2.9</td><td>0.8</td><td>3.5</td><td>3.9E-04</td><td>1.1E-03</td><td>BT20/MB468</td><td>MP</td></tr>
<tr><td>>hg38_MT_+_1621_3279_RNR2_16S_50ntFlanks@482.518.37|AAAGAGGAACAGCTCTTTGGACACTAGGAAAAAACCT</td><td>>hg38_MT_+_1621_3279_RNR2_16S_50ntFlanks</td><td>482</td><td>518</td><td>37</td><td>AAAGAGGAACAGCTCTTTGGACACTAGGAAAAAACCT</td><td>118.7</td><td>5.7</td><td>1.6</td><td>3.5</td><td>3.9E-04</td><td>1.1E-03</td><td>BT20/MB468</td><td>MP</td></tr>
<tr><td>>MI0000482|hsa-mir-185&WithFlank&22|+|20033133|20033226@21.42.22(+1A)&[MIMAT0000455&hsa-miR-185-5p&offsets|0|0(+1A);m-145&22|+|20033153|20033174&offsets|0|0(+1A)]|TGGAGAGAAAGGCAGTTCCTGAA</td><td>>MI0000482|hsa-mir-185&WithFlank&22|+|20033133|20033226</td><td>21</td><td>42</td><td>22(+1A)</td><td>TGGAGAGAAAGGCAGTTCCTGAA</td><td>303.9</td><td>2.4</td><td>0.6</td><td>3.8</td><td>1.4E-04</td><td>1.1E-03</td><td>BT20/MB231</td><td>NUC</td></tr>
<tr><td>>trna119_LysCTT_1_-_145395522_145395594@1.33.33,trna11_LysCTT_5_-_180648979_180649051@1.33.33,trna13_LysCTT_6_+_26556774_26556846@1.33.33,trna7_LysCTT_16_+_3225692_3225764@1.33.33,trna9_LysCTT_5_+_180634755_180634827@1.33.33|GCCCGGCTAGCTCAGTCGGTAGAGCATGAGACT</td><td>>trna119_LysCTT_1_-_145395522_145395594</td><td>1</td><td>33</td><td>33</td><td>GCCCGGCTAGCTCAGTCGGTAGAGCATGAGACT</td><td>8917.3</td><td>-4.3</td><td>1.1</td><td>-3.8</td><td>1.4E-04</td><td>1.1E-03</td><td>BT20/MB231</td><td>NUC</td></tr>
<tr><td>>hg38_21_+_8433172_8446622_RNA45SN1_45S_50ntFlanks@10318.10352.35|TGCAGATCTTGGTGGTAGTAGCAAATATTCAAACG</td><td>>hg38_21_+_8433172_8446622_RNA45SN1_45S_50ntFlanks</td><td>10318</td><td>10352</td><td>35</td><td>TGCAGATCTTGGTGGTAGTAGCAAATATTCAAACG</td><td>67.9</td><td>6.5</td><td>1.8</td><td>3.5</td><td>3.9E-04</td><td>1.1E-03</td><td>BT20/MB468</td><td>MP</td></tr>
<tr><td>>MI0000298|hsa-mir-221&WithFlank&X|-|45746151|45746272@31.55.25&[MIMAT0004568&hsa-miR-221-5p&offsets|0|+3;m-405&X|-|45746220|45746242&offsets|0|+2]|ACCTGGCATACAATGTAGATTTCTG</td><td>>MI0000298|hsa-mir-221&WithFlank&X|-|45746151|45746272</td><td>31</td><td>55</td><td>25</td><td>ACCTGGCATACAATGTAGATTTCTG</td><td>236.8</td><td>-2.4</td><td>0.6</td><td>-3.8</td><td>1.4E-04</td><td>1.1E-03</td><td>BT20/MB231</td><td>NUC</td></tr>
<tr><td>>hg38_21_+_8433172_8446622_RNA45SN1_45S_50ntFlanks@10345.10374.30|TTCAAACGAGAACTTTGAAGGCCGAAGTGG</td><td>>hg38_21_+_8433172_8446622_RNA45SN1_45S_50ntFlanks</td><td>10345</td><td>10374</td><td>30</td><td>TTCAAACGAGAACTTTGAAGGCCGAAGTGG</td><td>210.0</td><td>-4.2</td><td>1.2</td><td>-3.4</td><td>7.5E-04</td><td>1.1E-03</td><td>MB231/MB468</td><td>MP</td></tr>
<tr><td>>hg38_1_-_228634819_228635039_RNA5S12_5S_50ntFlanks@51.89.39|GTCTACGGCCATACCACCCTGAACGCGCCCGATCTCGTC</td><td>>hg38_1_-_228634819_228635039_RNA5S12_5S_50ntFlanks</td><td>51</td><td>89</td><td>39</td><td>GTCTACGGCCATACCACCCTGAACGCGCCCGATCTCGTC</td><td>26660.2</td><td>4.5</td><td>1.4</td><td>3.3</td><td>9.0E-04</td><td>1.1E-03</td><td>BT20/MB468</td><td>NUC</td></tr>
</table>

| | | | | | | | | | | | | | |
|---|---|---|---|---|---|---|---|---|---|---|---|---|---|
| >trna116_GluCTC_1_-_145399233_145399304@13.31.19,trna25_GluCTC_2_-_71273488_71273560@13.31.19,trna59_GluCTC_1_+_249168447_249168518@13.31.19,trna71_GluCTC_1_-_161439189_161439260@13.31.19,trna74_GluCTC_1_-_161431809_161431880@13.31.19,trna77_GluCTC_1_-_161424398_161424469@13.31.19,trna77_GluCTC_6_+_28949976_28950047@13.31.19,trna80_GluCTC_1_-_161417018_161417089@13.31.19,trna87_GluCTC_6_-_126101393_126101464@13.31.19\|TAGTGGTTAGGATTCGGCG | >trna116_GluCTC_1_-_145399233_145399304 | 13 | 31 | 19 | TAGTGGTTAGGATTCGGCG | 135.9 | -4.8 | 1.4 | -3.3 | 9.1E-04 | 1.1E-03 | MB231/MB468 | NUC |
| >am_trnaMT_HisGTG_MT_+_12138_12206@18.38.21\|AAAACATCAGATTGTGAATCT | >am_trnaMT_HisGTG_MT_+_12138_12206 | 18 | 38 | 21 | AAAACATCAGATTGTGAATCT | 243.2 | 5.1 | 1.4 | 3.5 | 3.9E-04 | 1.1E-03 | BT20/MB468 | MP |
| >MI0000457\|hsa-mir-141&WithFlank&12\|+\|6964091\|6964197@23.43.21&[MIMAT0004598&hsa-miR-141-5p&offsets\|0\|-1;m-221&12\|+\|6964113\|6964133&offsets\|0\|0]\|CATCTTCCAGTACAGTGTTGG | >MI0000457\|hsa-mir-141&WithFlank&12\|+\|6964091\|6964197 | 23 | 43 | 21 | CATCTTCCAGTACAGTGTTGG | 212.5 | 6.4 | 1.8 | 3.5 | 3.9E-04 | 1.1E-03 | BT20/MB468 | MP |
| >hg38_21_+_8433172_8446622_RNA45SN1_45S_50ntFlanks@10387.10407.21\|TGTGAACAGCAGTTGAACATG | >hg38_21_+_8433172_8446622_RNA45SN1_45S_50ntFlanks | 10387 | 10407 | 21 | TGTGAACAGCAGTTGAACATG | 361.3 | -6.2 | 1.7 | -3.5 | 3.9E-04 | 1.1E-03 | BT20/MB468 | MP |
| >hg38_21_+_8433172_8446622_RNA45SN1_45S_50ntFlanks@11573.11600.28\|CGACTTAGAACTGGTGCGGACCAGGGGA | >hg38_21_+_8433172_8446622_RNA45SN1_45S_50ntFlanks | 11573 | 11600 | 28 | CGACTTAGAACTGGTGCGGACCAGGGGA | 407.3 | -2.2 | 0.7 | -3.3 | 8.1E-04 | 1.1E-03 | BT20/MB468 | CYTO |
| >MI0000065\|hsa-let-7d&WithFlank&9\|+\|94178828\|94178926@14.34.21(+1C)&[MIMAT0000065&hsa-let-7d-5p&offsets\|0\|-1(+1C);m-90&9\|+\|94178841\|94178862&offsets\|0\|-1(+1C)]\|AGAGGTAGTAGGTTGCATAGTC | >MI0000065\|hsa-let-7d&WithFlank&9\|+\|94178828\|94178926 | 14 | 34 | 21(+1C) | AGAGGTAGTAGGTTGCATAGTC | 450.2 | 3.7 | 1.0 | 3.5 | 3.9E-04 | 1.1E-03 | BT20/MB468 | MP |
| >MI0000264\|hsa-mir-7-2&WithFlank&15\|+\|88611819\|88611940@38.60.23;MI0000265\|hsa-mir-7-3&WithFlank&19\|+\|4770664\|4770785@37.59.23;MI0000263\|hsa-mir-7-1&WithFlank&9\|-\|83969742\|83969863@30.52.23&[MIMAT0000252&hsa-miR-7-5p&offsets\|0\|-1;m-400&15\|+\|88611856\|88611879&offsets\|0\|-1];[MIMAT0000252_1&hsa-miR-7-5p&offsets\|0\|-1;m-402&19\|+\|4770700\|4770723&offsets\|0\|-1];[MIMAT0000252_2&hsa-miR-7-5p&offsets\|0\|-1;m-401&9\|-\|83969811\|83969834&offsets\|0\|-1]\|TGGAAGACTAGTGATTTTGTTGT | >MI0000264\|hsa-mir-7-2&WithFlank&15\|+\|88611819\|88611940 | 38 | 60 | 23 | TGGAAGACTAGTGATTTTGTTGT | 5802.7 | 3.6 | 1.0 | 3.5 | 3.9E-04 | 1.1E-03 | BT20/MB468 | MP |
| >hg38_21_+_8433172_8446622_RNA45SN1_45S_50ntFlanks@9169.9197.29\|GGCGGGTCGCGCCGTCGGGCCCGGGGGAG | >hg38_21_+_8433172_8446622_RNA45SN1_45S_50ntFlanks | 9169 | 9197 | 29 | GGCGGGTCGCGCCGTCGGGCCCGGGGGAG | 144.4 | -5.9 | 1.7 | -3.5 | 3.9E-04 | 1.1E-03 | BT20/MB468 | MP |
| >hg38_MT_+_1621_3279_RNR2_16S_50ntFlanks@1171.1215.45\|AACCTGCATTAAAAATTTCGGTTGGGGCGACCTCGGAGCAGAACC | >hg38_MT_+_1621_3279_RNR2_16S_50ntFlanks | 1171 | 1215 | 45 | AACCTGCATTAAAAATTTCGGTTGGGGCGACCTCGGAGCAGAACC | 59.5 | 9.6 | 2.7 | 3.5 | 3.9E-04 | 1.1E-03 | BT20/MB468 | MP |
| >hg38_21_+_8433172_8446622_RNA45SN1_45S_50ntFlanks@11540.11583.44\|CGCGGCGCCTCGCCTCGGCCGGCGCCTAGCAGCCGACTTAGAAC | >hg38_21_+_8433172_8446622_RNA45SN1_45S_50ntFlanks | 11540 | 11583 | 44 | CGCGGCGCCTCGCCTCGGCCGGCGCCTAGCAGCCGACTTAGAAC | 629.7 | 3.0 | 0.9 | 3.3 | 9.0E-04 | 1.1E-03 | MB231/MB468 | MITO |
| >MI0000085\|hsa-mir-27a&WithFlank&19\|-\|13836434\|13836523@57.75.19(+1C)&[MIMAT0000084&hsa-miR-27a-3p&offsets\|0\|-2(+1C);m-43&19\|-\|13836447\|13836467&offsets\|0\|-2(+1C)]\|TTCACAGTGGCTAAGTTCCC | >MI0000085\|hsa-mir-27a&WithFlank&19\|-\|13836434\|13836523 | 57 | 75 | 19(+1C) | TTCACAGTGGCTAAGTTCCC | 169.7 | -4.0 | 1.1 | -3.8 | 1.7E-04 | 1.1E-03 | BT20/MB231 | MITO |
| >hg38_21_+_8433172_8446622_RNA45SN1_45S_50ntFlanks@12946.12982.37\|TAGACGACCTGCTTCTGGGTCGGGGTTTCGTACGTAG | >hg38_21_+_8433172_8446622_RNA45SN1_45S_50ntFlanks | 12946 | 12982 | 37 | TAGACGACCTGCTTCTGGGTCGGGGTTTCGTACGTAG | 200.1 | -3.4 | 1.0 | -3.3 | 8.1E-04 | 1.1E-03 | BT20/MB468 | CYTO |
| >hg38_21_+_8433172_8446622_RNA45SN1_45S_50ntFlanks@7990.8007.18\|GACGTGGCGACCCGCTGA | >hg38_21_+_8433172_8446622_RNA45SN1_45S_50ntFlanks | 7990 | 8007 | 18 | GACGTGGCGACCCGCTGA | 632.4 | 7.6 | 2.0 | 3.8 | 1.4E-04 | 1.1E-03 | BT20/MB231 | MP |
| >hg38_MT_+_598_1651_RNR1_12S_50ntFlanks@965.1004.40\|GTAACATGGTAAGTGTACTGGAAAGTGCACTTGGACGAAC | >hg38_MT_+_598_1651_RNR1_12S_50ntFlanks | 965 | 1004 | 40 | GTAACATGGTAAGTGTACTGGAAAGTGCACTTGGACGAAC | 167.5 | 5.9 | 1.8 | 3.3 | 9.0E-04 | 1.1E-03 | MB231/MB468 | MITO |
| >hg38_21_+_8433172_8446622_RNA45SN1_45S_50ntFlanks@10727.10757.31\|GGAGAGGGTGTAAATCTCGCGCCGGGCCGTA | >hg38_21_+_8433172_8446622_RNA45SN1_45S_50ntFlanks | 10727 | 10757 | 31 | GGAGAGGGTGTAAATCTCGCGCCGGGCCGTA | 246.3 | -5.5 | 1.6 | -3.5 | 3.9E-04 | 1.1E-03 | BT20/MB468 | MP |
| >hg38_21_+_8433172_8446622_RNA45SN1_45S_50ntFlanks@4735.4767.33\|ACGAAAGTCGGAGGTTCGAAGACGATCAGATAC | >hg38_21_+_8433172_8446622_RNA45SN1_45S_50ntFlanks | 4735 | 4767 | 33 | ACGAAAGTCGGAGGTTCGAAGACGATCAGATAC | 50.1 | 8.9 | 2.6 | 3.4 | 7.5E-04 | 1.1E-03 | MB231/MB468 | MP |
| >MI0000070\|hsa-mir-16-1&WithFlank&13\|-\|50048967\|50049067@20.37.18;MI0000115\|hsa-mir-16-2&WithFlank&3\|+\|160404739\|160404831@16.33.18&[MIMAT0000069&hsa-miR-16-5p&offsets\|0\|-4;m-62&13\|-\|50049027\|50049048&offsets\|0\|-4];[MIMAT0000069_1&hsa-miR-16-5p&offsets\|0\|-4;m-61&3\|+\|160404754\|160404775&offsets\|0\|-4]\|TAGCAGCACGTAAATATT | >MI0000070\|hsa-mir-16-1&WithFlank&13\|-\|50048967\|50049067 | 20 | 37 | 18 | TAGCAGCACGTAAATATT | 259.6 | 5.7 | 1.7 | 3.4 | 7.5E-04 | 1.1E-03 | MB231/MB468 | MP |
| >hg38_21_+_8433172_8446622_RNA45SN1_45S_50ntFlanks@485.514.30\|GATGTGGTGACGTCGTGCTCTCCCGGGCCG | >hg38_21_+_8433172_8446622_RNA45SN1_45S_50ntFlanks | 485 | 514 | 30 | GATGTGGTGACGTCGTGCTCTCCCGGGCCG | 57.9 | 9.6 | 2.7 | 3.5 | 4.0E-04 | 1.1E-03 | BT20/MB468 | MP |
| >hg38_21_+_8433172_8446622_RNA45SN1_45S_50ntFlanks@10768.10785.18\|CAGCAGGTCTCCAAGGTG | >hg38_21_+_8433172_8446622_RNA45SN1_45S_50ntFlanks | 10768 | 10785 | 18 | CAGCAGGTCTCCAAGGTG | 1028.5 | -6.4 | 1.9 | -3.3 | 9.1E-04 | 1.1E-03 | BT20/MB468 | NUC |
| >hg38_21_+_8433172_8446622_RNA45SN1_45S_50ntFlanks@10353.10388.36\|AGAACTTTGAAGGCCGAAGTGGAGAAGGGTTCCATG | >hg38_21_+_8433172_8446622_RNA45SN1_45S_50ntFlanks | 10353 | 10388 | 36 | AGAACTTTGAAGGCCGAAGTGGAGAAGGGTTCCATG | 513.8 | -4.3 | 1.3 | -3.3 | 9.0E-04 | 1.1E-03 | MB231/MB468 | MITO |
| >hg38_21_+_8433172_8446622_RNA45SN1_45S_50ntFlanks@9833.9850.18\|CTTTTGGTAAGCAGAACT | >hg38_21_+_8433172_8446622_RNA45SN1_45S_50ntFlanks | 9833 | 9850 | 18 | CTTTTGGTAAGCAGAACT | 153.3 | -5.5 | 1.6 | -3.5 | 4.0E-04 | 1.1E-03 | BT20/MB468 | MP |
| >hg38_21_+_8433172_8446622_RNA45SN1_45S_50ntFlanks@11849.11872.24\|CGGGCTTGGCGGAATCAGCGGGGA | >hg38_21_+_8433172_8446622_RNA45SN1_45S_50ntFlanks | 11849 | 11872 | 24 | CGGGCTTGGCGGAATCAGCGGGGA | 1574.3 | -6.2 | 1.7 | -3.5 | 4.0E-04 | 1.1E-03 | BT20/MB468 | MP |
| >hg38_21_+_8433172_8446622_RNA45SN1_45S_50ntFlanks@6645.6666.22\|CTCGTACGACTCTTAGCGGTGG | >hg38_21_+_8433172_8446622_RNA45SN1_45S_50ntFlanks | 6645 | 6666 | 22 | CTCGTACGACTCTTAGCGGTGG | 4317.8 | 3.0 | 0.9 | 3.3 | 8.2E-04 | 1.1E-03 | BT20/MB468 | CYTO |
| >hg38_21_+_8433172_8446622_RNA45SN1_45S_50ntFlanks@3782.3805.24\|CAGTGAAACTGCGAATGGCTCATT | >hg38_21_+_8433172_8446622_RNA45SN1_45S_50ntFlanks | 3782 | 3805 | 24 | CAGTGAAACTGCGAATGGCTCATT | 501.1 | -3.5 | 1.0 | -3.5 | 4.0E-04 | 1.1E-03 | BT20/MB468 | MP |
| >hg38_21_+_8433172_8446622_RNA45SN1_45S_50ntFlanks@6661.6707.47\|CGGTGGATCACTCGGCTCGTGCGTCGATGAAGAACGCAGCTAGCTGC | >hg38_21_+_8433172_8446622_RNA45SN1_45S_50ntFlanks | 6661 | 6707 | 47 | CGGTGGATCACTCGGCTCGTGCGTCGATGAAGAACGCAGCTAGCTGC | 398.6 | 5.9 | 1.7 | 3.5 | 4.0E-04 | 1.1E-03 | BT20/MB468 | MP |
| >hg38_21_+_8433172_8446622_RNA45SN1_45S_50ntFlanks@10866.10890.25\|CTAAGGGCTGGGTCGGTCGGGCTGG | >hg38_21_+_8433172_8446622_RNA45SN1_45S_50ntFlanks | 10866 | 10890 | 25 | CTAAGGGCTGGGTCGGTCGGGCTGG | 762.5 | -4.1 | 1.2 | -3.3 | 8.2E-04 | 1.1E-03 | BT20/MB468 | CYTO |
| >hg38_21_+_8433172_8446622_RNA45SN1_45S_50ntFlanks@7970.8006.37\|CGAGACGCGACCTCAGATCAGACGTGGCGACCCGCTG | >hg38_21_+_8433172_8446622_RNA45SN1_45S_50ntFlanks | 7970 | 8006 | 37 | CGAGACGCGACCTCAGATCAGACGTGGCGACCCGCTG | 262.3 | 3.4 | 0.9 | 3.8 | 1.4E-04 | 1.1E-03 | BT20/MB231 | NUC |
| >hg38_21_+_8433172_8446622_RNA45SN1_45S_50ntFlanks@4005.4042.38\|AACCTCGGGCCGATCGCACGCCCCCCGTGGCGGCGACG | >hg38_21_+_8433172_8446622_RNA45SN1_45S_50ntFlanks | 4005 | 4042 | 38 | AACCTCGGGCCGATCGCACGCCCCCCGTGGCGGCGACG | 247.4 | -3.7 | 1.1 | -3.4 | 6.8E-04 | 1.1E-03 | BT20/MB468 | MITO |
| >hg38_21_+_8433172_8446622_RNA45SN1_45S_50ntFlanks@9034.9055.22\|TCGTGGGGGGGCCGGGCCACCC | >hg38_21_+_8433172_8446622_RNA45SN1_45S_50ntFlanks | 9034 | 9055 | 22 | TCGTGGGGGGGCCGGGCCACCC | 81.7 | -5.6 | 1.6 | -3.5 | 4.0E-04 | 1.1E-03 | BT20/MB468 | MP |
| >hg38_21_+_8433172_8446622_RNA45SN1_45S_50ntFlanks@12417.12444.28\|TCGATGTCGGCTCTTCCTATCATTGTGA | >hg38_21_+_8433172_8446622_RNA45SN1_45S_50ntFlanks | 12417 | 12444 | 28 | TCGATGTCGGCTCTTCCTATCATTGTGA | 272.0 | -5.5 | 1.6 | -3.4 | 7.6E-04 | 1.1E-03 | MB231/MB468 | MP |
| >hg38_21_+_8433172_8446622_RNA45SN1_45S_50ntFlanks@11907.11939.33\|ACGGTGAAGAGACATGAGAGGTGTAGAATAAGT | >hg38_21_+_8433172_8446622_RNA45SN1_45S_50ntFlanks | 11907 | 11939 | 33 | ACGGTGAAGAGACATGAGAGGTGTAGAATAAGT | 122.2 | 10.3 | 2.7 | 3.8 | 1.4E-04 | 1.1E-03 | BT20/MB231 | MP |
| >hg38_21_+_8433172_8446622_RNA45SN1_45S_50ntFlanks@11921.11948.28\|TGAGAGGTGTAGAATAAGTGGGAGGCCC | >hg38_21_+_8433172_8446622_RNA45SN1_45S_50ntFlanks | 11921 | 11948 | 28 | TGAGAGGTGTAGAATAAGTGGGAGGCCC | 125.3 | 10.3 | 2.7 | 3.8 | 1.4E-04 | 1.1E-03 | BT20/MB231 | MP |
| >hg38_21_+_8433172_8446622_RNA45SN1_45S_50ntFlanks@10479.10512.34\|CGATCGAAAGGGAGTCGGGTTCAGATCCCCGAAT | >hg38_21_+_8433172_8446622_RNA45SN1_45S_50ntFlanks | 10479 | 10512 | 34 | CGATCGAAAGGGAGTCGGGTTCAGATCCCCGAAT | 839.8 | -4.9 | 1.5 | -3.3 | 8.6E-04 | 1.1E-03 | MB231/MB468 | CYTO |
| >hg38_21_+_8433172_8446622_RNA45SN1_45S_50ntFlanks@11575.11607.33\|ACTTAGAACTGGTGCGGACCAGGGGAATCCGAC | >hg38_21_+_8433172_8446622_RNA45SN1_45S_50ntFlanks | 11575 | 11607 | 33 | ACTTAGAACTGGTGCGGACCAGGGGAATCCGAC | 630.0 | -2.1 | 0.6 | -3.3 | 8.2E-04 | 1.1E-03 | BT20/MB468 | CYTO |
| >MI0000542\|hsa-mir-320a&WithFlank&8\|-\|22244960\|22245043@48.69.22(+3U)&[MIMAT0000510&hsa-miR-320a-3p&offsets\|0\|0(+3U);m-60&8\|-\|22244975\|22244996&offsets\|0\|0(+3U)]\|AAAAGCTGGGTTGAGAGGGCGATTT | >MI0000542\|hsa-mir-320a&WithFlank&8\|-\|22244960\|22245043 | 48 | 69 | 22(+3U) | AAAAGCTGGGTTGAGAGGGCGATTT | 186.2 | 6.1 | 1.7 | 3.5 | 4.0E-04 | 1.1E-03 | BT20/MB468 | MP |
| >hg38_21_+_8433172_8446622_RNA45SN1_45S_50ntFlanks@10687.10709.23\|CGGCGTCCGGTGAGCTCTCGCTG | >hg38_21_+_8433172_8446622_RNA45SN1_45S_50ntFlanks | 10687 | 10709 | 23 | CGGCGTCCGGTGAGCTCTCGCTG | 1437.7 | -3.8 | 1.1 | -3.5 | 4.0E-04 | 1.1E-03 | BT20/MB468 | MP |
| >hg38_21_+_8433172_8446622_RNA45SN1_45S_50ntFlanks@475.494.20\|CTTCGTGATCGATGTGGTGA | >hg38_21_+_8433172_8446622_RNA45SN1_45S_50ntFlanks | 475 | 494 | 20 | CTTCGTGATCGATGTGGTGA | 179.2 | 7.1 | 2.0 | 3.5 | 4.0E-04 | 1.1E-03 | BT20/MB468 | MP |
| >hg38_21_+_8433172_8446622_RNA45SN1_45S_50ntFlanks@12474.12496.23\|ACCCACTAATAGGGAACGTGAGC | >hg38_21_+_8433172_8446622_RNA45SN1_45S_50ntFlanks | 12474 | 12496 | 23 | ACCCACTAATAGGGAACGTGAGC | 1605.4 | 2.5 | 0.7 | 3.3 | 8.6E-04 | 1.1E-03 | MB231/MB468 | CYTO |
| >MI0000101\|hsa-mir-99a&WithFlank&21\|+\|16539083\|16539175@19.40.22(+1A)&[MIMAT0000097&hsa-miR-99a-5p&offsets\|0\|0(+1A);m-64&21\|+\|16539101\|16539121&offsets\|0\|+1(+1A)]\|AACCCGTAGATCCGATCTTGTGA | >MI0000101\|hsa-mir-99a&WithFlank&21\|+\|16539083\|16539175 | 19 | 40 | 22(+1A) | AACCCGTAGATCCGATCTTGTGA | 379.1 | -5.8 | 1.5 | -3.8 | 1.5E-04 | 1.1E-03 | BT20/MB231 | NUC |
| >hg38_1_-_228634819_228635039_RNA5S12_5S_50ntFlanks@150.169.20\|AATACCGGGTGCTGTAGGCT | >hg38_1_-_228634819_228635039_RNA5S12_5S_50ntFlanks | 150 | 169 | 20 | AATACCGGGTGCTGTAGGCT | 1925.0 | 3.5 | 0.9 | 3.8 | 1.4E-04 | 1.1E-03 | BT20/MB231 | MP |
| >hg38_21_+_8433172_8446622_RNA45SN1_45S_50ntFlanks@11671.11704.34\|CCAGTGCTCTGAATGTCAAAGTGAAGAAATTCAA | >hg38_21_+_8433172_8446622_RNA45SN1_45S_50ntFlanks | 11671 | 11704 | 34 | CCAGTGCTCTGAATGTCAAAGTGAAGAAATTCAA | 207.6 | -5.8 | 1.7 | -3.4 | 7.6E-04 | 1.1E-03 | MB231/MB468 | MP |
| >hg38_MT_+_1621_3279_RNR2_16S_50ntFlanks@999.1018.20\|AGGGACCTGTATGAATGGCT | >hg38_MT_+_1621_3279_RNR2_16S_50ntFlanks | 999 | 1018 | 20 | AGGGACCTGTATGAATGGCT | 253.5 | 3.5 | 1.0 | 3.4 | 7.7E-04 | 1.1E-03 | MB231/MB468 | MP |
| >am_trnaMT_ValTAC_MT_+_1602_1670@42.68.27\|AGATTTCAACTTAACTTGACCGCTCTG | >am_trnaMT_ValTAC_MT_+_1602_1670 | 42 | 68 | 27 | AGATTTCAACTTAACTTGACCGCTCTG | 805.6 | -4.0 | 1.2 | -3.3 | 9.2E-04 | 1.1E-03 | BT20/MB468 | NUC |
| >hg38_21_+_8433172_8446622_RNA45SN1_45S_50ntFlanks@5188.5205.18\|ATAACAGGTCTGTGATGC | >hg38_21_+_8433172_8446622_RNA45SN1_45S_50ntFlanks | 5188 | 5205 | 18 | ATAACAGGTCTGTGATGC | 303.7 | 2.0 | 0.6 | 3.3 | 8.6E-04 | 1.1E-03 | MB231/MB468 | CYTO |
| >MI0000093\|hsa-mir-92a-1&WithFlank&13\|+\|91351308\|91351397@54.75.22(+3A);MI0000094\|hsa-mir-92a-2&WithFlank&X\|-\|134169532\|134169618@54.75.22(+3A)&[MIMAT0000092&hsa-miR-92a-3p&offsets\|0\|0(+3A);m-9&13\|+\|91351361\|91351382&offsets\|0\|0(+3A)];[MIMAT0000092_1&hsa-miR-92a-3p&offsets\|0\|0(+3A);m-11&X\|-\|134169544\|134169565&offsets\|0\|0(+3A)]\|TATTGCACTTGTCCCGGCCTGTAAA | >MI0000093\|hsa-mir-92a-1&WithFlank&13\|+\|91351308\|91351397 | 54 | 75 | 22(+3A) | TATTGCACTTGTCCCGGCCTGTAAA | 161.1 | 3.2 | 1.0 | 3.3 | 9.1E-04 | 1.1E-03 | MB231/MB468 | MITO |
| >hg38_21_+_8433172_8446622_RNA45SN1_45S_50ntFlanks@5485.5516.32\|AGCGCTGAGAAGACGGTCGAACTTGACTATCT | >hg38_21_+_8433172_8446622_RNA45SN1_45S_50ntFlanks | 5485 | 5516 | 32 | AGCGCTGAGAAGACGGTCGAACTTGACTATCT | 643.4 | -2.5 | 0.8 | -3.3 | 9.3E-04 | 1.1E-03 | MB231/MB468 | NUC |

<table>
<tr><td>>hg38_21_+_8433172_8446622_RNA45SN1_45S_50ntFlanks@6653.6688.36|ACTCTTAGCGGTGGATCACTCGGCTCGTGCGTCGAT</td><td>>hg38_21_+_8433172_8446622_RNA45SN1_45S_50ntFlanks</td><td>6653</td><td>6688</td><td>36</td><td>ACTCTTAGCGGTGGATCACTCGGCTCGTGCGTCGAT</td><td>943.3</td><td>-1.5</td><td>0.5</td><td>-3.3</td><td>9.2E-04</td><td>1.1E-03</td><td>BT20/MB468</td><td>NUC</td></tr>
<tr><td>>hg38_21_+_8433172_8446622_RNA45SN1_45S_50ntFlanks@3854.3872.19|ACTGTGGTAATTCTAGAGC</td><td>>hg38_21_+_8433172_8446622_RNA45SN1_45S_50ntFlanks</td><td>3854</td><td>3872</td><td>19</td><td>ACTGTGGTAATTCTAGAGC</td><td>215.1</td><td>-3.7</td><td>1.1</td><td>-3.3</td><td>9.3E-04</td><td>1.1E-03</td><td>MB231/MB468</td><td>NUC</td></tr>
<tr><td>>hg38_21_+_8433172_8446622_RNA45SN1_45S_50ntFlanks@10393.10411.19|CAGCAGTTGAACATGGGTC</td><td>>hg38_21_+_8433172_8446622_RNA45SN1_45S_50ntFlanks</td><td>10393</td><td>10411</td><td>19</td><td>CAGCAGTTGAACATGGGTC</td><td>130.9</td><td>-4.1</td><td>1.2</td><td>-3.4</td><td>7.7E-04</td><td>1.1E-03</td><td>MB231/MB468</td><td>MP</td></tr>
<tr><td>>hg38_21_+_8433172_8446622_RNA45SN1_45S_50ntFlanks@6660.6699.40|GCGGTGGATCACTCGGCTCGTGCGTCGATGAAGAACGCAG</td><td>>hg38_21_+_8433172_8446622_RNA45SN1_45S_50ntFlanks</td><td>6660</td><td>6699</td><td>40</td><td>GCGGTGGATCACTCGGCTCGTGCGTCGATGAAGAACGCAG</td><td>77.1</td><td>-4.3</td><td>1.3</td><td>-3.4</td><td>7.7E-04</td><td>1.1E-03</td><td>MB231/MB468</td><td>MP</td></tr>
<tr><td>>MI0000088|hsa-mir-30a&WithFlank&6|-|71403545|71403627@12.35.24(+1A)&[MIMAT0000087&hsa-miR-30a-5p&offsets|0|+2(+1A);m-15&6|-|71403595|71403616&offsets|0|+2(+1A)]|TGTAAACATCCTCGACTGGAAGCTA</td><td>>MI0000088|hsa-mir-30a&WithFlank&6|-|71403545|71403627</td><td>12</td><td>35</td><td>24(+1A)</td><td>TGTAAACATCCTCGACTGGAAGCTA</td><td>2707.4</td><td>-4.1</td><td>1.1</td><td>-3.8</td><td>1.4E-04</td><td>1.1E-03</td><td>BT20/MB231</td><td>MP</td></tr>
<tr><td>>hg38_21_+_8433172_8446622_RNA45SN1_45S_50ntFlanks@12617.12653.37|TGAGGAGCCAATGGGGCGAAGCTACCATCTGTGGGAT</td><td>>hg38_21_+_8433172_8446622_RNA45SN1_45S_50ntFlanks</td><td>12617</td><td>12653</td><td>37</td><td>TGAGGAGCCAATGGGGCGAAGCTACCATCTGTGGGAT</td><td>329.2</td><td>-3.8</td><td>1.1</td><td>-3.3</td><td>8.3E-04</td><td>1.1E-03</td><td>BT20/MB468</td><td>CYTO</td></tr>
<tr><td>>hg38_21_+_8433172_8446622_RNA45SN1_45S_50ntFlanks@6776.6802.27|CTCCCGGGGCTACGCCTGTCTGAGCGT</td><td>>hg38_21_+_8433172_8446622_RNA45SN1_45S_50ntFlanks</td><td>6776</td><td>6802</td><td>27</td><td>CTCCCGGGGCTACGCCTGTCTGAGCGT</td><td>381.3</td><td>2.2</td><td>0.6</td><td>3.3</td><td>8.3E-04</td><td>1.1E-03</td><td>BT20/MB468</td><td>CYTO</td></tr>
<tr><td>>MI0003820|hsa-mir-454&WithFlank&17|-|59137752|59137878@70.94.25&[MIMAT0003885&hsa-miR-454-3p&offsets|0|+2;m-272&17|-|59137787|59137809&offsets|0|+2]|TAGTGCAATATTGCTTATAGGGTTT</td><td>>MI0003820|hsa-mir-454&WithFlank&17|-|59137752|59137878</td><td>70</td><td>94</td><td>25</td><td>TAGTGCAATATTGCTTATAGGGTTT</td><td>125.7</td><td>3.5</td><td>1.1</td><td>3.3</td><td>9.3E-04</td><td>1.1E-03</td><td>BT20/MB468</td><td>NUC</td></tr>
<tr><td>>hg38_21_+_8433172_8446622_RNA45SN1_45S_50ntFlanks@12081.12107.27|CTCTCGCTTCTGGCGCCAAGCGCCCGG</td><td>>hg38_21_+_8433172_8446622_RNA45SN1_45S_50ntFlanks</td><td>12081</td><td>12107</td><td>27</td><td>CTCTCGCTTCTGGCGCCAAGCGCCCGG</td><td>201.1</td><td>-5.5</td><td>1.6</td><td>-3.5</td><td>4.0E-04</td><td>1.1E-03</td><td>BT20/MB468</td><td>MP</td></tr>
<tr><td>>hg38_21_+_8433172_8446622_RNA45SN1_45S_50ntFlanks@4230.4251.22|ACAATACAGGACTCTTTCGAGG</td><td>>hg38_21_+_8433172_8446622_RNA45SN1_45S_50ntFlanks</td><td>4230</td><td>4251</td><td>22</td><td>ACAATACAGGACTCTTTCGAGG</td><td>138.0</td><td>-2.2</td><td>0.7</td><td>-3.3</td><td>8.7E-04</td><td>1.1E-03</td><td>MB231/MB468</td><td>CYTO</td></tr>
<tr><td>>hg38_21_+_8433172_8446622_RNA45SN1_45S_50ntFlanks@10866.10889.24|CTAAGGGCTGGGTCGGTCGGGCTG</td><td>>hg38_21_+_8433172_8446622_RNA45SN1_45S_50ntFlanks</td><td>10866</td><td>10889</td><td>24</td><td>CTAAGGGCTGGGTCGGTCGGGCTG</td><td>333.9</td><td>-2.6</td><td>0.8</td><td>-3.4</td><td>6.9E-04</td><td>1.1E-03</td><td>BT20/MB468</td><td>MITO</td></tr>
<tr><td>>am_trna128_GlyGCC_6_-_27870686_27870756@3.31.29,trna133_GlyCCC_1_-_16872434_16872504@3.31.29,trna18_GlyGCC_16_+_70822597_70822667@3.31.29,trna19_GlyGCC_16_+_70823410_70823480@3.31.29,trna19_GlyGCC_2_-_157257659_157257729@3.31.29,trna24_GlyGCC_16_-_70812942_70813012@3.31.29,trna25_GlyGCC_16_-_70812114_70812184@3.31.29,trna4_GlyCCC_1_+_17188416_17188486@3.31.29,trna5_GlyGCC_17_+_8029064_8029134@3.31.29,trna68_GlyGCC_1_-_161493637_161493707@3.31.29|ATTGGTGGTTCAGTGGTAGAATTCTCGCC</td><td>>am_trna128_GlyGCC_6_-_27870686_27870756</td><td>3</td><td>31</td><td>29</td><td>ATTGGTGGTTCAGTGGTAGAATTCTCGCC</td><td>135.7</td><td>5.2</td><td>1.6</td><td>3.4</td><td>7.7E-04</td><td>1.1E-03</td><td>MB231/MB468</td><td>MP</td></tr>
<tr><td>>am_trnaMT_SerGCT_MT_+_12207_12265@14.32.19|AGAACTGCTAACTCATGCC</td><td>>am_trnaMT_SerGCT_MT_+_12207_12265</td><td>14</td><td>32</td><td>19</td><td>AGAACTGCTAACTCATGCC</td><td>622.2</td><td>3.0</td><td>0.8</td><td>3.8</td><td>1.5E-04</td><td>1.1E-03</td><td>BT20/MB231</td><td>CYTO</td></tr>
<tr><td>>am_trnaMT_HisGTG_MT_+_12138_12206@26.47.22|AGATTGTGAATCTGACAACAGA</td><td>>am_trnaMT_HisGTG_MT_+_12138_12206</td><td>26</td><td>47</td><td>22</td><td>AGATTGTGAATCTGACAACAGA</td><td>59.9</td><td>6.6</td><td>1.9</td><td>3.5</td><td>4.0E-04</td><td>1.1E-03</td><td>BT20/MB468</td><td>MP</td></tr>
<tr><td>>hg38_21_+_8433172_8446622_RNA45SN1_45S_50ntFlanks@11772.11804.33|TTAGTGACGCGCATGAATGGATGAACGAGATTC</td><td>>hg38_21_+_8433172_8446622_RNA45SN1_45S_50ntFlanks</td><td>11772</td><td>11804</td><td>33</td><td>TTAGTGACGCGCATGAATGGATGAACGAGATTC</td><td>84.0</td><td>-4.0</td><td>1.2</td><td>-3.4</td><td>7.7E-04</td><td>1.1E-03</td><td>MB231/MB468</td><td>MP</td></tr>
<tr><td>>MI0000439|hsa-mir-23b&WithFlank&9|+|95085202|95085310@67.85.19(+2U)&[MIMAT0000418&hsa-miR-23b-3p&offsets|+3|-1(+2U);m-42&9|+|95085265|95085285&offsets|+3|+1(+2U)]|ACATTGCCAGGGATTACCATT</td><td>>MI0000439|hsa-mir-23b&WithFlank&9|+|95085202|95085310</td><td>67</td><td>85</td><td>19(+2U)</td><td>ACATTGCCAGGGATTACCATT</td><td>449.2</td><td>12.0</td><td>3.6</td><td>3.3</td><td>9.2E-04</td><td>1.1E-03</td><td>MB231/MB468</td><td>MITO</td></tr>
<tr><td>>hg38_21_+_8433172_8446622_RNA45SN1_45S_50ntFlanks@10792.10816.25|CTCTGGCATGTTGGAACAATGTAGG</td><td>>hg38_21_+_8433172_8446622_RNA45SN1_45S_50ntFlanks</td><td>10792</td><td>10816</td><td>25</td><td>CTCTGGCATGTTGGAACAATGTAGG</td><td>230.4</td><td>-2.5</td><td>0.8</td><td>-3.3</td><td>9.3E-04</td><td>1.1E-03</td><td>MB231/MB468</td><td>NUC</td></tr>
<tr><td>>hg38_21_+_8433172_8446622_RNA45SN1_45S_50ntFlanks@9534.9564.31|AGGAAACTCTGGTGGAGGTCCGTAGCGGTCC</td><td>>hg38_21_+_8433172_8446622_RNA45SN1_45S_50ntFlanks</td><td>9534</td><td>9564</td><td>31</td><td>AGGAAACTCTGGTGGAGGTCCGTAGCGGTCC</td><td>1147.6</td><td>-3.8</td><td>1.1</td><td>-3.5</td><td>4.1E-04</td><td>1.1E-03</td><td>BT20/MB468</td><td>MP</td></tr>
<tr><td>>MI0001445|hsa-mir-423&WithFlank&17|+|30117073|30117178@58.80.23&[MIMAT0001340&hsa-miR-423-3p&offsets|-1|-1;m-100&17|+|30117131|30117153&offsets|-1|-1]|AAGCTCGGTCTGAGGCCCCTCAG</td><td>>MI0001445|hsa-mir-423&WithFlank&17|+|30117073|30117178</td><td>58</td><td>80</td><td>23</td><td>AAGCTCGGTCTGAGGCCCCTCAG</td><td>157.3</td><td>5.8</td><td>1.6</td><td>3.5</td><td>4.1E-04</td><td>1.1E-03</td><td>BT20/MB468</td><td>MP</td></tr>
<tr><td>>hg38_21_+_8433172_8446622_RNA45SN1_45S_50ntFlanks@6730.6772.43|ACACATTGATCATCGACACTTCGAACGCACTTGCGGCCCCGGG</td><td>>hg38_21_+_8433172_8446622_RNA45SN1_45S_50ntFlanks</td><td>6730</td><td>6772</td><td>43</td><td>ACACATTGATCATCGACACTTCGAACGCACTTGCGGCCCCGGG</td><td>207.9</td><td>-2.2</td><td>0.7</td><td>-3.4</td><td>7.0E-04</td><td>1.1E-03</td><td>BT20/MB468</td><td>MITO</td></tr>
<tr><td>>hg38_21_+_8433172_8446622_RNA45SN1_45S_50ntFlanks@11917.11940.24|GACATGAGAGGTGTAGAATAAGTG</td><td>>hg38_21_+_8433172_8446622_RNA45SN1_45S_50ntFlanks</td><td>11917</td><td>11940</td><td>24</td><td>GACATGAGAGGTGTAGAATAAGTG</td><td>336.1</td><td>7.2</td><td>1.9</td><td>3.8</td><td>1.4E-04</td><td>1.1E-03</td><td>BT20/MB231</td><td>MP</td></tr>
<tr><td>>hg38_21_+_8433172_8446622_RNA45SN1_45S_50ntFlanks@12081.12107.27|CTCTCGCTTCTGGCGCCAAGCGCCCGG</td><td>>hg38_21_+_8433172_8446622_RNA45SN1_45S_50ntFlanks</td><td>12081</td><td>12107</td><td>27</td><td>CTCTCGCTTCTGGCGCCAAGCGCCCGG</td><td>500.0</td><td>-3.9</td><td>1.2</td><td>-3.4</td><td>7.0E-04</td><td>1.1E-03</td><td>BT20/MB468</td><td>MITO</td></tr>
<tr><td>>hg38_21_+_8433172_8446622_RNA45SN1_45S_50ntFlanks@5482.5514.33|CGGAGCGCTGAGAAGACGGTCGAACTTGACTAT</td><td>>hg38_21_+_8433172_8446622_RNA45SN1_45S_50ntFlanks</td><td>5482</td><td>5514</td><td>33</td><td>CGGAGCGCTGAGAAGACGGTCGAACTTGACTAT</td><td>336.1</td><td>-2.3</td><td>0.7</td><td>-3.3</td><td>8.7E-04</td><td>1.1E-03</td><td>MB231/MB468</td><td>CYTO</td></tr>
<tr><td>>MI0000456|hsa-mir-140&WithFlank&16|+|69933075|69933186@69.87.19(+1A)&[MIMAT0004597&hsa-miR-140-3p&offsets|+1|-1(+1A);m-45&16|+|69933143|69933164&offsets|0|-3(+1A)]|ACCACAGGGTAGAACCACGA</td><td>>MI0000456|hsa-mir-140&WithFlank&16|+|69933075|69933186</td><td>69</td><td>87</td><td>19(+1A)</td><td>ACCACAGGGTAGAACCACGA</td><td>57.9</td><td>9.6</td><td>2.7</td><td>3.5</td><td>4.1E-04</td><td>1.1E-03</td><td>BT20/MB468</td><td>MP</td></tr>
<tr><td>>am_trna128_GlyGCC_6_-_27870686_27870756@1.28.28,trna133_GlyCCC_1_-_16872434_16872504@1.28.28,trna18_GlyGCC_16_+_70822597_70822667@1.28.28,trna19_GlyGCC_16_+_70823410_70823480@1.28.28,trna19_GlyGCC_2_-_157257659_157257729@1.28.28,trna24_GlyGCC_16_-_70812942_70813012@1.28.28,trna25_GlyGCC_16_-_70812114_70812184@1.28.28,trna4_GlyCCC_1_+_17188416_17188486@1.28.28,trna5_GlyGCC_17_+_8029064_8029134@1.28.28,trna68_GlyGCC_1_-_161493637_161493707@1.28.28|GCATTGGTGGTTCAGTGGTAGAATTCTC</td><td>>am_trna128_GlyGCC_6_-_27870686_27870756</td><td>1</td><td>28</td><td>28</td><td>GCATTGGTGGTTCAGTGGTAGAATTCTC</td><td>21370.8</td><td>-2.9</td><td>0.8</td><td>-3.5</td><td>4.1E-04</td><td>1.1E-03</td><td>BT20/MB468</td><td>MP</td></tr>
<tr><td>>hg38_21_+_8433172_8446622_RNA45SN1_45S_50ntFlanks@10238.10258.21|CGCTACGCCGCGACGAGTAGG</td><td>>hg38_21_+_8433172_8446622_RNA45SN1_45S_50ntFlanks</td><td>10238</td><td>10258</td><td>21</td><td>CGCTACGCCGCGACGAGTAGG</td><td>464.5</td><td>3.7</td><td>1.0</td><td>3.5</td><td>4.1E-04</td><td>1.1E-03</td><td>BT20/MB468</td><td>MP</td></tr>
<tr><td>>trna10_AspGTC_12_-_125424193_125424264@35.73.39,trna12_AspGTC_12_-_125411891_125411962@35.73.39,trna144_AspGTC_6_-_27551236_27551307@35.73.39,trna38_AspGTC_17_-_8125556_8125627@35.73.39,trna45_AspGTC_6_+_27447453_27447524@35.73.39,trna48_AspGTC_6_+_27471523_27471594@35.73.39,trna4_AspGTC_12_+_96429799_96429870@35.73.39,trna69_AspGTC_1_-_161440205_161440276@35.73.39,trna72_AspGTC_1_-_161432824_161432895@35.73.39,trna75_AspGTC_1_-_161425414_161425485@35.73.39,trna78_AspGTC_1_-_161418033_161418104@35.73.39,trna81_AspGTC_1_-_161410615_161410686@35.73.39|TCACGCGGGAGACCGGGGTTCGATTCCCCGACGGGGAGC</td><td>>trna10_AspGTC_12_-_125424193_125424264</td><td>35</td><td>73</td><td>39</td><td>TCACGCGGGAGACCGGGGTTCGATTCCCCGACGGGGAGC</td><td>325.7</td><td>2.8</td><td>0.8</td><td>3.3</td><td>8.4E-04</td><td>1.1E-03</td><td>BT20/MB468</td><td>CYTO</td></tr>
<tr><td>>hg38_21_+_8433172_8446622_RNA45SN1_45S_50ntFlanks@11920.11967.48|ATGAGAGGTGTAGAATAAGTGGGAGGCCCCCGGCGCCCCCCCGGTGTC</td><td>>hg38_21_+_8433172_8446622_RNA45SN1_45S_50ntFlanks</td><td>11920</td><td>11967</td><td>48</td><td>ATGAGAGGTGTAGAATAAGTGGGAGGCCCCCGGCGCCCCCCCGGTGTC</td><td>58.8</td><td>-9.0</td><td>2.5</td><td>-3.5</td><td>4.1E-04</td><td>1.1E-03</td><td>BT20/MB468</td><td>MP</td></tr>
<tr><td>>trnaMT_ProTGG_MT_-_15956_16023@40.58.19|AATGGTGGAGTTAAAGACT</td><td>>trnaMT_ProTGG_MT_-_15956_16023</td><td>40</td><td>58</td><td>19</td><td>AATGGTGGAGTTAAAGACT</td><td>838.9</td><td>2.9</td><td>0.9</td><td>3.4</td><td>7.8E-04</td><td>1.1E-03</td><td>MB231/MB468</td><td>MP</td></tr>
<tr><td>>hg38_21_+_8433172_8446622_RNA45SN1_45S_50ntFlanks@3855.3898.44|CTGTGGTAATTCTAGAGCTAATACATGCCGACGGGCGCTGACCC</td><td>>hg38_21_+_8433172_8446622_RNA45SN1_45S_50ntFlanks</td><td>3855</td><td>3898</td><td>44</td><td>CTGTGGTAATTCTAGAGCTAATACATGCCGACGGGCGCTGACCC</td><td>66.5</td><td>-9.9</td><td>3.0</td><td>-3.4</td><td>7.8E-04</td><td>1.1E-03</td><td>MB231/MB468</td><td>MP</td></tr>
<tr><td>>am_trnaMT_PheGAA_MT_+_577_647@40.60.21|TGTTTAGACGGGCTCACATCA</td><td>>am_trnaMT_PheGAA_MT_+_577_647</td><td>40</td><td>60</td><td>21</td><td>TGTTTAGACGGGCTCACATCA</td><td>109.5</td><td>6.9</td><td>1.9</td><td>3.5</td><td>4.1E-04</td><td>1.1E-03</td><td>BT20/MB468</td><td>MP</td></tr>
<tr><td>>hg38_21_+_8433172_8446622_RNA45SN1_45S_50ntFlanks@8359.8390.32|AAGAACTTTGAAGAGAGAGTTCAAGAGGGCGT</td><td>>hg38_21_+_8433172_8446622_RNA45SN1_45S_50ntFlanks</td><td>8359</td><td>8390</td><td>32</td><td>AAGAACTTTGAAGAGAGAGTTCAAGAGGGCGT</td><td>59.5</td><td>5.6</td><td>1.6</td><td>3.5</td><td>4.1E-04</td><td>1.1E-03</td><td>BT20/MB468</td><td>MP</td></tr>
<tr><td>>hg38_21_+_8433172_8446622_RNA45SN1_45S_50ntFlanks@3852.3874.23|TAACTGTGGTAATTCTAGAGCTA</td><td>>hg38_21_+_8433172_8446622_RNA45SN1_45S_50ntFlanks</td><td>3852</td><td>3874</td><td>23</td><td>TAACTGTGGTAATTCTAGAGCTA</td><td>199.1</td><td>-5.5</td><td>1.5</td><td>-3.5</td><td>4.1E-04</td><td>1.1E-03</td><td>BT20/MB468</td><td>MP</td></tr>
<tr><td>>hg38_21_+_8433172_8446622_RNA45SN1_45S_50ntFlanks@9494.9531.38|CCGAAAGATGGTGAACTATGCCTGGGCAGGGCGAAGCC</td><td>>hg38_21_+_8433172_8446622_RNA45SN1_45S_50ntFlanks</td><td>9494</td><td>9531</td><td>38</td><td>CCGAAAGATGGTGAACTATGCCTGGGCAGGGCGAAGCC</td><td>119.1</td><td>6.6</td><td>1.9</td><td>3.5</td><td>4.1E-04</td><td>1.1E-03</td><td>BT20/MB468</td><td>MP</td></tr>
<tr><td>>MI0005559|hsa-mir-744&WithFlank&17|+|12081893|12082002@17.36.20(+1A)&[MIMAT0004945&hsa-miR-744-5p&offsets|0|-2(+1A);m-172&17|+|12081909|12081929&offsets|0|-1(+1A)]|TGCGGGGCTAGGGCTAACAGA</td><td>>MI0005559|hsa-mir-744&WithFlank&17|+|12081893|12082002</td><td>17</td><td>36</td><td>20(+1A)</td><td>TGCGGGGCTAGGGCTAACAGA</td><td>137.4</td><td>6.6</td><td>1.9</td><td>3.5</td><td>4.1E-04</td><td>1.1E-03</td><td>BT20/MB468</td><td>MP</td></tr>
<tr><td>>trnaMT_ThrTGT_MT_+_15888_15953@37.63.27|ACCGGAGATGAAAACCTTTTTCCAAGG</td><td>>trnaMT_ThrTGT_MT_+_15888_15953</td><td>37</td><td>63</td><td>27</td><td>ACCGGAGATGAAAACCTTTTTCCAAGG</td><td>170.6</td><td>5.6</td><td>1.6</td><td>3.5</td><td>4.1E-04</td><td>1.1E-03</td><td>BT20/MB468</td><td>MP</td></tr>
<tr><td>>hg38_21_+_8433172_8446622_RNA45SN1_45S_50ntFlanks@11206.11232.27|CGGCGGCGACTCTGGACGCGAGCCGGG</td><td>>hg38_21_+_8433172_8446622_RNA45SN1_45S_50ntFlanks</td><td>11206</td><td>11232</td><td>27</td><td>CGGCGGCGACTCTGGACGCGAGCCGGG</td><td>931.3</td><td>-2.2</td><td>0.7</td><td>-3.3</td><td>8.4E-04</td><td>1.1E-03</td><td>BT20/MB468</td><td>CYTO</td></tr>
</table>

<table>
<tr><td>>trna10_AspGTC_12_-_125424193_125424264@38.69.32,trna12_AspGTC_12_-_125411891_125411962@38.69.32,trna144_AspGTC_6_-_27551236_27551307@38.69.32,trna38_AspGTC_17_-_8125556_8125627@38.69.32,trna45_AspGTC_6_+_27447453_27447524@38.69.32,trna48_AspGTC_6_+_27471523_27471594@38.69.32,trna4_AspGTC_12_+_96429799_96429870@38.69.32,trna69_AspGTC_1_-_161440205_161440276@38.69.32,trna72_AspGTC_1_-_161432824_161432895@38.69.32,trna75_AspGTC_1_-_161425414_161425485@38.69.32,trna78_AspGTC_1_-_161418033_161418104@38.69.32,trna81_AspGTC_1_-_161410615_161410686@38.69.32|CGCGGGAGACCGGGGTTCGATTCCCCGACGGG</td><td>>trna10_AspGTC_12_-_125424193_125424264</td><td>38</td><td>69</td><td>32</td><td>CGCGGGAGACCGGGGTTCGATTCCCCGACGGG</td><td>203.2</td><td>3.3</td><td>0.9</td><td>3.8</td><td>1.7E-04</td><td>1.1E-03</td><td>BT20/MB231</td><td>MITO</td></tr>
<tr><td>>hg38_21_+_8433172_8446622_RNA45SN1_45S_50ntFlanks@10395.10424.30|GCAGTTGAACATGGGTCAGTCGGTCCTGAG</td><td>>hg38_21_+_8433172_8446622_RNA45SN1_45S_50ntFlanks</td><td>10395</td><td>10424</td><td>30</td><td>GCAGTTGAACATGGGTCAGTCGGTCCTGAG</td><td>11468.3</td><td>3.5</td><td>0.9</td><td>3.8</td><td>1.7E-04</td><td>1.1E-03</td><td>BT20/MB231</td><td>MITO</td></tr>
<tr><td>>MI0000803|hsa-mir-330&WithFlank&19|-|45638988|45639093@64.87.24&[MIMAT0000751&hsa-miR-330-3p&offsets|+1|+2;m-347&19|-|45639009|45639030&offsets|0|+2]|CAAAGCACACGGCCTGCAGAGAGG</td><td>>MI0000803|hsa-mir-330&WithFlank&19|-|45638988|45639093</td><td>64</td><td>87</td><td>24</td><td>CAAAGCACACGGCCTGCAGAGAGG</td><td>212.8</td><td>5.2</td><td>1.5</td><td>3.4</td><td>7.8E-04</td><td>1.1E-03</td><td>MB231/MB468</td><td>MP</td></tr>
<tr><td>>MI0003184|hsa-mir-500a&WithFlank&X|+|50008425|50008520@58.78.21&[MIMAT0002871&hsa-miR-500a-3p&offsets|0|-1;m-111&X|+|50008482|50008504&offsets|0|-2]|ATGCACCTGGGCAAGGATTCT</td><td>>MI0003184|hsa-mir-500a&WithFlank&X|+|50008425|50008520</td><td>58</td><td>78</td><td>21</td><td>ATGCACCTGGGCAAGGATTCT</td><td>372.4</td><td>4.0</td><td>1.1</td><td>3.5</td><td>4.1E-04</td><td>1.1E-03</td><td>BT20/MB468</td><td>MP</td></tr>
<tr><td>>trna116_GluCTC_1_-_145399233_145399304@1.36.36,trna59_GluCTC_1_+_249168447_249168518@1.36.36,trna71_GluCTC_1_-_161439189_161439260@1.36.36,trna74_GluCTC_1_-_161431809_161431880@1.36.36,trna77_GluCTC_1_-_161424398_161424469@1.36.36,trna77_GluCTC_6_+_28949976_28950047@1.36.36,trna80_GluCTC_1_-_161417018_161417089@1.36.36,trna87_GluCTC_6_-_126101393_126101464@1.36.36|TCCCTGGTGGTCTAGTGGTTAGGATTCGGCGCTCTC</td><td>>trna116_GluCTC_1_-_145399233_145399304</td><td>1</td><td>36</td><td>36</td><td>TCCCTGGTGGTCTAGTGGTTAGGATTCGGCGCTCTC</td><td>868.1</td><td>4.7</td><td>1.4</td><td>3.4</td><td>7.8E-04</td><td>1.1E-03</td><td>MB231/MB468</td><td>MP</td></tr>
<tr><td>>MI0000749|hsa-mir-30e&WithFlank&1|+|40754349|40754452@65.85.21&[MIMAT0000693&hsa-miR-30e-3p&offsets|0|-1;m-25&1|+|40754413|40754434&offsets|0|-1]|CTTTCAGTCGGATGTTTACAG</td><td>>MI0000749|hsa-mir-30e&WithFlank&1|+|40754349|40754452</td><td>65</td><td>85</td><td>21</td><td>CTTTCAGTCGGATGTTTACAG</td><td>137.4</td><td>3.2</td><td>1.0</td><td>3.3</td><td>9.3E-04</td><td>1.1E-03</td><td>MB231/MB468</td><td>MITO</td></tr>
<tr><td>>hg38_21_+_8433172_8446622_RNA45SN1_45S_50ntFlanks@7989.8012.24|AGACGTGGCGACCCGCTGAATTTA</td><td>>hg38_21_+_8433172_8446622_RNA45SN1_45S_50ntFlanks</td><td>7989</td><td>8012</td><td>24</td><td>AGACGTGGCGACCCGCTGAATTTA</td><td>2198.3</td><td>-3.4</td><td>1.0</td><td>-3.3</td><td>9.4E-04</td><td>1.1E-03</td><td>BT20/MB468</td><td>NUC</td></tr>
<tr><td>>hg38_21_+_8433172_8446622_RNA45SN1_45S_50ntFlanks@5084.5115.32|CGAACGAGACTCTGGCATGCTAACTAGTTACG</td><td>>hg38_21_+_8433172_8446622_RNA45SN1_45S_50ntFlanks</td><td>5084</td><td>5115</td><td>32</td><td>CGAACGAGACTCTGGCATGCTAACTAGTTACG</td><td>460.6</td><td>-3.3</td><td>1.0</td><td>-3.4</td><td>7.0E-04</td><td>1.1E-03</td><td>BT20/MB468</td><td>MITO</td></tr>
<tr><td>>hg38_21_+_8433172_8446622_RNA45SN1_45S_50ntFlanks@10509.10532.24|GAATCCGGAGTGGCGGAGATGGGC</td><td>>hg38_21_+_8433172_8446622_RNA45SN1_45S_50ntFlanks</td><td>10509</td><td>10532</td><td>24</td><td>GAATCCGGAGTGGCGGAGATGGGC</td><td>2577.1</td><td>6.9</td><td>1.8</td><td>3.8</td><td>1.4E-04</td><td>1.1E-03</td><td>BT20/MB231</td><td>MP</td></tr>
<tr><td>>trna111_HisGTG_1_-_147774845_147774916@-1G.33.34,trna118_HisGTG_1_-_145396881_145396952@-1G.33.34,trna16_HisGTG_1_+_146544773_146544844@-1G.33.34,trna1_HisGTG_15_+_45493349_45493420@-1G.33.34,trna21_HisGTG_1_+_147753471_147753542@-1G.33.34,trna33_HisGTG_6_+_27125906_27125977@-1G.33.34,trna7_HisGTG_9_-_14433938_14434009@-1G.33.34,trna8_HisGTG_15_-_45492611_45492682@-1G.33.34,trna9_HisGTG_15_-_45490804_45490875@-1G.33.34|GGCCGTGATCGTATAGTGGTTAGTACTCTGCGTT</td><td>>trna111_HisGTG_1_-_147774845_147774916</td><td>-1G</td><td>33</td><td>34</td><td>GGCCGTGATCGTATAGTGGTTAGTACTCTGCGTT</td><td>421.6</td><td>2.3</td><td>0.7</td><td>3.3</td><td>9.4E-04</td><td>1.1E-03</td><td>BT20/MB468</td><td>NUC</td></tr>
<tr><td>>hg38_21_+_8433172_8446622_RNA45SN1_45S_50ntFlanks@6743.6770.28|CGACACTTCGAACGCACTTGCGGCCCCG</td><td>>hg38_21_+_8433172_8446622_RNA45SN1_45S_50ntFlanks</td><td>6743</td><td>6770</td><td>28</td><td>CGACACTTCGAACGCACTTGCGGCCCCG</td><td>2299.2</td><td>-6.3</td><td>1.8</td><td>-3.5</td><td>4.1E-04</td><td>1.1E-03</td><td>BT20/MB468</td><td>MP</td></tr>
<tr><td>>hg38_21_+_8433172_8446622_RNA45SN1_45S_50ntFlanks@4982.4999.18|ACGGACAGGATTGACAGA</td><td>>hg38_21_+_8433172_8446622_RNA45SN1_45S_50ntFlanks</td><td>4982</td><td>4999</td><td>18</td><td>ACGGACAGGATTGACAGA</td><td>398.0</td><td>-6.1</td><td>1.8</td><td>-3.4</td><td>7.0E-04</td><td>1.1E-03</td><td>BT20/MB468</td><td>MITO</td></tr>
<tr><td>>MI0000750|hsa-mir-26a-2&WithFlank&12|-|57824603|57824698@20.40.21(+1C);MI0000083|hsa-mir-26a-1&WithFlank&3|+|37969398|37969486@16.36.21(+1C)&[MIMAT0000082&hsa-miR-26a-5p&offsets|0|-1(+1C);m-33&12|-|57824658|57824679&offsets|0|-1(+1C)];[MIMAT0000082_1&hsa-miR-26a-5p&offsets|0|-1(+1C);m-34&3|+|37969413|37969434&offsets|0|-1(+1C)]|TTCAAGTAATCCAGGATAGGCC</td><td>>MI0000750|hsa-mir-26a-2&WithFlank&12|-|57824603|57824698</td><td>20</td><td>40</td><td>21(+1C)</td><td>TTCAAGTAATCCAGGATAGGCC</td><td>2773.1</td><td>1.6</td><td>0.5</td><td>3.3</td><td>9.4E-04</td><td>1.1E-03</td><td>BT20/MB468</td><td>NUC</td></tr>
<tr><td>>MI0000434|hsa-let-7i&WithFlank&12|+|62603680|62603775@12.30.19&[MIMAT0000415&hsa-let-7i-5p&offsets|0|-3;m-49&12|+|62603691|62603712&offsets|0|-3]|TGAGGTAGTAGTTTGTGCT</td><td>>MI0000434|hsa-let-7i&WithFlank&12|+|62603680|62603775</td><td>12</td><td>30</td><td>19</td><td>TGAGGTAGTAGTTTGTGCT</td><td>261.3</td><td>5.7</td><td>1.7</td><td>3.4</td><td>7.8E-04</td><td>1.1E-03</td><td>MB231/MB468</td><td>MP</td></tr>
<tr><td>>hg38_21_+_8433172_8446622_RNA45SN1_45S_50ntFlanks@13006.13039.34|CTATTGAAAGTCAGCCCTCGACACAAGGGTTTGT</td><td>>hg38_21_+_8433172_8446622_RNA45SN1_45S_50ntFlanks</td><td>13006</td><td>13039</td><td>34</td><td>CTATTGAAAGTCAGCCCTCGACACAAGGGTTTGT</td><td>798.0</td><td>2.8</td><td>0.8</td><td>3.3</td><td>9.4E-04</td><td>1.1E-03</td><td>MB231/MB468</td><td>NUC</td></tr>
<tr><td>>hg38_MT_+_1621_3279_RNR2_16S_50ntFlanks@1095.1120.26|AGACGAGAAGACCCTATGGAGCTTTA</td><td>>hg38_MT_+_1621_3279_RNR2_16S_50ntFlanks</td><td>1095</td><td>1120</td><td>26</td><td>AGACGAGAAGACCCTATGGAGCTTTA</td><td>505.8</td><td>3.5</td><td>1.0</td><td>3.5</td><td>4.1E-04</td><td>1.1E-03</td><td>BT20/MB468</td><td>MP</td></tr>
<tr><td>>hg38_21_+_8433172_8446622_RNA45SN1_45S_50ntFlanks@11553.11588.36|CTCGGCCGGCGCCTAGCAGCCGACTTAGAACTGGTG</td><td>>hg38_21_+_8433172_8446622_RNA45SN1_45S_50ntFlanks</td><td>11553</td><td>11588</td><td>36</td><td>CTCGGCCGGCGCCTAGCAGCCGACTTAGAACTGGTG</td><td>58.6</td><td>-4.0</td><td>1.2</td><td>-3.4</td><td>7.8E-04</td><td>1.1E-03</td><td>MB231/MB468</td><td>MP</td></tr>
<tr><td>>hg38_21_+_8433172_8446622_RNA45SN1_45S_50ntFlanks@11573.11605.33|CGACTTAGAACTGGTGCGGACCAGGGGAATCCG</td><td>>hg38_21_+_8433172_8446622_RNA45SN1_45S_50ntFlanks</td><td>11573</td><td>11605</td><td>33</td><td>CGACTTAGAACTGGTGCGGACCAGGGGAATCCG</td><td>254.9</td><td>7.3</td><td>1.9</td><td>3.8</td><td>1.4E-04</td><td>1.1E-03</td><td>BT20/MB231</td><td>MP</td></tr>
<tr><td>>hg38_21_+_8433172_8446622_RNA45SN1_45S_50ntFlanks@5485.5521.37|AGCGCTGAGAAGACGGTCGAACTTGACTATCTAGAGG</td><td>>hg38_21_+_8433172_8446622_RNA45SN1_45S_50ntFlanks</td><td>5485</td><td>5521</td><td>37</td><td>AGCGCTGAGAAGACGGTCGAACTTGACTATCTAGAGG</td><td>123.0</td><td>10.3</td><td>2.7</td><td>3.8</td><td>1.4E-04</td><td>1.1E-03</td><td>BT20/MB231</td><td>MP</td></tr>
<tr><td>>am_trnaMT_SerGCT_MT_+_12207_12265@14.35.22|AGAACTGCTAACTCATGCCCCC</td><td>>am_trnaMT_SerGCT_MT_+_12207_12265</td><td>14</td><td>35</td><td>22</td><td>AGAACTGCTAACTCATGCCCCC</td><td>299.9</td><td>6.4</td><td>1.8</td><td>3.5</td><td>4.1E-04</td><td>1.1E-03</td><td>BT20/MB468</td><td>MP</td></tr>
<tr><td>>hg38_21_+_8433172_8446622_RNA45SN1_45S_50ntFlanks@12926.12956.31|ACCTGGCGCTAAACCATTCGTAGACGACCTG</td><td>>hg38_21_+_8433172_8446622_RNA45SN1_45S_50ntFlanks</td><td>12926</td><td>12956</td><td>31</td><td>ACCTGGCGCTAAACCATTCGTAGACGACCTG</td><td>97.0</td><td>-4.6</td><td>1.4</td><td>-3.3</td><td>8.5E-04</td><td>1.1E-03</td><td>BT20/MB468</td><td>CYTO</td></tr>
<tr><td>>hg38_21_+_8433172_8446622_RNA45SN1_45S_50ntFlanks@12407.12448.42|TTTTGATCCTTCGATGTCGGCTCTTCCTATCATTGTGAAGCA</td><td>>hg38_21_+_8433172_8446622_RNA45SN1_45S_50ntFlanks</td><td>12407</td><td>12448</td><td>42</td><td>TTTTGATCCTTCGATGTCGGCTCTTCCTATCATTGTGAAGCA</td><td>317.1</td><td>3.4</td><td>1.0</td><td>3.3</td><td>9.5E-04</td><td>1.1E-03</td><td>MB231/MB468</td><td>NUC</td></tr>
<tr><td>>hg38_21_+_8433172_8446622_RNA45SN1_45S_50ntFlanks@4722.4748.27|TCATTAATCAAGAACGAAAGTCGGAGG</td><td>>hg38_21_+_8433172_8446622_RNA45SN1_45S_50ntFlanks</td><td>4722</td><td>4748</td><td>27</td><td>TCATTAATCAAGAACGAAAGTCGGAGG</td><td>557.8</td><td>-3.0</td><td>0.9</td><td>-3.3</td><td>8.5E-04</td><td>1.1E-03</td><td>BT20/MB468</td><td>CYTO</td></tr>
<tr><td>>MI0000239|hsa-mir-197&WithFlank&1|+|109598887|109598973@54.75.22(+1U)&[MIMAT0000227&hsa-miR-197-3p&offsets|0|0(+1U);m-99&1|+|109598940|109598961&offsets|0|0(+1U)]|TTCACCACCTTCTCCACCCAGCT</td><td>>MI0000239|hsa-mir-197&WithFlank&1|+|109598887|109598973</td><td>54</td><td>75</td><td>22(+1U)</td><td>TTCACCACCTTCTCCACCCAGCT</td><td>340.2</td><td>2.8</td><td>0.7</td><td>3.8</td><td>1.5E-04</td><td>1.1E-03</td><td>BT20/MB231</td><td>CYTO</td></tr>
<tr><td>>hg38_21_+_8433172_8446622_RNA45SN1_45S_50ntFlanks@4004.4024.21|TAACCTCGGGCCGATCGCACG</td><td>>hg38_21_+_8433172_8446622_RNA45SN1_45S_50ntFlanks</td><td>4004</td><td>4024</td><td>21</td><td>TAACCTCGGGCCGATCGCACG</td><td>258.7</td><td>-1.9</td><td>0.6</td><td>-3.3</td><td>8.8E-04</td><td>1.1E-03</td><td>MB231/MB468</td><td>CYTO</td></tr>
<tr><td>>hg38_21_+_8433172_8446622_RNA45SN1_45S_50ntFlanks@7983.8004.22|CAGATCAGACGTGGCGACCCGC</td><td>>hg38_21_+_8433172_8446622_RNA45SN1_45S_50ntFlanks</td><td>7983</td><td>8004</td><td>22</td><td>CAGATCAGACGTGGCGACCCGC</td><td>293.0</td><td>-2.5</td><td>0.7</td><td>-3.3</td><td>8.8E-04</td><td>1.1E-03</td><td>MB231/MB468</td><td>CYTO</td></tr>
<tr><td>>hg38_21_+_8433172_8446622_RNA45SN1_45S_50ntFlanks@10392.10429.38|ACAGCAGTTGAACATGGGTCAGTCGGTCCTGAGAGATG</td><td>>hg38_21_+_8433172_8446622_RNA45SN1_45S_50ntFlanks</td><td>10392</td><td>10429</td><td>38</td><td>ACAGCAGTTGAACATGGGTCAGTCGGTCCTGAGAGATG</td><td>1531.3</td><td>-3.7</td><td>1.1</td><td>-3.5</td><td>4.1E-04</td><td>1.1E-03</td><td>BT20/MB468</td><td>MP</td></tr>
<tr><td>>hg38_21_+_8433172_8446622_RNA45SN1_45S_50ntFlanks@3840.3862.23|CTCCTACTTGGATAACTGTGGTA</td><td>>hg38_21_+_8433172_8446622_RNA45SN1_45S_50ntFlanks</td><td>3840</td><td>3862</td><td>23</td><td>CTCCTACTTGGATAACTGTGGTA</td><td>106.5</td><td>-5.9</td><td>1.7</td><td>-3.4</td><td>7.9E-04</td><td>1.1E-03</td><td>MB231/MB468</td><td>MP</td></tr>
<tr><td>>hg38_21_+_8433172_8446622_RNA45SN1_45S_50ntFlanks@3783.3805.23|AGTGAAACTGCGAATGGCTCATT</td><td>>hg38_21_+_8433172_8446622_RNA45SN1_45S_50ntFlanks</td><td>3783</td><td>3805</td><td>23</td><td>AGTGAAACTGCGAATGGCTCATT</td><td>115.3</td><td>-4.1</td><td>1.2</td><td>-3.4</td><td>7.9E-04</td><td>1.2E-03</td><td>MB231/MB468</td><td>MP</td></tr>
<tr><td>>hg38_21_+_8433172_8446622_RNA45SN1_45S_50ntFlanks@11671.11697.27|CCAGTGCTCTGAATGTCAAAGTGAAGA</td><td>>hg38_21_+_8433172_8446622_RNA45SN1_45S_50ntFlanks</td><td>11671</td><td>11697</td><td>27</td><td>CCAGTGCTCTGAATGTCAAAGTGAAGA</td><td>106.5</td><td>-3.9</td><td>1.2</td><td>-3.4</td><td>7.9E-04</td><td>1.2E-03</td><td>MB231/MB468</td><td>MP</td></tr>
<tr><td>>hg38_21_+_8433172_8446622_RNA45SN1_45S_50ntFlanks@11582.11600.19|ACTGGTGCGGACCAGGGGA</td><td>>hg38_21_+_8433172_8446622_RNA45SN1_45S_50ntFlanks</td><td>11582</td><td>11600</td><td>19</td><td>ACTGGTGCGGACCAGGGGA</td><td>278.0</td><td>-4.5</td><td>1.3</td><td>-3.4</td><td>7.9E-04</td><td>1.2E-03</td><td>MB231/MB468</td><td>MP</td></tr>
<tr><td>>hg38_21_+_8433172_8446622_RNA45SN1_45S_50ntFlanks@10792.10815.24|CTCTGGCATGTTGGAACAATGTAG</td><td>>hg38_21_+_8433172_8446622_RNA45SN1_45S_50ntFlanks</td><td>10792</td><td>10815</td><td>24</td><td>CTCTGGCATGTTGGAACAATGTAG</td><td>215.7</td><td>-2.5</td><td>0.8</td><td>-3.3</td><td>8.9E-04</td><td>1.2E-03</td><td>MB231/MB468</td><td>CYTO</td></tr>
<tr><td>>hg38_21_+_8433172_8446622_RNA45SN1_45S_50ntFlanks@6651.6686.36|CGACTCTTAGCGGTGGATCACTCGGCTCGTGCGTCG</td><td>>hg38_21_+_8433172_8446622_RNA45SN1_45S_50ntFlanks</td><td>6651</td><td>6686</td><td>36</td><td>CGACTCTTAGCGGTGGATCACTCGGCTCGTGCGTCG</td><td>454.4</td><td>2.9</td><td>0.9</td><td>3.3</td><td>8.5E-04</td><td>1.2E-03</td><td>BT20/MB468</td><td>CYTO</td></tr>
<tr><td>>hg38_21_+_8433172_8446622_RNA45SN1_45S_50ntFlanks@10571.10596.26|GAGAAGCCGGCGGGAGCCCCGGGGAG</td><td>>hg38_21_+_8433172_8446622_RNA45SN1_45S_50ntFlanks</td><td>10571</td><td>10596</td><td>26</td><td>GAGAAGCCGGCGGGAGCCCCGGGGAG</td><td>165.4</td><td>-4.7</td><td>1.4</td><td>-3.3</td><td>8.5E-04</td><td>1.2E-03</td><td>BT20/MB468</td><td>CYTO</td></tr>
<tr><td>>hg38_21_+_8433172_8446622_RNA45SN1_45S_50ntFlanks@7975.8016.42|CGCGACCTCAGATCAGACGTGGCGACCCGCTGAATTTAAGCA</td><td>>hg38_21_+_8433172_8446622_RNA45SN1_45S_50ntFlanks</td><td>7975</td><td>8016</td><td>42</td><td>CGCGACCTCAGATCAGACGTGGCGACCCGCTGAATTTAAGCA</td><td>50510.2</td><td>2.9</td><td>0.9</td><td>3.3</td><td>9.4E-04</td><td>1.2E-03</td><td>MB231/MB468</td><td>MITO</td></tr>
<tr><td>>hg38_21_+_8433172_8446622_RNA45SN1_45S_50ntFlanks@9481.9511.31|CACGTGTTAGGACCCGAAAGATGGTGAACTA</td><td>>hg38_21_+_8433172_8446622_RNA45SN1_45S_50ntFlanks</td><td>9481</td><td>9511</td><td>31</td><td>CACGTGTTAGGACCCGAAAGATGGTGAACTA</td><td>124.7</td><td>10.3</td><td>2.7</td><td>3.8</td><td>1.5E-04</td><td>1.2E-03</td><td>BT20/MB231</td><td>MP</td></tr>
<tr><td>>trnaMT_LeuTAA_MT_+_3230_3304@18.44.27|CGGTAATCGCATAAAACTTAAAACTTT</td><td>>trnaMT_LeuTAA_MT_+_3230_3304</td><td>18</td><td>44</td><td>27</td><td>CGGTAATCGCATAAAACTTAAAACTTT</td><td>543.3</td><td>2.9</td><td>0.8</td><td>3.8</td><td>1.5E-04</td><td>1.2E-03</td><td>BT20/MB231</td><td>CYTO</td></tr>
<tr><td>>hg38_21_+_8433172_8446622_RNA45SN1_45S_50ntFlanks@5066.5093.28|TCTGGTTAATTCCGATAACGAACGAGAC</td><td>>hg38_21_+_8433172_8446622_RNA45SN1_45S_50ntFlanks</td><td>5066</td><td>5093</td><td>28</td><td>TCTGGTTAATTCCGATAACGAACGAGAC</td><td>66.6</td><td>-9.9</td><td>3.0</td><td>-3.4</td><td>7.9E-04</td><td>1.2E-03</td><td>MB231/MB468</td><td>MP</td></tr>
<tr><td>>hg38_21_+_8433172_8446622_RNA45SN1_45S_50ntFlanks@10408.10456.49|GGTCAGTCGGTCCTGAGAGATGGGCGAGCGCCGTTCCGAAGGGACGGGC</td><td>>hg38_21_+_8433172_8446622_RNA45SN1_45S_50ntFlanks</td><td>10408</td><td>10456</td><td>49</td><td>GGTCAGTCGGTCCTGAGAGATGGGCGAGCGCCGTTCCGAAGGGACGGGC</td><td>270.7</td><td>-4.4</td><td>1.3</td><td>-3.3</td><td>8.5E-04</td><td>1.2E-03</td><td>BT20/MB468</td><td>CYTO</td></tr>
<tr><td>>hg38_21_+_8433172_8446622_RNA45SN1_45S_50ntFlanks@11200.11223.24|CGGCGGCGGCGGCGACTCTGGACG</td><td>>hg38_21_+_8433172_8446622_RNA45SN1_45S_50ntFlanks</td><td>11200</td><td>11223</td><td>24</td><td>CGGCGGCGGCGGCGACTCTGGACG</td><td>297.9</td><td>-4.9</td><td>1.5</td><td>-3.3</td><td>8.5E-04</td><td>1.2E-03</td><td>BT20/MB468</td><td>CYTO</td></tr>
</table>

<table>
<thead>
<tr><th></th><th></th><th></th><th></th><th></th><th></th><th></th><th></th><th></th><th></th><th></th><th></th><th></th><th></th></tr>
</thead>
<tbody>
<tr><td>>hg38_21_+_8433172_8446622_RNA45SN1_45S_50ntFlanks@10476.10512.37|GGCCGATCGAAAGGGAGTCGGGTTCAGATCCCCGAAT</td><td>>hg38_21_+_8433172_8446622_RNA45SN1_45S_50ntFlanks</td><td>10476</td><td>10512</td><td>37</td><td>GGCCGATCGAAAGGGAGTCGGGTTCAGATCCCCGAAT</td><td>177.7</td><td>-3.6</td><td>1.1</td><td>-3.3</td><td>8.6E-04</td><td>1.2E-03</td><td>BT20/MB468</td><td>CYTO</td></tr>
<tr><td>>hg38_1_-_228634819_228635039_RNA5S12_5S_50ntFlanks@138.169.32|AGACCGCCTGGGAATACCGGGTGCTGTAGGCT</td><td>>hg38_1_-_228634819_228635039_RNA5S12_5S_50ntFlanks</td><td>138</td><td>169</td><td>32</td><td>AGACCGCCTGGGAATACCGGGTGCTGTAGGCT</td><td>462.6</td><td>3.0</td><td>0.9</td><td>3.3</td><td>9.4E-04</td><td>1.2E-03</td><td>MB231/MB468</td><td>MITO</td></tr>
<tr><td>>hg38_21_+_8433172_8446622_RNA45SN1_45S_50ntFlanks@5298.5333.36|ATTCGTGATGGGGATCGGGGATTGCAATTATTCCCC</td><td>>hg38_21_+_8433172_8446622_RNA45SN1_45S_50ntFlanks</td><td>5298</td><td>5333</td><td>36</td><td>ATTCGTGATGGGGATCGGGGATTGCAATTATTCCCC</td><td>67.0</td><td>-9.9</td><td>3.0</td><td>-3.4</td><td>7.9E-04</td><td>1.2E-03</td><td>MB231/MB468</td><td>MP</td></tr>
<tr><td>>trna11_SerAGA_8_-_96281885_96281966@1.24.24,trna145_SerAGA_6_-_27521192_27521273@1.24.24,trna147_SerAGA_6_-_27509554_27509635@1.24.24,trna148_SerTGA_6_-_27473607_27473688@1.24.24,trna172_SerTGA_6_-_26312824_26312905@1.24.24,trna35_SerAGA_17_-_8129928_8130009@1.24.24,trna44_SerAGA_6_+_27446591_27446672@1.24.24,trna46_SerAGA_6_+_27463593_27463674@1.24.24,trna47_SerAGA_6_+_27470818_27470899@1.24.24,trna50_SerAGA_6_+_27499987_27500068@1.24.24,trna51_SerTGA_6_+_27513468_27513549@1.24.24,trna5_SerAGA_6_+_26327817_26327898@1.24.24|GTAGTCGTGGCCGAGTGGTTAAGG</td><td>>trna11_SerAGA_8_-_96281885_96281966</td><td>1</td><td>24</td><td>24</td><td>GTAGTCGTGGCCGAGTGGTTAAGG</td><td>304.6</td><td>6.3</td><td>1.8</td><td>3.5</td><td>4.2E-04</td><td>1.2E-03</td><td>BT20/MB468</td><td>MP</td></tr>
<tr><td>>hg38_21_+_8433172_8446622_RNA45SN1_45S_50ntFlanks@11584.11612.29|TGGTGCGGACCAGGGGAATCCGACTGTTT</td><td>>hg38_21_+_8433172_8446622_RNA45SN1_45S_50ntFlanks</td><td>11584</td><td>11612</td><td>29</td><td>TGGTGCGGACCAGGGGAATCCGACTGTTT</td><td>356.3</td><td>2.9</td><td>0.8</td><td>3.7</td><td>1.8E-04</td><td>1.2E-03</td><td>BT20/MB231</td><td>MITO</td></tr>
<tr><td>>hg38_21_+_8433172_8446622_RNA45SN1_45S_50ntFlanks@11671.11697.27|CCAGTGCTCTGAATGTCAAAGTGAAGA</td><td>>hg38_21_+_8433172_8446622_RNA45SN1_45S_50ntFlanks</td><td>11671</td><td>11697</td><td>27</td><td>CCAGTGCTCTGAATGTCAAAGTGAAGA</td><td>276.3</td><td>-5.7</td><td>1.7</td><td>-3.3</td><td>9.6E-04</td><td>1.2E-03</td><td>BT20/MB468</td><td>NUC</td></tr>
<tr><td>>hg38_21_+_8433172_8446622_RNA45SN1_45S_50ntFlanks@9792.9812.21|GGCGTGGAGCCGGGCGTGGAA</td><td>>hg38_21_+_8433172_8446622_RNA45SN1_45S_50ntFlanks</td><td>9792</td><td>9812</td><td>21</td><td>GGCGTGGAGCCGGGCGTGGAA</td><td>53.6</td><td>9.5</td><td>2.7</td><td>3.5</td><td>4.2E-04</td><td>1.2E-03</td><td>BT20/MB468</td><td>MP</td></tr>
<tr><td>>hg38_21_+_8433172_8446622_RNA45SN1_45S_50ntFlanks@11664.11702.39|TTTCTGCCCAGTGCTCTGAATGTCAAAGTGAAGAAATTC</td><td>>hg38_21_+_8433172_8446622_RNA45SN1_45S_50ntFlanks</td><td>11664</td><td>11702</td><td>39</td><td>TTTCTGCCCAGTGCTCTGAATGTCAAAGTGAAGAAATTC</td><td>237.7</td><td>-3.1</td><td>0.8</td><td>-3.8</td><td>1.5E-04</td><td>1.2E-03</td><td>BT20/MB231</td><td>CYTO</td></tr>
<tr><td>>am_trna128_GlyGCC_6_-_27870686_27870756@1.33.33,trna18_GlyGCC_16_+_70822597_70822667@1.33.33,trna19_GlyGCC_16_+_70823410_70823480@1.33.33,trna19_GlyGCC_2_-_157257659_157257729@1.33.33,trna24_GlyGCC_16_-_70812942_70813012@1.33.33,trna25_GlyGCC_16_-_70812114_70812184@1.33.33,trna5_GlyGCC_17_+_8029064_8029134@1.33.33,trna68_GlyGCC_1_-_161493637_161493707@1.33.33|GCATTGGTGGTTCAGTGGTAGAATTCTCGCCTG</td><td>>am_trna128_GlyGCC_6_-_27870686_27870756</td><td>1</td><td>33</td><td>33</td><td>GCATTGGTGGTTCAGTGGTAGAATTCTCGCCTG</td><td>2042.4</td><td>4.8</td><td>1.4</td><td>3.5</td><td>4.2E-04</td><td>1.2E-03</td><td>BT20/MB468</td><td>MP</td></tr>
<tr><td>>MI0000803|hsa-mir-330&WithFlank&19|-|45638988|45639093@65.87.23&[MIMAT0000751&hsa-miR-330-3p&offsets|+2|+2;m-347&19|-|45639009|45639030&offsets|+1|+2]|AAAGCACACGGCCTGCAGAGAGG</td><td>>MI0000803|hsa-mir-330&WithFlank&19|-|45638988|45639093</td><td>65</td><td>87</td><td>23</td><td>AAAGCACACGGCCTGCAGAGAGG</td><td>60.7</td><td>5.9</td><td>1.8</td><td>3.4</td><td>8.0E-04</td><td>1.2E-03</td><td>MB231/MB468</td><td>MP</td></tr>
<tr><td>>MI0000095|hsa-mir-93&WithFlank&7|-|100093762|100093853@17.40.24&[MIMAT0000093&hsa-miR-93-5p&offsets|0|+1;m-29&7|-|100093815|100093837&offsets|0|+1]|CAAAGTGCTGTTCGTGCAGGTAGT</td><td>>MI0000095|hsa-mir-93&WithFlank&7|-|100093762|100093853</td><td>17</td><td>40</td><td>24</td><td>CAAAGTGCTGTTCGTGCAGGTAGT</td><td>1151.7</td><td>3.1</td><td>0.9</td><td>3.4</td><td>8.0E-04</td><td>1.2E-03</td><td>MB231/MB468</td><td>MP</td></tr>
<tr><td>>hg38_21_+_8433172_8446622_RNA45SN1_45S_50ntFlanks@6772.6800.29|GTTCCTCCCGGGGCTACGCCTGTCTGAGC</td><td>>hg38_21_+_8433172_8446622_RNA45SN1_45S_50ntFlanks</td><td>6772</td><td>6800</td><td>29</td><td>GTTCCTCCCGGGGCTACGCCTGTCTGAGC</td><td>551.7</td><td>2.4</td><td>0.7</td><td>3.4</td><td>7.2E-04</td><td>1.2E-03</td><td>BT20/MB468</td><td>MITO</td></tr>
<tr><td>>hg38_21_+_8433172_8446622_RNA45SN1_45S_50ntFlanks@5074.5108.35|ATTCCGATAACGAACGAGACTCTGGCATGCTAACT</td><td>>hg38_21_+_8433172_8446622_RNA45SN1_45S_50ntFlanks</td><td>5074</td><td>5108</td><td>35</td><td>ATTCCGATAACGAACGAGACTCTGGCATGCTAACT</td><td>68.3</td><td>-10.0</td><td>3.0</td><td>-3.4</td><td>8.0E-04</td><td>1.2E-03</td><td>MB231/MB468</td><td>MP</td></tr>
<tr><td>>hg38_21_+_8433172_8446622_RNA45SN1_45S_50ntFlanks@12502.12519.18|TTAGACCGTCGTGAGACA</td><td>>hg38_21_+_8433172_8446622_RNA45SN1_45S_50ntFlanks</td><td>12502</td><td>12519</td><td>18</td><td>TTAGACCGTCGTGAGACA</td><td>155.6</td><td>-4.3</td><td>1.3</td><td>-3.4</td><td>8.0E-04</td><td>1.2E-03</td><td>MB231/MB468</td><td>MP</td></tr>
<tr><td>>MI0001445|hsa-mir-423&WithFlank&17|+|30117073|30117178@58.78.21&[MIMAT0001340&hsa-miR-423-3p&offsets|-1|-3;m-100&17|+|30117131|30117153&offsets|-1|-3]|AAGCTCGGTCTGAGGCCCCTC</td><td>>MI0001445|hsa-mir-423&WithFlank&17|+|30117073|30117178</td><td>58</td><td>78</td><td>21</td><td>AAGCTCGGTCTGAGGCCCCTC</td><td>245.1</td><td>1.8</td><td>0.5</td><td>3.3</td><td>9.6E-04</td><td>1.2E-03</td><td>BT20/MB468</td><td>NUC</td></tr>
<tr><td>>hg38_1_-_228634819_228635039_RNA5S12_5S_50ntFlanks@51.80.30|GTCTACGGCCATACCACCCTGAACGCGCCC</td><td>>hg38_1_-_228634819_228635039_RNA5S12_5S_50ntFlanks</td><td>51</td><td>80</td><td>30</td><td>GTCTACGGCCATACCACCCTGAACGCGCCC</td><td>1354.7</td><td>5.4</td><td>1.6</td><td>3.3</td><td>9.0E-04</td><td>1.2E-03</td><td>MB231/MB468</td><td>CYTO</td></tr>
<tr><td>>hg38_21_+_8433172_8446622_RNA45SN1_45S_50ntFlanks@12394.12416.23|TAGCGACGTCGCTTTTTGATCCT</td><td>>hg38_21_+_8433172_8446622_RNA45SN1_45S_50ntFlanks</td><td>12394</td><td>12416</td><td>23</td><td>TAGCGACGTCGCTTTTTGATCCT</td><td>403.9</td><td>-3.1</td><td>0.9</td><td>-3.4</td><td>7.2E-04</td><td>1.2E-03</td><td>BT20/MB468</td><td>MITO</td></tr>
<tr><td>>trnaMT_ThrTGT_MT_+_15888_15953@31.69.39|TTGTAAACCGGAGATGAAAACCTTTTTCCAAGGACACCA</td><td>>trnaMT_ThrTGT_MT_+_15888_15953</td><td>31</td><td>69</td><td>39</td><td>TTGTAAACCGGAGATGAAAACCTTTTTCCAAGGACACCA</td><td>294.2</td><td>4.5</td><td>1.4</td><td>3.3</td><td>9.5E-04</td><td>1.2E-03</td><td>MB231/MB468</td><td>MITO</td></tr>
<tr><td>>MI0000469|hsa-mir-125a&WithFlank&19|+|51693248|51693345@60.79.20&[MIMAT0004602&hsa-miR-125a-3p&offsets|+1|-1;m-333&19|+|51693306|51693326&offsets|+1|0]|CAGGTGAGGTTCTTGGGAGC</td><td>>MI0000469|hsa-mir-125a&WithFlank&19|+|51693248|51693345</td><td>60</td><td>79</td><td>20</td><td>CAGGTGAGGTTCTTGGGAGC</td><td>112.0</td><td>7.0</td><td>2.0</td><td>3.5</td><td>4.2E-04</td><td>1.2E-03</td><td>BT20/MB468</td><td>MP</td></tr>
<tr><td>>hg38_21_+_8433172_8446622_RNA45SN1_45S_50ntFlanks@5125.5158.34|AGCGGTCGGCGTCCCCCAACTTCTTAGAGGGACA</td><td>>hg38_21_+_8433172_8446622_RNA45SN1_45S_50ntFlanks</td><td>5125</td><td>5158</td><td>34</td><td>AGCGGTCGGCGTCCCCCAACTTCTTAGAGGGACA</td><td>289.7</td><td>3.4</td><td>0.9</td><td>3.8</td><td>1.5E-04</td><td>1.2E-03</td><td>BT20/MB231</td><td>NUC</td></tr>
<tr><td>>MI0000446|hsa-mir-125b-1&WithFlank&11|-|122099751|122099850@21.41.21(+1U);MI0000470|hsa-mir-125b-2&WithFlank&21|+|16590231|16590331@23.43.21(+1U)&[MIMAT0000423&hsa-miR-125b-5p&offsets|0|-1(+1U);m-67&11|-|122099809|122099830&offsets|0|-1(+1U)];[MIMAT0000423_1&hsa-miR-125b-5p&offsets|0|-1(+1U);m-68&21|+|16590253|16590274&offsets|0|-1(+1U)]|TCCCTGAGACCCTAACTTGTGT</td><td>>MI0000446|hsa-mir-125b-1&WithFlank&11|-|122099751|122099850</td><td>21</td><td>41</td><td>21(+1U)</td><td>TCCCTGAGACCCTAACTTGTGT</td><td>661.7</td><td>-6.0</td><td>1.6</td><td>-3.8</td><td>1.5E-04</td><td>1.2E-03</td><td>BT20/MB231</td><td>NUC</td></tr>
<tr><td>>hg38_MT_+_1621_3279_RNR2_16S_50ntFlanks@51.81.31|GCTAAACCTAGCCCCAAACCCACTCCACCTT</td><td>>hg38_MT_+_1621_3279_RNR2_16S_50ntFlanks</td><td>51</td><td>81</td><td>31</td><td>GCTAAACCTAGCCCCAAACCCACTCCACCTT</td><td>238.5</td><td>2.4</td><td>0.7</td><td>3.3</td><td>9.6E-04</td><td>1.2E-03</td><td>BT20/MB468</td><td>NUC</td></tr>
<tr><td>>hg38_MT_+_598_1651_RNR1_12S_50ntFlanks@51.88.38|AATAGGTTTGGTCCTAGCCTTTCTATTAGCTCTTAGTA</td><td>>hg38_MT_+_598_1651_RNR1_12S_50ntFlanks</td><td>51</td><td>88</td><td>38</td><td>AATAGGTTTGGTCCTAGCCTTTCTATTAGCTCTTAGTA</td><td>859.9</td><td>3.0</td><td>0.9</td><td>3.3</td><td>9.5E-04</td><td>1.2E-03</td><td>MB231/MB468</td><td>MITO</td></tr>
<tr><td>>hg38_21_+_8433172_8446622_RNA45SN1_45S_50ntFlanks@6645.6675.31|CTCGTACGACTCTTAGCGGTGGATCACTCGG</td><td>>hg38_21_+_8433172_8446622_RNA45SN1_45S_50ntFlanks</td><td>6645</td><td>6675</td><td>31</td><td>CTCGTACGACTCTTAGCGGTGGATCACTCGG</td><td>2098.5</td><td>2.6</td><td>0.8</td><td>3.4</td><td>7.2E-04</td><td>1.2E-03</td><td>BT20/MB468</td><td>MITO</td></tr>
<tr><td>>MI0001445|hsa-mir-423&WithFlank&17|+|30117073|30117178@23.44.22(+1A)&[MIMAT0004748&hsa-miR-423-5p&offsets|0|-1(+1A);m-141&17|+|30117095|30117117&offsets|0|-1(+1A)]|TGAGGGGCAGAGAGCGAGACTTA</td><td>>MI0001445|hsa-mir-423&WithFlank&17|+|30117073|30117178</td><td>23</td><td>44</td><td>22(+1A)</td><td>TGAGGGGCAGAGAGCGAGACTTA</td><td>277.9</td><td>4.3</td><td>1.3</td><td>3.4</td><td>8.0E-04</td><td>1.2E-03</td><td>MB231/MB468</td><td>MP</td></tr>
<tr><td>>MI0003560|hsa-mir-92b&WithFlank&1|+|155195171|155195278@67.87.21(+1G)&[MIMAT0003218&hsa-miR-92b-3p&offsets|0|-1(+1G);m-115&1|+|155195237|155195258&offsets|0|-1(+1G)]|TATTGCACTCGTCCCGGCCTCG</td><td>>MI0003560|hsa-mir-92b&WithFlank&1|+|155195171|155195278</td><td>67</td><td>87</td><td>21(+1G)</td><td>TATTGCACTCGTCCCGGCCTCG</td><td>59.1</td><td>9.6</td><td>2.7</td><td>3.5</td><td>4.2E-04</td><td>1.2E-03</td><td>BT20/MB468</td><td>MP</td></tr>
<tr><td>>hg38_21_+_8433172_8446622_RNA45SN1_45S_50ntFlanks@9933.9955.23|TAGACAGCAGGACGGTGGCCATG</td><td>>hg38_21_+_8433172_8446622_RNA45SN1_45S_50ntFlanks</td><td>9933</td><td>9955</td><td>23</td><td>TAGACAGCAGGACGGTGGCCATG</td><td>132.6</td><td>5.7</td><td>1.6</td><td>3.5</td><td>4.2E-04</td><td>1.2E-03</td><td>BT20/MB468</td><td>MP</td></tr>
<tr><td>>hg38_21_+_8433172_8446622_RNA45SN1_45S_50ntFlanks@11200.11223.24|CGGCGGCGGCGGCGACTCTGGACG</td><td>>hg38_21_+_8433172_8446622_RNA45SN1_45S_50ntFlanks</td><td>11200</td><td>11223</td><td>24</td><td>CGGCGGCGGCGGCGACTCTGGACG</td><td>591.5</td><td>-5.6</td><td>1.7</td><td>-3.3</td><td>9.7E-04</td><td>1.2E-03</td><td>MB231/MB468</td><td>NUC</td></tr>
<tr><td>>MI0000809|hsa-mir-151a&WithFlank&8|-|140732558|140732659@53.74.22(+2A)&[MIMAT0000757&hsa-miR-151a-3p&offsets|0|+1(+2A);m-30&8|-|140732587|140732607&offsets|0|+1(+2A)]|CTAGACTGAAGCTCCTTGAGGAAA</td><td>>MI0000809|hsa-mir-151a&WithFlank&8|-|140732558|140732659</td><td>53</td><td>74</td><td>22(+2A)</td><td>CTAGACTGAAGCTCCTTGAGGAAA</td><td>1409.7</td><td>2.7</td><td>0.7</td><td>3.8</td><td>1.6E-04</td><td>1.2E-03</td><td>BT20/MB231</td><td>CYTO</td></tr>
<tr><td>>hg38_21_+_8433172_8446622_RNA45SN1_45S_50ntFlanks@11210.11254.45|GGCGACTCTGGACGCGAGCCGGGCCCTTCCCGTGGATCGCCCCAG</td><td>>hg38_21_+_8433172_8446622_RNA45SN1_45S_50ntFlanks</td><td>11210</td><td>11254</td><td>45</td><td>GGCGACTCTGGACGCGAGCCGGGCCCTTCCCGTGGATCGCCCCAG</td><td>689.3</td><td>-4.5</td><td>1.4</td><td>-3.3</td><td>8.7E-04</td><td>1.2E-03</td><td>BT20/MB468</td><td>CYTO</td></tr>
<tr><td>>hg38_21_+_8433172_8446622_RNA45SN1_45S_50ntFlanks@6650.6688.39|ACGACTCTTAGCGGTGGATCACTCGGCTCGTGCGTCGAT</td><td>>hg38_21_+_8433172_8446622_RNA45SN1_45S_50ntFlanks</td><td>6650</td><td>6688</td><td>39</td><td>ACGACTCTTAGCGGTGGATCACTCGGCTCGTGCGTCGAT</td><td>81.8</td><td>2.2</td><td>0.7</td><td>3.3</td><td>9.7E-04</td><td>1.2E-03</td><td>BT20/MB468</td><td>NUC</td></tr>
<tr><td>>hg38_21_+_8433172_8446622_RNA45SN1_45S_50ntFlanks@11847.11872.26|AACGGGCTTGGCGGAATCAGCGGGGA</td><td>>hg38_21_+_8433172_8446622_RNA45SN1_45S_50ntFlanks</td><td>11847</td><td>11872</td><td>26</td><td>AACGGGCTTGGCGGAATCAGCGGGGA</td><td>213.6</td><td>-3.4</td><td>1.0</td><td>-3.3</td><td>8.7E-04</td><td>1.2E-03</td><td>BT20/MB468</td><td>CYTO</td></tr>
<tr><td>>hg38_21_+_8433172_8446622_RNA45SN1_45S_50ntFlanks@8182.8203.22|ATCGAGGCCCAGCCCGTGGACG</td><td>>hg38_21_+_8433172_8446622_RNA45SN1_45S_50ntFlanks</td><td>8182</td><td>8203</td><td>22</td><td>ATCGAGGCCCAGCCCGTGGACG</td><td>854.7</td><td>-5.4</td><td>1.6</td><td>-3.3</td><td>9.6E-04</td><td>1.2E-03</td><td>MB231/MB468</td><td>MITO</td></tr>
<tr><td>>hg38_21_+_8433172_8446622_RNA45SN1_45S_50ntFlanks@11668.11692.25|TGCCCAGTGCTCTGAATGTCAAAGT</td><td>>hg38_21_+_8433172_8446622_RNA45SN1_45S_50ntFlanks</td><td>11668</td><td>11692</td><td>25</td><td>TGCCCAGTGCTCTGAATGTCAAAGT</td><td>134.6</td><td>7.0</td><td>2.0</td><td>3.5</td><td>4.2E-04</td><td>1.2E-03</td><td>BT20/MB468</td><td>MP</td></tr>
<tr><td>>hg38_21_+_8433172_8446622_RNA45SN1_45S_50ntFlanks@4494.4511.18|CCCGAAGCGTTTACTTTG</td><td>>hg38_21_+_8433172_8446622_RNA45SN1_45S_50ntFlanks</td><td>4494</td><td>4511</td><td>18</td><td>CCCGAAGCGTTTACTTTG</td><td>162.3</td><td>-5.2</td><td>1.5</td><td>-3.4</td><td>7.3E-04</td><td>1.2E-03</td><td>BT20/MB468</td><td>MITO</td></tr>
<tr><td>>hg38_21_+_8433172_8446622_RNA45SN1_45S_50ntFlanks@11710.11728.19|CGCGGGTAAACGGCGGGAG</td><td>>hg38_21_+_8433172_8446622_RNA45SN1_45S_50ntFlanks</td><td>11710</td><td>11728</td><td>19</td><td>CGCGGGTAAACGGCGGGAG</td><td>662.2</td><td>-5.9</td><td>1.7</td><td>-3.5</td><td>4.2E-04</td><td>1.2E-03</td><td>BT20/MB468</td><td>MP</td></tr>
<tr><td>>MI0000810|hsa-mir-135b&WithFlank&1|-|205448296|205448404@22.44.23&[MIMAT0000758&hsa-miR-135b-5p&offsets|0|0;m-321&1|-|205448361|205448383&offsets|0|0]|TATGGCTTTTCATTCCTATGTGA</td><td>>MI0000810|hsa-mir-135b&WithFlank&1|-|205448296|205448404</td><td>22</td><td>44</td><td>23</td><td>TATGGCTTTTCATTCCTATGTGA</td><td>847.4</td><td>3.3</td><td>1.0</td><td>3.3</td><td>9.6E-04</td><td>1.2E-03</td><td>MB231/MB468</td><td>MITO</td></tr>
<tr><td>>trna10_ValCAC_5_-_180649395_180649467@35.54.20,trna12_ValAAC_5_-_180645270_180645342@35.54.20,trna132_ValAAC_6_-_27721179_27721251@35.54.20,trna13_ValCAC_19_-_4724647_4724719@35.54.20,trna152_ValCAC_6_-_27248049_27248121@35.54.20,trna157_ValCAC_6_-_27173867_27173939@35.54.20,trna15_ValAAC_5_-_180615416_180615488@35.54.20,trna18_ValCAC_5_-_180529253_180529325@35.54.20,trna2_ValAAC_3_+_169490018_169490090@35.54.20,trna2_ValCAC_5_+_180524070_180524142@35.54.20,trna37_ValAAC_6_+_27203288_27203360@35.54.20,trna4_ValAAC_5_+_180591154_180591226@35.54.20,trna5_ValAAC_5_+_180596610_180596682@35.54.20,trna6_ValCAC_5_+_180600650_180600722@35.54.20,trna85_ValCAC_1_-_161369490_161369562@35.54.20,trna98_ValCAC_1_-_149298555_149298627@35.54.20,trna9_ValCAC_6_+_26538282_26538354@35.54.20|ACACGCGAAAGGTCCCCGGT</td><td>>trna10_ValCAC_5_-_180649395_180649467</td><td>35</td><td>54</td><td>20</td><td>ACACGCGAAAGGTCCCCGGT</td><td>641.3</td><td>-5.4</td><td>1.5</td><td>-3.5</td><td>4.3E-04</td><td>1.2E-03</td><td>BT20/MB468</td><td>MP</td></tr>
</tbody>
</table>

| | | | | | | | | | | | | | |
|---|---|---|---|---|---|---|---|---|---|---|---|---|---|
| >hg38_MT_+_1621_3279_RNR2_16S_50ntFlanks@1216.1262.47\|CAACCTCCGAGCAGTACATGCTAAGACTTCACCAGTCAAAGCGAACT | >hg38_MT_+_1621_3279_RNR2_16S_50ntFlanks | 1216 | 1262 | 47 | CAACCTCCGAGCAGTACATGCTAAGACTTCACCAGTCAAAGCGAACT | 1564.4 | 5.1 | 1.5 | 3.4 | 7.3E-04 | 1.2E-03 | BT20/MB468 | MITO |
| >MI0000447\|hsa-mir-128-1&WithFlank&2\|+\|135665391\|135665484@56.76.21;MI0000727\|hsa-mir-128-2&WithFlank&3\|+\|35744470\|35744565@58.78.21&[MIMAT0000424&hsa-miR-128-3p&offsets\|0\|0;m-108&2\|+\|135665446\|135665466&offsets\|0\|0];[MIMAT0000424_1&hsa-miR-128-3p&offsets\|0\|0;m-117&3\|+\|35744527\|35744547&offsets\|0\|0]\|TCACAGTGAACCGGTCTCTTT | >MI0000447\|hsa-mir-128-1&WithFlank&2\|+\|135665391\|135665484 | 56 | 76 | 21 | TCACAGTGAACCGGTCTCTTT | 7540.9 | 2.7 | 0.8 | 3.3 | 9.7E-04 | 1.2E-03 | MB231/MB468 | NUC |
| >hg38_21_+_8433172_8446622_RNA45SN1_45S_50ntFlanks@9546.9563.18\|TGGAGGTCCGTAGCGGTC | >hg38_21_+_8433172_8446622_RNA45SN1_45S_50ntFlanks | 9546 | 9563 | 18 | TGGAGGTCCGTAGCGGTC | 167.7 | -4.5 | 1.3 | -3.4 | 8.1E-04 | 1.2E-03 | MB231/MB468 | MP |
| >hg38_21_+_8433172_8446622_RNA45SN1_45S_50ntFlanks@11877.11898.22\|AGACCCTGTTGAGCTTGACTCT | >hg38_21_+_8433172_8446622_RNA45SN1_45S_50ntFlanks | 11877 | 11898 | 22 | AGACCCTGTTGAGCTTGACTCT | 88.1 | -5.2 | 1.6 | -3.3 | 8.1E-04 | 1.2E-03 | MB231/MB468 | MP |
| >hg38_21_+_8433172_8446622_RNA45SN1_45S_50ntFlanks@6650.6676.27\|ACGACTCTTAGCGGTGGATCACTCGGC | >hg38_21_+_8433172_8446622_RNA45SN1_45S_50ntFlanks | 6650 | 6676 | 27 | ACGACTCTTAGCGGTGGATCACTCGGC | 2137.2 | 3.2 | 1.0 | 3.3 | 9.6E-04 | 1.2E-03 | MB231/MB468 | MITO |
| >hg38_21_+_8433172_8446622_RNA45SN1_45S_50ntFlanks@5491.5514.24\|GAGAAGACGGTCGAACTTGACTAT | >hg38_21_+_8433172_8446622_RNA45SN1_45S_50ntFlanks | 5491 | 5514 | 24 | GAGAAGACGGTCGAACTTGACTAT | 321.8 | -3.6 | 1.1 | -3.3 | 9.6E-04 | 1.2E-03 | MB231/MB468 | MITO |
| >MI0000102\|hsa-mir-100&WithFlank&11\|-\|122152223\|122152314@22.40.19&[MIMAT0000098&hsa-miR-100-5p&offsets\|+3\|0;m-27&11\|-\|122152276\|122152296&offsets\|+3\|+1]\|CCGTAGATCCGAACTTGTG | >MI0000102\|hsa-mir-100&WithFlank&11\|-\|122152223\|122152314 | 22 | 40 | 19 | CCGTAGATCCGAACTTGTG | 577.8 | -8.2 | 2.2 | -3.7 | 1.8E-04 | 1.2E-03 | BT20/MB231 | MITO |
| >hg38_21_+_8433172_8446622_RNA45SN1_45S_50ntFlanks@5079.5120.42\|GATAACGAACGAGACTCTGGCATGCTAACTAGTTACGCGACC | >hg38_21_+_8433172_8446622_RNA45SN1_45S_50ntFlanks | 5079 | 5120 | 42 | GATAACGAACGAGACTCTGGCATGCTAACTAGTTACGCGACC | 179.8 | -3.1 | 0.8 | -3.8 | 1.6E-04 | 1.2E-03 | BT20/MB231 | CYTO |
| >MI0000650\|hsa-mir-200c&WithFlank&12\|+\|6963693\|6963772@50.72.23(+1U)&[MIMAT0000617&hsa-miR-200c-3p&offsets\|0\|0(+1U);m-54&12\|+\|6963742\|6963764&offsets\|0\|0(+1U)]\|TAATACTGCCGGGTAATGATGGAT | >MI0000650\|hsa-mir-200c&WithFlank&12\|+\|6963693\|6963772 | 50 | 72 | 23(+1U) | TAATACTGCCGGGTAATGATGGAT | 220.0 | -3.8 | 1.2 | -3.3 | 9.8E-04 | 1.2E-03 | MB231/MB468 | NUC |
| >MI0000285\|hsa-mir-205&WithFlank&1\|+\|209432127\|209432248@40.61.22(+1G)&[MIMAT0000266&hsa-miR-205-5p&offsets\|0\|0(+1G);m-351&1\|+\|209432166\|209432187&offsets\|0\|0(+1G)]\|TCCTTCATTCCACCGGAGTCTGG | >MI0000285\|hsa-mir-205&WithFlank&1\|+\|209432127\|209432248 | 40 | 61 | 22(+1G) | TCCTTCATTCCACCGGAGTCTGG | 781.7 | 2.5 | 0.8 | 3.3 | 9.8E-04 | 1.2E-03 | BT20/MB468 | NUC |
| >trnaMT_PheGAA_MT_+_577_647@33.63.31\|CTGAAAATGTTTAGACGGGCTCACATCACCC | >trnaMT_PheGAA_MT_+_577_647 | 33 | 63 | 31 | CTGAAAATGTTTAGACGGGCTCACATCACCC | 59.7 | 9.6 | 2.7 | 3.5 | 4.3E-04 | 1.2E-03 | BT20/MB468 | MP |
| >hg38_21_+_8433172_8446622_RNA45SN1_45S_50ntFlanks@5515.5535.21\|CTAGAGGAAGTAAAAGTCGTA | >hg38_21_+_8433172_8446622_RNA45SN1_45S_50ntFlanks | 5515 | 5535 | 21 | CTAGAGGAAGTAAAAGTCGTA | 480.9 | -2.4 | 0.7 | -3.3 | 8.8E-04 | 1.2E-03 | BT20/MB468 | CYTO |
| >hg38_21_+_8433172_8446622_RNA45SN1_45S_50ntFlanks@11663.11691.29\|ATTTCTGCCCAGTGCTCTGAATGTCAAAG | >hg38_21_+_8433172_8446622_RNA45SN1_45S_50ntFlanks | 11663 | 11691 | 29 | ATTTCTGCCCAGTGCTCTGAATGTCAAAG | 164.8 | 3.5 | 1.0 | 3.5 | 4.3E-04 | 1.2E-03 | BT20/MB468 | MP |
| >MI0000542\|hsa-mir-320a&WithFlank&8\|-\|22244960\|22245043@48.68.21(+1G)&[MIMAT0000510&hsa-miR-320a-3p&offsets\|0\|-1(+1G);m-60&8\|-\|22244975\|22244996&offsets\|0\|-1(+1G)]\|AAAAGCTGGGTTGAGAGGGCGG | >MI0000542\|hsa-mir-320a&WithFlank&8\|-\|22244960\|22245043 | 48 | 68 | 21(+1G) | AAAAGCTGGGTTGAGAGGGCGG | 430.8 | 2.8 | 0.7 | 3.8 | 1.6E-04 | 1.2E-03 | BT20/MB231 | NUC |
| >trnaMT_PheGAA_MT_+_577_647@38.59.22\|AATGTTTAGACGGGCTCACATC | >trnaMT_PheGAA_MT_+_577_647 | 38 | 59 | 22 | AATGTTTAGACGGGCTCACATC | 202.5 | 4.5 | 1.3 | 3.5 | 4.3E-04 | 1.2E-03 | BT20/MB468 | MP |
| >hg38_21_+_8433172_8446622_RNA45SN1_45S_50ntFlanks@11792.11821.30\|ATGAACGAGATTCCCACTGTCCCTACCTAC | >hg38_21_+_8433172_8446622_RNA45SN1_45S_50ntFlanks | 11792 | 11821 | 30 | ATGAACGAGATTCCCACTGTCCCTACCTAC | 193.7 | -4.9 | 1.4 | -3.4 | 7.3E-04 | 1.2E-03 | BT20/MB468 | MITO |
| >hg38_21_+_8433172_8446622_RNA45SN1_45S_50ntFlanks@5379.5398.20\|ATTAAGTCCCTGCCCTTTGT | >hg38_21_+_8433172_8446622_RNA45SN1_45S_50ntFlanks | 5379 | 5398 | 20 | ATTAAGTCCCTGCCCTTTGT | 372.7 | -5.0 | 1.5 | -3.4 | 7.3E-04 | 1.2E-03 | BT20/MB468 | MITO |
| >hg38_21_+_8433172_8446622_RNA45SN1_45S_50ntFlanks@11670.11693.24\|CCCAGTGCTCTGAATGTCAAAGTG | >hg38_21_+_8433172_8446622_RNA45SN1_45S_50ntFlanks | 11670 | 11693 | 24 | CCCAGTGCTCTGAATGTCAAAGTG | 2107.6 | -6.6 | 2.0 | -3.3 | 9.8E-04 | 1.2E-03 | BT20/MB468 | NUC |
| >MI0000255\|hsa-mir-30d&WithFlank&8\|-\|134804870\|134804951@12.35.24(+3U)&[MIMAT0000245&hsa-miR-30d-5p&offsets\|0\|+2(+3U);m-21&8\|-\|134804919\|134804940&offsets\|0\|+2(+3U)]\|TGTAAACATCCCCGACTGGAAGCTTTT | >MI0000255\|hsa-mir-30d&WithFlank&8\|-\|134804870\|134804951 | 12 | 35 | 24(+3U) | TGTAAACATCCCCGACTGGAAGCTTTT | 320.8 | -2.9 | 0.8 | -3.8 | 1.6E-04 | 1.2E-03 | BT20/MB231 | NUC |
| >hg38_21_+_8433172_8446622_RNA45SN1_45S_50ntFlanks@9552.9569.18\|TCCGTAGCGGTCCTGACG | >hg38_21_+_8433172_8446622_RNA45SN1_45S_50ntFlanks | 9552 | 9569 | 18 | TCCGTAGCGGTCCTGACG | 91.9 | -5.5 | 1.6 | -3.3 | 8.1E-04 | 1.2E-03 | MB231/MB468 | MP |
| >hg38_21_+_8433172_8446622_RNA45SN1_45S_50ntFlanks@12942.12961.20\|TTCGTAGACGACCTGCTTCT | >hg38_21_+_8433172_8446622_RNA45SN1_45S_50ntFlanks | 12942 | 12961 | 20 | TTCGTAGACGACCTGCTTCT | 210.7 | -4.2 | 1.2 | -3.3 | 8.2E-04 | 1.2E-03 | MB231/MB468 | MP |
| >hg38_21_+_8433172_8446622_RNA45SN1_45S_50ntFlanks@7974.7991.18\|ACGCGACCTCAGATCAGA | >hg38_21_+_8433172_8446622_RNA45SN1_45S_50ntFlanks | 7974 | 7991 | 18 | ACGCGACCTCAGATCAGA | 2350.0 | 3.5 | 1.0 | 3.5 | 4.3E-04 | 1.2E-03 | BT20/MB468 | MP |
| >hg38_21_+_8433172_8446622_RNA45SN1_45S_50ntFlanks@4673.4691.19\|TGGACCGGCGCAAGACGGA | >hg38_21_+_8433172_8446622_RNA45SN1_45S_50ntFlanks | 4673 | 4691 | 19 | TGGACCGGCGCAAGACGGA | 59.8 | -9.0 | 2.6 | -3.5 | 4.3E-04 | 1.2E-03 | BT20/MB468 | MP |
| >hg38_21_+_8433172_8446622_RNA45SN1_45S_50ntFlanks@5489.5516.28\|CTGAGAAGACGGTCGAACTTGACTATCT | >hg38_21_+_8433172_8446622_RNA45SN1_45S_50ntFlanks | 5489 | 5516 | 28 | CTGAGAAGACGGTCGAACTTGACTATCT | 3868.7 | 3.1 | 0.8 | 3.7 | 1.8E-04 | 1.2E-03 | BT20/MB231 | MITO |
| >hg38_21_+_8433172_8446622_RNA45SN1_45S_50ntFlanks@10726.10745.20\|GGGAGAGGGTGTAAATCTCG | >hg38_21_+_8433172_8446622_RNA45SN1_45S_50ntFlanks | 10726 | 10745 | 20 | GGGAGAGGGTGTAAATCTCG | 89.1 | -5.4 | 1.6 | -3.3 | 8.2E-04 | 1.2E-03 | MB231/MB468 | MP |
| >hg38_MT_+_1621_3279_RNR2_16S_50ntFlanks@644.676.33\|AATTGGACCAATCTATCACCCTATAGAAGAACT | >hg38_MT_+_1621_3279_RNR2_16S_50ntFlanks | 644 | 676 | 33 | AATTGGACCAATCTATCACCCTATAGAAGAACT | 139.6 | 5.8 | 1.8 | 3.3 | 9.7E-04 | 1.2E-03 | MB231/MB468 | MITO |
| >hg38_21_+_8433172_8446622_RNA45SN1_45S_50ntFlanks@12447.12486.40\|CAGAATTCACCAAGCGTTGGATTGTTCACCCACTAATAGG | >hg38_21_+_8433172_8446622_RNA45SN1_45S_50ntFlanks | 12447 | 12486 | 40 | CAGAATTCACCAAGCGTTGGATTGTTCACCCACTAATAGG | 207.6 | -4.6 | 1.4 | -3.3 | 9.7E-04 | 1.2E-03 | MB231/MB468 | MITO |
| >trnaMT_LeuTAA_MT_+_3230_3304@18.44.27\|CGGTAATCGCATAAAACTTAAAACTTT | >trnaMT_LeuTAA_MT_+_3230_3304 | 18 | 44 | 27 | CGGTAATCGCATAAAACTTAAAACTTT | 1016.4 | -3.4 | 1.0 | -3.3 | 8.2E-04 | 1.2E-03 | MB231/MB468 | MP |
| >MI0003820\|hsa-mir-454&WithFlank&17\|-\|59137752\|59137878@70.92.23&[MIMAT0003885&hsa-miR-454-3p&offsets\|0\|0;m-272&17\|-\|59137787\|59137809&offsets\|0\|0]\|TAGTGCAATATTGCTTATAGGGT | >MI0003820\|hsa-mir-454&WithFlank&17\|-\|59137752\|59137878 | 70 | 92 | 23 | TAGTGCAATATTGCTTATAGGGT | 379.2 | -3.0 | 0.8 | -3.7 | 1.8E-04 | 1.2E-03 | BT20/MB231 | MITO |
| >hg38_21_+_8433172_8446622_RNA45SN1_45S_50ntFlanks@10784.10818.35\|TGAACAGCCTCTGGCATGTTGGAACAATGTAGGTA | >hg38_21_+_8433172_8446622_RNA45SN1_45S_50ntFlanks | 10784 | 10818 | 35 | TGAACAGCCTCTGGCATGTTGGAACAATGTAGGTA | 359.5 | -3.3 | 0.9 | -3.5 | 4.3E-04 | 1.2E-03 | BT20/MB468 | MP |
| >hg38_21_+_8433172_8446622_RNA45SN1_45S_50ntFlanks@12476.12497.22\|CCACTAATAGGGAACGTGAGCT | >hg38_21_+_8433172_8446622_RNA45SN1_45S_50ntFlanks | 12476 | 12497 | 22 | CCACTAATAGGGAACGTGAGCT | 1909.8 | -2.1 | 0.6 | -3.3 | 9.2E-04 | 1.2E-03 | MB231/MB468 | CYTO |
| >hg38_21_+_8433172_8446622_RNA45SN1_45S_50ntFlanks@6651.6672.22\|CGACTCTTAGCGGTGGATCACT | >hg38_21_+_8433172_8446622_RNA45SN1_45S_50ntFlanks | 6651 | 6672 | 22 | CGACTCTTAGCGGTGGATCACT | 8826.4 | 2.8 | 0.8 | 3.3 | 9.7E-04 | 1.2E-03 | MB231/MB468 | MITO |
| >hg38_MT_+_598_1651_RNR1_12S_50ntFlanks@909.950.42\|TACTTCAAAGGACATTTAACTAAAACCCCTACGCATTTATAT | >hg38_MT_+_598_1651_RNR1_12S_50ntFlanks | 909 | 950 | 42 | TACTTCAAAGGACATTTAACTAAAACCCCTACGCATTTATAT | 58.7 | 9.6 | 2.7 | 3.5 | 4.3E-04 | 1.2E-03 | BT20/MB468 | MP |
| >hg38_21_+_8433172_8446622_RNA45SN1_45S_50ntFlanks@11568.11591.24\|GCAGCCGACTTAGAACTGGTGCGG | >hg38_21_+_8433172_8446622_RNA45SN1_45S_50ntFlanks | 11568 | 11591 | 24 | GCAGCCGACTTAGAACTGGTGCGG | 528.8 | -5.1 | 1.5 | -3.5 | 4.3E-04 | 1.2E-03 | BT20/MB468 | MP |
| >trnaMT_PheGAA_MT_+_577_647@32.65.34\|ACTGAAAATGTTTAGACGGGCTCACATCACCCCA | >trnaMT_PheGAA_MT_+_577_647 | 32 | 65 | 34 | ACTGAAAATGTTTAGACGGGCTCACATCACCCCA | 57.0 | 9.6 | 2.7 | 3.5 | 4.3E-04 | 1.2E-03 | BT20/MB468 | MP |
| >hg38_21_+_8433172_8446622_RNA45SN1_45S_50ntFlanks@9574.9605.32\|AATCGGTCGTCCGACCTGGGTATAGGGGCGAA | >hg38_21_+_8433172_8446622_RNA45SN1_45S_50ntFlanks | 9574 | 9605 | 32 | AATCGGTCGTCCGACCTGGGTATAGGGGCGAA | 1476.0 | -2.7 | 0.8 | -3.3 | 9.8E-04 | 1.2E-03 | MB231/MB468 | MITO |
| >hg38_21_+_8433172_8446622_RNA45SN1_45S_50ntFlanks@10405.10432.28\|ATGGGTCAGTCGGTCCTGAGAGATGGGC | >hg38_21_+_8433172_8446622_RNA45SN1_45S_50ntFlanks | 10405 | 10432 | 28 | ATGGGTCAGTCGGTCCTGAGAGATGGGC | 173.3 | 4.6 | 1.3 | 3.5 | 4.3E-04 | 1.2E-03 | BT20/MB468 | MP |
| >hg38_21_+_8433172_8446622_RNA45SN1_45S_50ntFlanks@8290.8308.19\|TAAACTCCATCTAAGGCTA | >hg38_21_+_8433172_8446622_RNA45SN1_45S_50ntFlanks | 8290 | 8308 | 19 | TAAACTCCATCTAAGGCTA | 56.1 | -8.9 | 2.5 | -3.5 | 4.3E-04 | 1.2E-03 | BT20/MB468 | MP |
| >hg38_21_+_8433172_8446622_RNA45SN1_45S_50ntFlanks@9572.9605.34\|CAAATCGGTCGTCCGACCTGGGTATAGGGGCGAA | >hg38_21_+_8433172_8446622_RNA45SN1_45S_50ntFlanks | 9572 | 9605 | 34 | CAAATCGGTCGTCCGACCTGGGTATAGGGGCGAA | 201.6 | -2.6 | 0.8 | -3.3 | 8.9E-04 | 1.2E-03 | BT20/MB468 | CYTO |
| >hg38_21_+_8433172_8446622_RNA45SN1_45S_50ntFlanks@6790.6807.18\|CCTGTCTGAGCGTCGCTT | >hg38_21_+_8433172_8446622_RNA45SN1_45S_50ntFlanks | 6790 | 6807 | 18 | CCTGTCTGAGCGTCGCTT | 1032.0 | 4.5 | 1.3 | 3.5 | 4.3E-04 | 1.2E-03 | BT20/MB468 | MP |
| >hg38_21_+_8433172_8446622_RNA45SN1_45S_50ntFlanks@7976.8011.36\|GCGACCTCAGATCAGACGTGGCGACCCGCTGAATTT | >hg38_21_+_8433172_8446622_RNA45SN1_45S_50ntFlanks | 7976 | 8011 | 36 | GCGACCTCAGATCAGACGTGGCGACCCGCTGAATTT | 394.7 | 2.4 | 0.7 | 3.3 | 9.8E-04 | 1.2E-03 | MB231/MB468 | MITO |
| >hg38_21_+_8433172_8446622_RNA45SN1_45S_50ntFlanks@3975.4005.31\|CGGGCGCCGGCGGCTTTGGTGACTCTAGATA | >hg38_21_+_8433172_8446622_RNA45SN1_45S_50ntFlanks | 3975 | 4005 | 31 | CGGGCGCCGGCGGCTTTGGTGACTCTAGATA | 229.7 | -2.8 | 0.8 | -3.4 | 7.4E-04 | 1.2E-03 | BT20/MB468 | MITO |
| >hg38_21_+_8433172_8446622_RNA45SN1_45S_50ntFlanks@12242.12273.32\|AGAAGGGCAAAAGCTCGCTTGATCTTGATTTT | >hg38_21_+_8433172_8446622_RNA45SN1_45S_50ntFlanks | 12242 | 12273 | 32 | AGAAGGGCAAAAGCTCGCTTGATCTTGATTTT | 402.4 | -2.3 | 0.7 | -3.4 | 7.4E-04 | 1.2E-03 | BT20/MB468 | MITO |
| >MI0000812\|hsa-mir-331&WithFlank&12\|+\|95308414\|95308519@32.52.21&[MIMAT0004700&hsa-miR-331-5p&offsets\|0\|-1;m-377&12\|+\|95308445\|95308465&offsets\|0\|0]\|CTAGGTATGGTCCCAGGGATC | >MI0000812\|hsa-mir-331&WithFlank&12\|+\|95308414\|95308519 | 32 | 52 | 21 | CTAGGTATGGTCCCAGGGATC | 263.7 | 7.4 | 2.1 | 3.5 | 4.3E-04 | 1.2E-03 | BT20/MB468 | MP |
| >hg38_21_+_8433172_8446622_RNA45SN1_45S_50ntFlanks@7989.8013.25\|AGACGTGGCGACCCGCTGAATTTAA | >hg38_21_+_8433172_8446622_RNA45SN1_45S_50ntFlanks | 7989 | 8013 | 25 | AGACGTGGCGACCCGCTGAATTTAA | 536.4 | -2.4 | 0.7 | -3.3 | 9.8E-04 | 1.2E-03 | MB231/MB468 | MITO |
| >trna10_LysCTT_16_+_3241501_3241573@4.34.31,trna13_LysCTT_14_-_58706613_58706685@4.34.31,trna2_LysCTT_15_+_79152904_79152976@4.34.31\|CGGCTAGCTCAGTCGGTAGAGCATGGGACTC | >trna10_LysCTT_16_+_3241501_3241573 | 4 | 34 | 31 | CGGCTAGCTCAGTCGGTAGAGCATGGGACTC | 59.0 | 9.6 | 2.7 | 3.5 | 4.3E-04 | 1.2E-03 | BT20/MB468 | MP |
| >hg38_21_+_8433172_8446622_RNA45SN1_45S_50ntFlanks@12419.12450.32\|GATGTCGGCTCTTCCTATCATTGTGAAGCAGA | >hg38_21_+_8433172_8446622_RNA45SN1_45S_50ntFlanks | 12419 | 12450 | 32 | GATGTCGGCTCTTCCTATCATTGTGAAGCAGA | 217.2 | 2.4 | 0.7 | 3.3 | 9.9E-04 | 1.2E-03 | MB231/MB468 | NUC |
| >hg38_21_+_8433172_8446622_RNA45SN1_45S_50ntFlanks@12632.12652.21\|GCGAAGCTACCATCTGTGGGA | >hg38_21_+_8433172_8446622_RNA45SN1_45S_50ntFlanks | 12632 | 12652 | 21 | GCGAAGCTACCATCTGTGGGA | 180.3 | -5.6 | 1.6 | -3.5 | 4.4E-04 | 1.2E-03 | BT20/MB468 | MP |
| >am_MI0006381\|hsa-mir-1246&WithFlank&2\|-\|176600974\|176601058@17.35.19(+1G)&[MIMAT0005898&hsa-miR-1246&offsets\|0\|0(+1G);m-338&2\|-\|176601023\|176601043&offsets\|+1\|-1(+1G)]\|AATGGATTTTTGGAGCAGGG | >am_MI0006381\|hsa-mir-1246&WithFlank&2\|-\|176600974\|176601058 | 17 | 35 | 19(+1G) | AATGGATTTTTGGAGCAGGG | 574.1 | 3.7 | 1.1 | 3.5 | 4.4E-04 | 1.2E-03 | BT20/MB468 | MP |
| >MI0000734\|hsa-mir-106b&WithFlank&7\|-\|100093987\|100094080@58.77.20&[MIMAT0004672&hsa-miR-106b-3p&offsets\|0\|-2;m-87&7\|-\|100094003\|100094023&offsets\|0\|-1]\|CCGCACTGTGGGTACTTGCT | >MI0000734\|hsa-mir-106b&WithFlank&7\|-\|100093987\|100094080 | 58 | 77 | 20 | CCGCACTGTGGGTACTTGCT | 546.8 | 3.6 | 1.1 | 3.3 | 8.3E-04 | 1.2E-03 | MB231/MB468 | MP |
| >trnaMT_ValTAC_MT_+_1602_1670@44.71.28\|ATTTCAACTTAACTTGACCGCTCTGACC | >trnaMT_ValTAC_MT_+_1602_1670 | 44 | 71 | 28 | ATTTCAACTTAACTTGACCGCTCTGACC | 3161.1 | 3.2 | 1.0 | 3.3 | 9.8E-04 | 1.2E-03 | MB231/MB468 | MITO |

<table>
<thead><tr><th></th><th></th><th></th><th></th><th></th><th></th><th></th><th></th><th></th><th></th><th></th><th></th><th></th><th></th></tr></thead>
<tbody>
<tr><td>>trna10_AspGTC_12_-_125424193_125424264@36.70.35,trna12_AspGTC_12_-_125411891_125411962@36.70.35,trna144_AspGTC_6_-_27551236_27551307@36.70.35,trna38_AspGTC_17_-_8125556_8125627@36.70.35,trna45_AspGTC_6_+_27447453_27447524@36.70.35,trna48_AspGTC_6_+_27471523_27471594@36.70.35,trna4_AspGTC_12_+_96429799_96429870@36.70.35,trna69_AspGTC_1_-_161440205_161440276@36.70.35,trna72_AspGTC_1_-_161432824_161432895@36.70.35,trna75_AspGTC_1_-_161425414_161425485@36.70.35,trna78_AspGTC_1_-_161418033_161418104@36.70.35,trna81_AspGTC_1_-_161410615_161410686@36.70.35|CACGCGGGAGACCGGGGTTCGATTCCCCGACGGGG</td><td>>trna10_AspGTC_12_-_125424193_125424264</td><td>36</td><td>70</td><td>35</td><td>CACGCGGGAGACCGGGGTTCGATTCCCCGACGGGG</td><td>503.5</td><td>-3.8</td><td>1.0</td><td>-3.8</td><td>1.6E-04</td><td>1.2E-03</td><td>BT20/MB231</td><td>CYTO</td></tr>
<tr><td>>hg38_21_+_8433172_8446622_RNA45SN1_45S_50ntFlanks@3927.3959.33|CAGATCAAAACCAACCCGGTCAGCCCCTCTCCG</td><td>>hg38_21_+_8433172_8446622_RNA45SN1_45S_50ntFlanks</td><td>3927</td><td>3959</td><td>33</td><td>CAGATCAAAACCAACCCGGTCAGCCCCTCTCCG</td><td>190.4</td><td>-4.7</td><td>1.4</td><td>-3.4</td><td>7.5E-04</td><td>1.2E-03</td><td>BT20/MB468</td><td>MITO</td></tr>
<tr><td>>hg38_21_+_8433172_8446622_RNA45SN1_45S_50ntFlanks@9926.9951.26|GTTGATATAGACAGCAGGACGGTGGC</td><td>>hg38_21_+_8433172_8446622_RNA45SN1_45S_50ntFlanks</td><td>9926</td><td>9951</td><td>26</td><td>GTTGATATAGACAGCAGGACGGTGGC</td><td>65.2</td><td>6.4</td><td>1.8</td><td>3.5</td><td>4.4E-04</td><td>1.2E-03</td><td>BT20/MB468</td><td>MP</td></tr>
<tr><td>>MI0000076|hsa-mir-20a&WithFlank&13|+|91351059|91351141@14.37.24&[MIMAT0000075&hsa-miR-20a-5p&offsets|0|+1;m-96&13|+|91351072|91351094&offsets|0|+1]|TAAAGTGCTTATAGTGCAGGTAGT</td><td>>MI0000076|hsa-mir-20a&WithFlank&13|+|91351059|91351141</td><td>14</td><td>37</td><td>24</td><td>TAAAGTGCTTATAGTGCAGGTAGT</td><td>504.2</td><td>2.7</td><td>0.8</td><td>3.3</td><td>8.3E-04</td><td>1.2E-03</td><td>MB231/MB468</td><td>MP</td></tr>
<tr><td>>hg38_MT_+_598_1651_RNR1_12S_50ntFlanks@568.611.44|CAAAGGACCTGGCGGTGCTTCATATCCCTCTAGAGGAGCCTGTT</td><td>>hg38_MT_+_598_1651_RNR1_12S_50ntFlanks</td><td>568</td><td>611</td><td>44</td><td>CAAAGGACCTGGCGGTGCTTCATATCCCTCTAGAGGAGCCTGTT</td><td>165.5</td><td>6.0</td><td>1.8</td><td>3.3</td><td>9.9E-04</td><td>1.2E-03</td><td>MB231/MB468</td><td>MITO</td></tr>
<tr><td>>hg38_21_+_8433172_8446622_RNA45SN1_45S_50ntFlanks@10357.10390.34|CTTTGAAGGCCGAAGTGGAGAAGGGTTCCATGTG</td><td>>hg38_21_+_8433172_8446622_RNA45SN1_45S_50ntFlanks</td><td>10357</td><td>10390</td><td>34</td><td>CTTTGAAGGCCGAAGTGGAGAAGGGTTCCATGTG</td><td>266.1</td><td>-2.9</td><td>0.8</td><td>-3.5</td><td>4.4E-04</td><td>1.2E-03</td><td>BT20/MB468</td><td>MP</td></tr>
<tr><td>>hg38_MT_+_1621_3279_RNR2_16S_50ntFlanks@51.73.23|GCTAAACCTAGCCCCAAACCCAC</td><td>>hg38_MT_+_1621_3279_RNR2_16S_50ntFlanks</td><td>51</td><td>73</td><td>23</td><td>GCTAAACCTAGCCCCAAACCCAC</td><td>2119.3</td><td>3.2</td><td>1.0</td><td>3.3</td><td>9.9E-04</td><td>1.2E-03</td><td>MB231/MB468</td><td>MITO</td></tr>
<tr><td>>hg38_21_+_8433172_8446622_RNA45SN1_45S_50ntFlanks@11209.11232.24|CGGCGACTCTGGACGCGAGCCGGG</td><td>>hg38_21_+_8433172_8446622_RNA45SN1_45S_50ntFlanks</td><td>11209</td><td>11232</td><td>24</td><td>CGGCGACTCTGGACGCGAGCCGGG</td><td>380.1</td><td>6.4</td><td>1.7</td><td>3.8</td><td>1.5E-04</td><td>1.2E-03</td><td>BT20/MB231</td><td>MP</td></tr>
<tr><td>>MI0000342|hsa-mir-200b&WithFlank&1|+|1167098|1167204@63.84.22&[MIMAT0000318&hsa-miR-200b-3p&offsets|0|0;m-93&1|+|1167160|1167182&offsets|0|-1]|TAATACTGCCTGGTAATGATGA</td><td>>MI0000342|hsa-mir-200b&WithFlank&1|+|1167098|1167204</td><td>63</td><td>84</td><td>22</td><td>TAATACTGCCTGGTAATGATGA</td><td>1132.5</td><td>-3.4</td><td>1.0</td><td>-3.3</td><td>9.9E-04</td><td>1.2E-03</td><td>MB231/MB468</td><td>MITO</td></tr>
<tr><td>>hg38_21_+_8433172_8446622_RNA45SN1_45S_50ntFlanks@10400.10424.25|TGAACATGGGTCAGTCGGTCCTGAG</td><td>>hg38_21_+_8433172_8446622_RNA45SN1_45S_50ntFlanks</td><td>10400</td><td>10424</td><td>25</td><td>TGAACATGGGTCAGTCGGTCCTGAG</td><td>441.7</td><td>3.3</td><td>1.0</td><td>3.3</td><td>8.3E-04</td><td>1.2E-03</td><td>MB231/MB468</td><td>MP</td></tr>
<tr><td>>hg38_1_-_228634819_228635039_RNA5S12_5S_50ntFlanks@122.170.49|TTAGTACTTGGATGGGAGACCGCCTGGGAATACCGGGTGCTGTAGGCTT</td><td>>hg38_1_-_228634819_228635039_RNA5S12_5S_50ntFlanks</td><td>122</td><td>170</td><td>49</td><td>TTAGTACTTGGATGGGAGACCGCCTGGGAATACCGGGTGCTGTAGGCTT</td><td>942.0</td><td>3.3</td><td>1.0</td><td>3.3</td><td>9.3E-04</td><td>1.2E-03</td><td>MB231/MB468</td><td>CYTO</td></tr>
<tr><td>>hg38_21_+_8433172_8446622_RNA45SN1_45S_50ntFlanks@4641.4663.23|CATTCGTATTGCGCCGCTAGAGG</td><td>>hg38_21_+_8433172_8446622_RNA45SN1_45S_50ntFlanks</td><td>4641</td><td>4663</td><td>23</td><td>CATTCGTATTGCGCCGCTAGAGG</td><td>157.6</td><td>-4.7</td><td>1.4</td><td>-3.4</td><td>7.5E-04</td><td>1.2E-03</td><td>BT20/MB468</td><td>MITO</td></tr>
<tr><td>>hg38_21_+_8433172_8446622_RNA45SN1_45S_50ntFlanks@3838.3860.23|CTCTCCTACTTGGATAACTGTGG</td><td>>hg38_21_+_8433172_8446622_RNA45SN1_45S_50ntFlanks</td><td>3838</td><td>3860</td><td>23</td><td>CTCTCCTACTTGGATAACTGTGG</td><td>1443.4</td><td>-1.8</td><td>0.6</td><td>-3.3</td><td>9.4E-04</td><td>1.2E-03</td><td>MB231/MB468</td><td>CYTO</td></tr>
<tr><td>>trnaMT_TyrGTA_MT_-_5826_5891@38.69.32|AAAGACAGGGGTTAGGCCTCTTTTTACCACCA</td><td>>trnaMT_TyrGTA_MT_-_5826_5891</td><td>38</td><td>69</td><td>32</td><td>AAAGACAGGGGTTAGGCCTCTTTTTACCACCA</td><td>302.0</td><td>4.3</td><td>1.3</td><td>3.3</td><td>8.3E-04</td><td>1.2E-03</td><td>MB231/MB468</td><td>MP</td></tr>
<tr><td>>MI0000103|hsa-mir-101-1&WithFlank&1|-|65058428|65058514@52.72.21;MI0000739|hsa-mir-101-2&WithFlank&9|+|4850291|4850381@54.74.21&[MIMAT0000099&hsa-miR-101-3p&offsets|-1|-1;m-17&1|-|65058443|65058462&offsets|-1|0];[MIMAT0000099_1&hsa-miR-101-3p&offsets|-1|-1;m-16&9|+|4850345|4850364&offsets|-1|0]|GTACAGTACTGTGATAACTGA</td><td>>MI0000103|hsa-mir-101-1&WithFlank&1|-|65058428|65058514</td><td>52</td><td>72</td><td>21</td><td>GTACAGTACTGTGATAACTGA</td><td>258.9</td><td>3.0</td><td>0.9</td><td>3.3</td><td>1.0E-03</td><td>1.2E-03</td><td>BT20/MB468</td><td>NUC</td></tr>
<tr><td>>hg38_21_+_8433172_8446622_RNA45SN1_45S_50ntFlanks@12933.12965.33|GCTAAACCATTCGTAGACGACCTGCTTCTGGGT</td><td>>hg38_21_+_8433172_8446622_RNA45SN1_45S_50ntFlanks</td><td>12933</td><td>12965</td><td>33</td><td>GCTAAACCATTCGTAGACGACCTGCTTCTGGGT</td><td>379.8</td><td>-2.2</td><td>0.7</td><td>-3.3</td><td>1.0E-03</td><td>1.2E-03</td><td>MB231/MB468</td><td>NUC</td></tr>
<tr><td>>hg38_21_+_8433172_8446622_RNA45SN1_45S_50ntFlanks@10408.10429.22|GGTCAGTCGGTCCTGAGAGATG</td><td>>hg38_21_+_8433172_8446622_RNA45SN1_45S_50ntFlanks</td><td>10408</td><td>10429</td><td>22</td><td>GGTCAGTCGGTCCTGAGAGATG</td><td>620.3</td><td>-4.2</td><td>1.2</td><td>-3.4</td><td>7.5E-04</td><td>1.2E-03</td><td>BT20/MB468</td><td>MITO</td></tr>
<tr><td>>hg38_21_+_8433172_8446622_RNA45SN1_45S_50ntFlanks@7980.8012.33|CCTCAGATCAGACGTGGCGACCCGCTGAATTTA</td><td>>hg38_21_+_8433172_8446622_RNA45SN1_45S_50ntFlanks</td><td>7980</td><td>8012</td><td>33</td><td>CCTCAGATCAGACGTGGCGACCCGCTGAATTTA</td><td>703.3</td><td>-4.1</td><td>1.2</td><td>-3.4</td><td>7.5E-04</td><td>1.2E-03</td><td>BT20/MB468</td><td>MITO</td></tr>
<tr><td>>hg38_21_+_8433172_8446622_RNA45SN1_45S_50ntFlanks@11921.11967.47|TGAGAGGTGTAGAATAAGTGGGAGGCCCCCGGCGCCCCCCCGGTGTC</td><td>>hg38_21_+_8433172_8446622_RNA45SN1_45S_50ntFlanks</td><td>11921</td><td>11967</td><td>47</td><td>TGAGAGGTGTAGAATAAGTGGGAGGCCCCCGGCGCCCCCCCGGTGTC</td><td>57.1</td><td>-8.9</td><td>2.5</td><td>-3.5</td><td>4.4E-04</td><td>1.2E-03</td><td>BT20/MB468</td><td>MP</td></tr>
<tr><td>>trnaMT_GluTTC_MT_-_14674_14742@24.67.44,trnalookalike8_GluTTC_5_-_93905172_93905240@24.67.44|ATGGTTTTTCATATCATTGGTCGTGGTTGTAGTCCGTGCGAGAA</td><td>>trnaMT_GluTTC_MT_-_14674_14742</td><td>24</td><td>67</td><td>44</td><td>ATGGTTTTTCATATCATTGGTCGTGGTTGTAGTCCGTGCGAGAA</td><td>57.1</td><td>9.6</td><td>2.7</td><td>3.5</td><td>4.4E-04</td><td>1.2E-03</td><td>BT20/MB468</td><td>MP</td></tr>
<tr><td>>hg38_21_+_8433172_8446622_RNA45SN1_45S_50ntFlanks@6703.6720.18|GCTGCGAGAATTAATGTG</td><td>>hg38_21_+_8433172_8446622_RNA45SN1_45S_50ntFlanks</td><td>6703</td><td>6720</td><td>18</td><td>GCTGCGAGAATTAATGTG</td><td>521.5</td><td>-4.5</td><td>1.3</td><td>-3.5</td><td>4.4E-04</td><td>1.2E-03</td><td>BT20/MB468</td><td>MP</td></tr>
<tr><td>>hg38_21_+_8433172_8446622_RNA45SN1_45S_50ntFlanks@6660.6707.48|GCGGTGGATCACTCGGCTCGTGCGTCGATGAAGAACGCAGCTAGCTGC</td><td>>hg38_21_+_8433172_8446622_RNA45SN1_45S_50ntFlanks</td><td>6660</td><td>6707</td><td>48</td><td>GCGGTGGATCACTCGGCTCGTGCGTCGATGAAGAACGCAGCTAGCTGC</td><td>1126.3</td><td>3.7</td><td>1.0</td><td>3.7</td><td>1.9E-04</td><td>1.2E-03</td><td>BT20/MB231</td><td>MITO</td></tr>
<tr><td>>hg38_21_+_8433172_8446622_RNA45SN1_45S_50ntFlanks@3705.3723.19|TACCTGGTTGATCCTGCCA</td><td>>hg38_21_+_8433172_8446622_RNA45SN1_45S_50ntFlanks</td><td>3705</td><td>3723</td><td>19</td><td>TACCTGGTTGATCCTGCCA</td><td>207.0</td><td>1.7</td><td>0.5</td><td>3.3</td><td>1.0E-03</td><td>1.2E-03</td><td>BT20/MB468</td><td>NUC</td></tr>
<tr><td>>MI0000079|hsa-mir-23a&WithFlank&19|-|13836581|13836665@54.74.21&[MIMAT0000078&hsa-miR-23a-3p&offsets|+3|+3;m-31&19|-|13836595|13836615&offsets|+3|+3]|ACATTGCCAGGGATTTCCAAC</td><td>>MI0000079|hsa-mir-23a&WithFlank&19|-|13836581|13836665</td><td>54</td><td>74</td><td>21</td><td>ACATTGCCAGGGATTTCCAAC</td><td>264.1</td><td>11.9</td><td>3.6</td><td>3.3</td><td>1.0E-03</td><td>1.2E-03</td><td>MB231/MB468</td><td>NUC</td></tr>
<tr><td>>hg38_21_+_8433172_8446622_RNA45SN1_45S_50ntFlanks@11666.11697.32|TCTGCCCAGTGCTCTGAATGTCAAAGTGAAGA</td><td>>hg38_21_+_8433172_8446622_RNA45SN1_45S_50ntFlanks</td><td>11666</td><td>11697</td><td>32</td><td>TCTGCCCAGTGCTCTGAATGTCAAAGTGAAGA</td><td>235.8</td><td>1.9</td><td>0.6</td><td>3.3</td><td>1.0E-03</td><td>1.2E-03</td><td>MB231/MB468</td><td>NUC</td></tr>
<tr><td>>trnaMT_PheGAA_MT_+_577_647@38.62.25|AATGTTTAGACGGGCTCACATCACC</td><td>>trnaMT_PheGAA_MT_+_577_647</td><td>38</td><td>62</td><td>25</td><td>AATGTTTAGACGGGCTCACATCACC</td><td>375.9</td><td>4.6</td><td>1.3</td><td>3.5</td><td>4.4E-04</td><td>1.2E-03</td><td>BT20/MB468</td><td>MP</td></tr>
<tr><td>>hg38_21_+_8433172_8446622_RNA45SN1_45S_50ntFlanks@10393.10411.19|CAGCAGTTGAACATGGGTC</td><td>>hg38_21_+_8433172_8446622_RNA45SN1_45S_50ntFlanks</td><td>10393</td><td>10411</td><td>19</td><td>CAGCAGTTGAACATGGGTC</td><td>187.8</td><td>-3.8</td><td>1.2</td><td>-3.3</td><td>1.0E-03</td><td>1.2E-03</td><td>MB231/MB468</td><td>NUC</td></tr>
<tr><td>>hg38_21_+_8433172_8446622_RNA45SN1_45S_50ntFlanks@12434.12455.22|TATCATTGTGAAGCAGAATTCA</td><td>>hg38_21_+_8433172_8446622_RNA45SN1_45S_50ntFlanks</td><td>12434</td><td>12455</td><td>22</td><td>TATCATTGTGAAGCAGAATTCA</td><td>160.2</td><td>-5.4</td><td>1.5</td><td>-3.5</td><td>4.4E-04</td><td>1.2E-03</td><td>BT20/MB468</td><td>MP</td></tr>
<tr><td>>hg38_21_+_8433172_8446622_RNA45SN1_45S_50ntFlanks@8636.8658.23|CGGCGACCGGCCGCCGCCGGGCG</td><td>>hg38_21_+_8433172_8446622_RNA45SN1_45S_50ntFlanks</td><td>8636</td><td>8658</td><td>23</td><td>CGGCGACCGGCCGCCGCCGGGCG</td><td>253.5</td><td>-5.7</td><td>1.6</td><td>-3.5</td><td>4.4E-04</td><td>1.2E-03</td><td>BT20/MB468</td><td>MP</td></tr>
<tr><td>>MI0017294|hsa-mir-4664&WithFlank&8|-|143733077|143733159@50.71.22(+1U)&[MIMAT0019738&hsa-miR-4664-3p&offsets|0|0(+1U);m-1025&8|-|143733094|143733110&offsets|0|+5(+1U)]|CTTCCGGTCTGTGAGCCCCGTCT</td><td>>MI0017294|hsa-mir-4664&WithFlank&8|-|143733077|143733159</td><td>50</td><td>71</td><td>22(+1U)</td><td>CTTCCGGTCTGTGAGCCCCGTCT</td><td>88.7</td><td>6.1</td><td>1.7</td><td>3.5</td><td>4.4E-04</td><td>1.2E-03</td><td>BT20/MB468</td><td>MP</td></tr>
<tr><td>>hg38_21_+_8433172_8446622_RNA45SN1_45S_50ntFlanks@11665.11693.29|TTCTGCCCAGTGCTCTGAATGTCAAAGTG</td><td>>hg38_21_+_8433172_8446622_RNA45SN1_45S_50ntFlanks</td><td>11665</td><td>11693</td><td>29</td><td>TTCTGCCCAGTGCTCTGAATGTCAAAGTG</td><td>50.2</td><td>6.5</td><td>1.8</td><td>3.5</td><td>4.4E-04</td><td>1.2E-03</td><td>BT20/MB468</td><td>MP</td></tr>
<tr><td>>hg38_21_+_8433172_8446622_RNA45SN1_45S_50ntFlanks@5488.5515.28|GCTGAGAAGACGGTCGAACTTGACTATC</td><td>>hg38_21_+_8433172_8446622_RNA45SN1_45S_50ntFlanks</td><td>5488</td><td>5515</td><td>28</td><td>GCTGAGAAGACGGTCGAACTTGACTATC</td><td>270.6</td><td>3.7</td><td>1.1</td><td>3.5</td><td>4.4E-04</td><td>1.2E-03</td><td>BT20/MB468</td><td>MP</td></tr>
<tr><td>>hg38_21_+_8433172_8446622_RNA45SN1_45S_50ntFlanks@4770.4793.24|TCGTAGTTCCGACCATAAACGATG</td><td>>hg38_21_+_8433172_8446622_RNA45SN1_45S_50ntFlanks</td><td>4770</td><td>4793</td><td>24</td><td>TCGTAGTTCCGACCATAAACGATG</td><td>164.9</td><td>-1.8</td><td>0.5</td><td>-3.4</td><td>7.5E-04</td><td>1.2E-03</td><td>BT20/MB468</td><td>MITO</td></tr>
<tr><td>>hg38_21_+_8433172_8446622_RNA45SN1_45S_50ntFlanks@11575.11612.38|ACTTAGAACTGGTGCGGACCAGGGGAATCCGACTGTTT</td><td>>hg38_21_+_8433172_8446622_RNA45SN1_45S_50ntFlanks</td><td>11575</td><td>11612</td><td>38</td><td>ACTTAGAACTGGTGCGGACCAGGGGAATCCGACTGTTT</td><td>265.3</td><td>-2.5</td><td>0.8</td><td>-3.3</td><td>9.4E-04</td><td>1.2E-03</td><td>MB231/MB468</td><td>CYTO</td></tr>
<tr><td>>MI0000239|hsa-mir-197&WithFlank&1|+|109598887|109598973@54.75.22&[MIMAT0000227&hsa-miR-197-3p&offsets|0|0;m-99&1|+|109598940|109598961&offsets|0|0]|TTCACCACCTTCTCCACCCAGC</td><td>>MI0000239|hsa-mir-197&WithFlank&1|+|109598887|109598973</td><td>54</td><td>75</td><td>22</td><td>TTCACCACCTTCTCCACCCAGC</td><td>16230.4</td><td>3.0</td><td>0.8</td><td>3.8</td><td>1.6E-04</td><td>1.2E-03</td><td>BT20/MB231</td><td>CYTO</td></tr>
<tr><td>>hg38_21_+_8433172_8446622_RNA45SN1_45S_50ntFlanks@7969.8010.42|CCGAGACGCGACCTCAGATCAGACGTGGCGACCCGCTGAATT</td><td>>hg38_21_+_8433172_8446622_RNA45SN1_45S_50ntFlanks</td><td>7969</td><td>8010</td><td>42</td><td>CCGAGACGCGACCTCAGATCAGACGTGGCGACCCGCTGAATT</td><td>384.9</td><td>3.8</td><td>1.2</td><td>3.3</td><td>1.0E-03</td><td>1.2E-03</td><td>BT20/MB468</td><td>NUC</td></tr>
<tr><td>>hg38_21_+_8433172_8446622_RNA45SN1_45S_50ntFlanks@10787.10816.30|ACAGCCTCTGGCATGTTGGAACAATGTAGG</td><td>>hg38_21_+_8433172_8446622_RNA45SN1_45S_50ntFlanks</td><td>10787</td><td>10816</td><td>30</td><td>ACAGCCTCTGGCATGTTGGAACAATGTAGG</td><td>513.5</td><td>-2.4</td><td>0.7</td><td>-3.3</td><td>9.1E-04</td><td>1.2E-03</td><td>BT20/MB468</td><td>CYTO</td></tr>
<tr><td>>MI0000082|hsa-mir-25&WithFlank&7|-|100093554|100093649@58.77.20&[MIMAT0000081&hsa-miR-25-3p&offsets|0|-2;m-23&7|-|100093571|100093592&offsets|0|-2]|CATTGCACTTGTCTCGGTCT</td><td>>MI0000082|hsa-mir-25&WithFlank&7|-|100093554|100093649</td><td>58</td><td>77</td><td>20</td><td>CATTGCACTTGTCTCGGTCT</td><td>310.3</td><td>5.1</td><td>1.4</td><td>3.5</td><td>4.4E-04</td><td>1.2E-03</td><td>BT20/MB468</td><td>MP</td></tr>
<tr><td>>am_trnaMT_GluTTC_MT_-_14674_14742@16.53.38,trnalookalike8_GluTTC_5_-_93905172_93905240@16.53.38|ATACAACGATGGTTTTTCATATCATTGGTCGTGGTTGT</td><td>>am_trnaMT_GluTTC_MT_-_14674_14742</td><td>16</td><td>53</td><td>38</td><td>ATACAACGATGGTTTTTCATATCATTGGTCGTGGTTGT</td><td>58.1</td><td>9.6</td><td>2.7</td><td>3.5</td><td>4.4E-04</td><td>1.2E-03</td><td>BT20/MB468</td><td>MP</td></tr>
<tr><td>>hg38_21_+_8433172_8446622_RNA45SN1_45S_50ntFlanks@9567.9595.29|ACGTGCAAATCGGTCGTCCGACCTGGGTA</td><td>>hg38_21_+_8433172_8446622_RNA45SN1_45S_50ntFlanks</td><td>9567</td><td>9595</td><td>29</td><td>ACGTGCAAATCGGTCGTCCGACCTGGGTA</td><td>1572.5</td><td>-5.2</td><td>1.5</td><td>-3.5</td><td>4.4E-04</td><td>1.2E-03</td><td>BT20/MB468</td><td>MP</td></tr>
<tr><td>>hg38_21_+_8433172_8446622_RNA45SN1_45S_50ntFlanks@10408.10456.49|GGTCAGTCGGTCCTGAGAGATGGGCGAGCGCCGTTCCGAAGGGACGGGC</td><td>>hg38_21_+_8433172_8446622_RNA45SN1_45S_50ntFlanks</td><td>10408</td><td>10456</td><td>49</td><td>GGTCAGTCGGTCCTGAGAGATGGGCGAGCGCCGTTCCGAAGGGACGGGC</td><td>350.6</td><td>-2.5</td><td>0.8</td><td>-3.3</td><td>9.4E-04</td><td>1.2E-03</td><td>MB231/MB468</td><td>CYTO</td></tr>
<tr><td>>MI0000750|hsa-mir-26a-2&WithFlank&12|-|57824603|57824698@23.41.19(+1U);MI0000083|hsa-mir-26a-1&WithFlank&3|+|37969398|37969486@19.37.19(+1U)&[MIMAT0000082&hsa-miR-26a-5p&offsets|+3|0(+1U);m-33&12|-|57824658|57824679&offsets|+3|0(+1U)];[MIMAT0000082_1&hsa-miR-26a-5p&offsets|+3|0(+1U);m-34&3|+|37969413|37969434&offsets|+3|0(+1U)]|AAGTAATCCAGGATAGGCTT</td><td>>MI0000750|hsa-mir-26a-2&WithFlank&12|-|57824603|57824698</td><td>23</td><td>41</td><td>19(+1U)</td><td>AAGTAATCCAGGATAGGCTT</td><td>52.8</td><td>8.9</td><td>2.7</td><td>3.3</td><td>8.4E-04</td><td>1.2E-03</td><td>MB231/MB468</td><td>MP</td></tr>
<tr><td>>hg38_21_+_8433172_8446622_RNA45SN1_45S_50ntFlanks@10829.10858.30|GCAAGCCGGATCCGTAACTTCGGGATAAGG</td><td>>hg38_21_+_8433172_8446622_RNA45SN1_45S_50ntFlanks</td><td>10829</td><td>10858</td><td>30</td><td>GCAAGCCGGATCCGTAACTTCGGGATAAGG</td><td>374.9</td><td>-6.0</td><td>1.7</td><td>-3.5</td><td>4.4E-04</td><td>1.2E-03</td><td>BT20/MB468</td><td>MP</td></tr>
<tr><td>>MI0000469|hsa-mir-125a&WithFlank&19|+|51693248|51693345@21.42.22(+1A)&[MIMAT0000443&hsa-miR-125a-5p&offsets|0|-2(+1A);m-63&19|+|51693268|51693289&offsets|0|0(+1A)]|TCCCTGAGACCCTTTAACCTGTA</td><td>>MI0000469|hsa-mir-125a&WithFlank&19|+|51693248|51693345</td><td>21</td><td>42</td><td>22(+1A)</td><td>TCCCTGAGACCCTTTAACCTGTA</td><td>2569.8</td><td>2.0</td><td>0.5</td><td>3.8</td><td>1.6E-04</td><td>1.2E-03</td><td>BT20/MB231</td><td>NUC</td></tr>
</tbody>
</table>

| | | | | | | | | | | | | | |
|---|---|---|---|---|---|---|---|---|---|---|---|---|---|
| >hg38_21_+_8433172_8446622_RNA45SN1_45S_50ntFlanks@6652.6683.32\|GACTCTTAGCGGTGGATCACTCGGCTCGTGCG | >hg38_21_+_8433172_8446622_RNA45SN1_45S_50ntFlanks | 6652 | 6683 | 32 | GACTCTTAGCGGTGGATCACTCGGCTCGTGCG | 16818.5 | -2.6 | 0.7 | -3.8 | 1.6E-04 | 1.2E-03 | BT20/MB231 | NUC |
| >hg38_21_+_8433172_8446622_RNA45SN1_45S_50ntFlanks@12418.12451.34\|CGATGTCGGCTCTTCCTATCATTGTGAAGCAGAA | >hg38_21_+_8433172_8446622_RNA45SN1_45S_50ntFlanks | 12418 | 12451 | 34 | CGATGTCGGCTCTTCCTATCATTGTGAAGCAGAA | 116.3 | -4.1 | 1.2 | -3.3 | 8.4E-04 | 1.2E-03 | MB231/MB468 | MP |
| >MI0033423\|hsa-mir-10399&WithFlank&7\|-\|139044094\|139044163@43.63.21(+1U)&[MIMAT0041630&hsa-miR-10399-3p&offsets\|-1\|-1(+1U);TJU_CMC_MD2.ID03061.3p-miR&offsets\|+3\|+2(+1U);m-2657&7\|-\|139044103\|139044124&offsets\|+3\|+2(+1U)]\|TCTCTCGGACAAGCTGTAGGTT | >MI0033423\|hsa-mir-10399&WithFlank&7\|-\|139044094\|139044163 | 43 | 63 | 21(+1U) | TCTCTCGGACAAGCTGTAGGTT | 203.0 | -3.8 | 1.0 | -3.7 | 1.9E-04 | 1.2E-03 | BT20/MB231 | MITO |
| >hg38_21_+_8433172_8446622_RNA45SN1_45S_50ntFlanks@11915.11940.26\|GAGACATGAGAGGTGTAGAATAAGTG | >hg38_21_+_8433172_8446622_RNA45SN1_45S_50ntFlanks | 11915 | 11940 | 26 | GAGACATGAGAGGTGTAGAATAAGTG | 295.1 | 7.0 | 1.9 | 3.8 | 1.5E-04 | 1.2E-03 | BT20/MB231 | MP |
| >hg38_21_+_8433172_8446622_RNA45SN1_45S_50ntFlanks@10798.10826.29\|CATGTTGGAACAATGTAGGTAAGGGAAGT | >hg38_21_+_8433172_8446622_RNA45SN1_45S_50ntFlanks | 10798 | 10826 | 29 | CATGTTGGAACAATGTAGGTAAGGGAAGT | 480.0 | -6.0 | 1.8 | -3.3 | 1.0E-03 | 1.2E-03 | BT20/MB468 | NUC |
| >hg38_1_-_228634819_228635039_RNA5S12_5S_50ntFlanks@133.170.38\|ATGGGAGACCGCCTGGGAATACCGGGTGCTGTAGGCTT | >hg38_1_-_228634819_228635039_RNA5S12_5S_50ntFlanks | 133 | 170 | 38 | ATGGGAGACCGCCTGGGAATACCGGGTGCTGTAGGCTT | 72464.8 | 4.9 | 1.3 | 3.8 | 1.6E-04 | 1.2E-03 | BT20/MB231 | MP |
| >hg38_21_+_8433172_8446622_RNA45SN1_45S_50ntFlanks@12477.12498.22\|CACTAATAGGGAACGTGAGCTG | >hg38_21_+_8433172_8446622_RNA45SN1_45S_50ntFlanks | 12477 | 12498 | 22 | CACTAATAGGGAACGTGAGCTG | 1866.3 | -2.1 | 0.6 | -3.3 | 9.5E-04 | 1.2E-03 | MB231/MB468 | CYTO |
| >hg38_21_+_8433172_8446622_RNA45SN1_45S_50ntFlanks@11808.11830.23\|CTGTCCCTACCTACTATCCAGCG | >hg38_21_+_8433172_8446622_RNA45SN1_45S_50ntFlanks | 11808 | 11830 | 23 | CTGTCCCTACCTACTATCCAGCG | 235.2 | -3.4 | 1.0 | -3.5 | 4.5E-04 | 1.2E-03 | BT20/MB468 | MP |
| >hg38_21_+_8433172_8446622_RNA45SN1_45S_50ntFlanks@11667.11693.27\|CTGCCCAGTGCTCTGAATGTCAAAGTG | >hg38_21_+_8433172_8446622_RNA45SN1_45S_50ntFlanks | 11667 | 11693 | 27 | CTGCCCAGTGCTCTGAATGTCAAAGTG | 248.1 | 4.6 | 1.4 | 3.3 | 9.1E-04 | 1.2E-03 | BT20/MB468 | CYTO |
| >hg38_21_+_8433172_8446622_RNA45SN1_45S_50ntFlanks@4357.4391.35\|AAAGTTGCTGCAGTTAAAAAGCTCGTAGTTGGATC | >hg38_21_+_8433172_8446622_RNA45SN1_45S_50ntFlanks | 4357 | 4391 | 35 | AAAGTTGCTGCAGTTAAAAAGCTCGTAGTTGGATC | 72.8 | -4.1 | 1.2 | -3.3 | 8.5E-04 | 1.2E-03 | MB231/MB468 | MP |
| >MI0000440\|hsa-mir-27b&WithFlank&9\|+\|95085439\|95085547@67.88.22(+1U)&[MIMAT0000419&hsa-miR-27b-3p&offsets\|0\|+1(+1U);m-39&9\|+\|95085505\|95085525&offsets\|0\|+1(+1U)]\|TTCACAGTGGCTAAGTTCTGCAT | >MI0000440\|hsa-mir-27b&WithFlank&9\|+\|95085439\|95085547 | 67 | 88 | 22(+1U) | TTCACAGTGGCTAAGTTCTGCAT | 111.8 | 4.1 | 1.2 | 3.3 | 8.5E-04 | 1.2E-03 | MB231/MB468 | MP |
| >hg38_21_+_8433172_8446622_RNA45SN1_45S_50ntFlanks@9563.9595.33\|CCTGACGTGCAAATCGGTCGTCCGACCTGGGTA | >hg38_21_+_8433172_8446622_RNA45SN1_45S_50ntFlanks | 9563 | 9595 | 33 | CCTGACGTGCAAATCGGTCGTCCGACCTGGGTA | 323.3 | -2.1 | 0.6 | -3.4 | 7.6E-04 | 1.2E-03 | BT20/MB468 | MITO |
| >hg38_21_+_8433172_8446622_RNA45SN1_45S_50ntFlanks@3996.4024.29\|ACTCTAGATAACCTCGGGCCGATCGCACG | >hg38_21_+_8433172_8446622_RNA45SN1_45S_50ntFlanks | 3996 | 4024 | 29 | ACTCTAGATAACCTCGGGCCGATCGCACG | 315.5 | -3.8 | 1.1 | -3.4 | 7.6E-04 | 1.2E-03 | BT20/MB468 | MITO |
| >hg38_1_-_228634819_228635039_RNA5S12_5S_50ntFlanks@139.171.33\|GACCGCCTGGGAATACCGGGTGCTGTAGGCTTT | >hg38_1_-_228634819_228635039_RNA5S12_5S_50ntFlanks | 139 | 171 | 33 | GACCGCCTGGGAATACCGGGTGCTGTAGGCTTT | 447.1 | 3.0 | 0.9 | 3.3 | 1.0E-03 | 1.2E-03 | MB231/MB468 | MITO |
| >hg38_21_+_8433172_8446622_RNA45SN1_45S_50ntFlanks@10916.10946.31\|CCGCGGCTGGACGAGGCGCCGCCGCCCCCCC | >hg38_21_+_8433172_8446622_RNA45SN1_45S_50ntFlanks | 10916 | 10946 | 31 | CCGCGGCTGGACGAGGCGCCGCCGCCCCCCC | 87.1 | -5.4 | 1.5 | -3.5 | 4.5E-04 | 1.2E-03 | BT20/MB468 | MP |
| >hg38_21_+_8433172_8446622_RNA45SN1_45S_50ntFlanks@12029.12057.29\|TCTGATCGTTTTTTCACTGACCCGGTGAG | >hg38_21_+_8433172_8446622_RNA45SN1_45S_50ntFlanks | 12029 | 12057 | 29 | TCTGATCGTTTTTTCACTGACCCGGTGAG | 123.6 | 10.3 | 2.7 | 3.8 | 1.6E-04 | 1.2E-03 | BT20/MB231 | MP |
| >trna10_AspGTC_12_-_125424193_125424264@1.35.35,trna12_AspGTC_12_-_125411891_125411962@1.35.35,trna38_AspGTC_17_-_8125556_8125627@1.35.35,trna45_AspGTC_6_+_27447453_27447524@1.35.35,trna48_AspGTC_6_+_27471523_27471594@1.35.35,trna4_AspGTC_12_+_96429799_96429870@1.35.35,trna69_AspGTC_1_-_161440205_161440276@1.35.35,trna72_AspGTC_1_-_161432824_161432895@1.35.35,trna75_AspGTC_1_-_161425414_161425485@1.35.35,trna78_AspGTC_1_-_161418033_161418104@1.35.35,trna81_AspGTC_1_-_161410615_161410686@1.35.35\|TCCTCGTTAGTATAGTGGTGAGTATCCCCGCCTGT | >trna10_AspGTC_12_-_125424193_125424264 | 1 | 35 | 35 | TCCTCGTTAGTATAGTGGTGAGTATCCCCGCCTGT | 342.8 | 8.5 | 2.6 | 3.3 | 9.5E-04 | 1.2E-03 | MB231/MB468 | CYTO |
| >hg38_21_+_8433172_8446622_RNA45SN1_45S_50ntFlanks@5074.5113.40\|ATTCCGATAACGAACGAGACTCTGGCATGCTAACTAGTTA | >hg38_21_+_8433172_8446622_RNA45SN1_45S_50ntFlanks | 5074 | 5113 | 40 | ATTCCGATAACGAACGAGACTCTGGCATGCTAACTAGTTA | 65.1 | -9.9 | 3.0 | -3.3 | 8.5E-04 | 1.2E-03 | MB231/MB468 | MP |
| >hg38_21_+_8433172_8446622_RNA45SN1_45S_50ntFlanks@10409.10429.21\|GTCAGTCGGTCCTGAGAGATG | >hg38_21_+_8433172_8446622_RNA45SN1_45S_50ntFlanks | 10409 | 10429 | 21 | GTCAGTCGGTCCTGAGAGATG | 231.2 | 3.2 | 0.9 | 3.7 | 1.9E-04 | 1.2E-03 | BT20/MB231 | MITO |
| >am_trnaMT_PheGAA_MT_+_577_647@41.65.25\|GTTTAGACGGGCTCACATCACCCCA | >am_trnaMT_PheGAA_MT_+_577_647 | 41 | 65 | 25 | GTTTAGACGGGCTCACATCACCCCA | 101.0 | 2.6 | 0.8 | 3.3 | 8.5E-04 | 1.2E-03 | MB231/MB468 | MP |
| >hg38_21_+_8433172_8446622_RNA45SN1_45S_50ntFlanks@9533.9569.37\|GAGGAAACTCTGGTGGAGGTCCGTAGCGGTCCTGACG | >hg38_21_+_8433172_8446622_RNA45SN1_45S_50ntFlanks | 9533 | 9569 | 37 | GAGGAAACTCTGGTGGAGGTCCGTAGCGGTCCTGACG | 247.7 | -3.1 | 0.9 | -3.3 | 1.0E-03 | 1.2E-03 | MB231/MB468 | MITO |
| >hg38_21_+_8433172_8446622_RNA45SN1_45S_50ntFlanks@10843.10867.25\|TAACTTCGGGATAAGGATTGGCTCT | >hg38_21_+_8433172_8446622_RNA45SN1_45S_50ntFlanks | 10843 | 10867 | 25 | TAACTTCGGGATAAGGATTGGCTCT | 477.3 | -2.3 | 0.7 | -3.3 | 9.6E-04 | 1.2E-03 | MB231/MB468 | CYTO |
| >hg38_21_+_8433172_8446622_RNA45SN1_45S_50ntFlanks@7983.8003.21\|CAGATCAGACGTGGCGACCCG | >hg38_21_+_8433172_8446622_RNA45SN1_45S_50ntFlanks | 7983 | 8003 | 21 | CAGATCAGACGTGGCGACCCG | 403.9 | -3.9 | 1.2 | -3.3 | 9.2E-04 | 1.2E-03 | BT20/MB468 | CYTO |
| >MI0000063\|hsa-let-7b&WithFlank&22\|+\|46113680\|46113774@12.33.22&[MIMAT0000063&hsa-let-7b-5p&offsets\|0\|0;m-14&22\|+\|46113691\|46113712&offsets\|0\|0]\|TGAGGTAGTAGGTTGTGTGGTT | >MI0000063\|hsa-let-7b&WithFlank&22\|+\|46113680\|46113774 | 12 | 33 | 22 | TGAGGTAGTAGGTTGTGTGGTT | 72936.3 | 3.0 | 0.8 | 3.8 | 1.7E-04 | 1.2E-03 | BT20/MB231 | CYTO |
| >MI0000065\|hsa-let-7d&WithFlank&9\|+\|94178828\|94178926@14.31.18&[MIMAT0000065&hsa-let-7d-5p&offsets\|0\|-4;m-90&9\|+\|94178841\|94178862&offsets\|0\|-4]\|AGAGGTAGTAGGTTGCAT | >MI0000065\|hsa-let-7d&WithFlank&9\|+\|94178828\|94178926 | 14 | 31 | 18 | AGAGGTAGTAGGTTGCAT | 209.5 | 5.7 | 1.6 | 3.5 | 4.5E-04 | 1.2E-03 | BT20/MB468 | MP |
| >hg38_21_+_8433172_8446622_RNA45SN1_45S_50ntFlanks@10396.10429.34\|CAGTTGAACATGGGTCAGTCGGTCCTGAGAGATG | >hg38_21_+_8433172_8446622_RNA45SN1_45S_50ntFlanks | 10396 | 10429 | 34 | CAGTTGAACATGGGTCAGTCGGTCCTGAGAGATG | 281.8 | -3.0 | 0.9 | -3.5 | 4.5E-04 | 1.2E-03 | BT20/MB468 | MP |
| >hg38_21_+_8433172_8446622_RNA45SN1_45S_50ntFlanks@12456.12485.30\|CCAAGCGTTGGATTGTTCACCCACTAATAG | >hg38_21_+_8433172_8446622_RNA45SN1_45S_50ntFlanks | 12456 | 12485 | 30 | CCAAGCGTTGGATTGTTCACCCACTAATAG | 1527.7 | -6.1 | 1.7 | -3.5 | 4.5E-04 | 1.2E-03 | BT20/MB468 | MP |
| >trna119_LysCTT_1_-_145395522_145395594@4.36.33,trna11_LysCTT_5_-_180648979_180649051@4.36.33,trna13_LysCTT_6_+_26556774_26556846@4.36.33,trna7_LysCTT_16_+_3225692_3225764@4.36.33,trna9_LysCTT_5_+_180634755_180634827@4.36.33\|CGGCTAGCTCAGTCGGTAGAGCATGAGACTCTT | >trna119_LysCTT_1_-_145395522_145395594 | 4 | 36 | 33 | CGGCTAGCTCAGTCGGTAGAGCATGAGACTCTT | 2238.0 | -6.9 | 1.8 | -3.7 | 1.9E-04 | 1.2E-03 | BT20/MB231 | MITO |
| >MI0000082\|hsa-mir-25&WithFlank&7\|-\|100093554\|100093649@58.77.20&[MIMAT0000081&hsa-miR-25-3p&offsets\|0\|-2;m-23&7\|-\|100093571\|100093592&offsets\|0\|-2]\|CATTGCACTTGTCTCGGTCT | >MI0000082\|hsa-mir-25&WithFlank&7\|-\|100093554\|100093649 | 58 | 77 | 20 | CATTGCACTTGTCTCGGTCT | 294.3 | 4.3 | 1.3 | 3.3 | 8.5E-04 | 1.2E-03 | MB231/MB468 | MP |
| >hg38_21_+_8433172_8446622_RNA45SN1_45S_50ntFlanks@4163.4186.24\|CATCCAAGGAAGGCAGCAGGCGCG | >hg38_21_+_8433172_8446622_RNA45SN1_45S_50ntFlanks | 4163 | 4186 | 24 | CATCCAAGGAAGGCAGCAGGCGCG | 407.1 | -1.9 | 0.6 | -3.3 | 9.2E-04 | 1.2E-03 | BT20/MB468 | CYTO |
| >MI0001445\|hsa-mir-423&WithFlank&17\|+\|30117073\|30117178@59.77.19(+2U)&[MIMAT0001340&hsa-miR-423-3p&offsets\|0\|-4(+2U);m-100&17\|+\|30117131\|30117153&offsets\|0\|-4(+2U)]\|AGCTCGGTCTGAGGCCCCTTT | >MI0001445\|hsa-mir-423&WithFlank&17\|+\|30117073\|30117178 | 59 | 77 | 19(+2U) | AGCTCGGTCTGAGGCCCCTTT | 66.5 | 6.9 | 2.0 | 3.5 | 4.5E-04 | 1.2E-03 | BT20/MB468 | MP |
| >hg38_21_+_8433172_8446622_RNA45SN1_45S_50ntFlanks@7984.8007.24\|AGATCAGACGTGGCGACCCGCTGA | >hg38_21_+_8433172_8446622_RNA45SN1_45S_50ntFlanks | 7984 | 8007 | 24 | AGATCAGACGTGGCGACCCGCTGA | 752.8 | -3.6 | 1.0 | -3.5 | 4.5E-04 | 1.2E-03 | BT20/MB468 | MP |
| >hg38_21_+_8433172_8446622_RNA45SN1_45S_50ntFlanks@11208.11225.18\|GCGGCGACTCTGGACGCG | >hg38_21_+_8433172_8446622_RNA45SN1_45S_50ntFlanks | 11208 | 11225 | 18 | GCGGCGACTCTGGACGCG | 1716.3 | -6.6 | 2.0 | -3.3 | 1.0E-03 | 1.2E-03 | BT20/MB468 | NUC |
| >hg38_21_+_8433172_8446622_RNA45SN1_45S_50ntFlanks@12412.12432.21\|ATCCTTCGATGTCGGCTCTTC | >hg38_21_+_8433172_8446622_RNA45SN1_45S_50ntFlanks | 12412 | 12432 | 21 | ATCCTTCGATGTCGGCTCTTC | 1943.6 | -5.7 | 1.8 | -3.3 | 1.0E-03 | 1.2E-03 | MB231/MB468 | NUC |
| >hg38_21_+_8433172_8446622_RNA45SN1_45S_50ntFlanks@3775.3798.24\|GCCGGTACAGTGAAACTGCGAATG | >hg38_21_+_8433172_8446622_RNA45SN1_45S_50ntFlanks | 3775 | 3798 | 24 | GCCGGTACAGTGAAACTGCGAATG | 297.2 | -2.3 | 0.7 | -3.3 | 1.0E-03 | 1.2E-03 | MB231/MB468 | NUC |
| >MI0003776\|hsa-mir-320b-1&WithFlank&1\|+\|116671740\|116671823@48.67.20;MI0003839\|hsa-mir-320b-2&WithFlank&1\|-\|224257034\|224257116@47.66.20;MI0000542\|hsa-mir-320a&WithFlank&8\|-\|22244960\|22245043@48.67.20&[MIMAT0005792&hsa-miR-320b&offsets\|0\|-2;m-183&1\|+\|116671787\|116671806&offsets\|0\|0];[MIMAT0005792_1&hsa-miR-320b&offsets\|0\|-2;m-183&1\|-\|224257051\|224257070&offsets\|0\|0];[MIMAT0000510&hsa-miR-320a-3p&offsets\|0\|-2;m-60&8\|-\|22244975\|22244996&offsets\|0\|-2]\|AAAAGCTGGGTTGAGAGGGC | >MI0003776\|hsa-mir-320b-1&WithFlank&1\|+\|116671740\|116671823 | 48 | 67 | 20 | AAAAGCTGGGTTGAGAGGGC | 680.8 | 3.1 | 0.9 | 3.3 | 8.6E-04 | 1.2E-03 | MB231/MB468 | MP |
| >trna111_HisGTG_1_-_147774845_147774916@-1A.29.30,trna118_HisGTG_1_-_145396881_145396952@-1A.29.30,trna16_HisGTG_1_+_146544773_146544844@-1A.29.30,trna1_HisGTG_15_+_45493349_45493420@-1A.29.30,trna21_HisGTG_1_+_147753471_147753542@-1A.29.30,trna33_HisGTG_6_+_27125906_27125977@-1A.29.30,trna7_HisGTG_9_-_14433938_14434009@-1A.29.30,trna8_HisGTG_15_-_45492611_45492682@-1A.29.30,trna9_HisGTG_15_-_45490804_45490875@-1A.29.30\|AGCCGTGATCGTATAGTGGTTAGTACTCTG | >trna111_HisGTG_1_-_147774845_147774916 | -1A | 29 | 30 | AGCCGTGATCGTATAGTGGTTAGTACTCTG | 972.9 | -5.5 | 1.7 | -3.3 | 9.3E-04 | 1.2E-03 | BT20/MB468 | CYTO |
| >hg38_21_+_8433172_8446622_RNA45SN1_45S_50ntFlanks@6652.6682.31\|GACTCTTAGCGGTGGATCACTCGGCTCGTGC | >hg38_21_+_8433172_8446622_RNA45SN1_45S_50ntFlanks | 6652 | 6682 | 31 | GACTCTTAGCGGTGGATCACTCGGCTCGTGC | 1485.0 | 4.4 | 1.2 | 3.8 | 1.6E-04 | 1.2E-03 | BT20/MB231 | MP |
| >hg38_MT_+_1621_3279_RNR2_16S_50ntFlanks@1416.1446.31\|GTTTGTTCAACGATTAAAGTCCTACGTGATC | >hg38_MT_+_1621_3279_RNR2_16S_50ntFlanks | 1416 | 1446 | 31 | GTTTGTTCAACGATTAAAGTCCTACGTGATC | 122.0 | 4.1 | 1.2 | 3.3 | 1.0E-03 | 1.2E-03 | MB231/MB468 | MITO |

| | | | | | | | | | | | | | |
|---|---|---|---|---|---|---|---|---|---|---|---|---|---|
| >trna10_LysCTT_16_+_3241501_3241573@14.32.19,trna13_LysCTT_14_-_58706613_58706685@14.32.19,trna2_LysCTT_15_+_79152904_79152976@14.32.19,trna4_LysCTT_18_-_43669271_43669343@14.32.19\|AGTCGGTAGAGCATGGGAC | >trna10_LysCTT_16_+_3241501_3241573 | 14 | 32 | 19 | AGTCGGTAGAGCATGGGAC | 166.0 | -4.8 | 1.5 | -3.3 | 1.0E-03 | 1.2E-03 | MB231/MB468 | NUC |
| >MI0005561\|hsa-mir-877&WithFlank&6\|+\|30584326\|30584423@7.25.19&[MIMAT0004949&hsa-miR-877-5p&offsets\|0\|-1;m-382&6\|+\|30584332\|30584354&offsets\|0\|-4]\|GTAGAGGAGATGGCGCAGG | >MI0005561\|hsa-mir-877&WithFlank&6\|+\|30584326\|30584423 | 7 | 25 | 19 | GTAGAGGAGATGGCGCAGG | 106.7 | 5.9 | 1.7 | 3.5 | 4.5E-04 | 1.2E-03 | BT20/MB468 | MP |
| >MI0000447\|hsa-mir-128-1&WithFlank&2\|+\|135665391\|135665484@56.76.21(+1A);MI0000727\|hsa-mir-128-2&WithFlank&3\|+\|35744470\|35744565@58.78.21(+1A)&[MIMAT0000424&hsa-miR-128-3p&offsets\|0\|0(+1A);m-108&2\|+\|135665446\|135665466&offsets\|0\|0(+1A)];[MIMAT0000424_1&hsa-miR-128-3p&offsets\|0\|0(+1A);m-117&3\|+\|35744527\|35744547&offsets\|0\|0(+1A)]\|TCACAGTGAACCGGTCTCTTTA | >MI0000447\|hsa-mir-128-1&WithFlank&2\|+\|135665391\|135665484 | 56 | 76 | 21(+1A) | TCACAGTGAACCGGTCTCTTTA | 389.9 | 2.4 | 0.7 | 3.3 | 9.6E-04 | 1.2E-03 | MB231/MB468 | CYTO |
| >hg38_21_+_8433172_8446622_RNA45SN1_45S_50ntFlanks@9481.9511.31\|CACGTGTTAGGACCCGAAAGATGGTGAACTA | >hg38_21_+_8433172_8446622_RNA45SN1_45S_50ntFlanks | 9481 | 9511 | 31 | CACGTGTTAGGACCCGAAAGATGGTGAACTA | 51.8 | 9.4 | 2.7 | 3.5 | 4.5E-04 | 1.3E-03 | BT20/MB468 | MP |
| >hg38_21_+_8433172_8446622_RNA45SN1_45S_50ntFlanks@10727.10756.30\|GGAGAGGGTGTAAATCTCGCGCCGGGCCGT | >hg38_21_+_8433172_8446622_RNA45SN1_45S_50ntFlanks | 10727 | 10756 | 30 | GGAGAGGGTGTAAATCTCGCGCCGGGCCGT | 168.7 | -5.4 | 1.5 | -3.5 | 4.5E-04 | 1.3E-03 | BT20/MB468 | MP |
| >hg38_21_+_8433172_8446622_RNA45SN1_45S_50ntFlanks@12690.12708.19\|GCGGAACGATACGGCAGCG | >hg38_21_+_8433172_8446622_RNA45SN1_45S_50ntFlanks | 12690 | 12708 | 19 | GCGGAACGATACGGCAGCG | 175.4 | -5.9 | 1.7 | -3.5 | 4.5E-04 | 1.3E-03 | BT20/MB468 | MP |
| >trna14_LysTTT_11_-_59327808_59327880@3.33.31,trna2_LysTTT_17_+_8022473_8022545@3.33.31,trna54_LysTTT_1_+_204475655_204475727@3.33.31,trna5_LysTTT_11_+_59323902_59323974@3.33.31,trna62_LysTTT_1_-_204476158_204476230@3.33.31,trna76_LysTTT_6_+_28918806_28918878@3.33.31\|CCGGATAGCTCAGTCGGTAGAGCATCAGACT | >trna14_LysTTT_11_-_59327808_59327880 | 3 | 33 | 31 | CCGGATAGCTCAGTCGGTAGAGCATCAGACT | 120.7 | 5.9 | 1.8 | 3.3 | 8.6E-04 | 1.3E-03 | MB231/MB468 | MP |
| >trnaMT_PheGAA_MT_+_577_647@22.62.41\|AAAGCAATACACTGAAAATGTTTAGACGGGCTCACATCACC | >trnaMT_PheGAA_MT_+_577_647 | 22 | 62 | 41 | AAAGCAATACACTGAAAATGTTTAGACGGGCTCACATCACC | 56.8 | 9.6 | 2.7 | 3.5 | 4.6E-04 | 1.3E-03 | BT20/MB468 | MP |
| >hg38_21_+_8433172_8446622_RNA45SN1_45S_50ntFlanks@12637.12659.23\|GCTACCATCTGTGGGATTATGAC | >hg38_21_+_8433172_8446622_RNA45SN1_45S_50ntFlanks | 12637 | 12659 | 23 | GCTACCATCTGTGGGATTATGAC | 1670.9 | -3.3 | 1.0 | -3.4 | 7.7E-04 | 1.3E-03 | BT20/MB468 | MITO |
| >MI0000446\|hsa-mir-125b-1&WithFlank&11\|-\|122099751\|122099850@21.39.19(+1C);MI0000470\|hsa-mir-125b-2&WithFlank&21\|+\|16590231\|16590331@23.41.19(+1C)&[MIMAT0000423&hsa-miR-125b-5p&offsets\|0\|-3(+1C);m-67&11\|-\|122099809\|122099830&offsets\|0\|-3(+1C)];[MIMAT0000423_1&hsa-miR-125b-5p&offsets\|0\|-3(+1C);m-68&21\|+\|16590253\|16590274&offsets\|0\|-3(+1C)]\|TCCCTGAGACCCTAACTTGC | >MI0000446\|hsa-mir-125b-1&WithFlank&11\|-\|122099751\|122099850 | 21 | 39 | 19(+1C) | TCCCTGAGACCCTAACTTGC | 103.7 | 6.3 | 1.9 | 3.3 | 8.6E-04 | 1.3E-03 | MB231/MB468 | MP |
| >hg38_21_+_8433172_8446622_RNA45SN1_45S_50ntFlanks@10392.10420.29\|ACAGCAGTTGAACATGGGTCAGTCGGTCC | >hg38_21_+_8433172_8446622_RNA45SN1_45S_50ntFlanks | 10392 | 10420 | 29 | ACAGCAGTTGAACATGGGTCAGTCGGTCC | 254.3 | -3.6 | 1.1 | -3.4 | 7.8E-04 | 1.3E-03 | BT20/MB468 | MITO |
| >hg38_21_+_8433172_8446622_RNA45SN1_45S_50ntFlanks@12655.12673.19\|ATGACTGAACGCCTCTAAG | >hg38_21_+_8433172_8446622_RNA45SN1_45S_50ntFlanks | 12655 | 12673 | 19 | ATGACTGAACGCCTCTAAG | 228.9 | -1.7 | 0.5 | -3.4 | 7.8E-04 | 1.3E-03 | BT20/MB468 | MITO |
| >hg38_21_+_8433172_8446622_RNA45SN1_45S_50ntFlanks@6735.6770.36\|TTGATCATCGACACTTCGAACGCACTTGCGGCCCCG | >hg38_21_+_8433172_8446622_RNA45SN1_45S_50ntFlanks | 6735 | 6770 | 36 | TTGATCATCGACACTTCGAACGCACTTGCGGCCCCG | 293.3 | -5.8 | 1.6 | -3.5 | 4.6E-04 | 1.3E-03 | BT20/MB468 | MP |
| >hg38_21_+_8433172_8446622_RNA45SN1_45S_50ntFlanks@4547.4565.19\|CTGGATACCGCAGCTAGGA | >hg38_21_+_8433172_8446622_RNA45SN1_45S_50ntFlanks | 4547 | 4565 | 19 | CTGGATACCGCAGCTAGGA | 146.8 | -4.7 | 1.4 | -3.3 | 1.0E-03 | 1.3E-03 | MB231/MB468 | NUC |
| >hg38_21_+_8433172_8446622_RNA45SN1_45S_50ntFlanks@8177.8203.27\|TTCTGATCGAGGCCCAGCCCGTGGACG | >hg38_21_+_8433172_8446622_RNA45SN1_45S_50ntFlanks | 8177 | 8203 | 27 | TTCTGATCGAGGCCCAGCCCGTGGACG | 457.6 | -4.4 | 1.3 | -3.3 | 8.6E-04 | 1.3E-03 | MB231/MB468 | MP |
| >MI0000255\|hsa-mir-30d&WithFlank&8\|-\|134804870\|134804951@12.33.22(+1U)&[MIMAT0000245&hsa-miR-30d-5p&offsets\|0\|0(+1U);m-21&8\|-\|134804919\|134804940&offsets\|0\|0(+1U)]\|TGTAAACATCCCCGACTGGAAGT | >MI0000255\|hsa-mir-30d&WithFlank&8\|-\|134804870\|134804951 | 12 | 33 | 22(+1U) | TGTAAACATCCCCGACTGGAAGT | 11110.1 | 2.0 | 0.5 | 3.8 | 1.7E-04 | 1.3E-03 | BT20/MB231 | NUC |
| >hg38_21_+_8433172_8446622_RNA45SN1_45S_50ntFlanks@7973.8012.40\|GACGCGACCTCAGATCAGACGTGGCGACCCGCTGAATTTA | >hg38_21_+_8433172_8446622_RNA45SN1_45S_50ntFlanks | 7973 | 8012 | 40 | GACGCGACCTCAGATCAGACGTGGCGACCCGCTGAATTTA | 418.7 | -3.3 | 0.9 | -3.8 | 1.7E-04 | 1.3E-03 | BT20/MB231 | CYTO |
| >MI0000072\|hsa-mir-18a&WithFlank&13\|+\|91350745\|91350827@12.34.23&[MIMAT0000072&hsa-miR-18a-5p&offsets\|0\|0;m-262&13\|+\|91350756\|91350778&offsets\|0\|0]\|TAAGGTGCATCTAGTGCAGATAG | >MI0000072\|hsa-mir-18a&WithFlank&13\|+\|91350745\|91350827 | 12 | 34 | 23 | TAAGGTGCATCTAGTGCAGATAG | 780.8 | 3.0 | 0.9 | 3.3 | 8.7E-04 | 1.3E-03 | MB231/MB468 | MP |
| >hg38_21_+_8433172_8446622_RNA45SN1_45S_50ntFlanks@3853.3880.28\|AACTGTGGTAATTCTAGAGCTAATACAT | >hg38_21_+_8433172_8446622_RNA45SN1_45S_50ntFlanks | 3853 | 3880 | 28 | AACTGTGGTAATTCTAGAGCTAATACAT | 136.5 | -3.1 | 0.9 | -3.3 | 8.7E-04 | 1.3E-03 | MB231/MB468 | MP |
| >hg38_21_+_8433172_8446622_RNA45SN1_45S_50ntFlanks@11602.11620.19\|TCCGACTGTTTAATTAAAA | >hg38_21_+_8433172_8446622_RNA45SN1_45S_50ntFlanks | 11602 | 11620 | 19 | TCCGACTGTTTAATTAAAA | 1274.3 | -3.0 | 0.9 | -3.3 | 9.7E-04 | 1.3E-03 | MB231/MB468 | CYTO |
| >MI0000061\|hsa-let-7a-2&WithFlank&11\|-\|122146516\|122146599@11.30.20(+1A);MI0000062\|hsa-let-7a-3&WithFlank&22\|+\|46112743\|46112828@10.29.20(+1A);MI0000060\|hsa-let-7a-1&WithFlank&9\|+\|94175951\|94176042@12.31.20(+1A)&[MIMAT0000062&hsa-let-7a-5p&offsets\|0\|-2(+1A);m-6&11\|-\|122146568\|122146589&offsets\|0\|-2(+1A)];[MIMAT0000062_1&hsa-let-7a-5p&offsets\|0\|-2(+1A);m-5&22\|+\|46112752\|46112773&offsets\|0\|-2(+1A)];[MIMAT0000062_2&hsa-let-7a-5p&offsets\|0\|-2(+1A);m-4&9\|+\|94175962\|94175983&offsets\|0\|-2(+1A)]\|TGAGGTAGTAGGTTGTATAGA | >MI0000061\|hsa-let-7a-2&WithFlank&11\|-\|122146516\|122146599 | 11 | 30 | 20(+1A) | TGAGGTAGTAGGTTGTATAGA | 152.4 | 5.3 | 1.6 | 3.3 | 8.7E-04 | 1.3E-03 | MB231/MB468 | MP |
| >hg38_21_+_8433172_8446622_RNA45SN1_45S_50ntFlanks@12474.12498.25\|ACCCACTAATAGGGAACGTGAGCTG | >hg38_21_+_8433172_8446622_RNA45SN1_45S_50ntFlanks | 12474 | 12498 | 25 | ACCCACTAATAGGGAACGTGAGCTG | 376.7 | -3.2 | 1.0 | -3.3 | 9.7E-04 | 1.3E-03 | MB231/MB468 | CYTO |
| >hg38_21_+_8433172_8446622_RNA45SN1_45S_50ntFlanks@6648.6689.42\|GTACGACTCTTAGCGGTGGATCACTCGGCTCGTGCGTCGATG | >hg38_21_+_8433172_8446622_RNA45SN1_45S_50ntFlanks | 6648 | 6689 | 42 | GTACGACTCTTAGCGGTGGATCACTCGGCTCGTGCGTCGATG | 2678.7 | -2.4 | 0.7 | -3.3 | 1.0E-03 | 1.3E-03 | BT20/MB468 | NUC |
| >MI0000088\|hsa-mir-30a&WithFlank&6\|-\|71403545\|71403627@12.31.20&[MIMAT0000087&hsa-miR-30a-5p&offsets\|0\|-2;m-15&6\|-\|71403595\|71403616&offsets\|0\|-2]\|TGTAAACATCCTCGACTGGA | >MI0000088\|hsa-mir-30a&WithFlank&6\|-\|71403545\|71403627 | 12 | 31 | 20 | TGTAAACATCCTCGACTGGA | 5473.6 | 4.3 | 1.2 | 3.5 | 4.6E-04 | 1.3E-03 | BT20/MB468 | MP |
| >hg38_21_+_8433172_8446622_RNA45SN1_45S_50ntFlanks@11666.11692.27\|TCTGCCCAGTGCTCTGAATGTCAAAGT | >hg38_21_+_8433172_8446622_RNA45SN1_45S_50ntFlanks | 11666 | 11692 | 27 | TCTGCCCAGTGCTCTGAATGTCAAAGT | 153.0 | 5.6 | 1.6 | 3.5 | 4.6E-04 | 1.3E-03 | BT20/MB468 | MP |
| >am_trnaMT_HisGTG_MT_+_12138_12206@19.50.32\|AAACATCAGATTGTGAATCTGACAACAGAGGC | >am_trnaMT_HisGTG_MT_+_12138_12206 | 19 | 50 | 32 | AAACATCAGATTGTGAATCTGACAACAGAGGC | 55.8 | 9.5 | 2.7 | 3.5 | 4.6E-04 | 1.3E-03 | BT20/MB468 | MP |
| >hg38_21_+_8433172_8446622_RNA45SN1_45S_50ntFlanks@6676.6696.21\|CTCGTGCGTCGATGAAGAACG | >hg38_21_+_8433172_8446622_RNA45SN1_45S_50ntFlanks | 6676 | 6696 | 21 | CTCGTGCGTCGATGAAGAACG | 237.7 | -3.7 | 1.1 | -3.3 | 9.4E-04 | 1.3E-03 | BT20/MB468 | CYTO |
| >MI0000750\|hsa-mir-26a-2&WithFlank&12\|-\|57824603\|57824698@20.43.24;MI0000083\|hsa-mir-26a-1&WithFlank&3\|+\|37969398\|37969486@16.39.24&[MIMAT0000082&hsa-miR-26a-5p&offsets\|0\|+2;m-33&12\|-\|57824658\|57824679&offsets\|0\|+2];[MIMAT0000082_1&hsa-miR-26a-5p&offsets\|0\|+2;m-34&3\|+\|37969413\|37969434&offsets\|0\|+2]\|TTCAAGTAATCCAGGATAGGCTGT | >MI0000750\|hsa-mir-26a-2&WithFlank&12\|-\|57824603\|57824698 | 20 | 43 | 24 | TTCAAGTAATCCAGGATAGGCTGT | 187.6 | 4.2 | 1.3 | 3.3 | 8.7E-04 | 1.3E-03 | MB231/MB468 | MP |
| >hg38_21_+_8433172_8446622_RNA45SN1_45S_50ntFlanks@474.505.32\|TCTTCGTGATCGATGTGGTGACGTCGTGCTCT | >hg38_21_+_8433172_8446622_RNA45SN1_45S_50ntFlanks | 474 | 505 | 32 | TCTTCGTGATCGATGTGGTGACGTCGTGCTCT | 61.4 | 6.6 | 1.9 | 3.5 | 4.6E-04 | 1.3E-03 | BT20/MB468 | MP |
| >hg38_21_+_8433172_8446622_RNA45SN1_45S_50ntFlanks@12800.12817.18\|AGGGCGCGTGCCCCGCCG | >hg38_21_+_8433172_8446622_RNA45SN1_45S_50ntFlanks | 12800 | 12817 | 18 | AGGGCGCGTGCCCCGCCG | 118.2 | -5.8 | 1.6 | -3.5 | 4.6E-04 | 1.3E-03 | BT20/MB468 | MP |
| >hg38_21_+_8433172_8446622_RNA45SN1_45S_50ntFlanks@5472.5499.28\|ACGGCCCTGGCGGAGCGCTGAGAAGACG | >hg38_21_+_8433172_8446622_RNA45SN1_45S_50ntFlanks | 5472 | 5499 | 28 | ACGGCCCTGGCGGAGCGCTGAGAAGACG | 3605.5 | -3.6 | 1.1 | -3.3 | 1.0E-03 | 1.3E-03 | BT20/MB468 | NUC |
| >MI0000071\|hsa-mir-17&WithFlank&13\|+\|91350599\|91350694@20.40.21&[MIMAT0000070&hsa-miR-17-5p&offsets\|0\|-2;m-88&13\|+\|91350618\|91350640&offsets\|0\|-2]\|CAAAGTGCTTACAGTGCAGGT | >MI0000071\|hsa-mir-17&WithFlank&13\|+\|91350599\|91350694 | 20 | 40 | 21 | CAAAGTGCTTACAGTGCAGGT | 1944.8 | 3.0 | 0.9 | 3.3 | 8.7E-04 | 1.3E-03 | MB231/MB468 | MP |
| >hg38_21_+_8433172_8446622_RNA45SN1_45S_50ntFlanks@6653.6676.24\|ACTCTTAGCGGTGGATCACTCGGC | >hg38_21_+_8433172_8446622_RNA45SN1_45S_50ntFlanks | 6653 | 6676 | 24 | ACTCTTAGCGGTGGATCACTCGGC | 201.6 | 4.7 | 1.4 | 3.5 | 4.6E-04 | 1.3E-03 | BT20/MB468 | MP |
| >hg38_21_+_8433172_8446622_RNA45SN1_45S_50ntFlanks@8381.8403.23\|AAGAGGGCGTGAAACCGTTAAGA | >hg38_21_+_8433172_8446622_RNA45SN1_45S_50ntFlanks | 8381 | 8403 | 23 | AAGAGGGCGTGAAACCGTTAAGA | 282.5 | -2.8 | 0.9 | -3.3 | 9.8E-04 | 1.3E-03 | MB231/MB468 | CYTO |
| >MI0000255\|hsa-mir-30d&WithFlank&8\|-\|134804870\|134804951@12.33.22(+1U)&[MIMAT0000245&hsa-miR-30d-5p&offsets\|0\|0(+1U);m-21&8\|-\|134804919\|134804940&offsets\|0\|0(+1U)]\|TGTAAACATCCCCGACTGGAAGT | >MI0000255\|hsa-mir-30d&WithFlank&8\|-\|134804870\|134804951 | 12 | 33 | 22(+1U) | TGTAAACATCCCCGACTGGAAGT | 11588.5 | 3.1 | 0.8 | 3.8 | 1.7E-04 | 1.3E-03 | BT20/MB231 | CYTO |
| >hg38_21_+_8433172_8446622_RNA45SN1_45S_50ntFlanks@12624.12660.37\|CCAATGGGGCGAAGCTACCATCTGTGGGATTATGACT | >hg38_21_+_8433172_8446622_RNA45SN1_45S_50ntFlanks | 12624 | 12660 | 37 | CCAATGGGGCGAAGCTACCATCTGTGGGATTATGACT | 166.1 | -4.0 | 1.2 | -3.4 | 7.8E-04 | 1.3E-03 | BT20/MB468 | MITO |
| >MI0000434\|hsa-let-7i&WithFlank&12\|+\|62603680\|62603775@12.31.20&[MIMAT0000415&hsa-let-7i-5p&offsets\|0\|-2;m-49&12\|+\|62603691\|62603712&offsets\|0\|-2]\|TGAGGTAGTAGTTTGTGCTG | >MI0000434\|hsa-let-7i&WithFlank&12\|+\|62603680\|62603775 | 12 | 31 | 20 | TGAGGTAGTAGTTTGTGCTG | 115.2 | 5.9 | 1.8 | 3.3 | 1.0E-03 | 1.3E-03 | MB231/MB468 | MITO |
| >MI0000284\|hsa-mir-204&WithFlank&9\|-\|70809969\|70810090@39.60.22&[MIMAT0000265&hsa-miR-204-5p&offsets\|0\|0;m-296&9\|-\|70810031\|70810052&offsets\|0\|0]\|TTCCCTTTGTCATCCTATGCCT | >MI0000284\|hsa-mir-204&WithFlank&9\|-\|70809969\|70810090 | 39 | 60 | 22 | TTCCCTTTGTCATCCTATGCCT | 102.9 | 5.9 | 1.8 | 3.3 | 8.7E-04 | 1.3E-03 | MB231/MB468 | MP |
| >MI0003560\|hsa-mir-92b&WithFlank&1\|+\|155195171\|155195278@67.86.20(+1A)&[MIMAT0003218&hsa-miR-92b-3p&offsets\|0\|-2(+1A);m-115&1\|+\|155195237\|155195258&offsets\|0\|-2(+1A)]\|TATTGCACTCGTCCCGGCCTA | >MI0003560\|hsa-mir-92b&WithFlank&1\|+\|155195171\|155195278 | 67 | 86 | 20(+1A) | TATTGCACTCGTCCCGGCCTA | 82.1 | 6.1 | 1.8 | 3.3 | 8.7E-04 | 1.3E-03 | MB231/MB468 | MP |
| >hg38_21_+_8433172_8446622_RNA45SN1_45S_50ntFlanks@10238.10261.24\|CGCTACGCCGCGACGAGTAGGAGG | >hg38_21_+_8433172_8446622_RNA45SN1_45S_50ntFlanks | 10238 | 10261 | 24 | CGCTACGCCGCGACGAGTAGGAGG | 397.4 | 3.5 | 1.0 | 3.5 | 4.6E-04 | 1.3E-03 | BT20/MB468 | MP |
| >hg38_21_+_8433172_8446622_RNA45SN1_45S_50ntFlanks@6776.6802.27\|CTCCCGGGGCTACGCCTGTCTGAGCGT | >hg38_21_+_8433172_8446622_RNA45SN1_45S_50ntFlanks | 6776 | 6802 | 27 | CTCCCGGGGCTACGCCTGTCTGAGCGT | 512.3 | -1.9 | 0.6 | -3.3 | 1.0E-03 | 1.3E-03 | BT20/MB468 | NUC |
| >hg38_21_+_8433172_8446622_RNA45SN1_45S_50ntFlanks@5103.5136.34\|CTAACTAGTTACGCGACCCCCGAGCGGTCGGCGT | >hg38_21_+_8433172_8446622_RNA45SN1_45S_50ntFlanks | 5103 | 5136 | 34 | CTAACTAGTTACGCGACCCCCGAGCGGTCGGCGT | 63.4 | -9.9 | 3.0 | -3.3 | 8.7E-04 | 1.3E-03 | MB231/MB468 | MP |
| >hg38_1_-_228634819_228635039_RNA5S12_5S_50ntFlanks@147.169.23\|GGGAATACCGGGTGCTGTAGGCT | >hg38_1_-_228634819_228635039_RNA5S12_5S_50ntFlanks | 147 | 169 | 23 | GGGAATACCGGGTGCTGTAGGCT | 6076.6 | 3.8 | 1.1 | 3.5 | 4.6E-04 | 1.3E-03 | BT20/MB468 | MP |

| | | | | | | | | | | | | | |
|---|---|---|---|---|---|---|---|---|---|---|---|---|---|
| >MI0000767\|hsa-mir-365a&WithFlank&16\|+\|14309279\|14309377@22.44.23&[MIMAT0009199&hsa-miR-365a-5p&offsets\|0\|0;m-595&16\|+\|14309300\|14309322&offsets\|0\|0]\|AGGGACTTTTGGGGGCAGATGTG | >MI0000767\|hsa-mir-365a&WithFlank&16\|+\|14309279\|14309377 | 22 | 44 | 23 | AGGGACTTTTGGGGGCAGATGTG | 109.1 | 5.7 | 1.6 | 3.5 | 4.6E-04 | 1.3E-03 | BT20/MB468 | MP |
| >hg38_MT_+_1621_3279_RNR2_16S_50ntFlanks@199.230.32\|TAATATAGCAAGGACTAACCCCTATACCTTCT | >hg38_MT_+_1621_3279_RNR2_16S_50ntFlanks | 199 | 230 | 32 | TAATATAGCAAGGACTAACCCCTATACCTTCT | 56.5 | 9.6 | 2.7 | 3.5 | 4.6E-04 | 1.3E-03 | BT20/MB468 | MP |
| >am_trna111_HisGTG_1_-_147774845_147774916@1.26.26,trna118_HisGTG_1_-_145396881_145396952@1.26.26,trna16_HisGTG_1_+_146544773_146544844@1.26.26,trna1_HisGTG_15_+_45493349_45493420@1.26.26,trna21_HisGTG_1_+_147753471_147753542@1.26.26,trna33_HisGTG_6_+_27125906_27125977@1.26.26,trna7_HisGTG_9_-_14433938_14434009@1.26.26,trna8_HisGTG_15_-_45492611_45492682@1.26.26,trna9_HisGTG_15_-_45490804_45490875@1.26.26\|GCCGTGATCGTATAGTGGTTAGTACT | >am_trna111_HisGTG_1_-_147774845_147774916 | 1 | 26 | 26 | GCCGTGATCGTATAGTGGTTAGTACT | 1775.1 | 3.8 | 1.2 | 3.3 | 9.8E-04 | 1.3E-03 | MB231/MB468 | CYTO |
| >hg38_21_+_8433172_8446622_RNA45SN1_45S_50ntFlanks@5487.5516.30\|CGCTGAGAAGACGGTCGAACTTGACTATCT | >hg38_21_+_8433172_8446622_RNA45SN1_45S_50ntFlanks | 5487 | 5516 | 30 | CGCTGAGAAGACGGTCGAACTTGACTATCT | 6073.9 | 2.8 | 0.8 | 3.3 | 9.8E-04 | 1.3E-03 | MB231/MB468 | CYTO |
| >hg38_21_+_8433172_8446622_RNA45SN1_45S_50ntFlanks@9352.9370.19\|CCGTGGCGCAATGAAGGTG | >hg38_21_+_8433172_8446622_RNA45SN1_45S_50ntFlanks | 9352 | 9370 | 19 | CCGTGGCGCAATGAAGGTG | 4215.7 | -6.5 | 1.9 | -3.5 | 4.6E-04 | 1.3E-03 | BT20/MB468 | MP |
| >hg38_21_+_8433172_8446622_RNA45SN1_45S_50ntFlanks@9929.9968.40\|GATATAGACAGCAGGACGGTGGCCATGGAAGTCGGAATCC | >hg38_21_+_8433172_8446622_RNA45SN1_45S_50ntFlanks | 9929 | 9968 | 40 | GATATAGACAGCAGGACGGTGGCCATGGAAGTCGGAATCC | 312.8 | 7.2 | 1.9 | 3.8 | 1.6E-04 | 1.3E-03 | BT20/MB231 | MP |
| >hg38_21_+_8433172_8446622_RNA45SN1_45S_50ntFlanks@12416.12450.35\|TTCGATGTCGGCTCTTCCTATCATTGTGAAGCAGA | >hg38_21_+_8433172_8446622_RNA45SN1_45S_50ntFlanks | 12416 | 12450 | 35 | TTCGATGTCGGCTCTTCCTATCATTGTGAAGCAGA | 164.4 | 3.9 | 1.2 | 3.3 | 1.0E-03 | 1.3E-03 | MB231/MB468 | MITO |
| >trnaMT_GluTTC_MT_-_14674_14742@47.70.24,trnalookalike8_GluTTC_5_-_93905172_93905240@47.70.24\|TGGTTGTAGTCCGTGCGAGAATAC | >trnaMT_GluTTC_MT_-_14674_14742 | 47 | 70 | 24 | TGGTTGTAGTCCGTGCGAGAATAC | 159.7 | 6.8 | 2.0 | 3.5 | 4.6E-04 | 1.3E-03 | BT20/MB468 | MP |
| >hg38_21_+_8433172_8446622_RNA45SN1_45S_50ntFlanks@12621.12654.34\|GAGCCAATGGGGCGAAGCTACCATCTGTGGGATT | >hg38_21_+_8433172_8446622_RNA45SN1_45S_50ntFlanks | 12621 | 12654 | 34 | GAGCCAATGGGGCGAAGCTACCATCTGTGGGATT | 192.9 | -4.2 | 1.2 | -3.5 | 4.6E-04 | 1.3E-03 | BT20/MB468 | MP |
| >Antisense_hg38_21_-_8433172_8446622_RNA45SN1_45S_50ntFlanks@7674.7692.19\|CTCTCTCTCTCTCCCCC | >Antisense_hg38_21_-_8433172_8446622_RNA45SN1_45S_50ntFlanks | 7674 | 7692 | 19 | CTCTCTCTCTCTCCCCC | 1083.8 | -6.6 | 2.0 | -3.3 | 1.1E-03 | 1.3E-03 | BT20/MB468 | NUC |
| >hg38_21_+_8433172_8446622_RNA45SN1_45S_50ntFlanks@11027.11045.19\|CTCTCTCTCTCTCCCCC | >hg38_21_+_8433172_8446622_RNA45SN1_45S_50ntFlanks | 11027 | 11045 | 19 | CTCTCTCTCTCTCCCCC | 1083.8 | -6.6 | 2.0 | -3.3 | 1.1E-03 | 1.3E-03 | BT20/MB468 | NUC |
| >MI0000782\|hsa-mir-374a&WithFlank&X\|-\|74287280\|74287363@18.38.21&[MIMAT0000727&hsa-miR-374a-5p&offsets\|0\|-1;m-184&X\|-\|74287325\|74287346&offsets\|0\|-1]\|TTATAATACAACCTGATAAGT | >MI0000782\|hsa-mir-374a&WithFlank&X\|-\|74287280\|74287363 | 18 | 38 | 21 | TTATAATACAACCTGATAAGT | 396.2 | -3.5 | 0.9 | -3.7 | 2.0E-04 | 1.3E-03 | BT20/MB231 | MITO |
| >MI0000101\|hsa-mir-99a&WithFlank&21\|+\|16539083\|16539175@19.39.21&[MIMAT0000097&hsa-miR-99a-5p&offsets\|0\|-1;m-64&21\|+\|16539101\|16539121&offsets\|0\|0]\|AACCCGTAGATCCGATCTTGT | >MI0000101\|hsa-mir-99a&WithFlank&21\|+\|16539083\|16539175 | 19 | 39 | 21 | AACCCGTAGATCCGATCTTGT | 1051.7 | -1.9 | 0.6 | -3.3 | 9.5E-04 | 1.3E-03 | BT20/MB468 | CYTO |
| >MI0000301\|hsa-mir-224&WithFlank&X\|-\|151958572\|151958664@14.34.21(+1C)&[MIMAT0000281&hsa-miR-224-5p&offsets\|+1\|-3(+1C);m-238&X\|-\|151958630\|151958651&offsets\|0\|-1(+1C)]\|CAAGTCACTAGTGGTTCCGTTC | >MI0000301\|hsa-mir-224&WithFlank&X\|-\|151958572\|151958664 | 14 | 34 | 21(+1C) | CAAGTCACTAGTGGTTCCGTTC | 93.6 | 6.2 | 1.9 | 3.3 | 8.8E-04 | 1.3E-03 | MB231/MB468 | MP |
| >am_trnaMT_GluTTC_MT_-_14674_14742@21.53.33,trnalookalike8_GluTTC_5_-_93905172_93905240@21.53.33\|ACGATGGTTTTTCATATCATTGGTCGTGGTTGT | >am_trnaMT_GluTTC_MT_-_14674_14742 | 21 | 53 | 33 | ACGATGGTTTTTCATATCATTGGTCGTGGTTGT | 263.7 | 6.4 | 1.8 | 3.5 | 4.6E-04 | 1.3E-03 | BT20/MB468 | MP |
| >hg38_21_+_8433172_8446622_RNA45SN1_45S_50ntFlanks@3843.3862.20\|CTACTTGGATAACTGTGGTA | >hg38_21_+_8433172_8446622_RNA45SN1_45S_50ntFlanks | 3843 | 3862 | 20 | CTACTTGGATAACTGTGGTA | 129.1 | -5.6 | 1.6 | -3.5 | 4.6E-04 | 1.3E-03 | BT20/MB468 | MP |
| >hg38_21_+_8433172_8446622_RNA45SN1_45S_50ntFlanks@11322.11342.21\|ACGTCTCGTCGCGCGCGCGTC | >hg38_21_+_8433172_8446622_RNA45SN1_45S_50ntFlanks | 11322 | 11342 | 21 | ACGTCTCGTCGCGCGCGCGTC | 2140.1 | -2.8 | 0.9 | -3.3 | 9.5E-04 | 1.3E-03 | BT20/MB468 | CYTO |
| >hg38_21_+_8433172_8446622_RNA45SN1_45S_50ntFlanks@12408.12443.36\|TTTGATCCTTCGATGTCGGCTCTTCCTATCATTGTG | >hg38_21_+_8433172_8446622_RNA45SN1_45S_50ntFlanks | 12408 | 12443 | 36 | TTTGATCCTTCGATGTCGGCTCTTCCTATCATTGTG | 316.1 | -2.6 | 0.8 | -3.3 | 1.0E-03 | 1.3E-03 | MB231/MB468 | MITO |
| >hg38_21_+_8433172_8446622_RNA45SN1_45S_50ntFlanks@8190.8213.24\|CCAGCCCGTGGACGGTGTGAGGCC | >hg38_21_+_8433172_8446622_RNA45SN1_45S_50ntFlanks | 8190 | 8213 | 24 | CCAGCCCGTGGACGGTGTGAGGCC | 345.4 | -2.8 | 0.8 | -3.3 | 1.0E-03 | 1.3E-03 | MB231/MB468 | MITO |
| >hg38_21_+_8433172_8446622_RNA45SN1_45S_50ntFlanks@11234.11254.21\|CCTTCCCGTGGATCGCCCCAG | >hg38_21_+_8433172_8446622_RNA45SN1_45S_50ntFlanks | 11234 | 11254 | 21 | CCTTCCCGTGGATCGCCCCAG | 196.7 | -5.6 | 1.6 | -3.5 | 4.7E-04 | 1.3E-03 | BT20/MB468 | MP |
| >hg38_21_+_8433172_8446622_RNA45SN1_45S_50ntFlanks@4225.4242.18\|AAATAACAATACAGGACT | >hg38_21_+_8433172_8446622_RNA45SN1_45S_50ntFlanks | 4225 | 4242 | 18 | AAATAACAATACAGGACT | 657.5 | -3.2 | 1.0 | -3.3 | 9.9E-04 | 1.3E-03 | MB231/MB468 | CYTO |
| >hg38_21_+_8433172_8446622_RNA45SN1_45S_50ntFlanks@10392.10432.41\|ACAGCAGTTGAACATGGGTCAGTCGGTCCTGAGAGATGGGC | >hg38_21_+_8433172_8446622_RNA45SN1_45S_50ntFlanks | 10392 | 10432 | 41 | ACAGCAGTTGAACATGGGTCAGTCGGTCCTGAGAGATGGGC | 237.5 | -3.4 | 1.0 | -3.3 | 9.9E-04 | 1.3E-03 | MB231/MB468 | CYTO |
| >trna119_LysCTT_1_-_145395522_145395594@5.36.32,trna11_LysCTT_5_-_180648979_180649051@5.36.32,trna13_LysCTT_6_+_26556774_26556846@5.36.32,trna7_LysCTT_16_+_3225692_3225764@5.36.32,trna9_LysCTT_5_+_180634755_180634827@5.36.32\|GGCTAGCTCAGTCGGTAGAGCATGAGACTCTT | >trna119_LysCTT_1_-_145395522_145395594 | 5 | 36 | 32 | GGCTAGCTCAGTCGGTAGAGCATGAGACTCTT | 226.6 | 4.4 | 1.3 | 3.3 | 8.8E-04 | 1.3E-03 | MB231/MB468 | MP |
| >hg38_21_+_8433172_8446622_RNA45SN1_45S_50ntFlanks@7975.7994.20\|CGCGACCTCAGATCAGACGT | >hg38_21_+_8433172_8446622_RNA45SN1_45S_50ntFlanks | 7975 | 7994 | 20 | CGCGACCTCAGATCAGACGT | 137816.8 | 2.9 | 0.8 | 3.5 | 4.7E-04 | 1.3E-03 | BT20/MB468 | MP |
| >MI0000456\|hsa-mir-140&WithFlank&16\|+\|69933075\|69933186@70.91.22&[MIMAT0004597&hsa-miR-140-3p&offsets\|+2\|+3;m-45&16\|+\|69933143\|69933164&offsets\|+1\|+1]\|CCACAGGGTAGAACCACGGACA | >MI0000456\|hsa-mir-140&WithFlank&16\|+\|69933075\|69933186 | 70 | 91 | 22 | CCACAGGGTAGAACCACGGACA | 77.9 | 6.9 | 2.0 | 3.5 | 4.7E-04 | 1.3E-03 | BT20/MB468 | MP |
| >hg38_21_+_8433172_8446622_RNA45SN1_45S_50ntFlanks@10907.10932.26\|GGGCGCGCGCCGCGGCTGGACGAGGC | >hg38_21_+_8433172_8446622_RNA45SN1_45S_50ntFlanks | 10907 | 10932 | 26 | GGGCGCGCGCCGCGGCTGGACGAGGC | 216.8 | 7.1 | 1.9 | 3.8 | 1.6E-04 | 1.3E-03 | BT20/MB231 | MP |
| >hg38_1_-_228634819_228635039_RNA5S12_5S_50ntFlanks@132.170.39\|GATGGGAGACCGCCTGGGAATACCGGGTGCTGTAGGCTT | >hg38_1_-_228634819_228635039_RNA5S12_5S_50ntFlanks | 132 | 170 | 39 | GATGGGAGACCGCCTGGGAATACCGGGTGCTGTAGGCTT | 68266.6 | 3.8 | 1.1 | 3.4 | 7.9E-04 | 1.3E-03 | BT20/MB468 | MITO |
| >hg38_MT_+_598_1651_RNR1_12S_50ntFlanks@428.452.25\|AAAGTGGCTTTAACATATCTGAACA | >hg38_MT_+_598_1651_RNR1_12S_50ntFlanks | 428 | 452 | 25 | AAAGTGGCTTTAACATATCTGAACA | 493.6 | 3.9 | 1.1 | 3.5 | 4.7E-04 | 1.3E-03 | BT20/MB468 | MP |
| >hg38_21_+_8433172_8446622_RNA45SN1_45S_50ntFlanks@11320.11344.25\|CCACGTCTCGTCGCGCGCGCGTCCG | >hg38_21_+_8433172_8446622_RNA45SN1_45S_50ntFlanks | 11320 | 11344 | 25 | CCACGTCTCGTCGCGCGCGCGTCCG | 400.4 | -5.7 | 1.6 | -3.5 | 4.7E-04 | 1.3E-03 | BT20/MB468 | MP |
| >hg38_21_+_8433172_8446622_RNA45SN1_45S_50ntFlanks@6652.6698.47\|GACTCTTAGCGGTGGATCACTCGGCTCGTGCGTCGATGAAGAACGCA | >hg38_21_+_8433172_8446622_RNA45SN1_45S_50ntFlanks | 6652 | 6698 | 47 | GACTCTTAGCGGTGGATCACTCGGCTCGTGCGTCGATGAAGAACGCA | 29357.6 | -2.9 | 0.9 | -3.3 | 1.1E-03 | 1.3E-03 | MB231/MB468 | MITO |
| >hg38_21_+_8433172_8446622_RNA45SN1_45S_50ntFlanks@10786.10820.35\|AACAGCCTCTGGCATGTTGGAACAATGTAGGTAAG | >hg38_21_+_8433172_8446622_RNA45SN1_45S_50ntFlanks | 10786 | 10820 | 35 | AACAGCCTCTGGCATGTTGGAACAATGTAGGTAAG | 284.2 | -3.8 | 1.2 | -3.3 | 9.5E-04 | 1.3E-03 | BT20/MB468 | CYTO |
| >hg38_21_+_8433172_8446622_RNA45SN1_45S_50ntFlanks@8380.8399.20\|CAAGAGGGCGTGAAACCGTT | >hg38_21_+_8433172_8446622_RNA45SN1_45S_50ntFlanks | 8380 | 8399 | 20 | CAAGAGGGCGTGAAACCGTT | 385.5 | -5.0 | 1.5 | -3.4 | 8.0E-04 | 1.3E-03 | BT20/MB468 | MITO |
| >MI0000286\|hsa-mir-210&WithFlank&11\|-\|568083\|568204@72.94.23&[MIMAT0000267&hsa-miR-210-3p&offsets\|0\|+1;m-102&11\|-\|568112\|568133&offsets\|0\|+1]\|CTGTGCGTGTGACAGCGGCTGAT | >MI0000286\|hsa-mir-210&WithFlank&11\|-\|568083\|568204 | 72 | 94 | 23 | CTGTGCGTGTGACAGCGGCTGAT | 500.6 | 4.5 | 1.4 | 3.3 | 8.8E-04 | 1.3E-03 | MB231/MB468 | MP |
| >hg38_21_+_8433172_8446622_RNA45SN1_45S_50ntFlanks@10771.10791.21\|CAGGTCTCCAAGGTGAACAGC | >hg38_21_+_8433172_8446622_RNA45SN1_45S_50ntFlanks | 10771 | 10791 | 21 | CAGGTCTCCAAGGTGAACAGC | 216.8 | -5.2 | 1.6 | -3.3 | 8.8E-04 | 1.3E-03 | MB231/MB468 | MP |
| >MI0000815\|hsa-mir-339&WithFlank&7\|-\|1022927\|1023032@56.76.21(+1U)&[MIMAT0004702&hsa-miR-339-3p&offsets\|0\|-2(+1U);m-235&7\|-\|1022957\|1022977&offsets\|0\|0(+1U)]\|TGAGCGCCTCGACGACAGAGCT | >MI0000815\|hsa-mir-339&WithFlank&7\|-\|1022927\|1023032 | 56 | 76 | 21(+1U) | TGAGCGCCTCGACGACAGAGCT | 85.0 | 6.0 | 1.8 | 3.3 | 8.8E-04 | 1.3E-03 | MB231/MB468 | MP |
| >hg38_21_+_8433172_8446622_RNA45SN1_45S_50ntFlanks@3992.4024.33\|GGTGACTCTAGATAACCTCGGGCCGATCGCACG | >hg38_21_+_8433172_8446622_RNA45SN1_45S_50ntFlanks | 3992 | 4024 | 33 | GGTGACTCTAGATAACCTCGGGCCGATCGCACG | 389.6 | -4.2 | 1.3 | -3.3 | 8.8E-04 | 1.3E-03 | MB231/MB468 | MP |
| >hg38_1_-_228634819_228635039_RNA5S12_5S_50ntFlanks@149.169.21\|GAATACCGGGTGCTGTAGGCT | >hg38_1_-_228634819_228635039_RNA5S12_5S_50ntFlanks | 149 | 169 | 21 | GAATACCGGGTGCTGTAGGCT | 2301.1 | 4.3 | 1.2 | 3.8 | 1.6E-04 | 1.3E-03 | BT20/MB231 | MP |
| >hg38_21_+_8433172_8446622_RNA45SN1_45S_50ntFlanks@9605.9622.18\|AAGACTAATCGAACCATC | >hg38_21_+_8433172_8446622_RNA45SN1_45S_50ntFlanks | 9605 | 9622 | 18 | AAGACTAATCGAACCATC | 224.8 | -2.2 | 0.7 | -3.4 | 8.0E-04 | 1.3E-03 | BT20/MB468 | MITO |
| >hg38_21_+_8433172_8446622_RNA45SN1_45S_50ntFlanks@12973.13000.28\|TCGTACGTAGCAGAGCAGCTCCCTCGCT | >hg38_21_+_8433172_8446622_RNA45SN1_45S_50ntFlanks | 12973 | 13000 | 28 | TCGTACGTAGCAGAGCAGCTCCCTCGCT | 117.5 | -1.8 | 0.5 | -3.4 | 8.0E-04 | 1.3E-03 | BT20/MB468 | MITO |
| >MI0033672\|hsa-mir-10525&WithFlank&7\|+\|74595962\|74596032@45.66.22&[MIMAT0041995&hsa-miR-10525-3p&offsets\|0\|+1]\|TGACTATGATGTGCACCTGATT | >MI0033672\|hsa-mir-10525&WithFlank&7\|+\|74595962\|74596032 | 45 | 66 | 22 | TGACTATGATGTGCACCTGATT | 316.7 | -3.0 | 0.8 | -3.8 | 1.7E-04 | 1.3E-03 | BT20/MB231 | NUC |
| >hg38_21_+_8433172_8446622_RNA45SN1_45S_50ntFlanks@4687.4707.21\|ACGGACCAGAGCGAAAGCATT | >hg38_21_+_8433172_8446622_RNA45SN1_45S_50ntFlanks | 4687 | 4707 | 21 | ACGGACCAGAGCGAAAGCATT | 429.3 | -3.1 | 0.9 | -3.4 | 8.0E-04 | 1.3E-03 | BT20/MB468 | MITO |
| >hg38_21_+_8433172_8446622_RNA45SN1_45S_50ntFlanks@7989.8016.28\|AGACGTGGCGACCCGCTGAATTTAAGCA | >hg38_21_+_8433172_8446622_RNA45SN1_45S_50ntFlanks | 7989 | 8016 | 28 | AGACGTGGCGACCCGCTGAATTTAAGCA | 529.3 | 2.5 | 0.8 | 3.3 | 1.1E-03 | 1.3E-03 | MB231/MB468 | NUC |
| >hg38_21_+_8433172_8446622_RNA45SN1_45S_50ntFlanks@485.504.20\|GATGTGGTGACGTCGTGCTC | >hg38_21_+_8433172_8446622_RNA45SN1_45S_50ntFlanks | 485 | 504 | 20 | GATGTGGTGACGTCGTGCTC | 461.1 | 12.1 | 3.6 | 3.3 | 8.9E-04 | 1.3E-03 | MB231/MB468 | MP |
| >am_trna111_HisGTG_1_-_147774845_147774916@2.29.28,trna118_HisGTG_1_-_145396881_145396952@2.29.28,trna16_HisGTG_1_+_146544773_146544844@2.29.28,trna1_HisGTG_15_+_45493349_45493420@2.29.28,trna21_HisGTG_1_+_147753471_147753542@2.29.28,trna33_HisGTG_6_+_27125906_27125977@2.29.28,trna7_HisGTG_9_-_14433938_14434009@2.29.28,trna8_HisGTG_15_-_45492611_45492682@2.29.28,trna9_HisGTG_15_-_45490804_45490875@2.29.28\|CCGTGATCGTATAGTGGTTAGTACTCTG | >am_trna111_HisGTG_1_-_147774845_147774916 | 2 | 29 | 28 | CCGTGATCGTATAGTGGTTAGTACTCTG | 281.7 | -1.8 | 0.5 | -3.3 | 9.6E-04 | 1.3E-03 | BT20/MB468 | CYTO |
| >hg38_21_+_8433172_8446622_RNA45SN1_45S_50ntFlanks@7974.7996.23\|ACGCGACCTCAGATCAGACGTGG | >hg38_21_+_8433172_8446622_RNA45SN1_45S_50ntFlanks | 7974 | 7996 | 23 | ACGCGACCTCAGATCAGACGTGG | 1907.6 | 4.0 | 1.2 | 3.3 | 9.6E-04 | 1.3E-03 | BT20/MB468 | CYTO |
| >hg38_21_+_8433172_8446622_RNA45SN1_45S_50ntFlanks@485.521.37\|GATGTGGTGACGTCGTGCTCTCCCGGGCCGGGTCCGA | >hg38_21_+_8433172_8446622_RNA45SN1_45S_50ntFlanks | 485 | 521 | 37 | GATGTGGTGACGTCGTGCTCTCCCGGGCCGGGTCCGA | 407.4 | 11.9 | 3.6 | 3.3 | 1.1E-03 | 1.3E-03 | MB231/MB468 | MITO |
| >hg38_MT_+_1621_3279_RNR2_16S_50ntFlanks@1349.1389.41\|ACAATAGGGTTTACGACCTCGATGTTGGATCAGGACATCCC | >hg38_MT_+_1621_3279_RNR2_16S_50ntFlanks | 1349 | 1389 | 41 | ACAATAGGGTTTACGACCTCGATGTTGGATCAGGACATCCC | 144.1 | 5.8 | 1.8 | 3.3 | 1.1E-03 | 1.3E-03 | MB231/MB468 | MITO |
| >hg38_21_+_8433172_8446622_RNA45SN1_45S_50ntFlanks@11317.11336.20\|ACCCCACGTCTCGTCGCGCG | >hg38_21_+_8433172_8446622_RNA45SN1_45S_50ntFlanks | 11317 | 11336 | 20 | ACCCCACGTCTCGTCGCGCG | 2805.7 | -6.8 | 2.1 | -3.3 | 1.1E-03 | 1.3E-03 | BT20/MB468 | NUC |
| >hg38_21_+_8433172_8446622_RNA45SN1_45S_50ntFlanks@10738.10756.19\|AAATCTCGCGCCGGGCCGT | >hg38_21_+_8433172_8446622_RNA45SN1_45S_50ntFlanks | 10738 | 10756 | 19 | AAATCTCGCGCCGGGCCGT | 851.8 | -3.5 | 0.9 | -3.8 | 1.7E-04 | 1.3E-03 | BT20/MB231 | CYTO |
| >MI0000809\|hsa-mir-151a&WithFlank&8\|-\|140732558\|140732659@53.74.22(+2U)&[MIMAT0000757&hsa-miR-151a-3p&offsets\|0\|+1(+2U);m-30&8\|-\|140732587\|140732607&offsets\|0\|+1(+2U)]\|CTAGACTGAAGCTCCTTGAGGATT | >MI0000809\|hsa-mir-151a&WithFlank&8\|-\|140732558\|140732659 | 53 | 74 | 22(+2U) | CTAGACTGAAGCTCCTTGAGGATT | 577.3 | 5.1 | 1.5 | 3.5 | 4.7E-04 | 1.3E-03 | BT20/MB468 | MP |

<table>
<tr><td>>trna5_AspGTC_12_+_98897281_98897352@37.70.34|ACGCGGGAGACCGGGGTTCAATTCCCCGACGGGG</td><td>>trna5_AspGTC_12_+_98897281_98897352</td><td>37</td><td>70</td><td>34</td><td>ACGCGGGAGACCGGGGTTCAATTCCCCGACGGGG</td><td>164.0</td><td>-3.1</td><td>0.9</td><td>-3.3</td><td>8.9E-04</td><td>1.3E-03</td><td>MB231/MB468</td><td>MP</td></tr>
<tr><td>>trnaMT_GluTTC_MT_-_14674_14742@39.71.33,trnalookalike8_GluTTC_5_-_93905172_93905240@39.71.33|ATTGGTCGTGGTTGTAGTCCGTGCGAGAATACC</td><td>>trnaMT_GluTTC_MT_-_14674_14742</td><td>39</td><td>71</td><td>33</td><td>ATTGGTCGTGGTTGTAGTCCGTGCGAGAATACC</td><td>1495.9</td><td>3.0</td><td>0.9</td><td>3.3</td><td>8.9E-04</td><td>1.3E-03</td><td>MB231/MB468</td><td>MP</td></tr>
<tr><td>>hg38_1_-_228634819_228635039_RNA5S12_5S_50ntFlanks@147.170.24|GGGAATACCGGGTGCTGTAGGCTT</td><td>>hg38_1_-_228634819_228635039_RNA5S12_5S_50ntFlanks</td><td>147</td><td>170</td><td>24</td><td>GGGAATACCGGGTGCTGTAGGCTT</td><td>21431.8</td><td>3.7</td><td>1.1</td><td>3.5</td><td>4.7E-04</td><td>1.3E-03</td><td>BT20/MB468</td><td>MP</td></tr>
<tr><td>>hg38_21_+_8433172_8446622_RNA45SN1_45S_50ntFlanks@11233.11255.23|CCCTTCCCGTGGATCGCCCCAGC</td><td>>hg38_21_+_8433172_8446622_RNA45SN1_45S_50ntFlanks</td><td>11233</td><td>11255</td><td>23</td><td>CCCTTCCCGTGGATCGCCCCAGC</td><td>97.8</td><td>-4.6</td><td>1.4</td><td>-3.4</td><td>8.0E-04</td><td>1.3E-03</td><td>BT20/MB468</td><td>MITO</td></tr>
<tr><td>>MI0000299|hsa-mir-222&WithFlank&X|-|45747009|45747130@75.99.25(+3U)&[MIMAT0000279&hsa-miR-222-3p&offsets|0|+4(+3U);m-140&X|-|45747033|45747056&offsets|0|+1(+3U)]|AGCTACATCTGGCTACTGGGTCTCTTTT</td><td>>MI0000299|hsa-mir-222&WithFlank&X|-|45747009|45747130</td><td>75</td><td>99</td><td>25(+3U)</td><td>AGCTACATCTGGCTACTGGGTCTCTTTT</td><td>2291.3</td><td>-2.9</td><td>0.8</td><td>-3.8</td><td>1.7E-04</td><td>1.3E-03</td><td>BT20/MB231</td><td>CYTO</td></tr>
<tr><td>>trna10_AspGTC_12_-_125424193_125424264@41.69.29,trna12_AspGTC_12_-_125411891_125411962@41.69.29,trna144_AspGTC_6_-_27551236_27551307@41.69.29,trna38_AspGTC_17_-_8125556_8125627@41.69.29,trna45_AspGTC_6_+_27447453_27447524@41.69.29,trna48_AspGTC_6_+_27471523_27471594@41.69.29,trna4_AspGTC_12_+_96429799_96429870@41.69.29,trna69_AspGTC_1_-_161440205_161440276@41.69.29,trna72_AspGTC_1_-_161432824_161432895@41.69.29,trna75_AspGTC_1_-_161425414_161425485@41.69.29,trna78_AspGTC_1_-_161418033_161418104@41.69.29,trna81_AspGTC_1_-_161410615_161410686@41.69.29|GGGAGACCGGGGTTCGATTCCCCGACGGG</td><td>>trna10_AspGTC_12_-_125424193_125424264</td><td>41</td><td>69</td><td>29</td><td>GGGAGACCGGGGTTCGATTCCCCGACGGG</td><td>83.6</td><td>6.5</td><td>1.9</td><td>3.5</td><td>4.7E-04</td><td>1.3E-03</td><td>BT20/MB468</td><td>MP</td></tr>
<tr><td>>MI0000299|hsa-mir-222&WithFlank&X|-|45747009|45747130@75.94.20(+1A)&[MIMAT0000279&hsa-miR-222-3p&offsets|0|-1(+1A);m-140&X|-|45747033|45747056&offsets|0|-4(+1A)]|AGCTACATCTGGCTACTGGGA</td><td>>MI0000299|hsa-mir-222&WithFlank&X|-|45747009|45747130</td><td>75</td><td>94</td><td>20(+1A)</td><td>AGCTACATCTGGCTACTGGGA</td><td>423.3</td><td>4.3</td><td>1.3</td><td>3.3</td><td>8.9E-04</td><td>1.3E-03</td><td>MB231/MB468</td><td>MP</td></tr>
<tr><td>>hg38_21_+_8433172_8446622_RNA45SN1_45S_50ntFlanks@12413.12432.20|TCCTTCGATGTCGGCTCTTC</td><td>>hg38_21_+_8433172_8446622_RNA45SN1_45S_50ntFlanks</td><td>12413</td><td>12432</td><td>20</td><td>TCCTTCGATGTCGGCTCTTC</td><td>866.9</td><td>-2.1</td><td>0.7</td><td>-3.3</td><td>1.0E-03</td><td>1.3E-03</td><td>MB231/MB468</td><td>CYTO</td></tr>
<tr><td>>hg38_21_+_8433172_8446622_RNA45SN1_45S_50ntFlanks@11595.11612.18|AGGGGAATCCGACTGTTT</td><td>>hg38_21_+_8433172_8446622_RNA45SN1_45S_50ntFlanks</td><td>11595</td><td>11612</td><td>18</td><td>AGGGGAATCCGACTGTTT</td><td>897.7</td><td>3.3</td><td>1.0</td><td>3.3</td><td>1.0E-03</td><td>1.3E-03</td><td>MB231/MB468</td><td>CYTO</td></tr>
<tr><td>>trnaMT_ThrTGT_MT_+_15888_15953@31.52.22|TTGTAAACCGGAGATGAAAACC</td><td>>trnaMT_ThrTGT_MT_+_15888_15953</td><td>31</td><td>52</td><td>22</td><td>TTGTAAACCGGAGATGAAAACC</td><td>153.0</td><td>6.5</td><td>1.9</td><td>3.5</td><td>4.7E-04</td><td>1.3E-03</td><td>BT20/MB468</td><td>MP</td></tr>
<tr><td>>trna143_LysTTT_6_-_27559593_27559665@14.31.18,trna149_LysTTT_6_-_27302769_27302841@14.31.18,trna14_LysTTT_11_-_59327808_59327880@14.31.18,trna2_LysTTT_17_+_8022473_8022545@14.31.18,trna54_LysTTT_1_+_204475655_204475727@14.31.18,trna5_LysTTT_11_+_59323902_59323974@14.31.18,trna62_LysTTT_1_-_204476158_204476230@14.31.18,trna76_LysTTT_6_+_28918806_28918878@14.31.18|AGTCGGTAGAGCATCAGA</td><td>>trna143_LysTTT_6_-_27559593_27559665</td><td>14</td><td>31</td><td>18</td><td>AGTCGGTAGAGCATCAGA</td><td>197.6</td><td>-4.0</td><td>1.2</td><td>-3.3</td><td>1.1E-03</td><td>1.3E-03</td><td>MB231/MB468</td><td>NUC</td></tr>
<tr><td>>hg38_21_+_8433172_8446622_RNA45SN1_45S_50ntFlanks@12597.12614.18|ATTTGGTGTATGTGCTTG</td><td>>hg38_21_+_8433172_8446622_RNA45SN1_45S_50ntFlanks</td><td>12597</td><td>12614</td><td>18</td><td>ATTTGGTGTATGTGCTTG</td><td>281.6</td><td>-2.6</td><td>0.8</td><td>-3.3</td><td>1.0E-03</td><td>1.3E-03</td><td>MB231/MB468</td><td>CYTO</td></tr>
<tr><td>>hg38_21_+_8433172_8446622_RNA45SN1_45S_50ntFlanks@12414.12434.21|CCTTCGATGTCGGCTCTTCCT</td><td>>hg38_21_+_8433172_8446622_RNA45SN1_45S_50ntFlanks</td><td>12414</td><td>12434</td><td>21</td><td>CCTTCGATGTCGGCTCTTCCT</td><td>450.0</td><td>3.5</td><td>1.1</td><td>3.3</td><td>9.6E-04</td><td>1.3E-03</td><td>BT20/MB468</td><td>CYTO</td></tr>
<tr><td>>hg38_21_+_8433172_8446622_RNA45SN1_45S_50ntFlanks@476.523.48|TTCGTGATCGATGTGGTGACGTCGTGCTCTCCCGGGCCGGGTCCGAGC</td><td>>hg38_21_+_8433172_8446622_RNA45SN1_45S_50ntFlanks</td><td>476</td><td>523</td><td>48</td><td>TTCGTGATCGATGTGGTGACGTCGTGCTCTCCCGGGCCGGGTCCGAGC</td><td>344.8</td><td>5.2</td><td>1.6</td><td>3.3</td><td>1.1E-03</td><td>1.3E-03</td><td>MB231/MB468</td><td>MITO</td></tr>
<tr><td>>hg38_21_+_8433172_8446622_RNA45SN1_45S_50ntFlanks@12447.12473.27|CAGAATTCACCAAGCGTTGGATTGTTC</td><td>>hg38_21_+_8433172_8446622_RNA45SN1_45S_50ntFlanks</td><td>12447</td><td>12473</td><td>27</td><td>CAGAATTCACCAAGCGTTGGATTGTTC</td><td>446.1</td><td>-4.7</td><td>1.4</td><td>-3.5</td><td>4.7E-04</td><td>1.3E-03</td><td>BT20/MB468</td><td>MP</td></tr>
<tr><td>>hg38_21_+_8433172_8446622_RNA45SN1_45S_50ntFlanks@6661.6685.25|CGGTGGATCACTCGGCTCGTGCGTC</td><td>>hg38_21_+_8433172_8446622_RNA45SN1_45S_50ntFlanks</td><td>6661</td><td>6685</td><td>25</td><td>CGGTGGATCACTCGGCTCGTGCGTC</td><td>1301.8</td><td>2.3</td><td>0.6</td><td>3.8</td><td>1.7E-04</td><td>1.3E-03</td><td>BT20/MB231</td><td>NUC</td></tr>
<tr><td>>trna119_LysCTT_1_-_145395522_145395594@2.33.32,trna11_LysCTT_5_-_180648979_180649051@2.33.32,trna13_LysCTT_6_+_26556774_26556846@2.33.32,trna7_LysCTT_16_+_3225692_3225764@2.33.32,trna9_LysCTT_5_+_180634755_180634827@2.33.32|CCCGGCTAGCTCAGTCGGTAGAGCATGAGACT</td><td>>trna119_LysCTT_1_-_145395522_145395594</td><td>2</td><td>33</td><td>32</td><td>CCCGGCTAGCTCAGTCGGTAGAGCATGAGACT</td><td>548.5</td><td>-5.3</td><td>1.4</td><td>-3.8</td><td>1.7E-04</td><td>1.3E-03</td><td>BT20/MB231</td><td>NUC</td></tr>
<tr><td>>hg38_21_+_8433172_8446622_RNA45SN1_45S_50ntFlanks@12995.13017.23|CTCGCTGCGATCTATTGAAAGTC</td><td>>hg38_21_+_8433172_8446622_RNA45SN1_45S_50ntFlanks</td><td>12995</td><td>13017</td><td>23</td><td>CTCGCTGCGATCTATTGAAAGTC</td><td>523.6</td><td>-2.5</td><td>0.7</td><td>-3.8</td><td>1.7E-04</td><td>1.3E-03</td><td>BT20/MB231</td><td>NUC</td></tr>
<tr><td>>am_trna2_GlyGCC_21_-_18827107_18827177@1.22.22,trna35_GlyGCC_1_+_161413094_161413164@1.22.22,trna37_GlyGCC_1_+_161420467_161420537@1.22.22,trna39_GlyGCC_1_+_161427898_161427968@1.22.22,trna41_GlyGCC_1_+_161435258_161435328@1.22.22|GCATGGGTGGTTCAGTGGTAGA</td><td>>am_trna2_GlyGCC_21_-_18827107_18827177</td><td>1</td><td>22</td><td>22</td><td>GCATGGGTGGTTCAGTGGTAGA</td><td>209.4</td><td>3.6</td><td>1.0</td><td>3.5</td><td>4.7E-04</td><td>1.3E-03</td><td>BT20/MB468</td><td>MP</td></tr>
<tr><td>>MI0000746|hsa-mir-99b&WithFlank&19|+|51692606|51692687@51.69.19&[MIMAT0004678&hsa-miR-99b-3p&offsets|0|-3;m-151&19|+|51692656|51692678&offsets|0|-4]|CAAGCTCGTGTCTGTGGGT</td><td>>MI0000746|hsa-mir-99b&WithFlank&19|+|51692606|51692687</td><td>51</td><td>69</td><td>19</td><td>CAAGCTCGTGTCTGTGGGT</td><td>338.1</td><td>5.3</td><td>1.5</td><td>3.5</td><td>4.7E-04</td><td>1.3E-03</td><td>BT20/MB468</td><td>MP</td></tr>
<tr><td>>hg38_21_+_8433172_8446622_RNA45SN1_45S_50ntFlanks@12397.12417.21|CGACGTCGCTTTTTGATCCTT</td><td>>hg38_21_+_8433172_8446622_RNA45SN1_45S_50ntFlanks</td><td>12397</td><td>12417</td><td>21</td><td>CGACGTCGCTTTTTGATCCTT</td><td>1322.5</td><td>-2.7</td><td>0.8</td><td>-3.4</td><td>8.0E-04</td><td>1.3E-03</td><td>BT20/MB468</td><td>MITO</td></tr>
<tr><td>>hg38_21_+_8433172_8446622_RNA45SN1_45S_50ntFlanks@7983.8011.29|CAGATCAGACGTGGCGACCCGCTGAATTT</td><td>>hg38_21_+_8433172_8446622_RNA45SN1_45S_50ntFlanks</td><td>7983</td><td>8011</td><td>29</td><td>CAGATCAGACGTGGCGACCCGCTGAATTT</td><td>236.4</td><td>-5.3</td><td>1.6</td><td>-3.3</td><td>8.9E-04</td><td>1.3E-03</td><td>MB231/MB468</td><td>MP</td></tr>
<tr><td>>trnaMT_PheGAA_MT_+_577_647@36.62.27|AAAATGTTTAGACGGGCTCACATCACC</td><td>>trnaMT_PheGAA_MT_+_577_647</td><td>36</td><td>62</td><td>27</td><td>AAAATGTTTAGACGGGCTCACATCACC</td><td>596.6</td><td>6.6</td><td>1.9</td><td>3.5</td><td>4.7E-04</td><td>1.3E-03</td><td>BT20/MB468</td><td>MP</td></tr>
<tr><td>>MI0001448|hsa-mir-425&WithFlank&3|-|49020142|49020240@64.81.18(+1U)&[MIMAT0001343&hsa-miR-425-3p&offsets|+3|-1(+1U);m-258&3|-|49020160|49020181&offsets|+4|0(+1U)]|GGGAATGTCGTGTCCGCCT</td><td>>MI0001448|hsa-mir-425&WithFlank&3|-|49020142|49020240</td><td>64</td><td>81</td><td>18(+1U)</td><td>GGGAATGTCGTGTCCGCCT</td><td>52.4</td><td>8.9</td><td>2.7</td><td>3.3</td><td>8.9E-04</td><td>1.3E-03</td><td>MB231/MB468</td><td>MP</td></tr>
<tr><td>>hg38_21_+_8433172_8446622_RNA45SN1_45S_50ntFlanks@10872.10890.19|GCTGGGTCGGTCGGGCTGG</td><td>>hg38_21_+_8433172_8446622_RNA45SN1_45S_50ntFlanks</td><td>10872</td><td>10890</td><td>19</td><td>GCTGGGTCGGTCGGGCTGG</td><td>796.1</td><td>6.9</td><td>1.8</td><td>3.8</td><td>1.6E-04</td><td>1.3E-03</td><td>BT20/MB231</td><td>MP</td></tr>
<tr><td>>hg38_21_+_8433172_8446622_RNA45SN1_45S_50ntFlanks@11714.11741.28|GGTAAACGGCGGGAGTAACTATGACTCT</td><td>>hg38_21_+_8433172_8446622_RNA45SN1_45S_50ntFlanks</td><td>11714</td><td>11741</td><td>28</td><td>GGTAAACGGCGGGAGTAACTATGACTCT</td><td>262.6</td><td>4.2</td><td>1.3</td><td>3.3</td><td>1.0E-03</td><td>1.3E-03</td><td>MB231/MB468</td><td>CYTO</td></tr>
<tr><td>>hg38_21_+_8433172_8446622_RNA45SN1_45S_50ntFlanks@12393.12414.22|ATAGCGACGTCGCTTTTTGATC</td><td>>hg38_21_+_8433172_8446622_RNA45SN1_45S_50ntFlanks</td><td>12393</td><td>12414</td><td>22</td><td>ATAGCGACGTCGCTTTTTGATC</td><td>1001.6</td><td>-3.4</td><td>1.1</td><td>-3.3</td><td>1.1E-03</td><td>1.3E-03</td><td>MB231/MB468</td><td>MITO</td></tr>
<tr><td>>MI0000086|hsa-mir-28&WithFlank&3|+|188688775|188688872@60.82.23&[MIMAT0004502&hsa-miR-28-3p&offsets|0|+1;m-28&3|+|188688834|188688855&offsets|0|+1]|CACTAGATTGTGAGCTCCTGGAG</td><td>>MI0000086|hsa-mir-28&WithFlank&3|+|188688775|188688872</td><td>60</td><td>82</td><td>23</td><td>CACTAGATTGTGAGCTCCTGGAG</td><td>175.7</td><td>5.1</td><td>1.5</td><td>3.3</td><td>8.9E-04</td><td>1.3E-03</td><td>MB231/MB468</td><td>MP</td></tr>
<tr><td>>hg38_21_+_8433172_8446622_RNA45SN1_45S_50ntFlanks@11561.11585.25|GCGCCTAGCAGCCGACTTAGAACTG</td><td>>hg38_21_+_8433172_8446622_RNA45SN1_45S_50ntFlanks</td><td>11561</td><td>11585</td><td>25</td><td>GCGCCTAGCAGCCGACTTAGAACTG</td><td>1103.2</td><td>-5.4</td><td>1.6</td><td>-3.3</td><td>9.7E-04</td><td>1.3E-03</td><td>BT20/MB468</td><td>CYTO</td></tr>
<tr><td>>hg38_21_+_8433172_8446622_RNA45SN1_45S_50ntFlanks@4008.4026.19|CTCGGGCCGATCGCACGCC</td><td>>hg38_21_+_8433172_8446622_RNA45SN1_45S_50ntFlanks</td><td>4008</td><td>4026</td><td>19</td><td>CTCGGGCCGATCGCACGCC</td><td>122.6</td><td>-3.2</td><td>1.0</td><td>-3.3</td><td>9.7E-04</td><td>1.3E-03</td><td>BT20/MB468</td><td>CYTO</td></tr>
<tr><td>>MI0000298|hsa-mir-221&WithFlank&X|-|45746151|45746272@71.93.23(+1C)&[MIMAT0000278&hsa-miR-221-3p&offsets|0|0(+1C);m-91&X|-|45746181|45746202&offsets|0|+1(+1C)]|AGCTACATTGTCTGCTGGGTTTCC</td><td>>MI0000298|hsa-mir-221&WithFlank&X|-|45746151|45746272</td><td>71</td><td>93</td><td>23(+1C)</td><td>AGCTACATTGTCTGCTGGGTTTCC</td><td>1152.5</td><td>-1.9</td><td>0.5</td><td>-3.8</td><td>1.8E-04</td><td>1.3E-03</td><td>BT20/MB231</td><td>NUC</td></tr>
<tr><td>>trnaMT_ValTAC_MT_+_1602_1670@49.72.24|AACTTAACTTGACCGCTCTGACCA</td><td>>trnaMT_ValTAC_MT_+_1602_1670</td><td>49</td><td>72</td><td>24</td><td>AACTTAACTTGACCGCTCTGACCA</td><td>8596.3</td><td>3.9</td><td>1.1</td><td>3.4</td><td>8.1E-04</td><td>1.3E-03</td><td>BT20/MB468</td><td>MITO</td></tr>
<tr><td>>hg38_21_+_8433172_8446622_RNA45SN1_45S_50ntFlanks@11875.11893.19|GAAGACCCTGTTGAGCTTG</td><td>>hg38_21_+_8433172_8446622_RNA45SN1_45S_50ntFlanks</td><td>11875</td><td>11893</td><td>19</td><td>GAAGACCCTGTTGAGCTTG</td><td>1537.5</td><td>-5.3</td><td>1.6</td><td>-3.3</td><td>1.1E-03</td><td>1.3E-03</td><td>BT20/MB468</td><td>NUC</td></tr>
<tr><td>>trna10_LysCTT_16_+_3241501_3241573@1.29.29,trna13_LysCTT_14_-_58706613_58706685@1.29.29,trna2_LysCTT_15_+_79152904_79152976@1.29.29|GCCCGGCTAGCTCAGTCGGTAGAGCATGG</td><td>>trna10_LysCTT_16_+_3241501_3241573</td><td>1</td><td>29</td><td>29</td><td>GCCCGGCTAGCTCAGTCGGTAGAGCATGG</td><td>260.2</td><td>-5.7</td><td>1.6</td><td>-3.5</td><td>4.8E-04</td><td>1.3E-03</td><td>BT20/MB468</td><td>MP</td></tr>
<tr><td>>hg38_21_+_8433172_8446622_RNA45SN1_45S_50ntFlanks@12008.12041.34|CCGCCGGTGAAATACCACTACTCTGATCGTTTTT</td><td>>hg38_21_+_8433172_8446622_RNA45SN1_45S_50ntFlanks</td><td>12008</td><td>12041</td><td>34</td><td>CCGCCGGTGAAATACCACTACTCTGATCGTTTTT</td><td>125.7</td><td>-5.9</td><td>1.7</td><td>-3.5</td><td>4.8E-04</td><td>1.3E-03</td><td>BT20/MB468</td><td>MP</td></tr>
<tr><td>>hg38_21_+_8433172_8446622_RNA45SN1_45S_50ntFlanks@4454.4473.20|CGGCGCCCCCTCGATGCTCT</td><td>>hg38_21_+_8433172_8446622_RNA45SN1_45S_50ntFlanks</td><td>4454</td><td>4473</td><td>20</td><td>CGGCGCCCCCTCGATGCTCT</td><td>216.9</td><td>-4.2</td><td>1.1</td><td>-3.8</td><td>1.8E-04</td><td>1.3E-03</td><td>BT20/MB231</td><td>CYTO</td></tr>
<tr><td>>MI0000077|hsa-mir-21&WithFlank&17|+|59841260|59841343@14.36.23(+1A)&[MIMAT0000076&hsa-miR-21-5p&offsets|0|+1(+1A);m-1&17|+|59841273|59841294&offsets|0|+1(+1A)]|TAGCTTATCAGACTGATGTTGACA</td><td>>MI0000077|hsa-mir-21&WithFlank&17|+|59841260|59841343</td><td>14</td><td>36</td><td>23(+1A)</td><td>TAGCTTATCAGACTGATGTTGACA</td><td>21219.9</td><td>2.8</td><td>0.8</td><td>3.3</td><td>9.0E-04</td><td>1.3E-03</td><td>MB231/MB468</td><td>MP</td></tr>
<tr><td>>MI0000088|hsa-mir-30a&WithFlank&6|-|71403545|71403627@12.36.25&[MIMAT0000087&hsa-miR-30a-5p&offsets|0|+3;m-15&6|-|71403595|71403616&offsets|0|+3]|TGTAAACATCCTCGACTGGAAGCTG</td><td>>MI0000088|hsa-mir-30a&WithFlank&6|-|71403545|71403627</td><td>12</td><td>36</td><td>25</td><td>TGTAAACATCCTCGACTGGAAGCTG</td><td>313.8</td><td>4.3</td><td>1.3</td><td>3.3</td><td>9.0E-04</td><td>1.3E-03</td><td>MB231/MB468</td><td>MP</td></tr>
<tr><td>>am_trnaMT_ValTAC_MT_+_1602_1670@48.65.18|CAACTTAACTTGACCGCT</td><td>>am_trnaMT_ValTAC_MT_+_1602_1670</td><td>48</td><td>65</td><td>18</td><td>CAACTTAACTTGACCGCT</td><td>145.8</td><td>5.3</td><td>1.6</td><td>3.3</td><td>1.1E-03</td><td>1.3E-03</td><td>MB231/MB468</td><td>MITO</td></tr>
<tr><td>>trnaMT_HisGTG_MT_+_12138_12206@25.58.34|CAGATTGTGAATCTGACAACAGAGGCTTACGACC</td><td>>trnaMT_HisGTG_MT_+_12138_12206</td><td>25</td><td>58</td><td>34</td><td>CAGATTGTGAATCTGACAACAGAGGCTTACGACC</td><td>55.7</td><td>9.5</td><td>2.7</td><td>3.5</td><td>4.8E-04</td><td>1.3E-03</td><td>BT20/MB468</td><td>MP</td></tr>
<tr><td>>hg38_21_+_8433172_8446622_RNA45SN1_45S_50ntFlanks@4975.5001.27|CCCGGACACGGACAGGATTGACAGATT</td><td>>hg38_21_+_8433172_8446622_RNA45SN1_45S_50ntFlanks</td><td>4975</td><td>5001</td><td>27</td><td>CCCGGACACGGACAGGATTGACAGATT</td><td>459.8</td><td>3.5</td><td>0.9</td><td>3.7</td><td>1.8E-04</td><td>1.3E-03</td><td>BT20/MB231</td><td>NUC</td></tr>
<tr><td>>hg38_21_+_8433172_8446622_RNA45SN1_45S_50ntFlanks@7976.8005.30|GCGACCTCAGATCAGACGTGGCGACCCGCT</td><td>>hg38_21_+_8433172_8446622_RNA45SN1_45S_50ntFlanks</td><td>7976</td><td>8005</td><td>30</td><td>GCGACCTCAGATCAGACGTGGCGACCCGCT</td><td>524.4</td><td>3.0</td><td>0.8</td><td>3.7</td><td>1.8E-04</td><td>1.3E-03</td><td>BT20/MB231</td><td>NUC</td></tr>
<tr><td>>MI0000446|hsa-mir-125b-1&WithFlank&11|-|122099751|122099850@21.42.22(+1A);MI0000470|hsa-mir-125b-2&WithFlank&21|+|16590231|16590331@23.44.22(+1A)&[MIMAT0000423&hsa-miR-125b-5p&offsets|0|0(+1A);m-67&11|-|122099809|122099830&offsets|0|0(+1A)];[MIMAT0000423_1&hsa-miR-125b-5p&offsets|0|0(+1A);m-68&21|+|16590253|16590274&offsets|0|0(+1A)]|TCCCTGAGACCCTAACTTGTGAA</td><td>>MI0000446|hsa-mir-125b-1&WithFlank&11|-|122099751|122099850</td><td>21</td><td>42</td><td>22(+1A)</td><td>TCCCTGAGACCCTAACTTGTGAA</td><td>3256.7</td><td>-5.0</td><td>1.3</td><td>-3.7</td><td>1.8E-04</td><td>1.3E-03</td><td>BT20/MB231</td><td>NUC</td></tr>
<tr><td>>hg38_21_+_8433172_8446622_RNA45SN1_45S_50ntFlanks@3851.3874.24|ATAACTGTGGTAATTCTAGAGCTA</td><td>>hg38_21_+_8433172_8446622_RNA45SN1_45S_50ntFlanks</td><td>3851</td><td>3874</td><td>24</td><td>ATAACTGTGGTAATTCTAGAGCTA</td><td>209.4</td><td>-5.4</td><td>1.6</td><td>-3.5</td><td>4.8E-04</td><td>1.3E-03</td><td>BT20/MB468</td><td>MP</td></tr>
</table>

| | | | | | | | | | | | | | |
|---|---|---|---|---|---|---|---|---|---|---|---|---|---|
| >hg38_21_+_8433172_8446622_RNA45SN1_45S_50ntFlanks@5527.5548.22\|AAAGTCGTAACAAGGTTTCCGT | >hg38_21_+_8433172_8446622_RNA45SN1_45S_50ntFlanks | 5527 | 5548 | 22 | AAAGTCGTAACAAGGTTTCCGT | 270.8 | -5.5 | 1.6 | -3.5 | 4.8E-04 | 1.3E-03 | BT20/MB468 | MP |
| >hg38_21_+_8433172_8446622_RNA45SN1_45S_50ntFlanks@12718.12742.25\|CTCGGTTGGCCTCGGATAGCCGGTC | >hg38_21_+_8433172_8446622_RNA45SN1_45S_50ntFlanks | 12718 | 12742 | 25 | CTCGGTTGGCCTCGGATAGCCGGTC | 224.9 | -4.8 | 1.4 | -3.5 | 4.8E-04 | 1.3E-03 | BT20/MB468 | MP |
| >hg38_21_+_8433172_8446622_RNA45SN1_45S_50ntFlanks@10783.10816.34\|GTGAACAGCCTCTGGCATGTTGGAACAATGTAGG | >hg38_21_+_8433172_8446622_RNA45SN1_45S_50ntFlanks | 10783 | 10816 | 34 | GTGAACAGCCTCTGGCATGTTGGAACAATGTAGG | 202.3 | -5.0 | 1.5 | -3.3 | 1.1E-03 | 1.3E-03 | MB231/MB468 | NUC |
| >hg38_21_+_8433172_8446622_RNA45SN1_45S_50ntFlanks@4555.4587.33\|CGCAGCTAGGAATAATGGAATAGGACCGCGGTT | >hg38_21_+_8433172_8446622_RNA45SN1_45S_50ntFlanks | 4555 | 4587 | 33 | CGCAGCTAGGAATAATGGAATAGGACCGCGGTT | 127.4 | -4.5 | 1.3 | -3.3 | 8.1E-04 | 1.3E-03 | BT20/MB468 | MITO |
| >hg38_21_+_8433172_8446622_RNA45SN1_45S_50ntFlanks@9029.9064.36\|TCTCCTCGTGGGGGGGCCGGGCCACCCCTCCCACGG | >hg38_21_+_8433172_8446622_RNA45SN1_45S_50ntFlanks | 9029 | 9064 | 36 | TCTCCTCGTGGGGGGGCCGGGCCACCCCTCCCACGG | 157.7 | -4.9 | 1.5 | -3.3 | 8.1E-04 | 1.3E-03 | BT20/MB468 | MITO |
| >hg38_21_+_8433172_8446622_RNA45SN1_45S_50ntFlanks@8191.8213.23\|CAGCCCGTGGACGGTGTGAGGCC | >hg38_21_+_8433172_8446622_RNA45SN1_45S_50ntFlanks | 8191 | 8213 | 23 | CAGCCCGTGGACGGTGTGAGGCC | 468.8 | -1.9 | 0.6 | -3.3 | 1.0E-03 | 1.3E-03 | MB231/MB468 | CYTO |
| >hg38_MT_+_1621_3279_RNR2_16S_50ntFlanks@1370.1398.29\|ATGTTGGATCAGGACATCCCGATGGTGCA | >hg38_MT_+_1621_3279_RNR2_16S_50ntFlanks | 1370 | 1398 | 29 | ATGTTGGATCAGGACATCCCGATGGTGCA | 164.5 | 5.1 | 1.4 | 3.5 | 4.8E-04 | 1.3E-03 | BT20/MB468 | MP |
| >hg38_21_+_8433172_8446622_RNA45SN1_45S_50ntFlanks@4590.4616.27\|ATTTTGTTGGTTTTCGGAACTGAGGCC | >hg38_21_+_8433172_8446622_RNA45SN1_45S_50ntFlanks | 4590 | 4616 | 27 | ATTTTGTTGGTTTTCGGAACTGAGGCC | 430.5 | -4.0 | 1.2 | -3.3 | 1.1E-03 | 1.3E-03 | MB231/MB468 | NUC |
| >hg38_21_+_8433172_8446622_RNA45SN1_45S_50ntFlanks@11561.11594.34\|GCGCCTAGCAGCCGACTTAGAACTGGTGCGGACC | >hg38_21_+_8433172_8446622_RNA45SN1_45S_50ntFlanks | 11561 | 11594 | 34 | GCGCCTAGCAGCCGACTTAGAACTGGTGCGGACC | 395.3 | -1.6 | 0.5 | -3.3 | 8.1E-04 | 1.3E-03 | BT20/MB468 | MITO |
| >hg38_21_+_8433172_8446622_RNA45SN1_45S_50ntFlanks@11582.11605.24\|ACTGGTGCGGACCAGGGGAATCCG | >hg38_21_+_8433172_8446622_RNA45SN1_45S_50ntFlanks | 11582 | 11605 | 24 | ACTGGTGCGGACCAGGGGAATCCG | 398.2 | -3.0 | 0.9 | -3.3 | 1.1E-03 | 1.3E-03 | MB231/MB468 | MITO |
| >MI0006444\|hsa-mir-1307&WithFlank&10\|-\|103394247\|103394407@87.107.21(+1U)&[MIMAT0005951&hsa-miR-1307-3p&offsets\|+1\|0(+1U);m-66&10\|-\|103394300\|103394322&offsets\|+1\|-1(+1U)]\|CTCGGCGTGGCGTCGGTCGTGT | >MI0006444\|hsa-mir-1307&WithFlank&10\|-\|103394247\|103394407 | 87 | 107 | 21(+1U) | CTCGGCGTGGCGTCGGTCGTGT | 234.1 | 3.5 | 1.0 | 3.5 | 4.8E-04 | 1.3E-03 | BT20/MB468 | MP |
| >hg38_21_+_8433172_8446622_RNA45SN1_45S_50ntFlanks@9862.9899.38\|ATGAACCGAACGCCGGGTTAAGGCGCCCGATGCCGACG | >hg38_21_+_8433172_8446622_RNA45SN1_45S_50ntFlanks | 9862 | 9899 | 38 | ATGAACCGAACGCCGGGTTAAGGCGCCCGATGCCGACG | 353.3 | -5.8 | 1.7 | -3.3 | 8.1E-04 | 1.3E-03 | BT20/MB468 | MITO |
| >hg38_21_+_8433172_8446622_RNA45SN1_45S_50ntFlanks@6646.6667.22\|TCGTACGACTCTTAGCGGTGGA | >hg38_21_+_8433172_8446622_RNA45SN1_45S_50ntFlanks | 6646 | 6667 | 22 | TCGTACGACTCTTAGCGGTGGA | 8919.7 | 2.4 | 0.7 | 3.3 | 1.1E-03 | 1.3E-03 | MB231/MB468 | MITO |
| >hg38_21_+_8433172_8446622_RNA45SN1_45S_50ntFlanks@10794.10822.29\|CTGGCATGTTGGAACAATGTAGGTAAGGG | >hg38_21_+_8433172_8446622_RNA45SN1_45S_50ntFlanks | 10794 | 10822 | 29 | CTGGCATGTTGGAACAATGTAGGTAAGGG | 296.6 | -3.8 | 1.2 | -3.3 | 9.8E-04 | 1.3E-03 | BT20/MB468 | CYTO |
| >hg38_21_+_8433172_8446622_RNA45SN1_45S_50ntFlanks@7991.8012.22\|ACGTGGCGACCCGCTGAATTTA | >hg38_21_+_8433172_8446622_RNA45SN1_45S_50ntFlanks | 7991 | 8012 | 22 | ACGTGGCGACCCGCTGAATTTA | 479.1 | -2.0 | 0.6 | -3.3 | 9.8E-04 | 1.3E-03 | BT20/MB468 | CYTO |
| >MI0000088\|hsa-mir-30a&WithFlank&6\|-\|71403545\|71403627@53.75.23&[MIMAT0000088&hsa-miR-30a-3p&offsets\|0\|+1;m-38&6\|-\|71403554\|71403575&offsets\|0\|+1]\|CTTTCAGTCGGATGTTTGCAGCT | >MI0000088\|hsa-mir-30a&WithFlank&6\|-\|71403545\|71403627 | 53 | 75 | 23 | CTTTCAGTCGGATGTTTGCAGCT | 184.8 | 5.1 | 1.5 | 3.3 | 9.0E-04 | 1.3E-03 | MB231/MB468 | MP |
| >hg38_21_+_8433172_8446622_RNA45SN1_45S_50ntFlanks@9569.9609.41\|GTGCAAATCGGTCGTCCGACCTGGGTATAGGGGCGAAAGAC | >hg38_21_+_8433172_8446622_RNA45SN1_45S_50ntFlanks | 9569 | 9609 | 41 | GTGCAAATCGGTCGTCCGACCTGGGTATAGGGGCGAAAGAC | 206.4 | 5.2 | 1.6 | 3.3 | 1.1E-03 | 1.3E-03 | MB231/MB468 | MITO |
| >MI0000077\|hsa-mir-21&WithFlank&17\|+\|59841260\|59841343@14.36.23&[MIMAT0000076&hsa-miR-21-5p&offsets\|0\|+1;m-1&17\|+\|59841273\|59841294&offsets\|0\|+1]\|TAGCTTATCAGACTGATGTTGAC | >MI0000077\|hsa-mir-21&WithFlank&17\|+\|59841260\|59841343 | 14 | 36 | 23 | TAGCTTATCAGACTGATGTTGAC | 1122078.4 | -2.3 | 0.6 | -3.7 | 2.0E-04 | 1.3E-03 | BT20/MB231 | MITO |
| >hg38_21_+_8433172_8446622_RNA45SN1_45S_50ntFlanks@4045.4063.19\|CCATTCGAACGTCTGCCCT | >hg38_21_+_8433172_8446622_RNA45SN1_45S_50ntFlanks | 4045 | 4063 | 19 | CCATTCGAACGTCTGCCCT | 550.8 | -5.7 | 1.6 | -3.5 | 4.8E-04 | 1.3E-03 | BT20/MB468 | MP |
| >hg38_21_+_8433172_8446622_RNA45SN1_45S_50ntFlanks@7975.7996.22\|CGCGACCTCAGATCAGACGTGG | >hg38_21_+_8433172_8446622_RNA45SN1_45S_50ntFlanks | 7975 | 7996 | 22 | CGCGACCTCAGATCAGACGTGG | 23088.5 | 4.1 | 1.2 | 3.3 | 9.8E-04 | 1.3E-03 | BT20/MB468 | CYTO |
| >MI0000286\|hsa-mir-210&WithFlank&11\|-\|568083\|568204@72.92.21(+1U)&[MIMAT0000267&hsa-miR-210-3p&offsets\|0\|-1(+1U);m-102&11\|-\|568112\|568133&offsets\|0\|-1(+1U)]\|CTGTGCGTGTGACAGCGGCTGT | >MI0000286\|hsa-mir-210&WithFlank&11\|-\|568083\|568204 | 72 | 92 | 21(+1U) | CTGTGCGTGTGACAGCGGCTGT | 206.8 | 4.2 | 1.1 | 3.7 | 1.8E-04 | 1.3E-03 | BT20/MB231 | CYTO |
| >hg38_21_+_8433172_8446622_RNA45SN1_45S_50ntFlanks@11667.11687.21\|CTGCCCAGTGCTCTGAATGTC | >hg38_21_+_8433172_8446622_RNA45SN1_45S_50ntFlanks | 11667 | 11687 | 21 | CTGCCCAGTGCTCTGAATGTC | 187.8 | 4.8 | 1.4 | 3.5 | 4.8E-04 | 1.3E-03 | BT20/MB468 | MP |
| >hg38_21_+_8433172_8446622_RNA45SN1_45S_50ntFlanks@9507.9524.18\|AACTATGCCTGGGCAGGG | >hg38_21_+_8433172_8446622_RNA45SN1_45S_50ntFlanks | 9507 | 9524 | 18 | AACTATGCCTGGGCAGGG | 54.9 | -8.9 | 2.5 | -3.5 | 4.8E-04 | 1.3E-03 | BT20/MB468 | MP |
| >hg38_MT_+_1621_3279_RNR2_16S_50ntFlanks@200.230.31\|AATATAGCAAGGACTAACCCCTATACCTTCT | >hg38_MT_+_1621_3279_RNR2_16S_50ntFlanks | 200 | 230 | 31 | AATATAGCAAGGACTAACCCCTATACCTTCT | 54.9 | 9.5 | 2.7 | 3.5 | 4.8E-04 | 1.3E-03 | BT20/MB468 | MP |
| >hg38_21_+_8433172_8446622_RNA45SN1_45S_50ntFlanks@3790.3810.21\|CTGCGAATGGCTCATTAAATC | >hg38_21_+_8433172_8446622_RNA45SN1_45S_50ntFlanks | 3790 | 3810 | 21 | CTGCGAATGGCTCATTAAATC | 171.3 | -4.9 | 1.5 | -3.3 | 8.2E-04 | 1.3E-03 | BT20/MB468 | MITO |
| >hg38_21_+_8433172_8446622_RNA45SN1_45S_50ntFlanks@8182.8208.27\|ATCGAGGCCCAGCCCGTGGACGGTGTG | >hg38_21_+_8433172_8446622_RNA45SN1_45S_50ntFlanks | 8182 | 8208 | 27 | ATCGAGGCCCAGCCCGTGGACGGTGTG | 696.0 | -5.3 | 1.6 | -3.3 | 1.1E-03 | 1.3E-03 | MB231/MB468 | NUC |
| >hg38_21_+_8433172_8446622_RNA45SN1_45S_50ntFlanks@12004.12036.33\|CGGGCCGCCGGTGAAATACCACTACTCTGATCG | >hg38_21_+_8433172_8446622_RNA45SN1_45S_50ntFlanks | 12004 | 12036 | 33 | CGGGCCGCCGGTGAAATACCACTACTCTGATCG | 661.2 | 3.7 | 1.0 | 3.7 | 2.1E-04 | 1.3E-03 | BT20/MB231 | MITO |
| >MI0000279\|hsa-mir-196a-2&WithFlank&12\|+\|53991732\|53991853@31.51.21(+1U);MI0000238\|hsa-mir-196a-1&WithFlank&17\|-\|48632484\|48632565@13.33.21(+1U)&[MIMAT0000226&hsa-miR-196a-5p&offsets\|0\|-1(+1U);m-261&12\|+\|53991762\|53991783&offsets\|0\|-1(+1U)];[MIMAT0000226_1&hsa-miR-196a-5p&offsets\|0\|-1(+1U);m-268&17\|-\|48632532\|48632553&offsets\|0\|-1(+1U)]\|TAGGTAGTTTCATGTTGTTGGT | >MI0000279\|hsa-mir-196a-2&WithFlank&12\|+\|53991732\|53991853 | 31 | 51 | 21(+1U) | TAGGTAGTTTCATGTTGTTGGT | 143.0 | 5.2 | 1.5 | 3.5 | 4.8E-04 | 1.3E-03 | BT20/MB468 | MP |
| >hg38_21_+_8433172_8446622_RNA45SN1_45S_50ntFlanks@12997.13036.40\|CGCTGCGATCTATTGAAAGTCAGCCCTCGACACAAGGGTT | >hg38_21_+_8433172_8446622_RNA45SN1_45S_50ntFlanks | 12997 | 13036 | 40 | CGCTGCGATCTATTGAAAGTCAGCCCTCGACACAAGGGTT | 56.7 | 5.8 | 1.7 | 3.3 | 9.1E-04 | 1.3E-03 | MB231/MB468 | MP |
| >hg38_21_+_8433172_8446622_RNA45SN1_45S_50ntFlanks@7989.8013.25\|AGACGTGGCGACCCGCTGAATTTAA | >hg38_21_+_8433172_8446622_RNA45SN1_45S_50ntFlanks | 7989 | 8013 | 25 | AGACGTGGCGACCCGCTGAATTTAA | 162.9 | 4.3 | 1.3 | 3.3 | 1.0E-03 | 1.3E-03 | MB231/MB468 | CYTO |
| >MI0000253\|hsa-mir-148a&WithFlank&7\|-\|25949913\|25949992@50.70.21&[MIMAT0000243&hsa-miR-148a-3p&offsets\|0\|-1;m-7&7\|-\|25949922\|25949943&offsets\|0\|-1]\|TCAGTGCACTACAGAACTTTG | >MI0000253\|hsa-mir-148a&WithFlank&7\|-\|25949913\|25949992 | 50 | 70 | 21 | TCAGTGCACTACAGAACTTTG | 234.8 | 6.8 | 1.8 | 3.7 | 1.8E-04 | 1.3E-03 | BT20/MB231 | CYTO |
| >hg38_21_+_8433172_8446622_RNA45SN1_45S_50ntFlanks@12150.12180.31\|GGTGGGGAGTTTGACTGGGGCGGTACACCTG | >hg38_21_+_8433172_8446622_RNA45SN1_45S_50ntFlanks | 12150 | 12180 | 31 | GGTGGGGAGTTTGACTGGGGCGGTACACCTG | 115.2 | -5.3 | 1.5 | -3.5 | 4.8E-04 | 1.3E-03 | BT20/MB468 | MP |
| >hg38_21_+_8433172_8446622_RNA45SN1_45S_50ntFlanks@4734.4752.19\|AACGAAAGTCGGAGGTTCG | >hg38_21_+_8433172_8446622_RNA45SN1_45S_50ntFlanks | 4734 | 4752 | 19 | AACGAAAGTCGGAGGTTCG | 204.1 | -3.6 | 1.1 | -3.3 | 8.2E-04 | 1.3E-03 | BT20/MB468 | MITO |
| >hg38_21_+_8433172_8446622_RNA45SN1_45S_50ntFlanks@9166.9195.30\|CCGGGCGGGTCGCGCCGTCGGGCCCGGGGG | >hg38_21_+_8433172_8446622_RNA45SN1_45S_50ntFlanks | 9166 | 9195 | 30 | CCGGGCGGGTCGCGCCGTCGGGCCCGGGGG | 244.1 | -5.1 | 1.5 | -3.3 | 8.2E-04 | 1.3E-03 | BT20/MB468 | MITO |
| >hg38_21_+_8433172_8446622_RNA45SN1_45S_50ntFlanks@11207.11228.22\|GGCGGCGACTCTGGACGCGAGC | >hg38_21_+_8433172_8446622_RNA45SN1_45S_50ntFlanks | 11207 | 11228 | 22 | GGCGGCGACTCTGGACGCGAGC | 238.5 | -3.1 | 0.9 | -3.3 | 9.1E-04 | 1.3E-03 | MB231/MB468 | MP |
| >hg38_21_+_8433172_8446622_RNA45SN1_45S_50ntFlanks@12074.12095.22\|CGAGGGGCTCTCGCTTCTGGCG | >hg38_21_+_8433172_8446622_RNA45SN1_45S_50ntFlanks | 12074 | 12095 | 22 | CGAGGGGCTCTCGCTTCTGGCG | 615.3 | -5.8 | 1.7 | -3.5 | 4.8E-04 | 1.3E-03 | BT20/MB468 | MP |
| >hg38_21_+_8433172_8446622_RNA45SN1_45S_50ntFlanks@5485.5513.29\|AGCGCTGAGAAGACGGTCGAACTTGACTA | >hg38_21_+_8433172_8446622_RNA45SN1_45S_50ntFlanks | 5485 | 5513 | 29 | AGCGCTGAGAAGACGGTCGAACTTGACTA | 1325.2 | -3.3 | 0.9 | -3.5 | 4.8E-04 | 1.3E-03 | BT20/MB468 | MP |
| >hg38_21_+_8433172_8446622_RNA45SN1_45S_50ntFlanks@11209.11251.43\|CGGCGACTCTGGACGCGAGCCGGGCCCTTCCCGTGGATCGCCC | >hg38_21_+_8433172_8446622_RNA45SN1_45S_50ntFlanks | 11209 | 11251 | 43 | CGGCGACTCTGGACGCGAGCCGGGCCCTTCCCGTGGATCGCCC | 356.0 | -5.2 | 1.6 | -3.3 | 1.1E-03 | 1.3E-03 | MB231/MB468 | NUC |
| >hg38_1_-_228634819_228635039_RNA5S12_5S_50ntFlanks@147.169.23\|GGGAATACCGGGTGCTGTAGGCT | >hg38_1_-_228634819_228635039_RNA5S12_5S_50ntFlanks | 147 | 169 | 23 | GGGAATACCGGGTGCTGTAGGCT | 14896.4 | 5.5 | 1.5 | 3.8 | 1.7E-04 | 1.3E-03 | BT20/MB231 | MP |
| >hg38_MT_+_598_1651_RNR1_12S_50ntFlanks@707.730.24\|CAAGTACCCACGTAAAGACGTTAG | >hg38_MT_+_598_1651_RNR1_12S_50ntFlanks | 707 | 730 | 24 | CAAGTACCCACGTAAAGACGTTAG | 189.3 | 3.6 | 1.1 | 3.3 | 9.1E-04 | 1.3E-03 | MB231/MB468 | MP |
| >hg38_21_+_8433172_8446622_RNA45SN1_45S_50ntFlanks@11665.11706.42\|TTCTGCCCAGTGCTCTGAATGTCAAAGTGAAGAAATTCAATG | >hg38_21_+_8433172_8446622_RNA45SN1_45S_50ntFlanks | 11665 | 11706 | 42 | TTCTGCCCAGTGCTCTGAATGTCAAAGTGAAGAAATTCAATG | 720.0 | -3.4 | 0.9 | -3.7 | 1.8E-04 | 1.3E-03 | BT20/MB231 | CYTO |
| >hg38_21_+_8433172_8446622_RNA45SN1_45S_50ntFlanks@7975.7995.21\|CGCGACCTCAGATCAGACGTG | >hg38_21_+_8433172_8446622_RNA45SN1_45S_50ntFlanks | 7975 | 7995 | 21 | CGCGACCTCAGATCAGACGTG | 16936.2 | 2.2 | 0.6 | 3.7 | 1.8E-04 | 1.3E-03 | BT20/MB231 | NUC |
| >MI0000088\|hsa-mir-30a&WithFlank&6\|-\|71403545\|71403627@54.74.21(+1C)&[MIMAT0000088&hsa-miR-30a-3p&offsets\|+1\|0(+1C);m-38&6\|-\|71403554\|71403575&offsets\|+1\|0(+1C)]\|TTTCAGTCGGATGTTTGCAGCC | >MI0000088\|hsa-mir-30a&WithFlank&6\|-\|71403545\|71403627 | 54 | 74 | 21(+1C) | TTTCAGTCGGATGTTTGCAGCC | 112.8 | 6.1 | 1.8 | 3.3 | 9.1E-04 | 1.3E-03 | MB231/MB468 | MP |
| >hg38_21_+_8433172_8446622_RNA45SN1_45S_50ntFlanks@6647.6680.34\|CGTACGACTCTTAGCGGTGGATCACTCGGCTCGT | >hg38_21_+_8433172_8446622_RNA45SN1_45S_50ntFlanks | 6647 | 6680 | 34 | CGTACGACTCTTAGCGGTGGATCACTCGGCTCGT | 1027.5 | 2.3 | 0.7 | 3.3 | 1.1E-03 | 1.3E-03 | MB231/MB468 | NUC |
| >hg38_21_+_8433172_8446622_RNA45SN1_45S_50ntFlanks@5044.5063.20\|TCTTAGTTGGTGGAGCGATT | >hg38_21_+_8433172_8446622_RNA45SN1_45S_50ntFlanks | 5044 | 5063 | 20 | TCTTAGTTGGTGGAGCGATT | 120.8 | 10.2 | 2.7 | 3.8 | 1.7E-04 | 1.3E-03 | BT20/MB231 | MP |
| >hg38_MT_+_598_1651_RNR1_12S_50ntFlanks@905.950.46\|AGTATACTTCAAAGGACATTTAACTAAAACCCCTACGCATTTATAT | >hg38_MT_+_598_1651_RNR1_12S_50ntFlanks | 905 | 950 | 46 | AGTATACTTCAAAGGACATTTAACTAAAACCCCTACGCATTTATAT | 55.0 | 9.5 | 2.7 | 3.5 | 4.9E-04 | 1.3E-03 | BT20/MB468 | MP |
| >hg38_21_+_8433172_8446622_RNA45SN1_45S_50ntFlanks@4217.4242.26\|GTGACGAAAAATAACAATACAGGACT | >hg38_21_+_8433172_8446622_RNA45SN1_45S_50ntFlanks | 4217 | 4242 | 26 | GTGACGAAAAATAACAATACAGGACT | 116.3 | -5.4 | 1.6 | -3.3 | 9.1E-04 | 1.3E-03 | MB231/MB468 | MP |
| >hg38_21_+_8433172_8446622_RNA45SN1_45S_50ntFlanks@12941.12961.21\|ATTCGTAGACGACCTGCTTCT | >hg38_21_+_8433172_8446622_RNA45SN1_45S_50ntFlanks | 12941 | 12961 | 21 | ATTCGTAGACGACCTGCTTCT | 530.0 | 2.2 | 0.7 | 3.3 | 1.0E-03 | 1.3E-03 | MB231/MB468 | CYTO |
| >hg38_21_+_8433172_8446622_RNA45SN1_45S_50ntFlanks@11583.11609.27\|CTGGTGCGGACCAGGGGAATCCGACTG | >hg38_21_+_8433172_8446622_RNA45SN1_45S_50ntFlanks | 11583 | 11609 | 27 | CTGGTGCGGACCAGGGGAATCCGACTG | 945.4 | -2.9 | 0.9 | -3.3 | 1.1E-03 | 1.3E-03 | BT20/MB468 | NUC |
| >MI0000103\|hsa-mir-101-1&WithFlank&1\|-\|65058428\|65058514@53.74.22;MI0000739\|hsa-mir-101-2&WithFlank&9\|+\|4850291\|4850381@55.76.22&[MIMAT0000099&hsa-miR-101-3p&offsets\|0\|+1;m-17&1\|-\|65058443\|65058462&offsets\|0\|+2];[MIMAT0000099_1&hsa-miR-101-3p&offsets\|0\|+1;m-16&9\|+\|4850345\|4850364&offsets\|0\|+2]\|TACAGTACTGTGATAACTGAAG | >MI0000103\|hsa-mir-101-1&WithFlank&1\|-\|65058428\|65058514 | 53 | 74 | 22 | TACAGTACTGTGATAACTGAAG | 774.5 | 3.1 | 0.8 | 3.7 | 1.8E-04 | 1.3E-03 | BT20/MB231 | CYTO |

| | | | | | | | | | | | | | |
|---|---|---|---|---|---|---|---|---|---|---|---|---|---|
| >hg38_21_+_8433172_8446622_RNA45SN1_45S_50ntFlanks@10362.10381.20\|AAGGCCGAAGTGGAGAAGGG | >hg38_21_+_8433172_8446622_RNA45SN1_45S_50ntFlanks | 10362 | 10381 | 20 | AAGGCCGAAGTGGAGAAGGG | 215.4 | -2.6 | 0.7 | -3.5 | 4.9E-04 | 1.3E-03 | BT20/MB468 | MP |
| >MI0000286\|hsa-mir-210&WithFlank&11\|-\|568083\|568204@72.93.22(+1C)&[MIMAT0000267&hsa-miR-210-3p&offsets\|0\|0(+1C);m-102&11\|-\|568112\|568133&offsets\|0\|0(+1C)]\|CTGTGCGTGTGACAGCGGCTGAC | >MI0000286\|hsa-mir-210&WithFlank&11\|-\|568083\|568204 | 72 | 93 | 22(+1C) | CTGTGCGTGTGACAGCGGCTGAC | 130.6 | 6.1 | 1.8 | 3.3 | 9.2E-04 | 1.3E-03 | MB231/MB468 | MP |
| >am_MI0016846\|hsa-mir-4485&WithFlank&11\|-\|10508264\|10508332@34.52.19&[MIMAT0019019&hsa-miR-4485-3p&offsets\|-3\|-4;m-604&11\|-\|10508277\|10508296&offsets\|-3\|-4]\|GTTTAACGGCCGCGGTACC | >am_MI0016846\|hsa-mir-4485&WithFlank&11\|-\|10508264\|10508332 | 34 | 52 | 19 | GTTTAACGGCCGCGGTACC | 68.5 | 6.1 | 1.8 | 3.3 | 9.2E-04 | 1.3E-03 | MB231/MB468 | MP |
| >hg38_MT_+_1621_3279_RNR2_16S_50ntFlanks@940.958.19\|GTTTAACGGCCGCGGTACC | >hg38_MT_+_1621_3279_RNR2_16S_50ntFlanks | 940 | 958 | 19 | GTTTAACGGCCGCGGTACC | 68.5 | 6.1 | 1.8 | 3.3 | 9.2E-04 | 1.3E-03 | MB231/MB468 | MP |
| >hg38_21_+_8433172_8446622_RNA45SN1_45S_50ntFlanks@476.521.46\|TTCGTGATCGATGTGGTGACGTCGTGCTCTCCCGGGCCGGGTCCGA | >hg38_21_+_8433172_8446622_RNA45SN1_45S_50ntFlanks | 476 | 521 | 46 | TTCGTGATCGATGTGGTGACGTCGTGCTCTCCCGGGCCGGGTCCGA | 356.3 | 5.2 | 1.6 | 3.3 | 1.1E-03 | 1.3E-03 | MB231/MB468 | MITO |
| >hg38_MT_+_1621_3279_RNR2_16S_50ntFlanks@284.310.27\|CCGAAACCAGACGAGCTACCTAAGAAC | >hg38_MT_+_1621_3279_RNR2_16S_50ntFlanks | 284 | 310 | 27 | CCGAAACCAGACGAGCTACCTAAGAAC | 73.0 | 6.6 | 1.9 | 3.5 | 4.9E-04 | 1.3E-03 | BT20/MB468 | MP |
| >trnaMT_MetCAT_MT_+_4402_4469@49.71.23\|GTTATACCCTTCCCGTACTACCA | >trnaMT_MetCAT_MT_+_4402_4469 | 49 | 71 | 23 | GTTATACCCTTCCCGTACTACCA | 236.2 | 3.5 | 1.1 | 3.3 | 1.1E-03 | 1.3E-03 | MB231/MB468 | MITO |
| >hg38_21_+_8433172_8446622_RNA45SN1_45S_50ntFlanks@3988.4026.39\|CTTTGGTGACTCTAGATAACCTCGGGCCGATCGCACGCC | >hg38_21_+_8433172_8446622_RNA45SN1_45S_50ntFlanks | 3988 | 4026 | 39 | CTTTGGTGACTCTAGATAACCTCGGGCCGATCGCACGCC | 90.4 | -4.7 | 1.4 | -3.3 | 8.3E-04 | 1.3E-03 | BT20/MB468 | MITO |
| >MI0000076\|hsa-mir-20a&WithFlank&13\|+\|91351059\|91351141@14.35.22(+1A)&[MIMAT0000075&hsa-miR-20a-5p&offsets\|0\|-1(+1A);m-96&13\|+\|91351072\|91351094&offsets\|0\|-1(+1A)]\|TAAAGTGCTTATAGTGCAGGTAA | >MI0000076\|hsa-mir-20a&WithFlank&13\|+\|91351059\|91351141 | 14 | 35 | 22(+1A) | TAAAGTGCTTATAGTGCAGGTAA | 326.3 | 3.8 | 1.2 | 3.3 | 1.0E-03 | 1.3E-03 | MB231/MB468 | CYTO |
| >hg38_21_+_8433172_8446622_RNA45SN1_45S_50ntFlanks@6644.6663.20\|CCTCGTACGACTCTTAGCGG | >hg38_21_+_8433172_8446622_RNA45SN1_45S_50ntFlanks | 6644 | 6663 | 20 | CCTCGTACGACTCTTAGCGG | 1241.8 | 3.6 | 1.1 | 3.3 | 1.1E-03 | 1.3E-03 | MB231/MB468 | MITO |
| >hg38_21_+_8433172_8446622_RNA45SN1_45S_50ntFlanks@4762.4780.19\|AGATACCGTCGTAGTTCCG | >hg38_21_+_8433172_8446622_RNA45SN1_45S_50ntFlanks | 4762 | 4780 | 19 | AGATACCGTCGTAGTTCCG | 349.4 | -2.0 | 0.6 | -3.3 | 8.3E-04 | 1.3E-03 | BT20/MB468 | MITO |
| >hg38_21_+_8433172_8446622_RNA45SN1_45S_50ntFlanks@12242.12272.31\|AGAAGGGCAAAAGCTCGCTTGATCTTGATTT | >hg38_21_+_8433172_8446622_RNA45SN1_45S_50ntFlanks | 12242 | 12272 | 31 | AGAAGGGCAAAAGCTCGCTTGATCTTGATTT | 221.2 | -5.5 | 1.7 | -3.3 | 9.2E-04 | 1.3E-03 | MB231/MB468 | MP |
| >MI0000542\|hsa-mir-320a&WithFlank&8\|-\|22244960\|22245043@49.69.21&[MIMAT0000510&hsa-miR-320a-3p&offsets\|+1\|0;m-60&8\|-\|22244975\|22244996&offsets\|+1\|0]\|AAAGCTGGGTTGAGAGGGCGA | >MI0000542\|hsa-mir-320a&WithFlank&8\|-\|22244960\|22245043 | 49 | 69 | 21 | AAAGCTGGGTTGAGAGGGCGA | 379.0 | 2.7 | 0.7 | 3.7 | 2.1E-04 | 1.3E-03 | BT20/MB231 | MITO |
| >hg38_21_+_8433172_8446622_RNA45SN1_45S_50ntFlanks@10794.10817.24\|CTGGCATGTTGGAACAATGTAGGT | >hg38_21_+_8433172_8446622_RNA45SN1_45S_50ntFlanks | 10794 | 10817 | 24 | CTGGCATGTTGGAACAATGTAGGT | 215.0 | -4.9 | 1.5 | -3.3 | 1.1E-03 | 1.3E-03 | MB231/MB468 | NUC |
| >hg38_21_+_8433172_8446622_RNA45SN1_45S_50ntFlanks@5072.5112.41\|TAATTCCGATAACGAACGAGACTCTGGCATGCTAACTAGTT | >hg38_21_+_8433172_8446622_RNA45SN1_45S_50ntFlanks | 5072 | 5112 | 41 | TAATTCCGATAACGAACGAGACTCTGGCATGCTAACTAGTT | 225.6 | -3.2 | 0.9 | -3.7 | 1.8E-04 | 1.3E-03 | BT20/MB231 | CYTO |
| >hg38_21_+_8433172_8446622_RNA45SN1_45S_50ntFlanks@9716.9737.22\|AATGATTAGAGGTCTTGGGGCC | >hg38_21_+_8433172_8446622_RNA45SN1_45S_50ntFlanks | 9716 | 9737 | 22 | AATGATTAGAGGTCTTGGGGCC | 526.8 | -4.7 | 1.3 | -3.5 | 4.9E-04 | 1.3E-03 | BT20/MB468 | MP |
| >am_trnaMT_ArgTCG_MT_+_10405_10469@29.48.20\|TTTCGACTCATTAAATTATG | >am_trnaMT_ArgTCG_MT_+_10405_10469 | 29 | 48 | 20 | TTTCGACTCATTAAATTATG | 165.1 | -5.3 | 1.6 | -3.3 | 9.2E-04 | 1.3E-03 | MB231/MB468 | MP |
| >hg38_1_-_228634819_228635039_RNA5S12_5S_50ntFlanks@84.102.19\|CTCGTCTGATCTCGGAAGC | >hg38_1_-_228634819_228635039_RNA5S12_5S_50ntFlanks | 84 | 102 | 19 | CTCGTCTGATCTCGGAAGC | 470.2 | -5.3 | 1.6 | -3.3 | 1.1E-03 | 1.3E-03 | MB231/MB468 | NUC |
| >hg38_21_+_8433172_8446622_RNA45SN1_45S_50ntFlanks@12415.12444.30\|CTTCGATGTCGGCTCTTCCTATCATTGTGA | >hg38_21_+_8433172_8446622_RNA45SN1_45S_50ntFlanks | 12415 | 12444 | 30 | CTTCGATGTCGGCTCTTCCTATCATTGTGA | 2348.5 | -2.8 | 0.8 | -3.3 | 1.1E-03 | 1.3E-03 | MB231/MB468 | MITO |
| >hg38_21_+_8433172_8446622_RNA45SN1_45S_50ntFlanks@8196.8213.18\|CGTGGACGGTGTGAGGCC | >hg38_21_+_8433172_8446622_RNA45SN1_45S_50ntFlanks | 8196 | 8213 | 18 | CGTGGACGGTGTGAGGCC | 3564.3 | 2.0 | 0.6 | 3.3 | 9.9E-04 | 1.3E-03 | BT20/MB468 | CYTO |
| >MI0003778\|hsa-mir-320c-1&WithFlank&18\|+\|21683512\|21683595@48.67.20;MI0008191\|hsa-mir-320c-2&WithFlank&18\|+\|24321669\|24321752@48.67.20&[MIMAT0005793&hsa-miR-320c&offsets\|0\|0;m-222&18\|+\|21683559\|21683578&offsets\|0\|0];[MIMAT0005793_1&hsa-miR-320c&offsets\|0\|0;m-219&18\|+\|24321716\|24321735&offsets\|0\|0]\|AAAAGCTGGGTTGAGAGGGT | >MI0003778\|hsa-mir-320c-1&WithFlank&18\|+\|21683512\|21683595 | 48 | 67 | 20 | AAAAGCTGGGTTGAGAGGGT | 273.2 | 2.6 | 0.7 | 3.7 | 2.1E-04 | 1.3E-03 | BT20/MB231 | MITO |
| >hg38_21_+_8433172_8446622_RNA45SN1_45S_50ntFlanks@12418.12454.37\|CGATGTCGGCTCTTCCTATCATTGTGAAGCAGAATTC | >hg38_21_+_8433172_8446622_RNA45SN1_45S_50ntFlanks | 12418 | 12454 | 37 | CGATGTCGGCTCTTCCTATCATTGTGAAGCAGAATTC | 244.5 | -2.7 | 0.7 | -3.7 | 1.8E-04 | 1.3E-03 | BT20/MB231 | CYTO |
| >hg38_21_+_8433172_8446622_RNA45SN1_45S_50ntFlanks@10351.10390.40\|CGAGAACTTTGAAGGCCGAAGTGGAGAAGGGTTCCATGTG | >hg38_21_+_8433172_8446622_RNA45SN1_45S_50ntFlanks | 10351 | 10390 | 40 | CGAGAACTTTGAAGGCCGAAGTGGAGAAGGGTTCCATGTG | 1229.3 | 5.4 | 1.4 | 3.8 | 1.7E-04 | 1.3E-03 | BT20/MB231 | MP |
| >hg38_21_+_8433172_8446622_RNA45SN1_45S_50ntFlanks@11671.11704.34\|CCAGTGCTCTGAATGTCAAAGTGAAGAAATTCAA | >hg38_21_+_8433172_8446622_RNA45SN1_45S_50ntFlanks | 11671 | 11704 | 34 | CCAGTGCTCTGAATGTCAAAGTGAAGAAATTCAA | 331.3 | -2.9 | 0.9 | -3.3 | 1.0E-03 | 1.3E-03 | BT20/MB468 | CYTO |
| >hg38_21_+_8433172_8446622_RNA45SN1_45S_50ntFlanks@483.505.23\|TCGATGTGGTGACGTCGTGCTCT | >hg38_21_+_8433172_8446622_RNA45SN1_45S_50ntFlanks | 483 | 505 | 23 | TCGATGTGGTGACGTCGTGCTCT | 526.3 | 2.5 | 0.8 | 3.3 | 1.1E-03 | 1.3E-03 | BT20/MB468 | NUC |
| >MI0000078\|hsa-mir-22&WithFlank&17\|-\|1713897\|1713993@59.79.21(+1G)&[MIMAT0000077&hsa-miR-22-3p&offsets\|0\|-1(+1G);m-2&17\|-\|1713914\|1713935&offsets\|0\|-1(+1G)]\|AAGCTGCCAGTTGAAGAACTGG | >MI0000078\|hsa-mir-22&WithFlank&17\|-\|1713897\|1713993 | 59 | 79 | 21(+1G) | AAGCTGCCAGTTGAAGAACTGG | 382.2 | 4.3 | 1.2 | 3.5 | 4.9E-04 | 1.3E-03 | BT20/MB468 | MP |
| >hg38_21_+_8433172_8446622_RNA45SN1_45S_50ntFlanks@3986.4023.38\|GGCTTTGGTGACTCTAGATAACCTCGGGCCGATCGCAC | >hg38_21_+_8433172_8446622_RNA45SN1_45S_50ntFlanks | 3986 | 4023 | 38 | GGCTTTGGTGACTCTAGATAACCTCGGGCCGATCGCAC | 118.9 | 3.5 | 1.1 | 3.3 | 1.1E-03 | 1.3E-03 | BT20/MB468 | NUC |
| >hg38_21_+_8433172_8446622_RNA45SN1_45S_50ntFlanks@5485.5522.38\|AGCGCTGAGAAGACGGTCGAACTTGACTATCTAGAGGA | >hg38_21_+_8433172_8446622_RNA45SN1_45S_50ntFlanks | 5485 | 5522 | 38 | AGCGCTGAGAAGACGGTCGAACTTGACTATCTAGAGGA | 174.1 | -2.5 | 0.8 | -3.3 | 1.0E-03 | 1.3E-03 | MB231/MB468 | CYTO |
| >hg38_MT_+_1621_3279_RNR2_16S_50ntFlanks@1369.1397.29\|GATGTTGGATCAGGACATCCCGATGGTGC | >hg38_MT_+_1621_3279_RNR2_16S_50ntFlanks | 1369 | 1397 | 29 | GATGTTGGATCAGGACATCCCGATGGTGC | 128.6 | 4.8 | 1.4 | 3.5 | 4.9E-04 | 1.3E-03 | BT20/MB468 | MP |
| >hg38_21_+_8433172_8446622_RNA45SN1_45S_50ntFlanks@11562.11592.31\|CGCCTAGCAGCCGACTTAGAACTGGTGCGGA | >hg38_21_+_8433172_8446622_RNA45SN1_45S_50ntFlanks | 11562 | 11592 | 31 | CGCCTAGCAGCCGACTTAGAACTGGTGCGGA | 318.7 | -2.5 | 0.8 | -3.3 | 1.1E-03 | 1.3E-03 | MB231/MB468 | NUC |
| >hg38_21_+_8433172_8446622_RNA45SN1_45S_50ntFlanks@12418.12448.31\|CGATGTCGGCTCTTCCTATCATTGTGAAGCA | >hg38_21_+_8433172_8446622_RNA45SN1_45S_50ntFlanks | 12418 | 12448 | 31 | CGATGTCGGCTCTTCCTATCATTGTGAAGCA | 2008.3 | -2.5 | 0.8 | -3.3 | 1.1E-03 | 1.3E-03 | MB231/MB468 | MITO |
| >hg38_21_+_8433172_8446622_RNA45SN1_45S_50ntFlanks@3785.3812.28\|TGAAACTGCGAATGGCTCATTAAATCAG | >hg38_21_+_8433172_8446622_RNA45SN1_45S_50ntFlanks | 3785 | 3812 | 28 | TGAAACTGCGAATGGCTCATTAAATCAG | 230.2 | -6.3 | 1.8 | -3.5 | 4.9E-04 | 1.3E-03 | BT20/MB468 | MP |
| >hg38_MT_+_598_1651_RNR1_12S_50ntFlanks@443.469.27\|TATCTGAACACACAATAGCTAAGACCC | >hg38_MT_+_598_1651_RNR1_12S_50ntFlanks | 443 | 469 | 27 | TATCTGAACACACAATAGCTAAGACCC | 264.7 | 4.8 | 1.5 | 3.3 | 9.3E-04 | 1.3E-03 | MB231/MB468 | MP |
| >hg38_21_+_8433172_8446622_RNA45SN1_45S_50ntFlanks@9035.9059.25\|CGTGGGGGGGCCGGGCCACCCCTCC | >hg38_21_+_8433172_8446622_RNA45SN1_45S_50ntFlanks | 9035 | 9059 | 25 | CGTGGGGGGGCCGGGCCACCCCTCC | 386.2 | -5.8 | 1.7 | -3.5 | 4.9E-04 | 1.3E-03 | BT20/MB468 | MP |
| >hg38_21_+_8433172_8446622_RNA45SN1_45S_50ntFlanks@11582.11613.32\|ACTGGTGCGGACCAGGGGAATCCGACTGTTTA | >hg38_21_+_8433172_8446622_RNA45SN1_45S_50ntFlanks | 11582 | 11613 | 32 | ACTGGTGCGGACCAGGGGAATCCGACTGTTTA | 140.9 | -2.5 | 0.8 | -3.3 | 9.3E-04 | 1.3E-03 | MB231/MB468 | MP |
| >hg38_21_+_8433172_8446622_RNA45SN1_45S_50ntFlanks@11565.11601.37\|CTAGCAGCCGACTTAGAACTGGTGCGGACCAGGGGAA | >hg38_21_+_8433172_8446622_RNA45SN1_45S_50ntFlanks | 11565 | 11601 | 37 | CTAGCAGCCGACTTAGAACTGGTGCGGACCAGGGGAA | 188.2 | 6.9 | 1.8 | 3.8 | 1.7E-04 | 1.3E-03 | BT20/MB231 | MP |
| >trnaMT_PheGAA_MT_+_577_647@27.60.34\|AATACACTGAAAATGTTTAGACGGGCTCACATCA | >trnaMT_PheGAA_MT_+_577_647 | 27 | 60 | 34 | AATACACTGAAAATGTTTAGACGGGCTCACATCA | 54.7 | 9.5 | 2.7 | 3.5 | 4.9E-04 | 1.3E-03 | BT20/MB468 | MP |
| >hg38_21_+_8433172_8446622_RNA45SN1_45S_50ntFlanks@3837.3860.24\|CCTCTCCTACTTGGATAACTGTGG | >hg38_21_+_8433172_8446622_RNA45SN1_45S_50ntFlanks | 3837 | 3860 | 24 | CCTCTCCTACTTGGATAACTGTGG | 158.4 | -5.8 | 1.7 | -3.5 | 4.9E-04 | 1.3E-03 | BT20/MB468 | MP |
| >hg38_21_+_8433172_8446622_RNA45SN1_45S_50ntFlanks@10458.10498.41\|ATGGCCTCCGTTGCCCTCGGCCGATCGAAAGGGAGTCGGGT | >hg38_21_+_8433172_8446622_RNA45SN1_45S_50ntFlanks | 10458 | 10498 | 41 | ATGGCCTCCGTTGCCCTCGGCCGATCGAAAGGGAGTCGGGT | 144.2 | -4.8 | 1.4 | -3.3 | 8.4E-04 | 1.3E-03 | BT20/MB468 | MITO |
| >hg38_21_+_8433172_8446622_RNA45SN1_45S_50ntFlanks@8114.8132.19\|GCGCGGGACATGTGGCGTA | >hg38_21_+_8433172_8446622_RNA45SN1_45S_50ntFlanks | 8114 | 8132 | 19 | GCGCGGGACATGTGGCGTA | 282.7 | 6.8 | 1.8 | 3.8 | 1.7E-04 | 1.3E-03 | BT20/MB231 | MP |
| >MI0000088\|hsa-mir-30a&WithFlank&6\|-\|71403545\|71403627@53.70.18(+1U)&[MIMAT0000088&hsa-miR-30a-3p&offsets\|0\|-4(+1U);m-38&6\|-\|71403554\|71403575&offsets\|0\|-4(+1U)]\|CTTTCAGTCGGATGTTTGT | >MI0000088\|hsa-mir-30a&WithFlank&6\|-\|71403545\|71403627 | 53 | 70 | 18(+1U) | CTTTCAGTCGGATGTTTGT | 1208.5 | -4.0 | 1.1 | -3.8 | 1.7E-04 | 1.3E-03 | BT20/MB231 | MP |
| >MI0000079\|hsa-mir-23a&WithFlank&19\|-\|13836581\|13836665@54.71.18(+1G)&[MIMAT0000078&hsa-miR-23a-3p&offsets\|+3\|0(+1G);m-31&19\|-\|13836595\|13836615&offsets\|+3\|0(+1G)]\|ACATTGCCAGGGATTTCCG | >MI0000079\|hsa-mir-23a&WithFlank&19\|-\|13836581\|13836665 | 54 | 71 | 18(+1G) | ACATTGCCAGGGATTTCCG | 50.8 | 8.9 | 2.7 | 3.3 | 9.3E-04 | 1.3E-03 | MB231/MB468 | MP |
| >hg38_21_+_8433172_8446622_RNA45SN1_45S_50ntFlanks@10061.10080.20\|CAGTCGAGAGTGGACGGGAG | >hg38_21_+_8433172_8446622_RNA45SN1_45S_50ntFlanks | 10061 | 10080 | 20 | CAGTCGAGAGTGGACGGGAG | 63.2 | -9.8 | 3.0 | -3.3 | 9.3E-04 | 1.3E-03 | MB231/MB468 | MP |
| >hg38_MT_+_1621_3279_RNR2_16S_50ntFlanks@1209.1231.23\|CAGAACCCAACCTCCGAGCAGTA | >hg38_MT_+_1621_3279_RNR2_16S_50ntFlanks | 1209 | 1231 | 23 | CAGAACCCAACCTCCGAGCAGTA | 131.8 | -6.0 | 1.7 | -3.5 | 5.0E-04 | 1.3E-03 | BT20/MB468 | MP |
| >am_MI0016800\|hsa-mir-4454&WithFlank&4\|-\|163093568\|163093634@12.29.18(+1U)&[MIMAT0018976&hsa-miR-4454&offsets\|+3\|+1(+1U);m-444&4\|-\|163093607\|163093623&offsets\|0\|+1(+1U)]\|TCCGAGTCACGGCACCAAT | >am_MI0016800\|hsa-mir-4454&WithFlank&4\|-\|163093568\|163093634 | 12 | 29 | 18(+1U) | TCCGAGTCACGGCACCAAT | 54.8 | 9.5 | 2.7 | 3.5 | 5.0E-04 | 1.3E-03 | BT20/MB468 | MP |
| >trna116_GluCTC_1_-_145399233_145399304@1.39.39,trna59_GluCTC_1_+_249168447_249168518@1.39.39,trna71_GluCTC_1_-_161439189_161439260@1.39.39,trna74_GluCTC_1_-_161431809_161431880@1.39.39,trna77_GluCTC_1_-_161424398_161424469@1.39.39,trna77_GluCTC_6_+_28949976_28950047@1.39.39,trna80_GluCTC_1_-_161417018_161417089@1.39.39,trna87_GluCTC_6_-_126101393_126101464@1.39.39\|TCCCTGGTGGTCTAGTGGTTAGGATTCGGCGCTCTCACC | >trna116_GluCTC_1_-_145399233_145399304 | 1 | 39 | 39 | TCCCTGGTGGTCTAGTGGTTAGGATTCGGCGCTCTCACC | 919.1 | 5.5 | 1.6 | 3.5 | 5.0E-04 | 1.3E-03 | BT20/MB468 | MP |
| >MI0014211\|hsa-mir-3177&WithFlank&16\|+\|1734979\|1735072@60.80.21&[MIMAT0015054&hsa-miR-3177-3p&offsets\|0\|0;m-745&16\|+\|1735038\|1735058&offsets\|0\|0]\|TGCACGGCACTGGGGACACGT | >MI0014211\|hsa-mir-3177&WithFlank&16\|+\|1734979\|1735072 | 60 | 80 | 21 | TGCACGGCACTGGGGACACGT | 171.3 | 5.6 | 1.6 | 3.5 | 5.0E-04 | 1.3E-03 | BT20/MB468 | MP |
| >hg38_MT_+_1621_3279_RNR2_16S_50ntFlanks@485.518.34\|GAGGAACAGCTCTTTGGACACTAGGAAAAAACCT | >hg38_MT_+_1621_3279_RNR2_16S_50ntFlanks | 485 | 518 | 34 | GAGGAACAGCTCTTTGGACACTAGGAAAAAACCT | 180.6 | 4.7 | 1.3 | 3.5 | 5.0E-04 | 1.3E-03 | BT20/MB468 | MP |
| >hg38_21_+_8433172_8446622_RNA45SN1_45S_50ntFlanks@9033.9075.43\|CTCGTGGGGGGGCCGGGCCACCCCTCCCACGGCGCGACCGCTC | >hg38_21_+_8433172_8446622_RNA45SN1_45S_50ntFlanks | 9033 | 9075 | 43 | CTCGTGGGGGGGCCGGGCCACCCCTCCCACGGCGCGACCGCTC | 61.9 | -9.8 | 3.0 | -3.3 | 9.3E-04 | 1.3E-03 | MB231/MB468 | MP |
| >hg38_21_+_8433172_8446622_RNA45SN1_45S_50ntFlanks@5073.5117.45\|AATTCCGATAACGAACGAGACTCTGGCATGCTAACTAGTTACGCG | >hg38_21_+_8433172_8446622_RNA45SN1_45S_50ntFlanks | 5073 | 5117 | 45 | AATTCCGATAACGAACGAGACTCTGGCATGCTAACTAGTTACGCG | 63.2 | -9.8 | 3.0 | -3.3 | 9.3E-04 | 1.3E-03 | MB231/MB468 | MP |

<table>
<tr><th></th><th></th><th></th><th></th><th></th><th></th><th></th><th></th><th></th><th></th><th></th><th></th><th></th><th></th></tr>
<tr><td>>MI0000279|hsa-mir-196a-2&WithFlank&12|+|53991732|53991853@31.51.21(+1A);MI0000238|hsa-mir-196a-1&WithFlank&17|-|48632484|48632565@13.33.21(+1A)&[MIMAT0000226&hsa-miR-196a-5p&offsets|0|-1(+1A);m-261&12|+|53991762|53991783&offsets|0|-1(+1A)];[MIMAT0000226_1&hsa-miR-196a-5p&offsets|0|-1(+1A);m-268&17|-|48632532|48632553&offsets|0|-1(+1A)]|TAGGTAGTTTCATGTTGTTGGA</td><td>>MI0000279|hsa-mir-196a-2&WithFlank&12|+|53991732|53991853</td><td>31</td><td>51</td><td>21(+1A)</td><td>TAGGTAGTTTCATGTTGTTGGA</td><td>483.7</td><td>5.1</td><td>1.5</td><td>3.5</td><td>5.0E-04</td><td>1.3E-03</td><td>BT20/MB468</td><td>MP</td></tr>
<tr><td>>hg38_21_+_8433172_8446622_RNA45SN1_45S_50ntFlanks@3835.3860.26|CTCCTCTCCTACTTGGATAACTGTGG</td><td>>hg38_21_+_8433172_8446622_RNA45SN1_45S_50ntFlanks</td><td>3835</td><td>3860</td><td>26</td><td>CTCCTCTCCTACTTGGATAACTGTGG</td><td>2578.7</td><td>-2.3</td><td>0.7</td><td>-3.3</td><td>1.0E-03</td><td>1.3E-03</td><td>MB231/MB468</td><td>CYTO</td></tr>
<tr><td>>MI0000734|hsa-mir-106b&WithFlank&7|-|100093987|100094080@58.78.21(+1A)&[MIMAT0004672&hsa-miR-106b-3p&offsets|0|-1(+1A);m-87&7|-|100094003|100094023&offsets|0|0(+1A)]|CCGCACTGTGGGTACTTGCTGA</td><td>>MI0000734|hsa-mir-106b&WithFlank&7|-|100093987|100094080</td><td>58</td><td>78</td><td>21(+1A)</td><td>CCGCACTGTGGGTACTTGCTGA</td><td>449.6</td><td>6.2</td><td>1.8</td><td>3.5</td><td>5.0E-04</td><td>1.3E-03</td><td>BT20/MB468</td><td>MP</td></tr>
<tr><td>>MI0000433|hsa-let-7g&WithFlank&3|-|52268272|52268367@11.31.21(+1A)&[MIMAT0000414&hsa-let-7g-5p&offsets|0|-1(+1A);m-58&3|-|52268336|52268357&offsets|0|-1(+1A)]|TGAGGTAGTAGTTTGTACAGTA</td><td>>MI0000433|hsa-let-7g&WithFlank&3|-|52268272|52268367</td><td>11</td><td>31</td><td>21(+1A)</td><td>TGAGGTAGTAGTTTGTACAGTA</td><td>507.6</td><td>3.5</td><td>1.0</td><td>3.5</td><td>5.0E-04</td><td>1.4E-03</td><td>BT20/MB468</td><td>MP</td></tr>
<tr><td>>MI0000299|hsa-mir-222&WithFlank&X|-|45747009|45747130@75.96.22(+1A)&[MIMAT0000279&hsa-miR-222-3p&offsets|0|+1(+1A);m-140&X|-|45747033|45747056&offsets|0|-2(+1A)]|AGCTACATCTGGCTACTGGGTCA</td><td>>MI0000299|hsa-mir-222&WithFlank&X|-|45747009|45747130</td><td>75</td><td>96</td><td>22(+1A)</td><td>AGCTACATCTGGCTACTGGGTCA</td><td>100.6</td><td>2.0</td><td>0.6</td><td>3.3</td><td>1.1E-03</td><td>1.4E-03</td><td>BT20/MB468</td><td>NUC</td></tr>
<tr><td>>hg38_21_+_8433172_8446622_RNA45SN1_45S_50ntFlanks@6652.6687.36|GACTCTTAGCGGTGGATCACTCGGCTCGTGCGTCGA</td><td>>hg38_21_+_8433172_8446622_RNA45SN1_45S_50ntFlanks</td><td>6652</td><td>6687</td><td>36</td><td>GACTCTTAGCGGTGGATCACTCGGCTCGTGCGTCGA</td><td>2506.4</td><td>2.6</td><td>0.8</td><td>3.3</td><td>8.4E-04</td><td>1.4E-03</td><td>BT20/MB468</td><td>MITO</td></tr>
<tr><td>>hg38_21_+_8433172_8446622_RNA45SN1_45S_50ntFlanks@10807.10829.23|ACAATGTAGGTAAGGGAAGTCGG</td><td>>hg38_21_+_8433172_8446622_RNA45SN1_45S_50ntFlanks</td><td>10807</td><td>10829</td><td>23</td><td>ACAATGTAGGTAAGGGAAGTCGG</td><td>145.1</td><td>-4.5</td><td>1.4</td><td>-3.3</td><td>8.4E-04</td><td>1.4E-03</td><td>BT20/MB468</td><td>MITO</td></tr>
<tr><td>>hg38_21_+_8433172_8446622_RNA45SN1_45S_50ntFlanks@7989.8010.22|AGACGTGGCGACCCGCTGAATT</td><td>>hg38_21_+_8433172_8446622_RNA45SN1_45S_50ntFlanks</td><td>7989</td><td>8010</td><td>22</td><td>AGACGTGGCGACCCGCTGAATT</td><td>1315.6</td><td>-2.3</td><td>0.7</td><td>-3.3</td><td>9.3E-04</td><td>1.4E-03</td><td>MB231/MB468</td><td>MP</td></tr>
<tr><td>>hg38_21_+_8433172_8446622_RNA45SN1_45S_50ntFlanks@10841.10865.25|CGTAACTTCGGGATAAGGATTGGCT</td><td>>hg38_21_+_8433172_8446622_RNA45SN1_45S_50ntFlanks</td><td>10841</td><td>10865</td><td>25</td><td>CGTAACTTCGGGATAAGGATTGGCT</td><td>4817.1</td><td>6.7</td><td>1.8</td><td>3.8</td><td>1.7E-04</td><td>1.4E-03</td><td>BT20/MB231</td><td>MP</td></tr>
<tr><td>>hg38_21_+_8433172_8446622_RNA45SN1_45S_50ntFlanks@4104.4121.18|CACGGGTGACGGGGAATC</td><td>>hg38_21_+_8433172_8446622_RNA45SN1_45S_50ntFlanks</td><td>4104</td><td>4121</td><td>18</td><td>CACGGGTGACGGGGAATC</td><td>211.1</td><td>-1.9</td><td>0.6</td><td>-3.3</td><td>1.0E-03</td><td>1.4E-03</td><td>MB231/MB468</td><td>CYTO</td></tr>
<tr><td>>hg38_21_+_8433172_8446622_RNA45SN1_45S_50ntFlanks@11641.11670.30|CGGCGGGTGTTGACGCGATGTGATTTCTGC</td><td>>hg38_21_+_8433172_8446622_RNA45SN1_45S_50ntFlanks</td><td>11641</td><td>11670</td><td>30</td><td>CGGCGGGTGTTGACGCGATGTGATTTCTGC</td><td>993.7</td><td>-4.4</td><td>1.4</td><td>-3.3</td><td>1.0E-03</td><td>1.4E-03</td><td>BT20/MB468</td><td>CYTO</td></tr>
<tr><td>>hg38_21_+_8433172_8446622_RNA45SN1_45S_50ntFlanks@11583.11623.41|CTGGTGCGGACCAGGGGAATCCGACTGTTTAATTAAAACAA</td><td>>hg38_21_+_8433172_8446622_RNA45SN1_45S_50ntFlanks</td><td>11583</td><td>11623</td><td>41</td><td>CTGGTGCGGACCAGGGGAATCCGACTGTTTAATTAAAACAA</td><td>416.8</td><td>6.0</td><td>1.7</td><td>3.5</td><td>5.0E-04</td><td>1.4E-03</td><td>BT20/MB468</td><td>MP</td></tr>
<tr><td>>hg38_21_+_8433172_8446622_RNA45SN1_45S_50ntFlanks@3777.3794.18|CGGTACAGTGAAACTGCG</td><td>>hg38_21_+_8433172_8446622_RNA45SN1_45S_50ntFlanks</td><td>3777</td><td>3794</td><td>18</td><td>CGGTACAGTGAAACTGCG</td><td>139.9</td><td>-2.9</td><td>0.9</td><td>-3.3</td><td>1.0E-03</td><td>1.4E-03</td><td>MB231/MB468</td><td>CYTO</td></tr>
<tr><td>>hg38_21_+_8433172_8446622_RNA45SN1_45S_50ntFlanks@11322.11342.21|ACGTCTCGTCGCGCGCGCGTC</td><td>>hg38_21_+_8433172_8446622_RNA45SN1_45S_50ntFlanks</td><td>11322</td><td>11342</td><td>21</td><td>ACGTCTCGTCGCGCGCGCGTC</td><td>5626.0</td><td>-4.5</td><td>1.4</td><td>-3.3</td><td>1.1E-03</td><td>1.4E-03</td><td>BT20/MB468</td><td>NUC</td></tr>
<tr><td>>hg38_21_+_8433172_8446622_RNA45SN1_45S_50ntFlanks@9711.9737.27|AAGCGAATGATTAGAGGTCTTGGGGCC</td><td>>hg38_21_+_8433172_8446622_RNA45SN1_45S_50ntFlanks</td><td>9711</td><td>9737</td><td>27</td><td>AAGCGAATGATTAGAGGTCTTGGGGCC</td><td>181.2</td><td>-5.0</td><td>1.5</td><td>-3.3</td><td>8.4E-04</td><td>1.4E-03</td><td>BT20/MB468</td><td>MITO</td></tr>
<tr><td>>MI0000471|hsa-mir-126&WithFlank&9|+|136670596|136670692@58.78.21(+1A)&[MIMAT0000445&hsa-miR-126-3p&offsets|0|-1(+1A);m-40&9|+|136670653|136670674&offsets|0|-1(+1A)]|TCGTACCGTGAGTAATAATGCA</td><td>>MI0000471|hsa-mir-126&WithFlank&9|+|136670596|136670692</td><td>58</td><td>78</td><td>21(+1A)</td><td>TCGTACCGTGAGTAATAATGCA</td><td>1063.4</td><td>3.5</td><td>0.9</td><td>3.7</td><td>1.9E-04</td><td>1.4E-03</td><td>BT20/MB231</td><td>CYTO</td></tr>
<tr><td>>hg38_21_+_8433172_8446622_RNA45SN1_45S_50ntFlanks@11210.11227.18|GGCGACTCTGGACGCGAG</td><td>>hg38_21_+_8433172_8446622_RNA45SN1_45S_50ntFlanks</td><td>11210</td><td>11227</td><td>18</td><td>GGCGACTCTGGACGCGAG</td><td>295.4</td><td>-5.5</td><td>1.6</td><td>-3.5</td><td>5.0E-04</td><td>1.4E-03</td><td>BT20/MB468</td><td>MP</td></tr>
<tr><td>>hg38_21_+_8433172_8446622_RNA45SN1_45S_50ntFlanks@12409.12433.25|TTGATCCTTCGATGTCGGCTCTTCC</td><td>>hg38_21_+_8433172_8446622_RNA45SN1_45S_50ntFlanks</td><td>12409</td><td>12433</td><td>25</td><td>TTGATCCTTCGATGTCGGCTCTTCC</td><td>331.8</td><td>1.6</td><td>0.5</td><td>3.3</td><td>1.1E-03</td><td>1.4E-03</td><td>MB231/MB468</td><td>NUC</td></tr>
<tr><td>>hg38_21_+_8433172_8446622_RNA45SN1_45S_50ntFlanks@11603.11626.24|CCGACTGTTTAATTAAAACAAAGC</td><td>>hg38_21_+_8433172_8446622_RNA45SN1_45S_50ntFlanks</td><td>11603</td><td>11626</td><td>24</td><td>CCGACTGTTTAATTAAAACAAAGC</td><td>105.6</td><td>5.7</td><td>1.6</td><td>3.5</td><td>5.0E-04</td><td>1.4E-03</td><td>BT20/MB468</td><td>MP</td></tr>
<tr><td>>hg38_21_+_8433172_8446622_RNA45SN1_45S_50ntFlanks@10238.10259.22|CGCTACGCCGCGACGAGTAGGA</td><td>>hg38_21_+_8433172_8446622_RNA45SN1_45S_50ntFlanks</td><td>10238</td><td>10259</td><td>22</td><td>CGCTACGCCGCGACGAGTAGGA</td><td>959.9</td><td>4.4</td><td>1.2</td><td>3.8</td><td>1.7E-04</td><td>1.4E-03</td><td>BT20/MB231</td><td>MP</td></tr>
<tr><td>>hg38_21_+_8433172_8446622_RNA45SN1_45S_50ntFlanks@4730.4752.23|CAAGAACGAAAGTCGGAGGTTCG</td><td>>hg38_21_+_8433172_8446622_RNA45SN1_45S_50ntFlanks</td><td>4730</td><td>4752</td><td>23</td><td>CAAGAACGAAAGTCGGAGGTTCG</td><td>279.9</td><td>-2.1</td><td>0.6</td><td>-3.3</td><td>1.0E-03</td><td>1.4E-03</td><td>BT20/MB468</td><td>CYTO</td></tr>
<tr><td>>MI0000438|hsa-mir-15b&WithFlank&3|+|160404582|160404691@26.47.22&[MIMAT0000417&hsa-miR-15b-5p&offsets|0|0;m-110&3|+|160404607|160404628&offsets|0|0]|TAGCAGCACATCATGGTTTACA</td><td>>MI0000438|hsa-mir-15b&WithFlank&3|+|160404582|160404691</td><td>26</td><td>47</td><td>22</td><td>TAGCAGCACATCATGGTTTACA</td><td>1220.0</td><td>1.8</td><td>0.6</td><td>3.3</td><td>1.1E-03</td><td>1.4E-03</td><td>BT20/MB468</td><td>NUC</td></tr>
<tr><td>>hg38_21_+_8433172_8446622_RNA45SN1_45S_50ntFlanks@10348.10372.25|AAACGAGAACTTTGAAGGCCGAAGT</td><td>>hg38_21_+_8433172_8446622_RNA45SN1_45S_50ntFlanks</td><td>10348</td><td>10372</td><td>25</td><td>AAACGAGAACTTTGAAGGCCGAAGT</td><td>928.2</td><td>2.1</td><td>0.6</td><td>3.3</td><td>1.1E-03</td><td>1.4E-03</td><td>MB231/MB468</td><td>NUC</td></tr>
<tr><td>>MI0000261|hsa-mir-139&WithFlank&11|-|72615057|72615136@13.34.22(+1C)&[MIMAT0000250&hsa-miR-139-5p&offsets|0|-1(+1C);m-186&11|-|72615102|72615124&offsets|0|-1(+1C)]|TCTACAGTGCACGTGTCTCCAGC</td><td>>MI0000261|hsa-mir-139&WithFlank&11|-|72615057|72615136</td><td>13</td><td>34</td><td>22(+1C)</td><td>TCTACAGTGCACGTGTCTCCAGC</td><td>339.8</td><td>-3.8</td><td>1.0</td><td>-3.8</td><td>1.7E-04</td><td>1.4E-03</td><td>BT20/MB231</td><td>MP</td></tr>
<tr><td>>trnaMT_PheGAA_MT_+_577_647@1.35.35|GTTTATGTAGCTTACCTCCTCAAAGCAATACACTG</td><td>>trnaMT_PheGAA_MT_+_577_647</td><td>1</td><td>35</td><td>35</td><td>GTTTATGTAGCTTACCTCCTCAAAGCAATACACTG</td><td>266.6</td><td>4.0</td><td>1.2</td><td>3.3</td><td>9.4E-04</td><td>1.4E-03</td><td>MB231/MB468</td><td>MP</td></tr>
<tr><td>>hg38_21_+_8433172_8446622_RNA45SN1_45S_50ntFlanks@10783.10816.34|GTGAACAGCCTCTGGCATGTTGGAACAATGTAGG</td><td>>hg38_21_+_8433172_8446622_RNA45SN1_45S_50ntFlanks</td><td>10783</td><td>10816</td><td>34</td><td>GTGAACAGCCTCTGGCATGTTGGAACAATGTAGG</td><td>285.3</td><td>-4.3</td><td>1.3</td><td>-3.3</td><td>1.0E-03</td><td>1.4E-03</td><td>BT20/MB468</td><td>CYTO</td></tr>
<tr><td>>trna134_GluTTC_1_-_16861774_16861845@1.32.32,trna5_GluTTC_1_+_17199078_17199149@1.32.32,trna84_GluTTC_1_-_161391883_161391954@1.32.32,trna94_GluTTC_1_-_149664355_149664427@1.32.32|TCCCTGGTGGTCTAGTGGCTAGGATTCGGCGC</td><td>>trna134_GluTTC_1_-_16861774_16861845</td><td>1</td><td>32</td><td>32</td><td>TCCCTGGTGGTCTAGTGGCTAGGATTCGGCGC</td><td>233.4</td><td>2.9</td><td>0.8</td><td>3.7</td><td>1.9E-04</td><td>1.4E-03</td><td>BT20/MB231</td><td>CYTO</td></tr>
<tr><td>>hg38_21_+_8433172_8446622_RNA45SN1_45S_50ntFlanks@12000.12032.33|CCTGCGGGCCGCCGGTGAAATACCACTACTCTG</td><td>>hg38_21_+_8433172_8446622_RNA45SN1_45S_50ntFlanks</td><td>12000</td><td>12032</td><td>33</td><td>CCTGCGGGCCGCCGGTGAAATACCACTACTCTG</td><td>192.7</td><td>-5.4</td><td>1.6</td><td>-3.5</td><td>5.0E-04</td><td>1.4E-03</td><td>BT20/MB468</td><td>MP</td></tr>
<tr><td>>MI0000815|hsa-mir-339&WithFlank&7|-|1022927|1023032@21.43.23&[MIMAT0000764&hsa-miR-339-5p&offsets|0|0;m-163&7|-|1022990|1023012&offsets|0|0]|TCCCTGTCCTCCAGGAGCTCACG</td><td>>MI0000815|hsa-mir-339&WithFlank&7|-|1022927|1023032</td><td>21</td><td>43</td><td>23</td><td>TCCCTGTCCTCCAGGAGCTCACG</td><td>217.0</td><td>2.3</td><td>0.7</td><td>3.3</td><td>1.1E-03</td><td>1.4E-03</td><td>BT20/MB468</td><td>NUC</td></tr>
<tr><td>>MI0000273|hsa-mir-183&WithFlank&7|-|129774899|129775020@33.54.22&[MIMAT0000261&hsa-miR-183-5p&offsets|0|0;m-41&7|-|129774966|129774987&offsets|-1|-1]|TATGGCACTGGTAGAATTCACT</td><td>>MI0000273|hsa-mir-183&WithFlank&7|-|129774899|129775020</td><td>33</td><td>54</td><td>22</td><td>TATGGCACTGGTAGAATTCACT</td><td>4722.0</td><td>1.2</td><td>0.4</td><td>3.3</td><td>1.1E-03</td><td>1.4E-03</td><td>BT20/MB468</td><td>NUC</td></tr>
<tr><td>>hg38_1_-_228634819_228635039_RNA5S12_5S_50ntFlanks@51.80.30|GTCTACGGCCATACCACCCTGAACGCGCCC</td><td>>hg38_1_-_228634819_228635039_RNA5S12_5S_50ntFlanks</td><td>51</td><td>80</td><td>30</td><td>GTCTACGGCCATACCACCCTGAACGCGCCC</td><td>2339.0</td><td>6.5</td><td>2.0</td><td>3.3</td><td>1.0E-03</td><td>1.4E-03</td><td>BT20/MB468</td><td>CYTO</td></tr>
<tr><td>>hg38_21_+_8433172_8446622_RNA45SN1_45S_50ntFlanks@12934.12966.33|CTAAACCATTCGTAGACGACCTGCTTCTGGGTC</td><td>>hg38_21_+_8433172_8446622_RNA45SN1_45S_50ntFlanks</td><td>12934</td><td>12966</td><td>33</td><td>CTAAACCATTCGTAGACGACCTGCTTCTGGGTC</td><td>163.2</td><td>-3.6</td><td>1.1</td><td>-3.3</td><td>8.5E-04</td><td>1.4E-03</td><td>BT20/MB468</td><td>MITO</td></tr>
<tr><td>>am_trna10_ValCAC_5_-_180649395_180649467@3.23.21,trna12_ValAAC_5_-_180645270_180645342@3.23.21,trna132_ValAAC_6_-_27721179_27721251@3.23.21,trna136_ValAAC_6_-_27648885_27648957@3.23.21,trna139_ValAAC_6_-_27618707_27618779@3.23.21,trna18_ValCAC_5_-_180529253_180529325@3.23.21,trna2_ValAAC_3_+_169490018_169490090@3.23.21,trna2_ValCAC_5_+_180524070_180524142@3.23.21,trna37_ValAAC_6_+_27203288_27203360@3.23.21,trna4_ValAAC_5_+_180591154_180591226@3.23.21,trna5_ValAAC_5_+_180596610_180596682@3.23.21,trna6_ValCAC_5_+_180600650_180600722@3.23.21,trna85_ValCAC_1_-_161369490_161369562@3.23.21,trna90_ValCAC_1_-_149684088_149684161@3.23.21,trna98_ValCAC_1_-_149298555_149298627@3.23.21,trna9_ValCAC_6_+_26538282_26538354@3.23.21|TTCCGTAGTGTAGTGGTTATC</td><td>>am_trna10_ValCAC_5_-_180649395_180649467</td><td>3</td><td>23</td><td>21</td><td>TTCCGTAGTGTAGTGGTTATC</td><td>81.7</td><td>6.6</td><td>1.9</td><td>3.5</td><td>5.0E-04</td><td>1.4E-03</td><td>BT20/MB468</td><td>MP</td></tr>
<tr><td>>hg38_21_+_8433172_8446622_RNA45SN1_45S_50ntFlanks@12991.13012.22|CTCCCTCGCTGCGATCTATTGA</td><td>>hg38_21_+_8433172_8446622_RNA45SN1_45S_50ntFlanks</td><td>12991</td><td>13012</td><td>22</td><td>CTCCCTCGCTGCGATCTATTGA</td><td>2144.5</td><td>-4.2</td><td>1.3</td><td>-3.3</td><td>1.0E-03</td><td>1.4E-03</td><td>BT20/MB468</td><td>CYTO</td></tr>
<tr><td>>hg38_21_+_8433172_8446622_RNA45SN1_45S_50ntFlanks@12997.13040.44|CGCTGCGATCTATTGAAAGTCAGCCCTCGACACAAGGGTTTGTC</td><td>>hg38_21_+_8433172_8446622_RNA45SN1_45S_50ntFlanks</td><td>12997</td><td>13040</td><td>44</td><td>CGCTGCGATCTATTGAAAGTCAGCCCTCGACACAAGGGTTTGTC</td><td>170.6</td><td>-5.9</td><td>1.6</td><td>-3.7</td><td>1.9E-04</td><td>1.4E-03</td><td>BT20/MB231</td><td>NUC</td></tr>
<tr><td>>hg38_21_+_8433172_8446622_RNA45SN1_45S_50ntFlanks@8357.8375.19|AAAAGAACTTTGAAGAGAG</td><td>>hg38_21_+_8433172_8446622_RNA45SN1_45S_50ntFlanks</td><td>8357</td><td>8375</td><td>19</td><td>AAAAGAACTTTGAAGAGAG</td><td>1618.2</td><td>-2.8</td><td>0.9</td><td>-3.3</td><td>1.0E-03</td><td>1.4E-03</td><td>BT20/MB468</td><td>CYTO</td></tr>
<tr><td>>MI0000289|hsa-mir-181a-1&WithFlank&1|-|198859038|198859159@70.90.21&[MIMAT0000270&hsa-miR-181a-3p&offsets|0|-1;m-125&1|-|198859069|198859090&offsets|0|-1]|ACCATCGACCGTTGATTGTAC</td><td>>MI0000289|hsa-mir-181a-1&WithFlank&1|-|198859038|198859159</td><td>70</td><td>90</td><td>21</td><td>ACCATCGACCGTTGATTGTAC</td><td>101.9</td><td>4.0</td><td>1.2</td><td>3.3</td><td>9.4E-04</td><td>1.4E-03</td><td>MB231/MB468</td><td>MP</td></tr>
<tr><td>>hg38_21_+_8433172_8446622_RNA45SN1_45S_50ntFlanks@10353.10388.36|AGAACTTTGAAGGCCGAAGTGGAGAAGGGTTCCATG</td><td>>hg38_21_+_8433172_8446622_RNA45SN1_45S_50ntFlanks</td><td>10353</td><td>10388</td><td>36</td><td>AGAACTTTGAAGGCCGAAGTGGAGAAGGGTTCCATG</td><td>476.6</td><td>-3.0</td><td>0.9</td><td>-3.3</td><td>1.1E-03</td><td>1.4E-03</td><td>MB231/MB468</td><td>NUC</td></tr>
<tr><td>>hg38_21_+_8433172_8446622_RNA45SN1_45S_50ntFlanks@3847.3879.33|TTGGATAACTGTGGTAATTCTAGAGCTAATACA</td><td>>hg38_21_+_8433172_8446622_RNA45SN1_45S_50ntFlanks</td><td>3847</td><td>3879</td><td>33</td><td>TTGGATAACTGTGGTAATTCTAGAGCTAATACA</td><td>49.0</td><td>6.4</td><td>1.8</td><td>3.5</td><td>5.0E-04</td><td>1.4E-03</td><td>BT20/MB468</td><td>MP</td></tr>
<tr><td>>hg38_21_+_8433172_8446622_RNA45SN1_45S_50ntFlanks@6648.6677.30|GTACGACTCTTAGCGGTGGATCACTCGGCT</td><td>>hg38_21_+_8433172_8446622_RNA45SN1_45S_50ntFlanks</td><td>6648</td><td>6677</td><td>30</td><td>GTACGACTCTTAGCGGTGGATCACTCGGCT</td><td>510.3</td><td>3.3</td><td>0.9</td><td>3.5</td><td>5.0E-04</td><td>1.4E-03</td><td>BT20/MB468</td><td>MP</td></tr>
<tr><td>>hg38_21_+_8433172_8446622_RNA45SN1_45S_50ntFlanks@12633.12651.19|CGAAGCTACCATCTGTGGG</td><td>>hg38_21_+_8433172_8446622_RNA45SN1_45S_50ntFlanks</td><td>12633</td><td>12651</td><td>19</td><td>CGAAGCTACCATCTGTGGG</td><td>461.3</td><td>5.3</td><td>1.4</td><td>3.8</td><td>1.8E-04</td><td>1.4E-03</td><td>BT20/MB231</td><td>MP</td></tr>
<tr><td>>hg38_21_+_8433172_8446622_RNA45SN1_45S_50ntFlanks@10351.10379.29|CGAGAACTTTGAAGGCCGAAGTGGAGAAG</td><td>>hg38_21_+_8433172_8446622_RNA45SN1_45S_50ntFlanks</td><td>10351</td><td>10379</td><td>29</td><td>CGAGAACTTTGAAGGCCGAAGTGGAGAAG</td><td>2243.9</td><td>-4.1</td><td>1.2</td><td>-3.5</td><td>5.1E-04</td><td>1.4E-03</td><td>BT20/MB468</td><td>MP</td></tr>
</table>

<table>
<tr><td>>hg38_21_+_8433172_8446622_RNA45SN1_45S_50ntFlanks@12539.12563.25|ATGATGTGTTGTTGCCATGGTAATC</td><td>>hg38_21_+_8433172_8446622_RNA45SN1_45S_50ntFlanks</td><td>12539</td><td>12563</td><td>25</td><td>ATGATGTGTTGTTGCCATGGTAATC</td><td>602.4</td><td>-5.8</td><td>1.7</td><td>-3.5</td><td>5.0E-04</td><td>1.4E-03</td><td>BT20/MB468</td><td>MP</td></tr>
<tr><td>>hg38_21_+_8433172_8446622_RNA45SN1_45S_50ntFlanks@9120.9152.33|TGCCGCGCGCGGGTCGGGGGGCGGGGCGGACTG</td><td>>hg38_21_+_8433172_8446622_RNA45SN1_45S_50ntFlanks</td><td>9120</td><td>9152</td><td>33</td><td>TGCCGCGCGCGGGTCGGGGGGCGGGGCGGACTG</td><td>206.7</td><td>-5.8</td><td>1.7</td><td>-3.5</td><td>5.1E-04</td><td>1.4E-03</td><td>BT20/MB468</td><td>MP</td></tr>
<tr><td>>hg38_21_+_8433172_8446622_RNA45SN1_45S_50ntFlanks@10829.10853.25|GCAAGCCGGATCCGTAACTTCGGGA</td><td>>hg38_21_+_8433172_8446622_RNA45SN1_45S_50ntFlanks</td><td>10829</td><td>10853</td><td>25</td><td>GCAAGCCGGATCCGTAACTTCGGGA</td><td>201.8</td><td>-3.6</td><td>1.1</td><td>-3.3</td><td>1.0E-03</td><td>1.4E-03</td><td>BT20/MB468</td><td>CYTO</td></tr>
<tr><td>>hg38_21_+_8433172_8446622_RNA45SN1_45S_50ntFlanks@12076.12095.20|AGGGGCTCTCGCTTCTGGCG</td><td>>hg38_21_+_8433172_8446622_RNA45SN1_45S_50ntFlanks</td><td>12076</td><td>12095</td><td>20</td><td>AGGGGCTCTCGCTTCTGGCG</td><td>1418.6</td><td>-4.2</td><td>1.2</td><td>-3.5</td><td>5.1E-04</td><td>1.4E-03</td><td>BT20/MB468</td><td>MP</td></tr>
<tr><td>>hg38_21_+_8433172_8446622_RNA45SN1_45S_50ntFlanks@5054.5082.29|TGGAGCGATTTGTCTGGTTAATTCCGATA</td><td>>hg38_21_+_8433172_8446622_RNA45SN1_45S_50ntFlanks</td><td>5054</td><td>5082</td><td>29</td><td>TGGAGCGATTTGTCTGGTTAATTCCGATA</td><td>62.6</td><td>-9.8</td><td>3.0</td><td>-3.3</td><td>9.5E-04</td><td>1.4E-03</td><td>MB231/MB468</td><td>MP</td></tr>
<tr><td>>MI0000482|hsa-mir-185&WithFlank&22|+|20033133|20033226@21.43.23&[MIMAT0000455&hsa-miR-185-5p&offsets|0|+1;m-145&22|+|20033153|20033174&offsets|0|+1]|TGGAGAGAAAGGCAGTTCCTGAT</td><td>>MI0000482|hsa-mir-185&WithFlank&22|+|20033133|20033226</td><td>21</td><td>43</td><td>23</td><td>TGGAGAGAAAGGCAGTTCCTGAT</td><td>537.2</td><td>3.2</td><td>0.9</td><td>3.7</td><td>1.9E-04</td><td>1.4E-03</td><td>BT20/MB231</td><td>CYTO</td></tr>
<tr><td>>hg38_21_+_8433172_8446622_RNA45SN1_45S_50ntFlanks@3916.3938.23|CGTGCATTTATCAGATCAAAACC</td><td>>hg38_21_+_8433172_8446622_RNA45SN1_45S_50ntFlanks</td><td>3916</td><td>3938</td><td>23</td><td>CGTGCATTTATCAGATCAAAACC</td><td>133.8</td><td>-5.4</td><td>1.6</td><td>-3.5</td><td>5.1E-04</td><td>1.4E-03</td><td>BT20/MB468</td><td>MP</td></tr>
<tr><td>>hg38_21_+_8433172_8446622_RNA45SN1_45S_50ntFlanks@10402.10432.31|AACATGGGTCAGTCGGTCCTGAGAGATGGGC</td><td>>hg38_21_+_8433172_8446622_RNA45SN1_45S_50ntFlanks</td><td>10402</td><td>10432</td><td>31</td><td>AACATGGGTCAGTCGGTCCTGAGAGATGGGC</td><td>214.7</td><td>-3.2</td><td>1.0</td><td>-3.3</td><td>1.0E-03</td><td>1.4E-03</td><td>BT20/MB468</td><td>CYTO</td></tr>
<tr><td>>MI0000093|hsa-mir-92a-1&WithFlank&13|+|91351308|91351397@54.76.23&[MIMAT0000092&hsa-miR-92a-3p&offsets|0|+1;m-9&13|+|91351361|91351382&offsets|0|+1]|TATTGCACTTGTCCCGGCCTGTT</td><td>>MI0000093|hsa-mir-92a-1&WithFlank&13|+|91351308|91351397</td><td>54</td><td>76</td><td>23</td><td>TATTGCACTTGTCCCGGCCTGTT</td><td>4125.3</td><td>-3.6</td><td>1.0</td><td>-3.8</td><td>1.8E-04</td><td>1.4E-03</td><td>BT20/MB231</td><td>MP</td></tr>
<tr><td>>hg38_21_+_8433172_8446622_RNA45SN1_45S_50ntFlanks@11655.11687.33|GCGATGTGATTTCTGCCCAGTGCTCTGAATGTC</td><td>>hg38_21_+_8433172_8446622_RNA45SN1_45S_50ntFlanks</td><td>11655</td><td>11687</td><td>33</td><td>GCGATGTGATTTCTGCCCAGTGCTCTGAATGTC</td><td>309.4</td><td>3.1</td><td>0.8</td><td>3.7</td><td>2.2E-04</td><td>1.4E-03</td><td>BT20/MB231</td><td>MITO</td></tr>
<tr><td>>hg38_21_+_8433172_8446622_RNA45SN1_45S_50ntFlanks@7976.8005.30|GCGACCTCAGATCAGACGTGGCGACCCGCT</td><td>>hg38_21_+_8433172_8446622_RNA45SN1_45S_50ntFlanks</td><td>7976</td><td>8005</td><td>30</td><td>GCGACCTCAGATCAGACGTGGCGACCCGCT</td><td>846.8</td><td>3.0</td><td>0.8</td><td>3.7</td><td>2.2E-04</td><td>1.4E-03</td><td>BT20/MB231</td><td>MITO</td></tr>
<tr><td>>hg38_21_+_8433172_8446622_RNA45SN1_45S_50ntFlanks@12446.12476.31|GCAGAATTCACCAAGCGTTGGATTGTTCACC</td><td>>hg38_21_+_8433172_8446622_RNA45SN1_45S_50ntFlanks</td><td>12446</td><td>12476</td><td>31</td><td>GCAGAATTCACCAAGCGTTGGATTGTTCACC</td><td>377.7</td><td>2.6</td><td>0.8</td><td>3.3</td><td>1.1E-03</td><td>1.4E-03</td><td>MB231/MB468</td><td>CYTO</td></tr>
<tr><td>>hg38_21_+_8433172_8446622_RNA45SN1_45S_50ntFlanks@9498.9523.26|AAGATGGTGAACTATGCCTGGGCAGG</td><td>>hg38_21_+_8433172_8446622_RNA45SN1_45S_50ntFlanks</td><td>9498</td><td>9523</td><td>26</td><td>AAGATGGTGAACTATGCCTGGGCAGG</td><td>117.3</td><td>10.2</td><td>2.7</td><td>3.8</td><td>1.8E-04</td><td>1.4E-03</td><td>BT20/MB231</td><td>MP</td></tr>
<tr><td>>hg38_21_+_8433172_8446622_RNA45SN1_45S_50ntFlanks@8105.8137.33|CGCGGCGGGGCGCGGGACATGTGGCGTACGGAA</td><td>>hg38_21_+_8433172_8446622_RNA45SN1_45S_50ntFlanks</td><td>8105</td><td>8137</td><td>33</td><td>CGCGGCGGGGCGCGGGACATGTGGCGTACGGAA</td><td>296.9</td><td>-5.6</td><td>1.7</td><td>-3.3</td><td>9.5E-04</td><td>1.4E-03</td><td>MB231/MB468</td><td>MP</td></tr>
<tr><td>>MI0000088|hsa-mir-30a&WithFlank&6|-|71403545|71403627@53.71.19&[MIMAT0000088&hsa-miR-30a-3p&offsets|0|-3;m-38&6|-|71403554|71403575&offsets|0|-3]|CTTTCAGTCGGATGTTTGC</td><td>>MI0000088|hsa-mir-30a&WithFlank&6|-|71403545|71403627</td><td>53</td><td>71</td><td>19</td><td>CTTTCAGTCGGATGTTTGC</td><td>3931.1</td><td>-3.5</td><td>0.9</td><td>-3.7</td><td>1.8E-04</td><td>1.4E-03</td><td>BT20/MB231</td><td>MP</td></tr>
<tr><td>>hg38_21_+_8433172_8446622_RNA45SN1_45S_50ntFlanks@11600.11621.22|AATCCGACTGTTTAATTAAAAC</td><td>>hg38_21_+_8433172_8446622_RNA45SN1_45S_50ntFlanks</td><td>11600</td><td>11621</td><td>22</td><td>AATCCGACTGTTTAATTAAAAC</td><td>203.6</td><td>7.0</td><td>1.9</td><td>3.7</td><td>1.8E-04</td><td>1.4E-03</td><td>BT20/MB231</td><td>MP</td></tr>
<tr><td>>am_trnaMT_PheGAA_MT_+_577_647@40.62.23|TGTTTAGACGGGCTCACATCACC</td><td>>am_trnaMT_PheGAA_MT_+_577_647</td><td>40</td><td>62</td><td>23</td><td>TGTTTAGACGGGCTCACATCACC</td><td>197.1</td><td>7.1</td><td>2.2</td><td>3.3</td><td>1.1E-03</td><td>1.4E-03</td><td>MB231/MB468</td><td>CYTO</td></tr>
<tr><td>>hg38_21_+_8433172_8446622_RNA45SN1_45S_50ntFlanks@6662.6695.34|GGTGGATCACTCGGCTCGTGCGTCGATGAAGAAC</td><td>>hg38_21_+_8433172_8446622_RNA45SN1_45S_50ntFlanks</td><td>6662</td><td>6695</td><td>34</td><td>GGTGGATCACTCGGCTCGTGCGTCGATGAAGAAC</td><td>164.2</td><td>5.5</td><td>1.6</td><td>3.5</td><td>5.1E-04</td><td>1.4E-03</td><td>BT20/MB468</td><td>MP</td></tr>
<tr><td>>MI0000299|hsa-mir-222&WithFlank&X|-|45747009|45747130@75.99.25(+1U)&[MIMAT0000279&hsa-miR-222-3p&offsets|0|+4(+1U);m-140&X|-|45747033|45747056&offsets|0|+1(+1U)]|AGCTACATCTGGCTACTGGGTCTCTT</td><td>>MI0000299|hsa-mir-222&WithFlank&X|-|45747009|45747130</td><td>75</td><td>99</td><td>25(+1U)</td><td>AGCTACATCTGGCTACTGGGTCTCTT</td><td>2825.4</td><td>-2.6</td><td>0.7</td><td>-3.7</td><td>1.9E-04</td><td>1.4E-03</td><td>BT20/MB231</td><td>NUC</td></tr>
<tr><td>>hg38_21_+_8433172_8446622_RNA45SN1_45S_50ntFlanks@3852.3881.30|TAACTGTGGTAATTCTAGAGCTAATACATG</td><td>>hg38_21_+_8433172_8446622_RNA45SN1_45S_50ntFlanks</td><td>3852</td><td>3881</td><td>30</td><td>TAACTGTGGTAATTCTAGAGCTAATACATG</td><td>116.4</td><td>-5.4</td><td>1.6</td><td>-3.5</td><td>5.1E-04</td><td>1.4E-03</td><td>BT20/MB468</td><td>MP</td></tr>
<tr><td>>hg38_1_-_228634819_228635039_RNA5S12_5S_50ntFlanks@151.169.19|ATACCGGGTGCTGTAGGCT</td><td>>hg38_1_-_228634819_228635039_RNA5S12_5S_50ntFlanks</td><td>151</td><td>169</td><td>19</td><td>ATACCGGGTGCTGTAGGCT</td><td>1860.5</td><td>3.1</td><td>0.8</td><td>3.7</td><td>1.8E-04</td><td>1.4E-03</td><td>BT20/MB231</td><td>MP</td></tr>
<tr><td>>hg38_21_+_8433172_8446622_RNA45SN1_45S_50ntFlanks@5078.5120.43|CGATAACGAACGAGACTCTGGCATGCTAACTAGTTACGCGACC</td><td>>hg38_21_+_8433172_8446622_RNA45SN1_45S_50ntFlanks</td><td>5078</td><td>5120</td><td>43</td><td>CGATAACGAACGAGACTCTGGCATGCTAACTAGTTACGCGACC</td><td>62.0</td><td>-9.8</td><td>3.0</td><td>-3.3</td><td>9.5E-04</td><td>1.4E-03</td><td>MB231/MB468</td><td>MP</td></tr>
<tr><td>>hg38_21_+_8433172_8446622_RNA45SN1_45S_50ntFlanks@474.514.41|TCTTCGTGATCGATGTGGTGACGTCGTGCTCTCCCGGGCCG</td><td>>hg38_21_+_8433172_8446622_RNA45SN1_45S_50ntFlanks</td><td>474</td><td>514</td><td>41</td><td>TCTTCGTGATCGATGTGGTGACGTCGTGCTCTCCCGGGCCG</td><td>305.2</td><td>3.4</td><td>1.1</td><td>3.3</td><td>1.1E-03</td><td>1.4E-03</td><td>MB231/MB468</td><td>NUC</td></tr>
<tr><td>>hg38_21_+_8433172_8446622_RNA45SN1_45S_50ntFlanks@8079.8103.25|AAGAGCCCAGCGCCGAATCCCCGCC</td><td>>hg38_21_+_8433172_8446622_RNA45SN1_45S_50ntFlanks</td><td>8079</td><td>8103</td><td>25</td><td>AAGAGCCCAGCGCCGAATCCCCGCC</td><td>1745.9</td><td>-4.0</td><td>1.1</td><td>-3.5</td><td>5.1E-04</td><td>1.4E-03</td><td>BT20/MB468</td><td>MP</td></tr>
<tr><td>>hg38_21_+_8433172_8446622_RNA45SN1_45S_50ntFlanks@6787.6806.20|ACGCCTGTCTGAGCGTCGCT</td><td>>hg38_21_+_8433172_8446622_RNA45SN1_45S_50ntFlanks</td><td>6787</td><td>6806</td><td>20</td><td>ACGCCTGTCTGAGCGTCGCT</td><td>940.5</td><td>3.1</td><td>0.9</td><td>3.5</td><td>5.1E-04</td><td>1.4E-03</td><td>BT20/MB468</td><td>MP</td></tr>
<tr><td>>hg38_21_+_8433172_8446622_RNA45SN1_45S_50ntFlanks@8358.8378.21|AAAGAACTTTGAAGAGAGAGT</td><td>>hg38_21_+_8433172_8446622_RNA45SN1_45S_50ntFlanks</td><td>8358</td><td>8378</td><td>21</td><td>AAAGAACTTTGAAGAGAGAGT</td><td>845.4</td><td>-3.9</td><td>1.1</td><td>-3.5</td><td>5.1E-04</td><td>1.4E-03</td><td>BT20/MB468</td><td>MP</td></tr>
<tr><td>>hg38_21_+_8433172_8446622_RNA45SN1_45S_50ntFlanks@9462.9503.42|GGGAGGTGGAGCACGAGCGCACGTGTTAGGACCCGAAAGATG</td><td>>hg38_21_+_8433172_8446622_RNA45SN1_45S_50ntFlanks</td><td>9462</td><td>9503</td><td>42</td><td>GGGAGGTGGAGCACGAGCGCACGTGTTAGGACCCGAAAGATG</td><td>212.0</td><td>-5.9</td><td>1.7</td><td>-3.5</td><td>5.1E-04</td><td>1.4E-03</td><td>BT20/MB468</td><td>MP</td></tr>
<tr><td>>hg38_21_+_8433172_8446622_RNA45SN1_45S_50ntFlanks@8105.8140.36|CGCGGCGGGGCGCGGGACATGTGGCGTACGGAAGAC</td><td>>hg38_21_+_8433172_8446622_RNA45SN1_45S_50ntFlanks</td><td>8105</td><td>8140</td><td>36</td><td>CGCGGCGGGGCGCGGGACATGTGGCGTACGGAAGAC</td><td>1543.4</td><td>-5.1</td><td>1.6</td><td>-3.3</td><td>1.1E-03</td><td>1.4E-03</td><td>BT20/MB468</td><td>NUC</td></tr>
<tr><td>>MI0000264|hsa-mir-7-2&WithFlank&15|+|88611819|88611940@38.59.22(+1C);MI0000265|hsa-mir-7-3&WithFlank&19|+|4770664|4770785@37.58.22(+1C);MI0000263|hsa-mir-7-1&WithFlank&9|-|83969742|83969863@30.51.22(+1C)&[MIMAT0000252&hsa-miR-7-5p&offsets|0|-2(+1C);m-400&15|+|88611856|88611879&offsets|0|-2(+1C)];[MIMAT0000252_1&hsa-miR-7-5p&offsets|0|-2(+1C);m-402&19|+|4770700|4770723&offsets|0|-2(+1C)];[MIMAT0000252_2&hsa-miR-7-5p&offsets|0|-2(+1C);m-401&9|-|83969811|83969834&offsets|0|-2(+1C)]|TGGAAGACTAGTGATTTTGTTGC</td><td>>MI0000264|hsa-mir-7-2&WithFlank&15|+|88611819|88611940</td><td>38</td><td>59</td><td>22(+1C)</td><td>TGGAAGACTAGTGATTTTGTTGC</td><td>863.9</td><td>3.7</td><td>1.1</td><td>3.5</td><td>5.1E-04</td><td>1.4E-03</td><td>BT20/MB468</td><td>MP</td></tr>
<tr><td>>hg38_21_+_8433172_8446622_RNA45SN1_45S_50ntFlanks@3838.3860.23|CTCTCCTACTTGGATAACTGTGG</td><td>>hg38_21_+_8433172_8446622_RNA45SN1_45S_50ntFlanks</td><td>3838</td><td>3860</td><td>23</td><td>CTCTCCTACTTGGATAACTGTGG</td><td>523.6</td><td>-2.5</td><td>0.8</td><td>-3.3</td><td>8.6E-04</td><td>1.4E-03</td><td>BT20/MB468</td><td>MITO</td></tr>
<tr><td>>hg38_21_+_8433172_8446622_RNA45SN1_45S_50ntFlanks@7978.8017.40|GACCTCAGATCAGACGTGGCGACCCGCTGAATTTAAGCAT</td><td>>hg38_21_+_8433172_8446622_RNA45SN1_45S_50ntFlanks</td><td>7978</td><td>8017</td><td>40</td><td>GACCTCAGATCAGACGTGGCGACCCGCTGAATTTAAGCAT</td><td>577.0</td><td>4.6</td><td>1.4</td><td>3.3</td><td>1.1E-03</td><td>1.4E-03</td><td>MB231/MB468</td><td>MITO</td></tr>
<tr><td>>hg38_21_+_8433172_8446622_RNA45SN1_45S_50ntFlanks@11718.11744.27|AACGGCGGGAGTAACTATGACTCTCTT</td><td>>hg38_21_+_8433172_8446622_RNA45SN1_45S_50ntFlanks</td><td>11718</td><td>11744</td><td>27</td><td>AACGGCGGGAGTAACTATGACTCTCTT</td><td>660.9</td><td>-3.4</td><td>1.0</td><td>-3.3</td><td>1.0E-03</td><td>1.4E-03</td><td>BT20/MB468</td><td>CYTO</td></tr>
<tr><td>>MI0000085|hsa-mir-27a&WithFlank&19|-|13836434|13836523@57.77.21(+1U)&[MIMAT0000084&hsa-miR-27a-3p&offsets|0|0(+1U);m-43&19|-|13836447|13836467&offsets|0|0(+1U)]|TTCACAGTGGCTAAGTTCCGCT</td><td>>MI0000085|hsa-mir-27a&WithFlank&19|-|13836434|13836523</td><td>57</td><td>77</td><td>21(+1U)</td><td>TTCACAGTGGCTAAGTTCCGCT</td><td>765.4</td><td>3.4</td><td>1.0</td><td>3.3</td><td>9.6E-04</td><td>1.4E-03</td><td>MB231/MB468</td><td>MP</td></tr>
<tr><td>>hg38_21_+_8433172_8446622_RNA45SN1_45S_50ntFlanks@9621.9650.30|TCTAGTAGCTGGTTCCCTCCGAAGTTTCCC</td><td>>hg38_21_+_8433172_8446622_RNA45SN1_45S_50ntFlanks</td><td>9621</td><td>9650</td><td>30</td><td>TCTAGTAGCTGGTTCCCTCCGAAGTTTCCC</td><td>156.4</td><td>-5.4</td><td>1.5</td><td>-3.5</td><td>5.1E-04</td><td>1.4E-03</td><td>BT20/MB468</td><td>MP</td></tr>
<tr><td>>hg38_21_+_8433172_8446622_RNA45SN1_45S_50ntFlanks@9855.9880.26|CTGCGGGATGAACCGAACGCCGGGTT</td><td>>hg38_21_+_8433172_8446622_RNA45SN1_45S_50ntFlanks</td><td>9855</td><td>9880</td><td>26</td><td>CTGCGGGATGAACCGAACGCCGGGTT</td><td>126.9</td><td>-3.5</td><td>1.1</td><td>-3.3</td><td>9.6E-04</td><td>1.4E-03</td><td>MB231/MB468</td><td>MP</td></tr>
<tr><td>>hg38_21_+_8433172_8446622_RNA45SN1_45S_50ntFlanks@4731.4753.23|AAGAACGAAAGTCGGAGGTTCGA</td><td>>hg38_21_+_8433172_8446622_RNA45SN1_45S_50ntFlanks</td><td>4731</td><td>4753</td><td>23</td><td>AAGAACGAAAGTCGGAGGTTCGA</td><td>130.1</td><td>-4.0</td><td>1.2</td><td>-3.3</td><td>9.6E-04</td><td>1.4E-03</td><td>MB231/MB468</td><td>MP</td></tr>
<tr><td>>hg38_21_+_8433172_8446622_RNA45SN1_45S_50ntFlanks@3831.3871.41|CTCGCTCCTCTCCTACTTGGATAACTGTGGTAATTCTAGAG</td><td>>hg38_21_+_8433172_8446622_RNA45SN1_45S_50ntFlanks</td><td>3831</td><td>3871</td><td>41</td><td>CTCGCTCCTCTCCTACTTGGATAACTGTGGTAATTCTAGAG</td><td>294.9</td><td>-4.3</td><td>1.3</td><td>-3.3</td><td>1.2E-03</td><td>1.4E-03</td><td>MB231/MB468</td><td>NUC</td></tr>
<tr><td>>trnaMT_ValTAC_MT_+_1602_1670@41.72.32|GAGATTTCAACTTAACTTGACCGCTCTGACCA</td><td>>trnaMT_ValTAC_MT_+_1602_1670</td><td>41</td><td>72</td><td>32</td><td>GAGATTTCAACTTAACTTGACCGCTCTGACCA</td><td>1865.8</td><td>3.6</td><td>1.1</td><td>3.3</td><td>8.6E-04</td><td>1.4E-03</td><td>BT20/MB468</td><td>MITO</td></tr>
<tr><td>>hg38_21_+_8433172_8446622_RNA45SN1_45S_50ntFlanks@11767.11790.24|TCTAATTAGTGACGCGCATGAATG</td><td>>hg38_21_+_8433172_8446622_RNA45SN1_45S_50ntFlanks</td><td>11767</td><td>11790</td><td>24</td><td>TCTAATTAGTGACGCGCATGAATG</td><td>286.1</td><td>-4.8</td><td>1.5</td><td>-3.3</td><td>1.0E-03</td><td>1.4E-03</td><td>BT20/MB468</td><td>CYTO</td></tr>
<tr><td>>hg38_21_+_8433172_8446622_RNA45SN1_45S_50ntFlanks@12418.12448.31|CGATGTCGGCTCTTCCTATCATTGTGAAGCA</td><td>>hg38_21_+_8433172_8446622_RNA45SN1_45S_50ntFlanks</td><td>12418</td><td>12448</td><td>31</td><td>CGATGTCGGCTCTTCCTATCATTGTGAAGCA</td><td>641.5</td><td>-3.4</td><td>1.1</td><td>-3.3</td><td>1.1E-03</td><td>1.4E-03</td><td>BT20/MB468</td><td>NUC</td></tr>
<tr><td>>hg38_21_+_8433172_8446622_RNA45SN1_45S_50ntFlanks@6651.6675.25|CGACTCTTAGCGGTGGATCACTCGG</td><td>>hg38_21_+_8433172_8446622_RNA45SN1_45S_50ntFlanks</td><td>6651</td><td>6675</td><td>25</td><td>CGACTCTTAGCGGTGGATCACTCGG</td><td>1501.1</td><td>-2.8</td><td>0.8</td><td>-3.3</td><td>9.6E-04</td><td>1.4E-03</td><td>MB231/MB468</td><td>MP</td></tr>
<tr><td>>hg38_21_+_8433172_8446622_RNA45SN1_45S_50ntFlanks@10421.10453.33|TGAGAGATGGGCGAGCGCCGTTCCGAAGGGACG</td><td>>hg38_21_+_8433172_8446622_RNA45SN1_45S_50ntFlanks</td><td>10421</td><td>10453</td><td>33</td><td>TGAGAGATGGGCGAGCGCCGTTCCGAAGGGACG</td><td>854.8</td><td>-2.4</td><td>0.7</td><td>-3.3</td><td>1.1E-03</td><td>1.4E-03</td><td>MB231/MB468</td><td>CYTO</td></tr>
<tr><td>>trnaMT_PheGAA_MT_+_577_647@28.64.37|ATACACTGAAAATGTTTAGACGGGCTCACATCACCCC</td><td>>trnaMT_PheGAA_MT_+_577_647</td><td>28</td><td>64</td><td>37</td><td>ATACACTGAAAATGTTTAGACGGGCTCACATCACCCC</td><td>55.9</td><td>9.6</td><td>2.8</td><td>3.5</td><td>5.1E-04</td><td>1.4E-03</td><td>BT20/MB468</td><td>MP</td></tr>
<tr><td>>hg38_21_+_8433172_8446622_RNA45SN1_45S_50ntFlanks@11598.11621.24|GGAATCCGACTGTTTAATTAAAAC</td><td>>hg38_21_+_8433172_8446622_RNA45SN1_45S_50ntFlanks</td><td>11598</td><td>11621</td><td>24</td><td>GGAATCCGACTGTTTAATTAAAAC</td><td>79.8</td><td>5.5</td><td>1.6</td><td>3.5</td><td>5.1E-04</td><td>1.4E-03</td><td>BT20/MB468</td><td>MP</td></tr>
<tr><td>>hg38_21_+_8433172_8446622_RNA45SN1_45S_50ntFlanks@11604.11627.24|CGACTGTTTAATTAAAACAAAGCA</td><td>>hg38_21_+_8433172_8446622_RNA45SN1_45S_50ntFlanks</td><td>11604</td><td>11627</td><td>24</td><td>CGACTGTTTAATTAAAACAAAGCA</td><td>265.4</td><td>6.9</td><td>1.8</td><td>3.7</td><td>1.8E-04</td><td>1.4E-03</td><td>BT20/MB231</td><td>MP</td></tr>
<tr><td>>hg38_21_+_8433172_8446622_RNA45SN1_45S_50ntFlanks@6786.6807.22|TACGCCTGTCTGAGCGTCGCTT</td><td>>hg38_21_+_8433172_8446622_RNA45SN1_45S_50ntFlanks</td><td>6786</td><td>6807</td><td>22</td><td>TACGCCTGTCTGAGCGTCGCTT</td><td>377.8</td><td>2.8</td><td>0.9</td><td>3.3</td><td>1.1E-03</td><td>1.4E-03</td><td>MB231/MB468</td><td>MITO</td></tr>
<tr><td>>hg38_21_+_8433172_8446622_RNA45SN1_45S_50ntFlanks@9574.9603.30|AATCGGTCGTCCGACCTGGGTATAGGGGCG</td><td>>hg38_21_+_8433172_8446622_RNA45SN1_45S_50ntFlanks</td><td>9574</td><td>9603</td><td>30</td><td>AATCGGTCGTCCGACCTGGGTATAGGGGCG</td><td>6713.8</td><td>-2.7</td><td>0.8</td><td>-3.3</td><td>1.1E-03</td><td>1.4E-03</td><td>MB231/MB468</td><td>MITO</td></tr>
<tr><td>>hg38_21_+_8433172_8446622_RNA45SN1_45S_50ntFlanks@12412.12441.30|ATCCTTCGATGTCGGCTCTTCCTATCATTG</td><td>>hg38_21_+_8433172_8446622_RNA45SN1_45S_50ntFlanks</td><td>12412</td><td>12441</td><td>30</td><td>ATCCTTCGATGTCGGCTCTTCCTATCATTG</td><td>836.0</td><td>-5.3</td><td>1.6</td><td>-3.2</td><td>1.2E-03</td><td>1.4E-03</td><td>MB231/MB468</td><td>NUC</td></tr>
</table>

<table>
<tr><td>>MI0000471|hsa-mir-126&WithFlank&9|+|136670596|136670692@58.77.20&[MIMAT0000445&hsa-miR-126-3p&offsets|0|-2;m-40&9|+|136670653|136670674&offsets|0|-2]|TCGTACCGTGAGTAATAATG</td><td>>MI0000471|hsa-mir-126&WithFlank&9|+|136670596|136670692</td><td>58</td><td>77</td><td>20</td><td>TCGTACCGTGAGTAATAATG</td><td>325.0</td><td>6.3</td><td>1.9</td><td>3.3</td><td>9.6E-04</td><td>1.4E-03</td><td>MB231/MB468</td><td>MP</td></tr>
<tr><td>>hg38_21_+_8433172_8446622_RNA45SN1_45S_50ntFlanks@3835.3869.35|CTCCTCTCCTACTTGGATAACTGTGGTAATTCTAG</td><td>>hg38_21_+_8433172_8446622_RNA45SN1_45S_50ntFlanks</td><td>3835</td><td>3869</td><td>35</td><td>CTCCTCTCCTACTTGGATAACTGTGGTAATTCTAG</td><td>254.5</td><td>-2.2</td><td>0.7</td><td>-3.3</td><td>1.1E-03</td><td>1.4E-03</td><td>MB231/MB468</td><td>CYTO</td></tr>
<tr><td>>hg38_21_+_8433172_8446622_RNA45SN1_45S_50ntFlanks@11873.11908.36|AAGAAGACCCTGTTGAGCTTGACTCTAGTCTGGCAC</td><td>>hg38_21_+_8433172_8446622_RNA45SN1_45S_50ntFlanks</td><td>11873</td><td>11908</td><td>36</td><td>AAGAAGACCCTGTTGAGCTTGACTCTAGTCTGGCAC</td><td>298.3</td><td>-5.1</td><td>1.6</td><td>-3.2</td><td>1.2E-03</td><td>1.4E-03</td><td>MB231/MB468</td><td>NUC</td></tr>
<tr><td>>hg38_21_+_8433172_8446622_RNA45SN1_45S_50ntFlanks@12914.12933.20|CGTTCGTGGGGAACCTGGCG</td><td>>hg38_21_+_8433172_8446622_RNA45SN1_45S_50ntFlanks</td><td>12914</td><td>12933</td><td>20</td><td>CGTTCGTGGGGAACCTGGCG</td><td>1135.0</td><td>-3.4</td><td>1.0</td><td>-3.5</td><td>5.2E-04</td><td>1.4E-03</td><td>BT20/MB468</td><td>MP</td></tr>
<tr><td>>hg38_21_+_8433172_8446622_RNA45SN1_45S_50ntFlanks@12444.12484.41|AAGCAGAATTCACCAAGCGTTGGATTGTTCACCCACTAATA</td><td>>hg38_21_+_8433172_8446622_RNA45SN1_45S_50ntFlanks</td><td>12444</td><td>12484</td><td>41</td><td>AAGCAGAATTCACCAAGCGTTGGATTGTTCACCCACTAATA</td><td>236.0</td><td>-5.1</td><td>1.5</td><td>-3.3</td><td>1.0E-03</td><td>1.4E-03</td><td>BT20/MB468</td><td>CYTO</td></tr>
<tr><td>>hg38_21_+_8433172_8446622_RNA45SN1_45S_50ntFlanks@6700.6719.20|CTAGCTGCGAGAATTAATGT</td><td>>hg38_21_+_8433172_8446622_RNA45SN1_45S_50ntFlanks</td><td>6700</td><td>6719</td><td>20</td><td>CTAGCTGCGAGAATTAATGT</td><td>524.2</td><td>-5.7</td><td>1.6</td><td>-3.5</td><td>5.2E-04</td><td>1.4E-03</td><td>BT20/MB468</td><td>MP</td></tr>
<tr><td>>hg38_21_+_8433172_8446622_RNA45SN1_45S_50ntFlanks@11565.11611.47|CTAGCAGCCGACTTAGAACTGGTGCGGACCAGGGGAATCCGACTGTT</td><td>>hg38_21_+_8433172_8446622_RNA45SN1_45S_50ntFlanks</td><td>11565</td><td>11611</td><td>47</td><td>CTAGCAGCCGACTTAGAACTGGTGCGGACCAGGGGAATCCGACTGTT</td><td>2887.9</td><td>3.9</td><td>1.2</td><td>3.3</td><td>8.7E-04</td><td>1.4E-03</td><td>BT20/MB468</td><td>MITO</td></tr>
<tr><td>>hg38_21_+_8433172_8446622_RNA45SN1_45S_50ntFlanks@11572.11607.36|CCGACTTAGAACTGGTGCGGACCAGGGGAATCCGAC</td><td>>hg38_21_+_8433172_8446622_RNA45SN1_45S_50ntFlanks</td><td>11572</td><td>11607</td><td>36</td><td>CCGACTTAGAACTGGTGCGGACCAGGGGAATCCGAC</td><td>307.5</td><td>-1.9</td><td>0.6</td><td>-3.3</td><td>1.1E-03</td><td>1.4E-03</td><td>MB231/MB468</td><td>CYTO</td></tr>
<tr><td>>hg38_21_+_8433172_8446622_RNA45SN1_45S_50ntFlanks@10348.10365.18|AAACGAGAACTTTGAAGG</td><td>>hg38_21_+_8433172_8446622_RNA45SN1_45S_50ntFlanks</td><td>10348</td><td>10365</td><td>18</td><td>AAACGAGAACTTTGAAGG</td><td>526.0</td><td>-4.4</td><td>1.3</td><td>-3.3</td><td>9.6E-04</td><td>1.4E-03</td><td>MB231/MB468</td><td>MP</td></tr>
<tr><td>>trnaMT_LysTTT_MT_+_8295_8364@56.73.18,trnalookalike7_LysTTT_1_+_568844_568913@56.73.18|ACCTCTTTACAGTGACCA</td><td>>trnaMT_LysTTT_MT_+_8295_8364</td><td>56</td><td>73</td><td>18</td><td>ACCTCTTTACAGTGACCA</td><td>120.6</td><td>4.9</td><td>1.5</td><td>3.3</td><td>9.6E-04</td><td>1.4E-03</td><td>MB231/MB468</td><td>MP</td></tr>
<tr><td>>hg38_21_+_8433172_8446622_RNA45SN1_45S_50ntFlanks@11582.11603.22|ACTGGTGCGGACCAGGGGAATC</td><td>>hg38_21_+_8433172_8446622_RNA45SN1_45S_50ntFlanks</td><td>11582</td><td>11603</td><td>22</td><td>ACTGGTGCGGACCAGGGGAATC</td><td>1411.2</td><td>-1.7</td><td>0.5</td><td>-3.3</td><td>1.0E-03</td><td>1.4E-03</td><td>BT20/MB468</td><td>CYTO</td></tr>
<tr><td>>hg38_21_+_8433172_8446622_RNA45SN1_45S_50ntFlanks@10854.10873.20|TAAGGATTGGCTCTAAGGGC</td><td>>hg38_21_+_8433172_8446622_RNA45SN1_45S_50ntFlanks</td><td>10854</td><td>10873</td><td>20</td><td>TAAGGATTGGCTCTAAGGGC</td><td>165.8</td><td>-2.4</td><td>0.7</td><td>-3.3</td><td>1.0E-03</td><td>1.4E-03</td><td>BT20/MB468</td><td>CYTO</td></tr>
<tr><td>>hg38_21_+_8433172_8446622_RNA45SN1_45S_50ntFlanks@11569.11597.29|CAGCCGACTTAGAACTGGTGCGGACCAGG</td><td>>hg38_21_+_8433172_8446622_RNA45SN1_45S_50ntFlanks</td><td>11569</td><td>11597</td><td>29</td><td>CAGCCGACTTAGAACTGGTGCGGACCAGG</td><td>307.4</td><td>-2.4</td><td>0.7</td><td>-3.2</td><td>1.2E-03</td><td>1.4E-03</td><td>MB231/MB468</td><td>NUC</td></tr>
<tr><td>>hg38_21_+_8433172_8446622_RNA45SN1_45S_50ntFlanks@11564.11603.40|CCTAGCAGCCGACTTAGAACTGGTGCGGACCAGGGGAATC</td><td>>hg38_21_+_8433172_8446622_RNA45SN1_45S_50ntFlanks</td><td>11564</td><td>11603</td><td>40</td><td>CCTAGCAGCCGACTTAGAACTGGTGCGGACCAGGGGAATC</td><td>748.0</td><td>-4.2</td><td>1.3</td><td>-3.3</td><td>1.0E-03</td><td>1.4E-03</td><td>BT20/MB468</td><td>CYTO</td></tr>
<tr><td>>MI0000811|hsa-mir-148b&WithFlank&12|+|54337210|54337320@69.90.22(+3U)&[MIMAT0000759&hsa-miR-148b-3p&offsets|0|0(+3U);m-77&12|+|54337278|54337299&offsets|0|0(+3U)]|TCAGTGCATCACAGAACTTTGTTTT</td><td>>MI0000811|hsa-mir-148b&WithFlank&12|+|54337210|54337320</td><td>69</td><td>90</td><td>22(+3U)</td><td>TCAGTGCATCACAGAACTTTGTTTT</td><td>184.2</td><td>4.1</td><td>1.2</td><td>3.3</td><td>9.7E-04</td><td>1.4E-03</td><td>MB231/MB468</td><td>MP</td></tr>
<tr><td>>hg38_21_+_8433172_8446622_RNA45SN1_45S_50ntFlanks@6659.6681.23|AGCGGTGGATCACTCGGCTCGTG</td><td>>hg38_21_+_8433172_8446622_RNA45SN1_45S_50ntFlanks</td><td>6659</td><td>6681</td><td>23</td><td>AGCGGTGGATCACTCGGCTCGTG</td><td>2004.7</td><td>-3.1</td><td>0.9</td><td>-3.5</td><td>5.2E-04</td><td>1.4E-03</td><td>BT20/MB468</td><td>MP</td></tr>
<tr><td>>hg38_1_-_228634819_228635039_RNA5S12_5S_50ntFlanks@133.170.38|ATGGGAGACCGCCTGGGAATACCGGGTGCTGTAGGCTT</td><td>>hg38_1_-_228634819_228635039_RNA5S12_5S_50ntFlanks</td><td>133</td><td>170</td><td>38</td><td>ATGGGAGACCGCCTGGGAATACCGGGTGCTGTAGGCTT</td><td>18782.6</td><td>3.3</td><td>0.9</td><td>3.7</td><td>2.2E-04</td><td>1.4E-03</td><td>BT20/MB231</td><td>MITO</td></tr>
<tr><td>>MI0000816|hsa-mir-335&WithFlank&7|+|130496105|130496210@58.78.21(+1U)&[MIMAT0004703&hsa-miR-335-3p&offsets|0|-1(+1U);m-159&7|+|130496162|130496183&offsets|0|-1(+1U)]|TTTTTCATTATTGCTCCTGACT</td><td>>MI0000816|hsa-mir-335&WithFlank&7|+|130496105|130496210</td><td>58</td><td>78</td><td>21(+1U)</td><td>TTTTTCATTATTGCTCCTGACT</td><td>1458.5</td><td>-3.3</td><td>0.9</td><td>-3.7</td><td>2.2E-04</td><td>1.4E-03</td><td>BT20/MB231</td><td>MITO</td></tr>
<tr><td>>hg38_21_+_8433172_8446622_RNA45SN1_45S_50ntFlanks@12418.12449.32|CGATGTCGGCTCTTCCTATCATTGTGAAGCAG</td><td>>hg38_21_+_8433172_8446622_RNA45SN1_45S_50ntFlanks</td><td>12418</td><td>12449</td><td>32</td><td>CGATGTCGGCTCTTCCTATCATTGTGAAGCAG</td><td>394.6</td><td>-3.0</td><td>0.8</td><td>-3.7</td><td>2.2E-04</td><td>1.4E-03</td><td>BT20/MB231</td><td>MITO</td></tr>
<tr><td>>hg38_21_+_8433172_8446622_RNA45SN1_45S_50ntFlanks@485.525.41|GATGTGGTGACGTCGTGCTCTCCCGGGCCGGGTCCGAGCCG</td><td>>hg38_21_+_8433172_8446622_RNA45SN1_45S_50ntFlanks</td><td>485</td><td>525</td><td>41</td><td>GATGTGGTGACGTCGTGCTCTCCCGGGCCGGGTCCGAGCCG</td><td>383.2</td><td>11.8</td><td>3.6</td><td>3.3</td><td>1.1E-03</td><td>1.4E-03</td><td>MB231/MB468</td><td>MITO</td></tr>
<tr><td>>hg38_MT_+_1621_3279_RNR2_16S_50ntFlanks@1182.1215.34|AAAATTTCGGTTGGGGCGACCTCGGAGCAGAACC</td><td>>hg38_MT_+_1621_3279_RNR2_16S_50ntFlanks</td><td>1182</td><td>1215</td><td>34</td><td>AAAATTTCGGTTGGGGCGACCTCGGAGCAGAACC</td><td>126.1</td><td>4.9</td><td>1.5</td><td>3.3</td><td>1.2E-03</td><td>1.4E-03</td><td>MB231/MB468</td><td>MITO</td></tr>
<tr><td>>hg38_21_+_8433172_8446622_RNA45SN1_45S_50ntFlanks@10786.10828.43|AACAGCCTCTGGCATGTTGGAACAATGTAGGTAAGGGAAGTCG</td><td>>hg38_21_+_8433172_8446622_RNA45SN1_45S_50ntFlanks</td><td>10786</td><td>10828</td><td>43</td><td>AACAGCCTCTGGCATGTTGGAACAATGTAGGTAAGGGAAGTCG</td><td>827.9</td><td>-4.3</td><td>1.3</td><td>-3.3</td><td>8.7E-04</td><td>1.4E-03</td><td>BT20/MB468</td><td>MITO</td></tr>
<tr><td>>hg38_21_+_8433172_8446622_RNA45SN1_45S_50ntFlanks@11563.11584.22|GCCTAGCAGCCGACTTAGAACT</td><td>>hg38_21_+_8433172_8446622_RNA45SN1_45S_50ntFlanks</td><td>11563</td><td>11584</td><td>22</td><td>GCCTAGCAGCCGACTTAGAACT</td><td>95.6</td><td>5.1</td><td>1.6</td><td>3.3</td><td>1.2E-03</td><td>1.4E-03</td><td>MB231/MB468</td><td>MITO</td></tr>
<tr><td>>hg38_21_+_8433172_8446622_RNA45SN1_45S_50ntFlanks@4991.5018.28|ATTGACAGATTGATAGCTCTTTCTCGAT</td><td>>hg38_21_+_8433172_8446622_RNA45SN1_45S_50ntFlanks</td><td>4991</td><td>5018</td><td>28</td><td>ATTGACAGATTGATAGCTCTTTCTCGAT</td><td>55.0</td><td>9.5</td><td>2.7</td><td>3.5</td><td>5.2E-04</td><td>1.4E-03</td><td>BT20/MB468</td><td>MP</td></tr>
<tr><td>>hg38_21_+_8433172_8446622_RNA45SN1_45S_50ntFlanks@10392.10419.28|ACAGCAGTTGAACATGGGTCAGTCGGTC</td><td>>hg38_21_+_8433172_8446622_RNA45SN1_45S_50ntFlanks</td><td>10392</td><td>10419</td><td>28</td><td>ACAGCAGTTGAACATGGGTCAGTCGGTC</td><td>150.8</td><td>-2.1</td><td>0.6</td><td>-3.3</td><td>8.8E-04</td><td>1.4E-03</td><td>BT20/MB468</td><td>MITO</td></tr>
<tr><td>>hg38_21_+_8433172_8446622_RNA45SN1_45S_50ntFlanks@6650.6679.30|ACGACTCTTAGCGGTGGATCACTCGGCTCG</td><td>>hg38_21_+_8433172_8446622_RNA45SN1_45S_50ntFlanks</td><td>6650</td><td>6679</td><td>30</td><td>ACGACTCTTAGCGGTGGATCACTCGGCTCG</td><td>2845.4</td><td>5.4</td><td>1.4</td><td>3.7</td><td>1.8E-04</td><td>1.4E-03</td><td>BT20/MB231</td><td>MP</td></tr>
<tr><td>>hg38_MT_+_1621_3279_RNR2_16S_50ntFlanks@826.855.30|AAGGAAAGGTTAAAAAAAGTAAAAGGAACT</td><td>>hg38_MT_+_1621_3279_RNR2_16S_50ntFlanks</td><td>826</td><td>855</td><td>30</td><td>AAGGAAAGGTTAAAAAAAGTAAAAGGAACT</td><td>263.3</td><td>4.1</td><td>1.3</td><td>3.3</td><td>9.7E-04</td><td>1.4E-03</td><td>MB231/MB468</td><td>MP</td></tr>
<tr><td>>hg38_21_+_8433172_8446622_RNA45SN1_45S_50ntFlanks@11641.11663.23|CGGCGGGTGTTGACGCGATGTGA</td><td>>hg38_21_+_8433172_8446622_RNA45SN1_45S_50ntFlanks</td><td>11641</td><td>11663</td><td>23</td><td>CGGCGGGTGTTGACGCGATGTGA</td><td>745.9</td><td>-5.3</td><td>1.6</td><td>-3.3</td><td>8.8E-04</td><td>1.4E-03</td><td>BT20/MB468</td><td>MITO</td></tr>
<tr><td>>hg38_21_+_8433172_8446622_RNA45SN1_45S_50ntFlanks@11319.11342.24|CCCACGTCTCGTCGCGCGCGCGTC</td><td>>hg38_21_+_8433172_8446622_RNA45SN1_45S_50ntFlanks</td><td>11319</td><td>11342</td><td>24</td><td>CCCACGTCTCGTCGCGCGCGCGTC</td><td>2056.2</td><td>-6.6</td><td>2.0</td><td>-3.2</td><td>1.2E-03</td><td>1.4E-03</td><td>BT20/MB468</td><td>NUC</td></tr>
<tr><td>>hg38_21_+_8433172_8446622_RNA45SN1_45S_50ntFlanks@7976.8011.36|GCGACCTCAGATCAGACGTGGCGACCCGCTGAATTT</td><td>>hg38_21_+_8433172_8446622_RNA45SN1_45S_50ntFlanks</td><td>7976</td><td>8011</td><td>36</td><td>GCGACCTCAGATCAGACGTGGCGACCCGCTGAATTT</td><td>491.5</td><td>2.5</td><td>0.8</td><td>3.2</td><td>1.2E-03</td><td>1.4E-03</td><td>BT20/MB468</td><td>NUC</td></tr>
<tr><td>>hg38_21_+_8433172_8446622_RNA45SN1_45S_50ntFlanks@3881.3898.18|GCCGACGGGCGCTGACCC</td><td>>hg38_21_+_8433172_8446622_RNA45SN1_45S_50ntFlanks</td><td>3881</td><td>3898</td><td>18</td><td>GCCGACGGGCGCTGACCC</td><td>394.0</td><td>-3.0</td><td>0.9</td><td>-3.3</td><td>1.1E-03</td><td>1.4E-03</td><td>BT20/MB468</td><td>CYTO</td></tr>
<tr><td>>hg38_21_+_8433172_8446622_RNA45SN1_45S_50ntFlanks@475.520.46|CTTCGTGATCGATGTGGTGACGTCGTGCTCTCCCGGGCCGGGTCCG</td><td>>hg38_21_+_8433172_8446622_RNA45SN1_45S_50ntFlanks</td><td>475</td><td>520</td><td>46</td><td>CTTCGTGATCGATGTGGTGACGTCGTGCTCTCCCGGGCCGGGTCCG</td><td>511.1</td><td>5.2</td><td>1.6</td><td>3.2</td><td>1.2E-03</td><td>1.4E-03</td><td>MB231/MB468</td><td>MITO</td></tr>
<tr><td>>MI0000434|hsa-let-7i&WithFlank&12|+|62603680|62603775@12.33.22(+1C)&[MIMAT0000415&hsa-let-7i-5p&offsets|0|0(+1C);m-49&12|+|62603691|62603712&offsets|0|0(+1C)]|TGAGGTAGTAGTTTGTGCTGTTC</td><td>>MI0000434|hsa-let-7i&WithFlank&12|+|62603680|62603775</td><td>12</td><td>33</td><td>22(+1C)</td><td>TGAGGTAGTAGTTTGTGCTGTTC</td><td>878.1</td><td>5.3</td><td>1.5</td><td>3.5</td><td>5.2E-04</td><td>1.4E-03</td><td>BT20/MB468</td><td>MP</td></tr>
<tr><td>>hg38_21_+_8433172_8446622_RNA45SN1_45S_50ntFlanks@8179.8210.32|CTGATCGAGGCCCAGCCCGTGGACGGTGTGAG</td><td>>hg38_21_+_8433172_8446622_RNA45SN1_45S_50ntFlanks</td><td>8179</td><td>8210</td><td>32</td><td>CTGATCGAGGCCCAGCCCGTGGACGGTGTGAG</td><td>218.0</td><td>-4.3</td><td>1.3</td><td>-3.5</td><td>5.2E-04</td><td>1.4E-03</td><td>BT20/MB468</td><td>MP</td></tr>
<tr><td>>hg38_21_+_8433172_8446622_RNA45SN1_45S_50ntFlanks@6647.6666.20|CGTACGACTCTTAGCGGTGG</td><td>>hg38_21_+_8433172_8446622_RNA45SN1_45S_50ntFlanks</td><td>6647</td><td>6666</td><td>20</td><td>CGTACGACTCTTAGCGGTGG</td><td>3335.4</td><td>2.8</td><td>0.8</td><td>3.5</td><td>5.2E-04</td><td>1.4E-03</td><td>BT20/MB468</td><td>MP</td></tr>
<tr><td>>trnaMT_PheGAA_MT_+_577_647@39.58.20|ATGTTTAGACGGGCTCACAT</td><td>>trnaMT_PheGAA_MT_+_577_647</td><td>39</td><td>58</td><td>20</td><td>ATGTTTAGACGGGCTCACAT</td><td>133.3</td><td>6.8</td><td>2.0</td><td>3.5</td><td>5.2E-04</td><td>1.4E-03</td><td>BT20/MB468</td><td>MP</td></tr>
<tr><td>>hg38_21_+_8433172_8446622_RNA45SN1_45S_50ntFlanks@6747.6770.24|ACTTCGAACGCACTTGCGGCCCCG</td><td>>hg38_21_+_8433172_8446622_RNA45SN1_45S_50ntFlanks</td><td>6747</td><td>6770</td><td>24</td><td>ACTTCGAACGCACTTGCGGCCCCG</td><td>1219.8</td><td>-6.0</td><td>1.7</td><td>-3.5</td><td>5.2E-04</td><td>1.4E-03</td><td>BT20/MB468</td><td>MP</td></tr>
<tr><td>>hg38_21_+_8433172_8446622_RNA45SN1_45S_50ntFlanks@9204.9222.19|TCGGGGCCACGCGCGCGTC</td><td>>hg38_21_+_8433172_8446622_RNA45SN1_45S_50ntFlanks</td><td>9204</td><td>9222</td><td>19</td><td>TCGGGGCCACGCGCGCGTC</td><td>1459.7</td><td>-6.1</td><td>1.8</td><td>-3.5</td><td>5.2E-04</td><td>1.4E-03</td><td>BT20/MB468</td><td>MP</td></tr>
<tr><td>>MI0000077|hsa-mir-21&WithFlank&17|+|59841260|59841343@14.35.22&[MIMAT0000076&hsa-miR-21-5p&offsets|0|0;m-1&17|+|59841273|59841294&offsets|0|0]|TAGCTTATCAGACTGATGTTGA</td><td>>MI0000077|hsa-mir-21&WithFlank&17|+|59841260|59841343</td><td>14</td><td>35</td><td>22</td><td>TAGCTTATCAGACTGATGTTGA</td><td>386483.9</td><td>2.6</td><td>0.8</td><td>3.3</td><td>9.8E-04</td><td>1.4E-03</td><td>MB231/MB468</td><td>MP</td></tr>
<tr><td>>hg38_MT_+_1621_3279_RNR2_16S_50ntFlanks@1370.1414.45|ATGTTGGATCAGGACATCCCGATGGTGCAGCCGCTATTAAAGGTT</td><td>>hg38_MT_+_1621_3279_RNR2_16S_50ntFlanks</td><td>1370</td><td>1414</td><td>45</td><td>ATGTTGGATCAGGACATCCCGATGGTGCAGCCGCTATTAAAGGTT</td><td>218.7</td><td>5.0</td><td>1.5</td><td>3.2</td><td>1.2E-03</td><td>1.4E-03</td><td>MB231/MB468</td><td>MITO</td></tr>
<tr><td>>trnaMT_ProTGG_MT_-_15956_16023@1.31.31|CAGAGAATAGTTTAAATTAGAATCTTAGCTT</td><td>>trnaMT_ProTGG_MT_-_15956_16023</td><td>1</td><td>31</td><td>31</td><td>CAGAGAATAGTTTAAATTAGAATCTTAGCTT</td><td>191.6</td><td>-6.6</td><td>1.8</td><td>-3.7</td><td>1.8E-04</td><td>1.4E-03</td><td>BT20/MB231</td><td>MP</td></tr>
<tr><td>>hg38_21_+_8433172_8446622_RNA45SN1_45S_50ntFlanks@3844.3868.25|TACTTGGATAACTGTGGTAATTCTA</td><td>>hg38_21_+_8433172_8446622_RNA45SN1_45S_50ntFlanks</td><td>3844</td><td>3868</td><td>25</td><td>TACTTGGATAACTGTGGTAATTCTA</td><td>359.7</td><td>-4.1</td><td>1.3</td><td>-3.2</td><td>1.2E-03</td><td>1.4E-03</td><td>MB231/MB468</td><td>MITO</td></tr>
<tr><td>>trnaMT_GluTTC_MT_-_14674_14742@26.61.36,trnalookalike8_GluTTC_5_-_93905172_93905240@26.61.36|GGTTTTTCATATCATTGGTCGTGGTTGTAGTCCGTG</td><td>>trnaMT_GluTTC_MT_-_14674_14742</td><td>26</td><td>61</td><td>36</td><td>GGTTTTTCATATCATTGGTCGTGGTTGTAGTCCGTG</td><td>170.3</td><td>3.3</td><td>1.0</td><td>3.2</td><td>1.2E-03</td><td>1.4E-03</td><td>MB231/MB468</td><td>MITO</td></tr>
<tr><td>>am_MI0000786|hsa-mir-378a&WithFlank&5|+|149732819|149732896@50.70.21(+1U)&[MIMAT0000732&hsa-miR-378a-3p&offsets|+1|0(+1U);m-44&5|+|149732867|149732888&offsets|+1|0(+1U)]|CTGGACTTGGAGTCAGAAGGCT</td><td>>am_MI0000786|hsa-mir-378a&WithFlank&5|+|149732819|149732896</td><td>50</td><td>70</td><td>21(+1U)</td><td>CTGGACTTGGAGTCAGAAGGCT</td><td>441.7</td><td>1.9</td><td>0.6</td><td>3.2</td><td>1.2E-03</td><td>1.4E-03</td><td>MB231/MB468</td><td>NUC</td></tr>
<tr><td>>trnaMT_GluTTC_MT_-_14674_14742@41.63.23,trnalookalike8_GluTTC_5_-_93905172_93905240@41.63.23|TGGTCGTGGTTGTAGTCCGTGCG</td><td>>trnaMT_GluTTC_MT_-_14674_14742</td><td>41</td><td>63</td><td>23</td><td>TGGTCGTGGTTGTAGTCCGTGCG</td><td>239.7</td><td>6.0</td><td>1.7</td><td>3.5</td><td>5.3E-04</td><td>1.4E-03</td><td>BT20/MB468</td><td>MP</td></tr>
<tr><td>>hg38_21_+_8433172_8446622_RNA45SN1_45S_50ntFlanks@10807.10828.22|ACAATGTAGGTAAGGGAAGTCG</td><td>>hg38_21_+_8433172_8446622_RNA45SN1_45S_50ntFlanks</td><td>10807</td><td>10828</td><td>22</td><td>ACAATGTAGGTAAGGGAAGTCG</td><td>134.5</td><td>-5.3</td><td>1.5</td><td>-3.5</td><td>5.3E-04</td><td>1.4E-03</td><td>BT20/MB468</td><td>MP</td></tr>
<tr><td>>hg38_21_+_8433172_8446622_RNA45SN1_45S_50ntFlanks@6649.6670.22|TACGACTCTTAGCGGTGGATCA</td><td>>hg38_21_+_8433172_8446622_RNA45SN1_45S_50ntFlanks</td><td>6649</td><td>6670</td><td>22</td><td>TACGACTCTTAGCGGTGGATCA</td><td>93.2</td><td>-5.4</td><td>1.6</td><td>-3.5</td><td>5.3E-04</td><td>1.4E-03</td><td>BT20/MB468</td><td>MP</td></tr>
<tr><td>>hg38_21_+_8433172_8446622_RNA45SN1_45S_50ntFlanks@6652.6681.30|GACTCTTAGCGGTGGATCACTCGGCTCGTG</td><td>>hg38_21_+_8433172_8446622_RNA45SN1_45S_50ntFlanks</td><td>6652</td><td>6681</td><td>30</td><td>GACTCTTAGCGGTGGATCACTCGGCTCGTG</td><td>41992.2</td><td>-3.5</td><td>1.1</td><td>-3.2</td><td>1.2E-03</td><td>1.4E-03</td><td>MB231/MB468</td><td>MITO</td></tr>
<tr><td>>am_trnaMT_ValTAC_MT_+_1602_1670@51.68.18|CTTAACTTGACCGCTCTG</td><td>>am_trnaMT_ValTAC_MT_+_1602_1670</td><td>51</td><td>68</td><td>18</td><td>CTTAACTTGACCGCTCTG</td><td>23753.2</td><td>-5.5</td><td>1.7</td><td>-3.3</td><td>1.1E-03</td><td>1.4E-03</td><td>MB231/MB468</td><td>CYTO</td></tr>
<tr><td>>hg38_21_+_8433172_8446622_RNA45SN1_45S_50ntFlanks@4732.4761.30|AGAACGAAAGTCGGAGGTTCGAAGACGATC</td><td>>hg38_21_+_8433172_8446622_RNA45SN1_45S_50ntFlanks</td><td>4732</td><td>4761</td><td>30</td><td>AGAACGAAAGTCGGAGGTTCGAAGACGATC</td><td>354.6</td><td>2.0</td><td>0.6</td><td>3.2</td><td>1.2E-03</td><td>1.4E-03</td><td>MB231/MB468</td><td>NUC</td></tr>
</table>

<table>
<tr><td>>hg38_21_+_8433172_8446622_RNA45SN1_45S_50ntFlanks@12281.12321.41|AATACAGACCGTGAAAGCGGGGCCTCACGATCCTTCTGACC</td><td>>hg38_21_+_8433172_8446622_RNA45SN1_45S_50ntFlanks</td><td>12281</td><td>12321</td><td>41</td><td>AATACAGACCGTGAAAGCGGGGCCTCACGATCCTTCTGACC</td><td>213.7</td><td>-3.7</td><td>1.0</td><td>-3.7</td><td>2.0E-04</td><td>1.4E-03</td><td>BT20/MB231</td><td>CYTO</td></tr>
<tr><td>>hg38_21_+_8433172_8446622_RNA45SN1_45S_50ntFlanks@12008.12042.35|CCGCCGGTGAAATACCACTACTCTGATCGTTTTTT</td><td>>hg38_21_+_8433172_8446622_RNA45SN1_45S_50ntFlanks</td><td>12008</td><td>12042</td><td>35</td><td>CCGCCGGTGAAATACCACTACTCTGATCGTTTTTT</td><td>584.3</td><td>-5.5</td><td>1.7</td><td>-3.3</td><td>1.1E-03</td><td>1.4E-03</td><td>BT20/MB468</td><td>CYTO</td></tr>
<tr><td>>hg38_21_+_8433172_8446622_RNA45SN1_45S_50ntFlanks@3840.3860.21|CTCCTACTTGGATAACTGTGG</td><td>>hg38_21_+_8433172_8446622_RNA45SN1_45S_50ntFlanks</td><td>3840</td><td>3860</td><td>21</td><td>CTCCTACTTGGATAACTGTGG</td><td>714.3</td><td>-3.1</td><td>1.0</td><td>-3.2</td><td>1.2E-03</td><td>1.4E-03</td><td>MB231/MB468</td><td>MITO</td></tr>
<tr><td>>MI0000289|hsa-mir-181a-1&WithFlank&1|-|198859038|198859159@30.53.24;MI0000269|hsa-mir-181a-2&WithFlank&9|+|124692436|124692557@45.68.24&[MIMAT0000256&hsa-miR-181a-5p&offsets|0|+1;m-32&1|-|198859108|198859130&offsets|0|+1];[MIMAT0000256_1&hsa-miR-181a-5p&offsets|0|+1;m-32&9|+|124692480|124692502&offsets|0|+1]|AACATTCAACGCTGTCGGTGAGTT</td><td>>MI0000289|hsa-mir-181a-1&WithFlank&1|-|198859038|198859159</td><td>30</td><td>53</td><td>24</td><td>AACATTCAACGCTGTCGGTGAGTT</td><td>5637.0</td><td>3.6</td><td>1.1</td><td>3.3</td><td>9.8E-04</td><td>1.4E-03</td><td>MB231/MB468</td><td>MP</td></tr>
<tr><td>>hg38_21_+_8433172_8446622_RNA45SN1_45S_50ntFlanks@3842.3873.32|CCTACTTGGATAACTGTGGTAATTCTAGAGCT</td><td>>hg38_21_+_8433172_8446622_RNA45SN1_45S_50ntFlanks</td><td>3842</td><td>3873</td><td>32</td><td>CCTACTTGGATAACTGTGGTAATTCTAGAGCT</td><td>128.4</td><td>-4.2</td><td>1.3</td><td>-3.3</td><td>9.8E-04</td><td>1.4E-03</td><td>MB231/MB468</td><td>MP</td></tr>
<tr><td>>hg38_21_+_8433172_8446622_RNA45SN1_45S_50ntFlanks@12029.12057.29|TCTGATCGTTTTTTCACTGACCCGGTGAG</td><td>>hg38_21_+_8433172_8446622_RNA45SN1_45S_50ntFlanks</td><td>12029</td><td>12057</td><td>29</td><td>TCTGATCGTTTTTTCACTGACCCGGTGAG</td><td>49.9</td><td>9.4</td><td>2.7</td><td>3.5</td><td>5.3E-04</td><td>1.4E-03</td><td>BT20/MB468</td><td>MP</td></tr>
<tr><td>>trnaMT_IleGAT_MT_+_4263_4331@42.61.20|AATAGGAGCTTAAACCCCCT</td><td>>trnaMT_IleGAT_MT_+_4263_4331</td><td>42</td><td>61</td><td>20</td><td>AATAGGAGCTTAAACCCCCT</td><td>53.7</td><td>9.5</td><td>2.7</td><td>3.5</td><td>5.3E-04</td><td>1.4E-03</td><td>BT20/MB468</td><td>MP</td></tr>
<tr><td>>trnaMT_PheGAA_MT_+_577_647@13.34.22|TACCTCCTCAAAGCAATACACT</td><td>>trnaMT_PheGAA_MT_+_577_647</td><td>13</td><td>34</td><td>22</td><td>TACCTCCTCAAAGCAATACACT</td><td>151.5</td><td>4.0</td><td>1.2</td><td>3.2</td><td>1.2E-03</td><td>1.4E-03</td><td>MB231/MB468</td><td>MITO</td></tr>
<tr><td>>hg38_21_+_8433172_8446622_RNA45SN1_45S_50ntFlanks@5074.5097.24|ATTCCGATAACGAACGAGACTCTG</td><td>>hg38_21_+_8433172_8446622_RNA45SN1_45S_50ntFlanks</td><td>5074</td><td>5097</td><td>24</td><td>ATTCCGATAACGAACGAGACTCTG</td><td>368.1</td><td>-3.2</td><td>1.0</td><td>-3.2</td><td>1.2E-03</td><td>1.4E-03</td><td>MB231/MB468</td><td>NUC</td></tr>
<tr><td>>hg38_21_+_8433172_8446622_RNA45SN1_45S_50ntFlanks@8347.8375.29|GGGAAAGTTGAAAAGAACTTTGAAGAGAG</td><td>>hg38_21_+_8433172_8446622_RNA45SN1_45S_50ntFlanks</td><td>8347</td><td>8375</td><td>29</td><td>GGGAAAGTTGAAAAGAACTTTGAAGAGAG</td><td>4006.7</td><td>5.3</td><td>1.4</td><td>3.7</td><td>1.8E-04</td><td>1.4E-03</td><td>BT20/MB231</td><td>MP</td></tr>
<tr><td>>hg38_21_+_8433172_8446622_RNA45SN1_45S_50ntFlanks@4380.4410.31|CGTAGTTGGATCTTGGGAGCGGGCGGGCGGT</td><td>>hg38_21_+_8433172_8446622_RNA45SN1_45S_50ntFlanks</td><td>4380</td><td>4410</td><td>31</td><td>CGTAGTTGGATCTTGGGAGCGGGCGGGCGGT</td><td>118.9</td><td>-2.7</td><td>0.8</td><td>-3.3</td><td>1.1E-03</td><td>1.4E-03</td><td>MB231/MB468</td><td>CYTO</td></tr>
<tr><td>>hg38_21_+_8433172_8446622_RNA45SN1_45S_50ntFlanks@7981.8006.26|CTCAGATCAGACGTGGCGACCCGCTG</td><td>>hg38_21_+_8433172_8446622_RNA45SN1_45S_50ntFlanks</td><td>7981</td><td>8006</td><td>26</td><td>CTCAGATCAGACGTGGCGACCCGCTG</td><td>1557.1</td><td>-4.2</td><td>1.3</td><td>-3.3</td><td>1.1E-03</td><td>1.4E-03</td><td>BT20/MB468</td><td>CYTO</td></tr>
<tr><td>>hg38_21_+_8433172_8446622_RNA45SN1_45S_50ntFlanks@9952.9979.28|CATGGAAGTCGGAATCCGCTAAGGAGTG</td><td>>hg38_21_+_8433172_8446622_RNA45SN1_45S_50ntFlanks</td><td>9952</td><td>9979</td><td>28</td><td>CATGGAAGTCGGAATCCGCTAAGGAGTG</td><td>499.8</td><td>7.5</td><td>2.0</td><td>3.7</td><td>1.8E-04</td><td>1.4E-03</td><td>BT20/MB231</td><td>MP</td></tr>
<tr><td>>hg38_21_+_8433172_8446622_RNA45SN1_45S_50ntFlanks@3852.3881.30|TAACTGTGGTAATTCTAGAGCTAATACATG</td><td>>hg38_21_+_8433172_8446622_RNA45SN1_45S_50ntFlanks</td><td>3852</td><td>3881</td><td>30</td><td>TAACTGTGGTAATTCTAGAGCTAATACATG</td><td>192.6</td><td>-3.6</td><td>1.1</td><td>-3.3</td><td>1.1E-03</td><td>1.4E-03</td><td>BT20/MB468</td><td>CYTO</td></tr>
<tr><td>>hg38_21_+_8433172_8446622_RNA45SN1_45S_50ntFlanks@9953.9979.27|ATGGAAGTCGGAATCCGCTAAGGAGTG</td><td>>hg38_21_+_8433172_8446622_RNA45SN1_45S_50ntFlanks</td><td>9953</td><td>9979</td><td>27</td><td>ATGGAAGTCGGAATCCGCTAAGGAGTG</td><td>427.8</td><td>6.4</td><td>1.7</td><td>3.7</td><td>1.9E-04</td><td>1.4E-03</td><td>BT20/MB231</td><td>MP</td></tr>
<tr><td>>hg38_21_+_8433172_8446622_RNA45SN1_45S_50ntFlanks@11317.11343.27|ACCCCACGTCTCGTCGCGCGCGCGTCC</td><td>>hg38_21_+_8433172_8446622_RNA45SN1_45S_50ntFlanks</td><td>11317</td><td>11343</td><td>27</td><td>ACCCCACGTCTCGTCGCGCGCGCGTCC</td><td>2553.4</td><td>-5.1</td><td>1.6</td><td>-3.2</td><td>1.2E-03</td><td>1.4E-03</td><td>MB231/MB468</td><td>NUC</td></tr>
<tr><td>>hg38_21_+_8433172_8446622_RNA45SN1_45S_50ntFlanks@12437.12473.37|CATTGTGAAGCAGAATTCACCAAGCGTTGGATTGTTC</td><td>>hg38_21_+_8433172_8446622_RNA45SN1_45S_50ntFlanks</td><td>12437</td><td>12473</td><td>37</td><td>CATTGTGAAGCAGAATTCACCAAGCGTTGGATTGTTC</td><td>598.7</td><td>-2.8</td><td>0.8</td><td>-3.3</td><td>8.9E-04</td><td>1.4E-03</td><td>BT20/MB468</td><td>MITO</td></tr>
<tr><td>>MI0000809|hsa-mir-151a&WithFlank&8|-|140732558|140732659@53.74.22(+1G)&[MIMAT0000757&hsa-miR-151a-3p&offsets|0|+1(+1G);m-30&8|-|140732587|140732607&offsets|0|+1(+1G)]|CTAGACTGAAGCTCCTTGAGGAG</td><td>>MI0000809|hsa-mir-151a&WithFlank&8|-|140732558|140732659</td><td>53</td><td>74</td><td>22(+1G)</td><td>CTAGACTGAAGCTCCTTGAGGAG</td><td>164.7</td><td>5.8</td><td>1.7</td><td>3.5</td><td>5.3E-04</td><td>1.4E-03</td><td>BT20/MB468</td><td>MP</td></tr>
<tr><td>>hg38_21_+_8433172_8446622_RNA45SN1_45S_50ntFlanks@9573.9605.33|AAATCGGTCGTCCGACCTGGGTATAGGGGCGAA</td><td>>hg38_21_+_8433172_8446622_RNA45SN1_45S_50ntFlanks</td><td>9573</td><td>9605</td><td>33</td><td>AAATCGGTCGTCCGACCTGGGTATAGGGGCGAA</td><td>494.2</td><td>-3.2</td><td>1.0</td><td>-3.3</td><td>9.9E-04</td><td>1.4E-03</td><td>MB231/MB468</td><td>MP</td></tr>
<tr><td>>hg38_21_+_8433172_8446622_RNA45SN1_45S_50ntFlanks@12273.12293.21|TCAGTACGAATACAGACCGTG</td><td>>hg38_21_+_8433172_8446622_RNA45SN1_45S_50ntFlanks</td><td>12273</td><td>12293</td><td>21</td><td>TCAGTACGAATACAGACCGTG</td><td>282.8</td><td>-5.6</td><td>1.6</td><td>-3.5</td><td>5.3E-04</td><td>1.4E-03</td><td>BT20/MB468</td><td>MP</td></tr>
<tr><td>>hg38_MT_+_598_1651_RNR1_12S_50ntFlanks@428.474.47|AAAGTGGCTTTAACATATCTGAACACACAATAGCTAAGACCCAAACT</td><td>>hg38_MT_+_598_1651_RNR1_12S_50ntFlanks</td><td>428</td><td>474</td><td>47</td><td>AAAGTGGCTTTAACATATCTGAACACACAATAGCTAAGACCCAAACT</td><td>53.4</td><td>9.5</td><td>2.7</td><td>3.5</td><td>5.3E-04</td><td>1.4E-03</td><td>BT20/MB468</td><td>MP</td></tr>
<tr><td>>hg38_21_+_8433172_8446622_RNA45SN1_45S_50ntFlanks@485.509.25|GATGTGGTGACGTCGTGCTCTCCCG</td><td>>hg38_21_+_8433172_8446622_RNA45SN1_45S_50ntFlanks</td><td>485</td><td>509</td><td>25</td><td>GATGTGGTGACGTCGTGCTCTCCCG</td><td>376.9</td><td>11.8</td><td>3.6</td><td>3.2</td><td>1.2E-03</td><td>1.4E-03</td><td>MB231/MB468</td><td>MITO</td></tr>
<tr><td>>hg38_21_+_8433172_8446622_RNA45SN1_45S_50ntFlanks@6677.6696.20|TCGTGCGTCGATGAAGAACG</td><td>>hg38_21_+_8433172_8446622_RNA45SN1_45S_50ntFlanks</td><td>6677</td><td>6696</td><td>20</td><td>TCGTGCGTCGATGAAGAACG</td><td>140.8</td><td>2.6</td><td>0.8</td><td>3.5</td><td>5.3E-04</td><td>1.4E-03</td><td>BT20/MB468</td><td>MP</td></tr>
<tr><td>>MI0000088|hsa-mir-30a&WithFlank&6|-|71403545|71403627@54.73.20(+2U)&[MIMAT0000088&hsa-miR-30a-3p&offsets|+1|-1(+2U);m-38&6|-|71403554|71403575&offsets|+1|-1(+2U)]|TTTCAGTCGGATGTTTGCAGTT</td><td>>MI0000088|hsa-mir-30a&WithFlank&6|-|71403545|71403627</td><td>54</td><td>73</td><td>20(+2U)</td><td>TTTCAGTCGGATGTTTGCAGTT</td><td>413.5</td><td>4.3</td><td>1.3</td><td>3.3</td><td>9.9E-04</td><td>1.4E-03</td><td>MB231/MB468</td><td>MP</td></tr>
<tr><td>>hg38_21_+_8433172_8446622_RNA45SN1_45S_50ntFlanks@5485.5518.34|AGCGCTGAGAAGACGGTCGAACTTGACTATCTAG</td><td>>hg38_21_+_8433172_8446622_RNA45SN1_45S_50ntFlanks</td><td>5485</td><td>5518</td><td>34</td><td>AGCGCTGAGAAGACGGTCGAACTTGACTATCTAG</td><td>1570.9</td><td>-3.3</td><td>1.0</td><td>-3.3</td><td>1.1E-03</td><td>1.4E-03</td><td>BT20/MB468</td><td>CYTO</td></tr>
<tr><td>>hg38_21_+_8433172_8446622_RNA45SN1_45S_50ntFlanks@7975.8017.43|CGCGACCTCAGATCAGACGTGGCGACCCGCTGAATTTAAGCAT</td><td>>hg38_21_+_8433172_8446622_RNA45SN1_45S_50ntFlanks</td><td>7975</td><td>8017</td><td>43</td><td>CGCGACCTCAGATCAGACGTGGCGACCCGCTGAATTTAAGCAT</td><td>11293.9</td><td>3.2</td><td>1.0</td><td>3.2</td><td>1.2E-03</td><td>1.4E-03</td><td>MB231/MB468</td><td>MITO</td></tr>
<tr><td>>MI0000088|hsa-mir-30a&WithFlank&6|-|71403545|71403627@12.34.23(+1G)&[MIMAT0000087&hsa-miR-30a-5p&offsets|0|+1(+1G);m-15&6|-|71403595|71403616&offsets|0|+1(+1G)]|TGTAAACATCCTCGACTGGAAGCG</td><td>>MI0000088|hsa-mir-30a&WithFlank&6|-|71403545|71403627</td><td>12</td><td>34</td><td>23(+1G)</td><td>TGTAAACATCCTCGACTGGAAGCG</td><td>464.3</td><td>3.5</td><td>1.1</td><td>3.2</td><td>1.2E-03</td><td>1.4E-03</td><td>BT20/MB468</td><td>NUC</td></tr>
<tr><td>>hg38_21_+_8433172_8446622_RNA45SN1_45S_50ntFlanks@10866.10892.27|CTAAGGGCTGGGTCGGTCGGGCTGGGG</td><td>>hg38_21_+_8433172_8446622_RNA45SN1_45S_50ntFlanks</td><td>10866</td><td>10892</td><td>27</td><td>CTAAGGGCTGGGTCGGTCGGGCTGGGG</td><td>533.3</td><td>-3.9</td><td>1.2</td><td>-3.3</td><td>9.0E-04</td><td>1.4E-03</td><td>BT20/MB468</td><td>MITO</td></tr>
<tr><td>>hg38_21_+_8433172_8446622_RNA45SN1_45S_50ntFlanks@5497.5525.29|ACGGTCGAACTTGACTATCTAGAGGAAGT</td><td>>hg38_21_+_8433172_8446622_RNA45SN1_45S_50ntFlanks</td><td>5497</td><td>5525</td><td>29</td><td>ACGGTCGAACTTGACTATCTAGAGGAAGT</td><td>111.1</td><td>-4.0</td><td>1.2</td><td>-3.3</td><td>9.9E-04</td><td>1.4E-03</td><td>MB231/MB468</td><td>MP</td></tr>
<tr><td>>hg38_21_+_8433172_8446622_RNA45SN1_45S_50ntFlanks@11641.11667.27|CGGCGGGTGTTGACGCGATGTGATTTC</td><td>>hg38_21_+_8433172_8446622_RNA45SN1_45S_50ntFlanks</td><td>11641</td><td>11667</td><td>27</td><td>CGGCGGGTGTTGACGCGATGTGATTTC</td><td>1608.7</td><td>-6.3</td><td>1.8</td><td>-3.5</td><td>5.3E-04</td><td>1.4E-03</td><td>BT20/MB468</td><td>MP</td></tr>
<tr><td>>hg38_21_+_8433172_8446622_RNA45SN1_45S_50ntFlanks@6744.6770.27|GACACTTCGAACGCACTTGCGGCCCCG</td><td>>hg38_21_+_8433172_8446622_RNA45SN1_45S_50ntFlanks</td><td>6744</td><td>6770</td><td>27</td><td>GACACTTCGAACGCACTTGCGGCCCCG</td><td>1696.0</td><td>-5.2</td><td>1.5</td><td>-3.5</td><td>5.3E-04</td><td>1.4E-03</td><td>BT20/MB468</td><td>MP</td></tr>
<tr><td>>hg38_21_+_8433172_8446622_RNA45SN1_45S_50ntFlanks@10248.10276.29|CGACGAGTAGGAGGGCCGCTGCGGTGAGC</td><td>>hg38_21_+_8433172_8446622_RNA45SN1_45S_50ntFlanks</td><td>10248</td><td>10276</td><td>29</td><td>CGACGAGTAGGAGGGCCGCTGCGGTGAGC</td><td>206.5</td><td>6.7</td><td>1.8</td><td>3.7</td><td>1.9E-04</td><td>1.4E-03</td><td>BT20/MB231</td><td>MP</td></tr>
<tr><td>>hg38_21_+_8433172_8446622_RNA45SN1_45S_50ntFlanks@6652.6679.28|GACTCTTAGCGGTGGATCACTCGGCTCG</td><td>>hg38_21_+_8433172_8446622_RNA45SN1_45S_50ntFlanks</td><td>6652</td><td>6679</td><td>28</td><td>GACTCTTAGCGGTGGATCACTCGGCTCG</td><td>3390.2</td><td>5.5</td><td>1.5</td><td>3.7</td><td>1.9E-04</td><td>1.4E-03</td><td>BT20/MB231</td><td>MP</td></tr>
<tr><td>>hg38_21_+_8433172_8446622_RNA45SN1_45S_50ntFlanks@12447.12486.40|CAGAATTCACCAAGCGTTGGATTGTTCACCCACTAATAGG</td><td>>hg38_21_+_8433172_8446622_RNA45SN1_45S_50ntFlanks</td><td>12447</td><td>12486</td><td>40</td><td>CAGAATTCACCAAGCGTTGGATTGTTCACCCACTAATAGG</td><td>48.4</td><td>-8.7</td><td>2.5</td><td>-3.5</td><td>5.3E-04</td><td>1.4E-03</td><td>BT20/MB468</td><td>MP</td></tr>
<tr><td>>hg38_21_+_8433172_8446622_RNA45SN1_45S_50ntFlanks@3860.3898.39|GTAATTCTAGAGCTAATACATGCCGACGGGCGCTGACCC</td><td>>hg38_21_+_8433172_8446622_RNA45SN1_45S_50ntFlanks</td><td>3860</td><td>3898</td><td>39</td><td>GTAATTCTAGAGCTAATACATGCCGACGGGCGCTGACCC</td><td>51.5</td><td>-8.8</td><td>2.5</td><td>-3.5</td><td>5.3E-04</td><td>1.4E-03</td><td>BT20/MB468</td><td>MP</td></tr>
<tr><td>>hg38_21_+_8433172_8446622_RNA45SN1_45S_50ntFlanks@11200.11234.35|CGGCGGCGGCGGCGACTCTGGACGCGAGCCGGGCC</td><td>>hg38_21_+_8433172_8446622_RNA45SN1_45S_50ntFlanks</td><td>11200</td><td>11234</td><td>35</td><td>CGGCGGCGGCGGCGACTCTGGACGCGAGCCGGGCC</td><td>359.7</td><td>-4.9</td><td>1.5</td><td>-3.3</td><td>1.1E-03</td><td>1.4E-03</td><td>BT20/MB468</td><td>CYTO</td></tr>
<tr><td>>am_trna130_GlnTTG_6_-_27763640_27763711@13.30.18,trna146_GlnCTG_6_-_27515531_27515602@13.30.18,trna16_GlnTTG_17_+_47269890_47269961@13.30.18,trna173_GlnTTG_6_-_26311975_26312046@13.30.18,trna174_GlnTTG_6_-_26311424_26311495@13.30.18,trna1_GlnCTG_6_+_18836402_18836473@13.30.18,trna3_GlnCTG_17_+_8023070_8023141@13.30.18,trna42_GlnCTG_6_+_27263212_27263283@13.30.18,trna49_GlnCTG_6_+_27487308_27487379@13.30.18,trna64_GlnTTG_6_+_28557156_28557227@13.30.18,trna7_GlnCTG_15_-_66161400_66161471@13.30.18,trna84_GlnTTG_6_+_145503859_145503930@13.30.18,trna99_GlnCTG_6_-_28909378_28909449@13.30.18|TAATGGTTAGCACTCTGG</td><td>>am_trna130_GlnTTG_6_-_27763640_27763711</td><td>13</td><td>30</td><td>18</td><td>TAATGGTTAGCACTCTGG</td><td>156.3</td><td>-3.1</td><td>1.0</td><td>-3.3</td><td>1.1E-03</td><td>1.4E-03</td><td>BT20/MB468</td><td>CYTO</td></tr>
<tr><td>>hg38_21_+_8433172_8446622_RNA45SN1_45S_50ntFlanks@3789.3810.22|ACTGCGAATGGCTCATTAAATC</td><td>>hg38_21_+_8433172_8446622_RNA45SN1_45S_50ntFlanks</td><td>3789</td><td>3810</td><td>22</td><td>ACTGCGAATGGCTCATTAAATC</td><td>371.0</td><td>-4.0</td><td>1.2</td><td>-3.3</td><td>9.0E-04</td><td>1.4E-03</td><td>BT20/MB468</td><td>MITO</td></tr>
<tr><td>>hg38_MT_+_1621_3279_RNR2_16S_50ntFlanks@1216.1261.46|CAACCTCCGAGCAGTACATGCTAAGACTTCACCAGTCAAAGCGAAC</td><td>>hg38_MT_+_1621_3279_RNR2_16S_50ntFlanks</td><td>1216</td><td>1261</td><td>46</td><td>CAACCTCCGAGCAGTACATGCTAAGACTTCACCAGTCAAAGCGAAC</td><td>357.2</td><td>5.2</td><td>1.6</td><td>3.2</td><td>1.2E-03</td><td>1.4E-03</td><td>MB231/MB468</td><td>MITO</td></tr>
<tr><td>>trnaMT_ValTAC_MT_+_1602_1670@39.72.34|AGGAGATTTCAACTTAACTTGACCGCTCTGACCA</td><td>>trnaMT_ValTAC_MT_+_1602_1670</td><td>39</td><td>72</td><td>34</td><td>AGGAGATTTCAACTTAACTTGACCGCTCTGACCA</td><td>686.9</td><td>4.9</td><td>1.4</td><td>3.5</td><td>5.4E-04</td><td>1.4E-03</td><td>BT20/MB468</td><td>MP</td></tr>
<tr><td>>hg38_21_+_8433172_8446622_RNA45SN1_45S_50ntFlanks@9498.9526.29|AAGATGGTGAACTATGCCTGGGCAGGGCG</td><td>>hg38_21_+_8433172_8446622_RNA45SN1_45S_50ntFlanks</td><td>9498</td><td>9526</td><td>29</td><td>AAGATGGTGAACTATGCCTGGGCAGGGCG</td><td>116.7</td><td>10.2</td><td>2.7</td><td>3.7</td><td>1.9E-04</td><td>1.4E-03</td><td>BT20/MB231</td><td>MP</td></tr>
<tr><td>>hg38_21_+_8433172_8446622_RNA45SN1_45S_50ntFlanks@7973.7991.19|GACGCGACCTCAGATCAGA</td><td>>hg38_21_+_8433172_8446622_RNA45SN1_45S_50ntFlanks</td><td>7973</td><td>7991</td><td>19</td><td>GACGCGACCTCAGATCAGA</td><td>646.7</td><td>3.5</td><td>1.0</td><td>3.5</td><td>5.4E-04</td><td>1.4E-03</td><td>BT20/MB468</td><td>MP</td></tr>
<tr><td>>hg38_21_+_8433172_8446622_RNA45SN1_45S_50ntFlanks@10105.10124.20|CGCGTGTGGTGTGCGTCGGA</td><td>>hg38_21_+_8433172_8446622_RNA45SN1_45S_50ntFlanks</td><td>10105</td><td>10124</td><td>20</td><td>CGCGTGTGGTGTGCGTCGGA</td><td>113.6</td><td>10.2</td><td>2.7</td><td>3.7</td><td>1.9E-04</td><td>1.4E-03</td><td>BT20/MB231</td><td>MP</td></tr>
<tr><td>>hg38_21_+_8433172_8446622_RNA45SN1_45S_50ntFlanks@4984.5018.35|GGACAGGATTGACAGATTGATAGCTCTTTCTCGAT</td><td>>hg38_21_+_8433172_8446622_RNA45SN1_45S_50ntFlanks</td><td>4984</td><td>5018</td><td>35</td><td>GGACAGGATTGACAGATTGATAGCTCTTTCTCGAT</td><td>63.6</td><td>6.4</td><td>1.8</td><td>3.5</td><td>5.4E-04</td><td>1.4E-03</td><td>BT20/MB468</td><td>MP</td></tr>
<tr><td>>hg38_21_+_8433172_8446622_RNA45SN1_45S_50ntFlanks@12324.12366.43|TTGGGTTTTAAGCAGGAGGTGTCAGAAAAGTTACCACAGGGAT</td><td>>hg38_21_+_8433172_8446622_RNA45SN1_45S_50ntFlanks</td><td>12324</td><td>12366</td><td>43</td><td>TTGGGTTTTAAGCAGGAGGTGTCAGAAAAGTTACCACAGGGAT</td><td>48.1</td><td>-8.7</td><td>2.5</td><td>-3.5</td><td>5.4E-04</td><td>1.4E-03</td><td>BT20/MB468</td><td>MP</td></tr>
<tr><td>>hg38_21_+_8433172_8446622_RNA45SN1_45S_50ntFlanks@4220.4243.24|ACGAAAAATAACAATACAGGACTC</td><td>>hg38_21_+_8433172_8446622_RNA45SN1_45S_50ntFlanks</td><td>4220</td><td>4243</td><td>24</td><td>ACGAAAAATAACAATACAGGACTC</td><td>228.2</td><td>-2.0</td><td>0.6</td><td>-3.3</td><td>1.1E-03</td><td>1.4E-03</td><td>BT20/MB468</td><td>CYTO</td></tr>
<tr><td>>hg38_21_+_8433172_8446622_RNA45SN1_45S_50ntFlanks@8359.8376.18|AAGAACTTTGAAGAGAGA</td><td>>hg38_21_+_8433172_8446622_RNA45SN1_45S_50ntFlanks</td><td>8359</td><td>8376</td><td>18</td><td>AAGAACTTTGAAGAGAGA</td><td>129.0</td><td>-4.1</td><td>1.2</td><td>-3.3</td><td>1.0E-03</td><td>1.4E-03</td><td>MB231/MB468</td><td>MP</td></tr>
<tr><td>>hg38_21_+_8433172_8446622_RNA45SN1_45S_50ntFlanks@8106.8138.33|GCGGCGGGGCGCGGGACATGTGGCGTACGGAAG</td><td>>hg38_21_+_8433172_8446622_RNA45SN1_45S_50ntFlanks</td><td>8106</td><td>8138</td><td>33</td><td>GCGGCGGGGCGCGGGACATGTGGCGTACGGAAG</td><td>705.4</td><td>-3.1</td><td>0.9</td><td>-3.5</td><td>5.4E-04</td><td>1.4E-03</td><td>BT20/MB468</td><td>MP</td></tr>
</table>

<table>
<tr><th></th><th></th><th></th><th></th><th></th><th></th><th></th><th></th><th></th><th></th><th></th><th></th><th></th><th></th></tr>
<tr><td>>hg38_21_+_8433172_8446622_RNA45SN1_45S_50ntFlanks@12444.12481.38|AAGCAGAATTCACCAAGCGTTGGATTGTTCACCCACTA</td><td>>hg38_21_+_8433172_8446622_RNA45SN1_45S_50ntFlanks</td><td>12444</td><td>12481</td><td>38</td><td>AAGCAGAATTCACCAAGCGTTGGATTGTTCACCCACTA</td><td>63.8</td><td>-5.3</td><td>1.6</td><td>-3.3</td><td>1.0E-03</td><td>1.4E-03</td><td>MB231/MB468</td><td>MP</td></tr>
<tr><td>>hg38_21_+_8433172_8446622_RNA45SN1_45S_50ntFlanks@9606.9623.18|AGACTAATCGAACCATCT</td><td>>hg38_21_+_8433172_8446622_RNA45SN1_45S_50ntFlanks</td><td>9606</td><td>9623</td><td>18</td><td>AGACTAATCGAACCATCT</td><td>201.0</td><td>-4.2</td><td>1.3</td><td>-3.3</td><td>1.0E-03</td><td>1.4E-03</td><td>MB231/MB468</td><td>MP</td></tr>
<tr><td>>hg38_21_+_8433172_8446622_RNA45SN1_45S_50ntFlanks@12934.12962.29|CTAAACCATTCGTAGACGACCTGCTTCTG</td><td>>hg38_21_+_8433172_8446622_RNA45SN1_45S_50ntFlanks</td><td>12934</td><td>12962</td><td>29</td><td>CTAAACCATTCGTAGACGACCTGCTTCTG</td><td>152.8</td><td>-5.3</td><td>1.5</td><td>-3.5</td><td>5.4E-04</td><td>1.4E-03</td><td>BT20/MB468</td><td>MP</td></tr>
<tr><td>>MI0000298|hsa-mir-221&WithFlank&X|-|45746151|45746272@71.91.21(+1C)&[MIMAT0000278&hsa-miR-221-3p&offsets|0|-2(+1C);m-91&X|-|45746181|45746202&offsets|0|-1(+1C)]|AGCTACATTGTCTGCTGGGTTC</td><td>>MI0000298|hsa-mir-221&WithFlank&X|-|45746151|45746272</td><td>71</td><td>91</td><td>21(+1C)</td><td>AGCTACATTGTCTGCTGGGTTC</td><td>2023.8</td><td>4.0</td><td>1.2</td><td>3.5</td><td>5.4E-04</td><td>1.4E-03</td><td>BT20/MB468</td><td>MP</td></tr>
<tr><td>>hg38_21_+_8433172_8446622_RNA45SN1_45S_50ntFlanks@10403.10429.27|ACATGGGTCAGTCGGTCCTGAGAGATG</td><td>>hg38_21_+_8433172_8446622_RNA45SN1_45S_50ntFlanks</td><td>10403</td><td>10429</td><td>27</td><td>ACATGGGTCAGTCGGTCCTGAGAGATG</td><td>474.7</td><td>-1.9</td><td>0.6</td><td>-3.3</td><td>1.1E-03</td><td>1.4E-03</td><td>BT20/MB468</td><td>CYTO</td></tr>
<tr><td>>am_MI0003639|hsa-mir-625&WithFlank&14|+|65471096|65471192@58.80.23&[MIMAT0004808&hsa-miR-625-3p&offsets|0|+1;m-137&14|+|65471153|65471174&offsets|0|+1]|GACTATAGAACTTTCCCCCTCAT</td><td>>am_MI0003639|hsa-mir-625&WithFlank&14|+|65471096|65471192</td><td>58</td><td>80</td><td>23</td><td>GACTATAGAACTTTCCCCCTCAT</td><td>677.9</td><td>-2.8</td><td>0.7</td><td>-3.7</td><td>2.0E-04</td><td>1.4E-03</td><td>BT20/MB231</td><td>CYTO</td></tr>
<tr><td>>hg38_21_+_8433172_8446622_RNA45SN1_45S_50ntFlanks@7974.7994.21|ACGCGACCTCAGATCAGACGT</td><td>>hg38_21_+_8433172_8446622_RNA45SN1_45S_50ntFlanks</td><td>7974</td><td>7994</td><td>21</td><td>ACGCGACCTCAGATCAGACGT</td><td>1222.5</td><td>2.3</td><td>0.7</td><td>3.3</td><td>1.1E-03</td><td>1.5E-03</td><td>BT20/MB468</td><td>CYTO</td></tr>
<tr><td>>MI0000077|hsa-mir-21&WithFlank&17|+|59841260|59841343@14.36.23&[MIMAT0000076&hsa-miR-21-5p&offsets|0|+1;m-1&17|+|59841273|59841294&offsets|0|+1]|TAGCTTATCAGACTGATGTTGAC</td><td>>MI0000077|hsa-mir-21&WithFlank&17|+|59841260|59841343</td><td>14</td><td>36</td><td>23</td><td>TAGCTTATCAGACTGATGTTGAC</td><td>775649.9</td><td>-3.3</td><td>0.9</td><td>-3.7</td><td>1.9E-04</td><td>1.5E-03</td><td>BT20/MB231</td><td>MP</td></tr>
<tr><td>>hg38_21_+_8433172_8446622_RNA45SN1_45S_50ntFlanks@6779.6803.25|CCGGGGCTACGCCTGTCTGAGCGTC</td><td>>hg38_21_+_8433172_8446622_RNA45SN1_45S_50ntFlanks</td><td>6779</td><td>6803</td><td>25</td><td>CCGGGGCTACGCCTGTCTGAGCGTC</td><td>995.6</td><td>-2.6</td><td>0.8</td><td>-3.3</td><td>9.1E-04</td><td>1.5E-03</td><td>BT20/MB468</td><td>MITO</td></tr>
<tr><td>>hg38_21_+_8433172_8446622_RNA45SN1_45S_50ntFlanks@6729.6771.43|GACACATTGATCATCGACACTTCGAACGCACTTGCGGCCCCGG</td><td>>hg38_21_+_8433172_8446622_RNA45SN1_45S_50ntFlanks</td><td>6729</td><td>6771</td><td>43</td><td>GACACATTGATCATCGACACTTCGAACGCACTTGCGGCCCCGG</td><td>575.0</td><td>-2.4</td><td>0.7</td><td>-3.3</td><td>9.1E-04</td><td>1.5E-03</td><td>BT20/MB468</td><td>MITO</td></tr>
<tr><td>>trnaMT_TyrGTA_MT_-_5826_5891@24.69.46|TGGACTGTAAATCTAAAGACAGGGGTTAGGCCTCTTTTTACCACCA</td><td>>trnaMT_TyrGTA_MT_-_5826_5891</td><td>24</td><td>69</td><td>46</td><td>TGGACTGTAAATCTAAAGACAGGGGTTAGGCCTCTTTTTACCACCA</td><td>53.3</td><td>9.5</td><td>2.7</td><td>3.5</td><td>5.4E-04</td><td>1.5E-03</td><td>BT20/MB468</td><td>MP</td></tr>
<tr><td>>hg38_21_+_8433172_8446622_RNA45SN1_45S_50ntFlanks@9483.9511.29|CGTGTTAGGACCCGAAAGATGGTGAACTA</td><td>>hg38_21_+_8433172_8446622_RNA45SN1_45S_50ntFlanks</td><td>9483</td><td>9511</td><td>29</td><td>CGTGTTAGGACCCGAAAGATGGTGAACTA</td><td>48.5</td><td>9.3</td><td>2.7</td><td>3.5</td><td>5.4E-04</td><td>1.5E-03</td><td>BT20/MB468</td><td>MP</td></tr>
<tr><td>>hg38_21_+_8433172_8446622_RNA45SN1_45S_50ntFlanks@12912.12932.21|CACGTTCGTGGGGAACCTGGC</td><td>>hg38_21_+_8433172_8446622_RNA45SN1_45S_50ntFlanks</td><td>12912</td><td>12932</td><td>21</td><td>CACGTTCGTGGGGAACCTGGC</td><td>206.7</td><td>-2.9</td><td>0.9</td><td>-3.2</td><td>1.2E-03</td><td>1.5E-03</td><td>MB231/MB468</td><td>NUC</td></tr>
<tr><td>>hg38_21_+_8433172_8446622_RNA45SN1_45S_50ntFlanks@12349.12375.27|AAAAGTTACCACAGGGATAACTGGCTT</td><td>>hg38_21_+_8433172_8446622_RNA45SN1_45S_50ntFlanks</td><td>12349</td><td>12375</td><td>27</td><td>AAAAGTTACCACAGGGATAACTGGCTT</td><td>59.5</td><td>-9.0</td><td>2.6</td><td>-3.5</td><td>5.4E-04</td><td>1.5E-03</td><td>BT20/MB468</td><td>MP</td></tr>
<tr><td>>hg38_21_+_8433172_8446622_RNA45SN1_45S_50ntFlanks@9604.9621.18|AAAGACTAATCGAACCAT</td><td>>hg38_21_+_8433172_8446622_RNA45SN1_45S_50ntFlanks</td><td>9604</td><td>9621</td><td>18</td><td>AAAGACTAATCGAACCAT</td><td>295.0</td><td>-3.9</td><td>1.2</td><td>-3.2</td><td>1.2E-03</td><td>1.5E-03</td><td>MB231/MB468</td><td>NUC</td></tr>
<tr><td>>hg38_21_+_8433172_8446622_RNA45SN1_45S_50ntFlanks@12251.12273.23|AAAGCTCGCTTGATCTTGATTTT</td><td>>hg38_21_+_8433172_8446622_RNA45SN1_45S_50ntFlanks</td><td>12251</td><td>12273</td><td>23</td><td>AAAGCTCGCTTGATCTTGATTTT</td><td>72.7</td><td>-5.2</td><td>1.6</td><td>-3.3</td><td>1.0E-03</td><td>1.5E-03</td><td>MB231/MB468</td><td>MP</td></tr>
<tr><td>>MI0000255|hsa-mir-30d&WithFlank&8|-|134804870|134804951@12.33.22(+1A)&[MIMAT0000245&hsa-miR-30d-5p&offsets|0|0(+1A);m-21&8|-|134804919|134804940&offsets|0|0(+1A)]|TGTAAACATCCCCGACTGGAAGA</td><td>>MI0000255|hsa-mir-30d&WithFlank&8|-|134804870|134804951</td><td>12</td><td>33</td><td>22(+1A)</td><td>TGTAAACATCCCCGACTGGAAGA</td><td>2352.4</td><td>2.0</td><td>0.5</td><td>3.7</td><td>2.0E-04</td><td>1.5E-03</td><td>BT20/MB231</td><td>NUC</td></tr>
<tr><td>>hg38_21_+_8433172_8446622_RNA45SN1_45S_50ntFlanks@9554.9584.31|CGTAGCGGTCCTGACGTGCAAATCGGTCGTC</td><td>>hg38_21_+_8433172_8446622_RNA45SN1_45S_50ntFlanks</td><td>9554</td><td>9584</td><td>31</td><td>CGTAGCGGTCCTGACGTGCAAATCGGTCGTC</td><td>988.7</td><td>-1.8</td><td>0.5</td><td>-3.3</td><td>9.1E-04</td><td>1.5E-03</td><td>BT20/MB468</td><td>MITO</td></tr>
<tr><td>>hg38_21_+_8433172_8446622_RNA45SN1_45S_50ntFlanks@11580.11602.23|GAACTGGTGCGGACCAGGGGAAT</td><td>>hg38_21_+_8433172_8446622_RNA45SN1_45S_50ntFlanks</td><td>11580</td><td>11602</td><td>23</td><td>GAACTGGTGCGGACCAGGGGAAT</td><td>308.6</td><td>-2.1</td><td>0.6</td><td>-3.3</td><td>9.1E-04</td><td>1.5E-03</td><td>BT20/MB468</td><td>MITO</td></tr>
<tr><td>>hg38_1_-_228634819_228635039_RNA5S12_5S_50ntFlanks@125.169.45|GTACTTGGATGGGAGACCGCCTGGGAATACCGGGTGCTGTAGGCT</td><td>>hg38_1_-_228634819_228635039_RNA5S12_5S_50ntFlanks</td><td>125</td><td>169</td><td>45</td><td>GTACTTGGATGGGAGACCGCCTGGGAATACCGGGTGCTGTAGGCT</td><td>281.1</td><td>4.3</td><td>1.2</td><td>3.7</td><td>2.3E-04</td><td>1.5E-03</td><td>BT20/MB231</td><td>MITO</td></tr>
<tr><td>>hg38_21_+_8433172_8446622_RNA45SN1_45S_50ntFlanks@7973.7992.20|GACGCGACCTCAGATCAGAC</td><td>>hg38_21_+_8433172_8446622_RNA45SN1_45S_50ntFlanks</td><td>7973</td><td>7992</td><td>20</td><td>GACGCGACCTCAGATCAGAC</td><td>1333.9</td><td>-3.8</td><td>1.0</td><td>-3.7</td><td>2.0E-04</td><td>1.5E-03</td><td>BT20/MB231</td><td>CYTO</td></tr>
<tr><td>>hg38_21_+_8433172_8446622_RNA45SN1_45S_50ntFlanks@8184.8203.20|CGAGGCCCAGCCCGTGGACG</td><td>>hg38_21_+_8433172_8446622_RNA45SN1_45S_50ntFlanks</td><td>8184</td><td>8203</td><td>20</td><td>CGAGGCCCAGCCCGTGGACG</td><td>173.6</td><td>-4.1</td><td>1.3</td><td>-3.3</td><td>1.0E-03</td><td>1.5E-03</td><td>MB231/MB468</td><td>MP</td></tr>
<tr><td>>hg38_21_+_8433172_8446622_RNA45SN1_45S_50ntFlanks@10345.10368.24|TTCAAACGAGAACTTTGAAGGCCG</td><td>>hg38_21_+_8433172_8446622_RNA45SN1_45S_50ntFlanks</td><td>10345</td><td>10368</td><td>24</td><td>TTCAAACGAGAACTTTGAAGGCCG</td><td>196.4</td><td>-4.4</td><td>1.3</td><td>-3.3</td><td>1.0E-03</td><td>1.5E-03</td><td>MB231/MB468</td><td>MP</td></tr>
<tr><td>>trnaMT_GluTTC_MT_-_14674_14742@40.63.24,trnalookalike8_GluTTC_5_-_93905172_93905240@40.63.24|TTGGTCGTGGTTGTAGTCCGTGCG</td><td>>trnaMT_GluTTC_MT_-_14674_14742</td><td>40</td><td>63</td><td>24</td><td>TTGGTCGTGGTTGTAGTCCGTGCG</td><td>224.8</td><td>2.9</td><td>0.9</td><td>3.3</td><td>1.0E-03</td><td>1.5E-03</td><td>MB231/MB468</td><td>MP</td></tr>
<tr><td>>hg38_21_+_8433172_8446622_RNA45SN1_45S_50ntFlanks@12627.12659.33|ATGGGGCGAAGCTACCATCTGTGGGATTATGAC</td><td>>hg38_21_+_8433172_8446622_RNA45SN1_45S_50ntFlanks</td><td>12627</td><td>12659</td><td>33</td><td>ATGGGGCGAAGCTACCATCTGTGGGATTATGAC</td><td>572.2</td><td>-3.5</td><td>1.1</td><td>-3.3</td><td>1.1E-03</td><td>1.5E-03</td><td>MB231/MB468</td><td>CYTO</td></tr>
<tr><td>>hg38_21_+_8433172_8446622_RNA45SN1_45S_50ntFlanks@10786.10821.36|AACAGCCTCTGGCATGTTGGAACAATGTAGGTAAGG</td><td>>hg38_21_+_8433172_8446622_RNA45SN1_45S_50ntFlanks</td><td>10786</td><td>10821</td><td>36</td><td>AACAGCCTCTGGCATGTTGGAACAATGTAGGTAAGG</td><td>413.9</td><td>-2.8</td><td>0.8</td><td>-3.5</td><td>5.4E-04</td><td>1.5E-03</td><td>BT20/MB468</td><td>MP</td></tr>
<tr><td>>hg38_21_+_8433172_8446622_RNA45SN1_45S_50ntFlanks@12416.12439.24|TTCGATGTCGGCTCTTCCTATCAT</td><td>>hg38_21_+_8433172_8446622_RNA45SN1_45S_50ntFlanks</td><td>12416</td><td>12439</td><td>24</td><td>TTCGATGTCGGCTCTTCCTATCAT</td><td>263.1</td><td>2.7</td><td>0.8</td><td>3.2</td><td>1.2E-03</td><td>1.5E-03</td><td>MB231/MB468</td><td>MITO</td></tr>
<tr><td>>hg38_21_+_8433172_8446622_RNA45SN1_45S_50ntFlanks@6651.6677.27|CGACTCTTAGCGGTGGATCACTCGGCT</td><td>>hg38_21_+_8433172_8446622_RNA45SN1_45S_50ntFlanks</td><td>6651</td><td>6677</td><td>27</td><td>CGACTCTTAGCGGTGGATCACTCGGCT</td><td>8832.8</td><td>3.2</td><td>1.0</td><td>3.2</td><td>1.2E-03</td><td>1.5E-03</td><td>MB231/MB468</td><td>NUC</td></tr>
<tr><td>>hg38_21_+_8433172_8446622_RNA45SN1_45S_50ntFlanks@10392.10409.18|ACAGCAGTTGAACATGGG</td><td>>hg38_21_+_8433172_8446622_RNA45SN1_45S_50ntFlanks</td><td>10392</td><td>10409</td><td>18</td><td>ACAGCAGTTGAACATGGG</td><td>2790.0</td><td>-5.9</td><td>1.8</td><td>-3.3</td><td>1.1E-03</td><td>1.5E-03</td><td>BT20/MB468</td><td>CYTO</td></tr>
<tr><td>>hg38_21_+_8433172_8446622_RNA45SN1_45S_50ntFlanks@12475.12498.24|CCCACTAATAGGGAACGTGAGCTG</td><td>>hg38_21_+_8433172_8446622_RNA45SN1_45S_50ntFlanks</td><td>12475</td><td>12498</td><td>24</td><td>CCCACTAATAGGGAACGTGAGCTG</td><td>478.2</td><td>-3.6</td><td>1.1</td><td>-3.3</td><td>1.1E-03</td><td>1.5E-03</td><td>BT20/MB468</td><td>CYTO</td></tr>
<tr><td>>hg38_21_+_8433172_8446622_RNA45SN1_45S_50ntFlanks@11666.11685.20|TCTGCCCAGTGCTCTGAATG</td><td>>hg38_21_+_8433172_8446622_RNA45SN1_45S_50ntFlanks</td><td>11666</td><td>11685</td><td>20</td><td>TCTGCCCAGTGCTCTGAATG</td><td>2618.4</td><td>-2.8</td><td>0.8</td><td>-3.5</td><td>5.4E-04</td><td>1.5E-03</td><td>BT20/MB468</td><td>MP</td></tr>
<tr><td>>hg38_21_+_8433172_8446622_RNA45SN1_45S_50ntFlanks@12487.12509.23|GAACGTGAGCTGGGTTTAGACCG</td><td>>hg38_21_+_8433172_8446622_RNA45SN1_45S_50ntFlanks</td><td>12487</td><td>12509</td><td>23</td><td>GAACGTGAGCTGGGTTTAGACCG</td><td>235.5</td><td>-2.1</td><td>0.6</td><td>-3.3</td><td>1.1E-03</td><td>1.5E-03</td><td>BT20/MB468</td><td>CYTO</td></tr>
<tr><td>>hg38_21_+_8433172_8446622_RNA45SN1_45S_50ntFlanks@5482.5499.18|CGGAGCGCTGAGAAGACG</td><td>>hg38_21_+_8433172_8446622_RNA45SN1_45S_50ntFlanks</td><td>5482</td><td>5499</td><td>18</td><td>CGGAGCGCTGAGAAGACG</td><td>195.0</td><td>-2.2</td><td>0.7</td><td>-3.3</td><td>9.2E-04</td><td>1.5E-03</td><td>BT20/MB468</td><td>MITO</td></tr>
<tr><td>>trnaMT_GluTTC_MT_-_14674_14742@41.72.32,trnalookalike8_GluTTC_5_-_93905172_93905240@41.72.32|TGGTCGTGGTTGTAGTCCGTGCGAGAATACCA</td><td>>trnaMT_GluTTC_MT_-_14674_14742</td><td>41</td><td>72</td><td>32</td><td>TGGTCGTGGTTGTAGTCCGTGCGAGAATACCA</td><td>8558.1</td><td>-3.6</td><td>1.0</td><td>-3.7</td><td>2.0E-04</td><td>1.5E-03</td><td>BT20/MB231</td><td>CYTO</td></tr>
<tr><td>>hg38_21_+_8433172_8446622_RNA45SN1_45S_50ntFlanks@8570.8593.24|CTCCGGCGGGTGCGGGGGTGGGCG</td><td>>hg38_21_+_8433172_8446622_RNA45SN1_45S_50ntFlanks</td><td>8570</td><td>8593</td><td>24</td><td>CTCCGGCGGGTGCGGGGGTGGGCG</td><td>250.6</td><td>-5.5</td><td>1.6</td><td>-3.5</td><td>5.5E-04</td><td>1.5E-03</td><td>BT20/MB468</td><td>MP</td></tr>
<tr><td>>hg38_21_+_8433172_8446622_RNA45SN1_45S_50ntFlanks@11885.11903.19|TTGAGCTTGACTCTAGTCT</td><td>>hg38_21_+_8433172_8446622_RNA45SN1_45S_50ntFlanks</td><td>11885</td><td>11903</td><td>19</td><td>TTGAGCTTGACTCTAGTCT</td><td>230.1</td><td>-2.4</td><td>0.7</td><td>-3.3</td><td>9.2E-04</td><td>1.5E-03</td><td>BT20/MB468</td><td>MITO</td></tr>
<tr><td>>trna111_HisGTG_1_-_147774845_147774916@1.36.36,trna118_HisGTG_1_-_145396881_145396952@1.36.36,trna16_HisGTG_1_+_146544773_146544844@1.36.36,trna1_HisGTG_15_+_45493349_45493420@1.36.36,trna21_HisGTG_1_+_147753471_147753542@1.36.36,trna33_HisGTG_6_+_27125906_27125977@1.36.36,trna7_HisGTG_9_-_14433938_14434009@1.36.36,trna8_HisGTG_15_-_45492611_45492682@1.36.36,trna9_HisGTG_15_-_45490804_45490875@1.36.36|GCCGTGATCGTATAGTGGTTAGTACTCTGCGTTGTG</td><td>>trna111_HisGTG_1_-_147774845_147774916</td><td>1</td><td>36</td><td>36</td><td>GCCGTGATCGTATAGTGGTTAGTACTCTGCGTTGTG</td><td>54.2</td><td>9.5</td><td>2.8</td><td>3.5</td><td>5.5E-04</td><td>1.5E-03</td><td>BT20/MB468</td><td>MP</td></tr>
<tr><td>>MI0000082|hsa-mir-25&WithFlank&7|-|100093554|100093649@58.77.20(+1A)&[MIMAT0000081&hsa-miR-25-3p&offsets|0|-2(+1A);m-23&7|-|100093571|100093592&offsets|0|-2(+1A)]|CATTGCACTTGTCTCGGTCTA</td><td>>MI0000082|hsa-mir-25&WithFlank&7|-|100093554|100093649</td><td>58</td><td>77</td><td>20(+1A)</td><td>CATTGCACTTGTCTCGGTCTA</td><td>729.7</td><td>5.0</td><td>1.4</td><td>3.5</td><td>5.5E-04</td><td>1.5E-03</td><td>BT20/MB468</td><td>MP</td></tr>
<tr><td>>hg38_21_+_8433172_8446622_RNA45SN1_45S_50ntFlanks@11913.11943.31|AAGAGACATGAGAGGTGTAGAATAAGTGGGA</td><td>>hg38_21_+_8433172_8446622_RNA45SN1_45S_50ntFlanks</td><td>11913</td><td>11943</td><td>31</td><td>AAGAGACATGAGAGGTGTAGAATAAGTGGGA</td><td>48.0</td><td>9.3</td><td>2.7</td><td>3.5</td><td>5.5E-04</td><td>1.5E-03</td><td>BT20/MB468</td><td>MP</td></tr>
<tr><td>>hg38_21_+_8433172_8446622_RNA45SN1_45S_50ntFlanks@7977.8010.34|CGACCTCAGATCAGACGTGGCGACCCGCTGAATT</td><td>>hg38_21_+_8433172_8446622_RNA45SN1_45S_50ntFlanks</td><td>7977</td><td>8010</td><td>34</td><td>CGACCTCAGATCAGACGTGGCGACCCGCTGAATT</td><td>465.5</td><td>4.3</td><td>1.2</td><td>3.5</td><td>5.5E-04</td><td>1.5E-03</td><td>BT20/MB468</td><td>MP</td></tr>
<tr><td>>hg38_21_+_8433172_8446622_RNA45SN1_45S_50ntFlanks@9498.9523.26|AAGATGGTGAACTATGCCTGGGCAGG</td><td>>hg38_21_+_8433172_8446622_RNA45SN1_45S_50ntFlanks</td><td>9498</td><td>9523</td><td>26</td><td>AAGATGGTGAACTATGCCTGGGCAGG</td><td>49.0</td><td>9.4</td><td>2.7</td><td>3.5</td><td>5.5E-04</td><td>1.5E-03</td><td>BT20/MB468</td><td>MP</td></tr>
<tr><td>>hg38_21_+_8433172_8446622_RNA45SN1_45S_50ntFlanks@11104.11123.20|AAGGGTCGGGGCGGCAGGGG</td><td>>hg38_21_+_8433172_8446622_RNA45SN1_45S_50ntFlanks</td><td>11104</td><td>11123</td><td>20</td><td>AAGGGTCGGGGCGGCAGGGG</td><td>683.6</td><td>-5.9</td><td>1.7</td><td>-3.5</td><td>5.5E-04</td><td>1.5E-03</td><td>BT20/MB468</td><td>MP</td></tr>
<tr><td>>hg38_21_+_8433172_8446622_RNA45SN1_45S_50ntFlanks@10846.10865.20|CTTCGGGATAAGGATTGGCT</td><td>>hg38_21_+_8433172_8446622_RNA45SN1_45S_50ntFlanks</td><td>10846</td><td>10865</td><td>20</td><td>CTTCGGGATAAGGATTGGCT</td><td>1911.8</td><td>-5.2</td><td>1.5</td><td>-3.5</td><td>5.5E-04</td><td>1.5E-03</td><td>BT20/MB468</td><td>MP</td></tr>
<tr><td>>hg38_21_+_8433172_8446622_RNA45SN1_45S_50ntFlanks@10393.10433.41|CAGCAGTTGAACATGGGTCAGTCGGTCCTGAGAGATGGGCG</td><td>>hg38_21_+_8433172_8446622_RNA45SN1_45S_50ntFlanks</td><td>10393</td><td>10433</td><td>41</td><td>CAGCAGTTGAACATGGGTCAGTCGGTCCTGAGAGATGGGCG</td><td>302.3</td><td>-2.6</td><td>0.8</td><td>-3.3</td><td>9.2E-04</td><td>1.5E-03</td><td>BT20/MB468</td><td>MITO</td></tr>
<tr><td>>hg38_21_+_8433172_8446622_RNA45SN1_45S_50ntFlanks@485.504.20|GATGTGGTGACGTCGTGCTC</td><td>>hg38_21_+_8433172_8446622_RNA45SN1_45S_50ntFlanks</td><td>485</td><td>504</td><td>20</td><td>GATGTGGTGACGTCGTGCTC</td><td>369.9</td><td>11.8</td><td>3.6</td><td>3.2</td><td>1.2E-03</td><td>1.5E-03</td><td>MB231/MB468</td><td>MITO</td></tr>
<tr><td>>hg38_21_+_8433172_8446622_RNA45SN1_45S_50ntFlanks@13007.13027.21|TATTGAAAGTCAGCCCTCGAC</td><td>>hg38_21_+_8433172_8446622_RNA45SN1_45S_50ntFlanks</td><td>13007</td><td>13027</td><td>21</td><td>TATTGAAAGTCAGCCCTCGAC</td><td>920.3</td><td>-3.3</td><td>1.0</td><td>-3.3</td><td>1.1E-03</td><td>1.5E-03</td><td>MB231/MB468</td><td>CYTO</td></tr>
<tr><td>>hg38_21_+_8433172_8446622_RNA45SN1_45S_50ntFlanks@11540.11584.45|CGCGGCGCCTCGCCTCGGCCGGCGCCTAGCAGCCGACTTAGAACT</td><td>>hg38_21_+_8433172_8446622_RNA45SN1_45S_50ntFlanks</td><td>11540</td><td>11584</td><td>45</td><td>CGCGGCGCCTCGCCTCGGCCGGCGCCTAGCAGCCGACTTAGAACT</td><td>1206.9</td><td>3.7</td><td>1.1</td><td>3.2</td><td>1.2E-03</td><td>1.5E-03</td><td>MB231/MB468</td><td>MITO</td></tr>
<tr><td>>MI0000079|hsa-mir-23a&WithFlank&19|-|13836581|13836665@54.73.20(+2U)&[MIMAT0000078&hsa-miR-23a-3p&offsets|+3|+2(+2U);m-31&19|-|13836595|13836615&offsets|+3|+2(+2U)]|ACATTGCCAGGGATTTCCAATT</td><td>>MI0000079|hsa-mir-23a&WithFlank&19|-|13836581|13836665</td><td>54</td><td>73</td><td>20(+2U)</td><td>ACATTGCCAGGGATTTCCAATT</td><td>369.4</td><td>11.8</td><td>3.6</td><td>3.2</td><td>1.2E-03</td><td>1.5E-03</td><td>MB231/MB468</td><td>MITO</td></tr>
<tr><td>>MI0000086|hsa-mir-28&WithFlank&3|+|188688775|188688872@20.41.22&[MIMAT0000085&hsa-miR-28-5p&offsets|0|0;m-133&3|+|188688794|188688815&offsets|0|0]|AAGGAGCTCACAGTCTATTGAG</td><td>>MI0000086|hsa-mir-28&WithFlank&3|+|188688775|188688872</td><td>20</td><td>41</td><td>22</td><td>AAGGAGCTCACAGTCTATTGAG</td><td>853.9</td><td>4.5</td><td>1.4</td><td>3.3</td><td>1.0E-03</td><td>1.5E-03</td><td>MB231/MB468</td><td>MP</td></tr>
</table>

<table>
<tr><td>>MI0000088|hsa-mir-30a&WithFlank&6|-|71403545|71403627@13.35.23(+1A)&[MIMAT0000087&hsa-miR-30a-5p&offsets|+1|+2(+1A);m-15&6|-|71403595|71403616&offsets|+1|+2(+1A)]|GTAAACATCCTCGACTGGAAGCTA</td><td>>MI0000088|hsa-mir-30a&WithFlank&6|-|71403545|71403627</td><td>13</td><td>35</td><td>23(+1A)</td><td>GTAAACATCCTCGACTGGAAGCTA</td><td>62.3</td><td>5.8</td><td>1.8</td><td>3.3</td><td>1.0E-03</td><td>1.5E-03</td><td>MB231/MB468</td><td>MP</td></tr>
<tr><td>>hg38_21_+_8433172_8446622_RNA45SN1_45S_50ntFlanks@4046.4064.19|CATTCGAACGTCTGCCCTA</td><td>>hg38_21_+_8433172_8446622_RNA45SN1_45S_50ntFlanks</td><td>4046</td><td>4064</td><td>19</td><td>CATTCGAACGTCTGCCCTA</td><td>214.1</td><td>-3.0</td><td>0.9</td><td>-3.3</td><td>1.1E-03</td><td>1.5E-03</td><td>MB231/MB468</td><td>CYTO</td></tr>
<tr><td>>hg38_21_+_8433172_8446622_RNA45SN1_45S_50ntFlanks@11580.11606.27|GAACTGGTGCGGACCAGGGGAATCCGA</td><td>>hg38_21_+_8433172_8446622_RNA45SN1_45S_50ntFlanks</td><td>11580</td><td>11606</td><td>27</td><td>GAACTGGTGCGGACCAGGGGAATCCGA</td><td>131.9</td><td>-1.7</td><td>0.5</td><td>-3.3</td><td>9.2E-04</td><td>1.5E-03</td><td>BT20/MB468</td><td>MITO</td></tr>
<tr><td>>MI0000750|hsa-mir-26a-2&WithFlank&12|-|57824603|57824698@20.41.22(+1A);MI0000083|hsa-mir-26a-1&WithFlank&3|+|37969398|37969486@16.37.22(+1A)&[MIMAT0000082&hsa-miR-26a-5p&offsets|0|0(+1A);m-33&12|-|57824658|57824679&offsets|0|0(+1A)];[MIMAT0000082_1&hsa-miR-26a-5p&offsets|0|0(+1A);m-34&3|+|37969413|37969434&offsets|0|0(+1A)]|TTCAAGTAATCCAGGATAGGCTA</td><td>>MI0000750|hsa-mir-26a-2&WithFlank&12|-|57824603|57824698</td><td>20</td><td>41</td><td>22(+1A)</td><td>TTCAAGTAATCCAGGATAGGCTA</td><td>1134.5</td><td>3.3</td><td>1.0</td><td>3.3</td><td>1.0E-03</td><td>1.5E-03</td><td>MB231/MB468</td><td>MP</td></tr>
<tr><td>>hg38_21_+_8433172_8446622_RNA45SN1_45S_50ntFlanks@6651.6680.30|CGACTCTTAGCGGTGGATCACTCGGCTCGT</td><td>>hg38_21_+_8433172_8446622_RNA45SN1_45S_50ntFlanks</td><td>6651</td><td>6680</td><td>30</td><td>CGACTCTTAGCGGTGGATCACTCGGCTCGT</td><td>3153.5</td><td>2.1</td><td>0.6</td><td>3.3</td><td>1.1E-03</td><td>1.5E-03</td><td>BT20/MB468</td><td>CYTO</td></tr>
<tr><td>>hg38_21_+_8433172_8446622_RNA45SN1_45S_50ntFlanks@5492.5516.25|AGAAGACGGTCGAACTTGACTATCT</td><td>>hg38_21_+_8433172_8446622_RNA45SN1_45S_50ntFlanks</td><td>5492</td><td>5516</td><td>25</td><td>AGAAGACGGTCGAACTTGACTATCT</td><td>6441.7</td><td>4.2</td><td>1.3</td><td>3.2</td><td>1.2E-03</td><td>1.5E-03</td><td>BT20/MB468</td><td>NUC</td></tr>
<tr><td>>hg38_21_+_8433172_8446622_RNA45SN1_45S_50ntFlanks@11999.12033.35|CCCTGCGGGCCGCCGGTGAAATACCACTACTCTGA</td><td>>hg38_21_+_8433172_8446622_RNA45SN1_45S_50ntFlanks</td><td>11999</td><td>12033</td><td>35</td><td>CCCTGCGGGCCGCCGGTGAAATACCACTACTCTGA</td><td>517.1</td><td>-3.9</td><td>1.2</td><td>-3.3</td><td>9.2E-04</td><td>1.5E-03</td><td>BT20/MB468</td><td>MITO</td></tr>
<tr><td>>trnaMT_HisGTG_MT_+_12138_12206@26.58.33|AGATTGTGAATCTGACAACAGAGGCTTACGACC</td><td>>trnaMT_HisGTG_MT_+_12138_12206</td><td>26</td><td>58</td><td>33</td><td>AGATTGTGAATCTGACAACAGAGGCTTACGACC</td><td>53.3</td><td>9.5</td><td>2.7</td><td>3.5</td><td>5.5E-04</td><td>1.5E-03</td><td>BT20/MB468</td><td>MP</td></tr>
<tr><td>>hg38_21_+_8433172_8446622_RNA45SN1_45S_50ntFlanks@9577.9595.19|CGGTCGTCCGACCTGGGTA</td><td>>hg38_21_+_8433172_8446622_RNA45SN1_45S_50ntFlanks</td><td>9577</td><td>9595</td><td>19</td><td>CGGTCGTCCGACCTGGGTA</td><td>51.8</td><td>-8.8</td><td>2.5</td><td>-3.5</td><td>5.5E-04</td><td>1.5E-03</td><td>BT20/MB468</td><td>MP</td></tr>
<tr><td>>MI0000077|hsa-mir-21&WithFlank&17|+|59841260|59841343@14.33.20&[MIMAT0000076&hsa-miR-21-5p&offsets|0|-2;m-1&17|+|59841273|59841294&offsets|0|-2]|TAGCTTATCAGACTGATGTT</td><td>>MI0000077|hsa-mir-21&WithFlank&17|+|59841260|59841343</td><td>14</td><td>33</td><td>20</td><td>TAGCTTATCAGACTGATGTT</td><td>1588.7</td><td>3.7</td><td>1.0</td><td>3.7</td><td>2.1E-04</td><td>1.5E-03</td><td>BT20/MB231</td><td>CYTO</td></tr>
<tr><td>>hg38_21_+_8433172_8446622_RNA45SN1_45S_50ntFlanks@12408.12436.29|TTTGATCCTTCGATGTCGGCTCTTCCTAT</td><td>>hg38_21_+_8433172_8446622_RNA45SN1_45S_50ntFlanks</td><td>12408</td><td>12436</td><td>29</td><td>TTTGATCCTTCGATGTCGGCTCTTCCTAT</td><td>1263.7</td><td>-6.4</td><td>2.0</td><td>-3.2</td><td>1.2E-03</td><td>1.5E-03</td><td>BT20/MB468</td><td>NUC</td></tr>
<tr><td>>hg38_21_+_8433172_8446622_RNA45SN1_45S_50ntFlanks@9533.9565.33|GAGGAAACTCTGGTGGAGGTCCGTAGCGGTCCT</td><td>>hg38_21_+_8433172_8446622_RNA45SN1_45S_50ntFlanks</td><td>9533</td><td>9565</td><td>33</td><td>GAGGAAACTCTGGTGGAGGTCCGTAGCGGTCCT</td><td>212.3</td><td>-2.4</td><td>0.8</td><td>-3.2</td><td>1.2E-03</td><td>1.5E-03</td><td>MB231/MB468</td><td>NUC</td></tr>
<tr><td>>MI0000298|hsa-mir-221&WithFlank&X|-|45746151|45746272@71.94.24&[MIMAT0000278&hsa-miR-221-3p&offsets|0|+1;m-91&X|-|45746181|45746202&offsets|0|+2]|AGCTACATTGTCTGCTGGGTTTCA</td><td>>MI0000298|hsa-mir-221&WithFlank&X|-|45746151|45746272</td><td>71</td><td>94</td><td>24</td><td>AGCTACATTGTCTGCTGGGTTTCA</td><td>3162.7</td><td>-2.7</td><td>0.7</td><td>-3.7</td><td>2.3E-04</td><td>1.5E-03</td><td>BT20/MB231</td><td>MITO</td></tr>
<tr><td>>MI0000299|hsa-mir-222&WithFlank&X|-|45747009|45747130@75.95.21&[MIMAT0000279&hsa-miR-222-3p&offsets|0|0;m-140&X|-|45747033|45747056&offsets|0|-3]|AGCTACATCTGGCTACTGGGT</td><td>>MI0000299|hsa-mir-222&WithFlank&X|-|45747009|45747130</td><td>75</td><td>95</td><td>21</td><td>AGCTACATCTGGCTACTGGGT</td><td>4284.4</td><td>3.9</td><td>1.1</td><td>3.5</td><td>5.5E-04</td><td>1.5E-03</td><td>BT20/MB468</td><td>MP</td></tr>
<tr><td>>hg38_21_+_8433172_8446622_RNA45SN1_45S_50ntFlanks@4547.4565.19|CTGGATACCGCAGCTAGGA</td><td>>hg38_21_+_8433172_8446622_RNA45SN1_45S_50ntFlanks</td><td>4547</td><td>4565</td><td>19</td><td>CTGGATACCGCAGCTAGGA</td><td>120.5</td><td>-4.6</td><td>1.4</td><td>-3.3</td><td>9.3E-04</td><td>1.5E-03</td><td>BT20/MB468</td><td>MITO</td></tr>
<tr><td>>hg38_21_+_8433172_8446622_RNA45SN1_45S_50ntFlanks@12151.12180.30|GTGGGGAGTTTGACTGGGGCGGTACACCTG</td><td>>hg38_21_+_8433172_8446622_RNA45SN1_45S_50ntFlanks</td><td>12151</td><td>12180</td><td>30</td><td>GTGGGGAGTTTGACTGGGGCGGTACACCTG</td><td>222.4</td><td>-4.6</td><td>1.4</td><td>-3.3</td><td>1.1E-03</td><td>1.5E-03</td><td>BT20/MB468</td><td>CYTO</td></tr>
<tr><td>>MI0000746|hsa-mir-99b&WithFlank&19|+|51692606|51692687@13.34.22(+1A)&[MIMAT0000689&hsa-miR-99b-5p&offsets|0|0(+1A);m-10&19|+|51692618|51692639&offsets|0|0(+1A)]|CACCCGTAGAACCGACCTTGCGA</td><td>>MI0000746|hsa-mir-99b&WithFlank&19|+|51692606|51692687</td><td>13</td><td>34</td><td>22(+1A)</td><td>CACCCGTAGAACCGACCTTGCGA</td><td>436.2</td><td>3.1</td><td>0.9</td><td>3.5</td><td>5.5E-04</td><td>1.5E-03</td><td>BT20/MB468</td><td>MP</td></tr>
<tr><td>>hg38_21_+_8433172_8446622_RNA45SN1_45S_50ntFlanks@9470.9500.31|GAGCACGAGCGCACGTGTTAGGACCCGAAAG</td><td>>hg38_21_+_8433172_8446622_RNA45SN1_45S_50ntFlanks</td><td>9470</td><td>9500</td><td>31</td><td>GAGCACGAGCGCACGTGTTAGGACCCGAAAG</td><td>121.4</td><td>-5.3</td><td>1.5</td><td>-3.5</td><td>5.5E-04</td><td>1.5E-03</td><td>BT20/MB468</td><td>MP</td></tr>
<tr><td>>hg38_21_+_8433172_8446622_RNA45SN1_45S_50ntFlanks@11029.11053.25|CTCTCTCTCTCTCCCCCGCTCCCCG</td><td>>hg38_21_+_8433172_8446622_RNA45SN1_45S_50ntFlanks</td><td>11029</td><td>11053</td><td>25</td><td>CTCTCTCTCTCTCCCCCGCTCCCCG</td><td>427.3</td><td>-5.7</td><td>1.8</td><td>-3.2</td><td>1.2E-03</td><td>1.5E-03</td><td>MB231/MB468</td><td>NUC</td></tr>
<tr><td>>hg38_MT_+_1621_3279_RNR2_16S_50ntFlanks@1367.1414.48|TCGATGTTGGATCAGGACATCCCGATGGTGCAGCCGCTATTAAAGGTT</td><td>>hg38_MT_+_1621_3279_RNR2_16S_50ntFlanks</td><td>1367</td><td>1414</td><td>48</td><td>TCGATGTTGGATCAGGACATCCCGATGGTGCAGCCGCTATTAAAGGTT</td><td>52.8</td><td>9.5</td><td>2.7</td><td>3.5</td><td>5.5E-04</td><td>1.5E-03</td><td>BT20/MB468</td><td>MP</td></tr>
<tr><td>>hg38_21_+_8433172_8446622_RNA45SN1_45S_50ntFlanks@4449.4466.18|CCTCTCGGCGCCCCCTCG</td><td>>hg38_21_+_8433172_8446622_RNA45SN1_45S_50ntFlanks</td><td>4449</td><td>4466</td><td>18</td><td>CCTCTCGGCGCCCCCTCG</td><td>502.5</td><td>-3.6</td><td>1.1</td><td>-3.2</td><td>1.2E-03</td><td>1.5E-03</td><td>MB231/MB468</td><td>NUC</td></tr>
<tr><td>>hg38_21_+_8433172_8446622_RNA45SN1_45S_50ntFlanks@8117.8145.29|CGGGACATGTGGCGTACGGAAGACCCGCT</td><td>>hg38_21_+_8433172_8446622_RNA45SN1_45S_50ntFlanks</td><td>8117</td><td>8145</td><td>29</td><td>CGGGACATGTGGCGTACGGAAGACCCGCT</td><td>266.8</td><td>6.0</td><td>1.6</td><td>3.7</td><td>1.9E-04</td><td>1.5E-03</td><td>BT20/MB231</td><td>MP</td></tr>
<tr><td>>MI0000650|hsa-mir-200c&WithFlank&12|+|6963693|6963772@49.70.22&[MIMAT0000617&hsa-miR-200c-3p&offsets|-1|-2;m-54&12|+|6963742|6963764&offsets|-1|-2]|CTAATACTGCCGGGTAATGATG</td><td>>MI0000650|hsa-mir-200c&WithFlank&12|+|6963693|6963772</td><td>49</td><td>70</td><td>22</td><td>CTAATACTGCCGGGTAATGATG</td><td>229.9</td><td>4.9</td><td>1.4</td><td>3.5</td><td>5.5E-04</td><td>1.5E-03</td><td>BT20/MB468</td><td>MP</td></tr>
<tr><td>>MI0000734|hsa-mir-106b&WithFlank&7|-|100093987|100094080@58.76.19&[MIMAT0004672&hsa-miR-106b-3p&offsets|0|-3;m-87&7|-|100094003|100094023&offsets|0|-2]|CCGCACTGTGGGTACTTGC</td><td>>MI0000734|hsa-mir-106b&WithFlank&7|-|100093987|100094080</td><td>58</td><td>76</td><td>19</td><td>CCGCACTGTGGGTACTTGC</td><td>370.9</td><td>7.0</td><td>2.0</td><td>3.5</td><td>5.5E-04</td><td>1.5E-03</td><td>BT20/MB468</td><td>MP</td></tr>
<tr><td>>hg38_21_+_8433172_8446622_RNA45SN1_45S_50ntFlanks@11579.11611.33|AGAACTGGTGCGGACCAGGGGAATCCGACTGTT</td><td>>hg38_21_+_8433172_8446622_RNA45SN1_45S_50ntFlanks</td><td>11579</td><td>11611</td><td>33</td><td>AGAACTGGTGCGGACCAGGGGAATCCGACTGTT</td><td>1285.7</td><td>-3.2</td><td>0.9</td><td>-3.5</td><td>5.5E-04</td><td>1.5E-03</td><td>BT20/MB468</td><td>MP</td></tr>
<tr><td>>hg38_21_+_8433172_8446622_RNA45SN1_45S_50ntFlanks@12947.12990.44|AGACGACCTGCTTCTGGGTCGGGGTTTCGTACGTAGCAGAGCAG</td><td>>hg38_21_+_8433172_8446622_RNA45SN1_45S_50ntFlanks</td><td>12947</td><td>12990</td><td>44</td><td>AGACGACCTGCTTCTGGGTCGGGGTTTCGTACGTAGCAGAGCAG</td><td>254.7</td><td>-4.4</td><td>1.4</td><td>-3.2</td><td>1.2E-03</td><td>1.5E-03</td><td>MB231/MB468</td><td>NUC</td></tr>
<tr><td>>hg38_21_+_8433172_8446622_RNA45SN1_45S_50ntFlanks@12933.12953.21|GCTAAACCATTCGTAGACGAC</td><td>>hg38_21_+_8433172_8446622_RNA45SN1_45S_50ntFlanks</td><td>12933</td><td>12953</td><td>21</td><td>GCTAAACCATTCGTAGACGAC</td><td>314.9</td><td>-3.4</td><td>1.0</td><td>-3.3</td><td>1.2E-03</td><td>1.5E-03</td><td>MB231/MB468</td><td>CYTO</td></tr>
<tr><td>>hg38_21_+_8433172_8446622_RNA45SN1_45S_50ntFlanks@6645.6662.18|CTCGTACGACTCTTAGCG</td><td>>hg38_21_+_8433172_8446622_RNA45SN1_45S_50ntFlanks</td><td>6645</td><td>6662</td><td>18</td><td>CTCGTACGACTCTTAGCG</td><td>8841.3</td><td>3.2</td><td>1.0</td><td>3.2</td><td>1.2E-03</td><td>1.5E-03</td><td>MB231/MB468</td><td>MITO</td></tr>
<tr><td>>hg38_21_+_8433172_8446622_RNA45SN1_45S_50ntFlanks@3705.3743.39|TACCTGGTTGATCCTGCCAGTAGCATATGCTTGTCTCAA</td><td>>hg38_21_+_8433172_8446622_RNA45SN1_45S_50ntFlanks</td><td>3705</td><td>3743</td><td>39</td><td>TACCTGGTTGATCCTGCCAGTAGCATATGCTTGTCTCAA</td><td>620.5</td><td>1.6</td><td>0.5</td><td>3.2</td><td>1.2E-03</td><td>1.5E-03</td><td>BT20/MB468</td><td>NUC</td></tr>
<tr><td>>hg38_21_+_8433172_8446622_RNA45SN1_45S_50ntFlanks@12419.12458.40|GATGTCGGCTCTTCCTATCATTGTGAAGCAGAATTCACCA</td><td>>hg38_21_+_8433172_8446622_RNA45SN1_45S_50ntFlanks</td><td>12419</td><td>12458</td><td>40</td><td>GATGTCGGCTCTTCCTATCATTGTGAAGCAGAATTCACCA</td><td>285.4</td><td>-3.2</td><td>0.9</td><td>-3.7</td><td>2.0E-04</td><td>1.5E-03</td><td>BT20/MB231</td><td>NUC</td></tr>
<tr><td>>hg38_21_+_8433172_8446622_RNA45SN1_45S_50ntFlanks@5485.5509.25|AGCGCTGAGAAGACGGTCGAACTTG</td><td>>hg38_21_+_8433172_8446622_RNA45SN1_45S_50ntFlanks</td><td>5485</td><td>5509</td><td>25</td><td>AGCGCTGAGAAGACGGTCGAACTTG</td><td>2799.2</td><td>-3.2</td><td>0.9</td><td>-3.5</td><td>5.6E-04</td><td>1.5E-03</td><td>BT20/MB468</td><td>MP</td></tr>
<tr><td>>hg38_21_+_8433172_8446622_RNA45SN1_45S_50ntFlanks@12637.12654.18|GCTACCATCTGTGGGATT</td><td>>hg38_21_+_8433172_8446622_RNA45SN1_45S_50ntFlanks</td><td>12637</td><td>12654</td><td>18</td><td>GCTACCATCTGTGGGATT</td><td>212.4</td><td>-4.6</td><td>1.4</td><td>-3.3</td><td>1.1E-03</td><td>1.5E-03</td><td>BT20/MB468</td><td>CYTO</td></tr>
<tr><td>>hg38_21_+_8433172_8446622_RNA45SN1_45S_50ntFlanks@6653.6698.46|ACTCTTAGCGGTGGATCACTCGGCTCGTGCGTCGATGAAGAACGCA</td><td>>hg38_21_+_8433172_8446622_RNA45SN1_45S_50ntFlanks</td><td>6653</td><td>6698</td><td>46</td><td>ACTCTTAGCGGTGGATCACTCGGCTCGTGCGTCGATGAAGAACGCA</td><td>251.5</td><td>2.3</td><td>0.7</td><td>3.2</td><td>1.2E-03</td><td>1.5E-03</td><td>MB231/MB468</td><td>CYTO</td></tr>
<tr><td>>hg38_21_+_8433172_8446622_RNA45SN1_45S_50ntFlanks@12331.12365.35|TTAAGCAGGAGGTGTCAGAAAAGTTACCACAGGGA</td><td>>hg38_21_+_8433172_8446622_RNA45SN1_45S_50ntFlanks</td><td>12331</td><td>12365</td><td>35</td><td>TTAAGCAGGAGGTGTCAGAAAAGTTACCACAGGGA</td><td>43.8</td><td>-8.5</td><td>2.5</td><td>-3.5</td><td>5.6E-04</td><td>1.5E-03</td><td>BT20/MB468</td><td>MP</td></tr>
<tr><td>>hg38_21_+_8433172_8446622_RNA45SN1_45S_50ntFlanks@12002.12030.29|TGCGGGCCGCCGGTGAAATACCACTACTC</td><td>>hg38_21_+_8433172_8446622_RNA45SN1_45S_50ntFlanks</td><td>12002</td><td>12030</td><td>29</td><td>TGCGGGCCGCCGGTGAAATACCACTACTC</td><td>319.2</td><td>-2.0</td><td>0.6</td><td>-3.3</td><td>9.3E-04</td><td>1.5E-03</td><td>BT20/MB468</td><td>MITO</td></tr>
<tr><td>>hg38_21_+_8433172_8446622_RNA45SN1_45S_50ntFlanks@9552.9572.21|TCCGTAGCGGTCCTGACGTGC</td><td>>hg38_21_+_8433172_8446622_RNA45SN1_45S_50ntFlanks</td><td>9552</td><td>9572</td><td>21</td><td>TCCGTAGCGGTCCTGACGTGC</td><td>315.2</td><td>-1.7</td><td>0.5</td><td>-3.2</td><td>1.2E-03</td><td>1.5E-03</td><td>MB231/MB468</td><td>CYTO</td></tr>
<tr><td>>hg38_21_+_8433172_8446622_RNA45SN1_45S_50ntFlanks@10574.10607.34|AAGCCGGCGGGAGCCCCGGGGAGAGTTCTCTTTT</td><td>>hg38_21_+_8433172_8446622_RNA45SN1_45S_50ntFlanks</td><td>10574</td><td>10607</td><td>34</td><td>AAGCCGGCGGGAGCCCCGGGGAGAGTTCTCTTTT</td><td>113.8</td><td>-4.5</td><td>1.4</td><td>-3.3</td><td>9.3E-04</td><td>1.5E-03</td><td>BT20/MB468</td><td>MITO</td></tr>
<tr><td>>hg38_21_+_8433172_8446622_RNA45SN1_45S_50ntFlanks@5471.5503.33|CACGGCCCTGGCGGAGCGCTGAGAAGACGGTCG</td><td>>hg38_21_+_8433172_8446622_RNA45SN1_45S_50ntFlanks</td><td>5471</td><td>5503</td><td>33</td><td>CACGGCCCTGGCGGAGCGCTGAGAAGACGGTCG</td><td>152.5</td><td>-2.2</td><td>0.7</td><td>-3.2</td><td>1.2E-03</td><td>1.5E-03</td><td>MB231/MB468</td><td>CYTO</td></tr>
<tr><td>>am_trna10_ValCAC_5_-_180649395_180649467@4.23.20,trna12_ValAAC_5_-_180645270_180645342@4.23.20,trna132_ValAAC_6_-_27721179_27721251@4.23.20,trna136_ValAAC_6_-_27648885_27648957@4.23.20,trna139_ValAAC_6_-_27618707_27618779@4.23.20,trna18_ValCAC_5_-_180529253_180529325@4.23.20,trna2_ValAAC_3_+_169490018_169490090@4.23.20,trna2_ValCAC_5_+_180524070_180524142@4.23.20,trna37_ValAAC_6_+_27203288_27203360@4.23.20,trna4_ValAAC_5_+_180591154_180591226@4.23.20,trna5_ValAAC_5_+_180596610_180596682@4.23.20,trna6_ValCAC_5_+_180600650_180600722@4.23.20,trna85_ValCAC_1_-_161369490_161369562@4.23.20,trna90_ValCAC_1_-_149684088_149684161@4.23.20,trna98_ValCAC_1_-_149298555_149298627@4.23.20,trna9_ValCAC_6_+_26538282_26538354@4.23.20|TCCGTAGTGTAGTGGTTATC</td><td>>am_trna10_ValCAC_5_-_180649395_180649467</td><td>4</td><td>23</td><td>20</td><td>TCCGTAGTGTAGTGGTTATC</td><td>209.2</td><td>3.8</td><td>1.2</td><td>3.2</td><td>1.2E-03</td><td>1.5E-03</td><td>MB231/MB468</td><td>MITO</td></tr>
<tr><td>>trnaMT_PheGAA_MT_+_577_647@35.53.19|GAAAATGTTTAGACGGGCT</td><td>>trnaMT_PheGAA_MT_+_577_647</td><td>35</td><td>53</td><td>19</td><td>GAAAATGTTTAGACGGGCT</td><td>280.4</td><td>2.6</td><td>0.8</td><td>3.3</td><td>1.0E-03</td><td>1.5E-03</td><td>MB231/MB468</td><td>MP</td></tr>
<tr><td>>hg38_21_+_8433172_8446622_RNA45SN1_45S_50ntFlanks@9470.9503.34|GAGCACGAGCGCACGTGTTAGGACCCGAAAGATG</td><td>>hg38_21_+_8433172_8446622_RNA45SN1_45S_50ntFlanks</td><td>9470</td><td>9503</td><td>34</td><td>GAGCACGAGCGCACGTGTTAGGACCCGAAAGATG</td><td>701.8</td><td>-2.1</td><td>0.6</td><td>-3.3</td><td>9.4E-04</td><td>1.5E-03</td><td>BT20/MB468</td><td>MITO</td></tr>
<tr><td>>hg38_21_+_8433172_8446622_RNA45SN1_45S_50ntFlanks@6649.6681.33|TACGACTCTTAGCGGTGGATCACTCGGCTCGTG</td><td>>hg38_21_+_8433172_8446622_RNA45SN1_45S_50ntFlanks</td><td>6649</td><td>6681</td><td>33</td><td>TACGACTCTTAGCGGTGGATCACTCGGCTCGTG</td><td>270.5</td><td>-2.0</td><td>0.6</td><td>-3.3</td><td>1.1E-03</td><td>1.5E-03</td><td>BT20/MB468</td><td>CYTO</td></tr>
<tr><td>>hg38_21_+_8433172_8446622_RNA45SN1_45S_50ntFlanks@6773.6799.27|TTCCTCCCGGGGCTACGCCTGTCTGAG</td><td>>hg38_21_+_8433172_8446622_RNA45SN1_45S_50ntFlanks</td><td>6773</td><td>6799</td><td>27</td><td>TTCCTCCCGGGGCTACGCCTGTCTGAG</td><td>312.1</td><td>-4.8</td><td>1.5</td><td>-3.3</td><td>1.1E-03</td><td>1.5E-03</td><td>BT20/MB468</td><td>CYTO</td></tr>
<tr><td>>trnaMT_HisGTG_MT_+_12138_12206@28.59.32|ATTGTGAATCTGACAACAGAGGCTTACGACCC</td><td>>trnaMT_HisGTG_MT_+_12138_12206</td><td>28</td><td>59</td><td>32</td><td>ATTGTGAATCTGACAACAGAGGCTTACGACCC</td><td>285.9</td><td>4.6</td><td>1.3</td><td>3.5</td><td>5.6E-04</td><td>1.5E-03</td><td>BT20/MB468</td><td>MP</td></tr>
<tr><td>>hg38_21_+_8433172_8446622_RNA45SN1_45S_50ntFlanks@6645.6663.19|CTCGTACGACTCTTAGCGG</td><td>>hg38_21_+_8433172_8446622_RNA45SN1_45S_50ntFlanks</td><td>6645</td><td>6663</td><td>19</td><td>CTCGTACGACTCTTAGCGG</td><td>4796.0</td><td>3.2</td><td>1.0</td><td>3.2</td><td>1.2E-03</td><td>1.5E-03</td><td>MB231/MB468</td><td>NUC</td></tr>
<tr><td>>MI0000082|hsa-mir-25&WithFlank&7|-|100093554|100093649@58.78.21(+1C)&[MIMAT0000081&hsa-miR-25-3p&offsets|0|-1(+1C);m-23&7|-|100093571|100093592&offsets|0|-1(+1C)]|CATTGCACTTGTCTCGGTCTGC</td><td>>MI0000082|hsa-mir-25&WithFlank&7|-|100093554|100093649</td><td>58</td><td>78</td><td>21(+1C)</td><td>CATTGCACTTGTCTCGGTCTGC</td><td>358.3</td><td>5.0</td><td>1.5</td><td>3.5</td><td>5.6E-04</td><td>1.5E-03</td><td>BT20/MB468</td><td>MP</td></tr>
<tr><td>>hg38_21_+_8433172_8446622_RNA45SN1_45S_50ntFlanks@10802.10826.25|TTGGAACAATGTAGGTAAGGGAAGT</td><td>>hg38_21_+_8433172_8446622_RNA45SN1_45S_50ntFlanks</td><td>10802</td><td>10826</td><td>25</td><td>TTGGAACAATGTAGGTAAGGGAAGT</td><td>121.6</td><td>-2.0</td><td>0.6</td><td>-3.3</td><td>9.4E-04</td><td>1.5E-03</td><td>BT20/MB468</td><td>MITO</td></tr>
</table>

<table>
<tr><td>>hg38_21_+_8433172_8446622_RNA45SN1_45S_50ntFlanks@12619.12653.35|AGGAGCCAATGGGGCGAAGCTACCATCTGTGGGAT</td><td>>hg38_21_+_8433172_8446622_RNA45SN1_45S_50ntFlanks</td><td>12619</td><td>12653</td><td>35</td><td>AGGAGCCAATGGGGCGAAGCTACCATCTGTGGGAT</td><td>494.0</td><td>-5.0</td><td>1.5</td><td>-3.3</td><td>1.1E-03</td><td>1.5E-03</td><td>BT20/MB468</td><td>CYTO</td></tr>
<tr><td>>hg38_21_+_8433172_8446622_RNA45SN1_45S_50ntFlanks@10425.10456.32|AGATGGGCGAGCGCCGTTCCGAAGGGACGGGC</td><td>>hg38_21_+_8433172_8446622_RNA45SN1_45S_50ntFlanks</td><td>10425</td><td>10456</td><td>32</td><td>AGATGGGCGAGCGCCGTTCCGAAGGGACGGGC</td><td>293.1</td><td>-3.0</td><td>0.8</td><td>-3.7</td><td>2.1E-04</td><td>1.5E-03</td><td>BT20/MB231</td><td>CYTO</td></tr>
<tr><td>>hg38_21_+_8433172_8446622_RNA45SN1_45S_50ntFlanks@5472.5519.48|ACGGCCCTGGCGGAGCGCTGAGAAGACGGTCGAACTTGACTATCTAGA</td><td>>hg38_21_+_8433172_8446622_RNA45SN1_45S_50ntFlanks</td><td>5472</td><td>5519</td><td>48</td><td>ACGGCCCTGGCGGAGCGCTGAGAAGACGGTCGAACTTGACTATCTAGA</td><td>587.5</td><td>-3.7</td><td>1.0</td><td>-3.7</td><td>2.1E-04</td><td>1.5E-03</td><td>BT20/MB231</td><td>CYTO</td></tr>
<tr><td>>hg38_21_+_8433172_8446622_RNA45SN1_45S_50ntFlanks@8340.8378.39|ACCGTAAGGGAAAGTTGAAAAGAACTTTGAAGAGAGAGT</td><td>>hg38_21_+_8433172_8446622_RNA45SN1_45S_50ntFlanks</td><td>8340</td><td>8378</td><td>39</td><td>ACCGTAAGGGAAAGTTGAAAAGAACTTTGAAGAGAGAGT</td><td>48.2</td><td>6.4</td><td>1.9</td><td>3.5</td><td>5.6E-04</td><td>1.5E-03</td><td>BT20/MB468</td><td>MP</td></tr>
<tr><td>>hg38_21_+_8433172_8446622_RNA45SN1_45S_50ntFlanks@8342.8365.24|CGTAAGGGAAAGTTGAAAAGAACT</td><td>>hg38_21_+_8433172_8446622_RNA45SN1_45S_50ntFlanks</td><td>8342</td><td>8365</td><td>24</td><td>CGTAAGGGAAAGTTGAAAAGAACT</td><td>395.8</td><td>4.7</td><td>1.4</td><td>3.4</td><td>5.6E-04</td><td>1.5E-03</td><td>BT20/MB468</td><td>MP</td></tr>
<tr><td>>hg38_21_+_8433172_8446622_RNA45SN1_45S_50ntFlanks@485.516.32|GATGTGGTGACGTCGTGCTCTCCCGGGCCGGG</td><td>>hg38_21_+_8433172_8446622_RNA45SN1_45S_50ntFlanks</td><td>485</td><td>516</td><td>32</td><td>GATGTGGTGACGTCGTGCTCTCCCGGGCCGGG</td><td>364.1</td><td>11.7</td><td>3.6</td><td>3.2</td><td>1.2E-03</td><td>1.5E-03</td><td>MB231/MB468</td><td>MITO</td></tr>
<tr><td>>hg38_21_+_8433172_8446622_RNA45SN1_45S_50ntFlanks@3854.3882.29|ACTGTGGTAATTCTAGAGCTAATACATGC</td><td>>hg38_21_+_8433172_8446622_RNA45SN1_45S_50ntFlanks</td><td>3854</td><td>3882</td><td>29</td><td>ACTGTGGTAATTCTAGAGCTAATACATGC</td><td>250.7</td><td>-2.5</td><td>0.8</td><td>-3.3</td><td>1.1E-03</td><td>1.5E-03</td><td>BT20/MB468</td><td>CYTO</td></tr>
<tr><td>>hg38_21_+_8433172_8446622_RNA45SN1_45S_50ntFlanks@9737.9754.18|CGAAACGATCTCAACCTA</td><td>>hg38_21_+_8433172_8446622_RNA45SN1_45S_50ntFlanks</td><td>9737</td><td>9754</td><td>18</td><td>CGAAACGATCTCAACCTA</td><td>134.2</td><td>-4.5</td><td>1.4</td><td>-3.3</td><td>1.1E-03</td><td>1.5E-03</td><td>BT20/MB468</td><td>CYTO</td></tr>
<tr><td>>hg38_21_+_8433172_8446622_RNA45SN1_45S_50ntFlanks@485.515.31|GATGTGGTGACGTCGTGCTCTCCCGGGCCGG</td><td>>hg38_21_+_8433172_8446622_RNA45SN1_45S_50ntFlanks</td><td>485</td><td>515</td><td>31</td><td>GATGTGGTGACGTCGTGCTCTCCCGGGCCGG</td><td>49.4</td><td>8.8</td><td>2.7</td><td>3.3</td><td>1.0E-03</td><td>1.5E-03</td><td>MB231/MB468</td><td>MP</td></tr>
<tr><td>>trnaMT_SerGCT_MT_+_12207_12265@27.62.36|CATGCCCCCATGTCTAACAACATGGCTTTCTCACCA</td><td>>trnaMT_SerGCT_MT_+_12207_12265</td><td>27</td><td>62</td><td>36</td><td>CATGCCCCCATGTCTAACAACATGGCTTTCTCACCA</td><td>260.5</td><td>3.5</td><td>1.1</td><td>3.2</td><td>1.2E-03</td><td>1.5E-03</td><td>MB231/MB468</td><td>MITO</td></tr>
<tr><td>>hg38_21_+_8433172_8446622_RNA45SN1_45S_50ntFlanks@485.520.36|GATGTGGTGACGTCGTGCTCTCCCGGGCCGGGTCCG</td><td>>hg38_21_+_8433172_8446622_RNA45SN1_45S_50ntFlanks</td><td>485</td><td>520</td><td>36</td><td>GATGTGGTGACGTCGTGCTCTCCCGGGCCGGGTCCG</td><td>363.6</td><td>11.7</td><td>3.6</td><td>3.2</td><td>1.2E-03</td><td>1.5E-03</td><td>MB231/MB468</td><td>MITO</td></tr>
<tr><td>>hg38_21_+_8433172_8446622_RNA45SN1_45S_50ntFlanks@12420.12454.35|ATGTCGGCTCTTCCTATCATTGTGAAGCAGAATTC</td><td>>hg38_21_+_8433172_8446622_RNA45SN1_45S_50ntFlanks</td><td>12420</td><td>12454</td><td>35</td><td>ATGTCGGCTCTTCCTATCATTGTGAAGCAGAATTC</td><td>367.6</td><td>-2.8</td><td>0.8</td><td>-3.7</td><td>2.4E-04</td><td>1.5E-03</td><td>BT20/MB231</td><td>MITO</td></tr>
<tr><td>>MI0000273|hsa-mir-183&WithFlank&7|-|129774899|129775020@34.54.21(+1A)&[MIMAT0000261&hsa-miR-183-5p&offsets|+1|0(+1A);m-41&7|-|129774966|129774987&offsets|0|-1(+1A)]|ATGGCACTGGTAGAATTCACTA</td><td>>MI0000273|hsa-mir-183&WithFlank&7|-|129774899|129775020</td><td>34</td><td>54</td><td>21(+1A)</td><td>ATGGCACTGGTAGAATTCACTA</td><td>421.5</td><td>2.8</td><td>0.8</td><td>3.7</td><td>2.4E-04</td><td>1.5E-03</td><td>BT20/MB231</td><td>MITO</td></tr>
<tr><td>>MI0000736|hsa-mir-30c-1&WithFlank&1|+|40757278|40757378@23.46.24;MI0000254|hsa-mir-30c-2&WithFlank&6|-|71376954|71377037@13.36.24&[MIMAT0000244&hsa-miR-30c-5p&offsets|0|+1;m-57&1|+|40757300|40757323&offsets|0|0];[MIMAT0000244_1&hsa-miR-30c-5p&offsets|0|+1;m-56&6|-|71377002|71377025&offsets|0|0]|TGTAAACATCCTACACTCTCAGCT</td><td>>MI0000736|hsa-mir-30c-1&WithFlank&1|+|40757278|40757378</td><td>23</td><td>46</td><td>24</td><td>TGTAAACATCCTACACTCTCAGCT</td><td>49904.7</td><td>-3.2</td><td>0.9</td><td>-3.7</td><td>2.4E-04</td><td>1.5E-03</td><td>BT20/MB231</td><td>MITO</td></tr>
<tr><td>>hg38_21_+_8433172_8446622_RNA45SN1_45S_50ntFlanks@10678.10709.32|CGGTTCCGGCGGCGTCCGGTGAGCTCTCGCTG</td><td>>hg38_21_+_8433172_8446622_RNA45SN1_45S_50ntFlanks</td><td>10678</td><td>10709</td><td>32</td><td>CGGTTCCGGCGGCGTCCGGTGAGCTCTCGCTG</td><td>738.3</td><td>3.9</td><td>1.1</td><td>3.7</td><td>2.4E-04</td><td>1.5E-03</td><td>BT20/MB231</td><td>MITO</td></tr>
<tr><td>>MI0000446|hsa-mir-125b-1&WithFlank&11|-|122099751|122099850@21.42.22(+1C);MI0000470|hsa-mir-125b-2&WithFlank&21|+|16590231|16590331@23.44.22(+1C)&[MIMAT0000423&hsa-miR-125b-5p&offsets|0|0(+1C);m-67&11|-|122099809|122099830&offsets|0|0(+1C)];[MIMAT0000423_1&hsa-miR-125b-5p&offsets|0|0(+1C);m-68&21|+|16590253|16590274&offsets|0|0(+1C)]|TCCCTGAGACCCTAACTTGTGAC</td><td>>MI0000446|hsa-mir-125b-1&WithFlank&11|-|122099751|122099850</td><td>21</td><td>42</td><td>22(+1C)</td><td>TCCCTGAGACCCTAACTTGTGAC</td><td>446.8</td><td>-7.5</td><td>2.0</td><td>-3.7</td><td>2.4E-04</td><td>1.5E-03</td><td>BT20/MB231</td><td>MITO</td></tr>
<tr><td>>MI0000483|hsa-mir-186&WithFlank&1|-|71067625|71067722@21.44.24&[MIMAT0000456&hsa-miR-186-5p&offsets|0|+2;m-84&1|-|71067681|71067702&offsets|0|+2]|CAAAGAATTCTCCTTTTGGGCTTT</td><td>>MI0000483|hsa-mir-186&WithFlank&1|-|71067625|71067722</td><td>21</td><td>44</td><td>24</td><td>CAAAGAATTCTCCTTTTGGGCTTT</td><td>569.9</td><td>-3.4</td><td>0.9</td><td>-3.7</td><td>2.4E-04</td><td>1.5E-03</td><td>BT20/MB231</td><td>MITO</td></tr>
<tr><td>>MI0000736|hsa-mir-30c-1&WithFlank&1|+|40757278|40757378@23.42.20;MI0000254|hsa-mir-30c-2&WithFlank&6|-|71376954|71377037@13.32.20&[MIMAT0000244&hsa-miR-30c-5p&offsets|0|-3;m-57&1|+|40757300|40757323&offsets|0|-4];[MIMAT0000244_1&hsa-miR-30c-5p&offsets|0|-3;m-56&6|-|71377002|71377025&offsets|0|-4]|TGTAAACATCCTACACTCTC</td><td>>MI0000736|hsa-mir-30c-1&WithFlank&1|+|40757278|40757378</td><td>23</td><td>42</td><td>20</td><td>TGTAAACATCCTACACTCTC</td><td>2628.7</td><td>-3.0</td><td>0.8</td><td>-3.7</td><td>2.4E-04</td><td>1.5E-03</td><td>BT20/MB231</td><td>MITO</td></tr>
<tr><td>>am_trna128_GlyGCC_6_-_27870686_27870756@8.35.28,trna18_GlyGCC_16_+_70822597_70822667@8.35.28,trna19_GlyGCC_16_+_70823410_70823480@8.35.28,trna19_GlyGCC_2_-_157257659_157257729@8.35.28,trna24_GlyGCC_16_-_70812942_70813012@8.35.28,trna25_GlyGCC_16_-_70812114_70812184@8.35.28,trna2_GlyGCC_21_-_18827107_18827177@8.35.28,trna35_GlyGCC_1_+_161413094_161413164@8.35.28,trna37_GlyGCC_1_+_161420467_161420537@8.35.28,trna39_GlyGCC_1_+_161427898_161427968@8.35.28,trna41_GlyGCC_1_+_161435258_161435328@8.35.28,trna5_GlyGCC_17_+_8029064_8029134@8.35.28,trna68_GlyGCC_1_-_161493637_161493707@8.35.28|TGGTTCAGTGGTAGAATTCTCGCCTGCC</td><td>>am_trna128_GlyGCC_6_-_27870686_27870756</td><td>8</td><td>35</td><td>28</td><td>TGGTTCAGTGGTAGAATTCTCGCCTGCC</td><td>286.0</td><td>-2.8</td><td>0.8</td><td>-3.7</td><td>2.4E-04</td><td>1.5E-03</td><td>BT20/MB231</td><td>MITO</td></tr>
<tr><td>>hg38_21_+_8433172_8446622_RNA45SN1_45S_50ntFlanks@9467.9503.37|GTGGAGCACGAGCGCACGTGTTAGGACCCGAAAGATG</td><td>>hg38_21_+_8433172_8446622_RNA45SN1_45S_50ntFlanks</td><td>9467</td><td>9503</td><td>37</td><td>GTGGAGCACGAGCGCACGTGTTAGGACCCGAAAGATG</td><td>853.3</td><td>-2.4</td><td>0.7</td><td>-3.3</td><td>9.4E-04</td><td>1.5E-03</td><td>BT20/MB468</td><td>MITO</td></tr>
<tr><td>>MI0000457|hsa-mir-141&WithFlank&12|+|6964091|6964197@65.85.21&[MIMAT0000432&hsa-miR-141-3p&offsets|0|-1;m-134&12|+|6964155|6964175&offsets|0|0]|TAACACTGTCTGGTAAAGATG</td><td>>MI0000457|hsa-mir-141&WithFlank&12|+|6964091|6964197</td><td>65</td><td>85</td><td>21</td><td>TAACACTGTCTGGTAAAGATG</td><td>1437.2</td><td>4.2</td><td>1.2</td><td>3.4</td><td>5.6E-04</td><td>1.5E-03</td><td>BT20/MB468</td><td>MP</td></tr>
<tr><td>>hg38_21_+_8433172_8446622_RNA45SN1_45S_50ntFlanks@9004.9026.23|TCTCCCCCGCGGGGGCGCGCCGG</td><td>>hg38_21_+_8433172_8446622_RNA45SN1_45S_50ntFlanks</td><td>9004</td><td>9026</td><td>23</td><td>TCTCCCCCGCGGGGGCGCGCCGG</td><td>126.5</td><td>-5.6</td><td>1.6</td><td>-3.4</td><td>5.6E-04</td><td>1.5E-03</td><td>BT20/MB468</td><td>MP</td></tr>
<tr><td>>hg38_21_+_8433172_8446622_RNA45SN1_45S_50ntFlanks@6687.6720.34|ATGAAGAACGCAGCTAGCTGCGAGAATTAATGTG</td><td>>hg38_21_+_8433172_8446622_RNA45SN1_45S_50ntFlanks</td><td>6687</td><td>6720</td><td>34</td><td>ATGAAGAACGCAGCTAGCTGCGAGAATTAATGTG</td><td>286.8</td><td>-4.9</td><td>1.5</td><td>-3.3</td><td>1.1E-03</td><td>1.5E-03</td><td>BT20/MB468</td><td>CYTO</td></tr>
<tr><td>>hg38_1_-_228634819_228635039_RNA5S12_5S_50ntFlanks@124.170.47|AGTACTTGGATGGGAGACCGCCTGGGAATACCGGGTGCTGTAGGCTT</td><td>>hg38_1_-_228634819_228635039_RNA5S12_5S_50ntFlanks</td><td>124</td><td>170</td><td>47</td><td>AGTACTTGGATGGGAGACCGCCTGGGAATACCGGGTGCTGTAGGCTT</td><td>933.6</td><td>5.0</td><td>1.5</td><td>3.3</td><td>9.4E-04</td><td>1.5E-03</td><td>BT20/MB468</td><td>MITO</td></tr>
<tr><td>>hg38_21_+_8433172_8446622_RNA45SN1_45S_50ntFlanks@7977.8018.42|CGACCTCAGATCAGACGTGGCGACCCGCTGAATTTAAGCATA</td><td>>hg38_21_+_8433172_8446622_RNA45SN1_45S_50ntFlanks</td><td>7977</td><td>8018</td><td>42</td><td>CGACCTCAGATCAGACGTGGCGACCCGCTGAATTTAAGCATA</td><td>512.0</td><td>-6.5</td><td>1.8</td><td>-3.7</td><td>2.1E-04</td><td>1.5E-03</td><td>BT20/MB231</td><td>CYTO</td></tr>
<tr><td>>hg38_21_+_8433172_8446622_RNA45SN1_45S_50ntFlanks@11774.11804.31|AGTGACGCGCATGAATGGATGAACGAGATTC</td><td>>hg38_21_+_8433172_8446622_RNA45SN1_45S_50ntFlanks</td><td>11774</td><td>11804</td><td>31</td><td>AGTGACGCGCATGAATGGATGAACGAGATTC</td><td>176.8</td><td>-4.1</td><td>1.2</td><td>-3.3</td><td>1.0E-03</td><td>1.5E-03</td><td>MB231/MB468</td><td>MP</td></tr>
<tr><td>>hg38_21_+_8433172_8446622_RNA45SN1_45S_50ntFlanks@10683.10708.26|CCGGCGGCGTCCGGTGAGCTCTCGCT</td><td>>hg38_21_+_8433172_8446622_RNA45SN1_45S_50ntFlanks</td><td>10683</td><td>10708</td><td>26</td><td>CCGGCGGCGTCCGGTGAGCTCTCGCT</td><td>269.1</td><td>-1.8</td><td>0.6</td><td>-3.3</td><td>9.4E-04</td><td>1.5E-03</td><td>BT20/MB468</td><td>MITO</td></tr>
<tr><td>>hg38_21_+_8433172_8446622_RNA45SN1_45S_50ntFlanks@6680.6718.39|TGCGTCGATGAAGAACGCAGCTAGCTGCGAGAATTAATG</td><td>>hg38_21_+_8433172_8446622_RNA45SN1_45S_50ntFlanks</td><td>6680</td><td>6718</td><td>39</td><td>TGCGTCGATGAAGAACGCAGCTAGCTGCGAGAATTAATG</td><td>171.7</td><td>-2.8</td><td>0.9</td><td>-3.2</td><td>1.2E-03</td><td>1.5E-03</td><td>MB231/MB468</td><td>CYTO</td></tr>
<tr><td>>trnaMT_GluTTC_MT_-_14674_14742@37.71.35,trnalookalike8_GluTTC_5_-_93905172_93905240@37.71.35|TCATTGGTCGTGGTTGTAGTCCGTGCGAGAATACC</td><td>>trnaMT_GluTTC_MT_-_14674_14742</td><td>37</td><td>71</td><td>35</td><td>TCATTGGTCGTGGTTGTAGTCCGTGCGAGAATACC</td><td>653.8</td><td>-3.9</td><td>1.0</td><td>-3.7</td><td>2.1E-04</td><td>1.5E-03</td><td>BT20/MB231</td><td>NUC</td></tr>
<tr><td>>hg38_21_+_8433172_8446622_RNA45SN1_45S_50ntFlanks@10355.10390.36|AACTTTGAAGGCCGAAGTGGAGAAGGGTTCCATGTG</td><td>>hg38_21_+_8433172_8446622_RNA45SN1_45S_50ntFlanks</td><td>10355</td><td>10390</td><td>36</td><td>AACTTTGAAGGCCGAAGTGGAGAAGGGTTCCATGTG</td><td>313.6</td><td>-5.0</td><td>1.5</td><td>-3.3</td><td>1.1E-03</td><td>1.5E-03</td><td>BT20/MB468</td><td>CYTO</td></tr>
<tr><td>>am_trnaMT_HisGTG_MT_+_12138_12206@27.44.18|GATTGTGAATCTGACAAC</td><td>>am_trnaMT_HisGTG_MT_+_12138_12206</td><td>27</td><td>44</td><td>18</td><td>GATTGTGAATCTGACAAC</td><td>424.2</td><td>3.0</td><td>0.9</td><td>3.3</td><td>9.5E-04</td><td>1.5E-03</td><td>BT20/MB468</td><td>MITO</td></tr>
<tr><td>>hg38_21_+_8433172_8446622_RNA45SN1_45S_50ntFlanks@12408.12448.41|TTTGATCCTTCGATGTCGGCTCTTCCTATCATTGTGAAGCA</td><td>>hg38_21_+_8433172_8446622_RNA45SN1_45S_50ntFlanks</td><td>12408</td><td>12448</td><td>41</td><td>TTTGATCCTTCGATGTCGGCTCTTCCTATCATTGTGAAGCA</td><td>353.8</td><td>3.4</td><td>1.0</td><td>3.3</td><td>9.5E-04</td><td>1.5E-03</td><td>BT20/MB468</td><td>MITO</td></tr>
<tr><td>>hg38_21_+_8433172_8446622_RNA45SN1_45S_50ntFlanks@10350.10377.28|ACGAGAACTTTGAAGGCCGAAGTGGAGA</td><td>>hg38_21_+_8433172_8446622_RNA45SN1_45S_50ntFlanks</td><td>10350</td><td>10377</td><td>28</td><td>ACGAGAACTTTGAAGGCCGAAGTGGAGA</td><td>805.2</td><td>-2.1</td><td>0.6</td><td>-3.3</td><td>9.5E-04</td><td>1.5E-03</td><td>BT20/MB468</td><td>MITO</td></tr>
<tr><td>>hg38_21_+_8433172_8446622_RNA45SN1_45S_50ntFlanks@3784.3801.18|GTGAAACTGCGAATGGCT</td><td>>hg38_21_+_8433172_8446622_RNA45SN1_45S_50ntFlanks</td><td>3784</td><td>3801</td><td>18</td><td>GTGAAACTGCGAATGGCT</td><td>224.9</td><td>-2.2</td><td>0.7</td><td>-3.3</td><td>9.5E-04</td><td>1.5E-03</td><td>BT20/MB468</td><td>MITO</td></tr>
<tr><td>>hg38_21_+_8433172_8446622_RNA45SN1_45S_50ntFlanks@7973.8008.36|GACGCGACCTCAGATCAGACGTGGCGACCCGCTGAA</td><td>>hg38_21_+_8433172_8446622_RNA45SN1_45S_50ntFlanks</td><td>7973</td><td>8008</td><td>36</td><td>GACGCGACCTCAGATCAGACGTGGCGACCCGCTGAA</td><td>2320.7</td><td>2.3</td><td>0.7</td><td>3.2</td><td>1.3E-03</td><td>1.5E-03</td><td>MB231/MB468</td><td>NUC</td></tr>
<tr><td>>hg38_21_+_8433172_8446622_RNA45SN1_45S_50ntFlanks@11571.11612.42|GCCGACTTAGAACTGGTGCGGACCAGGGGAATCCGACTGTTT</td><td>>hg38_21_+_8433172_8446622_RNA45SN1_45S_50ntFlanks</td><td>11571</td><td>11612</td><td>42</td><td>GCCGACTTAGAACTGGTGCGGACCAGGGGAATCCGACTGTTT</td><td>364.4</td><td>-3.6</td><td>1.0</td><td>-3.7</td><td>2.1E-04</td><td>1.5E-03</td><td>BT20/MB231</td><td>CYTO</td></tr>
<tr><td>>hg38_21_+_8433172_8446622_RNA45SN1_45S_50ntFlanks@8689.8706.18|GCTCCGGGACGGCTGGGA</td><td>>hg38_21_+_8433172_8446622_RNA45SN1_45S_50ntFlanks</td><td>8689</td><td>8706</td><td>18</td><td>GCTCCGGGACGGCTGGGA</td><td>354.7</td><td>-2.7</td><td>0.7</td><td>-3.7</td><td>2.1E-04</td><td>1.5E-03</td><td>BT20/MB231</td><td>CYTO</td></tr>
<tr><td>>MI0000298|hsa-mir-221&WithFlank&X|-|45746151|45746272@71.89.19(+1C)&[MIMAT0000278&hsa-miR-221-3p&offsets|0|-4(+1C);m-91&X|-|45746181|45746202&offsets|0|-3(+1C)]|AGCTACATTGTCTGCTGGGC</td><td>>MI0000298|hsa-mir-221&WithFlank&X|-|45746151|45746272</td><td>71</td><td>89</td><td>19(+1C)</td><td>AGCTACATTGTCTGCTGGGC</td><td>178.9</td><td>5.2</td><td>1.6</td><td>3.3</td><td>1.0E-03</td><td>1.5E-03</td><td>MB231/MB468</td><td>MP</td></tr>
<tr><td>>hg38_21_+_8433172_8446622_RNA45SN1_45S_50ntFlanks@7983.8005.23|CAGATCAGACGTGGCGACCCGCT</td><td>>hg38_21_+_8433172_8446622_RNA45SN1_45S_50ntFlanks</td><td>7983</td><td>8005</td><td>23</td><td>CAGATCAGACGTGGCGACCCGCT</td><td>329.2</td><td>-2.5</td><td>0.8</td><td>-3.2</td><td>1.2E-03</td><td>1.5E-03</td><td>MB231/MB468</td><td>CYTO</td></tr>
<tr><td>>hg38_21_+_8433172_8446622_RNA45SN1_45S_50ntFlanks@11588.11623.36|GCGGACCAGGGGAATCCGACTGTTTAATTAAAACAA</td><td>>hg38_21_+_8433172_8446622_RNA45SN1_45S_50ntFlanks</td><td>11588</td><td>11623</td><td>36</td><td>GCGGACCAGGGGAATCCGACTGTTTAATTAAAACAA</td><td>48.9</td><td>8.8</td><td>2.7</td><td>3.3</td><td>1.0E-03</td><td>1.5E-03</td><td>MB231/MB468</td><td>MP</td></tr>
<tr><td>>hg38_21_+_8433172_8446622_RNA45SN1_45S_50ntFlanks@7974.8011.38|ACGCGACCTCAGATCAGACGTGGCGACCCGCTGAATTT</td><td>>hg38_21_+_8433172_8446622_RNA45SN1_45S_50ntFlanks</td><td>7974</td><td>8011</td><td>38</td><td>ACGCGACCTCAGATCAGACGTGGCGACCCGCTGAATTT</td><td>15501.4</td><td>2.3</td><td>0.7</td><td>3.2</td><td>1.2E-03</td><td>1.5E-03</td><td>MB231/MB468</td><td>MITO</td></tr>
<tr><td>>hg38_21_+_8433172_8446622_RNA45SN1_45S_50ntFlanks@10427.10453.27|ATGGGCGAGCGCCGTTCCGAAGGGACG</td><td>>hg38_21_+_8433172_8446622_RNA45SN1_45S_50ntFlanks</td><td>10427</td><td>10453</td><td>27</td><td>ATGGGCGAGCGCCGTTCCGAAGGGACG</td><td>1255.3</td><td>-2.0</td><td>0.6</td><td>-3.2</td><td>1.2E-03</td><td>1.5E-03</td><td>MB231/MB468</td><td>CYTO</td></tr>
<tr><td>>MI0000439|hsa-mir-23b&WithFlank&9|+|95085202|95085310@67.85.19&[MIMAT0000418&hsa-miR-23b-3p&offsets|+3|-1;m-42&9|+|95085265|95085285&offsets|+3|+1]|ACATTGCCAGGGATTACCA</td><td>>MI0000439|hsa-mir-23b&WithFlank&9|+|95085202|95085310</td><td>67</td><td>85</td><td>19</td><td>ACATTGCCAGGGATTACCA</td><td>224.9</td><td>11.7</td><td>3.6</td><td>3.2</td><td>1.3E-03</td><td>1.5E-03</td><td>MB231/MB468</td><td>NUC</td></tr>
</table>

<table>
<tr><td></td><td></td><td></td><td></td><td></td><td></td><td></td><td></td><td></td><td></td><td></td><td></td><td></td><td></td></tr>
<tr><td>>MI0000736|hsa-mir-30c-1&WithFlank&1|+|40757278|40757378@23.45.23(+1G);MI0000254|hsa-mir-30c-2&WithFlank&6|-|71376954|71377037@13.35.23(+1G)&[MIMAT0000244&hsa-miR-30c-5p&offsets|0|0(+1G);m-57&1|+|40757300|40757323&offsets|0|-1(+1G)];[MIMAT0000244_1&hsa-miR-30c-5p&offsets|0|0(+1G);m-56&6|-|71377002|71377025&offsets|0|-1(+1G)]|TGTAAACATCCTACACTCTCAGCG</td><td>>MI0000736|hsa-mir-30c-1&WithFlank&1|+|40757278|40757378</td><td>23</td><td>45</td><td>23(+1G)</td><td>TGTAAACATCCTACACTCTCAGCG</td><td>1061.5</td><td>-3.5</td><td>0.9</td><td>-3.7</td><td>2.5E-04</td><td>1.5E-03</td><td>BT20/MB231</td><td>MITO</td></tr>
<tr><td>>am_trnaMT_MetCAT_MT_+_4402_4469@1.21.21|AGTAAGGTCAGCTAAATAAGC</td><td>>am_trnaMT_MetCAT_MT_+_4402_4469</td><td>1</td><td>21</td><td>21</td><td>AGTAAGGTCAGCTAAATAAGC</td><td>3192.7</td><td>2.7</td><td>0.8</td><td>3.3</td><td>1.1E-03</td><td>1.5E-03</td><td>MB231/MB468</td><td>MP</td></tr>
<tr><td>>hg38_1_-_228634819_228635039_RNA5S12_5S_50ntFlanks@51.84.34|GTCTACGGCCATACCACCCTGAACGCGCCCGATC</td><td>>hg38_1_-_228634819_228635039_RNA5S12_5S_50ntFlanks</td><td>51</td><td>84</td><td>34</td><td>GTCTACGGCCATACCACCCTGAACGCGCCCGATC</td><td>6890.1</td><td>2.0</td><td>0.6</td><td>3.2</td><td>1.3E-03</td><td>1.5E-03</td><td>MB231/MB468</td><td>NUC</td></tr>
<tr><td>>am_trnaMT_PheGAA_MT_+_577_647@44.62.19|TAGACGGGCTCACATCACC</td><td>>am_trnaMT_PheGAA_MT_+_577_647</td><td>44</td><td>62</td><td>19</td><td>TAGACGGGCTCACATCACC</td><td>433.3</td><td>3.9</td><td>1.2</td><td>3.2</td><td>1.2E-03</td><td>1.5E-03</td><td>MB231/MB468</td><td>CYTO</td></tr>
<tr><td>>hg38_21_+_8433172_8446622_RNA45SN1_45S_50ntFlanks@12475.12499.25|CCCACTAATAGGGAACGTGAGCTGG</td><td>>hg38_21_+_8433172_8446622_RNA45SN1_45S_50ntFlanks</td><td>12475</td><td>12499</td><td>25</td><td>CCCACTAATAGGGAACGTGAGCTGG</td><td>207.9</td><td>-3.3</td><td>1.0</td><td>-3.2</td><td>1.2E-03</td><td>1.5E-03</td><td>MB231/MB468</td><td>CYTO</td></tr>
<tr><td>>hg38_21_+_8433172_8446622_RNA45SN1_45S_50ntFlanks@5075.5110.36|TTCCGATAACGAACGAGACTCTGGCATGCTAACTAG</td><td>>hg38_21_+_8433172_8446622_RNA45SN1_45S_50ntFlanks</td><td>5075</td><td>5110</td><td>36</td><td>TTCCGATAACGAACGAGACTCTGGCATGCTAACTAG</td><td>59.8</td><td>-9.8</td><td>3.0</td><td>-3.3</td><td>1.1E-03</td><td>1.5E-03</td><td>MB231/MB468</td><td>MP</td></tr>
<tr><td>>hg38_21_+_8433172_8446622_RNA45SN1_45S_50ntFlanks@3815.3842.28|ATGGTTCCTTTGGTCGCTCGCTCCTCTC</td><td>>hg38_21_+_8433172_8446622_RNA45SN1_45S_50ntFlanks</td><td>3815</td><td>3842</td><td>28</td><td>ATGGTTCCTTTGGTCGCTCGCTCCTCTC</td><td>128.3</td><td>-3.6</td><td>1.1</td><td>-3.3</td><td>9.5E-04</td><td>1.5E-03</td><td>BT20/MB468</td><td>MITO</td></tr>
<tr><td>>hg38_21_+_8433172_8446622_RNA45SN1_45S_50ntFlanks@10420.10454.35|CTGAGAGATGGGCGAGCGCCGTTCCGAAGGGACGG</td><td>>hg38_21_+_8433172_8446622_RNA45SN1_45S_50ntFlanks</td><td>10420</td><td>10454</td><td>35</td><td>CTGAGAGATGGGCGAGCGCCGTTCCGAAGGGACGG</td><td>130.7</td><td>-4.7</td><td>1.4</td><td>-3.3</td><td>1.1E-03</td><td>1.5E-03</td><td>BT20/MB468</td><td>CYTO</td></tr>
<tr><td>>hg38_21_+_8433172_8446622_RNA45SN1_45S_50ntFlanks@6721.6757.37|AATTGCAGGACACATTGATCATCGACACTTCGAACGC</td><td>>hg38_21_+_8433172_8446622_RNA45SN1_45S_50ntFlanks</td><td>6721</td><td>6757</td><td>37</td><td>AATTGCAGGACACATTGATCATCGACACTTCGAACGC</td><td>280.0</td><td>-3.0</td><td>0.9</td><td>-3.2</td><td>1.2E-03</td><td>1.5E-03</td><td>MB231/MB468</td><td>CYTO</td></tr>
<tr><td>>hg38_21_+_8433172_8446622_RNA45SN1_45S_50ntFlanks@11208.11251.44|GCGGCGACTCTGGACGCGAGCCGGGCCCTTCCCGTGGATCGCCC</td><td>>hg38_21_+_8433172_8446622_RNA45SN1_45S_50ntFlanks</td><td>11208</td><td>11251</td><td>44</td><td>GCGGCGACTCTGGACGCGAGCCGGGCCCTTCCCGTGGATCGCCC</td><td>439.0</td><td>-5.2</td><td>1.6</td><td>-3.2</td><td>1.3E-03</td><td>1.5E-03</td><td>MB231/MB468</td><td>NUC</td></tr>
<tr><td>>hg38_MT_+_1621_3279_RNR2_16S_50ntFlanks@1415.1453.39|CGTTTGTTCAACGATTAAAGTCCTACGTGATCTGAGTTC</td><td>>hg38_MT_+_1621_3279_RNR2_16S_50ntFlanks</td><td>1415</td><td>1453</td><td>39</td><td>CGTTTGTTCAACGATTAAAGTCCTACGTGATCTGAGTTC</td><td>55.6</td><td>5.8</td><td>1.8</td><td>3.3</td><td>1.1E-03</td><td>1.5E-03</td><td>MB231/MB468</td><td>MP</td></tr>
<tr><td>>hg38_21_+_8433172_8446622_RNA45SN1_45S_50ntFlanks@476.521.46|TTCGTGATCGATGTGGTGACGTCGTGCTCTCCCGGGCCGGGTCCGA</td><td>>hg38_21_+_8433172_8446622_RNA45SN1_45S_50ntFlanks</td><td>476</td><td>521</td><td>46</td><td>TTCGTGATCGATGTGGTGACGTCGTGCTCTCCCGGGCCGGGTCCGA</td><td>3212.1</td><td>3.6</td><td>1.1</td><td>3.2</td><td>1.3E-03</td><td>1.5E-03</td><td>MB231/MB468</td><td>NUC</td></tr>
<tr><td>>hg38_21_+_8433172_8446622_RNA45SN1_45S_50ntFlanks@12437.12472.36|CATTGTGAAGCAGAATTCACCAAGCGTTGGATTGTT</td><td>>hg38_21_+_8433172_8446622_RNA45SN1_45S_50ntFlanks</td><td>12437</td><td>12472</td><td>36</td><td>CATTGTGAAGCAGAATTCACCAAGCGTTGGATTGTT</td><td>186.0</td><td>-2.4</td><td>0.8</td><td>-3.2</td><td>1.3E-03</td><td>1.5E-03</td><td>MB231/MB468</td><td>NUC</td></tr>
<tr><td>>MI0000270|hsa-mir-181b-1&WithFlank&1|-|198858867|198858988@42.64.23(+1C);MI0000683|hsa-mir-181b-2&WithFlank&9|+|124693704|124693804@22.44.23(+1C)&[MIMAT0000257&hsa-miR-181b-5p&offsets|0|0(+1C);m-73&1|-|198858925|198858947&offsets|0|0(+1C)];[MIMAT0000257_1&hsa-miR-181b-5p&offsets|0|0(+1C);m-72&9|+|124693725|124693747&offsets|0|0(+1C)]|AACATTCATTGCTGTCGGTGGGTC</td><td>>MI0000270|hsa-mir-181b-1&WithFlank&1|-|198858867|198858988</td><td>42</td><td>64</td><td>23(+1C)</td><td>AACATTCATTGCTGTCGGTGGGTC</td><td>560.1</td><td>4.3</td><td>1.3</td><td>3.3</td><td>1.1E-03</td><td>1.5E-03</td><td>MB231/MB468</td><td>MP</td></tr>
<tr><td>>hg38_21_+_8433172_8446622_RNA45SN1_45S_50ntFlanks@6647.6680.34|CGTACGACTCTTAGCGGTGGATCACTCGGCTCGT</td><td>>hg38_21_+_8433172_8446622_RNA45SN1_45S_50ntFlanks</td><td>6647</td><td>6680</td><td>34</td><td>CGTACGACTCTTAGCGGTGGATCACTCGGCTCGT</td><td>607.7</td><td>-2.6</td><td>0.8</td><td>-3.3</td><td>1.1E-03</td><td>1.5E-03</td><td>MB231/MB468</td><td>MP</td></tr>
<tr><td>>trna152_ValCAC_6_-_27248049_27248121@1.24.24|GCTTCTGTAGTGTAGTGGTTATCA</td><td>>trna152_ValCAC_6_-_27248049_27248121</td><td>1</td><td>24</td><td>24</td><td>GCTTCTGTAGTGTAGTGGTTATCA</td><td>282.5</td><td>-3.9</td><td>1.2</td><td>-3.3</td><td>1.1E-03</td><td>1.5E-03</td><td>BT20/MB468</td><td>CYTO</td></tr>
<tr><td>>MI0000063|hsa-let-7b&WithFlank&22|+|46113680|46113774@12.33.22(+1G)&[MIMAT0000063&hsa-let-7b-5p&offsets|0|0(+1G);m-14&22|+|46113691|46113712&offsets|0|0(+1G)]|TGAGGTAGTAGGTTGTGTGGTTG</td><td>>MI0000063|hsa-let-7b&WithFlank&22|+|46113680|46113774</td><td>12</td><td>33</td><td>22(+1G)</td><td>TGAGGTAGTAGGTTGTGTGGTTG</td><td>479.8</td><td>3.2</td><td>0.9</td><td>3.7</td><td>2.1E-04</td><td>1.5E-03</td><td>BT20/MB231</td><td>CYTO</td></tr>
<tr><td>>hg38_21_+_8433172_8446622_RNA45SN1_45S_50ntFlanks@11672.11707.36|CAGTGCTCTGAATGTCAAAGTGAAGAAATTCAATGA</td><td>>hg38_21_+_8433172_8446622_RNA45SN1_45S_50ntFlanks</td><td>11672</td><td>11707</td><td>36</td><td>CAGTGCTCTGAATGTCAAAGTGAAGAAATTCAATGA</td><td>705.1</td><td>3.0</td><td>0.9</td><td>3.2</td><td>1.2E-03</td><td>1.5E-03</td><td>MB231/MB468</td><td>CYTO</td></tr>
<tr><td>>trnaMT_GluTTC_MT_-_14674_14742@44.72.29,trnalookalike8_GluTTC_5_-_93905172_93905240@44.72.29|TCGTGGTTGTAGTCCGTGCGAGAATACCA</td><td>>trnaMT_GluTTC_MT_-_14674_14742</td><td>44</td><td>72</td><td>29</td><td>TCGTGGTTGTAGTCCGTGCGAGAATACCA</td><td>779.7</td><td>-3.1</td><td>0.8</td><td>-3.7</td><td>2.1E-04</td><td>1.5E-03</td><td>BT20/MB231</td><td>NUC</td></tr>
<tr><td>>MI0003591|hsa-mir-584&WithFlank&5|-|149062307|149062415@22.39.18(+1C)&[MIMAT0003249&hsa-miR-584-5p&offsets|0|-4(+1C);m-167&5|-|149062374|149062394&offsets|0|-3(+1C)]|TTATGGTTTGCCTGGGACC</td><td>>MI0003591|hsa-mir-584&WithFlank&5|-|149062307|149062415</td><td>22</td><td>39</td><td>18(+1C)</td><td>TTATGGTTTGCCTGGGACC</td><td>106.9</td><td>6.0</td><td>1.8</td><td>3.3</td><td>1.1E-03</td><td>1.5E-03</td><td>MB231/MB468</td><td>MP</td></tr>
<tr><td>>hg38_21_+_8433172_8446622_RNA45SN1_45S_50ntFlanks@10807.10828.22|ACAATGTAGGTAAGGGAAGTCG</td><td>>hg38_21_+_8433172_8446622_RNA45SN1_45S_50ntFlanks</td><td>10807</td><td>10828</td><td>22</td><td>ACAATGTAGGTAAGGGAAGTCG</td><td>210.7</td><td>-4.7</td><td>1.4</td><td>-3.3</td><td>1.1E-03</td><td>1.5E-03</td><td>BT20/MB468</td><td>CYTO</td></tr>
<tr><td>>hg38_MT_+_1621_3279_RNR2_16S_50ntFlanks@625.663.39|AACTGAACTCCTCACACCCAATTGGACCAATCTATCACC</td><td>>hg38_MT_+_1621_3279_RNR2_16S_50ntFlanks</td><td>625</td><td>663</td><td>39</td><td>AACTGAACTCCTCACACCCAATTGGACCAATCTATCACC</td><td>103.7</td><td>6.9</td><td>2.0</td><td>3.4</td><td>5.7E-04</td><td>1.5E-03</td><td>BT20/MB468</td><td>MP</td></tr>
<tr><td>>MI0000067|hsa-let-7f-1&WithFlank&9|+|94176341|94176439@13.35.23&[MIMAT0000067&hsa-let-7f-5p&offsets|0|+1;m-20&9|+|94176353|94176374&offsets|0|+1]|TGAGGTAGTAGATTGTATAGTTG</td><td>>MI0000067|hsa-let-7f-1&WithFlank&9|+|94176341|94176439</td><td>13</td><td>35</td><td>23</td><td>TGAGGTAGTAGATTGTATAGTTG</td><td>272.6</td><td>4.9</td><td>1.4</td><td>3.4</td><td>5.7E-04</td><td>1.5E-03</td><td>BT20/MB468</td><td>MP</td></tr>
<tr><td>>hg38_21_+_8433172_8446622_RNA45SN1_45S_50ntFlanks@10781.10817.37|AGGTGAACAGCCTCTGGCATGTTGGAACAATGTAGGT</td><td>>hg38_21_+_8433172_8446622_RNA45SN1_45S_50ntFlanks</td><td>10781</td><td>10817</td><td>37</td><td>AGGTGAACAGCCTCTGGCATGTTGGAACAATGTAGGT</td><td>277.0</td><td>-4.9</td><td>1.5</td><td>-3.2</td><td>1.3E-03</td><td>1.5E-03</td><td>MB231/MB468</td><td>NUC</td></tr>
<tr><td>>hg38_MT_+_1621_3279_RNR2_16S_50ntFlanks@1170.1209.40|AAACCTGCATTAAAAATTTCGGTTGGGGCGACCTCGGAGC</td><td>>hg38_MT_+_1621_3279_RNR2_16S_50ntFlanks</td><td>1170</td><td>1209</td><td>40</td><td>AAACCTGCATTAAAAATTTCGGTTGGGGCGACCTCGGAGC</td><td>81.1</td><td>5.7</td><td>1.8</td><td>3.2</td><td>1.3E-03</td><td>1.5E-03</td><td>MB231/MB468</td><td>MITO</td></tr>
<tr><td>>MI0000767|hsa-mir-365a&WithFlank&16|+|14309279|14309377@62.81.20;MI0000769|hsa-mir-365b&WithFlank&17|+|31575405|31575527@74.93.20&[MIMAT0000710&hsa-miR-365a-3p&offsets|0|-2;m-176&16|+|14309340|14309361&offsets|0|-2];[MIMAT0022834&hsa-miR-365b-3p&offsets|0|-2;m-175&17|+|31575478|31575499&offsets|0|-2]|TAATGCCCCTAAAAATCCTT</td><td>>MI0000767|hsa-mir-365a&WithFlank&16|+|14309279|14309377</td><td>62</td><td>81</td><td>20</td><td>TAATGCCCCTAAAAATCCTT</td><td>340.7</td><td>4.7</td><td>1.3</td><td>3.7</td><td>2.1E-04</td><td>1.5E-03</td><td>BT20/MB231</td><td>NUC</td></tr>
<tr><td>>hg38_21_+_8433172_8446622_RNA45SN1_45S_50ntFlanks@7975.8003.29|CGCGACCTCAGATCAGACGTGGCGACCCG</td><td>>hg38_21_+_8433172_8446622_RNA45SN1_45S_50ntFlanks</td><td>7975</td><td>8003</td><td>29</td><td>CGCGACCTCAGATCAGACGTGGCGACCCG</td><td>9602.5</td><td>2.7</td><td>0.8</td><td>3.3</td><td>9.6E-04</td><td>1.5E-03</td><td>BT20/MB468</td><td>MITO</td></tr>
<tr><td>>hg38_21_+_8433172_8446622_RNA45SN1_45S_50ntFlanks@11209.11229.21|CGGCGACTCTGGACGCGAGCC</td><td>>hg38_21_+_8433172_8446622_RNA45SN1_45S_50ntFlanks</td><td>11209</td><td>11229</td><td>21</td><td>CGGCGACTCTGGACGCGAGCC</td><td>545.2</td><td>3.5</td><td>1.0</td><td>3.7</td><td>2.5E-04</td><td>1.5E-03</td><td>BT20/MB231</td><td>MITO</td></tr>
<tr><td>>MI0001445|hsa-mir-423&WithFlank&17|+|30117073|30117178@58.77.20&[MIMAT0001340&hsa-miR-423-3p&offsets|-1|-4;m-100&17|+|30117131|30117153&offsets|-1|-4]|AAGCTCGGTCTGAGGCCCCT</td><td>>MI0001445|hsa-mir-423&WithFlank&17|+|30117073|30117178</td><td>58</td><td>77</td><td>20</td><td>AAGCTCGGTCTGAGGCCCCT</td><td>87.7</td><td>5.7</td><td>1.6</td><td>3.4</td><td>5.7E-04</td><td>1.5E-03</td><td>BT20/MB468</td><td>MP</td></tr>
<tr><td>>MI0000483|hsa-mir-186&WithFlank&1|-|71067625|71067722@21.40.20&[MIMAT0000456&hsa-miR-186-5p&offsets|0|-2;m-84&1|-|71067681|71067702&offsets|0|-2]|CAAAGAATTCTCCTTTTGGG</td><td>>MI0000483|hsa-mir-186&WithFlank&1|-|71067625|71067722</td><td>21</td><td>40</td><td>20</td><td>CAAAGAATTCTCCTTTTGGG</td><td>88.5</td><td>1.2</td><td>0.4</td><td>3.2</td><td>1.3E-03</td><td>1.5E-03</td><td>BT20/MB468</td><td>NUC</td></tr>
<tr><td>>hg38_21_+_8433172_8446622_RNA45SN1_45S_50ntFlanks@9935.9952.18|GACAGCAGGACGGTGGCC</td><td>>hg38_21_+_8433172_8446622_RNA45SN1_45S_50ntFlanks</td><td>9935</td><td>9952</td><td>18</td><td>GACAGCAGGACGGTGGCC</td><td>79.3</td><td>5.5</td><td>1.6</td><td>3.4</td><td>5.8E-04</td><td>1.5E-03</td><td>BT20/MB468</td><td>MP</td></tr>
<tr><td>>hg38_MT_+_1621_3279_RNR2_16S_50ntFlanks@644.663.20|AATTGGACCAATCTATCACC</td><td>>hg38_MT_+_1621_3279_RNR2_16S_50ntFlanks</td><td>644</td><td>663</td><td>20</td><td>AATTGGACCAATCTATCACC</td><td>165.7</td><td>4.1</td><td>1.2</td><td>3.4</td><td>5.8E-04</td><td>1.5E-03</td><td>BT20/MB468</td><td>MP</td></tr>
<tr><td>>am_trna128_GlyGCC_6_-_27870686_27870756@1.35.35,trna18_GlyGCC_16_+_70822597_70822667@1.35.35,trna19_GlyGCC_16_+_70823410_70823480@1.35.35,trna19_GlyGCC_2_-_157257659_157257729@1.35.35,trna24_GlyGCC_16_-_70812942_70813012@1.35.35,trna25_GlyGCC_16_-_70812114_70812184@1.35.35,trna5_GlyGCC_17_+_8029064_8029134@1.35.35,trna68_GlyGCC_1_-_161493637_161493707@1.35.35|GCATTGGTGGTTCAGTGGTAGAATTCTCGCCTGCC</td><td>>am_trna128_GlyGCC_6_-_27870686_27870756</td><td>1</td><td>35</td><td>35</td><td>GCATTGGTGGTTCAGTGGTAGAATTCTCGCCTGCC</td><td>11625.0</td><td>-3.2</td><td>0.9</td><td>-3.7</td><td>2.2E-04</td><td>1.5E-03</td><td>BT20/MB231</td><td>CYTO</td></tr>
<tr><td>>hg38_21_+_8433172_8446622_RNA45SN1_45S_50ntFlanks@10683.10709.27|CCGGCGGCGTCCGGTGAGCTCTCGCTG</td><td>>hg38_21_+_8433172_8446622_RNA45SN1_45S_50ntFlanks</td><td>10683</td><td>10709</td><td>27</td><td>CCGGCGGCGTCCGGTGAGCTCTCGCTG</td><td>772.7</td><td>-5.8</td><td>1.7</td><td>-3.4</td><td>5.8E-04</td><td>1.5E-03</td><td>BT20/MB468</td><td>MP</td></tr>
<tr><td>>hg38_21_+_8433172_8446622_RNA45SN1_45S_50ntFlanks@4884.4918.35|CAAAGCTGAAACTTAAAGGAATTGACGGAAGGGCA</td><td>>hg38_21_+_8433172_8446622_RNA45SN1_45S_50ntFlanks</td><td>4884</td><td>4918</td><td>35</td><td>CAAAGCTGAAACTTAAAGGAATTGACGGAAGGGCA</td><td>172.9</td><td>-4.6</td><td>1.4</td><td>-3.3</td><td>9.7E-04</td><td>1.5E-03</td><td>BT20/MB468</td><td>MITO</td></tr>
<tr><td>>hg38_21_+_8433172_8446622_RNA45SN1_45S_50ntFlanks@8191.8213.23|CAGCCCGTGGACGGTGTGAGGCC</td><td>>hg38_21_+_8433172_8446622_RNA45SN1_45S_50ntFlanks</td><td>8191</td><td>8213</td><td>23</td><td>CAGCCCGTGGACGGTGTGAGGCC</td><td>505.3</td><td>-2.0</td><td>0.6</td><td>-3.3</td><td>9.7E-04</td><td>1.5E-03</td><td>BT20/MB468</td><td>MITO</td></tr>
<tr><td>>hg38_21_+_8433172_8446622_RNA45SN1_45S_50ntFlanks@11670.11688.19|CCCAGTGCTCTGAATGTCA</td><td>>hg38_21_+_8433172_8446622_RNA45SN1_45S_50ntFlanks</td><td>11670</td><td>11688</td><td>19</td><td>CCCAGTGCTCTGAATGTCA</td><td>666.1</td><td>-3.2</td><td>1.0</td><td>-3.2</td><td>1.2E-03</td><td>1.5E-03</td><td>BT20/MB468</td><td>CYTO</td></tr>
<tr><td>>trnaMT_GluTTC_MT_-_14674_14742@45.71.27,trnalookalike8_GluTTC_5_-_93905172_93905240@45.71.27|CGTGGTTGTAGTCCGTGCGAGAATACC</td><td>>trnaMT_GluTTC_MT_-_14674_14742</td><td>45</td><td>71</td><td>27</td><td>CGTGGTTGTAGTCCGTGCGAGAATACC</td><td>111.2</td><td>3.0</td><td>0.9</td><td>3.2</td><td>1.3E-03</td><td>1.5E-03</td><td>MB231/MB468</td><td>MITO</td></tr>
<tr><td>>hg38_21_+_8433172_8446622_RNA45SN1_45S_50ntFlanks@4243.4271.29|CTTTCGAGGCCCTGTAATTGGAATGAGTC</td><td>>hg38_21_+_8433172_8446622_RNA45SN1_45S_50ntFlanks</td><td>4243</td><td>4271</td><td>29</td><td>CTTTCGAGGCCCTGTAATTGGAATGAGTC</td><td>352.0</td><td>-4.4</td><td>1.4</td><td>-3.2</td><td>1.3E-03</td><td>1.5E-03</td><td>BT20/MB468</td><td>NUC</td></tr>
<tr><td>>am_trna2_GlyGCC_21_-_18827107_18827177@1.26.26,trna35_GlyGCC_1_+_161413094_161413164@1.26.26,trna37_GlyGCC_1_+_161420467_161420537@1.26.26,trna39_GlyGCC_1_+_161427898_161427968@1.26.26,trna41_GlyGCC_1_+_161435258_161435328@1.26.26|GCATGGGTGGTTCAGTGGTAGAATTC</td><td>>am_trna2_GlyGCC_21_-_18827107_18827177</td><td>1</td><td>26</td><td>26</td><td>GCATGGGTGGTTCAGTGGTAGAATTC</td><td>1833.7</td><td>3.6</td><td>1.0</td><td>3.4</td><td>5.8E-04</td><td>1.5E-03</td><td>BT20/MB468</td><td>MP</td></tr>
<tr><td>>hg38_21_+_8433172_8446622_RNA45SN1_45S_50ntFlanks@10347.10383.37|CAAACGAGAACTTTGAAGGCCGAAGTGGAGAAGGGTT</td><td>>hg38_21_+_8433172_8446622_RNA45SN1_45S_50ntFlanks</td><td>10347</td><td>10383</td><td>37</td><td>CAAACGAGAACTTTGAAGGCCGAAGTGGAGAAGGGTT</td><td>332.8</td><td>-2.6</td><td>0.8</td><td>-3.2</td><td>1.3E-03</td><td>1.5E-03</td><td>MB231/MB468</td><td>MITO</td></tr>
<tr><td>>hg38_21_+_8433172_8446622_RNA45SN1_45S_50ntFlanks@10786.10816.31|AACAGCCTCTGGCATGTTGGAACAATGTAGG</td><td>>hg38_21_+_8433172_8446622_RNA45SN1_45S_50ntFlanks</td><td>10786</td><td>10816</td><td>31</td><td>AACAGCCTCTGGCATGTTGGAACAATGTAGG</td><td>475.2</td><td>-1.8</td><td>0.5</td><td>-3.3</td><td>9.7E-04</td><td>1.5E-03</td><td>BT20/MB468</td><td>MITO</td></tr>
<tr><td>>hg38_21_+_8433172_8446622_RNA45SN1_45S_50ntFlanks@7978.8011.34|GACCTCAGATCAGACGTGGCGACCCGCTGAATTT</td><td>>hg38_21_+_8433172_8446622_RNA45SN1_45S_50ntFlanks</td><td>7978</td><td>8011</td><td>34</td><td>GACCTCAGATCAGACGTGGCGACCCGCTGAATTT</td><td>357.8</td><td>5.1</td><td>1.5</td><td>3.4</td><td>5.8E-04</td><td>1.5E-03</td><td>BT20/MB468</td><td>MP</td></tr>
<tr><td>>hg38_21_+_8433172_8446622_RNA45SN1_45S_50ntFlanks@11876.11905.30|AAGACCCTGTTGAGCTTGACTCTAGTCTGG</td><td>>hg38_21_+_8433172_8446622_RNA45SN1_45S_50ntFlanks</td><td>11876</td><td>11905</td><td>30</td><td>AAGACCCTGTTGAGCTTGACTCTAGTCTGG</td><td>542.7</td><td>-5.7</td><td>1.7</td><td>-3.4</td><td>5.8E-04</td><td>1.5E-03</td><td>BT20/MB468</td><td>MP</td></tr>
<tr><td>>hg38_21_+_8433172_8446622_RNA45SN1_45S_50ntFlanks@485.517.33|GATGTGGTGACGTCGTGCTCTCCCGGGCCGGGT</td><td>>hg38_21_+_8433172_8446622_RNA45SN1_45S_50ntFlanks</td><td>485</td><td>517</td><td>33</td><td>GATGTGGTGACGTCGTGCTCTCCCGGGCCGGGT</td><td>355.8</td><td>11.7</td><td>3.6</td><td>3.2</td><td>1.3E-03</td><td>1.5E-03</td><td>MB231/MB468</td><td>MITO</td></tr>
</table>

| | | | | | | | | | | | | | |
|---|---|---|---|---|---|---|---|---|---|---|---|---|---|
| >hg38_21_+_8433172_8446622_RNA45SN1_45S_50ntFlanks@9939.9960.22\|GCAGGACGGTGGCCATGGAAGT | >hg38_21_+_8433172_8446622_RNA45SN1_45S_50ntFlanks | 9939 | 9960 | 22 | GCAGGACGGTGGCCATGGAAGT | 345.3 | 4.6 | 1.3 | 3.4 | 5.8E-04 | 1.5E-03 | BT20/MB468 | MP |
| >hg38_21_+_8433172_8446622_RNA45SN1_45S_50ntFlanks@5086.5120.35\|AACGAGACTCTGGCATGCTAACTAGTTACGCGACC | >hg38_21_+_8433172_8446622_RNA45SN1_45S_50ntFlanks | 5086 | 5120 | 35 | AACGAGACTCTGGCATGCTAACTAGTTACGCGACC | 303.4 | -3.8 | 1.2 | -3.3 | 1.1E-03 | 1.5E-03 | MB231/MB468 | MP |
| >hg38_21_+_8433172_8446622_RNA45SN1_45S_50ntFlanks@6659.6676.18\|AGCGGTGGATCACTCGGC | >hg38_21_+_8433172_8446622_RNA45SN1_45S_50ntFlanks | 6659 | 6676 | 18 | AGCGGTGGATCACTCGGC | 163.5 | 3.6 | 1.0 | 3.4 | 5.8E-04 | 1.5E-03 | BT20/MB468 | MP |
| >hg38_21_+_8433172_8446622_RNA45SN1_45S_50ntFlanks@8105.8136.32\|CGCGGCGGGGCGCGGGACATGTGGCGTACGGA | >hg38_21_+_8433172_8446622_RNA45SN1_45S_50ntFlanks | 8105 | 8136 | 32 | CGCGGCGGGGCGCGGGACATGTGGCGTACGGA | 687.6 | 6.0 | 1.6 | 3.7 | 2.0E-04 | 1.5E-03 | BT20/MB231 | MP |
| >hg38_21_+_8433172_8446622_RNA45SN1_45S_50ntFlanks@10882.10902.21\|TCGGGCTGGGGCGCGAAGCGG | >hg38_21_+_8433172_8446622_RNA45SN1_45S_50ntFlanks | 10882 | 10902 | 21 | TCGGGCTGGGGCGCGAAGCGG | 286.9 | -3.4 | 1.0 | -3.2 | 1.2E-03 | 1.5E-03 | BT20/MB468 | CYTO |
| >hg38_21_+_8433172_8446622_RNA45SN1_45S_50ntFlanks@9458.9503.46\|GCCGGGGAGGTGGAGCACGAGCGCACGTGTTAGGACCCGAAAGATG | >hg38_21_+_8433172_8446622_RNA45SN1_45S_50ntFlanks | 9458 | 9503 | 46 | GCCGGGGAGGTGGAGCACGAGCGCACGTGTTAGGACCCGAAAGATG | 58.6 | -9.7 | 3.0 | -3.3 | 1.1E-03 | 1.5E-03 | MB231/MB468 | MP |
| >hg38_21_+_8433172_8446622_RNA45SN1_45S_50ntFlanks@6719.6751.33\|TGAATTGCAGGACACATTGATCATCGACACTTC | >hg38_21_+_8433172_8446622_RNA45SN1_45S_50ntFlanks | 6719 | 6751 | 33 | TGAATTGCAGGACACATTGATCATCGACACTTC | 184.3 | -3.4 | 1.0 | -3.2 | 1.2E-03 | 1.5E-03 | BT20/MB468 | CYTO |
| >hg38_MT_+_1621_3279_RNR2_16S_50ntFlanks@341.378.38\|AAATAGTGGGAAGATTTATAGGTAGAGGCGACAAACCT | >hg38_MT_+_1621_3279_RNR2_16S_50ntFlanks | 341 | 378 | 38 | AAATAGTGGGAAGATTTATAGGTAGAGGCGACAAACCT | 52.0 | 9.4 | 2.7 | 3.4 | 5.8E-04 | 1.5E-03 | BT20/MB468 | MP |
| >hg38_1_-_228634819_228635039_RNA5S12_5S_50ntFlanks@51.75.25\|GTCTACGGCCATACCACCCTGAACG | >hg38_1_-_228634819_228635039_RNA5S12_5S_50ntFlanks | 51 | 75 | 25 | GTCTACGGCCATACCACCCTGAACG | 375.3 | -1.7 | 0.5 | -3.2 | 1.3E-03 | 1.6E-03 | BT20/MB468 | NUC |
| >hg38_21_+_8433172_8446622_RNA45SN1_45S_50ntFlanks@10440.10462.23\|GTTCCGAAGGGACGGGCGATGGC | >hg38_21_+_8433172_8446622_RNA45SN1_45S_50ntFlanks | 10440 | 10462 | 23 | GTTCCGAAGGGACGGGCGATGGC | 326.5 | -3.2 | 0.9 | -3.7 | 2.2E-04 | 1.6E-03 | BT20/MB231 | CYTO |
| >hg38_21_+_8433172_8446622_RNA45SN1_45S_50ntFlanks@10128.10147.20\|CGGCGGCGGCGGCGGCGGCG | >hg38_21_+_8433172_8446622_RNA45SN1_45S_50ntFlanks | 10128 | 10147 | 20 | CGGCGGCGGCGGCGGCGGCG | 1566.3 | -5.5 | 1.7 | -3.3 | 9.8E-04 | 1.6E-03 | BT20/MB468 | MITO |
| >hg38_21_+_8433172_8446622_RNA45SN1_45S_50ntFlanks@8845.8864.20\|CGGCGGCGGCGGCGGCGGCG | >hg38_21_+_8433172_8446622_RNA45SN1_45S_50ntFlanks | 8845 | 8864 | 20 | CGGCGGCGGCGGCGGCGGCG | 1566.3 | -5.5 | 1.7 | -3.3 | 9.8E-04 | 1.6E-03 | BT20/MB468 | MITO |
| >hg38_21_+_8433172_8446622_RNA45SN1_45S_50ntFlanks@6647.6678.32\|CGTACGACTCTTAGCGGTGGATCACTCGGCTC | >hg38_21_+_8433172_8446622_RNA45SN1_45S_50ntFlanks | 6647 | 6678 | 32 | CGTACGACTCTTAGCGGTGGATCACTCGGCTC | 520.2 | 2.7 | 0.8 | 3.2 | 1.3E-03 | 1.6E-03 | MB231/MB468 | NUC |
| >hg38_21_+_8433172_8446622_RNA45SN1_45S_50ntFlanks@3784.3801.18\|GTGAAACTGCGAATGGCT | >hg38_21_+_8433172_8446622_RNA45SN1_45S_50ntFlanks | 3784 | 3801 | 18 | GTGAAACTGCGAATGGCT | 417.4 | -5.6 | 1.7 | -3.3 | 1.1E-03 | 1.6E-03 | MB231/MB468 | MP |
| >hg38_21_+_8433172_8446622_RNA45SN1_45S_50ntFlanks@12003.12030.28\|GCGGGCCGCCGGTGAAATACCACTACTC | >hg38_21_+_8433172_8446622_RNA45SN1_45S_50ntFlanks | 12003 | 12030 | 28 | GCGGGCCGCCGGTGAAATACCACTACTC | 203.3 | -5.4 | 1.6 | -3.4 | 5.8E-04 | 1.6E-03 | BT20/MB468 | MP |
| >hg38_21_+_8433172_8446622_RNA45SN1_45S_50ntFlanks@4731.4761.31\|AAGAACGAAAGTCGGAGGTTCGAAGACGATC | >hg38_21_+_8433172_8446622_RNA45SN1_45S_50ntFlanks | 4731 | 4761 | 31 | AAGAACGAAAGTCGGAGGTTCGAAGACGATC | 667.9 | 2.1 | 0.6 | 3.3 | 9.8E-04 | 1.6E-03 | BT20/MB468 | MITO |
| >hg38_21_+_8433172_8446622_RNA45SN1_45S_50ntFlanks@4588.4608.21\|CTATTTTGTTGGTTTTCGGAA | >hg38_21_+_8433172_8446622_RNA45SN1_45S_50ntFlanks | 4588 | 4608 | 21 | CTATTTTGTTGGTTTTCGGAA | 442.8 | -4.3 | 1.3 | -3.2 | 1.2E-03 | 1.6E-03 | BT20/MB468 | CYTO |
| >hg38_21_+_8433172_8446622_RNA45SN1_45S_50ntFlanks@12991.13008.18\|CTCCCTCGCTGCGATCTA | >hg38_21_+_8433172_8446622_RNA45SN1_45S_50ntFlanks | 12991 | 13008 | 18 | CTCCCTCGCTGCGATCTA | 346.2 | -5.3 | 1.6 | -3.2 | 1.2E-03 | 1.6E-03 | BT20/MB468 | CYTO |
| >MI0000081\|hsa-mir-24-2&WithFlank&19\|-\|13836281\|13836365@18.40.23&[MIMAT0004497&hsa-miR-24-2-5p&offsets\|-1\|0;m-226&19\|-\|13836326\|13836348&offsets\|0\|0]\|GTGCCTACTGAGCTGAAACACAG | >MI0000081\|hsa-mir-24-2&WithFlank&19\|-\|13836281\|13836365 | 18 | 40 | 23 | GTGCCTACTGAGCTGAAACACAG | 134.0 | 3.3 | 1.0 | 3.2 | 1.3E-03 | 1.6E-03 | MB231/MB468 | MITO |
| >MI0000465\|hsa-mir-191&WithFlank&3\|-\|49020612\|49020715@22.43.22(+2U)&[MIMAT0000440&hsa-miR-191-5p&offsets\|0\|-1(+2U);m-48&3\|-\|49020672\|49020694&offsets\|0\|-1(+2U)]\|CAACGGAATCCCAAAAGCAGCTTT | >MI0000465\|hsa-mir-191&WithFlank&3\|-\|49020612\|49020715 | 22 | 43 | 22(+2U) | CAACGGAATCCCAAAAGCAGCTTT | 115.8 | 4.1 | 1.2 | 3.3 | 1.1E-03 | 1.6E-03 | MB231/MB468 | MP |
| >hg38_21_+_8433172_8446622_RNA45SN1_45S_50ntFlanks@8180.8208.29\|TGATCGAGGCCCAGCCCGTGGACGGTGTG | >hg38_21_+_8433172_8446622_RNA45SN1_45S_50ntFlanks | 8180 | 8208 | 29 | TGATCGAGGCCCAGCCCGTGGACGGTGTG | 69.3 | -5.1 | 1.6 | -3.3 | 1.1E-03 | 1.6E-03 | MB231/MB468 | MP |
| >hg38_21_+_8433172_8446622_RNA45SN1_45S_50ntFlanks@10411.10442.32\|CAGTCGGTCCTGAGAGATGGGCGAGCGCCGTT | >hg38_21_+_8433172_8446622_RNA45SN1_45S_50ntFlanks | 10411 | 10442 | 32 | CAGTCGGTCCTGAGAGATGGGCGAGCGCCGTT | 1186.3 | -3.2 | 0.9 | -3.4 | 5.8E-04 | 1.6E-03 | BT20/MB468 | MP |
| >hg38_21_+_8433172_8446622_RNA45SN1_45S_50ntFlanks@10685.10711.27\|GGCGGCGTCCGGTGAGCTCTCGCTGGC | >hg38_21_+_8433172_8446622_RNA45SN1_45S_50ntFlanks | 10685 | 10711 | 27 | GGCGGCGTCCGGTGAGCTCTCGCTGGC | 305.8 | -2.0 | 0.6 | -3.2 | 1.3E-03 | 1.6E-03 | MB231/MB468 | NUC |
| >MI0000465\|hsa-mir-191&WithFlank&3\|-\|49020612\|49020715@23.45.23&[MIMAT0000440&hsa-miR-191-5p&offsets\|+1\|+1;m-48&3\|-\|49020672\|49020694&offsets\|+1\|+1]\|AACGGAATCCCAAAAGCAGCTGT | >MI0000465\|hsa-mir-191&WithFlank&3\|-\|49020612\|49020715 | 23 | 45 | 23 | AACGGAATCCCAAAAGCAGCTGT | 146.3 | 6.0 | 1.8 | 3.3 | 1.1E-03 | 1.6E-03 | MB231/MB468 | MP |
| >hg38_21_+_8433172_8446622_RNA45SN1_45S_50ntFlanks@10786.10823.38\|AACAGCCTCTGGCATGTTGGAACAATGTAGGTAAGGGA | >hg38_21_+_8433172_8446622_RNA45SN1_45S_50ntFlanks | 10786 | 10823 | 38 | AACAGCCTCTGGCATGTTGGAACAATGTAGGTAAGGGA | 323.1 | -3.9 | 1.2 | -3.2 | 1.2E-03 | 1.6E-03 | BT20/MB468 | CYTO |
| >hg38_21_+_8433172_8446622_RNA45SN1_45S_50ntFlanks@9496.9523.28\|GAAAGATGGTGAACTATGCCTGGGCAGG | >hg38_21_+_8433172_8446622_RNA45SN1_45S_50ntFlanks | 9496 | 9523 | 28 | GAAAGATGGTGAACTATGCCTGGGCAGG | 194.8 | 6.1 | 1.8 | 3.4 | 5.9E-04 | 1.6E-03 | BT20/MB468 | MP |
| >hg38_21_+_8433172_8446622_RNA45SN1_45S_50ntFlanks@12157.12179.23\|AGTTTGACTGGGGCGGTACACCT | >hg38_21_+_8433172_8446622_RNA45SN1_45S_50ntFlanks | 12157 | 12179 | 23 | AGTTTGACTGGGGCGGTACACCT | 1923.2 | -6.1 | 1.8 | -3.4 | 5.9E-04 | 1.6E-03 | BT20/MB468 | MP |
| >hg38_21_+_8433172_8446622_RNA45SN1_45S_50ntFlanks@11571.11590.20\|GCCGACTTAGAACTGGTGCG | >hg38_21_+_8433172_8446622_RNA45SN1_45S_50ntFlanks | 11571 | 11590 | 20 | GCCGACTTAGAACTGGTGCG | 232.3 | 1.9 | 0.6 | 3.2 | 1.2E-03 | 1.6E-03 | MB231/MB468 | CYTO |
| >hg38_21_+_8433172_8446622_RNA45SN1_45S_50ntFlanks@11927.11950.24\|GTGTAGAATAAGTGGGAGGCCCCC | >hg38_21_+_8433172_8446622_RNA45SN1_45S_50ntFlanks | 11927 | 11950 | 24 | GTGTAGAATAAGTGGGAGGCCCCC | 178.5 | -2.4 | 0.7 | -3.2 | 1.2E-03 | 1.6E-03 | MB231/MB468 | CYTO |
| >trnaMT_ThrTGT_MT_+_15888_15953@36.69.34\|AACCGGAGATGAAAACCTTTTTCCAAGGACACCA | >trnaMT_ThrTGT_MT_+_15888_15953 | 36 | 69 | 34 | AACCGGAGATGAAAACCTTTTTCCAAGGACACCA | 463.4 | 4.6 | 1.3 | 3.4 | 5.9E-04 | 1.6E-03 | BT20/MB468 | MP |
| >hg38_21_+_8433172_8446622_RNA45SN1_45S_50ntFlanks@8673.8702.30\|CGGTGCGCCGCGACCGGCTCCGGGACGGCT | >hg38_21_+_8433172_8446622_RNA45SN1_45S_50ntFlanks | 8673 | 8702 | 30 | CGGTGCGCCGCGACCGGCTCCGGGACGGCT | 81.7 | 5.4 | 1.6 | 3.4 | 5.9E-04 | 1.6E-03 | BT20/MB468 | MP |
| >hg38_21_+_8433172_8446622_RNA45SN1_45S_50ntFlanks@3854.3883.30\|ACTGTGGTAATTCTAGAGCTAATACATGCC | >hg38_21_+_8433172_8446622_RNA45SN1_45S_50ntFlanks | 3854 | 3883 | 30 | ACTGTGGTAATTCTAGAGCTAATACATGCC | 256.1 | -2.1 | 0.6 | -3.2 | 1.3E-03 | 1.6E-03 | MB231/MB468 | NUC |
| >trnaMT_GluTTC_MT_-_14674_14742@26.61.36,trnalookalike8_GluTTC_5_-_93905172_93905240@26.61.36\|GGTTTTTCATATCATTGGTCGTGGTTGTAGTCCGTG | >trnaMT_GluTTC_MT_-_14674_14742 | 26 | 61 | 36 | GGTTTTTCATATCATTGGTCGTGGTTGTAGTCCGTG | 1149.6 | 3.1 | 1.0 | 3.2 | 1.3E-03 | 1.6E-03 | MB231/MB468 | NUC |
| >hg38_21_+_8433172_8446622_RNA45SN1_45S_50ntFlanks@6646.6694.49\|TCGTACGACTCTTAGCGGTGGATCACTCGGCTCGTGCGTCGATGAAGAA | >hg38_21_+_8433172_8446622_RNA45SN1_45S_50ntFlanks | 6646 | 6694 | 49 | TCGTACGACTCTTAGCGGTGGATCACTCGGCTCGTGCGTCGATGAAGAA | 704.1 | 3.8 | 1.1 | 3.3 | 9.9E-04 | 1.6E-03 | BT20/MB468 | MITO |
| >hg38_1_-_228634819_228635039_RNA5S12_5S_50ntFlanks@129.170.42\|TTGGATGGGAGACCGCCTGGGAATACCGGGTGCTGTAGGCTT | >hg38_1_-_228634819_228635039_RNA5S12_5S_50ntFlanks | 129 | 170 | 42 | TTGGATGGGAGACCGCCTGGGAATACCGGGTGCTGTAGGCTT | 2327.3 | 3.1 | 0.9 | 3.7 | 2.5E-04 | 1.6E-03 | BT20/MB231 | MITO |
| >trna119_LysCTT_1_-_145395522_145395594@1.36.36,trna11_LysCTT_5_-_180648979_180649051@1.36.36,trna13_LysCTT_6_+_26556774_26556846@1.36.36,trna7_LysCTT_16_+_3225692_3225764@1.36.36,trna9_LysCTT_5_+_180634755_180634827@1.36.36\|GCCCGGCTAGCTCAGTCGGTAGAGCATGAGACTCTT | >trna119_LysCTT_1_-_145395522_145395594 | 1 | 36 | 36 | GCCCGGCTAGCTCAGTCGGTAGAGCATGAGACTCTT | 1979.7 | -3.5 | 1.0 | -3.7 | 2.5E-04 | 1.6E-03 | BT20/MB231 | MITO |
| >hg38_21_+_8433172_8446622_RNA45SN1_45S_50ntFlanks@12912.12943.32\|CACGTTCGTGGGGAACCTGGCGCTAAACCATT | >hg38_21_+_8433172_8446622_RNA45SN1_45S_50ntFlanks | 12912 | 12943 | 32 | CACGTTCGTGGGGAACCTGGCGCTAAACCATT | 381.2 | -4.0 | 1.2 | -3.2 | 1.3E-03 | 1.6E-03 | MB231/MB468 | MITO |
| >hg38_21_+_8433172_8446622_RNA45SN1_45S_50ntFlanks@12449.12480.32\|GAATTCACCAAGCGTTGGATTGTTCACCCACT | >hg38_21_+_8433172_8446622_RNA45SN1_45S_50ntFlanks | 12449 | 12480 | 32 | GAATTCACCAAGCGTTGGATTGTTCACCCACT | 119.2 | 4.5 | 1.4 | 3.2 | 1.3E-03 | 1.6E-03 | BT20/MB468 | NUC |
| >MI0000077\|hsa-mir-21&WithFlank&17\|+\|59841260\|59841343@14.34.21(+2C)&[MIMAT0000076&hsa-miR-21-5p&offsets\|0\|-1(+2C);m-1&17\|+\|59841273\|59841294&offsets\|0\|-1(+2C)]\|TAGCTTATCAGACTGATGTTGCC | >MI0000077\|hsa-mir-21&WithFlank&17\|+\|59841260\|59841343 | 14 | 34 | 21(+2C) | TAGCTTATCAGACTGATGTTGCC | 1938.7 | -2.7 | 0.7 | -3.7 | 2.2E-04 | 1.6E-03 | BT20/MB231 | NUC |
| >MI0000255\|hsa-mir-30d&WithFlank&8\|-\|134804870\|134804951@12.32.21(+1A)&[MIMAT0000245&hsa-miR-30d-5p&offsets\|0\|-1(+1A);m-21&8\|-\|134804919\|134804940&offsets\|0\|-1(+1A)]\|TGTAAACATCCCCGACTGGAAA | >MI0000255\|hsa-mir-30d&WithFlank&8\|-\|134804870\|134804951 | 12 | 32 | 21(+1A) | TGTAAACATCCCCGACTGGAAA | 674.6 | 2.6 | 0.7 | 3.7 | 2.2E-04 | 1.6E-03 | BT20/MB231 | NUC |
| >hg38_21_+_8433172_8446622_RNA45SN1_45S_50ntFlanks@10440.10460.21\|GTTCCGAAGGGACGGGCGATGG | >hg38_21_+_8433172_8446622_RNA45SN1_45S_50ntFlanks | 10440 | 10460 | 21 | GTTCCGAAGGGACGGGCGATG | 236.2 | 2.8 | 0.9 | 3.2 | 1.2E-03 | 1.6E-03 | MB231/MB468 | CYTO |
| >hg38_MT_+_598_1651_RNR1_12S_50ntFlanks@52.84.33\|ATAGGTTTGGTCCTAGCCTTTCTATTAGCTCTT | >hg38_MT_+_598_1651_RNR1_12S_50ntFlanks | 52 | 84 | 33 | ATAGGTTTGGTCCTAGCCTTTCTATTAGCTCTT | 122.1 | 2.4 | 0.7 | 3.2 | 1.3E-03 | 1.6E-03 | BT20/MB468 | NUC |
| >trnaMT_SerGCT_MT_+_12207_12265@1.19.19\|GAGAAAGCTCACAAGAACT | >trnaMT_SerGCT_MT_+_12207_12265 | 1 | 19 | 19 | GAGAAAGCTCACAAGAACT | 5763.1 | 2.4 | 0.6 | 3.7 | 2.2E-04 | 1.6E-03 | BT20/MB231 | NUC |
| >hg38_21_+_8433172_8446622_RNA45SN1_45S_50ntFlanks@13000.13027.28\|TGCGATCTATTGAAAGTCAGCCCTCGAC | >hg38_21_+_8433172_8446622_RNA45SN1_45S_50ntFlanks | 13000 | 13027 | 28 | TGCGATCTATTGAAAGTCAGCCCTCGAC | 531.9 | -5.2 | 1.4 | -3.7 | 2.2E-04 | 1.6E-03 | BT20/MB231 | NUC |
| >hg38_21_+_8433172_8446622_RNA45SN1_45S_50ntFlanks@7971.8005.35\|GAGACGCGACCTCAGATCAGACGTGGCGACCCGCT | >hg38_21_+_8433172_8446622_RNA45SN1_45S_50ntFlanks | 7971 | 8005 | 35 | GAGACGCGACCTCAGATCAGACGTGGCGACCCGCT | 130.0 | 6.8 | 2.0 | 3.4 | 5.9E-04 | 1.6E-03 | BT20/MB468 | MP |
| >hg38_MT_+_1621_3279_RNR2_16S_50ntFlanks@622.657.36\|TATAACTGAACTCCTCACACCCAATTGGACCAATCT | >hg38_MT_+_1621_3279_RNR2_16S_50ntFlanks | 622 | 657 | 36 | TATAACTGAACTCCTCACACCCAATTGGACCAATCT | 52.2 | 9.5 | 2.8 | 3.4 | 5.9E-04 | 1.6E-03 | BT20/MB468 | MP |
| >MI0000439\|hsa-mir-23b&WithFlank&9\|+\|95085202\|95085310@67.85.19(+1U)&[MIMAT0000418&hsa-miR-23b-3p&offsets\|+3\|-1(+1U);m-42&9\|+\|95085265\|95085285&offsets\|+3\|+1(+1U)]\|ACATTGCCAGGGATTACCAT | >MI0000439\|hsa-mir-23b&WithFlank&9\|+\|95085202\|95085310 | 67 | 85 | 19(+1U) | ACATTGCCAGGGATTACCAT | 218.5 | 11.7 | 3.6 | 3.2 | 1.3E-03 | 1.6E-03 | MB231/MB468 | NUC |
| >hg38_21_+_8433172_8446622_RNA45SN1_45S_50ntFlanks@10420.10442.23\|CTGAGAGATGGGCGAGCGCCGTT | >hg38_21_+_8433172_8446622_RNA45SN1_45S_50ntFlanks | 10420 | 10442 | 23 | CTGAGAGATGGGCGAGCGCCGTT | 148.3 | -3.2 | 1.0 | -3.2 | 1.2E-03 | 1.6E-03 | BT20/MB468 | CYTO |
| >trnaMT_SerGCT_MT_+_12207_12265@20.61.42\|GCTAACTCATGCCCCCATGTCTAACAACATGGCTTTCTCACC | >trnaMT_SerGCT_MT_+_12207_12265 | 20 | 61 | 42 | GCTAACTCATGCCCCCATGTCTAACAACATGGCTTTCTCACC | 52.4 | 9.5 | 2.8 | 3.4 | 5.9E-04 | 1.6E-03 | BT20/MB468 | MP |
| >trnaMT_GlnTTG_MT_-_4329_4400@1.33.33,trnalookalike2_GlnTTG_1_-_564879_564950@1.33.33\|TAGGATGGGGTGTGATAGGTGGCACGGAGAATT | >trnaMT_GlnTTG_MT_-_4329_4400 | 1 | 33 | 33 | TAGGATGGGGTGTGATAGGTGGCACGGAGAATT | 893.4 | 4.0 | 1.1 | 3.7 | 2.2E-04 | 1.6E-03 | BT20/MB231 | CYTO |
| >hg38_21_+_8433172_8446622_RNA45SN1_45S_50ntFlanks@6646.6666.21\|TCGTACGACTCTTAGCGGTGG | >hg38_21_+_8433172_8446622_RNA45SN1_45S_50ntFlanks | 6646 | 6666 | 21 | TCGTACGACTCTTAGCGGTGG | 15204.8 | 2.9 | 0.8 | 3.4 | 5.9E-04 | 1.6E-03 | BT20/MB468 | MP |

| | | | | | | | | | | | | | |
|---|---|---|---|---|---|---|---|---|---|---|---|---|---|
| >MI0000746\|hsa-mir-99b&WithFlank&19\|+\|51692606\|51692687@13.30.18&[MIMAT0000689&hsa-miR-99b-5p&offsets\|0\|-4;m-10&19\|+\|51692618\|51692639&offsets\|0\|-4]\|CACCCGTAGAACCGACCT | >MI0000746\|hsa-mir-99b&WithFlank&19\|+\|51692606\|51692687 | 13 | 30 | 18 | CACCCGTAGAACCGACCT | 195.7 | 3.2 | 0.9 | 3.7 | 2.2E-04 | 1.6E-03 | BT20/MB231 | CYTO |
| >hg38_21_+_8433172_8446622_RNA45SN1_45S_50ntFlanks@4110.4131.22\|TGACGGGGAATCAGGGTTCGAT | >hg38_21_+_8433172_8446622_RNA45SN1_45S_50ntFlanks | 4110 | 4131 | 22 | TGACGGGGAATCAGGGTTCGAT | 199.9 | 6.8 | 2.0 | 3.4 | 5.9E-04 | 1.6E-03 | BT20/MB468 | MP |
| >hg38_21_+_8433172_8446622_RNA45SN1_45S_50ntFlanks@10822.10842.21\|GAAGTCGGCAAGCCGGATCCG | >hg38_21_+_8433172_8446622_RNA45SN1_45S_50ntFlanks | 10822 | 10842 | 21 | GAAGTCGGCAAGCCGGATCCG | 130.1 | -5.1 | 1.6 | -3.3 | 1.1E-03 | 1.6E-03 | MB231/MB468 | MP |
| >hg38_21_+_8433172_8446622_RNA45SN1_45S_50ntFlanks@11586.11609.24\|GTGCGGACCAGGGGAATCCGACTG | >hg38_21_+_8433172_8446622_RNA45SN1_45S_50ntFlanks | 11586 | 11609 | 24 | GTGCGGACCAGGGGAATCCGACTG | 705.1 | -2.5 | 0.8 | -3.2 | 1.2E-03 | 1.6E-03 | MB231/MB468 | CYTO |
| >hg38_21_+_8433172_8446622_RNA45SN1_45S_50ntFlanks@474.497.24\|TCTTCGTGATCGATGTGGTGACGT | >hg38_21_+_8433172_8446622_RNA45SN1_45S_50ntFlanks | 474 | 497 | 24 | TCTTCGTGATCGATGTGGTGACGT | 233.6 | 5.1 | 1.6 | 3.3 | 1.1E-03 | 1.6E-03 | MB231/MB468 | MP |
| >hg38_MT_+_598_1651_RNR1_12S_50ntFlanks@968.1004.37\|ACATGGTAAGTGTACTGGAAAGTGCACTTGGACGAAC | >hg38_MT_+_598_1651_RNR1_12S_50ntFlanks | 968 | 1004 | 37 | ACATGGTAAGTGTACTGGAAAGTGCACTTGGACGAAC | 307.9 | 4.1 | 1.3 | 3.2 | 1.3E-03 | 1.6E-03 | MB231/MB468 | MITO |
| >MI0000809\|hsa-mir-151a&WithFlank&8\|-\|140732558\|140732659@55.75.21(+1U)&[MIMAT0000757&hsa-miR-151a-3p&offsets\|+2\|+2(+1U);m-30&8\|-\|140732587\|140732607&offsets\|+2\|+2(+1U)]\|AGACTGAAGCTCCTTGAGGACT | >MI0000809\|hsa-mir-151a&WithFlank&8\|-\|140732558\|140732659 | 55 | 75 | 21(+1U) | AGACTGAAGCTCCTTGAGGACT | 135.7 | 6.9 | 2.0 | 3.4 | 5.9E-04 | 1.6E-03 | BT20/MB468 | MP |
| >hg38_21_+_8433172_8446622_RNA45SN1_45S_50ntFlanks@475.492.18\|CTTCGTGATCGATGTGGT | >hg38_21_+_8433172_8446622_RNA45SN1_45S_50ntFlanks | 475 | 492 | 18 | CTTCGTGATCGATGTGGT | 784.3 | 2.2 | 0.7 | 3.2 | 1.3E-03 | 1.6E-03 | MB231/MB468 | NUC |
| >hg38_21_+_8433172_8446622_RNA45SN1_45S_50ntFlanks@11190.11225.36\|CCCGGGGGGCCGGCGGCGGCGGCGACTCTGGACGCG | >hg38_21_+_8433172_8446622_RNA45SN1_45S_50ntFlanks | 11190 | 11225 | 36 | CCCGGGGGGCCGGCGGCGGCGGCGACTCTGGACGCG | 557.1 | -6.1 | 1.8 | -3.4 | 5.9E-04 | 1.6E-03 | BT20/MB468 | MP |
| >hg38_21_+_8433172_8446622_RNA45SN1_45S_50ntFlanks@8170.8200.31\|CAAGTCCTTCTGATCGAGGCCCAGCCCGTGG | >hg38_21_+_8433172_8446622_RNA45SN1_45S_50ntFlanks | 8170 | 8200 | 31 | CAAGTCCTTCTGATCGAGGCCCAGCCCGTGG | 80.2 | -5.4 | 1.6 | -3.4 | 5.9E-04 | 1.6E-03 | BT20/MB468 | MP |
| >trna20_GluTTC_2_-_131094701_131094772@1.31.31,trna5_GluTTC_13_-_41634874_41634945@1.31.31\|TCCCATATGGTCTAGCGGTTAGGATTCCTGG | >trna20_GluTTC_2_-_131094701_131094772 | 1 | 31 | 31 | TCCCATATGGTCTAGCGGTTAGGATTCCTGG | 471.7 | -1.9 | 0.6 | -3.2 | 1.2E-03 | 1.6E-03 | BT20/MB468 | CYTO |
| >hg38_21_+_8433172_8446622_RNA45SN1_45S_50ntFlanks@6651.6674.24\|CGACTCTTAGCGGTGGATCACTCG | >hg38_21_+_8433172_8446622_RNA45SN1_45S_50ntFlanks | 6651 | 6674 | 24 | CGACTCTTAGCGGTGGATCACTCG | 2202.8 | 3.3 | 1.0 | 3.4 | 5.9E-04 | 1.6E-03 | BT20/MB468 | MP |
| >hg38_MT_+_598_1651_RNR1_12S_50ntFlanks@52.77.26\|ATAGGTTTGGTCCTAGCCTTTCTATT | >hg38_MT_+_598_1651_RNR1_12S_50ntFlanks | 52 | 77 | 26 | ATAGGTTTGGTCCTAGCCTTTCTATT | 61542.7 | 4.9 | 1.4 | 3.4 | 5.9E-04 | 1.6E-03 | BT20/MB468 | MP |
| >hg38_21_+_8433172_8446622_RNA45SN1_45S_50ntFlanks@5489.5514.26\|CTGAGAAGACGGTCGAACTTGACTAT | >hg38_21_+_8433172_8446622_RNA45SN1_45S_50ntFlanks | 5489 | 5514 | 26 | CTGAGAAGACGGTCGAACTTGACTAT | 5171.7 | -6.4 | 1.9 | -3.4 | 5.9E-04 | 1.6E-03 | BT20/MB468 | MP |
| >hg38_21_+_8433172_8446622_RNA45SN1_45S_50ntFlanks@12912.12933.22\|CACGTTCGTGGGGAACCTGGCG | >hg38_21_+_8433172_8446622_RNA45SN1_45S_50ntFlanks | 12912 | 12933 | 22 | CACGTTCGTGGGGAACCTGGCG | 1075.9 | -5.0 | 1.4 | -3.4 | 5.9E-04 | 1.6E-03 | BT20/MB468 | MP |
| >MI0000067\|hsa-let-7f-1&WithFlank&9\|+\|94176341\|94176439@69.90.22&[MIMAT0004486&hsa-let-7f-1-3p&offsets\|0\|0;m-275&9\|+\|94176409\|94176430&offsets\|0\|0]\|CTATACAATCTATTGCCTTCCC | >MI0000067\|hsa-let-7f-1&WithFlank&9\|+\|94176341\|94176439 | 69 | 90 | 22 | CTATACAATCTATTGCCTTCCC | 64.2 | 6.0 | 1.8 | 3.3 | 1.1E-03 | 1.6E-03 | MB231/MB468 | MP |
| >hg38_21_+_8433172_8446622_RNA45SN1_45S_50ntFlanks@12456.12479.24\|CCAAGCGTTGGATTGTTCACCCAC | >hg38_21_+_8433172_8446622_RNA45SN1_45S_50ntFlanks | 12456 | 12479 | 24 | CCAAGCGTTGGATTGTTCACCCAC | 492.8 | -5.1 | 1.6 | -3.2 | 1.2E-03 | 1.6E-03 | BT20/MB468 | CYTO |
| >MI0000088\|hsa-mir-30a&WithFlank&6\|-\|71403545\|71403627@14.35.22&[MIMAT0000087&hsa-miR-30a-5p&offsets\|+2\|+2;m-15&6\|-\|71403595\|71403616&offsets\|+2\|+2]\|TAAACATCCTCGACTGGAAGCT | >MI0000088\|hsa-mir-30a&WithFlank&6\|-\|71403545\|71403627 | 14 | 35 | 22 | TAAACATCCTCGACTGGAAGCT | 87.2 | 4.9 | 1.5 | 3.3 | 1.1E-03 | 1.6E-03 | MB231/MB468 | MP |
| >MI0000255\|hsa-mir-30d&WithFlank&8\|-\|134804870\|134804951@12.33.22(+1A)&[MIMAT0000245&hsa-miR-30d-5p&offsets\|0\|0(+1A);m-21&8\|-\|134804919\|134804940&offsets\|0\|0(+1A)]\|TGTAAACATCCCCGACTGGAAGA | >MI0000255\|hsa-mir-30d&WithFlank&8\|-\|134804870\|134804951 | 12 | 33 | 22(+1A) | TGTAAACATCCCCGACTGGAAGA | 273.3 | 2.7 | 0.8 | 3.3 | 1.1E-03 | 1.6E-03 | MB231/MB468 | MP |
| >MI0001445\|hsa-mir-423&WithFlank&17\|+\|30117073\|30117178@59.79.21&[MIMAT0001340&hsa-miR-423-3p&offsets\|0\|-2;m-100&17\|+\|30117131\|30117153&offsets\|0\|-2]\|AGCTCGGTCTGAGGCCCCTCA | >MI0001445\|hsa-mir-423&WithFlank&17\|+\|30117073\|30117178 | 59 | 79 | 21 | AGCTCGGTCTGAGGCCCCTCA | 195.6 | 3.1 | 1.0 | 3.3 | 1.1E-03 | 1.6E-03 | MB231/MB468 | MP |
| >MI0000266\|hsa-mir-10a&WithFlank&17\|-\|48579832\|48579953@28.48.21(+1A)&[MIMAT0000253&hsa-miR-10a-5p&offsets\|0\|-2(+1A);m-18&17\|-\|48579905\|48579926&offsets\|0\|-1(+1A)]\|TACCCTGTAGATCCGAATTTGA | >MI0000266\|hsa-mir-10a&WithFlank&17\|-\|48579832\|48579953 | 28 | 48 | 21(+1A) | TACCCTGTAGATCCGAATTTGA | 1458.4 | 3.1 | 0.8 | 3.7 | 2.3E-04 | 1.6E-03 | BT20/MB231 | CYTO |
| >MI0000266\|hsa-mir-10a&WithFlank&17\|-\|48579832\|48579953@28.49.22(+1U)&[MIMAT0000253&hsa-miR-10a-5p&offsets\|0\|-1(+1U);m-18&17\|-\|48579905\|48579926&offsets\|0\|0(+1U)]\|TACCCTGTAGATCCGAATTTGTT | >MI0000266\|hsa-mir-10a&WithFlank&17\|-\|48579832\|48579953 | 28 | 49 | 22(+1U) | TACCCTGTAGATCCGAATTTGTT | 333.3 | 2.8 | 0.8 | 3.7 | 2.3E-04 | 1.6E-03 | BT20/MB231 | CYTO |
| >hg38_21_+_8433172_8446622_RNA45SN1_45S_50ntFlanks@6652.6669.18\|GACTCTTAGCGGTGGATC | >hg38_21_+_8433172_8446622_RNA45SN1_45S_50ntFlanks | 6652 | 6669 | 18 | GACTCTTAGCGGTGGATC | 500.3 | 3.0 | 0.8 | 3.7 | 2.2E-04 | 1.6E-03 | BT20/MB231 | NUC |
| >MI0005566\|hsa-mir-374b&WithFlank&X\|-\|74218541\|74218624@17.37.21&[MIMAT0004955&hsa-miR-374b-5p&offsets\|0\|-1;m-153&X\|-\|74218587\|74218608&offsets\|0\|-1]\|ATATAATACAACCTGCTAAGT | >MI0005566\|hsa-mir-374b&WithFlank&X\|-\|74218541\|74218624 | 17 | 37 | 21 | ATATAATACAACCTGCTAAGT | 449.3 | 3.3 | 1.0 | 3.3 | 1.1E-03 | 1.6E-03 | MB231/MB468 | MP |
| >hg38_21_+_8433172_8446622_RNA45SN1_45S_50ntFlanks@11575.11608.34\|ACTTAGAACTGGTGCGGACCAGGGGAATCCGACT | >hg38_21_+_8433172_8446622_RNA45SN1_45S_50ntFlanks | 11575 | 11608 | 34 | ACTTAGAACTGGTGCGGACCAGGGGAATCCGACT | 78.0 | -5.2 | 1.6 | -3.3 | 1.1E-03 | 1.6E-03 | MB231/MB468 | MP |
| >hg38_21_+_8433172_8446622_RNA45SN1_45S_50ntFlanks@5493.5516.24\|GAAGACGGTCGAACTTGACTATCT | >hg38_21_+_8433172_8446622_RNA45SN1_45S_50ntFlanks | 5493 | 5516 | 24 | GAAGACGGTCGAACTTGACTATCT | 217.8 | 3.1 | 1.0 | 3.2 | 1.3E-03 | 1.6E-03 | BT20/MB468 | NUC |
| >hg38_MT_+_1621_3279_RNR2_16S_50ntFlanks@1347.1388.42\|CAACAATAGGGTTTACGACCTCGATGTTGGATCAGGACATCC | >hg38_MT_+_1621_3279_RNR2_16S_50ntFlanks | 1347 | 1388 | 42 | CAACAATAGGGTTTACGACCTCGATGTTGGATCAGGACATCC | 101.4 | 5.7 | 1.8 | 3.2 | 1.3E-03 | 1.6E-03 | MB231/MB468 | MITO |
| >hg38_21_+_8433172_8446622_RNA45SN1_45S_50ntFlanks@12404.12434.31\|GCTTTTTGATCCTTCGATGTCGGCTCTTCCT | >hg38_21_+_8433172_8446622_RNA45SN1_45S_50ntFlanks | 12404 | 12434 | 31 | GCTTTTTGATCCTTCGATGTCGGCTCTTCCT | 284.8 | -2.4 | 0.7 | -3.2 | 1.3E-03 | 1.6E-03 | MB231/MB468 | MITO |
| >MI0000070\|hsa-mir-16-1&WithFlank&13\|-\|50048967\|50049067@20.40.21;MI0000115\|hsa-mir-16-2&WithFlank&3\|+\|160404739\|160404831@16.36.21&[MIMAT0000069&hsa-miR-16-5p&offsets\|0\|-1;m-62&13\|-\|50049027\|50049048&offsets\|0\|-1];[MIMAT0000069_1&hsa-miR-16-5p&offsets\|0\|-1;m-61&3\|+\|160404754\|160404775&offsets\|0\|-1]\|TAGCAGCACGTAAATATTGGC | >MI0000070\|hsa-mir-16-1&WithFlank&13\|-\|50048967\|50049067 | 20 | 40 | 21 | TAGCAGCACGTAAATATTGGC | 2317.4 | 2.8 | 0.8 | 3.3 | 1.1E-03 | 1.6E-03 | MB231/MB468 | MP |
| >MI0000298\|hsa-mir-221&WithFlank&X\|-\|45746151\|45746272@31.53.23&[MIMAT0004568&hsa-miR-221-5p&offsets\|0\|+1;m-405&X\|-\|45746220\|45746242&offsets\|0\|0]\|ACCTGGCATACAATGTAGATTTC | >MI0000298\|hsa-mir-221&WithFlank&X\|-\|45746151\|45746272 | 31 | 53 | 23 | ACCTGGCATACAATGTAGATTTC | 207.9 | 3.8 | 1.1 | 3.4 | 6.0E-04 | 1.6E-03 | BT20/MB468 | MP |
| >hg38_21_+_8433172_8446622_RNA45SN1_45S_50ntFlanks@11874.11905.32\|AGAAGACCCTGTTGAGCTTGACTCTAGTCTGG | >hg38_21_+_8433172_8446622_RNA45SN1_45S_50ntFlanks | 11874 | 11905 | 32 | AGAAGACCCTGTTGAGCTTGACTCTAGTCTGG | 1159.6 | -6.0 | 1.7 | -3.4 | 6.0E-04 | 1.6E-03 | BT20/MB468 | MP |
| >hg38_21_+_8433172_8446622_RNA45SN1_45S_50ntFlanks@10548.10574.27\|GTGCGGTAACGCGACCGATCCCGGAGA | >hg38_21_+_8433172_8446622_RNA45SN1_45S_50ntFlanks | 10548 | 10574 | 27 | GTGCGGTAACGCGACCGATCCCGGAGA | 250.6 | 3.0 | 0.8 | 3.7 | 2.6E-04 | 1.6E-03 | BT20/MB231 | MITO |
| >hg38_21_+_8433172_8446622_RNA45SN1_45S_50ntFlanks@11671.11692.22\|CCAGTGCTCTGAATGTCAAAGT | >hg38_21_+_8433172_8446622_RNA45SN1_45S_50ntFlanks | 11671 | 11692 | 22 | CCAGTGCTCTGAATGTCAAAGT | 452.7 | 3.0 | 0.8 | 3.7 | 2.6E-04 | 1.6E-03 | BT20/MB231 | MITO |
| >hg38_21_+_8433172_8446622_RNA45SN1_45S_50ntFlanks@12993.13017.25\|CCCTCGCTGCGATCTATTGAAAGTC | >hg38_21_+_8433172_8446622_RNA45SN1_45S_50ntFlanks | 12993 | 13017 | 25 | CCCTCGCTGCGATCTATTGAAAGTC | 279.4 | -3.1 | 0.8 | -3.7 | 2.3E-04 | 1.6E-03 | BT20/MB231 | CYTO |
| >hg38_21_+_8433172_8446622_RNA45SN1_45S_50ntFlanks@10798.10825.28\|CATGTTGGAACAATGTAGGTAAGGGAAG | >hg38_21_+_8433172_8446622_RNA45SN1_45S_50ntFlanks | 10798 | 10825 | 28 | CATGTTGGAACAATGTAGGTAAGGGAAG | 215.6 | -2.4 | 0.7 | -3.2 | 1.2E-03 | 1.6E-03 | BT20/MB468 | CYTO |
| >hg38_21_+_8433172_8446622_RNA45SN1_45S_50ntFlanks@9533.9551.19\|GAGGAAACTCTGGTGGAGG | >hg38_21_+_8433172_8446622_RNA45SN1_45S_50ntFlanks | 9533 | 9551 | 19 | GAGGAAACTCTGGTGGAGG | 574.3 | -2.4 | 0.7 | -3.2 | 1.2E-03 | 1.6E-03 | BT20/MB468 | CYTO |
| >hg38_1_-_228634819_228635039_RNA5S12_5S_50ntFlanks@89.123.35\|CTGATCTCGGAAGCTAAGCAGGGTCGGGCCTGGTT | >hg38_1_-_228634819_228635039_RNA5S12_5S_50ntFlanks | 89 | 123 | 35 | CTGATCTCGGAAGCTAAGCAGGGTCGGGCCTGGTT | 550.7 | -3.1 | 0.8 | -3.7 | 2.3E-04 | 1.6E-03 | BT20/MB231 | CYTO |
| >hg38_21_+_8433172_8446622_RNA45SN1_45S_50ntFlanks@8183.8223.41\|TCGAGGCCCAGCCCGTGGACGGTGTGAGGCCGGTAGCGGCC | >hg38_21_+_8433172_8446622_RNA45SN1_45S_50ntFlanks | 8183 | 8223 | 41 | TCGAGGCCCAGCCCGTGGACGGTGTGAGGCCGGTAGCGGCC | 107.5 | -4.8 | 1.4 | -3.3 | 1.0E-03 | 1.6E-03 | BT20/MB468 | MITO |
| >hg38_21_+_8433172_8446622_RNA45SN1_45S_50ntFlanks@4099.4127.29\|GTGACCACGGGTGACGGGGAATCAGGGTT | >hg38_21_+_8433172_8446622_RNA45SN1_45S_50ntFlanks | 4099 | 4127 | 29 | GTGACCACGGGTGACGGGGAATCAGGGTT | 90.3 | -5.3 | 1.5 | -3.4 | 6.0E-04 | 1.6E-03 | BT20/MB468 | MP |
| >hg38_21_+_8433172_8446622_RNA45SN1_45S_50ntFlanks@7980.8008.29\|CCTCAGATCAGACGTGGCGACCCGCTGAA | >hg38_21_+_8433172_8446622_RNA45SN1_45S_50ntFlanks | 7980 | 8008 | 29 | CCTCAGATCAGACGTGGCGACCCGCTGAA | 1040.9 | -2.9 | 0.9 | -3.2 | 1.3E-03 | 1.6E-03 | MB231/MB468 | NUC |
| >trnaMT_SerGCT_MT_+_12207_12265@13.54.42\|AAGAACTGCTAACTCATGCCCCCATGTCTAACAACATGGCTT | >trnaMT_SerGCT_MT_+_12207_12265 | 13 | 54 | 42 | AAGAACTGCTAACTCATGCCCCCATGTCTAACAACATGGCTT | 51.9 | 9.4 | 2.8 | 3.4 | 6.0E-04 | 1.6E-03 | BT20/MB468 | MP |
| >hg38_21_+_8433172_8446622_RNA45SN1_45S_50ntFlanks@12071.12106.36\|CCCCGAGGGGCTCTCGCTTCTGGCGCCAAGCGCCCG | >hg38_21_+_8433172_8446622_RNA45SN1_45S_50ntFlanks | 12071 | 12106 | 36 | CCCCGAGGGGCTCTCGCTTCTGGCGCCAAGCGCCCG | 953.1 | -6.3 | 2.0 | -3.2 | 1.3E-03 | 1.6E-03 | BT20/MB468 | NUC |
| >hg38_21_+_8433172_8446622_RNA45SN1_45S_50ntFlanks@12417.12446.30\|TCGATGTCGGCTCTTCCTATCATTGTGAAG | >hg38_21_+_8433172_8446622_RNA45SN1_45S_50ntFlanks | 12417 | 12446 | 30 | TCGATGTCGGCTCTTCCTATCATTGTGAAG | 2105.0 | -2.7 | 0.8 | -3.2 | 1.2E-03 | 1.6E-03 | BT20/MB468 | CYTO |
| >hg38_21_+_8433172_8446622_RNA45SN1_45S_50ntFlanks@12456.12491.36\|CCAAGCGTTGGATTGTTCACCCACTAATAGGGAACG | >hg38_21_+_8433172_8446622_RNA45SN1_45S_50ntFlanks | 12456 | 12491 | 36 | CCAAGCGTTGGATTGTTCACCCACTAATAGGGAACG | 275.1 | -4.3 | 1.3 | -3.2 | 1.3E-03 | 1.6E-03 | MB231/MB468 | NUC |
| >hg38_21_+_8433172_8446622_RNA45SN1_45S_50ntFlanks@8104.8146.43\|CCGCGGCGGGGCGCGGGACATGTGGCGTACGGAAGACCCGCTC | >hg38_21_+_8433172_8446622_RNA45SN1_45S_50ntFlanks | 8104 | 8146 | 43 | CCGCGGCGGGGCGCGGGACATGTGGCGTACGGAAGACCCGCTC | 58.8 | -9.7 | 3.0 | -3.3 | 1.1E-03 | 1.6E-03 | MB231/MB468 | MP |
| >hg38_21_+_8433172_8446622_RNA45SN1_45S_50ntFlanks@4884.4902.19\|CAAAGCTGAAACTTAAAGG | >hg38_21_+_8433172_8446622_RNA45SN1_45S_50ntFlanks | 4884 | 4902 | 19 | CAAAGCTGAAACTTAAAGG | 101.5 | -5.1 | 1.6 | -3.3 | 1.1E-03 | 1.6E-03 | MB231/MB468 | MP |
| >hg38_21_+_8433172_8446622_RNA45SN1_45S_50ntFlanks@4983.5001.19\|CGGACAGGATTGACAGATT | >hg38_21_+_8433172_8446622_RNA45SN1_45S_50ntFlanks | 4983 | 5001 | 19 | CGGACAGGATTGACAGATT | 329.9 | -5.1 | 1.6 | -3.2 | 1.3E-03 | 1.6E-03 | MB231/MB468 | MITO |
| >trnaMT_PheGAA_MT_+_577_647@36.66.31\|AAAATGTTTAGACGGGCTCACATCACCCCAT | >trnaMT_PheGAA_MT_+_577_647 | 36 | 66 | 31 | AAAATGTTTAGACGGGCTCACATCACCCCAT | 52.6 | 9.5 | 2.8 | 3.4 | 6.0E-04 | 1.6E-03 | BT20/MB468 | MP |
| >MI0000088\|hsa-mir-30a&WithFlank&6\|-\|71403545\|71403627@13.35.23(+1U)&[MIMAT0000087&hsa-miR-30a-5p&offsets\|+1\|+2(+1U);m-15&6\|-\|71403595\|71403616&offsets\|+1\|+2(+1U)]\|GTAAACATCCTCGACTGGAAGCTT | >MI0000088\|hsa-mir-30a&WithFlank&6\|-\|71403545\|71403627 | 13 | 35 | 23(+1U) | GTAAACATCCTCGACTGGAAGCTT | 314.1 | 5.3 | 1.6 | 3.4 | 6.0E-04 | 1.6E-03 | BT20/MB468 | MP |
| >trnaMT_PheGAA_MT_+_577_647@37.64.28\|AAATGTTTAGACGGGCTCACATCACCCC | >trnaMT_PheGAA_MT_+_577_647 | 37 | 64 | 28 | AAATGTTTAGACGGGCTCACATCACCCC | 52.8 | 9.5 | 2.8 | 3.4 | 6.0E-04 | 1.6E-03 | BT20/MB468 | MP |
| >hg38_21_+_8433172_8446622_RNA45SN1_45S_50ntFlanks@11873.11892.20\|AAGAAGACCCTGTTGAGCTT | >hg38_21_+_8433172_8446622_RNA45SN1_45S_50ntFlanks | 11873 | 11892 | 20 | AAGAAGACCCTGTTGAGCTT | 784.9 | -3.3 | 1.0 | -3.4 | 6.0E-04 | 1.6E-03 | BT20/MB468 | MP |

| | | | | | | | | | | | | | |
|---|---|---|---|---|---|---|---|---|---|---|---|---|---|
| >MI0001445\|hsa-mir-423&WithFlank&17\|+\|30117073\|30117178@58.78.21(+1U)&[MIMAT0001340&hsa-miR-423-3p&offsets\|-1\|-3(+1U);m-100&17\|+\|30117131\|30117153&offsets\|-1\|-3(+1U)]\|AAGCTCGGTCTGAGGCCCCTCT | >MI0001445\|hsa-mir-423&WithFlank&17\|+\|30117073\|30117178 | 58 | 78 | 21(+1U) | AAGCTCGGTCTGAGGCCCCTCT | 83.2 | 6.8 | 2.0 | 3.4 | 6.0E-04 | 1.6E-03 | BT20/MB468 | MP |
| >trnaMT_ProTGG_MT_-_15956_16023@44.70.27\|GTGGAGTTAAAGACTTTTTCTCTGACC | >trnaMT_ProTGG_MT_-_15956_16023 | 44 | 70 | 27 | GTGGAGTTAAAGACTTTTTCTCTGACC | 314.6 | 3.7 | 1.1 | 3.4 | 6.0E-04 | 1.6E-03 | BT20/MB468 | MP |
| >MI0000816\|hsa-mir-335&WithFlank&7\|+\|130496105\|130496210@22.43.22&[MIMAT0000765&hsa-miR-335-5p&offsets\|0\|-1;m-249&7\|+\|130496126\|130496147&offsets\|0\|0]\|TCAAGAGCAATAACGAAAAATG | >MI0000816\|hsa-mir-335&WithFlank&7\|+\|130496105\|130496210 | 22 | 43 | 22 | TCAAGAGCAATAACGAAAAATG | 314.7 | 2.7 | 0.8 | 3.3 | 1.1E-03 | 1.6E-03 | MB231/MB468 | MP |
| >MI0000289\|hsa-mir-181a-1&WithFlank&1\|-\|198859038\|198859159@70.88.19&[MIMAT0000270&hsa-miR-181a-3p&offsets\|0\|-3;m-125&1\|-\|198859069\|198859090&offsets\|0\|-3]\|ACCATCGACCGTTGATTGT | >MI0000289\|hsa-mir-181a-1&WithFlank&1\|-\|198859038\|198859159 | 70 | 88 | 19 | ACCATCGACCGTTGATTGT | 49.9 | 5.8 | 1.8 | 3.3 | 1.1E-03 | 1.6E-03 | MB231/MB468 | MP |
| >hg38_1_-_228634819_228635039_RNA5S12_5S_50ntFlanks@145.169.25\|CTGGGAATACCGGGTGCTGTAGGCT | >hg38_1_-_228634819_228635039_RNA5S12_5S_50ntFlanks | 145 | 169 | 25 | CTGGGAATACCGGGTGCTGTAGGCT | 254.9 | -2.7 | 0.8 | -3.3 | 1.1E-03 | 1.6E-03 | MB231/MB468 | MP |
| >trnaMT_LeuTAA_MT_+_3230_3304@25.44.20\|CGCATAAAACTTAAAACTTT | >trnaMT_LeuTAA_MT_+_3230_3304 | 25 | 44 | 20 | CGCATAAAACTTAAAACTTT | 134.5 | -5.1 | 1.6 | -3.3 | 1.1E-03 | 1.6E-03 | MB231/MB468 | MP |
| >trnaMT_PheGAA_MT_+_577_647@22.60.39\|AAAGCAATACACTGAAAATGTTTAGACGGGCTCACATCA | >trnaMT_PheGAA_MT_+_577_647 | 22 | 60 | 39 | AAAGCAATACACTGAAAATGTTTAGACGGGCTCACATCA | 51.8 | 9.4 | 2.8 | 3.4 | 6.0E-04 | 1.6E-03 | BT20/MB468 | MP |
| >hg38_21_+_8433172_8446622_RNA45SN1_45S_50ntFlanks@9544.9573.30\|GGTGGAGGTCCGTAGCGGTCCTGACGTGCA | >hg38_21_+_8433172_8446622_RNA45SN1_45S_50ntFlanks | 9544 | 9573 | 30 | GGTGGAGGTCCGTAGCGGTCCTGACGTGCA | 584.5 | 2.5 | 0.8 | 3.2 | 1.3E-03 | 1.6E-03 | MB231/MB468 | CYTO |
| >hg38_21_+_8433172_8446622_RNA45SN1_45S_50ntFlanks@9853.9874.22\|CGCTGCGGGATGAACCGAACGC | >hg38_21_+_8433172_8446622_RNA45SN1_45S_50ntFlanks | 9853 | 9874 | 22 | CGCTGCGGGATGAACCGAACGC | 80.9 | 4.5 | 1.3 | 3.4 | 6.1E-04 | 1.6E-03 | BT20/MB468 | MP |
| >trna10_LysCTT_16_+_3241501_3241573@4.34.31,trna13_LysCTT_14_-_58706613_58706685@4.34.31,trna2_LysCTT_15_+_79152904_79152976@4.34.31\|CGGCTAGCTCAGTCGGTAGAGCATGGGACTC | >trna10_LysCTT_16_+_3241501_3241573 | 4 | 34 | 31 | CGGCTAGCTCAGTCGGTAGAGCATGGGACTC | 437.8 | -3.9 | 1.1 | -3.6 | 2.6E-04 | 1.6E-03 | BT20/MB231 | MITO |
| >hg38_21_+_8433172_8446622_RNA45SN1_45S_50ntFlanks@11314.11336.23\|CCCACCCCACGTCTCGTCGCGCG | >hg38_21_+_8433172_8446622_RNA45SN1_45S_50ntFlanks | 11314 | 11336 | 23 | CCCACCCCACGTCTCGTCGCGCG | 434.4 | -6.0 | 1.9 | -3.2 | 1.3E-03 | 1.6E-03 | BT20/MB468 | NUC |
| >trnaMT_LeuTAA_MT_+_3230_3304@1.30.30\|GTTAAGATGGCAGAGCCCGGTAATCGCATA | >trnaMT_LeuTAA_MT_+_3230_3304 | 1 | 30 | 30 | GTTAAGATGGCAGAGCCCGGTAATCGCATA | 692.4 | 2.7 | 0.8 | 3.2 | 1.3E-03 | 1.6E-03 | MB231/MB468 | MITO |
| >am_trna128_GlyGCC_6_-_27870686_27870756@1.25.25,trna133_GlyCCC_1_-_16872434_16872504@1.25.25,trna18_GlyGCC_16_+_70822597_70822667@1.25.25,trna19_GlyGCC_16_+_70823410_70823480@1.25.25,trna19_GlyGCC_2_-_157257659_157257729@1.25.25,trna24_GlyGCC_16_-_70812942_70813012@1.25.25,trna25_GlyGCC_16_-_70812114_70812184@1.25.25,trna4_GlyCCC_1_+_17188416_17188486@1.25.25,trna5_GlyGCC_17_+_8029064_8029134@1.25.25,trna68_GlyGCC_1_-_161493637_161493707@1.25.25\|GCATTGGTGGTTCAGTGGTAGAATT | >am_trna128_GlyGCC_6_-_27870686_27870756 | 1 | 25 | 25 | GCATTGGTGGTTCAGTGGTAGAATT | 768.9 | -2.8 | 0.9 | -3.2 | 1.3E-03 | 1.6E-03 | MB231/MB468 | NUC |
| >am_MI0000809\|hsa-mir-151a&WithFlank&8\|-\|140732558\|140732659@53.72.20(+3A)&[MIMAT0000757&hsa-miR-151a-3p&offsets\|0\|-1(+3A);m-30&8\|-\|140732587\|140732607&offsets\|0\|-1(+3A)]\|CTAGACTGAAGCTCCTTGAGAAA | >am_MI0000809\|hsa-mir-151a&WithFlank&8\|-\|140732558\|140732659 | 53 | 72 | 20(+3A) | CTAGACTGAAGCTCCTTGAGAAA | 67.7 | 6.5 | 1.9 | 3.4 | 6.1E-04 | 1.6E-03 | BT20/MB468 | MP |
| >trnaMT_SerGCT_MT_+_12207_12265@23.61.39\|AACTCATGCCCCCATGTCTAACAACATGGCTTTCTCACC | >trnaMT_SerGCT_MT_+_12207_12265 | 23 | 61 | 39 | AACTCATGCCCCCATGTCTAACAACATGGCTTTCTCACC | 52.4 | 9.5 | 2.8 | 3.4 | 6.1E-04 | 1.6E-03 | BT20/MB468 | MP |
| >hg38_21_+_8433172_8446622_RNA45SN1_45S_50ntFlanks@11580.11609.30\|GAACTGGTGCGGACCAGGGGAATCCGACTG | >hg38_21_+_8433172_8446622_RNA45SN1_45S_50ntFlanks | 11580 | 11609 | 30 | GAACTGGTGCGGACCAGGGGAATCCGACTG | 1506.5 | -3.3 | 1.0 | -3.4 | 6.1E-04 | 1.6E-03 | BT20/MB468 | MP |
| >hg38_21_+_8433172_8446622_RNA45SN1_45S_50ntFlanks@10410.10428.19\|TCAGTCGGTCCTGAGAGAT | >hg38_21_+_8433172_8446622_RNA45SN1_45S_50ntFlanks | 10410 | 10428 | 19 | TCAGTCGGTCCTGAGAGAT | 245.6 | -2.3 | 0.7 | -3.3 | 1.0E-03 | 1.6E-03 | BT20/MB468 | MITO |
| >MI0000081\|hsa-mir-24-2&WithFlank&19\|-\|13836281\|13836365@56.76.21(+1C);MI0000080\|hsa-mir-24-1&WithFlank&9\|+\|95086015\|95086094@50.70.21(+1C)&[MIMAT0000080&hsa-miR-24-3p&offsets\|0\|-1(+1C);m-36&19\|-\|13836288\|13836310&offsets\|0\|-2(+1C)];[MIMAT0000080_1&hsa-miR-24-3p&offsets\|0\|-1(+1C);m-35&9\|+\|95086064\|95086086&offsets\|0\|-2(+1C)]\|TGGCTCAGTTCAGCAGGAACAC | >MI0000081\|hsa-mir-24-2&WithFlank&19\|-\|13836281\|13836365 | 56 | 76 | 21(+1C) | TGGCTCAGTTCAGCAGGAACAC | 324.4 | 4.3 | 1.3 | 3.3 | 1.1E-03 | 1.6E-03 | MB231/MB468 | MP |
| >MI0000736\|hsa-mir-30c-1&WithFlank&1\|+\|40757278\|40757378@23.45.23(+1C);MI0000254\|hsa-mir-30c-2&WithFlank&6\|-\|71376954\|71377037@13.35.23(+1C)&[MIMAT0000244&hsa-miR-30c-5p&offsets\|0\|0(+1C);m-57&1\|+\|40757300\|40757323&offsets\|0\|-1(+1C)];[MIMAT0000244_1&hsa-miR-30c-5p&offsets\|0\|0(+1C);m-56&6\|-\|71377002\|71377025&offsets\|0\|-1(+1C)]\|TGTAAACATCCTACACTCTCAGCC | >MI0000736\|hsa-mir-30c-1&WithFlank&1\|+\|40757278\|40757378 | 23 | 45 | 23(+1C) | TGTAAACATCCTACACTCTCAGCC | 1536.2 | 3.6 | 1.1 | 3.3 | 1.1E-03 | 1.6E-03 | MB231/MB468 | MP |
| >hg38_21_+_8433172_8446622_RNA45SN1_45S_50ntFlanks@4417.4438.22\|CGAGGCGAGCCACCGCCCGTCC | >hg38_21_+_8433172_8446622_RNA45SN1_45S_50ntFlanks | 4417 | 4438 | 22 | CGAGGCGAGCCACCGCCCGTCC | 52.0 | -8.8 | 2.6 | -3.4 | 6.1E-04 | 1.6E-03 | BT20/MB468 | MP |
| >hg38_21_+_8433172_8446622_RNA45SN1_45S_50ntFlanks@8198.8223.26\|TGGACGGTGTGAGGCCGGTAGCGGCC | >hg38_21_+_8433172_8446622_RNA45SN1_45S_50ntFlanks | 8198 | 8223 | 26 | TGGACGGTGTGAGGCCGGTAGCGGCC | 581.2 | -3.2 | 0.9 | -3.4 | 6.1E-04 | 1.6E-03 | BT20/MB468 | MP |
| >hg38_21_+_8433172_8446622_RNA45SN1_45S_50ntFlanks@4985.5018.34\|GACAGGATTGACAGATTGATAGCTCTTTCTCGAT | >hg38_21_+_8433172_8446622_RNA45SN1_45S_50ntFlanks | 4985 | 5018 | 34 | GACAGGATTGACAGATTGATAGCTCTTTCTCGAT | 50.1 | 6.5 | 1.9 | 3.4 | 6.1E-04 | 1.6E-03 | BT20/MB468 | MP |
| >hg38_21_+_8433172_8446622_RNA45SN1_45S_50ntFlanks@10686.10712.27\|GCGGCGTCCGGTGAGCTCTCGCTGGCC | >hg38_21_+_8433172_8446622_RNA45SN1_45S_50ntFlanks | 10686 | 10712 | 27 | GCGGCGTCCGGTGAGCTCTCGCTGGCC | 881.4 | -4.9 | 1.5 | -3.2 | 1.3E-03 | 1.6E-03 | BT20/MB468 | NUC |
| >hg38_21_+_8433172_8446622_RNA45SN1_45S_50ntFlanks@12635.12659.25\|AAGCTACCATCTGTGGGATTATGAC | >hg38_21_+_8433172_8446622_RNA45SN1_45S_50ntFlanks | 12635 | 12659 | 25 | AAGCTACCATCTGTGGGATTATGAC | 4464.0 | -4.5 | 1.3 | -3.4 | 6.1E-04 | 1.6E-03 | BT20/MB468 | MP |
| >hg38_21_+_8433172_8446622_RNA45SN1_45S_50ntFlanks@9805.9832.28\|GCGTGGAATGCGAGTGCCTAGTGGGCCA | >hg38_21_+_8433172_8446622_RNA45SN1_45S_50ntFlanks | 9805 | 9832 | 28 | GCGTGGAATGCGAGTGCCTAGTGGGCCA | 96.4 | -5.7 | 1.7 | -3.4 | 6.1E-04 | 1.6E-03 | BT20/MB468 | MP |
| >hg38_21_+_8433172_8446622_RNA45SN1_45S_50ntFlanks@10683.10707.25\|CCGGCGGCGTCCGGTGAGCTCTCGC | >hg38_21_+_8433172_8446622_RNA45SN1_45S_50ntFlanks | 10683 | 10707 | 25 | CCGGCGGCGTCCGGTGAGCTCTCGC | 69.0 | -5.0 | 1.5 | -3.3 | 1.1E-03 | 1.6E-03 | MB231/MB468 | MP |
| >hg38_21_+_8433172_8446622_RNA45SN1_45S_50ntFlanks@10806.10823.18\|AACAATGTAGGTAAGGGA | >hg38_21_+_8433172_8446622_RNA45SN1_45S_50ntFlanks | 10806 | 10823 | 18 | AACAATGTAGGTAAGGGA | 462.8 | -4.3 | 1.3 | -3.4 | 6.1E-04 | 1.6E-03 | BT20/MB468 | MP |
| >MI0003205\|hsa-mir-532&WithFlank&X\|+\|50003142\|50003244@26.46.21(+1C)&[MIMAT0002888&hsa-miR-532-5p&offsets\|0\|-1(+1C);m-65&X\|+\|50003167\|50003188&offsets\|0\|-1(+1C)]\|CATGCCTTGAGTGTAGGACCGC | >MI0003205\|hsa-mir-532&WithFlank&X\|+\|50003142\|50003244 | 26 | 46 | 21(+1C) | CATGCCTTGAGTGTAGGACCGC | 833.7 | 3.5 | 1.0 | 3.4 | 6.1E-04 | 1.6E-03 | BT20/MB468 | MP |
| >hg38_21_+_8433172_8446622_RNA45SN1_45S_50ntFlanks@9454.9502.49\|CCGCGCCGGGGAGGTGGAGCACGAGCGCACGTGTTAGGACCCGAAAGAT | >hg38_21_+_8433172_8446622_RNA45SN1_45S_50ntFlanks | 9454 | 9502 | 49 | CCGCGCCGGGGAGGTGGAGCACGAGCGCACGTGTTAGGACCCGAAAGAT | 57.6 | -9.7 | 3.0 | -3.3 | 1.1E-03 | 1.6E-03 | MB231/MB468 | MP |
| >MI0000456\|hsa-mir-140&WithFlank&16\|+\|69933075\|69933186@68.90.23&[MIMAT0004597&hsa-miR-140-3p&offsets\|0\|+2;m-45&16\|+\|69933143\|69933164&offsets\|-1\|0]\|TACCACAGGGTAGAACCACGGAC | >MI0000456\|hsa-mir-140&WithFlank&16\|+\|69933075\|69933186 | 68 | 90 | 23 | TACCACAGGGTAGAACCACGGAC | 192.5 | 4.9 | 1.4 | 3.4 | 6.1E-04 | 1.6E-03 | BT20/MB468 | MP |
| >hg38_MT_+_1621_3279_RNR2_16S_50ntFlanks@51.79.29\|GCTAAACCTAGCCCCAAACCCACTCCACC | >hg38_MT_+_1621_3279_RNR2_16S_50ntFlanks | 51 | 79 | 29 | GCTAAACCTAGCCCCAAACCCACTCCACC | 10841.2 | 4.3 | 1.3 | 3.4 | 6.1E-04 | 1.6E-03 | BT20/MB468 | MP |
| >hg38_21_+_8433172_8446622_RNA45SN1_45S_50ntFlanks@10376.10424.49\|GAAGGGTTCCATGTGAACAGCAGTTGAACATGGGTCAGTCGGTCCTGAG | >hg38_21_+_8433172_8446622_RNA45SN1_45S_50ntFlanks | 10376 | 10424 | 49 | GAAGGGTTCCATGTGAACAGCAGTTGAACATGGGTCAGTCGGTCCTGAG | 1250.0 | 4.1 | 1.1 | 3.6 | 2.7E-04 | 1.6E-03 | BT20/MB231 | MITO |
| >hg38_21_+_8433172_8446622_RNA45SN1_45S_50ntFlanks@7977.8011.35\|CGACCTCAGATCAGACGTGGCGACCCGCTGAATTT | >hg38_21_+_8433172_8446622_RNA45SN1_45S_50ntFlanks | 7977 | 8011 | 35 | CGACCTCAGATCAGACGTGGCGACCCGCTGAATTT | 4050.3 | -3.7 | 1.0 | -3.7 | 2.3E-04 | 1.6E-03 | BT20/MB231 | CYTO |
| >trna14_LysTTT_11_-_59327808_59327880@1.33.33,trna2_LysTTT_17_+_8022473_8022545@1.33.33,trna54_LysTTT_1_+_204475655_204475727@1.33.33,trna5_LysTTT_11_+_59323902_59323974@1.33.33,trna62_LysTTT_1_-_204476158_204476230@1.33.33,trna76_LysTTT_6_+_28918806_28918878@1.33.33\|GCCCGGATAGCTCAGTCGGTAGAGCATCAGACT | >trna14_LysTTT_11_-_59327808_59327880 | 1 | 33 | 33 | GCCCGGATAGCTCAGTCGGTAGAGCATCAGACT | 3557.2 | 4.7 | 1.4 | 3.4 | 6.1E-04 | 1.6E-03 | BT20/MB468 | MP |
| >MI0000081\|hsa-mir-24-2&WithFlank&19\|-\|13836281\|13836365@56.76.21;MI0000080\|hsa-mir-24-1&WithFlank&9\|+\|95086015\|95086094@50.70.21&[MIMAT0000080&hsa-miR-24-3p&offsets\|0\|-1;m-36&19\|-\|13836288\|13836310&offsets\|0\|-2];[MIMAT0000080_1&hsa-miR-24-3p&offsets\|0\|-1;m-35&9\|+\|95086064\|95086086&offsets\|0\|-2]\|TGGCTCAGTTCAGCAGGAACA | >MI0000081\|hsa-mir-24-2&WithFlank&19\|-\|13836281\|13836365 | 56 | 76 | 21 | TGGCTCAGTTCAGCAGGAACA | 6077.8 | 2.7 | 0.8 | 3.3 | 1.1E-03 | 1.6E-03 | MB231/MB468 | MP |
| >hg38_21_+_8433172_8446622_RNA45SN1_45S_50ntFlanks@10846.10872.27\|CTTCGGGATAAGGATTGGCTCTAAGGG | >hg38_21_+_8433172_8446622_RNA45SN1_45S_50ntFlanks | 10846 | 10872 | 27 | CTTCGGGATAAGGATTGGCTCTAAGGG | 211.5 | -3.0 | 0.9 | -3.2 | 1.3E-03 | 1.6E-03 | MB231/MB468 | CYTO |
| >hg38_21_+_8433172_8446622_RNA45SN1_45S_50ntFlanks@11209.11234.26\|CGGCGACTCTGGACGCGAGCCGGGCC | >hg38_21_+_8433172_8446622_RNA45SN1_45S_50ntFlanks | 11209 | 11234 | 26 | CGGCGACTCTGGACGCGAGCCGGGCC | 1560.9 | -6.0 | 1.8 | -3.4 | 6.1E-04 | 1.6E-03 | BT20/MB468 | MP |
| >am_trnaMT_TyrGTA_MT_-_5826_5891@13.32.20\|GAGTGAAGCATTGGACTGTA | >am_trnaMT_TyrGTA_MT_-_5826_5891 | 13 | 32 | 20 | GAGTGAAGCATTGGACTGTA | 137.0 | 4.0 | 1.3 | 3.2 | 1.3E-03 | 1.6E-03 | MB231/MB468 | MITO |
| >hg38_21_+_8433172_8446622_RNA45SN1_45S_50ntFlanks@10766.10797.32\|CGCAGCAGGTCTCCAAGGTGAACAGCCTCTGG | >hg38_21_+_8433172_8446622_RNA45SN1_45S_50ntFlanks | 10766 | 10797 | 32 | CGCAGCAGGTCTCCAAGGTGAACAGCCTCTGG | 163.8 | -5.6 | 1.6 | -3.4 | 6.1E-04 | 1.6E-03 | BT20/MB468 | MP |
| >hg38_21_+_8433172_8446622_RNA45SN1_45S_50ntFlanks@8171.8195.25\|AAGTCCTTCTGATCGAGGCCCAGCC | >hg38_21_+_8433172_8446622_RNA45SN1_45S_50ntFlanks | 8171 | 8195 | 25 | AAGTCCTTCTGATCGAGGCCCAGCC | 218.2 | -3.0 | 0.9 | -3.2 | 1.3E-03 | 1.6E-03 | MB231/MB468 | CYTO |
| >hg38_21_+_8433172_8446622_RNA45SN1_45S_50ntFlanks@4243.4264.22\|CTTTCGAGGCCCTGTAATTGGA | >hg38_21_+_8433172_8446622_RNA45SN1_45S_50ntFlanks | 4243 | 4264 | 22 | CTTTCGAGGCCCTGTAATTGGA | 174.9 | -2.3 | 0.7 | -3.3 | 1.0E-03 | 1.6E-03 | BT20/MB468 | MITO |
| >trna59_GluCTC_1_+_249168447_249168518@32.70.39\|CTCTCACCGCCGCGGCCCGGGTTCGATTCCCGGTCAGGA | >trna59_GluCTC_1_+_249168447_249168518 | 32 | 70 | 39 | CTCTCACCGCCGCGGCCCGGGTTCGATTCCCGGTCAGGA | 103.7 | -5.1 | 1.5 | -3.3 | 1.0E-03 | 1.6E-03 | BT20/MB468 | MITO |
| >hg38_21_+_8433172_8446622_RNA45SN1_45S_50ntFlanks@6649.6678.30\|TACGACTCTTAGCGGTGGATCACTCGGCTC | >hg38_21_+_8433172_8446622_RNA45SN1_45S_50ntFlanks | 6649 | 6678 | 30 | TACGACTCTTAGCGGTGGATCACTCGGCTC | 460.1 | 2.3 | 0.7 | 3.2 | 1.4E-03 | 1.6E-03 | MB231/MB468 | NUC |
| >hg38_21_+_8433172_8446622_RNA45SN1_45S_50ntFlanks@11729.11761.33\|TAACTATGACTCTCTTAAGGTAGCCAAATGCCT | >hg38_21_+_8433172_8446622_RNA45SN1_45S_50ntFlanks | 11729 | 11761 | 33 | TAACTATGACTCTCTTAAGGTAGCCAAATGCCT | 496.8 | -3.5 | 1.1 | -3.2 | 1.3E-03 | 1.6E-03 | MB231/MB468 | CYTO |

| | | | | | | | | | | | | | |
|---|---|---|---|---|---|---|---|---|---|---|---|---|---|
| >MI0005559\|hsa-mir-744&WithFlank&17\|+\|12081893\|12082002@17.38.22&[MIMAT0004945&hsa-miR-744-5p&offsets\|0\|0;m-172&17\|+\|12081909\|12081929&offsets\|0\|+1]\|TGCGGGGCTAGGGCTAACAGCA | >MI0005559\|hsa-mir-744&WithFlank&17\|+\|12081893\|12082002 | 17 | 38 | 22 | TGCGGGGCTAGGGCTAACAGCA | 418.5 | 3.0 | 0.9 | 3.3 | 1.1E-03 | 1.6E-03 | MB231/MB468 | MP |
| >hg38_21_+_8433172_8446622_RNA45SN1_45S_50ntFlanks@10350.10373.24\|ACGAGAACTTTGAAGGCCGAAGTG | >hg38_21_+_8433172_8446622_RNA45SN1_45S_50ntFlanks | 10350 | 10373 | 24 | ACGAGAACTTTGAAGGCCGAAGTG | 498.9 | -3.8 | 1.2 | -3.2 | 1.3E-03 | 1.6E-03 | MB231/MB468 | MITO |
| >hg38_21_+_8433172_8446622_RNA45SN1_45S_50ntFlanks@7988.8014.27\|CAGACGTGGCGACCCGCTGAATTTAAG | >hg38_21_+_8433172_8446622_RNA45SN1_45S_50ntFlanks | 7988 | 8014 | 27 | CAGACGTGGCGACCCGCTGAATTTAAG | 271.8 | -2.4 | 0.7 | -3.2 | 1.4E-03 | 1.6E-03 | MB231/MB468 | NUC |
| >MI0000746\|hsa-mir-99b&WithFlank&19\|+\|51692606\|51692687@51.68.18(+1C)&[MIMAT0004678&hsa-miR-99b-3p&offsets\|0\|-4(+1C);m-151&19\|+\|51692656\|51692678&offsets\|0\|-5(+1C)]\|CAAGCTCGTGTCTGTGGGC | >MI0000746\|hsa-mir-99b&WithFlank&19\|+\|51692606\|51692687 | 51 | 68 | 18(+1C) | CAAGCTCGTGTCTGTGGGC | 51.1 | 9.4 | 2.8 | 3.4 | 6.2E-04 | 1.6E-03 | BT20/MB468 | MP |
| >trna10_LysCTT_16_+_3241501_3241573@14.31.18,trna13_LysCTT_14_-_58706613_58706685@14.31.18,trna2_LysCTT_15_+_79152904_79152976@14.31.18,trna4_LysCTT_18_-_43669271_43669343@14.31.18\|AGTCGGTAGAGCATGGGA | >trna10_LysCTT_16_+_3241501_3241573 | 14 | 31 | 18 | AGTCGGTAGAGCATGGGA | 437.1 | -3.1 | 0.9 | -3.4 | 6.2E-04 | 1.6E-03 | BT20/MB468 | MP |
| >MI0000108\|hsa-mir-103a-2&WithFlank&20\|+\|3917488\|3917577@54.76.23(+2U);MI0000109\|hsa-mir-103a-1&WithFlank&5\|-\|168560890\|168560979@54.76.23(+2U)&[MIMAT0000101&hsa-miR-103a-3p&offsets\|0\|0(+2U);m-12&20\|+\|3917541\|3917563&offsets\|0\|0(+2U)];[MIMAT0000101_1&hsa-miR-103a-3p&offsets\|0\|0(+2U);m-13&5\|-\|168560904\|168560926&offsets\|0\|0(+2U);m-44220&5\|-\|168560908\|168560926&offsets\|0\|+4(+2U)]\|AGCAGCATTGTACAGGGCTATGATT | >MI0000108\|hsa-mir-103a-2&WithFlank&20\|+\|3917488\|3917577 | 54 | 76 | 23(+2U) | AGCAGCATTGTACAGGGCTATGATT | 306.5 | 4.1 | 1.3 | 3.3 | 1.1E-03 | 1.6E-03 | MB231/MB468 | MP |
| >hg38_21_+_8433172_8446622_RNA45SN1_45S_50ntFlanks@7989.8007.19\|AGACGTGGCGACCCGCTGA | >hg38_21_+_8433172_8446622_RNA45SN1_45S_50ntFlanks | 7989 | 8007 | 19 | AGACGTGGCGACCCGCTGA | 2371.1 | 5.3 | 1.4 | 3.7 | 2.1E-04 | 1.6E-03 | BT20/MB231 | MP |
| >hg38_21_+_8433172_8446622_RNA45SN1_45S_50ntFlanks@10351.10383.33\|CGAGAACTTTGAAGGCCGAAGTGGAGAAGGGTT | >hg38_21_+_8433172_8446622_RNA45SN1_45S_50ntFlanks | 10351 | 10383 | 33 | CGAGAACTTTGAAGGCCGAAGTGGAGAAGGGTT | 36109.1 | -4.7 | 1.4 | -3.4 | 6.2E-04 | 1.6E-03 | BT20/MB468 | MP |
| >hg38_21_+_8433172_8446622_RNA45SN1_45S_50ntFlanks@11579.11606.28\|AGAACTGGTGCGGACCAGGGGAATCCGA | >hg38_21_+_8433172_8446622_RNA45SN1_45S_50ntFlanks | 11579 | 11606 | 28 | AGAACTGGTGCGGACCAGGGGAATCCGA | 196.8 | -1.9 | 0.6 | -3.2 | 1.2E-03 | 1.6E-03 | BT20/MB468 | CYTO |
| >hg38_1_-_228634819_228635039_RNA5S12_5S_50ntFlanks@146.170.25\|TGGGAATACCGGGTGCTGTAGGCTT | >hg38_1_-_228634819_228635039_RNA5S12_5S_50ntFlanks | 146 | 170 | 25 | TGGGAATACCGGGTGCTGTAGGCTT | 4646.7 | 3.2 | 0.9 | 3.4 | 6.2E-04 | 1.6E-03 | BT20/MB468 | MP |
| >MI0000085\|hsa-mir-27a&WithFlank&19\|-\|13836434\|13836523@57.76.20&[MIMAT0000084&hsa-miR-27a-3p&offsets\|0\|-1;m-43&19\|-\|13836447\|13836467&offsets\|0\|-1]\|TTCACAGTGGCTAAGTTCCG | >MI0000085\|hsa-mir-27a&WithFlank&19\|-\|13836434\|13836523 | 57 | 76 | 20 | TTCACAGTGGCTAAGTTCCG | 17357.1 | -2.8 | 0.8 | -3.6 | 2.7E-04 | 1.6E-03 | BT20/MB231 | MITO |
| >MI0000088\|hsa-mir-30a&WithFlank&6\|-\|71403545\|71403627@53.73.21(+1A)&[MIMAT0000088&hsa-miR-30a-3p&offsets\|0\|-1(+1A);m-38&6\|-\|71403554\|71403575&offsets\|0\|-1(+1A)]\|CTTTCAGTCGGATGTTTGCAGA | >MI0000088\|hsa-mir-30a&WithFlank&6\|-\|71403545\|71403627 | 53 | 73 | 21(+1A) | CTTTCAGTCGGATGTTTGCAGA | 621.2 | -2.7 | 0.7 | -3.6 | 2.7E-04 | 1.6E-03 | BT20/MB231 | MITO |
| >hg38_21_+_8433172_8446622_RNA45SN1_45S_50ntFlanks@10349.10372.24\|AACGAGAACTTTGAAGGCCGAAGT | >hg38_21_+_8433172_8446622_RNA45SN1_45S_50ntFlanks | 10349 | 10372 | 24 | AACGAGAACTTTGAAGGCCGAAGT | 1320.9 | 2.5 | 0.8 | 3.2 | 1.4E-03 | 1.6E-03 | MB231/MB468 | NUC |
| >MI0001733\|hsa-mir-452&WithFlank&X\|-\|151959622\|151959718@20.42.23&[MIMAT0001635&hsa-miR-452-5p&offsets\|0\|+1;m-188&X\|-\|151959677\|151959699&offsets\|0\|0]\|AACTGTTTGCAGAGGAAACTGAG | >MI0001733\|hsa-mir-452&WithFlank&X\|-\|151959622\|151959718 | 20 | 42 | 23 | AACTGTTTGCAGAGGAAACTGAG | 799.3 | 2.7 | 0.7 | 3.6 | 2.7E-04 | 1.6E-03 | BT20/MB231 | MITO |
| >hg38_MT_+_1621_3279_RNR2_16S_50ntFlanks@1363.1388.26\|GACCTCGATGTTGGATCAGGACATCC | >hg38_MT_+_1621_3279_RNR2_16S_50ntFlanks | 1363 | 1388 | 26 | GACCTCGATGTTGGATCAGGACATCC | 190.2 | 4.0 | 1.2 | 3.2 | 1.3E-03 | 1.6E-03 | MB231/MB468 | MITO |
| >hg38_21_+_8433172_8446622_RNA45SN1_45S_50ntFlanks@12408.12446.39\|TTTGATCCTTCGATGTCGGCTCTTCCTATCATTGTGAAG | >hg38_21_+_8433172_8446622_RNA45SN1_45S_50ntFlanks | 12408 | 12446 | 39 | TTTGATCCTTCGATGTCGGCTCTTCCTATCATTGTGAAG | 267.7 | -5.7 | 1.8 | -3.2 | 1.4E-03 | 1.6E-03 | BT20/MB468 | NUC |
| >hg38_21_+_8433172_8446622_RNA45SN1_45S_50ntFlanks@11921.11956.36\|TGAGAGGTGTAGAATAAGTGGGAGGCCCCCGGCGCC | >hg38_21_+_8433172_8446622_RNA45SN1_45S_50ntFlanks | 11921 | 11956 | 36 | TGAGAGGTGTAGAATAAGTGGGAGGCCCCCGGCGCC | 216.2 | -5.0 | 1.6 | -3.2 | 1.4E-03 | 1.6E-03 | MB231/MB468 | NUC |
| >trnaMT_IleGAT_MT_+_4263_4331@44.68.25\|TAGGAGCTTAAACCCCCTTATTTCT | >trnaMT_IleGAT_MT_+_4263_4331 | 44 | 68 | 25 | TAGGAGCTTAAACCCCCTTATTTCT | 51.6 | 9.4 | 2.8 | 3.4 | 6.2E-04 | 1.6E-03 | BT20/MB468 | MP |
| >hg38_21_+_8433172_8446622_RNA45SN1_45S_50ntFlanks@6645.6664.20\|CTCGTACGACTCTTAGCGGT | >hg38_21_+_8433172_8446622_RNA45SN1_45S_50ntFlanks | 6645 | 6664 | 20 | CTCGTACGACTCTTAGCGGT | 4182.8 | 2.5 | 0.8 | 3.2 | 1.4E-03 | 1.6E-03 | MB231/MB468 | MITO |
| >hg38_MT_+_598_1651_RNR1_12S_50ntFlanks@844.873.30\|AGTAGAGTGCTTAGTTGAACAGGGCCCTGA | >hg38_MT_+_598_1651_RNR1_12S_50ntFlanks | 844 | 873 | 30 | AGTAGAGTGCTTAGTTGAACAGGGCCCTGA | 52.4 | 9.5 | 2.8 | 3.4 | 6.2E-04 | 1.6E-03 | BT20/MB468 | MP |
| >hg38_21_+_8433172_8446622_RNA45SN1_45S_50ntFlanks@8349.8379.31\|GAAAGTTGAAAAGAACTTTGAAGAGAGAGTT | >hg38_21_+_8433172_8446622_RNA45SN1_45S_50ntFlanks | 8349 | 8379 | 31 | GAAAGTTGAAAAGAACTTTGAAGAGAGAGTT | 77.6 | 5.5 | 1.6 | 3.4 | 6.2E-04 | 1.6E-03 | BT20/MB468 | MP |
| >hg38_21_+_8433172_8446622_RNA45SN1_45S_50ntFlanks@8169.8186.18\|CCAAGTCCTTCTGATCGA | >hg38_21_+_8433172_8446622_RNA45SN1_45S_50ntFlanks | 8169 | 8186 | 18 | CCAAGTCCTTCTGATCGA | 302.8 | -5.0 | 1.6 | -3.2 | 1.2E-03 | 1.6E-03 | BT20/MB468 | CYTO |
| >hg38_21_+_8433172_8446622_RNA45SN1_45S_50ntFlanks@10348.10368.21\|AAACGAGAACTTTGAAGGCCG | >hg38_21_+_8433172_8446622_RNA45SN1_45S_50ntFlanks | 10348 | 10368 | 21 | AAACGAGAACTTTGAAGGCCG | 560.4 | -2.1 | 0.7 | -3.3 | 1.0E-03 | 1.6E-03 | BT20/MB468 | MITO |
| >hg38_21_+_8433172_8446622_RNA45SN1_45S_50ntFlanks@485.527.43\|GATGTGGTGACGTCGTGCTCTCCCGGGCCGGGTCCGAGCCGCG | >hg38_21_+_8433172_8446622_RNA45SN1_45S_50ntFlanks | 485 | 527 | 43 | GATGTGGTGACGTCGTGCTCTCCCGGGCCGGGTCCGAGCCGCG | 340.1 | 11.6 | 3.6 | 3.2 | 1.4E-03 | 1.6E-03 | MB231/MB468 | MITO |
| >MI0000255\|hsa-mir-30d&WithFlank&8\|-\|134804870\|134804951@12.35.24(+2U)&[MIMAT0000245&hsa-miR-30d-5p&offsets\|0\|+2(+2U);m-21&8\|-\|134804919\|134804940&offsets\|0\|+2(+2U)]\|TGTAAACATCCCCGACTGGAAGCTTT | >MI0000255\|hsa-mir-30d&WithFlank&8\|-\|134804870\|134804951 | 12 | 35 | 24(+2U) | TGTAAACATCCCCGACTGGAAGCTTT | 265.2 | -3.8 | 1.0 | -3.7 | 2.1E-04 | 1.6E-03 | BT20/MB231 | MP |
| >MI0006273\|hsa-mir-1180&WithFlank&17\|-\|19344500\|19344580@47.67.21(+1C)&[MIMAT0005825&hsa-miR-1180-3p&offsets\|0\|-1(+1C);m-214&17\|-\|19344513\|19344534&offsets\|0\|-1(+1C)]\|TTTCCGGCTCGCGTGGGTGTGC | >MI0006273\|hsa-mir-1180&WithFlank&17\|-\|19344500\|19344580 | 47 | 67 | 21(+1C) | TTTCCGGCTCGCGTGGGTGTGC | 247.0 | 4.7 | 1.4 | 3.4 | 6.2E-04 | 1.6E-03 | BT20/MB468 | MP |
| >hg38_21_+_8433172_8446622_RNA45SN1_45S_50ntFlanks@8176.8194.19\|CTTCTGATCGAGGCCCAGC | >hg38_21_+_8433172_8446622_RNA45SN1_45S_50ntFlanks | 8176 | 8194 | 19 | CTTCTGATCGAGGCCCAGC | 250.0 | -2.7 | 0.8 | -3.4 | 6.2E-04 | 1.6E-03 | BT20/MB468 | MP |
| >hg38_21_+_8433172_8446622_RNA45SN1_45S_50ntFlanks@6646.6680.35\|TCGTACGACTCTTAGCGGTGGATCACTCGGCTCGT | >hg38_21_+_8433172_8446622_RNA45SN1_45S_50ntFlanks | 6646 | 6680 | 35 | TCGTACGACTCTTAGCGGTGGATCACTCGGCTCGT | 1789.6 | 1.7 | 0.5 | 3.2 | 1.3E-03 | 1.6E-03 | MB231/MB468 | CYTO |
| >hg38_21_+_8433172_8446622_RNA45SN1_45S_50ntFlanks@6760.6806.47\|TTGCGGCCCCGGGTTCCTCCCGGGGCTACGCCTGTCTGAGCGTCGCT | >hg38_21_+_8433172_8446622_RNA45SN1_45S_50ntFlanks | 6760 | 6806 | 47 | TTGCGGCCCCGGGTTCCTCCCGGGGCTACGCCTGTCTGAGCGTCGCT | 447.7 | 5.1 | 1.6 | 3.2 | 1.4E-03 | 1.6E-03 | MB231/MB468 | MITO |
| >hg38_21_+_8433172_8446622_RNA45SN1_45S_50ntFlanks@9548.9571.24\|GAGGTCCGTAGCGGTCCTGACGTG | >hg38_21_+_8433172_8446622_RNA45SN1_45S_50ntFlanks | 9548 | 9571 | 24 | GAGGTCCGTAGCGGTCCTGACGTG | 1159.2 | -6.3 | 2.0 | -3.2 | 1.4E-03 | 1.6E-03 | BT20/MB468 | NUC |
| >hg38_21_+_8433172_8446622_RNA45SN1_45S_50ntFlanks@9306.9335.30\|CAAGGAGTCTAACACGTGCGCGAGTCGGGG | >hg38_21_+_8433172_8446622_RNA45SN1_45S_50ntFlanks | 9306 | 9335 | 30 | CAAGGAGTCTAACACGTGCGCGAGTCGGGG | 390.8 | -2.5 | 0.8 | -3.3 | 1.0E-03 | 1.6E-03 | BT20/MB468 | MITO |
| >MI0003581\|hsa-mir-574&WithFlank&4\|+\|38868026\|38868133@67.87.21&[MIMAT0003239&hsa-miR-574-3p&offsets\|0\|-1;m-149&4\|+\|38868092\|38868113&offsets\|0\|-1]\|CACGCTCATGCACACACCCAC | >MI0003581\|hsa-mir-574&WithFlank&4\|+\|38868026\|38868133 | 67 | 87 | 21 | CACGCTCATGCACACACCCAC | 697.8 | 8.5 | 2.6 | 3.2 | 1.3E-03 | 1.6E-03 | MB231/MB468 | CYTO |
| >hg38_21_+_8433172_8446622_RNA45SN1_45S_50ntFlanks@11891.11910.20\|TTGACTCTAGTCTGGCACGG | >hg38_21_+_8433172_8446622_RNA45SN1_45S_50ntFlanks | 11891 | 11910 | 20 | TTGACTCTAGTCTGGCACGG | 530.7 | -2.5 | 0.8 | -3.2 | 1.3E-03 | 1.6E-03 | MB231/MB468 | CYTO |
| >hg38_21_+_8433172_8446622_RNA45SN1_45S_50ntFlanks@5236.5259.24\|CACTGACTGGCTCAGCGTGTGCCT | >hg38_21_+_8433172_8446622_RNA45SN1_45S_50ntFlanks | 5236 | 5259 | 24 | CACTGACTGGCTCAGCGTGTGCCT | 80.8 | -5.0 | 1.5 | -3.3 | 1.1E-03 | 1.6E-03 | MB231/MB468 | MP |
| >hg38_21_+_8433172_8446622_RNA45SN1_45S_50ntFlanks@6646.6663.18\|TCGTACGACTCTTAGCGG | >hg38_21_+_8433172_8446622_RNA45SN1_45S_50ntFlanks | 6646 | 6663 | 18 | TCGTACGACTCTTAGCGG | 22358.9 | 3.3 | 1.0 | 3.2 | 1.4E-03 | 1.6E-03 | MB231/MB468 | NUC |
| >hg38_21_+_8433172_8446622_RNA45SN1_45S_50ntFlanks@7973.8007.35\|GACGCGACCTCAGATCAGACGTGGCGACCCGCTGA | >hg38_21_+_8433172_8446622_RNA45SN1_45S_50ntFlanks | 7973 | 8007 | 35 | GACGCGACCTCAGATCAGACGTGGCGACCCGCTGA | 396.6 | 2.3 | 0.7 | 3.2 | 1.4E-03 | 1.6E-03 | MB231/MB468 | NUC |
| >hg38_21_+_8433172_8446622_RNA45SN1_45S_50ntFlanks@3835.3862.28\|CTCCTCTCCTACTTGGATAACTGTGGTA | >hg38_21_+_8433172_8446622_RNA45SN1_45S_50ntFlanks | 3835 | 3862 | 28 | CTCCTCTCCTACTTGGATAACTGTGGTA | 235.0 | -2.0 | 0.6 | -3.2 | 1.3E-03 | 1.6E-03 | MB231/MB468 | CYTO |
| >MI0000264\|hsa-mir-7-2&WithFlank&15\|+\|88611819\|88611940@38.61.24;MI0000265\|hsa-mir-7-3&WithFlank&19\|+\|4770664\|4770785@37.60.24;MI0000263\|hsa-mir-7-1&WithFlank&9\|-\|83969742\|83969863@30.53.24&[MIMAT0000252&hsa-miR-7-5p&offsets\|0\|0;m-400&15\|+\|88611856\|88611879&offsets\|0\|0];[MIMAT0000252_1&hsa-miR-7-5p&offsets\|0\|0;m-402&19\|+\|4770700\|4770723&offsets\|0\|0];[MIMAT0000252_2&hsa-miR-7-5p&offsets\|0\|0;m-401&9\|-\|83969811\|83969834&offsets\|0\|0]\|TGGAAGACTAGTGATTTTGTTGTT | >MI0000264\|hsa-mir-7-2&WithFlank&15\|+\|88611819\|88611940 | 38 | 61 | 24 | TGGAAGACTAGTGATTTTGTTGTT | 98014.0 | 3.1 | 0.9 | 3.7 | 2.4E-04 | 1.6E-03 | BT20/MB231 | CYTO |
| >hg38_21_+_8433172_8446622_RNA45SN1_45S_50ntFlanks@11572.11609.38\|CCGACTTAGAACTGGTGCGGACCAGGGGAATCCGACTG | >hg38_21_+_8433172_8446622_RNA45SN1_45S_50ntFlanks | 11572 | 11609 | 38 | CCGACTTAGAACTGGTGCGGACCAGGGGAATCCGACTG | 304.5 | -2.5 | 0.8 | -3.2 | 1.2E-03 | 1.7E-03 | BT20/MB468 | CYTO |
| >hg38_21_+_8433172_8446622_RNA45SN1_45S_50ntFlanks@11316.11340.25\|CACCCCACGTCTCGTCGCGCGCGCG | >hg38_21_+_8433172_8446622_RNA45SN1_45S_50ntFlanks | 11316 | 11340 | 25 | CACCCCACGTCTCGTCGCGCGCGCG | 2271.2 | -6.5 | 2.0 | -3.2 | 1.4E-03 | 1.7E-03 | BT20/MB468 | NUC |
| >hg38_21_+_8433172_8446622_RNA45SN1_45S_50ntFlanks@10869.10886.18\|AGGGCTGGGTCGGTCGGG | >hg38_21_+_8433172_8446622_RNA45SN1_45S_50ntFlanks | 10869 | 10886 | 18 | AGGGCTGGGTCGGTCGGG | 598.6 | -4.5 | 1.3 | -3.4 | 6.3E-04 | 1.7E-03 | BT20/MB468 | MP |
| >MI0000079\|hsa-mir-23a&WithFlank&19\|-\|13836581\|13836665@54.73.20(+1U)&[MIMAT0000078&hsa-miR-23a-3p&offsets\|+3\|+2(+1U);m-31&19\|-\|13836595\|13836615&offsets\|+3\|+2(+1U)]\|ACATTGCCAGGGATTTCCAAT | >MI0000079\|hsa-mir-23a&WithFlank&19\|-\|13836581\|13836665 | 54 | 73 | 20(+1U) | ACATTGCCAGGGATTTCCAAT | 383.7 | 11.8 | 3.6 | 3.3 | 1.1E-03 | 1.7E-03 | MB231/MB468 | MP |
| >hg38_21_+_8433172_8446622_RNA45SN1_45S_50ntFlanks@6661.6706.46\|CGGTGGATCACTCGGCTCGTGCGTCGATGAAGAACGCAGCTAGCTG | >hg38_21_+_8433172_8446622_RNA45SN1_45S_50ntFlanks | 6661 | 6706 | 46 | CGGTGGATCACTCGGCTCGTGCGTCGATGAAGAACGCAGCTAGCTG | 258.4 | 2.4 | 0.7 | 3.2 | 1.3E-03 | 1.7E-03 | MB231/MB468 | CYTO |
| >hg38_21_+_8433172_8446622_RNA45SN1_45S_50ntFlanks@11670.11693.24\|CCCAGTGCTCTGAATGTCAAAGTG | >hg38_21_+_8433172_8446622_RNA45SN1_45S_50ntFlanks | 11670 | 11693 | 24 | CCCAGTGCTCTGAATGTCAAAGTG | 735.1 | -2.3 | 0.7 | -3.2 | 1.3E-03 | 1.7E-03 | MB231/MB468 | CYTO |
| >hg38_21_+_8433172_8446622_RNA45SN1_45S_50ntFlanks@11570.11603.34\|AGCCGACTTAGAACTGGTGCGGACCAGGGGAATC | >hg38_21_+_8433172_8446622_RNA45SN1_45S_50ntFlanks | 11570 | 11603 | 34 | AGCCGACTTAGAACTGGTGCGGACCAGGGGAATC | 461.5 | -2.5 | 0.8 | -3.2 | 1.3E-03 | 1.7E-03 | MB231/MB468 | CYTO |
| >hg38_21_+_8433172_8446622_RNA45SN1_45S_50ntFlanks@11843.11872.30\|AGGGAACGGGCTTGGCGGAATCAGCGGGGA | >hg38_21_+_8433172_8446622_RNA45SN1_45S_50ntFlanks | 11843 | 11872 | 30 | AGGGAACGGGCTTGGCGGAATCAGCGGGGA | 1098.4 | -5.9 | 1.7 | -3.4 | 6.3E-04 | 1.7E-03 | BT20/MB468 | MP |
| >hg38_21_+_8433172_8446622_RNA45SN1_45S_50ntFlanks@7983.8003.21\|CAGATCAGACGTGGCGACCCG | >hg38_21_+_8433172_8446622_RNA45SN1_45S_50ntFlanks | 7983 | 8003 | 21 | CAGATCAGACGTGGCGACCCG | 429.8 | -1.9 | 0.6 | -3.3 | 1.0E-03 | 1.7E-03 | BT20/MB468 | MITO |
| >trnaMT_GluTTC_MT_-_14674_14742@39.69.31,trnalookalike8_GluTTC_5_-_93905172_93905240@39.69.31\|ATTGGTCGTGGTTGTAGTCCGTGCGAGAATA | >trnaMT_GluTTC_MT_-_14674_14742 | 39 | 69 | 31 | ATTGGTCGTGGTTGTAGTCCGTGCGAGAATA | 107.4 | 2.8 | 0.9 | 3.3 | 1.1E-03 | 1.7E-03 | MB231/MB468 | MP |

<table>
<tr><td>>hg38_21_+_8433172_8446622_RNA45SN1_45S_50ntFlanks@12417.12438.22|TCGATGTCGGCTCTTCCTATCA</td><td>>hg38_21_+_8433172_8446622_RNA45SN1_45S_50ntFlanks</td><td>12417</td><td>12438</td><td>22</td><td>TCGATGTCGGCTCTTCCTATCA</td><td>2217.8</td><td>-3.1</td><td>1.0</td><td>-3.2</td><td>1.4E-03</td><td>1.7E-03</td><td>MB231/MB468</td><td>MITO</td></tr>
<tr><td>>hg38_MT_+_1621_3279_RNR2_16S_50ntFlanks@518.543.26|TTGTAGAGAGAGTAAAAAATTTAACA</td><td>>hg38_MT_+_1621_3279_RNR2_16S_50ntFlanks</td><td>518</td><td>543</td><td>26</td><td>TTGTAGAGAGAGTAAAAAATTTAACA</td><td>119.5</td><td>5.3</td><td>1.6</td><td>3.3</td><td>1.2E-03</td><td>1.7E-03</td><td>MB231/MB468</td><td>MP</td></tr>
<tr><td>>MI0000266|hsa-mir-10a&WithFlank&17|-|48579832|48579953@69.88.20(+1C)&[MIMAT0004555&hsa-miR-10a-3p&offsets|0|-2(+1C);m-468&17|-|48579865|48579885&offsets|0|-1(+1C)]|CAAATTCGTATCTAGGGGAAC</td><td>>MI0000266|hsa-mir-10a&WithFlank&17|-|48579832|48579953</td><td>69</td><td>88</td><td>20(+1C)</td><td>CAAATTCGTATCTAGGGGAAC</td><td>51.7</td><td>9.4</td><td>2.8</td><td>3.4</td><td>6.3E-04</td><td>1.7E-03</td><td>BT20/MB468</td><td>MP</td></tr>
<tr><td>>trna10_AspGTC_12_-_125424193_125424264@36.75.40,trna12_AspGTC_12_-_125411891_125411962@36.75.40,trna144_AspGTC_6_-_27551236_27551307@36.75.40,trna38_AspGTC_17_-_8125556_8125627@36.75.40,trna45_AspGTC_6_+_27447453_27447524@36.75.40,trna48_AspGTC_6_+_27471523_27471594@36.75.40,trna4_AspGTC_12_+_96429799_96429870@36.75.40,trna69_AspGTC_1_-_161440205_161440276@36.75.40,trna72_AspGTC_1_-_161432824_161432895@36.75.40,trna75_AspGTC_1_-_161425414_161425485@36.75.40,trna78_AspGTC_1_-_161418033_161418104@36.75.40,trna81_AspGTC_1_-_161410615_161410686@36.75.40|CACGCGGGAGACCGGGGTTCGATTCCCCGACGGGGAGCCA</td><td>>trna10_AspGTC_12_-_125424193_125424264</td><td>36</td><td>75</td><td>40</td><td>CACGCGGGAGACCGGGGTTCGATTCCCCGACGGGGAGCCA</td><td>146.0</td><td>5.0</td><td>1.6</td><td>3.2</td><td>1.4E-03</td><td>1.7E-03</td><td>MB231/MB468</td><td>MITO</td></tr>
<tr><td>>hg38_21_+_8433172_8446622_RNA45SN1_45S_50ntFlanks@10689.10712.24|GCGTCCGGTGAGCTCTCGCTGGCC</td><td>>hg38_21_+_8433172_8446622_RNA45SN1_45S_50ntFlanks</td><td>10689</td><td>10712</td><td>24</td><td>GCGTCCGGTGAGCTCTCGCTGGCC</td><td>55.2</td><td>-8.9</td><td>2.6</td><td>-3.4</td><td>6.3E-04</td><td>1.7E-03</td><td>BT20/MB468</td><td>MP</td></tr>
<tr><td>>hg38_21_+_8433172_8446622_RNA45SN1_45S_50ntFlanks@12008.12036.29|CCGCCGGTGAAATACCACTACTCTGATCG</td><td>>hg38_21_+_8433172_8446622_RNA45SN1_45S_50ntFlanks</td><td>12008</td><td>12036</td><td>29</td><td>CCGCCGGTGAAATACCACTACTCTGATCG</td><td>470.2</td><td>-2.5</td><td>0.8</td><td>-3.3</td><td>1.0E-03</td><td>1.7E-03</td><td>BT20/MB468</td><td>MITO</td></tr>
<tr><td>>hg38_21_+_8433172_8446622_RNA45SN1_45S_50ntFlanks@12393.12419.27|ATAGCGACGTCGCTTTTTGATCCTTCG</td><td>>hg38_21_+_8433172_8446622_RNA45SN1_45S_50ntFlanks</td><td>12393</td><td>12419</td><td>27</td><td>ATAGCGACGTCGCTTTTTGATCCTTCG</td><td>214.0</td><td>-2.2</td><td>0.7</td><td>-3.3</td><td>1.0E-03</td><td>1.7E-03</td><td>BT20/MB468</td><td>MITO</td></tr>
<tr><td>>MI0000086|hsa-mir-28&WithFlank&3|+|188688775|188688872@62.82.21&[MIMAT0004502&hsa-miR-28-3p&offsets|+2|+1;m-28&3|+|188688834|188688855&offsets|+2|+1]|CTAGATTGTGAGCTCCTGGAG</td><td>>MI0000086|hsa-mir-28&WithFlank&3|+|188688775|188688872</td><td>62</td><td>82</td><td>21</td><td>CTAGATTGTGAGCTCCTGGAG</td><td>457.3</td><td>3.0</td><td>0.8</td><td>3.7</td><td>2.4E-04</td><td>1.7E-03</td><td>BT20/MB231</td><td>CYTO</td></tr>
<tr><td>>MI0000542|hsa-mir-320a&WithFlank&8|-|22244960|22245043@50.70.21(+1U)&[MIMAT0000510&hsa-miR-320a-3p&offsets|+2|+1(+1U);m-60&8|-|22244975|22244996&offsets|+2|+1(+1U)]|AAGCTGGGTTGAGAGGGCGAAT</td><td>>MI0000542|hsa-mir-320a&WithFlank&8|-|22244960|22245043</td><td>50</td><td>70</td><td>21(+1U)</td><td>AAGCTGGGTTGAGAGGGCGAAT</td><td>330.0</td><td>2.7</td><td>0.7</td><td>3.7</td><td>2.4E-04</td><td>1.7E-03</td><td>BT20/MB231</td><td>CYTO</td></tr>
<tr><td>>hg38_21_+_8433172_8446622_RNA45SN1_45S_50ntFlanks@11029.11051.23|CTCTCTCTCTCTCCCCCGCTCCC</td><td>>hg38_21_+_8433172_8446622_RNA45SN1_45S_50ntFlanks</td><td>11029</td><td>11051</td><td>23</td><td>CTCTCTCTCTCTCCCCCGCTCCC</td><td>302.6</td><td>-5.3</td><td>1.7</td><td>-3.2</td><td>1.4E-03</td><td>1.7E-03</td><td>MB231/MB468</td><td>NUC</td></tr>
<tr><td>>am_MI0015825|hsa-mir-378c&WithFlank&10|-|130962582|130962674@17.34.18;MI0016749|hsa-mir-378d-1&WithFlank&4|-|5923269|5923334@37.54.18;MI0000786|hsa-mir-378a&WithFlank&5|+|149732819|149732896@49.66.18;MI0003840|hsa-mir-378d-2&WithFlank&8|-|93916016|93916125@28.45.18&[MIMAT0016847&hsa-miR-378c&offsets|0|-7;m-130&10|-|130962639|130962658&offsets|0|-2];[MIMAT0018926&hsa-miR-378d&offsets|0|-2;m-228&4|-|5923281|5923298&offsets|0|0];[MIMAT0000732&hsa-miR-378a-3p&offsets|0|-4;m-44&5|+|149732867|149732888&offsets|0|-4];[MIMAT0018926_1&hsa-miR-378d&offsets|0|-2;m-227&8|-|93916081|93916098&offsets|0|0]|ACTGGACTTGGAGTCAGA</td><td>>am_MI0015825|hsa-mir-378c&WithFlank&10|-|130962582|130962674</td><td>17</td><td>34</td><td>18</td><td>ACTGGACTTGGAGTCAGA</td><td>194.6</td><td>3.4</td><td>1.1</td><td>3.2</td><td>1.4E-03</td><td>1.7E-03</td><td>MB231/MB468</td><td>MITO</td></tr>
<tr><td>>MI0003591|hsa-mir-584&WithFlank&5|-|149062307|149062415@22.43.22&[MIMAT0003249&hsa-miR-584-5p&offsets|0|0;m-167&5|-|149062374|149062394&offsets|0|+1]|TTATGGTTTGCCTGGGACTGAG</td><td>>MI0003591|hsa-mir-584&WithFlank&5|-|149062307|149062415</td><td>22</td><td>43</td><td>22</td><td>TTATGGTTTGCCTGGGACTGAG</td><td>201.9</td><td>4.0</td><td>1.2</td><td>3.3</td><td>1.2E-03</td><td>1.7E-03</td><td>MB231/MB468</td><td>MP</td></tr>
<tr><td>>hg38_21_+_8433172_8446622_RNA45SN1_45S_50ntFlanks@12658.12677.20|ACTGAACGCCTCTAAGTCAG</td><td>>hg38_21_+_8433172_8446622_RNA45SN1_45S_50ntFlanks</td><td>12658</td><td>12677</td><td>20</td><td>ACTGAACGCCTCTAAGTCAG</td><td>47.9</td><td>-8.7</td><td>2.5</td><td>-3.4</td><td>6.3E-04</td><td>1.7E-03</td><td>BT20/MB468</td><td>MP</td></tr>
<tr><td>>hg38_21_+_8433172_8446622_RNA45SN1_45S_50ntFlanks@4369.4391.23|GTTAAAAAGCTCGTAGTTGGATC</td><td>>hg38_21_+_8433172_8446622_RNA45SN1_45S_50ntFlanks</td><td>4369</td><td>4391</td><td>23</td><td>GTTAAAAAGCTCGTAGTTGGATC</td><td>282.3</td><td>-2.7</td><td>0.8</td><td>-3.4</td><td>6.3E-04</td><td>1.7E-03</td><td>BT20/MB468</td><td>MP</td></tr>
<tr><td>>hg38_21_+_8433172_8446622_RNA45SN1_45S_50ntFlanks@8359.8379.21|AAGAACTTTGAAGAGAGAGTT</td><td>>hg38_21_+_8433172_8446622_RNA45SN1_45S_50ntFlanks</td><td>8359</td><td>8379</td><td>21</td><td>AAGAACTTTGAAGAGAGAGTT</td><td>449.7</td><td>-3.5</td><td>1.1</td><td>-3.2</td><td>1.3E-03</td><td>1.7E-03</td><td>BT20/MB468</td><td>CYTO</td></tr>
<tr><td>>am_trnaMT_ValTAC_MT_+_1602_1670@48.68.21|CAACTTAACTTGACCGCTCTG</td><td>>am_trnaMT_ValTAC_MT_+_1602_1670</td><td>48</td><td>68</td><td>21</td><td>CAACTTAACTTGACCGCTCTG</td><td>1202.4</td><td>3.6</td><td>1.1</td><td>3.2</td><td>1.3E-03</td><td>1.7E-03</td><td>BT20/MB468</td><td>CYTO</td></tr>
<tr><td>>hg38_21_+_8433172_8446622_RNA45SN1_45S_50ntFlanks@5489.5516.28|CTGAGAAGACGGTCGAACTTGACTATCT</td><td>>hg38_21_+_8433172_8446622_RNA45SN1_45S_50ntFlanks</td><td>5489</td><td>5516</td><td>28</td><td>CTGAGAAGACGGTCGAACTTGACTATCT</td><td>10717.0</td><td>5.5</td><td>1.5</td><td>3.7</td><td>2.2E-04</td><td>1.7E-03</td><td>BT20/MB231</td><td>MP</td></tr>
<tr><td>>hg38_1_-_228634819_228635039_RNA5S12_5S_50ntFlanks@128.170.43|CTTGGATGGGAGACCGCCTGGGAATACCGGGTGCTGTAGGCTT</td><td>>hg38_1_-_228634819_228635039_RNA5S12_5S_50ntFlanks</td><td>128</td><td>170</td><td>43</td><td>CTTGGATGGGAGACCGCCTGGGAATACCGGGTGCTGTAGGCTT</td><td>905.7</td><td>5.0</td><td>1.5</td><td>3.4</td><td>6.3E-04</td><td>1.7E-03</td><td>BT20/MB468</td><td>MP</td></tr>
<tr><td>>hg38_21_+_8433172_8446622_RNA45SN1_45S_50ntFlanks@6646.6690.45|TCGTACGACTCTTAGCGGTGGATCACTCGGCTCGTGCGTCGATGA</td><td>>hg38_21_+_8433172_8446622_RNA45SN1_45S_50ntFlanks</td><td>6646</td><td>6690</td><td>45</td><td>TCGTACGACTCTTAGCGGTGGATCACTCGGCTCGTGCGTCGATGA</td><td>907.1</td><td>4.4</td><td>1.3</td><td>3.4</td><td>6.3E-04</td><td>1.7E-03</td><td>BT20/MB468</td><td>MP</td></tr>
<tr><td>>MI0000736|hsa-mir-30c-1&WithFlank&1|+|40757278|40757378@23.45.23(+1G);MI0000254|hsa-mir-30c-2&WithFlank&6|-|71376954|71377037@13.35.23(+1G)&[MIMAT0000244&hsa-miR-30c-5p&offsets|0|0(+1G);m-57&1|+|40757300|40757323&offsets|0|-1(+1G)];[MIMAT0000244_1&hsa-miR-30c-5p&offsets|0|0(+1G);m-56&6|-|71377002|71377025&offsets|0|-1(+1G)]|TGTAAACATCCTACACTCTCAGCG</td><td>>MI0000736|hsa-mir-30c-1&WithFlank&1|+|40757278|40757378</td><td>23</td><td>45</td><td>23(+1G)</td><td>TGTAAACATCCTACACTCTCAGCG</td><td>338.7</td><td>3.1</td><td>1.0</td><td>3.2</td><td>1.3E-03</td><td>1.7E-03</td><td>MB231/MB468</td><td>CYTO</td></tr>
<tr><td>>hg38_21_+_8433172_8446622_RNA45SN1_45S_50ntFlanks@11208.11239.32|GCGGCGACTCTGGACGCGAGCCGGGCCCTTCC</td><td>>hg38_21_+_8433172_8446622_RNA45SN1_45S_50ntFlanks</td><td>11208</td><td>11239</td><td>32</td><td>GCGGCGACTCTGGACGCGAGCCGGGCCCTTCC</td><td>628.0</td><td>-2.5</td><td>0.8</td><td>-3.2</td><td>1.3E-03</td><td>1.7E-03</td><td>BT20/MB468</td><td>CYTO</td></tr>
<tr><td>>hg38_21_+_8433172_8446622_RNA45SN1_45S_50ntFlanks@3939.3959.21|AACCCGGTCAGCCCCTCTCCG</td><td>>hg38_21_+_8433172_8446622_RNA45SN1_45S_50ntFlanks</td><td>3939</td><td>3959</td><td>21</td><td>AACCCGGTCAGCCCCTCTCCG</td><td>1448.0</td><td>-4.4</td><td>1.3</td><td>-3.3</td><td>1.1E-03</td><td>1.7E-03</td><td>BT20/MB468</td><td>MITO</td></tr>
<tr><td>>am_MI0000809|hsa-mir-151a&WithFlank&8|-|140732558|140732659@53.70.18(+3G)&[MIMAT0000757&hsa-miR-151a-3p&offsets|0|-3(+3G);m-30&8|-|140732587|140732607&offsets|0|-3(+3G)]|CTAGACTGAAGCTCCTTGGGG</td><td>>am_MI0000809|hsa-mir-151a&WithFlank&8|-|140732558|140732659</td><td>53</td><td>70</td><td>18(+3G)</td><td>CTAGACTGAAGCTCCTTGGGG</td><td>92.8</td><td>3.1</td><td>1.0</td><td>3.2</td><td>1.4E-03</td><td>1.7E-03</td><td>BT20/MB468</td><td>NUC</td></tr>
<tr><td>>hg38_21_+_8433172_8446622_RNA45SN1_45S_50ntFlanks@6658.6681.24|TAGCGGTGGATCACTCGGCTCGTG</td><td>>hg38_21_+_8433172_8446622_RNA45SN1_45S_50ntFlanks</td><td>6658</td><td>6681</td><td>24</td><td>TAGCGGTGGATCACTCGGCTCGTG</td><td>462.0</td><td>-4.2</td><td>1.3</td><td>-3.2</td><td>1.4E-03</td><td>1.7E-03</td><td>MB231/MB468</td><td>MITO</td></tr>
<tr><td>>hg38_21_+_8433172_8446622_RNA45SN1_45S_50ntFlanks@12163.12180.18|ACTGGGGCGGTACACCTG</td><td>>hg38_21_+_8433172_8446622_RNA45SN1_45S_50ntFlanks</td><td>12163</td><td>12180</td><td>18</td><td>ACTGGGGCGGTACACCTG</td><td>475.4</td><td>-5.0</td><td>1.6</td><td>-3.2</td><td>1.3E-03</td><td>1.7E-03</td><td>BT20/MB468</td><td>CYTO</td></tr>
<tr><td>>hg38_21_+_8433172_8446622_RNA45SN1_45S_50ntFlanks@12289.12316.28|CCGTGAAAGCGGGGCCTCACGATCCTTC</td><td>>hg38_21_+_8433172_8446622_RNA45SN1_45S_50ntFlanks</td><td>12289</td><td>12316</td><td>28</td><td>CCGTGAAAGCGGGGCCTCACGATCCTTC</td><td>269.4</td><td>-5.3</td><td>1.6</td><td>-3.3</td><td>1.1E-03</td><td>1.7E-03</td><td>BT20/MB468</td><td>MITO</td></tr>
<tr><td>>hg38_MT_+_598_1651_RNR1_12S_50ntFlanks@914.946.33|CAAAGGACATTTAACTAAAACCCCTACGCATTT</td><td>>hg38_MT_+_598_1651_RNR1_12S_50ntFlanks</td><td>914</td><td>946</td><td>33</td><td>CAAAGGACATTTAACTAAAACCCCTACGCATTT</td><td>781.2</td><td>5.3</td><td>1.5</td><td>3.4</td><td>6.3E-04</td><td>1.7E-03</td><td>BT20/MB468</td><td>MP</td></tr>
<tr><td>>hg38_21_+_8433172_8446622_RNA45SN1_45S_50ntFlanks@10905.10939.35|CTGGGCGCGCGCCGCGGCTGGACGAGGCGCCGCCG</td><td>>hg38_21_+_8433172_8446622_RNA45SN1_45S_50ntFlanks</td><td>10905</td><td>10939</td><td>35</td><td>CTGGGCGCGCGCCGCGGCTGGACGAGGCGCCGCCG</td><td>103.5</td><td>-4.5</td><td>1.4</td><td>-3.3</td><td>1.1E-03</td><td>1.7E-03</td><td>BT20/MB468</td><td>MITO</td></tr>
<tr><td>>trnaMT_SerGCT_MT_+_12207_12265@13.59.47|AAGAACTGCTAACTCATGCCCCCATGTCTAACAACATGGCTTTCTCA</td><td>>trnaMT_SerGCT_MT_+_12207_12265</td><td>13</td><td>59</td><td>47</td><td>AAGAACTGCTAACTCATGCCCCCATGTCTAACAACATGGCTTTCTCA</td><td>51.0</td><td>9.4</td><td>2.8</td><td>3.4</td><td>6.3E-04</td><td>1.7E-03</td><td>BT20/MB468</td><td>MP</td></tr>
<tr><td>>hg38_21_+_8433172_8446622_RNA45SN1_45S_50ntFlanks@11343.11362.20|CGCTGGGGGCGGGGAGCGGT</td><td>>hg38_21_+_8433172_8446622_RNA45SN1_45S_50ntFlanks</td><td>11343</td><td>11362</td><td>20</td><td>CGCTGGGGGCGGGGAGCGGT</td><td>344.9</td><td>-5.8</td><td>1.7</td><td>-3.4</td><td>6.3E-04</td><td>1.7E-03</td><td>BT20/MB468</td><td>MP</td></tr>
<tr><td>>hg38_21_+_8433172_8446622_RNA45SN1_45S_50ntFlanks@9926.9946.21|GTTGATATAGACAGCAGGACG</td><td>>hg38_21_+_8433172_8446622_RNA45SN1_45S_50ntFlanks</td><td>9926</td><td>9946</td><td>21</td><td>GTTGATATAGACAGCAGGACG</td><td>85.0</td><td>6.4</td><td>1.9</td><td>3.4</td><td>6.3E-04</td><td>1.7E-03</td><td>BT20/MB468</td><td>MP</td></tr>
<tr><td>>hg38_21_+_8433172_8446622_RNA45SN1_45S_50ntFlanks@8115.8140.26|CGCGGGACATGTGGCGTACGGAAGAC</td><td>>hg38_21_+_8433172_8446622_RNA45SN1_45S_50ntFlanks</td><td>8115</td><td>8140</td><td>26</td><td>CGCGGGACATGTGGCGTACGGAAGAC</td><td>4912.7</td><td>-3.7</td><td>1.1</td><td>-3.4</td><td>6.4E-04</td><td>1.7E-03</td><td>BT20/MB468</td><td>MP</td></tr>
<tr><td>>hg38_MT_+_598_1651_RNR1_12S_50ntFlanks@425.455.31|ACGAAAGTGGCTTTAACATATCTGAACACAC</td><td>>hg38_MT_+_598_1651_RNR1_12S_50ntFlanks</td><td>425</td><td>455</td><td>31</td><td>ACGAAAGTGGCTTTAACATATCTGAACACAC</td><td>500.0</td><td>5.2</td><td>1.5</td><td>3.4</td><td>6.4E-04</td><td>1.7E-03</td><td>BT20/MB468</td><td>MP</td></tr>
<tr><td>>hg38_21_+_8433172_8446622_RNA45SN1_45S_50ntFlanks@10687.10712.26|CGGCGTCCGGTGAGCTCTCGCTGGCC</td><td>>hg38_21_+_8433172_8446622_RNA45SN1_45S_50ntFlanks</td><td>10687</td><td>10712</td><td>26</td><td>CGGCGTCCGGTGAGCTCTCGCTGGCC</td><td>723.1</td><td>-4.7</td><td>1.4</td><td>-3.4</td><td>6.4E-04</td><td>1.7E-03</td><td>BT20/MB468</td><td>MP</td></tr>
<tr><td>>hg38_21_+_8433172_8446622_RNA45SN1_45S_50ntFlanks@11641.11670.30|CGGCGGGTGTTGACGCGATGTGATTTCTGC</td><td>>hg38_21_+_8433172_8446622_RNA45SN1_45S_50ntFlanks</td><td>11641</td><td>11670</td><td>30</td><td>CGGCGGGTGTTGACGCGATGTGATTTCTGC</td><td>6249.0</td><td>-6.7</td><td>2.0</td><td>-3.4</td><td>6.4E-04</td><td>1.7E-03</td><td>BT20/MB468</td><td>MP</td></tr>
<tr><td>>hg38_21_+_8433172_8446622_RNA45SN1_45S_50ntFlanks@474.498.25|TCTTCGTGATCGATGTGGTGACGTC</td><td>>hg38_21_+_8433172_8446622_RNA45SN1_45S_50ntFlanks</td><td>474</td><td>498</td><td>25</td><td>TCTTCGTGATCGATGTGGTGACGTC</td><td>253.5</td><td>7.1</td><td>2.1</td><td>3.4</td><td>6.4E-04</td><td>1.7E-03</td><td>BT20/MB468</td><td>MP</td></tr>
<tr><td>>MI0005561|hsa-mir-877&WithFlank&6|+|30584326|30584423@8.30.23&[MIMAT0004949&hsa-miR-877-5p&offsets|+1|+4;m-382&6|+|30584332|30584354&offsets|+1|+1]|TAGAGGAGATGGCGCAGGGGACA</td><td>>MI0005561|hsa-mir-877&WithFlank&6|+|30584326|30584423</td><td>8</td><td>30</td><td>23</td><td>TAGAGGAGATGGCGCAGGGGACA</td><td>51.4</td><td>9.4</td><td>2.8</td><td>3.4</td><td>6.4E-04</td><td>1.7E-03</td><td>BT20/MB468</td><td>MP</td></tr>
<tr><td>>am_trnaMT_ValTAC_MT_+_1602_1670@48.65.18|CAACTTAACTTGACCGCT</td><td>>am_trnaMT_ValTAC_MT_+_1602_1670</td><td>48</td><td>65</td><td>18</td><td>CAACTTAACTTGACCGCT</td><td>374.4</td><td>3.2</td><td>0.9</td><td>3.7</td><td>2.4E-04</td><td>1.7E-03</td><td>BT20/MB231</td><td>CYTO</td></tr>
<tr><td>>hg38_21_+_8433172_8446622_RNA45SN1_45S_50ntFlanks@12454.12496.43|CACCAAGCGTTGGATTGTTCACCCACTAATAGGGAACGTGAGC</td><td>>hg38_21_+_8433172_8446622_RNA45SN1_45S_50ntFlanks</td><td>12454</td><td>12496</td><td>43</td><td>CACCAAGCGTTGGATTGTTCACCCACTAATAGGGAACGTGAGC</td><td>235.5</td><td>-3.0</td><td>0.8</td><td>-3.7</td><td>2.4E-04</td><td>1.7E-03</td><td>BT20/MB231</td><td>CYTO</td></tr>
<tr><td>>hg38_21_+_8433172_8446622_RNA45SN1_45S_50ntFlanks@6659.6678.20|AGCGGTGGATCACTCGGCTC</td><td>>hg38_21_+_8433172_8446622_RNA45SN1_45S_50ntFlanks</td><td>6659</td><td>6678</td><td>20</td><td>AGCGGTGGATCACTCGGCTC</td><td>112.0</td><td>4.6</td><td>1.3</td><td>3.4</td><td>6.4E-04</td><td>1.7E-03</td><td>BT20/MB468</td><td>MP</td></tr>
<tr><td>>MI0000103|hsa-mir-101-1&WithFlank&1|-|65058428|65058514@53.72.20;MI0000739|hsa-mir-101-2&WithFlank&9|+|4850291|4850381@55.74.20&[MIMAT0000099&hsa-miR-101-3p&offsets|0|-1;m-17&1|-|65058443|65058462&offsets|0|0];[MIMAT0000099_1&hsa-miR-101-3p&offsets|0|-1;m-16&9|+|4850345|4850364&offsets|0|0]|TACAGTACTGTGATAACTGA</td><td>>MI0000103|hsa-mir-101-1&WithFlank&1|-|65058428|65058514</td><td>53</td><td>72</td><td>20</td><td>TACAGTACTGTGATAACTGA</td><td>321.7</td><td>2.7</td><td>0.8</td><td>3.2</td><td>1.4E-03</td><td>1.7E-03</td><td>BT20/MB468</td><td>NUC</td></tr>
<tr><td>>hg38_21_+_8433172_8446622_RNA45SN1_45S_50ntFlanks@8358.8379.22|AAAGAACTTTGAAGAGAGAGTT</td><td>>hg38_21_+_8433172_8446622_RNA45SN1_45S_50ntFlanks</td><td>8358</td><td>8379</td><td>22</td><td>AAAGAACTTTGAAGAGAGAGTT</td><td>453.9</td><td>-3.1</td><td>1.0</td><td>-3.2</td><td>1.3E-03</td><td>1.7E-03</td><td>BT20/MB468</td><td>CYTO</td></tr>
<tr><td>>hg38_21_+_8433172_8446622_RNA45SN1_45S_50ntFlanks@10229.10273.45|CCCCGCGGACGCTACGCCGCGACGAGTAGGAGGGCCGCTGCGGTG</td><td>>hg38_21_+_8433172_8446622_RNA45SN1_45S_50ntFlanks</td><td>10229</td><td>10273</td><td>45</td><td>CCCCGCGGACGCTACGCCGCGACGAGTAGGAGGGCCGCTGCGGTG</td><td>50.8</td><td>9.4</td><td>2.8</td><td>3.4</td><td>6.4E-04</td><td>1.7E-03</td><td>BT20/MB468</td><td>MP</td></tr>
</table>

| | | | | | | | | | | | | | |
|---|---|---|---|---|---|---|---|---|---|---|---|---|---|
| >MI0000086\|hsa-mir-28&WithFlank&3\|+\|188688775\|188688872@62.82.21(+1U)&[MIMAT0004502&hsa-miR-28-3p&offsets\|+2\|+1(+1U);m-28&3\|+\|188688834\|188688855&offsets\|+2\|+1(+1U)]\|CTAGATTGTGAGCTCCTGGAGT | >MI0000086\|hsa-mir-28&WithFlank&3\|+\|188688775\|188688872 | 62 | 82 | 21(+1U) | CTAGATTGTGAGCTCCTGGAGT | 291.6 | 5.2 | 1.6 | 3.2 | 1.2E-03 | 1.7E-03 | MB231/MB468 | MP |
| >hg38_21_+_8433172_8446622_RNA45SN1_45S_50ntFlanks@8184.8223.40\|CGAGGCCCAGCCCGTGGACGGTGTGAGGCCGGTAGCGGCC | >hg38_21_+_8433172_8446622_RNA45SN1_45S_50ntFlanks | 8184 | 8223 | 40 | CGAGGCCCAGCCCGTGGACGGTGTGAGGCCGGTAGCGGCC | 118.1 | -4.5 | 1.4 | -3.3 | 1.1E-03 | 1.7E-03 | BT20/MB468 | MITO |
| >MI0005561\|hsa-mir-877&WithFlank&6\|+\|30584326\|30584423@7.26.20(+1U)&[MIMAT0004949&hsa-miR-877-5p&offsets\|0\|0(+1U);m-382&6\|+\|30584332\|30584354&offsets\|0\|-3(+1U)]\|GTAGAGGAGATGGCGCAGGGT | >MI0005561\|hsa-mir-877&WithFlank&6\|+\|30584326\|30584423 | 7 | 26 | 20(+1U) | GTAGAGGAGATGGCGCAGGGT | 51.4 | 9.4 | 2.8 | 3.4 | 6.4E-04 | 1.7E-03 | BT20/MB468 | MP |
| >hg38_21_+_8433172_8446622_RNA45SN1_45S_50ntFlanks@6649.6695.47\|TACGACTCTTAGCGGTGGATCACTCGGCTCGTGCGTCGATGAAGAAC | >hg38_21_+_8433172_8446622_RNA45SN1_45S_50ntFlanks | 6649 | 6695 | 47 | TACGACTCTTAGCGGTGGATCACTCGGCTCGTGCGTCGATGAAGAAC | 480.0 | 3.6 | 1.1 | 3.2 | 1.4E-03 | 1.7E-03 | MB231/MB468 | MITO |
| >hg38_21_+_8433172_8446622_RNA45SN1_45S_50ntFlanks@12073.12108.36\|CCGAGGGGCTCTCGCTTCTGGCGCCAAGCGCCCGGC | >hg38_21_+_8433172_8446622_RNA45SN1_45S_50ntFlanks | 12073 | 12108 | 36 | CCGAGGGGCTCTCGCTTCTGGCGCCAAGCGCCCGGC | 256.1 | -5.5 | 1.6 | -3.4 | 6.4E-04 | 1.7E-03 | BT20/MB468 | MP |
| >hg38_21_+_8433172_8446622_RNA45SN1_45S_50ntFlanks@6649.6676.28\|TACGACTCTTAGCGGTGGATCACTCGGC | >hg38_21_+_8433172_8446622_RNA45SN1_45S_50ntFlanks | 6649 | 6676 | 28 | TACGACTCTTAGCGGTGGATCACTCGGC | 866.0 | 3.0 | 0.9 | 3.2 | 1.4E-03 | 1.7E-03 | MB231/MB468 | MITO |
| >MI0001445\|hsa-mir-423&WithFlank&17\|+\|30117073\|30117178@59.77.19&[MIMAT0001340&hsa-miR-423-3p&offsets\|0\|-4;m-100&17\|+\|30117131\|30117153&offsets\|0\|-4]\|AGCTCGGTCTGAGGCCCCT | >MI0001445\|hsa-mir-423&WithFlank&17\|+\|30117073\|30117178 | 59 | 77 | 19 | AGCTCGGTCTGAGGCCCCT | 114.5 | 5.5 | 1.6 | 3.4 | 6.4E-04 | 1.7E-03 | BT20/MB468 | MP |
| >MI0000065\|hsa-let-7d&WithFlank&9\|+\|94178828\|94178926@68.88.21(+1A)&[MIMAT0004484&hsa-let-7d-3p&offsets\|0\|-1(+1A);m-98&9\|+\|94178895\|94178916&offsets\|0\|-1(+1A)]\|CTATACGACCTGCTGCCTTTCA | >MI0000065\|hsa-let-7d&WithFlank&9\|+\|94178828\|94178926 | 68 | 88 | 21(+1A) | CTATACGACCTGCTGCCTTTCA | 299.4 | 5.2 | 1.5 | 3.4 | 6.4E-04 | 1.7E-03 | BT20/MB468 | MP |
| >hg38_21_+_8433172_8446622_RNA45SN1_45S_50ntFlanks@11665.11691.27\|TTCTGCCCAGTGCTCTGAATGTCAAAG | >hg38_21_+_8433172_8446622_RNA45SN1_45S_50ntFlanks | 11665 | 11691 | 27 | TTCTGCCCAGTGCTCTGAATGTCAAAG | 91.2 | 3.2 | 0.9 | 3.4 | 6.4E-04 | 1.7E-03 | BT20/MB468 | MP |
| >hg38_21_+_8433172_8446622_RNA45SN1_45S_50ntFlanks@11583.11613.31\|CTGGTGCGGACCAGGGGAATCCGACTGTTTA | >hg38_21_+_8433172_8446622_RNA45SN1_45S_50ntFlanks | 11583 | 11613 | 31 | CTGGTGCGGACCAGGGGAATCCGACTGTTTA | 281.0 | 2.7 | 0.8 | 3.6 | 2.8E-04 | 1.7E-03 | BT20/MB231 | MITO |
| >hg38_21_+_8433172_8446622_RNA45SN1_45S_50ntFlanks@6702.6720.19\|AGCTGCGAGAATTAATGTG | >hg38_21_+_8433172_8446622_RNA45SN1_45S_50ntFlanks | 6702 | 6720 | 19 | AGCTGCGAGAATTAATGTG | 1226.6 | -2.8 | 0.8 | -3.4 | 6.4E-04 | 1.7E-03 | BT20/MB468 | MP |
| >hg38_21_+_8433172_8446622_RNA45SN1_45S_50ntFlanks@12996.13013.18\|TCGCTGCGATCTATTGAA | >hg38_21_+_8433172_8446622_RNA45SN1_45S_50ntFlanks | 12996 | 13013 | 18 | TCGCTGCGATCTATTGAA | 2501.8 | -2.8 | 0.8 | -3.7 | 2.4E-04 | 1.7E-03 | BT20/MB231 | NUC |
| >hg38_21_+_8433172_8446622_RNA45SN1_45S_50ntFlanks@9546.9570.25\|TGGAGGTCCGTAGCGGTCCTGACGT | >hg38_21_+_8433172_8446622_RNA45SN1_45S_50ntFlanks | 9546 | 9570 | 25 | TGGAGGTCCGTAGCGGTCCTGACGT | 371.9 | 1.9 | 0.6 | 3.2 | 1.3E-03 | 1.7E-03 | MB231/MB468 | CYTO |
| >MI0000086\|hsa-mir-28&WithFlank&3\|+\|188688775\|188688872@20.40.21&[MIMAT0000085&hsa-miR-28-5p&offsets\|0\|-1;m-133&3\|+\|188688794\|188688815&offsets\|0\|-1]\|AAGGAGCTCACAGTCTATTGA | >MI0000086\|hsa-mir-28&WithFlank&3\|+\|188688775\|188688872 | 20 | 40 | 21 | AAGGAGCTCACAGTCTATTGA | 423.9 | 4.9 | 1.4 | 3.4 | 6.4E-04 | 1.7E-03 | BT20/MB468 | MP |
| >hg38_21_+_8433172_8446622_RNA45SN1_45S_50ntFlanks@6731.6770.40\|CACATTGATCATCGACACTTCGAACGCACTTGCGGCCCCG | >hg38_21_+_8433172_8446622_RNA45SN1_45S_50ntFlanks | 6731 | 6770 | 40 | CACATTGATCATCGACACTTCGAACGCACTTGCGGCCCCG | 726.7 | -6.2 | 1.8 | -3.4 | 6.4E-04 | 1.7E-03 | BT20/MB468 | MP |
| >MI0001448\|hsa-mir-425&WithFlank&3\|-\|49020142\|49020240@60.80.21&[MIMAT0001343&hsa-miR-425-3p&offsets\|-1\|-2;m-258&3\|-\|49020160\|49020181&offsets\|0\|-1]\|CATCGGGAATGTCGTGTCCGC | >MI0001448\|hsa-mir-425&WithFlank&3\|-\|49020142\|49020240 | 60 | 80 | 21 | CATCGGGAATGTCGTGTCCGC | 103.8 | 6.2 | 1.8 | 3.4 | 6.4E-04 | 1.7E-03 | BT20/MB468 | MP |
| >MI0000298\|hsa-mir-221&WithFlank&X\|-\|45746151\|45746272@31.55.25&[MIMAT0004568&hsa-miR-221-5p&offsets\|0\|+3;m-405&X\|-\|45746220\|45746242&offsets\|0\|+2]\|ACCTGGCATACAATGTAGATTTCTG | >MI0000298\|hsa-mir-221&WithFlank&X\|-\|45746151\|45746272 | 31 | 55 | 25 | ACCTGGCATACAATGTAGATTTCTG | 273.6 | -2.6 | 0.7 | -3.6 | 2.8E-04 | 1.7E-03 | BT20/MB231 | MITO |
| >hg38_21_+_8433172_8446622_RNA45SN1_45S_50ntFlanks@12640.12661.22\|ACCATCTGTGGGATTATGACTG | >hg38_21_+_8433172_8446622_RNA45SN1_45S_50ntFlanks | 12640 | 12661 | 22 | ACCATCTGTGGGATTATGACTG | 2518.9 | -3.9 | 1.2 | -3.4 | 6.4E-04 | 1.7E-03 | BT20/MB468 | MP |
| >hg38_21_+_8433172_8446622_RNA45SN1_45S_50ntFlanks@3854.3889.36\|ACTGTGGTAATTCTAGAGCTAATACATGCCGACGGG | >hg38_21_+_8433172_8446622_RNA45SN1_45S_50ntFlanks | 3854 | 3889 | 36 | ACTGTGGTAATTCTAGAGCTAATACATGCCGACGGG | 220.0 | -2.5 | 0.8 | -3.2 | 1.3E-03 | 1.7E-03 | MB231/MB468 | CYTO |
| >hg38_21_+_8433172_8446622_RNA45SN1_45S_50ntFlanks@11211.11231.21\|GCGACTCTGGACGCGAGCCGG | >hg38_21_+_8433172_8446622_RNA45SN1_45S_50ntFlanks | 11211 | 11231 | 21 | GCGACTCTGGACGCGAGCCGG | 156.3 | -2.0 | 0.6 | -3.2 | 1.3E-03 | 1.7E-03 | BT20/MB468 | CYTO |
| >hg38_21_+_8433172_8446622_RNA45SN1_45S_50ntFlanks@4258.4282.25\|AATTGGAATGAGTCCACTTTAAATC | >hg38_21_+_8433172_8446622_RNA45SN1_45S_50ntFlanks | 4258 | 4282 | 25 | AATTGGAATGAGTCCACTTTAAATC | 51.3 | -8.8 | 2.6 | -3.4 | 6.4E-04 | 1.7E-03 | BT20/MB468 | MP |
| >MI0000102\|hsa-mir-100&WithFlank&11\|-\|122152223\|122152314@22.39.18&[MIMAT0000098&hsa-miR-100-5p&offsets\|+3\|-1;m-27&11\|-\|122152276\|122152296&offsets\|+3\|0]\|CCGTAGATCCGAACTTGT | >MI0000102\|hsa-mir-100&WithFlank&11\|-\|122152223\|122152314 | 22 | 39 | 18 | CCGTAGATCCGAACTTGT | 207.5 | 11.6 | 3.6 | 3.2 | 1.4E-03 | 1.7E-03 | MB231/MB468 | NUC |
| >MI0000456\|hsa-mir-140&WithFlank&16\|+\|69933075\|69933186@68.90.23(+1U)&[MIMAT0004597&hsa-miR-140-3p&offsets\|0\|+2(+1U);m-45&16\|+\|69933143\|69933164&offsets\|-1\|0(+1U)]\|TACCACAGGGTAGAACCACGGACT | >MI0000456\|hsa-mir-140&WithFlank&16\|+\|69933075\|69933186 | 68 | 90 | 23(+1U) | TACCACAGGGTAGAACCACGGACT | 172.7 | 6.0 | 1.8 | 3.4 | 6.4E-04 | 1.7E-03 | BT20/MB468 | MP |
| >hg38_21_+_8433172_8446622_RNA45SN1_45S_50ntFlanks@485.519.35\|GATGTGGTGACGTCGTGCTCTCCCGGGCCGGGTCC | >hg38_21_+_8433172_8446622_RNA45SN1_45S_50ntFlanks | 485 | 519 | 35 | GATGTGGTGACGTCGTGCTCTCCCGGGCCGGGTCC | 47.6 | 8.8 | 2.7 | 3.2 | 1.2E-03 | 1.7E-03 | MB231/MB468 | MP |
| >hg38_21_+_8433172_8446622_RNA45SN1_45S_50ntFlanks@11567.11586.20\|AGCAGCCGACTTAGAACTGG | >hg38_21_+_8433172_8446622_RNA45SN1_45S_50ntFlanks | 11567 | 11586 | 20 | AGCAGCCGACTTAGAACTGG | 929.6 | 2.0 | 0.6 | 3.2 | 1.3E-03 | 1.7E-03 | MB231/MB468 | CYTO |
| >hg38_21_+_8433172_8446622_RNA45SN1_45S_50ntFlanks@6775.6803.29\|CCTCCCGGGGCTACGCCTGTCTGAGCGTC | >hg38_21_+_8433172_8446622_RNA45SN1_45S_50ntFlanks | 6775 | 6803 | 29 | CCTCCCGGGGCTACGCCTGTCTGAGCGTC | 811.3 | -2.4 | 0.7 | -3.3 | 1.1E-03 | 1.7E-03 | BT20/MB468 | MITO |
| >MI0000084\|hsa-mir-26b&WithFlank&2\|+\|218402640\|218402728@18.38.21(+1C)&[MIMAT0000083&hsa-miR-26b-5p&offsets\|0\|0(+1C);m-53&2\|+\|218402657\|218402678&offsets\|0\|-1(+1C)]\|TTCAAGTAATTCAGGATAGGTC | >MI0000084\|hsa-mir-26b&WithFlank&2\|+\|218402640\|218402728 | 18 | 38 | 21(+1C) | TTCAAGTAATTCAGGATAGGTC | 882.9 | 3.3 | 1.0 | 3.2 | 1.2E-03 | 1.7E-03 | MB231/MB468 | MP |
| >am_MI0000786\|hsa-mir-378a&WithFlank&5\|+\|149732819\|149732896@49.70.22(+1A)&[MIMAT0000732&hsa-miR-378a-3p&offsets\|0\|0(+1A);m-44&5\|+\|149732867\|149732888&offsets\|0\|0(+1A)]\|ACTGGACTTGGAGTCAGAAGGCA | >am_MI0000786\|hsa-mir-378a&WithFlank&5\|+\|149732819\|149732896 | 49 | 70 | 22(+1A) | ACTGGACTTGGAGTCAGAAGGCA | 912.2 | 1.9 | 0.6 | 3.2 | 1.4E-03 | 1.7E-03 | MB231/MB468 | NUC |
| >hg38_21_+_8433172_8446622_RNA45SN1_45S_50ntFlanks@6645.6682.38\|CTCGTACGACTCTTAGCGGTGGATCACTCGGCTCGTGC | >hg38_21_+_8433172_8446622_RNA45SN1_45S_50ntFlanks | 6645 | 6682 | 38 | CTCGTACGACTCTTAGCGGTGGATCACTCGGCTCGTGC | 200.3 | 5.3 | 1.6 | 3.2 | 1.4E-03 | 1.7E-03 | MB231/MB468 | NUC |
| >hg38_21_+_8433172_8446622_RNA45SN1_45S_50ntFlanks@12997.13014.18\|CGCTGCGATCTATTGAAA | >hg38_21_+_8433172_8446622_RNA45SN1_45S_50ntFlanks | 12997 | 13014 | 18 | CGCTGCGATCTATTGAAA | 2945.4 | -2.9 | 0.8 | -3.7 | 2.4E-04 | 1.7E-03 | BT20/MB231 | NUC |
| >hg38_21_+_8433172_8446622_RNA45SN1_45S_50ntFlanks@6648.6696.49\|GTACGACTCTTAGCGGTGGATCACTCGGCTCGTGCGTCGATGAAGAACG | >hg38_21_+_8433172_8446622_RNA45SN1_45S_50ntFlanks | 6648 | 6696 | 49 | GTACGACTCTTAGCGGTGGATCACTCGGCTCGTGCGTCGATGAAGAACG | 2905.8 | 4.1 | 1.3 | 3.2 | 1.4E-03 | 1.7E-03 | MB231/MB468 | NUC |
| >hg38_21_+_8433172_8446622_RNA45SN1_45S_50ntFlanks@4585.4616.32\|GTTCTATTTTGTTGGTTTTCGGAACTGAGGCC | >hg38_21_+_8433172_8446622_RNA45SN1_45S_50ntFlanks | 4585 | 4616 | 32 | GTTCTATTTTGTTGGTTTTCGGAACTGAGGCC | 262.0 | -2.9 | 0.8 | -3.7 | 2.5E-04 | 1.7E-03 | BT20/MB231 | CYTO |
| >hg38_21_+_8433172_8446622_RNA45SN1_45S_50ntFlanks@12394.12418.25\|TAGCGACGTCGCTTTTTGATCCTTC | >hg38_21_+_8433172_8446622_RNA45SN1_45S_50ntFlanks | 12394 | 12418 | 25 | TAGCGACGTCGCTTTTTGATCCTTC | 381.9 | 2.4 | 0.7 | 3.2 | 1.3E-03 | 1.7E-03 | MB231/MB468 | CYTO |
| >hg38_21_+_8433172_8446622_RNA45SN1_45S_50ntFlanks@10663.10709.47\|CCTTGGAAAGCGTCGCGGTTCCGGCGGCGTCCGGTGAGCTCTCGCTG | >hg38_21_+_8433172_8446622_RNA45SN1_45S_50ntFlanks | 10663 | 10709 | 47 | CCTTGGAAAGCGTCGCGGTTCCGGCGGCGTCCGGTGAGCTCTCGCTG | 181.7 | -5.4 | 1.6 | -3.3 | 1.1E-03 | 1.7E-03 | BT20/MB468 | MITO |
| >hg38_21_+_8433172_8446622_RNA45SN1_45S_50ntFlanks@4027.4059.33\|CCCCGTGGCGGCGACGACCCATTCGAACGTCTG | >hg38_21_+_8433172_8446622_RNA45SN1_45S_50ntFlanks | 4027 | 4059 | 33 | CCCCGTGGCGGCGACGACCCATTCGAACGTCTG | 109.9 | -4.4 | 1.3 | -3.3 | 1.1E-03 | 1.7E-03 | BT20/MB468 | MITO |
| >hg38_21_+_8433172_8446622_RNA45SN1_45S_50ntFlanks@10408.10437.30\|GGTCAGTCGGTCCTGAGAGATGGGCGAGCG | >hg38_21_+_8433172_8446622_RNA45SN1_45S_50ntFlanks | 10408 | 10437 | 30 | GGTCAGTCGGTCCTGAGAGATGGGCGAGCG | 172.0 | -5.3 | 1.6 | -3.4 | 6.5E-04 | 1.7E-03 | BT20/MB468 | MP |
| >hg38_21_+_8433172_8446622_RNA45SN1_45S_50ntFlanks@10317.10340.24\|GTGCAGATCTTGGTGGTAGTAGCA | >hg38_21_+_8433172_8446622_RNA45SN1_45S_50ntFlanks | 10317 | 10340 | 24 | GTGCAGATCTTGGTGGTAGTAGCA | 364.0 | 3.5 | 1.0 | 3.4 | 6.5E-04 | 1.7E-03 | BT20/MB468 | MP |
| >hg38_21_+_8433172_8446622_RNA45SN1_45S_50ntFlanks@7977.8007.31\|CGACCTCAGATCAGACGTGGCGACCCGCTGA | >hg38_21_+_8433172_8446622_RNA45SN1_45S_50ntFlanks | 7977 | 8007 | 31 | CGACCTCAGATCAGACGTGGCGACCCGCTGA | 454.6 | -2.8 | 0.9 | -3.2 | 1.4E-03 | 1.7E-03 | MB231/MB468 | MITO |
| >hg38_MT_+_1621_3279_RNR2_16S_50ntFlanks@1432.1458.27\|AAGTCCTACGTGATCTGAGTTCAGACC | >hg38_MT_+_1621_3279_RNR2_16S_50ntFlanks | 1432 | 1458 | 27 | AAGTCCTACGTGATCTGAGTTCAGACC | 227.8 | 3.2 | 1.0 | 3.2 | 1.2E-03 | 1.7E-03 | MB231/MB468 | MP |
| >hg38_21_+_8433172_8446622_RNA45SN1_45S_50ntFlanks@5492.5509.18\|AGAAGACGGTCGAACTTG | >hg38_21_+_8433172_8446622_RNA45SN1_45S_50ntFlanks | 5492 | 5509 | 18 | AGAAGACGGTCGAACTTG | 1173.2 | -3.4 | 1.0 | -3.4 | 6.5E-04 | 1.7E-03 | BT20/MB468 | MP |
| >hg38_21_+_8433172_8446622_RNA45SN1_45S_50ntFlanks@4243.4264.22\|CTTTCGAGGCCCTGTAATTGGA | >hg38_21_+_8433172_8446622_RNA45SN1_45S_50ntFlanks | 4243 | 4264 | 22 | CTTTCGAGGCCCTGTAATTGGA | 302.9 | -1.7 | 0.5 | -3.2 | 1.3E-03 | 1.7E-03 | MB231/MB468 | CYTO |
| >MI0000103\|hsa-mir-101-1&WithFlank&1\|-\|65058428\|65058514@53.73.21(+1A);MI0000739\|hsa-mir-101-2&WithFlank&9\|+\|4850291\|4850381@55.75.21(+1A)&[MIMAT0000099&hsa-miR-101-3p&offsets\|0\|0(+1A);m-17&1\|-\|65058443\|65058462&offsets\|0\|+1(+1A)];[MIMAT0000099_1&hsa-miR-101-3p&offsets\|0\|0(+1A);m-16&9\|+\|4850345\|4850364&offsets\|0\|+1(+1A)]\|TACAGTACTGTGATAACTGAAA | >MI0000103\|hsa-mir-101-1&WithFlank&1\|-\|65058428\|65058514 | 53 | 73 | 21(+1A) | TACAGTACTGTGATAACTGAAA | 137.0 | 3.5 | 1.1 | 3.2 | 1.4E-03 | 1.7E-03 | BT20/MB468 | NUC |
| >hg38_21_+_8433172_8446622_RNA45SN1_45S_50ntFlanks@10319.10340.22\|GCAGATCTTGGTGGTAGTAGCA | >hg38_21_+_8433172_8446622_RNA45SN1_45S_50ntFlanks | 10319 | 10340 | 22 | GCAGATCTTGGTGGTAGTAGCA | 302.0 | 4.9 | 1.4 | 3.4 | 6.5E-04 | 1.7E-03 | BT20/MB468 | MP |
| >hg38_21_+_8433172_8446622_RNA45SN1_45S_50ntFlanks@9534.9571.38\|AGGAAACTCTGGTGGAGGTCCGTAGCGGTCCTGACGTG | >hg38_21_+_8433172_8446622_RNA45SN1_45S_50ntFlanks | 9534 | 9571 | 38 | AGGAAACTCTGGTGGAGGTCCGTAGCGGTCCTGACGTG | 11776.4 | -4.4 | 1.3 | -3.4 | 6.5E-04 | 1.7E-03 | BT20/MB468 | MP |
| >hg38_21_+_8433172_8446622_RNA45SN1_45S_50ntFlanks@9565.9588.24\|TGACGTGCAAATCGGTCGTCCGAC | >hg38_21_+_8433172_8446622_RNA45SN1_45S_50ntFlanks | 9565 | 9588 | 24 | TGACGTGCAAATCGGTCGTCCGAC | 168.9 | -1.9 | 0.6 | -3.2 | 1.3E-03 | 1.7E-03 | BT20/MB468 | CYTO |
| >hg38_21_+_8433172_8446622_RNA45SN1_45S_50ntFlanks@10838.10865.28\|ATCCGTAACTTCGGGATAAGGATTGGCT | >hg38_21_+_8433172_8446622_RNA45SN1_45S_50ntFlanks | 10838 | 10865 | 28 | ATCCGTAACTTCGGGATAAGGATTGGCT | 673.1 | -4.9 | 1.5 | -3.2 | 1.3E-03 | 1.7E-03 | MB231/MB468 | CYTO |
| >hg38_21_+_8433172_8446622_RNA45SN1_45S_50ntFlanks@10392.10418.27\|ACAGCAGTTGAACATGGGTCAGTCGGT | >hg38_21_+_8433172_8446622_RNA45SN1_45S_50ntFlanks | 10392 | 10418 | 27 | ACAGCAGTTGAACATGGGTCAGTCGGT | 236.0 | -2.1 | 0.6 | -3.2 | 1.3E-03 | 1.7E-03 | MB231/MB468 | CYTO |

<table>
<tr><td>>MI0000255|hsa-mir-30d&WithFlank&8|-|134804870|134804951@12.35.24(+2U)&[MIMAT0000245&hsa-miR-30d-5p&offsets|0|+2(+2U);m-21&8|-|134804919|134804940&offsets|0|+2(+2U)]|TGTAAACATCCCCGACTGGAAGCTTT</td><td>>MI0000255|hsa-mir-30d&WithFlank&8|-|134804870|134804951</td><td>12</td><td>35</td><td>24(+2U)</td><td>TGTAAACATCCCCGACTGGAAGCTTT</td><td>143.0</td><td>4.1</td><td>1.3</td><td>3.2</td><td>1.2E-03</td><td>1.7E-03</td><td>MB231/MB468</td><td>MP</td></tr>
<tr><td>>hg38_21_+_8433172_8446622_RNA45SN1_45S_50ntFlanks@9736.9753.18|CCGAAACGATCTCAACCT</td><td>>hg38_21_+_8433172_8446622_RNA45SN1_45S_50ntFlanks</td><td>9736</td><td>9753</td><td>18</td><td>CCGAAACGATCTCAACCT</td><td>214.5</td><td>-2.8</td><td>0.9</td><td>-3.3</td><td>1.1E-03</td><td>1.7E-03</td><td>BT20/MB468</td><td>MITO</td></tr>
<tr><td>>hg38_21_+_8433172_8446622_RNA45SN1_45S_50ntFlanks@10060.10078.19|GCAGTCGAGAGTGGACGGG</td><td>>hg38_21_+_8433172_8446622_RNA45SN1_45S_50ntFlanks</td><td>10060</td><td>10078</td><td>19</td><td>GCAGTCGAGAGTGGACGGG</td><td>115.4</td><td>-4.2</td><td>1.3</td><td>-3.2</td><td>1.2E-03</td><td>1.7E-03</td><td>MB231/MB468</td><td>MP</td></tr>
<tr><td>>hg38_21_+_8433172_8446622_RNA45SN1_45S_50ntFlanks@9546.9571.26|TGGAGGTCCGTAGCGGTCCTGACGTG</td><td>>hg38_21_+_8433172_8446622_RNA45SN1_45S_50ntFlanks</td><td>9546</td><td>9571</td><td>26</td><td>TGGAGGTCCGTAGCGGTCCTGACGTG</td><td>609.1</td><td>-3.1</td><td>0.9</td><td>-3.2</td><td>1.2E-03</td><td>1.7E-03</td><td>MB231/MB468</td><td>MP</td></tr>
<tr><td>>hg38_21_+_8433172_8446622_RNA45SN1_45S_50ntFlanks@3880.3898.19|TGCCGACGGGCGCTGACCC</td><td>>hg38_21_+_8433172_8446622_RNA45SN1_45S_50ntFlanks</td><td>3880</td><td>3898</td><td>19</td><td>TGCCGACGGGCGCTGACCC</td><td>223.1</td><td>-2.7</td><td>0.8</td><td>-3.2</td><td>1.3E-03</td><td>1.7E-03</td><td>MB231/MB468</td><td>CYTO</td></tr>
<tr><td>>hg38_MT_+_1621_3279_RNR2_16S_50ntFlanks@51.75.25|GCTAAACCTAGCCCCAAACCCACTC</td><td>>hg38_MT_+_1621_3279_RNR2_16S_50ntFlanks</td><td>51</td><td>75</td><td>25</td><td>GCTAAACCTAGCCCCAAACCCACTC</td><td>395.0</td><td>3.0</td><td>0.9</td><td>3.2</td><td>1.4E-03</td><td>1.7E-03</td><td>MB231/MB468</td><td>MITO</td></tr>
<tr><td>>hg38_21_+_8433172_8446622_RNA45SN1_45S_50ntFlanks@11593.11616.24|CCAGGGGAATCCGACTGTTTAATT</td><td>>hg38_21_+_8433172_8446622_RNA45SN1_45S_50ntFlanks</td><td>11593</td><td>11616</td><td>24</td><td>CCAGGGGAATCCGACTGTTTAATT</td><td>374.9</td><td>-4.3</td><td>1.3</td><td>-3.2</td><td>1.4E-03</td><td>1.7E-03</td><td>BT20/MB468</td><td>NUC</td></tr>
<tr><td>>trna10_AspGTC_12_-_125424193_125424264@31.72.42,trna12_AspGTC_12_-_125411891_125411962@31.72.42,trna38_AspGTC_17_-_8125556_8125627@31.72.42,trna45_AspGTC_6_+_27447453_27447524@31.72.42,trna48_AspGTC_6_+_27471523_27471594@31.72.42,trna4_AspGTC_12_+_96429799_96429870@31.72.42,trna69_AspGTC_1_-_161440205_161440276@31.72.42,trna72_AspGTC_1_-_161432824_161432895@31.72.42,trna75_AspGTC_1_-_161425414_161425485@31.72.42,trna78_AspGTC_1_-_161418033_161418104@31.72.42,trna81_AspGTC_1_-_161410615_161410686@31.72.42|CCTGTCACGCGGGAGACCGGGGTTCGATTCCCCGACGGGGAG</td><td>>trna10_AspGTC_12_-_125424193_125424264</td><td>31</td><td>72</td><td>42</td><td>CCTGTCACGCGGGAGACCGGGGTTCGATTCCCCGACGGGGAG</td><td>1478.1</td><td>-5.5</td><td>1.7</td><td>-3.2</td><td>1.3E-03</td><td>1.7E-03</td><td>BT20/MB468</td><td>CYTO</td></tr>
<tr><td>>hg38_MT_+_1621_3279_RNR2_16S_50ntFlanks@827.860.34|AGGAAAGGTTAAAAAAAGTAAAAGGAACTCGGCA</td><td>>hg38_MT_+_1621_3279_RNR2_16S_50ntFlanks</td><td>827</td><td>860</td><td>34</td><td>AGGAAAGGTTAAAAAAAGTAAAAGGAACTCGGCA</td><td>77.8</td><td>6.7</td><td>2.0</td><td>3.4</td><td>6.5E-04</td><td>1.7E-03</td><td>BT20/MB468</td><td>MP</td></tr>
<tr><td>>hg38_21_+_8433172_8446622_RNA45SN1_45S_50ntFlanks@4102.4120.19|ACCACGGGTGACGGGGAAT</td><td>>hg38_21_+_8433172_8446622_RNA45SN1_45S_50ntFlanks</td><td>4102</td><td>4120</td><td>19</td><td>ACCACGGGTGACGGGGAAT</td><td>303.2</td><td>-2.9</td><td>0.9</td><td>-3.2</td><td>1.2E-03</td><td>1.7E-03</td><td>MB231/MB468</td><td>MP</td></tr>
<tr><td>>MI0000737|hsa-mir-200a&WithFlank&1|+|1167857|1167958@22.43.22&[MIMAT0001620&hsa-miR-200a-5p&offsets|0|0;m-154&1|+|1167878|1167899&offsets|0|0]|CATCTTACCGGACAGTGCTGGA</td><td>>MI0000737|hsa-mir-200a&WithFlank&1|+|1167857|1167958</td><td>22</td><td>43</td><td>22</td><td>CATCTTACCGGACAGTGCTGGA</td><td>841.8</td><td>-5.9</td><td>1.7</td><td>-3.4</td><td>6.5E-04</td><td>1.7E-03</td><td>BT20/MB468</td><td>MP</td></tr>
<tr><td>>hg38_21_+_8433172_8446622_RNA45SN1_45S_50ntFlanks@6788.6806.19|CGCCTGTCTGAGCGTCGCT</td><td>>hg38_21_+_8433172_8446622_RNA45SN1_45S_50ntFlanks</td><td>6788</td><td>6806</td><td>19</td><td>CGCCTGTCTGAGCGTCGCT</td><td>5540.0</td><td>2.2</td><td>0.7</td><td>3.2</td><td>1.4E-03</td><td>1.7E-03</td><td>MB231/MB468</td><td>MITO</td></tr>
<tr><td>>MI0000087|hsa-mir-29a&WithFlank&7|-|130876741|130876816@48.68.21(+2U)&[MIMAT0000086&hsa-miR-29a-3p&offsets|0|-1(+2U);m-26&7|-|130876749|130876769&offsets|0|0(+2U)]|TAGCACCATCTGAAATCGGTTTT</td><td>>MI0000087|hsa-mir-29a&WithFlank&7|-|130876741|130876816</td><td>48</td><td>68</td><td>21(+2U)</td><td>TAGCACCATCTGAAATCGGTTTT</td><td>51.3</td><td>9.4</td><td>2.8</td><td>3.4</td><td>6.5E-04</td><td>1.7E-03</td><td>BT20/MB468</td><td>MP</td></tr>
<tr><td>>hg38_21_+_8433172_8446622_RNA45SN1_45S_50ntFlanks@4673.4690.18|TGGACCGGCGCAAGACGG</td><td>>hg38_21_+_8433172_8446622_RNA45SN1_45S_50ntFlanks</td><td>4673</td><td>4690</td><td>18</td><td>TGGACCGGCGCAAGACGG</td><td>51.0</td><td>-8.8</td><td>2.6</td><td>-3.4</td><td>6.5E-04</td><td>1.7E-03</td><td>BT20/MB468</td><td>MP</td></tr>
<tr><td>>trnaMT_GluTTC_MT_-_14674_14742@46.63.18,trnalookalike8_GluTTC_5_-_93905172_93905240@46.63.18|GTGGTTGTAGTCCGTGCG</td><td>>trnaMT_GluTTC_MT_-_14674_14742</td><td>46</td><td>63</td><td>18</td><td>GTGGTTGTAGTCCGTGCG</td><td>106.4</td><td>5.7</td><td>1.7</td><td>3.4</td><td>6.5E-04</td><td>1.7E-03</td><td>BT20/MB468</td><td>MP</td></tr>
<tr><td>>trnaMT_SerGCT_MT_+_12207_12265@12.48.37|CAAGAACTGCTAACTCATGCCCCCATGTCTAACAACA</td><td>>trnaMT_SerGCT_MT_+_12207_12265</td><td>12</td><td>48</td><td>37</td><td>CAAGAACTGCTAACTCATGCCCCCATGTCTAACAACA</td><td>260.1</td><td>5.3</td><td>1.5</td><td>3.4</td><td>6.5E-04</td><td>1.7E-03</td><td>BT20/MB468</td><td>MP</td></tr>
<tr><td>>hg38_21_+_8433172_8446622_RNA45SN1_45S_50ntFlanks@4547.4565.19|CTGGATACCGCAGCTAGGA</td><td>>hg38_21_+_8433172_8446622_RNA45SN1_45S_50ntFlanks</td><td>4547</td><td>4565</td><td>19</td><td>CTGGATACCGCAGCTAGGA</td><td>588.6</td><td>-5.9</td><td>1.7</td><td>-3.4</td><td>6.5E-04</td><td>1.7E-03</td><td>BT20/MB468</td><td>MP</td></tr>
<tr><td>>hg38_21_+_8433172_8446622_RNA45SN1_45S_50ntFlanks@3853.3881.29|AACTGTGGTAATTCTAGAGCTAATACATG</td><td>>hg38_21_+_8433172_8446622_RNA45SN1_45S_50ntFlanks</td><td>3853</td><td>3881</td><td>29</td><td>AACTGTGGTAATTCTAGAGCTAATACATG</td><td>524.2</td><td>-5.6</td><td>1.7</td><td>-3.4</td><td>6.5E-04</td><td>1.7E-03</td><td>BT20/MB468</td><td>MP</td></tr>
<tr><td>>trnaMT_GluTTC_MT_-_14674_14742@31.72.42,trnalookalike8_GluTTC_5_-_93905172_93905240@31.72.42|TTCATATCATTGGTCGTGGTTGTAGTCCGTGCGAGAATACCA</td><td>>trnaMT_GluTTC_MT_-_14674_14742</td><td>31</td><td>72</td><td>42</td><td>TTCATATCATTGGTCGTGGTTGTAGTCCGTGCGAGAATACCA</td><td>924.6</td><td>-3.5</td><td>1.0</td><td>-3.7</td><td>2.4E-04</td><td>1.7E-03</td><td>BT20/MB231</td><td>NUC</td></tr>
<tr><td>>hg38_21_+_8433172_8446622_RNA45SN1_45S_50ntFlanks@12007.12039.33|GCCGCCGGTGAAATACCACTACTCTGATCGTTT</td><td>>hg38_21_+_8433172_8446622_RNA45SN1_45S_50ntFlanks</td><td>12007</td><td>12039</td><td>33</td><td>GCCGCCGGTGAAATACCACTACTCTGATCGTTT</td><td>94.4</td><td>-5.5</td><td>1.7</td><td>-3.2</td><td>1.2E-03</td><td>1.7E-03</td><td>MB231/MB468</td><td>MP</td></tr>
<tr><td>>hg38_21_+_8433172_8446622_RNA45SN1_45S_50ntFlanks@5084.5108.25|CGAACGAGACTCTGGCATGCTAACT</td><td>>hg38_21_+_8433172_8446622_RNA45SN1_45S_50ntFlanks</td><td>5084</td><td>5108</td><td>25</td><td>CGAACGAGACTCTGGCATGCTAACT</td><td>601.8</td><td>-4.0</td><td>1.2</td><td>-3.3</td><td>1.1E-03</td><td>1.7E-03</td><td>BT20/MB468</td><td>MITO</td></tr>
<tr><td>>MI0000748|hsa-mir-130b&WithFlank&22|+|21653298|21653391@57.76.20&[MIMAT0000691&hsa-miR-130b-3p&offsets|0|-2;m-205&22|+|21653354|21653375&offsets|0|-2]|CAGTGCAATGATGAAAGGGC</td><td>>MI0000748|hsa-mir-130b&WithFlank&22|+|21653298|21653391</td><td>57</td><td>76</td><td>20</td><td>CAGTGCAATGATGAAAGGGC</td><td>92.2</td><td>5.9</td><td>1.8</td><td>3.2</td><td>1.2E-03</td><td>1.7E-03</td><td>MB231/MB468</td><td>MP</td></tr>
<tr><td>>hg38_21_+_8433172_8446622_RNA45SN1_45S_50ntFlanks@9573.9591.19|AAATCGGTCGTCCGACCTG</td><td>>hg38_21_+_8433172_8446622_RNA45SN1_45S_50ntFlanks</td><td>9573</td><td>9591</td><td>19</td><td>AAATCGGTCGTCCGACCTG</td><td>47.4</td><td>-8.6</td><td>2.5</td><td>-3.4</td><td>6.6E-04</td><td>1.7E-03</td><td>BT20/MB468</td><td>MP</td></tr>
<tr><td>>MI0000299|hsa-mir-222&WithFlank&X|-|45747009|45747130@75.94.20&[MIMAT0000279&hsa-miR-222-3p&offsets|0|-1;m-140&X|-|45747033|45747056&offsets|0|-4]|AGCTACATCTGGCTACTGGG</td><td>>MI0000299|hsa-mir-222&WithFlank&X|-|45747009|45747130</td><td>75</td><td>94</td><td>20</td><td>AGCTACATCTGGCTACTGGG</td><td>528.7</td><td>4.3</td><td>1.3</td><td>3.2</td><td>1.2E-03</td><td>1.7E-03</td><td>MB231/MB468</td><td>MP</td></tr>
<tr><td>>trna15_ValAAC_5_-_180615416_180615488@1.32.32|GTTTCCGTAGTGTAGTGGTCATCACGTTCGCC</td><td>>trna15_ValAAC_5_-_180615416_180615488</td><td>1</td><td>32</td><td>32</td><td>GTTTCCGTAGTGTAGTGGTCATCACGTTCGCC</td><td>256.8</td><td>6.1</td><td>1.9</td><td>3.2</td><td>1.2E-03</td><td>1.7E-03</td><td>MB231/MB468</td><td>MP</td></tr>
<tr><td>>hg38_21_+_8433172_8446622_RNA45SN1_45S_50ntFlanks@6653.6683.31|ACTCTTAGCGGTGGATCACTCGGCTCGTGCG</td><td>>hg38_21_+_8433172_8446622_RNA45SN1_45S_50ntFlanks</td><td>6653</td><td>6683</td><td>31</td><td>ACTCTTAGCGGTGGATCACTCGGCTCGTGCG</td><td>621.7</td><td>2.7</td><td>0.8</td><td>3.3</td><td>1.1E-03</td><td>1.7E-03</td><td>BT20/MB468</td><td>MITO</td></tr>
<tr><td>>am_trnaMT_ValTAC_MT_+_1602_1670@50.68.19|ACTTAACTTGACCGCTCTG</td><td>>am_trnaMT_ValTAC_MT_+_1602_1670</td><td>50</td><td>68</td><td>19</td><td>ACTTAACTTGACCGCTCTG</td><td>3077.8</td><td>-3.1</td><td>0.9</td><td>-3.4</td><td>6.6E-04</td><td>1.7E-03</td><td>BT20/MB468</td><td>MP</td></tr>
<tr><td>>hg38_21_+_8433172_8446622_RNA45SN1_45S_50ntFlanks@6651.6669.19|CGACTCTTAGCGGTGGATC</td><td>>hg38_21_+_8433172_8446622_RNA45SN1_45S_50ntFlanks</td><td>6651</td><td>6669</td><td>19</td><td>CGACTCTTAGCGGTGGATC</td><td>55746.9</td><td>2.5</td><td>0.7</td><td>3.6</td><td>2.8E-04</td><td>1.7E-03</td><td>BT20/MB231</td><td>MITO</td></tr>
<tr><td>>MI0001445|hsa-mir-423&WithFlank&17|+|30117073|30117178@23.46.24(+1U)&[MIMAT0004748&hsa-miR-423-5p&offsets|0|+1(+1U);m-141&17|+|30117095|30117117&offsets|0|+1(+1U)]|TGAGGGGCAGAGAGCGAGACTTTTT</td><td>>MI0001445|hsa-mir-423&WithFlank&17|+|30117073|30117178</td><td>23</td><td>46</td><td>24(+1U)</td><td>TGAGGGGCAGAGAGCGAGACTTTTT</td><td>127.8</td><td>5.0</td><td>1.6</td><td>3.2</td><td>1.2E-03</td><td>1.7E-03</td><td>MB231/MB468</td><td>MP</td></tr>
<tr><td>>hg38_21_+_8433172_8446622_RNA45SN1_45S_50ntFlanks@12416.12449.34|TTCGATGTCGGCTCTTCCTATCATTGTGAAGCAG</td><td>>hg38_21_+_8433172_8446622_RNA45SN1_45S_50ntFlanks</td><td>12416</td><td>12449</td><td>34</td><td>TTCGATGTCGGCTCTTCCTATCATTGTGAAGCAG</td><td>661.6</td><td>-2.9</td><td>0.8</td><td>-3.6</td><td>2.9E-04</td><td>1.7E-03</td><td>BT20/MB231</td><td>MITO</td></tr>
<tr><td>>hg38_MT_+_598_1651_RNR1_12S_50ntFlanks@608.633.26|TGTTCTGTAATCGATAAACCCCGATC</td><td>>hg38_MT_+_598_1651_RNR1_12S_50ntFlanks</td><td>608</td><td>633</td><td>26</td><td>TGTTCTGTAATCGATAAACCCCGATC</td><td>237.2</td><td>4.0</td><td>1.2</td><td>3.4</td><td>6.6E-04</td><td>1.7E-03</td><td>BT20/MB468</td><td>MP</td></tr>
<tr><td>>hg38_MT_+_598_1651_RNR1_12S_50ntFlanks@368.404.37|CAATAAAGCTAAAACTCACCTGAGTTGTAAAAAACTC</td><td>>hg38_MT_+_598_1651_RNR1_12S_50ntFlanks</td><td>368</td><td>404</td><td>37</td><td>CAATAAAGCTAAAACTCACCTGAGTTGTAAAAAACTC</td><td>829.8</td><td>2.9</td><td>0.9</td><td>3.2</td><td>1.4E-03</td><td>1.7E-03</td><td>MB231/MB468</td><td>MITO</td></tr>
<tr><td>>hg38_21_+_8433172_8446622_RNA45SN1_45S_50ntFlanks@9574.9601.28|AATCGGTCGTCCGACCTGGGTATAGGGG</td><td>>hg38_21_+_8433172_8446622_RNA45SN1_45S_50ntFlanks</td><td>9574</td><td>9601</td><td>28</td><td>AATCGGTCGTCCGACCTGGGTATAGGGG</td><td>364.3</td><td>-3.2</td><td>1.0</td><td>-3.2</td><td>1.4E-03</td><td>1.7E-03</td><td>BT20/MB468</td><td>NUC</td></tr>
<tr><td>>hg38_21_+_8433172_8446622_RNA45SN1_45S_50ntFlanks@10421.10456.36|TGAGAGATGGGCGAGCGCCGTTCCGAAGGGACGGGC</td><td>>hg38_21_+_8433172_8446622_RNA45SN1_45S_50ntFlanks</td><td>10421</td><td>10456</td><td>36</td><td>TGAGAGATGGGCGAGCGCCGTTCCGAAGGGACGGGC</td><td>161.8</td><td>-3.6</td><td>1.1</td><td>-3.3</td><td>1.1E-03</td><td>1.7E-03</td><td>BT20/MB468</td><td>MITO</td></tr>
<tr><td>>MI0006273|hsa-mir-1180&WithFlank&17|-|19344500|19344580@47.66.20&[MIMAT0005825&hsa-miR-1180-3p&offsets|0|-2;m-214&17|-|19344513|19344534&offsets|0|-2]|TTTCCGGCTCGCGTGGGTGT</td><td>>MI0006273|hsa-mir-1180&WithFlank&17|-|19344500|19344580</td><td>47</td><td>66</td><td>20</td><td>TTTCCGGCTCGCGTGGGTGT</td><td>125.4</td><td>1.9</td><td>0.6</td><td>3.2</td><td>1.4E-03</td><td>1.7E-03</td><td>BT20/MB468</td><td>NUC</td></tr>
<tr><td>>hg38_21_+_8433172_8446622_RNA45SN1_45S_50ntFlanks@8104.8141.38|CCGCGGCGGGGCGCGGGACATGTGGCGTACGGAAGACC</td><td>>hg38_21_+_8433172_8446622_RNA45SN1_45S_50ntFlanks</td><td>8104</td><td>8141</td><td>38</td><td>CCGCGGCGGGGCGCGGGACATGTGGCGTACGGAAGACC</td><td>358.5</td><td>-4.1</td><td>1.3</td><td>-3.2</td><td>1.4E-03</td><td>1.7E-03</td><td>MB231/MB468</td><td>NUC</td></tr>
<tr><td>>hg38_21_+_8433172_8446622_RNA45SN1_45S_50ntFlanks@11212.11234.23|CGACTCTGGACGCGAGCCGGGCC</td><td>>hg38_21_+_8433172_8446622_RNA45SN1_45S_50ntFlanks</td><td>11212</td><td>11234</td><td>23</td><td>CGACTCTGGACGCGAGCCGGGCC</td><td>88.3</td><td>-5.3</td><td>1.6</td><td>-3.4</td><td>6.6E-04</td><td>1.7E-03</td><td>BT20/MB468</td><td>MP</td></tr>
<tr><td>>hg38_21_+_8433172_8446622_RNA45SN1_45S_50ntFlanks@9132.9152.21|GTCGGGGGGCGGGGCGGACTG</td><td>>hg38_21_+_8433172_8446622_RNA45SN1_45S_50ntFlanks</td><td>9132</td><td>9152</td><td>21</td><td>GTCGGGGGGCGGGGCGGACTG</td><td>218.4</td><td>-5.7</td><td>1.7</td><td>-3.4</td><td>6.6E-04</td><td>1.7E-03</td><td>BT20/MB468</td><td>MP</td></tr>
<tr><td>>hg38_21_+_8433172_8446622_RNA45SN1_45S_50ntFlanks@6651.6699.49|CGACTCTTAGCGGTGGATCACTCGGCTCGTGCGTCGATGAAGAACGCAG</td><td>>hg38_21_+_8433172_8446622_RNA45SN1_45S_50ntFlanks</td><td>6651</td><td>6699</td><td>49</td><td>CGACTCTTAGCGGTGGATCACTCGGCTCGTGCGTCGATGAAGAACGCAG</td><td>2806.4</td><td>-3.5</td><td>1.0</td><td>-3.7</td><td>2.5E-04</td><td>1.7E-03</td><td>BT20/MB231</td><td>CYTO</td></tr>
<tr><td>>hg38_MT_+_1621_3279_RNR2_16S_50ntFlanks@622.656.35|TATAACTGAACTCCTCACACCCAATTGGACCAATC</td><td>>hg38_MT_+_1621_3279_RNR2_16S_50ntFlanks</td><td>622</td><td>656</td><td>35</td><td>TATAACTGAACTCCTCACACCCAATTGGACCAATC</td><td>100.0</td><td>6.9</td><td>2.0</td><td>3.4</td><td>6.6E-04</td><td>1.7E-03</td><td>BT20/MB468</td><td>MP</td></tr>
<tr><td>>MI0005566|hsa-mir-374b&WithFlank&X|-|74218541|74218624@17.34.18&[MIMAT0004955&hsa-miR-374b-5p&offsets|0|-4;m-153&X|-|74218587|74218608&offsets|0|-4]|ATATAATACAACCTGCTA</td><td>>MI0005566|hsa-mir-374b&WithFlank&X|-|74218541|74218624</td><td>17</td><td>34</td><td>18</td><td>ATATAATACAACCTGCTA</td><td>319.3</td><td>-4.5</td><td>1.2</td><td>-3.7</td><td>2.3E-04</td><td>1.7E-03</td><td>BT20/MB231</td><td>MP</td></tr>
<tr><td>>hg38_21_+_8433172_8446622_RNA45SN1_45S_50ntFlanks@12399.12417.19|ACGTCGCTTTTTGATCCTT</td><td>>hg38_21_+_8433172_8446622_RNA45SN1_45S_50ntFlanks</td><td>12399</td><td>12417</td><td>19</td><td>ACGTCGCTTTTTGATCCTT</td><td>77.5</td><td>-5.2</td><td>1.5</td><td>-3.4</td><td>6.6E-04</td><td>1.7E-03</td><td>BT20/MB468</td><td>MP</td></tr>
<tr><td>>hg38_21_+_8433172_8446622_RNA45SN1_45S_50ntFlanks@12126.12148.23|CGACCCGCTCCGGGGACAGTGCC</td><td>>hg38_21_+_8433172_8446622_RNA45SN1_45S_50ntFlanks</td><td>12126</td><td>12148</td><td>23</td><td>CGACCCGCTCCGGGGACAGTGCC</td><td>406.3</td><td>-5.7</td><td>1.7</td><td>-3.4</td><td>6.6E-04</td><td>1.7E-03</td><td>BT20/MB468</td><td>MP</td></tr>
<tr><td>>hg38_21_+_8433172_8446622_RNA45SN1_45S_50ntFlanks@9738.9764.27|GAAACGATCTCAACCTATTCTCAAACT</td><td>>hg38_21_+_8433172_8446622_RNA45SN1_45S_50ntFlanks</td><td>9738</td><td>9764</td><td>27</td><td>GAAACGATCTCAACCTATTCTCAAACT</td><td>473.4</td><td>-2.1</td><td>0.7</td><td>-3.2</td><td>1.4E-03</td><td>1.7E-03</td><td>MB231/MB468</td><td>NUC</td></tr>
<tr><td>>MI0000542|hsa-mir-320a&WithFlank&8|-|22244960|22245043@48.68.21(+3U)&[MIMAT0000510&hsa-miR-320a-3p&offsets|0|-1(+3U);m-60&8|-|22244975|22244996&offsets|0|-1(+3U)]|AAAAGCTGGGTTGAGAGGGCGTTT</td><td>>MI0000542|hsa-mir-320a&WithFlank&8|-|22244960|22245043</td><td>48</td><td>68</td><td>21(+3U)</td><td>AAAAGCTGGGTTGAGAGGGCGTTT</td><td>257.5</td><td>5.3</td><td>1.6</td><td>3.2</td><td>1.2E-03</td><td>1.7E-03</td><td>MB231/MB468</td><td>MP</td></tr>
<tr><td>>MI0000469|hsa-mir-125a&WithFlank&19|+|51693248|51693345@21.43.23(+1U)&[MIMAT0000443&hsa-miR-125a-5p&offsets|0|-1(+1U);m-63&19|+|51693268|51693289&offsets|0|+1(+1U)]|TCCCTGAGACCCTTTAACCTGTGT</td><td>>MI0000469|hsa-mir-125a&WithFlank&19|+|51693248|51693345</td><td>21</td><td>43</td><td>23(+1U)</td><td>TCCCTGAGACCCTTTAACCTGTGT</td><td>188.5</td><td>3.0</td><td>0.9</td><td>3.4</td><td>6.6E-04</td><td>1.7E-03</td><td>BT20/MB468</td><td>MP</td></tr>
<tr><td>>hg38_21_+_8433172_8446622_RNA45SN1_45S_50ntFlanks@4731.4761.31|AAGAACGAAAGTCGGAGGTTCGAAGACGATC</td><td>>hg38_21_+_8433172_8446622_RNA45SN1_45S_50ntFlanks</td><td>4731</td><td>4761</td><td>31</td><td>AAGAACGAAAGTCGGAGGTTCGAAGACGATC</td><td>1164.5</td><td>2.4</td><td>0.7</td><td>3.2</td><td>1.4E-03</td><td>1.7E-03</td><td>MB231/MB468</td><td>NUC</td></tr>
<tr><td>>hg38_21_+_8433172_8446622_RNA45SN1_45S_50ntFlanks@11562.11588.27|CGCCTAGCAGCCGACTTAGAACTGGTG</td><td>>hg38_21_+_8433172_8446622_RNA45SN1_45S_50ntFlanks</td><td>11562</td><td>11588</td><td>27</td><td>CGCCTAGCAGCCGACTTAGAACTGGTG</td><td>647.5</td><td>-3.0</td><td>0.9</td><td>-3.4</td><td>6.6E-04</td><td>1.7E-03</td><td>BT20/MB468</td><td>MP</td></tr>
<tr><td>>hg38_1_-_228634819_228635039_RNA5S12_5S_50ntFlanks@51.84.34|GTCTACGGCCATACCACCCTGAACGCGCCCGATC</td><td>>hg38_1_-_228634819_228635039_RNA5S12_5S_50ntFlanks</td><td>51</td><td>84</td><td>34</td><td>GTCTACGGCCATACCACCCTGAACGCGCCCGATC</td><td>2113.9</td><td>2.3</td><td>0.7</td><td>3.2</td><td>1.4E-03</td><td>1.7E-03</td><td>MB231/MB468</td><td>MITO</td></tr>
<tr><td>>MI0000255|hsa-mir-30d&WithFlank&8|-|134804870|134804951@12.33.22(+1A)&[MIMAT0000245&hsa-miR-30d-5p&offsets|0|0(+1A);m-21&8|-|134804919|134804940&offsets|0|0(+1A)]|TGTAAACATCCCCGACTGGAAGA</td><td>>MI0000255|hsa-mir-30d&WithFlank&8|-|134804870|134804951</td><td>12</td><td>33</td><td>22(+1A)</td><td>TGTAAACATCCCCGACTGGAAGA</td><td>3293.3</td><td>3.2</td><td>0.9</td><td>3.7</td><td>2.5E-04</td><td>1.7E-03</td><td>BT20/MB231</td><td>CYTO</td></tr>
<tr><td>>trnaMT_HisGTG_MT_+_12138_12206@32.72.41|TGAATCTGACAACAGAGGCTTACGACCCCTTATTTACCCCA</td><td>>trnaMT_HisGTG_MT_+_12138_12206</td><td>32</td><td>72</td><td>41</td><td>TGAATCTGACAACAGAGGCTTACGACCCCTTATTTACCCCA</td><td>825.5</td><td>8.5</td><td>2.6</td><td>3.2</td><td>1.4E-03</td><td>1.7E-03</td><td>MB231/MB468</td><td>CYTO</td></tr>
</table>

<table>
<tr><td>>hg38_21_+_8433172_8446622_RNA45SN1_45S_50ntFlanks@11745.11768.24|AAGGTAGCCAAATGCCTCGTCATC</td><td>>hg38_21_+_8433172_8446622_RNA45SN1_45S_50ntFlanks</td><td>11745</td><td>11768</td><td>24</td><td>AAGGTAGCCAAATGCCTCGTCATC</td><td>47.7</td><td>-8.7</td><td>2.5</td><td>-3.4</td><td>6.6E-04</td><td>1.7E-03</td><td>BT20/MB468</td><td>MP</td></tr>
<tr><td>>hg38_21_+_8433172_8446622_RNA45SN1_45S_50ntFlanks@11779.11804.26|CGCGCATGAATGGATGAACGAGATTC</td><td>>hg38_21_+_8433172_8446622_RNA45SN1_45S_50ntFlanks</td><td>11779</td><td>11804</td><td>26</td><td>CGCGCATGAATGGATGAACGAGATTC</td><td>131.7</td><td>-5.4</td><td>1.7</td><td>-3.2</td><td>1.2E-03</td><td>1.7E-03</td><td>MB231/MB468</td><td>MP</td></tr>
<tr><td>>trnaMT_PheGAA_MT_+_577_647@53.74.22|TCACATCACCCCATAAACACCA</td><td>>trnaMT_PheGAA_MT_+_577_647</td><td>53</td><td>74</td><td>22</td><td>TCACATCACCCCATAAACACCA</td><td>306.3</td><td>4.3</td><td>1.3</td><td>3.2</td><td>1.3E-03</td><td>1.7E-03</td><td>BT20/MB468</td><td>CYTO</td></tr>
<tr><td>>hg38_21_+_8433172_8446622_RNA45SN1_45S_50ntFlanks@11672.11692.21|CAGTGCTCTGAATGTCAAAGT</td><td>>hg38_21_+_8433172_8446622_RNA45SN1_45S_50ntFlanks</td><td>11672</td><td>11692</td><td>21</td><td>CAGTGCTCTGAATGTCAAAGT</td><td>348.5</td><td>-2.3</td><td>0.7</td><td>-3.2</td><td>1.4E-03</td><td>1.7E-03</td><td>BT20/MB468</td><td>NUC</td></tr>
<tr><td>>hg38_21_+_8433172_8446622_RNA45SN1_45S_50ntFlanks@7971.8009.39|GAGACGCGACCTCAGATCAGACGTGGCGACCCGCTGAAT</td><td>>hg38_21_+_8433172_8446622_RNA45SN1_45S_50ntFlanks</td><td>7971</td><td>8009</td><td>39</td><td>GAGACGCGACCTCAGATCAGACGTGGCGACCCGCTGAAT</td><td>785.1</td><td>3.1</td><td>0.9</td><td>3.7</td><td>2.5E-04</td><td>1.7E-03</td><td>BT20/MB231</td><td>NUC</td></tr>
<tr><td>>am_trnaMT_ValTAC_MT_+_1602_1670@41.68.28|GAGATTTCAACTTAACTTGACCGCTCTG</td><td>>am_trnaMT_ValTAC_MT_+_1602_1670</td><td>41</td><td>68</td><td>28</td><td>GAGATTTCAACTTAACTTGACCGCTCTG</td><td>95.7</td><td>-2.6</td><td>0.8</td><td>-3.2</td><td>1.2E-03</td><td>1.7E-03</td><td>MB231/MB468</td><td>MP</td></tr>
<tr><td>>hg38_21_+_8433172_8446622_RNA45SN1_45S_50ntFlanks@7975.7994.20|CGCGACCTCAGATCAGACGT</td><td>>hg38_21_+_8433172_8446622_RNA45SN1_45S_50ntFlanks</td><td>7975</td><td>7994</td><td>20</td><td>CGCGACCTCAGATCAGACGT</td><td>497028.4</td><td>2.5</td><td>0.7</td><td>3.6</td><td>2.9E-04</td><td>1.7E-03</td><td>BT20/MB231</td><td>MITO</td></tr>
<tr><td>>trnaMT_GluTTC_MT_-_14674_14742@33.72.40,trnalookalike8_GluTTC_5_-_93905172_93905240@33.72.40|CATATCATTGGTCGTGGTTGTAGTCCGTGCGAGAATACCA</td><td>>trnaMT_GluTTC_MT_-_14674_14742</td><td>33</td><td>72</td><td>40</td><td>CATATCATTGGTCGTGGTTGTAGTCCGTGCGAGAATACCA</td><td>370.0</td><td>-3.0</td><td>0.9</td><td>-3.2</td><td>1.4E-03</td><td>1.7E-03</td><td>BT20/MB468</td><td>NUC</td></tr>
<tr><td>>MI0000825|hsa-mir-345&WithFlank&14|+|100307853|100307962@24.45.22&[MIMAT0000772&hsa-miR-345-5p&offsets|0|0;m-193&14|+|100307876|100307896&offsets|0|+1]|GCTGACTCCTAGTCCAGGGCTC</td><td>>MI0000825|hsa-mir-345&WithFlank&14|+|100307853|100307962</td><td>24</td><td>45</td><td>22</td><td>GCTGACTCCTAGTCCAGGGCTC</td><td>250.5</td><td>5.1</td><td>1.6</td><td>3.2</td><td>1.4E-03</td><td>1.7E-03</td><td>MB231/MB468</td><td>MITO</td></tr>
<tr><td>>hg38_21_+_8433172_8446622_RNA45SN1_45S_50ntFlanks@3990.4024.35|TTGGTGACTCTAGATAACCTCGGGCCGATCGCACG</td><td>>hg38_21_+_8433172_8446622_RNA45SN1_45S_50ntFlanks</td><td>3990</td><td>4024</td><td>35</td><td>TTGGTGACTCTAGATAACCTCGGGCCGATCGCACG</td><td>437.0</td><td>-2.4</td><td>0.8</td><td>-3.2</td><td>1.4E-03</td><td>1.7E-03</td><td>MB231/MB468</td><td>MITO</td></tr>
<tr><td>>hg38_21_+_8433172_8446622_RNA45SN1_45S_50ntFlanks@9933.9951.19|TAGACAGCAGGACGGTGGC</td><td>>hg38_21_+_8433172_8446622_RNA45SN1_45S_50ntFlanks</td><td>9933</td><td>9951</td><td>19</td><td>TAGACAGCAGGACGGTGGC</td><td>366.6</td><td>7.0</td><td>1.9</td><td>3.7</td><td>2.3E-04</td><td>1.7E-03</td><td>BT20/MB231</td><td>MP</td></tr>
<tr><td>>hg38_MT_+_1621_3279_RNR2_16S_50ntFlanks@847.869.23|AAAGGAACTCGGCAAATCTTACC</td><td>>hg38_MT_+_1621_3279_RNR2_16S_50ntFlanks</td><td>847</td><td>869</td><td>23</td><td>AAAGGAACTCGGCAAATCTTACC</td><td>50.1</td><td>6.5</td><td>1.9</td><td>3.4</td><td>6.6E-04</td><td>1.7E-03</td><td>BT20/MB468</td><td>MP</td></tr>
<tr><td>>MI0000088|hsa-mir-30a&WithFlank&6|-|71403545|71403627@53.73.21&[MIMAT0000088&hsa-miR-30a-3p&offsets|0|-1;m-38&6|-|71403554|71403575&offsets|0|-1]|CTTTCAGTCGGATGTTTGCAG</td><td>>MI0000088|hsa-mir-30a&WithFlank&6|-|71403545|71403627</td><td>53</td><td>73</td><td>21</td><td>CTTTCAGTCGGATGTTTGCAG</td><td>3165.2</td><td>-3.3</td><td>0.9</td><td>-3.7</td><td>2.3E-04</td><td>1.7E-03</td><td>BT20/MB231</td><td>MP</td></tr>
<tr><td>>trna146_GlnCTG_6_-_27515531_27515602@14.41.28,trna1_GlnCTG_6_+_18836402_18836473@14.41.28,trna3_GlnCTG_17_+_8023070_8023141@14.41.28,trna42_GlnCTG_6_+_27263212_27263283@14.41.28,trna49_GlnCTG_6_+_27487308_27487379@14.41.28,trna7_GlnCTG_15_-_66161400_66161471@14.41.28,trna99_GlnCTG_6_-_28909378_28909449@14.41.28|AATGGTTAGCACTCTGGACTCTGAATCC</td><td>>trna146_GlnCTG_6_-_27515531_27515602</td><td>14</td><td>41</td><td>28</td><td>AATGGTTAGCACTCTGGACTCTGAATCC</td><td>99.9</td><td>10.0</td><td>2.7</td><td>3.7</td><td>2.3E-04</td><td>1.7E-03</td><td>BT20/MB231</td><td>MP</td></tr>
<tr><td>>hg38_21_+_8433172_8446622_RNA45SN1_45S_50ntFlanks@3929.3959.31|GATCAAAACCAACCCGGTCAGCCCCTCTCCG</td><td>>hg38_21_+_8433172_8446622_RNA45SN1_45S_50ntFlanks</td><td>3929</td><td>3959</td><td>31</td><td>GATCAAAACCAACCCGGTCAGCCCCTCTCCG</td><td>122.8</td><td>-3.7</td><td>1.2</td><td>-3.2</td><td>1.4E-03</td><td>1.7E-03</td><td>MB231/MB468</td><td>CYTO</td></tr>
<tr><td>>hg38_1_-_228634819_228635039_RNA5S12_5S_50ntFlanks@51.71.21|GTCTACGGCCATACCACCCTG</td><td>>hg38_1_-_228634819_228635039_RNA5S12_5S_50ntFlanks</td><td>51</td><td>71</td><td>21</td><td>GTCTACGGCCATACCACCCTG</td><td>1199.0</td><td>-2.3</td><td>0.7</td><td>-3.2</td><td>1.5E-03</td><td>1.7E-03</td><td>MB231/MB468</td><td>NUC</td></tr>
<tr><td>>hg38_21_+_8433172_8446622_RNA45SN1_45S_50ntFlanks@12394.12414.21|TAGCGACGTCGCTTTTTGATC</td><td>>hg38_21_+_8433172_8446622_RNA45SN1_45S_50ntFlanks</td><td>12394</td><td>12414</td><td>21</td><td>TAGCGACGTCGCTTTTTGATC</td><td>1645.2</td><td>-2.7</td><td>0.8</td><td>-3.3</td><td>1.1E-03</td><td>1.7E-03</td><td>BT20/MB468</td><td>MITO</td></tr>
<tr><td>>trnaMT_HisGTG_MT_+_12138_12206@29.59.31|TTGTGAATCTGACAACAGAGGCTTACGACCC</td><td>>trnaMT_HisGTG_MT_+_12138_12206</td><td>29</td><td>59</td><td>31</td><td>TTGTGAATCTGACAACAGAGGCTTACGACCC</td><td>51.1</td><td>9.4</td><td>2.8</td><td>3.4</td><td>6.7E-04</td><td>1.7E-03</td><td>BT20/MB468</td><td>MP</td></tr>
<tr><td>>hg38_21_+_8433172_8446622_RNA45SN1_45S_50ntFlanks@7974.8018.45|ACGCGACCTCAGATCAGACGTGGCGACCCGCTGAATTTAAGCATA</td><td>>hg38_21_+_8433172_8446622_RNA45SN1_45S_50ntFlanks</td><td>7974</td><td>8018</td><td>45</td><td>ACGCGACCTCAGATCAGACGTGGCGACCCGCTGAATTTAAGCATA</td><td>2671.6</td><td>3.4</td><td>1.1</td><td>3.2</td><td>1.4E-03</td><td>1.7E-03</td><td>MB231/MB468</td><td>MITO</td></tr>
<tr><td>>hg38_21_+_8433172_8446622_RNA45SN1_45S_50ntFlanks@6661.6686.26|CGGTGGATCACTCGGCTCGTGCGTCG</td><td>>hg38_21_+_8433172_8446622_RNA45SN1_45S_50ntFlanks</td><td>6661</td><td>6686</td><td>26</td><td>CGGTGGATCACTCGGCTCGTGCGTCG</td><td>706.9</td><td>3.7</td><td>1.0</td><td>3.7</td><td>2.3E-04</td><td>1.7E-03</td><td>BT20/MB231</td><td>MP</td></tr>
<tr><td>>MI0000746|hsa-mir-99b&WithFlank&19|+|51692606|51692687@13.32.20&[MIMAT0000689&hsa-miR-99b-5p&offsets|0|-2;m-10&19|+|51692618|51692639&offsets|0|-2]|CACCCGTAGAACCGACCTTG</td><td>>MI0000746|hsa-mir-99b&WithFlank&19|+|51692606|51692687</td><td>13</td><td>32</td><td>20</td><td>CACCCGTAGAACCGACCTTG</td><td>343.7</td><td>3.0</td><td>0.9</td><td>3.2</td><td>1.2E-03</td><td>1.7E-03</td><td>MB231/MB468</td><td>MP</td></tr>
<tr><td>>hg38_21_+_8433172_8446622_RNA45SN1_45S_50ntFlanks@3835.3871.37|CTCCTCTCCTACTTGGATAACTGTGGTAATTCTAGAG</td><td>>hg38_21_+_8433172_8446622_RNA45SN1_45S_50ntFlanks</td><td>3835</td><td>3871</td><td>37</td><td>CTCCTCTCCTACTTGGATAACTGTGGTAATTCTAGAG</td><td>737.6</td><td>-3.1</td><td>1.0</td><td>-3.2</td><td>1.3E-03</td><td>1.7E-03</td><td>BT20/MB468</td><td>CYTO</td></tr>
<tr><td>>hg38_21_+_8433172_8446622_RNA45SN1_45S_50ntFlanks@4732.4768.37|AGAACGAAAGTCGGAGGTTCGAAGACGATCAGATACC</td><td>>hg38_21_+_8433172_8446622_RNA45SN1_45S_50ntFlanks</td><td>4732</td><td>4768</td><td>37</td><td>AGAACGAAAGTCGGAGGTTCGAAGACGATCAGATACC</td><td>58.8</td><td>5.8</td><td>1.8</td><td>3.2</td><td>1.2E-03</td><td>1.7E-03</td><td>MB231/MB468</td><td>MP</td></tr>
<tr><td>>hg38_21_+_8433172_8446622_RNA45SN1_45S_50ntFlanks@6745.6769.25|ACACTTCGAACGCACTTGCGGCCCC</td><td>>hg38_21_+_8433172_8446622_RNA45SN1_45S_50ntFlanks</td><td>6745</td><td>6769</td><td>25</td><td>ACACTTCGAACGCACTTGCGGCCCC</td><td>266.5</td><td>-3.7</td><td>1.1</td><td>-3.3</td><td>1.1E-03</td><td>1.7E-03</td><td>BT20/MB468</td><td>MITO</td></tr>
<tr><td>>hg38_21_+_8433172_8446622_RNA45SN1_45S_50ntFlanks@11566.11585.20|TAGCAGCCGACTTAGAACTG</td><td>>hg38_21_+_8433172_8446622_RNA45SN1_45S_50ntFlanks</td><td>11566</td><td>11585</td><td>20</td><td>TAGCAGCCGACTTAGAACTG</td><td>394.3</td><td>2.0</td><td>0.6</td><td>3.2</td><td>1.4E-03</td><td>1.7E-03</td><td>MB231/MB468</td><td>CYTO</td></tr>
<tr><td>>trna119_LysCTT_1_-_145395522_145395594@1.32.32,trna11_LysCTT_5_-_180648979_180649051@1.32.32,trna13_LysCTT_6_+_26556774_26556846@1.32.32,trna32_LysCTT_16_-_3207406_3207478@1.32.32,trna7_LysCTT_16_+_3225692_3225764@1.32.32,trna9_LysCTT_5_+_180634755_180634827@1.32.32|GCCCGGCTAGCTCAGTCGGTAGAGCATGAGAC</td><td>>trna119_LysCTT_1_-_145395522_145395594</td><td>1</td><td>32</td><td>32</td><td>GCCCGGCTAGCTCAGTCGGTAGAGCATGAGAC</td><td>1469.6</td><td>4.0</td><td>1.2</td><td>3.3</td><td>1.1E-03</td><td>1.7E-03</td><td>BT20/MB468</td><td>MITO</td></tr>
<tr><td>>hg38_21_+_8433172_8446622_RNA45SN1_45S_50ntFlanks@4724.4758.35|ATTAATCAAGAACGAAAGTCGGAGGTTCGAAGACG</td><td>>hg38_21_+_8433172_8446622_RNA45SN1_45S_50ntFlanks</td><td>4724</td><td>4758</td><td>35</td><td>ATTAATCAAGAACGAAAGTCGGAGGTTCGAAGACG</td><td>2193.0</td><td>6.2</td><td>1.7</td><td>3.7</td><td>2.3E-04</td><td>1.7E-03</td><td>BT20/MB231</td><td>MP</td></tr>
<tr><td>>MI0000434|hsa-let-7i&WithFlank&12|+|62603680|62603775@12.30.19&[MIMAT0000415&hsa-let-7i-5p&offsets|0|-3;m-49&12|+|62603691|62603712&offsets|0|-3]|TGAGGTAGTAGTTTGTGCT</td><td>>MI0000434|hsa-let-7i&WithFlank&12|+|62603680|62603775</td><td>12</td><td>30</td><td>19</td><td>TGAGGTAGTAGTTTGTGCT</td><td>170.1</td><td>5.9</td><td>1.7</td><td>3.4</td><td>6.7E-04</td><td>1.7E-03</td><td>BT20/MB468</td><td>MP</td></tr>
<tr><td>>am_MI0016800|hsa-mir-4454&WithFlank&4|-|163093568|163093634@12.29.18(+1G)&[MIMAT0018976&hsa-miR-4454&offsets|+3|+1(+1G);m-444&4|-|163093607|163093623&offsets|0|+1(+1G)]|TCCGAGTCACGGCACCAAG</td><td>>am_MI0016800|hsa-mir-4454&WithFlank&4|-|163093568|163093634</td><td>12</td><td>29</td><td>18(+1G)</td><td>TCCGAGTCACGGCACCAAG</td><td>50.7</td><td>9.4</td><td>2.8</td><td>3.4</td><td>6.7E-04</td><td>1.7E-03</td><td>BT20/MB468</td><td>MP</td></tr>
<tr><td>>MI0000102|hsa-mir-100&WithFlank&11|-|122152223|122152314@19.38.20(+1C)&[MIMAT0000098&hsa-miR-100-5p&offsets|0|-2(+1C);m-27&11|-|122152276|122152296&offsets|0|-1(+1C)]|AACCCGTAGATCCGAACTTGC</td><td>>MI0000102|hsa-mir-100&WithFlank&11|-|122152223|122152314</td><td>19</td><td>38</td><td>20(+1C)</td><td>AACCCGTAGATCCGAACTTGC</td><td>381.0</td><td>6.4</td><td>1.9</td><td>3.4</td><td>6.7E-04</td><td>1.7E-03</td><td>BT20/MB468</td><td>MP</td></tr>
<tr><td>>am_MI0008190|hsa-mir-320d-1&WithFlank&13|-|40727810|40727893@48.66.19;MI0008192|hsa-mir-320d-2&WithFlank&X|-|140926154|140926237@48.66.19&[MIMAT0006764&hsa-miR-320d&offsets|0|0;m-246&13|-|40727829|40727846&offsets|0|+1];[MIMAT0006764_1&hsa-miR-320d&offsets|0|0;m-253&X|-|140926173|140926190&offsets|0|+1]|AAAAGCTGGGTTGAGAGGA</td><td>>am_MI0008190|hsa-mir-320d-1&WithFlank&13|-|40727810|40727893</td><td>48</td><td>66</td><td>19</td><td>AAAAGCTGGGTTGAGAGGA</td><td>179.0</td><td>4.0</td><td>1.1</td><td>3.7</td><td>2.5E-04</td><td>1.7E-03</td><td>BT20/MB231</td><td>CYTO</td></tr>
<tr><td>>hg38_21_+_8433172_8446622_RNA45SN1_45S_50ntFlanks@4160.4186.27|CCACATCCAAGGAAGGCAGCAGGCGCG</td><td>>hg38_21_+_8433172_8446622_RNA45SN1_45S_50ntFlanks</td><td>4160</td><td>4186</td><td>27</td><td>CCACATCCAAGGAAGGCAGCAGGCGCG</td><td>111.1</td><td>-1.8</td><td>0.6</td><td>-3.2</td><td>1.3E-03</td><td>1.7E-03</td><td>BT20/MB468</td><td>CYTO</td></tr>
<tr><td>>hg38_21_+_8433172_8446622_RNA45SN1_45S_50ntFlanks@10247.10286.40|GCGACGAGTAGGAGGGCCGCTGCGGTGAGCCTTGAAGCCT</td><td>>hg38_21_+_8433172_8446622_RNA45SN1_45S_50ntFlanks</td><td>10247</td><td>10286</td><td>40</td><td>GCGACGAGTAGGAGGGCCGCTGCGGTGAGCCTTGAAGCCT</td><td>299.6</td><td>4.6</td><td>1.3</td><td>3.7</td><td>2.3E-04</td><td>1.7E-03</td><td>BT20/MB231</td><td>MP</td></tr>
<tr><td>>hg38_21_+_8433172_8446622_RNA45SN1_45S_50ntFlanks@10574.10591.18|AAGCCGGCGGGAGCCCCG</td><td>>hg38_21_+_8433172_8446622_RNA45SN1_45S_50ntFlanks</td><td>10574</td><td>10591</td><td>18</td><td>AAGCCGGCGGGAGCCCCG</td><td>359.2</td><td>-5.5</td><td>1.6</td><td>-3.4</td><td>6.7E-04</td><td>1.7E-03</td><td>BT20/MB468</td><td>MP</td></tr>
<tr><td>>hg38_21_+_8433172_8446622_RNA45SN1_45S_50ntFlanks@6661.6695.35|CGGTGGATCACTCGGCTCGTGCGTCGATGAAGAAC</td><td>>hg38_21_+_8433172_8446622_RNA45SN1_45S_50ntFlanks</td><td>6661</td><td>6695</td><td>35</td><td>CGGTGGATCACTCGGCTCGTGCGTCGATGAAGAAC</td><td>1624.3</td><td>2.4</td><td>0.6</td><td>3.7</td><td>2.5E-04</td><td>1.7E-03</td><td>BT20/MB231</td><td>NUC</td></tr>
<tr><td>>hg38_1_-_228634819_228635039_RNA5S12_5S_50ntFlanks@132.169.38|GATGGGAGACCGCCTGGGAATACCGGGTGCTGTAGGCT</td><td>>hg38_1_-_228634819_228635039_RNA5S12_5S_50ntFlanks</td><td>132</td><td>169</td><td>38</td><td>GATGGGAGACCGCCTGGGAATACCGGGTGCTGTAGGCT</td><td>6073.3</td><td>3.7</td><td>1.1</td><td>3.3</td><td>1.1E-03</td><td>1.7E-03</td><td>BT20/MB468</td><td>MITO</td></tr>
<tr><td>>MI0001445|hsa-mir-423&WithFlank&17|+|30117073|30117178@59.81.23(+1A)&[MIMAT0001340&hsa-miR-423-3p&offsets|0|0(+1A);m-100&17|+|30117131|30117153&offsets|0|0(+1A)]|AGCTCGGTCTGAGGCCCCTCAGTA</td><td>>MI0001445|hsa-mir-423&WithFlank&17|+|30117073|30117178</td><td>59</td><td>81</td><td>23(+1A)</td><td>AGCTCGGTCTGAGGCCCCTCAGTA</td><td>125.2</td><td>5.5</td><td>1.6</td><td>3.4</td><td>6.7E-04</td><td>1.7E-03</td><td>BT20/MB468</td><td>MP</td></tr>
<tr><td>>trna10_AspGTC_12_-_125424193_125424264@37.72.36,trna12_AspGTC_12_-_125411891_125411962@37.72.36,trna144_AspGTC_6_-_27551236_27551307@37.72.36,trna38_AspGTC_17_-_8125556_8125627@37.72.36,trna45_AspGTC_6_+_27447453_27447524@37.72.36,trna48_AspGTC_6_+_27471523_27471594@37.72.36,trna4_AspGTC_12_+_96429799_96429870@37.72.36,trna69_AspGTC_1_-_161440205_161440276@37.72.36,trna72_AspGTC_1_-_161432824_161432895@37.72.36,trna75_AspGTC_1_-_161425414_161425485@37.72.36,trna78_AspGTC_1_-_161418033_161418104@37.72.36,trna81_AspGTC_1_-_161410615_161410686@37.72.36|ACGCGGGAGACCGGGGTTCGATTCCCCGACGGGGAG</td><td>>trna10_AspGTC_12_-_125424193_125424264</td><td>37</td><td>72</td><td>36</td><td>ACGCGGGAGACCGGGGTTCGATTCCCCGACGGGGAG</td><td>879.3</td><td>-3.4</td><td>1.1</td><td>-3.2</td><td>1.2E-03</td><td>1.7E-03</td><td>MB231/MB468</td><td>MP</td></tr>
<tr><td>>hg38_21_+_8433172_8446622_RNA45SN1_45S_50ntFlanks@9503.9521.19|GGTGAACTATGCCTGGGCA</td><td>>hg38_21_+_8433172_8446622_RNA45SN1_45S_50ntFlanks</td><td>9503</td><td>9521</td><td>19</td><td>GGTGAACTATGCCTGGGCA</td><td>87.5</td><td>-5.1</td><td>1.6</td><td>-3.2</td><td>1.2E-03</td><td>1.8E-03</td><td>MB231/MB468</td><td>MP</td></tr>
<tr><td>>hg38_21_+_8433172_8446622_RNA45SN1_45S_50ntFlanks@9953.9981.29|ATGGAAGTCGGAATCCGCTAAGGAGTGTG</td><td>>hg38_21_+_8433172_8446622_RNA45SN1_45S_50ntFlanks</td><td>9953</td><td>9981</td><td>29</td><td>ATGGAAGTCGGAATCCGCTAAGGAGTGTG</td><td>79.4</td><td>5.4</td><td>1.6</td><td>3.4</td><td>6.7E-04</td><td>1.8E-03</td><td>BT20/MB468</td><td>MP</td></tr>
<tr><td>>hg38_MT_+_1621_3279_RNR2_16S_50ntFlanks@722.749.28|TGCGTCAGATTAAAACACTGAACTGACA</td><td>>hg38_MT_+_1621_3279_RNR2_16S_50ntFlanks</td><td>722</td><td>749</td><td>28</td><td>TGCGTCAGATTAAAACACTGAACTGACA</td><td>50.1</td><td>9.4</td><td>2.8</td><td>3.4</td><td>6.7E-04</td><td>1.8E-03</td><td>BT20/MB468</td><td>MP</td></tr>
<tr><td>>hg38_21_+_8433172_8446622_RNA45SN1_45S_50ntFlanks@6651.6669.19|CGACTCTTAGCGGTGGATC</td><td>>hg38_21_+_8433172_8446622_RNA45SN1_45S_50ntFlanks</td><td>6651</td><td>6669</td><td>19</td><td>CGACTCTTAGCGGTGGATC</td><td>5288.4</td><td>-3.0</td><td>0.9</td><td>-3.2</td><td>1.2E-03</td><td>1.8E-03</td><td>MB231/MB468</td><td>MP</td></tr>
<tr><td>>hg38_21_+_8433172_8446622_RNA45SN1_45S_50ntFlanks@12644.12669.26|TCTGTGGGATTATGACTGAACGCCTC</td><td>>hg38_21_+_8433172_8446622_RNA45SN1_45S_50ntFlanks</td><td>12644</td><td>12669</td><td>26</td><td>TCTGTGGGATTATGACTGAACGCCTC</td><td>325.4</td><td>3.6</td><td>1.1</td><td>3.2</td><td>1.4E-03</td><td>1.8E-03</td><td>MB231/MB468</td><td>CYTO</td></tr>
<tr><td>>hg38_21_+_8433172_8446622_RNA45SN1_45S_50ntFlanks@7989.8014.26|AGACGTGGCGACCCGCTGAATTTAAG</td><td>>hg38_21_+_8433172_8446622_RNA45SN1_45S_50ntFlanks</td><td>7989</td><td>8014</td><td>26</td><td>AGACGTGGCGACCCGCTGAATTTAAG</td><td>1075.0</td><td>-2.6</td><td>0.8</td><td>-3.4</td><td>6.7E-04</td><td>1.8E-03</td><td>BT20/MB468</td><td>MP</td></tr>
<tr><td>>MI0000087|hsa-mir-29a&WithFlank&7|-|130876741|130876816@48.67.20(+1G)&[MIMAT0000086&hsa-miR-29a-3p&offsets|0|-2(+1G);m-26&7|-|130876749|130876769&offsets|0|-1(+1G)]|TAGCACCATCTGAAATCGGTG</td><td>>MI0000087|hsa-mir-29a&WithFlank&7|-|130876741|130876816</td><td>48</td><td>67</td><td>20(+1G)</td><td>TAGCACCATCTGAAATCGGTG</td><td>311.8</td><td>5.2</td><td>1.6</td><td>3.2</td><td>1.2E-03</td><td>1.8E-03</td><td>MB231/MB468</td><td>MP</td></tr>
<tr><td>>trnaMT_GluTTC_MT_-_14674_14742@41.70.30,trnalookalike8_GluTTC_5_-_93905172_93905240@41.70.30|TGGTCGTGGTTGTAGTCCGTGCGAGAATAC</td><td>>trnaMT_GluTTC_MT_-_14674_14742</td><td>41</td><td>70</td><td>30</td><td>TGGTCGTGGTTGTAGTCCGTGCGAGAATAC</td><td>157.9</td><td>6.6</td><td>2.0</td><td>3.4</td><td>6.7E-04</td><td>1.8E-03</td><td>BT20/MB468</td><td>MP</td></tr>
</table>

| | | | | | | | | | | | | | |
|---|---|---|---|---|---|---|---|---|---|---|---|---|---|
| >hg38_21_+_8433172_8446622_RNA45SN1_45S_50ntFlanks@5472.5516.45\|ACGGCCCTGGCGGAGCGCTGAGAAGACGGTCGAACTTGACTATCT | >hg38_21_+_8433172_8446622_RNA45SN1_45S_50ntFlanks | 5472 | 5516 | 45 | ACGGCCCTGGCGGAGCGCTGAGAAGACGGTCGAACTTGACTATCT | 720.2 | 5.9 | 1.8 | 3.2 | 1.2E-03 | 1.8E-03 | MB231/MB468 | MP |
| >hg38_21_+_8433172_8446622_RNA45SN1_45S_50ntFlanks@12417.12459.43\|TCGATGTCGGCTCTTCCTATCATTGTGAAGCAGAATTCACCAA | >hg38_21_+_8433172_8446622_RNA45SN1_45S_50ntFlanks | 12417 | 12459 | 43 | TCGATGTCGGCTCTTCCTATCATTGTGAAGCAGAATTCACCAA | 227.2 | 5.0 | 1.6 | 3.2 | 1.5E-03 | 1.8E-03 | MB231/MB468 | MITO |
| >hg38_21_+_8433172_8446622_RNA45SN1_45S_50ntFlanks@6650.6675.26\|ACGACTCTTAGCGGTGGATCACTCGG | >hg38_21_+_8433172_8446622_RNA45SN1_45S_50ntFlanks | 6650 | 6675 | 26 | ACGACTCTTAGCGGTGGATCACTCGG | 1489.3 | 4.8 | 1.3 | 3.7 | 2.3E-04 | 1.8E-03 | BT20/MB231 | MP |
| >hg38_21_+_8433172_8446622_RNA45SN1_45S_50ntFlanks@5472.5521.50\|ACGGCCCTGGCGGAGCGCTGAGAAGACGGTCGAACTTGACTATCTAGAGG | >hg38_21_+_8433172_8446622_RNA45SN1_45S_50ntFlanks | 5472 | 5521 | 50 | ACGGCCCTGGCGGAGCGCTGAGAAGACGGTCGAACTTGACTATCTAGAGG | 395.7 | 7.7 | 2.4 | 3.2 | 1.4E-03 | 1.8E-03 | MB231/MB468 | CYTO |
| >hg38_21_+_8433172_8446622_RNA45SN1_45S_50ntFlanks@11577.11605.29\|TTAGAACTGGTGCGGACCAGGGGAATCCG | >hg38_21_+_8433172_8446622_RNA45SN1_45S_50ntFlanks | 11577 | 11605 | 29 | TTAGAACTGGTGCGGACCAGGGGAATCCG | 225.4 | -3.7 | 1.1 | -3.3 | 1.1E-03 | 1.8E-03 | BT20/MB468 | MITO |
| >hg38_21_+_8433172_8446622_RNA45SN1_45S_50ntFlanks@485.510.26\|GATGTGGTGACGTCGTGCTCTCCCGG | >hg38_21_+_8433172_8446622_RNA45SN1_45S_50ntFlanks | 485 | 510 | 26 | GATGTGGTGACGTCGTGCTCTCCCGG | 322.8 | 11.6 | 3.6 | 3.2 | 1.5E-03 | 1.8E-03 | MB231/MB468 | MITO |
| >MI0000298\|hsa-mir-221&WithFlank&X\|-\|45746151\|45746272@71.91.21(+1G)&[MIMAT0000278&hsa-miR-221-3p&offsets\|0\|-2(+1G);m-91&X\|-\|45746181\|45746202&offsets\|0\|-1(+1G)]\|AGCTACATTGTCTGCTGGGTTG | >MI0000298\|hsa-mir-221&WithFlank&X\|-\|45746151\|45746272 | 71 | 91 | 21(+1G) | AGCTACATTGTCTGCTGGGTTG | 352.3 | 4.3 | 1.3 | 3.2 | 1.2E-03 | 1.8E-03 | MB231/MB468 | MP |
| >trnaMT_GluTTC_MT_-_14674_14742@44.72.29,trnalookalike8_GluTTC_5_-_93905172_93905240@44.72.29\|TCGTGGTTGTAGTCCGTGCGAGAATACCA | >trnaMT_GluTTC_MT_-_14674_14742 | 44 | 72 | 29 | TCGTGGTTGTAGTCCGTGCGAGAATACCA | 2084.5 | 3.2 | 1.0 | 3.2 | 1.5E-03 | 1.8E-03 | MB231/MB468 | MITO |
| >hg38_21_+_8433172_8446622_RNA45SN1_45S_50ntFlanks@11666.11705.40\|TCTGCCCAGTGCTCTGAATGTCAAAGTGAAGAAATTCAAT | >hg38_21_+_8433172_8446622_RNA45SN1_45S_50ntFlanks | 11666 | 11705 | 40 | TCTGCCCAGTGCTCTGAATGTCAAAGTGAAGAAATTCAAT | 174.5 | -2.9 | 0.8 | -3.7 | 2.6E-04 | 1.8E-03 | BT20/MB231 | CYTO |
| >hg38_21_+_8433172_8446622_RNA45SN1_45S_50ntFlanks@11876.11908.33\|AAGACCCTGTTGAGCTTGACTCTAGTCTGGCAC | >hg38_21_+_8433172_8446622_RNA45SN1_45S_50ntFlanks | 11876 | 11908 | 33 | AAGACCCTGTTGAGCTTGACTCTAGTCTGGCAC | 399.0 | -1.8 | 0.5 | -3.3 | 1.1E-03 | 1.8E-03 | BT20/MB468 | MITO |
| >trna14_LysTTT_11_-_59327808_59327880@1.32.32,trna2_LysTTT_17_+_8022473_8022545@1.32.32,trna54_LysTTT_1_+_204475655_204475727@1.32.32,trna5_LysTTT_11_+_59323902_59323974@1.32.32,trna62_LysTTT_1_-_204476158_204476230@1.32.32,trna76_LysTTT_6_+_28918806_28918878@1.32.32\|GCCCGGATAGCTCAGTCGGTAGAGCATCAGAC | >trna14_LysTTT_11_-_59327808_59327880 | 1 | 32 | 32 | GCCCGGATAGCTCAGTCGGTAGAGCATCAGAC | 250.7 | 4.6 | 1.4 | 3.2 | 1.5E-03 | 1.8E-03 | MB231/MB468 | MITO |
| >MI0000736\|hsa-mir-30c-1&WithFlank&1\|+\|40757278\|40757378@23.45.23(+1A);MI0000254\|hsa-mir-30c-2&WithFlank&6\|-\|71376954\|71377037@13.35.23(+1A)&[MIMAT0000244&hsa-miR-30c-5p&offsets\|0\|0(+1A);m-57&1\|+\|40757300\|40757323&offsets\|0\|-1(+1A)];[MIMAT0000244_1&hsa-miR-30c-5p&offsets\|0\|0(+1A);m-56&6\|-\|71377002\|71377025&offsets\|0\|-1(+1A)]\|TGTAAACATCCTACACTCTCAGCA | >MI0000736\|hsa-mir-30c-1&WithFlank&1\|+\|40757278\|40757378 | 23 | 45 | 23(+1A) | TGTAAACATCCTACACTCTCAGCA | 1131.5 | 3.6 | 1.1 | 3.2 | 1.2E-03 | 1.8E-03 | MB231/MB468 | MP |
| >hg38_21_+_8433172_8446622_RNA45SN1_45S_50ntFlanks@6700.6717.18\|CTAGCTGCGAGAATTAAT | >hg38_21_+_8433172_8446622_RNA45SN1_45S_50ntFlanks | 6700 | 6717 | 18 | CTAGCTGCGAGAATTAAT | 286.1 | -3.0 | 1.0 | -3.2 | 1.4E-03 | 1.8E-03 | MB231/MB468 | CYTO |
| >hg38_21_+_8433172_8446622_RNA45SN1_45S_50ntFlanks@12488.12518.31\|AACGTGAGCTGGGTTTAGACCGTCGTGAGAC | >hg38_21_+_8433172_8446622_RNA45SN1_45S_50ntFlanks | 12488 | 12518 | 31 | AACGTGAGCTGGGTTTAGACCGTCGTGAGAC | 180.1 | -2.0 | 0.6 | -3.3 | 1.1E-03 | 1.8E-03 | BT20/MB468 | MITO |
| >hg38_21_+_8433172_8446622_RNA45SN1_45S_50ntFlanks@10348.10371.24\|AAACGAGAACTTTGAAGGCCGAAG | >hg38_21_+_8433172_8446622_RNA45SN1_45S_50ntFlanks | 10348 | 10371 | 24 | AAACGAGAACTTTGAAGGCCGAAG | 752.3 | 2.0 | 0.6 | 3.3 | 1.1E-03 | 1.8E-03 | BT20/MB468 | MITO |
| >hg38_21_+_8433172_8446622_RNA45SN1_45S_50ntFlanks@9934.9955.22\|AGACAGCAGGACGGTGGCCATG | >hg38_21_+_8433172_8446622_RNA45SN1_45S_50ntFlanks | 9934 | 9955 | 22 | AGACAGCAGGACGGTGGCCATG | 97.8 | 6.3 | 1.9 | 3.4 | 6.8E-04 | 1.8E-03 | BT20/MB468 | MP |
| >hg38_21_+_8433172_8446622_RNA45SN1_45S_50ntFlanks@11807.11827.21\|ACTGTCCCTACCTACTATCCA | >hg38_21_+_8433172_8446622_RNA45SN1_45S_50ntFlanks | 11807 | 11827 | 21 | ACTGTCCCTACCTACTATCCA | 54.5 | -8.8 | 2.6 | -3.4 | 6.8E-04 | 1.8E-03 | BT20/MB468 | MP |
| >hg38_21_+_8433172_8446622_RNA45SN1_45S_50ntFlanks@7988.8005.18\|CAGACGTGGCGACCCGCT | >hg38_21_+_8433172_8446622_RNA45SN1_45S_50ntFlanks | 7988 | 8005 | 18 | CAGACGTGGCGACCCGCT | 413.1 | -3.2 | 0.9 | -3.4 | 6.8E-04 | 1.8E-03 | BT20/MB468 | MP |
| >MI0000065\|hsa-let-7d&WithFlank&9\|+\|94178828\|94178926@14.36.23&[MIMAT0000065&hsa-let-7d-5p&offsets\|0\|+1;m-90&9\|+\|94178841\|94178862&offsets\|0\|+1]\|AGAGGTAGTAGGTTGCATAGTTT | >MI0000065\|hsa-let-7d&WithFlank&9\|+\|94178828\|94178926 | 14 | 36 | 23 | AGAGGTAGTAGGTTGCATAGTTT | 399.6 | 4.8 | 1.4 | 3.4 | 6.8E-04 | 1.8E-03 | BT20/MB468 | MP |
| >MI0000816\|hsa-mir-335&WithFlank&7\|+\|130496105\|130496210@58.78.21(+1A)&[MIMAT0004703&hsa-miR-335-3p&offsets\|0\|-1(+1A);m-159&7\|+\|130496162\|130496183&offsets\|0\|-1(+1A)]\|TTTTTCATTATTGCTCCTGACA | >MI0000816\|hsa-mir-335&WithFlank&7\|+\|130496105\|130496210 | 58 | 78 | 21(+1A) | TTTTTCATTATTGCTCCTGACA | 139.2 | 4.2 | 1.3 | 3.2 | 1.2E-03 | 1.8E-03 | MB231/MB468 | MP |
| >hg38_21_+_8433172_8446622_RNA45SN1_45S_50ntFlanks@3769.3792.24\|CGCACGGCCGGTACAGTGAAACTG | >hg38_21_+_8433172_8446622_RNA45SN1_45S_50ntFlanks | 3769 | 3792 | 24 | CGCACGGCCGGTACAGTGAAACTG | 158.2 | -3.0 | 0.9 | -3.2 | 1.3E-03 | 1.8E-03 | BT20/MB468 | CYTO |
| >hg38_MT_+_1621_3279_RNR2_16S_50ntFlanks@115.146.32\|AAATAAAGTATAGGCGATAGAAATTGAAACCT | >hg38_MT_+_1621_3279_RNR2_16S_50ntFlanks | 115 | 146 | 32 | AAATAAAGTATAGGCGATAGAAATTGAAACCT | 50.3 | 9.4 | 2.8 | 3.4 | 6.8E-04 | 1.8E-03 | BT20/MB468 | MP |
| >hg38_21_+_8433172_8446622_RNA45SN1_45S_50ntFlanks@8245.8269.25\|CTTCCCGGAGTCGGGTTGCTTGGGA | >hg38_21_+_8433172_8446622_RNA45SN1_45S_50ntFlanks | 8245 | 8269 | 25 | CTTCCCGGAGTCGGGTTGCTTGGGA | 94.1 | -4.0 | 1.2 | -3.2 | 1.2E-03 | 1.8E-03 | MB231/MB468 | MP |
| >hg38_21_+_8433172_8446622_RNA45SN1_45S_50ntFlanks@485.508.24\|GATGTGGTGACGTCGTGCTCTCCC | >hg38_21_+_8433172_8446622_RNA45SN1_45S_50ntFlanks | 485 | 508 | 24 | GATGTGGTGACGTCGTGCTCTCCC | 321.2 | 11.5 | 3.6 | 3.2 | 1.5E-03 | 1.8E-03 | MB231/MB468 | MITO |
| >trna10_AspGTC_12_-_125424193_125424264@1.31.31,trna12_AspGTC_12_-_125411891_125411962@1.31.31,trna38_AspGTC_17_-_8125556_8125627@1.31.31,trna45_AspGTC_6_+_27447453_27447524@1.31.31,trna48_AspGTC_6_+_27471523_27471594@1.31.31,trna4_AspGTC_12_+_96429799_96429870@1.31.31,trna69_AspGTC_1_-_161440205_161440276@1.31.31,trna72_AspGTC_1_-_161432824_161432895@1.31.31,trna75_AspGTC_1_-_161425414_161425485@1.31.31,trna78_AspGTC_1_-_161418033_161418104@1.31.31,trna81_AspGTC_1_-_161410615_161410686@1.31.31\|TCCTCGTTAGTATAGTGGTGAGTATCCCCGC | >trna10_AspGTC_12_-_125424193_125424264 | 1 | 31 | 31 | TCCTCGTTAGTATAGTGGTGAGTATCCCCGC | 380.6 | -2.9 | 0.9 | -3.2 | 1.4E-03 | 1.8E-03 | MB231/MB468 | CYTO |
| >hg38_21_+_8433172_8446622_RNA45SN1_45S_50ntFlanks@6722.6743.22\|ATTGCAGGACACATTGATCATC | >hg38_21_+_8433172_8446622_RNA45SN1_45S_50ntFlanks | 6722 | 6743 | 22 | ATTGCAGGACACATTGATCATC | 931.0 | -2.7 | 0.8 | -3.2 | 1.4E-03 | 1.8E-03 | MB231/MB468 | CYTO |
| >hg38_21_+_8433172_8446622_RNA45SN1_45S_50ntFlanks@12415.12433.19\|CTTCGATGTCGGCTCTTCC | >hg38_21_+_8433172_8446622_RNA45SN1_45S_50ntFlanks | 12415 | 12433 | 19 | CTTCGATGTCGGCTCTTCC | 806.5 | 1.4 | 0.4 | 3.3 | 1.1E-03 | 1.8E-03 | BT20/MB468 | MITO |
| >hg38_21_+_8433172_8446622_RNA45SN1_45S_50ntFlanks@10797.10823.27\|GCATGTTGGAACAATGTAGGTAAGGGA | >hg38_21_+_8433172_8446622_RNA45SN1_45S_50ntFlanks | 10797 | 10823 | 27 | GCATGTTGGAACAATGTAGGTAAGGGA | 594.8 | -2.6 | 0.8 | -3.4 | 6.8E-04 | 1.8E-03 | BT20/MB468 | MP |
| >hg38_21_+_8433172_8446622_RNA45SN1_45S_50ntFlanks@12957.12989.33\|CTTCTGGGTCGGGGTTTCGTACGTAGCAGAGCA | >hg38_21_+_8433172_8446622_RNA45SN1_45S_50ntFlanks | 12957 | 12989 | 33 | CTTCTGGGTCGGGGTTTCGTACGTAGCAGAGCA | 152.1 | -4.4 | 1.4 | -3.2 | 1.3E-03 | 1.8E-03 | BT20/MB468 | CYTO |
| >hg38_21_+_8433172_8446622_RNA45SN1_45S_50ntFlanks@12488.12518.31\|AACGTGAGCTGGGTTTAGACCGTCGTGAGAC | >hg38_21_+_8433172_8446622_RNA45SN1_45S_50ntFlanks | 12488 | 12518 | 31 | AACGTGAGCTGGGTTTAGACCGTCGTGAGAC | 296.4 | -4.4 | 1.3 | -3.4 | 6.8E-04 | 1.8E-03 | BT20/MB468 | MP |
| >hg38_1_-_228634819_228635039_RNA5S12_5S_50ntFlanks@124.169.46\|AGTACTTGGATGGGAGACCGCCTGGGAATACCGGGTGCTGTAGGCT | >hg38_1_-_228634819_228635039_RNA5S12_5S_50ntFlanks | 124 | 169 | 46 | AGTACTTGGATGGGAGACCGCCTGGGAATACCGGGTGCTGTAGGCT | 599.6 | 4.7 | 1.4 | 3.3 | 1.1E-03 | 1.8E-03 | BT20/MB468 | MITO |
| >hg38_21_+_8433172_8446622_RNA45SN1_45S_50ntFlanks@4380.4404.25\|CGTAGTTGGATCTTGGGAGCGGGCG | >hg38_21_+_8433172_8446622_RNA45SN1_45S_50ntFlanks | 4380 | 4404 | 25 | CGTAGTTGGATCTTGGGAGCGGGCG | 2940.1 | 5.8 | 1.7 | 3.4 | 6.8E-04 | 1.8E-03 | BT20/MB468 | MP |
| >MI0000456\|hsa-mir-140&WithFlank&16\|+\|69933075\|69933186@69.89.21&[MIMAT0004597&hsa-miR-140-3p&offsets\|+1\|+1;m-45&16\|+\|69933143\|69933164&offsets\|0\|-1]\|ACCACAGGGTAGAACCACGGA | >MI0000456\|hsa-mir-140&WithFlank&16\|+\|69933075\|69933186 | 69 | 89 | 21 | ACCACAGGGTAGAACCACGGA | 1010.7 | 5.4 | 1.6 | 3.4 | 6.8E-04 | 1.8E-03 | BT20/MB468 | MP |
| >hg38_21_+_8433172_8446622_RNA45SN1_45S_50ntFlanks@10384.10404.21\|CCATGTGAACAGCAGTTGAAC | >hg38_21_+_8433172_8446622_RNA45SN1_45S_50ntFlanks | 10384 | 10404 | 21 | CCATGTGAACAGCAGTTGAAC | 149.9 | -3.8 | 1.2 | -3.2 | 1.5E-03 | 1.8E-03 | MB231/MB468 | NUC |
| >hg38_21_+_8433172_8446622_RNA45SN1_45S_50ntFlanks@3984.4027.44\|GCGGCTTTGGTGACTCTAGATAACCTCGGGCCGATCGCACGCCC | >hg38_21_+_8433172_8446622_RNA45SN1_45S_50ntFlanks | 3984 | 4027 | 44 | GCGGCTTTGGTGACTCTAGATAACCTCGGGCCGATCGCACGCCC | 49.5 | -8.7 | 2.6 | -3.4 | 6.8E-04 | 1.8E-03 | BT20/MB468 | MP |
| >hg38_21_+_8433172_8446622_RNA45SN1_45S_50ntFlanks@4759.4777.19\|ATCAGATACCGTCGTAGTT | >hg38_21_+_8433172_8446622_RNA45SN1_45S_50ntFlanks | 4759 | 4777 | 19 | ATCAGATACCGTCGTAGTT | 478.4 | -5.4 | 1.7 | -3.2 | 1.2E-03 | 1.8E-03 | MB231/MB468 | MP |
| >hg38_1_-_228634819_228635039_RNA5S12_5S_50ntFlanks@138.170.33\|AGACCGCCTGGGAATACCGGGTGCTGTAGGCTT | >hg38_1_-_228634819_228635039_RNA5S12_5S_50ntFlanks | 138 | 170 | 33 | AGACCGCCTGGGAATACCGGGTGCTGTAGGCTT | 928.0 | 3.0 | 0.9 | 3.2 | 1.5E-03 | 1.8E-03 | MB231/MB468 | MITO |
| >hg38_21_+_8433172_8446622_RNA45SN1_45S_50ntFlanks@8171.8215.45\|AAGTCCTTCTGATCGAGGCCCAGCCCGTGGACGGTGTGAGGCCGG | >hg38_21_+_8433172_8446622_RNA45SN1_45S_50ntFlanks | 8171 | 8215 | 45 | AAGTCCTTCTGATCGAGGCCCAGCCCGTGGACGGTGTGAGGCCGG | 810.8 | -3.5 | 1.1 | -3.2 | 1.5E-03 | 1.8E-03 | MB231/MB468 | NUC |
| >MI0000264\|hsa-mir-7-2&WithFlank&15\|+\|88611819\|88611940@38.61.24(+1A);MI0000265\|hsa-mir-7-3&WithFlank&19\|+\|4770664\|4770785@37.60.24(+1A);MI0000263\|hsa-mir-7-1&WithFlank&9\|-\|83969742\|83969863@30.53.24(+1A)&[MIMAT0000252&hsa-miR-7-5p&offsets\|0\|0(+1A);m-400&15\|+\|88611856\|88611879&offsets\|0\|0(+1A)];[MIMAT0000252_1&hsa-miR-7-5p&offsets\|0\|0(+1A);m-402&19\|+\|4770700\|4770723&offsets\|0\|0(+1A)];[MIMAT0000252_2&hsa-miR-7-5p&offsets\|0\|0(+1A);m-401&9\|-\|83969811\|83969834&offsets\|0\|0(+1A)]\|TGGAAGACTAGTGATTTTGTTGTTA | >MI0000264\|hsa-mir-7-2&WithFlank&15\|+\|88611819\|88611940 | 38 | 61 | 24(+1A) | TGGAAGACTAGTGATTTTGTTGTTA | 447.1 | 4.2 | 1.3 | 3.2 | 1.2E-03 | 1.8E-03 | MB231/MB468 | MP |
| >hg38_21_+_8433172_8446622_RNA45SN1_45S_50ntFlanks@12003.12029.27\|GCGGGCCGCCGGTGAAATACCACTACT | >hg38_21_+_8433172_8446622_RNA45SN1_45S_50ntFlanks | 12003 | 12029 | 27 | GCGGGCCGCCGGTGAAATACCACTACT | 474.8 | -3.1 | 0.9 | -3.7 | 2.5E-04 | 1.8E-03 | BT20/MB231 | NUC |
| >MI0000086\|hsa-mir-28&WithFlank&3\|+\|188688775\|188688872@60.82.23(+1A)&[MIMAT0004502&hsa-miR-28-3p&offsets\|0\|+1(+1A);m-28&3\|+\|188688834\|188688855&offsets\|0\|+1(+1A)]\|CACTAGATTGTGAGCTCCTGGAGA | >MI0000086\|hsa-mir-28&WithFlank&3\|+\|188688775\|188688872 | 60 | 82 | 23(+1A) | CACTAGATTGTGAGCTCCTGGAGA | 211.4 | 3.0 | 0.8 | 3.7 | 2.6E-04 | 1.8E-03 | BT20/MB231 | CYTO |

<table>
<tr><th></th><th></th><th></th><th></th><th></th><th></th><th></th><th></th><th></th><th></th><th></th><th></th><th></th><th></th></tr>
<tr><td>>MI0000736|hsa-mir-30c-1&WithFlank&1|+|40757278|40757378@23.45.23(+1A);MI0000254|hsa-mir-30c-2&WithFlank&6|-|71376954|71377037@13.35.23(+1A)&[MIMAT0000244&hsa-miR-30c-5p&offsets|0|0(+1A);m-57&1|+|40757300|40757323&offsets|0|-1(+1A)];[MIMAT0000244_1&hsa-miR-30c-5p&offsets|0|0(+1A);m-56&6|-|71377002|71377025&offsets|0|-1(+1A)]|TGTAAACATCCTACACTCTCAGCA</td><td>>MI0000736|hsa-mir-30c-1&WithFlank&1|+|40757278|40757378</td><td>23</td><td>45</td><td>23(+1A)</td><td>TGTAAACATCCTACACTCTCAGCA</td><td>5673.1</td><td>-3.3</td><td>0.9</td><td>-3.6</td><td>3.0E-04</td><td>1.8E-03</td><td>BT20/MB231</td><td>MITO</td></tr>
<tr><td>>hg38_21_+_8433172_8446622_RNA45SN1_45S_50ntFlanks@5423.5443.21|ATTGGATGGTTTAGTGAGGCC</td><td>>hg38_21_+_8433172_8446622_RNA45SN1_45S_50ntFlanks</td><td>5423</td><td>5443</td><td>21</td><td>ATTGGATGGTTTAGTGAGGCC</td><td>354.0</td><td>-5.7</td><td>1.7</td><td>-3.4</td><td>6.8E-04</td><td>1.8E-03</td><td>BT20/MB468</td><td>MP</td></tr>
<tr><td>>trna129_MetCAT_6_-_27870271_27870342@16.33.18,trna142_MetCAT_6_-_27560600_27560671@16.33.18,trna150_MetCAT_6_-_27300764_27300835@16.33.18,trna151_ThrCGT_6_-_27271568_27271639@16.33.18,trna169_MetCAT_6_-_26330529_26330600@16.33.18,trna171_MetCAT_6_-_26313352_26313423@16.33.18,trna20_MetCAT_17_-_80452597_80452668@16.33.18,trna2_MetCAT_6_+_26286754_26286825@16.33.18,trna32_MetCAT_1_+_153643726_153643797@16.33.18,trna61_MetCAT_6_+_27745664_27745735@16.33.18|CGGAAGCGTGCTGGGCCC</td><td>>trna129_MetCAT_6_-_27870271_27870342</td><td>16</td><td>33</td><td>18</td><td>CGGAAGCGTGCTGGGCCC</td><td>179.5</td><td>-2.5</td><td>0.7</td><td>-3.6</td><td>3.0E-04</td><td>1.8E-03</td><td>BT20/MB231</td><td>MITO</td></tr>
<tr><td>>hg38_21_+_8433172_8446622_RNA45SN1_45S_50ntFlanks@10396.10435.40|CAGTTGAACATGGGTCAGTCGGTCCTGAGAGATGGGCGAG</td><td>>hg38_21_+_8433172_8446622_RNA45SN1_45S_50ntFlanks</td><td>10396</td><td>10435</td><td>40</td><td>CAGTTGAACATGGGTCAGTCGGTCCTGAGAGATGGGCGAG</td><td>148.2</td><td>-4.7</td><td>1.5</td><td>-3.2</td><td>1.3E-03</td><td>1.8E-03</td><td>BT20/MB468</td><td>CYTO</td></tr>
<tr><td>>MI0003628|hsa-mir-615&WithFlank&12|+|54033944|54034051@67.88.22&[MIMAT0003283&hsa-miR-615-3p&offsets|0|0;m-495&12|+|54034010|54034030&offsets|0|+1]|TCCGAGCCTGGGTCTCCCTCTT</td><td>>MI0003628|hsa-mir-615&WithFlank&12|+|54033944|54034051</td><td>67</td><td>88</td><td>22</td><td>TCCGAGCCTGGGTCTCCCTCTT</td><td>317.2</td><td>5.3</td><td>1.4</td><td>3.7</td><td>2.3E-04</td><td>1.8E-03</td><td>BT20/MB231</td><td>MP</td></tr>
<tr><td>>hg38_21_+_8433172_8446622_RNA45SN1_45S_50ntFlanks@6661.6679.19|CGGTGGATCACTCGGCTCG</td><td>>hg38_21_+_8433172_8446622_RNA45SN1_45S_50ntFlanks</td><td>6661</td><td>6679</td><td>19</td><td>CGGTGGATCACTCGGCTCG</td><td>477.6</td><td>4.1</td><td>1.1</td><td>3.7</td><td>2.4E-04</td><td>1.8E-03</td><td>BT20/MB231</td><td>MP</td></tr>
<tr><td>>hg38_21_+_8433172_8446622_RNA45SN1_45S_50ntFlanks@4002.4039.38|GATAACCTCGGGCCGATCGCACGCCCCCCGTGGCGGCG</td><td>>hg38_21_+_8433172_8446622_RNA45SN1_45S_50ntFlanks</td><td>4002</td><td>4039</td><td>38</td><td>GATAACCTCGGGCCGATCGCACGCCCCCCGTGGCGGCG</td><td>223.5</td><td>-2.6</td><td>0.8</td><td>-3.2</td><td>1.4E-03</td><td>1.8E-03</td><td>MB231/MB468</td><td>CYTO</td></tr>
<tr><td>>MI0000804|hsa-mir-328&WithFlank&16|-|67202315|67202401@54.75.22&[MIMAT0000752&hsa-miR-328-3p&offsets|0|0;m-160&16|-|67202327|67202348&offsets|0|0]|CTGGCCCTCTCTGCCCTTCCGT</td><td>>MI0000804|hsa-mir-328&WithFlank&16|-|67202315|67202401</td><td>54</td><td>75</td><td>22</td><td>CTGGCCCTCTCTGCCCTTCCGT</td><td>678.6</td><td>2.5</td><td>0.7</td><td>3.7</td><td>2.6E-04</td><td>1.8E-03</td><td>BT20/MB231</td><td>NUC</td></tr>
<tr><td>>hg38_21_+_8433172_8446622_RNA45SN1_45S_50ntFlanks@4008.4025.18|CTCGGGCCGATCGCACGC</td><td>>hg38_21_+_8433172_8446622_RNA45SN1_45S_50ntFlanks</td><td>4008</td><td>4025</td><td>18</td><td>CTCGGGCCGATCGCACGC</td><td>186.4</td><td>-2.4</td><td>0.7</td><td>-3.3</td><td>1.1E-03</td><td>1.8E-03</td><td>BT20/MB468</td><td>MITO</td></tr>
<tr><td>>hg38_21_+_8433172_8446622_RNA45SN1_45S_50ntFlanks@5473.5502.30|CGGCCCTGGCGGAGCGCTGAGAAGACGGTC</td><td>>hg38_21_+_8433172_8446622_RNA45SN1_45S_50ntFlanks</td><td>5473</td><td>5502</td><td>30</td><td>CGGCCCTGGCGGAGCGCTGAGAAGACGGTC</td><td>193.9</td><td>3.2</td><td>0.9</td><td>3.4</td><td>6.9E-04</td><td>1.8E-03</td><td>BT20/MB468</td><td>MP</td></tr>
<tr><td>>hg38_21_+_8433172_8446622_RNA45SN1_45S_50ntFlanks@10234.10272.39|CGGACGCTACGCCGCGACGAGTAGGAGGGCCGCTGCGGT</td><td>>hg38_21_+_8433172_8446622_RNA45SN1_45S_50ntFlanks</td><td>10234</td><td>10272</td><td>39</td><td>CGGACGCTACGCCGCGACGAGTAGGAGGGCCGCTGCGGT</td><td>169.3</td><td>3.5</td><td>1.0</td><td>3.4</td><td>6.9E-04</td><td>1.8E-03</td><td>BT20/MB468</td><td>MP</td></tr>
<tr><td>>hg38_21_+_8433172_8446622_RNA45SN1_45S_50ntFlanks@10681.10705.25|TTCCGGCGGCGTCCGGTGAGCTCTC</td><td>>hg38_21_+_8433172_8446622_RNA45SN1_45S_50ntFlanks</td><td>10681</td><td>10705</td><td>25</td><td>TTCCGGCGGCGTCCGGTGAGCTCTC</td><td>201.7</td><td>-2.3</td><td>0.7</td><td>-3.2</td><td>1.3E-03</td><td>1.8E-03</td><td>BT20/MB468</td><td>CYTO</td></tr>
<tr><td>>hg38_21_+_8433172_8446622_RNA45SN1_45S_50ntFlanks@11570.11590.21|AGCCGACTTAGAACTGGTGCG</td><td>>hg38_21_+_8433172_8446622_RNA45SN1_45S_50ntFlanks</td><td>11570</td><td>11590</td><td>21</td><td>AGCCGACTTAGAACTGGTGCG</td><td>187.8</td><td>-4.2</td><td>1.3</td><td>-3.2</td><td>1.2E-03</td><td>1.8E-03</td><td>MB231/MB468</td><td>MP</td></tr>
<tr><td>>hg38_21_+_8433172_8446622_RNA45SN1_45S_50ntFlanks@12996.13038.43|TCGCTGCGATCTATTGAAAGTCAGCCCTCGACACAAGGGTTTG</td><td>>hg38_21_+_8433172_8446622_RNA45SN1_45S_50ntFlanks</td><td>12996</td><td>13038</td><td>43</td><td>TCGCTGCGATCTATTGAAAGTCAGCCCTCGACACAAGGGTTTG</td><td>794.3</td><td>5.0</td><td>1.6</td><td>3.2</td><td>1.5E-03</td><td>1.8E-03</td><td>MB231/MB468</td><td>MITO</td></tr>
<tr><td>>hg38_MT_+_1621_3279_RNR2_16S_50ntFlanks@622.665.44|TATAACTGAACTCCTCACACCCAATTGGACCAATCTATCACCCT</td><td>>hg38_MT_+_1621_3279_RNR2_16S_50ntFlanks</td><td>622</td><td>665</td><td>44</td><td>TATAACTGAACTCCTCACACCCAATTGGACCAATCTATCACCCT</td><td>49.8</td><td>9.4</td><td>2.8</td><td>3.4</td><td>6.9E-04</td><td>1.8E-03</td><td>BT20/MB468</td><td>MP</td></tr>
<tr><td>>hg38_21_+_8433172_8446622_RNA45SN1_45S_50ntFlanks@10686.10711.26|GCGGCGTCCGGTGAGCTCTCGCTGGC</td><td>>hg38_21_+_8433172_8446622_RNA45SN1_45S_50ntFlanks</td><td>10686</td><td>10711</td><td>26</td><td>GCGGCGTCCGGTGAGCTCTCGCTGGC</td><td>590.7</td><td>-3.6</td><td>1.1</td><td>-3.2</td><td>1.4E-03</td><td>1.8E-03</td><td>BT20/MB468</td><td>CYTO</td></tr>
<tr><td>>hg38_21_+_8433172_8446622_RNA45SN1_45S_50ntFlanks@9171.9197.27|CGGGTCGCGCCGTCGGGCCCGGGGGAG</td><td>>hg38_21_+_8433172_8446622_RNA45SN1_45S_50ntFlanks</td><td>9171</td><td>9197</td><td>27</td><td>CGGGTCGCGCCGTCGGGCCCGGGGGAG</td><td>850.2</td><td>-5.9</td><td>1.7</td><td>-3.4</td><td>6.9E-04</td><td>1.8E-03</td><td>BT20/MB468</td><td>MP</td></tr>
<tr><td>>hg38_21_+_8433172_8446622_RNA45SN1_45S_50ntFlanks@12417.12439.23|TCGATGTCGGCTCTTCCTATCAT</td><td>>hg38_21_+_8433172_8446622_RNA45SN1_45S_50ntFlanks</td><td>12417</td><td>12439</td><td>23</td><td>TCGATGTCGGCTCTTCCTATCAT</td><td>1163.7</td><td>2.0</td><td>0.6</td><td>3.2</td><td>1.4E-03</td><td>1.8E-03</td><td>MB231/MB468</td><td>CYTO</td></tr>
<tr><td>>hg38_21_+_8433172_8446622_RNA45SN1_45S_50ntFlanks@11671.11688.18|CCAGTGCTCTGAATGTCA</td><td>>hg38_21_+_8433172_8446622_RNA45SN1_45S_50ntFlanks</td><td>11671</td><td>11688</td><td>18</td><td>CCAGTGCTCTGAATGTCA</td><td>935.0</td><td>-3.8</td><td>1.2</td><td>-3.2</td><td>1.4E-03</td><td>1.8E-03</td><td>MB231/MB468</td><td>CYTO</td></tr>
<tr><td>>trnaMT_PheGAA_MT_+_577_647@42.74.33|TTTAGACGGGCTCACATCACCCCATAAACACCA</td><td>>trnaMT_PheGAA_MT_+_577_647</td><td>42</td><td>74</td><td>33</td><td>TTTAGACGGGCTCACATCACCCCATAAACACCA</td><td>394.2</td><td>2.6</td><td>0.8</td><td>3.2</td><td>1.2E-03</td><td>1.8E-03</td><td>MB231/MB468</td><td>MP</td></tr>
<tr><td>>hg38_21_+_8433172_8446622_RNA45SN1_45S_50ntFlanks@12203.12240.38|TAAGGCGAGCTCAGGGAGGACAGAAACCTCCCGTGGAG</td><td>>hg38_21_+_8433172_8446622_RNA45SN1_45S_50ntFlanks</td><td>12203</td><td>12240</td><td>38</td><td>TAAGGCGAGCTCAGGGAGGACAGAAACCTCCCGTGGAG</td><td>55.5</td><td>-9.7</td><td>3.0</td><td>-3.2</td><td>1.2E-03</td><td>1.8E-03</td><td>MB231/MB468</td><td>MP</td></tr>
<tr><td>>hg38_21_+_8433172_8446622_RNA45SN1_45S_50ntFlanks@4730.4757.28|CAAGAACGAAAGTCGGAGGTTCGAAGAC</td><td>>hg38_21_+_8433172_8446622_RNA45SN1_45S_50ntFlanks</td><td>4730</td><td>4757</td><td>28</td><td>CAAGAACGAAAGTCGGAGGTTCGAAGAC</td><td>118.7</td><td>-4.1</td><td>1.3</td><td>-3.2</td><td>1.2E-03</td><td>1.8E-03</td><td>MB231/MB468</td><td>MP</td></tr>
<tr><td>>trnaMT_PheGAA_MT_+_577_647@45.74.30|AGACGGGCTCACATCACCCCATAAACACCA</td><td>>trnaMT_PheGAA_MT_+_577_647</td><td>45</td><td>74</td><td>30</td><td>AGACGGGCTCACATCACCCCATAAACACCA</td><td>1646.1</td><td>-3.2</td><td>1.0</td><td>-3.2</td><td>1.5E-03</td><td>1.8E-03</td><td>MB231/MB468</td><td>MITO</td></tr>
<tr><td>>MI0001445|hsa-mir-423&WithFlank&17|+|30117073|30117178@58.79.22&[MIMAT0001340&hsa-miR-423-3p&offsets|-1|-2;m-100&17|+|30117131|30117153&offsets|-1|-2]|AAGCTCGGTCTGAGGCCCCTCA</td><td>>MI0001445|hsa-mir-423&WithFlank&17|+|30117073|30117178</td><td>58</td><td>79</td><td>22</td><td>AAGCTCGGTCTGAGGCCCCTCA</td><td>207.5</td><td>5.6</td><td>1.7</td><td>3.4</td><td>6.9E-04</td><td>1.8E-03</td><td>BT20/MB468</td><td>MP</td></tr>
<tr><td>>hg38_21_+_8433172_8446622_RNA45SN1_45S_50ntFlanks@11566.11585.20|TAGCAGCCGACTTAGAACTG</td><td>>hg38_21_+_8433172_8446622_RNA45SN1_45S_50ntFlanks</td><td>11566</td><td>11585</td><td>20</td><td>TAGCAGCCGACTTAGAACTG</td><td>380.5</td><td>2.4</td><td>0.7</td><td>3.2</td><td>1.5E-03</td><td>1.8E-03</td><td>MB231/MB468</td><td>NUC</td></tr>
<tr><td>>hg38_21_+_8433172_8446622_RNA45SN1_45S_50ntFlanks@8171.8214.44|AAGTCCTTCTGATCGAGGCCCAGCCCGTGGACGGTGTGAGGCCG</td><td>>hg38_21_+_8433172_8446622_RNA45SN1_45S_50ntFlanks</td><td>8171</td><td>8214</td><td>44</td><td>AAGTCCTTCTGATCGAGGCCCAGCCCGTGGACGGTGTGAGGCCG</td><td>783.9</td><td>-3.2</td><td>1.0</td><td>-3.2</td><td>1.4E-03</td><td>1.8E-03</td><td>BT20/MB468</td><td>CYTO</td></tr>
<tr><td>>hg38_21_+_8433172_8446622_RNA45SN1_45S_50ntFlanks@6651.6678.28|CGACTCTTAGCGGTGGATCACTCGGCTC</td><td>>hg38_21_+_8433172_8446622_RNA45SN1_45S_50ntFlanks</td><td>6651</td><td>6678</td><td>28</td><td>CGACTCTTAGCGGTGGATCACTCGGCTC</td><td>9935.2</td><td>2.8</td><td>0.9</td><td>3.2</td><td>1.5E-03</td><td>1.8E-03</td><td>MB231/MB468</td><td>NUC</td></tr>
<tr><td>>hg38_21_+_8433172_8446622_RNA45SN1_45S_50ntFlanks@11668.11706.39|TGCCCAGTGCTCTGAATGTCAAAGTGAAGAAATTCAATG</td><td>>hg38_21_+_8433172_8446622_RNA45SN1_45S_50ntFlanks</td><td>11668</td><td>11706</td><td>39</td><td>TGCCCAGTGCTCTGAATGTCAAAGTGAAGAAATTCAATG</td><td>3457.6</td><td>-3.0</td><td>0.8</td><td>-3.6</td><td>2.6E-04</td><td>1.8E-03</td><td>BT20/MB231</td><td>CYTO</td></tr>
<tr><td>>MI0001445|hsa-mir-423&WithFlank&17|+|30117073|30117178@58.80.23(+1A)&[MIMAT0001340&hsa-miR-423-3p&offsets|-1|-1(+1A);m-100&17|+|30117131|30117153&offsets|-1|-1(+1A)]|AAGCTCGGTCTGAGGCCCCTCAGA</td><td>>MI0001445|hsa-mir-423&WithFlank&17|+|30117073|30117178</td><td>58</td><td>80</td><td>23(+1A)</td><td>AAGCTCGGTCTGAGGCCCCTCAGA</td><td>118.8</td><td>5.6</td><td>1.6</td><td>3.4</td><td>6.9E-04</td><td>1.8E-03</td><td>BT20/MB468</td><td>MP</td></tr>
<tr><td>>hg38_21_+_8433172_8446622_RNA45SN1_45S_50ntFlanks@3854.3872.19|ACTGTGGTAATTCTAGAGC</td><td>>hg38_21_+_8433172_8446622_RNA45SN1_45S_50ntFlanks</td><td>3854</td><td>3872</td><td>19</td><td>ACTGTGGTAATTCTAGAGC</td><td>262.6</td><td>-5.2</td><td>1.6</td><td>-3.3</td><td>1.1E-03</td><td>1.8E-03</td><td>BT20/MB468</td><td>MITO</td></tr>
<tr><td>>hg38_21_+_8433172_8446622_RNA45SN1_45S_50ntFlanks@10781.10803.23|AGGTGAACAGCCTCTGGCATGTT</td><td>>hg38_21_+_8433172_8446622_RNA45SN1_45S_50ntFlanks</td><td>10781</td><td>10803</td><td>23</td><td>AGGTGAACAGCCTCTGGCATGTT</td><td>188.1</td><td>-4.0</td><td>1.2</td><td>-3.4</td><td>6.9E-04</td><td>1.8E-03</td><td>BT20/MB468</td><td>MP</td></tr>
<tr><td>>hg38_21_+_8433172_8446622_RNA45SN1_45S_50ntFlanks@6670.6699.30|ACTCGGCTCGTGCGTCGATGAAGAACGCAG</td><td>>hg38_21_+_8433172_8446622_RNA45SN1_45S_50ntFlanks</td><td>6670</td><td>6699</td><td>30</td><td>ACTCGGCTCGTGCGTCGATGAAGAACGCAG</td><td>799.7</td><td>-3.3</td><td>1.0</td><td>-3.4</td><td>6.9E-04</td><td>1.8E-03</td><td>BT20/MB468</td><td>MP</td></tr>
<tr><td>>hg38_21_+_8433172_8446622_RNA45SN1_45S_50ntFlanks@7993.8010.18|GTGGCGACCCGCTGAATT</td><td>>hg38_21_+_8433172_8446622_RNA45SN1_45S_50ntFlanks</td><td>7993</td><td>8010</td><td>18</td><td>GTGGCGACCCGCTGAATT</td><td>298.4</td><td>2.6</td><td>0.7</td><td>3.7</td><td>2.6E-04</td><td>1.8E-03</td><td>BT20/MB231</td><td>NUC</td></tr>
<tr><td>>hg38_21_+_8433172_8446622_RNA45SN1_45S_50ntFlanks@9507.9529.23|AACTATGCCTGGGCAGGGCGAAG</td><td>>hg38_21_+_8433172_8446622_RNA45SN1_45S_50ntFlanks</td><td>9507</td><td>9529</td><td>23</td><td>AACTATGCCTGGGCAGGGCGAAG</td><td>47.3</td><td>-8.6</td><td>2.5</td><td>-3.4</td><td>6.9E-04</td><td>1.8E-03</td><td>BT20/MB468</td><td>MP</td></tr>
<tr><td>>MI0000079|hsa-mir-23a&WithFlank&19|-|13836581|13836665@54.73.20&[MIMAT0000078&hsa-miR-23a-3p&offsets|+3|+2;m-31&19|-|13836595|13836615&offsets|+3|+2]|ACATTGCCAGGGATTTCCAA</td><td>>MI0000079|hsa-mir-23a&WithFlank&19|-|13836581|13836665</td><td>54</td><td>73</td><td>20</td><td>ACATTGCCAGGGATTTCCAA</td><td>359.5</td><td>11.7</td><td>3.6</td><td>3.2</td><td>1.3E-03</td><td>1.8E-03</td><td>MB231/MB468</td><td>MP</td></tr>
<tr><td>>hg38_21_+_8433172_8446622_RNA45SN1_45S_50ntFlanks@11319.11344.26|CCCACGTCTCGTCGCGCGCGCGTCCG</td><td>>hg38_21_+_8433172_8446622_RNA45SN1_45S_50ntFlanks</td><td>11319</td><td>11344</td><td>26</td><td>CCCACGTCTCGTCGCGCGCGCGTCCG</td><td>2293.6</td><td>-3.6</td><td>1.1</td><td>-3.3</td><td>1.1E-03</td><td>1.8E-03</td><td>BT20/MB468</td><td>MITO</td></tr>
<tr><td>>hg38_21_+_8433172_8446622_RNA45SN1_45S_50ntFlanks@11783.11819.37|CATGAATGGATGAACGAGATTCCCACTGTCCCTACCT</td><td>>hg38_21_+_8433172_8446622_RNA45SN1_45S_50ntFlanks</td><td>11783</td><td>11819</td><td>37</td><td>CATGAATGGATGAACGAGATTCCCACTGTCCCTACCT</td><td>222.3</td><td>-2.7</td><td>0.8</td><td>-3.3</td><td>1.1E-03</td><td>1.8E-03</td><td>BT20/MB468</td><td>MITO</td></tr>
<tr><td>>hg38_21_+_8433172_8446622_RNA45SN1_45S_50ntFlanks@9539.9570.32|ACTCTGGTGGAGGTCCGTAGCGGTCCTGACGT</td><td>>hg38_21_+_8433172_8446622_RNA45SN1_45S_50ntFlanks</td><td>9539</td><td>9570</td><td>32</td><td>ACTCTGGTGGAGGTCCGTAGCGGTCCTGACGT</td><td>265.1</td><td>-2.1</td><td>0.7</td><td>-3.2</td><td>1.5E-03</td><td>1.8E-03</td><td>MB231/MB468</td><td>NUC</td></tr>
<tr><td>>trnaMT_SerGCT_MT_+_12207_12265@13.41.29|AAGAACTGCTAACTCATGCCCCCATGTCT</td><td>>trnaMT_SerGCT_MT_+_12207_12265</td><td>13</td><td>41</td><td>29</td><td>AAGAACTGCTAACTCATGCCCCCATGTCT</td><td>238.9</td><td>4.0</td><td>1.2</td><td>3.4</td><td>7.0E-04</td><td>1.8E-03</td><td>BT20/MB468</td><td>MP</td></tr>
<tr><td>>hg38_21_+_8433172_8446622_RNA45SN1_45S_50ntFlanks@11641.11684.44|CGGCGGGTGTTGACGCGATGTGATTTCTGCCCAGTGCTCTGAAT</td><td>>hg38_21_+_8433172_8446622_RNA45SN1_45S_50ntFlanks</td><td>11641</td><td>11684</td><td>44</td><td>CGGCGGGTGTTGACGCGATGTGATTTCTGCCCAGTGCTCTGAAT</td><td>55.6</td><td>-9.7</td><td>3.0</td><td>-3.2</td><td>1.3E-03</td><td>1.8E-03</td><td>MB231/MB468</td><td>MP</td></tr>
<tr><td>>trnaMT_GluTTC_MT_-_14674_14742@39.57.19,trnalookalike8_GluTTC_5_-_93905172_93905240@39.57.19|ATTGGTCGTGGTTGTAGTC</td><td>>trnaMT_GluTTC_MT_-_14674_14742</td><td>39</td><td>57</td><td>19</td><td>ATTGGTCGTGGTTGTAGTC</td><td>339.3</td><td>2.7</td><td>0.8</td><td>3.2</td><td>1.3E-03</td><td>1.8E-03</td><td>MB231/MB468</td><td>MP</td></tr>
<tr><td>>MI0003763|hsa-mir-767&WithFlank&X|-|152393415|152393535@33.52.20&[MIMAT0003882&hsa-miR-767-5p&offsets|0|-3;m-788&X|-|152393481|152393503&offsets|0|-3]|TGCACCATGGTTGTCTGAGC</td><td>>MI0003763|hsa-mir-767&WithFlank&X|-|152393415|152393535</td><td>33</td><td>52</td><td>20</td><td>TGCACCATGGTTGTCTGAGC</td><td>37.6</td><td>8.6</td><td>2.3</td><td>3.7</td><td>2.4E-04</td><td>1.8E-03</td><td>BT20/MB231</td><td>MP</td></tr>
<tr><td>>MI0000073|hsa-mir-19a&WithFlank&13|+|91350885|91350978@55.78.24&[MIMAT0000073&hsa-miR-19a-3p&offsets|0|+1;m-201&13|+|91350939|91350961&offsets|0|+1]|TGTGCAAATCTATGCAAAACTGAT</td><td>>MI0000073|hsa-mir-19a&WithFlank&13|+|91350885|91350978</td><td>55</td><td>78</td><td>24</td><td>TGTGCAAATCTATGCAAAACTGAT</td><td>884.3</td><td>-5.2</td><td>1.4</td><td>-3.7</td><td>2.4E-04</td><td>1.8E-03</td><td>BT20/MB231</td><td>MP</td></tr>
<tr><td>>trna116_GluCTC_1_-_145399233_145399304@37.75.39,trna134_GluTTC_1_-_16861774_16861845@37.75.39,trna71_GluCTC_1_-_161439189_161439260@37.75.39,trna74_GluCTC_1_-_161431809_161431880@37.75.39,trna77_GluCTC_1_-_161424398_161424469@37.75.39,trna77_GluCTC_6_+_28949976_28950047@37.75.39,trna80_GluCTC_1_-_161417018_161417089@37.75.39,trna84_GluTTC_1_-_161391883_161391954@37.75.39,trna87_GluCTC_6_-_126101393_126101464@37.75.39|ACCGCCGCGGCCCGGGTTCGATTCCCGGTCAGGGAACCA</td><td>>trna116_GluCTC_1_-_145399233_145399304</td><td>37</td><td>75</td><td>39</td><td>ACCGCCGCGGCCCGGGTTCGATTCCCGGTCAGGGAACCA</td><td>19048.3</td><td>-4.4</td><td>1.2</td><td>-3.6</td><td>2.7E-04</td><td>1.8E-03</td><td>BT20/MB231</td><td>CYTO</td></tr>
<tr><td>>trna11_GluTTC_15_-_26327381_26327452@1.26.26,trna3_GluTTC_13_-_45492062_45492133@1.26.26|TCCCACATGGTCTAGCGGTTAGGATT</td><td>>trna11_GluTTC_15_-_26327381_26327452</td><td>1</td><td>26</td><td>26</td><td>TCCCACATGGTCTAGCGGTTAGGATT</td><td>575.2</td><td>3.1</td><td>0.9</td><td>3.6</td><td>2.7E-04</td><td>1.8E-03</td><td>BT20/MB231</td><td>CYTO</td></tr>
</table>

<table>
<tr><td>>hg38_21_+_8433172_8446622_RNA45SN1_45S_50ntFlanks@4040.4063.24|ACGACCCATTCGAACGTCTGCCCT</td><td>>hg38_21_+_8433172_8446622_RNA45SN1_45S_50ntFlanks</td><td>4040</td><td>4063</td><td>24</td><td>ACGACCCATTCGAACGTCTGCCCT</td><td>172.2</td><td>-2.2</td><td>0.7</td><td>-3.2</td><td>1.4E-03</td><td>1.8E-03</td><td>MB231/MB468</td><td>CYTO</td></tr>
<tr><td>>am_trna128_GlyGCC_6_-_27870686_27870756@1.34.34,trna18_GlyGCC_16_+_70822597_70822667@1.34.34,trna19_GlyGCC_16_+_70823410_70823480@1.34.34,trna19_GlyGCC_2_-_157257659_157257729@1.34.34,trna24_GlyGCC_16_-_70812942_70813012@1.34.34,trna25_GlyGCC_16_-_70812114_70812184@1.34.34,trna5_GlyGCC_17_+_8029064_8029134@1.34.34,trna68_GlyGCC_1_-_161493637_161493707@1.34.34|GCATTGGTGGTTCAGTGGTAGAATTCTCGCCTGC</td><td>>am_trna128_GlyGCC_6_-_27870686_27870756</td><td>1</td><td>34</td><td>34</td><td>GCATTGGTGGTTCAGTGGTAGAATTCTCGCCTGC</td><td>3005.7</td><td>-4.1</td><td>1.1</td><td>-3.7</td><td>2.4E-04</td><td>1.8E-03</td><td>BT20/MB231</td><td>MP</td></tr>
<tr><td>>am_MI0016746|hsa-mir-548o-2&WithFlank&20|+|38516557|38516638@51.71.21(+1U);MI0006402|hsa-mir-548o&WithFlank&7|-|102405736|102405861@93.113.21(+1U)&[MIMAT0005919&hsa-miR-548o-3p&offsets|0|-1(+1U);m-431&20|+|38516607|38516628&offsets|0|-1(+1U)];[MIMAT0005919_1&hsa-miR-548o-3p&offsets|0|-1(+1U);m-430&7|-|102405748|102405769&offsets|0|-1(+1U)]|CCAAAACTGCAGTTACTTTTGT</td><td>>am_MI0016746|hsa-mir-548o-2&WithFlank&20|+|38516557|38516638</td><td>51</td><td>71</td><td>21(+1U)</td><td>CCAAAACTGCAGTTACTTTTGT</td><td>137.7</td><td>4.9</td><td>1.5</td><td>3.4</td><td>7.0E-04</td><td>1.8E-03</td><td>BT20/MB468</td><td>MP</td></tr>
<tr><td>>hg38_21_+_8433172_8446622_RNA45SN1_45S_50ntFlanks@4570.4589.20|TGGAATAGGACCGCGGTTCT</td><td>>hg38_21_+_8433172_8446622_RNA45SN1_45S_50ntFlanks</td><td>4570</td><td>4589</td><td>20</td><td>TGGAATAGGACCGCGGTTCT</td><td>180.6</td><td>-4.1</td><td>1.3</td><td>-3.2</td><td>1.3E-03</td><td>1.8E-03</td><td>MB231/MB468</td><td>MP</td></tr>
<tr><td>>MI0003186|hsa-mir-502&WithFlank&X|+|50014592|50014689@58.79.22&[MIMAT0004775&hsa-miR-502-3p&offsets|0|0;m-236&X|+|50014649|50014669&offsets|0|+1]|AATGCACCTGGGCAAGGATTCA</td><td>>MI0003186|hsa-mir-502&WithFlank&X|+|50014592|50014689</td><td>58</td><td>79</td><td>22</td><td>AATGCACCTGGGCAAGGATTCA</td><td>465.5</td><td>4.4</td><td>1.3</td><td>3.4</td><td>7.0E-04</td><td>1.8E-03</td><td>BT20/MB468</td><td>MP</td></tr>
<tr><td>>hg38_21_+_8433172_8446622_RNA45SN1_45S_50ntFlanks@11212.11252.41|CGACTCTGGACGCGAGCCGGGCCCTTCCCGTGGATCGCCCC</td><td>>hg38_21_+_8433172_8446622_RNA45SN1_45S_50ntFlanks</td><td>11212</td><td>11252</td><td>41</td><td>CGACTCTGGACGCGAGCCGGGCCCTTCCCGTGGATCGCCCC</td><td>687.6</td><td>-2.5</td><td>0.8</td><td>-3.2</td><td>1.4E-03</td><td>1.8E-03</td><td>MB231/MB468</td><td>CYTO</td></tr>
<tr><td>>hg38_21_+_8433172_8446622_RNA45SN1_45S_50ntFlanks@12931.12956.26|GCGCTAAACCATTCGTAGACGACCTG</td><td>>hg38_21_+_8433172_8446622_RNA45SN1_45S_50ntFlanks</td><td>12931</td><td>12956</td><td>26</td><td>GCGCTAAACCATTCGTAGACGACCTG</td><td>577.7</td><td>-3.6</td><td>1.1</td><td>-3.2</td><td>1.4E-03</td><td>1.8E-03</td><td>MB231/MB468</td><td>CYTO</td></tr>
<tr><td>>hg38_21_+_8433172_8446622_RNA45SN1_45S_50ntFlanks@4898.4916.19|AAAGGAATTGACGGAAGGG</td><td>>hg38_21_+_8433172_8446622_RNA45SN1_45S_50ntFlanks</td><td>4898</td><td>4916</td><td>19</td><td>AAAGGAATTGACGGAAGGG</td><td>138.3</td><td>-5.5</td><td>1.6</td><td>-3.4</td><td>7.0E-04</td><td>1.8E-03</td><td>BT20/MB468</td><td>MP</td></tr>
<tr><td>>trnaMT_PheGAA_MT_+_577_647@38.64.27|AATGTTTAGACGGGCTCACATCACCCC</td><td>>trnaMT_PheGAA_MT_+_577_647</td><td>38</td><td>64</td><td>27</td><td>AATGTTTAGACGGGCTCACATCACCCC</td><td>166.8</td><td>8.2</td><td>2.4</td><td>3.4</td><td>7.0E-04</td><td>1.8E-03</td><td>BT20/MB468</td><td>MP</td></tr>
<tr><td>>hg38_21_+_8433172_8446622_RNA45SN1_45S_50ntFlanks@3842.3862.21|CCTACTTGGATAACTGTGGTA</td><td>>hg38_21_+_8433172_8446622_RNA45SN1_45S_50ntFlanks</td><td>3842</td><td>3862</td><td>21</td><td>CCTACTTGGATAACTGTGGTA</td><td>393.0</td><td>3.0</td><td>0.8</td><td>3.6</td><td>2.7E-04</td><td>1.8E-03</td><td>BT20/MB231</td><td>CYTO</td></tr>
<tr><td>>hg38_21_+_8433172_8446622_RNA45SN1_45S_50ntFlanks@11561.11602.42|GCGCCTAGCAGCCGACTTAGAACTGGTGCGGACCAGGGGAAT</td><td>>hg38_21_+_8433172_8446622_RNA45SN1_45S_50ntFlanks</td><td>11561</td><td>11602</td><td>42</td><td>GCGCCTAGCAGCCGACTTAGAACTGGTGCGGACCAGGGGAAT</td><td>503.5</td><td>-3.8</td><td>1.2</td><td>-3.2</td><td>1.4E-03</td><td>1.8E-03</td><td>MB231/MB468</td><td>CYTO</td></tr>
<tr><td>>hg38_21_+_8433172_8446622_RNA45SN1_45S_50ntFlanks@7992.8016.25|CGTGGCGACCCGCTGAATTTAAGCA</td><td>>hg38_21_+_8433172_8446622_RNA45SN1_45S_50ntFlanks</td><td>7992</td><td>8016</td><td>25</td><td>CGTGGCGACCCGCTGAATTTAAGCA</td><td>204.1</td><td>-4.0</td><td>1.2</td><td>-3.2</td><td>1.3E-03</td><td>1.8E-03</td><td>MB231/MB468</td><td>MP</td></tr>
<tr><td>>hg38_MT_+_1621_3279_RNR2_16S_50ntFlanks@1438.1477.40|TACGTGATCTGAGTTCAGACCGGAGTAATCCAGGTCGGTT</td><td>>hg38_MT_+_1621_3279_RNR2_16S_50ntFlanks</td><td>1438</td><td>1477</td><td>40</td><td>TACGTGATCTGAGTTCAGACCGGAGTAATCCAGGTCGGTT</td><td>232.0</td><td>-2.6</td><td>0.7</td><td>-3.6</td><td>2.7E-04</td><td>1.8E-03</td><td>BT20/MB231</td><td>CYTO</td></tr>
<tr><td>>hg38_21_+_8433172_8446622_RNA45SN1_45S_50ntFlanks@6645.6676.32|CTCGTACGACTCTTAGCGGTGGATCACTCGGC</td><td>>hg38_21_+_8433172_8446622_RNA45SN1_45S_50ntFlanks</td><td>6645</td><td>6676</td><td>32</td><td>CTCGTACGACTCTTAGCGGTGGATCACTCGGC</td><td>1581.5</td><td>2.9</td><td>0.9</td><td>3.2</td><td>1.5E-03</td><td>1.8E-03</td><td>MB231/MB468</td><td>MITO</td></tr>
<tr><td>>hg38_21_+_8433172_8446622_RNA45SN1_45S_50ntFlanks@9622.9656.35|CTAGTAGCTGGTTCCCTCCGAAGTTTCCCTCAGGA</td><td>>hg38_21_+_8433172_8446622_RNA45SN1_45S_50ntFlanks</td><td>9622</td><td>9656</td><td>35</td><td>CTAGTAGCTGGTTCCCTCCGAAGTTTCCCTCAGGA</td><td>167.0</td><td>-2.1</td><td>0.6</td><td>-3.3</td><td>1.2E-03</td><td>1.8E-03</td><td>BT20/MB468</td><td>MITO</td></tr>
<tr><td>>hg38_21_+_8433172_8446622_RNA45SN1_45S_50ntFlanks@10796.10818.23|GGCATGTTGGAACAATGTAGGTA</td><td>>hg38_21_+_8433172_8446622_RNA45SN1_45S_50ntFlanks</td><td>10796</td><td>10818</td><td>23</td><td>GGCATGTTGGAACAATGTAGGTA</td><td>301.9</td><td>-2.0</td><td>0.6</td><td>-3.3</td><td>1.2E-03</td><td>1.8E-03</td><td>BT20/MB468</td><td>MITO</td></tr>
<tr><td>>hg38_21_+_8433172_8446622_RNA45SN1_45S_50ntFlanks@5486.5506.21|GCGCTGAGAAGACGGTCGAAC</td><td>>hg38_21_+_8433172_8446622_RNA45SN1_45S_50ntFlanks</td><td>5486</td><td>5506</td><td>21</td><td>GCGCTGAGAAGACGGTCGAAC</td><td>264.2</td><td>3.3</td><td>0.9</td><td>3.6</td><td>2.7E-04</td><td>1.8E-03</td><td>BT20/MB231</td><td>CYTO</td></tr>
<tr><td>>hg38_21_+_8433172_8446622_RNA45SN1_45S_50ntFlanks@7974.8017.44|ACGCGACCTCAGATCAGACGTGGCGACCCGCTGAATTTAAGCAT</td><td>>hg38_21_+_8433172_8446622_RNA45SN1_45S_50ntFlanks</td><td>7974</td><td>8017</td><td>44</td><td>ACGCGACCTCAGATCAGACGTGGCGACCCGCTGAATTTAAGCAT</td><td>1230.1</td><td>3.3</td><td>1.0</td><td>3.2</td><td>1.5E-03</td><td>1.8E-03</td><td>MB231/MB468</td><td>MITO</td></tr>
<tr><td>>hg38_21_+_8433172_8446622_RNA45SN1_45S_50ntFlanks@9930.9957.28|ATATAGACAGCAGGACGGTGGCCATGGA</td><td>>hg38_21_+_8433172_8446622_RNA45SN1_45S_50ntFlanks</td><td>9930</td><td>9957</td><td>28</td><td>ATATAGACAGCAGGACGGTGGCCATGGA</td><td>328.8</td><td>-3.2</td><td>1.0</td><td>-3.3</td><td>1.2E-03</td><td>1.8E-03</td><td>BT20/MB468</td><td>MITO</td></tr>
<tr><td>>hg38_21_+_8433172_8446622_RNA45SN1_45S_50ntFlanks@3839.3859.21|TCTCCTACTTGGATAACTGTG</td><td>>hg38_21_+_8433172_8446622_RNA45SN1_45S_50ntFlanks</td><td>3839</td><td>3859</td><td>21</td><td>TCTCCTACTTGGATAACTGTG</td><td>291.3</td><td>2.7</td><td>0.9</td><td>3.2</td><td>1.4E-03</td><td>1.8E-03</td><td>MB231/MB468</td><td>CYTO</td></tr>
<tr><td>>hg38_21_+_8433172_8446622_RNA45SN1_45S_50ntFlanks@10830.10853.24|CAAGCCGGATCCGTAACTTCGGGA</td><td>>hg38_21_+_8433172_8446622_RNA45SN1_45S_50ntFlanks</td><td>10830</td><td>10853</td><td>24</td><td>CAAGCCGGATCCGTAACTTCGGGA</td><td>259.3</td><td>-5.5</td><td>1.6</td><td>-3.4</td><td>7.0E-04</td><td>1.8E-03</td><td>BT20/MB468</td><td>MP</td></tr>
<tr><td>>hg38_21_+_8433172_8446622_RNA45SN1_45S_50ntFlanks@12916.12933.18|TTCGTGGGGAACCTGGCG</td><td>>hg38_21_+_8433172_8446622_RNA45SN1_45S_50ntFlanks</td><td>12916</td><td>12933</td><td>18</td><td>TTCGTGGGGAACCTGGCG</td><td>208.7</td><td>-2.3</td><td>0.7</td><td>-3.2</td><td>1.5E-03</td><td>1.8E-03</td><td>MB231/MB468</td><td>NUC</td></tr>
<tr><td>>trnaMT_PheGAA_MT_+_577_647@35.60.26|GAAAATGTTTAGACGGGCTCACATCA</td><td>>trnaMT_PheGAA_MT_+_577_647</td><td>35</td><td>60</td><td>26</td><td>GAAAATGTTTAGACGGGCTCACATCA</td><td>104.1</td><td>3.6</td><td>1.1</td><td>3.2</td><td>1.3E-03</td><td>1.8E-03</td><td>MB231/MB468</td><td>MP</td></tr>
<tr><td>>am_trna146_GlnCTG_6_-_27515531_27515602@1.29.29,trna1_GlnCTG_6_+_18836402_18836473@1.29.29,trna3_GlnCTG_17_+_8023070_8023141@1.29.29,trna42_GlnCTG_6_+_27263212_27263283@1.29.29,trna49_GlnCTG_6_+_27487308_27487379@1.29.29,trna7_GlnCTG_15_-_66161400_66161471@1.29.29,trna99_GlnCTG_6_-_28909378_28909449@1.29.29|GGTTCCATGGTGTAATGGTTAGCACTCTG</td><td>>am_trna146_GlnCTG_6_-_27515531_27515602</td><td>1</td><td>29</td><td>29</td><td>GGTTCCATGGTGTAATGGTTAGCACTCTG</td><td>16175.4</td><td>-5.3</td><td>1.7</td><td>-3.2</td><td>1.4E-03</td><td>1.8E-03</td><td>BT20/MB468</td><td>CYTO</td></tr>
<tr><td>>MI0006349|hsa-mir-1287&WithFlank&10|-|98395212|98395313@22.43.22&[MIMAT0005878&hsa-miR-1287-5p&offsets|0|0;m-244&10|-|98395272|98395292&offsets|0|+1]|TGCTGGATCAGTGGTTCGAGTC</td><td>>MI0006349|hsa-mir-1287&WithFlank&10|-|98395212|98395313</td><td>22</td><td>43</td><td>22</td><td>TGCTGGATCAGTGGTTCGAGTC</td><td>307.0</td><td>4.0</td><td>1.3</td><td>3.2</td><td>1.3E-03</td><td>1.8E-03</td><td>MB231/MB468</td><td>MP</td></tr>
<tr><td>>hg38_21_+_8433172_8446622_RNA45SN1_45S_50ntFlanks@9554.9581.28|CGTAGCGGTCCTGACGTGCAAATCGGTC</td><td>>hg38_21_+_8433172_8446622_RNA45SN1_45S_50ntFlanks</td><td>9554</td><td>9581</td><td>28</td><td>CGTAGCGGTCCTGACGTGCAAATCGGTC</td><td>305.7</td><td>1.2</td><td>0.4</td><td>3.2</td><td>1.2E-03</td><td>1.8E-03</td><td>BT20/MB468</td><td>MITO</td></tr>
<tr><td>>hg38_21_+_8433172_8446622_RNA45SN1_45S_50ntFlanks@11914.11943.30|AGAGACATGAGAGGTGTAGAATAAGTGGGA</td><td>>hg38_21_+_8433172_8446622_RNA45SN1_45S_50ntFlanks</td><td>11914</td><td>11943</td><td>30</td><td>AGAGACATGAGAGGTGTAGAATAAGTGGGA</td><td>54.8</td><td>6.4</td><td>1.9</td><td>3.4</td><td>7.0E-04</td><td>1.8E-03</td><td>BT20/MB468</td><td>MP</td></tr>
<tr><td>>MI0001445|hsa-mir-423&WithFlank&17|+|30117073|30117178@23.42.20(+1A)&[MIMAT0004748&hsa-miR-423-5p&offsets|0|-3(+1A);m-141&17|+|30117095|30117117&offsets|0|-3(+1A)]|TGAGGGGCAGAGAGCGAGACA</td><td>>MI0001445|hsa-mir-423&WithFlank&17|+|30117073|30117178</td><td>23</td><td>42</td><td>20(+1A)</td><td>TGAGGGGCAGAGAGCGAGACA</td><td>274.4</td><td>4.1</td><td>1.3</td><td>3.2</td><td>1.3E-03</td><td>1.8E-03</td><td>MB231/MB468</td><td>MP</td></tr>
<tr><td>>hg38_21_+_8433172_8446622_RNA45SN1_45S_50ntFlanks@12512.12538.27|GTGAGACAGGTTAGTTTTACCCTACTG</td><td>>hg38_21_+_8433172_8446622_RNA45SN1_45S_50ntFlanks</td><td>12512</td><td>12538</td><td>27</td><td>GTGAGACAGGTTAGTTTTACCCTACTG</td><td>71.5</td><td>-5.3</td><td>1.5</td><td>-3.4</td><td>7.0E-04</td><td>1.8E-03</td><td>BT20/MB468</td><td>MP</td></tr>
<tr><td>>hg38_21_+_8433172_8446622_RNA45SN1_45S_50ntFlanks@11848.11873.26|ACGGGCTTGGCGGAATCAGCGGGGAA</td><td>>hg38_21_+_8433172_8446622_RNA45SN1_45S_50ntFlanks</td><td>11848</td><td>11873</td><td>26</td><td>ACGGGCTTGGCGGAATCAGCGGGGAA</td><td>261.7</td><td>-4.2</td><td>1.3</td><td>-3.4</td><td>7.0E-04</td><td>1.8E-03</td><td>BT20/MB468</td><td>MP</td></tr>
<tr><td>>hg38_21_+_8433172_8446622_RNA45SN1_45S_50ntFlanks@12405.12443.39|CTTTTTGATCCTTCGATGTCGGCTCTTCCTATCATTGTG</td><td>>hg38_21_+_8433172_8446622_RNA45SN1_45S_50ntFlanks</td><td>12405</td><td>12443</td><td>39</td><td>CTTTTTGATCCTTCGATGTCGGCTCTTCCTATCATTGTG</td><td>5732.3</td><td>-3.3</td><td>1.0</td><td>-3.2</td><td>1.4E-03</td><td>1.8E-03</td><td>MB231/MB468</td><td>CYTO</td></tr>
<tr><td>>hg38_21_+_8433172_8446622_RNA45SN1_45S_50ntFlanks@11319.11343.25|CCCACGTCTCGTCGCGCGCGCGTCC</td><td>>hg38_21_+_8433172_8446622_RNA45SN1_45S_50ntFlanks</td><td>11319</td><td>11343</td><td>25</td><td>CCCACGTCTCGTCGCGCGCGCGTCC</td><td>226.0</td><td>-3.3</td><td>1.0</td><td>-3.2</td><td>1.4E-03</td><td>1.8E-03</td><td>BT20/MB468</td><td>CYTO</td></tr>
<tr><td>>trna111_HisGTG_1_-_147774845_147774916@2.30.29,trna118_HisGTG_1_-_145396881_145396952@2.30.29,trna16_HisGTG_1_+_146544773_146544844@2.30.29,trna1_HisGTG_15_+_45493349_45493420@2.30.29,trna21_HisGTG_1_+_147753471_147753542@2.30.29,trna33_HisGTG_6_+_27125906_27125977@2.30.29,trna7_HisGTG_9_-_14433938_14434009@2.30.29,trna8_HisGTG_15_-_45492611_45492682@2.30.29,trna9_HisGTG_15_-_45490804_45490875@2.30.29|CCGTGATCGTATAGTGGTTAGTACTCTGC</td><td>>trna111_HisGTG_1_-_147774845_147774916</td><td>2</td><td>30</td><td>29</td><td>CCGTGATCGTATAGTGGTTAGTACTCTGC</td><td>116.7</td><td>2.4</td><td>0.7</td><td>3.2</td><td>1.3E-03</td><td>1.8E-03</td><td>MB231/MB468</td><td>MP</td></tr>
<tr><td>>hg38_21_+_8433172_8446622_RNA45SN1_45S_50ntFlanks@11575.11613.39|ACTTAGAACTGGTGCGGACCAGGGGAATCCGACTGTTTA</td><td>>hg38_21_+_8433172_8446622_RNA45SN1_45S_50ntFlanks</td><td>11575</td><td>11613</td><td>39</td><td>ACTTAGAACTGGTGCGGACCAGGGGAATCCGACTGTTTA</td><td>1941.0</td><td>3.3</td><td>0.9</td><td>3.6</td><td>3.1E-04</td><td>1.8E-03</td><td>BT20/MB231</td><td>MITO</td></tr>
<tr><td>>hg38_21_+_8433172_8446622_RNA45SN1_45S_50ntFlanks@10822.10865.44|GAAGTCGGCAAGCCGGATCCGTAACTTCGGGATAAGGATTGGCT</td><td>>hg38_21_+_8433172_8446622_RNA45SN1_45S_50ntFlanks</td><td>10822</td><td>10865</td><td>44</td><td>GAAGTCGGCAAGCCGGATCCGTAACTTCGGGATAAGGATTGGCT</td><td>571.1</td><td>-2.8</td><td>0.9</td><td>-3.2</td><td>1.4E-03</td><td>1.8E-03</td><td>MB231/MB468</td><td>CYTO</td></tr>
<tr><td>>hg38_21_+_8433172_8446622_RNA45SN1_45S_50ntFlanks@5056.5097.42|GAGCGATTTGTCTGGTTAATTCCGATAACGAACGAGACTCTG</td><td>>hg38_21_+_8433172_8446622_RNA45SN1_45S_50ntFlanks</td><td>5056</td><td>5097</td><td>42</td><td>GAGCGATTTGTCTGGTTAATTCCGATAACGAACGAGACTCTG</td><td>54.8</td><td>-9.6</td><td>3.0</td><td>-3.2</td><td>1.3E-03</td><td>1.8E-03</td><td>MB231/MB468</td><td>MP</td></tr>
<tr><td>>hg38_21_+_8433172_8446622_RNA45SN1_45S_50ntFlanks@10663.10706.44|CCTTGGAAAGCGTCGCGGTTCCGGCGGCGTCCGGTGAGCTCTCG</td><td>>hg38_21_+_8433172_8446622_RNA45SN1_45S_50ntFlanks</td><td>10663</td><td>10706</td><td>44</td><td>CCTTGGAAAGCGTCGCGGTTCCGGCGGCGTCCGGTGAGCTCTCG</td><td>248.9</td><td>-4.9</td><td>1.5</td><td>-3.2</td><td>1.4E-03</td><td>1.8E-03</td><td>BT20/MB468</td><td>CYTO</td></tr>
<tr><td>>hg38_21_+_8433172_8446622_RNA45SN1_45S_50ntFlanks@475.523.49|CTTCGTGATCGATGTGGTGACGTCGTGCTCTCCCGGGCCGGGTCCGAGC</td><td>>hg38_21_+_8433172_8446622_RNA45SN1_45S_50ntFlanks</td><td>475</td><td>523</td><td>49</td><td>CTTCGTGATCGATGTGGTGACGTCGTGCTCTCCCGGGCCGGGTCCGAGC</td><td>387.2</td><td>6.1</td><td>1.9</td><td>3.2</td><td>1.5E-03</td><td>1.8E-03</td><td>MB231/MB468</td><td>MITO</td></tr>
<tr><td>>hg38_21_+_8433172_8446622_RNA45SN1_45S_50ntFlanks@3850.3874.25|GATAACTGTGGTAATTCTAGAGCTA</td><td>>hg38_21_+_8433172_8446622_RNA45SN1_45S_50ntFlanks</td><td>3850</td><td>3874</td><td>25</td><td>GATAACTGTGGTAATTCTAGAGCTA</td><td>232.7</td><td>-5.3</td><td>1.6</td><td>-3.4</td><td>7.1E-04</td><td>1.8E-03</td><td>BT20/MB468</td><td>MP</td></tr>
<tr><td>>hg38_21_+_8433172_8446622_RNA45SN1_45S_50ntFlanks@9564.9596.33|CTGACGTGCAAATCGGTCGTCCGACCTGGGTAT</td><td>>hg38_21_+_8433172_8446622_RNA45SN1_45S_50ntFlanks</td><td>9564</td><td>9596</td><td>33</td><td>CTGACGTGCAAATCGGTCGTCCGACCTGGGTAT</td><td>5359.0</td><td>-5.5</td><td>1.6</td><td>-3.4</td><td>7.1E-04</td><td>1.8E-03</td><td>BT20/MB468</td><td>MP</td></tr>
<tr><td>>MI0000746|hsa-mir-99b&WithFlank&19|+|51692606|51692687@51.72.22(+1C)&[MIMAT0004678&hsa-miR-99b-3p&offsets|0|0(+1C);m-151&19|+|51692656|51692678&offsets|0|-1(+1C)]|CAAGCTCGTGTCTGTGGGTCCGC</td><td>>MI0000746|hsa-mir-99b&WithFlank&19|+|51692606|51692687</td><td>51</td><td>72</td><td>22(+1C)</td><td>CAAGCTCGTGTCTGTGGGTCCGC</td><td>342.7</td><td>2.9</td><td>0.8</td><td>3.6</td><td>2.7E-04</td><td>1.8E-03</td><td>BT20/MB231</td><td>CYTO</td></tr>
<tr><td>>am_trnaMT_ValTAC_MT_+_1602_1670@28.62.35|AACTTACACTTAGGAGATTTCAACTTAACTTGACC</td><td>>am_trnaMT_ValTAC_MT_+_1602_1670</td><td>28</td><td>62</td><td>35</td><td>AACTTACACTTAGGAGATTTCAACTTAACTTGACC</td><td>115.1</td><td>4.8</td><td>1.5</td><td>3.2</td><td>1.5E-03</td><td>1.8E-03</td><td>MB231/MB468</td><td>MITO</td></tr>
<tr><td>>hg38_21_+_8433172_8446622_RNA45SN1_45S_50ntFlanks@12259.12286.28|CTTGATCTTGATTTTCAGTACGAATACA</td><td>>hg38_21_+_8433172_8446622_RNA45SN1_45S_50ntFlanks</td><td>12259</td><td>12286</td><td>28</td><td>CTTGATCTTGATTTTCAGTACGAATACA</td><td>64.3</td><td>-4.1</td><td>1.3</td><td>-3.2</td><td>1.3E-03</td><td>1.8E-03</td><td>MB231/MB468</td><td>MP</td></tr>
<tr><td>>MI0000086|hsa-mir-28&WithFlank&3|+|188688775|188688872@62.82.21&[MIMAT0004502&hsa-miR-28-3p&offsets|+2|+1;m-28&3|+|188688834|188688855&offsets|+2|+1]|CTAGATTGTGAGCTCCTGGAG</td><td>>MI0000086|hsa-mir-28&WithFlank&3|+|188688775|188688872</td><td>62</td><td>82</td><td>21</td><td>CTAGATTGTGAGCTCCTGGAG</td><td>56.4</td><td>5.8</td><td>1.8</td><td>3.2</td><td>1.3E-03</td><td>1.8E-03</td><td>MB231/MB468</td><td>MP</td></tr>
<tr><td>>hg38_21_+_8433172_8446622_RNA45SN1_45S_50ntFlanks@10552.10581.30|GGTAACGCGACCGATCCCGGAGAAGCCGGC</td><td>>hg38_21_+_8433172_8446622_RNA45SN1_45S_50ntFlanks</td><td>10552</td><td>10581</td><td>30</td><td>GGTAACGCGACCGATCCCGGAGAAGCCGGC</td><td>105.6</td><td>10.1</td><td>2.7</td><td>3.7</td><td>2.4E-04</td><td>1.8E-03</td><td>BT20/MB231</td><td>MP</td></tr>
<tr><td>>hg38_21_+_8433172_8446622_RNA45SN1_45S_50ntFlanks@11573.11602.30|CGACTTAGAACTGGTGCGGACCAGGGGAAT</td><td>>hg38_21_+_8433172_8446622_RNA45SN1_45S_50ntFlanks</td><td>11573</td><td>11602</td><td>30</td><td>CGACTTAGAACTGGTGCGGACCAGGGGAAT</td><td>662.3</td><td>-2.1</td><td>0.6</td><td>-3.2</td><td>1.4E-03</td><td>1.8E-03</td><td>MB231/MB468</td><td>CYTO</td></tr>
</table>

<table>
<tr><td>>hg38_MT_+_1621_3279_RNR2_16S_50ntFlanks@51.84.34|GCTAAACCTAGCCCCAAACCCACTCCACCTTACT</td><td>>hg38_MT_+_1621_3279_RNR2_16S_50ntFlanks</td><td>51</td><td>84</td><td>34</td><td>GCTAAACCTAGCCCCAAACCCACTCCACCTTACT</td><td>2901.3</td><td>3.2</td><td>1.0</td><td>3.2</td><td>1.5E-03</td><td>1.8E-03</td><td>MB231/MB468</td><td>MITO</td></tr>
<tr><td>>MI0000273|hsa-mir-183&WithFlank&7|-|129774899|129775020@34.54.21&[MIMAT0000261&hsa-miR-183-5p&offsets|+1|0;m-41&7|-|129774966|129774987&offsets|0|-1]|ATGGCACTGGTAGAATTCACT</td><td>>MI0000273|hsa-mir-183&WithFlank&7|-|129774899|129775020</td><td>34</td><td>54</td><td>21</td><td>ATGGCACTGGTAGAATTCACT</td><td>506.4</td><td>3.0</td><td>0.9</td><td>3.2</td><td>1.5E-03</td><td>1.8E-03</td><td>MB231/MB468</td><td>MITO</td></tr>
<tr><td>>trna20_GluTTC_2_-_131094701_131094772@1.36.36,trna5_GluTTC_13_-_41634874_41634945@1.36.36|TCCCATATGGTCTAGCGGTTAGGATTCCTGGTTTTC</td><td>>trna20_GluTTC_2_-_131094701_131094772</td><td>1</td><td>36</td><td>36</td><td>TCCCATATGGTCTAGCGGTTAGGATTCCTGGTTTTC</td><td>941.6</td><td>-4.0</td><td>1.1</td><td>-3.6</td><td>2.7E-04</td><td>1.8E-03</td><td>BT20/MB231</td><td>CYTO</td></tr>
<tr><td>>MI0000446|hsa-mir-125b-1&WithFlank&11|-|122099751|122099850@61.82.22&[MIMAT0004592&hsa-miR-125b-1-3p&offsets|0|0;m-485&11|-|122099769|122099790&offsets|0|0]|ACGGGTTAGGCTCTTGGGAGCT</td><td>>MI0000446|hsa-mir-125b-1&WithFlank&11|-|122099751|122099850</td><td>61</td><td>82</td><td>22</td><td>ACGGGTTAGGCTCTTGGGAGCT</td><td>462.8</td><td>-5.7</td><td>1.6</td><td>-3.6</td><td>2.6E-04</td><td>1.8E-03</td><td>BT20/MB231</td><td>NUC</td></tr>
<tr><td>>hg38_21_+_8433172_8446622_RNA45SN1_45S_50ntFlanks@4047.4073.27|ATTCGAACGTCTGCCCTATCAACTTTC</td><td>>hg38_21_+_8433172_8446622_RNA45SN1_45S_50ntFlanks</td><td>4047</td><td>4073</td><td>27</td><td>ATTCGAACGTCTGCCCTATCAACTTTC</td><td>189.3</td><td>-4.9</td><td>1.5</td><td>-3.2</td><td>1.2E-03</td><td>1.8E-03</td><td>BT20/MB468</td><td>MITO</td></tr>
<tr><td>>hg38_21_+_8433172_8446622_RNA45SN1_45S_50ntFlanks@7976.7993.18|GCGACCTCAGATCAGACG</td><td>>hg38_21_+_8433172_8446622_RNA45SN1_45S_50ntFlanks</td><td>7976</td><td>7993</td><td>18</td><td>GCGACCTCAGATCAGACG</td><td>151.2</td><td>5.6</td><td>1.6</td><td>3.4</td><td>7.1E-04</td><td>1.8E-03</td><td>BT20/MB468</td><td>MP</td></tr>
<tr><td>>hg38_21_+_8433172_8446622_RNA45SN1_45S_50ntFlanks@4353.4391.39|TATTAAAGTTGCTGCAGTTAAAAAGCTCGTAGTTGGATC</td><td>>hg38_21_+_8433172_8446622_RNA45SN1_45S_50ntFlanks</td><td>4353</td><td>4391</td><td>39</td><td>TATTAAAGTTGCTGCAGTTAAAAAGCTCGTAGTTGGATC</td><td>102.5</td><td>-4.5</td><td>1.4</td><td>-3.2</td><td>1.2E-03</td><td>1.8E-03</td><td>BT20/MB468</td><td>MITO</td></tr>
<tr><td>>hg38_MT_+_1621_3279_RNR2_16S_50ntFlanks@292.310.19|AGACGAGCTACCTAAGAAC</td><td>>hg38_MT_+_1621_3279_RNR2_16S_50ntFlanks</td><td>292</td><td>310</td><td>19</td><td>AGACGAGCTACCTAAGAAC</td><td>606.4</td><td>4.3</td><td>1.3</td><td>3.4</td><td>7.1E-04</td><td>1.8E-03</td><td>BT20/MB468</td><td>MP</td></tr>
<tr><td>>hg38_21_+_8433172_8446622_RNA45SN1_45S_50ntFlanks@6644.6668.25|CCTCGTACGACTCTTAGCGGTGGAT</td><td>>hg38_21_+_8433172_8446622_RNA45SN1_45S_50ntFlanks</td><td>6644</td><td>6668</td><td>25</td><td>CCTCGTACGACTCTTAGCGGTGGAT</td><td>94.3</td><td>2.9</td><td>0.8</td><td>3.4</td><td>7.1E-04</td><td>1.8E-03</td><td>BT20/MB468</td><td>MP</td></tr>
<tr><td>>trna111_HisGTG_1_-_147774845_147774916@-1T.20.21,trna118_HisGTG_1_-_145396881_145396952@-1T.20.21,trna16_HisGTG_1_+_146544773_146544844@-1T.20.21,trna1_HisGTG_15_+_45493349_45493420@-1T.20.21,trna21_HisGTG_1_+_147753471_147753542@-1T.20.21,trna33_HisGTG_6_+_27125906_27125977@-1T.20.21,trna7_HisGTG_9_-_14433938_14434009@-1T.20.21,trna8_HisGTG_15_-_45492611_45492682@-1T.20.21,trna9_HisGTG_15_-_45490804_45490875@-1T.20.21|TGCCGTGATCGTATAGTGGTT</td><td>>trna111_HisGTG_1_-_147774845_147774916</td><td>-1T</td><td>20</td><td>21</td><td>TGCCGTGATCGTATAGTGGTT</td><td>213.8</td><td>-3.9</td><td>1.1</td><td>-3.6</td><td>3.1E-04</td><td>1.8E-03</td><td>BT20/MB231</td><td>MITO</td></tr>
<tr><td>>hg38_21_+_8433172_8446622_RNA45SN1_45S_50ntFlanks@11552.11585.34|CCTCGGCCGGCGCCTAGCAGCCGACTTAGAACTG</td><td>>hg38_21_+_8433172_8446622_RNA45SN1_45S_50ntFlanks</td><td>11552</td><td>11585</td><td>34</td><td>CCTCGGCCGGCGCCTAGCAGCCGACTTAGAACTG</td><td>97.3</td><td>-5.0</td><td>1.6</td><td>-3.2</td><td>1.3E-03</td><td>1.8E-03</td><td>MB231/MB468</td><td>MP</td></tr>
<tr><td>>hg38_21_+_8433172_8446622_RNA45SN1_45S_50ntFlanks@475.512.38|CTTCGTGATCGATGTGGTGACGTCGTGCTCTCCCGGGC</td><td>>hg38_21_+_8433172_8446622_RNA45SN1_45S_50ntFlanks</td><td>475</td><td>512</td><td>38</td><td>CTTCGTGATCGATGTGGTGACGTCGTGCTCTCCCGGGC</td><td>170.2</td><td>5.8</td><td>1.8</td><td>3.2</td><td>1.5E-03</td><td>1.8E-03</td><td>MB231/MB468</td><td>MITO</td></tr>
<tr><td>>hg38_21_+_8433172_8446622_RNA45SN1_45S_50ntFlanks@6732.6774.43|ACATTGATCATCGACACTTCGAACGCACTTGCGGCCCCGGGTT</td><td>>hg38_21_+_8433172_8446622_RNA45SN1_45S_50ntFlanks</td><td>6732</td><td>6774</td><td>43</td><td>ACATTGATCATCGACACTTCGAACGCACTTGCGGCCCCGGGTT</td><td>134.1</td><td>2.3</td><td>0.7</td><td>3.2</td><td>1.5E-03</td><td>1.8E-03</td><td>BT20/MB468</td><td>NUC</td></tr>
<tr><td>>hg38_MT_+_1621_3279_RNR2_16S_50ntFlanks@1389.1413.25|CGATGGTGCAGCCGCTATTAAAGGT</td><td>>hg38_MT_+_1621_3279_RNR2_16S_50ntFlanks</td><td>1389</td><td>1413</td><td>25</td><td>CGATGGTGCAGCCGCTATTAAAGGT</td><td>45.5</td><td>8.7</td><td>2.7</td><td>3.2</td><td>1.3E-03</td><td>1.8E-03</td><td>MB231/MB468</td><td>MP</td></tr>
<tr><td>>hg38_21_+_8433172_8446622_RNA45SN1_45S_50ntFlanks@12158.12181.24|GTTTGACTGGGGCGGTACACCTGT</td><td>>hg38_21_+_8433172_8446622_RNA45SN1_45S_50ntFlanks</td><td>12158</td><td>12181</td><td>24</td><td>GTTTGACTGGGGCGGTACACCTGT</td><td>324.2</td><td>-5.4</td><td>1.6</td><td>-3.4</td><td>7.1E-04</td><td>1.8E-03</td><td>BT20/MB468</td><td>MP</td></tr>
<tr><td>>hg38_21_+_8433172_8446622_RNA45SN1_45S_50ntFlanks@11705.11736.32|TGAAGCGCGGGTAAACGGCGGGAGTAACTATG</td><td>>hg38_21_+_8433172_8446622_RNA45SN1_45S_50ntFlanks</td><td>11705</td><td>11736</td><td>32</td><td>TGAAGCGCGGGTAAACGGCGGGAGTAACTATG</td><td>135.7</td><td>-5.2</td><td>1.6</td><td>-3.2</td><td>1.3E-03</td><td>1.8E-03</td><td>MB231/MB468</td><td>MP</td></tr>
<tr><td>>hg38_21_+_8433172_8446622_RNA45SN1_45S_50ntFlanks@4498.4519.22|AAGCGTTTACTTTGAAAAAATT</td><td>>hg38_21_+_8433172_8446622_RNA45SN1_45S_50ntFlanks</td><td>4498</td><td>4519</td><td>22</td><td>AAGCGTTTACTTTGAAAAAATT</td><td>321.4</td><td>2.9</td><td>0.8</td><td>3.6</td><td>2.7E-04</td><td>1.8E-03</td><td>BT20/MB231</td><td>CYTO</td></tr>
<tr><td>>trna10_AspGTC_12_-_125424193_125424264@37.71.35,trna12_AspGTC_12_-_125411891_125411962@37.71.35,trna144_AspGTC_6_-_27551236_27551307@37.71.35,trna38_AspGTC_17_-_8125556_8125627@37.71.35,trna45_AspGTC_6_+_27447453_27447524@37.71.35,trna48_AspGTC_6_+_27471523_27471594@37.71.35,trna4_AspGTC_12_+_96429799_96429870@37.71.35,trna69_AspGTC_1_-_161440205_161440276@37.71.35,trna72_AspGTC_1_-_161432824_161432895@37.71.35,trna75_AspGTC_1_-_161425414_161425485@37.71.35,trna78_AspGTC_1_-_161418033_161418104@37.71.35,trna81_AspGTC_1_-_161410615_161410686@37.71.35|ACGCGGGAGACCGGGGTTCGATTCCCCGACGGGGA</td><td>>trna10_AspGTC_12_-_125424193_125424264</td><td>37</td><td>71</td><td>35</td><td>ACGCGGGAGACCGGGGTTCGATTCCCCGACGGGGA</td><td>501.3</td><td>-3.4</td><td>0.9</td><td>-3.6</td><td>2.7E-04</td><td>1.8E-03</td><td>BT20/MB231</td><td>CYTO</td></tr>
<tr><td>>MI0000286|hsa-mir-210&WithFlank&11|-|568083|568204@72.93.22&[MIMAT0000267&hsa-miR-210-3p&offsets|0|0;m-102&11|-|568112|568133&offsets|0|0]|CTGTGCGTGTGACAGCGGCTGA</td><td>>MI0000286|hsa-mir-210&WithFlank&11|-|568083|568204</td><td>72</td><td>93</td><td>22</td><td>CTGTGCGTGTGACAGCGGCTGA</td><td>765.9</td><td>3.0</td><td>0.8</td><td>3.6</td><td>2.7E-04</td><td>1.8E-03</td><td>BT20/MB231</td><td>NUC</td></tr>
<tr><td>>hg38_21_+_8433172_8446622_RNA45SN1_45S_50ntFlanks@3704.3725.22|CTACCTGGTTGATCCTGCCAGT</td><td>>hg38_21_+_8433172_8446622_RNA45SN1_45S_50ntFlanks</td><td>3704</td><td>3725</td><td>22</td><td>CTACCTGGTTGATCCTGCCAGT</td><td>205.8</td><td>5.2</td><td>1.4</td><td>3.6</td><td>2.7E-04</td><td>1.8E-03</td><td>BT20/MB231</td><td>NUC</td></tr>
<tr><td>>MI0001445|hsa-mir-423&WithFlank&17|+|30117073|30117178@59.78.20(+1U)&[MIMAT0001340&hsa-miR-423-3p&offsets|0|-3(+1U);m-100&17|+|30117131|30117153&offsets|0|-3(+1U)]|AGCTCGGTCTGAGGCCCCTCT</td><td>>MI0001445|hsa-mir-423&WithFlank&17|+|30117073|30117178</td><td>59</td><td>78</td><td>20(+1U)</td><td>AGCTCGGTCTGAGGCCCCTCT</td><td>68.0</td><td>6.0</td><td>1.9</td><td>3.2</td><td>1.3E-03</td><td>1.8E-03</td><td>MB231/MB468</td><td>MP</td></tr>
<tr><td>>trna10_AspGTC_12_-_125424193_125424264@31.72.42,trna12_AspGTC_12_-_125411891_125411962@31.72.42,trna38_AspGTC_17_-_8125556_8125627@31.72.42,trna45_AspGTC_6_+_27447453_27447524@31.72.42,trna48_AspGTC_6_+_27471523_27471594@31.72.42,trna4_AspGTC_12_+_96429799_96429870@31.72.42,trna69_AspGTC_1_-_161440205_161440276@31.72.42,trna72_AspGTC_1_-_161432824_161432895@31.72.42,trna75_AspGTC_1_-_161425414_161425485@31.72.42,trna78_AspGTC_1_-_161418033_161418104@31.72.42,trna81_AspGTC_1_-_161410615_161410686@31.72.42|CCTGTCACGCGGGAGACCGGGGTTCGATTCCCCGACGGGGAG</td><td>>trna10_AspGTC_12_-_125424193_125424264</td><td>31</td><td>72</td><td>42</td><td>CCTGTCACGCGGGAGACCGGGGTTCGATTCCCCGACGGGGAG</td><td>407.7</td><td>-6.0</td><td>1.8</td><td>-3.2</td><td>1.2E-03</td><td>1.8E-03</td><td>BT20/MB468</td><td>MITO</td></tr>
<tr><td>>hg38_21_+_8433172_8446622_RNA45SN1_45S_50ntFlanks@10391.10409.19|AACAGCAGTTGAACATGGG</td><td>>hg38_21_+_8433172_8446622_RNA45SN1_45S_50ntFlanks</td><td>10391</td><td>10409</td><td>19</td><td>AACAGCAGTTGAACATGGG</td><td>151.2</td><td>-3.5</td><td>1.1</td><td>-3.2</td><td>1.2E-03</td><td>1.8E-03</td><td>BT20/MB468</td><td>MITO</td></tr>
<tr><td>>hg38_21_+_8433172_8446622_RNA45SN1_45S_50ntFlanks@11670.11693.24|CCCAGTGCTCTGAATGTCAAAGTG</td><td>>hg38_21_+_8433172_8446622_RNA45SN1_45S_50ntFlanks</td><td>11670</td><td>11693</td><td>24</td><td>CCCAGTGCTCTGAATGTCAAAGTG</td><td>613.8</td><td>-2.5</td><td>0.8</td><td>-3.2</td><td>1.4E-03</td><td>1.8E-03</td><td>BT20/MB468</td><td>CYTO</td></tr>
<tr><td>>hg38_21_+_8433172_8446622_RNA45SN1_45S_50ntFlanks@9862.9899.38|ATGAACCGAACGCCGGGTTAAGGCGCCCGATGCCGACG</td><td>>hg38_21_+_8433172_8446622_RNA45SN1_45S_50ntFlanks</td><td>9862</td><td>9899</td><td>38</td><td>ATGAACCGAACGCCGGGTTAAGGCGCCCGATGCCGACG</td><td>375.6</td><td>-3.8</td><td>1.1</td><td>-3.4</td><td>7.2E-04</td><td>1.9E-03</td><td>BT20/MB468</td><td>MP</td></tr>
<tr><td>>hg38_21_+_8433172_8446622_RNA45SN1_45S_50ntFlanks@4045.4073.29|CCATTCGAACGTCTGCCCTATCAACTTTC</td><td>>hg38_21_+_8433172_8446622_RNA45SN1_45S_50ntFlanks</td><td>4045</td><td>4073</td><td>29</td><td>CCATTCGAACGTCTGCCCTATCAACTTTC</td><td>206.0</td><td>-3.0</td><td>0.8</td><td>-3.6</td><td>2.8E-04</td><td>1.9E-03</td><td>BT20/MB231</td><td>CYTO</td></tr>
<tr><td>>hg38_21_+_8433172_8446622_RNA45SN1_45S_50ntFlanks@8170.8213.44|CAAGTCCTTCTGATCGAGGCCCAGCCCGTGGACGGTGTGAGGCC</td><td>>hg38_21_+_8433172_8446622_RNA45SN1_45S_50ntFlanks</td><td>8170</td><td>8213</td><td>44</td><td>CAAGTCCTTCTGATCGAGGCCCAGCCCGTGGACGGTGTGAGGCC</td><td>442.8</td><td>-2.4</td><td>0.7</td><td>-3.2</td><td>1.5E-03</td><td>1.9E-03</td><td>MB231/MB468</td><td>CYTO</td></tr>
<tr><td>>hg38_21_+_8433172_8446622_RNA45SN1_45S_50ntFlanks@12307.12331.25|ACGATCCTTCTGACCTTTTGGGTTT</td><td>>hg38_21_+_8433172_8446622_RNA45SN1_45S_50ntFlanks</td><td>12307</td><td>12331</td><td>25</td><td>ACGATCCTTCTGACCTTTTGGGTTT</td><td>45.6</td><td>-8.6</td><td>2.5</td><td>-3.4</td><td>7.2E-04</td><td>1.9E-03</td><td>BT20/MB468</td><td>MP</td></tr>
<tr><td>>MI0000469|hsa-mir-125a&WithFlank&19|+|51693248|51693345@21.44.24&[MIMAT0000443&hsa-miR-125a-5p&offsets|0|0;m-63&19|+|51693268|51693289&offsets|0|+2]|TCCCTGAGACCCTTTAACCTGTGA</td><td>>MI0000469|hsa-mir-125a&WithFlank&19|+|51693248|51693345</td><td>21</td><td>44</td><td>24</td><td>TCCCTGAGACCCTTTAACCTGTGA</td><td>1748.8</td><td>1.7</td><td>0.5</td><td>3.2</td><td>1.5E-03</td><td>1.9E-03</td><td>BT20/MB468</td><td>NUC</td></tr>
<tr><td>>MI0000542|hsa-mir-320a&WithFlank&8|-|22244960|22245043@48.69.22(+1G)&[MIMAT0000510&hsa-miR-320a-3p&offsets|0|0(+1G);m-60&8|-|22244975|22244996&offsets|0|0(+1G)]|AAAAGCTGGGTTGAGAGGGCGAG</td><td>>MI0000542|hsa-mir-320a&WithFlank&8|-|22244960|22245043</td><td>48</td><td>69</td><td>22(+1G)</td><td>AAAAGCTGGGTTGAGAGGGCGAG</td><td>441.1</td><td>2.6</td><td>0.7</td><td>3.6</td><td>2.7E-04</td><td>1.9E-03</td><td>BT20/MB231</td><td>NUC</td></tr>
<tr><td>>am_trna162_MetCAT_6_-_26758550_26758622@31.54.24,trna164_MetCAT_6_-_26735574_26735646@31.54.24,trna20_MetCAT_16_+_71460396_71460468@31.54.24,trna21_MetCAT_6_+_26701712_26701784@31.54.24,trna27_MetCAT_6_+_26766444_26766516@31.54.24,trna75_MetCAT_6_+_28912352_28912424@31.54.24,trna97_MetCAT_6_-_28921042_28921114@31.54.24|TCTCATAATCTGAAGGTCCTGAGT</td><td>>am_trna162_MetCAT_6_-_26758550_26758622</td><td>31</td><td>54</td><td>24</td><td>TCTCATAATCTGAAGGTCCTGAGT</td><td>239.0</td><td>-3.7</td><td>1.2</td><td>-3.2</td><td>1.6E-03</td><td>1.9E-03</td><td>MB231/MB468</td><td>NUC</td></tr>
<tr><td>>MI0000737|hsa-mir-200a&WithFlank&1|+|1167857|1167958@60.82.23&[MIMAT0000682&hsa-miR-200a-3p&offsets|0|+1;m-142&1|+|1167916|1167938&offsets|0|0]|TAACACTGTCTGGTAACGATGTT</td><td>>MI0000737|hsa-mir-200a&WithFlank&1|+|1167857|1167958</td><td>60</td><td>82</td><td>23</td><td>TAACACTGTCTGGTAACGATGTT</td><td>435.0</td><td>-3.9</td><td>1.2</td><td>-3.2</td><td>1.2E-03</td><td>1.9E-03</td><td>BT20/MB468</td><td>MITO</td></tr>
<tr><td>>MI0003188|hsa-mir-503&WithFlank&X|-|134546322|134546404@12.32.21&[MIMAT0002874&hsa-miR-503-5p&offsets|0|-2;m-297&X|-|134546371|134546393&offsets|0|-2]|TAGCAGCGGGAACAGTTCTGC</td><td>>MI0003188|hsa-mir-503&WithFlank&X|-|134546322|134546404</td><td>12</td><td>32</td><td>21</td><td>TAGCAGCGGGAACAGTTCTGC</td><td>49.0</td><td>9.4</td><td>2.8</td><td>3.4</td><td>7.2E-04</td><td>1.9E-03</td><td>BT20/MB468</td><td>MP</td></tr>
<tr><td>>hg38_21_+_8433172_8446622_RNA45SN1_45S_50ntFlanks@6646.6679.34|TCGTACGACTCTTAGCGGTGGATCACTCGGCTCG</td><td>>hg38_21_+_8433172_8446622_RNA45SN1_45S_50ntFlanks</td><td>6646</td><td>6679</td><td>34</td><td>TCGTACGACTCTTAGCGGTGGATCACTCGGCTCG</td><td>2750.5</td><td>2.0</td><td>0.6</td><td>3.2</td><td>1.4E-03</td><td>1.9E-03</td><td>BT20/MB468</td><td>CYTO</td></tr>
<tr><td>>hg38_21_+_8433172_8446622_RNA45SN1_45S_50ntFlanks@12005.12026.22|GGGCCGCCGGTGAAATACCACT</td><td>>hg38_21_+_8433172_8446622_RNA45SN1_45S_50ntFlanks</td><td>12005</td><td>12026</td><td>22</td><td>GGGCCGCCGGTGAAATACCACT</td><td>725.3</td><td>7.4</td><td>2.2</td><td>3.4</td><td>7.2E-04</td><td>1.9E-03</td><td>BT20/MB468</td><td>MP</td></tr>
<tr><td>>hg38_21_+_8433172_8446622_RNA45SN1_45S_50ntFlanks@11208.11251.44|GCGGCGACTCTGGACGCGAGCCGGGCCCTTCCCGTGGATCGCCC</td><td>>hg38_21_+_8433172_8446622_RNA45SN1_45S_50ntFlanks</td><td>11208</td><td>11251</td><td>44</td><td>GCGGCGACTCTGGACGCGAGCCGGGCCCTTCCCGTGGATCGCCC</td><td>902.8</td><td>-6.2</td><td>2.0</td><td>-3.2</td><td>1.5E-03</td><td>1.9E-03</td><td>BT20/MB468</td><td>NUC</td></tr>
<tr><td>>hg38_21_+_8433172_8446622_RNA45SN1_45S_50ntFlanks@10393.10422.30|CAGCAGTTGAACATGGGTCAGTCGGTCCTG</td><td>>hg38_21_+_8433172_8446622_RNA45SN1_45S_50ntFlanks</td><td>10393</td><td>10422</td><td>30</td><td>CAGCAGTTGAACATGGGTCAGTCGGTCCTG</td><td>445.9</td><td>-3.5</td><td>1.1</td><td>-3.2</td><td>1.5E-03</td><td>1.9E-03</td><td>MB231/MB468</td><td>CYTO</td></tr>
<tr><td>>hg38_21_+_8433172_8446622_RNA45SN1_45S_50ntFlanks@4602.4619.18|TTCGGAACTGAGGCCATG</td><td>>hg38_21_+_8433172_8446622_RNA45SN1_45S_50ntFlanks</td><td>4602</td><td>4619</td><td>18</td><td>TTCGGAACTGAGGCCATG</td><td>194.3</td><td>-5.3</td><td>1.6</td><td>-3.4</td><td>7.2E-04</td><td>1.9E-03</td><td>BT20/MB468</td><td>MP</td></tr>
<tr><td>>hg38_21_+_8433172_8446622_RNA45SN1_45S_50ntFlanks@475.518.44|CTTCGTGATCGATGTGGTGACGTCGTGCTCTCCCGGGCCGGGTC</td><td>>hg38_21_+_8433172_8446622_RNA45SN1_45S_50ntFlanks</td><td>475</td><td>518</td><td>44</td><td>CTTCGTGATCGATGTGGTGACGTCGTGCTCTCCCGGGCCGGGTC</td><td>6717.1</td><td>3.4</td><td>1.1</td><td>3.2</td><td>1.6E-03</td><td>1.9E-03</td><td>MB231/MB468</td><td>NUC</td></tr>
<tr><td>>hg38_21_+_8433172_8446622_RNA45SN1_45S_50ntFlanks@10912.10932.21|CGCGCCGCGGCTGGACGAGGC</td><td>>hg38_21_+_8433172_8446622_RNA45SN1_45S_50ntFlanks</td><td>10912</td><td>10932</td><td>21</td><td>CGCGCCGCGGCTGGACGAGGC</td><td>444.1</td><td>-5.3</td><td>1.6</td><td>-3.2</td><td>1.4E-03</td><td>1.9E-03</td><td>BT20/MB468</td><td>CYTO</td></tr>
</table>

<table>
<tr><td>>trna16_TyrGTA_14_-_21131351_21131444@59.76.18,trna17_TyrGTA_14_-_21128117_21128210@59.76.18,trna19_TyrGTA_14_-_21121258_21121351@59.76.18,trna2_TyrGTA_2_+_27273650_27273738@59.76.18,trna4_TyrGTA_8_+_67025602_67025694@59.76.18,trna5_TyrGTA_14_+_21151432_21151520@59.76.18,trna5_TyrGTA_8_+_67026223_67026311@59.76.18|TTCCGGCTCGAAGGACCA</td><td>>trna16_TyrGTA_14_-_21131351_21131444</td><td>59</td><td>76</td><td>18</td><td>TTCCGGCTCGAAGGACCA</td><td>1946.0</td><td>4.3</td><td>1.3</td><td>3.4</td><td>7.2E-04</td><td>1.9E-03</td><td>BT20/MB468</td><td>MP</td></tr>
<tr><td>>hg38_21_+_8433172_8446622_RNA45SN1_45S_50ntFlanks@8077.8104.28|GGAAGAGCCCAGCGCCGAATCCCCGCCC</td><td>>hg38_21_+_8433172_8446622_RNA45SN1_45S_50ntFlanks</td><td>8077</td><td>8104</td><td>28</td><td>GGAAGAGCCCAGCGCCGAATCCCCGCCC</td><td>254.8</td><td>-6.0</td><td>1.8</td><td>-3.4</td><td>7.2E-04</td><td>1.9E-03</td><td>BT20/MB468</td><td>MP</td></tr>
<tr><td>>MI0000077|hsa-mir-21&WithFlank&17|+|59841260|59841343@52.73.22(+1C)&[MIMAT0004494&hsa-miR-21-3p&offsets|0|+1(+1C);m-52&17|+|59841311|59841332&offsets|0|0(+1C)]|CAACACCAGTCGATGGGCTGTCC</td><td>>MI0000077|hsa-mir-21&WithFlank&17|+|59841260|59841343</td><td>52</td><td>73</td><td>22(+1C)</td><td>CAACACCAGTCGATGGGCTGTCC</td><td>84.3</td><td>5.8</td><td>1.8</td><td>3.2</td><td>1.3E-03</td><td>1.9E-03</td><td>MB231/MB468</td><td>MP</td></tr>
<tr><td>>hg38_21_+_8433172_8446622_RNA45SN1_45S_50ntFlanks@4732.4771.40|AGAACGAAAGTCGGAGGTTCGAAGACGATCAGATACCGTC</td><td>>hg38_21_+_8433172_8446622_RNA45SN1_45S_50ntFlanks</td><td>4732</td><td>4771</td><td>40</td><td>AGAACGAAAGTCGGAGGTTCGAAGACGATCAGATACCGTC</td><td>58.4</td><td>6.3</td><td>1.9</td><td>3.4</td><td>7.2E-04</td><td>1.9E-03</td><td>BT20/MB468</td><td>MP</td></tr>
<tr><td>>MI0000746|hsa-mir-99b&WithFlank&19|+|51692606|51692687@13.31.19(+1A)&[MIMAT0000689&hsa-miR-99b-5p&offsets|0|-3(+1A);m-10&19|+|51692618|51692639&offsets|0|-3(+1A)]|CACCCGTAGAACCGACCTTA</td><td>>MI0000746|hsa-mir-99b&WithFlank&19|+|51692606|51692687</td><td>13</td><td>31</td><td>19(+1A)</td><td>CACCCGTAGAACCGACCTTA</td><td>203.6</td><td>2.7</td><td>0.8</td><td>3.6</td><td>2.7E-04</td><td>1.9E-03</td><td>BT20/MB231</td><td>NUC</td></tr>
<tr><td>>trnaMT_HisGTG_MT_+_12138_12206@20.52.33|AACATCAGATTGTGAATCTGACAACAGAGGCTT</td><td>>trnaMT_HisGTG_MT_+_12138_12206</td><td>20</td><td>52</td><td>33</td><td>AACATCAGATTGTGAATCTGACAACAGAGGCTT</td><td>84.0</td><td>6.5</td><td>1.9</td><td>3.4</td><td>7.2E-04</td><td>1.9E-03</td><td>BT20/MB468</td><td>MP</td></tr>
<tr><td>>MI0000815|hsa-mir-339&WithFlank&7|-|1022927|1023032@56.77.22&[MIMAT0004702&hsa-miR-339-3p&offsets|0|-1;m-235&7|-|1022957|1022977&offsets|0|+1]|TGAGCGCCTCGACGACAGAGCC</td><td>>MI0000815|hsa-mir-339&WithFlank&7|-|1022927|1023032</td><td>56</td><td>77</td><td>22</td><td>TGAGCGCCTCGACGACAGAGCC</td><td>305.4</td><td>6.0</td><td>1.8</td><td>3.4</td><td>7.2E-04</td><td>1.9E-03</td><td>BT20/MB468</td><td>MP</td></tr>
<tr><td>>hg38_21_+_8433172_8446622_RNA45SN1_45S_50ntFlanks@10392.10424.33|ACAGCAGTTGAACATGGGTCAGTCGGTCCTGAG</td><td>>hg38_21_+_8433172_8446622_RNA45SN1_45S_50ntFlanks</td><td>10392</td><td>10424</td><td>33</td><td>ACAGCAGTTGAACATGGGTCAGTCGGTCCTGAG</td><td>10235.5</td><td>3.4</td><td>0.9</td><td>3.6</td><td>3.1E-04</td><td>1.9E-03</td><td>BT20/MB231</td><td>MITO</td></tr>
<tr><td>>hg38_21_+_8433172_8446622_RNA45SN1_45S_50ntFlanks@9738.9766.29|GAAACGATCTCAACCTATTCTCAAACTTT</td><td>>hg38_21_+_8433172_8446622_RNA45SN1_45S_50ntFlanks</td><td>9738</td><td>9766</td><td>29</td><td>GAAACGATCTCAACCTATTCTCAAACTTT</td><td>290.4</td><td>-2.1</td><td>0.7</td><td>-3.2</td><td>1.6E-03</td><td>1.9E-03</td><td>MB231/MB468</td><td>NUC</td></tr>
<tr><td>>hg38_21_+_8433172_8446622_RNA45SN1_45S_50ntFlanks@6653.6677.25|ACTCTTAGCGGTGGATCACTCGGCT</td><td>>hg38_21_+_8433172_8446622_RNA45SN1_45S_50ntFlanks</td><td>6653</td><td>6677</td><td>25</td><td>ACTCTTAGCGGTGGATCACTCGGCT</td><td>294.0</td><td>3.2</td><td>0.9</td><td>3.4</td><td>7.2E-04</td><td>1.9E-03</td><td>BT20/MB468</td><td>MP</td></tr>
<tr><td>>am_trna116_GluCTC_1_-_145399233_145399304@1.23.23,trna59_GluCTC_1_+_249168447_249168518@1.23.23,trna71_GluCTC_1_-_161439189_161439260@1.23.23,trna74_GluCTC_1_-_161431809_161431880@1.23.23,trna77_GluCTC_1_-_161424398_161424469@1.23.23,trna77_GluCTC_6_+_28949976_28950047@1.23.23,trna80_GluCTC_1_-_161417018_161417089@1.23.23,trna87_GluCTC_6_-_126101393_126101464@1.23.23|TCCCTGGTGGTCTAGTGGTTAGG</td><td>>am_trna116_GluCTC_1_-_145399233_145399304</td><td>1</td><td>23</td><td>23</td><td>TCCCTGGTGGTCTAGTGGTTAGG</td><td>229.3</td><td>3.1</td><td>0.8</td><td>3.6</td><td>2.8E-04</td><td>1.9E-03</td><td>BT20/MB231</td><td>CYTO</td></tr>
<tr><td>>hg38_21_+_8433172_8446622_RNA45SN1_45S_50ntFlanks@12893.12934.42|CCTCGCCCGTCACGCACCGCACGTTCGTGGGGAACCTGGCGC</td><td>>hg38_21_+_8433172_8446622_RNA45SN1_45S_50ntFlanks</td><td>12893</td><td>12934</td><td>42</td><td>CCTCGCCCGTCACGCACCGCACGTTCGTGGGGAACCTGGCGC</td><td>186.7</td><td>-4.7</td><td>1.5</td><td>-3.2</td><td>1.2E-03</td><td>1.9E-03</td><td>BT20/MB468</td><td>MITO</td></tr>
<tr><td>>hg38_21_+_8433172_8446622_RNA45SN1_45S_50ntFlanks@6661.6688.28|CGGTGGATCACTCGGCTCGTGCGTCGAT</td><td>>hg38_21_+_8433172_8446622_RNA45SN1_45S_50ntFlanks</td><td>6661</td><td>6688</td><td>28</td><td>CGGTGGATCACTCGGCTCGTGCGTCGAT</td><td>3000.1</td><td>2.1</td><td>0.6</td><td>3.6</td><td>2.7E-04</td><td>1.9E-03</td><td>BT20/MB231</td><td>NUC</td></tr>
<tr><td>>hg38_21_+_8433172_8446622_RNA45SN1_45S_50ntFlanks@8194.8216.23|CCCGTGGACGGTGTGAGGCCGGT</td><td>>hg38_21_+_8433172_8446622_RNA45SN1_45S_50ntFlanks</td><td>8194</td><td>8216</td><td>23</td><td>CCCGTGGACGGTGTGAGGCCGGT</td><td>222.8</td><td>-3.7</td><td>1.2</td><td>-3.2</td><td>1.6E-03</td><td>1.9E-03</td><td>MB231/MB468</td><td>NUC</td></tr>
<tr><td>>hg38_21_+_8433172_8446622_RNA45SN1_45S_50ntFlanks@10248.10286.39|CGACGAGTAGGAGGGCCGCTGCGGTGAGCCTTGAAGCCT</td><td>>hg38_21_+_8433172_8446622_RNA45SN1_45S_50ntFlanks</td><td>10248</td><td>10286</td><td>39</td><td>CGACGAGTAGGAGGGCCGCTGCGGTGAGCCTTGAAGCCT</td><td>562.4</td><td>4.8</td><td>1.3</td><td>3.7</td><td>2.5E-04</td><td>1.9E-03</td><td>BT20/MB231</td><td>MP</td></tr>
<tr><td>>hg38_21_+_8433172_8446622_RNA45SN1_45S_50ntFlanks@6771.6803.33|GGTTCCTCCCGGGGCTACGCCTGTCTGAGCGTC</td><td>>hg38_21_+_8433172_8446622_RNA45SN1_45S_50ntFlanks</td><td>6771</td><td>6803</td><td>33</td><td>GGTTCCTCCCGGGGCTACGCCTGTCTGAGCGTC</td><td>493.7</td><td>-5.7</td><td>1.7</td><td>-3.4</td><td>7.2E-04</td><td>1.9E-03</td><td>BT20/MB468</td><td>MP</td></tr>
<tr><td>>MI0000078|hsa-mir-22&WithFlank&17|-|1713897|1713993@21.42.22&[MIMAT0004495&hsa-miR-22-5p&offsets|0|0;m-217&17|-|1713953|1713973&offsets|0|+1]|AGTTCTTCAGTGGCAAGCTTTA</td><td>>MI0000078|hsa-mir-22&WithFlank&17|-|1713897|1713993</td><td>21</td><td>42</td><td>22</td><td>AGTTCTTCAGTGGCAAGCTTTA</td><td>667.6</td><td>4.3</td><td>1.3</td><td>3.2</td><td>1.3E-03</td><td>1.9E-03</td><td>MB231/MB468</td><td>MP</td></tr>
<tr><td>>hg38_MT_+_1621_3279_RNR2_16S_50ntFlanks@1427.1448.22|GATTAAAGTCCTACGTGATCTG</td><td>>hg38_MT_+_1621_3279_RNR2_16S_50ntFlanks</td><td>1427</td><td>1448</td><td>22</td><td>GATTAAAGTCCTACGTGATCTG</td><td>236.8</td><td>4.1</td><td>1.3</td><td>3.2</td><td>1.3E-03</td><td>1.9E-03</td><td>MB231/MB468</td><td>MP</td></tr>
<tr><td>>MI0005561|hsa-mir-877&WithFlank&6|+|30584326|30584423@7.28.22(+1A)&[MIMAT0004949&hsa-miR-877-5p&offsets|0|+2(+1A);m-382&6|+|30584332|30584354&offsets|0|-1(+1A)]|GTAGAGGAGATGGCGCAGGGGAA</td><td>>MI0005561|hsa-mir-877&WithFlank&6|+|30584326|30584423</td><td>7</td><td>28</td><td>22(+1A)</td><td>GTAGAGGAGATGGCGCAGGGGAA</td><td>60.2</td><td>6.6</td><td>1.9</td><td>3.4</td><td>7.3E-04</td><td>1.9E-03</td><td>BT20/MB468</td><td>MP</td></tr>
<tr><td>>hg38_21_+_8433172_8446622_RNA45SN1_45S_50ntFlanks@10798.10822.25|CATGTTGGAACAATGTAGGTAAGGG</td><td>>hg38_21_+_8433172_8446622_RNA45SN1_45S_50ntFlanks</td><td>10798</td><td>10822</td><td>25</td><td>CATGTTGGAACAATGTAGGTAAGGG</td><td>243.8</td><td>-1.8</td><td>0.6</td><td>-3.2</td><td>1.4E-03</td><td>1.9E-03</td><td>BT20/MB468</td><td>CYTO</td></tr>
<tr><td>>hg38_21_+_8433172_8446622_RNA45SN1_45S_50ntFlanks@6773.6803.31|TTCCTCCCGGGGCTACGCCTGTCTGAGCGTC</td><td>>hg38_21_+_8433172_8446622_RNA45SN1_45S_50ntFlanks</td><td>6773</td><td>6803</td><td>31</td><td>TTCCTCCCGGGGCTACGCCTGTCTGAGCGTC</td><td>747.2</td><td>-4.0</td><td>1.3</td><td>-3.2</td><td>1.4E-03</td><td>1.9E-03</td><td>BT20/MB468</td><td>CYTO</td></tr>
<tr><td>>trnaMT_PheGAA_MT_+_577_647@54.74.21|CACATCACCCCATAAACACCA</td><td>>trnaMT_PheGAA_MT_+_577_647</td><td>54</td><td>74</td><td>21</td><td>CACATCACCCCATAAACACCA</td><td>267.4</td><td>-2.5</td><td>0.8</td><td>-3.2</td><td>1.3E-03</td><td>1.9E-03</td><td>MB231/MB468</td><td>MP</td></tr>
<tr><td>>hg38_21_+_8433172_8446622_RNA45SN1_45S_50ntFlanks@4732.4761.30|AGAACGAAAGTCGGAGGTTCGAAGACGATC</td><td>>hg38_21_+_8433172_8446622_RNA45SN1_45S_50ntFlanks</td><td>4732</td><td>4761</td><td>30</td><td>AGAACGAAAGTCGGAGGTTCGAAGACGATC</td><td>1649.9</td><td>3.8</td><td>1.1</td><td>3.4</td><td>7.3E-04</td><td>1.9E-03</td><td>BT20/MB468</td><td>MP</td></tr>
<tr><td>>hg38_21_+_8433172_8446622_RNA45SN1_45S_50ntFlanks@10353.10373.21|AGAACTTTGAAGGCCGAAGTG</td><td>>hg38_21_+_8433172_8446622_RNA45SN1_45S_50ntFlanks</td><td>10353</td><td>10373</td><td>21</td><td>AGAACTTTGAAGGCCGAAGTG</td><td>49.1</td><td>-8.7</td><td>2.6</td><td>-3.4</td><td>7.3E-04</td><td>1.9E-03</td><td>BT20/MB468</td><td>MP</td></tr>
<tr><td>>hg38_21_+_8433172_8446622_RNA45SN1_45S_50ntFlanks@6647.6668.22|CGTACGACTCTTAGCGGTGGAT</td><td>>hg38_21_+_8433172_8446622_RNA45SN1_45S_50ntFlanks</td><td>6647</td><td>6668</td><td>22</td><td>CGTACGACTCTTAGCGGTGGAT</td><td>1709.4</td><td>2.2</td><td>0.7</td><td>3.2</td><td>1.6E-03</td><td>1.9E-03</td><td>MB231/MB468</td><td>MITO</td></tr>
<tr><td>>hg38_21_+_8433172_8446622_RNA45SN1_45S_50ntFlanks@9549.9574.26|AGGTCCGTAGCGGTCCTGACGTGCAA</td><td>>hg38_21_+_8433172_8446622_RNA45SN1_45S_50ntFlanks</td><td>9549</td><td>9574</td><td>26</td><td>AGGTCCGTAGCGGTCCTGACGTGCAA</td><td>85.7</td><td>2.8</td><td>0.8</td><td>3.4</td><td>7.3E-04</td><td>1.9E-03</td><td>BT20/MB468</td><td>MP</td></tr>
<tr><td>>MI0000079|hsa-mir-23a&WithFlank&19|-|13836581|13836665@54.72.19(+1C)&[MIMAT0000078&hsa-miR-23a-3p&offsets|+3|+1(+1C);m-31&19|-|13836595|13836615&offsets|+3|+1(+1C)]|ACATTGCCAGGGATTTCCAC</td><td>>MI0000079|hsa-mir-23a&WithFlank&19|-|13836581|13836665</td><td>54</td><td>72</td><td>19(+1C)</td><td>ACATTGCCAGGGATTTCCAC</td><td>46.7</td><td>8.8</td><td>2.7</td><td>3.2</td><td>1.3E-03</td><td>1.9E-03</td><td>MB231/MB468</td><td>MP</td></tr>
<tr><td>>hg38_21_+_8433172_8446622_RNA45SN1_45S_50ntFlanks@5527.5549.23|AAAGTCGTAACAAGGTTTCCGTA</td><td>>hg38_21_+_8433172_8446622_RNA45SN1_45S_50ntFlanks</td><td>5527</td><td>5549</td><td>23</td><td>AAAGTCGTAACAAGGTTTCCGTA</td><td>511.7</td><td>-5.6</td><td>1.7</td><td>-3.4</td><td>7.3E-04</td><td>1.9E-03</td><td>BT20/MB468</td><td>MP</td></tr>
<tr><td>>hg38_21_+_8433172_8446622_RNA45SN1_45S_50ntFlanks@11209.11229.21|CGGCGACTCTGGACGCGAGCC</td><td>>hg38_21_+_8433172_8446622_RNA45SN1_45S_50ntFlanks</td><td>11209</td><td>11229</td><td>21</td><td>CGGCGACTCTGGACGCGAGCC</td><td>539.6</td><td>7.6</td><td>2.1</td><td>3.7</td><td>2.5E-04</td><td>1.9E-03</td><td>BT20/MB231</td><td>MP</td></tr>
<tr><td>>hg38_MT_+_1621_3279_RNR2_16S_50ntFlanks@498.521.24|TTGGACACTAGGAAAAAACCTTGT</td><td>>hg38_MT_+_1621_3279_RNR2_16S_50ntFlanks</td><td>498</td><td>521</td><td>24</td><td>TTGGACACTAGGAAAAAACCTTGT</td><td>88.0</td><td>6.5</td><td>1.9</td><td>3.4</td><td>7.3E-04</td><td>1.9E-03</td><td>BT20/MB468</td><td>MP</td></tr>
<tr><td>>hg38_21_+_8433172_8446622_RNA45SN1_45S_50ntFlanks@6698.6717.20|AGCTAGCTGCGAGAATTAAT</td><td>>hg38_21_+_8433172_8446622_RNA45SN1_45S_50ntFlanks</td><td>6698</td><td>6717</td><td>20</td><td>AGCTAGCTGCGAGAATTAAT</td><td>166.6</td><td>-4.5</td><td>1.4</td><td>-3.2</td><td>1.4E-03</td><td>1.9E-03</td><td>BT20/MB468</td><td>CYTO</td></tr>
<tr><td>>hg38_21_+_8433172_8446622_RNA45SN1_45S_50ntFlanks@12407.12446.40|TTTTGATCCTTCGATGTCGGCTCTTCCTATCATTGTGAAG</td><td>>hg38_21_+_8433172_8446622_RNA45SN1_45S_50ntFlanks</td><td>12407</td><td>12446</td><td>40</td><td>TTTTGATCCTTCGATGTCGGCTCTTCCTATCATTGTGAAG</td><td>348.4</td><td>-3.6</td><td>1.0</td><td>-3.6</td><td>2.8E-04</td><td>1.9E-03</td><td>BT20/MB231</td><td>CYTO</td></tr>
<tr><td>>hg38_21_+_8433172_8446622_RNA45SN1_45S_50ntFlanks@12454.12476.23|CACCAAGCGTTGGATTGTTCACC</td><td>>hg38_21_+_8433172_8446622_RNA45SN1_45S_50ntFlanks</td><td>12454</td><td>12476</td><td>23</td><td>CACCAAGCGTTGGATTGTTCACC</td><td>678.4</td><td>-3.0</td><td>0.8</td><td>-3.6</td><td>2.8E-04</td><td>1.9E-03</td><td>BT20/MB231</td><td>CYTO</td></tr>
<tr><td>>MI0005566|hsa-mir-374b&WithFlank&X|-|74218541|74218624@17.37.21&[MIMAT0004955&hsa-miR-374b-5p&offsets|0|-1;m-153&X|-|74218587|74218608&offsets|0|-1]|ATATAATACAACCTGCTAAGT</td><td>>MI0005566|hsa-mir-374b&WithFlank&X|-|74218541|74218624</td><td>17</td><td>37</td><td>21</td><td>ATATAATACAACCTGCTAAGT</td><td>226.5</td><td>1.9</td><td>0.6</td><td>3.2</td><td>1.6E-03</td><td>1.9E-03</td><td>BT20/MB468</td><td>NUC</td></tr>
<tr><td>>MI0000253|hsa-mir-148a&WithFlank&7|-|25949913|25949992@50.70.21(+1C)&[MIMAT0000243&hsa-miR-148a-3p&offsets|0|-1(+1C);m-7&7|-|25949922|25949943&offsets|0|-1(+1C)]|TCAGTGCACTACAGAACTTTGC</td><td>>MI0000253|hsa-mir-148a&WithFlank&7|-|25949913|25949992</td><td>50</td><td>70</td><td>21(+1C)</td><td>TCAGTGCACTACAGAACTTTGC</td><td>12633.2</td><td>3.1</td><td>0.9</td><td>3.6</td><td>3.2E-04</td><td>1.9E-03</td><td>BT20/MB231</td><td>MITO</td></tr>
<tr><td>>MI0000441|hsa-mir-30b&WithFlank&8|-|134800514|134800613@23.44.22&[MIMAT0000420&hsa-miR-30b-5p&offsets|0|0;m-76&8|-|134800570|134800591&offsets|0|0]|TGTAAACATCCTACACTCAGCT</td><td>>MI0000441|hsa-mir-30b&WithFlank&8|-|134800514|134800613</td><td>23</td><td>44</td><td>22</td><td>TGTAAACATCCTACACTCAGCT</td><td>9667.6</td><td>-3.7</td><td>1.0</td><td>-3.6</td><td>3.2E-04</td><td>1.9E-03</td><td>BT20/MB231</td><td>MITO</td></tr>
<tr><td>>hg38_21_+_8433172_8446622_RNA45SN1_45S_50ntFlanks@10380.10424.45|GGTTCCATGTGAACAGCAGTTGAACATGGGTCAGTCGGTCCTGAG</td><td>>hg38_21_+_8433172_8446622_RNA45SN1_45S_50ntFlanks</td><td>10380</td><td>10424</td><td>45</td><td>GGTTCCATGTGAACAGCAGTTGAACATGGGTCAGTCGGTCCTGAG</td><td>2730.8</td><td>3.2</td><td>0.9</td><td>3.6</td><td>3.2E-04</td><td>1.9E-03</td><td>BT20/MB231</td><td>MITO</td></tr>
<tr><td>>hg38_21_+_8433172_8446622_RNA45SN1_45S_50ntFlanks@7968.7986.19|TCCGAGACGCGACCTCAGA</td><td>>hg38_21_+_8433172_8446622_RNA45SN1_45S_50ntFlanks</td><td>7968</td><td>7986</td><td>19</td><td>TCCGAGACGCGACCTCAGA</td><td>110.5</td><td>2.2</td><td>0.7</td><td>3.2</td><td>1.6E-03</td><td>1.9E-03</td><td>BT20/MB468</td><td>NUC</td></tr>
<tr><td>>hg38_MT_+_1621_3279_RNR2_16S_50ntFlanks@622.660.39|TATAACTGAACTCCTCACACCCAATTGGACCAATCTATC</td><td>>hg38_MT_+_1621_3279_RNR2_16S_50ntFlanks</td><td>622</td><td>660</td><td>39</td><td>TATAACTGAACTCCTCACACCCAATTGGACCAATCTATC</td><td>49.1</td><td>9.4</td><td>2.8</td><td>3.4</td><td>7.3E-04</td><td>1.9E-03</td><td>BT20/MB468</td><td>MP</td></tr>
<tr><td>>hg38_21_+_8433172_8446622_RNA45SN1_45S_50ntFlanks@3705.3728.24|TACCTGGTTGATCCTGCCAGTAGC</td><td>>hg38_21_+_8433172_8446622_RNA45SN1_45S_50ntFlanks</td><td>3705</td><td>3728</td><td>24</td><td>TACCTGGTTGATCCTGCCAGTAGC</td><td>749.5</td><td>2.4</td><td>0.7</td><td>3.6</td><td>2.8E-04</td><td>1.9E-03</td><td>BT20/MB231</td><td>NUC</td></tr>
<tr><td>>hg38_21_+_8433172_8446622_RNA45SN1_45S_50ntFlanks@10846.10867.22|CTTCGGGATAAGGATTGGCTCT</td><td>>hg38_21_+_8433172_8446622_RNA45SN1_45S_50ntFlanks</td><td>10846</td><td>10867</td><td>22</td><td>CTTCGGGATAAGGATTGGCTCT</td><td>415.6</td><td>-2.4</td><td>0.8</td><td>-3.2</td><td>1.4E-03</td><td>1.9E-03</td><td>BT20/MB468</td><td>CYTO</td></tr>
<tr><td>>hg38_21_+_8433172_8446622_RNA45SN1_45S_50ntFlanks@9533.9554.22|GAGGAAACTCTGGTGGAGGTCC</td><td>>hg38_21_+_8433172_8446622_RNA45SN1_45S_50ntFlanks</td><td>9533</td><td>9554</td><td>22</td><td>GAGGAAACTCTGGTGGAGGTCC</td><td>214.0</td><td>-3.9</td><td>1.2</td><td>-3.2</td><td>1.6E-03</td><td>1.9E-03</td><td>MB231/MB468</td><td>NUC</td></tr>
<tr><td>>hg38_21_+_8433172_8446622_RNA45SN1_45S_50ntFlanks@12940.12965.26|CATTCGTAGACGACCTGCTTCTGGGT</td><td>>hg38_21_+_8433172_8446622_RNA45SN1_45S_50ntFlanks</td><td>12940</td><td>12965</td><td>26</td><td>CATTCGTAGACGACCTGCTTCTGGGT</td><td>195.2</td><td>-4.5</td><td>1.4</td><td>-3.2</td><td>1.4E-03</td><td>1.9E-03</td><td>BT20/MB468</td><td>CYTO</td></tr>
<tr><td>>hg38_21_+_8433172_8446622_RNA45SN1_45S_50ntFlanks@4390.4408.19|TCTTGGGAGCGGGCGGGCG</td><td>>hg38_21_+_8433172_8446622_RNA45SN1_45S_50ntFlanks</td><td>4390</td><td>4408</td><td>19</td><td>TCTTGGGAGCGGGCGGGCG</td><td>195.0</td><td>-4.7</td><td>1.5</td><td>-3.2</td><td>1.4E-03</td><td>1.9E-03</td><td>BT20/MB468</td><td>CYTO</td></tr>
<tr><td>>hg38_1_-_228634819_228635039_RNA5S12_5S_50ntFlanks@137.171.35|GAGACCGCCTGGGAATACCGGGTGCTGTAGGCTTT</td><td>>hg38_1_-_228634819_228635039_RNA5S12_5S_50ntFlanks</td><td>137</td><td>171</td><td>35</td><td>GAGACCGCCTGGGAATACCGGGTGCTGTAGGCTTT</td><td>557.7</td><td>3.0</td><td>1.0</td><td>3.2</td><td>1.6E-03</td><td>1.9E-03</td><td>MB231/MB468</td><td>MITO</td></tr>
<tr><td>>hg38_21_+_8433172_8446622_RNA45SN1_45S_50ntFlanks@12072.12089.18|CCCGAGGGGCTCTCGCTT</td><td>>hg38_21_+_8433172_8446622_RNA45SN1_45S_50ntFlanks</td><td>12072</td><td>12089</td><td>18</td><td>CCCGAGGGGCTCTCGCTT</td><td>117.6</td><td>-4.5</td><td>1.4</td><td>-3.2</td><td>1.2E-03</td><td>1.9E-03</td><td>BT20/MB468</td><td>MITO</td></tr>
<tr><td>>am_MI0000815|hsa-mir-339&WithFlank&7|-|1022927|1023032@21.40.20&[MIMAT0000764&hsa-miR-339-5p&offsets|0|-3;m-163&7|-|1022990|1023012&offsets|0|-3]|TCCCTGTCCTCCAGGAGCTC</td><td>>am_MI0000815|hsa-mir-339&WithFlank&7|-|1022927|1023032</td><td>21</td><td>40</td><td>20</td><td>TCCCTGTCCTCCAGGAGCTC</td><td>277.8</td><td>3.7</td><td>1.1</td><td>3.4</td><td>7.3E-04</td><td>1.9E-03</td><td>BT20/MB468</td><td>MP</td></tr>
</table>

| | | | | | | | | | | | | | |
|---|---|---|---|---|---|---|---|---|---|---|---|---|---|
| >trna111_HisGTG_1_-_147774845_147774916@-1G.32.33,trna118_HisGTG_1_-_145396881_145396952@-1G.32.33,trna16_HisGTG_1_+_146544773_146544844@-1G.32.33,trna1_HisGTG_15_+_45493349_45493420@-1G.32.33,trna21_HisGTG_1_+_147753471_147753542@-1G.32.33,trna33_HisGTG_6_+_27125906_27125977@-1G.32.33,trna7_HisGTG_9_-_14433938_14434009@-1G.32.33,trna8_HisGTG_15_-_45492611_45492682@-1G.32.33,trna9_HisGTG_15_-_45490804_45490875@-1G.32.33\|GGCCGTGATCGTATAGTGGTTAGTACTCTGCGT | >trna111_HisGTG_1_-_147774845_147774916 | -1G | 32 | 33 | GGCCGTGATCGTATAGTGGTTAGTACTCTGCGT | 5340.6 | -3.9 | 1.1 | -3.6 | 2.8E-04 | 1.9E-03 | BT20/MB231 | CYTO |
| >hg38_21_+_8433172_8446622_RNA45SN1_45S_50ntFlanks@8196.8222.27\|CGTGGACGGTGTGAGGCCGGTAGCGGC | >hg38_21_+_8433172_8446622_RNA45SN1_45S_50ntFlanks | 8196 | 8222 | 27 | CGTGGACGGTGTGAGGCCGGTAGCGGC | 494.3 | -3.6 | 1.1 | -3.2 | 1.5E-03 | 1.9E-03 | MB231/MB468 | CYTO |
| >hg38_21_+_8433172_8446622_RNA45SN1_45S_50ntFlanks@11190.11207.18\|CCCGGGGGGCCGGCGGCG | >hg38_21_+_8433172_8446622_RNA45SN1_45S_50ntFlanks | 11190 | 11207 | 18 | CCCGGGGGGCCGGCGGCG | 466.1 | -3.7 | 1.2 | -3.2 | 1.4E-03 | 1.9E-03 | BT20/MB468 | CYTO |
| >hg38_21_+_8433172_8446622_RNA45SN1_45S_50ntFlanks@8177.8208.32\|TTCTGATCGAGGCCCAGCCCGTGGACGGTGTG | >hg38_21_+_8433172_8446622_RNA45SN1_45S_50ntFlanks | 8177 | 8208 | 32 | TTCTGATCGAGGCCCAGCCCGTGGACGGTGTG | 254.3 | -2.1 | 0.7 | -3.2 | 1.6E-03 | 1.9E-03 | MB231/MB468 | NUC |
| >hg38_21_+_8433172_8446622_RNA45SN1_45S_50ntFlanks@12004.12026.23\|CGGGCCGCCGGTGAAATACCACT | >hg38_21_+_8433172_8446622_RNA45SN1_45S_50ntFlanks | 12004 | 12026 | 23 | CGGGCCGCCGGTGAAATACCACT | 303.2 | 6.4 | 1.8 | 3.7 | 2.5E-04 | 1.9E-03 | BT20/MB231 | MP |
| >hg38_MT_+_598_1651_RNR1_12S_50ntFlanks@854.871.18\|TTAGTTGAACAGGGCCCT | >hg38_MT_+_598_1651_RNR1_12S_50ntFlanks | 854 | 871 | 18 | TTAGTTGAACAGGGCCCT | 48.5 | 9.3 | 2.8 | 3.4 | 7.3E-04 | 1.9E-03 | BT20/MB468 | MP |
| >hg38_21_+_8433172_8446622_RNA45SN1_45S_50ntFlanks@11789.11810.22\|TGGATGAACGAGATTCCCACTG | >hg38_21_+_8433172_8446622_RNA45SN1_45S_50ntFlanks | 11789 | 11810 | 22 | TGGATGAACGAGATTCCCACTG | 46.3 | -8.6 | 2.6 | -3.4 | 7.3E-04 | 1.9E-03 | BT20/MB468 | MP |
| >hg38_21_+_8433172_8446622_RNA45SN1_45S_50ntFlanks@4725.4748.24\|TTAATCAAGAACGAAAGTCGGAGG | >hg38_21_+_8433172_8446622_RNA45SN1_45S_50ntFlanks | 4725 | 4748 | 24 | TTAATCAAGAACGAAAGTCGGAGG | 70.9 | -5.2 | 1.6 | -3.4 | 7.4E-04 | 1.9E-03 | BT20/MB468 | MP |
| >hg38_21_+_8433172_8446622_RNA45SN1_45S_50ntFlanks@11671.11700.30\|CCAGTGCTCTGAATGTCAAAGTGAAGAAAT | >hg38_21_+_8433172_8446622_RNA45SN1_45S_50ntFlanks | 11671 | 11700 | 30 | CCAGTGCTCTGAATGTCAAAGTGAAGAAAT | 165.3 | -3.2 | 1.0 | -3.2 | 1.4E-03 | 1.9E-03 | BT20/MB468 | CYTO |
| >hg38_21_+_8433172_8446622_RNA45SN1_45S_50ntFlanks@10910.10930.21\|CGCGCGCCGCGGCTGGACGAG | >hg38_21_+_8433172_8446622_RNA45SN1_45S_50ntFlanks | 10910 | 10930 | 21 | CGCGCGCCGCGGCTGGACGAG | 101.3 | 10.0 | 2.7 | 3.7 | 2.5E-04 | 1.9E-03 | BT20/MB231 | MP |
| >hg38_MT_+_1621_3279_RNR2_16S_50ntFlanks@1092.1119.28\|GCAAGACGAGAAGACCCTATGGAGCTTT | >hg38_MT_+_1621_3279_RNR2_16S_50ntFlanks | 1092 | 1119 | 28 | GCAAGACGAGAAGACCCTATGGAGCTTT | 105.3 | 3.4 | 1.0 | 3.4 | 7.4E-04 | 1.9E-03 | BT20/MB468 | MP |
| >trnaMT_GluTTC_MT_-_14674_14742@40.71.32,trnalookalike8_GluTTC_5_-_93905172_93905240@40.71.32\|TTGGTCGTGGTTGTAGTCCGTGCGAGAATACC | >trnaMT_GluTTC_MT_-_14674_14742 | 40 | 71 | 32 | TTGGTCGTGGTTGTAGTCCGTGCGAGAATACC | 1448.9 | 3.8 | 1.1 | 3.4 | 7.4E-04 | 1.9E-03 | BT20/MB468 | MP |
| >MI0000298\|hsa-mir-221&WithFlank&X\|-\|45746151\|45746272@71.92.22&[MIMAT0000278&hsa-miR-221-3p&offsets\|0\|-1;m-91&X\|-\|45746181\|45746202&offsets\|0\|0]\|AGCTACATTGTCTGCTGGGTTT | >MI0000298\|hsa-mir-221&WithFlank&X\|-\|45746151\|45746272 | 71 | 92 | 22 | AGCTACATTGTCTGCTGGGTTT | 15731.2 | 4.0 | 1.2 | 3.4 | 7.4E-04 | 1.9E-03 | BT20/MB468 | MP |
| >hg38_21_+_8433172_8446622_RNA45SN1_45S_50ntFlanks@12420.12448.29\|ATGTCGGCTCTTCCTATCATTGTGAAGCA | >hg38_21_+_8433172_8446622_RNA45SN1_45S_50ntFlanks | 12420 | 12448 | 29 | ATGTCGGCTCTTCCTATCATTGTGAAGCA | 1929.9 | -2.9 | 0.8 | -3.6 | 3.2E-04 | 1.9E-03 | BT20/MB231 | MITO |
| >hg38_21_+_8433172_8446622_RNA45SN1_45S_50ntFlanks@12162.12180.19\|GACTGGGGCGGTACACCTG | >hg38_21_+_8433172_8446622_RNA45SN1_45S_50ntFlanks | 12162 | 12180 | 19 | GACTGGGGCGGTACACCTG | 486.3 | -4.0 | 1.2 | -3.2 | 1.2E-03 | 1.9E-03 | BT20/MB468 | MITO |
| >trna116_GluCTC_1_-_145399233_145399304@38.74.37,trna134_GluTTC_1_-_16861774_16861845@38.74.37,trna71_GluCTC_1_-_161439189_161439260@38.74.37,trna74_GluCTC_1_-_161431809_161431880@38.74.37,trna77_GluCTC_1_-_161424398_161424469@38.74.37,trna77_GluCTC_6_+_28949976_28950047@38.74.37,trna80_GluCTC_1_-_161417018_161417089@38.74.37,trna84_GluTTC_1_-_161391883_161391954@38.74.37,trna87_GluCTC_6_-_126101393_126101464@38.74.37\|CCGCCGCGGCCCGGGTTCGATTCCCGGTCAGGGAACC | >trna116_GluCTC_1_-_145399233_145399304 | 38 | 74 | 37 | CCGCCGCGGCCCGGGTTCGATTCCCGGTCAGGGAACC | 269.1 | 4.2 | 1.3 | 3.2 | 1.4E-03 | 1.9E-03 | BT20/MB468 | CYTO |
| >hg38_21_+_8433172_8446622_RNA45SN1_45S_50ntFlanks@10679.10710.32\|GGTTCCGGCGGCGTCCGGTGAGCTCTCGCTGG | >hg38_21_+_8433172_8446622_RNA45SN1_45S_50ntFlanks | 10679 | 10710 | 32 | GGTTCCGGCGGCGTCCGGTGAGCTCTCGCTGG | 20281.6 | -4.5 | 1.3 | -3.4 | 7.4E-04 | 1.9E-03 | BT20/MB468 | MP |
| >hg38_21_+_8433172_8446622_RNA45SN1_45S_50ntFlanks@8176.8195.20\|CTTCTGATCGAGGCCCAGCC | >hg38_21_+_8433172_8446622_RNA45SN1_45S_50ntFlanks | 8176 | 8195 | 20 | CTTCTGATCGAGGCCCAGCC | 991.3 | -2.7 | 0.8 | -3.4 | 7.4E-04 | 1.9E-03 | BT20/MB468 | MP |
| >hg38_MT_+_598_1651_RNR1_12S_50ntFlanks@844.872.29\|AGTAGAGTGCTTAGTTGAACAGGGCCCTG | >hg38_MT_+_598_1651_RNR1_12S_50ntFlanks | 844 | 872 | 29 | AGTAGAGTGCTTAGTTGAACAGGGCCCTG | 78.9 | 7.0 | 2.1 | 3.4 | 7.4E-04 | 1.9E-03 | BT20/MB468 | MP |
| >hg38_21_+_8433172_8446622_RNA45SN1_45S_50ntFlanks@12416.12453.38\|TTCGATGTCGGCTCTTCCTATCATTGTGAAGCAGAATT | >hg38_21_+_8433172_8446622_RNA45SN1_45S_50ntFlanks | 12416 | 12453 | 38 | TTCGATGTCGGCTCTTCCTATCATTGTGAAGCAGAATT | 531.0 | -3.0 | 0.8 | -3.6 | 2.9E-04 | 1.9E-03 | BT20/MB231 | CYTO |
| >hg38_21_+_8433172_8446622_RNA45SN1_45S_50ntFlanks@12636.12669.34\|AGCTACCATCTGTGGGATTATGACTGAACGCCTC | >hg38_21_+_8433172_8446622_RNA45SN1_45S_50ntFlanks | 12636 | 12669 | 34 | AGCTACCATCTGTGGGATTATGACTGAACGCCTC | 59.5 | -4.3 | 1.3 | -3.2 | 1.3E-03 | 1.9E-03 | MB231/MB468 | MP |
| >am_MI0005763\|hsa-mir-941-1&WithFlank&20\|+\|63919443\|63919526@53.73.21(+1A);MI0005764\|hsa-mir-941-2&WithFlank&20\|+\|63919499\|63919582@53.73.21(+1A);MI0005765\|hsa-mir-941-3&WithFlank&20\|+\|63919555\|63919638@53.73.21(+1A);MI0005766\|hsa-mir-941-4&WithFlank&20\|+\|63919750\|63919833@53.73.21(+1A);MI0031520\|hsa-mir-941-5&WithFlank&20\|+\|63919862\|63919945@53.73.21(+1A)&[MIMAT0004984&hsa-miR-941&offsets\|0\|-2(+1A);m-263&20\|+\|63919495\|63919517&offsets\|0\|-2(+1A)];[MIMAT0004984_1&hsa-miR-941&offsets\|0\|-2(+1A);m-263&20\|+\|63919551\|63919573&offsets\|0\|-2(+1A)];[MIMAT0004984_2&hsa-miR-941&offsets\|0\|-2(+1A);m-263&20\|+\|63919607\|63919629&offsets\|0\|-2(+1A)];[MIMAT0004984_3&hsa-miR-941&offsets\|0\|-2(+1A);m-263&20\|+\|63919802\|63919824&offsets\|0\|-2(+1A)];[MIMAT0004984_4&hsa-miR-941&offsets\|0\|-2(+1A);m-263&20\|+\|63919914\|63919936&offsets\|0\|-2(+1A)]\|CACCCGGCTGTGTGCACATGTA | >am_MI0005763\|hsa-mir-941-1&WithFlank&20\|+\|63919443\|63919526 | 53 | 73 | 21(+1A) | CACCCGGCTGTGTGCACATGTA | 81.3 | 5.0 | 1.5 | 3.2 | 1.3E-03 | 1.9E-03 | MB231/MB468 | MP |
| >hg38_21_+_8433172_8446622_RNA45SN1_45S_50ntFlanks@9737.9758.22\|CGAAACGATCTCAACCTATTCT | >hg38_21_+_8433172_8446622_RNA45SN1_45S_50ntFlanks | 9737 | 9758 | 22 | CGAAACGATCTCAACCTATTCT | 106.3 | -4.1 | 1.3 | -3.2 | 1.3E-03 | 1.9E-03 | MB231/MB468 | MP |
| >hg38_21_+_8433172_8446622_RNA45SN1_45S_50ntFlanks@11208.11234.27\|GCGGCGACTCTGGACGCGAGCCGGGCC | >hg38_21_+_8433172_8446622_RNA45SN1_45S_50ntFlanks | 11208 | 11234 | 27 | GCGGCGACTCTGGACGCGAGCCGGGCC | 1334.0 | -2.3 | 0.7 | -3.2 | 1.2E-03 | 1.9E-03 | BT20/MB468 | MITO |
| >hg38_21_+_8433172_8446622_RNA45SN1_45S_50ntFlanks@10508.10533.26\|CGAATCCGGAGTGGCGGAGATGGGCG | >hg38_21_+_8433172_8446622_RNA45SN1_45S_50ntFlanks | 10508 | 10533 | 26 | CGAATCCGGAGTGGCGGAGATGGGCG | 281.7 | -3.6 | 1.1 | -3.2 | 1.2E-03 | 1.9E-03 | BT20/MB468 | MITO |
| >hg38_21_+_8433172_8446622_RNA45SN1_45S_50ntFlanks@8168.8185.18\|CCCAAGTCCTTCTGATCG | >hg38_21_+_8433172_8446622_RNA45SN1_45S_50ntFlanks | 8168 | 8185 | 18 | CCCAAGTCCTTCTGATCG | 303.7 | -2.3 | 0.7 | -3.2 | 1.2E-03 | 1.9E-03 | BT20/MB468 | MITO |
| >hg38_21_+_8433172_8446622_RNA45SN1_45S_50ntFlanks@11670.11690.21\|CCCAGTGCTCTGAATGTCAAA | >hg38_21_+_8433172_8446622_RNA45SN1_45S_50ntFlanks | 11670 | 11690 | 21 | CCCAGTGCTCTGAATGTCAAA | 90.5 | -5.0 | 1.5 | -3.2 | 1.3E-03 | 1.9E-03 | MB231/MB468 | MP |
| >hg38_21_+_8433172_8446622_RNA45SN1_45S_50ntFlanks@11881.11903.23\|CCTGTTGAGCTTGACTCTAGTCT | >hg38_21_+_8433172_8446622_RNA45SN1_45S_50ntFlanks | 11881 | 11903 | 23 | CCTGTTGAGCTTGACTCTAGTCT | 220.4 | -3.6 | 1.2 | -3.2 | 1.6E-03 | 1.9E-03 | MB231/MB468 | NUC |
| >MI0000746\|hsa-mir-99b&WithFlank&19\|+\|51692606\|51692687@51.72.22(+1A)&[MIMAT0004678&hsa-miR-99b-3p&offsets\|0\|0(+1A);m-151&19\|+\|51692656\|51692678&offsets\|0\|-1(+1A)]\|CAAGCTCGTGTCTGTGGGTCCGA | >MI0000746\|hsa-mir-99b&WithFlank&19\|+\|51692606\|51692687 | 51 | 72 | 22(+1A) | CAAGCTCGTGTCTGTGGGTCCGA | 531.3 | 2.6 | 0.7 | 3.6 | 2.8E-04 | 1.9E-03 | BT20/MB231 | NUC |
| >MI0000542\|hsa-mir-320a&WithFlank&8\|-\|22244960\|22245043@48.68.21(+3U)&[MIMAT0000510&hsa-miR-320a-3p&offsets\|0\|-1(+3U);m-60&8\|-\|22244975\|22244996&offsets\|0\|-1(+3U)]\|AAAAGCTGGGTTGAGAGGGCGTTT | >MI0000542\|hsa-mir-320a&WithFlank&8\|-\|22244960\|22245043 | 48 | 68 | 21(+3U) | AAAAGCTGGGTTGAGAGGGCGTTT | 610.0 | 2.8 | 0.8 | 3.6 | 2.8E-04 | 1.9E-03 | BT20/MB231 | NUC |
| >hg38_21_+_8433172_8446622_RNA45SN1_45S_50ntFlanks@3846.3874.29\|CTTGGATAACTGTGGTAATTCTAGAGCTA | >hg38_21_+_8433172_8446622_RNA45SN1_45S_50ntFlanks | 3846 | 3874 | 29 | CTTGGATAACTGTGGTAATTCTAGAGCTA | 167.7 | -5.4 | 1.6 | -3.4 | 7.4E-04 | 1.9E-03 | BT20/MB468 | MP |
| >hg38_21_+_8433172_8446622_RNA45SN1_45S_50ntFlanks@10685.10710.26\|GGCGGCGTCCGGTGAGCTCTCGCTGG | >hg38_21_+_8433172_8446622_RNA45SN1_45S_50ntFlanks | 10685 | 10710 | 26 | GGCGGCGTCCGGTGAGCTCTCGCTGG | 1722.0 | -5.1 | 1.6 | -3.2 | 1.6E-03 | 1.9E-03 | BT20/MB468 | NUC |
| >hg38_21_+_8433172_8446622_RNA45SN1_45S_50ntFlanks@11211.11254.44\|GCGACTCTGGACGCGAGCCGGGCCCTTCCCGTGGATCGCCCCAG | >hg38_21_+_8433172_8446622_RNA45SN1_45S_50ntFlanks | 11211 | 11254 | 44 | GCGACTCTGGACGCGAGCCGGGCCCTTCCCGTGGATCGCCCCAG | 292.2 | -3.8 | 1.2 | -3.2 | 1.4E-03 | 1.9E-03 | BT20/MB468 | CYTO |
| >hg38_21_+_8433172_8446622_RNA45SN1_45S_50ntFlanks@12419.12447.29\|GATGTCGGCTCTTCCTATCATTGTGAAGC | >hg38_21_+_8433172_8446622_RNA45SN1_45S_50ntFlanks | 12419 | 12447 | 29 | GATGTCGGCTCTTCCTATCATTGTGAAGC | 3580.5 | -3.0 | 0.8 | -3.6 | 2.9E-04 | 1.9E-03 | BT20/MB231 | CYTO |
| >trnaMT_ValTAC_MT_+_1602_1670@53.71.19\|TAACTTGACCGCTCTGACC | >trnaMT_ValTAC_MT_+_1602_1670 | 53 | 71 | 19 | TAACTTGACCGCTCTGACC | 386.3 | 2.3 | 0.7 | 3.2 | 1.6E-03 | 1.9E-03 | MB231/MB468 | MITO |
| >am_trnaMT_SerGCT_MT_+_12207_12265@14.34.21\|AGAACTGCTAACTCATGCCCC | >am_trnaMT_SerGCT_MT_+_12207_12265 | 14 | 34 | 21 | AGAACTGCTAACTCATGCCCC | 180.7 | 5.1 | 1.5 | 3.4 | 7.4E-04 | 1.9E-03 | BT20/MB468 | MP |
| >hg38_21_+_8433172_8446622_RNA45SN1_45S_50ntFlanks@12149.12182.34\|AGGTGGGGAGTTTGACTGGGGCGGTACACCTGTC | >hg38_21_+_8433172_8446622_RNA45SN1_45S_50ntFlanks | 12149 | 12182 | 34 | AGGTGGGGAGTTTGACTGGGGCGGTACACCTGTC | 72.9 | -5.3 | 1.6 | -3.4 | 7.4E-04 | 1.9E-03 | BT20/MB468 | MP |
| >am_MI0006370\|hsa-mir-1303&WithFlank&5\|+\|154685770\|154685867@57.79.23&[MIMAT0005891&hsa-miR-1303&offsets\|-1\|0;m-516&5\|+\|154685826\|154685848&offsets\|0\|0]\|TTTTAGAGACGGGGTCTTGCTCT | >am_MI0006370\|hsa-mir-1303&WithFlank&5\|+\|154685770\|154685867 | 57 | 79 | 23 | TTTTAGAGACGGGGTCTTGCTCT | 175.5 | 4.5 | 1.3 | 3.4 | 7.4E-04 | 1.9E-03 | BT20/MB468 | MP |
| >hg38_21_+_8433172_8446622_RNA45SN1_45S_50ntFlanks@10843.10867.25\|TAACTTCGGGATAAGGATTGGCTCT | >hg38_21_+_8433172_8446622_RNA45SN1_45S_50ntFlanks | 10843 | 10867 | 25 | TAACTTCGGGATAAGGATTGGCTCT | 403.8 | -2.4 | 0.8 | -3.2 | 1.5E-03 | 1.9E-03 | BT20/MB468 | CYTO |
| >MI0001446\|hsa-mir-424&WithFlank&X\|-\|134546608\|134546717@17.37.21&[MIMAT0001341&hsa-miR-424-5p&offsets\|0\|-1;m-152&X\|-\|134546681\|134546701&offsets\|0\|0]\|CAGCAGCAATTCATGTTTTGA | >MI0001446\|hsa-mir-424&WithFlank&X\|-\|134546608\|134546717 | 17 | 37 | 21 | CAGCAGCAATTCATGTTTTGA | 158.5 | 4.9 | 1.5 | 3.2 | 1.3E-03 | 1.9E-03 | MB231/MB468 | MP |
| >hg38_21_+_8433172_8446622_RNA45SN1_45S_50ntFlanks@7989.8011.23\|AGACGTGGCGACCCGCTGAATTT | >hg38_21_+_8433172_8446622_RNA45SN1_45S_50ntFlanks | 7989 | 8011 | 23 | AGACGTGGCGACCCGCTGAATTT | 2383.0 | -2.2 | 0.7 | -3.2 | 1.3E-03 | 1.9E-03 | MB231/MB468 | MP |
| >hg38_21_+_8433172_8446622_RNA45SN1_45S_50ntFlanks@8178.8207.30\|TCTGATCGAGGCCCAGCCCGTGGACGGTGT | >hg38_21_+_8433172_8446622_RNA45SN1_45S_50ntFlanks | 8178 | 8207 | 30 | TCTGATCGAGGCCCAGCCCGTGGACGGTGT | 191.9 | 8.2 | 2.4 | 3.4 | 7.4E-04 | 1.9E-03 | BT20/MB468 | MP |

<table>
<tr><td></td><td></td><td></td><td></td><td></td><td></td><td></td><td></td><td></td><td></td><td></td><td></td><td></td><td></td></tr>
<tr><td>>hg38_21_+_8433172_8446622_RNA45SN1_45S_50ntFlanks@10346.10388.43|TCAAACGAGAACTTTGAAGGCCGAAGTGGAGAAGGGTTCCATG</td><td>>hg38_21_+_8433172_8446622_RNA45SN1_45S_50ntFlanks</td><td>10346</td><td>10388</td><td>43</td><td>TCAAACGAGAACTTTGAAGGCCGAAGTGGAGAAGGGTTCCATG</td><td>193.0</td><td>-5.5</td><td>1.7</td><td>-3.2</td><td>1.3E-03</td><td>1.9E-03</td><td>MB231/MB468</td><td>MP</td></tr>
<tr><td>>hg38_21_+_8433172_8446622_RNA45SN1_45S_50ntFlanks@10389.10424.36|TGAACAGCAGTTGAACATGGGTCAGTCGGTCCTGAG</td><td>>hg38_21_+_8433172_8446622_RNA45SN1_45S_50ntFlanks</td><td>10389</td><td>10424</td><td>36</td><td>TGAACAGCAGTTGAACATGGGTCAGTCGGTCCTGAG</td><td>7513.4</td><td>3.8</td><td>1.1</td><td>3.4</td><td>7.5E-04</td><td>1.9E-03</td><td>BT20/MB468</td><td>MP</td></tr>
<tr><td>>MI0000299|hsa-mir-222&WithFlank&X|-|45747009|45747130@75.99.25(+1U)&[MIMAT0000279&hsa-miR-222-3p&offsets|0|+4(+1U);m-140&X|-|45747033|45747056&offsets|0|+1(+1U)]|AGCTACATCTGGCTACTGGGTCTCTT</td><td>>MI0000299|hsa-mir-222&WithFlank&X|-|45747009|45747130</td><td>75</td><td>99</td><td>25(+1U)</td><td>AGCTACATCTGGCTACTGGGTCTCTT</td><td>5602.8</td><td>-2.9</td><td>0.8</td><td>-3.6</td><td>3.3E-04</td><td>1.9E-03</td><td>BT20/MB231</td><td>MITO</td></tr>
<tr><td>>MI0000289|hsa-mir-181a-1&WithFlank&1|-|198859038|198859159@30.49.20;MI0000269|hsa-mir-181a-2&WithFlank&9|+|124692436|124692557@45.64.20&[MIMAT0000256&hsa-miR-181a-5p&offsets|0|-3;m-32&1|-|198859108|198859130&offsets|0|-3];[MIMAT0000256_1&hsa-miR-181a-5p&offsets|0|-3;m-32&9|+|124692480|124692502&offsets|0|-3]|AACATTCAACGCTGTCGGTG</td><td>>MI0000289|hsa-mir-181a-1&WithFlank&1|-|198859038|198859159</td><td>30</td><td>49</td><td>20</td><td>AACATTCAACGCTGTCGGTG</td><td>770.0</td><td>-2.7</td><td>0.7</td><td>-3.6</td><td>3.3E-04</td><td>1.9E-03</td><td>BT20/MB231</td><td>MITO</td></tr>
<tr><td>>MI0000456|hsa-mir-140&WithFlank&16|+|69933075|69933186@68.90.23(+1U)&[MIMAT0004597&hsa-miR-140-3p&offsets|0|+2(+1U);m-45&16|+|69933143|69933164&offsets|-1|0(+1U)]|TACCACAGGGTAGAACCACGGACT</td><td>>MI0000456|hsa-mir-140&WithFlank&16|+|69933075|69933186</td><td>68</td><td>90</td><td>23(+1U)</td><td>TACCACAGGGTAGAACCACGGACT</td><td>256.1</td><td>-2.7</td><td>0.8</td><td>-3.6</td><td>3.3E-04</td><td>1.9E-03</td><td>BT20/MB231</td><td>MITO</td></tr>
<tr><td>>trnaMT_MetCAT_MT_+_4402_4469@37.71.35|CCGAAAATGTTGGTTATACCCTTCCCGTACTACCA</td><td>>trnaMT_MetCAT_MT_+_4402_4469</td><td>37</td><td>71</td><td>35</td><td>CCGAAAATGTTGGTTATACCCTTCCCGTACTACCA</td><td>1075.1</td><td>5.6</td><td>1.7</td><td>3.4</td><td>7.5E-04</td><td>1.9E-03</td><td>BT20/MB468</td><td>MP</td></tr>
<tr><td>>hg38_21_+_8433172_8446622_RNA45SN1_45S_50ntFlanks@11874.11910.37|AGAAGACCCTGTTGAGCTTGACTCTAGTCTGGCACGG</td><td>>hg38_21_+_8433172_8446622_RNA45SN1_45S_50ntFlanks</td><td>11874</td><td>11910</td><td>37</td><td>AGAAGACCCTGTTGAGCTTGACTCTAGTCTGGCACGG</td><td>202.1</td><td>-4.6</td><td>1.4</td><td>-3.2</td><td>1.5E-03</td><td>1.9E-03</td><td>BT20/MB468</td><td>CYTO</td></tr>
<tr><td>>hg38_21_+_8433172_8446622_RNA45SN1_45S_50ntFlanks@10819.10836.18|AGGGAAGTCGGCAAGCCG</td><td>>hg38_21_+_8433172_8446622_RNA45SN1_45S_50ntFlanks</td><td>10819</td><td>10836</td><td>18</td><td>AGGGAAGTCGGCAAGCCG</td><td>397.6</td><td>-3.0</td><td>0.9</td><td>-3.4</td><td>7.5E-04</td><td>1.9E-03</td><td>BT20/MB468</td><td>MP</td></tr>
<tr><td>>hg38_21_+_8433172_8446622_RNA45SN1_45S_50ntFlanks@10823.10840.18|AAGTCGGCAAGCCGGATC</td><td>>hg38_21_+_8433172_8446622_RNA45SN1_45S_50ntFlanks</td><td>10823</td><td>10840</td><td>18</td><td>AAGTCGGCAAGCCGGATC</td><td>476.2</td><td>-6.1</td><td>1.8</td><td>-3.4</td><td>7.5E-04</td><td>1.9E-03</td><td>BT20/MB468</td><td>MP</td></tr>
<tr><td>>hg38_21_+_8433172_8446622_RNA45SN1_45S_50ntFlanks@4163.4184.22|CATCCAAGGAAGGCAGCAGGCG</td><td>>hg38_21_+_8433172_8446622_RNA45SN1_45S_50ntFlanks</td><td>4163</td><td>4184</td><td>22</td><td>CATCCAAGGAAGGCAGCAGGCG</td><td>885.0</td><td>-5.8</td><td>1.7</td><td>-3.4</td><td>7.5E-04</td><td>1.9E-03</td><td>BT20/MB468</td><td>MP</td></tr>
<tr><td>>hg38_21_+_8433172_8446622_RNA45SN1_45S_50ntFlanks@11564.11609.46|CCTAGCAGCCGACTTAGAACTGGTGCGGACCAGGGGAATCCGACTG</td><td>>hg38_21_+_8433172_8446622_RNA45SN1_45S_50ntFlanks</td><td>11564</td><td>11609</td><td>46</td><td>CCTAGCAGCCGACTTAGAACTGGTGCGGACCAGGGGAATCCGACTG</td><td>53.8</td><td>-9.6</td><td>3.0</td><td>-3.2</td><td>1.3E-03</td><td>1.9E-03</td><td>MB231/MB468</td><td>MP</td></tr>
<tr><td>>hg38_21_+_8433172_8446622_RNA45SN1_45S_50ntFlanks@6645.6668.24|CTCGTACGACTCTTAGCGGTGGAT</td><td>>hg38_21_+_8433172_8446622_RNA45SN1_45S_50ntFlanks</td><td>6645</td><td>6668</td><td>24</td><td>CTCGTACGACTCTTAGCGGTGGAT</td><td>1863.0</td><td>2.1</td><td>0.7</td><td>3.2</td><td>1.6E-03</td><td>1.9E-03</td><td>MB231/MB468</td><td>MITO</td></tr>
<tr><td>>trnaMT_GluTTC_MT_-_14674_14742@44.63.20,trnalookalike8_GluTTC_5_-_93905172_93905240@44.63.20|TCGTGGTTGTAGTCCGTGCG</td><td>>trnaMT_GluTTC_MT_-_14674_14742</td><td>44</td><td>63</td><td>20</td><td>TCGTGGTTGTAGTCCGTGCG</td><td>217.8</td><td>3.8</td><td>1.1</td><td>3.4</td><td>7.5E-04</td><td>1.9E-03</td><td>BT20/MB468</td><td>MP</td></tr>
<tr><td>>hg38_21_+_8433172_8446622_RNA45SN1_45S_50ntFlanks@3705.3748.44|TACCTGGTTGATCCTGCCAGTAGCATATGCTTGTCTCAAAGATT</td><td>>hg38_21_+_8433172_8446622_RNA45SN1_45S_50ntFlanks</td><td>3705</td><td>3748</td><td>44</td><td>TACCTGGTTGATCCTGCCAGTAGCATATGCTTGTCTCAAAGATT</td><td>1731.5</td><td>3.7</td><td>1.2</td><td>3.2</td><td>1.6E-03</td><td>1.9E-03</td><td>BT20/MB468</td><td>NUC</td></tr>
<tr><td>>hg38_21_+_8433172_8446622_RNA45SN1_45S_50ntFlanks@4727.4752.26|AATCAAGAACGAAAGTCGGAGGTTCG</td><td>>hg38_21_+_8433172_8446622_RNA45SN1_45S_50ntFlanks</td><td>4727</td><td>4752</td><td>26</td><td>AATCAAGAACGAAAGTCGGAGGTTCG</td><td>146.4</td><td>-3.7</td><td>1.2</td><td>-3.2</td><td>1.6E-03</td><td>1.9E-03</td><td>MB231/MB468</td><td>NUC</td></tr>
<tr><td>>hg38_21_+_8433172_8446622_RNA45SN1_45S_50ntFlanks@4685.4723.39|AGACGGACCAGAGCGAAAGCATTTGCCAAGAATGTTTTC</td><td>>hg38_21_+_8433172_8446622_RNA45SN1_45S_50ntFlanks</td><td>4685</td><td>4723</td><td>39</td><td>AGACGGACCAGAGCGAAAGCATTTGCCAAGAATGTTTTC</td><td>167.1</td><td>-5.6</td><td>1.7</td><td>-3.2</td><td>1.3E-03</td><td>1.9E-03</td><td>MB231/MB468</td><td>MP</td></tr>
<tr><td>>hg38_21_+_8433172_8446622_RNA45SN1_45S_50ntFlanks@8176.8197.22|CTTCTGATCGAGGCCCAGCCCG</td><td>>hg38_21_+_8433172_8446622_RNA45SN1_45S_50ntFlanks</td><td>8176</td><td>8197</td><td>22</td><td>CTTCTGATCGAGGCCCAGCCCG</td><td>693.8</td><td>-5.7</td><td>1.7</td><td>-3.4</td><td>7.5E-04</td><td>1.9E-03</td><td>BT20/MB468</td><td>MP</td></tr>
<tr><td>>trna10_LysCTT_16_+_3241501_3241573@1.36.36,trna13_LysCTT_14_-_58706613_58706685@1.36.36,trna2_LysCTT_15_+_79152904_79152976@1.36.36|GCCCGGCTAGCTCAGTCGGTAGAGCATGGGACTCTT</td><td>>trna10_LysCTT_16_+_3241501_3241573</td><td>1</td><td>36</td><td>36</td><td>GCCCGGCTAGCTCAGTCGGTAGAGCATGGGACTCTT</td><td>7454.7</td><td>3.5</td><td>1.0</td><td>3.6</td><td>2.9E-04</td><td>1.9E-03</td><td>BT20/MB231</td><td>CYTO</td></tr>
<tr><td>>hg38_21_+_8433172_8446622_RNA45SN1_45S_50ntFlanks@5073.5094.22|AATTCCGATAACGAACGAGACT</td><td>>hg38_21_+_8433172_8446622_RNA45SN1_45S_50ntFlanks</td><td>5073</td><td>5094</td><td>22</td><td>AATTCCGATAACGAACGAGACT</td><td>493.1</td><td>-4.0</td><td>1.2</td><td>-3.2</td><td>1.2E-03</td><td>1.9E-03</td><td>BT20/MB468</td><td>MITO</td></tr>
<tr><td>>hg38_21_+_8433172_8446622_RNA45SN1_45S_50ntFlanks@11921.11948.28|TGAGAGGTGTAGAATAAGTGGGAGGCCC</td><td>>hg38_21_+_8433172_8446622_RNA45SN1_45S_50ntFlanks</td><td>11921</td><td>11948</td><td>28</td><td>TGAGAGGTGTAGAATAAGTGGGAGGCCC</td><td>338.5</td><td>-3.3</td><td>1.0</td><td>-3.2</td><td>1.5E-03</td><td>1.9E-03</td><td>BT20/MB468</td><td>CYTO</td></tr>
<tr><td>>hg38_21_+_8433172_8446622_RNA45SN1_45S_50ntFlanks@13000.13027.28|TGCGATCTATTGAAAGTCAGCCCTCGAC</td><td>>hg38_21_+_8433172_8446622_RNA45SN1_45S_50ntFlanks</td><td>13000</td><td>13027</td><td>28</td><td>TGCGATCTATTGAAAGTCAGCCCTCGAC</td><td>302.9</td><td>-3.0</td><td>0.8</td><td>-3.6</td><td>2.9E-04</td><td>1.9E-03</td><td>BT20/MB231</td><td>CYTO</td></tr>
<tr><td>>hg38_21_+_8433172_8446622_RNA45SN1_45S_50ntFlanks@11876.11898.23|AAGACCCTGTTGAGCTTGACTCT</td><td>>hg38_21_+_8433172_8446622_RNA45SN1_45S_50ntFlanks</td><td>11876</td><td>11898</td><td>23</td><td>AAGACCCTGTTGAGCTTGACTCT</td><td>2649.8</td><td>-5.1</td><td>1.6</td><td>-3.2</td><td>1.6E-03</td><td>1.9E-03</td><td>BT20/MB468</td><td>NUC</td></tr>
<tr><td>>trnaMT_LeuTAA_MT_+_3230_3304@16.45.30|CCCGGTAATCGCATAAAACTTAAAACTTTA</td><td>>trnaMT_LeuTAA_MT_+_3230_3304</td><td>16</td><td>45</td><td>30</td><td>CCCGGTAATCGCATAAAACTTAAAACTTTA</td><td>118.1</td><td>-4.6</td><td>1.5</td><td>-3.2</td><td>1.5E-03</td><td>1.9E-03</td><td>BT20/MB468</td><td>CYTO</td></tr>
<tr><td>>hg38_21_+_8433172_8446622_RNA45SN1_45S_50ntFlanks@10771.10796.26|CAGGTCTCCAAGGTGAACAGCCTCTG</td><td>>hg38_21_+_8433172_8446622_RNA45SN1_45S_50ntFlanks</td><td>10771</td><td>10796</td><td>26</td><td>CAGGTCTCCAAGGTGAACAGCCTCTG</td><td>489.0</td><td>-3.9</td><td>1.2</td><td>-3.2</td><td>1.5E-03</td><td>1.9E-03</td><td>BT20/MB468</td><td>CYTO</td></tr>
<tr><td>>hg38_21_+_8433172_8446622_RNA45SN1_45S_50ntFlanks@4046.4065.20|CATTCGAACGTCTGCCCTAT</td><td>>hg38_21_+_8433172_8446622_RNA45SN1_45S_50ntFlanks</td><td>4046</td><td>4065</td><td>20</td><td>CATTCGAACGTCTGCCCTAT</td><td>280.7</td><td>-1.8</td><td>0.6</td><td>-3.2</td><td>1.5E-03</td><td>1.9E-03</td><td>MB231/MB468</td><td>CYTO</td></tr>
<tr><td>>hg38_21_+_8433172_8446622_RNA45SN1_45S_50ntFlanks@9939.9956.18|GCAGGACGGTGGCCATGG</td><td>>hg38_21_+_8433172_8446622_RNA45SN1_45S_50ntFlanks</td><td>9939</td><td>9956</td><td>18</td><td>GCAGGACGGTGGCCATGG</td><td>279.0</td><td>5.8</td><td>1.7</td><td>3.4</td><td>7.5E-04</td><td>1.9E-03</td><td>BT20/MB468</td><td>MP</td></tr>
<tr><td>>am_trnaMT_ValTAC_MT_+_1602_1670@38.66.29|TAGGAGATTTCAACTTAACTTGACCGCTC</td><td>>am_trnaMT_ValTAC_MT_+_1602_1670</td><td>38</td><td>66</td><td>29</td><td>TAGGAGATTTCAACTTAACTTGACCGCTC</td><td>205.9</td><td>8.3</td><td>2.5</td><td>3.4</td><td>7.5E-04</td><td>1.9E-03</td><td>BT20/MB468</td><td>MP</td></tr>
<tr><td>>MI0000088|hsa-mir-30a&WithFlank&6|-|71403545|71403627@54.74.21&[MIMAT0000088&hsa-miR-30a-3p&offsets|+1|0;m-38&6|-|71403554|71403575&offsets|+1|0]|TTTCAGTCGGATGTTTGCAGC</td><td>>MI0000088|hsa-mir-30a&WithFlank&6|-|71403545|71403627</td><td>54</td><td>74</td><td>21</td><td>TTTCAGTCGGATGTTTGCAGC</td><td>5051.4</td><td>-3.4</td><td>0.9</td><td>-3.7</td><td>2.6E-04</td><td>1.9E-03</td><td>BT20/MB231</td><td>MP</td></tr>
<tr><td>>hg38_21_+_8433172_8446622_RNA45SN1_45S_50ntFlanks@10114.10131.18|TGTGCGTCGGAGGGCGGC</td><td>>hg38_21_+_8433172_8446622_RNA45SN1_45S_50ntFlanks</td><td>10114</td><td>10131</td><td>18</td><td>TGTGCGTCGGAGGGCGGC</td><td>98.7</td><td>10.0</td><td>2.7</td><td>3.7</td><td>2.6E-04</td><td>1.9E-03</td><td>BT20/MB231</td><td>MP</td></tr>
<tr><td>>MI0000736|hsa-mir-30c-1&WithFlank&1|+|40757278|40757378@23.44.22(+1A);MI0000254|hsa-mir-30c-2&WithFlank&6|-|71376954|71377037@13.34.22(+1A)&[MIMAT0000244&hsa-miR-30c-5p&offsets|0|-1(+1A);m-57&1|+|40757300|40757323&offsets|0|-2(+1A)];[MIMAT0000244_1&hsa-miR-30c-5p&offsets|0|-1(+1A);m-56&6|-|71377002|71377025&offsets|0|-2(+1A)]|TGTAAACATCCTACACTCTCAGA</td><td>>MI0000736|hsa-mir-30c-1&WithFlank&1|+|40757278|40757378</td><td>23</td><td>44</td><td>22(+1A)</td><td>TGTAAACATCCTACACTCTCAGA</td><td>466.5</td><td>4.5</td><td>1.4</td><td>3.2</td><td>1.3E-03</td><td>1.9E-03</td><td>MB231/MB468</td><td>MP</td></tr>
<tr><td>>hg38_21_+_8433172_8446622_RNA45SN1_45S_50ntFlanks@11820.11844.25|ACTATCCAGCGAAACCACAGCCAAG</td><td>>hg38_21_+_8433172_8446622_RNA45SN1_45S_50ntFlanks</td><td>11820</td><td>11844</td><td>25</td><td>ACTATCCAGCGAAACCACAGCCAAG</td><td>143.3</td><td>-5.5</td><td>1.6</td><td>-3.4</td><td>7.5E-04</td><td>1.9E-03</td><td>BT20/MB468</td><td>MP</td></tr>
<tr><td>>hg38_21_+_8433172_8446622_RNA45SN1_45S_50ntFlanks@11552.11574.23|CCTCGGCCGGCGCCTAGCAGCCG</td><td>>hg38_21_+_8433172_8446622_RNA45SN1_45S_50ntFlanks</td><td>11552</td><td>11574</td><td>23</td><td>CCTCGGCCGGCGCCTAGCAGCCG</td><td>206.9</td><td>-3.8</td><td>1.2</td><td>-3.2</td><td>1.2E-03</td><td>1.9E-03</td><td>BT20/MB468</td><td>MITO</td></tr>
<tr><td>>hg38_MT_+_598_1651_RNR1_12S_50ntFlanks@372.404.33|AAAGCTAAAACTCACCTGAGTTGTAAAAAACTC</td><td>>hg38_MT_+_598_1651_RNR1_12S_50ntFlanks</td><td>372</td><td>404</td><td>33</td><td>AAAGCTAAAACTCACCTGAGTTGTAAAAAACTC</td><td>354.9</td><td>2.9</td><td>0.9</td><td>3.2</td><td>1.6E-03</td><td>1.9E-03</td><td>MB231/MB468</td><td>MITO</td></tr>
<tr><td>>MI0000088|hsa-mir-30a&WithFlank&6|-|71403545|71403627@53.74.22&[MIMAT0000088&hsa-miR-30a-3p&offsets|0|0;m-38&6|-|71403554|71403575&offsets|0|0]|CTTTCAGTCGGATGTTTGCAGC</td><td>>MI0000088|hsa-mir-30a&WithFlank&6|-|71403545|71403627</td><td>53</td><td>74</td><td>22</td><td>CTTTCAGTCGGATGTTTGCAGC</td><td>10391.1</td><td>-2.6</td><td>0.7</td><td>-3.6</td><td>3.3E-04</td><td>1.9E-03</td><td>BT20/MB231</td><td>MITO</td></tr>
<tr><td>>hg38_21_+_8433172_8446622_RNA45SN1_45S_50ntFlanks@11322.11343.22|ACGTCTCGTCGCGCGCGCGTCC</td><td>>hg38_21_+_8433172_8446622_RNA45SN1_45S_50ntFlanks</td><td>11322</td><td>11343</td><td>22</td><td>ACGTCTCGTCGCGCGCGCGTCC</td><td>238.8</td><td>-4.1</td><td>1.3</td><td>-3.2</td><td>1.4E-03</td><td>1.9E-03</td><td>MB231/MB468</td><td>MP</td></tr>
<tr><td>>hg38_21_+_8433172_8446622_RNA45SN1_45S_50ntFlanks@3900.3937.38|CTTCGCGGGGGGGATGCGTGCATTTATCAGATCAAAAC</td><td>>hg38_21_+_8433172_8446622_RNA45SN1_45S_50ntFlanks</td><td>3900</td><td>3937</td><td>38</td><td>CTTCGCGGGGGGGATGCGTGCATTTATCAGATCAAAAC</td><td>353.8</td><td>-6.1</td><td>1.8</td><td>-3.4</td><td>7.5E-04</td><td>1.9E-03</td><td>BT20/MB468</td><td>MP</td></tr>
<tr><td>>hg38_21_+_8433172_8446622_RNA45SN1_45S_50ntFlanks@4047.4074.28|ATTCGAACGTCTGCCCTATCAACTTTCG</td><td>>hg38_21_+_8433172_8446622_RNA45SN1_45S_50ntFlanks</td><td>4047</td><td>4074</td><td>28</td><td>ATTCGAACGTCTGCCCTATCAACTTTCG</td><td>91.5</td><td>-5.2</td><td>1.5</td><td>-3.4</td><td>7.5E-04</td><td>1.9E-03</td><td>BT20/MB468</td><td>MP</td></tr>
<tr><td>>hg38_21_+_8433172_8446622_RNA45SN1_45S_50ntFlanks@4090.4121.32|CCTACCATGGTGACCACGGGTGACGGGGAATC</td><td>>hg38_21_+_8433172_8446622_RNA45SN1_45S_50ntFlanks</td><td>4090</td><td>4121</td><td>32</td><td>CCTACCATGGTGACCACGGGTGACGGGGAATC</td><td>78.9</td><td>-5.2</td><td>1.5</td><td>-3.4</td><td>7.5E-04</td><td>1.9E-03</td><td>BT20/MB468</td><td>MP</td></tr>
<tr><td>>hg38_21_+_8433172_8446622_RNA45SN1_45S_50ntFlanks@12995.13017.23|CTCGCTGCGATCTATTGAAAGTC</td><td>>hg38_21_+_8433172_8446622_RNA45SN1_45S_50ntFlanks</td><td>12995</td><td>13017</td><td>23</td><td>CTCGCTGCGATCTATTGAAAGTC</td><td>122.8</td><td>2.8</td><td>0.9</td><td>3.2</td><td>1.6E-03</td><td>1.9E-03</td><td>MB231/MB468</td><td>MITO</td></tr>
<tr><td>>hg38_21_+_8433172_8446622_RNA45SN1_45S_50ntFlanks@10112.10131.20|GGTGTGCGTCGGAGGGCGGC</td><td>>hg38_21_+_8433172_8446622_RNA45SN1_45S_50ntFlanks</td><td>10112</td><td>10131</td><td>20</td><td>GGTGTGCGTCGGAGGGCGGC</td><td>43.9</td><td>9.2</td><td>2.7</td><td>3.4</td><td>7.5E-04</td><td>1.9E-03</td><td>BT20/MB468</td><td>MP</td></tr>
<tr><td>>hg38_21_+_8433172_8446622_RNA45SN1_45S_50ntFlanks@11570.11594.25|AGCCGACTTAGAACTGGTGCGGACC</td><td>>hg38_21_+_8433172_8446622_RNA45SN1_45S_50ntFlanks</td><td>11570</td><td>11594</td><td>25</td><td>AGCCGACTTAGAACTGGTGCGGACC</td><td>135.5</td><td>-2.2</td><td>0.7</td><td>-3.2</td><td>1.4E-03</td><td>1.9E-03</td><td>MB231/MB468</td><td>MP</td></tr>
<tr><td>>hg38_21_+_8433172_8446622_RNA45SN1_45S_50ntFlanks@10366.10383.18|CCGAAGTGGAGAAGGGTT</td><td>>hg38_21_+_8433172_8446622_RNA45SN1_45S_50ntFlanks</td><td>10366</td><td>10383</td><td>18</td><td>CCGAAGTGGAGAAGGGTT</td><td>81.2</td><td>-5.2</td><td>1.5</td><td>-3.4</td><td>7.6E-04</td><td>1.9E-03</td><td>BT20/MB468</td><td>MP</td></tr>
<tr><td>>MI0000439|hsa-mir-23b&WithFlank&9|+|95085202|95085310@67.85.19&[MIMAT0000418&hsa-miR-23b-3p&offsets|+3|-1;m-42&9|+|95085265|95085285&offsets|+3|+1]|ACATTGCCAGGGATTACCA</td><td>>MI0000439|hsa-mir-23b&WithFlank&9|+|95085202|95085310</td><td>67</td><td>85</td><td>19</td><td>ACATTGCCAGGGATTACCA</td><td>300.0</td><td>11.5</td><td>3.6</td><td>3.2</td><td>1.6E-03</td><td>1.9E-03</td><td>MB231/MB468</td><td>MITO</td></tr>
<tr><td>>hg38_21_+_8433172_8446622_RNA45SN1_45S_50ntFlanks@8381.8403.23|AAGAGGGCGTGAAACCGTTAAGA</td><td>>hg38_21_+_8433172_8446622_RNA45SN1_45S_50ntFlanks</td><td>8381</td><td>8403</td><td>23</td><td>AAGAGGGCGTGAAACCGTTAAGA</td><td>400.6</td><td>-2.5</td><td>0.8</td><td>-3.2</td><td>1.2E-03</td><td>1.9E-03</td><td>BT20/MB468</td><td>MITO</td></tr>
<tr><td>>hg38_21_+_8433172_8446622_RNA45SN1_45S_50ntFlanks@11561.11590.30|GCGCCTAGCAGCCGACTTAGAACTGGTGCG</td><td>>hg38_21_+_8433172_8446622_RNA45SN1_45S_50ntFlanks</td><td>11561</td><td>11590</td><td>30</td><td>GCGCCTAGCAGCCGACTTAGAACTGGTGCG</td><td>260.2</td><td>-1.9</td><td>0.6</td><td>-3.2</td><td>1.2E-03</td><td>1.9E-03</td><td>BT20/MB468</td><td>MITO</td></tr>
<tr><td>>hg38_21_+_8433172_8446622_RNA45SN1_45S_50ntFlanks@11029.11053.25|CTCTCTCTCTCTCCCCCGCTCCCCG</td><td>>hg38_21_+_8433172_8446622_RNA45SN1_45S_50ntFlanks</td><td>11029</td><td>11053</td><td>25</td><td>CTCTCTCTCTCTCCCCCGCTCCCCG</td><td>203.9</td><td>-2.7</td><td>0.9</td><td>-3.2</td><td>1.5E-03</td><td>1.9E-03</td><td>MB231/MB468</td><td>CYTO</td></tr>
<tr><td>>hg38_21_+_8433172_8446622_RNA45SN1_45S_50ntFlanks@8171.8215.45|AAGTCCTTCTGATCGAGGCCCAGCCCGTGGACGGTGTGAGGCCGG</td><td>>hg38_21_+_8433172_8446622_RNA45SN1_45S_50ntFlanks</td><td>8171</td><td>8215</td><td>45</td><td>AAGTCCTTCTGATCGAGGCCCAGCCCGTGGACGGTGTGAGGCCGG</td><td>408.9</td><td>-2.9</td><td>0.9</td><td>-3.2</td><td>1.5E-03</td><td>1.9E-03</td><td>MB231/MB468</td><td>CYTO</td></tr>
<tr><td>>hg38_21_+_8433172_8446622_RNA45SN1_45S_50ntFlanks@6645.6669.25|CTCGTACGACTCTTAGCGGTGGATC</td><td>>hg38_21_+_8433172_8446622_RNA45SN1_45S_50ntFlanks</td><td>6645</td><td>6669</td><td>25</td><td>CTCGTACGACTCTTAGCGGTGGATC</td><td>2275.3</td><td>2.7</td><td>0.9</td><td>3.2</td><td>1.5E-03</td><td>1.9E-03</td><td>MB231/MB468</td><td>CYTO</td></tr>
<tr><td>>MI0000298|hsa-mir-221&WithFlank&X|-|45746151|45746272@71.91.21(+1A)&[MIMAT0000278&hsa-miR-221-3p&offsets|0|-2(+1A);m-91&X|-|45746181|45746202&offsets|0|-1(+1A)]|AGCTACATTGTCTGCTGGGTTA</td><td>>MI0000298|hsa-mir-221&WithFlank&X|-|45746151|45746272</td><td>71</td><td>91</td><td>21(+1A)</td><td>AGCTACATTGTCTGCTGGGTTA</td><td>1228.4</td><td>-3.0</td><td>0.8</td><td>-3.6</td><td>3.3E-04</td><td>1.9E-03</td><td>BT20/MB231</td><td>MITO</td></tr>
<tr><td>>hg38_21_+_8433172_8446622_RNA45SN1_45S_50ntFlanks@10517.10534.18|AGTGGCGGAGATGGGCGC</td><td>>hg38_21_+_8433172_8446622_RNA45SN1_45S_50ntFlanks</td><td>10517</td><td>10534</td><td>18</td><td>AGTGGCGGAGATGGGCGC</td><td>148.8</td><td>-3.5</td><td>1.1</td><td>-3.2</td><td>1.2E-03</td><td>1.9E-03</td><td>BT20/MB468</td><td>MITO</td></tr>
</table>

<table>
<tr><td>>hg38_21_+_8433172_8446622_RNA45SN1_45S_50ntFlanks@10788.10827.40|CAGCCTCTGGCATGTTGGAACAATGTAGGTAAGGGAAGTC</td><td>>hg38_21_+_8433172_8446622_RNA45SN1_45S_50ntFlanks</td><td>10788</td><td>10827</td><td>40</td><td>CAGCCTCTGGCATGTTGGAACAATGTAGGTAAGGGAAGTC</td><td>57.0</td><td>-5.2</td><td>1.6</td><td>-3.2</td><td>1.4E-03</td><td>1.9E-03</td><td>MB231/MB468</td><td>MP</td></tr>
<tr><td>>hg38_21_+_8433172_8446622_RNA45SN1_45S_50ntFlanks@9035.9059.25|CGTGGGGGGGCCGGGCCACCCCTCC</td><td>>hg38_21_+_8433172_8446622_RNA45SN1_45S_50ntFlanks</td><td>9035</td><td>9059</td><td>25</td><td>CGTGGGGGGGCCGGGCCACCCCTCC</td><td>354.3</td><td>-4.0</td><td>1.2</td><td>-3.2</td><td>1.2E-03</td><td>1.9E-03</td><td>BT20/MB468</td><td>MITO</td></tr>
<tr><td>>hg38_21_+_8433172_8446622_RNA45SN1_45S_50ntFlanks@12244.12269.26|AAGGGCAAAAGCTCGCTTGATCTTGA</td><td>>hg38_21_+_8433172_8446622_RNA45SN1_45S_50ntFlanks</td><td>12244</td><td>12269</td><td>26</td><td>AAGGGCAAAAGCTCGCTTGATCTTGA</td><td>297.5</td><td>-3.1</td><td>1.0</td><td>-3.2</td><td>1.6E-03</td><td>1.9E-03</td><td>MB231/MB468</td><td>MITO</td></tr>
<tr><td>>MI0000342|hsa-mir-200b&WithFlank&1|+|1167098|1167204@63.85.23&[MIMAT0000318&hsa-miR-200b-3p&offsets|0|+1;m-93&1|+|1167160|1167182&offsets|0|0]|TAATACTGCCTGGTAATGATGAC</td><td>>MI0000342|hsa-mir-200b&WithFlank&1|+|1167098|1167204</td><td>63</td><td>85</td><td>23</td><td>TAATACTGCCTGGTAATGATGAC</td><td>2428.3</td><td>-3.0</td><td>1.0</td><td>-3.2</td><td>1.6E-03</td><td>1.9E-03</td><td>MB231/MB468</td><td>MITO</td></tr>
<tr><td>>hg38_21_+_8433172_8446622_RNA45SN1_45S_50ntFlanks@9122.9142.21|CCGCGCGCGGGTCGGGGGGCG</td><td>>hg38_21_+_8433172_8446622_RNA45SN1_45S_50ntFlanks</td><td>9122</td><td>9142</td><td>21</td><td>CCGCGCGCGGGTCGGGGGGCG</td><td>121.8</td><td>-5.4</td><td>1.6</td><td>-3.4</td><td>7.6E-04</td><td>1.9E-03</td><td>BT20/MB468</td><td>MP</td></tr>
<tr><td>>hg38_21_+_8433172_8446622_RNA45SN1_45S_50ntFlanks@11534.11580.47|GTTCCGCGCGGCGCCTCGCCTCGGCCGGCGCCTAGCAGCCGACTTAG</td><td>>hg38_21_+_8433172_8446622_RNA45SN1_45S_50ntFlanks</td><td>11534</td><td>11580</td><td>47</td><td>GTTCCGCGCGGCGCCTCGCCTCGGCCGGCGCCTAGCAGCCGACTTAG</td><td>182.7</td><td>-5.3</td><td>1.7</td><td>-3.2</td><td>1.6E-03</td><td>1.9E-03</td><td>MB231/MB468</td><td>MITO</td></tr>
<tr><td>>hg38_21_+_8433172_8446622_RNA45SN1_45S_50ntFlanks@6659.6681.23|AGCGGTGGATCACTCGGCTCGTG</td><td>>hg38_21_+_8433172_8446622_RNA45SN1_45S_50ntFlanks</td><td>6659</td><td>6681</td><td>23</td><td>AGCGGTGGATCACTCGGCTCGTG</td><td>292.2</td><td>5.7</td><td>1.6</td><td>3.7</td><td>2.6E-04</td><td>1.9E-03</td><td>BT20/MB231</td><td>MP</td></tr>
<tr><td>>hg38_21_+_8433172_8446622_RNA45SN1_45S_50ntFlanks@6661.6686.26|CGGTGGATCACTCGGCTCGTGCGTCG</td><td>>hg38_21_+_8433172_8446622_RNA45SN1_45S_50ntFlanks</td><td>6661</td><td>6686</td><td>26</td><td>CGGTGGATCACTCGGCTCGTGCGTCG</td><td>798.3</td><td>2.8</td><td>0.9</td><td>3.2</td><td>1.5E-03</td><td>1.9E-03</td><td>BT20/MB468</td><td>CYTO</td></tr>
<tr><td>>MI0000108|hsa-mir-103a-2&WithFlank&20|+|3917488|3917577@54.75.22(+1U);MI0000109|hsa-mir-103a-1&WithFlank&5|-|168560890|168560979@54.75.22(+1U)&[MIMAT0000101&hsa-miR-103a-3p&offsets|0|-1(+1U);m-12&20|+|3917541|3917563&offsets|0|-1(+1U)];[MIMAT0000101_1&hsa-miR-103a-3p&offsets|0|-1(+1U);m-13&5|-|168560904|168560926&offsets|0|-1(+1U);m-44220&5|-|168560908|168560926&offsets|0|+3(+1U)]|AGCAGCATTGTACAGGGCTATGT</td><td>>MI0000108|hsa-mir-103a-2&WithFlank&20|+|3917488|3917577</td><td>54</td><td>75</td><td>22(+1U)</td><td>AGCAGCATTGTACAGGGCTATGT</td><td>1967.2</td><td>3.0</td><td>1.0</td><td>3.2</td><td>1.4E-03</td><td>1.9E-03</td><td>MB231/MB468</td><td>MP</td></tr>
<tr><td>>hg38_21_+_8433172_8446622_RNA45SN1_45S_50ntFlanks@3787.3812.26|AAACTGCGAATGGCTCATTAAATCAG</td><td>>hg38_21_+_8433172_8446622_RNA45SN1_45S_50ntFlanks</td><td>3787</td><td>3812</td><td>26</td><td>AAACTGCGAATGGCTCATTAAATCAG</td><td>197.9</td><td>-5.8</td><td>1.7</td><td>-3.4</td><td>7.6E-04</td><td>1.9E-03</td><td>BT20/MB468</td><td>MP</td></tr>
<tr><td>>trna32_LysCTT_16_-_3207406_3207478@1.33.33|GCCCGGCTAGCTCAGTCGGTAGAGCATGAGACC</td><td>>trna32_LysCTT_16_-_3207406_3207478</td><td>1</td><td>33</td><td>33</td><td>GCCCGGCTAGCTCAGTCGGTAGAGCATGAGACC</td><td>1020.6</td><td>-4.0</td><td>1.1</td><td>-3.6</td><td>2.9E-04</td><td>1.9E-03</td><td>BT20/MB231</td><td>NUC</td></tr>
<tr><td>>hg38_21_+_8433172_8446622_RNA45SN1_45S_50ntFlanks@10778.10818.41|CCAAGGTGAACAGCCTCTGGCATGTTGGAACAATGTAGGTA</td><td>>hg38_21_+_8433172_8446622_RNA45SN1_45S_50ntFlanks</td><td>10778</td><td>10818</td><td>41</td><td>CCAAGGTGAACAGCCTCTGGCATGTTGGAACAATGTAGGTA</td><td>368.6</td><td>-4.1</td><td>1.3</td><td>-3.2</td><td>1.5E-03</td><td>1.9E-03</td><td>BT20/MB468</td><td>CYTO</td></tr>
<tr><td>>hg38_MT_+_1621_3279_RNR2_16S_50ntFlanks@1349.1384.36|ACAATAGGGTTTACGACCTCGATGTTGGATCAGGAC</td><td>>hg38_MT_+_1621_3279_RNR2_16S_50ntFlanks</td><td>1349</td><td>1384</td><td>36</td><td>ACAATAGGGTTTACGACCTCGATGTTGGATCAGGAC</td><td>54.8</td><td>5.7</td><td>1.8</td><td>3.2</td><td>1.4E-03</td><td>1.9E-03</td><td>MB231/MB468</td><td>MP</td></tr>
<tr><td>>hg38_21_+_8433172_8446622_RNA45SN1_45S_50ntFlanks@9739.9760.22|AAACGATCTCAACCTATTCTCA</td><td>>hg38_21_+_8433172_8446622_RNA45SN1_45S_50ntFlanks</td><td>9739</td><td>9760</td><td>22</td><td>AAACGATCTCAACCTATTCTCA</td><td>539.8</td><td>-3.1</td><td>1.0</td><td>-3.2</td><td>1.2E-03</td><td>1.9E-03</td><td>BT20/MB468</td><td>MITO</td></tr>
<tr><td>>MI0003513|hsa-mir-455&WithFlank&9|+|114209428|114209535@60.80.21(+1C)&[MIMAT0004784&hsa-miR-455-3p&offsets|0|0(+1C);m-135&9|+|114209487|114209508&offsets|0|-1(+1C)]|GCAGTCCATGGGCATATACACC</td><td>>MI0003513|hsa-mir-455&WithFlank&9|+|114209428|114209535</td><td>60</td><td>80</td><td>21(+1C)</td><td>GCAGTCCATGGGCATATACACC</td><td>161.2</td><td>-6.3</td><td>1.7</td><td>-3.7</td><td>2.6E-04</td><td>1.9E-03</td><td>BT20/MB231</td><td>MP</td></tr>
<tr><td>>MI0000734|hsa-mir-106b&WithFlank&7|-|100093987|100094080@58.77.20(+1A)&[MIMAT0004672&hsa-miR-106b-3p&offsets|0|-2(+1A);m-87&7|-|100094003|100094023&offsets|0|-1(+1A)]|CCGCACTGTGGGTACTTGCTA</td><td>>MI0000734|hsa-mir-106b&WithFlank&7|-|100093987|100094080</td><td>58</td><td>77</td><td>20(+1A)</td><td>CCGCACTGTGGGTACTTGCTA</td><td>254.6</td><td>6.7</td><td>2.0</td><td>3.4</td><td>7.6E-04</td><td>1.9E-03</td><td>BT20/MB468</td><td>MP</td></tr>
<tr><td>>hg38_21_+_8433172_8446622_RNA45SN1_45S_50ntFlanks@6649.6675.27|TACGACTCTTAGCGGTGGATCACTCGG</td><td>>hg38_21_+_8433172_8446622_RNA45SN1_45S_50ntFlanks</td><td>6649</td><td>6675</td><td>27</td><td>TACGACTCTTAGCGGTGGATCACTCGG</td><td>490.4</td><td>5.1</td><td>1.4</td><td>3.7</td><td>2.6E-04</td><td>1.9E-03</td><td>BT20/MB231</td><td>MP</td></tr>
<tr><td>>trnaMT_LeuTAA_MT_+_3230_3304@1.30.30|GTTAAGATGGCAGAGCCCGGTAATCGCATA</td><td>>trnaMT_LeuTAA_MT_+_3230_3304</td><td>1</td><td>30</td><td>30</td><td>GTTAAGATGGCAGAGCCCGGTAATCGCATA</td><td>465.5</td><td>2.4</td><td>0.7</td><td>3.6</td><td>2.9E-04</td><td>1.9E-03</td><td>BT20/MB231</td><td>NUC</td></tr>
<tr><td>>hg38_21_+_8433172_8446622_RNA45SN1_45S_50ntFlanks@12417.12444.28|TCGATGTCGGCTCTTCCTATCATTGTGA</td><td>>hg38_21_+_8433172_8446622_RNA45SN1_45S_50ntFlanks</td><td>12417</td><td>12444</td><td>28</td><td>TCGATGTCGGCTCTTCCTATCATTGTGA</td><td>1277.0</td><td>-1.9</td><td>0.6</td><td>-3.1</td><td>1.6E-03</td><td>2.0E-03</td><td>MB231/MB468</td><td>NUC</td></tr>
<tr><td>>MI0016746|hsa-mir-548o-2&WithFlank&20|+|38516557|38516638@51.72.22;MI0006402|hsa-mir-548o&WithFlank&7|-|102405736|102405861@93.114.22&[MIMAT0005919&hsa-miR-548o-3p&offsets|0|0;m-431&20|+|38516607|38516628&offsets|0|0];[MIMAT0005919_1&hsa-miR-548o-3p&offsets|0|0;m-430&7|-|102405748|102405769&offsets|0|0]|CCAAAACTGCAGTTACTTTTGC</td><td>>MI0016746|hsa-mir-548o-2&WithFlank&20|+|38516557|38516638</td><td>51</td><td>72</td><td>22</td><td>CCAAAACTGCAGTTACTTTTGC</td><td>123.3</td><td>2.0</td><td>0.6</td><td>3.2</td><td>1.6E-03</td><td>2.0E-03</td><td>BT20/MB468</td><td>NUC</td></tr>
<tr><td>>hg38_21_+_8433172_8446622_RNA45SN1_45S_50ntFlanks@12251.12284.34|AAAGCTCGCTTGATCTTGATTTTCAGTACGAATA</td><td>>hg38_21_+_8433172_8446622_RNA45SN1_45S_50ntFlanks</td><td>12251</td><td>12284</td><td>34</td><td>AAAGCTCGCTTGATCTTGATTTTCAGTACGAATA</td><td>53.5</td><td>-9.6</td><td>3.0</td><td>-3.2</td><td>1.4E-03</td><td>2.0E-03</td><td>MB231/MB468</td><td>MP</td></tr>
<tr><td>>hg38_21_+_8433172_8446622_RNA45SN1_45S_50ntFlanks@12656.12675.20|TGACTGAACGCCTCTAAGTC</td><td>>hg38_21_+_8433172_8446622_RNA45SN1_45S_50ntFlanks</td><td>12656</td><td>12675</td><td>20</td><td>TGACTGAACGCCTCTAAGTC</td><td>421.0</td><td>-5.1</td><td>1.6</td><td>-3.2</td><td>1.5E-03</td><td>2.0E-03</td><td>BT20/MB468</td><td>CYTO</td></tr>
<tr><td>>trnaMT_GluTTC_MT_-_14674_14742@38.63.26,trnalookalike8_GluTTC_5_-_93905172_93905240@38.63.26|CATTGGTCGTGGTTGTAGTCCGTGCG</td><td>>trnaMT_GluTTC_MT_-_14674_14742</td><td>38</td><td>63</td><td>26</td><td>CATTGGTCGTGGTTGTAGTCCGTGCG</td><td>253.5</td><td>5.9</td><td>1.8</td><td>3.4</td><td>7.6E-04</td><td>2.0E-03</td><td>BT20/MB468</td><td>MP</td></tr>
<tr><td>>hg38_21_+_8433172_8446622_RNA45SN1_45S_50ntFlanks@11672.11693.22|CAGTGCTCTGAATGTCAAAGTG</td><td>>hg38_21_+_8433172_8446622_RNA45SN1_45S_50ntFlanks</td><td>11672</td><td>11693</td><td>22</td><td>CAGTGCTCTGAATGTCAAAGTG</td><td>252.1</td><td>-4.0</td><td>1.3</td><td>-3.2</td><td>1.4E-03</td><td>2.0E-03</td><td>MB231/MB468</td><td>MP</td></tr>
<tr><td>>hg38_21_+_8433172_8446622_RNA45SN1_45S_50ntFlanks@12418.12439.22|CGATGTCGGCTCTTCCTATCAT</td><td>>hg38_21_+_8433172_8446622_RNA45SN1_45S_50ntFlanks</td><td>12418</td><td>12439</td><td>22</td><td>CGATGTCGGCTCTTCCTATCAT</td><td>81.4</td><td>-5.0</td><td>1.5</td><td>-3.2</td><td>1.4E-03</td><td>2.0E-03</td><td>MB231/MB468</td><td>MP</td></tr>
<tr><td>>hg38_21_+_8433172_8446622_RNA45SN1_45S_50ntFlanks@9554.9595.42|CGTAGCGGTCCTGACGTGCAAATCGGTCGTCCGACCTGGGTA</td><td>>hg38_21_+_8433172_8446622_RNA45SN1_45S_50ntFlanks</td><td>9554</td><td>9595</td><td>42</td><td>CGTAGCGGTCCTGACGTGCAAATCGGTCGTCCGACCTGGGTA</td><td>428.0</td><td>-2.8</td><td>0.9</td><td>-3.2</td><td>1.5E-03</td><td>2.0E-03</td><td>MB231/MB468</td><td>CYTO</td></tr>
<tr><td>>hg38_21_+_8433172_8446622_RNA45SN1_45S_50ntFlanks@12423.12457.35|TCGGCTCTTCCTATCATTGTGAAGCAGAATTCACC</td><td>>hg38_21_+_8433172_8446622_RNA45SN1_45S_50ntFlanks</td><td>12423</td><td>12457</td><td>35</td><td>TCGGCTCTTCCTATCATTGTGAAGCAGAATTCACC</td><td>768.1</td><td>-3.0</td><td>0.8</td><td>-3.6</td><td>3.0E-04</td><td>2.0E-03</td><td>BT20/MB231</td><td>CYTO</td></tr>
<tr><td>>hg38_21_+_8433172_8446622_RNA45SN1_45S_50ntFlanks@12635.12670.36|AAGCTACCATCTGTGGGATTATGACTGAACGCCTCT</td><td>>hg38_21_+_8433172_8446622_RNA45SN1_45S_50ntFlanks</td><td>12635</td><td>12670</td><td>36</td><td>AAGCTACCATCTGTGGGATTATGACTGAACGCCTCT</td><td>173.2</td><td>3.3</td><td>1.0</td><td>3.2</td><td>1.5E-03</td><td>2.0E-03</td><td>MB231/MB468</td><td>CYTO</td></tr>
<tr><td>>hg38_MT_+_598_1651_RNR1_12S_50ntFlanks@428.462.35|AAAGTGGCTTTAACATATCTGAACACACAATAGCT</td><td>>hg38_MT_+_598_1651_RNR1_12S_50ntFlanks</td><td>428</td><td>462</td><td>35</td><td>AAAGTGGCTTTAACATATCTGAACACACAATAGCT</td><td>257.0</td><td>5.7</td><td>1.8</td><td>3.2</td><td>1.6E-03</td><td>2.0E-03</td><td>MB231/MB468</td><td>MITO</td></tr>
<tr><td>>hg38_21_+_8433172_8446622_RNA45SN1_45S_50ntFlanks@6652.6677.26|GACTCTTAGCGGTGGATCACTCGGCT</td><td>>hg38_21_+_8433172_8446622_RNA45SN1_45S_50ntFlanks</td><td>6652</td><td>6677</td><td>26</td><td>GACTCTTAGCGGTGGATCACTCGGCT</td><td>1375.4</td><td>2.9</td><td>0.9</td><td>3.1</td><td>1.6E-03</td><td>2.0E-03</td><td>MB231/MB468</td><td>NUC</td></tr>
<tr><td>>MI0005531|hsa-mir-450b&WithFlank&X|-|134540179|134540268@17.37.21&[MIMAT0004909&hsa-miR-450b-5p&offsets|0|-1;m-266&X|-|134540232|134540252&offsets|0|0]|TTTTGCAATATGTTCCTGAAT</td><td>>MI0005531|hsa-mir-450b&WithFlank&X|-|134540179|134540268</td><td>17</td><td>37</td><td>21</td><td>TTTTGCAATATGTTCCTGAAT</td><td>590.9</td><td>3.2</td><td>0.9</td><td>3.6</td><td>3.0E-04</td><td>2.0E-03</td><td>BT20/MB231</td><td>CYTO</td></tr>
<tr><td>>trna10_AspGTC_12_-_125424193_125424264@31.53.23,trna12_AspGTC_12_-_125411891_125411962@31.53.23,trna38_AspGTC_17_-_8125556_8125627@31.53.23,trna45_AspGTC_6_+_27447453_27447524@31.53.23,trna48_AspGTC_6_+_27471523_27471594@31.53.23,trna4_AspGTC_12_+_96429799_96429870@31.53.23,trna5_AspGTC_12_+_98897281_98897352@31.53.23,trna69_AspGTC_1_-_161440205_161440276@31.53.23,trna6_AspGTC_9_-_77517990_77518061@31.53.23,trna72_AspGTC_1_-_161432824_161432895@31.53.23,trna75_AspGTC_1_-_161425414_161425485@31.53.23,trna78_AspGTC_1_-_161418033_161418104@31.53.23,trna81_AspGTC_1_-_161410615_161410686@31.53.23|CCTGTCACGCGGGAGACCGGGGT</td><td>>trna10_AspGTC_12_-_125424193_125424264</td><td>31</td><td>53</td><td>23</td><td>CCTGTCACGCGGGAGACCGGGGT</td><td>985.5</td><td>3.0</td><td>0.8</td><td>3.6</td><td>3.0E-04</td><td>2.0E-03</td><td>BT20/MB231</td><td>CYTO</td></tr>
<tr><td>>hg38_21_+_8433172_8446622_RNA45SN1_45S_50ntFlanks@7978.8012.35|GACCTCAGATCAGACGTGGCGACCCGCTGAATTTA</td><td>>hg38_21_+_8433172_8446622_RNA45SN1_45S_50ntFlanks</td><td>7978</td><td>8012</td><td>35</td><td>GACCTCAGATCAGACGTGGCGACCCGCTGAATTTA</td><td>487.9</td><td>-3.7</td><td>1.0</td><td>-3.6</td><td>3.0E-04</td><td>2.0E-03</td><td>BT20/MB231</td><td>CYTO</td></tr>
<tr><td>>hg38_21_+_8433172_8446622_RNA45SN1_45S_50ntFlanks@11581.11612.32|AACTGGTGCGGACCAGGGGAATCCGACTGTTT</td><td>>hg38_21_+_8433172_8446622_RNA45SN1_45S_50ntFlanks</td><td>11581</td><td>11612</td><td>32</td><td>AACTGGTGCGGACCAGGGGAATCCGACTGTTT</td><td>1988.8</td><td>-2.0</td><td>0.6</td><td>-3.2</td><td>1.5E-03</td><td>2.0E-03</td><td>MB231/MB468</td><td>CYTO</td></tr>
<tr><td>>hg38_MT_+_598_1651_RNR1_12S_50ntFlanks@51.77.27|AATAGGTTTGGTCCTAGCCTTTCTATT</td><td>>hg38_MT_+_598_1651_RNR1_12S_50ntFlanks</td><td>51</td><td>77</td><td>27</td><td>AATAGGTTTGGTCCTAGCCTTTCTATT</td><td>268.1</td><td>3.4</td><td>1.1</td><td>3.1</td><td>1.6E-03</td><td>2.0E-03</td><td>MB231/MB468</td><td>MITO</td></tr>
<tr><td>>MI0000816|hsa-mir-335&WithFlank&7|+|130496105|130496210@58.77.20&[MIMAT0004703&hsa-miR-335-3p&offsets|0|-2;m-159&7|+|130496162|130496183&offsets|0|-2]|TTTTTCATTATTGCTCCTGA</td><td>>MI0000816|hsa-mir-335&WithFlank&7|+|130496105|130496210</td><td>58</td><td>77</td><td>20</td><td>TTTTTCATTATTGCTCCTGA</td><td>84.7</td><td>5.9</td><td>1.9</td><td>3.2</td><td>1.4E-03</td><td>2.0E-03</td><td>MB231/MB468</td><td>MP</td></tr>
<tr><td>>hg38_21_+_8433172_8446622_RNA45SN1_45S_50ntFlanks@9934.9951.18|AGACAGCAGGACGGTGGC</td><td>>hg38_21_+_8433172_8446622_RNA45SN1_45S_50ntFlanks</td><td>9934</td><td>9951</td><td>18</td><td>AGACAGCAGGACGGTGGC</td><td>249.0</td><td>5.7</td><td>1.7</td><td>3.4</td><td>7.7E-04</td><td>2.0E-03</td><td>BT20/MB468</td><td>MP</td></tr>
<tr><td>>MI0000750|hsa-mir-26a-2&WithFlank&12|-|57824603|57824698@20.39.20;MI0000083|hsa-mir-26a-1&WithFlank&3|+|37969398|37969486@16.35.20&[MIMAT0000082&hsa-miR-26a-5p&offsets|0|-2;m-33&12|-|57824658|57824679&offsets|0|-2];[MIMAT0000082_1&hsa-miR-26a-5p&offsets|0|-2;m-34&3|+|37969413|37969434&offsets|0|-2]|TTCAAGTAATCCAGGATAGG</td><td>>MI0000750|hsa-mir-26a-2&WithFlank&12|-|57824603|57824698</td><td>20</td><td>39</td><td>20</td><td>TTCAAGTAATCCAGGATAGG</td><td>347.0</td><td>3.0</td><td>0.9</td><td>3.2</td><td>1.4E-03</td><td>2.0E-03</td><td>MB231/MB468</td><td>MP</td></tr>
<tr><td>>hg38_21_+_8433172_8446622_RNA45SN1_45S_50ntFlanks@11665.11702.38|TTCTGCCCAGTGCTCTGAATGTCAAAGTGAAGAAATTC</td><td>>hg38_21_+_8433172_8446622_RNA45SN1_45S_50ntFlanks</td><td>11665</td><td>11702</td><td>38</td><td>TTCTGCCCAGTGCTCTGAATGTCAAAGTGAAGAAATTC</td><td>435.9</td><td>2.5</td><td>0.8</td><td>3.1</td><td>1.7E-03</td><td>2.0E-03</td><td>MB231/MB468</td><td>NUC</td></tr>
<tr><td>>hg38_21_+_8433172_8446622_RNA45SN1_45S_50ntFlanks@6650.6669.20|ACGACTCTTAGCGGTGGATC</td><td>>hg38_21_+_8433172_8446622_RNA45SN1_45S_50ntFlanks</td><td>6650</td><td>6669</td><td>20</td><td>ACGACTCTTAGCGGTGGATC</td><td>7704.3</td><td>2.7</td><td>0.7</td><td>3.6</td><td>3.4E-04</td><td>2.0E-03</td><td>BT20/MB231</td><td>MITO</td></tr>
<tr><td>>hg38_21_+_8433172_8446622_RNA45SN1_45S_50ntFlanks@6655.6681.27|TCTTAGCGGTGGATCACTCGGCTCGTG</td><td>>hg38_21_+_8433172_8446622_RNA45SN1_45S_50ntFlanks</td><td>6655</td><td>6681</td><td>27</td><td>TCTTAGCGGTGGATCACTCGGCTCGTG</td><td>2266.3</td><td>-1.4</td><td>0.4</td><td>-3.2</td><td>1.3E-03</td><td>2.0E-03</td><td>BT20/MB468</td><td>MITO</td></tr>
<tr><td>>hg38_21_+_8433172_8446622_RNA45SN1_45S_50ntFlanks@6651.6700.50|CGACTCTTAGCGGTGGATCACTCGGCTCGTGCGTCGATGAAGAACGCAGC</td><td>>hg38_21_+_8433172_8446622_RNA45SN1_45S_50ntFlanks</td><td>6651</td><td>6700</td><td>50</td><td>CGACTCTTAGCGGTGGATCACTCGGCTCGTGCGTCGATGAAGAACGCAGC</td><td>2753.3</td><td>5.6</td><td>1.7</td><td>3.4</td><td>7.7E-04</td><td>2.0E-03</td><td>BT20/MB468</td><td>MP</td></tr>
<tr><td>>hg38_21_+_8433172_8446622_RNA45SN1_45S_50ntFlanks@10402.10424.23|AACATGGGTCAGTCGGTCCTGAG</td><td>>hg38_21_+_8433172_8446622_RNA45SN1_45S_50ntFlanks</td><td>10402</td><td>10424</td><td>23</td><td>AACATGGGTCAGTCGGTCCTGAG</td><td>7676.0</td><td>2.5</td><td>0.7</td><td>3.4</td><td>7.7E-04</td><td>2.0E-03</td><td>BT20/MB468</td><td>MP</td></tr>
</table>

<table>
<tr><td>>MI0000478|hsa-mir-149&WithFlank&2|+|240455995|240456095@21.41.21(+1U)&[MIMAT0000450&hsa-miR-149-5p&offsets|0|-2(+1U);m-195&2|+|240456015|240456037&offsets|0|-2(+1U)]|TCTGGCTCCGTGTCTTCACTCT</td><td>>MI0000478|hsa-mir-149&WithFlank&2|+|240455995|240456095</td><td>21</td><td>41</td><td>21(+1U)</td><td>TCTGGCTCCGTGTCTTCACTCT</td><td>56.7</td><td>5.9</td><td>1.9</td><td>3.2</td><td>1.4E-03</td><td>2.0E-03</td><td>MB231/MB468</td><td>MP</td></tr>
<tr><td>>hg38_21_+_8433172_8446622_RNA45SN1_45S_50ntFlanks@4983.5013.31|CGGACAGGATTGACAGATTGATAGCTCTTTC</td><td>>hg38_21_+_8433172_8446622_RNA45SN1_45S_50ntFlanks</td><td>4983</td><td>5013</td><td>31</td><td>CGGACAGGATTGACAGATTGATAGCTCTTTC</td><td>84.0</td><td>-5.1</td><td>1.6</td><td>-3.2</td><td>1.4E-03</td><td>2.0E-03</td><td>MB231/MB468</td><td>MP</td></tr>
<tr><td>>hg38_21_+_8433172_8446622_RNA45SN1_45S_50ntFlanks@4328.4346.19|CGGTAATTCCAGCTCCAAT</td><td>>hg38_21_+_8433172_8446622_RNA45SN1_45S_50ntFlanks</td><td>4328</td><td>4346</td><td>19</td><td>CGGTAATTCCAGCTCCAAT</td><td>67.8</td><td>-4.9</td><td>1.5</td><td>-3.2</td><td>1.4E-03</td><td>2.0E-03</td><td>MB231/MB468</td><td>MP</td></tr>
<tr><td>>hg38_21_+_8433172_8446622_RNA45SN1_45S_50ntFlanks@12937.12960.24|AACCATTCGTAGACGACCTGCTTC</td><td>>hg38_21_+_8433172_8446622_RNA45SN1_45S_50ntFlanks</td><td>12937</td><td>12960</td><td>24</td><td>AACCATTCGTAGACGACCTGCTTC</td><td>595.1</td><td>-2.1</td><td>0.7</td><td>-3.1</td><td>1.7E-03</td><td>2.0E-03</td><td>MB231/MB468</td><td>NUC</td></tr>
<tr><td>>hg38_21_+_8433172_8446622_RNA45SN1_45S_50ntFlanks@8380.8397.18|CAAGAGGGCGTGAAACCG</td><td>>hg38_21_+_8433172_8446622_RNA45SN1_45S_50ntFlanks</td><td>8380</td><td>8397</td><td>18</td><td>CAAGAGGGCGTGAAACCG</td><td>820.3</td><td>-5.1</td><td>1.6</td><td>-3.2</td><td>1.5E-03</td><td>2.0E-03</td><td>BT20/MB468</td><td>CYTO</td></tr>
<tr><td>>hg38_21_+_8433172_8446622_RNA45SN1_45S_50ntFlanks@6649.6677.29|TACGACTCTTAGCGGTGGATCACTCGGCT</td><td>>hg38_21_+_8433172_8446622_RNA45SN1_45S_50ntFlanks</td><td>6649</td><td>6677</td><td>29</td><td>TACGACTCTTAGCGGTGGATCACTCGGCT</td><td>511.1</td><td>2.4</td><td>0.8</td><td>3.1</td><td>1.7E-03</td><td>2.0E-03</td><td>MB231/MB468</td><td>NUC</td></tr>
<tr><td>>MI0000085|hsa-mir-27a&WithFlank&19|-|13836434|13836523@57.75.19(+3U)&[MIMAT0000084&hsa-miR-27a-3p&offsets|0|-2(+3U);m-43&19|-|13836447|13836467&offsets|0|-2(+3U)]|TTCACAGTGGCTAAGTTCCTTT</td><td>>MI0000085|hsa-mir-27a&WithFlank&19|-|13836434|13836523</td><td>57</td><td>75</td><td>19(+3U)</td><td>TTCACAGTGGCTAAGTTCCTTT</td><td>237.1</td><td>-3.1</td><td>0.9</td><td>-3.6</td><td>3.4E-04</td><td>2.0E-03</td><td>BT20/MB231</td><td>MITO</td></tr>
<tr><td>>hg38_21_+_8433172_8446622_RNA45SN1_45S_50ntFlanks@475.512.38|CTTCGTGATCGATGTGGTGACGTCGTGCTCTCCCGGGC</td><td>>hg38_21_+_8433172_8446622_RNA45SN1_45S_50ntFlanks</td><td>475</td><td>512</td><td>38</td><td>CTTCGTGATCGATGTGGTGACGTCGTGCTCTCCCGGGC</td><td>47.9</td><td>9.3</td><td>2.8</td><td>3.4</td><td>7.7E-04</td><td>2.0E-03</td><td>BT20/MB468</td><td>MP</td></tr>
<tr><td>>MI0000270|hsa-mir-181b-1&WithFlank&1|-|198858867|198858988@42.65.24(+1C);MI0000683|hsa-mir-181b-2&WithFlank&9|+|124693704|124693804@22.45.24(+1C)&[MIMAT0000257&hsa-miR-181b-5p&offsets|0|+1(+1C);m-73&1|-|198858925|198858947&offsets|0|+1(+1C)];[MIMAT0000257_1&hsa-miR-181b-5p&offsets|0|+1(+1C);m-72&9|+|124693725|124693747&offsets|0|+1(+1C)]|AACATTCATTGCTGTCGGTGGGTTC</td><td>>MI0000270|hsa-mir-181b-1&WithFlank&1|-|198858867|198858988</td><td>42</td><td>65</td><td>24(+1C)</td><td>AACATTCATTGCTGTCGGTGGGTTC</td><td>162.8</td><td>5.1</td><td>1.6</td><td>3.2</td><td>1.4E-03</td><td>2.0E-03</td><td>MB231/MB468</td><td>MP</td></tr>
<tr><td>>hg38_MT_+_1621_3279_RNR2_16S_50ntFlanks@617.652.36|AAACATATAACTGAACTCCTCACACCCAATTGGACC</td><td>>hg38_MT_+_1621_3279_RNR2_16S_50ntFlanks</td><td>617</td><td>652</td><td>36</td><td>AAACATATAACTGAACTCCTCACACCCAATTGGACC</td><td>103.8</td><td>4.8</td><td>1.5</td><td>3.1</td><td>1.6E-03</td><td>2.0E-03</td><td>MB231/MB468</td><td>MITO</td></tr>
<tr><td>>hg38_21_+_8433172_8446622_RNA45SN1_45S_50ntFlanks@12712.12752.41|CGGAGCCTCGGTTGGCCTCGGATAGCCGGTCCCCCGCCTGT</td><td>>hg38_21_+_8433172_8446622_RNA45SN1_45S_50ntFlanks</td><td>12712</td><td>12752</td><td>41</td><td>CGGAGCCTCGGTTGGCCTCGGATAGCCGGTCCCCCGCCTGT</td><td>45.6</td><td>-8.6</td><td>2.6</td><td>-3.4</td><td>7.7E-04</td><td>2.0E-03</td><td>BT20/MB468</td><td>MP</td></tr>
<tr><td>>hg38_21_+_8433172_8446622_RNA45SN1_45S_50ntFlanks@9498.9524.27|AAGATGGTGAACTATGCCTGGGCAGGG</td><td>>hg38_21_+_8433172_8446622_RNA45SN1_45S_50ntFlanks</td><td>9498</td><td>9524</td><td>27</td><td>AAGATGGTGAACTATGCCTGGGCAGGG</td><td>103.8</td><td>10.0</td><td>2.7</td><td>3.6</td><td>2.6E-04</td><td>2.0E-03</td><td>BT20/MB231</td><td>MP</td></tr>
<tr><td>>hg38_21_+_8433172_8446622_RNA45SN1_45S_50ntFlanks@12941.12959.19|ATTCGTAGACGACCTGCTT</td><td>>hg38_21_+_8433172_8446622_RNA45SN1_45S_50ntFlanks</td><td>12941</td><td>12959</td><td>19</td><td>ATTCGTAGACGACCTGCTT</td><td>511.1</td><td>1.8</td><td>0.6</td><td>3.2</td><td>1.5E-03</td><td>2.0E-03</td><td>BT20/MB468</td><td>CYTO</td></tr>
<tr><td>>hg38_21_+_8433172_8446622_RNA45SN1_45S_50ntFlanks@10686.10710.25|GCGGCGTCCGGTGAGCTCTCGCTGG</td><td>>hg38_21_+_8433172_8446622_RNA45SN1_45S_50ntFlanks</td><td>10686</td><td>10710</td><td>25</td><td>GCGGCGTCCGGTGAGCTCTCGCTGG</td><td>2339.0</td><td>-4.0</td><td>1.3</td><td>-3.2</td><td>1.5E-03</td><td>2.0E-03</td><td>BT20/MB468</td><td>CYTO</td></tr>
<tr><td>>hg38_21_+_8433172_8446622_RNA45SN1_45S_50ntFlanks@9929.9961.33|GATATAGACAGCAGGACGGTGGCCATGGAAGTC</td><td>>hg38_21_+_8433172_8446622_RNA45SN1_45S_50ntFlanks</td><td>9929</td><td>9961</td><td>33</td><td>GATATAGACAGCAGGACGGTGGCCATGGAAGTC</td><td>342.4</td><td>6.2</td><td>1.7</td><td>3.6</td><td>2.6E-04</td><td>2.0E-03</td><td>BT20/MB231</td><td>MP</td></tr>
<tr><td>>MI0003205|hsa-mir-532&WithFlank&X|+|50003142|50003244@26.47.22&[MIMAT0002888&hsa-miR-532-5p&offsets|0|0;m-65&X|+|50003167|50003188&offsets|0|0]|CATGCCTTGAGTGTAGGACCGT</td><td>>MI0003205|hsa-mir-532&WithFlank&X|+|50003142|50003244</td><td>26</td><td>47</td><td>22</td><td>CATGCCTTGAGTGTAGGACCGT</td><td>5943.8</td><td>3.3</td><td>1.0</td><td>3.4</td><td>7.7E-04</td><td>2.0E-03</td><td>BT20/MB468</td><td>MP</td></tr>
<tr><td>>hg38_21_+_8433172_8446622_RNA45SN1_45S_50ntFlanks@10681.10709.29|TTCCGGCGGCGTCCGGTGAGCTCTCGCTG</td><td>>hg38_21_+_8433172_8446622_RNA45SN1_45S_50ntFlanks</td><td>10681</td><td>10709</td><td>29</td><td>TTCCGGCGGCGTCCGGTGAGCTCTCGCTG</td><td>308.1</td><td>-3.5</td><td>1.1</td><td>-3.2</td><td>1.3E-03</td><td>2.0E-03</td><td>BT20/MB468</td><td>MITO</td></tr>
<tr><td>>hg38_21_+_8433172_8446622_RNA45SN1_45S_50ntFlanks@9574.9598.25|AATCGGTCGTCCGACCTGGGTATAG</td><td>>hg38_21_+_8433172_8446622_RNA45SN1_45S_50ntFlanks</td><td>9574</td><td>9598</td><td>25</td><td>AATCGGTCGTCCGACCTGGGTATAG</td><td>243.5</td><td>-4.4</td><td>1.3</td><td>-3.4</td><td>7.7E-04</td><td>2.0E-03</td><td>BT20/MB468</td><td>MP</td></tr>
<tr><td>>MI0006273|hsa-mir-1180&WithFlank&17|-|19344500|19344580@47.68.22&[MIMAT0005825&hsa-miR-1180-3p&offsets|0|0;m-214&17|-|19344513|19344534&offsets|0|0]|TTTCCGGCTCGCGTGGGTGTGT</td><td>>MI0006273|hsa-mir-1180&WithFlank&17|-|19344500|19344580</td><td>47</td><td>68</td><td>22</td><td>TTTCCGGCTCGCGTGGGTGTGT</td><td>3613.0</td><td>2.9</td><td>0.8</td><td>3.6</td><td>3.0E-04</td><td>2.0E-03</td><td>BT20/MB231</td><td>CYTO</td></tr>
<tr><td>>trna152_ValCAC_6_-_27248049_27248121@1.41.41|GCTTCTGTAGTGTAGTGGTTATCACGTTCGCCTCACACGCG</td><td>>trna152_ValCAC_6_-_27248049_27248121</td><td>1</td><td>41</td><td>41</td><td>GCTTCTGTAGTGTAGTGGTTATCACGTTCGCCTCACACGCG</td><td>242.0</td><td>-3.3</td><td>0.9</td><td>-3.6</td><td>3.0E-04</td><td>2.0E-03</td><td>BT20/MB231</td><td>CYTO</td></tr>
<tr><td>>hg38_21_+_8433172_8446622_RNA45SN1_45S_50ntFlanks@7977.8012.36|CGACCTCAGATCAGACGTGGCGACCCGCTGAATTTA</td><td>>hg38_21_+_8433172_8446622_RNA45SN1_45S_50ntFlanks</td><td>7977</td><td>8012</td><td>36</td><td>CGACCTCAGATCAGACGTGGCGACCCGCTGAATTTA</td><td>906.6</td><td>-3.9</td><td>1.1</td><td>-3.6</td><td>3.0E-04</td><td>2.0E-03</td><td>BT20/MB231</td><td>CYTO</td></tr>
<tr><td>>MI0000077|hsa-mir-21&WithFlank&17|+|59841260|59841343@13.34.22&[MIMAT0000076&hsa-miR-21-5p&offsets|-1|-1;m-1&17|+|59841273|59841294&offsets|-1|-1]|GTAGCTTATCAGACTGATGTTG</td><td>>MI0000077|hsa-mir-21&WithFlank&17|+|59841260|59841343</td><td>13</td><td>34</td><td>22</td><td>GTAGCTTATCAGACTGATGTTG</td><td>304.4</td><td>2.6</td><td>0.8</td><td>3.2</td><td>1.4E-03</td><td>2.0E-03</td><td>MB231/MB468</td><td>MP</td></tr>
<tr><td>>hg38_21_+_8433172_8446622_RNA45SN1_45S_50ntFlanks@11104.11121.18|AAGGGTCGGGGCGGCAGG</td><td>>hg38_21_+_8433172_8446622_RNA45SN1_45S_50ntFlanks</td><td>11104</td><td>11121</td><td>18</td><td>AAGGGTCGGGGCGGCAGG</td><td>96.2</td><td>9.9</td><td>2.7</td><td>3.6</td><td>2.7E-04</td><td>2.0E-03</td><td>BT20/MB231</td><td>MP</td></tr>
<tr><td>>hg38_21_+_8433172_8446622_RNA45SN1_45S_50ntFlanks@4380.4407.28|CGTAGTTGGATCTTGGGAGCGGGCGGGC</td><td>>hg38_21_+_8433172_8446622_RNA45SN1_45S_50ntFlanks</td><td>4380</td><td>4407</td><td>28</td><td>CGTAGTTGGATCTTGGGAGCGGGCGGGC</td><td>107.6</td><td>-5.0</td><td>1.6</td><td>-3.2</td><td>1.4E-03</td><td>2.0E-03</td><td>MB231/MB468</td><td>MP</td></tr>
<tr><td>>trnaMT_TrpTCA_MT_+_5512_5579@53.71.19,trnalookalike3_TrpTCA_1_+_566062_566129@53.71.19|CAATACTTAATTTCTGCCA</td><td>>trnaMT_TrpTCA_MT_+_5512_5579</td><td>53</td><td>71</td><td>19</td><td>CAATACTTAATTTCTGCCA</td><td>776.3</td><td>-3.2</td><td>1.0</td><td>-3.1</td><td>1.7E-03</td><td>2.0E-03</td><td>MB231/MB468</td><td>MITO</td></tr>
<tr><td>>MI0000433|hsa-let-7g&WithFlank&3|-|52268272|52268367@11.30.20(+1A)&[MIMAT0000414&hsa-let-7g-5p&offsets|0|-2(+1A);m-58&3|-|52268336|52268357&offsets|0|-2(+1A)]|TGAGGTAGTAGTTTGTACAGA</td><td>>MI0000433|hsa-let-7g&WithFlank&3|-|52268272|52268367</td><td>11</td><td>30</td><td>20(+1A)</td><td>TGAGGTAGTAGTTTGTACAGA</td><td>78.5</td><td>6.3</td><td>2.0</td><td>3.2</td><td>1.4E-03</td><td>2.0E-03</td><td>MB231/MB468</td><td>MP</td></tr>
<tr><td>>MI0000264|hsa-mir-7-2&WithFlank&15|+|88611819|88611940@38.59.22(+1C);MI0000265|hsa-mir-7-3&WithFlank&19|+|4770664|4770785@37.58.22(+1C);MI0000263|hsa-mir-7-1&WithFlank&9|-|83969742|83969863@30.51.22(+1C)&[MIMAT0000252&hsa-miR-7-5p&offsets|0|-2(+1C);m-400&15|+|88611856|88611879&offsets|0|-2(+1C)];[MIMAT0000252_1&hsa-miR-7-5p&offsets|0|-2(+1C);m-402&19|+|4770700|4770723&offsets|0|-2(+1C)];[MIMAT0000252_2&hsa-miR-7-5p&offsets|0|-2(+1C);m-401&9|-|83969811|83969834&offsets|0|-2(+1C)]|TGGAAGACTAGTGATTTTGTTGC</td><td>>MI0000264|hsa-mir-7-2&WithFlank&15|+|88611819|88611940</td><td>38</td><td>59</td><td>22(+1C)</td><td>TGGAAGACTAGTGATTTTGTTGC</td><td>1443.5</td><td>3.5</td><td>1.0</td><td>3.6</td><td>3.1E-04</td><td>2.0E-03</td><td>BT20/MB231</td><td>CYTO</td></tr>
<tr><td>>hg38_21_+_8433172_8446622_RNA45SN1_45S_50ntFlanks@11666.11697.32|TCTGCCCAGTGCTCTGAATGTCAAAGTGAAGA</td><td>>hg38_21_+_8433172_8446622_RNA45SN1_45S_50ntFlanks</td><td>11666</td><td>11697</td><td>32</td><td>TCTGCCCAGTGCTCTGAATGTCAAAGTGAAGA</td><td>165.6</td><td>4.7</td><td>1.4</td><td>3.4</td><td>7.8E-04</td><td>2.0E-03</td><td>BT20/MB468</td><td>MP</td></tr>
<tr><td>>hg38_21_+_8433172_8446622_RNA45SN1_45S_50ntFlanks@4161.4184.24|CACATCCAAGGAAGGCAGCAGGCG</td><td>>hg38_21_+_8433172_8446622_RNA45SN1_45S_50ntFlanks</td><td>4161</td><td>4184</td><td>24</td><td>CACATCCAAGGAAGGCAGCAGGCG</td><td>137.6</td><td>-5.4</td><td>1.6</td><td>-3.4</td><td>7.8E-04</td><td>2.0E-03</td><td>BT20/MB468</td><td>MP</td></tr>
<tr><td>>hg38_21_+_8433172_8446622_RNA45SN1_45S_50ntFlanks@5188.5206.19|ATAACAGGTCTGTGATGCC</td><td>>hg38_21_+_8433172_8446622_RNA45SN1_45S_50ntFlanks</td><td>5188</td><td>5206</td><td>19</td><td>ATAACAGGTCTGTGATGCC</td><td>150.2</td><td>-5.7</td><td>1.7</td><td>-3.4</td><td>7.8E-04</td><td>2.0E-03</td><td>BT20/MB468</td><td>MP</td></tr>
<tr><td>>hg38_21_+_8433172_8446622_RNA45SN1_45S_50ntFlanks@12417.12455.39|TCGATGTCGGCTCTTCCTATCATTGTGAAGCAGAATTCA</td><td>>hg38_21_+_8433172_8446622_RNA45SN1_45S_50ntFlanks</td><td>12417</td><td>12455</td><td>39</td><td>TCGATGTCGGCTCTTCCTATCATTGTGAAGCAGAATTCA</td><td>1807.7</td><td>-2.6</td><td>0.8</td><td>-3.2</td><td>1.6E-03</td><td>2.0E-03</td><td>MB231/MB468</td><td>CYTO</td></tr>
<tr><td>>hg38_21_+_8433172_8446622_RNA45SN1_45S_50ntFlanks@7979.8003.25|ACCTCAGATCAGACGTGGCGACCCG</td><td>>hg38_21_+_8433172_8446622_RNA45SN1_45S_50ntFlanks</td><td>7979</td><td>8003</td><td>25</td><td>ACCTCAGATCAGACGTGGCGACCCG</td><td>672.5</td><td>-2.5</td><td>0.8</td><td>-3.2</td><td>1.6E-03</td><td>2.0E-03</td><td>MB231/MB468</td><td>CYTO</td></tr>
<tr><td>>trna111_HisGTG_1_-_147774845_147774916@-1A.23.24,trna118_HisGTG_1_-_145396881_145396952@-1A.23.24,trna16_HisGTG_1_+_146544773_146544844@-1A.23.24,trna1_HisGTG_15_+_45493349_45493420@-1A.23.24,trna21_HisGTG_1_+_147753471_147753542@-1A.23.24,trna33_HisGTG_6_+_27125906_27125977@-1A.23.24,trna7_HisGTG_9_-_14433938_14434009@-1A.23.24,trna8_HisGTG_15_-_45492611_45492682@-1A.23.24,trna9_HisGTG_15_-_45490804_45490875@-1A.23.24|AGCCGTGATCGTATAGTGGTTAGT</td><td>>trna111_HisGTG_1_-_147774845_147774916</td><td>-1A</td><td>23</td><td>24</td><td>AGCCGTGATCGTATAGTGGTTAGT</td><td>486.2</td><td>-4.6</td><td>1.3</td><td>-3.6</td><td>2.9E-04</td><td>2.0E-03</td><td>BT20/MB231</td><td>NUC</td></tr>
<tr><td>>hg38_21_+_8433172_8446622_RNA45SN1_45S_50ntFlanks@12420.12451.32|ATGTCGGCTCTTCCTATCATTGTGAAGCAGAA</td><td>>hg38_21_+_8433172_8446622_RNA45SN1_45S_50ntFlanks</td><td>12420</td><td>12451</td><td>32</td><td>ATGTCGGCTCTTCCTATCATTGTGAAGCAGAA</td><td>315.0</td><td>-2.9</td><td>0.8</td><td>-3.6</td><td>3.4E-04</td><td>2.0E-03</td><td>BT20/MB231</td><td>MITO</td></tr>
<tr><td>>hg38_21_+_8433172_8446622_RNA45SN1_45S_50ntFlanks@4566.4587.22|ATAATGGAATAGGACCGCGGTT</td><td>>hg38_21_+_8433172_8446622_RNA45SN1_45S_50ntFlanks</td><td>4566</td><td>4587</td><td>22</td><td>ATAATGGAATAGGACCGCGGTT</td><td>184.5</td><td>-4.4</td><td>1.4</td><td>-3.2</td><td>1.4E-03</td><td>2.0E-03</td><td>MB231/MB468</td><td>MP</td></tr>
<tr><td>>hg38_21_+_8433172_8446622_RNA45SN1_45S_50ntFlanks@6662.6696.35|GGTGGATCACTCGGCTCGTGCGTCGATGAAGAACG</td><td>>hg38_21_+_8433172_8446622_RNA45SN1_45S_50ntFlanks</td><td>6662</td><td>6696</td><td>35</td><td>GGTGGATCACTCGGCTCGTGCGTCGATGAAGAACG</td><td>129.5</td><td>3.0</td><td>0.9</td><td>3.4</td><td>7.8E-04</td><td>2.0E-03</td><td>BT20/MB468</td><td>MP</td></tr>
<tr><td>>hg38_21_+_8433172_8446622_RNA45SN1_45S_50ntFlanks@10685.10709.25|GGCGGCGTCCGGTGAGCTCTCGCTG</td><td>>hg38_21_+_8433172_8446622_RNA45SN1_45S_50ntFlanks</td><td>10685</td><td>10709</td><td>25</td><td>GGCGGCGTCCGGTGAGCTCTCGCTG</td><td>331.7</td><td>-2.3</td><td>0.7</td><td>-3.1</td><td>1.7E-03</td><td>2.0E-03</td><td>MB231/MB468</td><td>MITO</td></tr>
<tr><td>>am_trnaMT_HisGTG_MT_+_12138_12206@18.37.20|AAAACATCAGATTGTGAATC</td><td>>am_trnaMT_HisGTG_MT_+_12138_12206</td><td>18</td><td>37</td><td>20</td><td>AAAACATCAGATTGTGAATC</td><td>41.8</td><td>8.6</td><td>2.7</td><td>3.2</td><td>1.4E-03</td><td>2.0E-03</td><td>MB231/MB468</td><td>MP</td></tr>
<tr><td>>MI0000093|hsa-mir-92a-1&WithFlank&13|+|91351308|91351397@54.75.22(+1C);MI0000094|hsa-mir-92a-2&WithFlank&X|-|134169532|134169618@54.75.22(+1C)&[MIMAT0000092&hsa-miR-92a-3p&offsets|0|0(+1C);m-9&13|+|91351361|91351382&offsets|0|0(+1C)];[MIMAT0000092_1&hsa-miR-92a-3p&offsets|0|0(+1C);m-11&X|-|134169544|134169565&offsets|0|0(+1C)]|TATTGCACTTGTCCCGGCCTGTC</td><td>>MI0000093|hsa-mir-92a-1&WithFlank&13|+|91351308|91351397</td><td>54</td><td>75</td><td>22(+1C)</td><td>TATTGCACTTGTCCCGGCCTGTC</td><td>705.1</td><td>-3.8</td><td>1.0</td><td>-3.6</td><td>2.7E-04</td><td>2.0E-03</td><td>BT20/MB231</td><td>MP</td></tr>
<tr><td>>MI0000283|hsa-mir-203a&WithFlank&14|+|104117399|104117520@71.92.22(+1U)&[MIMAT0000264&hsa-miR-203a-3p&offsets|0|0(+1U);m-79&14|+|104117469|104117490&offsets|0|0(+1U)]|GTGAAATGTTTAGGACCACTAGT</td><td>>MI0000283|hsa-mir-203a&WithFlank&14|+|104117399|104117520</td><td>71</td><td>92</td><td>22(+1U)</td><td>GTGAAATGTTTAGGACCACTAGT</td><td>36.6</td><td>8.5</td><td>2.3</td><td>3.6</td><td>2.7E-04</td><td>2.0E-03</td><td>BT20/MB231</td><td>MP</td></tr>
<tr><td>>hg38_21_+_8433172_8446622_RNA45SN1_45S_50ntFlanks@8381.8407.27|AAGAGGGCGTGAAACCGTTAAGAGGTA</td><td>>hg38_21_+_8433172_8446622_RNA45SN1_45S_50ntFlanks</td><td>8381</td><td>8407</td><td>27</td><td>AAGAGGGCGTGAAACCGTTAAGAGGTA</td><td>175.9</td><td>-5.1</td><td>1.6</td><td>-3.2</td><td>1.4E-03</td><td>2.0E-03</td><td>MB231/MB468</td><td>MP</td></tr>
<tr><td>>trnaMT_GluTTC_MT_-_14674_14742@26.72.47,trnalookalike8_GluTTC_5_-_93905172_93905240@26.72.47|GGTTTTTCATATCATTGGTCGTGGTTGTAGTCCGTGCGAGAATACCA</td><td>>trnaMT_GluTTC_MT_-_14674_14742</td><td>26</td><td>72</td><td>47</td><td>GGTTTTTCATATCATTGGTCGTGGTTGTAGTCCGTGCGAGAATACCA</td><td>346.6</td><td>6.2</td><td>2.0</td><td>3.1</td><td>1.7E-03</td><td>2.0E-03</td><td>MB231/MB468</td><td>MITO</td></tr>
</table>

<table>
<tr><td>>MI0000067|hsa-let-7f-1&WithFlank&9|+|94176341|94176439@13.34.22(+1C);MI0000068|hsa-let-7f-2&WithFlank&X|-|53557186|53557280@14.35.22(+1C)&[MIMAT0000067&hsa-let-7f-5p&offsets|0|0(+1C);m-20&9|+|94176353|94176374&offsets|0|0(+1C)];[MIMAT0000067_1&hsa-let-7f-5p&offsets|0|0(+1C);m-19&X|-|53557246|53557267&offsets|0|0(+1C)]|TGAGGTAGTAGATTGTATAGTTC</td><td>>MI0000067|hsa-let-7f-1&WithFlank&9|+|94176341|94176439</td><td>13</td><td>34</td><td>22(+1C)</td><td>TGAGGTAGTAGATTGTATAGTTC</td><td>349.3</td><td>5.0</td><td>1.5</td><td>3.4</td><td>7.8E-04</td><td>2.0E-03</td><td>BT20/MB468</td><td>MP</td></tr>
<tr><td>>hg38_21_+_8433172_8446622_RNA45SN1_45S_50ntFlanks@10059.10077.19|GGCAGTCGAGAGTGGACGG</td><td>>hg38_21_+_8433172_8446622_RNA45SN1_45S_50ntFlanks</td><td>10059</td><td>10077</td><td>19</td><td>GGCAGTCGAGAGTGGACGG</td><td>217.4</td><td>4.5</td><td>1.3</td><td>3.4</td><td>7.8E-04</td><td>2.0E-03</td><td>BT20/MB468</td><td>MP</td></tr>
<tr><td>>hg38_21_+_8433172_8446622_RNA45SN1_45S_50ntFlanks@10128.10157.30|CGGCGGCGGCGGCGGCGGCGGGGGTGTGGG</td><td>>hg38_21_+_8433172_8446622_RNA45SN1_45S_50ntFlanks</td><td>10128</td><td>10157</td><td>30</td><td>CGGCGGCGGCGGCGGCGGCGGGGGTGTGGG</td><td>91.4</td><td>-5.4</td><td>1.6</td><td>-3.4</td><td>7.8E-04</td><td>2.0E-03</td><td>BT20/MB468</td><td>MP</td></tr>
<tr><td>>hg38_21_+_8433172_8446622_RNA45SN1_45S_50ntFlanks@3843.3871.29|CTACTTGGATAACTGTGGTAATTCTAGAG</td><td>>hg38_21_+_8433172_8446622_RNA45SN1_45S_50ntFlanks</td><td>3843</td><td>3871</td><td>29</td><td>CTACTTGGATAACTGTGGTAATTCTAGAG</td><td>415.6</td><td>-4.5</td><td>1.3</td><td>-3.4</td><td>7.8E-04</td><td>2.0E-03</td><td>BT20/MB468</td><td>MP</td></tr>
<tr><td>>hg38_21_+_8433172_8446622_RNA45SN1_45S_50ntFlanks@7988.8011.24|CAGACGTGGCGACCCGCTGAATTT</td><td>>hg38_21_+_8433172_8446622_RNA45SN1_45S_50ntFlanks</td><td>7988</td><td>8011</td><td>24</td><td>CAGACGTGGCGACCCGCTGAATTT</td><td>352.1</td><td>2.1</td><td>0.7</td><td>3.2</td><td>1.6E-03</td><td>2.0E-03</td><td>MB231/MB468</td><td>CYTO</td></tr>
<tr><td>>hg38_21_+_8433172_8446622_RNA45SN1_45S_50ntFlanks@11570.11611.42|AGCCGACTTAGAACTGGTGCGGACCAGGGGAATCCGACTGTT</td><td>>hg38_21_+_8433172_8446622_RNA45SN1_45S_50ntFlanks</td><td>11570</td><td>11611</td><td>42</td><td>AGCCGACTTAGAACTGGTGCGGACCAGGGGAATCCGACTGTT</td><td>126.0</td><td>-4.3</td><td>1.4</td><td>-3.2</td><td>1.4E-03</td><td>2.0E-03</td><td>MB231/MB468</td><td>MP</td></tr>
<tr><td>>am_trnaMT_ValTAC_MT_+_1602_1670@48.68.21|CAACTTAACTTGACCGCTCTG</td><td>>am_trnaMT_ValTAC_MT_+_1602_1670</td><td>48</td><td>68</td><td>21</td><td>CAACTTAACTTGACCGCTCTG</td><td>864.3</td><td>-3.2</td><td>1.0</td><td>-3.1</td><td>1.7E-03</td><td>2.0E-03</td><td>MB231/MB468</td><td>MITO</td></tr>
<tr><td>>hg38_21_+_8433172_8446622_RNA45SN1_45S_50ntFlanks@12658.12675.18|ACTGAACGCCTCTAAGTC</td><td>>hg38_21_+_8433172_8446622_RNA45SN1_45S_50ntFlanks</td><td>12658</td><td>12675</td><td>18</td><td>ACTGAACGCCTCTAAGTC</td><td>660.6</td><td>-5.1</td><td>1.6</td><td>-3.2</td><td>1.5E-03</td><td>2.0E-03</td><td>BT20/MB468</td><td>CYTO</td></tr>
<tr><td>>hg38_21_+_8433172_8446622_RNA45SN1_45S_50ntFlanks@9576.9603.28|TCGGTCGTCCGACCTGGGTATAGGGGCG</td><td>>hg38_21_+_8433172_8446622_RNA45SN1_45S_50ntFlanks</td><td>9576</td><td>9603</td><td>28</td><td>TCGGTCGTCCGACCTGGGTATAGGGGCG</td><td>92.9</td><td>-5.3</td><td>1.6</td><td>-3.4</td><td>7.8E-04</td><td>2.0E-03</td><td>BT20/MB468</td><td>MP</td></tr>
<tr><td>>hg38_21_+_8433172_8446622_RNA45SN1_45S_50ntFlanks@6735.6771.37|TTGATCATCGACACTTCGAACGCACTTGCGGCCCCGG</td><td>>hg38_21_+_8433172_8446622_RNA45SN1_45S_50ntFlanks</td><td>6735</td><td>6771</td><td>37</td><td>TTGATCATCGACACTTCGAACGCACTTGCGGCCCCGG</td><td>580.6</td><td>-3.8</td><td>1.2</td><td>-3.2</td><td>1.3E-03</td><td>2.0E-03</td><td>BT20/MB468</td><td>MITO</td></tr>
<tr><td>>hg38_21_+_8433172_8446622_RNA45SN1_45S_50ntFlanks@11588.11611.24|GCGGACCAGGGGAATCCGACTGTT</td><td>>hg38_21_+_8433172_8446622_RNA45SN1_45S_50ntFlanks</td><td>11588</td><td>11611</td><td>24</td><td>GCGGACCAGGGGAATCCGACTGTT</td><td>383.7</td><td>2.0</td><td>0.6</td><td>3.1</td><td>1.7E-03</td><td>2.0E-03</td><td>MB231/MB468</td><td>NUC</td></tr>
<tr><td>>hg38_21_+_8433172_8446622_RNA45SN1_45S_50ntFlanks@12043.12060.18|CACTGACCCGGTGAGGCG</td><td>>hg38_21_+_8433172_8446622_RNA45SN1_45S_50ntFlanks</td><td>12043</td><td>12060</td><td>18</td><td>CACTGACCCGGTGAGGCG</td><td>111.6</td><td>-5.3</td><td>1.6</td><td>-3.4</td><td>7.8E-04</td><td>2.0E-03</td><td>BT20/MB468</td><td>MP</td></tr>
<tr><td>>hg38_21_+_8433172_8446622_RNA45SN1_45S_50ntFlanks@10572.10596.25|AGAAGCCGGCGGGAGCCCCGGGGAG</td><td>>hg38_21_+_8433172_8446622_RNA45SN1_45S_50ntFlanks</td><td>10572</td><td>10596</td><td>25</td><td>AGAAGCCGGCGGGAGCCCCGGGGAG</td><td>102.1</td><td>-5.2</td><td>1.6</td><td>-3.4</td><td>7.8E-04</td><td>2.0E-03</td><td>BT20/MB468</td><td>MP</td></tr>
<tr><td>>hg38_21_+_8433172_8446622_RNA45SN1_45S_50ntFlanks@8919.8936.18|TCCCGGGGCCGAGGGAGC</td><td>>hg38_21_+_8433172_8446622_RNA45SN1_45S_50ntFlanks</td><td>8919</td><td>8936</td><td>18</td><td>TCCCGGGGCCGAGGGAGC</td><td>71.6</td><td>5.4</td><td>1.6</td><td>3.4</td><td>7.9E-04</td><td>2.0E-03</td><td>BT20/MB468</td><td>MP</td></tr>
<tr><td>>hg38_21_+_8433172_8446622_RNA45SN1_45S_50ntFlanks@6661.6704.44|CGGTGGATCACTCGGCTCGTGCGTCGATGAAGAACGCAGCTAGC</td><td>>hg38_21_+_8433172_8446622_RNA45SN1_45S_50ntFlanks</td><td>6661</td><td>6704</td><td>44</td><td>CGGTGGATCACTCGGCTCGTGCGTCGATGAAGAACGCAGCTAGC</td><td>738.3</td><td>2.8</td><td>0.8</td><td>3.6</td><td>3.5E-04</td><td>2.0E-03</td><td>BT20/MB231</td><td>MITO</td></tr>
<tr><td>>hg38_MT_+_1621_3279_RNR2_16S_50ntFlanks@1243.1265.23|TTCACCAGTCAAAGCGAACTACT</td><td>>hg38_MT_+_1621_3279_RNR2_16S_50ntFlanks</td><td>1243</td><td>1265</td><td>23</td><td>TTCACCAGTCAAAGCGAACTACT</td><td>178.7</td><td>4.2</td><td>1.3</td><td>3.1</td><td>1.7E-03</td><td>2.0E-03</td><td>MB231/MB468</td><td>MITO</td></tr>
<tr><td>>MI0000487|hsa-mir-193a&WithFlank&17|+|31559990|31560089@27.46.20&[MIMAT0004614&hsa-miR-193a-5p&offsets|0|-2;m-123&17|+|31560016|31560037&offsets|0|-2]|TGGGTCTTTGCGGGCGAGAT</td><td>>MI0000487|hsa-mir-193a&WithFlank&17|+|31559990|31560089</td><td>27</td><td>46</td><td>20</td><td>TGGGTCTTTGCGGGCGAGAT</td><td>153.1</td><td>6.9</td><td>2.0</td><td>3.4</td><td>7.9E-04</td><td>2.0E-03</td><td>BT20/MB468</td><td>MP</td></tr>
<tr><td>>trnaMT_ThrTGT_MT_+_15888_15953@36.68.33|AACCGGAGATGAAAACCTTTTTCCAAGGACACC</td><td>>trnaMT_ThrTGT_MT_+_15888_15953</td><td>36</td><td>68</td><td>33</td><td>AACCGGAGATGAAAACCTTTTTCCAAGGACACC</td><td>47.7</td><td>9.3</td><td>2.8</td><td>3.4</td><td>7.9E-04</td><td>2.0E-03</td><td>BT20/MB468</td><td>MP</td></tr>
<tr><td>>hg38_21_+_8433172_8446622_RNA45SN1_45S_50ntFlanks@12488.12525.38|AACGTGAGCTGGGTTTAGACCGTCGTGAGACAGGTTAG</td><td>>hg38_21_+_8433172_8446622_RNA45SN1_45S_50ntFlanks</td><td>12488</td><td>12525</td><td>38</td><td>AACGTGAGCTGGGTTTAGACCGTCGTGAGACAGGTTAG</td><td>168.6</td><td>-4.6</td><td>1.4</td><td>-3.2</td><td>1.5E-03</td><td>2.0E-03</td><td>BT20/MB468</td><td>CYTO</td></tr>
<tr><td>>hg38_21_+_8433172_8446622_RNA45SN1_45S_50ntFlanks@8319.8356.38|CGAGACCGATAGTCAACAAGTACCGTAAGGGAAAGTTG</td><td>>hg38_21_+_8433172_8446622_RNA45SN1_45S_50ntFlanks</td><td>8319</td><td>8356</td><td>38</td><td>CGAGACCGATAGTCAACAAGTACCGTAAGGGAAAGTTG</td><td>444.6</td><td>-4.0</td><td>1.3</td><td>-3.2</td><td>1.5E-03</td><td>2.0E-03</td><td>BT20/MB468</td><td>CYTO</td></tr>
<tr><td>>hg38_21_+_8433172_8446622_RNA45SN1_45S_50ntFlanks@10346.10368.23|TCAAACGAGAACTTTGAAGGCCG</td><td>>hg38_21_+_8433172_8446622_RNA45SN1_45S_50ntFlanks</td><td>10346</td><td>10368</td><td>23</td><td>TCAAACGAGAACTTTGAAGGCCG</td><td>2839.5</td><td>-2.1</td><td>0.7</td><td>-3.1</td><td>1.7E-03</td><td>2.0E-03</td><td>MB231/MB468</td><td>NUC</td></tr>
<tr><td>>hg38_MT_+_1621_3279_RNR2_16S_50ntFlanks@440.459.20|ACAGAACCCTCTAAATCCCC</td><td>>hg38_MT_+_1621_3279_RNR2_16S_50ntFlanks</td><td>440</td><td>459</td><td>20</td><td>ACAGAACCCTCTAAATCCCC</td><td>104.5</td><td>4.9</td><td>1.6</td><td>3.1</td><td>1.7E-03</td><td>2.0E-03</td><td>MB231/MB468</td><td>MITO</td></tr>
<tr><td>>hg38_21_+_8433172_8446622_RNA45SN1_45S_50ntFlanks@11209.11234.26|CGGCGACTCTGGACGCGAGCCGGGCC</td><td>>hg38_21_+_8433172_8446622_RNA45SN1_45S_50ntFlanks</td><td>11209</td><td>11234</td><td>26</td><td>CGGCGACTCTGGACGCGAGCCGGGCC</td><td>2151.4</td><td>-6.4</td><td>2.0</td><td>-3.1</td><td>1.7E-03</td><td>2.0E-03</td><td>BT20/MB468</td><td>NUC</td></tr>
<tr><td>>hg38_MT_+_1621_3279_RNR2_16S_50ntFlanks@847.882.36|AAAGGAACTCGGCAAATCTTACCCCGCCTGTTTACC</td><td>>hg38_MT_+_1621_3279_RNR2_16S_50ntFlanks</td><td>847</td><td>882</td><td>36</td><td>AAAGGAACTCGGCAAATCTTACCCCGCCTGTTTACC</td><td>47.7</td><td>9.3</td><td>2.8</td><td>3.4</td><td>7.9E-04</td><td>2.0E-03</td><td>BT20/MB468</td><td>MP</td></tr>
<tr><td>>hg38_21_+_8433172_8446622_RNA45SN1_45S_50ntFlanks@6664.6688.25|TGGATCACTCGGCTCGTGCGTCGAT</td><td>>hg38_21_+_8433172_8446622_RNA45SN1_45S_50ntFlanks</td><td>6664</td><td>6688</td><td>25</td><td>TGGATCACTCGGCTCGTGCGTCGAT</td><td>276.9</td><td>3.8</td><td>1.1</td><td>3.6</td><td>3.5E-04</td><td>2.0E-03</td><td>BT20/MB231</td><td>MITO</td></tr>
<tr><td>>MI0000114|hsa-mir-107&WithFlank&10|-|89592741|89592833@56.75.20(+1C);MI0000108|hsa-mir-103a-2&WithFlank&20|+|3917488|3917577@54.73.20(+1C);MI0000109|hsa-mir-103a-1&WithFlank&5|-|168560890|168560979@54.73.20(+1C)&[MIMAT0000104&hsa-miR-107&offsets|0|-3(+1C);m-78&10|-|89592758|89592778&offsets|0|-1(+1C)];[MIMAT0000101&hsa-miR-103a-3p&offsets|0|-3(+1C);m-12&20|+|3917541|3917563&offsets|0|-3(+1C)];[MIMAT0000101_1&hsa-miR-103a-3p&offsets|0|-3(+1C);m-13&5|-|168560904|168560926&offsets|0|-3(+1C);m-44220&5|-|168560908|168560926&offsets|0|+1(+1C)]|AGCAGCATTGTACAGGGCTAC</td><td>>MI0000114|hsa-mir-107&WithFlank&10|-|89592741|89592833</td><td>56</td><td>75</td><td>20(+1C)</td><td>AGCAGCATTGTACAGGGCTAC</td><td>389.3</td><td>4.3</td><td>1.4</td><td>3.2</td><td>1.4E-03</td><td>2.0E-03</td><td>MB231/MB468</td><td>MP</td></tr>
<tr><td>>hg38_21_+_8433172_8446622_RNA45SN1_45S_50ntFlanks@4122.4144.23|AGGGTTCGATTCCGGAGAGGGAG</td><td>>hg38_21_+_8433172_8446622_RNA45SN1_45S_50ntFlanks</td><td>4122</td><td>4144</td><td>23</td><td>AGGGTTCGATTCCGGAGAGGGAG</td><td>281.4</td><td>-4.3</td><td>1.3</td><td>-3.4</td><td>7.9E-04</td><td>2.0E-03</td><td>BT20/MB468</td><td>MP</td></tr>
<tr><td>>hg38_21_+_8433172_8446622_RNA45SN1_45S_50ntFlanks@5495.5514.20|AGACGGTCGAACTTGACTAT</td><td>>hg38_21_+_8433172_8446622_RNA45SN1_45S_50ntFlanks</td><td>5495</td><td>5514</td><td>20</td><td>AGACGGTCGAACTTGACTAT</td><td>304.0</td><td>-5.6</td><td>1.7</td><td>-3.4</td><td>7.9E-04</td><td>2.0E-03</td><td>BT20/MB468</td><td>MP</td></tr>
<tr><td>>hg38_21_+_8433172_8446622_RNA45SN1_45S_50ntFlanks@5073.5115.43|AATTCCGATAACGAACGAGACTCTGGCATGCTAACTAGTTACG</td><td>>hg38_21_+_8433172_8446622_RNA45SN1_45S_50ntFlanks</td><td>5073</td><td>5115</td><td>43</td><td>AATTCCGATAACGAACGAGACTCTGGCATGCTAACTAGTTACG</td><td>52.7</td><td>-9.6</td><td>3.0</td><td>-3.2</td><td>1.4E-03</td><td>2.0E-03</td><td>MB231/MB468</td><td>MP</td></tr>
<tr><td>>hg38_21_+_8433172_8446622_RNA45SN1_45S_50ntFlanks@4445.4462.18|CTTGCCTCTCGGCGCCCC</td><td>>hg38_21_+_8433172_8446622_RNA45SN1_45S_50ntFlanks</td><td>4445</td><td>4462</td><td>18</td><td>CTTGCCTCTCGGCGCCCC</td><td>100.1</td><td>-5.4</td><td>1.6</td><td>-3.4</td><td>7.9E-04</td><td>2.0E-03</td><td>BT20/MB468</td><td>MP</td></tr>
<tr><td>>hg38_21_+_8433172_8446622_RNA45SN1_45S_50ntFlanks@11209.11250.42|CGGCGACTCTGGACGCGAGCCGGGCCCTTCCCGTGGATCGCC</td><td>>hg38_21_+_8433172_8446622_RNA45SN1_45S_50ntFlanks</td><td>11209</td><td>11250</td><td>42</td><td>CGGCGACTCTGGACGCGAGCCGGGCCCTTCCCGTGGATCGCC</td><td>332.1</td><td>-4.1</td><td>1.3</td><td>-3.2</td><td>1.5E-03</td><td>2.0E-03</td><td>BT20/MB468</td><td>CYTO</td></tr>
<tr><td>>hg38_21_+_8433172_8446622_RNA45SN1_45S_50ntFlanks@6646.6682.37|TCGTACGACTCTTAGCGGTGGATCACTCGGCTCGTGC</td><td>>hg38_21_+_8433172_8446622_RNA45SN1_45S_50ntFlanks</td><td>6646</td><td>6682</td><td>37</td><td>TCGTACGACTCTTAGCGGTGGATCACTCGGCTCGTGC</td><td>520.1</td><td>3.0</td><td>1.0</td><td>3.1</td><td>1.7E-03</td><td>2.0E-03</td><td>MB231/MB468</td><td>MITO</td></tr>
<tr><td>>hg38_21_+_8433172_8446622_RNA45SN1_45S_50ntFlanks@11579.11604.26|AGAACTGGTGCGGACCAGGGGAATCC</td><td>>hg38_21_+_8433172_8446622_RNA45SN1_45S_50ntFlanks</td><td>11579</td><td>11604</td><td>26</td><td>AGAACTGGTGCGGACCAGGGGAATCC</td><td>212.6</td><td>-3.7</td><td>1.2</td><td>-3.1</td><td>1.7E-03</td><td>2.0E-03</td><td>MB231/MB468</td><td>NUC</td></tr>
<tr><td>>hg38_21_+_8433172_8446622_RNA45SN1_45S_50ntFlanks@3849.3873.25|GGATAACTGTGGTAATTCTAGAGCT</td><td>>hg38_21_+_8433172_8446622_RNA45SN1_45S_50ntFlanks</td><td>3849</td><td>3873</td><td>25</td><td>GGATAACTGTGGTAATTCTAGAGCT</td><td>142.1</td><td>-3.9</td><td>1.2</td><td>-3.2</td><td>1.4E-03</td><td>2.0E-03</td><td>MB231/MB468</td><td>MP</td></tr>
<tr><td>>hg38_21_+_8433172_8446622_RNA45SN1_45S_50ntFlanks@8182.8204.23|ATCGAGGCCCAGCCCGTGGACGG</td><td>>hg38_21_+_8433172_8446622_RNA45SN1_45S_50ntFlanks</td><td>8182</td><td>8204</td><td>23</td><td>ATCGAGGCCCAGCCCGTGGACGG</td><td>396.2</td><td>-4.9</td><td>1.6</td><td>-3.1</td><td>1.7E-03</td><td>2.0E-03</td><td>MB231/MB468</td><td>NUC</td></tr>
<tr><td>>hg38_21_+_8433172_8446622_RNA45SN1_45S_50ntFlanks@4558.4587.30|AGCTAGGAATAATGGAATAGGACCGCGGTT</td><td>>hg38_21_+_8433172_8446622_RNA45SN1_45S_50ntFlanks</td><td>4558</td><td>4587</td><td>30</td><td>AGCTAGGAATAATGGAATAGGACCGCGGTT</td><td>131.8</td><td>-1.7</td><td>0.5</td><td>-3.2</td><td>1.5E-03</td><td>2.0E-03</td><td>BT20/MB468</td><td>CYTO</td></tr>
<tr><td>>MI0016887|hsa-mir-4521&WithFlank&17|+|8186939|8187010@14.34.21&[MIMAT0019058&hsa-miR-4521&offsets|+4|+3;m-765&17|+|8186948|8186969&offsets|+4|+3]|AGGAAGTCCTGTGCTCAGTTT</td><td>>MI0016887|hsa-mir-4521&WithFlank&17|+|8186939|8187010</td><td>14</td><td>34</td><td>21</td><td>AGGAAGTCCTGTGCTCAGTTT</td><td>496.5</td><td>2.9</td><td>0.8</td><td>3.6</td><td>3.1E-04</td><td>2.0E-03</td><td>BT20/MB231</td><td>CYTO</td></tr>
<tr><td>>hg38_21_+_8433172_8446622_RNA45SN1_45S_50ntFlanks@6650.6669.20|ACGACTCTTAGCGGTGGATC</td><td>>hg38_21_+_8433172_8446622_RNA45SN1_45S_50ntFlanks</td><td>6650</td><td>6669</td><td>20</td><td>ACGACTCTTAGCGGTGGATC</td><td>805.0</td><td>2.8</td><td>0.8</td><td>3.6</td><td>3.1E-04</td><td>2.0E-03</td><td>BT20/MB231</td><td>CYTO</td></tr>
<tr><td>>hg38_21_+_8433172_8446622_RNA45SN1_45S_50ntFlanks@12435.12454.20|ATCATTGTGAAGCAGAATTC</td><td>>hg38_21_+_8433172_8446622_RNA45SN1_45S_50ntFlanks</td><td>12435</td><td>12454</td><td>20</td><td>ATCATTGTGAAGCAGAATTC</td><td>486.8</td><td>-5.0</td><td>1.6</td><td>-3.1</td><td>1.7E-03</td><td>2.0E-03</td><td>MB231/MB468</td><td>NUC</td></tr>
<tr><td>>hg38_21_+_8433172_8446622_RNA45SN1_45S_50ntFlanks@12447.12476.30|CAGAATTCACCAAGCGTTGGATTGTTCACC</td><td>>hg38_21_+_8433172_8446622_RNA45SN1_45S_50ntFlanks</td><td>12447</td><td>12476</td><td>30</td><td>CAGAATTCACCAAGCGTTGGATTGTTCACC</td><td>677.7</td><td>-4.3</td><td>1.3</td><td>-3.2</td><td>1.4E-03</td><td>2.0E-03</td><td>MB231/MB468</td><td>MP</td></tr>
<tr><td>>hg38_21_+_8433172_8446622_RNA45SN1_45S_50ntFlanks@4283.4308.26|CTTTAACGAGGATCCATTGGAGGGCA</td><td>>hg38_21_+_8433172_8446622_RNA45SN1_45S_50ntFlanks</td><td>4283</td><td>4308</td><td>26</td><td>CTTTAACGAGGATCCATTGGAGGGCA</td><td>232.0</td><td>-5.4</td><td>1.6</td><td>-3.4</td><td>7.9E-04</td><td>2.0E-03</td><td>BT20/MB468</td><td>MP</td></tr>
<tr><td>>hg38_21_+_8433172_8446622_RNA45SN1_45S_50ntFlanks@4472.4496.25|CTTAGCTGAGTGTCCCGCGGGGCCC</td><td>>hg38_21_+_8433172_8446622_RNA45SN1_45S_50ntFlanks</td><td>4472</td><td>4496</td><td>25</td><td>CTTAGCTGAGTGTCCCGCGGGGCCC</td><td>467.6</td><td>-5.7</td><td>1.8</td><td>-3.1</td><td>1.7E-03</td><td>2.0E-03</td><td>BT20/MB468</td><td>NUC</td></tr>
<tr><td>>hg38_21_+_8433172_8446622_RNA45SN1_45S_50ntFlanks@9167.9195.29|CGGGCGGGTCGCGCCGTCGGGCCCGGGGG</td><td>>hg38_21_+_8433172_8446622_RNA45SN1_45S_50ntFlanks</td><td>9167</td><td>9195</td><td>29</td><td>CGGGCGGGTCGCGCCGTCGGGCCCGGGGG</td><td>159.3</td><td>-5.6</td><td>1.7</td><td>-3.4</td><td>7.9E-04</td><td>2.0E-03</td><td>BT20/MB468</td><td>MP</td></tr>
<tr><td>>trna119_LysCTT_1_-_145395522_145395594@3.32.30,trna11_LysCTT_5_-_180648979_180649051@3.32.30,trna13_LysCTT_6_+_26556774_26556846@3.32.30,trna32_LysCTT_16_-_3207406_3207478@3.32.30,trna7_LysCTT_16_+_3225692_3225764@3.32.30,trna9_LysCTT_5_+_180634755_180634827@3.32.30|CCGGCTAGCTCAGTCGGTAGAGCATGAGAC</td><td>>trna119_LysCTT_1_-_145395522_145395594</td><td>3</td><td>32</td><td>30</td><td>CCGGCTAGCTCAGTCGGTAGAGCATGAGAC</td><td>249.9</td><td>-3.8</td><td>1.1</td><td>-3.6</td><td>3.5E-04</td><td>2.0E-03</td><td>BT20/MB231</td><td>MITO</td></tr>
<tr><td>>hg38_21_+_8433172_8446622_RNA45SN1_45S_50ntFlanks@11719.11742.24|ACGGCGGGAGTAACTATGACTCTC</td><td>>hg38_21_+_8433172_8446622_RNA45SN1_45S_50ntFlanks</td><td>11719</td><td>11742</td><td>24</td><td>ACGGCGGGAGTAACTATGACTCTC</td><td>245.7</td><td>-2.3</td><td>0.7</td><td>-3.2</td><td>1.5E-03</td><td>2.0E-03</td><td>BT20/MB468</td><td>CYTO</td></tr>
<tr><td>>hg38_21_+_8433172_8446622_RNA45SN1_45S_50ntFlanks@7968.7988.21|TCCGAGACGCGACCTCAGATC</td><td>>hg38_21_+_8433172_8446622_RNA45SN1_45S_50ntFlanks</td><td>7968</td><td>7988</td><td>21</td><td>TCCGAGACGCGACCTCAGATC</td><td>144.3</td><td>4.6</td><td>1.4</td><td>3.4</td><td>8.0E-04</td><td>2.0E-03</td><td>BT20/MB468</td><td>MP</td></tr>
<tr><td>>hg38_21_+_8433172_8446622_RNA45SN1_45S_50ntFlanks@12419.12454.36|GATGTCGGCTCTTCCTATCATTGTGAAGCAGAATTC</td><td>>hg38_21_+_8433172_8446622_RNA45SN1_45S_50ntFlanks</td><td>12419</td><td>12454</td><td>36</td><td>GATGTCGGCTCTTCCTATCATTGTGAAGCAGAATTC</td><td>913.7</td><td>-2.8</td><td>0.8</td><td>-3.6</td><td>3.1E-04</td><td>2.0E-03</td><td>BT20/MB231</td><td>CYTO</td></tr>
<tr><td>>trnaMT_LysTTT_MT_+_8295_8364@56.73.18,trnalookalike7_LysTTT_1_+_568844_568913@56.73.18|ACCTCTTTACAGTGACCA</td><td>>trnaMT_LysTTT_MT_+_8295_8364</td><td>56</td><td>73</td><td>18</td><td>ACCTCTTTACAGTGACCA</td><td>205.3</td><td>6.2</td><td>1.8</td><td>3.4</td><td>8.0E-04</td><td>2.0E-03</td><td>BT20/MB468</td><td>MP</td></tr>
<tr><td>>hg38_21_+_8433172_8446622_RNA45SN1_45S_50ntFlanks@11719.11742.24|ACGGCGGGAGTAACTATGACTCTC</td><td>>hg38_21_+_8433172_8446622_RNA45SN1_45S_50ntFlanks</td><td>11719</td><td>11742</td><td>24</td><td>ACGGCGGGAGTAACTATGACTCTC</td><td>287.6</td><td>-2.3</td><td>0.7</td><td>-3.2</td><td>1.6E-03</td><td>2.0E-03</td><td>MB231/MB468</td><td>CYTO</td></tr>
<tr><td>>hg38_21_+_8433172_8446622_RNA45SN1_45S_50ntFlanks@9602.9622.21|CGAAAGACTAATCGAACCATC</td><td>>hg38_21_+_8433172_8446622_RNA45SN1_45S_50ntFlanks</td><td>9602</td><td>9622</td><td>21</td><td>CGAAAGACTAATCGAACCATC</td><td>119.2</td><td>-4.4</td><td>1.4</td><td>-3.2</td><td>1.3E-03</td><td>2.0E-03</td><td>BT20/MB468</td><td>MITO</td></tr>
<tr><td>>hg38_21_+_8433172_8446622_RNA45SN1_45S_50ntFlanks@13002.13039.38|CGATCTATTGAAAGTCAGCCCTCGACACAAGGGTTTGT</td><td>>hg38_21_+_8433172_8446622_RNA45SN1_45S_50ntFlanks</td><td>13002</td><td>13039</td><td>38</td><td>CGATCTATTGAAAGTCAGCCCTCGACACAAGGGTTTGT</td><td>1104.0</td><td>-5.6</td><td>1.8</td><td>-3.2</td><td>1.4E-03</td><td>2.0E-03</td><td>MB231/MB468</td><td>MP</td></tr>
</table>

<table>
<tr><td>>hg38_21_+_8433172_8446622_RNA45SN1_45S_50ntFlanks@8357.8391.35|AAAAGAACTTTGAAGAGAGAGTTCAAGAGGGCGTG</td><td>>hg38_21_+_8433172_8446622_RNA45SN1_45S_50ntFlanks</td><td>8357</td><td>8391</td><td>35</td><td>AAAAGAACTTTGAAGAGAGAGTTCAAGAGGGCGTG</td><td>626.5</td><td>-2.2</td><td>0.7</td><td>-3.2</td><td>1.6E-03</td><td>2.0E-03</td><td>MB231/MB468</td><td>CYTO</td></tr>
<tr><td>>trnaMT_PheGAA_MT_+_577_647@35.52.18|GAAAATGTTTAGACGGGC</td><td>>trnaMT_PheGAA_MT_+_577_647</td><td>35</td><td>52</td><td>18</td><td>GAAAATGTTTAGACGGGC</td><td>329.6</td><td>4.0</td><td>1.2</td><td>3.4</td><td>8.0E-04</td><td>2.0E-03</td><td>BT20/MB468</td><td>MP</td></tr>
<tr><td>>hg38_21_+_8433172_8446622_RNA45SN1_45S_50ntFlanks@9931.9952.22|TATAGACAGCAGGACGGTGGCC</td><td>>hg38_21_+_8433172_8446622_RNA45SN1_45S_50ntFlanks</td><td>9931</td><td>9952</td><td>22</td><td>TATAGACAGCAGGACGGTGGCC</td><td>105.6</td><td>6.4</td><td>1.9</td><td>3.4</td><td>8.0E-04</td><td>2.0E-03</td><td>BT20/MB468</td><td>MP</td></tr>
<tr><td>>hg38_21_+_8433172_8446622_RNA45SN1_45S_50ntFlanks@11569.11605.37|CAGCCGACTTAGAACTGGTGCGGACCAGGGGAATCCG</td><td>>hg38_21_+_8433172_8446622_RNA45SN1_45S_50ntFlanks</td><td>11569</td><td>11605</td><td>37</td><td>CAGCCGACTTAGAACTGGTGCGGACCAGGGGAATCCG</td><td>904.2</td><td>-3.6</td><td>1.1</td><td>-3.4</td><td>8.0E-04</td><td>2.0E-03</td><td>BT20/MB468</td><td>MP</td></tr>
<tr><td>>hg38_1_-_228634819_228635039_RNA5S12_5S_50ntFlanks@124.147.24|AGTACTTGGATGGGAGACCGCCTG</td><td>>hg38_1_-_228634819_228635039_RNA5S12_5S_50ntFlanks</td><td>124</td><td>147</td><td>24</td><td>AGTACTTGGATGGGAGACCGCCTG</td><td>158.6</td><td>-2.1</td><td>0.7</td><td>-3.2</td><td>1.6E-03</td><td>2.0E-03</td><td>BT20/MB468</td><td>CYTO</td></tr>
<tr><td>>hg38_21_+_8433172_8446622_RNA45SN1_45S_50ntFlanks@11875.11905.31|GAAGACCCTGTTGAGCTTGACTCTAGTCTGG</td><td>>hg38_21_+_8433172_8446622_RNA45SN1_45S_50ntFlanks</td><td>11875</td><td>11905</td><td>31</td><td>GAAGACCCTGTTGAGCTTGACTCTAGTCTGG</td><td>84.4</td><td>-5.4</td><td>1.6</td><td>-3.4</td><td>8.0E-04</td><td>2.0E-03</td><td>BT20/MB468</td><td>MP</td></tr>
<tr><td>>MI0000077|hsa-mir-21&WithFlank&17|+|59841260|59841343@14.32.19(+1C)&[MIMAT0000076&hsa-miR-21-5p&offsets|0|-3(+1C);m-1&17|+|59841273|59841294&offsets|0|-3(+1C)]|TAGCTTATCAGACTGATGTC</td><td>>MI0000077|hsa-mir-21&WithFlank&17|+|59841260|59841343</td><td>14</td><td>32</td><td>19(+1C)</td><td>TAGCTTATCAGACTGATGTC</td><td>209.2</td><td>4.1</td><td>1.1</td><td>3.6</td><td>3.2E-04</td><td>2.0E-03</td><td>BT20/MB231</td><td>CYTO</td></tr>
<tr><td>>hg38_21_+_8433172_8446622_RNA45SN1_45S_50ntFlanks@9554.9580.27|CGTAGCGGTCCTGACGTGCAAATCGGT</td><td>>hg38_21_+_8433172_8446622_RNA45SN1_45S_50ntFlanks</td><td>9554</td><td>9580</td><td>27</td><td>CGTAGCGGTCCTGACGTGCAAATCGGT</td><td>355.9</td><td>-2.8</td><td>0.8</td><td>-3.6</td><td>3.2E-04</td><td>2.0E-03</td><td>BT20/MB231</td><td>CYTO</td></tr>
<tr><td>>hg38_1_-_228634819_228635039_RNA5S12_5S_50ntFlanks@51.91.41|GTCTACGGCCATACCACCCTGAACGCGCCCGATCTCGTCTG</td><td>>hg38_1_-_228634819_228635039_RNA5S12_5S_50ntFlanks</td><td>51</td><td>91</td><td>41</td><td>GTCTACGGCCATACCACCCTGAACGCGCCCGATCTCGTCTG</td><td>796.6</td><td>2.8</td><td>0.9</td><td>3.1</td><td>1.7E-03</td><td>2.0E-03</td><td>MB231/MB468</td><td>NUC</td></tr>
<tr><td>>hg38_21_+_8433172_8446622_RNA45SN1_45S_50ntFlanks@9931.9951.21|TATAGACAGCAGGACGGTGGC</td><td>>hg38_21_+_8433172_8446622_RNA45SN1_45S_50ntFlanks</td><td>9931</td><td>9951</td><td>21</td><td>TATAGACAGCAGGACGGTGGC</td><td>413.1</td><td>5.9</td><td>1.6</td><td>3.6</td><td>2.7E-04</td><td>2.0E-03</td><td>BT20/MB231</td><td>MP</td></tr>
<tr><td>>hg38_21_+_8433172_8446622_RNA45SN1_45S_50ntFlanks@10806.10837.32|AACAATGTAGGTAAGGGAAGTCGGCAAGCCGG</td><td>>hg38_21_+_8433172_8446622_RNA45SN1_45S_50ntFlanks</td><td>10806</td><td>10837</td><td>32</td><td>AACAATGTAGGTAAGGGAAGTCGGCAAGCCGG</td><td>137.7</td><td>-5.0</td><td>1.6</td><td>-3.2</td><td>1.4E-03</td><td>2.0E-03</td><td>MB231/MB468</td><td>MP</td></tr>
<tr><td>>hg38_21_+_8433172_8446622_RNA45SN1_45S_50ntFlanks@11577.11605.29|TTAGAACTGGTGCGGACCAGGGGAATCCG</td><td>>hg38_21_+_8433172_8446622_RNA45SN1_45S_50ntFlanks</td><td>11577</td><td>11605</td><td>29</td><td>TTAGAACTGGTGCGGACCAGGGGAATCCG</td><td>242.5</td><td>-5.4</td><td>1.6</td><td>-3.4</td><td>8.0E-04</td><td>2.0E-03</td><td>BT20/MB468</td><td>MP</td></tr>
<tr><td>>trnaMT_PheGAA_MT_+_577_647@34.57.24|TGAAAATGTTTAGACGGGCTCACA</td><td>>trnaMT_PheGAA_MT_+_577_647</td><td>34</td><td>57</td><td>24</td><td>TGAAAATGTTTAGACGGGCTCACA</td><td>47.4</td><td>9.3</td><td>2.8</td><td>3.4</td><td>8.0E-04</td><td>2.0E-03</td><td>BT20/MB468</td><td>MP</td></tr>
<tr><td>>hg38_21_+_8433172_8446622_RNA45SN1_45S_50ntFlanks@4370.4398.29|TTAAAAAGCTCGTAGTTGGATCTTGGGAG</td><td>>hg38_21_+_8433172_8446622_RNA45SN1_45S_50ntFlanks</td><td>4370</td><td>4398</td><td>29</td><td>TTAAAAAGCTCGTAGTTGGATCTTGGGAG</td><td>113.7</td><td>-5.5</td><td>1.6</td><td>-3.4</td><td>8.0E-04</td><td>2.0E-03</td><td>BT20/MB468</td><td>MP</td></tr>
<tr><td>>MI0000746|hsa-mir-99b&WithFlank&19|+|51692606|51692687@51.69.19&[MIMAT0004678&hsa-miR-99b-3p&offsets|0|-3;m-151&19|+|51692656|51692678&offsets|0|-4]|CAAGCTCGTGTCTGTGGGT</td><td>>MI0000746|hsa-mir-99b&WithFlank&19|+|51692606|51692687</td><td>51</td><td>69</td><td>19</td><td>CAAGCTCGTGTCTGTGGGT</td><td>299.7</td><td>3.0</td><td>0.8</td><td>3.6</td><td>3.2E-04</td><td>2.0E-03</td><td>BT20/MB231</td><td>CYTO</td></tr>
<tr><td>>hg38_21_+_8433172_8446622_RNA45SN1_45S_50ntFlanks@11581.11611.31|AACTGGTGCGGACCAGGGGAATCCGACTGTT</td><td>>hg38_21_+_8433172_8446622_RNA45SN1_45S_50ntFlanks</td><td>11581</td><td>11611</td><td>31</td><td>AACTGGTGCGGACCAGGGGAATCCGACTGTT</td><td>2010.4</td><td>-2.2</td><td>0.7</td><td>-3.2</td><td>1.3E-03</td><td>2.0E-03</td><td>BT20/MB468</td><td>MITO</td></tr>
<tr><td>>hg38_MT_+_1621_3279_RNR2_16S_50ntFlanks@625.652.28|AACTGAACTCCTCACACCCAATTGGACC</td><td>>hg38_MT_+_1621_3279_RNR2_16S_50ntFlanks</td><td>625</td><td>652</td><td>28</td><td>AACTGAACTCCTCACACCCAATTGGACC</td><td>181.2</td><td>5.0</td><td>1.5</td><td>3.4</td><td>8.0E-04</td><td>2.0E-03</td><td>BT20/MB468</td><td>MP</td></tr>
<tr><td>>hg38_21_+_8433172_8446622_RNA45SN1_45S_50ntFlanks@10829.10855.27|GCAAGCCGGATCCGTAACTTCGGGATA</td><td>>hg38_21_+_8433172_8446622_RNA45SN1_45S_50ntFlanks</td><td>10829</td><td>10855</td><td>27</td><td>GCAAGCCGGATCCGTAACTTCGGGATA</td><td>261.8</td><td>-3.5</td><td>1.1</td><td>-3.2</td><td>1.6E-03</td><td>2.0E-03</td><td>MB231/MB468</td><td>CYTO</td></tr>
<tr><td>>trnaMT_ValTAC_MT_+_1602_1670@50.70.21|ACTTAACTTGACCGCTCTGAC</td><td>>trnaMT_ValTAC_MT_+_1602_1670</td><td>50</td><td>70</td><td>21</td><td>ACTTAACTTGACCGCTCTGAC</td><td>242.2</td><td>4.4</td><td>1.3</td><td>3.4</td><td>8.0E-04</td><td>2.0E-03</td><td>BT20/MB468</td><td>MP</td></tr>
<tr><td>>MI0000286|hsa-mir-210&WithFlank&11|-|568083|568204@72.94.23&[MIMAT0000267&hsa-miR-210-3p&offsets|0|+1;m-102&11|-|568112|568133&offsets|0|+1]|CTGTGCGTGTGACAGCGGCTGAT</td><td>>MI0000286|hsa-mir-210&WithFlank&11|-|568083|568204</td><td>72</td><td>94</td><td>23</td><td>CTGTGCGTGTGACAGCGGCTGAT</td><td>534.2</td><td>5.2</td><td>1.6</td><td>3.4</td><td>8.0E-04</td><td>2.0E-03</td><td>BT20/MB468</td><td>MP</td></tr>
<tr><td>>hg38_21_+_8433172_8446622_RNA45SN1_45S_50ntFlanks@8121.8142.22|ACATGTGGCGTACGGAAGACCC</td><td>>hg38_21_+_8433172_8446622_RNA45SN1_45S_50ntFlanks</td><td>8121</td><td>8142</td><td>22</td><td>ACATGTGGCGTACGGAAGACCC</td><td>123.6</td><td>5.7</td><td>1.7</td><td>3.4</td><td>8.0E-04</td><td>2.0E-03</td><td>BT20/MB468</td><td>MP</td></tr>
<tr><td>>hg38_21_+_8433172_8446622_RNA45SN1_45S_50ntFlanks@12081.12108.28|CTCTCGCTTCTGGCGCCAAGCGCCCGGC</td><td>>hg38_21_+_8433172_8446622_RNA45SN1_45S_50ntFlanks</td><td>12081</td><td>12108</td><td>28</td><td>CTCTCGCTTCTGGCGCCAAGCGCCCGGC</td><td>234.8</td><td>-5.3</td><td>1.6</td><td>-3.4</td><td>8.1E-04</td><td>2.0E-03</td><td>BT20/MB468</td><td>MP</td></tr>
<tr><td>>hg38_21_+_8433172_8446622_RNA45SN1_45S_50ntFlanks@11704.11728.25|ATGAAGCGCGGGTAAACGGCGGGAG</td><td>>hg38_21_+_8433172_8446622_RNA45SN1_45S_50ntFlanks</td><td>11704</td><td>11728</td><td>25</td><td>ATGAAGCGCGGGTAAACGGCGGGAG</td><td>1213.9</td><td>-5.9</td><td>1.8</td><td>-3.4</td><td>8.1E-04</td><td>2.0E-03</td><td>BT20/MB468</td><td>MP</td></tr>
<tr><td>>MI0000114|hsa-mir-107&WithFlank&10|-|89592741|89592833@56.76.21(+1A);MI0000108|hsa-mir-103a-2&WithFlank&20|+|3917488|3917577@54.74.21(+1A);MI0000109|hsa-mir-103a-1&WithFlank&5|-|168560890|168560979@54.74.21(+1A)&[MIMAT0000104&hsa-miR-107&offsets|0|-2(+1A);m-78&10|-|89592758|89592778&offsets|0|0(+1A)];[MIMAT0000101&hsa-miR-103a-3p&offsets|0|-2(+1A);m-12&20|+|3917541|3917563&offsets|0|-2(+1A)];[MIMAT0000101_1&hsa-miR-103a-3p&offsets|0|-2(+1A);m-13&5|-|168560904|168560926&offsets|0|-2(+1A);m-44220&5|-|168560908|168560926&offsets|0|+2(+1A)]|AGCAGCATTGTACAGGGCTATA</td><td>>MI0000114|hsa-mir-107&WithFlank&10|-|89592741|89592833</td><td>56</td><td>76</td><td>21(+1A)</td><td>AGCAGCATTGTACAGGGCTATA</td><td>1055.1</td><td>3.2</td><td>1.0</td><td>3.2</td><td>1.4E-03</td><td>2.0E-03</td><td>MB231/MB468</td><td>MP</td></tr>
<tr><td>>hg38_21_+_8433172_8446622_RNA45SN1_45S_50ntFlanks@8343.8375.33|GTAAGGGAAAGTTGAAAAGAACTTTGAAGAGAG</td><td>>hg38_21_+_8433172_8446622_RNA45SN1_45S_50ntFlanks</td><td>8343</td><td>8375</td><td>33</td><td>GTAAGGGAAAGTTGAAAAGAACTTTGAAGAGAG</td><td>2703.2</td><td>5.4</td><td>1.5</td><td>3.6</td><td>2.8E-04</td><td>2.0E-03</td><td>BT20/MB231</td><td>MP</td></tr>
<tr><td>>hg38_21_+_8433172_8446622_RNA45SN1_45S_50ntFlanks@6651.6695.45|CGACTCTTAGCGGTGGATCACTCGGCTCGTGCGTCGATGAAGAAC</td><td>>hg38_21_+_8433172_8446622_RNA45SN1_45S_50ntFlanks</td><td>6651</td><td>6695</td><td>45</td><td>CGACTCTTAGCGGTGGATCACTCGGCTCGTGCGTCGATGAAGAAC</td><td>8117.1</td><td>3.2</td><td>1.0</td><td>3.1</td><td>1.7E-03</td><td>2.1E-03</td><td>MB231/MB468</td><td>MITO</td></tr>
<tr><td>>hg38_21_+_8433172_8446622_RNA45SN1_45S_50ntFlanks@9847.9864.18|AACTGGCGCTGCGGGATG</td><td>>hg38_21_+_8433172_8446622_RNA45SN1_45S_50ntFlanks</td><td>9847</td><td>9864</td><td>18</td><td>AACTGGCGCTGCGGGATG</td><td>118.0</td><td>6.4</td><td>1.9</td><td>3.4</td><td>8.1E-04</td><td>2.1E-03</td><td>BT20/MB468</td><td>MP</td></tr>
<tr><td>>hg38_1_-_228634819_228635039_RNA5S12_5S_50ntFlanks@84.109.26|CTCGTCTGATCTCGGAAGCTAAGCAG</td><td>>hg38_1_-_228634819_228635039_RNA5S12_5S_50ntFlanks</td><td>84</td><td>109</td><td>26</td><td>CTCGTCTGATCTCGGAAGCTAAGCAG</td><td>248.5</td><td>-3.8</td><td>1.2</td><td>-3.1</td><td>1.7E-03</td><td>2.1E-03</td><td>MB231/MB468</td><td>NUC</td></tr>
<tr><td>>hg38_21_+_8433172_8446622_RNA45SN1_45S_50ntFlanks@11593.11616.24|CCAGGGGAATCCGACTGTTTAATT</td><td>>hg38_21_+_8433172_8446622_RNA45SN1_45S_50ntFlanks</td><td>11593</td><td>11616</td><td>24</td><td>CCAGGGGAATCCGACTGTTTAATT</td><td>110.3</td><td>-4.1</td><td>1.3</td><td>-3.2</td><td>1.4E-03</td><td>2.1E-03</td><td>MB231/MB468</td><td>MP</td></tr>
<tr><td>>hg38_MT_+_1621_3279_RNR2_16S_50ntFlanks@644.676.33|AATTGGACCAATCTATCACCCTATAGAAGAACT</td><td>>hg38_MT_+_1621_3279_RNR2_16S_50ntFlanks</td><td>644</td><td>676</td><td>33</td><td>AATTGGACCAATCTATCACCCTATAGAAGAACT</td><td>274.1</td><td>4.9</td><td>1.5</td><td>3.3</td><td>8.1E-04</td><td>2.1E-03</td><td>BT20/MB468</td><td>MP</td></tr>
<tr><td>>hg38_21_+_8433172_8446622_RNA45SN1_45S_50ntFlanks@10797.10831.35|GCATGTTGGAACAATGTAGGTAAGGGAAGTCGGCA</td><td>>hg38_21_+_8433172_8446622_RNA45SN1_45S_50ntFlanks</td><td>10797</td><td>10831</td><td>35</td><td>GCATGTTGGAACAATGTAGGTAAGGGAAGTCGGCA</td><td>80.8</td><td>-3.9</td><td>1.2</td><td>-3.2</td><td>1.4E-03</td><td>2.1E-03</td><td>MB231/MB468</td><td>MP</td></tr>
<tr><td>>hg38_21_+_8433172_8446622_RNA45SN1_45S_50ntFlanks@13000.13027.28|TGCGATCTATTGAAAGTCAGCCCTCGAC</td><td>>hg38_21_+_8433172_8446622_RNA45SN1_45S_50ntFlanks</td><td>13000</td><td>13027</td><td>28</td><td>TGCGATCTATTGAAAGTCAGCCCTCGAC</td><td>479.0</td><td>-2.5</td><td>0.8</td><td>-3.2</td><td>1.6E-03</td><td>2.1E-03</td><td>BT20/MB468</td><td>CYTO</td></tr>
<tr><td>>hg38_21_+_8433172_8446622_RNA45SN1_45S_50ntFlanks@10471.10495.25|CCCTCGGCCGATCGAAAGGGAGTCG</td><td>>hg38_21_+_8433172_8446622_RNA45SN1_45S_50ntFlanks</td><td>10471</td><td>10495</td><td>25</td><td>CCCTCGGCCGATCGAAAGGGAGTCG</td><td>136.4</td><td>-4.5</td><td>1.4</td><td>-3.2</td><td>1.3E-03</td><td>2.1E-03</td><td>BT20/MB468</td><td>MITO</td></tr>
<tr><td>>MI0003836|hsa-mir-766&WithFlank&X|-|119646732|119646854@71.92.22&[MIMAT0003888&hsa-miR-766-3p&offsets|0|0;m-286&X|-|119646763|119646784&offsets|0|0]|ACTCCAGCCCCACAGCCTCAGC</td><td>>MI0003836|hsa-mir-766&WithFlank&X|-|119646732|119646854</td><td>71</td><td>92</td><td>22</td><td>ACTCCAGCCCCACAGCCTCAGC</td><td>153.6</td><td>3.2</td><td>0.9</td><td>3.6</td><td>3.2E-04</td><td>2.1E-03</td><td>BT20/MB231</td><td>CYTO</td></tr>
<tr><td>>trna111_HisGTG_1_-_147774845_147774916@-1G.33.34,trna118_HisGTG_1_-_145396881_145396952@-1G.33.34,trna16_HisGTG_1_+_146544773_146544844@-1G.33.34,trna1_HisGTG_15_+_45493349_45493420@-1G.33.34,trna21_HisGTG_1_+_147753471_147753542@-1G.33.34,trna33_HisGTG_6_+_27125906_27125977@-1G.33.34,trna7_HisGTG_9_-_14433938_14434009@-1G.33.34,trna8_HisGTG_15_-_45492611_45492682@-1G.33.34,trna9_HisGTG_15_-_45490804_45490875@-1G.33.34|GGCCGTGATCGTATAGTGGTTAGTACTCTGCGTT</td><td>>trna111_HisGTG_1_-_147774845_147774916</td><td>-1G</td><td>33</td><td>34</td><td>GGCCGTGATCGTATAGTGGTTAGTACTCTGCGTT</td><td>69002.3</td><td>-3.8</td><td>1.0</td><td>-3.6</td><td>3.2E-04</td><td>2.1E-03</td><td>BT20/MB231</td><td>CYTO</td></tr>
<tr><td>>hg38_21_+_8433172_8446622_RNA45SN1_45S_50ntFlanks@11668.11707.40|TGCCCAGTGCTCTGAATGTCAAAGTGAAGAAATTCAATGA</td><td>>hg38_21_+_8433172_8446622_RNA45SN1_45S_50ntFlanks</td><td>11668</td><td>11707</td><td>40</td><td>TGCCCAGTGCTCTGAATGTCAAAGTGAAGAAATTCAATGA</td><td>2091.9</td><td>-2.9</td><td>0.8</td><td>-3.6</td><td>3.2E-04</td><td>2.1E-03</td><td>BT20/MB231</td><td>CYTO</td></tr>
<tr><td>>trnaMT_TyrGTA_MT_-_5826_5891@38.68.31|AAAGACAGGGGTTAGGCCTCTTTTTACCACC</td><td>>trnaMT_TyrGTA_MT_-_5826_5891</td><td>38</td><td>68</td><td>31</td><td>AAAGACAGGGGTTAGGCCTCTTTTTACCACC</td><td>47.3</td><td>9.3</td><td>2.8</td><td>3.3</td><td>8.1E-04</td><td>2.1E-03</td><td>BT20/MB468</td><td>MP</td></tr>
<tr><td>>hg38_21_+_8433172_8446622_RNA45SN1_45S_50ntFlanks@8380.8401.22|CAAGAGGGCGTGAAACCGTTAA</td><td>>hg38_21_+_8433172_8446622_RNA45SN1_45S_50ntFlanks</td><td>8380</td><td>8401</td><td>22</td><td>CAAGAGGGCGTGAAACCGTTAA</td><td>566.0</td><td>-5.7</td><td>1.7</td><td>-3.3</td><td>8.1E-04</td><td>2.1E-03</td><td>BT20/MB468</td><td>MP</td></tr>
<tr><td>>hg38_MT_+_1621_3279_RNR2_16S_50ntFlanks@1171.1215.45|AACCTGCATTAAAAATTTCGGTTGGGGCGACCTCGGAGCAGAACC</td><td>>hg38_MT_+_1621_3279_RNR2_16S_50ntFlanks</td><td>1171</td><td>1215</td><td>45</td><td>AACCTGCATTAAAAATTTCGGTTGGGGCGACCTCGGAGCAGAACC</td><td>123.5</td><td>5.8</td><td>1.8</td><td>3.1</td><td>1.7E-03</td><td>2.1E-03</td><td>MB231/MB468</td><td>MITO</td></tr>
<tr><td>>MI0000102|hsa-mir-100&WithFlank&11|-|122152223|122152314@19.38.20&[MIMAT0000098&hsa-miR-100-5p&offsets|0|-2;m-27&11|-|122152276|122152296&offsets|0|-1]|AACCCGTAGATCCGAACTTG</td><td>>MI0000102|hsa-mir-100&WithFlank&11|-|122152223|122152314</td><td>19</td><td>38</td><td>20</td><td>AACCCGTAGATCCGAACTTG</td><td>857.7</td><td>7.5</td><td>2.3</td><td>3.3</td><td>8.1E-04</td><td>2.1E-03</td><td>BT20/MB468</td><td>MP</td></tr>
<tr><td>>am_trna128_GlyGCC_6_-_27870686_27870756@19.37.19,trna19_GlyGCC_16_+_70823410_70823480@19.37.19,trna19_GlyGCC_2_-_157257659_157257729@19.37.19,trna24_GlyGCC_16_-_70812942_70813012@19.37.19,trna25_GlyGCC_16_-_70812114_70812184@19.37.19,trna2_GlyGCC_21_-_18827107_18827177@19.37.19,trna35_GlyGCC_1_+_161413094_161413164@19.37.19,trna37_GlyGCC_1_+_161420467_161420537@19.37.19,trna39_GlyGCC_1_+_161427898_161427968@19.37.19,trna41_GlyGCC_1_+_161435258_161435328@19.37.19,trna5_GlyGCC_17_+_8029064_8029134@19.37.19,trna68_GlyGCC_1_-_161493637_161493707@19.37.19|TAGAATTCTCGCCTGCCAC</td><td>>am_trna128_GlyGCC_6_-_27870686_27870756</td><td>19</td><td>37</td><td>19</td><td>TAGAATTCTCGCCTGCCAC</td><td>807.1</td><td>3.2</td><td>0.9</td><td>3.6</td><td>3.2E-04</td><td>2.1E-03</td><td>BT20/MB231</td><td>CYTO</td></tr>
<tr><td>>hg38_21_+_8433172_8446622_RNA45SN1_45S_50ntFlanks@5471.5513.43|CACGGCCCTGGCGGAGCGCTGAGAAGACGGTCGAACTTGACTA</td><td>>hg38_21_+_8433172_8446622_RNA45SN1_45S_50ntFlanks</td><td>5471</td><td>5513</td><td>43</td><td>CACGGCCCTGGCGGAGCGCTGAGAAGACGGTCGAACTTGACTA</td><td>230.4</td><td>-3.5</td><td>1.0</td><td>-3.6</td><td>3.2E-04</td><td>2.1E-03</td><td>BT20/MB231</td><td>CYTO</td></tr>
<tr><td>>hg38_21_+_8433172_8446622_RNA45SN1_45S_50ntFlanks@4005.4025.21|AACCTCGGGCCGATCGCACGC</td><td>>hg38_21_+_8433172_8446622_RNA45SN1_45S_50ntFlanks</td><td>4005</td><td>4025</td><td>21</td><td>AACCTCGGGCCGATCGCACGC</td><td>349.9</td><td>-2.1</td><td>0.7</td><td>-3.2</td><td>1.6E-03</td><td>2.1E-03</td><td>MB231/MB468</td><td>CYTO</td></tr>
<tr><td>>hg38_21_+_8433172_8446622_RNA45SN1_45S_50ntFlanks@8355.8375.21|TGAAAAGAACTTTGAAGAGAG</td><td>>hg38_21_+_8433172_8446622_RNA45SN1_45S_50ntFlanks</td><td>8355</td><td>8375</td><td>21</td><td>TGAAAAGAACTTTGAAGAGAG</td><td>1337.0</td><td>6.6</td><td>1.8</td><td>3.6</td><td>3.6E-04</td><td>2.1E-03</td><td>BT20/MB231</td><td>MITO</td></tr>
</table>

| | | | | | | | | | | | | | |
|---|---|---|---|---|---|---|---|---|---|---|---|---|---|
| >hg38_21_+_8433172_8446622_RNA45SN1_45S_50ntFlanks@483.532.50\|TCGATGTGGTGACGTCGTGCTCTCCCGGGCCGGGTCCGAGCCGCGACGGG | >hg38_21_+_8433172_8446622_RNA45SN1_45S_50ntFlanks | 483 | 532 | 50 | TCGATGTGGTGACGTCGTGCTCTCCCGGGCCGGGTCCGAGCCGCGACGGG | 152.3 | 5.9 | 1.9 | 3.1 | 1.7E-03 | 2.1E-03 | MB231/MB468 | MITO |
| >hg38_21_+_8433172_8446622_RNA45SN1_45S_50ntFlanks@12417.12444.28\|TCGATGTCGGCTCTTCCTATCATTGTGA | >hg38_21_+_8433172_8446622_RNA45SN1_45S_50ntFlanks | 12417 | 12444 | 28 | TCGATGTCGGCTCTTCCTATCATTGTGA | 2128.0 | -1.6 | 0.5 | -3.2 | 1.6E-03 | 2.1E-03 | BT20/MB468 | CYTO |
| >hg38_21_+_8433172_8446622_RNA45SN1_45S_50ntFlanks@10458.10486.29\|ATGGCCTCCGTTGCCCTCGGCCGATCGAA | >hg38_21_+_8433172_8446622_RNA45SN1_45S_50ntFlanks | 10458 | 10486 | 29 | ATGGCCTCCGTTGCCCTCGGCCGATCGAA | 159.5 | -3.5 | 1.1 | -3.2 | 1.6E-03 | 2.1E-03 | BT20/MB468 | CYTO |
| >trnaMT_GluTTC_MT_-_14674_14742@40.57.18,trnalookalike8_GluTTC_5_-_93905172_93905240@40.57.18\|TTGGTCGTGGTTGTAGTC | >trnaMT_GluTTC_MT_-_14674_14742 | 40 | 57 | 18 | TTGGTCGTGGTTGTAGTC | 254.5 | 2.7 | 0.8 | 3.2 | 1.4E-03 | 2.1E-03 | MB231/MB468 | MP |
| >hg38_21_+_8433172_8446622_RNA45SN1_45S_50ntFlanks@10341.10373.33\|AATATTCAAACGAGAACTTTGAAGGCCGAAGTG | >hg38_21_+_8433172_8446622_RNA45SN1_45S_50ntFlanks | 10341 | 10373 | 33 | AATATTCAAACGAGAACTTTGAAGGCCGAAGTG | 465.8 | 4.8 | 1.3 | 3.6 | 2.8E-04 | 2.1E-03 | BT20/MB231 | MP |
| >hg38_21_+_8433172_8446622_RNA45SN1_45S_50ntFlanks@11567.11602.36\|AGCAGCCGACTTAGAACTGGTGCGGACCAGGGGAAT | >hg38_21_+_8433172_8446622_RNA45SN1_45S_50ntFlanks | 11567 | 11602 | 36 | AGCAGCCGACTTAGAACTGGTGCGGACCAGGGGAAT | 521.8 | -2.8 | 0.9 | -3.1 | 1.7E-03 | 2.1E-03 | MB231/MB468 | MITO |
| >trnaMT_LysTTT_MT_+_8295_8364@1.43.43,trnalookalike7_LysTTT_1_+_568844_568913@1.43.43\|CACTGTAAAGCTAACTTAGCATTAACCTTTTAAGTTAAAGATT | >trnaMT_LysTTT_MT_+_8295_8364 | 1 | 43 | 43 | CACTGTAAAGCTAACTTAGCATTAACCTTTTAAGTTAAAGATT | 388.6 | -3.2 | 0.9 | -3.6 | 3.6E-04 | 2.1E-03 | BT20/MB231 | MITO |
| >MI0000273\|hsa-mir-183&WithFlank&7\|-\|129774899\|129775020@34.55.22&[MIMAT0000261&hsa-miR-183-5p&offsets\|+1\|+1;m-41&7\|-\|129774966\|129774987&offsets\|0\|0]\|ATGGCACTGGTAGAATTCACTG | >MI0000273\|hsa-mir-183&WithFlank&7\|-\|129774899\|129775020 | 34 | 55 | 22 | ATGGCACTGGTAGAATTCACTG | 2480.0 | 3.5 | 1.0 | 3.3 | 8.1E-04 | 2.1E-03 | BT20/MB468 | MP |
| >hg38_MT_+_1621_3279_RNR2_16S_50ntFlanks@1184.1215.32\|AATTTCGGTTGGGGCGACCTCGGAGCAGAACC | >hg38_MT_+_1621_3279_RNR2_16S_50ntFlanks | 1184 | 1215 | 32 | AATTTCGGTTGGGGCGACCTCGGAGCAGAACC | 429.5 | 4.8 | 1.4 | 3.3 | 8.1E-04 | 2.1E-03 | BT20/MB468 | MP |
| >hg38_1_-_228634819_228635039_RNA5S12_5S_50ntFlanks@51.85.35\|GTCTACGGCCATACCACCCTGAACGCGCCCGATCT | >hg38_1_-_228634819_228635039_RNA5S12_5S_50ntFlanks | 51 | 85 | 35 | GTCTACGGCCATACCACCCTGAACGCGCCCGATCT | 917.3 | -1.5 | 0.5 | -3.1 | 1.7E-03 | 2.1E-03 | BT20/MB468 | NUC |
| >trnaMT_MetCAT_MT_+_4402_4469@42.70.29\|AATGTTGGTTATACCCTTCCCGTACTACC | >trnaMT_MetCAT_MT_+_4402_4469 | 42 | 70 | 29 | AATGTTGGTTATACCCTTCCCGTACTACC | 230.9 | 2.2 | 0.7 | 3.2 | 1.5E-03 | 2.1E-03 | MB231/MB468 | MP |
| >am_MI0016846\|hsa-mir-4485&WithFlank&11\|-\|10508264\|10508332@34.53.20&[MIMAT0019019&hsa-miR-4485-3p&offsets\|-3\|-3;m-604&11\|-\|10508277\|10508296&offsets\|-3\|-3]\|GTTTAACGGCCGCGGTACCC | >am_MI0016846\|hsa-mir-4485&WithFlank&11\|-\|10508264\|10508332 | 34 | 53 | 20 | GTTTAACGGCCGCGGTACCC | 104.2 | 4.8 | 1.5 | 3.1 | 1.7E-03 | 2.1E-03 | MB231/MB468 | MITO |
| >hg38_MT_+_1621_3279_RNR2_16S_50ntFlanks@940.959.20\|GTTTAACGGCCGCGGTACCC | >hg38_MT_+_1621_3279_RNR2_16S_50ntFlanks | 940 | 959 | 20 | GTTTAACGGCCGCGGTACCC | 104.2 | 4.8 | 1.5 | 3.1 | 1.7E-03 | 2.1E-03 | MB231/MB468 | MITO |
| >hg38_21_+_8433172_8446622_RNA45SN1_45S_50ntFlanks@11670.11692.23\|CCCAGTGCTCTGAATGTCAAAGT | >hg38_21_+_8433172_8446622_RNA45SN1_45S_50ntFlanks | 11670 | 11692 | 23 | CCCAGTGCTCTGAATGTCAAAGT | 1162.3 | -4.6 | 1.5 | -3.1 | 1.7E-03 | 2.1E-03 | BT20/MB468 | NUC |
| >MI0003581\|hsa-mir-574&WithFlank&4\|+\|38868026\|38868133@67.87.21&[MIMAT0003239&hsa-miR-574-3p&offsets\|0\|-1;m-149&4\|+\|38868092\|38868113&offsets\|0\|-1]\|CACGCTCATGCACACACCCAC | >MI0003581\|hsa-mir-574&WithFlank&4\|+\|38868026\|38868133 | 67 | 87 | 21 | CACGCTCATGCACACACCCAC | 543.7 | -3.3 | 0.9 | -3.6 | 2.8E-04 | 2.1E-03 | BT20/MB231 | MP |
| >MI0000299\|hsa-mir-222&WithFlank&X\|-\|45747009\|45747130@75.98.24(+1G)&[MIMAT0000279&hsa-miR-222-3p&offsets\|0\|+3(+1G);m-140&X\|-\|45747033\|45747056&offsets\|0\|0(+1G)]\|AGCTACATCTGGCTACTGGGTCTCG | >MI0000299\|hsa-mir-222&WithFlank&X\|-\|45747009\|45747130 | 75 | 98 | 24(+1G) | AGCTACATCTGGCTACTGGGTCTCG | 257.7 | 4.1 | 1.3 | 3.2 | 1.5E-03 | 2.1E-03 | MB231/MB468 | MP |
| >MI0000809\|hsa-mir-151a&WithFlank&8\|-\|140732558\|140732659@51.71.21&[MIMAT0000757&hsa-miR-151a-3p&offsets\|-2\|-2;m-30&8\|-\|140732587\|140732607&offsets\|-2\|-2]\|TACTAGACTGAAGCTCCTTGA | >MI0000809\|hsa-mir-151a&WithFlank&8\|-\|140732558\|140732659 | 51 | 71 | 21 | TACTAGACTGAAGCTCCTTGA | 90.2 | 4.9 | 1.6 | 3.2 | 1.5E-03 | 2.1E-03 | MB231/MB468 | MP |
| >hg38_21_+_8433172_8446622_RNA45SN1_45S_50ntFlanks@4972.4999.28\|CGGCCCGGACACGGACAGGATTGACAGA | >hg38_21_+_8433172_8446622_RNA45SN1_45S_50ntFlanks | 4972 | 4999 | 28 | CGGCCCGGACACGGACAGGATTGACAGA | 210.1 | -2.7 | 0.8 | -3.2 | 1.3E-03 | 2.1E-03 | BT20/MB468 | MITO |
| >trna11_GluTTC_15_-_26327381_26327452@41.73.33,trna3_GluTTC_13_-_45492062_45492133@41.73.33\|AGGCGGCCCGGGTTCGACTCCCGGTGTGGGAAC | >trna11_GluTTC_15_-_26327381_26327452 | 41 | 73 | 33 | AGGCGGCCCGGGTTCGACTCCCGGTGTGGGAAC | 142.0 | 6.1 | 1.9 | 3.2 | 1.5E-03 | 2.1E-03 | MB231/MB468 | MP |
| >hg38_21_+_8433172_8446622_RNA45SN1_45S_50ntFlanks@11564.11605.42\|CCTAGCAGCCGACTTAGAACTGGTGCGGACCAGGGGAATCCG | >hg38_21_+_8433172_8446622_RNA45SN1_45S_50ntFlanks | 11564 | 11605 | 42 | CCTAGCAGCCGACTTAGAACTGGTGCGGACCAGGGGAATCCG | 850.0 | -4.0 | 1.3 | -3.2 | 1.3E-03 | 2.1E-03 | BT20/MB468 | MITO |
| >hg38_21_+_8433172_8446622_RNA45SN1_45S_50ntFlanks@9498.9524.27\|AAGATGGTGAACTATGCCTGGGCAGGG | >hg38_21_+_8433172_8446622_RNA45SN1_45S_50ntFlanks | 9498 | 9524 | 27 | AAGATGGTGAACTATGCCTGGGCAGGG | 43.5 | 9.2 | 2.7 | 3.3 | 8.2E-04 | 2.1E-03 | BT20/MB468 | MP |
| >trnaMT_GluTTC_MT_-_14674_14742@27.65.39,trnalookalike8_GluTTC_5_-_93905172_93905240@27.65.39\|GTTTTTCATATCATTGGTCGTGGTTGTAGTCCGTGCGAG | >trnaMT_GluTTC_MT_-_14674_14742 | 27 | 65 | 39 | GTTTTTCATATCATTGGTCGTGGTTGTAGTCCGTGCGAG | 231.6 | -4.3 | 1.4 | -3.1 | 1.7E-03 | 2.1E-03 | MB231/MB468 | NUC |
| >am_trna15_CysGCA_17_+_37023898_37023969@14.34.21,trna26_CysGCA_17_-_37310744_37310815@14.34.21,trna27_CysGCA_17_-_37309987_37310058@14.34.21,trna28_CysGCA_17_-_37025545_37025616@14.34.21,trna3_CysGCA_4_-_124430005_124430076@14.34.21,trna5_CysGCA_7_+_149007281_149007352@14.34.21,trna7_CysGCA_3_-_131947944_131948015@14.34.21\|AGTGGTAGAGCATTTGACTGC | >am_trna15_CysGCA_17_+_37023898_37023969 | 14 | 34 | 21 | AGTGGTAGAGCATTTGACTGC | 65.2 | 5.6 | 1.8 | 3.2 | 1.5E-03 | 2.1E-03 | MB231/MB468 | MP |
| >hg38_1_-_228634819_228635039_RNA5S12_5S_50ntFlanks@136.169.34\|GGAGACCGCCTGGGAATACCGGGTGCTGTAGGCT | >hg38_1_-_228634819_228635039_RNA5S12_5S_50ntFlanks | 136 | 169 | 34 | GGAGACCGCCTGGGAATACCGGGTGCTGTAGGCT | 1066.6 | 3.5 | 1.0 | 3.3 | 8.2E-04 | 2.1E-03 | BT20/MB468 | MP |
| >hg38_21_+_8433172_8446622_RNA45SN1_45S_50ntFlanks@7989.8018.30\|AGACGTGGCGACCCGCTGAATTTAAGCATA | >hg38_21_+_8433172_8446622_RNA45SN1_45S_50ntFlanks | 7989 | 8018 | 30 | AGACGTGGCGACCCGCTGAATTTAAGCATA | 209.3 | -2.5 | 0.7 | -3.3 | 8.2E-04 | 2.1E-03 | BT20/MB468 | MP |
| >hg38_MT_+_1621_3279_RNR2_16S_50ntFlanks@827.855.29\|AGGAAAGGTTAAAAAAAGTAAAAGGAACT | >hg38_MT_+_1621_3279_RNR2_16S_50ntFlanks | 827 | 855 | 29 | AGGAAAGGTTAAAAAAAGTAAAAGGAACT | 261.2 | 5.1 | 1.6 | 3.2 | 1.5E-03 | 2.1E-03 | MB231/MB468 | MP |
| >hg38_21_+_8433172_8446622_RNA45SN1_45S_50ntFlanks@10509.10532.24\|GAATCCGGAGTGGCGGAGATGGGC | >hg38_21_+_8433172_8446622_RNA45SN1_45S_50ntFlanks | 10509 | 10532 | 24 | GAATCCGGAGTGGCGGAGATGGGC | 1024.9 | 3.9 | 1.2 | 3.3 | 8.2E-04 | 2.1E-03 | BT20/MB468 | MP |
| >hg38_21_+_8433172_8446622_RNA45SN1_45S_50ntFlanks@10689.10709.21\|GCGTCCGGTGAGCTCTCGCTG | >hg38_21_+_8433172_8446622_RNA45SN1_45S_50ntFlanks | 10689 | 10709 | 21 | GCGTCCGGTGAGCTCTCGCTG | 400.8 | -3.7 | 1.2 | -3.2 | 1.3E-03 | 2.1E-03 | BT20/MB468 | MITO |
| >hg38_1_-_228634819_228635039_RNA5S12_5S_50ntFlanks@132.169.38\|GATGGGAGACCGCCTGGGAATACCGGGTGCTGTAGGCT | >hg38_1_-_228634819_228635039_RNA5S12_5S_50ntFlanks | 132 | 169 | 38 | GATGGGAGACCGCCTGGGAATACCGGGTGCTGTAGGCT | 10131.4 | 3.3 | 1.0 | 3.3 | 8.2E-04 | 2.1E-03 | BT20/MB468 | MP |
| >hg38_21_+_8433172_8446622_RNA45SN1_45S_50ntFlanks@3705.3740.36\|TACCTGGTTGATCCTGCCAGTAGCATATGCTTGTCT | >hg38_21_+_8433172_8446622_RNA45SN1_45S_50ntFlanks | 3705 | 3740 | 36 | TACCTGGTTGATCCTGCCAGTAGCATATGCTTGTCT | 158.7 | 4.9 | 1.6 | 3.1 | 1.7E-03 | 2.1E-03 | MB231/MB468 | MITO |
| >hg38_21_+_8433172_8446622_RNA45SN1_45S_50ntFlanks@3773.3791.19\|CGGCCGGTACAGTGAAACT | >hg38_21_+_8433172_8446622_RNA45SN1_45S_50ntFlanks | 3773 | 3791 | 19 | CGGCCGGTACAGTGAAACT | 274.6 | 2.6 | 0.8 | 3.1 | 1.6E-03 | 2.1E-03 | MB231/MB468 | CYTO |
| >hg38_21_+_8433172_8446622_RNA45SN1_45S_50ntFlanks@9940.9964.25\|CAGGACGGTGGCCATGGAAGTCGGA | >hg38_21_+_8433172_8446622_RNA45SN1_45S_50ntFlanks | 9940 | 9964 | 25 | CAGGACGGTGGCCATGGAAGTCGGA | 70.7 | 6.3 | 1.9 | 3.3 | 8.2E-04 | 2.1E-03 | BT20/MB468 | MP |
| >hg38_21_+_8433172_8446622_RNA45SN1_45S_50ntFlanks@10389.10429.41\|TGAACAGCAGTTGAACATGGGTCAGTCGGTCCTGAGAGATG | >hg38_21_+_8433172_8446622_RNA45SN1_45S_50ntFlanks | 10389 | 10429 | 41 | TGAACAGCAGTTGAACATGGGTCAGTCGGTCCTGAGAGATG | 840.3 | -2.7 | 0.9 | -3.1 | 1.6E-03 | 2.1E-03 | MB231/MB468 | CYTO |
| >MI0000298\|hsa-mir-221&WithFlank&X\|-\|45746151\|45746272@71.91.21(+1C)&[MIMAT0000278&hsa-miR-221-3p&offsets\|0\|-2(+1C);m-91&X\|-\|45746181\|45746202&offsets\|0\|-1(+1C)]\|AGCTACATTGTCTGCTGGGTTC | >MI0000298\|hsa-mir-221&WithFlank&X\|-\|45746151\|45746272 | 71 | 91 | 21(+1C) | AGCTACATTGTCTGCTGGGTTC | 5555.9 | -3.1 | 0.9 | -3.6 | 3.6E-04 | 2.1E-03 | BT20/MB231 | MITO |
| >hg38_21_+_8433172_8446622_RNA45SN1_45S_50ntFlanks@5237.5257.21\|ACTGACTGGCTCAGCGTGTGC | >hg38_21_+_8433172_8446622_RNA45SN1_45S_50ntFlanks | 5237 | 5257 | 21 | ACTGACTGGCTCAGCGTGTGC | 114.9 | -3.3 | 1.0 | -3.2 | 1.5E-03 | 2.1E-03 | MB231/MB468 | MP |
| >hg38_21_+_8433172_8446622_RNA45SN1_45S_50ntFlanks@9471.9504.34\|AGCACGAGCGCACGTGTTAGGACCCGAAAGATGG | >hg38_21_+_8433172_8446622_RNA45SN1_45S_50ntFlanks | 9471 | 9504 | 34 | AGCACGAGCGCACGTGTTAGGACCCGAAAGATGG | 56.9 | -5.3 | 1.7 | -3.2 | 1.5E-03 | 2.1E-03 | MB231/MB468 | MP |
| >MI0001445\|hsa-mir-423&WithFlank&17\|+\|30117073\|30117178@23.43.21(+1A)&[MIMAT0004748&hsa-miR-423-5p&offsets\|0\|-2(+1A);m-141&17\|+\|30117095\|30117117&offsets\|0\|-2(+1A)]\|TGAGGGGCAGAGAGCGAGACTA | >MI0001445\|hsa-mir-423&WithFlank&17\|+\|30117073\|30117178 | 23 | 43 | 21(+1A) | TGAGGGGCAGAGAGCGAGACTA | 91.0 | 5.1 | 1.6 | 3.2 | 1.5E-03 | 2.1E-03 | MB231/MB468 | MP |
| >MI0000299\|hsa-mir-222&WithFlank&X\|-\|45747009\|45747130@75.97.23(+2U)&[MIMAT0000279&hsa-miR-222-3p&offsets\|0\|+2(+2U);m-140&X\|-\|45747033\|45747056&offsets\|0\|-1(+2U)]\|AGCTACATCTGGCTACTGGGTCTTT | >MI0000299\|hsa-mir-222&WithFlank&X\|-\|45747009\|45747130 | 75 | 97 | 23(+2U) | AGCTACATCTGGCTACTGGGTCTTT | 1538.3 | -2.9 | 0.8 | -3.6 | 3.6E-04 | 2.1E-03 | BT20/MB231 | MITO |
| >MI0001445\|hsa-mir-423&WithFlank&17\|+\|30117073\|30117178@59.79.21(+1A)&[MIMAT0001340&hsa-miR-423-3p&offsets\|0\|-2(+1A);m-100&17\|+\|30117131\|30117153&offsets\|0\|-2(+1A)]\|AGCTCGGTCTGAGGCCCCTCAA | >MI0001445\|hsa-mir-423&WithFlank&17\|+\|30117073\|30117178 | 59 | 79 | 21(+1A) | AGCTCGGTCTGAGGCCCCTCAA | 62.9 | 5.9 | 1.9 | 3.2 | 1.5E-03 | 2.1E-03 | MB231/MB468 | MP |
| >hg38_21_+_8433172_8446622_RNA45SN1_45S_50ntFlanks@10458.10488.31\|ATGGCCTCCGTTGCCCTCGGCCGATCGAAAG | >hg38_21_+_8433172_8446622_RNA45SN1_45S_50ntFlanks | 10458 | 10488 | 31 | ATGGCCTCCGTTGCCCTCGGCCGATCGAAAG | 713.2 | -3.9 | 1.3 | -3.1 | 1.6E-03 | 2.1E-03 | MB231/MB468 | CYTO |
| >hg38_21_+_8433172_8446622_RNA45SN1_45S_50ntFlanks@4728.4757.30\|ATCAAGAACGAAAGTCGGAGGTTCGAAGAC | >hg38_21_+_8433172_8446622_RNA45SN1_45S_50ntFlanks | 4728 | 4757 | 30 | ATCAAGAACGAAAGTCGGAGGTTCGAAGAC | 140.0 | -5.0 | 1.6 | -3.2 | 1.5E-03 | 2.1E-03 | MB231/MB468 | MP |
| >hg38_21_+_8433172_8446622_RNA45SN1_45S_50ntFlanks@9252.9274.23\|CACGGGGTCGGCGGCGACGTCGG | >hg38_21_+_8433172_8446622_RNA45SN1_45S_50ntFlanks | 9252 | 9274 | 23 | CACGGGGTCGGCGGCGACGTCGG | 412.1 | -4.6 | 1.4 | -3.3 | 8.2E-04 | 2.1E-03 | BT20/MB468 | MP |
| >MI0000750\|hsa-mir-26a-2&WithFlank&12\|-\|57824603\|57824698@20.40.21(+1A);MI0000083\|hsa-mir-26a-1&WithFlank&3\|+\|37969398\|37969486@16.36.21(+1A)&[MIMAT0000082&hsa-miR-26a-5p&offsets\|0\|-1(+1A);m-33&12\|-\|57824658\|57824679&offsets\|0\|-1(+1A)];[MIMAT0000082_1&hsa-miR-26a-5p&offsets\|0\|-1(+1A);m-34&3\|+\|37969413\|37969434&offsets\|0\|-1(+1A)]\|TTCAAGTAATCCAGGATAGGCA | >MI0000750\|hsa-mir-26a-2&WithFlank&12\|-\|57824603\|57824698 | 20 | 40 | 21(+1A) | TTCAAGTAATCCAGGATAGGCA | 14756.5 | -3.9 | 1.1 | -3.6 | 2.8E-04 | 2.1E-03 | BT20/MB231 | MP |
| >hg38_21_+_8433172_8446622_RNA45SN1_45S_50ntFlanks@3789.3810.22\|ACTGCGAATGGCTCATTAAATC | >hg38_21_+_8433172_8446622_RNA45SN1_45S_50ntFlanks | 3789 | 3810 | 22 | ACTGCGAATGGCTCATTAAATC | 369.9 | -1.8 | 0.6 | -3.1 | 1.6E-03 | 2.1E-03 | MB231/MB468 | CYTO |
| >hg38_21_+_8433172_8446622_RNA45SN1_45S_50ntFlanks@11896.11933.38\|TCTAGTCTGGCACGGTGAAGAGACATGAGAGGTGTAGA | >hg38_21_+_8433172_8446622_RNA45SN1_45S_50ntFlanks | 11896 | 11933 | 38 | TCTAGTCTGGCACGGTGAAGAGACATGAGAGGTGTAGA | 101.9 | 10.0 | 2.8 | 3.6 | 2.8E-04 | 2.1E-03 | BT20/MB231 | MP |

<table>
<tr><td>>hg38_21_+_8433172_8446622_RNA45SN1_45S_50ntFlanks@12993.13039.47|CCCTCGCTGCGATCTATTGAAAGTCAGCCCTCGACACAAGGGTTTGT</td><td>>hg38_21_+_8433172_8446622_RNA45SN1_45S_50ntFlanks</td><td>12993</td><td>13039</td><td>47</td><td>CCCTCGCTGCGATCTATTGAAAGTCAGCCCTCGACACAAGGGTTTGT</td><td>9716.4</td><td>3.6</td><td>1.1</td><td>3.2</td><td>1.3E-03</td><td>2.1E-03</td><td>BT20/MB468</td><td>MITO</td></tr>
<tr><td>>hg38_21_+_8433172_8446622_RNA45SN1_45S_50ntFlanks@10116.10136.21|TGCGTCGGAGGGCGGCGGCGG</td><td>>hg38_21_+_8433172_8446622_RNA45SN1_45S_50ntFlanks</td><td>10116</td><td>10136</td><td>21</td><td>TGCGTCGGAGGGCGGCGGCGG</td><td>456.2</td><td>-5.8</td><td>1.7</td><td>-3.3</td><td>8.3E-04</td><td>2.1E-03</td><td>BT20/MB468</td><td>MP</td></tr>
<tr><td>>hg38_21_+_8433172_8446622_RNA45SN1_45S_50ntFlanks@3845.3872.28|ACTTGGATAACTGTGGTAATTCTAGAGC</td><td>>hg38_21_+_8433172_8446622_RNA45SN1_45S_50ntFlanks</td><td>3845</td><td>3872</td><td>28</td><td>ACTTGGATAACTGTGGTAATTCTAGAGC</td><td>85.5</td><td>-3.7</td><td>1.2</td><td>-3.2</td><td>1.5E-03</td><td>2.1E-03</td><td>MB231/MB468</td><td>MP</td></tr>
<tr><td>>hg38_21_+_8433172_8446622_RNA45SN1_45S_50ntFlanks@8122.8141.20|CATGTGGCGTACGGAAGACC</td><td>>hg38_21_+_8433172_8446622_RNA45SN1_45S_50ntFlanks</td><td>8122</td><td>8141</td><td>20</td><td>CATGTGGCGTACGGAAGACC</td><td>171.8</td><td>3.3</td><td>1.0</td><td>3.3</td><td>8.3E-04</td><td>2.1E-03</td><td>BT20/MB468</td><td>MP</td></tr>
<tr><td>>hg38_21_+_8433172_8446622_RNA45SN1_45S_50ntFlanks@3705.3728.24|TACCTGGTTGATCCTGCCAGTAGC</td><td>>hg38_21_+_8433172_8446622_RNA45SN1_45S_50ntFlanks</td><td>3705</td><td>3728</td><td>24</td><td>TACCTGGTTGATCCTGCCAGTAGC</td><td>7707.5</td><td>2.8</td><td>0.8</td><td>3.6</td><td>3.3E-04</td><td>2.1E-03</td><td>BT20/MB231</td><td>CYTO</td></tr>
<tr><td>>hg38_21_+_8433172_8446622_RNA45SN1_45S_50ntFlanks@8176.8203.28|CTTCTGATCGAGGCCCAGCCCGTGGACG</td><td>>hg38_21_+_8433172_8446622_RNA45SN1_45S_50ntFlanks</td><td>8176</td><td>8203</td><td>28</td><td>CTTCTGATCGAGGCCCAGCCCGTGGACG</td><td>1984.9</td><td>-2.8</td><td>0.9</td><td>-3.2</td><td>1.3E-03</td><td>2.1E-03</td><td>BT20/MB468</td><td>MITO</td></tr>
<tr><td>>hg38_21_+_8433172_8446622_RNA45SN1_45S_50ntFlanks@12444.12480.37|AAGCAGAATTCACCAAGCGTTGGATTGTTCACCCACT</td><td>>hg38_21_+_8433172_8446622_RNA45SN1_45S_50ntFlanks</td><td>12444</td><td>12480</td><td>37</td><td>AAGCAGAATTCACCAAGCGTTGGATTGTTCACCCACT</td><td>466.2</td><td>-2.7</td><td>0.9</td><td>-3.1</td><td>1.8E-03</td><td>2.1E-03</td><td>MB231/MB468</td><td>MITO</td></tr>
<tr><td>>hg38_21_+_8433172_8446622_RNA45SN1_45S_50ntFlanks@9556.9584.29|TAGCGGTCCTGACGTGCAAATCGGTCGTC</td><td>>hg38_21_+_8433172_8446622_RNA45SN1_45S_50ntFlanks</td><td>9556</td><td>9584</td><td>29</td><td>TAGCGGTCCTGACGTGCAAATCGGTCGTC</td><td>75.8</td><td>-5.2</td><td>1.6</td><td>-3.3</td><td>8.3E-04</td><td>2.1E-03</td><td>BT20/MB468</td><td>MP</td></tr>
<tr><td>>hg38_21_+_8433172_8446622_RNA45SN1_45S_50ntFlanks@9938.9961.24|AGCAGGACGGTGGCCATGGAAGTC</td><td>>hg38_21_+_8433172_8446622_RNA45SN1_45S_50ntFlanks</td><td>9938</td><td>9961</td><td>24</td><td>AGCAGGACGGTGGCCATGGAAGTC</td><td>655.7</td><td>5.7</td><td>1.6</td><td>3.6</td><td>2.8E-04</td><td>2.1E-03</td><td>BT20/MB231</td><td>MP</td></tr>
<tr><td>>hg38_21_+_8433172_8446622_RNA45SN1_45S_50ntFlanks@12456.12499.44|CCAAGCGTTGGATTGTTCACCCACTAATAGGGAACGTGAGCTGG</td><td>>hg38_21_+_8433172_8446622_RNA45SN1_45S_50ntFlanks</td><td>12456</td><td>12499</td><td>44</td><td>CCAAGCGTTGGATTGTTCACCCACTAATAGGGAACGTGAGCTGG</td><td>122.7</td><td>-5.8</td><td>1.7</td><td>-3.3</td><td>8.3E-04</td><td>2.1E-03</td><td>BT20/MB468</td><td>MP</td></tr>
<tr><td>>am_trnaMT_PheGAA_MT_+_577_647@45.65.21|AGACGGGCTCACATCACCCCA</td><td>>am_trnaMT_PheGAA_MT_+_577_647</td><td>45</td><td>65</td><td>21</td><td>AGACGGGCTCACATCACCCCA</td><td>106.6</td><td>-3.9</td><td>1.2</td><td>-3.2</td><td>1.5E-03</td><td>2.1E-03</td><td>MB231/MB468</td><td>MP</td></tr>
<tr><td>>hg38_21_+_8433172_8446622_RNA45SN1_45S_50ntFlanks@4723.4761.39|CATTAATCAAGAACGAAAGTCGGAGGTTCGAAGACGATC</td><td>>hg38_21_+_8433172_8446622_RNA45SN1_45S_50ntFlanks</td><td>4723</td><td>4761</td><td>39</td><td>CATTAATCAAGAACGAAAGTCGGAGGTTCGAAGACGATC</td><td>384.0</td><td>2.6</td><td>0.8</td><td>3.1</td><td>1.7E-03</td><td>2.1E-03</td><td>MB231/MB468</td><td>CYTO</td></tr>
<tr><td>>MI0000441|hsa-mir-30b&WithFlank&8|-|134800514|134800613@23.43.21(+1C)&[MIMAT0000420&hsa-miR-30b-5p&offsets|0|-1(+1C);m-76&8|-|134800570|134800591&offsets|0|-1(+1C)]|TGTAAACATCCTACACTCAGCC</td><td>>MI0000441|hsa-mir-30b&WithFlank&8|-|134800514|134800613</td><td>23</td><td>43</td><td>21(+1C)</td><td>TGTAAACATCCTACACTCAGCC</td><td>1684.0</td><td>-3.8</td><td>1.1</td><td>-3.6</td><td>3.7E-04</td><td>2.1E-03</td><td>BT20/MB231</td><td>MITO</td></tr>
<tr><td>>MI0000650|hsa-mir-200c&WithFlank&12|+|6963693|6963772@50.71.22(+1C)&[MIMAT0000617&hsa-miR-200c-3p&offsets|0|-1(+1C);m-54&12|+|6963742|6963764&offsets|0|-1(+1C)]|TAATACTGCCGGGTAATGATGGC</td><td>>MI0000650|hsa-mir-200c&WithFlank&12|+|6963693|6963772</td><td>50</td><td>71</td><td>22(+1C)</td><td>TAATACTGCCGGGTAATGATGGC</td><td>300.7</td><td>4.9</td><td>1.5</td><td>3.3</td><td>8.3E-04</td><td>2.1E-03</td><td>BT20/MB468</td><td>MP</td></tr>
<tr><td>>hg38_21_+_8433172_8446622_RNA45SN1_45S_50ntFlanks@11920.11949.30|ATGAGAGGTGTAGAATAAGTGGGAGGCCCC</td><td>>hg38_21_+_8433172_8446622_RNA45SN1_45S_50ntFlanks</td><td>11920</td><td>11949</td><td>30</td><td>ATGAGAGGTGTAGAATAAGTGGGAGGCCCC</td><td>165.2</td><td>-2.4</td><td>0.8</td><td>-3.2</td><td>1.6E-03</td><td>2.1E-03</td><td>BT20/MB468</td><td>CYTO</td></tr>
<tr><td>>hg38_21_+_8433172_8446622_RNA45SN1_45S_50ntFlanks@9740.9757.18|AACGATCTCAACCTATTC</td><td>>hg38_21_+_8433172_8446622_RNA45SN1_45S_50ntFlanks</td><td>9740</td><td>9757</td><td>18</td><td>AACGATCTCAACCTATTC</td><td>168.9</td><td>-3.2</td><td>1.0</td><td>-3.2</td><td>1.6E-03</td><td>2.1E-03</td><td>BT20/MB468</td><td>CYTO</td></tr>
<tr><td>>hg38_21_+_8433172_8446622_RNA45SN1_45S_50ntFlanks@10770.10805.36|GCAGGTCTCCAAGGTGAACAGCCTCTGGCATGTTGG</td><td>>hg38_21_+_8433172_8446622_RNA45SN1_45S_50ntFlanks</td><td>10770</td><td>10805</td><td>36</td><td>GCAGGTCTCCAAGGTGAACAGCCTCTGGCATGTTGG</td><td>70.4</td><td>-5.2</td><td>1.6</td><td>-3.3</td><td>8.3E-04</td><td>2.1E-03</td><td>BT20/MB468</td><td>MP</td></tr>
<tr><td>>hg38_21_+_8433172_8446622_RNA45SN1_45S_50ntFlanks@11845.11873.29|GGAACGGGCTTGGCGGAATCAGCGGGGAA</td><td>>hg38_21_+_8433172_8446622_RNA45SN1_45S_50ntFlanks</td><td>11845</td><td>11873</td><td>29</td><td>GGAACGGGCTTGGCGGAATCAGCGGGGAA</td><td>713.6</td><td>-5.6</td><td>1.7</td><td>-3.3</td><td>8.3E-04</td><td>2.1E-03</td><td>BT20/MB468</td><td>MP</td></tr>
<tr><td>>hg38_21_+_8433172_8446622_RNA45SN1_45S_50ntFlanks@10317.10340.24|GTGCAGATCTTGGTGGTAGTAGCA</td><td>>hg38_21_+_8433172_8446622_RNA45SN1_45S_50ntFlanks</td><td>10317</td><td>10340</td><td>24</td><td>GTGCAGATCTTGGTGGTAGTAGCA</td><td>506.8</td><td>3.0</td><td>0.9</td><td>3.2</td><td>1.3E-03</td><td>2.1E-03</td><td>BT20/MB468</td><td>MITO</td></tr>
<tr><td>>hg38_21_+_8433172_8446622_RNA45SN1_45S_50ntFlanks@7980.8005.26|CCTCAGATCAGACGTGGCGACCCGCT</td><td>>hg38_21_+_8433172_8446622_RNA45SN1_45S_50ntFlanks</td><td>7980</td><td>8005</td><td>26</td><td>CCTCAGATCAGACGTGGCGACCCGCT</td><td>180.1</td><td>-2.0</td><td>0.6</td><td>-3.1</td><td>1.7E-03</td><td>2.1E-03</td><td>MB231/MB468</td><td>CYTO</td></tr>
<tr><td>>MI0000542|hsa-mir-320a&WithFlank&8|-|22244960|22245043@50.69.20&[MIMAT0000510&hsa-miR-320a-3p&offsets|+2|0;m-60&8|-|22244975|22244996&offsets|+2|0]|AAGCTGGGTTGAGAGGGCGA</td><td>>MI0000542|hsa-mir-320a&WithFlank&8|-|22244960|22245043</td><td>50</td><td>69</td><td>20</td><td>AAGCTGGGTTGAGAGGGCGA</td><td>215.6</td><td>2.9</td><td>0.8</td><td>3.6</td><td>3.3E-04</td><td>2.1E-03</td><td>BT20/MB231</td><td>CYTO</td></tr>
<tr><td>>am_trnaMT_HisGTG_MT_+_12138_12206@10.51.42|GTTTAACCAAAACATCAGATTGTGAATCTGACAACAGAGGCT</td><td>>am_trnaMT_HisGTG_MT_+_12138_12206</td><td>10</td><td>51</td><td>42</td><td>GTTTAACCAAAACATCAGATTGTGAATCTGACAACAGAGGCT</td><td>46.9</td><td>9.3</td><td>2.8</td><td>3.3</td><td>8.3E-04</td><td>2.1E-03</td><td>BT20/MB468</td><td>MP</td></tr>
<tr><td>>hg38_21_+_8433172_8446622_RNA45SN1_45S_50ntFlanks@10387.10408.22|TGTGAACAGCAGTTGAACATGG</td><td>>hg38_21_+_8433172_8446622_RNA45SN1_45S_50ntFlanks</td><td>10387</td><td>10408</td><td>22</td><td>TGTGAACAGCAGTTGAACATGG</td><td>53.3</td><td>-9.6</td><td>3.0</td><td>-3.2</td><td>1.5E-03</td><td>2.1E-03</td><td>MB231/MB468</td><td>MP</td></tr>
<tr><td>>hg38_21_+_8433172_8446622_RNA45SN1_45S_50ntFlanks@11665.11707.43|TTCTGCCCAGTGCTCTGAATGTCAAAGTGAAGAAATTCAATGA</td><td>>hg38_21_+_8433172_8446622_RNA45SN1_45S_50ntFlanks</td><td>11665</td><td>11707</td><td>43</td><td>TTCTGCCCAGTGCTCTGAATGTCAAAGTGAAGAAATTCAATGA</td><td>262.8</td><td>-3.1</td><td>0.9</td><td>-3.6</td><td>3.3E-04</td><td>2.1E-03</td><td>BT20/MB231</td><td>CYTO</td></tr>
<tr><td>>hg38_21_+_8433172_8446622_RNA45SN1_45S_50ntFlanks@4369.4391.23|GTTAAAAAGCTCGTAGTTGGATC</td><td>>hg38_21_+_8433172_8446622_RNA45SN1_45S_50ntFlanks</td><td>4369</td><td>4391</td><td>23</td><td>GTTAAAAAGCTCGTAGTTGGATC</td><td>166.6</td><td>-2.6</td><td>0.8</td><td>-3.1</td><td>1.8E-03</td><td>2.1E-03</td><td>MB231/MB468</td><td>NUC</td></tr>
<tr><td>>am_trnaMT_ValTAC_MT_+_1602_1670@50.67.18|ACTTAACTTGACCGCTCT</td><td>>am_trnaMT_ValTAC_MT_+_1602_1670</td><td>50</td><td>67</td><td>18</td><td>ACTTAACTTGACCGCTCT</td><td>6363.5</td><td>3.2</td><td>0.9</td><td>3.6</td><td>3.3E-04</td><td>2.1E-03</td><td>BT20/MB231</td><td>CYTO</td></tr>
<tr><td>>hg38_21_+_8433172_8446622_RNA45SN1_45S_50ntFlanks@12909.12933.25|CCGCACGTTCGTGGGGAACCTGGCG</td><td>>hg38_21_+_8433172_8446622_RNA45SN1_45S_50ntFlanks</td><td>12909</td><td>12933</td><td>25</td><td>CCGCACGTTCGTGGGGAACCTGGCG</td><td>194.0</td><td>-4.5</td><td>1.4</td><td>-3.2</td><td>1.6E-03</td><td>2.1E-03</td><td>BT20/MB468</td><td>CYTO</td></tr>
<tr><td>>hg38_21_+_8433172_8446622_RNA45SN1_45S_50ntFlanks@11205.11227.23|GCGGCGGCGACTCTGGACGCGAG</td><td>>hg38_21_+_8433172_8446622_RNA45SN1_45S_50ntFlanks</td><td>11205</td><td>11227</td><td>23</td><td>GCGGCGGCGACTCTGGACGCGAG</td><td>145.3</td><td>-3.3</td><td>1.1</td><td>-3.2</td><td>1.6E-03</td><td>2.1E-03</td><td>BT20/MB468</td><td>CYTO</td></tr>
<tr><td>>trna116_GluCTC_1_-_145399233_145399304@1.35.35,trna59_GluCTC_1_+_249168447_249168518@1.35.35,trna71_GluCTC_1_-_161439189_161439260@1.35.35,trna74_GluCTC_1_-_161431809_161431880@1.35.35,trna77_GluCTC_1_-_161424398_161424469@1.35.35,trna77_GluCTC_6_+_28949976_28950047@1.35.35,trna80_GluCTC_1_-_161417018_161417089@1.35.35,trna87_GluCTC_6_-_126101393_126101464@1.35.35|TCCCTGGTGGTCTAGTGGTTAGGATTCGGCGCTCT</td><td>>trna116_GluCTC_1_-_145399233_145399304</td><td>1</td><td>35</td><td>35</td><td>TCCCTGGTGGTCTAGTGGTTAGGATTCGGCGCTCT</td><td>1573.9</td><td>3.2</td><td>1.0</td><td>3.1</td><td>1.8E-03</td><td>2.1E-03</td><td>BT20/MB468</td><td>NUC</td></tr>
<tr><td>>hg38_21_+_8433172_8446622_RNA45SN1_45S_50ntFlanks@5236.5260.25|CACTGACTGGCTCAGCGTGTGCCTA</td><td>>hg38_21_+_8433172_8446622_RNA45SN1_45S_50ntFlanks</td><td>5236</td><td>5260</td><td>25</td><td>CACTGACTGGCTCAGCGTGTGCCTA</td><td>81.8</td><td>-5.0</td><td>1.6</td><td>-3.2</td><td>1.5E-03</td><td>2.1E-03</td><td>MB231/MB468</td><td>MP</td></tr>
<tr><td>>hg38_21_+_8433172_8446622_RNA45SN1_45S_50ntFlanks@12164.12182.19|CTGGGGCGGTACACCTGTC</td><td>>hg38_21_+_8433172_8446622_RNA45SN1_45S_50ntFlanks</td><td>12164</td><td>12182</td><td>19</td><td>CTGGGGCGGTACACCTGTC</td><td>433.4</td><td>-3.2</td><td>1.0</td><td>-3.3</td><td>8.4E-04</td><td>2.1E-03</td><td>BT20/MB468</td><td>MP</td></tr>
<tr><td>>MI0003513|hsa-mir-455&WithFlank&9|+|114209428|114209535@60.80.21(+1C)&[MIMAT0004784&hsa-miR-455-3p&offsets|0|0(+1C);m-135&9|+|114209487|114209508&offsets|0|-1(+1C)]|GCAGTCCATGGGCATATACACC</td><td>>MI0003513|hsa-mir-455&WithFlank&9|+|114209428|114209535</td><td>60</td><td>80</td><td>21(+1C)</td><td>GCAGTCCATGGGCATATACACC</td><td>256.1</td><td>-2.7</td><td>0.8</td><td>-3.6</td><td>3.4E-04</td><td>2.1E-03</td><td>BT20/MB231</td><td>CYTO</td></tr>
<tr><td>>hg38_21_+_8433172_8446622_RNA45SN1_45S_50ntFlanks@6669.6699.31|CACTCGGCTCGTGCGTCGATGAAGAACGCAG</td><td>>hg38_21_+_8433172_8446622_RNA45SN1_45S_50ntFlanks</td><td>6669</td><td>6699</td><td>31</td><td>CACTCGGCTCGTGCGTCGATGAAGAACGCAG</td><td>467.5</td><td>-1.5</td><td>0.5</td><td>-3.2</td><td>1.4E-03</td><td>2.1E-03</td><td>BT20/MB468</td><td>MITO</td></tr>
<tr><td>>hg38_21_+_8433172_8446622_RNA45SN1_45S_50ntFlanks@11207.11229.23|GGCGGCGACTCTGGACGCGAGCC</td><td>>hg38_21_+_8433172_8446622_RNA45SN1_45S_50ntFlanks</td><td>11207</td><td>11229</td><td>23</td><td>GGCGGCGACTCTGGACGCGAGCC</td><td>106.0</td><td>-4.1</td><td>1.3</td><td>-3.2</td><td>1.5E-03</td><td>2.1E-03</td><td>MB231/MB468</td><td>MP</td></tr>
<tr><td>>trnaMT_GluTTC_MT_-_14674_14742@39.58.20,trnalookalike8_GluTTC_5_-_93905172_93905240@39.58.20|ATTGGTCGTGGTTGTAGTCC</td><td>>trnaMT_GluTTC_MT_-_14674_14742</td><td>39</td><td>58</td><td>20</td><td>ATTGGTCGTGGTTGTAGTCC</td><td>421.1</td><td>2.9</td><td>0.9</td><td>3.2</td><td>1.5E-03</td><td>2.1E-03</td><td>MB231/MB468</td><td>MP</td></tr>
<tr><td>>trnaMT_LysTTT_MT_+_8295_8364@34.72.39,trnalookalike7_LysTTT_1_+_568844_568913@34.72.39|GTTAAAGATTAAGAGAACCAACACCTCTTTACAGTGACC</td><td>>trnaMT_LysTTT_MT_+_8295_8364</td><td>34</td><td>72</td><td>39</td><td>GTTAAAGATTAAGAGAACCAACACCTCTTTACAGTGACC</td><td>121.8</td><td>5.9</td><td>1.9</td><td>3.2</td><td>1.5E-03</td><td>2.1E-03</td><td>MB231/MB468</td><td>MP</td></tr>
<tr><td>>hg38_21_+_8433172_8446622_RNA45SN1_45S_50ntFlanks@12408.12431.24|TTTGATCCTTCGATGTCGGCTCTT</td><td>>hg38_21_+_8433172_8446622_RNA45SN1_45S_50ntFlanks</td><td>12408</td><td>12431</td><td>24</td><td>TTTGATCCTTCGATGTCGGCTCTT</td><td>986.9</td><td>-2.8</td><td>0.9</td><td>-3.2</td><td>1.6E-03</td><td>2.1E-03</td><td>BT20/MB468</td><td>CYTO</td></tr>
<tr><td>>MI0003205|hsa-mir-532&WithFlank&X|+|50003142|50003244@26.46.21(+1A)&[MIMAT0002888&hsa-miR-532-5p&offsets|0|-1(+1A);m-65&X|+|50003167|50003188&offsets|0|-1(+1A)]|CATGCCTTGAGTGTAGGACCGA</td><td>>MI0003205|hsa-mir-532&WithFlank&X|+|50003142|50003244</td><td>26</td><td>46</td><td>21(+1A)</td><td>CATGCCTTGAGTGTAGGACCGA</td><td>233.3</td><td>2.8</td><td>0.8</td><td>3.6</td><td>3.4E-04</td><td>2.1E-03</td><td>BT20/MB231</td><td>CYTO</td></tr>
<tr><td>>hg38_21_+_8433172_8446622_RNA45SN1_45S_50ntFlanks@4574.4616.43|ATAGGACCGCGGTTCTATTTTGTTGGTTTTCGGAACTGAGGCC</td><td>>hg38_21_+_8433172_8446622_RNA45SN1_45S_50ntFlanks</td><td>4574</td><td>4616</td><td>43</td><td>ATAGGACCGCGGTTCTATTTTGTTGGTTTTCGGAACTGAGGCC</td><td>227.4</td><td>-3.8</td><td>1.2</td><td>-3.1</td><td>1.7E-03</td><td>2.1E-03</td><td>MB231/MB468</td><td>CYTO</td></tr>
<tr><td>>MI0000734|hsa-mir-106b&WithFlank&7|-|100093987|100094080@58.77.20(+1U)&[MIMAT0004672&hsa-miR-106b-3p&offsets|0|-2(+1U);m-87&7|-|100094003|100094023&offsets|0|-1(+1U)]|CCGCACTGTGGGTACTTGCTT</td><td>>MI0000734|hsa-mir-106b&WithFlank&7|-|100093987|100094080</td><td>58</td><td>77</td><td>20(+1U)</td><td>CCGCACTGTGGGTACTTGCTT</td><td>69.2</td><td>6.5</td><td>2.0</td><td>3.3</td><td>8.4E-04</td><td>2.1E-03</td><td>BT20/MB468</td><td>MP</td></tr>
<tr><td>>hg38_21_+_8433172_8446622_RNA45SN1_45S_50ntFlanks@6776.6803.28|CTCCCGGGGCTACGCCTGTCTGAGCGTC</td><td>>hg38_21_+_8433172_8446622_RNA45SN1_45S_50ntFlanks</td><td>6776</td><td>6803</td><td>28</td><td>CTCCCGGGGCTACGCCTGTCTGAGCGTC</td><td>2120.8</td><td>-2.4</td><td>0.8</td><td>-3.2</td><td>1.4E-03</td><td>2.1E-03</td><td>BT20/MB468</td><td>MITO</td></tr>
<tr><td>>hg38_21_+_8433172_8446622_RNA45SN1_45S_50ntFlanks@11805.11823.19|CCACTGTCCCTACCTACTA</td><td>>hg38_21_+_8433172_8446622_RNA45SN1_45S_50ntFlanks</td><td>11805</td><td>11823</td><td>19</td><td>CCACTGTCCCTACCTACTA</td><td>181.6</td><td>-3.4</td><td>1.1</td><td>-3.2</td><td>1.4E-03</td><td>2.1E-03</td><td>BT20/MB468</td><td>MITO</td></tr>
<tr><td>>hg38_21_+_8433172_8446622_RNA45SN1_45S_50ntFlanks@3705.3728.24|TACCTGGTTGATCCTGCCAGTAGC</td><td>>hg38_21_+_8433172_8446622_RNA45SN1_45S_50ntFlanks</td><td>3705</td><td>3728</td><td>24</td><td>TACCTGGTTGATCCTGCCAGTAGC</td><td>517.6</td><td>2.9</td><td>0.9</td><td>3.1</td><td>1.8E-03</td><td>2.1E-03</td><td>MB231/MB468</td><td>MITO</td></tr>
<tr><td>>MI0000299|hsa-mir-222&WithFlank&X|-|45747009|45747130@77.99.23&[MIMAT0000279&hsa-miR-222-3p&offsets|+2|+4;m-140&X|-|45747033|45747056&offsets|+2|+1]|CTACATCTGGCTACTGGGTCTCT</td><td>>MI0000299|hsa-mir-222&WithFlank&X|-|45747009|45747130</td><td>77</td><td>99</td><td>23</td><td>CTACATCTGGCTACTGGGTCTCT</td><td>48.4</td><td>5.7</td><td>1.8</td><td>3.2</td><td>1.5E-03</td><td>2.1E-03</td><td>MB231/MB468</td><td>MP</td></tr>
<tr><td>>hg38_21_+_8433172_8446622_RNA45SN1_45S_50ntFlanks@6652.6679.28|GACTCTTAGCGGTGGATCACTCGGCTCG</td><td>>hg38_21_+_8433172_8446622_RNA45SN1_45S_50ntFlanks</td><td>6652</td><td>6679</td><td>28</td><td>GACTCTTAGCGGTGGATCACTCGGCTCG</td><td>6031.5</td><td>4.2</td><td>1.3</td><td>3.2</td><td>1.6E-03</td><td>2.1E-03</td><td>BT20/MB468</td><td>CYTO</td></tr>
<tr><td>>hg38_21_+_8433172_8446622_RNA45SN1_45S_50ntFlanks@6698.6717.20|AGCTAGCTGCGAGAATTAAT</td><td>>hg38_21_+_8433172_8446622_RNA45SN1_45S_50ntFlanks</td><td>6698</td><td>6717</td><td>20</td><td>AGCTAGCTGCGAGAATTAAT</td><td>208.1</td><td>-3.1</td><td>1.0</td><td>-3.1</td><td>1.7E-03</td><td>2.1E-03</td><td>MB231/MB468</td><td>CYTO</td></tr>
<tr><td>>hg38_21_+_8433172_8446622_RNA45SN1_45S_50ntFlanks@11671.11709.39|CCAGTGCTCTGAATGTCAAAGTGAAGAAATTCAATGAAG</td><td>>hg38_21_+_8433172_8446622_RNA45SN1_45S_50ntFlanks</td><td>11671</td><td>11709</td><td>39</td><td>CCAGTGCTCTGAATGTCAAAGTGAAGAAATTCAATGAAG</td><td>195.8</td><td>-2.9</td><td>0.8</td><td>-3.6</td><td>3.4E-04</td><td>2.1E-03</td><td>BT20/MB231</td><td>CYTO</td></tr>
<tr><td>>hg38_21_+_8433172_8446622_RNA45SN1_45S_50ntFlanks@4726.4759.34|TAATCAAGAACGAAAGTCGGAGGTTCGAAGACGA</td><td>>hg38_21_+_8433172_8446622_RNA45SN1_45S_50ntFlanks</td><td>4726</td><td>4759</td><td>34</td><td>TAATCAAGAACGAAAGTCGGAGGTTCGAAGACGA</td><td>1196.5</td><td>-2.6</td><td>0.8</td><td>-3.1</td><td>1.7E-03</td><td>2.1E-03</td><td>MB231/MB468</td><td>CYTO</td></tr>
</table>

<table>
<tr><td>>hg38_MT_+_1621_3279_RNR2_16S_50ntFlanks@254.282.29|ACTTTGCAAGGAGAGCCAAAGCTAAGACC</td><td>>hg38_MT_+_1621_3279_RNR2_16S_50ntFlanks</td><td>254</td><td>282</td><td>29</td><td>ACTTTGCAAGGAGAGCCAAAGCTAAGACC</td><td>47.0</td><td>9.3</td><td>2.8</td><td>3.3</td><td>8.4E-04</td><td>2.1E-03</td><td>BT20/MB468</td><td>MP</td></tr>
<tr><td>>trnaMT_HisGTG_MT_+_12138_12206@46.71.26|GAGGCTTACGACCCCTTATTTACCCC</td><td>>trnaMT_HisGTG_MT_+_12138_12206</td><td>46</td><td>71</td><td>26</td><td>GAGGCTTACGACCCCTTATTTACCCC</td><td>204.5</td><td>6.1</td><td>1.8</td><td>3.3</td><td>8.4E-04</td><td>2.1E-03</td><td>BT20/MB468</td><td>MP</td></tr>
<tr><td>>hg38_21_+_8433172_8446622_RNA45SN1_45S_50ntFlanks@10394.10419.26|AGCAGTTGAACATGGGTCAGTCGGTC</td><td>>hg38_21_+_8433172_8446622_RNA45SN1_45S_50ntFlanks</td><td>10394</td><td>10419</td><td>26</td><td>AGCAGTTGAACATGGGTCAGTCGGTC</td><td>64.9</td><td>-5.2</td><td>1.6</td><td>-3.2</td><td>1.5E-03</td><td>2.1E-03</td><td>MB231/MB468</td><td>MP</td></tr>
<tr><td>>hg38_MT_+_598_1651_RNR1_12S_50ntFlanks@909.946.38|TACTTCAAAGGACATTTAACTAAAACCCCTACGCATTT</td><td>>hg38_MT_+_598_1651_RNR1_12S_50ntFlanks</td><td>909</td><td>946</td><td>38</td><td>TACTTCAAAGGACATTTAACTAAAACCCCTACGCATTT</td><td>314.4</td><td>5.9</td><td>1.8</td><td>3.3</td><td>8.4E-04</td><td>2.1E-03</td><td>BT20/MB468</td><td>MP</td></tr>
<tr><td>>hg38_21_+_8433172_8446622_RNA45SN1_45S_50ntFlanks@10731.10755.25|AGGGTGTAAATCTCGCGCCGGGCCG</td><td>>hg38_21_+_8433172_8446622_RNA45SN1_45S_50ntFlanks</td><td>10731</td><td>10755</td><td>25</td><td>AGGGTGTAAATCTCGCGCCGGGCCG</td><td>489.0</td><td>-5.5</td><td>1.6</td><td>-3.3</td><td>8.4E-04</td><td>2.1E-03</td><td>BT20/MB468</td><td>MP</td></tr>
<tr><td>>MI0000477|hsa-mir-146a&WithFlank&5|+|160485346|160485456@27.46.20(+1C)&[MIMAT0000449&hsa-miR-146a-5p&offsets|0|-2(+1C);m-129&5|+|160485372|160485393&offsets|0|-2(+1C)]|TGAGAACTGAATTCCATGGGC</td><td>>MI0000477|hsa-mir-146a&WithFlank&5|+|160485346|160485456</td><td>27</td><td>46</td><td>20(+1C)</td><td>TGAGAACTGAATTCCATGGGC</td><td>309.6</td><td>-7.6</td><td>2.1</td><td>-3.6</td><td>3.7E-04</td><td>2.1E-03</td><td>BT20/MB231</td><td>MITO</td></tr>
<tr><td>>hg38_21_+_8433172_8446622_RNA45SN1_45S_50ntFlanks@12627.12667.41|ATGGGGCGAAGCTACCATCTGTGGGATTATGACTGAACGCC</td><td>>hg38_21_+_8433172_8446622_RNA45SN1_45S_50ntFlanks</td><td>12627</td><td>12667</td><td>41</td><td>ATGGGGCGAAGCTACCATCTGTGGGATTATGACTGAACGCC</td><td>372.4</td><td>-3.0</td><td>0.8</td><td>-3.6</td><td>3.4E-04</td><td>2.1E-03</td><td>BT20/MB231</td><td>CYTO</td></tr>
<tr><td>>hg38_21_+_8433172_8446622_RNA45SN1_45S_50ntFlanks@6780.6803.24|CGGGGCTACGCCTGTCTGAGCGTC</td><td>>hg38_21_+_8433172_8446622_RNA45SN1_45S_50ntFlanks</td><td>6780</td><td>6803</td><td>24</td><td>CGGGGCTACGCCTGTCTGAGCGTC</td><td>541.7</td><td>-4.1</td><td>1.3</td><td>-3.1</td><td>1.8E-03</td><td>2.1E-03</td><td>MB231/MB468</td><td>MITO</td></tr>
<tr><td>>hg38_21_+_8433172_8446622_RNA45SN1_45S_50ntFlanks@11882.11905.24|CTGTTGAGCTTGACTCTAGTCTGG</td><td>>hg38_21_+_8433172_8446622_RNA45SN1_45S_50ntFlanks</td><td>11882</td><td>11905</td><td>24</td><td>CTGTTGAGCTTGACTCTAGTCTGG</td><td>132.2</td><td>-4.0</td><td>1.2</td><td>-3.3</td><td>8.4E-04</td><td>2.1E-03</td><td>BT20/MB468</td><td>MP</td></tr>
<tr><td>>hg38_21_+_8433172_8446622_RNA45SN1_45S_50ntFlanks@485.510.26|GATGTGGTGACGTCGTGCTCTCCCGG</td><td>>hg38_21_+_8433172_8446622_RNA45SN1_45S_50ntFlanks</td><td>485</td><td>510</td><td>26</td><td>GATGTGGTGACGTCGTGCTCTCCCGG</td><td>44.4</td><td>8.7</td><td>2.7</td><td>3.2</td><td>1.5E-03</td><td>2.1E-03</td><td>MB231/MB468</td><td>MP</td></tr>
<tr><td>>hg38_21_+_8433172_8446622_RNA45SN1_45S_50ntFlanks@10572.10608.37|AGAAGCCGGCGGGAGCCCCGGGGAGAGTTCTCTTTTC</td><td>>hg38_21_+_8433172_8446622_RNA45SN1_45S_50ntFlanks</td><td>10572</td><td>10608</td><td>37</td><td>AGAAGCCGGCGGGAGCCCCGGGGAGAGTTCTCTTTTC</td><td>1150.4</td><td>-5.3</td><td>1.7</td><td>-3.2</td><td>1.6E-03</td><td>2.1E-03</td><td>BT20/MB468</td><td>CYTO</td></tr>
<tr><td>>hg38_21_+_8433172_8446622_RNA45SN1_45S_50ntFlanks@10351.10392.42|CGAGAACTTTGAAGGCCGAAGTGGAGAAGGGTTCCATGTGAA</td><td>>hg38_21_+_8433172_8446622_RNA45SN1_45S_50ntFlanks</td><td>10351</td><td>10392</td><td>42</td><td>CGAGAACTTTGAAGGCCGAAGTGGAGAAGGGTTCCATGTGAA</td><td>239.7</td><td>-3.9</td><td>1.3</td><td>-3.1</td><td>1.8E-03</td><td>2.1E-03</td><td>MB231/MB468</td><td>NUC</td></tr>
<tr><td>>hg38_21_+_8433172_8446622_RNA45SN1_45S_50ntFlanks@10788.10825.38|CAGCCTCTGGCATGTTGGAACAATGTAGGTAAGGGAAG</td><td>>hg38_21_+_8433172_8446622_RNA45SN1_45S_50ntFlanks</td><td>10788</td><td>10825</td><td>38</td><td>CAGCCTCTGGCATGTTGGAACAATGTAGGTAAGGGAAG</td><td>115.1</td><td>-5.5</td><td>1.7</td><td>-3.3</td><td>8.5E-04</td><td>2.1E-03</td><td>BT20/MB468</td><td>MP</td></tr>
<tr><td>>hg38_21_+_8433172_8446622_RNA45SN1_45S_50ntFlanks@12159.12178.20|TTTGACTGGGGCGGTACACC</td><td>>hg38_21_+_8433172_8446622_RNA45SN1_45S_50ntFlanks</td><td>12159</td><td>12178</td><td>20</td><td>TTTGACTGGGGCGGTACACC</td><td>326.2</td><td>-2.0</td><td>0.6</td><td>-3.2</td><td>1.4E-03</td><td>2.1E-03</td><td>BT20/MB468</td><td>MITO</td></tr>
<tr><td>>hg38_21_+_8433172_8446622_RNA45SN1_45S_50ntFlanks@10471.10488.18|CCCTCGGCCGATCGAAAG</td><td>>hg38_21_+_8433172_8446622_RNA45SN1_45S_50ntFlanks</td><td>10471</td><td>10488</td><td>18</td><td>CCCTCGGCCGATCGAAAG</td><td>572.3</td><td>-3.9</td><td>1.2</td><td>-3.1</td><td>1.6E-03</td><td>2.1E-03</td><td>BT20/MB468</td><td>CYTO</td></tr>
<tr><td>>hg38_21_+_8433172_8446622_RNA45SN1_45S_50ntFlanks@11575.11612.38|ACTTAGAACTGGTGCGGACCAGGGGAATCCGACTGTTT</td><td>>hg38_21_+_8433172_8446622_RNA45SN1_45S_50ntFlanks</td><td>11575</td><td>11612</td><td>38</td><td>ACTTAGAACTGGTGCGGACCAGGGGAATCCGACTGTTT</td><td>244.7</td><td>-2.7</td><td>0.9</td><td>-3.1</td><td>1.8E-03</td><td>2.1E-03</td><td>MB231/MB468</td><td>NUC</td></tr>
<tr><td>>hg38_21_+_8433172_8446622_RNA45SN1_45S_50ntFlanks@11873.11909.37|AAGAAGACCCTGTTGAGCTTGACTCTAGTCTGGCACG</td><td>>hg38_21_+_8433172_8446622_RNA45SN1_45S_50ntFlanks</td><td>11873</td><td>11909</td><td>37</td><td>AAGAAGACCCTGTTGAGCTTGACTCTAGTCTGGCACG</td><td>196.5</td><td>-3.5</td><td>1.1</td><td>-3.1</td><td>1.6E-03</td><td>2.1E-03</td><td>BT20/MB468</td><td>CYTO</td></tr>
<tr><td>>hg38_21_+_8433172_8446622_RNA45SN1_45S_50ntFlanks@10392.10433.42|ACAGCAGTTGAACATGGGTCAGTCGGTCCTGAGAGATGGGCG</td><td>>hg38_21_+_8433172_8446622_RNA45SN1_45S_50ntFlanks</td><td>10392</td><td>10433</td><td>42</td><td>ACAGCAGTTGAACATGGGTCAGTCGGTCCTGAGAGATGGGCG</td><td>252.0</td><td>-4.8</td><td>1.4</td><td>-3.3</td><td>8.5E-04</td><td>2.1E-03</td><td>BT20/MB468</td><td>MP</td></tr>
<tr><td>>trnaMT_PheGAA_MT_+_577_647@1.34.34|GTTTATGTAGCTTACCTCCTCAAAGCAATACACT</td><td>>trnaMT_PheGAA_MT_+_577_647</td><td>1</td><td>34</td><td>34</td><td>GTTTATGTAGCTTACCTCCTCAAAGCAATACACT</td><td>538.3</td><td>3.0</td><td>0.9</td><td>3.2</td><td>1.5E-03</td><td>2.1E-03</td><td>MB231/MB468</td><td>MP</td></tr>
<tr><td>>hg38_MT_+_1621_3279_RNR2_16S_50ntFlanks@999.1029.31|AGGGACCTGTATGAATGGCTCCACGAGGGTT</td><td>>hg38_MT_+_1621_3279_RNR2_16S_50ntFlanks</td><td>999</td><td>1029</td><td>31</td><td>AGGGACCTGTATGAATGGCTCCACGAGGGTT</td><td>51.3</td><td>5.6</td><td>1.8</td><td>3.2</td><td>1.5E-03</td><td>2.1E-03</td><td>MB231/MB468</td><td>MP</td></tr>
<tr><td>>hg38_21_+_8433172_8446622_RNA45SN1_45S_50ntFlanks@12941.12960.20|ATTCGTAGACGACCTGCTTC</td><td>>hg38_21_+_8433172_8446622_RNA45SN1_45S_50ntFlanks</td><td>12941</td><td>12960</td><td>20</td><td>ATTCGTAGACGACCTGCTTC</td><td>158.3</td><td>-2.0</td><td>0.6</td><td>-3.2</td><td>1.5E-03</td><td>2.1E-03</td><td>MB231/MB468</td><td>MP</td></tr>
<tr><td>>hg38_21_+_8433172_8446622_RNA45SN1_45S_50ntFlanks@6645.6680.36|CTCGTACGACTCTTAGCGGTGGATCACTCGGCTCGT</td><td>>hg38_21_+_8433172_8446622_RNA45SN1_45S_50ntFlanks</td><td>6645</td><td>6680</td><td>36</td><td>CTCGTACGACTCTTAGCGGTGGATCACTCGGCTCGT</td><td>974.3</td><td>-1.6</td><td>0.5</td><td>-3.1</td><td>1.8E-03</td><td>2.1E-03</td><td>BT20/MB468</td><td>NUC</td></tr>
<tr><td>>hg38_21_+_8433172_8446622_RNA45SN1_45S_50ntFlanks@11602.11627.26|TCCGACTGTTTAATTAAAACAAAGCA</td><td>>hg38_21_+_8433172_8446622_RNA45SN1_45S_50ntFlanks</td><td>11602</td><td>11627</td><td>26</td><td>TCCGACTGTTTAATTAAAACAAAGCA</td><td>106.4</td><td>3.1</td><td>0.9</td><td>3.3</td><td>8.5E-04</td><td>2.1E-03</td><td>BT20/MB468</td><td>MP</td></tr>
<tr><td>>hg38_21_+_8433172_8446622_RNA45SN1_45S_50ntFlanks@480.498.19|TGATCGATGTGGTGACGTC</td><td>>hg38_21_+_8433172_8446622_RNA45SN1_45S_50ntFlanks</td><td>480</td><td>498</td><td>19</td><td>TGATCGATGTGGTGACGTC</td><td>58.7</td><td>5.7</td><td>1.8</td><td>3.2</td><td>1.5E-03</td><td>2.1E-03</td><td>MB231/MB468</td><td>MP</td></tr>
<tr><td>>hg38_21_+_8433172_8446622_RNA45SN1_45S_50ntFlanks@11326.11346.21|CTCGTCGCGCGCGCGTCCGCT</td><td>>hg38_21_+_8433172_8446622_RNA45SN1_45S_50ntFlanks</td><td>11326</td><td>11346</td><td>21</td><td>CTCGTCGCGCGCGCGTCCGCT</td><td>388.9</td><td>10.7</td><td>3.4</td><td>3.1</td><td>1.8E-03</td><td>2.1E-03</td><td>MB231/MB468</td><td>NUC</td></tr>
<tr><td>>hg38_21_+_8433172_8446622_RNA45SN1_45S_50ntFlanks@12893.12932.40|CCTCGCCCGTCACGCACCGCACGTTCGTGGGGAACCTGGC</td><td>>hg38_21_+_8433172_8446622_RNA45SN1_45S_50ntFlanks</td><td>12893</td><td>12932</td><td>40</td><td>CCTCGCCCGTCACGCACCGCACGTTCGTGGGGAACCTGGC</td><td>962.9</td><td>-6.1</td><td>1.9</td><td>-3.1</td><td>1.8E-03</td><td>2.1E-03</td><td>BT20/MB468</td><td>NUC</td></tr>
<tr><td>>hg38_21_+_8433172_8446622_RNA45SN1_45S_50ntFlanks@12636.12656.21|AGCTACCATCTGTGGGATTAT</td><td>>hg38_21_+_8433172_8446622_RNA45SN1_45S_50ntFlanks</td><td>12636</td><td>12656</td><td>21</td><td>AGCTACCATCTGTGGGATTAT</td><td>68.9</td><td>-5.2</td><td>1.6</td><td>-3.3</td><td>8.5E-04</td><td>2.1E-03</td><td>BT20/MB468</td><td>MP</td></tr>
<tr><td>>hg38_21_+_8433172_8446622_RNA45SN1_45S_50ntFlanks@10423.10456.34|AGAGATGGGCGAGCGCCGTTCCGAAGGGACGGGC</td><td>>hg38_21_+_8433172_8446622_RNA45SN1_45S_50ntFlanks</td><td>10423</td><td>10456</td><td>34</td><td>AGAGATGGGCGAGCGCCGTTCCGAAGGGACGGGC</td><td>1227.6</td><td>-3.1</td><td>0.9</td><td>-3.6</td><td>3.4E-04</td><td>2.1E-03</td><td>BT20/MB231</td><td>CYTO</td></tr>
<tr><td>>hg38_21_+_8433172_8446622_RNA45SN1_45S_50ntFlanks@12006.12032.27|GGCCGCCGGTGAAATACCACTACTCTG</td><td>>hg38_21_+_8433172_8446622_RNA45SN1_45S_50ntFlanks</td><td>12006</td><td>12032</td><td>27</td><td>GGCCGCCGGTGAAATACCACTACTCTG</td><td>415.3</td><td>-2.2</td><td>0.7</td><td>-3.1</td><td>1.6E-03</td><td>2.1E-03</td><td>BT20/MB468</td><td>CYTO</td></tr>
<tr><td>>hg38_21_+_8433172_8446622_RNA45SN1_45S_50ntFlanks@11591.11611.21|GACCAGGGGAATCCGACTGTT</td><td>>hg38_21_+_8433172_8446622_RNA45SN1_45S_50ntFlanks</td><td>11591</td><td>11611</td><td>21</td><td>GACCAGGGGAATCCGACTGTT</td><td>221.7</td><td>-4.5</td><td>1.3</td><td>-3.3</td><td>8.5E-04</td><td>2.1E-03</td><td>BT20/MB468</td><td>MP</td></tr>
<tr><td>>hg38_21_+_8433172_8446622_RNA45SN1_45S_50ntFlanks@9165.9183.19|CCCGGGCGGGTCGCGCCGT</td><td>>hg38_21_+_8433172_8446622_RNA45SN1_45S_50ntFlanks</td><td>9165</td><td>9183</td><td>19</td><td>CCCGGGCGGGTCGCGCCGT</td><td>79.8</td><td>-5.2</td><td>1.6</td><td>-3.3</td><td>8.5E-04</td><td>2.1E-03</td><td>BT20/MB468</td><td>MP</td></tr>
<tr><td>>hg38_21_+_8433172_8446622_RNA45SN1_45S_50ntFlanks@10826.10858.33|TCGGCAAGCCGGATCCGTAACTTCGGGATAAGG</td><td>>hg38_21_+_8433172_8446622_RNA45SN1_45S_50ntFlanks</td><td>10826</td><td>10858</td><td>33</td><td>TCGGCAAGCCGGATCCGTAACTTCGGGATAAGG</td><td>150.7</td><td>-2.2</td><td>0.7</td><td>-3.2</td><td>1.4E-03</td><td>2.1E-03</td><td>BT20/MB468</td><td>MITO</td></tr>
<tr><td>>hg38_21_+_8433172_8446622_RNA45SN1_45S_50ntFlanks@12415.12444.30|CTTCGATGTCGGCTCTTCCTATCATTGTGA</td><td>>hg38_21_+_8433172_8446622_RNA45SN1_45S_50ntFlanks</td><td>12415</td><td>12444</td><td>30</td><td>CTTCGATGTCGGCTCTTCCTATCATTGTGA</td><td>182.6</td><td>-5.2</td><td>1.6</td><td>-3.2</td><td>1.5E-03</td><td>2.1E-03</td><td>MB231/MB468</td><td>MP</td></tr>
<tr><td>>hg38_21_+_8433172_8446622_RNA45SN1_45S_50ntFlanks@4723.4748.26|CATTAATCAAGAACGAAAGTCGGAGG</td><td>>hg38_21_+_8433172_8446622_RNA45SN1_45S_50ntFlanks</td><td>4723</td><td>4748</td><td>26</td><td>CATTAATCAAGAACGAAAGTCGGAGG</td><td>626.6</td><td>-2.5</td><td>0.8</td><td>-3.1</td><td>1.7E-03</td><td>2.1E-03</td><td>MB231/MB468</td><td>CYTO</td></tr>
<tr><td>>am_trna128_GlyGCC_6_-_27870686_27870756@10.35.26,trna18_GlyGCC_16_+_70822597_70822667@10.35.26,trna19_GlyGCC_16_+_70823410_70823480@10.35.26,trna19_GlyGCC_2_-_157257659_157257729@10.35.26,trna24_GlyGCC_16_-_70812942_70813012@10.35.26,trna25_GlyGCC_16_-_70812114_70812184@10.35.26,trna2_GlyGCC_21_-_18827107_18827177@10.35.26,trna35_GlyGCC_1_+_161413094_161413164@10.35.26,trna37_GlyGCC_1_+_161420467_161420537@10.35.26,trna39_GlyGCC_1_+_161427898_161427968@10.35.26,trna41_GlyGCC_1_+_161435258_161435328@10.35.26,trna5_GlyGCC_17_+_8029064_8029134@10.35.26,trna68_GlyGCC_1_-_161493637_161493707@10.35.26|GTTCAGTGGTAGAATTCTCGCCTGCC</td><td>>am_trna128_GlyGCC_6_-_27870686_27870756</td><td>10</td><td>35</td><td>26</td><td>GTTCAGTGGTAGAATTCTCGCCTGCC</td><td>127.7</td><td>4.8</td><td>1.5</td><td>3.1</td><td>1.8E-03</td><td>2.2E-03</td><td>MB231/MB468</td><td>MITO</td></tr>
<tr><td>>hg38_21_+_8433172_8446622_RNA45SN1_45S_50ntFlanks@3988.4024.37|CTTTGGTGACTCTAGATAACCTCGGGCCGATCGCACG</td><td>>hg38_21_+_8433172_8446622_RNA45SN1_45S_50ntFlanks</td><td>3988</td><td>4024</td><td>37</td><td>CTTTGGTGACTCTAGATAACCTCGGGCCGATCGCACG</td><td>470.3</td><td>-1.6</td><td>0.5</td><td>-3.2</td><td>1.4E-03</td><td>2.2E-03</td><td>BT20/MB468</td><td>MITO</td></tr>
<tr><td>>trna111_HisGTG_1_-_147774845_147774916@-1T.34.35,trna118_HisGTG_1_-_145396881_145396952@-1T.34.35,trna16_HisGTG_1_+_146544773_146544844@-1T.34.35,trna1_HisGTG_15_+_45493349_45493420@-1T.34.35,trna21_HisGTG_1_+_147753471_147753542@-1T.34.35,trna33_HisGTG_6_+_27125906_27125977@-1T.34.35,trna7_HisGTG_9_-_14433938_14434009@-1T.34.35,trna8_HisGTG_15_-_45492611_45492682@-1T.34.35,trna9_HisGTG_15_-_45490804_45490875@-1T.34.35|TGCCGTGATCGTATAGTGGTTAGTACTCTGCGTTG</td><td>>trna111_HisGTG_1_-_147774845_147774916</td><td>-1T</td><td>34</td><td>35</td><td>TGCCGTGATCGTATAGTGGTTAGTACTCTGCGTTG</td><td>295.0</td><td>3.4</td><td>1.1</td><td>3.2</td><td>1.5E-03</td><td>2.2E-03</td><td>MB231/MB468</td><td>MP</td></tr>
<tr><td>>hg38_21_+_8433172_8446622_RNA45SN1_45S_50ntFlanks@7975.7992.18|CGCGACCTCAGATCAGAC</td><td>>hg38_21_+_8433172_8446622_RNA45SN1_45S_50ntFlanks</td><td>7975</td><td>7992</td><td>18</td><td>CGCGACCTCAGATCAGAC</td><td>44723.1</td><td>4.9</td><td>1.4</td><td>3.6</td><td>2.9E-04</td><td>2.2E-03</td><td>BT20/MB231</td><td>MP</td></tr>
<tr><td>>MI0000299|hsa-mir-222&WithFlank&X|-|45747009|45747130@75.99.25&[MIMAT0000279&hsa-miR-222-3p&offsets|0|+4;m-140&X|-|45747033|45747056&offsets|0|+1]|AGCTACATCTGGCTACTGGGTCTCT</td><td>>MI0000299|hsa-mir-222&WithFlank&X|-|45747009|45747130</td><td>75</td><td>99</td><td>25</td><td>AGCTACATCTGGCTACTGGGTCTCT</td><td>3082.7</td><td>3.1</td><td>0.9</td><td>3.3</td><td>8.5E-04</td><td>2.2E-03</td><td>BT20/MB468</td><td>MP</td></tr>
<tr><td>>hg38_21_+_8433172_8446622_RNA45SN1_45S_50ntFlanks@11918.11951.34|ACATGAGAGGTGTAGAATAAGTGGGAGGCCCCCG</td><td>>hg38_21_+_8433172_8446622_RNA45SN1_45S_50ntFlanks</td><td>11918</td><td>11951</td><td>34</td><td>ACATGAGAGGTGTAGAATAAGTGGGAGGCCCCCG</td><td>700.1</td><td>-4.6</td><td>1.4</td><td>-3.3</td><td>8.5E-04</td><td>2.2E-03</td><td>BT20/MB468</td><td>MP</td></tr>
<tr><td>>hg38_21_+_8433172_8446622_RNA45SN1_45S_50ntFlanks@10842.10864.23|GTAACTTCGGGATAAGGATTGGC</td><td>>hg38_21_+_8433172_8446622_RNA45SN1_45S_50ntFlanks</td><td>10842</td><td>10864</td><td>23</td><td>GTAACTTCGGGATAAGGATTGGC</td><td>235.8</td><td>-2.3</td><td>0.7</td><td>-3.1</td><td>1.7E-03</td><td>2.2E-03</td><td>MB231/MB468</td><td>CYTO</td></tr>
<tr><td>>trna2_GlyGCC_21_-_18827107_18827177@1.35.35,trna35_GlyGCC_1_+_161413094_161413164@1.35.35,trna37_GlyGCC_1_+_161420467_161420537@1.35.35,trna39_GlyGCC_1_+_161427898_161427968@1.35.35,trna41_GlyGCC_1_+_161435258_161435328@1.35.35|GCATGGGTGGTTCAGTGGTAGAATTCTCGCCTGCC</td><td>>trna2_GlyGCC_21_-_18827107_18827177</td><td>1</td><td>35</td><td>35</td><td>GCATGGGTGGTTCAGTGGTAGAATTCTCGCCTGCC</td><td>290.8</td><td>3.2</td><td>0.9</td><td>3.6</td><td>3.4E-04</td><td>2.2E-03</td><td>BT20/MB231</td><td>CYTO</td></tr>
<tr><td>>hg38_21_+_8433172_8446622_RNA45SN1_45S_50ntFlanks@12405.12440.36|CTTTTTGATCCTTCGATGTCGGCTCTTCCTATCATT</td><td>>hg38_21_+_8433172_8446622_RNA45SN1_45S_50ntFlanks</td><td>12405</td><td>12440</td><td>36</td><td>CTTTTTGATCCTTCGATGTCGGCTCTTCCTATCATT</td><td>299.2</td><td>-1.8</td><td>0.6</td><td>-3.2</td><td>1.4E-03</td><td>2.2E-03</td><td>BT20/MB468</td><td>MITO</td></tr>
<tr><td>>hg38_21_+_8433172_8446622_RNA45SN1_45S_50ntFlanks@11270.11293.24|CGGCCGCCCCCGGGGAGCCCGGCG</td><td>>hg38_21_+_8433172_8446622_RNA45SN1_45S_50ntFlanks</td><td>11270</td><td>11293</td><td>24</td><td>CGGCCGCCCCCGGGGAGCCCGGCG</td><td>592.3</td><td>-4.9</td><td>1.6</td><td>-3.1</td><td>1.6E-03</td><td>2.2E-03</td><td>BT20/MB468</td><td>CYTO</td></tr>
<tr><td>>hg38_21_+_8433172_8446622_RNA45SN1_45S_50ntFlanks@5074.5110.37|ATTCCGATAACGAACGAGACTCTGGCATGCTAACTAG</td><td>>hg38_21_+_8433172_8446622_RNA45SN1_45S_50ntFlanks</td><td>5074</td><td>5110</td><td>37</td><td>ATTCCGATAACGAACGAGACTCTGGCATGCTAACTAG</td><td>159.6</td><td>-6.8</td><td>2.2</td><td>-3.2</td><td>1.5E-03</td><td>2.2E-03</td><td>MB231/MB468</td><td>MP</td></tr>
<tr><td>>hg38_21_+_8433172_8446622_RNA45SN1_45S_50ntFlanks@8342.8379.38|CGTAAGGGAAAGTTGAAAAGAACTTTGAAGAGAGAGTT</td><td>>hg38_21_+_8433172_8446622_RNA45SN1_45S_50ntFlanks</td><td>8342</td><td>8379</td><td>38</td><td>CGTAAGGGAAAGTTGAAAAGAACTTTGAAGAGAGAGTT</td><td>161.3</td><td>4.5</td><td>1.3</td><td>3.3</td><td>8.6E-04</td><td>2.2E-03</td><td>BT20/MB468</td><td>MP</td></tr>
</table>

| | | | | | | | | | | | | | |
|---|---|---|---|---|---|---|---|---|---|---|---|---|---|
| >hg38_MT_+_1621_3279_RNR2_16S_50ntFlanks@1183.1215.33\|AAATTTCGGTTGGGGCGACCTCGGAGCAGAACC | >hg38_MT_+_1621_3279_RNR2_16S_50ntFlanks | 1183 | 1215 | 33 | AAATTTCGGTTGGGGCGACCTCGGAGCAGAACC | 234.7 | 4.9 | 1.5 | 3.3 | 8.6E-04 | 2.2E-03 | BT20/MB468 | MP |
| >hg38_21_+_8433172_8446622_RNA45SN1_45S_50ntFlanks@12324.12343.20\|TTGGGTTTTAAGCAGGAGGT | >hg38_21_+_8433172_8446622_RNA45SN1_45S_50ntFlanks | 12324 | 12343 | 20 | TTGGGTTTTAAGCAGGAGGT | 157.7 | -5.2 | 1.6 | -3.3 | 8.6E-04 | 2.2E-03 | BT20/MB468 | MP |
| >hg38_21_+_8433172_8446622_RNA45SN1_45S_50ntFlanks@4724.4751.28\|ATTAATCAAGAACGAAAGTCGGAGGTTC | >hg38_21_+_8433172_8446622_RNA45SN1_45S_50ntFlanks | 4724 | 4751 | 28 | ATTAATCAAGAACGAAAGTCGGAGGTTC | 1284.9 | -5.8 | 1.7 | -3.3 | 8.6E-04 | 2.2E-03 | BT20/MB468 | MP |
| >hg38_21_+_8433172_8446622_RNA45SN1_45S_50ntFlanks@10411.10440.30\|CAGTCGGTCCTGAGAGATGGGCGAGCGCCG | >hg38_21_+_8433172_8446622_RNA45SN1_45S_50ntFlanks | 10411 | 10440 | 30 | CAGTCGGTCCTGAGAGATGGGCGAGCGCCG | 358.0 | -5.6 | 1.7 | -3.3 | 8.6E-04 | 2.2E-03 | BT20/MB468 | MP |
| >hg38_21_+_8433172_8446622_RNA45SN1_45S_50ntFlanks@10771.10791.21\|CAGGTCTCCAAGGTGAACAGC | >hg38_21_+_8433172_8446622_RNA45SN1_45S_50ntFlanks | 10771 | 10791 | 21 | CAGGTCTCCAAGGTGAACAGC | 149.1 | -5.2 | 1.5 | -3.3 | 8.6E-04 | 2.2E-03 | BT20/MB468 | MP |
| >hg38_1_-_228634819_228635039_RNA5S12_5S_50ntFlanks@126.170.45\|TACTTGGATGGGAGACCGCCTGGGAATACCGGGTGCTGTAGGCTT | >hg38_1_-_228634819_228635039_RNA5S12_5S_50ntFlanks | 126 | 170 | 45 | TACTTGGATGGGAGACCGCCTGGGAATACCGGGTGCTGTAGGCTT | 512.2 | 5.9 | 1.8 | 3.3 | 8.6E-04 | 2.2E-03 | BT20/MB468 | MP |
| >hg38_21_+_8433172_8446622_RNA45SN1_45S_50ntFlanks@5485.5519.35\|AGCGCTGAGAAGACGGTCGAACTTGACTATCTAGA | >hg38_21_+_8433172_8446622_RNA45SN1_45S_50ntFlanks | 5485 | 5519 | 35 | AGCGCTGAGAAGACGGTCGAACTTGACTATCTAGA | 93.6 | 9.9 | 2.7 | 3.6 | 2.9E-04 | 2.2E-03 | BT20/MB231 | MP |
| >hg38_21_+_8433172_8446622_RNA45SN1_45S_50ntFlanks@11205.11225.21\|GCGGCGGCGACTCTGGACGCG | >hg38_21_+_8433172_8446622_RNA45SN1_45S_50ntFlanks | 11205 | 11225 | 21 | GCGGCGGCGACTCTGGACGCG | 627.4 | -4.3 | 1.3 | -3.3 | 8.6E-04 | 2.2E-03 | BT20/MB468 | MP |
| >hg38_21_+_8433172_8446622_RNA45SN1_45S_50ntFlanks@10391.10411.21\|AACAGCAGTTGAACATGGGTC | >hg38_21_+_8433172_8446622_RNA45SN1_45S_50ntFlanks | 10391 | 10411 | 21 | AACAGCAGTTGAACATGGGTC | 238.9 | -1.6 | 0.5 | -3.1 | 1.7E-03 | 2.2E-03 | BT20/MB468 | CYTO |
| >trnaMT_ValTAC_MT_+_1602_1670@50.71.22\|ACTTAACTTGACCGCTCTGACC | >trnaMT_ValTAC_MT_+_1602_1670 | 50 | 71 | 22 | ACTTAACTTGACCGCTCTGACC | 2669.1 | 3.7 | 1.1 | 3.2 | 1.4E-03 | 2.2E-03 | BT20/MB468 | MITO |
| >hg38_21_+_8433172_8446622_RNA45SN1_45S_50ntFlanks@12083.12110.28\|CTCGCTTCTGGCGCCAAGCGCCCGGCCG | >hg38_21_+_8433172_8446622_RNA45SN1_45S_50ntFlanks | 12083 | 12110 | 28 | CTCGCTTCTGGCGCCAAGCGCCCGGCCG | 194.2 | -5.2 | 1.6 | -3.3 | 8.6E-04 | 2.2E-03 | BT20/MB468 | MP |
| >hg38_21_+_8433172_8446622_RNA45SN1_45S_50ntFlanks@10238.10255.18\|CGCTACGCCGCGACGAGT | >hg38_21_+_8433172_8446622_RNA45SN1_45S_50ntFlanks | 10238 | 10255 | 18 | CGCTACGCCGCGACGAGT | 166.8 | 5.7 | 1.7 | 3.3 | 8.6E-04 | 2.2E-03 | BT20/MB468 | MP |
| >hg38_21_+_8433172_8446622_RNA45SN1_45S_50ntFlanks@10348.10390.43\|AAACGAGAACTTTGAAGGCCGAAGTGGAGAAGGGTTCCATGTG | >hg38_21_+_8433172_8446622_RNA45SN1_45S_50ntFlanks | 10348 | 10390 | 43 | AAACGAGAACTTTGAAGGCCGAAGTGGAGAAGGGTTCCATGTG | 549.2 | -3.3 | 1.1 | -3.1 | 1.8E-03 | 2.2E-03 | MB231/MB468 | MITO |
| >hg38_21_+_8433172_8446622_RNA45SN1_45S_50ntFlanks@12413.12443.31\|TCCTTCGATGTCGGCTCTTCCTATCATTGTG | >hg38_21_+_8433172_8446622_RNA45SN1_45S_50ntFlanks | 12413 | 12443 | 31 | TCCTTCGATGTCGGCTCTTCCTATCATTGTG | 320.9 | -2.9 | 0.9 | -3.1 | 1.8E-03 | 2.2E-03 | MB231/MB468 | MITO |
| >hg38_21_+_8433172_8446622_RNA45SN1_45S_50ntFlanks@12405.12426.22\|CTTTTTGATCCTTCGATGTCGG | >hg38_21_+_8433172_8446622_RNA45SN1_45S_50ntFlanks | 12405 | 12426 | 22 | CTTTTTGATCCTTCGATGTCGG | 2488.7 | -6.1 | 1.8 | -3.3 | 8.6E-04 | 2.2E-03 | BT20/MB468 | MP |
| >hg38_MT_+_598_1651_RNR1_12S_50ntFlanks@972.1004.33\|GGTAAGTGTACTGGAAAGTGCACTTGGACGAAC | >hg38_MT_+_598_1651_RNR1_12S_50ntFlanks | 972 | 1004 | 33 | GGTAAGTGTACTGGAAAGTGCACTTGGACGAAC | 273.3 | 3.9 | 1.2 | 3.2 | 1.5E-03 | 2.2E-03 | MB231/MB468 | MP |
| >MI0000298\|hsa-mir-221&WithFlank&X\|-\|45746151\|45746272@71.92.22(+1U)&[MIMAT0000278&hsa-miR-221-3p&offsets\|0\|-1(+1U);m-91&X\|-\|45746181\|45746202&offsets\|0\|0(+1U)]\|AGCTACATTGTCTGCTGGGTTTT | >MI0000298\|hsa-mir-221&WithFlank&X\|-\|45746151\|45746272 | 71 | 92 | 22(+1U) | AGCTACATTGTCTGCTGGGTTTT | 17207.1 | -3.6 | 1.0 | -3.6 | 2.9E-04 | 2.2E-03 | BT20/MB231 | MP |
| >MI0003177\|hsa-mir-522&WithFlank&19\|+\|53751205\|53751303@60.81.22&[MIMAT0002868&hsa-miR-522-3p&offsets\|0\|0;m-909&19\|+\|53751264\|53751285&offsets\|0\|0]\|AAAATGGTTCCCTTTAGAGTGT | >MI0003177\|hsa-mir-522&WithFlank&19\|+\|53751205\|53751303 | 60 | 81 | 22 | AAAATGGTTCCCTTTAGAGTGT | 246.3 | -5.1 | 1.4 | -3.6 | 2.9E-04 | 2.2E-03 | BT20/MB231 | MP |
| >hg38_21_+_8433172_8446622_RNA45SN1_45S_50ntFlanks@6651.6683.33\|CGACTCTTAGCGGTGGATCACTCGGCTCGTGCG | >hg38_21_+_8433172_8446622_RNA45SN1_45S_50ntFlanks | 6651 | 6683 | 33 | CGACTCTTAGCGGTGGATCACTCGGCTCGTGCG | 588.9 | 2.1 | 0.7 | 3.1 | 1.7E-03 | 2.2E-03 | MB231/MB468 | CYTO |
| >hg38_21_+_8433172_8446622_RNA45SN1_45S_50ntFlanks@9916.9948.33\|AAAGGTGTTGGTTGATATAGACAGCAGGACGGT | >hg38_21_+_8433172_8446622_RNA45SN1_45S_50ntFlanks | 9916 | 9948 | 33 | AAAGGTGTTGGTTGATATAGACAGCAGGACGGT | 56.9 | 6.2 | 1.9 | 3.3 | 8.6E-04 | 2.2E-03 | BT20/MB468 | MP |
| >hg38_MT_+_1621_3279_RNR2_16S_50ntFlanks@116.155.40\|AATAAAGTATAGGCGATAGAAATTGAAACCTGGCGCAATA | >hg38_MT_+_1621_3279_RNR2_16S_50ntFlanks | 116 | 155 | 40 | AATAAAGTATAGGCGATAGAAATTGAAACCTGGCGCAATA | 46.7 | 9.3 | 2.8 | 3.3 | 8.6E-04 | 2.2E-03 | BT20/MB468 | MP |
| >hg38_21_+_8433172_8446622_RNA45SN1_45S_50ntFlanks@5473.5514.42\|CGGCCCTGGCGGAGCGCTGAGAAGACGGTCGAACTTGACTAT | >hg38_21_+_8433172_8446622_RNA45SN1_45S_50ntFlanks | 5473 | 5514 | 42 | CGGCCCTGGCGGAGCGCTGAGAAGACGGTCGAACTTGACTAT | 666.3 | -2.5 | 0.8 | -3.1 | 1.8E-03 | 2.2E-03 | MB231/MB468 | NUC |
| >hg38_21_+_8433172_8446622_RNA45SN1_45S_50ntFlanks@12008.12032.25\|CCGCCGGTGAAATACCACTACTCTG | >hg38_21_+_8433172_8446622_RNA45SN1_45S_50ntFlanks | 12008 | 12032 | 25 | CCGCCGGTGAAATACCACTACTCTG | 15044.1 | -6.2 | 2.0 | -3.1 | 1.7E-03 | 2.2E-03 | BT20/MB468 | CYTO |
| >hg38_21_+_8433172_8446622_RNA45SN1_45S_50ntFlanks@4733.4759.27\|GAACGAAAGTCGGAGGTTCGAAGACGA | >hg38_21_+_8433172_8446622_RNA45SN1_45S_50ntFlanks | 4733 | 4759 | 27 | GAACGAAAGTCGGAGGTTCGAAGACGA | 3217.5 | 5.7 | 1.6 | 3.6 | 3.0E-04 | 2.2E-03 | BT20/MB231 | MP |
| >hg38_21_+_8433172_8446622_RNA45SN1_45S_50ntFlanks@4560.4587.28\|CTAGGAATAATGGAATAGGACCGCGGTT | >hg38_21_+_8433172_8446622_RNA45SN1_45S_50ntFlanks | 4560 | 4587 | 28 | CTAGGAATAATGGAATAGGACCGCGGTT | 490.8 | -5.5 | 1.7 | -3.3 | 8.7E-04 | 2.2E-03 | BT20/MB468 | MP |
| >hg38_21_+_8433172_8446622_RNA45SN1_45S_50ntFlanks@11565.11584.20\|CTAGCAGCCGACTTAGAACT | >hg38_21_+_8433172_8446622_RNA45SN1_45S_50ntFlanks | 11565 | 11584 | 20 | CTAGCAGCCGACTTAGAACT | 130.0 | 4.5 | 1.3 | 3.3 | 8.7E-04 | 2.2E-03 | BT20/MB468 | MP |
| >hg38_21_+_8433172_8446622_RNA45SN1_45S_50ntFlanks@11667.11704.38\|CTGCCCAGTGCTCTGAATGTCAAAGTGAAGAAATTCAA | >hg38_21_+_8433172_8446622_RNA45SN1_45S_50ntFlanks | 11667 | 11704 | 38 | CTGCCCAGTGCTCTGAATGTCAAAGTGAAGAAATTCAA | 297.5 | 3.0 | 0.8 | 3.6 | 3.8E-04 | 2.2E-03 | BT20/MB231 | MITO |
| >trna10_LysCTT_16_+_3241501_3241573@7.35.29,trna13_LysCTT_14_-_58706613_58706685@7.35.29,trna2_LysCTT_15_+_79152904_79152976@7.35.29,trna4_LysCTT_18_-_43669271_43669343@7.35.29\|CTAGCTCAGTCGGTAGAGCATGGGACTCT | >trna10_LysCTT_16_+_3241501_3241573 | 7 | 35 | 29 | CTAGCTCAGTCGGTAGAGCATGGGACTCT | 288.7 | -7.0 | 2.0 | -3.6 | 3.8E-04 | 2.2E-03 | BT20/MB231 | MITO |
| >hg38_21_+_8433172_8446622_RNA45SN1_45S_50ntFlanks@12420.12461.42\|ATGTCGGCTCTTCCTATCATTGTGAAGCAGAATTCACCAAGC | >hg38_21_+_8433172_8446622_RNA45SN1_45S_50ntFlanks | 12420 | 12461 | 42 | ATGTCGGCTCTTCCTATCATTGTGAAGCAGAATTCACCAAGC | 468.9 | -2.9 | 0.8 | -3.6 | 3.8E-04 | 2.2E-03 | BT20/MB231 | MITO |
| >trnaMT_HisGTG_MT_+_12138_12206@32.63.32\|TGAATCTGACAACAGAGGCTTACGACCCCTTA | >trnaMT_HisGTG_MT_+_12138_12206 | 32 | 63 | 32 | TGAATCTGACAACAGAGGCTTACGACCCCTTA | 4032.7 | 5.4 | 1.7 | 3.2 | 1.4E-03 | 2.2E-03 | BT20/MB468 | MITO |
| >hg38_21_+_8433172_8446622_RNA45SN1_45S_50ntFlanks@5473.5516.44\|CGGCCCTGGCGGAGCGCTGAGAAGACGGTCGAACTTGACTATCT | >hg38_21_+_8433172_8446622_RNA45SN1_45S_50ntFlanks | 5473 | 5516 | 44 | CGGCCCTGGCGGAGCGCTGAGAAGACGGTCGAACTTGACTATCT | 1189.1 | 6.3 | 1.9 | 3.3 | 8.7E-04 | 2.2E-03 | BT20/MB468 | MP |
| >hg38_21_+_8433172_8446622_RNA45SN1_45S_50ntFlanks@5473.5499.27\|CGGCCCTGGCGGAGCGCTGAGAAGACG | >hg38_21_+_8433172_8446622_RNA45SN1_45S_50ntFlanks | 5473 | 5499 | 27 | CGGCCCTGGCGGAGCGCTGAGAAGACG | 5700.8 | 2.9 | 0.9 | 3.1 | 1.7E-03 | 2.2E-03 | BT20/MB468 | CYTO |
| >hg38_21_+_8433172_8446622_RNA45SN1_45S_50ntFlanks@11578.11601.24\|TAGAACTGGTGCGGACCAGGGGAA | >hg38_21_+_8433172_8446622_RNA45SN1_45S_50ntFlanks | 11578 | 11601 | 24 | TAGAACTGGTGCGGACCAGGGGAA | 275.5 | -2.5 | 0.8 | -3.1 | 1.7E-03 | 2.2E-03 | BT20/MB468 | CYTO |
| >hg38_21_+_8433172_8446622_RNA45SN1_45S_50ntFlanks@7976.8016.41\|GCGACCTCAGATCAGACGTGGCGACCCGCTGAATTTAAGCA | >hg38_21_+_8433172_8446622_RNA45SN1_45S_50ntFlanks | 7976 | 8016 | 41 | GCGACCTCAGATCAGACGTGGCGACCCGCTGAATTTAAGCA | 212.0 | 2.9 | 0.9 | 3.1 | 1.8E-03 | 2.2E-03 | MB231/MB468 | MITO |
| >hg38_21_+_8433172_8446622_RNA45SN1_45S_50ntFlanks@7976.8005.30\|GCGACCTCAGATCAGACGTGGCGACCCGCT | >hg38_21_+_8433172_8446622_RNA45SN1_45S_50ntFlanks | 7976 | 8005 | 30 | GCGACCTCAGATCAGACGTGGCGACCCGCT | 426.6 | 5.4 | 1.5 | 3.6 | 3.0E-04 | 2.2E-03 | BT20/MB231 | MP |
| >hg38_21_+_8433172_8446622_RNA45SN1_45S_50ntFlanks@11663.11685.23\|ATTTCTGCCCAGTGCTCTGAATG | >hg38_21_+_8433172_8446622_RNA45SN1_45S_50ntFlanks | 11663 | 11685 | 23 | ATTTCTGCCCAGTGCTCTGAATG | 1135.9 | -3.7 | 1.1 | -3.3 | 8.7E-04 | 2.2E-03 | BT20/MB468 | MP |
| >hg38_21_+_8433172_8446622_RNA45SN1_45S_50ntFlanks@12418.12451.34\|CGATGTCGGCTCTTCCTATCATTGTGAAGCAGAA | >hg38_21_+_8433172_8446622_RNA45SN1_45S_50ntFlanks | 12418 | 12451 | 34 | CGATGTCGGCTCTTCCTATCATTGTGAAGCAGAA | 219.8 | -2.1 | 0.7 | -3.1 | 1.7E-03 | 2.2E-03 | BT20/MB468 | CYTO |
| >hg38_21_+_8433172_8446622_RNA45SN1_45S_50ntFlanks@11642.11660.19\|GGCGGGTGTTGACGCGATG | >hg38_21_+_8433172_8446622_RNA45SN1_45S_50ntFlanks | 11642 | 11660 | 19 | GGCGGGTGTTGACGCGATG | 174.2 | -3.7 | 1.2 | -3.2 | 1.4E-03 | 2.2E-03 | BT20/MB468 | MITO |
| >hg38_21_+_8433172_8446622_RNA45SN1_45S_50ntFlanks@12621.12654.34\|GAGCCAATGGGGCGAAGCTACCATCTGTGGGATT | >hg38_21_+_8433172_8446622_RNA45SN1_45S_50ntFlanks | 12621 | 12654 | 34 | GAGCCAATGGGGCGAAGCTACCATCTGTGGGATT | 120.3 | -2.0 | 0.6 | -3.1 | 1.7E-03 | 2.2E-03 | BT20/MB468 | CYTO |
| >hg38_21_+_8433172_8446622_RNA45SN1_45S_50ntFlanks@11843.11860.18\|AGGGAACGGGCTTGGCGG | >hg38_21_+_8433172_8446622_RNA45SN1_45S_50ntFlanks | 11843 | 11860 | 18 | AGGGAACGGGCTTGGCGG | 128.4 | -3.4 | 1.1 | -3.1 | 1.7E-03 | 2.2E-03 | BT20/MB468 | CYTO |
| >hg38_21_+_8433172_8446622_RNA45SN1_45S_50ntFlanks@8117.8137.21\|CGGGACATGTGGCGTACGGAA | >hg38_21_+_8433172_8446622_RNA45SN1_45S_50ntFlanks | 8117 | 8137 | 21 | CGGGACATGTGGCGTACGGAA | 276.6 | 4.0 | 1.2 | 3.3 | 8.7E-04 | 2.2E-03 | BT20/MB468 | MP |
| >am_trnaMT_PheGAA_MT_+_577_647@41.63.23\|GTTTAGACGGGCTCACATCACCC | >am_trnaMT_PheGAA_MT_+_577_647 | 41 | 63 | 23 | GTTTAGACGGGCTCACATCACCC | 194.5 | 4.7 | 1.4 | 3.3 | 8.7E-04 | 2.2E-03 | BT20/MB468 | MP |
| >hg38_21_+_8433172_8446622_RNA45SN1_45S_50ntFlanks@8170.8215.46\|CAAGTCCTTCTGATCGAGGCCCAGCCCGTGGACGGTGTGAGGCCGG | >hg38_21_+_8433172_8446622_RNA45SN1_45S_50ntFlanks | 8170 | 8215 | 46 | CAAGTCCTTCTGATCGAGGCCCAGCCCGTGGACGGTGTGAGGCCGG | 1675.9 | -4.3 | 1.3 | -3.2 | 1.4E-03 | 2.2E-03 | BT20/MB468 | MITO |
| >hg38_21_+_8433172_8446622_RNA45SN1_45S_50ntFlanks@4999.5016.18\|ATTGATAGCTCTTTCTCG | >hg38_21_+_8433172_8446622_RNA45SN1_45S_50ntFlanks | 4999 | 5016 | 18 | ATTGATAGCTCTTTCTCG | 295.6 | -6.1 | 1.9 | -3.2 | 1.4E-03 | 2.2E-03 | BT20/MB468 | MITO |
| >hg38_21_+_8433172_8446622_RNA45SN1_45S_50ntFlanks@4722.4755.34\|TCATTAATCAAGAACGAAAGTCGGAGGTTCGAAG | >hg38_21_+_8433172_8446622_RNA45SN1_45S_50ntFlanks | 4722 | 4755 | 34 | TCATTAATCAAGAACGAAAGTCGGAGGTTCGAAG | 109.7 | -4.6 | 1.5 | -3.1 | 1.7E-03 | 2.2E-03 | BT20/MB468 | CYTO |
| >hg38_21_+_8433172_8446622_RNA45SN1_45S_50ntFlanks@6647.6675.29\|CGTACGACTCTTAGCGGTGGATCACTCGG | >hg38_21_+_8433172_8446622_RNA45SN1_45S_50ntFlanks | 6647 | 6675 | 29 | CGTACGACTCTTAGCGGTGGATCACTCGG | 416.4 | -1.6 | 0.5 | -3.1 | 1.8E-03 | 2.2E-03 | BT20/MB468 | NUC |
| >hg38_21_+_8433172_8446622_RNA45SN1_45S_50ntFlanks@10818.10840.23\|AAGGGAAGTCGGCAAGCCGGATC | >hg38_21_+_8433172_8446622_RNA45SN1_45S_50ntFlanks | 10818 | 10840 | 23 | AAGGGAAGTCGGCAAGCCGGATC | 925.8 | 6.8 | 1.9 | 3.6 | 3.0E-04 | 2.2E-03 | BT20/MB231 | MP |
| >hg38_21_+_8433172_8446622_RNA45SN1_45S_50ntFlanks@4738.4759.22\|AAAGTCGGAGGTTCGAAGACGA | >hg38_21_+_8433172_8446622_RNA45SN1_45S_50ntFlanks | 4738 | 4759 | 22 | AAAGTCGGAGGTTCGAAGACGA | 314.3 | -2.2 | 0.7 | -3.2 | 1.4E-03 | 2.2E-03 | BT20/MB468 | MITO |
| >hg38_21_+_8433172_8446622_RNA45SN1_45S_50ntFlanks@10794.10825.32\|CTGGCATGTTGGAACAATGTAGGTAAGGGAAG | >hg38_21_+_8433172_8446622_RNA45SN1_45S_50ntFlanks | 10794 | 10825 | 32 | CTGGCATGTTGGAACAATGTAGGTAAGGGAAG | 424.9 | -5.5 | 1.6 | -3.3 | 8.7E-04 | 2.2E-03 | BT20/MB468 | MP |
| >hg38_21_+_8433172_8446622_RNA45SN1_45S_50ntFlanks@11199.11223.25\|CCGGCGGCGGCGGCGACTCTGGACG | >hg38_21_+_8433172_8446622_RNA45SN1_45S_50ntFlanks | 11199 | 11223 | 25 | CCGGCGGCGGCGGCGACTCTGGACG | 293.2 | -2.2 | 0.7 | -3.2 | 1.4E-03 | 2.2E-03 | BT20/MB468 | MITO |
| >hg38_21_+_8433172_8446622_RNA45SN1_45S_50ntFlanks@8191.8213.23\|CAGCCCGTGGACGGTGTGAGGCC | >hg38_21_+_8433172_8446622_RNA45SN1_45S_50ntFlanks | 8191 | 8213 | 23 | CAGCCCGTGGACGGTGTGAGGCC | 414.4 | -1.6 | 0.5 | -3.1 | 1.7E-03 | 2.2E-03 | BT20/MB468 | CYTO |
| >trnaMT_PheGAA_MT_+_577_647@35.56.22\|GAAAATGTTTAGACGGGCTCAC | >trnaMT_PheGAA_MT_+_577_647 | 35 | 56 | 22 | GAAAATGTTTAGACGGGCTCAC | 319.0 | 2.7 | 0.9 | 3.2 | 1.5E-03 | 2.2E-03 | MB231/MB468 | MP |

| | | | | | | | | | | | | |
|---|---|---|---|---|---|---|---|---|---|---|---|---|
| >hg38_21_+_8433172_8446622_RNA45SN1_45S_50ntFlanks@11917.11943.27\|GACATGAGAGGTGTAGAATAAGTGGGA | >hg38_21_+_8433172_8446622_RNA45SN1_45S_50ntFlanks | 11917 | 11943 | 27 | GACATGAGAGGTGTAGAATAAGTGGGA | 42.0 | 9.1 | 2.7 | 3.3 | 8.7E-04 | 2.2E-03 | BT20/MB468 | MP |
| >MI0000299\|hsa-mir-222&WithFlank&X\|-\|45747009\|45747130@75.98.24(+1C)&[MIMAT0000279&hsa-miR-222-3p&offsets\|0\|+3(+1C);m-140&X\|-\|45747033\|45747056&offsets\|0\|0(+1C)]\|AGCTACATCTGGCTACTGGGTCTCC | >MI0000299\|hsa-mir-222&WithFlank&X\|-\|45747009\|45747130 | 75 | 98 | 24(+1C) | AGCTACATCTGGCTACTGGGTCTCC | 5612.0 | -2.3 | 0.6 | -3.6 | 3.3E-04 | 2.2E-03 | BT20/MB231 | NUC |
| >trna119_LysCTT_1_-_145395522_145395594@4.33.30,trna11_LysCTT_5_-_180648979_180649051@4.33.30,trna13_LysCTT_6_+_26556774_26556846@4.33.30,trna7_LysCTT_16_+_3225692_3225764@4.33.30,trna9_LysCTT_5_+_180634755_180634827@4.33.30\|CGGCTAGCTCAGTCGGTAGAGCATGAGACT | >trna119_LysCTT_1_-_145395522_145395594 | 4 | 33 | 30 | CGGCTAGCTCAGTCGGTAGAGCATGAGACT | 1097.5 | -3.8 | 1.1 | -3.6 | 3.3E-04 | 2.2E-03 | BT20/MB231 | NUC |
| >hg38_21_+_8433172_8446622_RNA45SN1_45S_50ntFlanks@11319.11343.25\|CCCACGTCTCGTCGCGCGCGCGTCC | >hg38_21_+_8433172_8446622_RNA45SN1_45S_50ntFlanks | 11319 | 11343 | 25 | CCCACGTCTCGTCGCGCGCGCGTCC | 1385.8 | -6.3 | 2.0 | -3.1 | 1.8E-03 | 2.2E-03 | BT20/MB468 | NUC |
| >MI0000299\|hsa-mir-222&WithFlank&X\|-\|45747009\|45747130@75.99.25&[MIMAT0000279&hsa-miR-222-3p&offsets\|0\|+4;m-140&X\|-\|45747033\|45747056&offsets\|0\|+1]\|AGCTACATCTGGCTACTGGGTCTCT | >MI0000299\|hsa-mir-222&WithFlank&X\|-\|45747009\|45747130 | 75 | 99 | 25 | AGCTACATCTGGCTACTGGGTCTCT | 26336.2 | -2.3 | 0.6 | -3.6 | 3.3E-04 | 2.2E-03 | BT20/MB231 | NUC |
| >hg38_21_+_8433172_8446622_RNA45SN1_45S_50ntFlanks@10784.10816.33\|TGAACAGCCTCTGGCATGTTGGAACAATGTAGG | >hg38_21_+_8433172_8446622_RNA45SN1_45S_50ntFlanks | 10784 | 10816 | 33 | TGAACAGCCTCTGGCATGTTGGAACAATGTAGG | 218.8 | -2.4 | 0.8 | -3.1 | 1.7E-03 | 2.2E-03 | MB231/MB468 | CYTO |
| >hg38_21_+_8433172_8446622_RNA45SN1_45S_50ntFlanks@5481.5501.21\|GCGGAGCGCTGAGAAGACGGT | >hg38_21_+_8433172_8446622_RNA45SN1_45S_50ntFlanks | 5481 | 5501 | 21 | GCGGAGCGCTGAGAAGACGGT | 67.3 | 5.6 | 1.7 | 3.3 | 8.8E-04 | 2.2E-03 | BT20/MB468 | MP |
| >hg38_21_+_8433172_8446622_RNA45SN1_45S_50ntFlanks@12933.12951.19\|GCTAAACCATTCGTAGACG | >hg38_21_+_8433172_8446622_RNA45SN1_45S_50ntFlanks | 12933 | 12951 | 19 | GCTAAACCATTCGTAGACG | 90.1 | -4.9 | 1.6 | -3.2 | 1.6E-03 | 2.2E-03 | MB231/MB468 | MP |
| >hg38_21_+_8433172_8446622_RNA45SN1_45S_50ntFlanks@10735.10769.35\|TGTAAATCTCGCGCCGGGCCGTACCCATATCCGCA | >hg38_21_+_8433172_8446622_RNA45SN1_45S_50ntFlanks | 10735 | 10769 | 35 | TGTAAATCTCGCGCCGGGCCGTACCCATATCCGCA | 229.2 | -3.8 | 1.2 | -3.1 | 1.9E-03 | 2.2E-03 | MB231/MB468 | NUC |
| >hg38_21_+_8433172_8446622_RNA45SN1_45S_50ntFlanks@11541.11583.43\|GCGGCGCCTCGCCTCGGCCGGCGCCTAGCAGCCGACTTAGAAC | >hg38_21_+_8433172_8446622_RNA45SN1_45S_50ntFlanks | 11541 | 11583 | 43 | GCGGCGCCTCGCCTCGGCCGGCGCCTAGCAGCCGACTTAGAAC | 217.8 | -2.7 | 0.8 | -3.6 | 3.5E-04 | 2.2E-03 | BT20/MB231 | CYTO |
| >hg38_21_+_8433172_8446622_RNA45SN1_45S_50ntFlanks@10357.10377.21\|CTTTGAAGGCCGAAGTGGAGA | >hg38_21_+_8433172_8446622_RNA45SN1_45S_50ntFlanks | 10357 | 10377 | 21 | CTTTGAAGGCCGAAGTGGAGA | 167.0 | -1.9 | 0.6 | -3.1 | 1.7E-03 | 2.2E-03 | BT20/MB468 | CYTO |
| >hg38_21_+_8433172_8446622_RNA45SN1_45S_50ntFlanks@8116.8139.24\|GCGGGACATGTGGCGTACGGAAGA | >hg38_21_+_8433172_8446622_RNA45SN1_45S_50ntFlanks | 8116 | 8139 | 24 | GCGGGACATGTGGCGTACGGAAGA | 461.1 | 4.0 | 1.2 | 3.3 | 8.8E-04 | 2.2E-03 | BT20/MB468 | MP |
| >hg38_MT_+_1621_3279_RNR2_16S_50ntFlanks@1370.1414.45\|ATGTTGGATCAGGACATCCCGATGGTGCAGCCGCTATTAAAGGTT | >hg38_MT_+_1621_3279_RNR2_16S_50ntFlanks | 1370 | 1414 | 45 | ATGTTGGATCAGGACATCCCGATGGTGCAGCCGCTATTAAAGGTT | 289.1 | 6.0 | 1.8 | 3.3 | 8.8E-04 | 2.2E-03 | BT20/MB468 | MP |
| >hg38_21_+_8433172_8446622_RNA45SN1_45S_50ntFlanks@9570.9605.36\|TGCAAATCGGTCGTCCGACCTGGGTATAGGGGCGAA | >hg38_21_+_8433172_8446622_RNA45SN1_45S_50ntFlanks | 9570 | 9605 | 36 | TGCAAATCGGTCGTCCGACCTGGGTATAGGGGCGAA | 131.6 | -4.4 | 1.4 | -3.2 | 1.6E-03 | 2.2E-03 | MB231/MB468 | MP |
| >hg38_21_+_8433172_8446622_RNA45SN1_45S_50ntFlanks@12420.12443.24\|ATGTCGGCTCTTCCTATCATTGTG | >hg38_21_+_8433172_8446622_RNA45SN1_45S_50ntFlanks | 12420 | 12443 | 24 | ATGTCGGCTCTTCCTATCATTGTG | 6582.1 | -2.3 | 0.7 | -3.1 | 1.7E-03 | 2.2E-03 | MB231/MB468 | CYTO |
| >hg38_21_+_8433172_8446622_RNA45SN1_45S_50ntFlanks@11873.11898.26\|AAGAAGACCCTGTTGAGCTTGACTCT | >hg38_21_+_8433172_8446622_RNA45SN1_45S_50ntFlanks | 11873 | 11898 | 26 | AAGAAGACCCTGTTGAGCTTGACTCT | 735.1 | -1.8 | 0.6 | -3.2 | 1.4E-03 | 2.2E-03 | BT20/MB468 | MITO |
| >hg38_21_+_8433172_8446622_RNA45SN1_45S_50ntFlanks@8116.8137.22\|GCGGGACATGTGGCGTACGGAA | >hg38_21_+_8433172_8446622_RNA45SN1_45S_50ntFlanks | 8116 | 8137 | 22 | GCGGGACATGTGGCGTACGGAA | 760.5 | 7.6 | 2.3 | 3.3 | 8.8E-04 | 2.2E-03 | BT20/MB468 | MP |
| >hg38_21_+_8433172_8446622_RNA45SN1_45S_50ntFlanks@6646.6688.43\|TCGTACGACTCTTAGCGGTGGATCACTCGGCTCGTGCGTCGAT | >hg38_21_+_8433172_8446622_RNA45SN1_45S_50ntFlanks | 6646 | 6688 | 43 | TCGTACGACTCTTAGCGGTGGATCACTCGGCTCGTGCGTCGAT | 515.4 | 2.9 | 0.8 | 3.6 | 3.3E-04 | 2.2E-03 | BT20/MB231 | NUC |
| >hg38_21_+_8433172_8446622_RNA45SN1_45S_50ntFlanks@9533.9566.34\|GAGGAAACTCTGGTGGAGGTCCGTAGCGGTCCTG | >hg38_21_+_8433172_8446622_RNA45SN1_45S_50ntFlanks | 9533 | 9566 | 34 | GAGGAAACTCTGGTGGAGGTCCGTAGCGGTCCTG | 375.0 | -3.5 | 1.1 | -3.1 | 1.8E-03 | 2.2E-03 | MB231/MB468 | CYTO |
| >hg38_21_+_8433172_8446622_RNA45SN1_45S_50ntFlanks@10384.10430.47\|CCATGTGAACAGCAGTTGAACATGGGTCAGTCGGTCCTGAGAGATGG | >hg38_21_+_8433172_8446622_RNA45SN1_45S_50ntFlanks | 10384 | 10430 | 47 | CCATGTGAACAGCAGTTGAACATGGGTCAGTCGGTCCTGAGAGATGG | 159.5 | -2.8 | 0.9 | -3.1 | 1.8E-03 | 2.2E-03 | MB231/MB468 | CYTO |
| >hg38_MT_+_1621_3279_RNR2_16S_50ntFlanks@1176.1215.40\|GCATTAAAAATTTCGGTTGGGGCGACCTCGGAGCAGAACC | >hg38_MT_+_1621_3279_RNR2_16S_50ntFlanks | 1176 | 1215 | 40 | GCATTAAAAATTTCGGTTGGGGCGACCTCGGAGCAGAACC | 98.7 | 5.5 | 1.8 | 3.1 | 1.9E-03 | 2.2E-03 | MB231/MB468 | MITO |
| >hg38_21_+_8433172_8446622_RNA45SN1_45S_50ntFlanks@6670.6698.29\|ACTCGGCTCGTGCGTCGATGAAGAACGCA | >hg38_21_+_8433172_8446622_RNA45SN1_45S_50ntFlanks | 6670 | 6698 | 29 | ACTCGGCTCGTGCGTCGATGAAGAACGCA | 211.6 | -2.0 | 0.6 | -3.1 | 1.9E-03 | 2.2E-03 | MB231/MB468 | NUC |
| >trnaMT_SerGCT_MT_+_12207_12265@14.41.28\|AGAACTGCTAACTCATGCCCCCATGTCT | >trnaMT_SerGCT_MT_+_12207_12265 | 14 | 41 | 28 | AGAACTGCTAACTCATGCCCCCATGTCT | 184.3 | 3.2 | 1.0 | 3.2 | 1.6E-03 | 2.2E-03 | MB231/MB468 | MP |
| >hg38_21_+_8433172_8446622_RNA45SN1_45S_50ntFlanks@10350.10391.42\|ACGAGAACTTTGAAGGCCGAAGTGGAGAAGGGTTCCATGTGA | >hg38_21_+_8433172_8446622_RNA45SN1_45S_50ntFlanks | 10350 | 10391 | 42 | ACGAGAACTTTGAAGGCCGAAGTGGAGAAGGGTTCCATGTGA | 218.1 | -5.3 | 1.7 | -3.2 | 1.6E-03 | 2.2E-03 | MB231/MB468 | MP |
| >hg38_21_+_8433172_8446622_RNA45SN1_45S_50ntFlanks@5472.5499.28\|ACGGCCCTGGCGGAGCGCTGAGAAGACG | >hg38_21_+_8433172_8446622_RNA45SN1_45S_50ntFlanks | 5472 | 5499 | 28 | ACGGCCCTGGCGGAGCGCTGAGAAGACG | 10884.0 | 3.2 | 1.0 | 3.1 | 1.7E-03 | 2.2E-03 | BT20/MB468 | CYTO |
| >hg38_21_+_8433172_8446622_RNA45SN1_45S_50ntFlanks@5492.5525.34\|AGAAGACGGTCGAACTTGACTATCTAGAGGAAGT | >hg38_21_+_8433172_8446622_RNA45SN1_45S_50ntFlanks | 5492 | 5525 | 34 | AGAAGACGGTCGAACTTGACTATCTAGAGGAAGT | 646.8 | 4.0 | 1.1 | 3.6 | 3.3E-04 | 2.2E-03 | BT20/MB231 | NUC |
| >hg38_21_+_8433172_8446622_RNA45SN1_45S_50ntFlanks@11569.11605.37\|CAGCCGACTTAGAACTGGTGCGGACCAGGGGAATCCG | >hg38_21_+_8433172_8446622_RNA45SN1_45S_50ntFlanks | 11569 | 11605 | 37 | CAGCCGACTTAGAACTGGTGCGGACCAGGGGAATCCG | 348.2 | -4.7 | 1.5 | -3.1 | 1.7E-03 | 2.2E-03 | BT20/MB468 | CYTO |
| >hg38_21_+_8433172_8446622_RNA45SN1_45S_50ntFlanks@6661.6688.28\|CGGTGGATCACTCGGCTCGTGCGTCGAT | >hg38_21_+_8433172_8446622_RNA45SN1_45S_50ntFlanks | 6661 | 6688 | 28 | CGGTGGATCACTCGGCTCGTGCGTCGAT | 3481.4 | 2.6 | 0.8 | 3.3 | 8.9E-04 | 2.2E-03 | BT20/MB468 | MP |
| >hg38_21_+_8433172_8446622_RNA45SN1_45S_50ntFlanks@5485.5518.34\|AGCGCTGAGAAGACGGTCGAACTTGACTATCTAG | >hg38_21_+_8433172_8446622_RNA45SN1_45S_50ntFlanks | 5485 | 5518 | 34 | AGCGCTGAGAAGACGGTCGAACTTGACTATCTAG | 1419.6 | -5.9 | 1.8 | -3.3 | 8.9E-04 | 2.2E-03 | BT20/MB468 | MP |
| >hg38_21_+_8433172_8446622_RNA45SN1_45S_50ntFlanks@11568.11613.46\|GCAGCCGACTTAGAACTGGTGCGGACCAGGGGAATCCGACTGTTTA | >hg38_21_+_8433172_8446622_RNA45SN1_45S_50ntFlanks | 11568 | 11613 | 46 | GCAGCCGACTTAGAACTGGTGCGGACCAGGGGAATCCGACTGTTTA | 867.2 | 5.8 | 1.8 | 3.2 | 1.4E-03 | 2.2E-03 | BT20/MB468 | MITO |
| >MI0000253\|hsa-mir-148a&WithFlank&7\|-\|25949913\|25949992@50.72.23&[MIMAT0000243&hsa-miR-148a-3p&offsets\|0\|+1;m-7&7\|-\|25949922\|25949943&offsets\|0\|+1]\|TCAGTGCACTACAGAACTTTGTC | >MI0000253\|hsa-mir-148a&WithFlank&7\|-\|25949913\|25949992 | 50 | 72 | 23 | TCAGTGCACTACAGAACTTTGTC | 1103.2 | 2.9 | 0.8 | 3.5 | 3.9E-04 | 2.2E-03 | BT20/MB231 | MITO |
| >hg38_21_+_8433172_8446622_RNA45SN1_45S_50ntFlanks@10353.10377.25\|AGAACTTTGAAGGCCGAAGTGGAGA | >hg38_21_+_8433172_8446622_RNA45SN1_45S_50ntFlanks | 10353 | 10377 | 25 | AGAACTTTGAAGGCCGAAGTGGAGA | 241.6 | -4.2 | 1.3 | -3.2 | 1.6E-03 | 2.2E-03 | MB231/MB468 | MP |
| >hg38_21_+_8433172_8446622_RNA45SN1_45S_50ntFlanks@5469.5514.46\|CCCACGGCCCTGGCGGAGCGCTGAGAAGACGGTCGAACTTGACTAT | >hg38_21_+_8433172_8446622_RNA45SN1_45S_50ntFlanks | 5469 | 5514 | 46 | CCCACGGCCCTGGCGGAGCGCTGAGAAGACGGTCGAACTTGACTAT | 50.9 | -9.5 | 3.0 | -3.2 | 1.6E-03 | 2.2E-03 | MB231/MB468 | MP |
| >MI0000810\|hsa-mir-135b&WithFlank&1\|-\|205448296\|205448404@22.44.23&[MIMAT0000758&hsa-miR-135b-5p&offsets\|0\|0;m-321&1\|-\|205448361\|205448383&offsets\|0\|0]\|TATGGCTTTTCATTCCTATGTGA | >MI0000810\|hsa-mir-135b&WithFlank&1\|-\|205448296\|205448404 | 22 | 44 | 23 | TATGGCTTTTCATTCCTATGTGA | 740.8 | 3.3 | 1.1 | 3.1 | 1.9E-03 | 2.2E-03 | BT20/MB468 | NUC |
| >hg38_21_+_8433172_8446622_RNA45SN1_45S_50ntFlanks@11896.11933.38\|TCTAGTCTGGCACGGTGAAGAGACATGAGAGGTGTAGA | >hg38_21_+_8433172_8446622_RNA45SN1_45S_50ntFlanks | 11896 | 11933 | 38 | TCTAGTCTGGCACGGTGAAGAGACATGAGAGGTGTAGA | 42.5 | 9.2 | 2.8 | 3.3 | 8.9E-04 | 2.2E-03 | BT20/MB468 | MP |
| >hg38_21_+_8433172_8446622_RNA45SN1_45S_50ntFlanks@5420.5444.25\|CCGATTGGATGGTTTAGTGAGGCCC | >hg38_21_+_8433172_8446622_RNA45SN1_45S_50ntFlanks | 5420 | 5444 | 25 | CCGATTGGATGGTTTAGTGAGGCCC | 311.3 | -5.5 | 1.7 | -3.3 | 8.9E-04 | 2.2E-03 | BT20/MB468 | MP |
| >hg38_21_+_8433172_8446622_RNA45SN1_45S_50ntFlanks@8194.8214.21\|CCCGTGGACGGTGTGAGGCCG | >hg38_21_+_8433172_8446622_RNA45SN1_45S_50ntFlanks | 8194 | 8214 | 21 | CCCGTGGACGGTGTGAGGCCG | 987.4 | -4.9 | 1.5 | -3.3 | 8.9E-04 | 2.2E-03 | BT20/MB468 | MP |
| >hg38_21_+_8433172_8446622_RNA45SN1_45S_50ntFlanks@9573.9592.20\|AAATCGGTCGTCCGACCTGG | >hg38_21_+_8433172_8446622_RNA45SN1_45S_50ntFlanks | 9573 | 9592 | 20 | AAATCGGTCGTCCGACCTGG | 643.8 | -4.9 | 1.5 | -3.3 | 8.9E-04 | 2.2E-03 | BT20/MB468 | MP |
| >hg38_21_+_8433172_8446622_RNA45SN1_45S_50ntFlanks@12223.12247.25\|CAGAAACCTCCCGTGGAGCAGAAGG | >hg38_21_+_8433172_8446622_RNA45SN1_45S_50ntFlanks | 12223 | 12247 | 25 | CAGAAACCTCCCGTGGAGCAGAAGG | 42.3 | -8.5 | 2.6 | -3.3 | 8.9E-04 | 2.2E-03 | BT20/MB468 | MP |
| >hg38_21_+_8433172_8446622_RNA45SN1_45S_50ntFlanks@12715.12753.39\|AGCCTCGGTTGGCCTCGGATAGCCGGTCCCCCGCCTGTC | >hg38_21_+_8433172_8446622_RNA45SN1_45S_50ntFlanks | 12715 | 12753 | 39 | AGCCTCGGTTGGCCTCGGATAGCCGGTCCCCCGCCTGTC | 50.6 | -9.5 | 3.0 | -3.2 | 1.6E-03 | 2.2E-03 | MB231/MB468 | MP |
| >hg38_21_+_8433172_8446622_RNA45SN1_45S_50ntFlanks@9533.9574.42\|GAGGAAACTCTGGTGGAGGTCCGTAGCGGTCCTGACGTGCAA | >hg38_21_+_8433172_8446622_RNA45SN1_45S_50ntFlanks | 9533 | 9574 | 42 | GAGGAAACTCTGGTGGAGGTCCGTAGCGGTCCTGACGTGCAA | 366.2 | -3.1 | 1.0 | -3.1 | 1.9E-03 | 2.2E-03 | MB231/MB468 | NUC |
| >hg38_21_+_8433172_8446622_RNA45SN1_45S_50ntFlanks@4727.4765.39\|AATCAAGAACGAAAGTCGGAGGTTCGAAGACGATCAGAT | >hg38_21_+_8433172_8446622_RNA45SN1_45S_50ntFlanks | 4727 | 4765 | 39 | AATCAAGAACGAAAGTCGGAGGTTCGAAGACGATCAGAT | 123.7 | 4.8 | 1.5 | 3.2 | 1.6E-03 | 2.2E-03 | MB231/MB468 | MP |
| >hg38_21_+_8433172_8446622_RNA45SN1_45S_50ntFlanks@7973.8012.40\|GACGCGACCTCAGATCAGACGTGGCGACCCGCTGAATTTA | >hg38_21_+_8433172_8446622_RNA45SN1_45S_50ntFlanks | 7973 | 8012 | 40 | GACGCGACCTCAGATCAGACGTGGCGACCCGCTGAATTTA | 1291.0 | 2.3 | 0.8 | 3.1 | 1.9E-03 | 2.2E-03 | MB231/MB468 | NUC |
| >hg38_21_+_8433172_8446622_RNA45SN1_45S_50ntFlanks@10780.10818.39\|AAGGTGAACAGCCTCTGGCATGTTGGAACAATGTAGGTA | >hg38_21_+_8433172_8446622_RNA45SN1_45S_50ntFlanks | 10780 | 10818 | 39 | AAGGTGAACAGCCTCTGGCATGTTGGAACAATGTAGGTA | 71.7 | -5.1 | 1.6 | -3.2 | 1.6E-03 | 2.2E-03 | MB231/MB468 | MP |
| >hg38_21_+_8433172_8446622_RNA45SN1_45S_50ntFlanks@9542.9571.30\|CTGGTGGAGGTCCGTAGCGGTCCTGACGTG | >hg38_21_+_8433172_8446622_RNA45SN1_45S_50ntFlanks | 9542 | 9571 | 30 | CTGGTGGAGGTCCGTAGCGGTCCTGACGTG | 323.4 | -5.6 | 1.8 | -3.1 | 1.9E-03 | 2.2E-03 | BT20/MB468 | NUC |
| >MI0006273\|hsa-mir-1180&WithFlank&17\|-\|19344500\|19344580@47.68.22(+1U)&[MIMAT0005825&hsa-miR-1180-3p&offsets\|0\|0(+1U);m-214&17\|-\|19344513\|19344534&offsets\|0\|0(+1U)]\|TTTCCGGCTCGCGTGGGTGTGTT | >MI0006273\|hsa-mir-1180&WithFlank&17\|-\|19344500\|19344580 | 47 | 68 | 22(+1U) | TTTCCGGCTCGCGTGGGTGTGTT | 205.3 | 2.6 | 0.7 | 3.6 | 3.6E-04 | 2.2E-03 | BT20/MB231 | CYTO |
| >hg38_21_+_8433172_8446622_RNA45SN1_45S_50ntFlanks@6648.6696.49\|GTACGACTCTTAGCGGTGGATCACTCGGCTCGTGCGTCGATGAAGAACG | >hg38_21_+_8433172_8446622_RNA45SN1_45S_50ntFlanks | 6648 | 6696 | 49 | GTACGACTCTTAGCGGTGGATCACTCGGCTCGTGCGTCGATGAAGAACG | 588.4 | -3.5 | 1.0 | -3.6 | 3.6E-04 | 2.2E-03 | BT20/MB231 | CYTO |
| >hg38_21_+_8433172_8446622_RNA45SN1_45S_50ntFlanks@11569.11597.29\|CAGCCGACTTAGAACTGGTGCGGACCAGG | >hg38_21_+_8433172_8446622_RNA45SN1_45S_50ntFlanks | 11569 | 11597 | 29 | CAGCCGACTTAGAACTGGTGCGGACCAGG | 372.8 | -2.7 | 0.9 | -3.1 | 1.7E-03 | 2.2E-03 | BT20/MB468 | CYTO |
| >hg38_21_+_8433172_8446622_RNA45SN1_45S_50ntFlanks@11582.11601.20\|ACTGGTGCGGACCAGGGGAA | >hg38_21_+_8433172_8446622_RNA45SN1_45S_50ntFlanks | 11582 | 11601 | 20 | ACTGGTGCGGACCAGGGGAA | 334.1 | -4.7 | 1.4 | -3.3 | 8.9E-04 | 2.2E-03 | BT20/MB468 | MP |

| | | | | | | | | | | | | | |
|---|---|---|---|---|---|---|---|---|---|---|---|---|---|
| >hg38_21_+_8433172_8446622_RNA45SN1_45S_50ntFlanks@11671.11709.39\|CCAGTGCTCTGAATGTCAAAGTGAAGAAATTCAATGAAG | >hg38_21_+_8433172_8446622_RNA45SN1_45S_50ntFlanks | 11671 | 11709 | 39 | CCAGTGCTCTGAATGTCAAAGTGAAGAAATTCAATGAAG | 635.7 | -2.4 | 0.8 | -3.1 | 1.9E-03 | 2.2E-03 | MB231/MB468 | NUC |
| >hg38_MT_+_1621_3279_RNR2_16S_50ntFlanks@1188.1215.28\|TCGGTTGGGGCGACCTCGGAGCAGAACC | >hg38_MT_+_1621_3279_RNR2_16S_50ntFlanks | 1188 | 1215 | 28 | TCGGTTGGGGCGACCTCGGAGCAGAACC | 161.7 | 4.6 | 1.4 | 3.3 | 8.9E-04 | 2.2E-03 | BT20/MB468 | MP |
| >hg38_21_+_8433172_8446622_RNA45SN1_45S_50ntFlanks@3776.3800.25\|CCGGTACAGTGAAACTGCGAATGGC | >hg38_21_+_8433172_8446622_RNA45SN1_45S_50ntFlanks | 3776 | 3800 | 25 | CCGGTACAGTGAAACTGCGAATGGC | 175.3 | -5.3 | 1.6 | -3.3 | 8.9E-04 | 2.2E-03 | BT20/MB468 | MP |
| >hg38_21_+_8433172_8446622_RNA45SN1_45S_50ntFlanks@4730.4768.39\|CAAGAACGAAAGTCGGAGGTTCGAAGACGATCAGATACC | >hg38_21_+_8433172_8446622_RNA45SN1_45S_50ntFlanks | 4730 | 4768 | 39 | CAAGAACGAAAGTCGGAGGTTCGAAGACGATCAGATACC | 85.2 | 6.4 | 1.9 | 3.3 | 8.9E-04 | 2.2E-03 | BT20/MB468 | MP |
| >hg38_21_+_8433172_8446622_RNA45SN1_45S_50ntFlanks@12405.12444.40\|CTTTTTGATCCTTCGATGTCGGCTCTTCCTATCATTGTGA | >hg38_21_+_8433172_8446622_RNA45SN1_45S_50ntFlanks | 12405 | 12444 | 40 | CTTTTTGATCCTTCGATGTCGGCTCTTCCTATCATTGTGA | 750.8 | -2.6 | 0.8 | -3.1 | 1.8E-03 | 2.2E-03 | MB231/MB468 | CYTO |
| >hg38_21_+_8433172_8446622_RNA45SN1_45S_50ntFlanks@12420.12437.18\|ATGTCGGCTCTTCCTATC | >hg38_21_+_8433172_8446622_RNA45SN1_45S_50ntFlanks | 12420 | 12437 | 18 | ATGTCGGCTCTTCCTATC | 7168.1 | -2.2 | 0.7 | -3.1 | 1.7E-03 | 2.2E-03 | BT20/MB468 | CYTO |
| >hg38_21_+_8433172_8446622_RNA45SN1_45S_50ntFlanks@3705.3748.44\|TACCTGGTTGATCCTGCCAGTAGCATATGCTTGTCTCAAAGATT | >hg38_21_+_8433172_8446622_RNA45SN1_45S_50ntFlanks | 3705 | 3748 | 44 | TACCTGGTTGATCCTGCCAGTAGCATATGCTTGTCTCAAAGATT | 6010.0 | 4.4 | 1.2 | 3.6 | 3.4E-04 | 2.2E-03 | BT20/MB231 | NUC |
| >hg38_21_+_8433172_8446622_RNA45SN1_45S_50ntFlanks@6651.6668.18\|CGACTCTTAGCGGTGGAT | >hg38_21_+_8433172_8446622_RNA45SN1_45S_50ntFlanks | 6651 | 6668 | 18 | CGACTCTTAGCGGTGGAT | 16728.2 | 2.4 | 0.7 | 3.6 | 3.4E-04 | 2.2E-03 | BT20/MB231 | NUC |
| >hg38_MT_+_1621_3279_RNR2_16S_50ntFlanks@617.657.41\|AAACATATAACTGAACTCCTCACACCCAATTGGACCAATCT | >hg38_MT_+_1621_3279_RNR2_16S_50ntFlanks | 617 | 657 | 41 | AAACATATAACTGAACTCCTCACACCCAATTGGACCAATCT | 46.0 | 9.3 | 2.8 | 3.3 | 8.9E-04 | 2.2E-03 | BT20/MB468 | MP |
| >hg38_21_+_8433172_8446622_RNA45SN1_45S_50ntFlanks@8186.8212.27\|AGGCCCAGCCCGTGGACGGTGTGAGGC | >hg38_21_+_8433172_8446622_RNA45SN1_45S_50ntFlanks | 8186 | 8212 | 27 | AGGCCCAGCCCGTGGACGGTGTGAGGC | 52.6 | -8.8 | 2.6 | -3.3 | 8.9E-04 | 2.2E-03 | BT20/MB468 | MP |
| >hg38_21_+_8433172_8446622_RNA45SN1_45S_50ntFlanks@9556.9574.19\|TAGCGGTCCTGACGTGCAA | >hg38_21_+_8433172_8446622_RNA45SN1_45S_50ntFlanks | 9556 | 9574 | 19 | TAGCGGTCCTGACGTGCAA | 155.6 | -5.1 | 1.6 | -3.2 | 1.6E-03 | 2.2E-03 | MB231/MB468 | MP |
| >hg38_21_+_8433172_8446622_RNA45SN1_45S_50ntFlanks@12944.12962.19\|CGTAGACGACCTGCTTCTG | >hg38_21_+_8433172_8446622_RNA45SN1_45S_50ntFlanks | 12944 | 12962 | 19 | CGTAGACGACCTGCTTCTG | 296.5 | -1.9 | 0.6 | -3.1 | 1.8E-03 | 2.2E-03 | MB231/MB468 | CYTO |
| >hg38_21_+_8433172_8446622_RNA45SN1_45S_50ntFlanks@10808.10844.37\|CAATGTAGGTAAGGGAAGTCGGCAAGCCGGATCCGTA | >hg38_21_+_8433172_8446622_RNA45SN1_45S_50ntFlanks | 10808 | 10844 | 37 | CAATGTAGGTAAGGGAAGTCGGCAAGCCGGATCCGTA | 340.2 | -4.8 | 1.5 | -3.2 | 1.4E-03 | 2.2E-03 | BT20/MB468 | MITO |
| >hg38_21_+_8433172_8446622_RNA45SN1_45S_50ntFlanks@12632.12654.23\|GCGAAGCTACCATCTGTGGGATT | >hg38_21_+_8433172_8446622_RNA45SN1_45S_50ntFlanks | 12632 | 12654 | 23 | GCGAAGCTACCATCTGTGGGATT | 3163.2 | -5.0 | 1.5 | -3.3 | 8.9E-04 | 2.2E-03 | BT20/MB468 | MP |
| >hg38_21_+_8433172_8446622_RNA45SN1_45S_50ntFlanks@10061.10079.19\|CAGTCGAGAGTGGACGGGA | >hg38_21_+_8433172_8446622_RNA45SN1_45S_50ntFlanks | 10061 | 10079 | 19 | CAGTCGAGAGTGGACGGGA | 338.6 | 5.8 | 1.6 | 3.6 | 3.1E-04 | 2.2E-03 | BT20/MB231 | MP |
| >hg38_21_+_8433172_8446622_RNA45SN1_45S_50ntFlanks@11913.11943.31\|AAGAGACATGAGAGGTGTAGAATAAGTGGGA | >hg38_21_+_8433172_8446622_RNA45SN1_45S_50ntFlanks | 11913 | 11943 | 31 | AAGAGACATGAGAGGTGTAGAATAAGTGGGA | 93.6 | 9.9 | 2.7 | 3.6 | 3.1E-04 | 2.2E-03 | BT20/MB231 | MP |
| >hg38_21_+_8433172_8446622_RNA45SN1_45S_50ntFlanks@6650.6678.29\|ACGACTCTTAGCGGTGGATCACTCGGCTC | >hg38_21_+_8433172_8446622_RNA45SN1_45S_50ntFlanks | 6650 | 6678 | 29 | ACGACTCTTAGCGGTGGATCACTCGGCTC | 1793.8 | 4.7 | 1.3 | 3.6 | 3.1E-04 | 2.2E-03 | BT20/MB231 | MP |
| >hg38_21_+_8433172_8446622_RNA45SN1_45S_50ntFlanks@9941.9961.21\|AGGACGGTGGCCATGGAAGTC | >hg38_21_+_8433172_8446622_RNA45SN1_45S_50ntFlanks | 9941 | 9961 | 21 | AGGACGGTGGCCATGGAAGTC | 1334.0 | 6.5 | 1.8 | 3.6 | 3.1E-04 | 2.2E-03 | BT20/MB231 | MP |
| >MI0003186\|hsa-mir-502&WithFlank&X\|+\|50014592\|50014689@58.79.22&[MIMAT0004775&hsa-miR-502-3p&offsets\|0\|0;m-236&X\|+\|50014649\|50014669&offsets\|0\|+1]\|AATGCACCTGGGCAAGGATTCA | >MI0003186\|hsa-mir-502&WithFlank&X\|+\|50014592\|50014689 | 58 | 79 | 22 | AATGCACCTGGGCAAGGATTCA | 225.5 | 2.5 | 0.8 | 3.2 | 1.6E-03 | 2.2E-03 | MB231/MB468 | MP |
| >MI0000465\|hsa-mir-191&WithFlank&3\|-\|49020612\|49020715@22.44.23(+1C)&[MIMAT0000440&hsa-miR-191-5p&offsets\|0\|0(+1C);m-48&3\|-\|49020672\|49020694&offsets\|0\|0(+1C)]\|CAACGGAATCCCAAAAGCAGCTGC | >MI0000465\|hsa-mir-191&WithFlank&3\|-\|49020612\|49020715 | 22 | 44 | 23(+1C) | CAACGGAATCCCAAAAGCAGCTGC | 1094.9 | 4.6 | 1.4 | 3.2 | 1.6E-03 | 2.2E-03 | MB231/MB468 | MP |
| >trnaMT_GlnTTG_MT_-_4329_4400@1.21.21,trnalookalike2_GlnTTG_1_-_564879_564950@1.21.21\|TAGGATGGGGTGTGATAGGTG | >trnaMT_GlnTTG_MT_-_4329_4400 | 1 | 21 | 21 | TAGGATGGGGTGTGATAGGTG | 438.8 | 4.3 | 1.3 | 3.3 | 8.9E-04 | 2.2E-03 | BT20/MB468 | MP |
| >hg38_1_-_228634819_228635039_RNA5S12_5S_50ntFlanks@136.169.34\|GGAGACCGCCTGGGAATACCGGGTGCTGTAGGCT | >hg38_1_-_228634819_228635039_RNA5S12_5S_50ntFlanks | 136 | 169 | 34 | GGAGACCGCCTGGGAATACCGGGTGCTGTAGGCT | 185.7 | -2.2 | 0.7 | -3.2 | 1.6E-03 | 2.2E-03 | MB231/MB468 | MP |
| >hg38_21_+_8433172_8446622_RNA45SN1_45S_50ntFlanks@12957.12986.30\|CTTCTGGGTCGGGGTTTCGTACGTAGCAGA | >hg38_21_+_8433172_8446622_RNA45SN1_45S_50ntFlanks | 12957 | 12986 | 30 | CTTCTGGGTCGGGGTTTCGTACGTAGCAGA | 282.6 | -2.0 | 0.6 | -3.1 | 1.7E-03 | 2.2E-03 | BT20/MB468 | CYTO |
| >hg38_21_+_8433172_8446622_RNA45SN1_45S_50ntFlanks@11189.11225.37\|ACCCGGGGGGCCGGCGGCGGCGGCGACTCTGGACGCG | >hg38_21_+_8433172_8446622_RNA45SN1_45S_50ntFlanks | 11189 | 11225 | 37 | ACCCGGGGGGCCGGCGGCGGCGGCGACTCTGGACGCG | 332.3 | -3.6 | 1.2 | -3.1 | 1.7E-03 | 2.2E-03 | BT20/MB468 | CYTO |
| >hg38_21_+_8433172_8446622_RNA45SN1_45S_50ntFlanks@8168.8195.28\|CCCAAGTCCTTCTGATCGAGGCCCAGCC | >hg38_21_+_8433172_8446622_RNA45SN1_45S_50ntFlanks | 8168 | 8195 | 28 | CCCAAGTCCTTCTGATCGAGGCCCAGCC | 200.2 | -3.6 | 1.1 | -3.2 | 1.4E-03 | 2.2E-03 | BT20/MB468 | MITO |
| >hg38_21_+_8433172_8446622_RNA45SN1_45S_50ntFlanks@11881.11903.23\|CCTGTTGAGCTTGACTCTAGTCT | >hg38_21_+_8433172_8446622_RNA45SN1_45S_50ntFlanks | 11881 | 11903 | 23 | CCTGTTGAGCTTGACTCTAGTCT | 273.7 | -3.0 | 0.9 | -3.3 | 9.0E-04 | 2.2E-03 | BT20/MB468 | MP |
| >hg38_21_+_8433172_8446622_RNA45SN1_45S_50ntFlanks@4739.4758.20\|AAGTCGGAGGTTCGAAGACG | >hg38_21_+_8433172_8446622_RNA45SN1_45S_50ntFlanks | 4739 | 4758 | 20 | AAGTCGGAGGTTCGAAGACG | 184.7 | -2.9 | 0.9 | -3.1 | 1.8E-03 | 2.2E-03 | MB231/MB468 | CYTO |
| >hg38_21_+_8433172_8446622_RNA45SN1_45S_50ntFlanks@10786.10829.44\|AACAGCCTCTGGCATGTTGGAACAATGTAGGTAAGGGAAGTCGG | >hg38_21_+_8433172_8446622_RNA45SN1_45S_50ntFlanks | 10786 | 10829 | 44 | AACAGCCTCTGGCATGTTGGAACAATGTAGGTAAGGGAAGTCGG | 492.7 | -4.0 | 1.3 | -3.2 | 1.4E-03 | 2.2E-03 | BT20/MB468 | MITO |
| >hg38_21_+_8433172_8446622_RNA45SN1_45S_50ntFlanks@12418.12435.18\|CGATGTCGGCTCTTCCTA | >hg38_21_+_8433172_8446622_RNA45SN1_45S_50ntFlanks | 12418 | 12435 | 18 | CGATGTCGGCTCTTCCTA | 40027.6 | -6.2 | 2.0 | -3.1 | 1.9E-03 | 2.2E-03 | BT20/MB468 | NUC |
| >hg38_21_+_8433172_8446622_RNA45SN1_45S_50ntFlanks@11322.11340.19\|ACGTCTCGTCGCGCGCGCG | >hg38_21_+_8433172_8446622_RNA45SN1_45S_50ntFlanks | 11322 | 11340 | 19 | ACGTCTCGTCGCGCGCGCG | 498.6 | -3.1 | 1.0 | -3.1 | 1.8E-03 | 2.2E-03 | MB231/MB468 | CYTO |
| >hg38_21_+_8433172_8446622_RNA45SN1_45S_50ntFlanks@12605.12622.18\|TATGTGCTTGGCTGAGGA | >hg38_21_+_8433172_8446622_RNA45SN1_45S_50ntFlanks | 12605 | 12622 | 18 | TATGTGCTTGGCTGAGGA | 147.5 | -3.0 | 0.9 | -3.1 | 1.7E-03 | 2.2E-03 | BT20/MB468 | CYTO |
| >hg38_21_+_8433172_8446622_RNA45SN1_45S_50ntFlanks@11704.11736.33\|ATGAAGCGCGGGTAAACGGCGGGAGTAACTATG | >hg38_21_+_8433172_8446622_RNA45SN1_45S_50ntFlanks | 11704 | 11736 | 33 | ATGAAGCGCGGGTAAACGGCGGGAGTAACTATG | 510.7 | -5.8 | 1.9 | -3.1 | 1.9E-03 | 2.2E-03 | BT20/MB468 | NUC |
| >hg38_21_+_8433172_8446622_RNA45SN1_45S_50ntFlanks@11561.11587.27\|GCGCCTAGCAGCCGACTTAGAACTGGT | >hg38_21_+_8433172_8446622_RNA45SN1_45S_50ntFlanks | 11561 | 11587 | 27 | GCGCCTAGCAGCCGACTTAGAACTGGT | 295.3 | -3.1 | 0.9 | -3.3 | 9.0E-04 | 2.3E-03 | BT20/MB468 | MP |
| >hg38_21_+_8433172_8446622_RNA45SN1_45S_50ntFlanks@4685.4707.23\|AGACGGACCAGAGCGAAAGCATT | >hg38_21_+_8433172_8446622_RNA45SN1_45S_50ntFlanks | 4685 | 4707 | 23 | AGACGGACCAGAGCGAAAGCATT | 349.7 | -2.7 | 0.8 | -3.2 | 1.5E-03 | 2.3E-03 | BT20/MB468 | MITO |
| >hg38_21_+_8433172_8446622_RNA45SN1_45S_50ntFlanks@9933.9952.20\|TAGACAGCAGGACGGTGGCC | >hg38_21_+_8433172_8446622_RNA45SN1_45S_50ntFlanks | 9933 | 9952 | 20 | TAGACAGCAGGACGGTGGCC | 107.6 | 6.4 | 1.9 | 3.3 | 9.0E-04 | 2.3E-03 | BT20/MB468 | MP |
| >hg38_21_+_8433172_8446622_RNA45SN1_45S_50ntFlanks@11674.11697.24\|GTGCTCTGAATGTCAAAGTGAAGA | >hg38_21_+_8433172_8446622_RNA45SN1_45S_50ntFlanks | 11674 | 11697 | 24 | GTGCTCTGAATGTCAAAGTGAAGA | 41.7 | 9.1 | 2.7 | 3.3 | 9.0E-04 | 2.3E-03 | BT20/MB468 | MP |
| >hg38_MT_+_1621_3279_RNR2_16S_50ntFlanks@618.660.43\|AACATATAACTGAACTCCTCACACCCAATTGGACCAATCTATC | >hg38_MT_+_1621_3279_RNR2_16S_50ntFlanks | 618 | 660 | 43 | AACATATAACTGAACTCCTCACACCCAATTGGACCAATCTATC | 46.5 | 9.3 | 2.8 | 3.3 | 9.0E-04 | 2.3E-03 | BT20/MB468 | MP |
| >hg38_21_+_8433172_8446622_RNA45SN1_45S_50ntFlanks@9938.9982.45\|AGCAGGACGGTGGCCATGGAAGTCGGAATCCGCTAAGGAGTGTGT | >hg38_21_+_8433172_8446622_RNA45SN1_45S_50ntFlanks | 9938 | 9982 | 45 | AGCAGGACGGTGGCCATGGAAGTCGGAATCCGCTAAGGAGTGTGT | 988.8 | 7.4 | 2.2 | 3.3 | 9.0E-04 | 2.3E-03 | BT20/MB468 | MP |
| >hg38_21_+_8433172_8446622_RNA45SN1_45S_50ntFlanks@11913.11948.36\|AAGAGACATGAGAGGTGTAGAATAAGTGGGAGGCCC | >hg38_21_+_8433172_8446622_RNA45SN1_45S_50ntFlanks | 11913 | 11948 | 36 | AAGAGACATGAGAGGTGTAGAATAAGTGGGAGGCCC | 97.7 | 9.9 | 2.8 | 3.6 | 3.1E-04 | 2.3E-03 | BT20/MB231 | MP |
| >hg38_21_+_8433172_8446622_RNA45SN1_45S_50ntFlanks@11589.11611.23\|CGGACCAGGGGAATCCGACTGTT | >hg38_21_+_8433172_8446622_RNA45SN1_45S_50ntFlanks | 11589 | 11611 | 23 | CGGACCAGGGGAATCCGACTGTT | 419.6 | 2.5 | 0.7 | 3.5 | 4.0E-04 | 2.3E-03 | BT20/MB231 | MITO |
| >hg38_21_+_8433172_8446622_RNA45SN1_45S_50ntFlanks@11653.11691.39\|ACGCGATGTGATTTCTGCCCAGTGCTCTGAATGTCAAAG | >hg38_21_+_8433172_8446622_RNA45SN1_45S_50ntFlanks | 11653 | 11691 | 39 | ACGCGATGTGATTTCTGCCCAGTGCTCTGAATGTCAAAG | 290.8 | 3.2 | 0.9 | 3.5 | 4.0E-04 | 2.3E-03 | BT20/MB231 | MITO |
| >trna10_LysCTT_16_+_3241501_3241573@3.34.32,trna13_LysCTT_14_-_58706613_58706685@3.34.32,trna2_LysCTT_15_+_79152904_79152976@3.34.32\|CCGGCTAGCTCAGTCGGTAGAGCATGGGACTC | >trna10_LysCTT_16_+_3241501_3241573 | 3 | 34 | 32 | CCGGCTAGCTCAGTCGGTAGAGCATGGGACTC | 585.5 | -3.9 | 1.1 | -3.5 | 4.0E-04 | 2.3E-03 | BT20/MB231 | MITO |
| >trna116_GluCTC_1_-_145399233_145399304@41.74.34,trna134_GluTTC_1_-_16861774_16861845@41.74.34,trna71_GluCTC_1_-_161439189_161439260@41.74.34,trna74_GluCTC_1_-_161431809_161431880@41.74.34,trna77_GluCTC_1_-_161424398_161424469@41.74.34,trna77_GluCTC_6_+_28949976_28950047@41.74.34,trna80_GluCTC_1_-_161417018_161417089@41.74.34,trna84_GluTTC_1_-_161391883_161391954@41.74.34,trna87_GluCTC_6_-_126101393_126101464@41.74.34\|CCGCGGCCCGGGTTCGATTCCCGGTCAGGGAACC | >trna116_GluCTC_1_-_145399233_145399304 | 41 | 74 | 34 | CCGCGGCCCGGGTTCGATTCCCGGTCAGGGAACC | 304.4 | -3.2 | 0.9 | -3.5 | 4.0E-04 | 2.3E-03 | BT20/MB231 | MITO |
| >hg38_21_+_8433172_8446622_RNA45SN1_45S_50ntFlanks@11320.11344.25\|CCACGTCTCGTCGCGCGCGCGTCCG | >hg38_21_+_8433172_8446622_RNA45SN1_45S_50ntFlanks | 11320 | 11344 | 25 | CCACGTCTCGTCGCGCGCGCGTCCG | 4243.2 | -3.5 | 1.1 | -3.2 | 1.5E-03 | 2.3E-03 | BT20/MB468 | MITO |
| >hg38_21_+_8433172_8446622_RNA45SN1_45S_50ntFlanks@10866.10901.36\|CTAAGGGCTGGGTCGGTCGGGCTGGGGCGCGAAGCG | >hg38_21_+_8433172_8446622_RNA45SN1_45S_50ntFlanks | 10866 | 10901 | 36 | CTAAGGGCTGGGTCGGTCGGGCTGGGGCGCGAAGCG | 554.2 | -5.0 | 1.6 | -3.1 | 1.7E-03 | 2.3E-03 | BT20/MB468 | CYTO |
| >hg38_21_+_8433172_8446622_RNA45SN1_45S_50ntFlanks@4067.4089.23\|AACTTTCGATGGTAGTCGCCGTG | >hg38_21_+_8433172_8446622_RNA45SN1_45S_50ntFlanks | 4067 | 4089 | 23 | AACTTTCGATGGTAGTCGCCGTG | 126.4 | -3.2 | 1.0 | -3.1 | 1.7E-03 | 2.3E-03 | BT20/MB468 | CYTO |
| >hg38_21_+_8433172_8446622_RNA45SN1_45S_50ntFlanks@12638.12657.20\|CTACCATCTGTGGGATTATG | >hg38_21_+_8433172_8446622_RNA45SN1_45S_50ntFlanks | 12638 | 12657 | 20 | CTACCATCTGTGGGATTATG | 105.9 | -3.9 | 1.3 | -3.1 | 1.8E-03 | 2.3E-03 | MB231/MB468 | CYTO |
| >hg38_21_+_8433172_8446622_RNA45SN1_45S_50ntFlanks@12606.12624.19\|ATGTGCTTGGCTGAGGAGC | >hg38_21_+_8433172_8446622_RNA45SN1_45S_50ntFlanks | 12606 | 12624 | 19 | ATGTGCTTGGCTGAGGAGC | 99.9 | -3.7 | 1.2 | -3.2 | 1.6E-03 | 2.3E-03 | MB231/MB468 | MP |
| >MI0000299\|hsa-mir-222&WithFlank&X\|-\|45747009\|45747130@75.99.25&[MIMAT0000279&hsa-miR-222-3p&offsets\|0\|+4;m-140&X\|-\|45747033\|45747056&offsets\|0\|+1]\|AGCTACATCTGGCTACTGGGTCTCT | >MI0000299\|hsa-mir-222&WithFlank&X\|-\|45747009\|45747130 | 75 | 99 | 25 | AGCTACATCTGGCTACTGGGTCTCT | 45685.7 | -2.6 | 0.7 | -3.5 | 4.0E-04 | 2.3E-03 | BT20/MB231 | MITO |
| >hg38_21_+_8433172_8446622_RNA45SN1_45S_50ntFlanks@11564.11602.39\|CCTAGCAGCCGACTTAGAACTGGTGCGGACCAGGGGAAT | >hg38_21_+_8433172_8446622_RNA45SN1_45S_50ntFlanks | 11564 | 11602 | 39 | CCTAGCAGCCGACTTAGAACTGGTGCGGACCAGGGGAAT | 284.6 | 3.1 | 0.9 | 3.5 | 4.0E-04 | 2.3E-03 | BT20/MB231 | MITO |

<table>
<tr><td></td><td></td><td></td><td></td><td></td><td></td><td></td><td></td><td></td><td></td><td></td><td></td><td></td><td></td></tr>
<tr><td>>hg38_21_+_8433172_8446622_RNA45SN1_45S_50ntFlanks@11583.11609.27|CTGGTGCGGACCAGGGGAATCCGACTG</td><td>>hg38_21_+_8433172_8446622_RNA45SN1_45S_50ntFlanks</td><td>11583</td><td>11609</td><td>27</td><td>CTGGTGCGGACCAGGGGAATCCGACTG</td><td>391.2</td><td>2.9</td><td>0.8</td><td>3.5</td><td>4.0E-04</td><td>2.3E-03</td><td>BT20/MB231</td><td>MITO</td></tr>
<tr><td>>MI0000093|hsa-mir-92a-1&WithFlank&13|+|91351308|91351397@54.76.23(+1U)&[MIMAT0000092&hsa-miR-92a-3p&offsets|0|+1(+1U);m-9&13|+|91351361|91351382&offsets|0|+1(+1U)]|TATTGCACTTGTCCCGGCCTGTTT</td><td>>MI0000093|hsa-mir-92a-1&WithFlank&13|+|91351308|91351397</td><td>54</td><td>76</td><td>23(+1U)</td><td>TATTGCACTTGTCCCGGCCTGTTT</td><td>632.6</td><td>-2.6</td><td>0.7</td><td>-3.5</td><td>4.0E-04</td><td>2.3E-03</td><td>BT20/MB231</td><td>MITO</td></tr>
<tr><td>>hg38_MT_+_1621_3279_RNR2_16S_50ntFlanks@617.642.26|AAACATATAACTGAACTCCTCACACC</td><td>>hg38_MT_+_1621_3279_RNR2_16S_50ntFlanks</td><td>617</td><td>642</td><td>26</td><td>AAACATATAACTGAACTCCTCACACC</td><td>158.4</td><td>2.9</td><td>0.9</td><td>3.2</td><td>1.6E-03</td><td>2.3E-03</td><td>MB231/MB468</td><td>MP</td></tr>
<tr><td>>hg38_21_+_8433172_8446622_RNA45SN1_45S_50ntFlanks@6655.6689.35|TCTTAGCGGTGGATCACTCGGCTCGTGCGTCGATG</td><td>>hg38_21_+_8433172_8446622_RNA45SN1_45S_50ntFlanks</td><td>6655</td><td>6689</td><td>35</td><td>TCTTAGCGGTGGATCACTCGGCTCGTGCGTCGATG</td><td>1078.9</td><td>-1.9</td><td>0.6</td><td>-3.1</td><td>1.9E-03</td><td>2.3E-03</td><td>MB231/MB468</td><td>NUC</td></tr>
<tr><td>>trna111_HisGTG_1_-_147774845_147774916@-1G.30.31,trna118_HisGTG_1_-_145396881_145396952@-1G.30.31,trna16_HisGTG_1_+_146544773_146544844@-1G.30.31,trna1_HisGTG_15_+_45493349_45493420@-1G.30.31,trna21_HisGTG_1_+_147753471_147753542@-1G.30.31,trna33_HisGTG_6_+_27125906_27125977@-1G.30.31,trna7_HisGTG_9_-_14433938_14434009@-1G.30.31,trna8_HisGTG_15_-_45492611_45492682@-1G.30.31,trna9_HisGTG_15_-_45490804_45490875@-1G.30.31|GGCCGTGATCGTATAGTGGTTAGTACTCTGC</td><td>>trna111_HisGTG_1_-_147774845_147774916</td><td>-1G</td><td>30</td><td>31</td><td>GGCCGTGATCGTATAGTGGTTAGTACTCTGC</td><td>11677.4</td><td>2.8</td><td>0.9</td><td>3.1</td><td>1.8E-03</td><td>2.3E-03</td><td>MB231/MB468</td><td>CYTO</td></tr>
<tr><td>>MI0000440|hsa-mir-27b&WithFlank&9|+|95085439|95085547@66.87.22&[MIMAT0000419&hsa-miR-27b-3p&offsets|-1|0;m-39&9|+|95085505|95085525&offsets|-1|0]|GTTCACAGTGGCTAAGTTCTGC</td><td>>MI0000440|hsa-mir-27b&WithFlank&9|+|95085439|95085547</td><td>66</td><td>87</td><td>22</td><td>GTTCACAGTGGCTAAGTTCTGC</td><td>139.8</td><td>5.4</td><td>1.6</td><td>3.3</td><td>9.0E-04</td><td>2.3E-03</td><td>BT20/MB468</td><td>MP</td></tr>
<tr><td>>hg38_21_+_8433172_8446622_RNA45SN1_45S_50ntFlanks@4982.5016.35|ACGGACAGGATTGACAGATTGATAGCTCTTTCTCG</td><td>>hg38_21_+_8433172_8446622_RNA45SN1_45S_50ntFlanks</td><td>4982</td><td>5016</td><td>35</td><td>ACGGACAGGATTGACAGATTGATAGCTCTTTCTCG</td><td>278.2</td><td>-3.6</td><td>1.1</td><td>-3.2</td><td>1.5E-03</td><td>2.3E-03</td><td>BT20/MB468</td><td>MITO</td></tr>
<tr><td>>am_trna111_HisGTG_1_-_147774845_147774916@-1G.26.27,trna118_HisGTG_1_-_145396881_145396952@-1G.26.27,trna16_HisGTG_1_+_146544773_146544844@-1G.26.27,trna1_HisGTG_15_+_45493349_45493420@-1G.26.27,trna21_HisGTG_1_+_147753471_147753542@-1G.26.27,trna33_HisGTG_6_+_27125906_27125977@-1G.26.27,trna7_HisGTG_9_-_14433938_14434009@-1G.26.27,trna8_HisGTG_15_-_45492611_45492682@-1G.26.27,trna9_HisGTG_15_-_45490804_45490875@-1G.26.27|GGCCGTGATCGTATAGTGGTTAGTACT</td><td>>am_trna111_HisGTG_1_-_147774845_147774916</td><td>-1G</td><td>26</td><td>27</td><td>GGCCGTGATCGTATAGTGGTTAGTACT</td><td>1628.5</td><td>3.7</td><td>1.2</td><td>3.1</td><td>1.7E-03</td><td>2.3E-03</td><td>BT20/MB468</td><td>CYTO</td></tr>
<tr><td>>MI0000061|hsa-let-7a-2&WithFlank&11|-|122146516|122146599@11.30.20;MI0000062|hsa-let-7a-3&WithFlank&22|+|46112743|46112828@10.29.20;MI0000060|hsa-let-7a-1&WithFlank&9|+|94175951|94176042@12.31.20&[MIMAT0000062&hsa-let-7a-5p&offsets|0|-2;m-6&11|-|122146568|122146589&offsets|0|-2];[MIMAT0000062_1&hsa-let-7a-5p&offsets|0|-2;m-5&22|+|46112752|46112773&offsets|0|-2];[MIMAT0000062_2&hsa-let-7a-5p&offsets|0|-2;m-4&9|+|94175962|94175983&offsets|0|-2]|TGAGGTAGTAGGTTGTATAG</td><td>>MI0000061|hsa-let-7a-2&WithFlank&11|-|122146516|122146599</td><td>11</td><td>30</td><td>20</td><td>TGAGGTAGTAGGTTGTATAG</td><td>177.5</td><td>5.1</td><td>1.6</td><td>3.2</td><td>1.6E-03</td><td>2.3E-03</td><td>MB231/MB468</td><td>MP</td></tr>
<tr><td>>hg38_21_+_8433172_8446622_RNA45SN1_45S_50ntFlanks@8154.8183.30|CCGCTCGTGGGGGGCCCAAGTCCTTCTGAT</td><td>>hg38_21_+_8433172_8446622_RNA45SN1_45S_50ntFlanks</td><td>8154</td><td>8183</td><td>30</td><td>CCGCTCGTGGGGGGCCCAAGTCCTTCTGAT</td><td>228.7</td><td>-3.5</td><td>1.1</td><td>-3.2</td><td>1.5E-03</td><td>2.3E-03</td><td>BT20/MB468</td><td>MITO</td></tr>
<tr><td>>trnaMT_PheGAA_MT_+_577_647@36.59.24|AAAATGTTTAGACGGGCTCACATC</td><td>>trnaMT_PheGAA_MT_+_577_647</td><td>36</td><td>59</td><td>24</td><td>AAAATGTTTAGACGGGCTCACATC</td><td>331.7</td><td>6.1</td><td>1.8</td><td>3.3</td><td>9.1E-04</td><td>2.3E-03</td><td>BT20/MB468</td><td>MP</td></tr>
<tr><td>>am_trnaMT_HisGTG_MT_+_12138_12206@19.49.31|AAACATCAGATTGTGAATCTGACAACAGAGG</td><td>>am_trnaMT_HisGTG_MT_+_12138_12206</td><td>19</td><td>49</td><td>31</td><td>AAACATCAGATTGTGAATCTGACAACAGAGG</td><td>417.6</td><td>6.4</td><td>1.9</td><td>3.3</td><td>9.1E-04</td><td>2.3E-03</td><td>BT20/MB468</td><td>MP</td></tr>
<tr><td>>hg38_21_+_8433172_8446622_RNA45SN1_45S_50ntFlanks@12717.12745.29|CCTCGGTTGGCCTCGGATAGCCGGTCCCC</td><td>>hg38_21_+_8433172_8446622_RNA45SN1_45S_50ntFlanks</td><td>12717</td><td>12745</td><td>29</td><td>CCTCGGTTGGCCTCGGATAGCCGGTCCCC</td><td>352.1</td><td>-5.4</td><td>1.6</td><td>-3.3</td><td>9.1E-04</td><td>2.3E-03</td><td>BT20/MB468</td><td>MP</td></tr>
<tr><td>>trnaMT_ThrTGT_MT_+_15888_15953@32.69.38|TGTAAACCGGAGATGAAAACCTTTTTCCAAGGACACCA</td><td>>trnaMT_ThrTGT_MT_+_15888_15953</td><td>32</td><td>69</td><td>38</td><td>TGTAAACCGGAGATGAAAACCTTTTTCCAAGGACACCA</td><td>136.2</td><td>5.9</td><td>1.8</td><td>3.3</td><td>9.1E-04</td><td>2.3E-03</td><td>BT20/MB468</td><td>MP</td></tr>
<tr><td>>hg38_21_+_8433172_8446622_RNA45SN1_45S_50ntFlanks@10346.10367.22|TCAAACGAGAACTTTGAAGGCC</td><td>>hg38_21_+_8433172_8446622_RNA45SN1_45S_50ntFlanks</td><td>10346</td><td>10367</td><td>22</td><td>TCAAACGAGAACTTTGAAGGCC</td><td>286.1</td><td>-3.0</td><td>0.9</td><td>-3.3</td><td>9.1E-04</td><td>2.3E-03</td><td>BT20/MB468</td><td>MP</td></tr>
<tr><td>>hg38_21_+_8433172_8446622_RNA45SN1_45S_50ntFlanks@10678.10710.33|CGGTTCCGGCGGCGTCCGGTGAGCTCTCGCTGG</td><td>>hg38_21_+_8433172_8446622_RNA45SN1_45S_50ntFlanks</td><td>10678</td><td>10710</td><td>33</td><td>CGGTTCCGGCGGCGTCCGGTGAGCTCTCGCTGG</td><td>2225.0</td><td>3.6</td><td>1.0</td><td>3.5</td><td>4.1E-04</td><td>2.3E-03</td><td>BT20/MB231</td><td>MITO</td></tr>
<tr><td>>hg38_MT_+_1621_3279_RNR2_16S_50ntFlanks@1184.1216.33|AATTTCGGTTGGGGCGACCTCGGAGCAGAACCC</td><td>>hg38_MT_+_1621_3279_RNR2_16S_50ntFlanks</td><td>1184</td><td>1216</td><td>33</td><td>AATTTCGGTTGGGGCGACCTCGGAGCAGAACCC</td><td>348.0</td><td>4.6</td><td>1.4</td><td>3.3</td><td>9.1E-04</td><td>2.3E-03</td><td>BT20/MB468</td><td>MP</td></tr>
<tr><td>>hg38_21_+_8433172_8446622_RNA45SN1_45S_50ntFlanks@6728.6771.44|GGACACATTGATCATCGACACTTCGAACGCACTTGCGGCCCCGG</td><td>>hg38_21_+_8433172_8446622_RNA45SN1_45S_50ntFlanks</td><td>6728</td><td>6771</td><td>44</td><td>GGACACATTGATCATCGACACTTCGAACGCACTTGCGGCCCCGG</td><td>442.3</td><td>-2.9</td><td>0.9</td><td>-3.1</td><td>1.9E-03</td><td>2.3E-03</td><td>MB231/MB468</td><td>MITO</td></tr>
<tr><td>>MI0003591|hsa-mir-584&WithFlank&5|-|149062307|149062415@22.44.23&[MIMAT0003249&hsa-miR-584-5p&offsets|0|+1;m-167&5|-|149062374|149062394&offsets|0|+2]|TTATGGTTTGCCTGGGACTGAGG</td><td>>MI0003591|hsa-mir-584&WithFlank&5|-|149062307|149062415</td><td>22</td><td>44</td><td>23</td><td>TTATGGTTTGCCTGGGACTGAGG</td><td>178.3</td><td>4.9</td><td>1.6</td><td>3.2</td><td>1.6E-03</td><td>2.3E-03</td><td>MB231/MB468</td><td>MP</td></tr>
<tr><td>>MI0000079|hsa-mir-23a&WithFlank&19|-|13836581|13836665@54.72.19(+1U)&[MIMAT0000078&hsa-miR-23a-3p&offsets|+3|+1(+1U);m-31&19|-|13836595|13836615&offsets|+3|+1(+1U)]|ACATTGCCAGGGATTTCCAT</td><td>>MI0000079|hsa-mir-23a&WithFlank&19|-|13836581|13836665</td><td>54</td><td>72</td><td>19(+1U)</td><td>ACATTGCCAGGGATTTCCAT</td><td>301.4</td><td>11.5</td><td>3.6</td><td>3.2</td><td>1.6E-03</td><td>2.3E-03</td><td>MB231/MB468</td><td>MP</td></tr>
<tr><td>>MI0003834|hsa-mir-769&WithFlank&19|+|46018926|46019055@36.57.22(+2U)&[MIMAT0003886&hsa-miR-769-5p&offsets|0|0(+2U);m-216&19|+|46018961|46018982&offsets|0|0(+2U)]|TGAGACCTCTGGGTTCTGAGCTTT</td><td>>MI0003834|hsa-mir-769&WithFlank&19|+|46018926|46019055</td><td>36</td><td>57</td><td>22(+2U)</td><td>TGAGACCTCTGGGTTCTGAGCTTT</td><td>314.4</td><td>5.1</td><td>1.5</td><td>3.3</td><td>9.1E-04</td><td>2.3E-03</td><td>BT20/MB468</td><td>MP</td></tr>
<tr><td>>hg38_21_+_8433172_8446622_RNA45SN1_45S_50ntFlanks@6777.6806.30|TCCCGGGGCTACGCCTGTCTGAGCGTCGCT</td><td>>hg38_21_+_8433172_8446622_RNA45SN1_45S_50ntFlanks</td><td>6777</td><td>6806</td><td>30</td><td>TCCCGGGGCTACGCCTGTCTGAGCGTCGCT</td><td>772.7</td><td>2.1</td><td>0.7</td><td>3.1</td><td>1.9E-03</td><td>2.3E-03</td><td>MB231/MB468</td><td>MITO</td></tr>
<tr><td>>hg38_21_+_8433172_8446622_RNA45SN1_45S_50ntFlanks@475.519.45|CTTCGTGATCGATGTGGTGACGTCGTGCTCTCCCGGGCCGGGTCC</td><td>>hg38_21_+_8433172_8446622_RNA45SN1_45S_50ntFlanks</td><td>475</td><td>519</td><td>45</td><td>CTTCGTGATCGATGTGGTGACGTCGTGCTCTCCCGGGCCGGGTCC</td><td>225.2</td><td>5.7</td><td>1.9</td><td>3.1</td><td>1.9E-03</td><td>2.3E-03</td><td>MB231/MB468</td><td>MITO</td></tr>
<tr><td>>hg38_21_+_8433172_8446622_RNA45SN1_45S_50ntFlanks@8158.8188.31|TCGTGGGGGGCCCAAGTCCTTCTGATCGAGG</td><td>>hg38_21_+_8433172_8446622_RNA45SN1_45S_50ntFlanks</td><td>8158</td><td>8188</td><td>31</td><td>TCGTGGGGGGCCCAAGTCCTTCTGATCGAGG</td><td>272.8</td><td>-5.4</td><td>1.6</td><td>-3.3</td><td>9.1E-04</td><td>2.3E-03</td><td>BT20/MB468</td><td>MP</td></tr>
<tr><td>>hg38_21_+_8433172_8446622_RNA45SN1_45S_50ntFlanks@11552.11574.23|CCTCGGCCGGCGCCTAGCAGCCG</td><td>>hg38_21_+_8433172_8446622_RNA45SN1_45S_50ntFlanks</td><td>11552</td><td>11574</td><td>23</td><td>CCTCGGCCGGCGCCTAGCAGCCG</td><td>177.2</td><td>-3.9</td><td>1.2</td><td>-3.1</td><td>1.9E-03</td><td>2.3E-03</td><td>MB231/MB468</td><td>NUC</td></tr>
<tr><td>>hg38_21_+_8433172_8446622_RNA45SN1_45S_50ntFlanks@7965.8005.41|CCCTCCGAGACGCGACCTCAGATCAGACGTGGCGACCCGCT</td><td>>hg38_21_+_8433172_8446622_RNA45SN1_45S_50ntFlanks</td><td>7965</td><td>8005</td><td>41</td><td>CCCTCCGAGACGCGACCTCAGATCAGACGTGGCGACCCGCT</td><td>116.6</td><td>5.6</td><td>1.8</td><td>3.1</td><td>1.9E-03</td><td>2.3E-03</td><td>BT20/MB468</td><td>NUC</td></tr>
<tr><td>>hg38_21_+_8433172_8446622_RNA45SN1_45S_50ntFlanks@11771.11794.24|ATTAGTGACGCGCATGAATGGATG</td><td>>hg38_21_+_8433172_8446622_RNA45SN1_45S_50ntFlanks</td><td>11771</td><td>11794</td><td>24</td><td>ATTAGTGACGCGCATGAATGGATG</td><td>936.2</td><td>2.0</td><td>0.6</td><td>3.1</td><td>1.8E-03</td><td>2.3E-03</td><td>MB231/MB468</td><td>CYTO</td></tr>
<tr><td>>MI0000727|hsa-mir-128-2&WithFlank&3|+|35744470|35744565@58.79.22&[MIMAT0000424_1&hsa-miR-128-3p&offsets|0|+1;m-117&3|+|35744527|35744547&offsets|0|+1]|TCACAGTGAACCGGTCTCTTTC</td><td>>MI0000727|hsa-mir-128-2&WithFlank&3|+|35744470|35744565</td><td>58</td><td>79</td><td>22</td><td>TCACAGTGAACCGGTCTCTTTC</td><td>183.3</td><td>3.1</td><td>1.0</td><td>3.1</td><td>1.9E-03</td><td>2.3E-03</td><td>MB231/MB468</td><td>MITO</td></tr>
<tr><td>>trna116_GluCTC_1_-_145399233_145399304@39.75.37,trna134_GluTTC_1_-_16861774_16861845@39.75.37,trna71_GluCTC_1_-_161439189_161439260@39.75.37,trna74_GluCTC_1_-_161431809_161431880@39.75.37,trna77_GluCTC_1_-_161424398_161424469@39.75.37,trna77_GluCTC_6_+_28949976_28950047@39.75.37,trna80_GluCTC_1_-_161417018_161417089@39.75.37,trna84_GluTTC_1_-_161391883_161391954@39.75.37,trna87_GluCTC_6_-_126101393_126101464@39.75.37|CGCCGCGGCCCGGGTTCGATTCCCGGTCAGGGAACCA</td><td>>trna116_GluCTC_1_-_145399233_145399304</td><td>39</td><td>75</td><td>37</td><td>CGCCGCGGCCCGGGTTCGATTCCCGGTCAGGGAACCA</td><td>195.6</td><td>4.2</td><td>1.4</td><td>3.1</td><td>1.9E-03</td><td>2.3E-03</td><td>MB231/MB468</td><td>MITO</td></tr>
<tr><td>>hg38_21_+_8433172_8446622_RNA45SN1_45S_50ntFlanks@13000.13027.28|TGCGATCTATTGAAAGTCAGCCCTCGAC</td><td>>hg38_21_+_8433172_8446622_RNA45SN1_45S_50ntFlanks</td><td>13000</td><td>13027</td><td>28</td><td>TGCGATCTATTGAAAGTCAGCCCTCGAC</td><td>257.6</td><td>-2.9</td><td>0.8</td><td>-3.5</td><td>4.1E-04</td><td>2.3E-03</td><td>BT20/MB231</td><td>MITO</td></tr>
<tr><td>>hg38_MT_+_598_1651_RNR1_12S_50ntFlanks@608.633.26|TGTTCTGTAATCGATAAACCCCGATC</td><td>>hg38_MT_+_598_1651_RNR1_12S_50ntFlanks</td><td>608</td><td>633</td><td>26</td><td>TGTTCTGTAATCGATAAACCCCGATC</td><td>207.0</td><td>-2.5</td><td>0.7</td><td>-3.6</td><td>3.7E-04</td><td>2.3E-03</td><td>BT20/MB231</td><td>CYTO</td></tr>
<tr><td>>hg38_21_+_8433172_8446622_RNA45SN1_45S_50ntFlanks@6655.6680.26|TCTTAGCGGTGGATCACTCGGCTCGT</td><td>>hg38_21_+_8433172_8446622_RNA45SN1_45S_50ntFlanks</td><td>6655</td><td>6680</td><td>26</td><td>TCTTAGCGGTGGATCACTCGGCTCGT</td><td>344.3</td><td>1.5</td><td>0.5</td><td>3.2</td><td>1.5E-03</td><td>2.3E-03</td><td>BT20/MB468</td><td>MITO</td></tr>
<tr><td>>MI0000456|hsa-mir-140&WithFlank&16|+|69933075|69933186@29.49.21&[MIMAT0000431&hsa-miR-140-5p&offsets|0|-1;m-173&16|+|69933103|69933124&offsets|0|-1]|CAGTGGTTTTACCCTATGGTA</td><td>>MI0000456|hsa-mir-140&WithFlank&16|+|69933075|69933186</td><td>29</td><td>49</td><td>21</td><td>CAGTGGTTTTACCCTATGGTA</td><td>210.6</td><td>5.9</td><td>1.9</td><td>3.2</td><td>1.6E-03</td><td>2.3E-03</td><td>MB231/MB468</td><td>MP</td></tr>
<tr><td>>hg38_21_+_8433172_8446622_RNA45SN1_45S_50ntFlanks@5084.5121.38|CGAACGAGACTCTGGCATGCTAACTAGTTACGCGACCC</td><td>>hg38_21_+_8433172_8446622_RNA45SN1_45S_50ntFlanks</td><td>5084</td><td>5121</td><td>38</td><td>CGAACGAGACTCTGGCATGCTAACTAGTTACGCGACCC</td><td>228.5</td><td>-3.9</td><td>1.2</td><td>-3.2</td><td>1.5E-03</td><td>2.3E-03</td><td>BT20/MB468</td><td>MITO</td></tr>
<tr><td>>hg38_21_+_8433172_8446622_RNA45SN1_45S_50ntFlanks@4753.4783.31|AAGACGATCAGATACCGTCGTAGTTCCGACC</td><td>>hg38_21_+_8433172_8446622_RNA45SN1_45S_50ntFlanks</td><td>4753</td><td>4783</td><td>31</td><td>AAGACGATCAGATACCGTCGTAGTTCCGACC</td><td>738.4</td><td>-3.2</td><td>1.0</td><td>-3.2</td><td>1.5E-03</td><td>2.3E-03</td><td>BT20/MB468</td><td>MITO</td></tr>
<tr><td>>MI0000087|hsa-mir-29a&WithFlank&7|-|130876741|130876816@47.67.21(+1G)&[MIMAT0000086&hsa-miR-29a-3p&offsets|-1|-2(+1G);m-26&7|-|130876749|130876769&offsets|-1|-1(+1G)]|CTAGCACCATCTGAAATCGGTG</td><td>>MI0000087|hsa-mir-29a&WithFlank&7|-|130876741|130876816</td><td>47</td><td>67</td><td>21(+1G)</td><td>CTAGCACCATCTGAAATCGGTG</td><td>247.0</td><td>5.2</td><td>1.6</td><td>3.2</td><td>1.6E-03</td><td>2.3E-03</td><td>MB231/MB468</td><td>MP</td></tr>
<tr><td>>hg38_21_+_8433172_8446622_RNA45SN1_45S_50ntFlanks@11591.11612.22|GACCAGGGGAATCCGACTGTTT</td><td>>hg38_21_+_8433172_8446622_RNA45SN1_45S_50ntFlanks</td><td>11591</td><td>11612</td><td>22</td><td>GACCAGGGGAATCCGACTGTTT</td><td>149.2</td><td>-5.3</td><td>1.7</td><td>-3.2</td><td>1.6E-03</td><td>2.3E-03</td><td>MB231/MB468</td><td>MP</td></tr>
<tr><td>>MI0000439|hsa-mir-23b&WithFlank&9|+|95085202|95085310@67.85.19(+1U)&[MIMAT0000418&hsa-miR-23b-3p&offsets|+3|-1(+1U);m-42&9|+|95085265|95085285&offsets|+3|+1(+1U)]|ACATTGCCAGGGATTACCAT</td><td>>MI0000439|hsa-mir-23b&WithFlank&9|+|95085202|95085310</td><td>67</td><td>85</td><td>19(+1U)</td><td>ACATTGCCAGGGATTACCAT</td><td>264.4</td><td>11.3</td><td>3.6</td><td>3.1</td><td>1.9E-03</td><td>2.3E-03</td><td>MB231/MB468</td><td>MITO</td></tr>
<tr><td>>hg38_21_+_8433172_8446622_RNA45SN1_45S_50ntFlanks@3705.3748.44|TACCTGGTTGATCCTGCCAGTAGCATATGCTTGTCTCAAAGATT</td><td>>hg38_21_+_8433172_8446622_RNA45SN1_45S_50ntFlanks</td><td>3705</td><td>3748</td><td>44</td><td>TACCTGGTTGATCCTGCCAGTAGCATATGCTTGTCTCAAAGATT</td><td>813.3</td><td>3.6</td><td>1.2</td><td>3.1</td><td>1.9E-03</td><td>2.3E-03</td><td>MB231/MB468</td><td>MITO</td></tr>
</table>

<table>
<tr><td>>am_trnaMT_HisGTG_MT_+_12138_12206@19.38.20|AAACATCAGATTGTGAATCT</td><td>>am_trnaMT_HisGTG_MT_+_12138_12206</td><td>19</td><td>38</td><td>20</td><td>AAACATCAGATTGTGAATCT</td><td>172.1</td><td>3.9</td><td>1.2</td><td>3.3</td><td>9.1E-04</td><td>2.3E-03</td><td>BT20/MB468</td><td>MP</td></tr>
<tr><td>>hg38_21_+_8433172_8446622_RNA45SN1_45S_50ntFlanks@11628.11660.33|TCGCGAAGGCCCGCGGCGGGTGTTGACGCGATG</td><td>>hg38_21_+_8433172_8446622_RNA45SN1_45S_50ntFlanks</td><td>11628</td><td>11660</td><td>33</td><td>TCGCGAAGGCCCGCGGCGGGTGTTGACGCGATG</td><td>821.1</td><td>-4.9</td><td>1.5</td><td>-3.3</td><td>9.1E-04</td><td>2.3E-03</td><td>BT20/MB468</td><td>MP</td></tr>
<tr><td>>hg38_21_+_8433172_8446622_RNA45SN1_45S_50ntFlanks@10348.10388.41|AAACGAGAACTTTGAAGGCCGAAGTGGAGAAGGGTTCCATG</td><td>>hg38_21_+_8433172_8446622_RNA45SN1_45S_50ntFlanks</td><td>10348</td><td>10388</td><td>41</td><td>AAACGAGAACTTTGAAGGCCGAAGTGGAGAAGGGTTCCATG</td><td>450.5</td><td>-2.7</td><td>0.9</td><td>-3.1</td><td>1.9E-03</td><td>2.3E-03</td><td>MB231/MB468</td><td>NUC</td></tr>
<tr><td>>hg38_21_+_8433172_8446622_RNA45SN1_45S_50ntFlanks@11933.11967.35|AATAAGTGGGAGGCCCCCGGCGCCCCCCCGGTGTC</td><td>>hg38_21_+_8433172_8446622_RNA45SN1_45S_50ntFlanks</td><td>11933</td><td>11967</td><td>35</td><td>AATAAGTGGGAGGCCCCCGGCGCCCCCCCGGTGTC</td><td>179.0</td><td>-5.4</td><td>1.7</td><td>-3.2</td><td>1.5E-03</td><td>2.3E-03</td><td>BT20/MB468</td><td>MITO</td></tr>
<tr><td>>hg38_21_+_8433172_8446622_RNA45SN1_45S_50ntFlanks@11795.11820.26|AACGAGATTCCCACTGTCCCTACCTA</td><td>>hg38_21_+_8433172_8446622_RNA45SN1_45S_50ntFlanks</td><td>11795</td><td>11820</td><td>26</td><td>AACGAGATTCCCACTGTCCCTACCTA</td><td>192.0</td><td>-4.6</td><td>1.4</td><td>-3.2</td><td>1.5E-03</td><td>2.3E-03</td><td>BT20/MB468</td><td>MITO</td></tr>
<tr><td>>hg38_21_+_8433172_8446622_RNA45SN1_45S_50ntFlanks@12416.12451.36|TTCGATGTCGGCTCTTCCTATCATTGTGAAGCAGAA</td><td>>hg38_21_+_8433172_8446622_RNA45SN1_45S_50ntFlanks</td><td>12416</td><td>12451</td><td>36</td><td>TTCGATGTCGGCTCTTCCTATCATTGTGAAGCAGAA</td><td>106.3</td><td>2.6</td><td>0.8</td><td>3.1</td><td>1.9E-03</td><td>2.3E-03</td><td>MB231/MB468</td><td>MITO</td></tr>
<tr><td>>hg38_21_+_8433172_8446622_RNA45SN1_45S_50ntFlanks@5082.5102.21|AACGAACGAGACTCTGGCATG</td><td>>hg38_21_+_8433172_8446622_RNA45SN1_45S_50ntFlanks</td><td>5082</td><td>5102</td><td>21</td><td>AACGAACGAGACTCTGGCATG</td><td>352.2</td><td>-3.0</td><td>1.0</td><td>-3.2</td><td>1.6E-03</td><td>2.3E-03</td><td>MB231/MB468</td><td>MP</td></tr>
<tr><td>>MI0003560|hsa-mir-92b&WithFlank&1|+|155195171|155195278@67.87.21(+1G)&[MIMAT0003218&hsa-miR-92b-3p&offsets|0|-1(+1G);m-115&1|+|155195237|155195258&offsets|0|-1(+1G)]|TATTGCACTCGTCCCGGCCTCG</td><td>>MI0003560|hsa-mir-92b&WithFlank&1|+|155195171|155195278</td><td>67</td><td>87</td><td>21(+1G)</td><td>TATTGCACTCGTCCCGGCCTCG</td><td>42.7</td><td>8.6</td><td>2.7</td><td>3.2</td><td>1.6E-03</td><td>2.3E-03</td><td>MB231/MB468</td><td>MP</td></tr>
<tr><td>>am_trnaMT_PheGAA_MT_+_577_647@42.64.23|TTTAGACGGGCTCACATCACCCC</td><td>>am_trnaMT_PheGAA_MT_+_577_647</td><td>42</td><td>64</td><td>23</td><td>TTTAGACGGGCTCACATCACCCC</td><td>114.7</td><td>6.9</td><td>2.1</td><td>3.3</td><td>9.2E-04</td><td>2.3E-03</td><td>BT20/MB468</td><td>MP</td></tr>
<tr><td>>trnaMT_ValTAC_MT_+_1602_1670@28.72.45|AACTTACACTTAGGAGATTTCAACTTAACTTGACCGCTCTGACCA</td><td>>trnaMT_ValTAC_MT_+_1602_1670</td><td>28</td><td>72</td><td>45</td><td>AACTTACACTTAGGAGATTTCAACTTAACTTGACCGCTCTGACCA</td><td>1317.3</td><td>3.5</td><td>1.1</td><td>3.1</td><td>1.9E-03</td><td>2.3E-03</td><td>MB231/MB468</td><td>MITO</td></tr>
<tr><td>>hg38_21_+_8433172_8446622_RNA45SN1_45S_50ntFlanks@12914.12943.30|CGTTCGTGGGGAACCTGGCGCTAAACCATT</td><td>>hg38_21_+_8433172_8446622_RNA45SN1_45S_50ntFlanks</td><td>12914</td><td>12943</td><td>30</td><td>CGTTCGTGGGGAACCTGGCGCTAAACCATT</td><td>357.0</td><td>7.2</td><td>2.0</td><td>3.6</td><td>3.1E-04</td><td>2.3E-03</td><td>BT20/MB231</td><td>MP</td></tr>
<tr><td>>MI0000061|hsa-let-7a-2&WithFlank&11|-|122146516|122146599@11.30.20;MI0000062|hsa-let-7a-3&WithFlank&22|+|46112743|46112828@10.29.20;MI0000060|hsa-let-7a-1&WithFlank&9|+|94175951|94176042@12.31.20&[MIMAT0000062&hsa-let-7a-5p&offsets|0|-2;m-6&11|-|122146568|122146589&offsets|0|-2];[MIMAT0000062_1&hsa-let-7a-5p&offsets|0|-2;m-5&22|+|46112752|46112773&offsets|0|-2];[MIMAT0000062_2&hsa-let-7a-5p&offsets|0|-2;m-4&9|+|94175962|94175983&offsets|0|-2]|TGAGGTAGTAGGTTGTATAG</td><td>>MI0000061|hsa-let-7a-2&WithFlank&11|-|122146516|122146599</td><td>11</td><td>30</td><td>20</td><td>TGAGGTAGTAGGTTGTATAG</td><td>142.7</td><td>5.6</td><td>1.7</td><td>3.3</td><td>9.2E-04</td><td>2.3E-03</td><td>BT20/MB468</td><td>MP</td></tr>
<tr><td>>hg38_21_+_8433172_8446622_RNA45SN1_45S_50ntFlanks@12785.12805.21|CCCCGCGCGCGCGGGAGGGCG</td><td>>hg38_21_+_8433172_8446622_RNA45SN1_45S_50ntFlanks</td><td>12785</td><td>12805</td><td>21</td><td>CCCCGCGCGCGCGGGAGGGCG</td><td>637.6</td><td>-5.0</td><td>1.6</td><td>-3.1</td><td>1.8E-03</td><td>2.3E-03</td><td>BT20/MB468</td><td>CYTO</td></tr>
<tr><td>>hg38_21_+_8433172_8446622_RNA45SN1_45S_50ntFlanks@5418.5443.26|TACCGATTGGATGGTTTAGTGAGGCC</td><td>>hg38_21_+_8433172_8446622_RNA45SN1_45S_50ntFlanks</td><td>5418</td><td>5443</td><td>26</td><td>TACCGATTGGATGGTTTAGTGAGGCC</td><td>129.2</td><td>-1.7</td><td>0.5</td><td>-3.1</td><td>1.8E-03</td><td>2.3E-03</td><td>BT20/MB468</td><td>CYTO</td></tr>
<tr><td>>hg38_21_+_8433172_8446622_RNA45SN1_45S_50ntFlanks@9305.9335.31|CCAAGGAGTCTAACACGTGCGCGAGTCGGGG</td><td>>hg38_21_+_8433172_8446622_RNA45SN1_45S_50ntFlanks</td><td>9305</td><td>9335</td><td>31</td><td>CCAAGGAGTCTAACACGTGCGCGAGTCGGGG</td><td>326.9</td><td>-3.8</td><td>1.2</td><td>-3.2</td><td>1.5E-03</td><td>2.3E-03</td><td>BT20/MB468</td><td>MITO</td></tr>
<tr><td>>am_trna10_LysCTT_16_+_3241501_3241573@11.28.18,trna119_LysCTT_1_-_145395522_145395594@11.28.18,trna11_LysCTT_5_-_180648979_180649051@11.28.18,trna128_LysCTT_1_-_55423542_55423614@11.28.18,trna13_LysCTT_14_-_58706613_58706685@11.28.18,trna13_LysCTT_6_+_26556774_26556846@11.28.18,trna2_LysCTT_15_+_79152904_79152976@11.28.18,trna32_LysCTT_16_-_3207406_3207478@11.28.18,trna4_LysCTT_18_-_43669271_43669343@11.28.18,trna6_LysCTT_19_-_52425396_52425468@11.28.18,trna7_LysCTT_16_+_3225692_3225764@11.28.18,trna9_LysCTT_5_+_180634755_180634827@11.28.18|CTCAGTCGGTAGAGCATG</td><td>>am_trna10_LysCTT_16_+_3241501_3241573</td><td>11</td><td>28</td><td>18</td><td>CTCAGTCGGTAGAGCATG</td><td>365.5</td><td>-2.3</td><td>0.8</td><td>-3.1</td><td>1.9E-03</td><td>2.3E-03</td><td>MB231/MB468</td><td>NUC</td></tr>
<tr><td>>am_trna10_ValCAC_5_-_180649395_180649467@1.29.29,trna12_ValAAC_5_-_180645270_180645342@1.29.29,trna132_ValAAC_6_-_27721179_27721251@1.29.29,trna136_ValAAC_6_-_27648885_27648957@1.29.29,trna139_ValAAC_6_-_27618707_27618779@1.29.29,trna18_ValCAC_5_-_180529253_180529325@1.29.29,trna2_ValAAC_3_+_169490018_169490090@1.29.29,trna2_ValCAC_5_+_180524070_180524142@1.29.29,trna4_ValAAC_5_+_180591154_180591226@1.29.29,trna5_ValAAC_5_+_180596610_180596682@1.29.29,trna6_ValCAC_5_+_180600650_180600722@1.29.29,trna85_ValCAC_1_-_161369490_161369562@1.29.29,trna90_ValCAC_1_-_149684088_149684161@1.29.29,trna98_ValCAC_1_-_149298555_149298627@1.29.29,trna9_ValCAC_6_+_26538282_26538354@1.29.29|GTTTCCGTAGTGTAGTGGTTATCACGTTC</td><td>>am_trna10_ValCAC_5_-_180649395_180649467</td><td>1</td><td>29</td><td>29</td><td>GTTTCCGTAGTGTAGTGGTTATCACGTTC</td><td>4106.0</td><td>-3.5</td><td>1.1</td><td>-3.2</td><td>1.5E-03</td><td>2.3E-03</td><td>BT20/MB468</td><td>MITO</td></tr>
<tr><td>>am_trna10_ValCAC_5_-_180649395_180649467@1.23.23,trna12_ValAAC_5_-_180645270_180645342@1.23.23,trna132_ValAAC_6_-_27721179_27721251@1.23.23,trna136_ValAAC_6_-_27648885_27648957@1.23.23,trna139_ValAAC_6_-_27618707_27618779@1.23.23,trna18_ValCAC_5_-_180529253_180529325@1.23.23,trna2_ValAAC_3_+_169490018_169490090@1.23.23,trna2_ValCAC_5_+_180524070_180524142@1.23.23,trna37_ValAAC_6_+_27203288_27203360@1.23.23,trna4_ValAAC_5_+_180591154_180591226@1.23.23,trna5_ValAAC_5_+_180596610_180596682@1.23.23,trna6_ValCAC_5_+_180600650_180600722@1.23.23,trna85_ValCAC_1_-_161369490_161369562@1.23.23,trna90_ValCAC_1_-_149684088_149684161@1.23.23,trna98_ValCAC_1_-_149298555_149298627@1.23.23,trna9_ValCAC_6_+_26538282_26538354@1.23.23|GTTTCCGTAGTGTAGTGGTTATC</td><td>>am_trna10_ValCAC_5_-_180649395_180649467</td><td>1</td><td>23</td><td>23</td><td>GTTTCCGTAGTGTAGTGGTTATC</td><td>5005.4</td><td>4.5</td><td>1.4</td><td>3.3</td><td>9.2E-04</td><td>2.3E-03</td><td>BT20/MB468</td><td>MP</td></tr>
<tr><td>>hg38_21_+_8433172_8446622_RNA45SN1_45S_50ntFlanks@4882.4903.22|TGCAAAGCTGAAACTTAAAGGA</td><td>>hg38_21_+_8433172_8446622_RNA45SN1_45S_50ntFlanks</td><td>4882</td><td>4903</td><td>22</td><td>TGCAAAGCTGAAACTTAAAGGA</td><td>232.9</td><td>5.7</td><td>1.7</td><td>3.3</td><td>9.2E-04</td><td>2.3E-03</td><td>BT20/MB468</td><td>MP</td></tr>
<tr><td>>hg38_21_+_8433172_8446622_RNA45SN1_45S_50ntFlanks@11076.11103.28|CGCCGCGTGGGGGCGGCGGCGGGGGGAG</td><td>>hg38_21_+_8433172_8446622_RNA45SN1_45S_50ntFlanks</td><td>11076</td><td>11103</td><td>28</td><td>CGCCGCGTGGGGGCGGCGGCGGGGGGAG</td><td>159.4</td><td>-5.6</td><td>1.7</td><td>-3.3</td><td>9.2E-04</td><td>2.3E-03</td><td>BT20/MB468</td><td>MP</td></tr>
<tr><td>>hg38_21_+_8433172_8446622_RNA45SN1_45S_50ntFlanks@12597.12614.18|ATTTGGTGTATGTGCTTG</td><td>>hg38_21_+_8433172_8446622_RNA45SN1_45S_50ntFlanks</td><td>12597</td><td>12614</td><td>18</td><td>ATTTGGTGTATGTGCTTG</td><td>234.6</td><td>-2.6</td><td>0.8</td><td>-3.1</td><td>1.8E-03</td><td>2.3E-03</td><td>BT20/MB468</td><td>CYTO</td></tr>
<tr><td>>hg38_21_+_8433172_8446622_RNA45SN1_45S_50ntFlanks@7977.8016.40|CGACCTCAGATCAGACGTGGCGACCCGCTGAATTTAAGCA</td><td>>hg38_21_+_8433172_8446622_RNA45SN1_45S_50ntFlanks</td><td>7977</td><td>8016</td><td>40</td><td>CGACCTCAGATCAGACGTGGCGACCCGCTGAATTTAAGCA</td><td>961.2</td><td>3.3</td><td>1.1</td><td>3.1</td><td>1.9E-03</td><td>2.3E-03</td><td>MB231/MB468</td><td>MITO</td></tr>
<tr><td>>am_trnaMT_SerGCT_MT_+_12207_12265@18.38.21|CTGCTAACTCATGCCCCCATG</td><td>>am_trnaMT_SerGCT_MT_+_12207_12265</td><td>18</td><td>38</td><td>21</td><td>CTGCTAACTCATGCCCCCATG</td><td>86.2</td><td>-5.7</td><td>1.7</td><td>-3.3</td><td>9.2E-04</td><td>2.3E-03</td><td>BT20/MB468</td><td>MP</td></tr>
<tr><td>>hg38_21_+_8433172_8446622_RNA45SN1_45S_50ntFlanks@12417.12436.20|TCGATGTCGGCTCTTCCTAT</td><td>>hg38_21_+_8433172_8446622_RNA45SN1_45S_50ntFlanks</td><td>12417</td><td>12436</td><td>20</td><td>TCGATGTCGGCTCTTCCTAT</td><td>730.2</td><td>-5.6</td><td>1.7</td><td>-3.3</td><td>9.2E-04</td><td>2.3E-03</td><td>BT20/MB468</td><td>MP</td></tr>
<tr><td>>MI0003205|hsa-mir-532&WithFlank&X|+|50003142|50003244@26.46.21(+1C)&[MIMAT0002888&hsa-miR-532-5p&offsets|0|-1(+1C);m-65&X|+|50003167|50003188&offsets|0|-1(+1C)]|CATGCCTTGAGTGTAGGACCGC</td><td>>MI0003205|hsa-mir-532&WithFlank&X|+|50003142|50003244</td><td>26</td><td>46</td><td>21(+1C)</td><td>CATGCCTTGAGTGTAGGACCGC</td><td>587.6</td><td>2.5</td><td>0.7</td><td>3.6</td><td>3.7E-04</td><td>2.3E-03</td><td>BT20/MB231</td><td>CYTO</td></tr>
<tr><td>>hg38_21_+_8433172_8446622_RNA45SN1_45S_50ntFlanks@12259.12284.26|CTTGATCTTGATTTTCAGTACGAATA</td><td>>hg38_21_+_8433172_8446622_RNA45SN1_45S_50ntFlanks</td><td>12259</td><td>12284</td><td>26</td><td>CTTGATCTTGATTTTCAGTACGAATA</td><td>1737.2</td><td>-3.5</td><td>1.1</td><td>-3.3</td><td>9.2E-04</td><td>2.3E-03</td><td>BT20/MB468</td><td>MP</td></tr>
<tr><td>>trnaMT_GluTTC_MT_-_14674_14742@41.72.32,trnalookalike8_GluTTC_5_-_93905172_93905240@41.72.32|TGGTCGTGGTTGTAGTCCGTGCGAGAATACCA</td><td>>trnaMT_GluTTC_MT_-_14674_14742</td><td>41</td><td>72</td><td>32</td><td>TGGTCGTGGTTGTAGTCCGTGCGAGAATACCA</td><td>7138.2</td><td>-3.0</td><td>0.8</td><td>-3.6</td><td>3.5E-04</td><td>2.3E-03</td><td>BT20/MB231</td><td>NUC</td></tr>
<tr><td>>MI0000063|hsa-let-7b&WithFlank&22|+|46113680|46113774@12.32.21&[MIMAT0000063&hsa-let-7b-5p&offsets|0|-1;m-14&22|+|46113691|46113712&offsets|0|-1]|TGAGGTAGTAGGTTGTGTGGT</td><td>>MI0000063|hsa-let-7b&WithFlank&22|+|46113680|46113774</td><td>12</td><td>32</td><td>21</td><td>TGAGGTAGTAGGTTGTGTGGT</td><td>8883.3</td><td>2.9</td><td>0.8</td><td>3.6</td><td>3.7E-04</td><td>2.3E-03</td><td>BT20/MB231</td><td>CYTO</td></tr>
<tr><td>>hg38_21_+_8433172_8446622_RNA45SN1_45S_50ntFlanks@12477.12500.24|CACTAATAGGGAACGTGAGCTGGG</td><td>>hg38_21_+_8433172_8446622_RNA45SN1_45S_50ntFlanks</td><td>12477</td><td>12500</td><td>24</td><td>CACTAATAGGGAACGTGAGCTGGG</td><td>104.2</td><td>-3.8</td><td>1.2</td><td>-3.2</td><td>1.6E-03</td><td>2.3E-03</td><td>MB231/MB468</td><td>MP</td></tr>
<tr><td>>MI0000289|hsa-mir-181a-1&WithFlank&1|-|198859038|198859159@30.53.24(+1C);MI0000269|hsa-mir-181a-2&WithFlank&9|+|124692436|124692557@45.68.24(+1C)&[MIMAT0000256&hsa-miR-181a-5p&offsets|0|+1(+1C);m-32&1|-|198859108|198859130&offsets|0|+1(+1C)];[MIMAT0000256_1&hsa-miR-181a-5p&offsets|0|+1(+1C);m-32&9|+|124692480|124692502&offsets|0|+1(+1C)]|AACATTCAACGCTGTCGGTGAGTTC</td><td>>MI0000289|hsa-mir-181a-1&WithFlank&1|-|198859038|198859159</td><td>30</td><td>53</td><td>24(+1C)</td><td>AACATTCAACGCTGTCGGTGAGTTC</td><td>725.5</td><td>4.5</td><td>1.4</td><td>3.2</td><td>1.6E-03</td><td>2.3E-03</td><td>MB231/MB468</td><td>MP</td></tr>
<tr><td>>hg38_21_+_8433172_8446622_RNA45SN1_45S_50ntFlanks@9571.9595.25|GCAAATCGGTCGTCCGACCTGGGTA</td><td>>hg38_21_+_8433172_8446622_RNA45SN1_45S_50ntFlanks</td><td>9571</td><td>9595</td><td>25</td><td>GCAAATCGGTCGTCCGACCTGGGTA</td><td>205.1</td><td>-3.2</td><td>1.0</td><td>-3.1</td><td>1.8E-03</td><td>2.3E-03</td><td>BT20/MB468</td><td>CYTO</td></tr>
<tr><td>>hg38_21_+_8433172_8446622_RNA45SN1_45S_50ntFlanks@3984.4001.18|GCGGCTTTGGTGACTCTA</td><td>>hg38_21_+_8433172_8446622_RNA45SN1_45S_50ntFlanks</td><td>3984</td><td>4001</td><td>18</td><td>GCGGCTTTGGTGACTCTA</td><td>875.4</td><td>-4.1</td><td>1.3</td><td>-3.1</td><td>1.8E-03</td><td>2.3E-03</td><td>MB231/MB468</td><td>CYTO</td></tr>
<tr><td>>MI0000066|hsa-let-7e&WithFlank&19|+|51692780|51692870@14.35.22(+1A)&[MIMAT0000066&hsa-let-7e-5p&offsets|0|0(+1A);m-50&19|+|51692793|51692814&offsets|0|0(+1A)]|TGAGGTAGGAGGTTGTATAGTTA</td><td>>MI0000066|hsa-let-7e&WithFlank&19|+|51692780|51692870</td><td>14</td><td>35</td><td>22(+1A)</td><td>TGAGGTAGGAGGTTGTATAGTTA</td><td>139.9</td><td>1.2</td><td>0.4</td><td>3.1</td><td>1.9E-03</td><td>2.3E-03</td><td>BT20/MB468</td><td>NUC</td></tr>
<tr><td>>hg38_21_+_8433172_8446622_RNA45SN1_45S_50ntFlanks@11206.11232.27|CGGCGGCGACTCTGGACGCGAGCCGGG</td><td>>hg38_21_+_8433172_8446622_RNA45SN1_45S_50ntFlanks</td><td>11206</td><td>11232</td><td>27</td><td>CGGCGGCGACTCTGGACGCGAGCCGGG</td><td>652.6</td><td>-3.6</td><td>1.1</td><td>-3.1</td><td>1.9E-03</td><td>2.3E-03</td><td>BT20/MB468</td><td>NUC</td></tr>
<tr><td>>hg38_21_+_8433172_8446622_RNA45SN1_45S_50ntFlanks@11209.11226.18|CGGCGACTCTGGACGCGA</td><td>>hg38_21_+_8433172_8446622_RNA45SN1_45S_50ntFlanks</td><td>11209</td><td>11226</td><td>18</td><td>CGGCGACTCTGGACGCGA</td><td>91.2</td><td>-5.4</td><td>1.6</td><td>-3.3</td><td>9.3E-04</td><td>2.3E-03</td><td>BT20/MB468</td><td>MP</td></tr>
</table>

<table>
<tr><th></th><th></th><th></th><th></th><th></th><th></th><th></th><th></th><th></th><th></th><th></th><th></th><th></th><th></th></tr>
<tr><td>>hg38_21_+_8433172_8446622_RNA45SN1_45S_50ntFlanks@11755.11777.23|AATGCCTCGTCATCTAATTAGTG</td><td>>hg38_21_+_8433172_8446622_RNA45SN1_45S_50ntFlanks</td><td>11755</td><td>11777</td><td>23</td><td>AATGCCTCGTCATCTAATTAGTG</td><td>59.6</td><td>-5.1</td><td>1.5</td><td>-3.3</td><td>9.3E-04</td><td>2.3E-03</td><td>BT20/MB468</td><td>MP</td></tr>
<tr><td>>hg38_21_+_8433172_8446622_RNA45SN1_45S_50ntFlanks@7976.8004.29|GCGACCTCAGATCAGACGTGGCGACCCGC</td><td>>hg38_21_+_8433172_8446622_RNA45SN1_45S_50ntFlanks</td><td>7976</td><td>8004</td><td>29</td><td>GCGACCTCAGATCAGACGTGGCGACCCGC</td><td>68.1</td><td>5.5</td><td>1.7</td><td>3.3</td><td>9.3E-04</td><td>2.3E-03</td><td>BT20/MB468</td><td>MP</td></tr>
<tr><td>>MI0000465|hsa-mir-191&WithFlank&3|-|49020612|49020715@22.44.23(+1A)&[MIMAT0000440&hsa-miR-191-5p&offsets|0|0(+1A);m-48&3|-|49020672|49020694&offsets|0|0(+1A)]|CAACGGAATCCCAAAAGCAGCTGA</td><td>>MI0000465|hsa-mir-191&WithFlank&3|-|49020612|49020715</td><td>22</td><td>44</td><td>23(+1A)</td><td>CAACGGAATCCCAAAAGCAGCTGA</td><td>536.1</td><td>2.7</td><td>0.9</td><td>3.1</td><td>1.6E-03</td><td>2.3E-03</td><td>MB231/MB468</td><td>MP</td></tr>
<tr><td>>hg38_21_+_8433172_8446622_RNA45SN1_45S_50ntFlanks@7975.7995.21|CGCGACCTCAGATCAGACGTG</td><td>>hg38_21_+_8433172_8446622_RNA45SN1_45S_50ntFlanks</td><td>7975</td><td>7995</td><td>21</td><td>CGCGACCTCAGATCAGACGTG</td><td>30998.0</td><td>3.3</td><td>1.0</td><td>3.1</td><td>1.8E-03</td><td>2.3E-03</td><td>BT20/MB468</td><td>CYTO</td></tr>
<tr><td>>hg38_1_-_228634819_228635039_RNA5S12_5S_50ntFlanks@51.88.38|GTCTACGGCCATACCACCCTGAACGCGCCCGATCTCGT</td><td>>hg38_1_-_228634819_228635039_RNA5S12_5S_50ntFlanks</td><td>51</td><td>88</td><td>38</td><td>GTCTACGGCCATACCACCCTGAACGCGCCCGATCTCGT</td><td>681.4</td><td>3.7</td><td>1.1</td><td>3.3</td><td>9.3E-04</td><td>2.3E-03</td><td>BT20/MB468</td><td>MP</td></tr>
<tr><td>>MI0001448|hsa-mir-425&WithFlank&3|-|49020142|49020240@20.39.20&[MIMAT0003393&hsa-miR-425-5p&offsets|0|-3;m-95&3|-|49020200|49020221&offsets|0|-2]|AATGACACGATCACTCCCGT</td><td>>MI0001448|hsa-mir-425&WithFlank&3|-|49020142|49020240</td><td>20</td><td>39</td><td>20</td><td>AATGACACGATCACTCCCGT</td><td>127.5</td><td>5.8</td><td>1.9</td><td>3.1</td><td>1.9E-03</td><td>2.3E-03</td><td>MB231/MB468</td><td>MITO</td></tr>
<tr><td>>hg38_21_+_8433172_8446622_RNA45SN1_45S_50ntFlanks@12642.12659.18|CATCTGTGGGATTATGAC</td><td>>hg38_21_+_8433172_8446622_RNA45SN1_45S_50ntFlanks</td><td>12642</td><td>12659</td><td>18</td><td>CATCTGTGGGATTATGAC</td><td>911.8</td><td>-3.3</td><td>1.0</td><td>-3.2</td><td>1.5E-03</td><td>2.3E-03</td><td>BT20/MB468</td><td>MITO</td></tr>
<tr><td>>am_trnaMT_ArgTCG_MT_+_10405_10469@31.48.18|TCGACTCATTAAATTATG</td><td>>am_trnaMT_ArgTCG_MT_+_10405_10469</td><td>31</td><td>48</td><td>18</td><td>TCGACTCATTAAATTATG</td><td>178.5</td><td>-4.8</td><td>1.5</td><td>-3.1</td><td>1.8E-03</td><td>2.3E-03</td><td>BT20/MB468</td><td>CYTO</td></tr>
<tr><td>>trnaMT_GluTTC_MT_-_14674_14742@40.72.33,trnalookalike8_GluTTC_5_-_93905172_93905240@40.72.33|TTGGTCGTGGTTGTAGTCCGTGCGAGAATACCA</td><td>>trnaMT_GluTTC_MT_-_14674_14742</td><td>40</td><td>72</td><td>33</td><td>TTGGTCGTGGTTGTAGTCCGTGCGAGAATACCA</td><td>28129.1</td><td>3.9</td><td>1.2</td><td>3.3</td><td>9.3E-04</td><td>2.3E-03</td><td>BT20/MB468</td><td>MP</td></tr>
<tr><td>>MI0000087|hsa-mir-29a&WithFlank&7|-|130876741|130876816@47.69.23&[MIMAT0000086&hsa-miR-29a-3p&offsets|-1|0;m-26&7|-|130876749|130876769&offsets|-1|+1]|CTAGCACCATCTGAAATCGGTTA</td><td>>MI0000087|hsa-mir-29a&WithFlank&7|-|130876741|130876816</td><td>47</td><td>69</td><td>23</td><td>CTAGCACCATCTGAAATCGGTTA</td><td>3302.1</td><td>5.9</td><td>1.8</td><td>3.3</td><td>9.3E-04</td><td>2.3E-03</td><td>BT20/MB468</td><td>MP</td></tr>
<tr><td>>hg38_21_+_8433172_8446622_RNA45SN1_45S_50ntFlanks@8345.8365.21|AAGGGAAAGTTGAAAAGAACT</td><td>>hg38_21_+_8433172_8446622_RNA45SN1_45S_50ntFlanks</td><td>8345</td><td>8365</td><td>21</td><td>AAGGGAAAGTTGAAAAGAACT</td><td>74.5</td><td>6.4</td><td>1.9</td><td>3.3</td><td>9.3E-04</td><td>2.3E-03</td><td>BT20/MB468</td><td>MP</td></tr>
<tr><td>>hg38_21_+_8433172_8446622_RNA45SN1_45S_50ntFlanks@8366.8395.30|TTGAAGAGAGAGTTCAAGAGGGCGTGAAAC</td><td>>hg38_21_+_8433172_8446622_RNA45SN1_45S_50ntFlanks</td><td>8366</td><td>8395</td><td>30</td><td>TTGAAGAGAGAGTTCAAGAGGGCGTGAAAC</td><td>41.3</td><td>9.1</td><td>2.8</td><td>3.3</td><td>9.3E-04</td><td>2.3E-03</td><td>BT20/MB468</td><td>MP</td></tr>
<tr><td>>MI0003834|hsa-mir-769&WithFlank&19|+|46018926|46019055@36.57.22&[MIMAT0003886&hsa-miR-769-5p&offsets|0|0;m-216&19|+|46018961|46018982&offsets|0|0]|TGAGACCTCTGGGTTCTGAGCT</td><td>>MI0003834|hsa-mir-769&WithFlank&19|+|46018926|46019055</td><td>36</td><td>57</td><td>22</td><td>TGAGACCTCTGGGTTCTGAGCT</td><td>1622.3</td><td>2.5</td><td>0.8</td><td>3.1</td><td>1.6E-03</td><td>2.3E-03</td><td>MB231/MB468</td><td>MP</td></tr>
<tr><td>>hg38_21_+_8433172_8446622_RNA45SN1_45S_50ntFlanks@9533.9554.22|GAGGAAACTCTGGTGGAGGTCC</td><td>>hg38_21_+_8433172_8446622_RNA45SN1_45S_50ntFlanks</td><td>9533</td><td>9554</td><td>22</td><td>GAGGAAACTCTGGTGGAGGTCC</td><td>480.1</td><td>-4.2</td><td>1.3</td><td>-3.1</td><td>1.8E-03</td><td>2.3E-03</td><td>BT20/MB468</td><td>CYTO</td></tr>
<tr><td>>hg38_21_+_8433172_8446622_RNA45SN1_45S_50ntFlanks@12836.12871.36|CGGTGCGGAGTGCCCTTCGTCCTGGGAAACGGGGCG</td><td>>hg38_21_+_8433172_8446622_RNA45SN1_45S_50ntFlanks</td><td>12836</td><td>12871</td><td>36</td><td>CGGTGCGGAGTGCCCTTCGTCCTGGGAAACGGGGCG</td><td>504.1</td><td>-2.6</td><td>0.8</td><td>-3.2</td><td>1.5E-03</td><td>2.3E-03</td><td>BT20/MB468</td><td>MITO</td></tr>
<tr><td>>hg38_21_+_8433172_8446622_RNA45SN1_45S_50ntFlanks@6648.6673.26|GTACGACTCTTAGCGGTGGATCACTC</td><td>>hg38_21_+_8433172_8446622_RNA45SN1_45S_50ntFlanks</td><td>6648</td><td>6673</td><td>26</td><td>GTACGACTCTTAGCGGTGGATCACTC</td><td>618.3</td><td>2.9</td><td>0.9</td><td>3.1</td><td>1.9E-03</td><td>2.3E-03</td><td>MB231/MB468</td><td>MITO</td></tr>
<tr><td>>hg38_21_+_8433172_8446622_RNA45SN1_45S_50ntFlanks@11552.11591.40|CCTCGGCCGGCGCCTAGCAGCCGACTTAGAACTGGTGCGG</td><td>>hg38_21_+_8433172_8446622_RNA45SN1_45S_50ntFlanks</td><td>11552</td><td>11591</td><td>40</td><td>CCTCGGCCGGCGCCTAGCAGCCGACTTAGAACTGGTGCGG</td><td>595.6</td><td>-1.9</td><td>0.6</td><td>-3.2</td><td>1.5E-03</td><td>2.3E-03</td><td>BT20/MB468</td><td>MITO</td></tr>
<tr><td>>hg38_21_+_8433172_8446622_RNA45SN1_45S_50ntFlanks@7977.8019.43|CGACCTCAGATCAGACGTGGCGACCCGCTGAATTTAAGCATAT</td><td>>hg38_21_+_8433172_8446622_RNA45SN1_45S_50ntFlanks</td><td>7977</td><td>8019</td><td>43</td><td>CGACCTCAGATCAGACGTGGCGACCCGCTGAATTTAAGCATAT</td><td>333.7</td><td>6.0</td><td>1.9</td><td>3.1</td><td>1.9E-03</td><td>2.3E-03</td><td>MB231/MB468</td><td>MITO</td></tr>
<tr><td>>MI0006349|hsa-mir-1287&WithFlank&10|-|98395212|98395313@22.42.21(+1U)&[MIMAT0005878&hsa-miR-1287-5p&offsets|0|-1(+1U);m-244&10|-|98395272|98395292&offsets|0|0(+1U)]|TGCTGGATCAGTGGTTCGAGTT</td><td>>MI0006349|hsa-mir-1287&WithFlank&10|-|98395212|98395313</td><td>22</td><td>42</td><td>21(+1U)</td><td>TGCTGGATCAGTGGTTCGAGTT</td><td>65.8</td><td>5.8</td><td>1.8</td><td>3.1</td><td>1.6E-03</td><td>2.3E-03</td><td>MB231/MB468</td><td>MP</td></tr>
<tr><td>>hg38_21_+_8433172_8446622_RNA45SN1_45S_50ntFlanks@11202.11232.31|GCGGCGGCGGCGACTCTGGACGCGAGCCGGG</td><td>>hg38_21_+_8433172_8446622_RNA45SN1_45S_50ntFlanks</td><td>11202</td><td>11232</td><td>31</td><td>GCGGCGGCGGCGACTCTGGACGCGAGCCGGG</td><td>271.7</td><td>-3.5</td><td>1.1</td><td>-3.1</td><td>1.8E-03</td><td>2.3E-03</td><td>MB231/MB468</td><td>CYTO</td></tr>
<tr><td>>hg38_21_+_8433172_8446622_RNA45SN1_45S_50ntFlanks@12489.12509.21|ACGTGAGCTGGGTTTAGACCG</td><td>>hg38_21_+_8433172_8446622_RNA45SN1_45S_50ntFlanks</td><td>12489</td><td>12509</td><td>21</td><td>ACGTGAGCTGGGTTTAGACCG</td><td>1362.2</td><td>-6.2</td><td>2.0</td><td>-3.1</td><td>2.0E-03</td><td>2.3E-03</td><td>BT20/MB468</td><td>NUC</td></tr>
<tr><td>>hg38_21_+_8433172_8446622_RNA45SN1_45S_50ntFlanks@8191.8213.23|CAGCCCGTGGACGGTGTGAGGCC</td><td>>hg38_21_+_8433172_8446622_RNA45SN1_45S_50ntFlanks</td><td>8191</td><td>8213</td><td>23</td><td>CAGCCCGTGGACGGTGTGAGGCC</td><td>847.4</td><td>-3.2</td><td>1.0</td><td>-3.3</td><td>9.4E-04</td><td>2.3E-03</td><td>BT20/MB468</td><td>MP</td></tr>
<tr><td>>hg38_21_+_8433172_8446622_RNA45SN1_45S_50ntFlanks@11208.11252.45|GCGGCGACTCTGGACGCGAGCCGGGCCCTTCCCGTGGATCGCCCC</td><td>>hg38_21_+_8433172_8446622_RNA45SN1_45S_50ntFlanks</td><td>11208</td><td>11252</td><td>45</td><td>GCGGCGACTCTGGACGCGAGCCGGGCCCTTCCCGTGGATCGCCCC</td><td>778.4</td><td>-4.2</td><td>1.3</td><td>-3.1</td><td>1.8E-03</td><td>2.3E-03</td><td>BT20/MB468</td><td>CYTO</td></tr>
<tr><td>>hg38_21_+_8433172_8446622_RNA45SN1_45S_50ntFlanks@4587.4607.21|TCTATTTTGTTGGTTTTCGGA</td><td>>hg38_21_+_8433172_8446622_RNA45SN1_45S_50ntFlanks</td><td>4587</td><td>4607</td><td>21</td><td>TCTATTTTGTTGGTTTTCGGA</td><td>41.7</td><td>9.1</td><td>2.8</td><td>3.3</td><td>9.4E-04</td><td>2.3E-03</td><td>BT20/MB468</td><td>MP</td></tr>
<tr><td>>hg38_21_+_8433172_8446622_RNA45SN1_45S_50ntFlanks@10458.10487.30|ATGGCCTCCGTTGCCCTCGGCCGATCGAAA</td><td>>hg38_21_+_8433172_8446622_RNA45SN1_45S_50ntFlanks</td><td>10458</td><td>10487</td><td>30</td><td>ATGGCCTCCGTTGCCCTCGGCCGATCGAAA</td><td>550.5</td><td>-2.6</td><td>0.8</td><td>-3.1</td><td>1.8E-03</td><td>2.3E-03</td><td>MB231/MB468</td><td>CYTO</td></tr>
<tr><td>>hg38_21_+_8433172_8446622_RNA45SN1_45S_50ntFlanks@3990.4026.37|TTGGTGACTCTAGATAACCTCGGGCCGATCGCACGCC</td><td>>hg38_21_+_8433172_8446622_RNA45SN1_45S_50ntFlanks</td><td>3990</td><td>4026</td><td>37</td><td>TTGGTGACTCTAGATAACCTCGGGCCGATCGCACGCC</td><td>149.4</td><td>-1.7</td><td>0.5</td><td>-3.2</td><td>1.5E-03</td><td>2.3E-03</td><td>BT20/MB468</td><td>MITO</td></tr>
<tr><td>>hg38_21_+_8433172_8446622_RNA45SN1_45S_50ntFlanks@12841.12861.21|CGGAGTGCCCTTCGTCCTGGG</td><td>>hg38_21_+_8433172_8446622_RNA45SN1_45S_50ntFlanks</td><td>12841</td><td>12861</td><td>21</td><td>CGGAGTGCCCTTCGTCCTGGG</td><td>404.2</td><td>-5.9</td><td>1.8</td><td>-3.3</td><td>9.4E-04</td><td>2.3E-03</td><td>BT20/MB468</td><td>MP</td></tr>
<tr><td>>hg38_21_+_8433172_8446622_RNA45SN1_45S_50ntFlanks@4377.4405.29|GCTCGTAGTTGGATCTTGGGAGCGGGCGG</td><td>>hg38_21_+_8433172_8446622_RNA45SN1_45S_50ntFlanks</td><td>4377</td><td>4405</td><td>29</td><td>GCTCGTAGTTGGATCTTGGGAGCGGGCGG</td><td>858.4</td><td>8.8</td><td>2.7</td><td>3.3</td><td>9.4E-04</td><td>2.3E-03</td><td>BT20/MB468</td><td>MP</td></tr>
<tr><td>>hg38_21_+_8433172_8446622_RNA45SN1_45S_50ntFlanks@9180.9210.31|CCGTCGGGCCCGGGGGAGGTTCTCTCGGGGC</td><td>>hg38_21_+_8433172_8446622_RNA45SN1_45S_50ntFlanks</td><td>9180</td><td>9210</td><td>31</td><td>CCGTCGGGCCCGGGGGAGGTTCTCTCGGGGC</td><td>305.4</td><td>-3.1</td><td>0.9</td><td>-3.3</td><td>9.4E-04</td><td>2.3E-03</td><td>BT20/MB468</td><td>MP</td></tr>
<tr><td>>hg38_21_+_8433172_8446622_RNA45SN1_45S_50ntFlanks@10402.10428.27|AACATGGGTCAGTCGGTCCTGAGAGAT</td><td>>hg38_21_+_8433172_8446622_RNA45SN1_45S_50ntFlanks</td><td>10402</td><td>10428</td><td>27</td><td>AACATGGGTCAGTCGGTCCTGAGAGAT</td><td>230.6</td><td>-3.6</td><td>1.2</td><td>-3.1</td><td>2.0E-03</td><td>2.3E-03</td><td>MB231/MB468</td><td>MITO</td></tr>
<tr><td>>hg38_MT_+_1621_3279_RNR2_16S_50ntFlanks@1185.1215.31|ATTTCGGTTGGGGCGACCTCGGAGCAGAACC</td><td>>hg38_MT_+_1621_3279_RNR2_16S_50ntFlanks</td><td>1185</td><td>1215</td><td>31</td><td>ATTTCGGTTGGGGCGACCTCGGAGCAGAACC</td><td>301.7</td><td>4.6</td><td>1.4</td><td>3.3</td><td>9.4E-04</td><td>2.3E-03</td><td>BT20/MB468</td><td>MP</td></tr>
<tr><td>>hg38_21_+_8433172_8446622_RNA45SN1_45S_50ntFlanks@6719.6747.29|TGAATTGCAGGACACATTGATCATCGACA</td><td>>hg38_21_+_8433172_8446622_RNA45SN1_45S_50ntFlanks</td><td>6719</td><td>6747</td><td>29</td><td>TGAATTGCAGGACACATTGATCATCGACA</td><td>1137.8</td><td>-2.0</td><td>0.6</td><td>-3.2</td><td>1.5E-03</td><td>2.3E-03</td><td>BT20/MB468</td><td>MITO</td></tr>
<tr><td>>hg38_21_+_8433172_8446622_RNA45SN1_45S_50ntFlanks@11661.11685.25|TGATTTCTGCCCAGTGCTCTGAATG</td><td>>hg38_21_+_8433172_8446622_RNA45SN1_45S_50ntFlanks</td><td>11661</td><td>11685</td><td>25</td><td>TGATTTCTGCCCAGTGCTCTGAATG</td><td>4560.8</td><td>-4.0</td><td>1.2</td><td>-3.3</td><td>9.4E-04</td><td>2.3E-03</td><td>BT20/MB468</td><td>MP</td></tr>
<tr><td>>trna20_GluTTC_2_-_131094701_131094772@1.33.33,trna5_GluTTC_13_-_41634874_41634945@1.33.33|TCCCATATGGTCTAGCGGTTAGGATTCCTGGTT</td><td>>trna20_GluTTC_2_-_131094701_131094772</td><td>1</td><td>33</td><td>33</td><td>TCCCATATGGTCTAGCGGTTAGGATTCCTGGTT</td><td>2339.9</td><td>-3.6</td><td>1.0</td><td>-3.5</td><td>4.2E-04</td><td>2.3E-03</td><td>BT20/MB231</td><td>MITO</td></tr>
<tr><td>>hg38_21_+_8433172_8446622_RNA45SN1_45S_50ntFlanks@485.522.38|GATGTGGTGACGTCGTGCTCTCCCGGGCCGGGTCCGAG</td><td>>hg38_21_+_8433172_8446622_RNA45SN1_45S_50ntFlanks</td><td>485</td><td>522</td><td>38</td><td>GATGTGGTGACGTCGTGCTCTCCCGGGCCGGGTCCGAG</td><td>259.3</td><td>11.2</td><td>3.6</td><td>3.1</td><td>2.0E-03</td><td>2.3E-03</td><td>MB231/MB468</td><td>MITO</td></tr>
<tr><td>>MI0000301|hsa-mir-224&WithFlank&X|-|151958572|151958664@14.34.21&[MIMAT0000281&hsa-miR-224-5p&offsets|+1|-3;m-238&X|-|151958630|151958651&offsets|0|-1]|CAAGTCACTAGTGGTTCCGTT</td><td>>MI0000301|hsa-mir-224&WithFlank&X|-|151958572|151958664</td><td>14</td><td>34</td><td>21</td><td>CAAGTCACTAGTGGTTCCGTT</td><td>1308.5</td><td>2.2</td><td>0.6</td><td>3.6</td><td>3.6E-04</td><td>2.3E-03</td><td>BT20/MB231</td><td>NUC</td></tr>
<tr><td>>hg38_21_+_8433172_8446622_RNA45SN1_45S_50ntFlanks@11529.11546.18|CCGCGGTTCCGCGCGGCG</td><td>>hg38_21_+_8433172_8446622_RNA45SN1_45S_50ntFlanks</td><td>11529</td><td>11546</td><td>18</td><td>CCGCGGTTCCGCGCGGCG</td><td>51.0</td><td>-9.5</td><td>3.0</td><td>-3.1</td><td>1.7E-03</td><td>2.3E-03</td><td>MB231/MB468</td><td>MP</td></tr>
<tr><td>>trnaMT_GluTTC_MT_-_14674_14742@40.69.30,trnalookalike8_GluTTC_5_-_93905172_93905240@40.69.30|TTGGTCGTGGTTGTAGTCCGTGCGAGAATA</td><td>>trnaMT_GluTTC_MT_-_14674_14742</td><td>40</td><td>69</td><td>30</td><td>TTGGTCGTGGTTGTAGTCCGTGCGAGAATA</td><td>273.6</td><td>5.1</td><td>1.6</td><td>3.3</td><td>9.4E-04</td><td>2.3E-03</td><td>BT20/MB468</td><td>MP</td></tr>
<tr><td>>hg38_21_+_8433172_8446622_RNA45SN1_45S_50ntFlanks@6651.6696.46|CGACTCTTAGCGGTGGATCACTCGGCTCGTGCGTCGATGAAGAACG</td><td>>hg38_21_+_8433172_8446622_RNA45SN1_45S_50ntFlanks</td><td>6651</td><td>6696</td><td>46</td><td>CGACTCTTAGCGGTGGATCACTCGGCTCGTGCGTCGATGAAGAACG</td><td>1738.3</td><td>3.1</td><td>1.0</td><td>3.1</td><td>1.8E-03</td><td>2.3E-03</td><td>BT20/MB468</td><td>CYTO</td></tr>
<tr><td>>hg38_21_+_8433172_8446622_RNA45SN1_45S_50ntFlanks@4243.4264.22|CTTTCGAGGCCCTGTAATTGGA</td><td>>hg38_21_+_8433172_8446622_RNA45SN1_45S_50ntFlanks</td><td>4243</td><td>4264</td><td>22</td><td>CTTTCGAGGCCCTGTAATTGGA</td><td>266.6</td><td>-1.3</td><td>0.4</td><td>-3.1</td><td>1.8E-03</td><td>2.3E-03</td><td>BT20/MB468</td><td>CYTO</td></tr>
<tr><td>>hg38_21_+_8433172_8446622_RNA45SN1_45S_50ntFlanks@12073.12098.26|CCGAGGGGCTCTCGCTTCTGGCGCCA</td><td>>hg38_21_+_8433172_8446622_RNA45SN1_45S_50ntFlanks</td><td>12073</td><td>12098</td><td>26</td><td>CCGAGGGGCTCTCGCTTCTGGCGCCA</td><td>318.3</td><td>-3.6</td><td>1.1</td><td>-3.1</td><td>1.8E-03</td><td>2.3E-03</td><td>BT20/MB468</td><td>CYTO</td></tr>
<tr><td>>hg38_21_+_8433172_8446622_RNA45SN1_45S_50ntFlanks@11118.11145.28|CAGGGGCCGGCGGCGGCCCGCCGCGGGG</td><td>>hg38_21_+_8433172_8446622_RNA45SN1_45S_50ntFlanks</td><td>11118</td><td>11145</td><td>28</td><td>CAGGGGCCGGCGGCGGCCCGCCGCGGGG</td><td>261.6</td><td>-4.8</td><td>1.5</td><td>-3.1</td><td>1.8E-03</td><td>2.4E-03</td><td>BT20/MB468</td><td>CYTO</td></tr>
<tr><td>>hg38_21_+_8433172_8446622_RNA45SN1_45S_50ntFlanks@10420.10455.36|CTGAGAGATGGGCGAGCGCCGTTCCGAAGGGACGGG</td><td>>hg38_21_+_8433172_8446622_RNA45SN1_45S_50ntFlanks</td><td>10420</td><td>10455</td><td>36</td><td>CTGAGAGATGGGCGAGCGCCGTTCCGAAGGGACGGG</td><td>526.8</td><td>-2.2</td><td>0.7</td><td>-3.1</td><td>1.9E-03</td><td>2.4E-03</td><td>MB231/MB468</td><td>CYTO</td></tr>
<tr><td>>hg38_MT_+_1621_3279_RNR2_16S_50ntFlanks@1389.1430.42|CGATGGTGCAGCCGCTATTAAAGGTTCGTTTGTTCAACGATT</td><td>>hg38_MT_+_1621_3279_RNR2_16S_50ntFlanks</td><td>1389</td><td>1430</td><td>42</td><td>CGATGGTGCAGCCGCTATTAAAGGTTCGTTTGTTCAACGATT</td><td>45.4</td><td>9.3</td><td>2.8</td><td>3.3</td><td>9.5E-04</td><td>2.4E-03</td><td>BT20/MB468</td><td>MP</td></tr>
<tr><td>>am_MI0000809|hsa-mir-151a&WithFlank&8|-|140732558|140732659@52.72.21(+1A)&[MIMAT0000757&hsa-miR-151a-3p&offsets|-1|-1(+1A);m-30&8|-|140732587|140732607&offsets|-1|-1(+1A)]|ACTAGACTGAAGCTCCTTGAGA</td><td>>am_MI0000809|hsa-mir-151a&WithFlank&8|-|140732558|140732659</td><td>52</td><td>72</td><td>21(+1A)</td><td>ACTAGACTGAAGCTCCTTGAGA</td><td>114.4</td><td>4.4</td><td>1.3</td><td>3.3</td><td>9.5E-04</td><td>2.4E-03</td><td>BT20/MB468</td><td>MP</td></tr>
<tr><td>>MI0000279|hsa-mir-196a-2&WithFlank&12|+|53991732|53991853@31.50.20;MI0000238|hsa-mir-196a-1&WithFlank&17|-|48632484|48632565@13.32.20&[MIMAT0000226&hsa-miR-196a-5p&offsets|0|-2;m-261&12|+|53991762|53991783&offsets|0|-2];[MIMAT0000226_1&hsa-miR-196a-5p&offsets|0|-2;m-268&17|-|48632532|48632553&offsets|0|-2]|TAGGTAGTTTCATGTTGTTG</td><td>>MI0000279|hsa-mir-196a-2&WithFlank&12|+|53991732|53991853</td><td>31</td><td>50</td><td>20</td><td>TAGGTAGTTTCATGTTGTTG</td><td>215.5</td><td>4.6</td><td>1.4</td><td>3.3</td><td>9.5E-04</td><td>2.4E-03</td><td>BT20/MB468</td><td>MP</td></tr>
<tr><td>>hg38_21_+_8433172_8446622_RNA45SN1_45S_50ntFlanks@12407.12436.30|TTTTGATCCTTCGATGTCGGCTCTTCCTAT</td><td>>hg38_21_+_8433172_8446622_RNA45SN1_45S_50ntFlanks</td><td>12407</td><td>12436</td><td>30</td><td>TTTTGATCCTTCGATGTCGGCTCTTCCTAT</td><td>1243.7</td><td>-3.6</td><td>1.1</td><td>-3.1</td><td>1.9E-03</td><td>2.4E-03</td><td>MB231/MB468</td><td>CYTO</td></tr>
<tr><td>>hg38_21_+_8433172_8446622_RNA45SN1_45S_50ntFlanks@10402.10428.27|AACATGGGTCAGTCGGTCCTGAGAGAT</td><td>>hg38_21_+_8433172_8446622_RNA45SN1_45S_50ntFlanks</td><td>10402</td><td>10428</td><td>27</td><td>AACATGGGTCAGTCGGTCCTGAGAGAT</td><td>197.6</td><td>-2.8</td><td>0.9</td><td>-3.1</td><td>1.9E-03</td><td>2.4E-03</td><td>MB231/MB468</td><td>CYTO</td></tr>
</table>

| | | | | | | | | | | | | | |
|---|---|---|---|---|---|---|---|---|---|---|---|---|---|
| >trna116_GluCTC_1_-_145399233_145399304@38.75.38,trna134_GluTTC_1_-_16861774_16861845@38.75.38,trna71_GluCTC_1_-_161439189_161439260@38.75.38,trna74_GluCTC_1_-_161431809_161431880@38.75.38,trna77_GluCTC_1_-_161424398_161424469@38.75.38,trna77_GluCTC_6_+_28949976_28950047@38.75.38,trna80_GluCTC_1_-_161417018_161417089@38.75.38,trna84_GluTTC_1_-_161391883_161391954@38.75.38,trna87_GluCTC_6_-_126101393_126101464@38.75.38\|CCGCCGCGGCCCGGGTTCGATTCCCGGTCAGGGAACCA | >trna116_GluCTC_1_-_145399233_145399304 | 38 | 75 | 38 | CCGCCGCGGCCCGGGTTCGATTCCCGGTCAGGGAACCA | 559.0 | 2.9 | 0.9 | 3.1 | 1.8E-03 | 2.4E-03 | BT20/MB468 | CYTO |
| >hg38_21_+_8433172_8446622_RNA45SN1_45S_50ntFlanks@4163.4186.24\|CATCCAAGGAAGGCAGCAGGCGCG | >hg38_21_+_8433172_8446622_RNA45SN1_45S_50ntFlanks | 4163 | 4186 | 24 | CATCCAAGGAAGGCAGCAGGCGCG | 485.1 | -2.0 | 0.7 | -3.1 | 1.9E-03 | 2.4E-03 | MB231/MB468 | CYTO |
| >hg38_21_+_8433172_8446622_RNA45SN1_45S_50ntFlanks@10248.10278.31\|CGACGAGTAGGAGGGCCGCTGCGGTGAGCCT | >hg38_21_+_8433172_8446622_RNA45SN1_45S_50ntFlanks | 10248 | 10278 | 31 | CGACGAGTAGGAGGGCCGCTGCGGTGAGCCT | 127.8 | 6.3 | 1.9 | 3.3 | 9.5E-04 | 2.4E-03 | BT20/MB468 | MP |
| >hg38_1_-_228634819_228635039_RNA5S12_5S_50ntFlanks@148.169.22\|GGAATACCGGGTGCTGTAGGCT | >hg38_1_-_228634819_228635039_RNA5S12_5S_50ntFlanks | 148 | 169 | 22 | GGAATACCGGGTGCTGTAGGCT | 1419.9 | 2.0 | 0.7 | 3.1 | 1.9E-03 | 2.4E-03 | MB231/MB468 | CYTO |
| >hg38_21_+_8433172_8446622_RNA45SN1_45S_50ntFlanks@12071.12108.38\|CCCCGAGGGGCTCTCGCTTCTGGCGCCAAGCGCCCGGC | >hg38_21_+_8433172_8446622_RNA45SN1_45S_50ntFlanks | 12071 | 12108 | 38 | CCCCGAGGGGCTCTCGCTTCTGGCGCCAAGCGCCCGGC | 40.0 | -8.4 | 2.5 | -3.3 | 9.5E-04 | 2.4E-03 | BT20/MB468 | MP |
| >MI0000447\|hsa-mir-128-1&WithFlank&2\|+\|135665391\|135665484@56.78.23&[MIMAT0000424&hsa-miR-128-3p&offsets\|0\|+2;m-108&2\|+\|135665446\|135665466&offsets\|0\|+2]\|TCACAGTGAACCGGTCTCTTTTT | >MI0000447\|hsa-mir-128-1&WithFlank&2\|+\|135665391\|135665484 | 56 | 78 | 23 | TCACAGTGAACCGGTCTCTTTTT | 231.6 | 2.9 | 0.9 | 3.1 | 2.0E-03 | 2.4E-03 | MB231/MB468 | MITO |
| >trnaMT_ThrTGT_MT_+_15888_15953@34.69.36\|TAAACCGGAGATGAAAACCTTTTTCCAAGGACACCA | >trnaMT_ThrTGT_MT_+_15888_15953 | 34 | 69 | 36 | TAAACCGGAGATGAAAACCTTTTTCCAAGGACACCA | 193.5 | 5.0 | 1.6 | 3.1 | 2.0E-03 | 2.4E-03 | MB231/MB468 | MITO |
| >hg38_21_+_8433172_8446622_RNA45SN1_45S_50ntFlanks@6651.6682.32\|CGACTCTTAGCGGTGGATCACTCGGCTCGTGC | >hg38_21_+_8433172_8446622_RNA45SN1_45S_50ntFlanks | 6651 | 6682 | 32 | CGACTCTTAGCGGTGGATCACTCGGCTCGTGC | 8138.6 | 3.2 | 1.0 | 3.1 | 2.0E-03 | 2.4E-03 | MB231/MB468 | MITO |
| >hg38_21_+_8433172_8446622_RNA45SN1_45S_50ntFlanks@12415.12437.23\|CTTCGATGTCGGCTCTTCCTATC | >hg38_21_+_8433172_8446622_RNA45SN1_45S_50ntFlanks | 12415 | 12437 | 23 | CTTCGATGTCGGCTCTTCCTATC | 360.7 | -5.1 | 1.6 | -3.1 | 1.7E-03 | 2.4E-03 | MB231/MB468 | MP |
| >hg38_21_+_8433172_8446622_RNA45SN1_45S_50ntFlanks@10349.10376.28\|AACGAGAACTTTGAAGGCCGAAGTGGAG | >hg38_21_+_8433172_8446622_RNA45SN1_45S_50ntFlanks | 10349 | 10376 | 28 | AACGAGAACTTTGAAGGCCGAAGTGGAG | 721.9 | -1.9 | 0.6 | -3.1 | 1.9E-03 | 2.4E-03 | MB231/MB468 | CYTO |
| >hg38_21_+_8433172_8446622_RNA45SN1_45S_50ntFlanks@10922.10945.24\|CTGGACGAGGCGCCGCCGCCCCCC | >hg38_21_+_8433172_8446622_RNA45SN1_45S_50ntFlanks | 10922 | 10945 | 24 | CTGGACGAGGCGCCGCCGCCCCCC | 1043.4 | -4.6 | 1.4 | -3.3 | 9.5E-04 | 2.4E-03 | BT20/MB468 | MP |
| >hg38_21_+_8433172_8446622_RNA45SN1_45S_50ntFlanks@6671.6693.23\|CTCGGCTCGTGCGTCGATGAAGA | >hg38_21_+_8433172_8446622_RNA45SN1_45S_50ntFlanks | 6671 | 6693 | 23 | CTCGGCTCGTGCGTCGATGAAGA | 83.5 | -5.2 | 1.6 | -3.3 | 9.5E-04 | 2.4E-03 | BT20/MB468 | MP |
| >hg38_21_+_8433172_8446622_RNA45SN1_45S_50ntFlanks@12306.12328.23\|CACGATCCTTCTGACCTTTTGGG | >hg38_21_+_8433172_8446622_RNA45SN1_45S_50ntFlanks | 12306 | 12328 | 23 | CACGATCCTTCTGACCTTTTGGG | 1300.5 | -3.4 | 1.1 | -3.2 | 1.5E-03 | 2.4E-03 | BT20/MB468 | MITO |
| >hg38_21_+_8433172_8446622_RNA45SN1_45S_50ntFlanks@12539.12565.27\|ATGATGTGTTGTTGCCATGGTAATCCT | >hg38_21_+_8433172_8446622_RNA45SN1_45S_50ntFlanks | 12539 | 12565 | 27 | ATGATGTGTTGTTGCCATGGTAATCCT | 306.2 | -5.3 | 1.6 | -3.3 | 9.5E-04 | 2.4E-03 | BT20/MB468 | MP |
| >hg38_21_+_8433172_8446622_RNA45SN1_45S_50ntFlanks@11668.11701.34\|TGCCCAGTGCTCTGAATGTCAAAGTGAAGAAATT | >hg38_21_+_8433172_8446622_RNA45SN1_45S_50ntFlanks | 11668 | 11701 | 34 | TGCCCAGTGCTCTGAATGTCAAAGTGAAGAAATT | 970.0 | -2.1 | 0.7 | -3.1 | 2.0E-03 | 2.4E-03 | MB231/MB468 | NUC |
| >hg38_21_+_8433172_8446622_RNA45SN1_45S_50ntFlanks@9540.9570.31\|CTCTGGTGGAGGTCCGTAGCGGTCCTGACGT | >hg38_21_+_8433172_8446622_RNA45SN1_45S_50ntFlanks | 9540 | 9570 | 31 | CTCTGGTGGAGGTCCGTAGCGGTCCTGACGT | 219.8 | -2.1 | 0.7 | -3.1 | 2.0E-03 | 2.4E-03 | MB231/MB468 | NUC |
| >hg38_21_+_8433172_8446622_RNA45SN1_45S_50ntFlanks@10806.10827.22\|AACAATGTAGGTAAGGGAAGTC | >hg38_21_+_8433172_8446622_RNA45SN1_45S_50ntFlanks | 10806 | 10827 | 22 | AACAATGTAGGTAAGGGAAGTC | 153.0 | -4.5 | 1.4 | -3.1 | 1.8E-03 | 2.4E-03 | BT20/MB468 | CYTO |
| >hg38_21_+_8433172_8446622_RNA45SN1_45S_50ntFlanks@12072.12096.25\|CCCGAGGGGCTCTCGCTTCTGGCGC | >hg38_21_+_8433172_8446622_RNA45SN1_45S_50ntFlanks | 12072 | 12096 | 25 | CCCGAGGGGCTCTCGCTTCTGGCGC | 166.8 | -4.5 | 1.4 | -3.2 | 1.5E-03 | 2.4E-03 | BT20/MB468 | MITO |
| >am_trna128_GlyGCC_6_-_27870686_27870756@4.35.32,trna18_GlyGCC_16_+_70822597_70822667@4.35.32,trna19_GlyGCC_16_+_70823410_70823480@4.35.32,trna19_GlyGCC_2_-_157257659_157257729@4.35.32,trna24_GlyGCC_16_-_70812942_70813012@4.35.32,trna25_GlyGCC_16_-_70812114_70812184@4.35.32,trna5_GlyGCC_17_+_8029064_8029134@4.35.32,trna68_GlyGCC_1_-_161493637_161493707@4.35.32\|TTGGTGGTTCAGTGGTAGAATTCTCGCCTGCC | >am_trna128_GlyGCC_6_-_27870686_27870756 | 4 | 35 | 32 | TTGGTGGTTCAGTGGTAGAATTCTCGCCTGCC | 118.6 | 5.8 | 1.9 | 3.1 | 1.7E-03 | 2.4E-03 | MB231/MB468 | MP |
| >hg38_21_+_8433172_8446622_RNA45SN1_45S_50ntFlanks@6668.6686.19\|TCACTCGGCTCGTGCGTCG | >hg38_21_+_8433172_8446622_RNA45SN1_45S_50ntFlanks | 6668 | 6686 | 19 | TCACTCGGCTCGTGCGTCG | 109.8 | -5.1 | 1.5 | -3.3 | 9.6E-04 | 2.4E-03 | BT20/MB468 | MP |
| >hg38_MT_+_598_1651_RNR1_12S_50ntFlanks@442.469.28\|ATATCTGAACACACAATAGCTAAGACCC | >hg38_MT_+_598_1651_RNR1_12S_50ntFlanks | 442 | 469 | 28 | ATATCTGAACACACAATAGCTAAGACCC | 110.8 | 5.0 | 1.5 | 3.3 | 9.6E-04 | 2.4E-03 | BT20/MB468 | MP |
| >hg38_21_+_8433172_8446622_RNA45SN1_45S_50ntFlanks@4007.4024.18\|CCTCGGGCCGATCGCACG | >hg38_21_+_8433172_8446622_RNA45SN1_45S_50ntFlanks | 4007 | 4024 | 18 | CCTCGGGCCGATCGCACG | 456.5 | -3.8 | 1.2 | -3.2 | 1.5E-03 | 2.4E-03 | BT20/MB468 | MITO |
| >hg38_21_+_8433172_8446622_RNA45SN1_45S_50ntFlanks@10112.10131.20\|GGTGTGCGTCGGAGGGCGGC | >hg38_21_+_8433172_8446622_RNA45SN1_45S_50ntFlanks | 10112 | 10131 | 20 | GGTGTGCGTCGGAGGGCGGC | 94.1 | 9.9 | 2.8 | 3.6 | 3.3E-04 | 2.4E-03 | BT20/MB231 | MP |
| >hg38_21_+_8433172_8446622_RNA45SN1_45S_50ntFlanks@11896.11929.34\|TCTAGTCTGGCACGGTGAAGAGACATGAGAGGTG | >hg38_21_+_8433172_8446622_RNA45SN1_45S_50ntFlanks | 11896 | 11929 | 34 | TCTAGTCTGGCACGGTGAAGAGACATGAGAGGTG | 113.6 | 6.4 | 1.9 | 3.3 | 9.6E-04 | 2.4E-03 | BT20/MB468 | MP |
| >hg38_21_+_8433172_8446622_RNA45SN1_45S_50ntFlanks@4224.4242.19\|AAAATAACAATACAGGACT | >hg38_21_+_8433172_8446622_RNA45SN1_45S_50ntFlanks | 4224 | 4242 | 19 | AAAATAACAATACAGGACT | 802.9 | -3.2 | 1.0 | -3.1 | 1.8E-03 | 2.4E-03 | BT20/MB468 | CYTO |
| >hg38_21_+_8433172_8446622_RNA45SN1_45S_50ntFlanks@9929.9947.19\|GATATAGACAGCAGGACGG | >hg38_21_+_8433172_8446622_RNA45SN1_45S_50ntFlanks | 9929 | 9947 | 19 | GATATAGACAGCAGGACGG | 247.3 | 5.1 | 1.4 | 3.6 | 3.3E-04 | 2.4E-03 | BT20/MB231 | MP |
| >hg38_21_+_8433172_8446622_RNA45SN1_45S_50ntFlanks@4736.4758.23\|CGAAAGTCGGAGGTTCGAAGACG | >hg38_21_+_8433172_8446622_RNA45SN1_45S_50ntFlanks | 4736 | 4758 | 23 | CGAAAGTCGGAGGTTCGAAGACG | 266.3 | -2.7 | 0.9 | -3.2 | 1.5E-03 | 2.4E-03 | BT20/MB468 | MITO |
| >hg38_21_+_8433172_8446622_RNA45SN1_45S_50ntFlanks@9533.9572.40\|GAGGAAACTCTGGTGGAGGTCCGTAGCGGTCCTGACGTGC | >hg38_21_+_8433172_8446622_RNA45SN1_45S_50ntFlanks | 9533 | 9572 | 40 | GAGGAAACTCTGGTGGAGGTCCGTAGCGGTCCTGACGTGC | 267.0 | -3.0 | 1.0 | -3.1 | 2.0E-03 | 2.4E-03 | MB231/MB468 | MITO |
| >trnaMT_GlnTTG_MT_-_4329_4400@1.28.28,trnalookalike2_GlnTTG_1_-_564879_564950@1.28.28\|TAGGATGGGGTGTGATAGGTGGCACGGA | >trnaMT_GlnTTG_MT_-_4329_4400 | 1 | 28 | 28 | TAGGATGGGGTGTGATAGGTGGCACGGA | 425.6 | 4.4 | 1.3 | 3.3 | 9.6E-04 | 2.4E-03 | BT20/MB468 | MP |
| >hg38_21_+_8433172_8446622_RNA45SN1_45S_50ntFlanks@9937.9955.19\|CAGCAGGACGGTGGCCATG | >hg38_21_+_8433172_8446622_RNA45SN1_45S_50ntFlanks | 9937 | 9955 | 19 | CAGCAGGACGGTGGCCATG | 261.6 | 5.5 | 1.7 | 3.3 | 9.6E-04 | 2.4E-03 | BT20/MB468 | MP |
| >hg38_21_+_8433172_8446622_RNA45SN1_45S_50ntFlanks@11577.11602.26\|TTAGAACTGGTGCGGACCAGGGGAAT | >hg38_21_+_8433172_8446622_RNA45SN1_45S_50ntFlanks | 11577 | 11602 | 26 | TTAGAACTGGTGCGGACCAGGGGAAT | 95.9 | -5.3 | 1.6 | -3.3 | 9.6E-04 | 2.4E-03 | BT20/MB468 | MP |
| >hg38_21_+_8433172_8446622_RNA45SN1_45S_50ntFlanks@9554.9581.28\|CGTAGCGGTCCTGACGTGCAAATCGGTC | >hg38_21_+_8433172_8446622_RNA45SN1_45S_50ntFlanks | 9554 | 9581 | 28 | CGTAGCGGTCCTGACGTGCAAATCGGTC | 1706.1 | -2.8 | 0.8 | -3.6 | 3.8E-04 | 2.4E-03 | BT20/MB231 | CYTO |
| >hg38_21_+_8433172_8446622_RNA45SN1_45S_50ntFlanks@4657.4680.24\|CTAGAGGTGAAATTCTTGGACCGG | >hg38_21_+_8433172_8446622_RNA45SN1_45S_50ntFlanks | 4657 | 4680 | 24 | CTAGAGGTGAAATTCTTGGACCGG | 207.8 | -4.6 | 1.5 | -3.1 | 1.8E-03 | 2.4E-03 | BT20/MB468 | CYTO |
| >hg38_21_+_8433172_8446622_RNA45SN1_45S_50ntFlanks@4728.4758.31\|ATCAAGAACGAAAGTCGGAGGTTCGAAGACG | >hg38_21_+_8433172_8446622_RNA45SN1_45S_50ntFlanks | 4728 | 4758 | 31 | ATCAAGAACGAAAGTCGGAGGTTCGAAGACG | 6744.8 | -6.0 | 1.9 | -3.1 | 1.7E-03 | 2.4E-03 | MB231/MB468 | MP |
| >hg38_21_+_8433172_8446622_RNA45SN1_45S_50ntFlanks@6645.6665.21\|CTCGTACGACTCTTAGCGGTG | >hg38_21_+_8433172_8446622_RNA45SN1_45S_50ntFlanks | 6645 | 6665 | 21 | CTCGTACGACTCTTAGCGGTG | 8259.4 | 2.7 | 0.9 | 3.1 | 2.0E-03 | 2.4E-03 | MB231/MB468 | MITO |
| >MI0003820\|hsa-mir-454&WithFlank&17\|-\|59137752\|59137878@70.93.24&[MIMAT0003885&hsa-miR-454-3p&offsets\|0\|+1;m-272&17\|-\|59137787\|59137809&offsets\|0\|+1]\|TAGTGCAATATTGCTTATAGGGTT | >MI0003820\|hsa-mir-454&WithFlank&17\|-\|59137752\|59137878 | 70 | 93 | 24 | TAGTGCAATATTGCTTATAGGGTT | 1207.3 | -5.1 | 1.4 | -3.6 | 3.3E-04 | 2.4E-03 | BT20/MB231 | MP |
| >hg38_21_+_8433172_8446622_RNA45SN1_45S_50ntFlanks@9533.9573.41\|GAGGAAACTCTGGTGGAGGTCCGTAGCGGTCCTGACGTGCA | >hg38_21_+_8433172_8446622_RNA45SN1_45S_50ntFlanks | 9533 | 9573 | 41 | GAGGAAACTCTGGTGGAGGTCCGTAGCGGTCCTGACGTGCA | 618.5 | -6.1 | 2.0 | -3.1 | 2.0E-03 | 2.4E-03 | BT20/MB468 | NUC |
| >hg38_21_+_8433172_8446622_RNA45SN1_45S_50ntFlanks@9306.9336.31\|CAAGGAGTCTAACACGTGCGCGAGTCGGGGG | >hg38_21_+_8433172_8446622_RNA45SN1_45S_50ntFlanks | 9306 | 9336 | 31 | CAAGGAGTCTAACACGTGCGCGAGTCGGGGG | 259.4 | -2.3 | 0.7 | -3.2 | 1.5E-03 | 2.4E-03 | BT20/MB468 | MITO |
| >MI0000299\|hsa-mir-222&WithFlank&X\|-\|45747009\|45747130@75.98.24(+1C)&[MIMAT0000279&hsa-miR-222-3p&offsets\|0\|+3(+1C);m-140&X\|-\|45747033\|45747056&offsets\|0\|0(+1C)]\|AGCTACATCTGGCTACTGGGTCTCC | >MI0000299\|hsa-mir-222&WithFlank&X\|-\|45747009\|45747130 | 75 | 98 | 24(+1C) | AGCTACATCTGGCTACTGGGTCTCC | 535.8 | 3.0 | 0.9 | 3.3 | 9.6E-04 | 2.4E-03 | BT20/MB468 | MP |
| >MI0000465\|hsa-mir-191&WithFlank&3\|-\|49020612\|49020715@22.44.23&[MIMAT0000440&hsa-miR-191-5p&offsets\|0\|0;m-48&3\|-\|49020672\|49020694&offsets\|0\|0]\|CAACGGAATCCCAAAAGCAGCTG | >MI0000465\|hsa-mir-191&WithFlank&3\|-\|49020612\|49020715 | 22 | 44 | 23 | CAACGGAATCCCAAAAGCAGCTG | 4735.6 | 1.6 | 0.5 | 3.1 | 2.0E-03 | 2.4E-03 | BT20/MB468 | NUC |
| >am_trna111_HisGTG_1_-_147774845_147774916@3.28.26,trna118_HisGTG_1_-_145396881_145396952@3.28.26,trna16_HisGTG_1_+_146544773_146544844@3.28.26,trna1_HisGTG_15_+_45493349_45493420@3.28.26,trna21_HisGTG_1_+_147753471_147753542@3.28.26,trna33_HisGTG_6_+_27125906_27125977@3.28.26,trna7_HisGTG_9_-_14433938_14434009@3.28.26,trna8_HisGTG_15_-_45492611_45492682@3.28.26,trna9_HisGTG_15_-_45490804_45490875@3.28.26\|CGTGATCGTATAGTGGTTAGTACTCT | >am_trna111_HisGTG_1_-_147774845_147774916 | 3 | 28 | 26 | CGTGATCGTATAGTGGTTAGTACTCT | 321.6 | 3.9 | 1.2 | 3.3 | 9.6E-04 | 2.4E-03 | BT20/MB468 | MP |
| >hg38_21_+_8433172_8446622_RNA45SN1_45S_50ntFlanks@11895.11916.22\|CTCTAGTCTGGCACGGTGAAGA | >hg38_21_+_8433172_8446622_RNA45SN1_45S_50ntFlanks | 11895 | 11916 | 22 | CTCTAGTCTGGCACGGTGAAGA | 138.8 | -4.5 | 1.5 | -3.1 | 2.0E-03 | 2.4E-03 | MB231/MB468 | NUC |
| >MI0000299\|hsa-mir-222&WithFlank&X\|-\|45747009\|45747130@75.96.22&[MIMAT0000279&hsa-miR-222-3p&offsets\|0\|+1;m-140&X\|-\|45747033\|45747056&offsets\|0\|-2]\|AGCTACATCTGGCTACTGGGTC | >MI0000299\|hsa-mir-222&WithFlank&X\|-\|45747009\|45747130 | 75 | 96 | 22 | AGCTACATCTGGCTACTGGGTC | 946.8 | 3.7 | 1.1 | 3.3 | 9.7E-04 | 2.4E-03 | BT20/MB468 | MP |
| >hg38_21_+_8433172_8446622_RNA45SN1_45S_50ntFlanks@10240.10259.20\|CTACGCCGCGACGAGTAGGA | >hg38_21_+_8433172_8446622_RNA45SN1_45S_50ntFlanks | 10240 | 10259 | 20 | CTACGCCGCGACGAGTAGGA | 1175.1 | 2.4 | 0.7 | 3.5 | 4.3E-04 | 2.4E-03 | BT20/MB231 | MITO |
| >hg38_21_+_8433172_8446622_RNA45SN1_45S_50ntFlanks@9567.9586.20\|ACGTGCAAATCGGTCGTCCG | >hg38_21_+_8433172_8446622_RNA45SN1_45S_50ntFlanks | 9567 | 9586 | 20 | ACGTGCAAATCGGTCGTCCG | 218.3 | -4.3 | 1.3 | -3.3 | 9.7E-04 | 2.4E-03 | BT20/MB468 | MP |
| >hg38_MT_+_598_1651_RNR1_12S_50ntFlanks@907.929.23\|TATACTTCAAAGGACATTTAACT | >hg38_MT_+_598_1651_RNR1_12S_50ntFlanks | 907 | 929 | 23 | TATACTTCAAAGGACATTTAACT | 40.5 | 8.6 | 2.7 | 3.1 | 1.7E-03 | 2.4E-03 | MB231/MB468 | MP |
| >hg38_21_+_8433172_8446622_RNA45SN1_45S_50ntFlanks@5472.5500.29\|ACGGCCCTGGCGGAGCGCTGAGAAGACGG | >hg38_21_+_8433172_8446622_RNA45SN1_45S_50ntFlanks | 5472 | 5500 | 29 | ACGGCCCTGGCGGAGCGCTGAGAAGACGG | 252.6 | 3.3 | 1.0 | 3.3 | 9.7E-04 | 2.4E-03 | BT20/MB468 | MP |

<table>
<tr><th></th><th></th><th></th><th></th><th></th><th></th><th></th><th></th><th></th><th></th><th></th><th></th><th></th><th></th></tr>
<tr><td>>hg38_21_+_8433172_8446622_RNA45SN1_45S_50ntFlanks@11694.11716.23|AAGAAATTCAATGAAGCGCGGGT</td><td>>hg38_21_+_8433172_8446622_RNA45SN1_45S_50ntFlanks</td><td>11694</td><td>11716</td><td>23</td><td>AAGAAATTCAATGAAGCGCGGGT</td><td>101.8</td><td>-4.2</td><td>1.3</td><td>-3.1</td><td>1.7E-03</td><td>2.4E-03</td><td>MB231/MB468</td><td>MP</td></tr>
<tr><td>>MI0001445|hsa-mir-423&WithFlank&17|+|30117073|30117178@58.80.23(+1C)&[MIMAT0001340&hsa-miR-423-3p&offsets|-1|-1(+1C);m-100&17|+|30117131|30117153&offsets|-1|-1(+1C)]|AAGCTCGGTCTGAGGCCCCTCAGC</td><td>>MI0001445|hsa-mir-423&WithFlank&17|+|30117073|30117178</td><td>58</td><td>80</td><td>23(+1C)</td><td>AAGCTCGGTCTGAGGCCCCTCAGC</td><td>105.6</td><td>5.1</td><td>1.6</td><td>3.1</td><td>1.7E-03</td><td>2.4E-03</td><td>MB231/MB468</td><td>MP</td></tr>
<tr><td>>hg38_MT_+_1621_3279_RNR2_16S_50ntFlanks@1367.1401.35|TCGATGTTGGATCAGGACATCCCGATGGTGCAGCC</td><td>>hg38_MT_+_1621_3279_RNR2_16S_50ntFlanks</td><td>1367</td><td>1401</td><td>35</td><td>TCGATGTTGGATCAGGACATCCCGATGGTGCAGCC</td><td>279.6</td><td>4.7</td><td>1.4</td><td>3.3</td><td>9.7E-04</td><td>2.4E-03</td><td>BT20/MB468</td><td>MP</td></tr>
<tr><td>>trna112_GlnCTG_1_-_147737382_147737453@32.50.19,trna131_GlnCTG_6_-_27759135_27759206@32.50.19,trna146_GlnCTG_6_-_27515531_27515602@32.50.19,trna15_GlnCTG_1_+_145963304_145963375@32.50.19,trna19_GlnCTG_1_+_147505038_147505109@32.50.19,trna1_GlnCTG_6_+_18836402_18836473@32.50.19,trna28_GlnCTG_1_+_149186125_149186196@32.50.19,trna3_GlnCTG_17_+_8023070_8023141@32.50.19,trna49_GlnCTG_6_+_27487308_27487379@32.50.19,trna7_GlnCTG_15_-_66161400_66161471@32.50.19,trna99_GlnCTG_6_-_28909378_28909449@32.50.19|CTCTGAATCCAGCGATCCG</td><td>>trna112_GlnCTG_1_-_147737382_147737453</td><td>32</td><td>50</td><td>19</td><td>CTCTGAATCCAGCGATCCG</td><td>110.5</td><td>-5.1</td><td>1.6</td><td>-3.3</td><td>9.7E-04</td><td>2.4E-03</td><td>BT20/MB468</td><td>MP</td></tr>
<tr><td>>hg38_MT_+_1621_3279_RNR2_16S_50ntFlanks@1186.1215.30|TTTCGGTTGGGGCGACCTCGGAGCAGAACC</td><td>>hg38_MT_+_1621_3279_RNR2_16S_50ntFlanks</td><td>1186</td><td>1215</td><td>30</td><td>TTTCGGTTGGGGCGACCTCGGAGCAGAACC</td><td>143.6</td><td>5.6</td><td>1.7</td><td>3.3</td><td>9.7E-04</td><td>2.4E-03</td><td>BT20/MB468</td><td>MP</td></tr>
<tr><td>>hg38_21_+_8433172_8446622_RNA45SN1_45S_50ntFlanks@12417.12445.29|TCGATGTCGGCTCTTCCTATCATTGTGAA</td><td>>hg38_21_+_8433172_8446622_RNA45SN1_45S_50ntFlanks</td><td>12417</td><td>12445</td><td>29</td><td>TCGATGTCGGCTCTTCCTATCATTGTGAA</td><td>604.4</td><td>-2.2</td><td>0.7</td><td>-3.1</td><td>2.0E-03</td><td>2.4E-03</td><td>MB231/MB468</td><td>MITO</td></tr>
<tr><td>>hg38_21_+_8433172_8446622_RNA45SN1_45S_50ntFlanks@9313.9338.26|TCTAACACGTGCGCGAGTCGGGGGCT</td><td>>hg38_21_+_8433172_8446622_RNA45SN1_45S_50ntFlanks</td><td>9313</td><td>9338</td><td>26</td><td>TCTAACACGTGCGCGAGTCGGGGGCT</td><td>170.4</td><td>-3.6</td><td>1.2</td><td>-3.1</td><td>2.0E-03</td><td>2.4E-03</td><td>MB231/MB468</td><td>NUC</td></tr>
<tr><td>>hg38_21_+_8433172_8446622_RNA45SN1_45S_50ntFlanks@10351.10371.21|CGAGAACTTTGAAGGCCGAAG</td><td>>hg38_21_+_8433172_8446622_RNA45SN1_45S_50ntFlanks</td><td>10351</td><td>10371</td><td>21</td><td>CGAGAACTTTGAAGGCCGAAG</td><td>1955.8</td><td>-3.5</td><td>1.0</td><td>-3.3</td><td>9.7E-04</td><td>2.4E-03</td><td>BT20/MB468</td><td>MP</td></tr>
<tr><td>>hg38_21_+_8433172_8446622_RNA45SN1_45S_50ntFlanks@8195.8215.21|CCGTGGACGGTGTGAGGCCGG</td><td>>hg38_21_+_8433172_8446622_RNA45SN1_45S_50ntFlanks</td><td>8195</td><td>8215</td><td>21</td><td>CCGTGGACGGTGTGAGGCCGG</td><td>239.2</td><td>-2.8</td><td>0.9</td><td>-3.3</td><td>9.7E-04</td><td>2.4E-03</td><td>BT20/MB468</td><td>MP</td></tr>
<tr><td>>hg38_21_+_8433172_8446622_RNA45SN1_45S_50ntFlanks@10792.10818.27|CTCTGGCATGTTGGAACAATGTAGGTA</td><td>>hg38_21_+_8433172_8446622_RNA45SN1_45S_50ntFlanks</td><td>10792</td><td>10818</td><td>27</td><td>CTCTGGCATGTTGGAACAATGTAGGTA</td><td>237.4</td><td>-2.2</td><td>0.7</td><td>-3.2</td><td>1.6E-03</td><td>2.4E-03</td><td>BT20/MB468</td><td>MITO</td></tr>
<tr><td>>hg38_1_-_228634819_228635039_RNA5S12_5S_50ntFlanks@153.171.19|ACCGGGTGCTGTAGGCTTT</td><td>>hg38_1_-_228634819_228635039_RNA5S12_5S_50ntFlanks</td><td>153</td><td>171</td><td>19</td><td>ACCGGGTGCTGTAGGCTTT</td><td>3541.5</td><td>2.2</td><td>0.7</td><td>3.1</td><td>2.0E-03</td><td>2.4E-03</td><td>MB231/MB468</td><td>MITO</td></tr>
<tr><td>>hg38_21_+_8433172_8446622_RNA45SN1_45S_50ntFlanks@8380.8403.24|CAAGAGGGCGTGAAACCGTTAAGA</td><td>>hg38_21_+_8433172_8446622_RNA45SN1_45S_50ntFlanks</td><td>8380</td><td>8403</td><td>24</td><td>CAAGAGGGCGTGAAACCGTTAAGA</td><td>332.1</td><td>-3.8</td><td>1.2</td><td>-3.1</td><td>2.0E-03</td><td>2.4E-03</td><td>MB231/MB468</td><td>NUC</td></tr>
<tr><td>>trna111_HisGTG_1_-_147774845_147774916@1.31.31,trna118_HisGTG_1_-_145396881_145396952@1.31.31,trna16_HisGTG_1_+_146544773_146544844@1.31.31,trna1_HisGTG_15_+_45493349_45493420@1.31.31,trna21_HisGTG_1_+_147753471_147753542@1.31.31,trna33_HisGTG_6_+_27125906_27125977@1.31.31,trna7_HisGTG_9_-_14433938_14434009@1.31.31,trna8_HisGTG_15_-_45492611_45492682@1.31.31,trna9_HisGTG_15_-_45490804_45490875@1.31.31|GCCGTGATCGTATAGTGGTTAGTACTCTGCG</td><td>>trna111_HisGTG_1_-_147774845_147774916</td><td>1</td><td>31</td><td>31</td><td>GCCGTGATCGTATAGTGGTTAGTACTCTGCG</td><td>14297.8</td><td>-2.9</td><td>1.0</td><td>-3.1</td><td>2.0E-03</td><td>2.4E-03</td><td>MB231/MB468</td><td>NUC</td></tr>
<tr><td>>hg38_21_+_8433172_8446622_RNA45SN1_45S_50ntFlanks@8552.8569.18|CTCCCCGGAGGGGGCGGG</td><td>>hg38_21_+_8433172_8446622_RNA45SN1_45S_50ntFlanks</td><td>8552</td><td>8569</td><td>18</td><td>CTCCCCGGAGGGGGCGGG</td><td>648.0</td><td>-5.0</td><td>1.6</td><td>-3.1</td><td>1.9E-03</td><td>2.4E-03</td><td>BT20/MB468</td><td>CYTO</td></tr>
<tr><td>>hg38_MT_+_598_1651_RNR1_12S_50ntFlanks@634.665.32|AACCTCACCACCTCTTGCTCAGCCTATATACC</td><td>>hg38_MT_+_598_1651_RNR1_12S_50ntFlanks</td><td>634</td><td>665</td><td>32</td><td>AACCTCACCACCTCTTGCTCAGCCTATATACC</td><td>365.7</td><td>7.0</td><td>2.1</td><td>3.3</td><td>9.8E-04</td><td>2.4E-03</td><td>BT20/MB468</td><td>MP</td></tr>
<tr><td>>hg38_MT_+_598_1651_RNR1_12S_50ntFlanks@428.451.24|AAAGTGGCTTTAACATATCTGAAC</td><td>>hg38_MT_+_598_1651_RNR1_12S_50ntFlanks</td><td>428</td><td>451</td><td>24</td><td>AAAGTGGCTTTAACATATCTGAAC</td><td>189.4</td><td>5.9</td><td>1.8</td><td>3.3</td><td>9.8E-04</td><td>2.4E-03</td><td>BT20/MB468</td><td>MP</td></tr>
<tr><td>>hg38_21_+_8433172_8446622_RNA45SN1_45S_50ntFlanks@6660.6699.40|GCGGTGGATCACTCGGCTCGTGCGTCGATGAAGAACGCAG</td><td>>hg38_21_+_8433172_8446622_RNA45SN1_45S_50ntFlanks</td><td>6660</td><td>6699</td><td>40</td><td>GCGGTGGATCACTCGGCTCGTGCGTCGATGAAGAACGCAG</td><td>287.7</td><td>-4.0</td><td>1.3</td><td>-3.1</td><td>2.0E-03</td><td>2.4E-03</td><td>MB231/MB468</td><td>MITO</td></tr>
<tr><td>>hg38_21_+_8433172_8446622_RNA45SN1_45S_50ntFlanks@12161.12180.20|TGACTGGGGCGGTACACCTG</td><td>>hg38_21_+_8433172_8446622_RNA45SN1_45S_50ntFlanks</td><td>12161</td><td>12180</td><td>20</td><td>TGACTGGGGCGGTACACCTG</td><td>392.5</td><td>-3.8</td><td>1.2</td><td>-3.2</td><td>1.6E-03</td><td>2.4E-03</td><td>BT20/MB468</td><td>MITO</td></tr>
<tr><td>>hg38_21_+_8433172_8446622_RNA45SN1_45S_50ntFlanks@12934.12965.32|CTAAACCATTCGTAGACGACCTGCTTCTGGGT</td><td>>hg38_21_+_8433172_8446622_RNA45SN1_45S_50ntFlanks</td><td>12934</td><td>12965</td><td>32</td><td>CTAAACCATTCGTAGACGACCTGCTTCTGGGT</td><td>307.3</td><td>-1.7</td><td>0.5</td><td>-3.2</td><td>1.6E-03</td><td>2.4E-03</td><td>BT20/MB468</td><td>MITO</td></tr>
<tr><td>>hg38_21_+_8433172_8446622_RNA45SN1_45S_50ntFlanks@4726.4748.23|TAATCAAGAACGAAAGTCGGAGG</td><td>>hg38_21_+_8433172_8446622_RNA45SN1_45S_50ntFlanks</td><td>4726</td><td>4748</td><td>23</td><td>TAATCAAGAACGAAAGTCGGAGG</td><td>218.0</td><td>-2.3</td><td>0.8</td><td>-3.1</td><td>1.9E-03</td><td>2.4E-03</td><td>MB231/MB468</td><td>CYTO</td></tr>
<tr><td>>hg38_21_+_8433172_8446622_RNA45SN1_45S_50ntFlanks@4163.4186.24|CATCCAAGGAAGGCAGCAGGCGCG</td><td>>hg38_21_+_8433172_8446622_RNA45SN1_45S_50ntFlanks</td><td>4163</td><td>4186</td><td>24</td><td>CATCCAAGGAAGGCAGCAGGCGCG</td><td>359.4</td><td>-4.4</td><td>1.3</td><td>-3.3</td><td>9.8E-04</td><td>2.4E-03</td><td>BT20/MB468</td><td>MP</td></tr>
<tr><td>>hg38_21_+_8433172_8446622_RNA45SN1_45S_50ntFlanks@11569.11613.45|CAGCCGACTTAGAACTGGTGCGGACCAGGGGAATCCGACTGTTTA</td><td>>hg38_21_+_8433172_8446622_RNA45SN1_45S_50ntFlanks</td><td>11569</td><td>11613</td><td>45</td><td>CAGCCGACTTAGAACTGGTGCGGACCAGGGGAATCCGACTGTTTA</td><td>1043.9</td><td>3.7</td><td>1.1</td><td>3.5</td><td>4.4E-04</td><td>2.4E-03</td><td>BT20/MB231</td><td>MITO</td></tr>
<tr><td>>hg38_21_+_8433172_8446622_RNA45SN1_45S_50ntFlanks@4560.4578.19|CTAGGAATAATGGAATAGG</td><td>>hg38_21_+_8433172_8446622_RNA45SN1_45S_50ntFlanks</td><td>4560</td><td>4578</td><td>19</td><td>CTAGGAATAATGGAATAGG</td><td>104.6</td><td>-5.1</td><td>1.6</td><td>-3.3</td><td>9.8E-04</td><td>2.4E-03</td><td>BT20/MB468</td><td>MP</td></tr>
<tr><td>>hg38_21_+_8433172_8446622_RNA45SN1_45S_50ntFlanks@11569.11612.44|CAGCCGACTTAGAACTGGTGCGGACCAGGGGAATCCGACTGTTT</td><td>>hg38_21_+_8433172_8446622_RNA45SN1_45S_50ntFlanks</td><td>11569</td><td>11612</td><td>44</td><td>CAGCCGACTTAGAACTGGTGCGGACCAGGGGAATCCGACTGTTT</td><td>2110.1</td><td>-4.1</td><td>1.3</td><td>-3.1</td><td>2.0E-03</td><td>2.4E-03</td><td>MB231/MB468</td><td>NUC</td></tr>
<tr><td>>hg38_21_+_8433172_8446622_RNA45SN1_45S_50ntFlanks@6651.6676.26|CGACTCTTAGCGGTGGATCACTCGGC</td><td>>hg38_21_+_8433172_8446622_RNA45SN1_45S_50ntFlanks</td><td>6651</td><td>6676</td><td>26</td><td>CGACTCTTAGCGGTGGATCACTCGGC</td><td>24847.4</td><td>2.9</td><td>0.9</td><td>3.1</td><td>2.0E-03</td><td>2.4E-03</td><td>MB231/MB468</td><td>MITO</td></tr>
<tr><td>>MI0000076|hsa-mir-20a&WithFlank&13|+|91351059|91351141@14.34.21&[MIMAT0000075&hsa-miR-20a-5p&offsets|0|-2;m-96&13|+|91351072|91351094&offsets|0|-2]|TAAAGTGCTTATAGTGCAGGT</td><td>>MI0000076|hsa-mir-20a&WithFlank&13|+|91351059|91351141</td><td>14</td><td>34</td><td>21</td><td>TAAAGTGCTTATAGTGCAGGT</td><td>3257.7</td><td>-5.4</td><td>1.5</td><td>-3.6</td><td>3.3E-04</td><td>2.4E-03</td><td>BT20/MB231</td><td>MP</td></tr>
<tr><td>>MI0003513|hsa-mir-455&WithFlank&9|+|114209428|114209535@60.81.22&[MIMAT0004784&hsa-miR-455-3p&offsets|0|+1;m-135&9|+|114209487|114209508&offsets|0|0]|GCAGTCCATGGGCATATACACT</td><td>>MI0003513|hsa-mir-455&WithFlank&9|+|114209428|114209535</td><td>60</td><td>81</td><td>22</td><td>GCAGTCCATGGGCATATACACT</td><td>317.3</td><td>-2.9</td><td>0.8</td><td>-3.5</td><td>3.9E-04</td><td>2.4E-03</td><td>BT20/MB231</td><td>CYTO</td></tr>
<tr><td>>hg38_21_+_8433172_8446622_RNA45SN1_45S_50ntFlanks@8184.8209.26|CGAGGCCCAGCCCGTGGACGGTGTGA</td><td>>hg38_21_+_8433172_8446622_RNA45SN1_45S_50ntFlanks</td><td>8184</td><td>8209</td><td>26</td><td>CGAGGCCCAGCCCGTGGACGGTGTGA</td><td>107.3</td><td>-4.0</td><td>1.3</td><td>-3.1</td><td>1.7E-03</td><td>2.4E-03</td><td>MB231/MB468</td><td>MP</td></tr>
<tr><td>>hg38_21_+_8433172_8446622_RNA45SN1_45S_50ntFlanks@12936.12953.18|AAACCATTCGTAGACGAC</td><td>>hg38_21_+_8433172_8446622_RNA45SN1_45S_50ntFlanks</td><td>12936</td><td>12953</td><td>18</td><td>AAACCATTCGTAGACGAC</td><td>665.8</td><td>-5.1</td><td>1.6</td><td>-3.1</td><td>1.9E-03</td><td>2.4E-03</td><td>BT20/MB468</td><td>CYTO</td></tr>
<tr><td>>hg38_21_+_8433172_8446622_RNA45SN1_45S_50ntFlanks@8673.8700.28|CGGTGCGCCGCGACCGGCTCCGGGACGG</td><td>>hg38_21_+_8433172_8446622_RNA45SN1_45S_50ntFlanks</td><td>8673</td><td>8700</td><td>28</td><td>CGGTGCGCCGCGACCGGCTCCGGGACGG</td><td>157.3</td><td>5.6</td><td>1.7</td><td>3.3</td><td>9.8E-04</td><td>2.4E-03</td><td>BT20/MB468</td><td>MP</td></tr>
<tr><td>>hg38_21_+_8433172_8446622_RNA45SN1_45S_50ntFlanks@4591.4613.23|TTTTGTTGGTTTTCGGAACTGAG</td><td>>hg38_21_+_8433172_8446622_RNA45SN1_45S_50ntFlanks</td><td>4591</td><td>4613</td><td>23</td><td>TTTTGTTGGTTTTCGGAACTGAG</td><td>81.2</td><td>-5.2</td><td>1.6</td><td>-3.3</td><td>9.8E-04</td><td>2.4E-03</td><td>BT20/MB468</td><td>MP</td></tr>
<tr><td>>hg38_21_+_8433172_8446622_RNA45SN1_45S_50ntFlanks@4729.4765.37|TCAAGAACGAAAGTCGGAGGTTCGAAGACGATCAGAT</td><td>>hg38_21_+_8433172_8446622_RNA45SN1_45S_50ntFlanks</td><td>4729</td><td>4765</td><td>37</td><td>TCAAGAACGAAAGTCGGAGGTTCGAAGACGATCAGAT</td><td>175.8</td><td>4.4</td><td>1.3</td><td>3.3</td><td>9.8E-04</td><td>2.4E-03</td><td>BT20/MB468</td><td>MP</td></tr>
<tr><td>>hg38_21_+_8433172_8446622_RNA45SN1_45S_50ntFlanks@6652.6700.49|GACTCTTAGCGGTGGATCACTCGGCTCGTGCGTCGATGAAGAACGCAGC</td><td>>hg38_21_+_8433172_8446622_RNA45SN1_45S_50ntFlanks</td><td>6652</td><td>6700</td><td>49</td><td>GACTCTTAGCGGTGGATCACTCGGCTCGTGCGTCGATGAAGAACGCAGC</td><td>3647.9</td><td>6.5</td><td>2.0</td><td>3.3</td><td>9.8E-04</td><td>2.4E-03</td><td>BT20/MB468</td><td>MP</td></tr>
<tr><td>>hg38_21_+_8433172_8446622_RNA45SN1_45S_50ntFlanks@12158.12182.25|GTTTGACTGGGGCGGTACACCTGTC</td><td>>hg38_21_+_8433172_8446622_RNA45SN1_45S_50ntFlanks</td><td>12158</td><td>12182</td><td>25</td><td>GTTTGACTGGGGCGGTACACCTGTC</td><td>381.4</td><td>-5.6</td><td>1.7</td><td>-3.3</td><td>9.8E-04</td><td>2.4E-03</td><td>BT20/MB468</td><td>MP</td></tr>
<tr><td>>hg38_21_+_8433172_8446622_RNA45SN1_45S_50ntFlanks@4159.4184.26|ACCACATCCAAGGAAGGCAGCAGGCG</td><td>>hg38_21_+_8433172_8446622_RNA45SN1_45S_50ntFlanks</td><td>4159</td><td>4184</td><td>26</td><td>ACCACATCCAAGGAAGGCAGCAGGCG</td><td>117.6</td><td>-5.5</td><td>1.7</td><td>-3.3</td><td>9.8E-04</td><td>2.4E-03</td><td>BT20/MB468</td><td>MP</td></tr>
<tr><td>>hg38_21_+_8433172_8446622_RNA45SN1_45S_50ntFlanks@10677.10709.33|GCGGTTCCGGCGGCGTCCGGTGAGCTCTCGCTG</td><td>>hg38_21_+_8433172_8446622_RNA45SN1_45S_50ntFlanks</td><td>10677</td><td>10709</td><td>33</td><td>GCGGTTCCGGCGGCGTCCGGTGAGCTCTCGCTG</td><td>1681.0</td><td>3.8</td><td>1.1</td><td>3.5</td><td>4.4E-04</td><td>2.4E-03</td><td>BT20/MB231</td><td>MITO</td></tr>
<tr><td>>hg38_21_+_8433172_8446622_RNA45SN1_45S_50ntFlanks@6650.6674.25|ACGACTCTTAGCGGTGGATCACTCG</td><td>>hg38_21_+_8433172_8446622_RNA45SN1_45S_50ntFlanks</td><td>6650</td><td>6674</td><td>25</td><td>ACGACTCTTAGCGGTGGATCACTCG</td><td>2283.7</td><td>3.4</td><td>1.1</td><td>3.1</td><td>2.0E-03</td><td>2.4E-03</td><td>MB231/MB468</td><td>MITO</td></tr>
<tr><td>>MI0000816|hsa-mir-335&WithFlank&7|+|130496105|130496210@57.79.23&[MIMAT0004703&hsa-miR-335-3p&offsets|-1|0;m-159&7|+|130496162|130496183&offsets|-1|0]|GTTTTTCATTATTGCTCCTGACC</td><td>>MI0000816|hsa-mir-335&WithFlank&7|+|130496105|130496210</td><td>57</td><td>79</td><td>23</td><td>GTTTTTCATTATTGCTCCTGACC</td><td>290.8</td><td>-2.8</td><td>0.8</td><td>-3.5</td><td>4.4E-04</td><td>2.4E-03</td><td>BT20/MB231</td><td>MITO</td></tr>
<tr><td>>hg38_21_+_8433172_8446622_RNA45SN1_45S_50ntFlanks@7967.8009.43|CTCCGAGACGCGACCTCAGATCAGACGTGGCGACCCGCTGAAT</td><td>>hg38_21_+_8433172_8446622_RNA45SN1_45S_50ntFlanks</td><td>7967</td><td>8009</td><td>43</td><td>CTCCGAGACGCGACCTCAGATCAGACGTGGCGACCCGCTGAAT</td><td>58.3</td><td>4.2</td><td>1.4</td><td>3.1</td><td>2.0E-03</td><td>2.4E-03</td><td>BT20/MB468</td><td>NUC</td></tr>
<tr><td>>hg38_21_+_8433172_8446622_RNA45SN1_45S_50ntFlanks@12825.12871.47|GGACCGGGGTCCGGTGCGGAGTGCCCTTCGTCCTGGGAAACGGGGCG</td><td>>hg38_21_+_8433172_8446622_RNA45SN1_45S_50ntFlanks</td><td>12825</td><td>12871</td><td>47</td><td>GGACCGGGGTCCGGTGCGGAGTGCCCTTCGTCCTGGGAAACGGGGCG</td><td>48.8</td><td>-9.5</td><td>3.0</td><td>-3.1</td><td>1.7E-03</td><td>2.4E-03</td><td>MB231/MB468</td><td>MP</td></tr>
<tr><td>>am_trna128_GlyGCC_6_-_27870686_27870756@1.30.30,trna133_GlyCCC_1_-_16872434_16872504@1.30.30,trna18_GlyGCC_16_+_70822597_70822667@1.30.30,trna19_GlyGCC_16_+_70823410_70823480@1.30.30,trna19_GlyGCC_2_-_157257659_157257729@1.30.30,trna24_GlyGCC_16_-_70812942_70813012@1.30.30,trna25_GlyGCC_16_-_70812114_70812184@1.30.30,trna4_GlyCCC_1_+_17188416_17188486@1.30.30,trna5_GlyGCC_17_+_8029064_8029134@1.30.30,trna68_GlyGCC_1_-_161493637_161493707@1.30.30|GCATTGGTGGTTCAGTGGTAGAATTCTCGC</td><td>>am_trna128_GlyGCC_6_-_27870686_27870756</td><td>1</td><td>30</td><td>30</td><td>GCATTGGTGGTTCAGTGGTAGAATTCTCGC</td><td>1614.3</td><td>-3.4</td><td>0.9</td><td>-3.6</td><td>3.7E-04</td><td>2.4E-03</td><td>BT20/MB231</td><td>NUC</td></tr>
<tr><td>>MI0000081|hsa-mir-24-2&WithFlank&19|-|13836281|13836365@56.73.18(+2C);MI0000080|hsa-mir-24-1&WithFlank&9|+|95086015|95086094@50.67.18(+2C)&[MIMAT0000080&hsa-miR-24-3p&offsets|0|-4(+2C);m-36&19|-|13836288|13836310&offsets|0|-5(+2C)];[MIMAT0000080_1&hsa-miR-24-3p&offsets|0|-4(+2C);m-35&9|+|95086064|95086086&offsets|0|-5(+2C)]|TGGCTCAGTTCAGCAGGACC</td><td>>MI0000081|hsa-mir-24-2&WithFlank&19|-|13836281|13836365</td><td>56</td><td>73</td><td>18(+2C)</td><td>TGGCTCAGTTCAGCAGGACC</td><td>79.6</td><td>6.0</td><td>1.9</td><td>3.1</td><td>1.7E-03</td><td>2.4E-03</td><td>MB231/MB468</td><td>MP</td></tr>
<tr><td>>hg38_21_+_8433172_8446622_RNA45SN1_45S_50ntFlanks@6731.6758.28|CACATTGATCATCGACACTTCGAACGCA</td><td>>hg38_21_+_8433172_8446622_RNA45SN1_45S_50ntFlanks</td><td>6731</td><td>6758</td><td>28</td><td>CACATTGATCATCGACACTTCGAACGCA</td><td>241.2</td><td>-3.2</td><td>1.0</td><td>-3.1</td><td>1.9E-03</td><td>2.4E-03</td><td>BT20/MB468</td><td>CYTO</td></tr>
</table>

<table>
<tr><td>>hg38_21_+_8433172_8446622_RNA45SN1_45S_50ntFlanks@3900.3938.39|CTTCGCGGGGGGGATGCGTGCATTTATCAGATCAAAACC</td><td>>hg38_21_+_8433172_8446622_RNA45SN1_45S_50ntFlanks</td><td>3900</td><td>3938</td><td>39</td><td>CTTCGCGGGGGGGATGCGTGCATTTATCAGATCAAAACC</td><td>213.2</td><td>-3.1</td><td>1.0</td><td>-3.1</td><td>1.9E-03</td><td>2.4E-03</td><td>MB231/MB468</td><td>CYTO</td></tr>
<tr><td>>hg38_21_+_8433172_8446622_RNA45SN1_45S_50ntFlanks@6732.6770.39|ACATTGATCATCGACACTTCGAACGCACTTGCGGCCCCG</td><td>>hg38_21_+_8433172_8446622_RNA45SN1_45S_50ntFlanks</td><td>6732</td><td>6770</td><td>39</td><td>ACATTGATCATCGACACTTCGAACGCACTTGCGGCCCCG</td><td>974.2</td><td>-6.0</td><td>1.8</td><td>-3.3</td><td>9.9E-04</td><td>2.4E-03</td><td>BT20/MB468</td><td>MP</td></tr>
<tr><td>>hg38_21_+_8433172_8446622_RNA45SN1_45S_50ntFlanks@12395.12418.24|AGCGACGTCGCTTTTTGATCCTTC</td><td>>hg38_21_+_8433172_8446622_RNA45SN1_45S_50ntFlanks</td><td>12395</td><td>12418</td><td>24</td><td>AGCGACGTCGCTTTTTGATCCTTC</td><td>406.4</td><td>-2.5</td><td>0.7</td><td>-3.6</td><td>3.7E-04</td><td>2.4E-03</td><td>BT20/MB231</td><td>NUC</td></tr>
<tr><td>>hg38_21_+_8433172_8446622_RNA45SN1_45S_50ntFlanks@10291.10322.32|CGCGGGCCCGGGTGGAGCCGCCGCAGGTGCAG</td><td>>hg38_21_+_8433172_8446622_RNA45SN1_45S_50ntFlanks</td><td>10291</td><td>10322</td><td>32</td><td>CGCGGGCCCGGGTGGAGCCGCCGCAGGTGCAG</td><td>117.1</td><td>-5.5</td><td>1.7</td><td>-3.3</td><td>9.9E-04</td><td>2.4E-03</td><td>BT20/MB468</td><td>MP</td></tr>
<tr><td>>hg38_21_+_8433172_8446622_RNA45SN1_45S_50ntFlanks@10348.10377.30|AAACGAGAACTTTGAAGGCCGAAGTGGAGA</td><td>>hg38_21_+_8433172_8446622_RNA45SN1_45S_50ntFlanks</td><td>10348</td><td>10377</td><td>30</td><td>AAACGAGAACTTTGAAGGCCGAAGTGGAGA</td><td>317.8</td><td>1.3</td><td>0.4</td><td>3.1</td><td>1.9E-03</td><td>2.4E-03</td><td>BT20/MB468</td><td>CYTO</td></tr>
<tr><td>>hg38_21_+_8433172_8446622_RNA45SN1_45S_50ntFlanks@5443.5467.25|CCTCGGATCGGCCCCGCCGGGGTCG</td><td>>hg38_21_+_8433172_8446622_RNA45SN1_45S_50ntFlanks</td><td>5443</td><td>5467</td><td>25</td><td>CCTCGGATCGGCCCCGCCGGGGTCG</td><td>214.4</td><td>-5.3</td><td>1.6</td><td>-3.3</td><td>9.9E-04</td><td>2.4E-03</td><td>BT20/MB468</td><td>MP</td></tr>
<tr><td>>hg38_21_+_8433172_8446622_RNA45SN1_45S_50ntFlanks@12457.12474.18|CAAGCGTTGGATTGTTCA</td><td>>hg38_21_+_8433172_8446622_RNA45SN1_45S_50ntFlanks</td><td>12457</td><td>12474</td><td>18</td><td>CAAGCGTTGGATTGTTCA</td><td>388.9</td><td>-5.4</td><td>1.6</td><td>-3.3</td><td>9.9E-04</td><td>2.4E-03</td><td>BT20/MB468</td><td>MP</td></tr>
<tr><td>>MI0000093|hsa-mir-92a-1&WithFlank&13|+|91351308|91351397@54.76.23(+1U)&[MIMAT0000092&hsa-miR-92a-3p&offsets|0|+1(+1U);m-9&13|+|91351361|91351382&offsets|0|+1(+1U)]|TATTGCACTTGTCCCGGCCTGTTT</td><td>>MI0000093|hsa-mir-92a-1&WithFlank&13|+|91351308|91351397</td><td>54</td><td>76</td><td>23(+1U)</td><td>TATTGCACTTGTCCCGGCCTGTTT</td><td>193.0</td><td>4.1</td><td>1.3</td><td>3.1</td><td>1.7E-03</td><td>2.4E-03</td><td>MB231/MB468</td><td>MP</td></tr>
<tr><td>>hg38_21_+_8433172_8446622_RNA45SN1_45S_50ntFlanks@11570.11601.32|AGCCGACTTAGAACTGGTGCGGACCAGGGGAA</td><td>>hg38_21_+_8433172_8446622_RNA45SN1_45S_50ntFlanks</td><td>11570</td><td>11601</td><td>32</td><td>AGCCGACTTAGAACTGGTGCGGACCAGGGGAA</td><td>815.1</td><td>-2.6</td><td>0.8</td><td>-3.1</td><td>1.9E-03</td><td>2.4E-03</td><td>MB231/MB468</td><td>CYTO</td></tr>
<tr><td>>MI0000087|hsa-mir-29a&WithFlank&7|-|130876741|130876816@48.68.21(+1U)&[MIMAT0000086&hsa-miR-29a-3p&offsets|0|-1(+1U);m-26&7|-|130876749|130876769&offsets|0|0(+1U)]|TAGCACCATCTGAAATCGGTTT</td><td>>MI0000087|hsa-mir-29a&WithFlank&7|-|130876741|130876816</td><td>48</td><td>68</td><td>21(+1U)</td><td>TAGCACCATCTGAAATCGGTTT</td><td>1304.1</td><td>6.8</td><td>2.1</td><td>3.3</td><td>9.9E-04</td><td>2.4E-03</td><td>BT20/MB468</td><td>MP</td></tr>
<tr><td>>hg38_21_+_8433172_8446622_RNA45SN1_45S_50ntFlanks@10771.10817.47|CAGGTCTCCAAGGTGAACAGCCTCTGGCATGTTGGAACAATGTAGGT</td><td>>hg38_21_+_8433172_8446622_RNA45SN1_45S_50ntFlanks</td><td>10771</td><td>10817</td><td>47</td><td>CAGGTCTCCAAGGTGAACAGCCTCTGGCATGTTGGAACAATGTAGGT</td><td>183.8</td><td>-2.7</td><td>0.9</td><td>-3.1</td><td>1.9E-03</td><td>2.4E-03</td><td>MB231/MB468</td><td>CYTO</td></tr>
<tr><td>>hg38_21_+_8433172_8446622_RNA45SN1_45S_50ntFlanks@11718.11744.27|AACGGCGGGAGTAACTATGACTCTCTT</td><td>>hg38_21_+_8433172_8446622_RNA45SN1_45S_50ntFlanks</td><td>11718</td><td>11744</td><td>27</td><td>AACGGCGGGAGTAACTATGACTCTCTT</td><td>640.6</td><td>-4.5</td><td>1.4</td><td>-3.3</td><td>9.9E-04</td><td>2.4E-03</td><td>BT20/MB468</td><td>MP</td></tr>
<tr><td>>hg38_21_+_8433172_8446622_RNA45SN1_45S_50ntFlanks@4731.4752.22|AAGAACGAAAGTCGGAGGTTCG</td><td>>hg38_21_+_8433172_8446622_RNA45SN1_45S_50ntFlanks</td><td>4731</td><td>4752</td><td>22</td><td>AAGAACGAAAGTCGGAGGTTCG</td><td>321.2</td><td>-4.1</td><td>1.2</td><td>-3.3</td><td>9.9E-04</td><td>2.4E-03</td><td>BT20/MB468</td><td>MP</td></tr>
<tr><td>>hg38_21_+_8433172_8446622_RNA45SN1_45S_50ntFlanks@7982.8006.25|TCAGATCAGACGTGGCGACCCGCTG</td><td>>hg38_21_+_8433172_8446622_RNA45SN1_45S_50ntFlanks</td><td>7982</td><td>8006</td><td>25</td><td>TCAGATCAGACGTGGCGACCCGCTG</td><td>722.1</td><td>-3.8</td><td>1.2</td><td>-3.1</td><td>1.9E-03</td><td>2.4E-03</td><td>BT20/MB468</td><td>CYTO</td></tr>
<tr><td>>MI0000266|hsa-mir-10a&WithFlank&17|-|48579832|48579953@29.48.20(+1A)&[MIMAT0000253&hsa-miR-10a-5p&offsets|+1|-2(+1A);m-18&17|-|48579905|48579926&offsets|+1|-1(+1A)]|ACCCTGTAGATCCGAATTTGA</td><td>>MI0000266|hsa-mir-10a&WithFlank&17|-|48579832|48579953</td><td>29</td><td>48</td><td>20(+1A)</td><td>ACCCTGTAGATCCGAATTTGA</td><td>217.3</td><td>2.8</td><td>0.8</td><td>3.5</td><td>4.0E-04</td><td>2.4E-03</td><td>BT20/MB231</td><td>CYTO</td></tr>
<tr><td>>hg38_21_+_8433172_8446622_RNA45SN1_45S_50ntFlanks@3771.3794.24|CACGGCCGGTACAGTGAAACTGCG</td><td>>hg38_21_+_8433172_8446622_RNA45SN1_45S_50ntFlanks</td><td>3771</td><td>3794</td><td>24</td><td>CACGGCCGGTACAGTGAAACTGCG</td><td>314.9</td><td>-2.7</td><td>0.8</td><td>-3.2</td><td>1.6E-03</td><td>2.4E-03</td><td>BT20/MB468</td><td>MITO</td></tr>
<tr><td>>hg38_21_+_8433172_8446622_RNA45SN1_45S_50ntFlanks@12306.12325.20|CACGATCCTTCTGACCTTTT</td><td>>hg38_21_+_8433172_8446622_RNA45SN1_45S_50ntFlanks</td><td>12306</td><td>12325</td><td>20</td><td>CACGATCCTTCTGACCTTTT</td><td>1038.9</td><td>-3.1</td><td>1.0</td><td>-3.2</td><td>1.6E-03</td><td>2.4E-03</td><td>BT20/MB468</td><td>MITO</td></tr>
<tr><td>>hg38_21_+_8433172_8446622_RNA45SN1_45S_50ntFlanks@12615.12657.43|GCTGAGGAGCCAATGGGGCGAAGCTACCATCTGTGGGATTATG</td><td>>hg38_21_+_8433172_8446622_RNA45SN1_45S_50ntFlanks</td><td>12615</td><td>12657</td><td>43</td><td>GCTGAGGAGCCAATGGGGCGAAGCTACCATCTGTGGGATTATG</td><td>553.2</td><td>-4.0</td><td>1.3</td><td>-3.2</td><td>1.6E-03</td><td>2.4E-03</td><td>BT20/MB468</td><td>MITO</td></tr>
<tr><td>>hg38_MT_+_1621_3279_RNR2_16S_50ntFlanks@1431.1458.28|AAAGTCCTACGTGATCTGAGTTCAGACC</td><td>>hg38_MT_+_1621_3279_RNR2_16S_50ntFlanks</td><td>1431</td><td>1458</td><td>28</td><td>AAAGTCCTACGTGATCTGAGTTCAGACC</td><td>209.0</td><td>3.1</td><td>1.0</td><td>3.1</td><td>1.7E-03</td><td>2.4E-03</td><td>MB231/MB468</td><td>MP</td></tr>
<tr><td>>hg38_MT_+_1621_3279_RNR2_16S_50ntFlanks@999.1030.32|AGGGACCTGTATGAATGGCTCCACGAGGGTTC</td><td>>hg38_MT_+_1621_3279_RNR2_16S_50ntFlanks</td><td>999</td><td>1030</td><td>32</td><td>AGGGACCTGTATGAATGGCTCCACGAGGGTTC</td><td>191.0</td><td>4.9</td><td>1.5</td><td>3.3</td><td>9.9E-04</td><td>2.4E-03</td><td>BT20/MB468</td><td>MP</td></tr>
<tr><td>>hg38_21_+_8433172_8446622_RNA45SN1_45S_50ntFlanks@10391.10429.39|AACAGCAGTTGAACATGGGTCAGTCGGTCCTGAGAGATG</td><td>>hg38_21_+_8433172_8446622_RNA45SN1_45S_50ntFlanks</td><td>10391</td><td>10429</td><td>39</td><td>AACAGCAGTTGAACATGGGTCAGTCGGTCCTGAGAGATG</td><td>352.1</td><td>2.6</td><td>0.7</td><td>3.5</td><td>4.4E-04</td><td>2.5E-03</td><td>BT20/MB231</td><td>MITO</td></tr>
<tr><td>>hg38_21_+_8433172_8446622_RNA45SN1_45S_50ntFlanks@9566.9592.27|GACGTGCAAATCGGTCGTCCGACCTGG</td><td>>hg38_21_+_8433172_8446622_RNA45SN1_45S_50ntFlanks</td><td>9566</td><td>9592</td><td>27</td><td>GACGTGCAAATCGGTCGTCCGACCTGG</td><td>177.5</td><td>-1.7</td><td>0.6</td><td>-3.1</td><td>2.1E-03</td><td>2.5E-03</td><td>MB231/MB468</td><td>NUC</td></tr>
<tr><td>>hg38_21_+_8433172_8446622_RNA45SN1_45S_50ntFlanks@3773.3792.20|CGGCCGGTACAGTGAAACTG</td><td>>hg38_21_+_8433172_8446622_RNA45SN1_45S_50ntFlanks</td><td>3773</td><td>3792</td><td>20</td><td>CGGCCGGTACAGTGAAACTG</td><td>229.0</td><td>-3.0</td><td>1.0</td><td>-3.1</td><td>1.9E-03</td><td>2.5E-03</td><td>MB231/MB468</td><td>CYTO</td></tr>
<tr><td>>MI0000061|hsa-let-7a-2&WithFlank&11|-|122146516|122146599@11.32.22(+1G);MI0000062|hsa-let-7a-3&WithFlank&22|+|46112743|46112828@10.31.22(+1G);MI0000060|hsa-let-7a-1&WithFlank&9|+|94175951|94176042@12.33.22(+1G)&[MIMAT0000062&hsa-let-7a-5p&offsets|0|0(+1G);m-6&11|-|122146568|122146589&offsets|0|0(+1G)];[MIMAT0000062_1&hsa-let-7a-5p&offsets|0|0(+1G);m-5&22|+|46112752|46112773&offsets|0|0(+1G)];[MIMAT0000062_2&hsa-let-7a-5p&offsets|0|0(+1G);m-4&9|+|94175962|94175983&offsets|0|0(+1G)]|TGAGGTAGTAGGTTGTATAGTTG</td><td>>MI0000061|hsa-let-7a-2&WithFlank&11|-|122146516|122146599</td><td>11</td><td>32</td><td>22(+1G)</td><td>TGAGGTAGTAGGTTGTATAGTTG</td><td>165.6</td><td>4.7</td><td>1.4</td><td>3.3</td><td>9.9E-04</td><td>2.5E-03</td><td>BT20/MB468</td><td>MP</td></tr>
<tr><td>>hg38_21_+_8433172_8446622_RNA45SN1_45S_50ntFlanks@7974.8013.40|ACGCGACCTCAGATCAGACGTGGCGACCCGCTGAATTTAA</td><td>>hg38_21_+_8433172_8446622_RNA45SN1_45S_50ntFlanks</td><td>7974</td><td>8013</td><td>40</td><td>ACGCGACCTCAGATCAGACGTGGCGACCCGCTGAATTTAA</td><td>1753.1</td><td>2.8</td><td>0.9</td><td>3.1</td><td>2.1E-03</td><td>2.5E-03</td><td>MB231/MB468</td><td>MITO</td></tr>
<tr><td>>hg38_21_+_8433172_8446622_RNA45SN1_45S_50ntFlanks@6652.6688.37|GACTCTTAGCGGTGGATCACTCGGCTCGTGCGTCGAT</td><td>>hg38_21_+_8433172_8446622_RNA45SN1_45S_50ntFlanks</td><td>6652</td><td>6688</td><td>37</td><td>GACTCTTAGCGGTGGATCACTCGGCTCGTGCGTCGAT</td><td>5782.0</td><td>-1.9</td><td>0.6</td><td>-3.1</td><td>2.1E-03</td><td>2.5E-03</td><td>MB231/MB468</td><td>MITO</td></tr>
<tr><td>>MI0000734|hsa-mir-106b&WithFlank&7|-|100093987|100094080@58.76.19(+1C)&[MIMAT0004672&hsa-miR-106b-3p&offsets|0|-3(+1C);m-87&7|-|100094003|100094023&offsets|0|-2(+1C)]|CCGCACTGTGGGTACTTGCC</td><td>>MI0000734|hsa-mir-106b&WithFlank&7|-|100093987|100094080</td><td>58</td><td>76</td><td>19(+1C)</td><td>CCGCACTGTGGGTACTTGCC</td><td>152.7</td><td>6.5</td><td>2.0</td><td>3.3</td><td>1.0E-03</td><td>2.5E-03</td><td>BT20/MB468</td><td>MP</td></tr>
<tr><td>>hg38_21_+_8433172_8446622_RNA45SN1_45S_50ntFlanks@6730.6772.43|ACACATTGATCATCGACACTTCGAACGCACTTGCGGCCCCGGG</td><td>>hg38_21_+_8433172_8446622_RNA45SN1_45S_50ntFlanks</td><td>6730</td><td>6772</td><td>43</td><td>ACACATTGATCATCGACACTTCGAACGCACTTGCGGCCCCGGG</td><td>48.6</td><td>-9.5</td><td>3.0</td><td>-3.1</td><td>1.7E-03</td><td>2.5E-03</td><td>MB231/MB468</td><td>MP</td></tr>
<tr><td>>hg38_21_+_8433172_8446622_RNA45SN1_45S_50ntFlanks@11452.11477.26|CTCCTCGGCGCGCGGCGGCGGCGGCG</td><td>>hg38_21_+_8433172_8446622_RNA45SN1_45S_50ntFlanks</td><td>11452</td><td>11477</td><td>26</td><td>CTCCTCGGCGCGCGGCGGCGGCGG CG</td><td>446.2</td><td>-5.7</td><td>1.8</td><td>-3.1</td><td>2.1E-03</td><td>2.5E-03</td><td>BT20/MB468</td><td>NUC</td></tr>
<tr><td>>hg38_21_+_8433172_8446622_RNA45SN1_45S_50ntFlanks@4730.4748.19|CAAGAACGAAAGTCGGAGG</td><td>>hg38_21_+_8433172_8446622_RNA45SN1_45S_50ntFlanks</td><td>4730</td><td>4748</td><td>19</td><td>CAAGAACGAAAGTCGGAGG</td><td>431.2</td><td>-2.9</td><td>0.9</td><td>-3.3</td><td>1.0E-03</td><td>2.5E-03</td><td>BT20/MB468</td><td>MP</td></tr>
<tr><td>>MI0000085|hsa-mir-27a&WithFlank&19|-|13836434|13836523@57.77.21(+1U)&[MIMAT0000084&hsa-miR-27a-3p&offsets|0|0(+1U);m-43&19|-|13836447|13836467&offsets|0|0(+1U)]|TTCACAGTGGCTAAGTTCCGCT</td><td>>MI0000085|hsa-mir-27a&WithFlank&19|-|13836434|13836523</td><td>57</td><td>77</td><td>21(+1U)</td><td>TTCACAGTGGCTAAGTTCCGCT</td><td>1229.0</td><td>-5.3</td><td>1.5</td><td>-3.6</td><td>3.4E-04</td><td>2.5E-03</td><td>BT20/MB231</td><td>MP</td></tr>
<tr><td>>hg38_21_+_8433172_8446622_RNA45SN1_45S_50ntFlanks@10678.10706.29|CGGTTCCGGCGGCGTCCGGTGAGCTCTCG</td><td>>hg38_21_+_8433172_8446622_RNA45SN1_45S_50ntFlanks</td><td>10678</td><td>10706</td><td>29</td><td>CGGTTCCGGCGGCGTCCGGTGAGCTCTCG</td><td>276.9</td><td>5.7</td><td>1.6</td><td>3.6</td><td>3.4E-04</td><td>2.5E-03</td><td>BT20/MB231</td><td>MP</td></tr>
<tr><td>>hg38_MT_+_598_1651_RNR1_12S_50ntFlanks@425.456.32|ACGAAAGTGGCTTTAACATATCTGAACACACA</td><td>>hg38_MT_+_598_1651_RNR1_12S_50ntFlanks</td><td>425</td><td>456</td><td>32</td><td>ACGAAAGTGGCTTTAACATATCTGAACACACA</td><td>315.4</td><td>6.0</td><td>1.8</td><td>3.3</td><td>1.0E-03</td><td>2.5E-03</td><td>BT20/MB468</td><td>MP</td></tr>
<tr><td>>hg38_21_+_8433172_8446622_RNA45SN1_45S_50ntFlanks@12945.12982.38|GTAGACGACCTGCTTCTGGGTCGGGGTTTCGTACGTAG</td><td>>hg38_21_+_8433172_8446622_RNA45SN1_45S_50ntFlanks</td><td>12945</td><td>12982</td><td>38</td><td>GTAGACGACCTGCTTCTGGGTCGGGGTTTCGTACGTAG</td><td>118.0</td><td>-3.3</td><td>1.1</td><td>-3.1</td><td>1.9E-03</td><td>2.5E-03</td><td>BT20/MB468</td><td>CYTO</td></tr>
<tr><td>>hg38_21_+_8433172_8446622_RNA45SN1_45S_50ntFlanks@9549.9573.25|AGGTCCGTAGCGGTCCTGACGTGCA</td><td>>hg38_21_+_8433172_8446622_RNA45SN1_45S_50ntFlanks</td><td>9549</td><td>9573</td><td>25</td><td>AGGTCCGTAGCGGTCCTGACGTGCA</td><td>467.8</td><td>2.0</td><td>0.7</td><td>3.1</td><td>2.0E-03</td><td>2.5E-03</td><td>MB231/MB468</td><td>CYTO</td></tr>
<tr><td>>hg38_21_+_8433172_8446622_RNA45SN1_45S_50ntFlanks@11563.11587.25|GCCTAGCAGCCGACTTAGAACTGGT</td><td>>hg38_21_+_8433172_8446622_RNA45SN1_45S_50ntFlanks</td><td>11563</td><td>11587</td><td>25</td><td>GCCTAGCAGCCGACTTAGAACTGGT</td><td>63.3</td><td>6.2</td><td>1.9</td><td>3.3</td><td>1.0E-03</td><td>2.5E-03</td><td>BT20/MB468</td><td>MP</td></tr>
<tr><td>>hg38_21_+_8433172_8446622_RNA45SN1_45S_50ntFlanks@11874.11908.35|AGAAGACCCTGTTGAGCTTGACTCTAGTCTGGCAC</td><td>>hg38_21_+_8433172_8446622_RNA45SN1_45S_50ntFlanks</td><td>11874</td><td>11908</td><td>35</td><td>AGAAGACCCTGTTGAGCTTGACTCTAGTCTGGCAC</td><td>379.5</td><td>-4.9</td><td>1.6</td><td>-3.1</td><td>2.1E-03</td><td>2.5E-03</td><td>MB231/MB468</td><td>NUC</td></tr>
<tr><td>>MI0000438|hsa-mir-15b&WithFlank&3|+|160404582|160404691@64.84.21&[MIMAT0004586&hsa-miR-15b-3p&offsets|0|-1;m-204&3|+|160404645|160404665&offsets|0|0]|CGAATCATTATTTGCTGCTCT</td><td>>MI0000438|hsa-mir-15b&WithFlank&3|+|160404582|160404691</td><td>64</td><td>84</td><td>21</td><td>CGAATCATTATTTGCTGCTCT</td><td>388.4</td><td>3.6</td><td>1.1</td><td>3.3</td><td>1.0E-03</td><td>2.5E-03</td><td>BT20/MB468</td><td>MP</td></tr>
<tr><td>>trnaMT_PheGAA_MT_+_577_647@39.62.24|ATGTTTAGACGGGCTCACATCACC</td><td>>trnaMT_PheGAA_MT_+_577_647</td><td>39</td><td>62</td><td>24</td><td>ATGTTTAGACGGGCTCACATCACC</td><td>699.2</td><td>4.4</td><td>1.3</td><td>3.3</td><td>1.0E-03</td><td>2.5E-03</td><td>BT20/MB468</td><td>MP</td></tr>
<tr><td>>hg38_21_+_8433172_8446622_RNA45SN1_45S_50ntFlanks@6660.6706.47|GCGGTGGATCACTCGGCTCGTGCGTCGATGAAGAACGCAGCTAGCTG</td><td>>hg38_21_+_8433172_8446622_RNA45SN1_45S_50ntFlanks</td><td>6660</td><td>6706</td><td>47</td><td>GCGGTGGATCACTCGGCTCGTGCGTCGATGAAGAACGCAGCTAGCTG</td><td>800.8</td><td>3.3</td><td>0.9</td><td>3.5</td><td>4.5E-04</td><td>2.5E-03</td><td>BT20/MB231</td><td>MITO</td></tr>
<tr><td>>MI0000342|hsa-mir-200b&WithFlank&1|+|1167098|1167204@63.85.23(+1A)&[MIMAT0000318&hsa-miR-200b-3p&offsets|0|+1(+1A);m-93&1|+|1167160|1167182&offsets|0|0(+1A)]|TAATACTGCCTGGTAATGATGACA</td><td>>MI0000342|hsa-mir-200b&WithFlank&1|+|1167098|1167204</td><td>63</td><td>85</td><td>23(+1A)</td><td>TAATACTGCCTGGTAATGATGACA</td><td>329.6</td><td>-2.2</td><td>0.7</td><td>-3.2</td><td>1.6E-03</td><td>2.5E-03</td><td>BT20/MB468</td><td>MITO</td></tr>
<tr><td>>hg38_21_+_8433172_8446622_RNA45SN1_45S_50ntFlanks@8347.8379.33|GGGAAAGTTGAAAAGAACTTTGAAGAGAGAGTT</td><td>>hg38_21_+_8433172_8446622_RNA45SN1_45S_50ntFlanks</td><td>8347</td><td>8379</td><td>33</td><td>GGGAAAGTTGAAAAGAACTTTGAAGAGAGAGTT</td><td>111.2</td><td>4.4</td><td>1.3</td><td>3.3</td><td>1.0E-03</td><td>2.5E-03</td><td>BT20/MB468</td><td>MP</td></tr>
<tr><td>>hg38_21_+_8433172_8446622_RNA45SN1_45S_50ntFlanks@10393.10422.30|CAGCAGTTGAACATGGGTCAGTCGGTCCTG</td><td>>hg38_21_+_8433172_8446622_RNA45SN1_45S_50ntFlanks</td><td>10393</td><td>10422</td><td>30</td><td>CAGCAGTTGAACATGGGTCAGTCGGTCCTG</td><td>376.2</td><td>-3.8</td><td>1.2</td><td>-3.1</td><td>1.9E-03</td><td>2.5E-03</td><td>BT20/MB468</td><td>CYTO</td></tr>
<tr><td>>hg38_21_+_8433172_8446622_RNA45SN1_45S_50ntFlanks@8018.8046.29|ATTAGTCAGCGGAGGAGAAGAAACTAACC</td><td>>hg38_21_+_8433172_8446622_RNA45SN1_45S_50ntFlanks</td><td>8018</td><td>8046</td><td>29</td><td>ATTAGTCAGCGGAGGAGAAGAAACTAACC</td><td>147.4</td><td>6.8</td><td>2.2</td><td>3.1</td><td>2.0E-03</td><td>2.5E-03</td><td>MB231/MB468</td><td>CYTO</td></tr>
<tr><td>>hg38_21_+_8433172_8446622_RNA45SN1_45S_50ntFlanks@12456.12480.25|CCAAGCGTTGGATTGTTCACCCACT</td><td>>hg38_21_+_8433172_8446622_RNA45SN1_45S_50ntFlanks</td><td>12456</td><td>12480</td><td>25</td><td>CCAAGCGTTGGATTGTTCACCCACT</td><td>3447.6</td><td>-4.9</td><td>1.5</td><td>-3.3</td><td>1.0E-03</td><td>2.5E-03</td><td>BT20/MB468</td><td>MP</td></tr>
<tr><td>>trnaMT_GluTTC_MT_-_14674_14742@32.71.40,trnalookalike8_GluTTC_5_-_93905172_93905240@32.71.40|TCATATCATTGGTCGTGGTTGTAGTCCGTGCGAGAATACC</td><td>>trnaMT_GluTTC_MT_-_14674_14742</td><td>32</td><td>71</td><td>40</td><td>TCATATCATTGGTCGTGGTTGTAGTCCGTGCGAGAATACC</td><td>104.5</td><td>4.8</td><td>1.6</td><td>3.1</td><td>2.1E-03</td><td>2.5E-03</td><td>MB231/MB468</td><td>MITO</td></tr>
<tr><td>>MI0000750|hsa-mir-26a-2&WithFlank&12|-|57824603|57824698@20.41.22(+1U);MI0000083|hsa-mir-26a-1&WithFlank&3|+|37969398|37969486@16.37.22(+1U)&[MIMAT0000082&hsa-miR-26a-5p&offsets|0|0(+1U);m-33&12|-|57824658|57824679&offsets|0|0(+1U)];[MIMAT0000082_1&hsa-miR-26a-5p&offsets|0|0(+1U);m-34&3|+|37969413|37969434&offsets|0|0(+1U)]|TTCAAGTAATCCAGGATAGGCTT</td><td>>MI0000750|hsa-mir-26a-2&WithFlank&12|-|57824603|57824698</td><td>20</td><td>41</td><td>22(+1U)</td><td>TTCAAGTAATCCAGGATAGGCTT</td><td>1506.7</td><td>3.5</td><td>1.1</td><td>3.3</td><td>1.0E-03</td><td>2.5E-03</td><td>BT20/MB468</td><td>MP</td></tr>
</table>

| | | | | | | | | | | | | | |
|---|---|---|---|---|---|---|---|---|---|---|---|---|---|
| >hg38_21_+_8433172_8446622_RNA45SN1_45S_50ntFlanks@7992.8016.25\|CGTGGCGACCCGCTGAATTTAAGCA | >hg38_21_+_8433172_8446622_RNA45SN1_45S_50ntFlanks | 7992 | 8016 | 25 | CGTGGCGACCCGCTGAATTTAAGCA | 439.0 | 2.4 | 0.8 | 3.1 | 2.1E-03 | 2.5E-03 | MB231/MB468 | NUC |
| >hg38_21_+_8433172_8446622_RNA45SN1_45S_50ntFlanks@11576.11594.19\|CTTAGAACTGGTGCGGACC | >hg38_21_+_8433172_8446622_RNA45SN1_45S_50ntFlanks | 11576 | 11594 | 19 | CTTAGAACTGGTGCGGACC | 1724.9 | -1.8 | 0.6 | -3.1 | 2.0E-03 | 2.5E-03 | MB231/MB468 | CYTO |
| >hg38_21_+_8433172_8446622_RNA45SN1_45S_50ntFlanks@3845.3873.29\|ACTTGGATAACTGTGGTAATTCTAGAGCT | >hg38_21_+_8433172_8446622_RNA45SN1_45S_50ntFlanks | 3845 | 3873 | 29 | ACTTGGATAACTGTGGTAATTCTAGAGCT | 79.1 | -5.2 | 1.6 | -3.3 | 1.0E-03 | 2.5E-03 | BT20/MB468 | MP |
| >hg38_21_+_8433172_8446622_RNA45SN1_45S_50ntFlanks@12151.12179.29\|GTGGGGAGTTTGACTGGGGCGGTACACCT | >hg38_21_+_8433172_8446622_RNA45SN1_45S_50ntFlanks | 12151 | 12179 | 29 | GTGGGGAGTTTGACTGGGGCGGTACACCT | 164.8 | -4.4 | 1.4 | -3.1 | 1.9E-03 | 2.5E-03 | BT20/MB468 | CYTO |
| >hg38_21_+_8433172_8446622_RNA45SN1_45S_50ntFlanks@7990.8020.31\|GACGTGGCGACCCGCTGAATTTAAGCATATT | >hg38_21_+_8433172_8446622_RNA45SN1_45S_50ntFlanks | 7990 | 8020 | 31 | GACGTGGCGACCCGCTGAATTTAAGCATATT | 150.8 | 11.2 | 3.6 | 3.1 | 2.0E-03 | 2.5E-03 | MB231/MB468 | CYTO |
| >hg38_21_+_8433172_8446622_RNA45SN1_45S_50ntFlanks@7991.8010.20\|ACGTGGCGACCCGCTGAATT | >hg38_21_+_8433172_8446622_RNA45SN1_45S_50ntFlanks | 7991 | 8010 | 20 | ACGTGGCGACCCGCTGAATT | 350.5 | 2.2 | 0.7 | 3.1 | 2.0E-03 | 2.5E-03 | MB231/MB468 | CYTO |
| >hg38_21_+_8433172_8446622_RNA45SN1_45S_50ntFlanks@10248.10280.33\|CGACGAGTAGGAGGGCCGCTGCGGTGAGCCTTG | >hg38_21_+_8433172_8446622_RNA45SN1_45S_50ntFlanks | 10248 | 10280 | 33 | CGACGAGTAGGAGGGCCGCTGCGGTGAGCCTTG | 123.0 | 4.4 | 1.4 | 3.3 | 1.0E-03 | 2.5E-03 | BT20/MB468 | MP |
| >trna146_GlnCTG_6_-_27515531_27515602@1.36.36,trna1_GlnCTG_6_+_18836402_18836473@1.36.36,trna3_GlnCTG_17_+_8023070_8023141@1.36.36,trna42_GlnCTG_6_+_27263212_27263283@1.36.36,trna49_GlnCTG_6_+_27487308_27487379@1.36.36,trna7_GlnCTG_15_-_66161400_66161471@1.36.36,trna99_GlnCTG_6_-_28909378_28909449@1.36.36\|GGTTCCATGGTGTAATGGTTAGCACTCTGGACTCTG | >trna146_GlnCTG_6_-_27515531_27515602 | 1 | 36 | 36 | GGTTCCATGGTGTAATGGTTAGCACTCTGGACTCTG | 44.0 | 8.7 | 2.8 | 3.1 | 1.8E-03 | 2.5E-03 | MB231/MB468 | MP |
| >MI0000108\|hsa-mir-103a-2&WithFlank&20\|+\|3917488\|3917577@54.75.22;MI0000109\|hsa-mir-103a-1&WithFlank&5\|-\|168560890\|168560979@54.75.22&[MIMAT0000101&hsa-miR-103a-3p&offsets\|0\|-1;m-12&20\|+\|3917541\|3917563&offsets\|0\|-1];[MIMAT0000101_1&hsa-miR-103a-3p&offsets\|0\|-1;m-13&5\|-\|168560904\|168560926&offsets\|0\|-1;m-44220&5\|-\|168560908\|168560926&offsets\|0\|+3]\|AGCAGCATTGTACAGGGCTATG | >MI0000108\|hsa-mir-103a-2&WithFlank&20\|+\|3917488\|3917577 | 54 | 75 | 22 | AGCAGCATTGTACAGGGCTATG | 12822.9 | -3.2 | 0.9 | -3.6 | 3.4E-04 | 2.5E-03 | BT20/MB231 | MP |
| >hg38_21_+_8433172_8446622_RNA45SN1_45S_50ntFlanks@7989.8006.18\|AGACGTGGCGACCCGCTG | >hg38_21_+_8433172_8446622_RNA45SN1_45S_50ntFlanks | 7989 | 8006 | 18 | AGACGTGGCGACCCGCTG | 994.5 | 6.5 | 1.8 | 3.6 | 3.4E-04 | 2.5E-03 | BT20/MB231 | MP |
| >hg38_21_+_8433172_8446622_RNA45SN1_45S_50ntFlanks@9553.9571.19\|CCGTAGCGGTCCTGACGTG | >hg38_21_+_8433172_8446622_RNA45SN1_45S_50ntFlanks | 9553 | 9571 | 19 | CCGTAGCGGTCCTGACGTG | 305.8 | -1.6 | 0.5 | -3.1 | 2.0E-03 | 2.5E-03 | MB231/MB468 | CYTO |
| >trnaMT_SerGCT_MT_+_12207_12265@15.62.48\|GAACTGCTAACTCATGCCCCCATGTCTAACAACATGGCTTTCTCACCA | >trnaMT_SerGCT_MT_+_12207_12265 | 15 | 62 | 48 | GAACTGCTAACTCATGCCCCCATGTCTAACAACATGGCTTTCTCACCA | 587.0 | 5.6 | 1.8 | 3.2 | 1.6E-03 | 2.5E-03 | BT20/MB468 | MITO |
| >MI0000767\|hsa-mir-365a&WithFlank&16\|+\|14309279\|14309377@62.81.20(+2U);MI0000769\|hsa-mir-365b&WithFlank&17\|+\|31575405\|31575527@74.93.20(+2U)&[MIMAT0000710&hsa-miR-365a-3p&offsets\|0\|-2(+2U);m-176&16\|+\|14309340\|14309361&offsets\|0\|-2(+2U)];[MIMAT0022834&hsa-miR-365b-3p&offsets\|0\|-2(+2U);m-175&17\|+\|31575478\|31575499&offsets\|0\|-2(+2U)]\|TAATGCCCCTAAAAATCCTTTT | >MI0000767\|hsa-mir-365a&WithFlank&16\|+\|14309279\|14309377 | 62 | 81 | 20(+2U) | TAATGCCCCTAAAAATCCTTTT | 132.5 | 5.6 | 1.7 | 3.3 | 1.0E-03 | 2.5E-03 | BT20/MB468 | MP |
| >hg38_21_+_8433172_8446622_RNA45SN1_45S_50ntFlanks@3782.3805.24\|CAGTGAAACTGCGAATGGCTCATT | >hg38_21_+_8433172_8446622_RNA45SN1_45S_50ntFlanks | 3782 | 3805 | 24 | CAGTGAAACTGCGAATGGCTCATT | 319.9 | -1.6 | 0.5 | -3.1 | 1.9E-03 | 2.5E-03 | BT20/MB468 | CYTO |
| >hg38_21_+_8433172_8446622_RNA45SN1_45S_50ntFlanks@5188.5206.19\|ATAACAGGTCTGTGATGCC | >hg38_21_+_8433172_8446622_RNA45SN1_45S_50ntFlanks | 5188 | 5206 | 19 | ATAACAGGTCTGTGATGCC | 405.3 | -6.1 | 1.9 | -3.2 | 1.6E-03 | 2.5E-03 | BT20/MB468 | MITO |
| >MI0000093\|hsa-mir-92a-1&WithFlank&13\|+\|91351308\|91351397@54.74.21;MI0000094\|hsa-mir-92a-2&WithFlank&X\|-\|134169532\|134169618@54.74.21&[MIMAT0000092&hsa-miR-92a-3p&offsets\|0\|-1;m-9&13\|+\|91351361\|91351382&offsets\|0\|-1];[MIMAT0000092_1&hsa-miR-92a-3p&offsets\|0\|-1;m-11&X\|-\|134169544\|134169565&offsets\|0\|-1]\|TATTGCACTTGTCCCGGCCTG | >MI0000093\|hsa-mir-92a-1&WithFlank&13\|+\|91351308\|91351397 | 54 | 74 | 21 | TATTGCACTTGTCCCGGCCTG | 505.9 | 3.7 | 1.1 | 3.3 | 1.0E-03 | 2.5E-03 | BT20/MB468 | MP |
| >hg38_21_+_8433172_8446622_RNA45SN1_45S_50ntFlanks@11455.11477.23\|CTCGGCGCGCGGCGGCGGCGGCG | >hg38_21_+_8433172_8446622_RNA45SN1_45S_50ntFlanks | 11455 | 11477 | 23 | CTCGGCGCGCGGCGGCGGCGGCG | 250.6 | -5.5 | 1.7 | -3.3 | 1.0E-03 | 2.5E-03 | BT20/MB468 | MP |
| >hg38_21_+_8433172_8446622_RNA45SN1_45S_50ntFlanks@4370.4391.22\|TTAAAAAGCTCGTAGTTGGATC | >hg38_21_+_8433172_8446622_RNA45SN1_45S_50ntFlanks | 4370 | 4391 | 22 | TTAAAAAGCTCGTAGTTGGATC | 239.0 | -4.3 | 1.3 | -3.3 | 1.0E-03 | 2.5E-03 | BT20/MB468 | MP |
| >hg38_21_+_8433172_8446622_RNA45SN1_45S_50ntFlanks@11769.11806.38\|TAATTAGTGACGCGCATGAATGGATGAACGAGATTCCC | >hg38_21_+_8433172_8446622_RNA45SN1_45S_50ntFlanks | 11769 | 11806 | 38 | TAATTAGTGACGCGCATGAATGGATGAACGAGATTCCC | 250.5 | -2.0 | 0.7 | -3.1 | 2.0E-03 | 2.5E-03 | MB231/MB468 | CYTO |
| >MI0000298\|hsa-mir-221&WithFlank&X\|-\|45746151\|45746272@71.93.23(+1U)&[MIMAT0000278&hsa-miR-221-3p&offsets\|0\|0(+1U);m-91&X\|-\|45746181\|45746202&offsets\|0\|+1(+1U)]\|AGCTACATTGTCTGCTGGGTTTCT | >MI0000298\|hsa-mir-221&WithFlank&X\|-\|45746151\|45746272 | 71 | 93 | 23(+1U) | AGCTACATTGTCTGCTGGGTTTCT | 13229.4 | -2.6 | 0.8 | -3.5 | 4.5E-04 | 2.5E-03 | BT20/MB231 | MITO |
| >MI0000269\|hsa-mir-181a-2&WithFlank&9\|+\|124692436\|124692557@83.103.21(+1U)&[MIMAT0004558&hsa-miR-181a-2-3p&offsets\|0\|-1(+1U);m-71&9\|+\|124692518\|124692539&offsets\|0\|-1(+1U)]\|ACCACTGACCGTTGACTGTACT | >MI0000269\|hsa-mir-181a-2&WithFlank&9\|+\|124692436\|124692557 | 83 | 103 | 21(+1U) | ACCACTGACCGTTGACTGTACT | 539.7 | -2.7 | 0.8 | -3.5 | 4.5E-04 | 2.5E-03 | BT20/MB231 | MITO |
| >MI0000767\|hsa-mir-365a&WithFlank&16\|+\|14309279\|14309377@22.44.23&[MIMAT0009199&hsa-miR-365a-5p&offsets\|0\|0;m-595&16\|+\|14309300\|14309322&offsets\|0\|0]\|AGGGACTTTTGGGGGCAGATGTG | >MI0000767\|hsa-mir-365a&WithFlank&16\|+\|14309279\|14309377 | 22 | 44 | 23 | AGGGACTTTTGGGGGCAGATGTG | 280.3 | 2.8 | 0.8 | 3.5 | 4.0E-04 | 2.5E-03 | BT20/MB231 | CYTO |
| >hg38_21_+_8433172_8446622_RNA45SN1_45S_50ntFlanks@10467.10487.21\|GTTGCCCTCGGCCGATCGAAA | >hg38_21_+_8433172_8446622_RNA45SN1_45S_50ntFlanks | 10467 | 10487 | 21 | GTTGCCCTCGGCCGATCGAAA | 231.6 | -2.7 | 0.8 | -3.5 | 4.0E-04 | 2.5E-03 | BT20/MB231 | CYTO |
| >hg38_21_+_8433172_8446622_RNA45SN1_45S_50ntFlanks@9391.9411.21\|CCGAGGTGGGATCCCGAGGCC | >hg38_21_+_8433172_8446622_RNA45SN1_45S_50ntFlanks | 9391 | 9411 | 21 | CCGAGGTGGGATCCCGAGGCC | 121.1 | -5.2 | 1.6 | -3.3 | 1.0E-03 | 2.5E-03 | BT20/MB468 | MP |
| >trnaMT_MetCAT_MT_+_4402_4469@35.71.37\|CCCCGAAAATGTTGGTTATACCCTTCCCGTACTACCA | >trnaMT_MetCAT_MT_+_4402_4469 | 35 | 71 | 37 | CCCCGAAAATGTTGGTTATACCCTTCCCGTACTACCA | 414.3 | -4.8 | 1.4 | -3.6 | 3.8E-04 | 2.5E-03 | BT20/MB231 | NUC |
| >hg38_21_+_8433172_8446622_RNA45SN1_45S_50ntFlanks@10393.10431.39\|CAGCAGTTGAACATGGGTCAGTCGGTCCTGAGAGATGGG | >hg38_21_+_8433172_8446622_RNA45SN1_45S_50ntFlanks | 10393 | 10431 | 39 | CAGCAGTTGAACATGGGTCAGTCGGTCCTGAGAGATGGG | 165.0 | -4.6 | 1.5 | -3.1 | 1.9E-03 | 2.5E-03 | BT20/MB468 | CYTO |
| >hg38_21_+_8433172_8446622_RNA45SN1_45S_50ntFlanks@10393.10429.37\|CAGCAGTTGAACATGGGTCAGTCGGTCCTGAGAGATG | >hg38_21_+_8433172_8446622_RNA45SN1_45S_50ntFlanks | 10393 | 10429 | 37 | CAGCAGTTGAACATGGGTCAGTCGGTCCTGAGAGATG | 1634.4 | -2.4 | 0.8 | -3.1 | 2.0E-03 | 2.5E-03 | MB231/MB468 | CYTO |
| >hg38_MT_+_598_1651_RNR1_12S_50ntFlanks@843.871.29\|GAGTAGAGTGCTTAGTTGAACAGGGCCCT | >hg38_MT_+_598_1651_RNR1_12S_50ntFlanks | 843 | 871 | 29 | GAGTAGAGTGCTTAGTTGAACAGGGCCCT | 117.9 | 5.9 | 1.9 | 3.1 | 1.8E-03 | 2.5E-03 | MB231/MB468 | MP |
| >hg38_21_+_8433172_8446622_RNA45SN1_45S_50ntFlanks@11583.11611.29\|CTGGTGCGGACCAGGGGAATCCGACTGTT | >hg38_21_+_8433172_8446622_RNA45SN1_45S_50ntFlanks | 11583 | 11611 | 29 | CTGGTGCGGACCAGGGGAATCCGACTGTT | 1327.6 | 3.2 | 0.9 | 3.6 | 3.8E-04 | 2.5E-03 | BT20/MB231 | NUC |
| >MI0000298\|hsa-mir-221&WithFlank&X\|-\|45746151\|45746272@71.92.22&[MIMAT0000278&hsa-miR-221-3p&offsets\|0\|-1;m-91&X\|-\|45746181\|45746202&offsets\|0\|0]\|AGCTACATTGTCTGCTGGGTTT | >MI0000298\|hsa-mir-221&WithFlank&X\|-\|45746151\|45746272 | 71 | 92 | 22 | AGCTACATTGTCTGCTGGGTTT | 39033.2 | -2.9 | 0.8 | -3.5 | 4.6E-04 | 2.5E-03 | BT20/MB231 | MITO |
| >hg38_21_+_8433172_8446622_RNA45SN1_45S_50ntFlanks@11320.11338.19\|CCACGTCTCGTCGCGCGCG | >hg38_21_+_8433172_8446622_RNA45SN1_45S_50ntFlanks | 11320 | 11338 | 19 | CCACGTCTCGTCGCGCGCG | 324.0 | -5.5 | 1.8 | -3.1 | 2.1E-03 | 2.5E-03 | BT20/MB468 | NUC |
| >hg38_MT_+_598_1651_RNR1_12S_50ntFlanks@966.1004.39\|TAACATGGTAAGTGTACTGGAAAGTGCACTTGGACGAAC | >hg38_MT_+_598_1651_RNR1_12S_50ntFlanks | 966 | 1004 | 39 | TAACATGGTAAGTGTACTGGAAAGTGCACTTGGACGAAC | 297.6 | 3.1 | 1.0 | 3.1 | 2.1E-03 | 2.5E-03 | MB231/MB468 | MITO |
| >trna10_AspGTC_12_-_125424193_125424264@36.72.37,trna12_AspGTC_12_-_125411891_125411962@36.72.37,trna144_AspGTC_6_-_27551236_27551307@36.72.37,trna38_AspGTC_17_-_8125556_8125627@36.72.37,trna45_AspGTC_6_+_27447453_27447524@36.72.37,trna48_AspGTC_6_+_27471523_27471594@36.72.37,trna4_AspGTC_12_+_96429799_96429870@36.72.37,trna69_AspGTC_1_-_161440205_161440276@36.72.37,trna72_AspGTC_1_-_161432824_161432895@36.72.37,trna75_AspGTC_1_-_161425414_161425485@36.72.37,trna78_AspGTC_1_-_161418033_161418104@36.72.37,trna81_AspGTC_1_-_161410615_161410686@36.72.37\|CACGCGGGAGACCGGGGTTCGATTCCCCGACGGGGAG | >trna10_AspGTC_12_-_125424193_125424264 | 36 | 72 | 37 | CACGCGGGAGACCGGGGTTCGATTCCCCGACGGGGAG | 994.0 | 3.0 | 1.0 | 3.1 | 2.0E-03 | 2.5E-03 | MB231/MB468 | CYTO |
| >hg38_21_+_8433172_8446622_RNA45SN1_45S_50ntFlanks@4498.4517.20\|AAGCGTTTACTTTGAAAAAA | >hg38_21_+_8433172_8446622_RNA45SN1_45S_50ntFlanks | 4498 | 4517 | 20 | AAGCGTTTACTTTGAAAAAA | 584.9 | 2.8 | 0.9 | 3.1 | 1.9E-03 | 2.5E-03 | BT20/MB468 | CYTO |
| >hg38_21_+_8433172_8446622_RNA45SN1_45S_50ntFlanks@10786.10819.34\|AACAGCCTCTGGCATGTTGGAACAATGTAGGTAA | >hg38_21_+_8433172_8446622_RNA45SN1_45S_50ntFlanks | 10786 | 10819 | 34 | AACAGCCTCTGGCATGTTGGAACAATGTAGGTAA | 144.3 | -5.2 | 1.6 | -3.3 | 1.0E-03 | 2.5E-03 | BT20/MB468 | MP |
| >trnaMT_CysGCA_MT_-_5761_5826@38.69.32\|AAGAAGCAGCTTCAAACCTGCCGGGGCTTCCA | >trnaMT_CysGCA_MT_-_5761_5826 | 38 | 69 | 32 | AAGAAGCAGCTTCAAACCTGCCGGGGCTTCCA | 190.8 | 6.0 | 1.8 | 3.3 | 1.0E-03 | 2.5E-03 | BT20/MB468 | MP |
| >hg38_21_+_8433172_8446622_RNA45SN1_45S_50ntFlanks@4917.4940.24\|CACCACCAGGAGTGGAGCCTGCGG | >hg38_21_+_8433172_8446622_RNA45SN1_45S_50ntFlanks | 4917 | 4940 | 24 | CACCACCAGGAGTGGAGCCTGCGG | 484.2 | -4.5 | 1.4 | -3.3 | 1.0E-03 | 2.5E-03 | BT20/MB468 | MP |
| >hg38_21_+_8433172_8446622_RNA45SN1_45S_50ntFlanks@9932.9949.18\|ATAGACAGCAGGACGGTG | >hg38_21_+_8433172_8446622_RNA45SN1_45S_50ntFlanks | 9932 | 9949 | 18 | ATAGACAGCAGGACGGTG | 418.7 | 5.4 | 1.5 | 3.6 | 3.4E-04 | 2.5E-03 | BT20/MB231 | MP |
| >hg38_21_+_8433172_8446622_RNA45SN1_45S_50ntFlanks@12416.12457.42\|TTCGATGTCGGCTCTTCCTATCATTGTGAAGCAGAATTCACC | >hg38_21_+_8433172_8446622_RNA45SN1_45S_50ntFlanks | 12416 | 12457 | 42 | TTCGATGTCGGCTCTTCCTATCATTGTGAAGCAGAATTCACC | 198.6 | 4.7 | 1.5 | 3.1 | 2.1E-03 | 2.5E-03 | MB231/MB468 | MITO |
| >trnaMT_CysGCA_MT_-_5761_5826@47.69.23\|CTTCAAACCTGCCGGGGCTTCCA | >trnaMT_CysGCA_MT_-_5761_5826 | 47 | 69 | 23 | CTTCAAACCTGCCGGGGCTTCCA | 260.7 | -3.8 | 1.2 | -3.1 | 2.1E-03 | 2.5E-03 | MB231/MB468 | NUC |
| >hg38_21_+_8433172_8446622_RNA45SN1_45S_50ntFlanks@12539.12566.28\|ATGATGTGTTGTTGCCATGGTAATCCTG | >hg38_21_+_8433172_8446622_RNA45SN1_45S_50ntFlanks | 12539 | 12566 | 28 | ATGATGTGTTGTTGCCATGGTAATCCTG | 1496.3 | -5.8 | 1.8 | -3.3 | 1.0E-03 | 2.5E-03 | BT20/MB468 | MP |
| >hg38_21_+_8433172_8446622_RNA45SN1_45S_50ntFlanks@10353.10390.38\|AGAACTTTGAAGGCCGAAGTGGAGAAGGGTTCCATGTG | >hg38_21_+_8433172_8446622_RNA45SN1_45S_50ntFlanks | 10353 | 10390 | 38 | AGAACTTTGAAGGCCGAAGTGGAGAAGGGTTCCATGTG | 369.4 | -3.0 | 1.0 | -3.1 | 1.8E-03 | 2.5E-03 | MB231/MB468 | MP |
| >MI0000239\|hsa-mir-197&WithFlank&1\|+\|109598887\|109598973@54.76.23&[MIMAT0000227&hsa-miR-197-3p&offsets\|0\|+1;m-99&1\|+\|109598940\|109598961&offsets\|0\|+1]\|TTCACCACCTTCTCCACCCAGCA | >MI0000239\|hsa-mir-197&WithFlank&1\|+\|109598887\|109598973 | 54 | 76 | 23 | TTCACCACCTTCTCCACCCAGCA | 233.9 | 2.7 | 0.8 | 3.5 | 4.6E-04 | 2.5E-03 | BT20/MB231 | MITO |
| >hg38_21_+_8433172_8446622_RNA45SN1_45S_50ntFlanks@7979.8009.31\|ACCTCAGATCAGACGTGGCGACCCGCTGAAT | >hg38_21_+_8433172_8446622_RNA45SN1_45S_50ntFlanks | 7979 | 8009 | 31 | ACCTCAGATCAGACGTGGCGACCCGCTGAAT | 481.6 | 2.4 | 0.7 | 3.5 | 3.9E-04 | 2.5E-03 | BT20/MB231 | NUC |
| >hg38_21_+_8433172_8446622_RNA45SN1_45S_50ntFlanks@12072.12097.26\|CCCGAGGGGCTCTCGCTTCTGGCGCC | >hg38_21_+_8433172_8446622_RNA45SN1_45S_50ntFlanks | 12072 | 12097 | 26 | CCCGAGGGGCTCTCGCTTCTGGCGCC | 412.0 | -3.6 | 1.2 | -3.2 | 1.6E-03 | 2.5E-03 | BT20/MB468 | MITO |

| | | | | | | | | | | | | | |
|---|---|---|---|---|---|---|---|---|---|---|---|---|---|
| >hg38_21_+_8433172_8446622_RNA45SN1_45S_50ntFlanks@10679.10712.34\|GGTTCCGGCGGCGTCCGGTGAGCTCTCGCTGGCC | >hg38_21_+_8433172_8446622_RNA45SN1_45S_50ntFlanks | 10679 | 10712 | 34 | GGTTCCGGCGGCGTCCGGTGAGCTCTCGCTGGCC | 298.2 | -4.6 | 1.5 | -3.1 | 1.9E-03 | 2.5E-03 | BT20/MB468 | CYTO |
| >MI0000234\|hsa-mir-192&WithFlank&11\|-\|64891131\|64891252@30.49.20(+1U)&[MIMAT0000222&hsa-miR-192-5p&offsets\|0\|-1(+1U);m-59&11\|-\|64891203\|64891223&offsets\|0\|-1(+1U)]\|CTGACCTATGAATTGACAGCT | >MI0000234\|hsa-mir-192&WithFlank&11\|-\|64891131\|64891252 | 30 | 49 | 20(+1U) | CTGACCTATGAATTGACAGCT | 141.3 | 2.0 | 0.7 | 3.1 | 2.1E-03 | 2.5E-03 | BT20/MB468 | NUC |
| >hg38_21_+_8433172_8446622_RNA45SN1_45S_50ntFlanks@11666.11706.41\|TCTGCCCAGTGCTCTGAATGTCAAAGTGAAGAAATTCAATG | >hg38_21_+_8433172_8446622_RNA45SN1_45S_50ntFlanks | 11666 | 11706 | 41 | TCTGCCCAGTGCTCTGAATGTCAAAGTGAAGAAATTCAATG | 1505.6 | -3.2 | 0.9 | -3.5 | 4.1E-04 | 2.5E-03 | BT20/MB231 | CYTO |
| >hg38_21_+_8433172_8446622_RNA45SN1_45S_50ntFlanks@6646.6669.24\|TCGTACGACTCTTAGCGGTGGATC | >hg38_21_+_8433172_8446622_RNA45SN1_45S_50ntFlanks | 6646 | 6669 | 24 | TCGTACGACTCTTAGCGGTGGATC | 34081.8 | 2.9 | 0.8 | 3.5 | 3.9E-04 | 2.5E-03 | BT20/MB231 | NUC |
| >am_trna128_GlyGCC_6_-_27870686_27870756@1.27.27,trna133_GlyCCC_1_-_16872434_16872504@1.27.27,trna18_GlyGCC_16_+_70822597_70822667@1.27.27,trna19_GlyGCC_16_+_70823410_70823480@1.27.27,trna19_GlyGCC_2_-_157257659_157257729@1.27.27,trna24_GlyGCC_16_-_70812942_70813012@1.27.27,trna25_GlyGCC_16_-_70812114_70812184@1.27.27,trna4_GlyCCC_1_+_17188416_17188486@1.27.27,trna5_GlyGCC_17_+_8029064_8029134@1.27.27,trna68_GlyGCC_1_-_161493637_161493707@1.27.27\|GCATTGGTGGTTCAGTGGTAGAATTCT | >am_trna128_GlyGCC_6_-_27870686_27870756 | 1 | 27 | 27 | GCATTGGTGGTTCAGTGGTAGAATTCT | 848.1 | 3.2 | 1.0 | 3.2 | 1.6E-03 | 2.5E-03 | BT20/MB468 | MITO |
| >MI0000433\|hsa-let-7g&WithFlank&3\|-\|52268272\|52268367@11.30.20(+1C)&[MIMAT0000414&hsa-let-7g-5p&offsets\|0\|-2(+1C);m-58&3\|-\|52268336\|52268357&offsets\|0\|-2(+1C)]\|TGAGGTAGTAGTTTGTACAGC | >MI0000433\|hsa-let-7g&WithFlank&3\|-\|52268272\|52268367 | 11 | 30 | 20(+1C) | TGAGGTAGTAGTTTGTACAGC | 249.7 | 4.8 | 1.5 | 3.3 | 1.0E-03 | 2.5E-03 | BT20/MB468 | MP |
| >hg38_21_+_8433172_8446622_RNA45SN1_45S_50ntFlanks@3776.3814.39\|CCGGTACAGTGAAACTGCGAATGGCTCATTAAATCAGTT | >hg38_21_+_8433172_8446622_RNA45SN1_45S_50ntFlanks | 3776 | 3814 | 39 | CCGGTACAGTGAAACTGCGAATGGCTCATTAAATCAGTT | 48.4 | -9.5 | 3.0 | -3.1 | 1.8E-03 | 2.5E-03 | MB231/MB468 | MP |
| >hg38_21_+_8433172_8446622_RNA45SN1_45S_50ntFlanks@12162.12181.20\|GACTGGGGCGGTACACCTGT | >hg38_21_+_8433172_8446622_RNA45SN1_45S_50ntFlanks | 12162 | 12181 | 20 | GACTGGGGCGGTACACCTGT | 220.3 | -4.7 | 1.5 | -3.1 | 1.6E-03 | 2.5E-03 | BT20/MB468 | MITO |
| >hg38_21_+_8433172_8446622_RNA45SN1_45S_50ntFlanks@10389.10424.36\|TGAACAGCAGTTGAACATGGGTCAGTCGGTCCTGAG | >hg38_21_+_8433172_8446622_RNA45SN1_45S_50ntFlanks | 10389 | 10424 | 36 | TGAACAGCAGTTGAACATGGGTCAGTCGGTCCTGAG | 3424.4 | 2.0 | 0.7 | 3.1 | 2.0E-03 | 2.5E-03 | MB231/MB468 | CYTO |
| >hg38_21_+_8433172_8446622_RNA45SN1_45S_50ntFlanks@6778.6806.29\|CCCGGGGCTACGCCTGTCTGAGCGTCGCT | >hg38_21_+_8433172_8446622_RNA45SN1_45S_50ntFlanks | 6778 | 6806 | 29 | CCCGGGGCTACGCCTGTCTGAGCGTCGCT | 2027.2 | 2.2 | 0.7 | 3.1 | 2.1E-03 | 2.5E-03 | MB231/MB468 | MITO |
| >hg38_MT_+_1621_3279_RNR2_16S_50ntFlanks@1404.1421.18\|TATTAAAGGTTCGTTTGT | >hg38_MT_+_1621_3279_RNR2_16S_50ntFlanks | 1404 | 1421 | 18 | TATTAAAGGTTCGTTTGT | 634.5 | -5.0 | 1.4 | -3.6 | 3.5E-04 | 2.5E-03 | BT20/MB231 | MP |
| >hg38_21_+_8433172_8446622_RNA45SN1_45S_50ntFlanks@12419.12452.34\|GATGTCGGCTCTTCCTATCATTGTGAAGCAGAAT | >hg38_21_+_8433172_8446622_RNA45SN1_45S_50ntFlanks | 12419 | 12452 | 34 | GATGTCGGCTCTTCCTATCATTGTGAAGCAGAAT | 243.1 | 4.7 | 1.5 | 3.1 | 2.1E-03 | 2.5E-03 | MB231/MB468 | MITO |
| >hg38_21_+_8433172_8446622_RNA45SN1_45S_50ntFlanks@10344.10362.19\|ATTCAAACGAGAACTTTGA | >hg38_21_+_8433172_8446622_RNA45SN1_45S_50ntFlanks | 10344 | 10362 | 19 | ATTCAAACGAGAACTTTGA | 355.9 | 2.8 | 0.8 | 3.5 | 4.6E-04 | 2.5E-03 | BT20/MB231 | MITO |
| >hg38_21_+_8433172_8446622_RNA45SN1_45S_50ntFlanks@5476.5501.26\|CCCTGGCGGAGCGCTGAGAAGACGGT | >hg38_21_+_8433172_8446622_RNA45SN1_45S_50ntFlanks | 5476 | 5501 | 26 | CCCTGGCGGAGCGCTGAGAAGACGGT | 93.5 | 9.9 | 2.8 | 3.6 | 3.5E-04 | 2.5E-03 | BT20/MB231 | MP |
| >hg38_21_+_8433172_8446622_RNA45SN1_45S_50ntFlanks@8244.8269.26\|TCTTCCCGGAGTCGGGTTGCTTGGGA | >hg38_21_+_8433172_8446622_RNA45SN1_45S_50ntFlanks | 8244 | 8269 | 26 | TCTTCCCGGAGTCGGGTTGCTTGGGA | 92.2 | -4.0 | 1.3 | -3.1 | 1.8E-03 | 2.5E-03 | MB231/MB468 | MP |
| >hg38_21_+_8433172_8446622_RNA45SN1_45S_50ntFlanks@12420.12459.40\|ATGTCGGCTCTTCCTATCATTGTGAAGCAGAATTCACCAA | >hg38_21_+_8433172_8446622_RNA45SN1_45S_50ntFlanks | 12420 | 12459 | 40 | ATGTCGGCTCTTCCTATCATTGTGAAGCAGAATTCACCAA | 272.4 | 2.9 | 1.0 | 3.1 | 2.1E-03 | 2.5E-03 | MB231/MB468 | NUC |
| >hg38_21_+_8433172_8446622_RNA45SN1_45S_50ntFlanks@11573.11611.39\|CGACTTAGAACTGGTGCGGACCAGGGGAATCCGACTGTT | >hg38_21_+_8433172_8446622_RNA45SN1_45S_50ntFlanks | 11573 | 11611 | 39 | CGACTTAGAACTGGTGCGGACCAGGGGAATCCGACTGTT | 413.2 | -2.9 | 0.9 | -3.1 | 2.1E-03 | 2.5E-03 | MB231/MB468 | MITO |
| >hg38_21_+_8433172_8446622_RNA45SN1_45S_50ntFlanks@11845.11867.23\|GGAACGGGCTTGGCGGAATCAGC | >hg38_21_+_8433172_8446622_RNA45SN1_45S_50ntFlanks | 11845 | 11867 | 23 | GGAACGGGCTTGGCGGAATCAGC | 82.7 | -5.0 | 1.6 | -3.1 | 1.8E-03 | 2.5E-03 | MB231/MB468 | MP |
| >trnaMT_LysTTT_MT_+_8295_8364@43.72.30,trnalookalike7_LysTTT_1_+_568844_568913@43.72.30\|TAAGAGAACCAACACCTCTTTACAGTGACC | >trnaMT_LysTTT_MT_+_8295_8364 | 43 | 72 | 30 | TAAGAGAACCAACACCTCTTTACAGTGACC | 90.1 | 3.9 | 1.3 | 3.1 | 1.8E-03 | 2.5E-03 | MB231/MB468 | MP |
| >hg38_21_+_8433172_8446622_RNA45SN1_45S_50ntFlanks@10411.10428.18\|CAGTCGGTCCTGAGAGAT | >hg38_21_+_8433172_8446622_RNA45SN1_45S_50ntFlanks | 10411 | 10428 | 18 | CAGTCGGTCCTGAGAGAT | 1165.7 | -3.5 | 1.1 | -3.3 | 1.0E-03 | 2.5E-03 | BT20/MB468 | MP |
| >MI0000457\|hsa-mir-141&WithFlank&12\|+\|6964091\|6964197@23.44.22&[MIMAT0004598&hsa-miR-141-5p&offsets\|0\|0;m-221&12\|+\|6964113\|6964133&offsets\|0\|+1]\|CATCTTCCAGTACAGTGTTGGA | >MI0000457\|hsa-mir-141&WithFlank&12\|+\|6964091\|6964197 | 23 | 44 | 22 | CATCTTCCAGTACAGTGTTGGA | 435.0 | 4.0 | 1.2 | 3.3 | 1.0E-03 | 2.5E-03 | BT20/MB468 | MP |
| >MI0005559\|hsa-mir-744&WithFlank&17\|+\|12081893\|12082002@17.37.21&[MIMAT0004945&hsa-miR-744-5p&offsets\|0\|-1;m-172&17\|+\|12081909\|12081929&offsets\|0\|0]\|TGCGGGGCTAGGGCTAACAGC | >MI0005559\|hsa-mir-744&WithFlank&17\|+\|12081893\|12082002 | 17 | 37 | 21 | TGCGGGGCTAGGGCTAACAGC | 736.6 | 3.4 | 1.1 | 3.1 | 1.8E-03 | 2.5E-03 | MB231/MB468 | MP |
| >hg38_21_+_8433172_8446622_RNA45SN1_45S_50ntFlanks@8850.8867.18\|GCGGCGGCGGCGGCGGGA | >hg38_21_+_8433172_8446622_RNA45SN1_45S_50ntFlanks | 8850 | 8867 | 18 | GCGGCGGCGGCGGCGGGA | 210.2 | -4.8 | 1.5 | -3.1 | 1.6E-03 | 2.5E-03 | BT20/MB468 | MITO |
| >hg38_21_+_8433172_8446622_RNA45SN1_45S_50ntFlanks@11663.11702.40\|ATTTCTGCCCAGTGCTCTGAATGTCAAAGTGAAGAAATTC | >hg38_21_+_8433172_8446622_RNA45SN1_45S_50ntFlanks | 11663 | 11702 | 40 | ATTTCTGCCCAGTGCTCTGAATGTCAAAGTGAAGAAATTC | 309.5 | -2.9 | 0.8 | -3.5 | 4.1E-04 | 2.5E-03 | BT20/MB231 | CYTO |
| >hg38_MT_+_1621_3279_RNR2_16S_50ntFlanks@618.652.35\|AACATATAACTGAACTCCTCACACCCAATTGGACC | >hg38_MT_+_1621_3279_RNR2_16S_50ntFlanks | 618 | 652 | 35 | AACATATAACTGAACTCCTCACACCCAATTGGACC | 299.7 | 5.2 | 1.6 | 3.3 | 1.0E-03 | 2.5E-03 | BT20/MB468 | MP |
| >hg38_21_+_8433172_8446622_RNA45SN1_45S_50ntFlanks@11874.11892.19\|AGAAGACCCTGTTGAGCTT | >hg38_21_+_8433172_8446622_RNA45SN1_45S_50ntFlanks | 11874 | 11892 | 19 | AGAAGACCCTGTTGAGCTT | 291.3 | -1.8 | 0.6 | -3.1 | 1.6E-03 | 2.5E-03 | BT20/MB468 | MITO |
| >hg38_21_+_8433172_8446622_RNA45SN1_45S_50ntFlanks@9556.9582.27\|TAGCGGTCCTGACGTGCAAATCGGTCG | >hg38_21_+_8433172_8446622_RNA45SN1_45S_50ntFlanks | 9556 | 9582 | 27 | TAGCGGTCCTGACGTGCAAATCGGTCG | 170.1 | -1.8 | 0.6 | -3.1 | 1.6E-03 | 2.5E-03 | BT20/MB468 | MITO |
| >MI0003560\|hsa-mir-92b&WithFlank&1\|+\|155195171\|155195278@26.48.23&[MIMAT0004792&hsa-miR-92b-5p&offsets\|0\|+1;m-581&1\|+\|155195196\|155195219&offsets\|0\|-1]\|AGGGACGGGACGCGGTGCAGTGT | >MI0003560\|hsa-mir-92b&WithFlank&1\|+\|155195171\|155195278 | 26 | 48 | 23 | AGGGACGGGACGCGGTGCAGTGT | 151.2 | 6.5 | 2.0 | 3.3 | 1.0E-03 | 2.5E-03 | BT20/MB468 | MP |
| >hg38_21_+_8433172_8446622_RNA45SN1_45S_50ntFlanks@4453.4474.22\|TCGGCGCCCCCTCGATGCTCTT | >hg38_21_+_8433172_8446622_RNA45SN1_45S_50ntFlanks | 4453 | 4474 | 22 | TCGGCGCCCCCTCGATGCTCTT | 205.4 | -3.2 | 0.9 | -3.5 | 4.1E-04 | 2.5E-03 | BT20/MB231 | CYTO |
| >hg38_MT_+_598_1651_RNR1_12S_50ntFlanks@428.455.28\|AAAGTGGCTTTAACATATCTGAACACAC | >hg38_MT_+_598_1651_RNR1_12S_50ntFlanks | 428 | 455 | 28 | AAAGTGGCTTTAACATATCTGAACACAC | 572.4 | 3.2 | 1.0 | 3.1 | 1.8E-03 | 2.5E-03 | MB231/MB468 | MP |
| >hg38_21_+_8433172_8446622_RNA45SN1_45S_50ntFlanks@485.515.31\|GATGTGGTGACGTCGTGCTCTCCCGGGCCGG | >hg38_21_+_8433172_8446622_RNA45SN1_45S_50ntFlanks | 485 | 515 | 31 | GATGTGGTGACGTCGTGCTCTCCCGGGCCGG | 243.8 | 11.2 | 3.6 | 3.1 | 2.1E-03 | 2.5E-03 | MB231/MB468 | MITO |
| >MI0000299\|hsa-mir-222&WithFlank&X\|-\|45747009\|45747130@75.99.25(+4U)&[MIMAT0000279&hsa-miR-222-3p&offsets\|0\|+4(+4U);m-140&X\|-\|45747033\|45747056&offsets\|0\|+1(+4U)]\|AGCTACATCTGGCTACTGGGTCTCTTTTT | >MI0000299\|hsa-mir-222&WithFlank&X\|-\|45747009\|45747130 | 75 | 99 | 25(+4U) | AGCTACATCTGGCTACTGGGTCTCTTTTT | 245.4 | -4.4 | 1.3 | -3.5 | 3.9E-04 | 2.5E-03 | BT20/MB231 | NUC |
| >hg38_21_+_8433172_8446622_RNA45SN1_45S_50ntFlanks@5237.5257.21\|ACTGACTGGCTCAGCGTGTGC | >hg38_21_+_8433172_8446622_RNA45SN1_45S_50ntFlanks | 5237 | 5257 | 21 | ACTGACTGGCTCAGCGTGTGC | 388.6 | -4.0 | 1.3 | -3.1 | 2.1E-03 | 2.5E-03 | MB231/MB468 | MITO |
| >hg38_21_+_8433172_8446622_RNA45SN1_45S_50ntFlanks@11582.11600.19\|ACTGGTGCGGACCAGGGGA | >hg38_21_+_8433172_8446622_RNA45SN1_45S_50ntFlanks | 11582 | 11600 | 19 | ACTGGTGCGGACCAGGGGA | 294.3 | -2.2 | 0.7 | -3.1 | 2.0E-03 | 2.5E-03 | BT20/MB468 | CYTO |
| >hg38_21_+_8433172_8446622_RNA45SN1_45S_50ntFlanks@9955.9979.25\|GGAAGTCGGAATCCGCTAAGGAGTG | >hg38_21_+_8433172_8446622_RNA45SN1_45S_50ntFlanks | 9955 | 9979 | 25 | GGAAGTCGGAATCCGCTAAGGAGTG | 146.7 | 4.6 | 1.4 | 3.3 | 1.0E-03 | 2.5E-03 | BT20/MB468 | MP |
| >hg38_21_+_8433172_8446622_RNA45SN1_45S_50ntFlanks@10058.10077.20\|CGGCAGTCGAGAGTGGACGG | >hg38_21_+_8433172_8446622_RNA45SN1_45S_50ntFlanks | 10058 | 10077 | 20 | CGGCAGTCGAGAGTGGACGG | 248.4 | 5.6 | 1.6 | 3.6 | 3.5E-04 | 2.5E-03 | BT20/MB231 | MP |
| >hg38_21_+_8433172_8446622_RNA45SN1_45S_50ntFlanks@9496.9525.30\|GAAAGATGGTGAACTATGCCTGGGCAGGGC | >hg38_21_+_8433172_8446622_RNA45SN1_45S_50ntFlanks | 9496 | 9525 | 30 | GAAAGATGGTGAACTATGCCTGGGCAGGGC | 605.6 | 7.1 | 2.0 | 3.6 | 3.5E-04 | 2.5E-03 | BT20/MB231 | MP |
| >hg38_21_+_8433172_8446622_RNA45SN1_45S_50ntFlanks@5467.5494.28\|GGCCCACGGCCCTGGCGGAGCGCTGAGA | >hg38_21_+_8433172_8446622_RNA45SN1_45S_50ntFlanks | 5467 | 5494 | 28 | GGCCCACGGCCCTGGCGGAGCGCTGAGA | 1321.6 | 4.2 | 1.4 | 3.1 | 2.0E-03 | 2.5E-03 | BT20/MB468 | CYTO |
| >hg38_21_+_8433172_8446622_RNA45SN1_45S_50ntFlanks@7979.8009.31\|ACCTCAGATCAGACGTGGCGACCCGCTGAAT | >hg38_21_+_8433172_8446622_RNA45SN1_45S_50ntFlanks | 7979 | 8009 | 31 | ACCTCAGATCAGACGTGGCGACCCGCTGAAT | 400.3 | -1.2 | 0.4 | -3.1 | 2.1E-03 | 2.5E-03 | MB231/MB468 | NUC |
| >hg38_21_+_8433172_8446622_RNA45SN1_45S_50ntFlanks@6651.6698.48\|CGACTCTTAGCGGTGGATCACTCGGCTCGTGCGTCGATGAAGAACGCA | >hg38_21_+_8433172_8446622_RNA45SN1_45S_50ntFlanks | 6651 | 6698 | 48 | CGACTCTTAGCGGTGGATCACTCGGCTCGTGCGTCGATGAAGAACGCA | 55297.2 | -2.7 | 0.9 | -3.1 | 2.1E-03 | 2.5E-03 | MB231/MB468 | MITO |
| >trnaMT_ThrTGT_MT_+_15888_15953@37.69.33\|ACCGGAGATGAAAACCTTTTTCCAAGGACACCA | >trnaMT_ThrTGT_MT_+_15888_15953 | 37 | 69 | 33 | ACCGGAGATGAAAACCTTTTTCCAAGGACACCA | 918.7 | 4.5 | 1.4 | 3.3 | 1.0E-03 | 2.5E-03 | BT20/MB468 | MP |
| >hg38_21_+_8433172_8446622_RNA45SN1_45S_50ntFlanks@7972.7993.22\|AGACGCGACCTCAGATCAGACG | >hg38_21_+_8433172_8446622_RNA45SN1_45S_50ntFlanks | 7972 | 7993 | 22 | AGACGCGACCTCAGATCAGACG | 114.2 | 5.7 | 1.7 | 3.3 | 1.0E-03 | 2.5E-03 | BT20/MB468 | MP |
| >hg38_MT_+_1621_3279_RNR2_16S_50ntFlanks@826.860.35\|AAGGAAAGGTTAAAAAAAGTAAAAGGAACTCGGCA | >hg38_MT_+_1621_3279_RNR2_16S_50ntFlanks | 826 | 860 | 35 | AAGGAAAGGTTAAAAAAAGTAAAAGGAACTCGGCA | 62.6 | 6.4 | 1.9 | 3.3 | 1.0E-03 | 2.5E-03 | BT20/MB468 | MP |
| >hg38_MT_+_1621_3279_RNR2_16S_50ntFlanks@618.642.25\|AACATATAACTGAACTCCTCACACC | >hg38_MT_+_1621_3279_RNR2_16S_50ntFlanks | 618 | 642 | 25 | AACATATAACTGAACTCCTCACACC | 150.9 | 4.8 | 1.5 | 3.1 | 2.1E-03 | 2.5E-03 | MB231/MB468 | MITO |
| >MI0000253\|hsa-mir-148a&WithFlank&7\|-\|25949913\|25949992@50.70.21(+1G)&[MIMAT0000243&hsa-miR-148a-3p&offsets\|0\|-1(+1G);m-7&7\|-\|25949922\|25949943&offsets\|0\|-1(+1G)]\|TCAGTGCACTACAGAACTTTGG | >MI0000253\|hsa-mir-148a&WithFlank&7\|-\|25949913\|25949992 | 50 | 70 | 21(+1G) | TCAGTGCACTACAGAACTTTGG | 1328.9 | 2.8 | 0.9 | 3.1 | 2.1E-03 | 2.5E-03 | BT20/MB468 | NUC |
| >hg38_21_+_8433172_8446622_RNA45SN1_45S_50ntFlanks@3840.3860.21\|CTCCTACTTGGATAACTGTGG | >hg38_21_+_8433172_8446622_RNA45SN1_45S_50ntFlanks | 3840 | 3860 | 21 | CTCCTACTTGGATAACTGTGG | 830.5 | -2.1 | 0.7 | -3.1 | 2.1E-03 | 2.5E-03 | BT20/MB468 | NUC |
| >MI0000285\|hsa-mir-205&WithFlank&1\|+\|209432127\|209432248@40.59.20(+1G)&[MIMAT0000266&hsa-miR-205-5p&offsets\|0\|-2(+1G);m-351&1\|+\|209432166\|209432187&offsets\|0\|-2(+1G)]\|TCCTTCATTCCACCGGAGTCG | >MI0000285\|hsa-mir-205&WithFlank&1\|+\|209432127\|209432248 | 40 | 59 | 20(+1G) | TCCTTCATTCCACCGGAGTCG | 77.3 | 9.6 | 2.7 | 3.6 | 3.5E-04 | 2.5E-03 | BT20/MB231 | MP |
| >hg38_21_+_8433172_8446622_RNA45SN1_45S_50ntFlanks@8463.8497.35\|CCGTGTCGGCGGCCCGGCGGATCTTTCCCGCCCCC | >hg38_21_+_8433172_8446622_RNA45SN1_45S_50ntFlanks | 8463 | 8497 | 35 | CCGTGTCGGCGGCCCGGCGGATCTTTCCCGCCCCC | 183.4 | -4.6 | 1.5 | -3.1 | 2.2E-03 | 2.5E-03 | MB231/MB468 | NUC |
| >hg38_21_+_8433172_8446622_RNA45SN1_45S_50ntFlanks@11586.11603.18\|GTGCGGACCAGGGGAATC | >hg38_21_+_8433172_8446622_RNA45SN1_45S_50ntFlanks | 11586 | 11603 | 18 | GTGCGGACCAGGGGAATC | 2193.3 | -4.8 | 1.5 | -3.3 | 1.0E-03 | 2.5E-03 | BT20/MB468 | MP |

<table>
<tr><th></th><th></th><th></th><th></th><th></th><th></th><th></th><th></th><th></th><th></th><th></th><th></th><th></th><th></th></tr>
<tr><td>>hg38_21_+_8433172_8446622_RNA45SN1_45S_50ntFlanks@10237.10259.23|ACGCTACGCCGCGACGAGTAGGA</td><td>>hg38_21_+_8433172_8446622_RNA45SN1_45S_50ntFlanks</td><td>10237</td><td>10259</td><td>23</td><td>ACGCTACGCCGCGACGAGTAGGA</td><td>368.5</td><td>6.6</td><td>1.8</td><td>3.6</td><td>3.5E-04</td><td>2.5E-03</td><td>BT20/MB231</td><td>MP</td></tr>
<tr><td>>hg38_21_+_8433172_8446622_RNA45SN1_45S_50ntFlanks@9534.9572.39|AGGAAACTCTGGTGGAGGTCCGTAGCGGTCCTGACGTGC</td><td>>hg38_21_+_8433172_8446622_RNA45SN1_45S_50ntFlanks</td><td>9534</td><td>9572</td><td>39</td><td>AGGAAACTCTGGTGGAGGTCCGTAGCGGTCCTGACGTGC</td><td>784.0</td><td>6.4</td><td>1.8</td><td>3.6</td><td>3.5E-04</td><td>2.5E-03</td><td>BT20/MB231</td><td>MP</td></tr>
<tr><td>>hg38_21_+_8433172_8446622_RNA45SN1_45S_50ntFlanks@12633.12659.27|CGAAGCTACCATCTGTGGGATTATGAC</td><td>>hg38_21_+_8433172_8446622_RNA45SN1_45S_50ntFlanks</td><td>12633</td><td>12659</td><td>27</td><td>CGAAGCTACCATCTGTGGGATTATGAC</td><td>412.4</td><td>-2.9</td><td>0.9</td><td>-3.1</td><td>2.0E-03</td><td>2.5E-03</td><td>MB231/MB468</td><td>CYTO</td></tr>
<tr><td>>hg38_21_+_8433172_8446622_RNA45SN1_45S_50ntFlanks@7985.8003.19|GATCAGACGTGGCGACCCG</td><td>>hg38_21_+_8433172_8446622_RNA45SN1_45S_50ntFlanks</td><td>7985</td><td>8003</td><td>19</td><td>GATCAGACGTGGCGACCCG</td><td>262.8</td><td>-4.4</td><td>1.4</td><td>-3.1</td><td>2.0E-03</td><td>2.5E-03</td><td>MB231/MB468</td><td>CYTO</td></tr>
<tr><td>>hg38_21_+_8433172_8446622_RNA45SN1_45S_50ntFlanks@9549.9574.26|AGGTCCGTAGCGGTCCTGACGTGCAA</td><td>>hg38_21_+_8433172_8446622_RNA45SN1_45S_50ntFlanks</td><td>9549</td><td>9574</td><td>26</td><td>AGGTCCGTAGCGGTCCTGACGTGCAA</td><td>255.4</td><td>-4.3</td><td>1.4</td><td>-3.1</td><td>2.1E-03</td><td>2.5E-03</td><td>BT20/MB468</td><td>NUC</td></tr>
<tr><td>>MI0000088|hsa-mir-30a&WithFlank&6|-|71403545|71403627@53.75.23&[MIMAT0000088&hsa-miR-30a-3p&offsets|0|+1;m-38&6|-|71403554|71403575&offsets|0|+1]|CTTTCAGTCGGATGTTTGCAGCT</td><td>>MI0000088|hsa-mir-30a&WithFlank&6|-|71403545|71403627</td><td>53</td><td>75</td><td>23</td><td>CTTTCAGTCGGATGTTTGCAGCT</td><td>343.0</td><td>-4.0</td><td>1.1</td><td>-3.6</td><td>3.5E-04</td><td>2.5E-03</td><td>BT20/MB231</td><td>MP</td></tr>
<tr><td>>hg38_21_+_8433172_8446622_RNA45SN1_45S_50ntFlanks@3993.4031.39|GTGACTCTAGATAACCTCGGGCCGATCGCACGCCCCCCG</td><td>>hg38_21_+_8433172_8446622_RNA45SN1_45S_50ntFlanks</td><td>3993</td><td>4031</td><td>39</td><td>GTGACTCTAGATAACCTCGGGCCGATCGCACGCCCCCCG</td><td>697.7</td><td>-4.1</td><td>1.3</td><td>-3.1</td><td>2.1E-03</td><td>2.5E-03</td><td>MB231/MB468</td><td>MITO</td></tr>
<tr><td>>MI0000736|hsa-mir-30c-1&WithFlank&1|+|40757278|40757378@23.46.24;MI0000254|hsa-mir-30c-2&WithFlank&6|-|71376954|71377037@13.36.24&[MIMAT0000244&hsa-miR-30c-5p&offsets|0|+1;m-57&1|+|40757300|40757323&offsets|0|0];[MIMAT0000244_1&hsa-miR-30c-5p&offsets|0|+1;m-56&6|-|71377002|71377025&offsets|0|0]|TGTAAACATCCTACACTCTCAGCT</td><td>>MI0000736|hsa-mir-30c-1&WithFlank&1|+|40757278|40757378</td><td>23</td><td>46</td><td>24</td><td>TGTAAACATCCTACACTCTCAGCT</td><td>8833.2</td><td>3.3</td><td>1.1</td><td>3.1</td><td>1.8E-03</td><td>2.5E-03</td><td>MB231/MB468</td><td>MP</td></tr>
<tr><td>>hg38_21_+_8433172_8446622_RNA45SN1_45S_50ntFlanks@6646.6676.31|TCGTACGACTCTTAGCGGTGGATCACTCGGC</td><td>>hg38_21_+_8433172_8446622_RNA45SN1_45S_50ntFlanks</td><td>6646</td><td>6676</td><td>31</td><td>TCGTACGACTCTTAGCGGTGGATCACTCGGC</td><td>5217.5</td><td>2.6</td><td>0.8</td><td>3.1</td><td>2.1E-03</td><td>2.6E-03</td><td>MB231/MB468</td><td>MITO</td></tr>
<tr><td>>hg38_21_+_8433172_8446622_RNA45SN1_45S_50ntFlanks@4293.4317.25|GATCCATTGGAGGGCAAGTCTGGTG</td><td>>hg38_21_+_8433172_8446622_RNA45SN1_45S_50ntFlanks</td><td>4293</td><td>4317</td><td>25</td><td>GATCCATTGGAGGGCAAGTCTGGTG</td><td>172.2</td><td>-5.1</td><td>1.6</td><td>-3.1</td><td>1.8E-03</td><td>2.6E-03</td><td>MB231/MB468</td><td>MP</td></tr>
<tr><td>>am_MI0000809|hsa-mir-151a&WithFlank&8|-|140732558|140732659@53.71.19&[MIMAT0000757&hsa-miR-151a-3p&offsets|0|-2;m-30&8|-|140732587|140732607&offsets|0|-2]|CTAGACTGAAGCTCCTTGA</td><td>>am_MI0000809|hsa-mir-151a&WithFlank&8|-|140732558|140732659</td><td>53</td><td>71</td><td>19</td><td>CTAGACTGAAGCTCCTTGA</td><td>403.8</td><td>4.0</td><td>1.3</td><td>3.1</td><td>1.8E-03</td><td>2.6E-03</td><td>MB231/MB468</td><td>MP</td></tr>
<tr><td>>hg38_MT_+_1621_3279_RNR2_16S_50ntFlanks@114.145.32|CAAATAAAGTATAGGCGATAGAAATTGAAACC</td><td>>hg38_MT_+_1621_3279_RNR2_16S_50ntFlanks</td><td>114</td><td>145</td><td>32</td><td>CAAATAAAGTATAGGCGATAGAAATTGAAACC</td><td>95.1</td><td>5.8</td><td>1.9</td><td>3.1</td><td>1.8E-03</td><td>2.6E-03</td><td>MB231/MB468</td><td>MP</td></tr>
<tr><td>>hg38_21_+_8433172_8446622_RNA45SN1_45S_50ntFlanks@9471.9503.33|AGCACGAGCGCACGTGTTAGGACCCGAAAGATG</td><td>>hg38_21_+_8433172_8446622_RNA45SN1_45S_50ntFlanks</td><td>9471</td><td>9503</td><td>33</td><td>AGCACGAGCGCACGTGTTAGGACCCGAAAGATG</td><td>158.1</td><td>-4.6</td><td>1.5</td><td>-3.1</td><td>2.2E-03</td><td>2.6E-03</td><td>MB231/MB468</td><td>NUC</td></tr>
<tr><td>>MI0003591|hsa-mir-584&WithFlank&5|-|149062307|149062415@22.42.21(+1A)&[MIMAT0003249&hsa-miR-584-5p&offsets|0|-1(+1A);m-167&5|-|149062374|149062394&offsets|0|0(+1A)]|TTATGGTTTGCCTGGGACTGAA</td><td>>MI0003591|hsa-mir-584&WithFlank&5|-|149062307|149062415</td><td>22</td><td>42</td><td>21(+1A)</td><td>TTATGGTTTGCCTGGGACTGAA</td><td>196.5</td><td>4.6</td><td>1.4</td><td>3.3</td><td>1.0E-03</td><td>2.6E-03</td><td>BT20/MB468</td><td>MP</td></tr>
<tr><td>>hg38_21_+_8433172_8446622_RNA45SN1_45S_50ntFlanks@11200.11233.34|CGGCGGCGGCGGCGACTCTGGACGCGAGCCGGGC</td><td>>hg38_21_+_8433172_8446622_RNA45SN1_45S_50ntFlanks</td><td>11200</td><td>11233</td><td>34</td><td>CGGCGGCGGCGGCGACTCTGGACGCGAGCCGGGC</td><td>312.0</td><td>-3.6</td><td>1.2</td><td>-3.1</td><td>2.0E-03</td><td>2.6E-03</td><td>BT20/MB468</td><td>CYTO</td></tr>
<tr><td>>hg38_MT_+_1621_3279_RNR2_16S_50ntFlanks@1338.1366.29|AGTCCATATCAACAATAGGGTTTACGACC</td><td>>hg38_MT_+_1621_3279_RNR2_16S_50ntFlanks</td><td>1338</td><td>1366</td><td>29</td><td>AGTCCATATCAACAATAGGGTTTACGACC</td><td>44.2</td><td>9.2</td><td>2.8</td><td>3.3</td><td>1.0E-03</td><td>2.6E-03</td><td>BT20/MB468</td><td>MP</td></tr>
<tr><td>>hg38_21_+_8433172_8446622_RNA45SN1_45S_50ntFlanks@4731.4767.37|AAGAACGAAAGTCGGAGGTTCGAAGACGATCAGATAC</td><td>>hg38_21_+_8433172_8446622_RNA45SN1_45S_50ntFlanks</td><td>4731</td><td>4767</td><td>37</td><td>AAGAACGAAAGTCGGAGGTTCGAAGACGATCAGATAC</td><td>133.7</td><td>4.7</td><td>1.5</td><td>3.1</td><td>1.8E-03</td><td>2.6E-03</td><td>MB231/MB468</td><td>MP</td></tr>
<tr><td>>MI0000434|hsa-let-7i&WithFlank&12|+|62603680|62603775@12.34.23&[MIMAT0000415&hsa-let-7i-5p&offsets|0|+1;m-49&12|+|62603691|62603712&offsets|0|+1]|TGAGGTAGTAGTTTGTGCTGTTG</td><td>>MI0000434|hsa-let-7i&WithFlank&12|+|62603680|62603775</td><td>12</td><td>34</td><td>23</td><td>TGAGGTAGTAGTTTGTGCTGTTG</td><td>507.0</td><td>4.8</td><td>1.5</td><td>3.3</td><td>1.0E-03</td><td>2.6E-03</td><td>BT20/MB468</td><td>MP</td></tr>
<tr><td>>hg38_21_+_8433172_8446622_RNA45SN1_45S_50ntFlanks@10392.10426.35|ACAGCAGTTGAACATGGGTCAGTCGGTCCTGAGAG</td><td>>hg38_21_+_8433172_8446622_RNA45SN1_45S_50ntFlanks</td><td>10392</td><td>10426</td><td>35</td><td>ACAGCAGTTGAACATGGGTCAGTCGGTCCTGAGAG</td><td>398.5</td><td>-5.5</td><td>1.7</td><td>-3.3</td><td>1.0E-03</td><td>2.6E-03</td><td>BT20/MB468</td><td>MP</td></tr>
<tr><td>>hg38_21_+_8433172_8446622_RNA45SN1_45S_50ntFlanks@8357.8387.31|AAAAGAACTTTGAAGAGAGAGTTCAAGAGGG</td><td>>hg38_21_+_8433172_8446622_RNA45SN1_45S_50ntFlanks</td><td>8357</td><td>8387</td><td>31</td><td>AAAAGAACTTTGAAGAGAGAGTTCAAGAGGG</td><td>665.4</td><td>-3.4</td><td>1.0</td><td>-3.3</td><td>1.0E-03</td><td>2.6E-03</td><td>BT20/MB468</td><td>MP</td></tr>
<tr><td>>hg38_21_+_8433172_8446622_RNA45SN1_45S_50ntFlanks@9624.9648.25|AGTAGCTGGTTCCCTCCGAAGTTTC</td><td>>hg38_21_+_8433172_8446622_RNA45SN1_45S_50ntFlanks</td><td>9624</td><td>9648</td><td>25</td><td>AGTAGCTGGTTCCCTCCGAAGTTTC</td><td>169.6</td><td>-3.6</td><td>1.2</td><td>-3.1</td><td>1.7E-03</td><td>2.6E-03</td><td>BT20/MB468</td><td>MITO</td></tr>
<tr><td>>hg38_21_+_8433172_8446622_RNA45SN1_45S_50ntFlanks@10849.10866.18|CGGGATAAGGATTGGCTC</td><td>>hg38_21_+_8433172_8446622_RNA45SN1_45S_50ntFlanks</td><td>10849</td><td>10866</td><td>18</td><td>CGGGATAAGGATTGGCTC</td><td>312.1</td><td>-2.7</td><td>0.9</td><td>-3.1</td><td>2.0E-03</td><td>2.6E-03</td><td>BT20/MB468</td><td>CYTO</td></tr>
<tr><td>>hg38_21_+_8433172_8446622_RNA45SN1_45S_50ntFlanks@4660.4679.20|GAGGTGAAATTCTTGGACCG</td><td>>hg38_21_+_8433172_8446622_RNA45SN1_45S_50ntFlanks</td><td>4660</td><td>4679</td><td>20</td><td>GAGGTGAAATTCTTGGACCG</td><td>40.1</td><td>-8.4</td><td>2.6</td><td>-3.3</td><td>1.0E-03</td><td>2.6E-03</td><td>BT20/MB468</td><td>MP</td></tr>
<tr><td>>trnaMT_GluTTC_MT_-_14674_14742@40.72.33,trnalookalike8_GluTTC_5_-_93905172_93905240@40.72.33|TTGGTCGTGGTTGTAGTCCGTGCGAGAATACCA</td><td>>trnaMT_GluTTC_MT_-_14674_14742</td><td>40</td><td>72</td><td>33</td><td>TTGGTCGTGGTTGTAGTCCGTGCGAGAATACCA</td><td>15110.3</td><td>-3.1</td><td>0.9</td><td>-3.5</td><td>4.2E-04</td><td>2.6E-03</td><td>BT20/MB231</td><td>CYTO</td></tr>
<tr><td>>hg38_21_+_8433172_8446622_RNA45SN1_45S_50ntFlanks@5490.5533.44|TGAGAAGACGGTCGAACTTGACTATCTAGAGGAAGTAAAAGTCG</td><td>>hg38_21_+_8433172_8446622_RNA45SN1_45S_50ntFlanks</td><td>5490</td><td>5533</td><td>44</td><td>TGAGAAGACGGTCGAACTTGACTATCTAGAGGAAGTAAAAGTCG</td><td>225.7</td><td>-3.4</td><td>1.0</td><td>-3.5</td><td>4.2E-04</td><td>2.6E-03</td><td>BT20/MB231</td><td>CYTO</td></tr>
<tr><td>>hg38_21_+_8433172_8446622_RNA45SN1_45S_50ntFlanks@485.518.34|GATGTGGTGACGTCGTGCTCTCCCGGGCCGGGTC</td><td>>hg38_21_+_8433172_8446622_RNA45SN1_45S_50ntFlanks</td><td>485</td><td>518</td><td>34</td><td>GATGTGGTGACGTCGTGCTCTCCCGGGCCGGGTC</td><td>240.8</td><td>11.1</td><td>3.6</td><td>3.1</td><td>2.2E-03</td><td>2.6E-03</td><td>MB231/MB468</td><td>MITO</td></tr>
<tr><td>>hg38_21_+_8433172_8446622_RNA45SN1_45S_50ntFlanks@7984.8018.35|AGATCAGACGTGGCGACCCGCTGAATTTAAGCATA</td><td>>hg38_21_+_8433172_8446622_RNA45SN1_45S_50ntFlanks</td><td>7984</td><td>8018</td><td>35</td><td>AGATCAGACGTGGCGACCCGCTGAATTTAAGCATA</td><td>302.0</td><td>-3.1</td><td>1.0</td><td>-3.1</td><td>2.2E-03</td><td>2.6E-03</td><td>MB231/MB468</td><td>MITO</td></tr>
<tr><td>>MI0000255|hsa-mir-30d&WithFlank&8|-|134804870|134804951@12.32.21&[MIMAT0000245&hsa-miR-30d-5p&offsets|0|-1;m-21&8|-|134804919|134804940&offsets|0|-1]|TGTAAACATCCCCGACTGGAA</td><td>>MI0000255|hsa-mir-30d&WithFlank&8|-|134804870|134804951</td><td>12</td><td>32</td><td>21</td><td>TGTAAACATCCCCGACTGGAA</td><td>309.9</td><td>4.6</td><td>1.4</td><td>3.3</td><td>1.0E-03</td><td>2.6E-03</td><td>BT20/MB468</td><td>MP</td></tr>
<tr><td>>hg38_21_+_8433172_8446622_RNA45SN1_45S_50ntFlanks@9574.9602.29|AATCGGTCGTCCGACCTGGGTATAGGGGC</td><td>>hg38_21_+_8433172_8446622_RNA45SN1_45S_50ntFlanks</td><td>9574</td><td>9602</td><td>29</td><td>AATCGGTCGTCCGACCTGGGTATAGGGGC</td><td>584.6</td><td>2.3</td><td>0.7</td><td>3.1</td><td>2.2E-03</td><td>2.6E-03</td><td>MB231/MB468</td><td>NUC</td></tr>
<tr><td>>hg38_21_+_8433172_8446622_RNA45SN1_45S_50ntFlanks@10771.10817.47|CAGGTCTCCAAGGTGAACAGCCTCTGGCATGTTGGAACAATGTAGGT</td><td>>hg38_21_+_8433172_8446622_RNA45SN1_45S_50ntFlanks</td><td>10771</td><td>10817</td><td>47</td><td>CAGGTCTCCAAGGTGAACAGCCTCTGGCATGTTGGAACAATGTAGGT</td><td>279.3</td><td>-4.3</td><td>1.4</td><td>-3.1</td><td>2.2E-03</td><td>2.6E-03</td><td>MB231/MB468</td><td>NUC</td></tr>
<tr><td>>hg38_21_+_8433172_8446622_RNA45SN1_45S_50ntFlanks@12417.12435.19|TCGATGTCGGCTCTTCCTA</td><td>>hg38_21_+_8433172_8446622_RNA45SN1_45S_50ntFlanks</td><td>12417</td><td>12435</td><td>19</td><td>TCGATGTCGGCTCTTCCTA</td><td>784.8</td><td>-5.6</td><td>1.7</td><td>-3.3</td><td>1.1E-03</td><td>2.6E-03</td><td>BT20/MB468</td><td>MP</td></tr>
<tr><td>>hg38_21_+_8433172_8446622_RNA45SN1_45S_50ntFlanks@10922.10944.23|CTGGACGAGGCGCCGCCGCCCCC</td><td>>hg38_21_+_8433172_8446622_RNA45SN1_45S_50ntFlanks</td><td>10922</td><td>10944</td><td>23</td><td>CTGGACGAGGCGCCGCCGCCCCC</td><td>207.6</td><td>-3.2</td><td>1.0</td><td>-3.1</td><td>2.0E-03</td><td>2.6E-03</td><td>BT20/MB468</td><td>CYTO</td></tr>
<tr><td>>hg38_21_+_8433172_8446622_RNA45SN1_45S_50ntFlanks@6647.6687.41|CGTACGACTCTTAGCGGTGGATCACTCGGCTCGTGCGTCGA</td><td>>hg38_21_+_8433172_8446622_RNA45SN1_45S_50ntFlanks</td><td>6647</td><td>6687</td><td>41</td><td>CGTACGACTCTTAGCGGTGGATCACTCGGCTCGTGCGTCGA</td><td>265.2</td><td>3.0</td><td>1.0</td><td>3.1</td><td>2.2E-03</td><td>2.6E-03</td><td>MB231/MB468</td><td>MITO</td></tr>
<tr><td>>hg38_21_+_8433172_8446622_RNA45SN1_45S_50ntFlanks@10841.10862.22|CGTAACTTCGGGATAAGGATTG</td><td>>hg38_21_+_8433172_8446622_RNA45SN1_45S_50ntFlanks</td><td>10841</td><td>10862</td><td>22</td><td>CGTAACTTCGGGATAAGGATTG</td><td>2607.2</td><td>-6.1</td><td>1.9</td><td>-3.3</td><td>1.1E-03</td><td>2.6E-03</td><td>BT20/MB468</td><td>MP</td></tr>
<tr><td>>MI0000077|hsa-mir-21&WithFlank&17|+|59841260|59841343@14.37.24&[MIMAT0000076&hsa-miR-21-5p&offsets|0|+2;m-1&17|+|59841273|59841294&offsets|0|+2]|TAGCTTATCAGACTGATGTTGACT</td><td>>MI0000077|hsa-mir-21&WithFlank&17|+|59841260|59841343</td><td>14</td><td>37</td><td>24</td><td>TAGCTTATCAGACTGATGTTGACT</td><td>15614.0</td><td>2.9</td><td>0.9</td><td>3.3</td><td>1.1E-03</td><td>2.6E-03</td><td>BT20/MB468</td><td>MP</td></tr>
<tr><td>>hg38_21_+_8433172_8446622_RNA45SN1_45S_50ntFlanks@11674.11707.34|GTGCTCTGAATGTCAAAGTGAAGAAATTCAATGA</td><td>>hg38_21_+_8433172_8446622_RNA45SN1_45S_50ntFlanks</td><td>11674</td><td>11707</td><td>34</td><td>GTGCTCTGAATGTCAAAGTGAAGAAATTCAATGA</td><td>40.2</td><td>9.1</td><td>2.8</td><td>3.3</td><td>1.1E-03</td><td>2.6E-03</td><td>BT20/MB468</td><td>MP</td></tr>
<tr><td>>hg38_MT_+_598_1651_RNR1_12S_50ntFlanks@914.946.33|CAAAGGACATTTAACTAAAACCCCTACGCATTT</td><td>>hg38_MT_+_598_1651_RNR1_12S_50ntFlanks</td><td>914</td><td>946</td><td>33</td><td>CAAAGGACATTTAACTAAAACCCCTACGCATTT</td><td>336.6</td><td>2.7</td><td>0.8</td><td>3.5</td><td>4.7E-04</td><td>2.6E-03</td><td>BT20/MB231</td><td>MITO</td></tr>
<tr><td>>hg38_21_+_8433172_8446622_RNA45SN1_45S_50ntFlanks@11795.11821.27|AACGAGATTCCCACTGTCCCTACCTAC</td><td>>hg38_21_+_8433172_8446622_RNA45SN1_45S_50ntFlanks</td><td>11795</td><td>11821</td><td>27</td><td>AACGAGATTCCCACTGTCCCTACCTAC</td><td>242.8</td><td>-2.8</td><td>0.9</td><td>-3.1</td><td>1.7E-03</td><td>2.6E-03</td><td>BT20/MB468</td><td>MITO</td></tr>
<tr><td>>trna10_AspGTC_12_-_125424193_125424264@35.70.36,trna12_AspGTC_12_-_125411891_125411962@35.70.36,trna144_AspGTC_6_-_27551236_27551307@35.70.36,trna38_AspGTC_17_-_8125556_8125627@35.70.36,trna45_AspGTC_6_+_27447453_27447524@35.70.36,trna48_AspGTC_6_+_27471523_27471594@35.70.36,trna4_AspGTC_12_+_96429799_96429870@35.70.36,trna69_AspGTC_1_-_161440205_161440276@35.70.36,trna72_AspGTC_1_-_161432824_161432895@35.70.36,trna75_AspGTC_1_-_161425414_161425485@35.70.36,trna78_AspGTC_1_-_161418033_161418104@35.70.36,trna81_AspGTC_1_-_161410615_161410686@35.70.36|TCACGCGGGAGACCGGGGTTCGATTCCCCGACGGGG</td><td>>trna10_AspGTC_12_-_125424193_125424264</td><td>35</td><td>70</td><td>36</td><td>TCACGCGGGAGACCGGGGTTCGATTCCCCGACGGGG</td><td>1042.6</td><td>-2.7</td><td>0.9</td><td>-3.1</td><td>1.8E-03</td><td>2.6E-03</td><td>MB231/MB468</td><td>MP</td></tr>
<tr><td>>hg38_21_+_8433172_8446622_RNA45SN1_45S_50ntFlanks@10847.10865.19|TTCGGGATAAGGATTGGCT</td><td>>hg38_21_+_8433172_8446622_RNA45SN1_45S_50ntFlanks</td><td>10847</td><td>10865</td><td>19</td><td>TTCGGGATAAGGATTGGCT</td><td>192.0</td><td>-1.8</td><td>0.6</td><td>-3.1</td><td>2.0E-03</td><td>2.6E-03</td><td>BT20/MB468</td><td>CYTO</td></tr>
<tr><td>>hg38_21_+_8433172_8446622_RNA45SN1_45S_50ntFlanks@11710.11728.19|CGCGGGTAAACGGCGGGAG</td><td>>hg38_21_+_8433172_8446622_RNA45SN1_45S_50ntFlanks</td><td>11710</td><td>11728</td><td>19</td><td>CGCGGGTAAACGGCGGGAG</td><td>512.3</td><td>-3.6</td><td>1.2</td><td>-3.1</td><td>2.0E-03</td><td>2.6E-03</td><td>BT20/MB468</td><td>CYTO</td></tr>
<tr><td>>hg38_21_+_8433172_8446622_RNA45SN1_45S_50ntFlanks@12907.12940.34|CACCGCACGTTCGTGGGGAACCTGGCGCTAAACC</td><td>>hg38_21_+_8433172_8446622_RNA45SN1_45S_50ntFlanks</td><td>12907</td><td>12940</td><td>34</td><td>CACCGCACGTTCGTGGGGAACCTGGCGCTAAACC</td><td>484.3</td><td>-3.8</td><td>1.3</td><td>-3.1</td><td>2.2E-03</td><td>2.6E-03</td><td>MB231/MB468</td><td>MITO</td></tr>
<tr><td>>MI0000086|hsa-mir-28&WithFlank&3|+|188688775|188688872@60.79.20&[MIMAT0004502&hsa-miR-28-3p&offsets|0|-2;m-28&3|+|188688834|188688855&offsets|0|-2]|CACTAGATTGTGAGCTCCTG</td><td>>MI0000086|hsa-mir-28&WithFlank&3|+|188688775|188688872</td><td>60</td><td>79</td><td>20</td><td>CACTAGATTGTGAGCTCCTG</td><td>94.9</td><td>4.8</td><td>1.6</td><td>3.1</td><td>1.8E-03</td><td>2.6E-03</td><td>MB231/MB468</td><td>MP</td></tr>
<tr><td>>hg38_21_+_8433172_8446622_RNA45SN1_45S_50ntFlanks@5472.5507.36|ACGGCCCTGGCGGAGCGCTGAGAAGACGGTCGAACT</td><td>>hg38_21_+_8433172_8446622_RNA45SN1_45S_50ntFlanks</td><td>5472</td><td>5507</td><td>36</td><td>ACGGCCCTGGCGGAGCGCTGAGAAGACGGTCGAACT</td><td>141.7</td><td>3.8</td><td>1.3</td><td>3.1</td><td>2.2E-03</td><td>2.6E-03</td><td>BT20/MB468</td><td>NUC</td></tr>
<tr><td>>hg38_21_+_8433172_8446622_RNA45SN1_45S_50ntFlanks@12418.12444.27|CGATGTCGGCTCTTCCTATCATTGTGA</td><td>>hg38_21_+_8433172_8446622_RNA45SN1_45S_50ntFlanks</td><td>12418</td><td>12444</td><td>27</td><td>CGATGTCGGCTCTTCCTATCATTGTGA</td><td>1274.4</td><td>-1.8</td><td>0.6</td><td>-3.1</td><td>1.7E-03</td><td>2.6E-03</td><td>BT20/MB468</td><td>MITO</td></tr>
<tr><td>>trnaMT_GluTTC_MT_-_14674_14742@41.61.21,trnalookalike8_GluTTC_5_-_93905172_93905240@41.61.21|TGGTCGTGGTTGTAGTCCGTG</td><td>>trnaMT_GluTTC_MT_-_14674_14742</td><td>41</td><td>61</td><td>21</td><td>TGGTCGTGGTTGTAGTCCGTG</td><td>195.2</td><td>4.7</td><td>1.4</td><td>3.3</td><td>1.1E-03</td><td>2.6E-03</td><td>BT20/MB468</td><td>MP</td></tr>
<tr><td>>hg38_21_+_8433172_8446622_RNA45SN1_45S_50ntFlanks@4762.4780.19|AGATACCGTCGTAGTTCCG</td><td>>hg38_21_+_8433172_8446622_RNA45SN1_45S_50ntFlanks</td><td>4762</td><td>4780</td><td>19</td><td>AGATACCGTCGTAGTTCCG</td><td>446.2</td><td>-3.8</td><td>1.2</td><td>-3.1</td><td>2.2E-03</td><td>2.6E-03</td><td>MB231/MB468</td><td>NUC</td></tr>
<tr><td>>hg38_21_+_8433172_8446622_RNA45SN1_45S_50ntFlanks@12606.12625.20|ATGTGCTTGGCTGAGGAGCC</td><td>>hg38_21_+_8433172_8446622_RNA45SN1_45S_50ntFlanks</td><td>12606</td><td>12625</td><td>20</td><td>ATGTGCTTGGCTGAGGAGCC</td><td>675.8</td><td>-2.1</td><td>0.7</td><td>-3.1</td><td>2.1E-03</td><td>2.6E-03</td><td>MB231/MB468</td><td>CYTO</td></tr>
</table>

| | | | | | | | | | | | | | |
|---|---|---|---|---|---|---|---|---|---|---|---|---|---|
| >hg38_21_+_8433172_8446622_RNA45SN1_45S_50ntFlanks@9243.9274.32\|GAGCGAGCGCACGGGGTCGGCGGCGACGTCGG | >hg38_21_+_8433172_8446622_RNA45SN1_45S_50ntFlanks | 9243 | 9274 | 32 | GAGCGAGCGCACGGGGTCGGCGGCGACGTCGG | 167.5 | -5.3 | 1.6 | -3.3 | 1.1E-03 | 2.6E-03 | BT20/MB468 | MP |
| >trnaMT_GluTTC_MT_-_14674_14742@25.72.48,trnalookalike8_GluTTC_5_-_93905172_93905240@25.72.48\|TGGTTTTTCATATCATTGGTCGTGGTTGTAGTCCGTGCGAGAATACCA | >trnaMT_GluTTC_MT_-_14674_14742 | 25 | 72 | 48 | TGGTTTTTCATATCATTGGTCGTGGTTGTAGTCCGTGCGAGAATACCA | 370.4 | 4.3 | 1.4 | 3.1 | 2.2E-03 | 2.6E-03 | MB231/MB468 | MITO |
| >hg38_21_+_8433172_8446622_RNA45SN1_45S_50ntFlanks@12259.12288.30\|CTTGATCTTGATTTTCAGTACGAATACAGA | >hg38_21_+_8433172_8446622_RNA45SN1_45S_50ntFlanks | 12259 | 12288 | 30 | CTTGATCTTGATTTTCAGTACGAATACAGA | 232.3 | -3.2 | 1.0 | -3.3 | 1.1E-03 | 2.6E-03 | BT20/MB468 | MP |
| >trnaMT_ValTAC_MT_+_1602_1670@45.71.27\|TTTCAACTTAACTTGACCGCTCTGACC | >trnaMT_ValTAC_MT_+_1602_1670 | 45 | 71 | 27 | TTTCAACTTAACTTGACCGCTCTGACC | 342.6 | 3.0 | 1.0 | 3.1 | 2.2E-03 | 2.6E-03 | MB231/MB468 | MITO |
| >trnaMT_IleGAT_MT_+_4263_4331@45.63.19\|AGGAGCTTAAACCCCCTTA | >trnaMT_IleGAT_MT_+_4263_4331 | 45 | 63 | 19 | AGGAGCTTAAACCCCCTTA | 43.8 | 9.2 | 2.8 | 3.3 | 1.1E-03 | 2.6E-03 | BT20/MB468 | MP |
| >hg38_21_+_8433172_8446622_RNA45SN1_45S_50ntFlanks@9554.9597.44\|CGTAGCGGTCCTGACGTGCAAATCGGTCGTCCGACCTGGGTATA | >hg38_21_+_8433172_8446622_RNA45SN1_45S_50ntFlanks | 9554 | 9597 | 44 | CGTAGCGGTCCTGACGTGCAAATCGGTCGTCCGACCTGGGTATA | 210.6 | -6.2 | 1.9 | -3.3 | 1.1E-03 | 2.6E-03 | BT20/MB468 | MP |
| >hg38_MT_+_598_1651_RNR1_12S_50ntFlanks@608.633.26\|TGTTCTGTAATCGATAAACCCCGATC | >hg38_MT_+_598_1651_RNR1_12S_50ntFlanks | 608 | 633 | 26 | TGTTCTGTAATCGATAAACCCCGATC | 266.3 | 3.6 | 1.2 | 3.1 | 1.8E-03 | 2.6E-03 | MB231/MB468 | MP |
| >hg38_21_+_8433172_8446622_RNA45SN1_45S_50ntFlanks@11670.11699.30\|CCCAGTGCTCTGAATGTCAAAGTGAAGAAA | >hg38_21_+_8433172_8446622_RNA45SN1_45S_50ntFlanks | 11670 | 11699 | 30 | CCCAGTGCTCTGAATGTCAAAGTGAAGAAA | 265.3 | -3.3 | 1.1 | -3.1 | 2.1E-03 | 2.6E-03 | MB231/MB468 | CYTO |
| >hg38_21_+_8433172_8446622_RNA45SN1_45S_50ntFlanks@11671.11693.23\|CCAGTGCTCTGAATGTCAAAGTG | >hg38_21_+_8433172_8446622_RNA45SN1_45S_50ntFlanks | 11671 | 11693 | 23 | CCAGTGCTCTGAATGTCAAAGTG | 1386.9 | -3.4 | 1.1 | -3.1 | 2.1E-03 | 2.6E-03 | MB231/MB468 | CYTO |
| >hg38_21_+_8433172_8446622_RNA45SN1_45S_50ntFlanks@10384.10424.41\|CCATGTGAACAGCAGTTGAACATGGGTCAGTCGGTCCTGAG | >hg38_21_+_8433172_8446622_RNA45SN1_45S_50ntFlanks | 10384 | 10424 | 41 | CCATGTGAACAGCAGTTGAACATGGGTCAGTCGGTCCTGAG | 2072.2 | -2.9 | 0.9 | -3.1 | 2.2E-03 | 2.6E-03 | MB231/MB468 | MITO |
| >trnaMT_CysGCA_MT_-_5761_5826@47.69.23\|CTTCAAACCTGCCGGGGCTTCCA | >trnaMT_CysGCA_MT_-_5761_5826 | 47 | 69 | 23 | CTTCAAACCTGCCGGGGCTTCCA | 72.8 | -4.9 | 1.6 | -3.1 | 1.8E-03 | 2.6E-03 | MB231/MB468 | MP |
| >hg38_21_+_8433172_8446622_RNA45SN1_45S_50ntFlanks@12634.12661.28\|GAAGCTACCATCTGTGGGATTATGACTG | >hg38_21_+_8433172_8446622_RNA45SN1_45S_50ntFlanks | 12634 | 12661 | 28 | GAAGCTACCATCTGTGGGATTATGACTG | 551.4 | 7.5 | 2.1 | 3.6 | 3.6E-04 | 2.6E-03 | BT20/MB231 | MP |
| >hg38_21_+_8433172_8446622_RNA45SN1_45S_50ntFlanks@11315.11336.22\|CCACCCCACGTCTCGTCGCGCG | >hg38_21_+_8433172_8446622_RNA45SN1_45S_50ntFlanks | 11315 | 11336 | 22 | CCACCCCACGTCTCGTCGCGCG | 2485.9 | -6.6 | 2.2 | -3.1 | 2.2E-03 | 2.6E-03 | BT20/MB468 | NUC |
| >MI0000446\|hsa-mir-125b-1&WithFlank&11\|-\|122099751\|122099850@21.42.22(+1A);MI0000470\|hsa-mir-125b-2&WithFlank&21\|+\|16590231\|16590331@23.44.22(+1A)&[MIMAT0000423&hsa-miR-125b-5p&offsets\|0\|0(+1A);m-67&11\|-\|122099809\|122099830&offsets\|0\|0(+1A)];[MIMAT0000423_1&hsa-miR-125b-5p&offsets\|0\|0(+1A);m-68&21\|+\|16590253\|16590274&offsets\|0\|0(+1A)]\|TCCCTGAGACCCTAACTTGTGAA | >MI0000446\|hsa-mir-125b-1&WithFlank&11\|-\|122099751\|122099850 | 21 | 42 | 22(+1A) | TCCCTGAGACCCTAACTTGTGAA | 4121.7 | -3.8 | 1.1 | -3.5 | 4.3E-04 | 2.6E-03 | BT20/MB231 | CYTO |
| >hg38_21_+_8433172_8446622_RNA45SN1_45S_50ntFlanks@11666.11707.42\|TCTGCCCAGTGCTCTGAATGTCAAAGTGAAGAAATTCAATGA | >hg38_21_+_8433172_8446622_RNA45SN1_45S_50ntFlanks | 11666 | 11707 | 42 | TCTGCCCAGTGCTCTGAATGTCAAAGTGAAGAAATTCAATGA | 624.7 | -2.8 | 0.8 | -3.5 | 4.3E-04 | 2.6E-03 | BT20/MB231 | CYTO |
| >hg38_21_+_8433172_8446622_RNA45SN1_45S_50ntFlanks@8497.8514.18\|CCGTTCCTCCCGACCCCT | >hg38_21_+_8433172_8446622_RNA45SN1_45S_50ntFlanks | 8497 | 8514 | 18 | CCGTTCCTCCCGACCCCT | 114.8 | -4.4 | 1.4 | -3.1 | 1.7E-03 | 2.6E-03 | BT20/MB468 | MITO |
| >MI0000086\|hsa-mir-28&WithFlank&3\|+\|188688775\|188688872@61.82.22(+1C)&[MIMAT0004502&hsa-miR-28-3p&offsets\|+1\|+1(+1C);m-28&3\|+\|188688834\|188688855&offsets\|+1\|+1(+1C)]\|ACTAGATTGTGAGCTCCTGGAGC | >MI0000086\|hsa-mir-28&WithFlank&3\|+\|188688775\|188688872 | 61 | 82 | 22(+1C) | ACTAGATTGTGAGCTCCTGGAGC | 48.4 | 5.7 | 1.8 | 3.1 | 1.8E-03 | 2.6E-03 | MB231/MB468 | MP |
| >trna10_AspGTC_12_-_125424193_125424264@38.68.31,trna12_AspGTC_12_-_125411891_125411962@38.68.31,trna144_AspGTC_6_-_27551236_27551307@38.68.31,trna38_AspGTC_17_-_8125556_8125627@38.68.31,trna45_AspGTC_6_+_27447453_27447524@38.68.31,trna48_AspGTC_6_+_27471523_27471594@38.68.31,trna4_AspGTC_12_+_96429799_96429870@38.68.31,trna69_AspGTC_1_-_161440205_161440276@38.68.31,trna72_AspGTC_1_-_161432824_161432895@38.68.31,trna75_AspGTC_1_-_161425414_161425485@38.68.31,trna78_AspGTC_1_-_161418033_161418104@38.68.31,trna81_AspGTC_1_-_161410615_161410686@38.68.31\|CGCGGGAGACCGGGGTTCGATTCCCCGACGG | >trna10_AspGTC_12_-_125424193_125424264 | 38 | 68 | 31 | CGCGGGAGACCGGGGTTCGATTCCCCGACGG | 260.3 | -2.7 | 0.9 | -3.1 | 1.8E-03 | 2.6E-03 | MB231/MB468 | MP |
| >hg38_21_+_8433172_8446622_RNA45SN1_45S_50ntFlanks@10879.10901.23\|CGGTCGGGCTGGGGCGCGAAGCG | >hg38_21_+_8433172_8446622_RNA45SN1_45S_50ntFlanks | 10879 | 10901 | 23 | CGGTCGGGCTGGGGCGCGAAGCG | 362.9 | -5.6 | 1.8 | -3.1 | 2.2E-03 | 2.6E-03 | BT20/MB468 | NUC |
| >hg38_21_+_8433172_8446622_RNA45SN1_45S_50ntFlanks@11583.11612.30\|CTGGTGCGGACCAGGGGAATCCGACTGTTT | >hg38_21_+_8433172_8446622_RNA45SN1_45S_50ntFlanks | 11583 | 11612 | 30 | CTGGTGCGGACCAGGGGAATCCGACTGTTT | 1168.9 | 3.8 | 1.1 | 3.5 | 4.0E-04 | 2.6E-03 | BT20/MB231 | NUC |
| >hg38_21_+_8433172_8446622_RNA45SN1_45S_50ntFlanks@6650.6667.18\|ACGACTCTTAGCGGTGGA | >hg38_21_+_8433172_8446622_RNA45SN1_45S_50ntFlanks | 6650 | 6667 | 18 | ACGACTCTTAGCGGTGGA | 424.3 | 2.3 | 0.6 | 3.5 | 4.1E-04 | 2.6E-03 | BT20/MB231 | NUC |
| >hg38_21_+_8433172_8446622_RNA45SN1_45S_50ntFlanks@11318.11336.19\|CCCCACGTCTCGTCGCGCG | >hg38_21_+_8433172_8446622_RNA45SN1_45S_50ntFlanks | 11318 | 11336 | 19 | CCCCACGTCTCGTCGCGCG | 1277.7 | -6.2 | 2.0 | -3.1 | 2.2E-03 | 2.6E-03 | BT20/MB468 | NUC |
| >trna128_GlyGCC_6_-_27870686_27870756@19.38.20,trna19_GlyGCC_16_+_70823410_70823480@19.38.20,trna19_GlyGCC_2_-_157257659_157257729@19.38.20,trna24_GlyGCC_16_-_70812942_70813012@19.38.20,trna25_GlyGCC_16_-_70812114_70812184@19.38.20,trna2_GlyGCC_21_-_18827107_18827177@19.38.20,trna35_GlyGCC_1_+_161413094_161413164@19.38.20,trna37_GlyGCC_1_+_161420467_161420537@19.38.20,trna39_GlyGCC_1_+_161427898_161427968@19.38.20,trna41_GlyGCC_1_+_161435258_161435328@19.38.20,trna5_GlyGCC_17_+_8029064_8029134@19.38.20,trna68_GlyGCC_1_-_161493637_161493707@19.38.20\|TAGAATTCTCGCCTGCCACG | >trna128_GlyGCC_6_-_27870686_27870756 | 19 | 38 | 20 | TAGAATTCTCGCCTGCCACG | 229.1 | 6.2 | 1.9 | 3.3 | 1.1E-03 | 2.6E-03 | BT20/MB468 | MP |
| >MI0001448\|hsa-mir-425&WithFlank&3\|-\|49020142\|49020240@60.80.21&[MIMAT0001343&hsa-miR-425-3p&offsets\|-1\|-2;m-258&3\|-\|49020160\|49020181&offsets\|0\|-1]\|CATCGGGAATGTCGTGTCCGC | >MI0001448\|hsa-mir-425&WithFlank&3\|-\|49020142\|49020240 | 60 | 80 | 21 | CATCGGGAATGTCGTGTCCGC | 70.5 | 4.9 | 1.6 | 3.1 | 1.9E-03 | 2.6E-03 | MB231/MB468 | MP |
| >trna5_AspGTC_12_+_98897281_98897352@37.69.33\|ACGCGGGAGACCGGGGTTCAATTCCCCGACGGG | >trna5_AspGTC_12_+_98897281_98897352 | 37 | 69 | 33 | ACGCGGGAGACCGGGGTTCAATTCCCCGACGGG | 86.1 | -2.4 | 0.8 | -3.1 | 1.9E-03 | 2.6E-03 | MB231/MB468 | MP |
| >hg38_21_+_8433172_8446622_RNA45SN1_45S_50ntFlanks@3705.3723.19\|TACCTGGTTGATCCTGCCA | >hg38_21_+_8433172_8446622_RNA45SN1_45S_50ntFlanks | 3705 | 3723 | 19 | TACCTGGTTGATCCTGCCA | 185.4 | -2.2 | 0.7 | -3.1 | 1.9E-03 | 2.6E-03 | MB231/MB468 | MP |
| >MI0000446\|hsa-mir-125b-1&WithFlank&11\|-\|122099751\|122099850@21.39.19(+1C);MI0000470\|hsa-mir-125b-2&WithFlank&21\|+\|16590231\|16590331@23.41.19(+1C)&[MIMAT0000423&hsa-miR-125b-5p&offsets\|0\|-3(+1C);m-67&11\|-\|122099809\|122099830&offsets\|0\|-3(+1C)];[MIMAT0000423_1&hsa-miR-125b-5p&offsets\|0\|-3(+1C);m-68&21\|+\|16590253\|16590274&offsets\|0\|-3(+1C)]\|TCCCTGAGACCCTAACTTGC | >MI0000446\|hsa-mir-125b-1&WithFlank&11\|-\|122099751\|122099850 | 21 | 39 | 19(+1C) | TCCCTGAGACCCTAACTTGC | 319.8 | -6.9 | 2.0 | -3.5 | 4.8E-04 | 2.6E-03 | BT20/MB231 | MITO |
| >am_trnaMT_SerGCT_MT_+_12207_12265@12.32.21\|CAAGAACTGCTAACTCATGCC | >am_trnaMT_SerGCT_MT_+_12207_12265 | 12 | 32 | 21 | CAAGAACTGCTAACTCATGCC | 338.5 | 2.7 | 0.8 | 3.5 | 4.3E-04 | 2.6E-03 | BT20/MB231 | CYTO |
| >hg38_21_+_8433172_8446622_RNA45SN1_45S_50ntFlanks@5490.5532.43\|TGAGAAGACGGTCGAACTTGACTATCTAGAGGAAGTAAAAGTC | >hg38_21_+_8433172_8446622_RNA45SN1_45S_50ntFlanks | 5490 | 5532 | 43 | TGAGAAGACGGTCGAACTTGACTATCTAGAGGAAGTAAAAGTC | 2247.4 | -3.7 | 1.1 | -3.5 | 4.3E-04 | 2.6E-03 | BT20/MB231 | CYTO |
| >hg38_21_+_8433172_8446622_RNA45SN1_45S_50ntFlanks@10846.10864.19\|CTTCGGGATAAGGATTGGC | >hg38_21_+_8433172_8446622_RNA45SN1_45S_50ntFlanks | 10846 | 10864 | 19 | CTTCGGGATAAGGATTGGC | 299.3 | -5.5 | 1.7 | -3.3 | 1.1E-03 | 2.6E-03 | BT20/MB468 | MP |
| >MI0000077\|hsa-mir-21&WithFlank&17\|+\|59841260\|59841343@14.35.22(+1U)&[MIMAT0000076&hsa-miR-21-5p&offsets\|0\|0(+1U);m-1&17\|+\|59841273\|59841294&offsets\|0\|0(+1U)]\|TAGCTTATCAGACTGATGTTGAT | >MI0000077\|hsa-mir-21&WithFlank&17\|+\|59841260\|59841343 | 14 | 35 | 22(+1U) | TAGCTTATCAGACTGATGTTGAT | 132239.1 | -3.5 | 1.0 | -3.6 | 3.6E-04 | 2.6E-03 | BT20/MB231 | MP |
| >hg38_21_+_8433172_8446622_RNA45SN1_45S_50ntFlanks@11670.11692.23\|CCCAGTGCTCTGAATGTCAAAGT | >hg38_21_+_8433172_8446622_RNA45SN1_45S_50ntFlanks | 11670 | 11692 | 23 | CCCAGTGCTCTGAATGTCAAAGT | 657.5 | 5.2 | 1.4 | 3.6 | 3.6E-04 | 2.6E-03 | BT20/MB231 | MP |
| >hg38_21_+_8433172_8446622_RNA45SN1_45S_50ntFlanks@9504.9531.28\|GTGAACTATGCCTGGGCAGGGCGAAGCC | >hg38_21_+_8433172_8446622_RNA45SN1_45S_50ntFlanks | 9504 | 9531 | 28 | GTGAACTATGCCTGGGCAGGGCGAAGCC | 262.1 | -5.4 | 1.7 | -3.3 | 1.1E-03 | 2.6E-03 | BT20/MB468 | MP |
| >MI0000650\|hsa-mir-200c&WithFlank&12\|+\|6963693\|6963772@11.32.22&[MIMAT0004657&hsa-miR-200c-5p&offsets\|0\|0;m-483&12\|+\|6963703\|6963724&offsets\|0\|0]\|CGTCTTACCCAGCAGTGTTTGG | >MI0000650\|hsa-mir-200c&WithFlank&12\|+\|6963693\|6963772 | 11 | 32 | 22 | CGTCTTACCCAGCAGTGTTTGG | 36.5 | 8.5 | 2.4 | 3.6 | 3.6E-04 | 2.6E-03 | BT20/MB231 | MP |
| >hg38_21_+_8433172_8446622_RNA45SN1_45S_50ntFlanks@11918.11943.26\|ACATGAGAGGTGTAGAATAAGTGGGA | >hg38_21_+_8433172_8446622_RNA45SN1_45S_50ntFlanks | 11918 | 11943 | 26 | ACATGAGAGGTGTAGAATAAGTGGGA | 90.5 | 9.8 | 2.8 | 3.6 | 3.6E-04 | 2.6E-03 | BT20/MB231 | MP |
| >trnaMT_GluTTC_MT_-_14674_14742@45.70.26,trnalookalike8_GluTTC_5_-_93905172_93905240@45.70.26\|CGTGGTTGTAGTCCGTGCGAGAATAC | >trnaMT_GluTTC_MT_-_14674_14742 | 45 | 70 | 26 | CGTGGTTGTAGTCCGTGCGAGAATAC | 216.5 | 5.7 | 1.7 | 3.3 | 1.1E-03 | 2.6E-03 | BT20/MB468 | MP |
| >trna119_LysCTT_1_-_145395522_145395594@1.31.31,trna11_LysCTT_5_-_180648979_180649051@1.31.31,trna13_LysCTT_6_+_26556774_26556846@1.31.31,trna32_LysCTT_16_-_3207406_3207478@1.31.31,trna7_LysCTT_16_+_3225692_3225764@1.31.31,trna9_LysCTT_5_+_180634755_180634827@1.31.31\|GCCCGGCTAGCTCAGTCGGTAGAGCATGAGA | >trna119_LysCTT_1_-_145395522_145395594 | 1 | 31 | 31 | GCCCGGCTAGCTCAGTCGGTAGAGCATGAGA | 1204.8 | -4.6 | 1.4 | -3.3 | 1.1E-03 | 2.6E-03 | BT20/MB468 | MP |
| >hg38_21_+_8433172_8446622_RNA45SN1_45S_50ntFlanks@8104.8139.36\|CCGCGGCGGGGCGCGGGACATGTGGCGTACGGAAGA | >hg38_21_+_8433172_8446622_RNA45SN1_45S_50ntFlanks | 8104 | 8139 | 36 | CCGCGGCGGGGCGCGGGACATGTGGCGTACGGAAGA | 1520.6 | 6.3 | 1.8 | 3.6 | 3.6E-04 | 2.6E-03 | BT20/MB231 | MP |
| >hg38_21_+_8433172_8446622_RNA45SN1_45S_50ntFlanks@6783.6803.21\|GGCTACGCCTGTCTGAGCGTC | >hg38_21_+_8433172_8446622_RNA45SN1_45S_50ntFlanks | 6783 | 6803 | 21 | GGCTACGCCTGTCTGAGCGTC | 174.9 | -5.2 | 1.6 | -3.3 | 1.1E-03 | 2.6E-03 | BT20/MB468 | MP |
| >trnaMT_GluTTC_MT_-_14674_14742@45.63.19,trnalookalike8_GluTTC_5_-_93905172_93905240@45.63.19\|CGTGGTTGTAGTCCGTGCG | >trnaMT_GluTTC_MT_-_14674_14742 | 45 | 63 | 19 | CGTGGTTGTAGTCCGTGCG | 226.0 | 5.6 | 1.7 | 3.3 | 1.1E-03 | 2.6E-03 | BT20/MB468 | MP |
| >hg38_21_+_8433172_8446622_RNA45SN1_45S_50ntFlanks@485.524.40\|GATGTGGTGACGTCGTGCTCTCCCGGGCCGGGTCCGAGCC | >hg38_21_+_8433172_8446622_RNA45SN1_45S_50ntFlanks | 485 | 524 | 40 | GATGTGGTGACGTCGTGCTCTCCCGGGCCGGGTCCGAGCC | 237.1 | 11.1 | 3.6 | 3.1 | 2.2E-03 | 2.6E-03 | MB231/MB468 | MITO |
| >hg38_21_+_8433172_8446622_RNA45SN1_45S_50ntFlanks@10798.10826.29\|CATGTTGGAACAATGTAGGTAAGGGAAGT | >hg38_21_+_8433172_8446622_RNA45SN1_45S_50ntFlanks | 10798 | 10826 | 29 | CATGTTGGAACAATGTAGGTAAGGGAAGT | 516.4 | -4.2 | 1.3 | -3.3 | 1.1E-03 | 2.6E-03 | BT20/MB468 | MP |
| >MI0000734\|hsa-mir-106b&WithFlank&7\|-\|100093987\|100094080@56.77.22&[MIMAT0004672&hsa-miR-106b-3p&offsets\|-2\|-2;m-87&7\|-\|100094003\|100094023&offsets\|-2\|-1]\|TACCGCACTGTGGGTACTTGCT | >MI0000734\|hsa-mir-106b&WithFlank&7\|-\|100093987\|100094080 | 56 | 77 | 22 | TACCGCACTGTGGGTACTTGCT | 505.2 | 5.4 | 1.7 | 3.1 | 1.9E-03 | 2.6E-03 | MB231/MB468 | MP |

| | | | | | | | | | | | | | |
|---|---|---|---|---|---|---|---|---|---|---|---|---|---|
| >hg38_21_+_8433172_8446622_RNA45SN1_45S_50ntFlanks@5473.5498.26\|CGGCCCTGGCGGAGCGCTGAGAAGAC | >hg38_21_+_8433172_8446622_RNA45SN1_45S_50ntFlanks | 5473 | 5498 | 26 | CGGCCCTGGCGGAGCGCTGAGAAGAC | 107.6 | -5.0 | 1.6 | -3.1 | 1.9E-03 | 2.6E-03 | MB231/MB468 | MP |
| >hg38_21_+_8433172_8446622_RNA45SN1_45S_50ntFlanks@9308.9325.18\|AGGAGTCTAACACGTGCG | >hg38_21_+_8433172_8446622_RNA45SN1_45S_50ntFlanks | 9308 | 9325 | 18 | AGGAGTCTAACACGTGCG | 39.9 | -8.4 | 2.6 | -3.3 | 1.1E-03 | 2.6E-03 | BT20/MB468 | MP |
| >hg38_21_+_8433172_8446622_RNA45SN1_45S_50ntFlanks@4560.4589.30\|CTAGGAATAATGGAATAGGACCGCGGTTCT | >hg38_21_+_8433172_8446622_RNA45SN1_45S_50ntFlanks | 4560 | 4589 | 30 | CTAGGAATAATGGAATAGGACCGCGGTTCT | 276.6 | -2.2 | 0.7 | -3.1 | 2.0E-03 | 2.6E-03 | BT20/MB468 | CYTO |
| >am_trnaMT_ProTGG_MT_-_15956_16023@53.71.19\|AAGACTTTTTCTCTGACCA | >am_trnaMT_ProTGG_MT_-_15956_16023 | 53 | 71 | 19 | AAGACTTTTTCTCTGACCA | 2984.5 | 3.2 | 1.0 | 3.1 | 2.0E-03 | 2.6E-03 | BT20/MB468 | CYTO |
| >trnaMT_SerGCT_MT_+_12207_12265@10.30.21\|CACAAGAACTGCTAACTCATG | >trnaMT_SerGCT_MT_+_12207_12265 | 10 | 30 | 21 | CACAAGAACTGCTAACTCATG | 124.8 | -3.9 | 1.3 | -3.1 | 1.9E-03 | 2.6E-03 | MB231/MB468 | MP |
| >hg38_21_+_8433172_8446622_RNA45SN1_45S_50ntFlanks@6648.6675.28\|GTACGACTCTTAGCGGTGGATCACTCGG | >hg38_21_+_8433172_8446622_RNA45SN1_45S_50ntFlanks | 6648 | 6675 | 28 | GTACGACTCTTAGCGGTGGATCACTCGG | 683.1 | 2.6 | 0.8 | 3.1 | 2.2E-03 | 2.6E-03 | MB231/MB468 | NUC |
| >hg38_21_+_8433172_8446622_RNA45SN1_45S_50ntFlanks@5051.5081.31\|TGGTGGAGCGATTTGTCTGGTTAATTCCGAT | >hg38_21_+_8433172_8446622_RNA45SN1_45S_50ntFlanks | 5051 | 5081 | 31 | TGGTGGAGCGATTTGTCTGGTTAATTCCGAT | 490.6 | 4.1 | 1.1 | 3.6 | 3.7E-04 | 2.6E-03 | BT20/MB231 | MP |
| >hg38_1_-_228634819_228635039_RNA5S12_5S_50ntFlanks@152.170.19\|TACCGGGTGCTGTAGGCTT | >hg38_1_-_228634819_228635039_RNA5S12_5S_50ntFlanks | 152 | 170 | 19 | TACCGGGTGCTGTAGGCTT | 2009.7 | 1.7 | 0.6 | 3.1 | 1.7E-03 | 2.6E-03 | BT20/MB468 | MITO |
| >hg38_21_+_8433172_8446622_RNA45SN1_45S_50ntFlanks@11209.11255.47\|CGGCGACTCTGGACGCGAGCCGGGCCCTTCCCGTGGATCGCCCCAGC | >hg38_21_+_8433172_8446622_RNA45SN1_45S_50ntFlanks | 11209 | 11255 | 47 | CGGCGACTCTGGACGCGAGCCGGGCCCTTCCCGTGGATCGCCCCAGC | 310.0 | 3.8 | 1.1 | 3.5 | 4.9E-04 | 2.6E-03 | BT20/MB231 | MITO |
| >hg38_21_+_8433172_8446622_RNA45SN1_45S_50ntFlanks@11580.11613.34\|GAACTGGTGCGGACCAGGGGAATCCGACTGTTTA | >hg38_21_+_8433172_8446622_RNA45SN1_45S_50ntFlanks | 11580 | 11613 | 34 | GAACTGGTGCGGACCAGGGGAATCCGACTGTTTA | 2963.6 | 2.8 | 0.8 | 3.5 | 4.9E-04 | 2.6E-03 | BT20/MB231 | MITO |
| >hg38_21_+_8433172_8446622_RNA45SN1_45S_50ntFlanks@8342.8375.34\|CGTAAGGGAAAGTTGAAAAGAACTTTGAAGAGAG | >hg38_21_+_8433172_8446622_RNA45SN1_45S_50ntFlanks | 8342 | 8375 | 34 | CGTAAGGGAAAGTTGAAAAGAACTTTGAAGAGAG | 513.5 | 4.0 | 1.1 | 3.5 | 4.9E-04 | 2.6E-03 | BT20/MB231 | MITO |
| >MI0000446\|hsa-mir-125b-1&WithFlank&11\|-\|122099751\|122099850@21.41.21;MI0000470\|hsa-mir-125b-2&WithFlank&21\|+\|16590231\|16590331@23.43.21&[MIMAT0000423&hsa-miR-125b-5p&offsets\|0\|-1;m-67&11\|-\|122099809\|122099830&offsets\|0\|-1];[MIMAT0000423_1&hsa-miR-125b-5p&offsets\|0\|-1;m-68&21\|+\|16590253\|16590274&offsets\|0\|-1]\|TCCCTGAGACCCTAACTTGTG | >MI0000446\|hsa-mir-125b-1&WithFlank&11\|-\|122099751\|122099850 | 21 | 41 | 21 | TCCCTGAGACCCTAACTTGTG | 9988.8 | -6.7 | 1.9 | -3.5 | 4.9E-04 | 2.6E-03 | BT20/MB231 | MITO |
| >MI0000825\|hsa-mir-345&WithFlank&14\|+\|100307853\|100307962@24.45.22(+1U)&[MIMAT0000772&hsa-miR-345-5p&offsets\|0\|0(+1U);m-193&14\|+\|100307876\|100307896&offsets\|0\|+1(+1U)]\|GCTGACTCCTAGTCCAGGGCTCT | >MI0000825\|hsa-mir-345&WithFlank&14\|+\|100307853\|100307962 | 24 | 45 | 22(+1U) | GCTGACTCCTAGTCCAGGGCTCT | 379.2 | -3.1 | 0.9 | -3.5 | 4.9E-04 | 2.6E-03 | BT20/MB231 | MITO |
| >hg38_21_+_8433172_8446622_RNA45SN1_45S_50ntFlanks@11200.11225.26\|CGGCGGCGGCGGCGACTCTGGACGCG | >hg38_21_+_8433172_8446622_RNA45SN1_45S_50ntFlanks | 11200 | 11225 | 26 | CGGCGGCGGCGGCGACTCTGGACGCG | 1325.5 | -5.2 | 1.7 | -3.1 | 2.0E-03 | 2.6E-03 | BT20/MB468 | CYTO |
| >hg38_21_+_8433172_8446622_RNA45SN1_45S_50ntFlanks@11670.11701.32\|CCCAGTGCTCTGAATGTCAAAGTGAAGAAATT | >hg38_21_+_8433172_8446622_RNA45SN1_45S_50ntFlanks | 11670 | 11701 | 32 | CCCAGTGCTCTGAATGTCAAAGTGAAGAAATT | 462.5 | 6.4 | 1.8 | 3.6 | 3.7E-04 | 2.6E-03 | BT20/MB231 | MP |
| >hg38_21_+_8433172_8446622_RNA45SN1_45S_50ntFlanks@12455.12476.22\|ACCAAGCGTTGGATTGTTCACC | >hg38_21_+_8433172_8446622_RNA45SN1_45S_50ntFlanks | 12455 | 12476 | 22 | ACCAAGCGTTGGATTGTTCACC | 591.5 | -3.2 | 1.0 | -3.3 | 1.1E-03 | 2.6E-03 | BT20/MB468 | MP |
| >hg38_21_+_8433172_8446622_RNA45SN1_45S_50ntFlanks@9927.9946.20\|TTGATATAGACAGCAGGACG | >hg38_21_+_8433172_8446622_RNA45SN1_45S_50ntFlanks | 9927 | 9946 | 20 | TTGATATAGACAGCAGGACG | 98.3 | 6.3 | 1.9 | 3.3 | 1.1E-03 | 2.6E-03 | BT20/MB468 | MP |
| >hg38_21_+_8433172_8446622_RNA45SN1_45S_50ntFlanks@4160.4186.27\|CCACATCCAAGGAAGGCAGCAGGCGCG | >hg38_21_+_8433172_8446622_RNA45SN1_45S_50ntFlanks | 4160 | 4186 | 27 | CCACATCCAAGGAAGGCAGCAGGCGCG | 178.5 | -4.5 | 1.4 | -3.3 | 1.1E-03 | 2.6E-03 | BT20/MB468 | MP |
| >hg38_21_+_8433172_8446622_RNA45SN1_45S_50ntFlanks@12251.12272.22\|AAAGCTCGCTTGATCTTGATTT | >hg38_21_+_8433172_8446622_RNA45SN1_45S_50ntFlanks | 12251 | 12272 | 22 | AAAGCTCGCTTGATCTTGATTT | 105.3 | -5.0 | 1.6 | -3.1 | 1.9E-03 | 2.6E-03 | MB231/MB468 | MP |
| >hg38_21_+_8433172_8446622_RNA45SN1_45S_50ntFlanks@12184.12209.26\|AACGGTAACGCAGGTGTCCTAAGGCG | >hg38_21_+_8433172_8446622_RNA45SN1_45S_50ntFlanks | 12184 | 12209 | 26 | AACGGTAACGCAGGTGTCCTAAGGCG | 87.6 | -5.1 | 1.6 | -3.1 | 1.9E-03 | 2.6E-03 | MB231/MB468 | MP |
| >hg38_21_+_8433172_8446622_RNA45SN1_45S_50ntFlanks@10514.10532.19\|CGGAGTGGCGGAGATGGGC | >hg38_21_+_8433172_8446622_RNA45SN1_45S_50ntFlanks | 10514 | 10532 | 19 | CGGAGTGGCGGAGATGGGC | 3330.6 | 7.9 | 2.2 | 3.6 | 3.7E-04 | 2.6E-03 | BT20/MB231 | MP |
| >hg38_21_+_8433172_8446622_RNA45SN1_45S_50ntFlanks@6650.6674.25\|ACGACTCTTAGCGGTGGATCACTCG | >hg38_21_+_8433172_8446622_RNA45SN1_45S_50ntFlanks | 6650 | 6674 | 25 | ACGACTCTTAGCGGTGGATCACTCG | 1399.7 | 5.0 | 1.4 | 3.6 | 3.7E-04 | 2.6E-03 | BT20/MB231 | MP |
| >hg38_21_+_8433172_8446622_RNA45SN1_45S_50ntFlanks@9931.9949.19\|TATAGACAGCAGGACGGTG | >hg38_21_+_8433172_8446622_RNA45SN1_45S_50ntFlanks | 9931 | 9949 | 19 | TATAGACAGCAGGACGGTG | 226.6 | 3.8 | 1.1 | 3.6 | 3.7E-04 | 2.6E-03 | BT20/MB231 | MP |
| >MI0000273\|hsa-mir-183&WithFlank&7\|-\|129774899\|129775020@34.55.22&[MIMAT0000261&hsa-miR-183-5p&offsets\|+1\|+1;m-41&7\|-\|129774966\|129774987&offsets\|0\|0]\|ATGGCACTGGTAGAATTCACTG | >MI0000273\|hsa-mir-183&WithFlank&7\|-\|129774899\|129775020 | 34 | 55 | 22 | ATGGCACTGGTAGAATTCACTG | 345.9 | 2.6 | 0.8 | 3.1 | 2.2E-03 | 2.6E-03 | MB231/MB468 | MITO |
| >trnaMT_ValTAC_MT_+_1602_1670@35.72.38\|ACTTAGGAGATTTCAACTTAACTTGACCGCTCTGACCA | >trnaMT_ValTAC_MT_+_1602_1670 | 35 | 72 | 38 | ACTTAGGAGATTTCAACTTAACTTGACCGCTCTGACCA | 232.4 | 4.3 | 1.4 | 3.1 | 2.2E-03 | 2.6E-03 | MB231/MB468 | MITO |
| >hg38_21_+_8433172_8446622_RNA45SN1_45S_50ntFlanks@11580.11613.34\|GAACTGGTGCGGACCAGGGGAATCCGACTGTTTA | >hg38_21_+_8433172_8446622_RNA45SN1_45S_50ntFlanks | 11580 | 11613 | 34 | GAACTGGTGCGGACCAGGGGAATCCGACTGTTTA | 1787.1 | 1.8 | 0.6 | 3.1 | 2.0E-03 | 2.6E-03 | BT20/MB468 | CYTO |
| >hg38_21_+_8433172_8446622_RNA45SN1_45S_50ntFlanks@7979.8011.33\|ACCTCAGATCAGACGTGGCGACCCGCTGAATTT | >hg38_21_+_8433172_8446622_RNA45SN1_45S_50ntFlanks | 7979 | 8011 | 33 | ACCTCAGATCAGACGTGGCGACCCGCTGAATTT | 1256.2 | 2.3 | 0.7 | 3.1 | 1.7E-03 | 2.6E-03 | BT20/MB468 | MITO |
| >hg38_21_+_8433172_8446622_RNA45SN1_45S_50ntFlanks@12007.12031.25\|GCCGCCGGTGAAATACCACTACTCT | >hg38_21_+_8433172_8446622_RNA45SN1_45S_50ntFlanks | 12007 | 12031 | 25 | GCCGCCGGTGAAATACCACTACTCT | 1492.8 | -2.5 | 0.8 | -3.1 | 2.2E-03 | 2.6E-03 | MB231/MB468 | NUC |
| >trnaMT_GluTTC_MT_-_14674_14742@27.56.30,trnalookalike8_GluTTC_5_-_93905172_93905240@27.56.30\|GTTTTTCATATCATTGGTCGTGGTTGTAGT | >trnaMT_GluTTC_MT_-_14674_14742 | 27 | 56 | 30 | GTTTTTCATATCATTGGTCGTGGTTGTAGT | 95.3 | -3.9 | 1.3 | -3.1 | 1.9E-03 | 2.6E-03 | MB231/MB468 | MP |
| >hg38_21_+_8433172_8446622_RNA45SN1_45S_50ntFlanks@11235.11268.34\|CTTCCCGTGGATCGCCCCAGCTGCGGCGGGCGTC | >hg38_21_+_8433172_8446622_RNA45SN1_45S_50ntFlanks | 11235 | 11268 | 34 | CTTCCCGTGGATCGCCCCAGCTGCGGCGGGCGTC | 309.2 | -2.1 | 0.7 | -3.1 | 1.7E-03 | 2.6E-03 | BT20/MB468 | MITO |
| >hg38_21_+_8433172_8446622_RNA45SN1_45S_50ntFlanks@11804.11821.18\|CCCACTGTCCCTACCTAC | >hg38_21_+_8433172_8446622_RNA45SN1_45S_50ntFlanks | 11804 | 11821 | 18 | CCCACTGTCCCTACCTAC | 196.0 | -4.4 | 1.4 | -3.1 | 2.0E-03 | 2.6E-03 | BT20/MB468 | CYTO |
| >hg38_21_+_8433172_8446622_RNA45SN1_45S_50ntFlanks@11656.11687.32\|CGATGTGATTTCTGCCCAGTGCTCTGAATGTC | >hg38_21_+_8433172_8446622_RNA45SN1_45S_50ntFlanks | 11656 | 11687 | 32 | CGATGTGATTTCTGCCCAGTGCTCTGAATGTC | 627.5 | 6.2 | 1.8 | 3.6 | 3.7E-04 | 2.6E-03 | BT20/MB231 | MP |
| >hg38_21_+_8433172_8446622_RNA45SN1_45S_50ntFlanks@6776.6807.32\|CTCCCGGGGCTACGCCTGTCTGAGCGTCGCTT | >hg38_21_+_8433172_8446622_RNA45SN1_45S_50ntFlanks | 6776 | 6807 | 32 | CTCCCGGGGCTACGCCTGTCTGAGCGTCGCTT | 94.0 | -2.6 | 0.8 | -3.1 | 1.9E-03 | 2.6E-03 | MB231/MB468 | MP |
| >hg38_21_+_8433172_8446622_RNA45SN1_45S_50ntFlanks@7981.8003.23\|CTCAGATCAGACGTGGCGACCCG | >hg38_21_+_8433172_8446622_RNA45SN1_45S_50ntFlanks | 7981 | 8003 | 23 | CTCAGATCAGACGTGGCGACCCG | 341.1 | -4.9 | 1.6 | -3.1 | 2.2E-03 | 2.7E-03 | MB231/MB468 | NUC |
| >hg38_21_+_8433172_8446622_RNA45SN1_45S_50ntFlanks@6671.6695.25\|CTCGGCTCGTGCGTCGATGAAGAAC | >hg38_21_+_8433172_8446622_RNA45SN1_45S_50ntFlanks | 6671 | 6695 | 25 | CTCGGCTCGTGCGTCGATGAAGAAC | 129.4 | -2.6 | 0.8 | -3.3 | 1.1E-03 | 2.7E-03 | BT20/MB468 | MP |
| >hg38_21_+_8433172_8446622_RNA45SN1_45S_50ntFlanks@6650.6673.24\|ACGACTCTTAGCGGTGGATCACTC | >hg38_21_+_8433172_8446622_RNA45SN1_45S_50ntFlanks | 6650 | 6673 | 24 | ACGACTCTTAGCGGTGGATCACTC | 551.4 | 3.0 | 1.0 | 3.1 | 2.2E-03 | 2.7E-03 | MB231/MB468 | MITO |
| >hg38_21_+_8433172_8446622_RNA45SN1_45S_50ntFlanks@3777.3801.25\|CGGTACAGTGAAACTGCGAATGGCT | >hg38_21_+_8433172_8446622_RNA45SN1_45S_50ntFlanks | 3777 | 3801 | 25 | CGGTACAGTGAAACTGCGAATGGCT | 604.0 | -1.8 | 0.6 | -3.1 | 2.1E-03 | 2.7E-03 | BT20/MB468 | CYTO |
| >hg38_21_+_8433172_8446622_RNA45SN1_45S_50ntFlanks@475.511.37\|CTTCGTGATCGATGTGGTGACGTCGTGCTCTCCCGGG | >hg38_21_+_8433172_8446622_RNA45SN1_45S_50ntFlanks | 475 | 511 | 37 | CTTCGTGATCGATGTGGTGACGTCGTGCTCTCCCGGG | 119.9 | 6.5 | 2.0 | 3.3 | 1.1E-03 | 2.7E-03 | BT20/MB468 | MP |
| >hg38_21_+_8433172_8446622_RNA45SN1_45S_50ntFlanks@10353.10377.25\|AGAACTTTGAAGGCCGAAGTGGAGA | >hg38_21_+_8433172_8446622_RNA45SN1_45S_50ntFlanks | 10353 | 10377 | 25 | AGAACTTTGAAGGCCGAAGTGGAGA | 173.5 | -3.7 | 1.1 | -3.3 | 1.1E-03 | 2.7E-03 | BT20/MB468 | MP |
| >am_MI0003581\|hsa-mir-574&WithFlank&4\|+\|38868026\|38868133@31.52.22(+1A)&[MIMAT0004795&hsa-miR-574-5p&offsets\|0\|-1(+1A);m-213&4\|+\|38868056\|38868078&offsets\|0\|-1(+1A)]\|TGAGTGTGTGTGTGTGAGTGTGA | >am_MI0003581\|hsa-mir-574&WithFlank&4\|+\|38868026\|38868133 | 31 | 52 | 22(+1A) | TGAGTGTGTGTGTGTGAGTGTGA | 114.8 | 3.8 | 1.2 | 3.1 | 1.9E-03 | 2.7E-03 | MB231/MB468 | MP |
| >hg38_21_+_8433172_8446622_RNA45SN1_45S_50ntFlanks@10851.10879.29\|GGATAAGGATTGGCTCTAAGGGCTGGGTC | >hg38_21_+_8433172_8446622_RNA45SN1_45S_50ntFlanks | 10851 | 10879 | 29 | GGATAAGGATTGGCTCTAAGGGCTGGGTC | 39.3 | 9.0 | 2.8 | 3.3 | 1.1E-03 | 2.7E-03 | BT20/MB468 | MP |
| >trna32_LysCTT_16_-_3207406_3207478@4.33.30\|CGGCTAGCTCAGTCGGTAGAGCATGAGACC | >trna32_LysCTT_16_-_3207406_3207478 | 4 | 33 | 30 | CGGCTAGCTCAGTCGGTAGAGCATGAGACC | 86.4 | 5.8 | 1.9 | 3.1 | 1.9E-03 | 2.7E-03 | MB231/MB468 | MP |
| >trna10_AspGTC_12_-_125424193_125424264@31.73.43,trna12_AspGTC_12_-_125411891_125411962@31.73.43,trna38_AspGTC_17_-_8125556_8125627@31.73.43,trna45_AspGTC_6_+_27447453_27447524@31.73.43,trna48_AspGTC_6_+_27471523_27471594@31.73.43,trna4_AspGTC_12_+_96429799_96429870@31.73.43,trna69_AspGTC_1_-_161440205_161440276@31.73.43,trna72_AspGTC_1_-_161432824_161432895@31.73.43,trna75_AspGTC_1_-_161425414_161425485@31.73.43,trna78_AspGTC_1_-_161418033_161418104@31.73.43,trna81_AspGTC_1_-_161410615_161410686@31.73.43\|CCTGTCACGCGGGAGACCGGGGTTCGATTCCCCGACGGGGAGC | >trna10_AspGTC_12_-_125424193_125424264 | 31 | 73 | 43 | CCTGTCACGCGGGAGACCGGGGTTCGATTCCCCGACGGGGAGC | 208.0 | -3.4 | 1.1 | -3.1 | 2.1E-03 | 2.7E-03 | BT20/MB468 | CYTO |
| >hg38_21_+_8433172_8446622_RNA45SN1_45S_50ntFlanks@7983.8005.23\|CAGATCAGACGTGGCGACCCGCT | >hg38_21_+_8433172_8446622_RNA45SN1_45S_50ntFlanks | 7983 | 8005 | 23 | CAGATCAGACGTGGCGACCCGCT | 255.6 | -3.5 | 1.1 | -3.1 | 2.1E-03 | 2.7E-03 | BT20/MB468 | CYTO |
| >hg38_21_+_8433172_8446622_RNA45SN1_45S_50ntFlanks@12419.12454.36\|GATGTCGGCTCTTCCTATCATTGTGAAGCAGAATTC | >hg38_21_+_8433172_8446622_RNA45SN1_45S_50ntFlanks | 12419 | 12454 | 36 | GATGTCGGCTCTTCCTATCATTGTGAAGCAGAATTC | 413.6 | -2.9 | 0.8 | -3.5 | 4.1E-04 | 2.7E-03 | BT20/MB231 | NUC |
| >MI0000734\|hsa-mir-106b&WithFlank&7\|-\|100093987\|100094080@56.76.21&[MIMAT0004672&hsa-miR-106b-3p&offsets\|-2\|-3;m-87&7\|-\|100094003\|100094023&offsets\|-2\|-2]\|TACCGCACTGTGGGTACTTGC | >MI0000734\|hsa-mir-106b&WithFlank&7\|-\|100093987\|100094080 | 56 | 76 | 21 | TACCGCACTGTGGGTACTTGC | 55.2 | 5.7 | 1.8 | 3.1 | 1.9E-03 | 2.7E-03 | MB231/MB468 | MP |
| >hg38_MT_+_1621_3279_RNR2_16S_50ntFlanks@292.311.20\|AGACGAGCTACCTAAGAACA | >hg38_MT_+_1621_3279_RNR2_16S_50ntFlanks | 292 | 311 | 20 | AGACGAGCTACCTAAGAACA | 889.6 | 4.3 | 1.3 | 3.3 | 1.1E-03 | 2.7E-03 | BT20/MB468 | MP |
| >hg38_21_+_8433172_8446622_RNA45SN1_45S_50ntFlanks@11534.11551.18\|GTTCCGCGCGGCGCCTCG | >hg38_21_+_8433172_8446622_RNA45SN1_45S_50ntFlanks | 11534 | 11551 | 18 | GTTCCGCGCGGCGCCTCG | 101.0 | -5.1 | 1.6 | -3.3 | 1.1E-03 | 2.7E-03 | BT20/MB468 | MP |
| >hg38_21_+_8433172_8446622_RNA45SN1_45S_50ntFlanks@11876.11906.31\|AAGACCCTGTTGAGCTTGACTCTAGTCTGGC | >hg38_21_+_8433172_8446622_RNA45SN1_45S_50ntFlanks | 11876 | 11906 | 31 | AAGACCCTGTTGAGCTTGACTCTAGTCTGGC | 356.5 | -4.3 | 1.3 | -3.3 | 1.1E-03 | 2.7E-03 | BT20/MB468 | MP |

<table>
<tr><td>>hg38_21_+_8433172_8446622_RNA45SN1_45S_50ntFlanks@4728.4748.21|ATCAAGAACGAAAGTCGGAGG</td><td>>hg38_21_+_8433172_8446622_RNA45SN1_45S_50ntFlanks</td><td>4728</td><td>4748</td><td>21</td><td>ATCAAGAACGAAAGTCGGAGG</td><td>638.1</td><td>-5.4</td><td>1.7</td><td>-3.1</td><td>1.9E-03</td><td>2.7E-03</td><td>MB231/MB468</td><td>MP</td></tr>
<tr><td>>trnaMT_GluTTC_MT_-_14674_14742@26.71.46,trnalookalike8_GluTTC_5_-_93905172_93905240@26.71.46|GGTTTTTCATATCATTGGTCGTGGTTGTAGTCCGTGCGAGAATACC</td><td>>trnaMT_GluTTC_MT_-_14674_14742</td><td>26</td><td>71</td><td>46</td><td>GGTTTTTCATATCATTGGTCGTGGTTGTAGTCCGTGCGAGAATACC</td><td>147.0</td><td>5.8</td><td>1.9</td><td>3.1</td><td>2.2E-03</td><td>2.7E-03</td><td>MB231/MB468</td><td>MITO</td></tr>
<tr><td>>MI0000066|hsa-let-7e&WithFlank&19|+|51692780|51692870@14.31.18&[MIMAT0000066&hsa-let-7e-5p&offsets|0|-4;m-50&19|+|51692793|51692814&offsets|0|-4]|TGAGGTAGGAGGTTGTAT</td><td>>MI0000066|hsa-let-7e&WithFlank&19|+|51692780|51692870</td><td>14</td><td>31</td><td>18</td><td>TGAGGTAGGAGGTTGTAT</td><td>157.4</td><td>5.4</td><td>1.7</td><td>3.3</td><td>1.1E-03</td><td>2.7E-03</td><td>BT20/MB468</td><td>MP</td></tr>
<tr><td>>hg38_21_+_8433172_8446622_RNA45SN1_45S_50ntFlanks@8379.8402.24|TCAAGAGGGCGTGAAACCGTTAAG</td><td>>hg38_21_+_8433172_8446622_RNA45SN1_45S_50ntFlanks</td><td>8379</td><td>8402</td><td>24</td><td>TCAAGAGGGCGTGAAACCGTTAAG</td><td>202.1</td><td>-3.4</td><td>1.1</td><td>-3.1</td><td>1.7E-03</td><td>2.7E-03</td><td>BT20/MB468</td><td>MITO</td></tr>
<tr><td>>hg38_21_+_8433172_8446622_RNA45SN1_45S_50ntFlanks@6652.6697.46|GACTCTTAGCGGTGGATCACTCGGCTCGTGCGTCGATGAAGAACGC</td><td>>hg38_21_+_8433172_8446622_RNA45SN1_45S_50ntFlanks</td><td>6652</td><td>6697</td><td>46</td><td>GACTCTTAGCGGTGGATCACTCGGCTCGTGCGTCGATGAAGAACGC</td><td>1692.3</td><td>-3.0</td><td>1.0</td><td>-3.1</td><td>2.2E-03</td><td>2.7E-03</td><td>BT20/MB468</td><td>NUC</td></tr>
<tr><td>>MI0003581|hsa-mir-574&WithFlank&4|+|38868026|38868133@67.87.21&[MIMAT0003239&hsa-miR-574-3p&offsets|0|-1;m-149&4|+|38868092|38868113&offsets|0|-1]|CACGCTCATGCACACACCCAC</td><td>>MI0003581|hsa-mir-574&WithFlank&4|+|38868026|38868133</td><td>67</td><td>87</td><td>21</td><td>CACGCTCATGCACACACCCAC</td><td>243.6</td><td>-2.6</td><td>0.7</td><td>-3.5</td><td>5.0E-04</td><td>2.7E-03</td><td>BT20/MB231</td><td>MITO</td></tr>
<tr><td>>am_trna128_GlyGCC_6_-_27870686_27870756@3.31.29,trna133_GlyCCC_1_-_16872434_16872504@3.31.29,trna18_GlyGCC_16_+_70822597_70822667@3.31.29,trna19_GlyGCC_16_+_70823410_70823480@3.31.29,trna19_GlyGCC_2_-_157257659_157257729@3.31.29,trna24_GlyGCC_16_-_70812942_70813012@3.31.29,trna25_GlyGCC_16_-_70812114_70812184@3.31.29,trna4_GlyCCC_1_+_17188416_17188486@3.31.29,trna5_GlyGCC_17_+_8029064_8029134@3.31.29,trna68_GlyGCC_1_-_161493637_161493707@3.31.29|ATTGGTGGTTCAGTGGTAGAATTCTCGCC</td><td>>am_trna128_GlyGCC_6_-_27870686_27870756</td><td>3</td><td>31</td><td>29</td><td>ATTGGTGGTTCAGTGGTAGAATTCTCGCC</td><td>291.2</td><td>-3.7</td><td>1.1</td><td>-3.5</td><td>5.0E-04</td><td>2.7E-03</td><td>BT20/MB231</td><td>MITO</td></tr>
<tr><td>>hg38_21_+_8433172_8446622_RNA45SN1_45S_50ntFlanks@10735.10757.23|TGTAAATCTCGCGCCGGGCCGTA</td><td>>hg38_21_+_8433172_8446622_RNA45SN1_45S_50ntFlanks</td><td>10735</td><td>10757</td><td>23</td><td>TGTAAATCTCGCGCCGGGCCGTA</td><td>208.6</td><td>-3.6</td><td>1.2</td><td>-3.1</td><td>2.1E-03</td><td>2.7E-03</td><td>BT20/MB468</td><td>CYTO</td></tr>
<tr><td>>hg38_21_+_8433172_8446622_RNA45SN1_45S_50ntFlanks@9198.9218.21|GTTCTCTCGGGGCCACGCGCG</td><td>>hg38_21_+_8433172_8446622_RNA45SN1_45S_50ntFlanks</td><td>9198</td><td>9218</td><td>21</td><td>GTTCTCTCGGGGCCACGCGCG</td><td>522.4</td><td>-6.1</td><td>2.0</td><td>-3.1</td><td>2.2E-03</td><td>2.7E-03</td><td>BT20/MB468</td><td>NUC</td></tr>
<tr><td>>hg38_21_+_8433172_8446622_RNA45SN1_45S_50ntFlanks@11161.11178.18|CACGGTCCCCCGCGAGGG</td><td>>hg38_21_+_8433172_8446622_RNA45SN1_45S_50ntFlanks</td><td>11161</td><td>11178</td><td>18</td><td>CACGGTCCCCCGCGAGGG</td><td>1335.8</td><td>-5.8</td><td>1.8</td><td>-3.3</td><td>1.1E-03</td><td>2.7E-03</td><td>BT20/MB468</td><td>MP</td></tr>
<tr><td>>hg38_21_+_8433172_8446622_RNA45SN1_45S_50ntFlanks@11575.11594.20|ACTTAGAACTGGTGCGGACC</td><td>>hg38_21_+_8433172_8446622_RNA45SN1_45S_50ntFlanks</td><td>11575</td><td>11594</td><td>20</td><td>ACTTAGAACTGGTGCGGACC</td><td>2042.0</td><td>-1.9</td><td>0.6</td><td>-3.1</td><td>2.1E-03</td><td>2.7E-03</td><td>MB231/MB468</td><td>CYTO</td></tr>
<tr><td>>hg38_21_+_8433172_8446622_RNA45SN1_45S_50ntFlanks@10797.10817.21|GCATGTTGGAACAATGTAGGT</td><td>>hg38_21_+_8433172_8446622_RNA45SN1_45S_50ntFlanks</td><td>10797</td><td>10817</td><td>21</td><td>GCATGTTGGAACAATGTAGGT</td><td>701.5</td><td>-4.3</td><td>1.3</td><td>-3.3</td><td>1.1E-03</td><td>2.7E-03</td><td>BT20/MB468</td><td>MP</td></tr>
<tr><td>>hg38_21_+_8433172_8446622_RNA45SN1_45S_50ntFlanks@11670.11707.38|CCCAGTGCTCTGAATGTCAAAGTGAAGAAATTCAATGA</td><td>>hg38_21_+_8433172_8446622_RNA45SN1_45S_50ntFlanks</td><td>11670</td><td>11707</td><td>38</td><td>CCCAGTGCTCTGAATGTCAAAGTGAAGAAATTCAATGA</td><td>298.8</td><td>2.6</td><td>0.8</td><td>3.1</td><td>2.1E-03</td><td>2.7E-03</td><td>MB231/MB468</td><td>CYTO</td></tr>
<tr><td>>hg38_21_+_8433172_8446622_RNA45SN1_45S_50ntFlanks@8176.8203.28|CTTCTGATCGAGGCCCAGCCCGTGGACG</td><td>>hg38_21_+_8433172_8446622_RNA45SN1_45S_50ntFlanks</td><td>8176</td><td>8203</td><td>28</td><td>CTTCTGATCGAGGCCCAGCCCGTGGACG</td><td>3286.9</td><td>-2.4</td><td>0.8</td><td>-3.1</td><td>2.1E-03</td><td>2.7E-03</td><td>MB231/MB468</td><td>CYTO</td></tr>
<tr><td>>hg38_21_+_8433172_8446622_RNA45SN1_45S_50ntFlanks@13007.13024.18|TATTGAAAGTCAGCCCTC</td><td>>hg38_21_+_8433172_8446622_RNA45SN1_45S_50ntFlanks</td><td>13007</td><td>13024</td><td>18</td><td>TATTGAAAGTCAGCCCTC</td><td>190.1</td><td>-2.8</td><td>0.9</td><td>-3.1</td><td>2.1E-03</td><td>2.7E-03</td><td>MB231/MB468</td><td>CYTO</td></tr>
<tr><td>>hg38_21_+_8433172_8446622_RNA45SN1_45S_50ntFlanks@7975.8009.35|CGCGACCTCAGATCAGACGTGGCGACCCGCTGAAT</td><td>>hg38_21_+_8433172_8446622_RNA45SN1_45S_50ntFlanks</td><td>7975</td><td>8009</td><td>35</td><td>CGCGACCTCAGATCAGACGTGGCGACCCGCTGAAT</td><td>364130.0</td><td>3.2</td><td>0.9</td><td>3.5</td><td>4.2E-04</td><td>2.7E-03</td><td>BT20/MB231</td><td>NUC</td></tr>
<tr><td>>hg38_21_+_8433172_8446622_RNA45SN1_45S_50ntFlanks@5071.5102.32|TTAATTCCGATAACGAACGAGACTCTGGCATG</td><td>>hg38_21_+_8433172_8446622_RNA45SN1_45S_50ntFlanks</td><td>5071</td><td>5102</td><td>32</td><td>TTAATTCCGATAACGAACGAGACTCTGGCATG</td><td>193.2</td><td>-3.3</td><td>1.0</td><td>-3.1</td><td>1.7E-03</td><td>2.7E-03</td><td>BT20/MB468</td><td>MITO</td></tr>
<tr><td>>MI0000081|hsa-mir-24-2&WithFlank&19|-|13836281|13836365@56.76.21(+1C);MI0000080|hsa-mir-24-1&WithFlank&9|+|95086015|95086094@50.70.21(+1C)&[MIMAT0000080&hsa-miR-24-3p&offsets|0|-1(+1C);m-36&19|-|13836288|13836310&offsets|0|-2(+1C)];[MIMAT0000080_1&hsa-miR-24-3p&offsets|0|-1(+1C);m-35&9|+|95086064|95086086&offsets|0|-2(+1C)]|TGGCTCAGTTCAGCAGGAACAC</td><td>>MI0000081|hsa-mir-24-2&WithFlank&19|-|13836281|13836365</td><td>56</td><td>76</td><td>21(+1C)</td><td>TGGCTCAGTTCAGCAGGAACAC</td><td>310.8</td><td>4.9</td><td>1.5</td><td>3.3</td><td>1.1E-03</td><td>2.7E-03</td><td>BT20/MB468</td><td>MP</td></tr>
<tr><td>>hg38_21_+_8433172_8446622_RNA45SN1_45S_50ntFlanks@5237.5254.18|ACTGACTGGCTCAGCGTG</td><td>>hg38_21_+_8433172_8446622_RNA45SN1_45S_50ntFlanks</td><td>5237</td><td>5254</td><td>18</td><td>ACTGACTGGCTCAGCGTG</td><td>1435.2</td><td>-4.1</td><td>1.3</td><td>-3.1</td><td>2.3E-03</td><td>2.7E-03</td><td>MB231/MB468</td><td>MITO</td></tr>
<tr><td>>hg38_21_+_8433172_8446622_RNA45SN1_45S_50ntFlanks@12160.12180.21|TTGACTGGGGCGGTACACCTG</td><td>>hg38_21_+_8433172_8446622_RNA45SN1_45S_50ntFlanks</td><td>12160</td><td>12180</td><td>21</td><td>TTGACTGGGGCGGTACACCTG</td><td>407.0</td><td>-5.3</td><td>1.6</td><td>-3.3</td><td>1.1E-03</td><td>2.7E-03</td><td>BT20/MB468</td><td>MP</td></tr>
<tr><td>>trna104_AlaTGC_6_-_28785012_28785083@58.75.18,trna107_AlaTGC_6_-_28770577_28770647@57.74.18|TCCCCGACACCTCCACCA</td><td>>trna104_AlaTGC_6_-_28785012_28785083</td><td>58</td><td>75</td><td>18</td><td>TCCCCGACACCTCCACCA</td><td>215.4</td><td>3.0</td><td>0.9</td><td>3.5</td><td>4.4E-04</td><td>2.7E-03</td><td>BT20/MB231</td><td>CYTO</td></tr>
<tr><td>>hg38_21_+_8433172_8446622_RNA45SN1_45S_50ntFlanks@6649.6669.21|TACGACTCTTAGCGGTGGATC</td><td>>hg38_21_+_8433172_8446622_RNA45SN1_45S_50ntFlanks</td><td>6649</td><td>6669</td><td>21</td><td>TACGACTCTTAGCGGTGGATC</td><td>420.3</td><td>2.6</td><td>0.7</td><td>3.5</td><td>4.4E-04</td><td>2.7E-03</td><td>BT20/MB231</td><td>CYTO</td></tr>
<tr><td>>hg38_MT_+_1621_3279_RNR2_16S_50ntFlanks@1349.1385.37|ACAATAGGGTTTACGACCTCGATGTTGGATCAGGACA</td><td>>hg38_MT_+_1621_3279_RNR2_16S_50ntFlanks</td><td>1349</td><td>1385</td><td>37</td><td>ACAATAGGGTTTACGACCTCGATGTTGGATCAGGACA</td><td>230.4</td><td>3.7</td><td>1.2</td><td>3.1</td><td>1.9E-03</td><td>2.7E-03</td><td>MB231/MB468</td><td>MP</td></tr>
<tr><td>>hg38_21_+_8433172_8446622_RNA45SN1_45S_50ntFlanks@8687.8706.20|CGGCTCCGGGACGGCTGGGA</td><td>>hg38_21_+_8433172_8446622_RNA45SN1_45S_50ntFlanks</td><td>8687</td><td>8706</td><td>20</td><td>CGGCTCCGGGACGGCTGGGA</td><td>1676.8</td><td>-2.9</td><td>0.8</td><td>-3.5</td><td>4.4E-04</td><td>2.7E-03</td><td>BT20/MB231</td><td>CYTO</td></tr>
<tr><td>>hg38_21_+_8433172_8446622_RNA45SN1_45S_50ntFlanks@6789.6807.19|GCCTGTCTGAGCGTCGCTT</td><td>>hg38_21_+_8433172_8446622_RNA45SN1_45S_50ntFlanks</td><td>6789</td><td>6807</td><td>19</td><td>GCCTGTCTGAGCGTCGCTT</td><td>290.4</td><td>5.1</td><td>1.6</td><td>3.3</td><td>1.1E-03</td><td>2.7E-03</td><td>BT20/MB468</td><td>MP</td></tr>
<tr><td>>hg38_21_+_8433172_8446622_RNA45SN1_45S_50ntFlanks@12003.12034.32|GCGGGCCGCCGGTGAAATACCACTACTCTGAT</td><td>>hg38_21_+_8433172_8446622_RNA45SN1_45S_50ntFlanks</td><td>12003</td><td>12034</td><td>32</td><td>GCGGGCCGCCGGTGAAATACCACTACTCTGAT</td><td>614.8</td><td>-2.9</td><td>0.8</td><td>-3.5</td><td>4.2E-04</td><td>2.7E-03</td><td>BT20/MB231</td><td>NUC</td></tr>
<tr><td>>hg38_21_+_8433172_8446622_RNA45SN1_45S_50ntFlanks@10059.10084.26|GGCAGTCGAGAGTGGACGGGAGCGGC</td><td>>hg38_21_+_8433172_8446622_RNA45SN1_45S_50ntFlanks</td><td>10059</td><td>10084</td><td>26</td><td>GGCAGTCGAGAGTGGACGGGAGCGGC</td><td>92.8</td><td>9.9</td><td>2.8</td><td>3.6</td><td>3.8E-04</td><td>2.7E-03</td><td>BT20/MB231</td><td>MP</td></tr>
<tr><td>>trna2_GlyGCC_21_-_18827107_18827177@1.32.32,trna35_GlyGCC_1_+_161413094_161413164@1.32.32,trna37_GlyGCC_1_+_161420467_161420537@1.32.32,trna39_GlyGCC_1_+_161427898_161427968@1.32.32,trna41_GlyGCC_1_+_161435258_161435328@1.32.32|GCATGGGTGGTTCAGTGGTAGAATTCTCGCCT</td><td>>trna2_GlyGCC_21_-_18827107_18827177</td><td>1</td><td>32</td><td>32</td><td>GCATGGGTGGTTCAGTGGTAGAATTCTCGCCT</td><td>19789.9</td><td>5.2</td><td>1.4</td><td>3.6</td><td>3.8E-04</td><td>2.7E-03</td><td>BT20/MB231</td><td>MP</td></tr>
<tr><td>>trna152_ValCAC_6_-_27248049_27248121@1.31.31|GCTTCTGTAGTGTAGTGGTTATCACGTTCGC</td><td>>trna152_ValCAC_6_-_27248049_27248121</td><td>1</td><td>31</td><td>31</td><td>GCTTCTGTAGTGTAGTGGTTATCACGTTCGC</td><td>701.0</td><td>-4.3</td><td>1.2</td><td>-3.6</td><td>3.8E-04</td><td>2.7E-03</td><td>BT20/MB231</td><td>MP</td></tr>
<tr><td>>hg38_21_+_8433172_8446622_RNA45SN1_45S_50ntFlanks@9952.9978.27|CATGGAAGTCGGAATCCGCTAAGGAGT</td><td>>hg38_21_+_8433172_8446622_RNA45SN1_45S_50ntFlanks</td><td>9952</td><td>9978</td><td>27</td><td>CATGGAAGTCGGAATCCGCTAAGGAGT</td><td>132.6</td><td>4.4</td><td>1.3</td><td>3.3</td><td>1.1E-03</td><td>2.7E-03</td><td>BT20/MB468</td><td>MP</td></tr>
<tr><td>>hg38_21_+_8433172_8446622_RNA45SN1_45S_50ntFlanks@9738.9760.23|GAAACGATCTCAACCTATTCTCA</td><td>>hg38_21_+_8433172_8446622_RNA45SN1_45S_50ntFlanks</td><td>9738</td><td>9760</td><td>23</td><td>GAAACGATCTCAACCTATTCTCA</td><td>130.3</td><td>-2.8</td><td>0.9</td><td>-3.1</td><td>2.1E-03</td><td>2.7E-03</td><td>MB231/MB468</td><td>CYTO</td></tr>
<tr><td>>hg38_21_+_8433172_8446622_RNA45SN1_45S_50ntFlanks@4049.4069.21|TCGAACGTCTGCCCTATCAAC</td><td>>hg38_21_+_8433172_8446622_RNA45SN1_45S_50ntFlanks</td><td>4049</td><td>4069</td><td>21</td><td>TCGAACGTCTGCCCTATCAAC</td><td>290.2</td><td>-2.7</td><td>0.8</td><td>-3.5</td><td>4.5E-04</td><td>2.7E-03</td><td>BT20/MB231</td><td>CYTO</td></tr>
<tr><td>>MI0000434|hsa-let-7i&WithFlank&12|+|62603680|62603775@12.31.20(+1C)&[MIMAT0000415&hsa-let-7i-5p&offsets|0|-2(+1C);m-49&12|+|62603691|62603712&offsets|0|-2(+1C)]|TGAGGTAGTAGTTTGTGCTGC</td><td>>MI0000434|hsa-let-7i&WithFlank&12|+|62603680|62603775</td><td>12</td><td>31</td><td>20(+1C)</td><td>TGAGGTAGTAGTTTGTGCTGC</td><td>525.0</td><td>5.0</td><td>1.5</td><td>3.3</td><td>1.1E-03</td><td>2.7E-03</td><td>BT20/MB468</td><td>MP</td></tr>
<tr><td>>hg38_21_+_8433172_8446622_RNA45SN1_45S_50ntFlanks@11849.11872.24|CGGGCTTGGCGGAATCAGCGGGGA</td><td>>hg38_21_+_8433172_8446622_RNA45SN1_45S_50ntFlanks</td><td>11849</td><td>11872</td><td>24</td><td>CGGGCTTGGCGGAATCAGCGGGGA</td><td>257.4</td><td>-3.3</td><td>1.1</td><td>-3.1</td><td>2.1E-03</td><td>2.7E-03</td><td>BT20/MB468</td><td>CYTO</td></tr>
<tr><td>>hg38_21_+_8433172_8446622_RNA45SN1_45S_50ntFlanks@11568.11587.20|GCAGCCGACTTAGAACTGGT</td><td>>hg38_21_+_8433172_8446622_RNA45SN1_45S_50ntFlanks</td><td>11568</td><td>11587</td><td>20</td><td>GCAGCCGACTTAGAACTGGT</td><td>337.5</td><td>-2.2</td><td>0.7</td><td>-3.1</td><td>1.9E-03</td><td>2.7E-03</td><td>MB231/MB468</td><td>MP</td></tr>
<tr><td>>trnaMT_GluTTC_MT_-_14674_14742@28.72.45,trnalookalike8_GluTTC_5_-_93905172_93905240@28.72.45|TTTTTCATATCATTGGTCGTGGTTGTAGTCCGTGCGAGAATACCA</td><td>>trnaMT_GluTTC_MT_-_14674_14742</td><td>28</td><td>72</td><td>45</td><td>TTTTTCATATCATTGGTCGTGGTTGTAGTCCGTGCGAGAATACCA</td><td>495.4</td><td>4.4</td><td>1.4</td><td>3.1</td><td>2.3E-03</td><td>2.7E-03</td><td>MB231/MB468</td><td>MITO</td></tr>
<tr><td>>hg38_1_-_228634819_228635039_RNA5S12_5S_50ntFlanks@152.169.18|TACCGGGTGCTGTAGGCT</td><td>>hg38_1_-_228634819_228635039_RNA5S12_5S_50ntFlanks</td><td>152</td><td>169</td><td>18</td><td>TACCGGGTGCTGTAGGCT</td><td>394.3</td><td>-2.4</td><td>0.8</td><td>-3.1</td><td>2.3E-03</td><td>2.7E-03</td><td>MB231/MB468</td><td>MITO</td></tr>
<tr><td>>hg38_21_+_8433172_8446622_RNA45SN1_45S_50ntFlanks@5019.5044.26|TCCGTGGGTGGTGGTGCATGGCCGTT</td><td>>hg38_21_+_8433172_8446622_RNA45SN1_45S_50ntFlanks</td><td>5019</td><td>5044</td><td>26</td><td>TCCGTGGGTGGTGGTGCATGGCCGTT</td><td>58.8</td><td>6.5</td><td>2.0</td><td>3.3</td><td>1.1E-03</td><td>2.7E-03</td><td>BT20/MB468</td><td>MP</td></tr>
<tr><td>>hg38_21_+_8433172_8446622_RNA45SN1_45S_50ntFlanks@11572.11600.29|CCGACTTAGAACTGGTGCGGACCAGGGGA</td><td>>hg38_21_+_8433172_8446622_RNA45SN1_45S_50ntFlanks</td><td>11572</td><td>11600</td><td>29</td><td>CCGACTTAGAACTGGTGCGGACCAGGGGA</td><td>375.9</td><td>3.0</td><td>0.9</td><td>3.5</td><td>5.0E-04</td><td>2.7E-03</td><td>BT20/MB231</td><td>MITO</td></tr>
<tr><td>>hg38_21_+_8433172_8446622_RNA45SN1_45S_50ntFlanks@11873.11903.31|AAGAAGACCCTGTTGAGCTTGACTCTAGTCT</td><td>>hg38_21_+_8433172_8446622_RNA45SN1_45S_50ntFlanks</td><td>11873</td><td>11903</td><td>31</td><td>AAGAAGACCCTGTTGAGCTTGACTCTAGTCT</td><td>380.0</td><td>-2.1</td><td>0.7</td><td>-3.1</td><td>1.8E-03</td><td>2.7E-03</td><td>BT20/MB468</td><td>MITO</td></tr>
<tr><td>>hg38_21_+_8433172_8446622_RNA45SN1_45S_50ntFlanks@6670.6696.27|ACTCGGCTCGTGCGTCGATGAAGAACG</td><td>>hg38_21_+_8433172_8446622_RNA45SN1_45S_50ntFlanks</td><td>6670</td><td>6696</td><td>27</td><td>ACTCGGCTCGTGCGTCGATGAAGAACG</td><td>642.1</td><td>-2.7</td><td>0.8</td><td>-3.3</td><td>1.1E-03</td><td>2.7E-03</td><td>BT20/MB468</td><td>MP</td></tr>
<tr><td>>trnaMT_GlnTTG_MT_-_4329_4400@57.75.19,trnalookalike2_GlnTTG_1_-_564879_564950@57.75.19|ATTCTCATAGTCCTAGCCA</td><td>>trnaMT_GlnTTG_MT_-_4329_4400</td><td>57</td><td>75</td><td>19</td><td>ATTCTCATAGTCCTAGCCA</td><td>141.3</td><td>2.9</td><td>0.9</td><td>3.1</td><td>1.9E-03</td><td>2.7E-03</td><td>MB231/MB468</td><td>MP</td></tr>
<tr><td>>hg38_21_+_8433172_8446622_RNA45SN1_45S_50ntFlanks@11564.11586.23|CCTAGCAGCCGACTTAGAACTGG</td><td>>hg38_21_+_8433172_8446622_RNA45SN1_45S_50ntFlanks</td><td>11564</td><td>11586</td><td>23</td><td>CCTAGCAGCCGACTTAGAACTGG</td><td>764.3</td><td>-2.2</td><td>0.7</td><td>-3.1</td><td>1.8E-03</td><td>2.7E-03</td><td>BT20/MB468</td><td>MITO</td></tr>
<tr><td>>hg38_MT_+_598_1651_RNR1_12S_50ntFlanks@972.1004.33|GGTAAGTGTACTGGAAAGTGCACTTGGACGAAC</td><td>>hg38_MT_+_598_1651_RNR1_12S_50ntFlanks</td><td>972</td><td>1004</td><td>33</td><td>GGTAAGTGTACTGGAAAGTGCACTTGGACGAAC</td><td>548.5</td><td>-5.1</td><td>1.4</td><td>-3.6</td><td>3.8E-04</td><td>2.7E-03</td><td>BT20/MB231</td><td>MP</td></tr>
<tr><td>>hg38_21_+_8433172_8446622_RNA45SN1_45S_50ntFlanks@11190.11223.34|CCCGGGGGGCCGGCGGCGGCGGCGACTCTGGACG</td><td>>hg38_21_+_8433172_8446622_RNA45SN1_45S_50ntFlanks</td><td>11190</td><td>11223</td><td>34</td><td>CCCGGGGGGCCGGCGGCGGCGGCGACTCTGGACG</td><td>115.0</td><td>-5.5</td><td>1.7</td><td>-3.3</td><td>1.1E-03</td><td>2.7E-03</td><td>BT20/MB468</td><td>MP</td></tr>
<tr><td>>hg38_21_+_8433172_8446622_RNA45SN1_45S_50ntFlanks@6661.6687.27|CGGTGGATCACTCGGCTCGTGCGTCGA</td><td>>hg38_21_+_8433172_8446622_RNA45SN1_45S_50ntFlanks</td><td>6661</td><td>6687</td><td>27</td><td>CGGTGGATCACTCGGCTCGTGCGTCGA</td><td>986.7</td><td>3.9</td><td>1.1</td><td>3.6</td><td>3.8E-04</td><td>2.7E-03</td><td>BT20/MB231</td><td>MP</td></tr>
<tr><td>>MI0000299|hsa-mir-222&WithFlank&X|-|45747009|45747130@75.99.25(+2U)&[MIMAT0000279&hsa-miR-222-3p&offsets|0|+4(+2U);m-140&X|-|45747033|45747056&offsets|0|+1(+2U)]|AGCTACATCTGGCTACTGGGTCTCTTT</td><td>>MI0000299|hsa-mir-222&WithFlank&X|-|45747009|45747130</td><td>75</td><td>99</td><td>25(+2U)</td><td>AGCTACATCTGGCTACTGGGTCTCTTT</td><td>310.0</td><td>5.2</td><td>1.7</td><td>3.1</td><td>1.9E-03</td><td>2.7E-03</td><td>MB231/MB468</td><td>MP</td></tr>
<tr><td>>hg38_21_+_8433172_8446622_RNA45SN1_45S_50ntFlanks@6645.6663.19|CTCGTACGACTCTTAGCGG</td><td>>hg38_21_+_8433172_8446622_RNA45SN1_45S_50ntFlanks</td><td>6645</td><td>6663</td><td>19</td><td>CTCGTACGACTCTTAGCGG</td><td>4079.1</td><td>2.5</td><td>0.8</td><td>3.3</td><td>1.1E-03</td><td>2.7E-03</td><td>BT20/MB468</td><td>MP</td></tr>
</table>

| | | | | | | | | | | | | | |
|---|---|---|---|---|---|---|---|---|---|---|---|---|---|
| >hg38_21_+_8433172_8446622_RNA45SN1_45S_50ntFlanks@4969.4999.31\|ACCCGGCCCGGACACGGACAGGATTGACAGA | >hg38_21_+_8433172_8446622_RNA45SN1_45S_50ntFlanks | 4969 | 4999 | 31 | ACCCGGCCCGGACACGGACAGGATTGACAGA | 114.4 | -3.1 | 1.0 | -3.1 | 1.9E-03 | 2.7E-03 | MB231/MB468 | MP |
| >hg38_MT_+_1621_3279_RNR2_16S_50ntFlanks@1363.1388.26\|GACCTCGATGTTGGATCAGGACATCC | >hg38_MT_+_1621_3279_RNR2_16S_50ntFlanks | 1363 | 1388 | 26 | GACCTCGATGTTGGATCAGGACATCC | 448.8 | 3.3 | 1.1 | 3.1 | 1.9E-03 | 2.7E-03 | MB231/MB468 | MP |
| >trna111_HisGTG_1_-_147774845_147774916@1.30.30,trna118_HisGTG_1_-_145396881_145396952@1.30.30,trna16_HisGTG_1_+_146544773_146544844@1.30.30,trna1_HisGTG_15_+_45493349_45493420@1.30.30,trna21_HisGTG_1_+_147753471_147753542@1.30.30,trna33_HisGTG_6_+_27125906_27125977@1.30.30,trna7_HisGTG_9_-_14433938_14434009@1.30.30,trna8_HisGTG_15_-_45492611_45492682@1.30.30,trna9_HisGTG_15_-_45490804_45490875@1.30.30\|GCCGTGATCGTATAGTGGTTAGTACTCTGC | >trna111_HisGTG_1_-_147774845_147774916 | 1 | 30 | 30 | GCCGTGATCGTATAGTGGTTAGTACTCTGC | 357.2 | 2.3 | 0.7 | 3.1 | 1.8E-03 | 2.7E-03 | BT20/MB468 | MITO |
| >hg38_21_+_8433172_8446622_RNA45SN1_45S_50ntFlanks@10822.10865.44\|GAAGTCGGCAAGCCGGATCCGTAACTTCGGGATAAGGATTGGCT | >hg38_21_+_8433172_8446622_RNA45SN1_45S_50ntFlanks | 10822 | 10865 | 44 | GAAGTCGGCAAGCCGGATCCGTAACTTCGGGATAAGGATTGGCT | 449.4 | -5.3 | 1.7 | -3.1 | 2.1E-03 | 2.7E-03 | BT20/MB468 | CYTO |
| >MI0000342\|hsa-mir-200b&WithFlank&1\|+\|1167098\|1167204@63.85.23(+1U)&[MIMAT0000318&hsa-miR-200b-3p&offsets\|0\|+1(+1U);m-93&1\|+\|1167160\|1167182&offsets\|0\|0(+1U)]\|TAATACTGCCTGGTAATGATGACT | >MI0000342\|hsa-mir-200b&WithFlank&1\|+\|1167098\|1167204 | 63 | 85 | 23(+1U) | TAATACTGCCTGGTAATGATGACT | 454.7 | 3.2 | 0.9 | 3.5 | 4.5E-04 | 2.7E-03 | BT20/MB231 | CYTO |
| >hg38_21_+_8433172_8446622_RNA45SN1_45S_50ntFlanks@9198.9220.23\|GTTCTCTCGGGGCCACGCGCGCG | >hg38_21_+_8433172_8446622_RNA45SN1_45S_50ntFlanks | 9198 | 9220 | 23 | GTTCTCTCGGGGCCACGCGCGCG | 499.9 | -6.0 | 2.0 | -3.1 | 2.3E-03 | 2.7E-03 | BT20/MB468 | NUC |
| >hg38_MT_+_1621_3279_RNR2_16S_50ntFlanks@1183.1216.34\|AAATTTCGGTTGGGGCGACCTCGGAGCAGAACCC | >hg38_MT_+_1621_3279_RNR2_16S_50ntFlanks | 1183 | 1216 | 34 | AAATTTCGGTTGGGGCGACCTCGGAGCAGAACCC | 135.5 | 4.5 | 1.4 | 3.3 | 1.1E-03 | 2.7E-03 | BT20/MB468 | MP |
| >hg38_21_+_8433172_8446622_RNA45SN1_45S_50ntFlanks@3902.3938.37\|TCGCGGGGGGGATGCGTGCATTTATCAGATCAAAACC | >hg38_21_+_8433172_8446622_RNA45SN1_45S_50ntFlanks | 3902 | 3938 | 37 | TCGCGGGGGGGATGCGTGCATTTATCAGATCAAAACC | 39.7 | -8.4 | 2.6 | -3.3 | 1.1E-03 | 2.7E-03 | BT20/MB468 | MP |
| >hg38_21_+_8433172_8446622_RNA45SN1_45S_50ntFlanks@4489.4511.23\|CGGGGCCCGAAGCGTTTACTTTG | >hg38_21_+_8433172_8446622_RNA45SN1_45S_50ntFlanks | 4489 | 4511 | 23 | CGGGGCCCGAAGCGTTTACTTTG | 196.2 | -4.9 | 1.6 | -3.1 | 1.8E-03 | 2.7E-03 | BT20/MB468 | MITO |
| >hg38_21_+_8433172_8446622_RNA45SN1_45S_50ntFlanks@9534.9567.34\|AGGAAACTCTGGTGGAGGTCCGTAGCGGTCCTGA | >hg38_21_+_8433172_8446622_RNA45SN1_45S_50ntFlanks | 9534 | 9567 | 34 | AGGAAACTCTGGTGGAGGTCCGTAGCGGTCCTGA | 237.1 | -1.8 | 0.6 | -3.1 | 1.8E-03 | 2.7E-03 | BT20/MB468 | MITO |
| >MI0006273\|hsa-mir-1180&WithFlank&17\|-\|19344500\|19344580@47.68.22&[MIMAT0005825&hsa-miR-1180-3p&offsets\|0\|0;m-214&17\|-\|19344513\|19344534&offsets\|0\|0]\|TTTCCGGCTCGCGTGGGTGTGT | >MI0006273\|hsa-mir-1180&WithFlank&17\|-\|19344500\|19344580 | 47 | 68 | 22 | TTTCCGGCTCGCGTGGGTGTGT | 1468.6 | 1.9 | 0.5 | 3.5 | 4.3E-04 | 2.7E-03 | BT20/MB231 | NUC |
| >trna130_GlnTTG_6_-_27763640_27763711@1.29.29,trna173_GlnTTG_6_-_26311975_26312046@1.29.29,trna174_GlnTTG_6_-_26311424_26311495@1.29.29\|GGCCCCATGGTGTAATGGTTAGCACTCTG | >trna130_GlnTTG_6_-_27763640_27763711 | 1 | 29 | 29 | GGCCCCATGGTGTAATGGTTAGCACTCTG | 1562.1 | -5.4 | 1.8 | -3.1 | 2.1E-03 | 2.7E-03 | BT20/MB468 | CYTO |
| >MI0000108\|hsa-mir-103a-2&WithFlank&20\|+\|3917488\|3917577@54.76.23(+2U);MI0000109\|hsa-mir-103a-1&WithFlank&5\|-\|168560890\|168560979@54.76.23(+2U)&[MIMAT0000101&hsa-miR-103a-3p&offsets\|0\|0(+2U);m-12&20\|+\|3917541\|3917563&offsets\|0\|0(+2U)];[MIMAT0000101_1&hsa-miR-103a-3p&offsets\|0\|0(+2U);m-13&5\|-\|168560904\|168560926&offsets\|0\|0(+2U);m-44220&5\|-\|168560908\|168560926&offsets\|0\|+4(+2U)]\|AGCAGCATTGTACAGGGCTATGATT | >MI0000108\|hsa-mir-103a-2&WithFlank&20\|+\|3917488\|3917577 | 54 | 76 | 23(+2U) | AGCAGCATTGTACAGGGCTATGATT | 574.8 | -3.5 | 1.0 | -3.6 | 3.8E-04 | 2.7E-03 | BT20/MB231 | MP |
| >hg38_21_+_8433172_8446622_RNA45SN1_45S_50ntFlanks@12417.12445.29\|TCGATGTCGGCTCTTCCTATCATTGTGAA | >hg38_21_+_8433172_8446622_RNA45SN1_45S_50ntFlanks | 12417 | 12445 | 29 | TCGATGTCGGCTCTTCCTATCATTGTGAA | 526.7 | -1.6 | 0.5 | -3.0 | 2.3E-03 | 2.7E-03 | MB231/MB468 | NUC |
| >hg38_21_+_8433172_8446622_RNA45SN1_45S_50ntFlanks@12420.12445.26\|ATGTCGGCTCTTCCTATCATTGTGAA | >hg38_21_+_8433172_8446622_RNA45SN1_45S_50ntFlanks | 12420 | 12445 | 26 | ATGTCGGCTCTTCCTATCATTGTGAA | 870.0 | -2.4 | 0.8 | -3.0 | 2.3E-03 | 2.7E-03 | MB231/MB468 | MITO |
| >hg38_21_+_8433172_8446622_RNA45SN1_45S_50ntFlanks@11323.11344.22\|CGTCTCGTCGCGCGCGCGTCCG | >hg38_21_+_8433172_8446622_RNA45SN1_45S_50ntFlanks | 11323 | 11344 | 22 | CGTCTCGTCGCGCGCGCGTCCG | 15009.0 | -4.2 | 1.3 | -3.1 | 1.8E-03 | 2.7E-03 | BT20/MB468 | MITO |
| >am_MI0000086\|hsa-mir-28&WithFlank&3\|+\|188688775\|188688872@20.38.19(+1A)&[MIMAT0000085&hsa-miR-28-5p&offsets\|0\|-3(+1A);m-133&3\|+\|188688794\|188688815&offsets\|0\|-3(+1A)]\|AAGGAGCTCACAGTCTATTA | >am_MI0000086\|hsa-mir-28&WithFlank&3\|+\|188688775\|188688872 | 20 | 38 | 19(+1A) | AAGGAGCTCACAGTCTATTA | 77.3 | 5.9 | 1.9 | 3.1 | 1.9E-03 | 2.7E-03 | MB231/MB468 | MP |
| >hg38_21_+_8433172_8446622_RNA45SN1_45S_50ntFlanks@12416.12436.21\|TTCGATGTCGGCTCTTCCTAT | >hg38_21_+_8433172_8446622_RNA45SN1_45S_50ntFlanks | 12416 | 12436 | 21 | TTCGATGTCGGCTCTTCCTAT | 154.4 | -4.9 | 1.6 | -3.1 | 1.9E-03 | 2.7E-03 | MB231/MB468 | MP |
| >hg38_21_+_8433172_8446622_RNA45SN1_45S_50ntFlanks@10059.10084.26\|GGCAGTCGAGAGTGGACGGGAGCGGC | >hg38_21_+_8433172_8446622_RNA45SN1_45S_50ntFlanks | 10059 | 10084 | 26 | GGCAGTCGAGAGTGGACGGGAGCGGC | 39.5 | 9.1 | 2.8 | 3.3 | 1.1E-03 | 2.7E-03 | BT20/MB468 | MP |
| >am_trna128_GlyGCC_6_-_27870686_27870756@4.28.25,trna133_GlyCCC_1_-_16872434_16872504@4.28.25,trna18_GlyGCC_16_+_70822597_70822667@4.28.25,trna19_GlyGCC_16_+_70823410_70823480@4.28.25,trna19_GlyGCC_2_-_157257659_157257729@4.28.25,trna24_GlyGCC_16_-_70812942_70813012@4.28.25,trna25_GlyGCC_16_-_70812114_70812184@4.28.25,trna4_GlyCCC_1_+_17188416_17188486@4.28.25,trna5_GlyGCC_17_+_8029064_8029134@4.28.25,trna68_GlyGCC_1_-_161493637_161493707@4.28.25\|TTGGTGGTTCAGTGGTAGAATTCTC | >am_trna128_GlyGCC_6_-_27870686_27870756 | 4 | 28 | 25 | TTGGTGGTTCAGTGGTAGAATTCTC | 460.5 | -5.7 | 1.9 | -3.1 | 2.3E-03 | 2.7E-03 | BT20/MB468 | NUC |
| >hg38_21_+_8433172_8446622_RNA45SN1_45S_50ntFlanks@11594.11613.20\|CAGGGGAATCCGACTGTTTA | >hg38_21_+_8433172_8446622_RNA45SN1_45S_50ntFlanks | 11594 | 11613 | 20 | CAGGGGAATCCGACTGTTTA | 269.9 | -1.7 | 0.6 | -3.1 | 2.1E-03 | 2.7E-03 | BT20/MB468 | CYTO |
| >hg38_21_+_8433172_8446622_RNA45SN1_45S_50ntFlanks@7980.7998.19\|CCTCAGATCAGACGTGGCG | >hg38_21_+_8433172_8446622_RNA45SN1_45S_50ntFlanks | 7980 | 7998 | 19 | CCTCAGATCAGACGTGGCG | 154.4 | -4.4 | 1.4 | -3.1 | 2.1E-03 | 2.7E-03 | BT20/MB468 | CYTO |
| >MI0000085\|hsa-mir-27a&WithFlank&19\|-\|13836434\|13836523@57.74.18(+1A);MI0000440\|hsa-mir-27b&WithFlank&9\|+\|95085439\|95085547@67.84.18(+1A)&[MIMAT0000084&hsa-miR-27a-3p&offsets\|0\|-3(+1A);m-43&19\|-\|13836447\|13836467&offsets\|0\|-3(+1A)];[MIMAT0000419&hsa-miR-27b-3p&offsets\|0\|-3(+1A);m-39&9\|+\|95085505\|95085525&offsets\|0\|-3(+1A)]\|TTCACAGTGGCTAAGTTCA | >MI0000085\|hsa-mir-27a&WithFlank&19\|-\|13836434\|13836523 | 57 | 74 | 18(+1A) | TTCACAGTGGCTAAGTTCA | 261.9 | 4.2 | 1.3 | 3.1 | 1.9E-03 | 2.7E-03 | MB231/MB468 | MP |
| >hg38_21_+_8433172_8446622_RNA45SN1_45S_50ntFlanks@11709.11736.28\|GCGCGGGTAAACGGCGGGAGTAACTATG | >hg38_21_+_8433172_8446622_RNA45SN1_45S_50ntFlanks | 11709 | 11736 | 28 | GCGCGGGTAAACGGCGGGAGTAACTATG | 156.2 | 2.9 | 1.0 | 3.1 | 2.2E-03 | 2.7E-03 | MB231/MB468 | CYTO |
| >am_trna116_GluCTC_1_-_145399233_145399304@13.30.18,trna25_GluCTC_2_-_71273488_71273560@13.30.18,trna59_GluCTC_1_+_249168447_249168518@13.30.18,trna71_GluCTC_1_-_161439189_161439260@13.30.18,trna74_GluCTC_1_-_161431809_161431880@13.30.18,trna77_GluCTC_1_-_161424398_161424469@13.30.18,trna77_GluCTC_6_+_28949976_28950047@13.30.18,trna80_GluCTC_1_-_161417018_161417089@13.30.18,trna87_GluCTC_6_-_126101393_126101464@13.30.18\|TAGTGGTTAGGATTCGGC | >am_trna116_GluCTC_1_-_145399233_145399304 | 13 | 30 | 18 | TAGTGGTTAGGATTCGGC | 420.5 | 2.4 | 0.8 | 3.1 | 2.2E-03 | 2.7E-03 | MB231/MB468 | CYTO |
| >MI0000088\|hsa-mir-30a&WithFlank&6\|-\|71403545\|71403627@53.73.21(+1U)&[MIMAT0000088&hsa-miR-30a-3p&offsets\|0\|-1(+1U);m-38&6\|-\|71403554\|71403575&offsets\|0\|-1(+1U)]\|CTTTCAGTCGGATGTTTGCAGT | >MI0000088\|hsa-mir-30a&WithFlank&6\|-\|71403545\|71403627 | 53 | 73 | 21(+1U) | CTTTCAGTCGGATGTTTGCAGT | 8551.1 | -2.5 | 0.7 | -3.5 | 5.1E-04 | 2.7E-03 | BT20/MB231 | MITO |
| >hg38_1_-_228634819_228635039_RNA5S12_5S_50ntFlanks@125.170.46\|GTACTTGGATGGGAGACCGCCTGGGAATACCGGGTGCTGTAGGCTT | >hg38_1_-_228634819_228635039_RNA5S12_5S_50ntFlanks | 125 | 170 | 46 | GTACTTGGATGGGAGACCGCCTGGGAATACCGGGTGCTGTAGGCTT | 860.4 | 3.7 | 1.1 | 3.5 | 5.1E-04 | 2.7E-03 | BT20/MB231 | MITO |
| >hg38_21_+_8433172_8446622_RNA45SN1_45S_50ntFlanks@10349.10381.33\|AACGAGAACTTTGAAGGCCGAAGTGGAGAAGGG | >hg38_21_+_8433172_8446622_RNA45SN1_45S_50ntFlanks | 10349 | 10381 | 33 | AACGAGAACTTTGAAGGCCGAAGTGGAGAAGGG | 327.7 | -4.0 | 1.3 | -3.1 | 1.9E-03 | 2.7E-03 | MB231/MB468 | MP |
| >hg38_21_+_8433172_8446622_RNA45SN1_45S_50ntFlanks@10767.10785.19\|GCAGCAGGTCTCCAAGGTG | >hg38_21_+_8433172_8446622_RNA45SN1_45S_50ntFlanks | 10767 | 10785 | 19 | GCAGCAGGTCTCCAAGGTG | 236.9 | -3.2 | 1.0 | -3.1 | 2.2E-03 | 2.7E-03 | MB231/MB468 | CYTO |
| >MI0000273\|hsa-mir-183&WithFlank&7\|-\|129774899\|129775020@33.54.22&[MIMAT0000261&hsa-miR-183-5p&offsets\|0\|0;m-41&7\|-\|129774966\|129774987&offsets\|-1\|-1]\|TATGGCACTGGTAGAATTCACT | >MI0000273\|hsa-mir-183&WithFlank&7\|-\|129774899\|129775020 | 33 | 54 | 22 | TATGGCACTGGTAGAATTCACT | 3171.7 | 2.5 | 0.8 | 3.3 | 1.1E-03 | 2.7E-03 | BT20/MB468 | MP |
| >hg38_21_+_8433172_8446622_RNA45SN1_45S_50ntFlanks@5074.5102.29\|ATTCCGATAACGAACGAGACTCTGGCATG | >hg38_21_+_8433172_8446622_RNA45SN1_45S_50ntFlanks | 5074 | 5102 | 29 | ATTCCGATAACGAACGAGACTCTGGCATG | 337.3 | -3.3 | 1.0 | -3.1 | 1.8E-03 | 2.7E-03 | BT20/MB468 | MITO |
| >hg38_21_+_8433172_8446622_RNA45SN1_45S_50ntFlanks@10394.10424.31\|AGCAGTTGAACATGGGTCAGTCGGTCCTGAG | >hg38_21_+_8433172_8446622_RNA45SN1_45S_50ntFlanks | 10394 | 10424 | 31 | AGCAGTTGAACATGGGTCAGTCGGTCCTGAG | 9669.2 | 3.4 | 1.0 | 3.5 | 5.1E-04 | 2.7E-03 | BT20/MB231 | MITO |
| >hg38_21_+_8433172_8446622_RNA45SN1_45S_50ntFlanks@8168.8185.18\|CCCAAGTCCTTCTGATCG | >hg38_21_+_8433172_8446622_RNA45SN1_45S_50ntFlanks | 8168 | 8185 | 18 | CCCAAGTCCTTCTGATCG | 502.2 | -3.4 | 1.1 | -3.1 | 2.2E-03 | 2.7E-03 | MB231/MB468 | CYTO |
| >MI0000264\|hsa-mir-7-2&WithFlank&15\|+\|88611819\|88611940@38.57.20(+1A);MI0000265\|hsa-mir-7-3&WithFlank&19\|+\|4770664\|4770785@37.56.20(+1A);MI0000263\|hsa-mir-7-1&WithFlank&9\|-\|83969742\|83969863@30.49.20(+1A)&[MIMAT0000252&hsa-miR-7-5p&offsets\|0\|-4(+1A);m-400&15\|+\|88611856\|88611879&offsets\|0\|-4(+1A)];[MIMAT0000252_1&hsa-miR-7-5p&offsets\|0\|-4(+1A);m-402&19\|+\|4770700\|4770723&offsets\|0\|-4(+1A)];[MIMAT0000252_2&hsa-miR-7-5p&offsets\|0\|-4(+1A);m-401&9\|-\|83969811\|83969834&offsets\|0\|-4(+1A)]\|TGGAAGACTAGTGATTTTGTA | >MI0000264\|hsa-mir-7-2&WithFlank&15\|+\|88611819\|88611940 | 38 | 57 | 20(+1A) | TGGAAGACTAGTGATTTTGTA | 62.4 | 3.9 | 1.3 | 3.1 | 1.9E-03 | 2.7E-03 | MB231/MB468 | MP |
| >am_MI0003583\|hsa-mir-576&WithFlank&4\|+\|109488692\|109488801@61.83.23&[MIMAT0004796&hsa-miR-576-3p&offsets\|0\|+1;m-524&4\|+\|109488752\|109488772&offsets\|0\|+2]\|AAGATGTGGAAAAATTGGAATCC | >am_MI0003583\|hsa-mir-576&WithFlank&4\|+\|109488692\|109488801 | 61 | 83 | 23 | AAGATGTGGAAAAATTGGAATCC | 292.0 | 2.8 | 0.8 | 3.5 | 4.6E-04 | 2.7E-03 | BT20/MB231 | CYTO |
| >hg38_21_+_8433172_8446622_RNA45SN1_45S_50ntFlanks@3769.3786.18\|CGCACGGCCGGTACAGTG | >hg38_21_+_8433172_8446622_RNA45SN1_45S_50ntFlanks | 3769 | 3786 | 18 | CGCACGGCCGGTACAGTG | 218.7 | -4.0 | 1.3 | -3.1 | 1.8E-03 | 2.7E-03 | BT20/MB468 | MITO |

| | | | | | | | | | | | | | |
|---|---|---|---|---|---|---|---|---|---|---|---|---|---|
| >hg38_21_+_8433172_8446622_RNA45SN1_45S_50ntFlanks@8194.8214.21\|CCCGTGGACGGTGTGAGGCCG | >hg38_21_+_8433172_8446622_RNA45SN1_45S_50ntFlanks | 8194 | 8214 | 21 | CCCGTGGACGGTGTGAGGCCG | 884.0 | -5.0 | 1.6 | -3.1 | 2.1E-03 | 2.7E-03 | BT20/MB468 | CYTO |
| >hg38_21_+_8433172_8446622_RNA45SN1_45S_50ntFlanks@7984.8003.20\|AGATCAGACGTGGCGACCCG | >hg38_21_+_8433172_8446622_RNA45SN1_45S_50ntFlanks | 7984 | 8003 | 20 | AGATCAGACGTGGCGACCCG | 292.5 | -1.9 | 0.6 | -3.1 | 1.8E-03 | 2.7E-03 | BT20/MB468 | MITO |
| >hg38_21_+_8433172_8446622_RNA45SN1_45S_50ntFlanks@12324.12355.32\|TTGGGTTTTAAGCAGGAGGTGTCAGAAAAGTT | >hg38_21_+_8433172_8446622_RNA45SN1_45S_50ntFlanks | 12324 | 12355 | 32 | TTGGGTTTTAAGCAGGAGGTGTCAGAAAAGTT | 251.8 | -5.4 | 1.6 | -3.3 | 1.1E-03 | 2.7E-03 | BT20/MB468 | MP |
| >trna111_HisGTG_1_-_147774845_147774916@1.31.31,trna118_HisGTG_1_-_145396881_145396952@1.31.31,trna16_HisGTG_1_+_146544773_146544844@1.31.31,trna1_HisGTG_15_+_45493349_45493420@1.31.31,trna21_HisGTG_1_+_147753471_147753542@1.31.31,trna33_HisGTG_6_+_27125906_27125977@1.31.31,trna7_HisGTG_9_-_14433938_14434009@1.31.31,trna8_HisGTG_15_-_45492611_45492682@1.31.31,trna9_HisGTG_15_-_45490804_45490875@1.31.31\|GCCGTGATCGTATAGTGGTTAGTACTCTGCG | >trna111_HisGTG_1_-_147774845_147774916 | 1 | 31 | 31 | GCCGTGATCGTATAGTGGTTAGTACTCTGCG | 2003.2 | -2.8 | 0.8 | -3.5 | 5.2E-04 | 2.7E-03 | BT20/MB231 | MITO |
| >MI0000750\|hsa-mir-26a-2&WithFlank&12\|-\|57824603\|57824698@20.39.20(+1U);MI0000083\|hsa-mir-26a-1&WithFlank&3\|+\|37969398\|37969486@16.35.20(+1U)&[MIMAT0000082&hsa-miR-26a-5p&offsets\|0\|-2(+1U);m-33&12\|-\|57824658\|57824679&offsets\|0\|-2(+1U)];[MIMAT0000082_1&hsa-miR-26a-5p&offsets\|0\|-2(+1U);m-34&3\|+\|37969413\|37969434&offsets\|0\|-2(+1U)]\|TTCAAGTAATCCAGGATAGGT | >MI0000750\|hsa-mir-26a-2&WithFlank&12\|-\|57824603\|57824698 | 20 | 39 | 20(+1U) | TTCAAGTAATCCAGGATAGGT | 478.5 | -3.1 | 0.9 | -3.5 | 5.2E-04 | 2.7E-03 | BT20/MB231 | MITO |
| >MI0003513\|hsa-mir-455&WithFlank&9\|+\|114209428\|114209535@24.44.21&[MIMAT0003150&hsa-miR-455-5p&offsets\|+2\|+1;m-232&9\|+\|114209449\|114209470&offsets\|+2\|+1]\|TGTGCCTTTGGACTACATCGT | >MI0003513\|hsa-mir-455&WithFlank&9\|+\|114209428\|114209535 | 24 | 44 | 21 | TGTGCCTTTGGACTACATCGT | 156.5 | -6.1 | 1.7 | -3.5 | 3.9E-04 | 2.7E-03 | BT20/MB231 | MP |
| >hg38_21_+_8433172_8446622_RNA45SN1_45S_50ntFlanks@6728.6770.43\|GGACACATTGATCATCGACACTTCGAACGCACTTGCGGCCCCG | >hg38_21_+_8433172_8446622_RNA45SN1_45S_50ntFlanks | 6728 | 6770 | 43 | GGACACATTGATCATCGACACTTCGAACGCACTTGCGGCCCCG | 539.3 | -2.7 | 0.9 | -3.0 | 2.3E-03 | 2.7E-03 | MB231/MB468 | MITO |
| >hg38_21_+_8433172_8446622_RNA45SN1_45S_50ntFlanks@12420.12452.33\|ATGTCGGCTCTTCCTATCATTGTGAAGCAGAAT | >hg38_21_+_8433172_8446622_RNA45SN1_45S_50ntFlanks | 12420 | 12452 | 33 | ATGTCGGCTCTTCCTATCATTGTGAAGCAGAAT | 276.2 | 2.2 | 0.7 | 3.1 | 2.2E-03 | 2.7E-03 | MB231/MB468 | CYTO |
| >hg38_21_+_8433172_8446622_RNA45SN1_45S_50ntFlanks@10784.10816.33\|TGAACAGCCTCTGGCATGTTGGAACAATGTAGG | >hg38_21_+_8433172_8446622_RNA45SN1_45S_50ntFlanks | 10784 | 10816 | 33 | TGAACAGCCTCTGGCATGTTGGAACAATGTAGG | 187.4 | -2.3 | 0.7 | -3.1 | 2.1E-03 | 2.7E-03 | BT20/MB468 | CYTO |
| >hg38_21_+_8433172_8446622_RNA45SN1_45S_50ntFlanks@9482.9520.39\|ACGTGTTAGGACCCGAAAGATGGTGAACTATGCCTGGGC | >hg38_21_+_8433172_8446622_RNA45SN1_45S_50ntFlanks | 9482 | 9520 | 39 | ACGTGTTAGGACCCGAAAGATGGTGAACTATGCCTGGGC | 38.9 | 9.0 | 2.8 | 3.3 | 1.1E-03 | 2.7E-03 | BT20/MB468 | MP |
| >MI0000440\|hsa-mir-27b&WithFlank&9\|+\|95085439\|95085547@66.86.21(+1U)&[MIMAT0000419&hsa-miR-27b-3p&offsets\|-1\|-1(+1U);m-39&9\|+\|95085505\|95085525&offsets\|-1\|-1(+1U)]\|GTTCACAGTGGCTAAGTTCTGT | >MI0000440\|hsa-mir-27b&WithFlank&9\|+\|95085439\|95085547 | 66 | 86 | 21(+1U) | GTTCACAGTGGCTAAGTTCTGT | 212.2 | 2.9 | 0.8 | 3.5 | 4.6E-04 | 2.7E-03 | BT20/MB231 | CYTO |
| >hg38_21_+_8433172_8446622_RNA45SN1_45S_50ntFlanks@11768.11803.36\|CTAATTAGTGACGCGCATGAATGGATGAACGAGATT | >hg38_21_+_8433172_8446622_RNA45SN1_45S_50ntFlanks | 11768 | 11803 | 36 | CTAATTAGTGACGCGCATGAATGGATGAACGAGATT | 1107.2 | -3.5 | 1.1 | -3.3 | 1.1E-03 | 2.7E-03 | BT20/MB468 | MP |
| >hg38_21_+_8433172_8446622_RNA45SN1_45S_50ntFlanks@10355.10376.22\|AACTTTGAAGGCCGAAGTGGAG | >hg38_21_+_8433172_8446622_RNA45SN1_45S_50ntFlanks | 10355 | 10376 | 22 | AACTTTGAAGGCCGAAGTGGAG | 43.1 | -8.5 | 2.6 | -3.3 | 1.1E-03 | 2.7E-03 | BT20/MB468 | MP |
| >hg38_21_+_8433172_8446622_RNA45SN1_45S_50ntFlanks@4557.4575.19\|CAGCTAGGAATAATGGAAT | >hg38_21_+_8433172_8446622_RNA45SN1_45S_50ntFlanks | 4557 | 4575 | 19 | CAGCTAGGAATAATGGAAT | 132.9 | -2.2 | 0.7 | -3.0 | 2.3E-03 | 2.7E-03 | MB231/MB468 | NUC |
| >hg38_1_-_228634819_228635039_RNA5S12_5S_50ntFlanks@51.76.26\|GTCTACGGCCATACCACCCTGAACGC | >hg38_1_-_228634819_228635039_RNA5S12_5S_50ntFlanks | 51 | 76 | 26 | GTCTACGGCCATACCACCCTGAACGC | 1813.5 | 3.7 | 1.1 | 3.3 | 1.1E-03 | 2.8E-03 | BT20/MB468 | MP |
| >hg38_21_+_8433172_8446622_RNA45SN1_45S_50ntFlanks@6732.6770.39\|ACATTGATCATCGACACTTCGAACGCACTTGCGGCCCCG | >hg38_21_+_8433172_8446622_RNA45SN1_45S_50ntFlanks | 6732 | 6770 | 39 | ACATTGATCATCGACACTTCGAACGCACTTGCGGCCCCG | 732.8 | -2.3 | 0.8 | -3.1 | 2.2E-03 | 2.8E-03 | MB231/MB468 | CYTO |
| >hg38_MT_+_1621_3279_RNR2_16S_50ntFlanks@930.958.29\|AGTGACACATGTTTAACGGCCGCGGTACC | >hg38_MT_+_1621_3279_RNR2_16S_50ntFlanks | 930 | 958 | 29 | AGTGACACATGTTTAACGGCCGCGGTACC | 314.7 | 3.8 | 1.2 | 3.1 | 2.0E-03 | 2.8E-03 | MB231/MB468 | MP |
| >hg38_21_+_8433172_8446622_RNA45SN1_45S_50ntFlanks@11571.11600.30\|GCCGACTTAGAACTGGTGCGGACCAGGGGA | >hg38_21_+_8433172_8446622_RNA45SN1_45S_50ntFlanks | 11571 | 11600 | 30 | GCCGACTTAGAACTGGTGCGGACCAGGGGA | 776.9 | -1.7 | 0.6 | -3.0 | 2.3E-03 | 2.8E-03 | MB231/MB468 | NUC |
| >hg38_21_+_8433172_8446622_RNA45SN1_45S_50ntFlanks@10059.10078.20\|GGCAGTCGAGAGTGGACGGG | >hg38_21_+_8433172_8446622_RNA45SN1_45S_50ntFlanks | 10059 | 10078 | 20 | GGCAGTCGAGAGTGGACGGG | 360.2 | 5.5 | 1.6 | 3.5 | 3.9E-04 | 2.8E-03 | BT20/MB231 | MP |
| >hg38_21_+_8433172_8446622_RNA45SN1_45S_50ntFlanks@11783.11819.37\|CATGAATGGATGAACGAGATTCCCACTGTCCCTACCT | >hg38_21_+_8433172_8446622_RNA45SN1_45S_50ntFlanks | 11783 | 11819 | 37 | CATGAATGGATGAACGAGATTCCCACTGTCCCTACCT | 268.5 | -3.4 | 1.0 | -3.3 | 1.1E-03 | 2.8E-03 | BT20/MB468 | MP |
| >hg38_21_+_8433172_8446622_RNA45SN1_45S_50ntFlanks@5073.5102.30\|AATTCCGATAACGAACGAGACTCTGGCATG | >hg38_21_+_8433172_8446622_RNA45SN1_45S_50ntFlanks | 5073 | 5102 | 30 | AATTCCGATAACGAACGAGACTCTGGCATG | 436.0 | -4.3 | 1.4 | -3.1 | 1.8E-03 | 2.8E-03 | BT20/MB468 | MITO |
| >hg38_21_+_8433172_8446622_RNA45SN1_45S_50ntFlanks@10876.10902.27\|GGTCGGTCGGGCTGGGGCGCGAAGCGG | >hg38_21_+_8433172_8446622_RNA45SN1_45S_50ntFlanks | 10876 | 10902 | 27 | GGTCGGTCGGGCTGGGGCGCGAAGCGG | 124.6 | -5.2 | 1.6 | -3.3 | 1.1E-03 | 2.8E-03 | BT20/MB468 | MP |
| >hg38_21_+_8433172_8446622_RNA45SN1_45S_50ntFlanks@5421.5438.18\|CGATTGGATGGTTTAGTG | >hg38_21_+_8433172_8446622_RNA45SN1_45S_50ntFlanks | 5421 | 5438 | 18 | CGATTGGATGGTTTAGTG | 172.2 | -5.3 | 1.6 | -3.3 | 1.1E-03 | 2.8E-03 | BT20/MB468 | MP |
| >trna116_GluCTC_1_-_145399233_145399304@2.33.32,trna59_GluCTC_1_+_249168447_249168518@2.33.32,trna71_GluCTC_1_-_161439189_161439260@2.33.32,trna74_GluCTC_1_-_161431809_161431880@2.33.32,trna77_GluCTC_1_-_161424398_161424469@2.33.32,trna77_GluCTC_6_+_28949976_28950047@2.33.32,trna80_GluCTC_1_-_161417018_161417089@2.33.32,trna87_GluCTC_6_-_126101393_126101464@2.33.32\|CCCTGGTGGTCTAGTGGTTAGGATTCGGCGCT | >trna116_GluCTC_1_-_145399233_145399304 | 2 | 33 | 32 | CCCTGGTGGTCTAGTGGTTAGGATTCGGCGCT | 618.0 | 3.5 | 1.0 | 3.5 | 5.2E-04 | 2.8E-03 | BT20/MB231 | MITO |
| >MI0000456\|hsa-mir-140&WithFlank&16\|+\|69933075\|69933186@69.90.22(+1U)&[MIMAT0004597&hsa-miR-140-3p&offsets\|+1\|+2(+1U);m-45&16\|+\|69933143\|69933164&offsets\|0\|0(+1U)]\|ACCACAGGGTAGAACCACGGACT | >MI0000456\|hsa-mir-140&WithFlank&16\|+\|69933075\|69933186 | 69 | 90 | 22(+1U) | ACCACAGGGTAGAACCACGGACT | 836.0 | -2.9 | 0.8 | -3.5 | 5.2E-04 | 2.8E-03 | BT20/MB231 | MITO |
| >hg38_21_+_8433172_8446622_RNA45SN1_45S_50ntFlanks@12728.12747.20\|CTCGGATAGCCGGTCCCCCG | >hg38_21_+_8433172_8446622_RNA45SN1_45S_50ntFlanks | 12728 | 12747 | 20 | CTCGGATAGCCGGTCCCCCG | 95.3 | -5.3 | 1.7 | -3.1 | 2.1E-03 | 2.8E-03 | BT20/MB468 | CYTO |
| >MI0000299\|hsa-mir-222&WithFlank&X\|-\|45747009\|45747130@75.98.24(+1A)&[MIMAT0000279&hsa-miR-222-3p&offsets\|0\|+3(+1A);m-140&X\|-\|45747033\|45747056&offsets\|0\|0(+1A)]\|AGCTACATCTGGCTACTGGGTCTCA | >MI0000299\|hsa-mir-222&WithFlank&X\|-\|45747009\|45747130 | 75 | 98 | 24(+1A) | AGCTACATCTGGCTACTGGGTCTCA | 2528.8 | -3.3 | 0.9 | -3.5 | 3.9E-04 | 2.8E-03 | BT20/MB231 | MP |
| >hg38_21_+_8433172_8446622_RNA45SN1_45S_50ntFlanks@12415.12442.28\|CTTCGATGTCGGCTCTTCCTATCATTGT | >hg38_21_+_8433172_8446622_RNA45SN1_45S_50ntFlanks | 12415 | 12442 | 28 | CTTCGATGTCGGCTCTTCCTATCATTGT | 475.7 | 1.9 | 0.6 | 3.1 | 1.8E-03 | 2.8E-03 | BT20/MB468 | MITO |
| >trnaMT_ThrTGT_MT_+_15888_15953@31.69.39\|TTGTAAACCGGAGATGAAAACCTTTTTCCAAGGACACCA | >trnaMT_ThrTGT_MT_+_15888_15953 | 31 | 69 | 39 | TTGTAAACCGGAGATGAAAACCTTTTTCCAAGGACACCA | 277.9 | 4.0 | 1.3 | 3.1 | 2.0E-03 | 2.8E-03 | MB231/MB468 | MP |
| >hg38_21_+_8433172_8446622_RNA45SN1_45S_50ntFlanks@3900.3938.39\|CTTCGCGGGGGGGATGCGTGCATTTATCAGATCAAAACC | >hg38_21_+_8433172_8446622_RNA45SN1_45S_50ntFlanks | 3900 | 3938 | 39 | CTTCGCGGGGGGGATGCGTGCATTTATCAGATCAAAACC | 276.8 | -4.8 | 1.5 | -3.1 | 1.8E-03 | 2.8E-03 | BT20/MB468 | MITO |
| >hg38_21_+_8433172_8446622_RNA45SN1_45S_50ntFlanks@3854.3898.45\|ACTGTGGTAATTCTAGAGCTAATACATGCCGACGGGCGCTGACCC | >hg38_21_+_8433172_8446622_RNA45SN1_45S_50ntFlanks | 3854 | 3898 | 45 | ACTGTGGTAATTCTAGAGCTAATACATGCCGACGGGCGCTGACCC | 229.7 | -6.4 | 2.0 | -3.3 | 1.1E-03 | 2.8E-03 | BT20/MB468 | MP |
| >hg38_21_+_8433172_8446622_RNA45SN1_45S_50ntFlanks@9953.9978.26\|ATGGAAGTCGGAATCCGCTAAGGAGT | >hg38_21_+_8433172_8446622_RNA45SN1_45S_50ntFlanks | 9953 | 9978 | 26 | ATGGAAGTCGGAATCCGCTAAGGAGT | 202.5 | 4.3 | 1.3 | 3.3 | 1.1E-03 | 2.8E-03 | BT20/MB468 | MP |
| >hg38_21_+_8433172_8446622_RNA45SN1_45S_50ntFlanks@4591.4614.24\|TTTTGTTGGTTTTCGGAACTGAGG | >hg38_21_+_8433172_8446622_RNA45SN1_45S_50ntFlanks | 4591 | 4614 | 24 | TTTTGTTGGTTTTCGGAACTGAGG | 80.0 | -5.1 | 1.6 | -3.3 | 1.1E-03 | 2.8E-03 | BT20/MB468 | MP |
| >hg38_21_+_8433172_8446622_RNA45SN1_45S_50ntFlanks@4723.4752.30\|CATTAATCAAGAACGAAAGTCGGAGGTTCG | >hg38_21_+_8433172_8446622_RNA45SN1_45S_50ntFlanks | 4723 | 4752 | 30 | CATTAATCAAGAACGAAAGTCGGAGGTTCG | 884.4 | -5.6 | 1.7 | -3.3 | 1.1E-03 | 2.8E-03 | BT20/MB468 | MP |
| >hg38_21_+_8433172_8446622_RNA45SN1_45S_50ntFlanks@12404.12443.40\|GCTTTTTGATCCTTCGATGTCGGCTCTTCCTATCATTGTG | >hg38_21_+_8433172_8446622_RNA45SN1_45S_50ntFlanks | 12404 | 12443 | 40 | GCTTTTTGATCCTTCGATGTCGGCTCTTCCTATCATTGTG | 1130.8 | -3.1 | 1.0 | -3.0 | 2.3E-03 | 2.8E-03 | MB231/MB468 | NUC |
| >am_trnaMT_SerGCT_MT_+_12207_12265@13.34.22\|AAGAACTGCTAACTCATGCCCC | >am_trnaMT_SerGCT_MT_+_12207_12265 | 13 | 34 | 22 | AAGAACTGCTAACTCATGCCCC | 168.9 | 5.1 | 1.6 | 3.3 | 1.1E-03 | 2.8E-03 | BT20/MB468 | MP |
| >hg38_21_+_8433172_8446622_RNA45SN1_45S_50ntFlanks@11891.11910.20\|TTGACTCTAGTCTGGCACGG | >hg38_21_+_8433172_8446622_RNA45SN1_45S_50ntFlanks | 11891 | 11910 | 20 | TTGACTCTAGTCTGGCACGG | 293.5 | -2.4 | 0.8 | -3.0 | 2.4E-03 | 2.8E-03 | MB231/MB468 | NUC |
| >hg38_21_+_8433172_8446622_RNA45SN1_45S_50ntFlanks@8196.8230.35\|CGTGGACGGTGTGAGGCCGGTAGCGGCCCCCGGCG | >hg38_21_+_8433172_8446622_RNA45SN1_45S_50ntFlanks | 8196 | 8230 | 35 | CGTGGACGGTGTGAGGCCGGTAGCGGCCCCCGGCG | 140.9 | -4.0 | 1.3 | -3.1 | 2.0E-03 | 2.8E-03 | MB231/MB468 | MP |
| >MI0000088\|hsa-mir-30a&WithFlank&6\|-\|71403545\|71403627@12.35.24&[MIMAT0000087&hsa-miR-30a-5p&offsets\|0\|+2;m-15&6\|-\|71403595\|71403616&offsets\|0\|+2]\|TGTAAACATCCTCGACTGGAAGCT | >MI0000088\|hsa-mir-30a&WithFlank&6\|-\|71403545\|71403627 | 12 | 35 | 24 | TGTAAACATCCTCGACTGGAAGCT | 11627.3 | 3.2 | 1.0 | 3.0 | 2.3E-03 | 2.8E-03 | BT20/MB468 | NUC |
| >am_MI0000786\|hsa-mir-378a&WithFlank&5\|+\|149732819\|149732896@49.70.22(+1U)&[MIMAT0000732&hsa-miR-378a-3p&offsets\|0\|0(+1U);m-44&5\|+\|149732867\|149732888&offsets\|0\|0(+1U)]\|ACTGGACTTGGAGTCAGAAGGCT | >am_MI0000786\|hsa-mir-378a&WithFlank&5\|+\|149732819\|149732896 | 49 | 70 | 22(+1U) | ACTGGACTTGGAGTCAGAAGGCT | 777.8 | -1.3 | 0.4 | -3.0 | 2.3E-03 | 2.8E-03 | BT20/MB468 | NUC |
| >hg38_21_+_8433172_8446622_RNA45SN1_45S_50ntFlanks@11208.11254.47\|GCGGCGACTCTGGACGCGAGCCGGGCCCTTCCCGTGGATCGCCCCAG | >hg38_21_+_8433172_8446622_RNA45SN1_45S_50ntFlanks | 11208 | 11254 | 47 | GCGGCGACTCTGGACGCGAGCCGGGCCCTTCCCGTGGATCGCCCCAG | 5933.0 | -3.5 | 1.1 | -3.1 | 1.8E-03 | 2.8E-03 | BT20/MB468 | MITO |
| >hg38_21_+_8433172_8446622_RNA45SN1_45S_50ntFlanks@5487.5516.30\|CGCTGAGAAGACGGTCGAACTTGACTATCT | >hg38_21_+_8433172_8446622_RNA45SN1_45S_50ntFlanks | 5487 | 5516 | 30 | CGCTGAGAAGACGGTCGAACTTGACTATCT | 1302.9 | -3.2 | 1.0 | -3.1 | 2.0E-03 | 2.8E-03 | MB231/MB468 | MP |
| >hg38_21_+_8433172_8446622_RNA45SN1_45S_50ntFlanks@12022.12055.34\|CCACTACTCTGATCGTTTTTTCACTGACCCGGTG | >hg38_21_+_8433172_8446622_RNA45SN1_45S_50ntFlanks | 12022 | 12055 | 34 | CCACTACTCTGATCGTTTTTTCACTGACCCGGTG | 203.5 | 5.6 | 1.6 | 3.5 | 3.9E-04 | 2.8E-03 | BT20/MB231 | MP |
| >hg38_21_+_8433172_8446622_RNA45SN1_45S_50ntFlanks@11199.11220.22\|CCGGCGGCGGCGGCGACTCTGG | >hg38_21_+_8433172_8446622_RNA45SN1_45S_50ntFlanks | 11199 | 11220 | 22 | CCGGCGGCGGCGGCGACTCTGG | 481.7 | -3.7 | 1.2 | -3.1 | 2.2E-03 | 2.8E-03 | BT20/MB468 | CYTO |
| >trnaMT_GluTTC_MT_-_14674_14742@38.71.34,trnalookalike8_GluTTC_5_-_93905172_93905240@38.71.34\|CATTGGTCGTGGTTGTAGTCCGTGCGAGAATACC | >trnaMT_GluTTC_MT_-_14674_14742 | 38 | 71 | 34 | CATTGGTCGTGGTTGTAGTCCGTGCGAGAATACC | 351.4 | -2.7 | 0.8 | -3.5 | 4.6E-04 | 2.8E-03 | BT20/MB231 | CYTO |
| >am_trnaMT_MetCAT_MT_+_4402_4469@1.19.19\|AGTAAGGTCAGCTAAATAA | >am_trnaMT_MetCAT_MT_+_4402_4469 | 1 | 19 | 19 | AGTAAGGTCAGCTAAATAA | 327.1 | 4.3 | 1.4 | 3.0 | 2.3E-03 | 2.8E-03 | MB231/MB468 | MITO |

| | | | | | | | | | | | |
|---|---|---|---|---|---|---|---|---|---|---|---|
| >hg38_21_+_8433172_8446622_RNA45SN1_45S_50ntFlanks@11209.11251.43\|CGGCGACTCTGGACGCGAGCCGGGCCCTTCCCGTGGATCGCCC | >hg38_21_+_8433172_8446622_RNA45SN1_45S_50ntFlanks | 11209 | 11251 | 43 | CGGCGACTCTGGACGCGAGCCGGGCCCTTCCCGTGGATCGCCC | 733.8 | -5.9 | 1.9 | -3.0 | 2.3E-03 | 2.8E-03 | BT20/MB468 | NUC |
| >hg38_21_+_8433172_8446622_RNA45SN1_45S_50ntFlanks@7975.8006.32\|CGCGACCTCAGATCAGACGTGGCGACCCGCTG | >hg38_21_+_8433172_8446622_RNA45SN1_45S_50ntFlanks | 7975 | 8006 | 32 | CGCGACCTCAGATCAGACGTGGCGACCCGCTG | 25007.0 | 4.6 | 1.3 | 3.5 | 3.9E-04 | 2.8E-03 | BT20/MB231 | MP |
| >MI0000088\|hsa-mir-30a&WithFlank&6\|-\|71403545\|71403627@54.73.20(+2U)&[MIMAT0000088&hsa-miR-30a-3p&offsets\|+1\|-1(+2U);m-38&6\|-\|71403554\|71403575&offsets\|+1\|-1(+2U)]\|TTTCAGTCGGATGTTTGCAGTT | >MI0000088\|hsa-mir-30a&WithFlank&6\|-\|71403545\|71403627 | 54 | 73 | 20(+2U) | TTTCAGTCGGATGTTTGCAGTT | 763.5 | -3.5 | 1.0 | -3.5 | 3.9E-04 | 2.8E-03 | BT20/MB231 | MP |
| >MI0000108\|hsa-mir-103a-2&WithFlank&20\|+\|3917488\|3917577@17.40.24&[MIMAT0009196&hsa-miR-103a-2-5p&offsets\|0\|+1;m-481&20\|+\|3917504\|3917526&offsets\|0\|+1]\|AGCTTCTTTACAGTGCTGCCTTGT | >MI0000108\|hsa-mir-103a-2&WithFlank&20\|+\|3917488\|3917577 | 17 | 40 | 24 | AGCTTCTTTACAGTGCTGCCTTGT | 55.9 | 4.8 | 1.5 | 3.1 | 2.0E-03 | 2.8E-03 | MB231/MB468 | MP |
| >hg38_21_+_8433172_8446622_RNA45SN1_45S_50ntFlanks@3987.4019.33\|GCTTTGGTGACTCTAGATAACCTCGGGCCGATC | >hg38_21_+_8433172_8446622_RNA45SN1_45S_50ntFlanks | 3987 | 4019 | 33 | GCTTTGGTGACTCTAGATAACCTCGGGCCGATC | 575.4 | 7.5 | 2.1 | 3.5 | 3.9E-04 | 2.8E-03 | BT20/MB231 | MP |
| >MI0000095\|hsa-mir-93&WithFlank&7\|-\|100093762\|100093853@17.37.21&[MIMAT0000093&hsa-miR-93-5p&offsets\|0\|-2;m-29&7\|-\|100093815\|100093837&offsets\|0\|-2]\|CAAAGTGCTGTTCGTGCAGGT | >MI0000095\|hsa-mir-93&WithFlank&7\|-\|100093762\|100093853 | 17 | 37 | 21 | CAAAGTGCTGTTCGTGCAGGT | 326.7 | 2.0 | 0.7 | 3.0 | 2.3E-03 | 2.8E-03 | BT20/MB468 | NUC |
| >hg38_21_+_8433172_8446622_RNA45SN1_45S_50ntFlanks@10351.10392.42\|CGAGAACTTTGAAGGCCGAAGTGGAGAAGGGTTCCATGTGAA | >hg38_21_+_8433172_8446622_RNA45SN1_45S_50ntFlanks | 10351 | 10392 | 42 | CGAGAACTTTGAAGGCCGAAGTGGAGAAGGGTTCCATGTGAA | 60.9 | -5.2 | 1.7 | -3.1 | 2.0E-03 | 2.8E-03 | MB231/MB468 | MP |
| >hg38_1_-_228634819_228635039_RNA5S12_5S_50ntFlanks@146.169.24\|TGGGAATACCGGGTGCTGTAGGCT | >hg38_1_-_228634819_228635039_RNA5S12_5S_50ntFlanks | 146 | 169 | 24 | TGGGAATACCGGGTGCTGTAGGCT | 2185.1 | 4.5 | 1.3 | 3.5 | 4.0E-04 | 2.8E-03 | BT20/MB231 | MP |
| >hg38_21_+_8433172_8446622_RNA45SN1_45S_50ntFlanks@8366.8395.30\|TTGAAGAGAGAGTTCAAGAGGGCGTGAAAC | >hg38_21_+_8433172_8446622_RNA45SN1_45S_50ntFlanks | 8366 | 8395 | 30 | TTGAAGAGAGAGTTCAAGAGGGCGTGAAAC | 88.9 | 9.8 | 2.8 | 3.5 | 4.0E-04 | 2.8E-03 | BT20/MB231 | MP |
| >hg38_21_+_8433172_8446622_RNA45SN1_45S_50ntFlanks@9932.9952.21\|ATAGACAGCAGGACGGTGGCC | >hg38_21_+_8433172_8446622_RNA45SN1_45S_50ntFlanks | 9932 | 9952 | 21 | ATAGACAGCAGGACGGTGGCC | 318.1 | 6.8 | 1.9 | 3.5 | 4.0E-04 | 2.8E-03 | BT20/MB231 | MP |
| >hg38_21_+_8433172_8446622_RNA45SN1_45S_50ntFlanks@11582.11605.24\|ACTGGTGCGGACCAGGGGAATCCG | >hg38_21_+_8433172_8446622_RNA45SN1_45S_50ntFlanks | 11582 | 11605 | 24 | ACTGGTGCGGACCAGGGGAATCCG | 430.5 | -2.5 | 0.8 | -3.1 | 2.2E-03 | 2.8E-03 | MB231/MB468 | CYTO |
| >MI0000465\|hsa-mir-191&WithFlank&3\|-\|49020612\|49020715@22.39.18&[MIMAT0000440&hsa-miR-191-5p&offsets\|0\|-5;m-48&3\|-\|49020672\|49020694&offsets\|0\|-5]\|CAACGGAATCCCAAAAGC | >MI0000465\|hsa-mir-191&WithFlank&3\|-\|49020612\|49020715 | 22 | 39 | 18 | CAACGGAATCCCAAAAGC | 258.1 | 2.9 | 0.9 | 3.1 | 2.0E-03 | 2.8E-03 | MB231/MB468 | MP |
| >hg38_MT_+_1621_3279_RNR2_16S_50ntFlanks@1095.1120.26\|AGACGAGAAGACCCTATGGAGCTTTA | >hg38_MT_+_1621_3279_RNR2_16S_50ntFlanks | 1095 | 1120 | 26 | AGACGAGAAGACCCTATGGAGCTTTA | 386.8 | 2.5 | 0.8 | 3.1 | 2.0E-03 | 2.8E-03 | MB231/MB468 | MP |
| >hg38_21_+_8433172_8446622_RNA45SN1_45S_50ntFlanks@7975.7994.20\|CGCGACCTCAGATCAGACGT | >hg38_21_+_8433172_8446622_RNA45SN1_45S_50ntFlanks | 7975 | 7994 | 20 | CGCGACCTCAGATCAGACGT | 76425.6 | 2.5 | 0.8 | 3.1 | 2.2E-03 | 2.8E-03 | MB231/MB468 | CYTO |
| >hg38_1_-_228634819_228635039_RNA5S12_5S_50ntFlanks@144.169.26\|CCTGGGAATACCGGGTGCTGTAGGCT | >hg38_1_-_228634819_228635039_RNA5S12_5S_50ntFlanks | 144 | 169 | 26 | CCTGGGAATACCGGGTGCTGTAGGCT | 264.6 | -2.8 | 0.9 | -3.1 | 2.0E-03 | 2.8E-03 | MB231/MB468 | MP |
| >hg38_21_+_8433172_8446622_RNA45SN1_45S_50ntFlanks@9203.9223.21\|CTCGGGGCCACGCGCGCGTCC | >hg38_21_+_8433172_8446622_RNA45SN1_45S_50ntFlanks | 9203 | 9223 | 21 | CTCGGGGCCACGCGCGCGTCC | 478.4 | -5.5 | 1.7 | -3.3 | 1.2E-03 | 2.8E-03 | BT20/MB468 | MP |
| >hg38_MT_+_1621_3279_RNR2_16S_50ntFlanks@469.491.23\|TAACTGTTAGTCCAAAGAGGAAC | >hg38_MT_+_1621_3279_RNR2_16S_50ntFlanks | 469 | 491 | 23 | TAACTGTTAGTCCAAAGAGGAAC | 55.5 | 6.2 | 1.9 | 3.3 | 1.2E-03 | 2.8E-03 | BT20/MB468 | MP |
| >MI0000255\|hsa-mir-30d&WithFlank&8\|-\|134804870\|134804951@12.32.21&[MIMAT0000245&hsa-miR-30d-5p&offsets\|0\|-1;m-21&8\|-\|134804919\|134804940&offsets\|0\|-1]\|TGTAAACATCCCCGACTGGAA | >MI0000255\|hsa-mir-30d&WithFlank&8\|-\|134804870\|134804951 | 12 | 32 | 21 | TGTAAACATCCCCGACTGGAA | 1161.0 | 3.2 | 0.9 | 3.5 | 4.7E-04 | 2.8E-03 | BT20/MB231 | CYTO |
| >hg38_21_+_8433172_8446622_RNA45SN1_45S_50ntFlanks@8168.8186.19\|CCCAAGTCCTTCTGATCGA | >hg38_21_+_8433172_8446622_RNA45SN1_45S_50ntFlanks | 8168 | 8186 | 19 | CCCAAGTCCTTCTGATCGA | 986.1 | -4.4 | 1.4 | -3.1 | 2.2E-03 | 2.8E-03 | BT20/MB468 | CYTO |
| >hg38_21_+_8433172_8446622_RNA45SN1_45S_50ntFlanks@10798.10816.19\|CATGTTGGAACAATGTAGG | >hg38_21_+_8433172_8446622_RNA45SN1_45S_50ntFlanks | 10798 | 10816 | 19 | CATGTTGGAACAATGTAGG | 242.8 | -5.2 | 1.6 | -3.3 | 1.2E-03 | 2.8E-03 | BT20/MB468 | MP |
| >hg38_21_+_8433172_8446622_RNA45SN1_45S_50ntFlanks@12406.12444.39\|TTTTTGATCCTTCGATGTCGGCTCTTCCTATCATTGTGA | >hg38_21_+_8433172_8446622_RNA45SN1_45S_50ntFlanks | 12406 | 12444 | 39 | TTTTTGATCCTTCGATGTCGGCTCTTCCTATCATTGTGA | 354.8 | -2.6 | 0.8 | -3.0 | 2.4E-03 | 2.8E-03 | MB231/MB468 | NUC |
| >hg38_21_+_8433172_8446622_RNA45SN1_45S_50ntFlanks@12399.12435.37\|ACGTCGCTTTTTGATCCTTCGATGTCGGCTCTTCCTA | >hg38_21_+_8433172_8446622_RNA45SN1_45S_50ntFlanks | 12399 | 12435 | 37 | ACGTCGCTTTTTGATCCTTCGATGTCGGCTCTTCCTA | 126.0 | -4.4 | 1.4 | -3.1 | 1.8E-03 | 2.8E-03 | BT20/MB468 | MITO |
| >hg38_21_+_8433172_8446622_RNA45SN1_45S_50ntFlanks@12423.12441.19\|TCGGCTCTTCCTATCATTG | >hg38_21_+_8433172_8446622_RNA45SN1_45S_50ntFlanks | 12423 | 12441 | 19 | TCGGCTCTTCCTATCATTG | 177.1 | -3.2 | 1.0 | -3.1 | 2.2E-03 | 2.8E-03 | BT20/MB468 | CYTO |
| >trnaMT_TyrGTA_MT_-_5826_5891@32.69.38\|AAATCTAAAGACAGGGGTTAGGCCTCTTTTTACCACCA | >trnaMT_TyrGTA_MT_-_5826_5891 | 32 | 69 | 38 | AAATCTAAAGACAGGGGTTAGGCCTCTTTTTACCACCA | 216.7 | 5.2 | 1.7 | 3.1 | 2.0E-03 | 2.8E-03 | MB231/MB468 | MP |
| >MI0000433\|hsa-let-7g&WithFlank&3\|-\|52268272\|52268367@11.31.21(+1G)&[MIMAT0000414&hsa-let-7g-5p&offsets\|0\|-1(+1G);m-58&3\|-\|52268336\|52268357&offsets\|0\|-1(+1G)]\|TGAGGTAGTAGTTTGTACAGTG | >MI0000433\|hsa-let-7g&WithFlank&3\|-\|52268272\|52268367 | 11 | 31 | 21(+1G) | TGAGGTAGTAGTTTGTACAGTG | 502.3 | 2.4 | 0.8 | 3.0 | 2.4E-03 | 2.8E-03 | BT20/MB468 | NUC |
| >hg38_21_+_8433172_8446622_RNA45SN1_45S_50ntFlanks@12417.12435.19\|TCGATGTCGGCTCTTCCTA | >hg38_21_+_8433172_8446622_RNA45SN1_45S_50ntFlanks | 12417 | 12435 | 19 | TCGATGTCGGCTCTTCCTA | 13304.2 | -6.9 | 2.3 | -3.0 | 2.4E-03 | 2.8E-03 | BT20/MB468 | NUC |
| >hg38_21_+_8433172_8446622_RNA45SN1_45S_50ntFlanks@8194.8212.19\|CCCGTGGACGGTGTGAGGC | >hg38_21_+_8433172_8446622_RNA45SN1_45S_50ntFlanks | 8194 | 8212 | 19 | CCCGTGGACGGTGTGAGGC | 641.8 | -2.2 | 0.7 | -3.1 | 2.2E-03 | 2.8E-03 | MB231/MB468 | CYTO |
| >Antisense_hg38_21_-_8433172_8446622_RNA45SN1_45S_50ntFlanks@7674.7691.18\|CTCTCTCTCTCTCTCCCC | >Antisense_hg38_21_-_8433172_8446622_RNA45SN1_45S_50ntFlanks | 7674 | 7691 | 18 | CTCTCTCTCTCTCTCCCC | 188.7 | -3.4 | 1.1 | -3.1 | 2.2E-03 | 2.8E-03 | BT20/MB468 | CYTO |
| >hg38_21_+_8433172_8446622_RNA45SN1_45S_50ntFlanks@11027.11044.18\|CTCTCTCTCTCTCTCCCC | >hg38_21_+_8433172_8446622_RNA45SN1_45S_50ntFlanks | 11027 | 11044 | 18 | CTCTCTCTCTCTCTCCCC | 188.7 | -3.4 | 1.1 | -3.1 | 2.2E-03 | 2.8E-03 | BT20/MB468 | CYTO |
| >hg38_21_+_8433172_8446622_RNA45SN1_45S_50ntFlanks@10840.10864.25\|CCGTAACTTCGGGATAAGGATTGGC | >hg38_21_+_8433172_8446622_RNA45SN1_45S_50ntFlanks | 10840 | 10864 | 25 | CCGTAACTTCGGGATAAGGATTGGC | 207.9 | -3.6 | 1.2 | -3.1 | 2.2E-03 | 2.8E-03 | BT20/MB468 | CYTO |
| >MI0000298\|hsa-mir-221&WithFlank&X\|-\|45746151\|45746272@71.91.21&[MIMAT0000278&hsa-miR-221-3p&offsets\|0\|-2;m-91&X\|-\|45746181\|45746202&offsets\|0\|-1]\|AGCTACATTGTCTGCTGGGTT | >MI0000298\|hsa-mir-221&WithFlank&X\|-\|45746151\|45746272 | 71 | 91 | 21 | AGCTACATTGTCTGCTGGGTT | 2227.2 | 4.0 | 1.2 | 3.2 | 1.2E-03 | 2.8E-03 | BT20/MB468 | MP |
| >hg38_21_+_8433172_8446622_RNA45SN1_45S_50ntFlanks@9457.9488.32\|CGCCGGGGAGGTGGAGCACGAGCGCACGTGTT | >hg38_21_+_8433172_8446622_RNA45SN1_45S_50ntFlanks | 9457 | 9488 | 32 | CGCCGGGGAGGTGGAGCACGAGCGCACGTGTT | 163.1 | -3.3 | 1.0 | -3.1 | 1.8E-03 | 2.8E-03 | BT20/MB468 | MITO |
| >trnaMT_SerGCT_MT_+_12207_12265@1.21.21\|GAGAAAGCTCACAAGAACTGC | >trnaMT_SerGCT_MT_+_12207_12265 | 1 | 21 | 21 | GAGAAAGCTCACAAGAACTGC | 20443.9 | -3.3 | 0.9 | -3.5 | 4.0E-04 | 2.8E-03 | BT20/MB231 | MP |
| >hg38_21_+_8433172_8446622_RNA45SN1_45S_50ntFlanks@476.520.45\|TTCGTGATCGATGTGGTGACGTCGTGCTCTCCCGGGCCGGGTCCG | >hg38_21_+_8433172_8446622_RNA45SN1_45S_50ntFlanks | 476 | 520 | 45 | TTCGTGATCGATGTGGTGACGTCGTGCTCTCCCGGGCCGGGTCCG | 234.8 | 4.7 | 1.6 | 3.0 | 2.4E-03 | 2.8E-03 | MB231/MB468 | MITO |
| >hg38_21_+_8433172_8446622_RNA45SN1_45S_50ntFlanks@4353.4371.19\|TATTAAAGTTGCTGCAGTT | >hg38_21_+_8433172_8446622_RNA45SN1_45S_50ntFlanks | 4353 | 4371 | 19 | TATTAAAGTTGCTGCAGTT | 391.0 | -1.9 | 0.6 | -3.1 | 2.2E-03 | 2.8E-03 | MB231/MB468 | CYTO |
| >hg38_21_+_8433172_8446622_RNA45SN1_45S_50ntFlanks@5477.5499.23\|CCTGGCGGAGCGCTGAGAAGACG | >hg38_21_+_8433172_8446622_RNA45SN1_45S_50ntFlanks | 5477 | 5499 | 23 | CCTGGCGGAGCGCTGAGAAGACG | 84.0 | -3.8 | 1.2 | -3.1 | 2.0E-03 | 2.8E-03 | MB231/MB468 | MP |
| >hg38_21_+_8433172_8446622_RNA45SN1_45S_50ntFlanks@11917.11943.27\|GACATGAGAGGTGTAGAATAAGTGGGA | >hg38_21_+_8433172_8446622_RNA45SN1_45S_50ntFlanks | 11917 | 11943 | 27 | GACATGAGAGGTGTAGAATAAGTGGGA | 88.0 | 9.8 | 2.8 | 3.5 | 4.0E-04 | 2.8E-03 | BT20/MB231 | MP |
| >hg38_21_+_8433172_8446622_RNA45SN1_45S_50ntFlanks@6719.6758.40\|TGAATTGCAGGACACATTGATCATCGACACTTCGAACGCA | >hg38_21_+_8433172_8446622_RNA45SN1_45S_50ntFlanks | 6719 | 6758 | 40 | TGAATTGCAGGACACATTGATCATCGACACTTCGAACGCA | 288.5 | -1.8 | 0.6 | -3.1 | 2.3E-03 | 2.8E-03 | MB231/MB468 | CYTO |
| >MI0000253\|hsa-mir-148a&WithFlank&7\|-\|25949913\|25949992@50.70.21&[MIMAT0000243&hsa-miR-148a-3p&offsets\|0\|-1;m-7&7\|-\|25949922\|25949943&offsets\|0\|-1]\|TCAGTGCACTACAGAACTTTG | >MI0000253\|hsa-mir-148a&WithFlank&7\|-\|25949913\|25949992 | 50 | 70 | 21 | TCAGTGCACTACAGAACTTTG | 482.2 | 4.0 | 1.2 | 3.2 | 1.2E-03 | 2.8E-03 | BT20/MB468 | MP |
| >hg38_21_+_8433172_8446622_RNA45SN1_45S_50ntFlanks@12456.12484.29\|CCAAGCGTTGGATTGTTCACCCACTAATA | >hg38_21_+_8433172_8446622_RNA45SN1_45S_50ntFlanks | 12456 | 12484 | 29 | CCAAGCGTTGGATTGTTCACCCACTAATA | 3886.6 | -6.1 | 1.9 | -3.2 | 1.2E-03 | 2.8E-03 | BT20/MB468 | MP |
| >MI0000067\|hsa-let-7f-1&WithFlank&9\|+\|94176341\|94176439@13.31.19;MI0000068\|hsa-let-7f-2&WithFlank&X\|-\|53557186\|53557280@14.32.19&[MIMAT0000067&hsa-let-7f-5p&offsets\|0\|-3;m-20&9\|+\|94176353\|94176374&offsets\|0\|-3];[MIMAT0000067_1&hsa-let-7f-5p&offsets\|0\|-3;m-19&X\|-\|53557246\|53557267&offsets\|0\|-3]\|TGAGGTAGTAGATTGTATA | >MI0000067\|hsa-let-7f-1&WithFlank&9\|+\|94176341\|94176439 | 13 | 31 | 19 | TGAGGTAGTAGATTGTATA | 2859.2 | 3.5 | 1.0 | 3.5 | 4.7E-04 | 2.8E-03 | BT20/MB231 | CYTO |
| >MI0000264\|hsa-mir-7-2&WithFlank&15\|+\|88611819\|88611940@38.61.24(+1A);MI0000265\|hsa-mir-7-3&WithFlank&19\|+\|4770664\|4770785@37.60.24(+1A);MI0000263\|hsa-mir-7-1&WithFlank&9\|-\|83969742\|83969863@30.53.24(+1A)&[MIMAT0000252&hsa-miR-7-5p&offsets\|0\|0(+1A);m-400&15\|+\|88611856\|88611879&offsets\|0\|0(+1A)];[MIMAT0000252_1&hsa-miR-7-5p&offsets\|0\|0(+1A);m-402&19\|+\|4770700\|4770723&offsets\|0\|0(+1A)];[MIMAT0000252_2&hsa-miR-7-5p&offsets\|0\|0(+1A);m-401&9\|-\|83969811\|83969834&offsets\|0\|0(+1A)]\|TGGAAGACTAGTGATTTTGTTGTTA | >MI0000264\|hsa-mir-7-2&WithFlank&15\|+\|88611819\|88611940 | 38 | 61 | 24(+1A) | TGGAAGACTAGTGATTTTGTTGTTA | 918.9 | 3.1 | 0.9 | 3.5 | 4.8E-04 | 2.8E-03 | BT20/MB231 | CYTO |
| >hg38_21_+_8433172_8446622_RNA45SN1_45S_50ntFlanks@10427.10453.27\|ATGGGCGAGCGCCGTTCCGAAGGGACG | >hg38_21_+_8433172_8446622_RNA45SN1_45S_50ntFlanks | 10427 | 10453 | 27 | ATGGGCGAGCGCCGTTCCGAAGGGACG | 217.5 | -2.9 | 0.8 | -3.5 | 4.7E-04 | 2.8E-03 | BT20/MB231 | CYTO |
| >hg38_21_+_8433172_8446622_RNA45SN1_45S_50ntFlanks@6650.6698.49\|ACGACTCTTAGCGGTGGATCACTCGGCTCGTGCGTCGATGAAGAACGCA | >hg38_21_+_8433172_8446622_RNA45SN1_45S_50ntFlanks | 6650 | 6698 | 49 | ACGACTCTTAGCGGTGGATCACTCGGCTCGTGCGTCGATGAAGAACGCA | 598.7 | -3.6 | 1.0 | -3.5 | 4.8E-04 | 2.8E-03 | BT20/MB231 | CYTO |
| >hg38_21_+_8433172_8446622_RNA45SN1_45S_50ntFlanks@6649.6685.37\|TACGACTCTTAGCGGTGGATCACTCGGCTCGTGCGTC | >hg38_21_+_8433172_8446622_RNA45SN1_45S_50ntFlanks | 6649 | 6685 | 37 | TACGACTCTTAGCGGTGGATCACTCGGCTCGTGCGTC | 334.8 | 3.1 | 1.0 | 3.2 | 1.2E-03 | 2.8E-03 | BT20/MB468 | MP |
| >hg38_21_+_8433172_8446622_RNA45SN1_45S_50ntFlanks@8182.8205.24\|ATCGAGGCCCAGCCCGTGGACGGT | >hg38_21_+_8433172_8446622_RNA45SN1_45S_50ntFlanks | 8182 | 8205 | 24 | ATCGAGGCCCAGCCCGTGGACGGT | 183.1 | -5.0 | 1.6 | -3.1 | 2.0E-03 | 2.8E-03 | MB231/MB468 | MP |
| >hg38_21_+_8433172_8446622_RNA45SN1_45S_50ntFlanks@11318.11344.27\|CCCCACGTCTCGTCGCGCGCGCGTCCG | >hg38_21_+_8433172_8446622_RNA45SN1_45S_50ntFlanks | 11318 | 11344 | 27 | CCCCACGTCTCGTCGCGCGCGCGTCCG | 7556.5 | -5.8 | 1.9 | -3.1 | 1.8E-03 | 2.8E-03 | BT20/MB468 | MITO |
| >hg38_21_+_8433172_8446622_RNA45SN1_45S_50ntFlanks@3900.3935.36\|CTTCGCGGGGGGGATGCGTGCATTTATCAGATCAAA | >hg38_21_+_8433172_8446622_RNA45SN1_45S_50ntFlanks | 3900 | 3935 | 36 | CTTCGCGGGGGGGATGCGTGCATTTATCAGATCAAA | 161.0 | -5.5 | 1.7 | -3.2 | 1.2E-03 | 2.8E-03 | BT20/MB468 | MP |
| >MI0000469\|hsa-mir-125a&WithFlank&19\|+\|51693248\|51693345@21.42.22(+1A)&[MIMAT0000443&hsa-miR-125a-5p&offsets\|0\|-2(+1A);m-63&19\|+\|51693268\|51693289&offsets\|0\|0(+1A)]\|TCCCTGAGACCCTTTAACCTGTA | >MI0000469\|hsa-mir-125a&WithFlank&19\|+\|51693248\|51693345 | 21 | 42 | 22(+1A) | TCCCTGAGACCCTTTAACCTGTA | 307.4 | 3.0 | 0.9 | 3.2 | 1.2E-03 | 2.8E-03 | BT20/MB468 | MP |
| >hg38_21_+_8433172_8446622_RNA45SN1_45S_50ntFlanks@11575.11601.27\|ACTTAGAACTGGTGCGGACCAGGGGAA | >hg38_21_+_8433172_8446622_RNA45SN1_45S_50ntFlanks | 11575 | 11601 | 27 | ACTTAGAACTGGTGCGGACCAGGGGAA | 337.8 | -3.5 | 1.1 | -3.1 | 1.9E-03 | 2.8E-03 | BT20/MB468 | MITO |

| | | | | | | | | | | | | |
|---|---|---|---|---|---|---|---|---|---|---|---|---|
| >hg38_21_+_8433172_8446622_RNA45SN1_45S_50ntFlanks@12418.12454.37\|CGATGTCGGCTCTTCCTATCATTGTGAAGCAGAATTC | >hg38_21_+_8433172_8446622_RNA45SN1_45S_50ntFlanks | 12418 | 12454 | 37 | CGATGTCGGCTCTTCCTATCATTGTGAAGCAGAATTC | 717.0 | -2.5 | 0.8 | -3.0 | 2.4E-03 | 2.8E-03 | MB231/MB468 | MITO |
| >hg38_21_+_8433172_8446622_RNA45SN1_45S_50ntFlanks@5061.5098.38\|ATTTGTCTGGTTAATTCCGATAACGAACGAGACTCTGG | >hg38_21_+_8433172_8446622_RNA45SN1_45S_50ntFlanks | 5061 | 5098 | 38 | ATTTGTCTGGTTAATTCCGATAACGAACGAGACTCTGG | 46.2 | -9.4 | 3.0 | -3.1 | 2.0E-03 | 2.8E-03 | MB231/MB468 | MP |
| >hg38_21_+_8433172_8446622_RNA45SN1_45S_50ntFlanks@8358.8375.18\|AAAGAACTTTGAAGAGAG | >hg38_21_+_8433172_8446622_RNA45SN1_45S_50ntFlanks | 8358 | 8375 | 18 | AAAGAACTTTGAAGAGAG | 1090.7 | 6.3 | 1.8 | 3.5 | 5.4E-04 | 2.8E-03 | BT20/MB231 | MITO |
| >hg38_21_+_8433172_8446622_RNA45SN1_45S_50ntFlanks@9553.9570.18\|CCGTAGCGGTCCTGACGT | >hg38_21_+_8433172_8446622_RNA45SN1_45S_50ntFlanks | 9553 | 9570 | 18 | CCGTAGCGGTCCTGACGT | 225.8 | 1.4 | 0.5 | 3.0 | 2.4E-03 | 2.8E-03 | MB231/MB468 | NUC |
| >MI0000067\|hsa-let-7f-1&WithFlank&9\|+\|94176341\|94176439@13.32.20(+1A);MI0000068\|hsa-let-7f-2&WithFlank&X\|-\|53557186\|53557280@14.33.20(+1A)&[MIMAT0000067&hsa-let-7f-5p&offsets\|0\|-2(+1A);m-20&9\|+\|94176353\|94176374&offsets\|0\|-2(+1A)];[MIMAT0000067_1&hsa-let-7f-5p&offsets\|0\|-2(+1A);m-19&X\|-\|53557246\|53557267&offsets\|0\|-2(+1A)]\|TGAGGTAGTAGATTGTATAGA | >MI0000067\|hsa-let-7f-1&WithFlank&9\|+\|94176341\|94176439 | 13 | 32 | 20(+1A) | TGAGGTAGTAGATTGTATAGA | 135.4 | 5.0 | 1.6 | 3.1 | 2.0E-03 | 2.8E-03 | MB231/MB468 | MP |
| >MI0000289\|hsa-mir-181a-1&WithFlank&1\|-\|198859038\|198859159@30.54.25(+1U);MI0000269\|hsa-mir-181a-2&WithFlank&9\|+\|124692436\|124692557@45.69.25(+1U)&[MIMAT0000256&hsa-miR-181a-5p&offsets\|0\|+2(+1U);m-32&1\|-\|198859108\|198859130&offsets\|0\|+2(+1U)];[MIMAT0000256_1&hsa-miR-181a-5p&offsets\|0\|+2(+1U);m-32&9\|+\|124692480\|124692502&offsets\|0\|+2(+1U)]\|AACATTCAACGCTGTCGGTGAGTTTT | >MI0000289\|hsa-mir-181a-1&WithFlank&1\|-\|198859038\|198859159 | 30 | 54 | 25(+1U) | AACATTCAACGCTGTCGGTGAGTTTT | 806.8 | 4.3 | 1.4 | 3.1 | 2.0E-03 | 2.8E-03 | MB231/MB468 | MP |
| >MI0000088\|hsa-mir-30a&WithFlank&6\|-\|71403545\|71403627@12.37.26&[MIMAT0000087&hsa-miR-30a-5p&offsets\|0\|+4;m-15&6\|-\|71403595\|71403616&offsets\|0\|+4]\|TGTAAACATCCTCGACTGGAAGCTGT | >MI0000088\|hsa-mir-30a&WithFlank&6\|-\|71403545\|71403627 | 12 | 37 | 26 | TGTAAACATCCTCGACTGGAAGCTGT | 64.8 | 5.6 | 1.8 | 3.1 | 2.0E-03 | 2.8E-03 | MB231/MB468 | MP |
| >MI0000108\|hsa-mir-103a-2&WithFlank&20\|+\|3917488\|3917577@54.78.25&[MIMAT0000101&hsa-miR-103a-3p&offsets\|0\|+2;m-12&20\|+\|3917541\|3917563&offsets\|0\|+2]\|AGCAGCATTGTACAGGGCTATGAAA | >MI0000108\|hsa-mir-103a-2&WithFlank&20\|+\|3917488\|3917577 | 54 | 78 | 25 | AGCAGCATTGTACAGGGCTATGAAA | 69.3 | 1.9 | 0.6 | 3.0 | 2.4E-03 | 2.8E-03 | BT20/MB468 | NUC |
| >hg38_21_+_8433172_8446622_RNA45SN1_45S_50ntFlanks@9858.9899.42\|CGGGATGAACCGAACGCCGGGTTAAGGCGCCCGATGCCGACG | >hg38_21_+_8433172_8446622_RNA45SN1_45S_50ntFlanks | 9858 | 9899 | 42 | CGGGATGAACCGAACGCCGGGTTAAGGCGCCCGATGCCGACG | 1495.6 | -3.8 | 1.2 | -3.1 | 1.9E-03 | 2.8E-03 | BT20/MB468 | MITO |
| >hg38_MT_+_1621_3279_RNR2_16S_50ntFlanks@51.88.38\|GCTAAACCTAGCCCCAAACCCACTCCACCTTACTACCA | >hg38_MT_+_1621_3279_RNR2_16S_50ntFlanks | 51 | 88 | 38 | GCTAAACCTAGCCCCAAACCCACTCCACCTTACTACCA | 4676.3 | 4.5 | 1.4 | 3.2 | 1.2E-03 | 2.8E-03 | BT20/MB468 | MP |
| >hg38_21_+_8433172_8446622_RNA45SN1_45S_50ntFlanks@7982.8003.22\|TCAGATCAGACGTGGCGACCCG | >hg38_21_+_8433172_8446622_RNA45SN1_45S_50ntFlanks | 7982 | 8003 | 22 | TCAGATCAGACGTGGCGACCCG | 158.2 | -3.7 | 1.2 | -3.0 | 2.4E-03 | 2.8E-03 | MB231/MB468 | NUC |
| >hg38_21_+_8433172_8446622_RNA45SN1_45S_50ntFlanks@3781.3801.21\|ACAGTGAAACTGCGAATGGCT | >hg38_21_+_8433172_8446622_RNA45SN1_45S_50ntFlanks | 3781 | 3801 | 21 | ACAGTGAAACTGCGAATGGCT | 373.0 | -3.3 | 1.1 | -3.1 | 2.0E-03 | 2.8E-03 | MB231/MB468 | MP |
| >hg38_21_+_8433172_8446622_RNA45SN1_45S_50ntFlanks@9569.9597.29\|GTGCAAATCGGTCGTCCGACCTGGGTATA | >hg38_21_+_8433172_8446622_RNA45SN1_45S_50ntFlanks | 9569 | 9597 | 29 | GTGCAAATCGGTCGTCCGACCTGGGTATA | 562.7 | -5.0 | 1.6 | -3.1 | 2.2E-03 | 2.8E-03 | BT20/MB468 | CYTO |
| >MI0003591\|hsa-mir-584&WithFlank&5\|-\|149062307\|149062415@22.41.20(+1G)&[MIMAT0003249&hsa-miR-584-5p&offsets\|0\|-2(+1G);m-167&5\|-\|149062374\|149062394&offsets\|0\|-1(+1G)]\|TTATGGTTTGCCTGGGACTGG | >MI0003591\|hsa-mir-584&WithFlank&5\|-\|149062307\|149062415 | 22 | 41 | 20(+1G) | TTATGGTTTGCCTGGGACTGG | 163.4 | 5.8 | 1.8 | 3.2 | 1.2E-03 | 2.8E-03 | BT20/MB468 | MP |
| >hg38_MT_+_1621_3279_RNR2_16S_50ntFlanks@1343.1385.43\|ATATCAACAATAGGGTTTACGACCTCGATGTTGGATCAGGACA | >hg38_MT_+_1621_3279_RNR2_16S_50ntFlanks | 1343 | 1385 | 43 | ATATCAACAATAGGGTTTACGACCTCGATGTTGGATCAGGACA | 42.5 | 9.2 | 2.8 | 3.2 | 1.2E-03 | 2.8E-03 | BT20/MB468 | MP |
| >hg38_21_+_8433172_8446622_RNA45SN1_45S_50ntFlanks@12324.12342.19\|TTGGGTTTTAAGCAGGAGG | >hg38_21_+_8433172_8446622_RNA45SN1_45S_50ntFlanks | 12324 | 12342 | 19 | TTGGGTTTTAAGCAGGAGG | 241.2 | -5.3 | 1.6 | -3.2 | 1.2E-03 | 2.8E-03 | BT20/MB468 | MP |
| >hg38_21_+_8433172_8446622_RNA45SN1_45S_50ntFlanks@5524.5550.27\|GTAAAAGTCGTAACAAGGTTTCCGTAG | >hg38_21_+_8433172_8446622_RNA45SN1_45S_50ntFlanks | 5524 | 5550 | 27 | GTAAAAGTCGTAACAAGGTTTCCGTAG | 126.7 | -5.1 | 1.6 | -3.2 | 1.2E-03 | 2.8E-03 | BT20/MB468 | MP |
| >hg38_21_+_8433172_8446622_RNA45SN1_45S_50ntFlanks@12416.12436.21\|TTCGATGTCGGCTCTTCCTAT | >hg38_21_+_8433172_8446622_RNA45SN1_45S_50ntFlanks | 12416 | 12436 | 21 | TTCGATGTCGGCTCTTCCTAT | 1923.8 | -2.0 | 0.7 | -3.0 | 2.4E-03 | 2.8E-03 | MB231/MB468 | NUC |
| >hg38_21_+_8433172_8446622_RNA45SN1_45S_50ntFlanks@475.524.50\|CTTCGTGATCGATGTGGTGACGTCGTGCTCTCCCGGGCCGGGTCCGAGCC | >hg38_21_+_8433172_8446622_RNA45SN1_45S_50ntFlanks | 475 | 524 | 50 | CTTCGTGATCGATGTGGTGACGTCGTGCTCTCCCGGGCCGGGTCCGAGCC | 6789.1 | 4.5 | 1.5 | 3.0 | 2.4E-03 | 2.8E-03 | MB231/MB468 | NUC |
| >MI0000542\|hsa-mir-320a&WithFlank&8\|-\|22244960\|22245043@48.69.22(+1C)&[MIMAT0000510&hsa-miR-320a-3p&offsets\|0\|0(+1C);m-60&8\|-\|22244975\|22244996&offsets\|0\|0(+1C)]\|AAAAGCTGGGTTGAGAGGGCGAC | >MI0000542\|hsa-mir-320a&WithFlank&8\|-\|22244960\|22245043 | 48 | 69 | 22(+1C) | AAAAGCTGGGTTGAGAGGGCGAC | 355.1 | 4.3 | 1.4 | 3.1 | 2.0E-03 | 2.8E-03 | MB231/MB468 | MP |
| >hg38_MT_+_1621_3279_RNR2_16S_50ntFlanks@486.518.33\|AGGAACAGCTCTTTGGACACTAGGAAAAAACCT | >hg38_MT_+_1621_3279_RNR2_16S_50ntFlanks | 486 | 518 | 33 | AGGAACAGCTCTTTGGACACTAGGAAAAAACCT | 289.1 | 4.0 | 1.2 | 3.2 | 1.2E-03 | 2.8E-03 | BT20/MB468 | MP |
| >hg38_21_+_8433172_8446622_RNA45SN1_45S_50ntFlanks@8342.8375.34\|CGTAAGGGAAAGTTGAAAAGAACTTTGAAGAGAG | >hg38_21_+_8433172_8446622_RNA45SN1_45S_50ntFlanks | 8342 | 8375 | 34 | CGTAAGGGAAAGTTGAAAAGAACTTTGAAGAGAG | 4726.0 | 4.7 | 1.3 | 3.5 | 4.1E-04 | 2.8E-03 | BT20/MB231 | MP |
| >hg38_21_+_8433172_8446622_RNA45SN1_45S_50ntFlanks@12999.13027.29\|CTGCGATCTATTGAAAGTCAGCCCTCGAC | >hg38_21_+_8433172_8446622_RNA45SN1_45S_50ntFlanks | 12999 | 13027 | 29 | CTGCGATCTATTGAAAGTCAGCCCTCGAC | 555.4 | -3.0 | 0.9 | -3.5 | 4.8E-04 | 2.8E-03 | BT20/MB231 | CYTO |
| >am_trnaMT_ValTAC_MT_+_1602_1670@42.68.27\|AGATTTCAACTTAACTTGACCGCTCTG | >am_trnaMT_ValTAC_MT_+_1602_1670 | 42 | 68 | 27 | AGATTTCAACTTAACTTGACCGCTCTG | 378.8 | -2.8 | 0.9 | -3.0 | 2.4E-03 | 2.8E-03 | MB231/MB468 | NUC |
| >hg38_21_+_8433172_8446622_RNA45SN1_45S_50ntFlanks@7978.8011.34\|GACCTCAGATCAGACGTGGCGACCCGCTGAATTT | >hg38_21_+_8433172_8446622_RNA45SN1_45S_50ntFlanks | 7978 | 8011 | 34 | GACCTCAGATCAGACGTGGCGACCCGCTGAATTT | 2087.4 | -3.3 | 0.9 | -3.5 | 4.8E-04 | 2.8E-03 | BT20/MB231 | CYTO |
| >MI0005561\|hsa-mir-877&WithFlank&6\|+\|30584326\|30584423@7.29.23(+1U)&[MIMAT0004949&hsa-miR-877-5p&offsets\|0\|+3(+1U);m-382&6\|+\|30584332\|30584354&offsets\|0\|0(+1U)]\|GTAGAGGAGATGGCGCAGGGGACT | >MI0005561\|hsa-mir-877&WithFlank&6\|+\|30584326\|30584423 | 7 | 29 | 23(+1U) | GTAGAGGAGATGGCGCAGGGGACT | 209.2 | 4.6 | 1.4 | 3.2 | 1.2E-03 | 2.8E-03 | BT20/MB468 | MP |
| >hg38_21_+_8433172_8446622_RNA45SN1_45S_50ntFlanks@5494.5513.20\|AAGACGGTCGAACTTGACTA | >hg38_21_+_8433172_8446622_RNA45SN1_45S_50ntFlanks | 5494 | 5513 | 20 | AAGACGGTCGAACTTGACTA | 348.5 | -4.1 | 1.3 | -3.2 | 1.2E-03 | 2.8E-03 | BT20/MB468 | MP |
| >hg38_21_+_8433172_8446622_RNA45SN1_45S_50ntFlanks@10513.10531.19\|CCGGAGTGGCGGAGATGGG | >hg38_21_+_8433172_8446622_RNA45SN1_45S_50ntFlanks | 10513 | 10531 | 19 | CCGGAGTGGCGGAGATGGG | 163.3 | 6.7 | 1.9 | 3.5 | 4.1E-04 | 2.9E-03 | BT20/MB231 | MP |
| >MI0000736\|hsa-mir-30c-1&WithFlank&1\|+\|40757278\|40757378@23.46.24(+1U);MI0000254\|hsa-mir-30c-2&WithFlank&6\|-\|71376954\|71377037@13.36.24(+1U)&[MIMAT0000244&hsa-miR-30c-5p&offsets\|0\|+1(+1U);m-57&1\|+\|40757300\|40757323&offsets\|0\|0(+1U)];[MIMAT0000244_1&hsa-miR-30c-5p&offsets\|0\|+1(+1U);m-56&6\|-\|71377002\|71377025&offsets\|0\|0(+1U)]\|TGTAAACATCCTACACTCTCAGCTT | >MI0000736\|hsa-mir-30c-1&WithFlank&1\|+\|40757278\|40757378 | 23 | 46 | 24(+1U) | TGTAAACATCCTACACTCTCAGCTT | 474.6 | 4.4 | 1.4 | 3.1 | 2.0E-03 | 2.9E-03 | MB231/MB468 | MP |
| >hg38_21_+_8433172_8446622_RNA45SN1_45S_50ntFlanks@6649.6696.48\|TACGACTCTTAGCGGTGGATCACTCGGCTCGTGCGTCGATGAAGAACG | >hg38_21_+_8433172_8446622_RNA45SN1_45S_50ntFlanks | 6649 | 6696 | 48 | TACGACTCTTAGCGGTGGATCACTCGGCTCGTGCGTCGATGAAGAACG | 752.3 | 3.9 | 1.3 | 3.0 | 2.4E-03 | 2.9E-03 | MB231/MB468 | NUC |
| >trnaMT_GluTTC_MT_-_14674_14742@17.57.41,trnalookalike8_GluTTC_5_-_93905172_93905240@17.57.41\|TACAACGATGGTTTTTCATATCATTGGTCGTGGTTGTAGTC | >trnaMT_GluTTC_MT_-_14674_14742 | 17 | 57 | 41 | TACAACGATGGTTTTTCATATCATTGGTCGTGGTTGTAGTC | 42.3 | 9.1 | 2.8 | 3.2 | 1.2E-03 | 2.9E-03 | BT20/MB468 | MP |
| >hg38_21_+_8433172_8446622_RNA45SN1_45S_50ntFlanks@9591.9610.20\|GGGTATAGGGGCGAAAGACT | >hg38_21_+_8433172_8446622_RNA45SN1_45S_50ntFlanks | 9591 | 9610 | 20 | GGGTATAGGGGCGAAAGACT | 38.6 | 9.0 | 2.8 | 3.2 | 1.2E-03 | 2.9E-03 | BT20/MB468 | MP |
| >hg38_21_+_8433172_8446622_RNA45SN1_45S_50ntFlanks@11881.11900.20\|CCTGTTGAGCTTGACTCTAG | >hg38_21_+_8433172_8446622_RNA45SN1_45S_50ntFlanks | 11881 | 11900 | 20 | CCTGTTGAGCTTGACTCTAG | 250.9 | -5.4 | 1.7 | -3.2 | 1.2E-03 | 2.9E-03 | BT20/MB468 | MP |
| >hg38_21_+_8433172_8446622_RNA45SN1_45S_50ntFlanks@9202.9226.25\|TCTCGGGGCCACGCGCGCGTCCCCC | >hg38_21_+_8433172_8446622_RNA45SN1_45S_50ntFlanks | 9202 | 9226 | 25 | TCTCGGGGCCACGCGCGCGTCCCCC | 228.5 | -5.3 | 1.6 | -3.2 | 1.2E-03 | 2.9E-03 | BT20/MB468 | MP |
| >trnaMT_LysTTT_MT_+_8295_8364@40.72.33,trnalookalike7_LysTTT_1_+_568844_568913@40.72.33\|GATTAAGAGAACCAACACCTCTTTACAGTGACC | >trnaMT_LysTTT_MT_+_8295_8364 | 40 | 72 | 33 | GATTAAGAGAACCAACACCTCTTTACAGTGACC | 463.2 | -5.2 | 1.5 | -3.5 | 4.1E-04 | 2.9E-03 | BT20/MB231 | MP |
| >hg38_21_+_8433172_8446622_RNA45SN1_45S_50ntFlanks@11568.11590.23\|GCAGCCGACTTAGAACTGGTGCG | >hg38_21_+_8433172_8446622_RNA45SN1_45S_50ntFlanks | 11568 | 11590 | 23 | GCAGCCGACTTAGAACTGGTGCG | 242.4 | -4.2 | 1.4 | -3.1 | 2.0E-03 | 2.9E-03 | MB231/MB468 | MP |
| >hg38_21_+_8433172_8446622_RNA45SN1_45S_50ntFlanks@8194.8213.20\|CCCGTGGACGGTGTGAGGCC | >hg38_21_+_8433172_8446622_RNA45SN1_45S_50ntFlanks | 8194 | 8213 | 20 | CCCGTGGACGGTGTGAGGCC | 1448.2 | -3.1 | 1.0 | -3.2 | 1.2E-03 | 2.9E-03 | BT20/MB468 | MP |
| >MI0003815\|hsa-mir-1301&WithFlank&2\|-\|25328634\|25328727@54.77.24&[MIMAT0005797&hsa-miR-1301-3p&offsets\|0\|0;m-256&2\|-\|25328650\|25328674&offsets\|0\|-1]\|TTGCAGCTGCCTGGGAGTGACTTC | >MI0003815\|hsa-mir-1301&WithFlank&2\|-\|25328634\|25328727 | 54 | 77 | 24 | TTGCAGCTGCCTGGGAGTGACTTC | 222.2 | 4.4 | 1.4 | 3.2 | 1.2E-03 | 2.9E-03 | BT20/MB468 | MP |
| >hg38_21_+_8433172_8446622_RNA45SN1_45S_50ntFlanks@10402.10419.18\|AACATGGGTCAGTCGGTC | >hg38_21_+_8433172_8446622_RNA45SN1_45S_50ntFlanks | 10402 | 10419 | 18 | AACATGGGTCAGTCGGTC | 213.4 | -3.4 | 1.1 | -3.1 | 1.9E-03 | 2.9E-03 | BT20/MB468 | MITO |
| >hg38_21_+_8433172_8446622_RNA45SN1_45S_50ntFlanks@6653.6688.36\|ACTCTTAGCGGTGGATCACTCGGCTCGTGCGTCGAT | >hg38_21_+_8433172_8446622_RNA45SN1_45S_50ntFlanks | 6653 | 6688 | 36 | ACTCTTAGCGGTGGATCACTCGGCTCGTGCGTCGAT | 983.7 | 5.2 | 1.5 | 3.5 | 4.1E-04 | 2.9E-03 | BT20/MB231 | MP |
| >MI0000088\|hsa-mir-30a&WithFlank&6\|-\|71403545\|71403627@13.35.23(+2U)&[MIMAT0000087&hsa-miR-30a-5p&offsets\|+1\|+2(+2U);m-15&6\|-\|71403595\|71403616&offsets\|+1\|+2(+2U)]\|GTAAACATCCTCGACTGGAAGCTTT | >MI0000088\|hsa-mir-30a&WithFlank&6\|-\|71403545\|71403627 | 13 | 35 | 23(+2U) | GTAAACATCCTCGACTGGAAGCTTT | 194.8 | -6.1 | 1.7 | -3.5 | 4.1E-04 | 2.9E-03 | BT20/MB231 | MP |
| >hg38_21_+_8433172_8446622_RNA45SN1_45S_50ntFlanks@10841.10858.18\|CGTAACTTCGGGATAAGG | >hg38_21_+_8433172_8446622_RNA45SN1_45S_50ntFlanks | 10841 | 10858 | 18 | CGTAACTTCGGGATAAGG | 156.0 | -2.2 | 0.7 | -3.1 | 1.9E-03 | 2.9E-03 | BT20/MB468 | MITO |
| >hg38_21_+_8433172_8446622_RNA45SN1_45S_50ntFlanks@5084.5117.34\|CGAACGAGACTCTGGCATGCTAACTAGTTACGCG | >hg38_21_+_8433172_8446622_RNA45SN1_45S_50ntFlanks | 5084 | 5117 | 34 | CGAACGAGACTCTGGCATGCTAACTAGTTACGCG | 186.1 | -2.7 | 0.9 | -3.1 | 1.9E-03 | 2.9E-03 | BT20/MB468 | MITO |
| >hg38_21_+_8433172_8446622_RNA45SN1_45S_50ntFlanks@11592.11613.22\|ACCAGGGGAATCCGACTGTTTA | >hg38_21_+_8433172_8446622_RNA45SN1_45S_50ntFlanks | 11592 | 11613 | 22 | ACCAGGGGAATCCGACTGTTTA | 331.2 | -2.0 | 0.6 | -3.1 | 1.9E-03 | 2.9E-03 | BT20/MB468 | MITO |
| >hg38_MT_+_1621_3279_RNR2_16S_50ntFlanks@1185.1216.32\|ATTTCGGTTGGGGCGACCTCGGAGCAGAACCC | >hg38_MT_+_1621_3279_RNR2_16S_50ntFlanks | 1185 | 1216 | 32 | ATTTCGGTTGGGGCGACCTCGGAGCAGAACCC | 256.7 | 4.4 | 1.4 | 3.2 | 1.2E-03 | 2.9E-03 | BT20/MB468 | MP |
| >MI0000255\|hsa-mir-30d&WithFlank&8\|-\|134804870\|134804951@12.35.24(+2U)&[MIMAT0000245&hsa-miR-30d-5p&offsets\|0\|+2(+2U);m-21&8\|-\|134804919\|134804940&offsets\|0\|+2(+2U)]\|TGTAAACATCCCCGACTGGAAGCTTT | >MI0000255\|hsa-mir-30d&WithFlank&8\|-\|134804870\|134804951 | 12 | 35 | 24(+2U) | TGTAAACATCCCCGACTGGAAGCTTT | 878.7 | -2.8 | 0.8 | -3.5 | 5.4E-04 | 2.9E-03 | BT20/MB231 | MITO |

| | | | | | | | | | | | | | |
|---|---|---|---|---|---|---|---|---|---|---|---|---|---|
| >hg38_21_+_8433172_8446622_RNA45SN1_45S_50ntFlanks@11876.11900.25\|AAGACCCTGTTGAGCTTGACTCTAG | >hg38_21_+_8433172_8446622_RNA45SN1_45S_50ntFlanks | 11876 | 11900 | 25 | AAGACCCTGTTGAGCTTGACTCTAG | 1024.4 | -6.0 | 2.0 | -3.0 | 2.4E-03 | 2.9E-03 | BT20/MB468 | NUC |
| >hg38_21_+_8433172_8446622_RNA45SN1_45S_50ntFlanks@4569.4587.19\|ATGGAATAGGACCGCGGTT | >hg38_21_+_8433172_8446622_RNA45SN1_45S_50ntFlanks | 4569 | 4587 | 19 | ATGGAATAGGACCGCGGTT | 273.5 | -1.8 | 0.6 | -3.1 | 2.2E-03 | 2.9E-03 | BT20/MB468 | CYTO |
| >hg38_21_+_8433172_8446622_RNA45SN1_45S_50ntFlanks@6773.6793.21\|TTCCTCCCGGGGCTACGCCTG | >hg38_21_+_8433172_8446622_RNA45SN1_45S_50ntFlanks | 6773 | 6793 | 21 | TTCCTCCCGGGGCTACGCCTG | 335.2 | -4.7 | 1.5 | -3.1 | 2.2E-03 | 2.9E-03 | BT20/MB468 | CYTO |
| >hg38_21_+_8433172_8446622_RNA45SN1_45S_50ntFlanks@10349.10374.26\|AACGAGAACTTTGAAGGCCGAAGTGG | >hg38_21_+_8433172_8446622_RNA45SN1_45S_50ntFlanks | 10349 | 10374 | 26 | AACGAGAACTTTGAAGGCCGAAGTGG | 935.9 | -1.9 | 0.6 | -3.1 | 2.3E-03 | 2.9E-03 | MB231/MB468 | CYTO |
| >hg38_21_+_8433172_8446622_RNA45SN1_45S_50ntFlanks@9549.9573.25\|AGGTCCGTAGCGGTCCTGACGTGCA | >hg38_21_+_8433172_8446622_RNA45SN1_45S_50ntFlanks | 9549 | 9573 | 25 | AGGTCCGTAGCGGTCCTGACGTGCA | 131.6 | 2.7 | 0.8 | 3.2 | 1.2E-03 | 2.9E-03 | BT20/MB468 | MP |
| >hg38_21_+_8433172_8446622_RNA45SN1_45S_50ntFlanks@12632.12661.30\|GCGAAGCTACCATCTGTGGGATTATGACTG | >hg38_21_+_8433172_8446622_RNA45SN1_45S_50ntFlanks | 12632 | 12661 | 30 | GCGAAGCTACCATCTGTGGGATTATGACTG | 510.2 | 7.4 | 2.1 | 3.5 | 4.1E-04 | 2.9E-03 | BT20/MB231 | MP |
| >MI0000088\|hsa-mir-30a&WithFlank&6\|-\|71403545\|71403627@54.73.20(+1U)&[MIMAT0000088&hsa-miR-30a-3p&offsets\|+1\|-1(+1U);m-38&6\|-\|71403554\|71403575&offsets\|+1\|-1(+1U)]\|TTTCAGTCGGATGTTTGCAGT | >MI0000088\|hsa-mir-30a&WithFlank&6\|-\|71403545\|71403627 | 54 | 73 | 20(+1U) | TTTCAGTCGGATGTTTGCAGT | 4204.3 | -2.5 | 0.7 | -3.5 | 5.4E-04 | 2.9E-03 | BT20/MB231 | MITO |
| >MI0000749\|hsa-mir-30e&WithFlank&1\|+\|40754349\|40754452@65.86.22&[MIMAT0000693&hsa-miR-30e-3p&offsets\|0\|0;m-25&1\|+\|40754413\|40754434&offsets\|0\|0]\|CTTTCAGTCGGATGTTTACAGC | >MI0000749\|hsa-mir-30e&WithFlank&1\|+\|40754349\|40754452 | 65 | 86 | 22 | CTTTCAGTCGGATGTTTACAGC | 1291.7 | -1.2 | 0.4 | -3.0 | 2.4E-03 | 2.9E-03 | BT20/MB468 | NUC |
| >hg38_21_+_8433172_8446622_RNA45SN1_45S_50ntFlanks@4723.4751.29\|CATTAATCAAGAACGAAAGTCGGAGGTTC | >hg38_21_+_8433172_8446622_RNA45SN1_45S_50ntFlanks | 4723 | 4751 | 29 | CATTAATCAAGAACGAAAGTCGGAGGTTC | 923.1 | -5.6 | 1.7 | -3.2 | 1.2E-03 | 2.9E-03 | BT20/MB468 | MP |
| >hg38_21_+_8433172_8446622_RNA45SN1_45S_50ntFlanks@12288.12323.36\|ACCGTGAAAGCGGGGCCTCACGATCCTTCTGACCTT | >hg38_21_+_8433172_8446622_RNA45SN1_45S_50ntFlanks | 12288 | 12323 | 36 | ACCGTGAAAGCGGGGCCTCACGATCCTTCTGACCTT | 313.3 | -4.9 | 1.6 | -3.0 | 2.4E-03 | 2.9E-03 | MB231/MB468 | NUC |
| >hg38_21_+_8433172_8446622_RNA45SN1_45S_50ntFlanks@6648.6681.34\|GTACGACTCTTAGCGGTGGATCACTCGGCTCGTG | >hg38_21_+_8433172_8446622_RNA45SN1_45S_50ntFlanks | 6648 | 6681 | 34 | GTACGACTCTTAGCGGTGGATCACTCGGCTCGTG | 9616.4 | 2.3 | 0.8 | 3.0 | 2.4E-03 | 2.9E-03 | MB231/MB468 | NUC |
| >trna10_LysCTT_16_+_3241501_3241573@4.35.32,trna13_LysCTT_14_-_58706613_58706685@4.35.32,trna2_LysCTT_15_+_79152904_79152976@4.35.32\|CGGCTAGCTCAGTCGGTAGAGCATGGGACTCT | >trna10_LysCTT_16_+_3241501_3241573 | 4 | 35 | 32 | CGGCTAGCTCAGTCGGTAGAGCATGGGACTCT | 137.8 | 4.8 | 1.5 | 3.2 | 1.2E-03 | 2.9E-03 | BT20/MB468 | MP |
| >hg38_21_+_8433172_8446622_RNA45SN1_45S_50ntFlanks@7988.8007.20\|CAGACGTGGCGACCCGCTGA | >hg38_21_+_8433172_8446622_RNA45SN1_45S_50ntFlanks | 7988 | 8007 | 20 | CAGACGTGGCGACCCGCTGA | 2079.7 | -3.5 | 1.1 | -3.2 | 1.2E-03 | 2.9E-03 | BT20/MB468 | MP |
| >hg38_21_+_8433172_8446622_RNA45SN1_45S_50ntFlanks@8170.8212.43\|CAAGTCCTTCTGATCGAGGCCCAGCCCGTGGACGGTGTGAGGC | >hg38_21_+_8433172_8446622_RNA45SN1_45S_50ntFlanks | 8170 | 8212 | 43 | CAAGTCCTTCTGATCGAGGCCCAGCCCGTGGACGGTGTGAGGC | 1039.1 | -3.7 | 1.2 | -3.0 | 2.3E-03 | 2.9E-03 | MB231/MB468 | CYTO |
| >hg38_21_+_8433172_8446622_RNA45SN1_45S_50ntFlanks@8170.8214.45\|CAAGTCCTTCTGATCGAGGCCCAGCCCGTGGACGGTGTGAGGCCG | >hg38_21_+_8433172_8446622_RNA45SN1_45S_50ntFlanks | 8170 | 8214 | 45 | CAAGTCCTTCTGATCGAGGCCCAGCCCGTGGACGGTGTGAGGCCG | 2176.0 | -3.3 | 1.1 | -3.0 | 2.3E-03 | 2.9E-03 | MB231/MB468 | CYTO |
| >hg38_21_+_8433172_8446622_RNA45SN1_45S_50ntFlanks@6650.6680.31\|ACGACTCTTAGCGGTGGATCACTCGGCTCGT | >hg38_21_+_8433172_8446622_RNA45SN1_45S_50ntFlanks | 6650 | 6680 | 31 | ACGACTCTTAGCGGTGGATCACTCGGCTCGT | 236.9 | 2.3 | 0.8 | 3.0 | 2.4E-03 | 2.9E-03 | MB231/MB468 | NUC |
| >hg38_21_+_8433172_8446622_RNA45SN1_45S_50ntFlanks@6650.6675.26\|ACGACTCTTAGCGGTGGATCACTCGG | >hg38_21_+_8433172_8446622_RNA45SN1_45S_50ntFlanks | 6650 | 6675 | 26 | ACGACTCTTAGCGGTGGATCACTCGG | 1179.9 | 2.5 | 0.8 | 3.0 | 2.4E-03 | 2.9E-03 | MB231/MB468 | NUC |
| >hg38_21_+_8433172_8446622_RNA45SN1_45S_50ntFlanks@11583.11611.29\|CTGGTGCGGACCAGGGGAATCCGACTGTT | >hg38_21_+_8433172_8446622_RNA45SN1_45S_50ntFlanks | 11583 | 11611 | 29 | CTGGTGCGGACCAGGGGAATCCGACTGTT | 1010.6 | 2.6 | 0.7 | 3.5 | 5.5E-04 | 2.9E-03 | BT20/MB231 | MITO |
| >hg38_21_+_8433172_8446622_RNA45SN1_45S_50ntFlanks@12992.13013.22\|TCCCTCGCTGCGATCTATTGAA | >hg38_21_+_8433172_8446622_RNA45SN1_45S_50ntFlanks | 12992 | 13013 | 22 | TCCCTCGCTGCGATCTATTGAA | 1047.7 | -2.5 | 0.8 | -3.1 | 2.2E-03 | 2.9E-03 | BT20/MB468 | CYTO |
| >hg38_21_+_8433172_8446622_RNA45SN1_45S_50ntFlanks@5073.5098.26\|AATTCCGATAACGAACGAGACTCTGG | >hg38_21_+_8433172_8446622_RNA45SN1_45S_50ntFlanks | 5073 | 5098 | 26 | AATTCCGATAACGAACGAGACTCTGG | 299.8 | -2.0 | 0.7 | -3.1 | 2.2E-03 | 2.9E-03 | BT20/MB468 | CYTO |
| >hg38_21_+_8433172_8446622_RNA45SN1_45S_50ntFlanks@11709.11728.20\|GCGCGGGTAAACGGCGGGAG | >hg38_21_+_8433172_8446622_RNA45SN1_45S_50ntFlanks | 11709 | 11728 | 20 | GCGCGGGTAAACGGCGGGAG | 339.0 | -5.4 | 1.8 | -3.0 | 2.4E-03 | 2.9E-03 | BT20/MB468 | NUC |
| >hg38_21_+_8433172_8446622_RNA45SN1_45S_50ntFlanks@11602.11622.21\|TCCGACTGTTTAATTAAAACA | >hg38_21_+_8433172_8446622_RNA45SN1_45S_50ntFlanks | 11602 | 11622 | 21 | TCCGACTGTTTAATTAAAACA | 123.1 | -4.9 | 1.6 | -3.1 | 2.1E-03 | 2.9E-03 | MB231/MB468 | MP |
| >hg38_21_+_8433172_8446622_RNA45SN1_45S_50ntFlanks@3842.3861.20\|CCTACTTGGATAACTGTGGT | >hg38_21_+_8433172_8446622_RNA45SN1_45S_50ntFlanks | 3842 | 3861 | 20 | CCTACTTGGATAACTGTGGT | 688.4 | 2.0 | 0.7 | 3.1 | 2.2E-03 | 2.9E-03 | BT20/MB468 | CYTO |
| >hg38_21_+_8433172_8446622_RNA45SN1_45S_50ntFlanks@11906.11927.22\|CACGGTGAAGAGACATGAGAGG | >hg38_21_+_8433172_8446622_RNA45SN1_45S_50ntFlanks | 11906 | 11927 | 22 | CACGGTGAAGAGACATGAGAGG | 88.2 | 9.8 | 2.8 | 3.5 | 4.1E-04 | 2.9E-03 | BT20/MB231 | MP |
| >trnaMT_SerGCT_MT_+_12207_12265@13.62.50\|AAGAACTGCTAACTCATGCCCCCATGTCTAACAACATGGCTTTCTCACCA | >trnaMT_SerGCT_MT_+_12207_12265 | 13 | 62 | 50 | AAGAACTGCTAACTCATGCCCCCATGTCTAACAACATGGCTTTCTCACCA | 579.0 | 4.6 | 1.5 | 3.0 | 2.4E-03 | 2.9E-03 | MB231/MB468 | MITO |
| >hg38_21_+_8433172_8446622_RNA45SN1_45S_50ntFlanks@12007.12029.23\|GCCGCCGGTGAAATACCACTACT | >hg38_21_+_8433172_8446622_RNA45SN1_45S_50ntFlanks | 12007 | 12029 | 23 | GCCGCCGGTGAAATACCACTACT | 1017.7 | 3.6 | 1.0 | 3.5 | 5.5E-04 | 2.9E-03 | BT20/MB231 | MITO |
| >trnaMT_ThrTGT_MT_+_15888_15953@37.69.33\|ACCGGAGATGAAAACCTTTTTCCAAGGACACCA | >trnaMT_ThrTGT_MT_+_15888_15953 | 37 | 69 | 33 | ACCGGAGATGAAAACCTTTTTCCAAGGACACCA | 502.5 | 3.1 | 1.0 | 3.1 | 2.1E-03 | 2.9E-03 | MB231/MB468 | MP |
| >hg38_21_+_8433172_8446622_RNA45SN1_45S_50ntFlanks@4726.4758.33\|TAATCAAGAACGAAAGTCGGAGGTTCGAAGACG | >hg38_21_+_8433172_8446622_RNA45SN1_45S_50ntFlanks | 4726 | 4758 | 33 | TAATCAAGAACGAAAGTCGGAGGTTCGAAGACG | 3004.8 | -3.7 | 1.1 | -3.2 | 1.2E-03 | 2.9E-03 | BT20/MB468 | MP |
| >MI0000433\|hsa-let-7g&WithFlank&3\|-\|52268272\|52268367@11.30.20&[MIMAT0000414&hsa-let-7g-5p&offsets\|0\|-2;m-58&3\|-\|52268336\|52268357&offsets\|0\|-2]\|TGAGGTAGTAGTTTGTACAG | >MI0000433\|hsa-let-7g&WithFlank&3\|-\|52268272\|52268367 | 11 | 30 | 20 | TGAGGTAGTAGTTTGTACAG | 164.9 | 6.2 | 2.0 | 3.1 | 2.1E-03 | 2.9E-03 | MB231/MB468 | MP |
| >hg38_MT_+_1621_3279_RNR2_16S_50ntFlanks@1369.1407.39\|GATGTTGGATCAGGACATCCCGATGGTGCAGCCGCTATT | >hg38_MT_+_1621_3279_RNR2_16S_50ntFlanks | 1369 | 1407 | 39 | GATGTTGGATCAGGACATCCCGATGGTGCAGCCGCTATT | 62.5 | 6.4 | 2.0 | 3.2 | 1.2E-03 | 2.9E-03 | BT20/MB468 | MP |
| >hg38_21_+_8433172_8446622_RNA45SN1_45S_50ntFlanks@12616.12653.38\|CTGAGGAGCCAATGGGGCGAAGCTACCATCTGTGGGAT | >hg38_21_+_8433172_8446622_RNA45SN1_45S_50ntFlanks | 12616 | 12653 | 38 | CTGAGGAGCCAATGGGGCGAAGCTACCATCTGTGGGAT | 427.8 | -5.5 | 1.7 | -3.2 | 1.2E-03 | 2.9E-03 | BT20/MB468 | MP |
| >hg38_21_+_8433172_8446622_RNA45SN1_45S_50ntFlanks@12405.12427.23\|CTTTTTGATCCTTCGATGTCGGC | >hg38_21_+_8433172_8446622_RNA45SN1_45S_50ntFlanks | 12405 | 12427 | 23 | CTTTTTGATCCTTCGATGTCGGC | 107.3 | -5.1 | 1.6 | -3.2 | 1.2E-03 | 2.9E-03 | BT20/MB468 | MP |
| >hg38_21_+_8433172_8446622_RNA45SN1_45S_50ntFlanks@7989.8012.24\|AGACGTGGCGACCCGCTGAATTTA | >hg38_21_+_8433172_8446622_RNA45SN1_45S_50ntFlanks | 7989 | 8012 | 24 | AGACGTGGCGACCCGCTGAATTTA | 5207.9 | -3.0 | 0.9 | -3.2 | 1.2E-03 | 2.9E-03 | BT20/MB468 | MP |
| >MI0000077\|hsa-mir-21&WithFlank&17\|+\|59841260\|59841343@13.34.22&[MIMAT0000076&hsa-miR-21-5p&offsets\|-1\|-1;m-1&17\|+\|59841273\|59841294&offsets\|-1\|-1]\|GTAGCTTATCAGACTGATGTTG | >MI0000077\|hsa-mir-21&WithFlank&17\|+\|59841260\|59841343 | 13 | 34 | 22 | GTAGCTTATCAGACTGATGTTG | 344.4 | 3.4 | 1.1 | 3.2 | 1.2E-03 | 2.9E-03 | BT20/MB468 | MP |
| >MI0000438\|hsa-mir-15b&WithFlank&3\|+\|160404582\|160404691@26.46.21&[MIMAT0000417&hsa-miR-15b-5p&offsets\|0\|-1;m-110&3\|+\|160404607\|160404628&offsets\|0\|-1]\|TAGCAGCACATCATGGTTTAC | >MI0000438\|hsa-mir-15b&WithFlank&3\|+\|160404582\|160404691 | 26 | 46 | 21 | TAGCAGCACATCATGGTTTAC | 468.6 | 5.2 | 1.6 | 3.2 | 1.2E-03 | 2.9E-03 | BT20/MB468 | MP |
| >trna16_ValTAC_11_-_59318460_59318532@37.75.39,trna17_ValTAC_11_-_59318102_59318174@37.75.39,trna4_ValTAC_X_-_18693029_18693101@37.75.39\|ACGCAGAAGGTCCTGGGTTCGAGCCCCAGTGGAACCACC | >trna16_ValTAC_11_-_59318460_59318532 | 37 | 75 | 39 | ACGCAGAAGGTCCTGGGTTCGAGCCCCAGTGGAACCACC | 643.1 | -6.7 | 1.9 | -3.5 | 4.9E-04 | 2.9E-03 | BT20/MB231 | CYTO |
| >hg38_21_+_8433172_8446622_RNA45SN1_45S_50ntFlanks@12244.12280.37\|AAGGGCAAAAGCTCGCTTGATCTTGATTTTCAGTACG | >hg38_21_+_8433172_8446622_RNA45SN1_45S_50ntFlanks | 12244 | 12280 | 37 | AAGGGCAAAAGCTCGCTTGATCTTGATTTTCAGTACG | 149.0 | -3.3 | 0.9 | -3.5 | 4.9E-04 | 2.9E-03 | BT20/MB231 | CYTO |
| >hg38_21_+_8433172_8446622_RNA45SN1_45S_50ntFlanks@9570.9605.36\|TGCAAATCGGTCGTCCGACCTGGGTATAGGGGCGAA | >hg38_21_+_8433172_8446622_RNA45SN1_45S_50ntFlanks | 9570 | 9605 | 36 | TGCAAATCGGTCGTCCGACCTGGGTATAGGGGCGAA | 405.8 | -3.9 | 1.3 | -3.0 | 2.4E-03 | 2.9E-03 | MB231/MB468 | MITO |
| >hg38_1_-_228634819_228635039_RNA5S12_5S_50ntFlanks@149.169.21\|GAATACCGGGTGCTGTAGGCT | >hg38_1_-_228634819_228635039_RNA5S12_5S_50ntFlanks | 149 | 169 | 21 | GAATACCGGGTGCTGTAGGCT | 234.3 | 2.3 | 0.8 | 3.0 | 2.5E-03 | 2.9E-03 | MB231/MB468 | NUC |
| >hg38_21_+_8433172_8446622_RNA45SN1_45S_50ntFlanks@12002.12030.29\|TGCGGGCCGCCGGTGAAATACCACTACTC | >hg38_21_+_8433172_8446622_RNA45SN1_45S_50ntFlanks | 12002 | 12030 | 29 | TGCGGGCCGCCGGTGAAATACCACTACTC | 418.8 | -3.3 | 1.1 | -3.1 | 2.3E-03 | 2.9E-03 | BT20/MB468 | CYTO |
| >hg38_21_+_8433172_8446622_RNA45SN1_45S_50ntFlanks@12420.12450.31\|ATGTCGGCTCTTCCTATCATTGTGAAGCAGA | >hg38_21_+_8433172_8446622_RNA45SN1_45S_50ntFlanks | 12420 | 12450 | 31 | ATGTCGGCTCTTCCTATCATTGTGAAGCAGA | 345.8 | -1.6 | 0.5 | -3.1 | 2.3E-03 | 2.9E-03 | BT20/MB468 | CYTO |
| >hg38_21_+_8433172_8446622_RNA45SN1_45S_50ntFlanks@10681.10708.28\|TTCCGGCGGCGTCCGGTGAGCTCTCGCT | >hg38_21_+_8433172_8446622_RNA45SN1_45S_50ntFlanks | 10681 | 10708 | 28 | TTCCGGCGGCGTCCGGTGAGCTCTCGCT | 299.3 | -3.4 | 1.1 | -3.1 | 1.9E-03 | 2.9E-03 | BT20/MB468 | MITO |
| >MI0000299\|hsa-mir-222&WithFlank&X\|-\|45747009\|45747130@75.97.23(+3U)&[MIMAT0000279&hsa-miR-222-3p&offsets\|0\|+2(+3U);m-140&X\|-\|45747033\|45747056&offsets\|0\|-1(+3U)]\|AGCTACATCTGGCTACTGGGTCTTTT | >MI0000299\|hsa-mir-222&WithFlank&X\|-\|45747009\|45747130 | 75 | 97 | 23(+3U) | AGCTACATCTGGCTACTGGGTCTTTT | 86.0 | 5.7 | 1.8 | 3.1 | 2.1E-03 | 2.9E-03 | MB231/MB468 | MP |
| >hg38_21_+_8433172_8446622_RNA45SN1_45S_50ntFlanks@10400.10424.25\|TGAACATGGGTCAGTCGGTCCTGAG | >hg38_21_+_8433172_8446622_RNA45SN1_45S_50ntFlanks | 10400 | 10424 | 25 | TGAACATGGGTCAGTCGGTCCTGAG | 394.7 | 1.6 | 0.5 | 3.0 | 2.4E-03 | 2.9E-03 | BT20/MB468 | NUC |
| >trnaMT_GluTTC_MT_-_14674_14742@44.71.28,trnalookalike8_GluTTC_5_-_93905172_93905240@44.71.28\|TCGTGGTTGTAGTCCGTGCGAGAATACC | >trnaMT_GluTTC_MT_-_14674_14742 | 44 | 71 | 28 | TCGTGGTTGTAGTCCGTGCGAGAATACC | 204.0 | 3.1 | 1.0 | 3.0 | 2.5E-03 | 2.9E-03 | MB231/MB468 | MITO |
| >hg38_21_+_8433172_8446622_RNA45SN1_45S_50ntFlanks@11672.11709.38\|CAGTGCTCTGAATGTCAAAGTGAAGAAATTCAATGAAG | >hg38_21_+_8433172_8446622_RNA45SN1_45S_50ntFlanks | 11672 | 11709 | 38 | CAGTGCTCTGAATGTCAAAGTGAAGAAATTCAATGAAG | 681.2 | 2.8 | 0.9 | 3.0 | 2.3E-03 | 2.9E-03 | MB231/MB468 | CYTO |
| >hg38_21_+_8433172_8446622_RNA45SN1_45S_50ntFlanks@9569.9603.35\|GTGCAAATCGGTCGTCCGACCTGGGTATAGGGGCG | >hg38_21_+_8433172_8446622_RNA45SN1_45S_50ntFlanks | 9569 | 9603 | 35 | GTGCAAATCGGTCGTCCGACCTGGGTATAGGGGCG | 516.5 | 2.2 | 0.7 | 3.1 | 1.9E-03 | 2.9E-03 | BT20/MB468 | MITO |
| >hg38_1_-_228634819_228635039_RNA5S12_5S_50ntFlanks@136.170.35\|GGAGACCGCCTGGGAATACCGGGTGCTGTAGGCTT | >hg38_1_-_228634819_228635039_RNA5S12_5S_50ntFlanks | 136 | 170 | 35 | GGAGACCGCCTGGGAATACCGGGTGCTGTAGGCTT | 21317.2 | 4.9 | 1.4 | 3.5 | 4.2E-04 | 2.9E-03 | BT20/MB231 | MP |
| >am_trna162_MetCAT_6_-_26758550_26758622@31.54.24,trna164_MetCAT_6_-_26735574_26735646@31.54.24,trna20_MetCAT_16_+_71460396_71460468@31.54.24,trna21_MetCAT_6_+_26701712_26701784@31.54.24,trna27_MetCAT_6_+_26766444_26766516@31.54.24,trna75_MetCAT_6_+_28912352_28912424@31.54.24,trna97_MetCAT_6_-_28921042_28921114@31.54.24\|TCTCATAATCTGAAGGTCCTGAGT | >am_trna162_MetCAT_6_-_26758550_26758622 | 31 | 54 | 24 | TCTCATAATCTGAAGGTCCTGAGT | 262.7 | -5.4 | 1.7 | -3.2 | 1.2E-03 | 2.9E-03 | BT20/MB468 | MP |
| >hg38_21_+_8433172_8446622_RNA45SN1_45S_50ntFlanks@4008.4026.19\|CTCGGGCCGATCGCACGCC | >hg38_21_+_8433172_8446622_RNA45SN1_45S_50ntFlanks | 4008 | 4026 | 19 | CTCGGGCCGATCGCACGCC | 262.9 | -5.3 | 1.6 | -3.2 | 1.2E-03 | 2.9E-03 | BT20/MB468 | MP |

<table>
<tr><th></th><th></th><th></th><th></th><th></th><th></th><th></th><th></th><th></th><th></th><th></th><th></th><th></th><th></th></tr>
<tr><td>>hg38_21_+_8433172_8446622_RNA45SN1_45S_50ntFlanks@11604.11629.26|CGACTGTTTAATTAAAACAAAGCATC</td><td>>hg38_21_+_8433172_8446622_RNA45SN1_45S_50ntFlanks</td><td>11604</td><td>11629</td><td>26</td><td>CGACTGTTTAATTAAAACAAAGCATC</td><td>76.8</td><td>6.2</td><td>1.9</td><td>3.2</td><td>1.2E-03</td><td>2.9E-03</td><td>BT20/MB468</td><td>MP</td></tr>
<tr><td>>hg38_21_+_8433172_8446622_RNA45SN1_45S_50ntFlanks@7981.8009.29|CTCAGATCAGACGTGGCGACCCGCTGAAT</td><td>>hg38_21_+_8433172_8446622_RNA45SN1_45S_50ntFlanks</td><td>7981</td><td>8009</td><td>29</td><td>CTCAGATCAGACGTGGCGACCCGCTGAAT</td><td>709.5</td><td>1.4</td><td>0.5</td><td>3.1</td><td>1.9E-03</td><td>2.9E-03</td><td>BT20/MB468</td><td>MITO</td></tr>
<tr><td>>hg38_21_+_8433172_8446622_RNA45SN1_45S_50ntFlanks@4982.5002.21|ACGGACAGGATTGACAGATTG</td><td>>hg38_21_+_8433172_8446622_RNA45SN1_45S_50ntFlanks</td><td>4982</td><td>5002</td><td>21</td><td>ACGGACAGGATTGACAGATTG</td><td>352.2</td><td>-4.7</td><td>1.5</td><td>-3.1</td><td>1.9E-03</td><td>2.9E-03</td><td>BT20/MB468</td><td>MITO</td></tr>
<tr><td>>trnaMT_GluTTC_MT_-_14674_14742@26.69.44,trnalookalike8_GluTTC_5_-_93905172_93905240@26.69.44|GGTTTTTCATATCATTGGTCGTGGTTGTAGTCCGTGCGAGAATA</td><td>>trnaMT_GluTTC_MT_-_14674_14742</td><td>26</td><td>69</td><td>44</td><td>GGTTTTTCATATCATTGGTCGTGGTTGTAGTCCGTGCGAGAATA</td><td>41.9</td><td>9.1</td><td>2.8</td><td>3.2</td><td>1.2E-03</td><td>2.9E-03</td><td>BT20/MB468</td><td>MP</td></tr>
<tr><td>>hg38_1_-_228634819_228635039_RNA5S12_5S_50ntFlanks@51.84.34|GTCTACGGCCATACCACCCTGAACGCGCCCGATC</td><td>>hg38_1_-_228634819_228635039_RNA5S12_5S_50ntFlanks</td><td>51</td><td>84</td><td>34</td><td>GTCTACGGCCATACCACCCTGAACGCGCCCGATC</td><td>7580.7</td><td>3.6</td><td>1.1</td><td>3.2</td><td>1.2E-03</td><td>2.9E-03</td><td>BT20/MB468</td><td>MP</td></tr>
<tr><td>>trnaMT_ProTGG_MT_-_15956_16023@39.58.20|TAATGGTGGAGTTAAAGACT</td><td>>trnaMT_ProTGG_MT_-_15956_16023</td><td>39</td><td>58</td><td>20</td><td>TAATGGTGGAGTTAAAGACT</td><td>135.3</td><td>4.4</td><td>1.4</td><td>3.2</td><td>1.2E-03</td><td>2.9E-03</td><td>BT20/MB468</td><td>MP</td></tr>
<tr><td>>hg38_21_+_8433172_8446622_RNA45SN1_45S_50ntFlanks@7991.8012.22|ACGTGGCGACCCGCTGAATTTA</td><td>>hg38_21_+_8433172_8446622_RNA45SN1_45S_50ntFlanks</td><td>7991</td><td>8012</td><td>22</td><td>ACGTGGCGACCCGCTGAATTTA</td><td>580.5</td><td>-2.0</td><td>0.6</td><td>-3.0</td><td>2.3E-03</td><td>2.9E-03</td><td>MB231/MB468</td><td>CYTO</td></tr>
<tr><td>>MI0001641|hsa-mir-429&WithFlank&1|+|1168999|1169093@57.78.22&[MIMAT0001536&hsa-miR-429&offsets|0|0;m-207&1|+|1169055|1169076&offsets|0|0]|TAATACTGTCTGGTAAAACCGT</td><td>>MI0001641|hsa-mir-429&WithFlank&1|+|1168999|1169093</td><td>57</td><td>78</td><td>22</td><td>TAATACTGTCTGGTAAAACCGT</td><td>575.5</td><td>-4.1</td><td>1.4</td><td>-3.0</td><td>2.5E-03</td><td>2.9E-03</td><td>MB231/MB468</td><td>MITO</td></tr>
<tr><td>>hg38_21_+_8433172_8446622_RNA45SN1_45S_50ntFlanks@12635.12660.26|AAGCTACCATCTGTGGGATTATGACT</td><td>>hg38_21_+_8433172_8446622_RNA45SN1_45S_50ntFlanks</td><td>12635</td><td>12660</td><td>26</td><td>AAGCTACCATCTGTGGGATTATGACT</td><td>562.9</td><td>-4.3</td><td>1.3</td><td>-3.2</td><td>1.2E-03</td><td>2.9E-03</td><td>BT20/MB468</td><td>MP</td></tr>
<tr><td>>hg38_21_+_8433172_8446622_RNA45SN1_45S_50ntFlanks@6774.6802.29|TCCTCCCGGGGCTACGCCTGTCTGAGCGT</td><td>>hg38_21_+_8433172_8446622_RNA45SN1_45S_50ntFlanks</td><td>6774</td><td>6802</td><td>29</td><td>TCCTCCCGGGGCTACGCCTGTCTGAGCGT</td><td>132.6</td><td>2.9</td><td>0.9</td><td>3.0</td><td>2.5E-03</td><td>2.9E-03</td><td>MB231/MB468</td><td>MITO</td></tr>
<tr><td>>MI0003776|hsa-mir-320b-1&WithFlank&1|+|116671740|116671823@48.67.20;MI0003839|hsa-mir-320b-2&WithFlank&1|-|224257034|224257116@47.66.20;MI0000542|hsa-mir-320a&WithFlank&8|-|22244960|22245043@48.67.20&[MIMAT0005792&hsa-miR-320b&offsets|0|-2;m-183&1|+|116671787|116671806&offsets|0|0];[MIMAT0005792_1&hsa-miR-320b&offsets|0|-2;m-183&1|-|224257051|224257070&offsets|0|0];[MIMAT0000510&hsa-miR-320a-3p&offsets|0|-2;m-60&8|-|22244975|22244996&offsets|0|-2]|AAAAGCTGGGTTGAGAGGGC</td><td>>MI0003776|hsa-mir-320b-1&WithFlank&1|+|116671740|116671823</td><td>48</td><td>67</td><td>20</td><td>AAAAGCTGGGTTGAGAGGGC</td><td>10429.8</td><td>3.1</td><td>0.9</td><td>3.5</td><td>4.2E-04</td><td>2.9E-03</td><td>BT20/MB231</td><td>MP</td></tr>
<tr><td>>hg38_21_+_8433172_8446622_RNA45SN1_45S_50ntFlanks@11849.11887.39|CGGGCTTGGCGGAATCAGCGGGGAAAGAAGACCCTGTTG</td><td>>hg38_21_+_8433172_8446622_RNA45SN1_45S_50ntFlanks</td><td>11849</td><td>11887</td><td>39</td><td>CGGGCTTGGCGGAATCAGCGGGGAAAGAAGACCCTGTTG</td><td>85.2</td><td>9.7</td><td>2.8</td><td>3.5</td><td>4.2E-04</td><td>2.9E-03</td><td>BT20/MB231</td><td>MP</td></tr>
<tr><td>>hg38_21_+_8433172_8446622_RNA45SN1_45S_50ntFlanks@8105.8143.39|CGCGGCGGGGCGCGGGACATGTGGCGTACGGAAGACCCG</td><td>>hg38_21_+_8433172_8446622_RNA45SN1_45S_50ntFlanks</td><td>8105</td><td>8143</td><td>39</td><td>CGCGGCGGGGCGCGGGACATGTGGCGTACGGAAGACCCG</td><td>410.0</td><td>4.9</td><td>1.4</td><td>3.5</td><td>4.2E-04</td><td>2.9E-03</td><td>BT20/MB231</td><td>MP</td></tr>
<tr><td>>MI0000088|hsa-mir-30a&WithFlank&6|-|71403545|71403627@12.35.24(+1U)&[MIMAT0000087&hsa-miR-30a-5p&offsets|0|+2(+1U);m-15&6|-|71403595|71403616&offsets|0|+2(+1U)]|TGTAAACATCCTCGACTGGAAGCTT</td><td>>MI0000088|hsa-mir-30a&WithFlank&6|-|71403545|71403627</td><td>12</td><td>35</td><td>24(+1U)</td><td>TGTAAACATCCTCGACTGGAAGCTT</td><td>313.9</td><td>3.2</td><td>1.1</td><td>3.0</td><td>2.5E-03</td><td>2.9E-03</td><td>BT20/MB468</td><td>NUC</td></tr>
<tr><td>>trnaMT_GluTTC_MT_-_14674_14742@28.71.44,trnalookalike8_GluTTC_5_-_93905172_93905240@28.71.44|TTTTTCATATCATTGGTCGTGGTTGTAGTCCGTGCGAGAATACC</td><td>>trnaMT_GluTTC_MT_-_14674_14742</td><td>28</td><td>71</td><td>44</td><td>TTTTTCATATCATTGGTCGTGGTTGTAGTCCGTGCGAGAATACC</td><td>117.3</td><td>5.8</td><td>1.9</td><td>3.0</td><td>2.5E-03</td><td>2.9E-03</td><td>MB231/MB468</td><td>MITO</td></tr>
<tr><td>>hg38_21_+_8433172_8446622_RNA45SN1_45S_50ntFlanks@8357.8405.49|AAAAGAACTTTGAAGAGAGAGTTCAAGAGGGCGTGAAACCGTTAAGAGG</td><td>>hg38_21_+_8433172_8446622_RNA45SN1_45S_50ntFlanks</td><td>8357</td><td>8405</td><td>49</td><td>AAAAGAACTTTGAAGAGAGAGTTCAAGAGGGCGTGAAACCGTTAAGAGG</td><td>45.7</td><td>-9.4</td><td>3.0</td><td>-3.1</td><td>2.1E-03</td><td>2.9E-03</td><td>MB231/MB468</td><td>MP</td></tr>
<tr><td>>hg38_21_+_8433172_8446622_RNA45SN1_45S_50ntFlanks@12241.12273.33|CAGAAGGGCAAAAGCTCGCTTGATCTTGATTTT</td><td>>hg38_21_+_8433172_8446622_RNA45SN1_45S_50ntFlanks</td><td>12241</td><td>12273</td><td>33</td><td>CAGAAGGGCAAAAGCTCGCTTGATCTTGATTTT</td><td>187.1</td><td>-2.5</td><td>0.8</td><td>-3.0</td><td>2.5E-03</td><td>2.9E-03</td><td>MB231/MB468</td><td>NUC</td></tr>
<tr><td>>hg38_21_+_8433172_8446622_RNA45SN1_45S_50ntFlanks@11537.11580.44|CCGCGCGGCGCCTCGCCTCGGCCGGCGCCTAGCAGCCGACTTAG</td><td>>hg38_21_+_8433172_8446622_RNA45SN1_45S_50ntFlanks</td><td>11537</td><td>11580</td><td>44</td><td>CCGCGCGGCGCCTCGCCTCGGCCGGCGCCTAGCAGCCGACTTAG</td><td>464.6</td><td>-2.9</td><td>0.9</td><td>-3.1</td><td>1.9E-03</td><td>2.9E-03</td><td>BT20/MB468</td><td>MITO</td></tr>
<tr><td>>hg38_21_+_8433172_8446622_RNA45SN1_45S_50ntFlanks@12641.12664.24|CCATCTGTGGGATTATGACTGAAC</td><td>>hg38_21_+_8433172_8446622_RNA45SN1_45S_50ntFlanks</td><td>12641</td><td>12664</td><td>24</td><td>CCATCTGTGGGATTATGACTGAAC</td><td>187.6</td><td>6.8</td><td>1.9</td><td>3.5</td><td>4.2E-04</td><td>2.9E-03</td><td>BT20/MB231</td><td>MP</td></tr>
<tr><td>>MI0000298|hsa-mir-221&WithFlank&X|-|45746151|45746272@71.91.21&[MIMAT0000278&hsa-miR-221-3p&offsets|0|-2;m-91&X|-|45746181|45746202&offsets|0|-1]|AGCTACATTGTCTGCTGGGTT</td><td>>MI0000298|hsa-mir-221&WithFlank&X|-|45746151|45746272</td><td>71</td><td>91</td><td>21</td><td>AGCTACATTGTCTGCTGGGTT</td><td>2921.0</td><td>-2.6</td><td>0.7</td><td>-3.5</td><td>5.6E-04</td><td>2.9E-03</td><td>BT20/MB231</td><td>MITO</td></tr>
<tr><td>>MI0000478|hsa-mir-149&WithFlank&2|+|240455995|240456095@21.41.21(+1U)&[MIMAT0000450&hsa-miR-149-5p&offsets|0|-2(+1U);m-195&2|+|240456015|240456037&offsets|0|-2(+1U)]|TCTGGCTCCGTGTCTTCACTCT</td><td>>MI0000478|hsa-mir-149&WithFlank&2|+|240455995|240456095</td><td>21</td><td>41</td><td>21(+1U)</td><td>TCTGGCTCCGTGTCTTCACTCT</td><td>490.9</td><td>2.6</td><td>0.8</td><td>3.5</td><td>5.6E-04</td><td>2.9E-03</td><td>BT20/MB231</td><td>MITO</td></tr>
<tr><td>>MI0000542|hsa-mir-320a&WithFlank&8|-|22244960|22245043@48.68.21(+2U)&[MIMAT0000510&hsa-miR-320a-3p&offsets|0|-1(+2U);m-60&8|-|22244975|22244996&offsets|0|-1(+2U)]|AAAAGCTGGGTTGAGAGGGCGTT</td><td>>MI0000542|hsa-mir-320a&WithFlank&8|-|22244960|22245043</td><td>48</td><td>68</td><td>21(+2U)</td><td>AAAAGCTGGGTTGAGAGGGCGTT</td><td>1396.6</td><td>2.5</td><td>0.7</td><td>3.5</td><td>5.6E-04</td><td>2.9E-03</td><td>BT20/MB231</td><td>MITO</td></tr>
<tr><td>>hg38_MT_+_1621_3279_RNR2_16S_50ntFlanks@1432.1451.20|AAGTCCTACGTGATCTGAGT</td><td>>hg38_MT_+_1621_3279_RNR2_16S_50ntFlanks</td><td>1432</td><td>1451</td><td>20</td><td>AAGTCCTACGTGATCTGAGT</td><td>98.4</td><td>2.9</td><td>1.0</td><td>3.1</td><td>2.1E-03</td><td>2.9E-03</td><td>MB231/MB468</td><td>MP</td></tr>
<tr><td>>hg38_21_+_8433172_8446622_RNA45SN1_45S_50ntFlanks@8319.8356.38|CGAGACCGATAGTCAACAAGTACCGTAAGGGAAAGTTG</td><td>>hg38_21_+_8433172_8446622_RNA45SN1_45S_50ntFlanks</td><td>8319</td><td>8356</td><td>38</td><td>CGAGACCGATAGTCAACAAGTACCGTAAGGGAAAGTTG</td><td>306.4</td><td>-1.7</td><td>0.5</td><td>-3.1</td><td>1.9E-03</td><td>2.9E-03</td><td>BT20/MB468</td><td>MITO</td></tr>
<tr><td>>hg38_21_+_8433172_8446622_RNA45SN1_45S_50ntFlanks@4006.4039.34|ACCTCGGGCCGATCGCACGCCCCCCGTGGCGGCG</td><td>>hg38_21_+_8433172_8446622_RNA45SN1_45S_50ntFlanks</td><td>4006</td><td>4039</td><td>34</td><td>ACCTCGGGCCGATCGCACGCCCCCCGTGGCGGCG</td><td>292.5</td><td>-4.7</td><td>1.6</td><td>-3.0</td><td>2.5E-03</td><td>2.9E-03</td><td>MB231/MB468</td><td>NUC</td></tr>
<tr><td>>hg38_21_+_8433172_8446622_RNA45SN1_45S_50ntFlanks@12957.12987.31|CTTCTGGGTCGGGGTTTCGTACGTAGCAGAG</td><td>>hg38_21_+_8433172_8446622_RNA45SN1_45S_50ntFlanks</td><td>12957</td><td>12987</td><td>31</td><td>CTTCTGGGTCGGGGTTTCGTACGTAGCAGAG</td><td>175.0</td><td>-3.2</td><td>1.0</td><td>-3.1</td><td>1.9E-03</td><td>2.9E-03</td><td>BT20/MB468</td><td>MITO</td></tr>
<tr><td>>MI0000283|hsa-mir-203a&WithFlank&14|+|104117399|104117520@71.90.20&[MIMAT0000264&hsa-miR-203a-3p&offsets|0|-2;m-79&14|+|104117469|104117490&offsets|0|-2]|GTGAAATGTTTAGGACCACT</td><td>>MI0000283|hsa-mir-203a&WithFlank&14|+|104117399|104117520</td><td>71</td><td>90</td><td>20</td><td>GTGAAATGTTTAGGACCACT</td><td>430.1</td><td>5.8</td><td>1.8</td><td>3.2</td><td>1.2E-03</td><td>2.9E-03</td><td>BT20/MB468</td><td>MP</td></tr>
<tr><td>>hg38_MT_+_1621_3279_RNR2_16S_50ntFlanks@112.146.35|CCCAAATAAAGTATAGGCGATAGAAATTGAAACCT</td><td>>hg38_MT_+_1621_3279_RNR2_16S_50ntFlanks</td><td>112</td><td>146</td><td>35</td><td>CCCAAATAAAGTATAGGCGATAGAAATTGAAACCT</td><td>41.7</td><td>9.1</td><td>2.8</td><td>3.2</td><td>1.2E-03</td><td>2.9E-03</td><td>BT20/MB468</td><td>MP</td></tr>
<tr><td>>hg38_21_+_8433172_8446622_RNA45SN1_45S_50ntFlanks@6652.6680.29|GACTCTTAGCGGTGGATCACTCGGCTCGT</td><td>>hg38_21_+_8433172_8446622_RNA45SN1_45S_50ntFlanks</td><td>6652</td><td>6680</td><td>29</td><td>GACTCTTAGCGGTGGATCACTCGGCTCGT</td><td>418.6</td><td>2.0</td><td>0.6</td><td>3.1</td><td>2.3E-03</td><td>2.9E-03</td><td>BT20/MB468</td><td>CYTO</td></tr>
<tr><td>>hg38_21_+_8433172_8446622_RNA45SN1_45S_50ntFlanks@11029.11050.22|CTCTCTCTCTCTCCCCCGCTCC</td><td>>hg38_21_+_8433172_8446622_RNA45SN1_45S_50ntFlanks</td><td>11029</td><td>11050</td><td>22</td><td>CTCTCTCTCTCTCCCCCGCTCC</td><td>172.4</td><td>-3.3</td><td>1.1</td><td>-3.1</td><td>2.3E-03</td><td>2.9E-03</td><td>BT20/MB468</td><td>CYTO</td></tr>
<tr><td>>hg38_21_+_8433172_8446622_RNA45SN1_45S_50ntFlanks@12992.13016.25|TCCCTCGCTGCGATCTATTGAAAGT</td><td>>hg38_21_+_8433172_8446622_RNA45SN1_45S_50ntFlanks</td><td>12992</td><td>13016</td><td>25</td><td>TCCCTCGCTGCGATCTATTGAAAGT</td><td>420.2</td><td>2.0</td><td>0.7</td><td>3.0</td><td>2.3E-03</td><td>2.9E-03</td><td>MB231/MB468</td><td>CYTO</td></tr>
<tr><td>>hg38_MT_+_1621_3279_RNR2_16S_50ntFlanks@1431.1452.22|AAAGTCCTACGTGATCTGAGTT</td><td>>hg38_MT_+_1621_3279_RNR2_16S_50ntFlanks</td><td>1431</td><td>1452</td><td>22</td><td>AAAGTCCTACGTGATCTGAGTT</td><td>160.1</td><td>3.2</td><td>1.0</td><td>3.1</td><td>2.1E-03</td><td>2.9E-03</td><td>MB231/MB468</td><td>MP</td></tr>
<tr><td>>hg38_21_+_8433172_8446622_RNA45SN1_45S_50ntFlanks@10879.10902.24|CGGTCGGGCTGGGGCGCGAAGCGG</td><td>>hg38_21_+_8433172_8446622_RNA45SN1_45S_50ntFlanks</td><td>10879</td><td>10902</td><td>24</td><td>CGGTCGGGCTGGGGCGCGAAGCGG</td><td>643.8</td><td>-5.8</td><td>1.9</td><td>-3.0</td><td>2.5E-03</td><td>2.9E-03</td><td>BT20/MB468</td><td>NUC</td></tr>
<tr><td>>hg38_21_+_8433172_8446622_RNA45SN1_45S_50ntFlanks@3985.4005.21|CGGCTTTGGTGACTCTAGATA</td><td>>hg38_21_+_8433172_8446622_RNA45SN1_45S_50ntFlanks</td><td>3985</td><td>4005</td><td>21</td><td>CGGCTTTGGTGACTCTAGATA</td><td>267.9</td><td>-3.5</td><td>1.2</td><td>-3.0</td><td>2.4E-03</td><td>2.9E-03</td><td>MB231/MB468</td><td>CYTO</td></tr>
<tr><td>>hg38_21_+_8433172_8446622_RNA45SN1_45S_50ntFlanks@8114.8139.26|GCGCGGGACATGTGGCGTACGGAAGA</td><td>>hg38_21_+_8433172_8446622_RNA45SN1_45S_50ntFlanks</td><td>8114</td><td>8139</td><td>26</td><td>GCGCGGGACATGTGGCGTACGGAAGA</td><td>279.7</td><td>-4.2</td><td>1.4</td><td>-3.0</td><td>2.5E-03</td><td>2.9E-03</td><td>BT20/MB468</td><td>NUC</td></tr>
<tr><td>>hg38_21_+_8433172_8446622_RNA45SN1_45S_50ntFlanks@9604.9630.27|AAAGACTAATCGAACCATCTAGTAGCT</td><td>>hg38_21_+_8433172_8446622_RNA45SN1_45S_50ntFlanks</td><td>9604</td><td>9630</td><td>27</td><td>AAAGACTAATCGAACCATCTAGTAGCT</td><td>242.5</td><td>2.6</td><td>0.9</td><td>3.0</td><td>2.5E-03</td><td>2.9E-03</td><td>MB231/MB468</td><td>NUC</td></tr>
<tr><td>>hg38_21_+_8433172_8446622_RNA45SN1_45S_50ntFlanks@3705.3742.38|TACCTGGTTGATCCTGCCAGTAGCATATGCTTGTCTCA</td><td>>hg38_21_+_8433172_8446622_RNA45SN1_45S_50ntFlanks</td><td>3705</td><td>3742</td><td>38</td><td>TACCTGGTTGATCCTGCCAGTAGCATATGCTTGTCTCA</td><td>1907.6</td><td>3.0</td><td>0.9</td><td>3.5</td><td>4.6E-04</td><td>2.9E-03</td><td>BT20/MB231</td><td>NUC</td></tr>
<tr><td>>hg38_21_+_8433172_8446622_RNA45SN1_45S_50ntFlanks@11593.11621.29|CCAGGGGAATCCGACTGTTTAATTAAAAC</td><td>>hg38_21_+_8433172_8446622_RNA45SN1_45S_50ntFlanks</td><td>11593</td><td>11621</td><td>29</td><td>CCAGGGGAATCCGACTGTTTAATTAAAAC</td><td>202.2</td><td>-1.9</td><td>0.6</td><td>-3.0</td><td>2.5E-03</td><td>2.9E-03</td><td>MB231/MB468</td><td>NUC</td></tr>
<tr><td>>hg38_21_+_8433172_8446622_RNA45SN1_45S_50ntFlanks@12716.12751.36|GCCTCGGTTGGCCTCGGATAGCCGGTCCCCCGCCTG</td><td>>hg38_21_+_8433172_8446622_RNA45SN1_45S_50ntFlanks</td><td>12716</td><td>12751</td><td>36</td><td>GCCTCGGTTGGCCTCGGATAGCCGGTCCCCCGCCTG</td><td>45.8</td><td>-8.6</td><td>2.7</td><td>-3.2</td><td>1.2E-03</td><td>2.9E-03</td><td>BT20/MB468</td><td>MP</td></tr>
<tr><td>>am_MI0000809|hsa-mir-151a&WithFlank&8|-|140732558|140732659@53.72.20(+2U)&[MIMAT0000757&hsa-miR-151a-3p&offsets|0|-1(+2U);m-30&8|-|140732587|140732607&offsets|0|-1(+2U)]|CTAGACTGAAGCTCCTTGAGTT</td><td>>am_MI0000809|hsa-mir-151a&WithFlank&8|-|140732558|140732659</td><td>53</td><td>72</td><td>20(+2U)</td><td>CTAGACTGAAGCTCCTTGAGTT</td><td>112.9</td><td>6.4</td><td>2.0</td><td>3.2</td><td>1.2E-03</td><td>2.9E-03</td><td>BT20/MB468</td><td>MP</td></tr>
<tr><td>>hg38_21_+_8433172_8446622_RNA45SN1_45S_50ntFlanks@11197.11232.36|GGCCGGCGGCGGCGGCGACTCTGGACGCGAGCCGGG</td><td>>hg38_21_+_8433172_8446622_RNA45SN1_45S_50ntFlanks</td><td>11197</td><td>11232</td><td>36</td><td>GGCCGGCGGCGGCGGCGACTCTGGACGCGAGCCGGG</td><td>71.9</td><td>-5.1</td><td>1.6</td><td>-3.2</td><td>1.2E-03</td><td>2.9E-03</td><td>BT20/MB468</td><td>MP</td></tr>
<tr><td>>hg38_21_+_8433172_8446622_RNA45SN1_45S_50ntFlanks@9574.9610.37|AATCGGTCGTCCGACCTGGGTATAGGGGCGAAAGACT</td><td>>hg38_21_+_8433172_8446622_RNA45SN1_45S_50ntFlanks</td><td>9574</td><td>9610</td><td>37</td><td>AATCGGTCGTCCGACCTGGGTATAGGGGCGAAAGACT</td><td>563.0</td><td>2.8</td><td>0.9</td><td>3.0</td><td>2.5E-03</td><td>2.9E-03</td><td>MB231/MB468</td><td>MITO</td></tr>
<tr><td>>hg38_21_+_8433172_8446622_RNA45SN1_45S_50ntFlanks@8169.8195.27|CCAAGTCCTTCTGATCGAGGCCCAGCC</td><td>>hg38_21_+_8433172_8446622_RNA45SN1_45S_50ntFlanks</td><td>8169</td><td>8195</td><td>27</td><td>CCAAGTCCTTCTGATCGAGGCCCAGCC</td><td>113.4</td><td>-3.4</td><td>1.1</td><td>-3.1</td><td>1.9E-03</td><td>2.9E-03</td><td>BT20/MB468</td><td>MITO</td></tr>
<tr><td>>hg38_21_+_8433172_8446622_RNA45SN1_45S_50ntFlanks@11873.11900.28|AAGAAGACCCTGTTGAGCTTGACTCTAG</td><td>>hg38_21_+_8433172_8446622_RNA45SN1_45S_50ntFlanks</td><td>11873</td><td>11900</td><td>28</td><td>AAGAAGACCCTGTTGAGCTTGACTCTAG</td><td>625.3</td><td>-5.5</td><td>1.7</td><td>-3.2</td><td>1.2E-03</td><td>3.0E-03</td><td>BT20/MB468</td><td>MP</td></tr>
<tr><td>>hg38_21_+_8433172_8446622_RNA45SN1_45S_50ntFlanks@6654.6698.45|CTCTTAGCGGTGGATCACTCGGCTCGTGCGTCGATGAAGAACGCA</td><td>>hg38_21_+_8433172_8446622_RNA45SN1_45S_50ntFlanks</td><td>6654</td><td>6698</td><td>45</td><td>CTCTTAGCGGTGGATCACTCGGCTCGTGCGTCGATGAAGAACGCA</td><td>394.1</td><td>-2.5</td><td>0.8</td><td>-3.0</td><td>2.5E-03</td><td>3.0E-03</td><td>MB231/MB468</td><td>MITO</td></tr>
<tr><td>>MI0003560|hsa-mir-92b&WithFlank&1|+|155195171|155195278@67.88.22(+1C)&[MIMAT0003218&hsa-miR-92b-3p&offsets|0|0(+1C);m-115&1|+|155195237|155195258&offsets|0|0(+1C)]|TATTGCACTCGTCCCGGCCTCCC</td><td>>MI0003560|hsa-mir-92b&WithFlank&1|+|155195171|155195278</td><td>67</td><td>88</td><td>22(+1C)</td><td>TATTGCACTCGTCCCGGCCTCCC</td><td>39.4</td><td>8.5</td><td>2.8</td><td>3.1</td><td>2.1E-03</td><td>3.0E-03</td><td>MB231/MB468</td><td>MP</td></tr>
</table>

| | | | | | | | | | | | | | |
|---|---|---|---|---|---|---|---|---|---|---|---|---|---|
| >trnaMT_ValTAC_MT_+_1602_1670@35.72.38\|ACTTAGGAGATTTCAACTTAACTTGACCGCTCTGACCA | >trnaMT_ValTAC_MT_+_1602_1670 | 35 | 72 | 38 | ACTTAGGAGATTTCAACTTAACTTGACCGCTCTGACCA | 257.4 | 4.1 | 1.3 | 3.1 | 2.1E-03 | 3.0E-03 | MB231/MB468 | MP |
| >hg38_21_+_8433172_8446622_RNA45SN1_45S_50ntFlanks@6650.6668.19\|ACGACTCTTAGCGGTGGAT | >hg38_21_+_8433172_8446622_RNA45SN1_45S_50ntFlanks | 6650 | 6668 | 19 | ACGACTCTTAGCGGTGGAT | 1944.2 | 2.5 | 0.7 | 3.5 | 4.7E-04 | 3.0E-03 | BT20/MB231 | NUC |
| >hg38_21_+_8433172_8446622_RNA45SN1_45S_50ntFlanks@8197.8224.28\|GTGGACGGTGTGAGGCCGGTAGCGGCCC | >hg38_21_+_8433172_8446622_RNA45SN1_45S_50ntFlanks | 8197 | 8224 | 28 | GTGGACGGTGTGAGGCCGGTAGCGGCCC | 89.1 | -3.9 | 1.3 | -3.1 | 2.1E-03 | 3.0E-03 | MB231/MB468 | MP |
| >hg38_21_+_8433172_8446622_RNA45SN1_45S_50ntFlanks@9527.9570.44\|AAGCCAGAGGAAACTCTGGTGGAGGTCCGTAGCGGTCCTGACGT | >hg38_21_+_8433172_8446622_RNA45SN1_45S_50ntFlanks | 9527 | 9570 | 44 | AAGCCAGAGGAAACTCTGGTGGAGGTCCGTAGCGGTCCTGACGT | 254.2 | 2.7 | 0.9 | 3.0 | 2.4E-03 | 3.0E-03 | MB231/MB468 | CYTO |
| >hg38_21_+_8433172_8446622_RNA45SN1_45S_50ntFlanks@5075.5112.38\|TTCCGATAACGAACGAGACTCTGGCATGCTAACTAGTT | >hg38_21_+_8433172_8446622_RNA45SN1_45S_50ntFlanks | 5075 | 5112 | 38 | TTCCGATAACGAACGAGACTCTGGCATGCTAACTAGTT | 45.6 | -9.4 | 3.0 | -3.1 | 2.1E-03 | 3.0E-03 | MB231/MB468 | MP |
| >hg38_1_-_228634819_228635039_RNA5S12_5S_50ntFlanks@126.170.45\|TACTTGGATGGGAGACCGCCTGGGAATACCGGGTGCTGTAGGCTT | >hg38_1_-_228634819_228635039_RNA5S12_5S_50ntFlanks | 126 | 170 | 45 | TACTTGGATGGGAGACCGCCTGGGAATACCGGGTGCTGTAGGCTT | 1003.1 | 4.9 | 1.6 | 3.0 | 2.5E-03 | 3.0E-03 | MB231/MB468 | NUC |
| >hg38_21_+_8433172_8446622_RNA45SN1_45S_50ntFlanks@12413.12435.23\|TCCTTCGATGTCGGCTCTTCCTA | >hg38_21_+_8433172_8446622_RNA45SN1_45S_50ntFlanks | 12413 | 12435 | 23 | TCCTTCGATGTCGGCTCTTCCTA | 551.4 | -4.2 | 1.4 | -3.0 | 2.5E-03 | 3.0E-03 | MB231/MB468 | NUC |
| >hg38_21_+_8433172_8446622_RNA45SN1_45S_50ntFlanks@12640.12661.22\|ACCATCTGTGGGATTATGACTG | >hg38_21_+_8433172_8446622_RNA45SN1_45S_50ntFlanks | 12640 | 12661 | 22 | ACCATCTGTGGGATTATGACTG | 303.7 | 7.1 | 2.0 | 3.5 | 4.3E-04 | 3.0E-03 | BT20/MB231 | MP |
| >hg38_21_+_8433172_8446622_RNA45SN1_45S_50ntFlanks@483.505.23\|TCGATGTGGTGACGTCGTGCTCT | >hg38_21_+_8433172_8446622_RNA45SN1_45S_50ntFlanks | 483 | 505 | 23 | TCGATGTGGTGACGTCGTGCTCT | 184.3 | 3.6 | 1.1 | 3.2 | 1.2E-03 | 3.0E-03 | BT20/MB468 | MP |
| >am_trna128_GlyGCC_6_-_27870686_27870756@12.29.18,trna133_GlyCCC_1_-_16872434_16872504@12.29.18,trna17_GluTTC_2_-_204229131_204229201@12.29.18,trna18_GlyGCC_16_+_70822597_70822667@12.29.18,trna19_GlyGCC_16_+_70823410_70823480@12.29.18,trna19_GlyGCC_2_-_157257659_157257729@12.29.18,trna24_GlyGCC_16_-_70812942_70813012@12.29.18,trna25_GlyGCC_16_-_70812114_70812184@12.29.18,trna2_GlyGCC_21_-_18827107_18827177@12.29.18,trna35_GlyGCC_1_+_161413094_161413164@12.29.18,trna37_GlyGCC_1_+_161420467_161420537@12.29.18,trna39_GlyGCC_1_+_161427898_161427968@12.29.18,trna41_GlyGCC_1_+_161435258_161435328@12.29.18,trna4_GlyCCC_1_+_17188416_17188486@12.29.18,trna5_GlyGCC_17_+_8029064_8029134@12.29.18,trna68_GlyGCC_1_-_161493637_161493707@12.29.18,trna91_GlyCCC_1_-_149680210_149680280@12.29.18,trna99_ValCAC_1_-_149294666_149294736@12.29.18\|TCAGTGGTAGAATTCTCG | >am_trna128_GlyGCC_6_-_27870686_27870756 | 12 | 29 | 18 | TCAGTGGTAGAATTCTCG | 39.7 | -8.4 | 2.6 | -3.2 | 1.2E-03 | 3.0E-03 | BT20/MB468 | MP |
| >hg38_21_+_8433172_8446622_RNA45SN1_45S_50ntFlanks@10794.10821.28\|CTGGCATGTTGGAACAATGTAGGTAAGG | >hg38_21_+_8433172_8446622_RNA45SN1_45S_50ntFlanks | 10794 | 10821 | 28 | CTGGCATGTTGGAACAATGTAGGTAAGG | 184.9 | -4.0 | 1.2 | -3.2 | 1.2E-03 | 3.0E-03 | BT20/MB468 | MP |
| >hg38_21_+_8433172_8446622_RNA45SN1_45S_50ntFlanks@10392.10429.38\|ACAGCAGTTGAACATGGGTCAGTCGGTCCTGAGAGATG | >hg38_21_+_8433172_8446622_RNA45SN1_45S_50ntFlanks | 10392 | 10429 | 38 | ACAGCAGTTGAACATGGGTCAGTCGGTCCTGAGAGATG | 1341.4 | -2.1 | 0.7 | -3.1 | 1.9E-03 | 3.0E-03 | BT20/MB468 | MITO |
| >hg38_21_+_8433172_8446622_RNA45SN1_45S_50ntFlanks@9939.9957.19\|GCAGGACGGTGGCCATGGA | >hg38_21_+_8433172_8446622_RNA45SN1_45S_50ntFlanks | 9939 | 9957 | 19 | GCAGGACGGTGGCCATGGA | 307.7 | 4.7 | 1.5 | 3.2 | 1.2E-03 | 3.0E-03 | BT20/MB468 | MP |
| >hg38_21_+_8433172_8446622_RNA45SN1_45S_50ntFlanks@12406.12446.41\|TTTTTGATCCTTCGATGTCGGCTCTTCCTATCATTGTGAAG | >hg38_21_+_8433172_8446622_RNA45SN1_45S_50ntFlanks | 12406 | 12446 | 41 | TTTTTGATCCTTCGATGTCGGCTCTTCCTATCATTGTGAAG | 332.2 | -2.9 | 0.9 | -3.0 | 2.5E-03 | 3.0E-03 | MB231/MB468 | NUC |
| >hg38_21_+_8433172_8446622_RNA45SN1_45S_50ntFlanks@3899.3927.29\|CCTTCGCGGGGGGGATGCGTGCATTTATC | >hg38_21_+_8433172_8446622_RNA45SN1_45S_50ntFlanks | 3899 | 3927 | 29 | CCTTCGCGGGGGGGATGCGTGCATTTATC | 756.9 | -6.0 | 2.0 | -3.0 | 2.5E-03 | 3.0E-03 | BT20/MB468 | NUC |
| >hg38_21_+_8433172_8446622_RNA45SN1_45S_50ntFlanks@10411.10441.31\|CAGTCGGTCCTGAGAGATGGGCGAGCGCCGT | >hg38_21_+_8433172_8446622_RNA45SN1_45S_50ntFlanks | 10411 | 10441 | 31 | CAGTCGGTCCTGAGAGATGGGCGAGCGCCGT | 123.7 | -4.3 | 1.4 | -3.1 | 1.9E-03 | 3.0E-03 | BT20/MB468 | MITO |
| >hg38_21_+_8433172_8446622_RNA45SN1_45S_50ntFlanks@5445.5466.22\|TCGGATCGGCCCCGCCGGGGTC | >hg38_21_+_8433172_8446622_RNA45SN1_45S_50ntFlanks | 5445 | 5466 | 22 | TCGGATCGGCCCCGCCGGGGTC | 7834.7 | -6.7 | 2.2 | -3.0 | 2.5E-03 | 3.0E-03 | BT20/MB468 | NUC |
| >hg38_21_+_8433172_8446622_RNA45SN1_45S_50ntFlanks@4981.5001.21\|CACGGACAGGATTGACAGATT | >hg38_21_+_8433172_8446622_RNA45SN1_45S_50ntFlanks | 4981 | 5001 | 21 | CACGGACAGGATTGACAGATT | 167.7 | 2.9 | 0.9 | 3.2 | 1.2E-03 | 3.0E-03 | BT20/MB468 | MP |
| >hg38_MT_+_1621_3279_RNR2_16S_50ntFlanks@1182.1216.35\|AAAATTTCGGTTGGGGCGACCTCGGAGCAGAACCC | >hg38_MT_+_1621_3279_RNR2_16S_50ntFlanks | 1182 | 1216 | 35 | AAAATTTCGGTTGGGGCGACCTCGGAGCAGAACCC | 210.5 | 4.5 | 1.4 | 3.2 | 1.2E-03 | 3.0E-03 | BT20/MB468 | MP |
| >trnaMT_GluTTC_MT_-_14674_14742@26.66.41,trnalookalike8_GluTTC_5_-_93905172_93905240@26.66.41\|GGTTTTTCATATCATTGGTCGTGGTTGTAGTCCGTGCGAGA | >trnaMT_GluTTC_MT_-_14674_14742 | 26 | 66 | 41 | GGTTTTTCATATCATTGGTCGTGGTTGTAGTCCGTGCGAGA | 661.6 | 3.3 | 1.1 | 3.0 | 2.5E-03 | 3.0E-03 | MB231/MB468 | NUC |
| >MI0000298\|hsa-mir-221&WithFlank&X\|-\|45746151\|45746272@71.93.23(+1C)&[MIMAT0000278&hsa-miR-221-3p&offsets\|0\|0(+1C);m-91&X\|-\|45746181\|45746202&offsets\|0\|+1(+1C)]\|AGCTACATTGTCTGCTGGGTTTCC | >MI0000298\|hsa-mir-221&WithFlank&X\|-\|45746151\|45746272 | 71 | 93 | 23(+1C) | AGCTACATTGTCTGCTGGGTTTCC | 2078.5 | -2.6 | 0.8 | -3.4 | 5.7E-04 | 3.0E-03 | BT20/MB231 | MITO |
| >trnaMT_ProTGG_MT_-_15956_16023@50.71.22\|TTAAAGACTTTTTCTCTGACCA | >trnaMT_ProTGG_MT_-_15956_16023 | 50 | 71 | 22 | TTAAAGACTTTTTCTCTGACCA | 443.2 | 2.9 | 1.0 | 3.0 | 2.5E-03 | 3.0E-03 | MB231/MB468 | MITO |
| >hg38_21_+_8433172_8446622_RNA45SN1_45S_50ntFlanks@12420.12459.40\|ATGTCGGCTCTTCCTATCATTGTGAAGCAGAATTCACCAA | >hg38_21_+_8433172_8446622_RNA45SN1_45S_50ntFlanks | 12420 | 12459 | 40 | ATGTCGGCTCTTCCTATCATTGTGAAGCAGAATTCACCAA | 309.7 | -2.8 | 0.8 | -3.5 | 5.0E-04 | 3.0E-03 | BT20/MB231 | CYTO |
| >hg38_21_+_8433172_8446622_RNA45SN1_45S_50ntFlanks@10778.10818.41\|CCAAGGTGAACAGCCTCTGGCATGTTGGAACAATGTAGGTA | >hg38_21_+_8433172_8446622_RNA45SN1_45S_50ntFlanks | 10778 | 10818 | 41 | CCAAGGTGAACAGCCTCTGGCATGTTGGAACAATGTAGGTA | 226.4 | -3.6 | 1.2 | -3.1 | 2.0E-03 | 3.0E-03 | BT20/MB468 | MITO |
| >hg38_21_+_8433172_8446622_RNA45SN1_45S_50ntFlanks@11894.11916.23\|ACTCTAGTCTGGCACGGTGAAGA | >hg38_21_+_8433172_8446622_RNA45SN1_45S_50ntFlanks | 11894 | 11916 | 23 | ACTCTAGTCTGGCACGGTGAAGA | 221.5 | -2.1 | 0.7 | -3.0 | 2.3E-03 | 3.0E-03 | BT20/MB468 | CYTO |
| >hg38_21_+_8433172_8446622_RNA45SN1_45S_50ntFlanks@12416.12452.37\|TTCGATGTCGGCTCTTCCTATCATTGTGAAGCAGAAT | >hg38_21_+_8433172_8446622_RNA45SN1_45S_50ntFlanks | 12416 | 12452 | 37 | TTCGATGTCGGCTCTTCCTATCATTGTGAAGCAGAAT | 560.3 | -2.9 | 0.8 | -3.5 | 5.1E-04 | 3.0E-03 | BT20/MB231 | CYTO |
| >trna111_HisGTG_1_-_147774845_147774916@-1T.20.21,trna118_HisGTG_1_-_145396881_145396952@-1T.20.21,trna16_HisGTG_1_+_146544773_146544844@-1T.20.21,trna1_HisGTG_15_+_45493349_45493420@-1T.20.21,trna21_HisGTG_1_+_147753471_147753542@-1T.20.21,trna33_HisGTG_6_+_27125906_27125977@-1T.20.21,trna7_HisGTG_9_-_14433938_14434009@-1T.20.21,trna8_HisGTG_15_-_45492611_45492682@-1T.20.21,trna9_HisGTG_15_-_45490804_45490875@-1T.20.21\|TGCCGTGATCGTATAGTGGTT | >trna111_HisGTG_1_-_147774845_147774916 | -1T | 20 | 21 | TGCCGTGATCGTATAGTGGTT | 148.3 | -5.7 | 1.6 | -3.5 | 4.7E-04 | 3.0E-03 | BT20/MB231 | NUC |
| >hg38_21_+_8433172_8446622_RNA45SN1_45S_50ntFlanks@12995.13040.46\|CTCGCTGCGATCTATTGAAAGTCAGCCCTCGACACAAGGGTTTGTC | >hg38_21_+_8433172_8446622_RNA45SN1_45S_50ntFlanks | 12995 | 13040 | 46 | CTCGCTGCGATCTATTGAAAGTCAGCCCTCGACACAAGGGTTTGTC | 548.7 | -5.8 | 1.7 | -3.5 | 4.7E-04 | 3.0E-03 | BT20/MB231 | NUC |
| >MI0006444\|hsa-mir-1307&WithFlank&10\|-\|103394247\|103394407@86.107.22&[MIMAT0005951&hsa-miR-1307-3p&offsets\|0\|0;m-66&10\|-\|103394300\|103394322&offsets\|0\|-1]\|ACTCGGCGTGGCGTCGGTCGTG | >MI0006444\|hsa-mir-1307&WithFlank&10\|-\|103394247\|103394407 | 86 | 107 | 22 | ACTCGGCGTGGCGTCGGTCGTG | 323.9 | 3.0 | 0.9 | 3.2 | 1.2E-03 | 3.0E-03 | BT20/MB468 | MP |
| >am_trnaMT_ValTAC_MT_+_1602_1670@36.66.31\|CTTAGGAGATTTCAACTTAACTTGACCGCTC | >am_trnaMT_ValTAC_MT_+_1602_1670 | 36 | 66 | 31 | CTTAGGAGATTTCAACTTAACTTGACCGCTC | 291.8 | 8.1 | 2.5 | 3.2 | 1.2E-03 | 3.0E-03 | BT20/MB468 | MP |
| >hg38_21_+_8433172_8446622_RNA45SN1_45S_50ntFlanks@10350.10381.32\|ACGAGAACTTTGAAGGCCGAAGTGGAGAAGGG | >hg38_21_+_8433172_8446622_RNA45SN1_45S_50ntFlanks | 10350 | 10381 | 32 | ACGAGAACTTTGAAGGCCGAAGTGGAGAAGGG | 674.5 | -3.0 | 0.9 | -3.2 | 1.2E-03 | 3.0E-03 | BT20/MB468 | MP |
| >hg38_21_+_8433172_8446622_RNA45SN1_45S_50ntFlanks@8186.8206.21\|AGGCCCAGCCCGTGGACGGTG | >hg38_21_+_8433172_8446622_RNA45SN1_45S_50ntFlanks | 8186 | 8206 | 21 | AGGCCCAGCCCGTGGACGGTG | 91.9 | -5.0 | 1.6 | -3.1 | 2.1E-03 | 3.0E-03 | MB231/MB468 | MP |
| >hg38_21_+_8433172_8446622_RNA45SN1_45S_50ntFlanks@6778.6803.26\|CCCGGGGCTACGCCTGTCTGAGCGTC | >hg38_21_+_8433172_8446622_RNA45SN1_45S_50ntFlanks | 6778 | 6803 | 26 | CCCGGGGCTACGCCTGTCTGAGCGTC | 238.5 | -4.1 | 1.3 | -3.1 | 2.1E-03 | 3.0E-03 | MB231/MB468 | MP |
| >hg38_21_+_8433172_8446622_RNA45SN1_45S_50ntFlanks@11755.11777.23\|AATGCCTCGTCATCTAATTAGTG | >hg38_21_+_8433172_8446622_RNA45SN1_45S_50ntFlanks | 11755 | 11777 | 23 | AATGCCTCGTCATCTAATTAGTG | 279.5 | -3.2 | 1.1 | -3.0 | 2.4E-03 | 3.0E-03 | MB231/MB468 | CYTO |
| >hg38_21_+_8433172_8446622_RNA45SN1_45S_50ntFlanks@10787.10826.40\|ACAGCCTCTGGCATGTTGGAACAATGTAGGTAAGGGAAGT | >hg38_21_+_8433172_8446622_RNA45SN1_45S_50ntFlanks | 10787 | 10826 | 40 | ACAGCCTCTGGCATGTTGGAACAATGTAGGTAAGGGAAGT | 178.9 | -3.7 | 1.2 | -3.0 | 2.5E-03 | 3.0E-03 | MB231/MB468 | NUC |
| >hg38_21_+_8433172_8446622_RNA45SN1_45S_50ntFlanks@12413.12437.25\|TCCTTCGATGTCGGCTCTTCCTATC | >hg38_21_+_8433172_8446622_RNA45SN1_45S_50ntFlanks | 12413 | 12437 | 25 | TCCTTCGATGTCGGCTCTTCCTATC | 374.5 | -2.0 | 0.7 | -3.0 | 2.5E-03 | 3.0E-03 | MB231/MB468 | NUC |
| >MI0000433\|hsa-let-7g&WithFlank&3\|-\|52268272\|52268367@11.29.19&[MIMAT0000414&hsa-let-7g-5p&offsets\|0\|-3;m-58&3\|-\|52268336\|52268357&offsets\|0\|-3]\|TGAGGTAGTAGTTTGTACA | >MI0000433\|hsa-let-7g&WithFlank&3\|-\|52268272\|52268367 | 11 | 29 | 19 | TGAGGTAGTAGTTTGTACA | 82.2 | 3.2 | 1.1 | 3.0 | 2.5E-03 | 3.0E-03 | BT20/MB468 | NUC |
| >am_trna119_LysCTT_1_-_145395522_145395594@13.30.18,trna11_LysCTT_5_-_180648979_180649051@13.30.18,trna128_LysCTT_1_-_55423542_55423614@13.30.18,trna13_LysCTT_6_+_26556774_26556846@13.30.18,trna32_LysCTT_16_-_3207406_3207478@13.30.18,trna6_LysCTT_19_-_52425396_52425468@13.30.18,trna7_LysCTT_16_+_3225692_3225764@13.30.18,trna9_LysCTT_5_+_180634755_180634827@13.30.18\|CAGTCGGTAGAGCATGAG | >am_trna119_LysCTT_1_-_145395522_145395594 | 13 | 30 | 18 | CAGTCGGTAGAGCATGAG | 246.2 | -3.6 | 1.2 | -3.0 | 2.5E-03 | 3.0E-03 | MB231/MB468 | NUC |
| >MI0000289\|hsa-mir-181a-1&WithFlank&1\|-\|198859038\|198859159@30.52.23(+1C);MI0000269\|hsa-mir-181a-2&WithFlank&9\|+\|124692436\|124692557@45.67.23(+1C)&[MIMAT0000256&hsa-miR-181a-5p&offsets\|0\|0(+1C);m-32&1\|-\|198859108\|198859130&offsets\|0\|0(+1C)];[MIMAT0000256_1&hsa-miR-181a-5p&offsets\|0\|0(+1C);m-32&9\|+\|124692480\|124692502&offsets\|0\|0(+1C)]\|AACATTCAACGCTGTCGGTGAGTC | >MI0000289\|hsa-mir-181a-1&WithFlank&1\|-\|198859038\|198859159 | 30 | 52 | 23(+1C) | AACATTCAACGCTGTCGGTGAGTC | 1603.3 | -5.3 | 1.5 | -3.5 | 4.3E-04 | 3.0E-03 | BT20/MB231 | MP |
| >hg38_21_+_8433172_8446622_RNA45SN1_45S_50ntFlanks@4160.4186.27\|CCACATCCAAGGAAGGCAGCAGGCGCG | >hg38_21_+_8433172_8446622_RNA45SN1_45S_50ntFlanks | 4160 | 4186 | 27 | CCACATCCAAGGAAGGCAGCAGGCGCG | 134.1 | -1.8 | 0.6 | -3.0 | 2.4E-03 | 3.0E-03 | MB231/MB468 | CYTO |

| | | | | | | | | | | | | | |
|---|---|---|---|---|---|---|---|---|---|---|---|---|---|
| >hg38_21_+_8433172_8446622_RNA45SN1_45S_50ntFlanks@12082.12110.29\|TCTCGCTTCTGGCGCCAAGCGCCCGGCCG | >hg38_21_+_8433172_8446622_RNA45SN1_45S_50ntFlanks | 12082 | 12110 | 29 | TCTCGCTTCTGGCGCCAAGCGCCCGGCCG | 66.2 | -5.0 | 1.6 | -3.2 | 1.2E-03 | 3.0E-03 | BT20/MB468 | MP |
| >hg38_21_+_8433172_8446622_RNA45SN1_45S_50ntFlanks@12912.12933.22\|CACGTTCGTGGGGAACCTGGCG | >hg38_21_+_8433172_8446622_RNA45SN1_45S_50ntFlanks | 12912 | 12933 | 22 | CACGTTCGTGGGGAACCTGGCG | 1297.8 | -4.4 | 1.5 | -3.0 | 2.4E-03 | 3.0E-03 | MB231/MB468 | CYTO |
| >MI0001641\|hsa-mir-429&WithFlank&1\|+\|1168999\|1169093@57.78.22&[MIMAT0001536&hsa-miR-429&offsets\|0\|0;m-207&1\|+\|1169055\|1169076&offsets\|0\|0]\|TAATACTGTCTGGTAAAACCGT | >MI0001641\|hsa-mir-429&WithFlank&1\|+\|1168999\|1169093 | 57 | 78 | 22 | TAATACTGTCTGGTAAAACCGT | 237.5 | -4.5 | 1.5 | -3.1 | 2.0E-03 | 3.0E-03 | BT20/MB468 | MITO |
| >hg38_21_+_8433172_8446622_RNA45SN1_45S_50ntFlanks@6654.6699.46\|CTCTTAGCGGTGGATCACTCGGCTCGTGCGTCGATGAAGAACGCAG | >hg38_21_+_8433172_8446622_RNA45SN1_45S_50ntFlanks | 6654 | 6699 | 46 | CTCTTAGCGGTGGATCACTCGGCTCGTGCGTCGATGAAGAACGCAG | 234.9 | -2.8 | 0.9 | -3.0 | 2.5E-03 | 3.0E-03 | MB231/MB468 | MITO |
| >hg38_21_+_8433172_8446622_RNA45SN1_45S_50ntFlanks@476.498.23\|TTCGTGATCGATGTGGTGACGTC | >hg38_21_+_8433172_8446622_RNA45SN1_45S_50ntFlanks | 476 | 498 | 23 | TTCGTGATCGATGTGGTGACGTC | 419.1 | 4.3 | 1.4 | 3.1 | 2.1E-03 | 3.0E-03 | MB231/MB468 | MP |
| >hg38_21_+_8433172_8446622_RNA45SN1_45S_50ntFlanks@11729.11748.20\|TAACTATGACTCTCTTAAGG | >hg38_21_+_8433172_8446622_RNA45SN1_45S_50ntFlanks | 11729 | 11748 | 20 | TAACTATGACTCTCTTAAGG | 4252.0 | -2.6 | 0.9 | -3.0 | 2.3E-03 | 3.0E-03 | BT20/MB468 | CYTO |
| >trnaMT_ValTAC_MT_+_1602_1670@33.71.39\|ACACTTAGGAGATTTCAACTTAACTTGACCGCTCTGACC | >trnaMT_ValTAC_MT_+_1602_1670 | 33 | 71 | 39 | ACACTTAGGAGATTTCAACTTAACTTGACCGCTCTGACC | 244.1 | 4.0 | 1.3 | 3.1 | 2.1E-03 | 3.0E-03 | MB231/MB468 | MP |
| >hg38_21_+_8433172_8446622_RNA45SN1_45S_50ntFlanks@9941.9979.39\|AGGACGGTGGCCATGGAAGTCGGAATCCGCTAAGGAGTG | >hg38_21_+_8433172_8446622_RNA45SN1_45S_50ntFlanks | 9941 | 9979 | 39 | AGGACGGTGGCCATGGAAGTCGGAATCCGCTAAGGAGTG | 411.2 | 7.3 | 2.1 | 3.5 | 4.3E-04 | 3.0E-03 | BT20/MB231 | MP |
| >hg38_21_+_8433172_8446622_RNA45SN1_45S_50ntFlanks@6651.6688.38\|CGACTCTTAGCGGTGGATCACTCGGCTCGTGCGTCGAT | >hg38_21_+_8433172_8446622_RNA45SN1_45S_50ntFlanks | 6651 | 6688 | 38 | CGACTCTTAGCGGTGGATCACTCGGCTCGTGCGTCGAT | 2246.6 | -2.3 | 0.8 | -3.1 | 2.1E-03 | 3.0E-03 | MB231/MB468 | MP |
| >hg38_21_+_8433172_8446622_RNA45SN1_45S_50ntFlanks@11208.11229.22\|GCGGCGACTCTGGACGCGAGCC | >hg38_21_+_8433172_8446622_RNA45SN1_45S_50ntFlanks | 11208 | 11229 | 22 | GCGGCGACTCTGGACGCGAGCC | 566.7 | 7.4 | 2.1 | 3.5 | 4.3E-04 | 3.0E-03 | BT20/MB231 | MP |
| >MI0000542\|hsa-mir-320a&WithFlank&8\|-\|22244960\|22245043@49.72.24&[MIMAT0000510&hsa-miR-320a-3p&offsets\|+1\|+3;m-60&8\|-\|22244975\|22244996&offsets\|+1\|+3]\|AAAGCTGGGTTGAGAGGGCGAAAA | >MI0000542\|hsa-mir-320a&WithFlank&8\|-\|22244960\|22245043 | 49 | 72 | 24 | AAAGCTGGGTTGAGAGGGCGAAAA | 67.4 | 6.2 | 1.9 | 3.2 | 1.2E-03 | 3.0E-03 | BT20/MB468 | MP |
| >trna10_AspGTC_12_-_125424193_125424264@37.70.34,trna12_AspGTC_12_-_125411891_125411962@37.70.34,trna144_AspGTC_6_-_27551236_27551307@37.70.34,trna38_AspGTC_17_-_8125556_8125627@37.70.34,trna45_AspGTC_6_+_27447453_27447524@37.70.34,trna48_AspGTC_6_+_27471523_27471594@37.70.34,trna4_AspGTC_12_+_96429799_96429870@37.70.34,trna69_AspGTC_1_-_161440205_161440276@37.70.34,trna72_AspGTC_1_-_161432824_161432895@37.70.34,trna75_AspGTC_1_-_161425414_161425485@37.70.34,trna78_AspGTC_1_-_161418033_161418104@37.70.34,trna81_AspGTC_1_-_161410615_161410686@37.70.34\|ACGCGGGAGACCGGGGTTCGATTCCCCGACGGGG | >trna10_AspGTC_12_-_125424193_125424264 | 37 | 70 | 34 | ACGCGGGAGACCGGGGTTCGATTCCCCGACGGGG | 327.5 | -2.9 | 1.0 | -3.0 | 2.5E-03 | 3.0E-03 | MB231/MB468 | MITO |
| >hg38_1_-_228634819_228635039_RNA5S12_5S_50ntFlanks@127.171.45\|ACTTGGATGGGAGACCGCCTGGGAATACCGGGTGCTGTAGGCTTT | >hg38_1_-_228634819_228635039_RNA5S12_5S_50ntFlanks | 127 | 171 | 45 | ACTTGGATGGGAGACCGCCTGGGAATACCGGGTGCTGTAGGCTTT | 39.7 | 8.5 | 2.8 | 3.1 | 2.1E-03 | 3.0E-03 | MB231/MB468 | MP |
| >MI0000081\|hsa-mir-24-2&WithFlank&19\|-\|13836281\|13836365@56.75.20(+1G);MI0000080\|hsa-mir-24-1&WithFlank&9\|+\|95086015\|95086094@50.69.20(+1G)&[MIMAT0000080&hsa-miR-24-3p&offsets\|0\|-2(+1G);m-36&19\|-\|13836288\|13836310&offsets\|0\|-3(+1G)];[MIMAT0000080_1&hsa-miR-24-3p&offsets\|0\|-2(+1G);m-35&9\|+\|95086064\|95086086&offsets\|0\|-3(+1G)]\|TGGCTCAGTTCAGCAGGAACG | >MI0000081\|hsa-mir-24-2&WithFlank&19\|-\|13836281\|13836365 | 56 | 75 | 20(+1G) | TGGCTCAGTTCAGCAGGAACG | 166.2 | 3.3 | 1.0 | 3.2 | 1.2E-03 | 3.0E-03 | BT20/MB468 | MP |
| >MI0000095\|hsa-mir-93&WithFlank&7\|-\|100093762\|100093853@17.38.22(+1A)&[MIMAT0000093&hsa-miR-93-5p&offsets\|0\|-1(+1A);m-29&7\|-\|100093815\|100093837&offsets\|0\|-1(+1A)]\|CAAAGTGCTGTTCGTGCAGGTAA | >MI0000095\|hsa-mir-93&WithFlank&7\|-\|100093762\|100093853 | 17 | 38 | 22(+1A) | CAAAGTGCTGTTCGTGCAGGTAA | 2533.3 | 4.0 | 1.2 | 3.2 | 1.3E-03 | 3.0E-03 | BT20/MB468 | MP |
| >MI0000434\|hsa-let-7i&WithFlank&12\|+\|62603680\|62603775@12.31.20&[MIMAT0000415&hsa-let-7i-5p&offsets\|0\|-2;m-49&12\|+\|62603691\|62603712&offsets\|0\|-2]\|TGAGGTAGTAGTTTGTGCTG | >MI0000434\|hsa-let-7i&WithFlank&12\|+\|62603680\|62603775 | 12 | 31 | 20 | TGAGGTAGTAGTTTGTGCTG | 224.3 | 4.8 | 1.5 | 3.2 | 1.3E-03 | 3.0E-03 | BT20/MB468 | MP |
| >hg38_MT_+_1621_3279_RNR2_16S_50ntFlanks@624.652.29\|TAACTGAACTCCTCACACCCAATTGGACC | >hg38_MT_+_1621_3279_RNR2_16S_50ntFlanks | 624 | 652 | 29 | TAACTGAACTCCTCACACCCAATTGGACC | 75.3 | 6.3 | 2.0 | 3.2 | 1.3E-03 | 3.0E-03 | BT20/MB468 | MP |
| >hg38_21_+_8433172_8446622_RNA45SN1_45S_50ntFlanks@10800.10818.19\|TGTTGGAACAATGTAGGTA | >hg38_21_+_8433172_8446622_RNA45SN1_45S_50ntFlanks | 10800 | 10818 | 19 | TGTTGGAACAATGTAGGTA | 391.3 | -3.1 | 1.0 | -3.0 | 2.3E-03 | 3.0E-03 | BT20/MB468 | CYTO |
| >hg38_21_+_8433172_8446622_RNA45SN1_45S_50ntFlanks@11199.11233.35\|CCGGCGGCGGCGGCGACTCTGGACGCGAGCCGGGC | >hg38_21_+_8433172_8446622_RNA45SN1_45S_50ntFlanks | 11199 | 11233 | 35 | CCGGCGGCGGCGGCGACTCTGGACGCGAGCCGGGC | 163.6 | -5.3 | 1.6 | -3.2 | 1.3E-03 | 3.0E-03 | BT20/MB468 | MP |
| >hg38_21_+_8433172_8446622_RNA45SN1_45S_50ntFlanks@6648.6669.22\|GTACGACTCTTAGCGGTGGATC | >hg38_21_+_8433172_8446622_RNA45SN1_45S_50ntFlanks | 6648 | 6669 | 22 | GTACGACTCTTAGCGGTGGATC | 616.2 | 1.2 | 0.4 | 3.0 | 2.5E-03 | 3.0E-03 | BT20/MB468 | NUC |
| >hg38_21_+_8433172_8446622_RNA45SN1_45S_50ntFlanks@10234.10260.27\|CGGACGCTACGCCGCGACGAGTAGGAG | >hg38_21_+_8433172_8446622_RNA45SN1_45S_50ntFlanks | 10234 | 10260 | 27 | CGGACGCTACGCCGCGACGAGTAGGAG | 364.0 | -2.8 | 0.9 | -3.2 | 1.3E-03 | 3.0E-03 | BT20/MB468 | MP |
| >hg38_21_+_8433172_8446622_RNA45SN1_45S_50ntFlanks@8171.8201.31\|AAGTCCTTCTGATCGAGGCCCAGCCCGTGGA | >hg38_21_+_8433172_8446622_RNA45SN1_45S_50ntFlanks | 8171 | 8201 | 31 | AAGTCCTTCTGATCGAGGCCCAGCCCGTGGA | 57.0 | -5.1 | 1.6 | -3.1 | 2.1E-03 | 3.0E-03 | MB231/MB468 | MP |
| >trnaMT_GluTTC_MT_-_14674_14742@27.58.32,trnalookalike8_GluTTC_5_-_93905172_93905240@27.58.32\|GTTTTTCATATCATTGGTCGTGGTTGTAGTCC | >trnaMT_GluTTC_MT_-_14674_14742 | 27 | 58 | 32 | GTTTTTCATATCATTGGTCGTGGTTGTAGTCC | 198.2 | 4.7 | 1.4 | 3.2 | 1.3E-03 | 3.0E-03 | BT20/MB468 | MP |
| >am_trnaMT_SerGCT_MT_+_12207_12265@12.38.27\|CAAGAACTGCTAACTCATGCCCCCATG | >am_trnaMT_SerGCT_MT_+_12207_12265 | 12 | 38 | 27 | CAAGAACTGCTAACTCATGCCCCCATG | 205.4 | -3.7 | 1.2 | -3.1 | 2.2E-03 | 3.0E-03 | MB231/MB468 | MP |
| >MI0000086\|hsa-mir-28&WithFlank&3\|+\|188688775\|188688872@20.41.22(+1A)&[MIMAT0000085&hsa-miR-28-5p&offsets\|0\|0(+1A);m-133&3\|+\|188688794\|188688815&offsets\|0\|0(+1A)]\|AAGGAGCTCACAGTCTATTGAGA | >MI0000086\|hsa-mir-28&WithFlank&3\|+\|188688775\|188688872 | 20 | 41 | 22(+1A) | AAGGAGCTCACAGTCTATTGAGA | 381.0 | 2.2 | 0.6 | 3.5 | 4.8E-04 | 3.0E-03 | BT20/MB231 | NUC |
| >hg38_21_+_8433172_8446622_RNA45SN1_45S_50ntFlanks@6722.6770.49\|ATTGCAGGACACATTGATCATCGACACTTCGAACGCACTTGCGGCCCCG | >hg38_21_+_8433172_8446622_RNA45SN1_45S_50ntFlanks | 6722 | 6770 | 49 | ATTGCAGGACACATTGATCATCGACACTTCGAACGCACTTGCGGCCCCG | 191.8 | -4.9 | 1.6 | -3.1 | 2.0E-03 | 3.0E-03 | BT20/MB468 | MITO |
| >MI0000090\|hsa-mir-32&WithFlank&9\|-\|109046223\|109046304@12.33.22&[MIMAT0000090&hsa-miR-32-5p&offsets\|0\|0;m-208&9\|-\|109046273\|109046293&offsets\|0\|+1]\|TATTGCACATTACTAAGTTGCA | >MI0000090\|hsa-mir-32&WithFlank&9\|-\|109046223\|109046304 | 12 | 33 | 22 | TATTGCACATTACTAAGTTGCA | 646.5 | 2.5 | 0.8 | 3.1 | 2.2E-03 | 3.0E-03 | MB231/MB468 | MP |
| >MI0000298\|hsa-mir-221&WithFlank&X\|-\|45746151\|45746272@71.90.20&[MIMAT0000278&hsa-miR-221-3p&offsets\|0\|-3;m-91&X\|-\|45746181\|45746202&offsets\|0\|-2]\|AGCTACATTGTCTGCTGGGT | >MI0000298\|hsa-mir-221&WithFlank&X\|-\|45746151\|45746272 | 71 | 90 | 20 | AGCTACATTGTCTGCTGGGT | 659.6 | 3.9 | 1.2 | 3.2 | 1.3E-03 | 3.0E-03 | BT20/MB468 | MP |
| >hg38_21_+_8433172_8446622_RNA45SN1_45S_50ntFlanks@6669.6697.29\|CACTCGGCTCGTGCGTCGATGAAGAACGC | >hg38_21_+_8433172_8446622_RNA45SN1_45S_50ntFlanks | 6669 | 6697 | 29 | CACTCGGCTCGTGCGTCGATGAAGAACGC | 81.6 | -4.8 | 1.6 | -3.1 | 2.2E-03 | 3.0E-03 | MB231/MB468 | MP |
| >MI0003820\|hsa-mir-454&WithFlank&17\|-\|59137752\|59137878@70.92.23(+1C)&[MIMAT0003885&hsa-miR-454-3p&offsets\|0\|0(+1C);m-272&17\|-\|59137787\|59137809&offsets\|0\|0(+1C)]\|TAGTGCAATATTGCTTATAGGGTC | >MI0003820\|hsa-mir-454&WithFlank&17\|-\|59137752\|59137878 | 70 | 92 | 23(+1C) | TAGTGCAATATTGCTTATAGGGTC | 122.5 | 3.9 | 1.3 | 3.1 | 2.2E-03 | 3.0E-03 | MB231/MB468 | MP |
| >hg38_21_+_8433172_8446622_RNA45SN1_45S_50ntFlanks@6648.6688.41\|GTACGACTCTTAGCGGTGGATCACTCGGCTCGTGCGTCGAT | >hg38_21_+_8433172_8446622_RNA45SN1_45S_50ntFlanks | 6648 | 6688 | 41 | GTACGACTCTTAGCGGTGGATCACTCGGCTCGTGCGTCGAT | 789.8 | 2.2 | 0.7 | 3.1 | 2.0E-03 | 3.0E-03 | BT20/MB468 | MITO |
| >trnaMT_ValTAC_MT_+_1602_1670@45.72.28\|TTTCAACTTAACTTGACCGCTCTGACCA | >trnaMT_ValTAC_MT_+_1602_1670 | 45 | 72 | 28 | TTTCAACTTAACTTGACCGCTCTGACCA | 743.4 | 2.9 | 1.0 | 3.0 | 2.6E-03 | 3.0E-03 | MB231/MB468 | MITO |
| >hg38_21_+_8433172_8446622_RNA45SN1_45S_50ntFlanks@10343.10362.20\|TATTCAAACGAGAACTTTGA | >hg38_21_+_8433172_8446622_RNA45SN1_45S_50ntFlanks | 10343 | 10362 | 20 | TATTCAAACGAGAACTTTGA | 506.9 | 2.9 | 0.8 | 3.4 | 5.8E-04 | 3.0E-03 | BT20/MB231 | MITO |
| >hg38_21_+_8433172_8446622_RNA45SN1_45S_50ntFlanks@10319.10340.22\|GCAGATCTTGGTGGTAGTAGCA | >hg38_21_+_8433172_8446622_RNA45SN1_45S_50ntFlanks | 10319 | 10340 | 22 | GCAGATCTTGGTGGTAGTAGCA | 664.9 | 4.7 | 1.3 | 3.5 | 4.4E-04 | 3.0E-03 | BT20/MB231 | MP |
| >hg38_21_+_8433172_8446622_RNA45SN1_45S_50ntFlanks@4006.4023.18\|ACCTCGGGCCGATCGCAC | >hg38_21_+_8433172_8446622_RNA45SN1_45S_50ntFlanks | 4006 | 4023 | 18 | ACCTCGGGCCGATCGCAC | 1218.4 | -2.3 | 0.8 | -3.0 | 2.6E-03 | 3.0E-03 | MB231/MB468 | NUC |
| >MI0000465\|hsa-mir-191&WithFlank&3\|-\|49020612\|49020715@22.43.22(+1U)&[MIMAT0000440&hsa-miR-191-5p&offsets\|0\|-1(+1U);m-48&3\|-\|49020672\|49020694&offsets\|0\|-1(+1U)]\|CAACGGAATCCCAAAAGCAGCTT | >MI0000465\|hsa-mir-191&WithFlank&3\|-\|49020612\|49020715 | 22 | 43 | 22(+1U) | CAACGGAATCCCAAAAGCAGCTT | 380.3 | 3.8 | 1.2 | 3.2 | 1.3E-03 | 3.0E-03 | BT20/MB468 | MP |
| >hg38_21_+_8433172_8446622_RNA45SN1_45S_50ntFlanks@4732.4767.36\|AGAACGAAAGTCGGAGGTTCGAAGACGATCAGATAC | >hg38_21_+_8433172_8446622_RNA45SN1_45S_50ntFlanks | 4732 | 4767 | 36 | AGAACGAAAGTCGGAGGTTCGAAGACGATCAGATAC | 110.2 | 4.7 | 1.5 | 3.1 | 2.2E-03 | 3.0E-03 | MB231/MB468 | MP |
| >hg38_21_+_8433172_8446622_RNA45SN1_45S_50ntFlanks@10389.10406.18\|TGAACAGCAGTTGAACAT | >hg38_21_+_8433172_8446622_RNA45SN1_45S_50ntFlanks | 10389 | 10406 | 18 | TGAACAGCAGTTGAACAT | 72.2 | -5.0 | 1.6 | -3.2 | 1.3E-03 | 3.0E-03 | BT20/MB468 | MP |
| >hg38_21_+_8433172_8446622_RNA45SN1_45S_50ntFlanks@6661.6708.48\|CGGTGGATCACTCGGCTCGTGCGTCGATGAAGAACGCAGCTAGCTGCG | >hg38_21_+_8433172_8446622_RNA45SN1_45S_50ntFlanks | 6661 | 6708 | 48 | CGGTGGATCACTCGGCTCGTGCGTCGATGAAGAACGCAGCTAGCTGCG | 295.4 | 2.6 | 0.8 | 3.0 | 2.4E-03 | 3.0E-03 | MB231/MB468 | CYTO |
| >hg38_21_+_8433172_8446622_RNA45SN1_45S_50ntFlanks@11452.11472.21\|CTCCTCGGCGCGCGGCGGCGG | >hg38_21_+_8433172_8446622_RNA45SN1_45S_50ntFlanks | 11452 | 11472 | 21 | CTCCTCGGCGCGCGGCGGCGG | 145.3 | -5.1 | 1.6 | -3.2 | 1.3E-03 | 3.0E-03 | BT20/MB468 | MP |
| >hg38_MT_+_598_1651_RNR1_12S_50ntFlanks@52.93.42\|ATAGGTTTGGTCCTAGCCTTTCTATTAGCTCTTAGTAAGATT | >hg38_MT_+_598_1651_RNR1_12S_50ntFlanks | 52 | 93 | 42 | ATAGGTTTGGTCCTAGCCTTTCTATTAGCTCTTAGTAAGATT | 1135.6 | 3.5 | 1.2 | 3.0 | 2.6E-03 | 3.0E-03 | MB231/MB468 | MITO |
| >hg38_21_+_8433172_8446622_RNA45SN1_45S_50ntFlanks@6698.6719.22\|AGCTAGCTGCGAGAATTAATGT | >hg38_21_+_8433172_8446622_RNA45SN1_45S_50ntFlanks | 6698 | 6719 | 22 | AGCTAGCTGCGAGAATTAATGT | 319.9 | -3.5 | 1.1 | -3.1 | 2.0E-03 | 3.0E-03 | BT20/MB468 | MITO |
| >hg38_MT_+_1621_3279_RNR2_16S_50ntFlanks@1338.1373.36\|AGTCCATATCAACAATAGGGTTTACGACCTCGATGT | >hg38_MT_+_1621_3279_RNR2_16S_50ntFlanks | 1338 | 1373 | 36 | AGTCCATATCAACAATAGGGTTTACGACCTCGATGT | 72.3 | 5.5 | 1.8 | 3.1 | 2.2E-03 | 3.0E-03 | MB231/MB468 | MP |
| >MI0000088\|hsa-mir-30a&WithFlank&6\|-\|71403545\|71403627@54.74.21&[MIMAT0000088&hsa-miR-30a-3p&offsets\|+1\|0;m-38&6\|-\|71403554\|71403575&offsets\|+1\|0]\|TTTCAGTCGGATGTTTGCAGC | >MI0000088\|hsa-mir-30a&WithFlank&6\|-\|71403545\|71403627 | 54 | 74 | 21 | TTTCAGTCGGATGTTTGCAGC | 535.6 | 1.8 | 0.6 | 3.0 | 2.6E-03 | 3.0E-03 | BT20/MB468 | NUC |
| >hg38_21_+_8433172_8446622_RNA45SN1_45S_50ntFlanks@4068.4089.22\|ACTTTCGATGGTAGTCGCCGTG | >hg38_21_+_8433172_8446622_RNA45SN1_45S_50ntFlanks | 4068 | 4089 | 22 | ACTTTCGATGGTAGTCGCCGTG | 193.7 | -5.1 | 1.6 | -3.2 | 1.3E-03 | 3.0E-03 | BT20/MB468 | MP |

| | | | | | | | | | | | | | |
|---|---|---|---|---|---|---|---|---|---|---|---|---|---|
| >MI0000746\|hsa-mir-99b&WithFlank&19\|+\|51692606\|51692687@13.32.20(+1G)&[MIMAT0000689&hsa-miR-99b-5p&offsets\|0\|-2(+1G);m-10&19\|+\|51692618\|51692639&offsets\|0\|-2(+1G)]\|CACCCGTAGAACCGACCTTGG | >MI0000746\|hsa-mir-99b&WithFlank&19\|+\|51692606\|51692687 | 13 | 32 | 20(+1G) | CACCCGTAGAACCGACCTTGG | 218.2 | 4.3 | 1.2 | 3.5 | 4.8E-04 | 3.0E-03 | BT20/MB231 | NUC |
| >MI0000750\|hsa-mir-26a-2&WithFlank&12\|-\|57824603\|57824698@20.41.22(+3U);MI0000083\|hsa-mir-26a-1&WithFlank&3\|+\|37969398\|37969486@16.37.22(+3U)&[MIMAT0000082&hsa-miR-26a-5p&offsets\|0\|0(+3U);m-33&12\|-\|57824658\|57824679&offsets\|0\|0(+3U)];[MIMAT0000082_1&hsa-miR-26a-5p&offsets\|0\|0(+3U);m-34&3\|+\|37969413\|37969434&offsets\|0\|0(+3U)]\|TTCAAGTAATCCAGGATAGGCTTTT | >MI0000750\|hsa-mir-26a-2&WithFlank&12\|-\|57824603\|57824698 | 20 | 41 | 22(+3U) | TTCAAGTAATCCAGGATAGGCTTTT | 366.6 | -4.9 | 1.4 | -3.5 | 4.4E-04 | 3.0E-03 | BT20/MB231 | MP |
| >hg38_21_+_8433172_8446622_RNA45SN1_45S_50ntFlanks@9562.9584.23\|TCCTGACGTGCAAATCGGTCGTC | >hg38_21_+_8433172_8446622_RNA45SN1_45S_50ntFlanks | 9562 | 9584 | 23 | TCCTGACGTGCAAATCGGTCGTC | 76.8 | -5.0 | 1.6 | -3.2 | 1.3E-03 | 3.0E-03 | BT20/MB468 | MP |
| >MI0000542\|hsa-mir-320a&WithFlank&8\|-\|22244960\|22245043@48.69.22(+1G)&[MIMAT0000510&hsa-miR-320a-3p&offsets\|0\|0(+1G);m-60&8\|-\|22244975\|22244996&offsets\|0\|0(+1G)]\|AAAAGCTGGGTTGAGAGGGCGAG | >MI0000542\|hsa-mir-320a&WithFlank&8\|-\|22244960\|22245043 | 48 | 69 | 22(+1G) | AAAAGCTGGGTTGAGAGGGCGAG | 123.0 | 4.1 | 1.4 | 3.0 | 2.6E-03 | 3.0E-03 | BT20/MB468 | NUC |
| >MI0000095\|hsa-mir-93&WithFlank&7\|-\|100093762\|100093853@17.38.22&[MIMAT0000093&hsa-miR-93-5p&offsets\|0\|-1;m-29&7\|-\|100093815\|100093837&offsets\|0\|-1]\|CAAAGTGCTGTTCGTGCAGGTA | >MI0000095\|hsa-mir-93&WithFlank&7\|-\|100093762\|100093853 | 17 | 38 | 22 | CAAAGTGCTGTTCGTGCAGGTA | 1440.3 | 2.8 | 0.9 | 3.1 | 2.2E-03 | 3.0E-03 | MB231/MB468 | MP |
| >hg38_21_+_8433172_8446622_RNA45SN1_45S_50ntFlanks@9454.9498.45\|CCGCGCCGGGGAGGTGGAGCACGAGCGCACGTGTTAGGACCCGAA | >hg38_21_+_8433172_8446622_RNA45SN1_45S_50ntFlanks | 9454 | 9498 | 45 | CCGCGCCGGGGAGGTGGAGCACGAGCGCACGTGTTAGGACCCGAA | 44.9 | -9.4 | 3.1 | -3.1 | 2.2E-03 | 3.0E-03 | MB231/MB468 | MP |
| >MI0000085\|hsa-mir-27a&WithFlank&19\|-\|13836434\|13836523@57.74.18;MI0000440\|hsa-mir-27b&WithFlank&9\|+\|95085439\|95085547@67.84.18&[MIMAT0000084&hsa-miR-27a-3p&offsets\|0\|-3;m-43&19\|-\|13836447\|13836467&offsets\|0\|-3];[MIMAT0000419&hsa-miR-27b-3p&offsets\|0\|-3;m-39&9\|+\|95085505\|95085525&offsets\|0\|-3]\|TTCACAGTGGCTAAGTTC | >MI0000085\|hsa-mir-27a&WithFlank&19\|-\|13836434\|13836523 | 57 | 74 | 18 | TTCACAGTGGCTAAGTTC | 900.1 | -2.7 | 0.8 | -3.4 | 5.8E-04 | 3.0E-03 | BT20/MB231 | MITO |
| >trna11_SerAGA_8_-_96281885_96281966@1.24.24,trna145_SerAGA_6_-_27521192_27521273@1.24.24,trna147_SerAGA_6_-_27509554_27509635@1.24.24,trna148_SerTGA_6_-_27473607_27473688@1.24.24,trna172_SerTGA_6_-_26312824_26312905@1.24.24,trna35_SerAGA_17_-_8129928_8130009@1.24.24,trna44_SerAGA_6_+_27446591_27446672@1.24.24,trna46_SerAGA_6_+_27463593_27463674@1.24.24,trna47_SerAGA_6_+_27470818_27470899@1.24.24,trna50_SerAGA_6_+_27499987_27500068@1.24.24,trna51_SerTGA_6_+_27513468_27513549@1.24.24,trna5_SerAGA_6_+_26327817_26327898@1.24.24\|GTAGTCGTGGCCGAGTGGTTAAGG | >trna11_SerAGA_8_-_96281885_96281966 | 1 | 24 | 24 | GTAGTCGTGGCCGAGTGGTTAAGG | 694.1 | 5.0 | 1.4 | 3.5 | 4.4E-04 | 3.0E-03 | BT20/MB231 | MP |
| >hg38_21_+_8433172_8446622_RNA45SN1_45S_50ntFlanks@11875.11893.19\|GAAGACCCTGTTGAGCTTG | >hg38_21_+_8433172_8446622_RNA45SN1_45S_50ntFlanks | 11875 | 11893 | 19 | GAAGACCCTGTTGAGCTTG | 210.9 | 6.9 | 2.0 | 3.5 | 4.4E-04 | 3.0E-03 | BT20/MB231 | MP |
| >am_trna117_GlyTCC_1_-_145397864_145397935@1.32.32,trna45_GlyTCC_1_+_161500903_161500974@1.32.32,trna70_GlyTCC_1_-_161439547_161439618@1.32.32,trna73_GlyTCC_1_-_161432166_161432237@1.32.32,trna76_GlyTCC_1_-_161424756_161424827@1.32.32,trna79_GlyTCC_1_-_161417375_161417446@1.32.32\|GCGTTGGTGGTATAGTGGTGAGCATAGCTGCC | >am_trna117_GlyTCC_1_-_145397864_145397935 | 1 | 32 | 32 | GCGTTGGTGGTATAGTGGTGAGCATAGCTGCC | 151.3 | 2.6 | 0.8 | 3.1 | 2.2E-03 | 3.0E-03 | MB231/MB468 | MP |
| >hg38_21_+_8433172_8446622_RNA45SN1_45S_50ntFlanks@11583.11602.20\|CTGGTGCGGACCAGGGGAAT | >hg38_21_+_8433172_8446622_RNA45SN1_45S_50ntFlanks | 11583 | 11602 | 20 | CTGGTGCGGACCAGGGGAAT | 464.8 | -2.2 | 0.7 | -3.1 | 2.0E-03 | 3.0E-03 | BT20/MB468 | MITO |
| >MI0001445\|hsa-mir-423&WithFlank&17\|+\|30117073\|30117178@23.45.23(+1A)&[MIMAT0004748&hsa-miR-423-5p&offsets\|0\|0(+1A);m-141&17\|+\|30117095\|30117117&offsets\|0\|0(+1A)]\|TGAGGGGCAGAGAGCGAGACTTTA | >MI0001445\|hsa-mir-423&WithFlank&17\|+\|30117073\|30117178 | 23 | 45 | 23(+1A) | TGAGGGGCAGAGAGCGAGACTTTA | 130.2 | 3.9 | 1.3 | 3.1 | 2.2E-03 | 3.0E-03 | MB231/MB468 | MP |
| >MI0000266\|hsa-mir-10a&WithFlank&17\|-\|48579832\|48579953@29.49.21(+1A)&[MIMAT0000253&hsa-miR-10a-5p&offsets\|+1\|-1(+1A);m-18&17\|-\|48579905\|48579926&offsets\|+1\|0(+1A)]\|ACCCTGTAGATCCGAATTTGTA | >MI0000266\|hsa-mir-10a&WithFlank&17\|-\|48579832\|48579953 | 29 | 49 | 21(+1A) | ACCCTGTAGATCCGAATTTGTA | 2235.5 | 2.7 | 0.8 | 3.5 | 5.2E-04 | 3.0E-03 | BT20/MB231 | CYTO |
| >hg38_21_+_8433172_8446622_RNA45SN1_45S_50ntFlanks@10678.10707.30\|CGGTTCCGGCGGCGTCCGGTGAGCTCTCGC | >hg38_21_+_8433172_8446622_RNA45SN1_45S_50ntFlanks | 10678 | 10707 | 30 | CGGTTCCGGCGGCGTCCGGTGAGCTCTCGC | 409.0 | 7.1 | 2.0 | 3.5 | 4.4E-04 | 3.0E-03 | BT20/MB231 | MP |
| >trna119_LysCTT_1_-_145395522_145395594@1.31.31,trna11_LysCTT_5_-_180648979_180649051@1.31.31,trna13_LysCTT_6_+_26556774_26556846@1.31.31,trna32_LysCTT_16_-_3207406_3207478@1.31.31,trna7_LysCTT_16_+_3225692_3225764@1.31.31,trna9_LysCTT_5_+_180634755_180634827@1.31.31\|GCCCGGCTAGCTCAGTCGGTAGAGCATGAGA | >trna119_LysCTT_1_-_145395522_145395594 | 1 | 31 | 31 | GCCCGGCTAGCTCAGTCGGTAGAGCATGAGA | 2721.6 | -2.9 | 1.0 | -3.0 | 2.6E-03 | 3.0E-03 | MB231/MB468 | NUC |
| >hg38_21_+_8433172_8446622_RNA45SN1_45S_50ntFlanks@12988.13013.26\|CAGCTCCCTCGCTGCGATCTATTGAA | >hg38_21_+_8433172_8446622_RNA45SN1_45S_50ntFlanks | 12988 | 13013 | 26 | CAGCTCCCTCGCTGCGATCTATTGAA | 149.6 | -3.8 | 1.2 | -3.1 | 2.2E-03 | 3.0E-03 | MB231/MB468 | MP |
| >hg38_21_+_8433172_8446622_RNA45SN1_45S_50ntFlanks@7974.7997.24\|ACGCGACCTCAGATCAGACGTGGC | >hg38_21_+_8433172_8446622_RNA45SN1_45S_50ntFlanks | 7974 | 7997 | 24 | ACGCGACCTCAGATCAGACGTGGC | 2657.1 | 2.4 | 0.8 | 3.0 | 2.6E-03 | 3.0E-03 | MB231/MB468 | MITO |
| >hg38_21_+_8433172_8446622_RNA45SN1_45S_50ntFlanks@12478.12500.23\|ACTAATAGGGAACGTGAGCTGGG | >hg38_21_+_8433172_8446622_RNA45SN1_45S_50ntFlanks | 12478 | 12500 | 23 | ACTAATAGGGAACGTGAGCTGGG | 135.6 | -3.8 | 1.2 | -3.1 | 2.2E-03 | 3.0E-03 | MB231/MB468 | MP |
| >hg38_21_+_8433172_8446622_RNA45SN1_45S_50ntFlanks@4759.4783.25\|ATCAGATACCGTCGTAGTTCCGACC | >hg38_21_+_8433172_8446622_RNA45SN1_45S_50ntFlanks | 4759 | 4783 | 25 | ATCAGATACCGTCGTAGTTCCGACC | 745.9 | -5.0 | 1.6 | -3.0 | 2.6E-03 | 3.0E-03 | MB231/MB468 | NUC |
| >hg38_21_+_8433172_8446622_RNA45SN1_45S_50ntFlanks@11561.11600.40\|GCGCCTAGCAGCCGACTTAGAACTGGTGCGGACCAGGGGA | >hg38_21_+_8433172_8446622_RNA45SN1_45S_50ntFlanks | 11561 | 11600 | 40 | GCGCCTAGCAGCCGACTTAGAACTGGTGCGGACCAGGGGA | 325.3 | -4.7 | 1.5 | -3.0 | 2.4E-03 | 3.0E-03 | BT20/MB468 | CYTO |
| >hg38_21_+_8433172_8446622_RNA45SN1_45S_50ntFlanks@6655.6695.41\|TCTTAGCGGTGGATCACTCGGCTCGTGCGTCGATGAAGAAC | >hg38_21_+_8433172_8446622_RNA45SN1_45S_50ntFlanks | 6655 | 6695 | 41 | TCTTAGCGGTGGATCACTCGGCTCGTGCGTCGATGAAGAAC | 111.8 | 3.7 | 1.2 | 3.0 | 2.6E-03 | 3.0E-03 | MB231/MB468 | MITO |
| >hg38_21_+_8433172_8446622_RNA45SN1_45S_50ntFlanks@7973.8009.37\|GACGCGACCTCAGATCAGACGTGGCGACCCGCTGAAT | >hg38_21_+_8433172_8446622_RNA45SN1_45S_50ntFlanks | 7973 | 8009 | 37 | GACGCGACCTCAGATCAGACGTGGCGACCCGCTGAAT | 1247.8 | 3.0 | 1.0 | 3.1 | 2.2E-03 | 3.0E-03 | MB231/MB468 | MP |
| >hg38_21_+_8433172_8446622_RNA45SN1_45S_50ntFlanks@9533.9552.20\|GAGGAAACTCTGGTGGAGGT | >hg38_21_+_8433172_8446622_RNA45SN1_45S_50ntFlanks | 9533 | 9552 | 20 | GAGGAAACTCTGGTGGAGGT | 279.0 | -4.2 | 1.4 | -3.1 | 2.2E-03 | 3.0E-03 | MB231/MB468 | MP |
| >trnaMT_GluTTC_MT_-_14674_14742@44.72.29,trnalookalike8_GluTTC_5_-_93905172_93905240@44.72.29\|TCGTGGTTGTAGTCCGTGCGAGAATACCA | >trnaMT_GluTTC_MT_-_14674_14742 | 44 | 72 | 29 | TCGTGGTTGTAGTCCGTGCGAGAATACCA | 1229.7 | -3.5 | 1.0 | -3.5 | 5.2E-04 | 3.0E-03 | BT20/MB231 | CYTO |
| >MI0000063\|hsa-let-7b&WithFlank&22\|+\|46113680\|46113774@12.35.24&[MIMAT0000063&hsa-let-7b-5p&offsets\|0\|+2;m-14&22\|+\|46113691\|46113712&offsets\|0\|+2]\|TGAGGTAGTAGGTTGTGTGGTTTC | >MI0000063\|hsa-let-7b&WithFlank&22\|+\|46113680\|46113774 | 12 | 35 | 24 | TGAGGTAGTAGGTTGTGTGGTTTC | 77.3 | 5.7 | 1.9 | 3.1 | 2.2E-03 | 3.0E-03 | MB231/MB468 | MP |
| >MI0000446\|hsa-mir-125b-1&WithFlank&11\|-\|122099751\|122099850@21.41.21;MI0000470\|hsa-mir-125b-2&WithFlank&21\|+\|16590231\|16590331@23.43.21&[MIMAT0000423&hsa-miR-125b-5p&offsets\|0\|-1;m-67&11\|-\|122099809\|122099830&offsets\|0\|-1];[MIMAT0000423_1&hsa-miR-125b-5p&offsets\|0\|-1;m-68&21\|+\|16590253\|16590274&offsets\|0\|-1]\|TCCCTGAGACCCTAACTTGTG | >MI0000446\|hsa-mir-125b-1&WithFlank&11\|-\|122099751\|122099850 | 21 | 41 | 21 | TCCCTGAGACCCTAACTTGTG | 10197.7 | -3.7 | 1.1 | -3.5 | 5.2E-04 | 3.0E-03 | BT20/MB231 | CYTO |
| >hg38_21_+_8433172_8446622_RNA45SN1_45S_50ntFlanks@5478.5512.35\|CTGGCGGAGCGCTGAGAAGACGGTCGAACTTGACT | >hg38_21_+_8433172_8446622_RNA45SN1_45S_50ntFlanks | 5478 | 5512 | 35 | CTGGCGGAGCGCTGAGAAGACGGTCGAACTTGACT | 77.0 | 5.3 | 1.6 | 3.2 | 1.3E-03 | 3.1E-03 | BT20/MB468 | MP |
| >am_trnaMT_ProTGG_MT_-_15956_16023@53.70.18\|AAGACTTTTTCTCTGACC | >am_trnaMT_ProTGG_MT_-_15956_16023 | 53 | 70 | 18 | AAGACTTTTTCTCTGACC | 1808.3 | 3.4 | 1.1 | 3.1 | 2.0E-03 | 3.1E-03 | BT20/MB468 | MITO |
| >trnaMT_HisGTG_MT_+_12138_12206@24.55.32\|TCAGATTGTGAATCTGACAACAGAGGCTTACG | >trnaMT_HisGTG_MT_+_12138_12206 | 24 | 55 | 32 | TCAGATTGTGAATCTGACAACAGAGGCTTACG | 41.2 | 9.1 | 2.8 | 3.2 | 1.3E-03 | 3.1E-03 | BT20/MB468 | MP |
| >trnaMT_HisGTG_MT_+_12138_12206@26.61.36\|AGATTGTGAATCTGACAACAGAGGCTTACGACCCCT | >trnaMT_HisGTG_MT_+_12138_12206 | 26 | 61 | 36 | AGATTGTGAATCTGACAACAGAGGCTTACGACCCCT | 264.6 | 6.5 | 2.0 | 3.2 | 1.3E-03 | 3.1E-03 | BT20/MB468 | MP |
| >trnaMT_ProTGG_MT_-_15956_16023@40.58.19\|AATGGTGGAGTTAAAGACT | >trnaMT_ProTGG_MT_-_15956_16023 | 40 | 58 | 19 | AATGGTGGAGTTAAAGACT | 769.3 | 3.4 | 1.0 | 3.2 | 1.3E-03 | 3.1E-03 | BT20/MB468 | MP |
| >hg38_21_+_8433172_8446622_RNA45SN1_45S_50ntFlanks@11643.11665.23\|GCGGGTGTTGACGCGATGTGATT | >hg38_21_+_8433172_8446622_RNA45SN1_45S_50ntFlanks | 11643 | 11665 | 23 | GCGGGTGTTGACGCGATGTGATT | 73.9 | -5.1 | 1.6 | -3.2 | 1.3E-03 | 3.1E-03 | BT20/MB468 | MP |
| >hg38_21_+_8433172_8446622_RNA45SN1_45S_50ntFlanks@8345.8375.31\|AAGGGAAAGTTGAAAAGAACTTTGAAGAGAG | >hg38_21_+_8433172_8446622_RNA45SN1_45S_50ntFlanks | 8345 | 8375 | 31 | AAGGGAAAGTTGAAAAGAACTTTGAAGAGAG | 178.8 | 6.2 | 2.0 | 3.1 | 2.2E-03 | 3.1E-03 | MB231/MB468 | MP |
| >hg38_21_+_8433172_8446622_RNA45SN1_45S_50ntFlanks@10783.10825.43\|GTGAACAGCCTCTGGCATGTTGGAACAATGTAGGTAAGGGAAG | >hg38_21_+_8433172_8446622_RNA45SN1_45S_50ntFlanks | 10783 | 10825 | 43 | GTGAACAGCCTCTGGCATGTTGGAACAATGTAGGTAAGGGAAG | 35.6 | -8.2 | 2.6 | -3.2 | 1.3E-03 | 3.1E-03 | BT20/MB468 | MP |
| >trna111_HisGTG_1_-_147774845_147774916@1.31.31,trna118_HisGTG_1_-_145396881_145396952@1.31.31,trna16_HisGTG_1_+_146544773_146544844@1.31.31,trna1_HisGTG_15_+_45493349_45493420@1.31.31,trna21_HisGTG_1_+_147753471_147753542@1.31.31,trna33_HisGTG_6_+_27125906_27125977@1.31.31,trna7_HisGTG_9_-_14433938_14434009@1.31.31,trna8_HisGTG_15_-_45492611_45492682@1.31.31,trna9_HisGTG_15_-_45490804_45490875@1.31.31\|GCCGTGATCGTATAGTGGTTAGTACTCTGCG | >trna111_HisGTG_1_-_147774845_147774916 | 1 | 31 | 31 | GCCGTGATCGTATAGTGGTTAGTACTCTGCG | 3320.7 | -3.6 | 1.0 | -3.5 | 4.4E-04 | 3.1E-03 | BT20/MB231 | MP |
| >MI0003780\|hsa-mir-1296&WithFlank&10\|-\|63372951\|63373054@22.42.21(+1A)&[MIMAT0005794&hsa-miR-1296-5p&offsets\|0\|-1(+1A);m-304&10\|-\|63373012\|63373033&offsets\|0\|-1(+1A)]\|TTAGGGCCCTGGCTCCATCTCA | >MI0003780\|hsa-mir-1296&WithFlank&10\|-\|63372951\|63373054 | 22 | 42 | 21(+1A) | TTAGGGCCCTGGCTCCATCTCA | 48.9 | 5.7 | 1.9 | 3.1 | 2.2E-03 | 3.1E-03 | MB231/MB468 | MP |
| >MI0015866\|hsa-mir-4326&WithFlank&20\|+\|63286802\|63286872@17.40.24&[MIMAT0016888&hsa-miR-4326&offsets\|0\|+4;m-339&20\|+\|63286818\|63286841&offsets\|0\|0]\|TGTTCCTCTGTCTCCCAGACTCTG | >MI0015866\|hsa-mir-4326&WithFlank&20\|+\|63286802\|63286872 | 17 | 40 | 24 | TGTTCCTCTGTCTCCCAGACTCTG | 335.3 | 3.9 | 1.3 | 3.1 | 2.2E-03 | 3.1E-03 | MB231/MB468 | MP |
| >hg38_21_+_8433172_8446622_RNA45SN1_45S_50ntFlanks@9861.9885.25\|GATGAACCGAACGCCGGGTTAAGGC | >hg38_21_+_8433172_8446622_RNA45SN1_45S_50ntFlanks | 9861 | 9885 | 25 | GATGAACCGAACGCCGGGTTAAGGC | 308.7 | 4.3 | 1.2 | 3.5 | 4.4E-04 | 3.1E-03 | BT20/MB231 | MP |
| >MI0000456\|hsa-mir-140&WithFlank&16\|+\|69933075\|69933186@68.89.22(+1A)&[MIMAT0004597&hsa-miR-140-3p&offsets\|0\|+1(+1A);m-45&16\|+\|69933143\|69933164&offsets\|-1\|-1(+1A)]\|TACCACAGGGTAGAACCACGGAA | >MI0000456\|hsa-mir-140&WithFlank&16\|+\|69933075\|69933186 | 68 | 89 | 22(+1A) | TACCACAGGGTAGAACCACGGAA | 43.3 | 5.5 | 1.8 | 3.1 | 2.2E-03 | 3.1E-03 | MB231/MB468 | MP |

| | | | | | | | | | | | | | |
|---|---|---|---|---|---|---|---|---|---|---|---|---|---|
| >hg38_21_+_8433172_8446622_RNA45SN1_45S_50ntFlanks@6648.6669.22\|GTACGACTCTTAGCGGTGGATC | >hg38_21_+_8433172_8446622_RNA45SN1_45S_50ntFlanks | 6648 | 6669 | 22 | GTACGACTCTTAGCGGTGGATC | 353.7 | 1.6 | 0.5 | 3.0 | 2.6E-03 | 3.1E-03 | MB231/MB468 | NUC |
| >hg38_21_+_8433172_8446622_RNA45SN1_45S_50ntFlanks@9956.9983.28\|GAAGTCGGAATCCGCTAAGGAGTGTGTA | >hg38_21_+_8433172_8446622_RNA45SN1_45S_50ntFlanks | 9956 | 9983 | 28 | GAAGTCGGAATCCGCTAAGGAGTGTGTA | 79.8 | 5.3 | 1.6 | 3.2 | 1.3E-03 | 3.1E-03 | BT20/MB468 | MP |
| >hg38_MT_+_1621_3279_RNR2_16S_50ntFlanks@1090.1120.31\|CAGCAAGACGAGAAGACCCTATGGAGCTTTA | >hg38_MT_+_1621_3279_RNR2_16S_50ntFlanks | 1090 | 1120 | 31 | CAGCAAGACGAGAAGACCCTATGGAGCTTTA | 176.9 | 3.6 | 1.1 | 3.2 | 1.3E-03 | 3.1E-03 | BT20/MB468 | MP |
| >hg38_21_+_8433172_8446622_RNA45SN1_45S_50ntFlanks@6648.6667.20\|GTACGACTCTTAGCGGTGGA | >hg38_21_+_8433172_8446622_RNA45SN1_45S_50ntFlanks | 6648 | 6667 | 20 | GTACGACTCTTAGCGGTGGA | 481.6 | 2.5 | 0.8 | 3.0 | 2.6E-03 | 3.1E-03 | MB231/MB468 | MITO |
| >hg38_21_+_8433172_8446622_RNA45SN1_45S_50ntFlanks@8019.8046.28\|TTAGTCAGCGGAGGAGAAGAAACTAACC | >hg38_21_+_8433172_8446622_RNA45SN1_45S_50ntFlanks | 8019 | 8046 | 28 | TTAGTCAGCGGAGGAGAAGAAACTAACC | 305.9 | 7.3 | 2.4 | 3.0 | 2.5E-03 | 3.1E-03 | MB231/MB468 | CYTO |
| >MI0000542\|hsa-mir-320a&WithFlank&8\|-\|22244960\|22245043@49.70.22(+1U)&[MIMAT0000510&hsa-miR-320a-3p&offsets\|+1\|+1(+1U);m-60&8\|-\|22244975\|22244996&offsets\|+1\|+1(+1U)]\|AAAGCTGGGTTGAGAGGGCGAAT | >MI0000542\|hsa-mir-320a&WithFlank&8\|-\|22244960\|22245043 | 49 | 70 | 22(+1U) | AAAGCTGGGTTGAGAGGGCGAAT | 316.7 | 2.5 | 0.7 | 3.4 | 5.9E-04 | 3.1E-03 | BT20/MB231 | MITO |
| >hg38_21_+_8433172_8446622_RNA45SN1_45S_50ntFlanks@9305.9334.30\|CCAAGGAGTCTAACACGTGCGCGAGTCGGG | >hg38_21_+_8433172_8446622_RNA45SN1_45S_50ntFlanks | 9305 | 9334 | 30 | CCAAGGAGTCTAACACGTGCGCGAGTCGGG | 86.8 | 9.8 | 2.8 | 3.5 | 4.5E-04 | 3.1E-03 | BT20/MB231 | MP |
| >hg38_21_+_8433172_8446622_RNA45SN1_45S_50ntFlanks@5493.5525.33\|GAAGACGGTCGAACTTGACTATCTAGAGGAAGT | >hg38_21_+_8433172_8446622_RNA45SN1_45S_50ntFlanks | 5493 | 5525 | 33 | GAAGACGGTCGAACTTGACTATCTAGAGGAAGT | 229.4 | 3.6 | 1.1 | 3.2 | 1.3E-03 | 3.1E-03 | BT20/MB468 | MP |
| >hg38_21_+_8433172_8446622_RNA45SN1_45S_50ntFlanks@12420.12456.37\|ATGTCGGCTCTTCCTATCATTGTGAAGCAGAATTCAC | >hg38_21_+_8433172_8446622_RNA45SN1_45S_50ntFlanks | 12420 | 12456 | 37 | ATGTCGGCTCTTCCTATCATTGTGAAGCAGAATTCAC | 377.3 | -2.8 | 0.8 | -3.5 | 5.3E-04 | 3.1E-03 | BT20/MB231 | CYTO |
| >hg38_21_+_8433172_8446622_RNA45SN1_45S_50ntFlanks@7991.8014.24\|ACGTGGCGACCCGCTGAATTTAAG | >hg38_21_+_8433172_8446622_RNA45SN1_45S_50ntFlanks | 7991 | 8014 | 24 | ACGTGGCGACCCGCTGAATTTAAG | 439.0 | -2.2 | 0.7 | -3.0 | 2.6E-03 | 3.1E-03 | MB231/MB468 | NUC |
| >hg38_21_+_8433172_8446622_RNA45SN1_45S_50ntFlanks@12907.12933.27\|CACCGCACGTTCGTGGGGAACCTGGCG | >hg38_21_+_8433172_8446622_RNA45SN1_45S_50ntFlanks | 12907 | 12933 | 27 | CACCGCACGTTCGTGGGGAACCTGGCG | 864.5 | -4.9 | 1.6 | -3.0 | 2.4E-03 | 3.1E-03 | BT20/MB468 | CYTO |
| >trnaMT_GluTTC_MT_-_14674_14742@45.71.27,trnalookalike8_GluTTC_5_-_93905172_93905240@45.71.27\|CGTGGTTGTAGTCCGTGCGAGAATACC | >trnaMT_GluTTC_MT_-_14674_14742 | 45 | 71 | 27 | CGTGGTTGTAGTCCGTGCGAGAATACC | 154.6 | 3.4 | 1.0 | 3.2 | 1.3E-03 | 3.1E-03 | BT20/MB468 | MP |
| >hg38_21_+_8433172_8446622_RNA45SN1_45S_50ntFlanks@7980.8002.23\|CCTCAGATCAGACGTGGCGACCC | >hg38_21_+_8433172_8446622_RNA45SN1_45S_50ntFlanks | 7980 | 8002 | 23 | CCTCAGATCAGACGTGGCGACCC | 87.3 | 9.8 | 2.8 | 3.5 | 4.5E-04 | 3.1E-03 | BT20/MB231 | MP |
| >hg38_21_+_8433172_8446622_RNA45SN1_45S_50ntFlanks@6661.6689.29\|CGGTGGATCACTCGGCTCGTGCGTCGATG | >hg38_21_+_8433172_8446622_RNA45SN1_45S_50ntFlanks | 6661 | 6689 | 29 | CGGTGGATCACTCGGCTCGTGCGTCGATG | 10374.4 | 4.5 | 1.3 | 3.5 | 4.5E-04 | 3.1E-03 | BT20/MB231 | MP |
| >MI0003513\|hsa-mir-455&WithFlank&9\|+\|114209428\|114209535@60.81.22&[MIMAT0004784&hsa-miR-455-3p&offsets\|0\|+1;m-135&9\|+\|114209487\|114209508&offsets\|0\|0]\|GCAGTCCATGGGCATATACACT | >MI0003513\|hsa-mir-455&WithFlank&9\|+\|114209428\|114209535 | 60 | 81 | 22 | GCAGTCCATGGGCATATACACT | 205.1 | -6.3 | 1.8 | -3.5 | 4.5E-04 | 3.1E-03 | BT20/MB231 | MP |
| >hg38_21_+_8433172_8446622_RNA45SN1_45S_50ntFlanks@4046.4063.18\|CATTCGAACGTCTGCCCT | >hg38_21_+_8433172_8446622_RNA45SN1_45S_50ntFlanks | 4046 | 4063 | 18 | CATTCGAACGTCTGCCCT | 1170.7 | -5.7 | 1.8 | -3.2 | 1.3E-03 | 3.1E-03 | BT20/MB468 | MP |
| >hg38_21_+_8433172_8446622_RNA45SN1_45S_50ntFlanks@11586.11606.21\|GTGCGGACCAGGGGAATCCGA | >hg38_21_+_8433172_8446622_RNA45SN1_45S_50ntFlanks | 11586 | 11606 | 21 | GTGCGGACCAGGGGAATCCGA | 87.8 | -5.3 | 1.7 | -3.2 | 1.3E-03 | 3.1E-03 | BT20/MB468 | MP |
| >hg38_21_+_8433172_8446622_RNA45SN1_45S_50ntFlanks@12161.12179.19\|TGACTGGGGCGGTACACCT | >hg38_21_+_8433172_8446622_RNA45SN1_45S_50ntFlanks | 12161 | 12179 | 19 | TGACTGGGGCGGTACACCT | 1157.2 | -5.7 | 1.8 | -3.2 | 1.3E-03 | 3.1E-03 | BT20/MB468 | MP |
| >hg38_21_+_8433172_8446622_RNA45SN1_45S_50ntFlanks@10357.10382.26\|CTTTGAAGGCCGAAGTGGAGAAGGGT | >hg38_21_+_8433172_8446622_RNA45SN1_45S_50ntFlanks | 10357 | 10382 | 26 | CTTTGAAGGCCGAAGTGGAGAAGGGT | 136.0 | -3.7 | 1.2 | -3.1 | 2.2E-03 | 3.1E-03 | MB231/MB468 | MP |
| >hg38_MT_+_1621_3279_RNR2_16S_50ntFlanks@929.958.30\|CAGTGACACATGTTTAACGGCCGCGGTACC | >hg38_MT_+_1621_3279_RNR2_16S_50ntFlanks | 929 | 958 | 30 | CAGTGACACATGTTTAACGGCCGCGGTACC | 204.0 | 3.7 | 1.2 | 3.1 | 2.2E-03 | 3.1E-03 | MB231/MB468 | MP |
| >hg38_21_+_8433172_8446622_RNA45SN1_45S_50ntFlanks@11874.11898.25\|AGAAGACCCTGTTGAGCTTGACTCT | >hg38_21_+_8433172_8446622_RNA45SN1_45S_50ntFlanks | 11874 | 11898 | 25 | AGAAGACCCTGTTGAGCTTGACTCT | 1376.9 | -1.9 | 0.6 | -3.1 | 2.0E-03 | 3.1E-03 | BT20/MB468 | MITO |
| >hg38_MT_+_1621_3279_RNR2_16S_50ntFlanks@1170.1215.46\|AAACCTGCATTAAAAATTTCGGTTGGGGCGACCTCGGAGCAGAACC | >hg38_MT_+_1621_3279_RNR2_16S_50ntFlanks | 1170 | 1215 | 46 | AAACCTGCATTAAAAATTTCGGTTGGGGCGACCTCGGAGCAGAACC | 245.5 | 5.7 | 1.9 | 3.0 | 2.6E-03 | 3.1E-03 | MB231/MB468 | MITO |
| >hg38_21_+_8433172_8446622_RNA45SN1_45S_50ntFlanks@4297.4316.20\|CATTGGAGGGCAAGTCTGGT | >hg38_21_+_8433172_8446622_RNA45SN1_45S_50ntFlanks | 4297 | 4316 | 20 | CATTGGAGGGCAAGTCTGGT | 291.3 | -3.6 | 1.2 | -3.0 | 2.6E-03 | 3.1E-03 | MB231/MB468 | NUC |
| >hg38_21_+_8433172_8446622_RNA45SN1_45S_50ntFlanks@11920.11948.29\|ATGAGAGGTGTAGAATAAGTGGGAGGCCC | >hg38_21_+_8433172_8446622_RNA45SN1_45S_50ntFlanks | 11920 | 11948 | 29 | ATGAGAGGTGTAGAATAAGTGGGAGGCCC | 336.1 | -2.6 | 0.9 | -3.0 | 2.4E-03 | 3.1E-03 | BT20/MB468 | CYTO |
| >hg38_21_+_8433172_8446622_RNA45SN1_45S_50ntFlanks@6647.6665.19\|CGTACGACTCTTAGCGGTG | >hg38_21_+_8433172_8446622_RNA45SN1_45S_50ntFlanks | 6647 | 6665 | 19 | CGTACGACTCTTAGCGGTG | 2555.9 | 2.2 | 0.7 | 3.0 | 2.6E-03 | 3.1E-03 | BT20/MB468 | NUC |
| >MI0000750\|hsa-mir-26a-2&WithFlank&12\|-\|57824603\|57824698@23.40.18(+1C);MI0000083\|hsa-mir-26a-1&WithFlank&3\|+\|37969398\|37969486@19.36.18(+1C)&[MIMAT0000082&hsa-miR-26a-5p&offsets\|+3\|-1(+1C);m-33&12\|-\|57824658\|57824679&offsets\|+3\|-1(+1C)];[MIMAT0000082_1&hsa-miR-26a-5p&offsets\|+3\|-1(+1C);m-34&3\|+\|37969413\|37969434&offsets\|+3\|-1(+1C)]\|AAGTAATCCAGGATAGGCC | >MI0000750\|hsa-mir-26a-2&WithFlank&12\|-\|57824603\|57824698 | 23 | 40 | 18(+1C) | AAGTAATCCAGGATAGGCC | 208.2 | 10.9 | 3.6 | 3.0 | 2.6E-03 | 3.1E-03 | MB231/MB468 | MITO |
| >hg38_21_+_8433172_8446622_RNA45SN1_45S_50ntFlanks@11808.11826.19\|CTGTCCCTACCTACTATCC | >hg38_21_+_8433172_8446622_RNA45SN1_45S_50ntFlanks | 11808 | 11826 | 19 | CTGTCCCTACCTACTATCC | 76.1 | -4.0 | 1.3 | -3.1 | 2.2E-03 | 3.1E-03 | MB231/MB468 | MP |
| >hg38_21_+_8433172_8446622_RNA45SN1_45S_50ntFlanks@12458.12478.21\|AAGCGTTGGATTGTTCACCCA | >hg38_21_+_8433172_8446622_RNA45SN1_45S_50ntFlanks | 12458 | 12478 | 21 | AAGCGTTGGATTGTTCACCCA | 1078.3 | -2.2 | 0.7 | -3.0 | 2.6E-03 | 3.1E-03 | MB231/MB468 | NUC |
| >MI0000095\|hsa-mir-93&WithFlank&7\|-\|100093762\|100093853@17.39.23&[MIMAT0000093&hsa-miR-93-5p&offsets\|0\|0;m-29&7\|-\|100093815\|100093837&offsets\|0\|0]\|CAAAGTGCTGTTCGTGCAGGTAG | >MI0000095\|hsa-mir-93&WithFlank&7\|-\|100093762\|100093853 | 17 | 39 | 23 | CAAAGTGCTGTTCGTGCAGGTAG | 25753.8 | 3.9 | 1.2 | 3.2 | 1.3E-03 | 3.1E-03 | BT20/MB468 | MP |
| >hg38_1_-_228634819_228635039_RNA5S12_5S_50ntFlanks@125.170.46\|GTACTTGGATGGGAGACCGCCTGGGAATACCGGGTGCTGTAGGCTT | >hg38_1_-_228634819_228635039_RNA5S12_5S_50ntFlanks | 125 | 170 | 46 | GTACTTGGATGGGAGACCGCCTGGGAATACCGGGTGCTGTAGGCTT | 362.7 | 6.7 | 2.1 | 3.2 | 1.3E-03 | 3.1E-03 | BT20/MB468 | MP |
| >hg38_21_+_8433172_8446622_RNA45SN1_45S_50ntFlanks@3840.3868.29\|CTCCTACTTGGATAACTGTGGTAATTCTA | >hg38_21_+_8433172_8446622_RNA45SN1_45S_50ntFlanks | 3840 | 3868 | 29 | CTCCTACTTGGATAACTGTGGTAATTCTA | 637.7 | 2.7 | 0.8 | 3.5 | 4.9E-04 | 3.1E-03 | BT20/MB231 | NUC |
| >hg38_21_+_8433172_8446622_RNA45SN1_45S_50ntFlanks@5493.5516.24\|GAAGACGGTCGAACTTGACTATCT | >hg38_21_+_8433172_8446622_RNA45SN1_45S_50ntFlanks | 5493 | 5516 | 24 | GAAGACGGTCGAACTTGACTATCT | 470.0 | 6.6 | 2.1 | 3.2 | 1.3E-03 | 3.1E-03 | BT20/MB468 | MP |
| >hg38_21_+_8433172_8446622_RNA45SN1_45S_50ntFlanks@12894.12933.40\|CTCGCCCGTCACGCACCGCACGTTCGTGGGGAACCTGGCG | >hg38_21_+_8433172_8446622_RNA45SN1_45S_50ntFlanks | 12894 | 12933 | 40 | CTCGCCCGTCACGCACCGCACGTTCGTGGGGAACCTGGCG | 162.1 | -5.3 | 1.6 | -3.2 | 1.3E-03 | 3.1E-03 | BT20/MB468 | MP |
| >hg38_21_+_8433172_8446622_RNA45SN1_45S_50ntFlanks@10574.10608.35\|AAGCCGGCGGGAGCCCCGGGGAGAGTTCTCTTTTC | >hg38_21_+_8433172_8446622_RNA45SN1_45S_50ntFlanks | 10574 | 10608 | 35 | AAGCCGGCGGGAGCCCCGGGGAGAGTTCTCTTTTC | 1449.8 | -5.3 | 1.7 | -3.0 | 2.4E-03 | 3.1E-03 | BT20/MB468 | CYTO |
| >trnaMT_TyrGTA_MT_-_5826_5891@39.69.31\|AAGACAGGGGTTAGGCCTCTTTTTACCACCA | >trnaMT_TyrGTA_MT_-_5826_5891 | 39 | 69 | 31 | AAGACAGGGGTTAGGCCTCTTTTTACCACCA | 154.9 | 2.8 | 0.9 | 3.1 | 2.2E-03 | 3.1E-03 | MB231/MB468 | MP |
| >hg38_21_+_8433172_8446622_RNA45SN1_45S_50ntFlanks@10409.10443.35\|GTCAGTCGGTCCTGAGAGATGGGCGAGCGCCGTTC | >hg38_21_+_8433172_8446622_RNA45SN1_45S_50ntFlanks | 10409 | 10443 | 35 | GTCAGTCGGTCCTGAGAGATGGGCGAGCGCCGTTC | 203.6 | -3.8 | 1.3 | -3.1 | 2.2E-03 | 3.1E-03 | MB231/MB468 | MP |
| >hg38_21_+_8433172_8446622_RNA45SN1_45S_50ntFlanks@9495.9526.32\|CGAAAGATGGTGAACTATGCCTGGGCAGGGCG | >hg38_21_+_8433172_8446622_RNA45SN1_45S_50ntFlanks | 9495 | 9526 | 32 | CGAAAGATGGTGAACTATGCCTGGGCAGGGCG | 206.4 | 7.0 | 2.0 | 3.5 | 4.5E-04 | 3.1E-03 | BT20/MB231 | MP |
| >hg38_21_+_8433172_8446622_RNA45SN1_45S_50ntFlanks@11317.11340.24\|ACCCCACGTCTCGTCGCGCGCGCG | >hg38_21_+_8433172_8446622_RNA45SN1_45S_50ntFlanks | 11317 | 11340 | 24 | ACCCCACGTCTCGTCGCGCGCGCG | 186.8 | -3.2 | 1.0 | -3.0 | 2.4E-03 | 3.1E-03 | BT20/MB468 | CYTO |
| >hg38_21_+_8433172_8446622_RNA45SN1_45S_50ntFlanks@8170.8187.18\|CAAGTCCTTCTGATCGAG | >hg38_21_+_8433172_8446622_RNA45SN1_45S_50ntFlanks | 8170 | 8187 | 18 | CAAGTCCTTCTGATCGAG | 456.8 | -5.3 | 1.7 | -3.2 | 1.3E-03 | 3.1E-03 | BT20/MB468 | MP |
| >hg38_21_+_8433172_8446622_RNA45SN1_45S_50ntFlanks@9131.9152.22\|GGTCGGGGGGCGGGGCGGACTG | >hg38_21_+_8433172_8446622_RNA45SN1_45S_50ntFlanks | 9131 | 9152 | 22 | GGTCGGGGGGCGGGGCGGACTG | 270.0 | -4.6 | 1.5 | -3.1 | 2.0E-03 | 3.1E-03 | BT20/MB468 | MITO |
| >hg38_MT_+_598_1651_RNR1_12S_50ntFlanks@969.1004.36\|CATGGTAAGTGTACTGGAAAGTGCACTTGGACGAAC | >hg38_MT_+_598_1651_RNR1_12S_50ntFlanks | 969 | 1004 | 36 | CATGGTAAGTGTACTGGAAAGTGCACTTGGACGAAC | 155.6 | 3.9 | 1.3 | 3.0 | 2.6E-03 | 3.1E-03 | MB231/MB468 | MITO |
| >hg38_21_+_8433172_8446622_RNA45SN1_45S_50ntFlanks@6649.6675.27\|TACGACTCTTAGCGGTGGATCACTCGG | >hg38_21_+_8433172_8446622_RNA45SN1_45S_50ntFlanks | 6649 | 6675 | 27 | TACGACTCTTAGCGGTGGATCACTCGG | 88.0 | -2.9 | 1.0 | -3.1 | 2.2E-03 | 3.1E-03 | MB231/MB468 | MP |
| >hg38_21_+_8433172_8446622_RNA45SN1_45S_50ntFlanks@8344.8375.32\|TAAGGGAAAGTTGAAAAGAACTTTGAAGAGAG | >hg38_21_+_8433172_8446622_RNA45SN1_45S_50ntFlanks | 8344 | 8375 | 32 | TAAGGGAAAGTTGAAAAGAACTTTGAAGAGAG | 2728.2 | 5.4 | 1.5 | 3.5 | 4.5E-04 | 3.1E-03 | BT20/MB231 | MP |
| >trnaMT_SerGCT_MT_+_12207_12265@12.41.30\|CAAGAACTGCTAACTCATGCCCCCATGTCT | >trnaMT_SerGCT_MT_+_12207_12265 | 12 | 41 | 30 | CAAGAACTGCTAACTCATGCCCCCATGTCT | 172.5 | 4.1 | 1.3 | 3.2 | 1.3E-03 | 3.1E-03 | BT20/MB468 | MP |
| >hg38_21_+_8433172_8446622_RNA45SN1_45S_50ntFlanks@9129.9151.23\|CGGGTCGGGGGGCGGGGCGGACT | >hg38_21_+_8433172_8446622_RNA45SN1_45S_50ntFlanks | 9129 | 9151 | 23 | CGGGTCGGGGGGCGGGGCGGACT | 192.9 | -4.5 | 1.5 | -3.1 | 2.1E-03 | 3.1E-03 | BT20/MB468 | MITO |
| >hg38_21_+_8433172_8446622_RNA45SN1_45S_50ntFlanks@11571.11602.32\|GCCGACTTAGAACTGGTGCGGACCAGGGGAAT | >hg38_21_+_8433172_8446622_RNA45SN1_45S_50ntFlanks | 11571 | 11602 | 32 | GCCGACTTAGAACTGGTGCGGACCAGGGGAAT | 573.8 | -2.8 | 0.9 | -3.0 | 2.6E-03 | 3.1E-03 | MB231/MB468 | MITO |
| >hg38_21_+_8433172_8446622_RNA45SN1_45S_50ntFlanks@12420.12444.25\|ATGTCGGCTCTTCCTATCATTGTGA | >hg38_21_+_8433172_8446622_RNA45SN1_45S_50ntFlanks | 12420 | 12444 | 25 | ATGTCGGCTCTTCCTATCATTGTGA | 1838.9 | -2.0 | 0.6 | -3.0 | 2.4E-03 | 3.1E-03 | BT20/MB468 | CYTO |
| >hg38_21_+_8433172_8446622_RNA45SN1_45S_50ntFlanks@3840.3871.32\|CTCCTACTTGGATAACTGTGGTAATTCTAGAG | >hg38_21_+_8433172_8446622_RNA45SN1_45S_50ntFlanks | 3840 | 3871 | 32 | CTCCTACTTGGATAACTGTGGTAATTCTAGAG | 75.9 | -5.1 | 1.7 | -3.1 | 2.2E-03 | 3.1E-03 | MB231/MB468 | MP |
| >hg38_21_+_8433172_8446622_RNA45SN1_45S_50ntFlanks@12397.12416.20\|CGACGTCGCTTTTTGATCCT | >hg38_21_+_8433172_8446622_RNA45SN1_45S_50ntFlanks | 12397 | 12416 | 20 | CGACGTCGCTTTTTGATCCT | 231.5 | -1.8 | 0.6 | -3.0 | 2.7E-03 | 3.1E-03 | MB231/MB468 | NUC |
| >MI0000087\|hsa-mir-29a&WithFlank&7\|-\|130876741\|130876816@47.68.22(+1G)&[MIMAT0000086&hsa-miR-29a-3p&offsets\|-1\|-1(+1G);m-26&7\|-\|130876749\|130876769&offsets\|-1\|0(+1G)]\|CTAGCACCATCTGAAATCGGTTG | >MI0000087\|hsa-mir-29a&WithFlank&7\|-\|130876741\|130876816 | 47 | 68 | 22(+1G) | CTAGCACCATCTGAAATCGGTTG | 59.7 | 5.7 | 1.9 | 3.1 | 2.2E-03 | 3.1E-03 | MB231/MB468 | MP |
| >hg38_21_+_8433172_8446622_RNA45SN1_45S_50ntFlanks@4660.4690.31\|GAGGTGAAATTCTTGGACCGGCGCAAGACGG | >hg38_21_+_8433172_8446622_RNA45SN1_45S_50ntFlanks | 4660 | 4690 | 31 | GAGGTGAAATTCTTGGACCGGCGCAAGACGG | 61.9 | -5.2 | 1.6 | -3.2 | 1.3E-03 | 3.1E-03 | BT20/MB468 | MP |
| >hg38_21_+_8433172_8446622_RNA45SN1_45S_50ntFlanks@4726.4768.43\|TAATCAAGAACGAAAGTCGGAGGTTCGAAGACGATCAGATACC | >hg38_21_+_8433172_8446622_RNA45SN1_45S_50ntFlanks | 4726 | 4768 | 43 | TAATCAAGAACGAAAGTCGGAGGTTCGAAGACGATCAGATACC | 524.9 | -3.3 | 0.9 | -3.5 | 5.4E-04 | 3.1E-03 | BT20/MB231 | CYTO |

<table>
<tr><td>>hg38_21_+_8433172_8446622_RNA45SN1_45S_50ntFlanks@12420.12444.25|ATGTCGGCTCTTCCTATCATTGTGA</td><td>>hg38_21_+_8433172_8446622_RNA45SN1_45S_50ntFlanks</td><td>12420</td><td>12444</td><td>25</td><td>ATGTCGGCTCTTCCTATCATTGTGA</td><td>469.7</td><td>-5.1</td><td>1.7</td><td>-3.1</td><td>2.2E-03</td><td>3.1E-03</td><td>MB231/MB468</td><td>MP</td></tr>
<tr><td>>MI0000253|hsa-mir-148a&WithFlank&7|-|25949913|25949992@50.71.22&[MIMAT0000243&hsa-miR-148a-3p&offsets|0|0;m-7&7|-|25949922|25949943&offsets|0|0]|TCAGTGCACTACAGAACTTTGT</td><td>>MI0000253|hsa-mir-148a&WithFlank&7|-|25949913|25949992</td><td>50</td><td>71</td><td>22</td><td>TCAGTGCACTACAGAACTTTGT</td><td>56692.0</td><td>2.2</td><td>0.7</td><td>3.0</td><td>2.6E-03</td><td>3.1E-03</td><td>BT20/MB468</td><td>NUC</td></tr>
<tr><td>>hg38_21_+_8433172_8446622_RNA45SN1_45S_50ntFlanks@10856.10873.18|AGGATTGGCTCTAAGGGC</td><td>>hg38_21_+_8433172_8446622_RNA45SN1_45S_50ntFlanks</td><td>10856</td><td>10873</td><td>18</td><td>AGGATTGGCTCTAAGGGC</td><td>316.2</td><td>-4.2</td><td>1.4</td><td>-3.0</td><td>2.6E-03</td><td>3.1E-03</td><td>BT20/MB468</td><td>NUC</td></tr>
<tr><td>>hg38_21_+_8433172_8446622_RNA45SN1_45S_50ntFlanks@4145.4175.31|CCTGAGAAACGGCTACCACATCCAAGGAAGG</td><td>>hg38_21_+_8433172_8446622_RNA45SN1_45S_50ntFlanks</td><td>4145</td><td>4175</td><td>31</td><td>CCTGAGAAACGGCTACCACATCCAAGGAAGG</td><td>42.3</td><td>-8.5</td><td>2.6</td><td>-3.2</td><td>1.3E-03</td><td>3.1E-03</td><td>BT20/MB468</td><td>MP</td></tr>
<tr><td>>am_MI0003581|hsa-mir-574&WithFlank&4|+|38868026|38868133@31.55.25&[MIMAT0004795&hsa-miR-574-5p&offsets|0|+2;m-213&4|+|38868056|38868078&offsets|0|+2]|TGAGTGTGTGTGTGTGAGTGTGTGT</td><td>>am_MI0003581|hsa-mir-574&WithFlank&4|+|38868026|38868133</td><td>31</td><td>55</td><td>25</td><td>TGAGTGTGTGTGTGTGAGTGTGTGT</td><td>674.1</td><td>4.1</td><td>1.3</td><td>3.1</td><td>2.2E-03</td><td>3.1E-03</td><td>MB231/MB468</td><td>MP</td></tr>
<tr><td>>hg38_21_+_8433172_8446622_RNA45SN1_45S_50ntFlanks@7984.8016.33|AGATCAGACGTGGCGACCCGCTGAATTTAAGCA</td><td>>hg38_21_+_8433172_8446622_RNA45SN1_45S_50ntFlanks</td><td>7984</td><td>8016</td><td>33</td><td>AGATCAGACGTGGCGACCCGCTGAATTTAAGCA</td><td>100.6</td><td>-5.1</td><td>1.6</td><td>-3.2</td><td>1.3E-03</td><td>3.1E-03</td><td>BT20/MB468</td><td>MP</td></tr>
<tr><td>>MI0000102|hsa-mir-100&WithFlank&11|-|122152223|122152314@19.39.21&[MIMAT0000098&hsa-miR-100-5p&offsets|0|-1;m-27&11|-|122152276|122152296&offsets|0|0]|AACCCGTAGATCCGAACTTGT</td><td>>MI0000102|hsa-mir-100&WithFlank&11|-|122152223|122152314</td><td>19</td><td>39</td><td>21</td><td>AACCCGTAGATCCGAACTTGT</td><td>167790.4</td><td>-4.1</td><td>1.2</td><td>-3.5</td><td>5.4E-04</td><td>3.1E-03</td><td>BT20/MB231</td><td>CYTO</td></tr>
<tr><td>>hg38_21_+_8433172_8446622_RNA45SN1_45S_50ntFlanks@11200.11232.33|CGGCGGCGGCGGCGACTCTGGACGCGAGCCGGG</td><td>>hg38_21_+_8433172_8446622_RNA45SN1_45S_50ntFlanks</td><td>11200</td><td>11232</td><td>33</td><td>CGGCGGCGGCGGCGACTCTGGACGCGAGCCGGG</td><td>2251.8</td><td>-4.3</td><td>1.4</td><td>-3.0</td><td>2.7E-03</td><td>3.1E-03</td><td>MB231/MB468</td><td>NUC</td></tr>
<tr><td>>hg38_21_+_8433172_8446622_RNA45SN1_45S_50ntFlanks@6648.6674.27|GTACGACTCTTAGCGGTGGATCACTCG</td><td>>hg38_21_+_8433172_8446622_RNA45SN1_45S_50ntFlanks</td><td>6648</td><td>6674</td><td>27</td><td>GTACGACTCTTAGCGGTGGATCACTCG</td><td>1416.9</td><td>3.3</td><td>1.1</td><td>3.0</td><td>2.7E-03</td><td>3.1E-03</td><td>MB231/MB468</td><td>MITO</td></tr>
<tr><td>>hg38_21_+_8433172_8446622_RNA45SN1_45S_50ntFlanks@11670.11698.29|CCCAGTGCTCTGAATGTCAAAGTGAAGAA</td><td>>hg38_21_+_8433172_8446622_RNA45SN1_45S_50ntFlanks</td><td>11670</td><td>11698</td><td>29</td><td>CCCAGTGCTCTGAATGTCAAAGTGAAGAA</td><td>211.7</td><td>-3.1</td><td>1.0</td><td>-3.0</td><td>2.4E-03</td><td>3.1E-03</td><td>BT20/MB468</td><td>CYTO</td></tr>
<tr><td>>hg38_21_+_8433172_8446622_RNA45SN1_45S_50ntFlanks@6647.6669.23|CGTACGACTCTTAGCGGTGGATC</td><td>>hg38_21_+_8433172_8446622_RNA45SN1_45S_50ntFlanks</td><td>6647</td><td>6669</td><td>23</td><td>CGTACGACTCTTAGCGGTGGATC</td><td>5324.6</td><td>2.8</td><td>0.8</td><td>3.5</td><td>5.0E-04</td><td>3.1E-03</td><td>BT20/MB231</td><td>NUC</td></tr>
<tr><td>>hg38_21_+_8433172_8446622_RNA45SN1_45S_50ntFlanks@12249.12284.36|CAAAAGCTCGCTTGATCTTGATTTTCAGTACGAATA</td><td>>hg38_21_+_8433172_8446622_RNA45SN1_45S_50ntFlanks</td><td>12249</td><td>12284</td><td>36</td><td>CAAAAGCTCGCTTGATCTTGATTTTCAGTACGAATA</td><td>44.1</td><td>-9.3</td><td>3.1</td><td>-3.1</td><td>2.2E-03</td><td>3.1E-03</td><td>MB231/MB468</td><td>MP</td></tr>
<tr><td>>hg38_21_+_8433172_8446622_RNA45SN1_45S_50ntFlanks@5473.5510.38|CGGCCCTGGCGGAGCGCTGAGAAGACGGTCGAACTTGA</td><td>>hg38_21_+_8433172_8446622_RNA45SN1_45S_50ntFlanks</td><td>5473</td><td>5510</td><td>38</td><td>CGGCCCTGGCGGAGCGCTGAGAAGACGGTCGAACTTGA</td><td>60.6</td><td>-4.2</td><td>1.4</td><td>-3.1</td><td>2.2E-03</td><td>3.1E-03</td><td>MB231/MB468</td><td>MP</td></tr>
<tr><td>>hg38_21_+_8433172_8446622_RNA45SN1_45S_50ntFlanks@9938.9979.42|AGCAGGACGGTGGCCATGGAAGTCGGAATCCGCTAAGGAGTG</td><td>>hg38_21_+_8433172_8446622_RNA45SN1_45S_50ntFlanks</td><td>9938</td><td>9979</td><td>42</td><td>AGCAGGACGGTGGCCATGGAAGTCGGAATCCGCTAAGGAGTG</td><td>159.4</td><td>6.3</td><td>2.0</td><td>3.2</td><td>1.3E-03</td><td>3.1E-03</td><td>BT20/MB468</td><td>MP</td></tr>
<tr><td>>hg38_21_+_8433172_8446622_RNA45SN1_45S_50ntFlanks@476.525.50|TTCGTGATCGATGTGGTGACGTCGTGCTCTCCCGGGCCGGGTCCGAGCCG</td><td>>hg38_21_+_8433172_8446622_RNA45SN1_45S_50ntFlanks</td><td>476</td><td>525</td><td>50</td><td>TTCGTGATCGATGTGGTGACGTCGTGCTCTCCCGGGCCGGGTCCGAGCCG</td><td>232.9</td><td>5.8</td><td>1.9</td><td>3.0</td><td>2.7E-03</td><td>3.1E-03</td><td>MB231/MB468</td><td>MITO</td></tr>
<tr><td>>hg38_21_+_8433172_8446622_RNA45SN1_45S_50ntFlanks@11573.11593.21|CGACTTAGAACTGGTGCGGAC</td><td>>hg38_21_+_8433172_8446622_RNA45SN1_45S_50ntFlanks</td><td>11573</td><td>11593</td><td>21</td><td>CGACTTAGAACTGGTGCGGAC</td><td>375.7</td><td>-1.8</td><td>0.6</td><td>-3.0</td><td>2.5E-03</td><td>3.1E-03</td><td>MB231/MB468</td><td>CYTO</td></tr>
<tr><td>>hg38_21_+_8433172_8446622_RNA45SN1_45S_50ntFlanks@4588.4619.32|CTATTTTGTTGGTTTTCGGAACTGAGGCCATG</td><td>>hg38_21_+_8433172_8446622_RNA45SN1_45S_50ntFlanks</td><td>4588</td><td>4619</td><td>32</td><td>CTATTTTGTTGGTTTTCGGAACTGAGGCCATG</td><td>441.0</td><td>-3.0</td><td>0.9</td><td>-3.2</td><td>1.3E-03</td><td>3.1E-03</td><td>BT20/MB468</td><td>MP</td></tr>
<tr><td>>trnaMT_GluTTC_MT_-_14674_14742@39.61.23,trnalookalike8_GluTTC_5_-_93905172_93905240@39.61.23|ATTGGTCGTGGTTGTAGTCCGTG</td><td>>trnaMT_GluTTC_MT_-_14674_14742</td><td>39</td><td>61</td><td>23</td><td>ATTGGTCGTGGTTGTAGTCCGTG</td><td>278.4</td><td>2.4</td><td>0.8</td><td>3.1</td><td>2.2E-03</td><td>3.1E-03</td><td>MB231/MB468</td><td>MP</td></tr>
<tr><td>>hg38_21_+_8433172_8446622_RNA45SN1_45S_50ntFlanks@6728.6756.29|GGACACATTGATCATCGACACTTCGAACG</td><td>>hg38_21_+_8433172_8446622_RNA45SN1_45S_50ntFlanks</td><td>6728</td><td>6756</td><td>29</td><td>GGACACATTGATCATCGACACTTCGAACG</td><td>342.2</td><td>-3.1</td><td>0.9</td><td>-3.5</td><td>5.4E-04</td><td>3.1E-03</td><td>BT20/MB231</td><td>CYTO</td></tr>
<tr><td>>MI0000272|hsa-mir-182&WithFlank&7|-|129770377|129770498@29.51.23(+1A)&[MIMAT0000259&hsa-miR-182-5p&offsets|0|-1(+1A);m-22&7|-|129770449|129770470&offsets|0|+1(+1A)]|TTTGGCAATGGTAGAACTCACACA</td><td>>MI0000272|hsa-mir-182&WithFlank&7|-|129770377|129770498</td><td>29</td><td>51</td><td>23(+1A)</td><td>TTTGGCAATGGTAGAACTCACACA</td><td>306.0</td><td>4.1</td><td>1.4</td><td>3.0</td><td>2.7E-03</td><td>3.1E-03</td><td>MB231/MB468</td><td>MITO</td></tr>
<tr><td>>hg38_21_+_8433172_8446622_RNA45SN1_45S_50ntFlanks@6650.6679.30|ACGACTCTTAGCGGTGGATCACTCGGCTCG</td><td>>hg38_21_+_8433172_8446622_RNA45SN1_45S_50ntFlanks</td><td>6650</td><td>6679</td><td>30</td><td>ACGACTCTTAGCGGTGGATCACTCGGCTCG</td><td>2820.7</td><td>3.5</td><td>1.2</td><td>3.0</td><td>2.5E-03</td><td>3.1E-03</td><td>BT20/MB468</td><td>CYTO</td></tr>
<tr><td>>MI0000087|hsa-mir-29a&WithFlank&7|-|130876741|130876816@48.68.21(+1G)&[MIMAT0000086&hsa-miR-29a-3p&offsets|0|-1(+1G);m-26&7|-|130876749|130876769&offsets|0|0(+1G)]|TAGCACCATCTGAAATCGGTTG</td><td>>MI0000087|hsa-mir-29a&WithFlank&7|-|130876741|130876816</td><td>48</td><td>68</td><td>21(+1G)</td><td>TAGCACCATCTGAAATCGGTTG</td><td>228.3</td><td>4.9</td><td>1.6</td><td>3.1</td><td>2.3E-03</td><td>3.1E-03</td><td>MB231/MB468</td><td>MP</td></tr>
<tr><td>>trna10_AspGTC_12_-_125424193_125424264@35.68.34,trna12_AspGTC_12_-_125411891_125411962@35.68.34,trna144_AspGTC_6_-_27551236_27551307@35.68.34,trna38_AspGTC_17_-_8125556_8125627@35.68.34,trna45_AspGTC_6_+_27447453_27447524@35.68.34,trna48_AspGTC_6_+_27471523_27471594@35.68.34,trna4_AspGTC_12_+_96429799_96429870@35.68.34,trna69_AspGTC_1_-_161440205_161440276@35.68.34,trna72_AspGTC_1_-_161432824_161432895@35.68.34,trna75_AspGTC_1_-_161425414_161425485@35.68.34,trna78_AspGTC_1_-_161418033_161418104@35.68.34,trna81_AspGTC_1_-_161410615_161410686@35.68.34|TCACGCGGGAGACCGGGGTTCGATTCCCCGACGG</td><td>>trna10_AspGTC_12_-_125424193_125424264</td><td>35</td><td>68</td><td>34</td><td>TCACGCGGGAGACCGGGGTTCGATTCCCCGACGG</td><td>139.5</td><td>-2.7</td><td>0.9</td><td>-3.1</td><td>2.3E-03</td><td>3.1E-03</td><td>MB231/MB468</td><td>MP</td></tr>
<tr><td>>hg38_21_+_8433172_8446622_RNA45SN1_45S_50ntFlanks@9788.9820.33|CGCTGGCGTGGAGCCGGGCGTGGAATGCGAGTG</td><td>>hg38_21_+_8433172_8446622_RNA45SN1_45S_50ntFlanks</td><td>9788</td><td>9820</td><td>33</td><td>CGCTGGCGTGGAGCCGGGCGTGGAATGCGAGTG</td><td>158.6</td><td>-3.5</td><td>1.2</td><td>-3.0</td><td>2.7E-03</td><td>3.1E-03</td><td>MB231/MB468</td><td>NUC</td></tr>
<tr><td>>hg38_21_+_8433172_8446622_RNA45SN1_45S_50ntFlanks@10795.10817.23|TGGCATGTTGGAACAATGTAGGT</td><td>>hg38_21_+_8433172_8446622_RNA45SN1_45S_50ntFlanks</td><td>10795</td><td>10817</td><td>23</td><td>TGGCATGTTGGAACAATGTAGGT</td><td>203.7</td><td>-2.9</td><td>0.9</td><td>-3.2</td><td>1.3E-03</td><td>3.1E-03</td><td>BT20/MB468</td><td>MP</td></tr>
<tr><td>>am_trnaMT_ValTAC_MT_+_1602_1670@29.57.29|ACTTACACTTAGGAGATTTCAACTTAACT</td><td>>am_trnaMT_ValTAC_MT_+_1602_1670</td><td>29</td><td>57</td><td>29</td><td>ACTTACACTTAGGAGATTTCAACTTAACT</td><td>167.6</td><td>4.3</td><td>1.4</td><td>3.1</td><td>2.3E-03</td><td>3.1E-03</td><td>MB231/MB468</td><td>MP</td></tr>
<tr><td>>hg38_1_-_228634819_228635039_RNA5S12_5S_50ntFlanks@133.169.37|ATGGGAGACCGCCTGGGAATACCGGGTGCTGTAGGCT</td><td>>hg38_1_-_228634819_228635039_RNA5S12_5S_50ntFlanks</td><td>133</td><td>169</td><td>37</td><td>ATGGGAGACCGCCTGGGAATACCGGGTGCTGTAGGCT</td><td>781.1</td><td>-2.3</td><td>0.8</td><td>-3.1</td><td>2.3E-03</td><td>3.1E-03</td><td>MB231/MB468</td><td>MP</td></tr>
<tr><td>>hg38_21_+_8433172_8446622_RNA45SN1_45S_50ntFlanks@8379.8400.22|TCAAGAGGGCGTGAAACCGTTA</td><td>>hg38_21_+_8433172_8446622_RNA45SN1_45S_50ntFlanks</td><td>8379</td><td>8400</td><td>22</td><td>TCAAGAGGGCGTGAAACCGTTA</td><td>288.9</td><td>-5.4</td><td>1.7</td><td>-3.2</td><td>1.3E-03</td><td>3.1E-03</td><td>BT20/MB468</td><td>MP</td></tr>
<tr><td>>hg38_21_+_8433172_8446622_RNA45SN1_45S_50ntFlanks@11315.11334.20|CCACCCCACGTCTCGTCGCG</td><td>>hg38_21_+_8433172_8446622_RNA45SN1_45S_50ntFlanks</td><td>11315</td><td>11334</td><td>20</td><td>CCACCCCACGTCTCGTCGCG</td><td>432.5</td><td>-4.0</td><td>1.3</td><td>-3.0</td><td>2.5E-03</td><td>3.1E-03</td><td>BT20/MB468</td><td>CYTO</td></tr>
<tr><td>>trna103_AsnGTT_1_-_149230570_149230643@29.55.27,trna107_AsnGTT_1_-_148760356_148760429@29.55.27,trna108_AsnGTT_1_-_148598314_148598387@29.55.27,trna1_AsnGTT_19_+_1383562_1383635@29.55.27,trna26_AsnGTT_1_+_148248115_148248188@29.55.27,trna31_AsnGTT_17_-_36908034_36908107@29.55.27,trna47_AsnGTT_1_+_161510031_161510104@29.55.27,trna4_AsnGTT_10_-_22518438_22518511@29.55.27,trna7_AsnGTT_13_-_31248101_31248174@29.55.27,trna83_AsnGTT_1_-_161397867_161397940@29.55.27|TCGGCTGTTAACCGAAAGGTTGGTGGT</td><td>>trna103_AsnGTT_1_-_149230570_149230643</td><td>29</td><td>55</td><td>27</td><td>TCGGCTGTTAACCGAAAGGTTGGTGGT</td><td>386.9</td><td>3.6</td><td>1.0</td><td>3.5</td><td>5.4E-04</td><td>3.1E-03</td><td>BT20/MB231</td><td>CYTO</td></tr>
<tr><td>>hg38_21_+_8433172_8446622_RNA45SN1_45S_50ntFlanks@11667.11706.40|CTGCCCAGTGCTCTGAATGTCAAAGTGAAGAAATTCAATG</td><td>>hg38_21_+_8433172_8446622_RNA45SN1_45S_50ntFlanks</td><td>11667</td><td>11706</td><td>40</td><td>CTGCCCAGTGCTCTGAATGTCAAAGTGAAGAAATTCAATG</td><td>1167.2</td><td>-2.9</td><td>0.8</td><td>-3.5</td><td>5.4E-04</td><td>3.1E-03</td><td>BT20/MB231</td><td>CYTO</td></tr>
<tr><td>>hg38_21_+_8433172_8446622_RNA45SN1_45S_50ntFlanks@5469.5514.46|CCCACGGCCCTGGCGGAGCGCTGAGAAGACGGTCGAACTTGACTAT</td><td>>hg38_21_+_8433172_8446622_RNA45SN1_45S_50ntFlanks</td><td>5469</td><td>5514</td><td>46</td><td>CCCACGGCCCTGGCGGAGCGCTGAGAAGACGGTCGAACTTGACTAT</td><td>226.1</td><td>-3.6</td><td>1.2</td><td>-3.0</td><td>2.5E-03</td><td>3.1E-03</td><td>MB231/MB468</td><td>CYTO</td></tr>
<tr><td>>hg38_21_+_8433172_8446622_RNA45SN1_45S_50ntFlanks@5472.5506.35|ACGGCCCTGGCGGAGCGCTGAGAAGACGGTCGAAC</td><td>>hg38_21_+_8433172_8446622_RNA45SN1_45S_50ntFlanks</td><td>5472</td><td>5506</td><td>35</td><td>ACGGCCCTGGCGGAGCGCTGAGAAGACGGTCGAAC</td><td>167.6</td><td>3.9</td><td>1.3</td><td>3.0</td><td>2.7E-03</td><td>3.2E-03</td><td>MB231/MB468</td><td>MITO</td></tr>
<tr><td>>hg38_21_+_8433172_8446622_RNA45SN1_45S_50ntFlanks@10914.10946.33|CGCCGCGGCTGGACGAGGCGCCGCCGCCCCCCC</td><td>>hg38_21_+_8433172_8446622_RNA45SN1_45S_50ntFlanks</td><td>10914</td><td>10946</td><td>33</td><td>CGCCGCGGCTGGACGAGGCGCCGCCGCCCCCCC</td><td>306.0</td><td>-4.5</td><td>1.5</td><td>-3.0</td><td>2.5E-03</td><td>3.2E-03</td><td>BT20/MB468</td><td>CYTO</td></tr>
<tr><td>>hg38_21_+_8433172_8446622_RNA45SN1_45S_50ntFlanks@4817.4835.19|ATTCCCATGACCCGCCGGG</td><td>>hg38_21_+_8433172_8446622_RNA45SN1_45S_50ntFlanks</td><td>4817</td><td>4835</td><td>19</td><td>ATTCCCATGACCCGCCGGG</td><td>77.7</td><td>-5.0</td><td>1.6</td><td>-3.2</td><td>1.3E-03</td><td>3.2E-03</td><td>BT20/MB468</td><td>MP</td></tr>
<tr><td>>hg38_21_+_8433172_8446622_RNA45SN1_45S_50ntFlanks@12456.12490.35|CCAAGCGTTGGATTGTTCACCCACTAATAGGGAAC</td><td>>hg38_21_+_8433172_8446622_RNA45SN1_45S_50ntFlanks</td><td>12456</td><td>12490</td><td>35</td><td>CCAAGCGTTGGATTGTTCACCCACTAATAGGGAAC</td><td>1071.0</td><td>-3.5</td><td>1.1</td><td>-3.2</td><td>1.3E-03</td><td>3.2E-03</td><td>BT20/MB468</td><td>MP</td></tr>
<tr><td>>hg38_21_+_8433172_8446622_RNA45SN1_45S_50ntFlanks@10869.10902.34|AGGGCTGGGTCGGTCGGGCTGGGGCGCGAAGCGG</td><td>>hg38_21_+_8433172_8446622_RNA45SN1_45S_50ntFlanks</td><td>10869</td><td>10902</td><td>34</td><td>AGGGCTGGGTCGGTCGGGCTGGGGCGCGAAGCGG</td><td>1024.2</td><td>-4.1</td><td>1.3</td><td>-3.1</td><td>2.1E-03</td><td>3.2E-03</td><td>BT20/MB468</td><td>MITO</td></tr>
<tr><td>>MI0000285|hsa-mir-205&WithFlank&1|+|209432127|209432248@40.62.23(+1U)&[MIMAT0000266&hsa-miR-205-5p&offsets|0|+1(+1U);m-351&1|+|209432166|209432187&offsets|0|+1(+1U)]|TCCTTCATTCCACCGGAGTCTGTT</td><td>>MI0000285|hsa-mir-205&WithFlank&1|+|209432127|209432248</td><td>40</td><td>62</td><td>23(+1U)</td><td>TCCTTCATTCCACCGGAGTCTGTT</td><td>426.2</td><td>2.2</td><td>0.7</td><td>3.0</td><td>2.7E-03</td><td>3.2E-03</td><td>BT20/MB468</td><td>NUC</td></tr>
<tr><td>>hg38_21_+_8433172_8446622_RNA45SN1_45S_50ntFlanks@6647.6675.29|CGTACGACTCTTAGCGGTGGATCACTCGG</td><td>>hg38_21_+_8433172_8446622_RNA45SN1_45S_50ntFlanks</td><td>6647</td><td>6675</td><td>29</td><td>CGTACGACTCTTAGCGGTGGATCACTCGG</td><td>1142.0</td><td>2.4</td><td>0.8</td><td>3.1</td><td>2.1E-03</td><td>3.2E-03</td><td>BT20/MB468</td><td>MITO</td></tr>
<tr><td>>MI0000298|hsa-mir-221&WithFlank&X|-|45746151|45746272@74.93.20&[MIMAT0000278&hsa-miR-221-3p&offsets|+3|0;m-91&X|-|45746181|45746202&offsets|+3|+1]|TACATTGTCTGCTGGGTTTC</td><td>>MI0000298|hsa-mir-221&WithFlank&X|-|45746151|45746272</td><td>74</td><td>93</td><td>20</td><td>TACATTGTCTGCTGGGTTTC</td><td>39.2</td><td>8.5</td><td>2.8</td><td>3.1</td><td>2.3E-03</td><td>3.2E-03</td><td>MB231/MB468</td><td>MP</td></tr>
<tr><td>>hg38_21_+_8433172_8446622_RNA45SN1_45S_50ntFlanks@11575.11602.28|ACTTAGAACTGGTGCGGACCAGGGGAAT</td><td>>hg38_21_+_8433172_8446622_RNA45SN1_45S_50ntFlanks</td><td>11575</td><td>11602</td><td>28</td><td>ACTTAGAACTGGTGCGGACCAGGGGAAT</td><td>376.6</td><td>-1.9</td><td>0.6</td><td>-3.1</td><td>2.1E-03</td><td>3.2E-03</td><td>BT20/MB468</td><td>MITO</td></tr>
<tr><td>>MI0000272|hsa-mir-182&WithFlank&7|-|129770377|129770498@29.53.25(+1U)&[MIMAT0000259&hsa-miR-182-5p&offsets|0|+1(+1U);m-22&7|-|129770449|129770470&offsets|0|+3(+1U)]|TTTGGCAATGGTAGAACTCACACTGT</td><td>>MI0000272|hsa-mir-182&WithFlank&7|-|129770377|129770498</td><td>29</td><td>53</td><td>25(+1U)</td><td>TTTGGCAATGGTAGAACTCACACTGT</td><td>2986.4</td><td>2.8</td><td>0.8</td><td>3.5</td><td>5.5E-04</td><td>3.2E-03</td><td>BT20/MB231</td><td>CYTO</td></tr>
</table>

<table>
<tr><td>>hg38_21_+_8433172_8446622_RNA45SN1_45S_50ntFlanks@12397.12420.24|CGACGTCGCTTTTTGATCCTTCGA</td><td>>hg38_21_+_8433172_8446622_RNA45SN1_45S_50ntFlanks</td><td>12397</td><td>12420</td><td>24</td><td>CGACGTCGCTTTTTGATCCTTCGA</td><td>588.7</td><td>-3.1</td><td>1.0</td><td>-3.0</td><td>2.7E-03</td><td>3.2E-03</td><td>MB231/MB468</td><td>MITO</td></tr>
<tr><td>>hg38_21_+_8433172_8446622_RNA45SN1_45S_50ntFlanks@11570.11599.30|AGCCGACTTAGAACTGGTGCGGACCAGGGG</td><td>>hg38_21_+_8433172_8446622_RNA45SN1_45S_50ntFlanks</td><td>11570</td><td>11599</td><td>30</td><td>AGCCGACTTAGAACTGGTGCGGACCAGGGG</td><td>98.8</td><td>-3.9</td><td>1.3</td><td>-3.1</td><td>2.3E-03</td><td>3.2E-03</td><td>MB231/MB468</td><td>MP</td></tr>
<tr><td>>hg38_21_+_8433172_8446622_RNA45SN1_45S_50ntFlanks@475.503.29|CTTCGTGATCGATGTGGTGACGTCGTGCT</td><td>>hg38_21_+_8433172_8446622_RNA45SN1_45S_50ntFlanks</td><td>475</td><td>503</td><td>29</td><td>CTTCGTGATCGATGTGGTGACGTCGTGCT</td><td>624.9</td><td>4.8</td><td>1.5</td><td>3.2</td><td>1.3E-03</td><td>3.2E-03</td><td>BT20/MB468</td><td>MP</td></tr>
<tr><td>>trna14_LysTTT_11_-_59327808_59327880@1.28.28,trna2_LysTTT_17_+_8022473_8022545@1.28.28,trna54_LysTTT_1_+_204475655_204475727@1.28.28,trna5_LysTTT_11_+_59323902_59323974@1.28.28,trna62_LysTTT_1_-_204476158_204476230@1.28.28,trna76_LysTTT_6_+_28918806_28918878@1.28.28|GCCCGGATAGCTCAGTCGGTAGAGCATC</td><td>>trna14_LysTTT_11_-_59327808_59327880</td><td>1</td><td>28</td><td>28</td><td>GCCCGGATAGCTCAGTCGGTAGAGCATC</td><td>261.3</td><td>-2.5</td><td>0.8</td><td>-3.0</td><td>2.5E-03</td><td>3.2E-03</td><td>MB231/MB468</td><td>CYTO</td></tr>
<tr><td>>MI0000067|hsa-let-7f-1&WithFlank&9|+|94176341|94176439@13.34.22;MI0000068|hsa-let-7f-2&WithFlank&X|-|53557186|53557280@14.35.22&[MIMAT0000067&hsa-let-7f-5p&offsets|0|0;m-20&9|+|94176353|94176374&offsets|0|0];[MIMAT0000067_1&hsa-let-7f-5p&offsets|0|0;m-19&X|-|53557246|53557267&offsets|0|0]|TGAGGTAGTAGATTGTATAGTT</td><td>>MI0000067|hsa-let-7f-1&WithFlank&9|+|94176341|94176439</td><td>13</td><td>34</td><td>22</td><td>TGAGGTAGTAGATTGTATAGTT</td><td>55707.5</td><td>1.4</td><td>0.5</td><td>3.0</td><td>2.7E-03</td><td>3.2E-03</td><td>BT20/MB468</td><td>NUC</td></tr>
<tr><td>>hg38_21_+_8433172_8446622_RNA45SN1_45S_50ntFlanks@7974.7994.21|ACGCGACCTCAGATCAGACGT</td><td>>hg38_21_+_8433172_8446622_RNA45SN1_45S_50ntFlanks</td><td>7974</td><td>7994</td><td>21</td><td>ACGCGACCTCAGATCAGACGT</td><td>2183.0</td><td>2.3</td><td>0.8</td><td>3.0</td><td>2.7E-03</td><td>3.2E-03</td><td>BT20/MB468</td><td>NUC</td></tr>
<tr><td>>hg38_21_+_8433172_8446622_RNA45SN1_45S_50ntFlanks@10318.10336.19|TGCAGATCTTGGTGGTAGT</td><td>>hg38_21_+_8433172_8446622_RNA45SN1_45S_50ntFlanks</td><td>10318</td><td>10336</td><td>19</td><td>TGCAGATCTTGGTGGTAGT</td><td>182.3</td><td>6.7</td><td>1.9</td><td>3.5</td><td>4.6E-04</td><td>3.2E-03</td><td>BT20/MB231</td><td>MP</td></tr>
<tr><td>>MI0000077|hsa-mir-21&WithFlank&17|+|59841260|59841343@14.31.18(+1A)&[MIMAT0000076&hsa-miR-21-5p&offsets|0|-4(+1A);m-1&17|+|59841273|59841294&offsets|0|-4(+1A)]|TAGCTTATCAGACTGATGA</td><td>>MI0000077|hsa-mir-21&WithFlank&17|+|59841260|59841343</td><td>14</td><td>31</td><td>18(+1A)</td><td>TAGCTTATCAGACTGATGA</td><td>106.6</td><td>4.9</td><td>1.6</td><td>3.1</td><td>2.3E-03</td><td>3.2E-03</td><td>MB231/MB468</td><td>MP</td></tr>
<tr><td>>hg38_21_+_8433172_8446622_RNA45SN1_45S_50ntFlanks@9575.9603.29|ATCGGTCGTCCGACCTGGGTATAGGGGCG</td><td>>hg38_21_+_8433172_8446622_RNA45SN1_45S_50ntFlanks</td><td>9575</td><td>9603</td><td>29</td><td>ATCGGTCGTCCGACCTGGGTATAGGGGCG</td><td>2601.2</td><td>-5.3</td><td>1.8</td><td>-3.0</td><td>2.7E-03</td><td>3.2E-03</td><td>MB231/MB468</td><td>MITO</td></tr>
<tr><td>>hg38_21_+_8433172_8446622_RNA45SN1_45S_50ntFlanks@4103.4120.18|CCACGGGTGACGGGGAAT</td><td>>hg38_21_+_8433172_8446622_RNA45SN1_45S_50ntFlanks</td><td>4103</td><td>4120</td><td>18</td><td>CCACGGGTGACGGGGAAT</td><td>142.5</td><td>-4.3</td><td>1.3</td><td>-3.2</td><td>1.3E-03</td><td>3.2E-03</td><td>BT20/MB468</td><td>MP</td></tr>
<tr><td>>MI0000272|hsa-mir-182&WithFlank&7|-|129770377|129770498@29.48.20(+1U)&[MIMAT0000259&hsa-miR-182-5p&offsets|0|-4(+1U);m-22&7|-|129770449|129770470&offsets|0|-2(+1U)]|TTTGGCAATGGTAGAACTCAT</td><td>>MI0000272|hsa-mir-182&WithFlank&7|-|129770377|129770498</td><td>29</td><td>48</td><td>20(+1U)</td><td>TTTGGCAATGGTAGAACTCAT</td><td>149.6</td><td>5.8</td><td>1.8</td><td>3.2</td><td>1.3E-03</td><td>3.2E-03</td><td>BT20/MB468</td><td>MP</td></tr>
<tr><td>>hg38_21_+_8433172_8446622_RNA45SN1_45S_50ntFlanks@10414.10443.30|TCGGTCCTGAGAGATGGGCGAGCGCCGTTC</td><td>>hg38_21_+_8433172_8446622_RNA45SN1_45S_50ntFlanks</td><td>10414</td><td>10443</td><td>30</td><td>TCGGTCCTGAGAGATGGGCGAGCGCCGTTC</td><td>90.3</td><td>-5.2</td><td>1.6</td><td>-3.2</td><td>1.3E-03</td><td>3.2E-03</td><td>BT20/MB468</td><td>MP</td></tr>
<tr><td>>hg38_21_+_8433172_8446622_RNA45SN1_45S_50ntFlanks@6771.6807.37|GGTTCCTCCCGGGGCTACGCCTGTCTGAGCGTCGCTT</td><td>>hg38_21_+_8433172_8446622_RNA45SN1_45S_50ntFlanks</td><td>6771</td><td>6807</td><td>37</td><td>GGTTCCTCCCGGGGCTACGCCTGTCTGAGCGTCGCTT</td><td>603.1</td><td>-1.4</td><td>0.5</td><td>-3.1</td><td>2.1E-03</td><td>3.2E-03</td><td>BT20/MB468</td><td>MITO</td></tr>
<tr><td>>trnaMT_ValTAC_MT_+_1602_1670@29.72.44|ACTTACACTTAGGAGATTTCAACTTAACTTGACCGCTCTGACCA</td><td>>trnaMT_ValTAC_MT_+_1602_1670</td><td>29</td><td>72</td><td>44</td><td>ACTTACACTTAGGAGATTTCAACTTAACTTGACCGCTCTGACCA</td><td>1458.7</td><td>3.6</td><td>1.2</td><td>3.0</td><td>2.7E-03</td><td>3.2E-03</td><td>MB231/MB468</td><td>MITO</td></tr>
<tr><td>>trnaMT_ProTGG_MT_-_15956_16023@50.70.21|TTAAAGACTTTTTCTCTGACC</td><td>>trnaMT_ProTGG_MT_-_15956_16023</td><td>50</td><td>70</td><td>21</td><td>TTAAAGACTTTTTCTCTGACC</td><td>170.2</td><td>2.7</td><td>0.9</td><td>3.1</td><td>2.3E-03</td><td>3.2E-03</td><td>MB231/MB468</td><td>MP</td></tr>
<tr><td>>hg38_21_+_8433172_8446622_RNA45SN1_45S_50ntFlanks@3846.3869.24|CTTGGATAACTGTGGTAATTCTAG</td><td>>hg38_21_+_8433172_8446622_RNA45SN1_45S_50ntFlanks</td><td>3846</td><td>3869</td><td>24</td><td>CTTGGATAACTGTGGTAATTCTAG</td><td>441.3</td><td>-3.9</td><td>1.3</td><td>-3.1</td><td>2.1E-03</td><td>3.2E-03</td><td>BT20/MB468</td><td>MITO</td></tr>
<tr><td>>hg38_21_+_8433172_8446622_RNA45SN1_45S_50ntFlanks@11920.11967.48|ATGAGAGGTGTAGAATAAGTGGGAGGCCCCCGGCGCCCCCCCGGTGTC</td><td>>hg38_21_+_8433172_8446622_RNA45SN1_45S_50ntFlanks</td><td>11920</td><td>11967</td><td>48</td><td>ATGAGAGGTGTAGAATAAGTGGGAGGCCCCCGGCGCCCCCCCGGTGTC</td><td>325.6</td><td>-4.3</td><td>1.4</td><td>-3.0</td><td>2.6E-03</td><td>3.2E-03</td><td>MB231/MB468</td><td>CYTO</td></tr>
<tr><td>>hg38_21_+_8433172_8446622_RNA45SN1_45S_50ntFlanks@7975.8018.44|CGCGACCTCAGATCAGACGTGGCGACCCGCTGAATTTAAGCATA</td><td>>hg38_21_+_8433172_8446622_RNA45SN1_45S_50ntFlanks</td><td>7975</td><td>8018</td><td>44</td><td>CGCGACCTCAGATCAGACGTGGCGACCCGCTGAATTTAAGCATA</td><td>2056.5</td><td>4.0</td><td>1.3</td><td>3.0</td><td>2.7E-03</td><td>3.2E-03</td><td>MB231/MB468</td><td>NUC</td></tr>
<tr><td>>hg38_21_+_8433172_8446622_RNA45SN1_45S_50ntFlanks@7979.8009.31|ACCTCAGATCAGACGTGGCGACCCGCTGAAT</td><td>>hg38_21_+_8433172_8446622_RNA45SN1_45S_50ntFlanks</td><td>7979</td><td>8009</td><td>31</td><td>ACCTCAGATCAGACGTGGCGACCCGCTGAAT</td><td>258.0</td><td>1.8</td><td>0.6</td><td>3.0</td><td>2.6E-03</td><td>3.2E-03</td><td>MB231/MB468</td><td>CYTO</td></tr>
<tr><td>>hg38_21_+_8433172_8446622_RNA45SN1_45S_50ntFlanks@6652.6690.39|GACTCTTAGCGGTGGATCACTCGGCTCGTGCGTCGATGA</td><td>>hg38_21_+_8433172_8446622_RNA45SN1_45S_50ntFlanks</td><td>6652</td><td>6690</td><td>39</td><td>GACTCTTAGCGGTGGATCACTCGGCTCGTGCGTCGATGA</td><td>1598.3</td><td>4.2</td><td>1.3</td><td>3.2</td><td>1.3E-03</td><td>3.2E-03</td><td>BT20/MB468</td><td>MP</td></tr>
<tr><td>>hg38_21_+_8433172_8446622_RNA45SN1_45S_50ntFlanks@6646.6665.20|TCGTACGACTCTTAGCGGTG</td><td>>hg38_21_+_8433172_8446622_RNA45SN1_45S_50ntFlanks</td><td>6646</td><td>6665</td><td>20</td><td>TCGTACGACTCTTAGCGGTG</td><td>15597.7</td><td>2.3</td><td>0.7</td><td>3.0</td><td>2.7E-03</td><td>3.2E-03</td><td>BT20/MB468</td><td>NUC</td></tr>
<tr><td>>hg38_21_+_8433172_8446622_RNA45SN1_45S_50ntFlanks@12008.12030.23|CCGCCGGTGAAATACCACTACTC</td><td>>hg38_21_+_8433172_8446622_RNA45SN1_45S_50ntFlanks</td><td>12008</td><td>12030</td><td>23</td><td>CCGCCGGTGAAATACCACTACTC</td><td>930.8</td><td>-5.1</td><td>1.7</td><td>-3.0</td><td>2.5E-03</td><td>3.2E-03</td><td>BT20/MB468</td><td>CYTO</td></tr>
<tr><td>>hg38_MT_+_1621_3279_RNR2_16S_50ntFlanks@1367.1397.31|TCGATGTTGGATCAGGACATCCCGATGGTGC</td><td>>hg38_MT_+_1621_3279_RNR2_16S_50ntFlanks</td><td>1367</td><td>1397</td><td>31</td><td>TCGATGTTGGATCAGGACATCCCGATGGTGC</td><td>152.2</td><td>5.5</td><td>1.7</td><td>3.2</td><td>1.3E-03</td><td>3.2E-03</td><td>BT20/MB468</td><td>MP</td></tr>
<tr><td>>hg38_21_+_8433172_8446622_RNA45SN1_45S_50ntFlanks@6652.6680.29|GACTCTTAGCGGTGGATCACTCGGCTCGT</td><td>>hg38_21_+_8433172_8446622_RNA45SN1_45S_50ntFlanks</td><td>6652</td><td>6680</td><td>29</td><td>GACTCTTAGCGGTGGATCACTCGGCTCGT</td><td>555.9</td><td>-2.7</td><td>0.9</td><td>-3.1</td><td>2.3E-03</td><td>3.2E-03</td><td>MB231/MB468</td><td>MP</td></tr>
<tr><td>>hg38_21_+_8433172_8446622_RNA45SN1_45S_50ntFlanks@51.74.24|GCTGACACGCTGTCCTCTGGCGAC</td><td>>hg38_21_+_8433172_8446622_RNA45SN1_45S_50ntFlanks</td><td>51</td><td>74</td><td>24</td><td>GCTGACACGCTGTCCTCTGGCGAC</td><td>1229.8</td><td>3.0</td><td>1.0</td><td>3.0</td><td>2.7E-03</td><td>3.2E-03</td><td>MB231/MB468</td><td>NUC</td></tr>
<tr><td>>hg38_21_+_8433172_8446622_RNA45SN1_45S_50ntFlanks@11710.11728.19|CGCGGGTAAACGGCGGGAG</td><td>>hg38_21_+_8433172_8446622_RNA45SN1_45S_50ntFlanks</td><td>11710</td><td>11728</td><td>19</td><td>CGCGGGTAAACGGCGGGAG</td><td>1379.2</td><td>-6.0</td><td>2.0</td><td>-3.0</td><td>2.7E-03</td><td>3.2E-03</td><td>BT20/MB468</td><td>NUC</td></tr>
<tr><td>>trnaMT_GluTTC_MT_-_14674_14742@35.66.32,trnalookalike8_GluTTC_5_-_93905172_93905240@35.66.32|TATCATTGGTCGTGGTTGTAGTCCGTGCGAGA</td><td>>trnaMT_GluTTC_MT_-_14674_14742</td><td>35</td><td>66</td><td>32</td><td>TATCATTGGTCGTGGTTGTAGTCCGTGCGAGA</td><td>175.8</td><td>4.3</td><td>1.4</td><td>3.2</td><td>1.3E-03</td><td>3.2E-03</td><td>BT20/MB468</td><td>MP</td></tr>
<tr><td>>hg38_21_+_8433172_8446622_RNA45SN1_45S_50ntFlanks@9554.9582.29|CGTAGCGGTCCTGACGTGCAAATCGGTCG</td><td>>hg38_21_+_8433172_8446622_RNA45SN1_45S_50ntFlanks</td><td>9554</td><td>9582</td><td>29</td><td>CGTAGCGGTCCTGACGTGCAAATCGGTCG</td><td>465.1</td><td>-4.9</td><td>1.6</td><td>-3.0</td><td>2.5E-03</td><td>3.2E-03</td><td>BT20/MB468</td><td>CYTO</td></tr>
<tr><td>>hg38_21_+_8433172_8446622_RNA45SN1_45S_50ntFlanks@10346.10382.37|TCAAACGAGAACTTTGAAGGCCGAAGTGGAGAAGGGT</td><td>>hg38_21_+_8433172_8446622_RNA45SN1_45S_50ntFlanks</td><td>10346</td><td>10382</td><td>37</td><td>TCAAACGAGAACTTTGAAGGCCGAAGTGGAGAAGGGT</td><td>309.1</td><td>1.6</td><td>0.5</td><td>3.1</td><td>2.1E-03</td><td>3.2E-03</td><td>BT20/MB468</td><td>MITO</td></tr>
<tr><td>>hg38_21_+_8433172_8446622_RNA45SN1_45S_50ntFlanks@8342.8365.24|CGTAAGGGAAAGTTGAAAAGAACT</td><td>>hg38_21_+_8433172_8446622_RNA45SN1_45S_50ntFlanks</td><td>8342</td><td>8365</td><td>24</td><td>CGTAAGGGAAAGTTGAAAAGAACT</td><td>383.6</td><td>4.1</td><td>1.3</td><td>3.0</td><td>2.3E-03</td><td>3.2E-03</td><td>MB231/MB468</td><td>MP</td></tr>
<tr><td>>hg38_1_-_228634819_228635039_RNA5S12_5S_50ntFlanks@124.170.47|AGTACTTGGATGGGAGACCGCCTGGGAATACCGGGTGCTGTAGGCTT</td><td>>hg38_1_-_228634819_228635039_RNA5S12_5S_50ntFlanks</td><td>124</td><td>170</td><td>47</td><td>AGTACTTGGATGGGAGACCGCCTGGGAATACCGGGTGCTGTAGGCTT</td><td>807.2</td><td>3.8</td><td>1.1</td><td>3.4</td><td>6.2E-04</td><td>3.2E-03</td><td>BT20/MB231</td><td>MITO</td></tr>
<tr><td>>hg38_21_+_8433172_8446622_RNA45SN1_45S_50ntFlanks@11038.11060.23|TCTCCCCCGCTCCCCGTCCTCCC</td><td>>hg38_21_+_8433172_8446622_RNA45SN1_45S_50ntFlanks</td><td>11038</td><td>11060</td><td>23</td><td>TCTCCCCCGCTCCCCGTCCTCCC</td><td>2533.2</td><td>-6.5</td><td>2.2</td><td>-3.0</td><td>2.7E-03</td><td>3.2E-03</td><td>BT20/MB468</td><td>NUC</td></tr>
<tr><td>>hg38_21_+_8433172_8446622_RNA45SN1_45S_50ntFlanks@12414.12432.19|CCTTCGATGTCGGCTCTTC</td><td>>hg38_21_+_8433172_8446622_RNA45SN1_45S_50ntFlanks</td><td>12414</td><td>12432</td><td>19</td><td>CCTTCGATGTCGGCTCTTC</td><td>514.7</td><td>1.9</td><td>0.6</td><td>3.0</td><td>2.6E-03</td><td>3.2E-03</td><td>MB231/MB468</td><td>CYTO</td></tr>
<tr><td>>hg38_21_+_8433172_8446622_RNA45SN1_45S_50ntFlanks@5467.5501.35|GGCCCACGGCCCTGGCGGAGCGCTGAGAAGACGGT</td><td>>hg38_21_+_8433172_8446622_RNA45SN1_45S_50ntFlanks</td><td>5467</td><td>5501</td><td>35</td><td>GGCCCACGGCCCTGGCGGAGCGCTGAGAAGACGGT</td><td>83.7</td><td>9.7</td><td>2.8</td><td>3.5</td><td>4.7E-04</td><td>3.2E-03</td><td>BT20/MB231</td><td>MP</td></tr>
<tr><td>>hg38_21_+_8433172_8446622_RNA45SN1_45S_50ntFlanks@9931.9951.21|TATAGACAGCAGGACGGTGGC</td><td>>hg38_21_+_8433172_8446622_RNA45SN1_45S_50ntFlanks</td><td>9931</td><td>9951</td><td>21</td><td>TATAGACAGCAGGACGGTGGC</td><td>208.1</td><td>5.6</td><td>1.8</td><td>3.2</td><td>1.3E-03</td><td>3.2E-03</td><td>BT20/MB468</td><td>MP</td></tr>
<tr><td>>MI0000299|hsa-mir-222&WithFlank&X|-|45747009|45747130@75.99.25(+1C)&[MIMAT0000279&hsa-miR-222-3p&offsets|0|+4(+1C);m-140&X|-|45747033|45747056&offsets|0|+1(+1C)]|AGCTACATCTGGCTACTGGGTCTCTC</td><td>>MI0000299|hsa-mir-222&WithFlank&X|-|45747009|45747130</td><td>75</td><td>99</td><td>25(+1C)</td><td>AGCTACATCTGGCTACTGGGTCTCTC</td><td>764.1</td><td>-2.8</td><td>0.8</td><td>-3.4</td><td>6.2E-04</td><td>3.2E-03</td><td>BT20/MB231</td><td>MITO</td></tr>
<tr><td>>hg38_21_+_8433172_8446622_RNA45SN1_45S_50ntFlanks@10788.10817.30|CAGCCTCTGGCATGTTGGAACAATGTAGGT</td><td>>hg38_21_+_8433172_8446622_RNA45SN1_45S_50ntFlanks</td><td>10788</td><td>10817</td><td>30</td><td>CAGCCTCTGGCATGTTGGAACAATGTAGGT</td><td>419.7</td><td>-2.7</td><td>0.8</td><td>-3.2</td><td>1.3E-03</td><td>3.2E-03</td><td>BT20/MB468</td><td>MP</td></tr>
<tr><td>>hg38_MT_+_1621_3279_RNR2_16S_50ntFlanks@622.642.21|TATAACTGAACTCCTCACACC</td><td>>hg38_MT_+_1621_3279_RNR2_16S_50ntFlanks</td><td>622</td><td>642</td><td>21</td><td>TATAACTGAACTCCTCACACC</td><td>105.2</td><td>4.7</td><td>1.6</td><td>3.0</td><td>2.7E-03</td><td>3.2E-03</td><td>MB231/MB468</td><td>MITO</td></tr>
<tr><td>>am_trnaMT_HisGTG_MT_+_12138_12206@26.45.20|AGATTGTGAATCTGACAACA</td><td>>am_trnaMT_HisGTG_MT_+_12138_12206</td><td>26</td><td>45</td><td>20</td><td>AGATTGTGAATCTGACAACA</td><td>446.7</td><td>3.5</td><td>1.2</td><td>3.0</td><td>2.7E-03</td><td>3.2E-03</td><td>MB231/MB468</td><td>MITO</td></tr>
<tr><td>>hg38_21_+_8433172_8446622_RNA45SN1_45S_50ntFlanks@6730.6770.41|ACACATTGATCATCGACACTTCGAACGCACTTGCGGCCCCG</td><td>>hg38_21_+_8433172_8446622_RNA45SN1_45S_50ntFlanks</td><td>6730</td><td>6770</td><td>41</td><td>ACACATTGATCATCGACACTTCGAACGCACTTGCGGCCCCG</td><td>231.7</td><td>-5.7</td><td>1.8</td><td>-3.2</td><td>1.3E-03</td><td>3.2E-03</td><td>BT20/MB468</td><td>MP</td></tr>
<tr><td>>MI0000457|hsa-mir-141&WithFlank&12|+|6964091|6964197@65.87.23&[MIMAT0000432&hsa-miR-141-3p&offsets|0|+1;m-134&12|+|6964155|6964175&offsets|0|+2]|TAACACTGTCTGGTAAAGATGGC</td><td>>MI0000457|hsa-mir-141&WithFlank&12|+|6964091|6964197</td><td>65</td><td>87</td><td>23</td><td>TAACACTGTCTGGTAAAGATGGC</td><td>71.0</td><td>9.5</td><td>2.7</td><td>3.5</td><td>4.7E-04</td><td>3.2E-03</td><td>BT20/MB231</td><td>MP</td></tr>
<tr><td>>am_trnaMT_ProTGG_MT_-_15956_16023@53.70.18|AAGACTTTTTCTCTGACC</td><td>>am_trnaMT_ProTGG_MT_-_15956_16023</td><td>53</td><td>70</td><td>18</td><td>AAGACTTTTTCTCTGACC</td><td>709.4</td><td>4.3</td><td>1.4</td><td>3.0</td><td>2.5E-03</td><td>3.2E-03</td><td>BT20/MB468</td><td>CYTO</td></tr>
<tr><td>>MI0000088|hsa-mir-30a&WithFlank&6|-|71403545|71403627@12.32.21(+1A)&[MIMAT0000087&hsa-miR-30a-5p&offsets|0|-1(+1A);m-15&6|-|71403595|71403616&offsets|0|-1(+1A)]|TGTAAACATCCTCGACTGGAAA</td><td>>MI0000088|hsa-mir-30a&WithFlank&6|-|71403545|71403627</td><td>12</td><td>32</td><td>21(+1A)</td><td>TGTAAACATCCTCGACTGGAAA</td><td>815.5</td><td>2.7</td><td>0.9</td><td>3.0</td><td>2.3E-03</td><td>3.2E-03</td><td>MB231/MB468</td><td>MP</td></tr>
<tr><td>>MI0000298|hsa-mir-221&WithFlank&X|-|45746151|45746272@71.91.21(+1A)&[MIMAT0000278&hsa-miR-221-3p&offsets|0|-2(+1A);m-91&X|-|45746181|45746202&offsets|0|-1(+1A)]|AGCTACATTGTCTGCTGGGTTA</td><td>>MI0000298|hsa-mir-221&WithFlank&X|-|45746151|45746272</td><td>71</td><td>91</td><td>21(+1A)</td><td>AGCTACATTGTCTGCTGGGTTA</td><td>541.6</td><td>3.9</td><td>1.2</td><td>3.2</td><td>1.4E-03</td><td>3.2E-03</td><td>BT20/MB468</td><td>MP</td></tr>
<tr><td>>MI0000254|hsa-mir-30c-2&WithFlank&6|-|71376954|71377037@53.73.21(+1C)&[MIMAT0004550&hsa-miR-30c-2-3p&offsets|0|-1(+1C);m-206&6|-|71376964|71376985&offsets|0|-1(+1C)]|CTGGGAGAAGGCTGTTTACTCC</td><td>>MI0000254|hsa-mir-30c-2&WithFlank&6|-|71376954|71377037</td><td>53</td><td>73</td><td>21(+1C)</td><td>CTGGGAGAAGGCTGTTTACTCC</td><td>60.0</td><td>4.8</td><td>1.6</td><td>3.0</td><td>2.3E-03</td><td>3.2E-03</td><td>MB231/MB468</td><td>MP</td></tr>
<tr><td>>hg38_MT_+_598_1651_RNR1_12S_50ntFlanks@994.1016.23|CTTGGACGAACCAGAGTGTAGCT</td><td>>hg38_MT_+_598_1651_RNR1_12S_50ntFlanks</td><td>994</td><td>1016</td><td>23</td><td>CTTGGACGAACCAGAGTGTAGCT</td><td>47.3</td><td>5.5</td><td>1.8</td><td>3.0</td><td>2.3E-03</td><td>3.2E-03</td><td>MB231/MB468</td><td>MP</td></tr>
<tr><td>>hg38_21_+_8433172_8446622_RNA45SN1_45S_50ntFlanks@8184.8208.25|CGAGGCCCAGCCCGTGGACGGTGTG</td><td>>hg38_21_+_8433172_8446622_RNA45SN1_45S_50ntFlanks</td><td>8184</td><td>8208</td><td>25</td><td>CGAGGCCCAGCCCGTGGACGGTGTG</td><td>564.6</td><td>-5.7</td><td>1.9</td><td>-3.0</td><td>2.7E-03</td><td>3.2E-03</td><td>BT20/MB468</td><td>NUC</td></tr>
<tr><td>>hg38_MT_+_1621_3279_RNR2_16S_50ntFlanks@1175.1216.42|TGCATTAAAAATTTCGGTTGGGGCGACCTCGGAGCAGAACCC</td><td>>hg38_MT_+_1621_3279_RNR2_16S_50ntFlanks</td><td>1175</td><td>1216</td><td>42</td><td>TGCATTAAAAATTTCGGTTGGGGCGACCTCGGAGCAGAACCC</td><td>98.6</td><td>4.6</td><td>1.5</td><td>3.0</td><td>2.7E-03</td><td>3.2E-03</td><td>MB231/MB468</td><td>MITO</td></tr>
<tr><td>>hg38_21_+_8433172_8446622_RNA45SN1_45S_50ntFlanks@10845.10863.19|ACTTCGGGATAAGGATTGG</td><td>>hg38_21_+_8433172_8446622_RNA45SN1_45S_50ntFlanks</td><td>10845</td><td>10863</td><td>19</td><td>ACTTCGGGATAAGGATTGG</td><td>317.0</td><td>-5.3</td><td>1.7</td><td>-3.2</td><td>1.4E-03</td><td>3.2E-03</td><td>BT20/MB468</td><td>MP</td></tr>
</table>

<table>
<tr><th></th><th></th><th></th><th></th><th></th><th></th><th></th><th></th><th></th><th></th><th></th><th></th><th></th><th></th></tr>
<tr><td>>MI0000071|hsa-mir-17&WithFlank&13|+|91350599|91350694@20.41.22(+1A)&[MIMAT0000070&hsa-miR-17-5p&offsets|0|-1(+1A);m-88&13|+|91350618|91350640&offsets|0|-1(+1A)]|CAAAGTGCTTACAGTGCAGGTAA</td><td>>MI0000071|hsa-mir-17&WithFlank&13|+|91350599|91350694</td><td>20</td><td>41</td><td>22(+1A)</td><td>CAAAGTGCTTACAGTGCAGGTAA</td><td>1877.5</td><td>2.9</td><td>0.9</td><td>3.0</td><td>2.3E-03</td><td>3.2E-03</td><td>MB231/MB468</td><td>MP</td></tr>
<tr><td>>MI0000264|hsa-mir-7-2&WithFlank&15|+|88611819|88611940@38.59.22;MI0000265|hsa-mir-7-3&WithFlank&19|+|4770664|4770785@37.58.22;MI0000263|hsa-mir-7-1&WithFlank&9|-|83969742|83969863@30.51.22&[MIMAT0000252&hsa-miR-7-5p&offsets|0|-2;m-400&15|+|88611856|88611879&offsets|0|-2];[MIMAT0000252_1&hsa-miR-7-5p&offsets|0|-2;m-402&19|+|4770700|4770723&offsets|0|-2];[MIMAT0000252_2&hsa-miR-7-5p&offsets|0|-2;m-401&9|-|83969811|83969834&offsets|0|-2]|TGGAAGACTAGTGATTTTGTTG</td><td>>MI0000264|hsa-mir-7-2&WithFlank&15|+|88611819|88611940</td><td>38</td><td>59</td><td>22</td><td>TGGAAGACTAGTGATTTTGTTG</td><td>805.3</td><td>3.2</td><td>1.0</td><td>3.2</td><td>1.4E-03</td><td>3.2E-03</td><td>BT20/MB468</td><td>MP</td></tr>
<tr><td>>hg38_21_+_8433172_8446622_RNA45SN1_45S_50ntFlanks@12073.12104.32|CCGAGGGGCTCTCGCTTCTGGCGCCAAGCGCC</td><td>>hg38_21_+_8433172_8446622_RNA45SN1_45S_50ntFlanks</td><td>12073</td><td>12104</td><td>32</td><td>CCGAGGGGCTCTCGCTTCTGGCGCCAAGCGCC</td><td>180.6</td><td>-3.1</td><td>1.0</td><td>-3.0</td><td>2.5E-03</td><td>3.2E-03</td><td>BT20/MB468</td><td>CYTO</td></tr>
<tr><td>>trnaMT_MetCAT_MT_+_4402_4469@43.71.29|ATGTTGGTTATACCCTTCCCGTACTACCA</td><td>>trnaMT_MetCAT_MT_+_4402_4469</td><td>43</td><td>71</td><td>29</td><td>ATGTTGGTTATACCCTTCCCGTACTACCA</td><td>343.9</td><td>4.0</td><td>1.3</td><td>3.2</td><td>1.4E-03</td><td>3.2E-03</td><td>BT20/MB468</td><td>MP</td></tr>
<tr><td>>hg38_21_+_8433172_8446622_RNA45SN1_45S_50ntFlanks@12417.12452.36|TCGATGTCGGCTCTTCCTATCATTGTGAAGCAGAAT</td><td>>hg38_21_+_8433172_8446622_RNA45SN1_45S_50ntFlanks</td><td>12417</td><td>12452</td><td>36</td><td>TCGATGTCGGCTCTTCCTATCATTGTGAAGCAGAAT</td><td>182.1</td><td>2.7</td><td>0.9</td><td>3.0</td><td>2.7E-03</td><td>3.2E-03</td><td>MB231/MB468</td><td>MITO</td></tr>
<tr><td>>hg38_21_+_8433172_8446622_RNA45SN1_45S_50ntFlanks@11795.11832.38|AACGAGATTCCCACTGTCCCTACCTACTATCCAGCGAA</td><td>>hg38_21_+_8433172_8446622_RNA45SN1_45S_50ntFlanks</td><td>11795</td><td>11832</td><td>38</td><td>AACGAGATTCCCACTGTCCCTACCTACTATCCAGCGAA</td><td>40.5</td><td>9.1</td><td>2.8</td><td>3.2</td><td>1.4E-03</td><td>3.2E-03</td><td>BT20/MB468</td><td>MP</td></tr>
<tr><td>>hg38_21_+_8433172_8446622_RNA45SN1_45S_50ntFlanks@9836.9875.40|TTGGTAAGCAGAACTGGCGCTGCGGGATGAACCGAACGCC</td><td>>hg38_21_+_8433172_8446622_RNA45SN1_45S_50ntFlanks</td><td>9836</td><td>9875</td><td>40</td><td>TTGGTAAGCAGAACTGGCGCTGCGGGATGAACCGAACGCC</td><td>37.1</td><td>9.0</td><td>2.8</td><td>3.2</td><td>1.4E-03</td><td>3.2E-03</td><td>BT20/MB468</td><td>MP</td></tr>
<tr><td>>hg38_21_+_8433172_8446622_RNA45SN1_45S_50ntFlanks@4069.4098.30|CTTTCGATGGTAGTCGCCGTGCCTACCATG</td><td>>hg38_21_+_8433172_8446622_RNA45SN1_45S_50ntFlanks</td><td>4069</td><td>4098</td><td>30</td><td>CTTTCGATGGTAGTCGCCGTGCCTACCATG</td><td>129.0</td><td>-5.0</td><td>1.6</td><td>-3.2</td><td>1.4E-03</td><td>3.2E-03</td><td>BT20/MB468</td><td>MP</td></tr>
<tr><td>>hg38_21_+_8433172_8446622_RNA45SN1_45S_50ntFlanks@474.498.25|TCTTCGTGATCGATGTGGTGACGTC</td><td>>hg38_21_+_8433172_8446622_RNA45SN1_45S_50ntFlanks</td><td>474</td><td>498</td><td>25</td><td>TCTTCGTGATCGATGTGGTGACGTC</td><td>140.1</td><td>5.7</td><td>1.9</td><td>3.0</td><td>2.3E-03</td><td>3.2E-03</td><td>MB231/MB468</td><td>MP</td></tr>
<tr><td>>hg38_21_+_8433172_8446622_RNA45SN1_45S_50ntFlanks@7979.8011.33|ACCTCAGATCAGACGTGGCGACCCGCTGAATTT</td><td>>hg38_21_+_8433172_8446622_RNA45SN1_45S_50ntFlanks</td><td>7979</td><td>8011</td><td>33</td><td>ACCTCAGATCAGACGTGGCGACCCGCTGAATTT</td><td>252.2</td><td>-2.6</td><td>0.8</td><td>-3.0</td><td>2.3E-03</td><td>3.2E-03</td><td>MB231/MB468</td><td>MP</td></tr>
<tr><td>>hg38_21_+_8433172_8446622_RNA45SN1_45S_50ntFlanks@3993.4020.28|GTGACTCTAGATAACCTCGGGCCGATCG</td><td>>hg38_21_+_8433172_8446622_RNA45SN1_45S_50ntFlanks</td><td>3993</td><td>4020</td><td>28</td><td>GTGACTCTAGATAACCTCGGGCCGATCG</td><td>330.6</td><td>-2.9</td><td>0.9</td><td>-3.1</td><td>2.1E-03</td><td>3.2E-03</td><td>BT20/MB468</td><td>MITO</td></tr>
<tr><td>>hg38_21_+_8433172_8446622_RNA45SN1_45S_50ntFlanks@9503.9533.31|GGTGAACTATGCCTGGGCAGGGCGAAGCCAG</td><td>>hg38_21_+_8433172_8446622_RNA45SN1_45S_50ntFlanks</td><td>9503</td><td>9533</td><td>31</td><td>GGTGAACTATGCCTGGGCAGGGCGAAGCCAG</td><td>44.3</td><td>-9.3</td><td>3.1</td><td>-3.0</td><td>2.3E-03</td><td>3.2E-03</td><td>MB231/MB468</td><td>MP</td></tr>
<tr><td>>MI0000082|hsa-mir-25&WithFlank&7|-|100093554|100093649@59.79.21&[MIMAT0000081&hsa-miR-25-3p&offsets|+1|0;m-23&7|-|100093571|100093592&offsets|+1|0]|ATTGCACTTGTCTCGGTCTGA</td><td>>MI0000082|hsa-mir-25&WithFlank&7|-|100093554|100093649</td><td>59</td><td>79</td><td>21</td><td>ATTGCACTTGTCTCGGTCTGA</td><td>77.5</td><td>5.4</td><td>1.7</td><td>3.2</td><td>1.4E-03</td><td>3.2E-03</td><td>BT20/MB468</td><td>MP</td></tr>
<tr><td>>hg38_21_+_8433172_8446622_RNA45SN1_45S_50ntFlanks@8358.8392.35|AAAGAACTTTGAAGAGAGAGTTCAAGAGGGCGTGA</td><td>>hg38_21_+_8433172_8446622_RNA45SN1_45S_50ntFlanks</td><td>8358</td><td>8392</td><td>35</td><td>AAAGAACTTTGAAGAGAGAGTTCAAGAGGGCGTGA</td><td>75.1</td><td>6.0</td><td>1.9</td><td>3.2</td><td>1.4E-03</td><td>3.2E-03</td><td>BT20/MB468</td><td>MP</td></tr>
<tr><td>>hg38_21_+_8433172_8446622_RNA45SN1_45S_50ntFlanks@9204.9223.20|TCGGGGCCACGCGCGCGTCC</td><td>>hg38_21_+_8433172_8446622_RNA45SN1_45S_50ntFlanks</td><td>9204</td><td>9223</td><td>20</td><td>TCGGGGCCACGCGCGCGTCC</td><td>482.7</td><td>-5.4</td><td>1.7</td><td>-3.2</td><td>1.4E-03</td><td>3.2E-03</td><td>BT20/MB468</td><td>MP</td></tr>
<tr><td>>hg38_21_+_8433172_8446622_RNA45SN1_45S_50ntFlanks@11921.11967.47|TGAGAGGTGTAGAATAAGTGGGAGGCCCCCGGCGCCCCCCCGGTGTC</td><td>>hg38_21_+_8433172_8446622_RNA45SN1_45S_50ntFlanks</td><td>11921</td><td>11967</td><td>47</td><td>TGAGAGGTGTAGAATAAGTGGGAGGCCCCCGGCGCCCCCCCGGTGTC</td><td>296.0</td><td>-4.2</td><td>1.4</td><td>-3.0</td><td>2.6E-03</td><td>3.2E-03</td><td>MB231/MB468</td><td>CYTO</td></tr>
<tr><td>>hg38_21_+_8433172_8446622_RNA45SN1_45S_50ntFlanks@4753.4780.28|AAGACGATCAGATACCGTCGTAGTTCCG</td><td>>hg38_21_+_8433172_8446622_RNA45SN1_45S_50ntFlanks</td><td>4753</td><td>4780</td><td>28</td><td>AAGACGATCAGATACCGTCGTAGTTCCG</td><td>64.1</td><td>-5.1</td><td>1.6</td><td>-3.2</td><td>1.4E-03</td><td>3.2E-03</td><td>BT20/MB468</td><td>MP</td></tr>
<tr><td>>MI0000447|hsa-mir-128-1&WithFlank&2|+|135665391|135665484@56.75.20;MI0000727|hsa-mir-128-2&WithFlank&3|+|35744470|35744565@58.77.20&[MIMAT0000424&hsa-miR-128-3p&offsets|0|-1;m-108&2|+|135665446|135665466&offsets|0|-1];[MIMAT0000424_1&hsa-miR-128-3p&offsets|0|-1;m-117&3|+|35744527|35744547&offsets|0|-1]|TCACAGTGAACCGGTCTCTT</td><td>>MI0000447|hsa-mir-128-1&WithFlank&2|+|135665391|135665484</td><td>56</td><td>75</td><td>20</td><td>TCACAGTGAACCGGTCTCTT</td><td>590.2</td><td>3.0</td><td>0.9</td><td>3.2</td><td>1.4E-03</td><td>3.2E-03</td><td>BT20/MB468</td><td>MP</td></tr>
<tr><td>>hg38_21_+_8433172_8446622_RNA45SN1_45S_50ntFlanks@12711.12747.37|GCGGAGCCTCGGTTGGCCTCGGATAGCCGGTCCCCCG</td><td>>hg38_21_+_8433172_8446622_RNA45SN1_45S_50ntFlanks</td><td>12711</td><td>12747</td><td>37</td><td>GCGGAGCCTCGGTTGGCCTCGGATAGCCGGTCCCCCG</td><td>159.5</td><td>-5.3</td><td>1.7</td><td>-3.2</td><td>1.4E-03</td><td>3.2E-03</td><td>BT20/MB468</td><td>MP</td></tr>
<tr><td>>hg38_21_+_8433172_8446622_RNA45SN1_45S_50ntFlanks@10780.10801.22|AAGGTGAACAGCCTCTGGCATG</td><td>>hg38_21_+_8433172_8446622_RNA45SN1_45S_50ntFlanks</td><td>10780</td><td>10801</td><td>22</td><td>AAGGTGAACAGCCTCTGGCATG</td><td>208.1</td><td>-1.6</td><td>0.5</td><td>-3.0</td><td>2.5E-03</td><td>3.2E-03</td><td>BT20/MB468</td><td>CYTO</td></tr>
<tr><td>>hg38_21_+_8433172_8446622_RNA45SN1_45S_50ntFlanks@4006.4024.19|ACCTCGGGCCGATCGCACG</td><td>>hg38_21_+_8433172_8446622_RNA45SN1_45S_50ntFlanks</td><td>4006</td><td>4024</td><td>19</td><td>ACCTCGGGCCGATCGCACG</td><td>521.0</td><td>-5.4</td><td>1.7</td><td>-3.2</td><td>1.4E-03</td><td>3.2E-03</td><td>BT20/MB468</td><td>MP</td></tr>
<tr><td>>hg38_21_+_8433172_8446622_RNA45SN1_45S_50ntFlanks@10351.10383.33|CGAGAACTTTGAAGGCCGAAGTGGAGAAGGGTT</td><td>>hg38_21_+_8433172_8446622_RNA45SN1_45S_50ntFlanks</td><td>10351</td><td>10383</td><td>33</td><td>CGAGAACTTTGAAGGCCGAAGTGGAGAAGGGTT</td><td>3059.6</td><td>5.7</td><td>1.6</td><td>3.5</td><td>4.8E-04</td><td>3.2E-03</td><td>BT20/MB231</td><td>MP</td></tr>
<tr><td>>hg38_21_+_8433172_8446622_RNA45SN1_45S_50ntFlanks@11670.11707.38|CCCAGTGCTCTGAATGTCAAAGTGAAGAAATTCAATGA</td><td>>hg38_21_+_8433172_8446622_RNA45SN1_45S_50ntFlanks</td><td>11670</td><td>11707</td><td>38</td><td>CCCAGTGCTCTGAATGTCAAAGTGAAGAAATTCAATGA</td><td>258.5</td><td>-2.8</td><td>0.8</td><td>-3.4</td><td>5.6E-04</td><td>3.3E-03</td><td>BT20/MB231</td><td>CYTO</td></tr>
<tr><td>>hg38_21_+_8433172_8446622_RNA45SN1_45S_50ntFlanks@6780.6803.24|CGGGGCTACGCCTGTCTGAGCGTC</td><td>>hg38_21_+_8433172_8446622_RNA45SN1_45S_50ntFlanks</td><td>6780</td><td>6803</td><td>24</td><td>CGGGGCTACGCCTGTCTGAGCGTC</td><td>383.0</td><td>-2.8</td><td>0.9</td><td>-3.2</td><td>1.4E-03</td><td>3.3E-03</td><td>BT20/MB468</td><td>MP</td></tr>
<tr><td>>trna111_HisGTG_1_-_147774845_147774916@-1A.34.35,trna118_HisGTG_1_-_145396881_145396952@-1A.34.35,trna16_HisGTG_1_+_146544773_146544844@-1A.34.35,trna1_HisGTG_15_+_45493349_45493420@-1A.34.35,trna21_HisGTG_1_+_147753471_147753542@-1A.34.35,trna33_HisGTG_6_+_27125906_27125977@-1A.34.35,trna7_HisGTG_9_-_14433938_14434009@-1A.34.35,trna8_HisGTG_15_-_45492611_45492682@-1A.34.35,trna9_HisGTG_15_-_45490804_45490875@-1A.34.35|AGCCGTGATCGTATAGTGGTTAGTACTCTGCGTTG</td><td>>trna111_HisGTG_1_-_147774845_147774916</td><td>-1A</td><td>34</td><td>35</td><td>AGCCGTGATCGTATAGTGGTTAGTACTCTGCGTTG</td><td>125.6</td><td>4.2</td><td>1.4</td><td>3.0</td><td>2.8E-03</td><td>3.3E-03</td><td>MB231/MB468</td><td>MITO</td></tr>
<tr><td>>hg38_21_+_8433172_8446622_RNA45SN1_45S_50ntFlanks@4129.4151.23|GATTCCGGAGAGGGAGCCTGAGA</td><td>>hg38_21_+_8433172_8446622_RNA45SN1_45S_50ntFlanks</td><td>4129</td><td>4151</td><td>23</td><td>GATTCCGGAGAGGGAGCCTGAGA</td><td>86.1</td><td>-3.7</td><td>1.2</td><td>-3.0</td><td>2.3E-03</td><td>3.3E-03</td><td>MB231/MB468</td><td>MP</td></tr>
<tr><td>>trnaMT_HisGTG_MT_+_12138_12206@43.72.30|ACAGAGGCTTACGACCCCTTATTTACCCCA</td><td>>trnaMT_HisGTG_MT_+_12138_12206</td><td>43</td><td>72</td><td>30</td><td>ACAGAGGCTTACGACCCCTTATTTACCCCA</td><td>58.8</td><td>3.7</td><td>1.2</td><td>3.0</td><td>2.3E-03</td><td>3.3E-03</td><td>MB231/MB468</td><td>MP</td></tr>
<tr><td>>hg38_21_+_8433172_8446622_RNA45SN1_45S_50ntFlanks@5473.5514.42|CGGCCCTGGCGGAGCGCTGAGAAGACGGTCGAACTTGACTAT</td><td>>hg38_21_+_8433172_8446622_RNA45SN1_45S_50ntFlanks</td><td>5473</td><td>5514</td><td>42</td><td>CGGCCCTGGCGGAGCGCTGAGAAGACGGTCGAACTTGACTAT</td><td>1006.3</td><td>-2.9</td><td>0.8</td><td>-3.4</td><td>5.6E-04</td><td>3.3E-03</td><td>BT20/MB231</td><td>CYTO</td></tr>
<tr><td>>hg38_21_+_8433172_8446622_RNA45SN1_45S_50ntFlanks@10393.10424.32|CAGCAGTTGAACATGGGTCAGTCGGTCCTGAG</td><td>>hg38_21_+_8433172_8446622_RNA45SN1_45S_50ntFlanks</td><td>10393</td><td>10424</td><td>32</td><td>CAGCAGTTGAACATGGGTCAGTCGGTCCTGAG</td><td>6670.2</td><td>3.7</td><td>1.1</td><td>3.2</td><td>1.4E-03</td><td>3.3E-03</td><td>BT20/MB468</td><td>MP</td></tr>
<tr><td>>hg38_21_+_8433172_8446622_RNA45SN1_45S_50ntFlanks@10832.10852.21|AGCCGGATCCGTAACTTCGGG</td><td>>hg38_21_+_8433172_8446622_RNA45SN1_45S_50ntFlanks</td><td>10832</td><td>10852</td><td>21</td><td>AGCCGGATCCGTAACTTCGGG</td><td>70.6</td><td>-5.0</td><td>1.6</td><td>-3.2</td><td>1.4E-03</td><td>3.3E-03</td><td>BT20/MB468</td><td>MP</td></tr>
<tr><td>>hg38_21_+_8433172_8446622_RNA45SN1_45S_50ntFlanks@11785.11803.19|TGAATGGATGAACGAGATT</td><td>>hg38_21_+_8433172_8446622_RNA45SN1_45S_50ntFlanks</td><td>11785</td><td>11803</td><td>19</td><td>TGAATGGATGAACGAGATT</td><td>301.6</td><td>-1.7</td><td>0.5</td><td>-3.1</td><td>2.2E-03</td><td>3.3E-03</td><td>BT20/MB468</td><td>MITO</td></tr>
<tr><td>>am_trna128_GlyGCC_6_-_27870686_27870756@1.26.26,trna133_GlyCCC_1_-_16872434_16872504@1.26.26,trna18_GlyGCC_16_+_70822597_70822667@1.26.26,trna19_GlyGCC_16_+_70823410_70823480@1.26.26,trna19_GlyGCC_2_-_157257659_157257729@1.26.26,trna24_GlyGCC_16_-_70812942_70813012@1.26.26,trna25_GlyGCC_16_-_70812114_70812184@1.26.26,trna4_GlyCCC_1_+_17188416_17188486@1.26.26,trna5_GlyGCC_17_+_8029064_8029134@1.26.26,trna68_GlyGCC_1_-_161493637_161493707@1.26.26|GCATTGGTGGTTCAGTGGTAGAATTC</td><td>>am_trna128_GlyGCC_6_-_27870686_27870756</td><td>1</td><td>26</td><td>26</td><td>GCATTGGTGGTTCAGTGGTAGAATTC</td><td>1148.4</td><td>2.3</td><td>0.8</td><td>3.0</td><td>2.6E-03</td><td>3.3E-03</td><td>MB231/MB468</td><td>CYTO</td></tr>
<tr><td>>hg38_21_+_8433172_8446622_RNA45SN1_45S_50ntFlanks@8353.8379.27|GTTGAAAAGAACTTTGAAGAGAGAGTT</td><td>>hg38_21_+_8433172_8446622_RNA45SN1_45S_50ntFlanks</td><td>8353</td><td>8379</td><td>27</td><td>GTTGAAAAGAACTTTGAAGAGAGAGTT</td><td>69.6</td><td>-5.0</td><td>1.6</td><td>-3.0</td><td>2.3E-03</td><td>3.3E-03</td><td>MB231/MB468</td><td>MP</td></tr>
<tr><td>>hg38_21_+_8433172_8446622_RNA45SN1_45S_50ntFlanks@9622.9655.34|CTAGTAGCTGGTTCCCTCCGAAGTTTCCCTCAGG</td><td>>hg38_21_+_8433172_8446622_RNA45SN1_45S_50ntFlanks</td><td>9622</td><td>9655</td><td>34</td><td>CTAGTAGCTGGTTCCCTCCGAAGTTTCCCTCAGG</td><td>172.6</td><td>-3.3</td><td>1.1</td><td>-3.1</td><td>2.2E-03</td><td>3.3E-03</td><td>BT20/MB468</td><td>MITO</td></tr>
<tr><td>>trnaMT_GluTTC_MT_-_14674_14742@27.69.43,trnalookalike8_GluTTC_5_-_93905172_93905240@27.69.43|GTTTTTCATATCATTGGTCGTGGTTGTAGTCCGTGCGAGAATA</td><td>>trnaMT_GluTTC_MT_-_14674_14742</td><td>27</td><td>69</td><td>43</td><td>GTTTTTCATATCATTGGTCGTGGTTGTAGTCCGTGCGAGAATA</td><td>555.8</td><td>-4.8</td><td>1.6</td><td>-3.0</td><td>2.8E-03</td><td>3.3E-03</td><td>MB231/MB468</td><td>NUC</td></tr>
<tr><td>>trna10_AspGTC_12_-_125424193_125424264@38.75.38,trna12_AspGTC_12_-_125411891_125411962@38.75.38,trna144_AspGTC_6_-_27551236_27551307@38.75.38,trna38_AspGTC_17_-_8125556_8125627@38.75.38,trna45_AspGTC_6_+_27447453_27447524@38.75.38,trna48_AspGTC_6_+_27471523_27471594@38.75.38,trna4_AspGTC_12_+_96429799_96429870@38.75.38,trna69_AspGTC_1_-_161440205_161440276@38.75.38,trna72_AspGTC_1_-_161432824_161432895@38.75.38,trna75_AspGTC_1_-_161425414_161425485@38.75.38,trna78_AspGTC_1_-_161418033_161418104@38.75.38,trna81_AspGTC_1_-_161410615_161410686@38.75.38|CGCGGGAGACCGGGGTTCGATTCCCCGACGGGGAGCCA</td><td>>trna10_AspGTC_12_-_125424193_125424264</td><td>38</td><td>75</td><td>38</td><td>CGCGGGAGACCGGGGTTCGATTCCCCGACGGGGAGCCA</td><td>212.0</td><td>4.2</td><td>1.3</td><td>3.2</td><td>1.4E-03</td><td>3.3E-03</td><td>BT20/MB468</td><td>MP</td></tr>
<tr><td>>MI0001448|hsa-mir-425&WithFlank&3|-|49020142|49020240@20.40.21&[MIMAT0003393&hsa-miR-425-5p&offsets|0|-2;m-95&3|-|49020200|49020221&offsets|0|-1]|AATGACACGATCACTCCCGTT</td><td>>MI0001448|hsa-mir-425&WithFlank&3|-|49020142|49020240</td><td>20</td><td>40</td><td>21</td><td>AATGACACGATCACTCCCGTT</td><td>148.9</td><td>5.7</td><td>1.8</td><td>3.2</td><td>1.4E-03</td><td>3.3E-03</td><td>BT20/MB468</td><td>MP</td></tr>
<tr><td>>hg38_21_+_8433172_8446622_RNA45SN1_45S_50ntFlanks@6651.6695.45|CGACTCTTAGCGGTGGATCACTCGGCTCGTGCGTCGATGAAGAAC</td><td>>hg38_21_+_8433172_8446622_RNA45SN1_45S_50ntFlanks</td><td>6651</td><td>6695</td><td>45</td><td>CGACTCTTAGCGGTGGATCACTCGGCTCGTGCGTCGATGAAGAAC</td><td>1917.9</td><td>3.9</td><td>1.2</td><td>3.2</td><td>1.4E-03</td><td>3.3E-03</td><td>BT20/MB468</td><td>MP</td></tr>
<tr><td>>hg38_21_+_8433172_8446622_RNA45SN1_45S_50ntFlanks@9200.9222.23|TCTCTCGGGGCCACGCGCGCGTC</td><td>>hg38_21_+_8433172_8446622_RNA45SN1_45S_50ntFlanks</td><td>9200</td><td>9222</td><td>23</td><td>TCTCTCGGGGCCACGCGCGCGTC</td><td>78.2</td><td>-5.2</td><td>1.6</td><td>-3.2</td><td>1.4E-03</td><td>3.3E-03</td><td>BT20/MB468</td><td>MP</td></tr>
</table>

<table>
<tr><td>>MI0000299|hsa-mir-222&WithFlank&X|-|45747009|45747130@75.98.24(+1A)&[MIMAT0000279&hsa-miR-222-3p&offsets|0|+3(+1A);m-140&X|-|45747033|45747056&offsets|0|0(+1A)]|AGCTACATCTGGCTACTGGGTCTCA</td><td>>MI0000299|hsa-mir-222&WithFlank&X|-|45747009|45747130</td><td>75</td><td>98</td><td>24(+1A)</td><td>AGCTACATCTGGCTACTGGGTCTCA</td><td>5740.2</td><td>-2.6</td><td>0.8</td><td>-3.4</td><td>6.3E-04</td><td>3.3E-03</td><td>BT20/MB231</td><td>MITO</td></tr>
<tr><td>>hg38_21_+_8433172_8446622_RNA45SN1_45S_50ntFlanks@4003.4039.37|ATAACCTCGGGCCGATCGCACGCCCCCCGTGGCGGCG</td><td>>hg38_21_+_8433172_8446622_RNA45SN1_45S_50ntFlanks</td><td>4003</td><td>4039</td><td>37</td><td>ATAACCTCGGGCCGATCGCACGCCCCCCGTGGCGGCG</td><td>279.3</td><td>-2.7</td><td>0.9</td><td>-3.0</td><td>2.6E-03</td><td>3.3E-03</td><td>MB231/MB468</td><td>CYTO</td></tr>
<tr><td>>hg38_MT_+_1621_3279_RNR2_16S_50ntFlanks@1369.1396.28|GATGTTGGATCAGGACATCCCGATGGTG</td><td>>hg38_MT_+_1621_3279_RNR2_16S_50ntFlanks</td><td>1369</td><td>1396</td><td>28</td><td>GATGTTGGATCAGGACATCCCGATGGTG</td><td>388.9</td><td>3.9</td><td>1.2</td><td>3.2</td><td>1.4E-03</td><td>3.3E-03</td><td>BT20/MB468</td><td>MP</td></tr>
<tr><td>>MI0000542|hsa-mir-320a&WithFlank&8|-|22244960|22245043@48.71.24(+1U)&[MIMAT0000510&hsa-miR-320a-3p&offsets|0|+2(+1U);m-60&8|-|22244975|22244996&offsets|0|+2(+1U)]|AAAAGCTGGGTTGAGAGGGCGAAAT</td><td>>MI0000542|hsa-mir-320a&WithFlank&8|-|22244960|22245043</td><td>48</td><td>71</td><td>24(+1U)</td><td>AAAAGCTGGGTTGAGAGGGCGAAAT</td><td>229.9</td><td>4.0</td><td>1.3</td><td>3.0</td><td>2.4E-03</td><td>3.3E-03</td><td>MB231/MB468</td><td>MP</td></tr>
<tr><td>>hg38_21_+_8433172_8446622_RNA45SN1_45S_50ntFlanks@5487.5519.33|CGCTGAGAAGACGGTCGAACTTGACTATCTAGA</td><td>>hg38_21_+_8433172_8446622_RNA45SN1_45S_50ntFlanks</td><td>5487</td><td>5519</td><td>33</td><td>CGCTGAGAAGACGGTCGAACTTGACTATCTAGA</td><td>202.1</td><td>-4.4</td><td>1.4</td><td>-3.2</td><td>1.4E-03</td><td>3.3E-03</td><td>BT20/MB468</td><td>MP</td></tr>
<tr><td>>trnaMT_ThrTGT_MT_+_15888_15953@36.69.34|AACCGGAGATGAAAACCTTTTTCCAAGGACACCA</td><td>>trnaMT_ThrTGT_MT_+_15888_15953</td><td>36</td><td>69</td><td>34</td><td>AACCGGAGATGAAAACCTTTTTCCAAGGACACCA</td><td>136.1</td><td>5.5</td><td>1.8</td><td>3.0</td><td>2.8E-03</td><td>3.3E-03</td><td>MB231/MB468</td><td>MITO</td></tr>
<tr><td>>hg38_21_+_8433172_8446622_RNA45SN1_45S_50ntFlanks@11677.11702.26|CTCTGAATGTCAAAGTGAAGAAATTC</td><td>>hg38_21_+_8433172_8446622_RNA45SN1_45S_50ntFlanks</td><td>11677</td><td>11702</td><td>26</td><td>CTCTGAATGTCAAAGTGAAGAAATTC</td><td>410.1</td><td>-2.2</td><td>0.7</td><td>-3.0</td><td>2.8E-03</td><td>3.3E-03</td><td>MB231/MB468</td><td>NUC</td></tr>
<tr><td>>hg38_21_+_8433172_8446622_RNA45SN1_45S_50ntFlanks@3854.3877.24|ACTGTGGTAATTCTAGAGCTAATA</td><td>>hg38_21_+_8433172_8446622_RNA45SN1_45S_50ntFlanks</td><td>3854</td><td>3877</td><td>24</td><td>ACTGTGGTAATTCTAGAGCTAATA</td><td>83.3</td><td>-5.1</td><td>1.6</td><td>-3.2</td><td>1.4E-03</td><td>3.3E-03</td><td>BT20/MB468</td><td>MP</td></tr>
<tr><td>>MI0000076|hsa-mir-20a&WithFlank&13|+|91351059|91351141@14.36.23&[MIMAT0000075&hsa-miR-20a-5p&offsets|0|0;m-96&13|+|91351072|91351094&offsets|0|0]|TAAAGTGCTTATAGTGCAGGTAG</td><td>>MI0000076|hsa-mir-20a&WithFlank&13|+|91351059|91351141</td><td>14</td><td>36</td><td>23</td><td>TAAAGTGCTTATAGTGCAGGTAG</td><td>32801.5</td><td>-4.1</td><td>1.2</td><td>-3.5</td><td>4.8E-04</td><td>3.3E-03</td><td>BT20/MB231</td><td>MP</td></tr>
<tr><td>>hg38_21_+_8433172_8446622_RNA45SN1_45S_50ntFlanks@5467.5494.28|GGCCCACGGCCCTGGCGGAGCGCTGAGA</td><td>>hg38_21_+_8433172_8446622_RNA45SN1_45S_50ntFlanks</td><td>5467</td><td>5494</td><td>28</td><td>GGCCCACGGCCCTGGCGGAGCGCTGAGA</td><td>85.5</td><td>9.8</td><td>2.8</td><td>3.5</td><td>4.8E-04</td><td>3.3E-03</td><td>BT20/MB231</td><td>MP</td></tr>
<tr><td>>hg38_21_+_8433172_8446622_RNA45SN1_45S_50ntFlanks@11535.11552.18|TTCCGCGCGGCGCCTCGC</td><td>>hg38_21_+_8433172_8446622_RNA45SN1_45S_50ntFlanks</td><td>11535</td><td>11552</td><td>18</td><td>TTCCGCGCGGCGCCTCGC</td><td>144.7</td><td>-4.3</td><td>1.4</td><td>-3.1</td><td>2.2E-03</td><td>3.3E-03</td><td>BT20/MB468</td><td>MITO</td></tr>
<tr><td>>MI0000440|hsa-mir-27b&WithFlank&9|+|95085439|95085547@67.85.19(+1U)&[MIMAT0000419&hsa-miR-27b-3p&offsets|0|-2(+1U);m-39&9|+|95085505|95085525&offsets|0|-2(+1U)]|TTCACAGTGGCTAAGTTCTT</td><td>>MI0000440|hsa-mir-27b&WithFlank&9|+|95085439|95085547</td><td>67</td><td>85</td><td>19(+1U)</td><td>TTCACAGTGGCTAAGTTCTT</td><td>74.2</td><td>4.0</td><td>1.3</td><td>3.0</td><td>2.4E-03</td><td>3.3E-03</td><td>MB231/MB468</td><td>MP</td></tr>
<tr><td>>hg38_21_+_8433172_8446622_RNA45SN1_45S_50ntFlanks@12406.12429.24|TTTTTGATCCTTCGATGTCGGCTC</td><td>>hg38_21_+_8433172_8446622_RNA45SN1_45S_50ntFlanks</td><td>12406</td><td>12429</td><td>24</td><td>TTTTTGATCCTTCGATGTCGGCTC</td><td>229.7</td><td>-1.7</td><td>0.6</td><td>-3.0</td><td>2.6E-03</td><td>3.3E-03</td><td>MB231/MB468</td><td>CYTO</td></tr>
<tr><td>>hg38_21_+_8433172_8446622_RNA45SN1_45S_50ntFlanks@10411.10456.46|CAGTCGGTCCTGAGAGATGGGCGAGCGCCGTTCCGAAGGGACGGGC</td><td>>hg38_21_+_8433172_8446622_RNA45SN1_45S_50ntFlanks</td><td>10411</td><td>10456</td><td>46</td><td>CAGTCGGTCCTGAGAGATGGGCGAGCGCCGTTCCGAAGGGACGGGC</td><td>203.8</td><td>-4.5</td><td>1.4</td><td>-3.2</td><td>1.4E-03</td><td>3.3E-03</td><td>BT20/MB468</td><td>MP</td></tr>
<tr><td>>hg38_21_+_8433172_8446622_RNA45SN1_45S_50ntFlanks@12336.12365.30|CAGGAGGTGTCAGAAAAGTTACCACAGGGA</td><td>>hg38_21_+_8433172_8446622_RNA45SN1_45S_50ntFlanks</td><td>12336</td><td>12365</td><td>30</td><td>CAGGAGGTGTCAGAAAAGTTACCACAGGGA</td><td>71.1</td><td>-5.0</td><td>1.6</td><td>-3.2</td><td>1.4E-03</td><td>3.3E-03</td><td>BT20/MB468</td><td>MP</td></tr>
<tr><td>>hg38_21_+_8433172_8446622_RNA45SN1_45S_50ntFlanks@6651.6673.23|CGACTCTTAGCGGTGGATCACTC</td><td>>hg38_21_+_8433172_8446622_RNA45SN1_45S_50ntFlanks</td><td>6651</td><td>6673</td><td>23</td><td>CGACTCTTAGCGGTGGATCACTC</td><td>6011.4</td><td>2.7</td><td>0.9</td><td>3.0</td><td>2.8E-03</td><td>3.3E-03</td><td>MB231/MB468</td><td>MITO</td></tr>
<tr><td>>MI0001445|hsa-mir-423&WithFlank&17|+|30117073|30117178@58.79.22&[MIMAT0001340&hsa-miR-423-3p&offsets|-1|-2;m-100&17|+|30117131|30117153&offsets|-1|-2]|AAGCTCGGTCTGAGGCCCCTCA</td><td>>MI0001445|hsa-mir-423&WithFlank&17|+|30117073|30117178</td><td>58</td><td>79</td><td>22</td><td>AAGCTCGGTCTGAGGCCCCTCA</td><td>80.9</td><td>1.8</td><td>0.6</td><td>3.0</td><td>2.8E-03</td><td>3.3E-03</td><td>BT20/MB468</td><td>NUC</td></tr>
<tr><td>>hg38_21_+_8433172_8446622_RNA45SN1_45S_50ntFlanks@4722.4759.38|TCATTAATCAAGAACGAAAGTCGGAGGTTCGAAGACGA</td><td>>hg38_21_+_8433172_8446622_RNA45SN1_45S_50ntFlanks</td><td>4722</td><td>4759</td><td>38</td><td>TCATTAATCAAGAACGAAAGTCGGAGGTTCGAAGACGA</td><td>440.0</td><td>-2.8</td><td>0.8</td><td>-3.4</td><td>5.7E-04</td><td>3.3E-03</td><td>BT20/MB231</td><td>CYTO</td></tr>
<tr><td>>am_trnaMT_ValTAC_MT_+_1602_1670@29.67.39|ACTTACACTTAGGAGATTTCAACTTAACTTGACCGCTCT</td><td>>am_trnaMT_ValTAC_MT_+_1602_1670</td><td>29</td><td>67</td><td>39</td><td>ACTTACACTTAGGAGATTTCAACTTAACTTGACCGCTCT</td><td>356.9</td><td>6.1</td><td>1.9</td><td>3.2</td><td>1.4E-03</td><td>3.3E-03</td><td>BT20/MB468</td><td>MP</td></tr>
<tr><td>>hg38_1_-_228634819_228635039_RNA5S12_5S_50ntFlanks@149.170.22|GAATACCGGGTGCTGTAGGCTT</td><td>>hg38_1_-_228634819_228635039_RNA5S12_5S_50ntFlanks</td><td>149</td><td>170</td><td>22</td><td>GAATACCGGGTGCTGTAGGCTT</td><td>1058.3</td><td>-2.0</td><td>0.7</td><td>-3.0</td><td>2.8E-03</td><td>3.3E-03</td><td>MB231/MB468</td><td>MITO</td></tr>
<tr><td>>hg38_MT_+_1621_3279_RNR2_16S_50ntFlanks@1431.1458.28|AAAGTCCTACGTGATCTGAGTTCAGACC</td><td>>hg38_MT_+_1621_3279_RNR2_16S_50ntFlanks</td><td>1431</td><td>1458</td><td>28</td><td>AAAGTCCTACGTGATCTGAGTTCAGACC</td><td>126.2</td><td>3.8</td><td>1.3</td><td>3.0</td><td>2.8E-03</td><td>3.3E-03</td><td>MB231/MB468</td><td>MITO</td></tr>
<tr><td>>trnaMT_ValTAC_MT_+_1602_1670@32.72.41|TACACTTAGGAGATTTCAACTTAACTTGACCGCTCTGACCA</td><td>>trnaMT_ValTAC_MT_+_1602_1670</td><td>32</td><td>72</td><td>41</td><td>TACACTTAGGAGATTTCAACTTAACTTGACCGCTCTGACCA</td><td>330.5</td><td>3.6</td><td>1.2</td><td>3.0</td><td>2.8E-03</td><td>3.3E-03</td><td>MB231/MB468</td><td>MITO</td></tr>
<tr><td>>hg38_MT_+_1621_3279_RNR2_16S_50ntFlanks@644.676.33|AATTGGACCAATCTATCACCCTATAGAAGAACT</td><td>>hg38_MT_+_1621_3279_RNR2_16S_50ntFlanks</td><td>644</td><td>676</td><td>33</td><td>AATTGGACCAATCTATCACCCTATAGAAGAACT</td><td>165.9</td><td>3.6</td><td>1.2</td><td>3.0</td><td>2.4E-03</td><td>3.3E-03</td><td>MB231/MB468</td><td>MP</td></tr>
<tr><td>>hg38_21_+_8433172_8446622_RNA45SN1_45S_50ntFlanks@3777.3799.23|CGGTACAGTGAAACTGCGAATGG</td><td>>hg38_21_+_8433172_8446622_RNA45SN1_45S_50ntFlanks</td><td>3777</td><td>3799</td><td>23</td><td>CGGTACAGTGAAACTGCGAATGG</td><td>205.2</td><td>-5.1</td><td>1.6</td><td>-3.2</td><td>1.4E-03</td><td>3.3E-03</td><td>BT20/MB468</td><td>MP</td></tr>
<tr><td>>trna10_AspGTC_12_-_125424193_125424264@37.70.34,trna12_AspGTC_12_-_125411891_125411962@37.70.34,trna144_AspGTC_6_-_27551236_27551307@37.70.34,trna38_AspGTC_17_-_8125556_8125627@37.70.34,trna45_AspGTC_6_+_27447453_27447524@37.70.34,trna48_AspGTC_6_+_27471523_27471594@37.70.34,trna4_AspGTC_12_+_96429799_96429870@37.70.34,trna69_AspGTC_1_-_161440205_161440276@37.70.34,trna72_AspGTC_1_-_161432824_161432895@37.70.34,trna75_AspGTC_1_-_161425414_161425485@37.70.34,trna78_AspGTC_1_-_161418033_161418104@37.70.34,trna81_AspGTC_1_-_161410615_161410686@37.70.34|ACGCGGGAGACCGGGGTTCGATTCCCCGACGGGG</td><td>>trna10_AspGTC_12_-_125424193_125424264</td><td>37</td><td>70</td><td>34</td><td>ACGCGGGAGACCGGGGTTCGATTCCCCGACGGGG</td><td>443.5</td><td>-1.3</td><td>0.4</td><td>-3.0</td><td>2.6E-03</td><td>3.3E-03</td><td>BT20/MB468</td><td>CYTO</td></tr>
<tr><td>>hg38_21_+_8433172_8446622_RNA45SN1_45S_50ntFlanks@10905.10939.35|CTGGGCGCGCGCCGCGGCTGGACGAGGCGCCGCCG</td><td>>hg38_21_+_8433172_8446622_RNA45SN1_45S_50ntFlanks</td><td>10905</td><td>10939</td><td>35</td><td>CTGGGCGCGCGCCGCGGCTGGACGAGGCGCCGCCG</td><td>219.2</td><td>-4.8</td><td>1.6</td><td>-3.0</td><td>2.6E-03</td><td>3.3E-03</td><td>BT20/MB468</td><td>CYTO</td></tr>
<tr><td>>am_trna10_ValCAC_5_-_180649395_180649467@1.19.19,trna12_ValAAC_5_-_180645270_180645342@1.19.19,trna132_ValAAC_6_-_27721179_27721251@1.19.19,trna133_ValCAC_6_-_27696327_27696399@1.19.19,trna136_ValAAC_6_-_27648885_27648957@1.19.19,trna139_ValAAC_6_-_27618707_27618779@1.19.19,trna15_ValAAC_5_-_180615416_180615488@1.19.19,trna18_ValCAC_5_-_180529253_180529325@1.19.19,trna2_ValAAC_3_+_169490018_169490090@1.19.19,trna2_ValCAC_5_+_180524070_180524142@1.19.19,trna37_ValAAC_6_+_27203288_27203360@1.19.19,trna4_ValAAC_5_+_180591154_180591226@1.19.19,trna5_ValAAC_5_+_180596610_180596682@1.19.19,trna6_ValCAC_5_+_180600650_180600722@1.19.19,trna85_ValCAC_1_-_161369490_161369562@1.19.19,trna90_ValCAC_1_-_149684088_149684161@1.19.19,trna98_ValCAC_1_-_149298555_149298627@1.19.19,trna9_ValCAC_6_+_26538282_26538354@1.19.19|GTTTCCGTAGTGTAGTGGT</td><td>>am_trna10_ValCAC_5_-_180649395_180649467</td><td>1</td><td>19</td><td>19</td><td>GTTTCCGTAGTGTAGTGGT</td><td>212.7</td><td>3.9</td><td>1.3</td><td>3.0</td><td>2.4E-03</td><td>3.3E-03</td><td>MB231/MB468</td><td>MP</td></tr>
<tr><td>>hg38_21_+_8433172_8446622_RNA45SN1_45S_50ntFlanks@11569.11588.20|CAGCCGACTTAGAACTGGTG</td><td>>hg38_21_+_8433172_8446622_RNA45SN1_45S_50ntFlanks</td><td>11569</td><td>11588</td><td>20</td><td>CAGCCGACTTAGAACTGGTG</td><td>400.2</td><td>-5.6</td><td>1.7</td><td>-3.2</td><td>1.4E-03</td><td>3.3E-03</td><td>BT20/MB468</td><td>MP</td></tr>
<tr><td>>MI0000650|hsa-mir-200c&WithFlank&12|+|6963693|6963772@50.70.21(+1C)&[MIMAT0000617&hsa-miR-200c-3p&offsets|0|-2(+1C);m-54&12|+|6963742|6963764&offsets|0|-2(+1C)]|TAATACTGCCGGGTAATGATGC</td><td>>MI0000650|hsa-mir-200c&WithFlank&12|+|6963693|6963772</td><td>50</td><td>70</td><td>21(+1C)</td><td>TAATACTGCCGGGTAATGATGC</td><td>179.4</td><td>5.6</td><td>1.7</td><td>3.2</td><td>1.4E-03</td><td>3.3E-03</td><td>BT20/MB468</td><td>MP</td></tr>
<tr><td>>trna10_LysCTT_16_+_3241501_3241573@1.34.34,trna13_LysCTT_14_-_58706613_58706685@1.34.34,trna2_LysCTT_15_+_79152904_79152976@1.34.34|GCCCGGCTAGCTCAGTCGGTAGAGCATGGGACTC</td><td>>trna10_LysCTT_16_+_3241501_3241573</td><td>1</td><td>34</td><td>34</td><td>GCCCGGCTAGCTCAGTCGGTAGAGCATGGGACTC</td><td>5942.1</td><td>4.9</td><td>1.5</td><td>3.2</td><td>1.4E-03</td><td>3.3E-03</td><td>BT20/MB468</td><td>MP</td></tr>
<tr><td>>hg38_21_+_8433172_8446622_RNA45SN1_45S_50ntFlanks@7984.8002.19|AGATCAGACGTGGCGACCC</td><td>>hg38_21_+_8433172_8446622_RNA45SN1_45S_50ntFlanks</td><td>7984</td><td>8002</td><td>19</td><td>AGATCAGACGTGGCGACCC</td><td>289.5</td><td>6.6</td><td>1.9</td><td>3.5</td><td>4.9E-04</td><td>3.3E-03</td><td>BT20/MB231</td><td>MP</td></tr>
<tr><td>>hg38_21_+_8433172_8446622_RNA45SN1_45S_50ntFlanks@9593.9612.20|GTATAGGGGCGAAAGACTAA</td><td>>hg38_21_+_8433172_8446622_RNA45SN1_45S_50ntFlanks</td><td>9593</td><td>9612</td><td>20</td><td>GTATAGGGGCGAAAGACTAA</td><td>82.4</td><td>9.7</td><td>2.8</td><td>3.5</td><td>4.9E-04</td><td>3.3E-03</td><td>BT20/MB231</td><td>MP</td></tr>
<tr><td>>MI0000088|hsa-mir-30a&WithFlank&6|-|71403545|71403627@53.72.20&[MIMAT0000088&hsa-miR-30a-3p&offsets|0|-2;m-38&6|-|71403554|71403575&offsets|0|-2]|CTTTCAGTCGGATGTTTGCA</td><td>>MI0000088|hsa-mir-30a&WithFlank&6|-|71403545|71403627</td><td>53</td><td>72</td><td>20</td><td>CTTTCAGTCGGATGTTTGCA</td><td>503.1</td><td>-3.3</td><td>1.0</td><td>-3.5</td><td>4.9E-04</td><td>3.3E-03</td><td>BT20/MB231</td><td>MP</td></tr>
<tr><td>>hg38_21_+_8433172_8446622_RNA45SN1_45S_50ntFlanks@11312.11344.33|CCCCCACCCCACGTCTCGTCGCGCGCGCGTCCG</td><td>>hg38_21_+_8433172_8446622_RNA45SN1_45S_50ntFlanks</td><td>11312</td><td>11344</td><td>33</td><td>CCCCCACCCCACGTCTCGTCGCGCGCGCGTCCG</td><td>111.1</td><td>-5.4</td><td>1.7</td><td>-3.2</td><td>1.4E-03</td><td>3.3E-03</td><td>BT20/MB468</td><td>MP</td></tr>
<tr><td>>hg38_21_+_8433172_8446622_RNA45SN1_45S_50ntFlanks@12636.12660.25|AGCTACCATCTGTGGGATTATGACT</td><td>>hg38_21_+_8433172_8446622_RNA45SN1_45S_50ntFlanks</td><td>12636</td><td>12660</td><td>25</td><td>AGCTACCATCTGTGGGATTATGACT</td><td>169.4</td><td>-1.9</td><td>0.6</td><td>-3.1</td><td>2.2E-03</td><td>3.3E-03</td><td>BT20/MB468</td><td>MITO</td></tr>
<tr><td>>am_trnaMT_ValTAC_MT_+_1602_1670@30.54.25|CTTACACTTAGGAGATTTCAACTTA</td><td>>am_trnaMT_ValTAC_MT_+_1602_1670</td><td>30</td><td>54</td><td>25</td><td>CTTACACTTAGGAGATTTCAACTTA</td><td>102.0</td><td>-4.9</td><td>1.6</td><td>-3.0</td><td>2.4E-03</td><td>3.3E-03</td><td>MB231/MB468</td><td>MP</td></tr>
<tr><td>>hg38_21_+_8433172_8446622_RNA45SN1_45S_50ntFlanks@9454.9503.50|CCGCGCCGGGGAGGTGGAGCACGAGCGCACGTGTTAGGACCCGAAAGATG</td><td>>hg38_21_+_8433172_8446622_RNA45SN1_45S_50ntFlanks</td><td>9454</td><td>9503</td><td>50</td><td>CCGCGCCGGGGAGGTGGAGCACGAGCGCACGTGTTAGGACCCGAAAGATG</td><td>309.7</td><td>-4.3</td><td>1.4</td><td>-3.0</td><td>2.7E-03</td><td>3.3E-03</td><td>MB231/MB468</td><td>CYTO</td></tr>
<tr><td>>hg38_21_+_8433172_8446622_RNA45SN1_45S_50ntFlanks@11668.11702.35|TGCCCAGTGCTCTGAATGTCAAAGTGAAGAAATTC</td><td>>hg38_21_+_8433172_8446622_RNA45SN1_45S_50ntFlanks</td><td>11668</td><td>11702</td><td>35</td><td>TGCCCAGTGCTCTGAATGTCAAAGTGAAGAAATTC</td><td>1506.5</td><td>-2.3</td><td>0.8</td><td>-3.0</td><td>2.8E-03</td><td>3.3E-03</td><td>MB231/MB468</td><td>MITO</td></tr>
<tr><td>>hg38_21_+_8433172_8446622_RNA45SN1_45S_50ntFlanks@12927.12956.30|CCTGGCGCTAAACCATTCGTAGACGACCTG</td><td>>hg38_21_+_8433172_8446622_RNA45SN1_45S_50ntFlanks</td><td>12927</td><td>12956</td><td>30</td><td>CCTGGCGCTAAACCATTCGTAGACGACCTG</td><td>367.9</td><td>-5.3</td><td>1.6</td><td>-3.2</td><td>1.4E-03</td><td>3.3E-03</td><td>BT20/MB468</td><td>MP</td></tr>
<tr><td>>hg38_21_+_8433172_8446622_RNA45SN1_45S_50ntFlanks@10468.10488.21|TTGCCCTCGGCCGATCGAAAG</td><td>>hg38_21_+_8433172_8446622_RNA45SN1_45S_50ntFlanks</td><td>10468</td><td>10488</td><td>21</td><td>TTGCCCTCGGCCGATCGAAAG</td><td>895.7</td><td>-2.7</td><td>0.9</td><td>-3.0</td><td>2.8E-03</td><td>3.3E-03</td><td>MB231/MB468</td><td>NUC</td></tr>
<tr><td>>hg38_21_+_8433172_8446622_RNA45SN1_45S_50ntFlanks@10352.10369.18|GAGAACTTTGAAGGCCGA</td><td>>hg38_21_+_8433172_8446622_RNA45SN1_45S_50ntFlanks</td><td>10352</td><td>10369</td><td>18</td><td>GAGAACTTTGAAGGCCGA</td><td>279.0</td><td>1.9</td><td>0.6</td><td>3.0</td><td>2.4E-03</td><td>3.3E-03</td><td>MB231/MB468</td><td>MP</td></tr>
<tr><td>>hg38_21_+_8433172_8446622_RNA45SN1_45S_50ntFlanks@11843.11860.18|AGGGAACGGGCTTGGCGG</td><td>>hg38_21_+_8433172_8446622_RNA45SN1_45S_50ntFlanks</td><td>11843</td><td>11860</td><td>18</td><td>AGGGAACGGGCTTGGCGG</td><td>771.0</td><td>-3.2</td><td>1.0</td><td>-3.2</td><td>1.4E-03</td><td>3.3E-03</td><td>BT20/MB468</td><td>MP</td></tr>
<tr><td>>hg38_21_+_8433172_8446622_RNA45SN1_45S_50ntFlanks@6778.6806.29|CCCGGGGCTACGCCTGTCTGAGCGTCGCT</td><td>>hg38_21_+_8433172_8446622_RNA45SN1_45S_50ntFlanks</td><td>6778</td><td>6806</td><td>29</td><td>CCCGGGGCTACGCCTGTCTGAGCGTCGCT</td><td>493.9</td><td>2.9</td><td>0.9</td><td>3.2</td><td>1.4E-03</td><td>3.3E-03</td><td>BT20/MB468</td><td>MP</td></tr>
<tr><td>>trnaMT_ThrTGT_MT_+_15888_15953@38.69.32|CCGGAGATGAAAACCTTTTTCCAAGGACACCA</td><td>>trnaMT_ThrTGT_MT_+_15888_15953</td><td>38</td><td>69</td><td>32</td><td>CCGGAGATGAAAACCTTTTTCCAAGGACACCA</td><td>138.0</td><td>4.9</td><td>1.7</td><td>3.0</td><td>2.8E-03</td><td>3.3E-03</td><td>MB231/MB468</td><td>MITO</td></tr>
</table>

<table>
<tr><td>>hg38_21_+_8433172_8446622_RNA45SN1_45S_50ntFlanks@6745.6771.27|ACACTTCGAACGCACTTGCGGCCCCGG</td><td>>hg38_21_+_8433172_8446622_RNA45SN1_45S_50ntFlanks</td><td>6745</td><td>6771</td><td>27</td><td>ACACTTCGAACGCACTTGCGGCCCCGG</td><td>4735.3</td><td>-5.2</td><td>1.6</td><td>-3.2</td><td>1.4E-03</td><td>3.3E-03</td><td>BT20/MB468</td><td>MP</td></tr>
<tr><td>>MI0000466|hsa-mir-9-1&WithFlank&1|-|156420335|156420435@22.44.23;MI0000468|hsa-mir-9-3&WithFlank&15|+|89368011|89368112@22.44.23;MI0000467|hsa-mir-9-2&WithFlank&5|-|88666847|88666945@22.44.23&[MIMAT0000441&hsa-miR-9-5p&offsets|0|0;m-104&1|-|156420392|156420414&offsets|0|0];[MIMAT0000441_1&hsa-miR-9-5p&offsets|0|0;m-105&15|+|89368032|89368054&offsets|0|0];[MIMAT0000441_2&hsa-miR-9-5p&offsets|0|0;m-104&5|-|88666902|88666924&offsets|0|0]|TCTTTGGTTATCTAGCTGTATGA</td><td>>MI0000466|hsa-mir-9-1&WithFlank&1|-|156420335|156420435</td><td>22</td><td>44</td><td>23</td><td>TCTTTGGTTATCTAGCTGTATGA</td><td>1834.3</td><td>2.7</td><td>0.8</td><td>3.5</td><td>5.3E-04</td><td>3.3E-03</td><td>BT20/MB231</td><td>NUC</td></tr>
<tr><td>>hg38_21_+_8433172_8446622_RNA45SN1_45S_50ntFlanks@10391.10418.28|AACAGCAGTTGAACATGGGTCAGTCGGT</td><td>>hg38_21_+_8433172_8446622_RNA45SN1_45S_50ntFlanks</td><td>10391</td><td>10418</td><td>28</td><td>AACAGCAGTTGAACATGGGTCAGTCGGT</td><td>154.5</td><td>-2.9</td><td>1.0</td><td>-3.0</td><td>2.7E-03</td><td>3.3E-03</td><td>MB231/MB468</td><td>CYTO</td></tr>
<tr><td>>hg38_21_+_8433172_8446622_RNA45SN1_45S_50ntFlanks@6662.6685.24|GGTGGATCACTCGGCTCGTGCGTC</td><td>>hg38_21_+_8433172_8446622_RNA45SN1_45S_50ntFlanks</td><td>6662</td><td>6685</td><td>24</td><td>GGTGGATCACTCGGCTCGTGCGTC</td><td>178.4</td><td>5.3</td><td>1.7</td><td>3.2</td><td>1.4E-03</td><td>3.3E-03</td><td>BT20/MB468</td><td>MP</td></tr>
<tr><td>>hg38_21_+_8433172_8446622_RNA45SN1_45S_50ntFlanks@10344.10369.26|ATTCAAACGAGAACTTTGAAGGCCGA</td><td>>hg38_21_+_8433172_8446622_RNA45SN1_45S_50ntFlanks</td><td>10344</td><td>10369</td><td>26</td><td>ATTCAAACGAGAACTTTGAAGGCCGA</td><td>167.1</td><td>5.3</td><td>1.7</td><td>3.2</td><td>1.4E-03</td><td>3.3E-03</td><td>BT20/MB468</td><td>MP</td></tr>
<tr><td>>MI0000811|hsa-mir-148b&WithFlank&12|+|54337210|54337320@69.90.22(+2U)&[MIMAT0000759&hsa-miR-148b-3p&offsets|0|0(+2U);m-77&12|+|54337278|54337299&offsets|0|0(+2U)]|TCAGTGCATCACAGAACTTTGTTT</td><td>>MI0000811|hsa-mir-148b&WithFlank&12|+|54337210|54337320</td><td>69</td><td>90</td><td>22(+2U)</td><td>TCAGTGCATCACAGAACTTTGTTT</td><td>718.3</td><td>-4.0</td><td>1.1</td><td>-3.5</td><td>4.9E-04</td><td>3.3E-03</td><td>BT20/MB231</td><td>MP</td></tr>
<tr><td>>hg38_21_+_8433172_8446622_RNA45SN1_45S_50ntFlanks@6779.6803.25|CCGGGGCTACGCCTGTCTGAGCGTC</td><td>>hg38_21_+_8433172_8446622_RNA45SN1_45S_50ntFlanks</td><td>6779</td><td>6803</td><td>25</td><td>CCGGGGCTACGCCTGTCTGAGCGTC</td><td>350.8</td><td>-1.7</td><td>0.6</td><td>-3.0</td><td>2.7E-03</td><td>3.3E-03</td><td>MB231/MB468</td><td>CYTO</td></tr>
<tr><td>>hg38_21_+_8433172_8446622_RNA45SN1_45S_50ntFlanks@11572.11593.22|CCGACTTAGAACTGGTGCGGAC</td><td>>hg38_21_+_8433172_8446622_RNA45SN1_45S_50ntFlanks</td><td>11572</td><td>11593</td><td>22</td><td>CCGACTTAGAACTGGTGCGGAC</td><td>392.1</td><td>-2.4</td><td>0.8</td><td>-3.1</td><td>2.2E-03</td><td>3.3E-03</td><td>BT20/MB468</td><td>MITO</td></tr>
<tr><td>>trnaMT_GluTTC_MT_-_14674_14742@39.63.25,trnalookalike8_GluTTC_5_-_93905172_93905240@39.63.25|ATTGGTCGTGGTTGTAGTCCGTGCG</td><td>>trnaMT_GluTTC_MT_-_14674_14742</td><td>39</td><td>63</td><td>25</td><td>ATTGGTCGTGGTTGTAGTCCGTGCG</td><td>237.7</td><td>2.8</td><td>0.9</td><td>3.0</td><td>2.4E-03</td><td>3.3E-03</td><td>MB231/MB468</td><td>MP</td></tr>
<tr><td>>trnaMT_ArgTCG_MT_+_10405_10469@28.68.41|ATTTCGACTCATTAAATTATGATAATCATATTTACCAACCA</td><td>>trnaMT_ArgTCG_MT_+_10405_10469</td><td>28</td><td>68</td><td>41</td><td>ATTTCGACTCATTAAATTATGATAATCATATTTACCAACCA</td><td>345.6</td><td>-3.3</td><td>1.1</td><td>-3.0</td><td>2.8E-03</td><td>3.3E-03</td><td>MB231/MB468</td><td>MITO</td></tr>
<tr><td>>hg38_MT_+_1621_3279_RNR2_16S_50ntFlanks@1368.1401.34|CGATGTTGGATCAGGACATCCCGATGGTGCAGCC</td><td>>hg38_MT_+_1621_3279_RNR2_16S_50ntFlanks</td><td>1368</td><td>1401</td><td>34</td><td>CGATGTTGGATCAGGACATCCCGATGGTGCAGCC</td><td>166.2</td><td>5.5</td><td>1.7</td><td>3.2</td><td>1.4E-03</td><td>3.3E-03</td><td>BT20/MB468</td><td>MP</td></tr>
<tr><td>>hg38_21_+_8433172_8446622_RNA45SN1_45S_50ntFlanks@4415.4438.24|CGCGAGGCGAGCCACCGCCCGTCC</td><td>>hg38_21_+_8433172_8446622_RNA45SN1_45S_50ntFlanks</td><td>4415</td><td>4438</td><td>24</td><td>CGCGAGGCGAGCCACCGCCCGTCC</td><td>43.1</td><td>-8.5</td><td>2.7</td><td>-3.2</td><td>1.4E-03</td><td>3.3E-03</td><td>BT20/MB468</td><td>MP</td></tr>
<tr><td>>hg38_21_+_8433172_8446622_RNA45SN1_45S_50ntFlanks@3769.3794.26|CGCACGGCCGGTACAGTGAAACTGCG</td><td>>hg38_21_+_8433172_8446622_RNA45SN1_45S_50ntFlanks</td><td>3769</td><td>3794</td><td>26</td><td>CGCACGGCCGGTACAGTGAAACTGCG</td><td>298.2</td><td>-3.9</td><td>1.3</td><td>-3.1</td><td>2.2E-03</td><td>3.3E-03</td><td>BT20/MB468</td><td>MITO</td></tr>
<tr><td>>hg38_21_+_8433172_8446622_RNA45SN1_45S_50ntFlanks@7979.8003.25|ACCTCAGATCAGACGTGGCGACCCG</td><td>>hg38_21_+_8433172_8446622_RNA45SN1_45S_50ntFlanks</td><td>7979</td><td>8003</td><td>25</td><td>ACCTCAGATCAGACGTGGCGACCCG</td><td>393.8</td><td>-2.0</td><td>0.7</td><td>-3.1</td><td>2.2E-03</td><td>3.3E-03</td><td>BT20/MB468</td><td>MITO</td></tr>
<tr><td>>hg38_21_+_8433172_8446622_RNA45SN1_45S_50ntFlanks@6649.6683.35|TACGACTCTTAGCGGTGGATCACTCGGCTCGTGCG</td><td>>hg38_21_+_8433172_8446622_RNA45SN1_45S_50ntFlanks</td><td>6649</td><td>6683</td><td>35</td><td>TACGACTCTTAGCGGTGGATCACTCGGCTCGTGCG</td><td>4453.1</td><td>-2.7</td><td>0.8</td><td>-3.5</td><td>5.4E-04</td><td>3.3E-03</td><td>BT20/MB231</td><td>NUC</td></tr>
<tr><td>>hg38_21_+_8433172_8446622_RNA45SN1_45S_50ntFlanks@5525.5550.26|TAAAAGTCGTAACAAGGTTTCCGTAG</td><td>>hg38_21_+_8433172_8446622_RNA45SN1_45S_50ntFlanks</td><td>5525</td><td>5550</td><td>26</td><td>TAAAAGTCGTAACAAGGTTTCCGTAG</td><td>187.1</td><td>-5.1</td><td>1.6</td><td>-3.2</td><td>1.4E-03</td><td>3.3E-03</td><td>BT20/MB468</td><td>MP</td></tr>
<tr><td>>MI0000746|hsa-mir-99b&WithFlank&19|+|51692606|51692687@51.72.22&[MIMAT0004678&hsa-miR-99b-3p&offsets|0|0;m-151&19|+|51692656|51692678&offsets|0|-1]|CAAGCTCGTGTCTGTGGGTCCG</td><td>>MI0000746|hsa-mir-99b&WithFlank&19|+|51692606|51692687</td><td>51</td><td>72</td><td>22</td><td>CAAGCTCGTGTCTGTGGGTCCG</td><td>300.1</td><td>2.6</td><td>0.9</td><td>3.0</td><td>2.8E-03</td><td>3.3E-03</td><td>MB231/MB468</td><td>MITO</td></tr>
<tr><td>>MI0000746|hsa-mir-99b&WithFlank&19|+|51692606|51692687@13.33.21(+1U)&[MIMAT0000689&hsa-miR-99b-5p&offsets|0|-1(+1U);m-10&19|+|51692618|51692639&offsets|0|-1(+1U)]|CACCCGTAGAACCGACCTTGCT</td><td>>MI0000746|hsa-mir-99b&WithFlank&19|+|51692606|51692687</td><td>13</td><td>33</td><td>21(+1U)</td><td>CACCCGTAGAACCGACCTTGCT</td><td>230.5</td><td>2.6</td><td>0.8</td><td>3.0</td><td>2.4E-03</td><td>3.3E-03</td><td>MB231/MB468</td><td>MP</td></tr>
<tr><td>>hg38_21_+_8433172_8446622_RNA45SN1_45S_50ntFlanks@8175.8216.42|CCTTCTGATCGAGGCCCAGCCCGTGGACGGTGTGAGGCCGGT</td><td>>hg38_21_+_8433172_8446622_RNA45SN1_45S_50ntFlanks</td><td>8175</td><td>8216</td><td>42</td><td>CCTTCTGATCGAGGCCCAGCCCGTGGACGGTGTGAGGCCGGT</td><td>551.9</td><td>4.1</td><td>1.4</td><td>3.0</td><td>2.8E-03</td><td>3.3E-03</td><td>MB231/MB468</td><td>MITO</td></tr>
<tr><td>>hg38_21_+_8433172_8446622_RNA45SN1_45S_50ntFlanks@12002.12029.28|TGCGGGCCGCCGGTGAAATACCACTACT</td><td>>hg38_21_+_8433172_8446622_RNA45SN1_45S_50ntFlanks</td><td>12002</td><td>12029</td><td>28</td><td>TGCGGGCCGCCGGTGAAATACCACTACT</td><td>693.2</td><td>-5.7</td><td>1.9</td><td>-3.0</td><td>2.8E-03</td><td>3.3E-03</td><td>BT20/MB468</td><td>NUC</td></tr>
<tr><td>>MI0000299|hsa-mir-222&WithFlank&X|-|45747009|45747130@75.97.23&[MIMAT0000279&hsa-miR-222-3p&offsets|0|+2;m-140&X|-|45747033|45747056&offsets|0|-1]|AGCTACATCTGGCTACTGGGTCT</td><td>>MI0000299|hsa-mir-222&WithFlank&X|-|45747009|45747130</td><td>75</td><td>97</td><td>23</td><td>AGCTACATCTGGCTACTGGGTCT</td><td>2145.0</td><td>3.5</td><td>1.1</td><td>3.2</td><td>1.4E-03</td><td>3.3E-03</td><td>BT20/MB468</td><td>MP</td></tr>
<tr><td>>hg38_21_+_8433172_8446622_RNA45SN1_45S_50ntFlanks@10349.10366.18|AACGAGAACTTTGAAGGC</td><td>>hg38_21_+_8433172_8446622_RNA45SN1_45S_50ntFlanks</td><td>10349</td><td>10366</td><td>18</td><td>AACGAGAACTTTGAAGGC</td><td>1858.1</td><td>-4.3</td><td>1.4</td><td>-3.0</td><td>2.4E-03</td><td>3.3E-03</td><td>MB231/MB468</td><td>MP</td></tr>
<tr><td>>hg38_MT_+_1621_3279_RNR2_16S_50ntFlanks@1427.1450.24|GATTAAAGTCCTACGTGATCTGAG</td><td>>hg38_MT_+_1621_3279_RNR2_16S_50ntFlanks</td><td>1427</td><td>1450</td><td>24</td><td>GATTAAAGTCCTACGTGATCTGAG</td><td>165.4</td><td>4.9</td><td>1.6</td><td>3.0</td><td>2.4E-03</td><td>3.3E-03</td><td>MB231/MB468</td><td>MP</td></tr>
<tr><td>>hg38_21_+_8433172_8446622_RNA45SN1_45S_50ntFlanks@9567.9592.26|ACGTGCAAATCGGTCGTCCGACCTGG</td><td>>hg38_21_+_8433172_8446622_RNA45SN1_45S_50ntFlanks</td><td>9567</td><td>9592</td><td>26</td><td>ACGTGCAAATCGGTCGTCCGACCTGG</td><td>342.7</td><td>-1.9</td><td>0.6</td><td>-3.1</td><td>2.2E-03</td><td>3.3E-03</td><td>BT20/MB468</td><td>MITO</td></tr>
<tr><td>>hg38_21_+_8433172_8446622_RNA45SN1_45S_50ntFlanks@9927.9950.24|TTGATATAGACAGCAGGACGGTGG</td><td>>hg38_21_+_8433172_8446622_RNA45SN1_45S_50ntFlanks</td><td>9927</td><td>9950</td><td>24</td><td>TTGATATAGACAGCAGGACGGTGG</td><td>132.0</td><td>6.3</td><td>2.0</td><td>3.2</td><td>1.4E-03</td><td>3.3E-03</td><td>BT20/MB468</td><td>MP</td></tr>
<tr><td>>hg38_MT_+_598_1651_RNR1_12S_50ntFlanks@853.875.23|CTTAGTTGAACAGGGCCCTGAAG</td><td>>hg38_MT_+_598_1651_RNR1_12S_50ntFlanks</td><td>853</td><td>875</td><td>23</td><td>CTTAGTTGAACAGGGCCCTGAAG</td><td>53.5</td><td>6.4</td><td>2.0</td><td>3.2</td><td>1.4E-03</td><td>3.3E-03</td><td>BT20/MB468</td><td>MP</td></tr>
<tr><td>>hg38_21_+_8433172_8446622_RNA45SN1_45S_50ntFlanks@10841.10863.23|CGTAACTTCGGGATAAGGATTGG</td><td>>hg38_21_+_8433172_8446622_RNA45SN1_45S_50ntFlanks</td><td>10841</td><td>10863</td><td>23</td><td>CGTAACTTCGGGATAAGGATTGG</td><td>3281.3</td><td>-4.0</td><td>1.3</td><td>-3.2</td><td>1.4E-03</td><td>3.3E-03</td><td>BT20/MB468</td><td>MP</td></tr>
<tr><td>>hg38_21_+_8433172_8446622_RNA45SN1_45S_50ntFlanks@7992.8017.26|CGTGGCGACCCGCTGAATTTAAGCAT</td><td>>hg38_21_+_8433172_8446622_RNA45SN1_45S_50ntFlanks</td><td>7992</td><td>8017</td><td>26</td><td>CGTGGCGACCCGCTGAATTTAAGCAT</td><td>393.5</td><td>-2.1</td><td>0.7</td><td>-3.0</td><td>2.9E-03</td><td>3.3E-03</td><td>MB231/MB468</td><td>MITO</td></tr>
<tr><td>>hg38_21_+_8433172_8446622_RNA45SN1_45S_50ntFlanks@11553.11588.36|CTCGGCCGGCGCCTAGCAGCCGACTTAGAACTGGTG</td><td>>hg38_21_+_8433172_8446622_RNA45SN1_45S_50ntFlanks</td><td>11553</td><td>11588</td><td>36</td><td>CTCGGCCGGCGCCTAGCAGCCGACTTAGAACTGGTG</td><td>246.2</td><td>-2.4</td><td>0.8</td><td>-3.0</td><td>2.9E-03</td><td>3.3E-03</td><td>MB231/MB468</td><td>MITO</td></tr>
<tr><td>>hg38_21_+_8433172_8446622_RNA45SN1_45S_50ntFlanks@6651.6680.30|CGACTCTTAGCGGTGGATCACTCGGCTCGT</td><td>>hg38_21_+_8433172_8446622_RNA45SN1_45S_50ntFlanks</td><td>6651</td><td>6680</td><td>30</td><td>CGACTCTTAGCGGTGGATCACTCGGCTCGT</td><td>4941.3</td><td>-1.7</td><td>0.6</td><td>-3.0</td><td>2.8E-03</td><td>3.3E-03</td><td>BT20/MB468</td><td>NUC</td></tr>
<tr><td>>hg38_21_+_8433172_8446622_RNA45SN1_45S_50ntFlanks@9175.9197.23|TCGCGCCGTCGGGCCCGGGGGAG</td><td>>hg38_21_+_8433172_8446622_RNA45SN1_45S_50ntFlanks</td><td>9175</td><td>9197</td><td>23</td><td>TCGCGCCGTCGGGCCCGGGGGAG</td><td>165.8</td><td>-5.2</td><td>1.6</td><td>-3.2</td><td>1.4E-03</td><td>3.4E-03</td><td>BT20/MB468</td><td>MP</td></tr>
<tr><td>>hg38_21_+_8433172_8446622_RNA45SN1_45S_50ntFlanks@11207.11232.26|GGCGGCGACTCTGGACGCGAGCCGGG</td><td>>hg38_21_+_8433172_8446622_RNA45SN1_45S_50ntFlanks</td><td>11207</td><td>11232</td><td>26</td><td>GGCGGCGACTCTGGACGCGAGCCGGG</td><td>1223.7</td><td>-6.0</td><td>2.0</td><td>-3.0</td><td>2.8E-03</td><td>3.4E-03</td><td>BT20/MB468</td><td>NUC</td></tr>
<tr><td>>hg38_21_+_8433172_8446622_RNA45SN1_45S_50ntFlanks@11778.11800.23|ACGCGCATGAATGGATGAACGAG</td><td>>hg38_21_+_8433172_8446622_RNA45SN1_45S_50ntFlanks</td><td>11778</td><td>11800</td><td>23</td><td>ACGCGCATGAATGGATGAACGAG</td><td>95.5</td><td>-5.2</td><td>1.6</td><td>-3.2</td><td>1.4E-03</td><td>3.4E-03</td><td>BT20/MB468</td><td>MP</td></tr>
<tr><td>>hg38_21_+_8433172_8446622_RNA45SN1_45S_50ntFlanks@10783.10817.35|GTGAACAGCCTCTGGCATGTTGGAACAATGTAGGT</td><td>>hg38_21_+_8433172_8446622_RNA45SN1_45S_50ntFlanks</td><td>10783</td><td>10817</td><td>35</td><td>GTGAACAGCCTCTGGCATGTTGGAACAATGTAGGT</td><td>381.4</td><td>-2.7</td><td>0.9</td><td>-3.0</td><td>2.6E-03</td><td>3.4E-03</td><td>BT20/MB468</td><td>CYTO</td></tr>
<tr><td>>trnaMT_ValTAC_MT_+_1602_1670@49.71.23|AACTTAACTTGACCGCTCTGACC</td><td>>trnaMT_ValTAC_MT_+_1602_1670</td><td>49</td><td>71</td><td>23</td><td>AACTTAACTTGACCGCTCTGACC</td><td>2092.2</td><td>3.3</td><td>1.1</td><td>3.1</td><td>2.2E-03</td><td>3.4E-03</td><td>BT20/MB468</td><td>MITO</td></tr>
<tr><td>>hg38_21_+_8433172_8446622_RNA45SN1_45S_50ntFlanks@8380.8397.18|CAAGAGGGCGTGAAACCG</td><td>>hg38_21_+_8433172_8446622_RNA45SN1_45S_50ntFlanks</td><td>8380</td><td>8397</td><td>18</td><td>CAAGAGGGCGTGAAACCG</td><td>332.6</td><td>-3.6</td><td>1.2</td><td>-3.1</td><td>2.2E-03</td><td>3.4E-03</td><td>BT20/MB468</td><td>MITO</td></tr>
<tr><td>>hg38_21_+_8433172_8446622_RNA45SN1_45S_50ntFlanks@3850.3871.22|GATAACTGTGGTAATTCTAGAG</td><td>>hg38_21_+_8433172_8446622_RNA45SN1_45S_50ntFlanks</td><td>3850</td><td>3871</td><td>22</td><td>GATAACTGTGGTAATTCTAGAG</td><td>242.6</td><td>-2.2</td><td>0.7</td><td>-3.0</td><td>2.7E-03</td><td>3.4E-03</td><td>MB231/MB468</td><td>CYTO</td></tr>
<tr><td>>MI0000542|hsa-mir-320a&WithFlank&8|-|22244960|22245043@49.68.20(+1U)&[MIMAT0000510&hsa-miR-320a-3p&offsets|+1|-1(+1U);m-60&8|-|22244975|22244996&offsets|+1|-1(+1U)]|AAAGCTGGGTTGAGAGGGCGT</td><td>>MI0000542|hsa-mir-320a&WithFlank&8|-|22244960|22245043</td><td>49</td><td>68</td><td>20(+1U)</td><td>AAAGCTGGGTTGAGAGGGCGT</td><td>179.7</td><td>4.0</td><td>1.3</td><td>3.0</td><td>2.4E-03</td><td>3.4E-03</td><td>MB231/MB468</td><td>MP</td></tr>
<tr><td>>hg38_MT_+_1621_3279_RNR2_16S_50ntFlanks@824.855.32|ATAAGGAAAGGTTAAAAAAAGTAAAAGGAACT</td><td>>hg38_MT_+_1621_3279_RNR2_16S_50ntFlanks</td><td>824</td><td>855</td><td>32</td><td>ATAAGGAAAGGTTAAAAAAAGTAAAAGGAACT</td><td>81.3</td><td>4.6</td><td>1.5</td><td>3.0</td><td>2.4E-03</td><td>3.4E-03</td><td>MB231/MB468</td><td>MP</td></tr>
<tr><td>>MI0000077|hsa-mir-21&WithFlank&17|+|59841260|59841343@13.35.23&[MIMAT0000076&hsa-miR-21-5p&offsets|-1|0;m-1&17|+|59841273|59841294&offsets|-1|0]|GTAGCTTATCAGACTGATGTTGA</td><td>>MI0000077|hsa-mir-21&WithFlank&17|+|59841260|59841343</td><td>13</td><td>35</td><td>23</td><td>GTAGCTTATCAGACTGATGTTGA</td><td>2340.9</td><td>3.8</td><td>1.2</td><td>3.2</td><td>1.4E-03</td><td>3.4E-03</td><td>BT20/MB468</td><td>MP</td></tr>
<tr><td>>hg38_21_+_8433172_8446622_RNA45SN1_45S_50ntFlanks@12973.13000.28|TCGTACGTAGCAGAGCAGCTCCCTCGCT</td><td>>hg38_21_+_8433172_8446622_RNA45SN1_45S_50ntFlanks</td><td>12973</td><td>13000</td><td>28</td><td>TCGTACGTAGCAGAGCAGCTCCCTCGCT</td><td>152.3</td><td>-2.9</td><td>1.0</td><td>-3.0</td><td>2.6E-03</td><td>3.4E-03</td><td>BT20/MB468</td><td>CYTO</td></tr>
<tr><td>>hg38_21_+_8433172_8446622_RNA45SN1_45S_50ntFlanks@11208.11229.22|GCGGCGACTCTGGACGCGAGCC</td><td>>hg38_21_+_8433172_8446622_RNA45SN1_45S_50ntFlanks</td><td>11208</td><td>11229</td><td>22</td><td>GCGGCGACTCTGGACGCGAGCC</td><td>1123.9</td><td>-1.9</td><td>0.6</td><td>-3.0</td><td>2.6E-03</td><td>3.4E-03</td><td>BT20/MB468</td><td>CYTO</td></tr>
<tr><td>>hg38_21_+_8433172_8446622_RNA45SN1_45S_50ntFlanks@12444.12479.36|AAGCAGAATTCACCAAGCGTTGGATTGTTCACCCAC</td><td>>hg38_21_+_8433172_8446622_RNA45SN1_45S_50ntFlanks</td><td>12444</td><td>12479</td><td>36</td><td>AAGCAGAATTCACCAAGCGTTGGATTGTTCACCCAC</td><td>100.1</td><td>-3.9</td><td>1.3</td><td>-3.0</td><td>2.4E-03</td><td>3.4E-03</td><td>MB231/MB468</td><td>MP</td></tr>
<tr><td>>hg38_21_+_8433172_8446622_RNA45SN1_45S_50ntFlanks@9570.9594.25|TGCAAATCGGTCGTCCGACCTGGGT</td><td>>hg38_21_+_8433172_8446622_RNA45SN1_45S_50ntFlanks</td><td>9570</td><td>9594</td><td>25</td><td>TGCAAATCGGTCGTCCGACCTGGGT</td><td>400.4</td><td>-2.6</td><td>0.8</td><td>-3.1</td><td>2.2E-03</td><td>3.4E-03</td><td>BT20/MB468</td><td>MITO</td></tr>
<tr><td>>hg38_21_+_8433172_8446622_RNA45SN1_45S_50ntFlanks@12476.12497.22|CCACTAATAGGGAACGTGAGCT</td><td>>hg38_21_+_8433172_8446622_RNA45SN1_45S_50ntFlanks</td><td>12476</td><td>12497</td><td>22</td><td>CCACTAATAGGGAACGTGAGCT</td><td>1557.4</td><td>-2.3</td><td>0.7</td><td>-3.0</td><td>2.6E-03</td><td>3.4E-03</td><td>BT20/MB468</td><td>CYTO</td></tr>
<tr><td>>hg38_21_+_8433172_8446622_RNA45SN1_45S_50ntFlanks@12406.12434.29|TTTTTGATCCTTCGATGTCGGCTCTTCCT</td><td>>hg38_21_+_8433172_8446622_RNA45SN1_45S_50ntFlanks</td><td>12406</td><td>12434</td><td>29</td><td>TTTTTGATCCTTCGATGTCGGCTCTTCCT</td><td>627.8</td><td>-1.8</td><td>0.6</td><td>-3.0</td><td>2.6E-03</td><td>3.4E-03</td><td>BT20/MB468</td><td>CYTO</td></tr>
<tr><td>>MI0003180|hsa-mir-516a-1&WithFlank&19|+|53756735|53756836@22.43.22;MI0003181|hsa-mir-516a-2&WithFlank&19|+|53761127|53761228@22.43.22&[MIMAT0004770&hsa-miR-516a-5p&offsets|0|-1;m-852&19|+|53756756|53756777&offsets|0|0];[MIMAT0004770_1&hsa-miR-516a-5p&offsets|0|-1;m-852&19|+|53761148|53761169&offsets|0|0]|TTCTCGAGGAAAGAAGCACTTT</td><td>>MI0003180|hsa-mir-516a-1&WithFlank&19|+|53756735|53756836</td><td>22</td><td>43</td><td>22</td><td>TTCTCGAGGAAAGAAGCACTTT</td><td>203.0</td><td>-4.7</td><td>1.4</td><td>-3.5</td><td>5.0E-04</td><td>3.4E-03</td><td>BT20/MB231</td><td>MP</td></tr>
<tr><td>>hg38_21_+_8433172_8446622_RNA45SN1_45S_50ntFlanks@11799.11820.22|AGATTCCCACTGTCCCTACCTA</td><td>>hg38_21_+_8433172_8446622_RNA45SN1_45S_50ntFlanks</td><td>11799</td><td>11820</td><td>22</td><td>AGATTCCCACTGTCCCTACCTA</td><td>100.9</td><td>-4.4</td><td>1.4</td><td>-3.1</td><td>2.2E-03</td><td>3.4E-03</td><td>BT20/MB468</td><td>MITO</td></tr>
</table>

<table>
<tr><td>>MI0000298|hsa-mir-221&WithFlank&X|-|45746151|45746272@70.93.24&[MIMAT0000278&hsa-miR-221-3p&offsets|-1|0;m-91&X|-|45746181|45746202&offsets|-1|+1]|CAGCTACATTGTCTGCTGGGTTTC</td><td>>MI0000298|hsa-mir-221&WithFlank&X|-|45746151|45746272</td><td>70</td><td>93</td><td>24</td><td>CAGCTACATTGTCTGCTGGGTTTC</td><td>219.9</td><td>-4.7</td><td>1.4</td><td>-3.5</td><td>5.0E-04</td><td>3.4E-03</td><td>BT20/MB231</td><td>MP</td></tr>
<tr><td>>hg38_21_+_8433172_8446622_RNA45SN1_45S_50ntFlanks@9387.9409.23|CCGGCCGAGGTGGGATCCCGAGG</td><td>>hg38_21_+_8433172_8446622_RNA45SN1_45S_50ntFlanks</td><td>9387</td><td>9409</td><td>23</td><td>CCGGCCGAGGTGGGATCCCGAGG</td><td>103.7</td><td>-5.3</td><td>1.7</td><td>-3.2</td><td>1.4E-03</td><td>3.4E-03</td><td>BT20/MB468</td><td>MP</td></tr>
<tr><td>>MI0000086|hsa-mir-28&WithFlank&3|+|188688775|188688872@61.81.21(+1A)&[MIMAT0004502&hsa-miR-28-3p&offsets|+1|0(+1A);m-28&3|+|188688834|188688855&offsets|+1|0(+1A)]|ACTAGATTGTGAGCTCCTGGAA</td><td>>MI0000086|hsa-mir-28&WithFlank&3|+|188688775|188688872</td><td>61</td><td>81</td><td>21(+1A)</td><td>ACTAGATTGTGAGCTCCTGGAA</td><td>251.5</td><td>3.4</td><td>1.0</td><td>3.4</td><td>5.9E-04</td><td>3.4E-03</td><td>BT20/MB231</td><td>CYTO</td></tr>
<tr><td>>MI0000809|hsa-mir-151a&WithFlank&8|-|140732558|140732659@53.74.22(+1A)&[MIMAT0000757&hsa-miR-151a-3p&offsets|0|+1(+1A);m-30&8|-|140732587|140732607&offsets|0|+1(+1A)]|CTAGACTGAAGCTCCTTGAGGAA</td><td>>MI0000809|hsa-mir-151a&WithFlank&8|-|140732558|140732659</td><td>53</td><td>74</td><td>22(+1A)</td><td>CTAGACTGAAGCTCCTTGAGGAA</td><td>7813.8</td><td>2.5</td><td>0.7</td><td>3.4</td><td>5.9E-04</td><td>3.4E-03</td><td>BT20/MB231</td><td>CYTO</td></tr>
<tr><td>>hg38_21_+_8433172_8446622_RNA45SN1_45S_50ntFlanks@11667.11707.41|CTGCCCAGTGCTCTGAATGTCAAAGTGAAGAAATTCAATGA</td><td>>hg38_21_+_8433172_8446622_RNA45SN1_45S_50ntFlanks</td><td>11667</td><td>11707</td><td>41</td><td>CTGCCCAGTGCTCTGAATGTCAAAGTGAAGAAATTCAATGA</td><td>506.9</td><td>-2.7</td><td>0.8</td><td>-3.4</td><td>5.9E-04</td><td>3.4E-03</td><td>BT20/MB231</td><td>CYTO</td></tr>
<tr><td>>MI0000478|hsa-mir-149&WithFlank&2|+|240455995|240456095@21.42.22(+1U)&[MIMAT0000450&hsa-miR-149-5p&offsets|0|-1(+1U);m-195&2|+|240456015|240456037&offsets|0|-1(+1U)]|TCTGGCTCCGTGTCTTCACTCCT</td><td>>MI0000478|hsa-mir-149&WithFlank&2|+|240455995|240456095</td><td>21</td><td>42</td><td>22(+1U)</td><td>TCTGGCTCCGTGTCTTCACTCCT</td><td>48.6</td><td>5.7</td><td>1.9</td><td>3.0</td><td>2.4E-03</td><td>3.4E-03</td><td>MB231/MB468</td><td>MP</td></tr>
<tr><td>>hg38_21_+_8433172_8446622_RNA45SN1_45S_50ntFlanks@7982.8010.29|TCAGATCAGACGTGGCGACCCGCTGAATT</td><td>>hg38_21_+_8433172_8446622_RNA45SN1_45S_50ntFlanks</td><td>7982</td><td>8010</td><td>29</td><td>TCAGATCAGACGTGGCGACCCGCTGAATT</td><td>67.1</td><td>-3.7</td><td>1.2</td><td>-3.0</td><td>2.4E-03</td><td>3.4E-03</td><td>MB231/MB468</td><td>MP</td></tr>
<tr><td>>trna116_GluCTC_1_-_145399233_145399304@32.72.41,trna71_GluCTC_1_-_161439189_161439260@32.72.41,trna74_GluCTC_1_-_161431809_161431880@32.72.41,trna77_GluCTC_1_-_161424398_161424469@32.72.41,trna77_GluCTC_6_+_28949976_28950047@32.72.41,trna80_GluCTC_1_-_161417018_161417089@32.72.41,trna87_GluCTC_6_-_126101393_126101464@32.72.41|CTCTCACCGCCGCGGCCCGGGTTCGATTCCCGGTCAGGGAA</td><td>>trna116_GluCTC_1_-_145399233_145399304</td><td>32</td><td>72</td><td>41</td><td>CTCTCACCGCCGCGGCCCGGGTTCGATTCCCGGTCAGGGAA</td><td>112.8</td><td>-5.0</td><td>1.6</td><td>-3.1</td><td>2.2E-03</td><td>3.4E-03</td><td>BT20/MB468</td><td>MITO</td></tr>
<tr><td>>hg38_21_+_8433172_8446622_RNA45SN1_45S_50ntFlanks@12417.12448.32|TCGATGTCGGCTCTTCCTATCATTGTGAAGCA</td><td>>hg38_21_+_8433172_8446622_RNA45SN1_45S_50ntFlanks</td><td>12417</td><td>12448</td><td>32</td><td>TCGATGTCGGCTCTTCCTATCATTGTGAAGCA</td><td>378.2</td><td>-2.5</td><td>0.8</td><td>-3.0</td><td>2.9E-03</td><td>3.4E-03</td><td>BT20/MB468</td><td>NUC</td></tr>
<tr><td>>hg38_21_+_8433172_8446622_RNA45SN1_45S_50ntFlanks@4560.4582.23|CTAGGAATAATGGAATAGGACCG</td><td>>hg38_21_+_8433172_8446622_RNA45SN1_45S_50ntFlanks</td><td>4560</td><td>4582</td><td>23</td><td>CTAGGAATAATGGAATAGGACCG</td><td>106.5</td><td>-4.1</td><td>1.4</td><td>-3.0</td><td>2.6E-03</td><td>3.4E-03</td><td>BT20/MB468</td><td>CYTO</td></tr>
<tr><td>>hg38_21_+_8433172_8446622_RNA45SN1_45S_50ntFlanks@11674.11706.33|GTGCTCTGAATGTCAAAGTGAAGAAATTCAATG</td><td>>hg38_21_+_8433172_8446622_RNA45SN1_45S_50ntFlanks</td><td>11674</td><td>11706</td><td>33</td><td>GTGCTCTGAATGTCAAAGTGAAGAAATTCAATG</td><td>48.5</td><td>6.2</td><td>1.9</td><td>3.2</td><td>1.4E-03</td><td>3.4E-03</td><td>BT20/MB468</td><td>MP</td></tr>
<tr><td>>trna10_ValCAC_5_-_180649395_180649467@35.53.19,trna12_ValAAC_5_-_180645270_180645342@35.53.19,trna132_ValAAC_6_-_27721179_27721251@35.53.19,trna13_ValCAC_19_-_4724647_4724719@35.53.19,trna152_ValCAC_6_-_27248049_27248121@35.53.19,trna157_ValCAC_6_-_27173867_27173939@35.53.19,trna15_ValAAC_5_-_180615416_180615488@35.53.19,trna18_ValCAC_5_-_180529253_180529325@35.53.19,trna2_ValAAC_3_+_169490018_169490090@35.53.19,trna2_ValCAC_5_+_180524070_180524142@35.53.19,trna37_ValAAC_6_+_27203288_27203360@35.53.19,trna4_ValAAC_5_+_180591154_180591226@35.53.19,trna5_ValAAC_5_+_180596610_180596682@35.53.19,trna6_ValCAC_5_+_180600650_180600722@35.53.19,trna85_ValCAC_1_-_161369490_161369562@35.53.19,trna98_ValCAC_1_-_149298555_149298627@35.53.19,trna9_ValCAC_6_+_26538282_26538354@35.53.19|ACACGCGAAAGGTCCCCGG</td><td>>trna10_ValCAC_5_-_180649395_180649467</td><td>35</td><td>53</td><td>19</td><td>ACACGCGAAAGGTCCCCGG</td><td>37.1</td><td>-8.3</td><td>2.6</td><td>-3.2</td><td>1.4E-03</td><td>3.4E-03</td><td>BT20/MB468</td><td>MP</td></tr>
<tr><td>>MI0000065|hsa-let-7d&WithFlank&9|+|94178828|94178926@68.87.20&[MIMAT0004484&hsa-let-7d-3p&offsets|0|-2;m-98&9|+|94178895|94178916&offsets|0|-2]|CTATACGACCTGCTGCCTTT</td><td>>MI0000065|hsa-let-7d&WithFlank&9|+|94178828|94178926</td><td>68</td><td>87</td><td>20</td><td>CTATACGACCTGCTGCCTTT</td><td>49.6</td><td>5.6</td><td>1.8</td><td>3.0</td><td>2.4E-03</td><td>3.4E-03</td><td>MB231/MB468</td><td>MP</td></tr>
<tr><td>>trna116_GluCTC_1_-_145399233_145399304@3.35.33,trna59_GluCTC_1_+_249168447_249168518@3.35.33,trna71_GluCTC_1_-_161439189_161439260@3.35.33,trna74_GluCTC_1_-_161431809_161431880@3.35.33,trna77_GluCTC_1_-_161424398_161424469@3.35.33,trna77_GluCTC_6_+_28949976_28950047@3.35.33,trna80_GluCTC_1_-_161417018_161417089@3.35.33,trna87_GluCTC_6_-_126101393_126101464@3.35.33|CCTGGTGGTCTAGTGGTTAGGATTCGGCGCTCT</td><td>>trna116_GluCTC_1_-_145399233_145399304</td><td>3</td><td>35</td><td>33</td><td>CCTGGTGGTCTAGTGGTTAGGATTCGGCGCTCT</td><td>139.3</td><td>3.2</td><td>1.1</td><td>3.0</td><td>2.4E-03</td><td>3.4E-03</td><td>MB231/MB468</td><td>MP</td></tr>
<tr><td>>hg38_21_+_8433172_8446622_RNA45SN1_45S_50ntFlanks@10420.10443.24|CTGAGAGATGGGCGAGCGCCGTTC</td><td>>hg38_21_+_8433172_8446622_RNA45SN1_45S_50ntFlanks</td><td>10420</td><td>10443</td><td>24</td><td>CTGAGAGATGGGCGAGCGCCGTTC</td><td>762.5</td><td>-6.0</td><td>1.9</td><td>-3.2</td><td>1.4E-03</td><td>3.4E-03</td><td>BT20/MB468</td><td>MP</td></tr>
<tr><td>>hg38_21_+_8433172_8446622_RNA45SN1_45S_50ntFlanks@9033.9077.45|CTCGTGGGGGGGCCGGGCCACCCCTCCCACGGCGCGACCGCTCTC</td><td>>hg38_21_+_8433172_8446622_RNA45SN1_45S_50ntFlanks</td><td>9033</td><td>9077</td><td>45</td><td>CTCGTGGGGGGGCCGGGCCACCCCTCCCACGGCGCGACCGCTCTC</td><td>42.6</td><td>-9.3</td><td>3.1</td><td>-3.0</td><td>2.4E-03</td><td>3.4E-03</td><td>MB231/MB468</td><td>MP</td></tr>
<tr><td>>MI0000298|hsa-mir-221&WithFlank&X|-|45746151|45746272@71.93.23&[MIMAT0000278&hsa-miR-221-3p&offsets|0|0;m-91&X|-|45746181|45746202&offsets|0|+1]|AGCTACATTGTCTGCTGGGTTTC</td><td>>MI0000298|hsa-mir-221&WithFlank&X|-|45746151|45746272</td><td>71</td><td>93</td><td>23</td><td>AGCTACATTGTCTGCTGGGTTTC</td><td>43451.4</td><td>-2.3</td><td>0.7</td><td>-3.4</td><td>6.6E-04</td><td>3.4E-03</td><td>BT20/MB231</td><td>MITO</td></tr>
<tr><td>>hg38_21_+_8433172_8446622_RNA45SN1_45S_50ntFlanks@7968.8010.43|TCCGAGACGCGACCTCAGATCAGACGTGGCGACCCGCTGAATT</td><td>>hg38_21_+_8433172_8446622_RNA45SN1_45S_50ntFlanks</td><td>7968</td><td>8010</td><td>43</td><td>TCCGAGACGCGACCTCAGATCAGACGTGGCGACCCGCTGAATT</td><td>444.0</td><td>3.6</td><td>1.2</td><td>3.0</td><td>2.9E-03</td><td>3.4E-03</td><td>BT20/MB468</td><td>NUC</td></tr>
<tr><td>>hg38_21_+_8433172_8446622_RNA45SN1_45S_50ntFlanks@11208.11232.25|GCGGCGACTCTGGACGCGAGCCGGG</td><td>>hg38_21_+_8433172_8446622_RNA45SN1_45S_50ntFlanks</td><td>11208</td><td>11232</td><td>25</td><td>GCGGCGACTCTGGACGCGAGCCGGG</td><td>3982.3</td><td>-2.7</td><td>0.9</td><td>-3.1</td><td>2.2E-03</td><td>3.4E-03</td><td>BT20/MB468</td><td>MITO</td></tr>
<tr><td>>trnaMT_ThrTGT_MT_+_15888_15953@38.69.32|CCGGAGATGAAAACCTTTTTCCAAGGACACCA</td><td>>trnaMT_ThrTGT_MT_+_15888_15953</td><td>38</td><td>69</td><td>32</td><td>CCGGAGATGAAAACCTTTTTCCAAGGACACCA</td><td>356.9</td><td>4.1</td><td>1.3</td><td>3.2</td><td>1.4E-03</td><td>3.4E-03</td><td>BT20/MB468</td><td>MP</td></tr>
<tr><td>>MI0000299|hsa-mir-222&WithFlank&X|-|45747009|45747130@75.98.24(+1C)&[MIMAT0000279&hsa-miR-222-3p&offsets|0|+3(+1C);m-140&X|-|45747033|45747056&offsets|0|0(+1C)]|AGCTACATCTGGCTACTGGGTCTCC</td><td>>MI0000299|hsa-mir-222&WithFlank&X|-|45747009|45747130</td><td>75</td><td>98</td><td>24(+1C)</td><td>AGCTACATCTGGCTACTGGGTCTCC</td><td>11304.5</td><td>-2.6</td><td>0.8</td><td>-3.4</td><td>6.6E-04</td><td>3.4E-03</td><td>BT20/MB231</td><td>MITO</td></tr>
<tr><td>>MI0000301|hsa-mir-224&WithFlank&X|-|151958572|151958664@14.36.23&[MIMAT0000281&hsa-miR-224-5p&offsets|+1|-1;m-238&X|-|151958630|151958651&offsets|0|+1]|CAAGTCACTAGTGGTTCCGTTTA</td><td>>MI0000301|hsa-mir-224&WithFlank&X|-|151958572|151958664</td><td>14</td><td>36</td><td>23</td><td>CAAGTCACTAGTGGTTCCGTTTA</td><td>2357.0</td><td>3.2</td><td>0.9</td><td>3.4</td><td>5.9E-04</td><td>3.4E-03</td><td>BT20/MB231</td><td>CYTO</td></tr>
<tr><td>>hg38_21_+_8433172_8446622_RNA45SN1_45S_50ntFlanks@12417.12441.25|TCGATGTCGGCTCTTCCTATCATTG</td><td>>hg38_21_+_8433172_8446622_RNA45SN1_45S_50ntFlanks</td><td>12417</td><td>12441</td><td>25</td><td>TCGATGTCGGCTCTTCCTATCATTG</td><td>2119.3</td><td>-2.1</td><td>0.7</td><td>-3.0</td><td>2.7E-03</td><td>3.4E-03</td><td>MB231/MB468</td><td>CYTO</td></tr>
<tr><td>>MI0000066|hsa-let-7e&WithFlank&19|+|51692780|51692870@14.36.23&[MIMAT0000066&hsa-let-7e-5p&offsets|0|+1;m-50&19|+|51692793|51692814&offsets|0|+1]|TGAGGTAGGAGGTTGTATAGTTG</td><td>>MI0000066|hsa-let-7e&WithFlank&19|+|51692780|51692870</td><td>14</td><td>36</td><td>23</td><td>TGAGGTAGGAGGTTGTATAGTTG</td><td>243.0</td><td>4.4</td><td>1.4</td><td>3.2</td><td>1.4E-03</td><td>3.4E-03</td><td>BT20/MB468</td><td>MP</td></tr>
<tr><td>>hg38_21_+_8433172_8446622_RNA45SN1_45S_50ntFlanks@10237.10264.28|ACGCTACGCCGCGACGAGTAGGAGGGCC</td><td>>hg38_21_+_8433172_8446622_RNA45SN1_45S_50ntFlanks</td><td>10237</td><td>10264</td><td>28</td><td>ACGCTACGCCGCGACGAGTAGGAGGGCC</td><td>134.6</td><td>4.6</td><td>1.4</td><td>3.2</td><td>1.4E-03</td><td>3.4E-03</td><td>BT20/MB468</td><td>MP</td></tr>
<tr><td>>hg38_21_+_8433172_8446622_RNA45SN1_45S_50ntFlanks@10678.10711.34|CGGTTCCGGCGGCGTCCGGTGAGCTCTCGCTGGC</td><td>>hg38_21_+_8433172_8446622_RNA45SN1_45S_50ntFlanks</td><td>10678</td><td>10711</td><td>34</td><td>CGGTTCCGGCGGCGTCCGGTGAGCTCTCGCTGGC</td><td>14751.5</td><td>-5.3</td><td>1.7</td><td>-3.2</td><td>1.4E-03</td><td>3.4E-03</td><td>BT20/MB468</td><td>MP</td></tr>
<tr><td>>trna119_LysCTT_1_-_145395522_145395594@1.31.31,trna11_LysCTT_5_-_180648979_180649051@1.31.31,trna13_LysCTT_6_+_26556774_26556846@1.31.31,trna32_LysCTT_16_-_3207406_3207478@1.31.31,trna7_LysCTT_16_+_3225692_3225764@1.31.31,trna9_LysCTT_5_+_180634755_180634827@1.31.31|GCCCGGCTAGCTCAGTCGGTAGAGCATGAGA</td><td>>trna119_LysCTT_1_-_145395522_145395594</td><td>1</td><td>31</td><td>31</td><td>GCCCGGCTAGCTCAGTCGGTAGAGCATGAGA</td><td>563.3</td><td>-3.8</td><td>1.1</td><td>-3.4</td><td>6.6E-04</td><td>3.4E-03</td><td>BT20/MB231</td><td>MITO</td></tr>
<tr><td>>hg38_21_+_8433172_8446622_RNA45SN1_45S_50ntFlanks@12408.12438.31|TTTGATCCTTCGATGTCGGCTCTTCCTATCA</td><td>>hg38_21_+_8433172_8446622_RNA45SN1_45S_50ntFlanks</td><td>12408</td><td>12438</td><td>31</td><td>TTTGATCCTTCGATGTCGGCTCTTCCTATCA</td><td>210.7</td><td>1.9</td><td>0.6</td><td>3.0</td><td>2.9E-03</td><td>3.4E-03</td><td>MB231/MB468</td><td>NUC</td></tr>
<tr><td>>hg38_21_+_8433172_8446622_RNA45SN1_45S_50ntFlanks@12909.12933.25|CCGCACGTTCGTGGGGAACCTGGCG</td><td>>hg38_21_+_8433172_8446622_RNA45SN1_45S_50ntFlanks</td><td>12909</td><td>12933</td><td>25</td><td>CCGCACGTTCGTGGGGAACCTGGCG</td><td>349.7</td><td>-3.0</td><td>1.0</td><td>-3.2</td><td>1.4E-03</td><td>3.4E-03</td><td>BT20/MB468</td><td>MP</td></tr>
<tr><td>>hg38_21_+_8433172_8446622_RNA45SN1_45S_50ntFlanks@10514.10533.20|CGGAGTGGCGGAGATGGGCG</td><td>>hg38_21_+_8433172_8446622_RNA45SN1_45S_50ntFlanks</td><td>10514</td><td>10533</td><td>20</td><td>CGGAGTGGCGGAGATGGGCG</td><td>185.9</td><td>-3.2</td><td>1.1</td><td>-3.0</td><td>2.7E-03</td><td>3.4E-03</td><td>BT20/MB468</td><td>CYTO</td></tr>
<tr><td>>hg38_21_+_8433172_8446622_RNA45SN1_45S_50ntFlanks@12957.12981.25|CTTCTGGGTCGGGGTTTCGTACGTA</td><td>>hg38_21_+_8433172_8446622_RNA45SN1_45S_50ntFlanks</td><td>12957</td><td>12981</td><td>25</td><td>CTTCTGGGTCGGGGTTTCGTACGTA</td><td>194.6</td><td>-1.9</td><td>0.6</td><td>-3.1</td><td>2.3E-03</td><td>3.4E-03</td><td>BT20/MB468</td><td>MITO</td></tr>
<tr><td>>hg38_21_+_8433172_8446622_RNA45SN1_45S_50ntFlanks@8380.8419.40|CAAGAGGGCGTGAAACCGTTAAGAGGTAAACGGGTGGGGT</td><td>>hg38_21_+_8433172_8446622_RNA45SN1_45S_50ntFlanks</td><td>8380</td><td>8419</td><td>40</td><td>CAAGAGGGCGTGAAACCGTTAAGAGGTAAACGGGTGGGGT</td><td>35.6</td><td>-8.2</td><td>2.6</td><td>-3.2</td><td>1.4E-03</td><td>3.4E-03</td><td>BT20/MB468</td><td>MP</td></tr>
<tr><td>>MI0003591|hsa-mir-584&WithFlank&5|-|149062307|149062415@22.42.21(+2U)&[MIMAT0003249&hsa-miR-584-5p&offsets|0|-1(+2U);m-167&5|-|149062374|149062394&offsets|0|0(+2U)]|TTATGGTTTGCCTGGGACTGATT</td><td>>MI0003591|hsa-mir-584&WithFlank&5|-|149062307|149062415</td><td>22</td><td>42</td><td>21(+2U)</td><td>TTATGGTTTGCCTGGGACTGATT</td><td>74.5</td><td>6.4</td><td>2.0</td><td>3.2</td><td>1.4E-03</td><td>3.4E-03</td><td>BT20/MB468</td><td>MP</td></tr>
<tr><td>>hg38_MT_+_598_1651_RNR1_12S_50ntFlanks@428.461.34|AAAGTGGCTTTAACATATCTGAACACACAATAGC</td><td>>hg38_MT_+_598_1651_RNR1_12S_50ntFlanks</td><td>428</td><td>461</td><td>34</td><td>AAAGTGGCTTTAACATATCTGAACACACAATAGC</td><td>94.8</td><td>4.6</td><td>1.5</td><td>3.0</td><td>2.9E-03</td><td>3.4E-03</td><td>MB231/MB468</td><td>MITO</td></tr>
<tr><td>>hg38_21_+_8433172_8446622_RNA45SN1_45S_50ntFlanks@12892.12935.44|CCCTCGCCCGTCACGCACCGCACGTTCGTGGGGAACCTGGCGCT</td><td>>hg38_21_+_8433172_8446622_RNA45SN1_45S_50ntFlanks</td><td>12892</td><td>12935</td><td>44</td><td>CCCTCGCCCGTCACGCACCGCACGTTCGTGGGGAACCTGGCGCT</td><td>449.4</td><td>-4.7</td><td>1.5</td><td>-3.1</td><td>2.3E-03</td><td>3.4E-03</td><td>BT20/MB468</td><td>MITO</td></tr>
<tr><td>>hg38_21_+_8433172_8446622_RNA45SN1_45S_50ntFlanks@12420.12453.34|ATGTCGGCTCTTCCTATCATTGTGAAGCAGAATT</td><td>>hg38_21_+_8433172_8446622_RNA45SN1_45S_50ntFlanks</td><td>12420</td><td>12453</td><td>34</td><td>ATGTCGGCTCTTCCTATCATTGTGAAGCAGAATT</td><td>247.9</td><td>-2.6</td><td>0.8</td><td>-3.4</td><td>6.0E-04</td><td>3.4E-03</td><td>BT20/MB231</td><td>CYTO</td></tr>
<tr><td>>MI0000734|hsa-mir-106b&WithFlank&7|-|100093987|100094080@58.76.19(+1A)&[MIMAT0004672&hsa-miR-106b-3p&offsets|0|-3(+1A);m-87&7|-|100094003|100094023&offsets|0|-2(+1A)]|CCGCACTGTGGGTACTTGCA</td><td>>MI0000734|hsa-mir-106b&WithFlank&7|-|100093987|100094080</td><td>58</td><td>76</td><td>19(+1A)</td><td>CCGCACTGTGGGTACTTGCA</td><td>96.5</td><td>6.3</td><td>2.0</td><td>3.2</td><td>1.4E-03</td><td>3.4E-03</td><td>BT20/MB468</td><td>MP</td></tr>
</table>

| | | | | | | | | | | | | | |
|---|---|---|---|---|---|---|---|---|---|---|---|---|---|
| >trnaMT_AsnGTT_MT_-_5657_5729@1.40.40,trnalookalike5_AsnGTT_1_-_566207_566279@1.40.40\|TAGATTGAAGCCAGTTGATTAGGGTGCTTAGCTGTTAACT | >trnaMT_AsnGTT_MT_-_5657_5729 | 1 | 40 | 40 | TAGATTGAAGCCAGTTGATTAGGGTGCTTAGCTGTTAACT | 133.8 | 2.9 | 1.0 | 3.0 | 2.9E-03 | 3.4E-03 | MB231/MB468 | MITO |
| >hg38_21_+_8433172_8446622_RNA45SN1_45S_50ntFlanks@10389.10433.45\|TGAACAGCAGTTGAACATGGGTCAGTCGGTCCTGAGAGATGGGCG | >hg38_21_+_8433172_8446622_RNA45SN1_45S_50ntFlanks | 10389 | 10433 | 45 | TGAACAGCAGTTGAACATGGGTCAGTCGGTCCTGAGAGATGGGCG | 37.5 | -8.3 | 2.6 | -3.2 | 1.4E-03 | 3.4E-03 | BT20/MB468 | MP |
| >hg38_21_+_8433172_8446622_RNA45SN1_45S_50ntFlanks@6719.6752.34\|TGAATTGCAGGACACATTGATCATCGACACTTCG | >hg38_21_+_8433172_8446622_RNA45SN1_45S_50ntFlanks | 6719 | 6752 | 34 | TGAATTGCAGGACACATTGATCATCGACACTTCG | 256.0 | -4.5 | 1.5 | -3.1 | 2.3E-03 | 3.4E-03 | BT20/MB468 | MITO |
| >hg38_21_+_8433172_8446622_RNA45SN1_45S_50ntFlanks@8189.8213.25\|CCCAGCCCGTGGACGGTGTGAGGCC | >hg38_21_+_8433172_8446622_RNA45SN1_45S_50ntFlanks | 8189 | 8213 | 25 | CCCAGCCCGTGGACGGTGTGAGGCC | 215.3 | -4.3 | 1.3 | -3.2 | 1.5E-03 | 3.4E-03 | BT20/MB468 | MP |
| >MI0000809\|hsa-mir-151a&WithFlank&8\|-\|140732558\|140732659@51.70.20&[MIMAT0000757&hsa-miR-151a-3p&offsets\|-2\|-3;m-30&8\|-\|140732587\|140732607&offsets\|-2\|-3]\|TACTAGACTGAAGCTCCTTG | >MI0000809\|hsa-mir-151a&WithFlank&8\|-\|140732558\|140732659 | 51 | 70 | 20 | TACTAGACTGAAGCTCCTTG | 129.1 | 4.7 | 1.6 | 3.0 | 2.5E-03 | 3.4E-03 | MB231/MB468 | MP |
| >hg38_21_+_8433172_8446622_RNA45SN1_45S_50ntFlanks@12415.12432.18\|CTTCGATGTCGGCTCTTC | >hg38_21_+_8433172_8446622_RNA45SN1_45S_50ntFlanks | 12415 | 12432 | 18 | CTTCGATGTCGGCTCTTC | 8386.1 | -1.7 | 0.6 | -3.0 | 2.7E-03 | 3.4E-03 | MB231/MB468 | CYTO |
| >MI0000447\|hsa-mir-128-1&WithFlank&2\|+\|135665391\|135665484@56.76.21(+1A);MI0000727\|hsa-mir-128-2&WithFlank&3\|+\|35744470\|35744565@58.78.21(+1A)&[MIMAT0000424&hsa-miR-128-3p&offsets\|0\|0(+1A);m-108&2\|+\|135665446\|135665466&offsets\|0\|0(+1A)];[MIMAT0000424_1&hsa-miR-128-3p&offsets\|0\|0(+1A);m-117&3\|+\|35744527\|35744547&offsets\|0\|0(+1A)]\|TCACAGTGAACCGGTCTCTTTA | >MI0000447\|hsa-mir-128-1&WithFlank&2\|+\|135665391\|135665484 | 56 | 76 | 21(+1A) | TCACAGTGAACCGGTCTCTTTA | 1503.7 | 2.5 | 0.7 | 3.4 | 6.0E-04 | 3.4E-03 | BT20/MB231 | CYTO |
| >hg38_21_+_8433172_8446622_RNA45SN1_45S_50ntFlanks@7988.8016.29\|CAGACGTGGCGACCCGCTGAATTTAAGCA | >hg38_21_+_8433172_8446622_RNA45SN1_45S_50ntFlanks | 7988 | 8016 | 29 | CAGACGTGGCGACCCGCTGAATTTAAGCA | 537.1 | -3.1 | 1.0 | -3.0 | 2.9E-03 | 3.4E-03 | BT20/MB468 | NUC |
| >hg38_1_-_228634819_228635039_RNA5S12_5S_50ntFlanks@51.85.35\|GTCTACGGCCATACCACCCTGAACGCGCCCGATCT | >hg38_1_-_228634819_228635039_RNA5S12_5S_50ntFlanks | 51 | 85 | 35 | GTCTACGGCCATACCACCCTGAACGCGCCCGATCT | 1556.1 | 2.2 | 0.8 | 3.0 | 2.9E-03 | 3.4E-03 | MB231/MB468 | NUC |
| >hg38_21_+_8433172_8446622_RNA45SN1_45S_50ntFlanks@7987.8008.22\|TCAGACGTGGCGACCCGCTGAA | >hg38_21_+_8433172_8446622_RNA45SN1_45S_50ntFlanks | 7987 | 8008 | 22 | TCAGACGTGGCGACCCGCTGAA | 255.3 | -3.8 | 1.2 | -3.1 | 2.3E-03 | 3.4E-03 | BT20/MB468 | MITO |
| >hg38_21_+_8433172_8446622_RNA45SN1_45S_50ntFlanks@11580.11611.32\|GAACTGGTGCGGACCAGGGGAATCCGACTGTT | >hg38_21_+_8433172_8446622_RNA45SN1_45S_50ntFlanks | 11580 | 11611 | 32 | GAACTGGTGCGGACCAGGGGAATCCGACTGTT | 1279.0 | -2.0 | 0.7 | -3.0 | 2.7E-03 | 3.4E-03 | BT20/MB468 | CYTO |
| >hg38_21_+_8433172_8446622_RNA45SN1_45S_50ntFlanks@6721.6753.33\|AATTGCAGGACACATTGATCATCGACACTTCGA | >hg38_21_+_8433172_8446622_RNA45SN1_45S_50ntFlanks | 6721 | 6753 | 33 | AATTGCAGGACACATTGATCATCGACACTTCGA | 105.3 | -4.0 | 1.3 | -3.0 | 2.7E-03 | 3.4E-03 | BT20/MB468 | CYTO |
| >trna10_AspGTC_12_-_125424193_125424264@36.75.40,trna12_AspGTC_12_-_125411891_125411962@36.75.40,trna144_AspGTC_6_-_27551236_27551307@36.75.40,trna38_AspGTC_17_-_8125556_8125627@36.75.40,trna45_AspGTC_6_+_27447453_27447524@36.75.40,trna48_AspGTC_6_+_27471523_27471594@36.75.40,trna4_AspGTC_12_+_96429799_96429870@36.75.40,trna69_AspGTC_1_-_161440205_161440276@36.75.40,trna72_AspGTC_1_-_161432824_161432895@36.75.40,trna75_AspGTC_1_-_161425414_161425485@36.75.40,trna78_AspGTC_1_-_161418033_161418104@36.75.40,trna81_AspGTC_1_-_161410615_161410686@36.75.40\|CACGCGGGAGACCGGGGTTCGATTCCCCGACGGGGAGCCA | >trna10_AspGTC_12_-_125424193_125424264 | 36 | 75 | 40 | CACGCGGGAGACCGGGGTTCGATTCCCCGACGGGGAGCCA | 182.0 | 4.9 | 1.6 | 3.0 | 2.5E-03 | 3.4E-03 | MB231/MB468 | MP |
| >hg38_21_+_8433172_8446622_RNA45SN1_45S_50ntFlanks@10463.10480.18\|CTCCGTTGCCCTCGGCCG | >hg38_21_+_8433172_8446622_RNA45SN1_45S_50ntFlanks | 10463 | 10480 | 18 | CTCCGTTGCCCTCGGCCG | 97.3 | -5.1 | 1.6 | -3.2 | 1.5E-03 | 3.4E-03 | BT20/MB468 | MP |
| >hg38_21_+_8433172_8446622_RNA45SN1_45S_50ntFlanks@4995.5020.26\|ACAGATTGATAGCTCTTTCTCGATTC | >hg38_21_+_8433172_8446622_RNA45SN1_45S_50ntFlanks | 4995 | 5020 | 26 | ACAGATTGATAGCTCTTTCTCGATTC | 169.3 | -4.3 | 1.3 | -3.2 | 1.5E-03 | 3.4E-03 | BT20/MB468 | MP |
| >hg38_21_+_8433172_8446622_RNA45SN1_45S_50ntFlanks@10237.10261.25\|ACGCTACGCCGCGACGAGTAGGAGG | >hg38_21_+_8433172_8446622_RNA45SN1_45S_50ntFlanks | 10237 | 10261 | 25 | ACGCTACGCCGCGACGAGTAGGAGG | 59.8 | -4.9 | 1.6 | -3.0 | 2.5E-03 | 3.4E-03 | MB231/MB468 | MP |
| >hg38_21_+_8433172_8446622_RNA45SN1_45S_50ntFlanks@485.508.24\|GATGTGGTGACGTCGTGCTCTCCC | >hg38_21_+_8433172_8446622_RNA45SN1_45S_50ntFlanks | 485 | 508 | 24 | GATGTGGTGACGTCGTGCTCTCCC | 38.9 | 8.5 | 2.8 | 3.0 | 2.5E-03 | 3.4E-03 | MB231/MB468 | MP |
| >MI0000746\|hsa-mir-99b&WithFlank&19\|+\|51692606\|51692687@13.33.21(+1U)&[MIMAT0000689&hsa-miR-99b-5p&offsets\|0\|-1(+1U);m-10&19\|+\|51692618\|51692639&offsets\|0\|-1(+1U)]\|CACCCGTAGAACCGACCTTGCT | >MI0000746\|hsa-mir-99b&WithFlank&19\|+\|51692606\|51692687 | 13 | 33 | 21(+1U) | CACCCGTAGAACCGACCTTGCT | 187.0 | 2.8 | 0.9 | 3.2 | 1.5E-03 | 3.4E-03 | BT20/MB468 | MP |
| >trnaMT_SerGCT_MT_+_12207_12265@14.44.31\|AGAACTGCTAACTCATGCCCCCATGTCTAAC | >trnaMT_SerGCT_MT_+_12207_12265 | 14 | 44 | 31 | AGAACTGCTAACTCATGCCCCCATGTCTAAC | 269.2 | 4.0 | 1.3 | 3.2 | 1.5E-03 | 3.4E-03 | BT20/MB468 | MP |
| >hg38_21_+_8433172_8446622_RNA45SN1_45S_50ntFlanks@9491.9515.25\|GACCCGAAAGATGGTGAACTATGCC | >hg38_21_+_8433172_8446622_RNA45SN1_45S_50ntFlanks | 9491 | 9515 | 25 | GACCCGAAAGATGGTGAACTATGCC | 74.2 | 5.3 | 1.7 | 3.2 | 1.5E-03 | 3.4E-03 | BT20/MB468 | MP |
| >hg38_21_+_8433172_8446622_RNA45SN1_45S_50ntFlanks@3843.3869.27\|CTACTTGGATAACTGTGGTAATTCTAG | >hg38_21_+_8433172_8446622_RNA45SN1_45S_50ntFlanks | 3843 | 3869 | 27 | CTACTTGGATAACTGTGGTAATTCTAG | 286.5 | -2.1 | 0.7 | -3.0 | 2.8E-03 | 3.4E-03 | MB231/MB468 | CYTO |
| >MI0000115\|hsa-mir-16-2&WithFlank&3\|+\|160404739\|160404831@58.79.22(+1U)&[MIMAT0004518&hsa-miR-16-2-3p&offsets\|-1\|-1(+1U);m-252&3\|+\|160404796\|160404816&offsets\|0\|+1(+1U)]\|ACCAATATTACTGTGCTGCTTTT | >MI0000115\|hsa-mir-16-2&WithFlank&3\|+\|160404739\|160404831 | 58 | 79 | 22(+1U) | ACCAATATTACTGTGCTGCTTTT | 55.1 | 5.7 | 1.9 | 3.0 | 2.5E-03 | 3.4E-03 | MB231/MB468 | MP |
| >MI0000465\|hsa-mir-191&WithFlank&3\|-\|49020612\|49020715@23.43.21(+1A)&[MIMAT0000440&hsa-miR-191-5p&offsets\|+1\|-1(+1A);m-48&3\|-\|49020672\|49020694&offsets\|+1\|-1(+1A)]\|AACGGAATCCCAAAAGCAGCTA | >MI0000465\|hsa-mir-191&WithFlank&3\|-\|49020612\|49020715 | 23 | 43 | 21(+1A) | AACGGAATCCCAAAAGCAGCTA | 52.0 | 5.7 | 1.9 | 3.0 | 2.5E-03 | 3.4E-03 | MB231/MB468 | MP |
| >MI0000077\|hsa-mir-21&WithFlank&17\|+\|59841260\|59841343@14.35.22(+1A)&[MIMAT0000076&hsa-miR-21-5p&offsets\|0\|0(+1A);m-1&17\|+\|59841273\|59841294&offsets\|0\|0(+1A)]\|TAGCTTATCAGACTGATGTTGAA | >MI0000077\|hsa-mir-21&WithFlank&17\|+\|59841260\|59841343 | 14 | 35 | 22(+1A) | TAGCTTATCAGACTGATGTTGAA | 22917.7 | -3.3 | 1.0 | -3.5 | 5.1E-04 | 3.4E-03 | BT20/MB231 | MP |
| >MI0000088\|hsa-mir-30a&WithFlank&6\|-\|71403545\|71403627@54.73.20(+1U)&[MIMAT0000088&hsa-miR-30a-3p&offsets\|+1\|-1(+1U);m-38&6\|-\|71403554\|71403575&offsets\|+1\|-1(+1U)]\|TTTCAGTCGGATGTTTGCAGT | >MI0000088\|hsa-mir-30a&WithFlank&6\|-\|71403545\|71403627 | 54 | 73 | 20(+1U) | TTTCAGTCGGATGTTTGCAGT | 4623.8 | -3.4 | 1.0 | -3.5 | 5.1E-04 | 3.4E-03 | BT20/MB231 | MP |
| >hg38_21_+_8433172_8446622_RNA45SN1_45S_50ntFlanks@10404.10430.27\|CATGGGTCAGTCGGTCCTGAGAGATGG | >hg38_21_+_8433172_8446622_RNA45SN1_45S_50ntFlanks | 10404 | 10430 | 27 | CATGGGTCAGTCGGTCCTGAGAGATGG | 209.4 | -4.4 | 1.4 | -3.1 | 2.3E-03 | 3.4E-03 | BT20/MB468 | MITO |
| >hg38_21_+_8433172_8446622_RNA45SN1_45S_50ntFlanks@3842.3860.19\|CCTACTTGGATAACTGTGG | >hg38_21_+_8433172_8446622_RNA45SN1_45S_50ntFlanks | 3842 | 3860 | 19 | CCTACTTGGATAACTGTGG | 214.6 | -2.1 | 0.7 | -3.0 | 2.8E-03 | 3.4E-03 | MB231/MB468 | CYTO |
| >hg38_1_-_228634819_228635039_RNA5S12_5S_50ntFlanks@146.169.24\|TGGGAATACCGGGTGCTGTAGGCT | >hg38_1_-_228634819_228635039_RNA5S12_5S_50ntFlanks | 146 | 169 | 24 | TGGGAATACCGGGTGCTGTAGGCT | 1112.5 | 3.0 | 1.0 | 3.2 | 1.5E-03 | 3.4E-03 | BT20/MB468 | MP |
| >MI0000804\|hsa-mir-328&WithFlank&16\|-\|67202315\|67202401@54.75.22&[MIMAT0000752&hsa-miR-328-3p&offsets\|0\|0;m-160&16\|-\|67202327\|67202348&offsets\|0\|0]\|CTGGCCCTCTCTGCCCTTCCGT | >MI0000804\|hsa-mir-328&WithFlank&16\|-\|67202315\|67202401 | 54 | 75 | 22 | CTGGCCCTCTCTGCCCTTCCGT | 189.6 | 5.5 | 1.7 | 3.2 | 1.5E-03 | 3.4E-03 | BT20/MB468 | MP |
| >hg38_21_+_8433172_8446622_RNA45SN1_45S_50ntFlanks@5082.5108.27\|AACGAACGAGACTCTGGCATGCTAACT | >hg38_21_+_8433172_8446622_RNA45SN1_45S_50ntFlanks | 5082 | 5108 | 27 | AACGAACGAGACTCTGGCATGCTAACT | 837.7 | 5.2 | 1.5 | 3.5 | 5.1E-04 | 3.5E-03 | BT20/MB231 | MP |
| >trnaMT_HisGTG_MT_+_12138_12206@32.72.41\|TGAATCTGACAACAGAGGCTTACGACCCCTTATTTACCCCA | >trnaMT_HisGTG_MT_+_12138_12206 | 32 | 72 | 41 | TGAATCTGACAACAGAGGCTTACGACCCCTTATTTACCCCA | 5488.4 | 4.1 | 1.3 | 3.1 | 2.3E-03 | 3.5E-03 | BT20/MB468 | MITO |
| >trnaMT_LeuTAA_MT_+_3230_3304@16.44.29\|CCCGGTAATCGCATAAAACTTAAAACTTT | >trnaMT_LeuTAA_MT_+_3230_3304 | 16 | 44 | 29 | CCCGGTAATCGCATAAAACTTAAAACTTT | 940.3 | -3.1 | 1.0 | -3.1 | 2.3E-03 | 3.5E-03 | BT20/MB468 | MITO |
| >MI0005561\|hsa-mir-877&WithFlank&6\|+\|30584326\|30584423@7.25.19(+1A)&[MIMAT0004949&hsa-miR-877-5p&offsets\|0\|-1(+1A);m-382&6\|+\|30584332\|30584354&offsets\|0\|-4(+1A)]\|GTAGAGGAGATGGCGCAGGA | >MI0005561\|hsa-mir-877&WithFlank&6\|+\|30584326\|30584423 | 7 | 25 | 19(+1A) | GTAGAGGAGATGGCGCAGGA | 86.5 | 5.7 | 1.8 | 3.2 | 1.5E-03 | 3.5E-03 | BT20/MB468 | MP |
| >hg38_21_+_8433172_8446622_RNA45SN1_45S_50ntFlanks@9555.9581.27\|GTAGCGGTCCTGACGTGCAAATCGGTC | >hg38_21_+_8433172_8446622_RNA45SN1_45S_50ntFlanks | 9555 | 9581 | 27 | GTAGCGGTCCTGACGTGCAAATCGGTC | 296.5 | -3.1 | 0.9 | -3.4 | 6.1E-04 | 3.5E-03 | BT20/MB231 | CYTO |
| >hg38_21_+_8433172_8446622_RNA45SN1_45S_50ntFlanks@8176.8220.45\|CTTCTGATCGAGGCCCAGCCCGTGGACGGTGTGAGGCCGGTAGCG | >hg38_21_+_8433172_8446622_RNA45SN1_45S_50ntFlanks | 8176 | 8220 | 45 | CTTCTGATCGAGGCCCAGCCCGTGGACGGTGTGAGGCCGGTAGCG | 510.9 | -2.7 | 0.9 | -3.0 | 3.0E-03 | 3.5E-03 | MB231/MB468 | NUC |
| >hg38_21_+_8433172_8446622_RNA45SN1_45S_50ntFlanks@9927.9951.25\|TTGATATAGACAGCAGGACGGTGGC | >hg38_21_+_8433172_8446622_RNA45SN1_45S_50ntFlanks | 9927 | 9951 | 25 | TTGATATAGACAGCAGGACGGTGGC | 129.4 | 6.2 | 1.9 | 3.2 | 1.5E-03 | 3.5E-03 | BT20/MB468 | MP |
| >hg38_21_+_8433172_8446622_RNA45SN1_45S_50ntFlanks@12990.13012.23\|GCTCCCTCGCTGCGATCTATTGA | >hg38_21_+_8433172_8446622_RNA45SN1_45S_50ntFlanks | 12990 | 13012 | 23 | GCTCCCTCGCTGCGATCTATTGA | 392.3 | -3.6 | 1.2 | -3.0 | 2.9E-03 | 3.5E-03 | MB231/MB468 | MITO |
| >hg38_21_+_8433172_8446622_RNA45SN1_45S_50ntFlanks@11326.11345.20\|CTCGTCGCGCGCGCGTCCGC | >hg38_21_+_8433172_8446622_RNA45SN1_45S_50ntFlanks | 11326 | 11345 | 20 | CTCGTCGCGCGCGCGTCCGC | 37.8 | 8.5 | 2.8 | 3.0 | 2.5E-03 | 3.5E-03 | MB231/MB468 | MP |
| >MI0000086\|hsa-mir-28&WithFlank&3\|+\|188688775\|188688872@20.41.22&[MIMAT0000085&hsa-miR-28-5p&offsets\|0\|0;m-133&3\|+\|188688794\|188688815&offsets\|0\|0]\|AAGGAGCTCACAGTCTATTGAG | >MI0000086\|hsa-mir-28&WithFlank&3\|+\|188688775\|188688872 | 20 | 41 | 22 | AAGGAGCTCACAGTCTATTGAG | 635.0 | 4.7 | 1.5 | 3.2 | 1.5E-03 | 3.5E-03 | BT20/MB468 | MP |
| >MI0003778\|hsa-mir-320c-1&WithFlank&18\|+\|21683512\|21683595@48.67.20(+1U);MI0008191\|hsa-mir-320c-2&WithFlank&18\|+\|24321669\|24321752@48.67.20(+1U)&[MIMAT0005793&hsa-miR-320c&offsets\|0\|0(+1U);m-222&18\|+\|21683559\|21683578&offsets\|0\|0(+1U)];[MIMAT0005793_1&hsa-miR-320c&offsets\|0\|0(+1U);m-219&18\|+\|24321716\|24321735&offsets\|0\|0(+1U)]\|AAAAGCTGGGTTGAGAGGGTT | >MI0003778\|hsa-mir-320c-1&WithFlank&18\|+\|21683512\|21683595 | 48 | 67 | 20(+1U) | AAAAGCTGGGTTGAGAGGGTT | 104.0 | 5.9 | 1.8 | 3.2 | 1.5E-03 | 3.5E-03 | BT20/MB468 | MP |
| >hg38_21_+_8433172_8446622_RNA45SN1_45S_50ntFlanks@5053.5081.29\|GTGGAGCGATTTGTCTGGTTAATTCCGAT | >hg38_21_+_8433172_8446622_RNA45SN1_45S_50ntFlanks | 5053 | 5081 | 29 | GTGGAGCGATTTGTCTGGTTAATTCCGAT | 488.0 | 7.0 | 2.0 | 3.5 | 5.1E-04 | 3.5E-03 | BT20/MB231 | MP |
| >hg38_21_+_8433172_8446622_RNA45SN1_45S_50ntFlanks@3705.3723.19\|TACCTGGTTGATCCTGCCA | >hg38_21_+_8433172_8446622_RNA45SN1_45S_50ntFlanks | 3705 | 3723 | 19 | TACCTGGTTGATCCTGCCA | 595.1 | 2.3 | 0.7 | 3.5 | 5.6E-04 | 3.5E-03 | BT20/MB231 | NUC |
| >hg38_21_+_8433172_8446622_RNA45SN1_45S_50ntFlanks@5488.5516.29\|GCTGAGAAGACGGTCGAACTTGACTATCT | >hg38_21_+_8433172_8446622_RNA45SN1_45S_50ntFlanks | 5488 | 5516 | 29 | GCTGAGAAGACGGTCGAACTTGACTATCT | 396.2 | 2.4 | 0.7 | 3.5 | 5.6E-04 | 3.5E-03 | BT20/MB231 | NUC |

| | | | | | | | | | | | | | |
|---|---|---|---|---|---|---|---|---|---|---|---|---|---|
| >hg38_21_+_8433172_8446622_RNA45SN1_45S_50ntFlanks@9572.9593.22\|CAAATCGGTCGTCCGACCTGGG | >hg38_21_+_8433172_8446622_RNA45SN1_45S_50ntFlanks | 9572 | 9593 | 22 | CAAATCGGTCGTCCGACCTGGG | 261.8 | -3.3 | 1.0 | -3.2 | 1.5E-03 | 3.5E-03 | BT20/MB468 | MP |
| >hg38_21_+_8433172_8446622_RNA45SN1_45S_50ntFlanks@6731.6756.26\|CACATTGATCATCGACACTTCGAACG | >hg38_21_+_8433172_8446622_RNA45SN1_45S_50ntFlanks | 6731 | 6756 | 26 | CACATTGATCATCGACACTTCGAACG | 571.3 | -4.4 | 1.4 | -3.2 | 1.5E-03 | 3.5E-03 | BT20/MB468 | MP |
| >hg38_21_+_8433172_8446622_RNA45SN1_45S_50ntFlanks@11577.11594.18\|TTAGAACTGGTGCGGACC | >hg38_21_+_8433172_8446622_RNA45SN1_45S_50ntFlanks | 11577 | 11594 | 18 | TTAGAACTGGTGCGGACC | 414.0 | -1.8 | 0.6 | -3.0 | 2.7E-03 | 3.5E-03 | BT20/MB468 | CYTO |
| >hg38_21_+_8433172_8446622_RNA45SN1_45S_50ntFlanks@10788.10816.29\|CAGCCTCTGGCATGTTGGAACAATGTAGG | >hg38_21_+_8433172_8446622_RNA45SN1_45S_50ntFlanks | 10788 | 10816 | 29 | CAGCCTCTGGCATGTTGGAACAATGTAGG | 189.3 | 6.5 | 1.9 | 3.5 | 5.1E-04 | 3.5E-03 | BT20/MB231 | MP |
| >hg38_21_+_8433172_8446622_RNA45SN1_45S_50ntFlanks@9928.9951.24\|TGATATAGACAGCAGGACGGTGGC | >hg38_21_+_8433172_8446622_RNA45SN1_45S_50ntFlanks | 9928 | 9951 | 24 | TGATATAGACAGCAGGACGGTGGC | 241.8 | 6.3 | 1.8 | 3.5 | 5.1E-04 | 3.5E-03 | BT20/MB231 | MP |
| >trnaMT_PheGAA_MT_+_577_647@39.65.27\|ATGTTTAGACGGGCTCACATCACCCCA | >trnaMT_PheGAA_MT_+_577_647 | 39 | 65 | 27 | ATGTTTAGACGGGCTCACATCACCCCA | 251.7 | 4.4 | 1.4 | 3.2 | 1.5E-03 | 3.5E-03 | BT20/MB468 | MP |
| >hg38_21_+_8433172_8446622_RNA45SN1_45S_50ntFlanks@11323.11340.18\|CGTCTCGTCGCGCGCGCG | >hg38_21_+_8433172_8446622_RNA45SN1_45S_50ntFlanks | 11323 | 11340 | 18 | CGTCTCGTCGCGCGCGCG | 550.3 | -4.3 | 1.4 | -3.2 | 1.5E-03 | 3.5E-03 | BT20/MB468 | MP |
| >hg38_21_+_8433172_8446622_RNA45SN1_45S_50ntFlanks@9623.9649.27\|TAGTAGCTGGTTCCCTCCGAAGTTTCC | >hg38_21_+_8433172_8446622_RNA45SN1_45S_50ntFlanks | 9623 | 9649 | 27 | TAGTAGCTGGTTCCCTCCGAAGTTTCC | 133.2 | -3.2 | 1.1 | -3.0 | 3.0E-03 | 3.5E-03 | MB231/MB468 | NUC |
| >hg38_21_+_8433172_8446622_RNA45SN1_45S_50ntFlanks@6651.6698.48\|CGACTCTTAGCGGTGGATCACTCGGCTCGTGCGTCGATGAAGAACGCA | >hg38_21_+_8433172_8446622_RNA45SN1_45S_50ntFlanks | 6651 | 6698 | 48 | CGACTCTTAGCGGTGGATCACTCGGCTCGTGCGTCGATGAAGAACGCA | 3571.6 | -4.5 | 1.5 | -3.0 | 2.5E-03 | 3.5E-03 | MB231/MB468 | MP |
| >MI0000064\|hsa-let-7c&WithFlank&21\|+\|16539822\|16539917@17.38.22&[MIMAT0000064&hsa-let-7c-5p&offsets\|0\|0;m-37&21\|+\|16539838\|16539859&offsets\|0\|0]\|TGAGGTAGTAGGTTGTATGGTT | >MI0000064\|hsa-let-7c&WithFlank&21\|+\|16539822\|16539917 | 17 | 38 | 22 | TGAGGTAGTAGGTTGTATGGTT | 860.7 | -2.9 | 0.9 | -3.2 | 1.5E-03 | 3.5E-03 | BT20/MB468 | MP |
| >hg38_MT_+_1621_3279_RNR2_16S_50ntFlanks@1427.1451.25\|GATTAAAGTCCTACGTGATCTGAGT | >hg38_MT_+_1621_3279_RNR2_16S_50ntFlanks | 1427 | 1451 | 25 | GATTAAAGTCCTACGTGATCTGAGT | 179.0 | 3.5 | 1.1 | 3.2 | 1.5E-03 | 3.5E-03 | BT20/MB468 | MP |
| >hg38_21_+_8433172_8446622_RNA45SN1_45S_50ntFlanks@475.522.48\|CTTCGTGATCGATGTGGTGACGTCGTGCTCTCCCGGGCCGGGTCCGAG | >hg38_21_+_8433172_8446622_RNA45SN1_45S_50ntFlanks | 475 | 522 | 48 | CTTCGTGATCGATGTGGTGACGTCGTGCTCTCCCGGGCCGGGTCCGAG | 251.5 | 4.9 | 1.6 | 3.0 | 3.0E-03 | 3.5E-03 | MB231/MB468 | MITO |
| >hg38_21_+_8433172_8446622_RNA45SN1_45S_50ntFlanks@485.524.40\|GATGTGGTGACGTCGTGCTCTCCCGGGCCGGGTCCGAGCC | >hg38_21_+_8433172_8446622_RNA45SN1_45S_50ntFlanks | 485 | 524 | 40 | GATGTGGTGACGTCGTGCTCTCCCGGGCCGGGTCCGAGCC | 38.5 | 8.5 | 2.8 | 3.0 | 2.5E-03 | 3.5E-03 | MB231/MB468 | MP |
| >hg38_MT_+_598_1651_RNR1_12S_50ntFlanks@52.85.34\|ATAGGTTTGGTCCTAGCCTTTCTATTAGCTCTTA | >hg38_MT_+_598_1651_RNR1_12S_50ntFlanks | 52 | 85 | 34 | ATAGGTTTGGTCCTAGCCTTTCTATTAGCTCTTA | 19303.8 | 4.3 | 1.4 | 3.2 | 1.5E-03 | 3.5E-03 | BT20/MB468 | MP |
| >hg38_21_+_8433172_8446622_RNA45SN1_45S_50ntFlanks@12714.12751.38\|GAGCCTCGGTTGGCCTCGGATAGCCGGTCCCCCGCCTG | >hg38_21_+_8433172_8446622_RNA45SN1_45S_50ntFlanks | 12714 | 12751 | 38 | GAGCCTCGGTTGGCCTCGGATAGCCGGTCCCCCGCCTG | 515.3 | -4.7 | 1.5 | -3.0 | 2.3E-03 | 3.5E-03 | BT20/MB468 | MITO |
| >hg38_21_+_8433172_8446622_RNA45SN1_45S_50ntFlanks@12289.12325.37\|CCGTGAAAGCGGGGCCTCACGATCCTTCTGACCTTTT | >hg38_21_+_8433172_8446622_RNA45SN1_45S_50ntFlanks | 12289 | 12325 | 37 | CCGTGAAAGCGGGGCCTCACGATCCTTCTGACCTTTT | 196.4 | -2.3 | 0.8 | -3.0 | 2.3E-03 | 3.5E-03 | BT20/MB468 | MITO |
| >hg38_21_+_8433172_8446622_RNA45SN1_45S_50ntFlanks@10410.10428.19\|TCAGTCGGTCCTGAGAGAT | >hg38_21_+_8433172_8446622_RNA45SN1_45S_50ntFlanks | 10410 | 10428 | 19 | TCAGTCGGTCCTGAGAGAT | 308.9 | -4.1 | 1.3 | -3.2 | 1.5E-03 | 3.5E-03 | BT20/MB468 | MP |
| >hg38_21_+_8433172_8446622_RNA45SN1_45S_50ntFlanks@6773.6797.25\|TTCCTCCCGGGGCTACGCCTGTCTG | >hg38_21_+_8433172_8446622_RNA45SN1_45S_50ntFlanks | 6773 | 6797 | 25 | TTCCTCCCGGGGCTACGCCTGTCTG | 113.7 | -5.1 | 1.6 | -3.2 | 1.5E-03 | 3.5E-03 | BT20/MB468 | MP |
| >hg38_21_+_8433172_8446622_RNA45SN1_45S_50ntFlanks@10392.10431.40\|ACAGCAGTTGAACATGGGTCAGTCGGTCCTGAGAGATGGG | >hg38_21_+_8433172_8446622_RNA45SN1_45S_50ntFlanks | 10392 | 10431 | 40 | ACAGCAGTTGAACATGGGTCAGTCGGTCCTGAGAGATGGG | 276.1 | -4.7 | 1.6 | -3.0 | 2.7E-03 | 3.5E-03 | BT20/MB468 | CYTO |
| >hg38_21_+_8433172_8446622_RNA45SN1_45S_50ntFlanks@3899.3935.37\|CCTTCGCGGGGGGGATGCGTGCATTTATCAGATCAAA | >hg38_21_+_8433172_8446622_RNA45SN1_45S_50ntFlanks | 3899 | 3935 | 37 | CCTTCGCGGGGGGGATGCGTGCATTTATCAGATCAAA | 227.0 | -4.5 | 1.5 | -3.0 | 2.3E-03 | 3.5E-03 | BT20/MB468 | MITO |
| >hg38_21_+_8433172_8446622_RNA45SN1_45S_50ntFlanks@9953.9980.28\|ATGGAAGTCGGAATCCGCTAAGGAGTGT | >hg38_21_+_8433172_8446622_RNA45SN1_45S_50ntFlanks | 9953 | 9980 | 28 | ATGGAAGTCGGAATCCGCTAAGGAGTGT | 99.0 | 6.2 | 2.0 | 3.2 | 1.5E-03 | 3.5E-03 | BT20/MB468 | MP |
| >trna143_LysTTT_6_-_27559593_27559665@11.28.18,trna149_LysTTT_6_-_27302769_27302841@11.28.18,trna14_LysTTT_11_-_59327808_59327880@11.28.18,trna2_LysTTT_17_+_8022473_8022545@11.28.18,trna54_LysTTT_1_+_204475655_204475727@11.28.18,trna5_LysTTT_11_+_59323902_59323974@11.28.18,trna62_LysTTT_1_-_204476158_204476230@11.28.18,trna76_LysTTT_6_+_28918806_28918878@11.28.18\|CTCAGTCGGTAGAGCATC | >trna143_LysTTT_6_-_27559593_27559665 | 11 | 28 | 18 | CTCAGTCGGTAGAGCATC | 36.8 | -8.3 | 2.6 | -3.2 | 1.5E-03 | 3.5E-03 | BT20/MB468 | MP |
| >hg38_21_+_8433172_8446622_RNA45SN1_45S_50ntFlanks@12628.12651.24\|TGGGGCGAAGCTACCATCTGTGGG | >hg38_21_+_8433172_8446622_RNA45SN1_45S_50ntFlanks | 12628 | 12651 | 24 | TGGGGCGAAGCTACCATCTGTGGG | 64.6 | -5.1 | 1.6 | -3.2 | 1.5E-03 | 3.5E-03 | BT20/MB468 | MP |
| >hg38_21_+_8433172_8446622_RNA45SN1_45S_50ntFlanks@10912.10931.20\|CGCGCCGCGGCTGGACGAGG | >hg38_21_+_8433172_8446622_RNA45SN1_45S_50ntFlanks | 10912 | 10931 | 20 | CGCGCCGCGGCTGGACGAGG | 1176.9 | -6.1 | 2.1 | -3.0 | 3.0E-03 | 3.5E-03 | BT20/MB468 | NUC |
| >hg38_21_+_8433172_8446622_RNA45SN1_45S_50ntFlanks@3993.4011.19\|GTGACTCTAGATAACCTCG | >hg38_21_+_8433172_8446622_RNA45SN1_45S_50ntFlanks | 3993 | 4011 | 19 | GTGACTCTAGATAACCTCG | 118.3 | -3.7 | 1.2 | -3.0 | 2.5E-03 | 3.5E-03 | MB231/MB468 | MP |
| >trnaMT_PheGAA_MT_+_577_647@1.18.18\|GTTTATGTAGCTTACCTC | >trnaMT_PheGAA_MT_+_577_647 | 1 | 18 | 18 | GTTTATGTAGCTTACCTC | 670.3 | 2.6 | 0.9 | 3.0 | 3.0E-03 | 3.5E-03 | MB231/MB468 | MITO |
| >MI0000078\|hsa-mir-22&WithFlank&17\|-\|1713897\|1713993@21.40.20(+1C)&[MIMAT0004495&hsa-miR-22-5p&offsets\|0\|-2(+1C);m-217&17\|-\|1713953\|1713973&offsets\|0\|-1(+1C)]\|AGTTCTTCAGTGGCAAGCTTC | >MI0000078\|hsa-mir-22&WithFlank&17\|-\|1713897\|1713993 | 21 | 40 | 20(+1C) | AGTTCTTCAGTGGCAAGCTTC | 95.4 | 4.5 | 1.4 | 3.2 | 1.5E-03 | 3.5E-03 | BT20/MB468 | MP |
| >hg38_21_+_8433172_8446622_RNA45SN1_45S_50ntFlanks@11923.11947.25\|AGAGGTGTAGAATAAGTGGGAGGCC | >hg38_21_+_8433172_8446622_RNA45SN1_45S_50ntFlanks | 11923 | 11947 | 25 | AGAGGTGTAGAATAAGTGGGAGGCC | 57.4 | 6.2 | 2.0 | 3.2 | 1.5E-03 | 3.5E-03 | BT20/MB468 | MP |
| >hg38_MT_+_1621_3279_RNR2_16S_50ntFlanks@1095.1118.24\|AGACGAGAAGACCCTATGGAGCTT | >hg38_MT_+_1621_3279_RNR2_16S_50ntFlanks | 1095 | 1118 | 24 | AGACGAGAAGACCCTATGGAGCTT | 121.8 | 3.1 | 1.0 | 3.2 | 1.5E-03 | 3.5E-03 | BT20/MB468 | MP |
| >hg38_1_-_228634819_228635039_RNA5S12_5S_50ntFlanks@51.79.29\|GTCTACGGCCATACCACCCTGAACGCGCC | >hg38_1_-_228634819_228635039_RNA5S12_5S_50ntFlanks | 51 | 79 | 29 | GTCTACGGCCATACCACCCTGAACGCGCC | 602.8 | 3.5 | 1.1 | 3.0 | 2.3E-03 | 3.5E-03 | BT20/MB468 | MITO |
| >hg38_21_+_8433172_8446622_RNA45SN1_45S_50ntFlanks@5484.5526.43\|GAGCGCTGAGAAGACGGTCGAACTTGACTATCTAGAGGAAGTA | >hg38_21_+_8433172_8446622_RNA45SN1_45S_50ntFlanks | 5484 | 5526 | 43 | GAGCGCTGAGAAGACGGTCGAACTTGACTATCTAGAGGAAGTA | 239.2 | -3.3 | 1.1 | -3.0 | 2.3E-03 | 3.5E-03 | BT20/MB468 | MITO |
| >hg38_21_+_8433172_8446622_RNA45SN1_45S_50ntFlanks@7968.7994.27\|TCCGAGACGCGACCTCAGATCAGACGT | >hg38_21_+_8433172_8446622_RNA45SN1_45S_50ntFlanks | 7968 | 7994 | 27 | TCCGAGACGCGACCTCAGATCAGACGT | 97.7 | 6.2 | 2.0 | 3.2 | 1.5E-03 | 3.5E-03 | BT20/MB468 | MP |
| >hg38_21_+_8433172_8446622_RNA45SN1_45S_50ntFlanks@6644.6683.40\|CCTCGTACGACTCTTAGCGGTGGATCACTCGGCTCGTGCG | >hg38_21_+_8433172_8446622_RNA45SN1_45S_50ntFlanks | 6644 | 6683 | 40 | CCTCGTACGACTCTTAGCGGTGGATCACTCGGCTCGTGCG | 890.2 | -2.5 | 0.7 | -3.4 | 5.7E-04 | 3.5E-03 | BT20/MB231 | NUC |
| >hg38_21_+_8433172_8446622_RNA45SN1_45S_50ntFlanks@9561.9597.37\|GTCCTGACGTGCAAATCGGTCGTCCGACCTGGGTATA | >hg38_21_+_8433172_8446622_RNA45SN1_45S_50ntFlanks | 9561 | 9597 | 37 | GTCCTGACGTGCAAATCGGTCGTCCGACCTGGGTATA | 166.0 | -4.5 | 1.5 | -3.0 | 2.8E-03 | 3.5E-03 | BT20/MB468 | CYTO |
| >hg38_MT_+_598_1651_RNR1_12S_50ntFlanks@425.454.30\|ACGAAAGTGGCTTTAACATATCTGAACACA | >hg38_MT_+_598_1651_RNR1_12S_50ntFlanks | 425 | 454 | 30 | ACGAAAGTGGCTTTAACATATCTGAACACA | 224.9 | 5.6 | 1.8 | 3.2 | 1.5E-03 | 3.5E-03 | BT20/MB468 | MP |
| >MI0000077\|hsa-mir-21&WithFlank&17\|+\|59841260\|59841343@14.34.21(+1U)&[MIMAT0000076&hsa-miR-21-5p&offsets\|0\|-1(+1U);m-1&17\|+\|59841273\|59841294&offsets\|0\|-1(+1U)]\|TAGCTTATCAGACTGATGTTGT | >MI0000077\|hsa-mir-21&WithFlank&17\|+\|59841260\|59841343 | 14 | 34 | 21(+1U) | TAGCTTATCAGACTGATGTTGT | 6314.8 | 2.8 | 0.9 | 3.0 | 2.5E-03 | 3.5E-03 | MB231/MB468 | MP |
| >MI0000103\|hsa-mir-101-1&WithFlank&1\|-\|65058428\|65058514@53.74.22(+1U);MI0000739\|hsa-mir-101-2&WithFlank&9\|+\|4850291\|4850381@55.76.22(+1U)&[MIMAT0000099&hsa-miR-101-3p&offsets\|0\|+1(+1U);m-17&1\|-\|65058443\|65058462&offsets\|0\|+2(+1U)];[MIMAT0000099_1&hsa-miR-101-3p&offsets\|0\|+1(+1U);m-16&9\|+\|4850345\|4850364&offsets\|0\|+2(+1U)]\|TACAGTACTGTGATAACTGAAGT | >MI0000103\|hsa-mir-101-1&WithFlank&1\|-\|65058428\|65058514 | 53 | 74 | 22(+1U) | TACAGTACTGTGATAACTGAAGT | 594.9 | -3.1 | 0.9 | -3.4 | 6.9E-04 | 3.5E-03 | BT20/MB231 | MITO |
| >am_trna128_GlyGCC_6_-_27870686_27870756@10.35.26,trna18_GlyGCC_16_+_70822597_70822667@10.35.26,trna19_GlyGCC_16_+_70823410_70823480@10.35.26,trna19_GlyGCC_2_-_157257659_157257729@10.35.26,trna24_GlyGCC_16_-_70812942_70813012@10.35.26,trna25_GlyGCC_16_-_70812114_70812184@10.35.26,trna2_GlyGCC_21_-_18827107_18827177@10.35.26,trna35_GlyGCC_1_+_161413094_161413164@10.35.26,trna37_GlyGCC_1_+_161420467_161420537@10.35.26,trna39_GlyGCC_1_+_161427898_161427968@10.35.26,trna41_GlyGCC_1_+_161435258_161435328@10.35.26,trna5_GlyGCC_17_+_8029064_8029134@10.35.26,trna68_GlyGCC_1_-_161493637_161493707@10.35.26\|GTTCAGTGGTAGAATTCTCGCCTGCC | >am_trna128_GlyGCC_6_-_27870686_27870756 | 10 | 35 | 26 | GTTCAGTGGTAGAATTCTCGCCTGCC | 164.8 | -3.4 | 1.0 | -3.4 | 6.9E-04 | 3.5E-03 | BT20/MB231 | MITO |
| >MI0000736\|hsa-mir-30c-1&WithFlank&1\|+\|40757278\|40757378@23.46.24(+1A);MI0000254\|hsa-mir-30c-2&WithFlank&6\|-\|71376954\|71377037@13.36.24(+1A)&[MIMAT0000244&hsa-miR-30c-5p&offsets\|0\|+1(+1A);m-57&1\|+\|40757300\|40757323&offsets\|0\|0(+1A)];[MIMAT0000244_1&hsa-miR-30c-5p&offsets\|0\|+1(+1A);m-56&6\|-\|71377002\|71377025&offsets\|0\|0(+1A)]\|TGTAAACATCCTACACTCTCAGCTA | >MI0000736\|hsa-mir-30c-1&WithFlank&1\|+\|40757278\|40757378 | 23 | 46 | 24(+1A) | TGTAAACATCCTACACTCTCAGCTA | 506.8 | -3.1 | 0.9 | -3.4 | 6.9E-04 | 3.5E-03 | BT20/MB231 | MITO |
| >hg38_21_+_8433172_8446622_RNA45SN1_45S_50ntFlanks@4983.5016.34\|CGGACAGGATTGACAGATTGATAGCTCTTTCTCG | >hg38_21_+_8433172_8446622_RNA45SN1_45S_50ntFlanks | 4983 | 5016 | 34 | CGGACAGGATTGACAGATTGATAGCTCTTTCTCG | 144.5 | -4.5 | 1.5 | -3.0 | 2.5E-03 | 3.5E-03 | MB231/MB468 | MP |
| >hg38_MT_+_1621_3279_RNR2_16S_50ntFlanks@1349.1382.34\|ACAATAGGGTTTACGACCTCGATGTTGGATCAGG | >hg38_MT_+_1621_3279_RNR2_16S_50ntFlanks | 1349 | 1382 | 34 | ACAATAGGGTTTACGACCTCGATGTTGGATCAGG | 90.8 | 4.6 | 1.5 | 3.0 | 3.0E-03 | 3.5E-03 | MB231/MB468 | MITO |
| >hg38_21_+_8433172_8446622_RNA45SN1_45S_50ntFlanks@6785.6807.23\|CTACGCCTGTCTGAGCGTCGCTT | >hg38_21_+_8433172_8446622_RNA45SN1_45S_50ntFlanks | 6785 | 6807 | 23 | CTACGCCTGTCTGAGCGTCGCTT | 796.9 | 2.6 | 0.9 | 3.0 | 2.3E-03 | 3.5E-03 | BT20/MB468 | MITO |
| >hg38_21_+_8433172_8446622_RNA45SN1_45S_50ntFlanks@11585.11603.19\|GGTGCGGACCAGGGGAATC | >hg38_21_+_8433172_8446622_RNA45SN1_45S_50ntFlanks | 11585 | 11603 | 19 | GGTGCGGACCAGGGGAATC | 78.6 | -3.8 | 1.3 | -3.0 | 2.5E-03 | 3.5E-03 | MB231/MB468 | MP |
| >trna119_LysCTT_1_-_145395522_145395594@3.34.32,trna11_LysCTT_5_-_180648979_180649051@3.34.32,trna13_LysCTT_6_+_26556774_26556846@3.34.32,trna7_LysCTT_16_+_3225692_3225764@3.34.32,trna9_LysCTT_5_+_180634755_180634827@3.34.32\|CCGGCTAGCTCAGTCGGTAGAGCATGAGACTC | >trna119_LysCTT_1_-_145395522_145395594 | 3 | 34 | 32 | CCGGCTAGCTCAGTCGGTAGAGCATGAGACTC | 93.5 | 5.5 | 1.8 | 3.0 | 2.5E-03 | 3.5E-03 | MB231/MB468 | MP |

| | | | | | | | | | | | | | |
|---|---|---|---|---|---|---|---|---|---|---|---|---|---|
| >trna128_GlyGCC_6_-_27870686_27870756@19.38.20,trna19_GlyGCC_16_+_70823410_70823480@19.38.20,trna19_GlyGCC_2_-_157257659_157257729@19.38.20,trna24_GlyGCC_16_-_70812942_70813012@19.38.20,trna25_GlyGCC_16_-_70812114_70812184@19.38.20,trna2_GlyGCC_21_-_18827107_18827177@19.38.20,trna35_GlyGCC_1_+_161413094_161413164@19.38.20,trna37_GlyGCC_1_+_161420467_161420537@19.38.20,trna39_GlyGCC_1_+_161427898_161427968@19.38.20,trna41_GlyGCC_1_+_161435258_161435328@19.38.20,trna5_GlyGCC_17_+_8029064_8029134@19.38.20,trna68_GlyGCC_1_-_161493637_161493707@19.38.20\|TAGAATTCTCGCCTGCCACG | >trna128_GlyGCC_6_-_27870686_27870756 | 19 | 38 | 20 | TAGAATTCTCGCCTGCCACG | 295.3 | 3.6 | 1.2 | 3.0 | 2.8E-03 | 3.5E-03 | MB231/MB468 | CYTO |
| >MI0000114\|hsa-mir-107&WithFlank&10\|-\|89592741\|89592833@56.76.21(+1A);MI0000108\|hsa-mir-103a-2&WithFlank&20\|+\|3917488\|3917577@54.74.21(+1A);MI0000109\|hsa-mir-103a-1&WithFlank&5\|-\|168560890\|168560979@54.74.21(+1A)&[MIMAT0000104&hsa-miR-107&offsets\|0\|-2(+1A);m-78&10\|-\|89592758\|89592778&offsets\|0\|0(+1A)];[MIMAT0000101&hsa-miR-103a-3p&offsets\|0\|-2(+1A);m-12&20\|+\|3917541\|3917563&offsets\|0\|-2(+1A)];[MIMAT0000101_1&hsa-miR-103a-3p&offsets\|0\|-2(+1A);m-13&5\|-\|168560904\|168560926&offsets\|0\|-2(+1A);m-44220&5\|-\|168560908\|168560926&offsets\|0\|+2(+1A)]\|AGCAGCATTGTACAGGGCTATA | >MI0000114\|hsa-mir-107&WithFlank&10\|-\|89592741\|89592833 | 56 | 76 | 21(+1A) | AGCAGCATTGTACAGGGCTATA | 1814.8 | -3.5 | 1.0 | -3.5 | 5.2E-04 | 3.5E-03 | BT20/MB231 | MP |
| >hg38_21_+_8433172_8446622_RNA45SN1_45S_50ntFlanks@8171.8197.27\|AAGTCCTTCTGATCGAGGCCCAGCCCG | >hg38_21_+_8433172_8446622_RNA45SN1_45S_50ntFlanks | 8171 | 8197 | 27 | AAGTCCTTCTGATCGAGGCCCAGCCCG | 314.0 | -2.2 | 0.7 | -3.0 | 2.8E-03 | 3.5E-03 | MB231/MB468 | CYTO |
| >hg38_21_+_8433172_8446622_RNA45SN1_45S_50ntFlanks@11579.11609.31\|AGAACTGGTGCGGACCAGGGGAATCCGACTG | >hg38_21_+_8433172_8446622_RNA45SN1_45S_50ntFlanks | 11579 | 11609 | 31 | AGAACTGGTGCGGACCAGGGGAATCCGACTG | 1037.1 | -3.3 | 1.0 | -3.2 | 1.5E-03 | 3.5E-03 | BT20/MB468 | MP |
| >hg38_21_+_8433172_8446622_RNA45SN1_45S_50ntFlanks@10678.10705.28\|CGGTTCCGGCGGCGTCCGGTGAGCTCTC | >hg38_21_+_8433172_8446622_RNA45SN1_45S_50ntFlanks | 10678 | 10705 | 28 | CGGTTCCGGCGGCGTCCGGTGAGCTCTC | 1427.7 | -6.0 | 2.0 | -3.0 | 3.0E-03 | 3.5E-03 | BT20/MB468 | NUC |
| >hg38_1_-_228634819_228635039_RNA5S12_5S_50ntFlanks@134.172.39\|TGGGAGACCGCCTGGGAATACCGGGTGCTGTAGGCTTTT | >hg38_1_-_228634819_228635039_RNA5S12_5S_50ntFlanks | 134 | 172 | 39 | TGGGAGACCGCCTGGGAATACCGGGTGCTGTAGGCTTTT | 1844.7 | -3.6 | 1.1 | -3.4 | 6.2E-04 | 3.5E-03 | BT20/MB231 | CYTO |
| >MI0000103\|hsa-mir-101-1&WithFlank&1\|-\|65058428\|65058514@53.73.21;MI0000739\|hsa-mir-101-2&WithFlank&9\|+\|4850291\|4850381@55.75.21&[MIMAT0000099&hsa-miR-101-3p&offsets\|0\|0;m-17&1\|-\|65058443\|65058462&offsets\|0\|+1];[MIMAT0000099_1&hsa-miR-101-3p&offsets\|0\|0;m-16&9\|+\|4850345\|4850364&offsets\|0\|+1]\|TACAGTACTGTGATAACTGAA | >MI0000103\|hsa-mir-101-1&WithFlank&1\|-\|65058428\|65058514 | 53 | 73 | 21 | TACAGTACTGTGATAACTGAA | 244.2 | 3.0 | 0.9 | 3.4 | 6.2E-04 | 3.5E-03 | BT20/MB231 | CYTO |
| >hg38_21_+_8433172_8446622_RNA45SN1_45S_50ntFlanks@10794.10828.35\|CTGGCATGTTGGAACAATGTAGGTAAGGGAAGTCG | >hg38_21_+_8433172_8446622_RNA45SN1_45S_50ntFlanks | 10794 | 10828 | 35 | CTGGCATGTTGGAACAATGTAGGTAAGGGAAGTCG | 200.1 | -3.3 | 1.1 | -3.0 | 2.4E-03 | 3.5E-03 | BT20/MB468 | MITO |
| >hg38_21_+_8433172_8446622_RNA45SN1_45S_50ntFlanks@9554.9595.42\|CGTAGCGGTCCTGACGTGCAAATCGGTCGTCCGACCTGGGTA | >hg38_21_+_8433172_8446622_RNA45SN1_45S_50ntFlanks | 9554 | 9595 | 42 | CGTAGCGGTCCTGACGTGCAAATCGGTCGTCCGACCTGGGTA | 922.3 | -4.5 | 1.5 | -3.0 | 2.4E-03 | 3.5E-03 | BT20/MB468 | MITO |
| >hg38_21_+_8433172_8446622_RNA45SN1_45S_50ntFlanks@4687.4723.37\|ACGGACCAGAGCGAAAGCATTTGCCAAGAATGTTTTC | >hg38_21_+_8433172_8446622_RNA45SN1_45S_50ntFlanks | 4687 | 4723 | 37 | ACGGACCAGAGCGAAAGCATTTGCCAAGAATGTTTTC | 568.4 | -3.1 | 1.0 | -3.0 | 2.4E-03 | 3.5E-03 | BT20/MB468 | MITO |
| >hg38_21_+_8433172_8446622_RNA45SN1_45S_50ntFlanks@8381.8404.24\|AAGAGGGCGTGAAACCGTTAAGAG | >hg38_21_+_8433172_8446622_RNA45SN1_45S_50ntFlanks | 8381 | 8404 | 24 | AAGAGGGCGTGAAACCGTTAAGAG | 200.7 | -2.7 | 0.9 | -3.0 | 2.4E-03 | 3.5E-03 | BT20/MB468 | MITO |
| >hg38_21_+_8433172_8446622_RNA45SN1_45S_50ntFlanks@6661.6697.37\|CGGTGGATCACTCGGCTCGTGCGTCGATGAAGAACGC | >hg38_21_+_8433172_8446622_RNA45SN1_45S_50ntFlanks | 6661 | 6697 | 37 | CGGTGGATCACTCGGCTCGTGCGTCGATGAAGAACGC | 3355.3 | 1.9 | 0.6 | 3.0 | 2.4E-03 | 3.5E-03 | BT20/MB468 | MITO |
| >am_trnaMT_MetCAT_MT_+_4402_4469@19.36.18\|AGCTATCGGGCCCATACC | >am_trnaMT_MetCAT_MT_+_4402_4469 | 19 | 36 | 18 | AGCTATCGGGCCCATACC | 231.4 | 2.4 | 0.7 | 3.4 | 6.3E-04 | 3.5E-03 | BT20/MB231 | CYTO |
| >hg38_21_+_8433172_8446622_RNA45SN1_45S_50ntFlanks@11707.11736.30\|AAGCGCGGGTAAACGGCGGGAGTAACTATG | >hg38_21_+_8433172_8446622_RNA45SN1_45S_50ntFlanks | 11707 | 11736 | 30 | AAGCGCGGGTAAACGGCGGGAGTAACTATG | 173.5 | -4.1 | 1.4 | -3.0 | 2.6E-03 | 3.5E-03 | MB231/MB468 | MP |
| >hg38_21_+_8433172_8446622_RNA45SN1_45S_50ntFlanks@6646.6665.20\|TCGTACGACTCTTAGCGGTG | >hg38_21_+_8433172_8446622_RNA45SN1_45S_50ntFlanks | 6646 | 6665 | 20 | TCGTACGACTCTTAGCGGTG | 42645.1 | 2.5 | 0.7 | 3.4 | 6.3E-04 | 3.5E-03 | BT20/MB231 | CYTO |
| >hg38_21_+_8433172_8446622_RNA45SN1_45S_50ntFlanks@8018.8046.29\|ATTAGTCAGCGGAGGAGAAGAAACTAACC | >hg38_21_+_8433172_8446622_RNA45SN1_45S_50ntFlanks | 8018 | 8046 | 29 | ATTAGTCAGCGGAGGAGAAGAAACTAACC | 156.7 | -3.1 | 0.9 | -3.4 | 6.3E-04 | 3.6E-03 | BT20/MB231 | CYTO |
| >hg38_21_+_8433172_8446622_RNA45SN1_45S_50ntFlanks@12937.12960.24\|AACCATTCGTAGACGACCTGCTTC | >hg38_21_+_8433172_8446622_RNA45SN1_45S_50ntFlanks | 12937 | 12960 | 24 | AACCATTCGTAGACGACCTGCTTC | 456.3 | 2.0 | 0.7 | 3.0 | 2.9E-03 | 3.6E-03 | MB231/MB468 | CYTO |
| >am_MI0003581\|hsa-mir-574&WithFlank&4\|+\|38868026\|38868133@31.51.21&[MIMAT0004795&hsa-miR-574-5p&offsets\|0\|-2;m-213&4\|+\|38868056\|38868078&offsets\|0\|-2]\|TGAGTGTGTGTGTGTGAGTGT | >am_MI0003581\|hsa-mir-574&WithFlank&4\|+\|38868026\|38868133 | 31 | 51 | 21 | TGAGTGTGTGTGTGTGAGTGT | 403.9 | 3.9 | 1.3 | 3.0 | 2.6E-03 | 3.6E-03 | MB231/MB468 | MP |
| >hg38_21_+_8433172_8446622_RNA45SN1_45S_50ntFlanks@6657.6689.33\|TTAGCGGTGGATCACTCGGCTCGTGCGTCGATG | >hg38_21_+_8433172_8446622_RNA45SN1_45S_50ntFlanks | 6657 | 6689 | 33 | TTAGCGGTGGATCACTCGGCTCGTGCGTCGATG | 181.3 | -1.6 | 0.5 | -3.0 | 3.0E-03 | 3.6E-03 | MB231/MB468 | NUC |
| >hg38_MT_+_1621_3279_RNR2_16S_50ntFlanks@1380.1401.22\|AGGACATCCCGATGGTGCAGCC | >hg38_MT_+_1621_3279_RNR2_16S_50ntFlanks | 1380 | 1401 | 22 | AGGACATCCCGATGGTGCAGCC | 175.5 | 5.0 | 1.6 | 3.0 | 2.6E-03 | 3.6E-03 | MB231/MB468 | MP |
| >hg38_21_+_8433172_8446622_RNA45SN1_45S_50ntFlanks@11573.11592.20\|CGACTTAGAACTGGTGCGGA | >hg38_21_+_8433172_8446622_RNA45SN1_45S_50ntFlanks | 11573 | 11592 | 20 | CGACTTAGAACTGGTGCGGA | 1235.6 | -4.6 | 1.5 | -3.2 | 1.5E-03 | 3.6E-03 | BT20/MB468 | MP |
| >hg38_21_+_8433172_8446622_RNA45SN1_45S_50ntFlanks@12365.12383.19\|ATAACTGGCTTGTGGCGGC | >hg38_21_+_8433172_8446622_RNA45SN1_45S_50ntFlanks | 12365 | 12383 | 19 | ATAACTGGCTTGTGGCGGC | 140.9 | -2.1 | 0.7 | -3.0 | 2.4E-03 | 3.6E-03 | BT20/MB468 | MITO |
| >hg38_21_+_8433172_8446622_RNA45SN1_45S_50ntFlanks@9806.9831.26\|CGTGGAATGCGAGTGCCTAGTGGGCC | >hg38_21_+_8433172_8446622_RNA45SN1_45S_50ntFlanks | 9806 | 9831 | 26 | CGTGGAATGCGAGTGCCTAGTGGGCC | 152.0 | -1.8 | 0.6 | -3.0 | 2.4E-03 | 3.6E-03 | BT20/MB468 | MITO |
| >MI0000433\|hsa-let-7g&WithFlank&3\|-\|52268272\|52268367@11.31.21(+1G)&[MIMAT0000414&hsa-let-7g-5p&offsets\|0\|-1(+1G);m-58&3\|-\|52268336\|52268357&offsets\|0\|-1(+1G)]\|TGAGGTAGTAGTTTGTACAGTG | >MI0000433\|hsa-let-7g&WithFlank&3\|-\|52268272\|52268367 | 11 | 31 | 21(+1G) | TGAGGTAGTAGTTTGTACAGTG | 367.2 | 2.9 | 1.0 | 3.0 | 2.9E-03 | 3.6E-03 | MB231/MB468 | CYTO |
| >hg38_21_+_8433172_8446622_RNA45SN1_45S_50ntFlanks@9736.9755.20\|CCGAAACGATCTCAACCTAT | >hg38_21_+_8433172_8446622_RNA45SN1_45S_50ntFlanks | 9736 | 9755 | 20 | CCGAAACGATCTCAACCTAT | 117.1 | -5.0 | 1.6 | -3.2 | 1.5E-03 | 3.6E-03 | BT20/MB468 | MP |
| >MI0000465\|hsa-mir-191&WithFlank&3\|-\|49020612\|49020715@22.42.21(+1A)&[MIMAT0000440&hsa-miR-191-5p&offsets\|0\|-2(+1A);m-48&3\|-\|49020672\|49020694&offsets\|0\|-2(+1A)]\|CAACGGAATCCCAAAAGCAGCA | >MI0000465\|hsa-mir-191&WithFlank&3\|-\|49020612\|49020715 | 22 | 42 | 21(+1A) | CAACGGAATCCCAAAAGCAGCA | 741.3 | 2.9 | 1.0 | 3.0 | 2.6E-03 | 3.6E-03 | MB231/MB468 | MP |
| >MI0000446\|hsa-mir-125b-1&WithFlank&11\|-\|122099751\|122099850@21.42.22(+2A);MI0000470\|hsa-mir-125b-2&WithFlank&21\|+\|16590231\|16590331@23.44.22(+2A)&[MIMAT0000423&hsa-miR-125b-5p&offsets\|0\|0(+2A);m-67&11\|-\|122099809\|122099830&offsets\|0\|0(+2A)];[MIMAT0000423_1&hsa-miR-125b-5p&offsets\|0\|0(+2A);m-68&21\|+\|16590253\|16590274&offsets\|0\|0(+2A)]\|TCCCTGAGACCCTAACTTGTGAAA | >MI0000446\|hsa-mir-125b-1&WithFlank&11\|-\|122099751\|122099850 | 21 | 42 | 22(+2A) | TCCCTGAGACCCTAACTTGTGAAA | 81.8 | 3.8 | 1.2 | 3.0 | 2.6E-03 | 3.6E-03 | MB231/MB468 | MP |
| >hg38_21_+_8433172_8446622_RNA45SN1_45S_50ntFlanks@8348.8375.28\|GGAAAGTTGAAAAGAACTTTGAAGAGAG | >hg38_21_+_8433172_8446622_RNA45SN1_45S_50ntFlanks | 8348 | 8375 | 28 | GGAAAGTTGAAAAGAACTTTGAAGAGAG | 4677.6 | 4.6 | 1.3 | 3.5 | 5.3E-04 | 3.6E-03 | BT20/MB231 | MP |
| >hg38_21_+_8433172_8446622_RNA45SN1_45S_50ntFlanks@8117.8145.29\|CGGGACATGTGGCGTACGGAAGACCCGCT | >hg38_21_+_8433172_8446622_RNA45SN1_45S_50ntFlanks | 8117 | 8145 | 29 | CGGGACATGTGGCGTACGGAAGACCCGCT | 142.0 | 5.3 | 1.7 | 3.2 | 1.5E-03 | 3.6E-03 | BT20/MB468 | MP |
| >hg38_21_+_8433172_8446622_RNA45SN1_45S_50ntFlanks@12007.12035.29\|GCCGCCGGTGAAATACCACTACTCTGATC | >hg38_21_+_8433172_8446622_RNA45SN1_45S_50ntFlanks | 12007 | 12035 | 29 | GCCGCCGGTGAAATACCACTACTCTGATC | 322.9 | -2.1 | 0.7 | -3.0 | 2.6E-03 | 3.6E-03 | MB231/MB468 | MP |
| >hg38_21_+_8433172_8446622_RNA45SN1_45S_50ntFlanks@4160.4184.25\|CCACATCCAAGGAAGGCAGCAGGCG | >hg38_21_+_8433172_8446622_RNA45SN1_45S_50ntFlanks | 4160 | 4184 | 25 | CCACATCCAAGGAAGGCAGCAGGCG | 119.3 | -4.1 | 1.4 | -3.0 | 2.8E-03 | 3.6E-03 | BT20/MB468 | CYTO |
| >hg38_21_+_8433172_8446622_RNA45SN1_45S_50ntFlanks@6699.6717.19\|GCTAGCTGCGAGAATTAAT | >hg38_21_+_8433172_8446622_RNA45SN1_45S_50ntFlanks | 6699 | 6717 | 19 | GCTAGCTGCGAGAATTAAT | 358.2 | -4.6 | 1.5 | -3.0 | 2.4E-03 | 3.6E-03 | BT20/MB468 | MITO |
| >hg38_21_+_8433172_8446622_RNA45SN1_45S_50ntFlanks@5210.5240.31\|AGATGTCCGGGGCTGCACGCGCGCTACACTG | >hg38_21_+_8433172_8446622_RNA45SN1_45S_50ntFlanks | 5210 | 5240 | 31 | AGATGTCCGGGGCTGCACGCGCGCTACACTG | 412.2 | -4.6 | 1.5 | -3.0 | 2.4E-03 | 3.6E-03 | BT20/MB468 | MITO |
| >hg38_21_+_8433172_8446622_RNA45SN1_45S_50ntFlanks@9602.9620.19\|CGAAAGACTAATCGAACCA | >hg38_21_+_8433172_8446622_RNA45SN1_45S_50ntFlanks | 9602 | 9620 | 19 | CGAAAGACTAATCGAACCA | 37.0 | -8.3 | 2.6 | -3.2 | 1.5E-03 | 3.6E-03 | BT20/MB468 | MP |
| >hg38_21_+_8433172_8446622_RNA45SN1_45S_50ntFlanks@8357.8390.34\|AAAAGAACTTTGAAGAGAGAGTTCAAGAGGGCGT | >hg38_21_+_8433172_8446622_RNA45SN1_45S_50ntFlanks | 8357 | 8390 | 34 | AAAAGAACTTTGAAGAGAGAGTTCAAGAGGGCGT | 148.2 | -3.8 | 1.3 | -3.0 | 2.6E-03 | 3.6E-03 | MB231/MB468 | MP |
| >hg38_21_+_8433172_8446622_RNA45SN1_45S_50ntFlanks@11575.11613.39\|ACTTAGAACTGGTGCGGACCAGGGGAATCCGACTGTTTA | >hg38_21_+_8433172_8446622_RNA45SN1_45S_50ntFlanks | 11575 | 11613 | 39 | ACTTAGAACTGGTGCGGACCAGGGGAATCCGACTGTTTA | 318.5 | -3.2 | 1.1 | -3.0 | 2.6E-03 | 3.6E-03 | MB231/MB468 | MP |
| >hg38_21_+_8433172_8446622_RNA45SN1_45S_50ntFlanks@7992.8014.23\|CGTGGCGACCCGCTGAATTTAAG | >hg38_21_+_8433172_8446622_RNA45SN1_45S_50ntFlanks | 7992 | 8014 | 23 | CGTGGCGACCCGCTGAATTTAAG | 491.6 | -1.8 | 0.6 | -3.0 | 2.4E-03 | 3.6E-03 | BT20/MB468 | MITO |
| >hg38_21_+_8433172_8446622_RNA45SN1_45S_50ntFlanks@5472.5506.35\|ACGGCCCTGGCGGAGCGCTGAGAAGACGGTCGAAC | >hg38_21_+_8433172_8446622_RNA45SN1_45S_50ntFlanks | 5472 | 5506 | 35 | ACGGCCCTGGCGGAGCGCTGAGAAGACGGTCGAAC | 241.7 | 3.6 | 1.1 | 3.2 | 1.5E-03 | 3.6E-03 | BT20/MB468 | MP |
| >hg38_21_+_8433172_8446622_RNA45SN1_45S_50ntFlanks@3846.3871.26\|CTTGGATAACTGTGGTAATTCTAGAG | >hg38_21_+_8433172_8446622_RNA45SN1_45S_50ntFlanks | 3846 | 3871 | 26 | CTTGGATAACTGTGGTAATTCTAGAG | 124.0 | -4.1 | 1.3 | -3.2 | 1.5E-03 | 3.6E-03 | BT20/MB468 | MP |
| >hg38_21_+_8433172_8446622_RNA45SN1_45S_50ntFlanks@12259.12289.31\|CTTGATCTTGATTTTCAGTACGAATACAGAC | >hg38_21_+_8433172_8446622_RNA45SN1_45S_50ntFlanks | 12259 | 12289 | 31 | CTTGATCTTGATTTTCAGTACGAATACAGAC | 62.4 | -5.2 | 1.6 | -3.2 | 1.5E-03 | 3.6E-03 | BT20/MB468 | MP |
| >trnaMT_SerGCT_MT_+_12207_12265@14.48.35\|AGAACTGCTAACTCATGCCCCCATGTCTAACAACA | >trnaMT_SerGCT_MT_+_12207_12265 | 14 | 48 | 35 | AGAACTGCTAACTCATGCCCCCATGTCTAACAACA | 369.3 | 4.9 | 1.5 | 3.2 | 1.5E-03 | 3.6E-03 | BT20/MB468 | MP |
| >hg38_21_+_8433172_8446622_RNA45SN1_45S_50ntFlanks@11694.11717.24\|AAGAAATTCAATGAAGCGCGGGTA | >hg38_21_+_8433172_8446622_RNA45SN1_45S_50ntFlanks | 11694 | 11717 | 24 | AAGAAATTCAATGAAGCGCGGGTA | 338.8 | -3.8 | 1.3 | -3.0 | 2.8E-03 | 3.6E-03 | BT20/MB468 | CYTO |
| >hg38_21_+_8433172_8446622_RNA45SN1_45S_50ntFlanks@11583.11611.29\|CTGGTGCGGACCAGGGGAATCCGACTGTT | >hg38_21_+_8433172_8446622_RNA45SN1_45S_50ntFlanks | 11583 | 11611 | 29 | CTGGTGCGGACCAGGGGAATCCGACTGTT | 2447.3 | -1.7 | 0.6 | -3.0 | 2.9E-03 | 3.6E-03 | MB231/MB468 | CYTO |
| >hg38_21_+_8433172_8446622_RNA45SN1_45S_50ntFlanks@6719.6745.27\|TGAATTGCAGGACACATTGATCATCGA | >hg38_21_+_8433172_8446622_RNA45SN1_45S_50ntFlanks | 6719 | 6745 | 27 | TGAATTGCAGGACACATTGATCATCGA | 149.4 | -3.5 | 1.1 | -3.0 | 2.4E-03 | 3.6E-03 | BT20/MB468 | MITO |

<table>
<tr><th></th><th></th><th></th><th></th><th></th><th></th><th></th><th></th><th></th><th></th><th></th><th></th><th></th><th></th></tr>
<tr><td>>hg38_21_+_8433172_8446622_RNA45SN1_45S_50ntFlanks@6646.6670.25|TCGTACGACTCTTAGCGGTGGATCA</td><td>>hg38_21_+_8433172_8446622_RNA45SN1_45S_50ntFlanks</td><td>6646</td><td>6670</td><td>25</td><td>TCGTACGACTCTTAGCGGTGGATCA</td><td>1738.1</td><td>-1.4</td><td>0.5</td><td>-3.0</td><td>2.9E-03</td><td>3.6E-03</td><td>MB231/MB468</td><td>CYTO</td></tr>
<tr><td>>MI0000115|hsa-mir-16-2&WithFlank&3|+|160404739|160404831@58.79.22&[MIMAT0004518&hsa-miR-16-2-3p&offsets|-1|-1;m-252&3|+|160404796|160404816&offsets|0|+1]|ACCAATATTACTGTGCTGCTTT</td><td>>MI0000115|hsa-mir-16-2&WithFlank&3|+|160404739|160404831</td><td>58</td><td>79</td><td>22</td><td>ACCAATATTACTGTGCTGCTTT</td><td>445.1</td><td>2.9</td><td>0.9</td><td>3.2</td><td>1.5E-03</td><td>3.6E-03</td><td>BT20/MB468</td><td>MP</td></tr>
<tr><td>>MI0000750|hsa-mir-26a-2&WithFlank&12|-|57824603|57824698@20.40.21(+1G);MI0000083|hsa-mir-26a-1&WithFlank&3|+|37969398|37969486@16.36.21(+1G)&[MIMAT0000082&hsa-miR-26a-5p&offsets|0|-1(+1G);m-33&12|-|57824658|57824679&offsets|0|-1(+1G)];[MIMAT0000082_1&hsa-miR-26a-5p&offsets|0|-1(+1G);m-34&3|+|37969413|37969434&offsets|0|-1(+1G)]|TTCAAGTAATCCAGGATAGGCG</td><td>>MI0000750|hsa-mir-26a-2&WithFlank&12|-|57824603|57824698</td><td>20</td><td>40</td><td>21(+1G)</td><td>TTCAAGTAATCCAGGATAGGCG</td><td>1740.0</td><td>3.0</td><td>1.0</td><td>3.0</td><td>2.6E-03</td><td>3.6E-03</td><td>MB231/MB468</td><td>MP</td></tr>
<tr><td>>MI0000065|hsa-let-7d&WithFlank&9|+|94178828|94178926@14.35.22(+1C)&[MIMAT0000065&hsa-let-7d-5p&offsets|0|0(+1C);m-90&9|+|94178841|94178862&offsets|0|0(+1C)]|AGAGGTAGTAGGTTGCATAGTTC</td><td>>MI0000065|hsa-let-7d&WithFlank&9|+|94178828|94178926</td><td>14</td><td>35</td><td>22(+1C)</td><td>AGAGGTAGTAGGTTGCATAGTTC</td><td>126.1</td><td>4.9</td><td>1.6</td><td>3.0</td><td>2.6E-03</td><td>3.6E-03</td><td>MB231/MB468</td><td>MP</td></tr>
<tr><td>>hg38_21_+_8433172_8446622_RNA45SN1_45S_50ntFlanks@11882.11904.23|CTGTTGAGCTTGACTCTAGTCTG</td><td>>hg38_21_+_8433172_8446622_RNA45SN1_45S_50ntFlanks</td><td>11882</td><td>11904</td><td>23</td><td>CTGTTGAGCTTGACTCTAGTCTG</td><td>653.0</td><td>-2.4</td><td>0.8</td><td>-3.0</td><td>2.4E-03</td><td>3.6E-03</td><td>BT20/MB468</td><td>MITO</td></tr>
<tr><td>>hg38_21_+_8433172_8446622_RNA45SN1_45S_50ntFlanks@5444.5465.22|CTCGGATCGGCCCCGCCGGGGT</td><td>>hg38_21_+_8433172_8446622_RNA45SN1_45S_50ntFlanks</td><td>5444</td><td>5465</td><td>22</td><td>CTCGGATCGGCCCCGCCGGGGT</td><td>278.9</td><td>-2.3</td><td>0.8</td><td>-3.0</td><td>2.9E-03</td><td>3.6E-03</td><td>MB231/MB468</td><td>CYTO</td></tr>
<tr><td>>hg38_MT_+_1621_3279_RNR2_16S_50ntFlanks@930.959.30|AGTGACACATGTTTAACGGCCGCGGTACCC</td><td>>hg38_MT_+_1621_3279_RNR2_16S_50ntFlanks</td><td>930</td><td>959</td><td>30</td><td>AGTGACACATGTTTAACGGCCGCGGTACCC</td><td>172.0</td><td>3.7</td><td>1.2</td><td>3.0</td><td>2.6E-03</td><td>3.6E-03</td><td>MB231/MB468</td><td>MP</td></tr>
<tr><td>>hg38_21_+_8433172_8446622_RNA45SN1_45S_50ntFlanks@12204.12240.37|AAGGCGAGCTCAGGGAGGACAGAAACCTCCCGTGGAG</td><td>>hg38_21_+_8433172_8446622_RNA45SN1_45S_50ntFlanks</td><td>12204</td><td>12240</td><td>37</td><td>AAGGCGAGCTCAGGGAGGACAGAAACCTCCCGTGGAG</td><td>41.7</td><td>-9.2</td><td>3.1</td><td>-3.0</td><td>2.6E-03</td><td>3.6E-03</td><td>MB231/MB468</td><td>MP</td></tr>
<tr><td>>hg38_21_+_8433172_8446622_RNA45SN1_45S_50ntFlanks@10347.10383.37|CAAACGAGAACTTTGAAGGCCGAAGTGGAGAAGGGTT</td><td>>hg38_21_+_8433172_8446622_RNA45SN1_45S_50ntFlanks</td><td>10347</td><td>10383</td><td>37</td><td>CAAACGAGAACTTTGAAGGCCGAAGTGGAGAAGGGTT</td><td>295.5</td><td>-2.3</td><td>0.8</td><td>-3.0</td><td>3.1E-03</td><td>3.6E-03</td><td>MB231/MB468</td><td>NUC</td></tr>
<tr><td>>hg38_21_+_8433172_8446622_RNA45SN1_45S_50ntFlanks@3999.4024.26|CTAGATAACCTCGGGCCGATCGCACG</td><td>>hg38_21_+_8433172_8446622_RNA45SN1_45S_50ntFlanks</td><td>3999</td><td>4024</td><td>26</td><td>CTAGATAACCTCGGGCCGATCGCACG</td><td>135.3</td><td>-3.3</td><td>1.1</td><td>-3.0</td><td>2.4E-03</td><td>3.6E-03</td><td>BT20/MB468</td><td>MITO</td></tr>
<tr><td>>hg38_21_+_8433172_8446622_RNA45SN1_45S_50ntFlanks@10343.10361.19|TATTCAAACGAGAACTTTG</td><td>>hg38_21_+_8433172_8446622_RNA45SN1_45S_50ntFlanks</td><td>10343</td><td>10361</td><td>19</td><td>TATTCAAACGAGAACTTTG</td><td>270.0</td><td>-2.9</td><td>1.0</td><td>-3.0</td><td>2.8E-03</td><td>3.6E-03</td><td>BT20/MB468</td><td>CYTO</td></tr>
<tr><td>>hg38_21_+_8433172_8446622_RNA45SN1_45S_50ntFlanks@10361.10390.30|GAAGGCCGAAGTGGAGAAGGGTTCCATGTG</td><td>>hg38_21_+_8433172_8446622_RNA45SN1_45S_50ntFlanks</td><td>10361</td><td>10390</td><td>30</td><td>GAAGGCCGAAGTGGAGAAGGGTTCCATGTG</td><td>395.3</td><td>-5.3</td><td>1.7</td><td>-3.2</td><td>1.5E-03</td><td>3.6E-03</td><td>BT20/MB468</td><td>MP</td></tr>
<tr><td>>hg38_21_+_8433172_8446622_RNA45SN1_45S_50ntFlanks@9545.9564.20|GTGGAGGTCCGTAGCGGTCC</td><td>>hg38_21_+_8433172_8446622_RNA45SN1_45S_50ntFlanks</td><td>9545</td><td>9564</td><td>20</td><td>GTGGAGGTCCGTAGCGGTCC</td><td>120.2</td><td>-3.2</td><td>1.1</td><td>-3.0</td><td>2.4E-03</td><td>3.6E-03</td><td>BT20/MB468</td><td>MITO</td></tr>
<tr><td>>MI0000289|hsa-mir-181a-1&WithFlank&1|-|198859038|198859159@30.51.22;MI0000269|hsa-mir-181a-2&WithFlank&9|+|124692436|124692557@45.66.22&[MIMAT0000256&hsa-miR-181a-5p&offsets|0|-1;m-32&1|-|198859108|198859130&offsets|0|-1];[MIMAT0000256_1&hsa-miR-181a-5p&offsets|0|-1;m-32&9|+|124692480|124692502&offsets|0|-1]|AACATTCAACGCTGTCGGTGAG</td><td>>MI0000289|hsa-mir-181a-1&WithFlank&1|-|198859038|198859159</td><td>30</td><td>51</td><td>22</td><td>AACATTCAACGCTGTCGGTGAG</td><td>1149.9</td><td>-3.4</td><td>1.0</td><td>-3.5</td><td>5.4E-04</td><td>3.6E-03</td><td>BT20/MB231</td><td>MP</td></tr>
<tr><td>>hg38_21_+_8433172_8446622_RNA45SN1_45S_50ntFlanks@10236.10263.28|GACGCTACGCCGCGACGAGTAGGAGGGC</td><td>>hg38_21_+_8433172_8446622_RNA45SN1_45S_50ntFlanks</td><td>10236</td><td>10263</td><td>28</td><td>GACGCTACGCCGCGACGAGTAGGAGGGC</td><td>281.1</td><td>4.1</td><td>1.2</td><td>3.5</td><td>5.4E-04</td><td>3.6E-03</td><td>BT20/MB231</td><td>MP</td></tr>
<tr><td>>hg38_21_+_8433172_8446622_RNA45SN1_45S_50ntFlanks@11581.11611.31|AACTGGTGCGGACCAGGGGAATCCGACTGTT</td><td>>hg38_21_+_8433172_8446622_RNA45SN1_45S_50ntFlanks</td><td>11581</td><td>11611</td><td>31</td><td>AACTGGTGCGGACCAGGGGAATCCGACTGTT</td><td>1135.7</td><td>-2.3</td><td>0.8</td><td>-3.0</td><td>2.8E-03</td><td>3.6E-03</td><td>BT20/MB468</td><td>CYTO</td></tr>
<tr><td>>hg38_21_+_8433172_8446622_RNA45SN1_45S_50ntFlanks@12141.12158.18|ACAGTGCCAGGTGGGGAG</td><td>>hg38_21_+_8433172_8446622_RNA45SN1_45S_50ntFlanks</td><td>12141</td><td>12158</td><td>18</td><td>ACAGTGCCAGGTGGGGAG</td><td>134.4</td><td>-4.2</td><td>1.4</td><td>-3.0</td><td>2.8E-03</td><td>3.6E-03</td><td>BT20/MB468</td><td>CYTO</td></tr>
<tr><td>>hg38_21_+_8433172_8446622_RNA45SN1_45S_50ntFlanks@4163.4184.22|CATCCAAGGAAGGCAGCAGGCG</td><td>>hg38_21_+_8433172_8446622_RNA45SN1_45S_50ntFlanks</td><td>4163</td><td>4184</td><td>22</td><td>CATCCAAGGAAGGCAGCAGGCG</td><td>514.7</td><td>-4.7</td><td>1.6</td><td>-3.0</td><td>2.8E-03</td><td>3.6E-03</td><td>BT20/MB468</td><td>CYTO</td></tr>
<tr><td>>hg38_21_+_8433172_8446622_RNA45SN1_45S_50ntFlanks@12420.12442.23|ATGTCGGCTCTTCCTATCATTGT</td><td>>hg38_21_+_8433172_8446622_RNA45SN1_45S_50ntFlanks</td><td>12420</td><td>12442</td><td>23</td><td>ATGTCGGCTCTTCCTATCATTGT</td><td>1258.4</td><td>2.0</td><td>0.7</td><td>3.0</td><td>2.9E-03</td><td>3.6E-03</td><td>MB231/MB468</td><td>CYTO</td></tr>
<tr><td>>hg38_21_+_8433172_8446622_RNA45SN1_45S_50ntFlanks@11768.11790.23|CTAATTAGTGACGCGCATGAATG</td><td>>hg38_21_+_8433172_8446622_RNA45SN1_45S_50ntFlanks</td><td>11768</td><td>11790</td><td>23</td><td>CTAATTAGTGACGCGCATGAATG</td><td>509.4</td><td>-4.1</td><td>1.3</td><td>-3.2</td><td>1.5E-03</td><td>3.6E-03</td><td>BT20/MB468</td><td>MP</td></tr>
<tr><td>>hg38_21_+_8433172_8446622_RNA45SN1_45S_50ntFlanks@12152.12182.31|TGGGGAGTTTGACTGGGGCGGTACACCTGTC</td><td>>hg38_21_+_8433172_8446622_RNA45SN1_45S_50ntFlanks</td><td>12152</td><td>12182</td><td>31</td><td>TGGGGAGTTTGACTGGGGCGGTACACCTGTC</td><td>398.6</td><td>-4.0</td><td>1.3</td><td>-3.2</td><td>1.5E-03</td><td>3.6E-03</td><td>BT20/MB468</td><td>MP</td></tr>
<tr><td>>hg38_21_+_8433172_8446622_RNA45SN1_45S_50ntFlanks@12223.12248.26|CAGAAACCTCCCGTGGAGCAGAAGGG</td><td>>hg38_21_+_8433172_8446622_RNA45SN1_45S_50ntFlanks</td><td>12223</td><td>12248</td><td>26</td><td>CAGAAACCTCCCGTGGAGCAGAAGGG</td><td>113.3</td><td>-5.0</td><td>1.6</td><td>-3.2</td><td>1.5E-03</td><td>3.6E-03</td><td>BT20/MB468</td><td>MP</td></tr>
<tr><td>>hg38_21_+_8433172_8446622_RNA45SN1_45S_50ntFlanks@12569.12587.19|CAGTACGAGAGGAACCGCA</td><td>>hg38_21_+_8433172_8446622_RNA45SN1_45S_50ntFlanks</td><td>12569</td><td>12587</td><td>19</td><td>CAGTACGAGAGGAACCGCA</td><td>179.6</td><td>-1.8</td><td>0.6</td><td>-3.0</td><td>2.4E-03</td><td>3.6E-03</td><td>BT20/MB468</td><td>MITO</td></tr>
<tr><td>>hg38_21_+_8433172_8446622_RNA45SN1_45S_50ntFlanks@7978.8010.33|GACCTCAGATCAGACGTGGCGACCCGCTGAATT</td><td>>hg38_21_+_8433172_8446622_RNA45SN1_45S_50ntFlanks</td><td>7978</td><td>8010</td><td>33</td><td>GACCTCAGATCAGACGTGGCGACCCGCTGAATT</td><td>138.8</td><td>6.3</td><td>2.0</td><td>3.2</td><td>1.5E-03</td><td>3.6E-03</td><td>BT20/MB468</td><td>MP</td></tr>
<tr><td>>hg38_21_+_8433172_8446622_RNA45SN1_45S_50ntFlanks@12154.12183.30|GGGAGTTTGACTGGGGCGGTACACCTGTCA</td><td>>hg38_21_+_8433172_8446622_RNA45SN1_45S_50ntFlanks</td><td>12154</td><td>12183</td><td>30</td><td>GGGAGTTTGACTGGGGCGGTACACCTGTCA</td><td>232.6</td><td>-4.1</td><td>1.3</td><td>-3.2</td><td>1.5E-03</td><td>3.6E-03</td><td>BT20/MB468</td><td>MP</td></tr>
<tr><td>>hg38_21_+_8433172_8446622_RNA45SN1_45S_50ntFlanks@5431.5466.36|GTTTAGTGAGGCCCTCGGATCGGCCCCGCCGGGGTC</td><td>>hg38_21_+_8433172_8446622_RNA45SN1_45S_50ntFlanks</td><td>5431</td><td>5466</td><td>36</td><td>GTTTAGTGAGGCCCTCGGATCGGCCCCGCCGGGGTC</td><td>59.1</td><td>-5.0</td><td>1.6</td><td>-3.2</td><td>1.5E-03</td><td>3.6E-03</td><td>BT20/MB468</td><td>MP</td></tr>
<tr><td>>hg38_21_+_8433172_8446622_RNA45SN1_45S_50ntFlanks@9738.9760.23|GAAACGATCTCAACCTATTCTCA</td><td>>hg38_21_+_8433172_8446622_RNA45SN1_45S_50ntFlanks</td><td>9738</td><td>9760</td><td>23</td><td>GAAACGATCTCAACCTATTCTCA</td><td>166.4</td><td>-3.4</td><td>1.2</td><td>-3.0</td><td>3.1E-03</td><td>3.6E-03</td><td>MB231/MB468</td><td>NUC</td></tr>
<tr><td>>hg38_21_+_8433172_8446622_RNA45SN1_45S_50ntFlanks@9837.9856.20|TGGTAAGCAGAACTGGCGCT</td><td>>hg38_21_+_8433172_8446622_RNA45SN1_45S_50ntFlanks</td><td>9837</td><td>9856</td><td>20</td><td>TGGTAAGCAGAACTGGCGCT</td><td>35.5</td><td>8.9</td><td>2.8</td><td>3.2</td><td>1.5E-03</td><td>3.6E-03</td><td>BT20/MB468</td><td>MP</td></tr>
<tr><td>>MI0000298|hsa-mir-221&WithFlank&X|-|45746151|45746272@71.93.23&[MIMAT0000278&hsa-miR-221-3p&offsets|0|0;m-91&X|-|45746181|45746202&offsets|0|+1]|AGCTACATTGTCTGCTGGGTTTC</td><td>>MI0000298|hsa-mir-221&WithFlank&X|-|45746151|45746272</td><td>71</td><td>93</td><td>23</td><td>AGCTACATTGTCTGCTGGGTTTC</td><td>94264.4</td><td>-3.1</td><td>0.9</td><td>-3.5</td><td>5.4E-04</td><td>3.6E-03</td><td>BT20/MB231</td><td>MP</td></tr>
<tr><td>>am_trna128_GlyGCC_6_-_27870686_27870756@2.30.29,trna133_GlyCCC_1_-_16872434_16872504@2.30.29,trna18_GlyGCC_16_+_70822597_70822667@2.30.29,trna19_GlyGCC_16_+_70823410_70823480@2.30.29,trna19_GlyGCC_2_-_157257659_157257729@2.30.29,trna24_GlyGCC_16_-_70812942_70813012@2.30.29,trna25_GlyGCC_16_-_70812114_70812184@2.30.29,trna4_GlyCCC_1_+_17188416_17188486@2.30.29,trna5_GlyGCC_17_+_8029064_8029134@2.30.29,trna68_GlyGCC_1_-_161493637_161493707@2.30.29|CATTGGTGGTTCAGTGGTAGAATTCTCGC</td><td>>am_trna128_GlyGCC_6_-_27870686_27870756</td><td>2</td><td>30</td><td>29</td><td>CATTGGTGGTTCAGTGGTAGAATTCTCGC</td><td>527.5</td><td>-5.6</td><td>1.6</td><td>-3.5</td><td>5.4E-04</td><td>3.6E-03</td><td>BT20/MB231</td><td>MP</td></tr>
<tr><td>>hg38_21_+_8433172_8446622_RNA45SN1_45S_50ntFlanks@3843.3873.31|CTACTTGGATAACTGTGGTAATTCTAGAGCT</td><td>>hg38_21_+_8433172_8446622_RNA45SN1_45S_50ntFlanks</td><td>3843</td><td>3873</td><td>31</td><td>CTACTTGGATAACTGTGGTAATTCTAGAGCT</td><td>177.3</td><td>-4.3</td><td>1.4</td><td>-3.2</td><td>1.5E-03</td><td>3.6E-03</td><td>BT20/MB468</td><td>MP</td></tr>
<tr><td>>trna143_LysTTT_6_-_27559593_27559665@14.32.19,trna149_LysTTT_6_-_27302769_27302841@14.32.19,trna14_LysTTT_11_-_59327808_59327880@14.32.19,trna2_LysTTT_17_+_8022473_8022545@14.32.19,trna54_LysTTT_1_+_204475655_204475727@14.32.19,trna5_LysTTT_11_+_59323902_59323974@14.32.19,trna62_LysTTT_1_-_204476158_204476230@14.32.19,trna76_LysTTT_6_+_28918806_28918878@14.32.19|AGTCGGTAGAGCATCAGAC</td><td>>trna143_LysTTT_6_-_27559593_27559665</td><td>14</td><td>32</td><td>19</td><td>AGTCGGTAGAGCATCAGAC</td><td>332.0</td><td>2.5</td><td>0.8</td><td>3.0</td><td>2.9E-03</td><td>3.6E-03</td><td>MB231/MB468</td><td>CYTO</td></tr>
<tr><td>>hg38_21_+_8433172_8446622_RNA45SN1_45S_50ntFlanks@11569.11600.32|CAGCCGACTTAGAACTGGTGCGGACCAGGGGA</td><td>>hg38_21_+_8433172_8446622_RNA45SN1_45S_50ntFlanks</td><td>11569</td><td>11600</td><td>32</td><td>CAGCCGACTTAGAACTGGTGCGGACCAGGGGA</td><td>1114.4</td><td>-1.7</td><td>0.6</td><td>-3.0</td><td>2.4E-03</td><td>3.6E-03</td><td>BT20/MB468</td><td>MITO</td></tr>
<tr><td>>hg38_21_+_8433172_8446622_RNA45SN1_45S_50ntFlanks@8380.8406.27|CAAGAGGGCGTGAAACCGTTAAGAGGT</td><td>>hg38_21_+_8433172_8446622_RNA45SN1_45S_50ntFlanks</td><td>8380</td><td>8406</td><td>27</td><td>CAAGAGGGCGTGAAACCGTTAAGAGGT</td><td>478.8</td><td>-2.2</td><td>0.7</td><td>-3.0</td><td>2.8E-03</td><td>3.6E-03</td><td>BT20/MB468</td><td>CYTO</td></tr>
<tr><td>>MI0000478|hsa-mir-149&WithFlank&2|+|240455995|240456095@21.41.21&[MIMAT0000450&hsa-miR-149-5p&offsets|0|-2;m-195&2|+|240456015|240456037&offsets|0|-2]|TCTGGCTCCGTGTCTTCACTC</td><td>>MI0000478|hsa-mir-149&WithFlank&2|+|240455995|240456095</td><td>21</td><td>41</td><td>21</td><td>TCTGGCTCCGTGTCTTCACTC</td><td>490.1</td><td>2.6</td><td>0.8</td><td>3.4</td><td>6.4E-04</td><td>3.6E-03</td><td>BT20/MB231</td><td>CYTO</td></tr>
<tr><td>>hg38_21_+_8433172_8446622_RNA45SN1_45S_50ntFlanks@9716.9735.20|AATGATTAGAGGTCTTGGGG</td><td>>hg38_21_+_8433172_8446622_RNA45SN1_45S_50ntFlanks</td><td>9716</td><td>9735</td><td>20</td><td>AATGATTAGAGGTCTTGGGG</td><td>93.8</td><td>-3.1</td><td>1.0</td><td>-3.0</td><td>2.8E-03</td><td>3.6E-03</td><td>BT20/MB468</td><td>CYTO</td></tr>
<tr><td>>hg38_21_+_8433172_8446622_RNA45SN1_45S_50ntFlanks@12407.12426.20|TTTTGATCCTTCGATGTCGG</td><td>>hg38_21_+_8433172_8446622_RNA45SN1_45S_50ntFlanks</td><td>12407</td><td>12426</td><td>20</td><td>TTTTGATCCTTCGATGTCGG</td><td>332.4</td><td>-5.3</td><td>1.7</td><td>-3.2</td><td>1.6E-03</td><td>3.6E-03</td><td>BT20/MB468</td><td>MP</td></tr>
<tr><td>>hg38_MT_+_598_1651_RNR1_12S_50ntFlanks@416.456.41|AAATAGACTACGAAAGTGGCTTTAACATATCTGAACACACA</td><td>>hg38_MT_+_598_1651_RNR1_12S_50ntFlanks</td><td>416</td><td>456</td><td>41</td><td>AAATAGACTACGAAAGTGGCTTTAACATATCTGAACACACA</td><td>121.8</td><td>7.8</td><td>2.5</td><td>3.2</td><td>1.6E-03</td><td>3.6E-03</td><td>BT20/MB468</td><td>MP</td></tr>
<tr><td>>hg38_MT_+_1621_3279_RNR2_16S_50ntFlanks@503.530.28|CACTAGGAAAAAACCTTGTAGAGAGAGT</td><td>>hg38_MT_+_1621_3279_RNR2_16S_50ntFlanks</td><td>503</td><td>530</td><td>28</td><td>CACTAGGAAAAAACCTTGTAGAGAGAGT</td><td>83.2</td><td>6.1</td><td>1.9</td><td>3.2</td><td>1.6E-03</td><td>3.6E-03</td><td>BT20/MB468</td><td>MP</td></tr>
<tr><td>>hg38_21_+_8433172_8446622_RNA45SN1_45S_50ntFlanks@7972.8009.38|AGACGCGACCTCAGATCAGACGTGGCGACCCGCTGAAT</td><td>>hg38_21_+_8433172_8446622_RNA45SN1_45S_50ntFlanks</td><td>7972</td><td>8009</td><td>38</td><td>AGACGCGACCTCAGATCAGACGTGGCGACCCGCTGAAT</td><td>84.3</td><td>6.2</td><td>2.0</td><td>3.2</td><td>1.6E-03</td><td>3.6E-03</td><td>BT20/MB468</td><td>MP</td></tr>
<tr><td>>hg38_21_+_8433172_8446622_RNA45SN1_45S_50ntFlanks@5487.5526.40|CGCTGAGAAGACGGTCGAACTTGACTATCTAGAGGAAGTA</td><td>>hg38_21_+_8433172_8446622_RNA45SN1_45S_50ntFlanks</td><td>5487</td><td>5526</td><td>40</td><td>CGCTGAGAAGACGGTCGAACTTGACTATCTAGAGGAAGTA</td><td>685.4</td><td>-4.5</td><td>1.4</td><td>-3.2</td><td>1.6E-03</td><td>3.6E-03</td><td>BT20/MB468</td><td>MP</td></tr>
<tr><td>>hg38_21_+_8433172_8446622_RNA45SN1_45S_50ntFlanks@5071.5110.40|TTAATTCCGATAACGAACGAGACTCTGGCATGCTAACTAG</td><td>>hg38_21_+_8433172_8446622_RNA45SN1_45S_50ntFlanks</td><td>5071</td><td>5110</td><td>40</td><td>TTAATTCCGATAACGAACGAGACTCTGGCATGCTAACTAG</td><td>225.1</td><td>-3.4</td><td>1.1</td><td>-3.0</td><td>2.4E-03</td><td>3.6E-03</td><td>BT20/MB468</td><td>MITO</td></tr>
<tr><td>>hg38_21_+_8433172_8446622_RNA45SN1_45S_50ntFlanks@12003.12030.28|GCGGGCCGCCGGTGAAATACCACTACTC</td><td>>hg38_21_+_8433172_8446622_RNA45SN1_45S_50ntFlanks</td><td>12003</td><td>12030</td><td>28</td><td>GCGGGCCGCCGGTGAAATACCACTACTC</td><td>362.6</td><td>-1.6</td><td>0.5</td><td>-3.0</td><td>2.4E-03</td><td>3.6E-03</td><td>BT20/MB468</td><td>MITO</td></tr>
<tr><td>>hg38_MT_+_1621_3279_RNR2_16S_50ntFlanks@1249.1275.27|AGTCAAAGCGAACTACTATACTCAATT</td><td>>hg38_MT_+_1621_3279_RNR2_16S_50ntFlanks</td><td>1249</td><td>1275</td><td>27</td><td>AGTCAAAGCGAACTACTATACTCAATT</td><td>340.8</td><td>4.2</td><td>1.4</td><td>3.0</td><td>2.6E-03</td><td>3.6E-03</td><td>MB231/MB468</td><td>MP</td></tr>
<tr><td>>MI0000095|hsa-mir-93&WithFlank&7|-|100093762|100093853@17.38.22(+1C)&[MIMAT0000093&hsa-miR-93-5p&offsets|0|-1(+1C);m-29&7|-|100093815|100093837&offsets|0|-1(+1C)]|CAAAGTGCTGTTCGTGCAGGTAC</td><td>>MI0000095|hsa-mir-93&WithFlank&7|-|100093762|100093853</td><td>17</td><td>38</td><td>22(+1C)</td><td>CAAAGTGCTGTTCGTGCAGGTAC</td><td>160.1</td><td>4.0</td><td>1.3</td><td>3.0</td><td>2.6E-03</td><td>3.6E-03</td><td>MB231/MB468</td><td>MP</td></tr>
</table>

<table>
<tr><td>>hg38_21_+_8433172_8446622_RNA45SN1_45S_50ntFlanks@11770.11792.23|AATTAGTGACGCGCATGAATGGA</td><td>>hg38_21_+_8433172_8446622_RNA45SN1_45S_50ntFlanks</td><td>11770</td><td>11792</td><td>23</td><td>AATTAGTGACGCGCATGAATGGA</td><td>167.0</td><td>-3.3</td><td>1.1</td><td>-3.0</td><td>3.1E-03</td><td>3.6E-03</td><td>MB231/MB468</td><td>NUC</td></tr>
<tr><td>>hg38_21_+_8433172_8446622_RNA45SN1_45S_50ntFlanks@5485.5516.32|AGCGCTGAGAAGACGGTCGAACTTGACTATCT</td><td>>hg38_21_+_8433172_8446622_RNA45SN1_45S_50ntFlanks</td><td>5485</td><td>5516</td><td>32</td><td>AGCGCTGAGAAGACGGTCGAACTTGACTATCT</td><td>1539.8</td><td>-3.3</td><td>1.1</td><td>-3.0</td><td>2.6E-03</td><td>3.6E-03</td><td>MB231/MB468</td><td>MP</td></tr>
<tr><td>>hg38_21_+_8433172_8446622_RNA45SN1_45S_50ntFlanks@10717.10736.20|AAAATCCGGGGGAGAGGGTG</td><td>>hg38_21_+_8433172_8446622_RNA45SN1_45S_50ntFlanks</td><td>10717</td><td>10736</td><td>20</td><td>AAAATCCGGGGGAGAGGGTG</td><td>424.6</td><td>-3.9</td><td>1.3</td><td>-3.0</td><td>2.6E-03</td><td>3.6E-03</td><td>MB231/MB468</td><td>MP</td></tr>
<tr><td>>trnaMT_LeuTAG_MT_+_12266_12336@38.74.37|CCCAAAAATTTTGGTGCAACTCCAAATAAAAGTACCA</td><td>>trnaMT_LeuTAG_MT_+_12266_12336</td><td>38</td><td>74</td><td>37</td><td>CCCAAAAATTTTGGTGCAACTCCAAATAAAAGTACCA</td><td>204.7</td><td>4.5</td><td>1.5</td><td>3.0</td><td>3.1E-03</td><td>3.6E-03</td><td>MB231/MB468</td><td>MITO</td></tr>
<tr><td>>trna10_ValCAC_5_-_180649395_180649467@31.54.24,trna13_ValCAC_19_-_4724647_4724719@31.54.24,trna152_ValCAC_6_-_27248049_27248121@31.54.24,trna157_ValCAC_6_-_27173867_27173939@31.54.24,trna18_ValCAC_5_-_180529253_180529325@31.54.24,trna2_ValCAC_5_+_180524070_180524142@31.54.24,trna6_ValCAC_5_+_180600650_180600722@31.54.24,trna85_ValCAC_1_-_161369490_161369562@31.54.24,trna98_ValCAC_1_-_149298555_149298627@31.54.24,trna9_ValCAC_6_+_26538282_26538354@31.54.24|CCTCACACGCGAAAGGTCCCCGGT</td><td>>trna10_ValCAC_5_-_180649395_180649467</td><td>31</td><td>54</td><td>24</td><td>CCTCACACGCGAAAGGTCCCCGGT</td><td>37.8</td><td>-8.3</td><td>2.6</td><td>-3.2</td><td>1.6E-03</td><td>3.6E-03</td><td>BT20/MB468</td><td>MP</td></tr>
<tr><td>>hg38_MT_+_1621_3279_RNR2_16S_50ntFlanks@1427.1457.31|GATTAAAGTCCTACGTGATCTGAGTTCAGAC</td><td>>hg38_MT_+_1621_3279_RNR2_16S_50ntFlanks</td><td>1427</td><td>1457</td><td>31</td><td>GATTAAAGTCCTACGTGATCTGAGTTCAGAC</td><td>111.2</td><td>3.8</td><td>1.3</td><td>3.0</td><td>3.1E-03</td><td>3.6E-03</td><td>MB231/MB468</td><td>MITO</td></tr>
<tr><td>>am_trna128_GlyGCC_6_-_27870686_27870756@1.36.36,trna18_GlyGCC_16_+_70822597_70822667@1.36.36,trna19_GlyGCC_16_+_70823410_70823480@1.36.36,trna19_GlyGCC_2_-_157257659_157257729@1.36.36,trna24_GlyGCC_16_-_70812942_70813012@1.36.36,trna25_GlyGCC_16_-_70812114_70812184@1.36.36,trna5_GlyGCC_17_+_8029064_8029134@1.36.36,trna68_GlyGCC_1_-_161493637_161493707@1.36.36|GCATTGGTGGTTCAGTGGTAGAATTCTCGCCTGCCA</td><td>>am_trna128_GlyGCC_6_-_27870686_27870756</td><td>1</td><td>36</td><td>36</td><td>GCATTGGTGGTTCAGTGGTAGAATTCTCGCCTGCCA</td><td>1236.5</td><td>-2.5</td><td>0.7</td><td>-3.4</td><td>6.5E-04</td><td>3.6E-03</td><td>BT20/MB231</td><td>CYTO</td></tr>
<tr><td>>hg38_21_+_8433172_8446622_RNA45SN1_45S_50ntFlanks@11665.11703.39|TTCTGCCCAGTGCTCTGAATGTCAAAGTGAAGAAATTCA</td><td>>hg38_21_+_8433172_8446622_RNA45SN1_45S_50ntFlanks</td><td>11665</td><td>11703</td><td>39</td><td>TTCTGCCCAGTGCTCTGAATGTCAAAGTGAAGAAATTCA</td><td>353.7</td><td>-2.8</td><td>0.8</td><td>-3.4</td><td>6.5E-04</td><td>3.6E-03</td><td>BT20/MB231</td><td>CYTO</td></tr>
<tr><td>>MI0000071|hsa-mir-17&WithFlank&13|+|91350599|91350694@20.42.23&[MIMAT0000070&hsa-miR-17-5p&offsets|0|0;m-88&13|+|91350618|91350640&offsets|0|0]|CAAAGTGCTTACAGTGCAGGTAG</td><td>>MI0000071|hsa-mir-17&WithFlank&13|+|91350599|91350694</td><td>20</td><td>42</td><td>23</td><td>CAAAGTGCTTACAGTGCAGGTAG</td><td>13998.2</td><td>2.4</td><td>0.8</td><td>3.0</td><td>2.6E-03</td><td>3.6E-03</td><td>MB231/MB468</td><td>MP</td></tr>
<tr><td>>hg38_21_+_8433172_8446622_RNA45SN1_45S_50ntFlanks@11537.11581.45|CCGCGCGGCGCCTCGCCTCGGCCGGCGCCTAGCAGCCGACTTAGA</td><td>>hg38_21_+_8433172_8446622_RNA45SN1_45S_50ntFlanks</td><td>11537</td><td>11581</td><td>45</td><td>CCGCGCGGCGCCTCGCCTCGGCCGGCGCCTAGCAGCCGACTTAGA</td><td>257.4</td><td>-3.1</td><td>1.1</td><td>-3.0</td><td>3.1E-03</td><td>3.6E-03</td><td>MB231/MB468</td><td>MITO</td></tr>
<tr><td>>hg38_21_+_8433172_8446622_RNA45SN1_45S_50ntFlanks@5237.5282.46|ACTGACTGGCTCAGCGTGTGCCTACCCTACGCCGGCAGGCGCGGGT</td><td>>hg38_21_+_8433172_8446622_RNA45SN1_45S_50ntFlanks</td><td>5237</td><td>5282</td><td>46</td><td>ACTGACTGGCTCAGCGTGTGCCTACCCTACGCCGGCAGGCGCGGGT</td><td>81.7</td><td>5.7</td><td>1.9</td><td>3.0</td><td>2.6E-03</td><td>3.6E-03</td><td>MB231/MB468</td><td>MP</td></tr>
<tr><td>>MI0000108|hsa-mir-103a-2&WithFlank&20|+|3917488|3917577@54.75.22;MI0000109|hsa-mir-103a-1&WithFlank&5|-|168560890|168560979@54.75.22&[MIMAT0000101&hsa-miR-103a-3p&offsets|0|-1;m-12&20|+|3917541|3917563&offsets|0|-1];[MIMAT0000101_1&hsa-miR-103a-3p&offsets|0|-1;m-13&5|-|168560904|168560926&offsets|0|-1;m-44220&5|-|168560908|168560926&offsets|0|+3]|AGCAGCATTGTACAGGGCTATG</td><td>>MI0000108|hsa-mir-103a-2&WithFlank&20|+|3917488|3917577</td><td>54</td><td>75</td><td>22</td><td>AGCAGCATTGTACAGGGCTATG</td><td>7474.1</td><td>2.7</td><td>0.9</td><td>3.0</td><td>2.6E-03</td><td>3.6E-03</td><td>MB231/MB468</td><td>MP</td></tr>
<tr><td>>hg38_21_+_8433172_8446622_RNA45SN1_45S_50ntFlanks@6648.6669.22|GTACGACTCTTAGCGGTGGATC</td><td>>hg38_21_+_8433172_8446622_RNA45SN1_45S_50ntFlanks</td><td>6648</td><td>6669</td><td>22</td><td>GTACGACTCTTAGCGGTGGATC</td><td>924.3</td><td>2.9</td><td>0.8</td><td>3.4</td><td>6.5E-04</td><td>3.6E-03</td><td>BT20/MB231</td><td>CYTO</td></tr>
<tr><td>>hg38_21_+_8433172_8446622_RNA45SN1_45S_50ntFlanks@11203.11225.23|CGGCGGCGGCGACTCTGGACGCG</td><td>>hg38_21_+_8433172_8446622_RNA45SN1_45S_50ntFlanks</td><td>11203</td><td>11225</td><td>23</td><td>CGGCGGCGGCGACTCTGGACGCG</td><td>81.7</td><td>9.7</td><td>2.8</td><td>3.5</td><td>5.4E-04</td><td>3.6E-03</td><td>BT20/MB231</td><td>MP</td></tr>
<tr><td>>hg38_21_+_8433172_8446622_RNA45SN1_45S_50ntFlanks@3861.3899.39|TAATTCTAGAGCTAATACATGCCGACGGGCGCTGACCCC</td><td>>hg38_21_+_8433172_8446622_RNA45SN1_45S_50ntFlanks</td><td>3861</td><td>3899</td><td>39</td><td>TAATTCTAGAGCTAATACATGCCGACGGGCGCTGACCCC</td><td>160.7</td><td>-2.2</td><td>0.7</td><td>-3.0</td><td>2.9E-03</td><td>3.7E-03</td><td>MB231/MB468</td><td>CYTO</td></tr>
<tr><td>>hg38_21_+_8433172_8446622_RNA45SN1_45S_50ntFlanks@9951.9981.31|CCATGGAAGTCGGAATCCGCTAAGGAGTGTG</td><td>>hg38_21_+_8433172_8446622_RNA45SN1_45S_50ntFlanks</td><td>9951</td><td>9981</td><td>31</td><td>CCATGGAAGTCGGAATCCGCTAAGGAGTGTG</td><td>77.3</td><td>2.8</td><td>0.9</td><td>3.2</td><td>1.6E-03</td><td>3.7E-03</td><td>BT20/MB468</td><td>MP</td></tr>
<tr><td>>hg38_21_+_8433172_8446622_RNA45SN1_45S_50ntFlanks@8192.8211.20|AGCCCGTGGACGGTGTGAGG</td><td>>hg38_21_+_8433172_8446622_RNA45SN1_45S_50ntFlanks</td><td>8192</td><td>8211</td><td>20</td><td>AGCCCGTGGACGGTGTGAGG</td><td>300.7</td><td>-3.9</td><td>1.3</td><td>-3.0</td><td>3.1E-03</td><td>3.7E-03</td><td>MB231/MB468</td><td>NUC</td></tr>
<tr><td>>trnaMT_ValTAC_MT_+_1602_1670@54.72.19|AACTTGACCGCTCTGACCA</td><td>>trnaMT_ValTAC_MT_+_1602_1670</td><td>54</td><td>72</td><td>19</td><td>AACTTGACCGCTCTGACCA</td><td>985.8</td><td>2.9</td><td>0.9</td><td>3.2</td><td>1.6E-03</td><td>3.7E-03</td><td>BT20/MB468</td><td>MP</td></tr>
<tr><td>>hg38_MT_+_598_1651_RNR1_12S_50ntFlanks@428.455.28|AAAGTGGCTTTAACATATCTGAACACAC</td><td>>hg38_MT_+_598_1651_RNR1_12S_50ntFlanks</td><td>428</td><td>455</td><td>28</td><td>AAAGTGGCTTTAACATATCTGAACACAC</td><td>592.2</td><td>3.8</td><td>1.2</td><td>3.2</td><td>1.6E-03</td><td>3.7E-03</td><td>BT20/MB468</td><td>MP</td></tr>
<tr><td>>hg38_21_+_8433172_8446622_RNA45SN1_45S_50ntFlanks@10377.10401.25|AAGGGTTCCATGTGAACAGCAGTTG</td><td>>hg38_21_+_8433172_8446622_RNA45SN1_45S_50ntFlanks</td><td>10377</td><td>10401</td><td>25</td><td>AAGGGTTCCATGTGAACAGCAGTTG</td><td>37.1</td><td>-8.3</td><td>2.6</td><td>-3.2</td><td>1.6E-03</td><td>3.7E-03</td><td>BT20/MB468</td><td>MP</td></tr>
<tr><td>>MI0000469|hsa-mir-125a&WithFlank&19|+|51693248|51693345@21.41.21(+1C)&[MIMAT0000443&hsa-miR-125a-5p&offsets|0|-3(+1C);m-63&19|+|51693268|51693289&offsets|0|-1(+1C)]|TCCCTGAGACCCTTTAACCTGC</td><td>>MI0000469|hsa-mir-125a&WithFlank&19|+|51693248|51693345</td><td>21</td><td>41</td><td>21(+1C)</td><td>TCCCTGAGACCCTTTAACCTGC</td><td>724.4</td><td>2.1</td><td>0.6</td><td>3.4</td><td>5.9E-04</td><td>3.7E-03</td><td>BT20/MB231</td><td>NUC</td></tr>
<tr><td>>hg38_21_+_8433172_8446622_RNA45SN1_45S_50ntFlanks@5467.5491.25|GGCCCACGGCCCTGGCGGAGCGCTG</td><td>>hg38_21_+_8433172_8446622_RNA45SN1_45S_50ntFlanks</td><td>5467</td><td>5491</td><td>25</td><td>GGCCCACGGCCCTGGCGGAGCGCTG</td><td>562.9</td><td>-5.6</td><td>1.9</td><td>-3.0</td><td>3.1E-03</td><td>3.7E-03</td><td>BT20/MB468</td><td>NUC</td></tr>
<tr><td>>am_trna128_GlyGCC_6_-_27870686_27870756@4.28.25,trna133_GlyCCC_1_-_16872434_16872504@4.28.25,trna18_GlyGCC_16_+_70822597_70822667@4.28.25,trna19_GlyGCC_16_+_70823410_70823480@4.28.25,trna19_GlyGCC_2_-_157257659_157257729@4.28.25,trna24_GlyGCC_16_-_70812942_70813012@4.28.25,trna25_GlyGCC_16_-_70812114_70812184@4.28.25,trna4_GlyCCC_1_+_17188416_17188486@4.28.25,trna5_GlyGCC_17_+_8029064_8029134@4.28.25,trna68_GlyGCC_1_-_161493637_161493707@4.28.25|TTGGTGGTTCAGTGGTAGAATTCTC</td><td>>am_trna128_GlyGCC_6_-_27870686_27870756</td><td>4</td><td>28</td><td>25</td><td>TTGGTGGTTCAGTGGTAGAATTCTC</td><td>262.6</td><td>-2.3</td><td>0.8</td><td>-3.0</td><td>2.9E-03</td><td>3.7E-03</td><td>MB231/MB468</td><td>CYTO</td></tr>
<tr><td>>hg38_21_+_8433172_8446622_RNA45SN1_45S_50ntFlanks@12996.13014.19|TCGCTGCGATCTATTGAAA</td><td>>hg38_21_+_8433172_8446622_RNA45SN1_45S_50ntFlanks</td><td>12996</td><td>13014</td><td>19</td><td>TCGCTGCGATCTATTGAAA</td><td>1425.6</td><td>-2.7</td><td>0.8</td><td>-3.4</td><td>5.9E-04</td><td>3.7E-03</td><td>BT20/MB231</td><td>NUC</td></tr>
<tr><td>>hg38_21_+_8433172_8446622_RNA45SN1_45S_50ntFlanks@12458.12487.30|AAGCGTTGGATTGTTCACCCACTAATAGGG</td><td>>hg38_21_+_8433172_8446622_RNA45SN1_45S_50ntFlanks</td><td>12458</td><td>12487</td><td>30</td><td>AAGCGTTGGATTGTTCACCCACTAATAGGG</td><td>613.6</td><td>-4.8</td><td>1.6</td><td>-3.0</td><td>2.9E-03</td><td>3.7E-03</td><td>BT20/MB468</td><td>CYTO</td></tr>
<tr><td>>hg38_21_+_8433172_8446622_RNA45SN1_45S_50ntFlanks@11200.11233.34|CGGCGGCGGCGGCGACTCTGGACGCGAGCCGGGC</td><td>>hg38_21_+_8433172_8446622_RNA45SN1_45S_50ntFlanks</td><td>11200</td><td>11233</td><td>34</td><td>CGGCGGCGGCGGCGACTCTGGACGCGAGCCGGGC</td><td>170.8</td><td>-3.5</td><td>1.2</td><td>-3.0</td><td>3.1E-03</td><td>3.7E-03</td><td>MB231/MB468</td><td>NUC</td></tr>
<tr><td>>MI0000093|hsa-mir-92a-1&WithFlank&13|+|91351308|91351397@54.75.22(+1C);MI0000094|hsa-mir-92a-2&WithFlank&X|-|134169532|134169618@54.75.22(+1C)&[MIMAT0000092&hsa-miR-92a-3p&offsets|0|0(+1C);m-9&13|+|91351361|91351382&offsets|0|0(+1C)];[MIMAT0000092_1&hsa-miR-92a-3p&offsets|0|0(+1C);m-11&X|-|134169544|134169565&offsets|0|0(+1C)]|TATTGCACTTGTCCCGGCCTGTC</td><td>>MI0000093|hsa-mir-92a-1&WithFlank&13|+|91351308|91351397</td><td>54</td><td>75</td><td>22(+1C)</td><td>TATTGCACTTGTCCCGGCCTGTC</td><td>1164.0</td><td>-2.6</td><td>0.8</td><td>-3.4</td><td>7.2E-04</td><td>3.7E-03</td><td>BT20/MB231</td><td>MITO</td></tr>
<tr><td>>hg38_MT_+_598_1651_RNR1_12S_50ntFlanks@966.1004.39|TAACATGGTAAGTGTACTGGAAAGTGCACTTGGACGAAC</td><td>>hg38_MT_+_598_1651_RNR1_12S_50ntFlanks</td><td>966</td><td>1004</td><td>39</td><td>TAACATGGTAAGTGTACTGGAAAGTGCACTTGGACGAAC</td><td>140.1</td><td>-5.4</td><td>1.6</td><td>-3.4</td><td>6.0E-04</td><td>3.7E-03</td><td>BT20/MB231</td><td>NUC</td></tr>
<tr><td>>hg38_21_+_8433172_8446622_RNA45SN1_45S_50ntFlanks@12075.12093.19|GAGGGGCTCTCGCTTCTGG</td><td>>hg38_21_+_8433172_8446622_RNA45SN1_45S_50ntFlanks</td><td>12075</td><td>12093</td><td>19</td><td>GAGGGGCTCTCGCTTCTGG</td><td>41.1</td><td>-8.4</td><td>2.7</td><td>-3.2</td><td>1.6E-03</td><td>3.7E-03</td><td>BT20/MB468</td><td>MP</td></tr>
<tr><td>>MI0000266|hsa-mir-10a&WithFlank&17|-|48579832|48579953@28.48.21(+1G)&[MIMAT0000253&hsa-miR-10a-5p&offsets|0|-2(+1G);m-18&17|-|48579905|48579926&offsets|0|-1(+1G)]|TACCCTGTAGATCCGAATTTGG</td><td>>MI0000266|hsa-mir-10a&WithFlank&17|-|48579832|48579953</td><td>28</td><td>48</td><td>21(+1G)</td><td>TACCCTGTAGATCCGAATTTGG</td><td>786.3</td><td>3.0</td><td>0.9</td><td>3.4</td><td>6.5E-04</td><td>3.7E-03</td><td>BT20/MB231</td><td>CYTO</td></tr>
<tr><td>>hg38_21_+_8433172_8446622_RNA45SN1_45S_50ntFlanks@6661.6678.18|CGGTGGATCACTCGGCTC</td><td>>hg38_21_+_8433172_8446622_RNA45SN1_45S_50ntFlanks</td><td>6661</td><td>6678</td><td>18</td><td>CGGTGGATCACTCGGCTC</td><td>1653.5</td><td>4.2</td><td>1.2</td><td>3.5</td><td>5.5E-04</td><td>3.7E-03</td><td>BT20/MB231</td><td>MP</td></tr>
<tr><td>>trnaMT_GluTTC_MT_-_14674_14742@40.64.25,trnalookalike8_GluTTC_5_-_93905172_93905240@40.64.25|TTGGTCGTGGTTGTAGTCCGTGCGA</td><td>>trnaMT_GluTTC_MT_-_14674_14742</td><td>40</td><td>64</td><td>25</td><td>TTGGTCGTGGTTGTAGTCCGTGCGA</td><td>105.5</td><td>5.4</td><td>1.7</td><td>3.2</td><td>1.6E-03</td><td>3.7E-03</td><td>BT20/MB468</td><td>MP</td></tr>
<tr><td>>MI0000298|hsa-mir-221&WithFlank&X|-|45746151|45746272@71.92.22(+1A)&[MIMAT0000278&hsa-miR-221-3p&offsets|0|-1(+1A);m-91&X|-|45746181|45746202&offsets|0|0(+1A)]|AGCTACATTGTCTGCTGGGTTTA</td><td>>MI0000298|hsa-mir-221&WithFlank&X|-|45746151|45746272</td><td>71</td><td>92</td><td>22(+1A)</td><td>AGCTACATTGTCTGCTGGGTTTA</td><td>1316.6</td><td>-2.4</td><td>0.7</td><td>-3.4</td><td>7.2E-04</td><td>3.7E-03</td><td>BT20/MB231</td><td>MITO</td></tr>
<tr><td>>hg38_21_+_8433172_8446622_RNA45SN1_45S_50ntFlanks@7977.7995.19|CGACCTCAGATCAGACGTG</td><td>>hg38_21_+_8433172_8446622_RNA45SN1_45S_50ntFlanks</td><td>7977</td><td>7995</td><td>19</td><td>CGACCTCAGATCAGACGTG</td><td>311.3</td><td>-4.2</td><td>1.3</td><td>-3.2</td><td>1.6E-03</td><td>3.7E-03</td><td>BT20/MB468</td><td>MP</td></tr>
<tr><td>>hg38_21_+_8433172_8446622_RNA45SN1_45S_50ntFlanks@12616.12634.19|CTGAGGAGCCAATGGGGCG</td><td>>hg38_21_+_8433172_8446622_RNA45SN1_45S_50ntFlanks</td><td>12616</td><td>12634</td><td>19</td><td>CTGAGGAGCCAATGGGGCG</td><td>81.7</td><td>-5.1</td><td>1.6</td><td>-3.2</td><td>1.6E-03</td><td>3.7E-03</td><td>BT20/MB468</td><td>MP</td></tr>
<tr><td>>hg38_21_+_8433172_8446622_RNA45SN1_45S_50ntFlanks@10234.10272.39|CGGACGCTACGCCGCGACGAGTAGGAGGGCCGCTGCGGT</td><td>>hg38_21_+_8433172_8446622_RNA45SN1_45S_50ntFlanks</td><td>10234</td><td>10272</td><td>39</td><td>CGGACGCTACGCCGCGACGAGTAGGAGGGCCGCTGCGGT</td><td>237.4</td><td>4.3</td><td>1.2</td><td>3.5</td><td>5.5E-04</td><td>3.7E-03</td><td>BT20/MB231</td><td>MP</td></tr>
<tr><td>>hg38_21_+_8433172_8446622_RNA45SN1_45S_50ntFlanks@9308.9336.29|AGGAGTCTAACACGTGCGCGAGTCGGGGG</td><td>>hg38_21_+_8433172_8446622_RNA45SN1_45S_50ntFlanks</td><td>9308</td><td>9336</td><td>29</td><td>AGGAGTCTAACACGTGCGCGAGTCGGGGG</td><td>2071.0</td><td>-4.7</td><td>1.5</td><td>-3.2</td><td>1.6E-03</td><td>3.7E-03</td><td>BT20/MB468</td><td>MP</td></tr>
<tr><td>>hg38_21_+_8433172_8446622_RNA45SN1_45S_50ntFlanks@10680.10705.26|GTTCCGGCGGCGTCCGGTGAGCTCTC</td><td>>hg38_21_+_8433172_8446622_RNA45SN1_45S_50ntFlanks</td><td>10680</td><td>10705</td><td>26</td><td>GTTCCGGCGGCGTCCGGTGAGCTCTC</td><td>512.0</td><td>-2.7</td><td>0.9</td><td>-3.2</td><td>1.6E-03</td><td>3.7E-03</td><td>BT20/MB468</td><td>MP</td></tr>
<tr><td>>hg38_21_+_8433172_8446622_RNA45SN1_45S_50ntFlanks@10458.10490.33|ATGGCCTCCGTTGCCCTCGGCCGATCGAAAGGG</td><td>>hg38_21_+_8433172_8446622_RNA45SN1_45S_50ntFlanks</td><td>10458</td><td>10490</td><td>33</td><td>ATGGCCTCCGTTGCCCTCGGCCGATCGAAAGGG</td><td>170.1</td><td>-5.0</td><td>1.6</td><td>-3.2</td><td>1.6E-03</td><td>3.7E-03</td><td>BT20/MB468</td><td>MP</td></tr>
</table>

<table>
<tr><td>>MI0000750|hsa-mir-26a-2&WithFlank&12|-|57824603|57824698@20.40.21(+1C);MI0000083|hsa-mir-26a-1&WithFlank&3|+|37969398|37969486@16.36.21(+1C)&[MIMAT0000082&hsa-miR-26a-5p&offsets|0|-1(+1C);m-33&12|-|57824658|57824679&offsets|0|-1(+1C)];[MIMAT0000082_1&hsa-miR-26a-5p&offsets|0|-1(+1C);m-34&3|+|37969413|37969434&offsets|0|-1(+1C)]|TTCAAGTAATCCAGGATAGGCC</td><td>>MI0000750|hsa-mir-26a-2&WithFlank&12|-|57824603|57824698</td><td>20</td><td>40</td><td>21(+1C)</td><td>TTCAAGTAATCCAGGATAGGCC</td><td>21535.0</td><td>-3.6</td><td>1.1</td><td>-3.5</td><td>5.5E-04</td><td>3.7E-03</td><td>BT20/MB231</td><td>MP</td></tr>
<tr><td>>hg38_MT_+_1621_3279_RNR2_16S_50ntFlanks@1370.1396.27|ATGTTGGATCAGGACATCCCGATGGTG</td><td>>hg38_MT_+_1621_3279_RNR2_16S_50ntFlanks</td><td>1370</td><td>1396</td><td>27</td><td>ATGTTGGATCAGGACATCCCGATGGTG</td><td>1313.8</td><td>3.4</td><td>1.1</td><td>3.2</td><td>1.6E-03</td><td>3.7E-03</td><td>BT20/MB468</td><td>MP</td></tr>
<tr><td>>hg38_21_+_8433172_8446622_RNA45SN1_45S_50ntFlanks@12995.13040.46|CTCGCTGCGATCTATTGAAAGTCAGCCCTCGACACAAGGGTTTGTC</td><td>>hg38_21_+_8433172_8446622_RNA45SN1_45S_50ntFlanks</td><td>12995</td><td>13040</td><td>46</td><td>CTCGCTGCGATCTATTGAAAGTCAGCCCTCGACACAAGGGTTTGTC</td><td>1338.5</td><td>-3.0</td><td>0.9</td><td>-3.4</td><td>7.2E-04</td><td>3.7E-03</td><td>BT20/MB231</td><td>MITO</td></tr>
<tr><td>>hg38_21_+_8433172_8446622_RNA45SN1_45S_50ntFlanks@6661.6700.40|CGGTGGATCACTCGGCTCGTGCGTCGATGAAGAACGCAGC</td><td>>hg38_21_+_8433172_8446622_RNA45SN1_45S_50ntFlanks</td><td>6661</td><td>6700</td><td>40</td><td>CGGTGGATCACTCGGCTCGTGCGTCGATGAAGAACGCAGC</td><td>1468.0</td><td>2.2</td><td>0.7</td><td>3.4</td><td>6.0E-04</td><td>3.7E-03</td><td>BT20/MB231</td><td>NUC</td></tr>
<tr><td>>am_MI0000815|hsa-mir-339&WithFlank&7|-|1022927|1023032@21.39.19(+1U)&[MIMAT0000764&hsa-miR-339-5p&offsets|0|-4(+1U);m-163&7|-|1022990|1023012&offsets|0|-4(+1U)]|TCCCTGTCCTCCAGGAGCTT</td><td>>am_MI0000815|hsa-mir-339&WithFlank&7|-|1022927|1023032</td><td>21</td><td>39</td><td>19(+1U)</td><td>TCCCTGTCCTCCAGGAGCTT</td><td>55.9</td><td>6.5</td><td>2.0</td><td>3.2</td><td>1.6E-03</td><td>3.7E-03</td><td>BT20/MB468</td><td>MP</td></tr>
<tr><td>>hg38_21_+_8433172_8446622_RNA45SN1_45S_50ntFlanks@8174.8212.39|TCCTTCTGATCGAGGCCCAGCCCGTGGACGGTGTGAGGC</td><td>>hg38_21_+_8433172_8446622_RNA45SN1_45S_50ntFlanks</td><td>8174</td><td>8212</td><td>39</td><td>TCCTTCTGATCGAGGCCCAGCCCGTGGACGGTGTGAGGC</td><td>144.6</td><td>-5.2</td><td>1.7</td><td>-3.0</td><td>2.7E-03</td><td>3.7E-03</td><td>MB231/MB468</td><td>MP</td></tr>
<tr><td>>hg38_21_+_8433172_8446622_RNA45SN1_45S_50ntFlanks@10458.10487.30|ATGGCCTCCGTTGCCCTCGGCCGATCGAAA</td><td>>hg38_21_+_8433172_8446622_RNA45SN1_45S_50ntFlanks</td><td>10458</td><td>10487</td><td>30</td><td>ATGGCCTCCGTTGCCCTCGGCCGATCGAAA</td><td>441.7</td><td>-3.8</td><td>1.3</td><td>-3.0</td><td>2.9E-03</td><td>3.7E-03</td><td>BT20/MB468</td><td>CYTO</td></tr>
<tr><td>>hg38_21_+_8433172_8446622_RNA45SN1_45S_50ntFlanks@4241.4263.23|CTCTTTCGAGGCCCTGTAATTGG</td><td>>hg38_21_+_8433172_8446622_RNA45SN1_45S_50ntFlanks</td><td>4241</td><td>4263</td><td>23</td><td>CTCTTTCGAGGCCCTGTAATTGG</td><td>59.9</td><td>-5.0</td><td>1.6</td><td>-3.2</td><td>1.6E-03</td><td>3.7E-03</td><td>BT20/MB468</td><td>MP</td></tr>
<tr><td>>hg38_21_+_8433172_8446622_RNA45SN1_45S_50ntFlanks@11661.11687.27|TGATTTCTGCCCAGTGCTCTGAATGTC</td><td>>hg38_21_+_8433172_8446622_RNA45SN1_45S_50ntFlanks</td><td>11661</td><td>11687</td><td>27</td><td>TGATTTCTGCCCAGTGCTCTGAATGTC</td><td>231.6</td><td>-1.8</td><td>0.6</td><td>-3.0</td><td>3.2E-03</td><td>3.7E-03</td><td>MB231/MB468</td><td>NUC</td></tr>
<tr><td>>MI0000088|hsa-mir-30a&WithFlank&6|-|71403545|71403627@53.73.21(+1U)&[MIMAT0000088&hsa-miR-30a-3p&offsets|0|-1(+1U);m-38&6|-|71403554|71403575&offsets|0|-1(+1U)]|CTTTCAGTCGGATGTTTGCAGT</td><td>>MI0000088|hsa-mir-30a&WithFlank&6|-|71403545|71403627</td><td>53</td><td>73</td><td>21(+1U)</td><td>CTTTCAGTCGGATGTTTGCAGT</td><td>1513.9</td><td>1.3</td><td>0.4</td><td>3.0</td><td>3.1E-03</td><td>3.7E-03</td><td>BT20/MB468</td><td>NUC</td></tr>
<tr><td>>hg38_21_+_8433172_8446622_RNA45SN1_45S_50ntFlanks@9339.9378.40|CGCACGAAAGCCGCCGTGGCGCAATGAAGGTGAAGGCCGG</td><td>>hg38_21_+_8433172_8446622_RNA45SN1_45S_50ntFlanks</td><td>9339</td><td>9378</td><td>40</td><td>CGCACGAAAGCCGCCGTGGCGCAATGAAGGTGAAGGCCGG</td><td>104.7</td><td>-4.7</td><td>1.6</td><td>-3.0</td><td>2.5E-03</td><td>3.7E-03</td><td>BT20/MB468</td><td>MITO</td></tr>
<tr><td>>hg38_21_+_8433172_8446622_RNA45SN1_45S_50ntFlanks@8343.8379.37|GTAAGGGAAAGTTGAAAAGAACTTTGAAGAGAGAGTT</td><td>>hg38_21_+_8433172_8446622_RNA45SN1_45S_50ntFlanks</td><td>8343</td><td>8379</td><td>37</td><td>GTAAGGGAAAGTTGAAAAGAACTTTGAAGAGAGAGTT</td><td>44.0</td><td>6.0</td><td>1.9</td><td>3.2</td><td>1.6E-03</td><td>3.7E-03</td><td>BT20/MB468</td><td>MP</td></tr>
<tr><td>>hg38_21_+_8433172_8446622_RNA45SN1_45S_50ntFlanks@12994.13015.22|CCTCGCTGCGATCTATTGAAAG</td><td>>hg38_21_+_8433172_8446622_RNA45SN1_45S_50ntFlanks</td><td>12994</td><td>13015</td><td>22</td><td>CCTCGCTGCGATCTATTGAAAG</td><td>107.7</td><td>-4.8</td><td>1.6</td><td>-3.0</td><td>2.7E-03</td><td>3.7E-03</td><td>MB231/MB468</td><td>MP</td></tr>
<tr><td>>hg38_21_+_8433172_8446622_RNA45SN1_45S_50ntFlanks@8353.8375.23|GTTGAAAAGAACTTTGAAGAGAG</td><td>>hg38_21_+_8433172_8446622_RNA45SN1_45S_50ntFlanks</td><td>8353</td><td>8375</td><td>23</td><td>GTTGAAAAGAACTTTGAAGAGAG</td><td>7962.2</td><td>4.7</td><td>1.4</td><td>3.5</td><td>5.5E-04</td><td>3.7E-03</td><td>BT20/MB231</td><td>MP</td></tr>
<tr><td>>hg38_21_+_8433172_8446622_RNA45SN1_45S_50ntFlanks@12249.12284.36|CAAAAGCTCGCTTGATCTTGATTTTCAGTACGAATA</td><td>>hg38_21_+_8433172_8446622_RNA45SN1_45S_50ntFlanks</td><td>12249</td><td>12284</td><td>36</td><td>CAAAAGCTCGCTTGATCTTGATTTTCAGTACGAATA</td><td>351.6</td><td>-2.2</td><td>0.7</td><td>-3.0</td><td>2.5E-03</td><td>3.7E-03</td><td>BT20/MB468</td><td>MITO</td></tr>
<tr><td>>am_trna119_LysCTT_1_-_145395522_145395594@13.35.23,trna11_LysCTT_5_-_180648979_180649051@13.35.23,trna128_LysCTT_1_-_55423542_55423614@13.35.23,trna13_LysCTT_6_+_26556774_26556846@13.35.23,trna6_LysCTT_19_-_52425396_52425468@13.35.23,trna7_LysCTT_16_+_3225692_3225764@13.35.23,trna9_LysCTT_5_+_180634755_180634827@13.35.23|CAGTCGGTAGAGCATGAGACTCT</td><td>>am_trna119_LysCTT_1_-_145395522_145395594</td><td>13</td><td>35</td><td>23</td><td>CAGTCGGTAGAGCATGAGACTCT</td><td>70.9</td><td>5.5</td><td>1.8</td><td>3.0</td><td>2.7E-03</td><td>3.7E-03</td><td>MB231/MB468</td><td>MP</td></tr>
<tr><td>>trna10_AspGTC_12_-_125424193_125424264@1.30.30,trna12_AspGTC_12_-_125411891_125411962@1.30.30,trna38_AspGTC_17_-_8125556_8125627@1.30.30,trna45_AspGTC_6_+_27447453_27447524@1.30.30,trna48_AspGTC_6_+_27471523_27471594@1.30.30,trna4_AspGTC_12_+_96429799_96429870@1.30.30,trna69_AspGTC_1_-_161440205_161440276@1.30.30,trna72_AspGTC_1_-_161432824_161432895@1.30.30,trna75_AspGTC_1_-_161425414_161425485@1.30.30,trna78_AspGTC_1_-_161418033_161418104@1.30.30,trna81_AspGTC_1_-_161410615_161410686@1.30.30|TCCTCGTTAGTATAGTGGTGAGTATCCCCG</td><td>>trna10_AspGTC_12_-_125424193_125424264</td><td>1</td><td>30</td><td>30</td><td>TCCTCGTTAGTATAGTGGTGAGTATCCCCG</td><td>818.0</td><td>-4.9</td><td>1.6</td><td>-3.0</td><td>2.9E-03</td><td>3.7E-03</td><td>BT20/MB468</td><td>CYTO</td></tr>
<tr><td>>hg38_21_+_8433172_8446622_RNA45SN1_45S_50ntFlanks@10440.10460.21|GTTCCGAAGGGACGGGCGATG</td><td>>hg38_21_+_8433172_8446622_RNA45SN1_45S_50ntFlanks</td><td>10440</td><td>10460</td><td>21</td><td>GTTCCGAAGGGACGGGCGATG</td><td>199.4</td><td>2.3</td><td>0.8</td><td>3.0</td><td>3.2E-03</td><td>3.7E-03</td><td>MB231/MB468</td><td>NUC</td></tr>
<tr><td>>trna10_AspGTC_12_-_125424193_125424264@37.71.35,trna12_AspGTC_12_-_125411891_125411962@37.71.35,trna144_AspGTC_6_-_27551236_27551307@37.71.35,trna38_AspGTC_17_-_8125556_8125627@37.71.35,trna45_AspGTC_6_+_27447453_27447524@37.71.35,trna48_AspGTC_6_+_27471523_27471594@37.71.35,trna4_AspGTC_12_+_96429799_96429870@37.71.35,trna69_AspGTC_1_-_161440205_161440276@37.71.35,trna72_AspGTC_1_-_161432824_161432895@37.71.35,trna75_AspGTC_1_-_161425414_161425485@37.71.35,trna78_AspGTC_1_-_161418033_161418104@37.71.35,trna81_AspGTC_1_-_161410615_161410686@37.71.35|ACGCGGGAGACCGGGGTTCGATTCCCCGACGGGGA</td><td>>trna10_AspGTC_12_-_125424193_125424264</td><td>37</td><td>71</td><td>35</td><td>ACGCGGGAGACCGGGGTTCGATTCCCCGACGGGGA</td><td>888.4</td><td>2.9</td><td>1.0</td><td>3.0</td><td>3.0E-03</td><td>3.7E-03</td><td>MB231/MB468</td><td>CYTO</td></tr>
<tr><td>>hg38_21_+_8433172_8446622_RNA45SN1_45S_50ntFlanks@10792.10820.29|CTCTGGCATGTTGGAACAATGTAGGTAAG</td><td>>hg38_21_+_8433172_8446622_RNA45SN1_45S_50ntFlanks</td><td>10792</td><td>10820</td><td>29</td><td>CTCTGGCATGTTGGAACAATGTAGGTAAG</td><td>104.7</td><td>-3.5</td><td>1.2</td><td>-3.0</td><td>2.7E-03</td><td>3.7E-03</td><td>MB231/MB468</td><td>MP</td></tr>
<tr><td>>hg38_21_+_8433172_8446622_RNA45SN1_45S_50ntFlanks@12957.12975.19|CTTCTGGGTCGGGGTTTCG</td><td>>hg38_21_+_8433172_8446622_RNA45SN1_45S_50ntFlanks</td><td>12957</td><td>12975</td><td>19</td><td>CTTCTGGGTCGGGGTTTCG</td><td>206.3</td><td>-2.7</td><td>0.9</td><td>-3.0</td><td>3.0E-03</td><td>3.7E-03</td><td>MB231/MB468</td><td>CYTO</td></tr>
<tr><td>>hg38_21_+_8433172_8446622_RNA45SN1_45S_50ntFlanks@3854.3889.36|ACTGTGGTAATTCTAGAGCTAATACATGCCGACGGG</td><td>>hg38_21_+_8433172_8446622_RNA45SN1_45S_50ntFlanks</td><td>3854</td><td>3889</td><td>36</td><td>ACTGTGGTAATTCTAGAGCTAATACATGCCGACGGG</td><td>168.5</td><td>-4.5</td><td>1.5</td><td>-3.0</td><td>2.9E-03</td><td>3.7E-03</td><td>BT20/MB468</td><td>CYTO</td></tr>
<tr><td>>MI0000065|hsa-let-7d&WithFlank&9|+|94178828|94178926@68.88.21(+1A)&[MIMAT0004484&hsa-let-7d-3p&offsets|0|-1(+1A);m-98&9|+|94178895|94178916&offsets|0|-1(+1A)]|CTATACGACCTGCTGCCTTTCA</td><td>>MI0000065|hsa-let-7d&WithFlank&9|+|94178828|94178926</td><td>68</td><td>88</td><td>21(+1A)</td><td>CTATACGACCTGCTGCCTTTCA</td><td>214.3</td><td>4.1</td><td>1.4</td><td>3.0</td><td>2.7E-03</td><td>3.7E-03</td><td>MB231/MB468</td><td>MP</td></tr>
<tr><td>>hg38_21_+_8433172_8446622_RNA45SN1_45S_50ntFlanks@6777.6802.26|TCCCGGGGCTACGCCTGTCTGAGCGT</td><td>>hg38_21_+_8433172_8446622_RNA45SN1_45S_50ntFlanks</td><td>6777</td><td>6802</td><td>26</td><td>TCCCGGGGCTACGCCTGTCTGAGCGT</td><td>257.5</td><td>2.7</td><td>0.9</td><td>3.0</td><td>3.2E-03</td><td>3.7E-03</td><td>MB231/MB468</td><td>MITO</td></tr>
<tr><td>>hg38_21_+_8433172_8446622_RNA45SN1_45S_50ntFlanks@11270.11295.26|CGGCCGCCCCCGGGGAGCCCGGCGGG</td><td>>hg38_21_+_8433172_8446622_RNA45SN1_45S_50ntFlanks</td><td>11270</td><td>11295</td><td>26</td><td>CGGCCGCCCCCGGGGAGCCCGGCGGG</td><td>968.1</td><td>-5.0</td><td>1.7</td><td>-3.0</td><td>2.9E-03</td><td>3.7E-03</td><td>BT20/MB468</td><td>CYTO</td></tr>
<tr><td>>trnaMT_SerGCT_MT_+_12207_12265@1.38.38|GAGAAAGCTCACAAGAACTGCTAACTCATGCCCCCATG</td><td>>trnaMT_SerGCT_MT_+_12207_12265</td><td>1</td><td>38</td><td>38</td><td>GAGAAAGCTCACAAGAACTGCTAACTCATGCCCCCATG</td><td>363.1</td><td>3.8</td><td>1.3</td><td>3.0</td><td>3.0E-03</td><td>3.7E-03</td><td>MB231/MB468</td><td>CYTO</td></tr>
<tr><td>>hg38_21_+_8433172_8446622_RNA45SN1_45S_50ntFlanks@8182.8210.29|ATCGAGGCCCAGCCCGTGGACGGTGTGAG</td><td>>hg38_21_+_8433172_8446622_RNA45SN1_45S_50ntFlanks</td><td>8182</td><td>8210</td><td>29</td><td>ATCGAGGCCCAGCCCGTGGACGGTGTGAG</td><td>1012.5</td><td>-5.1</td><td>1.7</td><td>-3.0</td><td>3.2E-03</td><td>3.7E-03</td><td>MB231/MB468</td><td>NUC</td></tr>
<tr><td>>hg38_21_+_8433172_8446622_RNA45SN1_45S_50ntFlanks@12336.12370.35|CAGGAGGTGTCAGAAAAGTTACCACAGGGATAACT</td><td>>hg38_21_+_8433172_8446622_RNA45SN1_45S_50ntFlanks</td><td>12336</td><td>12370</td><td>35</td><td>CAGGAGGTGTCAGAAAAGTTACCACAGGGATAACT</td><td>63.6</td><td>6.2</td><td>2.0</td><td>3.2</td><td>1.6E-03</td><td>3.7E-03</td><td>BT20/MB468</td><td>MP</td></tr>
<tr><td>>hg38_21_+_8433172_8446622_RNA45SN1_45S_50ntFlanks@12407.12443.37|TTTTGATCCTTCGATGTCGGCTCTTCCTATCATTGTG</td><td>>hg38_21_+_8433172_8446622_RNA45SN1_45S_50ntFlanks</td><td>12407</td><td>12443</td><td>37</td><td>TTTTGATCCTTCGATGTCGGCTCTTCCTATCATTGTG</td><td>1260.0</td><td>-1.8</td><td>0.6</td><td>-3.0</td><td>2.5E-03</td><td>3.7E-03</td><td>BT20/MB468</td><td>MITO</td></tr>
<tr><td>>hg38_21_+_8433172_8446622_RNA45SN1_45S_50ntFlanks@11566.11586.21|TAGCAGCCGACTTAGAACTGG</td><td>>hg38_21_+_8433172_8446622_RNA45SN1_45S_50ntFlanks</td><td>11566</td><td>11586</td><td>21</td><td>TAGCAGCCGACTTAGAACTGG</td><td>283.2</td><td>-3.2</td><td>0.9</td><td>-3.4</td><td>6.1E-04</td><td>3.7E-03</td><td>BT20/MB231</td><td>NUC</td></tr>
<tr><td>>hg38_21_+_8433172_8446622_RNA45SN1_45S_50ntFlanks@11672.11702.31|CAGTGCTCTGAATGTCAAAGTGAAGAAATTC</td><td>>hg38_21_+_8433172_8446622_RNA45SN1_45S_50ntFlanks</td><td>11672</td><td>11702</td><td>31</td><td>CAGTGCTCTGAATGTCAAAGTGAAGAAATTC</td><td>551.7</td><td>-2.6</td><td>0.8</td><td>-3.4</td><td>6.1E-04</td><td>3.7E-03</td><td>BT20/MB231</td><td>NUC</td></tr>
<tr><td>>hg38_21_+_8433172_8446622_RNA45SN1_45S_50ntFlanks@11029.11051.23|CTCTCTCTCTCTCCCCCGCTCCC</td><td>>hg38_21_+_8433172_8446622_RNA45SN1_45S_50ntFlanks</td><td>11029</td><td>11051</td><td>23</td><td>CTCTCTCTCTCTCCCCCGCTCCC</td><td>259.0</td><td>-4.2</td><td>1.4</td><td>-3.0</td><td>3.0E-03</td><td>3.7E-03</td><td>MB231/MB468</td><td>CYTO</td></tr>
<tr><td>>MI0000811|hsa-mir-148b&WithFlank&12|+|54337210|54337320@69.91.23&[MIMAT0000759&hsa-miR-148b-3p&offsets|0|+1;m-77&12|+|54337278|54337299&offsets|0|+1]|TCAGTGCATCACAGAACTTTGTC</td><td>>MI0000811|hsa-mir-148b&WithFlank&12|+|54337210|54337320</td><td>69</td><td>91</td><td>23</td><td>TCAGTGCATCACAGAACTTTGTC</td><td>412.9</td><td>2.7</td><td>0.9</td><td>3.0</td><td>2.7E-03</td><td>3.7E-03</td><td>MB231/MB468</td><td>MP</td></tr>
<tr><td>>MI0001445|hsa-mir-423&WithFlank&17|+|30117073|30117178@59.81.23(+1U)&[MIMAT0001340&hsa-miR-423-3p&offsets|0|0(+1U);m-100&17|+|30117131|30117153&offsets|0|0(+1U)]|AGCTCGGTCTGAGGCCCCTCAGTT</td><td>>MI0001445|hsa-mir-423&WithFlank&17|+|30117073|30117178</td><td>59</td><td>81</td><td>23(+1U)</td><td>AGCTCGGTCTGAGGCCCCTCAGTT</td><td>234.8</td><td>3.9</td><td>1.3</td><td>3.0</td><td>2.7E-03</td><td>3.7E-03</td><td>MB231/MB468</td><td>MP</td></tr>
<tr><td>>hg38_21_+_8433172_8446622_RNA45SN1_45S_50ntFlanks@11847.11871.25|AACGGGCTTGGCGGAATCAGCGGGG</td><td>>hg38_21_+_8433172_8446622_RNA45SN1_45S_50ntFlanks</td><td>11847</td><td>11871</td><td>25</td><td>AACGGGCTTGGCGGAATCAGCGGGG</td><td>481.5</td><td>-2.3</td><td>0.7</td><td>-3.2</td><td>1.6E-03</td><td>3.7E-03</td><td>BT20/MB468</td><td>MP</td></tr>
<tr><td>>MI0000803|hsa-mir-330&WithFlank&19|-|45638988|45639093@64.87.24(+1U)&[MIMAT0000751&hsa-miR-330-3p&offsets|+1|+2(+1U);m-347&19|-|45639009|45639030&offsets|0|+2(+1U)]|CAAAGCACACGGCCTGCAGAGAGGT</td><td>>MI0000803|hsa-mir-330&WithFlank&19|-|45638988|45639093</td><td>64</td><td>87</td><td>24(+1U)</td><td>CAAAGCACACGGCCTGCAGAGAGGT</td><td>99.4</td><td>5.6</td><td>1.9</td><td>3.0</td><td>2.7E-03</td><td>3.7E-03</td><td>MB231/MB468</td><td>MP</td></tr>
<tr><td>>hg38_21_+_8433172_8446622_RNA45SN1_45S_50ntFlanks@3862.3898.37|AATTCTAGAGCTAATACATGCCGACGGGCGCTGACCC</td><td>>hg38_21_+_8433172_8446622_RNA45SN1_45S_50ntFlanks</td><td>3862</td><td>3898</td><td>37</td><td>AATTCTAGAGCTAATACATGCCGACGGGCGCTGACCC</td><td>149.7</td><td>-4.4</td><td>1.5</td><td>-3.0</td><td>2.9E-03</td><td>3.7E-03</td><td>BT20/MB468</td><td>CYTO</td></tr>
<tr><td>>hg38_21_+_8433172_8446622_RNA45SN1_45S_50ntFlanks@4588.4617.30|CTATTTTGTTGGTTTTCGGAACTGAGGCCA</td><td>>hg38_21_+_8433172_8446622_RNA45SN1_45S_50ntFlanks</td><td>4588</td><td>4617</td><td>30</td><td>CTATTTTGTTGGTTTTCGGAACTGAGGCCA</td><td>152.8</td><td>-5.1</td><td>1.6</td><td>-3.2</td><td>1.6E-03</td><td>3.7E-03</td><td>BT20/MB468</td><td>MP</td></tr>
</table>

<table>
<tr><th></th><th></th><th></th><th></th><th></th><th></th><th></th><th></th><th></th><th></th><th></th><th></th><th></th><th></th></tr>
<tr><td>>MI0000067|hsa-let-7f-1&WithFlank&9|+|94176341|94176439@13.33.21;MI0000068|hsa-let-7f-2&WithFlank&X|-|53557186|53557280@14.34.21&[MIMAT0000067&hsa-let-7f-5p&offsets|0|-1;m-20&9|+|94176353|94176374&offsets|0|-1];[MIMAT0000067_1&hsa-let-7f-5p&offsets|0|-1;m-19&X|-|53557246|53557267&offsets|0|-1]|TGAGGTAGTAGATTGTATAGT</td><td>>MI0000067|hsa-let-7f-1&WithFlank&9|+|94176341|94176439</td><td>13</td><td>33</td><td>21</td><td>TGAGGTAGTAGATTGTATAGT</td><td>3913.3</td><td>1.6</td><td>0.5</td><td>3.0</td><td>3.2E-03</td><td>3.7E-03</td><td>BT20/MB468</td><td>NUC</td></tr>
<tr><td>>hg38_21_+_8433172_8446622_RNA45SN1_45S_50ntFlanks@6700.6717.18|CTAGCTGCGAGAATTAAT</td><td>>hg38_21_+_8433172_8446622_RNA45SN1_45S_50ntFlanks</td><td>6700</td><td>6717</td><td>18</td><td>CTAGCTGCGAGAATTAAT</td><td>177.5</td><td>-4.4</td><td>1.5</td><td>-3.0</td><td>2.5E-03</td><td>3.7E-03</td><td>BT20/MB468</td><td>MITO</td></tr>
<tr><td>>hg38_1_-_228634819_228635039_RNA5S12_5S_50ntFlanks@51.68.18|GTCTACGGCCATACCACC</td><td>>hg38_1_-_228634819_228635039_RNA5S12_5S_50ntFlanks</td><td>51</td><td>68</td><td>18</td><td>GTCTACGGCCATACCACC</td><td>1292.3</td><td>4.9</td><td>1.4</td><td>3.5</td><td>5.6E-04</td><td>3.7E-03</td><td>BT20/MB231</td><td>MP</td></tr>
<tr><td>>MI0000082|hsa-mir-25&WithFlank&7|-|100093554|100093649@58.78.21(+1C)&[MIMAT0000081&hsa-miR-25-3p&offsets|0|-1(+1C);m-23&7|-|100093571|100093592&offsets|0|-1(+1C)]|CATTGCACTTGTCTCGGTCTGC</td><td>>MI0000082|hsa-mir-25&WithFlank&7|-|100093554|100093649</td><td>58</td><td>78</td><td>21(+1C)</td><td>CATTGCACTTGTCTCGGTCTGC</td><td>289.9</td><td>4.1</td><td>1.4</td><td>3.0</td><td>2.7E-03</td><td>3.7E-03</td><td>MB231/MB468</td><td>MP</td></tr>
<tr><td>>hg38_21_+_8433172_8446622_RNA45SN1_45S_50ntFlanks@11582.11601.20|ACTGGTGCGGACCAGGGGAA</td><td>>hg38_21_+_8433172_8446622_RNA45SN1_45S_50ntFlanks</td><td>11582</td><td>11601</td><td>20</td><td>ACTGGTGCGGACCAGGGGAA</td><td>527.9</td><td>-2.8</td><td>0.9</td><td>-3.0</td><td>3.0E-03</td><td>3.7E-03</td><td>MB231/MB468</td><td>CYTO</td></tr>
<tr><td>>hg38_MT_+_598_1651_RNR1_12S_50ntFlanks@721.744.24|AAGACGTTAGGTCAAGGTGTAGCC</td><td>>hg38_MT_+_598_1651_RNR1_12S_50ntFlanks</td><td>721</td><td>744</td><td>24</td><td>AAGACGTTAGGTCAAGGTGTAGCC</td><td>116.1</td><td>4.0</td><td>1.3</td><td>3.0</td><td>2.7E-03</td><td>3.7E-03</td><td>MB231/MB468</td><td>MP</td></tr>
<tr><td>>hg38_21_+_8433172_8446622_RNA45SN1_45S_50ntFlanks@5074.5092.19|ATTCCGATAACGAACGAGA</td><td>>hg38_21_+_8433172_8446622_RNA45SN1_45S_50ntFlanks</td><td>5074</td><td>5092</td><td>19</td><td>ATTCCGATAACGAACGAGA</td><td>113.7</td><td>-5.4</td><td>1.7</td><td>-3.2</td><td>1.6E-03</td><td>3.7E-03</td><td>BT20/MB468</td><td>MP</td></tr>
<tr><td>>MI0000093|hsa-mir-92a-1&WithFlank&13|+|91351308|91351397@54.76.23&[MIMAT0000092&hsa-miR-92a-3p&offsets|0|+1;m-9&13|+|91351361|91351382&offsets|0|+1]|TATTGCACTTGTCCCGGCCTGTT</td><td>>MI0000093|hsa-mir-92a-1&WithFlank&13|+|91351308|91351397</td><td>54</td><td>76</td><td>23</td><td>TATTGCACTTGTCCCGGCCTGTT</td><td>7940.6</td><td>-2.5</td><td>0.7</td><td>-3.4</td><td>7.4E-04</td><td>3.7E-03</td><td>BT20/MB231</td><td>MITO</td></tr>
<tr><td>>MI0000272|hsa-mir-182&WithFlank&7|-|129770377|129770498@29.51.23&[MIMAT0000259&hsa-miR-182-5p&offsets|0|-1;m-22&7|-|129770449|129770470&offsets|0|+1]|TTTGGCAATGGTAGAACTCACAC</td><td>>MI0000272|hsa-mir-182&WithFlank&7|-|129770377|129770498</td><td>29</td><td>51</td><td>23</td><td>TTTGGCAATGGTAGAACTCACAC</td><td>1249.7</td><td>2.4</td><td>0.7</td><td>3.4</td><td>7.4E-04</td><td>3.7E-03</td><td>BT20/MB231</td><td>MITO</td></tr>
<tr><td>>MI0000542|hsa-mir-320a&WithFlank&8|-|22244960|22245043@48.71.24&[MIMAT0000510&hsa-miR-320a-3p&offsets|0|+2;m-60&8|-|22244975|22244996&offsets|0|+2]|AAAAGCTGGGTTGAGAGGGCGAAA</td><td>>MI0000542|hsa-mir-320a&WithFlank&8|-|22244960|22245043</td><td>48</td><td>71</td><td>24</td><td>AAAAGCTGGGTTGAGAGGGCGAAA</td><td>952.7</td><td>2.5</td><td>0.7</td><td>3.4</td><td>7.4E-04</td><td>3.7E-03</td><td>BT20/MB231</td><td>MITO</td></tr>
<tr><td>>MI0000736|hsa-mir-30c-1&WithFlank&1|+|40757278|40757378@23.45.23(+1C);MI0000254|hsa-mir-30c-2&WithFlank&6|-|71376954|71377037@13.35.23(+1C)&[MIMAT0000244&hsa-miR-30c-5p&offsets|0|0(+1C);m-57&1|+|40757300|40757323&offsets|0|-1(+1C)];[MIMAT0000244_1&hsa-miR-30c-5p&offsets|0|0(+1C);m-56&6|-|71377002|71377025&offsets|0|-1(+1C)]|TGTAAACATCCTACACTCTCAGCC</td><td>>MI0000736|hsa-mir-30c-1&WithFlank&1|+|40757278|40757378</td><td>23</td><td>45</td><td>23(+1C)</td><td>TGTAAACATCCTACACTCTCAGCC</td><td>8184.2</td><td>-3.2</td><td>0.9</td><td>-3.4</td><td>7.4E-04</td><td>3.7E-03</td><td>BT20/MB231</td><td>MITO</td></tr>
<tr><td>>hg38_21_+_8433172_8446622_RNA45SN1_45S_50ntFlanks@10341.10373.33|AATATTCAAACGAGAACTTTGAAGGCCGAAGTG</td><td>>hg38_21_+_8433172_8446622_RNA45SN1_45S_50ntFlanks</td><td>10341</td><td>10373</td><td>33</td><td>AATATTCAAACGAGAACTTTGAAGGCCGAAGTG</td><td>305.0</td><td>3.0</td><td>0.9</td><td>3.4</td><td>7.4E-04</td><td>3.7E-03</td><td>BT20/MB231</td><td>MITO</td></tr>
<tr><td>>hg38_21_+_8433172_8446622_RNA45SN1_45S_50ntFlanks@6772.6800.29|GTTCCTCCCGGGGCTACGCCTGTCTGAGC</td><td>>hg38_21_+_8433172_8446622_RNA45SN1_45S_50ntFlanks</td><td>6772</td><td>6800</td><td>29</td><td>GTTCCTCCCGGGGCTACGCCTGTCTGAGC</td><td>255.7</td><td>2.7</td><td>0.8</td><td>3.4</td><td>7.4E-04</td><td>3.7E-03</td><td>BT20/MB231</td><td>MITO</td></tr>
<tr><td>>MI0000811|hsa-mir-148b&WithFlank&12|+|54337210|54337320@69.90.22(+2U)&[MIMAT0000759&hsa-miR-148b-3p&offsets|0|0(+2U);m-77&12|+|54337278|54337299&offsets|0|0(+2U)]|TCAGTGCATCACAGAACTTTGTTT</td><td>>MI0000811|hsa-mir-148b&WithFlank&12|+|54337210|54337320</td><td>69</td><td>90</td><td>22(+2U)</td><td>TCAGTGCATCACAGAACTTTGTTT</td><td>687.7</td><td>-3.0</td><td>0.9</td><td>-3.4</td><td>7.4E-04</td><td>3.7E-03</td><td>BT20/MB231</td><td>MITO</td></tr>
<tr><td>>hg38_21_+_8433172_8446622_RNA45SN1_45S_50ntFlanks@11317.11334.18|ACCCCACGTCTCGTCGCG</td><td>>hg38_21_+_8433172_8446622_RNA45SN1_45S_50ntFlanks</td><td>11317</td><td>11334</td><td>18</td><td>ACCCCACGTCTCGTCGCG</td><td>618.0</td><td>-5.8</td><td>2.0</td><td>-3.0</td><td>3.2E-03</td><td>3.7E-03</td><td>BT20/MB468</td><td>NUC</td></tr>
<tr><td>>hg38_21_+_8433172_8446622_RNA45SN1_45S_50ntFlanks@11583.11613.31|CTGGTGCGGACCAGGGGAATCCGACTGTTTA</td><td>>hg38_21_+_8433172_8446622_RNA45SN1_45S_50ntFlanks</td><td>11583</td><td>11613</td><td>31</td><td>CTGGTGCGGACCAGGGGAATCCGACTGTTTA</td><td>566.2</td><td>-2.3</td><td>0.8</td><td>-3.0</td><td>3.2E-03</td><td>3.7E-03</td><td>BT20/MB468</td><td>NUC</td></tr>
<tr><td>>hg38_21_+_8433172_8446622_RNA45SN1_45S_50ntFlanks@11784.11806.23|ATGAATGGATGAACGAGATTCCC</td><td>>hg38_21_+_8433172_8446622_RNA45SN1_45S_50ntFlanks</td><td>11784</td><td>11806</td><td>23</td><td>ATGAATGGATGAACGAGATTCCC</td><td>406.4</td><td>-2.0</td><td>0.7</td><td>-3.0</td><td>2.5E-03</td><td>3.7E-03</td><td>BT20/MB468</td><td>MITO</td></tr>
<tr><td>>hg38_21_+_8433172_8446622_RNA45SN1_45S_50ntFlanks@12606.12626.21|ATGTGCTTGGCTGAGGAGCCA</td><td>>hg38_21_+_8433172_8446622_RNA45SN1_45S_50ntFlanks</td><td>12606</td><td>12626</td><td>21</td><td>ATGTGCTTGGCTGAGGAGCCA</td><td>292.9</td><td>-5.2</td><td>1.6</td><td>-3.2</td><td>1.6E-03</td><td>3.7E-03</td><td>BT20/MB468</td><td>MP</td></tr>
<tr><td>>hg38_21_+_8433172_8446622_RNA45SN1_45S_50ntFlanks@10874.10892.19|TGGGTCGGTCGGGCTGGGG</td><td>>hg38_21_+_8433172_8446622_RNA45SN1_45S_50ntFlanks</td><td>10874</td><td>10892</td><td>19</td><td>TGGGTCGGTCGGGCTGGGG</td><td>103.2</td><td>-5.1</td><td>1.6</td><td>-3.2</td><td>1.6E-03</td><td>3.7E-03</td><td>BT20/MB468</td><td>MP</td></tr>
<tr><td>>hg38_21_+_8433172_8446622_RNA45SN1_45S_50ntFlanks@6784.6806.23|GCTACGCCTGTCTGAGCGTCGCT</td><td>>hg38_21_+_8433172_8446622_RNA45SN1_45S_50ntFlanks</td><td>6784</td><td>6806</td><td>23</td><td>GCTACGCCTGTCTGAGCGTCGCT</td><td>176.1</td><td>-3.4</td><td>1.1</td><td>-3.0</td><td>2.7E-03</td><td>3.7E-03</td><td>MB231/MB468</td><td>MP</td></tr>
<tr><td>>MI0003667|hsa-mir-652&WithFlank&X|+|110055323|110055432@67.85.19(+1C)&[MIMAT0003322&hsa-miR-652-3p&offsets|0|-2(+1C);m-148&X|+|110055389|110055410&offsets|0|-3(+1C)]|AATGGCGCCACTAGGGTTGC</td><td>>MI0003667|hsa-mir-652&WithFlank&X|+|110055323|110055432</td><td>67</td><td>85</td><td>19(+1C)</td><td>AATGGCGCCACTAGGGTTGC</td><td>127.5</td><td>6.6</td><td>2.1</td><td>3.2</td><td>1.6E-03</td><td>3.7E-03</td><td>BT20/MB468</td><td>MP</td></tr>
<tr><td>>hg38_21_+_8433172_8446622_RNA45SN1_45S_50ntFlanks@11567.11593.27|AGCAGCCGACTTAGAACTGGTGCGGAC</td><td>>hg38_21_+_8433172_8446622_RNA45SN1_45S_50ntFlanks</td><td>11567</td><td>11593</td><td>27</td><td>AGCAGCCGACTTAGAACTGGTGCGGAC</td><td>176.7</td><td>-4.1</td><td>1.4</td><td>-3.0</td><td>2.7E-03</td><td>3.7E-03</td><td>MB231/MB468</td><td>MP</td></tr>
<tr><td>>hg38_21_+_8433172_8446622_RNA45SN1_45S_50ntFlanks@11603.11621.19|CCGACTGTTTAATTAAAAC</td><td>>hg38_21_+_8433172_8446622_RNA45SN1_45S_50ntFlanks</td><td>11603</td><td>11621</td><td>19</td><td>CCGACTGTTTAATTAAAAC</td><td>199.3</td><td>-4.9</td><td>1.6</td><td>-3.0</td><td>2.7E-03</td><td>3.7E-03</td><td>MB231/MB468</td><td>MP</td></tr>
<tr><td>>hg38_21_+_8433172_8446622_RNA45SN1_45S_50ntFlanks@10771.10818.48|CAGGTCTCCAAGGTGAACAGCCTCTGGCATGTTGGAACAATGTAGGTA</td><td>>hg38_21_+_8433172_8446622_RNA45SN1_45S_50ntFlanks</td><td>10771</td><td>10818</td><td>48</td><td>CAGGTCTCCAAGGTGAACAGCCTCTGGCATGTTGGAACAATGTAGGTA</td><td>443.0</td><td>-4.1</td><td>1.4</td><td>-3.0</td><td>2.5E-03</td><td>3.7E-03</td><td>BT20/MB468</td><td>MITO</td></tr>
<tr><td>>MI0000103|hsa-mir-101-1&WithFlank&1|-|65058428|65058514@53.73.21(+2U);MI0000739|hsa-mir-101-2&WithFlank&9|+|4850291|4850381@55.75.21(+2U)&[MIMAT0000099&hsa-miR-101-3p&offsets|0|0(+2U);m-17&1|-|65058443|65058462&offsets|0|+1(+2U)];[MIMAT0000099_1&hsa-miR-101-3p&offsets|0|0(+2U);m-16&9|+|4850345|4850364&offsets|0|+1(+2U)]|TACAGTACTGTGATAACTGAATT</td><td>>MI0000103|hsa-mir-101-1&WithFlank&1|-|65058428|65058514</td><td>53</td><td>73</td><td>21(+2U)</td><td>TACAGTACTGTGATAACTGAATT</td><td>135.4</td><td>-6.1</td><td>1.8</td><td>-3.4</td><td>5.6E-04</td><td>3.7E-03</td><td>BT20/MB231</td><td>MP</td></tr>
<tr><td>>hg38_21_+_8433172_8446622_RNA45SN1_45S_50ntFlanks@12388.12406.19|CGTTCATAGCGACGTCGCT</td><td>>hg38_21_+_8433172_8446622_RNA45SN1_45S_50ntFlanks</td><td>12388</td><td>12406</td><td>19</td><td>CGTTCATAGCGACGTCGCT</td><td>301.7</td><td>1.7</td><td>0.6</td><td>3.0</td><td>3.0E-03</td><td>3.7E-03</td><td>MB231/MB468</td><td>CYTO</td></tr>
<tr><td>>hg38_21_+_8433172_8446622_RNA45SN1_45S_50ntFlanks@10841.10874.34|CGTAACTTCGGGATAAGGATTGGCTCTAAGGGCT</td><td>>hg38_21_+_8433172_8446622_RNA45SN1_45S_50ntFlanks</td><td>10841</td><td>10874</td><td>34</td><td>CGTAACTTCGGGATAAGGATTGGCTCTAAGGGCT</td><td>1042.3</td><td>-2.2</td><td>0.7</td><td>-3.0</td><td>3.0E-03</td><td>3.7E-03</td><td>MB231/MB468</td><td>CYTO</td></tr>
<tr><td>>hg38_21_+_8433172_8446622_RNA45SN1_45S_50ntFlanks@10786.10826.41|AACAGCCTCTGGCATGTTGGAACAATGTAGGTAAGGGAAGT</td><td>>hg38_21_+_8433172_8446622_RNA45SN1_45S_50ntFlanks</td><td>10786</td><td>10826</td><td>41</td><td>AACAGCCTCTGGCATGTTGGAACAATGTAGGTAAGGGAAGT</td><td>554.8</td><td>-4.7</td><td>1.5</td><td>-3.2</td><td>1.6E-03</td><td>3.8E-03</td><td>BT20/MB468</td><td>MP</td></tr>
<tr><td>>hg38_21_+_8433172_8446622_RNA45SN1_45S_50ntFlanks@12934.12951.18|CTAAACCATTCGTAGACG</td><td>>hg38_21_+_8433172_8446622_RNA45SN1_45S_50ntFlanks</td><td>12934</td><td>12951</td><td>18</td><td>CTAAACCATTCGTAGACG</td><td>813.0</td><td>-5.4</td><td>1.7</td><td>-3.2</td><td>1.6E-03</td><td>3.8E-03</td><td>BT20/MB468</td><td>MP</td></tr>
<tr><td>>am_trnaMT_LysTTT_MT_+_8295_8364@31.52.22,trnalookalike7_LysTTT_1_+_568844_568913@31.52.22|TAAGTTAAAGATTAAGAGAACC</td><td>>am_trnaMT_LysTTT_MT_+_8295_8364</td><td>31</td><td>52</td><td>22</td><td>TAAGTTAAAGATTAAGAGAACC</td><td>280.6</td><td>3.0</td><td>0.9</td><td>3.4</td><td>6.7E-04</td><td>3.8E-03</td><td>BT20/MB231</td><td>CYTO</td></tr>
<tr><td>>MI0003560|hsa-mir-92b&WithFlank&1|+|155195171|155195278@67.88.22(+2U)&[MIMAT0003218&hsa-miR-92b-3p&offsets|0|0(+2U);m-115&1|+|155195237|155195258&offsets|0|0(+2U)]|TATTGCACTCGTCCCGGCCTCCTT</td><td>>MI0003560|hsa-mir-92b&WithFlank&1|+|155195171|155195278</td><td>67</td><td>88</td><td>22(+2U)</td><td>TATTGCACTCGTCCCGGCCTCCTT</td><td>79.5</td><td>5.7</td><td>1.9</td><td>3.0</td><td>2.7E-03</td><td>3.8E-03</td><td>MB231/MB468</td><td>MP</td></tr>
<tr><td>>hg38_21_+_8433172_8446622_RNA45SN1_45S_50ntFlanks@4724.4748.25|ATTAATCAAGAACGAAAGTCGGAGG</td><td>>hg38_21_+_8433172_8446622_RNA45SN1_45S_50ntFlanks</td><td>4724</td><td>4748</td><td>25</td><td>ATTAATCAAGAACGAAAGTCGGAGG</td><td>621.2</td><td>-2.6</td><td>0.9</td><td>-3.0</td><td>3.0E-03</td><td>3.8E-03</td><td>MB231/MB468</td><td>CYTO</td></tr>
<tr><td>>hg38_21_+_8433172_8446622_RNA45SN1_45S_50ntFlanks@11587.11623.37|TGCGGACCAGGGGAATCCGACTGTTTAATTAAAACAA</td><td>>hg38_21_+_8433172_8446622_RNA45SN1_45S_50ntFlanks</td><td>11587</td><td>11623</td><td>37</td><td>TGCGGACCAGGGGAATCCGACTGTTTAATTAAAACAA</td><td>36.4</td><td>8.4</td><td>2.8</td><td>3.0</td><td>2.7E-03</td><td>3.8E-03</td><td>MB231/MB468</td><td>MP</td></tr>
<tr><td>>hg38_21_+_8433172_8446622_RNA45SN1_45S_50ntFlanks@7974.7997.24|ACGCGACCTCAGATCAGACGTGGC</td><td>>hg38_21_+_8433172_8446622_RNA45SN1_45S_50ntFlanks</td><td>7974</td><td>7997</td><td>24</td><td>ACGCGACCTCAGATCAGACGTGGC</td><td>11317.7</td><td>3.1</td><td>0.9</td><td>3.4</td><td>6.2E-04</td><td>3.8E-03</td><td>BT20/MB231</td><td>NUC</td></tr>
<tr><td>>trna152_ValCAC_6_-_27248049_27248121@1.36.36|GCTTCTGTAGTGTAGTGGTTATCACGTTCGCCTCAC</td><td>>trna152_ValCAC_6_-_27248049_27248121</td><td>1</td><td>36</td><td>36</td><td>GCTTCTGTAGTGTAGTGGTTATCACGTTCGCCTCAC</td><td>476.0</td><td>-4.7</td><td>1.4</td><td>-3.4</td><td>6.2E-04</td><td>3.8E-03</td><td>BT20/MB231</td><td>NUC</td></tr>
<tr><td>>hg38_21_+_8433172_8446622_RNA45SN1_45S_50ntFlanks@12634.12654.21|GAAGCTACCATCTGTGGGATT</td><td>>hg38_21_+_8433172_8446622_RNA45SN1_45S_50ntFlanks</td><td>12634</td><td>12654</td><td>21</td><td>GAAGCTACCATCTGTGGGATT</td><td>391.1</td><td>-3.9</td><td>1.3</td><td>-3.0</td><td>2.5E-03</td><td>3.8E-03</td><td>BT20/MB468</td><td>MITO</td></tr>
<tr><td>>hg38_21_+_8433172_8446622_RNA45SN1_45S_50ntFlanks@5477.5501.25|CCTGGCGGAGCGCTGAGAAGACGGT</td><td>>hg38_21_+_8433172_8446622_RNA45SN1_45S_50ntFlanks</td><td>5477</td><td>5501</td><td>25</td><td>CCTGGCGGAGCGCTGAGAAGACGGT</td><td>53.2</td><td>6.1</td><td>1.9</td><td>3.2</td><td>1.6E-03</td><td>3.8E-03</td><td>BT20/MB468</td><td>MP</td></tr>
<tr><td>>hg38_21_+_8433172_8446622_RNA45SN1_45S_50ntFlanks@12913.12933.21|ACGTTCGTGGGGAACCTGGCG</td><td>>hg38_21_+_8433172_8446622_RNA45SN1_45S_50ntFlanks</td><td>12913</td><td>12933</td><td>21</td><td>ACGTTCGTGGGGAACCTGGCG</td><td>310.6</td><td>-3.6</td><td>1.2</td><td>-3.0</td><td>3.0E-03</td><td>3.8E-03</td><td>BT20/MB468</td><td>CYTO</td></tr>
<tr><td>>hg38_21_+_8433172_8446622_RNA45SN1_45S_50ntFlanks@9621.9658.38|TCTAGTAGCTGGTTCCCTCCGAAGTTTCCCTCAGGATA</td><td>>hg38_21_+_8433172_8446622_RNA45SN1_45S_50ntFlanks</td><td>9621</td><td>9658</td><td>38</td><td>TCTAGTAGCTGGTTCCCTCCGAAGTTTCCCTCAGGATA</td><td>204.7</td><td>-3.6</td><td>1.2</td><td>-2.9</td><td>3.2E-03</td><td>3.8E-03</td><td>MB231/MB468</td><td>NUC</td></tr>
<tr><td>>hg38_21_+_8433172_8446622_RNA45SN1_45S_50ntFlanks@3775.3794.20|GCCGGTACAGTGAAACTGCG</td><td>>hg38_21_+_8433172_8446622_RNA45SN1_45S_50ntFlanks</td><td>3775</td><td>3794</td><td>20</td><td>GCCGGTACAGTGAAACTGCG</td><td>262.0</td><td>-5.3</td><td>1.7</td><td>-3.2</td><td>1.6E-03</td><td>3.8E-03</td><td>BT20/MB468</td><td>MP</td></tr>
<tr><td>>hg38_21_+_8433172_8446622_RNA45SN1_45S_50ntFlanks@11200.11232.33|CGGCGGCGGCGGCGACTCTGGACGCGAGCCGGG</td><td>>hg38_21_+_8433172_8446622_RNA45SN1_45S_50ntFlanks</td><td>11200</td><td>11232</td><td>33</td><td>CGGCGGCGGCGGCGACTCTGGACGCGAGCCGGG</td><td>778.9</td><td>-2.2</td><td>0.7</td><td>-3.0</td><td>2.5E-03</td><td>3.8E-03</td><td>BT20/MB468</td><td>MITO</td></tr>
<tr><td>>hg38_21_+_8433172_8446622_RNA45SN1_45S_50ntFlanks@11210.11231.22|GGCGACTCTGGACGCGAGCCGG</td><td>>hg38_21_+_8433172_8446622_RNA45SN1_45S_50ntFlanks</td><td>11210</td><td>11231</td><td>22</td><td>GGCGACTCTGGACGCGAGCCGG</td><td>204.8</td><td>-2.2</td><td>0.7</td><td>-3.0</td><td>3.0E-03</td><td>3.8E-03</td><td>BT20/MB468</td><td>CYTO</td></tr>
<tr><td>>hg38_21_+_8433172_8446622_RNA45SN1_45S_50ntFlanks@12001.12030.30|CTGCGGGCCGCCGGTGAAATACCACTACTC</td><td>>hg38_21_+_8433172_8446622_RNA45SN1_45S_50ntFlanks</td><td>12001</td><td>12030</td><td>30</td><td>CTGCGGGCCGCCGGTGAAATACCACTACTC</td><td>106.9</td><td>-4.9</td><td>1.6</td><td>-3.2</td><td>1.6E-03</td><td>3.8E-03</td><td>BT20/MB468</td><td>MP</td></tr>
<tr><td>>trnaMT_ValTAC_MT_+_1602_1670@40.72.33|GGAGATTTCAACTTAACTTGACCGCTCTGACCA</td><td>>trnaMT_ValTAC_MT_+_1602_1670</td><td>40</td><td>72</td><td>33</td><td>GGAGATTTCAACTTAACTTGACCGCTCTGACCA</td><td>363.8</td><td>4.2</td><td>1.3</td><td>3.2</td><td>1.6E-03</td><td>3.8E-03</td><td>BT20/MB468</td><td>MP</td></tr>
<tr><td>>hg38_21_+_8433172_8446622_RNA45SN1_45S_50ntFlanks@4008.4042.35|CTCGGGCCGATCGCACGCCCCCCGTGGCGGCGACG</td><td>>hg38_21_+_8433172_8446622_RNA45SN1_45S_50ntFlanks</td><td>4008</td><td>4042</td><td>35</td><td>CTCGGGCCGATCGCACGCCCCCCGTGGCGGCGACG</td><td>439.2</td><td>-2.3</td><td>0.8</td><td>-3.0</td><td>3.0E-03</td><td>3.8E-03</td><td>MB231/MB468</td><td>CYTO</td></tr>
<tr><td>>hg38_21_+_8433172_8446622_RNA45SN1_45S_50ntFlanks@9564.9605.42|CTGACGTGCAAATCGGTCGTCCGACCTGGGTATAGGGGCGAA</td><td>>hg38_21_+_8433172_8446622_RNA45SN1_45S_50ntFlanks</td><td>9564</td><td>9605</td><td>42</td><td>CTGACGTGCAAATCGGTCGTCCGACCTGGGTATAGGGGCGAA</td><td>383.9</td><td>-3.9</td><td>1.3</td><td>-2.9</td><td>3.2E-03</td><td>3.8E-03</td><td>MB231/MB468</td><td>MITO</td></tr>
<tr><td>>hg38_21_+_8433172_8446622_RNA45SN1_45S_50ntFlanks@12894.12933.40|CTCGCCCGTCACGCACCGCACGTTCGTGGGGAACCTGGCG</td><td>>hg38_21_+_8433172_8446622_RNA45SN1_45S_50ntFlanks</td><td>12894</td><td>12933</td><td>40</td><td>CTCGCCCGTCACGCACCGCACGTTCGTGGGGAACCTGGCG</td><td>146.3</td><td>-4.4</td><td>1.4</td><td>-3.0</td><td>2.5E-03</td><td>3.8E-03</td><td>BT20/MB468</td><td>MITO</td></tr>
<tr><td>>MI0000272|hsa-mir-182&WithFlank&7|-|129770377|129770498@29.52.24&[MIMAT0000259&hsa-miR-182-5p&offsets|0|0;m-22&7|-|129770449|129770470&offsets|0|+2]|TTTGGCAATGGTAGAACTCACACT</td><td>>MI0000272|hsa-mir-182&WithFlank&7|-|129770377|129770498</td><td>29</td><td>52</td><td>24</td><td>TTTGGCAATGGTAGAACTCACACT</td><td>8556.2</td><td>3.4</td><td>1.1</td><td>3.2</td><td>1.6E-03</td><td>3.8E-03</td><td>BT20/MB468</td><td>MP</td></tr>
</table>

<table>
<tr><td>>trnaMT_HisGTG_MT_+_12138_12206@42.72.31|AACAGAGGCTTACGACCCCTTATTTACCCCA</td><td>>trnaMT_HisGTG_MT_+_12138_12206</td><td>42</td><td>72</td><td>31</td><td>AACAGAGGCTTACGACCCCTTATTTACCCCA</td><td>345.0</td><td>5.1</td><td>1.6</td><td>3.2</td><td>1.6E-03</td><td>3.8E-03</td><td>BT20/MB468</td><td>MP</td></tr>
<tr><td>>hg38_21_+_8433172_8446622_RNA45SN1_45S_50ntFlanks@11874.11903.30|AGAAGACCCTGTTGAGCTTGACTCTAGTCT</td><td>>hg38_21_+_8433172_8446622_RNA45SN1_45S_50ntFlanks</td><td>11874</td><td>11903</td><td>30</td><td>AGAAGACCCTGTTGAGCTTGACTCTAGTCT</td><td>428.9</td><td>-2.1</td><td>0.7</td><td>-3.0</td><td>2.5E-03</td><td>3.8E-03</td><td>BT20/MB468</td><td>MITO</td></tr>
<tr><td>>hg38_21_+_8433172_8446622_RNA45SN1_45S_50ntFlanks@5125.5145.21|AGCGGTCGGCGTCCCCCAACT</td><td>>hg38_21_+_8433172_8446622_RNA45SN1_45S_50ntFlanks</td><td>5125</td><td>5145</td><td>21</td><td>AGCGGTCGGCGTCCCCCAACT</td><td>692.4</td><td>5.1</td><td>1.5</td><td>3.4</td><td>6.2E-04</td><td>3.8E-03</td><td>BT20/MB231</td><td>NUC</td></tr>
<tr><td>>hg38_21_+_8433172_8446622_RNA45SN1_45S_50ntFlanks@3785.3810.26|TGAAACTGCGAATGGCTCATTAAATC</td><td>>hg38_21_+_8433172_8446622_RNA45SN1_45S_50ntFlanks</td><td>3785</td><td>3810</td><td>26</td><td>TGAAACTGCGAATGGCTCATTAAATC</td><td>249.4</td><td>-1.6</td><td>0.5</td><td>-3.0</td><td>3.0E-03</td><td>3.8E-03</td><td>BT20/MB468</td><td>CYTO</td></tr>
<tr><td>>hg38_21_+_8433172_8446622_RNA45SN1_45S_50ntFlanks@3931.3959.29|TCAAAACCAACCCGGTCAGCCCCTCTCCG</td><td>>hg38_21_+_8433172_8446622_RNA45SN1_45S_50ntFlanks</td><td>3931</td><td>3959</td><td>29</td><td>TCAAAACCAACCCGGTCAGCCCCTCTCCG</td><td>445.6</td><td>-3.3</td><td>1.1</td><td>-3.0</td><td>3.0E-03</td><td>3.8E-03</td><td>BT20/MB468</td><td>CYTO</td></tr>
<tr><td>>hg38_21_+_8433172_8446622_RNA45SN1_45S_50ntFlanks@9533.9550.18|GAGGAAACTCTGGTGGAG</td><td>>hg38_21_+_8433172_8446622_RNA45SN1_45S_50ntFlanks</td><td>9533</td><td>9550</td><td>18</td><td>GAGGAAACTCTGGTGGAG</td><td>421.3</td><td>-2.3</td><td>0.8</td><td>-2.9</td><td>3.2E-03</td><td>3.8E-03</td><td>MB231/MB468</td><td>NUC</td></tr>
<tr><td>>hg38_21_+_8433172_8446622_RNA45SN1_45S_50ntFlanks@6687.6706.20|ATGAAGAACGCAGCTAGCTG</td><td>>hg38_21_+_8433172_8446622_RNA45SN1_45S_50ntFlanks</td><td>6687</td><td>6706</td><td>20</td><td>ATGAAGAACGCAGCTAGCTG</td><td>127.3</td><td>-5.0</td><td>1.6</td><td>-3.2</td><td>1.6E-03</td><td>3.8E-03</td><td>BT20/MB468</td><td>MP</td></tr>
<tr><td>>hg38_21_+_8433172_8446622_RNA45SN1_45S_50ntFlanks@11564.11583.20|CCTAGCAGCCGACTTAGAAC</td><td>>hg38_21_+_8433172_8446622_RNA45SN1_45S_50ntFlanks</td><td>11564</td><td>11583</td><td>20</td><td>CCTAGCAGCCGACTTAGAAC</td><td>296.7</td><td>-2.0</td><td>0.7</td><td>-2.9</td><td>3.2E-03</td><td>3.8E-03</td><td>MB231/MB468</td><td>NUC</td></tr>
<tr><td>>hg38_21_+_8433172_8446622_RNA45SN1_45S_50ntFlanks@12412.12443.32|ATCCTTCGATGTCGGCTCTTCCTATCATTGTG</td><td>>hg38_21_+_8433172_8446622_RNA45SN1_45S_50ntFlanks</td><td>12412</td><td>12443</td><td>32</td><td>ATCCTTCGATGTCGGCTCTTCCTATCATTGTG</td><td>2003.0</td><td>-5.3</td><td>1.8</td><td>-2.9</td><td>3.3E-03</td><td>3.8E-03</td><td>MB231/MB468</td><td>NUC</td></tr>
<tr><td>>hg38_21_+_8433172_8446622_RNA45SN1_45S_50ntFlanks@10546.10568.23|CAGTGCGGTAACGCGACCGATCC</td><td>>hg38_21_+_8433172_8446622_RNA45SN1_45S_50ntFlanks</td><td>10546</td><td>10568</td><td>23</td><td>CAGTGCGGTAACGCGACCGATCC</td><td>96.4</td><td>-5.0</td><td>1.7</td><td>-3.0</td><td>2.7E-03</td><td>3.8E-03</td><td>MB231/MB468</td><td>MP</td></tr>
<tr><td>>hg38_21_+_8433172_8446622_RNA45SN1_45S_50ntFlanks@10427.10456.30|ATGGGCGAGCGCCGTTCCGAAGGGACGGGC</td><td>>hg38_21_+_8433172_8446622_RNA45SN1_45S_50ntFlanks</td><td>10427</td><td>10456</td><td>30</td><td>ATGGGCGAGCGCCGTTCCGAAGGGACGGGC</td><td>279.3</td><td>-2.2</td><td>0.7</td><td>-3.0</td><td>2.5E-03</td><td>3.8E-03</td><td>BT20/MB468</td><td>MITO</td></tr>
<tr><td>>hg38_21_+_8433172_8446622_RNA45SN1_45S_50ntFlanks@12419.12457.39|GATGTCGGCTCTTCCTATCATTGTGAAGCAGAATTCACC</td><td>>hg38_21_+_8433172_8446622_RNA45SN1_45S_50ntFlanks</td><td>12419</td><td>12457</td><td>39</td><td>GATGTCGGCTCTTCCTATCATTGTGAAGCAGAATTCACC</td><td>181.2</td><td>3.7</td><td>1.3</td><td>2.9</td><td>3.2E-03</td><td>3.8E-03</td><td>MB231/MB468</td><td>MITO</td></tr>
<tr><td>>hg38_21_+_8433172_8446622_RNA45SN1_45S_50ntFlanks@10342.10373.32|ATATTCAAACGAGAACTTTGAAGGCCGAAGTG</td><td>>hg38_21_+_8433172_8446622_RNA45SN1_45S_50ntFlanks</td><td>10342</td><td>10373</td><td>32</td><td>ATATTCAAACGAGAACTTTGAAGGCCGAAGTG</td><td>197.2</td><td>1.9</td><td>0.7</td><td>2.9</td><td>3.3E-03</td><td>3.8E-03</td><td>MB231/MB468</td><td>NUC</td></tr>
<tr><td>>hg38_21_+_8433172_8446622_RNA45SN1_45S_50ntFlanks@12003.12031.29|GCGGGCCGCCGGTGAAATACCACTACTCT</td><td>>hg38_21_+_8433172_8446622_RNA45SN1_45S_50ntFlanks</td><td>12003</td><td>12031</td><td>29</td><td>GCGGGCCGCCGGTGAAATACCACTACTCT</td><td>486.1</td><td>-4.3</td><td>1.4</td><td>-3.1</td><td>1.6E-03</td><td>3.8E-03</td><td>BT20/MB468</td><td>MP</td></tr>
<tr><td>>am_trnaMT_HisGTG_MT_+_12138_12206@27.51.25|GATTGTGAATCTGACAACAGAGGCT</td><td>>am_trnaMT_HisGTG_MT_+_12138_12206</td><td>27</td><td>51</td><td>25</td><td>GATTGTGAATCTGACAACAGAGGCT</td><td>89.5</td><td>4.8</td><td>1.6</td><td>3.0</td><td>2.7E-03</td><td>3.8E-03</td><td>MB231/MB468</td><td>MP</td></tr>
<tr><td>>hg38_21_+_8433172_8446622_RNA45SN1_45S_50ntFlanks@3839.3862.24|TCTCCTACTTGGATAACTGTGGTA</td><td>>hg38_21_+_8433172_8446622_RNA45SN1_45S_50ntFlanks</td><td>3839</td><td>3862</td><td>24</td><td>TCTCCTACTTGGATAACTGTGGTA</td><td>1191.6</td><td>3.5</td><td>1.0</td><td>3.4</td><td>6.8E-04</td><td>3.8E-03</td><td>BT20/MB231</td><td>CYTO</td></tr>
<tr><td>>hg38_21_+_8433172_8446622_RNA45SN1_45S_50ntFlanks@6646.6668.23|TCGTACGACTCTTAGCGGTGGAT</td><td>>hg38_21_+_8433172_8446622_RNA45SN1_45S_50ntFlanks</td><td>6646</td><td>6668</td><td>23</td><td>TCGTACGACTCTTAGCGGTGGAT</td><td>9323.1</td><td>2.4</td><td>0.7</td><td>3.4</td><td>6.8E-04</td><td>3.8E-03</td><td>BT20/MB231</td><td>CYTO</td></tr>
<tr><td>>MI0000084|hsa-mir-26b&WithFlank&2|+|218402640|218402728@18.39.22(+1C)&[MIMAT0000083&hsa-miR-26b-5p&offsets|0|+1(+1C);m-53&2|+|218402657|218402678&offsets|0|0(+1C)]|TTCAAGTAATTCAGGATAGGTTC</td><td>>MI0000084|hsa-mir-26b&WithFlank&2|+|218402640|218402728</td><td>18</td><td>39</td><td>22(+1C)</td><td>TTCAAGTAATTCAGGATAGGTTC</td><td>143.2</td><td>4.0</td><td>1.3</td><td>3.0</td><td>2.8E-03</td><td>3.8E-03</td><td>MB231/MB468</td><td>MP</td></tr>
<tr><td>>MI0000269|hsa-mir-181a-2&WithFlank&9|+|124692436|124692557@83.104.22&[MIMAT0004558&hsa-miR-181a-2-3p&offsets|0|0;m-71&9|+|124692518|124692539&offsets|0|0]|ACCACTGACCGTTGACTGTACC</td><td>>MI0000269|hsa-mir-181a-2&WithFlank&9|+|124692436|124692557</td><td>83</td><td>104</td><td>22</td><td>ACCACTGACCGTTGACTGTACC</td><td>1375.8</td><td>-2.4</td><td>0.7</td><td>-3.4</td><td>7.6E-04</td><td>3.8E-03</td><td>BT20/MB231</td><td>MITO</td></tr>
<tr><td>>hg38_21_+_8433172_8446622_RNA45SN1_45S_50ntFlanks@12458.12478.21|AAGCGTTGGATTGTTCACCCA</td><td>>hg38_21_+_8433172_8446622_RNA45SN1_45S_50ntFlanks</td><td>12458</td><td>12478</td><td>21</td><td>AAGCGTTGGATTGTTCACCCA</td><td>1436.4</td><td>-2.3</td><td>0.8</td><td>-3.0</td><td>3.1E-03</td><td>3.8E-03</td><td>MB231/MB468</td><td>CYTO</td></tr>
<tr><td>>hg38_MT_+_598_1651_RNR1_12S_50ntFlanks@718.744.27|GTAAAGACGTTAGGTCAAGGTGTAGCC</td><td>>hg38_MT_+_598_1651_RNR1_12S_50ntFlanks</td><td>718</td><td>744</td><td>27</td><td>GTAAAGACGTTAGGTCAAGGTGTAGCC</td><td>168.3</td><td>3.8</td><td>1.3</td><td>2.9</td><td>3.3E-03</td><td>3.8E-03</td><td>MB231/MB468</td><td>MITO</td></tr>
<tr><td>>hg38_21_+_8433172_8446622_RNA45SN1_45S_50ntFlanks@5237.5282.46|ACTGACTGGCTCAGCGTGTGCCTACCCTACGCCGGCAGGCGCGGGT</td><td>>hg38_21_+_8433172_8446622_RNA45SN1_45S_50ntFlanks</td><td>5237</td><td>5282</td><td>46</td><td>ACTGACTGGCTCAGCGTGTGCCTACCCTACGCCGGCAGGCGCGGGT</td><td>153.7</td><td>3.3</td><td>1.1</td><td>2.9</td><td>3.3E-03</td><td>3.8E-03</td><td>MB231/MB468</td><td>MITO</td></tr>
<tr><td>>hg38_21_+_8433172_8446622_RNA45SN1_45S_50ntFlanks@13006.13028.23|CTATTGAAAGTCAGCCCTCGACA</td><td>>hg38_21_+_8433172_8446622_RNA45SN1_45S_50ntFlanks</td><td>13006</td><td>13028</td><td>23</td><td>CTATTGAAAGTCAGCCCTCGACA</td><td>133.6</td><td>-4.3</td><td>1.4</td><td>-3.0</td><td>2.6E-03</td><td>3.8E-03</td><td>BT20/MB468</td><td>MITO</td></tr>
<tr><td>>hg38_21_+_8433172_8446622_RNA45SN1_45S_50ntFlanks@475.521.47|CTTCGTGATCGATGTGGTGACGTCGTGCTCTCCCGGGCCGGGTCCGA</td><td>>hg38_21_+_8433172_8446622_RNA45SN1_45S_50ntFlanks</td><td>475</td><td>521</td><td>47</td><td>CTTCGTGATCGATGTGGTGACGTCGTGCTCTCCCGGGCCGGGTCCGA</td><td>1080.9</td><td>5.1</td><td>1.7</td><td>2.9</td><td>3.3E-03</td><td>3.8E-03</td><td>MB231/MB468</td><td>MITO</td></tr>
<tr><td>>hg38_21_+_8433172_8446622_RNA45SN1_45S_50ntFlanks@8104.8140.37|CCGCGGCGGGGCGCGGGACATGTGGCGTACGGAAGAC</td><td>>hg38_21_+_8433172_8446622_RNA45SN1_45S_50ntFlanks</td><td>8104</td><td>8140</td><td>37</td><td>CCGCGGCGGGGCGCGGGACATGTGGCGTACGGAAGAC</td><td>1588.1</td><td>6.1</td><td>1.8</td><td>3.4</td><td>5.7E-04</td><td>3.8E-03</td><td>BT20/MB231</td><td>MP</td></tr>
<tr><td>>hg38_MT_+_598_1651_RNR1_12S_50ntFlanks@912.946.35|TTCAAAGGACATTTAACTAAAACCCCTACGCATTT</td><td>>hg38_MT_+_598_1651_RNR1_12S_50ntFlanks</td><td>912</td><td>946</td><td>35</td><td>TTCAAAGGACATTTAACTAAAACCCCTACGCATTT</td><td>307.0</td><td>5.5</td><td>1.7</td><td>3.1</td><td>1.6E-03</td><td>3.8E-03</td><td>BT20/MB468</td><td>MP</td></tr>
<tr><td>>hg38_21_+_8433172_8446622_RNA45SN1_45S_50ntFlanks@10107.10125.19|CGTGTGGTGTGCGTCGGAG</td><td>>hg38_21_+_8433172_8446622_RNA45SN1_45S_50ntFlanks</td><td>10107</td><td>10125</td><td>19</td><td>CGTGTGGTGTGCGTCGGAG</td><td>495.6</td><td>-3.6</td><td>1.2</td><td>-3.0</td><td>3.0E-03</td><td>3.8E-03</td><td>BT20/MB468</td><td>CYTO</td></tr>
<tr><td>>MI0000264|hsa-mir-7-2&WithFlank&15|+|88611819|88611940@38.60.23;MI0000265|hsa-mir-7-3&WithFlank&19|+|4770664|4770785@37.59.23;MI0000263|hsa-mir-7-1&WithFlank&9|-|83969742|83969863@30.52.23&[MIMAT0000252&hsa-miR-7-5p&offsets|0|-1;m-400&15|+|88611856|88611879&offsets|0|-1];[MIMAT0000252_1&hsa-miR-7-5p&offsets|0|-1;m-402&19|+|4770700|4770723&offsets|0|-1];[MIMAT0000252_2&hsa-miR-7-5p&offsets|0|-1;m-401&9|-|83969811|83969834&offsets|0|-1]|TGGAAGACTAGTGATTTTGTTGT</td><td>>MI0000264|hsa-mir-7-2&WithFlank&15|+|88611819|88611940</td><td>38</td><td>60</td><td>23</td><td>TGGAAGACTAGTGATTTTGTTGT</td><td>10515.6</td><td>3.1</td><td>0.9</td><td>3.4</td><td>6.9E-04</td><td>3.8E-03</td><td>BT20/MB231</td><td>CYTO</td></tr>
<tr><td>>MI0000102|hsa-mir-100&WithFlank&11|-|122152223|122152314@19.40.22(+3A)&[MIMAT0000098&hsa-miR-100-5p&offsets|0|0(+3A);m-27&11|-|122152276|122152296&offsets|0|+1(+3A)]|AACCCGTAGATCCGAACTTGTGAAA</td><td>>MI0000102|hsa-mir-100&WithFlank&11|-|122152223|122152314</td><td>19</td><td>40</td><td>22(+3A)</td><td>AACCCGTAGATCCGAACTTGTGAAA</td><td>215.5</td><td>-3.8</td><td>1.1</td><td>-3.4</td><td>6.9E-04</td><td>3.8E-03</td><td>BT20/MB231</td><td>CYTO</td></tr>
<tr><td>>hg38_21_+_8433172_8446622_RNA45SN1_45S_50ntFlanks@6661.6681.21|CGGTGGATCACTCGGCTCGTG</td><td>>hg38_21_+_8433172_8446622_RNA45SN1_45S_50ntFlanks</td><td>6661</td><td>6681</td><td>21</td><td>CGGTGGATCACTCGGCTCGTG</td><td>4765.1</td><td>-3.1</td><td>1.1</td><td>-2.9</td><td>3.3E-03</td><td>3.8E-03</td><td>MB231/MB468</td><td>MITO</td></tr>
<tr><td>>hg38_21_+_8433172_8446622_RNA45SN1_45S_50ntFlanks@9305.9338.34|CCAAGGAGTCTAACACGTGCGCGAGTCGGGGGCT</td><td>>hg38_21_+_8433172_8446622_RNA45SN1_45S_50ntFlanks</td><td>9305</td><td>9338</td><td>34</td><td>CCAAGGAGTCTAACACGTGCGCGAGTCGGGGGCT</td><td>145.4</td><td>-2.3</td><td>0.8</td><td>-3.0</td><td>3.1E-03</td><td>3.8E-03</td><td>MB231/MB468</td><td>CYTO</td></tr>
<tr><td>>hg38_21_+_8433172_8446622_RNA45SN1_45S_50ntFlanks@11659.11692.34|TGTGATTTCTGCCCAGTGCTCTGAATGTCAAAGT</td><td>>hg38_21_+_8433172_8446622_RNA45SN1_45S_50ntFlanks</td><td>11659</td><td>11692</td><td>34</td><td>TGTGATTTCTGCCCAGTGCTCTGAATGTCAAAGT</td><td>34.7</td><td>8.9</td><td>2.8</td><td>3.1</td><td>1.7E-03</td><td>3.8E-03</td><td>BT20/MB468</td><td>MP</td></tr>
<tr><td>>hg38_21_+_8433172_8446622_RNA45SN1_45S_50ntFlanks@11755.11777.23|AATGCCTCGTCATCTAATTAGTG</td><td>>hg38_21_+_8433172_8446622_RNA45SN1_45S_50ntFlanks</td><td>11755</td><td>11777</td><td>23</td><td>AATGCCTCGTCATCTAATTAGTG</td><td>226.2</td><td>-4.5</td><td>1.5</td><td>-3.0</td><td>3.0E-03</td><td>3.8E-03</td><td>BT20/MB468</td><td>CYTO</td></tr>
<tr><td>>hg38_21_+_8433172_8446622_RNA45SN1_45S_50ntFlanks@6648.6669.22|GTACGACTCTTAGCGGTGGATC</td><td>>hg38_21_+_8433172_8446622_RNA45SN1_45S_50ntFlanks</td><td>6648</td><td>6669</td><td>22</td><td>GTACGACTCTTAGCGGTGGATC</td><td>1572.2</td><td>2.7</td><td>0.8</td><td>3.4</td><td>6.3E-04</td><td>3.8E-03</td><td>BT20/MB231</td><td>NUC</td></tr>
<tr><td>>am_trnaMT_PheGAA_MT_+_577_647@40.64.25|TGTTTAGACGGGCTCACATCACCCC</td><td>>am_trnaMT_PheGAA_MT_+_577_647</td><td>40</td><td>64</td><td>25</td><td>TGTTTAGACGGGCTCACATCACCCC</td><td>125.1</td><td>7.8</td><td>2.5</td><td>3.1</td><td>1.7E-03</td><td>3.8E-03</td><td>BT20/MB468</td><td>MP</td></tr>
<tr><td>>hg38_21_+_8433172_8446622_RNA45SN1_45S_50ntFlanks@11660.11692.33|GTGATTTCTGCCCAGTGCTCTGAATGTCAAAGT</td><td>>hg38_21_+_8433172_8446622_RNA45SN1_45S_50ntFlanks</td><td>11660</td><td>11692</td><td>33</td><td>GTGATTTCTGCCCAGTGCTCTGAATGTCAAAGT</td><td>34.9</td><td>8.9</td><td>2.8</td><td>3.1</td><td>1.7E-03</td><td>3.8E-03</td><td>BT20/MB468</td><td>MP</td></tr>
<tr><td>>hg38_21_+_8433172_8446622_RNA45SN1_45S_50ntFlanks@8358.8390.33|AAAGAACTTTGAAGAGAGAGTTCAAGAGGGCGT</td><td>>hg38_21_+_8433172_8446622_RNA45SN1_45S_50ntFlanks</td><td>8358</td><td>8390</td><td>33</td><td>AAAGAACTTTGAAGAGAGAGTTCAAGAGGGCGT</td><td>82.1</td><td>5.1</td><td>1.6</td><td>3.1</td><td>1.7E-03</td><td>3.8E-03</td><td>BT20/MB468</td><td>MP</td></tr>
<tr><td>>hg38_21_+_8433172_8446622_RNA45SN1_45S_50ntFlanks@12896.12921.26|CGCCCGTCACGCACCGCACGTTCGTG</td><td>>hg38_21_+_8433172_8446622_RNA45SN1_45S_50ntFlanks</td><td>12896</td><td>12921</td><td>26</td><td>CGCCCGTCACGCACCGCACGTTCGTG</td><td>38.9</td><td>-8.4</td><td>2.7</td><td>-3.1</td><td>1.7E-03</td><td>3.8E-03</td><td>BT20/MB468</td><td>MP</td></tr>
<tr><td>>hg38_21_+_8433172_8446622_RNA45SN1_45S_50ntFlanks@10458.10488.31|ATGGCCTCCGTTGCCCTCGGCCGATCGAAAG</td><td>>hg38_21_+_8433172_8446622_RNA45SN1_45S_50ntFlanks</td><td>10458</td><td>10488</td><td>31</td><td>ATGGCCTCCGTTGCCCTCGGCCGATCGAAAG</td><td>250.0</td><td>-5.1</td><td>1.6</td><td>-3.1</td><td>1.7E-03</td><td>3.8E-03</td><td>BT20/MB468</td><td>MP</td></tr>
<tr><td>>hg38_21_+_8433172_8446622_RNA45SN1_45S_50ntFlanks@11570.11604.35|AGCCGACTTAGAACTGGTGCGGACCAGGGGAATCC</td><td>>hg38_21_+_8433172_8446622_RNA45SN1_45S_50ntFlanks</td><td>11570</td><td>11604</td><td>35</td><td>AGCCGACTTAGAACTGGTGCGGACCAGGGGAATCC</td><td>257.5</td><td>5.9</td><td>1.7</td><td>3.4</td><td>5.8E-04</td><td>3.8E-03</td><td>BT20/MB231</td><td>MP</td></tr>
<tr><td>>hg38_21_+_8433172_8446622_RNA45SN1_45S_50ntFlanks@9494.9525.32|CCGAAAGATGGTGAACTATGCCTGGGCAGGGC</td><td>>hg38_21_+_8433172_8446622_RNA45SN1_45S_50ntFlanks</td><td>9494</td><td>9525</td><td>32</td><td>CCGAAAGATGGTGAACTATGCCTGGGCAGGGC</td><td>291.8</td><td>6.9</td><td>2.0</td><td>3.4</td><td>5.8E-04</td><td>3.8E-03</td><td>BT20/MB231</td><td>MP</td></tr>
<tr><td>>hg38_21_+_8433172_8446622_RNA45SN1_45S_50ntFlanks@10664.10706.43|CTTGGAAAGCGTCGCGGTTCCGGCGGCGTCCGGTGAGCTCTCG</td><td>>hg38_21_+_8433172_8446622_RNA45SN1_45S_50ntFlanks</td><td>10664</td><td>10706</td><td>43</td><td>CTTGGAAAGCGTCGCGGTTCCGGCGGCGTCCGGTGAGCTCTCG</td><td>580.4</td><td>-4.1</td><td>1.4</td><td>-3.0</td><td>3.0E-03</td><td>3.8E-03</td><td>BT20/MB468</td><td>CYTO</td></tr>
<tr><td>>hg38_21_+_8433172_8446622_RNA45SN1_45S_50ntFlanks@4735.4768.34|ACGAAAGTCGGAGGTTCGAAGACGATCAGATACC</td><td>>hg38_21_+_8433172_8446622_RNA45SN1_45S_50ntFlanks</td><td>4735</td><td>4768</td><td>34</td><td>ACGAAAGTCGGAGGTTCGAAGACGATCAGATACC</td><td>34.1</td><td>8.3</td><td>2.8</td><td>3.0</td><td>2.8E-03</td><td>3.8E-03</td><td>MB231/MB468</td><td>MP</td></tr>
<tr><td>>trnaMT_PheGAA_MT_+_577_647@1.44.44|GTTTATGTAGCTTACCTCCTCAAAGCAATACACTGAAAATGTTT</td><td>>trnaMT_PheGAA_MT_+_577_647</td><td>1</td><td>44</td><td>44</td><td>GTTTATGTAGCTTACCTCCTCAAAGCAATACACTGAAAATGTTT</td><td>3275.3</td><td>7.0</td><td>2.2</td><td>3.1</td><td>1.7E-03</td><td>3.8E-03</td><td>BT20/MB468</td><td>MP</td></tr>
<tr><td>>MI0000446|hsa-mir-125b-1&WithFlank&11|-|122099751|122099850@21.42.22;MI0000470|hsa-mir-125b-2&WithFlank&21|+|16590231|16590331@23.44.22&[MIMAT0000423&hsa-miR-125b-5p&offsets|0|0;m-67&11|-|122099809|122099830&offsets|0|0];[MIMAT0000423_1&hsa-miR-125b-5p&offsets|0|0;m-68&21|+|16590253|16590274&offsets|0|0]|TCCCTGAGACCCTAACTTGTGA</td><td>>MI0000446|hsa-mir-125b-1&WithFlank&11|-|122099751|122099850</td><td>21</td><td>42</td><td>22</td><td>TCCCTGAGACCCTAACTTGTGA</td><td>11440.2</td><td>-3.7</td><td>1.1</td><td>-3.4</td><td>6.9E-04</td><td>3.8E-03</td><td>BT20/MB231</td><td>CYTO</td></tr>
<tr><td>>hg38_21_+_8433172_8446622_RNA45SN1_45S_50ntFlanks@11570.11600.31|AGCCGACTTAGAACTGGTGCGGACCAGGGGA</td><td>>hg38_21_+_8433172_8446622_RNA45SN1_45S_50ntFlanks</td><td>11570</td><td>11600</td><td>31</td><td>AGCCGACTTAGAACTGGTGCGGACCAGGGGA</td><td>576.9</td><td>4.3</td><td>1.3</td><td>3.4</td><td>5.8E-04</td><td>3.9E-03</td><td>BT20/MB231</td><td>MP</td></tr>
<tr><td>>hg38_21_+_8433172_8446622_RNA45SN1_45S_50ntFlanks@12191.12209.19|ACGCAGGTGTCCTAAGGCG</td><td>>hg38_21_+_8433172_8446622_RNA45SN1_45S_50ntFlanks</td><td>12191</td><td>12209</td><td>19</td><td>ACGCAGGTGTCCTAAGGCG</td><td>75.0</td><td>-5.2</td><td>1.7</td><td>-3.1</td><td>1.7E-03</td><td>3.9E-03</td><td>BT20/MB468</td><td>MP</td></tr>
<tr><td>>hg38_21_+_8433172_8446622_RNA45SN1_45S_50ntFlanks@7971.7988.18|GAGACGCGACCTCAGATC</td><td>>hg38_21_+_8433172_8446622_RNA45SN1_45S_50ntFlanks</td><td>7971</td><td>7988</td><td>18</td><td>GAGACGCGACCTCAGATC</td><td>510.2</td><td>2.7</td><td>0.8</td><td>3.4</td><td>7.7E-04</td><td>3.9E-03</td><td>BT20/MB231</td><td>MITO</td></tr>
<tr><td>>hg38_21_+_8433172_8446622_RNA45SN1_45S_50ntFlanks@6655.6688.34|TCTTAGCGGTGGATCACTCGGCTCGTGCGTCGAT</td><td>>hg38_21_+_8433172_8446622_RNA45SN1_45S_50ntFlanks</td><td>6655</td><td>6688</td><td>34</td><td>TCTTAGCGGTGGATCACTCGGCTCGTGCGTCGAT</td><td>69.3</td><td>-3.8</td><td>1.3</td><td>-3.0</td><td>2.8E-03</td><td>3.9E-03</td><td>MB231/MB468</td><td>MP</td></tr>
<tr><td>>hg38_21_+_8433172_8446622_RNA45SN1_45S_50ntFlanks@6664.6685.22|TGGATCACTCGGCTCGTGCGTC</td><td>>hg38_21_+_8433172_8446622_RNA45SN1_45S_50ntFlanks</td><td>6664</td><td>6685</td><td>22</td><td>TGGATCACTCGGCTCGTGCGTC</td><td>84.2</td><td>-4.7</td><td>1.6</td><td>-3.0</td><td>2.8E-03</td><td>3.9E-03</td><td>MB231/MB468</td><td>MP</td></tr>
</table>

| | | | | | | | | | | | | | |
|---|---|---|---|---|---|---|---|---|---|---|---|---|---|
| >MI0000102\|hsa-mir-100&WithFlank&11\|-\|122152223\|122152314@19.40.22(+1A)&[MIMAT0000098&hsa-miR-100-5p&offsets\|0\|0(+1A);m-27&11\|-\|122152276\|122152296&offsets\|0\|+1(+1A)]\|AACCCGTAGATCCGAACTTGTGA | >MI0000102\|hsa-mir-100&WithFlank&11\|-\|122152223\|122152314 | 19 | 40 | 22(+1A) | AACCCGTAGATCCGAACTTGTGA | 22967.0 | -3.7 | 1.1 | -3.4 | 7.0E-04 | 3.9E-03 | BT20/MB231 | CYTO |
| >MI0000088\|hsa-mir-30a&WithFlank&6\|-\|71403545\|71403627@13.33.21(+1U)&[MIMAT0000087&hsa-miR-30a-5p&offsets\|+1\|0(+1U);m-15&6\|-\|71403595\|71403616&offsets\|+1\|0(+1U)]\|GTAAACATCCTCGACTGGAAGT | >MI0000088\|hsa-mir-30a&WithFlank&6\|-\|71403545\|71403627 | 13 | 33 | 21(+1U) | GTAAACATCCTCGACTGGAAGT | 83.0 | 4.6 | 1.5 | 3.0 | 2.8E-03 | 3.9E-03 | MB231/MB468 | MP |
| >hg38_21_+_8433172_8446622_RNA45SN1_45S_50ntFlanks@12008.12038.31\|CCGCCGGTGAAATACCACTACTCTGATCGTT | >hg38_21_+_8433172_8446622_RNA45SN1_45S_50ntFlanks | 12008 | 12038 | 31 | CCGCCGGTGAAATACCACTACTCTGATCGTT | 148.2 | -3.3 | 1.1 | -3.0 | 3.0E-03 | 3.9E-03 | BT20/MB468 | CYTO |
| >hg38_21_+_8433172_8446622_RNA45SN1_45S_50ntFlanks@6661.6698.38\|CGGTGGATCACTCGGCTCGTGCGTCGATGAAGAACGCA | >hg38_21_+_8433172_8446622_RNA45SN1_45S_50ntFlanks | 6661 | 6698 | 38 | CGGTGGATCACTCGGCTCGTGCGTCGATGAAGAACGCA | 1790.1 | 1.7 | 0.5 | 3.0 | 2.6E-03 | 3.9E-03 | BT20/MB468 | MITO |
| >hg38_21_+_8433172_8446622_RNA45SN1_45S_50ntFlanks@12008.12036.29\|CCGCCGGTGAAATACCACTACTCTGATCG | >hg38_21_+_8433172_8446622_RNA45SN1_45S_50ntFlanks | 12008 | 12036 | 29 | CCGCCGGTGAAATACCACTACTCTGATCG | 175.3 | -3.3 | 1.1 | -3.0 | 3.0E-03 | 3.9E-03 | BT20/MB468 | CYTO |
| >MI0000253\|hsa-mir-148a&WithFlank&7\|-\|25949913\|25949992@50.70.21(+1G)&[MIMAT0000243&hsa-miR-148a-3p&offsets\|0\|-1(+1G);m-7&7\|-\|25949922\|25949943&offsets\|0\|-1(+1G)]\|TCAGTGCACTACAGAACTTTGG | >MI0000253\|hsa-mir-148a&WithFlank&7\|-\|25949913\|25949992 | 50 | 70 | 21(+1G) | TCAGTGCACTACAGAACTTTGG | 896.0 | 2.8 | 0.8 | 3.4 | 7.7E-04 | 3.9E-03 | BT20/MB231 | MITO |
| >hg38_21_+_8433172_8446622_RNA45SN1_45S_50ntFlanks@5492.5509.18\|AGAAGACGGTCGAACTTG | >hg38_21_+_8433172_8446622_RNA45SN1_45S_50ntFlanks | 5492 | 5509 | 18 | AGAAGACGGTCGAACTTG | 188.2 | 6.8 | 2.0 | 3.4 | 5.8E-04 | 3.9E-03 | BT20/MB231 | MP |
| >hg38_21_+_8433172_8446622_RNA45SN1_45S_50ntFlanks@10240.10262.23\|CTACGCCGCGACGAGTAGGAGGG | >hg38_21_+_8433172_8446622_RNA45SN1_45S_50ntFlanks | 10240 | 10262 | 23 | CTACGCCGCGACGAGTAGGAGGG | 157.7 | 6.3 | 1.8 | 3.4 | 5.8E-04 | 3.9E-03 | BT20/MB231 | MP |
| >hg38_21_+_8433172_8446622_RNA45SN1_45S_50ntFlanks@10510.10532.23\|AATCCGGAGTGGCGGAGATGGGC | >hg38_21_+_8433172_8446622_RNA45SN1_45S_50ntFlanks | 10510 | 10532 | 23 | AATCCGGAGTGGCGGAGATGGGC | 2588.4 | 6.6 | 1.9 | 3.4 | 5.8E-04 | 3.9E-03 | BT20/MB231 | MP |
| >hg38_21_+_8433172_8446622_RNA45SN1_45S_50ntFlanks@10851.10879.29\|GGATAAGGATTGGCTCTAAGGGCTGGGTC | >hg38_21_+_8433172_8446622_RNA45SN1_45S_50ntFlanks | 10851 | 10879 | 29 | GGATAAGGATTGGCTCTAAGGGCTGGGTC | 77.7 | 9.6 | 2.8 | 3.4 | 5.8E-04 | 3.9E-03 | BT20/MB231 | MP |
| >hg38_21_+_8433172_8446622_RNA45SN1_45S_50ntFlanks@9929.9960.32\|GATATAGACAGCAGGACGGTGGCCATGGAAGT | >hg38_21_+_8433172_8446622_RNA45SN1_45S_50ntFlanks | 9929 | 9960 | 32 | GATATAGACAGCAGGACGGTGGCCATGGAAGT | 713.1 | -2.2 | 0.7 | -3.1 | 1.7E-03 | 3.9E-03 | BT20/MB468 | MP |
| >trnaMT_ThrTGT_MT_+_15888_15953@36.69.34\|AACCGGAGATGAAAACCTTTTTCCAAGGACACCA | >trnaMT_ThrTGT_MT_+_15888_15953 | 36 | 69 | 34 | AACCGGAGATGAAAACCTTTTTCCAAGGACACCA | 218.3 | 2.9 | 1.0 | 3.0 | 2.8E-03 | 3.9E-03 | MB231/MB468 | MP |
| >hg38_21_+_8433172_8446622_RNA45SN1_45S_50ntFlanks@476.512.37\|TTCGTGATCGATGTGGTGACGTCGTGCTCTCCCGGGC | >hg38_21_+_8433172_8446622_RNA45SN1_45S_50ntFlanks | 476 | 512 | 37 | TTCGTGATCGATGTGGTGACGTCGTGCTCTCCCGGGC | 149.8 | 4.6 | 1.6 | 2.9 | 3.3E-03 | 3.9E-03 | MB231/MB468 | MITO |
| >MI0000746\|hsa-mir-99b&WithFlank&19\|+\|51692606\|51692687@51.72.22(+1C)&[MIMAT0004678&hsa-miR-99b-3p&offsets\|0\|0(+1C);m-151&19\|+\|51692656\|51692678&offsets\|0\|-1(+1C)]\|CAAGCTCGTGTCTGTGGGTCCGC | >MI0000746\|hsa-mir-99b&WithFlank&19\|+\|51692606\|51692687 | 51 | 72 | 22(+1C) | CAAGCTCGTGTCTGTGGGTCCGC | 73.8 | 5.1 | 1.6 | 3.1 | 1.7E-03 | 3.9E-03 | BT20/MB468 | MP |
| >hg38_21_+_8433172_8446622_RNA45SN1_45S_50ntFlanks@10829.10851.23\|GCAAGCCGGATCCGTAACTTCGG | >hg38_21_+_8433172_8446622_RNA45SN1_45S_50ntFlanks | 10829 | 10851 | 23 | GCAAGCCGGATCCGTAACTTCGG | 172.6 | -1.6 | 0.6 | -3.0 | 3.1E-03 | 3.9E-03 | BT20/MB468 | CYTO |
| >MI0000298\|hsa-mir-221&WithFlank&X\|-\|45746151\|45746272@71.89.19(+1A)&[MIMAT0000278&hsa-miR-221-3p&offsets\|0\|-4(+1A);m-91&X\|-\|45746181\|45746202&offsets\|0\|-3(+1A)]\|AGCTACATTGTCTGCTGGGA | >MI0000298\|hsa-mir-221&WithFlank&X\|-\|45746151\|45746272 | 71 | 89 | 19(+1A) | AGCTACATTGTCTGCTGGGA | 75.3 | 3.7 | 1.2 | 3.0 | 2.8E-03 | 3.9E-03 | MB231/MB468 | MP |
| >hg38_MT_+_1621_3279_RNR2_16S_50ntFlanks@470.491.22\|AACTGTTAGTCCAAAGAGGAAC | >hg38_MT_+_1621_3279_RNR2_16S_50ntFlanks | 470 | 491 | 22 | AACTGTTAGTCCAAAGAGGAAC | 97.5 | 2.7 | 0.9 | 3.0 | 2.8E-03 | 3.9E-03 | MB231/MB468 | MP |
| >hg38_21_+_8433172_8446622_RNA45SN1_45S_50ntFlanks@3802.3829.28\|CATTAAATCAGTTATGGTTCCTTTGGTC | >hg38_21_+_8433172_8446622_RNA45SN1_45S_50ntFlanks | 3802 | 3829 | 28 | CATTAAATCAGTTATGGTTCCTTTGGTC | 98.5 | -5.1 | 1.6 | -3.1 | 1.7E-03 | 3.9E-03 | BT20/MB468 | MP |
| >hg38_21_+_8433172_8446622_RNA45SN1_45S_50ntFlanks@12405.12433.29\|CTTTTTGATCCTTCGATGTCGGCTCTTCC | >hg38_21_+_8433172_8446622_RNA45SN1_45S_50ntFlanks | 12405 | 12433 | 29 | CTTTTTGATCCTTCGATGTCGGCTCTTCC | 5020.6 | -2.9 | 1.0 | -3.0 | 3.1E-03 | 3.9E-03 | MB231/MB468 | CYTO |
| >hg38_21_+_8433172_8446622_RNA45SN1_45S_50ntFlanks@12623.12654.32\|GCCAATGGGGCGAAGCTACCATCTGTGGGATT | >hg38_21_+_8433172_8446622_RNA45SN1_45S_50ntFlanks | 12623 | 12654 | 32 | GCCAATGGGGCGAAGCTACCATCTGTGGGATT | 141.7 | -2.3 | 0.8 | -3.0 | 3.1E-03 | 3.9E-03 | BT20/MB468 | CYTO |
| >hg38_21_+_8433172_8446622_RNA45SN1_45S_50ntFlanks@4723.4767.45\|CATTAATCAAGAACGAAAGTCGGAGGTTCGAAGACGATCAGATAC | >hg38_21_+_8433172_8446622_RNA45SN1_45S_50ntFlanks | 4723 | 4767 | 45 | CATTAATCAAGAACGAAAGTCGGAGGTTCGAAGACGATCAGATAC | 709.2 | -3.6 | 1.1 | -3.4 | 7.0E-04 | 3.9E-03 | BT20/MB231 | CYTO |
| >hg38_21_+_8433172_8446622_RNA45SN1_45S_50ntFlanks@5487.5510.24\|CGCTGAGAAGACGGTCGAACTTGA | >hg38_21_+_8433172_8446622_RNA45SN1_45S_50ntFlanks | 5487 | 5510 | 24 | CGCTGAGAAGACGGTCGAACTTGA | 784.8 | -4.3 | 1.4 | -3.1 | 1.7E-03 | 3.9E-03 | BT20/MB468 | MP |
| >hg38_21_+_8433172_8446622_RNA45SN1_45S_50ntFlanks@12626.12657.32\|AATGGGGCGAAGCTACCATCTGTGGGATTATG | >hg38_21_+_8433172_8446622_RNA45SN1_45S_50ntFlanks | 12626 | 12657 | 32 | AATGGGGCGAAGCTACCATCTGTGGGATTATG | 752.7 | -5.4 | 1.7 | -3.1 | 1.7E-03 | 3.9E-03 | BT20/MB468 | MP |
| >hg38_21_+_8433172_8446622_RNA45SN1_45S_50ntFlanks@6651.6675.25\|CGACTCTTAGCGGTGGATCACTCGG | >hg38_21_+_8433172_8446622_RNA45SN1_45S_50ntFlanks | 6651 | 6675 | 25 | CGACTCTTAGCGGTGGATCACTCGG | 4307.5 | -1.9 | 0.6 | -2.9 | 3.3E-03 | 3.9E-03 | BT20/MB468 | NUC |
| >am_trnaMT_PheGAA_MT_+_577_647@45.62.18\|AGACGGGCTCACATCACC | >am_trnaMT_PheGAA_MT_+_577_647 | 45 | 62 | 18 | AGACGGGCTCACATCACC | 958.2 | 4.2 | 1.4 | 3.0 | 3.1E-03 | 3.9E-03 | BT20/MB468 | CYTO |
| >MI0000279\|hsa-mir-196a-2&WithFlank&12\|+\|53991732\|53991853@31.51.21;MI0000238\|hsa-mir-196a-1&WithFlank&17\|-\|48632484\|48632565@13.33.21&[MIMAT0000226&hsa-miR-196a-5p&offsets\|0\|-1;m-261&12\|+\|53991762\|53991783&offsets\|0\|-1];[MIMAT0000226_1&hsa-miR-196a-5p&offsets\|0\|-1;m-268&17\|-\|48632532\|48632553&offsets\|0\|-1]\|TAGGTAGTTTCATGTTGTTGG | >MI0000279\|hsa-mir-196a-2&WithFlank&12\|+\|53991732\|53991853 | 31 | 51 | 21 | TAGGTAGTTTCATGTTGTTGG | 523.7 | 4.5 | 1.4 | 3.1 | 1.7E-03 | 3.9E-03 | BT20/MB468 | MP |
| >hg38_21_+_8433172_8446622_RNA45SN1_45S_50ntFlanks@12412.12435.24\|ATCCTTCGATGTCGGCTCTTCCTA | >hg38_21_+_8433172_8446622_RNA45SN1_45S_50ntFlanks | 12412 | 12435 | 24 | ATCCTTCGATGTCGGCTCTTCCTA | 279.0 | -4.2 | 1.4 | -3.0 | 3.1E-03 | 3.9E-03 | MB231/MB468 | CYTO |
| >hg38_21_+_8433172_8446622_RNA45SN1_45S_50ntFlanks@9792.9819.28\|GGCGTGGAGCCGGGCGTGGAATGCGAGT | >hg38_21_+_8433172_8446622_RNA45SN1_45S_50ntFlanks | 9792 | 9819 | 28 | GGCGTGGAGCCGGGCGTGGAATGCGAGT | 310.5 | 6.4 | 1.9 | 3.4 | 5.9E-04 | 3.9E-03 | BT20/MB231 | MP |
| >hg38_1_-_228634819_228635039_RNA5S12_5S_50ntFlanks@127.170.44\|ACTTGGATGGGAGACCGCCTGGGAATACCGGGTGCTGTAGGCTT | >hg38_1_-_228634819_228635039_RNA5S12_5S_50ntFlanks | 127 | 170 | 44 | ACTTGGATGGGAGACCGCCTGGGAATACCGGGTGCTGTAGGCTT | 1569.4 | 3.1 | 0.9 | 3.4 | 7.8E-04 | 3.9E-03 | BT20/MB231 | MITO |
| >hg38_21_+_8433172_8446622_RNA45SN1_45S_50ntFlanks@12996.13040.45\|TCGCTGCGATCTATTGAAAGTCAGCCCTCGACACAAGGGTTTGTC | >hg38_21_+_8433172_8446622_RNA45SN1_45S_50ntFlanks | 12996 | 13040 | 45 | TCGCTGCGATCTATTGAAAGTCAGCCCTCGACACAAGGGTTTGTC | 290.3 | -4.5 | 1.3 | -3.4 | 6.5E-04 | 3.9E-03 | BT20/MB231 | NUC |
| >trnaMT_TyrGTA_MT_-_5826_5891@39.69.31\|AAGACAGGGGTTAGGCCTCTTTTTACCACCA | >trnaMT_TyrGTA_MT_-_5826_5891 | 39 | 69 | 31 | AAGACAGGGGTTAGGCCTCTTTTTACCACCA | 208.0 | 3.9 | 1.2 | 3.1 | 1.7E-03 | 3.9E-03 | BT20/MB468 | MP |
| >hg38_21_+_8433172_8446622_RNA45SN1_45S_50ntFlanks@7980.8012.33\|CCTCAGATCAGACGTGGCGACCCGCTGAATTTA | >hg38_21_+_8433172_8446622_RNA45SN1_45S_50ntFlanks | 7980 | 8012 | 33 | CCTCAGATCAGACGTGGCGACCCGCTGAATTTA | 366.7 | -4.4 | 1.4 | -3.1 | 1.7E-03 | 3.9E-03 | BT20/MB468 | MP |
| >MI0000294\|hsa-mir-218-1&WithFlank&4\|+\|20528269\|20528390@31.51.21;MI0000295\|hsa-mir-218-2&WithFlank&5\|-\|168768140\|168768261@31.51.21&[MIMAT0000275&hsa-miR-218-5p&offsets\|0\|0;m-185&4\|+\|20528299\|20528319&offsets\|0\|0];[MIMAT0000275_1&hsa-miR-218-5p&offsets\|0\|0;m-187&5\|-\|168768211\|168768231&offsets\|0\|0]\|TTGTGCTTGATCTAACCATGT | >MI0000294\|hsa-mir-218-1&WithFlank&4\|+\|20528269\|20528390 | 31 | 51 | 21 | TTGTGCTTGATCTAACCATGT | 1045.4 | -6.6 | 1.9 | -3.4 | 5.9E-04 | 3.9E-03 | BT20/MB231 | MP |
| >hg38_21_+_8433172_8446622_RNA45SN1_45S_50ntFlanks@10393.10433.41\|CAGCAGTTGAACATGGGTCAGTCGGTCCTGAGAGATGGGCG | >hg38_21_+_8433172_8446622_RNA45SN1_45S_50ntFlanks | 10393 | 10433 | 41 | CAGCAGTTGAACATGGGTCAGTCGGTCCTGAGAGATGGGCG | 285.5 | -4.7 | 1.6 | -3.0 | 3.1E-03 | 3.9E-03 | BT20/MB468 | CYTO |
| >hg38_21_+_8433172_8446622_RNA45SN1_45S_50ntFlanks@12416.12437.22\|TTCGATGTCGGCTCTTCCTATC | >hg38_21_+_8433172_8446622_RNA45SN1_45S_50ntFlanks | 12416 | 12437 | 22 | TTCGATGTCGGCTCTTCCTATC | 1914.4 | 1.9 | 0.6 | 2.9 | 3.3E-03 | 3.9E-03 | MB231/MB468 | NUC |
| >MI0000434\|hsa-let-7i&WithFlank&12\|+\|62603680\|62603775@12.29.18(+1A)&[MIMAT0000415&hsa-let-7i-5p&offsets\|0\|-4(+1A);m-49&12\|+\|62603691\|62603712&offsets\|0\|-4(+1A)]\|TGAGGTAGTAGTTTGTGCA | >MI0000434\|hsa-let-7i&WithFlank&12\|+\|62603680\|62603775 | 12 | 29 | 18(+1A) | TGAGGTAGTAGTTTGTGCA | 36.5 | 8.4 | 2.8 | 3.0 | 2.8E-03 | 3.9E-03 | MB231/MB468 | MP |
| >hg38_21_+_8433172_8446622_RNA45SN1_45S_50ntFlanks@5073.5112.40\|AATTCCGATAACGAACGAGACTCTGGCATGCTAACTAGTT | >hg38_21_+_8433172_8446622_RNA45SN1_45S_50ntFlanks | 5073 | 5112 | 40 | AATTCCGATAACGAACGAGACTCTGGCATGCTAACTAGTT | 318.9 | -2.8 | 0.9 | -2.9 | 3.3E-03 | 3.9E-03 | MB231/MB468 | NUC |
| >am_trna10_LysCTT_16_+_3241501_3241573@11.28.18,trna119_LysCTT_1_-_145395522_145395594@11.28.18,trna11_LysCTT_5_-_180648979_180649051@11.28.18,trna128_LysCTT_1_-_55423542_55423614@11.28.18,trna13_LysCTT_14_-_58706613_58706685@11.28.18,trna13_LysCTT_6_+_26556774_26556846@11.28.18,trna2_LysCTT_15_+_79152904_79152976@11.28.18,trna32_LysCTT_16_-_3207406_3207478@11.28.18,trna4_LysCTT_18_-_43669271_43669343@11.28.18,trna6_LysCTT_19_-_52425396_52425468@11.28.18,trna7_LysCTT_16_+_3225692_3225764@11.28.18,trna9_LysCTT_5_+_180634755_180634827@11.28.18\|CTCAGTCGGTAGAGCATG | >am_trna10_LysCTT_16_+_3241501_3241573 | 11 | 28 | 18 | CTCAGTCGGTAGAGCATG | 532.4 | -4.3 | 1.4 | -3.1 | 1.7E-03 | 3.9E-03 | BT20/MB468 | MP |
| >MI0000077\|hsa-mir-21&WithFlank&17\|+\|59841260\|59841343@14.35.22(+1U)&[MIMAT0000076&hsa-miR-21-5p&offsets\|0\|0(+1U);m-1&17\|+\|59841273\|59841294&offsets\|0\|0(+1U)]\|TAGCTTATCAGACTGATGTTGAT | >MI0000077\|hsa-mir-21&WithFlank&17\|+\|59841260\|59841343 | 14 | 35 | 22(+1U) | TAGCTTATCAGACTGATGTTGAT | 138949.8 | -2.5 | 0.7 | -3.4 | 7.8E-04 | 3.9E-03 | BT20/MB231 | MITO |
| >hg38_21_+_8433172_8446622_RNA45SN1_45S_50ntFlanks@7974.8010.37\|ACGCGACCTCAGATCAGACGTGGCGACCCGCTGAATT | >hg38_21_+_8433172_8446622_RNA45SN1_45S_50ntFlanks | 7974 | 8010 | 37 | ACGCGACCTCAGATCAGACGTGGCGACCCGCTGAATT | 67102.9 | 2.9 | 0.9 | 3.4 | 6.5E-04 | 3.9E-03 | BT20/MB231 | NUC |
| >hg38_21_+_8433172_8446622_RNA45SN1_45S_50ntFlanks@12474.12496.23\|ACCCACTAATAGGGAACGTGAGC | >hg38_21_+_8433172_8446622_RNA45SN1_45S_50ntFlanks | 12474 | 12496 | 23 | ACCCACTAATAGGGAACGTGAGC | 647.7 | -3.1 | 0.9 | -3.4 | 6.5E-04 | 3.9E-03 | BT20/MB231 | NUC |
| >hg38_21_+_8433172_8446622_RNA45SN1_45S_50ntFlanks@12003.12034.32\|GCGGGCCGCCGGTGAAATACCACTACTCTGAT | >hg38_21_+_8433172_8446622_RNA45SN1_45S_50ntFlanks | 12003 | 12034 | 32 | GCGGGCCGCCGGTGAAATACCACTACTCTGAT | 1663.9 | -3.3 | 1.0 | -3.4 | 7.0E-04 | 3.9E-03 | BT20/MB231 | CYTO |
| >MI0000063\|hsa-let-7b&WithFlank&22\|+\|46113680\|46113774@12.34.23(+1A)&[MIMAT0000063&hsa-let-7b-5p&offsets\|0\|+1(+1A);m-14&22\|+\|46113691\|46113712&offsets\|0\|+1(+1A)]\|TGAGGTAGTAGGTTGTGTGGTTTA | >MI0000063\|hsa-let-7b&WithFlank&22\|+\|46113680\|46113774 | 12 | 34 | 23(+1A) | TGAGGTAGTAGGTTGTGTGGTTTA | 122.8 | 4.5 | 1.4 | 3.1 | 1.7E-03 | 3.9E-03 | BT20/MB468 | MP |

| | | | | | | | | | | | | | |
|---|---|---|---|---|---|---|---|---|---|---|---|---|---|
| >MI0000650\|hsa-mir-200c&WithFlank&12\|+\|6963693\|6963772@50.72.23(+2U)&[MIMAT0000617&hsa-miR-200c-3p&offsets\|0\|0(+2U);m-54&12\|+\|6963742\|6963764&offsets\|0\|0(+2U)]\|TAATACTGCCGGGTAATGATGGATT | >MI0000650\|hsa-mir-200c&WithFlank&12\|+\|6963693\|6963772 | 50 | 72 | 23(+2U) | TAATACTGCCGGGTAATGATGGATT | 89.4 | 4.2 | 1.3 | 3.1 | 1.7E-03 | 3.9E-03 | BT20/MB468 | MP |
| >hg38_21_+_8433172_8446622_RNA45SN1_45S_50ntFlanks@10843.10872.30\|TAACTTCGGGATAAGGATTGGCTCTAAGGG | >hg38_21_+_8433172_8446622_RNA45SN1_45S_50ntFlanks | 10843 | 10872 | 30 | TAACTTCGGGATAAGGATTGGCTCTAAGGG | 148.1 | -4.4 | 1.5 | -2.9 | 3.4E-03 | 3.9E-03 | MB231/MB468 | NUC |
| >hg38_21_+_8433172_8446622_RNA45SN1_45S_50ntFlanks@6646.6675.30\|TCGTACGACTCTTAGCGGTGGATCACTCGG | >hg38_21_+_8433172_8446622_RNA45SN1_45S_50ntFlanks | 6646 | 6675 | 30 | TCGTACGACTCTTAGCGGTGGATCACTCGG | 1091.2 | -2.5 | 0.8 | -3.0 | 2.8E-03 | 3.9E-03 | MB231/MB468 | MP |
| >am_trna128_GlyGCC_6_-_27870686_27870756@8.35.28,trna18_GlyGCC_16_+_70822597_70822667@8.35.28,trna19_GlyGCC_16_+_70823410_70823480@8.35.28,trna19_GlyGCC_2_-_157257659_157257729@8.35.28,trna24_GlyGCC_16_-_70812942_70813012@8.35.28,trna25_GlyGCC_16_-_70812114_70812184@8.35.28,trna2_GlyGCC_21_-_18827107_18827177@8.35.28,trna35_GlyGCC_1_+_161413094_161413164@8.35.28,trna37_GlyGCC_1_+_161420467_161420537@8.35.28,trna39_GlyGCC_1_+_161427898_161427968@8.35.28,trna41_GlyGCC_1_+_161435258_161435328@8.35.28,trna5_GlyGCC_17_+_8029064_8029134@8.35.28,trna68_GlyGCC_1_-_161493637_161493707@8.35.28\|TGGTTCAGTGGTAGAATTCTCGCCTGCC | >am_trna128_GlyGCC_6_-_27870686_27870756 | 8 | 35 | 28 | TGGTTCAGTGGTAGAATTCTCGCCTGCC | 63.1 | 5.5 | 1.8 | 3.0 | 2.8E-03 | 3.9E-03 | MB231/MB468 | MP |
| >hg38_21_+_8433172_8446622_RNA45SN1_45S_50ntFlanks@5471.5514.44\|CACGGCCCTGGCGGAGCGCTGAGAAGACGGTCGAACTTGACTAT | >hg38_21_+_8433172_8446622_RNA45SN1_45S_50ntFlanks | 5471 | 5514 | 44 | CACGGCCCTGGCGGAGCGCTGAGAAGACGGTCGAACTTGACTAT | 533.3 | -4.2 | 1.3 | -3.1 | 1.7E-03 | 3.9E-03 | BT20/MB468 | MP |
| >hg38_21_+_8433172_8446622_RNA45SN1_45S_50ntFlanks@10458.10486.29\|ATGGCCTCCGTTGCCCTCGGCCGATCGAA | >hg38_21_+_8433172_8446622_RNA45SN1_45S_50ntFlanks | 10458 | 10486 | 29 | ATGGCCTCCGTTGCCCTCGGCCGATCGAA | 192.1 | -2.9 | 1.0 | -3.0 | 3.2E-03 | 3.9E-03 | MB231/MB468 | CYTO |
| >hg38_21_+_8433172_8446622_RNA45SN1_45S_50ntFlanks@10420.10456.37\|CTGAGAGATGGGCGAGCGCCGTTCCGAAGGGACGGGC | >hg38_21_+_8433172_8446622_RNA45SN1_45S_50ntFlanks | 10420 | 10456 | 37 | CTGAGAGATGGGCGAGCGCCGTTCCGAAGGGACGGGC | 976.5 | -3.0 | 0.9 | -3.4 | 7.1E-04 | 3.9E-03 | BT20/MB231 | CYTO |
| >hg38_21_+_8433172_8446622_RNA45SN1_45S_50ntFlanks@12616.12661.46\|CTGAGGAGCCAATGGGGCGAAGCTACCATCTGTGGGATTATGACTG | >hg38_21_+_8433172_8446622_RNA45SN1_45S_50ntFlanks | 12616 | 12661 | 46 | CTGAGGAGCCAATGGGGCGAAGCTACCATCTGTGGGATTATGACTG | 262.5 | -4.3 | 1.5 | -2.9 | 3.4E-03 | 3.9E-03 | MB231/MB468 | NUC |
| >hg38_21_+_8433172_8446622_RNA45SN1_45S_50ntFlanks@10916.10935.20\|CCGCGGCTGGACGAGGCGCC | >hg38_21_+_8433172_8446622_RNA45SN1_45S_50ntFlanks | 10916 | 10935 | 20 | CCGCGGCTGGACGAGGCGCC | 253.2 | -4.6 | 1.5 | -3.0 | 3.1E-03 | 3.9E-03 | BT20/MB468 | CYTO |
| >hg38_MT_+_598_1651_RNR1_12S_50ntFlanks@916.946.31\|AAGGACATTTAACTAAAACCCCTACGCATTT | >hg38_MT_+_598_1651_RNR1_12S_50ntFlanks | 916 | 946 | 31 | AAGGACATTTAACTAAAACCCCTACGCATTT | 694.8 | 2.5 | 0.8 | 3.4 | 7.9E-04 | 3.9E-03 | BT20/MB231 | MITO |
| >hg38_21_+_8433172_8446622_RNA45SN1_45S_50ntFlanks@11161.11178.18\|CACGGTCCCCCGCGAGGG | >hg38_21_+_8433172_8446622_RNA45SN1_45S_50ntFlanks | 11161 | 11178 | 18 | CACGGTCCCCCGCGAGGG | 2327.9 | -5.3 | 1.8 | -3.0 | 3.1E-03 | 3.9E-03 | BT20/MB468 | CYTO |
| >am_trnaMT_ValTAC_MT_+_1602_1670@48.68.21\|CAACTTAACTTGACCGCTCTG | >am_trnaMT_ValTAC_MT_+_1602_1670 | 48 | 68 | 21 | CAACTTAACTTGACCGCTCTG | 163.2 | -3.5 | 1.2 | -3.0 | 2.8E-03 | 3.9E-03 | MB231/MB468 | MP |
| >hg38_1_-_228634819_228635039_RNA5S12_5S_50ntFlanks@136.171.36\|GGAGACCGCCTGGGAATACCGGGTGCTGTAGGCTTT | >hg38_1_-_228634819_228635039_RNA5S12_5S_50ntFlanks | 136 | 171 | 36 | GGAGACCGCCTGGGAATACCGGGTGCTGTAGGCTTT | 10430.7 | 4.1 | 1.3 | 3.1 | 1.7E-03 | 3.9E-03 | BT20/MB468 | MP |
| >hg38_21_+_8433172_8446622_RNA45SN1_45S_50ntFlanks@10901.10928.28\|GGGGCTGGGCGCGCGCCGCGGCTGGACG | >hg38_21_+_8433172_8446622_RNA45SN1_45S_50ntFlanks | 10901 | 10928 | 28 | GGGGCTGGGCGCGCGCCGCGGCTGGACG | 34.4 | 8.9 | 2.8 | 3.1 | 1.7E-03 | 3.9E-03 | BT20/MB468 | MP |
| >am_trnaMT_PheGAA_MT_+_577_647@41.67.27\|GTTTAGACGGGCTCACATCACCCCATA | >am_trnaMT_PheGAA_MT_+_577_647 | 41 | 67 | 27 | GTTTAGACGGGCTCACATCACCCCATA | 297.6 | 5.2 | 1.7 | 3.1 | 1.7E-03 | 3.9E-03 | BT20/MB468 | MP |
| >hg38_21_+_8433172_8446622_RNA45SN1_45S_50ntFlanks@5083.5105.23\|ACGAACGAGACTCTGGCATGCTA | >hg38_21_+_8433172_8446622_RNA45SN1_45S_50ntFlanks | 5083 | 5105 | 23 | ACGAACGAGACTCTGGCATGCTA | 202.7 | -4.2 | 1.4 | -3.0 | 2.6E-03 | 3.9E-03 | BT20/MB468 | MITO |
| >MI0006442\|hsa-mir-664a&WithFlank&1\|-\|220200532\|220200625@17.38.22&[MIMAT0005948&hsa-miR-664a-5p&offsets\|0\|-2;m-353&1\|-\|220200588\|220200609&offsets\|0\|0]\|ACTGGCTAGGGAAAATGATTGG | >MI0006442\|hsa-mir-664a&WithFlank&1\|-\|220200532\|220200625 | 17 | 38 | 22 | ACTGGCTAGGGAAAATGATTGG | 84.5 | 6.4 | 2.0 | 3.1 | 1.7E-03 | 3.9E-03 | BT20/MB468 | MP |
| >hg38_21_+_8433172_8446622_RNA45SN1_45S_50ntFlanks@10827.10858.32\|CGGCAAGCCGGATCCGTAACTTCGGGATAAGG | >hg38_21_+_8433172_8446622_RNA45SN1_45S_50ntFlanks | 10827 | 10858 | 32 | CGGCAAGCCGGATCCGTAACTTCGGGATAAGG | 253.7 | -4.6 | 1.5 | -3.1 | 1.7E-03 | 3.9E-03 | BT20/MB468 | MP |
| >MI0000750\|hsa-mir-26a-2&WithFlank&12\|-\|57824603\|57824698@20.41.22(+2A);MI0000083\|hsa-mir-26a-1&WithFlank&3\|+\|37969398\|37969486@16.37.22(+2A)&[MIMAT0000082&hsa-miR-26a-5p&offsets\|0\|0(+2A);m-33&12\|-\|57824658\|57824679&offsets\|0\|0(+2A)];[MIMAT0000082_1&hsa-miR-26a-5p&offsets\|0\|0(+2A);m-34&3\|+\|37969413\|37969434&offsets\|0\|0(+2A)]\|TTCAAGTAATCCAGGATAGGCTAA | >MI0000750\|hsa-mir-26a-2&WithFlank&12\|-\|57824603\|57824698 | 20 | 41 | 22(+2A) | TTCAAGTAATCCAGGATAGGCTAA | 860.3 | -3.8 | 1.1 | -3.4 | 5.9E-04 | 3.9E-03 | BT20/MB231 | MP |
| >MI0000273\|hsa-mir-183&WithFlank&7\|-\|129774899\|129775020@33.55.23&[MIMAT0000261&hsa-miR-183-5p&offsets\|0\|+1;m-41&7\|-\|129774966\|129774987&offsets\|-1\|0]\|TATGGCACTGGTAGAATTCACTG | >MI0000273\|hsa-mir-183&WithFlank&7\|-\|129774899\|129775020 | 33 | 55 | 23 | TATGGCACTGGTAGAATTCACTG | 2904.4 | 2.2 | 0.7 | 3.4 | 7.9E-04 | 3.9E-03 | BT20/MB231 | MITO |
| >hg38_21_+_8433172_8446622_RNA45SN1_45S_50ntFlanks@10107.10134.28\|CGTGTGGTGTGCGTCGGAGGGCGGCGGC | >hg38_21_+_8433172_8446622_RNA45SN1_45S_50ntFlanks | 10107 | 10134 | 28 | CGTGTGGTGTGCGTCGGAGGGCGGCGGC | 75.8 | 9.6 | 2.8 | 3.4 | 5.9E-04 | 3.9E-03 | BT20/MB231 | MP |
| >hg38_21_+_8433172_8446622_RNA45SN1_45S_50ntFlanks@8189.8213.25\|CCCAGCCCGTGGACGGTGTGAGGCC | >hg38_21_+_8433172_8446622_RNA45SN1_45S_50ntFlanks | 8189 | 8213 | 25 | CCCAGCCCGTGGACGGTGTGAGGCC | 332.1 | -4.0 | 1.3 | -3.0 | 2.8E-03 | 3.9E-03 | MB231/MB468 | MP |
| >hg38_21_+_8433172_8446622_RNA45SN1_45S_50ntFlanks@12415.12455.41\|CTTCGATGTCGGCTCTTCCTATCATTGTGAAGCAGAATTCA | >hg38_21_+_8433172_8446622_RNA45SN1_45S_50ntFlanks | 12415 | 12455 | 41 | CTTCGATGTCGGCTCTTCCTATCATTGTGAAGCAGAATTCA | 422.6 | -4.2 | 1.4 | -3.0 | 3.1E-03 | 3.9E-03 | BT20/MB468 | CYTO |
| >hg38_21_+_8433172_8446622_RNA45SN1_45S_50ntFlanks@10348.10384.37\|AAACGAGAACTTTGAAGGCCGAAGTGGAGAAGGGTTC | >hg38_21_+_8433172_8446622_RNA45SN1_45S_50ntFlanks | 10348 | 10384 | 37 | AAACGAGAACTTTGAAGGCCGAAGTGGAGAAGGGTTC | 263.9 | -2.7 | 0.9 | -2.9 | 3.4E-03 | 3.9E-03 | MB231/MB468 | MITO |
| >MI0000285\|hsa-mir-205&WithFlank&1\|+\|209432127\|209432248@40.61.22(+2A)&[MIMAT0000266&hsa-miR-205-5p&offsets\|0\|0(+2A);m-351&1\|+\|209432166\|209432187&offsets\|0\|0(+2A)]\|TCCTTCATTCCACCGGAGTCTGAA | >MI0000285\|hsa-mir-205&WithFlank&1\|+\|209432127\|209432248 | 40 | 61 | 22(+2A) | TCCTTCATTCCACCGGAGTCTGAA | 421.1 | 1.9 | 0.6 | 2.9 | 3.3E-03 | 3.9E-03 | BT20/MB468 | NUC |
| >hg38_21_+_8433172_8446622_RNA45SN1_45S_50ntFlanks@7971.8009.39\|GAGACGCGACCTCAGATCAGACGTGGCGACCCGCTGAAT | >hg38_21_+_8433172_8446622_RNA45SN1_45S_50ntFlanks | 7971 | 8009 | 39 | GAGACGCGACCTCAGATCAGACGTGGCGACCCGCTGAAT | 354.0 | 2.5 | 0.7 | 3.4 | 7.9E-04 | 3.9E-03 | BT20/MB231 | MITO |
| >hg38_21_+_8433172_8446622_RNA45SN1_45S_50ntFlanks@3785.3810.26\|TGAAACTGCGAATGGCTCATTAAATC | >hg38_21_+_8433172_8446622_RNA45SN1_45S_50ntFlanks | 3785 | 3810 | 26 | TGAAACTGCGAATGGCTCATTAAATC | 191.0 | -3.6 | 1.2 | -3.0 | 2.6E-03 | 3.9E-03 | BT20/MB468 | MITO |
| >hg38_21_+_8433172_8446622_RNA45SN1_45S_50ntFlanks@7989.8012.24\|AGACGTGGCGACCCGCTGAATTTA | >hg38_21_+_8433172_8446622_RNA45SN1_45S_50ntFlanks | 7989 | 8012 | 24 | AGACGTGGCGACCCGCTGAATTTA | 1071.7 | 4.6 | 1.3 | 3.4 | 6.0E-04 | 3.9E-03 | BT20/MB231 | MP |
| >hg38_21_+_8433172_8446622_RNA45SN1_45S_50ntFlanks@11197.11225.29\|GGCCGGCGGCGGCGGCGACTCTGGACGCG | >hg38_21_+_8433172_8446622_RNA45SN1_45S_50ntFlanks | 11197 | 11225 | 29 | GGCCGGCGGCGGCGGCGACTCTGGACGCG | 268.7 | -5.2 | 1.7 | -3.1 | 1.7E-03 | 3.9E-03 | BT20/MB468 | MP |
| >hg38_21_+_8433172_8446622_RNA45SN1_45S_50ntFlanks@10845.10868.24\|ACTTCGGGATAAGGATTGGCTCTA | >hg38_21_+_8433172_8446622_RNA45SN1_45S_50ntFlanks | 10845 | 10868 | 24 | ACTTCGGGATAAGGATTGGCTCTA | 592.5 | -5.4 | 1.7 | -3.1 | 1.7E-03 | 3.9E-03 | BT20/MB468 | MP |
| >MI0000299\|hsa-mir-222&WithFlank&X\|-\|45747009\|45747130@75.98.24(+1A)&[MIMAT0000279&hsa-miR-222-3p&offsets\|0\|+3(+1A);m-140&X\|-\|45747033\|45747056&offsets\|0\|0(+1A)]\|AGCTACATCTGGCTACTGGGTCTCA | >MI0000299\|hsa-mir-222&WithFlank&X\|-\|45747009\|45747130 | 75 | 98 | 24(+1A) | AGCTACATCTGGCTACTGGGTCTCA | 320.5 | 3.0 | 0.9 | 3.1 | 1.7E-03 | 3.9E-03 | BT20/MB468 | MP |
| >hg38_21_+_8433172_8446622_RNA45SN1_45S_50ntFlanks@7990.8008.19\|GACGTGGCGACCCGCTGAA | >hg38_21_+_8433172_8446622_RNA45SN1_45S_50ntFlanks | 7990 | 8008 | 19 | GACGTGGCGACCCGCTGAA | 578.1 | -2.8 | 1.0 | -2.9 | 3.3E-03 | 3.9E-03 | BT20/MB468 | NUC |
| >hg38_21_+_8433172_8446622_RNA45SN1_45S_50ntFlanks@4884.4917.34\|CAAAGCTGAAACTTAAAGGAATTGACGGAAGGGC | >hg38_21_+_8433172_8446622_RNA45SN1_45S_50ntFlanks | 4884 | 4917 | 34 | CAAAGCTGAAACTTAAAGGAATTGACGGAAGGGC | 203.5 | -1.8 | 0.6 | -3.0 | 2.6E-03 | 3.9E-03 | BT20/MB468 | MITO |
| >am_trnaMT_GluTTC_MT_-_14674_14742@27.54.28,trnalookalike8_GluTTC_5_-_93905172_93905240@27.54.28\|GTTTTTCATATCATTGGTCGTGGTTGTA | >am_trnaMT_GluTTC_MT_-_14674_14742 | 27 | 54 | 28 | GTTTTTCATATCATTGGTCGTGGTTGTA | 223.5 | -3.9 | 1.3 | -3.0 | 2.9E-03 | 3.9E-03 | MB231/MB468 | MP |
| >hg38_21_+_8433172_8446622_RNA45SN1_45S_50ntFlanks@8176.8194.19\|CTTCTGATCGAGGCCCAGC | >hg38_21_+_8433172_8446622_RNA45SN1_45S_50ntFlanks | 8176 | 8194 | 19 | CTTCTGATCGAGGCCCAGC | 333.7 | -4.0 | 1.4 | -2.9 | 3.3E-03 | 3.9E-03 | BT20/MB468 | NUC |
| >hg38_21_+_8433172_8446622_RNA45SN1_45S_50ntFlanks@12616.12654.39\|CTGAGGAGCCAATGGGGCGAAGCTACCATCTGTGGGATT | >hg38_21_+_8433172_8446622_RNA45SN1_45S_50ntFlanks | 12616 | 12654 | 39 | CTGAGGAGCCAATGGGGCGAAGCTACCATCTGTGGGATT | 659.2 | -3.2 | 1.0 | -3.1 | 1.7E-03 | 3.9E-03 | BT20/MB468 | MP |
| >hg38_MT_+_1621_3279_RNR2_16S_50ntFlanks@1368.1398.31\|CGATGTTGGATCAGGACATCCCGATGGTGCA | >hg38_MT_+_1621_3279_RNR2_16S_50ntFlanks | 1368 | 1398 | 31 | CGATGTTGGATCAGGACATCCCGATGGTGCA | 90.8 | 4.7 | 1.5 | 3.1 | 1.7E-03 | 3.9E-03 | BT20/MB468 | MP |
| >hg38_21_+_8433172_8446622_RNA45SN1_45S_50ntFlanks@12918.12940.23\|CGTGGGGAACCTGGCGCTAAACC | >hg38_21_+_8433172_8446622_RNA45SN1_45S_50ntFlanks | 12918 | 12940 | 23 | CGTGGGGAACCTGGCGCTAAACC | 204.5 | 3.4 | 1.2 | 2.9 | 3.2E-03 | 3.9E-03 | MB231/MB468 | CYTO |
| >hg38_21_+_8433172_8446622_RNA45SN1_45S_50ntFlanks@3783.3801.19\|AGTGAAACTGCGAATGGCT | >hg38_21_+_8433172_8446622_RNA45SN1_45S_50ntFlanks | 3783 | 3801 | 19 | AGTGAAACTGCGAATGGCT | 249.0 | -1.8 | 0.6 | -2.9 | 3.2E-03 | 3.9E-03 | MB231/MB468 | CYTO |
| >am_trna2_GlyGCC_21_-_18827107_18827177@1.27.27,trna35_GlyGCC_1_+_161413094_161413164@1.27.27,trna37_GlyGCC_1_+_161420467_161420537@1.27.27,trna39_GlyGCC_1_+_161427898_161427968@1.27.27,trna41_GlyGCC_1_+_161435258_161435328@1.27.27\|GCATGGTGGTTCAGTGGTAGAATTCT | >am_trna2_GlyGCC_21_-_18827107_18827177 | 1 | 27 | 27 | GCATGGTGGTTCAGTGGTAGAATTCT | 3801.2 | 3.9 | 1.2 | 3.1 | 1.7E-03 | 3.9E-03 | BT20/MB468 | MP |
| >hg38_21_+_8433172_8446622_RNA45SN1_45S_50ntFlanks@6647.6681.35\|CGTACGACTCTTAGCGGTGGATCACTCGGCTCGTG | >hg38_21_+_8433172_8446622_RNA45SN1_45S_50ntFlanks | 6647 | 6681 | 35 | CGTACGACTCTTAGCGGTGGATCACTCGGCTCGTG | 11677.8 | 2.1 | 0.7 | 2.9 | 3.4E-03 | 3.9E-03 | MB231/MB468 | NUC |
| >hg38_21_+_8433172_8446622_RNA45SN1_45S_50ntFlanks@11729.11750.22\|TAACTATGACTCTCTTAAGGTA | >hg38_21_+_8433172_8446622_RNA45SN1_45S_50ntFlanks | 11729 | 11750 | 22 | TAACTATGACTCTCTTAAGGTA | 375.9 | -4.2 | 1.3 | -3.1 | 1.7E-03 | 3.9E-03 | BT20/MB468 | MP |
| >hg38_MT_+_1621_3279_RNR2_16S_50ntFlanks@1367.1385.19\|TCGATGTTGGATCAGGACA | >hg38_MT_+_1621_3279_RNR2_16S_50ntFlanks | 1367 | 1385 | 19 | TCGATGTTGGATCAGGACA | 186.4 | 3.6 | 1.1 | 3.1 | 1.7E-03 | 4.0E-03 | BT20/MB468 | MP |
| >hg38_21_+_8433172_8446622_RNA45SN1_45S_50ntFlanks@9489.9525.37\|AGGACCCGAAAGATGGTGAACTATGCCTGGGCAGGGC | >hg38_21_+_8433172_8446622_RNA45SN1_45S_50ntFlanks | 9489 | 9525 | 37 | AGGACCCGAAAGATGGTGAACTATGCCTGGGCAGGGC | 97.8 | 5.2 | 1.7 | 3.1 | 1.7E-03 | 4.0E-03 | BT20/MB468 | MP |
| >hg38_21_+_8433172_8446622_RNA45SN1_45S_50ntFlanks@7975.8020.46\|CGCGACCTCAGATCAGACGTGGCGACCCGCTGAATTTAAGCATATT | >hg38_21_+_8433172_8446622_RNA45SN1_45S_50ntFlanks | 7975 | 8020 | 46 | CGCGACCTCAGATCAGACGTGGCGACCCGCTGAATTTAAGCATATT | 311.9 | 7.2 | 2.4 | 2.9 | 3.2E-03 | 4.0E-03 | MB231/MB468 | CYTO |
| >hg38_21_+_8433172_8446622_RNA45SN1_45S_50ntFlanks@4565.4587.23\|AATAATGGAATAGGACCGCGGTT | >hg38_21_+_8433172_8446622_RNA45SN1_45S_50ntFlanks | 4565 | 4587 | 23 | AATAATGGAATAGGACCGCGGTT | 420.3 | -3.0 | 1.0 | -3.0 | 2.9E-03 | 4.0E-03 | MB231/MB468 | MP |

<table>
<tr><td>>hg38_21_+_8433172_8446622_RNA45SN1_45S_50ntFlanks@4730.4759.30|CAAGAACGAAAGTCGGAGGTTCGAAGACGA</td><td>>hg38_21_+_8433172_8446622_RNA45SN1_45S_50ntFlanks</td><td>4730</td><td>4759</td><td>30</td><td>CAAGAACGAAAGTCGGAGGTTCGAAGACGA</td><td>1068.0</td><td>5.4</td><td>1.6</td><td>3.4</td><td>6.0E-04</td><td>4.0E-03</td><td>BT20/MB231</td><td>MP</td></tr>
<tr><td>>hg38_21_+_8433172_8446622_RNA45SN1_45S_50ntFlanks@9573.9604.32|AAATCGGTCGTCCGACCTGGGTATAGGGGCGA</td><td>>hg38_21_+_8433172_8446622_RNA45SN1_45S_50ntFlanks</td><td>9573</td><td>9604</td><td>32</td><td>AAATCGGTCGTCCGACCTGGGTATAGGGGCGA</td><td>177.0</td><td>-4.0</td><td>1.3</td><td>-3.0</td><td>2.9E-03</td><td>4.0E-03</td><td>MB231/MB468</td><td>MP</td></tr>
<tr><td>>hg38_21_+_8433172_8446622_RNA45SN1_45S_50ntFlanks@6709.6731.23|AGAATTAATGTGAATTGCAGGAC</td><td>>hg38_21_+_8433172_8446622_RNA45SN1_45S_50ntFlanks</td><td>6709</td><td>6731</td><td>23</td><td>AGAATTAATGTGAATTGCAGGAC</td><td>83.7</td><td>-3.7</td><td>1.2</td><td>-3.0</td><td>2.9E-03</td><td>4.0E-03</td><td>MB231/MB468</td><td>MP</td></tr>
<tr><td>>hg38_21_+_8433172_8446622_RNA45SN1_45S_50ntFlanks@5071.5112.42|TTAATTCCGATAACGAACGAGACTCTGGCATGCTAACTAGTT</td><td>>hg38_21_+_8433172_8446622_RNA45SN1_45S_50ntFlanks</td><td>5071</td><td>5112</td><td>42</td><td>TTAATTCCGATAACGAACGAGACTCTGGCATGCTAACTAGTT</td><td>40.0</td><td>-9.2</td><td>3.1</td><td>-3.0</td><td>2.9E-03</td><td>4.0E-03</td><td>MB231/MB468</td><td>MP</td></tr>
<tr><td>>hg38_21_+_8433172_8446622_RNA45SN1_45S_50ntFlanks@10414.10431.18|TCGGTCCTGAGAGATGGG</td><td>>hg38_21_+_8433172_8446622_RNA45SN1_45S_50ntFlanks</td><td>10414</td><td>10431</td><td>18</td><td>TCGGTCCTGAGAGATGGG</td><td>386.4</td><td>-3.9</td><td>1.3</td><td>-3.1</td><td>1.7E-03</td><td>4.0E-03</td><td>BT20/MB468</td><td>MP</td></tr>
<tr><td>>hg38_21_+_8433172_8446622_RNA45SN1_45S_50ntFlanks@10727.10753.27|GGAGAGGGTGTAAATCTCGCGCCGGGC</td><td>>hg38_21_+_8433172_8446622_RNA45SN1_45S_50ntFlanks</td><td>10727</td><td>10753</td><td>27</td><td>GGAGAGGGTGTAAATCTCGCGCCGGGC</td><td>177.5</td><td>-5.1</td><td>1.6</td><td>-3.1</td><td>1.7E-03</td><td>4.0E-03</td><td>BT20/MB468</td><td>MP</td></tr>
<tr><td>>hg38_21_+_8433172_8446622_RNA45SN1_45S_50ntFlanks@11568.11600.33|GCAGCCGACTTAGAACTGGTGCGGACCAGGGGA</td><td>>hg38_21_+_8433172_8446622_RNA45SN1_45S_50ntFlanks</td><td>11568</td><td>11600</td><td>33</td><td>GCAGCCGACTTAGAACTGGTGCGGACCAGGGGA</td><td>365.1</td><td>-3.6</td><td>1.2</td><td>-2.9</td><td>3.4E-03</td><td>4.0E-03</td><td>MB231/MB468</td><td>MITO</td></tr>
<tr><td>>MI0000100|hsa-mir-98&WithFlank&X|-|53556217|53556347@28.49.22&[MIMAT0000096&hsa-miR-98-5p&offsets|0|0;m-143&X|-|53556299|53556320&offsets|0|0]|TGAGGTAGTAAGTTGTATTGTT</td><td>>MI0000100|hsa-mir-98&WithFlank&X|-|53556217|53556347</td><td>28</td><td>49</td><td>22</td><td>TGAGGTAGTAAGTTGTATTGTT</td><td>1973.2</td><td>2.3</td><td>0.8</td><td>2.9</td><td>3.2E-03</td><td>4.0E-03</td><td>MB231/MB468</td><td>CYTO</td></tr>
<tr><td>>am_trna111_HisGTG_1_-_147774845_147774916@1.29.29,trna118_HisGTG_1_-_145396881_145396952@1.29.29,trna16_HisGTG_1_+_146544773_146544844@1.29.29,trna1_HisGTG_15_+_45493349_45493420@1.29.29,trna21_HisGTG_1_+_147753471_147753542@1.29.29,trna33_HisGTG_6_+_27125906_27125977@1.29.29,trna7_HisGTG_9_-_14433938_14434009@1.29.29,trna8_HisGTG_15_-_45492611_45492682@1.29.29,trna9_HisGTG_15_-_45490804_45490875@1.29.29|GCCGTGATCGTATAGTGGTTAGTACTCTG</td><td>>am_trna111_HisGTG_1_-_147774845_147774916</td><td>1</td><td>29</td><td>29</td><td>GCCGTGATCGTATAGTGGTTAGTACTCTG</td><td>1302.6</td><td>-2.5</td><td>0.8</td><td>-3.0</td><td>2.7E-03</td><td>4.0E-03</td><td>BT20/MB468</td><td>MITO</td></tr>
<tr><td>>hg38_21_+_8433172_8446622_RNA45SN1_45S_50ntFlanks@5492.5526.35|AGAAGACGGTCGAACTTGACTATCTAGAGGAAGTA</td><td>>hg38_21_+_8433172_8446622_RNA45SN1_45S_50ntFlanks</td><td>5492</td><td>5526</td><td>35</td><td>AGAAGACGGTCGAACTTGACTATCTAGAGGAAGTA</td><td>1218.1</td><td>-3.2</td><td>1.0</td><td>-3.1</td><td>1.7E-03</td><td>4.0E-03</td><td>BT20/MB468</td><td>MP</td></tr>
<tr><td>>hg38_21_+_8433172_8446622_RNA45SN1_45S_50ntFlanks@12892.12932.41|CCCTCGCCCGTCACGCACCGCACGTTCGTGGGGAACCTGGC</td><td>>hg38_21_+_8433172_8446622_RNA45SN1_45S_50ntFlanks</td><td>12892</td><td>12932</td><td>41</td><td>CCCTCGCCCGTCACGCACCGCACGTTCGTGGGGAACCTGGC</td><td>151.3</td><td>-4.3</td><td>1.5</td><td>-3.0</td><td>3.1E-03</td><td>4.0E-03</td><td>BT20/MB468</td><td>CYTO</td></tr>
<tr><td>>hg38_21_+_8433172_8446622_RNA45SN1_45S_50ntFlanks@6646.6687.42|TCGTACGACTCTTAGCGGTGGATCACTCGGCTCGTGCGTCGA</td><td>>hg38_21_+_8433172_8446622_RNA45SN1_45S_50ntFlanks</td><td>6646</td><td>6687</td><td>42</td><td>TCGTACGACTCTTAGCGGTGGATCACTCGGCTCGTGCGTCGA</td><td>142.2</td><td>3.6</td><td>1.1</td><td>3.1</td><td>1.7E-03</td><td>4.0E-03</td><td>BT20/MB468</td><td>MP</td></tr>
<tr><td>>hg38_21_+_8433172_8446622_RNA45SN1_45S_50ntFlanks@12893.12929.37|CCTCGCCCGTCACGCACCGCACGTTCGTGGGGAACCT</td><td>>hg38_21_+_8433172_8446622_RNA45SN1_45S_50ntFlanks</td><td>12893</td><td>12929</td><td>37</td><td>CCTCGCCCGTCACGCACCGCACGTTCGTGGGGAACCT</td><td>103.3</td><td>-5.2</td><td>1.7</td><td>-3.1</td><td>1.7E-03</td><td>4.0E-03</td><td>BT20/MB468</td><td>MP</td></tr>
<tr><td>>hg38_21_+_8433172_8446622_RNA45SN1_45S_50ntFlanks@12420.12445.26|ATGTCGGCTCTTCCTATCATTGTGAA</td><td>>hg38_21_+_8433172_8446622_RNA45SN1_45S_50ntFlanks</td><td>12420</td><td>12445</td><td>26</td><td>ATGTCGGCTCTTCCTATCATTGTGAA</td><td>1169.3</td><td>-2.0</td><td>0.7</td><td>-3.0</td><td>3.1E-03</td><td>4.0E-03</td><td>BT20/MB468</td><td>CYTO</td></tr>
<tr><td>>hg38_MT_+_1621_3279_RNR2_16S_50ntFlanks@1094.1120.27|AAGACGAGAAGACCCTATGGAGCTTTA</td><td>>hg38_MT_+_1621_3279_RNR2_16S_50ntFlanks</td><td>1094</td><td>1120</td><td>27</td><td>AAGACGAGAAGACCCTATGGAGCTTTA</td><td>231.7</td><td>3.4</td><td>1.1</td><td>3.1</td><td>1.7E-03</td><td>4.0E-03</td><td>BT20/MB468</td><td>MP</td></tr>
<tr><td>>hg38_21_+_8433172_8446622_RNA45SN1_45S_50ntFlanks@6736.6771.36|TGATCATCGACACTTCGAACGCACTTGCGGCCCCGG</td><td>>hg38_21_+_8433172_8446622_RNA45SN1_45S_50ntFlanks</td><td>6736</td><td>6771</td><td>36</td><td>TGATCATCGACACTTCGAACGCACTTGCGGCCCCGG</td><td>192.2</td><td>-3.3</td><td>1.1</td><td>-3.0</td><td>2.7E-03</td><td>4.0E-03</td><td>BT20/MB468</td><td>MITO</td></tr>
<tr><td>>hg38_21_+_8433172_8446622_RNA45SN1_45S_50ntFlanks@5086.5105.20|AACGAGACTCTGGCATGCTA</td><td>>hg38_21_+_8433172_8446622_RNA45SN1_45S_50ntFlanks</td><td>5086</td><td>5105</td><td>20</td><td>AACGAGACTCTGGCATGCTA</td><td>181.4</td><td>-4.3</td><td>1.4</td><td>-3.0</td><td>2.9E-03</td><td>4.0E-03</td><td>MB231/MB468</td><td>MP</td></tr>
<tr><td>>hg38_21_+_8433172_8446622_RNA45SN1_45S_50ntFlanks@10319.10341.23|GCAGATCTTGGTGGTAGTAGCAA</td><td>>hg38_21_+_8433172_8446622_RNA45SN1_45S_50ntFlanks</td><td>10319</td><td>10341</td><td>23</td><td>GCAGATCTTGGTGGTAGTAGCAA</td><td>169.5</td><td>6.7</td><td>1.9</td><td>3.4</td><td>6.0E-04</td><td>4.0E-03</td><td>BT20/MB231</td><td>MP</td></tr>
<tr><td>>hg38_21_+_8433172_8446622_RNA45SN1_45S_50ntFlanks@11927.11950.24|GTGTAGAATAAGTGGGAGGCCCCC</td><td>>hg38_21_+_8433172_8446622_RNA45SN1_45S_50ntFlanks</td><td>11927</td><td>11950</td><td>24</td><td>GTGTAGAATAAGTGGGAGGCCCCC</td><td>155.7</td><td>-2.0</td><td>0.7</td><td>-3.0</td><td>3.1E-03</td><td>4.0E-03</td><td>BT20/MB468</td><td>CYTO</td></tr>
<tr><td>>hg38_21_+_8433172_8446622_RNA45SN1_45S_50ntFlanks@12841.12862.22|CGGAGTGCCCTTCGTCCTGGGA</td><td>>hg38_21_+_8433172_8446622_RNA45SN1_45S_50ntFlanks</td><td>12841</td><td>12862</td><td>22</td><td>CGGAGTGCCCTTCGTCCTGGGA</td><td>86.3</td><td>-6.6</td><td>2.2</td><td>-3.0</td><td>3.1E-03</td><td>4.0E-03</td><td>BT20/MB468</td><td>CYTO</td></tr>
<tr><td>>hg38_21_+_8433172_8446622_RNA45SN1_45S_50ntFlanks@7980.8006.27|CCTCAGATCAGACGTGGCGACCCGCTG</td><td>>hg38_21_+_8433172_8446622_RNA45SN1_45S_50ntFlanks</td><td>7980</td><td>8006</td><td>27</td><td>CCTCAGATCAGACGTGGCGACCCGCTG</td><td>1759.3</td><td>-4.0</td><td>1.3</td><td>-3.0</td><td>3.1E-03</td><td>4.0E-03</td><td>BT20/MB468</td><td>CYTO</td></tr>
<tr><td>>hg38_21_+_8433172_8446622_RNA45SN1_45S_50ntFlanks@11873.11890.18|AAGAAGACCCTGTTGAGC</td><td>>hg38_21_+_8433172_8446622_RNA45SN1_45S_50ntFlanks</td><td>11873</td><td>11890</td><td>18</td><td>AAGAAGACCCTGTTGAGC</td><td>1401.1</td><td>-3.7</td><td>1.2</td><td>-3.1</td><td>1.7E-03</td><td>4.0E-03</td><td>BT20/MB468</td><td>MP</td></tr>
<tr><td>>trna10_AspGTC_12_-_125424193_125424264@38.73.36,trna12_AspGTC_12_-_125411891_125411962@38.73.36,trna144_AspGTC_6_-_27551236_27551307@38.73.36,trna38_AspGTC_17_-_8125556_8125627@38.73.36,trna45_AspGTC_6_+_27447453_27447524@38.73.36,trna48_AspGTC_6_+_27471523_27471594@38.73.36,trna4_AspGTC_12_+_96429799_96429870@38.73.36,trna69_AspGTC_1_-_161440205_161440276@38.73.36,trna72_AspGTC_1_-_161432824_161432895@38.73.36,trna75_AspGTC_1_-_161425414_161425485@38.73.36,trna78_AspGTC_1_-_161418033_161418104@38.73.36,trna81_AspGTC_1_-_161410615_161410686@38.73.36|CGCGGGAGACCGGGGTTCGATTCCCCGACGGGGAGC</td><td>>trna10_AspGTC_12_-_125424193_125424264</td><td>38</td><td>73</td><td>36</td><td>CGCGGGAGACCGGGGTTCGATTCCCCGACGGGGAGC</td><td>796.9</td><td>2.9</td><td>1.0</td><td>3.0</td><td>3.1E-03</td><td>4.0E-03</td><td>BT20/MB468</td><td>CYTO</td></tr>
<tr><td>>hg38_21_+_8433172_8446622_RNA45SN1_45S_50ntFlanks@4734.4758.25|AACGAAAGTCGGAGGTTCGAAGACG</td><td>>hg38_21_+_8433172_8446622_RNA45SN1_45S_50ntFlanks</td><td>4734</td><td>4758</td><td>25</td><td>AACGAAAGTCGGAGGTTCGAAGACG</td><td>2374.9</td><td>-2.6</td><td>0.9</td><td>-2.9</td><td>3.4E-03</td><td>4.0E-03</td><td>MB231/MB468</td><td>MITO</td></tr>
<tr><td>>hg38_21_+_8433172_8446622_RNA45SN1_45S_50ntFlanks@7974.8007.34|ACGCGACCTCAGATCAGACGTGGCGACCCGCTGA</td><td>>hg38_21_+_8433172_8446622_RNA45SN1_45S_50ntFlanks</td><td>7974</td><td>8007</td><td>34</td><td>ACGCGACCTCAGATCAGACGTGGCGACCCGCTGA</td><td>6129.0</td><td>2.9</td><td>0.9</td><td>3.4</td><td>6.7E-04</td><td>4.0E-03</td><td>BT20/MB231</td><td>NUC</td></tr>
<tr><td>>hg38_21_+_8433172_8446622_RNA45SN1_45S_50ntFlanks@11568.11601.34|GCAGCCGACTTAGAACTGGTGCGGACCAGGGGAA</td><td>>hg38_21_+_8433172_8446622_RNA45SN1_45S_50ntFlanks</td><td>11568</td><td>11601</td><td>34</td><td>GCAGCCGACTTAGAACTGGTGCGGACCAGGGGAA</td><td>413.5</td><td>-1.6</td><td>0.6</td><td>-3.0</td><td>3.2E-03</td><td>4.0E-03</td><td>BT20/MB468</td><td>CYTO</td></tr>
<tr><td>>hg38_21_+_8433172_8446622_RNA45SN1_45S_50ntFlanks@8350.8378.29|AAAGTTGAAAAGAACTTTGAAGAGAGAGT</td><td>>hg38_21_+_8433172_8446622_RNA45SN1_45S_50ntFlanks</td><td>8350</td><td>8378</td><td>29</td><td>AAAGTTGAAAAGAACTTTGAAGAGAGAGT</td><td>193.8</td><td>3.8</td><td>1.1</td><td>3.4</td><td>6.0E-04</td><td>4.0E-03</td><td>BT20/MB231</td><td>MP</td></tr>
<tr><td>>hg38_21_+_8433172_8446622_RNA45SN1_45S_50ntFlanks@10500.10523.24|CAGATCCCCGAATCCGGAGTGGCG</td><td>>hg38_21_+_8433172_8446622_RNA45SN1_45S_50ntFlanks</td><td>10500</td><td>10523</td><td>24</td><td>CAGATCCCCGAATCCGGAGTGGCG</td><td>36.3</td><td>-8.3</td><td>2.6</td><td>-3.1</td><td>1.7E-03</td><td>4.0E-03</td><td>BT20/MB468</td><td>MP</td></tr>
<tr><td>>hg38_21_+_8433172_8446622_RNA45SN1_45S_50ntFlanks@12416.12435.20|TTCGATGTCGGCTCTTCCTA</td><td>>hg38_21_+_8433172_8446622_RNA45SN1_45S_50ntFlanks</td><td>12416</td><td>12435</td><td>20</td><td>TTCGATGTCGGCTCTTCCTA</td><td>1026.1</td><td>-1.9</td><td>0.6</td><td>-3.0</td><td>3.2E-03</td><td>4.0E-03</td><td>BT20/MB468</td><td>CYTO</td></tr>
<tr><td>>hg38_21_+_8433172_8446622_RNA45SN1_45S_50ntFlanks@5057.5094.38|AGCGATTTGTCTGGTTAATTCCGATAACGAACGAGACT</td><td>>hg38_21_+_8433172_8446622_RNA45SN1_45S_50ntFlanks</td><td>5057</td><td>5094</td><td>38</td><td>AGCGATTTGTCTGGTTAATTCCGATAACGAACGAGACT</td><td>39.9</td><td>-9.2</td><td>3.1</td><td>-3.0</td><td>2.9E-03</td><td>4.0E-03</td><td>MB231/MB468</td><td>MP</td></tr>
<tr><td>>hg38_21_+_8433172_8446622_RNA45SN1_45S_50ntFlanks@10682.10712.31|TCCGGCGGCGTCCGGTGAGCTCTCGCTGGCC</td><td>>hg38_21_+_8433172_8446622_RNA45SN1_45S_50ntFlanks</td><td>10682</td><td>10712</td><td>31</td><td>TCCGGCGGCGTCCGGTGAGCTCTCGCTGGCC</td><td>341.2</td><td>-3.5</td><td>1.2</td><td>-3.0</td><td>3.2E-03</td><td>4.0E-03</td><td>BT20/MB468</td><td>CYTO</td></tr>
<tr><td>>hg38_21_+_8433172_8446622_RNA45SN1_45S_50ntFlanks@6652.6684.33|GACTCTTAGCGGTGGATCACTCGGCTCGTGCGT</td><td>>hg38_21_+_8433172_8446622_RNA45SN1_45S_50ntFlanks</td><td>6652</td><td>6684</td><td>33</td><td>GACTCTTAGCGGTGGATCACTCGGCTCGTGCGT</td><td>588.4</td><td>3.1</td><td>1.0</td><td>3.1</td><td>1.7E-03</td><td>4.0E-03</td><td>BT20/MB468</td><td>MP</td></tr>
<tr><td>>hg38_21_+_8433172_8446622_RNA45SN1_45S_50ntFlanks@3900.3938.39|CTTCGCGGGGGGGATGCGTGCATTTATCAGATCAAAACC</td><td>>hg38_21_+_8433172_8446622_RNA45SN1_45S_50ntFlanks</td><td>3900</td><td>3938</td><td>39</td><td>CTTCGCGGGGGGGATGCGTGCATTTATCAGATCAAAACC</td><td>176.8</td><td>-4.0</td><td>1.4</td><td>-3.0</td><td>3.2E-03</td><td>4.0E-03</td><td>BT20/MB468</td><td>CYTO</td></tr>
<tr><td>>hg38_21_+_8433172_8446622_RNA45SN1_45S_50ntFlanks@7980.8009.30|CCTCAGATCAGACGTGGCGACCCGCTGAAT</td><td>>hg38_21_+_8433172_8446622_RNA45SN1_45S_50ntFlanks</td><td>7980</td><td>8009</td><td>30</td><td>CCTCAGATCAGACGTGGCGACCCGCTGAAT</td><td>298.5</td><td>2.3</td><td>0.7</td><td>3.4</td><td>8.1E-04</td><td>4.0E-03</td><td>BT20/MB231</td><td>MITO</td></tr>
<tr><td>>MI0000477|hsa-mir-146a&WithFlank&5|+|160485346|160485456@27.47.21&[MIMAT0000449&hsa-miR-146a-5p&offsets|0|-1;m-129&5|+|160485372|160485393&offsets|0|-1]|TGAGAACTGAATTCCATGGGT</td><td>>MI0000477|hsa-mir-146a&WithFlank&5|+|160485346|160485456</td><td>27</td><td>47</td><td>21</td><td>TGAGAACTGAATTCCATGGGT</td><td>2351.5</td><td>-8.4</td><td>2.5</td><td>-3.4</td><td>8.1E-04</td><td>4.0E-03</td><td>BT20/MB231</td><td>MITO</td></tr>
<tr><td>>hg38_21_+_8433172_8446622_RNA45SN1_45S_50ntFlanks@3992.4024.33|GGTGACTCTAGATAACCTCGGGCCGATCGCACG</td><td>>hg38_21_+_8433172_8446622_RNA45SN1_45S_50ntFlanks</td><td>3992</td><td>4024</td><td>33</td><td>GGTGACTCTAGATAACCTCGGGCCGATCGCACG</td><td>680.1</td><td>-2.6</td><td>0.9</td><td>-2.9</td><td>3.4E-03</td><td>4.0E-03</td><td>MB231/MB468</td><td>MITO</td></tr>
<tr><td>>hg38_21_+_8433172_8446622_RNA45SN1_45S_50ntFlanks@4983.5013.31|CGGACAGGATTGACAGATTGATAGCTCTTTC</td><td>>hg38_21_+_8433172_8446622_RNA45SN1_45S_50ntFlanks</td><td>4983</td><td>5013</td><td>31</td><td>CGGACAGGATTGACAGATTGATAGCTCTTTC</td><td>198.1</td><td>-4.1</td><td>1.4</td><td>-3.0</td><td>2.7E-03</td><td>4.0E-03</td><td>BT20/MB468</td><td>MITO</td></tr>
<tr><td>>hg38_21_+_8433172_8446622_RNA45SN1_45S_50ntFlanks@11564.11590.27|CCTAGCAGCCGACTTAGAACTGGTGCG</td><td>>hg38_21_+_8433172_8446622_RNA45SN1_45S_50ntFlanks</td><td>11564</td><td>11590</td><td>27</td><td>CCTAGCAGCCGACTTAGAACTGGTGCG</td><td>271.2</td><td>-1.8</td><td>0.6</td><td>-3.0</td><td>2.7E-03</td><td>4.0E-03</td><td>BT20/MB468</td><td>MITO</td></tr>
<tr><td>>hg38_21_+_8433172_8446622_RNA45SN1_45S_50ntFlanks@9495.9525.31|CGAAAGATGGTGAACTATGCCTGGGCAGGGC</td><td>>hg38_21_+_8433172_8446622_RNA45SN1_45S_50ntFlanks</td><td>9495</td><td>9525</td><td>31</td><td>CGAAAGATGGTGAACTATGCCTGGGCAGGGC</td><td>598.0</td><td>6.0</td><td>1.7</td><td>3.4</td><td>6.1E-04</td><td>4.0E-03</td><td>BT20/MB231</td><td>MP</td></tr>
<tr><td>>hg38_21_+_8433172_8446622_RNA45SN1_45S_50ntFlanks@9564.9592.29|CTGACGTGCAAATCGGTCGTCCGACCTGG</td><td>>hg38_21_+_8433172_8446622_RNA45SN1_45S_50ntFlanks</td><td>9564</td><td>9592</td><td>29</td><td>CTGACGTGCAAATCGGTCGTCCGACCTGG</td><td>891.1</td><td>-3.1</td><td>1.0</td><td>-3.1</td><td>1.7E-03</td><td>4.0E-03</td><td>BT20/MB468</td><td>MP</td></tr>
<tr><td>>hg38_21_+_8433172_8446622_RNA45SN1_45S_50ntFlanks@9737.9765.29|CGAAACGATCTCAACCTATTCTCAAACTT</td><td>>hg38_21_+_8433172_8446622_RNA45SN1_45S_50ntFlanks</td><td>9737</td><td>9765</td><td>29</td><td>CGAAACGATCTCAACCTATTCTCAAACTT</td><td>799.8</td><td>-2.5</td><td>0.8</td><td>-3.0</td><td>2.7E-03</td><td>4.0E-03</td><td>BT20/MB468</td><td>MITO</td></tr>
<tr><td>>hg38_21_+_8433172_8446622_RNA45SN1_45S_50ntFlanks@12419.12446.28|GATGTCGGCTCTTCCTATCATTGTGAAG</td><td>>hg38_21_+_8433172_8446622_RNA45SN1_45S_50ntFlanks</td><td>12419</td><td>12446</td><td>28</td><td>GATGTCGGCTCTTCCTATCATTGTGAAG</td><td>1991.2</td><td>-3.0</td><td>0.9</td><td>-3.4</td><td>7.3E-04</td><td>4.0E-03</td><td>BT20/MB231</td><td>CYTO</td></tr>
<tr><td>>MI0000750|hsa-mir-26a-2&WithFlank&12|-|57824603|57824698@20.40.21;MI0000083|hsa-mir-26a-1&WithFlank&3|+|37969398|37969486@16.36.21&[MIMAT0000082&hsa-miR-26a-5p&offsets|0|-1;m-33&12|-|57824658|57824679&offsets|0|-1];[MIMAT0000082_1&hsa-miR-26a-5p&offsets|0|-1;m-34&3|+|37969413|37969434&offsets|0|-1]|TTCAAGTAATCCAGGATAGGC</td><td>>MI0000750|hsa-mir-26a-2&WithFlank&12|-|57824603|57824698</td><td>20</td><td>40</td><td>21</td><td>TTCAAGTAATCCAGGATAGGC</td><td>1439.0</td><td>3.0</td><td>1.0</td><td>3.0</td><td>2.9E-03</td><td>4.0E-03</td><td>MB231/MB468</td><td>MP</td></tr>
<tr><td>>hg38_21_+_8433172_8446622_RNA45SN1_45S_50ntFlanks@9500.9520.21|GATGGTGAACTATGCCTGGGC</td><td>>hg38_21_+_8433172_8446622_RNA45SN1_45S_50ntFlanks</td><td>9500</td><td>9520</td><td>21</td><td>GATGGTGAACTATGCCTGGGC</td><td>187.0</td><td>6.6</td><td>1.9</td><td>3.4</td><td>6.1E-04</td><td>4.0E-03</td><td>BT20/MB231</td><td>MP</td></tr>
</table>

<table>
<tr><td>>MI0000816|hsa-mir-335&WithFlank&7|+|130496105|130496210@58.78.21(+1A)&[MIMAT0004703&hsa-miR-335-3p&offsets|0|-1(+1A);m-159&7|+|130496162|130496183&offsets|0|-1(+1A)]|TTTTTCATTATTGCTCCTGACA</td><td>>MI0000816|hsa-mir-335&WithFlank&7|+|130496105|130496210</td><td>58</td><td>78</td><td>21(+1A)</td><td>TTTTTCATTATTGCTCCTGACA</td><td>621.6</td><td>-3.1</td><td>0.9</td><td>-3.3</td><td>8.1E-04</td><td>4.0E-03</td><td>BT20/MB231</td><td>MITO</td></tr>
<tr><td>>hg38_21_+_8433172_8446622_RNA45SN1_45S_50ntFlanks@8352.8375.24|AGTTGAAAAGAACTTTGAAGAGAG</td><td>>hg38_21_+_8433172_8446622_RNA45SN1_45S_50ntFlanks</td><td>8352</td><td>8375</td><td>24</td><td>AGTTGAAAAGAACTTTGAAGAGAG</td><td>8644.2</td><td>4.9</td><td>1.4</td><td>3.4</td><td>6.1E-04</td><td>4.0E-03</td><td>BT20/MB231</td><td>MP</td></tr>
<tr><td>>hg38_21_+_8433172_8446622_RNA45SN1_45S_50ntFlanks@9534.9568.35|AGGAAACTCTGGTGGAGGTCCGTAGCGGTCCTGAC</td><td>>hg38_21_+_8433172_8446622_RNA45SN1_45S_50ntFlanks</td><td>9534</td><td>9568</td><td>35</td><td>AGGAAACTCTGGTGGAGGTCCGTAGCGGTCCTGAC</td><td>545.2</td><td>-2.7</td><td>0.9</td><td>-2.9</td><td>3.2E-03</td><td>4.0E-03</td><td>BT20/MB468</td><td>CYTO</td></tr>
<tr><td>>hg38_21_+_8433172_8446622_RNA45SN1_45S_50ntFlanks@12459.12478.20|AGCGTTGGATTGTTCACCCA</td><td>>hg38_21_+_8433172_8446622_RNA45SN1_45S_50ntFlanks</td><td>12459</td><td>12478</td><td>20</td><td>AGCGTTGGATTGTTCACCCA</td><td>245.4</td><td>-4.5</td><td>1.5</td><td>-3.0</td><td>2.7E-03</td><td>4.0E-03</td><td>BT20/MB468</td><td>MITO</td></tr>
<tr><td>>hg38_21_+_8433172_8446622_RNA45SN1_45S_50ntFlanks@11576.11611.36|CTTAGAACTGGTGCGGACCAGGGGAATCCGACTGTT</td><td>>hg38_21_+_8433172_8446622_RNA45SN1_45S_50ntFlanks</td><td>11576</td><td>11611</td><td>36</td><td>CTTAGAACTGGTGCGGACCAGGGGAATCCGACTGTT</td><td>371.9</td><td>-2.7</td><td>0.9</td><td>-2.9</td><td>3.2E-03</td><td>4.0E-03</td><td>BT20/MB468</td><td>CYTO</td></tr>
<tr><td>>hg38_21_+_8433172_8446622_RNA45SN1_45S_50ntFlanks@12405.12442.38|CTTTTTGATCCTTCGATGTCGGCTCTTCCTATCATTGT</td><td>>hg38_21_+_8433172_8446622_RNA45SN1_45S_50ntFlanks</td><td>12405</td><td>12442</td><td>38</td><td>CTTTTTGATCCTTCGATGTCGGCTCTTCCTATCATTGT</td><td>167.6</td><td>-4.2</td><td>1.4</td><td>-2.9</td><td>3.2E-03</td><td>4.0E-03</td><td>BT20/MB468</td><td>CYTO</td></tr>
<tr><td>>hg38_21_+_8433172_8446622_RNA45SN1_45S_50ntFlanks@10685.10709.25|GGCGGCGTCCGGTGAGCTCTCGCTG</td><td>>hg38_21_+_8433172_8446622_RNA45SN1_45S_50ntFlanks</td><td>10685</td><td>10709</td><td>25</td><td>GGCGGCGTCCGGTGAGCTCTCGCTG</td><td>857.5</td><td>3.0</td><td>1.0</td><td>3.0</td><td>2.7E-03</td><td>4.0E-03</td><td>BT20/MB468</td><td>MITO</td></tr>
<tr><td>>hg38_21_+_8433172_8446622_RNA45SN1_45S_50ntFlanks@6650.6680.31|ACGACTCTTAGCGGTGGATCACTCGGCTCGT</td><td>>hg38_21_+_8433172_8446622_RNA45SN1_45S_50ntFlanks</td><td>6650</td><td>6680</td><td>31</td><td>ACGACTCTTAGCGGTGGATCACTCGGCTCGT</td><td>979.4</td><td>1.7</td><td>0.6</td><td>3.0</td><td>2.7E-03</td><td>4.0E-03</td><td>BT20/MB468</td><td>MITO</td></tr>
<tr><td>>hg38_MT_+_598_1651_RNR1_12S_50ntFlanks@965.1004.40|GTAACATGGTAAGTGTACTGGAAAGTGCACTTGGACGAAC</td><td>>hg38_MT_+_598_1651_RNR1_12S_50ntFlanks</td><td>965</td><td>1004</td><td>40</td><td>GTAACATGGTAAGTGTACTGGAAAGTGCACTTGGACGAAC</td><td>80.1</td><td>5.4</td><td>1.8</td><td>3.0</td><td>2.9E-03</td><td>4.0E-03</td><td>MB231/MB468</td><td>MP</td></tr>
<tr><td>>hg38_21_+_8433172_8446622_RNA45SN1_45S_50ntFlanks@10869.10901.33|AGGGCTGGGTCGGTCGGGCTGGGGCGCGAAGCG</td><td>>hg38_21_+_8433172_8446622_RNA45SN1_45S_50ntFlanks</td><td>10869</td><td>10901</td><td>33</td><td>AGGGCTGGGTCGGTCGGGCTGGGGCGCGAAGCG</td><td>805.4</td><td>-4.4</td><td>1.4</td><td>-3.1</td><td>1.7E-03</td><td>4.0E-03</td><td>BT20/MB468</td><td>MP</td></tr>
<tr><td>>trna10_GluCTC_3_-_103879524_103879595@16.42.27,trna116_GluCTC_1_-_145399233_145399304@16.42.27,trna59_GluCTC_1_+_249168447_249168518@16.42.27,trna71_GluCTC_1_-_161439189_161439260@16.42.27,trna74_GluCTC_1_-_161431809_161431880@16.42.27,trna77_GluCTC_1_-_161424398_161424469@16.42.27,trna77_GluCTC_6_+_28949976_28950047@16.42.27,trna80_GluCTC_1_-_161417018_161417089@16.42.27,trna87_GluCTC_6_-_126101393_126101464@16.42.27|TGGTTAGGATTCGGCGCTCTCACCGCC</td><td>>trna10_GluCTC_3_-_103879524_103879595</td><td>16</td><td>42</td><td>27</td><td>TGGTTAGGATTCGGCGCTCTCACCGCC</td><td>483.8</td><td>4.3</td><td>1.4</td><td>3.1</td><td>1.7E-03</td><td>4.0E-03</td><td>BT20/MB468</td><td>MP</td></tr>
<tr><td>>hg38_21_+_8433172_8446622_RNA45SN1_45S_50ntFlanks@6648.6667.20|GTACGACTCTTAGCGGTGGA</td><td>>hg38_21_+_8433172_8446622_RNA45SN1_45S_50ntFlanks</td><td>6648</td><td>6667</td><td>20</td><td>GTACGACTCTTAGCGGTGGA</td><td>275.0</td><td>2.6</td><td>0.8</td><td>3.4</td><td>7.3E-04</td><td>4.0E-03</td><td>BT20/MB231</td><td>CYTO</td></tr>
<tr><td>>hg38_21_+_8433172_8446622_RNA45SN1_45S_50ntFlanks@6661.6680.20|CGGTGGATCACTCGGCTCGT</td><td>>hg38_21_+_8433172_8446622_RNA45SN1_45S_50ntFlanks</td><td>6661</td><td>6680</td><td>20</td><td>CGGTGGATCACTCGGCTCGT</td><td>409.3</td><td>-4.0</td><td>1.4</td><td>-3.0</td><td>2.9E-03</td><td>4.0E-03</td><td>MB231/MB468</td><td>MP</td></tr>
<tr><td>>trnaMT_ValTAC_MT_+_1602_1670@28.72.45|AACTTACACTTAGGAGATTTCAACTTAACTTGACCGCTCTGACCA</td><td>>trnaMT_ValTAC_MT_+_1602_1670</td><td>28</td><td>72</td><td>45</td><td>AACTTACACTTAGGAGATTTCAACTTAACTTGACCGCTCTGACCA</td><td>1590.6</td><td>5.8</td><td>1.9</td><td>3.1</td><td>1.7E-03</td><td>4.0E-03</td><td>BT20/MB468</td><td>MP</td></tr>
<tr><td>>hg38_21_+_8433172_8446622_RNA45SN1_45S_50ntFlanks@4220.4242.23|ACGAAAAATAACAATACAGGACT</td><td>>hg38_21_+_8433172_8446622_RNA45SN1_45S_50ntFlanks</td><td>4220</td><td>4242</td><td>23</td><td>ACGAAAAATAACAATACAGGACT</td><td>945.0</td><td>-2.5</td><td>0.9</td><td>-2.9</td><td>3.2E-03</td><td>4.0E-03</td><td>BT20/MB468</td><td>CYTO</td></tr>
<tr><td>>am_trna111_HisGTG_1_-_147774845_147774916@1.28.28,trna118_HisGTG_1_-_145396881_145396952@1.28.28,trna16_HisGTG_1_+_146544773_146544844@1.28.28,trna1_HisGTG_15_+_45493349_45493420@1.28.28,trna21_HisGTG_1_+_147753471_147753542@1.28.28,trna33_HisGTG_6_+_27125906_27125977@1.28.28,trna7_HisGTG_9_-_14433938_14434009@1.28.28,trna8_HisGTG_15_-_45492611_45492682@1.28.28,trna9_HisGTG_15_-_45490804_45490875@1.28.28|GCCGTGATCGTATAGTGGTTAGTACTCT</td><td>>am_trna111_HisGTG_1_-_147774845_147774916</td><td>1</td><td>28</td><td>28</td><td>GCCGTGATCGTATAGTGGTTAGTACTCT</td><td>974.0</td><td>-2.5</td><td>0.9</td><td>-2.9</td><td>3.5E-03</td><td>4.0E-03</td><td>MB231/MB468</td><td>NUC</td></tr>
<tr><td>>trnaMT_SerGCT_MT_+_12207_12265@1.48.48|GAGAAAGCTCACAAGAACTGCTAACTCATGCCCCCATGTCTAACAACA</td><td>>trnaMT_SerGCT_MT_+_12207_12265</td><td>1</td><td>48</td><td>48</td><td>GAGAAAGCTCACAAGAACTGCTAACTCATGCCCCCATGTCTAACAACA</td><td>1788.1</td><td>-3.9</td><td>1.2</td><td>-3.3</td><td>8.2E-04</td><td>4.0E-03</td><td>BT20/MB231</td><td>MITO</td></tr>
<tr><td>>hg38_21_+_8433172_8446622_RNA45SN1_45S_50ntFlanks@9311.9330.20|AGTCTAACACGTGCGCGAGT</td><td>>hg38_21_+_8433172_8446622_RNA45SN1_45S_50ntFlanks</td><td>9311</td><td>9330</td><td>20</td><td>AGTCTAACACGTGCGCGAGT</td><td>185.6</td><td>5.5</td><td>1.8</td><td>3.1</td><td>1.8E-03</td><td>4.0E-03</td><td>BT20/MB468</td><td>MP</td></tr>
<tr><td>>hg38_21_+_8433172_8446622_RNA45SN1_45S_50ntFlanks@12133.12150.18|CTCCGGGGACAGTGCCAG</td><td>>hg38_21_+_8433172_8446622_RNA45SN1_45S_50ntFlanks</td><td>12133</td><td>12150</td><td>18</td><td>CTCCGGGGACAGTGCCAG</td><td>1527.7</td><td>-5.2</td><td>1.8</td><td>-2.9</td><td>3.2E-03</td><td>4.0E-03</td><td>BT20/MB468</td><td>CYTO</td></tr>
<tr><td>>hg38_21_+_8433172_8446622_RNA45SN1_45S_50ntFlanks@11570.11601.32|AGCCGACTTAGAACTGGTGCGGACCAGGGGAA</td><td>>hg38_21_+_8433172_8446622_RNA45SN1_45S_50ntFlanks</td><td>11570</td><td>11601</td><td>32</td><td>AGCCGACTTAGAACTGGTGCGGACCAGGGGAA</td><td>1573.8</td><td>-2.6</td><td>0.8</td><td>-3.1</td><td>1.8E-03</td><td>4.0E-03</td><td>BT20/MB468</td><td>MP</td></tr>
<tr><td>>hg38_21_+_8433172_8446622_RNA45SN1_45S_50ntFlanks@8381.8404.24|AAGAGGGCGTGAAACCGTTAAGAG</td><td>>hg38_21_+_8433172_8446622_RNA45SN1_45S_50ntFlanks</td><td>8381</td><td>8404</td><td>24</td><td>AAGAGGGCGTGAAACCGTTAAGAG</td><td>168.6</td><td>-4.4</td><td>1.5</td><td>-2.9</td><td>3.5E-03</td><td>4.0E-03</td><td>MB231/MB468</td><td>NUC</td></tr>
<tr><td>>hg38_21_+_8433172_8446622_RNA45SN1_45S_50ntFlanks@12416.12436.21|TTCGATGTCGGCTCTTCCTAT</td><td>>hg38_21_+_8433172_8446622_RNA45SN1_45S_50ntFlanks</td><td>12416</td><td>12436</td><td>21</td><td>TTCGATGTCGGCTCTTCCTAT</td><td>687.0</td><td>-1.8</td><td>0.6</td><td>-3.0</td><td>2.7E-03</td><td>4.0E-03</td><td>BT20/MB468</td><td>MITO</td></tr>
<tr><td>>hg38_21_+_8433172_8446622_RNA45SN1_45S_50ntFlanks@11568.11603.36|GCAGCCGACTTAGAACTGGTGCGGACCAGGGGAATC</td><td>>hg38_21_+_8433172_8446622_RNA45SN1_45S_50ntFlanks</td><td>11568</td><td>11603</td><td>36</td><td>GCAGCCGACTTAGAACTGGTGCGGACCAGGGGAATC</td><td>444.5</td><td>-2.3</td><td>0.8</td><td>-2.9</td><td>3.3E-03</td><td>4.0E-03</td><td>MB231/MB468</td><td>CYTO</td></tr>
<tr><td>>MI0000811|hsa-mir-148b&WithFlank&12|+|54337210|54337320@69.90.22(+1U)&[MIMAT0000759&hsa-miR-148b-3p&offsets|0|0(+1U);m-77&12|+|54337278|54337299&offsets|0|0(+1U)]|TCAGTGCATCACAGAACTTTGTT</td><td>>MI0000811|hsa-mir-148b&WithFlank&12|+|54337210|54337320</td><td>69</td><td>90</td><td>22(+1U)</td><td>TCAGTGCATCACAGAACTTTGTT</td><td>2643.0</td><td>2.5</td><td>0.9</td><td>3.0</td><td>2.9E-03</td><td>4.0E-03</td><td>MB231/MB468</td><td>MP</td></tr>
<tr><td>>hg38_21_+_8433172_8446622_RNA45SN1_45S_50ntFlanks@12407.12430.24|TTTTGATCCTTCGATGTCGGCTCT</td><td>>hg38_21_+_8433172_8446622_RNA45SN1_45S_50ntFlanks</td><td>12407</td><td>12430</td><td>24</td><td>TTTTGATCCTTCGATGTCGGCTCT</td><td>82.1</td><td>-4.8</td><td>1.6</td><td>-3.0</td><td>2.9E-03</td><td>4.0E-03</td><td>MB231/MB468</td><td>MP</td></tr>
<tr><td>>hg38_21_+_8433172_8446622_RNA45SN1_45S_50ntFlanks@10402.10428.27|AACATGGGTCAGTCGGTCCTGAGAGAT</td><td>>hg38_21_+_8433172_8446622_RNA45SN1_45S_50ntFlanks</td><td>10402</td><td>10428</td><td>27</td><td>AACATGGGTCAGTCGGTCCTGAGAGAT</td><td>184.0</td><td>-5.0</td><td>1.6</td><td>-3.1</td><td>1.8E-03</td><td>4.0E-03</td><td>BT20/MB468</td><td>MP</td></tr>
<tr><td>>hg38_21_+_8433172_8446622_RNA45SN1_45S_50ntFlanks@10677.10707.31|GCGGTTCCGGCGGCGTCCGGTGAGCTCTCGC</td><td>>hg38_21_+_8433172_8446622_RNA45SN1_45S_50ntFlanks</td><td>10677</td><td>10707</td><td>31</td><td>GCGGTTCCGGCGGCGTCCGGTGAGCTCTCGC</td><td>806.7</td><td>3.7</td><td>1.1</td><td>3.3</td><td>8.2E-04</td><td>4.0E-03</td><td>BT20/MB231</td><td>MITO</td></tr>
<tr><td>>hg38_21_+_8433172_8446622_RNA45SN1_45S_50ntFlanks@3783.3801.19|AGTGAAACTGCGAATGGCT</td><td>>hg38_21_+_8433172_8446622_RNA45SN1_45S_50ntFlanks</td><td>3783</td><td>3801</td><td>19</td><td>AGTGAAACTGCGAATGGCT</td><td>456.3</td><td>-3.9</td><td>1.3</td><td>-2.9</td><td>3.5E-03</td><td>4.0E-03</td><td>MB231/MB468</td><td>NUC</td></tr>
<tr><td>>hg38_21_+_8433172_8446622_RNA45SN1_45S_50ntFlanks@10408.10426.19|GGTCAGTCGGTCCTGAGAG</td><td>>hg38_21_+_8433172_8446622_RNA45SN1_45S_50ntFlanks</td><td>10408</td><td>10426</td><td>19</td><td>GGTCAGTCGGTCCTGAGAG</td><td>353.9</td><td>-5.2</td><td>1.7</td><td>-3.1</td><td>1.8E-03</td><td>4.1E-03</td><td>BT20/MB468</td><td>MP</td></tr>
<tr><td>>hg38_21_+_8433172_8446622_RNA45SN1_45S_50ntFlanks@7990.8009.20|GACGTGGCGACCCGCTGAAT</td><td>>hg38_21_+_8433172_8446622_RNA45SN1_45S_50ntFlanks</td><td>7990</td><td>8009</td><td>20</td><td>GACGTGGCGACCCGCTGAAT</td><td>435.6</td><td>2.7</td><td>0.9</td><td>2.9</td><td>3.5E-03</td><td>4.1E-03</td><td>MB231/MB468</td><td>NUC</td></tr>
<tr><td>>hg38_21_+_8433172_8446622_RNA45SN1_45S_50ntFlanks@12418.12450.33|CGATGTCGGCTCTTCCTATCATTGTGAAGCAGA</td><td>>hg38_21_+_8433172_8446622_RNA45SN1_45S_50ntFlanks</td><td>12418</td><td>12450</td><td>33</td><td>CGATGTCGGCTCTTCCTATCATTGTGAAGCAGA</td><td>128.9</td><td>-3.7</td><td>1.2</td><td>-3.0</td><td>2.9E-03</td><td>4.1E-03</td><td>MB231/MB468</td><td>MP</td></tr>
<tr><td>>hg38_21_+_8433172_8446622_RNA45SN1_45S_50ntFlanks@11918.11967.50|ACATGAGAGGTGTAGAATAAGTGGGAGGCCCCCGGCGCCCCCCCGGTGTC</td><td>>hg38_21_+_8433172_8446622_RNA45SN1_45S_50ntFlanks</td><td>11918</td><td>11967</td><td>50</td><td>ACATGAGAGGTGTAGAATAAGTGGGAGGCCCCCGGCGCCCCCCCGGTGTC</td><td>34.6</td><td>-8.2</td><td>2.6</td><td>-3.1</td><td>1.8E-03</td><td>4.1E-03</td><td>BT20/MB468</td><td>MP</td></tr>
<tr><td>>hg38_21_+_8433172_8446622_RNA45SN1_45S_50ntFlanks@12944.12966.23|CGTAGACGACCTGCTTCTGGGTC</td><td>>hg38_21_+_8433172_8446622_RNA45SN1_45S_50ntFlanks</td><td>12944</td><td>12966</td><td>23</td><td>CGTAGACGACCTGCTTCTGGGTC</td><td>86.1</td><td>-5.1</td><td>1.6</td><td>-3.1</td><td>1.8E-03</td><td>4.1E-03</td><td>BT20/MB468</td><td>MP</td></tr>
<tr><td>>MI0000456|hsa-mir-140&WithFlank&16|+|69933075|69933186@68.88.21&[MIMAT0004597&hsa-miR-140-3p&offsets|0|0;m-45&16|+|69933143|69933164&offsets|-1|-2]|TACCACAGGGTAGAACCACGG</td><td>>MI0000456|hsa-mir-140&WithFlank&16|+|69933075|69933186</td><td>68</td><td>88</td><td>21</td><td>TACCACAGGGTAGAACCACGG</td><td>217.3</td><td>6.5</td><td>2.1</td><td>3.1</td><td>1.8E-03</td><td>4.1E-03</td><td>BT20/MB468</td><td>MP</td></tr>
<tr><td>>MI0000748|hsa-mir-130b&WithFlank&22|+|21653298|21653391@57.76.20&[MIMAT0000691&hsa-miR-130b-3p&offsets|0|-2;m-205&22|+|21653354|21653375&offsets|0|-2]|CAGTGCAATGATGAAAGGGC</td><td>>MI0000748|hsa-mir-130b&WithFlank&22|+|21653298|21653391</td><td>57</td><td>76</td><td>20</td><td>CAGTGCAATGATGAAAGGGC</td><td>93.8</td><td>6.5</td><td>2.1</td><td>3.1</td><td>1.8E-03</td><td>4.1E-03</td><td>BT20/MB468</td><td>MP</td></tr>
<tr><td>>hg38_MT_+_598_1651_RNR1_12S_50ntFlanks@967.1004.38|AACATGGTAAGTGTACTGGAAAGTGCACTTGGACGAAC</td><td>>hg38_MT_+_598_1651_RNR1_12S_50ntFlanks</td><td>967</td><td>1004</td><td>38</td><td>AACATGGTAAGTGTACTGGAAAGTGCACTTGGACGAAC</td><td>201.0</td><td>3.8</td><td>1.3</td><td>2.9</td><td>3.5E-03</td><td>4.1E-03</td><td>MB231/MB468</td><td>MITO</td></tr>
<tr><td>>trnaMT_GluTTC_MT_-_14674_14742@37.72.36,trnalookalike8_GluTTC_5_-_93905172_93905240@37.72.36|TCATTGGTCGTGGTTGTAGTCCGTGCGAGAATACCA</td><td>>trnaMT_GluTTC_MT_-_14674_14742</td><td>37</td><td>72</td><td>36</td><td>TCATTGGTCGTGGTTGTAGTCCGTGCGAGAATACCA</td><td>6958.0</td><td>-3.5</td><td>1.0</td><td>-3.4</td><td>6.8E-04</td><td>4.1E-03</td><td>BT20/MB231</td><td>NUC</td></tr>
<tr><td>>hg38_21_+_8433172_8446622_RNA45SN1_45S_50ntFlanks@11553.11589.37|CTCGGCCGGCGCCTAGCAGCCGACTTAGAACTGGTGC</td><td>>hg38_21_+_8433172_8446622_RNA45SN1_45S_50ntFlanks</td><td>11553</td><td>11589</td><td>37</td><td>CTCGGCCGGCGCCTAGCAGCCGACTTAGAACTGGTGC</td><td>66.6</td><td>5.1</td><td>1.6</td><td>3.1</td><td>1.8E-03</td><td>4.1E-03</td><td>BT20/MB468</td><td>MP</td></tr>
<tr><td>>hg38_21_+_8433172_8446622_RNA45SN1_45S_50ntFlanks@10808.10830.23|CAATGTAGGTAAGGGAAGTCGGC</td><td>>hg38_21_+_8433172_8446622_RNA45SN1_45S_50ntFlanks</td><td>10808</td><td>10830</td><td>23</td><td>CAATGTAGGTAAGGGAAGTCGGC</td><td>381.4</td><td>-2.6</td><td>0.8</td><td>-3.1</td><td>1.8E-03</td><td>4.1E-03</td><td>BT20/MB468</td><td>MP</td></tr>
<tr><td>>hg38_MT_+_598_1651_RNR1_12S_50ntFlanks@699.730.32|AGTAAGCGCAAGTACCCACGTAAAGACGTTAG</td><td>>hg38_MT_+_598_1651_RNR1_12S_50ntFlanks</td><td>699</td><td>730</td><td>32</td><td>AGTAAGCGCAAGTACCCACGTAAAGACGTTAG</td><td>33.8</td><td>8.3</td><td>2.8</td><td>3.0</td><td>2.9E-03</td><td>4.1E-03</td><td>MB231/MB468</td><td>MP</td></tr>
<tr><td>>trnaMT_SerGCT_MT_+_12207_12265@16.62.47|AACTGCTAACTCATGCCCCCATGTCTAACAACATGGCTTTCTCACCA</td><td>>trnaMT_SerGCT_MT_+_12207_12265</td><td>16</td><td>62</td><td>47</td><td>AACTGCTAACTCATGCCCCCATGTCTAACAACATGGCTTTCTCACCA</td><td>448.0</td><td>3.1</td><td>1.0</td><td>3.0</td><td>2.7E-03</td><td>4.1E-03</td><td>BT20/MB468</td><td>MITO</td></tr>
<tr><td>>am_trnaMT_SerGCT_MT_+_12207_12265@12.34.23|CAAGAACTGCTAACTCATGCCCC</td><td>>am_trnaMT_SerGCT_MT_+_12207_12265</td><td>12</td><td>34</td><td>23</td><td>CAAGAACTGCTAACTCATGCCCC</td><td>167.0</td><td>5.6</td><td>1.8</td><td>3.1</td><td>1.8E-03</td><td>4.1E-03</td><td>BT20/MB468</td><td>MP</td></tr>
<tr><td>>hg38_21_+_8433172_8446622_RNA45SN1_45S_50ntFlanks@12407.12432.26|TTTTGATCCTTCGATGTCGGCTCTTC</td><td>>hg38_21_+_8433172_8446622_RNA45SN1_45S_50ntFlanks</td><td>12407</td><td>12432</td><td>26</td><td>TTTTGATCCTTCGATGTCGGCTCTTC</td><td>1150.6</td><td>-2.2</td><td>0.8</td><td>-2.9</td><td>3.3E-03</td><td>4.1E-03</td><td>MB231/MB468</td><td>CYTO</td></tr>
<tr><td>>MI0005561|hsa-mir-877&WithFlank&6|+|30584326|30584423@7.26.20(+1A)&[MIMAT0004949&hsa-miR-877-5p&offsets|0|0(+1A);m-382&6|+|30584332|30584354&offsets|0|-3(+1A)]|GTAGAGGAGATGGCGCAGGGA</td><td>>MI0005561|hsa-mir-877&WithFlank&6|+|30584326|30584423</td><td>7</td><td>26</td><td>20(+1A)</td><td>GTAGAGGAGATGGCGCAGGGA</td><td>93.0</td><td>4.6</td><td>1.5</td><td>3.1</td><td>1.8E-03</td><td>4.1E-03</td><td>BT20/MB468</td><td>MP</td></tr>
<tr><td>>hg38_21_+_8433172_8446622_RNA45SN1_45S_50ntFlanks@10350.10379.30|ACGAGAACTTTGAAGGCCGAAGTGGAGAAG</td><td>>hg38_21_+_8433172_8446622_RNA45SN1_45S_50ntFlanks</td><td>10350</td><td>10379</td><td>30</td><td>ACGAGAACTTTGAAGGCCGAAGTGGAGAAG</td><td>946.0</td><td>-3.0</td><td>1.0</td><td>-3.1</td><td>1.8E-03</td><td>4.1E-03</td><td>BT20/MB468</td><td>MP</td></tr>
<tr><td>>hg38_21_+_8433172_8446622_RNA45SN1_45S_50ntFlanks@12274.12306.33|CAGTACGAATACAGACCGTGAAAGCGGGGCCTC</td><td>>hg38_21_+_8433172_8446622_RNA45SN1_45S_50ntFlanks</td><td>12274</td><td>12306</td><td>33</td><td>CAGTACGAATACAGACCGTGAAAGCGGGGCCTC</td><td>83.5</td><td>-4.4</td><td>1.5</td><td>-3.0</td><td>2.7E-03</td><td>4.1E-03</td><td>BT20/MB468</td><td>MITO</td></tr>
<tr><td>>MI0000650|hsa-mir-200c&WithFlank&12|+|6963693|6963772@50.73.24&[MIMAT0000617&hsa-miR-200c-3p&offsets|0|+1;m-54&12|+|6963742|6963764&offsets|0|+1]|TAATACTGCCGGGTAATGATGGAG</td><td>>MI0000650|hsa-mir-200c&WithFlank&12|+|6963693|6963772</td><td>50</td><td>73</td><td>24</td><td>TAATACTGCCGGGTAATGATGGAG</td><td>292.4</td><td>5.7</td><td>1.8</td><td>3.1</td><td>1.8E-03</td><td>4.1E-03</td><td>BT20/MB468</td><td>MP</td></tr>
</table>

<table>
<tr><td>>hg38_21_+_8433172_8446622_RNA45SN1_45S_50ntFlanks@11595.11621.27|AGGGGAATCCGACTGTTTAATTAAAAC</td><td>>hg38_21_+_8433172_8446622_RNA45SN1_45S_50ntFlanks</td><td>11595</td><td>11621</td><td>27</td><td>AGGGGAATCCGACTGTTTAATTAAAAC</td><td>68.3</td><td>6.0</td><td>1.9</td><td>3.1</td><td>1.8E-03</td><td>4.1E-03</td><td>BT20/MB468</td><td>MP</td></tr>
<tr><td>>hg38_21_+_8433172_8446622_RNA45SN1_45S_50ntFlanks@6697.6720.24|CAGCTAGCTGCGAGAATTAATGTG</td><td>>hg38_21_+_8433172_8446622_RNA45SN1_45S_50ntFlanks</td><td>6697</td><td>6720</td><td>24</td><td>CAGCTAGCTGCGAGAATTAATGTG</td><td>1199.1</td><td>-2.5</td><td>0.8</td><td>-3.1</td><td>1.8E-03</td><td>4.1E-03</td><td>BT20/MB468</td><td>MP</td></tr>
<tr><td>>hg38_21_+_8433172_8446622_RNA45SN1_45S_50ntFlanks@3870.3898.29|AGCTAATACATGCCGACGGGCGCTGACCC</td><td>>hg38_21_+_8433172_8446622_RNA45SN1_45S_50ntFlanks</td><td>3870</td><td>3898</td><td>29</td><td>AGCTAATACATGCCGACGGGCGCTGACCC</td><td>225.9</td><td>-5.2</td><td>1.7</td><td>-3.1</td><td>1.8E-03</td><td>4.1E-03</td><td>BT20/MB468</td><td>MP</td></tr>
<tr><td>>hg38_21_+_8433172_8446622_RNA45SN1_45S_50ntFlanks@4297.4318.22|CATTGGAGGGCAAGTCTGGTGC</td><td>>hg38_21_+_8433172_8446622_RNA45SN1_45S_50ntFlanks</td><td>4297</td><td>4318</td><td>22</td><td>CATTGGAGGGCAAGTCTGGTGC</td><td>174.0</td><td>-1.8</td><td>0.6</td><td>-2.9</td><td>3.2E-03</td><td>4.1E-03</td><td>BT20/MB468</td><td>CYTO</td></tr>
<tr><td>>am_trnaMT_GlyTCC_MT_+_9991_10058@28.48.21|ACTTCCAATTAACTAGTTTTG</td><td>>am_trnaMT_GlyTCC_MT_+_9991_10058</td><td>28</td><td>48</td><td>21</td><td>ACTTCCAATTAACTAGTTTTG</td><td>89.7</td><td>-6.2</td><td>2.1</td><td>-2.9</td><td>3.2E-03</td><td>4.1E-03</td><td>BT20/MB468</td><td>CYTO</td></tr>
<tr><td>>hg38_21_+_8433172_8446622_RNA45SN1_45S_50ntFlanks@9554.9592.39|CGTAGCGGTCCTGACGTGCAAATCGGTCGTCCGACCTGG</td><td>>hg38_21_+_8433172_8446622_RNA45SN1_45S_50ntFlanks</td><td>9554</td><td>9592</td><td>39</td><td>CGTAGCGGTCCTGACGTGCAAATCGGTCGTCCGACCTGG</td><td>111.0</td><td>-4.8</td><td>1.6</td><td>-3.0</td><td>2.7E-03</td><td>4.1E-03</td><td>BT20/MB468</td><td>MITO</td></tr>
<tr><td>>MI0000446|hsa-mir-125b-1&WithFlank&11|-|122099751|122099850@21.39.19;MI0000470|hsa-mir-125b-2&WithFlank&21|+|16590231|16590331@23.41.19&[MIMAT0000423&hsa-miR-125b-5p&offsets|0|-3;m-67&11|-|122099809|122099830&offsets|0|-3];[MIMAT0000423_1&hsa-miR-125b-5p&offsets|0|-3;m-68&21|+|16590253|16590274&offsets|0|-3]|TCCCTGAGACCCTAACTTG</td><td>>MI0000446|hsa-mir-125b-1&WithFlank&11|-|122099751|122099850</td><td>21</td><td>39</td><td>19</td><td>TCCCTGAGACCCTAACTTG</td><td>788.1</td><td>-6.5</td><td>1.9</td><td>-3.4</td><td>6.2E-04</td><td>4.1E-03</td><td>BT20/MB231</td><td>MP</td></tr>
<tr><td>>hg38_21_+_8433172_8446622_RNA45SN1_45S_50ntFlanks@12954.12972.19|CTGCTTCTGGGTCGGGGTT</td><td>>hg38_21_+_8433172_8446622_RNA45SN1_45S_50ntFlanks</td><td>12954</td><td>12972</td><td>19</td><td>CTGCTTCTGGGTCGGGGTT</td><td>225.6</td><td>-3.3</td><td>1.1</td><td>-3.0</td><td>2.7E-03</td><td>4.1E-03</td><td>BT20/MB468</td><td>MITO</td></tr>
<tr><td>>MI0000093|hsa-mir-92a-1&WithFlank&13|+|91351308|91351397@54.75.22(+1A);MI0000094|hsa-mir-92a-2&WithFlank&X|-|134169532|134169618@54.75.22(+1A)&[MIMAT0000092&hsa-miR-92a-3p&offsets|0|0(+1A);m-9&13|+|91351361|91351382&offsets|0|0(+1A)];[MIMAT0000092_1&hsa-miR-92a-3p&offsets|0|0(+1A);m-11&X|-|134169544|134169565&offsets|0|0(+1A)]|TATTGCACTTGTCCCGGCCTGTA</td><td>>MI0000093|hsa-mir-92a-1&WithFlank&13|+|91351308|91351397</td><td>54</td><td>75</td><td>22(+1A)</td><td>TATTGCACTTGTCCCGGCCTGTA</td><td>3238.3</td><td>2.0</td><td>0.7</td><td>3.0</td><td>3.0E-03</td><td>4.1E-03</td><td>MB231/MB468</td><td>MP</td></tr>
<tr><td>>hg38_21_+_8433172_8446622_RNA45SN1_45S_50ntFlanks@11584.11611.28|TGGTGCGGACCAGGGGAATCCGACTGTT</td><td>>hg38_21_+_8433172_8446622_RNA45SN1_45S_50ntFlanks</td><td>11584</td><td>11611</td><td>28</td><td>TGGTGCGGACCAGGGGAATCCGACTGTT</td><td>339.4</td><td>2.3</td><td>0.8</td><td>2.9</td><td>3.5E-03</td><td>4.1E-03</td><td>BT20/MB468</td><td>NUC</td></tr>
<tr><td>>hg38_21_+_8433172_8446622_RNA45SN1_45S_50ntFlanks@5106.5136.31|ACTAGTTACGCGACCCCCGAGCGGTCGGCGT</td><td>>hg38_21_+_8433172_8446622_RNA45SN1_45S_50ntFlanks</td><td>5106</td><td>5136</td><td>31</td><td>ACTAGTTACGCGACCCCCGAGCGGTCGGCGT</td><td>35.7</td><td>-8.2</td><td>2.6</td><td>-3.1</td><td>1.8E-03</td><td>4.1E-03</td><td>BT20/MB468</td><td>MP</td></tr>
<tr><td>>hg38_21_+_8433172_8446622_RNA45SN1_45S_50ntFlanks@5078.5098.21|CGATAACGAACGAGACTCTGG</td><td>>hg38_21_+_8433172_8446622_RNA45SN1_45S_50ntFlanks</td><td>5078</td><td>5098</td><td>21</td><td>CGATAACGAACGAGACTCTGG</td><td>366.6</td><td>-2.7</td><td>0.9</td><td>-3.0</td><td>2.8E-03</td><td>4.1E-03</td><td>BT20/MB468</td><td>MITO</td></tr>
<tr><td>>MI0016887|hsa-mir-4521&WithFlank&17|+|8186939|8187010@10.35.26&[MIMAT0019058&hsa-miR-4521&offsets|0|+4;m-765&17|+|8186948|8186969&offsets|0|+4]|GCTAAGGAAGTCCTGTGCTCAGTTTT</td><td>>MI0016887|hsa-mir-4521&WithFlank&17|+|8186939|8187010</td><td>10</td><td>35</td><td>26</td><td>GCTAAGGAAGTCCTGTGCTCAGTTTT</td><td>77.7</td><td>4.5</td><td>1.5</td><td>3.0</td><td>3.0E-03</td><td>4.1E-03</td><td>MB231/MB468</td><td>MP</td></tr>
<tr><td>>MI0000253|hsa-mir-148a&WithFlank&7|-|25949913|25949992@50.70.21(+1G)&[MIMAT0000243&hsa-miR-148a-3p&offsets|0|-1(+1G);m-7&7|-|25949922|25949943&offsets|0|-1(+1G)]|TCAGTGCACTACAGAACTTTGG</td><td>>MI0000253|hsa-mir-148a&WithFlank&7|-|25949913|25949992</td><td>50</td><td>70</td><td>21(+1G)</td><td>TCAGTGCACTACAGAACTTTGG</td><td>8152.3</td><td>4.4</td><td>1.3</td><td>3.4</td><td>6.2E-04</td><td>4.1E-03</td><td>BT20/MB231</td><td>MP</td></tr>
<tr><td>>MI0000438|hsa-mir-15b&WithFlank&3|+|160404582|160404691@64.85.22(+1U)&[MIMAT0004586&hsa-miR-15b-3p&offsets|0|0(+1U);m-204&3|+|160404645|160404665&offsets|0|+1(+1U)]|CGAATCATTATTTGCTGCTCTAT</td><td>>MI0000438|hsa-mir-15b&WithFlank&3|+|160404582|160404691</td><td>64</td><td>85</td><td>22(+1U)</td><td>CGAATCATTATTTGCTGCTCTAT</td><td>78.2</td><td>4.9</td><td>1.7</td><td>3.0</td><td>3.0E-03</td><td>4.1E-03</td><td>MB231/MB468</td><td>MP</td></tr>
<tr><td>>hg38_21_+_8433172_8446622_RNA45SN1_45S_50ntFlanks@9482.9511.30|ACGTGTTAGGACCCGAAAGATGGTGAACTA</td><td>>hg38_21_+_8433172_8446622_RNA45SN1_45S_50ntFlanks</td><td>9482</td><td>9511</td><td>30</td><td>ACGTGTTAGGACCCGAAAGATGGTGAACTA</td><td>102.3</td><td>7.5</td><td>2.4</td><td>3.1</td><td>1.8E-03</td><td>4.1E-03</td><td>BT20/MB468</td><td>MP</td></tr>
<tr><td>>hg38_21_+_8433172_8446622_RNA45SN1_45S_50ntFlanks@8168.8186.19|CCCAAGTCCTTCTGATCGA</td><td>>hg38_21_+_8433172_8446622_RNA45SN1_45S_50ntFlanks</td><td>8168</td><td>8186</td><td>19</td><td>CCCAAGTCCTTCTGATCGA</td><td>747.0</td><td>-5.7</td><td>1.8</td><td>-3.1</td><td>1.8E-03</td><td>4.1E-03</td><td>BT20/MB468</td><td>MP</td></tr>
<tr><td>>hg38_21_+_8433172_8446622_RNA45SN1_45S_50ntFlanks@5492.5525.34|AGAAGACGGTCGAACTTGACTATCTAGAGGAAGT</td><td>>hg38_21_+_8433172_8446622_RNA45SN1_45S_50ntFlanks</td><td>5492</td><td>5525</td><td>34</td><td>AGAAGACGGTCGAACTTGACTATCTAGAGGAAGT</td><td>1503.6</td><td>3.1</td><td>1.1</td><td>2.9</td><td>3.2E-03</td><td>4.1E-03</td><td>BT20/MB468</td><td>CYTO</td></tr>
<tr><td>>hg38_21_+_8433172_8446622_RNA45SN1_45S_50ntFlanks@12626.12667.42|AATGGGGCGAAGCTACCATCTGTGGGATTATGACTGAACGCC</td><td>>hg38_21_+_8433172_8446622_RNA45SN1_45S_50ntFlanks</td><td>12626</td><td>12667</td><td>42</td><td>AATGGGGCGAAGCTACCATCTGTGGGATTATGACTGAACGCC</td><td>1294.3</td><td>-2.8</td><td>0.8</td><td>-3.4</td><td>7.5E-04</td><td>4.1E-03</td><td>BT20/MB231</td><td>CYTO</td></tr>
<tr><td>>hg38_21_+_8433172_8446622_RNA45SN1_45S_50ntFlanks@3840.3862.23|CTCCTACTTGGATAACTGTGGTA</td><td>>hg38_21_+_8433172_8446622_RNA45SN1_45S_50ntFlanks</td><td>3840</td><td>3862</td><td>23</td><td>CTCCTACTTGGATAACTGTGGTA</td><td>182.4</td><td>1.8</td><td>0.6</td><td>2.9</td><td>3.5E-03</td><td>4.1E-03</td><td>BT20/MB468</td><td>NUC</td></tr>
<tr><td>>hg38_21_+_8433172_8446622_RNA45SN1_45S_50ntFlanks@12419.12452.34|GATGTCGGCTCTTCCTATCATTGTGAAGCAGAAT</td><td>>hg38_21_+_8433172_8446622_RNA45SN1_45S_50ntFlanks</td><td>12419</td><td>12452</td><td>34</td><td>GATGTCGGCTCTTCCTATCATTGTGAAGCAGAAT</td><td>1550.6</td><td>-2.8</td><td>0.8</td><td>-3.4</td><td>7.5E-04</td><td>4.1E-03</td><td>BT20/MB231</td><td>CYTO</td></tr>
<tr><td>>am_trnaMT_ValTAC_MT_+_1602_1670@28.57.30|AACTTACACTTAGGAGATTTCAACTTAACT</td><td>>am_trnaMT_ValTAC_MT_+_1602_1670</td><td>28</td><td>57</td><td>30</td><td>AACTTACACTTAGGAGATTTCAACTTAACT</td><td>401.1</td><td>6.1</td><td>1.9</td><td>3.1</td><td>1.8E-03</td><td>4.1E-03</td><td>BT20/MB468</td><td>MP</td></tr>
<tr><td>>MI0003599|hsa-mir-589&WithFlank&7|-|5495813|5495923@30.52.23&[MIMAT0004799&hsa-miR-589-5p&offsets|0|+1;m-162&7|-|5495874|5495894&offsets|0|+2]|TGAGAACCACGTCTGCTCTGAGC</td><td>>MI0003599|hsa-mir-589&WithFlank&7|-|5495813|5495923</td><td>30</td><td>52</td><td>23</td><td>TGAGAACCACGTCTGCTCTGAGC</td><td>194.8</td><td>4.8</td><td>1.6</td><td>3.0</td><td>3.0E-03</td><td>4.1E-03</td><td>MB231/MB468</td><td>MP</td></tr>
<tr><td>>hg38_21_+_8433172_8446622_RNA45SN1_45S_50ntFlanks@10350.10390.41|ACGAGAACTTTGAAGGCCGAAGTGGAGAAGGGTTCCATGTG</td><td>>hg38_21_+_8433172_8446622_RNA45SN1_45S_50ntFlanks</td><td>10350</td><td>10390</td><td>41</td><td>ACGAGAACTTTGAAGGCCGAAGTGGAGAAGGGTTCCATGTG</td><td>1725.2</td><td>-3.0</td><td>1.0</td><td>-3.1</td><td>1.8E-03</td><td>4.1E-03</td><td>BT20/MB468</td><td>MP</td></tr>
<tr><td>>hg38_21_+_8433172_8446622_RNA45SN1_45S_50ntFlanks@5044.5081.38|TCTTAGTTGGTGGAGCGATTTGTCTGGTTAATTCCGAT</td><td>>hg38_21_+_8433172_8446622_RNA45SN1_45S_50ntFlanks</td><td>5044</td><td>5081</td><td>38</td><td>TCTTAGTTGGTGGAGCGATTTGTCTGGTTAATTCCGAT</td><td>54.6</td><td>6.0</td><td>1.9</td><td>3.1</td><td>1.8E-03</td><td>4.1E-03</td><td>BT20/MB468</td><td>MP</td></tr>
<tr><td>>hg38_21_+_8433172_8446622_RNA45SN1_45S_50ntFlanks@8347.8365.19|GGGAAAGTTGAAAAGAACT</td><td>>hg38_21_+_8433172_8446622_RNA45SN1_45S_50ntFlanks</td><td>8347</td><td>8365</td><td>19</td><td>GGGAAAGTTGAAAAGAACT</td><td>67.7</td><td>6.0</td><td>1.9</td><td>3.1</td><td>1.8E-03</td><td>4.1E-03</td><td>BT20/MB468</td><td>MP</td></tr>
<tr><td>>hg38_21_+_8433172_8446622_RNA45SN1_45S_50ntFlanks@6645.6674.30|CTCGTACGACTCTTAGCGGTGGATCACTCG</td><td>>hg38_21_+_8433172_8446622_RNA45SN1_45S_50ntFlanks</td><td>6645</td><td>6674</td><td>30</td><td>CTCGTACGACTCTTAGCGGTGGATCACTCG</td><td>940.0</td><td>3.3</td><td>1.0</td><td>3.1</td><td>1.8E-03</td><td>4.1E-03</td><td>BT20/MB468</td><td>MP</td></tr>
<tr><td>>hg38_21_+_8433172_8446622_RNA45SN1_45S_50ntFlanks@4683.4723.41|CAAGACGGACCAGAGCGAAAGCATTTGCCAAGAATGTTTTC</td><td>>hg38_21_+_8433172_8446622_RNA45SN1_45S_50ntFlanks</td><td>4683</td><td>4723</td><td>41</td><td>CAAGACGGACCAGAGCGAAAGCATTTGCCAAGAATGTTTTC</td><td>180.4</td><td>-6.3</td><td>2.1</td><td>-3.0</td><td>3.0E-03</td><td>4.1E-03</td><td>MB231/MB468</td><td>MP</td></tr>
<tr><td>>hg38_21_+_8433172_8446622_RNA45SN1_45S_50ntFlanks@10830.10851.22|CAAGCCGGATCCGTAACTTCGG</td><td>>hg38_21_+_8433172_8446622_RNA45SN1_45S_50ntFlanks</td><td>10830</td><td>10851</td><td>22</td><td>CAAGCCGGATCCGTAACTTCGG</td><td>308.5</td><td>-3.1</td><td>1.1</td><td>-2.9</td><td>3.3E-03</td><td>4.1E-03</td><td>MB231/MB468</td><td>CYTO</td></tr>
<tr><td>>hg38_21_+_8433172_8446622_RNA45SN1_45S_50ntFlanks@6651.6669.19|CGACTCTTAGCGGTGGATC</td><td>>hg38_21_+_8433172_8446622_RNA45SN1_45S_50ntFlanks</td><td>6651</td><td>6669</td><td>19</td><td>CGACTCTTAGCGGTGGATC</td><td>7978.8</td><td>2.4</td><td>0.7</td><td>3.4</td><td>7.5E-04</td><td>4.1E-03</td><td>BT20/MB231</td><td>CYTO</td></tr>
<tr><td>>hg38_21_+_8433172_8446622_RNA45SN1_45S_50ntFlanks@10809.10830.22|AATGTAGGTAAGGGAAGTCGGC</td><td>>hg38_21_+_8433172_8446622_RNA45SN1_45S_50ntFlanks</td><td>10809</td><td>10830</td><td>22</td><td>AATGTAGGTAAGGGAAGTCGGC</td><td>995.5</td><td>-3.6</td><td>1.1</td><td>-3.4</td><td>7.5E-04</td><td>4.1E-03</td><td>BT20/MB231</td><td>CYTO</td></tr>
<tr><td>>hg38_21_+_8433172_8446622_RNA45SN1_45S_50ntFlanks@476.505.30|TTCGTGATCGATGTGGTGACGTCGTGCTCT</td><td>>hg38_21_+_8433172_8446622_RNA45SN1_45S_50ntFlanks</td><td>476</td><td>505</td><td>30</td><td>TTCGTGATCGATGTGGTGACGTCGTGCTCT</td><td>197.4</td><td>3.8</td><td>1.3</td><td>3.0</td><td>3.0E-03</td><td>4.1E-03</td><td>MB231/MB468</td><td>MP</td></tr>
<tr><td>>hg38_21_+_8433172_8446622_RNA45SN1_45S_50ntFlanks@12943.12960.18|TCGTAGACGACCTGCTTC</td><td>>hg38_21_+_8433172_8446622_RNA45SN1_45S_50ntFlanks</td><td>12943</td><td>12960</td><td>18</td><td>TCGTAGACGACCTGCTTC</td><td>78.1</td><td>-2.2</td><td>0.7</td><td>-3.0</td><td>3.0E-03</td><td>4.1E-03</td><td>MB231/MB468</td><td>MP</td></tr>
<tr><td>>am_trna12_ArgCCT_16_+_3243918_3243990@36.53.18,trna18_ArgCCT_17_+_73030001_73030073@36.53.18,trna21_ArgCCT_17_-_73030526_73030598@36.53.18,trna2_ArgCCT_16_+_3202901_3202973@36.53.18,trna3_ArgCCT_7_+_139025446_139025518@36.53.18|TAAGCCAGGGATTGTGGG</td><td>>am_trna12_ArgCCT_16_+_3243918_3243990</td><td>36</td><td>53</td><td>18</td><td>TAAGCCAGGGATTGTGGG</td><td>216.3</td><td>-2.4</td><td>0.8</td><td>-2.9</td><td>3.3E-03</td><td>4.1E-03</td><td>MB231/MB468</td><td>CYTO</td></tr>
<tr><td>>hg38_21_+_8433172_8446622_RNA45SN1_45S_50ntFlanks@6649.6677.29|TACGACTCTTAGCGGTGGATCACTCGGCT</td><td>>hg38_21_+_8433172_8446622_RNA45SN1_45S_50ntFlanks</td><td>6649</td><td>6677</td><td>29</td><td>TACGACTCTTAGCGGTGGATCACTCGGCT</td><td>588.9</td><td>4.5</td><td>1.3</td><td>3.4</td><td>6.3E-04</td><td>4.1E-03</td><td>BT20/MB231</td><td>MP</td></tr>
<tr><td>>MI0000433|hsa-let-7g&WithFlank&3|-|52268272|52268367@11.32.22(+1C)&[MIMAT0000414&hsa-let-7g-5p&offsets|0|0(+1C);m-58&3|-|52268336|52268357&offsets|0|0(+1C)]|TGAGGTAGTAGTTTGTACAGTTC</td><td>>MI0000433|hsa-let-7g&WithFlank&3|-|52268272|52268367</td><td>11</td><td>32</td><td>22(+1C)</td><td>TGAGGTAGTAGTTTGTACAGTTC</td><td>156.7</td><td>5.6</td><td>1.8</td><td>3.1</td><td>1.8E-03</td><td>4.1E-03</td><td>BT20/MB468</td><td>MP</td></tr>
<tr><td>>hg38_21_+_8433172_8446622_RNA45SN1_45S_50ntFlanks@12155.12182.28|GGAGTTTGACTGGGGCGGTACACCTGTC</td><td>>hg38_21_+_8433172_8446622_RNA45SN1_45S_50ntFlanks</td><td>12155</td><td>12182</td><td>28</td><td>GGAGTTTGACTGGGGCGGTACACCTGTC</td><td>367.9</td><td>-4.2</td><td>1.3</td><td>-3.1</td><td>1.8E-03</td><td>4.1E-03</td><td>BT20/MB468</td><td>MP</td></tr>
<tr><td>>hg38_21_+_8433172_8446622_RNA45SN1_45S_50ntFlanks@11671.11690.20|CCAGTGCTCTGAATGTCAAA</td><td>>hg38_21_+_8433172_8446622_RNA45SN1_45S_50ntFlanks</td><td>11671</td><td>11690</td><td>20</td><td>CCAGTGCTCTGAATGTCAAA</td><td>80.0</td><td>-4.8</td><td>1.6</td><td>-3.0</td><td>3.0E-03</td><td>4.1E-03</td><td>MB231/MB468</td><td>MP</td></tr>
<tr><td>>hg38_21_+_8433172_8446622_RNA45SN1_45S_50ntFlanks@10386.10407.22|ATGTGAACAGCAGTTGAACATG</td><td>>hg38_21_+_8433172_8446622_RNA45SN1_45S_50ntFlanks</td><td>10386</td><td>10407</td><td>22</td><td>ATGTGAACAGCAGTTGAACATG</td><td>35.5</td><td>-8.2</td><td>2.6</td><td>-3.1</td><td>1.8E-03</td><td>4.1E-03</td><td>BT20/MB468</td><td>MP</td></tr>
<tr><td>>hg38_1_-_228634819_228635039_RNA5S12_5S_50ntFlanks@129.170.42|TTGGATGGGAGACCGCCTGGGAATACCGGGTGCTGTAGGCTT</td><td>>hg38_1_-_228634819_228635039_RNA5S12_5S_50ntFlanks</td><td>129</td><td>170</td><td>42</td><td>TTGGATGGGAGACCGCCTGGGAATACCGGGTGCTGTAGGCTT</td><td>321.3</td><td>3.3</td><td>1.1</td><td>3.0</td><td>3.0E-03</td><td>4.1E-03</td><td>MB231/MB468</td><td>MP</td></tr>
<tr><td>>hg38_MT_+_1621_3279_RNR2_16S_50ntFlanks@1370.1393.24|ATGTTGGATCAGGACATCCCGATG</td><td>>hg38_MT_+_1621_3279_RNR2_16S_50ntFlanks</td><td>1370</td><td>1393</td><td>24</td><td>ATGTTGGATCAGGACATCCCGATG</td><td>368.8</td><td>2.7</td><td>0.9</td><td>3.0</td><td>3.0E-03</td><td>4.1E-03</td><td>MB231/MB468</td><td>MP</td></tr>
<tr><td>>hg38_21_+_8433172_8446622_RNA45SN1_45S_50ntFlanks@9568.9592.25|CGTGCAAATCGGTCGTCCGACCTGG</td><td>>hg38_21_+_8433172_8446622_RNA45SN1_45S_50ntFlanks</td><td>9568</td><td>9592</td><td>25</td><td>CGTGCAAATCGGTCGTCCGACCTGG</td><td>722.5</td><td>-5.4</td><td>1.7</td><td>-3.1</td><td>1.8E-03</td><td>4.1E-03</td><td>BT20/MB468</td><td>MP</td></tr>
<tr><td>>hg38_21_+_8433172_8446622_RNA45SN1_45S_50ntFlanks@5056.5082.27|GAGCGATTTGTCTGGTTAATTCCGATA</td><td>>hg38_21_+_8433172_8446622_RNA45SN1_45S_50ntFlanks</td><td>5056</td><td>5082</td><td>27</td><td>GAGCGATTTGTCTGGTTAATTCCGATA</td><td>1209.8</td><td>-4.3</td><td>1.4</td><td>-3.1</td><td>1.8E-03</td><td>4.1E-03</td><td>BT20/MB468</td><td>MP</td></tr>
<tr><td>>hg38_21_+_8433172_8446622_RNA45SN1_45S_50ntFlanks@12942.12959.18|TTCGTAGACGACCTGCTT</td><td>>hg38_21_+_8433172_8446622_RNA45SN1_45S_50ntFlanks</td><td>12942</td><td>12959</td><td>18</td><td>TTCGTAGACGACCTGCTT</td><td>197.8</td><td>-3.9</td><td>1.3</td><td>-3.1</td><td>1.8E-03</td><td>4.1E-03</td><td>BT20/MB468</td><td>MP</td></tr>
<tr><td>>MI0000093|hsa-mir-92a-1&WithFlank&13|+|91351308|91351397@54.75.22(+1C);MI0000094|hsa-mir-92a-2&WithFlank&X|-|134169532|134169618@54.75.22(+1C)&[MIMAT0000092&hsa-miR-92a-3p&offsets|0|0(+1C);m-9&13|+|91351361|91351382&offsets|0|0(+1C)];[MIMAT0000092_1&hsa-miR-92a-3p&offsets|0|0(+1C);m-11&X|-|134169544|134169565&offsets|0|0(+1C)]|TATTGCACTTGTCCCGGCCTGTC</td><td>>MI0000093|hsa-mir-92a-1&WithFlank&13|+|91351308|91351397</td><td>54</td><td>75</td><td>22(+1C)</td><td>TATTGCACTTGTCCCGGCCTGTC</td><td>403.7</td><td>4.0</td><td>1.3</td><td>3.0</td><td>3.0E-03</td><td>4.1E-03</td><td>MB231/MB468</td><td>MP</td></tr>
<tr><td>>hg38_21_+_8433172_8446622_RNA45SN1_45S_50ntFlanks@7980.8007.28|CCTCAGATCAGACGTGGCGACCCGCTGA</td><td>>hg38_21_+_8433172_8446622_RNA45SN1_45S_50ntFlanks</td><td>7980</td><td>8007</td><td>28</td><td>CCTCAGATCAGACGTGGCGACCCGCTGA</td><td>401.0</td><td>-2.1</td><td>0.7</td><td>-3.0</td><td>2.8E-03</td><td>4.1E-03</td><td>BT20/MB468</td><td>MITO</td></tr>
<tr><td>>hg38_21_+_8433172_8446622_RNA45SN1_45S_50ntFlanks@10686.10709.24|GCGGCGTCCGGTGAGCTCTCGCTG</td><td>>hg38_21_+_8433172_8446622_RNA45SN1_45S_50ntFlanks</td><td>10686</td><td>10709</td><td>24</td><td>GCGGCGTCCGGTGAGCTCTCGCTG</td><td>1302.8</td><td>3.4</td><td>1.0</td><td>3.3</td><td>8.4E-04</td><td>4.1E-03</td><td>BT20/MB231</td><td>MITO</td></tr>
</table>

| | | | | | | | | | | | | | |
|---|---|---|---|---|---|---|---|---|---|---|---|---|---|
| >trnaMT_AlaTGC_MT_-_5587_5655@35.72.38,trnalookalike1_AlaTGC_14_+_32954252_32954320@35.72.38,trnalookalike4_AlaTGC_1_-_566137_566205@35.72.38\|TTCAGTTGATGCAGAGTGGGGTTTTGCAGTCCTTACCA | >trnaMT_AlaTGC_MT_-_5587_5655 | 35 | 72 | 38 | TTCAGTTGATGCAGAGTGGGGTTTTGCAGTCCTTACCA | 335.3 | 3.3 | 1.1 | 2.9 | 3.5E-03 | 4.1E-03 | MB231/MB468 | MITO |
| >hg38_21_+_8433172_8446622_RNA45SN1_45S_50ntFlanks@7977.8010.34\|CGACCTCAGATCAGACGTGGCGACCCGCTGAATT | >hg38_21_+_8433172_8446622_RNA45SN1_45S_50ntFlanks | 7977 | 8010 | 34 | CGACCTCAGATCAGACGTGGCGACCCGCTGAATT | 499.4 | 2.2 | 0.7 | 2.9 | 3.5E-03 | 4.1E-03 | MB231/MB468 | MITO |
| >hg38_21_+_8433172_8446622_RNA45SN1_45S_50ntFlanks@11565.11611.47\|CTAGCAGCCGACTTAGAACTGGTGCGGACCAGGGGAATCCGACTGTT | >hg38_21_+_8433172_8446622_RNA45SN1_45S_50ntFlanks | 11565 | 11611 | 47 | CTAGCAGCCGACTTAGAACTGGTGCGGACCAGGGGAATCCGACTGTT | 277.5 | 4.3 | 1.4 | 3.1 | 1.8E-03 | 4.1E-03 | BT20/MB468 | MP |
| >hg38_21_+_8433172_8446622_RNA45SN1_45S_50ntFlanks@10066.10085.20\|GAGAGTGGACGGGAGCGGCG | >hg38_21_+_8433172_8446622_RNA45SN1_45S_50ntFlanks | 10066 | 10085 | 20 | GAGAGTGGACGGGAGCGGCG | 42.5 | 6.1 | 1.9 | 3.1 | 1.8E-03 | 4.1E-03 | BT20/MB468 | MP |
| >hg38_21_+_8433172_8446622_RNA45SN1_45S_50ntFlanks@6732.6769.38\|ACATTGATCATCGACACTTCGAACGCACTTGCGGCCCC | >hg38_21_+_8433172_8446622_RNA45SN1_45S_50ntFlanks | 6732 | 6769 | 38 | ACATTGATCATCGACACTTCGAACGCACTTGCGGCCCC | 67.1 | -5.2 | 1.8 | -3.0 | 3.0E-03 | 4.2E-03 | MB231/MB468 | MP |
| >hg38_MT_+_1621_3279_RNR2_16S_50ntFlanks@486.517.32\|AGGAACAGCTCTTTGGACACTAGGAAAAAACC | >hg38_MT_+_1621_3279_RNR2_16S_50ntFlanks | 486 | 517 | 32 | AGGAACAGCTCTTTGGACACTAGGAAAAAACC | 141.6 | 2.8 | 0.9 | 3.0 | 3.0E-03 | 4.2E-03 | MB231/MB468 | MP |
| >hg38_21_+_8433172_8446622_RNA45SN1_45S_50ntFlanks@10686.10706.21\|GCGGCGTCCGGTGAGCTCTCG | >hg38_21_+_8433172_8446622_RNA45SN1_45S_50ntFlanks | 10686 | 10706 | 21 | GCGGCGTCCGGTGAGCTCTCG | 3326.1 | -6.3 | 2.2 | -2.9 | 3.5E-03 | 4.2E-03 | BT20/MB468 | NUC |
| >hg38_21_+_8433172_8446622_RNA45SN1_45S_50ntFlanks@12936.12955.20\|AAACCATTCGTAGACGACCT | >hg38_21_+_8433172_8446622_RNA45SN1_45S_50ntFlanks | 12936 | 12955 | 20 | AAACCATTCGTAGACGACCT | 314.3 | -4.4 | 1.5 | -2.9 | 3.3E-03 | 4.2E-03 | BT20/MB468 | CYTO |
| >hg38_21_+_8433172_8446622_RNA45SN1_45S_50ntFlanks@6648.6692.45\|GTACGACTCTTAGCGGTGGATCACTCGGCTCGTGCGTCGATGAAG | >hg38_21_+_8433172_8446622_RNA45SN1_45S_50ntFlanks | 6648 | 6692 | 45 | GTACGACTCTTAGCGGTGGATCACTCGGCTCGTGCGTCGATGAAG | 774.6 | 2.8 | 1.0 | 2.9 | 3.6E-03 | 4.2E-03 | MB231/MB468 | NUC |
| >hg38_21_+_8433172_8446622_RNA45SN1_45S_50ntFlanks@12459.12487.29\|AGCGTTGGATTGTTCACCCACTAATAGGG | >hg38_21_+_8433172_8446622_RNA45SN1_45S_50ntFlanks | 12459 | 12487 | 29 | AGCGTTGGATTGTTCACCCACTAATAGGG | 170.4 | -5.2 | 1.7 | -3.1 | 1.8E-03 | 4.2E-03 | BT20/MB468 | MP |
| >hg38_21_+_8433172_8446622_RNA45SN1_45S_50ntFlanks@9157.9187.31\|CAGTGCGCCCCGGGCGGGTCGCGCCGTCGGG | >hg38_21_+_8433172_8446622_RNA45SN1_45S_50ntFlanks | 9157 | 9187 | 31 | CAGTGCGCCCCGGGCGGGTCGCGCCGTCGGG | 152.2 | -5.0 | 1.6 | -3.1 | 1.8E-03 | 4.2E-03 | BT20/MB468 | MP |
| >hg38_21_+_8433172_8446622_RNA45SN1_45S_50ntFlanks@10690.10707.18\|CGTCCGGTGAGCTCTCGC | >hg38_21_+_8433172_8446622_RNA45SN1_45S_50ntFlanks | 10690 | 10707 | 18 | CGTCCGGTGAGCTCTCGC | 159.8 | -1.7 | 0.6 | -2.9 | 3.3E-03 | 4.2E-03 | BT20/MB468 | CYTO |
| >hg38_21_+_8433172_8446622_RNA45SN1_45S_50ntFlanks@5073.5097.25\|AATTCCGATAACGAACGAGACTCTG | >hg38_21_+_8433172_8446622_RNA45SN1_45S_50ntFlanks | 5073 | 5097 | 25 | AATTCCGATAACGAACGAGACTCTG | 580.0 | -3.1 | 1.0 | -3.0 | 2.8E-03 | 4.2E-03 | BT20/MB468 | MITO |
| >hg38_21_+_8433172_8446622_RNA45SN1_45S_50ntFlanks@11208.11230.23\|GCGGCGACTCTGGACGCGAGCCG | >hg38_21_+_8433172_8446622_RNA45SN1_45S_50ntFlanks | 11208 | 11230 | 23 | GCGGCGACTCTGGACGCGAGCCG | 3095.2 | -6.2 | 2.1 | -2.9 | 3.5E-03 | 4.2E-03 | BT20/MB468 | NUC |
| >hg38_21_+_8433172_8446622_RNA45SN1_45S_50ntFlanks@8346.8365.20\|AGGGAAAGTTGAAAAGAACT | >hg38_21_+_8433172_8446622_RNA45SN1_45S_50ntFlanks | 8346 | 8365 | 20 | AGGGAAAGTTGAAAAGAACT | 143.7 | 4.2 | 1.4 | 3.1 | 1.8E-03 | 4.2E-03 | BT20/MB468 | MP |
| >hg38_21_+_8433172_8446622_RNA45SN1_45S_50ntFlanks@12974.12992.19\|CGTACGTAGCAGAGCAGCT | >hg38_21_+_8433172_8446622_RNA45SN1_45S_50ntFlanks | 12974 | 12992 | 19 | CGTACGTAGCAGAGCAGCT | 114.6 | -4.2 | 1.4 | -3.0 | 2.8E-03 | 4.2E-03 | BT20/MB468 | MITO |
| >hg38_21_+_8433172_8446622_RNA45SN1_45S_50ntFlanks@12420.12457.38\|ATGTCGGCTCTTCCTATCATTGTGAAGCAGAATTCACC | >hg38_21_+_8433172_8446622_RNA45SN1_45S_50ntFlanks | 12420 | 12457 | 38 | ATGTCGGCTCTTCCTATCATTGTGAAGCAGAATTCACC | 762.2 | 2.6 | 0.9 | 2.9 | 3.6E-03 | 4.2E-03 | MB231/MB468 | MITO |
| >hg38_MT_+_598_1651_RNR1_12S_50ntFlanks@974.1004.31\|TAAGTGTACTGGAAAGTGCACTTGGACGAAC | >hg38_MT_+_598_1651_RNR1_12S_50ntFlanks | 974 | 1004 | 31 | TAAGTGTACTGGAAAGTGCACTTGGACGAAC | 136.1 | 4.6 | 1.6 | 2.9 | 3.6E-03 | 4.2E-03 | MB231/MB468 | MITO |
| >hg38_21_+_8433172_8446622_RNA45SN1_45S_50ntFlanks@9569.9610.42\|GTGCAAATCGGTCGTCCGACCTGGGTATAGGGGCGAAAGACT | >hg38_21_+_8433172_8446622_RNA45SN1_45S_50ntFlanks | 9569 | 9610 | 42 | GTGCAAATCGGTCGTCCGACCTGGGTATAGGGGCGAAAGACT | 214.6 | 4.6 | 1.6 | 2.9 | 3.6E-03 | 4.2E-03 | MB231/MB468 | MITO |
| >MI0000542\|hsa-mir-320a&WithFlank&8\|-\|22244960\|22245043@48.70.23(+1C)&[MIMAT0000510&hsa-miR-320a-3p&offsets\|0\|+1(+1C);m-60&8\|-\|22244975\|22244996&offsets\|0\|+1(+1C)]\|AAAAGCTGGGTTGAGAGGGCGAAC | >MI0000542\|hsa-mir-320a&WithFlank&8\|-\|22244960\|22245043 | 48 | 70 | 23(+1C) | AAAAGCTGGGTTGAGAGGGCGAAC | 647.2 | 2.3 | 0.7 | 3.4 | 7.0E-04 | 4.2E-03 | BT20/MB231 | NUC |
| >hg38_1_-_228634819_228635039_RNA5S12_5S_50ntFlanks@133.169.37\|ATGGGAGACCGCCTGGGAATACCGGGTGCTGTAGGCT | >hg38_1_-_228634819_228635039_RNA5S12_5S_50ntFlanks | 133 | 169 | 37 | ATGGGAGACCGCCTGGGAATACCGGGTGCTGTAGGCT | 3806.5 | 3.2 | 1.0 | 3.1 | 1.8E-03 | 4.2E-03 | BT20/MB468 | MP |
| >hg38_MT_+_598_1651_RNR1_12S_50ntFlanks@781.809.29\|CAGAAAACTACGATAGCCCTTATGAAACT | >hg38_MT_+_598_1651_RNR1_12S_50ntFlanks | 781 | 809 | 29 | CAGAAAACTACGATAGCCCTTATGAAACT | 118.3 | 3.7 | 1.3 | 2.9 | 3.6E-03 | 4.2E-03 | MB231/MB468 | MITO |
| >hg38_21_+_8433172_8446622_RNA45SN1_45S_50ntFlanks@7985.8011.27\|GATCAGACGTGGCGACCCGCTGAATTT | >hg38_21_+_8433172_8446622_RNA45SN1_45S_50ntFlanks | 7985 | 8011 | 27 | GATCAGACGTGGCGACCCGCTGAATTT | 269.0 | -4.7 | 1.6 | -2.9 | 3.6E-03 | 4.2E-03 | MB231/MB468 | MITO |
| >hg38_21_+_8433172_8446622_RNA45SN1_45S_50ntFlanks@6759.6807.49\|CTTGCGGCCCCGGGTTCCTCCCGGGGCTACGCCTGTCTGAGCGTCGCTT | >hg38_21_+_8433172_8446622_RNA45SN1_45S_50ntFlanks | 6759 | 6807 | 49 | CTTGCGGCCCCGGGTTCCTCCCGGGGCTACGCCTGTCTGAGCGTCGCTT | 610.2 | 3.4 | 1.0 | 3.3 | 8.5E-04 | 4.2E-03 | BT20/MB231 | MITO |
| >hg38_21_+_8433172_8446622_RNA45SN1_45S_50ntFlanks@11578.11611.34\|TAGAACTGGTGCGGACCAGGGGAATCCGACTGTT | >hg38_21_+_8433172_8446622_RNA45SN1_45S_50ntFlanks | 11578 | 11611 | 34 | TAGAACTGGTGCGGACCAGGGGAATCCGACTGTT | 885.3 | 1.7 | 0.6 | 3.0 | 2.8E-03 | 4.2E-03 | BT20/MB468 | MITO |
| >hg38_21_+_8433172_8446622_RNA45SN1_45S_50ntFlanks@12717.12755.39\|CCTCGGTTGGCCTCGGATAGCCGGTCCCCCGCCTGTCCC | >hg38_21_+_8433172_8446622_RNA45SN1_45S_50ntFlanks | 12717 | 12755 | 39 | CCTCGGTTGGCCTCGGATAGCCGGTCCCCCGCCTGTCCC | 303.1 | -2.1 | 0.7 | -3.0 | 2.8E-03 | 4.2E-03 | BT20/MB468 | MITO |
| >hg38_21_+_8433172_8446622_RNA45SN1_45S_50ntFlanks@6646.6675.30\|TCGTACGACTCTTAGCGGTGGATCACTCGG | >hg38_21_+_8433172_8446622_RNA45SN1_45S_50ntFlanks | 6646 | 6675 | 30 | TCGTACGACTCTTAGCGGTGGATCACTCGG | 5836.3 | 2.3 | 0.8 | 2.9 | 3.6E-03 | 4.2E-03 | MB231/MB468 | NUC |
| >hg38_21_+_8433172_8446622_RNA45SN1_45S_50ntFlanks@4983.5000.18\|CGGACAGGATTGACAGAT | >hg38_21_+_8433172_8446622_RNA45SN1_45S_50ntFlanks | 4983 | 5000 | 18 | CGGACAGGATTGACAGAT | 55.9 | -4.9 | 1.7 | -3.0 | 3.0E-03 | 4.2E-03 | MB231/MB468 | MP |
| >hg38_21_+_8433172_8446622_RNA45SN1_45S_50ntFlanks@12043.12070.28\|CACTGACCCGGTGAGGCGGGGGGGCGAG | >hg38_21_+_8433172_8446622_RNA45SN1_45S_50ntFlanks | 12043 | 12070 | 28 | CACTGACCCGGTGAGGCGGGGGGGCGAG | 132.0 | -4.3 | 1.4 | -3.0 | 2.8E-03 | 4.2E-03 | BT20/MB468 | MITO |
| >hg38_21_+_8433172_8446622_RNA45SN1_45S_50ntFlanks@5125.5158.34\|AGCGGTCGGCGTCCCCCAACTTCTTAGAGGGACA | >hg38_21_+_8433172_8446622_RNA45SN1_45S_50ntFlanks | 5125 | 5158 | 34 | AGCGGTCGGCGTCCCCCAACTTCTTAGAGGGACA | 99.9 | 2.2 | 0.7 | 2.9 | 3.6E-03 | 4.2E-03 | BT20/MB468 | NUC |
| >hg38_21_+_8433172_8446622_RNA45SN1_45S_50ntFlanks@11566.11611.46\|TAGCAGCCGACTTAGAACTGGTGCGGACCAGGGGAATCCGACTGTT | >hg38_21_+_8433172_8446622_RNA45SN1_45S_50ntFlanks | 11566 | 11611 | 46 | TAGCAGCCGACTTAGAACTGGTGCGGACCAGGGGAATCCGACTGTT | 458.0 | 5.8 | 1.9 | 3.1 | 1.8E-03 | 4.2E-03 | BT20/MB468 | MP |
| >trna111_HisGTG_1_-_147774845_147774916@1.32.32,trna118_HisGTG_1_-_145396881_145396952@1.32.32,trna16_HisGTG_1_+_146544773_146544844@1.32.32,trna1_HisGTG_15_+_45493349_45493420@1.32.32,trna21_HisGTG_1_+_147753471_147753542@1.32.32,trna33_HisGTG_6_+_27125906_27125977@1.32.32,trna7_HisGTG_9_-_14433938_14434009@1.32.32,trna8_HisGTG_15_-_45492611_45492682@1.32.32,trna9_HisGTG_15_-_45490804_45490875@1.32.32\|GCCGTGATCGTATAGTGGTTAGTACTCTGCGT | >trna111_HisGTG_1_-_147774845_147774916 | 1 | 32 | 32 | GCCGTGATCGTATAGTGGTTAGTACTCTGCGT | 143.6 | 6.4 | 2.1 | 3.1 | 1.8E-03 | 4.2E-03 | BT20/MB468 | MP |
| >hg38_21_+_8433172_8446622_RNA45SN1_45S_50ntFlanks@9568.9597.30\|CGTGCAAATCGGTCGTCCGACCTGGGTATA | >hg38_21_+_8433172_8446622_RNA45SN1_45S_50ntFlanks | 9568 | 9597 | 30 | CGTGCAAATCGGTCGTCCGACCTGGGTATA | 3137.6 | -5.8 | 1.9 | -3.1 | 1.8E-03 | 4.2E-03 | BT20/MB468 | MP |
| >hg38_21_+_8433172_8446622_RNA45SN1_45S_50ntFlanks@6651.6677.27\|CGACTCTTAGCGGTGGATCACTCGGCT | >hg38_21_+_8433172_8446622_RNA45SN1_45S_50ntFlanks | 6651 | 6677 | 27 | CGACTCTTAGCGGTGGATCACTCGGCT | 2071.3 | -2.2 | 0.7 | -3.0 | 3.0E-03 | 4.2E-03 | MB231/MB468 | MP |
| >hg38_21_+_8433172_8446622_RNA45SN1_45S_50ntFlanks@8528.8551.24\|CTTCCCCCGCCGCCCCTCCTCCTC | >hg38_21_+_8433172_8446622_RNA45SN1_45S_50ntFlanks | 8528 | 8551 | 24 | CTTCCCCCGCCGCCCCTCCTCCTC | 436.6 | -4.1 | 1.4 | -2.9 | 3.3E-03 | 4.2E-03 | BT20/MB468 | CYTO |
| >hg38_21_+_8433172_8446622_RNA45SN1_45S_50ntFlanks@10514.10532.19\|CGGAGTGGCGGAGATGGGC | >hg38_21_+_8433172_8446622_RNA45SN1_45S_50ntFlanks | 10514 | 10532 | 19 | CGGAGTGGCGGAGATGGGC | 1244.6 | 3.9 | 1.3 | 3.1 | 1.8E-03 | 4.2E-03 | BT20/MB468 | MP |
| >hg38_21_+_8433172_8446622_RNA45SN1_45S_50ntFlanks@12157.12176.20\|AGTTTGACTGGGGCGGTACA | >hg38_21_+_8433172_8446622_RNA45SN1_45S_50ntFlanks | 12157 | 12176 | 20 | AGTTTGACTGGGGCGGTACA | 106.1 | -5.0 | 1.6 | -3.1 | 1.8E-03 | 4.2E-03 | BT20/MB468 | MP |
| >hg38_21_+_8433172_8446622_RNA45SN1_45S_50ntFlanks@9933.9956.24\|TAGACAGCAGGACGGTGGCCATGG | >hg38_21_+_8433172_8446622_RNA45SN1_45S_50ntFlanks | 9933 | 9956 | 24 | TAGACAGCAGGACGGTGGCCATGG | 57.0 | 6.1 | 1.9 | 3.1 | 1.8E-03 | 4.2E-03 | BT20/MB468 | MP |
| >hg38_21_+_8433172_8446622_RNA45SN1_45S_50ntFlanks@4128.4156.29\|CGATTCCGGAGAGGGAGCCTGAGAAACGG | >hg38_21_+_8433172_8446622_RNA45SN1_45S_50ntFlanks | 4128 | 4156 | 29 | CGATTCCGGAGAGGGAGCCTGAGAAACGG | 121.2 | -4.7 | 1.6 | -3.0 | 3.1E-03 | 4.2E-03 | MB231/MB468 | MP |
| >hg38_21_+_8433172_8446622_RNA45SN1_45S_50ntFlanks@11767.11794.28\|TCTAATTAGTGACGCGCATGAATGGATG | >hg38_21_+_8433172_8446622_RNA45SN1_45S_50ntFlanks | 11767 | 11794 | 28 | TCTAATTAGTGACGCGCATGAATGGATG | 1358.9 | -1.9 | 0.6 | -2.9 | 3.4E-03 | 4.2E-03 | MB231/MB468 | CYTO |
| >hg38_21_+_8433172_8446622_RNA45SN1_45S_50ntFlanks@11569.11609.41\|CAGCCGACTTAGAACTGGTGCGGACCAGGGGAATCCGACTG | >hg38_21_+_8433172_8446622_RNA45SN1_45S_50ntFlanks | 11569 | 11609 | 41 | CAGCCGACTTAGAACTGGTGCGGACCAGGGGAATCCGACTG | 315.0 | -2.8 | 0.9 | -2.9 | 3.3E-03 | 4.2E-03 | BT20/MB468 | CYTO |
| >trna111_HisGTG_1_-_147774845_147774916@-1G.31.32,trna118_HisGTG_1_-_145396881_145396952@-1G.31.32,trna16_HisGTG_1_+_146544773_146544844@-1G.31.32,trna1_HisGTG_15_+_45493349_45493420@-1G.31.32,trna21_HisGTG_1_+_147753471_147753542@-1G.31.32,trna33_HisGTG_6_+_27125906_27125977@-1G.31.32,trna7_HisGTG_9_-_14433938_14434009@-1G.31.32,trna8_HisGTG_15_-_45492611_45492682@-1G.31.32,trna9_HisGTG_15_-_45490804_45490875@-1G.31.32\|GGCCGTGATCGTATAGTGGTTAGTACTCTGCG | >trna111_HisGTG_1_-_147774845_147774916 | -1G | 31 | 32 | GGCCGTGATCGTATAGTGGTTAGTACTCTGCG | 25851.9 | -4.0 | 1.2 | -3.4 | 7.7E-04 | 4.2E-03 | BT20/MB231 | CYTO |
| >trna5_AspGTC_12_+_98897281_98897352@39.74.36\|GCGGGAGACCGGGGTTCAATTCCCCGACGGGGAGCC | >trna5_AspGTC_12_+_98897281_98897352 | 39 | 74 | 36 | GCGGGAGACCGGGGTTCAATTCCCCGACGGGGAGCC | 430.0 | -3.4 | 1.0 | -3.4 | 7.7E-04 | 4.2E-03 | BT20/MB231 | CYTO |
| >hg38_21_+_8433172_8446622_RNA45SN1_45S_50ntFlanks@8238.8262.25\|CCCGGGTCTTCCCGGAGTCGGGTTG | >hg38_21_+_8433172_8446622_RNA45SN1_45S_50ntFlanks | 8238 | 8262 | 25 | CCCGGGTCTTCCCGGAGTCGGGTTG | 473.9 | -2.3 | 0.8 | -3.0 | 2.8E-03 | 4.2E-03 | BT20/MB468 | MITO |
| >hg38_21_+_8433172_8446622_RNA45SN1_45S_50ntFlanks@12995.13015.21\|CTCGCTGCGATCTATTGAAAG | >hg38_21_+_8433172_8446622_RNA45SN1_45S_50ntFlanks | 12995 | 13015 | 21 | CTCGCTGCGATCTATTGAAAG | 821.0 | 2.7 | 0.9 | 2.9 | 3.3E-03 | 4.2E-03 | BT20/MB468 | CYTO |

<table>
<tr><td>>hg38_21_+_8433172_8446622_RNA45SN1_45S_50ntFlanks@4734.4759.26|AACGAAAGTCGGAGGTTCGAAGACGA</td><td>>hg38_21_+_8433172_8446622_RNA45SN1_45S_50ntFlanks</td><td>4734</td><td>4759</td><td>26</td><td>AACGAAAGTCGGAGGTTCGAAGACGA</td><td>2899.8</td><td>4.2</td><td>1.2</td><td>3.4</td><td>6.4E-04</td><td>4.2E-03</td><td>BT20/MB231</td><td>MP</td></tr>
<tr><td>>hg38_21_+_8433172_8446622_RNA45SN1_45S_50ntFlanks@7980.8007.28|CCTCAGATCAGACGTGGCGACCCGCTGA</td><td>>hg38_21_+_8433172_8446622_RNA45SN1_45S_50ntFlanks</td><td>7980</td><td>8007</td><td>28</td><td>CCTCAGATCAGACGTGGCGACCCGCTGA</td><td>323.7</td><td>-2.8</td><td>0.9</td><td>-3.1</td><td>1.8E-03</td><td>4.2E-03</td><td>BT20/MB468</td><td>MP</td></tr>
<tr><td>>hg38_21_+_8433172_8446622_RNA45SN1_45S_50ntFlanks@12003.12035.33|GCGGGCCGCCGGTGAAATACCACTACTCTGATC</td><td>>hg38_21_+_8433172_8446622_RNA45SN1_45S_50ntFlanks</td><td>12003</td><td>12035</td><td>33</td><td>GCGGGCCGCCGGTGAAATACCACTACTCTGATC</td><td>1236.3</td><td>2.9</td><td>0.9</td><td>3.3</td><td>8.6E-04</td><td>4.2E-03</td><td>BT20/MB231</td><td>MITO</td></tr>
<tr><td>>am_MI0003583|hsa-mir-576&WithFlank&4|+|109488692|109488801@61.83.23&[MIMAT0004796&hsa-miR-576-3p&offsets|0|+1;m-524&4|+|109488752|109488772&offsets|0|+2]|AAGATGTGGAAAAATTGGAATCC</td><td>>am_MI0003583|hsa-mir-576&WithFlank&4|+|109488692|109488801</td><td>61</td><td>83</td><td>23</td><td>AAGATGTGGAAAAATTGGAATCC</td><td>1016.8</td><td>4.9</td><td>1.6</td><td>3.1</td><td>1.8E-03</td><td>4.2E-03</td><td>BT20/MB468</td><td>MP</td></tr>
<tr><td>>hg38_21_+_8433172_8446622_RNA45SN1_45S_50ntFlanks@7977.8015.39|CGACCTCAGATCAGACGTGGCGACCCGCTGAATTTAAGC</td><td>>hg38_21_+_8433172_8446622_RNA45SN1_45S_50ntFlanks</td><td>7977</td><td>8015</td><td>39</td><td>CGACCTCAGATCAGACGTGGCGACCCGCTGAATTTAAGC</td><td>170.2</td><td>2.9</td><td>1.0</td><td>2.9</td><td>3.6E-03</td><td>4.2E-03</td><td>MB231/MB468</td><td>MITO</td></tr>
<tr><td>>hg38_21_+_8433172_8446622_RNA45SN1_45S_50ntFlanks@5478.5516.39|CTGGCGGAGCGCTGAGAAGACGGTCGAACTTGACTATCT</td><td>>hg38_21_+_8433172_8446622_RNA45SN1_45S_50ntFlanks</td><td>5478</td><td>5516</td><td>39</td><td>CTGGCGGAGCGCTGAGAAGACGGTCGAACTTGACTATCT</td><td>384.1</td><td>3.0</td><td>0.9</td><td>3.3</td><td>8.6E-04</td><td>4.2E-03</td><td>BT20/MB231</td><td>MITO</td></tr>
<tr><td>>hg38_21_+_8433172_8446622_RNA45SN1_45S_50ntFlanks@10869.10894.26|AGGGCTGGGTCGGTCGGGCTGGGGCG</td><td>>hg38_21_+_8433172_8446622_RNA45SN1_45S_50ntFlanks</td><td>10869</td><td>10894</td><td>26</td><td>AGGGCTGGGTCGGTCGGGCTGGGGCG</td><td>303.3</td><td>-2.4</td><td>0.8</td><td>-3.0</td><td>2.8E-03</td><td>4.2E-03</td><td>BT20/MB468</td><td>MITO</td></tr>
<tr><td>>MI0000289|hsa-mir-181a-1&WithFlank&1|-|198859038|198859159@30.49.20;MI0000269|hsa-mir-181a-2&WithFlank&9|+|124692436|124692557@45.64.20&[MIMAT0000256&hsa-miR-181a-5p&offsets|0|-3;m-32&1|-|198859108|198859130&offsets|0|-3];[MIMAT0000256_1&hsa-miR-181a-5p&offsets|0|-3;m-32&9|+|124692480|124692502&offsets|0|-3]|AACATTCAACGCTGTCGGTG</td><td>>MI0000289|hsa-mir-181a-1&WithFlank&1|-|198859038|198859159</td><td>30</td><td>49</td><td>20</td><td>AACATTCAACGCTGTCGGTG</td><td>1488.7</td><td>-3.8</td><td>1.1</td><td>-3.4</td><td>6.4E-04</td><td>4.2E-03</td><td>BT20/MB231</td><td>MP</td></tr>
<tr><td>>MI0000816|hsa-mir-335&WithFlank&7|+|130496105|130496210@58.79.22&[MIMAT0004703&hsa-miR-335-3p&offsets|0|0;m-159&7|+|130496162|130496183&offsets|0|0]|TTTTTCATTATTGCTCCTGACC</td><td>>MI0000816|hsa-mir-335&WithFlank&7|+|130496105|130496210</td><td>58</td><td>79</td><td>22</td><td>TTTTTCATTATTGCTCCTGACC</td><td>3330.9</td><td>-2.9</td><td>0.9</td><td>-3.3</td><td>8.7E-04</td><td>4.2E-03</td><td>BT20/MB231</td><td>MITO</td></tr>
<tr><td>>hg38_21_+_8433172_8446622_RNA45SN1_45S_50ntFlanks@12411.12435.25|GATCCTTCGATGTCGGCTCTTCCTA</td><td>>hg38_21_+_8433172_8446622_RNA45SN1_45S_50ntFlanks</td><td>12411</td><td>12435</td><td>25</td><td>GATCCTTCGATGTCGGCTCTTCCTA</td><td>222.0</td><td>-4.6</td><td>1.6</td><td>-2.9</td><td>3.6E-03</td><td>4.2E-03</td><td>MB231/MB468</td><td>NUC</td></tr>
<tr><td>>hg38_21_+_8433172_8446622_RNA45SN1_45S_50ntFlanks@4047.4067.21|ATTCGAACGTCTGCCCTATCA</td><td>>hg38_21_+_8433172_8446622_RNA45SN1_45S_50ntFlanks</td><td>4047</td><td>4067</td><td>21</td><td>ATTCGAACGTCTGCCCTATCA</td><td>153.1</td><td>-4.9</td><td>1.6</td><td>-3.1</td><td>1.9E-03</td><td>4.2E-03</td><td>BT20/MB468</td><td>MP</td></tr>
<tr><td>>hg38_21_+_8433172_8446622_RNA45SN1_45S_50ntFlanks@8528.8551.24|CTTCCCCCGCCGCCCCTCCTCCTC</td><td>>hg38_21_+_8433172_8446622_RNA45SN1_45S_50ntFlanks</td><td>8528</td><td>8551</td><td>24</td><td>CTTCCCCCGCCGCCCCTCCTCCTC</td><td>87.4</td><td>-5.2</td><td>1.7</td><td>-3.1</td><td>1.9E-03</td><td>4.2E-03</td><td>BT20/MB468</td><td>MP</td></tr>
<tr><td>>hg38_21_+_8433172_8446622_RNA45SN1_45S_50ntFlanks@12004.12029.26|CGGGCCGCCGGTGAAATACCACTACT</td><td>>hg38_21_+_8433172_8446622_RNA45SN1_45S_50ntFlanks</td><td>12004</td><td>12029</td><td>26</td><td>CGGGCCGCCGGTGAAATACCACTACT</td><td>350.2</td><td>2.5</td><td>0.8</td><td>3.3</td><td>8.7E-04</td><td>4.2E-03</td><td>BT20/MB231</td><td>MITO</td></tr>
<tr><td>>hg38_21_+_8433172_8446622_RNA45SN1_45S_50ntFlanks@6653.6688.36|ACTCTTAGCGGTGGATCACTCGGCTCGTGCGTCGAT</td><td>>hg38_21_+_8433172_8446622_RNA45SN1_45S_50ntFlanks</td><td>6653</td><td>6688</td><td>36</td><td>ACTCTTAGCGGTGGATCACTCGGCTCGTGCGTCGAT</td><td>1277.0</td><td>-2.2</td><td>0.7</td><td>-2.9</td><td>3.6E-03</td><td>4.2E-03</td><td>MB231/MB468</td><td>MITO</td></tr>
<tr><td>>hg38_21_+_8433172_8446622_RNA45SN1_45S_50ntFlanks@10353.10383.31|AGAACTTTGAAGGCCGAAGTGGAGAAGGGTT</td><td>>hg38_21_+_8433172_8446622_RNA45SN1_45S_50ntFlanks</td><td>10353</td><td>10383</td><td>31</td><td>AGAACTTTGAAGGCCGAAGTGGAGAAGGGTT</td><td>432.3</td><td>-2.7</td><td>0.9</td><td>-3.1</td><td>1.9E-03</td><td>4.2E-03</td><td>BT20/MB468</td><td>MP</td></tr>
<tr><td>>hg38_21_+_8433172_8446622_RNA45SN1_45S_50ntFlanks@3813.3834.22|TTATGGTTCCTTTGGTCGCTCG</td><td>>hg38_21_+_8433172_8446622_RNA45SN1_45S_50ntFlanks</td><td>3813</td><td>3834</td><td>22</td><td>TTATGGTTCCTTTGGTCGCTCG</td><td>108.7</td><td>-4.9</td><td>1.6</td><td>-3.1</td><td>1.9E-03</td><td>4.2E-03</td><td>BT20/MB468</td><td>MP</td></tr>
<tr><td>>hg38_21_+_8433172_8446622_RNA45SN1_45S_50ntFlanks@3837.3868.32|CCTCTCCTACTTGGATAACTGTGGTAATTCTA</td><td>>hg38_21_+_8433172_8446622_RNA45SN1_45S_50ntFlanks</td><td>3837</td><td>3868</td><td>32</td><td>CCTCTCCTACTTGGATAACTGTGGTAATTCTA</td><td>181.5</td><td>-2.3</td><td>0.8</td><td>-3.0</td><td>2.9E-03</td><td>4.2E-03</td><td>BT20/MB468</td><td>MITO</td></tr>
<tr><td>>hg38_21_+_8433172_8446622_RNA45SN1_45S_50ntFlanks@11660.11692.33|GTGATTTCTGCCCAGTGCTCTGAATGTCAAAGT</td><td>>hg38_21_+_8433172_8446622_RNA45SN1_45S_50ntFlanks</td><td>11660</td><td>11692</td><td>33</td><td>GTGATTTCTGCCCAGTGCTCTGAATGTCAAAGT</td><td>77.4</td><td>9.6</td><td>2.8</td><td>3.4</td><td>6.5E-04</td><td>4.2E-03</td><td>BT20/MB231</td><td>MP</td></tr>
<tr><td>>hg38_21_+_8433172_8446622_RNA45SN1_45S_50ntFlanks@8242.8269.28|GGTCTTCCCGGAGTCGGGTTGCTTGGGA</td><td>>hg38_21_+_8433172_8446622_RNA45SN1_45S_50ntFlanks</td><td>8242</td><td>8269</td><td>28</td><td>GGTCTTCCCGGAGTCGGGTTGCTTGGGA</td><td>292.6</td><td>6.7</td><td>2.0</td><td>3.4</td><td>6.5E-04</td><td>4.2E-03</td><td>BT20/MB231</td><td>MP</td></tr>
<tr><td>>MI0000095|hsa-mir-93&WithFlank&7|-|100093762|100093853@17.39.23(+1C)&[MIMAT0000093&hsa-miR-93-5p&offsets|0|0(+1C);m-29&7|-|100093815|100093837&offsets|0|0(+1C)]|CAAAGTGCTGTTCGTGCAGGTAGC</td><td>>MI0000095|hsa-mir-93&WithFlank&7|-|100093762|100093853</td><td>17</td><td>39</td><td>23(+1C)</td><td>CAAAGTGCTGTTCGTGCAGGTAGC</td><td>303.5</td><td>-3.8</td><td>1.1</td><td>-3.4</td><td>6.5E-04</td><td>4.2E-03</td><td>BT20/MB231</td><td>MP</td></tr>
<tr><td>>hg38_21_+_8433172_8446622_RNA45SN1_45S_50ntFlanks@11768.11804.37|CTAATTAGTGACGCGCATGAATGGATGAACGAGATTC</td><td>>hg38_21_+_8433172_8446622_RNA45SN1_45S_50ntFlanks</td><td>11768</td><td>11804</td><td>37</td><td>CTAATTAGTGACGCGCATGAATGGATGAACGAGATTC</td><td>472.5</td><td>-4.3</td><td>1.4</td><td>-3.1</td><td>1.9E-03</td><td>4.2E-03</td><td>BT20/MB468</td><td>MP</td></tr>
<tr><td>>MI0000263|hsa-mir-7-1&WithFlank&9|-|83969742|83969863@30.55.26&[MIMAT0000252_2&hsa-miR-7-5p&offsets|0|+2;m-401&9|-|83969811|83969834&offsets|0|+2]|TGGAAGACTAGTGATTTTGTTGTTTT</td><td>>MI0000263|hsa-mir-7-1&WithFlank&9|-|83969742|83969863</td><td>30</td><td>55</td><td>26</td><td>TGGAAGACTAGTGATTTTGTTGTTTT</td><td>122.3</td><td>4.7</td><td>1.6</td><td>3.0</td><td>3.1E-03</td><td>4.2E-03</td><td>MB231/MB468</td><td>MP</td></tr>
<tr><td>>hg38_21_+_8433172_8446622_RNA45SN1_45S_50ntFlanks@9932.9960.29|ATAGACAGCAGGACGGTGGCCATGGAAGT</td><td>>hg38_21_+_8433172_8446622_RNA45SN1_45S_50ntFlanks</td><td>9932</td><td>9960</td><td>29</td><td>ATAGACAGCAGGACGGTGGCCATGGAAGT</td><td>83.9</td><td>6.1</td><td>2.0</td><td>3.1</td><td>1.9E-03</td><td>4.2E-03</td><td>BT20/MB468</td><td>MP</td></tr>
<tr><td>>trnaMT_GluTTC_MT_-_14674_14742@46.71.26,trnalookalike8_GluTTC_5_-_93905172_93905240@46.71.26|GTGGTTGTAGTCCGTGCGAGAATACC</td><td>>trnaMT_GluTTC_MT_-_14674_14742</td><td>46</td><td>71</td><td>26</td><td>GTGGTTGTAGTCCGTGCGAGAATACC</td><td>258.2</td><td>3.7</td><td>1.2</td><td>3.1</td><td>1.9E-03</td><td>4.2E-03</td><td>BT20/MB468</td><td>MP</td></tr>
<tr><td>>hg38_21_+_8433172_8446622_RNA45SN1_45S_50ntFlanks@11581.11612.32|AACTGGTGCGGACCAGGGGAATCCGACTGTTT</td><td>>hg38_21_+_8433172_8446622_RNA45SN1_45S_50ntFlanks</td><td>11581</td><td>11612</td><td>32</td><td>AACTGGTGCGGACCAGGGGAATCCGACTGTTT</td><td>2019.2</td><td>-3.1</td><td>1.0</td><td>-3.1</td><td>1.9E-03</td><td>4.2E-03</td><td>BT20/MB468</td><td>MP</td></tr>
<tr><td>>MI0000063|hsa-let-7b&WithFlank&22|+|46113680|46113774@12.33.22(+2A)&[MIMAT0000063&hsa-let-7b-5p&offsets|0|0(+2A);m-14&22|+|46113691|46113712&offsets|0|0(+2A)]|TGAGGTAGTAGGTTGTGTGGTTAA</td><td>>MI0000063|hsa-let-7b&WithFlank&22|+|46113680|46113774</td><td>12</td><td>33</td><td>22(+2A)</td><td>TGAGGTAGTAGGTTGTGTGGTTAA</td><td>338.8</td><td>2.6</td><td>0.8</td><td>3.4</td><td>7.8E-04</td><td>4.2E-03</td><td>BT20/MB231</td><td>CYTO</td></tr>
<tr><td>>MI0000103|hsa-mir-101-1&WithFlank&1|-|65058428|65058514@52.72.21;MI0000739|hsa-mir-101-2&WithFlank&9|+|4850291|4850381@54.74.21&[MIMAT0000099&hsa-miR-101-3p&offsets|-1|-1;m-17&1|-|65058443|65058462&offsets|-1|0];[MIMAT0000099_1&hsa-miR-101-3p&offsets|-1|-1;m-16&9|+|4850345|4850364&offsets|-1|0]|GTACAGTACTGTGATAACTGA</td><td>>MI0000103|hsa-mir-101-1&WithFlank&1|-|65058428|65058514</td><td>52</td><td>72</td><td>21</td><td>GTACAGTACTGTGATAACTGA</td><td>233.6</td><td>2.6</td><td>0.8</td><td>3.4</td><td>7.8E-04</td><td>4.2E-03</td><td>BT20/MB231</td><td>CYTO</td></tr>
<tr><td>>hg38_21_+_8433172_8446622_RNA45SN1_45S_50ntFlanks@12488.12520.33|AACGTGAGCTGGGTTTAGACCGTCGTGAGACAG</td><td>>hg38_21_+_8433172_8446622_RNA45SN1_45S_50ntFlanks</td><td>12488</td><td>12520</td><td>33</td><td>AACGTGAGCTGGGTTTAGACCGTCGTGAGACAG</td><td>211.6</td><td>-4.4</td><td>1.5</td><td>-3.0</td><td>2.9E-03</td><td>4.2E-03</td><td>BT20/MB468</td><td>MITO</td></tr>
<tr><td>>hg38_21_+_8433172_8446622_RNA45SN1_45S_50ntFlanks@12408.12436.29|TTTGATCCTTCGATGTCGGCTCTTCCTAT</td><td>>hg38_21_+_8433172_8446622_RNA45SN1_45S_50ntFlanks</td><td>12408</td><td>12436</td><td>29</td><td>TTTGATCCTTCGATGTCGGCTCTTCCTAT</td><td>135.9</td><td>-3.1</td><td>1.0</td><td>-3.0</td><td>2.9E-03</td><td>4.2E-03</td><td>BT20/MB468</td><td>MITO</td></tr>
<tr><td>>MI0000234|hsa-mir-192&WithFlank&11|-|64891131|64891252@30.51.22(+1U)&[MIMAT0000222&hsa-miR-192-5p&offsets|0|+1(+1U);m-59&11|-|64891203|64891223&offsets|0|+1(+1U)]|CTGACCTATGAATTGACAGCCAT</td><td>>MI0000234|hsa-mir-192&WithFlank&11|-|64891131|64891252</td><td>30</td><td>51</td><td>22(+1U)</td><td>CTGACCTATGAATTGACAGCCAT</td><td>1017.5</td><td>2.0</td><td>0.7</td><td>3.0</td><td>3.1E-03</td><td>4.2E-03</td><td>MB231/MB468</td><td>MP</td></tr>
<tr><td>>trna14_LysTTT_11_-_59327808_59327880@1.33.33,trna2_LysTTT_17_+_8022473_8022545@1.33.33,trna54_LysTTT_1_+_204475655_204475727@1.33.33,trna5_LysTTT_11_+_59323902_59323974@1.33.33,trna62_LysTTT_1_-_204476158_204476230@1.33.33,trna76_LysTTT_6_+_28918806_28918878@1.33.33|GCCCGGATAGCTCAGTCGGTAGAGCATCAGACT</td><td>>trna14_LysTTT_11_-_59327808_59327880</td><td>1</td><td>33</td><td>33</td><td>GCCCGGATAGCTCAGTCGGTAGAGCATCAGACT</td><td>12089.2</td><td>-3.4</td><td>1.0</td><td>-3.3</td><td>8.7E-04</td><td>4.2E-03</td><td>BT20/MB231</td><td>MITO</td></tr>
<tr><td>>trna119_LysCTT_1_-_145395522_145395594@1.32.32,trna11_LysCTT_5_-_180648979_180649051@1.32.32,trna13_LysCTT_6_+_26556774_26556846@1.32.32,trna32_LysCTT_16_-_3207406_3207478@1.32.32,trna7_LysCTT_16_+_3225692_3225764@1.32.32,trna9_LysCTT_5_+_180634755_180634827@1.32.32|GCCCGGCTAGCTCAGTCGGTAGAGCATGAGAC</td><td>>trna119_LysCTT_1_-_145395522_145395594</td><td>1</td><td>32</td><td>32</td><td>GCCCGGCTAGCTCAGTCGGTAGAGCATGAGAC</td><td>916.7</td><td>-3.3</td><td>1.0</td><td>-3.4</td><td>7.2E-04</td><td>4.2E-03</td><td>BT20/MB231</td><td>NUC</td></tr>
<tr><td>>hg38_21_+_8433172_8446622_RNA45SN1_45S_50ntFlanks@12419.12457.39|GATGTCGGCTCTTCCTATCATTGTGAAGCAGAATTCACC</td><td>>hg38_21_+_8433172_8446622_RNA45SN1_45S_50ntFlanks</td><td>12419</td><td>12457</td><td>39</td><td>GATGTCGGCTCTTCCTATCATTGTGAAGCAGAATTCACC</td><td>715.5</td><td>-2.9</td><td>0.9</td><td>-3.4</td><td>7.2E-04</td><td>4.2E-03</td><td>BT20/MB231</td><td>NUC</td></tr>
<tr><td>>hg38_21_+_8433172_8446622_RNA45SN1_45S_50ntFlanks@12162.12179.18|GACTGGGGCGGTACACCT</td><td>>hg38_21_+_8433172_8446622_RNA45SN1_45S_50ntFlanks</td><td>12162</td><td>12179</td><td>18</td><td>GACTGGGGCGGTACACCT</td><td>631.5</td><td>-5.5</td><td>1.8</td><td>-3.1</td><td>1.9E-03</td><td>4.2E-03</td><td>BT20/MB468</td><td>MP</td></tr>
<tr><td>>hg38_21_+_8433172_8446622_RNA45SN1_45S_50ntFlanks@12202.12240.39|CTAAGGCGAGCTCAGGGAGGACAGAAACCTCCCGTGGAG</td><td>>hg38_21_+_8433172_8446622_RNA45SN1_45S_50ntFlanks</td><td>12202</td><td>12240</td><td>39</td><td>CTAAGGCGAGCTCAGGGAGGACAGAAACCTCCCGTGGAG</td><td>104.1</td><td>-5.8</td><td>1.9</td><td>-3.1</td><td>1.9E-03</td><td>4.3E-03</td><td>BT20/MB468</td><td>MP</td></tr>
<tr><td>>hg38_MT_+_1621_3279_RNR2_16S_50ntFlanks@732.749.18|TAAAACACTGAACTGACA</td><td>>hg38_MT_+_1621_3279_RNR2_16S_50ntFlanks</td><td>732</td><td>749</td><td>18</td><td>TAAAACACTGAACTGACA</td><td>32.9</td><td>8.3</td><td>2.8</td><td>3.0</td><td>3.1E-03</td><td>4.3E-03</td><td>MB231/MB468</td><td>MP</td></tr>
<tr><td>>trnaMT_ValTAC_MT_+_1602_1670@35.72.38|ACTTAGGAGATTTCAACTTAACTTGACCGCTCTGACCA</td><td>>trnaMT_ValTAC_MT_+_1602_1670</td><td>35</td><td>72</td><td>38</td><td>ACTTAGGAGATTTCAACTTAACTTGACCGCTCTGACCA</td><td>378.6</td><td>5.0</td><td>1.6</td><td>3.1</td><td>1.9E-03</td><td>4.3E-03</td><td>BT20/MB468</td><td>MP</td></tr>
<tr><td>>hg38_21_+_8433172_8446622_RNA45SN1_45S_50ntFlanks@11913.11947.35|AAGAGACATGAGAGGTGTAGAATAAGTGGGAGGCC</td><td>>hg38_21_+_8433172_8446622_RNA45SN1_45S_50ntFlanks</td><td>11913</td><td>11947</td><td>35</td><td>AAGAGACATGAGAGGTGTAGAATAAGTGGGAGGCC</td><td>33.5</td><td>8.8</td><td>2.8</td><td>3.1</td><td>1.9E-03</td><td>4.3E-03</td><td>BT20/MB468</td><td>MP</td></tr>
<tr><td>>hg38_21_+_8433172_8446622_RNA45SN1_45S_50ntFlanks@11913.11946.34|AAGAGACATGAGAGGTGTAGAATAAGTGGGAGGC</td><td>>hg38_21_+_8433172_8446622_RNA45SN1_45S_50ntFlanks</td><td>11913</td><td>11946</td><td>34</td><td>AAGAGACATGAGAGGTGTAGAATAAGTGGGAGGC</td><td>33.6</td><td>8.8</td><td>2.8</td><td>3.1</td><td>1.9E-03</td><td>4.3E-03</td><td>BT20/MB468</td><td>MP</td></tr>
<tr><td>>MI0000070|hsa-mir-16-1&WithFlank&13|-|50048967|50049067@20.42.23;MI0000115|hsa-mir-16-2&WithFlank&3|+|160404739|160404831@16.38.23&[MIMAT0000069&hsa-miR-16-5p&offsets|0|+1;m-62&13|-|50049027|50049048&offsets|0|+1];[MIMAT0000069_1&hsa-miR-16-5p&offsets|0|+1;m-61&3|+|160404754|160404775&offsets|0|+1]|TAGCAGCACGTAAATATTGGCGT</td><td>>MI0000070|hsa-mir-16-1&WithFlank&13|-|50048967|50049067</td><td>20</td><td>42</td><td>23</td><td>TAGCAGCACGTAAATATTGGCGT</td><td>3727.9</td><td>2.5</td><td>0.8</td><td>3.0</td><td>3.1E-03</td><td>4.3E-03</td><td>MB231/MB468</td><td>MP</td></tr>
<tr><td>>hg38_21_+_8433172_8446622_RNA45SN1_45S_50ntFlanks@7975.8018.44|CGCGACCTCAGATCAGACGTGGCGACCCGCTGAATTTAAGCATA</td><td>>hg38_21_+_8433172_8446622_RNA45SN1_45S_50ntFlanks</td><td>7975</td><td>8018</td><td>44</td><td>CGCGACCTCAGATCAGACGTGGCGACCCGCTGAATTTAAGCATA</td><td>24918.8</td><td>3.1</td><td>1.1</td><td>2.9</td><td>3.6E-03</td><td>4.3E-03</td><td>MB231/MB468</td><td>MITO</td></tr>
<tr><td>>hg38_21_+_8433172_8446622_RNA45SN1_45S_50ntFlanks@12628.12661.34|TGGGGCGAAGCTACCATCTGTGGGATTATGACTG</td><td>>hg38_21_+_8433172_8446622_RNA45SN1_45S_50ntFlanks</td><td>12628</td><td>12661</td><td>34</td><td>TGGGGCGAAGCTACCATCTGTGGGATTATGACTG</td><td>226.7</td><td>-2.8</td><td>1.0</td><td>-2.9</td><td>3.5E-03</td><td>4.3E-03</td><td>MB231/MB468</td><td>CYTO</td></tr>
<tr><td>>hg38_21_+_8433172_8446622_RNA45SN1_45S_50ntFlanks@8915.8942.28|CGAATCCCGGGGCCGAGGGAGCGAGACC</td><td>>hg38_21_+_8433172_8446622_RNA45SN1_45S_50ntFlanks</td><td>8915</td><td>8942</td><td>28</td><td>CGAATCCCGGGGCCGAGGGAGCGAGACC</td><td>100.8</td><td>5.3</td><td>1.7</td><td>3.1</td><td>1.9E-03</td><td>4.3E-03</td><td>BT20/MB468</td><td>MP</td></tr>
<tr><td>>hg38_21_+_8433172_8446622_RNA45SN1_45S_50ntFlanks@9534.9559.26|AGGAAACTCTGGTGGAGGTCCGTAGC</td><td>>hg38_21_+_8433172_8446622_RNA45SN1_45S_50ntFlanks</td><td>9534</td><td>9559</td><td>26</td><td>AGGAAACTCTGGTGGAGGTCCGTAGC</td><td>235.6</td><td>-1.9</td><td>0.6</td><td>-2.9</td><td>3.4E-03</td><td>4.3E-03</td><td>BT20/MB468</td><td>CYTO</td></tr>
<tr><td>>hg38_21_+_8433172_8446622_RNA45SN1_45S_50ntFlanks@6776.6797.22|CTCCCGGGGCTACGCCTGTCTG</td><td>>hg38_21_+_8433172_8446622_RNA45SN1_45S_50ntFlanks</td><td>6776</td><td>6797</td><td>22</td><td>CTCCCGGGGCTACGCCTGTCTG</td><td>64.4</td><td>-5.0</td><td>1.6</td><td>-3.1</td><td>1.9E-03</td><td>4.3E-03</td><td>BT20/MB468</td><td>MP</td></tr>
</table>

<table>
<tr><th></th><th></th><th></th><th></th><th></th><th></th><th></th><th></th><th></th><th></th><th></th><th></th><th></th><th></th></tr>
<tr><td>>hg38_21_+_8433172_8446622_RNA45SN1_45S_50ntFlanks@10353.10384.32|AGAACTTTGAAGGCCGAAGTGGAGAAGGGTTC</td><td>>hg38_21_+_8433172_8446622_RNA45SN1_45S_50ntFlanks</td><td>10353</td><td>10384</td><td>32</td><td>AGAACTTTGAAGGCCGAAGTGGAGAAGGGTTC</td><td>38.4</td><td>-8.3</td><td>2.7</td><td>-3.1</td><td>1.9E-03</td><td>4.3E-03</td><td>BT20/MB468</td><td>MP</td></tr>
<tr><td>>trnaMT_SerGCT_MT_+_12207_12265@14.48.35|AGAACTGCTAACTCATGCCCCCATGTCTAACAACA</td><td>>trnaMT_SerGCT_MT_+_12207_12265</td><td>14</td><td>48</td><td>35</td><td>AGAACTGCTAACTCATGCCCCCATGTCTAACAACA</td><td>252.1</td><td>3.9</td><td>1.3</td><td>3.0</td><td>3.1E-03</td><td>4.3E-03</td><td>MB231/MB468</td><td>MP</td></tr>
<tr><td>>hg38_21_+_8433172_8446622_RNA45SN1_45S_50ntFlanks@10393.10424.32|CAGCAGTTGAACATGGGTCAGTCGGTCCTGAG</td><td>>hg38_21_+_8433172_8446622_RNA45SN1_45S_50ntFlanks</td><td>10393</td><td>10424</td><td>32</td><td>CAGCAGTTGAACATGGGTCAGTCGGTCCTGAG</td><td>10803.0</td><td>3.1</td><td>0.9</td><td>3.3</td><td>8.8E-04</td><td>4.3E-03</td><td>BT20/MB231</td><td>MITO</td></tr>
<tr><td>>hg38_21_+_8433172_8446622_RNA45SN1_45S_50ntFlanks@12971.13010.40|TTTCGTACGTAGCAGAGCAGCTCCCTCGCTGCGATCTATT</td><td>>hg38_21_+_8433172_8446622_RNA45SN1_45S_50ntFlanks</td><td>12971</td><td>13010</td><td>40</td><td>TTTCGTACGTAGCAGAGCAGCTCCCTCGCTGCGATCTATT</td><td>296.0</td><td>3.9</td><td>1.2</td><td>3.3</td><td>8.8E-04</td><td>4.3E-03</td><td>BT20/MB231</td><td>MITO</td></tr>
<tr><td>>hg38_21_+_8433172_8446622_RNA45SN1_45S_50ntFlanks@8174.8216.43|TCCTTCTGATCGAGGCCCAGCCCGTGGACGGTGTGAGGCCGGT</td><td>>hg38_21_+_8433172_8446622_RNA45SN1_45S_50ntFlanks</td><td>8174</td><td>8216</td><td>43</td><td>TCCTTCTGATCGAGGCCCAGCCCGTGGACGGTGTGAGGCCGGT</td><td>116.3</td><td>6.0</td><td>1.9</td><td>3.1</td><td>1.9E-03</td><td>4.3E-03</td><td>BT20/MB468</td><td>MP</td></tr>
<tr><td>>hg38_21_+_8433172_8446622_RNA45SN1_45S_50ntFlanks@12418.12440.23|CGATGTCGGCTCTTCCTATCATT</td><td>>hg38_21_+_8433172_8446622_RNA45SN1_45S_50ntFlanks</td><td>12418</td><td>12440</td><td>23</td><td>CGATGTCGGCTCTTCCTATCATT</td><td>615.4</td><td>-2.4</td><td>0.8</td><td>-2.9</td><td>3.7E-03</td><td>4.3E-03</td><td>MB231/MB468</td><td>MITO</td></tr>
<tr><td>>hg38_21_+_8433172_8446622_RNA45SN1_45S_50ntFlanks@9482.9520.39|ACGTGTTAGGACCCGAAAGATGGTGAACTATGCCTGGGC</td><td>>hg38_21_+_8433172_8446622_RNA45SN1_45S_50ntFlanks</td><td>9482</td><td>9520</td><td>39</td><td>ACGTGTTAGGACCCGAAAGATGGTGAACTATGCCTGGGC</td><td>74.7</td><td>9.6</td><td>2.8</td><td>3.4</td><td>6.5E-04</td><td>4.3E-03</td><td>BT20/MB231</td><td>MP</td></tr>
<tr><td>>hg38_MT_+_1621_3279_RNR2_16S_50ntFlanks@1367.1398.32|TCGATGTTGGATCAGGACATCCCGATGGTGCA</td><td>>hg38_MT_+_1621_3279_RNR2_16S_50ntFlanks</td><td>1367</td><td>1398</td><td>32</td><td>TCGATGTTGGATCAGGACATCCCGATGGTGCA</td><td>140.3</td><td>6.2</td><td>2.0</td><td>3.1</td><td>1.9E-03</td><td>4.3E-03</td><td>BT20/MB468</td><td>MP</td></tr>
<tr><td>>hg38_21_+_8433172_8446622_RNA45SN1_45S_50ntFlanks@7977.7994.18|CGACCTCAGATCAGACGT</td><td>>hg38_21_+_8433172_8446622_RNA45SN1_45S_50ntFlanks</td><td>7977</td><td>7994</td><td>18</td><td>CGACCTCAGATCAGACGT</td><td>296.7</td><td>-1.9</td><td>0.6</td><td>-2.9</td><td>3.7E-03</td><td>4.3E-03</td><td>MB231/MB468</td><td>NUC</td></tr>
<tr><td>>trna116_GluCTC_1_-_145399233_145399304@32.73.42,trna71_GluCTC_1_-_161439189_161439260@32.73.42,trna74_GluCTC_1_-_161431809_161431880@32.73.42,trna77_GluCTC_1_-_161424398_161424469@32.73.42,trna77_GluCTC_6_+_28949976_28950047@32.73.42,trna80_GluCTC_1_-_161417018_161417089@32.73.42,trna87_GluCTC_6_-_126101393_126101464@32.73.42|CTCTCACCGCCGCGGCCCGGGTTCGATTCCCGGTCAGGGAAC</td><td>>trna116_GluCTC_1_-_145399233_145399304</td><td>32</td><td>73</td><td>42</td><td>CTCTCACCGCCGCGGCCCGGGTTCGATTCCCGGTCAGGGAAC</td><td>1380.7</td><td>-5.2</td><td>1.8</td><td>-2.9</td><td>3.4E-03</td><td>4.3E-03</td><td>BT20/MB468</td><td>CYTO</td></tr>
<tr><td>>hg38_21_+_8433172_8446622_RNA45SN1_45S_50ntFlanks@9545.9573.29|GTGGAGGTCCGTAGCGGTCCTGACGTGCA</td><td>>hg38_21_+_8433172_8446622_RNA45SN1_45S_50ntFlanks</td><td>9545</td><td>9573</td><td>29</td><td>GTGGAGGTCCGTAGCGGTCCTGACGTGCA</td><td>2051.7</td><td>-1.8</td><td>0.6</td><td>-2.9</td><td>3.5E-03</td><td>4.3E-03</td><td>MB231/MB468</td><td>CYTO</td></tr>
<tr><td>>MI0000815|hsa-mir-339&WithFlank&7|-|1022927|1023032@56.75.20(+1A)&[MIMAT0004702&hsa-miR-339-3p&offsets|0|-3(+1A);m-235&7|-|1022957|1022977&offsets|0|-1(+1A)]|TGAGCGCCTCGACGACAGAGA</td><td>>MI0000815|hsa-mir-339&WithFlank&7|-|1022927|1023032</td><td>56</td><td>75</td><td>20(+1A)</td><td>TGAGCGCCTCGACGACAGAGA</td><td>72.4</td><td>6.3</td><td>2.0</td><td>3.1</td><td>1.9E-03</td><td>4.3E-03</td><td>BT20/MB468</td><td>MP</td></tr>
<tr><td>>am_trnaMT_LysTTT_MT_+_8295_8364@34.52.19,trnalookalike7_LysTTT_1_+_568844_568913@34.52.19|GTTAAAGATTAAGAGAACC</td><td>>am_trnaMT_LysTTT_MT_+_8295_8364</td><td>34</td><td>52</td><td>19</td><td>GTTAAAGATTAAGAGAACC</td><td>137.3</td><td>3.7</td><td>1.3</td><td>3.0</td><td>3.1E-03</td><td>4.3E-03</td><td>MB231/MB468</td><td>MP</td></tr>
<tr><td>>hg38_21_+_8433172_8446622_RNA45SN1_45S_50ntFlanks@8154.8181.28|CCGCTCGTGGGGGGCCCAAGTCCTTCTG</td><td>>hg38_21_+_8433172_8446622_RNA45SN1_45S_50ntFlanks</td><td>8154</td><td>8181</td><td>28</td><td>CCGCTCGTGGGGGGCCCAAGTCCTTCTG</td><td>281.7</td><td>-3.0</td><td>1.0</td><td>-2.9</td><td>3.5E-03</td><td>4.3E-03</td><td>MB231/MB468</td><td>CYTO</td></tr>
<tr><td>>MI0000650|hsa-mir-200c&WithFlank&12|+|6963693|6963772@50.69.20(+1U)&[MIMAT0000617&hsa-miR-200c-3p&offsets|0|-3(+1U);m-54&12|+|6963742|6963764&offsets|0|-3(+1U)]|TAATACTGCCGGGTAATGATT</td><td>>MI0000650|hsa-mir-200c&WithFlank&12|+|6963693|6963772</td><td>50</td><td>69</td><td>20(+1U)</td><td>TAATACTGCCGGGTAATGATT</td><td>58.5</td><td>6.2</td><td>2.0</td><td>3.1</td><td>1.9E-03</td><td>4.3E-03</td><td>BT20/MB468</td><td>MP</td></tr>
<tr><td>>hg38_21_+_8433172_8446622_RNA45SN1_45S_50ntFlanks@4353.4371.19|TATTAAAGTTGCTGCAGTT</td><td>>hg38_21_+_8433172_8446622_RNA45SN1_45S_50ntFlanks</td><td>4353</td><td>4371</td><td>19</td><td>TATTAAAGTTGCTGCAGTT</td><td>123.7</td><td>-2.6</td><td>0.8</td><td>-3.1</td><td>1.9E-03</td><td>4.3E-03</td><td>BT20/MB468</td><td>MP</td></tr>
<tr><td>>hg38_21_+_8433172_8446622_RNA45SN1_45S_50ntFlanks@11027.11061.35|CTCTCTCTCTCTCTCCCCCGCTCCCCGTCCTCCCC</td><td>>hg38_21_+_8433172_8446622_RNA45SN1_45S_50ntFlanks</td><td>11027</td><td>11061</td><td>35</td><td>CTCTCTCTCTCTCTCCCCCGCTCCCCGTCCTCCCC</td><td>335.8</td><td>-3.4</td><td>1.2</td><td>-2.9</td><td>3.4E-03</td><td>4.3E-03</td><td>BT20/MB468</td><td>CYTO</td></tr>
<tr><td>>hg38_21_+_8433172_8446622_RNA45SN1_45S_50ntFlanks@12416.12446.31|TTCGATGTCGGCTCTTCCTATCATTGTGAAG</td><td>>hg38_21_+_8433172_8446622_RNA45SN1_45S_50ntFlanks</td><td>12416</td><td>12446</td><td>31</td><td>TTCGATGTCGGCTCTTCCTATCATTGTGAAG</td><td>649.2</td><td>-4.4</td><td>1.5</td><td>-2.9</td><td>3.6E-03</td><td>4.3E-03</td><td>BT20/MB468</td><td>NUC</td></tr>
<tr><td>>trnaMT_ValTAC_MT_+_1602_1670@44.72.29|ATTTCAACTTAACTTGACCGCTCTGACCA</td><td>>trnaMT_ValTAC_MT_+_1602_1670</td><td>44</td><td>72</td><td>29</td><td>ATTTCAACTTAACTTGACCGCTCTGACCA</td><td>5611.2</td><td>3.1</td><td>1.1</td><td>2.9</td><td>3.7E-03</td><td>4.3E-03</td><td>MB231/MB468</td><td>MITO</td></tr>
<tr><td>>hg38_21_+_8433172_8446622_RNA45SN1_45S_50ntFlanks@12417.12452.36|TCGATGTCGGCTCTTCCTATCATTGTGAAGCAGAAT</td><td>>hg38_21_+_8433172_8446622_RNA45SN1_45S_50ntFlanks</td><td>12417</td><td>12452</td><td>36</td><td>TCGATGTCGGCTCTTCCTATCATTGTGAAGCAGAAT</td><td>394.4</td><td>-2.6</td><td>0.8</td><td>-3.4</td><td>7.9E-04</td><td>4.3E-03</td><td>BT20/MB231</td><td>CYTO</td></tr>
<tr><td>>MI0000253|hsa-mir-148a&WithFlank&7|-|25949913|25949992@50.70.21(+1G)&[MIMAT0000243&hsa-miR-148a-3p&offsets|0|-1(+1G);m-7&7|-|25949922|25949943&offsets|0|-1(+1G)]|TCAGTGCACTACAGAACTTTGG</td><td>>MI0000253|hsa-mir-148a&WithFlank&7|-|25949913|25949992</td><td>50</td><td>70</td><td>21(+1G)</td><td>TCAGTGCACTACAGAACTTTGG</td><td>9978.7</td><td>4.1</td><td>1.3</td><td>3.1</td><td>1.9E-03</td><td>4.3E-03</td><td>BT20/MB468</td><td>MP</td></tr>
<tr><td>>hg38_21_+_8433172_8446622_RNA45SN1_45S_50ntFlanks@11920.11951.32|ATGAGAGGTGTAGAATAAGTGGGAGGCCCCCG</td><td>>hg38_21_+_8433172_8446622_RNA45SN1_45S_50ntFlanks</td><td>11920</td><td>11951</td><td>32</td><td>ATGAGAGGTGTAGAATAAGTGGGAGGCCCCCG</td><td>583.3</td><td>-4.1</td><td>1.3</td><td>-3.1</td><td>1.9E-03</td><td>4.3E-03</td><td>BT20/MB468</td><td>MP</td></tr>
<tr><td>>hg38_21_+_8433172_8446622_RNA45SN1_45S_50ntFlanks@9551.9571.21|GTCCGTAGCGGTCCTGACGTG</td><td>>hg38_21_+_8433172_8446622_RNA45SN1_45S_50ntFlanks</td><td>9551</td><td>9571</td><td>21</td><td>GTCCGTAGCGGTCCTGACGTG</td><td>403.7</td><td>-4.1</td><td>1.4</td><td>-2.9</td><td>3.7E-03</td><td>4.3E-03</td><td>BT20/MB468</td><td>NUC</td></tr>
<tr><td>>MI0000269|hsa-mir-181a-2&WithFlank&9|+|124692436|124692557@83.103.21&[MIMAT0004558&hsa-miR-181a-2-3p&offsets|0|-1;m-71&9|+|124692518|124692539&offsets|0|-1]|ACCACTGACCGTTGACTGTAC</td><td>>MI0000269|hsa-mir-181a-2&WithFlank&9|+|124692436|124692557</td><td>83</td><td>103</td><td>21</td><td>ACCACTGACCGTTGACTGTAC</td><td>139.6</td><td>3.7</td><td>1.2</td><td>3.0</td><td>3.1E-03</td><td>4.3E-03</td><td>MB231/MB468</td><td>MP</td></tr>
<tr><td>>hg38_21_+_8433172_8446622_RNA45SN1_45S_50ntFlanks@10351.10378.28|CGAGAACTTTGAAGGCCGAAGTGGAGAA</td><td>>hg38_21_+_8433172_8446622_RNA45SN1_45S_50ntFlanks</td><td>10351</td><td>10378</td><td>28</td><td>CGAGAACTTTGAAGGCCGAAGTGGAGAA</td><td>2501.3</td><td>-4.9</td><td>1.6</td><td>-3.1</td><td>1.9E-03</td><td>4.3E-03</td><td>BT20/MB468</td><td>MP</td></tr>
<tr><td>>hg38_21_+_8433172_8446622_RNA45SN1_45S_50ntFlanks@12324.12354.31|TTGGGTTTTAAGCAGGAGGTGTCAGAAAAGT</td><td>>hg38_21_+_8433172_8446622_RNA45SN1_45S_50ntFlanks</td><td>12324</td><td>12354</td><td>31</td><td>TTGGGTTTTAAGCAGGAGGTGTCAGAAAAGT</td><td>62.2</td><td>-5.1</td><td>1.6</td><td>-3.1</td><td>1.9E-03</td><td>4.3E-03</td><td>BT20/MB468</td><td>MP</td></tr>
<tr><td>>hg38_21_+_8433172_8446622_RNA45SN1_45S_50ntFlanks@12902.12933.32|TCACGCACCGCACGTTCGTGGGGAACCTGGCG</td><td>>hg38_21_+_8433172_8446622_RNA45SN1_45S_50ntFlanks</td><td>12902</td><td>12933</td><td>32</td><td>TCACGCACCGCACGTTCGTGGGGAACCTGGCG</td><td>90.8</td><td>-5.0</td><td>1.6</td><td>-3.1</td><td>1.9E-03</td><td>4.3E-03</td><td>BT20/MB468</td><td>MP</td></tr>
<tr><td>>hg38_21_+_8433172_8446622_RNA45SN1_45S_50ntFlanks@10392.10411.20|ACAGCAGTTGAACATGGGTC</td><td>>hg38_21_+_8433172_8446622_RNA45SN1_45S_50ntFlanks</td><td>10392</td><td>10411</td><td>20</td><td>ACAGCAGTTGAACATGGGTC</td><td>290.4</td><td>-2.0</td><td>0.7</td><td>-2.9</td><td>3.4E-03</td><td>4.3E-03</td><td>BT20/MB468</td><td>CYTO</td></tr>
<tr><td>>hg38_21_+_8433172_8446622_RNA45SN1_45S_50ntFlanks@10412.10430.19|AGTCGGTCCTGAGAGATGG</td><td>>hg38_21_+_8433172_8446622_RNA45SN1_45S_50ntFlanks</td><td>10412</td><td>10430</td><td>19</td><td>AGTCGGTCCTGAGAGATGG</td><td>271.6</td><td>-5.2</td><td>1.7</td><td>-3.1</td><td>1.9E-03</td><td>4.3E-03</td><td>BT20/MB468</td><td>MP</td></tr>
<tr><td>>hg38_21_+_8433172_8446622_RNA45SN1_45S_50ntFlanks@9847.9864.18|AACTGGCGCTGCGGGATG</td><td>>hg38_21_+_8433172_8446622_RNA45SN1_45S_50ntFlanks</td><td>9847</td><td>9864</td><td>18</td><td>AACTGGCGCTGCGGGATG</td><td>193.5</td><td>4.2</td><td>1.2</td><td>3.4</td><td>6.6E-04</td><td>4.3E-03</td><td>BT20/MB231</td><td>MP</td></tr>
<tr><td>>MI0000542|hsa-mir-320a&WithFlank&8|-|22244960|22245043@48.69.22&[MIMAT0000510&hsa-miR-320a-3p&offsets|0|0;m-60&8|-|22244975|22244996&offsets|0|0]|AAAAGCTGGGTTGAGAGGGCGA</td><td>>MI0000542|hsa-mir-320a&WithFlank&8|-|22244960|22245043</td><td>48</td><td>69</td><td>22</td><td>AAAAGCTGGGTTGAGAGGGCGA</td><td>6313.5</td><td>2.4</td><td>0.7</td><td>3.3</td><td>8.9E-04</td><td>4.3E-03</td><td>BT20/MB231</td><td>MITO</td></tr>
<tr><td>>hg38_21_+_8433172_8446622_RNA45SN1_45S_50ntFlanks@10351.10382.32|CGAGAACTTTGAAGGCCGAAGTGGAGAAGGGT</td><td>>hg38_21_+_8433172_8446622_RNA45SN1_45S_50ntFlanks</td><td>10351</td><td>10382</td><td>32</td><td>CGAGAACTTTGAAGGCCGAAGTGGAGAAGGGT</td><td>930.9</td><td>-1.8</td><td>0.6</td><td>-3.0</td><td>2.9E-03</td><td>4.3E-03</td><td>BT20/MB468</td><td>MITO</td></tr>
<tr><td>>hg38_21_+_8433172_8446622_RNA45SN1_45S_50ntFlanks@7975.7992.18|CGCGACCTCAGATCAGAC</td><td>>hg38_21_+_8433172_8446622_RNA45SN1_45S_50ntFlanks</td><td>7975</td><td>7992</td><td>18</td><td>CGCGACCTCAGATCAGAC</td><td>128166.7</td><td>2.9</td><td>0.9</td><td>3.3</td><td>8.9E-04</td><td>4.3E-03</td><td>BT20/MB231</td><td>MITO</td></tr>
<tr><td>>am_trna111_HisGTG_1_-_147774845_147774916@2.29.28,trna118_HisGTG_1_-_145396881_145396952@2.29.28,trna16_HisGTG_1_+_146544773_146544844@2.29.28,trna1_HisGTG_15_+_45493349_45493420@2.29.28,trna21_HisGTG_1_+_147753471_147753542@2.29.28,trna33_HisGTG_6_+_27125906_27125977@2.29.28,trna7_HisGTG_9_-_14433938_14434009@2.29.28,trna8_HisGTG_15_-_45492611_45492682@2.29.28,trna9_HisGTG_15_-_45490804_45490875@2.29.28|CCGTGATCGTATAGTGGTTAGTACTCTG</td><td>>am_trna111_HisGTG_1_-_147774845_147774916</td><td>2</td><td>29</td><td>28</td><td>CCGTGATCGTATAGTGGTTAGTACTCTG</td><td>323.9</td><td>-3.8</td><td>1.3</td><td>-2.9</td><td>3.7E-03</td><td>4.3E-03</td><td>MB231/MB468</td><td>MITO</td></tr>
<tr><td>>MI0000087|hsa-mir-29a&WithFlank&7|-|130876741|130876816@47.67.21&[MIMAT0000086&hsa-miR-29a-3p&offsets|-1|-2;m-26&7|-|130876749|130876769&offsets|-1|-1]|CTAGCACCATCTGAAATCGGT</td><td>>MI0000087|hsa-mir-29a&WithFlank&7|-|130876741|130876816</td><td>47</td><td>67</td><td>21</td><td>CTAGCACCATCTGAAATCGGT</td><td>165.5</td><td>6.6</td><td>2.1</td><td>3.1</td><td>1.9E-03</td><td>4.3E-03</td><td>BT20/MB468</td><td>MP</td></tr>
<tr><td>>trnaMT_HisGTG_MT_+_12138_12206@43.71.29|ACAGAGGCTTACGACCCCTTATTTACCCC</td><td>>trnaMT_HisGTG_MT_+_12138_12206</td><td>43</td><td>71</td><td>29</td><td>ACAGAGGCTTACGACCCCTTATTTACCCC</td><td>198.5</td><td>7.7</td><td>2.5</td><td>3.1</td><td>1.9E-03</td><td>4.3E-03</td><td>BT20/MB468</td><td>MP</td></tr>
<tr><td>>hg38_21_+_8433172_8446622_RNA45SN1_45S_50ntFlanks@3705.3744.40|TACCTGGTTGATCCTGCCAGTAGCATATGCTTGTCTCAAA</td><td>>hg38_21_+_8433172_8446622_RNA45SN1_45S_50ntFlanks</td><td>3705</td><td>3744</td><td>40</td><td>TACCTGGTTGATCCTGCCAGTAGCATATGCTTGTCTCAAA</td><td>69.7</td><td>6.2</td><td>2.0</td><td>3.1</td><td>1.9E-03</td><td>4.3E-03</td><td>BT20/MB468</td><td>MP</td></tr>
<tr><td>>MI0000061|hsa-let-7a-2&WithFlank&11|-|122146516|122146599@11.32.22(+1G);MI0000062|hsa-let-7a-3&WithFlank&22|+|46112743|46112828@10.31.22(+1G);MI0000060|hsa-let-7a-1&WithFlank&9|+|94175951|94176042@12.33.22(+1G)&[MIMAT0000062&hsa-let-7a-5p&offsets|0|0(+1G);m-6&11|-|122146568|122146589&offsets|0|0(+1G)];[MIMAT0000062_1&hsa-let-7a-5p&offsets|0|0(+1G);m-5&22|+|46112752|46112773&offsets|0|0(+1G)];[MIMAT0000062_2&hsa-let-7a-5p&offsets|0|0(+1G);m-4&9|+|94175962|94175983&offsets|0|0(+1G)]|TGAGGTAGTAGGTTGTATAGTTG</td><td>>MI0000061|hsa-let-7a-2&WithFlank&11|-|122146516|122146599</td><td>11</td><td>32</td><td>22(+1G)</td><td>TGAGGTAGTAGGTTGTATAGTTG</td><td>139.6</td><td>3.8</td><td>1.3</td><td>3.0</td><td>3.1E-03</td><td>4.3E-03</td><td>MB231/MB468</td><td>MP</td></tr>
<tr><td>>hg38_21_+_8433172_8446622_RNA45SN1_45S_50ntFlanks@12455.12480.26|ACCAAGCGTTGGATTGTTCACCCACT</td><td>>hg38_21_+_8433172_8446622_RNA45SN1_45S_50ntFlanks</td><td>12455</td><td>12480</td><td>26</td><td>ACCAAGCGTTGGATTGTTCACCCACT</td><td>140.8</td><td>-2.5</td><td>0.8</td><td>-3.0</td><td>3.1E-03</td><td>4.3E-03</td><td>MB231/MB468</td><td>MP</td></tr>
<tr><td>>hg38_21_+_8433172_8446622_RNA45SN1_45S_50ntFlanks@10682.10705.24|TCCGGCGGCGTCCGGTGAGCTCTC</td><td>>hg38_21_+_8433172_8446622_RNA45SN1_45S_50ntFlanks</td><td>10682</td><td>10705</td><td>24</td><td>TCCGGCGGCGTCCGGTGAGCTCTC</td><td>139.7</td><td>-2.0</td><td>0.7</td><td>-2.9</td><td>3.4E-03</td><td>4.3E-03</td><td>BT20/MB468</td><td>CYTO</td></tr>
<tr><td>>hg38_21_+_8433172_8446622_RNA45SN1_45S_50ntFlanks@10781.10817.37|AGGTGAACAGCCTCTGGCATGTTGGAACAATGTAGGT</td><td>>hg38_21_+_8433172_8446622_RNA45SN1_45S_50ntFlanks</td><td>10781</td><td>10817</td><td>37</td><td>AGGTGAACAGCCTCTGGCATGTTGGAACAATGTAGGT</td><td>253.2</td><td>-2.0</td><td>0.7</td><td>-3.0</td><td>2.9E-03</td><td>4.3E-03</td><td>BT20/MB468</td><td>MITO</td></tr>
<tr><td>>hg38_21_+_8433172_8446622_RNA45SN1_45S_50ntFlanks@11641.11681.41|CGGCGGGTGTTGACGCGATGTGATTTCTGCCCAGTGCTCTG</td><td>>hg38_21_+_8433172_8446622_RNA45SN1_45S_50ntFlanks</td><td>11641</td><td>11681</td><td>41</td><td>CGGCGGGTGTTGACGCGATGTGATTTCTGCCCAGTGCTCTG</td><td>294.3</td><td>-4.4</td><td>1.4</td><td>-3.1</td><td>1.9E-03</td><td>4.3E-03</td><td>BT20/MB468</td><td>MP</td></tr>
<tr><td>>trna152_ValCAC_6_-_27248049_27248121@1.32.32|GCTTCTGTAGTGTAGTGGTTATCACGTTCGCC</td><td>>trna152_ValCAC_6_-_27248049_27248121</td><td>1</td><td>32</td><td>32</td><td>GCTTCTGTAGTGTAGTGGTTATCACGTTCGCC</td><td>7726.2</td><td>-3.4</td><td>1.0</td><td>-3.4</td><td>8.0E-04</td><td>4.3E-03</td><td>BT20/MB231</td><td>CYTO</td></tr>
<tr><td>>hg38_21_+_8433172_8446622_RNA45SN1_45S_50ntFlanks@12417.12453.37|TCGATGTCGGCTCTTCCTATCATTGTGAAGCAGAATT</td><td>>hg38_21_+_8433172_8446622_RNA45SN1_45S_50ntFlanks</td><td>12417</td><td>12453</td><td>37</td><td>TCGATGTCGGCTCTTCCTATCATTGTGAAGCAGAATT</td><td>319.7</td><td>-2.7</td><td>0.8</td><td>-3.4</td><td>8.0E-04</td><td>4.3E-03</td><td>BT20/MB231</td><td>CYTO</td></tr>
<tr><td>>MI0000077|hsa-mir-21&WithFlank&17|+|59841260|59841343@14.34.21(+1C)&[MIMAT0000076&hsa-miR-21-5p&offsets|0|-1(+1C);m-1&17|+|59841273|59841294&offsets|0|-1(+1C)]|TAGCTTATCAGACTGATGTTGC</td><td>>MI0000077|hsa-mir-21&WithFlank&17|+|59841260|59841343</td><td>14</td><td>34</td><td>21(+1C)</td><td>TAGCTTATCAGACTGATGTTGC</td><td>776.6</td><td>2.9</td><td>1.0</td><td>3.0</td><td>3.1E-03</td><td>4.3E-03</td><td>MB231/MB468</td><td>MP</td></tr>
</table>

| | | | | | | | | | | | | | |
|---|---|---|---|---|---|---|---|---|---|---|---|---|---|
| >MI0000542\|hsa-mir-320a&WithFlank&8\|-\|22244960\|22245043@48.70.23(+2U)&[MIMAT0000510&hsa-miR-320a-3p&offsets\|0\|+1(+2U);m-60&8\|-\|22244975\|22244996&offsets\|0\|+1(+2U)]\|AAAAGCTGGGTTGAGAGGGCGAATT | >MI0000542\|hsa-mir-320a&WithFlank&8\|-\|22244960\|22245043 | 48 | 70 | 23(+2U) | AAAAGCTGGGTTGAGAGGGCGAATT | 185.7 | 3.9 | 1.3 | 3.0 | 3.1E-03 | 4.3E-03 | MB231/MB468 | MP |
| >hg38_21_+_8433172_8446622_RNA45SN1_45S_50ntFlanks@12479.12496.18\|CTAATAGGGAACGTGAGC | >hg38_21_+_8433172_8446622_RNA45SN1_45S_50ntFlanks | 12479 | 12496 | 18 | CTAATAGGGAACGTGAGC | 2655.1 | 2.1 | 0.7 | 2.9 | 3.4E-03 | 4.3E-03 | BT20/MB468 | CYTO |
| >hg38_21_+_8433172_8446622_RNA45SN1_45S_50ntFlanks@9381.9409.29\|CGCTCGCCGGCCGAGGTGGGATCCCGAGG | >hg38_21_+_8433172_8446622_RNA45SN1_45S_50ntFlanks | 9381 | 9409 | 29 | CGCTCGCCGGCCGAGGTGGGATCCCGAGG | 1148.6 | -5.9 | 2.0 | -2.9 | 3.7E-03 | 4.3E-03 | BT20/MB468 | NUC |
| >hg38_21_+_8433172_8446622_RNA45SN1_45S_50ntFlanks@10797.10840.44\|GCATGTTGGAACAATGTAGGTAAGGGAAGTCGGCAAGCCGGATC | >hg38_21_+_8433172_8446622_RNA45SN1_45S_50ntFlanks | 10797 | 10840 | 44 | GCATGTTGGAACAATGTAGGTAAGGGAAGTCGGCAAGCCGGATC | 255.1 | -3.8 | 1.1 | -3.4 | 8.0E-04 | 4.3E-03 | BT20/MB231 | CYTO |
| >hg38_21_+_8433172_8446622_RNA45SN1_45S_50ntFlanks@9622.9648.27\|CTAGTAGCTGGTTCCCTCCGAAGTTTC | >hg38_21_+_8433172_8446622_RNA45SN1_45S_50ntFlanks | 9622 | 9648 | 27 | CTAGTAGCTGGTTCCCTCCGAAGTTTC | 399.7 | -5.4 | 1.8 | -3.1 | 1.9E-03 | 4.3E-03 | BT20/MB468 | MP |
| >hg38_21_+_8433172_8446622_RNA45SN1_45S_50ntFlanks@10410.10453.44\|TCAGTCGGTCCTGAGAGATGGGCGAGCGCCGTTCCGAAGGGACG | >hg38_21_+_8433172_8446622_RNA45SN1_45S_50ntFlanks | 10410 | 10453 | 44 | TCAGTCGGTCCTGAGAGATGGGCGAGCGCCGTTCCGAAGGGACG | 531.5 | -2.9 | 1.0 | -3.0 | 2.9E-03 | 4.3E-03 | BT20/MB468 | MITO |
| >hg38_MT_+_1621_3279_RNR2_16S_50ntFlanks@1370.1414.45\|ATGTTGGATCAGGACATCCCGATGGTGCAGCCGCTATTAAAGGTT | >hg38_MT_+_1621_3279_RNR2_16S_50ntFlanks | 1370 | 1414 | 45 | ATGTTGGATCAGGACATCCCGATGGTGCAGCCGCTATTAAAGGTT | 186.3 | 4.9 | 1.7 | 3.0 | 3.1E-03 | 4.3E-03 | MB231/MB468 | MP |
| >hg38_21_+_8433172_8446622_RNA45SN1_45S_50ntFlanks@8168.8212.45\|CCCAAGTCCTTCTGATCGAGGCCCAGCCCGTGGACGGTGTGAGGC | >hg38_21_+_8433172_8446622_RNA45SN1_45S_50ntFlanks | 8168 | 8212 | 45 | CCCAAGTCCTTCTGATCGAGGCCCAGCCCGTGGACGGTGTGAGGC | 533.5 | -2.4 | 0.8 | -3.0 | 2.9E-03 | 4.3E-03 | BT20/MB468 | MITO |
| >hg38_MT_+_1621_3279_RNR2_16S_50ntFlanks@507.530.24\|AGGAAAAAACCTTGTAGAGAGAGT | >hg38_MT_+_1621_3279_RNR2_16S_50ntFlanks | 507 | 530 | 24 | AGGAAAAAACCTTGTAGAGAGAGT | 100.8 | 4.5 | 1.4 | 3.1 | 1.9E-03 | 4.3E-03 | BT20/MB468 | MP |
| >MI0001445\|hsa-mir-423&WithFlank&17\|+\|30117073\|30117178@59.77.19(+1U)&[MIMAT0001340&hsa-miR-423-3p&offsets\|0\|-4(+1U);m-100&17\|+\|30117131\|30117153&offsets\|0\|-4(+1U)]\|AGCTCGGTCTGAGGCCCCTT | >MI0001445\|hsa-mir-423&WithFlank&17\|+\|30117073\|30117178 | 59 | 77 | 19(+1U) | AGCTCGGTCTGAGGCCCCTT | 200.7 | 4.9 | 1.7 | 3.0 | 3.1E-03 | 4.3E-03 | MB231/MB468 | MP |
| >hg38_21_+_8433172_8446622_RNA45SN1_45S_50ntFlanks@10347.10366.20\|CAAACGAGAACTTTGAAGGC | >hg38_21_+_8433172_8446622_RNA45SN1_45S_50ntFlanks | 10347 | 10366 | 20 | CAAACGAGAACTTTGAAGGC | 137.5 | -3.3 | 1.0 | -3.1 | 1.9E-03 | 4.3E-03 | BT20/MB468 | MP |
| >hg38_21_+_8433172_8446622_RNA45SN1_45S_50ntFlanks@11769.11792.24\|TAATTAGTGACGCGCATGAATGGA | >hg38_21_+_8433172_8446622_RNA45SN1_45S_50ntFlanks | 11769 | 11792 | 24 | TAATTAGTGACGCGCATGAATGGA | 181.1 | -5.1 | 1.6 | -3.1 | 1.9E-03 | 4.3E-03 | BT20/MB468 | MP |
| >hg38_21_+_8433172_8446622_RNA45SN1_45S_50ntFlanks@12419.12449.31\|GATGTCGGCTCTTCCTATCATTGTGAAGCAG | >hg38_21_+_8433172_8446622_RNA45SN1_45S_50ntFlanks | 12419 | 12449 | 31 | GATGTCGGCTCTTCCTATCATTGTGAAGCAG | 2957.3 | -2.8 | 0.8 | -3.4 | 8.0E-04 | 4.3E-03 | BT20/MB231 | CYTO |
| >hg38_21_+_8433172_8446622_RNA45SN1_45S_50ntFlanks@12419.12453.35\|GATGTCGGCTCTTCCTATCATTGTGAAGCAGAATT | >hg38_21_+_8433172_8446622_RNA45SN1_45S_50ntFlanks | 12419 | 12453 | 35 | GATGTCGGCTCTTCCTATCATTGTGAAGCAGAATT | 889.3 | -2.7 | 0.8 | -3.4 | 8.0E-04 | 4.3E-03 | BT20/MB231 | CYTO |
| >hg38_21_+_8433172_8446622_RNA45SN1_45S_50ntFlanks@11517.11539.23\|CCGCCCCCGGGGCCGCGGTTCCG | >hg38_21_+_8433172_8446622_RNA45SN1_45S_50ntFlanks | 11517 | 11539 | 23 | CCGCCCCCGGGGCCGCGGTTCCG | 253.7 | -4.4 | 1.5 | -2.9 | 3.4E-03 | 4.3E-03 | BT20/MB468 | CYTO |
| >hg38_21_+_8433172_8446622_RNA45SN1_45S_50ntFlanks@12633.12669.37\|CGAAGCTACCATCTGTGGGATTATGACTGAACGCCTC | >hg38_21_+_8433172_8446622_RNA45SN1_45S_50ntFlanks | 12633 | 12669 | 37 | CGAAGCTACCATCTGTGGGATTATGACTGAACGCCTC | 68.5 | -4.0 | 1.4 | -3.0 | 3.1E-03 | 4.3E-03 | MB231/MB468 | MP |
| >trnaMT_GluTTC_MT_-_14674_14742@38.62.25,trnalookalike8_GluTTC_5_-_93905172_93905240@38.62.25\|CATTGGTCGTGGTTGTAGTCCGTGC | >trnaMT_GluTTC_MT_-_14674_14742 | 38 | 62 | 25 | CATTGGTCGTGGTTGTAGTCCGTGC | 92.2 | 6.2 | 2.0 | 3.1 | 1.9E-03 | 4.3E-03 | BT20/MB468 | MP |
| >hg38_21_+_8433172_8446622_RNA45SN1_45S_50ntFlanks@6644.6667.24\|CCTCGTACGACTCTTAGCGGTGGA | >hg38_21_+_8433172_8446622_RNA45SN1_45S_50ntFlanks | 6644 | 6667 | 24 | CCTCGTACGACTCTTAGCGGTGGA | 70.9 | 4.3 | 1.4 | 3.1 | 1.9E-03 | 4.3E-03 | BT20/MB468 | MP |
| >hg38_21_+_8433172_8446622_RNA45SN1_45S_50ntFlanks@10466.10497.32\|CGTTGCCCTCGGCCGATCGAAAGGGAGTCGGG | >hg38_21_+_8433172_8446622_RNA45SN1_45S_50ntFlanks | 10466 | 10497 | 32 | CGTTGCCCTCGGCCGATCGAAAGGGAGTCGGG | 628.4 | -5.3 | 1.7 | -3.1 | 1.9E-03 | 4.3E-03 | BT20/MB468 | MP |
| >trnaMT_LeuTAA_MT_+_3230_3304@16.44.29\|CCCGGTAATCGCATAAAACTTAAAACTTT | >trnaMT_LeuTAA_MT_+_3230_3304 | 16 | 44 | 29 | CCCGGTAATCGCATAAAACTTAAAACTTT | 1135.0 | -4.9 | 1.6 | -3.1 | 1.9E-03 | 4.3E-03 | BT20/MB468 | MP |
| >hg38_21_+_8433172_8446622_RNA45SN1_45S_50ntFlanks@12418.12448.31\|CGATGTCGGCTCTTCCTATCATTGTGAAGCA | >hg38_21_+_8433172_8446622_RNA45SN1_45S_50ntFlanks | 12418 | 12448 | 31 | CGATGTCGGCTCTTCCTATCATTGTGAAGCA | 914.4 | 2.6 | 0.9 | 2.9 | 3.4E-03 | 4.3E-03 | BT20/MB468 | CYTO |
| >hg38_21_+_8433172_8446622_RNA45SN1_45S_50ntFlanks@8117.8141.25\|CGGGACATGTGGCGTACGGAAGACC | >hg38_21_+_8433172_8446622_RNA45SN1_45S_50ntFlanks | 8117 | 8141 | 25 | CGGGACATGTGGCGTACGGAAGACC | 546.5 | 6.4 | 1.9 | 3.4 | 6.6E-04 | 4.3E-03 | BT20/MB231 | MP |
| >hg38_21_+_8433172_8446622_RNA45SN1_45S_50ntFlanks@4731.4767.37\|AAGAACGAAAGTCGGAGGTTCGAAGACGATCAGATAC | >hg38_21_+_8433172_8446622_RNA45SN1_45S_50ntFlanks | 4731 | 4767 | 37 | AAGAACGAAAGTCGGAGGTTCGAAGACGATCAGATAC | 83.5 | 2.0 | 0.7 | 2.9 | 3.7E-03 | 4.3E-03 | BT20/MB468 | NUC |
| >hg38_21_+_8433172_8446622_RNA45SN1_45S_50ntFlanks@11999.12030.32\|CCCTGCGGGCCGCCGGTGAAATACCACTACTC | >hg38_21_+_8433172_8446622_RNA45SN1_45S_50ntFlanks | 11999 | 12030 | 32 | CCCTGCGGGCCGCCGGTGAAATACCACTACTC | 57.3 | -4.9 | 1.6 | -3.1 | 1.9E-03 | 4.3E-03 | BT20/MB468 | MP |
| >hg38_21_+_8433172_8446622_RNA45SN1_45S_50ntFlanks@12141.12179.39\|ACAGTGCCAGGTGGGGAGTTTGACTGGGGCGGTACACCT | >hg38_21_+_8433172_8446622_RNA45SN1_45S_50ntFlanks | 12141 | 12179 | 39 | ACAGTGCCAGGTGGGGAGTTTGACTGGGGCGGTACACCT | 176.7 | -5.3 | 1.7 | -3.1 | 1.9E-03 | 4.3E-03 | BT20/MB468 | MP |
| >hg38_21_+_8433172_8446622_RNA45SN1_45S_50ntFlanks@7981.8008.28\|CTCAGATCAGACGTGGCGACCCGCTGAA | >hg38_21_+_8433172_8446622_RNA45SN1_45S_50ntFlanks | 7981 | 8008 | 28 | CTCAGATCAGACGTGGCGACCCGCTGAA | 1264.8 | -4.5 | 1.4 | -3.1 | 1.9E-03 | 4.3E-03 | BT20/MB468 | MP |
| >MI0000298\|hsa-mir-221&WithFlank&X\|-\|45746151\|45746272@71.92.22(+1A)&[MIMAT0000278&hsa-miR-221-3p&offsets\|0\|-1(+1A);m-91&X\|-\|45746181\|45746202&offsets\|0\|0(+1A)]\|AGCTACATTGTCTGCTGGGTTTA | >MI0000298\|hsa-mir-221&WithFlank&X\|-\|45746151\|45746272 | 71 | 92 | 22(+1A) | AGCTACATTGTCTGCTGGGTTTA | 3654.9 | -3.5 | 1.0 | -3.4 | 6.7E-04 | 4.3E-03 | BT20/MB231 | MP |
| >hg38_21_+_8433172_8446622_RNA45SN1_45S_50ntFlanks@7977.8011.35\|CGACCTCAGATCAGACGTGGCGACCCGCTGAATTT | >hg38_21_+_8433172_8446622_RNA45SN1_45S_50ntFlanks | 7977 | 8011 | 35 | CGACCTCAGATCAGACGTGGCGACCCGCTGAATTT | 1134.7 | 4.5 | 1.5 | 3.1 | 1.9E-03 | 4.3E-03 | BT20/MB468 | MP |
| >hg38_21_+_8433172_8446622_RNA45SN1_45S_50ntFlanks@3783.3800.18\|AGTGAAACTGCGAATGGC | >hg38_21_+_8433172_8446622_RNA45SN1_45S_50ntFlanks | 3783 | 3800 | 18 | AGTGAAACTGCGAATGGC | 375.2 | 1.5 | 0.5 | 2.9 | 3.4E-03 | 4.3E-03 | BT20/MB468 | CYTO |
| >MI0000072\|hsa-mir-18a&WithFlank&13\|+\|91350745\|91350827@12.34.23&[MIMAT0000072&hsa-miR-18a-5p&offsets\|0\|0;m-262&13\|+\|91350756\|91350778&offsets\|0\|0]\|TAAGGTGCATCTAGTGCAGATAG | >MI0000072\|hsa-mir-18a&WithFlank&13\|+\|91350745\|91350827 | 12 | 34 | 23 | TAAGGTGCATCTAGTGCAGATAG | 1280.3 | -3.9 | 1.1 | -3.4 | 6.7E-04 | 4.3E-03 | BT20/MB231 | MP |
| >MI0000750\|hsa-mir-26a-2&WithFlank&12\|-\|57824603\|57824698@20.41.22;MI0000083\|hsa-mir-26a-1&WithFlank&3\|+\|37969398\|37969486@16.37.22&[MIMAT0000082&hsa-miR-26a-5p&offsets\|0\|0;m-33&12\|-\|57824658\|57824679&offsets\|0\|0];[MIMAT0000082_1&hsa-miR-26a-5p&offsets\|0\|0;m-34&3\|+\|37969413\|37969434&offsets\|0\|0]\|TTCAAGTAATCCAGGATAGGCT | >MI0000750\|hsa-mir-26a-2&WithFlank&12\|-\|57824603\|57824698 | 20 | 41 | 22 | TTCAAGTAATCCAGGATAGGCT | 128321.4 | -3.4 | 1.0 | -3.4 | 6.7E-04 | 4.3E-03 | BT20/MB231 | MP |
| >hg38_21_+_8433172_8446622_RNA45SN1_45S_50ntFlanks@11570.11597.28\|AGCCGACTTAGAACTGGTGCGGACCAGG | >hg38_21_+_8433172_8446622_RNA45SN1_45S_50ntFlanks | 11570 | 11597 | 28 | AGCCGACTTAGAACTGGTGCGGACCAGG | 277.8 | 6.5 | 1.9 | 3.4 | 6.7E-04 | 4.3E-03 | BT20/MB231 | MP |
| >hg38_21_+_8433172_8446622_RNA45SN1_45S_50ntFlanks@4724.4759.36\|ATTAATCAAGAACGAAAGTCGGAGGTTCGAAGACGA | >hg38_21_+_8433172_8446622_RNA45SN1_45S_50ntFlanks | 4724 | 4759 | 36 | ATTAATCAAGAACGAAAGTCGGAGGTTCGAAGACGA | 1214.3 | 7.5 | 2.2 | 3.4 | 6.7E-04 | 4.3E-03 | BT20/MB231 | MP |
| >hg38_21_+_8433172_8446622_RNA45SN1_45S_50ntFlanks@12480.12497.18\|TAATAGGGAACGTGAGCT | >hg38_21_+_8433172_8446622_RNA45SN1_45S_50ntFlanks | 12480 | 12497 | 18 | TAATAGGGAACGTGAGCT | 231.6 | 2.9 | 0.9 | 3.1 | 1.9E-03 | 4.3E-03 | BT20/MB468 | MP |
| >hg38_21_+_8433172_8446622_RNA45SN1_45S_50ntFlanks@12415.12448.34\|CTTCGATGTCGGCTCTTCCTATCATTGTGAAGCA | >hg38_21_+_8433172_8446622_RNA45SN1_45S_50ntFlanks | 12415 | 12448 | 34 | CTTCGATGTCGGCTCTTCCTATCATTGTGAAGCA | 1918.8 | 1.8 | 0.6 | 3.0 | 2.9E-03 | 4.3E-03 | BT20/MB468 | MITO |
| >hg38_21_+_8433172_8446622_RNA45SN1_45S_50ntFlanks@11805.11823.19\|CCACTGTCCCTACCTACTA | >hg38_21_+_8433172_8446622_RNA45SN1_45S_50ntFlanks | 11805 | 11823 | 19 | CCACTGTCCCTACCTACTA | 147.1 | -3.0 | 1.0 | -2.9 | 3.5E-03 | 4.3E-03 | MB231/MB468 | CYTO |
| >am_trna128_GlyGCC_6_-_27870686_27870756@14.31.18,trna131_GlyCCC_1_-_17004766_17004836@14.31.18,trna133_GlyCCC_1_-_16872434_16872504@14.31.18,trna17_GluTTC_2_-_204229131_204229201@14.31.18,trna18_GlyGCC_16_+_70822597_70822667@14.31.18,trna19_GlyGCC_16_+_70823410_70823480@14.31.18,trna19_GlyGCC_2_-_157257659_157257729@14.31.18,trna24_GlyGCC_16_-_70812942_70813012@14.31.18,trna25_GlyGCC_16_-_70812114_70812184@14.31.18,trna2_GlyCCC_1_+_17053780_17053850@14.31.18,trna2_GlyGCC_21_-_18827107_18827177@14.31.18,trna35_GlyGCC_1_+_161413094_161413164@14.31.18,trna37_GlyGCC_1_+_161420467_161420537@14.31.18,trna39_GlyGCC_1_+_161427898_161427968@14.31.18,trna41_GlyGCC_1_+_161435258_161435328@14.31.18,trna4_GlyCCC_1_+_17188416_17188486@14.31.18,trna5_GlyGCC_17_+_8029064_8029134@14.31.18,trna68_GlyGCC_1_-_161493637_161493707@14.31.18,trna91_GlyCCC_1_-_149680210_149680280@14.31.18,trna99_ValCAC_1_-_149294666_149294736@14.31.18\|AGTGGTAGAATTCTCGCC | >am_trna128_GlyGCC_6_-_27870686_27870756 | 14 | 31 | 18 | AGTGGTAGAATTCTCGCC | 133.4 | 4.3 | 1.4 | 3.1 | 1.9E-03 | 4.3E-03 | BT20/MB468 | MP |
| >hg38_21_+_8433172_8446622_RNA45SN1_45S_50ntFlanks@7985.8016.32\|GATCAGACGTGGCGACCCGCTGAATTTAAGCA | >hg38_21_+_8433172_8446622_RNA45SN1_45S_50ntFlanks | 7985 | 8016 | 32 | GATCAGACGTGGCGACCCGCTGAATTTAAGCA | 239.6 | -4.9 | 1.7 | -3.0 | 3.2E-03 | 4.3E-03 | MB231/MB468 | MP |
| >MI0003685\|hsa-mir-421&WithFlank&X\|-\|74218371\|74218467@57.75.19&[MIMAT0003339&hsa-miR-421&offsets\|+3\|-1;m-309&X\|-\|74218392\|74218414&offsets\|+3\|-1]\|AACAGACATTAATTGGGCG | >MI0003685\|hsa-mir-421&WithFlank&X\|-\|74218371\|74218467 | 57 | 75 | 19 | AACAGACATTAATTGGGCG | 35.0 | 8.3 | 2.8 | 3.0 | 3.2E-03 | 4.4E-03 | MB231/MB468 | MP |
| >hg38_21_+_8433172_8446622_RNA45SN1_45S_50ntFlanks@9548.9569.22\|GAGGTCCGTAGCGGTCCTGACG | >hg38_21_+_8433172_8446622_RNA45SN1_45S_50ntFlanks | 9548 | 9569 | 22 | GAGGTCCGTAGCGGTCCTGACG | 103.5 | -3.7 | 1.2 | -3.0 | 3.2E-03 | 4.4E-03 | MB231/MB468 | MP |
| >trna111_HisGTG_1_-_147774845_147774916@-1T.31.32,trna118_HisGTG_1_-_145396881_145396952@-1T.31.32,trna16_HisGTG_1_+_146544773_146544844@-1T.31.32,trna1_HisGTG_15_+_45493349_45493420@-1T.31.32,trna21_HisGTG_1_+_147753471_147753542@-1T.31.32,trna33_HisGTG_6_+_27125906_27125977@-1T.31.32,trna7_HisGTG_9_-_14433938_14434009@-1T.31.32,trna8_HisGTG_15_-_45492611_45492682@-1T.31.32,trna9_HisGTG_15_-_45490804_45490875@-1T.31.32\|TGCCGTGATCGTATAGTGGTTAGTACTCTGCG | >trna111_HisGTG_1_-_147774845_147774916 | -1T | 31 | 32 | TGCCGTGATCGTATAGTGGTTAGTACTCTGCG | 3939.4 | -2.7 | 0.9 | -2.9 | 3.7E-03 | 4.4E-03 | MB231/MB468 | MITO |
| >hg38_21_+_8433172_8446622_RNA45SN1_45S_50ntFlanks@12420.12451.32\|ATGTCGGCTCTTCCTATCATTGTGAAGCAGAA | >hg38_21_+_8433172_8446622_RNA45SN1_45S_50ntFlanks | 12420 | 12451 | 32 | ATGTCGGCTCTTCCTATCATTGTGAAGCAGAA | 138.6 | -2.8 | 0.9 | -2.9 | 3.5E-03 | 4.4E-03 | BT20/MB468 | CYTO |

<table>
<tr><td></td><td></td><td></td><td></td><td></td><td></td><td></td><td></td><td></td><td></td><td></td><td></td><td></td><td></td></tr>
<tr><td>>hg38_21_+_8433172_8446622_RNA45SN1_45S_50ntFlanks@11918.11936.19|ACATGAGAGGTGTAGAATA</td><td>>hg38_21_+_8433172_8446622_RNA45SN1_45S_50ntFlanks</td><td>11918</td><td>11936</td><td>19</td><td>ACATGAGAGGTGTAGAATA</td><td>69.7</td><td>-4.9</td><td>1.6</td><td>-3.0</td><td>3.2E-03</td><td>4.4E-03</td><td>MB231/MB468</td><td>MP</td></tr>
<tr><td>>hg38_21_+_8433172_8446622_RNA45SN1_45S_50ntFlanks@9837.9856.20|TGGTAAGCAGAACTGGCGCT</td><td>>hg38_21_+_8433172_8446622_RNA45SN1_45S_50ntFlanks</td><td>9837</td><td>9856</td><td>20</td><td>TGGTAAGCAGAACTGGCGCT</td><td>75.8</td><td>9.6</td><td>2.8</td><td>3.4</td><td>6.7E-04</td><td>4.4E-03</td><td>BT20/MB231</td><td>MP</td></tr>
<tr><td>>hg38_MT_+_1621_3279_RNR2_16S_50ntFlanks@1216.1262.47|CAACCTCCGAGCAGTACATGCTAAGACTTCACCAGTCAAAGCGAACT</td><td>>hg38_MT_+_1621_3279_RNR2_16S_50ntFlanks</td><td>1216</td><td>1262</td><td>47</td><td>CAACCTCCGAGCAGTACATGCTAAGACTTCACCAGTCAAAGCGAACT</td><td>2282.4</td><td>4.4</td><td>1.5</td><td>2.9</td><td>3.7E-03</td><td>4.4E-03</td><td>MB231/MB468</td><td>MITO</td></tr>
<tr><td>>hg38_21_+_8433172_8446622_RNA45SN1_45S_50ntFlanks@4889.4917.29|CTGAAACTTAAAGGAATTGACGGAAGGGC</td><td>>hg38_21_+_8433172_8446622_RNA45SN1_45S_50ntFlanks</td><td>4889</td><td>4917</td><td>29</td><td>CTGAAACTTAAAGGAATTGACGGAAGGGC</td><td>77.5</td><td>-4.9</td><td>1.6</td><td>-3.1</td><td>1.9E-03</td><td>4.4E-03</td><td>BT20/MB468</td><td>MP</td></tr>
<tr><td>>hg38_1_-_228634819_228635039_RNA5S12_5S_50ntFlanks@138.170.33|AGACCGCCTGGGAATACCGGGTGCTGTAGGCTT</td><td>>hg38_1_-_228634819_228635039_RNA5S12_5S_50ntFlanks</td><td>138</td><td>170</td><td>33</td><td>AGACCGCCTGGGAATACCGGGTGCTGTAGGCTT</td><td>1637.1</td><td>2.5</td><td>0.8</td><td>3.1</td><td>1.9E-03</td><td>4.4E-03</td><td>BT20/MB468</td><td>MP</td></tr>
<tr><td>>trna15_ValAAC_5_-_180615416_180615488@1.29.29|GTTTCCGTAGTGTAGTGGTCATCACGTTC</td><td>>trna15_ValAAC_5_-_180615416_180615488</td><td>1</td><td>29</td><td>29</td><td>GTTTCCGTAGTGTAGTGGTCATCACGTTC</td><td>3842.7</td><td>-4.8</td><td>1.5</td><td>-3.1</td><td>1.9E-03</td><td>4.4E-03</td><td>BT20/MB468</td><td>MP</td></tr>
<tr><td>>hg38_21_+_8433172_8446622_RNA45SN1_45S_50ntFlanks@11912.11943.32|GAAGAGACATGAGAGGTGTAGAATAAGTGGGA</td><td>>hg38_21_+_8433172_8446622_RNA45SN1_45S_50ntFlanks</td><td>11912</td><td>11943</td><td>32</td><td>GAAGAGACATGAGAGGTGTAGAATAAGTGGGA</td><td>33.1</td><td>8.8</td><td>2.8</td><td>3.1</td><td>1.9E-03</td><td>4.4E-03</td><td>BT20/MB468</td><td>MP</td></tr>
<tr><td>>hg38_MT_+_1621_3279_RNR2_16S_50ntFlanks@1347.1389.43|CAACAATAGGGTTTACGACCTCGATGTTGGATCAGGACATCCC</td><td>>hg38_MT_+_1621_3279_RNR2_16S_50ntFlanks</td><td>1347</td><td>1389</td><td>43</td><td>CAACAATAGGGTTTACGACCTCGATGTTGGATCAGGACATCCC</td><td>208.0</td><td>4.5</td><td>1.6</td><td>2.9</td><td>3.7E-03</td><td>4.4E-03</td><td>MB231/MB468</td><td>MITO</td></tr>
<tr><td>>MI0000299|hsa-mir-222&WithFlank&X|-|45747009|45747130@75.97.23(+2U)&[MIMAT0000279&hsa-miR-222-3p&offsets|0|+2(+2U);m-140&X|-|45747033|45747056&offsets|0|-1(+2U)]|AGCTACATCTGGCTACTGGGTCTTT</td><td>>MI0000299|hsa-mir-222&WithFlank&X|-|45747009|45747130</td><td>75</td><td>97</td><td>23(+2U)</td><td>AGCTACATCTGGCTACTGGGTCTTT</td><td>855.6</td><td>-2.5</td><td>0.7</td><td>-3.4</td><td>7.4E-04</td><td>4.4E-03</td><td>BT20/MB231</td><td>NUC</td></tr>
<tr><td>>hg38_1_-_228634819_228635039_RNA5S12_5S_50ntFlanks@147.169.23|GGGAATACCGGGTGCTGTAGGCT</td><td>>hg38_1_-_228634819_228635039_RNA5S12_5S_50ntFlanks</td><td>147</td><td>169</td><td>23</td><td>GGGAATACCGGGTGCTGTAGGCT</td><td>305.9</td><td>1.9</td><td>0.6</td><td>2.9</td><td>3.8E-03</td><td>4.4E-03</td><td>MB231/MB468</td><td>NUC</td></tr>
<tr><td>>hg38_21_+_8433172_8446622_RNA45SN1_45S_50ntFlanks@8183.8212.30|TCGAGGCCCAGCCCGTGGACGGTGTGAGGC</td><td>>hg38_21_+_8433172_8446622_RNA45SN1_45S_50ntFlanks</td><td>8183</td><td>8212</td><td>30</td><td>TCGAGGCCCAGCCCGTGGACGGTGTGAGGC</td><td>129.0</td><td>-5.3</td><td>1.7</td><td>-3.1</td><td>1.9E-03</td><td>4.4E-03</td><td>BT20/MB468</td><td>MP</td></tr>
<tr><td>>MI0000440|hsa-mir-27b&WithFlank&9|+|95085439|95085547@67.87.21(+1C)&[MIMAT0000419&hsa-miR-27b-3p&offsets|0|0(+1C);m-39&9|+|95085505|95085525&offsets|0|0(+1C)]|TTCACAGTGGCTAAGTTCTGCC</td><td>>MI0000440|hsa-mir-27b&WithFlank&9|+|95085439|95085547</td><td>67</td><td>87</td><td>21(+1C)</td><td>TTCACAGTGGCTAAGTTCTGCC</td><td>257.9</td><td>4.1</td><td>1.4</td><td>2.9</td><td>3.2E-03</td><td>4.4E-03</td><td>MB231/MB468</td><td>MP</td></tr>
<tr><td>>hg38_21_+_8433172_8446622_RNA45SN1_45S_50ntFlanks@5175.5208.34|CCGAGATTGAGCAATAACAGGTCTGTGATGCCCT</td><td>>hg38_21_+_8433172_8446622_RNA45SN1_45S_50ntFlanks</td><td>5175</td><td>5208</td><td>34</td><td>CCGAGATTGAGCAATAACAGGTCTGTGATGCCCT</td><td>38.4</td><td>-9.1</td><td>3.1</td><td>-2.9</td><td>3.2E-03</td><td>4.4E-03</td><td>MB231/MB468</td><td>MP</td></tr>
<tr><td>>hg38_21_+_8433172_8446622_RNA45SN1_45S_50ntFlanks@11876.11908.33|AAGACCCTGTTGAGCTTGACTCTAGTCTGGCAC</td><td>>hg38_21_+_8433172_8446622_RNA45SN1_45S_50ntFlanks</td><td>11876</td><td>11908</td><td>33</td><td>AAGACCCTGTTGAGCTTGACTCTAGTCTGGCAC</td><td>151.5</td><td>-3.3</td><td>1.1</td><td>-2.9</td><td>3.5E-03</td><td>4.4E-03</td><td>BT20/MB468</td><td>CYTO</td></tr>
<tr><td>>hg38_21_+_8433172_8446622_RNA45SN1_45S_50ntFlanks@10882.10901.20|TCGGGCTGGGGCGCGAAGCG</td><td>>hg38_21_+_8433172_8446622_RNA45SN1_45S_50ntFlanks</td><td>10882</td><td>10901</td><td>20</td><td>TCGGGCTGGGGCGCGAAGCG</td><td>129.2</td><td>-2.8</td><td>1.0</td><td>-2.9</td><td>3.5E-03</td><td>4.4E-03</td><td>BT20/MB468</td><td>CYTO</td></tr>
<tr><td>>MI0000070|hsa-mir-16-1&WithFlank&13|-|50048967|50049067@20.41.22(+1C);MI0000115|hsa-mir-16-2&WithFlank&3|+|160404739|160404831@16.37.22(+1C)&[MIMAT0000069&hsa-miR-16-5p&offsets|0|0(+1C);m-62&13|-|50049027|50049048&offsets|0|0(+1C)];[MIMAT0000069_1&hsa-miR-16-5p&offsets|0|0(+1C);m-61&3|+|160404754|160404775&offsets|0|0(+1C)]|TAGCAGCACGTAAATATTGGCGC</td><td>>MI0000070|hsa-mir-16-1&WithFlank&13|-|50048967|50049067</td><td>20</td><td>41</td><td>22(+1C)</td><td>TAGCAGCACGTAAATATTGGCGC</td><td>583.2</td><td>2.8</td><td>0.9</td><td>2.9</td><td>3.2E-03</td><td>4.4E-03</td><td>MB231/MB468</td><td>MP</td></tr>
<tr><td>>MI0000268|hsa-mir-34a&WithFlank&1|-|9151662|9151783@28.49.22&[MIMAT0000255&hsa-miR-34a-5p&offsets|0|0;m-119&1|-|9151735|9151756&offsets|0|0]|TGGCAGTGTCTTAGCTGGTTGT</td><td>>MI0000268|hsa-mir-34a&WithFlank&1|-|9151662|9151783</td><td>28</td><td>49</td><td>22</td><td>TGGCAGTGTCTTAGCTGGTTGT</td><td>891.2</td><td>-2.5</td><td>0.9</td><td>-2.9</td><td>3.2E-03</td><td>4.4E-03</td><td>MB231/MB468</td><td>MP</td></tr>
<tr><td>>hg38_1_-_228634819_228635039_RNA5S12_5S_50ntFlanks@152.170.19|TACCGGGTGCTGTAGGCTT</td><td>>hg38_1_-_228634819_228635039_RNA5S12_5S_50ntFlanks</td><td>152</td><td>170</td><td>19</td><td>TACCGGGTGCTGTAGGCTT</td><td>1589.3</td><td>-2.0</td><td>0.7</td><td>-2.9</td><td>3.8E-03</td><td>4.4E-03</td><td>MB231/MB468</td><td>MITO</td></tr>
<tr><td>>trna10_AspGTC_12_-_125424193_125424264@35.74.40,trna12_AspGTC_12_-_125411891_125411962@35.74.40,trna144_AspGTC_6_-_27551236_27551307@35.74.40,trna38_AspGTC_17_-_8125556_8125627@35.74.40,trna45_AspGTC_6_+_27447453_27447524@35.74.40,trna48_AspGTC_6_+_27471523_27471594@35.74.40,trna4_AspGTC_12_+_96429799_96429870@35.74.40,trna69_AspGTC_1_-_161440205_161440276@35.74.40,trna72_AspGTC_1_-_161432824_161432895@35.74.40,trna75_AspGTC_1_-_161425414_161425485@35.74.40,trna78_AspGTC_1_-_161418033_161418104@35.74.40,trna81_AspGTC_1_-_161410615_161410686@35.74.40|TCACGCGGGAGACCGGGGTTCGATTCCCCGACGGGGAGCC</td><td>>trna10_AspGTC_12_-_125424193_125424264</td><td>35</td><td>74</td><td>40</td><td>TCACGCGGGAGACCGGGGTTCGATTCCCCGACGGGGAGCC</td><td>145.3</td><td>5.8</td><td>2.0</td><td>2.9</td><td>3.2E-03</td><td>4.4E-03</td><td>MB231/MB468</td><td>MP</td></tr>
<tr><td>>hg38_MT_+_1621_3279_RNR2_16S_50ntFlanks@1370.1397.28|ATGTTGGATCAGGACATCCCGATGGTGC</td><td>>hg38_MT_+_1621_3279_RNR2_16S_50ntFlanks</td><td>1370</td><td>1397</td><td>28</td><td>ATGTTGGATCAGGACATCCCGATGGTGC</td><td>98.2</td><td>3.8</td><td>1.3</td><td>2.9</td><td>3.2E-03</td><td>4.4E-03</td><td>MB231/MB468</td><td>MP</td></tr>
<tr><td>>hg38_21_+_8433172_8446622_RNA45SN1_45S_50ntFlanks@8196.8219.24|CGTGGACGGTGTGAGGCCGGTAGC</td><td>>hg38_21_+_8433172_8446622_RNA45SN1_45S_50ntFlanks</td><td>8196</td><td>8219</td><td>24</td><td>CGTGGACGGTGTGAGGCCGGTAGC</td><td>522.6</td><td>2.6</td><td>0.9</td><td>2.9</td><td>3.6E-03</td><td>4.4E-03</td><td>MB231/MB468</td><td>CYTO</td></tr>
<tr><td>>trnaMT_GluTTC_MT_-_14674_14742@30.72.43,trnalookalike8_GluTTC_5_-_93905172_93905240@30.72.43|TTTCATATCATTGGTCGTGGTTGTAGTCCGTGCGAGAATACCA</td><td>>trnaMT_GluTTC_MT_-_14674_14742</td><td>30</td><td>72</td><td>43</td><td>TTTCATATCATTGGTCGTGGTTGTAGTCCGTGCGAGAATACCA</td><td>245.0</td><td>-2.9</td><td>0.9</td><td>-3.3</td><td>8.2E-04</td><td>4.4E-03</td><td>BT20/MB231</td><td>CYTO</td></tr>
<tr><td>>hg38_21_+_8433172_8446622_RNA45SN1_45S_50ntFlanks@6656.6689.34|CTTAGCGGTGGATCACTCGGCTCGTGCGTCGATG</td><td>>hg38_21_+_8433172_8446622_RNA45SN1_45S_50ntFlanks</td><td>6656</td><td>6689</td><td>34</td><td>CTTAGCGGTGGATCACTCGGCTCGTGCGTCGATG</td><td>902.5</td><td>-1.6</td><td>0.6</td><td>-3.0</td><td>3.0E-03</td><td>4.4E-03</td><td>BT20/MB468</td><td>MITO</td></tr>
<tr><td>>hg38_21_+_8433172_8446622_RNA45SN1_45S_50ntFlanks@12234.12262.29|CGTGGAGCAGAAGGGCAAAAGCTCGCTTG</td><td>>hg38_21_+_8433172_8446622_RNA45SN1_45S_50ntFlanks</td><td>12234</td><td>12262</td><td>29</td><td>CGTGGAGCAGAAGGGCAAAAGCTCGCTTG</td><td>51.8</td><td>-4.9</td><td>1.6</td><td>-3.1</td><td>1.9E-03</td><td>4.4E-03</td><td>BT20/MB468</td><td>MP</td></tr>
<tr><td>>hg38_21_+_8433172_8446622_RNA45SN1_45S_50ntFlanks@12632.12661.30|GCGAAGCTACCATCTGTGGGATTATGACTG</td><td>>hg38_21_+_8433172_8446622_RNA45SN1_45S_50ntFlanks</td><td>12632</td><td>12661</td><td>30</td><td>GCGAAGCTACCATCTGTGGGATTATGACTG</td><td>1552.3</td><td>-2.2</td><td>0.7</td><td>-3.0</td><td>3.0E-03</td><td>4.4E-03</td><td>BT20/MB468</td><td>MITO</td></tr>
<tr><td>>hg38_21_+_8433172_8446622_RNA45SN1_45S_50ntFlanks@6661.6681.21|CGGTGGATCACTCGGCTCGTG</td><td>>hg38_21_+_8433172_8446622_RNA45SN1_45S_50ntFlanks</td><td>6661</td><td>6681</td><td>21</td><td>CGGTGGATCACTCGGCTCGTG</td><td>1536.1</td><td>4.0</td><td>1.2</td><td>3.4</td><td>6.8E-04</td><td>4.4E-03</td><td>BT20/MB231</td><td>MP</td></tr>
<tr><td>>MI0000102|hsa-mir-100&WithFlank&11|-|122152223|122152314@19.40.22(+1A)&[MIMAT0000098&hsa-miR-100-5p&offsets|0|0(+1A);m-27&11|-|122152276|122152296&offsets|0|+1(+1A)]|AACCCGTAGATCCGAACTTGTGA</td><td>>MI0000102|hsa-mir-100&WithFlank&11|-|122152223|122152314</td><td>19</td><td>40</td><td>22(+1A)</td><td>AACCCGTAGATCCGAACTTGTGA</td><td>347.2</td><td>4.9</td><td>1.6</td><td>3.1</td><td>1.9E-03</td><td>4.4E-03</td><td>BT20/MB468</td><td>MP</td></tr>
<tr><td>>trna152_ValCAC_6_-_27248049_27248121@1.32.32|GCTTCTGTAGTGTAGTGGTTATCACGTTCGCC</td><td>>trna152_ValCAC_6_-_27248049_27248121</td><td>1</td><td>32</td><td>32</td><td>GCTTCTGTAGTGTAGTGGTTATCACGTTCGCC</td><td>1101.4</td><td>-3.6</td><td>1.1</td><td>-3.4</td><td>7.5E-04</td><td>4.4E-03</td><td>BT20/MB231</td><td>NUC</td></tr>
<tr><td>>hg38_21_+_8433172_8446622_RNA45SN1_45S_50ntFlanks@11564.11588.25|CCTAGCAGCCGACTTAGAACTGGTG</td><td>>hg38_21_+_8433172_8446622_RNA45SN1_45S_50ntFlanks</td><td>11564</td><td>11588</td><td>25</td><td>CCTAGCAGCCGACTTAGAACTGGTG</td><td>763.0</td><td>-1.8</td><td>0.6</td><td>-3.0</td><td>3.0E-03</td><td>4.4E-03</td><td>BT20/MB468</td><td>MITO</td></tr>
<tr><td>>hg38_21_+_8433172_8446622_RNA45SN1_45S_50ntFlanks@12437.12467.31|CATTGTGAAGCAGAATTCACCAAGCGTTGGA</td><td>>hg38_21_+_8433172_8446622_RNA45SN1_45S_50ntFlanks</td><td>12437</td><td>12467</td><td>31</td><td>CATTGTGAAGCAGAATTCACCAAGCGTTGGA</td><td>229.2</td><td>-5.4</td><td>1.8</td><td>-3.1</td><td>1.9E-03</td><td>4.4E-03</td><td>BT20/MB468</td><td>MP</td></tr>
<tr><td>>hg38_MT_+_598_1651_RNR1_12S_50ntFlanks@558.580.23|GCTTAAAACTCAAAGGACCTGGC</td><td>>hg38_MT_+_598_1651_RNR1_12S_50ntFlanks</td><td>558</td><td>580</td><td>23</td><td>GCTTAAAACTCAAAGGACCTGGC</td><td>87.8</td><td>5.3</td><td>1.7</td><td>3.1</td><td>1.9E-03</td><td>4.4E-03</td><td>BT20/MB468</td><td>MP</td></tr>
<tr><td>>hg38_21_+_8433172_8446622_RNA45SN1_45S_50ntFlanks@9819.9838.20|TGCCTAGTGGGCCACTTTTG</td><td>>hg38_21_+_8433172_8446622_RNA45SN1_45S_50ntFlanks</td><td>9819</td><td>9838</td><td>20</td><td>TGCCTAGTGGGCCACTTTTG</td><td>35.2</td><td>-8.2</td><td>2.7</td><td>-3.1</td><td>1.9E-03</td><td>4.4E-03</td><td>BT20/MB468</td><td>MP</td></tr>
<tr><td>>hg38_21_+_8433172_8446622_RNA45SN1_45S_50ntFlanks@4066.4101.36|CAACTTTCGATGGTAGTCGCCGTGCCTACCATGGTG</td><td>>hg38_21_+_8433172_8446622_RNA45SN1_45S_50ntFlanks</td><td>4066</td><td>4101</td><td>36</td><td>CAACTTTCGATGGTAGTCGCCGTGCCTACCATGGTG</td><td>199.2</td><td>-1.5</td><td>0.5</td><td>-2.9</td><td>3.6E-03</td><td>4.4E-03</td><td>MB231/MB468</td><td>CYTO</td></tr>
<tr><td>>MI0003581|hsa-mir-574&WithFlank&4|+|38868026|38868133@67.87.21(+1U)&[MIMAT0003239&hsa-miR-574-3p&offsets|0|-1(+1U);m-149&4|+|38868092|38868113&offsets|0|-1(+1U)]|CACGCTCATGCACACACCCACT</td><td>>MI0003581|hsa-mir-574&WithFlank&4|+|38868026|38868133</td><td>67</td><td>87</td><td>21(+1U)</td><td>CACGCTCATGCACACACCCACT</td><td>199.4</td><td>-3.7</td><td>1.1</td><td>-3.4</td><td>6.8E-04</td><td>4.4E-03</td><td>BT20/MB231</td><td>MP</td></tr>
<tr><td>>MI0000299|hsa-mir-222&WithFlank&X|-|45747009|45747130@75.99.25(+2U)&[MIMAT0000279&hsa-miR-222-3p&offsets|0|+4(+2U);m-140&X|-|45747033|45747056&offsets|0|+1(+2U)]|AGCTACATCTGGCTACTGGGTCTCTTT</td><td>>MI0000299|hsa-mir-222&WithFlank&X|-|45747009|45747130</td><td>75</td><td>99</td><td>25(+2U)</td><td>AGCTACATCTGGCTACTGGGTCTCTTT</td><td>3257.6</td><td>-2.7</td><td>0.8</td><td>-3.3</td><td>8.2E-04</td><td>4.4E-03</td><td>BT20/MB231</td><td>CYTO</td></tr>
<tr><td>>hg38_21_+_8433172_8446622_RNA45SN1_45S_50ntFlanks@10783.10817.35|GTGAACAGCCTCTGGCATGTTGGAACAATGTAGGT</td><td>>hg38_21_+_8433172_8446622_RNA45SN1_45S_50ntFlanks</td><td>10783</td><td>10817</td><td>35</td><td>GTGAACAGCCTCTGGCATGTTGGAACAATGTAGGT</td><td>311.9</td><td>-2.8</td><td>0.9</td><td>-3.1</td><td>2.0E-03</td><td>4.4E-03</td><td>BT20/MB468</td><td>MP</td></tr>
<tr><td>>MI0000264|hsa-mir-7-2&WithFlank&15|+|88611819|88611940@38.59.22(+1A);MI0000265|hsa-mir-7-3&WithFlank&19|+|4770664|4770785@37.58.22(+1A);MI0000263|hsa-mir-7-1&WithFlank&9|-|83969742|83969863@30.51.22(+1A)&[MIMAT0000252&hsa-miR-7-5p&offsets|0|-2(+1A);m-400&15|+|88611856|88611879&offsets|0|-2(+1A)];[MIMAT0000252_1&hsa-miR-7-5p&offsets|0|-2(+1A);m-402&19|+|4770700|4770723&offsets|0|-2(+1A)];[MIMAT0000252_2&hsa-miR-7-5p&offsets|0|-2(+1A);m-401&9|-|83969811|83969834&offsets|0|-2(+1A)]|TGGAAGACTAGTGATTTTGTTGA</td><td>>MI0000264|hsa-mir-7-2&WithFlank&15|+|88611819|88611940</td><td>38</td><td>59</td><td>22(+1A)</td><td>TGGAAGACTAGTGATTTTGTTGA</td><td>338.1</td><td>3.7</td><td>1.3</td><td>2.9</td><td>3.2E-03</td><td>4.4E-03</td><td>MB231/MB468</td><td>MP</td></tr>
<tr><td>>hg38_MT_+_1621_3279_RNR2_16S_50ntFlanks@1095.1119.25|AGACGAGAAGACCCTATGGAGCTTT</td><td>>hg38_MT_+_1621_3279_RNR2_16S_50ntFlanks</td><td>1095</td><td>1119</td><td>25</td><td>AGACGAGAAGACCCTATGGAGCTTT</td><td>203.4</td><td>3.3</td><td>1.1</td><td>3.1</td><td>2.0E-03</td><td>4.4E-03</td><td>BT20/MB468</td><td>MP</td></tr>
<tr><td>>hg38_21_+_8433172_8446622_RNA45SN1_45S_50ntFlanks@8155.8178.24|CGCTCGTGGGGGGCCCAAGTCCTT</td><td>>hg38_21_+_8433172_8446622_RNA45SN1_45S_50ntFlanks</td><td>8155</td><td>8178</td><td>24</td><td>CGCTCGTGGGGGGCCCAAGTCCTT</td><td>119.7</td><td>-5.0</td><td>1.6</td><td>-3.1</td><td>2.0E-03</td><td>4.4E-03</td><td>BT20/MB468</td><td>MP</td></tr>
<tr><td>>MI0000289|hsa-mir-181a-1&WithFlank&1|-|198859038|198859159@30.52.23(+1A);MI0000269|hsa-mir-181a-2&WithFlank&9|+|124692436|124692557@45.67.23(+1A)&[MIMAT0000256&hsa-miR-181a-5p&offsets|0|0(+1A);m-32&1|-|198859108|198859130&offsets|0|0(+1A)];[MIMAT0000256_1&hsa-miR-181a-5p&offsets|0|0(+1A);m-32&9|+|124692480|124692502&offsets|0|0(+1A)]|AACATTCAACGCTGTCGGTGAGTA</td><td>>MI0000289|hsa-mir-181a-1&WithFlank&1|-|198859038|198859159</td><td>30</td><td>52</td><td>23(+1A)</td><td>AACATTCAACGCTGTCGGTGAGTA</td><td>452.2</td><td>-4.8</td><td>1.4</td><td>-3.4</td><td>6.8E-04</td><td>4.4E-03</td><td>BT20/MB231</td><td>MP</td></tr>
<tr><td>>hg38_21_+_8433172_8446622_RNA45SN1_45S_50ntFlanks@11539.11584.46|GCGCGGCGCCTCGCCTCGGCCGGCGCCTAGCAGCCGACTTAGAACT</td><td>>hg38_21_+_8433172_8446622_RNA45SN1_45S_50ntFlanks</td><td>11539</td><td>11584</td><td>46</td><td>GCGCGGCGCCTCGCCTCGGCCGGCGCCTAGCAGCCGACTTAGAACT</td><td>272.1</td><td>4.6</td><td>1.6</td><td>2.9</td><td>3.8E-03</td><td>4.4E-03</td><td>MB231/MB468</td><td>MITO</td></tr>
<tr><td>>trnaMT_SerGCT_MT_+_12207_12265@14.40.27|AGAACTGCTAACTCATGCCCCCATGTC</td><td>>trnaMT_SerGCT_MT_+_12207_12265</td><td>14</td><td>40</td><td>27</td><td>AGAACTGCTAACTCATGCCCCCATGTC</td><td>172.0</td><td>2.4</td><td>0.8</td><td>2.9</td><td>3.2E-03</td><td>4.4E-03</td><td>MB231/MB468</td><td>MP</td></tr>
<tr><td>>hg38_21_+_8433172_8446622_RNA45SN1_45S_50ntFlanks@8176.8194.19|CTTCTGATCGAGGCCCAGC</td><td>>hg38_21_+_8433172_8446622_RNA45SN1_45S_50ntFlanks</td><td>8176</td><td>8194</td><td>19</td><td>CTTCTGATCGAGGCCCAGC</td><td>355.0</td><td>-3.7</td><td>1.3</td><td>-2.9</td><td>3.2E-03</td><td>4.4E-03</td><td>MB231/MB468</td><td>MP</td></tr>
<tr><td>>trnaMT_ValTAC_MT_+_1602_1670@50.71.22|ACTTAACTTGACCGCTCTGACC</td><td>>trnaMT_ValTAC_MT_+_1602_1670</td><td>50</td><td>71</td><td>22</td><td>ACTTAACTTGACCGCTCTGACC</td><td>913.0</td><td>-3.4</td><td>1.2</td><td>-2.9</td><td>3.8E-03</td><td>4.4E-03</td><td>BT20/MB468</td><td>NUC</td></tr>
</table>

<table>
<tr><td>>MI0000077|hsa-mir-21&WithFlank&17|+|59841260|59841343@52.71.20(+1C)&[MIMAT0004494&hsa-miR-21-3p&offsets|0|-1(+1C);m-52&17|+|59841311|59841332&offsets|0|-2(+1C)]|CAACACCAGTCGATGGGCTGC</td><td>>MI0000077|hsa-mir-21&WithFlank&17|+|59841260|59841343</td><td>52</td><td>71</td><td>20(+1C)</td><td>CAACACCAGTCGATGGGCTGC</td><td>316.5</td><td>5.9</td><td>1.9</td><td>3.1</td><td>2.0E-03</td><td>4.4E-03</td><td>BT20/MB468</td><td>MP</td></tr>
<tr><td>>MI0003205|hsa-mir-532&WithFlank&X|+|50003142|50003244@27.47.21&[MIMAT0002888&hsa-miR-532-5p&offsets|+1|0;m-65&X|+|50003167|50003188&offsets|+1|0]|ATGCCTTGAGTGTAGGACCGT</td><td>>MI0003205|hsa-mir-532&WithFlank&X|+|50003142|50003244</td><td>27</td><td>47</td><td>21</td><td>ATGCCTTGAGTGTAGGACCGT</td><td>404.0</td><td>3.2</td><td>1.0</td><td>3.1</td><td>2.0E-03</td><td>4.4E-03</td><td>BT20/MB468</td><td>MP</td></tr>
<tr><td>>MI0000077|hsa-mir-21&WithFlank&17|+|59841260|59841343@14.33.20&[MIMAT0000076&hsa-miR-21-5p&offsets|0|-2;m-1&17|+|59841273|59841294&offsets|0|-2]|TAGCTTATCAGACTGATGTT</td><td>>MI0000077|hsa-mir-21&WithFlank&17|+|59841260|59841343</td><td>14</td><td>33</td><td>20</td><td>TAGCTTATCAGACTGATGTT</td><td>1672.7</td><td>3.5</td><td>1.1</td><td>3.1</td><td>2.0E-03</td><td>4.4E-03</td><td>BT20/MB468</td><td>MP</td></tr>
<tr><td>>am_MI0000803|hsa-mir-330&WithFlank&19|-|45638988|45639093@63.84.22(+1U)&[MIMAT0000751&hsa-miR-330-3p&offsets|0|-1(+1U);m-347&19|-|45639009|45639030&offsets|-1|-1(+1U)]|GCAAAGCACACGGCCTGCAGAGT</td><td>>am_MI0000803|hsa-mir-330&WithFlank&19|-|45638988|45639093</td><td>63</td><td>84</td><td>22(+1U)</td><td>GCAAAGCACACGGCCTGCAGAGT</td><td>80.4</td><td>6.3</td><td>2.0</td><td>3.1</td><td>2.0E-03</td><td>4.4E-03</td><td>BT20/MB468</td><td>MP</td></tr>
<tr><td>>hg38_21_+_8433172_8446622_RNA45SN1_45S_50ntFlanks@3838.3874.37|CTCTCCTACTTGGATAACTGTGGTAATTCTAGAGCTA</td><td>>hg38_21_+_8433172_8446622_RNA45SN1_45S_50ntFlanks</td><td>3838</td><td>3874</td><td>37</td><td>CTCTCCTACTTGGATAACTGTGGTAATTCTAGAGCTA</td><td>32.8</td><td>-8.1</td><td>2.6</td><td>-3.1</td><td>2.0E-03</td><td>4.4E-03</td><td>BT20/MB468</td><td>MP</td></tr>
<tr><td>>hg38_21_+_8433172_8446622_RNA45SN1_45S_50ntFlanks@5486.5514.29|GCGCTGAGAAGACGGTCGAACTTGACTAT</td><td>>hg38_21_+_8433172_8446622_RNA45SN1_45S_50ntFlanks</td><td>5486</td><td>5514</td><td>29</td><td>GCGCTGAGAAGACGGTCGAACTTGACTAT</td><td>3141.1</td><td>-3.7</td><td>1.2</td><td>-3.1</td><td>2.0E-03</td><td>4.4E-03</td><td>BT20/MB468</td><td>MP</td></tr>
<tr><td>>hg38_MT_+_1621_3279_RNR2_16S_50ntFlanks@883.906.24|AAAAACATCACCTCTAGCATCACC</td><td>>hg38_MT_+_1621_3279_RNR2_16S_50ntFlanks</td><td>883</td><td>906</td><td>24</td><td>AAAAACATCACCTCTAGCATCACC</td><td>109.2</td><td>2.6</td><td>0.9</td><td>2.9</td><td>3.2E-03</td><td>4.4E-03</td><td>MB231/MB468</td><td>MP</td></tr>
<tr><td>>trnaMT_GluTTC_MT_-_14674_14742@41.71.31,trnalookalike8_GluTTC_5_-_93905172_93905240@41.71.31|TGGTCGTGGTTGTAGTCCGTGCGAGAATACC</td><td>>trnaMT_GluTTC_MT_-_14674_14742</td><td>41</td><td>71</td><td>31</td><td>TGGTCGTGGTTGTAGTCCGTGCGAGAATACC</td><td>419.7</td><td>-2.9</td><td>0.9</td><td>-3.4</td><td>7.5E-04</td><td>4.4E-03</td><td>BT20/MB231</td><td>NUC</td></tr>
<tr><td>>hg38_21_+_8433172_8446622_RNA45SN1_45S_50ntFlanks@11769.11790.22|TAATTAGTGACGCGCATGAATG</td><td>>hg38_21_+_8433172_8446622_RNA45SN1_45S_50ntFlanks</td><td>11769</td><td>11790</td><td>22</td><td>TAATTAGTGACGCGCATGAATG</td><td>234.4</td><td>-4.9</td><td>1.7</td><td>-2.9</td><td>3.2E-03</td><td>4.4E-03</td><td>MB231/MB468</td><td>MP</td></tr>
<tr><td>>MI0000749|hsa-mir-30e&WithFlank&1|+|40754349|40754452@65.85.21(+2U)&[MIMAT0000693&hsa-miR-30e-3p&offsets|0|-1(+2U);m-25&1|+|40754413|40754434&offsets|0|-1(+2U)]|CTTTCAGTCGGATGTTTACAGTT</td><td>>MI0000749|hsa-mir-30e&WithFlank&1|+|40754349|40754452</td><td>65</td><td>85</td><td>21(+2U)</td><td>CTTTCAGTCGGATGTTTACAGTT</td><td>602.4</td><td>-3.2</td><td>1.0</td><td>-3.4</td><td>6.9E-04</td><td>4.4E-03</td><td>BT20/MB231</td><td>MP</td></tr>
<tr><td>>hg38_21_+_8433172_8446622_RNA45SN1_45S_50ntFlanks@4739.4758.20|AAGTCGGAGGTTCGAAGACG</td><td>>hg38_21_+_8433172_8446622_RNA45SN1_45S_50ntFlanks</td><td>4739</td><td>4758</td><td>20</td><td>AAGTCGGAGGTTCGAAGACG</td><td>173.3</td><td>-1.7</td><td>0.6</td><td>-2.9</td><td>3.5E-03</td><td>4.4E-03</td><td>BT20/MB468</td><td>CYTO</td></tr>
<tr><td>>hg38_21_+_8433172_8446622_RNA45SN1_45S_50ntFlanks@6650.6696.47|ACGACTCTTAGCGGTGGATCACTCGGCTCGTGCGTCGATGAAGAACG</td><td>>hg38_21_+_8433172_8446622_RNA45SN1_45S_50ntFlanks</td><td>6650</td><td>6696</td><td>47</td><td>ACGACTCTTAGCGGTGGATCACTCGGCTCGTGCGTCGATGAAGAACG</td><td>460.0</td><td>-2.9</td><td>0.9</td><td>-3.3</td><td>8.3E-04</td><td>4.4E-03</td><td>BT20/MB231</td><td>CYTO</td></tr>
<tr><td>>hg38_1_-_228634819_228635039_RNA5S12_5S_50ntFlanks@133.171.39|ATGGGAGACCGCCTGGGAATACCGGGTGCTGTAGGCTTT</td><td>>hg38_1_-_228634819_228635039_RNA5S12_5S_50ntFlanks</td><td>133</td><td>171</td><td>39</td><td>ATGGGAGACCGCCTGGGAATACCGGGTGCTGTAGGCTTT</td><td>6093.6</td><td>3.4</td><td>1.1</td><td>3.0</td><td>3.0E-03</td><td>4.4E-03</td><td>BT20/MB468</td><td>MITO</td></tr>
<tr><td>>hg38_21_+_8433172_8446622_RNA45SN1_45S_50ntFlanks@10826.10865.40|TCGGCAAGCCGGATCCGTAACTTCGGGATAAGGATTGGCT</td><td>>hg38_21_+_8433172_8446622_RNA45SN1_45S_50ntFlanks</td><td>10826</td><td>10865</td><td>40</td><td>TCGGCAAGCCGGATCCGTAACTTCGGGATAAGGATTGGCT</td><td>72.1</td><td>-5.3</td><td>1.7</td><td>-3.1</td><td>2.0E-03</td><td>4.4E-03</td><td>BT20/MB468</td><td>MP</td></tr>
<tr><td>>MI0000760|hsa-mir-361&WithFlank&X|-|85903630|85903713@51.75.25&[MIMAT0004682&hsa-miR-361-3p&offsets|0|+2;m-124&X|-|85903642|85903663&offsets|0|+3]|TCCCCCAGGTGTGATTCTGATTTGC</td><td>>MI0000760|hsa-mir-361&WithFlank&X|-|85903630|85903713</td><td>51</td><td>75</td><td>25</td><td>TCCCCCAGGTGTGATTCTGATTTGC</td><td>162.5</td><td>-2.5</td><td>0.7</td><td>-3.4</td><td>7.6E-04</td><td>4.4E-03</td><td>BT20/MB231</td><td>NUC</td></tr>
<tr><td>>hg38_21_+_8433172_8446622_RNA45SN1_45S_50ntFlanks@8318.8356.39|ACGAGACCGATAGTCAACAAGTACCGTAAGGGAAAGTTG</td><td>>hg38_21_+_8433172_8446622_RNA45SN1_45S_50ntFlanks</td><td>8318</td><td>8356</td><td>39</td><td>ACGAGACCGATAGTCAACAAGTACCGTAAGGGAAAGTTG</td><td>156.3</td><td>-2.2</td><td>0.8</td><td>-2.9</td><td>3.6E-03</td><td>4.4E-03</td><td>MB231/MB468</td><td>CYTO</td></tr>
<tr><td>>hg38_MT_+_1621_3279_RNR2_16S_50ntFlanks@826.860.35|AAGGAAAGGTTAAAAAAAGTAAAAGGAACTCGGCA</td><td>>hg38_MT_+_1621_3279_RNR2_16S_50ntFlanks</td><td>826</td><td>860</td><td>35</td><td>AAGGAAAGGTTAAAAAAAGTAAAAGGAACTCGGCA</td><td>57.1</td><td>5.4</td><td>1.8</td><td>2.9</td><td>3.2E-03</td><td>4.4E-03</td><td>MB231/MB468</td><td>MP</td></tr>
<tr><td>>hg38_21_+_8433172_8446622_RNA45SN1_45S_50ntFlanks@8911.8936.26|TCGCCGAATCCCGGGGCCGAGGGAGC</td><td>>hg38_21_+_8433172_8446622_RNA45SN1_45S_50ntFlanks</td><td>8911</td><td>8936</td><td>26</td><td>TCGCCGAATCCCGGGGCCGAGGGAGC</td><td>102.8</td><td>6.2</td><td>2.0</td><td>3.1</td><td>2.0E-03</td><td>4.4E-03</td><td>BT20/MB468</td><td>MP</td></tr>
<tr><td>>hg38_21_+_8433172_8446622_RNA45SN1_45S_50ntFlanks@12007.12030.24|GCCGCCGGTGAAATACCACTACTC</td><td>>hg38_21_+_8433172_8446622_RNA45SN1_45S_50ntFlanks</td><td>12007</td><td>12030</td><td>24</td><td>GCCGCCGGTGAAATACCACTACTC</td><td>489.1</td><td>-3.8</td><td>1.3</td><td>-2.9</td><td>3.8E-03</td><td>4.5E-03</td><td>MB231/MB468</td><td>NUC</td></tr>
<tr><td>>hg38_21_+_8433172_8446622_RNA45SN1_45S_50ntFlanks@4131.4151.21|TTCCGGAGAGGGAGCCTGAGA</td><td>>hg38_21_+_8433172_8446622_RNA45SN1_45S_50ntFlanks</td><td>4131</td><td>4151</td><td>21</td><td>TTCCGGAGAGGGAGCCTGAGA</td><td>143.6</td><td>5.3</td><td>1.7</td><td>3.1</td><td>2.0E-03</td><td>4.5E-03</td><td>BT20/MB468</td><td>MP</td></tr>
<tr><td>>hg38_21_+_8433172_8446622_RNA45SN1_45S_50ntFlanks@9198.9222.25|GTTCTCTCGGGGCCACGCGCGCGTC</td><td>>hg38_21_+_8433172_8446622_RNA45SN1_45S_50ntFlanks</td><td>9198</td><td>9222</td><td>25</td><td>GTTCTCTCGGGGCCACGCGCGCGTC</td><td>1309.8</td><td>-6.0</td><td>2.1</td><td>-2.9</td><td>3.8E-03</td><td>4.5E-03</td><td>BT20/MB468</td><td>NUC</td></tr>
<tr><td>>hg38_21_+_8433172_8446622_RNA45SN1_45S_50ntFlanks@4557.4587.31|CAGCTAGGAATAATGGAATAGGACCGCGGTT</td><td>>hg38_21_+_8433172_8446622_RNA45SN1_45S_50ntFlanks</td><td>4557</td><td>4587</td><td>31</td><td>CAGCTAGGAATAATGGAATAGGACCGCGGTT</td><td>165.7</td><td>-3.9</td><td>1.3</td><td>-3.1</td><td>2.0E-03</td><td>4.5E-03</td><td>BT20/MB468</td><td>MP</td></tr>
<tr><td>>hg38_21_+_8433172_8446622_RNA45SN1_45S_50ntFlanks@12994.13017.24|CCTCGCTGCGATCTATTGAAAGTC</td><td>>hg38_21_+_8433172_8446622_RNA45SN1_45S_50ntFlanks</td><td>12994</td><td>13017</td><td>24</td><td>CCTCGCTGCGATCTATTGAAAGTC</td><td>172.2</td><td>-2.7</td><td>0.8</td><td>-3.3</td><td>8.3E-04</td><td>4.5E-03</td><td>BT20/MB231</td><td>CYTO</td></tr>
<tr><td>>hg38_21_+_8433172_8446622_RNA45SN1_45S_50ntFlanks@12404.12432.29|GCTTTTTGATCCTTCGATGTCGGCTCTTC</td><td>>hg38_21_+_8433172_8446622_RNA45SN1_45S_50ntFlanks</td><td>12404</td><td>12432</td><td>29</td><td>GCTTTTTGATCCTTCGATGTCGGCTCTTC</td><td>483.2</td><td>-1.9</td><td>0.6</td><td>-2.9</td><td>3.6E-03</td><td>4.5E-03</td><td>MB231/MB468</td><td>CYTO</td></tr>
<tr><td>>hg38_21_+_8433172_8446622_RNA45SN1_45S_50ntFlanks@12658.12683.26|ACTGAACGCCTCTAAGTCAGAATCCC</td><td>>hg38_21_+_8433172_8446622_RNA45SN1_45S_50ntFlanks</td><td>12658</td><td>12683</td><td>26</td><td>ACTGAACGCCTCTAAGTCAGAATCCC</td><td>58.9</td><td>-5.0</td><td>1.6</td><td>-3.1</td><td>2.0E-03</td><td>4.5E-03</td><td>BT20/MB468</td><td>MP</td></tr>
<tr><td>>hg38_21_+_8433172_8446622_RNA45SN1_45S_50ntFlanks@10318.10354.37|TGCAGATCTTGGTGGTAGTAGCAAATATTCAAACGAG</td><td>>hg38_21_+_8433172_8446622_RNA45SN1_45S_50ntFlanks</td><td>10318</td><td>10354</td><td>37</td><td>TGCAGATCTTGGTGGTAGTAGCAAATATTCAAACGAG</td><td>52.1</td><td>5.9</td><td>1.9</td><td>3.1</td><td>2.0E-03</td><td>4.5E-03</td><td>BT20/MB468</td><td>MP</td></tr>
<tr><td>>hg38_21_+_8433172_8446622_RNA45SN1_45S_50ntFlanks@10910.10932.23|CGCGCGCCGCGGCTGGACGAGGC</td><td>>hg38_21_+_8433172_8446622_RNA45SN1_45S_50ntFlanks</td><td>10910</td><td>10932</td><td>23</td><td>CGCGCGCCGCGGCTGGACGAGGC</td><td>258.5</td><td>-4.7</td><td>1.6</td><td>-2.9</td><td>3.5E-03</td><td>4.5E-03</td><td>BT20/MB468</td><td>CYTO</td></tr>
<tr><td>>MI0000750|hsa-mir-26a-2&WithFlank&12|-|57824603|57824698@20.41.22(+2U);MI0000083|hsa-mir-26a-1&WithFlank&3|+|37969398|37969486@16.37.22(+2U)&[MIMAT0000082&hsa-miR-26a-5p&offsets|0|0(+2U);m-33&12|-|57824658|57824679&offsets|0|0(+2U)];[MIMAT0000082_1&hsa-miR-26a-5p&offsets|0|0(+2U);m-34&3|+|37969413|37969434&offsets|0|0(+2U)]|TTCAAGTAATCCAGGATAGGCTTT</td><td>>MI0000750|hsa-mir-26a-2&WithFlank&12|-|57824603|57824698</td><td>20</td><td>41</td><td>22(+2U)</td><td>TTCAAGTAATCCAGGATAGGCTTT</td><td>737.9</td><td>3.5</td><td>1.1</td><td>3.1</td><td>2.0E-03</td><td>4.5E-03</td><td>BT20/MB468</td><td>MP</td></tr>
<tr><td>>MI0000086|hsa-mir-28&WithFlank&3|+|188688775|188688872@60.81.22(+2A)&[MIMAT0004502&hsa-miR-28-3p&offsets|0|0(+2A);m-28&3|+|188688834|188688855&offsets|0|0(+2A)]|CACTAGATTGTGAGCTCCTGGAAA</td><td>>MI0000086|hsa-mir-28&WithFlank&3|+|188688775|188688872</td><td>60</td><td>81</td><td>22(+2A)</td><td>CACTAGATTGTGAGCTCCTGGAAA</td><td>483.6</td><td>2.8</td><td>0.8</td><td>3.3</td><td>8.3E-04</td><td>4.5E-03</td><td>BT20/MB231</td><td>CYTO</td></tr>
<tr><td>>hg38_21_+_8433172_8446622_RNA45SN1_45S_50ntFlanks@4200.4219.20|CCCGACCCGGGGAGGTAGTG</td><td>>hg38_21_+_8433172_8446622_RNA45SN1_45S_50ntFlanks</td><td>4200</td><td>4219</td><td>20</td><td>CCCGACCCGGGGAGGTAGTG</td><td>117.5</td><td>-2.6</td><td>0.8</td><td>-3.1</td><td>2.0E-03</td><td>4.5E-03</td><td>BT20/MB468</td><td>MP</td></tr>
<tr><td>>hg38_21_+_8433172_8446622_RNA45SN1_45S_50ntFlanks@10677.10710.34|GCGGTTCCGGCGGCGTCCGGTGAGCTCTCGCTGG</td><td>>hg38_21_+_8433172_8446622_RNA45SN1_45S_50ntFlanks</td><td>10677</td><td>10710</td><td>34</td><td>GCGGTTCCGGCGGCGTCCGGTGAGCTCTCGCTGG</td><td>10905.0</td><td>-4.2</td><td>1.5</td><td>-2.9</td><td>3.8E-03</td><td>4.5E-03</td><td>BT20/MB468</td><td>NUC</td></tr>
<tr><td>>hg38_21_+_8433172_8446622_RNA45SN1_45S_50ntFlanks@11541.11584.44|GCGGCGCCTCGCCTCGGCCGGCGCCTAGCAGCCGACTTAGAACT</td><td>>hg38_21_+_8433172_8446622_RNA45SN1_45S_50ntFlanks</td><td>11541</td><td>11584</td><td>44</td><td>GCGGCGCCTCGCCTCGGCCGGCGCCTAGCAGCCGACTTAGAACT</td><td>142.8</td><td>5.3</td><td>1.8</td><td>2.9</td><td>3.8E-03</td><td>4.5E-03</td><td>MB231/MB468</td><td>MITO</td></tr>
<tr><td>>hg38_21_+_8433172_8446622_RNA45SN1_45S_50ntFlanks@10348.10366.19|AAACGAGAACTTTGAAGGC</td><td>>hg38_21_+_8433172_8446622_RNA45SN1_45S_50ntFlanks</td><td>10348</td><td>10366</td><td>19</td><td>AAACGAGAACTTTGAAGGC</td><td>311.2</td><td>-2.1</td><td>0.7</td><td>-2.9</td><td>3.8E-03</td><td>4.5E-03</td><td>MB231/MB468</td><td>MITO</td></tr>
<tr><td>>trna129_MetCAT_6_-_27870271_27870342@14.31.18,trna142_MetCAT_6_-_27560600_27560671@14.31.18,trna150_MetCAT_6_-_27300764_27300835@14.31.18,trna151_ThrCGT_6_-_27271568_27271639@14.31.18,trna169_MetCAT_6_-_26330529_26330600@14.31.18,trna171_MetCAT_6_-_26313352_26313423@14.31.18,trna20_MetCAT_17_-_80452597_80452668@14.31.18,trna2_MetCAT_6_+_26286754_26286825@14.31.18,trna32_MetCAT_1_+_153643726_153643797@14.31.18,trna61_MetCAT_6_+_27745664_27745735@14.31.18|AGCGGAAGCGTGCTGGGC</td><td>>trna129_MetCAT_6_-_27870271_27870342</td><td>14</td><td>31</td><td>18</td><td>AGCGGAAGCGTGCTGGGC</td><td>399.4</td><td>-3.8</td><td>1.3</td><td>-2.9</td><td>3.9E-03</td><td>4.5E-03</td><td>MB231/MB468</td><td>NUC</td></tr>
<tr><td>>trna111_HisGTG_1_-_147774845_147774916@-1A.31.32,trna118_HisGTG_1_-_145396881_145396952@-1A.31.32,trna16_HisGTG_1_+_146544773_146544844@-1A.31.32,trna1_HisGTG_15_+_45493349_45493420@-1A.31.32,trna21_HisGTG_1_+_147753471_147753542@-1A.31.32,trna33_HisGTG_6_+_27125906_27125977@-1A.31.32,trna7_HisGTG_9_-_14433938_14434009@-1A.31.32,trna8_HisGTG_15_-_45492611_45492682@-1A.31.32,trna9_HisGTG_15_-_45490804_45490875@-1A.31.32|AGCCGTGATCGTATAGTGGTTAGTACTCTGCG</td><td>>trna111_HisGTG_1_-_147774845_147774916</td><td>-1A</td><td>31</td><td>32</td><td>AGCCGTGATCGTATAGTGGTTAGTACTCTGCG</td><td>540.1</td><td>-2.8</td><td>1.0</td><td>-2.9</td><td>3.8E-03</td><td>4.5E-03</td><td>MB231/MB468</td><td>MITO</td></tr>
<tr><td>>trna116_GluCTC_1_-_145399233_145399304@1.33.33,trna59_GluCTC_1_+_249168447_249168518@1.33.33,trna71_GluCTC_1_-_161439189_161439260@1.33.33,trna74_GluCTC_1_-_161431809_161431880@1.33.33,trna77_GluCTC_1_-_161424398_161424469@1.33.33,trna77_GluCTC_6_+_28949976_28950047@1.33.33,trna80_GluCTC_1_-_161417018_161417089@1.33.33,trna87_GluCTC_6_-_126101393_126101464@1.33.33|TCCCTGGTGGTCTAGTGGTTAGGATTCGGCGCT</td><td>>trna116_GluCTC_1_-_145399233_145399304</td><td>1</td><td>33</td><td>33</td><td>TCCCTGGTGGTCTAGTGGTTAGGATTCGGCGCT</td><td>3853.9</td><td>3.4</td><td>1.1</td><td>2.9</td><td>3.3E-03</td><td>4.5E-03</td><td>MB231/MB468</td><td>MP</td></tr>
<tr><td>>hg38_21_+_8433172_8446622_RNA45SN1_45S_50ntFlanks@11874.11909.36|AGAAGACCCTGTTGAGCTTGACTCTAGTCTGGCACG</td><td>>hg38_21_+_8433172_8446622_RNA45SN1_45S_50ntFlanks</td><td>11874</td><td>11909</td><td>36</td><td>AGAAGACCCTGTTGAGCTTGACTCTAGTCTGGCACG</td><td>255.9</td><td>-3.4</td><td>1.2</td><td>-2.9</td><td>3.6E-03</td><td>4.5E-03</td><td>BT20/MB468</td><td>CYTO</td></tr>
<tr><td>>hg38_MT_+_1621_3279_RNR2_16S_50ntFlanks@1089.1120.32|ACAGCAAGACGAGAAGACCCTATGGAGCTTTA</td><td>>hg38_MT_+_1621_3279_RNR2_16S_50ntFlanks</td><td>1089</td><td>1120</td><td>32</td><td>ACAGCAAGACGAGAAGACCCTATGGAGCTTTA</td><td>183.4</td><td>4.3</td><td>1.4</td><td>3.1</td><td>2.0E-03</td><td>4.5E-03</td><td>BT20/MB468</td><td>MP</td></tr>
<tr><td>>hg38_21_+_8433172_8446622_RNA45SN1_45S_50ntFlanks@4588.4619.32|CTATTTTGTTGGTTTTCGGAACTGAGGCCATG</td><td>>hg38_21_+_8433172_8446622_RNA45SN1_45S_50ntFlanks</td><td>4588</td><td>4619</td><td>32</td><td>CTATTTTGTTGGTTTTCGGAACTGAGGCCATG</td><td>328.7</td><td>-2.2</td><td>0.7</td><td>-2.9</td><td>3.6E-03</td><td>4.5E-03</td><td>BT20/MB468</td><td>CYTO</td></tr>
</table>

| | | | | | | | | | | | | | |
|---|---|---|---|---|---|---|---|---|---|---|---|---|---|
| >hg38_21_+_8433172_8446622_RNA45SN1_45S_50ntFlanks@7974.8001.28\|ACGCGACCTCAGATCAGACGTGGCGACC | >hg38_21_+_8433172_8446622_RNA45SN1_45S_50ntFlanks | 7974 | 8001 | 28 | ACGCGACCTCAGATCAGACGTGGCGACC | 151.3 | 2.7 | 0.9 | 2.9 | 3.8E-03 | 4.5E-03 | MB231/MB468 | MITO |
| >hg38_21_+_8433172_8446622_RNA45SN1_45S_50ntFlanks@9939.9980.42\|GCAGGACGGTGGCCATGGAAGTCGGAATCCGCTAAGGAGTGT | >hg38_21_+_8433172_8446622_RNA45SN1_45S_50ntFlanks | 9939 | 9980 | 42 | GCAGGACGGTGGCCATGGAAGTCGGAATCCGCTAAGGAGTGT | 731.7 | 7.1 | 2.1 | 3.4 | 6.9E-04 | 4.5E-03 | BT20/MB231 | MP |
| >hg38_21_+_8433172_8446622_RNA45SN1_45S_50ntFlanks@4130.4148.19\|ATTCCGGAGAGGGAGCCTG | >hg38_21_+_8433172_8446622_RNA45SN1_45S_50ntFlanks | 4130 | 4148 | 19 | ATTCCGGAGAGGGAGCCTG | 192.9 | -3.3 | 1.1 | -2.9 | 3.9E-03 | 4.5E-03 | MB231/MB468 | NUC |
| >hg38_21_+_8433172_8446622_RNA45SN1_45S_50ntFlanks@6652.6696.45\|GACTCTTAGCGGTGGATCACTCGGCTCGTGCGTCGATGAAGAACG | >hg38_21_+_8433172_8446622_RNA45SN1_45S_50ntFlanks | 6652 | 6696 | 45 | GACTCTTAGCGGTGGATCACTCGGCTCGTGCGTCGATGAAGAACG | 1570.7 | -2.9 | 0.9 | -3.3 | 8.4E-04 | 4.5E-03 | BT20/MB231 | CYTO |
| >trnaMT_ValTAC_MT_+_1602_1670@31.72.42\|TTACACTTAGGAGATTTCAACTTAACTTGACCGCTCTGACCA | >trnaMT_ValTAC_MT_+_1602_1670 | 31 | 72 | 42 | TTACACTTAGGAGATTTCAACTTAACTTGACCGCTCTGACCA | 366.5 | 4.9 | 1.7 | 2.9 | 3.8E-03 | 4.5E-03 | MB231/MB468 | MITO |
| >hg38_21_+_8433172_8446622_RNA45SN1_45S_50ntFlanks@9939.9959.21\|GCAGGACGGTGGCCATGGAAG | >hg38_21_+_8433172_8446622_RNA45SN1_45S_50ntFlanks | 9939 | 9959 | 21 | GCAGGACGGTGGCCATGGAAG | 69.9 | 5.0 | 1.6 | 3.1 | 2.0E-03 | 4.5E-03 | BT20/MB468 | MP |
| >MI0000077\|hsa-mir-21&WithFlank&17\|+\|59841260\|59841343@14.34.21(+1G)&[MIMAT0000076&hsa-miR-21-5p&offsets\|0\|-1(+1G);m-1&17\|+\|59841273\|59841294&offsets\|0\|-1(+1G)]\|TAGCTTATCAGACTGATGTTGG | >MI0000077\|hsa-mir-21&WithFlank&17\|+\|59841260\|59841343 | 14 | 34 | 21(+1G) | TAGCTTATCAGACTGATGTTGG | 3462.4 | 3.5 | 1.1 | 3.3 | 8.4E-04 | 4.5E-03 | BT20/MB231 | CYTO |
| >hg38_21_+_8433172_8446622_RNA45SN1_45S_50ntFlanks@12410.12434.25\|TGATCCTTCGATGTCGGCTCTTCCT | >hg38_21_+_8433172_8446622_RNA45SN1_45S_50ntFlanks | 12410 | 12434 | 25 | TGATCCTTCGATGTCGGCTCTTCCT | 549.7 | 3.2 | 1.1 | 2.9 | 3.6E-03 | 4.5E-03 | BT20/MB468 | CYTO |
| >hg38_21_+_8433172_8446622_RNA45SN1_45S_50ntFlanks@6733.6756.24\|CATTGATCATCGACACTTCGAACG | >hg38_21_+_8433172_8446622_RNA45SN1_45S_50ntFlanks | 6733 | 6756 | 24 | CATTGATCATCGACACTTCGAACG | 1620.4 | -3.2 | 1.0 | -3.1 | 2.0E-03 | 4.5E-03 | BT20/MB468 | MP |
| >hg38_21_+_8433172_8446622_RNA45SN1_45S_50ntFlanks@9005.9026.22\|CTCCCCCGCGGGGGCGCGCCGG | >hg38_21_+_8433172_8446622_RNA45SN1_45S_50ntFlanks | 9005 | 9026 | 22 | CTCCCCCGCGGGGGCGCGCCGG | 181.7 | -4.5 | 1.6 | -2.9 | 3.9E-03 | 4.5E-03 | MB231/MB468 | NUC |
| >hg38_21_+_8433172_8446622_RNA45SN1_45S_50ntFlanks@12658.12675.18\|ACTGAACGCCTCTAAGTC | >hg38_21_+_8433172_8446622_RNA45SN1_45S_50ntFlanks | 12658 | 12675 | 18 | ACTGAACGCCTCTAAGTC | 437.0 | -4.3 | 1.4 | -3.1 | 2.0E-03 | 4.5E-03 | BT20/MB468 | MP |
| >hg38_21_+_8433172_8446622_RNA45SN1_45S_50ntFlanks@10808.10838.31\|CAATGTAGGTAAGGGAAGTCGGCAAGCCGGA | >hg38_21_+_8433172_8446622_RNA45SN1_45S_50ntFlanks | 10808 | 10838 | 31 | CAATGTAGGTAAGGGAAGTCGGCAAGCCGGA | 184.0 | -4.3 | 1.4 | -3.0 | 3.0E-03 | 4.5E-03 | BT20/MB468 | MITO |
| >hg38_21_+_8433172_8446622_RNA45SN1_45S_50ntFlanks@12487.12509.23\|GAACGTGAGCTGGGTTTAGACCG | >hg38_21_+_8433172_8446622_RNA45SN1_45S_50ntFlanks | 12487 | 12509 | 23 | GAACGTGAGCTGGGTTTAGACCG | 273.6 | -2.3 | 0.8 | -2.9 | 3.6E-03 | 4.5E-03 | MB231/MB468 | CYTO |
| >hg38_21_+_8433172_8446622_RNA45SN1_45S_50ntFlanks@7991.8015.25\|ACGTGGCGACCCGCTGAATTTAAGC | >hg38_21_+_8433172_8446622_RNA45SN1_45S_50ntFlanks | 7991 | 8015 | 25 | ACGTGGCGACCCGCTGAATTTAAGC | 492.0 | -2.4 | 0.8 | -2.9 | 3.9E-03 | 4.5E-03 | MB231/MB468 | MITO |
| >MI0000082\|hsa-mir-25&WithFlank&7\|-\|100093554\|100093649@58.79.22(+1U)&[MIMAT0000081&hsa-miR-25-3p&offsets\|0\|0(+1U);m-23&7\|-\|100093571\|100093592&offsets\|0\|0(+1U)]\|CATTGCACTTGTCTCGGTCTGAT | >MI0000082\|hsa-mir-25&WithFlank&7\|-\|100093554\|100093649 | 58 | 79 | 22(+1U) | CATTGCACTTGTCTCGGTCTGAT | 219.5 | 1.2 | 0.4 | 2.9 | 3.8E-03 | 4.5E-03 | BT20/MB468 | NUC |
| >hg38_21_+_8433172_8446622_RNA45SN1_45S_50ntFlanks@12243.12262.20\|GAAGGGCAAAAGCTCGCTTG | >hg38_21_+_8433172_8446622_RNA45SN1_45S_50ntFlanks | 12243 | 12262 | 20 | GAAGGGCAAAAGCTCGCTTG | 61.5 | -5.0 | 1.6 | -3.1 | 2.0E-03 | 4.5E-03 | BT20/MB468 | MP |
| >am_MI0000809\|hsa-mir-151a&WithFlank&8\|-\|140732558\|140732659@53.71.19(+1A)&[MIMAT0000757&hsa-miR-151a-3p&offsets\|0\|-2(+1A);m-30&8\|-\|140732587\|140732607&offsets\|0\|-2(+1A)]\|CTAGACTGAAGCTCCTTGAA | >am_MI0000809\|hsa-mir-151a&WithFlank&8\|-\|140732558\|140732659 | 53 | 71 | 19(+1A) | CTAGACTGAAGCTCCTTGAA | 213.4 | 3.7 | 1.3 | 2.9 | 3.3E-03 | 4.5E-03 | MB231/MB468 | MP |
| >trna10_AspGTC_12_-_125424193_125424264@39.73.35,trna12_AspGTC_12_-_125411891_125411962@39.73.35,trna144_AspGTC_6_-_27551236_27551307@39.73.35,trna38_AspGTC_17_-_8125556_8125627@39.73.35,trna45_AspGTC_6_+_27447453_27447524@39.73.35,trna48_AspGTC_6_+_27471523_27471594@39.73.35,trna4_AspGTC_12_+_96429799_96429870@39.73.35,trna69_AspGTC_1_-_161440205_161440276@39.73.35,trna72_AspGTC_1_-_161432824_161432895@39.73.35,trna75_AspGTC_1_-_161425414_161425485@39.73.35,trna78_AspGTC_1_-_161418033_161418104@39.73.35,trna81_AspGTC_1_-_161410615_161410686@39.73.35\|GCGGGAGACCGGGGTTCGATTCCCCGACGGGGAGC | >trna10_AspGTC_12_-_125424193_125424264 | 39 | 73 | 35 | GCGGGAGACCGGGGTTCGATTCCCCGACGGGGAGC | 730.9 | 2.4 | 0.8 | 2.9 | 3.6E-03 | 4.5E-03 | BT20/MB468 | CYTO |
| >hg38_21_+_8433172_8446622_RNA45SN1_45S_50ntFlanks@11577.11602.26\|TTAGAACTGGTGCGGACCAGGGGAAT | >hg38_21_+_8433172_8446622_RNA45SN1_45S_50ntFlanks | 11577 | 11602 | 26 | TTAGAACTGGTGCGGACCAGGGGAAT | 300.3 | -2.7 | 0.9 | -2.9 | 3.6E-03 | 4.5E-03 | BT20/MB468 | CYTO |
| >trnaMT_PheGAA_MT_+_577_647@27.56.30\|AATACACTGAAAATGTTTAGACGGGCTCAC | >trnaMT_PheGAA_MT_+_577_647 | 27 | 56 | 30 | AATACACTGAAAATGTTTAGACGGGCTCAC | 32.0 | 8.3 | 2.5 | 3.4 | 7.0E-04 | 4.5E-03 | BT20/MB231 | MP |
| >hg38_21_+_8433172_8446622_RNA45SN1_45S_50ntFlanks@9573.9602.30\|AAATCGGTCGTCCGACCTGGGTATAGGGGC | >hg38_21_+_8433172_8446622_RNA45SN1_45S_50ntFlanks | 9573 | 9602 | 30 | AAATCGGTCGTCCGACCTGGGTATAGGGGC | 403.4 | 2.0 | 0.7 | 3.0 | 3.0E-03 | 4.5E-03 | BT20/MB468 | MITO |
| >MI0000088\|hsa-mir-30a&WithFlank&6\|-\|71403545\|71403627@54.74.21(+1A)&[MIMAT0000088&hsa-miR-30a-3p&offsets\|+1\|0(+1A);m-38&6\|-\|71403554\|71403575&offsets\|+1\|0(+1A)]\|TTTCAGTCGGATGTTTGCAGCA | >MI0000088\|hsa-mir-30a&WithFlank&6\|-\|71403545\|71403627 | 54 | 74 | 21(+1A) | TTTCAGTCGGATGTTTGCAGCA | 212.0 | -2.6 | 0.8 | -3.3 | 9.3E-04 | 4.5E-03 | BT20/MB231 | MITO |
| >hg38_21_+_8433172_8446622_RNA45SN1_45S_50ntFlanks@13003.13027.25\|GATCTATTGAAAGTCAGCCCTCGAC | >hg38_21_+_8433172_8446622_RNA45SN1_45S_50ntFlanks | 13003 | 13027 | 25 | GATCTATTGAAAGTCAGCCCTCGAC | 258.9 | -4.5 | 1.6 | -2.9 | 3.9E-03 | 4.5E-03 | MB231/MB468 | NUC |
| >hg38_21_+_8433172_8446622_RNA45SN1_45S_50ntFlanks@10771.10788.18\|CAGGTCTCCAAGGTGAAC | >hg38_21_+_8433172_8446622_RNA45SN1_45S_50ntFlanks | 10771 | 10788 | 18 | CAGGTCTCCAAGGTGAAC | 57.1 | -4.9 | 1.6 | -3.1 | 2.0E-03 | 4.5E-03 | BT20/MB468 | MP |
| >hg38_21_+_8433172_8446622_RNA45SN1_45S_50ntFlanks@12004.12031.28\|CGGGCCGCCGGTGAAATACCACTACTCT | >hg38_21_+_8433172_8446622_RNA45SN1_45S_50ntFlanks | 12004 | 12031 | 28 | CGGGCCGCCGGTGAAATACCACTACTCT | 463.5 | 1.9 | 0.7 | 3.0 | 3.0E-03 | 4.5E-03 | BT20/MB468 | MITO |
| >hg38_21_+_8433172_8446622_RNA45SN1_45S_50ntFlanks@9937.9960.24\|CAGCAGGACGGTGGCCATGGAAGT | >hg38_21_+_8433172_8446622_RNA45SN1_45S_50ntFlanks | 9937 | 9960 | 24 | CAGCAGGACGGTGGCCATGGAAGT | 55.4 | 5.1 | 1.6 | 3.1 | 2.0E-03 | 4.5E-03 | BT20/MB468 | MP |
| >hg38_21_+_8433172_8446622_RNA45SN1_45S_50ntFlanks@13000.13039.40\|TGCGATCTATTGAAAGTCAGCCCTCGACACAAGGGTTTGT | >hg38_21_+_8433172_8446622_RNA45SN1_45S_50ntFlanks | 13000 | 13039 | 40 | TGCGATCTATTGAAAGTCAGCCCTCGACACAAGGGTTTGT | 2617.4 | -2.3 | 0.7 | -3.3 | 9.4E-04 | 4.5E-03 | BT20/MB231 | MITO |
| >hg38_21_+_8433172_8446622_RNA45SN1_45S_50ntFlanks@6655.6688.34\|TCTTAGCGGTGGATCACTCGGCTCGTGCGTCGAT | >hg38_21_+_8433172_8446622_RNA45SN1_45S_50ntFlanks | 6655 | 6688 | 34 | TCTTAGCGGTGGATCACTCGGCTCGTGCGTCGAT | 380.1 | 2.3 | 0.7 | 3.3 | 9.4E-04 | 4.5E-03 | BT20/MB231 | MITO |
| >MI0000088\|hsa-mir-30a&WithFlank&6\|-\|71403545\|71403627@54.73.20(+1A)&[MIMAT0000088&hsa-miR-30a-3p&offsets\|+1\|-1(+1A);m-38&6\|-\|71403554\|71403575&offsets\|+1\|-1(+1A)]\|TTTCAGTCGGATGTTTGCAGA | >MI0000088\|hsa-mir-30a&WithFlank&6\|-\|71403545\|71403627 | 54 | 73 | 20(+1A) | TTTCAGTCGGATGTTTGCAGA | 244.6 | -2.5 | 0.8 | -3.3 | 9.4E-04 | 4.5E-03 | BT20/MB231 | MITO |
| >MI0000088\|hsa-mir-30a&WithFlank&6\|-\|71403545\|71403627@54.75.22&[MIMAT0000088&hsa-miR-30a-3p&offsets\|+1\|+1;m-38&6\|-\|71403554\|71403575&offsets\|+1\|+1]\|TTTCAGTCGGATGTTTGCAGCT | >MI0000088\|hsa-mir-30a&WithFlank&6\|-\|71403545\|71403627 | 54 | 75 | 22 | TTTCAGTCGGATGTTTGCAGCT | 938.1 | -2.4 | 0.7 | -3.3 | 9.4E-04 | 4.5E-03 | BT20/MB231 | MITO |
| >hg38_21_+_8433172_8446622_RNA45SN1_45S_50ntFlanks@9171.9195.25\|CGGGTCGCGCCGTCGGGCCCGGGGG | >hg38_21_+_8433172_8446622_RNA45SN1_45S_50ntFlanks | 9171 | 9195 | 25 | CGGGTCGCGCCGTCGGGCCCGGGGG | 207.9 | -5.3 | 1.7 | -3.1 | 2.0E-03 | 4.5E-03 | BT20/MB468 | MP |
| >hg38_21_+_8433172_8446622_RNA45SN1_45S_50ntFlanks@4732.4758.27\|AGAACGAAAGTCGGAGGTTCGAAGACG | >hg38_21_+_8433172_8446622_RNA45SN1_45S_50ntFlanks | 4732 | 4758 | 27 | AGAACGAAAGTCGGAGGTTCGAAGACG | 1077.4 | -2.4 | 0.8 | -3.0 | 3.1E-03 | 4.5E-03 | BT20/MB468 | MITO |
| >hg38_21_+_8433172_8446622_RNA45SN1_45S_50ntFlanks@10841.10860.20\|CGTAACTTCGGGATAAGGAT | >hg38_21_+_8433172_8446622_RNA45SN1_45S_50ntFlanks | 10841 | 10860 | 20 | CGTAACTTCGGGATAAGGAT | 469.0 | -2.6 | 0.9 | -2.9 | 3.6E-03 | 4.5E-03 | BT20/MB468 | CYTO |
| >MI0000103\|hsa-mir-101-1&WithFlank&1\|-\|65058428\|65058514@52.73.22(+1A);MI0000739\|hsa-mir-101-2&WithFlank&9\|+\|4850291\|4850381@54.75.22(+1A)&[MIMAT0000099&hsa-miR-101-3p&offsets\|-1\|0(+1A);m-17&1\|-\|65058443\|65058462&offsets\|-1\|+1(+1A)];[MIMAT0000099_1&hsa-miR-101-3p&offsets\|-1\|0(+1A);m-16&9\|+\|4850345\|4850364&offsets\|-1\|+1(+1A)]\|GTACAGTACTGTGATAACTGAAA | >MI0000103\|hsa-mir-101-1&WithFlank&1\|-\|65058428\|65058514 | 52 | 73 | 22(+1A) | GTACAGTACTGTGATAACTGAAA | 117.3 | 3.0 | 1.0 | 2.9 | 3.9E-03 | 4.5E-03 | BT20/MB468 | NUC |
| >hg38_21_+_8433172_8446622_RNA45SN1_45S_50ntFlanks@3984.4002.19\|GCGGCTTTGGTGACTCTAG | >hg38_21_+_8433172_8446622_RNA45SN1_45S_50ntFlanks | 3984 | 4002 | 19 | GCGGCTTTGGTGACTCTAG | 479.1 | -4.6 | 1.6 | -2.9 | 3.6E-03 | 4.5E-03 | BT20/MB468 | CYTO |
| >hg38_21_+_8433172_8446622_RNA45SN1_45S_50ntFlanks@12715.12751.37\|AGCCTCGGTTGGCCTCGGATAGCCGGTCCCCCGCCTG | >hg38_21_+_8433172_8446622_RNA45SN1_45S_50ntFlanks | 12715 | 12751 | 37 | AGCCTCGGTTGGCCTCGGATAGCCGGTCCCCCGCCTG | 275.5 | -4.4 | 1.5 | -3.0 | 3.1E-03 | 4.5E-03 | BT20/MB468 | MITO |
| >MI0000816\|hsa-mir-335&WithFlank&7\|+\|130496105\|130496210@58.77.20(+1U)&[MIMAT0004703&hsa-miR-335-3p&offsets\|0\|-2(+1U);m-159&7\|+\|130496162\|130496183&offsets\|0\|-2(+1U)]\|TTTTTCATTATTGCTCCTGAT | >MI0000816\|hsa-mir-335&WithFlank&7\|+\|130496105\|130496210 | 58 | 77 | 20(+1U) | TTTTTCATTATTGCTCCTGAT | 34.8 | 8.3 | 2.8 | 2.9 | 3.3E-03 | 4.5E-03 | MB231/MB468 | MP |
| >hg38_21_+_8433172_8446622_RNA45SN1_45S_50ntFlanks@10385.10424.40\|CATGTGAACAGCAGTTGAACATGGGTCAGTCGGTCCTGAG | >hg38_21_+_8433172_8446622_RNA45SN1_45S_50ntFlanks | 10385 | 10424 | 40 | CATGTGAACAGCAGTTGAACATGGGTCAGTCGGTCCTGAG | 427.9 | 3.4 | 1.1 | 3.1 | 2.0E-03 | 4.5E-03 | BT20/MB468 | MP |
| >hg38_21_+_8433172_8446622_RNA45SN1_45S_50ntFlanks@9122.9152.31\|CCGCGCGCGGGTCGGGGGGCGGGGCGGACTG | >hg38_21_+_8433172_8446622_RNA45SN1_45S_50ntFlanks | 9122 | 9152 | 31 | CCGCGCGCGGGTCGGGGGGCGGGGCGGACTG | 299.5 | -5.3 | 1.7 | -3.1 | 2.0E-03 | 4.5E-03 | BT20/MB468 | MP |
| >MI0025750\|hsa-mir-7974&WithFlank&19\|-\|11495538\|11495628@62.84.23(+1U)&[MIMAT0031177&hsa-miR-7974&offsets\|0\|-1(+1U);m-867&19\|-\|11495544\|11495567&offsets\|0\|-1(+1U)]\|AGGCTGTGATGCTCTCCTGAGCCT | >MI0025750\|hsa-mir-7974&WithFlank&19\|-\|11495538\|11495628 | 62 | 84 | 23(+1U) | AGGCTGTGATGCTCTCCTGAGCCT | 413.8 | -2.8 | 1.0 | -2.9 | 3.9E-03 | 4.5E-03 | BT20/MB468 | NUC |
| >hg38_21_+_8433172_8446622_RNA45SN1_45S_50ntFlanks@4727.4759.33\|AATCAAGAACGAAAGTCGGAGGTTCGAAGACGA | >hg38_21_+_8433172_8446622_RNA45SN1_45S_50ntFlanks | 4727 | 4759 | 33 | AATCAAGAACGAAAGTCGGAGGTTCGAAGACGA | 295.4 | -1.9 | 0.7 | -2.9 | 3.9E-03 | 4.5E-03 | MB231/MB468 | NUC |
| >hg38_21_+_8433172_8446622_RNA45SN1_45S_50ntFlanks@9454.9495.42\|CCGCGCCGGGGAGGTGGAGCACGAGCGCACGTGTTAGGACCC | >hg38_21_+_8433172_8446622_RNA45SN1_45S_50ntFlanks | 9454 | 9495 | 42 | CCGCGCCGGGGAGGTGGAGCACGAGCGCACGTGTTAGGACCC | 90.8 | -5.3 | 1.7 | -3.1 | 2.0E-03 | 4.5E-03 | BT20/MB468 | MP |
| >hg38_21_+_8433172_8446622_RNA45SN1_45S_50ntFlanks@8196.8223.28\|CGTGGACGGTGTGAGGCCGGTAGCGGCC | >hg38_21_+_8433172_8446622_RNA45SN1_45S_50ntFlanks | 8196 | 8223 | 28 | CGTGGACGGTGTGAGGCCGGTAGCGGCC | 1029.7 | -2.2 | 0.8 | -2.9 | 3.7E-03 | 4.5E-03 | MB231/MB468 | CYTO |
| >hg38_21_+_8433172_8446622_RNA45SN1_45S_50ntFlanks@6649.6678.30\|TACGACTCTTAGCGGTGGATCACTCGGCTC | >hg38_21_+_8433172_8446622_RNA45SN1_45S_50ntFlanks | 6649 | 6678 | 30 | TACGACTCTTAGCGGTGGATCACTCGGCTC | 461.3 | 4.6 | 1.4 | 3.4 | 7.0E-04 | 4.5E-03 | BT20/MB231 | MP |

<table>
<tr><td>>hg38_21_+_8433172_8446622_RNA45SN1_45S_50ntFlanks@9954.9979.26|TGGAAGTCGGAATCCGCTAAGGAGTG</td><td>>hg38_21_+_8433172_8446622_RNA45SN1_45S_50ntFlanks</td><td>9954</td><td>9979</td><td>26</td><td>TGGAAGTCGGAATCCGCTAAGGAGTG</td><td>648.8</td><td>7.0</td><td>2.1</td><td>3.4</td><td>7.1E-04</td><td>4.5E-03</td><td>BT20/MB231</td><td>MP</td></tr>
<tr><td>>hg38_21_+_8433172_8446622_RNA45SN1_45S_50ntFlanks@9536.9563.28|GAAACTCTGGTGGAGGTCCGTAGCGGTC</td><td>>hg38_21_+_8433172_8446622_RNA45SN1_45S_50ntFlanks</td><td>9536</td><td>9563</td><td>28</td><td>GAAACTCTGGTGGAGGTCCGTAGCGGTC</td><td>226.3</td><td>-3.6</td><td>1.2</td><td>-2.9</td><td>3.6E-03</td><td>4.6E-03</td><td>BT20/MB468</td><td>CYTO</td></tr>
<tr><td>>am_trnaMT_HisGTG_MT_+_12138_12206@19.51.33|AAACATCAGATTGTGAATCTGACAACAGAGGCT</td><td>>am_trnaMT_HisGTG_MT_+_12138_12206</td><td>19</td><td>51</td><td>33</td><td>AAACATCAGATTGTGAATCTGACAACAGAGGCT</td><td>197.9</td><td>3.7</td><td>1.3</td><td>2.9</td><td>3.9E-03</td><td>4.6E-03</td><td>MB231/MB468</td><td>MITO</td></tr>
<tr><td>>hg38_21_+_8433172_8446622_RNA45SN1_45S_50ntFlanks@10571.10596.26|GAGAAGCCGGCGGGAGCCCCGGGGAG</td><td>>hg38_21_+_8433172_8446622_RNA45SN1_45S_50ntFlanks</td><td>10571</td><td>10596</td><td>26</td><td>GAGAAGCCGGCGGGAGCCCCGGGGAG</td><td>163.9</td><td>-5.0</td><td>1.6</td><td>-3.1</td><td>2.0E-03</td><td>4.6E-03</td><td>BT20/MB468</td><td>MP</td></tr>
<tr><td>>hg38_21_+_8433172_8446622_RNA45SN1_45S_50ntFlanks@11552.11569.18|CCTCGGCCGGCGCCTAGC</td><td>>hg38_21_+_8433172_8446622_RNA45SN1_45S_50ntFlanks</td><td>11552</td><td>11569</td><td>18</td><td>CCTCGGCCGGCGCCTAGC</td><td>801.9</td><td>-2.0</td><td>0.7</td><td>-2.9</td><td>3.9E-03</td><td>4.6E-03</td><td>MB231/MB468</td><td>NUC</td></tr>
<tr><td>>hg38_21_+_8433172_8446622_RNA45SN1_45S_50ntFlanks@7984.8005.22|AGATCAGACGTGGCGACCCGCT</td><td>>hg38_21_+_8433172_8446622_RNA45SN1_45S_50ntFlanks</td><td>7984</td><td>8005</td><td>22</td><td>AGATCAGACGTGGCGACCCGCT</td><td>422.7</td><td>6.3</td><td>1.9</td><td>3.4</td><td>7.1E-04</td><td>4.6E-03</td><td>BT20/MB231</td><td>MP</td></tr>
<tr><td>>hg38_21_+_8433172_8446622_RNA45SN1_45S_50ntFlanks@3705.3743.39|TACCTGGTTGATCCTGCCAGTAGCATATGCTTGTCTCAA</td><td>>hg38_21_+_8433172_8446622_RNA45SN1_45S_50ntFlanks</td><td>3705</td><td>3743</td><td>39</td><td>TACCTGGTTGATCCTGCCAGTAGCATATGCTTGTCTCAA</td><td>363.2</td><td>5.5</td><td>1.8</td><td>3.1</td><td>2.0E-03</td><td>4.6E-03</td><td>BT20/MB468</td><td>MP</td></tr>
<tr><td>>hg38_21_+_8433172_8446622_RNA45SN1_45S_50ntFlanks@12489.12509.21|ACGTGAGCTGGGTTTAGACCG</td><td>>hg38_21_+_8433172_8446622_RNA45SN1_45S_50ntFlanks</td><td>12489</td><td>12509</td><td>21</td><td>ACGTGAGCTGGGTTTAGACCG</td><td>380.5</td><td>-5.2</td><td>1.7</td><td>-3.1</td><td>2.0E-03</td><td>4.6E-03</td><td>BT20/MB468</td><td>MP</td></tr>
<tr><td>>hg38_21_+_8433172_8446622_RNA45SN1_45S_50ntFlanks@6659.6681.23|AGCGGTGGATCACTCGGCTCGTG</td><td>>hg38_21_+_8433172_8446622_RNA45SN1_45S_50ntFlanks</td><td>6659</td><td>6681</td><td>23</td><td>AGCGGTGGATCACTCGGCTCGTG</td><td>438.7</td><td>-3.3</td><td>1.1</td><td>-2.9</td><td>3.6E-03</td><td>4.6E-03</td><td>BT20/MB468</td><td>CYTO</td></tr>
<tr><td>>hg38_21_+_8433172_8446622_RNA45SN1_45S_50ntFlanks@11576.11613.38|CTTAGAACTGGTGCGGACCAGGGGAATCCGACTGTTTA</td><td>>hg38_21_+_8433172_8446622_RNA45SN1_45S_50ntFlanks</td><td>11576</td><td>11613</td><td>38</td><td>CTTAGAACTGGTGCGGACCAGGGGAATCCGACTGTTTA</td><td>1697.9</td><td>2.2</td><td>0.8</td><td>3.0</td><td>3.1E-03</td><td>4.6E-03</td><td>BT20/MB468</td><td>MITO</td></tr>
<tr><td>>hg38_21_+_8433172_8446622_RNA45SN1_45S_50ntFlanks@474.497.24|TCTTCGTGATCGATGTGGTGACGT</td><td>>hg38_21_+_8433172_8446622_RNA45SN1_45S_50ntFlanks</td><td>474</td><td>497</td><td>24</td><td>TCTTCGTGATCGATGTGGTGACGT</td><td>764.2</td><td>-2.7</td><td>0.8</td><td>-3.3</td><td>9.5E-04</td><td>4.6E-03</td><td>BT20/MB231</td><td>MITO</td></tr>
<tr><td>>MI0005566|hsa-mir-374b&WithFlank&X|-|74218541|74218624@17.38.22&[MIMAT0004955&hsa-miR-374b-5p&offsets|0|0;m-153&X|-|74218587|74218608&offsets|0|0]|ATATAATACAACCTGCTAAGTG</td><td>>MI0005566|hsa-mir-374b&WithFlank&X|-|74218541|74218624</td><td>17</td><td>38</td><td>22</td><td>ATATAATACAACCTGCTAAGTG</td><td>1501.6</td><td>-3.6</td><td>1.1</td><td>-3.4</td><td>7.1E-04</td><td>4.6E-03</td><td>BT20/MB231</td><td>MP</td></tr>
<tr><td>>am_trnaMT_GluTTC_MT_-_14674_14742@26.54.29,trnalookalike8_GluTTC_5_-_93905172_93905240@26.54.29|GGTTTTTCATATCATTGGTCGTGGTTGTA</td><td>>am_trnaMT_GluTTC_MT_-_14674_14742</td><td>26</td><td>54</td><td>29</td><td>GGTTTTTCATATCATTGGTCGTGGTTGTA</td><td>300.5</td><td>3.1</td><td>0.9</td><td>3.4</td><td>7.1E-04</td><td>4.6E-03</td><td>BT20/MB231</td><td>MP</td></tr>
<tr><td>>trna119_LysCTT_1_-_145395522_145395594@3.32.30,trna11_LysCTT_5_-_180648979_180649051@3.32.30,trna13_LysCTT_6_+_26556774_26556846@3.32.30,trna32_LysCTT_16_-_3207406_3207478@3.32.30,trna7_LysCTT_16_+_3225692_3225764@3.32.30,trna9_LysCTT_5_+_180634755_180634827@3.32.30|CCGGCTAGCTCAGTCGGTAGAGCATGAGAC</td><td>>trna119_LysCTT_1_-_145395522_145395594</td><td>3</td><td>32</td><td>30</td><td>CCGGCTAGCTCAGTCGGTAGAGCATGAGAC</td><td>52.2</td><td>5.4</td><td>1.8</td><td>2.9</td><td>3.3E-03</td><td>4.6E-03</td><td>MB231/MB468</td><td>MP</td></tr>
<tr><td>>hg38_21_+_8433172_8446622_RNA45SN1_45S_50ntFlanks@10238.10258.21|CGCTACGCCGCGACGAGTAGG</td><td>>hg38_21_+_8433172_8446622_RNA45SN1_45S_50ntFlanks</td><td>10238</td><td>10258</td><td>21</td><td>CGCTACGCCGCGACGAGTAGG</td><td>383.5</td><td>3.7</td><td>1.1</td><td>3.4</td><td>7.1E-04</td><td>4.6E-03</td><td>BT20/MB231</td><td>MP</td></tr>
<tr><td>>hg38_21_+_8433172_8446622_RNA45SN1_45S_50ntFlanks@8185.8209.25|GAGGCCCAGCCCGTGGACGGTGTGA</td><td>>hg38_21_+_8433172_8446622_RNA45SN1_45S_50ntFlanks</td><td>8185</td><td>8209</td><td>25</td><td>GAGGCCCAGCCCGTGGACGGTGTGA</td><td>189.5</td><td>5.2</td><td>1.7</td><td>3.1</td><td>2.0E-03</td><td>4.6E-03</td><td>BT20/MB468</td><td>MP</td></tr>
<tr><td>>hg38_21_+_8433172_8446622_RNA45SN1_45S_50ntFlanks@4297.4315.19|CATTGGAGGGCAAGTCTGG</td><td>>hg38_21_+_8433172_8446622_RNA45SN1_45S_50ntFlanks</td><td>4297</td><td>4315</td><td>19</td><td>CATTGGAGGGCAAGTCTGG</td><td>883.5</td><td>6.0</td><td>1.8</td><td>3.4</td><td>7.1E-04</td><td>4.6E-03</td><td>BT20/MB231</td><td>MP</td></tr>
<tr><td>>MI0000298|hsa-mir-221&WithFlank&X|-|45746151|45746272@74.93.20&[MIMAT0000278&hsa-miR-221-3p&offsets|+3|0;m-91&X|-|45746181|45746202&offsets|+3|+1]|TACATTGTCTGCTGGGTTTC</td><td>>MI0000298|hsa-mir-221&WithFlank&X|-|45746151|45746272</td><td>74</td><td>93</td><td>20</td><td>TACATTGTCTGCTGGGTTTC</td><td>67.4</td><td>-9.5</td><td>2.8</td><td>-3.4</td><td>7.1E-04</td><td>4.6E-03</td><td>BT20/MB231</td><td>MP</td></tr>
<tr><td>>hg38_21_+_8433172_8446622_RNA45SN1_45S_50ntFlanks@7973.7997.25|GACGCGACCTCAGATCAGACGTGGC</td><td>>hg38_21_+_8433172_8446622_RNA45SN1_45S_50ntFlanks</td><td>7973</td><td>7997</td><td>25</td><td>GACGCGACCTCAGATCAGACGTGGC</td><td>295.2</td><td>3.3</td><td>1.1</td><td>3.1</td><td>2.0E-03</td><td>4.6E-03</td><td>BT20/MB468</td><td>MP</td></tr>
<tr><td>>hg38_21_+_8433172_8446622_RNA45SN1_45S_50ntFlanks@11569.11611.43|CAGCCGACTTAGAACTGGTGCGGACCAGGGGAATCCGACTGTT</td><td>>hg38_21_+_8433172_8446622_RNA45SN1_45S_50ntFlanks</td><td>11569</td><td>11611</td><td>43</td><td>CAGCCGACTTAGAACTGGTGCGGACCAGGGGAATCCGACTGTT</td><td>431.2</td><td>-2.5</td><td>0.9</td><td>-2.9</td><td>3.7E-03</td><td>4.6E-03</td><td>MB231/MB468</td><td>CYTO</td></tr>
<tr><td>>hg38_21_+_8433172_8446622_RNA45SN1_45S_50ntFlanks@11567.11586.20|AGCAGCCGACTTAGAACTGG</td><td>>hg38_21_+_8433172_8446622_RNA45SN1_45S_50ntFlanks</td><td>11567</td><td>11586</td><td>20</td><td>AGCAGCCGACTTAGAACTGG</td><td>1109.6</td><td>-2.9</td><td>0.9</td><td>-3.4</td><td>7.8E-04</td><td>4.6E-03</td><td>BT20/MB231</td><td>NUC</td></tr>
<tr><td>>hg38_21_+_8433172_8446622_RNA45SN1_45S_50ntFlanks@11671.11706.36|CCAGTGCTCTGAATGTCAAAGTGAAGAAATTCAATG</td><td>>hg38_21_+_8433172_8446622_RNA45SN1_45S_50ntFlanks</td><td>11671</td><td>11706</td><td>36</td><td>CCAGTGCTCTGAATGTCAAAGTGAAGAAATTCAATG</td><td>164.4</td><td>-4.3</td><td>1.5</td><td>-2.9</td><td>3.3E-03</td><td>4.6E-03</td><td>MB231/MB468</td><td>MP</td></tr>
<tr><td>>MI0000434|hsa-let-7i&WithFlank&12|+|62603680|62603775@12.29.18&[MIMAT0000415&hsa-let-7i-5p&offsets|0|-4;m-49&12|+|62603691|62603712&offsets|0|-4]|TGAGGTAGTAGTTTGTGC</td><td>>MI0000434|hsa-let-7i&WithFlank&12|+|62603680|62603775</td><td>12</td><td>29</td><td>18</td><td>TGAGGTAGTAGTTTGTGC</td><td>104.6</td><td>5.8</td><td>2.0</td><td>2.9</td><td>3.3E-03</td><td>4.6E-03</td><td>MB231/MB468</td><td>MP</td></tr>
<tr><td>>hg38_21_+_8433172_8446622_RNA45SN1_45S_50ntFlanks@4297.4316.20|CATTGGAGGGCAAGTCTGGT</td><td>>hg38_21_+_8433172_8446622_RNA45SN1_45S_50ntFlanks</td><td>4297</td><td>4316</td><td>20</td><td>CATTGGAGGGCAAGTCTGGT</td><td>6137.2</td><td>6.1</td><td>1.8</td><td>3.4</td><td>7.2E-04</td><td>4.6E-03</td><td>BT20/MB231</td><td>MP</td></tr>
<tr><td>>MI0000085|hsa-mir-27a&WithFlank&19|-|13836434|13836523@57.76.20(+4U)&[MIMAT0000084&hsa-miR-27a-3p&offsets|0|-1(+4U);m-43&19|-|13836447|13836467&offsets|0|-1(+4U)]|TTCACAGTGGCTAAGTTCCGTTTT</td><td>>MI0000085|hsa-mir-27a&WithFlank&19|-|13836434|13836523</td><td>57</td><td>76</td><td>20(+4U)</td><td>TTCACAGTGGCTAAGTTCCGTTTT</td><td>282.0</td><td>-6.3</td><td>1.9</td><td>-3.4</td><td>7.1E-04</td><td>4.6E-03</td><td>BT20/MB231</td><td>MP</td></tr>
<tr><td>>hg38_21_+_8433172_8446622_RNA45SN1_45S_50ntFlanks@6661.6698.38|CGGTGGATCACTCGGCTCGTGCGTCGATGAAGAACGCA</td><td>>hg38_21_+_8433172_8446622_RNA45SN1_45S_50ntFlanks</td><td>6661</td><td>6698</td><td>38</td><td>CGGTGGATCACTCGGCTCGTGCGTCGATGAAGAACGCA</td><td>1831.5</td><td>3.0</td><td>1.0</td><td>3.1</td><td>2.0E-03</td><td>4.6E-03</td><td>BT20/MB468</td><td>MP</td></tr>
<tr><td>>hg38_21_+_8433172_8446622_RNA45SN1_45S_50ntFlanks@12003.12031.29|GCGGGCCGCCGGTGAAATACCACTACTCT</td><td>>hg38_21_+_8433172_8446622_RNA45SN1_45S_50ntFlanks</td><td>12003</td><td>12031</td><td>29</td><td>GCGGGCCGCCGGTGAAATACCACTACTCT</td><td>723.8</td><td>-1.3</td><td>0.4</td><td>-3.0</td><td>3.1E-03</td><td>4.6E-03</td><td>BT20/MB468</td><td>MITO</td></tr>
<tr><td>>hg38_21_+_8433172_8446622_RNA45SN1_45S_50ntFlanks@11580.11606.27|GAACTGGTGCGGACCAGGGGAATCCGA</td><td>>hg38_21_+_8433172_8446622_RNA45SN1_45S_50ntFlanks</td><td>11580</td><td>11606</td><td>27</td><td>GAACTGGTGCGGACCAGGGGAATCCGA</td><td>278.3</td><td>-2.8</td><td>0.9</td><td>-3.1</td><td>2.0E-03</td><td>4.6E-03</td><td>BT20/MB468</td><td>MP</td></tr>
<tr><td>>hg38_21_+_8433172_8446622_RNA45SN1_45S_50ntFlanks@8343.8373.31|GTAAGGGAAAGTTGAAAAGAACTTTGAAGAG</td><td>>hg38_21_+_8433172_8446622_RNA45SN1_45S_50ntFlanks</td><td>8343</td><td>8373</td><td>31</td><td>GTAAGGGAAAGTTGAAAAGAACTTTGAAGAG</td><td>1057.1</td><td>6.0</td><td>1.8</td><td>3.4</td><td>7.2E-04</td><td>4.6E-03</td><td>BT20/MB231</td><td>MP</td></tr>
<tr><td>>hg38_21_+_8433172_8446622_RNA45SN1_45S_50ntFlanks@6645.6680.36|CTCGTACGACTCTTAGCGGTGGATCACTCGGCTCGT</td><td>>hg38_21_+_8433172_8446622_RNA45SN1_45S_50ntFlanks</td><td>6645</td><td>6680</td><td>36</td><td>CTCGTACGACTCTTAGCGGTGGATCACTCGGCTCGT</td><td>509.4</td><td>-2.7</td><td>0.9</td><td>-2.9</td><td>3.4E-03</td><td>4.6E-03</td><td>MB231/MB468</td><td>MP</td></tr>
<tr><td>>MI0003581|hsa-mir-574&WithFlank&4|+|38868026|38868133@67.88.22&[MIMAT0003239&hsa-miR-574-3p&offsets|0|0;m-149&4|+|38868092|38868113&offsets|0|0]|CACGCTCATGCACACACCCACA</td><td>>MI0003581|hsa-mir-574&WithFlank&4|+|38868026|38868133</td><td>67</td><td>88</td><td>22</td><td>CACGCTCATGCACACACCCACA</td><td>1327.1</td><td>-2.5</td><td>0.7</td><td>-3.3</td><td>9.6E-04</td><td>4.6E-03</td><td>BT20/MB231</td><td>MITO</td></tr>
<tr><td>>am_trnaMT_ValTAC_MT_+_1602_1670@38.65.28|TAGGAGATTTCAACTTAACTTGACCGCT</td><td>>am_trnaMT_ValTAC_MT_+_1602_1670</td><td>38</td><td>65</td><td>28</td><td>TAGGAGATTTCAACTTAACTTGACCGCT</td><td>279.1</td><td>6.1</td><td>2.0</td><td>3.1</td><td>2.0E-03</td><td>4.6E-03</td><td>BT20/MB468</td><td>MP</td></tr>
<tr><td>>hg38_MT_+_598_1651_RNR1_12S_50ntFlanks@428.452.25|AAAGTGGCTTTAACATATCTGAACA</td><td>>hg38_MT_+_598_1651_RNR1_12S_50ntFlanks</td><td>428</td><td>452</td><td>25</td><td>AAAGTGGCTTTAACATATCTGAACA</td><td>225.8</td><td>2.6</td><td>0.9</td><td>2.9</td><td>4.0E-03</td><td>4.6E-03</td><td>MB231/MB468</td><td>MITO</td></tr>
<tr><td>>hg38_21_+_8433172_8446622_RNA45SN1_45S_50ntFlanks@12072.12097.26|CCCGAGGGGCTCTCGCTTCTGGCGCC</td><td>>hg38_21_+_8433172_8446622_RNA45SN1_45S_50ntFlanks</td><td>12072</td><td>12097</td><td>26</td><td>CCCGAGGGGCTCTCGCTTCTGGCGCC</td><td>873.6</td><td>-2.1</td><td>0.7</td><td>-2.9</td><td>3.7E-03</td><td>4.6E-03</td><td>MB231/MB468</td><td>CYTO</td></tr>
<tr><td>>hg38_21_+_8433172_8446622_RNA45SN1_45S_50ntFlanks@10574.10608.35|AAGCCGGCGGGAGCCCCGGGGAGAGTTCTCTTTTC</td><td>>hg38_21_+_8433172_8446622_RNA45SN1_45S_50ntFlanks</td><td>10574</td><td>10608</td><td>35</td><td>AAGCCGGCGGGAGCCCCGGGGAGAGTTCTCTTTTC</td><td>178.3</td><td>-5.0</td><td>1.6</td><td>-3.1</td><td>2.0E-03</td><td>4.6E-03</td><td>BT20/MB468</td><td>MP</td></tr>
<tr><td>>hg38_21_+_8433172_8446622_RNA45SN1_45S_50ntFlanks@8359.8378.20|AAGAACTTTGAAGAGAGAGT</td><td>>hg38_21_+_8433172_8446622_RNA45SN1_45S_50ntFlanks</td><td>8359</td><td>8378</td><td>20</td><td>AAGAACTTTGAAGAGAGAGT</td><td>651.5</td><td>-3.2</td><td>1.0</td><td>-3.1</td><td>2.0E-03</td><td>4.6E-03</td><td>BT20/MB468</td><td>MP</td></tr>
<tr><td>>hg38_21_+_8433172_8446622_RNA45SN1_45S_50ntFlanks@8117.8134.18|CGGGACATGTGGCGTACG</td><td>>hg38_21_+_8433172_8446622_RNA45SN1_45S_50ntFlanks</td><td>8117</td><td>8134</td><td>18</td><td>CGGGACATGTGGCGTACG</td><td>375.8</td><td>2.4</td><td>0.8</td><td>3.1</td><td>2.0E-03</td><td>4.6E-03</td><td>BT20/MB468</td><td>MP</td></tr>
<tr><td>>hg38_21_+_8433172_8446622_RNA45SN1_45S_50ntFlanks@10410.10442.33|TCAGTCGGTCCTGAGAGATGGGCGAGCGCCGTT</td><td>>hg38_21_+_8433172_8446622_RNA45SN1_45S_50ntFlanks</td><td>10410</td><td>10442</td><td>33</td><td>TCAGTCGGTCCTGAGAGATGGGCGAGCGCCGTT</td><td>272.4</td><td>-2.7</td><td>0.9</td><td>-3.1</td><td>2.0E-03</td><td>4.6E-03</td><td>BT20/MB468</td><td>MP</td></tr>
<tr><td>>MI0000289|hsa-mir-181a-1&WithFlank&1|-|198859038|198859159@30.53.24(+1A);MI0000269|hsa-mir-181a-2&WithFlank&9|+|124692436|124692557@45.68.24(+1A)&[MIMAT0000256&hsa-miR-181a-5p&offsets|0|+1(+1A);m-32&1|-|198859108|198859130&offsets|0|+1(+1A)];[MIMAT0000256_1&hsa-miR-181a-5p&offsets|0|+1(+1A);m-32&9|+|124692480|124692502&offsets|0|+1(+1A)]|AACATTCAACGCTGTCGGTGAGTTA</td><td>>MI0000289|hsa-mir-181a-1&WithFlank&1|-|198859038|198859159</td><td>30</td><td>53</td><td>24(+1A)</td><td>AACATTCAACGCTGTCGGTGAGTTA</td><td>157.1</td><td>3.9</td><td>1.3</td><td>2.9</td><td>3.4E-03</td><td>4.6E-03</td><td>MB231/MB468</td><td>MP</td></tr>
<tr><td>>hg38_21_+_8433172_8446622_RNA45SN1_45S_50ntFlanks@4588.4613.26|CTATTTTGTTGGTTTTCGGAACTGAG</td><td>>hg38_21_+_8433172_8446622_RNA45SN1_45S_50ntFlanks</td><td>4588</td><td>4613</td><td>26</td><td>CTATTTTGTTGGTTTTCGGAACTGAG</td><td>661.9</td><td>-5.3</td><td>1.7</td><td>-3.1</td><td>2.1E-03</td><td>4.6E-03</td><td>BT20/MB468</td><td>MP</td></tr>
<tr><td>>hg38_21_+_8433172_8446622_RNA45SN1_45S_50ntFlanks@11210.11252.43|GGCGACTCTGGACGCGAGCCGGGCCCTTCCCGTGGATCGCCCC</td><td>>hg38_21_+_8433172_8446622_RNA45SN1_45S_50ntFlanks</td><td>11210</td><td>11252</td><td>43</td><td>GGCGACTCTGGACGCGAGCCGGGCCCTTCCCGTGGATCGCCCC</td><td>240.9</td><td>-4.7</td><td>1.6</td><td>-2.9</td><td>4.0E-03</td><td>4.6E-03</td><td>MB231/MB468</td><td>NUC</td></tr>
<tr><td>>hg38_21_+_8433172_8446622_RNA45SN1_45S_50ntFlanks@11918.11940.23|ACATGAGAGGTGTAGAATAAGTG</td><td>>hg38_21_+_8433172_8446622_RNA45SN1_45S_50ntFlanks</td><td>11918</td><td>11940</td><td>23</td><td>ACATGAGAGGTGTAGAATAAGTG</td><td>651.6</td><td>4.4</td><td>1.3</td><td>3.4</td><td>7.2E-04</td><td>4.6E-03</td><td>BT20/MB231</td><td>MP</td></tr>
<tr><td>>hg38_21_+_8433172_8446622_RNA45SN1_45S_50ntFlanks@12008.12038.31|CCGCCGGTGAAATACCACTACTCTGATCGTT</td><td>>hg38_21_+_8433172_8446622_RNA45SN1_45S_50ntFlanks</td><td>12008</td><td>12038</td><td>31</td><td>CCGCCGGTGAAATACCACTACTCTGATCGTT</td><td>433.1</td><td>-5.3</td><td>1.7</td><td>-3.1</td><td>2.1E-03</td><td>4.6E-03</td><td>BT20/MB468</td><td>MP</td></tr>
<tr><td>>hg38_21_+_8433172_8446622_RNA45SN1_45S_50ntFlanks@11321.11338.18|CACGTCTCGTCGCGCGCG</td><td>>hg38_21_+_8433172_8446622_RNA45SN1_45S_50ntFlanks</td><td>11321</td><td>11338</td><td>18</td><td>CACGTCTCGTCGCGCGCG</td><td>227.8</td><td>-3.8</td><td>1.2</td><td>-3.1</td><td>2.1E-03</td><td>4.6E-03</td><td>BT20/MB468</td><td>MP</td></tr>
<tr><td>>hg38_21_+_8433172_8446622_RNA45SN1_45S_50ntFlanks@3874.3891.18|AATACATGCCGACGGGCG</td><td>>hg38_21_+_8433172_8446622_RNA45SN1_45S_50ntFlanks</td><td>3874</td><td>3891</td><td>18</td><td>AATACATGCCGACGGGCG</td><td>33.5</td><td>-8.1</td><td>2.6</td><td>-3.1</td><td>2.1E-03</td><td>4.6E-03</td><td>BT20/MB468</td><td>MP</td></tr>
<tr><td>>trnaMT_GluTTC_MT_-_14674_14742@21.58.38,trnalookalike8_GluTTC_5_-_93905172_93905240@21.58.38|ACGATGGTTTTTCATATCATTGGTCGTGGTTGTAGTCC</td><td>>trnaMT_GluTTC_MT_-_14674_14742</td><td>21</td><td>58</td><td>38</td><td>ACGATGGTTTTTCATATCATTGGTCGTGGTTGTAGTCC</td><td>87.7</td><td>5.4</td><td>1.8</td><td>2.9</td><td>3.4E-03</td><td>4.6E-03</td><td>MB231/MB468</td><td>MP</td></tr>
<tr><td>>hg38_21_+_8433172_8446622_RNA45SN1_45S_50ntFlanks@11774.11792.19|AGTGACGCGCATGAATGGA</td><td>>hg38_21_+_8433172_8446622_RNA45SN1_45S_50ntFlanks</td><td>11774</td><td>11792</td><td>19</td><td>AGTGACGCGCATGAATGGA</td><td>95.9</td><td>-4.7</td><td>1.6</td><td>-2.9</td><td>3.4E-03</td><td>4.6E-03</td><td>MB231/MB468</td><td>MP</td></tr>
<tr><td>>hg38_21_+_8433172_8446622_RNA45SN1_45S_50ntFlanks@9203.9221.19|CTCGGGGCCACGCGCGCGT</td><td>>hg38_21_+_8433172_8446622_RNA45SN1_45S_50ntFlanks</td><td>9203</td><td>9221</td><td>19</td><td>CTCGGGGCCACGCGCGCGT</td><td>90.8</td><td>-4.9</td><td>1.6</td><td>-3.1</td><td>2.1E-03</td><td>4.6E-03</td><td>BT20/MB468</td><td>MP</td></tr>
<tr><td>>trnaMT_SerGCT_MT_+_12207_12265@1.38.38|GAGAAAGCTCACAAGAACTGCTAACTCATGCCCCCATG</td><td>>trnaMT_SerGCT_MT_+_12207_12265</td><td>1</td><td>38</td><td>38</td><td>GAGAAAGCTCACAAGAACTGCTAACTCATGCCCCCATG</td><td>2522.2</td><td>3.1</td><td>1.0</td><td>3.0</td><td>3.1E-03</td><td>4.6E-03</td><td>BT20/MB468</td><td>MITO</td></tr>
<tr><td>>hg38_21_+_8433172_8446622_RNA45SN1_45S_50ntFlanks@11847.11873.27|AACGGGCTTGGCGGAATCAGCGGGGAA</td><td>>hg38_21_+_8433172_8446622_RNA45SN1_45S_50ntFlanks</td><td>11847</td><td>11873</td><td>27</td><td>AACGGGCTTGGCGGAATCAGCGGGGAA</td><td>141.1</td><td>-1.7</td><td>0.6</td><td>-3.0</td><td>3.1E-03</td><td>4.6E-03</td><td>BT20/MB468</td><td>MITO</td></tr>
<tr><td>>hg38_21_+_8433172_8446622_RNA45SN1_45S_50ntFlanks@10389.10429.41|TGAACAGCAGTTGAACATGGGTCAGTCGGTCCTGAGAGATG</td><td>>hg38_21_+_8433172_8446622_RNA45SN1_45S_50ntFlanks</td><td>10389</td><td>10429</td><td>41</td><td>TGAACAGCAGTTGAACATGGGTCAGTCGGTCCTGAGAGATG</td><td>707.8</td><td>-3.0</td><td>1.0</td><td>-2.9</td><td>3.7E-03</td><td>4.6E-03</td><td>BT20/MB468</td><td>CYTO</td></tr>
<tr><td>>hg38_21_+_8433172_8446622_RNA45SN1_45S_50ntFlanks@9555.9584.30|GTAGCGGTCCTGACGTGCAAATCGGTCGTC</td><td>>hg38_21_+_8433172_8446622_RNA45SN1_45S_50ntFlanks</td><td>9555</td><td>9584</td><td>30</td><td>GTAGCGGTCCTGACGTGCAAATCGGTCGTC</td><td>95.1</td><td>-5.0</td><td>1.7</td><td>-2.9</td><td>3.4E-03</td><td>4.6E-03</td><td>MB231/MB468</td><td>MP</td></tr>
</table>

<table>
<tr><td>>trna10_AspGTC_12_-_125424193_125424264@38.72.35,trna12_AspGTC_12_-_125411891_125411962@38.72.35,trna144_AspGTC_6_-_27551236_27551307@38.72.35,trna38_AspGTC_17_-_8125556_8125627@38.72.35,trna45_AspGTC_6_+_27447453_27447524@38.72.35,trna48_AspGTC_6_+_27471523_27471594@38.72.35,trna4_AspGTC_12_+_96429799_96429870@38.72.35,trna69_AspGTC_1_-_161440205_161440276@38.72.35,trna72_AspGTC_1_-_161432824_161432895@38.72.35,trna75_AspGTC_1_-_161425414_161425485@38.72.35,trna78_AspGTC_1_-_161418033_161418104@38.72.35,trna81_AspGTC_1_-_161410615_161410686@38.72.35|CGCGGGAGACCGGGGTTCGATTCCCCGACGGGGAG</td><td>>trna10_AspGTC_12_-_125424193_125424264</td><td>38</td><td>72</td><td>35</td><td>CGCGGGAGACCGGGGTTCGATTCCCCGACGGGGAG</td><td>542.9</td><td>-1.9</td><td>0.7</td><td>-2.9</td><td>3.7E-03</td><td>4.6E-03</td><td>BT20/MB468</td><td>CYTO</td></tr>
<tr><td>>MI0000749|hsa-mir-30e&WithFlank&1|+|40754349|40754452@23.46.24(+1U)&[MIMAT0000692&hsa-miR-30e-5p&offsets|0|+2(+1U);m-24&1|+|40754371|40754394&offsets|0|0(+1U)]|TGTAAACATCCTTGACTGGAAGCTT</td><td>>MI0000749|hsa-mir-30e&WithFlank&1|+|40754349|40754452</td><td>23</td><td>46</td><td>24(+1U)</td><td>TGTAAACATCCTTGACTGGAAGCTT</td><td>1572.1</td><td>2.3</td><td>0.8</td><td>2.9</td><td>3.4E-03</td><td>4.6E-03</td><td>MB231/MB468</td><td>MP</td></tr>
<tr><td>>MI0000272|hsa-mir-182&WithFlank&7|-|129770377|129770498@29.51.23&[MIMAT0000259&hsa-miR-182-5p&offsets|0|-1;m-22&7|-|129770449|129770470&offsets|0|+1]|TTTGGCAATGGTAGAACTCACAC</td><td>>MI0000272|hsa-mir-182&WithFlank&7|-|129770377|129770498</td><td>29</td><td>51</td><td>23</td><td>TTTGGCAATGGTAGAACTCACAC</td><td>867.2</td><td>3.5</td><td>1.1</td><td>3.1</td><td>2.1E-03</td><td>4.6E-03</td><td>BT20/MB468</td><td>MP</td></tr>
<tr><td>>hg38_21_+_8433172_8446622_RNA45SN1_45S_50ntFlanks@8113.8141.29|GGCGCGGGACATGTGGCGTACGGAAGACC</td><td>>hg38_21_+_8433172_8446622_RNA45SN1_45S_50ntFlanks</td><td>8113</td><td>8141</td><td>29</td><td>GGCGCGGGACATGTGGCGTACGGAAGACC</td><td>1479.2</td><td>-3.0</td><td>1.0</td><td>-3.1</td><td>2.1E-03</td><td>4.6E-03</td><td>BT20/MB468</td><td>MP</td></tr>
<tr><td>>hg38_21_+_8433172_8446622_RNA45SN1_45S_50ntFlanks@11968.11985.18|CCCGCGAGGGGCCCGGGG</td><td>>hg38_21_+_8433172_8446622_RNA45SN1_45S_50ntFlanks</td><td>11968</td><td>11985</td><td>18</td><td>CCCGCGAGGGGCCCGGGG</td><td>112.4</td><td>-4.2</td><td>1.4</td><td>-3.0</td><td>3.1E-03</td><td>4.6E-03</td><td>BT20/MB468</td><td>MITO</td></tr>
<tr><td>>hg38_21_+_8433172_8446622_RNA45SN1_45S_50ntFlanks@6649.6679.31|TACGACTCTTAGCGGTGGATCACTCGGCTCG</td><td>>hg38_21_+_8433172_8446622_RNA45SN1_45S_50ntFlanks</td><td>6649</td><td>6679</td><td>31</td><td>TACGACTCTTAGCGGTGGATCACTCGGCTCG</td><td>920.6</td><td>5.1</td><td>1.5</td><td>3.4</td><td>7.2E-04</td><td>4.6E-03</td><td>BT20/MB231</td><td>MP</td></tr>
<tr><td>>hg38_21_+_8433172_8446622_RNA45SN1_45S_50ntFlanks@9268.9285.18|ACGTCGGCTACCCACCCG</td><td>>hg38_21_+_8433172_8446622_RNA45SN1_45S_50ntFlanks</td><td>9268</td><td>9285</td><td>18</td><td>ACGTCGGCTACCCACCCG</td><td>85.0</td><td>-4.7</td><td>1.6</td><td>-2.9</td><td>3.7E-03</td><td>4.6E-03</td><td>BT20/MB468</td><td>CYTO</td></tr>
<tr><td>>hg38_21_+_8433172_8446622_RNA45SN1_45S_50ntFlanks@11552.11588.37|CCTCGGCCGGCGCCTAGCAGCCGACTTAGAACTGGTG</td><td>>hg38_21_+_8433172_8446622_RNA45SN1_45S_50ntFlanks</td><td>11552</td><td>11588</td><td>37</td><td>CCTCGGCCGGCGCCTAGCAGCCGACTTAGAACTGGTG</td><td>187.8</td><td>-2.0</td><td>0.7</td><td>-2.9</td><td>3.7E-03</td><td>4.6E-03</td><td>BT20/MB468</td><td>CYTO</td></tr>
<tr><td>>hg38_MT_+_1621_3279_RNR2_16S_50ntFlanks@1380.1400.21|AGGACATCCCGATGGTGCAGC</td><td>>hg38_MT_+_1621_3279_RNR2_16S_50ntFlanks</td><td>1380</td><td>1400</td><td>21</td><td>AGGACATCCCGATGGTGCAGC</td><td>98.9</td><td>4.8</td><td>1.6</td><td>2.9</td><td>3.4E-03</td><td>4.6E-03</td><td>MB231/MB468</td><td>MP</td></tr>
<tr><td>>MI0000803|hsa-mir-330&WithFlank&19|-|45638988|45639093@24.43.20&[MIMAT0004693&hsa-miR-330-5p&offsets|0|-2;m-200&19|-|45639050|45639070&offsets|0|-1]|TCTCTGGGCCTGTGTCTTAG</td><td>>MI0000803|hsa-mir-330&WithFlank&19|-|45638988|45639093</td><td>24</td><td>43</td><td>20</td><td>TCTCTGGGCCTGTGTCTTAG</td><td>130.3</td><td>5.4</td><td>1.8</td><td>3.1</td><td>2.1E-03</td><td>4.6E-03</td><td>BT20/MB468</td><td>MP</td></tr>
<tr><td>>hg38_21_+_8433172_8446622_RNA45SN1_45S_50ntFlanks@11571.11599.29|GCCGACTTAGAACTGGTGCGGACCAGGGG</td><td>>hg38_21_+_8433172_8446622_RNA45SN1_45S_50ntFlanks</td><td>11571</td><td>11599</td><td>29</td><td>GCCGACTTAGAACTGGTGCGGACCAGGGG</td><td>472.9</td><td>5.7</td><td>1.7</td><td>3.4</td><td>7.3E-04</td><td>4.6E-03</td><td>BT20/MB231</td><td>MP</td></tr>
<tr><td>>trna119_LysCTT_1_-_145395522_145395594@1.33.33,trna11_LysCTT_5_-_180648979_180649051@1.33.33,trna13_LysCTT_6_+_26556774_26556846@1.33.33,trna7_LysCTT_16_+_3225692_3225764@1.33.33,trna9_LysCTT_5_+_180634755_180634827@1.33.33|GCCCGGCTAGCTCAGTCGGTAGAGCATGAGACT</td><td>>trna119_LysCTT_1_-_145395522_145395594</td><td>1</td><td>33</td><td>33</td><td>GCCCGGCTAGCTCAGTCGGTAGAGCATGAGACT</td><td>38208.6</td><td>-3.3</td><td>1.0</td><td>-3.3</td><td>8.7E-04</td><td>4.6E-03</td><td>BT20/MB231</td><td>CYTO</td></tr>
<tr><td>>hg38_21_+_8433172_8446622_RNA45SN1_45S_50ntFlanks@11669.11707.39|GCCCAGTGCTCTGAATGTCAAAGTGAAGAAATTCAATGA</td><td>>hg38_21_+_8433172_8446622_RNA45SN1_45S_50ntFlanks</td><td>11669</td><td>11707</td><td>39</td><td>GCCCAGTGCTCTGAATGTCAAAGTGAAGAAATTCAATGA</td><td>1898.2</td><td>-2.6</td><td>0.8</td><td>-3.3</td><td>8.7E-04</td><td>4.6E-03</td><td>BT20/MB231</td><td>CYTO</td></tr>
<tr><td>>hg38_21_+_8433172_8446622_RNA45SN1_45S_50ntFlanks@6645.6672.28|CTCGTACGACTCTTAGCGGTGGATCACT</td><td>>hg38_21_+_8433172_8446622_RNA45SN1_45S_50ntFlanks</td><td>6645</td><td>6672</td><td>28</td><td>CTCGTACGACTCTTAGCGGTGGATCACT</td><td>472.1</td><td>1.8</td><td>0.6</td><td>2.9</td><td>4.0E-03</td><td>4.6E-03</td><td>BT20/MB468</td><td>NUC</td></tr>
<tr><td>>hg38_21_+_8433172_8446622_RNA45SN1_45S_50ntFlanks@8693.8715.23|CGGGACGGCTGGGAAGGCCCGGC</td><td>>hg38_21_+_8433172_8446622_RNA45SN1_45S_50ntFlanks</td><td>8693</td><td>8715</td><td>23</td><td>CGGGACGGCTGGGAAGGCCCGGC</td><td>73.8</td><td>5.1</td><td>1.6</td><td>3.1</td><td>2.1E-03</td><td>4.7E-03</td><td>BT20/MB468</td><td>MP</td></tr>
<tr><td>>trna27_GlyCCC_2_-_70476123_70476193@57.74.18,trna34_GlyCCC_16_-_686736_686806@57.74.18|TTCCCGGGCGGCGCACCA</td><td>>trna27_GlyCCC_2_-_70476123_70476193</td><td>57</td><td>74</td><td>18</td><td>TTCCCGGGCGGCGCACCA</td><td>295.4</td><td>2.9</td><td>0.9</td><td>3.3</td><td>8.8E-04</td><td>4.7E-03</td><td>BT20/MB231</td><td>CYTO</td></tr>
<tr><td>>trnaMT_MetCAT_MT_+_4402_4469@32.71.40|ATACCCCGAAAATGTTGGTTATACCCTTCCCGTACTACCA</td><td>>trnaMT_MetCAT_MT_+_4402_4469</td><td>32</td><td>71</td><td>40</td><td>ATACCCCGAAAATGTTGGTTATACCCTTCCCGTACTACCA</td><td>245.2</td><td>-4.8</td><td>1.4</td><td>-3.4</td><td>8.0E-04</td><td>4.7E-03</td><td>BT20/MB231</td><td>NUC</td></tr>
<tr><td>>hg38_21_+_8433172_8446622_RNA45SN1_45S_50ntFlanks@6650.6699.50|ACGACTCTTAGCGGTGGATCACTCGGCTCGTGCGTCGATGAAGAACGCAG</td><td>>hg38_21_+_8433172_8446622_RNA45SN1_45S_50ntFlanks</td><td>6650</td><td>6699</td><td>50</td><td>ACGACTCTTAGCGGTGGATCACTCGGCTCGTGCGTCGATGAAGAACGCAG</td><td>189.5</td><td>-3.9</td><td>1.2</td><td>-3.3</td><td>8.8E-04</td><td>4.7E-03</td><td>BT20/MB231</td><td>CYTO</td></tr>
<tr><td>>am_trnaMT_GluTTC_MT_-_14674_14742@49.69.21,trnalookalike8_GluTTC_5_-_93905172_93905240@49.69.21|GTTGTAGTCCGTGCGAGAATA</td><td>>am_trnaMT_GluTTC_MT_-_14674_14742</td><td>49</td><td>69</td><td>21</td><td>GTTGTAGTCCGTGCGAGAATA</td><td>129.6</td><td>5.3</td><td>1.7</td><td>3.1</td><td>2.1E-03</td><td>4.7E-03</td><td>BT20/MB468</td><td>MP</td></tr>
<tr><td>>hg38_21_+_8433172_8446622_RNA45SN1_45S_50ntFlanks@9198.9218.21|GTTCTCTCGGGGCCACGCGCG</td><td>>hg38_21_+_8433172_8446622_RNA45SN1_45S_50ntFlanks</td><td>9198</td><td>9218</td><td>21</td><td>GTTCTCTCGGGGCCACGCGCG</td><td>887.0</td><td>-4.9</td><td>1.7</td><td>-3.0</td><td>3.2E-03</td><td>4.7E-03</td><td>BT20/MB468</td><td>MITO</td></tr>
<tr><td>>hg38_21_+_8433172_8446622_RNA45SN1_45S_50ntFlanks@11705.11728.24|TGAAGCGCGGGTAAACGGCGGGAG</td><td>>hg38_21_+_8433172_8446622_RNA45SN1_45S_50ntFlanks</td><td>11705</td><td>11728</td><td>24</td><td>TGAAGCGCGGGTAAACGGCGGGAG</td><td>697.8</td><td>-5.4</td><td>1.7</td><td>-3.1</td><td>2.1E-03</td><td>4.7E-03</td><td>BT20/MB468</td><td>MP</td></tr>
<tr><td>>MI0000542|hsa-mir-320a&WithFlank&8|-|22244960|22245043@48.71.24&[MIMAT0000510&hsa-miR-320a-3p&offsets|0|+2;m-60&8|-|22244975|22244996&offsets|0|+2]|AAAAGCTGGGTTGAGAGGGCGAAA</td><td>>MI0000542|hsa-mir-320a&WithFlank&8|-|22244960|22245043</td><td>48</td><td>71</td><td>24</td><td>AAAAGCTGGGTTGAGAGGGCGAAA</td><td>2077.2</td><td>2.3</td><td>0.7</td><td>3.4</td><td>8.0E-04</td><td>4.7E-03</td><td>BT20/MB231</td><td>NUC</td></tr>
<tr><td>>hg38_21_+_8433172_8446622_RNA45SN1_45S_50ntFlanks@10574.10598.25|AAGCCGGCGGGAGCCCCGGGGAGAG</td><td>>hg38_21_+_8433172_8446622_RNA45SN1_45S_50ntFlanks</td><td>10574</td><td>10598</td><td>25</td><td>AAGCCGGCGGGAGCCCCGGGGAGAG</td><td>53.8</td><td>-5.0</td><td>1.6</td><td>-3.1</td><td>2.1E-03</td><td>4.7E-03</td><td>BT20/MB468</td><td>MP</td></tr>
<tr><td>>trnaMT_GluTTC_MT_-_14674_14742@40.71.32,trnalookalike8_GluTTC_5_-_93905172_93905240@40.71.32|TTGGTCGTGGTTGTAGTCCGTGCGAGAATACC</td><td>>trnaMT_GluTTC_MT_-_14674_14742</td><td>40</td><td>71</td><td>32</td><td>TTGGTCGTGGTTGTAGTCCGTGCGAGAATACC</td><td>1111.3</td><td>2.8</td><td>0.9</td><td>2.9</td><td>3.4E-03</td><td>4.7E-03</td><td>MB231/MB468</td><td>MP</td></tr>
<tr><td>>hg38_21_+_8433172_8446622_RNA45SN1_45S_50ntFlanks@4297.4319.23|CATTGGAGGGCAAGTCTGGTGCC</td><td>>hg38_21_+_8433172_8446622_RNA45SN1_45S_50ntFlanks</td><td>4297</td><td>4319</td><td>23</td><td>CATTGGAGGGCAAGTCTGGTGCC</td><td>169.2</td><td>-3.6</td><td>1.2</td><td>-3.0</td><td>3.2E-03</td><td>4.7E-03</td><td>BT20/MB468</td><td>MITO</td></tr>
<tr><td>>hg38_21_+_8433172_8446622_RNA45SN1_45S_50ntFlanks@4403.4423.21|CGGGCGGTCCGCCGCGAGGCG</td><td>>hg38_21_+_8433172_8446622_RNA45SN1_45S_50ntFlanks</td><td>4403</td><td>4423</td><td>21</td><td>CGGGCGGTCCGCCGCGAGGCG</td><td>108.0</td><td>-5.2</td><td>1.7</td><td>-3.1</td><td>2.1E-03</td><td>4.7E-03</td><td>BT20/MB468</td><td>MP</td></tr>
<tr><td>>MI0000088|hsa-mir-30a&WithFlank&6|-|71403545|71403627@53.71.19&[MIMAT0000088&hsa-miR-30a-3p&offsets|0|-3;m-38&6|-|71403554|71403575&offsets|0|-3]|CTTTCAGTCGGATGTTTGC</td><td>>MI0000088|hsa-mir-30a&WithFlank&6|-|71403545|71403627</td><td>53</td><td>71</td><td>19</td><td>CTTTCAGTCGGATGTTTGC</td><td>2601.7</td><td>-2.0</td><td>0.6</td><td>-3.4</td><td>8.1E-04</td><td>4.7E-03</td><td>BT20/MB231</td><td>NUC</td></tr>
<tr><td>>MI0000298|hsa-mir-221&WithFlank&X|-|45746151|45746272@71.93.23(+1G)&[MIMAT0000278&hsa-miR-221-3p&offsets|0|0(+1G);m-91&X|-|45746181|45746202&offsets|0|+1(+1G)]|AGCTACATTGTCTGCTGGGTTTCG</td><td>>MI0000298|hsa-mir-221&WithFlank&X|-|45746151|45746272</td><td>71</td><td>93</td><td>23(+1G)</td><td>AGCTACATTGTCTGCTGGGTTTCG</td><td>363.2</td><td>-1.9</td><td>0.6</td><td>-3.4</td><td>8.1E-04</td><td>4.7E-03</td><td>BT20/MB231</td><td>NUC</td></tr>
<tr><td>>hg38_21_+_8433172_8446622_RNA45SN1_45S_50ntFlanks@12244.12275.32|AAGGGCAAAAGCTCGCTTGATCTTGATTTTCA</td><td>>hg38_21_+_8433172_8446622_RNA45SN1_45S_50ntFlanks</td><td>12244</td><td>12275</td><td>32</td><td>AAGGGCAAAAGCTCGCTTGATCTTGATTTTCA</td><td>237.6</td><td>2.3</td><td>0.7</td><td>3.3</td><td>9.8E-04</td><td>4.7E-03</td><td>BT20/MB231</td><td>MITO</td></tr>
<tr><td>>am_trna128_GlyGCC_6_-_27870686_27870756@7.28.22,trna133_GlyCCC_1_-_16872434_16872504@7.28.22,trna17_GluTTC_2_-_204229131_204229201@7.28.22,trna18_GlyGCC_16_+_70822597_70822667@7.28.22,trna19_GlyGCC_16_+_70823410_70823480@7.28.22,trna19_GlyGCC_2_-_157257659_157257729@7.28.22,trna24_GlyGCC_16_-_70812942_70813012@7.28.22,trna25_GlyGCC_16_-_70812114_70812184@7.28.22,trna2_GlyGCC_21_-_18827107_18827177@7.28.22,trna35_GlyGCC_1_+_161413094_161413164@7.28.22,trna37_GlyGCC_1_+_161420467_161420537@7.28.22,trna39_GlyGCC_1_+_161427898_161427968@7.28.22,trna41_GlyGCC_1_+_161435258_161435328@7.28.22,trna4_GlyCCC_1_+_17188416_17188486@7.28.22,trna5_GlyGCC_17_+_8029064_8029134@7.28.22,trna68_GlyGCC_1_-_161493637_161493707@7.28.22,trna91_GlyCCC_1_-_149680210_149680280@7.28.22,trna99_ValCAC_1_-_149294666_149294736@7.28.22|GTGGTTCAGTGGTAGAATTCTC</td><td>>am_trna128_GlyGCC_6_-_27870686_27870756</td><td>7</td><td>28</td><td>22</td><td>GTGGTTCAGTGGTAGAATTCTC</td><td>214.6</td><td>-2.6</td><td>0.9</td><td>-2.9</td><td>3.7E-03</td><td>4.7E-03</td><td>BT20/MB468</td><td>CYTO</td></tr>
<tr><td>>hg38_21_+_8433172_8446622_RNA45SN1_45S_50ntFlanks@11581.11606.26|AACTGGTGCGGACCAGGGGAATCCGA</td><td>>hg38_21_+_8433172_8446622_RNA45SN1_45S_50ntFlanks</td><td>11581</td><td>11606</td><td>26</td><td>AACTGGTGCGGACCAGGGGAATCCGA</td><td>300.0</td><td>-1.9</td><td>0.6</td><td>-2.9</td><td>3.7E-03</td><td>4.7E-03</td><td>BT20/MB468</td><td>CYTO</td></tr>
<tr><td>>hg38_21_+_8433172_8446622_RNA45SN1_45S_50ntFlanks@10107.10125.19|CGTGTGGTGTGCGTCGGAG</td><td>>hg38_21_+_8433172_8446622_RNA45SN1_45S_50ntFlanks</td><td>10107</td><td>10125</td><td>19</td><td>CGTGTGGTGTGCGTCGGAG</td><td>73.4</td><td>9.5</td><td>2.8</td><td>3.4</td><td>7.3E-04</td><td>4.7E-03</td><td>BT20/MB231</td><td>MP</td></tr>
<tr><td>>hg38_21_+_8433172_8446622_RNA45SN1_45S_50ntFlanks@12642.12661.20|CATCTGTGGGATTATGACTG</td><td>>hg38_21_+_8433172_8446622_RNA45SN1_45S_50ntFlanks</td><td>12642</td><td>12661</td><td>20</td><td>CATCTGTGGGATTATGACTG</td><td>441.3</td><td>8.2</td><td>2.4</td><td>3.4</td><td>7.3E-04</td><td>4.7E-03</td><td>BT20/MB231</td><td>MP</td></tr>
<tr><td>>trnaMT_HisGTG_MT_+_12138_12206@27.60.34|GATTGTGAATCTGACAACAGAGGCTTACGACCCC</td><td>>trnaMT_HisGTG_MT_+_12138_12206</td><td>27</td><td>60</td><td>34</td><td>GATTGTGAATCTGACAACAGAGGCTTACGACCCC</td><td>101.5</td><td>5.4</td><td>1.8</td><td>2.9</td><td>3.4E-03</td><td>4.7E-03</td><td>MB231/MB468</td><td>MP</td></tr>
<tr><td>>MI0000272|hsa-mir-182&WithFlank&7|-|129770377|129770498@29.51.23(+1C)&[MIMAT0000259&hsa-miR-182-5p&offsets|0|-1(+1C);m-22&7|-|129770449|129770470&offsets|0|+1(+1C)]|TTTGGCAATGGTAGAACTCACACC</td><td>>MI0000272|hsa-mir-182&WithFlank&7|-|129770377|129770498</td><td>29</td><td>51</td><td>23(+1C)</td><td>TTTGGCAATGGTAGAACTCACACC</td><td>1245.2</td><td>3.4</td><td>1.1</td><td>3.1</td><td>2.1E-03</td><td>4.7E-03</td><td>BT20/MB468</td><td>MP</td></tr>
<tr><td>>trna119_LysCTT_1_-_145395522_145395594@5.36.32,trna11_LysCTT_5_-_180648979_180649051@5.36.32,trna13_LysCTT_6_+_26556774_26556846@5.36.32,trna7_LysCTT_16_+_3225692_3225764@5.36.32,trna9_LysCTT_5_+_180634755_180634827@5.36.32|GGCTAGCTCAGTCGGTAGAGCATGAGACTCTT</td><td>>trna119_LysCTT_1_-_145395522_145395594</td><td>5</td><td>36</td><td>32</td><td>GGCTAGCTCAGTCGGTAGAGCATGAGACTCTT</td><td>236.3</td><td>-4.4</td><td>1.3</td><td>-3.3</td><td>8.1E-04</td><td>4.7E-03</td><td>BT20/MB231</td><td>NUC</td></tr>
<tr><td>>hg38_21_+_8433172_8446622_RNA45SN1_45S_50ntFlanks@475.522.48|CTTCGTGATCGATGTGGTGACGTCGTGCTCTCCCGGGCCGGGTCCGAG</td><td>>hg38_21_+_8433172_8446622_RNA45SN1_45S_50ntFlanks</td><td>475</td><td>522</td><td>48</td><td>CTTCGTGATCGATGTGGTGACGTCGTGCTCTCCCGGGCCGGGTCCGAG</td><td>2963.5</td><td>3.5</td><td>1.2</td><td>2.9</td><td>4.0E-03</td><td>4.7E-03</td><td>MB231/MB468</td><td>NUC</td></tr>
<tr><td>>hg38_21_+_8433172_8446622_RNA45SN1_45S_50ntFlanks@12914.12933.20|CGTTCGTGGGGAACCTGGCG</td><td>>hg38_21_+_8433172_8446622_RNA45SN1_45S_50ntFlanks</td><td>12914</td><td>12933</td><td>20</td><td>CGTTCGTGGGGAACCTGGCG</td><td>354.0</td><td>-3.4</td><td>1.2</td><td>-2.9</td><td>3.7E-03</td><td>4.7E-03</td><td>BT20/MB468</td><td>CYTO</td></tr>
<tr><td>>MI0000086|hsa-mir-28&WithFlank&3|+|188688775|188688872@20.41.22(+1A)&[MIMAT0000085&hsa-miR-28-5p&offsets|0|0(+1A);m-133&3|+|188688794|188688815&offsets|0|0(+1A)]|AAGGAGCTCACAGTCTATTGAGA</td><td>>MI0000086|hsa-mir-28&WithFlank&3|+|188688775|188688872</td><td>20</td><td>41</td><td>22(+1A)</td><td>AAGGAGCTCACAGTCTATTGAGA</td><td>254.4</td><td>3.1</td><td>0.9</td><td>3.3</td><td>8.9E-04</td><td>4.7E-03</td><td>BT20/MB231</td><td>CYTO</td></tr>
<tr><td>>hg38_21_+_8433172_8446622_RNA45SN1_45S_50ntFlanks@12418.12438.21|CGATGTCGGCTCTTCCTATCA</td><td>>hg38_21_+_8433172_8446622_RNA45SN1_45S_50ntFlanks</td><td>12418</td><td>12438</td><td>21</td><td>CGATGTCGGCTCTTCCTATCA</td><td>155.9</td><td>-5.0</td><td>1.7</td><td>-2.9</td><td>3.4E-03</td><td>4.7E-03</td><td>MB231/MB468</td><td>MP</td></tr>
</table>

<table>
<tr><th></th><th></th><th></th><th></th><th></th><th></th><th></th><th></th><th></th><th></th><th></th><th></th><th></th><th></th></tr>
<tr><td>>hg38_21_+_8433172_8446622_RNA45SN1_45S_50ntFlanks@11666.11690.25|TCTGCCCAGTGCTCTGAATGTCAAA</td><td>>hg38_21_+_8433172_8446622_RNA45SN1_45S_50ntFlanks</td><td>11666</td><td>11690</td><td>25</td><td>TCTGCCCAGTGCTCTGAATGTCAAA</td><td>48.8</td><td>5.2</td><td>1.7</td><td>3.1</td><td>2.1E-03</td><td>4.7E-03</td><td>BT20/MB468</td><td>MP</td></tr>
<tr><td>>hg38_21_+_8433172_8446622_RNA45SN1_45S_50ntFlanks@11674.11704.31|GTGCTCTGAATGTCAAAGTGAAGAAATTCAA</td><td>>hg38_21_+_8433172_8446622_RNA45SN1_45S_50ntFlanks</td><td>11674</td><td>11704</td><td>31</td><td>GTGCTCTGAATGTCAAAGTGAAGAAATTCAA</td><td>73.1</td><td>9.5</td><td>2.8</td><td>3.4</td><td>7.4E-04</td><td>4.7E-03</td><td>BT20/MB231</td><td>MP</td></tr>
<tr><td>>hg38_21_+_8433172_8446622_RNA45SN1_45S_50ntFlanks@11923.11949.27|AGAGGTGTAGAATAAGTGGGAGGCCCC</td><td>>hg38_21_+_8433172_8446622_RNA45SN1_45S_50ntFlanks</td><td>11923</td><td>11949</td><td>27</td><td>AGAGGTGTAGAATAAGTGGGAGGCCCC</td><td>203.1</td><td>6.3</td><td>1.9</td><td>3.4</td><td>7.4E-04</td><td>4.7E-03</td><td>BT20/MB231</td><td>MP</td></tr>
<tr><td>>hg38_21_+_8433172_8446622_RNA45SN1_45S_50ntFlanks@11323.11343.21|CGTCTCGTCGCGCGCGCGTCC</td><td>>hg38_21_+_8433172_8446622_RNA45SN1_45S_50ntFlanks</td><td>11323</td><td>11343</td><td>21</td><td>CGTCTCGTCGCGCGCGCGTCC</td><td>889.8</td><td>-4.2</td><td>1.4</td><td>-3.1</td><td>2.1E-03</td><td>4.7E-03</td><td>BT20/MB468</td><td>MP</td></tr>
<tr><td>>hg38_MT_+_1621_3279_RNR2_16S_50ntFlanks@827.860.34|AGGAAAGGTTAAAAAAAGTAAAAGGAACTCGGCA</td><td>>hg38_MT_+_1621_3279_RNR2_16S_50ntFlanks</td><td>827</td><td>860</td><td>34</td><td>AGGAAAGGTTAAAAAAAGTAAAAGGAACTCGGCA</td><td>57.1</td><td>5.4</td><td>1.8</td><td>2.9</td><td>3.4E-03</td><td>4.7E-03</td><td>MB231/MB468</td><td>MP</td></tr>
<tr><td>>MI0000434|hsa-let-7i&WithFlank&12|+|62603680|62603775@12.32.21(+1G)&[MIMAT0000415&hsa-let-7i-5p&offsets|0|-1(+1G);m-49&12|+|62603691|62603712&offsets|0|-1(+1G)]|TGAGGTAGTAGTTTGTGCTGTG</td><td>>MI0000434|hsa-let-7i&WithFlank&12|+|62603680|62603775</td><td>12</td><td>32</td><td>21(+1G)</td><td>TGAGGTAGTAGTTTGTGCTGTG</td><td>1168.9</td><td>3.7</td><td>1.2</td><td>3.1</td><td>2.1E-03</td><td>4.7E-03</td><td>BT20/MB468</td><td>MP</td></tr>
<tr><td>>hg38_21_+_8433172_8446622_RNA45SN1_45S_50ntFlanks@11571.11594.24|GCCGACTTAGAACTGGTGCGGACC</td><td>>hg38_21_+_8433172_8446622_RNA45SN1_45S_50ntFlanks</td><td>11571</td><td>11594</td><td>24</td><td>GCCGACTTAGAACTGGTGCGGACC</td><td>382.0</td><td>2.1</td><td>0.7</td><td>2.9</td><td>3.8E-03</td><td>4.7E-03</td><td>MB231/MB468</td><td>CYTO</td></tr>
<tr><td>>hg38_21_+_8433172_8446622_RNA45SN1_45S_50ntFlanks@10686.10711.26|GCGGCGTCCGGTGAGCTCTCGCTGGC</td><td>>hg38_21_+_8433172_8446622_RNA45SN1_45S_50ntFlanks</td><td>10686</td><td>10711</td><td>26</td><td>GCGGCGTCCGGTGAGCTCTCGCTGGC</td><td>1335.9</td><td>-5.9</td><td>2.0</td><td>-2.9</td><td>4.0E-03</td><td>4.7E-03</td><td>BT20/MB468</td><td>NUC</td></tr>
<tr><td>>hg38_21_+_8433172_8446622_RNA45SN1_45S_50ntFlanks@12687.12704.18|CAGGCGGAACGATACGGC</td><td>>hg38_21_+_8433172_8446622_RNA45SN1_45S_50ntFlanks</td><td>12687</td><td>12704</td><td>18</td><td>CAGGCGGAACGATACGGC</td><td>89.2</td><td>-4.7</td><td>1.6</td><td>-2.9</td><td>3.4E-03</td><td>4.7E-03</td><td>MB231/MB468</td><td>MP</td></tr>
<tr><td>>MI0003137|hsa-mir-193b&WithFlank&16|+|14303961|14304055@57.78.22(+1A)&[MIMAT0002819&hsa-miR-193b-3p&offsets|0|0(+1A);m-166&16|+|14304017|14304038&offsets|0|0(+1A)]|AACTGGCCCTCAAAGTCCCGCTA</td><td>>MI0003137|hsa-mir-193b&WithFlank&16|+|14303961|14304055</td><td>57</td><td>78</td><td>22(+1A)</td><td>AACTGGCCCTCAAAGTCCCGCTA</td><td>210.0</td><td>4.6</td><td>1.5</td><td>3.1</td><td>2.1E-03</td><td>4.7E-03</td><td>BT20/MB468</td><td>MP</td></tr>
<tr><td>>MI0000261|hsa-mir-139&WithFlank&11|-|72615057|72615136@13.35.23&[MIMAT0000250&hsa-miR-139-5p&offsets|0|0;m-186&11|-|72615102|72615124&offsets|0|0]|TCTACAGTGCACGTGTCTCCAGT</td><td>>MI0000261|hsa-mir-139&WithFlank&11|-|72615057|72615136</td><td>13</td><td>35</td><td>23</td><td>TCTACAGTGCACGTGTCTCCAGT</td><td>2483.0</td><td>-3.1</td><td>0.9</td><td>-3.4</td><td>7.4E-04</td><td>4.7E-03</td><td>BT20/MB231</td><td>MP</td></tr>
<tr><td>>MI0000071|hsa-mir-17&WithFlank&13|+|91350599|91350694@19.42.24&[MIMAT0000070&hsa-miR-17-5p&offsets|-1|0;m-88&13|+|91350618|91350640&offsets|-1|0]|TCAAAGTGCTTACAGTGCAGGTAG</td><td>>MI0000071|hsa-mir-17&WithFlank&13|+|91350599|91350694</td><td>19</td><td>42</td><td>24</td><td>TCAAAGTGCTTACAGTGCAGGTAG</td><td>146.0</td><td>3.8</td><td>1.3</td><td>2.9</td><td>3.4E-03</td><td>4.7E-03</td><td>MB231/MB468</td><td>MP</td></tr>
<tr><td>>hg38_21_+_8433172_8446622_RNA45SN1_45S_50ntFlanks@12709.12743.35|CCGCGGAGCCTCGGTTGGCCTCGGATAGCCGGTCC</td><td>>hg38_21_+_8433172_8446622_RNA45SN1_45S_50ntFlanks</td><td>12709</td><td>12743</td><td>35</td><td>CCGCGGAGCCTCGGTTGGCCTCGGATAGCCGGTCC</td><td>156.6</td><td>-4.3</td><td>1.4</td><td>-2.9</td><td>3.2E-03</td><td>4.7E-03</td><td>BT20/MB468</td><td>MITO</td></tr>
<tr><td>>hg38_21_+_8433172_8446622_RNA45SN1_45S_50ntFlanks@6646.6684.39|TCGTACGACTCTTAGCGGTGGATCACTCGGCTCGTGCGT</td><td>>hg38_21_+_8433172_8446622_RNA45SN1_45S_50ntFlanks</td><td>6646</td><td>6684</td><td>39</td><td>TCGTACGACTCTTAGCGGTGGATCACTCGGCTCGTGCGT</td><td>196.2</td><td>3.9</td><td>1.3</td><td>3.1</td><td>2.1E-03</td><td>4.7E-03</td><td>BT20/MB468</td><td>MP</td></tr>
<tr><td>>hg38_21_+_8433172_8446622_RNA45SN1_45S_50ntFlanks@10772.10806.35|AGGTCTCCAAGGTGAACAGCCTCTGGCATGTTGGA</td><td>>hg38_21_+_8433172_8446622_RNA45SN1_45S_50ntFlanks</td><td>10772</td><td>10806</td><td>35</td><td>AGGTCTCCAAGGTGAACAGCCTCTGGCATGTTGGA</td><td>59.5</td><td>-5.0</td><td>1.6</td><td>-3.1</td><td>2.1E-03</td><td>4.7E-03</td><td>BT20/MB468</td><td>MP</td></tr>
<tr><td>>hg38_21_+_8433172_8446622_RNA45SN1_45S_50ntFlanks@6645.6674.30|CTCGTACGACTCTTAGCGGTGGATCACTCG</td><td>>hg38_21_+_8433172_8446622_RNA45SN1_45S_50ntFlanks</td><td>6645</td><td>6674</td><td>30</td><td>CTCGTACGACTCTTAGCGGTGGATCACTCG</td><td>3883.0</td><td>2.8</td><td>1.0</td><td>2.9</td><td>4.1E-03</td><td>4.7E-03</td><td>MB231/MB468</td><td>MITO</td></tr>
<tr><td>>hg38_21_+_8433172_8446622_RNA45SN1_45S_50ntFlanks@6678.6695.18|CGTGCGTCGATGAAGAAC</td><td>>hg38_21_+_8433172_8446622_RNA45SN1_45S_50ntFlanks</td><td>6678</td><td>6695</td><td>18</td><td>CGTGCGTCGATGAAGAAC</td><td>85.8</td><td>4.2</td><td>1.4</td><td>3.1</td><td>2.1E-03</td><td>4.7E-03</td><td>BT20/MB468</td><td>MP</td></tr>
<tr><td>>MI0000298|hsa-mir-221&WithFlank&X|-|45746151|45746272@70.93.24&[MIMAT0000278&hsa-miR-221-3p&offsets|-1|0;m-91&X|-|45746181|45746202&offsets|-1|+1]|CAGCTACATTGTCTGCTGGGTTTC</td><td>>MI0000298|hsa-mir-221&WithFlank&X|-|45746151|45746272</td><td>70</td><td>93</td><td>24</td><td>CAGCTACATTGTCTGCTGGGTTTC</td><td>116.2</td><td>5.4</td><td>1.9</td><td>2.9</td><td>3.4E-03</td><td>4.7E-03</td><td>MB231/MB468</td><td>MP</td></tr>
<tr><td>>hg38_21_+_8433172_8446622_RNA45SN1_45S_50ntFlanks@10814.10840.27|AGGTAAGGGAAGTCGGCAAGCCGGATC</td><td>>hg38_21_+_8433172_8446622_RNA45SN1_45S_50ntFlanks</td><td>10814</td><td>10840</td><td>27</td><td>AGGTAAGGGAAGTCGGCAAGCCGGATC</td><td>241.8</td><td>5.8</td><td>1.9</td><td>3.1</td><td>2.1E-03</td><td>4.7E-03</td><td>BT20/MB468</td><td>MP</td></tr>
<tr><td>>hg38_21_+_8433172_8446622_RNA45SN1_45S_50ntFlanks@12944.12962.19|CGTAGACGACCTGCTTCTG</td><td>>hg38_21_+_8433172_8446622_RNA45SN1_45S_50ntFlanks</td><td>12944</td><td>12962</td><td>19</td><td>CGTAGACGACCTGCTTCTG</td><td>244.4</td><td>-2.1</td><td>0.7</td><td>-2.9</td><td>3.8E-03</td><td>4.7E-03</td><td>BT20/MB468</td><td>CYTO</td></tr>
<tr><td>>am_trnaMT_LysTTT_MT_+_8295_8364@34.52.19,trnalookalike7_LysTTT_1_+_568844_568913@34.52.19|GTTAAAGATTAAGAGAACC</td><td>>am_trnaMT_LysTTT_MT_+_8295_8364</td><td>34</td><td>52</td><td>19</td><td>GTTAAAGATTAAGAGAACC</td><td>709.3</td><td>2.8</td><td>0.8</td><td>3.3</td><td>8.9E-04</td><td>4.7E-03</td><td>BT20/MB231</td><td>CYTO</td></tr>
<tr><td>>trnaMT_ValTAC_MT_+_1602_1670@49.72.24|AACTTAACTTGACCGCTCTGACCA</td><td>>trnaMT_ValTAC_MT_+_1602_1670</td><td>49</td><td>72</td><td>24</td><td>AACTTAACTTGACCGCTCTGACCA</td><td>1214.7</td><td>3.2</td><td>1.0</td><td>3.1</td><td>2.1E-03</td><td>4.7E-03</td><td>BT20/MB468</td><td>MP</td></tr>
<tr><td>>hg38_21_+_8433172_8446622_RNA45SN1_45S_50ntFlanks@4592.4614.23|TTTGTTGGTTTTCGGAACTGAGG</td><td>>hg38_21_+_8433172_8446622_RNA45SN1_45S_50ntFlanks</td><td>4592</td><td>4614</td><td>23</td><td>TTTGTTGGTTTTCGGAACTGAGG</td><td>77.3</td><td>-4.8</td><td>1.6</td><td>-3.1</td><td>2.1E-03</td><td>4.7E-03</td><td>BT20/MB468</td><td>MP</td></tr>
<tr><td>>hg38_21_+_8433172_8446622_RNA45SN1_45S_50ntFlanks@3939.3958.20|AACCCGGTCAGCCCCTCTCC</td><td>>hg38_21_+_8433172_8446622_RNA45SN1_45S_50ntFlanks</td><td>3939</td><td>3958</td><td>20</td><td>AACCCGGTCAGCCCCTCTCC</td><td>760.7</td><td>-4.6</td><td>1.6</td><td>-2.9</td><td>4.0E-03</td><td>4.7E-03</td><td>BT20/MB468</td><td>NUC</td></tr>
<tr><td>>hg38_21_+_8433172_8446622_RNA45SN1_45S_50ntFlanks@7975.7995.21|CGCGACCTCAGATCAGACGTG</td><td>>hg38_21_+_8433172_8446622_RNA45SN1_45S_50ntFlanks</td><td>7975</td><td>7995</td><td>21</td><td>CGCGACCTCAGATCAGACGTG</td><td>5528.0</td><td>1.6</td><td>0.5</td><td>2.9</td><td>4.1E-03</td><td>4.7E-03</td><td>MB231/MB468</td><td>NUC</td></tr>
<tr><td>>hg38_MT_+_1621_3279_RNR2_16S_50ntFlanks@486.518.33|AGGAACAGCTCTTTGGACACTAGGAAAAAACCT</td><td>>hg38_MT_+_1621_3279_RNR2_16S_50ntFlanks</td><td>486</td><td>518</td><td>33</td><td>AGGAACAGCTCTTTGGACACTAGGAAAAAACCT</td><td>214.8</td><td>3.0</td><td>1.0</td><td>2.9</td><td>3.5E-03</td><td>4.7E-03</td><td>MB231/MB468</td><td>MP</td></tr>
<tr><td>>hg38_21_+_8433172_8446622_RNA45SN1_45S_50ntFlanks@12957.12986.30|CTTCTGGGTCGGGGTTTCGTACGTAGCAGA</td><td>>hg38_21_+_8433172_8446622_RNA45SN1_45S_50ntFlanks</td><td>12957</td><td>12986</td><td>30</td><td>CTTCTGGGTCGGGGTTTCGTACGTAGCAGA</td><td>333.8</td><td>-2.2</td><td>0.8</td><td>-2.9</td><td>3.8E-03</td><td>4.7E-03</td><td>MB231/MB468</td><td>CYTO</td></tr>
<tr><td>>hg38_21_+_8433172_8446622_RNA45SN1_45S_50ntFlanks@12893.12930.38|CCTCGCCCGTCACGCACCGCACGTTCGTGGGGAACCTG</td><td>>hg38_21_+_8433172_8446622_RNA45SN1_45S_50ntFlanks</td><td>12893</td><td>12930</td><td>38</td><td>CCTCGCCCGTCACGCACCGCACGTTCGTGGGGAACCTG</td><td>1899.1</td><td>-2.9</td><td>1.0</td><td>-2.9</td><td>3.2E-03</td><td>4.7E-03</td><td>BT20/MB468</td><td>MITO</td></tr>
<tr><td>>hg38_21_+_8433172_8446622_RNA45SN1_45S_50ntFlanks@12685.12708.24|CCCAGGCGGAACGATACGGCAGCG</td><td>>hg38_21_+_8433172_8446622_RNA45SN1_45S_50ntFlanks</td><td>12685</td><td>12708</td><td>24</td><td>CCCAGGCGGAACGATACGGCAGCG</td><td>73.4</td><td>-4.9</td><td>1.6</td><td>-3.1</td><td>2.1E-03</td><td>4.7E-03</td><td>BT20/MB468</td><td>MP</td></tr>
<tr><td>>hg38_1_-_228634819_228635039_RNA5S12_5S_50ntFlanks@51.91.41|GTCTACGGCCATACCACCCTGAACGCGCCCGATCTCGTCTG</td><td>>hg38_1_-_228634819_228635039_RNA5S12_5S_50ntFlanks</td><td>51</td><td>91</td><td>41</td><td>GTCTACGGCCATACCACCCTGAACGCGCCCGATCTCGTCTG</td><td>185.7</td><td>2.3</td><td>0.8</td><td>2.9</td><td>4.1E-03</td><td>4.7E-03</td><td>MB231/MB468</td><td>MITO</td></tr>
<tr><td>>hg38_21_+_8433172_8446622_RNA45SN1_45S_50ntFlanks@11565.11600.36|CTAGCAGCCGACTTAGAACTGGTGCGGACCAGGGGA</td><td>>hg38_21_+_8433172_8446622_RNA45SN1_45S_50ntFlanks</td><td>11565</td><td>11600</td><td>36</td><td>CTAGCAGCCGACTTAGAACTGGTGCGGACCAGGGGA</td><td>248.5</td><td>5.6</td><td>1.7</td><td>3.4</td><td>7.5E-04</td><td>4.7E-03</td><td>BT20/MB231</td><td>MP</td></tr>
<tr><td>>hg38_21_+_8433172_8446622_RNA45SN1_45S_50ntFlanks@9861.9892.32|GATGAACCGAACGCCGGGTTAAGGCGCCCGAT</td><td>>hg38_21_+_8433172_8446622_RNA45SN1_45S_50ntFlanks</td><td>9861</td><td>9892</td><td>32</td><td>GATGAACCGAACGCCGGGTTAAGGCGCCCGAT</td><td>209.5</td><td>5.4</td><td>1.6</td><td>3.4</td><td>7.5E-04</td><td>4.7E-03</td><td>BT20/MB231</td><td>MP</td></tr>
<tr><td>>MI0000746|hsa-mir-99b&WithFlank&19|+|51692606|51692687@51.71.21&[MIMAT0004678&hsa-miR-99b-3p&offsets|0|-1;m-151&19|+|51692656|51692678&offsets|0|-2]|CAAGCTCGTGTCTGTGGGTCC</td><td>>MI0000746|hsa-mir-99b&WithFlank&19|+|51692606|51692687</td><td>51</td><td>71</td><td>21</td><td>CAAGCTCGTGTCTGTGGGTCC</td><td>766.7</td><td>2.7</td><td>0.8</td><td>3.3</td><td>9.0E-04</td><td>4.7E-03</td><td>BT20/MB231</td><td>CYTO</td></tr>
<tr><td>>hg38_21_+_8433172_8446622_RNA45SN1_45S_50ntFlanks@9571.9603.33|GCAAATCGGTCGTCCGACCTGGGTATAGGGGCG</td><td>>hg38_21_+_8433172_8446622_RNA45SN1_45S_50ntFlanks</td><td>9571</td><td>9603</td><td>33</td><td>GCAAATCGGTCGTCCGACCTGGGTATAGGGGCG</td><td>146.1</td><td>-4.1</td><td>1.4</td><td>-2.9</td><td>3.5E-03</td><td>4.7E-03</td><td>MB231/MB468</td><td>MP</td></tr>
<tr><td>>hg38_21_+_8433172_8446622_RNA45SN1_45S_50ntFlanks@12940.12966.27|CATTCGTAGACGACCTGCTTCTGGGTC</td><td>>hg38_21_+_8433172_8446622_RNA45SN1_45S_50ntFlanks</td><td>12940</td><td>12966</td><td>27</td><td>CATTCGTAGACGACCTGCTTCTGGGTC</td><td>195.1</td><td>-2.3</td><td>0.8</td><td>-2.9</td><td>3.2E-03</td><td>4.8E-03</td><td>BT20/MB468</td><td>MITO</td></tr>
<tr><td>>hg38_21_+_8433172_8446622_RNA45SN1_45S_50ntFlanks@4979.5000.22|GACACGGACAGGATTGACAGAT</td><td>>hg38_21_+_8433172_8446622_RNA45SN1_45S_50ntFlanks</td><td>4979</td><td>5000</td><td>22</td><td>GACACGGACAGGATTGACAGAT</td><td>122.9</td><td>-5.0</td><td>1.6</td><td>-3.1</td><td>2.1E-03</td><td>4.8E-03</td><td>BT20/MB468</td><td>MP</td></tr>
<tr><td>>hg38_21_+_8433172_8446622_RNA45SN1_45S_50ntFlanks@4918.4940.23|ACCACCAGGAGTGGAGCCTGCGG</td><td>>hg38_21_+_8433172_8446622_RNA45SN1_45S_50ntFlanks</td><td>4918</td><td>4940</td><td>23</td><td>ACCACCAGGAGTGGAGCCTGCGG</td><td>246.7</td><td>-5.1</td><td>1.6</td><td>-3.1</td><td>2.1E-03</td><td>4.8E-03</td><td>BT20/MB468</td><td>MP</td></tr>
<tr><td>>am_MI0000809|hsa-mir-151a&WithFlank&8|-|140732558|140732659@53.73.21(+1G)&[MIMAT0000757&hsa-miR-151a-3p&offsets|0|0(+1G);m-30&8|-|140732587|140732607&offsets|0|0(+1G)]|CTAGACTGAAGCTCCTTGAGGG</td><td>>am_MI0000809|hsa-mir-151a&WithFlank&8|-|140732558|140732659</td><td>53</td><td>73</td><td>21(+1G)</td><td>CTAGACTGAAGCTCCTTGAGGG</td><td>172.9</td><td>3.7</td><td>1.3</td><td>2.9</td><td>3.5E-03</td><td>4.8E-03</td><td>MB231/MB468</td><td>MP</td></tr>
<tr><td>>hg38_21_+_8433172_8446622_RNA45SN1_45S_50ntFlanks@12445.12473.29|AGCAGAATTCACCAAGCGTTGGATTGTTC</td><td>>hg38_21_+_8433172_8446622_RNA45SN1_45S_50ntFlanks</td><td>12445</td><td>12473</td><td>29</td><td>AGCAGAATTCACCAAGCGTTGGATTGTTC</td><td>291.0</td><td>-3.7</td><td>1.3</td><td>-2.9</td><td>3.5E-03</td><td>4.8E-03</td><td>MB231/MB468</td><td>MP</td></tr>
<tr><td>>hg38_21_+_8433172_8446622_RNA45SN1_45S_50ntFlanks@8171.8214.44|AAGTCCTTCTGATCGAGGCCCAGCCCGTGGACGGTGTGAGGCCG</td><td>>hg38_21_+_8433172_8446622_RNA45SN1_45S_50ntFlanks</td><td>8171</td><td>8214</td><td>44</td><td>AAGTCCTTCTGATCGAGGCCCAGCCCGTGGACGGTGTGAGGCCG</td><td>973.5</td><td>-2.6</td><td>0.9</td><td>-2.9</td><td>3.9E-03</td><td>4.8E-03</td><td>MB231/MB468</td><td>CYTO</td></tr>
<tr><td>>hg38_21_+_8433172_8446622_RNA45SN1_45S_50ntFlanks@6645.6673.29|CTCGTACGACTCTTAGCGGTGGATCACTC</td><td>>hg38_21_+_8433172_8446622_RNA45SN1_45S_50ntFlanks</td><td>6645</td><td>6673</td><td>29</td><td>CTCGTACGACTCTTAGCGGTGGATCACTC</td><td>560.0</td><td>2.7</td><td>1.0</td><td>2.9</td><td>4.1E-03</td><td>4.8E-03</td><td>MB231/MB468</td><td>MITO</td></tr>
<tr><td>>am_trnaMT_PheGAA_MT_+_577_647@45.62.18|AGACGGGCTCACATCACC</td><td>>am_trnaMT_PheGAA_MT_+_577_647</td><td>45</td><td>62</td><td>18</td><td>AGACGGGCTCACATCACC</td><td>754.4</td><td>-3.0</td><td>1.1</td><td>-2.9</td><td>4.1E-03</td><td>4.8E-03</td><td>BT20/MB468</td><td>NUC</td></tr>
<tr><td>>trna2_GlyGCC_21_-_18827107_18827177@1.30.30,trna35_GlyGCC_1_+_161413094_161413164@1.30.30,trna37_GlyGCC_1_+_161420467_161420537@1.30.30,trna39_GlyGCC_1_+_161427898_161427968@1.30.30,trna41_GlyGCC_1_+_161435258_161435328@1.30.30|GCATGGGTGGTTCAGTGGTAGAATTCTCGC</td><td>>trna2_GlyGCC_21_-_18827107_18827177</td><td>1</td><td>30</td><td>30</td><td>GCATGGGTGGTTCAGTGGTAGAATTCTCGC</td><td>755.5</td><td>3.2</td><td>1.0</td><td>3.3</td><td>1.0E-03</td><td>4.8E-03</td><td>BT20/MB231</td><td>MITO</td></tr>
<tr><td>>MI0000447|hsa-mir-128-1&WithFlank&2|+|135665391|135665484@56.75.20(+1C);MI0000727|hsa-mir-128-2&WithFlank&3|+|35744470|35744565@58.77.20(+1C)&[MIMAT0000424&hsa-miR-128-3p&offsets|0|-1(+1C);m-108&2|+|135665446|135665466&offsets|0|-1(+1C)];[MIMAT0000424_1&hsa-miR-128-3p&offsets|0|-1(+1C);m-117&3|+|35744527|35744547&offsets|0|-1(+1C)]|TCACAGTGAACCGGTCTCTTC</td><td>>MI0000447|hsa-mir-128-1&WithFlank&2|+|135665391|135665484</td><td>56</td><td>75</td><td>20(+1C)</td><td>TCACAGTGAACCGGTCTCTTC</td><td>716.2</td><td>2.8</td><td>0.9</td><td>3.1</td><td>2.1E-03</td><td>4.8E-03</td><td>BT20/MB468</td><td>MP</td></tr>
<tr><td>>hg38_21_+_8433172_8446622_RNA45SN1_45S_50ntFlanks@12423.12458.36|TCGGCTCTTCCTATCATTGTGAAGCAGAATTCACCA</td><td>>hg38_21_+_8433172_8446622_RNA45SN1_45S_50ntFlanks</td><td>12423</td><td>12458</td><td>36</td><td>TCGGCTCTTCCTATCATTGTGAAGCAGAATTCACCA</td><td>230.0</td><td>-2.7</td><td>0.8</td><td>-3.3</td><td>9.0E-04</td><td>4.8E-03</td><td>BT20/MB231</td><td>CYTO</td></tr>
<tr><td>>hg38_21_+_8433172_8446622_RNA45SN1_45S_50ntFlanks@4731.4748.18|AAGAACGAAAGTCGGAGG</td><td>>hg38_21_+_8433172_8446622_RNA45SN1_45S_50ntFlanks</td><td>4731</td><td>4748</td><td>18</td><td>AAGAACGAAAGTCGGAGG</td><td>311.0</td><td>-2.9</td><td>1.0</td><td>-2.9</td><td>4.1E-03</td><td>4.8E-03</td><td>MB231/MB468</td><td>MITO</td></tr>
<tr><td>>hg38_21_+_8433172_8446622_RNA45SN1_45S_50ntFlanks@5071.5098.28|TTAATTCCGATAACGAACGAGACTCTGG</td><td>>hg38_21_+_8433172_8446622_RNA45SN1_45S_50ntFlanks</td><td>5071</td><td>5098</td><td>28</td><td>TTAATTCCGATAACGAACGAGACTCTGG</td><td>738.5</td><td>-4.3</td><td>1.5</td><td>-2.9</td><td>3.2E-03</td><td>4.8E-03</td><td>BT20/MB468</td><td>MITO</td></tr>
<tr><td>>hg38_21_+_8433172_8446622_RNA45SN1_45S_50ntFlanks@3853.3884.32|AACTGTGGTAATTCTAGAGCTAATACATGCCG</td><td>>hg38_21_+_8433172_8446622_RNA45SN1_45S_50ntFlanks</td><td>3853</td><td>3884</td><td>32</td><td>AACTGTGGTAATTCTAGAGCTAATACATGCCG</td><td>152.7</td><td>-4.1</td><td>1.4</td><td>-2.9</td><td>3.5E-03</td><td>4.8E-03</td><td>MB231/MB468</td><td>MP</td></tr>
<tr><td>>hg38_21_+_8433172_8446622_RNA45SN1_45S_50ntFlanks@12635.12656.22|AAGCTACCATCTGTGGGATTAT</td><td>>hg38_21_+_8433172_8446622_RNA45SN1_45S_50ntFlanks</td><td>12635</td><td>12656</td><td>22</td><td>AAGCTACCATCTGTGGGATTAT</td><td>136.4</td><td>-4.6</td><td>1.6</td><td>-2.9</td><td>3.2E-03</td><td>4.8E-03</td><td>BT20/MB468</td><td>MITO</td></tr>
</table>

<table>
<tr><td></td><td></td><td></td><td></td><td></td><td></td><td></td><td></td><td></td><td></td><td></td><td></td><td></td><td></td></tr>
<tr><td>>hg38_21_+_8433172_8446622_RNA45SN1_45S_50ntFlanks@12994.13014.21|CCTCGCTGCGATCTATTGAAA</td><td>>hg38_21_+_8433172_8446622_RNA45SN1_45S_50ntFlanks</td><td>12994</td><td>13014</td><td>21</td><td>CCTCGCTGCGATCTATTGAAA</td><td>266.9</td><td>-2.3</td><td>0.8</td><td>-2.9</td><td>3.8E-03</td><td>4.8E-03</td><td>BT20/MB468</td><td>CYTO</td></tr>
<tr><td>>hg38_21_+_8433172_8446622_RNA45SN1_45S_50ntFlanks@10059.10079.21|GGCAGTCGAGAGTGGACGGGA</td><td>>hg38_21_+_8433172_8446622_RNA45SN1_45S_50ntFlanks</td><td>10059</td><td>10079</td><td>21</td><td>GGCAGTCGAGAGTGGACGGGA</td><td>188.7</td><td>6.1</td><td>1.8</td><td>3.4</td><td>7.5E-04</td><td>4.8E-03</td><td>BT20/MB231</td><td>MP</td></tr>
<tr><td>>hg38_21_+_8433172_8446622_RNA45SN1_45S_50ntFlanks@12484.12508.25|AGGGAACGTGAGCTGGGTTTAGACC</td><td>>hg38_21_+_8433172_8446622_RNA45SN1_45S_50ntFlanks</td><td>12484</td><td>12508</td><td>25</td><td>AGGGAACGTGAGCTGGGTTTAGACC</td><td>68.5</td><td>6.0</td><td>2.0</td><td>3.1</td><td>2.1E-03</td><td>4.8E-03</td><td>BT20/MB468</td><td>MP</td></tr>
<tr><td>>hg38_21_+_8433172_8446622_RNA45SN1_45S_50ntFlanks@6655.6677.23|TCTTAGCGGTGGATCACTCGGCT</td><td>>hg38_21_+_8433172_8446622_RNA45SN1_45S_50ntFlanks</td><td>6655</td><td>6677</td><td>23</td><td>TCTTAGCGGTGGATCACTCGGCT</td><td>107.0</td><td>2.5</td><td>0.8</td><td>3.1</td><td>2.1E-03</td><td>4.8E-03</td><td>BT20/MB468</td><td>MP</td></tr>
<tr><td>>trna116_GluCTC_1_-_145399233_145399304@1.35.35,trna59_GluCTC_1_+_249168447_249168518@1.35.35,trna71_GluCTC_1_-_161439189_161439260@1.35.35,trna74_GluCTC_1_-_161431809_161431880@1.35.35,trna77_GluCTC_1_-_161424398_161424469@1.35.35,trna77_GluCTC_6_+_28949976_28950047@1.35.35,trna80_GluCTC_1_-_161417018_161417089@1.35.35,trna87_GluCTC_6_-_126101393_126101464@1.35.35|TCCCTGGTGGTCTAGTGGTTAGGATTCGGCGCTCT</td><td>>trna116_GluCTC_1_-_145399233_145399304</td><td>1</td><td>35</td><td>35</td><td>TCCCTGGTGGTCTAGTGGTTAGGATTCGGCGCTCT</td><td>821.5</td><td>3.4</td><td>1.2</td><td>2.9</td><td>3.5E-03</td><td>4.8E-03</td><td>MB231/MB468</td><td>MP</td></tr>
<tr><td>>hg38_21_+_8433172_8446622_RNA45SN1_45S_50ntFlanks@11670.11690.21|CCCAGTGCTCTGAATGTCAAA</td><td>>hg38_21_+_8433172_8446622_RNA45SN1_45S_50ntFlanks</td><td>11670</td><td>11690</td><td>21</td><td>CCCAGTGCTCTGAATGTCAAA</td><td>213.4</td><td>-3.3</td><td>1.2</td><td>-2.9</td><td>4.1E-03</td><td>4.8E-03</td><td>MB231/MB468</td><td>NUC</td></tr>
<tr><td>>hg38_21_+_8433172_8446622_RNA45SN1_45S_50ntFlanks@4230.4251.22|ACAATACAGGACTCTTTCGAGG</td><td>>hg38_21_+_8433172_8446622_RNA45SN1_45S_50ntFlanks</td><td>4230</td><td>4251</td><td>22</td><td>ACAATACAGGACTCTTTCGAGG</td><td>108.7</td><td>-3.0</td><td>1.1</td><td>-2.9</td><td>3.8E-03</td><td>4.8E-03</td><td>BT20/MB468</td><td>CYTO</td></tr>
<tr><td>>hg38_21_+_8433172_8446622_RNA45SN1_45S_50ntFlanks@3844.3861.18|TACTTGGATAACTGTGGT</td><td>>hg38_21_+_8433172_8446622_RNA45SN1_45S_50ntFlanks</td><td>3844</td><td>3861</td><td>18</td><td>TACTTGGATAACTGTGGT</td><td>475.1</td><td>1.9</td><td>0.7</td><td>2.9</td><td>3.8E-03</td><td>4.8E-03</td><td>BT20/MB468</td><td>CYTO</td></tr>
<tr><td>>hg38_21_+_8433172_8446622_RNA45SN1_45S_50ntFlanks@12415.12443.29|CTTCGATGTCGGCTCTTCCTATCATTGTG</td><td>>hg38_21_+_8433172_8446622_RNA45SN1_45S_50ntFlanks</td><td>12415</td><td>12443</td><td>29</td><td>CTTCGATGTCGGCTCTTCCTATCATTGTG</td><td>3865.7</td><td>-2.5</td><td>0.9</td><td>-2.9</td><td>4.1E-03</td><td>4.8E-03</td><td>MB231/MB468</td><td>NUC</td></tr>
<tr><td>>hg38_21_+_8433172_8446622_RNA45SN1_45S_50ntFlanks@11589.11621.33|CGGACCAGGGGAATCCGACTGTTTAATTAAAAC</td><td>>hg38_21_+_8433172_8446622_RNA45SN1_45S_50ntFlanks</td><td>11589</td><td>11621</td><td>33</td><td>CGGACCAGGGGAATCCGACTGTTTAATTAAAAC</td><td>75.7</td><td>5.9</td><td>1.9</td><td>3.1</td><td>2.1E-03</td><td>4.8E-03</td><td>BT20/MB468</td><td>MP</td></tr>
<tr><td>>hg38_21_+_8433172_8446622_RNA45SN1_45S_50ntFlanks@6661.6693.33|CGGTGGATCACTCGGCTCGTGCGTCGATGAAGA</td><td>>hg38_21_+_8433172_8446622_RNA45SN1_45S_50ntFlanks</td><td>6661</td><td>6693</td><td>33</td><td>CGGTGGATCACTCGGCTCGTGCGTCGATGAAGA</td><td>557.2</td><td>1.9</td><td>0.6</td><td>3.3</td><td>8.3E-04</td><td>4.8E-03</td><td>BT20/MB231</td><td>NUC</td></tr>
<tr><td>>hg38_21_+_8433172_8446622_RNA45SN1_45S_50ntFlanks@6645.6681.37|CTCGTACGACTCTTAGCGGTGGATCACTCGGCTCGTG</td><td>>hg38_21_+_8433172_8446622_RNA45SN1_45S_50ntFlanks</td><td>6645</td><td>6681</td><td>37</td><td>CTCGTACGACTCTTAGCGGTGGATCACTCGGCTCGTG</td><td>8132.8</td><td>-1.6</td><td>0.5</td><td>-2.9</td><td>3.3E-03</td><td>4.8E-03</td><td>BT20/MB468</td><td>MITO</td></tr>
<tr><td>>trna111_HisGTG_1_-_147774845_147774916@-1A.26.27,trna118_HisGTG_1_-_145396881_145396952@-1A.26.27,trna16_HisGTG_1_+_146544773_146544844@-1A.26.27,trna1_HisGTG_15_+_45493349_45493420@-1A.26.27,trna21_HisGTG_1_+_147753471_147753542@-1A.26.27,trna33_HisGTG_6_+_27125906_27125977@-1A.26.27,trna7_HisGTG_9_-_14433938_14434009@-1A.26.27,trna8_HisGTG_15_-_45492611_45492682@-1A.26.27,trna9_HisGTG_15_-_45490804_45490875@-1A.26.27|AGCCGTGATCGTATAGTGGTTAGTACT</td><td>>trna111_HisGTG_1_-_147774845_147774916</td><td>-1A</td><td>26</td><td>27</td><td>AGCCGTGATCGTATAGTGGTTAGTACT</td><td>55.9</td><td>4.8</td><td>1.6</td><td>2.9</td><td>3.5E-03</td><td>4.8E-03</td><td>MB231/MB468</td><td>MP</td></tr>
<tr><td>>hg38_MT_+_1621_3279_RNR2_16S_50ntFlanks@482.518.37|AAAGAGGAACAGCTCTTTGGACACTAGGAAAAAACCT</td><td>>hg38_MT_+_1621_3279_RNR2_16S_50ntFlanks</td><td>482</td><td>518</td><td>37</td><td>AAAGAGGAACAGCTCTTTGGACACTAGGAAAAAACCT</td><td>87.3</td><td>4.6</td><td>1.6</td><td>2.9</td><td>3.5E-03</td><td>4.8E-03</td><td>MB231/MB468</td><td>MP</td></tr>
<tr><td>>am_trnaMT_GluTTC_MT_-_14674_14742@24.54.31,trnalookalike8_GluTTC_5_-_93905172_93905240@24.54.31|ATGGTTTTTCATATCATTGGTCGTGGTTGTA</td><td>>am_trnaMT_GluTTC_MT_-_14674_14742</td><td>24</td><td>54</td><td>31</td><td>ATGGTTTTTCATATCATTGGTCGTGGTTGTA</td><td>351.2</td><td>4.7</td><td>1.4</td><td>3.4</td><td>7.6E-04</td><td>4.8E-03</td><td>BT20/MB231</td><td>MP</td></tr>
<tr><td>>MI0005531|hsa-mir-450b&WithFlank&X|-|134540179|134540268@17.38.22&[MIMAT0004909&hsa-miR-450b-5p&offsets|0|0;m-266&X|-|134540232|134540252&offsets|0|+1]|TTTTGCAATATGTTCCTGAATA</td><td>>MI0005531|hsa-mir-450b&WithFlank&X|-|134540179|134540268</td><td>17</td><td>38</td><td>22</td><td>TTTTGCAATATGTTCCTGAATA</td><td>55.2</td><td>5.4</td><td>1.8</td><td>2.9</td><td>3.5E-03</td><td>4.8E-03</td><td>MB231/MB468</td><td>MP</td></tr>
<tr><td>>hg38_21_+_8433172_8446622_RNA45SN1_45S_50ntFlanks@4734.4764.31|AACGAAAGTCGGAGGTTCGAAGACGATCAGA</td><td>>hg38_21_+_8433172_8446622_RNA45SN1_45S_50ntFlanks</td><td>4734</td><td>4764</td><td>31</td><td>AACGAAAGTCGGAGGTTCGAAGACGATCAGA</td><td>138.6</td><td>4.3</td><td>1.4</td><td>3.1</td><td>2.2E-03</td><td>4.8E-03</td><td>BT20/MB468</td><td>MP</td></tr>
<tr><td>>hg38_21_+_8433172_8446622_RNA45SN1_45S_50ntFlanks@3823.3850.28|TTTGGTCGCTCGCTCCTCTCCTACTTGG</td><td>>hg38_21_+_8433172_8446622_RNA45SN1_45S_50ntFlanks</td><td>3823</td><td>3850</td><td>28</td><td>TTTGGTCGCTCGCTCCTCTCCTACTTGG</td><td>248.7</td><td>2.3</td><td>0.8</td><td>2.9</td><td>3.9E-03</td><td>4.8E-03</td><td>MB231/MB468</td><td>CYTO</td></tr>
<tr><td>>trnaMT_GluTTC_MT_-_14674_14742@35.63.29,trnalookalike8_GluTTC_5_-_93905172_93905240@35.63.29|TATCATTGGTCGTGGTTGTAGTCCGTGCG</td><td>>trnaMT_GluTTC_MT_-_14674_14742</td><td>35</td><td>63</td><td>29</td><td>TATCATTGGTCGTGGTTGTAGTCCGTGCG</td><td>141.3</td><td>6.2</td><td>2.0</td><td>3.1</td><td>2.2E-03</td><td>4.8E-03</td><td>BT20/MB468</td><td>MP</td></tr>
<tr><td>>hg38_21_+_8433172_8446622_RNA45SN1_45S_50ntFlanks@10822.10844.23|GAAGTCGGCAAGCCGGATCCGTA</td><td>>hg38_21_+_8433172_8446622_RNA45SN1_45S_50ntFlanks</td><td>10822</td><td>10844</td><td>23</td><td>GAAGTCGGCAAGCCGGATCCGTA</td><td>261.0</td><td>-1.9</td><td>0.7</td><td>-2.9</td><td>3.9E-03</td><td>4.8E-03</td><td>MB231/MB468</td><td>CYTO</td></tr>
<tr><td>>trnaMT_ThrTGT_MT_+_15888_15953@34.69.36|TAAACCGGAGATGAAAACCTTTTTCCAAGGACACCA</td><td>>trnaMT_ThrTGT_MT_+_15888_15953</td><td>34</td><td>69</td><td>36</td><td>TAAACCGGAGATGAAAACCTTTTTCCAAGGACACCA</td><td>168.4</td><td>3.6</td><td>1.2</td><td>2.9</td><td>3.5E-03</td><td>4.8E-03</td><td>MB231/MB468</td><td>MP</td></tr>
<tr><td>>MI0000746|hsa-mir-99b&WithFlank&19|+|51692606|51692687@13.34.22&[MIMAT0000689&hsa-miR-99b-5p&offsets|0|0;m-10&19|+|51692618|51692639&offsets|0|0]|CACCCGTAGAACCGACCTTGCG</td><td>>MI0000746|hsa-mir-99b&WithFlank&19|+|51692606|51692687</td><td>13</td><td>34</td><td>22</td><td>CACCCGTAGAACCGACCTTGCG</td><td>8487.9</td><td>1.9</td><td>0.7</td><td>2.9</td><td>3.5E-03</td><td>4.8E-03</td><td>MB231/MB468</td><td>MP</td></tr>
<tr><td>>hg38_MT_+_1621_3279_RNR2_16S_50ntFlanks@1348.1382.35|AACAATAGGGTTTACGACCTCGATGTTGGATCAGG</td><td>>hg38_MT_+_1621_3279_RNR2_16S_50ntFlanks</td><td>1348</td><td>1382</td><td>35</td><td>AACAATAGGGTTTACGACCTCGATGTTGGATCAGG</td><td>125.0</td><td>3.6</td><td>1.2</td><td>2.9</td><td>3.5E-03</td><td>4.8E-03</td><td>MB231/MB468</td><td>MP</td></tr>
<tr><td>>hg38_21_+_8433172_8446622_RNA45SN1_45S_50ntFlanks@7974.7991.18|ACGCGACCTCAGATCAGA</td><td>>hg38_21_+_8433172_8446622_RNA45SN1_45S_50ntFlanks</td><td>7974</td><td>7991</td><td>18</td><td>ACGCGACCTCAGATCAGA</td><td>3084.1</td><td>2.7</td><td>0.9</td><td>2.9</td><td>4.2E-03</td><td>4.8E-03</td><td>MB231/MB468</td><td>NUC</td></tr>
<tr><td>>MI0000286|hsa-mir-210&WithFlank&11|-|568083|568204@72.93.22(+1A)&[MIMAT0000267&hsa-miR-210-3p&offsets|0|0(+1A);m-102&11|-|568112|568133&offsets|0|0(+1A)]|CTGTGCGTGTGACAGCGGCTGAA</td><td>>MI0000286|hsa-mir-210&WithFlank&11|-|568083|568204</td><td>72</td><td>93</td><td>22(+1A)</td><td>CTGTGCGTGTGACAGCGGCTGAA</td><td>430.6</td><td>4.0</td><td>1.4</td><td>2.9</td><td>3.5E-03</td><td>4.8E-03</td><td>MB231/MB468</td><td>MP</td></tr>
<tr><td>>hg38_21_+_8433172_8446622_RNA45SN1_45S_50ntFlanks@11671.11701.31|CCAGTGCTCTGAATGTCAAAGTGAAGAAATT</td><td>>hg38_21_+_8433172_8446622_RNA45SN1_45S_50ntFlanks</td><td>11671</td><td>11701</td><td>31</td><td>CCAGTGCTCTGAATGTCAAAGTGAAGAAATT</td><td>257.7</td><td>6.6</td><td>2.0</td><td>3.4</td><td>7.6E-04</td><td>4.8E-03</td><td>BT20/MB231</td><td>MP</td></tr>
<tr><td>>hg38_21_+_8433172_8446622_RNA45SN1_45S_50ntFlanks@11874.11893.20|AGAAGACCCTGTTGAGCTTG</td><td>>hg38_21_+_8433172_8446622_RNA45SN1_45S_50ntFlanks</td><td>11874</td><td>11893</td><td>20</td><td>AGAAGACCCTGTTGAGCTTG</td><td>287.2</td><td>6.8</td><td>2.0</td><td>3.4</td><td>7.6E-04</td><td>4.8E-03</td><td>BT20/MB231</td><td>MP</td></tr>
<tr><td>>hg38_21_+_8433172_8446622_RNA45SN1_45S_50ntFlanks@11913.11946.34|AAGAGACATGAGAGGTGTAGAATAAGTGGGAGGC</td><td>>hg38_21_+_8433172_8446622_RNA45SN1_45S_50ntFlanks</td><td>11913</td><td>11946</td><td>34</td><td>AAGAGACATGAGAGGTGTAGAATAAGTGGGAGGC</td><td>73.4</td><td>9.5</td><td>2.8</td><td>3.4</td><td>7.6E-04</td><td>4.8E-03</td><td>BT20/MB231</td><td>MP</td></tr>
<tr><td>>hg38_21_+_8433172_8446622_RNA45SN1_45S_50ntFlanks@12419.12452.34|GATGTCGGCTCTTCCTATCATTGTGAAGCAGAAT</td><td>>hg38_21_+_8433172_8446622_RNA45SN1_45S_50ntFlanks</td><td>12419</td><td>12452</td><td>34</td><td>GATGTCGGCTCTTCCTATCATTGTGAAGCAGAAT</td><td>492.9</td><td>-2.7</td><td>0.8</td><td>-3.3</td><td>8.4E-04</td><td>4.8E-03</td><td>BT20/MB231</td><td>NUC</td></tr>
<tr><td>>hg38_21_+_8433172_8446622_RNA45SN1_45S_50ntFlanks@11190.11208.19|CCCGGGGGGCCGGCGGCGG</td><td>>hg38_21_+_8433172_8446622_RNA45SN1_45S_50ntFlanks</td><td>11190</td><td>11208</td><td>19</td><td>CCCGGGGGGCCGGCGGCGG</td><td>207.8</td><td>-3.2</td><td>1.1</td><td>-2.9</td><td>3.9E-03</td><td>4.8E-03</td><td>BT20/MB468</td><td>CYTO</td></tr>
<tr><td>>hg38_21_+_8433172_8446622_RNA45SN1_45S_50ntFlanks@10116.10135.20|TGCGTCGGAGGGCGGCGGCG</td><td>>hg38_21_+_8433172_8446622_RNA45SN1_45S_50ntFlanks</td><td>10116</td><td>10135</td><td>20</td><td>TGCGTCGGAGGGCGGCGGCG</td><td>258.0</td><td>-5.4</td><td>1.8</td><td>-3.1</td><td>2.2E-03</td><td>4.8E-03</td><td>BT20/MB468</td><td>MP</td></tr>
<tr><td>>MI0000255|hsa-mir-30d&WithFlank&8|-|134804870|134804951@12.30.19&[MIMAT0000245&hsa-miR-30d-5p&offsets|0|-3;m-21&8|-|134804919|134804940&offsets|0|-3]|TGTAAACATCCCCGACTGG</td><td>>MI0000255|hsa-mir-30d&WithFlank&8|-|134804870|134804951</td><td>12</td><td>30</td><td>19</td><td>TGTAAACATCCCCGACTGG</td><td>94.8</td><td>5.4</td><td>1.8</td><td>3.1</td><td>2.2E-03</td><td>4.9E-03</td><td>BT20/MB468</td><td>MP</td></tr>
<tr><td>>hg38_21_+_8433172_8446622_RNA45SN1_45S_50ntFlanks@11920.11952.33|ATGAGAGGTGTAGAATAAGTGGGAGGCCCCCGG</td><td>>hg38_21_+_8433172_8446622_RNA45SN1_45S_50ntFlanks</td><td>11920</td><td>11952</td><td>33</td><td>ATGAGAGGTGTAGAATAAGTGGGAGGCCCCCGG</td><td>126.9</td><td>-4.1</td><td>1.4</td><td>-2.9</td><td>3.9E-03</td><td>4.9E-03</td><td>BT20/MB468</td><td>CYTO</td></tr>
<tr><td>>hg38_21_+_8433172_8446622_RNA45SN1_45S_50ntFlanks@9504.9524.21|GTGAACTATGCCTGGGCAGGG</td><td>>hg38_21_+_8433172_8446622_RNA45SN1_45S_50ntFlanks</td><td>9504</td><td>9524</td><td>21</td><td>GTGAACTATGCCTGGGCAGGG</td><td>159.1</td><td>-3.2</td><td>1.1</td><td>-2.9</td><td>3.3E-03</td><td>4.9E-03</td><td>BT20/MB468</td><td>MITO</td></tr>
<tr><td>>hg38_21_+_8433172_8446622_RNA45SN1_45S_50ntFlanks@10798.10817.20|CATGTTGGAACAATGTAGGT</td><td>>hg38_21_+_8433172_8446622_RNA45SN1_45S_50ntFlanks</td><td>10798</td><td>10817</td><td>20</td><td>CATGTTGGAACAATGTAGGT</td><td>635.7</td><td>-4.1</td><td>1.3</td><td>-3.1</td><td>2.2E-03</td><td>4.9E-03</td><td>BT20/MB468</td><td>MP</td></tr>
<tr><td>>hg38_21_+_8433172_8446622_RNA45SN1_45S_50ntFlanks@11782.11804.23|GCATGAATGGATGAACGAGATTC</td><td>>hg38_21_+_8433172_8446622_RNA45SN1_45S_50ntFlanks</td><td>11782</td><td>11804</td><td>23</td><td>GCATGAATGGATGAACGAGATTC</td><td>312.1</td><td>5.7</td><td>1.7</td><td>3.4</td><td>7.7E-04</td><td>4.9E-03</td><td>BT20/MB231</td><td>MP</td></tr>
<tr><td>>hg38_21_+_8433172_8446622_RNA45SN1_45S_50ntFlanks@8971.8990.20|CCCACCCCCGCGGGGAATCC</td><td>>hg38_21_+_8433172_8446622_RNA45SN1_45S_50ntFlanks</td><td>8971</td><td>8990</td><td>20</td><td>CCCACCCCCGCGGGGAATCC</td><td>387.5</td><td>-4.7</td><td>1.6</td><td>-2.9</td><td>3.3E-03</td><td>4.9E-03</td><td>BT20/MB468</td><td>MITO</td></tr>
<tr><td>>trna111_HisGTG_1_-_147774845_147774916@-1G.30.31,trna118_HisGTG_1_-_145396881_145396952@-1G.30.31,trna16_HisGTG_1_+_146544773_146544844@-1G.30.31,trna1_HisGTG_15_+_45493349_45493420@-1G.30.31,trna21_HisGTG_1_+_147753471_147753542@-1G.30.31,trna33_HisGTG_6_+_27125906_27125977@-1G.30.31,trna7_HisGTG_9_-_14433938_14434009@-1G.30.31,trna8_HisGTG_15_-_45492611_45492682@-1G.30.31,trna9_HisGTG_15_-_45490804_45490875@-1G.30.31|GGCCGTGATCGTATAGTGGTTAGTACTCTGC</td><td>>trna111_HisGTG_1_-_147774845_147774916</td><td>-1G</td><td>30</td><td>31</td><td>GGCCGTGATCGTATAGTGGTTAGTACTCTGC</td><td>7740.3</td><td>-3.4</td><td>1.0</td><td>-3.3</td><td>9.2E-04</td><td>4.9E-03</td><td>BT20/MB231</td><td>CYTO</td></tr>
<tr><td>>hg38_21_+_8433172_8446622_RNA45SN1_45S_50ntFlanks@10678.10709.32|CGGTTCCGGCGGCGTCCGGTGAGCTCTCGCTG</td><td>>hg38_21_+_8433172_8446622_RNA45SN1_45S_50ntFlanks</td><td>10678</td><td>10709</td><td>32</td><td>CGGTTCCGGCGGCGTCCGGTGAGCTCTCGCTG</td><td>415.6</td><td>6.0</td><td>1.8</td><td>3.4</td><td>7.7E-04</td><td>4.9E-03</td><td>BT20/MB231</td><td>MP</td></tr>
<tr><td>>hg38_21_+_8433172_8446622_RNA45SN1_45S_50ntFlanks@6646.6686.41|TCGTACGACTCTTAGCGGTGGATCACTCGGCTCGTGCGTCG</td><td>>hg38_21_+_8433172_8446622_RNA45SN1_45S_50ntFlanks</td><td>6646</td><td>6686</td><td>41</td><td>TCGTACGACTCTTAGCGGTGGATCACTCGGCTCGTGCGTCG</td><td>92.4</td><td>2.0</td><td>0.7</td><td>2.9</td><td>4.2E-03</td><td>4.9E-03</td><td>BT20/MB468</td><td>NUC</td></tr>
<tr><td>>hg38_21_+_8433172_8446622_RNA45SN1_45S_50ntFlanks@11641.11668.28|CGGCGGGTGTTGACGCGATGTGATTTCT</td><td>>hg38_21_+_8433172_8446622_RNA45SN1_45S_50ntFlanks</td><td>11641</td><td>11668</td><td>28</td><td>CGGCGGGTGTTGACGCGATGTGATTTCT</td><td>144.4</td><td>-4.2</td><td>1.5</td><td>-2.9</td><td>3.9E-03</td><td>4.9E-03</td><td>BT20/MB468</td><td>CYTO</td></tr>
<tr><td>>am_MI0003581|hsa-mir-574&WithFlank&4|+|38868026|38868133@31.52.22&[MIMAT0004795&hsa-miR-574-5p&offsets|0|-1;m-213&4|+|38868056|38868078&offsets|0|-1]|TGAGTGTGTGTGTGTGAGTGTG</td><td>>am_MI0003581|hsa-mir-574&WithFlank&4|+|38868026|38868133</td><td>31</td><td>52</td><td>22</td><td>TGAGTGTGTGTGTGTGAGTGTG</td><td>256.0</td><td>3.7</td><td>1.3</td><td>2.9</td><td>3.6E-03</td><td>4.9E-03</td><td>MB231/MB468</td><td>MP</td></tr>
<tr><td>>MI0005531|hsa-mir-450b&WithFlank&X|-|134540179|134540268@17.38.22&[MIMAT0004909&hsa-miR-450b-5p&offsets|0|0;m-266&X|-|134540232|134540252&offsets|0|+1]|TTTTGCAATATGTTCCTGAATA</td><td>>MI0005531|hsa-mir-450b&WithFlank&X|-|134540179|134540268</td><td>17</td><td>38</td><td>22</td><td>TTTTGCAATATGTTCCTGAATA</td><td>289.2</td><td>2.9</td><td>0.9</td><td>3.3</td><td>9.3E-04</td><td>4.9E-03</td><td>BT20/MB231</td><td>CYTO</td></tr>
</table>

| | | | | | | | | | | | | | |
|---|---|---|---|---|---|---|---|---|---|---|---|---|---|
| >MI0000465\|hsa-mir-191&WithFlank&3\|-\|49020612\|49020715@22.42.21(+1C)&[MIMAT0000440&hsa-miR-191-5p&offsets\|0\|-2(+1C);m-48&3\|-\|49020672\|49020694&offsets\|0\|-2(+1C)]\|CAACGGAATCCCAAAAGCAGCC | >MI0000465\|hsa-mir-191&WithFlank&3\|-\|49020612\|49020715 | 22 | 42 | 21(+1C) | CAACGGAATCCCAAAAGCAGCC | 1140.7 | 2.9 | 1.0 | 2.9 | 3.6E-03 | 4.9E-03 | MB231/MB468 | MP |
| >hg38_21_+_8433172_8446622_RNA45SN1_45S_50ntFlanks@11568.11585.18\|GCAGCCGACTTAGAACTG | >hg38_21_+_8433172_8446622_RNA45SN1_45S_50ntFlanks | 11568 | 11585 | 18 | GCAGCCGACTTAGAACTG | 1882.0 | -2.6 | 0.8 | -3.1 | 2.2E-03 | 4.9E-03 | BT20/MB468 | MP |
| >MI0000273\|hsa-mir-183&WithFlank&7\|-\|129774899\|129775020@33.52.20&[MIMAT0000261&hsa-miR-183-5p&offsets\|0\|-2;m-41&7\|-\|129774966\|129774987&offsets\|-1\|-3]\|TATGGCACTGGTAGAATTCA | >MI0000273\|hsa-mir-183&WithFlank&7\|-\|129774899\|129775020 | 33 | 52 | 20 | TATGGCACTGGTAGAATTCA | 127.9 | 6.3 | 2.1 | 3.1 | 2.2E-03 | 4.9E-03 | BT20/MB468 | MP |
| >hg38_MT_+_1621_3279_RNR2_16S_50ntFlanks@285.313.29\|CGAAACCAGACGAGCTACCTAAGAACAGC | >hg38_MT_+_1621_3279_RNR2_16S_50ntFlanks | 285 | 313 | 29 | CGAAACCAGACGAGCTACCTAAGAACAGC | 79.8 | 5.3 | 1.7 | 3.1 | 2.2E-03 | 4.9E-03 | BT20/MB468 | MP |
| >hg38_21_+_8433172_8446622_RNA45SN1_45S_50ntFlanks@9598.9615.18\|GGGGCGAAAGACTAATCG | >hg38_21_+_8433172_8446622_RNA45SN1_45S_50ntFlanks | 9598 | 9615 | 18 | GGGGCGAAAGACTAATCG | 123.3 | -4.9 | 1.6 | -3.1 | 2.2E-03 | 4.9E-03 | BT20/MB468 | MP |
| >hg38_21_+_8433172_8446622_RNA45SN1_45S_50ntFlanks@5470.5514.45\|CCACGGCCCTGGCGGAGCGCTGAGAAGACGGTCGAACTTGACTAT | >hg38_21_+_8433172_8446622_RNA45SN1_45S_50ntFlanks | 5470 | 5514 | 45 | CCACGGCCCTGGCGGAGCGCTGAGAAGACGGTCGAACTTGACTAT | 868.9 | -2.7 | 0.9 | -2.9 | 4.0E-03 | 4.9E-03 | MB231/MB468 | CYTO |
| >hg38_21_+_8433172_8446622_RNA45SN1_45S_50ntFlanks@10806.10826.21\|AACAATGTAGGTAAGGGAAGT | >hg38_21_+_8433172_8446622_RNA45SN1_45S_50ntFlanks | 10806 | 10826 | 21 | AACAATGTAGGTAAGGGAAGT | 285.8 | -5.0 | 1.6 | -3.1 | 2.2E-03 | 4.9E-03 | BT20/MB468 | MP |
| >trnaMT_MetCAT_MT_+_4402_4469@38.71.34\|CGAAAATGTTGGTTATACCCTTCCCGTACTACCA | >trnaMT_MetCAT_MT_+_4402_4469 | 38 | 71 | 34 | CGAAAATGTTGGTTATACCCTTCCCGTACTACCA | 560.1 | 5.8 | 1.9 | 3.1 | 2.2E-03 | 4.9E-03 | BT20/MB468 | MP |
| >hg38_21_+_8433172_8446622_RNA45SN1_45S_50ntFlanks@6777.6803.27\|TCCCGGGGCTACGCCTGTCTGAGCGTC | >hg38_21_+_8433172_8446622_RNA45SN1_45S_50ntFlanks | 6777 | 6803 | 27 | TCCCGGGGCTACGCCTGTCTGAGCGTC | 83.4 | -4.7 | 1.6 | -2.9 | 3.6E-03 | 4.9E-03 | MB231/MB468 | MP |
| >trnaMT_ValTAC_MT_+_1602_1670@34.72.39\|CACTTAGGAGATTTCAACTTAACTTGACCGCTCTGACCA | >trnaMT_ValTAC_MT_+_1602_1670 | 34 | 72 | 39 | CACTTAGGAGATTTCAACTTAACTTGACCGCTCTGACCA | 341.9 | 5.1 | 1.6 | 3.1 | 2.2E-03 | 4.9E-03 | BT20/MB468 | MP |
| >hg38_21_+_8433172_8446622_RNA45SN1_45S_50ntFlanks@3776.3801.26\|CCGGTACAGTGAAACTGCGAATGGCT | >hg38_21_+_8433172_8446622_RNA45SN1_45S_50ntFlanks | 3776 | 3801 | 26 | CCGGTACAGTGAAACTGCGAATGGCT | 780.3 | -5.3 | 1.7 | -3.1 | 2.2E-03 | 4.9E-03 | BT20/MB468 | MP |
| >hg38_MT_+_598_1651_RNR1_12S_50ntFlanks@440.480.41\|ACATATCTGAACACACAATAGCTAAGACCCAAACTGGGATT | >hg38_MT_+_598_1651_RNR1_12S_50ntFlanks | 440 | 480 | 41 | ACATATCTGAACACACAATAGCTAAGACCCAAACTGGGATT | 167.1 | 3.1 | 1.1 | 2.9 | 4.2E-03 | 4.9E-03 | MB231/MB468 | MITO |
| >hg38_21_+_8433172_8446622_RNA45SN1_45S_50ntFlanks@6675.6696.22\|GCTCGTGCGTCGATGAAGAACG | >hg38_21_+_8433172_8446622_RNA45SN1_45S_50ntFlanks | 6675 | 6696 | 22 | GCTCGTGCGTCGATGAAGAACG | 158.7 | -3.7 | 1.3 | -2.9 | 3.6E-03 | 4.9E-03 | MB231/MB468 | MP |
| >hg38_21_+_8433172_8446622_RNA45SN1_45S_50ntFlanks@12004.12026.23\|CGGGCCGCCGGTGAAATACCACT | >hg38_21_+_8433172_8446622_RNA45SN1_45S_50ntFlanks | 12004 | 12026 | 23 | CGGGCCGCCGGTGAAATACCACT | 465.1 | 7.3 | 2.2 | 3.3 | 1.0E-03 | 4.9E-03 | BT20/MB231 | MITO |
| >hg38_21_+_8433172_8446622_RNA45SN1_45S_50ntFlanks@12997.13014.18\|CGCTGCGATCTATTGAAA | >hg38_21_+_8433172_8446622_RNA45SN1_45S_50ntFlanks | 12997 | 13014 | 18 | CGCTGCGATCTATTGAAA | 1225.9 | -2.5 | 0.8 | -3.3 | 1.0E-03 | 4.9E-03 | BT20/MB231 | MITO |
| >hg38_21_+_8433172_8446622_RNA45SN1_45S_50ntFlanks@8159.8191.33\|CGTGGGGGGCCCAAGTCCTTCTGATCGAGGCCC | >hg38_21_+_8433172_8446622_RNA45SN1_45S_50ntFlanks | 8159 | 8191 | 33 | CGTGGGGGGCCCAAGTCCTTCTGATCGAGGCCC | 74.8 | 4.9 | 1.7 | 2.9 | 3.6E-03 | 4.9E-03 | MB231/MB468 | MP |
| >hg38_21_+_8433172_8446622_RNA45SN1_45S_50ntFlanks@11662.11681.20\|GATTTCTGCCCAGTGCTCTG | >hg38_21_+_8433172_8446622_RNA45SN1_45S_50ntFlanks | 11662 | 11681 | 20 | GATTTCTGCCCAGTGCTCTG | 177.5 | -1.5 | 0.5 | -2.9 | 3.9E-03 | 4.9E-03 | BT20/MB468 | CYTO |
| >hg38_21_+_8433172_8446622_RNA45SN1_45S_50ntFlanks@10711.10736.26\|CCCTTGAAAATCCGGGGGAGAGGGTG | >hg38_21_+_8433172_8446622_RNA45SN1_45S_50ntFlanks | 10711 | 10736 | 26 | CCCTTGAAAATCCGGGGGAGAGGGTG | 56.3 | -5.0 | 1.6 | -3.1 | 2.2E-03 | 4.9E-03 | BT20/MB468 | MP |
| >hg38_21_+_8433172_8446622_RNA45SN1_45S_50ntFlanks@5515.5534.20\|CTAGAGGAAGTAAAAGTCGT | >hg38_21_+_8433172_8446622_RNA45SN1_45S_50ntFlanks | 5515 | 5534 | 20 | CTAGAGGAAGTAAAAGTCGT | 270.2 | -5.0 | 1.6 | -3.1 | 2.2E-03 | 4.9E-03 | BT20/MB468 | MP |
| >MI0002468\|hsa-mir-484&WithFlank&16\|+\|15643288\|15643378@14.35.22(+1U)&[MIMAT0002174&hsa-miR-484&offsets\|0\|0(+1U);m-127&16\|+\|15643301\|15643321&offsets\|0\|+1(+1U)]\|TCAGGCTCAGTCCCCTCCCGATT | >MI0002468\|hsa-mir-484&WithFlank&16\|+\|15643288\|15643378 | 14 | 35 | 22(+1U) | TCAGGCTCAGTCCCCTCCCGATT | 177.4 | 2.3 | 0.8 | 2.9 | 3.6E-03 | 4.9E-03 | MB231/MB468 | MP |
| >hg38_21_+_8433172_8446622_RNA45SN1_45S_50ntFlanks@12163.12188.26\|ACTGGGGCGGTACACCTGTCAAACGG | >hg38_21_+_8433172_8446622_RNA45SN1_45S_50ntFlanks | 12163 | 12188 | 26 | ACTGGGGCGGTACACCTGTCAAACGG | 229.9 | -3.3 | 1.1 | -2.9 | 3.3E-03 | 4.9E-03 | BT20/MB468 | MITO |
| >hg38_21_+_8433172_8446622_RNA45SN1_45S_50ntFlanks@11677.11702.26\|CTCTGAATGTCAAAGTGAAGAAATTC | >hg38_21_+_8433172_8446622_RNA45SN1_45S_50ntFlanks | 11677 | 11702 | 26 | CTCTGAATGTCAAAGTGAAGAAATTC | 552.5 | -3.2 | 1.1 | -2.9 | 4.0E-03 | 4.9E-03 | MB231/MB468 | CYTO |
| >trnaMT_ProTGG_MT_-_15956_16023@44.70.27\|GTGGAGTTAAAGACTTTTTCTCTGACC | >trnaMT_ProTGG_MT_-_15956_16023 | 44 | 70 | 27 | GTGGAGTTAAAGACTTTTTCTCTGACC | 357.8 | 3.9 | 1.4 | 2.9 | 4.2E-03 | 4.9E-03 | MB231/MB468 | MITO |
| >hg38_21_+_8433172_8446622_RNA45SN1_45S_50ntFlanks@10401.10439.39\|GAACATGGGTCAGTCGGTCCTGAGAGATGGGCGAGCGCC | >hg38_21_+_8433172_8446622_RNA45SN1_45S_50ntFlanks | 10401 | 10439 | 39 | GAACATGGGTCAGTCGGTCCTGAGAGATGGGCGAGCGCC | 222.0 | 3.5 | 1.2 | 2.9 | 4.2E-03 | 4.9E-03 | MB231/MB468 | MITO |
| >hg38_21_+_8433172_8446622_RNA45SN1_45S_50ntFlanks@9345.9370.26\|AAAGCCGCCGTGGCGCAATGAAGGTG | >hg38_21_+_8433172_8446622_RNA45SN1_45S_50ntFlanks | 9345 | 9370 | 26 | AAAGCCGCCGTGGCGCAATGAAGGTG | 307.1 | -4.4 | 1.5 | -2.9 | 3.3E-03 | 4.9E-03 | BT20/MB468 | MITO |
| >hg38_MT_+_1621_3279_RNR2_16S_50ntFlanks@486.517.32\|AGGAACAGCTCTTTGGACACTAGGAAAAAACC | >hg38_MT_+_1621_3279_RNR2_16S_50ntFlanks | 486 | 517 | 32 | AGGAACAGCTCTTTGGACACTAGGAAAAAACC | 130.8 | 3.4 | 1.2 | 2.9 | 4.2E-03 | 4.9E-03 | MB231/MB468 | MITO |
| >hg38_21_+_8433172_8446622_RNA45SN1_45S_50ntFlanks@9534.9552.19\|AGGAAACTCTGGTGGAGGT | >hg38_21_+_8433172_8446622_RNA45SN1_45S_50ntFlanks | 9534 | 9552 | 19 | AGGAAACTCTGGTGGAGGT | 381.1 | -1.6 | 0.5 | -2.9 | 3.3E-03 | 4.9E-03 | BT20/MB468 | MITO |
| >hg38_21_+_8433172_8446622_RNA45SN1_45S_50ntFlanks@10347.10373.27\|CAAACGAGAACTTTGAAGGCCGAAGTG | >hg38_21_+_8433172_8446622_RNA45SN1_45S_50ntFlanks | 10347 | 10373 | 27 | CAAACGAGAACTTTGAAGGCCGAAGTG | 199.3 | -4.0 | 1.4 | -2.9 | 3.6E-03 | 4.9E-03 | MB231/MB468 | MP |
| >hg38_21_+_8433172_8446622_RNA45SN1_45S_50ntFlanks@8104.8142.39\|CCGCGGCGGGGCGCGGGACATGTGGCGTACGGAAGACCC | >hg38_21_+_8433172_8446622_RNA45SN1_45S_50ntFlanks | 8104 | 8142 | 39 | CCGCGGCGGGGCGCGGGACATGTGGCGTACGGAAGACCC | 654.6 | -4.7 | 1.5 | -3.1 | 2.2E-03 | 4.9E-03 | BT20/MB468 | MP |
| >hg38_21_+_8433172_8446622_RNA45SN1_45S_50ntFlanks@12045.12071.27\|CTGACCCGGTGAGGCGGGGGGGCGAGC | >hg38_21_+_8433172_8446622_RNA45SN1_45S_50ntFlanks | 12045 | 12071 | 27 | CTGACCCGGTGAGGCGGGGGGGCGAGC | 195.0 | -5.0 | 1.6 | -3.1 | 2.2E-03 | 4.9E-03 | BT20/MB468 | MP |
| >trna152_ValCAC_6_-_27248049_27248121@1.36.36\|GCTTCTGTAGTGTAGTGGTTATCACGTTCGCCTCAC | >trna152_ValCAC_6_-_27248049_27248121 | 1 | 36 | 36 | GCTTCTGTAGTGTAGTGGTTATCACGTTCGCCTCAC | 809.2 | -2.6 | 0.8 | -3.3 | 1.0E-03 | 4.9E-03 | BT20/MB231 | MITO |
| >hg38_21_+_8433172_8446622_RNA45SN1_45S_50ntFlanks@10420.10453.34\|CTGAGAGATGGGCGAGCGCCGTTCCGAAGGGACG | >hg38_21_+_8433172_8446622_RNA45SN1_45S_50ntFlanks | 10420 | 10453 | 34 | CTGAGAGATGGGCGAGCGCCGTTCCGAAGGGACG | 283.6 | -5.1 | 1.7 | -3.1 | 2.2E-03 | 4.9E-03 | BT20/MB468 | MP |
| >hg38_21_+_8433172_8446622_RNA45SN1_45S_50ntFlanks@6689.6706.18\|GAAGAACGCAGCTAGCTG | >hg38_21_+_8433172_8446622_RNA45SN1_45S_50ntFlanks | 6689 | 6706 | 18 | GAAGAACGCAGCTAGCTG | 78.2 | -4.9 | 1.6 | -3.1 | 2.2E-03 | 4.9E-03 | BT20/MB468 | MP |
| >hg38_21_+_8433172_8446622_RNA45SN1_45S_50ntFlanks@10109.10127.19\|TGTGGTGTGCGTCGGAGGG | >hg38_21_+_8433172_8446622_RNA45SN1_45S_50ntFlanks | 10109 | 10127 | 19 | TGTGGTGTGCGTCGGAGGG | 159.4 | -3.4 | 1.2 | -2.9 | 3.9E-03 | 4.9E-03 | BT20/MB468 | CYTO |
| >MI0000082\|hsa-mir-25&WithFlank&7\|-\|100093554\|100093649@58.78.21(+1G)&[MIMAT0000081&hsa-miR-25-3p&offsets\|0\|-1(+1G);m-23&7\|-\|100093571\|100093592&offsets\|0\|-1(+1G)]\|CATTGCACTTGTCTCGGTCTGG | >MI0000082\|hsa-mir-25&WithFlank&7\|-\|100093554\|100093649 | 58 | 78 | 21(+1G) | CATTGCACTTGTCTCGGTCTGG | 394.8 | 2.6 | 0.9 | 2.9 | 3.6E-03 | 4.9E-03 | MB231/MB468 | MP |
| >MI0000298\|hsa-mir-221&WithFlank&X\|-\|45746151\|45746272@31.54.24&[MIMAT0004568&hsa-miR-221-5p&offsets\|0\|+2;m-405&X\|-\|45746220\|45746242&offsets\|0\|+1]\|ACCTGGCATACAATGTAGATTTCT | >MI0000298\|hsa-mir-221&WithFlank&X\|-\|45746151\|45746272 | 31 | 54 | 24 | ACCTGGCATACAATGTAGATTTCT | 286.1 | 3.9 | 1.3 | 2.9 | 3.6E-03 | 4.9E-03 | MB231/MB468 | MP |
| >hg38_21_+_8433172_8446622_RNA45SN1_45S_50ntFlanks@7994.8011.18\|TGGCGACCCGCTGAATTT | >hg38_21_+_8433172_8446622_RNA45SN1_45S_50ntFlanks | 7994 | 8011 | 18 | TGGCGACCCGCTGAATTT | 93.0 | 3.8 | 1.3 | 2.9 | 4.2E-03 | 4.9E-03 | BT20/MB468 | NUC |
| >hg38_21_+_8433172_8446622_RNA45SN1_45S_50ntFlanks@6655.6678.24\|TCTTAGCGGTGGATCACTCGGCTC | >hg38_21_+_8433172_8446622_RNA45SN1_45S_50ntFlanks | 6655 | 6678 | 24 | TCTTAGCGGTGGATCACTCGGCTC | 112.4 | 5.2 | 1.7 | 3.1 | 2.2E-03 | 4.9E-03 | BT20/MB468 | MP |
| >hg38_21_+_8433172_8446622_RNA45SN1_45S_50ntFlanks@9932.9955.24\|ATAGACAGCAGGACGGTGGCCATG | >hg38_21_+_8433172_8446622_RNA45SN1_45S_50ntFlanks | 9932 | 9955 | 24 | ATAGACAGCAGGACGGTGGCCATG | 150.0 | 5.1 | 1.7 | 3.1 | 2.2E-03 | 4.9E-03 | BT20/MB468 | MP |
| >MI0000441\|hsa-mir-30b&WithFlank&8\|-\|134800514\|134800613@23.42.20(+1U)&[MIMAT0000420&hsa-miR-30b-5p&offsets\|0\|-2(+1U);m-76&8\|-\|134800570\|134800591&offsets\|0\|-2(+1U)]\|TGTAAACATCCTACACTCAGT | >MI0000441\|hsa-mir-30b&WithFlank&8\|-\|134800514\|134800613 | 23 | 42 | 20(+1U) | TGTAAACATCCTACACTCAGT | 102.6 | 4.8 | 1.6 | 2.9 | 3.6E-03 | 4.9E-03 | MB231/MB468 | MP |
| >MI0000782\|hsa-mir-374a&WithFlank&X\|-\|74287280\|74287363@48.69.22&[MIMAT0004688&hsa-miR-374a-3p&offsets\|0\|0;m-74&X\|-\|74287295\|74287316&offsets\|0\|0]\|CTTATCAGATTGTATTGTAATT | >MI0000782\|hsa-mir-374a&WithFlank&X\|-\|74287280\|74287363 | 48 | 69 | 22 | CTTATCAGATTGTATTGTAATT | 243.0 | -5.8 | 1.7 | -3.4 | 7.8E-04 | 4.9E-03 | BT20/MB231 | MP |
| >MI0000084\|hsa-mir-26b&WithFlank&2\|+\|218402640\|218402728@18.39.22&[MIMAT0000083&hsa-miR-26b-5p&offsets\|0\|+1;m-53&2\|+\|218402657\|218402678&offsets\|0\|0]\|TTCAAGTAATTCAGGATAGGTT | >MI0000084\|hsa-mir-26b&WithFlank&2\|+\|218402640\|218402728 | 18 | 39 | 22 | TTCAAGTAATTCAGGATAGGTT | 8711.6 | -3.6 | 1.1 | -3.4 | 7.8E-04 | 5.0E-03 | BT20/MB231 | MP |
| >hg38_21_+_8433172_8446622_RNA45SN1_45S_50ntFlanks@9720.9748.29\|ATTAGAGGTCTTGGGGCCGAAACGATCTC | >hg38_21_+_8433172_8446622_RNA45SN1_45S_50ntFlanks | 9720 | 9748 | 29 | ATTAGAGGTCTTGGGGCCGAAACGATCTC | 142.8 | -3.4 | 1.2 | -2.9 | 3.4E-03 | 5.0E-03 | BT20/MB468 | MITO |
| >hg38_21_+_8433172_8446622_RNA45SN1_45S_50ntFlanks@6718.6743.26\|GTGAATTGCAGGACACATTGATCATC | >hg38_21_+_8433172_8446622_RNA45SN1_45S_50ntFlanks | 6718 | 6743 | 26 | GTGAATTGCAGGACACATTGATCATC | 173.8 | -3.7 | 1.3 | -2.9 | 3.6E-03 | 5.0E-03 | MB231/MB468 | MP |
| >hg38_21_+_8433172_8446622_RNA45SN1_45S_50ntFlanks@11208.11234.27\|GCGGCGACTCTGGACGCGAGCCGGGCC | >hg38_21_+_8433172_8446622_RNA45SN1_45S_50ntFlanks | 11208 | 11234 | 27 | GCGGCGACTCTGGACGCGAGCCGGGCC | 1887.4 | -4.5 | 1.5 | -3.1 | 2.2E-03 | 5.0E-03 | BT20/MB468 | MP |
| >hg38_21_+_8433172_8446622_RNA45SN1_45S_50ntFlanks@11886.11903.18\|TGAGCTTGACTCTAGTCT | >hg38_21_+_8433172_8446622_RNA45SN1_45S_50ntFlanks | 11886 | 11903 | 18 | TGAGCTTGACTCTAGTCT | 166.5 | -4.9 | 1.6 | -3.1 | 2.2E-03 | 5.0E-03 | BT20/MB468 | MP |
| >hg38_21_+_8433172_8446622_RNA45SN1_45S_50ntFlanks@475.501.27\|CTTCGTGATCGATGTGGTGACGTCGTG | >hg38_21_+_8433172_8446622_RNA45SN1_45S_50ntFlanks | 475 | 501 | 27 | CTTCGTGATCGATGTGGTGACGTCGTG | 573.1 | 1.6 | 0.6 | 2.9 | 4.2E-03 | 5.0E-03 | BT20/MB468 | NUC |
| >hg38_21_+_8433172_8446622_RNA45SN1_45S_50ntFlanks@10346.10390.45\|TCAAACGAGAACTTTGAAGGCCGAAGTGGAGAAGGGTTCCATGTG | >hg38_21_+_8433172_8446622_RNA45SN1_45S_50ntFlanks | 10346 | 10390 | 45 | TCAAACGAGAACTTTGAAGGCCGAAGTGGAGAAGGGTTCCATGTG | 559.4 | -3.1 | 1.1 | -2.9 | 4.3E-03 | 5.0E-03 | MB231/MB468 | NUC |
| >hg38_21_+_8433172_8446622_RNA45SN1_45S_50ntFlanks@11670.11700.31\|CCCAGTGCTCTGAATGTCAAAGTGAAGAAAT | >hg38_21_+_8433172_8446622_RNA45SN1_45S_50ntFlanks | 11670 | 11700 | 31 | CCCAGTGCTCTGAATGTCAAAGTGAAGAAAT | 172.3 | -2.5 | 0.9 | -2.9 | 4.0E-03 | 5.0E-03 | MB231/MB468 | CYTO |
| >hg38_MT_+_1621_3279_RNR2_16S_50ntFlanks@1175.1215.41\|TGCATTAAAAATTTCGGTTGGGGCGACCTCGGAGCAGAACC | >hg38_MT_+_1621_3279_RNR2_16S_50ntFlanks | 1175 | 1215 | 41 | TGCATTAAAAATTTCGGTTGGGGCGACCTCGGAGCAGAACC | 158.4 | 5.2 | 1.8 | 2.9 | 4.3E-03 | 5.0E-03 | MB231/MB468 | MITO |
| >hg38_21_+_8433172_8446622_RNA45SN1_45S_50ntFlanks@3835.3868.34\|CTCCTCTCCTACTTGGATAACTGTGGTAATTCTA | >hg38_21_+_8433172_8446622_RNA45SN1_45S_50ntFlanks | 3835 | 3868 | 34 | CTCCTCTCCTACTTGGATAACTGTGGTAATTCTA | 218.6 | -2.2 | 0.8 | -2.9 | 3.4E-03 | 5.0E-03 | BT20/MB468 | MITO |
| >hg38_21_+_8433172_8446622_RNA45SN1_45S_50ntFlanks@10239.10273.35\|GCTACGCCGCGACGAGTAGGAGGGCCGCTGCGGTG | >hg38_21_+_8433172_8446622_RNA45SN1_45S_50ntFlanks | 10239 | 10273 | 35 | GCTACGCCGCGACGAGTAGGAGGGCCGCTGCGGTG | 296.1 | 4.7 | 1.5 | 3.1 | 2.2E-03 | 5.0E-03 | BT20/MB468 | MP |
| >MI0000066\|hsa-let-7e&WithFlank&19\|+\|51692780\|51692870@14.34.21&[MIMAT0000066&hsa-let-7e-5p&offsets\|0\|-1;m-50&19\|+\|51692793\|51692814&offsets\|0\|-1]\|TGAGGTAGGAGGTTGTATAGT | >MI0000066\|hsa-let-7e&WithFlank&19\|+\|51692780\|51692870 | 14 | 34 | 21 | TGAGGTAGGAGGTTGTATAGT | 470.2 | 2.5 | 0.8 | 2.9 | 3.6E-03 | 5.0E-03 | MB231/MB468 | MP |
| >hg38_21_+_8433172_8446622_RNA45SN1_45S_50ntFlanks@4721.4758.38\|TTCATTAATCAAGAACGAAAGTCGGAGGTTCGAAGACG | >hg38_21_+_8433172_8446622_RNA45SN1_45S_50ntFlanks | 4721 | 4758 | 38 | TTCATTAATCAAGAACGAAAGTCGGAGGTTCGAAGACG | 150.9 | -4.0 | 1.4 | -2.9 | 3.6E-03 | 5.0E-03 | MB231/MB468 | MP |

<table>
<tr><td>>MI0000077|hsa-mir-21&WithFlank&17|+|59841260|59841343@14.32.19(+1C)&[MIMAT0000076&hsa-miR-21-5p&offsets|0|-3(+1C);m-1&17|+|59841273|59841294&offsets|0|-3(+1C)]|TAGCTTATCAGACTGATGTC</td><td>>MI0000077|hsa-mir-21&WithFlank&17|+|59841260|59841343</td><td>14</td><td>32</td><td>19(+1C)</td><td>TAGCTTATCAGACTGATGTC</td><td>232.1</td><td>3.7</td><td>1.2</td><td>3.1</td><td>2.2E-03</td><td>5.0E-03</td><td>BT20/MB468</td><td>MP</td></tr>
<tr><td>>hg38_MT_+_1621_3279_RNR2_16S_50ntFlanks@126.150.25|AGGCGATAGAAATTGAAACCTGGCG</td><td>>hg38_MT_+_1621_3279_RNR2_16S_50ntFlanks</td><td>126</td><td>150</td><td>25</td><td>AGGCGATAGAAATTGAAACCTGGCG</td><td>62.1</td><td>6.0</td><td>2.0</td><td>3.1</td><td>2.2E-03</td><td>5.0E-03</td><td>BT20/MB468</td><td>MP</td></tr>
<tr><td>>hg38_21_+_8433172_8446622_RNA45SN1_45S_50ntFlanks@6650.6675.26|ACGACTCTTAGCGGTGGATCACTCGG</td><td>>hg38_21_+_8433172_8446622_RNA45SN1_45S_50ntFlanks</td><td>6650</td><td>6675</td><td>26</td><td>ACGACTCTTAGCGGTGGATCACTCGG</td><td>839.5</td><td>3.2</td><td>1.0</td><td>3.1</td><td>2.2E-03</td><td>5.0E-03</td><td>BT20/MB468</td><td>MP</td></tr>
<tr><td>>MI0005566|hsa-mir-374b&WithFlank&X|-|74218541|74218624@17.37.21(+1A)&[MIMAT0004955&hsa-miR-374b-5p&offsets|0|-1(+1A);m-153&X|-|74218587|74218608&offsets|0|-1(+1A)]|ATATAATACAACCTGCTAAGTA</td><td>>MI0005566|hsa-mir-374b&WithFlank&X|-|74218541|74218624</td><td>17</td><td>37</td><td>21(+1A)</td><td>ATATAATACAACCTGCTAAGTA</td><td>375.2</td><td>-2.9</td><td>0.9</td><td>-3.3</td><td>1.1E-03</td><td>5.0E-03</td><td>BT20/MB231</td><td>MITO</td></tr>
<tr><td>>hg38_21_+_8433172_8446622_RNA45SN1_45S_50ntFlanks@12635.12661.27|AAGCTACCATCTGTGGGATTATGACTG</td><td>>hg38_21_+_8433172_8446622_RNA45SN1_45S_50ntFlanks</td><td>12635</td><td>12661</td><td>27</td><td>AAGCTACCATCTGTGGGATTATGACTG</td><td>875.6</td><td>6.2</td><td>1.8</td><td>3.4</td><td>7.9E-04</td><td>5.0E-03</td><td>BT20/MB231</td><td>MP</td></tr>
<tr><td>>hg38_21_+_8433172_8446622_RNA45SN1_45S_50ntFlanks@4297.4317.21|CATTGGAGGGCAAGTCTGGTG</td><td>>hg38_21_+_8433172_8446622_RNA45SN1_45S_50ntFlanks</td><td>4297</td><td>4317</td><td>21</td><td>CATTGGAGGGCAAGTCTGGTG</td><td>453.7</td><td>-2.4</td><td>0.8</td><td>-2.9</td><td>3.4E-03</td><td>5.0E-03</td><td>BT20/MB468</td><td>MITO</td></tr>
<tr><td>>hg38_21_+_8433172_8446622_RNA45SN1_45S_50ntFlanks@9656.9675.20|ATAGCTGGCGCTCTCGCAGA</td><td>>hg38_21_+_8433172_8446622_RNA45SN1_45S_50ntFlanks</td><td>9656</td><td>9675</td><td>20</td><td>ATAGCTGGCGCTCTCGCAGA</td><td>45.4</td><td>6.2</td><td>2.0</td><td>3.1</td><td>2.2E-03</td><td>5.0E-03</td><td>BT20/MB468</td><td>MP</td></tr>
<tr><td>>hg38_21_+_8433172_8446622_RNA45SN1_45S_50ntFlanks@6646.6683.38|TCGTACGACTCTTAGCGGTGGATCACTCGGCTCGTGCG</td><td>>hg38_21_+_8433172_8446622_RNA45SN1_45S_50ntFlanks</td><td>6646</td><td>6683</td><td>38</td><td>TCGTACGACTCTTAGCGGTGGATCACTCGGCTCGTGCG</td><td>824.5</td><td>2.9</td><td>0.9</td><td>3.3</td><td>9.5E-04</td><td>5.0E-03</td><td>BT20/MB231</td><td>CYTO</td></tr>
<tr><td>>am_trna10_ValCAC_5_-_180649395_180649467@1.34.34,trna18_ValCAC_5_-_180529253_180529325@1.34.34,trna2_ValCAC_5_+_180524070_180524142@1.34.34,trna6_ValCAC_5_+_180600650_180600722@1.34.34,trna85_ValCAC_1_-_161369490_161369562@1.34.34,trna90_ValCAC_1_-_149684088_149684161@1.34.34,trna98_ValCAC_1_-_149298555_149298627@1.34.34,trna9_ValCAC_6_+_26538282_26538354@1.34.34|GTTTCCGTAGTGTAGTGGTTATCACGTTCGCCTC</td><td>>am_trna10_ValCAC_5_-_180649395_180649467</td><td>1</td><td>34</td><td>34</td><td>GTTTCCGTAGTGTAGTGGTTATCACGTTCGCCTC</td><td>6410.3</td><td>-4.1</td><td>1.2</td><td>-3.3</td><td>9.5E-04</td><td>5.0E-03</td><td>BT20/MB231</td><td>CYTO</td></tr>
<tr><td>>hg38_21_+_8433172_8446622_RNA45SN1_45S_50ntFlanks@12990.13016.27|GCTCCCTCGCTGCGATCTATTGAAAGT</td><td>>hg38_21_+_8433172_8446622_RNA45SN1_45S_50ntFlanks</td><td>12990</td><td>13016</td><td>27</td><td>GCTCCCTCGCTGCGATCTATTGAAAGT</td><td>147.3</td><td>-3.6</td><td>1.1</td><td>-3.3</td><td>9.5E-04</td><td>5.0E-03</td><td>BT20/MB231</td><td>CYTO</td></tr>
<tr><td>>MI0001448|hsa-mir-425&WithFlank&3|-|49020142|49020240@20.44.25&[MIMAT0003393&hsa-miR-425-5p&offsets|0|+2;m-95&3|-|49020200|49020221&offsets|0|+3]|AATGACACGATCACTCCCGTTGAGT</td><td>>MI0001448|hsa-mir-425&WithFlank&3|-|49020142|49020240</td><td>20</td><td>44</td><td>25</td><td>AATGACACGATCACTCCCGTTGAGT</td><td>6571.4</td><td>-2.7</td><td>0.8</td><td>-3.3</td><td>1.1E-03</td><td>5.0E-03</td><td>BT20/MB231</td><td>MITO</td></tr>
<tr><td>>MI0000746|hsa-mir-99b&WithFlank&19|+|51692606|51692687@51.71.21(+1A)&[MIMAT0004678&hsa-miR-99b-3p&offsets|0|-1(+1A);m-151&19|+|51692656|51692678&offsets|0|-2(+1A)]|CAAGCTCGTGTCTGTGGGTCCA</td><td>>MI0000746|hsa-mir-99b&WithFlank&19|+|51692606|51692687</td><td>51</td><td>71</td><td>21(+1A)</td><td>CAAGCTCGTGTCTGTGGGTCCA</td><td>249.3</td><td>5.2</td><td>1.7</td><td>3.1</td><td>2.2E-03</td><td>5.0E-03</td><td>BT20/MB468</td><td>MP</td></tr>
<tr><td>>hg38_MT_+_1621_3279_RNR2_16S_50ntFlanks@1368.1397.30|CGATGTTGGATCAGGACATCCCGATGGTGC</td><td>>hg38_MT_+_1621_3279_RNR2_16S_50ntFlanks</td><td>1368</td><td>1397</td><td>30</td><td>CGATGTTGGATCAGGACATCCCGATGGTGC</td><td>80.6</td><td>6.0</td><td>2.0</td><td>3.1</td><td>2.2E-03</td><td>5.0E-03</td><td>BT20/MB468</td><td>MP</td></tr>
<tr><td>>hg38_21_+_8433172_8446622_RNA45SN1_45S_50ntFlanks@5470.5514.45|CCACGGCCCTGGCGGAGCGCTGAGAAGACGGTCGAACTTGACTAT</td><td>>hg38_21_+_8433172_8446622_RNA45SN1_45S_50ntFlanks</td><td>5470</td><td>5514</td><td>45</td><td>CCACGGCCCTGGCGGAGCGCTGAGAAGACGGTCGAACTTGACTAT</td><td>247.2</td><td>-3.7</td><td>1.3</td><td>-2.9</td><td>3.4E-03</td><td>5.0E-03</td><td>BT20/MB468</td><td>MITO</td></tr>
<tr><td>>am_trnaMT_ValTAC_MT_+_1602_1670@29.68.40|ACTTACACTTAGGAGATTTCAACTTAACTTGACCGCTCTG</td><td>>am_trnaMT_ValTAC_MT_+_1602_1670</td><td>29</td><td>68</td><td>40</td><td>ACTTACACTTAGGAGATTTCAACTTAACTTGACCGCTCTG</td><td>170.4</td><td>-3.7</td><td>1.3</td><td>-2.9</td><td>3.7E-03</td><td>5.0E-03</td><td>MB231/MB468</td><td>MP</td></tr>
<tr><td>>MI0000081|hsa-mir-24-2&WithFlank&19|-|13836281|13836365@56.77.22(+1C);MI0000080|hsa-mir-24-1&WithFlank&9|+|95086015|95086094@50.71.22(+1C)&[MIMAT0000080&hsa-miR-24-3p&offsets|0|0(+1C);m-36&19|-|13836288|13836310&offsets|0|-1(+1C)];[MIMAT0000080_1&hsa-miR-24-3p&offsets|0|0(+1C);m-35&9|+|95086064|95086086&offsets|0|-1(+1C)]|TGGCTCAGTTCAGCAGGAACAGC</td><td>>MI0000081|hsa-mir-24-2&WithFlank&19|-|13836281|13836365</td><td>56</td><td>77</td><td>22(+1C)</td><td>TGGCTCAGTTCAGCAGGAACAGC</td><td>1659.0</td><td>3.1</td><td>1.1</td><td>2.9</td><td>3.7E-03</td><td>5.0E-03</td><td>MB231/MB468</td><td>MP</td></tr>
<tr><td>>hg38_21_+_8433172_8446622_RNA45SN1_45S_50ntFlanks@11877.11906.30|AGACCCTGTTGAGCTTGACTCTAGTCTGGC</td><td>>hg38_21_+_8433172_8446622_RNA45SN1_45S_50ntFlanks</td><td>11877</td><td>11906</td><td>30</td><td>AGACCCTGTTGAGCTTGACTCTAGTCTGGC</td><td>159.4</td><td>-1.7</td><td>0.6</td><td>-2.9</td><td>3.4E-03</td><td>5.0E-03</td><td>BT20/MB468</td><td>MITO</td></tr>
<tr><td>>MI0000434|hsa-let-7i&WithFlank&12|+|62603680|62603775@12.31.20(+1A)&[MIMAT0000415&hsa-let-7i-5p&offsets|0|-2(+1A);m-49&12|+|62603691|62603712&offsets|0|-2(+1A)]|TGAGGTAGTAGTTTGTGCTGA</td><td>>MI0000434|hsa-let-7i&WithFlank&12|+|62603680|62603775</td><td>12</td><td>31</td><td>20(+1A)</td><td>TGAGGTAGTAGTTTGTGCTGA</td><td>118.5</td><td>5.5</td><td>1.8</td><td>3.1</td><td>2.2E-03</td><td>5.0E-03</td><td>BT20/MB468</td><td>MP</td></tr>
<tr><td>>trna119_LysCTT_1_-_145395522_145395594@3.33.31,trna11_LysCTT_5_-_180648979_180649051@3.33.31,trna13_LysCTT_6_+_26556774_26556846@3.33.31,trna7_LysCTT_16_+_3225692_3225764@3.33.31,trna9_LysCTT_5_+_180634755_180634827@3.33.31|CCGGCTAGCTCAGTCGGTAGAGCATGAGACT</td><td>>trna119_LysCTT_1_-_145395522_145395594</td><td>3</td><td>33</td><td>31</td><td>CCGGCTAGCTCAGTCGGTAGAGCATGAGACT</td><td>454.1</td><td>-4.8</td><td>1.4</td><td>-3.4</td><td>7.9E-04</td><td>5.0E-03</td><td>BT20/MB231</td><td>MP</td></tr>
<tr><td>>hg38_MT_+_1621_3279_RNR2_16S_50ntFlanks@1347.1385.39|CAACAATAGGGTTTACGACCTCGATGTTGGATCAGGACA</td><td>>hg38_MT_+_1621_3279_RNR2_16S_50ntFlanks</td><td>1347</td><td>1385</td><td>39</td><td>CAACAATAGGGTTTACGACCTCGATGTTGGATCAGGACA</td><td>48.9</td><td>5.3</td><td>1.8</td><td>2.9</td><td>3.7E-03</td><td>5.0E-03</td><td>MB231/MB468</td><td>MP</td></tr>
<tr><td>>hg38_21_+_8433172_8446622_RNA45SN1_45S_50ntFlanks@12233.12265.33|CCGTGGAGCAGAAGGGCAAAAGCTCGCTTGATC</td><td>>hg38_21_+_8433172_8446622_RNA45SN1_45S_50ntFlanks</td><td>12233</td><td>12265</td><td>33</td><td>CCGTGGAGCAGAAGGGCAAAAGCTCGCTTGATC</td><td>234.0</td><td>6.7</td><td>2.0</td><td>3.4</td><td>7.9E-04</td><td>5.0E-03</td><td>BT20/MB231</td><td>MP</td></tr>
<tr><td>>hg38_21_+_8433172_8446622_RNA45SN1_45S_50ntFlanks@12416.12458.43|TTCGATGTCGGCTCTTCCTATCATTGTGAAGCAGAATTCACCA</td><td>>hg38_21_+_8433172_8446622_RNA45SN1_45S_50ntFlanks</td><td>12416</td><td>12458</td><td>43</td><td>TTCGATGTCGGCTCTTCCTATCATTGTGAAGCAGAATTCACCA</td><td>142.0</td><td>4.5</td><td>1.6</td><td>2.9</td><td>4.3E-03</td><td>5.0E-03</td><td>MB231/MB468</td><td>MITO</td></tr>
<tr><td>>MI0000071|hsa-mir-17&WithFlank&13|+|91350599|91350694@20.39.20(+1C)&[MIMAT0000070&hsa-miR-17-5p&offsets|0|-3(+1C);m-88&13|+|91350618|91350640&offsets|0|-3(+1C)]|CAAAGTGCTTACAGTGCAGGC</td><td>>MI0000071|hsa-mir-17&WithFlank&13|+|91350599|91350694</td><td>20</td><td>39</td><td>20(+1C)</td><td>CAAAGTGCTTACAGTGCAGGC</td><td>559.1</td><td>-4.9</td><td>1.5</td><td>-3.4</td><td>8.0E-04</td><td>5.0E-03</td><td>BT20/MB231</td><td>MP</td></tr>
<tr><td>>MI0000469|hsa-mir-125a&WithFlank&19|+|51693248|51693345@21.43.23(+1U)&[MIMAT0000443&hsa-miR-125a-5p&offsets|0|-1(+1U);m-63&19|+|51693268|51693289&offsets|0|+1(+1U)]|TCCCTGAGACCCTTTAACCTGTGT</td><td>>MI0000469|hsa-mir-125a&WithFlank&19|+|51693248|51693345</td><td>21</td><td>43</td><td>23(+1U)</td><td>TCCCTGAGACCCTTTAACCTGTGT</td><td>191.0</td><td>2.4</td><td>0.8</td><td>2.9</td><td>3.7E-03</td><td>5.0E-03</td><td>MB231/MB468</td><td>MP</td></tr>
<tr><td>>hg38_21_+_8433172_8446622_RNA45SN1_45S_50ntFlanks@11582.11609.28|ACTGGTGCGGACCAGGGGAATCCGACTG</td><td>>hg38_21_+_8433172_8446622_RNA45SN1_45S_50ntFlanks</td><td>11582</td><td>11609</td><td>28</td><td>ACTGGTGCGGACCAGGGGAATCCGACTG</td><td>401.6</td><td>1.4</td><td>0.5</td><td>2.9</td><td>3.4E-03</td><td>5.0E-03</td><td>BT20/MB468</td><td>MITO</td></tr>
<tr><td>>hg38_21_+_8433172_8446622_RNA45SN1_45S_50ntFlanks@8194.8211.18|CCCGTGGACGGTGTGAGG</td><td>>hg38_21_+_8433172_8446622_RNA45SN1_45S_50ntFlanks</td><td>8194</td><td>8211</td><td>18</td><td>CCCGTGGACGGTGTGAGG</td><td>418.3</td><td>-3.5</td><td>1.2</td><td>-2.9</td><td>3.4E-03</td><td>5.0E-03</td><td>BT20/MB468</td><td>MITO</td></tr>
<tr><td>>hg38_21_+_8433172_8446622_RNA45SN1_45S_50ntFlanks@10677.10706.30|GCGGTTCCGGCGGCGTCCGGTGAGCTCTCG</td><td>>hg38_21_+_8433172_8446622_RNA45SN1_45S_50ntFlanks</td><td>10677</td><td>10706</td><td>30</td><td>GCGGTTCCGGCGGCGTCCGGTGAGCTCTCG</td><td>299.7</td><td>-3.8</td><td>1.3</td><td>-2.9</td><td>4.3E-03</td><td>5.0E-03</td><td>MB231/MB468</td><td>MITO</td></tr>
<tr><td>>trnaMT_SerGCT_MT_+_12207_12265@28.62.35|ATGCCCCCATGTCTAACAACATGGCTTTCTCACCA</td><td>>trnaMT_SerGCT_MT_+_12207_12265</td><td>28</td><td>62</td><td>35</td><td>ATGCCCCCATGTCTAACAACATGGCTTTCTCACCA</td><td>141.8</td><td>3.0</td><td>1.1</td><td>2.9</td><td>4.3E-03</td><td>5.0E-03</td><td>MB231/MB468</td><td>MITO</td></tr>
<tr><td>>trna10_LysCTT_16_+_3241501_3241573@3.33.31,trna13_LysCTT_14_-_58706613_58706685@3.33.31,trna2_LysCTT_15_+_79152904_79152976@3.33.31|CCGGCTAGCTCAGTCGGTAGAGCATGGGACT</td><td>>trna10_LysCTT_16_+_3241501_3241573</td><td>3</td><td>33</td><td>31</td><td>CCGGCTAGCTCAGTCGGTAGAGCATGGGACT</td><td>2140.4</td><td>-2.6</td><td>0.8</td><td>-3.3</td><td>1.1E-03</td><td>5.0E-03</td><td>BT20/MB231</td><td>MITO</td></tr>
<tr><td>>hg38_21_+_8433172_8446622_RNA45SN1_45S_50ntFlanks@9739.9765.27|AAACGATCTCAACCTATTCTCAAACTT</td><td>>hg38_21_+_8433172_8446622_RNA45SN1_45S_50ntFlanks</td><td>9739</td><td>9765</td><td>27</td><td>AAACGATCTCAACCTATTCTCAAACTT</td><td>349.7</td><td>-2.3</td><td>0.8</td><td>-2.9</td><td>4.0E-03</td><td>5.0E-03</td><td>BT20/MB468</td><td>CYTO</td></tr>
<tr><td>>MI0000456|hsa-mir-140&WithFlank&16|+|69933075|69933186@69.89.21(+1A)&[MIMAT0004597&hsa-miR-140-3p&offsets|+1|+1(+1A);m-45&16|+|69933143|69933164&offsets|0|-1(+1A)]|ACCACAGGGTAGAACCACGGAA</td><td>>MI0000456|hsa-mir-140&WithFlank&16|+|69933075|69933186</td><td>69</td><td>89</td><td>21(+1A)</td><td>ACCACAGGGTAGAACCACGGAA</td><td>106.7</td><td>6.4</td><td>2.1</td><td>3.1</td><td>2.3E-03</td><td>5.0E-03</td><td>BT20/MB468</td><td>MP</td></tr>
<tr><td>>hg38_MT_+_598_1651_RNR1_12S_50ntFlanks@424.455.32|TACGAAAGTGGCTTTAACATATCTGAACACAC</td><td>>hg38_MT_+_598_1651_RNR1_12S_50ntFlanks</td><td>424</td><td>455</td><td>32</td><td>TACGAAAGTGGCTTTAACATATCTGAACACAC</td><td>322.2</td><td>4.7</td><td>1.5</td><td>3.1</td><td>2.3E-03</td><td>5.0E-03</td><td>BT20/MB468</td><td>MP</td></tr>
<tr><td>>hg38_MT_+_1621_3279_RNR2_16S_50ntFlanks@1090.1117.28|CAGCAAGACGAGAAGACCCTATGGAGCT</td><td>>hg38_MT_+_1621_3279_RNR2_16S_50ntFlanks</td><td>1090</td><td>1117</td><td>28</td><td>CAGCAAGACGAGAAGACCCTATGGAGCT</td><td>103.9</td><td>4.2</td><td>1.4</td><td>3.1</td><td>2.3E-03</td><td>5.0E-03</td><td>BT20/MB468</td><td>MP</td></tr>
<tr><td>>hg38_21_+_8433172_8446622_RNA45SN1_45S_50ntFlanks@9470.9503.34|GAGCACGAGCGCACGTGTTAGGACCCGAAAGATG</td><td>>hg38_21_+_8433172_8446622_RNA45SN1_45S_50ntFlanks</td><td>9470</td><td>9503</td><td>34</td><td>GAGCACGAGCGCACGTGTTAGGACCCGAAAGATG</td><td>1090.8</td><td>-3.7</td><td>1.2</td><td>-3.1</td><td>2.3E-03</td><td>5.0E-03</td><td>BT20/MB468</td><td>MP</td></tr>
<tr><td>>hg38_21_+_8433172_8446622_RNA45SN1_45S_50ntFlanks@9503.9520.18|GGTGAACTATGCCTGGGC</td><td>>hg38_21_+_8433172_8446622_RNA45SN1_45S_50ntFlanks</td><td>9503</td><td>9520</td><td>18</td><td>GGTGAACTATGCCTGGGC</td><td>397.0</td><td>5.9</td><td>1.7</td><td>3.4</td><td>8.0E-04</td><td>5.0E-03</td><td>BT20/MB231</td><td>MP</td></tr>
<tr><td>>hg38_21_+_8433172_8446622_RNA45SN1_45S_50ntFlanks@12995.13037.43|CTCGCTGCGATCTATTGAAAGTCAGCCCTCGACACAAGGGTTT</td><td>>hg38_21_+_8433172_8446622_RNA45SN1_45S_50ntFlanks</td><td>12995</td><td>13037</td><td>43</td><td>CTCGCTGCGATCTATTGAAAGTCAGCCCTCGACACAAGGGTTT</td><td>3320.5</td><td>-3.7</td><td>1.1</td><td>-3.3</td><td>9.6E-04</td><td>5.0E-03</td><td>BT20/MB231</td><td>CYTO</td></tr>
<tr><td>>MI0000273|hsa-mir-183&WithFlank&7|-|129774899|129775020@33.56.24&[MIMAT0000261&hsa-miR-183-5p&offsets|0|+2;m-41&7|-|129774966|129774987&offsets|-1|+1]|TATGGCACTGGTAGAATTCACTGT</td><td>>MI0000273|hsa-mir-183&WithFlank&7|-|129774899|129775020</td><td>33</td><td>56</td><td>24</td><td>TATGGCACTGGTAGAATTCACTGT</td><td>1511.7</td><td>2.6</td><td>0.8</td><td>3.3</td><td>9.6E-04</td><td>5.0E-03</td><td>BT20/MB231</td><td>CYTO</td></tr>
<tr><td>>trna10_AspGTC_12_-_125424193_125424264@36.73.38,trna12_AspGTC_12_-_125411891_125411962@36.73.38,trna144_AspGTC_6_-_27551236_27551307@36.73.38,trna38_AspGTC_17_-_8125556_8125627@36.73.38,trna45_AspGTC_6_+_27447453_27447524@36.73.38,trna48_AspGTC_6_+_27471523_27471594@36.73.38,trna4_AspGTC_12_+_96429799_96429870@36.73.38,trna69_AspGTC_1_-_161440205_161440276@36.73.38,trna72_AspGTC_1_-_161432824_161432895@36.73.38,trna75_AspGTC_1_-_161425414_161425485@36.73.38,trna78_AspGTC_1_-_161418033_161418104@36.73.38,trna81_AspGTC_1_-_161410615_161410686@36.73.38|CACGCGGGAGACCGGGGTTCGATTCCCCGACGGGGAGC</td><td>>trna10_AspGTC_12_-_125424193_125424264</td><td>36</td><td>73</td><td>38</td><td>CACGCGGGAGACCGGGGTTCGATTCCCCGACGGGGAGC</td><td>5464.1</td><td>-3.5</td><td>1.1</td><td>-3.3</td><td>9.7E-04</td><td>5.0E-03</td><td>BT20/MB231</td><td>CYTO</td></tr>
<tr><td>>hg38_21_+_8433172_8446622_RNA45SN1_45S_50ntFlanks@5152.5186.35|AGGGACAAGTGGCGTTCAGCCACCCGAGATTGAGC</td><td>>hg38_21_+_8433172_8446622_RNA45SN1_45S_50ntFlanks</td><td>5152</td><td>5186</td><td>35</td><td>AGGGACAAGTGGCGTTCAGCCACCCGAGATTGAGC</td><td>57.6</td><td>-5.1</td><td>1.8</td><td>-2.9</td><td>3.7E-03</td><td>5.0E-03</td><td>MB231/MB468</td><td>MP</td></tr>
<tr><td>>hg38_21_+_8433172_8446622_RNA45SN1_45S_50ntFlanks@6663.6699.37|GTGGATCACTCGGCTCGTGCGTCGATGAAGAACGCAG</td><td>>hg38_21_+_8433172_8446622_RNA45SN1_45S_50ntFlanks</td><td>6663</td><td>6699</td><td>37</td><td>GTGGATCACTCGGCTCGTGCGTCGATGAAGAACGCAG</td><td>521.4</td><td>-3.0</td><td>1.0</td><td>-3.1</td><td>2.3E-03</td><td>5.0E-03</td><td>BT20/MB468</td><td>MP</td></tr>
<tr><td>>hg38_21_+_8433172_8446622_RNA45SN1_45S_50ntFlanks@11555.11588.34|CGGCCGGCGCCTAGCAGCCGACTTAGAACTGGTG</td><td>>hg38_21_+_8433172_8446622_RNA45SN1_45S_50ntFlanks</td><td>11555</td><td>11588</td><td>34</td><td>CGGCCGGCGCCTAGCAGCCGACTTAGAACTGGTG</td><td>49.6</td><td>6.0</td><td>2.0</td><td>3.1</td><td>2.3E-03</td><td>5.0E-03</td><td>BT20/MB468</td><td>MP</td></tr>
</table>

| | | | | | | | | | | | | | |
|---|---|---|---|---|---|---|---|---|---|---|---|---|---|
| >MI0000077\|hsa-mir-21&WithFlank&17\|+\|59841260\|59841343@52.71.20(+1C)&[MIMAT0004494&hsa-miR-21-3p&offsets\|0\|-1(+1C);m-52&17\|+\|59841311\|59841332&offsets\|0\|-2(+1C)]\|CAACACCAGTCGATGGGCTGC | >MI0000077\|hsa-mir-21&WithFlank&17\|+\|59841260\|59841343 | 52 | 71 | 20(+1C) | CAACACCAGTCGATGGGCTGC | 204.0 | 4.7 | 1.6 | 2.9 | 3.7E-03 | 5.0E-03 | MB231/MB468 | MP |
| >hg38_21_+_8433172_8446622_RNA45SN1_45S_50ntFlanks@11845.11875.31\|GGAACGGGCTTGGCGGAATCAGCGGGGAAAG | >hg38_21_+_8433172_8446622_RNA45SN1_45S_50ntFlanks | 11845 | 11875 | 31 | GGAACGGGCTTGGCGGAATCAGCGGGGAAAG | 109.5 | -5.0 | 1.6 | -3.1 | 2.3E-03 | 5.0E-03 | BT20/MB468 | MP |
| >MI0000286\|hsa-mir-210&WithFlank&11\|-\|568083\|568204@72.93.22(+2A)&[MIMAT0000267&hsa-miR-210-3p&offsets\|0\|0(+2A);m-102&11\|-\|568112\|568133&offsets\|0\|0(+2A)]\|CTGTGCGTGTGACAGCGGCTGAAA | >MI0000286\|hsa-mir-210&WithFlank&11\|-\|568083\|568204 | 72 | 93 | 22(+2A) | CTGTGCGTGTGACAGCGGCTGAAA | 3411.5 | 3.3 | 1.0 | 3.3 | 8.8E-04 | 5.0E-03 | BT20/MB231 | NUC |
| >hg38_21_+_8433172_8446622_RNA45SN1_45S_50ntFlanks@12932.12973.42\|CGCTAAACCATTCGTAGACGACCTGCTTCTGGGTCGGGGTTT | >hg38_21_+_8433172_8446622_RNA45SN1_45S_50ntFlanks | 12932 | 12973 | 42 | CGCTAAACCATTCGTAGACGACCTGCTTCTGGGTCGGGGTTT | 347.4 | -2.4 | 0.8 | -2.9 | 4.1E-03 | 5.0E-03 | MB231/MB468 | CYTO |
| >hg38_1_-_228634819_228635039_RNA5S12_5S_50ntFlanks@135.172.38\|GGGAGACCGCCTGGGAATACCGGGTGCTGTAGGCTTTT | >hg38_1_-_228634819_228635039_RNA5S12_5S_50ntFlanks | 135 | 172 | 38 | GGGAGACCGCCTGGGAATACCGGGTGCTGTAGGCTTTT | 51.8 | 3.6 | 1.3 | 2.9 | 4.3E-03 | 5.0E-03 | BT20/MB468 | NUC |
| >trna119_LysCTT_1_-_145395522_145395594@1.33.33,trna11_LysCTT_5_-_180648979_180649051@1.33.33,trna13_LysCTT_6_+_26556774_26556846@1.33.33,trna7_LysCTT_16_+_3225692_3225764@1.33.33,trna9_LysCTT_5_+_180634755_180634827@1.33.33\|GCCCGGCTAGCTCAGTCGGTAGAGCATGAGACT | >trna119_LysCTT_1_-_145395522_145395594 | 1 | 33 | 33 | GCCCGGCTAGCTCAGTCGGTAGAGCATGAGACT | 2014.0 | 4.4 | 1.4 | 3.1 | 2.3E-03 | 5.0E-03 | BT20/MB468 | MP |
| >hg38_21_+_8433172_8446622_RNA45SN1_45S_50ntFlanks@6772.6799.28\|GTTCCTCCCGGGGCTACGCCTGTCTGAG | >hg38_21_+_8433172_8446622_RNA45SN1_45S_50ntFlanks | 6772 | 6799 | 28 | GTTCCTCCCGGGGCTACGCCTGTCTGAG | 597.5 | -4.6 | 1.6 | -2.9 | 4.0E-03 | 5.0E-03 | BT20/MB468 | CYTO |
| >hg38_21_+_8433172_8446622_RNA45SN1_45S_50ntFlanks@9353.9370.18\|CGTGGCGCAATGAAGGTG | >hg38_21_+_8433172_8446622_RNA45SN1_45S_50ntFlanks | 9353 | 9370 | 18 | CGTGGCGCAATGAAGGTG | 176.2 | -3.4 | 1.2 | -2.9 | 3.4E-03 | 5.0E-03 | BT20/MB468 | MITO |
| >hg38_21_+_8433172_8446622_RNA45SN1_45S_50ntFlanks@11659.11692.34\|TGTGATTTCTGCCCAGTGCTCTGAATGTCAAAGT | >hg38_21_+_8433172_8446622_RNA45SN1_45S_50ntFlanks | 11659 | 11692 | 34 | TGTGATTTCTGCCCAGTGCTCTGAATGTCAAAGT | 71.2 | 9.5 | 2.8 | 3.4 | 8.0E-04 | 5.0E-03 | BT20/MB231 | MP |
| >hg38_21_+_8433172_8446622_RNA45SN1_45S_50ntFlanks@10895.10913.19\|CGAAGCGGGGCTGGGCGCG | >hg38_21_+_8433172_8446622_RNA45SN1_45S_50ntFlanks | 10895 | 10913 | 19 | CGAAGCGGGGCTGGGCGCG | 71.8 | -4.8 | 1.6 | -3.1 | 2.3E-03 | 5.1E-03 | BT20/MB468 | MP |
| >hg38_21_+_8433172_8446622_RNA45SN1_45S_50ntFlanks@8112.8142.31\|GGGCGCGGGACATGTGGCGTACGGAAGACCC | >hg38_21_+_8433172_8446622_RNA45SN1_45S_50ntFlanks | 8112 | 8142 | 31 | GGGCGCGGGACATGTGGCGTACGGAAGACCC | 51.5 | -4.9 | 1.6 | -3.1 | 2.3E-03 | 5.1E-03 | BT20/MB468 | MP |
| >MI0000081\|hsa-mir-24-2&WithFlank&19\|-\|13836281\|13836365@56.73.18;MI0000080\|hsa-mir-24-1&WithFlank&9\|+\|95086015\|95086094@50.67.18&[MIMAT0000080&hsa-miR-24-3p&offsets\|0\|-4;m-36&19\|-\|13836288\|13836310&offsets\|0\|-5];[MIMAT0000080_1&hsa-miR-24-3p&offsets\|0\|-4;m-35&9\|+\|95086064\|95086086&offsets\|0\|-5]\|TGGCTCAGTTCAGCAGGA | >MI0000081\|hsa-mir-24-2&WithFlank&19\|-\|13836281\|13836365 | 56 | 73 | 18 | TGGCTCAGTTCAGCAGGA | 221.5 | 5.5 | 1.8 | 3.1 | 2.3E-03 | 5.1E-03 | BT20/MB468 | MP |
| >hg38_21_+_8433172_8446622_RNA45SN1_45S_50ntFlanks@6645.6681.37\|CTCGTACGACTCTTAGCGGTGGATCACTCGGCTCGTG | >hg38_21_+_8433172_8446622_RNA45SN1_45S_50ntFlanks | 6645 | 6681 | 37 | CTCGTACGACTCTTAGCGGTGGATCACTCGGCTCGTG | 17580.5 | -2.9 | 1.0 | -2.9 | 4.4E-03 | 5.1E-03 | MB231/MB468 | MITO |
| >hg38_21_+_8433172_8446622_RNA45SN1_45S_50ntFlanks@11534.11580.47\|GTTCCGCGCGGCGCCTCGCCTCGGCCGGCGCCTAGCAGCCGACTTAG | >hg38_21_+_8433172_8446622_RNA45SN1_45S_50ntFlanks | 11534 | 11580 | 47 | GTTCCGCGCGGCGCCTCGCCTCGGCCGGCGCCTAGCAGCCGACTTAG | 82.2 | -4.7 | 1.6 | -2.9 | 3.4E-03 | 5.1E-03 | BT20/MB468 | MITO |
| >hg38_21_+_8433172_8446622_RNA45SN1_45S_50ntFlanks@10694.10711.18\|CGGTGAGCTCTCGCTGGC | >hg38_21_+_8433172_8446622_RNA45SN1_45S_50ntFlanks | 10694 | 10711 | 18 | CGGTGAGCTCTCGCTGGC | 795.8 | -3.3 | 1.2 | -2.9 | 4.1E-03 | 5.1E-03 | MB231/MB468 | CYTO |
| >MI0001445\|hsa-mir-423&WithFlank&17\|+\|30117073\|30117178@23.43.21(+2A)&[MIMAT0004748&hsa-miR-423-5p&offsets\|0\|-2(+2A);m-141&17\|+\|30117095\|30117117&offsets\|0\|-2(+2A)]\|TGAGGGGCAGAGAGCGAGACTAA | >MI0001445\|hsa-mir-423&WithFlank&17\|+\|30117073\|30117178 | 23 | 43 | 21(+2A) | TGAGGGGCAGAGAGCGAGACTAA | 94.6 | 6.2 | 2.0 | 3.1 | 2.3E-03 | 5.1E-03 | BT20/MB468 | MP |
| >MI0025242\|hsa-mir-7706&WithFlank&15\|+\|85380590\|85380668@50.71.22&[MIMAT0030021&hsa-miR-7706&offsets\|0\|-2;m-390&15\|+\|85380639\|85380662&offsets\|0\|-2]\|TGAAGCGCCTGTGCTCTGCCGA | >MI0025242\|hsa-mir-7706&WithFlank&15\|+\|85380590\|85380668 | 50 | 71 | 22 | TGAAGCGCCTGTGCTCTGCCGA | 200.0 | 4.1 | 1.3 | 3.1 | 2.3E-03 | 5.1E-03 | BT20/MB468 | MP |
| >hg38_21_+_8433172_8446622_RNA45SN1_45S_50ntFlanks@12715.12742.28\|AGCCTCGGTTGGCCTCGGATAGCCGGTC | >hg38_21_+_8433172_8446622_RNA45SN1_45S_50ntFlanks | 12715 | 12742 | 28 | AGCCTCGGTTGGCCTCGGATAGCCGGTC | 294.7 | -5.1 | 1.7 | -3.1 | 2.3E-03 | 5.1E-03 | BT20/MB468 | MP |
| >hg38_21_+_8433172_8446622_RNA45SN1_45S_50ntFlanks@11029.11060.32\|CTCTCTCTCTCTCCCCCGCTCCCCGTCCTCCC | >hg38_21_+_8433172_8446622_RNA45SN1_45S_50ntFlanks | 11029 | 11060 | 32 | CTCTCTCTCTCTCCCCCGCTCCCCGTCCTCCC | 1078.8 | -6.1 | 2.1 | -2.9 | 4.3E-03 | 5.1E-03 | BT20/MB468 | NUC |
| >hg38_21_+_8433172_8446622_RNA45SN1_45S_50ntFlanks@3823.3861.39\|TTTGGTCGCTCGCTCCTCTCCTACTTGGATAACTGTGGT | >hg38_21_+_8433172_8446622_RNA45SN1_45S_50ntFlanks | 3823 | 3861 | 39 | TTTGGTCGCTCGCTCCTCTCCTACTTGGATAACTGTGGT | 319.8 | -3.0 | 0.9 | -3.3 | 9.7E-04 | 5.1E-03 | BT20/MB231 | CYTO |
| >hg38_21_+_8433172_8446622_RNA45SN1_45S_50ntFlanks@11027.11060.34\|CTCTCTCTCTCTCTCCCCCGCTCCCCGTCCTCCC | >hg38_21_+_8433172_8446622_RNA45SN1_45S_50ntFlanks | 11027 | 11060 | 34 | CTCTCTCTCTCTCTCCCCCGCTCCCCGTCCTCCC | 1074.0 | -3.7 | 1.3 | -2.9 | 4.0E-03 | 5.1E-03 | BT20/MB468 | CYTO |
| >hg38_21_+_8433172_8446622_RNA45SN1_45S_50ntFlanks@9941.9975.35\|AGGACGGTGGCCATGGAAGTCGGAATCCGCTAAGG | >hg38_21_+_8433172_8446622_RNA45SN1_45S_50ntFlanks | 9941 | 9975 | 35 | AGGACGGTGGCCATGGAAGTCGGAATCCGCTAAGG | 305.3 | 5.5 | 1.6 | 3.4 | 8.1E-04 | 5.1E-03 | BT20/MB231 | MP |
| >hg38_21_+_8433172_8446622_RNA45SN1_45S_50ntFlanks@6780.6802.23\|CGGGGCTACGCCTGTCTGAGCGT | >hg38_21_+_8433172_8446622_RNA45SN1_45S_50ntFlanks | 6780 | 6802 | 23 | CGGGGCTACGCCTGTCTGAGCGT | 226.3 | 2.2 | 0.8 | 2.9 | 4.4E-03 | 5.1E-03 | MB231/MB468 | MITO |
| >trnaMT_PheGAA_MT_+_577_647@31.53.23\|CACTGAAAATGTTTAGACGGGCT | >trnaMT_PheGAA_MT_+_577_647 | 31 | 53 | 23 | CACTGAAAATGTTTAGACGGGCT | 93.5 | -2.5 | 0.9 | -2.9 | 3.7E-03 | 5.1E-03 | MB231/MB468 | MP |
| >MI0000065\|hsa-let-7d&WithFlank&9\|+\|94178828\|94178926@14.35.22(+1A)&[MIMAT0000065&hsa-let-7d-5p&offsets\|0\|0(+1A);m-90&9\|+\|94178841\|94178862&offsets\|0\|0(+1A)]\|AGAGGTAGTAGGTTGCATAGTTA | >MI0000065\|hsa-let-7d&WithFlank&9\|+\|94178828\|94178926 | 14 | 35 | 22(+1A) | AGAGGTAGTAGGTTGCATAGTTA | 202.7 | 4.4 | 1.5 | 3.1 | 2.3E-03 | 5.1E-03 | BT20/MB468 | MP |
| >hg38_21_+_8433172_8446622_RNA45SN1_45S_50ntFlanks@11209.11230.22\|CGGCGACTCTGGACGCGAGCCG | >hg38_21_+_8433172_8446622_RNA45SN1_45S_50ntFlanks | 11209 | 11230 | 22 | CGGCGACTCTGGACGCGAGCCG | 1448.5 | -5.8 | 2.0 | -2.9 | 4.3E-03 | 5.1E-03 | BT20/MB468 | NUC |
| >MI0001445\|hsa-mir-423&WithFlank&17\|+\|30117073\|30117178@59.81.23(+1A)&[MIMAT0001340&hsa-miR-423-3p&offsets\|0\|0(+1A);m-100&17\|+\|30117131\|30117153&offsets\|0\|0(+1A)]\|AGCTCGGTCTGAGGCCCCTCAGTA | >MI0001445\|hsa-mir-423&WithFlank&17\|+\|30117073\|30117178 | 59 | 81 | 23(+1A) | AGCTCGGTCTGAGGCCCCTCAGTA | 323.8 | 1.6 | 0.6 | 2.9 | 4.3E-03 | 5.1E-03 | BT20/MB468 | NUC |
| >hg38_21_+_8433172_8446622_RNA45SN1_45S_50ntFlanks@4555.4572.18\|CGCAGCTAGGAATAATGG | >hg38_21_+_8433172_8446622_RNA45SN1_45S_50ntFlanks | 4555 | 4572 | 18 | CGCAGCTAGGAATAATGG | 220.4 | -3.4 | 1.2 | -2.9 | 3.4E-03 | 5.1E-03 | BT20/MB468 | MITO |
| >hg38_21_+_8433172_8446622_RNA45SN1_45S_50ntFlanks@11188.11207.20\|CACCCGGGGGGCCGGCGGCG | >hg38_21_+_8433172_8446622_RNA45SN1_45S_50ntFlanks | 11188 | 11207 | 20 | CACCCGGGGGGCCGGCGGCG | 167.0 | -4.2 | 1.5 | -2.9 | 4.1E-03 | 5.1E-03 | BT20/MB468 | CYTO |
| >trna16_ValTAC_11_-_59318460_59318532@37.76.40,trna17_ValTAC_11_-_59318102_59318174@37.76.40,trna4_ValTAC_X_-_18693029_18693101@37.76.40\|ACGCAGAAGGTCCTGGGTTCGAGCCCCAGTGGAACCACCA | >trna16_ValTAC_11_-_59318460_59318532 | 37 | 76 | 40 | ACGCAGAAGGTCCTGGGTTCGAGCCCCAGTGGAACCACCA | 335.4 | -7.8 | 2.4 | -3.3 | 9.7E-04 | 5.1E-03 | BT20/MB231 | CYTO |
| >hg38_21_+_8433172_8446622_RNA45SN1_45S_50ntFlanks@12407.12444.38\|TTTTGATCCTTCGATGTCGGCTCTTCCTATCATTGTGA | >hg38_21_+_8433172_8446622_RNA45SN1_45S_50ntFlanks | 12407 | 12444 | 38 | TTTTGATCCTTCGATGTCGGCTCTTCCTATCATTGTGA | 985.9 | -2.1 | 0.7 | -2.8 | 4.4E-03 | 5.1E-03 | MB231/MB468 | NUC |
| >MI0000101\|hsa-mir-99a&WithFlank&21\|+\|16539083\|16539175@19.40.22&[MIMAT0000097&hsa-miR-99a-5p&offsets\|0\|0;m-64&21\|+\|16539101\|16539121&offsets\|0\|+1]\|AACCCGTAGATCCGATCTTGTG | >MI0000101\|hsa-mir-99a&WithFlank&21\|+\|16539083\|16539175 | 19 | 40 | 22 | AACCCGTAGATCCGATCTTGTG | 385.5 | -4.5 | 1.5 | -2.9 | 3.5E-03 | 5.1E-03 | BT20/MB468 | MITO |
| >hg38_21_+_8433172_8446622_RNA45SN1_45S_50ntFlanks@11538.11583.46\|CGCGCGGCGCCTCGCCTCGGCCGGCGCCTAGCAGCCGACTTAGAAC | >hg38_21_+_8433172_8446622_RNA45SN1_45S_50ntFlanks | 11538 | 11583 | 46 | CGCGCGGCGCCTCGCCTCGGCCGGCGCCTAGCAGCCGACTTAGAAC | 255.9 | -3.7 | 1.1 | -3.3 | 9.8E-04 | 5.1E-03 | BT20/MB231 | CYTO |
| >hg38_21_+_8433172_8446622_RNA45SN1_45S_50ntFlanks@4555.4574.20\|CGCAGCTAGGAATAATGGAA | >hg38_21_+_8433172_8446622_RNA45SN1_45S_50ntFlanks | 4555 | 4574 | 20 | CGCAGCTAGGAATAATGGAA | 191.8 | -4.9 | 1.6 | -3.1 | 2.3E-03 | 5.1E-03 | BT20/MB468 | MP |
| >hg38_21_+_8433172_8446622_RNA45SN1_45S_50ntFlanks@6647.6667.21\|CGTACGACTCTTAGCGGTGGA | >hg38_21_+_8433172_8446622_RNA45SN1_45S_50ntFlanks | 6647 | 6667 | 21 | CGTACGACTCTTAGCGGTGGA | 1749.0 | 2.3 | 0.7 | 3.3 | 9.8E-04 | 5.1E-03 | BT20/MB231 | CYTO |
| >hg38_21_+_8433172_8446622_RNA45SN1_45S_50ntFlanks@12625.12667.43\|CAATGGGGCGAAGCTACCATCTGTGGGATTATGACTGAACGCC | >hg38_21_+_8433172_8446622_RNA45SN1_45S_50ntFlanks | 12625 | 12667 | 43 | CAATGGGGCGAAGCTACCATCTGTGGGATTATGACTGAACGCC | 580.0 | -2.9 | 0.9 | -3.3 | 9.8E-04 | 5.1E-03 | BT20/MB231 | CYTO |
| >hg38_21_+_8433172_8446622_RNA45SN1_45S_50ntFlanks@12219.12241.23\|AGGACAGAAACCTCCCGTGGAGC | >hg38_21_+_8433172_8446622_RNA45SN1_45S_50ntFlanks | 12219 | 12241 | 23 | AGGACAGAAACCTCCCGTGGAGC | 376.0 | 5.8 | 1.7 | 3.3 | 8.1E-04 | 5.1E-03 | BT20/MB231 | MP |
| >MI0000298\|hsa-mir-221&WithFlank&X\|-\|45746151\|45746272@74.92.19&[MIMAT0000278&hsa-miR-221-3p&offsets\|+3\|-1;m-91&X\|-\|45746181\|45746202&offsets\|+3\|0]\|TACATTGTCTGCTGGGTTT | >MI0000298\|hsa-mir-221&WithFlank&X\|-\|45746151\|45746272 | 74 | 92 | 19 | TACATTGTCTGCTGGGTTT | 33.2 | 8.3 | 2.9 | 2.9 | 3.7E-03 | 5.1E-03 | MB231/MB468 | MP |
| >hg38_21_+_8433172_8446622_RNA45SN1_45S_50ntFlanks@7972.7992.21\|AGACGCGACCTCAGATCAGAC | >hg38_21_+_8433172_8446622_RNA45SN1_45S_50ntFlanks | 7972 | 7992 | 21 | AGACGCGACCTCAGATCAGAC | 64.1 | 6.1 | 2.0 | 3.0 | 2.3E-03 | 5.1E-03 | BT20/MB468 | MP |
| >hg38_21_+_8433172_8446622_RNA45SN1_45S_50ntFlanks@11672.11704.33\|CAGTGCTCTGAATGTCAAAGTGAAGAAATTCAA | >hg38_21_+_8433172_8446622_RNA45SN1_45S_50ntFlanks | 11672 | 11704 | 33 | CAGTGCTCTGAATGTCAAAGTGAAGAAATTCAA | 158.0 | -3.9 | 1.3 | -2.9 | 3.7E-03 | 5.1E-03 | MB231/MB468 | MP |
| >hg38_21_+_8433172_8446622_RNA45SN1_45S_50ntFlanks@4735.4752.18\|ACGAAAGTCGGAGGTTCG | >hg38_21_+_8433172_8446622_RNA45SN1_45S_50ntFlanks | 4735 | 4752 | 18 | ACGAAAGTCGGAGGTTCG | 90.3 | -4.8 | 1.6 | -3.0 | 2.3E-03 | 5.1E-03 | BT20/MB468 | MP |
| >hg38_21_+_8433172_8446622_RNA45SN1_45S_50ntFlanks@6650.6677.28\|ACGACTCTTAGCGGTGGATCACTCGGCT | >hg38_21_+_8433172_8446622_RNA45SN1_45S_50ntFlanks | 6650 | 6677 | 28 | ACGACTCTTAGCGGTGGATCACTCGGCT | 1990.8 | 4.6 | 1.4 | 3.3 | 8.1E-04 | 5.1E-03 | BT20/MB231 | MP |
| >trnaMT_ValTAC_MT_+_1602_1670@49.72.24\|AACTTAACTTGACCGCTCTGACCA | >trnaMT_ValTAC_MT_+_1602_1670 | 49 | 72 | 24 | AACTTAACTTGACCGCTCTGACCA | 1231.1 | 2.6 | 0.8 | 3.3 | 9.9E-04 | 5.1E-03 | BT20/MB231 | CYTO |
| >hg38_21_+_8433172_8446622_RNA45SN1_45S_50ntFlanks@13008.13039.32\|ATTGAAAGTCAGCCCTCGACACAAGGGTTTGT | >hg38_21_+_8433172_8446622_RNA45SN1_45S_50ntFlanks | 13008 | 13039 | 32 | ATTGAAAGTCAGCCCTCGACACAAGGGTTTGT | 8139.8 | 2.7 | 0.8 | 3.3 | 1.1E-03 | 5.1E-03 | BT20/MB231 | MITO |
| >trnaMT_TrpTCA_MT_+_5512_5579@18.35.18,trnalookalike3_TrpTCA_1_+_566062_566129@18.35.18\|ACAGACCAAGAGCCTTCA | >trnaMT_TrpTCA_MT_+_5512_5579 | 18 | 35 | 18 | ACAGACCAAGAGCCTTCA | 242.6 | 3.9 | 1.3 | 3.0 | 2.3E-03 | 5.1E-03 | BT20/MB468 | MP |
| >hg38_21_+_8433172_8446622_RNA45SN1_45S_50ntFlanks@10922.10943.22\|CTGGACGAGGCGCCGCCGCCCC | >hg38_21_+_8433172_8446622_RNA45SN1_45S_50ntFlanks | 10922 | 10943 | 22 | CTGGACGAGGCGCCGCCGCCCC | 325.9 | -5.3 | 1.8 | -2.9 | 4.4E-03 | 5.1E-03 | BT20/MB468 | NUC |
| >am_trnaMT_ValTAC_MT_+_1602_1670@38.59.22\|TAGGAGATTTCAACTTAACTTG | >am_trnaMT_ValTAC_MT_+_1602_1670 | 38 | 59 | 22 | TAGGAGATTTCAACTTAACTTG | 143.0 | 3.8 | 1.3 | 3.0 | 2.3E-03 | 5.1E-03 | BT20/MB468 | MP |
| >MI0000746\|hsa-mir-99b&WithFlank&19\|+\|51692606\|51692687@51.73.23&[MIMAT0004678&hsa-miR-99b-3p&offsets\|0\|+1;m-151&19\|+\|51692656\|51692678&offsets\|0\|0]\|CAAGCTCGTGTCTGTGGGTCCGT | >MI0000746\|hsa-mir-99b&WithFlank&19\|+\|51692606\|51692687 | 51 | 73 | 23 | CAAGCTCGTGTCTGTGGGTCCGT | 237.7 | 2.6 | 0.9 | 2.9 | 3.7E-03 | 5.1E-03 | MB231/MB468 | MP |
| >hg38_21_+_8433172_8446622_RNA45SN1_45S_50ntFlanks@11568.11603.36\|GCAGCCGACTTAGAACTGGTGCGGACCAGGGGAATC | >hg38_21_+_8433172_8446622_RNA45SN1_45S_50ntFlanks | 11568 | 11603 | 36 | GCAGCCGACTTAGAACTGGTGCGGACCAGGGGAATC | 1543.5 | -1.9 | 0.7 | -2.9 | 3.5E-03 | 5.1E-03 | BT20/MB468 | MITO |
| >hg38_21_+_8433172_8446622_RNA45SN1_45S_50ntFlanks@9489.9506.18\|AGGACCCGAAAGATGGTG | >hg38_21_+_8433172_8446622_RNA45SN1_45S_50ntFlanks | 9489 | 9506 | 18 | AGGACCCGAAAGATGGTG | 387.3 | -2.9 | 1.0 | -3.0 | 2.3E-03 | 5.1E-03 | BT20/MB468 | MP |

<table>
<tr><th></th><th></th><th></th><th></th><th></th><th></th><th></th><th></th><th></th><th></th><th></th><th></th><th></th><th></th></tr>
<tr><td>>hg38_21_+_8433172_8446622_RNA45SN1_45S_50ntFlanks@11668.11707.40|TGCCCAGTGCTCTGAATGTCAAAGTGAAGAAATTCAATGA</td><td>>hg38_21_+_8433172_8446622_RNA45SN1_45S_50ntFlanks</td><td>11668</td><td>11707</td><td>40</td><td>TGCCCAGTGCTCTGAATGTCAAAGTGAAGAAATTCAATGA</td><td>487.2</td><td>-3.5</td><td>1.0</td><td>-3.3</td><td>8.9E-04</td><td>5.1E-03</td><td>BT20/MB231</td><td>NUC</td></tr>
<tr><td>>hg38_21_+_8433172_8446622_RNA45SN1_45S_50ntFlanks@11677.11702.26|CTCTGAATGTCAAAGTGAAGAAATTC</td><td>>hg38_21_+_8433172_8446622_RNA45SN1_45S_50ntFlanks</td><td>11677</td><td>11702</td><td>26</td><td>CTCTGAATGTCAAAGTGAAGAAATTC</td><td>442.5</td><td>-4.6</td><td>1.6</td><td>-2.9</td><td>4.1E-03</td><td>5.1E-03</td><td>BT20/MB468</td><td>CYTO</td></tr>
<tr><td>>MI0000270|hsa-mir-181b-1&WithFlank&1|-|198858867|198858988@42.65.24;MI0000683|hsa-mir-181b-2&WithFlank&9|+|124693704|124693804@22.45.24&[MIMAT0000257&hsa-miR-181b-5p&offsets|0|+1;m-73&1|-|198858925|198858947&offsets|0|+1];[MIMAT0000257_1&hsa-miR-181b-5p&offsets|0|+1;m-72&9|+|124693725|124693747&offsets|0|+1]|AACATTCATTGCTGTCGGTGGGTT</td><td>>MI0000270|hsa-mir-181b-1&WithFlank&1|-|198858867|198858988</td><td>42</td><td>65</td><td>24</td><td>AACATTCATTGCTGTCGGTGGGTT</td><td>3321.6</td><td>-2.6</td><td>0.8</td><td>-3.3</td><td>1.1E-03</td><td>5.1E-03</td><td>BT20/MB231</td><td>MITO</td></tr>
<tr><td>>hg38_MT_+_598_1651_RNR1_12S_50ntFlanks@911.946.36|CTTCAAAGGACATTTAACTAAAACCCCTACGCATTT</td><td>>hg38_MT_+_598_1651_RNR1_12S_50ntFlanks</td><td>911</td><td>946</td><td>36</td><td>CTTCAAAGGACATTTAACTAAAACCCCTACGCATTT</td><td>210.4</td><td>2.9</td><td>0.9</td><td>3.3</td><td>1.1E-03</td><td>5.1E-03</td><td>BT20/MB231</td><td>MITO</td></tr>
<tr><td>>MI0000073|hsa-mir-19a&WithFlank&13|+|91350885|91350978@55.76.22(+1U)&[MIMAT0000073&hsa-miR-19a-3p&offsets|0|-1(+1U);m-201&13|+|91350939|91350961&offsets|0|-1(+1U)]|TGTGCAAATCTATGCAAAACTGT</td><td>>MI0000073|hsa-mir-19a&WithFlank&13|+|91350885|91350978</td><td>55</td><td>76</td><td>22(+1U)</td><td>TGTGCAAATCTATGCAAAACTGT</td><td>345.0</td><td>-3.0</td><td>0.9</td><td>-3.3</td><td>1.1E-03</td><td>5.1E-03</td><td>BT20/MB231</td><td>MITO</td></tr>
<tr><td>>hg38_21_+_8433172_8446622_RNA45SN1_45S_50ntFlanks@12418.12448.31|CGATGTCGGCTCTTCCTATCATTGTGAAGCA</td><td>>hg38_21_+_8433172_8446622_RNA45SN1_45S_50ntFlanks</td><td>12418</td><td>12448</td><td>31</td><td>CGATGTCGGCTCTTCCTATCATTGTGAAGCA</td><td>151.1</td><td>-2.7</td><td>0.9</td><td>-2.9</td><td>3.7E-03</td><td>5.1E-03</td><td>MB231/MB468</td><td>MP</td></tr>
<tr><td>>hg38_1_-_228634819_228635039_RNA5S12_5S_50ntFlanks@132.169.38|GATGGGAGACCGCCTGGGAATACCGGGTGCTGTAGGCT</td><td>>hg38_1_-_228634819_228635039_RNA5S12_5S_50ntFlanks</td><td>132</td><td>169</td><td>38</td><td>GATGGGAGACCGCCTGGGAATACCGGGTGCTGTAGGCT</td><td>1423.0</td><td>-2.9</td><td>1.0</td><td>-2.8</td><td>4.4E-03</td><td>5.1E-03</td><td>MB231/MB468</td><td>MITO</td></tr>
<tr><td>>hg38_1_-_228634819_228635039_RNA5S12_5S_50ntFlanks@51.75.25|GTCTACGGCCATACCACCCTGAACG</td><td>>hg38_1_-_228634819_228635039_RNA5S12_5S_50ntFlanks</td><td>51</td><td>75</td><td>25</td><td>GTCTACGGCCATACCACCCTGAACG</td><td>687.6</td><td>2.3</td><td>0.8</td><td>2.8</td><td>4.4E-03</td><td>5.1E-03</td><td>MB231/MB468</td><td>NUC</td></tr>
<tr><td>>hg38_21_+_8433172_8446622_RNA45SN1_45S_50ntFlanks@4005.4026.22|AACCTCGGGCCGATCGCACGCC</td><td>>hg38_21_+_8433172_8446622_RNA45SN1_45S_50ntFlanks</td><td>4005</td><td>4026</td><td>22</td><td>AACCTCGGGCCGATCGCACGCC</td><td>406.2</td><td>-4.8</td><td>1.7</td><td>-2.8</td><td>4.4E-03</td><td>5.1E-03</td><td>MB231/MB468</td><td>NUC</td></tr>
<tr><td>>trnaMT_TyrGTA_MT_-_5826_5891@24.65.42|TGGACTGTAAATCTAAAGACAGGGGTTAGGCCTCTTTTTACC</td><td>>trnaMT_TyrGTA_MT_-_5826_5891</td><td>24</td><td>65</td><td>42</td><td>TGGACTGTAAATCTAAAGACAGGGGTTAGGCCTCTTTTTACC</td><td>183.6</td><td>4.0</td><td>1.4</td><td>2.8</td><td>4.4E-03</td><td>5.1E-03</td><td>MB231/MB468</td><td>MITO</td></tr>
<tr><td>>trna10_AspGTC_12_-_125424193_125424264@32.72.41,trna12_AspGTC_12_-_125411891_125411962@32.72.41,trna144_AspGTC_6_-_27551236_27551307@32.72.41,trna38_AspGTC_17_-_8125556_8125627@32.72.41,trna45_AspGTC_6_+_27447453_27447524@32.72.41,trna48_AspGTC_6_+_27471523_27471594@32.72.41,trna4_AspGTC_12_+_96429799_96429870@32.72.41,trna69_AspGTC_1_-_161440205_161440276@32.72.41,trna72_AspGTC_1_-_161432824_161432895@32.72.41,trna75_AspGTC_1_-_161425414_161425485@32.72.41,trna78_AspGTC_1_-_161418033_161418104@32.72.41,trna81_AspGTC_1_-_161410615_161410686@32.72.41|CTGTCACGCGGGAGACCGGGGTTCGATTCCCCGACGGGGAG</td><td>>trna10_AspGTC_12_-_125424193_125424264</td><td>32</td><td>72</td><td>41</td><td>CTGTCACGCGGGAGACCGGGGTTCGATTCCCCGACGGGGAG</td><td>249.2</td><td>-5.0</td><td>1.7</td><td>-2.9</td><td>3.5E-03</td><td>5.1E-03</td><td>BT20/MB468</td><td>MITO</td></tr>
<tr><td>>hg38_21_+_8433172_8446622_RNA45SN1_45S_50ntFlanks@12399.12443.45|ACGTCGCTTTTTGATCCTTCGATGTCGGCTCTTCCTATCATTGTG</td><td>>hg38_21_+_8433172_8446622_RNA45SN1_45S_50ntFlanks</td><td>12399</td><td>12443</td><td>45</td><td>ACGTCGCTTTTTGATCCTTCGATGTCGGCTCTTCCTATCATTGTG</td><td>129.4</td><td>-4.4</td><td>1.5</td><td>-2.9</td><td>3.5E-03</td><td>5.1E-03</td><td>BT20/MB468</td><td>MITO</td></tr>
<tr><td>>hg38_21_+_8433172_8446622_RNA45SN1_45S_50ntFlanks@4657.4684.28|CTAGAGGTGAAATTCTTGGACCGGCGCA</td><td>>hg38_21_+_8433172_8446622_RNA45SN1_45S_50ntFlanks</td><td>4657</td><td>4684</td><td>28</td><td>CTAGAGGTGAAATTCTTGGACCGGCGCA</td><td>275.8</td><td>-2.6</td><td>0.9</td><td>-2.9</td><td>4.2E-03</td><td>5.1E-03</td><td>MB231/MB468</td><td>CYTO</td></tr>
<tr><td>>hg38_21_+_8433172_8446622_RNA45SN1_45S_50ntFlanks@10116.10139.24|TGCGTCGGAGGGCGGCGGCGGCGG</td><td>>hg38_21_+_8433172_8446622_RNA45SN1_45S_50ntFlanks</td><td>10116</td><td>10139</td><td>24</td><td>TGCGTCGGAGGGCGGCGGCGGCGG</td><td>192.3</td><td>-5.2</td><td>1.7</td><td>-3.0</td><td>2.3E-03</td><td>5.1E-03</td><td>BT20/MB468</td><td>MP</td></tr>
<tr><td>>hg38_21_+_8433172_8446622_RNA45SN1_45S_50ntFlanks@5470.5516.47|CCACGGCCCTGGCGGAGCGCTGAGAAGACGGTCGAACTTGACTATCT</td><td>>hg38_21_+_8433172_8446622_RNA45SN1_45S_50ntFlanks</td><td>5470</td><td>5516</td><td>47</td><td>CCACGGCCCTGGCGGAGCGCTGAGAAGACGGTCGAACTTGACTATCT</td><td>318.0</td><td>3.5</td><td>1.1</td><td>3.3</td><td>1.1E-03</td><td>5.1E-03</td><td>BT20/MB231</td><td>MITO</td></tr>
<tr><td>>hg38_21_+_8433172_8446622_RNA45SN1_45S_50ntFlanks@11572.11601.30|CCGACTTAGAACTGGTGCGGACCAGGGGAA</td><td>>hg38_21_+_8433172_8446622_RNA45SN1_45S_50ntFlanks</td><td>11572</td><td>11601</td><td>30</td><td>CCGACTTAGAACTGGTGCGGACCAGGGGAA</td><td>516.9</td><td>4.3</td><td>1.3</td><td>3.3</td><td>8.2E-04</td><td>5.1E-03</td><td>BT20/MB231</td><td>MP</td></tr>
<tr><td>>MI0000086|hsa-mir-28&WithFlank&3|+|188688775|188688872@20.42.23&[MIMAT0000085&hsa-miR-28-5p&offsets|0|+1;m-133&3|+|188688794|188688815&offsets|0|+1]|AAGGAGCTCACAGTCTATTGAGT</td><td>>MI0000086|hsa-mir-28&WithFlank&3|+|188688775|188688872</td><td>20</td><td>42</td><td>23</td><td>AAGGAGCTCACAGTCTATTGAGT</td><td>229.1</td><td>-3.3</td><td>1.0</td><td>-3.3</td><td>8.2E-04</td><td>5.1E-03</td><td>BT20/MB231</td><td>MP</td></tr>
<tr><td>>MI0000272|hsa-mir-182&WithFlank&7|-|129770377|129770498@29.49.21&[MIMAT0000259&hsa-miR-182-5p&offsets|0|-3;m-22&7|-|129770449|129770470&offsets|0|-1]|TTTGGCAATGGTAGAACTCAC</td><td>>MI0000272|hsa-mir-182&WithFlank&7|-|129770377|129770498</td><td>29</td><td>49</td><td>21</td><td>TTTGGCAATGGTAGAACTCAC</td><td>910.4</td><td>4.7</td><td>1.6</td><td>3.0</td><td>2.3E-03</td><td>5.1E-03</td><td>BT20/MB468</td><td>MP</td></tr>
<tr><td>>hg38_21_+_8433172_8446622_RNA45SN1_45S_50ntFlanks@6719.6736.18|TGAATTGCAGGACACATT</td><td>>hg38_21_+_8433172_8446622_RNA45SN1_45S_50ntFlanks</td><td>6719</td><td>6736</td><td>18</td><td>TGAATTGCAGGACACATT</td><td>771.8</td><td>-1.7</td><td>0.6</td><td>-2.9</td><td>3.5E-03</td><td>5.1E-03</td><td>BT20/MB468</td><td>MITO</td></tr>
<tr><td>>hg38_21_+_8433172_8446622_RNA45SN1_45S_50ntFlanks@11582.11605.24|ACTGGTGCGGACCAGGGGAATCCG</td><td>>hg38_21_+_8433172_8446622_RNA45SN1_45S_50ntFlanks</td><td>11582</td><td>11605</td><td>24</td><td>ACTGGTGCGGACCAGGGGAATCCG</td><td>497.5</td><td>-2.6</td><td>0.9</td><td>-2.9</td><td>3.8E-03</td><td>5.1E-03</td><td>MB231/MB468</td><td>MP</td></tr>
<tr><td>>hg38_21_+_8433172_8446622_RNA45SN1_45S_50ntFlanks@11209.11253.45|CGGCGACTCTGGACGCGAGCCGGGCCCTTCCCGTGGATCGCCCCA</td><td>>hg38_21_+_8433172_8446622_RNA45SN1_45S_50ntFlanks</td><td>11209</td><td>11253</td><td>45</td><td>CGGCGACTCTGGACGCGAGCCGGGCCCTTCCCGTGGATCGCCCCA</td><td>170.0</td><td>-3.8</td><td>1.3</td><td>-2.9</td><td>4.1E-03</td><td>5.1E-03</td><td>BT20/MB468</td><td>CYTO</td></tr>
<tr><td>>hg38_21_+_8433172_8446622_RNA45SN1_45S_50ntFlanks@11563.11611.49|GCCTAGCAGCCGACTTAGAACTGGTGCGGACCAGGGGAATCCGACTGTT</td><td>>hg38_21_+_8433172_8446622_RNA45SN1_45S_50ntFlanks</td><td>11563</td><td>11611</td><td>49</td><td>GCCTAGCAGCCGACTTAGAACTGGTGCGGACCAGGGGAATCCGACTGTT</td><td>884.9</td><td>4.0</td><td>1.2</td><td>3.3</td><td>1.1E-03</td><td>5.1E-03</td><td>BT20/MB231</td><td>MITO</td></tr>
<tr><td>>hg38_21_+_8433172_8446622_RNA45SN1_45S_50ntFlanks@12639.12661.23|TACCATCTGTGGGATTATGACTG</td><td>>hg38_21_+_8433172_8446622_RNA45SN1_45S_50ntFlanks</td><td>12639</td><td>12661</td><td>23</td><td>TACCATCTGTGGGATTATGACTG</td><td>685.3</td><td>-3.1</td><td>1.0</td><td>-3.0</td><td>2.3E-03</td><td>5.1E-03</td><td>BT20/MB468</td><td>MP</td></tr>
<tr><td>>am_MI0005763|hsa-mir-941-1&WithFlank&20|+|63919443|63919526@53.74.22;MI0005764|hsa-mir-941-2&WithFlank&20|+|63919499|63919582@53.74.22;MI0005765|hsa-mir-941-3&WithFlank&20|+|63919555|63919638@53.74.22;MI0005766|hsa-mir-941-4&WithFlank&20|+|63919750|63919833@53.74.22;MI0031520|hsa-mir-941-5&WithFlank&20|+|63919862|63919945@53.74.22&[MIMAT0004984&hsa-miR-941&offsets|0|-1;m-263&20|+|63919495|63919517&offsets|0|-1];[MIMAT0004984_1&hsa-miR-941&offsets|0|-1;m-263&20|+|63919551|63919573&offsets|0|-1];[MIMAT0004984_2&hsa-miR-941&offsets|0|-1;m-263&20|+|63919607|63919629&offsets|0|-1];[MIMAT0004984_3&hsa-miR-941&offsets|0|-1;m-263&20|+|63919802|63919824&offsets|0|-1];[MIMAT0004984_4&hsa-miR-941&offsets|0|-1;m-263&20|+|63919914|63919936&offsets|0|-1]|CACCCGGCTGTGTGCACATGTG</td><td>>am_MI0005763|hsa-mir-941-1&WithFlank&20|+|63919443|63919526</td><td>53</td><td>74</td><td>22</td><td>CACCCGGCTGTGTGCACATGTG</td><td>517.0</td><td>3.8</td><td>1.3</td><td>2.9</td><td>3.8E-03</td><td>5.1E-03</td><td>MB231/MB468</td><td>MP</td></tr>
<tr><td>>MI0000769|hsa-mir-365b&WithFlank&17|+|31575405|31575527@35.56.22&[MIMAT0022833&hsa-miR-365b-5p&offsets|0|0;m-644&17|+|31575439|31575461&offsets|0|-1]|AGGGACTTTCAGGGGCAGCTGT</td><td>>MI0000769|hsa-mir-365b&WithFlank&17|+|31575405|31575527</td><td>35</td><td>56</td><td>22</td><td>AGGGACTTTCAGGGGCAGCTGT</td><td>47.0</td><td>6.2</td><td>2.0</td><td>3.0</td><td>2.3E-03</td><td>5.1E-03</td><td>BT20/MB468</td><td>MP</td></tr>
<tr><td>>hg38_21_+_8433172_8446622_RNA45SN1_45S_50ntFlanks@10548.10577.30|GTGCGGTAACGCGACCGATCCCGGAGAAGC</td><td>>hg38_21_+_8433172_8446622_RNA45SN1_45S_50ntFlanks</td><td>10548</td><td>10577</td><td>30</td><td>GTGCGGTAACGCGACCGATCCCGGAGAAGC</td><td>266.4</td><td>3.0</td><td>0.9</td><td>3.3</td><td>1.1E-03</td><td>5.1E-03</td><td>BT20/MB231</td><td>MITO</td></tr>
<tr><td>>MI0000750|hsa-mir-26a-2&WithFlank&12|-|57824603|57824698@20.39.20(+2U);MI0000083|hsa-mir-26a-1&WithFlank&3|+|37969398|37969486@16.35.20(+2U)&[MIMAT0000082&hsa-miR-26a-5p&offsets|0|-2(+2U);m-33&12|-|57824658|57824679&offsets|0|-2(+2U)];[MIMAT0000082_1&hsa-miR-26a-5p&offsets|0|-2(+2U);m-34&3|+|37969413|37969434&offsets|0|-2(+2U)]|TTCAAGTAATCCAGGATAGGTT</td><td>>MI0000750|hsa-mir-26a-2&WithFlank&12|-|57824603|57824698</td><td>20</td><td>39</td><td>20(+2U)</td><td>TTCAAGTAATCCAGGATAGGTT</td><td>361.0</td><td>-2.9</td><td>0.9</td><td>-3.3</td><td>1.1E-03</td><td>5.1E-03</td><td>BT20/MB231</td><td>MITO</td></tr>
<tr><td>>MI0000238|hsa-mir-196a-1&WithFlank&17|-|48632484|48632565@13.35.23&[MIMAT0000226_1&hsa-miR-196a-5p&offsets|0|+1;m-268&17|-|48632532|48632553&offsets|0|+1]|TAGGTAGTTTCATGTTGTTGGGC</td><td>>MI0000238|hsa-mir-196a-1&WithFlank&17|-|48632484|48632565</td><td>13</td><td>35</td><td>23</td><td>TAGGTAGTTTCATGTTGTTGGGC</td><td>90.9</td><td>4.5</td><td>1.5</td><td>3.0</td><td>2.3E-03</td><td>5.2E-03</td><td>BT20/MB468</td><td>MP</td></tr>
<tr><td>>hg38_21_+_8433172_8446622_RNA45SN1_45S_50ntFlanks@12995.13013.19|CTCGCTGCGATCTATTGAA</td><td>>hg38_21_+_8433172_8446622_RNA45SN1_45S_50ntFlanks</td><td>12995</td><td>13013</td><td>19</td><td>CTCGCTGCGATCTATTGAA</td><td>146.7</td><td>-2.5</td><td>0.9</td><td>-2.9</td><td>3.8E-03</td><td>5.2E-03</td><td>MB231/MB468</td><td>MP</td></tr>
<tr><td>>hg38_21_+_8433172_8446622_RNA45SN1_45S_50ntFlanks@9202.9225.24|TCTCGGGGCCACGCGCGCGTCCCC</td><td>>hg38_21_+_8433172_8446622_RNA45SN1_45S_50ntFlanks</td><td>9202</td><td>9225</td><td>24</td><td>TCTCGGGGCCACGCGCGCGTCCCC</td><td>122.1</td><td>-4.2</td><td>1.4</td><td>-2.9</td><td>3.5E-03</td><td>5.2E-03</td><td>BT20/MB468</td><td>MITO</td></tr>
<tr><td>>hg38_1_-_228634819_228635039_RNA5S12_5S_50ntFlanks@135.171.37|GGGAGACCGCCTGGGAATACCGGGTGCTGTAGGCTTT</td><td>>hg38_1_-_228634819_228635039_RNA5S12_5S_50ntFlanks</td><td>135</td><td>171</td><td>37</td><td>GGGAGACCGCCTGGGAATACCGGGTGCTGTAGGCTTT</td><td>4085.4</td><td>2.8</td><td>0.9</td><td>3.0</td><td>2.3E-03</td><td>5.2E-03</td><td>BT20/MB468</td><td>MP</td></tr>
<tr><td>>hg38_21_+_8433172_8446622_RNA45SN1_45S_50ntFlanks@12419.12443.25|GATGTCGGCTCTTCCTATCATTGTG</td><td>>hg38_21_+_8433172_8446622_RNA45SN1_45S_50ntFlanks</td><td>12419</td><td>12443</td><td>25</td><td>GATGTCGGCTCTTCCTATCATTGTG</td><td>497.7</td><td>1.7</td><td>0.6</td><td>2.9</td><td>3.5E-03</td><td>5.2E-03</td><td>BT20/MB468</td><td>MITO</td></tr>
<tr><td>>MI0000750|hsa-mir-26a-2&WithFlank&12|-|57824603|57824698@20.40.21;MI0000083|hsa-mir-26a-1&WithFlank&3|+|37969398|37969486@16.36.21&[MIMAT0000082&hsa-miR-26a-5p&offsets|0|-1;m-33&12|-|57824658|57824679&offsets|0|-1];[MIMAT0000082_1&hsa-miR-26a-5p&offsets|0|-1;m-34&3|+|37969413|37969434&offsets|0|-1]|TTCAAGTAATCCAGGATAGGC</td><td>>MI0000750|hsa-mir-26a-2&WithFlank&12|-|57824603|57824698</td><td>20</td><td>40</td><td>21</td><td>TTCAAGTAATCCAGGATAGGC</td><td>1656.7</td><td>-2.5</td><td>0.8</td><td>-3.3</td><td>1.1E-03</td><td>5.2E-03</td><td>BT20/MB231</td><td>MITO</td></tr>
<tr><td>>hg38_MT_+_598_1651_RNR1_12S_50ntFlanks@915.946.32|AAAGGACATTTAACTAAAACCCCTACGCATTT</td><td>>hg38_MT_+_598_1651_RNR1_12S_50ntFlanks</td><td>915</td><td>946</td><td>32</td><td>AAAGGACATTTAACTAAAACCCCTACGCATTT</td><td>382.6</td><td>2.6</td><td>0.8</td><td>3.3</td><td>1.1E-03</td><td>5.2E-03</td><td>BT20/MB231</td><td>MITO</td></tr>
<tr><td>>hg38_21_+_8433172_8446622_RNA45SN1_45S_50ntFlanks@9484.9520.37|GTGTTAGGACCCGAAAGATGGTGAACTATGCCTGGGC</td><td>>hg38_21_+_8433172_8446622_RNA45SN1_45S_50ntFlanks</td><td>9484</td><td>9520</td><td>37</td><td>GTGTTAGGACCCGAAAGATGGTGAACTATGCCTGGGC</td><td>31.3</td><td>8.7</td><td>2.9</td><td>3.0</td><td>2.3E-03</td><td>5.2E-03</td><td>BT20/MB468</td><td>MP</td></tr>
<tr><td>>hg38_21_+_8433172_8446622_RNA45SN1_45S_50ntFlanks@12907.12932.26|CACCGCACGTTCGTGGGGAACCTGGC</td><td>>hg38_21_+_8433172_8446622_RNA45SN1_45S_50ntFlanks</td><td>12907</td><td>12932</td><td>26</td><td>CACCGCACGTTCGTGGGGAACCTGGC</td><td>235.0</td><td>-3.8</td><td>1.3</td><td>-2.8</td><td>4.5E-03</td><td>5.2E-03</td><td>MB231/MB468</td><td>NUC</td></tr>
<tr><td>>hg38_21_+_8433172_8446622_RNA45SN1_45S_50ntFlanks@9544.9571.28|GGTGGAGGTCCGTAGCGGTCCTGACGTG</td><td>>hg38_21_+_8433172_8446622_RNA45SN1_45S_50ntFlanks</td><td>9544</td><td>9571</td><td>28</td><td>GGTGGAGGTCCGTAGCGGTCCTGACGTG</td><td>2235.3</td><td>-2.1</td><td>0.7</td><td>-2.9</td><td>4.1E-03</td><td>5.2E-03</td><td>BT20/MB468</td><td>CYTO</td></tr>
<tr><td>>hg38_21_+_8433172_8446622_RNA45SN1_45S_50ntFlanks@10802.10823.22|TTGGAACAATGTAGGTAAGGGA</td><td>>hg38_21_+_8433172_8446622_RNA45SN1_45S_50ntFlanks</td><td>10802</td><td>10823</td><td>22</td><td>TTGGAACAATGTAGGTAAGGGA</td><td>129.4</td><td>-3.2</td><td>1.1</td><td>-2.9</td><td>4.1E-03</td><td>5.2E-03</td><td>BT20/MB468</td><td>CYTO</td></tr>
<tr><td>>hg38_21_+_8433172_8446622_RNA45SN1_45S_50ntFlanks@12444.12476.33|AAGCAGAATTCACCAAGCGTTGGATTGTTCACC</td><td>>hg38_21_+_8433172_8446622_RNA45SN1_45S_50ntFlanks</td><td>12444</td><td>12476</td><td>33</td><td>AAGCAGAATTCACCAAGCGTTGGATTGTTCACC</td><td>659.3</td><td>-2.9</td><td>1.0</td><td>-3.0</td><td>2.3E-03</td><td>5.2E-03</td><td>BT20/MB468</td><td>MP</td></tr>
</table>

| | | | | | | | | | | | | | |
|---|---|---|---|---|---|---|---|---|---|---|---|---|---|
| >MI0000077\|hsa-mir-21&WithFlank&17\|+\|59841260\|59841343@13.37.25&[MIMAT0000076&hsa-miR-21-5p&offsets\|-1\|+2;m-1&17\|+\|59841273\|59841294&offsets\|-1\|+2]\|GTAGCTTATCAGACTGATGTTGACT | >MI0000077\|hsa-mir-21&WithFlank&17\|+\|59841260\|59841343 | 13 | 37 | 25 | GTAGCTTATCAGACTGATGTTGACT | 348.3 | -2.4 | 0.7 | -3.3 | 1.1E-03 | 5.2E-03 | BT20/MB231 | MITO |
| >MI0000289\|hsa-mir-181a-1&WithFlank&1\|-\|198859038\|198859159@30.51.22;MI0000269\|hsa-mir-181a-2&WithFlank&9\|+\|124692436\|124692557@45.66.22&[MIMAT0000256&hsa-miR-181a-5p&offsets\|0\|-1;m-32&1\|-\|198859108\|198859130&offsets\|0\|-1];[MIMAT0000256_1&hsa-miR-181a-5p&offsets\|0\|-1;m-32&9\|+\|124692480\|124692502&offsets\|0\|-1]\|AACATTCAACGCTGTCGGTGAG | >MI0000289\|hsa-mir-181a-1&WithFlank&1\|-\|198859038\|198859159 | 30 | 51 | 22 | AACATTCAACGCTGTCGGTGAG | 607.3 | -2.4 | 0.7 | -3.3 | 1.1E-03 | 5.2E-03 | BT20/MB231 | MITO |
| >hg38_21_+_8433172_8446622_RNA45SN1_45S_50ntFlanks@11571.11599.29\|GCCGACTTAGAACTGGTGCGGACCAGGGG | >hg38_21_+_8433172_8446622_RNA45SN1_45S_50ntFlanks | 11571 | 11599 | 29 | GCCGACTTAGAACTGGTGCGGACCAGGGG | 240.7 | 2.8 | 0.9 | 3.0 | 2.3E-03 | 5.2E-03 | BT20/MB468 | MP |
| >hg38_21_+_8433172_8446622_RNA45SN1_45S_50ntFlanks@11675.11702.28\|TGCTCTGAATGTCAAAGTGAAGAAATTC | >hg38_21_+_8433172_8446622_RNA45SN1_45S_50ntFlanks | 11675 | 11702 | 28 | TGCTCTGAATGTCAAAGTGAAGAAATTC | 946.5 | -2.9 | 0.9 | -3.3 | 9.0E-04 | 5.2E-03 | BT20/MB231 | NUC |
| >MI0000542\|hsa-mir-320a&WithFlank&8\|-\|22244960\|22245043@48.70.23&[MIMAT0000510&hsa-miR-320a-3p&offsets\|0\|+1;m-60&8\|-\|22244975\|22244996&offsets\|0\|+1]\|AAAAGCTGGGTTGAGAGGGCGAA | >MI0000542\|hsa-mir-320a&WithFlank&8\|-\|22244960\|22245043 | 48 | 70 | 23 | AAAAGCTGGGTTGAGAGGGCGAA | 3851.5 | 2.3 | 0.7 | 3.3 | 1.1E-03 | 5.2E-03 | BT20/MB231 | MITO |
| >hg38_21_+_8433172_8446622_RNA45SN1_45S_50ntFlanks@11667.11692.26\|CTGCCCAGTGCTCTGAATGTCAAAGT | >hg38_21_+_8433172_8446622_RNA45SN1_45S_50ntFlanks | 11667 | 11692 | 26 | CTGCCCAGTGCTCTGAATGTCAAAGT | 551.4 | 2.6 | 0.8 | 3.3 | 1.1E-03 | 5.2E-03 | BT20/MB231 | MITO |
| >trna119_LysCTT_1_-_145395522_145395594@1.34.34,trna11_LysCTT_5_-_180648979_180649051@1.34.34,trna13_LysCTT_6_+_26556774_26556846@1.34.34,trna7_LysCTT_16_+_3225692_3225764@1.34.34,trna9_LysCTT_5_+_180634755_180634827@1.34.34\|GCCCGGCTAGCTCAGTCGGTAGAGCATGAGACTC | >trna119_LysCTT_1_-_145395522_145395594 | 1 | 34 | 34 | GCCCGGCTAGCTCAGTCGGTAGAGCATGAGACTC | 3851.0 | -3.4 | 1.0 | -3.3 | 9.1E-04 | 5.2E-03 | BT20/MB231 | NUC |
| >MI0000483\|hsa-mir-186&WithFlank&1\|-\|71067625\|71067722@21.43.23(+1A)&[MIMAT0000456&hsa-miR-186-5p&offsets\|0\|+1(+1A);m-84&1\|-\|71067681\|71067702&offsets\|0\|+1(+1A)]\|CAAAGAATTCTCCTTTTGGGCTTA | >MI0000483\|hsa-mir-186&WithFlank&1\|-\|71067625\|71067722 | 21 | 43 | 23(+1A) | CAAAGAATTCTCCTTTTGGGCTTA | 178.2 | -3.5 | 1.0 | -3.3 | 8.3E-04 | 5.2E-03 | BT20/MB231 | MP |
| >hg38_21_+_8433172_8446622_RNA45SN1_45S_50ntFlanks@5491.5512.22\|GAGAAGACGGTCGAACTTGACT | >hg38_21_+_8433172_8446622_RNA45SN1_45S_50ntFlanks | 5491 | 5512 | 22 | GAGAAGACGGTCGAACTTGACT | 4126.3 | 3.7 | 1.1 | 3.3 | 1.0E-03 | 5.2E-03 | BT20/MB231 | CYTO |
| >MI0003560\|hsa-mir-92b&WithFlank&1\|+\|155195171\|155195278@67.88.22(+1A)&[MIMAT0003218&hsa-miR-92b-3p&offsets\|0\|0(+1A);m-115&1\|+\|155195237\|155195258&offsets\|0\|0(+1A)]\|TATTGCACTCGTCCCGGCCTCCA | >MI0003560\|hsa-mir-92b&WithFlank&1\|+\|155195171\|155195278 | 67 | 88 | 22(+1A) | TATTGCACTCGTCCCGGCCTCCA | 52.8 | 5.7 | 2.0 | 2.9 | 3.8E-03 | 5.2E-03 | MB231/MB468 | MP |
| >hg38_21_+_8433172_8446622_RNA45SN1_45S_50ntFlanks@9929.9968.40\|GATATAGACAGCAGGACGGTGGCCATGGAAGTCGGAATCC | >hg38_21_+_8433172_8446622_RNA45SN1_45S_50ntFlanks | 9929 | 9968 | 40 | GATATAGACAGCAGGACGGTGGCCATGGAAGTCGGAATCC | 58.2 | -5.1 | 1.8 | -2.9 | 3.8E-03 | 5.2E-03 | MB231/MB468 | MP |
| >hg38_21_+_8433172_8446622_RNA45SN1_45S_50ntFlanks@9604.9624.21\|AAAGACTAATCGAACCATCTA | >hg38_21_+_8433172_8446622_RNA45SN1_45S_50ntFlanks | 9604 | 9624 | 21 | AAAGACTAATCGAACCATCTA | 124.4 | -5.3 | 1.7 | -3.0 | 2.3E-03 | 5.2E-03 | BT20/MB468 | MP |
| >hg38_21_+_8433172_8446622_RNA45SN1_45S_50ntFlanks@6663.6699.37\|GTGGATCACTCGGCTCGTGCGTCGATGAAGAACGCAG | >hg38_21_+_8433172_8446622_RNA45SN1_45S_50ntFlanks | 6663 | 6699 | 37 | GTGGATCACTCGGCTCGTGCGTCGATGAAGAACGCAG | 151.9 | -1.7 | 0.6 | -2.9 | 4.2E-03 | 5.2E-03 | BT20/MB468 | CYTO |
| >MI0000088\|hsa-mir-30a&WithFlank&6\|-\|71403545\|71403627@12.31.20&[MIMAT0000087&hsa-miR-30a-5p&offsets\|0\|-2;m-15&6\|-\|71403595\|71403616&offsets\|0\|-2]\|TGTAAACATCCTCGACTGGA | >MI0000088\|hsa-mir-30a&WithFlank&6\|-\|71403545\|71403627 | 12 | 31 | 20 | TGTAAACATCCTCGACTGGA | 3244.4 | -2.6 | 0.8 | -3.3 | 1.1E-03 | 5.2E-03 | BT20/MB231 | MITO |
| >hg38_21_+_8433172_8446622_RNA45SN1_45S_50ntFlanks@10348.10365.18\|AAACGAGAACTTTGAAGG | >hg38_21_+_8433172_8446622_RNA45SN1_45S_50ntFlanks | 10348 | 10365 | 18 | AAACGAGAACTTTGAAGG | 204.8 | -1.7 | 0.6 | -2.9 | 4.2E-03 | 5.2E-03 | BT20/MB468 | CYTO |
| >hg38_21_+_8433172_8446622_RNA45SN1_45S_50ntFlanks@11670.11688.19\|CCCAGTGCTCTGAATGTCA | >hg38_21_+_8433172_8446622_RNA45SN1_45S_50ntFlanks | 11670 | 11688 | 19 | CCCAGTGCTCTGAATGTCA | 1796.3 | -5.9 | 2.1 | -2.8 | 4.4E-03 | 5.2E-03 | BT20/MB468 | NUC |
| >hg38_21_+_8433172_8446622_RNA45SN1_45S_50ntFlanks@6645.6675.31\|CTCGTACGACTCTTAGCGGTGGATCACTCGG | >hg38_21_+_8433172_8446622_RNA45SN1_45S_50ntFlanks | 6645 | 6675 | 31 | CTCGTACGACTCTTAGCGGTGGATCACTCGG | 1123.5 | 4.7 | 1.4 | 3.3 | 8.3E-04 | 5.2E-03 | BT20/MB231 | MP |
| >hg38_21_+_8433172_8446622_RNA45SN1_45S_50ntFlanks@7974.8009.36\|ACGCGACCTCAGATCAGACGTGGCGACCCGCTGAAT | >hg38_21_+_8433172_8446622_RNA45SN1_45S_50ntFlanks | 7974 | 8009 | 36 | ACGCGACCTCAGATCAGACGTGGCGACCCGCTGAAT | 6381.3 | 2.5 | 0.8 | 2.9 | 3.8E-03 | 5.2E-03 | MB231/MB468 | MP |
| >hg38_MT_+_1621_3279_RNR2_16S_50ntFlanks@275.305.31\|CTAAGACCCCCGAAACCAGACGAGCTACCTA | >hg38_MT_+_1621_3279_RNR2_16S_50ntFlanks | 275 | 305 | 31 | CTAAGACCCCCGAAACCAGACGAGCTACCTA | 88.1 | 6.2 | 2.0 | 3.0 | 2.3E-03 | 5.2E-03 | BT20/MB468 | MP |
| >hg38_21_+_8433172_8446622_RNA45SN1_45S_50ntFlanks@6645.6678.34\|CTCGTACGACTCTTAGCGGTGGATCACTCGGCTC | >hg38_21_+_8433172_8446622_RNA45SN1_45S_50ntFlanks | 6645 | 6678 | 34 | CTCGTACGACTCTTAGCGGTGGATCACTCGGCTC | 516.6 | 2.3 | 0.8 | 2.8 | 4.5E-03 | 5.2E-03 | MB231/MB468 | NUC |
| >hg38_21_+_8433172_8446622_RNA45SN1_45S_50ntFlanks@8347.8373.27\|GGGAAAGTTGAAAAGAACTTTGAAGAG | >hg38_21_+_8433172_8446622_RNA45SN1_45S_50ntFlanks | 8347 | 8373 | 27 | GGGAAAGTTGAAAAGAACTTTGAAGAG | 1589.0 | 5.6 | 1.7 | 3.3 | 8.3E-04 | 5.2E-03 | BT20/MB231 | MP |
| >hg38_21_+_8433172_8446622_RNA45SN1_45S_50ntFlanks@9158.9187.30\|AGTGCGCCCCGGGCGGGTCGCGCCGTCGGG | >hg38_21_+_8433172_8446622_RNA45SN1_45S_50ntFlanks | 9158 | 9187 | 30 | AGTGCGCCCCGGGCGGGTCGCGCCGTCGGG | 198.1 | -4.9 | 1.6 | -3.0 | 2.3E-03 | 5.2E-03 | BT20/MB468 | MP |
| >trna10_AspGTC_12_-_125424193_125424264@32.53.22,trna12_AspGTC_12_-_125411891_125411962@32.53.22,trna144_AspGTC_6_-_27551236_27551307@32.53.22,trna38_AspGTC_17_-_8125556_8125627@32.53.22,trna45_AspGTC_6_+_27447453_27447524@32.53.22,trna48_AspGTC_6_+_27471523_27471594@32.53.22,trna4_AspGTC_12_+_96429799_96429870@32.53.22,trna5_AspGTC_12_+_98897281_98897352@32.53.22,trna69_AspGTC_1_-_161440205_161440276@32.53.22,trna6_AspGTC_9_-_77517990_77518061@32.53.22,trna72_AspGTC_1_-_161432824_161432895@32.53.22,trna75_AspGTC_1_-_161425414_161425485@32.53.22,trna78_AspGTC_1_-_161418033_161418104@32.53.22,trna81_AspGTC_1_-_161410615_161410686@32.53.22\|CTGTCACGCGGGAGACCGGGGT | >trna10_AspGTC_12_-_125424193_125424264 | 32 | 53 | 22 | CTGTCACGCGGGAGACCGGGGT | 458.6 | 2.9 | 0.9 | 3.3 | 1.0E-03 | 5.2E-03 | BT20/MB231 | CYTO |
| >hg38_21_+_8433172_8446622_RNA45SN1_45S_50ntFlanks@11233.11252.20\|CCCTTCCCGTGGATCGCCCC | >hg38_21_+_8433172_8446622_RNA45SN1_45S_50ntFlanks | 11233 | 11252 | 20 | CCCTTCCCGTGGATCGCCCC | 213.2 | -5.1 | 1.7 | -3.0 | 2.4E-03 | 5.2E-03 | BT20/MB468 | MP |
| >hg38_21_+_8433172_8446622_RNA45SN1_45S_50ntFlanks@11629.11665.37\|CGCGAAGGCCCGCGGCGGGTGTTGACGCGATGTGATT | >hg38_21_+_8433172_8446622_RNA45SN1_45S_50ntFlanks | 11629 | 11665 | 37 | CGCGAAGGCCCGCGGCGGGTGTTGACGCGATGTGATT | 147.9 | -5.0 | 1.7 | -3.0 | 2.4E-03 | 5.2E-03 | BT20/MB468 | MP |
| >hg38_21_+_8433172_8446622_RNA45SN1_45S_50ntFlanks@3840.3858.19\|CTCCTACTTGGATAACTGT | >hg38_21_+_8433172_8446622_RNA45SN1_45S_50ntFlanks | 3840 | 3858 | 19 | CTCCTACTTGGATAACTGT | 351.1 | 3.3 | 1.1 | 2.9 | 4.2E-03 | 5.2E-03 | BT20/MB468 | CYTO |
| >hg38_21_+_8433172_8446622_RNA45SN1_45S_50ntFlanks@10237.10263.27\|ACGCTACGCCGCGACGAGTAGGAGGGC | >hg38_21_+_8433172_8446622_RNA45SN1_45S_50ntFlanks | 10237 | 10263 | 27 | ACGCTACGCCGCGACGAGTAGGAGGGC | 335.4 | 2.8 | 0.9 | 3.0 | 2.4E-03 | 5.2E-03 | BT20/MB468 | MP |
| >hg38_21_+_8433172_8446622_RNA45SN1_45S_50ntFlanks@10353.10383.31\|AGAACTTTGAAGGCCGAAGTGGAGAAGGGTT | >hg38_21_+_8433172_8446622_RNA45SN1_45S_50ntFlanks | 10353 | 10383 | 31 | AGAACTTTGAAGGCCGAAGTGGAGAAGGGTT | 260.3 | -2.4 | 0.8 | -2.8 | 4.5E-03 | 5.2E-03 | MB231/MB468 | MITO |
| >hg38_21_+_8433172_8446622_RNA45SN1_45S_50ntFlanks@12626.12657.32\|AATGGGGCGAAGCTACCATCTGTGGGATTATG | >hg38_21_+_8433172_8446622_RNA45SN1_45S_50ntFlanks | 12626 | 12657 | 32 | AATGGGGCGAAGCTACCATCTGTGGGATTATG | 253.5 | -3.2 | 1.1 | -2.9 | 4.2E-03 | 5.2E-03 | BT20/MB468 | CYTO |
| >hg38_21_+_8433172_8446622_RNA45SN1_45S_50ntFlanks@12420.12441.22\|ATGTCGGCTCTTCCTATCATTG | >hg38_21_+_8433172_8446622_RNA45SN1_45S_50ntFlanks | 12420 | 12441 | 22 | ATGTCGGCTCTTCCTATCATTG | 3871.9 | -1.9 | 0.7 | -2.9 | 4.2E-03 | 5.2E-03 | MB231/MB468 | CYTO |
| >hg38_21_+_8433172_8446622_RNA45SN1_45S_50ntFlanks@4102.4120.19\|ACCACGGGTGACGGGGAAT | >hg38_21_+_8433172_8446622_RNA45SN1_45S_50ntFlanks | 4102 | 4120 | 19 | ACCACGGGTGACGGGGAAT | 305.2 | -2.1 | 0.7 | -2.9 | 4.2E-03 | 5.2E-03 | BT20/MB468 | CYTO |
| >hg38_21_+_8433172_8446622_RNA45SN1_45S_50ntFlanks@11552.11569.18\|CCTCGGCCGGCGCCTAGC | >hg38_21_+_8433172_8446622_RNA45SN1_45S_50ntFlanks | 11552 | 11569 | 18 | CCTCGGCCGGCGCCTAGC | 377.2 | -2.5 | 0.9 | -2.9 | 4.2E-03 | 5.2E-03 | BT20/MB468 | CYTO |
| >MI0000088\|hsa-mir-30a&WithFlank&6\|-\|71403545\|71403627@12.35.24(+2U)&[MIMAT0000087&hsa-miR-30a-5p&offsets\|0\|+2(+2U);m-15&6\|-\|71403595\|71403616&offsets\|0\|+2(+2U)]\|TGTAAACATCCTCGACTGGAAGCTTT | >MI0000088\|hsa-mir-30a&WithFlank&6\|-\|71403545\|71403627 | 12 | 35 | 24(+2U) | TGTAAACATCCTCGACTGGAAGCTTT | 1366.8 | -3.4 | 1.0 | -3.3 | 9.1E-04 | 5.2E-03 | BT20/MB231 | NUC |
| >hg38_21_+_8433172_8446622_RNA45SN1_45S_50ntFlanks@9457.9493.37\|CGCCGGGGAGGTGGAGCACGAGCGCACGTGTTAGGAC | >hg38_21_+_8433172_8446622_RNA45SN1_45S_50ntFlanks | 9457 | 9493 | 37 | CGCCGGGGAGGTGGAGCACGAGCGCACGTGTTAGGAC | 65.2 | -5.1 | 1.7 | -3.0 | 2.4E-03 | 5.2E-03 | BT20/MB468 | MP |
| >hg38_1_-_228634819_228635039_RNA5S12_5S_50ntFlanks@132.171.40\|GATGGGAGACCGCCTGGGAATACCGGGTGCTGTAGGCTTT | >hg38_1_-_228634819_228635039_RNA5S12_5S_50ntFlanks | 132 | 171 | 40 | GATGGGAGACCGCCTGGGAATACCGGGTGCTGTAGGCTTT | 16534.9 | 3.3 | 1.1 | 3.0 | 2.4E-03 | 5.2E-03 | BT20/MB468 | MP |
| >hg38_21_+_8433172_8446622_RNA45SN1_45S_50ntFlanks@10683.10708.26\|CCGGCGGCGTCCGGTGAGCTCTCGCT | >hg38_21_+_8433172_8446622_RNA45SN1_45S_50ntFlanks | 10683 | 10708 | 26 | CCGGCGGCGTCCGGTGAGCTCTCGCT | 178.8 | -3.2 | 1.1 | -2.9 | 4.2E-03 | 5.2E-03 | BT20/MB468 | CYTO |
| >hg38_21_+_8433172_8446622_RNA45SN1_45S_50ntFlanks@6650.6689.40\|ACGACTCTTAGCGGTGGATCACTCGGCTCGTGCGTCGATG | >hg38_21_+_8433172_8446622_RNA45SN1_45S_50ntFlanks | 6650 | 6689 | 40 | ACGACTCTTAGCGGTGGATCACTCGGCTCGTGCGTCGATG | 1787.0 | -1.7 | 0.6 | -2.8 | 4.5E-03 | 5.2E-03 | BT20/MB468 | NUC |
| >MI0000254\|hsa-mir-30c-2&WithFlank&6\|-\|71376954\|71377037@53.74.22&[MIMAT0004550&hsa-miR-30c-2-3p&offsets\|0\|0;m-206&6\|-\|71376964\|71376985&offsets\|0\|0]\|CTGGGAGAAGGCTGTTTACTCT | >MI0000254\|hsa-mir-30c-2&WithFlank&6\|-\|71376954\|71377037 | 53 | 74 | 22 | CTGGGAGAAGGCTGTTTACTCT | 169.0 | 2.5 | 0.8 | 3.0 | 2.4E-03 | 5.2E-03 | BT20/MB468 | MP |
| >MI0003780\|hsa-mir-1296&WithFlank&10\|-\|63372951\|63373054@22.43.22&[MIMAT0005794&hsa-miR-1296-5p&offsets\|0\|0;m-304&10\|-\|63373012\|63373033&offsets\|0\|0]\|TTAGGGCCCTGGCTCCATCTCC | >MI0003780\|hsa-mir-1296&WithFlank&10\|-\|63372951\|63373054 | 22 | 43 | 22 | TTAGGGCCCTGGCTCCATCTCC | 375.1 | 4.5 | 1.5 | 3.0 | 2.4E-03 | 5.2E-03 | BT20/MB468 | MP |
| >hg38_21_+_8433172_8446622_RNA45SN1_45S_50ntFlanks@10229.10259.31\|CCCCGCGGACGCTACGCCGCGACGAGTAGGA | >hg38_21_+_8433172_8446622_RNA45SN1_45S_50ntFlanks | 10229 | 10259 | 31 | CCCCGCGGACGCTACGCCGCGACGAGTAGGA | 189.4 | 3.8 | 1.3 | 3.0 | 2.4E-03 | 5.2E-03 | BT20/MB468 | MP |
| >hg38_21_+_8433172_8446622_RNA45SN1_45S_50ntFlanks@5494.5516.23\|AAGACGGTCGAACTTGACTATCT | >hg38_21_+_8433172_8446622_RNA45SN1_45S_50ntFlanks | 5494 | 5516 | 23 | AAGACGGTCGAACTTGACTATCT | 897.0 | 4.7 | 1.5 | 3.0 | 2.4E-03 | 5.2E-03 | BT20/MB468 | MP |
| >hg38_21_+_8433172_8446622_RNA45SN1_45S_50ntFlanks@4859.4876.18\|TTGGGTTCCGGGGGGAGT | >hg38_21_+_8433172_8446622_RNA45SN1_45S_50ntFlanks | 4859 | 4876 | 18 | TTGGGTTCCGGGGGGAGT | 32.5 | 8.2 | 2.9 | 2.9 | 3.8E-03 | 5.2E-03 | MB231/MB468 | MP |
| >hg38_21_+_8433172_8446622_RNA45SN1_45S_50ntFlanks@10357.10390.34\|CTTTGAAGGCCGAAGTGGAGAAGGGTTCCATGTG | >hg38_21_+_8433172_8446622_RNA45SN1_45S_50ntFlanks | 10357 | 10390 | 34 | CTTTGAAGGCCGAAGTGGAGAAGGGTTCCATGTG | 480.4 | -3.1 | 1.1 | -2.9 | 4.3E-03 | 5.2E-03 | MB231/MB468 | CYTO |
| >hg38_21_+_8433172_8446622_RNA45SN1_45S_50ntFlanks@4367.4400.34\|CAGTTAAAAAGCTCGTAGTTGGATCTTGGGAGCG | >hg38_21_+_8433172_8446622_RNA45SN1_45S_50ntFlanks | 4367 | 4400 | 34 | CAGTTAAAAAGCTCGTAGTTGGATCTTGGGAGCG | 107.6 | -5.1 | 1.7 | -3.0 | 2.4E-03 | 5.2E-03 | BT20/MB468 | MP |
| >MI0002468\|hsa-mir-484&WithFlank&16\|+\|15643288\|15643378@14.33.20(+2U)&[MIMAT0002174&hsa-miR-484&offsets\|0\|-2(+2U);m-127&16\|+\|15643301\|15643321&offsets\|0\|-1(+2U)]\|TCAGGCTCAGTCCCCTCCCGTT | >MI0002468\|hsa-mir-484&WithFlank&16\|+\|15643288\|15643378 | 14 | 33 | 20(+2U) | TCAGGCTCAGTCCCCTCCCGTT | 75.0 | 5.2 | 1.7 | 3.0 | 2.4E-03 | 5.2E-03 | BT20/MB468 | MP |

<table>
<tr><td>>hg38_MT_+_1621_3279_RNR2_16S_50ntFlanks@1366.1401.36|CTCGATGTTGGATCAGGACATCCCGATGGTGCAGCC</td><td>>hg38_MT_+_1621_3279_RNR2_16S_50ntFlanks</td><td>1366</td><td>1401</td><td>36</td><td>CTCGATGTTGGATCAGGACATCCCGATGGTGCAGCC</td><td>59.1</td><td>6.1</td><td>2.0</td><td>3.0</td><td>2.4E-03</td><td>5.2E-03</td><td>BT20/MB468</td><td>MP</td></tr>
<tr><td>>hg38_21_+_8433172_8446622_RNA45SN1_45S_50ntFlanks@13001.13027.27|GCGATCTATTGAAAGTCAGCCCTCGAC</td><td>>hg38_21_+_8433172_8446622_RNA45SN1_45S_50ntFlanks</td><td>13001</td><td>13027</td><td>27</td><td>GCGATCTATTGAAAGTCAGCCCTCGAC</td><td>473.9</td><td>-2.4</td><td>0.9</td><td>-2.8</td><td>4.5E-03</td><td>5.2E-03</td><td>MB231/MB468</td><td>NUC</td></tr>
<tr><td>>hg38_21_+_8433172_8446622_RNA45SN1_45S_50ntFlanks@8687.8707.21|CGGCTCCGGGACGGCTGGGAA</td><td>>hg38_21_+_8433172_8446622_RNA45SN1_45S_50ntFlanks</td><td>8687</td><td>8707</td><td>21</td><td>CGGCTCCGGGACGGCTGGGAA</td><td>608.2</td><td>-3.0</td><td>0.9</td><td>-3.3</td><td>1.0E-03</td><td>5.2E-03</td><td>BT20/MB231</td><td>CYTO</td></tr>
<tr><td>>hg38_21_+_8433172_8446622_RNA45SN1_45S_50ntFlanks@3999.4024.26|CTAGATAACCTCGGGCCGATCGCACG</td><td>>hg38_21_+_8433172_8446622_RNA45SN1_45S_50ntFlanks</td><td>3999</td><td>4024</td><td>26</td><td>CTAGATAACCTCGGGCCGATCGCACG</td><td>216.0</td><td>-3.2</td><td>1.1</td><td>-2.9</td><td>4.2E-03</td><td>5.2E-03</td><td>BT20/MB468</td><td>CYTO</td></tr>
<tr><td>>hg38_21_+_8433172_8446622_RNA45SN1_45S_50ntFlanks@12397.12419.23|CGACGTCGCTTTTTGATCCTTCG</td><td>>hg38_21_+_8433172_8446622_RNA45SN1_45S_50ntFlanks</td><td>12397</td><td>12419</td><td>23</td><td>CGACGTCGCTTTTTGATCCTTCG</td><td>186.9</td><td>-2.2</td><td>0.8</td><td>-2.9</td><td>3.6E-03</td><td>5.2E-03</td><td>BT20/MB468</td><td>MITO</td></tr>
<tr><td>>MI0000471|hsa-mir-126&WithFlank&9|+|136670596|136670692@21.41.21&[MIMAT0000444&hsa-miR-126-5p&offsets|0|0;m-114&9|+|136670616|136670636&offsets|0|0]|CATTATTACTTTTGGTACGCG</td><td>>MI0000471|hsa-mir-126&WithFlank&9|+|136670596|136670692</td><td>21</td><td>41</td><td>21</td><td>CATTATTACTTTTGGTACGCG</td><td>252.3</td><td>3.1</td><td>0.9</td><td>3.3</td><td>1.0E-03</td><td>5.2E-03</td><td>BT20/MB231</td><td>CYTO</td></tr>
<tr><td>>MI0000102|hsa-mir-100&WithFlank&11|-|122152223|122152314@19.36.18&[MIMAT0000098&hsa-miR-100-5p&offsets|0|-4;m-27&11|-|122152276|122152296&offsets|0|-3]|AACCCGTAGATCCGAACT</td><td>>MI0000102|hsa-mir-100&WithFlank&11|-|122152223|122152314</td><td>19</td><td>36</td><td>18</td><td>AACCCGTAGATCCGAACT</td><td>740.8</td><td>-6.1</td><td>1.8</td><td>-3.3</td><td>8.4E-04</td><td>5.2E-03</td><td>BT20/MB231</td><td>MP</td></tr>
<tr><td>>hg38_21_+_8433172_8446622_RNA45SN1_45S_50ntFlanks@9566.9595.30|GACGTGCAAATCGGTCGTCCGACCTGGGTA</td><td>>hg38_21_+_8433172_8446622_RNA45SN1_45S_50ntFlanks</td><td>9566</td><td>9595</td><td>30</td><td>GACGTGCAAATCGGTCGTCCGACCTGGGTA</td><td>477.5</td><td>-1.9</td><td>0.7</td><td>-2.9</td><td>4.3E-03</td><td>5.2E-03</td><td>MB231/MB468</td><td>CYTO</td></tr>
<tr><td>>hg38_21_+_8433172_8446622_RNA45SN1_45S_50ntFlanks@10315.10340.26|AGGTGCAGATCTTGGTGGTAGTAGCA</td><td>>hg38_21_+_8433172_8446622_RNA45SN1_45S_50ntFlanks</td><td>10315</td><td>10340</td><td>26</td><td>AGGTGCAGATCTTGGTGGTAGTAGCA</td><td>828.2</td><td>6.2</td><td>1.9</td><td>3.3</td><td>8.4E-04</td><td>5.2E-03</td><td>BT20/MB231</td><td>MP</td></tr>
<tr><td>>hg38_21_+_8433172_8446622_RNA45SN1_45S_50ntFlanks@10408.10433.26|GGTCAGTCGGTCCTGAGAGATGGGCG</td><td>>hg38_21_+_8433172_8446622_RNA45SN1_45S_50ntFlanks</td><td>10408</td><td>10433</td><td>26</td><td>GGTCAGTCGGTCCTGAGAGATGGGCG</td><td>415.6</td><td>-4.8</td><td>1.6</td><td>-2.9</td><td>3.6E-03</td><td>5.3E-03</td><td>BT20/MB468</td><td>MITO</td></tr>
<tr><td>>hg38_21_+_8433172_8446622_RNA45SN1_45S_50ntFlanks@5083.5102.20|ACGAACGAGACTCTGGCATG</td><td>>hg38_21_+_8433172_8446622_RNA45SN1_45S_50ntFlanks</td><td>5083</td><td>5102</td><td>20</td><td>ACGAACGAGACTCTGGCATG</td><td>322.4</td><td>-2.9</td><td>1.0</td><td>-2.9</td><td>3.6E-03</td><td>5.3E-03</td><td>BT20/MB468</td><td>MITO</td></tr>
<tr><td>>hg38_MT_+_1621_3279_RNR2_16S_50ntFlanks@1349.1388.40|ACAATAGGGTTTACGACCTCGATGTTGGATCAGGACATCC</td><td>>hg38_MT_+_1621_3279_RNR2_16S_50ntFlanks</td><td>1349</td><td>1388</td><td>40</td><td>ACAATAGGGTTTACGACCTCGATGTTGGATCAGGACATCC</td><td>396.7</td><td>4.9</td><td>1.6</td><td>3.0</td><td>2.4E-03</td><td>5.3E-03</td><td>BT20/MB468</td><td>MP</td></tr>
<tr><td>>hg38_MT_+_1621_3279_RNR2_16S_50ntFlanks@51.92.42|GCTAAACCTAGCCCCAAACCCACTCCACCTTACTACCAGACA</td><td>>hg38_MT_+_1621_3279_RNR2_16S_50ntFlanks</td><td>51</td><td>92</td><td>42</td><td>GCTAAACCTAGCCCCAAACCCACTCCACCTTACTACCAGACA</td><td>1692.9</td><td>3.9</td><td>1.4</td><td>2.9</td><td>3.9E-03</td><td>5.3E-03</td><td>MB231/MB468</td><td>MP</td></tr>
<tr><td>>hg38_21_+_8433172_8446622_RNA45SN1_45S_50ntFlanks@11594.11613.20|CAGGGGAATCCGACTGTTTA</td><td>>hg38_21_+_8433172_8446622_RNA45SN1_45S_50ntFlanks</td><td>11594</td><td>11613</td><td>20</td><td>CAGGGGAATCCGACTGTTTA</td><td>288.4</td><td>-2.8</td><td>1.0</td><td>-2.9</td><td>4.3E-03</td><td>5.3E-03</td><td>MB231/MB468</td><td>CYTO</td></tr>
<tr><td>>MI0000809|hsa-mir-151a&WithFlank&8|-|140732558|140732659@53.75.23(+1U)&[MIMAT0000757&hsa-miR-151a-3p&offsets|0|+2(+1U);m-30&8|-|140732587|140732607&offsets|0|+2(+1U)]|CTAGACTGAAGCTCCTTGAGGACT</td><td>>MI0000809|hsa-mir-151a&WithFlank&8|-|140732558|140732659</td><td>53</td><td>75</td><td>23(+1U)</td><td>CTAGACTGAAGCTCCTTGAGGACT</td><td>118.5</td><td>3.7</td><td>1.3</td><td>2.9</td><td>3.9E-03</td><td>5.3E-03</td><td>MB231/MB468</td><td>MP</td></tr>
<tr><td>>hg38_1_-_228634819_228635039_RNA5S12_5S_50ntFlanks@51.92.42|GTCTACGGCCATACCACCCTGAACGCGCCCGATCTCGTCTGA</td><td>>hg38_1_-_228634819_228635039_RNA5S12_5S_50ntFlanks</td><td>51</td><td>92</td><td>42</td><td>GTCTACGGCCATACCACCCTGAACGCGCCCGATCTCGTCTGA</td><td>589.2</td><td>2.6</td><td>0.9</td><td>2.8</td><td>4.6E-03</td><td>5.3E-03</td><td>MB231/MB468</td><td>NUC</td></tr>
<tr><td>>hg38_1_-_228634819_228635039_RNA5S12_5S_50ntFlanks@128.169.42|CTTGGATGGGAGACCGCCTGGGAATACCGGGTGCTGTAGGCT</td><td>>hg38_1_-_228634819_228635039_RNA5S12_5S_50ntFlanks</td><td>128</td><td>169</td><td>42</td><td>CTTGGATGGGAGACCGCCTGGGAATACCGGGTGCTGTAGGCT</td><td>295.3</td><td>5.2</td><td>1.7</td><td>3.0</td><td>2.4E-03</td><td>5.3E-03</td><td>BT20/MB468</td><td>MP</td></tr>
<tr><td>>MI0000085|hsa-mir-27a&WithFlank&19|-|13836434|13836523@57.76.20(+4U)&[MIMAT0000084&hsa-miR-27a-3p&offsets|0|-1(+4U);m-43&19|-|13836447|13836467&offsets|0|-1(+4U)]|TTCACAGTGGCTAAGTTCCGTTTT</td><td>>MI0000085|hsa-mir-27a&WithFlank&19|-|13836434|13836523</td><td>57</td><td>76</td><td>20(+4U)</td><td>TTCACAGTGGCTAAGTTCCGTTTT</td><td>237.9</td><td>-6.5</td><td>2.0</td><td>-3.3</td><td>1.1E-03</td><td>5.3E-03</td><td>BT20/MB231</td><td>MITO</td></tr>
<tr><td>>hg38_21_+_8433172_8446622_RNA45SN1_45S_50ntFlanks@4367.4396.30|CAGTTAAAAAGCTCGTAGTTGGATCTTGGG</td><td>>hg38_21_+_8433172_8446622_RNA45SN1_45S_50ntFlanks</td><td>4367</td><td>4396</td><td>30</td><td>CAGTTAAAAAGCTCGTAGTTGGATCTTGGG</td><td>149.8</td><td>-2.1</td><td>0.7</td><td>-2.9</td><td>4.3E-03</td><td>5.3E-03</td><td>MB231/MB468</td><td>CYTO</td></tr>
<tr><td>>MI0000301|hsa-mir-224&WithFlank&X|-|151958572|151958664@14.36.23(+1A)&[MIMAT0000281&hsa-miR-224-5p&offsets|+1|-1(+1A);m-238&X|-|151958630|151958651&offsets|0|+1(+1A)]|CAAGTCACTAGTGGTTCCGTTTAA</td><td>>MI0000301|hsa-mir-224&WithFlank&X|-|151958572|151958664</td><td>14</td><td>36</td><td>23(+1A)</td><td>CAAGTCACTAGTGGTTCCGTTTAA</td><td>453.5</td><td>2.7</td><td>0.8</td><td>3.3</td><td>1.0E-03</td><td>5.3E-03</td><td>BT20/MB231</td><td>CYTO</td></tr>
<tr><td>>MI0000746|hsa-mir-99b&WithFlank&19|+|51692606|51692687@13.31.19&[MIMAT0000689&hsa-miR-99b-5p&offsets|0|-3;m-10&19|+|51692618|51692639&offsets|0|-3]|CACCCGTAGAACCGACCTT</td><td>>MI0000746|hsa-mir-99b&WithFlank&19|+|51692606|51692687</td><td>13</td><td>31</td><td>19</td><td>CACCCGTAGAACCGACCTT</td><td>337.7</td><td>2.5</td><td>0.8</td><td>3.3</td><td>1.0E-03</td><td>5.3E-03</td><td>BT20/MB231</td><td>CYTO</td></tr>
<tr><td>>am_trnaMT_ValTAC_MT_+_1602_1670@33.52.20|ACACTTAGGAGATTTCAACT</td><td>>am_trnaMT_ValTAC_MT_+_1602_1670</td><td>33</td><td>52</td><td>20</td><td>ACACTTAGGAGATTTCAACT</td><td>334.2</td><td>3.0</td><td>0.9</td><td>3.3</td><td>1.0E-03</td><td>5.3E-03</td><td>BT20/MB231</td><td>CYTO</td></tr>
<tr><td>>trna116_GluCTC_1_-_145399233_145399304@37.73.37,trna134_GluTTC_1_-_16861774_16861845@37.73.37,trna71_GluCTC_1_-_161439189_161439260@37.73.37,trna74_GluCTC_1_-_161431809_161431880@37.73.37,trna77_GluCTC_1_-_161424398_161424469@37.73.37,trna77_GluCTC_6_+_28949976_28950047@37.73.37,trna80_GluCTC_1_-_161417018_161417089@37.73.37,trna84_GluTTC_1_-_161391883_161391954@37.73.37,trna87_GluCTC_6_-_126101393_126101464@37.73.37|ACCGCCGCGGCCCGGGTTCGATTCCCGGTCAGGGAAC</td><td>>trna116_GluCTC_1_-_145399233_145399304</td><td>37</td><td>73</td><td>37</td><td>ACCGCCGCGGCCCGGGTTCGATTCCCGGTCAGGGAAC</td><td>2599.5</td><td>-3.8</td><td>1.1</td><td>-3.3</td><td>1.0E-03</td><td>5.3E-03</td><td>BT20/MB231</td><td>CYTO</td></tr>
<tr><td>>hg38_21_+_8433172_8446622_RNA45SN1_45S_50ntFlanks@11672.11708.37|CAGTGCTCTGAATGTCAAAGTGAAGAAATTCAATGAA</td><td>>hg38_21_+_8433172_8446622_RNA45SN1_45S_50ntFlanks</td><td>11672</td><td>11708</td><td>37</td><td>CAGTGCTCTGAATGTCAAAGTGAAGAAATTCAATGAA</td><td>252.5</td><td>-2.7</td><td>0.8</td><td>-3.3</td><td>1.0E-03</td><td>5.3E-03</td><td>BT20/MB231</td><td>CYTO</td></tr>
<tr><td>>hg38_21_+_8433172_8446622_RNA45SN1_45S_50ntFlanks@3823.3859.37|TTTGGTCGCTCGCTCCTCTCCTACTTGGATAACTGTG</td><td>>hg38_21_+_8433172_8446622_RNA45SN1_45S_50ntFlanks</td><td>3823</td><td>3859</td><td>37</td><td>TTTGGTCGCTCGCTCCTCTCCTACTTGGATAACTGTG</td><td>261.9</td><td>-2.9</td><td>0.9</td><td>-3.3</td><td>1.0E-03</td><td>5.3E-03</td><td>BT20/MB231</td><td>CYTO</td></tr>
<tr><td>>hg38_21_+_8433172_8446622_RNA45SN1_45S_50ntFlanks@6721.6758.38|AATTGCAGGACACATTGATCATCGACACTTCGAACGCA</td><td>>hg38_21_+_8433172_8446622_RNA45SN1_45S_50ntFlanks</td><td>6721</td><td>6758</td><td>38</td><td>AATTGCAGGACACATTGATCATCGACACTTCGAACGCA</td><td>184.1</td><td>-2.8</td><td>0.8</td><td>-3.3</td><td>1.0E-03</td><td>5.3E-03</td><td>BT20/MB231</td><td>CYTO</td></tr>
<tr><td>>hg38_21_+_8433172_8446622_RNA45SN1_45S_50ntFlanks@11571.11597.27|GCCGACTTAGAACTGGTGCGGACCAGG</td><td>>hg38_21_+_8433172_8446622_RNA45SN1_45S_50ntFlanks</td><td>11571</td><td>11597</td><td>27</td><td>GCCGACTTAGAACTGGTGCGGACCAGG</td><td>118.0</td><td>2.6</td><td>0.9</td><td>3.0</td><td>2.4E-03</td><td>5.3E-03</td><td>BT20/MB468</td><td>MP</td></tr>
<tr><td>>hg38_21_+_8433172_8446622_RNA45SN1_45S_50ntFlanks@6650.6672.23|ACGACTCTTAGCGGTGGATCACT</td><td>>hg38_21_+_8433172_8446622_RNA45SN1_45S_50ntFlanks</td><td>6650</td><td>6672</td><td>23</td><td>ACGACTCTTAGCGGTGGATCACT</td><td>268.2</td><td>5.3</td><td>1.7</td><td>3.0</td><td>2.4E-03</td><td>5.3E-03</td><td>BT20/MB468</td><td>MP</td></tr>
<tr><td>>hg38_21_+_8433172_8446622_RNA45SN1_45S_50ntFlanks@12243.12266.24|GAAGGGCAAAAGCTCGCTTGATCT</td><td>>hg38_21_+_8433172_8446622_RNA45SN1_45S_50ntFlanks</td><td>12243</td><td>12266</td><td>24</td><td>GAAGGGCAAAAGCTCGCTTGATCT</td><td>147.2</td><td>-4.6</td><td>1.6</td><td>-2.9</td><td>3.9E-03</td><td>5.3E-03</td><td>MB231/MB468</td><td>MP</td></tr>
<tr><td>>trnaMT_HisGTG_MT_+_12138_12206@32.65.34|TGAATCTGACAACAGAGGCTTACGACCCCTTATT</td><td>>trnaMT_HisGTG_MT_+_12138_12206</td><td>32</td><td>65</td><td>34</td><td>TGAATCTGACAACAGAGGCTTACGACCCCTTATT</td><td>428.8</td><td>2.6</td><td>0.9</td><td>2.8</td><td>4.6E-03</td><td>5.3E-03</td><td>MB231/MB468</td><td>MITO</td></tr>
<tr><td>>MI0003780|hsa-mir-1296&WithFlank&10|-|63372951|63373054@22.42.21&[MIMAT0005794&hsa-miR-1296-5p&offsets|0|-1;m-304&10|-|63373012|63373033&offsets|0|-1]|TTAGGGCCCTGGCTCCATCTC</td><td>>MI0003780|hsa-mir-1296&WithFlank&10|-|63372951|63373054</td><td>22</td><td>42</td><td>21</td><td>TTAGGGCCCTGGCTCCATCTC</td><td>158.2</td><td>5.5</td><td>1.8</td><td>3.0</td><td>2.4E-03</td><td>5.3E-03</td><td>BT20/MB468</td><td>MP</td></tr>
<tr><td>>MI0000805|hsa-mir-342&WithFlank&14|+|100109649|100109759@67.89.23&[MIMAT0000753&hsa-miR-342-3p&offsets|0|0;m-128&14|+|100109715|100109737&offsets|0|0]|TCTCACACAGAAATCGCACCCGT</td><td>>MI0000805|hsa-mir-342&WithFlank&14|+|100109649|100109759</td><td>67</td><td>89</td><td>23</td><td>TCTCACACAGAAATCGCACCCGT</td><td>253.2</td><td>2.8</td><td>0.8</td><td>3.3</td><td>1.0E-03</td><td>5.3E-03</td><td>BT20/MB231</td><td>CYTO</td></tr>
<tr><td>>MI0000809|hsa-mir-151a&WithFlank&8|-|140732558|140732659@53.74.22(+2A)&[MIMAT0000757&hsa-miR-151a-3p&offsets|0|+1(+2A);m-30&8|-|140732587|140732607&offsets|0|+1(+2A)]|CTAGACTGAAGCTCCTTGAGGAAA</td><td>>MI0000809|hsa-mir-151a&WithFlank&8|-|140732558|140732659</td><td>53</td><td>74</td><td>22(+2A)</td><td>CTAGACTGAAGCTCCTTGAGGAAA</td><td>272.4</td><td>3.6</td><td>1.2</td><td>2.9</td><td>3.9E-03</td><td>5.3E-03</td><td>MB231/MB468</td><td>MP</td></tr>
<tr><td>>MI0005756|hsa-mir-934&WithFlank&X|+|136550872|136550966@21.43.23(+1U)&[MIMAT0004977&hsa-miR-934&offsets|0|+1(+1U);m-943&X|+|136550892|136550913&offsets|0|+1(+1U)]|TGTCTACTACTGGAGACACTGGTT</td><td>>MI0005756|hsa-mir-934&WithFlank&X|+|136550872|136550966</td><td>21</td><td>43</td><td>23(+1U)</td><td>TGTCTACTACTGGAGACACTGGTT</td><td>60.5</td><td>9.3</td><td>2.8</td><td>3.3</td><td>8.5E-04</td><td>5.3E-03</td><td>BT20/MB231</td><td>MP</td></tr>
<tr><td>>trna152_ValCAC_6_-_27248049_27248121@1.23.23|GCTTCTGTAGTGTAGTGGTTATC</td><td>>trna152_ValCAC_6_-_27248049_27248121</td><td>1</td><td>23</td><td>23</td><td>GCTTCTGTAGTGTAGTGGTTATC</td><td>203.0</td><td>-2.1</td><td>0.6</td><td>-3.3</td><td>9.3E-04</td><td>5.3E-03</td><td>BT20/MB231</td><td>NUC</td></tr>
<tr><td>>trnaMT_GluTTC_MT_-_14674_14742@46.71.26,trnalookalike8_GluTTC_5_-_93905172_93905240@46.71.26|GTGGTTGTAGTCCGTGCGAGAATACC</td><td>>trnaMT_GluTTC_MT_-_14674_14742</td><td>46</td><td>71</td><td>26</td><td>GTGGTTGTAGTCCGTGCGAGAATACC</td><td>233.9</td><td>-2.8</td><td>0.9</td><td>-3.3</td><td>1.0E-03</td><td>5.3E-03</td><td>BT20/MB231</td><td>CYTO</td></tr>
<tr><td>>trnaMT_GluTTC_MT_-_14674_14742@45.66.22,trnalookalike8_GluTTC_5_-_93905172_93905240@45.66.22|CGTGGTTGTAGTCCGTGCGAGA</td><td>>trnaMT_GluTTC_MT_-_14674_14742</td><td>45</td><td>66</td><td>22</td><td>CGTGGTTGTAGTCCGTGCGAGA</td><td>80.1</td><td>5.2</td><td>1.7</td><td>3.0</td><td>2.4E-03</td><td>5.3E-03</td><td>BT20/MB468</td><td>MP</td></tr>
<tr><td>>am_MI0003581|hsa-mir-574&WithFlank&4|+|38868026|38868133@31.50.20(+1C)&[MIMAT0004795&hsa-miR-574-5p&offsets|0|-3(+1C);m-213&4|+|38868056|38868078&offsets|0|-3(+1C)]|TGAGTGTGTGTGTGTGAGTGC</td><td>>am_MI0003581|hsa-mir-574&WithFlank&4|+|38868026|38868133</td><td>31</td><td>50</td><td>20(+1C)</td><td>TGAGTGTGTGTGTGTGAGTGC</td><td>60.6</td><td>4.5</td><td>1.6</td><td>2.9</td><td>3.9E-03</td><td>5.3E-03</td><td>MB231/MB468</td><td>MP</td></tr>
<tr><td>>hg38_21_+_8433172_8446622_RNA45SN1_45S_50ntFlanks@12407.12438.32|TTTTGATCCTTCGATGTCGGCTCTTCCTATCA</td><td>>hg38_21_+_8433172_8446622_RNA45SN1_45S_50ntFlanks</td><td>12407</td><td>12438</td><td>32</td><td>TTTTGATCCTTCGATGTCGGCTCTTCCTATCA</td><td>519.5</td><td>-2.3</td><td>0.8</td><td>-2.8</td><td>4.6E-03</td><td>5.3E-03</td><td>MB231/MB468</td><td>MITO</td></tr>
<tr><td>>hg38_21_+_8433172_8446622_RNA45SN1_45S_50ntFlanks@12644.12661.18|TCTGTGGGATTATGACTG</td><td>>hg38_21_+_8433172_8446622_RNA45SN1_45S_50ntFlanks</td><td>12644</td><td>12661</td><td>18</td><td>TCTGTGGGATTATGACTG</td><td>274.8</td><td>-3.4</td><td>1.2</td><td>-2.8</td><td>4.6E-03</td><td>5.3E-03</td><td>MB231/MB468</td><td>NUC</td></tr>
<tr><td>>MI0001445|hsa-mir-423&WithFlank&17|+|30117073|30117178@58.79.22&[MIMAT0001340&hsa-miR-423-3p&offsets|-1|-2;m-100&17|+|30117131|30117153&offsets|-1|-2]|AAGCTCGGTCTGAGGCCCCTCA</td><td>>MI0001445|hsa-mir-423&WithFlank&17|+|30117073|30117178</td><td>58</td><td>79</td><td>22</td><td>AAGCTCGGTCTGAGGCCCCTCA</td><td>129.5</td><td>3.9</td><td>1.4</td><td>2.8</td><td>4.6E-03</td><td>5.3E-03</td><td>MB231/MB468</td><td>MITO</td></tr>
<tr><td>>hg38_21_+_8433172_8446622_RNA45SN1_45S_50ntFlanks@8238.8261.24|CCCGGGTCTTCCCGGAGTCGGGTT</td><td>>hg38_21_+_8433172_8446622_RNA45SN1_45S_50ntFlanks</td><td>8238</td><td>8261</td><td>24</td><td>CCCGGGTCTTCCCGGAGTCGGGTT</td><td>133.8</td><td>-3.9</td><td>1.3</td><td>-3.0</td><td>2.4E-03</td><td>5.3E-03</td><td>BT20/MB468</td><td>MP</td></tr>
<tr><td>>hg38_21_+_8433172_8446622_RNA45SN1_45S_50ntFlanks@9489.9515.27|AGGACCCGAAAGATGGTGAACTATGCC</td><td>>hg38_21_+_8433172_8446622_RNA45SN1_45S_50ntFlanks</td><td>9489</td><td>9515</td><td>27</td><td>AGGACCCGAAAGATGGTGAACTATGCC</td><td>67.0</td><td>5.0</td><td>1.7</td><td>3.0</td><td>2.4E-03</td><td>5.3E-03</td><td>BT20/MB468</td><td>MP</td></tr>
<tr><td>>hg38_21_+_8433172_8446622_RNA45SN1_45S_50ntFlanks@9454.9493.40|CCGCGCCGGGGAGGTGGAGCACGAGCGCACGTGTTAGGAC</td><td>>hg38_21_+_8433172_8446622_RNA45SN1_45S_50ntFlanks</td><td>9454</td><td>9493</td><td>40</td><td>CCGCGCCGGGGAGGTGGAGCACGAGCGCACGTGTTAGGAC</td><td>343.5</td><td>-5.2</td><td>1.7</td><td>-3.0</td><td>2.4E-03</td><td>5.3E-03</td><td>BT20/MB468</td><td>MP</td></tr>
<tr><td>>hg38_21_+_8433172_8446622_RNA45SN1_45S_50ntFlanks@11208.11225.18|GCGGCGACTCTGGACGCG</td><td>>hg38_21_+_8433172_8446622_RNA45SN1_45S_50ntFlanks</td><td>11208</td><td>11225</td><td>18</td><td>GCGGCGACTCTGGACGCG</td><td>139.6</td><td>6.4</td><td>1.9</td><td>3.3</td><td>8.5E-04</td><td>5.3E-03</td><td>BT20/MB231</td><td>MP</td></tr>
</table>

<table>
<tr><td>>hg38_21_+_8433172_8446622_RNA45SN1_45S_50ntFlanks@4596.4616.21|TTGGTTTTCGGAACTGAGGCC</td><td>>hg38_21_+_8433172_8446622_RNA45SN1_45S_50ntFlanks</td><td>4596</td><td>4616</td><td>21</td><td>TTGGTTTTCGGAACTGAGGCC</td><td>96.7</td><td>-2.6</td><td>0.8</td><td>-3.0</td><td>2.4E-03</td><td>5.3E-03</td><td>BT20/MB468</td><td>MP</td></tr>
<tr><td>>hg38_21_+_8433172_8446622_RNA45SN1_45S_50ntFlanks@9623.9650.28|TAGTAGCTGGTTCCCTCCGAAGTTTCCC</td><td>>hg38_21_+_8433172_8446622_RNA45SN1_45S_50ntFlanks</td><td>9623</td><td>9650</td><td>28</td><td>TAGTAGCTGGTTCCCTCCGAAGTTTCCC</td><td>241.6</td><td>-5.0</td><td>1.6</td><td>-3.0</td><td>2.4E-03</td><td>5.3E-03</td><td>BT20/MB468</td><td>MP</td></tr>
<tr><td>>MI0000088|hsa-mir-30a&WithFlank&6|-|71403545|71403627@53.70.18(+1U)&[MIMAT0000088&hsa-miR-30a-3p&offsets|0|-4(+1U);m-38&6|-|71403554|71403575&offsets|0|-4(+1U)]|CTTTCAGTCGGATGTTTGT</td><td>>MI0000088|hsa-mir-30a&WithFlank&6|-|71403545|71403627</td><td>53</td><td>70</td><td>18(+1U)</td><td>CTTTCAGTCGGATGTTTGT</td><td>399.2</td><td>-2.4</td><td>0.7</td><td>-3.3</td><td>1.2E-03</td><td>5.3E-03</td><td>BT20/MB231</td><td>MITO</td></tr>
<tr><td>>hg38_21_+_8433172_8446622_RNA45SN1_45S_50ntFlanks@6776.6807.32|CTCCCGGGGCTACGCCTGTCTGAGCGTCGCTT</td><td>>hg38_21_+_8433172_8446622_RNA45SN1_45S_50ntFlanks</td><td>6776</td><td>6807</td><td>32</td><td>CTCCCGGGGCTACGCCTGTCTGAGCGTCGCTT</td><td>1233.6</td><td>-2.2</td><td>0.8</td><td>-2.8</td><td>4.6E-03</td><td>5.3E-03</td><td>MB231/MB468</td><td>MITO</td></tr>
<tr><td>>hg38_21_+_8433172_8446622_RNA45SN1_45S_50ntFlanks@6779.6807.29|CCGGGGCTACGCCTGTCTGAGCGTCGCTT</td><td>>hg38_21_+_8433172_8446622_RNA45SN1_45S_50ntFlanks</td><td>6779</td><td>6807</td><td>29</td><td>CCGGGGCTACGCCTGTCTGAGCGTCGCTT</td><td>586.2</td><td>2.7</td><td>0.9</td><td>2.9</td><td>3.6E-03</td><td>5.3E-03</td><td>BT20/MB468</td><td>MITO</td></tr>
<tr><td>>hg38_21_+_8433172_8446622_RNA45SN1_45S_50ntFlanks@4043.4063.21|ACCCATTCGAACGTCTGCCCT</td><td>>hg38_21_+_8433172_8446622_RNA45SN1_45S_50ntFlanks</td><td>4043</td><td>4063</td><td>21</td><td>ACCCATTCGAACGTCTGCCCT</td><td>215.9</td><td>-4.3</td><td>1.5</td><td>-2.9</td><td>4.3E-03</td><td>5.3E-03</td><td>BT20/MB468</td><td>CYTO</td></tr>
<tr><td>>hg38_21_+_8433172_8446622_RNA45SN1_45S_50ntFlanks@12893.12935.43|CCTCGCCCGTCACGCACCGCACGTTCGTGGGGAACCTGGCGCT</td><td>>hg38_21_+_8433172_8446622_RNA45SN1_45S_50ntFlanks</td><td>12893</td><td>12935</td><td>43</td><td>CCTCGCCCGTCACGCACCGCACGTTCGTGGGGAACCTGGCGCT</td><td>117.2</td><td>-5.6</td><td>1.8</td><td>-3.0</td><td>2.4E-03</td><td>5.3E-03</td><td>BT20/MB468</td><td>MP</td></tr>
<tr><td>>MI0000299|hsa-mir-222&WithFlank&X|-|45747009|45747130@75.98.24&[MIMAT0000279&hsa-miR-222-3p&offsets|0|+3;m-140&X|-|45747033|45747056&offsets|0|0]|AGCTACATCTGGCTACTGGGTCTC</td><td>>MI0000299|hsa-mir-222&WithFlank&X|-|45747009|45747130</td><td>75</td><td>98</td><td>24</td><td>AGCTACATCTGGCTACTGGGTCTC</td><td>16102.6</td><td>-2.1</td><td>0.6</td><td>-3.3</td><td>9.4E-04</td><td>5.3E-03</td><td>BT20/MB231</td><td>NUC</td></tr>
<tr><td>>trnaMT_PheGAA_MT_+_577_647@35.57.23|GAAAATGTTTAGACGGGCTCACA</td><td>>trnaMT_PheGAA_MT_+_577_647</td><td>35</td><td>57</td><td>23</td><td>GAAAATGTTTAGACGGGCTCACA</td><td>2160.9</td><td>3.4</td><td>1.0</td><td>3.3</td><td>9.4E-04</td><td>5.3E-03</td><td>BT20/MB231</td><td>NUC</td></tr>
<tr><td>>hg38_21_+_8433172_8446622_RNA45SN1_45S_50ntFlanks@10687.10708.22|CGGCGTCCGGTGAGCTCTCGCT</td><td>>hg38_21_+_8433172_8446622_RNA45SN1_45S_50ntFlanks</td><td>10687</td><td>10708</td><td>22</td><td>CGGCGTCCGGTGAGCTCTCGCT</td><td>590.6</td><td>2.9</td><td>0.9</td><td>3.2</td><td>1.2E-03</td><td>5.3E-03</td><td>BT20/MB231</td><td>MITO</td></tr>
<tr><td>>MI0000298|hsa-mir-221&WithFlank&X|-|45746151|45746272@71.90.20(+1C)&[MIMAT0000278&hsa-miR-221-3p&offsets|0|-3(+1C);m-91&X|-|45746181|45746202&offsets|0|-2(+1C)]|AGCTACATTGTCTGCTGGGTC</td><td>>MI0000298|hsa-mir-221&WithFlank&X|-|45746151|45746272</td><td>71</td><td>90</td><td>20(+1C)</td><td>AGCTACATTGTCTGCTGGGTC</td><td>420.6</td><td>-2.7</td><td>0.8</td><td>-3.2</td><td>1.2E-03</td><td>5.3E-03</td><td>BT20/MB231</td><td>MITO</td></tr>
<tr><td>>MI0000299|hsa-mir-222&WithFlank&X|-|45747009|45747130@75.99.25(+1A)&[MIMAT0000279&hsa-miR-222-3p&offsets|0|+4(+1A);m-140&X|-|45747033|45747056&offsets|0|+1(+1A)]|AGCTACATCTGGCTACTGGGTCTCTA</td><td>>MI0000299|hsa-mir-222&WithFlank&X|-|45747009|45747130</td><td>75</td><td>99</td><td>25(+1A)</td><td>AGCTACATCTGGCTACTGGGTCTCTA</td><td>377.0</td><td>-2.6</td><td>0.8</td><td>-3.2</td><td>1.2E-03</td><td>5.3E-03</td><td>BT20/MB231</td><td>MITO</td></tr>
<tr><td>>hg38_21_+_8433172_8446622_RNA45SN1_45S_50ntFlanks@7978.8015.38|GACCTCAGATCAGACGTGGCGACCCGCTGAATTTAAGC</td><td>>hg38_21_+_8433172_8446622_RNA45SN1_45S_50ntFlanks</td><td>7978</td><td>8015</td><td>38</td><td>GACCTCAGATCAGACGTGGCGACCCGCTGAATTTAAGC</td><td>118.2</td><td>4.5</td><td>1.6</td><td>2.8</td><td>4.6E-03</td><td>5.3E-03</td><td>MB231/MB468</td><td>MITO</td></tr>
<tr><td>>hg38_21_+_8433172_8446622_RNA45SN1_45S_50ntFlanks@3885.3915.31|ACGGGCGCTGACCCCCTTCGCGGGGGGGATG</td><td>>hg38_21_+_8433172_8446622_RNA45SN1_45S_50ntFlanks</td><td>3885</td><td>3915</td><td>31</td><td>ACGGGCGCTGACCCCCTTCGCGGGGGGGATG</td><td>696.2</td><td>-2.5</td><td>0.9</td><td>-2.9</td><td>4.3E-03</td><td>5.3E-03</td><td>MB231/MB468</td><td>CYTO</td></tr>
<tr><td>>hg38_21_+_8433172_8446622_RNA45SN1_45S_50ntFlanks@10351.10370.20|CGAGAACTTTGAAGGCCGAA</td><td>>hg38_21_+_8433172_8446622_RNA45SN1_45S_50ntFlanks</td><td>10351</td><td>10370</td><td>20</td><td>CGAGAACTTTGAAGGCCGAA</td><td>1114.8</td><td>-3.1</td><td>1.0</td><td>-3.0</td><td>2.4E-03</td><td>5.3E-03</td><td>BT20/MB468</td><td>MP</td></tr>
<tr><td>>hg38_21_+_8433172_8446622_RNA45SN1_45S_50ntFlanks@6646.6688.43|TCGTACGACTCTTAGCGGTGGATCACTCGGCTCGTGCGTCGAT</td><td>>hg38_21_+_8433172_8446622_RNA45SN1_45S_50ntFlanks</td><td>6646</td><td>6688</td><td>43</td><td>TCGTACGACTCTTAGCGGTGGATCACTCGGCTCGTGCGTCGAT</td><td>164.5</td><td>2.3</td><td>0.8</td><td>2.8</td><td>4.6E-03</td><td>5.3E-03</td><td>BT20/MB468</td><td>NUC</td></tr>
<tr><td>>hg38_21_+_8433172_8446622_RNA45SN1_45S_50ntFlanks@5472.5514.43|ACGGCCCTGGCGGAGCGCTGAGAAGACGGTCGAACTTGACTAT</td><td>>hg38_21_+_8433172_8446622_RNA45SN1_45S_50ntFlanks</td><td>5472</td><td>5514</td><td>43</td><td>ACGGCCCTGGCGGAGCGCTGAGAAGACGGTCGAACTTGACTAT</td><td>7025.0</td><td>2.4</td><td>0.8</td><td>2.9</td><td>4.3E-03</td><td>5.3E-03</td><td>MB231/MB468</td><td>CYTO</td></tr>
<tr><td>>hg38_21_+_8433172_8446622_RNA45SN1_45S_50ntFlanks@3853.3883.31|AACTGTGGTAATTCTAGAGCTAATACATGCC</td><td>>hg38_21_+_8433172_8446622_RNA45SN1_45S_50ntFlanks</td><td>3853</td><td>3883</td><td>31</td><td>AACTGTGGTAATTCTAGAGCTAATACATGCC</td><td>360.0</td><td>-4.0</td><td>1.4</td><td>-2.8</td><td>4.6E-03</td><td>5.3E-03</td><td>BT20/MB468</td><td>NUC</td></tr>
<tr><td>>trnaMT_GluTTC_MT_-_14674_14742@27.61.35,trnalookalike8_GluTTC_5_-_93905172_93905240@27.61.35|GTTTTTCATATCATTGGTCGTGGTTGTAGTCCGTG</td><td>>trnaMT_GluTTC_MT_-_14674_14742</td><td>27</td><td>61</td><td>35</td><td>GTTTTTCATATCATTGGTCGTGGTTGTAGTCCGTG</td><td>129.7</td><td>-4.8</td><td>1.7</td><td>-2.9</td><td>4.3E-03</td><td>5.3E-03</td><td>BT20/MB468</td><td>CYTO</td></tr>
<tr><td>>hg38_21_+_8433172_8446622_RNA45SN1_45S_50ntFlanks@10466.10487.22|CGTTGCCCTCGGCCGATCGAAA</td><td>>hg38_21_+_8433172_8446622_RNA45SN1_45S_50ntFlanks</td><td>10466</td><td>10487</td><td>22</td><td>CGTTGCCCTCGGCCGATCGAAA</td><td>260.1</td><td>-2.1</td><td>0.7</td><td>-2.9</td><td>4.4E-03</td><td>5.3E-03</td><td>MB231/MB468</td><td>CYTO</td></tr>
<tr><td>>hg38_MT_+_598_1651_RNR1_12S_50ntFlanks@52.77.26|ATAGGTTTGGTCCTAGCCTTTCTATT</td><td>>hg38_MT_+_598_1651_RNR1_12S_50ntFlanks</td><td>52</td><td>77</td><td>26</td><td>ATAGGTTTGGTCCTAGCCTTTCTATT</td><td>2923.6</td><td>2.9</td><td>1.0</td><td>2.8</td><td>4.6E-03</td><td>5.3E-03</td><td>MB231/MB468</td><td>MITO</td></tr>
<tr><td>>hg38_21_+_8433172_8446622_RNA45SN1_45S_50ntFlanks@10393.10426.34|CAGCAGTTGAACATGGGTCAGTCGGTCCTGAGAG</td><td>>hg38_21_+_8433172_8446622_RNA45SN1_45S_50ntFlanks</td><td>10393</td><td>10426</td><td>34</td><td>CAGCAGTTGAACATGGGTCAGTCGGTCCTGAGAG</td><td>265.7</td><td>-3.5</td><td>1.2</td><td>-2.9</td><td>4.3E-03</td><td>5.3E-03</td><td>BT20/MB468</td><td>CYTO</td></tr>
<tr><td>>hg38_21_+_8433172_8446622_RNA45SN1_45S_50ntFlanks@3835.3859.25|CTCCTCTCCTACTTGGATAACTGTG</td><td>>hg38_21_+_8433172_8446622_RNA45SN1_45S_50ntFlanks</td><td>3835</td><td>3859</td><td>25</td><td>CTCCTCTCCTACTTGGATAACTGTG</td><td>65.9</td><td>-4.9</td><td>1.6</td><td>-3.0</td><td>2.4E-03</td><td>5.3E-03</td><td>BT20/MB468</td><td>MP</td></tr>
<tr><td>>hg38_21_+_8433172_8446622_RNA45SN1_45S_50ntFlanks@11873.11910.38|AAGAAGACCCTGTTGAGCTTGACTCTAGTCTGGCACGG</td><td>>hg38_21_+_8433172_8446622_RNA45SN1_45S_50ntFlanks</td><td>11873</td><td>11910</td><td>38</td><td>AAGAAGACCCTGTTGAGCTTGACTCTAGTCTGGCACGG</td><td>155.2</td><td>-5.1</td><td>1.7</td><td>-3.0</td><td>2.4E-03</td><td>5.3E-03</td><td>BT20/MB468</td><td>MP</td></tr>
<tr><td>>am_trnaMT_MetCAT_MT_+_4402_4469@1.22.22|AGTAAGGTCAGCTAAATAAGCT</td><td>>am_trnaMT_MetCAT_MT_+_4402_4469</td><td>1</td><td>22</td><td>22</td><td>AGTAAGGTCAGCTAAATAAGCT</td><td>924.0</td><td>3.3</td><td>1.2</td><td>2.8</td><td>4.6E-03</td><td>5.3E-03</td><td>MB231/MB468</td><td>MITO</td></tr>
<tr><td>>hg38_21_+_8433172_8446622_RNA45SN1_45S_50ntFlanks@12892.12930.39|CCCTCGCCCGTCACGCACCGCACGTTCGTGGGGAACCTG</td><td>>hg38_21_+_8433172_8446622_RNA45SN1_45S_50ntFlanks</td><td>12892</td><td>12930</td><td>39</td><td>CCCTCGCCCGTCACGCACCGCACGTTCGTGGGGAACCTG</td><td>590.0</td><td>-4.6</td><td>1.6</td><td>-2.9</td><td>3.7E-03</td><td>5.4E-03</td><td>BT20/MB468</td><td>MITO</td></tr>
<tr><td>>hg38_21_+_8433172_8446622_RNA45SN1_45S_50ntFlanks@10711.10736.26|CCCTTGAAAATCCGGGGGAGAGGGTG</td><td>>hg38_21_+_8433172_8446622_RNA45SN1_45S_50ntFlanks</td><td>10711</td><td>10736</td><td>26</td><td>CCCTTGAAAATCCGGGGGAGAGGGTG</td><td>146.9</td><td>-4.2</td><td>1.5</td><td>-2.9</td><td>4.3E-03</td><td>5.4E-03</td><td>BT20/MB468</td><td>CYTO</td></tr>
<tr><td>>MI0000255|hsa-mir-30d&WithFlank&8|-|134804870|134804951@12.33.22&[MIMAT0000245&hsa-miR-30d-5p&offsets|0|0;m-21&8|-|134804919|134804940&offsets|0|0]|TGTAAACATCCCCGACTGGAAG</td><td>>MI0000255|hsa-mir-30d&WithFlank&8|-|134804870|134804951</td><td>12</td><td>33</td><td>22</td><td>TGTAAACATCCCCGACTGGAAG</td><td>5120.1</td><td>3.0</td><td>0.9</td><td>3.3</td><td>1.1E-03</td><td>5.4E-03</td><td>BT20/MB231</td><td>CYTO</td></tr>
<tr><td>>hg38_21_+_8433172_8446622_RNA45SN1_45S_50ntFlanks@9936.9955.20|ACAGCAGGACGGTGGCCATG</td><td>>hg38_21_+_8433172_8446622_RNA45SN1_45S_50ntFlanks</td><td>9936</td><td>9955</td><td>20</td><td>ACAGCAGGACGGTGGCCATG</td><td>143.9</td><td>4.0</td><td>1.3</td><td>3.0</td><td>2.4E-03</td><td>5.4E-03</td><td>BT20/MB468</td><td>MP</td></tr>
<tr><td>>MI0000067|hsa-let-7f-1&WithFlank&9|+|94176341|94176439@13.35.23&[MIMAT0000067&hsa-let-7f-5p&offsets|0|+1;m-20&9|+|94176353|94176374&offsets|0|+1]|TGAGGTAGTAGATTGTATAGTTG</td><td>>MI0000067|hsa-let-7f-1&WithFlank&9|+|94176341|94176439</td><td>13</td><td>35</td><td>23</td><td>TGAGGTAGTAGATTGTATAGTTG</td><td>436.6</td><td>2.6</td><td>0.8</td><td>3.3</td><td>1.1E-03</td><td>5.4E-03</td><td>BT20/MB231</td><td>CYTO</td></tr>
<tr><td>>hg38_21_+_8433172_8446622_RNA45SN1_45S_50ntFlanks@12407.12448.42|TTTTGATCCTTCGATGTCGGCTCTTCCTATCATTGTGAAGCA</td><td>>hg38_21_+_8433172_8446622_RNA45SN1_45S_50ntFlanks</td><td>12407</td><td>12448</td><td>42</td><td>TTTTGATCCTTCGATGTCGGCTCTTCCTATCATTGTGAAGCA</td><td>711.4</td><td>-3.5</td><td>1.1</td><td>-3.3</td><td>1.1E-03</td><td>5.4E-03</td><td>BT20/MB231</td><td>CYTO</td></tr>
<tr><td>>am_trnaMT_LysTTT_MT_+_8295_8364@29.52.24,trnalookalike7_LysTTT_1_+_568844_568913@29.52.24|TTTAAGTTAAAGATTAAGAGAACC</td><td>>am_trnaMT_LysTTT_MT_+_8295_8364</td><td>29</td><td>52</td><td>24</td><td>TTTAAGTTAAAGATTAAGAGAACC</td><td>489.5</td><td>2.9</td><td>0.9</td><td>3.3</td><td>1.1E-03</td><td>5.4E-03</td><td>BT20/MB231</td><td>CYTO</td></tr>
<tr><td>>hg38_21_+_8433172_8446622_RNA45SN1_45S_50ntFlanks@12605.12623.19|TATGTGCTTGGCTGAGGAG</td><td>>hg38_21_+_8433172_8446622_RNA45SN1_45S_50ntFlanks</td><td>12605</td><td>12623</td><td>19</td><td>TATGTGCTTGGCTGAGGAG</td><td>122.1</td><td>-3.0</td><td>1.0</td><td>-2.9</td><td>3.7E-03</td><td>5.4E-03</td><td>BT20/MB468</td><td>MITO</td></tr>
<tr><td>>MI0000298|hsa-mir-221&WithFlank&X|-|45746151|45746272@31.53.23(+1C)&[MIMAT0004568&hsa-miR-221-5p&offsets|0|+1(+1C);m-405&X|-|45746220|45746242&offsets|0|0(+1C)]|ACCTGGCATACAATGTAGATTTCC</td><td>>MI0000298|hsa-mir-221&WithFlank&X|-|45746151|45746272</td><td>31</td><td>53</td><td>23(+1C)</td><td>ACCTGGCATACAATGTAGATTTCC</td><td>46.2</td><td>5.4</td><td>1.9</td><td>2.9</td><td>3.9E-03</td><td>5.4E-03</td><td>MB231/MB468</td><td>MP</td></tr>
<tr><td>>hg38_21_+_8433172_8446622_RNA45SN1_45S_50ntFlanks@9941.9960.20|AGGACGGTGGCCATGGAAGT</td><td>>hg38_21_+_8433172_8446622_RNA45SN1_45S_50ntFlanks</td><td>9941</td><td>9960</td><td>20</td><td>AGGACGGTGGCCATGGAAGT</td><td>482.6</td><td>4.5</td><td>1.3</td><td>3.3</td><td>8.6E-04</td><td>5.4E-03</td><td>BT20/MB231</td><td>MP</td></tr>
<tr><td>>hg38_MT_+_1621_3279_RNR2_16S_50ntFlanks@1346.1385.40|TCAACAATAGGGTTTACGACCTCGATGTTGGATCAGGACA</td><td>>hg38_MT_+_1621_3279_RNR2_16S_50ntFlanks</td><td>1346</td><td>1385</td><td>40</td><td>TCAACAATAGGGTTTACGACCTCGATGTTGGATCAGGACA</td><td>41.6</td><td>5.3</td><td>1.8</td><td>2.9</td><td>3.9E-03</td><td>5.4E-03</td><td>MB231/MB468</td><td>MP</td></tr>
<tr><td>>hg38_MT_+_1621_3279_RNR2_16S_50ntFlanks@760.777.18|CAATATCTACAATCAACC</td><td>>hg38_MT_+_1621_3279_RNR2_16S_50ntFlanks</td><td>760</td><td>777</td><td>18</td><td>CAATATCTACAATCAACC</td><td>97.8</td><td>5.3</td><td>1.8</td><td>2.9</td><td>3.9E-03</td><td>5.4E-03</td><td>MB231/MB468</td><td>MP</td></tr>
<tr><td>>am_MI0000748|hsa-mir-130b&WithFlank&22|+|21653298|21653391@19.39.21&[MIMAT0004680&hsa-miR-130b-5p&offsets|0|0;m-234&22|+|21653316|21653337&offsets|0|-1]|ACTCTTTCCCTGTTGCACTAC</td><td>>am_MI0000748|hsa-mir-130b&WithFlank&22|+|21653298|21653391</td><td>19</td><td>39</td><td>21</td><td>ACTCTTTCCCTGTTGCACTAC</td><td>441.7</td><td>-2.2</td><td>0.7</td><td>-3.2</td><td>1.2E-03</td><td>5.4E-03</td><td>BT20/MB231</td><td>MITO</td></tr>
<tr><td>>hg38_21_+_8433172_8446622_RNA45SN1_45S_50ntFlanks@6646.6668.23|TCGTACGACTCTTAGCGGTGGAT</td><td>>hg38_21_+_8433172_8446622_RNA45SN1_45S_50ntFlanks</td><td>6646</td><td>6668</td><td>23</td><td>TCGTACGACTCTTAGCGGTGGAT</td><td>3601.9</td><td>1.7</td><td>0.6</td><td>2.8</td><td>4.6E-03</td><td>5.4E-03</td><td>BT20/MB468</td><td>NUC</td></tr>
<tr><td>>hg38_21_+_8433172_8446622_RNA45SN1_45S_50ntFlanks@6777.6803.27|TCCCGGGGCTACGCCTGTCTGAGCGTC</td><td>>hg38_21_+_8433172_8446622_RNA45SN1_45S_50ntFlanks</td><td>6777</td><td>6803</td><td>27</td><td>TCCCGGGGCTACGCCTGTCTGAGCGTC</td><td>268.4</td><td>-3.5</td><td>1.2</td><td>-2.8</td><td>4.6E-03</td><td>5.4E-03</td><td>MB231/MB468</td><td>MITO</td></tr>
<tr><td>>hg38_21_+_8433172_8446622_RNA45SN1_45S_50ntFlanks@3738.3760.23|TCTCAAAGATTAAGCCATGCATG</td><td>>hg38_21_+_8433172_8446622_RNA45SN1_45S_50ntFlanks</td><td>3738</td><td>3760</td><td>23</td><td>TCTCAAAGATTAAGCCATGCATG</td><td>173.0</td><td>-4.9</td><td>1.6</td><td>-3.0</td><td>2.4E-03</td><td>5.4E-03</td><td>BT20/MB468</td><td>MP</td></tr>
<tr><td>>hg38_21_+_8433172_8446622_RNA45SN1_45S_50ntFlanks@4132.4150.19|TCCGGAGAGGGAGCCTGAG</td><td>>hg38_21_+_8433172_8446622_RNA45SN1_45S_50ntFlanks</td><td>4132</td><td>4150</td><td>19</td><td>TCCGGAGAGGGAGCCTGAG</td><td>117.4</td><td>6.0</td><td>2.0</td><td>3.0</td><td>2.4E-03</td><td>5.4E-03</td><td>BT20/MB468</td><td>MP</td></tr>
<tr><td>>hg38_21_+_8433172_8446622_RNA45SN1_45S_50ntFlanks@6648.6679.32|GTACGACTCTTAGCGGTGGATCACTCGGCTCG</td><td>>hg38_21_+_8433172_8446622_RNA45SN1_45S_50ntFlanks</td><td>6648</td><td>6679</td><td>32</td><td>GTACGACTCTTAGCGGTGGATCACTCGGCTCG</td><td>1657.5</td><td>5.2</td><td>1.6</td><td>3.3</td><td>8.7E-04</td><td>5.4E-03</td><td>BT20/MB231</td><td>MP</td></tr>
<tr><td>>hg38_21_+_8433172_8446622_RNA45SN1_45S_50ntFlanks@8351.8375.25|AAGTTGAAAAGAACTTTGAAGAGAG</td><td>>hg38_21_+_8433172_8446622_RNA45SN1_45S_50ntFlanks</td><td>8351</td><td>8375</td><td>25</td><td>AAGTTGAAAAGAACTTTGAAGAGAG</td><td>7199.8</td><td>4.6</td><td>1.4</td><td>3.3</td><td>8.7E-04</td><td>5.4E-03</td><td>BT20/MB231</td><td>MP</td></tr>
<tr><td>>hg38_21_+_8433172_8446622_RNA45SN1_45S_50ntFlanks@8183.8203.21|TCGAGGCCCAGCCCGTGGACG</td><td>>hg38_21_+_8433172_8446622_RNA45SN1_45S_50ntFlanks</td><td>8183</td><td>8203</td><td>21</td><td>TCGAGGCCCAGCCCGTGGACG</td><td>126.1</td><td>-3.6</td><td>1.2</td><td>-2.9</td><td>4.0E-03</td><td>5.4E-03</td><td>MB231/MB468</td><td>MP</td></tr>
<tr><td>>hg38_21_+_8433172_8446622_RNA45SN1_45S_50ntFlanks@8195.8213.19|CCGTGGACGGTGTGAGGCC</td><td>>hg38_21_+_8433172_8446622_RNA45SN1_45S_50ntFlanks</td><td>8195</td><td>8213</td><td>19</td><td>CCGTGGACGGTGTGAGGCC</td><td>927.6</td><td>2.6</td><td>0.8</td><td>3.0</td><td>2.4E-03</td><td>5.4E-03</td><td>BT20/MB468</td><td>MP</td></tr>
<tr><td>>trna111_HisGTG_1_-_147774845_147774916@-1G.30.31,trna118_HisGTG_1_-_145396881_145396952@-1G.30.31,trna16_HisGTG_1_+_146544773_146544844@-1G.30.31,trna1_HisGTG_15_+_45493349_45493420@-1G.30.31,trna21_HisGTG_1_+_147753471_147753542@-1G.30.31,trna33_HisGTG_6_+_27125906_27125977@-1G.30.31,trna7_HisGTG_9_-_14433938_14434009@-1G.30.31,trna8_HisGTG_15_-_45492611_45492682@-1G.30.31,trna9_HisGTG_15_-_45490804_45490875@-1G.30.31|GGCCGTGATCGTATAGTGGTTAGTACTCTGC</td><td>>trna111_HisGTG_1_-_147774845_147774916</td><td>-1G</td><td>30</td><td>31</td><td>GGCCGTGATCGTATAGTGGTTAGTACTCTGC</td><td>4099.8</td><td>-3.9</td><td>1.2</td><td>-3.3</td><td>8.7E-04</td><td>5.4E-03</td><td>BT20/MB231</td><td>MP</td></tr>
<tr><td>>MI0000086|hsa-mir-28&WithFlank&3|+|188688775|188688872@20.41.22(+1A)&[MIMAT0000085&hsa-miR-28-5p&offsets|0|0(+1A);m-133&3|+|188688794|188688815&offsets|0|0(+1A)]|AAGGAGCTCACAGTCTATTGAGA</td><td>>MI0000086|hsa-mir-28&WithFlank&3|+|188688775|188688872</td><td>20</td><td>41</td><td>22(+1A)</td><td>AAGGAGCTCACAGTCTATTGAGA</td><td>160.5</td><td>4.3</td><td>1.4</td><td>3.0</td><td>2.4E-03</td><td>5.4E-03</td><td>BT20/MB468</td><td>MP</td></tr>
</table>

<table>
<tr><td>>MI0000825|hsa-mir-345&WithFlank&14|+|100307853|100307962@24.44.21(+1A)&[MIMAT0000772&hsa-miR-345-5p&offsets|0|-1(+1A);m-193&14|+|100307876|100307896&offsets|0|0(+1A)]|GCTGACTCCTAGTCCAGGGCTA</td><td>>MI0000825|hsa-mir-345&WithFlank&14|+|100307853|100307962</td><td>24</td><td>44</td><td>21(+1A)</td><td>GCTGACTCCTAGTCCAGGGCTA</td><td>60.3</td><td>4.6</td><td>1.6</td><td>2.9</td><td>4.0E-03</td><td>5.4E-03</td><td>MB231/MB468</td><td>MP</td></tr>
<tr><td>>hg38_MT_+_1621_3279_RNR2_16S_50ntFlanks@825.855.31|TAAGGAAAGGTTAAAAAAAGTAAAAGGAACT</td><td>>hg38_MT_+_1621_3279_RNR2_16S_50ntFlanks</td><td>825</td><td>855</td><td>31</td><td>TAAGGAAAGGTTAAAAAAAGTAAAAGGAACT</td><td>171.5</td><td>4.5</td><td>1.6</td><td>2.9</td><td>4.0E-03</td><td>5.4E-03</td><td>MB231/MB468</td><td>MP</td></tr>
<tr><td>>hg38_MT_+_1621_3279_RNR2_16S_50ntFlanks@1096.1120.25|GACGAGAAGACCCTATGGAGCTTTA</td><td>>hg38_MT_+_1621_3279_RNR2_16S_50ntFlanks</td><td>1096</td><td>1120</td><td>25</td><td>GACGAGAAGACCCTATGGAGCTTTA</td><td>125.0</td><td>3.1</td><td>1.0</td><td>3.0</td><td>2.5E-03</td><td>5.4E-03</td><td>BT20/MB468</td><td>MP</td></tr>
<tr><td>>hg38_21_+_8433172_8446622_RNA45SN1_45S_50ntFlanks@11209.11227.19|CGGCGACTCTGGACGCGAG</td><td>>hg38_21_+_8433172_8446622_RNA45SN1_45S_50ntFlanks</td><td>11209</td><td>11227</td><td>19</td><td>CGGCGACTCTGGACGCGAG</td><td>930.8</td><td>-5.6</td><td>2.0</td><td>-2.8</td><td>4.6E-03</td><td>5.4E-03</td><td>BT20/MB468</td><td>NUC</td></tr>
<tr><td>>hg38_MT_+_598_1651_RNR1_12S_50ntFlanks@428.462.35|AAAGTGGCTTTAACATATCTGAACACACAATAGCT</td><td>>hg38_MT_+_598_1651_RNR1_12S_50ntFlanks</td><td>428</td><td>462</td><td>35</td><td>AAAGTGGCTTTAACATATCTGAACACACAATAGCT</td><td>171.3</td><td>3.6</td><td>1.3</td><td>2.9</td><td>4.0E-03</td><td>5.4E-03</td><td>MB231/MB468</td><td>MP</td></tr>
<tr><td>>hg38_21_+_8433172_8446622_RNA45SN1_45S_50ntFlanks@12490.12521.32|CGTGAGCTGGGTTTAGACCGTCGTGAGACAGG</td><td>>hg38_21_+_8433172_8446622_RNA45SN1_45S_50ntFlanks</td><td>12490</td><td>12521</td><td>32</td><td>CGTGAGCTGGGTTTAGACCGTCGTGAGACAGG</td><td>116.3</td><td>-4.9</td><td>1.6</td><td>-3.0</td><td>2.5E-03</td><td>5.4E-03</td><td>BT20/MB468</td><td>MP</td></tr>
<tr><td>>hg38_21_+_8433172_8446622_RNA45SN1_45S_50ntFlanks@8189.8213.25|CCCAGCCCGTGGACGGTGTGAGGCC</td><td>>hg38_21_+_8433172_8446622_RNA45SN1_45S_50ntFlanks</td><td>8189</td><td>8213</td><td>25</td><td>CCCAGCCCGTGGACGGTGTGAGGCC</td><td>318.9</td><td>-3.5</td><td>1.2</td><td>-2.8</td><td>4.7E-03</td><td>5.4E-03</td><td>MB231/MB468</td><td>MITO</td></tr>
<tr><td>>hg38_21_+_8433172_8446622_RNA45SN1_45S_50ntFlanks@9969.9989.21|GCTAAGGAGTGTGTAACAACT</td><td>>hg38_21_+_8433172_8446622_RNA45SN1_45S_50ntFlanks</td><td>9969</td><td>9989</td><td>21</td><td>GCTAAGGAGTGTGTAACAACT</td><td>115.3</td><td>6.0</td><td>2.0</td><td>3.0</td><td>2.5E-03</td><td>5.4E-03</td><td>BT20/MB468</td><td>MP</td></tr>
<tr><td>>hg38_21_+_8433172_8446622_RNA45SN1_45S_50ntFlanks@11317.11340.24|ACCCCACGTCTCGTCGCGCGCGCG</td><td>>hg38_21_+_8433172_8446622_RNA45SN1_45S_50ntFlanks</td><td>11317</td><td>11340</td><td>24</td><td>ACCCCACGTCTCGTCGCGCGCGCG</td><td>1183.1</td><td>-5.7</td><td>2.0</td><td>-2.8</td><td>4.6E-03</td><td>5.4E-03</td><td>BT20/MB468</td><td>NUC</td></tr>
<tr><td>>hg38_21_+_8433172_8446622_RNA45SN1_45S_50ntFlanks@7975.7998.24|CGCGACCTCAGATCAGACGTGGCG</td><td>>hg38_21_+_8433172_8446622_RNA45SN1_45S_50ntFlanks</td><td>7975</td><td>7998</td><td>24</td><td>CGCGACCTCAGATCAGACGTGGCG</td><td>423.7</td><td>2.6</td><td>0.8</td><td>3.3</td><td>1.1E-03</td><td>5.4E-03</td><td>BT20/MB231</td><td>CYTO</td></tr>
<tr><td>>hg38_21_+_8433172_8446622_RNA45SN1_45S_50ntFlanks@5236.5257.22|CACTGACTGGCTCAGCGTGTGC</td><td>>hg38_21_+_8433172_8446622_RNA45SN1_45S_50ntFlanks</td><td>5236</td><td>5257</td><td>22</td><td>CACTGACTGGCTCAGCGTGTGC</td><td>626.7</td><td>-3.8</td><td>1.3</td><td>-2.9</td><td>3.7E-03</td><td>5.4E-03</td><td>BT20/MB468</td><td>MITO</td></tr>
<tr><td>>trna37_ValAAC_6_+_27203288_27203360@1.29.29|GTTTCCGTAGTGTAGTGGTTATCACGTTT</td><td>>trna37_ValAAC_6_+_27203288_27203360</td><td>1</td><td>29</td><td>29</td><td>GTTTCCGTAGTGTAGTGGTTATCACGTTT</td><td>1195.3</td><td>-5.1</td><td>1.8</td><td>-2.9</td><td>4.3E-03</td><td>5.4E-03</td><td>BT20/MB468</td><td>CYTO</td></tr>
<tr><td>>hg38_21_+_8433172_8446622_RNA45SN1_45S_50ntFlanks@10866.10886.21|CTAAGGGCTGGGTCGGTCGGG</td><td>>hg38_21_+_8433172_8446622_RNA45SN1_45S_50ntFlanks</td><td>10866</td><td>10886</td><td>21</td><td>CTAAGGGCTGGGTCGGTCGGG</td><td>472.3</td><td>-4.5</td><td>1.6</td><td>-2.9</td><td>4.3E-03</td><td>5.4E-03</td><td>BT20/MB468</td><td>CYTO</td></tr>
<tr><td>>trna10_AspGTC_12_-_125424193_125424264@38.72.35,trna12_AspGTC_12_-_125411891_125411962@38.72.35,trna144_AspGTC_6_-_27551236_27551307@38.72.35,trna38_AspGTC_17_-_8125556_8125627@38.72.35,trna45_AspGTC_6_+_27447453_27447524@38.72.35,trna48_AspGTC_6_+_27471523_27471594@38.72.35,trna4_AspGTC_12_+_96429799_96429870@38.72.35,trna69_AspGTC_1_-_161440205_161440276@38.72.35,trna72_AspGTC_1_-_161432824_161432895@38.72.35,trna75_AspGTC_1_-_161425414_161425485@38.72.35,trna78_AspGTC_1_-_161418033_161418104@38.72.35,trna81_AspGTC_1_-_161410615_161410686@38.72.35|CGCGGGAGACCGGGGTTCGATTCCCCGACGGGGAG</td><td>>trna10_AspGTC_12_-_125424193_125424264</td><td>38</td><td>72</td><td>35</td><td>CGCGGGAGACCGGGGTTCGATTCCCCGACGGGGAG</td><td>552.4</td><td>-4.4</td><td>1.4</td><td>-3.0</td><td>2.5E-03</td><td>5.4E-03</td><td>BT20/MB468</td><td>MP</td></tr>
<tr><td>>hg38_21_+_8433172_8446622_RNA45SN1_45S_50ntFlanks@3855.3881.27|CTGTGGTAATTCTAGAGCTAATACATG</td><td>>hg38_21_+_8433172_8446622_RNA45SN1_45S_50ntFlanks</td><td>3855</td><td>3881</td><td>27</td><td>CTGTGGTAATTCTAGAGCTAATACATG</td><td>196.5</td><td>-2.6</td><td>0.9</td><td>-2.9</td><td>4.3E-03</td><td>5.4E-03</td><td>BT20/MB468</td><td>CYTO</td></tr>
<tr><td>>MI0000811|hsa-mir-148b&WithFlank&12|+|54337210|54337320@69.90.22(+3U)&[MIMAT0000759&hsa-miR-148b-3p&offsets|0|0(+3U);m-77&12|+|54337278|54337299&offsets|0|0(+3U)]|TCAGTGCATCACAGAACTTTGTTTT</td><td>>MI0000811|hsa-mir-148b&WithFlank&12|+|54337210|54337320</td><td>69</td><td>90</td><td>22(+3U)</td><td>TCAGTGCATCACAGAACTTTGTTTT</td><td>169.6</td><td>-2.1</td><td>0.6</td><td>-3.3</td><td>9.6E-04</td><td>5.4E-03</td><td>BT20/MB231</td><td>NUC</td></tr>
<tr><td>>trna152_ValCAC_6_-_27248049_27248121@1.33.33|GCTTCTGTAGTGTAGTGGTTATCACGTTCGCCT</td><td>>trna152_ValCAC_6_-_27248049_27248121</td><td>1</td><td>33</td><td>33</td><td>GCTTCTGTAGTGTAGTGGTTATCACGTTCGCCT</td><td>6261.7</td><td>-3.8</td><td>1.2</td><td>-3.3</td><td>1.1E-03</td><td>5.4E-03</td><td>BT20/MB231</td><td>CYTO</td></tr>
<tr><td>>hg38_21_+_8433172_8446622_RNA45SN1_45S_50ntFlanks@12638.12659.22|CTACCATCTGTGGGATTATGAC</td><td>>hg38_21_+_8433172_8446622_RNA45SN1_45S_50ntFlanks</td><td>12638</td><td>12659</td><td>22</td><td>CTACCATCTGTGGGATTATGAC</td><td>258.9</td><td>-3.4</td><td>1.2</td><td>-2.8</td><td>4.4E-03</td><td>5.4E-03</td><td>MB231/MB468</td><td>CYTO</td></tr>
<tr><td>>hg38_21_+_8433172_8446622_RNA45SN1_45S_50ntFlanks@7981.8003.23|CTCAGATCAGACGTGGCGACCCG</td><td>>hg38_21_+_8433172_8446622_RNA45SN1_45S_50ntFlanks</td><td>7981</td><td>8003</td><td>23</td><td>CTCAGATCAGACGTGGCGACCCG</td><td>144.8</td><td>-3.2</td><td>1.1</td><td>-2.9</td><td>4.4E-03</td><td>5.4E-03</td><td>BT20/MB468</td><td>CYTO</td></tr>
<tr><td>>hg38_21_+_8433172_8446622_RNA45SN1_45S_50ntFlanks@3787.3805.19|AAACTGCGAATGGCTCATT</td><td>>hg38_21_+_8433172_8446622_RNA45SN1_45S_50ntFlanks</td><td>3787</td><td>3805</td><td>19</td><td>AAACTGCGAATGGCTCATT</td><td>227.1</td><td>-2.6</td><td>0.9</td><td>-2.9</td><td>4.0E-03</td><td>5.4E-03</td><td>MB231/MB468</td><td>MP</td></tr>
<tr><td>>hg38_21_+_8433172_8446622_RNA45SN1_45S_50ntFlanks@8184.8210.27|CGAGGCCCAGCCCGTGGACGGTGTGAG</td><td>>hg38_21_+_8433172_8446622_RNA45SN1_45S_50ntFlanks</td><td>8184</td><td>8210</td><td>27</td><td>CGAGGCCCAGCCCGTGGACGGTGTGAG</td><td>241.1</td><td>-2.7</td><td>0.9</td><td>-3.0</td><td>2.5E-03</td><td>5.4E-03</td><td>BT20/MB468</td><td>MP</td></tr>
<tr><td>>MI0000088|hsa-mir-30a&WithFlank&6|-|71403545|71403627@13.34.22(+1C)&[MIMAT0000087&hsa-miR-30a-5p&offsets|+1|+1(+1C);m-15&6|-|71403595|71403616&offsets|+1|+1(+1C)]|GTAAACATCCTCGACTGGAAGCC</td><td>>MI0000088|hsa-mir-30a&WithFlank&6|-|71403545|71403627</td><td>13</td><td>34</td><td>22(+1C)</td><td>GTAAACATCCTCGACTGGAAGCC</td><td>382.8</td><td>4.7</td><td>1.5</td><td>3.0</td><td>2.5E-03</td><td>5.4E-03</td><td>BT20/MB468</td><td>MP</td></tr>
<tr><td>>am_trnaMT_ValTAC_MT_+_1602_1670@28.52.25|AACTTACACTTAGGAGATTTCAACT</td><td>>am_trnaMT_ValTAC_MT_+_1602_1670</td><td>28</td><td>52</td><td>25</td><td>AACTTACACTTAGGAGATTTCAACT</td><td>314.6</td><td>2.7</td><td>0.8</td><td>3.3</td><td>1.1E-03</td><td>5.4E-03</td><td>BT20/MB231</td><td>CYTO</td></tr>
<tr><td>>hg38_21_+_8433172_8446622_RNA45SN1_45S_50ntFlanks@12434.12455.22|TATCATTGTGAAGCAGAATTCA</td><td>>hg38_21_+_8433172_8446622_RNA45SN1_45S_50ntFlanks</td><td>12434</td><td>12455</td><td>22</td><td>TATCATTGTGAAGCAGAATTCA</td><td>164.8</td><td>-4.2</td><td>1.5</td><td>-2.9</td><td>3.7E-03</td><td>5.4E-03</td><td>BT20/MB468</td><td>MITO</td></tr>
<tr><td>>hg38_21_+_8433172_8446622_RNA45SN1_45S_50ntFlanks@11665.11702.38|TTCTGCCCAGTGCTCTGAATGTCAAAGTGAAGAAATTC</td><td>>hg38_21_+_8433172_8446622_RNA45SN1_45S_50ntFlanks</td><td>11665</td><td>11702</td><td>38</td><td>TTCTGCCCAGTGCTCTGAATGTCAAAGTGAAGAAATTC</td><td>1742.0</td><td>-2.8</td><td>0.9</td><td>-3.3</td><td>1.1E-03</td><td>5.4E-03</td><td>BT20/MB231</td><td>CYTO</td></tr>
<tr><td>>hg38_21_+_8433172_8446622_RNA45SN1_45S_50ntFlanks@6645.6685.41|CTCGTACGACTCTTAGCGGTGGATCACTCGGCTCGTGCGTC</td><td>>hg38_21_+_8433172_8446622_RNA45SN1_45S_50ntFlanks</td><td>6645</td><td>6685</td><td>41</td><td>CTCGTACGACTCTTAGCGGTGGATCACTCGGCTCGTGCGTC</td><td>530.3</td><td>3.1</td><td>0.9</td><td>3.3</td><td>1.1E-03</td><td>5.4E-03</td><td>BT20/MB231</td><td>CYTO</td></tr>
<tr><td>>MI0000760|hsa-mir-361&WithFlank&X|-|85903630|85903713@12.32.21(+1U)&[MIMAT0000703&hsa-miR-361-5p&offsets|0|-1(+1U);m-89&X|-|85903681|85903702&offsets|0|-1(+1U)]|TTATCAGAATCTCCAGGGGTAT</td><td>>MI0000760|hsa-mir-361&WithFlank&X|-|85903630|85903713</td><td>12</td><td>32</td><td>21(+1U)</td><td>TTATCAGAATCTCCAGGGGTAT</td><td>108.6</td><td>3.9</td><td>1.4</td><td>2.9</td><td>4.0E-03</td><td>5.4E-03</td><td>MB231/MB468</td><td>MP</td></tr>
<tr><td>>hg38_21_+_8433172_8446622_RNA45SN1_45S_50ntFlanks@12181.12211.31|TCAAACGGTAACGCAGGTGTCCTAAGGCGAG</td><td>>hg38_21_+_8433172_8446622_RNA45SN1_45S_50ntFlanks</td><td>12181</td><td>12211</td><td>31</td><td>TCAAACGGTAACGCAGGTGTCCTAAGGCGAG</td><td>84.5</td><td>-4.8</td><td>1.6</td><td>-3.0</td><td>2.5E-03</td><td>5.4E-03</td><td>BT20/MB468</td><td>MP</td></tr>
<tr><td>>hg38_21_+_8433172_8446622_RNA45SN1_45S_50ntFlanks@9934.9951.18|AGACAGCAGGACGGTGGC</td><td>>hg38_21_+_8433172_8446622_RNA45SN1_45S_50ntFlanks</td><td>9934</td><td>9951</td><td>18</td><td>AGACAGCAGGACGGTGGC</td><td>550.6</td><td>6.6</td><td>2.0</td><td>3.3</td><td>8.7E-04</td><td>5.4E-03</td><td>BT20/MB231</td><td>MP</td></tr>
<tr><td>>am_trna146_GlnCTG_6_-_27515531_27515602@1.29.29,trna1_GlnCTG_6_+_18836402_18836473@1.29.29,trna3_GlnCTG_17_+_8023070_8023141@1.29.29,trna42_GlnCTG_6_+_27263212_27263283@1.29.29,trna49_GlnCTG_6_+_27487308_27487379@1.29.29,trna7_GlnCTG_15_-_66161400_66161471@1.29.29,trna99_GlnCTG_6_-_28909378_28909449@1.29.29|GGTTCCATGGTGTAATGGTTAGCACTCTG</td><td>>am_trna146_GlnCTG_6_-_27515531_27515602</td><td>1</td><td>29</td><td>29</td><td>GGTTCCATGGTGTAATGGTTAGCACTCTG</td><td>1429.5</td><td>-4.1</td><td>1.4</td><td>-2.9</td><td>3.7E-03</td><td>5.4E-03</td><td>BT20/MB468</td><td>MITO</td></tr>
<tr><td>>hg38_MT_+_1621_3279_RNR2_16S_50ntFlanks@1091.1117.27|AGCAAGACGAGAAGACCCTATGGAGCT</td><td>>hg38_MT_+_1621_3279_RNR2_16S_50ntFlanks</td><td>1091</td><td>1117</td><td>27</td><td>AGCAAGACGAGAAGACCCTATGGAGCT</td><td>73.2</td><td>5.1</td><td>1.7</td><td>3.0</td><td>2.5E-03</td><td>5.4E-03</td><td>BT20/MB468</td><td>MP</td></tr>
<tr><td>>MI0000811|hsa-mir-148b&WithFlank&12|+|54337210|54337320@69.89.21&[MIMAT0000759&hsa-miR-148b-3p&offsets|0|-1;m-77&12|+|54337278|54337299&offsets|0|-1]|TCAGTGCATCACAGAACTTTG</td><td>>MI0000811|hsa-mir-148b&WithFlank&12|+|54337210|54337320</td><td>69</td><td>89</td><td>21</td><td>TCAGTGCATCACAGAACTTTG</td><td>695.8</td><td>3.5</td><td>1.2</td><td>3.0</td><td>2.5E-03</td><td>5.4E-03</td><td>BT20/MB468</td><td>MP</td></tr>
<tr><td>>hg38_21_+_8433172_8446622_RNA45SN1_45S_50ntFlanks@6657.6681.25|TTAGCGGTGGATCACTCGGCTCGTG</td><td>>hg38_21_+_8433172_8446622_RNA45SN1_45S_50ntFlanks</td><td>6657</td><td>6681</td><td>25</td><td>TTAGCGGTGGATCACTCGGCTCGTG</td><td>434.8</td><td>-3.1</td><td>1.1</td><td>-2.9</td><td>4.4E-03</td><td>5.4E-03</td><td>BT20/MB468</td><td>CYTO</td></tr>
<tr><td>>hg38_21_+_8433172_8446622_RNA45SN1_45S_50ntFlanks@7975.8011.37|CGCGACCTCAGATCAGACGTGGCGACCCGCTGAATTT</td><td>>hg38_21_+_8433172_8446622_RNA45SN1_45S_50ntFlanks</td><td>7975</td><td>8011</td><td>37</td><td>CGCGACCTCAGATCAGACGTGGCGACCCGCTGAATTT</td><td>25711.2</td><td>3.8</td><td>1.3</td><td>3.0</td><td>2.5E-03</td><td>5.4E-03</td><td>BT20/MB468</td><td>MP</td></tr>
<tr><td>>hg38_21_+_8433172_8446622_RNA45SN1_45S_50ntFlanks@4390.4410.21|TCTTGGGAGCGGGCGGGCGGT</td><td>>hg38_21_+_8433172_8446622_RNA45SN1_45S_50ntFlanks</td><td>4390</td><td>4410</td><td>21</td><td>TCTTGGGAGCGGGCGGGCGGT</td><td>103.7</td><td>-4.0</td><td>1.4</td><td>-2.9</td><td>4.4E-03</td><td>5.4E-03</td><td>BT20/MB468</td><td>CYTO</td></tr>
<tr><td>>hg38_21_+_8433172_8446622_RNA45SN1_45S_50ntFlanks@12418.12451.34|CGATGTCGGCTCTTCCTATCATTGTGAAGCAGAA</td><td>>hg38_21_+_8433172_8446622_RNA45SN1_45S_50ntFlanks</td><td>12418</td><td>12451</td><td>34</td><td>CGATGTCGGCTCTTCCTATCATTGTGAAGCAGAA</td><td>562.7</td><td>-2.0</td><td>0.7</td><td>-2.8</td><td>4.7E-03</td><td>5.4E-03</td><td>MB231/MB468</td><td>MITO</td></tr>
<tr><td>>MI0000077|hsa-mir-21&WithFlank&17|+|59841260|59841343@14.35.22(+1G)&[MIMAT0000076&hsa-miR-21-5p&offsets|0|0(+1G);m-1&17|+|59841273|59841294&offsets|0|0(+1G)]|TAGCTTATCAGACTGATGTTGAG</td><td>>MI0000077|hsa-mir-21&WithFlank&17|+|59841260|59841343</td><td>14</td><td>35</td><td>22(+1G)</td><td>TAGCTTATCAGACTGATGTTGAG</td><td>2198.2</td><td>-3.8</td><td>1.1</td><td>-3.3</td><td>8.8E-04</td><td>5.4E-03</td><td>BT20/MB231</td><td>MP</td></tr>
<tr><td>>hg38_21_+_8433172_8446622_RNA45SN1_45S_50ntFlanks@10393.10410.18|CAGCAGTTGAACATGGGT</td><td>>hg38_21_+_8433172_8446622_RNA45SN1_45S_50ntFlanks</td><td>10393</td><td>10410</td><td>18</td><td>CAGCAGTTGAACATGGGT</td><td>1413.4</td><td>-4.3</td><td>1.4</td><td>-3.0</td><td>2.5E-03</td><td>5.4E-03</td><td>BT20/MB468</td><td>MP</td></tr>
<tr><td>>hg38_21_+_8433172_8446622_RNA45SN1_45S_50ntFlanks@12997.13039.43|CGCTGCGATCTATTGAAAGTCAGCCCTCGACACAAGGGTTTGT</td><td>>hg38_21_+_8433172_8446622_RNA45SN1_45S_50ntFlanks</td><td>12997</td><td>13039</td><td>43</td><td>CGCTGCGATCTATTGAAAGTCAGCCCTCGACACAAGGGTTTGT</td><td>163063.4</td><td>-3.1</td><td>0.9</td><td>-3.3</td><td>9.6E-04</td><td>5.4E-03</td><td>BT20/MB231</td><td>NUC</td></tr>
<tr><td>>hg38_21_+_8433172_8446622_RNA45SN1_45S_50ntFlanks@8185.8205.21|GAGGCCCAGCCCGTGGACGGT</td><td>>hg38_21_+_8433172_8446622_RNA45SN1_45S_50ntFlanks</td><td>8185</td><td>8205</td><td>21</td><td>GAGGCCCAGCCCGTGGACGGT</td><td>126.4</td><td>4.0</td><td>1.3</td><td>3.0</td><td>2.5E-03</td><td>5.4E-03</td><td>BT20/MB468</td><td>MP</td></tr>
<tr><td>>MI0000289|hsa-mir-181a-1&WithFlank&1|-|198859038|198859159@70.90.21(+1A)&[MIMAT0000270&hsa-miR-181a-3p&offsets|0|-1(+1A);m-125&1|-|198859069|198859090&offsets|0|-1(+1A)]|ACCATCGACCGTTGATTGTACA</td><td>>MI0000289|hsa-mir-181a-1&WithFlank&1|-|198859038|198859159</td><td>70</td><td>90</td><td>21(+1A)</td><td>ACCATCGACCGTTGATTGTACA</td><td>102.4</td><td>4.5</td><td>1.6</td><td>2.9</td><td>4.0E-03</td><td>5.4E-03</td><td>MB231/MB468</td><td>MP</td></tr>
<tr><td>>trnaMT_HisGTG_MT_+_12138_12206@32.72.41|TGAATCTGACAACAGAGGCTTACGACCCCTTATTTACCCCA</td><td>>trnaMT_HisGTG_MT_+_12138_12206</td><td>32</td><td>72</td><td>41</td><td>TGAATCTGACAACAGAGGCTTACGACCCCTTATTTACCCCA</td><td>7782.3</td><td>3.3</td><td>1.2</td><td>2.8</td><td>4.7E-03</td><td>5.4E-03</td><td>MB231/MB468</td><td>MITO</td></tr>
<tr><td>>hg38_21_+_8433172_8446622_RNA45SN1_45S_50ntFlanks@9533.9573.41|GAGGAAACTCTGGTGGAGGTCCGTAGCGGTCCTGACGTGCA</td><td>>hg38_21_+_8433172_8446622_RNA45SN1_45S_50ntFlanks</td><td>9533</td><td>9573</td><td>41</td><td>GAGGAAACTCTGGTGGAGGTCCGTAGCGGTCCTGACGTGCA</td><td>373.2</td><td>-2.5</td><td>0.8</td><td>-3.3</td><td>1.1E-03</td><td>5.4E-03</td><td>BT20/MB231</td><td>CYTO</td></tr>
<tr><td>>MI0003778|hsa-mir-320c-1&WithFlank&18|+|21683512|21683595@48.67.20;MI0008191|hsa-mir-320c-2&WithFlank&18|+|24321669|24321752@48.67.20&[MIMAT0005793&hsa-miR-320c&offsets|0|0;m-222&18|+|21683559|21683578&offsets|0|0];[MIMAT0005793_1&hsa-miR-320c&offsets|0|0;m-219&18|+|24321716|24321735&offsets|0|0]|AAAAGCTGGGTTGAGAGGGT</td><td>>MI0003778|hsa-mir-320c-1&WithFlank&18|+|21683512|21683595</td><td>48</td><td>67</td><td>20</td><td>AAAAGCTGGGTTGAGAGGGT</td><td>349.2</td><td>3.9</td><td>1.4</td><td>2.9</td><td>4.0E-03</td><td>5.4E-03</td><td>MB231/MB468</td><td>MP</td></tr>
<tr><td>>hg38_21_+_8433172_8446622_RNA45SN1_45S_50ntFlanks@4723.4759.37|CATTAATCAAGAACGAAAGTCGGAGGTTCGAAGACGA</td><td>>hg38_21_+_8433172_8446622_RNA45SN1_45S_50ntFlanks</td><td>4723</td><td>4759</td><td>37</td><td>CATTAATCAAGAACGAAAGTCGGAGGTTCGAAGACGA</td><td>1719.7</td><td>-2.5</td><td>0.9</td><td>-2.8</td><td>4.4E-03</td><td>5.4E-03</td><td>MB231/MB468</td><td>CYTO</td></tr>
</table>

| | | | | | | | | | | | | | |
|---|---|---|---|---|---|---|---|---|---|---|---|---|---|
| >hg38_21_+_8433172_8446622_RNA45SN1_45S_50ntFlanks@11843.11872.30\|AGGGAACGGGCTTGGCGGAATCAGCGGGGA | >hg38_21_+_8433172_8446622_RNA45SN1_45S_50ntFlanks | 11843 | 11872 | 30 | AGGGAACGGGCTTGGCGGAATCAGCGGGGA | 239.8 | -3.3 | 1.2 | -2.9 | 3.7E-03 | 5.4E-03 | BT20/MB468 | MITO |
| >hg38_1_-_228634819_228635039_RNA5S12_5S_50ntFlanks@148.172.25\|GGAATACCGGGTGCTGTAGGCTTTT | >hg38_1_-_228634819_228635039_RNA5S12_5S_50ntFlanks | 148 | 172 | 25 | GGAATACCGGGTGCTGTAGGCTTTT | 169.3 | -5.5 | 1.9 | -2.9 | 4.0E-03 | 5.5E-03 | MB231/MB468 | MP |
| >hg38_21_+_8433172_8446622_RNA45SN1_45S_50ntFlanks@11710.11742.33\|CGCGGGTAAACGGCGGGAGTAACTATGACTCTC | >hg38_21_+_8433172_8446622_RNA45SN1_45S_50ntFlanks | 11710 | 11742 | 33 | CGCGGGTAAACGGCGGGAGTAACTATGACTCTC | 120.1 | -4.0 | 1.4 | -2.9 | 4.0E-03 | 5.5E-03 | MB231/MB468 | MP |
| >MI0000088\|hsa-mir-30a&WithFlank&6\|-\|71403545\|71403627@12.35.24(+4U)&[MIMAT0000087&hsa-miR-30a-5p&offsets\|0\|+2(+4U);m-15&6\|-\|71403595\|71403616&offsets\|0\|+2(+4U)]\|TGTAAACATCCTCGACTGGAAGCTTTTT | >MI0000088\|hsa-mir-30a&WithFlank&6\|-\|71403545\|71403627 | 12 | 35 | 24(+4U) | TGTAAACATCCTCGACTGGAAGCTTTTT | 47.9 | 5.4 | 1.9 | 2.9 | 4.0E-03 | 5.5E-03 | MB231/MB468 | MP |
| >hg38_21_+_8433172_8446622_RNA45SN1_45S_50ntFlanks@11584.11616.33\|TGGTGCGGACCAGGGGAATCCGACTGTTTAATT | >hg38_21_+_8433172_8446622_RNA45SN1_45S_50ntFlanks | 11584 | 11616 | 33 | TGGTGCGGACCAGGGGAATCCGACTGTTTAATT | 1145.7 | 3.9 | 1.2 | 3.3 | 9.7E-04 | 5.5E-03 | BT20/MB231 | NUC |
| >hg38_MT_+_598_1651_RNR1_12S_50ntFlanks@52.88.37\|ATAGGTTTGGTCCTAGCCTTTCTATTAGCTCTTAGTA | >hg38_MT_+_598_1651_RNR1_12S_50ntFlanks | 52 | 88 | 37 | ATAGGTTTGGTCCTAGCCTTTCTATTAGCTCTTAGTA | 203588.9 | 4.8 | 1.6 | 3.0 | 2.5E-03 | 5.5E-03 | BT20/MB468 | MP |
| >hg38_21_+_8433172_8446622_RNA45SN1_45S_50ntFlanks@10679.10708.30\|GGTTCCGGCGGCGTCCGGTGAGCTCTCGCT | >hg38_21_+_8433172_8446622_RNA45SN1_45S_50ntFlanks | 10679 | 10708 | 30 | GGTTCCGGCGGCGTCCGGTGAGCTCTCGCT | 623.5 | 5.2 | 1.6 | 3.3 | 8.8E-04 | 5.5E-03 | BT20/MB231 | MP |
| >hg38_21_+_8433172_8446622_RNA45SN1_45S_50ntFlanks@6646.6669.24\|TCGTACGACTCTTAGCGGTGGATC | >hg38_21_+_8433172_8446622_RNA45SN1_45S_50ntFlanks | 6646 | 6669 | 24 | TCGTACGACTCTTAGCGGTGGATC | 8710.7 | -2.7 | 0.9 | -2.9 | 4.0E-03 | 5.5E-03 | MB231/MB468 | MP |
| >am_trnaMT_ValTAC_MT_+_1602_1670@49.67.19\|AACTTAACTTGACCGCTCT | >am_trnaMT_ValTAC_MT_+_1602_1670 | 49 | 67 | 19 | AACTTAACTTGACCGCTCT | 188.6 | 3.7 | 1.2 | 3.0 | 2.5E-03 | 5.5E-03 | BT20/MB468 | MP |
| >hg38_21_+_8433172_8446622_RNA45SN1_45S_50ntFlanks@6651.6677.27\|CGACTCTTAGCGGTGGATCACTCGGCT | >hg38_21_+_8433172_8446622_RNA45SN1_45S_50ntFlanks | 6651 | 6677 | 27 | CGACTCTTAGCGGTGGATCACTCGGCT | 14214.1 | 4.5 | 1.3 | 3.3 | 8.8E-04 | 5.5E-03 | BT20/MB231 | MP |
| >hg38_21_+_8433172_8446622_RNA45SN1_45S_50ntFlanks@8196.8226.31\|CGTGGACGGTGTGAGGCCGGTAGCGGCCCCC | >hg38_21_+_8433172_8446622_RNA45SN1_45S_50ntFlanks | 8196 | 8226 | 31 | CGTGGACGGTGTGAGGCCGGTAGCGGCCCCC | 269.9 | -3.3 | 1.0 | -3.3 | 1.1E-03 | 5.5E-03 | BT20/MB231 | CYTO |
| >hg38_21_+_8433172_8446622_RNA45SN1_45S_50ntFlanks@8356.8375.20\|GAAAAGAACTTTGAAGAGAG | >hg38_21_+_8433172_8446622_RNA45SN1_45S_50ntFlanks | 8356 | 8375 | 20 | GAAAAGAACTTTGAAGAGAG | 918.5 | 2.5 | 0.9 | 2.9 | 3.7E-03 | 5.5E-03 | BT20/MB468 | MITO |
| >hg38_21_+_8433172_8446622_RNA45SN1_45S_50ntFlanks@11778.11795.18\|ACGCGCATGAATGGATGA | >hg38_21_+_8433172_8446622_RNA45SN1_45S_50ntFlanks | 11778 | 11795 | 18 | ACGCGCATGAATGGATGA | 102.3 | -4.9 | 1.6 | -3.0 | 2.5E-03 | 5.5E-03 | BT20/MB468 | MP |
| >hg38_21_+_8433172_8446622_RNA45SN1_45S_50ntFlanks@6670.6689.20\|ACTCGGCTCGTGCGTCGATG | >hg38_21_+_8433172_8446622_RNA45SN1_45S_50ntFlanks | 6670 | 6689 | 20 | ACTCGGCTCGTGCGTCGATG | 247.7 | -3.7 | 1.2 | -3.0 | 2.5E-03 | 5.5E-03 | BT20/MB468 | MP |
| >hg38_MT_+_1621_3279_RNR2_16S_50ntFlanks@1181.1215.35\|AAAAATTTCGGTTGGGGCGACCTCGGAGCAGAACC | >hg38_MT_+_1621_3279_RNR2_16S_50ntFlanks | 1181 | 1215 | 35 | AAAAATTTCGGTTGGGGCGACCTCGGAGCAGAACC | 71.0 | 5.3 | 1.9 | 2.9 | 4.0E-03 | 5.5E-03 | MB231/MB468 | MP |
| >hg38_21_+_8433172_8446622_RNA45SN1_45S_50ntFlanks@10355.10388.34\|AACTTTGAAGGCCGAAGTGGAGAAGGGTTCCATG | >hg38_21_+_8433172_8446622_RNA45SN1_45S_50ntFlanks | 10355 | 10388 | 34 | AACTTTGAAGGCCGAAGTGGAGAAGGGTTCCATG | 236.4 | -3.5 | 1.2 | -2.9 | 3.8E-03 | 5.5E-03 | BT20/MB468 | MITO |
| >hg38_21_+_8433172_8446622_RNA45SN1_45S_50ntFlanks@12637.12657.21\|GCTACCATCTGTGGGATTATG | >hg38_21_+_8433172_8446622_RNA45SN1_45S_50ntFlanks | 12637 | 12657 | 21 | GCTACCATCTGTGGGATTATG | 292.5 | -2.4 | 0.8 | -2.9 | 3.8E-03 | 5.5E-03 | BT20/MB468 | MITO |
| >hg38_21_+_8433172_8446622_RNA45SN1_45S_50ntFlanks@3769.3792.24\|CGCACGGCCGGTACAGTGAAACTG | >hg38_21_+_8433172_8446622_RNA45SN1_45S_50ntFlanks | 3769 | 3792 | 24 | CGCACGGCCGGTACAGTGAAACTG | 195.7 | -2.5 | 0.9 | -2.8 | 4.5E-03 | 5.5E-03 | MB231/MB468 | CYTO |
| >am_MI0000809\|hsa-mir-151a&WithFlank&8\|-\|140732558\|140732659@53.73.21(+1G)&[MIMAT0000757&hsa-miR-151a-3p&offsets\|0\|0(+1G);m-30&8\|-\|140732587\|140732607&offsets\|0\|0(+1G)]\|CTAGACTGAAGCTCCTTGAGGG | >am_MI0000809\|hsa-mir-151a&WithFlank&8\|-\|140732558\|140732659 | 53 | 73 | 21(+1G) | CTAGACTGAAGCTCCTTGAGGG | 880.6 | 2.5 | 0.8 | 3.3 | 1.1E-03 | 5.5E-03 | BT20/MB231 | CYTO |
| >trnaMT_ProTGG_MT_-_15956_16023@41.67.27\|ATGGTGGAGTTAAAGACTTTTTCTCTG | >trnaMT_ProTGG_MT_-_15956_16023 | 41 | 67 | 27 | ATGGTGGAGTTAAAGACTTTTTCTCTG | 210.7 | -4.1 | 1.3 | -3.0 | 2.5E-03 | 5.5E-03 | BT20/MB468 | MP |
| >hg38_21_+_8433172_8446622_RNA45SN1_45S_50ntFlanks@6646.6679.34\|TCGTACGACTCTTAGCGGTGGATCACTCGGCTCG | >hg38_21_+_8433172_8446622_RNA45SN1_45S_50ntFlanks | 6646 | 6679 | 34 | TCGTACGACTCTTAGCGGTGGATCACTCGGCTCG | 8661.9 | 5.0 | 1.5 | 3.3 | 8.9E-04 | 5.5E-03 | BT20/MB231 | MP |
| >hg38_21_+_8433172_8446622_RNA45SN1_45S_50ntFlanks@10693.10712.20\|CCGGTGAGCTCTCGCTGGCC | >hg38_21_+_8433172_8446622_RNA45SN1_45S_50ntFlanks | 10693 | 10712 | 20 | CCGGTGAGCTCTCGCTGGCC | 117.1 | -4.0 | 1.4 | -2.9 | 4.0E-03 | 5.5E-03 | MB231/MB468 | MP |
| >hg38_21_+_8433172_8446622_RNA45SN1_45S_50ntFlanks@9548.9570.23\|GAGGTCCGTAGCGGTCCTGACGT | >hg38_21_+_8433172_8446622_RNA45SN1_45S_50ntFlanks | 9548 | 9570 | 23 | GAGGTCCGTAGCGGTCCTGACGT | 95.6 | -2.2 | 0.8 | -2.9 | 4.0E-03 | 5.5E-03 | MB231/MB468 | MP |
| >hg38_21_+_8433172_8446622_RNA45SN1_45S_50ntFlanks@8115.8138.24\|CGCGGGACATGTGGCGTACGGAAG | >hg38_21_+_8433172_8446622_RNA45SN1_45S_50ntFlanks | 8115 | 8138 | 24 | CGCGGGACATGTGGCGTACGGAAG | 505.2 | -4.3 | 1.5 | -2.8 | 4.7E-03 | 5.5E-03 | BT20/MB468 | NUC |
| >am_trnaMT_ValTAC_MT_+_1602_1670@29.68.40\|ACTTACACTTAGGAGATTTCAACTTAACTTGACCGCTCTG | >am_trnaMT_ValTAC_MT_+_1602_1670 | 29 | 68 | 40 | ACTTACACTTAGGAGATTTCAACTTAACTTGACCGCTCTG | 204.0 | -3.0 | 1.1 | -2.8 | 4.8E-03 | 5.5E-03 | MB231/MB468 | MITO |
| >hg38_21_+_8433172_8446622_RNA45SN1_45S_50ntFlanks@12408.12437.30\|TTTGATCCTTCGATGTCGGCTCTTCCTATC | >hg38_21_+_8433172_8446622_RNA45SN1_45S_50ntFlanks | 12408 | 12437 | 30 | TTTGATCCTTCGATGTCGGCTCTTCCTATC | 298.7 | 2.5 | 0.9 | 2.8 | 4.5E-03 | 5.5E-03 | MB231/MB468 | CYTO |
| >am_MI0003639\|hsa-mir-625&WithFlank&14\|+\|65471096\|65471192@58.79.22&[MIMAT0004808&hsa-miR-625-3p&offsets\|0\|0;m-137&14\|+\|65471153\|65471174&offsets\|0\|0]\|GACTATAGAACTTTCCCCCTCA | >am_MI0003639\|hsa-mir-625&WithFlank&14\|+\|65471096\|65471192 | 58 | 79 | 22 | GACTATAGAACTTTCCCCCTCA | 98.8 | 5.4 | 1.8 | 3.0 | 2.5E-03 | 5.5E-03 | BT20/MB468 | MP |
| >MI0001448\|hsa-mir-425&WithFlank&3\|-\|49020142\|49020240@60.80.21(+1A)&[MIMAT0001343&hsa-miR-425-3p&offsets\|-1\|-2(+1A);m-258&3\|-\|49020160\|49020181&offsets\|0\|-1(+1A)]\|CATCGGGAATGTCGTGTCCGCA | >MI0001448\|hsa-mir-425&WithFlank&3\|-\|49020142\|49020240 | 60 | 80 | 21(+1A) | CATCGGGAATGTCGTGTCCGCA | 75.8 | 4.8 | 1.7 | 2.9 | 4.1E-03 | 5.5E-03 | MB231/MB468 | MP |
| >hg38_21_+_8433172_8446622_RNA45SN1_45S_50ntFlanks@9243.9273.31\|GAGCGAGCGCACGGGGTCGGCGGCGACGTCG | >hg38_21_+_8433172_8446622_RNA45SN1_45S_50ntFlanks | 9243 | 9273 | 31 | GAGCGAGCGCACGGGGTCGGCGGCGACGTCG | 97.4 | -4.9 | 1.6 | -3.0 | 2.5E-03 | 5.5E-03 | BT20/MB468 | MP |
| >trnaMT_GluTTC_MT_-_14674_14742@46.72.27,trnalookalike8_GluTTC_5_-_93905172_93905240@46.72.27\|GTGGTTGTAGTCCGTGCGAGAATACCA | >trnaMT_GluTTC_MT_-_14674_14742 | 46 | 72 | 27 | GTGGTTGTAGTCCGTGCGAGAATACCA | 839.8 | 2.4 | 0.8 | 2.9 | 4.1E-03 | 5.5E-03 | MB231/MB468 | MP |
| >hg38_21_+_8433172_8446622_RNA45SN1_45S_50ntFlanks@6772.6802.31\|GTTCCTCCCGGGGCTACGCCTGTCTGAGCGT | >hg38_21_+_8433172_8446622_RNA45SN1_45S_50ntFlanks | 6772 | 6802 | 31 | GTTCCTCCCGGGGCTACGCCTGTCTGAGCGT | 102.8 | -3.7 | 1.3 | -2.9 | 4.1E-03 | 5.5E-03 | MB231/MB468 | MP |
| >trnaMT_ValTAC_MT_+_1602_1670@53.72.20\|TAACTTGACCGCTCTGACCA | >trnaMT_ValTAC_MT_+_1602_1670 | 53 | 72 | 20 | TAACTTGACCGCTCTGACCA | 1182.9 | 2.0 | 0.7 | 2.8 | 4.8E-03 | 5.5E-03 | MB231/MB468 | MITO |
| >trna119_LysCTT_1_-_145395522_145395594@4.35.32,trna11_LysCTT_5_-_180648979_180649051@4.35.32,trna13_LysCTT_6_+_26556774_26556846@4.35.32,trna7_LysCTT_16_+_3225692_3225764@4.35.32,trna9_LysCTT_5_+_180634755_180634827@4.35.32\|CGGCTAGCTCAGTCGGTAGAGCATGAGACTCT | >trna119_LysCTT_1_-_145395522_145395594 | 4 | 35 | 32 | CGGCTAGCTCAGTCGGTAGAGCATGAGACTCT | 1217.1 | -3.7 | 1.1 | -3.3 | 8.9E-04 | 5.5E-03 | BT20/MB231 | MP |
| >hg38_21_+_8433172_8446622_RNA45SN1_45S_50ntFlanks@10351.10390.40\|CGAGAACTTTGAAGGCCGAAGTGGAGAAGGGTTCCATGTG | >hg38_21_+_8433172_8446622_RNA45SN1_45S_50ntFlanks | 10351 | 10390 | 40 | CGAGAACTTTGAAGGCCGAAGTGGAGAAGGGTTCCATGTG | 7094.1 | -3.4 | 1.1 | -3.0 | 2.5E-03 | 5.5E-03 | BT20/MB468 | MP |
| >hg38_21_+_8433172_8446622_RNA45SN1_45S_50ntFlanks@11913.11947.35\|AAGAGACATGAGAGGTGTAGAATAAGTGGGAGGCC | >hg38_21_+_8433172_8446622_RNA45SN1_45S_50ntFlanks | 11913 | 11947 | 35 | AAGAGACATGAGAGGTGTAGAATAAGTGGGAGGCC | 69.1 | 9.4 | 2.8 | 3.3 | 8.9E-04 | 5.5E-03 | BT20/MB231 | MP |
| >hg38_21_+_8433172_8446622_RNA45SN1_45S_50ntFlanks@8351.8378.28\|AAGTTGAAAAGAACTTTGAAGAGAGAGT | >hg38_21_+_8433172_8446622_RNA45SN1_45S_50ntFlanks | 8351 | 8378 | 28 | AAGTTGAAAAGAACTTTGAAGAGAGAGT | 301.3 | 4.2 | 1.3 | 3.3 | 9.0E-04 | 5.5E-03 | BT20/MB231 | MP |
| >hg38_21_+_8433172_8446622_RNA45SN1_45S_50ntFlanks@9929.9951.23\|GATATAGACAGCAGGACGGTGGC | >hg38_21_+_8433172_8446622_RNA45SN1_45S_50ntFlanks | 9929 | 9951 | 23 | GATATAGACAGCAGGACGGTGGC | 585.9 | 6.3 | 1.9 | 3.3 | 9.0E-04 | 5.5E-03 | BT20/MB231 | MP |
| >hg38_21_+_8433172_8446622_RNA45SN1_45S_50ntFlanks@5082.5120.39\|AACGAACGAGACTCTGGCATGCTAACTAGTTACGCGACC | >hg38_21_+_8433172_8446622_RNA45SN1_45S_50ntFlanks | 5082 | 5120 | 39 | AACGAACGAGACTCTGGCATGCTAACTAGTTACGCGACC | 116.5 | -5.3 | 1.8 | -2.9 | 4.1E-03 | 5.5E-03 | MB231/MB468 | MP |
| >hg38_21_+_8433172_8446622_RNA45SN1_45S_50ntFlanks@10248.10285.38\|CGACGAGTAGGAGGGCCGCTGCGGTGAGCCTTGAAGCC | >hg38_21_+_8433172_8446622_RNA45SN1_45S_50ntFlanks | 10248 | 10285 | 38 | CGACGAGTAGGAGGGCCGCTGCGGTGAGCCTTGAAGCC | 254.1 | 6.4 | 1.9 | 3.3 | 9.0E-04 | 5.5E-03 | BT20/MB231 | MP |
| >trnaMT_HisGTG_MT_+_12138_12206@26.63.38\|AGATTGTGAATCTGACAACAGAGGCTTACGACCCCTTA | >trnaMT_HisGTG_MT_+_12138_12206 | 26 | 63 | 38 | AGATTGTGAATCTGACAACAGAGGCTTACGACCCCTTA | 173.5 | 3.8 | 1.3 | 2.9 | 4.1E-03 | 5.5E-03 | MB231/MB468 | MP |
| >MI0000286\|hsa-mir-210&WithFlank&11\|-\|568083\|568204@72.93.22&[MIMAT0000267&hsa-miR-210-3p&offsets\|0\|0;m-102&11\|-\|568112\|568133&offsets\|0\|0]\|CTGTGCGTGTGACAGCGGCTGA | >MI0000286\|hsa-mir-210&WithFlank&11\|-\|568083\|568204 | 72 | 93 | 22 | CTGTGCGTGTGACAGCGGCTGA | 2503.1 | 5.2 | 1.7 | 3.0 | 2.5E-03 | 5.5E-03 | BT20/MB468 | MP |
| >hg38_21_+_8433172_8446622_RNA45SN1_45S_50ntFlanks@12481.12499.19\|AATAGGGAACGTGAGCTGG | >hg38_21_+_8433172_8446622_RNA45SN1_45S_50ntFlanks | 12481 | 12499 | 19 | AATAGGGAACGTGAGCTGG | 140.8 | -3.6 | 1.3 | -2.9 | 4.1E-03 | 5.5E-03 | MB231/MB468 | MP |
| >MI0000760\|hsa-mir-361&WithFlank&X\|-\|85903630\|85903713@12.32.21&[MIMAT0000703&hsa-miR-361-5p&offsets\|0\|-1;m-89&X\|-\|85903681\|85903702&offsets\|0\|-1]\|TTATCAGAATCTCCAGGGGTA | >MI0000760\|hsa-mir-361&WithFlank&X\|-\|85903630\|85903713 | 12 | 32 | 21 | TTATCAGAATCTCCAGGGGTA | 83.6 | 3.6 | 1.3 | 2.9 | 4.1E-03 | 5.5E-03 | MB231/MB468 | MP |
| >hg38_1_-_228634819_228635039_RNA5S12_5S_50ntFlanks@151.169.19\|ATACCGGGTGCTGTAGGCT | >hg38_1_-_228634819_228635039_RNA5S12_5S_50ntFlanks | 151 | 169 | 19 | ATACCGGGTGCTGTAGGCT | 271.0 | 2.4 | 0.9 | 2.8 | 4.8E-03 | 5.5E-03 | MB231/MB468 | NUC |
| >hg38_1_-_228634819_228635039_RNA5S12_5S_50ntFlanks@153.170.18\|ACCGGGTGCTGTAGGCTT | >hg38_1_-_228634819_228635039_RNA5S12_5S_50ntFlanks | 153 | 170 | 18 | ACCGGGTGCTGTAGGCTT | 3489.2 | 2.1 | 0.7 | 2.8 | 4.8E-03 | 5.5E-03 | MB231/MB468 | MITO |
| >hg38_21_+_8433172_8446622_RNA45SN1_45S_50ntFlanks@6651.6679.29\|CGACTCTTAGCGGTGGATCACTCGGCTCG | >hg38_21_+_8433172_8446622_RNA45SN1_45S_50ntFlanks | 6651 | 6679 | 29 | CGACTCTTAGCGGTGGATCACTCGGCTCG | 18998.8 | 3.5 | 1.2 | 2.8 | 4.4E-03 | 5.5E-03 | BT20/MB468 | CYTO |
| >MI0000286\|hsa-mir-210&WithFlank&11\|-\|568083\|568204@72.93.22(+3A)&[MIMAT0000267&hsa-miR-210-3p&offsets\|0\|0(+3A);m-102&11\|-\|568112\|568133&offsets\|0\|0(+3A)]\|CTGTGCGTGTGACAGCGGCTGAAAA | >MI0000286\|hsa-mir-210&WithFlank&11\|-\|568083\|568204 | 72 | 93 | 22(+3A) | CTGTGCGTGTGACAGCGGCTGAAAA | 194.1 | 5.3 | 1.7 | 3.0 | 2.5E-03 | 5.5E-03 | BT20/MB468 | MP |
| >hg38_21_+_8433172_8446622_RNA45SN1_45S_50ntFlanks@6650.6667.18\|ACGACTCTTAGCGGTGGA | >hg38_21_+_8433172_8446622_RNA45SN1_45S_50ntFlanks | 6650 | 6667 | 18 | ACGACTCTTAGCGGTGGA | 418.0 | 2.6 | 0.9 | 2.8 | 4.8E-03 | 5.5E-03 | MB231/MB468 | MITO |
| >hg38_21_+_8433172_8446622_RNA45SN1_45S_50ntFlanks@10350.10370.21\|ACGAGAACTTTGAAGGCCGAA | >hg38_21_+_8433172_8446622_RNA45SN1_45S_50ntFlanks | 10350 | 10370 | 21 | ACGAGAACTTTGAAGGCCGAA | 223.3 | -1.4 | 0.5 | -2.8 | 4.5E-03 | 5.5E-03 | MB231/MB468 | CYTO |
| >hg38_21_+_8433172_8446622_RNA45SN1_45S_50ntFlanks@11583.11616.34\|CTGGTGCGGACCAGGGGAATCCGACTGTTTAATT | >hg38_21_+_8433172_8446622_RNA45SN1_45S_50ntFlanks | 11583 | 11616 | 34 | CTGGTGCGGACCAGGGGAATCCGACTGTTTAATT | 396.4 | 2.0 | 0.7 | 2.9 | 3.8E-03 | 5.5E-03 | BT20/MB468 | MITO |
| >hg38_MT_+_1621_3279_RNR2_16S_50ntFlanks@51.85.35\|GCTAAACCTAGCCCCAAACCCACTCCACCTTACTA | >hg38_MT_+_1621_3279_RNR2_16S_50ntFlanks | 51 | 85 | 35 | GCTAAACCTAGCCCCAAACCCACTCCACCTTACTA | 1134.4 | 4.1 | 1.4 | 3.0 | 2.5E-03 | 5.5E-03 | BT20/MB468 | MP |
| >hg38_21_+_8433172_8446622_RNA45SN1_45S_50ntFlanks@12568.12587.20\|TCAGTACGAGAGGAACCGCA | >hg38_21_+_8433172_8446622_RNA45SN1_45S_50ntFlanks | 12568 | 12587 | 20 | TCAGTACGAGAGGAACCGCA | 141.2 | -4.9 | 1.6 | -3.0 | 2.5E-03 | 5.5E-03 | BT20/MB468 | MP |

<table>
<tr><td>>trna10_LysCTT_16_+_3241501_3241573@3.35.33,trna13_LysCTT_14_-_58706613_58706685@3.35.33,trna2_LysCTT_15_+_79152904_79152976@3.35.33|CCGGCTAGCTCAGTCGGTAGAGCATGGGACTCT</td><td>>trna10_LysCTT_16_+_3241501_3241573</td><td>3</td><td>35</td><td>33</td><td>CCGGCTAGCTCAGTCGGTAGAGCATGGGACTCT</td><td>269.8</td><td>4.8</td><td>1.6</td><td>3.0</td><td>2.5E-03</td><td>5.5E-03</td><td>BT20/MB468</td><td>MP</td></tr>
<tr><td>>trnaMT_TyrGTA_MT_-_5826_5891@38.69.32|AAAGACAGGGGTTAGGCCTCTTTTTACCACCA</td><td>>trnaMT_TyrGTA_MT_-_5826_5891</td><td>38</td><td>69</td><td>32</td><td>AAAGACAGGGGTTAGGCCTCTTTTTACCACCA</td><td>331.3</td><td>4.9</td><td>1.6</td><td>3.0</td><td>2.5E-03</td><td>5.5E-03</td><td>BT20/MB468</td><td>MP</td></tr>
<tr><td>>hg38_21_+_8433172_8446622_RNA45SN1_45S_50ntFlanks@11922.11948.27|GAGAGGTGTAGAATAAGTGGGAGGCCC</td><td>>hg38_21_+_8433172_8446622_RNA45SN1_45S_50ntFlanks</td><td>11922</td><td>11948</td><td>27</td><td>GAGAGGTGTAGAATAAGTGGGAGGCCC</td><td>153.1</td><td>-3.9</td><td>1.4</td><td>-2.9</td><td>4.1E-03</td><td>5.5E-03</td><td>MB231/MB468</td><td>MP</td></tr>
<tr><td>>hg38_1_-_228634819_228635039_RNA5S12_5S_50ntFlanks@51.83.33|GTCTACGGCCATACCACCCTGAACGCGCCCGAT</td><td>>hg38_1_-_228634819_228635039_RNA5S12_5S_50ntFlanks</td><td>51</td><td>83</td><td>33</td><td>GTCTACGGCCATACCACCCTGAACGCGCCCGAT</td><td>4831.9</td><td>2.2</td><td>0.8</td><td>2.8</td><td>4.5E-03</td><td>5.5E-03</td><td>BT20/MB468</td><td>CYTO</td></tr>
<tr><td>>hg38_21_+_8433172_8446622_RNA45SN1_45S_50ntFlanks@7991.8019.29|ACGTGGCGACCCGCTGAATTTAAGCATAT</td><td>>hg38_21_+_8433172_8446622_RNA45SN1_45S_50ntFlanks</td><td>7991</td><td>8019</td><td>29</td><td>ACGTGGCGACCCGCTGAATTTAAGCATAT</td><td>348.8</td><td>-3.3</td><td>1.0</td><td>-3.3</td><td>1.1E-03</td><td>5.5E-03</td><td>BT20/MB231</td><td>CYTO</td></tr>
<tr><td>>hg38_21_+_8433172_8446622_RNA45SN1_45S_50ntFlanks@12645.12666.22|CTGTGGGATTATGACTGAACGC</td><td>>hg38_21_+_8433172_8446622_RNA45SN1_45S_50ntFlanks</td><td>12645</td><td>12666</td><td>22</td><td>CTGTGGGATTATGACTGAACGC</td><td>146.4</td><td>6.1</td><td>2.0</td><td>3.0</td><td>2.5E-03</td><td>5.6E-03</td><td>BT20/MB468</td><td>MP</td></tr>
<tr><td>>hg38_MT_+_1621_3279_RNR2_16S_50ntFlanks@292.322.31|AGACGAGCTACCTAAGAACAGCTAAAAGAGC</td><td>>hg38_MT_+_1621_3279_RNR2_16S_50ntFlanks</td><td>292</td><td>322</td><td>31</td><td>AGACGAGCTACCTAAGAACAGCTAAAAGAGC</td><td>67.3</td><td>4.3</td><td>1.4</td><td>3.0</td><td>2.5E-03</td><td>5.6E-03</td><td>BT20/MB468</td><td>MP</td></tr>
<tr><td>>hg38_MT_+_1621_3279_RNR2_16S_50ntFlanks@930.959.30|AGTGACACATGTTTAACGGCCGCGGTACCC</td><td>>hg38_MT_+_1621_3279_RNR2_16S_50ntFlanks</td><td>930</td><td>959</td><td>30</td><td>AGTGACACATGTTTAACGGCCGCGGTACCC</td><td>242.3</td><td>4.7</td><td>1.6</td><td>3.0</td><td>2.5E-03</td><td>5.6E-03</td><td>BT20/MB468</td><td>MP</td></tr>
<tr><td>>hg38_21_+_8433172_8446622_RNA45SN1_45S_50ntFlanks@11668.11688.21|TGCCCAGTGCTCTGAATGTCA</td><td>>hg38_21_+_8433172_8446622_RNA45SN1_45S_50ntFlanks</td><td>11668</td><td>11688</td><td>21</td><td>TGCCCAGTGCTCTGAATGTCA</td><td>116.8</td><td>3.5</td><td>1.2</td><td>2.8</td><td>4.8E-03</td><td>5.6E-03</td><td>MB231/MB468</td><td>MITO</td></tr>
<tr><td>>MI0000062|hsa-let-7a-3&WithFlank&22|+|46112743|46112828@58.78.21;MI0000060|hsa-let-7a-1&WithFlank&9|+|94175951|94176042@63.83.21&[MIMAT0004481&hsa-let-7a-3p&offsets|0|0;m-180&22|+|46112800|46112821&offsets|0|-1];[MIMAT0004481_1&hsa-let-7a-3p&offsets|0|0;m-181&9|+|94176013|94176034&offsets|0|-1]|CTATACAATCTACTGTCTTTC</td><td>>MI0000062|hsa-let-7a-3&WithFlank&22|+|46112743|46112828</td><td>58</td><td>78</td><td>21</td><td>CTATACAATCTACTGTCTTTC</td><td>83.1</td><td>3.7</td><td>1.3</td><td>2.9</td><td>4.1E-03</td><td>5.6E-03</td><td>MB231/MB468</td><td>MP</td></tr>
<tr><td>>hg38_21_+_8433172_8446622_RNA45SN1_45S_50ntFlanks@9917.9948.32|AAGGTGTTGGTTGATATAGACAGCAGGACGGT</td><td>>hg38_21_+_8433172_8446622_RNA45SN1_45S_50ntFlanks</td><td>9917</td><td>9948</td><td>32</td><td>AAGGTGTTGGTTGATATAGACAGCAGGACGGT</td><td>65.0</td><td>5.8</td><td>1.9</td><td>3.0</td><td>2.5E-03</td><td>5.6E-03</td><td>BT20/MB468</td><td>MP</td></tr>
<tr><td>>hg38_21_+_8433172_8446622_RNA45SN1_45S_50ntFlanks@10802.10827.26|TTGGAACAATGTAGGTAAGGGAAGTC</td><td>>hg38_21_+_8433172_8446622_RNA45SN1_45S_50ntFlanks</td><td>10802</td><td>10827</td><td>26</td><td>TTGGAACAATGTAGGTAAGGGAAGTC</td><td>171.6</td><td>-2.9</td><td>1.0</td><td>-3.0</td><td>2.5E-03</td><td>5.6E-03</td><td>BT20/MB468</td><td>MP</td></tr>
<tr><td>>hg38_21_+_8433172_8446622_RNA45SN1_45S_50ntFlanks@6649.6674.26|TACGACTCTTAGCGGTGGATCACTCG</td><td>>hg38_21_+_8433172_8446622_RNA45SN1_45S_50ntFlanks</td><td>6649</td><td>6674</td><td>26</td><td>TACGACTCTTAGCGGTGGATCACTCG</td><td>147.5</td><td>3.6</td><td>1.2</td><td>3.0</td><td>2.5E-03</td><td>5.6E-03</td><td>BT20/MB468</td><td>MP</td></tr>
<tr><td>>hg38_21_+_8433172_8446622_RNA45SN1_45S_50ntFlanks@10901.10928.28|GGGGCTGGGCGCGCGCCGCGGCTGGACG</td><td>>hg38_21_+_8433172_8446622_RNA45SN1_45S_50ntFlanks</td><td>10901</td><td>10928</td><td>28</td><td>GGGGCTGGGCGCGCGCCGCGGCTGGACG</td><td>68.3</td><td>9.4</td><td>2.8</td><td>3.3</td><td>9.0E-04</td><td>5.6E-03</td><td>BT20/MB231</td><td>MP</td></tr>
<tr><td>>hg38_21_+_8433172_8446622_RNA45SN1_45S_50ntFlanks@10846.10867.22|CTTCGGGATAAGGATTGGCTCT</td><td>>hg38_21_+_8433172_8446622_RNA45SN1_45S_50ntFlanks</td><td>10846</td><td>10867</td><td>22</td><td>CTTCGGGATAAGGATTGGCTCT</td><td>503.1</td><td>-2.1</td><td>0.7</td><td>-2.8</td><td>4.5E-03</td><td>5.6E-03</td><td>MB231/MB468</td><td>CYTO</td></tr>
<tr><td>>MI0000103|hsa-mir-101-1&WithFlank&1|-|65058428|65058514@53.74.22(+1U);MI0000739|hsa-mir-101-2&WithFlank&9|+|4850291|4850381@55.76.22(+1U)&[MIMAT0000099&hsa-miR-101-3p&offsets|0|+1(+1U);m-17&1|-|65058443|65058462&offsets|0|+2(+1U)];[MIMAT0000099_1&hsa-miR-101-3p&offsets|0|+1(+1U);m-16&9|+|4850345|4850364&offsets|0|+2(+1U)]|TACAGTACTGTGATAACTGAAGT</td><td>>MI0000103|hsa-mir-101-1&WithFlank&1|-|65058428|65058514</td><td>53</td><td>74</td><td>22(+1U)</td><td>TACAGTACTGTGATAACTGAAGT</td><td>491.1</td><td>2.6</td><td>0.9</td><td>2.9</td><td>4.1E-03</td><td>5.6E-03</td><td>MB231/MB468</td><td>MP</td></tr>
<tr><td>>MI0000750|hsa-mir-26a-2&WithFlank&12|-|57824603|57824698@21.41.21;MI0000083|hsa-mir-26a-1&WithFlank&3|+|37969398|37969486@17.37.21&[MIMAT0000082&hsa-miR-26a-5p&offsets|+1|0;m-33&12|-|57824658|57824679&offsets|+1|0];[MIMAT0000082_1&hsa-miR-26a-5p&offsets|+1|0;m-34&3|+|37969413|37969434&offsets|+1|0]|TCAAGTAATCCAGGATAGGCT</td><td>>MI0000750|hsa-mir-26a-2&WithFlank&12|-|57824603|57824698</td><td>21</td><td>41</td><td>21</td><td>TCAAGTAATCCAGGATAGGCT</td><td>114.5</td><td>4.7</td><td>1.6</td><td>2.9</td><td>4.1E-03</td><td>5.6E-03</td><td>MB231/MB468</td><td>MP</td></tr>
<tr><td>>hg38_21_+_8433172_8446622_RNA45SN1_45S_50ntFlanks@8342.8378.37|CGTAAGGGAAAGTTGAAAAGAACTTTGAAGAGAGAGT</td><td>>hg38_21_+_8433172_8446622_RNA45SN1_45S_50ntFlanks</td><td>8342</td><td>8378</td><td>37</td><td>CGTAAGGGAAAGTTGAAAAGAACTTTGAAGAGAGAGT</td><td>160.7</td><td>4.6</td><td>1.6</td><td>2.9</td><td>4.1E-03</td><td>5.6E-03</td><td>MB231/MB468</td><td>MP</td></tr>
<tr><td>>am_MI0000809|hsa-mir-151a&WithFlank&8|-|140732558|140732659@52.72.21&[MIMAT0000757&hsa-miR-151a-3p&offsets|-1|-1;m-30&8|-|140732587|140732607&offsets|-1|-1]|ACTAGACTGAAGCTCCTTGAG</td><td>>am_MI0000809|hsa-mir-151a&WithFlank&8|-|140732558|140732659</td><td>52</td><td>72</td><td>21</td><td>ACTAGACTGAAGCTCCTTGAG</td><td>132.6</td><td>5.2</td><td>1.7</td><td>3.0</td><td>2.5E-03</td><td>5.6E-03</td><td>BT20/MB468</td><td>MP</td></tr>
<tr><td>>hg38_21_+_8433172_8446622_RNA45SN1_45S_50ntFlanks@7981.8012.32|CTCAGATCAGACGTGGCGACCCGCTGAATTTA</td><td>>hg38_21_+_8433172_8446622_RNA45SN1_45S_50ntFlanks</td><td>7981</td><td>8012</td><td>32</td><td>CTCAGATCAGACGTGGCGACCCGCTGAATTTA</td><td>270.9</td><td>-5.0</td><td>1.7</td><td>-3.0</td><td>2.5E-03</td><td>5.6E-03</td><td>BT20/MB468</td><td>MP</td></tr>
<tr><td>>trna2_GlyGCC_21_-_18827107_18827177@1.29.29,trna35_GlyGCC_1_+_161413094_161413164@1.29.29,trna37_GlyGCC_1_+_161420467_161420537@1.29.29,trna39_GlyGCC_1_+_161427898_161427968@1.29.29,trna41_GlyGCC_1_+_161435258_161435328@1.29.29|GCATGGGTGGTTCAGTGGTAGAATTCTCG</td><td>>trna2_GlyGCC_21_-_18827107_18827177</td><td>1</td><td>29</td><td>29</td><td>GCATGGGTGGTTCAGTGGTAGAATTCTCG</td><td>1021.9</td><td>5.3</td><td>1.6</td><td>3.3</td><td>9.1E-04</td><td>5.6E-03</td><td>BT20/MB231</td><td>MP</td></tr>
<tr><td>>hg38_1_-_228634819_228635039_RNA5S12_5S_50ntFlanks@149.170.22|GAATACCGGGTGCTGTAGGCTT</td><td>>hg38_1_-_228634819_228635039_RNA5S12_5S_50ntFlanks</td><td>149</td><td>170</td><td>22</td><td>GAATACCGGGTGCTGTAGGCTT</td><td>7205.0</td><td>3.6</td><td>1.1</td><td>3.3</td><td>9.1E-04</td><td>5.6E-03</td><td>BT20/MB231</td><td>MP</td></tr>
<tr><td>>hg38_MT_+_1621_3279_RNR2_16S_50ntFlanks@518.539.22|TTGTAGAGAGAGTAAAAAATTT</td><td>>hg38_MT_+_1621_3279_RNR2_16S_50ntFlanks</td><td>518</td><td>539</td><td>22</td><td>TTGTAGAGAGAGTAAAAAATTT</td><td>92.0</td><td>4.8</td><td>1.7</td><td>2.9</td><td>4.1E-03</td><td>5.6E-03</td><td>MB231/MB468</td><td>MP</td></tr>
<tr><td>>hg38_21_+_8433172_8446622_RNA45SN1_45S_50ntFlanks@6649.6674.26|TACGACTCTTAGCGGTGGATCACTCG</td><td>>hg38_21_+_8433172_8446622_RNA45SN1_45S_50ntFlanks</td><td>6649</td><td>6674</td><td>26</td><td>TACGACTCTTAGCGGTGGATCACTCG</td><td>806.8</td><td>3.0</td><td>1.1</td><td>2.8</td><td>4.8E-03</td><td>5.6E-03</td><td>MB231/MB468</td><td>MITO</td></tr>
<tr><td>>hg38_21_+_8433172_8446622_RNA45SN1_45S_50ntFlanks@3782.3812.31|CAGTGAAACTGCGAATGGCTCATTAAATCAG</td><td>>hg38_21_+_8433172_8446622_RNA45SN1_45S_50ntFlanks</td><td>3782</td><td>3812</td><td>31</td><td>CAGTGAAACTGCGAATGGCTCATTAAATCAG</td><td>153.6</td><td>-4.1</td><td>1.5</td><td>-2.8</td><td>4.5E-03</td><td>5.6E-03</td><td>BT20/MB468</td><td>CYTO</td></tr>
<tr><td>>hg38_21_+_8433172_8446622_RNA45SN1_45S_50ntFlanks@10238.10262.25|CGCTACGCCGCGACGAGTAGGAGGG</td><td>>hg38_21_+_8433172_8446622_RNA45SN1_45S_50ntFlanks</td><td>10238</td><td>10262</td><td>25</td><td>CGCTACGCCGCGACGAGTAGGAGGG</td><td>226.7</td><td>6.1</td><td>1.8</td><td>3.3</td><td>9.1E-04</td><td>5.6E-03</td><td>BT20/MB231</td><td>MP</td></tr>
<tr><td>>am_trnaMT_TyrGTA_MT_-_5826_5891@13.33.21|GAGTGAAGCATTGGACTGTAA</td><td>>am_trnaMT_TyrGTA_MT_-_5826_5891</td><td>13</td><td>33</td><td>21</td><td>GAGTGAAGCATTGGACTGTAA</td><td>29.9</td><td>8.1</td><td>2.8</td><td>2.9</td><td>4.1E-03</td><td>5.6E-03</td><td>MB231/MB468</td><td>MP</td></tr>
<tr><td>>hg38_21_+_8433172_8446622_RNA45SN1_45S_50ntFlanks@10681.10709.29|TTCCGGCGGCGTCCGGTGAGCTCTCGCTG</td><td>>hg38_21_+_8433172_8446622_RNA45SN1_45S_50ntFlanks</td><td>10681</td><td>10709</td><td>29</td><td>TTCCGGCGGCGTCCGGTGAGCTCTCGCTG</td><td>183.4</td><td>-4.9</td><td>1.6</td><td>-3.0</td><td>2.6E-03</td><td>5.6E-03</td><td>BT20/MB468</td><td>MP</td></tr>
<tr><td>>hg38_MT_+_1621_3279_RNR2_16S_50ntFlanks@1370.1398.29|ATGTTGGATCAGGACATCCCGATGGTGCA</td><td>>hg38_MT_+_1621_3279_RNR2_16S_50ntFlanks</td><td>1370</td><td>1398</td><td>29</td><td>ATGTTGGATCAGGACATCCCGATGGTGCA</td><td>108.4</td><td>3.8</td><td>1.3</td><td>2.9</td><td>4.1E-03</td><td>5.6E-03</td><td>MB231/MB468</td><td>MP</td></tr>
<tr><td>>hg38_21_+_8433172_8446622_RNA45SN1_45S_50ntFlanks@10357.10383.27|CTTTGAAGGCCGAAGTGGAGAAGGGTT</td><td>>hg38_21_+_8433172_8446622_RNA45SN1_45S_50ntFlanks</td><td>10357</td><td>10383</td><td>27</td><td>CTTTGAAGGCCGAAGTGGAGAAGGGTT</td><td>187.0</td><td>-1.7</td><td>0.6</td><td>-2.9</td><td>3.8E-03</td><td>5.6E-03</td><td>BT20/MB468</td><td>MITO</td></tr>
<tr><td>>hg38_21_+_8433172_8446622_RNA45SN1_45S_50ntFlanks@4898.4918.21|AAAGGAATTGACGGAAGGGCA</td><td>>hg38_21_+_8433172_8446622_RNA45SN1_45S_50ntFlanks</td><td>4898</td><td>4918</td><td>21</td><td>AAAGGAATTGACGGAAGGGCA</td><td>126.4</td><td>-5.4</td><td>1.8</td><td>-3.0</td><td>2.6E-03</td><td>5.6E-03</td><td>BT20/MB468</td><td>MP</td></tr>
<tr><td>>hg38_21_+_8433172_8446622_RNA45SN1_45S_50ntFlanks@4283.4307.25|CTTTAACGAGGATCCATTGGAGGGC</td><td>>hg38_21_+_8433172_8446622_RNA45SN1_45S_50ntFlanks</td><td>4283</td><td>4307</td><td>25</td><td>CTTTAACGAGGATCCATTGGAGGGC</td><td>440.0</td><td>-5.1</td><td>1.7</td><td>-3.0</td><td>2.6E-03</td><td>5.6E-03</td><td>BT20/MB468</td><td>MP</td></tr>
<tr><td>>hg38_21_+_8433172_8446622_RNA45SN1_45S_50ntFlanks@4728.4751.24|ATCAAGAACGAAAGTCGGAGGTTC</td><td>>hg38_21_+_8433172_8446622_RNA45SN1_45S_50ntFlanks</td><td>4728</td><td>4751</td><td>24</td><td>ATCAAGAACGAAAGTCGGAGGTTC</td><td>699.1</td><td>-4.7</td><td>1.7</td><td>-2.8</td><td>4.9E-03</td><td>5.6E-03</td><td>MB231/MB468</td><td>NUC</td></tr>
<tr><td>>hg38_21_+_8433172_8446622_RNA45SN1_45S_50ntFlanks@5238.5257.20|CTGACTGGCTCAGCGTGTGC</td><td>>hg38_21_+_8433172_8446622_RNA45SN1_45S_50ntFlanks</td><td>5238</td><td>5257</td><td>20</td><td>CTGACTGGCTCAGCGTGTGC</td><td>81.3</td><td>-3.9</td><td>1.3</td><td>-2.9</td><td>4.1E-03</td><td>5.6E-03</td><td>MB231/MB468</td><td>MP</td></tr>
<tr><td>>hg38_21_+_8433172_8446622_RNA45SN1_45S_50ntFlanks@4511.4528.18|GAAAAAATTAGAGTGTTC</td><td>>hg38_21_+_8433172_8446622_RNA45SN1_45S_50ntFlanks</td><td>4511</td><td>4528</td><td>18</td><td>GAAAAAATTAGAGTGTTC</td><td>147.2</td><td>-5.3</td><td>1.7</td><td>-3.0</td><td>2.6E-03</td><td>5.6E-03</td><td>BT20/MB468</td><td>MP</td></tr>
<tr><td>>MI0000086|hsa-mir-28&WithFlank&3|+|188688775|188688872@20.40.21(+1A)&[MIMAT0000085&hsa-miR-28-5p&offsets|0|-1(+1A);m-133&3|+|188688794|188688815&offsets|0|-1(+1A)]|AAGGAGCTCACAGTCTATTGAA</td><td>>MI0000086|hsa-mir-28&WithFlank&3|+|188688775|188688872</td><td>20</td><td>40</td><td>21(+1A)</td><td>AAGGAGCTCACAGTCTATTGAA</td><td>87.4</td><td>5.2</td><td>1.7</td><td>3.0</td><td>2.6E-03</td><td>5.6E-03</td><td>BT20/MB468</td><td>MP</td></tr>
<tr><td>>hg38_21_+_8433172_8446622_RNA45SN1_45S_50ntFlanks@3837.3862.26|CCTCTCCTACTTGGATAACTGTGGTA</td><td>>hg38_21_+_8433172_8446622_RNA45SN1_45S_50ntFlanks</td><td>3837</td><td>3862</td><td>26</td><td>CCTCTCCTACTTGGATAACTGTGGTA</td><td>152.5</td><td>-3.8</td><td>1.3</td><td>-2.9</td><td>3.8E-03</td><td>5.6E-03</td><td>BT20/MB468</td><td>MITO</td></tr>
<tr><td>>hg38_21_+_8433172_8446622_RNA45SN1_45S_50ntFlanks@10350.10378.29|ACGAGAACTTTGAAGGCCGAAGTGGAGAA</td><td>>hg38_21_+_8433172_8446622_RNA45SN1_45S_50ntFlanks</td><td>10350</td><td>10378</td><td>29</td><td>ACGAGAACTTTGAAGGCCGAAGTGGAGAA</td><td>253.2</td><td>-2.7</td><td>1.0</td><td>-2.8</td><td>4.6E-03</td><td>5.6E-03</td><td>MB231/MB468</td><td>CYTO</td></tr>
<tr><td>>MI0000077|hsa-mir-21&WithFlank&17|+|59841260|59841343@14.34.21(+1U)&[MIMAT0000076&hsa-miR-21-5p&offsets|0|-1(+1U);m-1&17|+|59841273|59841294&offsets|0|-1(+1U)]|TAGCTTATCAGACTGATGTTGT</td><td>>MI0000077|hsa-mir-21&WithFlank&17|+|59841260|59841343</td><td>14</td><td>34</td><td>21(+1U)</td><td>TAGCTTATCAGACTGATGTTGT</td><td>2910.9</td><td>3.4</td><td>1.1</td><td>3.3</td><td>1.1E-03</td><td>5.6E-03</td><td>BT20/MB231</td><td>CYTO</td></tr>
<tr><td>>hg38_21_+_8433172_8446622_RNA45SN1_45S_50ntFlanks@12420.12446.27|ATGTCGGCTCTTCCTATCATTGTGAAG</td><td>>hg38_21_+_8433172_8446622_RNA45SN1_45S_50ntFlanks</td><td>12420</td><td>12446</td><td>27</td><td>ATGTCGGCTCTTCCTATCATTGTGAAG</td><td>1104.6</td><td>-2.9</td><td>0.9</td><td>-3.3</td><td>1.1E-03</td><td>5.6E-03</td><td>BT20/MB231</td><td>CYTO</td></tr>
<tr><td>>MI0000087|hsa-mir-29a&WithFlank&7|-|130876741|130876816@48.70.23&[MIMAT0000086&hsa-miR-29a-3p&offsets|0|+1;m-26&7|-|130876749|130876769&offsets|0|+2]|TAGCACCATCTGAAATCGGTTAT</td><td>>MI0000087|hsa-mir-29a&WithFlank&7|-|130876741|130876816</td><td>48</td><td>70</td><td>23</td><td>TAGCACCATCTGAAATCGGTTAT</td><td>451.0</td><td>6.0</td><td>2.0</td><td>3.0</td><td>2.6E-03</td><td>5.6E-03</td><td>BT20/MB468</td><td>MP</td></tr>
<tr><td>>hg38_21_+_8433172_8446622_RNA45SN1_45S_50ntFlanks@4169.4186.18|AGGAAGGCAGCAGGCGCG</td><td>>hg38_21_+_8433172_8446622_RNA45SN1_45S_50ntFlanks</td><td>4169</td><td>4186</td><td>18</td><td>AGGAAGGCAGCAGGCGCG</td><td>75.0</td><td>-4.8</td><td>1.6</td><td>-3.0</td><td>2.6E-03</td><td>5.6E-03</td><td>BT20/MB468</td><td>MP</td></tr>
<tr><td>>MI0003126|hsa-mir-491&WithFlank&9|+|20716099|20716194@22.44.23&[MIMAT0002807&hsa-miR-491-5p&offsets|0|+1;m-451&9|+|20716120|20716141&offsets|0|+1]|AGTGGGGAACCCTTCCATGAGGA</td><td>>MI0003126|hsa-mir-491&WithFlank&9|+|20716099|20716194</td><td>22</td><td>44</td><td>23</td><td>AGTGGGGAACCCTTCCATGAGGA</td><td>157.5</td><td>-5.9</td><td>1.8</td><td>-3.3</td><td>9.2E-04</td><td>5.6E-03</td><td>BT20/MB231</td><td>MP</td></tr>
<tr><td>>trna14_LysTTT_11_-_59327808_59327880@1.31.31,trna2_LysTTT_17_+_8022473_8022545@1.31.31,trna54_LysTTT_1_+_204475655_204475727@1.31.31,trna5_LysTTT_11_+_59323902_59323974@1.31.31,trna62_LysTTT_1_-_204476158_204476230@1.31.31,trna76_LysTTT_6_+_28918806_28918878@1.31.31|GCCCGGATAGCTCAGTCGGTAGAGCATCAGA</td><td>>trna14_LysTTT_11_-_59327808_59327880</td><td>1</td><td>31</td><td>31</td><td>GCCCGGATAGCTCAGTCGGTAGAGCATCAGA</td><td>232.5</td><td>-2.4</td><td>0.8</td><td>-2.8</td><td>4.6E-03</td><td>5.6E-03</td><td>MB231/MB468</td><td>CYTO</td></tr>
<tr><td>>hg38_21_+_8433172_8446622_RNA45SN1_45S_50ntFlanks@12632.12659.28|GCGAAGCTACCATCTGTGGGATTATGAC</td><td>>hg38_21_+_8433172_8446622_RNA45SN1_45S_50ntFlanks</td><td>12632</td><td>12659</td><td>28</td><td>GCGAAGCTACCATCTGTGGGATTATGAC</td><td>1446.6</td><td>-2.5</td><td>0.9</td><td>-2.9</td><td>3.9E-03</td><td>5.6E-03</td><td>BT20/MB468</td><td>MITO</td></tr>
<tr><td>>hg38_21_+_8433172_8446622_RNA45SN1_45S_50ntFlanks@10512.10532.21|TCCGGAGTGGCGGAGATGGGC</td><td>>hg38_21_+_8433172_8446622_RNA45SN1_45S_50ntFlanks</td><td>10512</td><td>10532</td><td>21</td><td>TCCGGAGTGGCGGAGATGGGC</td><td>789.1</td><td>6.1</td><td>1.8</td><td>3.3</td><td>9.2E-04</td><td>5.6E-03</td><td>BT20/MB231</td><td>MP</td></tr>
</table>

<table>
<tr><td>>trna119_LysCTT_1_-_145395522_145395594@1.32.32,trna11_LysCTT_5_-_180648979_180649051@1.32.32,trna13_LysCTT_6_+_26556774_26556846@1.32.32,trna32_LysCTT_16_-_3207406_3207478@1.32.32,trna7_LysCTT_16_+_3225692_3225764@1.32.32,trna9_LysCTT_5_+_180634755_180634827@1.32.32|GCCCGGCTAGCTCAGTCGGTAGAGCATGAGAC</td><td>>trna119_LysCTT_1_-_145395522_145395594</td><td>1</td><td>32</td><td>32</td><td>GCCCGGCTAGCTCAGTCGGTAGAGCATGAGAC</td><td>8288.8</td><td>2.9</td><td>1.0</td><td>2.8</td><td>4.6E-03</td><td>5.6E-03</td><td>MB231/MB468</td><td>CYTO</td></tr>
<tr><td>>MI0000084|hsa-mir-26b&WithFlank&2|+|218402640|218402728@18.39.22&[MIMAT0000083&hsa-miR-26b-5p&offsets|0|+1;m-53&2|+|218402657|218402678&offsets|0|0]|TTCAAGTAATTCAGGATAGGTT</td><td>>MI0000084|hsa-mir-26b&WithFlank&2|+|218402640|218402728</td><td>18</td><td>39</td><td>22</td><td>TTCAAGTAATTCAGGATAGGTT</td><td>5432.4</td><td>2.8</td><td>1.0</td><td>2.9</td><td>4.2E-03</td><td>5.6E-03</td><td>MB231/MB468</td><td>MP</td></tr>
<tr><td>>hg38_21_+_8433172_8446622_RNA45SN1_45S_50ntFlanks@11210.11232.23|GGCGACTCTGGACGCGAGCCGGG</td><td>>hg38_21_+_8433172_8446622_RNA45SN1_45S_50ntFlanks</td><td>11210</td><td>11232</td><td>23</td><td>GGCGACTCTGGACGCGAGCCGGG</td><td>1090.0</td><td>-5.0</td><td>1.8</td><td>-2.8</td><td>4.5E-03</td><td>5.6E-03</td><td>BT20/MB468</td><td>CYTO</td></tr>
<tr><td>>MI0000088|hsa-mir-30a&WithFlank&6|-|71403545|71403627@53.72.20&[MIMAT0000088&hsa-miR-30a-3p&offsets|0|-2;m-38&6|-|71403554|71403575&offsets|0|-2]|CTTTCAGTCGGATGTTTGCA</td><td>>MI0000088|hsa-mir-30a&WithFlank&6|-|71403545|71403627</td><td>53</td><td>72</td><td>20</td><td>CTTTCAGTCGGATGTTTGCA</td><td>267.8</td><td>3.7</td><td>1.3</td><td>2.9</td><td>4.2E-03</td><td>5.6E-03</td><td>MB231/MB468</td><td>MP</td></tr>
<tr><td>>hg38_21_+_8433172_8446622_RNA45SN1_45S_50ntFlanks@10822.10839.18|GAAGTCGGCAAGCCGGAT</td><td>>hg38_21_+_8433172_8446622_RNA45SN1_45S_50ntFlanks</td><td>10822</td><td>10839</td><td>18</td><td>GAAGTCGGCAAGCCGGAT</td><td>114.6</td><td>-6.2</td><td>2.1</td><td>-2.9</td><td>4.2E-03</td><td>5.6E-03</td><td>MB231/MB468</td><td>MP</td></tr>
<tr><td>>hg38_21_+_8433172_8446622_RNA45SN1_45S_50ntFlanks@9534.9570.37|AGGAAACTCTGGTGGAGGTCCGTAGCGGTCCTGACGT</td><td>>hg38_21_+_8433172_8446622_RNA45SN1_45S_50ntFlanks</td><td>9534</td><td>9570</td><td>37</td><td>AGGAAACTCTGGTGGAGGTCCGTAGCGGTCCTGACGT</td><td>2824.9</td><td>-1.8</td><td>0.6</td><td>-2.8</td><td>4.6E-03</td><td>5.6E-03</td><td>MB231/MB468</td><td>CYTO</td></tr>
<tr><td>>hg38_21_+_8433172_8446622_RNA45SN1_45S_50ntFlanks@7975.7996.22|CGCGACCTCAGATCAGACGTGG</td><td>>hg38_21_+_8433172_8446622_RNA45SN1_45S_50ntFlanks</td><td>7975</td><td>7996</td><td>22</td><td>CGCGACCTCAGATCAGACGTGG</td><td>4858.4</td><td>1.9</td><td>0.7</td><td>2.8</td><td>4.9E-03</td><td>5.6E-03</td><td>MB231/MB468</td><td>NUC</td></tr>
<tr><td>>hg38_21_+_8433172_8446622_RNA45SN1_45S_50ntFlanks@3785.3812.28|TGAAACTGCGAATGGCTCATTAAATCAG</td><td>>hg38_21_+_8433172_8446622_RNA45SN1_45S_50ntFlanks</td><td>3785</td><td>3812</td><td>28</td><td>TGAAACTGCGAATGGCTCATTAAATCAG</td><td>164.1</td><td>-4.1</td><td>1.5</td><td>-2.8</td><td>4.5E-03</td><td>5.7E-03</td><td>BT20/MB468</td><td>CYTO</td></tr>
<tr><td>>hg38_21_+_8433172_8446622_RNA45SN1_45S_50ntFlanks@10781.10809.29|AGGTGAACAGCCTCTGGCATGTTGGAACA</td><td>>hg38_21_+_8433172_8446622_RNA45SN1_45S_50ntFlanks</td><td>10781</td><td>10809</td><td>29</td><td>AGGTGAACAGCCTCTGGCATGTTGGAACA</td><td>195.6</td><td>-3.3</td><td>1.2</td><td>-2.8</td><td>4.9E-03</td><td>5.7E-03</td><td>MB231/MB468</td><td>NUC</td></tr>
<tr><td>>hg38_21_+_8433172_8446622_RNA45SN1_45S_50ntFlanks@8178.8203.26|TCTGATCGAGGCCCAGCCCGTGGACG</td><td>>hg38_21_+_8433172_8446622_RNA45SN1_45S_50ntFlanks</td><td>8178</td><td>8203</td><td>26</td><td>TCTGATCGAGGCCCAGCCCGTGGACG</td><td>340.3</td><td>1.3</td><td>0.5</td><td>2.9</td><td>3.9E-03</td><td>5.7E-03</td><td>BT20/MB468</td><td>MITO</td></tr>
<tr><td>>hg38_21_+_8433172_8446622_RNA45SN1_45S_50ntFlanks@10392.10409.18|ACAGCAGTTGAACATGGG</td><td>>hg38_21_+_8433172_8446622_RNA45SN1_45S_50ntFlanks</td><td>10392</td><td>10409</td><td>18</td><td>ACAGCAGTTGAACATGGG</td><td>668.4</td><td>-5.3</td><td>1.7</td><td>-3.0</td><td>2.6E-03</td><td>5.7E-03</td><td>BT20/MB468</td><td>MP</td></tr>
<tr><td>>hg38_21_+_8433172_8446622_RNA45SN1_45S_50ntFlanks@8192.8211.20|AGCCCGTGGACGGTGTGAGG</td><td>>hg38_21_+_8433172_8446622_RNA45SN1_45S_50ntFlanks</td><td>8192</td><td>8211</td><td>20</td><td>AGCCCGTGGACGGTGTGAGG</td><td>514.4</td><td>-2.6</td><td>0.8</td><td>-3.0</td><td>2.6E-03</td><td>5.7E-03</td><td>BT20/MB468</td><td>MP</td></tr>
<tr><td>>hg38_21_+_8433172_8446622_RNA45SN1_45S_50ntFlanks@11672.11701.30|CAGTGCTCTGAATGTCAAAGTGAAGAAATT</td><td>>hg38_21_+_8433172_8446622_RNA45SN1_45S_50ntFlanks</td><td>11672</td><td>11701</td><td>30</td><td>CAGTGCTCTGAATGTCAAAGTGAAGAAATT</td><td>118.7</td><td>-2.4</td><td>0.8</td><td>-2.9</td><td>4.2E-03</td><td>5.7E-03</td><td>MB231/MB468</td><td>MP</td></tr>
<tr><td>>hg38_21_+_8433172_8446622_RNA45SN1_45S_50ntFlanks@9533.9562.30|GAGGAAACTCTGGTGGAGGTCCGTAGCGGT</td><td>>hg38_21_+_8433172_8446622_RNA45SN1_45S_50ntFlanks</td><td>9533</td><td>9562</td><td>30</td><td>GAGGAAACTCTGGTGGAGGTCCGTAGCGGT</td><td>321.2</td><td>-2.5</td><td>0.8</td><td>-3.0</td><td>2.6E-03</td><td>5.7E-03</td><td>BT20/MB468</td><td>MP</td></tr>
<tr><td>>hg38_21_+_8433172_8446622_RNA45SN1_45S_50ntFlanks@6734.6773.40|ATTGATCATCGACACTTCGAACGCACTTGCGGCCCCGGGT</td><td>>hg38_21_+_8433172_8446622_RNA45SN1_45S_50ntFlanks</td><td>6734</td><td>6773</td><td>40</td><td>ATTGATCATCGACACTTCGAACGCACTTGCGGCCCCGGGT</td><td>80.9</td><td>5.7</td><td>2.0</td><td>2.9</td><td>4.2E-03</td><td>5.7E-03</td><td>MB231/MB468</td><td>MP</td></tr>
<tr><td>>hg38_21_+_8433172_8446622_RNA45SN1_45S_50ntFlanks@10905.10933.29|CTGGGCGCGCGCCGCGGCTGGACGAGGCG</td><td>>hg38_21_+_8433172_8446622_RNA45SN1_45S_50ntFlanks</td><td>10905</td><td>10933</td><td>29</td><td>CTGGGCGCGCGCCGCGGCTGGACGAGGCG</td><td>250.8</td><td>-4.4</td><td>1.5</td><td>-2.8</td><td>4.6E-03</td><td>5.7E-03</td><td>BT20/MB468</td><td>CYTO</td></tr>
<tr><td>>hg38_21_+_8433172_8446622_RNA45SN1_45S_50ntFlanks@5446.5465.20|CGGATCGGCCCCGCCGGGGT</td><td>>hg38_21_+_8433172_8446622_RNA45SN1_45S_50ntFlanks</td><td>5446</td><td>5465</td><td>20</td><td>CGGATCGGCCCCGCCGGGGT</td><td>370.2</td><td>-5.1</td><td>1.7</td><td>-3.0</td><td>2.6E-03</td><td>5.7E-03</td><td>BT20/MB468</td><td>MP</td></tr>
<tr><td>>hg38_21_+_8433172_8446622_RNA45SN1_45S_50ntFlanks@10856.10873.18|AGGATTGGCTCTAAGGGC</td><td>>hg38_21_+_8433172_8446622_RNA45SN1_45S_50ntFlanks</td><td>10856</td><td>10873</td><td>18</td><td>AGGATTGGCTCTAAGGGC</td><td>177.9</td><td>-1.8</td><td>0.6</td><td>-2.8</td><td>4.9E-03</td><td>5.7E-03</td><td>MB231/MB468</td><td>NUC</td></tr>
<tr><td>>hg38_21_+_8433172_8446622_RNA45SN1_45S_50ntFlanks@12913.12933.21|ACGTTCGTGGGGAACCTGGCG</td><td>>hg38_21_+_8433172_8446622_RNA45SN1_45S_50ntFlanks</td><td>12913</td><td>12933</td><td>21</td><td>ACGTTCGTGGGGAACCTGGCG</td><td>384.6</td><td>-3.3</td><td>1.2</td><td>-2.8</td><td>4.6E-03</td><td>5.7E-03</td><td>MB231/MB468</td><td>CYTO</td></tr>
<tr><td>>hg38_21_+_8433172_8446622_RNA45SN1_45S_50ntFlanks@8357.8391.35|AAAAGAACTTTGAAGAGAGAGTTCAAGAGGGCGTG</td><td>>hg38_21_+_8433172_8446622_RNA45SN1_45S_50ntFlanks</td><td>8357</td><td>8391</td><td>35</td><td>AAAAGAACTTTGAAGAGAGAGTTCAAGAGGGCGTG</td><td>353.2</td><td>4.4</td><td>1.3</td><td>3.3</td><td>9.3E-04</td><td>5.7E-03</td><td>BT20/MB231</td><td>MP</td></tr>
<tr><td>>hg38_21_+_8433172_8446622_RNA45SN1_45S_50ntFlanks@6787.6807.21|ACGCCTGTCTGAGCGTCGCTT</td><td>>hg38_21_+_8433172_8446622_RNA45SN1_45S_50ntFlanks</td><td>6787</td><td>6807</td><td>21</td><td>ACGCCTGTCTGAGCGTCGCTT</td><td>142.1</td><td>4.1</td><td>1.4</td><td>3.0</td><td>2.6E-03</td><td>5.7E-03</td><td>BT20/MB468</td><td>MP</td></tr>
<tr><td>>am_trnaMT_HisGTG_MT_+_12138_12206@19.51.33|AAACATCAGATTGTGAATCTGACAACAGAGGCT</td><td>>am_trnaMT_HisGTG_MT_+_12138_12206</td><td>19</td><td>51</td><td>33</td><td>AAACATCAGATTGTGAATCTGACAACAGAGGCT</td><td>244.3</td><td>3.6</td><td>1.2</td><td>2.9</td><td>4.2E-03</td><td>5.7E-03</td><td>MB231/MB468</td><td>MP</td></tr>
<tr><td>>am_trna117_GlyTCC_1_-_145397864_145397935@1.31.31,trna45_GlyTCC_1_+_161500903_161500974@1.31.31,trna70_GlyTCC_1_-_161439547_161439618@1.31.31,trna73_GlyTCC_1_-_161432166_161432237@1.31.31,trna76_GlyTCC_1_-_161424756_161424827@1.31.31,trna79_GlyTCC_1_-_161417375_161417446@1.31.31|GCGTTGGTGGTATAGTGGTGAGCATAGCTGC</td><td>>am_trna117_GlyTCC_1_-_145397864_145397935</td><td>1</td><td>31</td><td>31</td><td>GCGTTGGTGGTATAGTGGTGAGCATAGCTGC</td><td>116.2</td><td>-2.5</td><td>0.9</td><td>-2.9</td><td>4.2E-03</td><td>5.7E-03</td><td>MB231/MB468</td><td>MP</td></tr>
<tr><td>>hg38_21_+_8433172_8446622_RNA45SN1_45S_50ntFlanks@11666.11691.26|TCTGCCCAGTGCTCTGAATGTCAAAG</td><td>>hg38_21_+_8433172_8446622_RNA45SN1_45S_50ntFlanks</td><td>11666</td><td>11691</td><td>26</td><td>TCTGCCCAGTGCTCTGAATGTCAAAG</td><td>483.3</td><td>3.0</td><td>1.0</td><td>2.9</td><td>3.9E-03</td><td>5.7E-03</td><td>BT20/MB468</td><td>MITO</td></tr>
<tr><td>>hg38_21_+_8433172_8446622_RNA45SN1_45S_50ntFlanks@12250.12279.30|AAAAGCTCGCTTGATCTTGATTTTCAGTAC</td><td>>hg38_21_+_8433172_8446622_RNA45SN1_45S_50ntFlanks</td><td>12250</td><td>12279</td><td>30</td><td>AAAAGCTCGCTTGATCTTGATTTTCAGTAC</td><td>689.1</td><td>3.3</td><td>1.2</td><td>2.8</td><td>4.6E-03</td><td>5.7E-03</td><td>MB231/MB468</td><td>CYTO</td></tr>
<tr><td>>hg38_MT_+_598_1651_RNR1_12S_50ntFlanks@917.935.19|AGGACATTTAACTAAAACC</td><td>>hg38_MT_+_598_1651_RNR1_12S_50ntFlanks</td><td>917</td><td>935</td><td>19</td><td>AGGACATTTAACTAAAACC</td><td>379.6</td><td>3.0</td><td>1.0</td><td>2.9</td><td>4.2E-03</td><td>5.7E-03</td><td>MB231/MB468</td><td>MP</td></tr>
<tr><td>>hg38_21_+_8433172_8446622_RNA45SN1_45S_50ntFlanks@10350.10367.18|ACGAGAACTTTGAAGGCC</td><td>>hg38_21_+_8433172_8446622_RNA45SN1_45S_50ntFlanks</td><td>10350</td><td>10367</td><td>18</td><td>ACGAGAACTTTGAAGGCC</td><td>981.7</td><td>-2.6</td><td>0.9</td><td>-2.8</td><td>4.9E-03</td><td>5.7E-03</td><td>MB231/MB468</td><td>NUC</td></tr>
<tr><td>>hg38_21_+_8433172_8446622_RNA45SN1_45S_50ntFlanks@12607.12626.20|TGTGCTTGGCTGAGGAGCCA</td><td>>hg38_21_+_8433172_8446622_RNA45SN1_45S_50ntFlanks</td><td>12607</td><td>12626</td><td>20</td><td>TGTGCTTGGCTGAGGAGCCA</td><td>174.2</td><td>-4.9</td><td>1.6</td><td>-3.0</td><td>2.6E-03</td><td>5.7E-03</td><td>BT20/MB468</td><td>MP</td></tr>
<tr><td>>hg38_1_-_228634819_228635039_RNA5S12_5S_50ntFlanks@51.74.24|GTCTACGGCCATACCACCCTGAAC</td><td>>hg38_1_-_228634819_228635039_RNA5S12_5S_50ntFlanks</td><td>51</td><td>74</td><td>24</td><td>GTCTACGGCCATACCACCCTGAAC</td><td>1091.0</td><td>3.8</td><td>1.3</td><td>3.0</td><td>2.6E-03</td><td>5.7E-03</td><td>BT20/MB468</td><td>MP</td></tr>
<tr><td>>hg38_21_+_8433172_8446622_RNA45SN1_45S_50ntFlanks@9597.9615.19|AGGGGCGAAAGACTAATCG</td><td>>hg38_21_+_8433172_8446622_RNA45SN1_45S_50ntFlanks</td><td>9597</td><td>9615</td><td>19</td><td>AGGGGCGAAAGACTAATCG</td><td>199.4</td><td>-3.8</td><td>1.3</td><td>-3.0</td><td>2.6E-03</td><td>5.7E-03</td><td>BT20/MB468</td><td>MP</td></tr>
<tr><td>>hg38_21_+_8433172_8446622_RNA45SN1_45S_50ntFlanks@6646.6675.30|TCGTACGACTCTTAGCGGTGGATCACTCGG</td><td>>hg38_21_+_8433172_8446622_RNA45SN1_45S_50ntFlanks</td><td>6646</td><td>6675</td><td>30</td><td>TCGTACGACTCTTAGCGGTGGATCACTCGG</td><td>6074.2</td><td>4.6</td><td>1.4</td><td>3.3</td><td>9.3E-04</td><td>5.7E-03</td><td>BT20/MB231</td><td>MP</td></tr>
<tr><td>>hg38_21_+_8433172_8446622_RNA45SN1_45S_50ntFlanks@6745.6766.22|ACACTTCGAACGCACTTGCGGC</td><td>>hg38_21_+_8433172_8446622_RNA45SN1_45S_50ntFlanks</td><td>6745</td><td>6766</td><td>22</td><td>ACACTTCGAACGCACTTGCGGC</td><td>227.1</td><td>-5.0</td><td>1.7</td><td>-3.0</td><td>2.6E-03</td><td>5.7E-03</td><td>BT20/MB468</td><td>MP</td></tr>
<tr><td>>hg38_21_+_8433172_8446622_RNA45SN1_45S_50ntFlanks@10786.10815.30|AACAGCCTCTGGCATGTTGGAACAATGTAG</td><td>>hg38_21_+_8433172_8446622_RNA45SN1_45S_50ntFlanks</td><td>10786</td><td>10815</td><td>30</td><td>AACAGCCTCTGGCATGTTGGAACAATGTAG</td><td>164.8</td><td>-1.6</td><td>0.5</td><td>-2.9</td><td>3.9E-03</td><td>5.7E-03</td><td>BT20/MB468</td><td>MITO</td></tr>
<tr><td>>hg38_21_+_8433172_8446622_RNA45SN1_45S_50ntFlanks@12416.12444.29|TTCGATGTCGGCTCTTCCTATCATTGTGA</td><td>>hg38_21_+_8433172_8446622_RNA45SN1_45S_50ntFlanks</td><td>12416</td><td>12444</td><td>29</td><td>TTCGATGTCGGCTCTTCCTATCATTGTGA</td><td>946.9</td><td>2.3</td><td>0.8</td><td>2.8</td><td>4.9E-03</td><td>5.7E-03</td><td>MB231/MB468</td><td>NUC</td></tr>
<tr><td>>hg38_21_+_8433172_8446622_RNA45SN1_45S_50ntFlanks@10391.10426.36|AACAGCAGTTGAACATGGGTCAGTCGGTCCTGAGAG</td><td>>hg38_21_+_8433172_8446622_RNA45SN1_45S_50ntFlanks</td><td>10391</td><td>10426</td><td>36</td><td>AACAGCAGTTGAACATGGGTCAGTCGGTCCTGAGAG</td><td>292.8</td><td>-3.5</td><td>1.2</td><td>-2.8</td><td>4.6E-03</td><td>5.7E-03</td><td>BT20/MB468</td><td>CYTO</td></tr>
<tr><td>>hg38_MT_+_1621_3279_RNR2_16S_50ntFlanks@292.313.22|AGACGAGCTACCTAAGAACAGC</td><td>>hg38_MT_+_1621_3279_RNR2_16S_50ntFlanks</td><td>292</td><td>313</td><td>22</td><td>AGACGAGCTACCTAAGAACAGC</td><td>90.4</td><td>4.6</td><td>1.5</td><td>3.0</td><td>2.6E-03</td><td>5.7E-03</td><td>BT20/MB468</td><td>MP</td></tr>
<tr><td>>hg38_21_+_8433172_8446622_RNA45SN1_45S_50ntFlanks@6647.6664.18|CGTACGACTCTTAGCGGT</td><td>>hg38_21_+_8433172_8446622_RNA45SN1_45S_50ntFlanks</td><td>6647</td><td>6664</td><td>18</td><td>CGTACGACTCTTAGCGGT</td><td>1684.7</td><td>1.6</td><td>0.6</td><td>2.8</td><td>4.9E-03</td><td>5.7E-03</td><td>BT20/MB468</td><td>NUC</td></tr>
<tr><td>>hg38_21_+_8433172_8446622_RNA45SN1_45S_50ntFlanks@11855.11872.18|TGGCGGAATCAGCGGGGA</td><td>>hg38_21_+_8433172_8446622_RNA45SN1_45S_50ntFlanks</td><td>11855</td><td>11872</td><td>18</td><td>TGGCGGAATCAGCGGGGA</td><td>206.1</td><td>-3.9</td><td>1.3</td><td>-3.0</td><td>2.6E-03</td><td>5.7E-03</td><td>BT20/MB468</td><td>MP</td></tr>
<tr><td>>hg38_21_+_8433172_8446622_RNA45SN1_45S_50ntFlanks@12001.12032.32|CTGCGGGCCGCCGGTGAAATACCACTACTCTG</td><td>>hg38_21_+_8433172_8446622_RNA45SN1_45S_50ntFlanks</td><td>12001</td><td>12032</td><td>32</td><td>CTGCGGGCCGCCGGTGAAATACCACTACTCTG</td><td>3244.6</td><td>-5.2</td><td>1.8</td><td>-2.8</td><td>4.6E-03</td><td>5.7E-03</td><td>BT20/MB468</td><td>CYTO</td></tr>
<tr><td>>am_trnaMT_ValTAC_MT_+_1602_1670@47.67.21|TCAACTTAACTTGACCGCTCT</td><td>>am_trnaMT_ValTAC_MT_+_1602_1670</td><td>47</td><td>67</td><td>21</td><td>TCAACTTAACTTGACCGCTCT</td><td>321.4</td><td>3.0</td><td>0.9</td><td>3.3</td><td>1.2E-03</td><td>5.7E-03</td><td>BT20/MB231</td><td>CYTO</td></tr>
<tr><td>>hg38_21_+_8433172_8446622_RNA45SN1_45S_50ntFlanks@9927.9948.22|TTGATATAGACAGCAGGACGGT</td><td>>hg38_21_+_8433172_8446622_RNA45SN1_45S_50ntFlanks</td><td>9927</td><td>9948</td><td>22</td><td>TTGATATAGACAGCAGGACGGT</td><td>288.5</td><td>3.2</td><td>1.0</td><td>3.3</td><td>9.3E-04</td><td>5.7E-03</td><td>BT20/MB231</td><td>MP</td></tr>
<tr><td>>hg38_21_+_8433172_8446622_RNA45SN1_45S_50ntFlanks@10510.10533.24|AATCCGGAGTGGCGGAGATGGGCG</td><td>>hg38_21_+_8433172_8446622_RNA45SN1_45S_50ntFlanks</td><td>10510</td><td>10533</td><td>24</td><td>AATCCGGAGTGGCGGAGATGGGCG</td><td>615.3</td><td>7.0</td><td>2.1</td><td>3.3</td><td>9.4E-04</td><td>5.7E-03</td><td>BT20/MB231</td><td>MP</td></tr>
<tr><td>>MI0000088|hsa-mir-30a&WithFlank&6|-|71403545|71403627@53.70.18(+1A)&[MIMAT0000088&hsa-miR-30a-3p&offsets|0|-4(+1A);m-38&6|-|71403554|71403575&offsets|0|-4(+1A)]|CTTTCAGTCGGATGTTTGA</td><td>>MI0000088|hsa-mir-30a&WithFlank&6|-|71403545|71403627</td><td>53</td><td>70</td><td>18(+1A)</td><td>CTTTCAGTCGGATGTTTGA</td><td>348.4</td><td>-3.8</td><td>1.1</td><td>-3.3</td><td>9.4E-04</td><td>5.7E-03</td><td>BT20/MB231</td><td>MP</td></tr>
<tr><td>>hg38_21_+_8433172_8446622_RNA45SN1_45S_50ntFlanks@11670.11688.19|CCCAGTGCTCTGAATGTCA</td><td>>hg38_21_+_8433172_8446622_RNA45SN1_45S_50ntFlanks</td><td>11670</td><td>11688</td><td>19</td><td>CCCAGTGCTCTGAATGTCA</td><td>226.6</td><td>-2.5</td><td>0.8</td><td>-3.0</td><td>2.6E-03</td><td>5.7E-03</td><td>BT20/MB468</td><td>MP</td></tr>
<tr><td>>hg38_21_+_8433172_8446622_RNA45SN1_45S_50ntFlanks@11874.11902.29|AGAAGACCCTGTTGAGCTTGACTCTAGTC</td><td>>hg38_21_+_8433172_8446622_RNA45SN1_45S_50ntFlanks</td><td>11874</td><td>11902</td><td>29</td><td>AGAAGACCCTGTTGAGCTTGACTCTAGTC</td><td>226.7</td><td>-2.6</td><td>0.8</td><td>-3.0</td><td>2.6E-03</td><td>5.7E-03</td><td>BT20/MB468</td><td>MP</td></tr>
<tr><td>>MI0000298|hsa-mir-221&WithFlank&X|-|45746151|45746272@71.92.22(+1U)&[MIMAT0000278&hsa-miR-221-3p&offsets|0|-1(+1U);m-91&X|-|45746181|45746202&offsets|0|0(+1U)]|AGCTACATTGTCTGCTGGGTTTT</td><td>>MI0000298|hsa-mir-221&WithFlank&X|-|45746151|45746272</td><td>71</td><td>92</td><td>22(+1U)</td><td>AGCTACATTGTCTGCTGGGTTTT</td><td>2364.6</td><td>3.4</td><td>1.1</td><td>3.0</td><td>2.6E-03</td><td>5.7E-03</td><td>BT20/MB468</td><td>MP</td></tr>
<tr><td>>hg38_21_+_8433172_8446622_RNA45SN1_45S_50ntFlanks@5493.5514.22|GAAGACGGTCGAACTTGACTAT</td><td>>hg38_21_+_8433172_8446622_RNA45SN1_45S_50ntFlanks</td><td>5493</td><td>5514</td><td>22</td><td>GAAGACGGTCGAACTTGACTAT</td><td>409.6</td><td>-5.2</td><td>1.7</td><td>-3.0</td><td>2.6E-03</td><td>5.7E-03</td><td>BT20/MB468</td><td>MP</td></tr>
<tr><td>>hg38_21_+_8433172_8446622_RNA45SN1_45S_50ntFlanks@7974.7995.22|ACGCGACCTCAGATCAGACGTG</td><td>>hg38_21_+_8433172_8446622_RNA45SN1_45S_50ntFlanks</td><td>7974</td><td>7995</td><td>22</td><td>ACGCGACCTCAGATCAGACGTG</td><td>7984.7</td><td>3.3</td><td>1.2</td><td>2.8</td><td>4.6E-03</td><td>5.7E-03</td><td>BT20/MB468</td><td>CYTO</td></tr>
<tr><td>>hg38_21_+_8433172_8446622_RNA45SN1_45S_50ntFlanks@10840.10867.28|CCGTAACTTCGGGATAAGGATTGGCTCT</td><td>>hg38_21_+_8433172_8446622_RNA45SN1_45S_50ntFlanks</td><td>10840</td><td>10867</td><td>28</td><td>CCGTAACTTCGGGATAAGGATTGGCTCT</td><td>129.4</td><td>-4.7</td><td>1.7</td><td>-2.9</td><td>4.2E-03</td><td>5.7E-03</td><td>MB231/MB468</td><td>MP</td></tr>
<tr><td>>hg38_21_+_8433172_8446622_RNA45SN1_45S_50ntFlanks@12408.12435.28|TTTGATCCTTCGATGTCGGCTCTTCCTA</td><td>>hg38_21_+_8433172_8446622_RNA45SN1_45S_50ntFlanks</td><td>12408</td><td>12435</td><td>28</td><td>TTTGATCCTTCGATGTCGGCTCTTCCTA</td><td>507.2</td><td>-5.4</td><td>1.9</td><td>-2.8</td><td>4.9E-03</td><td>5.7E-03</td><td>BT20/MB468</td><td>NUC</td></tr>
<tr><td>>MI0000066|hsa-let-7e&WithFlank&19|+|51692780|51692870@14.35.22(+1U)&[MIMAT0000066&hsa-let-7e-5p&offsets|0|0(+1U);m-50&19|+|51692793|51692814&offsets|0|0(+1U)]|TGAGGTAGGAGGTTGTATAGTTT</td><td>>MI0000066|hsa-let-7e&WithFlank&19|+|51692780|51692870</td><td>14</td><td>35</td><td>22(+1U)</td><td>TGAGGTAGGAGGTTGTATAGTTT</td><td>449.7</td><td>2.6</td><td>0.9</td><td>2.9</td><td>4.2E-03</td><td>5.7E-03</td><td>MB231/MB468</td><td>MP</td></tr>
<tr><td>>hg38_21_+_8433172_8446622_RNA45SN1_45S_50ntFlanks@5487.5515.29|CGCTGAGAAGACGGTCGAACTTGACTATC</td><td>>hg38_21_+_8433172_8446622_RNA45SN1_45S_50ntFlanks</td><td>5487</td><td>5515</td><td>29</td><td>CGCTGAGAAGACGGTCGAACTTGACTATC</td><td>1167.8</td><td>-3.0</td><td>1.0</td><td>-3.0</td><td>2.6E-03</td><td>5.7E-03</td><td>BT20/MB468</td><td>MP</td></tr>
<tr><td>>hg38_MT_+_598_1651_RNR1_12S_50ntFlanks@822.840.19|GTGGATTTAGCAGTAAACT</td><td>>hg38_MT_+_598_1651_RNR1_12S_50ntFlanks</td><td>822</td><td>840</td><td>19</td><td>GTGGATTTAGCAGTAAACT</td><td>613.1</td><td>4.8</td><td>1.6</td><td>3.0</td><td>2.6E-03</td><td>5.7E-03</td><td>BT20/MB468</td><td>MP</td></tr>
</table>

<table>
<tr><td></td><td></td><td></td><td></td><td></td><td></td><td></td><td></td><td></td><td></td><td></td><td></td><td></td><td></td></tr>
<tr><td>>MI0000298|hsa-mir-221&WithFlank&X|-|45746151|45746272@71.90.20&[MIMAT0000278&hsa-miR-221-3p&offsets|0|-3;m-91&X|-|45746181|45746202&offsets|0|-2]|AGCTACATTGTCTGCTGGGT</td><td>>MI0000298|hsa-mir-221&WithFlank&X|-|45746151|45746272</td><td>71</td><td>90</td><td>20</td><td>AGCTACATTGTCTGCTGGGT</td><td>866.1</td><td>-2.4</td><td>0.7</td><td>-3.2</td><td>1.3E-03</td><td>5.7E-03</td><td>BT20/MB231</td><td>MITO</td></tr>
<tr><td>>hg38_21_+_8433172_8446622_RNA45SN1_45S_50ntFlanks@3854.3882.29|ACTGTGGTAATTCTAGAGCTAATACATGC</td><td>>hg38_21_+_8433172_8446622_RNA45SN1_45S_50ntFlanks</td><td>3854</td><td>3882</td><td>29</td><td>ACTGTGGTAATTCTAGAGCTAATACATGC</td><td>299.6</td><td>-2.3</td><td>0.8</td><td>-2.8</td><td>4.7E-03</td><td>5.7E-03</td><td>MB231/MB468</td><td>CYTO</td></tr>
<tr><td>>MI0000070|hsa-mir-16-1&WithFlank&13|-|50048967|50049067@20.41.22(+1A);MI0000115|hsa-mir-16-2&WithFlank&3|+|160404739|160404831@16.37.22(+1A)&[MIMAT0000069&hsa-miR-16-5p&offsets|0|0(+1A);m-62&13|-|50049027|50049048&offsets|0|0(+1A)];[MIMAT0000069_1&hsa-miR-16-5p&offsets|0|0(+1A);m-61&3|+|160404754|160404775&offsets|0|0(+1A)]|TAGCAGCACGTAAATATTGGCGA</td><td>>MI0000070|hsa-mir-16-1&WithFlank&13|-|50048967|50049067</td><td>20</td><td>41</td><td>22(+1A)</td><td>TAGCAGCACGTAAATATTGGCGA</td><td>186.1</td><td>3.2</td><td>1.0</td><td>3.2</td><td>1.2E-03</td><td>5.7E-03</td><td>BT20/MB231</td><td>CYTO</td></tr>
<tr><td>>hg38_21_+_8433172_8446622_RNA45SN1_45S_50ntFlanks@12644.12669.26|TCTGTGGGATTATGACTGAACGCCTC</td><td>>hg38_21_+_8433172_8446622_RNA45SN1_45S_50ntFlanks</td><td>12644</td><td>12669</td><td>26</td><td>TCTGTGGGATTATGACTGAACGCCTC</td><td>378.4</td><td>5.2</td><td>1.6</td><td>3.3</td><td>9.4E-04</td><td>5.8E-03</td><td>BT20/MB231</td><td>MP</td></tr>
<tr><td>>MI0000086|hsa-mir-28&WithFlank&3|+|188688775|188688872@20.41.22(+1A)&[MIMAT0000085&hsa-miR-28-5p&offsets|0|0(+1A);m-133&3|+|188688794|188688815&offsets|0|0(+1A)]|AAGGAGCTCACAGTCTATTGAGA</td><td>>MI0000086|hsa-mir-28&WithFlank&3|+|188688775|188688872</td><td>20</td><td>41</td><td>22(+1A)</td><td>AAGGAGCTCACAGTCTATTGAGA</td><td>170.7</td><td>3.8</td><td>1.3</td><td>2.9</td><td>4.2E-03</td><td>5.8E-03</td><td>MB231/MB468</td><td>MP</td></tr>
<tr><td>>hg38_21_+_8433172_8446622_RNA45SN1_45S_50ntFlanks@4722.4757.36|TCATTAATCAAGAACGAAAGTCGGAGGTTCGAAGAC</td><td>>hg38_21_+_8433172_8446622_RNA45SN1_45S_50ntFlanks</td><td>4722</td><td>4757</td><td>36</td><td>TCATTAATCAAGAACGAAAGTCGGAGGTTCGAAGAC</td><td>109.8</td><td>-2.5</td><td>0.9</td><td>-2.8</td><td>4.6E-03</td><td>5.8E-03</td><td>BT20/MB468</td><td>CYTO</td></tr>
<tr><td>>hg38_1_-_228634819_228635039_RNA5S12_5S_50ntFlanks@51.93.43|GTCTACGGCCATACCACCCTGAACGCGCCCGATCTCGTCTGAT</td><td>>hg38_1_-_228634819_228635039_RNA5S12_5S_50ntFlanks</td><td>51</td><td>93</td><td>43</td><td>GTCTACGGCCATACCACCCTGAACGCGCCCGATCTCGTCTGAT</td><td>512.0</td><td>7.3</td><td>2.6</td><td>2.8</td><td>4.7E-03</td><td>5.8E-03</td><td>MB231/MB468</td><td>CYTO</td></tr>
<tr><td>>hg38_21_+_8433172_8446622_RNA45SN1_45S_50ntFlanks@10769.10796.28|AGCAGGTCTCCAAGGTGAACAGCCTCTG</td><td>>hg38_21_+_8433172_8446622_RNA45SN1_45S_50ntFlanks</td><td>10769</td><td>10796</td><td>28</td><td>AGCAGGTCTCCAAGGTGAACAGCCTCTG</td><td>630.0</td><td>-2.5</td><td>0.9</td><td>-2.8</td><td>5.0E-03</td><td>5.8E-03</td><td>MB231/MB468</td><td>MITO</td></tr>
<tr><td>>hg38_21_+_8433172_8446622_RNA45SN1_45S_50ntFlanks@5498.5526.29|CGGTCGAACTTGACTATCTAGAGGAAGTA</td><td>>hg38_21_+_8433172_8446622_RNA45SN1_45S_50ntFlanks</td><td>5498</td><td>5526</td><td>29</td><td>CGGTCGAACTTGACTATCTAGAGGAAGTA</td><td>320.1</td><td>-3.9</td><td>1.3</td><td>-2.9</td><td>3.9E-03</td><td>5.8E-03</td><td>BT20/MB468</td><td>MITO</td></tr>
<tr><td>>hg38_21_+_8433172_8446622_RNA45SN1_45S_50ntFlanks@4924.4943.20|AGGAGTGGAGCCTGCGGCTT</td><td>>hg38_21_+_8433172_8446622_RNA45SN1_45S_50ntFlanks</td><td>4924</td><td>4943</td><td>20</td><td>AGGAGTGGAGCCTGCGGCTT</td><td>176.5</td><td>3.7</td><td>1.2</td><td>3.0</td><td>2.6E-03</td><td>5.8E-03</td><td>BT20/MB468</td><td>MP</td></tr>
<tr><td>>MI0000736|hsa-mir-30c-1&WithFlank&1|+|40757278|40757378@23.46.24(+1C);MI0000254|hsa-mir-30c-2&WithFlank&6|-|71376954|71377037@13.36.24(+1C)&[MIMAT0000244&hsa-miR-30c-5p&offsets|0|+1(+1C);m-57&1|+|40757300|40757323&offsets|0|0(+1C)];[MIMAT0000244_1&hsa-miR-30c-5p&offsets|0|+1(+1C);m-56&6|-|71377002|71377025&offsets|0|0(+1C)]|TGTAAACATCCTACACTCTCAGCTC</td><td>>MI0000736|hsa-mir-30c-1&WithFlank&1|+|40757278|40757378</td><td>23</td><td>46</td><td>24(+1C)</td><td>TGTAAACATCCTACACTCTCAGCTC</td><td>65.6</td><td>5.6</td><td>2.0</td><td>2.9</td><td>4.3E-03</td><td>5.8E-03</td><td>MB231/MB468</td><td>MP</td></tr>
<tr><td>>hg38_21_+_8433172_8446622_RNA45SN1_45S_50ntFlanks@10343.10373.31|TATTCAAACGAGAACTTTGAAGGCCGAAGTG</td><td>>hg38_21_+_8433172_8446622_RNA45SN1_45S_50ntFlanks</td><td>10343</td><td>10373</td><td>31</td><td>TATTCAAACGAGAACTTTGAAGGCCGAAGTG</td><td>155.0</td><td>4.1</td><td>1.4</td><td>3.0</td><td>2.6E-03</td><td>5.8E-03</td><td>BT20/MB468</td><td>MP</td></tr>
<tr><td>>MI0000102|hsa-mir-100&WithFlank&11|-|122152223|122152314@19.36.18(+2G)&[MIMAT0000098&hsa-miR-100-5p&offsets|0|-4(+2G);m-27&11|-|122152276|122152296&offsets|0|-3(+2G)]|AACCCGTAGATCCGAACTGG</td><td>>MI0000102|hsa-mir-100&WithFlank&11|-|122152223|122152314</td><td>19</td><td>36</td><td>18(+2G)</td><td>AACCCGTAGATCCGAACTGG</td><td>31.6</td><td>8.2</td><td>2.9</td><td>2.9</td><td>4.3E-03</td><td>5.8E-03</td><td>MB231/MB468</td><td>MP</td></tr>
<tr><td>>am_trna116_GluCTC_1_-_145399233_145399304@1.27.27,trna59_GluCTC_1_+_249168447_249168518@1.27.27,trna71_GluCTC_1_-_161439189_161439260@1.27.27,trna74_GluCTC_1_-_161431809_161431880@1.27.27,trna77_GluCTC_1_-_161424398_161424469@1.27.27,trna77_GluCTC_6_+_28949976_28950047@1.27.27,trna80_GluCTC_1_-_161417018_161417089@1.27.27,trna87_GluCTC_6_-_126101393_126101464@1.27.27|TCCCTGGTGGTCTAGTGGTTAGGATTC</td><td>>am_trna116_GluCTC_1_-_145399233_145399304</td><td>1</td><td>27</td><td>27</td><td>TCCCTGGTGGTCTAGTGGTTAGGATTC</td><td>1428.5</td><td>2.3</td><td>0.8</td><td>2.9</td><td>4.0E-03</td><td>5.8E-03</td><td>BT20/MB468</td><td>MITO</td></tr>
<tr><td>>MI0000542|hsa-mir-320a&WithFlank&8|-|22244960|22245043@49.69.21&[MIMAT0000510&hsa-miR-320a-3p&offsets|+1|0;m-60&8|-|22244975|22244996&offsets|+1|0]|AAAGCTGGGTTGAGAGGGCGA</td><td>>MI0000542|hsa-mir-320a&WithFlank&8|-|22244960|22245043</td><td>49</td><td>69</td><td>21</td><td>AAAGCTGGGTTGAGAGGGCGA</td><td>613.3</td><td>2.2</td><td>0.7</td><td>3.3</td><td>1.0E-03</td><td>5.8E-03</td><td>BT20/MB231</td><td>NUC</td></tr>
<tr><td>>hg38_21_+_8433172_8446622_RNA45SN1_45S_50ntFlanks@9822.9850.29|CTAGTGGGCCACTTTTGGTAAGCAGAACT</td><td>>hg38_21_+_8433172_8446622_RNA45SN1_45S_50ntFlanks</td><td>9822</td><td>9850</td><td>29</td><td>CTAGTGGGCCACTTTTGGTAAGCAGAACT</td><td>245.9</td><td>-2.6</td><td>0.9</td><td>-2.8</td><td>4.6E-03</td><td>5.8E-03</td><td>BT20/MB468</td><td>CYTO</td></tr>
<tr><td>>hg38_21_+_8433172_8446622_RNA45SN1_45S_50ntFlanks@12476.12497.22|CCACTAATAGGGAACGTGAGCT</td><td>>hg38_21_+_8433172_8446622_RNA45SN1_45S_50ntFlanks</td><td>12476</td><td>12497</td><td>22</td><td>CCACTAATAGGGAACGTGAGCT</td><td>1356.4</td><td>-2.4</td><td>0.8</td><td>-3.0</td><td>2.6E-03</td><td>5.8E-03</td><td>BT20/MB468</td><td>MP</td></tr>
<tr><td>>hg38_21_+_8433172_8446622_RNA45SN1_45S_50ntFlanks@6657.6681.25|TTAGCGGTGGATCACTCGGCTCGTG</td><td>>hg38_21_+_8433172_8446622_RNA45SN1_45S_50ntFlanks</td><td>6657</td><td>6681</td><td>25</td><td>TTAGCGGTGGATCACTCGGCTCGTG</td><td>1202.4</td><td>-4.4</td><td>1.5</td><td>-3.0</td><td>2.6E-03</td><td>5.8E-03</td><td>BT20/MB468</td><td>MP</td></tr>
<tr><td>>hg38_21_+_8433172_8446622_RNA45SN1_45S_50ntFlanks@5190.5220.31|AACAGGTCTGTGATGCCCTTAGATGTCCGGG</td><td>>hg38_21_+_8433172_8446622_RNA45SN1_45S_50ntFlanks</td><td>5190</td><td>5220</td><td>31</td><td>AACAGGTCTGTGATGCCCTTAGATGTCCGGG</td><td>86.3</td><td>6.1</td><td>2.0</td><td>3.0</td><td>2.7E-03</td><td>5.8E-03</td><td>BT20/MB468</td><td>MP</td></tr>
<tr><td>>hg38_21_+_8433172_8446622_RNA45SN1_45S_50ntFlanks@5488.5506.19|GCTGAGAAGACGGTCGAAC</td><td>>hg38_21_+_8433172_8446622_RNA45SN1_45S_50ntFlanks</td><td>5488</td><td>5506</td><td>19</td><td>GCTGAGAAGACGGTCGAAC</td><td>129.8</td><td>5.3</td><td>1.8</td><td>3.0</td><td>2.7E-03</td><td>5.8E-03</td><td>BT20/MB468</td><td>MP</td></tr>
<tr><td>>hg38_1_-_228634819_228635039_RNA5S12_5S_50ntFlanks@147.171.25|GGGAATACCGGGTGCTGTAGGCTTT</td><td>>hg38_1_-_228634819_228635039_RNA5S12_5S_50ntFlanks</td><td>147</td><td>171</td><td>25</td><td>GGGAATACCGGGTGCTGTAGGCTTT</td><td>24827.1</td><td>5.2</td><td>1.6</td><td>3.3</td><td>9.5E-04</td><td>5.8E-03</td><td>BT20/MB231</td><td>MP</td></tr>
<tr><td>>hg38_21_+_8433172_8446622_RNA45SN1_45S_50ntFlanks@4005.4039.35|AACCTCGGGCCGATCGCACGCCCCCCGTGGCGGCG</td><td>>hg38_21_+_8433172_8446622_RNA45SN1_45S_50ntFlanks</td><td>4005</td><td>4039</td><td>35</td><td>AACCTCGGGCCGATCGCACGCCCCCCGTGGCGGCG</td><td>120.4</td><td>-2.0</td><td>0.7</td><td>-2.8</td><td>4.7E-03</td><td>5.8E-03</td><td>MB231/MB468</td><td>CYTO</td></tr>
<tr><td>>MI0000433|hsa-let-7g&WithFlank&3|-|52268272|52268367@11.32.22(+1G)&[MIMAT0000414&hsa-let-7g-5p&offsets|0|0(+1G);m-58&3|-|52268336|52268357&offsets|0|0(+1G)]|TGAGGTAGTAGTTTGTACAGTTG</td><td>>MI0000433|hsa-let-7g&WithFlank&3|-|52268272|52268367</td><td>11</td><td>32</td><td>22(+1G)</td><td>TGAGGTAGTAGTTTGTACAGTTG</td><td>78.9</td><td>4.7</td><td>1.7</td><td>2.9</td><td>4.3E-03</td><td>5.8E-03</td><td>MB231/MB468</td><td>MP</td></tr>
<tr><td>>MI0000542|hsa-mir-320a&WithFlank&8|-|22244960|22245043@48.68.21&[MIMAT0000510&hsa-miR-320a-3p&offsets|0|-1;m-60&8|-|22244975|22244996&offsets|0|-1]|AAAAGCTGGGTTGAGAGGGCG</td><td>>MI0000542|hsa-mir-320a&WithFlank&8|-|22244960|22245043</td><td>48</td><td>68</td><td>21</td><td>AAAAGCTGGGTTGAGAGGGCG</td><td>1648.5</td><td>2.8</td><td>1.0</td><td>2.9</td><td>4.3E-03</td><td>5.8E-03</td><td>MB231/MB468</td><td>MP</td></tr>
<tr><td>>MI0000084|hsa-mir-26b&WithFlank&2|+|218402640|218402728@18.39.22(+1U)&[MIMAT0000083&hsa-miR-26b-5p&offsets|0|+1(+1U);m-53&2|+|218402657|218402678&offsets|0|0(+1U)]|TTCAAGTAATTCAGGATAGGTTT</td><td>>MI0000084|hsa-mir-26b&WithFlank&2|+|218402640|218402728</td><td>18</td><td>39</td><td>22(+1U)</td><td>TTCAAGTAATTCAGGATAGGTTT</td><td>1539.3</td><td>-3.6</td><td>1.1</td><td>-3.3</td><td>9.5E-04</td><td>5.8E-03</td><td>BT20/MB231</td><td>MP</td></tr>
<tr><td>>hg38_21_+_8433172_8446622_RNA45SN1_45S_50ntFlanks@11999.12031.33|CCCTGCGGGCCGCCGGTGAAATACCACTACTCT</td><td>>hg38_21_+_8433172_8446622_RNA45SN1_45S_50ntFlanks</td><td>11999</td><td>12031</td><td>33</td><td>CCCTGCGGGCCGCCGGTGAAATACCACTACTCT</td><td>183.7</td><td>-4.9</td><td>1.6</td><td>-3.0</td><td>2.7E-03</td><td>5.8E-03</td><td>BT20/MB468</td><td>MP</td></tr>
<tr><td>>MI0000760|hsa-mir-361&WithFlank&X|-|85903630|85903713@12.33.22&[MIMAT0000703&hsa-miR-361-5p&offsets|0|0;m-89&X|-|85903681|85903702&offsets|0|0]|TTATCAGAATCTCCAGGGGTAC</td><td>>MI0000760|hsa-mir-361&WithFlank&X|-|85903630|85903713</td><td>12</td><td>33</td><td>22</td><td>TTATCAGAATCTCCAGGGGTAC</td><td>298.7</td><td>3.7</td><td>1.2</td><td>3.0</td><td>2.7E-03</td><td>5.8E-03</td><td>BT20/MB468</td><td>MP</td></tr>
<tr><td>>hg38_MT_+_1621_3279_RNR2_16S_50ntFlanks@1427.1449.23|GATTAAAGTCCTACGTGATCTGA</td><td>>hg38_MT_+_1621_3279_RNR2_16S_50ntFlanks</td><td>1427</td><td>1449</td><td>23</td><td>GATTAAAGTCCTACGTGATCTGA</td><td>92.4</td><td>5.4</td><td>1.9</td><td>2.9</td><td>4.3E-03</td><td>5.8E-03</td><td>MB231/MB468</td><td>MP</td></tr>
<tr><td>>trna116_GluCTC_1_-_145399233_145399304@3.36.34,trna59_GluCTC_1_+_249168447_249168518@3.36.34,trna71_GluCTC_1_-_161439189_161439260@3.36.34,trna74_GluCTC_1_-_161431809_161431880@3.36.34,trna77_GluCTC_1_-_161424398_161424469@3.36.34,trna77_GluCTC_6_+_28949976_28950047@3.36.34,trna80_GluCTC_1_-_161417018_161417089@3.36.34,trna87_GluCTC_6_-_126101393_126101464@3.36.34|CCTGGTGGTCTAGTGGTTAGGATTCGGCGCTCTC</td><td>>trna116_GluCTC_1_-_145399233_145399304</td><td>3</td><td>36</td><td>34</td><td>CCTGGTGGTCTAGTGGTTAGGATTCGGCGCTCTC</td><td>158.5</td><td>4.7</td><td>1.6</td><td>2.9</td><td>4.3E-03</td><td>5.8E-03</td><td>MB231/MB468</td><td>MP</td></tr>
<tr><td>>hg38_21_+_8433172_8446622_RNA45SN1_45S_50ntFlanks@11667.11701.35|CTGCCCAGTGCTCTGAATGTCAAAGTGAAGAAATT</td><td>>hg38_21_+_8433172_8446622_RNA45SN1_45S_50ntFlanks</td><td>11667</td><td>11701</td><td>35</td><td>CTGCCCAGTGCTCTGAATGTCAAAGTGAAGAAATT</td><td>233.0</td><td>5.6</td><td>1.7</td><td>3.3</td><td>9.5E-04</td><td>5.8E-03</td><td>BT20/MB231</td><td>MP</td></tr>
<tr><td>>MI0000088|hsa-mir-30a&WithFlank&6|-|71403545|71403627@53.72.20(+1A)&[MIMAT0000088&hsa-miR-30a-3p&offsets|0|-2(+1A);m-38&6|-|71403554|71403575&offsets|0|-2(+1A)]|CTTTCAGTCGGATGTTTGCAA</td><td>>MI0000088|hsa-mir-30a&WithFlank&6|-|71403545|71403627</td><td>53</td><td>72</td><td>20(+1A)</td><td>CTTTCAGTCGGATGTTTGCAA</td><td>419.8</td><td>-3.6</td><td>1.1</td><td>-3.3</td><td>9.5E-04</td><td>5.8E-03</td><td>BT20/MB231</td><td>MP</td></tr>
<tr><td>>hg38_21_+_8433172_8446622_RNA45SN1_45S_50ntFlanks@6650.6696.47|ACGACTCTTAGCGGTGGATCACTCGGCTCGTGCGTCGATGAAGAACG</td><td>>hg38_21_+_8433172_8446622_RNA45SN1_45S_50ntFlanks</td><td>6650</td><td>6696</td><td>47</td><td>ACGACTCTTAGCGGTGGATCACTCGGCTCGTGCGTCGATGAAGAACG</td><td>1669.5</td><td>2.5</td><td>0.9</td><td>2.9</td><td>4.0E-03</td><td>5.8E-03</td><td>BT20/MB468</td><td>MITO</td></tr>
<tr><td>>MI0000542|hsa-mir-320a&WithFlank&8|-|22244960|22245043@48.71.24&[MIMAT0000510&hsa-miR-320a-3p&offsets|0|+2;m-60&8|-|22244975|22244996&offsets|0|+2]|AAAAGCTGGGTTGAGAGGGCGAAA</td><td>>MI0000542|hsa-mir-320a&WithFlank&8|-|22244960|22245043</td><td>48</td><td>71</td><td>24</td><td>AAAAGCTGGGTTGAGAGGGCGAAA</td><td>418.5</td><td>3.8</td><td>1.3</td><td>2.9</td><td>4.3E-03</td><td>5.8E-03</td><td>MB231/MB468</td><td>MP</td></tr>
<tr><td>>hg38_21_+_8433172_8446622_RNA45SN1_45S_50ntFlanks@11787.11804.18|AATGGATGAACGAGATTC</td><td>>hg38_21_+_8433172_8446622_RNA45SN1_45S_50ntFlanks</td><td>11787</td><td>11804</td><td>18</td><td>AATGGATGAACGAGATTC</td><td>1261.0</td><td>-2.1</td><td>0.7</td><td>-2.9</td><td>4.0E-03</td><td>5.8E-03</td><td>BT20/MB468</td><td>MITO</td></tr>
<tr><td>>hg38_MT_+_598_1651_RNR1_12S_50ntFlanks@401.427.27|ACTCCAGTTGACACAAAATAGACTACG</td><td>>hg38_MT_+_598_1651_RNR1_12S_50ntFlanks</td><td>401</td><td>427</td><td>27</td><td>ACTCCAGTTGACACAAAATAGACTACG</td><td>153.7</td><td>4.8</td><td>1.7</td><td>2.9</td><td>4.3E-03</td><td>5.8E-03</td><td>MB231/MB468</td><td>MP</td></tr>
<tr><td>>hg38_21_+_8433172_8446622_RNA45SN1_45S_50ntFlanks@11536.11553.18|TCCGCGCGGCGCCTCGCC</td><td>>hg38_21_+_8433172_8446622_RNA45SN1_45S_50ntFlanks</td><td>11536</td><td>11553</td><td>18</td><td>TCCGCGCGGCGCCTCGCC</td><td>169.0</td><td>-4.2</td><td>1.5</td><td>-2.8</td><td>4.7E-03</td><td>5.8E-03</td><td>BT20/MB468</td><td>CYTO</td></tr>
<tr><td>>hg38_21_+_8433172_8446622_RNA45SN1_45S_50ntFlanks@5472.5512.41|ACGGCCCTGGCGGAGCGCTGAGAAGACGGTCGAACTTGACT</td><td>>hg38_21_+_8433172_8446622_RNA45SN1_45S_50ntFlanks</td><td>5472</td><td>5512</td><td>41</td><td>ACGGCCCTGGCGGAGCGCTGAGAAGACGGTCGAACTTGACT</td><td>396.2</td><td>3.4</td><td>1.2</td><td>2.8</td><td>5.0E-03</td><td>5.8E-03</td><td>MB231/MB468</td><td>MITO</td></tr>
<tr><td>>trnaMT_SerGCT_MT_+_12207_12265@12.40.29|CAAGAACTGCTAACTCATGCCCCCATGTC</td><td>>trnaMT_SerGCT_MT_+_12207_12265</td><td>12</td><td>40</td><td>29</td><td>CAAGAACTGCTAACTCATGCCCCCATGTC</td><td>260.6</td><td>3.7</td><td>1.2</td><td>3.0</td><td>2.7E-03</td><td>5.8E-03</td><td>BT20/MB468</td><td>MP</td></tr>
<tr><td>>hg38_21_+_8433172_8446622_RNA45SN1_45S_50ntFlanks@6719.6751.33|TGAATTGCAGGACACATTGATCATCGACACTTC</td><td>>hg38_21_+_8433172_8446622_RNA45SN1_45S_50ntFlanks</td><td>6719</td><td>6751</td><td>33</td><td>TGAATTGCAGGACACATTGATCATCGACACTTC</td><td>428.9</td><td>-4.1</td><td>1.4</td><td>-3.0</td><td>2.7E-03</td><td>5.8E-03</td><td>BT20/MB468</td><td>MP</td></tr>
<tr><td>>hg38_21_+_8433172_8446622_RNA45SN1_45S_50ntFlanks@12934.12969.36|CTAAACCATTCGTAGACGACCTGCTTCTGGGTCGGG</td><td>>hg38_21_+_8433172_8446622_RNA45SN1_45S_50ntFlanks</td><td>12934</td><td>12969</td><td>36</td><td>CTAAACCATTCGTAGACGACCTGCTTCTGGGTCGGG</td><td>226.8</td><td>-5.2</td><td>1.7</td><td>-3.0</td><td>2.7E-03</td><td>5.8E-03</td><td>BT20/MB468</td><td>MP</td></tr>
<tr><td>>hg38_21_+_8433172_8446622_RNA45SN1_45S_50ntFlanks@12407.12433.27|TTTTGATCCTTCGATGTCGGCTCTTCC</td><td>>hg38_21_+_8433172_8446622_RNA45SN1_45S_50ntFlanks</td><td>12407</td><td>12433</td><td>27</td><td>TTTTGATCCTTCGATGTCGGCTCTTCC</td><td>511.6</td><td>-1.8</td><td>0.6</td><td>-2.8</td><td>5.0E-03</td><td>5.8E-03</td><td>MB231/MB468</td><td>MITO</td></tr>
<tr><td>>hg38_21_+_8433172_8446622_RNA45SN1_45S_50ntFlanks@11579.11601.23|AGAACTGGTGCGGACCAGGGGAA</td><td>>hg38_21_+_8433172_8446622_RNA45SN1_45S_50ntFlanks</td><td>11579</td><td>11601</td><td>23</td><td>AGAACTGGTGCGGACCAGGGGAA</td><td>246.1</td><td>-3.2</td><td>1.1</td><td>-2.9</td><td>4.0E-03</td><td>5.8E-03</td><td>BT20/MB468</td><td>MITO</td></tr>
</table>

| | | | | | | | | | | | | | |
|---|---|---|---|---|---|---|---|---|---|---|---|---|---|
| >hg38_21_+_8433172_8446622_RNA45SN1_45S_50ntFlanks@10248.10281.34\|CGACGAGTAGGAGGGCCGCTGCGGTGAGCCTTGA | >hg38_21_+_8433172_8446622_RNA45SN1_45S_50ntFlanks | 10248 | 10281 | 34 | CGACGAGTAGGAGGGCCGCTGCGGTGAGCCTTGA | 385.1 | 6.3 | 1.9 | 3.3 | 9.6E-04 | 5.8E-03 | BT20/MB231 | MP |
| >hg38_21_+_8433172_8446622_RNA45SN1_45S_50ntFlanks@9929.9952.24\|GATATAGACAGCAGGACGGTGGCC | >hg38_21_+_8433172_8446622_RNA45SN1_45S_50ntFlanks | 9929 | 9952 | 24 | GATATAGACAGCAGGACGGTGGCC | 362.8 | 4.9 | 1.5 | 3.3 | 9.6E-04 | 5.8E-03 | BT20/MB231 | MP |
| >MI0000465\|hsa-mir-191&WithFlank&3\|-\|49020612\|49020715@22.43.22&[MIMAT0000440&hsa-miR-191-5p&offsets\|0\|-1;m-48&3\|-\|49020672\|49020694&offsets\|0\|-1]\|CAACGGAATCCCAAAAGCAGCT | >MI0000465\|hsa-mir-191&WithFlank&3\|-\|49020612\|49020715 | 22 | 43 | 22 | CAACGGAATCCCAAAAGCAGCT | 6705.1 | 2.6 | 0.9 | 2.9 | 4.3E-03 | 5.8E-03 | MB231/MB468 | MP |
| >hg38_21_+_8433172_8446622_RNA45SN1_45S_50ntFlanks@9352.9369.18\|CCGTGGCGCAATGAAGGT | >hg38_21_+_8433172_8446622_RNA45SN1_45S_50ntFlanks | 9352 | 9369 | 18 | CCGTGGCGCAATGAAGGT | 152.1 | -3.2 | 1.1 | -2.9 | 4.0E-03 | 5.8E-03 | BT20/MB468 | MITO |
| >hg38_21_+_8433172_8446622_RNA45SN1_45S_50ntFlanks@11912.11943.32\|GAAGAGACATGAGAGGTGTAGAATAAGTGGGA | >hg38_21_+_8433172_8446622_RNA45SN1_45S_50ntFlanks | 11912 | 11943 | 32 | GAAGAGACATGAGAGGTGTAGAATAAGTGGGA | 67.5 | 9.4 | 2.8 | 3.3 | 9.6E-04 | 5.8E-03 | BT20/MB231 | MP |
| >hg38_21_+_8433172_8446622_RNA45SN1_45S_50ntFlanks@8191.8216.26\|CAGCCCGTGGACGGTGTGAGGCCGGT | >hg38_21_+_8433172_8446622_RNA45SN1_45S_50ntFlanks | 8191 | 8216 | 26 | CAGCCCGTGGACGGTGTGAGGCCGGT | 211.7 | -5.2 | 1.7 | -3.0 | 2.7E-03 | 5.8E-03 | BT20/MB468 | MP |
| >trnaMT_ProTGG_MT_-_15956_16023@40.57.18\|AATGGTGGAGTTAAAGAC | >trnaMT_ProTGG_MT_-_15956_16023 | 40 | 57 | 18 | AATGGTGGAGTTAAAGAC | 100.1 | 5.0 | 1.7 | 3.0 | 2.7E-03 | 5.8E-03 | BT20/MB468 | MP |
| >MI0000438\|hsa-mir-15b&WithFlank&3\|+\|160404582\|160404691@64.83.20(+1A)&[MIMAT0004586&hsa-miR-15b-3p&offsets\|0\|-2(+1A);m-204&3\|+\|160404645\|160404665&offsets\|0\|-1(+1A)]\|CGAATCATTATTTGCTGCTCA | >MI0000438\|hsa-mir-15b&WithFlank&3\|+\|160404582\|160404691 | 64 | 83 | 20(+1A) | CGAATCATTATTTGCTGCTCA | 47.2 | 5.7 | 2.0 | 2.9 | 4.3E-03 | 5.8E-03 | MB231/MB468 | MP |
| >hg38_21_+_8433172_8446622_RNA45SN1_45S_50ntFlanks@8381.8406.26\|AAGAGGGCGTGAAACCGTTAAGAGGT | >hg38_21_+_8433172_8446622_RNA45SN1_45S_50ntFlanks | 8381 | 8406 | 26 | AAGAGGGCGTGAAACCGTTAAGAGGT | 966.9 | 2.5 | 0.9 | 2.8 | 4.8E-03 | 5.8E-03 | MB231/MB468 | CYTO |
| >hg38_21_+_8433172_8446622_RNA45SN1_45S_50ntFlanks@4554.4574.21\|CCGCAGCTAGGAATAATGGAA | >hg38_21_+_8433172_8446622_RNA45SN1_45S_50ntFlanks | 4554 | 4574 | 21 | CCGCAGCTAGGAATAATGGAA | 86.5 | -4.8 | 1.6 | -3.0 | 2.7E-03 | 5.8E-03 | BT20/MB468 | MP |
| >hg38_21_+_8433172_8446622_RNA45SN1_45S_50ntFlanks@4995.5024.30\|ACAGATTGATAGCTCTTTCTCGATTCCGTG | >hg38_21_+_8433172_8446622_RNA45SN1_45S_50ntFlanks | 4995 | 5024 | 30 | ACAGATTGATAGCTCTTTCTCGATTCCGTG | 236.5 | -2.1 | 0.7 | -2.8 | 4.7E-03 | 5.8E-03 | BT20/MB468 | CYTO |
| >hg38_21_+_8433172_8446622_RNA45SN1_45S_50ntFlanks@12631.12653.23\|GGCGAAGCTACCATCTGTGGGAT | >hg38_21_+_8433172_8446622_RNA45SN1_45S_50ntFlanks | 12631 | 12653 | 23 | GGCGAAGCTACCATCTGTGGGAT | 164.7 | -4.8 | 1.6 | -3.0 | 2.7E-03 | 5.8E-03 | BT20/MB468 | MP |
| >hg38_21_+_8433172_8446622_RNA45SN1_45S_50ntFlanks@5087.5110.24\|ACGAGACTCTGGCATGCTAACTAG | >hg38_21_+_8433172_8446622_RNA45SN1_45S_50ntFlanks | 5087 | 5110 | 24 | ACGAGACTCTGGCATGCTAACTAG | 140.7 | -3.1 | 1.1 | -2.9 | 4.3E-03 | 5.8E-03 | MB231/MB468 | MP |
| >MI0000255\|hsa-mir-30d&WithFlank&8\|-\|134804870\|134804951@12.35.24(+1C)&[MIMAT0000245&hsa-miR-30d-5p&offsets\|0\|+2(+1C);m-21&8\|-\|134804919\|134804940&offsets\|0\|+2(+1C)]\|TGTAAACATCCCCGACTGGAAGCTC | >MI0000255\|hsa-mir-30d&WithFlank&8\|-\|134804870\|134804951 | 12 | 35 | 24(+1C) | TGTAAACATCCCCGACTGGAAGCTC | 110.5 | 4.3 | 1.4 | 3.0 | 2.7E-03 | 5.8E-03 | BT20/MB468 | MP |
| >hg38_21_+_8433172_8446622_RNA45SN1_45S_50ntFlanks@11917.11942.26\|GACATGAGAGGTGTAGAATAAGTGGG | >hg38_21_+_8433172_8446622_RNA45SN1_45S_50ntFlanks | 11917 | 11942 | 26 | GACATGAGAGGTGTAGAATAAGTGGG | 30.0 | 8.7 | 2.9 | 3.0 | 2.7E-03 | 5.8E-03 | BT20/MB468 | MP |
| >hg38_MT_+_1621_3279_RNR2_16S_50ntFlanks@488.518.31\|GAACAGCTCTTTGGACACTAGGAAAAAACCT | >hg38_MT_+_1621_3279_RNR2_16S_50ntFlanks | 488 | 518 | 31 | GAACAGCTCTTTGGACACTAGGAAAAAACCT | 191.9 | 4.2 | 1.4 | 3.0 | 2.7E-03 | 5.8E-03 | BT20/MB468 | MP |
| >hg38_21_+_8433172_8446622_RNA45SN1_45S_50ntFlanks@12154.12182.29\|GGGAGTTTGACTGGGGCGGTACACCTGTC | >hg38_21_+_8433172_8446622_RNA45SN1_45S_50ntFlanks | 12154 | 12182 | 29 | GGGAGTTTGACTGGGGCGGTACACCTGTC | 125.0 | -3.1 | 1.1 | -2.9 | 4.0E-03 | 5.8E-03 | BT20/MB468 | MITO |
| >hg38_21_+_8433172_8446622_RNA45SN1_45S_50ntFlanks@9861.9892.32\|GATGAACCGAACGCCGGGTTAAGGCGCCCGAT | >hg38_21_+_8433172_8446622_RNA45SN1_45S_50ntFlanks | 9861 | 9892 | 32 | GATGAACCGAACGCCGGGTTAAGGCGCCCGAT | 128.2 | 2.8 | 0.9 | 3.0 | 2.7E-03 | 5.8E-03 | BT20/MB468 | MP |
| >hg38_21_+_8433172_8446622_RNA45SN1_45S_50ntFlanks@8068.8104.37\|AGTGAACAGGGAAGAGCCCAGCGCCGAATCCCCGCCC | >hg38_21_+_8433172_8446622_RNA45SN1_45S_50ntFlanks | 8068 | 8104 | 37 | AGTGAACAGGGAAGAGCCCAGCGCCGAATCCCCGCCC | 123.8 | -3.6 | 1.3 | -2.9 | 4.3E-03 | 5.8E-03 | MB231/MB468 | MP |
| >hg38_21_+_8433172_8446622_RNA45SN1_45S_50ntFlanks@13000.13039.40\|TGCGATCTATTGAAAGTCAGCCCTCGACACAAGGGTTTGT | >hg38_21_+_8433172_8446622_RNA45SN1_45S_50ntFlanks | 13000 | 13039 | 40 | TGCGATCTATTGAAAGTCAGCCCTCGACACAAGGGTTTGT | 138.0 | 3.4 | 1.2 | 2.9 | 4.3E-03 | 5.8E-03 | MB231/MB468 | MP |
| >hg38_21_+_8433172_8446622_RNA45SN1_45S_50ntFlanks@4226.4243.18\|AATAACAATACAGGACTC | >hg38_21_+_8433172_8446622_RNA45SN1_45S_50ntFlanks | 4226 | 4243 | 18 | AATAACAATACAGGACTC | 90.1 | -4.9 | 1.6 | -3.0 | 2.7E-03 | 5.8E-03 | BT20/MB468 | MP |
| >MI0000078\|hsa-mir-22&WithFlank&17\|-\|1713897\|1713993@59.79.21&[MIMAT0000077&hsa-miR-22-3p&offsets\|0\|-1;m-2&17\|-\|1713914\|1713935&offsets\|0\|-1]\|AAGCTGCCAGTTGAAGAACTG | >MI0000078\|hsa-mir-22&WithFlank&17\|-\|1713897\|1713993 | 59 | 79 | 21 | AAGCTGCCAGTTGAAGAACTG | 159.4 | 5.7 | 2.0 | 2.9 | 4.3E-03 | 5.8E-03 | MB231/MB468 | MP |
| >hg38_MT_+_1621_3279_RNR2_16S_50ntFlanks@482.504.23\|AAAGAGGAACAGCTCTTTGGACA | >hg38_MT_+_1621_3279_RNR2_16S_50ntFlanks | 482 | 504 | 23 | AAAGAGGAACAGCTCTTTGGACA | 148.5 | 1.9 | 0.7 | 2.9 | 4.3E-03 | 5.8E-03 | MB231/MB468 | MP |
| >hg38_21_+_8433172_8446622_RNA45SN1_45S_50ntFlanks@6655.6688.34\|TCTTAGCGGTGGATCACTCGGCTCGTGCGTCGAT | >hg38_21_+_8433172_8446622_RNA45SN1_45S_50ntFlanks | 6655 | 6688 | 34 | TCTTAGCGGTGGATCACTCGGCTCGTGCGTCGAT | 410.1 | -1.4 | 0.5 | -2.8 | 5.0E-03 | 5.8E-03 | BT20/MB468 | NUC |
| >MI0000065\|hsa-let-7d&WithFlank&9\|+\|94178828\|94178926@69.89.21&[MIMAT0004484&hsa-let-7d-3p&offsets\|+1\|0;m-98&9\|+\|94178895\|94178916&offsets\|+1\|0]\|TATACGACCTGCTGCCTTTCT | >MI0000065\|hsa-let-7d&WithFlank&9\|+\|94178828\|94178926 | 69 | 89 | 21 | TATACGACCTGCTGCCTTTCT | 95.5 | 3.6 | 1.3 | 2.9 | 4.3E-03 | 5.8E-03 | MB231/MB468 | MP |
| >MI0000542\|hsa-mir-320a&WithFlank&8\|-\|22244960\|22245043@48.70.23(+1G)&[MIMAT0000510&hsa-miR-320a-3p&offsets\|0\|+1(+1G);m-60&8\|-\|22244975\|22244996&offsets\|0\|+1(+1G)]\|AAAAGCTGGGTTGAGAGGGCGAAG | >MI0000542\|hsa-mir-320a&WithFlank&8\|-\|22244960\|22245043 | 48 | 70 | 23(+1G) | AAAAGCTGGGTTGAGAGGGCGAAG | 104.8 | 3.7 | 1.3 | 2.9 | 4.3E-03 | 5.9E-03 | MB231/MB468 | MP |
| >hg38_21_+_8433172_8446622_RNA45SN1_45S_50ntFlanks@5492.5526.35\|AGAAGACGGTCGAACTTGACTATCTAGAGGAAGTA | >hg38_21_+_8433172_8446622_RNA45SN1_45S_50ntFlanks | 5492 | 5526 | 35 | AGAAGACGGTCGAACTTGACTATCTAGAGGAAGTA | 801.1 | -4.3 | 1.5 | -2.8 | 5.0E-03 | 5.9E-03 | BT20/MB468 | NUC |
| >trna152_ValCAC_6_-_27248049_27248121@1.42.42\|GCTTCTGTAGTGTAGTGGTTATCACGTTCGCCTCACACGCGA | >trna152_ValCAC_6_-_27248049_27248121 | 1 | 42 | 42 | GCTTCTGTAGTGTAGTGGTTATCACGTTCGCCTCACACGCGA | 786.7 | -3.5 | 1.1 | -3.2 | 1.2E-03 | 5.9E-03 | BT20/MB231 | CYTO |
| >trna14_LysTTT_11_-_59327808_59327880@1.34.34,trna2_LysTTT_17_+_8022473_8022545@1.34.34,trna54_LysTTT_1_+_204475655_204475727@1.34.34,trna5_LysTTT_11_+_59323902_59323974@1.34.34,trna62_LysTTT_1_-_204476158_204476230@1.34.34,trna76_LysTTT_6_+_28918806_28918878@1.34.34\|GCCCGGATAGCTCAGTCGGTAGAGCATCAGACTT | >trna14_LysTTT_11_-_59327808_59327880 | 1 | 34 | 34 | GCCCGGATAGCTCAGTCGGTAGAGCATCAGACTT | 77.7 | 4.7 | 1.6 | 2.9 | 4.3E-03 | 5.9E-03 | MB231/MB468 | MP |
| >hg38_MT_+_1621_3279_RNR2_16S_50ntFlanks@778.807.30\|AACAAGTCATTATTACCCTCACTGTCAACC | >hg38_MT_+_1621_3279_RNR2_16S_50ntFlanks | 778 | 807 | 30 | AACAAGTCATTATTACCCTCACTGTCAACC | 217.6 | 2.9 | 1.0 | 2.9 | 4.3E-03 | 5.9E-03 | MB231/MB468 | MP |
| >hg38_21_+_8433172_8446622_RNA45SN1_45S_50ntFlanks@8176.8201.26\|CTTCTGATCGAGGCCCAGCCCGTGGA | >hg38_21_+_8433172_8446622_RNA45SN1_45S_50ntFlanks | 8176 | 8201 | 26 | CTTCTGATCGAGGCCCAGCCCGTGGA | 253.4 | -1.7 | 0.6 | -2.9 | 4.0E-03 | 5.9E-03 | BT20/MB468 | MITO |
| >MI0000746\|hsa-mir-99b&WithFlank&19\|+\|51692606\|51692687@51.70.20&[MIMAT0004678&hsa-miR-99b-3p&offsets\|0\|-2;m-151&19\|+\|51692656\|51692678&offsets\|0\|-3]\|CAAGCTCGTGTCTGTGGGTC | >MI0000746\|hsa-mir-99b&WithFlank&19\|+\|51692606\|51692687 | 51 | 70 | 20 | CAAGCTCGTGTCTGTGGGTC | 735.2 | 2.7 | 0.8 | 3.2 | 1.2E-03 | 5.9E-03 | BT20/MB231 | CYTO |
| >hg38_21_+_8433172_8446622_RNA45SN1_45S_50ntFlanks@11324.11347.24\|GTCTCGTCGCGCGCGCGTCCGCTG | >hg38_21_+_8433172_8446622_RNA45SN1_45S_50ntFlanks | 11324 | 11347 | 24 | GTCTCGTCGCGCGCGCGTCCGCTG | 2658.9 | 7.1 | 2.2 | 3.2 | 1.3E-03 | 5.9E-03 | BT20/MB231 | MITO |
| >trna11_GluTTC_15_-_26327381_26327452@1.34.34,trna3_GluTTC_13_-_45492062_45492133@1.34.34\|TCCCACATGGTCTAGCGGTTAGGATTCCTGGTTT | >trna11_GluTTC_15_-_26327381_26327452 | 1 | 34 | 34 | TCCCACATGGTCTAGCGGTTAGGATTCCTGGTTT | 53.0 | 5.4 | 1.9 | 2.9 | 4.3E-03 | 5.9E-03 | MB231/MB468 | MP |
| >hg38_21_+_8433172_8446622_RNA45SN1_45S_50ntFlanks@12274.12293.20\|CAGTACGAATACAGACCGTG | >hg38_21_+_8433172_8446622_RNA45SN1_45S_50ntFlanks | 12274 | 12293 | 20 | CAGTACGAATACAGACCGTG | 187.4 | -4.3 | 1.5 | -2.9 | 4.0E-03 | 5.9E-03 | BT20/MB468 | MITO |
| >MI0000483\|hsa-mir-186&WithFlank&1\|-\|71067625\|71067722@21.41.21(+1C)&[MIMAT0000456&hsa-miR-186-5p&offsets\|0\|-1(+1C);m-84&1\|-\|71067681\|71067702&offsets\|0\|-1(+1C)]\|CAAAGAATTCTCCTTTTGGGCC | >MI0000483\|hsa-mir-186&WithFlank&1\|-\|71067625\|71067722 | 21 | 41 | 21(+1C) | CAAAGAATTCTCCTTTTGGGCC | 234.5 | 2.9 | 0.9 | 3.2 | 1.2E-03 | 5.9E-03 | BT20/MB231 | CYTO |
| >MI0000088\|hsa-mir-30a&WithFlank&6\|-\|71403545\|71403627@13.34.22&[MIMAT0000087&hsa-miR-30a-5p&offsets\|+1\|+1;m-15&6\|-\|71403595\|71403616&offsets\|+1\|+1]\|GTAAACATCCTCGACTGGAAGC | >MI0000088\|hsa-mir-30a&WithFlank&6\|-\|71403545\|71403627 | 13 | 34 | 22 | GTAAACATCCTCGACTGGAAGC | 246.1 | 3.6 | 1.3 | 2.9 | 4.3E-03 | 5.9E-03 | MB231/MB468 | MP |
| >hg38_21_+_8433172_8446622_RNA45SN1_45S_50ntFlanks@6647.6673.27\|CGTACGACTCTTAGCGGTGGATCACTC | >hg38_21_+_8433172_8446622_RNA45SN1_45S_50ntFlanks | 6647 | 6673 | 27 | CGTACGACTCTTAGCGGTGGATCACTC | 469.4 | 2.6 | 0.9 | 2.8 | 5.1E-03 | 5.9E-03 | MB231/MB468 | MITO |
| >hg38_MT_+_1621_3279_RNR2_16S_50ntFlanks@51.82.32\|GCTAAACCTAGCCCCAAACCCACTCCACCTTA | >hg38_MT_+_1621_3279_RNR2_16S_50ntFlanks | 51 | 82 | 32 | GCTAAACCTAGCCCCAAACCCACTCCACCTTA | 1009.3 | 2.7 | 1.0 | 2.8 | 5.1E-03 | 5.9E-03 | MB231/MB468 | MITO |
| >hg38_21_+_8433172_8446622_RNA45SN1_45S_50ntFlanks@12453.12496.44\|TCACCAAGCGTTGGATTGTTCACCCACTAATAGGGAACGTGAGC | >hg38_21_+_8433172_8446622_RNA45SN1_45S_50ntFlanks | 12453 | 12496 | 44 | TCACCAAGCGTTGGATTGTTCACCCACTAATAGGGAACGTGAGC | 257.1 | -2.9 | 0.9 | -3.2 | 1.2E-03 | 5.9E-03 | BT20/MB231 | CYTO |
| >hg38_21_+_8433172_8446622_RNA45SN1_45S_50ntFlanks@10807.10829.23\|ACAATGTAGGTAAGGGAAGTCGG | >hg38_21_+_8433172_8446622_RNA45SN1_45S_50ntFlanks | 10807 | 10829 | 23 | ACAATGTAGGTAAGGGAAGTCGG | 126.0 | -3.7 | 1.2 | -3.0 | 2.7E-03 | 5.9E-03 | BT20/MB468 | MP |
| >hg38_21_+_8433172_8446622_RNA45SN1_45S_50ntFlanks@3793.3810.18\|CGAATGGCTCATTAAATC | >hg38_21_+_8433172_8446622_RNA45SN1_45S_50ntFlanks | 3793 | 3810 | 18 | CGAATGGCTCATTAAATC | 154.0 | -4.9 | 1.6 | -3.0 | 2.7E-03 | 5.9E-03 | BT20/MB468 | MP |
| >hg38_21_+_8433172_8446622_RNA45SN1_45S_50ntFlanks@9939.9959.21\|GCAGGACGGTGGCCATGGAAG | >hg38_21_+_8433172_8446622_RNA45SN1_45S_50ntFlanks | 9939 | 9959 | 21 | GCAGGACGGTGGCCATGGAAG | 112.0 | 6.2 | 1.9 | 3.3 | 9.7E-04 | 5.9E-03 | BT20/MB231 | MP |
| >trnaMT_HisGTG_MT_+_12138_12206@25.59.35\|CAGATTGTGAATCTGACAACAGAGGCTTACGACCC | >trnaMT_HisGTG_MT_+_12138_12206 | 25 | 59 | 35 | CAGATTGTGAATCTGACAACAGAGGCTTACGACCC | 29.4 | 8.1 | 2.8 | 2.9 | 4.3E-03 | 5.9E-03 | MB231/MB468 | MP |
| >hg38_21_+_8433172_8446622_RNA45SN1_45S_50ntFlanks@10842.10864.23\|GTAACTTCGGGATAAGGATTGGC | >hg38_21_+_8433172_8446622_RNA45SN1_45S_50ntFlanks | 10842 | 10864 | 23 | GTAACTTCGGGATAAGGATTGGC | 188.0 | -3.3 | 1.2 | -2.8 | 4.7E-03 | 5.9E-03 | BT20/MB468 | CYTO |
| >hg38_21_+_8433172_8446622_RNA45SN1_45S_50ntFlanks@11580.11604.25\|GAACTGGTGCGGACCAGGGGAATCC | >hg38_21_+_8433172_8446622_RNA45SN1_45S_50ntFlanks | 11580 | 11604 | 25 | GAACTGGTGCGGACCAGGGGAATCC | 198.6 | -1.7 | 0.6 | -2.8 | 5.1E-03 | 5.9E-03 | MB231/MB468 | NUC |
| >hg38_21_+_8433172_8446622_RNA45SN1_45S_50ntFlanks@9482.9506.25\|ACGTGTTAGGACCCGAAAGATGGTG | >hg38_21_+_8433172_8446622_RNA45SN1_45S_50ntFlanks | 9482 | 9506 | 25 | ACGTGTTAGGACCCGAAAGATGGTG | 78.4 | 5.2 | 1.7 | 3.0 | 2.7E-03 | 5.9E-03 | BT20/MB468 | MP |
| >am_trna2_GlyGCC_21_-_18827107_18827177@1.26.26,trna35_GlyGCC_1_+_161413094_161413164@1.26.26,trna37_GlyGCC_1_+_161420467_161420537@1.26.26,trna39_GlyGCC_1_+_161427898_161427968@1.26.26,trna41_GlyGCC_1_+_161435258_161435328@1.26.26\|GCATGGGTGGTTCAGTGGTAGAATTC | >am_trna2_GlyGCC_21_-_18827107_18827177 | 1 | 26 | 26 | GCATGGGTGGTTCAGTGGTAGAATTC | 275.0 | -4.2 | 1.5 | -2.9 | 4.4E-03 | 5.9E-03 | MB231/MB468 | MP |
| >hg38_21_+_8433172_8446622_RNA45SN1_45S_50ntFlanks@5078.5110.33\|CGATAACGAACGAGACTCTGGCATGCTAACTAG | >hg38_21_+_8433172_8446622_RNA45SN1_45S_50ntFlanks | 5078 | 5110 | 33 | CGATAACGAACGAGACTCTGGCATGCTAACTAG | 308.2 | -3.8 | 1.3 | -2.9 | 4.0E-03 | 5.9E-03 | BT20/MB468 | MITO |
| >hg38_21_+_8433172_8446622_RNA45SN1_45S_50ntFlanks@11729.11752.24\|TAACTATGACTCTCTTAAGGTAGC | >hg38_21_+_8433172_8446622_RNA45SN1_45S_50ntFlanks | 11729 | 11752 | 24 | TAACTATGACTCTCTTAAGGTAGC | 382.9 | -5.0 | 1.7 | -3.0 | 2.7E-03 | 5.9E-03 | BT20/MB468 | MP |

| | | | | | | | | | | | | | |
|---|---|---|---|---|---|---|---|---|---|---|---|---|---|
| >hg38_21_+_8433172_8446622_RNA45SN1_45S_50ntFlanks@9578.9605.28\|GGTCGTCCGACCTGGGTATAGGGGCGAA | >hg38_21_+_8433172_8446622_RNA45SN1_45S_50ntFlanks | 9578 | 9605 | 28 | GGTCGTCCGACCTGGGTATAGGGGCGAA | 31.1 | 8.2 | 2.9 | 2.9 | 4.4E-03 | 5.9E-03 | MB231/MB468 | MP |
| >hg38_21_+_8433172_8446622_RNA45SN1_45S_50ntFlanks@7984.8005.22\|AGATCAGACGTGGCGACCCGCT | >hg38_21_+_8433172_8446622_RNA45SN1_45S_50ntFlanks | 7984 | 8005 | 22 | AGATCAGACGTGGCGACCCGCT | 596.8 | -2.0 | 0.7 | -2.9 | 4.1E-03 | 5.9E-03 | BT20/MB468 | MITO |
| >MI0003628\|hsa-mir-615&WithFlank&12\|+\|54033944\|54034051@67.86.20(+1A)&[MIMAT0003283&hsa-miR-615-3p&offsets\|0\|-2(+1A);m-495&12\|+\|54034010\|54034030&offsets\|0\|-1(+1A)]\|TCCGAGCCTGGGTCTCCCTCA | >MI0003628\|hsa-mir-615&WithFlank&12\|+\|54033944\|54034051 | 67 | 86 | 20(+1A) | TCCGAGCCTGGGTCTCCCTCA | 88.5 | 1.6 | 0.6 | 2.8 | 5.1E-03 | 5.9E-03 | BT20/MB468 | NUC |
| >hg38_1_-_228634819_228635039_RNA5S12_5S_50ntFlanks@132.167.36\|GATGGGAGACCGCCTGGGAATACCGGGTGCTGTAGG | >hg38_1_-_228634819_228635039_RNA5S12_5S_50ntFlanks | 132 | 167 | 36 | GATGGGAGACCGCCTGGGAATACCGGGTGCTGTAGG | 284.1 | 5.7 | 1.7 | 3.3 | 9.7E-04 | 5.9E-03 | BT20/MB231 | MP |
| >hg38_21_+_8433172_8446622_RNA45SN1_45S_50ntFlanks@10773.10796.24\|GGTCTCCAAGGTGAACAGCCTCTG | >hg38_21_+_8433172_8446622_RNA45SN1_45S_50ntFlanks | 10773 | 10796 | 24 | GGTCTCCAAGGTGAACAGCCTCTG | 101.1 | -3.8 | 1.3 | -2.9 | 4.4E-03 | 5.9E-03 | MB231/MB468 | MP |
| >hg38_21_+_8433172_8446622_RNA45SN1_45S_50ntFlanks@9554.9580.27\|CGTAGCGGTCCTGACGTGCAAATCGGT | >hg38_21_+_8433172_8446622_RNA45SN1_45S_50ntFlanks | 9554 | 9580 | 27 | CGTAGCGGTCCTGACGTGCAAATCGGT | 377.4 | -1.6 | 0.6 | -2.8 | 4.8E-03 | 5.9E-03 | BT20/MB468 | CYTO |
| >am_trnaMT_PheGAA_MT_+_577_647@40.62.23\|TGTTTAGACGGGCTCACATCACC | >am_trnaMT_PheGAA_MT_+_577_647 | 40 | 62 | 23 | TGTTTAGACGGGCTCACATCACC | 84.6 | 2.3 | 0.8 | 2.9 | 4.4E-03 | 5.9E-03 | MB231/MB468 | MP |
| >hg38_21_+_8433172_8446622_RNA45SN1_45S_50ntFlanks@4557.4576.20\|CAGCTAGGAATAATGGAATA | >hg38_21_+_8433172_8446622_RNA45SN1_45S_50ntFlanks | 4557 | 4576 | 20 | CAGCTAGGAATAATGGAATA | 68.0 | -4.7 | 1.6 | -3.0 | 2.7E-03 | 5.9E-03 | BT20/MB468 | MP |
| >hg38_21_+_8433172_8446622_RNA45SN1_45S_50ntFlanks@6661.6692.32\|CGGTGGATCACTCGGCTCGTGCGTCGATGAAG | >hg38_21_+_8433172_8446622_RNA45SN1_45S_50ntFlanks | 6661 | 6692 | 32 | CGGTGGATCACTCGGCTCGTGCGTCGATGAAG | 849.1 | -2.4 | 0.9 | -2.8 | 5.1E-03 | 5.9E-03 | MB231/MB468 | MITO |
| >trnaMT_PheGAA_MT_+_577_647@37.56.20\|AAATGTTTAGACGGGCTCAC | >trnaMT_PheGAA_MT_+_577_647 | 37 | 56 | 20 | AAATGTTTAGACGGGCTCAC | 68.2 | 3.6 | 1.3 | 2.8 | 4.4E-03 | 5.9E-03 | MB231/MB468 | MP |
| >hg38_21_+_8433172_8446622_RNA45SN1_45S_50ntFlanks@9454.9494.41\|CCGCGCCGGGGAGGTGGAGCACGAGCGCACGTGTTAGGACC | >hg38_21_+_8433172_8446622_RNA45SN1_45S_50ntFlanks | 9454 | 9494 | 41 | CCGCGCCGGGGAGGTGGAGCACGAGCGCACGTGTTAGGACC | 339.1 | -4.4 | 1.5 | -2.9 | 4.1E-03 | 5.9E-03 | BT20/MB468 | MITO |
| >hg38_21_+_8433172_8446622_RNA45SN1_45S_50ntFlanks@6774.6806.33\|TCCTCCCGGGGCTACGCCTGTCTGAGCGTCGCT | >hg38_21_+_8433172_8446622_RNA45SN1_45S_50ntFlanks | 6774 | 6806 | 33 | TCCTCCCGGGGCTACGCCTGTCTGAGCGTCGCT | 154.8 | 2.7 | 0.9 | 2.8 | 4.4E-03 | 5.9E-03 | MB231/MB468 | MP |
| >MI0000299\|hsa-mir-222&WithFlank&X\|-\|45747009\|45747130@75.99.25&[MIMAT0000279&hsa-miR-222-3p&offsets\|0\|+4;m-140&X\|-\|45747033\|45747056&offsets\|0\|+1]\|AGCTACATCTGGCTACTGGGTCTCT | >MI0000299\|hsa-mir-222&WithFlank&X\|-\|45747009\|45747130 | 75 | 99 | 25 | AGCTACATCTGGCTACTGGGTCTCT | 20677.8 | -3.0 | 0.9 | -3.3 | 9.8E-04 | 5.9E-03 | BT20/MB231 | MP |
| >hg38_21_+_8433172_8446622_RNA45SN1_45S_50ntFlanks@12645.12666.22\|CTGTGGGATTATGACTGAACGC | >hg38_21_+_8433172_8446622_RNA45SN1_45S_50ntFlanks | 12645 | 12666 | 22 | CTGTGGGATTATGACTGAACGC | 368.7 | 6.6 | 2.0 | 3.3 | 9.8E-04 | 5.9E-03 | BT20/MB231 | MP |
| >hg38_21_+_8433172_8446622_RNA45SN1_45S_50ntFlanks@6650.6686.37\|ACGACTCTTAGCGGTGGATCACTCGGCTCGTGCGTCG | >hg38_21_+_8433172_8446622_RNA45SN1_45S_50ntFlanks | 6650 | 6686 | 37 | ACGACTCTTAGCGGTGGATCACTCGGCTCGTGCGTCG | 114.1 | -2.7 | 0.9 | -2.8 | 4.4E-03 | 5.9E-03 | MB231/MB468 | MP |
| >hg38_21_+_8433172_8446622_RNA45SN1_45S_50ntFlanks@5491.5511.21\|GAGAAGACGGTCGAACTTGAC | >hg38_21_+_8433172_8446622_RNA45SN1_45S_50ntFlanks | 5491 | 5511 | 21 | GAGAAGACGGTCGAACTTGAC | 80.6 | 5.0 | 1.7 | 3.0 | 2.7E-03 | 5.9E-03 | BT20/MB468 | MP |
| >hg38_21_+_8433172_8446622_RNA45SN1_45S_50ntFlanks@10830.10852.23\|CAAGCCGGATCCGTAACTTCGGG | >hg38_21_+_8433172_8446622_RNA45SN1_45S_50ntFlanks | 10830 | 10852 | 23 | CAAGCCGGATCCGTAACTTCGGG | 212.0 | -4.9 | 1.6 | -3.0 | 2.7E-03 | 5.9E-03 | BT20/MB468 | MP |
| >hg38_21_+_8433172_8446622_RNA45SN1_45S_50ntFlanks@7981.8011.31\|CTCAGATCAGACGTGGCGACCCGCTGAATTT | >hg38_21_+_8433172_8446622_RNA45SN1_45S_50ntFlanks | 7981 | 8011 | 31 | CTCAGATCAGACGTGGCGACCCGCTGAATTT | 158.5 | -4.6 | 1.6 | -2.8 | 4.4E-03 | 5.9E-03 | MB231/MB468 | MP |
| >hg38_21_+_8433172_8446622_RNA45SN1_45S_50ntFlanks@5073.5105.33\|AATTCCGATAACGAACGAGACTCTGGCATGCTA | >hg38_21_+_8433172_8446622_RNA45SN1_45S_50ntFlanks | 5073 | 5105 | 33 | AATTCCGATAACGAACGAGACTCTGGCATGCTA | 201.5 | -4.7 | 1.6 | -2.9 | 4.1E-03 | 5.9E-03 | BT20/MB468 | MITO |
| >hg38_21_+_8433172_8446622_RNA45SN1_45S_50ntFlanks@8356.8391.36\|GAAAAGAACTTTGAAGAGAGAGTTCAAGAGGGCGTG | >hg38_21_+_8433172_8446622_RNA45SN1_45S_50ntFlanks | 8356 | 8391 | 36 | GAAAAGAACTTTGAAGAGAGAGTTCAAGAGGGCGTG | 421.2 | 2.1 | 0.7 | 2.9 | 4.1E-03 | 5.9E-03 | BT20/MB468 | MITO |
| >hg38_21_+_8433172_8446622_RNA45SN1_45S_50ntFlanks@8914.8936.23\|CCGAATCCCGGGGCCGAGGGAGC | >hg38_21_+_8433172_8446622_RNA45SN1_45S_50ntFlanks | 8914 | 8936 | 23 | CCGAATCCCGGGGCCGAGGGAGC | 102.4 | 4.3 | 1.4 | 3.0 | 2.7E-03 | 5.9E-03 | BT20/MB468 | MP |
| >hg38_21_+_8433172_8446622_RNA45SN1_45S_50ntFlanks@3845.3873.29\|ACTTGGATAACTGTGGTAATTCTAGAGCT | >hg38_21_+_8433172_8446622_RNA45SN1_45S_50ntFlanks | 3845 | 3873 | 29 | ACTTGGATAACTGTGGTAATTCTAGAGCT | 118.7 | -3.5 | 1.2 | -2.8 | 4.4E-03 | 5.9E-03 | MB231/MB468 | MP |
| >hg38_21_+_8433172_8446622_RNA45SN1_45S_50ntFlanks@7975.8017.43\|CGCGACCTCAGATCAGACGTGGCGACCCGCTGAATTTAAGCAT | >hg38_21_+_8433172_8446622_RNA45SN1_45S_50ntFlanks | 7975 | 8017 | 43 | CGCGACCTCAGATCAGACGTGGCGACCCGCTGAATTTAAGCAT | 2011.1 | 3.4 | 1.2 | 2.8 | 5.1E-03 | 5.9E-03 | MB231/MB468 | NUC |
| >hg38_21_+_8433172_8446622_RNA45SN1_45S_50ntFlanks@9503.9525.23\|GGTGAACTATGCCTGGGCAGGGC | >hg38_21_+_8433172_8446622_RNA45SN1_45S_50ntFlanks | 9503 | 9525 | 23 | GGTGAACTATGCCTGGGCAGGGC | 166.4 | -4.8 | 1.7 | -2.8 | 4.4E-03 | 5.9E-03 | MB231/MB468 | MP |
| >hg38_21_+_8433172_8446622_RNA45SN1_45S_50ntFlanks@9939.9961.23\|GCAGGACGGTGGCCATGGAAGTC | >hg38_21_+_8433172_8446622_RNA45SN1_45S_50ntFlanks | 9939 | 9961 | 23 | GCAGGACGGTGGCCATGGAAGTC | 626.3 | 6.7 | 2.0 | 3.3 | 9.8E-04 | 5.9E-03 | BT20/MB231 | MP |
| >MI0000077\|hsa-mir-21&WithFlank&17\|+\|59841260\|59841343@14.31.18(+1C)&[MIMAT0000076&hsa-miR-21-5p&offsets\|0\|-4(+1C);m-1&17\|+\|59841273\|59841294&offsets\|0\|-4(+1C)]\|TAGCTTATCAGACTGATGC | >MI0000077\|hsa-mir-21&WithFlank&17\|+\|59841260\|59841343 | 14 | 31 | 18(+1C) | TAGCTTATCAGACTGATGC | 203.2 | 3.7 | 1.2 | 3.2 | 1.2E-03 | 5.9E-03 | BT20/MB231 | CYTO |
| >hg38_21_+_8433172_8446622_RNA45SN1_45S_50ntFlanks@12437.12472.36\|CATTGTGAAGCAGAATTCACCAAGCGTTGGATTGTT | >hg38_21_+_8433172_8446622_RNA45SN1_45S_50ntFlanks | 12437 | 12472 | 36 | CATTGTGAAGCAGAATTCACCAAGCGTTGGATTGTT | 444.8 | -2.3 | 0.8 | -2.8 | 4.9E-03 | 5.9E-03 | MB231/MB468 | CYTO |
| >MI0000456\|hsa-mir-140&WithFlank&16\|+\|69933075\|69933186@29.48.20&[MIMAT0000431&hsa-miR-140-5p&offsets\|0\|-2;m-173&16\|+\|69933103\|69933124&offsets\|0\|-2]\|CAGTGGTTTTACCCTATGGT | >MI0000456\|hsa-mir-140&WithFlank&16\|+\|69933075\|69933186 | 29 | 48 | 20 | CAGTGGTTTTACCCTATGGT | 113.6 | 5.6 | 1.9 | 3.0 | 2.7E-03 | 5.9E-03 | BT20/MB468 | MP |
| >hg38_21_+_8433172_8446622_RNA45SN1_45S_50ntFlanks@10406.10424.19\|TGGGTCAGTCGGTCCTGAG | >hg38_21_+_8433172_8446622_RNA45SN1_45S_50ntFlanks | 10406 | 10424 | 19 | TGGGTCAGTCGGTCCTGAG | 499.3 | 3.0 | 1.1 | 2.8 | 4.4E-03 | 5.9E-03 | MB231/MB468 | MP |
| >hg38_21_+_8433172_8446622_RNA45SN1_45S_50ntFlanks@6779.6802.24\|CCGGGGCTACGCCTGTCTGAGCGT | >hg38_21_+_8433172_8446622_RNA45SN1_45S_50ntFlanks | 6779 | 6802 | 24 | CCGGGGCTACGCCTGTCTGAGCGT | 437.5 | 2.1 | 0.8 | 2.8 | 5.2E-03 | 5.9E-03 | MB231/MB468 | NUC |
| >hg38_21_+_8433172_8446622_RNA45SN1_45S_50ntFlanks@3775.3792.18\|GCCGGTACAGTGAAACTG | >hg38_21_+_8433172_8446622_RNA45SN1_45S_50ntFlanks | 3775 | 3792 | 18 | GCCGGTACAGTGAAACTG | 2181.3 | -5.6 | 1.9 | -3.0 | 2.7E-03 | 5.9E-03 | BT20/MB468 | MP |
| >hg38_21_+_8433172_8446622_RNA45SN1_45S_50ntFlanks@4760.4786.27\|TCAGATACCGTCGTAGTTCCGACCATA | >hg38_21_+_8433172_8446622_RNA45SN1_45S_50ntFlanks | 4760 | 4786 | 27 | TCAGATACCGTCGTAGTTCCGACCATA | 507.0 | -4.2 | 1.4 | -3.0 | 2.7E-03 | 5.9E-03 | BT20/MB468 | MP |
| >trna5_AspGTC_12_+_98897281_98897352@35.73.39\|TCACGCGGGAGACCGGGGTTCAATTCCCCGACGGGGAGC | >trna5_AspGTC_12_+_98897281_98897352 | 35 | 73 | 39 | TCACGCGGGAGACCGGGGTTCAATTCCCCGACGGGGAGC | 864.4 | -6.1 | 1.9 | -3.2 | 1.2E-03 | 6.0E-03 | BT20/MB231 | CYTO |
| >hg38_21_+_8433172_8446622_RNA45SN1_45S_50ntFlanks@6647.6696.50\|CGTACGACTCTTAGCGGTGGATCACTCGGCTCGTGCGTCGATGAAGAACG | >hg38_21_+_8433172_8446622_RNA45SN1_45S_50ntFlanks | 6647 | 6696 | 50 | CGTACGACTCTTAGCGGTGGATCACTCGGCTCGTGCGTCGATGAAGAACG | 639.1 | 3.6 | 1.3 | 2.8 | 5.2E-03 | 6.0E-03 | MB231/MB468 | NUC |
| >hg38_21_+_8433172_8446622_RNA45SN1_45S_50ntFlanks@10388.10407.20\|GTGAACAGCAGTTGAACATG | >hg38_21_+_8433172_8446622_RNA45SN1_45S_50ntFlanks | 10388 | 10407 | 20 | GTGAACAGCAGTTGAACATG | 140.7 | -4.8 | 1.6 | -3.0 | 2.7E-03 | 6.0E-03 | BT20/MB468 | MP |
| >hg38_MT_+_1621_3279_RNR2_16S_50ntFlanks@1425.1450.26\|ACGATTAAAGTCCTACGTGATCTGAG | >hg38_MT_+_1621_3279_RNR2_16S_50ntFlanks | 1425 | 1450 | 26 | ACGATTAAAGTCCTACGTGATCTGAG | 110.9 | 4.7 | 1.6 | 2.8 | 4.4E-03 | 6.0E-03 | MB231/MB468 | MP |
| >trnaMT_ValTAC_MT_+_1602_1670@39.71.33\|AGGAGATTTCAACTTAACTTGACCGCTCTGACC | >trnaMT_ValTAC_MT_+_1602_1670 | 39 | 71 | 33 | AGGAGATTTCAACTTAACTTGACCGCTCTGACC | 322.1 | 4.1 | 1.4 | 3.0 | 2.7E-03 | 6.0E-03 | BT20/MB468 | MP |
| >hg38_21_+_8433172_8446622_RNA45SN1_45S_50ntFlanks@10845.10864.20\|ACTTCGGGATAAGGATTGGC | >hg38_21_+_8433172_8446622_RNA45SN1_45S_50ntFlanks | 10845 | 10864 | 20 | ACTTCGGGATAAGGATTGGC | 351.8 | -5.2 | 1.7 | -3.0 | 2.7E-03 | 6.0E-03 | BT20/MB468 | MP |
| >hg38_21_+_8433172_8446622_RNA45SN1_45S_50ntFlanks@9300.9334.35\|ACGGACCAAGGAGTCTAACACGTGCGCGAGTCGGG | >hg38_21_+_8433172_8446622_RNA45SN1_45S_50ntFlanks | 9300 | 9334 | 35 | ACGGACCAAGGAGTCTAACACGTGCGCGAGTCGGG | 80.6 | 4.0 | 1.3 | 3.0 | 2.7E-03 | 6.0E-03 | BT20/MB468 | MP |
| >hg38_21_+_8433172_8446622_RNA45SN1_45S_50ntFlanks@6661.6707.47\|CGGTGGATCACTCGGCTCGTGCGTCGATGAAGAACGCAGCTAGCTGC | >hg38_21_+_8433172_8446622_RNA45SN1_45S_50ntFlanks | 6661 | 6707 | 47 | CGGTGGATCACTCGGCTCGTGCGTCGATGAAGAACGCAGCTAGCTGC | 465.6 | 3.4 | 1.2 | 2.8 | 5.1E-03 | 6.0E-03 | BT20/MB468 | NUC |
| >hg38_21_+_8433172_8446622_RNA45SN1_45S_50ntFlanks@4727.4759.33\|AATCAAGAACGAAAGTCGGAGGTTCGAAGACGA | >hg38_21_+_8433172_8446622_RNA45SN1_45S_50ntFlanks | 4727 | 4759 | 33 | AATCAAGAACGAAAGTCGGAGGTTCGAAGACGA | 248.3 | -1.6 | 0.6 | -2.9 | 4.1E-03 | 6.0E-03 | BT20/MB468 | MITO |
| >hg38_21_+_8433172_8446622_RNA45SN1_45S_50ntFlanks@12625.12657.33\|CAATGGGGCGAAGCTACCATCTGTGGGATTATG | >hg38_21_+_8433172_8446622_RNA45SN1_45S_50ntFlanks | 12625 | 12657 | 33 | CAATGGGGCGAAGCTACCATCTGTGGGATTATG | 294.5 | -5.0 | 1.7 | -3.0 | 2.8E-03 | 6.0E-03 | BT20/MB468 | MP |
| >hg38_21_+_8433172_8446622_RNA45SN1_45S_50ntFlanks@12626.12667.42\|AATGGGGCGAAGCTACCATCTGTGGGATTATGACTGAACGCC | >hg38_21_+_8433172_8446622_RNA45SN1_45S_50ntFlanks | 12626 | 12667 | 42 | AATGGGGCGAAGCTACCATCTGTGGGATTATGACTGAACGCC | 108.2 | 4.0 | 1.3 | 3.0 | 2.8E-03 | 6.0E-03 | BT20/MB468 | MP |
| >hg38_21_+_8433172_8446622_RNA45SN1_45S_50ntFlanks@4727.4748.22\|AATCAAGAACGAAAGTCGGAGG | >hg38_21_+_8433172_8446622_RNA45SN1_45S_50ntFlanks | 4727 | 4748 | 22 | AATCAAGAACGAAAGTCGGAGG | 33.6 | -8.2 | 2.7 | -3.0 | 2.8E-03 | 6.0E-03 | BT20/MB468 | MP |
| >MI0000093\|hsa-mir-92a-1&WithFlank&13\|+\|91351308\|91351397@54.73.20;MI0000094\|hsa-mir-92a-2&WithFlank&X\|-\|134169532\|134169618@54.73.20&[MIMAT0000092&hsa-miR-92a-3p&offsets\|0\|-2;m-9&13\|+\|91351361\|91351382&offsets\|0\|-2];[MIMAT0000092_1&hsa-miR-92a-3p&offsets\|0\|-2;m-11&X\|-\|134169544\|134169565&offsets\|0\|-2]\|TATTGCACTTGTCCCGGCCT | >MI0000093\|hsa-mir-92a-1&WithFlank&13\|+\|91351308\|91351397 | 54 | 73 | 20 | TATTGCACTTGTCCCGGCCT | 366.0 | -2.5 | 0.8 | -3.2 | 1.3E-03 | 6.0E-03 | BT20/MB231 | MITO |
| >hg38_21_+_8433172_8446622_RNA45SN1_45S_50ntFlanks@12994.13015.22\|CCTCGCTGCGATCTATTGAAAG | >hg38_21_+_8433172_8446622_RNA45SN1_45S_50ntFlanks | 12994 | 13015 | 22 | CCTCGCTGCGATCTATTGAAAG | 345.1 | -2.2 | 0.8 | -2.8 | 4.8E-03 | 6.0E-03 | BT20/MB468 | CYTO |
| >hg38_21_+_8433172_8446622_RNA45SN1_45S_50ntFlanks@11873.11902.30\|AAGAAGACCCTGTTGAGCTTGACTCTAGTC | >hg38_21_+_8433172_8446622_RNA45SN1_45S_50ntFlanks | 11873 | 11902 | 30 | AAGAAGACCCTGTTGAGCTTGACTCTAGTC | 205.1 | -2.5 | 0.8 | -3.0 | 2.8E-03 | 6.0E-03 | BT20/MB468 | MP |
| >hg38_MT_+_598_1651_RNR1_12S_50ntFlanks@716.744.29\|ACGTAAAGACGTTAGGTCAAGGTGTAGCC | >hg38_MT_+_598_1651_RNR1_12S_50ntFlanks | 716 | 744 | 29 | ACGTAAAGACGTTAGGTCAAGGTGTAGCC | 127.7 | 4.7 | 1.6 | 2.8 | 4.4E-03 | 6.0E-03 | MB231/MB468 | MP |
| >hg38_21_+_8433172_8446622_RNA45SN1_45S_50ntFlanks@10787.10817.31\|ACAGCCTCTGGCATGTTGGAACAATGTAGGT | >hg38_21_+_8433172_8446622_RNA45SN1_45S_50ntFlanks | 10787 | 10817 | 31 | ACAGCCTCTGGCATGTTGGAACAATGTAGGT | 372.0 | -2.3 | 0.8 | -3.0 | 2.8E-03 | 6.0E-03 | BT20/MB468 | MP |
| >hg38_MT_+_1621_3279_RNR2_16S_50ntFlanks@51.80.30\|GCTAAACCTAGCCCCAAACCCACTCCACCT | >hg38_MT_+_1621_3279_RNR2_16S_50ntFlanks | 51 | 80 | 30 | GCTAAACCTAGCCCCAAACCCACTCCACCT | 1531.9 | 3.6 | 1.2 | 3.0 | 2.8E-03 | 6.0E-03 | BT20/MB468 | MP |
| >hg38_21_+_8433172_8446622_RNA45SN1_45S_50ntFlanks@6663.6699.37\|GTGGATCACTCGGCTCGTGCGTCGATGAAGAACGCAG | >hg38_21_+_8433172_8446622_RNA45SN1_45S_50ntFlanks | 6663 | 6699 | 37 | GTGGATCACTCGGCTCGTGCGTCGATGAAGAACGCAG | 235.1 | -1.8 | 0.7 | -2.8 | 5.2E-03 | 6.0E-03 | MB231/MB468 | NUC |

| | | | | | | | | | | | | | |
|---|---|---|---|---|---|---|---|---|---|---|---|---|---|
| >MI0000483\|hsa-mir-186&WithFlank&1\|-\|71067625\|71067722@21.41.21(+1A)&[MIMAT0000456&hsa-miR-186-5p&offsets\|0\|-1(+1A);m-84&1\|-\|71067681\|71067702&offsets\|0\|-1(+1A)]\|CAAAGAATTCTCCTTTTGGGCA | >MI0000483\|hsa-mir-186&WithFlank&1\|-\|71067625\|71067722 | 21 | 41 | 21(+1A) | CAAAGAATTCTCCTTTTGGGCA | 278.2 | 3.1 | 1.0 | 3.2 | 1.2E-03 | 6.0E-03 | BT20/MB231 | CYTO |
| >trnaMT_PheGAA_MT_+_577_647@35.64.30\|GAAAATGTTTAGACGGGCTCACATCACCCC | >trnaMT_PheGAA_MT_+_577_647 | 35 | 64 | 30 | GAAAATGTTTAGACGGGCTCACATCACCCC | 664.7 | 6.9 | 2.3 | 3.0 | 2.8E-03 | 6.0E-03 | BT20/MB468 | MP |
| >am_trna128_GlyGCC_6_-_27870686_27870756@13.30.18,trna133_GlyCCC_1_-_16872434_16872504@13.30.18,trna17_GluTTC_2_-_204229131_204229201@13.30.18,trna18_GlyGCC_16_+_70822597_70822667@13.30.18,trna19_GlyGCC_16_+_70823410_70823480@13.30.18,trna19_GlyGCC_2_-_157257659_157257729@13.30.18,trna24_GlyGCC_16_-_70812942_70813012@13.30.18,trna25_GlyGCC_16_-_70812114_70812184@13.30.18,trna2_GlyCCC_1_+_17053780_17053850@13.30.18,trna2_GlyGCC_21_-_18827107_18827177@13.30.18,trna35_GlyGCC_1_+_161413094_161413164@13.30.18,trna37_GlyGCC_1_+_161420467_161420537@13.30.18,trna39_GlyGCC_1_+_161427898_161427968@13.30.18,trna41_GlyGCC_1_+_161435258_161435328@13.30.18,trna4_GlyCCC_1_+_17188416_17188486@13.30.18,trna5_GlyGCC_17_+_8029064_8029134@13.30.18,trna68_GlyGCC_1_-_161493637_161493707@13.30.18,trna91_GlyCCC_1_-_149680210_149680280@13.30.18,trna99_ValCAC_1_-_149294666_149294736@13.30.18\|CAGTGGTAGAATTCTCGC | >am_trna128_GlyGCC_6_-_27870686_27870756 | 13 | 30 | 18 | CAGTGGTAGAATTCTCGC | 116.0 | -4.8 | 1.7 | -2.8 | 4.4E-03 | 6.0E-03 | MB231/MB468 | MP |
| >hg38_21_+_8433172_8446622_RNA45SN1_45S_50ntFlanks@11568.11611.44\|GCAGCCGACTTAGAACTGGTGCGGACCAGGGGAATCCGACTGTT | >hg38_21_+_8433172_8446622_RNA45SN1_45S_50ntFlanks | 11568 | 11611 | 44 | GCAGCCGACTTAGAACTGGTGCGGACCAGGGGAATCCGACTGTT | 875.5 | 3.9 | 1.3 | 3.0 | 2.8E-03 | 6.0E-03 | BT20/MB468 | MP |
| >hg38_21_+_8433172_8446622_RNA45SN1_45S_50ntFlanks@10349.10383.35\|AACGAGAACTTTGAAGGCCGAAGTGGAGAAGGGTT | >hg38_21_+_8433172_8446622_RNA45SN1_45S_50ntFlanks | 10349 | 10383 | 35 | AACGAGAACTTTGAAGGCCGAAGTGGAGAAGGGTT | 4990.3 | -1.8 | 0.6 | -2.8 | 4.9E-03 | 6.0E-03 | MB231/MB468 | CYTO |
| >hg38_21_+_8433172_8446622_RNA45SN1_45S_50ntFlanks@4735.4758.24\|ACGAAAGTCGGAGGTTCGAAGACG | >hg38_21_+_8433172_8446622_RNA45SN1_45S_50ntFlanks | 4735 | 4758 | 24 | ACGAAAGTCGGAGGTTCGAAGACG | 1371.6 | 2.1 | 0.8 | 2.8 | 4.9E-03 | 6.0E-03 | MB231/MB468 | CYTO |
| >MI0000101\|hsa-mir-99a&WithFlank&21\|+\|16539083\|16539175@19.40.22(+1A)&[MIMAT0000097&hsa-miR-99a-5p&offsets\|0\|0(+1A);m-64&21\|+\|16539101\|16539121&offsets\|0\|+1(+1A)]\|AACCCGTAGATCCGATCTTGTGA | >MI0000101\|hsa-mir-99a&WithFlank&21\|+\|16539083\|16539175 | 19 | 40 | 22(+1A) | AACCCGTAGATCCGATCTTGTGA | 699.8 | -1.7 | 0.6 | -2.8 | 4.8E-03 | 6.0E-03 | BT20/MB468 | CYTO |
| >hg38_21_+_8433172_8446622_RNA45SN1_45S_50ntFlanks@10787.10815.29\|ACAGCCTCTGGCATGTTGGAACAATGTAG | >hg38_21_+_8433172_8446622_RNA45SN1_45S_50ntFlanks | 10787 | 10815 | 29 | ACAGCCTCTGGCATGTTGGAACAATGTAG | 207.4 | -3.5 | 1.2 | -2.8 | 4.8E-03 | 6.0E-03 | BT20/MB468 | CYTO |
| >MI0000270\|hsa-mir-181b-1&WithFlank&1\|-\|198858867\|198858988@43.65.23;MI0000683\|hsa-mir-181b-2&WithFlank&9\|+\|124693704\|124693804@23.45.23&[MIMAT0000257&hsa-miR-181b-5p&offsets\|+1\|+1;m-73&1\|-\|198858925\|198858947&offsets\|+1\|+1];[MIMAT0000257_1&hsa-miR-181b-5p&offsets\|+1\|+1;m-72&9\|+\|124693725\|124693747&offsets\|+1\|+1]\|ACATTCATTGCTGTCGGTGGGTT | >MI0000270\|hsa-mir-181b-1&WithFlank&1\|-\|198858867\|198858988 | 43 | 65 | 23 | ACATTCATTGCTGTCGGTGGGTT | 56.5 | 4.5 | 1.6 | 2.8 | 4.4E-03 | 6.0E-03 | MB231/MB468 | MP |
| >hg38_21_+_8433172_8446622_RNA45SN1_45S_50ntFlanks@11577.11611.35\|TTAGAACTGGTGCGGACCAGGGGAATCCGACTGTT | >hg38_21_+_8433172_8446622_RNA45SN1_45S_50ntFlanks | 11577 | 11611 | 35 | TTAGAACTGGTGCGGACCAGGGGAATCCGACTGTT | 243.8 | -1.8 | 0.7 | -2.8 | 4.8E-03 | 6.0E-03 | BT20/MB468 | CYTO |
| >hg38_MT_+_1621_3279_RNR2_16S_50ntFlanks@493.518.26\|GCTCTTTGGACACTAGGAAAAAACCT | >hg38_MT_+_1621_3279_RNR2_16S_50ntFlanks | 493 | 518 | 26 | GCTCTTTGGACACTAGGAAAAAACCT | 285.5 | 2.4 | 0.8 | 2.8 | 4.5E-03 | 6.0E-03 | MB231/MB468 | MP |
| >hg38_21_+_8433172_8446622_RNA45SN1_45S_50ntFlanks@12934.12960.27\|CTAAACCATTCGTAGACGACCTGCTTC | >hg38_21_+_8433172_8446622_RNA45SN1_45S_50ntFlanks | 12934 | 12960 | 27 | CTAAACCATTCGTAGACGACCTGCTTC | 317.2 | -2.1 | 0.7 | -2.8 | 4.9E-03 | 6.0E-03 | BT20/MB468 | CYTO |
| >hg38_21_+_8433172_8446622_RNA45SN1_45S_50ntFlanks@9605.9622.18\|AAGACTAATCGAACCATC | >hg38_21_+_8433172_8446622_RNA45SN1_45S_50ntFlanks | 9605 | 9622 | 18 | AAGACTAATCGAACCATC | 632.1 | -2.5 | 0.9 | -2.8 | 4.9E-03 | 6.0E-03 | BT20/MB468 | CYTO |
| >MI0000542\|hsa-mir-320a&WithFlank&8\|-\|22244960\|22245043@48.69.22&[MIMAT0000510&hsa-miR-320a-3p&offsets\|0\|0;m-60&8\|-\|22244975\|22244996&offsets\|0\|0]\|AAAAGCTGGGTTGAGAGGGCGA | >MI0000542\|hsa-mir-320a&WithFlank&8\|-\|22244960\|22245043 | 48 | 69 | 22 | AAAAGCTGGGTTGAGAGGGCGA | 12988.8 | 2.3 | 0.7 | 3.3 | 1.1E-03 | 6.0E-03 | BT20/MB231 | NUC |
| >MI0006273\|hsa-mir-1180&WithFlank&17\|-\|19344500\|19344580@47.66.20&[MIMAT0005825&hsa-miR-1180-3p&offsets\|0\|-2;m-214&17\|-\|19344513\|19344534&offsets\|0\|-2]\|TTTCCGGCTCGCGTGGGTGT | >MI0006273\|hsa-mir-1180&WithFlank&17\|-\|19344500\|19344580 | 47 | 66 | 20 | TTTCCGGCTCGCGTGGGTGT | 378.5 | 2.3 | 0.7 | 3.3 | 1.1E-03 | 6.0E-03 | BT20/MB231 | NUC |
| >hg38_21_+_8433172_8446622_RNA45SN1_45S_50ntFlanks@4729.4751.23\|TCAAGAACGAAAGTCGGAGGTTC | >hg38_21_+_8433172_8446622_RNA45SN1_45S_50ntFlanks | 4729 | 4751 | 23 | TCAAGAACGAAAGTCGGAGGTTC | 104.4 | -4.1 | 1.4 | -2.9 | 4.1E-03 | 6.0E-03 | BT20/MB468 | MITO |
| >hg38_21_+_8433172_8446622_RNA45SN1_45S_50ntFlanks@6647.6675.29\|CGTACGACTCTTAGCGGTGGATCACTCGG | >hg38_21_+_8433172_8446622_RNA45SN1_45S_50ntFlanks | 6647 | 6675 | 29 | CGTACGACTCTTAGCGGTGGATCACTCGG | 153.8 | -2.5 | 0.9 | -2.8 | 4.5E-03 | 6.0E-03 | MB231/MB468 | MP |
| >hg38_21_+_8433172_8446622_RNA45SN1_45S_50ntFlanks@12152.12183.32\|TGGGGAGTTTGACTGGGGCGGTACACCTGTCA | >hg38_21_+_8433172_8446622_RNA45SN1_45S_50ntFlanks | 12152 | 12183 | 32 | TGGGGAGTTTGACTGGGGCGGTACACCTGTCA | 153.8 | -4.8 | 1.6 | -3.0 | 2.8E-03 | 6.0E-03 | BT20/MB468 | MP |
| >hg38_21_+_8433172_8446622_RNA45SN1_45S_50ntFlanks@3981.4001.21\|CCGGCGGCTTTGGTGACTCTA | >hg38_21_+_8433172_8446622_RNA45SN1_45S_50ntFlanks | 3981 | 4001 | 21 | CCGGCGGCTTTGGTGACTCTA | 144.5 | -3.6 | 1.3 | -2.9 | 4.2E-03 | 6.0E-03 | BT20/MB468 | MITO |
| >hg38_21_+_8433172_8446622_RNA45SN1_45S_50ntFlanks@7989.8008.20\|AGACGTGGCGACCCGCTGAA | >hg38_21_+_8433172_8446622_RNA45SN1_45S_50ntFlanks | 7989 | 8008 | 20 | AGACGTGGCGACCCGCTGAA | 2441.9 | 2.8 | 1.0 | 2.8 | 4.9E-03 | 6.0E-03 | MB231/MB468 | CYTO |
| >hg38_21_+_8433172_8446622_RNA45SN1_45S_50ntFlanks@8527.8549.23\|CCTTCCCCCGCCGCCCCTCCTCC | >hg38_21_+_8433172_8446622_RNA45SN1_45S_50ntFlanks | 8527 | 8549 | 23 | CCTTCCCCCGCCGCCCCTCCTCC | 251.1 | -4.3 | 1.5 | -2.8 | 4.9E-03 | 6.0E-03 | MB231/MB468 | CYTO |
| >hg38_21_+_8433172_8446622_RNA45SN1_45S_50ntFlanks@11568.11599.32\|GCAGCCGACTTAGAACTGGTGCGGACCAGGGG | >hg38_21_+_8433172_8446622_RNA45SN1_45S_50ntFlanks | 11568 | 11599 | 32 | GCAGCCGACTTAGAACTGGTGCGGACCAGGGG | 294.9 | -4.0 | 1.4 | -2.8 | 5.2E-03 | 6.0E-03 | BT20/MB468 | NUC |
| >hg38_21_+_8433172_8446622_RNA45SN1_45S_50ntFlanks@12456.12498.43\|CCAAGCGTTGGATTGTTCACCCACTAATAGGGAACGTGAGCTG | >hg38_21_+_8433172_8446622_RNA45SN1_45S_50ntFlanks | 12456 | 12498 | 43 | CCAAGCGTTGGATTGTTCACCCACTAATAGGGAACGTGAGCTG | 130.8 | -5.0 | 1.8 | -2.9 | 4.2E-03 | 6.0E-03 | BT20/MB468 | MITO |
| >hg38_21_+_8433172_8446622_RNA45SN1_45S_50ntFlanks@7989.8017.29\|AGACGTGGCGACCCGCTGAATTTAAGCAT | >hg38_21_+_8433172_8446622_RNA45SN1_45S_50ntFlanks | 7989 | 8017 | 29 | AGACGTGGCGACCCGCTGAATTTAAGCAT | 419.5 | 2.2 | 0.8 | 2.8 | 5.2E-03 | 6.0E-03 | MB231/MB468 | NUC |
| >hg38_21_+_8433172_8446622_RNA45SN1_45S_50ntFlanks@5486.5512.27\|GCGCTGAGAAGACGGTCGAACTTGACT | >hg38_21_+_8433172_8446622_RNA45SN1_45S_50ntFlanks | 5486 | 5512 | 27 | GCGCTGAGAAGACGGTCGAACTTGACT | 216.3 | -4.8 | 1.7 | -2.8 | 4.5E-03 | 6.0E-03 | MB231/MB468 | MP |
| >hg38_21_+_8433172_8446622_RNA45SN1_45S_50ntFlanks@12385.12414.30\|AAGCGTTCATAGCGACGTCGCTTTTTGATC | >hg38_21_+_8433172_8446622_RNA45SN1_45S_50ntFlanks | 12385 | 12414 | 30 | AAGCGTTCATAGCGACGTCGCTTTTTGATC | 234.2 | -3.8 | 1.3 | -2.9 | 4.2E-03 | 6.0E-03 | BT20/MB468 | MITO |
| >hg38_21_+_8433172_8446622_RNA45SN1_45S_50ntFlanks@12475.12496.22\|CCCACTAATAGGGAACGTGAGC | >hg38_21_+_8433172_8446622_RNA45SN1_45S_50ntFlanks | 12475 | 12496 | 22 | CCCACTAATAGGGAACGTGAGC | 269.0 | -3.5 | 1.2 | -2.8 | 4.5E-03 | 6.0E-03 | MB231/MB468 | MP |
| >hg38_MT_+_1621_3279_RNR2_16S_50ntFlanks@485.518.34\|GAGGAACAGCTCTTTGGACACTAGGAAAAAACCT | >hg38_MT_+_1621_3279_RNR2_16S_50ntFlanks | 485 | 518 | 34 | GAGGAACAGCTCTTTGGACACTAGGAAAAAACCT | 120.1 | 3.6 | 1.3 | 2.8 | 4.5E-03 | 6.0E-03 | MB231/MB468 | MP |
| >hg38_21_+_8433172_8446622_RNA45SN1_45S_50ntFlanks@4574.4611.38\|ATAGGACCGCGGTTCTATTTTGTTGGTTTTCGGAACTG | >hg38_21_+_8433172_8446622_RNA45SN1_45S_50ntFlanks | 4574 | 4611 | 38 | ATAGGACCGCGGTTCTATTTTGTTGGTTTTCGGAACTG | 81.9 | -5.9 | 2.1 | -2.8 | 4.5E-03 | 6.0E-03 | MB231/MB468 | MP |
| >hg38_21_+_8433172_8446622_RNA45SN1_45S_50ntFlanks@7973.8018.46\|GACGCGACCTCAGATCAGACGTGGCGACCCGCTGAATTTAAGCATA | >hg38_21_+_8433172_8446622_RNA45SN1_45S_50ntFlanks | 7973 | 8018 | 46 | GACGCGACCTCAGATCAGACGTGGCGACCCGCTGAATTTAAGCATA | 307.6 | 4.0 | 1.4 | 2.8 | 5.2E-03 | 6.0E-03 | MB231/MB468 | MITO |
| >MI0003591\|hsa-mir-584&WithFlank&5\|-\|149062307\|149062415@22.42.21(+1U)&[MIMAT0003249&hsa-miR-584-5p&offsets\|0\|-1(+1U);m-167&5\|-\|149062374\|149062394&offsets\|0\|0(+1U)]\|TTATGGTTTGCCTGGGACTGAT | >MI0003591\|hsa-mir-584&WithFlank&5\|-\|149062307\|149062415 | 22 | 42 | 21(+1U) | TTATGGTTTGCCTGGGACTGAT | 169.6 | 4.5 | 1.6 | 2.8 | 4.5E-03 | 6.0E-03 | MB231/MB468 | MP |
| >MI0000813\|hsa-mir-324&WithFlank&17\|-\|7223291\|7223385@59.78.20(+1U)&[MIMAT0000762&hsa-miR-324-3p&offsets\|+3\|0(+1U);m-203&17\|-\|7223308\|7223329&offsets\|+2\|0(+1U)]\|ACTGCCCCAGGTGCTGCTGGT | >MI0000813\|hsa-mir-324&WithFlank&17\|-\|7223291\|7223385 | 59 | 78 | 20(+1U) | ACTGCCCCAGGTGCTGCTGGT | 290.1 | 3.8 | 1.3 | 2.8 | 4.5E-03 | 6.0E-03 | MB231/MB468 | MP |
| >hg38_21_+_8433172_8446622_RNA45SN1_45S_50ntFlanks@10866.10891.26\|CTAAGGGCTGGGTCGGTCGGGCTGGG | >hg38_21_+_8433172_8446622_RNA45SN1_45S_50ntFlanks | 10866 | 10891 | 26 | CTAAGGGCTGGGTCGGTCGGGCTGGG | 847.1 | -4.2 | 1.4 | -3.0 | 2.8E-03 | 6.0E-03 | BT20/MB468 | MP |
| >hg38_21_+_8433172_8446622_RNA45SN1_45S_50ntFlanks@4686.4720.35\|GACGGACCAGAGCGAAAGCATTTGCCAAGAATGTT | >hg38_21_+_8433172_8446622_RNA45SN1_45S_50ntFlanks | 4686 | 4720 | 35 | GACGGACCAGAGCGAAAGCATTTGCCAAGAATGTT | 92.1 | 5.1 | 1.7 | 3.0 | 2.8E-03 | 6.1E-03 | BT20/MB468 | MP |
| >hg38_21_+_8433172_8446622_RNA45SN1_45S_50ntFlanks@9306.9338.33\|CAAGGAGTCTAACACGTGCGCGAGTCGGGGGCT | >hg38_21_+_8433172_8446622_RNA45SN1_45S_50ntFlanks | 9306 | 9338 | 33 | CAAGGAGTCTAACACGTGCGCGAGTCGGGGGCT | 1577.4 | -4.5 | 1.5 | -3.0 | 2.8E-03 | 6.1E-03 | BT20/MB468 | MP |
| >am_trnaMT_PheGAA_MT_+_577_647@40.57.18\|TGTTTAGACGGGCTCACA | >am_trnaMT_PheGAA_MT_+_577_647 | 40 | 57 | 18 | TGTTTAGACGGGCTCACA | 186.1 | 1.9 | 0.7 | 2.8 | 4.5E-03 | 6.1E-03 | MB231/MB468 | MP |
| >hg38_21_+_8433172_8446622_RNA45SN1_45S_50ntFlanks@6728.6770.43\|GGACACATTGATCATCGACACTTCGAACGCACTTGCGGCCCCG | >hg38_21_+_8433172_8446622_RNA45SN1_45S_50ntFlanks | 6728 | 6770 | 43 | GGACACATTGATCATCGACACTTCGAACGCACTTGCGGCCCCG | 99.2 | -4.3 | 1.5 | -2.8 | 4.5E-03 | 6.1E-03 | MB231/MB468 | MP |
| >hg38_21_+_8433172_8446622_RNA45SN1_45S_50ntFlanks@3782.3814.33\|CAGTGAAACTGCGAATGGCTCATTAAATCAGTT | >hg38_21_+_8433172_8446622_RNA45SN1_45S_50ntFlanks | 3782 | 3814 | 33 | CAGTGAAACTGCGAATGGCTCATTAAATCAGTT | 167.1 | -2.5 | 0.9 | -2.8 | 5.0E-03 | 6.1E-03 | MB231/MB468 | CYTO |
| >hg38_21_+_8433172_8446622_RNA45SN1_45S_50ntFlanks@11578.11611.34\|TAGAACTGGTGCGGACCAGGGGAATCCGACTGTT | >hg38_21_+_8433172_8446622_RNA45SN1_45S_50ntFlanks | 11578 | 11611 | 34 | TAGAACTGGTGCGGACCAGGGGAATCCGACTGTT | 367.3 | -1.1 | 0.4 | -2.8 | 5.2E-03 | 6.1E-03 | BT20/MB468 | NUC |
| >hg38_21_+_8433172_8446622_RNA45SN1_45S_50ntFlanks@9736.9766.31\|CCGAAACGATCTCAACCTATTCTCAAACTTT | >hg38_21_+_8433172_8446622_RNA45SN1_45S_50ntFlanks | 9736 | 9766 | 31 | CCGAAACGATCTCAACCTATTCTCAAACTTT | 166.2 | -3.5 | 1.2 | -2.9 | 4.2E-03 | 6.1E-03 | BT20/MB468 | MITO |
| >hg38_21_+_8433172_8446622_RNA45SN1_45S_50ntFlanks@12913.12943.31\|ACGTTCGTGGGGAACCTGGCGCTAAACCATT | >hg38_21_+_8433172_8446622_RNA45SN1_45S_50ntFlanks | 12913 | 12943 | 31 | ACGTTCGTGGGGAACCTGGCGCTAAACCATT | 244.1 | 5.7 | 1.7 | 3.3 | 1.0E-03 | 6.1E-03 | BT20/MB231 | MP |
| >hg38_21_+_8433172_8446622_RNA45SN1_45S_50ntFlanks@4995.5013.19\|ACAGATTGATAGCTCTTTC | >hg38_21_+_8433172_8446622_RNA45SN1_45S_50ntFlanks | 4995 | 5013 | 19 | ACAGATTGATAGCTCTTTC | 197.2 | -2.6 | 0.9 | -2.8 | 5.0E-03 | 6.1E-03 | MB231/MB468 | CYTO |
| >MI0001445\|hsa-mir-423&WithFlank&17\|+\|30117073\|30117178@59.77.19(+1A)&[MIMAT0001340&hsa-miR-423-3p&offsets\|0\|-4(+1A);m-100&17\|+\|30117131\|30117153&offsets\|0\|-4(+1A)]\|AGCTCGGTCTGAGGCCCCTA | >MI0001445\|hsa-mir-423&WithFlank&17\|+\|30117073\|30117178 | 59 | 77 | 19(+1A) | AGCTCGGTCTGAGGCCCCTA | 67.7 | 4.7 | 1.7 | 2.8 | 4.5E-03 | 6.1E-03 | MB231/MB468 | MP |
| >hg38_21_+_8433172_8446622_RNA45SN1_45S_50ntFlanks@11768.11790.23\|CTAATTAGTGACGCGCATGAATG | >hg38_21_+_8433172_8446622_RNA45SN1_45S_50ntFlanks | 11768 | 11790 | 23 | CTAATTAGTGACGCGCATGAATG | 330.1 | -1.6 | 0.6 | -2.9 | 4.2E-03 | 6.1E-03 | BT20/MB468 | MITO |
| >hg38_21_+_8433172_8446622_RNA45SN1_45S_50ntFlanks@4730.4747.18\|CAAGAACGAAAGTCGGAG | >hg38_21_+_8433172_8446622_RNA45SN1_45S_50ntFlanks | 4730 | 4747 | 18 | CAAGAACGAAAGTCGGAG | 131.9 | -3.8 | 1.3 | -2.8 | 4.5E-03 | 6.1E-03 | MB231/MB468 | MP |

<table>
<tr><td>>hg38_21_+_8433172_8446622_RNA45SN1_45S_50ntFlanks@8082.8100.19|AGCCCAGCGCCGAATCCCC</td><td>>hg38_21_+_8433172_8446622_RNA45SN1_45S_50ntFlanks</td><td>8082</td><td>8100</td><td>19</td><td>AGCCCAGCGCCGAATCCCC</td><td>157.9</td><td>7.4</td><td>2.5</td><td>3.0</td><td>2.8E-03</td><td>6.1E-03</td><td>BT20/MB468</td><td>MP</td></tr>
<tr><td>>hg38_21_+_8433172_8446622_RNA45SN1_45S_50ntFlanks@10821.10865.45|GGAAGTCGGCAAGCCGGATCCGTAACTTCGGGATAAGGATTGGCT</td><td>>hg38_21_+_8433172_8446622_RNA45SN1_45S_50ntFlanks</td><td>10821</td><td>10865</td><td>45</td><td>GGAAGTCGGCAAGCCGGATCCGTAACTTCGGGATAAGGATTGGCT</td><td>656.0</td><td>-3.4</td><td>1.2</td><td>-2.8</td><td>5.0E-03</td><td>6.1E-03</td><td>MB231/MB468</td><td>CYTO</td></tr>
<tr><td>>hg38_MT_+_1621_3279_RNR2_16S_50ntFlanks@1362.1388.27|CGACCTCGATGTTGGATCAGGACATCC</td><td>>hg38_MT_+_1621_3279_RNR2_16S_50ntFlanks</td><td>1362</td><td>1388</td><td>27</td><td>CGACCTCGATGTTGGATCAGGACATCC</td><td>107.5</td><td>3.6</td><td>1.3</td><td>2.8</td><td>5.3E-03</td><td>6.1E-03</td><td>MB231/MB468</td><td>MITO</td></tr>
<tr><td>>hg38_21_+_8433172_8446622_RNA45SN1_45S_50ntFlanks@6721.6741.21|AATTGCAGGACACATTGATCA</td><td>>hg38_21_+_8433172_8446622_RNA45SN1_45S_50ntFlanks</td><td>6721</td><td>6741</td><td>21</td><td>AATTGCAGGACACATTGATCA</td><td>203.2</td><td>-4.9</td><td>1.6</td><td>-3.0</td><td>2.8E-03</td><td>6.1E-03</td><td>BT20/MB468</td><td>MP</td></tr>
<tr><td>>hg38_21_+_8433172_8446622_RNA45SN1_45S_50ntFlanks@6662.6698.37|GGTGGATCACTCGGCTCGTGCGTCGATGAAGAACGCA</td><td>>hg38_21_+_8433172_8446622_RNA45SN1_45S_50ntFlanks</td><td>6662</td><td>6698</td><td>37</td><td>GGTGGATCACTCGGCTCGTGCGTCGATGAAGAACGCA</td><td>201.6</td><td>3.4</td><td>1.1</td><td>3.0</td><td>2.8E-03</td><td>6.1E-03</td><td>BT20/MB468</td><td>MP</td></tr>
<tr><td>>hg38_21_+_8433172_8446622_RNA45SN1_45S_50ntFlanks@10781.10809.29|AGGTGAACAGCCTCTGGCATGTTGGAACA</td><td>>hg38_21_+_8433172_8446622_RNA45SN1_45S_50ntFlanks</td><td>10781</td><td>10809</td><td>29</td><td>AGGTGAACAGCCTCTGGCATGTTGGAACA</td><td>236.0</td><td>-2.5</td><td>0.9</td><td>-2.8</td><td>5.0E-03</td><td>6.1E-03</td><td>MB231/MB468</td><td>CYTO</td></tr>
<tr><td>>hg38_MT_+_1621_3279_RNR2_16S_50ntFlanks@1370.1401.32|ATGTTGGATCAGGACATCCCGATGGTGCAGCC</td><td>>hg38_MT_+_1621_3279_RNR2_16S_50ntFlanks</td><td>1370</td><td>1401</td><td>32</td><td>ATGTTGGATCAGGACATCCCGATGGTGCAGCC</td><td>158.7</td><td>3.5</td><td>1.2</td><td>2.8</td><td>4.5E-03</td><td>6.1E-03</td><td>MB231/MB468</td><td>MP</td></tr>
<tr><td>>hg38_21_+_8433172_8446622_RNA45SN1_45S_50ntFlanks@4882.4903.22|TGCAAAGCTGAAACTTAAAGGA</td><td>>hg38_21_+_8433172_8446622_RNA45SN1_45S_50ntFlanks</td><td>4882</td><td>4903</td><td>22</td><td>TGCAAAGCTGAAACTTAAAGGA</td><td>303.1</td><td>2.7</td><td>0.8</td><td>3.2</td><td>1.3E-03</td><td>6.1E-03</td><td>BT20/MB231</td><td>MITO</td></tr>
<tr><td>>hg38_21_+_8433172_8446622_RNA45SN1_45S_50ntFlanks@5443.5467.25|CCTCGGATCGGCCCCGCCGGGGTCG</td><td>>hg38_21_+_8433172_8446622_RNA45SN1_45S_50ntFlanks</td><td>5443</td><td>5467</td><td>25</td><td>CCTCGGATCGGCCCCGCCGGGGTCG</td><td>107.8</td><td>-4.1</td><td>1.4</td><td>-2.9</td><td>4.2E-03</td><td>6.1E-03</td><td>BT20/MB468</td><td>MITO</td></tr>
<tr><td>>hg38_MT_+_1621_3279_RNR2_16S_50ntFlanks@1096.1117.22|GACGAGAAGACCCTATGGAGCT</td><td>>hg38_MT_+_1621_3279_RNR2_16S_50ntFlanks</td><td>1096</td><td>1117</td><td>22</td><td>GACGAGAAGACCCTATGGAGCT</td><td>81.7</td><td>5.8</td><td>2.0</td><td>3.0</td><td>2.8E-03</td><td>6.1E-03</td><td>BT20/MB468</td><td>MP</td></tr>
<tr><td>>hg38_21_+_8433172_8446622_RNA45SN1_45S_50ntFlanks@5490.5526.37|TGAGAAGACGGTCGAACTTGACTATCTAGAGGAAGTA</td><td>>hg38_21_+_8433172_8446622_RNA45SN1_45S_50ntFlanks</td><td>5490</td><td>5526</td><td>37</td><td>TGAGAAGACGGTCGAACTTGACTATCTAGAGGAAGTA</td><td>1017.8</td><td>5.1</td><td>1.6</td><td>3.3</td><td>1.0E-03</td><td>6.1E-03</td><td>BT20/MB231</td><td>MP</td></tr>
<tr><td>>MI0000061|hsa-let-7a-2&WithFlank&11|-|122146516|122146599@11.32.22(+1C);MI0000062|hsa-let-7a-3&WithFlank&22|+|46112743|46112828@10.31.22(+1C);MI0000060|hsa-let-7a-1&WithFlank&9|+|94175951|94176042@12.33.22(+1C)&[MIMAT0000062&hsa-let-7a-5p&offsets|0|0(+1C);m-6&11|-|122146568|122146589&offsets|0|0(+1C)];[MIMAT0000062_1&hsa-let-7a-5p&offsets|0|0(+1C);m-5&22|+|46112752|46112773&offsets|0|0(+1C)];[MIMAT0000062_2&hsa-let-7a-5p&offsets|0|0(+1C);m-4&9|+|94175962|94175983&offsets|0|0(+1C)]|TGAGGTAGTAGGTTGTATAGTTC</td><td>>MI0000061|hsa-let-7a-2&WithFlank&11|-|122146516|122146599</td><td>11</td><td>32</td><td>22(+1C)</td><td>TGAGGTAGTAGGTTGTATAGTTC</td><td>202.1</td><td>4.2</td><td>1.4</td><td>3.0</td><td>2.8E-03</td><td>6.1E-03</td><td>BT20/MB468</td><td>MP</td></tr>
<tr><td>>hg38_21_+_8433172_8446622_RNA45SN1_45S_50ntFlanks@11703.11727.25|AATGAAGCGCGGGTAAACGGCGGGA</td><td>>hg38_21_+_8433172_8446622_RNA45SN1_45S_50ntFlanks</td><td>11703</td><td>11727</td><td>25</td><td>AATGAAGCGCGGGTAAACGGCGGGA</td><td>173.4</td><td>-3.0</td><td>1.1</td><td>-2.8</td><td>5.0E-03</td><td>6.1E-03</td><td>MB231/MB468</td><td>CYTO</td></tr>
<tr><td>>hg38_21_+_8433172_8446622_RNA45SN1_45S_50ntFlanks@7988.8009.22|CAGACGTGGCGACCCGCTGAAT</td><td>>hg38_21_+_8433172_8446622_RNA45SN1_45S_50ntFlanks</td><td>7988</td><td>8009</td><td>22</td><td>CAGACGTGGCGACCCGCTGAAT</td><td>2994.0</td><td>-2.7</td><td>0.9</td><td>-3.0</td><td>2.8E-03</td><td>6.1E-03</td><td>BT20/MB468</td><td>MP</td></tr>
<tr><td>>hg38_21_+_8433172_8446622_RNA45SN1_45S_50ntFlanks@8180.8203.24|TGATCGAGGCCCAGCCCGTGGACG</td><td>>hg38_21_+_8433172_8446622_RNA45SN1_45S_50ntFlanks</td><td>8180</td><td>8203</td><td>24</td><td>TGATCGAGGCCCAGCCCGTGGACG</td><td>111.9</td><td>-3.7</td><td>1.3</td><td>-2.8</td><td>4.5E-03</td><td>6.1E-03</td><td>MB231/MB468</td><td>MP</td></tr>
<tr><td>>hg38_21_+_8433172_8446622_RNA45SN1_45S_50ntFlanks@9307.9336.30|AAGGAGTCTAACACGTGCGCGAGTCGGGGG</td><td>>hg38_21_+_8433172_8446622_RNA45SN1_45S_50ntFlanks</td><td>9307</td><td>9336</td><td>30</td><td>AAGGAGTCTAACACGTGCGCGAGTCGGGGG</td><td>1192.3</td><td>-2.7</td><td>0.9</td><td>-3.0</td><td>2.8E-03</td><td>6.1E-03</td><td>BT20/MB468</td><td>MP</td></tr>
<tr><td>>hg38_21_+_8433172_8446622_RNA45SN1_45S_50ntFlanks@4122.4156.35|AGGGTTCGATTCCGGAGAGGGAGCCTGAGAAACGG</td><td>>hg38_21_+_8433172_8446622_RNA45SN1_45S_50ntFlanks</td><td>4122</td><td>4156</td><td>35</td><td>AGGGTTCGATTCCGGAGAGGGAGCCTGAGAAACGG</td><td>65.7</td><td>-4.8</td><td>1.7</td><td>-2.8</td><td>4.5E-03</td><td>6.1E-03</td><td>MB231/MB468</td><td>MP</td></tr>
<tr><td>>hg38_21_+_8433172_8446622_RNA45SN1_45S_50ntFlanks@10355.10388.34|AACTTTGAAGGCCGAAGTGGAGAAGGGTTCCATG</td><td>>hg38_21_+_8433172_8446622_RNA45SN1_45S_50ntFlanks</td><td>10355</td><td>10388</td><td>34</td><td>AACTTTGAAGGCCGAAGTGGAGAAGGGTTCCATG</td><td>340.7</td><td>-2.8</td><td>0.9</td><td>-3.0</td><td>2.8E-03</td><td>6.1E-03</td><td>BT20/MB468</td><td>MP</td></tr>
<tr><td>>hg38_21_+_8433172_8446622_RNA45SN1_45S_50ntFlanks@7975.8004.30|CGCGACCTCAGATCAGACGTGGCGACCCGC</td><td>>hg38_21_+_8433172_8446622_RNA45SN1_45S_50ntFlanks</td><td>7975</td><td>8004</td><td>30</td><td>CGCGACCTCAGATCAGACGTGGCGACCCGC</td><td>48245.2</td><td>2.4</td><td>0.7</td><td>3.2</td><td>1.3E-03</td><td>6.1E-03</td><td>BT20/MB231</td><td>MITO</td></tr>
<tr><td>>hg38_21_+_8433172_8446622_RNA45SN1_45S_50ntFlanks@8105.8140.36|CGCGGCGGGGCGCGGGACATGTGGCGTACGGAAGAC</td><td>>hg38_21_+_8433172_8446622_RNA45SN1_45S_50ntFlanks</td><td>8105</td><td>8140</td><td>36</td><td>CGCGGCGGGGCGCGGGACATGTGGCGTACGGAAGAC</td><td>1144.8</td><td>5.6</td><td>1.7</td><td>3.3</td><td>1.0E-03</td><td>6.1E-03</td><td>BT20/MB231</td><td>MP</td></tr>
<tr><td>>hg38_21_+_8433172_8446622_RNA45SN1_45S_50ntFlanks@4727.4767.41|AATCAAGAACGAAAGTCGGAGGTTCGAAGACGATCAGATAC</td><td>>hg38_21_+_8433172_8446622_RNA45SN1_45S_50ntFlanks</td><td>4727</td><td>4767</td><td>41</td><td>AATCAAGAACGAAAGTCGGAGGTTCGAAGACGATCAGATAC</td><td>69.5</td><td>4.5</td><td>1.6</td><td>2.8</td><td>5.3E-03</td><td>6.1E-03</td><td>MB231/MB468</td><td>MITO</td></tr>
<tr><td>>hg38_21_+_8433172_8446622_RNA45SN1_45S_50ntFlanks@6644.6670.27|CCTCGTACGACTCTTAGCGGTGGATCA</td><td>>hg38_21_+_8433172_8446622_RNA45SN1_45S_50ntFlanks</td><td>6644</td><td>6670</td><td>27</td><td>CCTCGTACGACTCTTAGCGGTGGATCA</td><td>256.5</td><td>-2.4</td><td>0.9</td><td>-2.8</td><td>5.3E-03</td><td>6.1E-03</td><td>MB231/MB468</td><td>MITO</td></tr>
<tr><td>>hg38_21_+_8433172_8446622_RNA45SN1_45S_50ntFlanks@4002.4039.38|GATAACCTCGGGCCGATCGCACGCCCCCCGTGGCGGCG</td><td>>hg38_21_+_8433172_8446622_RNA45SN1_45S_50ntFlanks</td><td>4002</td><td>4039</td><td>38</td><td>GATAACCTCGGGCCGATCGCACGCCCCCCGTGGCGGCG</td><td>264.5</td><td>-4.5</td><td>1.6</td><td>-2.8</td><td>5.3E-03</td><td>6.1E-03</td><td>MB231/MB468</td><td>NUC</td></tr>
<tr><td>>hg38_21_+_8433172_8446622_RNA45SN1_45S_50ntFlanks@10903.10939.37|GGCTGGGCGCGCGCCGCGGCTGGACGAGGCGCCGCCG</td><td>>hg38_21_+_8433172_8446622_RNA45SN1_45S_50ntFlanks</td><td>10903</td><td>10939</td><td>37</td><td>GGCTGGGCGCGCGCCGCGGCTGGACGAGGCGCCGCCG</td><td>124.0</td><td>-4.1</td><td>1.4</td><td>-2.9</td><td>4.2E-03</td><td>6.1E-03</td><td>BT20/MB468</td><td>MITO</td></tr>
<tr><td>>trna14_LysTTT_11_-_59327808_59327880@1.35.35,trna2_LysTTT_17_+_8022473_8022545@1.35.35,trna54_LysTTT_1_+_204475655_204475727@1.35.35,trna5_LysTTT_11_+_59323902_59323974@1.35.35,trna62_LysTTT_1_-_204476158_204476230@1.35.35,trna76_LysTTT_6_+_28918806_28918878@1.35.35|GCCCGGATAGCTCAGTCGGTAGAGCATCAGACTTT</td><td>>trna14_LysTTT_11_-_59327808_59327880</td><td>1</td><td>35</td><td>35</td><td>GCCCGGATAGCTCAGTCGGTAGAGCATCAGACTTT</td><td>307.5</td><td>2.8</td><td>0.9</td><td>3.2</td><td>1.3E-03</td><td>6.1E-03</td><td>BT20/MB231</td><td>CYTO</td></tr>
<tr><td>>hg38_21_+_8433172_8446622_RNA45SN1_45S_50ntFlanks@11668.11705.38|TGCCCAGTGCTCTGAATGTCAAAGTGAAGAAATTCAAT</td><td>>hg38_21_+_8433172_8446622_RNA45SN1_45S_50ntFlanks</td><td>11668</td><td>11705</td><td>38</td><td>TGCCCAGTGCTCTGAATGTCAAAGTGAAGAAATTCAAT</td><td>547.8</td><td>-2.6</td><td>0.8</td><td>-3.2</td><td>1.3E-03</td><td>6.1E-03</td><td>BT20/MB231</td><td>CYTO</td></tr>
<tr><td>>hg38_21_+_8433172_8446622_RNA45SN1_45S_50ntFlanks@12909.12933.25|CCGCACGTTCGTGGGGAACCTGGCG</td><td>>hg38_21_+_8433172_8446622_RNA45SN1_45S_50ntFlanks</td><td>12909</td><td>12933</td><td>25</td><td>CCGCACGTTCGTGGGGAACCTGGCG</td><td>300.5</td><td>-1.5</td><td>0.5</td><td>-2.9</td><td>4.2E-03</td><td>6.1E-03</td><td>BT20/MB468</td><td>MITO</td></tr>
<tr><td>>hg38_21_+_8433172_8446622_RNA45SN1_45S_50ntFlanks@6646.6665.20|TCGTACGACTCTTAGCGGTG</td><td>>hg38_21_+_8433172_8446622_RNA45SN1_45S_50ntFlanks</td><td>6646</td><td>6665</td><td>20</td><td>TCGTACGACTCTTAGCGGTG</td><td>72276.7</td><td>4.0</td><td>1.2</td><td>3.3</td><td>1.0E-03</td><td>6.1E-03</td><td>BT20/MB231</td><td>MP</td></tr>
<tr><td>>MI0000239|hsa-mir-197&WithFlank&1|+|109598887|109598973@54.72.19&[MIMAT0000227&hsa-miR-197-3p&offsets|0|-3;m-99&1|+|109598940|109598961&offsets|0|-3]|TTCACCACCTTCTCCACCC</td><td>>MI0000239|hsa-mir-197&WithFlank&1|+|109598887|109598973</td><td>54</td><td>72</td><td>19</td><td>TTCACCACCTTCTCCACCC</td><td>524.8</td><td>2.6</td><td>0.8</td><td>3.2</td><td>1.3E-03</td><td>6.1E-03</td><td>BT20/MB231</td><td>CYTO</td></tr>
<tr><td>>MI0000266|hsa-mir-10a&WithFlank&17|-|48579832|48579953@28.50.23&[MIMAT0000253&hsa-miR-10a-5p&offsets|0|0;m-18&17|-|48579905|48579926&offsets|0|+1]|TACCCTGTAGATCCGAATTTGTG</td><td>>MI0000266|hsa-mir-10a&WithFlank&17|-|48579832|48579953</td><td>28</td><td>50</td><td>23</td><td>TACCCTGTAGATCCGAATTTGTG</td><td>18886.1</td><td>2.5</td><td>0.8</td><td>3.2</td><td>1.3E-03</td><td>6.1E-03</td><td>BT20/MB231</td><td>CYTO</td></tr>
<tr><td>>hg38_21_+_8433172_8446622_RNA45SN1_45S_50ntFlanks@6651.6696.46|CGACTCTTAGCGGTGGATCACTCGGCTCGTGCGTCGATGAAGAACG</td><td>>hg38_21_+_8433172_8446622_RNA45SN1_45S_50ntFlanks</td><td>6651</td><td>6696</td><td>46</td><td>CGACTCTTAGCGGTGGATCACTCGGCTCGTGCGTCGATGAAGAACG</td><td>25982.7</td><td>2.2</td><td>0.8</td><td>2.9</td><td>4.2E-03</td><td>6.1E-03</td><td>BT20/MB468</td><td>MITO</td></tr>
<tr><td>>trnaMT_SerGCT_MT_+_12207_12265@1.21.21|GAGAAAGCTCACAAGAACTGC</td><td>>trnaMT_SerGCT_MT_+_12207_12265</td><td>1</td><td>21</td><td>21</td><td>GAGAAAGCTCACAAGAACTGC</td><td>12143.7</td><td>2.3</td><td>0.8</td><td>2.8</td><td>4.6E-03</td><td>6.1E-03</td><td>MB231/MB468</td><td>MP</td></tr>
<tr><td>>hg38_21_+_8433172_8446622_RNA45SN1_45S_50ntFlanks@8168.8212.45|CCCAAGTCCTTCTGATCGAGGCCCAGCCCGTGGACGGTGTGAGGC</td><td>>hg38_21_+_8433172_8446622_RNA45SN1_45S_50ntFlanks</td><td>8168</td><td>8212</td><td>45</td><td>CCCAAGTCCTTCTGATCGAGGCCCAGCCCGTGGACGGTGTGAGGC</td><td>146.9</td><td>-3.5</td><td>1.2</td><td>-2.8</td><td>5.0E-03</td><td>6.1E-03</td><td>BT20/MB468</td><td>CYTO</td></tr>
<tr><td>>hg38_21_+_8433172_8446622_RNA45SN1_45S_50ntFlanks@6650.6668.19|ACGACTCTTAGCGGTGGAT</td><td>>hg38_21_+_8433172_8446622_RNA45SN1_45S_50ntFlanks</td><td>6650</td><td>6668</td><td>19</td><td>ACGACTCTTAGCGGTGGAT</td><td>754.2</td><td>2.4</td><td>0.8</td><td>3.2</td><td>1.3E-03</td><td>6.1E-03</td><td>BT20/MB231</td><td>CYTO</td></tr>
<tr><td>>hg38_21_+_8433172_8446622_RNA45SN1_45S_50ntFlanks@10391.10424.34|AACAGCAGTTGAACATGGGTCAGTCGGTCCTGAG</td><td>>hg38_21_+_8433172_8446622_RNA45SN1_45S_50ntFlanks</td><td>10391</td><td>10424</td><td>34</td><td>AACAGCAGTTGAACATGGGTCAGTCGGTCCTGAG</td><td>13909.1</td><td>4.3</td><td>1.3</td><td>3.3</td><td>1.0E-03</td><td>6.1E-03</td><td>BT20/MB231</td><td>MP</td></tr>
<tr><td>>hg38_21_+_8433172_8446622_RNA45SN1_45S_50ntFlanks@11920.11943.24|ATGAGAGGTGTAGAATAAGTGGGA</td><td>>hg38_21_+_8433172_8446622_RNA45SN1_45S_50ntFlanks</td><td>11920</td><td>11943</td><td>24</td><td>ATGAGAGGTGTAGAATAAGTGGGA</td><td>65.6</td><td>9.4</td><td>2.9</td><td>3.3</td><td>1.0E-03</td><td>6.1E-03</td><td>BT20/MB231</td><td>MP</td></tr>
<tr><td>>MI0000266|hsa-mir-10a&WithFlank&17|-|48579832|48579953@28.49.22(+1A)&[MIMAT0000253&hsa-miR-10a-5p&offsets|0|-1(+1A);m-18&17|-|48579905|48579926&offsets|0|0(+1A)]|TACCCTGTAGATCCGAATTTGTA</td><td>>MI0000266|hsa-mir-10a&WithFlank&17|-|48579832|48579953</td><td>28</td><td>49</td><td>22(+1A)</td><td>TACCCTGTAGATCCGAATTTGTA</td><td>1202.3</td><td>1.8</td><td>0.6</td><td>3.3</td><td>1.1E-03</td><td>6.1E-03</td><td>BT20/MB231</td><td>NUC</td></tr>
<tr><td>>MI0000746|hsa-mir-99b&WithFlank&19|+|51692606|51692687@51.70.20(+1U)&[MIMAT0004678&hsa-miR-99b-3p&offsets|0|-2(+1U);m-151&19|+|51692656|51692678&offsets|0|-3(+1U)]|CAAGCTCGTGTCTGTGGGTCT</td><td>>MI0000746|hsa-mir-99b&WithFlank&19|+|51692606|51692687</td><td>51</td><td>70</td><td>20(+1U)</td><td>CAAGCTCGTGTCTGTGGGTCT</td><td>240.5</td><td>2.4</td><td>0.7</td><td>3.3</td><td>1.1E-03</td><td>6.1E-03</td><td>BT20/MB231</td><td>NUC</td></tr>
<tr><td>>hg38_21_+_8433172_8446622_RNA45SN1_45S_50ntFlanks@6649.6669.21|TACGACTCTTAGCGGTGGATC</td><td>>hg38_21_+_8433172_8446622_RNA45SN1_45S_50ntFlanks</td><td>6649</td><td>6669</td><td>21</td><td>TACGACTCTTAGCGGTGGATC</td><td>958.4</td><td>2.6</td><td>0.8</td><td>3.3</td><td>1.1E-03</td><td>6.1E-03</td><td>BT20/MB231</td><td>NUC</td></tr>
<tr><td>>hg38_21_+_8433172_8446622_RNA45SN1_45S_50ntFlanks@4242.4264.23|TCTTTCGAGGCCCTGTAATTGGA</td><td>>hg38_21_+_8433172_8446622_RNA45SN1_45S_50ntFlanks</td><td>4242</td><td>4264</td><td>23</td><td>TCTTTCGAGGCCCTGTAATTGGA</td><td>134.8</td><td>-1.3</td><td>0.5</td><td>-2.8</td><td>5.0E-03</td><td>6.1E-03</td><td>BT20/MB468</td><td>CYTO</td></tr>
<tr><td>>hg38_21_+_8433172_8446622_RNA45SN1_45S_50ntFlanks@9575.9602.28|ATCGGTCGTCCGACCTGGGTATAGGGGC</td><td>>hg38_21_+_8433172_8446622_RNA45SN1_45S_50ntFlanks</td><td>9575</td><td>9602</td><td>28</td><td>ATCGGTCGTCCGACCTGGGTATAGGGGC</td><td>325.8</td><td>-4.9</td><td>1.7</td><td>-2.8</td><td>4.6E-03</td><td>6.1E-03</td><td>MB231/MB468</td><td>MP</td></tr>
<tr><td>>hg38_21_+_8433172_8446622_RNA45SN1_45S_50ntFlanks@4687.4720.34|ACGGACCAGAGCGAAAGCATTTGCCAAGAATGTT</td><td>>hg38_21_+_8433172_8446622_RNA45SN1_45S_50ntFlanks</td><td>4687</td><td>4720</td><td>34</td><td>ACGGACCAGAGCGAAAGCATTTGCCAAGAATGTT</td><td>451.1</td><td>3.4</td><td>1.2</td><td>2.8</td><td>5.0E-03</td><td>6.1E-03</td><td>MB231/MB468</td><td>CYTO</td></tr>
<tr><td>>trna10_GluCTC_3_-_103879524_103879595@16.42.27,trna116_GluCTC_1_-_145399233_145399304@16.42.27,trna59_GluCTC_1_+_249168447_249168518@16.42.27,trna71_GluCTC_1_-_161439189_161439260@16.42.27,trna74_GluCTC_1_-_161431809_161431880@16.42.27,trna77_GluCTC_1_-_161424398_161424469@16.42.27,trna77_GluCTC_6_+_28949976_28950047@16.42.27,trna80_GluCTC_1_-_161417018_161417089@16.42.27,trna87_GluCTC_6_-_126101393_126101464@16.42.27|TGGTTAGGATTCGGCGCTCTCACCGCC</td><td>>trna10_GluCTC_3_-_103879524_103879595</td><td>16</td><td>42</td><td>27</td><td>TGGTTAGGATTCGGCGCTCTCACCGCC</td><td>1002.2</td><td>2.6</td><td>0.9</td><td>2.8</td><td>5.0E-03</td><td>6.1E-03</td><td>MB231/MB468</td><td>CYTO</td></tr>
<tr><td>>hg38_21_+_8433172_8446622_RNA45SN1_45S_50ntFlanks@12934.12961.28|CTAAACCATTCGTAGACGACCTGCTTCT</td><td>>hg38_21_+_8433172_8446622_RNA45SN1_45S_50ntFlanks</td><td>12934</td><td>12961</td><td>28</td><td>CTAAACCATTCGTAGACGACCTGCTTCT</td><td>498.0</td><td>-3.1</td><td>1.1</td><td>-2.8</td><td>5.0E-03</td><td>6.1E-03</td><td>MB231/MB468</td><td>CYTO</td></tr>
</table>

<table>
<tr><td>>am_trna111_HisGTG_1_-_147774845_147774916@-1G.26.27,trna118_HisGTG_1_-_145396881_145396952@-1G.26.27,trna16_HisGTG_1_+_146544773_146544844@-1G.26.27,trna1_HisGTG_15_+_45493349_45493420@-1G.26.27,trna21_HisGTG_1_+_147753471_147753542@-1G.26.27,trna33_HisGTG_6_+_27125906_27125977@-1G.26.27,trna7_HisGTG_9_-_14433938_14434009@-1G.26.27,trna8_HisGTG_15_-_45492611_45492682@-1G.26.27,trna9_HisGTG_15_-_45490804_45490875@-1G.26.27|GGCCGTGATCGTATAGTGGTTAGTACT</td><td>>am_trna111_HisGTG_1_-_147774845_147774916</td><td>-1G</td><td>26</td><td>27</td><td>GGCCGTGATCGTATAGTGGTTAGTACT</td><td>1766.1</td><td>-3.4</td><td>1.0</td><td>-3.3</td><td>1.1E-03</td><td>6.2E-03</td><td>BT20/MB231</td><td>NUC</td></tr>
<tr><td>>hg38_21_+_8433172_8446622_RNA45SN1_45S_50ntFlanks@4731.4759.29|AAGAACGAAAGTCGGAGGTTCGAAGACGA</td><td>>hg38_21_+_8433172_8446622_RNA45SN1_45S_50ntFlanks</td><td>4731</td><td>4759</td><td>29</td><td>AAGAACGAAAGTCGGAGGTTCGAAGACGA</td><td>3462.4</td><td>5.1</td><td>1.5</td><td>3.3</td><td>1.0E-03</td><td>6.2E-03</td><td>BT20/MB231</td><td>MP</td></tr>
<tr><td>>hg38_21_+_8433172_8446622_RNA45SN1_45S_50ntFlanks@4723.4761.39|CATTAATCAAGAACGAAAGTCGGAGGTTCGAAGACGATC</td><td>>hg38_21_+_8433172_8446622_RNA45SN1_45S_50ntFlanks</td><td>4723</td><td>4761</td><td>39</td><td>CATTAATCAAGAACGAAAGTCGGAGGTTCGAAGACGATC</td><td>82.4</td><td>-4.1</td><td>1.5</td><td>-2.8</td><td>4.6E-03</td><td>6.2E-03</td><td>MB231/MB468</td><td>MP</td></tr>
<tr><td>>hg38_21_+_8433172_8446622_RNA45SN1_45S_50ntFlanks@10351.10380.30|CGAGAACTTTGAAGGCCGAAGTGGAGAAGG</td><td>>hg38_21_+_8433172_8446622_RNA45SN1_45S_50ntFlanks</td><td>10351</td><td>10380</td><td>30</td><td>CGAGAACTTTGAAGGCCGAAGTGGAGAAGG</td><td>687.5</td><td>5.9</td><td>1.8</td><td>3.3</td><td>1.0E-03</td><td>6.2E-03</td><td>BT20/MB231</td><td>MP</td></tr>
<tr><td>>hg38_21_+_8433172_8446622_RNA45SN1_45S_50ntFlanks@9484.9506.23|GTGTTAGGACCCGAAAGATGGTG</td><td>>hg38_21_+_8433172_8446622_RNA45SN1_45S_50ntFlanks</td><td>9484</td><td>9506</td><td>23</td><td>GTGTTAGGACCCGAAAGATGGTG</td><td>75.7</td><td>5.9</td><td>2.0</td><td>3.0</td><td>2.9E-03</td><td>6.2E-03</td><td>BT20/MB468</td><td>MP</td></tr>
<tr><td>>MI0000299|hsa-mir-222&WithFlank&X|-|45747009|45747130@75.99.25(+1C)&[MIMAT0000279&hsa-miR-222-3p&offsets|0|+4(+1C);m-140&X|-|45747033|45747056&offsets|0|+1(+1C)]|AGCTACATCTGGCTACTGGGTCTCTC</td><td>>MI0000299|hsa-mir-222&WithFlank&X|-|45747009|45747130</td><td>75</td><td>99</td><td>25(+1C)</td><td>AGCTACATCTGGCTACTGGGTCTCTC</td><td>97.2</td><td>4.7</td><td>1.6</td><td>2.8</td><td>4.6E-03</td><td>6.2E-03</td><td>MB231/MB468</td><td>MP</td></tr>
<tr><td>>hg38_21_+_8433172_8446622_RNA45SN1_45S_50ntFlanks@12420.12451.32|ATGTCGGCTCTTCCTATCATTGTGAAGCAGAA</td><td>>hg38_21_+_8433172_8446622_RNA45SN1_45S_50ntFlanks</td><td>12420</td><td>12451</td><td>32</td><td>ATGTCGGCTCTTCCTATCATTGTGAAGCAGAA</td><td>211.2</td><td>-1.7</td><td>0.6</td><td>-2.8</td><td>5.3E-03</td><td>6.2E-03</td><td>MB231/MB468</td><td>NUC</td></tr>
<tr><td>>hg38_21_+_8433172_8446622_RNA45SN1_45S_50ntFlanks@6653.6695.43|ACTCTTAGCGGTGGATCACTCGGCTCGTGCGTCGATGAAGAAC</td><td>>hg38_21_+_8433172_8446622_RNA45SN1_45S_50ntFlanks</td><td>6653</td><td>6695</td><td>43</td><td>ACTCTTAGCGGTGGATCACTCGGCTCGTGCGTCGATGAAGAAC</td><td>176.9</td><td>4.3</td><td>1.6</td><td>2.8</td><td>5.3E-03</td><td>6.2E-03</td><td>MB231/MB468</td><td>MITO</td></tr>
<tr><td>>hg38_21_+_8433172_8446622_RNA45SN1_45S_50ntFlanks@11573.11601.29|CGACTTAGAACTGGTGCGGACCAGGGGAA</td><td>>hg38_21_+_8433172_8446622_RNA45SN1_45S_50ntFlanks</td><td>11573</td><td>11601</td><td>29</td><td>CGACTTAGAACTGGTGCGGACCAGGGGAA</td><td>1739.3</td><td>-2.7</td><td>0.9</td><td>-3.0</td><td>2.9E-03</td><td>6.2E-03</td><td>BT20/MB468</td><td>MP</td></tr>
<tr><td>>hg38_21_+_8433172_8446622_RNA45SN1_45S_50ntFlanks@9928.9946.19|TGATATAGACAGCAGGACG</td><td>>hg38_21_+_8433172_8446622_RNA45SN1_45S_50ntFlanks</td><td>9928</td><td>9946</td><td>19</td><td>TGATATAGACAGCAGGACG</td><td>114.8</td><td>5.9</td><td>2.0</td><td>3.0</td><td>2.9E-03</td><td>6.2E-03</td><td>BT20/MB468</td><td>MP</td></tr>
<tr><td>>trnaMT_HisGTG_MT_+_12138_12206@32.70.39|TGAATCTGACAACAGAGGCTTACGACCCCTTATTTACCC</td><td>>trnaMT_HisGTG_MT_+_12138_12206</td><td>32</td><td>70</td><td>39</td><td>TGAATCTGACAACAGAGGCTTACGACCCCTTATTTACCC</td><td>1078.8</td><td>-4.5</td><td>1.4</td><td>-3.3</td><td>1.1E-03</td><td>6.2E-03</td><td>BT20/MB231</td><td>NUC</td></tr>
<tr><td>>hg38_MT_+_1621_3279_RNR2_16S_50ntFlanks@1380.1401.22|AGGACATCCCGATGGTGCAGCC</td><td>>hg38_MT_+_1621_3279_RNR2_16S_50ntFlanks</td><td>1380</td><td>1401</td><td>22</td><td>AGGACATCCCGATGGTGCAGCC</td><td>168.0</td><td>5.5</td><td>1.8</td><td>3.0</td><td>2.9E-03</td><td>6.2E-03</td><td>BT20/MB468</td><td>MP</td></tr>
<tr><td>>hg38_21_+_8433172_8446622_RNA45SN1_45S_50ntFlanks@5485.5508.24|AGCGCTGAGAAGACGGTCGAACTT</td><td>>hg38_21_+_8433172_8446622_RNA45SN1_45S_50ntFlanks</td><td>5485</td><td>5508</td><td>24</td><td>AGCGCTGAGAAGACGGTCGAACTT</td><td>97.5</td><td>-4.7</td><td>1.6</td><td>-3.0</td><td>2.9E-03</td><td>6.2E-03</td><td>BT20/MB468</td><td>MP</td></tr>
<tr><td>>hg38_MT_+_1621_3279_RNR2_16S_50ntFlanks@1000.1029.30|GGGACCTGTATGAATGGCTCCACGAGGGTT</td><td>>hg38_MT_+_1621_3279_RNR2_16S_50ntFlanks</td><td>1000</td><td>1029</td><td>30</td><td>GGGACCTGTATGAATGGCTCCACGAGGGTT</td><td>53.6</td><td>6.1</td><td>2.1</td><td>3.0</td><td>2.9E-03</td><td>6.2E-03</td><td>BT20/MB468</td><td>MP</td></tr>
<tr><td>>hg38_21_+_8433172_8446622_RNA45SN1_45S_50ntFlanks@5419.5443.25|ACCGATTGGATGGTTTAGTGAGGCC</td><td>>hg38_21_+_8433172_8446622_RNA45SN1_45S_50ntFlanks</td><td>5419</td><td>5443</td><td>25</td><td>ACCGATTGGATGGTTTAGTGAGGCC</td><td>82.3</td><td>-4.8</td><td>1.6</td><td>-3.0</td><td>2.9E-03</td><td>6.2E-03</td><td>BT20/MB468</td><td>MP</td></tr>
<tr><td>>hg38_21_+_8433172_8446622_RNA45SN1_45S_50ntFlanks@5485.5511.27|AGCGCTGAGAAGACGGTCGAACTTGAC</td><td>>hg38_21_+_8433172_8446622_RNA45SN1_45S_50ntFlanks</td><td>5485</td><td>5511</td><td>27</td><td>AGCGCTGAGAAGACGGTCGAACTTGAC</td><td>66.4</td><td>9.4</td><td>2.9</td><td>3.3</td><td>1.0E-03</td><td>6.2E-03</td><td>BT20/MB231</td><td>MP</td></tr>
<tr><td>>hg38_21_+_8433172_8446622_RNA45SN1_45S_50ntFlanks@9821.9850.30|CCTAGTGGGCCACTTTTGGTAAGCAGAACT</td><td>>hg38_21_+_8433172_8446622_RNA45SN1_45S_50ntFlanks</td><td>9821</td><td>9850</td><td>30</td><td>CCTAGTGGGCCACTTTTGGTAAGCAGAACT</td><td>268.6</td><td>-2.6</td><td>0.9</td><td>-2.8</td><td>5.0E-03</td><td>6.2E-03</td><td>BT20/MB468</td><td>CYTO</td></tr>
<tr><td>>hg38_21_+_8433172_8446622_RNA45SN1_45S_50ntFlanks@11583.11611.29|CTGGTGCGGACCAGGGGAATCCGACTGTT</td><td>>hg38_21_+_8433172_8446622_RNA45SN1_45S_50ntFlanks</td><td>11583</td><td>11611</td><td>29</td><td>CTGGTGCGGACCAGGGGAATCCGACTGTT</td><td>2127.4</td><td>-1.5</td><td>0.5</td><td>-2.8</td><td>5.0E-03</td><td>6.2E-03</td><td>BT20/MB468</td><td>CYTO</td></tr>
<tr><td>>am_trnaMT_PheGAA_MT_+_577_647@43.62.20|TTAGACGGGCTCACATCACC</td><td>>am_trnaMT_PheGAA_MT_+_577_647</td><td>43</td><td>62</td><td>20</td><td>TTAGACGGGCTCACATCACC</td><td>219.7</td><td>6.7</td><td>2.4</td><td>2.8</td><td>5.1E-03</td><td>6.2E-03</td><td>MB231/MB468</td><td>CYTO</td></tr>
<tr><td>>hg38_21_+_8433172_8446622_RNA45SN1_45S_50ntFlanks@10408.10453.46|GGTCAGTCGGTCCTGAGAGATGGGCGAGCGCCGTTCCGAAGGGACG</td><td>>hg38_21_+_8433172_8446622_RNA45SN1_45S_50ntFlanks</td><td>10408</td><td>10453</td><td>46</td><td>GGTCAGTCGGTCCTGAGAGATGGGCGAGCGCCGTTCCGAAGGGACG</td><td>93.0</td><td>-5.0</td><td>1.8</td><td>-2.8</td><td>4.6E-03</td><td>6.2E-03</td><td>MB231/MB468</td><td>MP</td></tr>
<tr><td>>hg38_MT_+_1621_3279_RNR2_16S_50ntFlanks@643.676.34|CAATTGGACCAATCTATCACCCTATAGAAGAACT</td><td>>hg38_MT_+_1621_3279_RNR2_16S_50ntFlanks</td><td>643</td><td>676</td><td>34</td><td>CAATTGGACCAATCTATCACCCTATAGAAGAACT</td><td>122.6</td><td>5.2</td><td>1.8</td><td>3.0</td><td>2.9E-03</td><td>6.2E-03</td><td>BT20/MB468</td><td>MP</td></tr>
<tr><td>>trnaMT_SerGCT_MT_+_12207_12265@1.25.25|GAGAAAGCTCACAAGAACTGCTAAC</td><td>>trnaMT_SerGCT_MT_+_12207_12265</td><td>1</td><td>25</td><td>25</td><td>GAGAAAGCTCACAAGAACTGCTAAC</td><td>1297.3</td><td>3.2</td><td>1.1</td><td>2.8</td><td>5.3E-03</td><td>6.2E-03</td><td>BT20/MB468</td><td>NUC</td></tr>
<tr><td>>am_trna111_HisGTG_1_-_147774845_147774916@2.28.27,trna118_HisGTG_1_-_145396881_145396952@2.28.27,trna16_HisGTG_1_+_146544773_146544844@2.28.27,trna1_HisGTG_15_+_45493349_45493420@2.28.27,trna21_HisGTG_1_+_147753471_147753542@2.28.27,trna33_HisGTG_6_+_27125906_27125977@2.28.27,trna7_HisGTG_9_-_14433938_14434009@2.28.27,trna8_HisGTG_15_-_45492611_45492682@2.28.27,trna9_HisGTG_15_-_45490804_45490875@2.28.27|CCGTGATCGTATAGTGGTTAGTACTCT</td><td>>am_trna111_HisGTG_1_-_147774845_147774916</td><td>2</td><td>28</td><td>27</td><td>CCGTGATCGTATAGTGGTTAGTACTCT</td><td>84.6</td><td>2.8</td><td>0.9</td><td>3.0</td><td>2.9E-03</td><td>6.2E-03</td><td>BT20/MB468</td><td>MP</td></tr>
<tr><td>>hg38_21_+_8433172_8446622_RNA45SN1_45S_50ntFlanks@7987.8017.31|TCAGACGTGGCGACCCGCTGAATTTAAGCAT</td><td>>hg38_21_+_8433172_8446622_RNA45SN1_45S_50ntFlanks</td><td>7987</td><td>8017</td><td>31</td><td>TCAGACGTGGCGACCCGCTGAATTTAAGCAT</td><td>53.5</td><td>-4.8</td><td>1.6</td><td>-3.0</td><td>2.9E-03</td><td>6.2E-03</td><td>BT20/MB468</td><td>MP</td></tr>
<tr><td>>MI0000085|hsa-mir-27a&WithFlank&19|-|13836434|13836523@57.76.20(+2U)&[MIMAT0000084&hsa-miR-27a-3p&offsets|0|-1(+2U);m-43&19|-|13836447|13836467&offsets|0|-1(+2U)]|TTCACAGTGGCTAAGTTCCGTT</td><td>>MI0000085|hsa-mir-27a&WithFlank&19|-|13836434|13836523</td><td>57</td><td>76</td><td>20(+2U)</td><td>TTCACAGTGGCTAAGTTCCGTT</td><td>4179.4</td><td>2.2</td><td>0.8</td><td>2.8</td><td>4.6E-03</td><td>6.2E-03</td><td>MB231/MB468</td><td>MP</td></tr>
<tr><td>>hg38_21_+_8433172_8446622_RNA45SN1_45S_50ntFlanks@12994.13039.46|CCTCGCTGCGATCTATTGAAAGTCAGCCCTCGACACAAGGGTTTGT</td><td>>hg38_21_+_8433172_8446622_RNA45SN1_45S_50ntFlanks</td><td>12994</td><td>13039</td><td>46</td><td>CCTCGCTGCGATCTATTGAAAGTCAGCCCTCGACACAAGGGTTTGT</td><td>12000.4</td><td>2.9</td><td>1.0</td><td>2.8</td><td>5.4E-03</td><td>6.2E-03</td><td>MB231/MB468</td><td>MITO</td></tr>
<tr><td>>hg38_21_+_8433172_8446622_RNA45SN1_45S_50ntFlanks@6649.6692.44|TACGACTCTTAGCGGTGGATCACTCGGCTCGTGCGTCGATGAAG</td><td>>hg38_21_+_8433172_8446622_RNA45SN1_45S_50ntFlanks</td><td>6649</td><td>6692</td><td>44</td><td>TACGACTCTTAGCGGTGGATCACTCGGCTCGTGCGTCGATGAAG</td><td>359.9</td><td>2.7</td><td>1.0</td><td>2.8</td><td>5.4E-03</td><td>6.2E-03</td><td>MB231/MB468</td><td>NUC</td></tr>
<tr><td>>hg38_21_+_8433172_8446622_RNA45SN1_45S_50ntFlanks@8346.8373.28|AGGGAAAGTTGAAAAGAACTTTGAAGAG</td><td>>hg38_21_+_8433172_8446622_RNA45SN1_45S_50ntFlanks</td><td>8346</td><td>8373</td><td>28</td><td>AGGGAAAGTTGAAAAGAACTTTGAAGAG</td><td>1464.0</td><td>5.6</td><td>1.7</td><td>3.3</td><td>1.0E-03</td><td>6.2E-03</td><td>BT20/MB231</td><td>MP</td></tr>
<tr><td>>hg38_21_+_8433172_8446622_RNA45SN1_45S_50ntFlanks@12413.12446.34|TCCTTCGATGTCGGCTCTTCCTATCATTGTGAAG</td><td>>hg38_21_+_8433172_8446622_RNA45SN1_45S_50ntFlanks</td><td>12413</td><td>12446</td><td>34</td><td>TCCTTCGATGTCGGCTCTTCCTATCATTGTGAAG</td><td>437.2</td><td>-5.4</td><td>1.9</td><td>-2.8</td><td>5.3E-03</td><td>6.2E-03</td><td>BT20/MB468</td><td>NUC</td></tr>
<tr><td>>hg38_21_+_8433172_8446622_RNA45SN1_45S_50ntFlanks@10780.10801.22|AAGGTGAACAGCCTCTGGCATG</td><td>>hg38_21_+_8433172_8446622_RNA45SN1_45S_50ntFlanks</td><td>10780</td><td>10801</td><td>22</td><td>AAGGTGAACAGCCTCTGGCATG</td><td>184.0</td><td>-1.4</td><td>0.5</td><td>-2.8</td><td>5.4E-03</td><td>6.2E-03</td><td>MB231/MB468</td><td>NUC</td></tr>
<tr><td>>hg38_MT_+_1621_3279_RNR2_16S_50ntFlanks@1095.1119.25|AGACGAGAAGACCCTATGGAGCTTT</td><td>>hg38_MT_+_1621_3279_RNR2_16S_50ntFlanks</td><td>1095</td><td>1119</td><td>25</td><td>AGACGAGAAGACCCTATGGAGCTTT</td><td>226.9</td><td>2.9</td><td>1.0</td><td>2.8</td><td>4.6E-03</td><td>6.2E-03</td><td>MB231/MB468</td><td>MP</td></tr>
<tr><td>>hg38_21_+_8433172_8446622_RNA45SN1_45S_50ntFlanks@3815.3834.20|ATGGTTCCTTTGGTCGCTCG</td><td>>hg38_21_+_8433172_8446622_RNA45SN1_45S_50ntFlanks</td><td>3815</td><td>3834</td><td>20</td><td>ATGGTTCCTTTGGTCGCTCG</td><td>369.6</td><td>-4.1</td><td>1.5</td><td>-2.8</td><td>5.3E-03</td><td>6.2E-03</td><td>BT20/MB468</td><td>NUC</td></tr>
<tr><td>>hg38_21_+_8433172_8446622_RNA45SN1_45S_50ntFlanks@10786.10827.42|AACAGCCTCTGGCATGTTGGAACAATGTAGGTAAGGGAAGTC</td><td>>hg38_21_+_8433172_8446622_RNA45SN1_45S_50ntFlanks</td><td>10786</td><td>10827</td><td>42</td><td>AACAGCCTCTGGCATGTTGGAACAATGTAGGTAAGGGAAGTC</td><td>427.3</td><td>-2.4</td><td>0.8</td><td>-2.9</td><td>4.3E-03</td><td>6.2E-03</td><td>BT20/MB468</td><td>MITO</td></tr>
<tr><td>>hg38_MT_+_1621_3279_RNR2_16S_50ntFlanks@507.530.24|AGGAAAAAACCTTGTAGAGAGAGT</td><td>>hg38_MT_+_1621_3279_RNR2_16S_50ntFlanks</td><td>507</td><td>530</td><td>24</td><td>AGGAAAAAACCTTGTAGAGAGAGT</td><td>97.2</td><td>3.8</td><td>1.3</td><td>2.8</td><td>4.6E-03</td><td>6.2E-03</td><td>MB231/MB468</td><td>MP</td></tr>
<tr><td>>hg38_21_+_8433172_8446622_RNA45SN1_45S_50ntFlanks@4594.4614.21|TGTTGGTTTTCGGAACTGAGG</td><td>>hg38_21_+_8433172_8446622_RNA45SN1_45S_50ntFlanks</td><td>4594</td><td>4614</td><td>21</td><td>TGTTGGTTTTCGGAACTGAGG</td><td>53.8</td><td>-4.8</td><td>1.6</td><td>-3.0</td><td>2.9E-03</td><td>6.2E-03</td><td>BT20/MB468</td><td>MP</td></tr>
<tr><td>>hg38_21_+_8433172_8446622_RNA45SN1_45S_50ntFlanks@9598.9617.20|GGGGCGAAAGACTAATCGAA</td><td>>hg38_21_+_8433172_8446622_RNA45SN1_45S_50ntFlanks</td><td>9598</td><td>9617</td><td>20</td><td>GGGGCGAAAGACTAATCGAA</td><td>110.0</td><td>-4.8</td><td>1.6</td><td>-3.0</td><td>2.9E-03</td><td>6.2E-03</td><td>BT20/MB468</td><td>MP</td></tr>
<tr><td>>trnaMT_LysTTT_MT_+_8295_8364@35.72.38,trnalookalike7_LysTTT_1_+_568844_568913@35.72.38|TTAAAGATTAAGAGAACCAACACCTCTTTACAGTGACC</td><td>>trnaMT_LysTTT_MT_+_8295_8364</td><td>35</td><td>72</td><td>38</td><td>TTAAAGATTAAGAGAACCAACACCTCTTTACAGTGACC</td><td>69.9</td><td>5.3</td><td>1.9</td><td>2.8</td><td>4.6E-03</td><td>6.2E-03</td><td>MB231/MB468</td><td>MP</td></tr>
<tr><td>>hg38_21_+_8433172_8446622_RNA45SN1_45S_50ntFlanks@10677.10709.33|GCGGTTCCGGCGGCGTCCGGTGAGCTCTCGCTG</td><td>>hg38_21_+_8433172_8446622_RNA45SN1_45S_50ntFlanks</td><td>10677</td><td>10709</td><td>33</td><td>GCGGTTCCGGCGGCGTCCGGTGAGCTCTCGCTG</td><td>2613.1</td><td>3.3</td><td>1.2</td><td>2.9</td><td>4.3E-03</td><td>6.2E-03</td><td>BT20/MB468</td><td>MITO</td></tr>
<tr><td>>hg38_21_+_8433172_8446622_RNA45SN1_45S_50ntFlanks@12407.12433.27|TTTTGATCCTTCGATGTCGGCTCTTCC</td><td>>hg38_21_+_8433172_8446622_RNA45SN1_45S_50ntFlanks</td><td>12407</td><td>12433</td><td>27</td><td>TTTTGATCCTTCGATGTCGGCTCTTCC</td><td>557.9</td><td>1.5</td><td>0.5</td><td>2.9</td><td>4.3E-03</td><td>6.2E-03</td><td>BT20/MB468</td><td>MITO</td></tr>
<tr><td>>hg38_21_+_8433172_8446622_RNA45SN1_45S_50ntFlanks@8353.8375.23|GTTGAAAAGAACTTTGAAGAGAG</td><td>>hg38_21_+_8433172_8446622_RNA45SN1_45S_50ntFlanks</td><td>8353</td><td>8375</td><td>23</td><td>GTTGAAAAGAACTTTGAAGAGAG</td><td>1005.7</td><td>7.9</td><td>2.5</td><td>3.2</td><td>1.4E-03</td><td>6.2E-03</td><td>BT20/MB231</td><td>MITO</td></tr>
<tr><td>>hg38_21_+_8433172_8446622_RNA45SN1_45S_50ntFlanks@475.514.40|CTTCGTGATCGATGTGGTGACGTCGTGCTCTCCCGGGCCG</td><td>>hg38_21_+_8433172_8446622_RNA45SN1_45S_50ntFlanks</td><td>475</td><td>514</td><td>40</td><td>CTTCGTGATCGATGTGGTGACGTCGTGCTCTCCCGGGCCG</td><td>511.7</td><td>4.6</td><td>1.6</td><td>2.8</td><td>5.4E-03</td><td>6.2E-03</td><td>MB231/MB468</td><td>MITO</td></tr>
<tr><td>>hg38_21_+_8433172_8446622_RNA45SN1_45S_50ntFlanks@3766.3798.33|GTACGCACGGCCGGTACAGTGAAACTGCGAATG</td><td>>hg38_21_+_8433172_8446622_RNA45SN1_45S_50ntFlanks</td><td>3766</td><td>3798</td><td>33</td><td>GTACGCACGGCCGGTACAGTGAAACTGCGAATG</td><td>451.4</td><td>-2.9</td><td>0.9</td><td>-3.2</td><td>1.3E-03</td><td>6.2E-03</td><td>BT20/MB231</td><td>CYTO</td></tr>
<tr><td>>MI0000088|hsa-mir-30a&WithFlank&6|-|71403545|71403627@54.71.18&[MIMAT0000088&hsa-miR-30a-3p&offsets|+1|-3;m-38&6|-|71403554|71403575&offsets|+1|-3]|TTTCAGTCGGATGTTTGC</td><td>>MI0000088|hsa-mir-30a&WithFlank&6|-|71403545|71403627</td><td>54</td><td>71</td><td>18</td><td>TTTCAGTCGGATGTTTGC</td><td>217.3</td><td>4.1</td><td>1.4</td><td>3.0</td><td>2.9E-03</td><td>6.2E-03</td><td>BT20/MB468</td><td>MP</td></tr>
<tr><td>>hg38_21_+_8433172_8446622_RNA45SN1_45S_50ntFlanks@8357.8386.30|AAAAGAACTTTGAAGAGAGAGTTCAAGAGG</td><td>>hg38_21_+_8433172_8446622_RNA45SN1_45S_50ntFlanks</td><td>8357</td><td>8386</td><td>30</td><td>AAAAGAACTTTGAAGAGAGAGTTCAAGAGG</td><td>413.9</td><td>-2.7</td><td>0.9</td><td>-3.0</td><td>2.9E-03</td><td>6.2E-03</td><td>BT20/MB468</td><td>MP</td></tr>
<tr><td>>hg38_21_+_8433172_8446622_RNA45SN1_45S_50ntFlanks@10866.10904.39|CTAAGGGCTGGGTCGGTCGGGCTGGGGCGCGAAGCGGGG</td><td>>hg38_21_+_8433172_8446622_RNA45SN1_45S_50ntFlanks</td><td>10866</td><td>10904</td><td>39</td><td>CTAAGGGCTGGGTCGGTCGGGCTGGGGCGCGAAGCGGGG</td><td>207.8</td><td>-4.3</td><td>1.5</td><td>-2.9</td><td>4.3E-03</td><td>6.2E-03</td><td>BT20/MB468</td><td>MITO</td></tr>
<tr><td>>MI0000273|hsa-mir-183&WithFlank&7|-|129774899|129775020@33.56.24&[MIMAT0000261&hsa-miR-183-5p&offsets|0|+2;m-41&7|-|129774966|129774987&offsets|-1|+1]|TATGGCACTGGTAGAATTCACTGT</td><td>>MI0000273|hsa-mir-183&WithFlank&7|-|129774899|129775020</td><td>33</td><td>56</td><td>24</td><td>TATGGCACTGGTAGAATTCACTGT</td><td>197.9</td><td>1.4</td><td>0.5</td><td>2.8</td><td>5.3E-03</td><td>6.2E-03</td><td>BT20/MB468</td><td>NUC</td></tr>
<tr><td>>hg38_21_+_8433172_8446622_RNA45SN1_45S_50ntFlanks@11669.11706.38|GCCCAGTGCTCTGAATGTCAAAGTGAAGAAATTCAATG</td><td>>hg38_21_+_8433172_8446622_RNA45SN1_45S_50ntFlanks</td><td>11669</td><td>11706</td><td>38</td><td>GCCCAGTGCTCTGAATGTCAAAGTGAAGAAATTCAATG</td><td>3424.0</td><td>-2.7</td><td>0.8</td><td>-3.2</td><td>1.3E-03</td><td>6.2E-03</td><td>BT20/MB231</td><td>CYTO</td></tr>
<tr><td>>hg38_21_+_8433172_8446622_RNA45SN1_45S_50ntFlanks@3854.3887.34|ACTGTGGTAATTCTAGAGCTAATACATGCCGACG</td><td>>hg38_21_+_8433172_8446622_RNA45SN1_45S_50ntFlanks</td><td>3854</td><td>3887</td><td>34</td><td>ACTGTGGTAATTCTAGAGCTAATACATGCCGACG</td><td>216.9</td><td>-1.8</td><td>0.7</td><td>-2.8</td><td>5.4E-03</td><td>6.2E-03</td><td>MB231/MB468</td><td>NUC</td></tr>
</table>

| | | | | | | | | | | | | | |
|---|---|---|---|---|---|---|---|---|---|---|---|---|---|
| >hg38_21_+_8433172_8446622_RNA45SN1_45S_50ntFlanks@7975.7998.24\|CGCGACCTCAGATCAGACGTGGCG | >hg38_21_+_8433172_8446622_RNA45SN1_45S_50ntFlanks | 7975 | 7998 | 24 | CGCGACCTCAGATCAGACGTGGCG | 316.3 | -3.7 | 1.3 | -2.8 | 4.6E-03 | 6.2E-03 | MB231/MB468 | MP |
| >hg38_21_+_8433172_8446622_RNA45SN1_45S_50ntFlanks@11668.11701.34\|TGCCCAGTGCTCTGAATGTCAAAGTGAAGAAATT | >hg38_21_+_8433172_8446622_RNA45SN1_45S_50ntFlanks | 11668 | 11701 | 34 | TGCCCAGTGCTCTGAATGTCAAAGTGAAGAAATT | 277.8 | 1.3 | 0.5 | 2.8 | 5.0E-03 | 6.2E-03 | BT20/MB468 | CYTO |
| >hg38_21_+_8433172_8446622_RNA45SN1_45S_50ntFlanks@10475.10512.38\|CGGCCGATCGAAAGGGAGTCGGGTTCAGATCCCCGAAT | >hg38_21_+_8433172_8446622_RNA45SN1_45S_50ntFlanks | 10475 | 10512 | 38 | CGGCCGATCGAAAGGGAGTCGGGTTCAGATCCCCGAAT | 114.7 | -2.8 | 1.0 | -2.8 | 5.0E-03 | 6.2E-03 | BT20/MB468 | CYTO |
| >hg38_21_+_8433172_8446622_RNA45SN1_45S_50ntFlanks@10794.10820.27\|CTGGCATGTTGGAACAATGTAGGTAAG | >hg38_21_+_8433172_8446622_RNA45SN1_45S_50ntFlanks | 10794 | 10820 | 27 | CTGGCATGTTGGAACAATGTAGGTAAG | 171.2 | -3.7 | 1.2 | -3.0 | 2.9E-03 | 6.2E-03 | BT20/MB468 | MP |
| >hg38_21_+_8433172_8446622_RNA45SN1_45S_50ntFlanks@12626.12667.42\|AATGGGGCGAAGCTACCATCTGTGGGATTATGACTGAACGCC | >hg38_21_+_8433172_8446622_RNA45SN1_45S_50ntFlanks | 12626 | 12667 | 42 | AATGGGGCGAAGCTACCATCTGTGGGATTATGACTGAACGCC | 456.2 | 2.8 | 1.0 | 2.8 | 5.4E-03 | 6.2E-03 | MB231/MB468 | MITO |
| >hg38_21_+_8433172_8446622_RNA45SN1_45S_50ntFlanks@12419.12449.31\|GATGTCGGCTCTTCCTATCATTGTGAAGCAG | >hg38_21_+_8433172_8446622_RNA45SN1_45S_50ntFlanks | 12419 | 12449 | 31 | GATGTCGGCTCTTCCTATCATTGTGAAGCAG | 553.5 | 2.1 | 0.8 | 2.8 | 5.4E-03 | 6.2E-03 | MB231/MB468 | NUC |
| >hg38_MT_+_1621_3279_RNR2_16S_50ntFlanks@617.652.36\|AAACATATAACTGAACTCCTCACACCCAATTGGACC | >hg38_MT_+_1621_3279_RNR2_16S_50ntFlanks | 617 | 652 | 36 | AAACATATAACTGAACTCCTCACACCCAATTGGACC | 111.4 | 3.4 | 1.2 | 2.8 | 4.6E-03 | 6.2E-03 | MB231/MB468 | MP |
| >hg38_21_+_8433172_8446622_RNA45SN1_45S_50ntFlanks@12617.12654.38\|TGAGGAGCCAATGGGGCGAAGCTACCATCTGTGGGATT | >hg38_21_+_8433172_8446622_RNA45SN1_45S_50ntFlanks | 12617 | 12654 | 38 | TGAGGAGCCAATGGGGCGAAGCTACCATCTGTGGGATT | 960.1 | -3.4 | 1.2 | -2.8 | 5.1E-03 | 6.2E-03 | MB231/MB468 | CYTO |
| >hg38_21_+_8433172_8446622_RNA45SN1_45S_50ntFlanks@4046.4065.20\|CATTCGAACGTCTGCCCTAT | >hg38_21_+_8433172_8446622_RNA45SN1_45S_50ntFlanks | 4046 | 4065 | 20 | CATTCGAACGTCTGCCCTAT | 247.1 | -1.5 | 0.5 | -2.8 | 5.0E-03 | 6.2E-03 | BT20/MB468 | CYTO |
| >hg38_21_+_8433172_8446622_RNA45SN1_45S_50ntFlanks@12395.12420.26\|AGCGACGTCGCTTTTTGATCCTTCGA | >hg38_21_+_8433172_8446622_RNA45SN1_45S_50ntFlanks | 12395 | 12420 | 26 | AGCGACGTCGCTTTTTGATCCTTCGA | 222.8 | -2.2 | 0.7 | -3.3 | 1.1E-03 | 6.2E-03 | BT20/MB231 | NUC |
| >am_trna146_GlnCTG_6_-_27515531_27515602@1.28.28,trna1_GlnCTG_6_+_18836402_18836473@1.28.28,trna3_GlnCTG_17_+_8023070_8023141@1.28.28,trna42_GlnCTG_6_+_27263212_27263283@1.28.28,trna49_GlnCTG_6_+_27487308_27487379@1.28.28,trna7_GlnCTG_15_-_66161400_66161471@1.28.28,trna99_GlnCTG_6_-_28909378_28909449@1.28.28\|GGTTCCATGGTGTAATGGTTAGCACTCT | >am_trna146_GlnCTG_6_-_27515531_27515602 | 1 | 28 | 28 | GGTTCCATGGTGTAATGGTTAGCACTCT | 295.0 | -3.5 | 1.3 | -2.8 | 5.1E-03 | 6.2E-03 | MB231/MB468 | CYTO |
| >hg38_21_+_8433172_8446622_RNA45SN1_45S_50ntFlanks@12457.12493.37\|CAAGCGTTGGATTGTTCACCCACTAATAGGGAACGTG | >hg38_21_+_8433172_8446622_RNA45SN1_45S_50ntFlanks | 12457 | 12493 | 37 | CAAGCGTTGGATTGTTCACCCACTAATAGGGAACGTG | 76.3 | -3.9 | 1.4 | -2.8 | 4.6E-03 | 6.2E-03 | MB231/MB468 | MP |
| >hg38_21_+_8433172_8446622_RNA45SN1_45S_50ntFlanks@12417.12449.33\|TCGATGTCGGCTCTTCCTATCATTGTGAAGCAG | >hg38_21_+_8433172_8446622_RNA45SN1_45S_50ntFlanks | 12417 | 12449 | 33 | TCGATGTCGGCTCTTCCTATCATTGTGAAGCAG | 170.3 | -3.4 | 1.1 | -3.2 | 1.4E-03 | 6.3E-03 | BT20/MB231 | MITO |
| >trna143_LysTTT_6_-_27559593_27559665@14.31.18,trna149_LysTTT_6_-_27302769_27302841@14.31.18,trna14_LysTTT_11_-_59327808_59327880@14.31.18,trna2_LysTTT_17_+_8022473_8022545@14.31.18,trna54_LysTTT_1_+_204475655_204475727@14.31.18,trna5_LysTTT_11_+_59323902_59323974@14.31.18,trna62_LysTTT_1_-_204476158_204476230@14.31.18,trna76_LysTTT_6_+_28918806_28918878@14.31.18\|AGTCGGTAGAGCATCAGA | >trna143_LysTTT_6_-_27559593_27559665 | 14 | 31 | 18 | AGTCGGTAGAGCATCAGA | 389.5 | -3.4 | 1.0 | -3.3 | 1.0E-03 | 6.3E-03 | BT20/MB231 | MP |
| >MI0000090\|hsa-mir-32&WithFlank&9\|-\|109046223\|109046304@12.31.20(+1U)&[MIMAT0000090&hsa-miR-32-5p&offsets\|0\|-2(+1U);m-208&9\|-\|109046273\|109046293&offsets\|0\|-1(+1U)]\|TATTGCACATTACTAAGTTGT | >MI0000090\|hsa-mir-32&WithFlank&9\|-\|109046223\|109046304 | 12 | 31 | 20(+1U) | TATTGCACATTACTAAGTTGT | 77.8 | 4.6 | 1.6 | 2.8 | 4.6E-03 | 6.3E-03 | MB231/MB468 | MP |
| >hg38_MT_+_1621_3279_RNR2_16S_50ntFlanks@1370.1396.27\|ATGTTGGATCAGGACATCCCGATGGTG | >hg38_MT_+_1621_3279_RNR2_16S_50ntFlanks | 1370 | 1396 | 27 | ATGTTGGATCAGGACATCCCGATGGTG | 992.7 | 2.4 | 0.8 | 2.8 | 4.6E-03 | 6.3E-03 | MB231/MB468 | MP |
| >hg38_21_+_8433172_8446622_RNA45SN1_45S_50ntFlanks@8121.8141.21\|ACATGTGGCGTACGGAAGACC | >hg38_21_+_8433172_8446622_RNA45SN1_45S_50ntFlanks | 8121 | 8141 | 21 | ACATGTGGCGTACGGAAGACC | 174.9 | 4.8 | 1.5 | 3.3 | 1.0E-03 | 6.3E-03 | BT20/MB231 | MP |
| >hg38_21_+_8433172_8446622_RNA45SN1_45S_50ntFlanks@3771.3798.28\|CACGGCCGGTACAGTGAAACTGCGAATG | >hg38_21_+_8433172_8446622_RNA45SN1_45S_50ntFlanks | 3771 | 3798 | 28 | CACGGCCGGTACAGTGAAACTGCGAATG | 227.4 | -2.1 | 0.7 | -2.8 | 5.4E-03 | 6.3E-03 | MB231/MB468 | NUC |
| >MI0000273\|hsa-mir-183&WithFlank&7\|-\|129774899\|129775020@33.53.21(+1C)&[MIMAT0000261&hsa-miR-183-5p&offsets\|0\|-1(+1C);m-41&7\|-\|129774966\|129774987&offsets\|-1\|-2(+1C)]\|TATGGCACTGGTAGAATTCACC | >MI0000273\|hsa-mir-183&WithFlank&7\|-\|129774899\|129775020 | 33 | 53 | 21(+1C) | TATGGCACTGGTAGAATTCACC | 470.4 | 2.5 | 0.8 | 3.0 | 2.9E-03 | 6.3E-03 | BT20/MB468 | MP |
| >MI0000085\|hsa-mir-27a&WithFlank&19\|-\|13836434\|13836523@16.37.22&[MIMAT0004501&hsa-miR-27a-5p&offsets\|0\|0;m-270&19\|-\|13836487\|13836508&offsets\|0\|0]\|AGGGCTTAGCTGCTTGTGAGCA | >MI0000085\|hsa-mir-27a&WithFlank&19\|-\|13836434\|13836523 | 16 | 37 | 22 | AGGGCTTAGCTGCTTGTGAGCA | 104.0 | 4.5 | 1.6 | 2.8 | 4.7E-03 | 6.3E-03 | MB231/MB468 | MP |
| >hg38_21_+_8433172_8446622_RNA45SN1_45S_50ntFlanks@3854.3874.21\|ACTGTGGTAATTCTAGAGCTA | >hg38_21_+_8433172_8446622_RNA45SN1_45S_50ntFlanks | 3854 | 3874 | 21 | ACTGTGGTAATTCTAGAGCTA | 344.5 | -5.2 | 1.9 | -2.8 | 5.4E-03 | 6.3E-03 | BT20/MB468 | NUC |
| >trna10_AspGTC_12_-_125424193_125424264@35.72.38,trna12_AspGTC_12_-_125411891_125411962@35.72.38,trna144_AspGTC_6_-_27551236_27551307@35.72.38,trna38_AspGTC_17_-_8125556_8125627@35.72.38,trna45_AspGTC_6_+_27447453_27447524@35.72.38,trna48_AspGTC_6_+_27471523_27471594@35.72.38,trna4_AspGTC_12_+_96429799_96429870@35.72.38,trna69_AspGTC_1_-_161440205_161440276@35.72.38,trna72_AspGTC_1_-_161432824_161432895@35.72.38,trna75_AspGTC_1_-_161425414_161425485@35.72.38,trna78_AspGTC_1_-_161418033_161418104@35.72.38,trna81_AspGTC_1_-_161410615_161410686@35.72.38\|TCACGCGGGAGACCGGGGTTCGATTCCCCGACGGGGAG | >trna10_AspGTC_12_-_125424193_125424264 | 35 | 72 | 38 | TCACGCGGGAGACCGGGGTTCGATTCCCCGACGGGGAG | 665.3 | -5.5 | 1.8 | -3.0 | 2.9E-03 | 6.3E-03 | BT20/MB468 | MP |
| >MI0000270\|hsa-mir-181b-1&WithFlank&1\|-\|198858867\|198858988@42.65.24(+1C);MI0000683\|hsa-mir-181b-2&WithFlank&9\|+\|124693704\|124693804@22.45.24(+1C)&[MIMAT0000257&hsa-miR-181b-5p&offsets\|0\|+1(+1C);m-73&1\|-\|198858925\|198858947&offsets\|0\|+1(+1C)];[MIMAT0000257_1&hsa-miR-181b-5p&offsets\|0\|+1(+1C);m-72&9\|+\|124693725\|124693747&offsets\|0\|+1(+1C)]\|AACATTCATTGCTGTCGGTGGGTTC | >MI0000270\|hsa-mir-181b-1&WithFlank&1\|-\|198858867\|198858988 | 42 | 65 | 24(+1C) | AACATTCATTGCTGTCGGTGGGTTC | 290.3 | -4.6 | 1.4 | -3.3 | 1.0E-03 | 6.3E-03 | BT20/MB231 | MP |
| >hg38_21_+_8433172_8446622_RNA45SN1_45S_50ntFlanks@9928.9948.21\|TGATATAGACAGCAGGACGGT | >hg38_21_+_8433172_8446622_RNA45SN1_45S_50ntFlanks | 9928 | 9948 | 21 | TGATATAGACAGCAGGACGGT | 344.5 | 6.2 | 2.1 | 3.0 | 2.9E-03 | 6.3E-03 | BT20/MB468 | MP |
| >hg38_21_+_8433172_8446622_RNA45SN1_45S_50ntFlanks@3771.3799.29\|CACGGCCGGTACAGTGAAACTGCGAATGG | >hg38_21_+_8433172_8446622_RNA45SN1_45S_50ntFlanks | 3771 | 3799 | 29 | CACGGCCGGTACAGTGAAACTGCGAATGG | 134.0 | -2.4 | 0.9 | -2.8 | 5.1E-03 | 6.3E-03 | MB231/MB468 | CYTO |
| >hg38_21_+_8433172_8446622_RNA45SN1_45S_50ntFlanks@11674.11705.32\|GTGCTCTGAATGTCAAAGTGAAGAAATTCAAT | >hg38_21_+_8433172_8446622_RNA45SN1_45S_50ntFlanks | 11674 | 11705 | 32 | GTGCTCTGAATGTCAAAGTGAAGAAATTCAAT | 29.2 | 8.6 | 2.9 | 3.0 | 2.9E-03 | 6.3E-03 | BT20/MB468 | MP |
| >MI0000342\|hsa-mir-200b&WithFlank&1\|+\|1167098\|1167204@63.85.23(+1A)&[MIMAT0000318&hsa-miR-200b-3p&offsets\|0\|+1(+1A);m-93&1\|+\|1167160\|1167182&offsets\|0\|0(+1A)]\|TAATACTGCCTGGTAATGATGACA | >MI0000342\|hsa-mir-200b&WithFlank&1\|+\|1167098\|1167204 | 63 | 85 | 23(+1A) | TAATACTGCCTGGTAATGATGACA | 708.1 | -3.9 | 1.4 | -2.8 | 5.4E-03 | 6.3E-03 | MB231/MB468 | MITO |
| >hg38_21_+_8433172_8446622_RNA45SN1_45S_50ntFlanks@3844.3874.31\|TACTTGGATAACTGTGGTAATTCTAGAGCTA | >hg38_21_+_8433172_8446622_RNA45SN1_45S_50ntFlanks | 3844 | 3874 | 31 | TACTTGGATAACTGTGGTAATTCTAGAGCTA | 91.4 | -4.8 | 1.7 | -2.8 | 4.7E-03 | 6.3E-03 | MB231/MB468 | MP |
| >hg38_21_+_8433172_8446622_RNA45SN1_45S_50ntFlanks@8176.8215.40\|CTTCTGATCGAGGCCCAGCCCGTGGACGGTGTGAGGCCGG | >hg38_21_+_8433172_8446622_RNA45SN1_45S_50ntFlanks | 8176 | 8215 | 40 | CTTCTGATCGAGGCCCAGCCCGTGGACGGTGTGAGGCCGG | 172.8 | -3.3 | 1.2 | -2.8 | 5.5E-03 | 6.3E-03 | MB231/MB468 | NUC |
| >am_MI0005763\|hsa-mir-941-1&WithFlank&20\|+\|63919443\|63919526@53.75.23;MI0005764\|hsa-mir-941-2&WithFlank&20\|+\|63919499\|63919582@53.75.23;MI0005765\|hsa-mir-941-3&WithFlank&20\|+\|63919555\|63919638@53.75.23;MI0005766\|hsa-mir-941-4&WithFlank&20\|+\|63919750\|63919833@53.75.23;MI0031520\|hsa-mir-941-5&WithFlank&20\|+\|63919862\|63919945@53.75.23&[MIMAT0004984&hsa-miR-941&offsets\|0\|0;m-263&20\|+\|63919495\|63919517&offsets\|0\|0];[MIMAT0004984_1&hsa-miR-941&offsets\|0\|0;m-263&20\|+\|63919551\|63919573&offsets\|0\|0];[MIMAT0004984_2&hsa-miR-941&offsets\|0\|0;m-263&20\|+\|63919607\|63919629&offsets\|0\|0];[MIMAT0004984_3&hsa-miR-941&offsets\|0\|0;m-263&20\|+\|63919802\|63919824&offsets\|0\|0];[MIMAT0004984_4&hsa-miR-941&offsets\|0\|0;m-263&20\|+\|63919914\|63919936&offsets\|0\|0]\|CACCCGGCTGTGTGCACATGTGC | >am_MI0005763\|hsa-mir-941-1&WithFlank&20\|+\|63919443\|63919526 | 53 | 75 | 23 | CACCCGGCTGTGTGCACATGTGC | 2649.4 | 1.3 | 0.4 | 2.8 | 5.4E-03 | 6.3E-03 | BT20/MB468 | NUC |
| >hg38_21_+_8433172_8446622_RNA45SN1_45S_50ntFlanks@12941.12959.19\|ATTCGTAGACGACCTGCTT | >hg38_21_+_8433172_8446622_RNA45SN1_45S_50ntFlanks | 12941 | 12959 | 19 | ATTCGTAGACGACCTGCTT | 584.9 | 1.7 | 0.6 | 2.8 | 5.2E-03 | 6.3E-03 | MB231/MB468 | CYTO |
| >hg38_21_+_8433172_8446622_RNA45SN1_45S_50ntFlanks@11209.11255.47\|CGGCGACTCTGGACGCGAGCCGGGCCCTTCCCGTGGATCGCCCCAGC | >hg38_21_+_8433172_8446622_RNA45SN1_45S_50ntFlanks | 11209 | 11255 | 47 | CGGCGACTCTGGACGCGAGCCGGGCCCTTCCCGTGGATCGCCCCAGC | 416.7 | 3.4 | 1.2 | 2.9 | 4.3E-03 | 6.3E-03 | BT20/MB468 | MITO |
| >hg38_21_+_8433172_8446622_RNA45SN1_45S_50ntFlanks@7975.7994.20\|CGCGACCTCAGATCAGACGT | >hg38_21_+_8433172_8446622_RNA45SN1_45S_50ntFlanks | 7975 | 7994 | 20 | CGCGACCTCAGATCAGACGT | 56456.4 | 2.3 | 0.8 | 2.8 | 5.1E-03 | 6.3E-03 | BT20/MB468 | CYTO |
| >trnaMT_SerGCT_MT_+_12207_12265@14.62.49\|AGAACTGCTAACTCATGCCCCCATGTCTAACAACATGGCTTTCTCACCA | >trnaMT_SerGCT_MT_+_12207_12265 | 14 | 62 | 49 | AGAACTGCTAACTCATGCCCCCATGTCTAACAACATGGCTTTCTCACCA | 684.7 | 4.9 | 1.8 | 2.8 | 5.5E-03 | 6.3E-03 | MB231/MB468 | MITO |
| >hg38_MT_+_1621_3279_RNR2_16S_50ntFlanks@507.531.25\|AGGAAAAAAACCTTGTAGAGAGAGTA | >hg38_MT_+_1621_3279_RNR2_16S_50ntFlanks | 507 | 531 | 25 | AGGAAAAAAACCTTGTAGAGAGAGTA | 166.1 | 4.7 | 1.6 | 3.0 | 2.9E-03 | 6.3E-03 | BT20/MB468 | MP |
| >hg38_21_+_8433172_8446622_RNA45SN1_45S_50ntFlanks@9737.9765.29\|CGAAACGATCTCAACCTATTCTCAAACTT | >hg38_21_+_8433172_8446622_RNA45SN1_45S_50ntFlanks | 9737 | 9765 | 29 | CGAAACGATCTCAACCTATTCTCAAACTT | 398.8 | -3.0 | 0.9 | -3.2 | 1.3E-03 | 6.3E-03 | BT20/MB231 | CYTO |
| >hg38_21_+_8433172_8446622_RNA45SN1_45S_50ntFlanks@6723.6743.21\|TTGCAGGACACATTGATCATC | >hg38_21_+_8433172_8446622_RNA45SN1_45S_50ntFlanks | 6723 | 6743 | 21 | TTGCAGGACACATTGATCATC | 393.7 | -2.6 | 0.9 | -2.8 | 5.1E-03 | 6.3E-03 | BT20/MB468 | CYTO |
| >hg38_21_+_8433172_8446622_RNA45SN1_45S_50ntFlanks@6650.6675.26\|ACGACTCTTAGCGGTGGATCACTCGG | >hg38_21_+_8433172_8446622_RNA45SN1_45S_50ntFlanks | 6650 | 6675 | 26 | ACGACTCTTAGCGGTGGATCACTCGG | 189.9 | -2.3 | 0.8 | -2.8 | 4.7E-03 | 6.3E-03 | MB231/MB468 | MP |
| >hg38_21_+_8433172_8446622_RNA45SN1_45S_50ntFlanks@8157.8179.23\|CTCGTGGGGGGCCCAAGTCCTTC | >hg38_21_+_8433172_8446622_RNA45SN1_45S_50ntFlanks | 8157 | 8179 | 23 | CTCGTGGGGGGCCCAAGTCCTTC | 58.2 | -5.0 | 1.7 | -3.0 | 2.9E-03 | 6.3E-03 | BT20/MB468 | MP |

| | | | | | | | | | | | | | |
|---|---|---|---|---|---|---|---|---|---|---|---|---|---|
| >MI0000542\|hsa-mir-320a&WithFlank&8\|-\|22244960\|22245043@48.69.22(+2U)&[MIMAT0000510&hsa-miR-320a-3p&offsets\|0\|0(+2U);m-60&8\|-\|22244975\|22244996&offsets\|0\|0(+2U)]\|AAAAGCTGGGTTGAGAGGGCGATT | >MI0000542\|hsa-mir-320a&WithFlank&8\|-\|22244960\|22245043 | 48 | 69 | 22(+2U) | AAAAGCTGGGTTGAGAGGGCGATT | 817.3 | 4.0 | 1.4 | 2.8 | 4.7E-03 | 6.3E-03 | MB231/MB468 | MP |
| >hg38_21_+_8433172_8446622_RNA45SN1_45S_50ntFlanks@8356.8375.20\|GAAAAGAACTTTGAAGAGAG | >hg38_21_+_8433172_8446622_RNA45SN1_45S_50ntFlanks | 8356 | 8375 | 20 | GAAAAGAACTTTGAAGAGAG | 249.0 | -1.9 | 0.7 | -2.8 | 5.2E-03 | 6.3E-03 | MB231/MB468 | CYTO |
| >hg38_21_+_8433172_8446622_RNA45SN1_45S_50ntFlanks@12912.12943.32\|CACGTTCGTGGGGAACCTGGCGCTAAACCATT | >hg38_21_+_8433172_8446622_RNA45SN1_45S_50ntFlanks | 12912 | 12943 | 32 | CACGTTCGTGGGGAACCTGGCGCTAAACCATT | 107.3 | -4.6 | 1.6 | -2.8 | 4.7E-03 | 6.3E-03 | MB231/MB468 | MP |
| >hg38_21_+_8433172_8446622_RNA45SN1_45S_50ntFlanks@12937.12954.18\|AACCATTCGTAGACGACC | >hg38_21_+_8433172_8446622_RNA45SN1_45S_50ntFlanks | 12937 | 12954 | 18 | AACCATTCGTAGACGACC | 208.3 | -1.8 | 0.7 | -2.8 | 5.5E-03 | 6.3E-03 | MB231/MB468 | NUC |
| >hg38_21_+_8433172_8446622_RNA45SN1_45S_50ntFlanks@11582.11606.25\|ACTGGTGCGGACCAGGGGAATCCGA | >hg38_21_+_8433172_8446622_RNA45SN1_45S_50ntFlanks | 11582 | 11606 | 25 | ACTGGTGCGGACCAGGGGAATCCGA | 369.2 | 3.3 | 1.1 | 3.0 | 2.9E-03 | 6.3E-03 | BT20/MB468 | MP |
| >hg38_21_+_8433172_8446622_RNA45SN1_45S_50ntFlanks@12008.12031.24\|CCGCCGGTGAAATACCACTACTCT | >hg38_21_+_8433172_8446622_RNA45SN1_45S_50ntFlanks | 12008 | 12031 | 24 | CCGCCGGTGAAATACCACTACTCT | 6909.0 | -5.9 | 2.0 | -3.0 | 2.9E-03 | 6.3E-03 | BT20/MB468 | MP |
| >hg38_21_+_8433172_8446622_RNA45SN1_45S_50ntFlanks@11663.11702.40\|ATTTCTGCCCAGTGCTCTGAATGTCAAAGTGAAGAAATTC | >hg38_21_+_8433172_8446622_RNA45SN1_45S_50ntFlanks | 11663 | 11702 | 40 | ATTTCTGCCCAGTGCTCTGAATGTCAAAGTGAAGAAATTC | 326.4 | 3.2 | 1.2 | 2.8 | 5.2E-03 | 6.3E-03 | MB231/MB468 | CYTO |
| >MI0000078\|hsa-mir-22&WithFlank&17\|-\|1713897\|1713993@59.79.21(+1A)&[MIMAT0000077&hsa-miR-22-3p&offsets\|0\|-1(+1A);m-2&17\|-\|1713914\|1713935&offsets\|0\|-1(+1A)]\|AAGCTGCCAGTTGAAGAACTGA | >MI0000078\|hsa-mir-22&WithFlank&17\|-\|1713897\|1713993 | 59 | 79 | 21(+1A) | AAGCTGCCAGTTGAAGAACTGA | 656.6 | 3.3 | 1.0 | 3.2 | 1.3E-03 | 6.3E-03 | BT20/MB231 | CYTO |
| >hg38_MT_+_1621_3279_RNR2_16S_50ntFlanks@194.214.21\|AAGCATAATATAGCAAGGACT | >hg38_MT_+_1621_3279_RNR2_16S_50ntFlanks | 194 | 214 | 21 | AAGCATAATATAGCAAGGACT | 236.7 | 3.5 | 1.2 | 3.0 | 2.9E-03 | 6.3E-03 | BT20/MB468 | MP |
| >hg38_21_+_8433172_8446622_RNA45SN1_45S_50ntFlanks@12940.12966.27\|CATTCGTAGACGACCTGCTTCTGGGTC | >hg38_21_+_8433172_8446622_RNA45SN1_45S_50ntFlanks | 12940 | 12966 | 27 | CATTCGTAGACGACCTGCTTCTGGGTC | 445.0 | -5.1 | 1.7 | -3.0 | 2.9E-03 | 6.3E-03 | BT20/MB468 | MP |
| >hg38_21_+_8433172_8446622_RNA45SN1_45S_50ntFlanks@7973.8002.30\|GACGCGACCTCAGATCAGACGTGGCGACCC | >hg38_21_+_8433172_8446622_RNA45SN1_45S_50ntFlanks | 7973 | 8002 | 30 | GACGCGACCTCAGATCAGACGTGGCGACCC | 163.5 | 4.5 | 1.5 | 3.0 | 2.9E-03 | 6.3E-03 | BT20/MB468 | MP |
| >hg38_21_+_8433172_8446622_RNA45SN1_45S_50ntFlanks@4139.4156.18\|AGGGAGCCTGAGAAACGG | >hg38_21_+_8433172_8446622_RNA45SN1_45S_50ntFlanks | 4139 | 4156 | 18 | AGGGAGCCTGAGAAACGG | 122.4 | -3.5 | 1.2 | -2.8 | 4.7E-03 | 6.3E-03 | MB231/MB468 | MP |
| >trna10_AspGTC_12_-_125424193_125424264@37.68.32,trna12_AspGTC_12_-_125411891_125411962@37.68.32,trna144_AspGTC_6_-_27551236_27551307@37.68.32,trna38_AspGTC_17_-_8125556_8125627@37.68.32,trna45_AspGTC_6_+_27447453_27447524@37.68.32,trna48_AspGTC_6_+_27471523_27471594@37.68.32,trna4_AspGTC_12_+_96429799_96429870@37.68.32,trna69_AspGTC_1_-_161440205_161440276@37.68.32,trna72_AspGTC_1_-_161432824_161432895@37.68.32,trna75_AspGTC_1_-_161425414_161425485@37.68.32,trna78_AspGTC_1_-_161418033_161418104@37.68.32,trna81_AspGTC_1_-_161410615_161410686@37.68.32\|ACGCGGGAGACCGGGGTTCGATTCCCCGACGG | >trna10_AspGTC_12_-_125424193_125424264 | 37 | 68 | 32 | ACGCGGGAGACCGGGGTTCGATTCCCCGACGG | 319.2 | -2.4 | 0.8 | -2.8 | 4.7E-03 | 6.3E-03 | MB231/MB468 | MP |
| >hg38_21_+_8433172_8446622_RNA45SN1_45S_50ntFlanks@4657.4684.28\|CTAGAGGTGAAATTCTTGGACCGGCGCA | >hg38_21_+_8433172_8446622_RNA45SN1_45S_50ntFlanks | 4657 | 4684 | 28 | CTAGAGGTGAAATTCTTGGACCGGCGCA | 235.0 | -2.2 | 0.8 | -2.8 | 5.1E-03 | 6.3E-03 | BT20/MB468 | CYTO |
| >hg38_21_+_8433172_8446622_RNA45SN1_45S_50ntFlanks@9533.9572.40\|GAGGAAACTCTGGTGGAGGTCCGTAGCGGTCCTGACGTGC | >hg38_21_+_8433172_8446622_RNA45SN1_45S_50ntFlanks | 9533 | 9572 | 40 | GAGGAAACTCTGGTGGAGGTCCGTAGCGGTCCTGACGTGC | 549.1 | 2.5 | 0.9 | 2.8 | 4.4E-03 | 6.3E-03 | BT20/MB468 | MITO |
| >trna10_AspGTC_12_-_125424193_125424264@35.72.38,trna12_AspGTC_12_-_125411891_125411962@35.72.38,trna144_AspGTC_6_-_27551236_27551307@35.72.38,trna38_AspGTC_17_-_8125556_8125627@35.72.38,trna45_AspGTC_6_+_27447453_27447524@35.72.38,trna48_AspGTC_6_+_27471523_27471594@35.72.38,trna4_AspGTC_12_+_96429799_96429870@35.72.38,trna69_AspGTC_1_-_161440205_161440276@35.72.38,trna72_AspGTC_1_-_161432824_161432895@35.72.38,trna75_AspGTC_1_-_161425414_161425485@35.72.38,trna78_AspGTC_1_-_161418033_161418104@35.72.38,trna81_AspGTC_1_-_161410615_161410686@35.72.38\|TCACGCGGGAGACCGGGGTTCGATTCCCCGACGGGGAG | >trna10_AspGTC_12_-_125424193_125424264 | 35 | 72 | 38 | TCACGCGGGAGACCGGGGTTCGATTCCCCGACGGGGAG | 540.3 | -2.7 | 1.0 | -2.8 | 5.5E-03 | 6.3E-03 | MB231/MB468 | MITO |
| >am_MI0039740\|hsa-mir-12136&WithFlank&1\|-\|632609\|632691@7.24.18&[MIMAT0049032&hsa-miR-12136&offsets\|0\|0]\|GAAAAAGTCATGGAGGCC | >am_MI0039740\|hsa-mir-12136&WithFlank&1\|-\|632609\|632691 | 7 | 24 | 18 | GAAAAAGTCATGGAGGCC | 1012.4 | 3.5 | 1.3 | 2.8 | 5.5E-03 | 6.3E-03 | MB231/MB468 | MITO |
| >trnaMT_SerTGA_MT_-_7446_7514@1.18.18,trnalookalike6_SerTGA_1_-_567997_568065@1.18.18\|GAAAAAGTCATGGAGGCC | >trnaMT_SerTGA_MT_-_7446_7514 | 1 | 18 | 18 | GAAAAAGTCATGGAGGCC | 1012.4 | 3.5 | 1.3 | 2.8 | 5.5E-03 | 6.3E-03 | MB231/MB468 | MITO |
| >hg38_21_+_8433172_8446622_RNA45SN1_45S_50ntFlanks@3782.3805.24\|CAGTGAAACTGCGAATGGCTCATT | >hg38_21_+_8433172_8446622_RNA45SN1_45S_50ntFlanks | 3782 | 3805 | 24 | CAGTGAAACTGCGAATGGCTCATT | 571.8 | -3.8 | 1.3 | -2.8 | 4.4E-03 | 6.3E-03 | BT20/MB468 | MITO |
| >hg38_21_+_8433172_8446622_RNA45SN1_45S_50ntFlanks@11568.11599.32\|GCAGCCGACTTAGAACTGGTGCGGACCAGGGG | >hg38_21_+_8433172_8446622_RNA45SN1_45S_50ntFlanks | 11568 | 11599 | 32 | GCAGCCGACTTAGAACTGGTGCGGACCAGGGG | 159.1 | -1.8 | 0.6 | -2.8 | 5.5E-03 | 6.3E-03 | MB231/MB468 | NUC |
| >hg38_21_+_8433172_8446622_RNA45SN1_45S_50ntFlanks@12404.12433.30\|GCTTTTTGATCCTTCGATGTCGGCTCTTCC | >hg38_21_+_8433172_8446622_RNA45SN1_45S_50ntFlanks | 12404 | 12433 | 30 | GCTTTTTGATCCTTCGATGTCGGCTCTTCC | 524.2 | 2.8 | 1.0 | 2.8 | 5.2E-03 | 6.3E-03 | MB231/MB468 | CYTO |
| >MI0001445\|hsa-mir-423&WithFlank&17\|+\|30117073\|30117178@59.82.24&[MIMAT0001340&hsa-miR-423-3p&offsets\|0\|+1;m-100&17\|+\|30117131\|30117153&offsets\|0\|+1]\|AGCTCGGTCTGAGGCCCCTCAGTC | >MI0001445\|hsa-mir-423&WithFlank&17\|+\|30117073\|30117178 | 59 | 82 | 24 | AGCTCGGTCTGAGGCCCCTCAGTC | 43.4 | 5.5 | 2.0 | 2.8 | 4.7E-03 | 6.3E-03 | MB231/MB468 | MP |
| >hg38_21_+_8433172_8446622_RNA45SN1_45S_50ntFlanks@10922.10941.20\|CTGGACGAGGCGCCGCCGCC | >hg38_21_+_8433172_8446622_RNA45SN1_45S_50ntFlanks | 10922 | 10941 | 20 | CTGGACGAGGCGCCGCCGCC | 1873.6 | -5.9 | 2.1 | -2.8 | 5.5E-03 | 6.4E-03 | BT20/MB468 | NUC |
| >hg38_21_+_8433172_8446622_RNA45SN1_45S_50ntFlanks@12394.12414.21\|TAGCGACGTCGCTTTTTGATC | >hg38_21_+_8433172_8446622_RNA45SN1_45S_50ntFlanks | 12394 | 12414 | 21 | TAGCGACGTCGCTTTTTGATC | 1409.6 | -1.8 | 0.6 | -2.8 | 5.1E-03 | 6.4E-03 | BT20/MB468 | CYTO |
| >hg38_21_+_8433172_8446622_RNA45SN1_45S_50ntFlanks@9786.9803.18\|CTCGCTGGCGTGGAGCCG | >hg38_21_+_8433172_8446622_RNA45SN1_45S_50ntFlanks | 9786 | 9803 | 18 | CTCGCTGGCGTGGAGCCG | 79.0 | -4.9 | 1.6 | -3.0 | 3.0E-03 | 6.4E-03 | BT20/MB468 | MP |
| >MI0000066\|hsa-let-7e&WithFlank&19\|+\|51692780\|51692870@14.36.23&[MIMAT0000066&hsa-let-7e-5p&offsets\|0\|+1;m-50&19\|+\|51692793\|51692814&offsets\|0\|+1]\|TGAGGTAGGAGGTTGTATAGTTG | >MI0000066\|hsa-let-7e&WithFlank&19\|+\|51692780\|51692870 | 14 | 36 | 23 | TGAGGTAGGAGGTTGTATAGTTG | 508.8 | 1.8 | 0.5 | 3.2 | 1.2E-03 | 6.4E-03 | BT20/MB231 | NUC |
| >hg38_21_+_8433172_8446622_RNA45SN1_45S_50ntFlanks@7966.8005.40\|CCTCCGAGACGCGACCTCAGATCAGACGTGGCGACCCGCT | >hg38_21_+_8433172_8446622_RNA45SN1_45S_50ntFlanks | 7966 | 8005 | 40 | CCTCCGAGACGCGACCTCAGATCAGACGTGGCGACCCGCT | 651.1 | 4.1 | 1.3 | 3.2 | 1.2E-03 | 6.4E-03 | BT20/MB231 | NUC |
| >hg38_21_+_8433172_8446622_RNA45SN1_45S_50ntFlanks@6647.6665.19\|CGTACGACTCTTAGCGGTG | >hg38_21_+_8433172_8446622_RNA45SN1_45S_50ntFlanks | 6647 | 6665 | 19 | CGTACGACTCTTAGCGGTG | 4981.3 | 2.4 | 0.9 | 2.8 | 5.5E-03 | 6.4E-03 | MB231/MB468 | MITO |
| >MI0000482\|hsa-mir-185&WithFlank&22\|+\|20033133\|20033226@56.76.21&[MIMAT0004611&hsa-miR-185-3p&offsets\|0\|-1;m-306&22\|+\|20033188\|20033208&offsets\|0\|0]\|AGGGGCTGGCTTTCCTCTGGT | >MI0000482\|hsa-mir-185&WithFlank&22\|+\|20033133\|20033226 | 56 | 76 | 21 | AGGGGCTGGCTTTCCTCTGGT | 139.8 | 4.3 | 1.4 | 3.0 | 3.0E-03 | 6.4E-03 | BT20/MB468 | MP |
| >hg38_21_+_8433172_8446622_RNA45SN1_45S_50ntFlanks@10350.10369.20\|ACGAGAACTTTGAAGGCCGA | >hg38_21_+_8433172_8446622_RNA45SN1_45S_50ntFlanks | 10350 | 10369 | 20 | ACGAGAACTTTGAAGGCCGA | 771.7 | 1.8 | 0.6 | 2.8 | 4.4E-03 | 6.4E-03 | BT20/MB468 | MITO |
| >hg38_21_+_8433172_8446622_RNA45SN1_45S_50ntFlanks@12007.12033.27\|GCCGCCGGTGAAATACCACTACTCTGA | >hg38_21_+_8433172_8446622_RNA45SN1_45S_50ntFlanks | 12007 | 12033 | 27 | GCCGCCGGTGAAATACCACTACTCTGA | 166.5 | -3.5 | 1.3 | -2.8 | 4.7E-03 | 6.4E-03 | MB231/MB468 | MP |
| >hg38_21_+_8433172_8446622_RNA45SN1_45S_50ntFlanks@6649.6666.18\|TACGACTCTTAGCGGTGG | >hg38_21_+_8433172_8446622_RNA45SN1_45S_50ntFlanks | 6649 | 6666 | 18 | TACGACTCTTAGCGGTGG | 730.9 | 2.8 | 1.0 | 2.8 | 4.4E-03 | 6.4E-03 | BT20/MB468 | MITO |
| >hg38_MT_+_1621_3279_RNR2_16S_50ntFlanks@962.983.22\|ACCGTGCAAAGGTAGCATAATC | >hg38_MT_+_1621_3279_RNR2_16S_50ntFlanks | 962 | 983 | 22 | ACCGTGCAAAGGTAGCATAATC | 144.1 | -5.9 | 1.8 | -3.3 | 1.1E-03 | 6.4E-03 | BT20/MB231 | MP |
| >am_trna128_GlyGCC_6_-_27870686_27870756@16.35.20,trna18_GlyGCC_16_+_70822597_70822667@16.35.20,trna19_GlyGCC_16_+_70823410_70823480@16.35.20,trna19_GlyGCC_2_-_157257659_157257729@16.35.20,trna24_GlyGCC_16_-_70812942_70813012@16.35.20,trna25_GlyGCC_16_-_70812114_70812184@16.35.20,trna2_GlyGCC_21_-_18827107_18827177@16.35.20,trna35_GlyGCC_1_+_161413094_161413164@16.35.20,trna37_GlyGCC_1_+_161420467_161420537@16.35.20,trna39_GlyGCC_1_+_161427898_161427968@16.35.20,trna41_GlyGCC_1_+_161435258_161435328@16.35.20,trna5_GlyGCC_17_+_8029064_8029134@16.35.20,trna68_GlyGCC_1_-_161493637_161493707@16.35.20\|TGGTAGAATTCTCGCCTGCC | >am_trna128_GlyGCC_6_-_27870686_27870756 | 16 | 35 | 20 | TGGTAGAATTCTCGCCTGCC | 162.4 | 4.8 | 1.6 | 3.0 | 3.0E-03 | 6.4E-03 | BT20/MB468 | MP |
| >MI0000438\|hsa-mir-15b&WithFlank&3\|+\|160404582\|160404691@64.85.22&[MIMAT0004586&hsa-miR-15b-3p&offsets\|0\|0;m-204&3\|+\|160404645\|160404665&offsets\|0\|+1]\|CGAATCATTATTTGCTGCTCTA | >MI0000438\|hsa-mir-15b&WithFlank&3\|+\|160404582\|160404691 | 64 | 85 | 22 | CGAATCATTATTTGCTGCTCTA | 183.9 | 4.5 | 1.5 | 3.0 | 3.0E-03 | 6.4E-03 | BT20/MB468 | MP |
| >hg38_21_+_8433172_8446622_RNA45SN1_45S_50ntFlanks@10840.10868.29\|CCGTAACTTCGGGATAAGGATTGGCTCTA | >hg38_21_+_8433172_8446622_RNA45SN1_45S_50ntFlanks | 10840 | 10868 | 29 | CCGTAACTTCGGGATAAGGATTGGCTCTA | 432.4 | -5.1 | 1.7 | -3.0 | 3.0E-03 | 6.4E-03 | BT20/MB468 | MP |
| >trnaMT_LysTTT_MT_+_8295_8364@1.36.36,trnalookalike7_LysTTT_1_+_568844_568913@1.36.36\|CACTGTAAAGCTAACTTAGCATTAACCTTTTAAGTT | >trnaMT_LysTTT_MT_+_8295_8364 | 1 | 36 | 36 | CACTGTAAAGCTAACTTAGCATTAACCTTTTAAGTT | 1372.0 | 3.5 | 1.2 | 2.8 | 4.4E-03 | 6.4E-03 | BT20/MB468 | MITO |
| >hg38_21_+_8433172_8446622_RNA45SN1_45S_50ntFlanks@12007.12033.27\|GCCGCCGGTGAAATACCACTACTCTGA | >hg38_21_+_8433172_8446622_RNA45SN1_45S_50ntFlanks | 12007 | 12033 | 27 | GCCGCCGGTGAAATACCACTACTCTGA | 1083.3 | 2.4 | 0.8 | 2.8 | 4.4E-03 | 6.4E-03 | BT20/MB468 | MITO |
| >hg38_21_+_8433172_8446622_RNA45SN1_45S_50ntFlanks@3843.3860.18\|CTACTTGGATAACTGTGG | >hg38_21_+_8433172_8446622_RNA45SN1_45S_50ntFlanks | 3843 | 3860 | 18 | CTACTTGGATAACTGTGG | 208.4 | -4.6 | 1.6 | -2.8 | 4.4E-03 | 6.4E-03 | BT20/MB468 | MITO |
| >MI0017385\|hsa-mir-4746&WithFlank&19\|+\|4445972\|4446054@16.38.23&[MIMAT0019880&hsa-miR-4746-5p&offsets\|0\|0;m-432&19\|+\|4445987\|4446009&offsets\|0\|0]\|CCGGTCCCAGGAGAACCTGCAGA | >MI0017385\|hsa-mir-4746&WithFlank&19\|+\|4445972\|4446054 | 16 | 38 | 23 | CCGGTCCCAGGAGAACCTGCAGA | 36.8 | 5.3 | 1.9 | 2.8 | 4.8E-03 | 6.4E-03 | MB231/MB468 | MP |
| >MI0000088\|hsa-mir-30a&WithFlank&6\|-\|71403545\|71403627@54.74.21&[MIMAT0000088&hsa-miR-30a-3p&offsets\|+1\|0;m-38&6\|-\|71403554\|71403575&offsets\|+1\|0]\|TTTCAGTCGGATGTTTGCAGC | >MI0000088\|hsa-mir-30a&WithFlank&6\|-\|71403545\|71403627 | 54 | 74 | 21 | TTTCAGTCGGATGTTTGCAGC | 3942.7 | -2.3 | 0.7 | -3.2 | 1.4E-03 | 6.4E-03 | BT20/MB231 | MITO |

<table>
<tr><td></td><td></td><td></td><td></td><td></td><td></td><td></td><td></td><td></td><td></td><td></td><td></td><td></td><td></td></tr>
<tr><td>>hg38_21_+_8433172_8446622_RNA45SN1_45S_50ntFlanks@3813.3836.24|TTATGGTTCCTTTGGTCGCTCGCT</td><td>>hg38_21_+_8433172_8446622_RNA45SN1_45S_50ntFlanks</td><td>3813</td><td>3836</td><td>24</td><td>TTATGGTTCCTTTGGTCGCTCGCT</td><td>521.4</td><td>-2.0</td><td>0.7</td><td>-2.8</td><td>5.6E-03</td><td>6.4E-03</td><td>MB231/MB468</td><td>NUC</td></tr>
<tr><td>>hg38_21_+_8433172_8446622_RNA45SN1_45S_50ntFlanks@5485.5506.22|AGCGCTGAGAAGACGGTCGAAC</td><td>>hg38_21_+_8433172_8446622_RNA45SN1_45S_50ntFlanks</td><td>5485</td><td>5506</td><td>22</td><td>AGCGCTGAGAAGACGGTCGAAC</td><td>435.8</td><td>6.3</td><td>1.9</td><td>3.3</td><td>1.1E-03</td><td>6.4E-03</td><td>BT20/MB231</td><td>MP</td></tr>
<tr><td>>hg38_21_+_8433172_8446622_RNA45SN1_45S_50ntFlanks@10291.10319.29|CGCGGGCCCGGGTGGAGCCGCCGCAGGTG</td><td>>hg38_21_+_8433172_8446622_RNA45SN1_45S_50ntFlanks</td><td>10291</td><td>10319</td><td>29</td><td>CGCGGGCCCGGGTGGAGCCGCCGCAGGTG</td><td>1054.7</td><td>-4.3</td><td>1.4</td><td>-3.0</td><td>3.0E-03</td><td>6.4E-03</td><td>BT20/MB468</td><td>MP</td></tr>
<tr><td>>hg38_MT_+_598_1651_RNR1_12S_50ntFlanks@52.87.36|ATAGGTTTGGTCCTAGCCTTTCTATTAGCTCTTAGT</td><td>>hg38_MT_+_598_1651_RNR1_12S_50ntFlanks</td><td>52</td><td>87</td><td>36</td><td>ATAGGTTTGGTCCTAGCCTTTCTATTAGCTCTTAGT</td><td>18166.7</td><td>4.0</td><td>1.4</td><td>3.0</td><td>3.0E-03</td><td>6.4E-03</td><td>BT20/MB468</td><td>MP</td></tr>
<tr><td>>hg38_21_+_8433172_8446622_RNA45SN1_45S_50ntFlanks@6652.6678.27|GACTCTTAGCGGTGGATCACTCGGCTC</td><td>>hg38_21_+_8433172_8446622_RNA45SN1_45S_50ntFlanks</td><td>6652</td><td>6678</td><td>27</td><td>GACTCTTAGCGGTGGATCACTCGGCTC</td><td>2397.7</td><td>2.3</td><td>0.8</td><td>2.8</td><td>5.6E-03</td><td>6.4E-03</td><td>MB231/MB468</td><td>NUC</td></tr>
<tr><td>>hg38_21_+_8433172_8446622_RNA45SN1_45S_50ntFlanks@4730.4752.23|CAAGAACGAAAGTCGGAGGTTCG</td><td>>hg38_21_+_8433172_8446622_RNA45SN1_45S_50ntFlanks</td><td>4730</td><td>4752</td><td>23</td><td>CAAGAACGAAAGTCGGAGGTTCG</td><td>307.9</td><td>-4.9</td><td>1.7</td><td>-3.0</td><td>3.0E-03</td><td>6.4E-03</td><td>BT20/MB468</td><td>MP</td></tr>
<tr><td>>hg38_21_+_8433172_8446622_RNA45SN1_45S_50ntFlanks@12479.12496.18|CTAATAGGGAACGTGAGC</td><td>>hg38_21_+_8433172_8446622_RNA45SN1_45S_50ntFlanks</td><td>12479</td><td>12496</td><td>18</td><td>CTAATAGGGAACGTGAGC</td><td>538.2</td><td>-2.1</td><td>0.7</td><td>-2.8</td><td>4.8E-03</td><td>6.4E-03</td><td>MB231/MB468</td><td>MP</td></tr>
<tr><td>>hg38_21_+_8433172_8446622_RNA45SN1_45S_50ntFlanks@10729.10752.24|AGAGGGTGTAAATCTCGCGCCGGG</td><td>>hg38_21_+_8433172_8446622_RNA45SN1_45S_50ntFlanks</td><td>10729</td><td>10752</td><td>24</td><td>AGAGGGTGTAAATCTCGCGCCGGG</td><td>225.3</td><td>-3.9</td><td>1.3</td><td>-3.0</td><td>3.0E-03</td><td>6.4E-03</td><td>BT20/MB468</td><td>MP</td></tr>
<tr><td>>trnaMT_ValTAC_MT_+_1602_1670@45.72.28|TTTCAACTTAACTTGACCGCTCTGACCA</td><td>>trnaMT_ValTAC_MT_+_1602_1670</td><td>45</td><td>72</td><td>28</td><td>TTTCAACTTAACTTGACCGCTCTGACCA</td><td>735.2</td><td>-3.0</td><td>0.9</td><td>-3.2</td><td>1.2E-03</td><td>6.4E-03</td><td>BT20/MB231</td><td>NUC</td></tr>
<tr><td>>hg38_21_+_8433172_8446622_RNA45SN1_45S_50ntFlanks@10247.10273.27|GCGACGAGTAGGAGGGCCGCTGCGGTG</td><td>>hg38_21_+_8433172_8446622_RNA45SN1_45S_50ntFlanks</td><td>10247</td><td>10273</td><td>27</td><td>GCGACGAGTAGGAGGGCCGCTGCGGTG</td><td>72.8</td><td>3.9</td><td>1.3</td><td>3.0</td><td>3.0E-03</td><td>6.4E-03</td><td>BT20/MB468</td><td>MP</td></tr>
<tr><td>>hg38_21_+_8433172_8446622_RNA45SN1_45S_50ntFlanks@10905.10935.31|CTGGGCGCGCGCCGCGGCTGGACGAGGCGCC</td><td>>hg38_21_+_8433172_8446622_RNA45SN1_45S_50ntFlanks</td><td>10905</td><td>10935</td><td>31</td><td>CTGGGCGCGCGCCGCGGCTGGACGAGGCGCC</td><td>143.9</td><td>-3.7</td><td>1.2</td><td>-3.0</td><td>3.0E-03</td><td>6.4E-03</td><td>BT20/MB468</td><td>MP</td></tr>
<tr><td>>hg38_21_+_8433172_8446622_RNA45SN1_45S_50ntFlanks@9225.9247.23|CCGAAGAGGGGGACGGCGGAGCG</td><td>>hg38_21_+_8433172_8446622_RNA45SN1_45S_50ntFlanks</td><td>9225</td><td>9247</td><td>23</td><td>CCGAAGAGGGGGACGGCGGAGCG</td><td>60.2</td><td>-4.8</td><td>1.6</td><td>-3.0</td><td>3.0E-03</td><td>6.4E-03</td><td>BT20/MB468</td><td>MP</td></tr>
<tr><td>>hg38_21_+_8433172_8446622_RNA45SN1_45S_50ntFlanks@12626.12652.27|AATGGGGCGAAGCTACCATCTGTGGGA</td><td>>hg38_21_+_8433172_8446622_RNA45SN1_45S_50ntFlanks</td><td>12626</td><td>12652</td><td>27</td><td>AATGGGGCGAAGCTACCATCTGTGGGA</td><td>241.5</td><td>-2.9</td><td>1.0</td><td>-2.8</td><td>5.2E-03</td><td>6.4E-03</td><td>BT20/MB468</td><td>CYTO</td></tr>
<tr><td>>am_trna2_LeuTAA_X_-_55207755_55207829@1.18.18,trna4_LeuTAA_20_-_55933428_55933502@1.18.18,trna60_LeuTAA_1_-_238106956_238107030@1.18.18,trna9_LeuTAA_2_+_117781417_117781491@1.18.18,trnaMT_LeuTAA_MT_+_3230_3304@1.18.18|GTTAAGATGGCAGAGCCC</td><td>>am_trna2_LeuTAA_X_-_55207755_55207829</td><td>1</td><td>18</td><td>18</td><td>GTTAAGATGGCAGAGCCC</td><td>359.9</td><td>4.1</td><td>1.4</td><td>3.0</td><td>3.0E-03</td><td>6.4E-03</td><td>BT20/MB468</td><td>MP</td></tr>
<tr><td>>trnaMT_PheGAA_MT_+_577_647@38.67.30|AATGTTTAGACGGGCTCACATCACCCCATA</td><td>>trnaMT_PheGAA_MT_+_577_647</td><td>38</td><td>67</td><td>30</td><td>AATGTTTAGACGGGCTCACATCACCCCATA</td><td>107.7</td><td>7.4</td><td>2.5</td><td>3.0</td><td>3.0E-03</td><td>6.4E-03</td><td>BT20/MB468</td><td>MP</td></tr>
<tr><td>>hg38_MT_+_1621_3279_RNR2_16S_50ntFlanks@731.759.29|TTAAAACACTGAACTGACAATTAACAGCC</td><td>>hg38_MT_+_1621_3279_RNR2_16S_50ntFlanks</td><td>731</td><td>759</td><td>29</td><td>TTAAAACACTGAACTGACAATTAACAGCC</td><td>28.6</td><td>8.1</td><td>2.9</td><td>2.8</td><td>4.8E-03</td><td>6.4E-03</td><td>MB231/MB468</td><td>MP</td></tr>
<tr><td>>hg38_21_+_8433172_8446622_RNA45SN1_45S_50ntFlanks@11209.11238.30|CGGCGACTCTGGACGCGAGCCGGGCCCTTC</td><td>>hg38_21_+_8433172_8446622_RNA45SN1_45S_50ntFlanks</td><td>11209</td><td>11238</td><td>30</td><td>CGGCGACTCTGGACGCGAGCCGGGCCCTTC</td><td>64.1</td><td>9.3</td><td>2.9</td><td>3.3</td><td>1.1E-03</td><td>6.4E-03</td><td>BT20/MB231</td><td>MP</td></tr>
<tr><td>>hg38_21_+_8433172_8446622_RNA45SN1_45S_50ntFlanks@12420.12457.38|ATGTCGGCTCTTCCTATCATTGTGAAGCAGAATTCACC</td><td>>hg38_21_+_8433172_8446622_RNA45SN1_45S_50ntFlanks</td><td>12420</td><td>12457</td><td>38</td><td>ATGTCGGCTCTTCCTATCATTGTGAAGCAGAATTCACC</td><td>648.8</td><td>-3.0</td><td>0.9</td><td>-3.2</td><td>1.2E-03</td><td>6.4E-03</td><td>BT20/MB231</td><td>NUC</td></tr>
<tr><td>>hg38_21_+_8433172_8446622_RNA45SN1_45S_50ntFlanks@10398.10432.35|GTTGAACATGGGTCAGTCGGTCCTGAGAGATGGGC</td><td>>hg38_21_+_8433172_8446622_RNA45SN1_45S_50ntFlanks</td><td>10398</td><td>10432</td><td>35</td><td>GTTGAACATGGGTCAGTCGGTCCTGAGAGATGGGC</td><td>45.1</td><td>5.9</td><td>2.0</td><td>3.0</td><td>3.0E-03</td><td>6.4E-03</td><td>BT20/MB468</td><td>MP</td></tr>
<tr><td>>trna119_LysCTT_1_-_145395522_145395594@1.40.40,trna11_LysCTT_5_-_180648979_180649051@1.40.40,trna13_LysCTT_6_+_26556774_26556846@1.40.40,trna7_LysCTT_16_+_3225692_3225764@1.40.40,trna9_LysCTT_5_+_180634755_180634827@1.40.40|GCCCGGCTAGCTCAGTCGGTAGAGCATGAGACTCTTAATC</td><td>>trna119_LysCTT_1_-_145395522_145395594</td><td>1</td><td>40</td><td>40</td><td>GCCCGGCTAGCTCAGTCGGTAGAGCATGAGACTCTTAATC</td><td>461.1</td><td>4.7</td><td>1.7</td><td>2.8</td><td>5.3E-03</td><td>6.4E-03</td><td>MB231/MB468</td><td>CYTO</td></tr>
<tr><td>>hg38_1_-_228634819_228635039_RNA5S12_5S_50ntFlanks@84.106.23|CTCGTCTGATCTCGGAAGCTAAG</td><td>>hg38_1_-_228634819_228635039_RNA5S12_5S_50ntFlanks</td><td>84</td><td>106</td><td>23</td><td>CTCGTCTGATCTCGGAAGCTAAG</td><td>141.2</td><td>-3.4</td><td>1.2</td><td>-2.8</td><td>5.6E-03</td><td>6.4E-03</td><td>MB231/MB468</td><td>NUC</td></tr>
<tr><td>>MI0000264|hsa-mir-7-2&WithFlank&15|+|88611819|88611940@38.61.24(+1A);MI0000265|hsa-mir-7-3&WithFlank&19|+|4770664|4770785@37.60.24(+1A);MI0000263|hsa-mir-7-1&WithFlank&9|-|83969742|83969863@30.53.24(+1A)&[MIMAT0000252&hsa-miR-7-5p&offsets|0|0(+1A);m-400&15|+|88611856|88611879&offsets|0|0(+1A)];[MIMAT0000252_1&hsa-miR-7-5p&offsets|0|0(+1A);m-402&19|+|4770700|4770723&offsets|0|0(+1A)];[MIMAT0000252_2&hsa-miR-7-5p&offsets|0|0(+1A);m-401&9|-|83969811|83969834&offsets|0|0(+1A)]|TGGAAGACTAGTGATTTTGTTGTTA</td><td>>MI0000264|hsa-mir-7-2&WithFlank&15|+|88611819|88611940</td><td>38</td><td>61</td><td>24(+1A)</td><td>TGGAAGACTAGTGATTTTGTTGTTA</td><td>875.8</td><td>-3.3</td><td>1.0</td><td>-3.3</td><td>1.1E-03</td><td>6.4E-03</td><td>BT20/MB231</td><td>MP</td></tr>
<tr><td>>hg38_21_+_8433172_8446622_RNA45SN1_45S_50ntFlanks@12633.12660.28|CGAAGCTACCATCTGTGGGATTATGACT</td><td>>hg38_21_+_8433172_8446622_RNA45SN1_45S_50ntFlanks</td><td>12633</td><td>12660</td><td>28</td><td>CGAAGCTACCATCTGTGGGATTATGACT</td><td>189.3</td><td>-1.9</td><td>0.7</td><td>-2.8</td><td>4.5E-03</td><td>6.4E-03</td><td>BT20/MB468</td><td>MITO</td></tr>
<tr><td>>hg38_21_+_8433172_8446622_RNA45SN1_45S_50ntFlanks@11576.11611.36|CTTAGAACTGGTGCGGACCAGGGGAATCCGACTGTT</td><td>>hg38_21_+_8433172_8446622_RNA45SN1_45S_50ntFlanks</td><td>11576</td><td>11611</td><td>36</td><td>CTTAGAACTGGTGCGGACCAGGGGAATCCGACTGTT</td><td>531.6</td><td>-3.2</td><td>1.1</td><td>-3.0</td><td>3.0E-03</td><td>6.4E-03</td><td>BT20/MB468</td><td>MP</td></tr>
<tr><td>>hg38_21_+_8433172_8446622_RNA45SN1_45S_50ntFlanks@12828.12847.20|CCGGGGTCCGGTGCGGAGTG</td><td>>hg38_21_+_8433172_8446622_RNA45SN1_45S_50ntFlanks</td><td>12828</td><td>12847</td><td>20</td><td>CCGGGGTCCGGTGCGGAGTG</td><td>62.4</td><td>-5.2</td><td>1.8</td><td>-3.0</td><td>3.0E-03</td><td>6.4E-03</td><td>BT20/MB468</td><td>MP</td></tr>
<tr><td>>hg38_21_+_8433172_8446622_RNA45SN1_45S_50ntFlanks@4728.4748.21|ATCAAGAACGAAAGTCGGAGG</td><td>>hg38_21_+_8433172_8446622_RNA45SN1_45S_50ntFlanks</td><td>4728</td><td>4748</td><td>21</td><td>ATCAAGAACGAAAGTCGGAGG</td><td>131.1</td><td>-4.1</td><td>1.5</td><td>-2.8</td><td>5.6E-03</td><td>6.4E-03</td><td>MB231/MB468</td><td>NUC</td></tr>
<tr><td>>hg38_21_+_8433172_8446622_RNA45SN1_45S_50ntFlanks@6661.6685.25|CGGTGGATCACTCGGCTCGTGCGTC</td><td>>hg38_21_+_8433172_8446622_RNA45SN1_45S_50ntFlanks</td><td>6661</td><td>6685</td><td>25</td><td>CGGTGGATCACTCGGCTCGTGCGTC</td><td>490.5</td><td>2.7</td><td>0.8</td><td>3.2</td><td>1.3E-03</td><td>6.4E-03</td><td>BT20/MB231</td><td>CYTO</td></tr>
<tr><td>>hg38_21_+_8433172_8446622_RNA45SN1_45S_50ntFlanks@9535.9570.36|GGAAACTCTGGTGGAGGTCCGTAGCGGTCCTGACGT</td><td>>hg38_21_+_8433172_8446622_RNA45SN1_45S_50ntFlanks</td><td>9535</td><td>9570</td><td>36</td><td>GGAAACTCTGGTGGAGGTCCGTAGCGGTCCTGACGT</td><td>559.3</td><td>-2.6</td><td>0.8</td><td>-3.2</td><td>1.3E-03</td><td>6.4E-03</td><td>BT20/MB231</td><td>CYTO</td></tr>
<tr><td>>MI0000542|hsa-mir-320a&WithFlank&8|-|22244960|22245043@49.68.20&[MIMAT0000510&hsa-miR-320a-3p&offsets|+1|-1;m-60&8|-|22244975|22244996&offsets|+1|-1]|AAAGCTGGGTTGAGAGGGCG</td><td>>MI0000542|hsa-mir-320a&WithFlank&8|-|22244960|22245043</td><td>49</td><td>68</td><td>20</td><td>AAAGCTGGGTTGAGAGGGCG</td><td>220.7</td><td>2.3</td><td>0.7</td><td>3.2</td><td>1.2E-03</td><td>6.4E-03</td><td>BT20/MB231</td><td>NUC</td></tr>
<tr><td>>am_MI0000748|hsa-mir-130b&WithFlank&22|+|21653298|21653391@19.39.21&[MIMAT0004680&hsa-miR-130b-5p&offsets|0|0;m-234&22|+|21653316|21653337&offsets|0|-1]|ACTCTTTCCCTGTTGCACTAC</td><td>>am_MI0000748|hsa-mir-130b&WithFlank&22|+|21653298|21653391</td><td>19</td><td>39</td><td>21</td><td>ACTCTTTCCCTGTTGCACTAC</td><td>568.6</td><td>-2.1</td><td>0.7</td><td>-3.2</td><td>1.2E-03</td><td>6.4E-03</td><td>BT20/MB231</td><td>NUC</td></tr>
<tr><td>>hg38_21_+_8433172_8446622_RNA45SN1_45S_50ntFlanks@5472.5514.43|ACGGCCCTGGCGGAGCGCTGAGAAGACGGTCGAACTTGACTAT</td><td>>hg38_21_+_8433172_8446622_RNA45SN1_45S_50ntFlanks</td><td>5472</td><td>5514</td><td>43</td><td>ACGGCCCTGGCGGAGCGCTGAGAAGACGGTCGAACTTGACTAT</td><td>738.2</td><td>-3.0</td><td>0.9</td><td>-3.2</td><td>1.2E-03</td><td>6.4E-03</td><td>BT20/MB231</td><td>NUC</td></tr>
<tr><td>>hg38_21_+_8433172_8446622_RNA45SN1_45S_50ntFlanks@6653.6678.26|ACTCTTAGCGGTGGATCACTCGGCTC</td><td>>hg38_21_+_8433172_8446622_RNA45SN1_45S_50ntFlanks</td><td>6653</td><td>6678</td><td>26</td><td>ACTCTTAGCGGTGGATCACTCGGCTC</td><td>112.3</td><td>5.1</td><td>1.7</td><td>3.0</td><td>3.0E-03</td><td>6.4E-03</td><td>BT20/MB468</td><td>MP</td></tr>
<tr><td>>hg38_21_+_8433172_8446622_RNA45SN1_45S_50ntFlanks@12407.12425.19|TTTTGATCCTTCGATGTCG</td><td>>hg38_21_+_8433172_8446622_RNA45SN1_45S_50ntFlanks</td><td>12407</td><td>12425</td><td>19</td><td>TTTTGATCCTTCGATGTCG</td><td>72.1</td><td>-4.7</td><td>1.6</td><td>-3.0</td><td>3.0E-03</td><td>6.4E-03</td><td>BT20/MB468</td><td>MP</td></tr>
<tr><td>>am_trna128_GlyGCC_6_-_27870686_27870756@7.35.29,trna18_GlyGCC_16_+_70822597_70822667@7.35.29,trna19_GlyGCC_16_+_70823410_70823480@7.35.29,trna19_GlyGCC_2_-_157257659_157257729@7.35.29,trna24_GlyGCC_16_-_70812942_70813012@7.35.29,trna25_GlyGCC_16_-_70812114_70812184@7.35.29,trna2_GlyGCC_21_-_18827107_18827177@7.35.29,trna35_GlyGCC_1_+_161413094_161413164@7.35.29,trna37_GlyGCC_1_+_161420467_161420537@7.35.29,trna39_GlyGCC_1_+_161427898_161427968@7.35.29,trna41_GlyGCC_1_+_161435258_161435328@7.35.29,trna5_GlyGCC_17_+_8029064_8029134@7.35.29,trna68_GlyGCC_1_-_161493637_161493707@7.35.29|GTGGTTCAGTGGTAGAATTCTCGCCTGCC</td><td>>am_trna128_GlyGCC_6_-_27870686_27870756</td><td>7</td><td>35</td><td>29</td><td>GTGGTTCAGTGGTAGAATTCTCGCCTGCC</td><td>138.0</td><td>-5.2</td><td>1.6</td><td>-3.2</td><td>1.2E-03</td><td>6.5E-03</td><td>BT20/MB231</td><td>NUC</td></tr>
<tr><td>>hg38_21_+_8433172_8446622_RNA45SN1_45S_50ntFlanks@6643.6681.39|ACCTCGTACGACTCTTAGCGGTGGATCACTCGGCTCGTG</td><td>>hg38_21_+_8433172_8446622_RNA45SN1_45S_50ntFlanks</td><td>6643</td><td>6681</td><td>39</td><td>ACCTCGTACGACTCTTAGCGGTGGATCACTCGGCTCGTG</td><td>413.5</td><td>-2.7</td><td>0.8</td><td>-3.2</td><td>1.2E-03</td><td>6.5E-03</td><td>BT20/MB231</td><td>NUC</td></tr>
<tr><td>>hg38_21_+_8433172_8446622_RNA45SN1_45S_50ntFlanks@6772.6797.26|GTTCCTCCCGGGGCTACGCCTGTCTG</td><td>>hg38_21_+_8433172_8446622_RNA45SN1_45S_50ntFlanks</td><td>6772</td><td>6797</td><td>26</td><td>GTTCCTCCCGGGGCTACGCCTGTCTG</td><td>422.7</td><td>-3.5</td><td>1.2</td><td>-2.8</td><td>4.5E-03</td><td>6.5E-03</td><td>BT20/MB468</td><td>MITO</td></tr>
<tr><td>>hg38_21_+_8433172_8446622_RNA45SN1_45S_50ntFlanks@10856.10873.18|AGGATTGGCTCTAAGGGC</td><td>>hg38_21_+_8433172_8446622_RNA45SN1_45S_50ntFlanks</td><td>10856</td><td>10873</td><td>18</td><td>AGGATTGGCTCTAAGGGC</td><td>249.8</td><td>5.3</td><td>1.6</td><td>3.3</td><td>1.1E-03</td><td>6.5E-03</td><td>BT20/MB231</td><td>MP</td></tr>
<tr><td>>MI0000071|hsa-mir-17&WithFlank&13|+|91350599|91350694@19.42.24&[MIMAT0000070&hsa-miR-17-5p&offsets|-1|0;m-88&13|+|91350618|91350640&offsets|-1|0]|TCAAAGTGCTTACAGTGCAGGTAG</td><td>>MI0000071|hsa-mir-17&WithFlank&13|+|91350599|91350694</td><td>19</td><td>42</td><td>24</td><td>TCAAAGTGCTTACAGTGCAGGTAG</td><td>252.6</td><td>-4.5</td><td>1.4</td><td>-3.3</td><td>1.1E-03</td><td>6.5E-03</td><td>BT20/MB231</td><td>MP</td></tr>
<tr><td>>MI0000440|hsa-mir-27b&WithFlank&9|+|95085439|95085547@25.46.22&[MIMAT0004588&hsa-miR-27b-5p&offsets|0|0;m-240&9|+|95085463|95085484&offsets|0|0]|AGAGCTTAGCTGATTGGTGAAC</td><td>>MI0000440|hsa-mir-27b&WithFlank&9|+|95085439|95085547</td><td>25</td><td>46</td><td>22</td><td>AGAGCTTAGCTGATTGGTGAAC</td><td>51.3</td><td>5.8</td><td>1.9</td><td>3.0</td><td>3.0E-03</td><td>6.5E-03</td><td>BT20/MB468</td><td>MP</td></tr>
<tr><td>>hg38_21_+_8433172_8446622_RNA45SN1_45S_50ntFlanks@5086.5112.27|AACGAGACTCTGGCATGCTAACTAGTT</td><td>>hg38_21_+_8433172_8446622_RNA45SN1_45S_50ntFlanks</td><td>5086</td><td>5112</td><td>27</td><td>AACGAGACTCTGGCATGCTAACTAGTT</td><td>435.8</td><td>-3.0</td><td>1.1</td><td>-2.8</td><td>4.8E-03</td><td>6.5E-03</td><td>MB231/MB468</td><td>MP</td></tr>
<tr><td>>hg38_MT_+_598_1651_RNR1_12S_50ntFlanks@416.456.41|AAATAGACTACGAAAGTGGCTTTAACATATCTGAACACACA</td><td>>hg38_MT_+_598_1651_RNR1_12S_50ntFlanks</td><td>416</td><td>456</td><td>41</td><td>AAATAGACTACGAAAGTGGCTTTAACATATCTGAACACACA</td><td>41.1</td><td>5.3</td><td>1.9</td><td>2.8</td><td>4.8E-03</td><td>6.5E-03</td><td>MB231/MB468</td><td>MP</td></tr>
<tr><td>>hg38_21_+_8433172_8446622_RNA45SN1_45S_50ntFlanks@475.510.36|CTTCGTGATCGATGTGGTGACGTCGTGCTCTCCCGG</td><td>>hg38_21_+_8433172_8446622_RNA45SN1_45S_50ntFlanks</td><td>475</td><td>510</td><td>36</td><td>CTTCGTGATCGATGTGGTGACGTCGTGCTCTCCCGG</td><td>173.4</td><td>5.3</td><td>1.8</td><td>3.0</td><td>3.0E-03</td><td>6.5E-03</td><td>BT20/MB468</td><td>MP</td></tr>
<tr><td>>hg38_21_+_8433172_8446622_RNA45SN1_45S_50ntFlanks@9562.9596.35|TCCTGACGTGCAAATCGGTCGTCCGACCTGGGTAT</td><td>>hg38_21_+_8433172_8446622_RNA45SN1_45S_50ntFlanks</td><td>9562</td><td>9596</td><td>35</td><td>TCCTGACGTGCAAATCGGTCGTCCGACCTGGGTAT</td><td>332.9</td><td>-5.2</td><td>1.8</td><td>-3.0</td><td>3.0E-03</td><td>6.5E-03</td><td>BT20/MB468</td><td>MP</td></tr>
<tr><td>>hg38_21_+_8433172_8446622_RNA45SN1_45S_50ntFlanks@4588.4607.20|CTATTTTGTTGGTTTTCGGA</td><td>>hg38_21_+_8433172_8446622_RNA45SN1_45S_50ntFlanks</td><td>4588</td><td>4607</td><td>20</td><td>CTATTTTGTTGGTTTTCGGA</td><td>181.0</td><td>-1.9</td><td>0.7</td><td>-2.8</td><td>5.2E-03</td><td>6.5E-03</td><td>BT20/MB468</td><td>CYTO</td></tr>
<tr><td>>hg38_21_+_8433172_8446622_RNA45SN1_45S_50ntFlanks@10347.10383.37|CAAACGAGAACTTTGAAGGCCGAAGTGGAGAAGGGTT</td><td>>hg38_21_+_8433172_8446622_RNA45SN1_45S_50ntFlanks</td><td>10347</td><td>10383</td><td>37</td><td>CAAACGAGAACTTTGAAGGCCGAAGTGGAGAAGGGTT</td><td>252.6</td><td>-2.5</td><td>0.9</td><td>-2.8</td><td>4.8E-03</td><td>6.5E-03</td><td>MB231/MB468</td><td>MP</td></tr>
<tr><td>>MI0000438|hsa-mir-15b&WithFlank&3|+|160404582|160404691@64.85.22&[MIMAT0004586&hsa-miR-15b-3p&offsets|0|0;m-204&3|+|160404645|160404665&offsets|0|+1]|CGAATCATTATTTGCTGCTCTA</td><td>>MI0000438|hsa-mir-15b&WithFlank&3|+|160404582|160404691</td><td>64</td><td>85</td><td>22</td><td>CGAATCATTATTTGCTGCTCTA</td><td>196.2</td><td>3.9</td><td>1.4</td><td>2.8</td><td>4.8E-03</td><td>6.5E-03</td><td>MB231/MB468</td><td>MP</td></tr>
<tr><td>>hg38_21_+_8433172_8446622_RNA45SN1_45S_50ntFlanks@475.492.18|CTTCGTGATCGATGTGGT</td><td>>hg38_21_+_8433172_8446622_RNA45SN1_45S_50ntFlanks</td><td>475</td><td>492</td><td>18</td><td>CTTCGTGATCGATGTGGT</td><td>966.6</td><td>1.1</td><td>0.4</td><td>2.8</td><td>5.6E-03</td><td>6.5E-03</td><td>BT20/MB468</td><td>NUC</td></tr>
</table>

<table>
<thead>
<tr><th></th><th></th><th></th><th></th><th></th><th></th><th></th><th></th><th></th><th></th><th></th><th></th><th></th><th></th></tr>
</thead>
<tbody>
<tr><td>>hg38_21_+_8433172_8446622_RNA45SN1_45S_50ntFlanks@11710.11736.27|CGCGGGTAAACGGCGGGAGTAACTATG</td><td>>hg38_21_+_8433172_8446622_RNA45SN1_45S_50ntFlanks</td><td>11710</td><td>11736</td><td>27</td><td>CGCGGGTAAACGGCGGGAGTAACTATG</td><td>244.3</td><td>-2.6</td><td>0.9</td><td>-3.0</td><td>3.0E-03</td><td>6.5E-03</td><td>BT20/MB468</td><td>MP</td></tr>
<tr><td>>MI0000086|hsa-mir-28&WithFlank&3|+|188688775|188688872@60.80.21(+1G)&[MIMAT0004502&hsa-miR-28-3p&offsets|0|-1(+1G);m-28&3|+|188688834|188688855&offsets|0|-1(+1G)]|CACTAGATTGTGAGCTCCTGGG</td><td>>MI0000086|hsa-mir-28&WithFlank&3|+|188688775|188688872</td><td>60</td><td>80</td><td>21(+1G)</td><td>CACTAGATTGTGAGCTCCTGGG</td><td>830.4</td><td>3.2</td><td>1.0</td><td>3.2</td><td>1.3E-03</td><td>6.5E-03</td><td>BT20/MB231</td><td>CYTO</td></tr>
<tr><td>>hg38_21_+_8433172_8446622_RNA45SN1_45S_50ntFlanks@4497.4520.24|GAAGCGTTTACTTTGAAAAAATTA</td><td>>hg38_21_+_8433172_8446622_RNA45SN1_45S_50ntFlanks</td><td>4497</td><td>4520</td><td>24</td><td>GAAGCGTTTACTTTGAAAAAATTA</td><td>267.7</td><td>3.5</td><td>1.1</td><td>3.2</td><td>1.3E-03</td><td>6.5E-03</td><td>BT20/MB231</td><td>CYTO</td></tr>
<tr><td>>hg38_21_+_8433172_8446622_RNA45SN1_45S_50ntFlanks@12415.12432.18|CTTCGATGTCGGCTCTTC</td><td>>hg38_21_+_8433172_8446622_RNA45SN1_45S_50ntFlanks</td><td>12415</td><td>12432</td><td>18</td><td>CTTCGATGTCGGCTCTTC</td><td>1387.4</td><td>-3.1</td><td>1.0</td><td>-3.0</td><td>3.0E-03</td><td>6.5E-03</td><td>BT20/MB468</td><td>MP</td></tr>
<tr><td>>hg38_21_+_8433172_8446622_RNA45SN1_45S_50ntFlanks@10852.10878.27|GATAAGGATTGGCTCTAAGGGCTGGGT</td><td>>hg38_21_+_8433172_8446622_RNA45SN1_45S_50ntFlanks</td><td>10852</td><td>10878</td><td>27</td><td>GATAAGGATTGGCTCTAAGGGCTGGGT</td><td>28.9</td><td>8.6</td><td>2.9</td><td>3.0</td><td>3.0E-03</td><td>6.5E-03</td><td>BT20/MB468</td><td>MP</td></tr>
<tr><td>>hg38_21_+_8433172_8446622_RNA45SN1_45S_50ntFlanks@12632.12657.26|GCGAAGCTACCATCTGTGGGATTATG</td><td>>hg38_21_+_8433172_8446622_RNA45SN1_45S_50ntFlanks</td><td>12632</td><td>12657</td><td>26</td><td>GCGAAGCTACCATCTGTGGGATTATG</td><td>405.9</td><td>-2.0</td><td>0.7</td><td>-2.8</td><td>4.5E-03</td><td>6.5E-03</td><td>BT20/MB468</td><td>MITO</td></tr>
<tr><td>>hg38_21_+_8433172_8446622_RNA45SN1_45S_50ntFlanks@9534.9559.26|AGGAAACTCTGGTGGAGGTCCGTAGC</td><td>>hg38_21_+_8433172_8446622_RNA45SN1_45S_50ntFlanks</td><td>9534</td><td>9559</td><td>26</td><td>AGGAAACTCTGGTGGAGGTCCGTAGC</td><td>144.0</td><td>-3.4</td><td>1.2</td><td>-2.8</td><td>4.5E-03</td><td>6.5E-03</td><td>BT20/MB468</td><td>MITO</td></tr>
<tr><td>>hg38_21_+_8433172_8446622_RNA45SN1_45S_50ntFlanks@12456.12500.45|CCAAGCGTTGGATTGTTCACCCACTAATAGGGAACGTGAGCTGGG</td><td>>hg38_21_+_8433172_8446622_RNA45SN1_45S_50ntFlanks</td><td>12456</td><td>12500</td><td>45</td><td>CCAAGCGTTGGATTGTTCACCCACTAATAGGGAACGTGAGCTGGG</td><td>137.5</td><td>-4.6</td><td>1.6</td><td>-2.8</td><td>4.5E-03</td><td>6.5E-03</td><td>BT20/MB468</td><td>MITO</td></tr>
<tr><td>>hg38_MT_+_1621_3279_RNR2_16S_50ntFlanks@51.90.40|GCTAAACCTAGCCCCAAACCCACTCCACCTTACTACCAGA</td><td>>hg38_MT_+_1621_3279_RNR2_16S_50ntFlanks</td><td>51</td><td>90</td><td>40</td><td>GCTAAACCTAGCCCCAAACCCACTCCACCTTACTACCAGA</td><td>208.1</td><td>-3.7</td><td>1.2</td><td>-3.2</td><td>1.3E-03</td><td>6.5E-03</td><td>BT20/MB231</td><td>CYTO</td></tr>
<tr><td>>hg38_21_+_8433172_8446622_RNA45SN1_45S_50ntFlanks@11566.11601.36|TAGCAGCCGACTTAGAACTGGTGCGGACCAGGGGAA</td><td>>hg38_21_+_8433172_8446622_RNA45SN1_45S_50ntFlanks</td><td>11566</td><td>11601</td><td>36</td><td>TAGCAGCCGACTTAGAACTGGTGCGGACCAGGGGAA</td><td>88.3</td><td>-4.8</td><td>1.7</td><td>-2.8</td><td>4.8E-03</td><td>6.5E-03</td><td>MB231/MB468</td><td>MP</td></tr>
<tr><td>>hg38_MT_+_1621_3279_RNR2_16S_50ntFlanks@645.676.32|ATTGGACCAATCTATCACCCTATAGAAGAACT</td><td>>hg38_MT_+_1621_3279_RNR2_16S_50ntFlanks</td><td>645</td><td>676</td><td>32</td><td>ATTGGACCAATCTATCACCCTATAGAAGAACT</td><td>93.4</td><td>5.0</td><td>1.7</td><td>3.0</td><td>3.0E-03</td><td>6.5E-03</td><td>BT20/MB468</td><td>MP</td></tr>
<tr><td>>hg38_21_+_8433172_8446622_RNA45SN1_45S_50ntFlanks@8659.8677.19|CATTTCCACCGCGGCGGTG</td><td>>hg38_21_+_8433172_8446622_RNA45SN1_45S_50ntFlanks</td><td>8659</td><td>8677</td><td>19</td><td>CATTTCCACCGCGGCGGTG</td><td>32.0</td><td>-8.1</td><td>2.7</td><td>-3.0</td><td>3.0E-03</td><td>6.5E-03</td><td>BT20/MB468</td><td>MP</td></tr>
<tr><td>>hg38_21_+_8433172_8446622_RNA45SN1_45S_50ntFlanks@11452.11471.20|CTCCTCGGCGCGCGGCGGCG</td><td>>hg38_21_+_8433172_8446622_RNA45SN1_45S_50ntFlanks</td><td>11452</td><td>11471</td><td>20</td><td>CTCCTCGGCGCGCGGCGGCG</td><td>117.0</td><td>-4.8</td><td>1.6</td><td>-3.0</td><td>3.0E-03</td><td>6.5E-03</td><td>BT20/MB468</td><td>MP</td></tr>
<tr><td>>am_MI0005763|hsa-mir-941-1&WithFlank&20|+|63919443|63919526@53.74.22;MI0005764|hsa-mir-941-2&WithFlank&20|+|63919499|63919582@53.74.22;MI0005765|hsa-mir-941-3&WithFlank&20|+|63919555|63919638@53.74.22;MI0005766|hsa-mir-941-4&WithFlank&20|+|63919750|63919833@53.74.22;MI0031520|hsa-mir-941-5&WithFlank&20|+|63919862|63919945@53.74.22&[MIMAT0004984&hsa-miR-941&offsets|0|-1;m-263&20|+|63919495|63919517&offsets|0|-1];[MIMAT0004984_1&hsa-miR-941&offsets|0|-1;m-263&20|+|63919551|63919573&offsets|0|-1];[MIMAT0004984_2&hsa-miR-941&offsets|0|-1;m-263&20|+|63919607|63919629&offsets|0|-1];[MIMAT0004984_3&hsa-miR-941&offsets|0|-1;m-263&20|+|63919802|63919824&offsets|0|-1];[MIMAT0004984_4&hsa-miR-941&offsets|0|-1;m-263&20|+|63919914|63919936&offsets|0|-1]|CACCCGGCTGTGTGCACATGTG</td><td>>am_MI0005763|hsa-mir-941-1&WithFlank&20|+|63919443|63919526</td><td>53</td><td>74</td><td>22</td><td>CACCCGGCTGTGTGCACATGTG</td><td>437.3</td><td>4.2</td><td>1.4</td><td>3.0</td><td>3.0E-03</td><td>6.5E-03</td><td>BT20/MB468</td><td>MP</td></tr>
<tr><td>>trna111_HisGTG_1_-_147774845_147774916@1.34.34,trna118_HisGTG_1_-_145396881_145396952@1.34.34,trna16_HisGTG_1_+_146544773_146544844@1.34.34,trna1_HisGTG_15_+_45493349_45493420@1.34.34,trna21_HisGTG_1_+_147753471_147753542@1.34.34,trna33_HisGTG_6_+_27125906_27125977@1.34.34,trna7_HisGTG_9_-_14433938_14434009@1.34.34,trna8_HisGTG_15_-_45492611_45492682@1.34.34,trna9_HisGTG_15_-_45490804_45490875@1.34.34|GCCGTGATCGTATAGTGGTTAGTACTCTGCGTTG</td><td>>trna111_HisGTG_1_-_147774845_147774916</td><td>1</td><td>34</td><td>34</td><td>GCCGTGATCGTATAGTGGTTAGTACTCTGCGTTG</td><td>450.2</td><td>3.8</td><td>1.3</td><td>3.0</td><td>3.0E-03</td><td>6.5E-03</td><td>BT20/MB468</td><td>MP</td></tr>
<tr><td>>hg38_21_+_8433172_8446622_RNA45SN1_45S_50ntFlanks@12405.12436.32|CTTTTTGATCCTTCGATGTCGGCTCTTCCTAT</td><td>>hg38_21_+_8433172_8446622_RNA45SN1_45S_50ntFlanks</td><td>12405</td><td>12436</td><td>32</td><td>CTTTTTGATCCTTCGATGTCGGCTCTTCCTAT</td><td>1236.0</td><td>-5.4</td><td>1.8</td><td>-3.0</td><td>3.0E-03</td><td>6.5E-03</td><td>BT20/MB468</td><td>MP</td></tr>
<tr><td>>hg38_21_+_8433172_8446622_RNA45SN1_45S_50ntFlanks@4685.4720.36|AGACGGACCAGAGCGAAAGCATTTGCCAAGAATGTT</td><td>>hg38_21_+_8433172_8446622_RNA45SN1_45S_50ntFlanks</td><td>4685</td><td>4720</td><td>36</td><td>AGACGGACCAGAGCGAAAGCATTTGCCAAGAATGTT</td><td>312.3</td><td>3.6</td><td>1.3</td><td>2.8</td><td>5.3E-03</td><td>6.5E-03</td><td>BT20/MB468</td><td>CYTO</td></tr>
<tr><td>>hg38_21_+_8433172_8446622_RNA45SN1_45S_50ntFlanks@12457.12490.34|CAAGCGTTGGATTGTTCACCCACTAATAGGGAAC</td><td>>hg38_21_+_8433172_8446622_RNA45SN1_45S_50ntFlanks</td><td>12457</td><td>12490</td><td>34</td><td>CAAGCGTTGGATTGTTCACCCACTAATAGGGAAC</td><td>215.1</td><td>-1.7</td><td>0.6</td><td>-2.8</td><td>5.7E-03</td><td>6.5E-03</td><td>MB231/MB468</td><td>NUC</td></tr>
<tr><td>>trna17_ValTAC_11_-_59318102_59318174@1.36.36,trna4_ValTAC_X_-_18693029_18693101@1.36.36|GGTTCCATAGTGTAGTGGTTATCACGTCTGCTTTAC</td><td>>trna17_ValTAC_11_-_59318102_59318174</td><td>1</td><td>36</td><td>36</td><td>GGTTCCATAGTGTAGTGGTTATCACGTCTGCTTTAC</td><td>197.6</td><td>-3.4</td><td>1.1</td><td>-3.2</td><td>1.4E-03</td><td>6.5E-03</td><td>BT20/MB231</td><td>MITO</td></tr>
<tr><td>>hg38_21_+_8433172_8446622_RNA45SN1_45S_50ntFlanks@12182.12209.28|CAAACGGTAACGCAGGTGTCCTAAGGCG</td><td>>hg38_21_+_8433172_8446622_RNA45SN1_45S_50ntFlanks</td><td>12182</td><td>12209</td><td>28</td><td>CAAACGGTAACGCAGGTGTCCTAAGGCG</td><td>62.6</td><td>-5.1</td><td>1.7</td><td>-3.0</td><td>3.0E-03</td><td>6.5E-03</td><td>BT20/MB468</td><td>MP</td></tr>
<tr><td>>hg38_21_+_8433172_8446622_RNA45SN1_45S_50ntFlanks@11910.11933.24|GTGAAGAGACATGAGAGGTGTAGA</td><td>>hg38_21_+_8433172_8446622_RNA45SN1_45S_50ntFlanks</td><td>11910</td><td>11933</td><td>24</td><td>GTGAAGAGACATGAGAGGTGTAGA</td><td>64.3</td><td>9.3</td><td>2.9</td><td>3.3</td><td>1.1E-03</td><td>6.5E-03</td><td>BT20/MB231</td><td>MP</td></tr>
<tr><td>>trna111_HisGTG_1_-_147774845_147774916@-1G.34.35,trna118_HisGTG_1_-_145396881_145396952@-1G.34.35,trna16_HisGTG_1_+_146544773_146544844@-1G.34.35,trna1_HisGTG_15_+_45493349_45493420@-1G.34.35,trna21_HisGTG_1_+_147753471_147753542@-1G.34.35,trna33_HisGTG_6_+_27125906_27125977@-1G.34.35,trna7_HisGTG_9_-_14433938_14434009@-1G.34.35,trna8_HisGTG_15_-_45492611_45492682@-1G.34.35,trna9_HisGTG_15_-_45490804_45490875@-1G.34.35|GGCCGTGATCGTATAGTGGTTAGTACTCTGCGTTG</td><td>>trna111_HisGTG_1_-_147774845_147774916</td><td>-1G</td><td>34</td><td>35</td><td>GGCCGTGATCGTATAGTGGTTAGTACTCTGCGTTG</td><td>1431.6</td><td>4.9</td><td>1.7</td><td>3.0</td><td>3.0E-03</td><td>6.5E-03</td><td>BT20/MB468</td><td>MP</td></tr>
<tr><td>>hg38_21_+_8433172_8446622_RNA45SN1_45S_50ntFlanks@6647.6666.20|CGTACGACTCTTAGCGGTGG</td><td>>hg38_21_+_8433172_8446622_RNA45SN1_45S_50ntFlanks</td><td>6647</td><td>6666</td><td>20</td><td>CGTACGACTCTTAGCGGTGG</td><td>2277.6</td><td>2.5</td><td>0.9</td><td>2.8</td><td>5.3E-03</td><td>6.5E-03</td><td>BT20/MB468</td><td>CYTO</td></tr>
<tr><td>>hg38_21_+_8433172_8446622_RNA45SN1_45S_50ntFlanks@7981.8004.24|CTCAGATCAGACGTGGCGACCCGC</td><td>>hg38_21_+_8433172_8446622_RNA45SN1_45S_50ntFlanks</td><td>7981</td><td>8004</td><td>24</td><td>CTCAGATCAGACGTGGCGACCCGC</td><td>331.0</td><td>-1.7</td><td>0.6</td><td>-2.8</td><td>4.5E-03</td><td>6.5E-03</td><td>BT20/MB468</td><td>MITO</td></tr>
<tr><td>>am_MI0000809|hsa-mir-151a&WithFlank&8|-|140732558|140732659@52.72.21(+1A)&[MIMAT0000757&hsa-miR-151a-3p&offsets|-1|-1(+1A);m-30&8|-|140732587|140732607&offsets|-1|-1(+1A)]|ACTAGACTGAAGCTCCTTGAGA</td><td>>am_MI0000809|hsa-mir-151a&WithFlank&8|-|140732558|140732659</td><td>52</td><td>72</td><td>21(+1A)</td><td>ACTAGACTGAAGCTCCTTGAGA</td><td>97.1</td><td>3.4</td><td>1.2</td><td>2.8</td><td>4.9E-03</td><td>6.5E-03</td><td>MB231/MB468</td><td>MP</td></tr>
<tr><td>>hg38_21_+_8433172_8446622_RNA45SN1_45S_50ntFlanks@7991.8011.21|ACGTGGCGACCCGCTGAATTT</td><td>>hg38_21_+_8433172_8446622_RNA45SN1_45S_50ntFlanks</td><td>7991</td><td>8011</td><td>21</td><td>ACGTGGCGACCCGCTGAATTT</td><td>8923.2</td><td>-2.5</td><td>0.9</td><td>-2.8</td><td>5.7E-03</td><td>6.5E-03</td><td>MB231/MB468</td><td>MITO</td></tr>
<tr><td>>hg38_21_+_8433172_8446622_RNA45SN1_45S_50ntFlanks@3853.3872.20|AACTGTGGTAATTCTAGAGC</td><td>>hg38_21_+_8433172_8446622_RNA45SN1_45S_50ntFlanks</td><td>3853</td><td>3872</td><td>20</td><td>AACTGTGGTAATTCTAGAGC</td><td>198.2</td><td>-1.7</td><td>0.6</td><td>-2.8</td><td>5.3E-03</td><td>6.5E-03</td><td>BT20/MB468</td><td>CYTO</td></tr>
<tr><td>>hg38_21_+_8433172_8446622_RNA45SN1_45S_50ntFlanks@11320.11342.23|CCACGTCTCGTCGCGCGCGCGTC</td><td>>hg38_21_+_8433172_8446622_RNA45SN1_45S_50ntFlanks</td><td>11320</td><td>11342</td><td>23</td><td>CCACGTCTCGTCGCGCGCGCGTC</td><td>3753.9</td><td>-6.1</td><td>2.2</td><td>-2.8</td><td>5.6E-03</td><td>6.5E-03</td><td>BT20/MB468</td><td>NUC</td></tr>
<tr><td>>am_MI0000809|hsa-mir-151a&WithFlank&8|-|140732558|140732659@53.71.19&[MIMAT0000757&hsa-miR-151a-3p&offsets|0|-2;m-30&8|-|140732587|140732607&offsets|0|-2]|CTAGACTGAAGCTCCTTGA</td><td>>am_MI0000809|hsa-mir-151a&WithFlank&8|-|140732558|140732659</td><td>53</td><td>71</td><td>19</td><td>CTAGACTGAAGCTCCTTGA</td><td>1216.0</td><td>2.5</td><td>0.8</td><td>3.2</td><td>1.4E-03</td><td>6.5E-03</td><td>BT20/MB231</td><td>CYTO</td></tr>
<tr><td>>hg38_21_+_8433172_8446622_RNA45SN1_45S_50ntFlanks@12202.12220.19|CTAAGGCGAGCTCAGGGAG</td><td>>hg38_21_+_8433172_8446622_RNA45SN1_45S_50ntFlanks</td><td>12202</td><td>12220</td><td>19</td><td>CTAAGGCGAGCTCAGGGAG</td><td>176.2</td><td>-2.5</td><td>0.9</td><td>-2.8</td><td>5.4E-03</td><td>6.5E-03</td><td>MB231/MB468</td><td>CYTO</td></tr>
<tr><td>>trnaMT_ProTGG_MT_-_15956_16023@37.70.34|GCTAATGGTGGAGTTAAAGACTTTTTCTCTGACC</td><td>>trnaMT_ProTGG_MT_-_15956_16023</td><td>37</td><td>70</td><td>34</td><td>GCTAATGGTGGAGTTAAAGACTTTTTCTCTGACC</td><td>53.8</td><td>5.3</td><td>1.9</td><td>2.8</td><td>4.9E-03</td><td>6.5E-03</td><td>MB231/MB468</td><td>MP</td></tr>
<tr><td>>hg38_21_+_8433172_8446622_RNA45SN1_45S_50ntFlanks@8184.8203.20|CGAGGCCCAGCCCGTGGACG</td><td>>hg38_21_+_8433172_8446622_RNA45SN1_45S_50ntFlanks</td><td>8184</td><td>8203</td><td>20</td><td>CGAGGCCCAGCCCGTGGACG</td><td>127.4</td><td>-2.5</td><td>0.9</td><td>-3.0</td><td>3.1E-03</td><td>6.5E-03</td><td>BT20/MB468</td><td>MP</td></tr>
<tr><td>>hg38_21_+_8433172_8446622_RNA45SN1_45S_50ntFlanks@12420.12449.30|ATGTCGGCTCTTCCTATCATTGTGAAGCAG</td><td>>hg38_21_+_8433172_8446622_RNA45SN1_45S_50ntFlanks</td><td>12420</td><td>12449</td><td>30</td><td>ATGTCGGCTCTTCCTATCATTGTGAAGCAG</td><td>258.4</td><td>-2.4</td><td>0.8</td><td>-3.0</td><td>3.1E-03</td><td>6.5E-03</td><td>BT20/MB468</td><td>MP</td></tr>
<tr><td>>hg38_21_+_8433172_8446622_RNA45SN1_45S_50ntFlanks@10344.10372.29|ATTCAAACGAGAACTTTGAAGGCCGAAGT</td><td>>hg38_21_+_8433172_8446622_RNA45SN1_45S_50ntFlanks</td><td>10344</td><td>10372</td><td>29</td><td>ATTCAAACGAGAACTTTGAAGGCCGAAGT</td><td>63.5</td><td>4.9</td><td>1.7</td><td>3.0</td><td>3.1E-03</td><td>6.5E-03</td><td>BT20/MB468</td><td>MP</td></tr>
<tr><td>>hg38_21_+_8433172_8446622_RNA45SN1_45S_50ntFlanks@4385.4423.39|TTGGATCTTGGGAGCGGGCGGGCGGTCCGCCGCGAGGCG</td><td>>hg38_21_+_8433172_8446622_RNA45SN1_45S_50ntFlanks</td><td>4385</td><td>4423</td><td>39</td><td>TTGGATCTTGGGAGCGGGCGGGCGGTCCGCCGCGAGGCG</td><td>236.7</td><td>-2.1</td><td>0.8</td><td>-2.8</td><td>5.4E-03</td><td>6.5E-03</td><td>MB231/MB468</td><td>CYTO</td></tr>
<tr><td>>hg38_21_+_8433172_8446622_RNA45SN1_45S_50ntFlanks@12992.13011.20|TCCCTCGCTGCGATCTATTG</td><td>>hg38_21_+_8433172_8446622_RNA45SN1_45S_50ntFlanks</td><td>12992</td><td>13011</td><td>20</td><td>TCCCTCGCTGCGATCTATTG</td><td>242.8</td><td>-4.4</td><td>1.6</td><td>-2.8</td><td>5.3E-03</td><td>6.5E-03</td><td>BT20/MB468</td><td>CYTO</td></tr>
<tr><td>>hg38_21_+_8433172_8446622_RNA45SN1_45S_50ntFlanks@9501.9526.26|ATGGTGAACTATGCCTGGGCAGGGCG</td><td>>hg38_21_+_8433172_8446622_RNA45SN1_45S_50ntFlanks</td><td>9501</td><td>9526</td><td>26</td><td>ATGGTGAACTATGCCTGGGCAGGGCG</td><td>65.4</td><td>9.4</td><td>2.9</td><td>3.3</td><td>1.1E-03</td><td>6.5E-03</td><td>BT20/MB231</td><td>MP</td></tr>
<tr><td>>hg38_MT_+_1621_3279_RNR2_16S_50ntFlanks@441.459.19|CAGAACCCTCTAAATCCCC</td><td>>hg38_MT_+_1621_3279_RNR2_16S_50ntFlanks</td><td>441</td><td>459</td><td>19</td><td>CAGAACCCTCTAAATCCCC</td><td>99.3</td><td>5.1</td><td>1.9</td><td>2.8</td><td>5.7E-03</td><td>6.6E-03</td><td>MB231/MB468</td><td>MITO</td></tr>
<tr><td>>hg38_21_+_8433172_8446622_RNA45SN1_45S_50ntFlanks@12419.12443.25|GATGTCGGCTCTTCCTATCATTGTG</td><td>>hg38_21_+_8433172_8446622_RNA45SN1_45S_50ntFlanks</td><td>12419</td><td>12443</td><td>25</td><td>GATGTCGGCTCTTCCTATCATTGTG</td><td>3186.3</td><td>-2.7</td><td>0.8</td><td>-3.2</td><td>1.4E-03</td><td>6.6E-03</td><td>BT20/MB231</td><td>CYTO</td></tr>
<tr><td>>hg38_21_+_8433172_8446622_RNA45SN1_45S_50ntFlanks@9544.9566.23|GGTGGAGGTCCGTAGCGGTCCTG</td><td>>hg38_21_+_8433172_8446622_RNA45SN1_45S_50ntFlanks</td><td>9544</td><td>9566</td><td>23</td><td>GGTGGAGGTCCGTAGCGGTCCTG</td><td>176.2</td><td>-4.1</td><td>1.5</td><td>-2.8</td><td>5.3E-03</td><td>6.6E-03</td><td>BT20/MB468</td><td>CYTO</td></tr>
<tr><td>>hg38_21_+_8433172_8446622_RNA45SN1_45S_50ntFlanks@4068.4101.34|ACTTTCGATGGTAGTCGCCGTGCCTACCATGGTG</td><td>>hg38_21_+_8433172_8446622_RNA45SN1_45S_50ntFlanks</td><td>4068</td><td>4101</td><td>34</td><td>ACTTTCGATGGTAGTCGCCGTGCCTACCATGGTG</td><td>118.4</td><td>-1.9</td><td>0.7</td><td>-2.8</td><td>4.9E-03</td><td>6.6E-03</td><td>MB231/MB468</td><td>MP</td></tr>
<tr><td>>trnaMT_ThrTGT_MT_+_15888_15953@33.69.37|GTAAACCGGAGATGAAAACCTTTTTCCAAGGACACCA</td><td>>trnaMT_ThrTGT_MT_+_15888_15953</td><td>33</td><td>69</td><td>37</td><td>GTAAACCGGAGATGAAAACCTTTTTCCAAGGACACCA</td><td>104.9</td><td>4.4</td><td>1.6</td><td>2.8</td><td>4.9E-03</td><td>6.6E-03</td><td>MB231/MB468</td><td>MP</td></tr>
<tr><td>>hg38_MT_+_598_1651_RNR1_12S_50ntFlanks@697.730.34|AAAGTAAGCGCAAGTACCCACGTAAAGACGTTAG</td><td>>hg38_MT_+_598_1651_RNR1_12S_50ntFlanks</td><td>697</td><td>730</td><td>34</td><td>AAAGTAAGCGCAAGTACCCACGTAAAGACGTTAG</td><td>28.4</td><td>8.1</td><td>2.9</td><td>2.8</td><td>4.9E-03</td><td>6.6E-03</td><td>MB231/MB468</td><td>MP</td></tr>
<tr><td>>hg38_21_+_8433172_8446622_RNA45SN1_45S_50ntFlanks@8192.8213.22|AGCCCGTGGACGGTGTGAGGCC</td><td>>hg38_21_+_8433172_8446622_RNA45SN1_45S_50ntFlanks</td><td>8192</td><td>8213</td><td>22</td><td>AGCCCGTGGACGGTGTGAGGCC</td><td>321.0</td><td>-2.0</td><td>0.7</td><td>-2.8</td><td>5.7E-03</td><td>6.6E-03</td><td>MB231/MB468</td><td>NUC</td></tr>
</tbody>
</table>

<table>
<tr><td>>hg38_21_+_8433172_8446622_RNA45SN1_45S_50ntFlanks@9526.9573.48|GAAGCCAGAGGAAACTCTGGTGGAGGTCCGTAGCGGTCCTGACGTGCA</td><td>>hg38_21_+_8433172_8446622_RNA45SN1_45S_50ntFlanks</td><td>9526</td><td>9573</td><td>48</td><td>GAAGCCAGAGGAAACTCTGGTGGAGGTCCGTAGCGGTCCTGACGTGCA</td><td>321.1</td><td>-5.5</td><td>1.7</td><td>-3.2</td><td>1.4E-03</td><td>6.6E-03</td><td>BT20/MB231</td><td>CYTO</td></tr>
<tr><td>>hg38_21_+_8433172_8446622_RNA45SN1_45S_50ntFlanks@6649.6668.20|TACGACTCTTAGCGGTGGAT</td><td>>hg38_21_+_8433172_8446622_RNA45SN1_45S_50ntFlanks</td><td>6649</td><td>6668</td><td>20</td><td>TACGACTCTTAGCGGTGGAT</td><td>322.1</td><td>2.3</td><td>0.8</td><td>3.0</td><td>3.1E-03</td><td>6.6E-03</td><td>BT20/MB468</td><td>MP</td></tr>
<tr><td>>am_MI0000809|hsa-mir-151a&WithFlank&8|-|140732558|140732659@53.72.20(+1A)&[MIMAT0000757&hsa-miR-151a-3p&offsets|0|-1(+1A);m-30&8|-|140732587|140732607&offsets|0|-1(+1A)]|CTAGACTGAAGCTCCTTGAGA</td><td>>am_MI0000809|hsa-mir-151a&WithFlank&8|-|140732558|140732659</td><td>53</td><td>72</td><td>20(+1A)</td><td>CTAGACTGAAGCTCCTTGAGA</td><td>22719.1</td><td>2.4</td><td>0.8</td><td>3.2</td><td>1.4E-03</td><td>6.6E-03</td><td>BT20/MB231</td><td>CYTO</td></tr>
<tr><td>>hg38_21_+_8433172_8446622_RNA45SN1_45S_50ntFlanks@5473.5515.43|CGGCCCTGGCGGAGCGCTGAGAAGACGGTCGAACTTGACTATC</td><td>>hg38_21_+_8433172_8446622_RNA45SN1_45S_50ntFlanks</td><td>5473</td><td>5515</td><td>43</td><td>CGGCCCTGGCGGAGCGCTGAGAAGACGGTCGAACTTGACTATC</td><td>929.6</td><td>-3.0</td><td>0.9</td><td>-3.2</td><td>1.4E-03</td><td>6.6E-03</td><td>BT20/MB231</td><td>CYTO</td></tr>
<tr><td>>hg38_21_+_8433172_8446622_RNA45SN1_45S_50ntFlanks@12002.12029.28|TGCGGGCCGCCGGTGAAATACCACTACT</td><td>>hg38_21_+_8433172_8446622_RNA45SN1_45S_50ntFlanks</td><td>12002</td><td>12029</td><td>28</td><td>TGCGGGCCGCCGGTGAAATACCACTACT</td><td>378.5</td><td>-2.3</td><td>0.8</td><td>-2.8</td><td>5.7E-03</td><td>6.6E-03</td><td>MB231/MB468</td><td>NUC</td></tr>
<tr><td>>hg38_21_+_8433172_8446622_RNA45SN1_45S_50ntFlanks@4738.4759.22|AAAGTCGGAGGTTCGAAGACGA</td><td>>hg38_21_+_8433172_8446622_RNA45SN1_45S_50ntFlanks</td><td>4738</td><td>4759</td><td>22</td><td>AAAGTCGGAGGTTCGAAGACGA</td><td>305.1</td><td>-3.3</td><td>1.2</td><td>-2.8</td><td>5.7E-03</td><td>6.6E-03</td><td>MB231/MB468</td><td>NUC</td></tr>
<tr><td>>hg38_21_+_8433172_8446622_RNA45SN1_45S_50ntFlanks@9573.9595.23|AAATCGGTCGTCCGACCTGGGTA</td><td>>hg38_21_+_8433172_8446622_RNA45SN1_45S_50ntFlanks</td><td>9573</td><td>9595</td><td>23</td><td>AAATCGGTCGTCCGACCTGGGTA</td><td>281.4</td><td>-2.3</td><td>0.8</td><td>-2.8</td><td>5.7E-03</td><td>6.6E-03</td><td>MB231/MB468</td><td>NUC</td></tr>
<tr><td>>hg38_21_+_8433172_8446622_RNA45SN1_45S_50ntFlanks@8169.8195.27|CCAAGTCCTTCTGATCGAGGCCCAGCC</td><td>>hg38_21_+_8433172_8446622_RNA45SN1_45S_50ntFlanks</td><td>8169</td><td>8195</td><td>27</td><td>CCAAGTCCTTCTGATCGAGGCCCAGCC</td><td>212.6</td><td>-3.9</td><td>1.3</td><td>-3.0</td><td>3.1E-03</td><td>6.6E-03</td><td>BT20/MB468</td><td>MP</td></tr>
<tr><td>>hg38_21_+_8433172_8446622_RNA45SN1_45S_50ntFlanks@11603.11620.18|CCGACTGTTTAATTAAAA</td><td>>hg38_21_+_8433172_8446622_RNA45SN1_45S_50ntFlanks</td><td>11603</td><td>11620</td><td>18</td><td>CCGACTGTTTAATTAAAA</td><td>251.8</td><td>-2.5</td><td>0.9</td><td>-3.0</td><td>3.1E-03</td><td>6.6E-03</td><td>BT20/MB468</td><td>MP</td></tr>
<tr><td>>MI0000825|hsa-mir-345&WithFlank&14|+|100307853|100307962@24.45.22(+1U)&[MIMAT0000772&hsa-miR-345-5p&offsets|0|0(+1U);m-193&14|+|100307876|100307896&offsets|0|+1(+1U)]|GCTGACTCCTAGTCCAGGGCTCT</td><td>>MI0000825|hsa-mir-345&WithFlank&14|+|100307853|100307962</td><td>24</td><td>45</td><td>22(+1U)</td><td>GCTGACTCCTAGTCCAGGGCTCT</td><td>193.0</td><td>3.8</td><td>1.3</td><td>3.0</td><td>3.1E-03</td><td>6.6E-03</td><td>BT20/MB468</td><td>MP</td></tr>
<tr><td>>hg38_21_+_8433172_8446622_RNA45SN1_45S_50ntFlanks@475.497.23|CTTCGTGATCGATGTGGTGACGT</td><td>>hg38_21_+_8433172_8446622_RNA45SN1_45S_50ntFlanks</td><td>475</td><td>497</td><td>23</td><td>CTTCGTGATCGATGTGGTGACGT</td><td>2656.1</td><td>4.4</td><td>1.5</td><td>3.0</td><td>3.1E-03</td><td>6.6E-03</td><td>BT20/MB468</td><td>MP</td></tr>
<tr><td>>Antisense_hg38_21_-_8433172_8446622_RNA45SN1_45S_50ntFlanks@7123.7140.18|GCGGCGGCGGCGGCGGCG</td><td>>Antisense_hg38_21_-_8433172_8446622_RNA45SN1_45S_50ntFlanks</td><td>7123</td><td>7140</td><td>18</td><td>GCGGCGGCGGCGGCGGCG</td><td>150.0</td><td>-4.9</td><td>1.6</td><td>-3.0</td><td>3.1E-03</td><td>6.6E-03</td><td>BT20/MB468</td><td>MP</td></tr>
<tr><td>>hg38_21_+_8433172_8446622_RNA45SN1_45S_50ntFlanks@10127.10144.18|GCGGCGGCGGCGGCGGCG</td><td>>hg38_21_+_8433172_8446622_RNA45SN1_45S_50ntFlanks</td><td>10127</td><td>10144</td><td>18</td><td>GCGGCGGCGGCGGCGGCG</td><td>150.0</td><td>-4.9</td><td>1.6</td><td>-3.0</td><td>3.1E-03</td><td>6.6E-03</td><td>BT20/MB468</td><td>MP</td></tr>
<tr><td>>hg38_21_+_8433172_8446622_RNA45SN1_45S_50ntFlanks@10130.10147.18|GCGGCGGCGGCGGCGGCG</td><td>>hg38_21_+_8433172_8446622_RNA45SN1_45S_50ntFlanks</td><td>10130</td><td>10147</td><td>18</td><td>GCGGCGGCGGCGGCGGCG</td><td>150.0</td><td>-4.9</td><td>1.6</td><td>-3.0</td><td>3.1E-03</td><td>6.6E-03</td><td>BT20/MB468</td><td>MP</td></tr>
<tr><td>>hg38_21_+_8433172_8446622_RNA45SN1_45S_50ntFlanks@8823.8840.18|GCGGCGGCGGCGGCGGCG</td><td>>hg38_21_+_8433172_8446622_RNA45SN1_45S_50ntFlanks</td><td>8823</td><td>8840</td><td>18</td><td>GCGGCGGCGGCGGCGGCG</td><td>150.0</td><td>-4.9</td><td>1.6</td><td>-3.0</td><td>3.1E-03</td><td>6.6E-03</td><td>BT20/MB468</td><td>MP</td></tr>
<tr><td>>hg38_21_+_8433172_8446622_RNA45SN1_45S_50ntFlanks@8844.8861.18|GCGGCGGCGGCGGCGGCG</td><td>>hg38_21_+_8433172_8446622_RNA45SN1_45S_50ntFlanks</td><td>8844</td><td>8861</td><td>18</td><td>GCGGCGGCGGCGGCGGCG</td><td>150.0</td><td>-4.9</td><td>1.6</td><td>-3.0</td><td>3.1E-03</td><td>6.6E-03</td><td>BT20/MB468</td><td>MP</td></tr>
<tr><td>>hg38_21_+_8433172_8446622_RNA45SN1_45S_50ntFlanks@8847.8864.18|GCGGCGGCGGCGGCGGCG</td><td>>hg38_21_+_8433172_8446622_RNA45SN1_45S_50ntFlanks</td><td>8847</td><td>8864</td><td>18</td><td>GCGGCGGCGGCGGCGGCG</td><td>150.0</td><td>-4.9</td><td>1.6</td><td>-3.0</td><td>3.1E-03</td><td>6.6E-03</td><td>BT20/MB468</td><td>MP</td></tr>
<tr><td>>hg38_21_+_8433172_8446622_RNA45SN1_45S_50ntFlanks@4743.4765.23|CGGAGGTTCGAAGACGATCAGAT</td><td>>hg38_21_+_8433172_8446622_RNA45SN1_45S_50ntFlanks</td><td>4743</td><td>4765</td><td>23</td><td>CGGAGGTTCGAAGACGATCAGAT</td><td>108.7</td><td>5.0</td><td>1.7</td><td>3.0</td><td>3.1E-03</td><td>6.6E-03</td><td>BT20/MB468</td><td>MP</td></tr>
<tr><td>>trna14_LysTTT_11_-_59327808_59327880@1.36.36,trna2_LysTTT_17_+_8022473_8022545@1.36.36,trna54_LysTTT_1_+_204475655_204475727@1.36.36,trna5_LysTTT_11_+_59323902_59323974@1.36.36,trna62_LysTTT_1_-_204476158_204476230@1.36.36,trna76_LysTTT_6_+_28918806_28918878@1.36.36|GCCCGGATAGCTCAGTCGGTAGAGCATCAGACTTTT</td><td>>trna14_LysTTT_11_-_59327808_59327880</td><td>1</td><td>36</td><td>36</td><td>GCCCGGATAGCTCAGTCGGTAGAGCATCAGACTTTT</td><td>281.0</td><td>4.6</td><td>1.7</td><td>2.8</td><td>5.7E-03</td><td>6.6E-03</td><td>MB231/MB468</td><td>MITO</td></tr>
<tr><td>>trnaMT_GluTTC_MT_-_14674_14742@34.72.39,trnalookalike8_GluTTC_5_-_93905172_93905240@34.72.39|ATATCATTGGTCGTGGTTGTAGTCCGTGCGAGAATACCA</td><td>>trnaMT_GluTTC_MT_-_14674_14742</td><td>34</td><td>72</td><td>39</td><td>ATATCATTGGTCGTGGTTGTAGTCCGTGCGAGAATACCA</td><td>2937.1</td><td>-2.9</td><td>0.9</td><td>-3.2</td><td>1.4E-03</td><td>6.6E-03</td><td>BT20/MB231</td><td>CYTO</td></tr>
<tr><td>>hg38_21_+_8433172_8446622_RNA45SN1_45S_50ntFlanks@6651.6668.18|CGACTCTTAGCGGTGGAT</td><td>>hg38_21_+_8433172_8446622_RNA45SN1_45S_50ntFlanks</td><td>6651</td><td>6668</td><td>18</td><td>CGACTCTTAGCGGTGGAT</td><td>9493.9</td><td>2.2</td><td>0.7</td><td>3.2</td><td>1.4E-03</td><td>6.6E-03</td><td>BT20/MB231</td><td>CYTO</td></tr>
<tr><td>>hg38_21_+_8433172_8446622_RNA45SN1_45S_50ntFlanks@4684.4726.43|AAGACGGACCAGAGCGAAAGCATTTGCCAAGAATGTTTTCATT</td><td>>hg38_21_+_8433172_8446622_RNA45SN1_45S_50ntFlanks</td><td>4684</td><td>4726</td><td>43</td><td>AAGACGGACCAGAGCGAAAGCATTTGCCAAGAATGTTTTCATT</td><td>459.3</td><td>-3.2</td><td>1.0</td><td>-3.2</td><td>1.4E-03</td><td>6.6E-03</td><td>BT20/MB231</td><td>CYTO</td></tr>
<tr><td>>hg38_21_+_8433172_8446622_RNA45SN1_45S_50ntFlanks@6722.6770.49|ATTGCAGGACACATTGATCATCGACACTTCGAACGCACTTGCGGCCCCG</td><td>>hg38_21_+_8433172_8446622_RNA45SN1_45S_50ntFlanks</td><td>6722</td><td>6770</td><td>49</td><td>ATTGCAGGACACATTGATCATCGACACTTCGAACGCACTTGCGGCCCCG</td><td>67.9</td><td>-9.2</td><td>3.1</td><td>-3.0</td><td>3.1E-03</td><td>6.6E-03</td><td>BT20/MB468</td><td>MP</td></tr>
<tr><td>>am_trnaMT_ValTAC_MT_+_1602_1670@29.52.24|ACTTACACTTAGGAGATTTCAACT</td><td>>am_trnaMT_ValTAC_MT_+_1602_1670</td><td>29</td><td>52</td><td>24</td><td>ACTTACACTTAGGAGATTTCAACT</td><td>435.9</td><td>2.7</td><td>0.8</td><td>3.2</td><td>1.4E-03</td><td>6.6E-03</td><td>BT20/MB231</td><td>CYTO</td></tr>
<tr><td>>hg38_21_+_8433172_8446622_RNA45SN1_45S_50ntFlanks@6661.6689.29|CGGTGGATCACTCGGCTCGTGCGTCGATG</td><td>>hg38_21_+_8433172_8446622_RNA45SN1_45S_50ntFlanks</td><td>6661</td><td>6689</td><td>29</td><td>CGGTGGATCACTCGGCTCGTGCGTCGATG</td><td>3772.7</td><td>2.3</td><td>0.7</td><td>3.2</td><td>1.2E-03</td><td>6.6E-03</td><td>BT20/MB231</td><td>NUC</td></tr>
<tr><td>>hg38_MT_+_1621_3279_RNR2_16S_50ntFlanks@621.652.32|ATATAACTGAACTCCTCACACCCAATTGGACC</td><td>>hg38_MT_+_1621_3279_RNR2_16S_50ntFlanks</td><td>621</td><td>652</td><td>32</td><td>ATATAACTGAACTCCTCACACCCAATTGGACC</td><td>79.2</td><td>4.5</td><td>1.6</td><td>2.8</td><td>4.9E-03</td><td>6.6E-03</td><td>MB231/MB468</td><td>MP</td></tr>
<tr><td>>MI0000487|hsa-mir-193a&WithFlank&17|+|31559990|31560089@27.46.20(+1A)&[MIMAT0004614&hsa-miR-193a-5p&offsets|0|-2(+1A);m-123&17|+|31560016|31560037&offsets|0|-2(+1A)]|TGGGTCTTTGCGGGCGAGATA</td><td>>MI0000487|hsa-mir-193a&WithFlank&17|+|31559990|31560089</td><td>27</td><td>46</td><td>20(+1A)</td><td>TGGGTCTTTGCGGGCGAGATA</td><td>115.9</td><td>5.9</td><td>1.8</td><td>3.3</td><td>1.1E-03</td><td>6.6E-03</td><td>BT20/MB231</td><td>MP</td></tr>
<tr><td>>hg38_21_+_8433172_8446622_RNA45SN1_45S_50ntFlanks@3872.3898.27|CTAATACATGCCGACGGGCGCTGACCC</td><td>>hg38_21_+_8433172_8446622_RNA45SN1_45S_50ntFlanks</td><td>3872</td><td>3898</td><td>27</td><td>CTAATACATGCCGACGGGCGCTGACCC</td><td>333.8</td><td>-5.0</td><td>1.7</td><td>-3.0</td><td>3.1E-03</td><td>6.6E-03</td><td>BT20/MB468</td><td>MP</td></tr>
<tr><td>>hg38_MT_+_1621_3279_RNR2_16S_50ntFlanks@1427.1453.27|GATTAAAGTCCTACGTGATCTGAGTTC</td><td>>hg38_MT_+_1621_3279_RNR2_16S_50ntFlanks</td><td>1427</td><td>1453</td><td>27</td><td>GATTAAAGTCCTACGTGATCTGAGTTC</td><td>182.7</td><td>3.8</td><td>1.3</td><td>3.0</td><td>3.1E-03</td><td>6.6E-03</td><td>BT20/MB468</td><td>MP</td></tr>
<tr><td>>hg38_21_+_8433172_8446622_RNA45SN1_45S_50ntFlanks@11866.11893.28|GCGGGGAAAGAAGACCCTGTTGAGCTTG</td><td>>hg38_21_+_8433172_8446622_RNA45SN1_45S_50ntFlanks</td><td>11866</td><td>11893</td><td>28</td><td>GCGGGGAAAGAAGACCCTGTTGAGCTTG</td><td>64.8</td><td>9.3</td><td>2.9</td><td>3.3</td><td>1.1E-03</td><td>6.6E-03</td><td>BT20/MB231</td><td>MP</td></tr>
<tr><td>>hg38_21_+_8433172_8446622_RNA45SN1_45S_50ntFlanks@7976.8008.33|GCGACCTCAGATCAGACGTGGCGACCCGCTGAA</td><td>>hg38_21_+_8433172_8446622_RNA45SN1_45S_50ntFlanks</td><td>7976</td><td>8008</td><td>33</td><td>GCGACCTCAGATCAGACGTGGCGACCCGCTGAA</td><td>697.4</td><td>3.9</td><td>1.3</td><td>3.0</td><td>3.1E-03</td><td>6.6E-03</td><td>BT20/MB468</td><td>MP</td></tr>
<tr><td>>hg38_21_+_8433172_8446622_RNA45SN1_45S_50ntFlanks@8192.8215.24|AGCCCGTGGACGGTGTGAGGCCGG</td><td>>hg38_21_+_8433172_8446622_RNA45SN1_45S_50ntFlanks</td><td>8192</td><td>8215</td><td>24</td><td>AGCCCGTGGACGGTGTGAGGCCGG</td><td>154.9</td><td>-3.9</td><td>1.3</td><td>-3.0</td><td>3.1E-03</td><td>6.6E-03</td><td>BT20/MB468</td><td>MP</td></tr>
<tr><td>>hg38_21_+_8433172_8446622_RNA45SN1_45S_50ntFlanks@6653.6673.21|ACTCTTAGCGGTGGATCACTC</td><td>>hg38_21_+_8433172_8446622_RNA45SN1_45S_50ntFlanks</td><td>6653</td><td>6673</td><td>21</td><td>ACTCTTAGCGGTGGATCACTC</td><td>63.5</td><td>9.3</td><td>2.9</td><td>3.3</td><td>1.1E-03</td><td>6.6E-03</td><td>BT20/MB231</td><td>MP</td></tr>
<tr><td>>trnaMT_GluTTC_MT_-_14674_14742@34.69.36,trnalookalike8_GluTTC_5_-_93905172_93905240@34.69.36|ATATCATTGGTCGTGGTTGTAGTCCGTGCGAGAATA</td><td>>trnaMT_GluTTC_MT_-_14674_14742</td><td>34</td><td>69</td><td>36</td><td>ATATCATTGGTCGTGGTTGTAGTCCGTGCGAGAATA</td><td>83.2</td><td>5.3</td><td>1.8</td><td>3.0</td><td>3.1E-03</td><td>6.6E-03</td><td>BT20/MB468</td><td>MP</td></tr>
<tr><td>>MI0000825|hsa-mir-345&WithFlank&14|+|100307853|100307962@24.44.21(+1U)&[MIMAT0000772&hsa-miR-345-5p&offsets|0|-1(+1U);m-193&14|+|100307876|100307896&offsets|0|0(+1U)]|GCTGACTCCTAGTCCAGGGCTT</td><td>>MI0000825|hsa-mir-345&WithFlank&14|+|100307853|100307962</td><td>24</td><td>44</td><td>21(+1U)</td><td>GCTGACTCCTAGTCCAGGGCTT</td><td>159.4</td><td>4.6</td><td>1.6</td><td>2.8</td><td>4.9E-03</td><td>6.6E-03</td><td>MB231/MB468</td><td>MP</td></tr>
<tr><td>>MI0000272|hsa-mir-182&WithFlank&7|-|129770377|129770498@29.54.26(+1U)&[MIMAT0000259&hsa-miR-182-5p&offsets|0|+2(+1U);m-22&7|-|129770449|129770470&offsets|0|+4(+1U)]|TTTGGCAATGGTAGAACTCACACTGGT</td><td>>MI0000272|hsa-mir-182&WithFlank&7|-|129770377|129770498</td><td>29</td><td>54</td><td>26(+1U)</td><td>TTTGGCAATGGTAGAACTCACACTGGT</td><td>630.9</td><td>1.9</td><td>0.6</td><td>3.2</td><td>1.2E-03</td><td>6.6E-03</td><td>BT20/MB231</td><td>NUC</td></tr>
<tr><td>>trnaMT_GlnTTG_MT_-_4329_4400@1.35.35,trnalookalike2_GlnTTG_1_-_564879_564950@1.35.35|TAGGATGGGGTGTGATAGGTGGCACGGAGAATTTT</td><td>>trnaMT_GlnTTG_MT_-_4329_4400</td><td>1</td><td>35</td><td>35</td><td>TAGGATGGGGTGTGATAGGTGGCACGGAGAATTTT</td><td>176.0</td><td>4.5</td><td>1.5</td><td>3.0</td><td>3.1E-03</td><td>6.6E-03</td><td>BT20/MB468</td><td>MP</td></tr>
<tr><td>>hg38_21_+_8433172_8446622_RNA45SN1_45S_50ntFlanks@12202.12232.31|CTAAGGCGAGCTCAGGGAGGACAGAAACCTC</td><td>>hg38_21_+_8433172_8446622_RNA45SN1_45S_50ntFlanks</td><td>12202</td><td>12232</td><td>31</td><td>CTAAGGCGAGCTCAGGGAGGACAGAAACCTC</td><td>312.9</td><td>-3.9</td><td>1.4</td><td>-2.8</td><td>4.6E-03</td><td>6.6E-03</td><td>BT20/MB468</td><td>MITO</td></tr>
<tr><td>>trnaMT_PheGAA_MT_+_577_647@42.74.33|TTTAGACGGGCTCACATCACCCCATAAACACCA</td><td>>trnaMT_PheGAA_MT_+_577_647</td><td>42</td><td>74</td><td>33</td><td>TTTAGACGGGCTCACATCACCCCATAAACACCA</td><td>64.4</td><td>4.4</td><td>1.6</td><td>2.8</td><td>5.7E-03</td><td>6.6E-03</td><td>BT20/MB468</td><td>NUC</td></tr>
<tr><td>>hg38_MT_+_1621_3279_RNR2_16S_50ntFlanks@285.310.26|CGAAACCAGACGAGCTACCTAAGAAC</td><td>>hg38_MT_+_1621_3279_RNR2_16S_50ntFlanks</td><td>285</td><td>310</td><td>26</td><td>CGAAACCAGACGAGCTACCTAAGAAC</td><td>141.3</td><td>5.2</td><td>1.8</td><td>3.0</td><td>3.1E-03</td><td>6.6E-03</td><td>BT20/MB468</td><td>MP</td></tr>
<tr><td>>hg38_21_+_8433172_8446622_RNA45SN1_45S_50ntFlanks@10392.10424.33|ACAGCAGTTGAACATGGGTCAGTCGGTCCTGAG</td><td>>hg38_21_+_8433172_8446622_RNA45SN1_45S_50ntFlanks</td><td>10392</td><td>10424</td><td>33</td><td>ACAGCAGTTGAACATGGGTCAGTCGGTCCTGAG</td><td>4527.4</td><td>-2.6</td><td>1.0</td><td>-2.8</td><td>5.8E-03</td><td>6.6E-03</td><td>MB231/MB468</td><td>MITO</td></tr>
<tr><td>>trnaMT_PheGAA_MT_+_577_647@30.57.28|ACACTGAAAATGTTTAGACGGGCTCACA</td><td>>trnaMT_PheGAA_MT_+_577_647</td><td>30</td><td>57</td><td>28</td><td>ACACTGAAAATGTTTAGACGGGCTCACA</td><td>48.5</td><td>6.0</td><td>2.0</td><td>3.0</td><td>3.1E-03</td><td>6.6E-03</td><td>BT20/MB468</td><td>MP</td></tr>
<tr><td>>hg38_21_+_8433172_8446622_RNA45SN1_45S_50ntFlanks@11569.11597.29|CAGCCGACTTAGAACTGGTGCGGACCAGG</td><td>>hg38_21_+_8433172_8446622_RNA45SN1_45S_50ntFlanks</td><td>11569</td><td>11597</td><td>29</td><td>CAGCCGACTTAGAACTGGTGCGGACCAGG</td><td>437.5</td><td>-2.7</td><td>1.0</td><td>-2.8</td><td>5.5E-03</td><td>6.6E-03</td><td>MB231/MB468</td><td>CYTO</td></tr>
<tr><td>>trnaMT_HisGTG_MT_+_12138_12206@28.59.32|ATTGTGAATCTGACAACAGAGGCTTACGACCC</td><td>>trnaMT_HisGTG_MT_+_12138_12206</td><td>28</td><td>59</td><td>32</td><td>ATTGTGAATCTGACAACAGAGGCTTACGACCC</td><td>128.0</td><td>2.9</td><td>1.0</td><td>2.8</td><td>4.9E-03</td><td>6.6E-03</td><td>MB231/MB468</td><td>MP</td></tr>
<tr><td>>hg38_21_+_8433172_8446622_RNA45SN1_45S_50ntFlanks@10766.10785.20|CGCAGCAGGTCTCCAAGGTG</td><td>>hg38_21_+_8433172_8446622_RNA45SN1_45S_50ntFlanks</td><td>10766</td><td>10785</td><td>20</td><td>CGCAGCAGGTCTCCAAGGTG</td><td>222.8</td><td>-4.9</td><td>1.7</td><td>-3.0</td><td>3.1E-03</td><td>6.6E-03</td><td>BT20/MB468</td><td>MP</td></tr>
<tr><td>>hg38_21_+_8433172_8446622_RNA45SN1_45S_50ntFlanks@8194.8225.32|CCCGTGGACGGTGTGAGGCCGGTAGCGGCCCC</td><td>>hg38_21_+_8433172_8446622_RNA45SN1_45S_50ntFlanks</td><td>8194</td><td>8225</td><td>32</td><td>CCCGTGGACGGTGTGAGGCCGGTAGCGGCCCC</td><td>150.2</td><td>-4.9</td><td>1.6</td><td>-3.0</td><td>3.1E-03</td><td>6.6E-03</td><td>BT20/MB468</td><td>MP</td></tr>
<tr><td>>trnaMT_GluTTC_MT_-_14674_14742@53.72.20,trnalookalike8_GluTTC_5_-_93905172_93905240@53.72.20|TAGTCCGTGCGAGAATACCA</td><td>>trnaMT_GluTTC_MT_-_14674_14742</td><td>53</td><td>72</td><td>20</td><td>TAGTCCGTGCGAGAATACCA</td><td>138.3</td><td>-1.8</td><td>0.7</td><td>-2.8</td><td>5.0E-03</td><td>6.7E-03</td><td>MB231/MB468</td><td>MP</td></tr>
<tr><td>>hg38_21_+_8433172_8446622_RNA45SN1_45S_50ntFlanks@12419.12446.28|GATGTCGGCTCTTCCTATCATTGTGAAG</td><td>>hg38_21_+_8433172_8446622_RNA45SN1_45S_50ntFlanks</td><td>12419</td><td>12446</td><td>28</td><td>GATGTCGGCTCTTCCTATCATTGTGAAG</td><td>166.5</td><td>-2.1</td><td>0.7</td><td>-2.8</td><td>5.0E-03</td><td>6.7E-03</td><td>MB231/MB468</td><td>MP</td></tr>
<tr><td>>trnaMT_PheGAA_MT_+_577_647@43.74.32|TTAGACGGGCTCACATCACCCCATAAACACCA</td><td>>trnaMT_PheGAA_MT_+_577_647</td><td>43</td><td>74</td><td>32</td><td>TTAGACGGGCTCACATCACCCCATAAACACCA</td><td>102.4</td><td>4.8</td><td>1.8</td><td>2.8</td><td>5.7E-03</td><td>6.7E-03</td><td>BT20/MB468</td><td>NUC</td></tr>
<tr><td>>hg38_21_+_8433172_8446622_RNA45SN1_45S_50ntFlanks@10375.10407.33|AGAAGGGTTCCATGTGAACAGCAGTTGAACATG</td><td>>hg38_21_+_8433172_8446622_RNA45SN1_45S_50ntFlanks</td><td>10375</td><td>10407</td><td>33</td><td>AGAAGGGTTCCATGTGAACAGCAGTTGAACATG</td><td>186.9</td><td>-4.9</td><td>1.7</td><td>-3.0</td><td>3.1E-03</td><td>6.7E-03</td><td>BT20/MB468</td><td>MP</td></tr>
<tr><td>>hg38_21_+_8433172_8446622_RNA45SN1_45S_50ntFlanks@12082.12107.26|TCTCGCTTCTGGCGCCAAGCGCCCGG</td><td>>hg38_21_+_8433172_8446622_RNA45SN1_45S_50ntFlanks</td><td>12082</td><td>12107</td><td>26</td><td>TCTCGCTTCTGGCGCCAAGCGCCCGG</td><td>66.7</td><td>-4.7</td><td>1.6</td><td>-3.0</td><td>3.1E-03</td><td>6.7E-03</td><td>BT20/MB468</td><td>MP</td></tr>
<tr><td>>trnaMT_GluTTC_MT_-_14674_14742@39.60.22,trnalookalike8_GluTTC_5_-_93905172_93905240@39.60.22|ATTGGTCGTGGTTGTAGTCCGT</td><td>>trnaMT_GluTTC_MT_-_14674_14742</td><td>39</td><td>60</td><td>22</td><td>ATTGGTCGTGGTTGTAGTCCGT</td><td>112.3</td><td>4.9</td><td>1.7</td><td>3.0</td><td>3.1E-03</td><td>6.7E-03</td><td>BT20/MB468</td><td>MP</td></tr>
</table>

<table>
<tr><td>>hg38_21_+_8433172_8446622_RNA45SN1_45S_50ntFlanks@9955.9978.24|GGAAGTCGGAATCCGCTAAGGAGT</td><td>>hg38_21_+_8433172_8446622_RNA45SN1_45S_50ntFlanks</td><td>9955</td><td>9978</td><td>24</td><td>GGAAGTCGGAATCCGCTAAGGAGT</td><td>105.9</td><td>5.0</td><td>1.7</td><td>3.0</td><td>3.1E-03</td><td>6.7E-03</td><td>BT20/MB468</td><td>MP</td></tr>
<tr><td>>trna11_GluTTC_15_-_26327381_26327452@1.30.30,trna3_GluTTC_13_-_45492062_45492133@1.30.30|TCCCACATGGTCTAGCGGTTAGGATTCCTG</td><td>>trna11_GluTTC_15_-_26327381_26327452</td><td>1</td><td>30</td><td>30</td><td>TCCCACATGGTCTAGCGGTTAGGATTCCTG</td><td>2634.2</td><td>-2.5</td><td>0.9</td><td>-2.8</td><td>5.5E-03</td><td>6.7E-03</td><td>MB231/MB468</td><td>CYTO</td></tr>
<tr><td>>hg38_21_+_8433172_8446622_RNA45SN1_45S_50ntFlanks@9561.9597.37|GTCCTGACGTGCAAATCGGTCGTCCGACCTGGGTATA</td><td>>hg38_21_+_8433172_8446622_RNA45SN1_45S_50ntFlanks</td><td>9561</td><td>9597</td><td>37</td><td>GTCCTGACGTGCAAATCGGTCGTCCGACCTGGGTATA</td><td>383.9</td><td>-2.2</td><td>0.8</td><td>-2.8</td><td>5.8E-03</td><td>6.7E-03</td><td>MB231/MB468</td><td>NUC</td></tr>
<tr><td>>MI0000085|hsa-mir-27a&WithFlank&19|-|13836434|13836523@57.75.19&[MIMAT0000084&hsa-miR-27a-3p&offsets|0|-2;m-43&19|-|13836447|13836467&offsets|0|-2]|TTCACAGTGGCTAAGTTCC</td><td>>MI0000085|hsa-mir-27a&WithFlank&19|-|13836434|13836523</td><td>57</td><td>75</td><td>19</td><td>TTCACAGTGGCTAAGTTCC</td><td>142.4</td><td>1.4</td><td>0.5</td><td>2.8</td><td>5.7E-03</td><td>6.7E-03</td><td>BT20/MB468</td><td>NUC</td></tr>
<tr><td>>trna111_HisGTG_1_-_147774845_147774916@-1T.26.27,trna118_HisGTG_1_-_145396881_145396952@-1T.26.27,trna16_HisGTG_1_+_146544773_146544844@-1T.26.27,trna1_HisGTG_15_+_45493349_45493420@-1T.26.27,trna21_HisGTG_1_+_147753471_147753542@-1T.26.27,trna33_HisGTG_6_+_27125906_27125977@-1T.26.27,trna7_HisGTG_9_-_14433938_14434009@-1T.26.27,trna8_HisGTG_15_-_45492611_45492682@-1T.26.27,trna9_HisGTG_15_-_45490804_45490875@-1T.26.27|TGCCGTGATCGTATAGTGGTTAGTACT</td><td>>trna111_HisGTG_1_-_147774845_147774916</td><td>-1T</td><td>26</td><td>27</td><td>TGCCGTGATCGTATAGTGGTTAGTACT</td><td>4133.8</td><td>-3.0</td><td>0.9</td><td>-3.2</td><td>1.5E-03</td><td>6.7E-03</td><td>BT20/MB231</td><td>MITO</td></tr>
<tr><td>>hg38_1_-_228634819_228635039_RNA5S12_5S_50ntFlanks@150.169.20|AATACCGGGTGCTGTAGGCT</td><td>>hg38_1_-_228634819_228635039_RNA5S12_5S_50ntFlanks</td><td>150</td><td>169</td><td>20</td><td>AATACCGGGTGCTGTAGGCT</td><td>295.7</td><td>2.2</td><td>0.7</td><td>3.2</td><td>1.5E-03</td><td>6.7E-03</td><td>BT20/MB231</td><td>MITO</td></tr>
<tr><td>>hg38_21_+_8433172_8446622_RNA45SN1_45S_50ntFlanks@476.504.29|TTCGTGATCGATGTGGTGACGTCGTGCTC</td><td>>hg38_21_+_8433172_8446622_RNA45SN1_45S_50ntFlanks</td><td>476</td><td>504</td><td>29</td><td>TTCGTGATCGATGTGGTGACGTCGTGCTC</td><td>270.6</td><td>5.2</td><td>1.8</td><td>3.0</td><td>3.1E-03</td><td>6.7E-03</td><td>BT20/MB468</td><td>MP</td></tr>
<tr><td>>hg38_21_+_8433172_8446622_RNA45SN1_45S_50ntFlanks@11594.11613.20|CAGGGGAATCCGACTGTTTA</td><td>>hg38_21_+_8433172_8446622_RNA45SN1_45S_50ntFlanks</td><td>11594</td><td>11613</td><td>20</td><td>CAGGGGAATCCGACTGTTTA</td><td>491.1</td><td>-1.9</td><td>0.7</td><td>-2.8</td><td>4.6E-03</td><td>6.7E-03</td><td>BT20/MB468</td><td>MITO</td></tr>
<tr><td>>hg38_21_+_8433172_8446622_RNA45SN1_45S_50ntFlanks@11718.11736.19|AACGGCGGGAGTAACTATG</td><td>>hg38_21_+_8433172_8446622_RNA45SN1_45S_50ntFlanks</td><td>11718</td><td>11736</td><td>19</td><td>AACGGCGGGAGTAACTATG</td><td>561.6</td><td>-4.1</td><td>1.4</td><td>-3.0</td><td>3.1E-03</td><td>6.7E-03</td><td>BT20/MB468</td><td>MP</td></tr>
<tr><td>>hg38_21_+_8433172_8446622_RNA45SN1_45S_50ntFlanks@12481.12498.18|AATAGGGAACGTGAGCTG</td><td>>hg38_21_+_8433172_8446622_RNA45SN1_45S_50ntFlanks</td><td>12481</td><td>12498</td><td>18</td><td>AATAGGGAACGTGAGCTG</td><td>903.4</td><td>-3.2</td><td>1.2</td><td>-2.8</td><td>5.7E-03</td><td>6.7E-03</td><td>BT20/MB468</td><td>NUC</td></tr>
<tr><td>>hg38_21_+_8433172_8446622_RNA45SN1_45S_50ntFlanks@12479.12498.20|CTAATAGGGAACGTGAGCTG</td><td>>hg38_21_+_8433172_8446622_RNA45SN1_45S_50ntFlanks</td><td>12479</td><td>12498</td><td>20</td><td>CTAATAGGGAACGTGAGCTG</td><td>1276.9</td><td>-1.8</td><td>0.6</td><td>-2.8</td><td>5.4E-03</td><td>6.7E-03</td><td>BT20/MB468</td><td>CYTO</td></tr>
<tr><td>>MI0000065|hsa-let-7d&WithFlank&9|+|94178828|94178926@14.35.22(+1A)&[MIMAT0000065&hsa-let-7d-5p&offsets|0|0(+1A);m-90&9|+|94178841|94178862&offsets|0|0(+1A)]|AGAGGTAGTAGGTTGCATAGTTA</td><td>>MI0000065|hsa-let-7d&WithFlank&9|+|94178828|94178926</td><td>14</td><td>35</td><td>22(+1A)</td><td>AGAGGTAGTAGGTTGCATAGTTA</td><td>193.6</td><td>3.7</td><td>1.3</td><td>2.8</td><td>5.0E-03</td><td>6.7E-03</td><td>MB231/MB468</td><td>MP</td></tr>
<tr><td>>hg38_21_+_8433172_8446622_RNA45SN1_45S_50ntFlanks@10362.10379.18|AAGGCCGAAGTGGAGAAG</td><td>>hg38_21_+_8433172_8446622_RNA45SN1_45S_50ntFlanks</td><td>10362</td><td>10379</td><td>18</td><td>AAGGCCGAAGTGGAGAAG</td><td>179.9</td><td>-2.1</td><td>0.7</td><td>-2.8</td><td>5.4E-03</td><td>6.7E-03</td><td>BT20/MB468</td><td>CYTO</td></tr>
<tr><td>>MI0000440|hsa-mir-27b&WithFlank&9|+|95085439|95085547@67.88.22(+1A)&[MIMAT0000419&hsa-miR-27b-3p&offsets|0|+1(+1A);m-39&9|+|95085505|95085525&offsets|0|+1(+1A)]|TTCACAGTGGCTAAGTTCTGCAA</td><td>>MI0000440|hsa-mir-27b&WithFlank&9|+|95085439|95085547</td><td>67</td><td>88</td><td>22(+1A)</td><td>TTCACAGTGGCTAAGTTCTGCAA</td><td>146.9</td><td>2.4</td><td>0.9</td><td>2.8</td><td>5.0E-03</td><td>6.7E-03</td><td>MB231/MB468</td><td>MP</td></tr>
<tr><td>>hg38_21_+_8433172_8446622_RNA45SN1_45S_50ntFlanks@11591.11609.19|GACCAGGGGAATCCGACTG</td><td>>hg38_21_+_8433172_8446622_RNA45SN1_45S_50ntFlanks</td><td>11591</td><td>11609</td><td>19</td><td>GACCAGGGGAATCCGACTG</td><td>48.6</td><td>-4.8</td><td>1.6</td><td>-3.0</td><td>3.1E-03</td><td>6.7E-03</td><td>BT20/MB468</td><td>MP</td></tr>
<tr><td>>MI0000066|hsa-let-7e&WithFlank&19|+|51692780|51692870@14.35.22(+1A)&[MIMAT0000066&hsa-let-7e-5p&offsets|0|0(+1A);m-50&19|+|51692793|51692814&offsets|0|0(+1A)]|TGAGGTAGGAGGTTGTATAGTTA</td><td>>MI0000066|hsa-let-7e&WithFlank&19|+|51692780|51692870</td><td>14</td><td>35</td><td>22(+1A)</td><td>TGAGGTAGGAGGTTGTATAGTTA</td><td>206.4</td><td>4.1</td><td>1.4</td><td>3.0</td><td>3.1E-03</td><td>6.7E-03</td><td>BT20/MB468</td><td>MP</td></tr>
<tr><td>>hg38_21_+_8433172_8446622_RNA45SN1_45S_50ntFlanks@11778.11806.29|ACGCGCATGAATGGATGAACGAGATTCCC</td><td>>hg38_21_+_8433172_8446622_RNA45SN1_45S_50ntFlanks</td><td>11778</td><td>11806</td><td>29</td><td>ACGCGCATGAATGGATGAACGAGATTCCC</td><td>168.1</td><td>-4.0</td><td>1.5</td><td>-2.8</td><td>5.4E-03</td><td>6.7E-03</td><td>BT20/MB468</td><td>CYTO</td></tr>
<tr><td>>MI0000272|hsa-mir-182&WithFlank&7|-|129770377|129770498@29.54.26(+1U)&[MIMAT0000259&hsa-miR-182-5p&offsets|0|+2(+1U);m-22&7|-|129770449|129770470&offsets|0|+4(+1U)]|TTTGGCAATGGTAGAACTCACACTGGT</td><td>>MI0000272|hsa-mir-182&WithFlank&7|-|129770377|129770498</td><td>29</td><td>54</td><td>26(+1U)</td><td>TTTGGCAATGGTAGAACTCACACTGGT</td><td>895.3</td><td>2.4</td><td>0.8</td><td>3.2</td><td>1.4E-03</td><td>6.7E-03</td><td>BT20/MB231</td><td>CYTO</td></tr>
<tr><td>>hg38_21_+_8433172_8446622_RNA45SN1_45S_50ntFlanks@12445.12480.36|AGCAGAATTCACCAAGCGTTGGATTGTTCACCCACT</td><td>>hg38_21_+_8433172_8446622_RNA45SN1_45S_50ntFlanks</td><td>12445</td><td>12480</td><td>36</td><td>AGCAGAATTCACCAAGCGTTGGATTGTTCACCCACT</td><td>324.9</td><td>2.0</td><td>0.7</td><td>2.8</td><td>5.5E-03</td><td>6.7E-03</td><td>MB231/MB468</td><td>CYTO</td></tr>
<tr><td>>MI0000108|hsa-mir-103a-2&WithFlank&20|+|3917488|3917577@54.76.23(+1U);MI0000109|hsa-mir-103a-1&WithFlank&5|-|168560890|168560979@54.76.23(+1U)&[MIMAT0000101&hsa-miR-103a-3p&offsets|0|0(+1U);m-12&20|+|3917541|3917563&offsets|0|0(+1U)];[MIMAT0000101_1&hsa-miR-103a-3p&offsets|0|0(+1U);m-13&5|-|168560904|168560926&offsets|0|0(+1U);m-44220&5|-|168560908|168560926&offsets|0|+4(+1U)]|AGCAGCATTGTACAGGGCTATGAT</td><td>>MI0000108|hsa-mir-103a-2&WithFlank&20|+|3917488|3917577</td><td>54</td><td>76</td><td>23(+1U)</td><td>AGCAGCATTGTACAGGGCTATGAT</td><td>5908.7</td><td>-2.6</td><td>0.8</td><td>-3.2</td><td>1.5E-03</td><td>6.7E-03</td><td>BT20/MB231</td><td>MITO</td></tr>
<tr><td>>hg38_21_+_8433172_8446622_RNA45SN1_45S_50ntFlanks@9548.9571.24|GAGGTCCGTAGCGGTCCTGACGTG</td><td>>hg38_21_+_8433172_8446622_RNA45SN1_45S_50ntFlanks</td><td>9548</td><td>9571</td><td>24</td><td>GAGGTCCGTAGCGGTCCTGACGTG</td><td>1179.5</td><td>-1.4</td><td>0.5</td><td>-2.8</td><td>5.4E-03</td><td>6.7E-03</td><td>BT20/MB468</td><td>CYTO</td></tr>
<tr><td>>hg38_21_+_8433172_8446622_RNA45SN1_45S_50ntFlanks@10315.10340.26|AGGTGCAGATCTTGGTGGTAGTAGCA</td><td>>hg38_21_+_8433172_8446622_RNA45SN1_45S_50ntFlanks</td><td>10315</td><td>10340</td><td>26</td><td>AGGTGCAGATCTTGGTGGTAGTAGCA</td><td>309.7</td><td>4.8</td><td>1.6</td><td>3.0</td><td>3.1E-03</td><td>6.7E-03</td><td>BT20/MB468</td><td>MP</td></tr>
<tr><td>>MI0000255|hsa-mir-30d&WithFlank&8|-|134804870|134804951@12.33.22(+1G)&[MIMAT0000245&hsa-miR-30d-5p&offsets|0|0(+1G);m-21&8|-|134804919|134804940&offsets|0|0(+1G)]|TGTAAACATCCCCGACTGGAAGG</td><td>>MI0000255|hsa-mir-30d&WithFlank&8|-|134804870|134804951</td><td>12</td><td>33</td><td>22(+1G)</td><td>TGTAAACATCCCCGACTGGAAGG</td><td>193.2</td><td>3.5</td><td>1.1</td><td>3.2</td><td>1.4E-03</td><td>6.7E-03</td><td>BT20/MB231</td><td>CYTO</td></tr>
<tr><td>>hg38_21_+_8433172_8446622_RNA45SN1_45S_50ntFlanks@10468.10497.30|TTGCCCTCGGCCGATCGAAAGGGAGTCGGG</td><td>>hg38_21_+_8433172_8446622_RNA45SN1_45S_50ntFlanks</td><td>10468</td><td>10497</td><td>30</td><td>TTGCCCTCGGCCGATCGAAAGGGAGTCGGG</td><td>154.7</td><td>-3.0</td><td>1.1</td><td>-2.8</td><td>4.6E-03</td><td>6.7E-03</td><td>BT20/MB468</td><td>MITO</td></tr>
<tr><td>>trnaMT_ProTGG_MT_-_15956_16023@40.60.21|AATGGTGGAGTTAAAGACTTT</td><td>>trnaMT_ProTGG_MT_-_15956_16023</td><td>40</td><td>60</td><td>21</td><td>AATGGTGGAGTTAAAGACTTT</td><td>100.6</td><td>3.5</td><td>1.3</td><td>2.8</td><td>5.0E-03</td><td>6.7E-03</td><td>MB231/MB468</td><td>MP</td></tr>
<tr><td>>hg38_21_+_8433172_8446622_RNA45SN1_45S_50ntFlanks@12999.13039.41|CTGCGATCTATTGAAAGTCAGCCCTCGACACAAGGGTTTGT</td><td>>hg38_21_+_8433172_8446622_RNA45SN1_45S_50ntFlanks</td><td>12999</td><td>13039</td><td>41</td><td>CTGCGATCTATTGAAAGTCAGCCCTCGACACAAGGGTTTGT</td><td>771.5</td><td>3.5</td><td>1.2</td><td>3.0</td><td>3.1E-03</td><td>6.7E-03</td><td>BT20/MB468</td><td>MP</td></tr>
<tr><td>>hg38_21_+_8433172_8446622_RNA45SN1_45S_50ntFlanks@11703.11728.26|AATGAAGCGCGGGTAAACGGCGGGAG</td><td>>hg38_21_+_8433172_8446622_RNA45SN1_45S_50ntFlanks</td><td>11703</td><td>11728</td><td>26</td><td>AATGAAGCGCGGGTAAACGGCGGGAG</td><td>2680.4</td><td>-5.6</td><td>1.9</td><td>-3.0</td><td>3.1E-03</td><td>6.7E-03</td><td>BT20/MB468</td><td>MP</td></tr>
<tr><td>>hg38_21_+_8433172_8446622_RNA45SN1_45S_50ntFlanks@12419.12455.37|GATGTCGGCTCTTCCTATCATTGTGAAGCAGAATTCA</td><td>>hg38_21_+_8433172_8446622_RNA45SN1_45S_50ntFlanks</td><td>12419</td><td>12455</td><td>37</td><td>GATGTCGGCTCTTCCTATCATTGTGAAGCAGAATTCA</td><td>447.5</td><td>-2.5</td><td>0.8</td><td>-3.2</td><td>1.4E-03</td><td>6.7E-03</td><td>BT20/MB231</td><td>CYTO</td></tr>
<tr><td>>hg38_21_+_8433172_8446622_RNA45SN1_45S_50ntFlanks@9572.9601.30|CAAATCGGTCGTCCGACCTGGGTATAGGGG</td><td>>hg38_21_+_8433172_8446622_RNA45SN1_45S_50ntFlanks</td><td>9572</td><td>9601</td><td>30</td><td>CAAATCGGTCGTCCGACCTGGGTATAGGGG</td><td>194.9</td><td>-1.9</td><td>0.7</td><td>-2.8</td><td>5.5E-03</td><td>6.7E-03</td><td>MB231/MB468</td><td>CYTO</td></tr>
<tr><td>>hg38_21_+_8433172_8446622_RNA45SN1_45S_50ntFlanks@11907.11928.22|ACGGTGAAGAGACATGAGAGGT</td><td>>hg38_21_+_8433172_8446622_RNA45SN1_45S_50ntFlanks</td><td>11907</td><td>11928</td><td>22</td><td>ACGGTGAAGAGACATGAGAGGT</td><td>139.3</td><td>6.4</td><td>2.0</td><td>3.3</td><td>1.1E-03</td><td>6.7E-03</td><td>BT20/MB231</td><td>MP</td></tr>
<tr><td>>hg38_21_+_8433172_8446622_RNA45SN1_45S_50ntFlanks@7979.8012.34|ACCTCAGATCAGACGTGGCGACCCGCTGAATTTA</td><td>>hg38_21_+_8433172_8446622_RNA45SN1_45S_50ntFlanks</td><td>7979</td><td>8012</td><td>34</td><td>ACCTCAGATCAGACGTGGCGACCCGCTGAATTTA</td><td>399.6</td><td>-4.2</td><td>1.4</td><td>-3.0</td><td>3.2E-03</td><td>6.7E-03</td><td>BT20/MB468</td><td>MP</td></tr>
<tr><td>>hg38_MT_+_1621_3279_RNR2_16S_50ntFlanks@1427.1457.31|GATTAAAGTCCTACGTGATCTGAGTTCAGAC</td><td>>hg38_MT_+_1621_3279_RNR2_16S_50ntFlanks</td><td>1427</td><td>1457</td><td>31</td><td>GATTAAAGTCCTACGTGATCTGAGTTCAGAC</td><td>213.3</td><td>3.5</td><td>1.2</td><td>2.8</td><td>5.0E-03</td><td>6.7E-03</td><td>MB231/MB468</td><td>MP</td></tr>
<tr><td>>hg38_21_+_8433172_8446622_RNA45SN1_45S_50ntFlanks@3853.3874.22|AACTGTGGTAATTCTAGAGCTA</td><td>>hg38_21_+_8433172_8446622_RNA45SN1_45S_50ntFlanks</td><td>3853</td><td>3874</td><td>22</td><td>AACTGTGGTAATTCTAGAGCTA</td><td>147.3</td><td>-3.6</td><td>1.3</td><td>-2.8</td><td>4.7E-03</td><td>6.7E-03</td><td>BT20/MB468</td><td>MITO</td></tr>
<tr><td>>hg38_21_+_8433172_8446622_RNA45SN1_45S_50ntFlanks@6658.6681.24|TAGCGGTGGATCACTCGGCTCGTG</td><td>>hg38_21_+_8433172_8446622_RNA45SN1_45S_50ntFlanks</td><td>6658</td><td>6681</td><td>24</td><td>TAGCGGTGGATCACTCGGCTCGTG</td><td>240.5</td><td>-1.5</td><td>0.5</td><td>-2.8</td><td>4.7E-03</td><td>6.7E-03</td><td>BT20/MB468</td><td>MITO</td></tr>
<tr><td>>hg38_21_+_8433172_8446622_RNA45SN1_45S_50ntFlanks@8172.8212.41|AGTCCTTCTGATCGAGGCCCAGCCCGTGGACGGTGTGAGGC</td><td>>hg38_21_+_8433172_8446622_RNA45SN1_45S_50ntFlanks</td><td>8172</td><td>8212</td><td>41</td><td>AGTCCTTCTGATCGAGGCCCAGCCCGTGGACGGTGTGAGGC</td><td>184.2</td><td>-1.7</td><td>0.6</td><td>-2.8</td><td>5.5E-03</td><td>6.7E-03</td><td>BT20/MB468</td><td>CYTO</td></tr>
<tr><td>>MI0001446|hsa-mir-424&WithFlank&X|-|134546608|134546717@17.38.22&[MIMAT0001341&hsa-miR-424-5p&offsets|0|0;m-152&X|-|134546681|134546701&offsets|0|+1]|CAGCAGCAATTCATGTTTTGAA</td><td>>MI0001446|hsa-mir-424&WithFlank&X|-|134546608|134546717</td><td>17</td><td>38</td><td>22</td><td>CAGCAGCAATTCATGTTTTGAA</td><td>76.6</td><td>4.4</td><td>1.6</td><td>2.8</td><td>5.0E-03</td><td>6.7E-03</td><td>MB231/MB468</td><td>MP</td></tr>
<tr><td>>hg38_21_+_8433172_8446622_RNA45SN1_45S_50ntFlanks@11999.12035.37|CCCTGCGGGCCGCCGGTGAAATACCACTACTCTGATC</td><td>>hg38_21_+_8433172_8446622_RNA45SN1_45S_50ntFlanks</td><td>11999</td><td>12035</td><td>37</td><td>CCCTGCGGGCCGCCGGTGAAATACCACTACTCTGATC</td><td>659.6</td><td>-2.6</td><td>0.9</td><td>-2.8</td><td>4.7E-03</td><td>6.7E-03</td><td>BT20/MB468</td><td>MITO</td></tr>
<tr><td>>am_trna10_LysCTT_16_+_3241501_3241573@31.51.21,trna119_LysCTT_1_-_145395522_145395594@31.51.21,trna11_LysCTT_5_-_180648979_180649051@31.51.21,trna13_LysCTT_16_+_3246153_3246225@31.51.21,trna13_LysCTT_6_+_26556774_26556846@31.51.21,trna30_LysCTT_16_-_3230555_3230627@31.51.21,trna5_LysCTT_16_+_3214939_3215011@31.51.21,trna7_LysCTT_16_+_3225692_3225764@31.51.21,trna8_LysCTT_1_+_39970195_39970267@31.51.21,trna9_LysCTT_5_+_180634755_180634827@31.51.21|ACTCTTAATCTCAGGGTCGTG</td><td>>am_trna10_LysCTT_16_+_3241501_3241573</td><td>31</td><td>51</td><td>21</td><td>ACTCTTAATCTCAGGGTCGTG</td><td>324.2</td><td>-4.9</td><td>1.7</td><td>-3.0</td><td>3.2E-03</td><td>6.7E-03</td><td>BT20/MB468</td><td>MP</td></tr>
<tr><td>>hg38_21_+_8433172_8446622_RNA45SN1_45S_50ntFlanks@8917.8941.25|AATCCCGGGGCCGAGGGAGCGAGAC</td><td>>hg38_21_+_8433172_8446622_RNA45SN1_45S_50ntFlanks</td><td>8917</td><td>8941</td><td>25</td><td>AATCCCGGGGCCGAGGGAGCGAGAC</td><td>142.9</td><td>2.5</td><td>0.8</td><td>3.0</td><td>3.2E-03</td><td>6.8E-03</td><td>BT20/MB468</td><td>MP</td></tr>
<tr><td>>hg38_21_+_8433172_8446622_RNA45SN1_45S_50ntFlanks@9738.9755.18|GAAACGATCTCAACCTAT</td><td>>hg38_21_+_8433172_8446622_RNA45SN1_45S_50ntFlanks</td><td>9738</td><td>9755</td><td>18</td><td>GAAACGATCTCAACCTAT</td><td>644.1</td><td>-5.1</td><td>1.7</td><td>-3.0</td><td>3.2E-03</td><td>6.8E-03</td><td>BT20/MB468</td><td>MP</td></tr>
<tr><td>>am_trnaMT_LysTTT_MT_+_8295_8364@34.52.19,trnalookalike7_LysTTT_1_+_568844_568913@34.52.19|GTTAAAGATTAAGAGAACC</td><td>>am_trnaMT_LysTTT_MT_+_8295_8364</td><td>34</td><td>52</td><td>19</td><td>GTTAAAGATTAAGAGAACC</td><td>146.1</td><td>4.4</td><td>1.5</td><td>3.0</td><td>3.2E-03</td><td>6.8E-03</td><td>BT20/MB468</td><td>MP</td></tr>
<tr><td>>am_trnaMT_HisGTG_MT_+_12138_12206@20.49.30|AACATCAGATTGTGAATCTGACAACAGAGG</td><td>>am_trnaMT_HisGTG_MT_+_12138_12206</td><td>20</td><td>49</td><td>30</td><td>AACATCAGATTGTGAATCTGACAACAGAGG</td><td>441.7</td><td>6.5</td><td>2.2</td><td>3.0</td><td>3.2E-03</td><td>6.8E-03</td><td>BT20/MB468</td><td>MP</td></tr>
<tr><td>>hg38_21_+_8433172_8446622_RNA45SN1_45S_50ntFlanks@11674.11708.35|GTGCTCTGAATGTCAAAGTGAAGAAATTCAATGAA</td><td>>hg38_21_+_8433172_8446622_RNA45SN1_45S_50ntFlanks</td><td>11674</td><td>11708</td><td>35</td><td>GTGCTCTGAATGTCAAAGTGAAGAAATTCAATGAA</td><td>28.4</td><td>8.6</td><td>2.9</td><td>3.0</td><td>3.2E-03</td><td>6.8E-03</td><td>BT20/MB468</td><td>MP</td></tr>
<tr><td>>trnaMT_LysTTT_MT_+_8295_8364@34.72.39,trnalookalike7_LysTTT_1_+_568844_568913@34.72.39|GTTAAAGATTAAGAGAACCAACACCTCTTTACAGTGACC</td><td>>trnaMT_LysTTT_MT_+_8295_8364</td><td>34</td><td>72</td><td>39</td><td>GTTAAAGATTAAGAGAACCAACACCTCTTTACAGTGACC</td><td>279.2</td><td>-5.0</td><td>1.6</td><td>-3.3</td><td>1.1E-03</td><td>6.8E-03</td><td>BT20/MB231</td><td>MP</td></tr>
</table>

<table>
<tr><td>>MI0000466|hsa-mir-9-1&WithFlank&1|-|156420335|156420435@22.43.22;MI0000468|hsa-mir-9-3&WithFlank&15|+|89368011|89368112@22.43.22;MI0000467|hsa-mir-9-2&WithFlank&5|-|88666847|88666945@22.43.22&[MIMAT0000441&hsa-miR-9-5p&offsets|0|-1;m-104&1|-|156420392|156420414&offsets|0|-1];[MIMAT0000441_1&hsa-miR-9-5p&offsets|0|-1;m-105&15|+|89368032|89368054&offsets|0|-1];[MIMAT0000441_2&hsa-miR-9-5p&offsets|0|-1;m-104&5|-|88666902|88666924&offsets|0|-1]|TCTTTGGTTATCTAGCTGTATG</td><td>>MI0000466|hsa-mir-9-1&WithFlank&1|-|156420335|156420435</td><td>22</td><td>43</td><td>22</td><td>TCTTTGGTTATCTAGCTGTATG</td><td>1214.1</td><td>2.4</td><td>0.7</td><td>3.2</td><td>1.3E-03</td><td>6.8E-03</td><td>BT20/MB231</td><td>NUC</td></tr>
<tr><td>>hg38_21_+_8433172_8446622_RNA45SN1_45S_50ntFlanks@6774.6807.34|TCCTCCCGGGGCTACGCCTGTCTGAGCGTCGCTT</td><td>>hg38_21_+_8433172_8446622_RNA45SN1_45S_50ntFlanks</td><td>6774</td><td>6807</td><td>34</td><td>TCCTCCCGGGGCTACGCCTGTCTGAGCGTCGCTT</td><td>287.1</td><td>2.1</td><td>0.6</td><td>3.2</td><td>1.3E-03</td><td>6.8E-03</td><td>BT20/MB231</td><td>NUC</td></tr>
<tr><td>>hg38_21_+_8433172_8446622_RNA45SN1_45S_50ntFlanks@9938.9958.21|AGCAGGACGGTGGCCATGGAA</td><td>>hg38_21_+_8433172_8446622_RNA45SN1_45S_50ntFlanks</td><td>9938</td><td>9958</td><td>21</td><td>AGCAGGACGGTGGCCATGGAA</td><td>70.9</td><td>4.9</td><td>1.7</td><td>3.0</td><td>3.2E-03</td><td>6.8E-03</td><td>BT20/MB468</td><td>MP</td></tr>
<tr><td>>MI0000750|hsa-mir-26a-2&WithFlank&12|-|57824603|57824698@20.42.23(+1A);MI0000083|hsa-mir-26a-1&WithFlank&3|+|37969398|37969486@16.38.23(+1A)&[MIMAT0000082&hsa-miR-26a-5p&offsets|0|+1(+1A);m-33&12|-|57824658|57824679&offsets|0|+1(+1A)];[MIMAT0000082_1&hsa-miR-26a-5p&offsets|0|+1(+1A);m-34&3|+|37969413|37969434&offsets|0|+1(+1A)]|TTCAAGTAATCCAGGATAGGCTGA</td><td>>MI0000750|hsa-mir-26a-2&WithFlank&12|-|57824603|57824698</td><td>20</td><td>42</td><td>23(+1A)</td><td>TTCAAGTAATCCAGGATAGGCTGA</td><td>153.3</td><td>3.6</td><td>1.3</td><td>2.8</td><td>5.0E-03</td><td>6.8E-03</td><td>MB231/MB468</td><td>MP</td></tr>
<tr><td>>hg38_21_+_8433172_8446622_RNA45SN1_45S_50ntFlanks@6649.6696.48|TACGACTCTTAGCGGTGGATCACTCGGCTCGTGCGTCGATGAAGAACG</td><td>>hg38_21_+_8433172_8446622_RNA45SN1_45S_50ntFlanks</td><td>6649</td><td>6696</td><td>48</td><td>TACGACTCTTAGCGGTGGATCACTCGGCTCGTGCGTCGATGAAGAACG</td><td>176.4</td><td>-3.6</td><td>1.1</td><td>-3.2</td><td>1.4E-03</td><td>6.8E-03</td><td>BT20/MB231</td><td>CYTO</td></tr>
<tr><td>>hg38_21_+_8433172_8446622_RNA45SN1_45S_50ntFlanks@8174.8203.30|TCCTTCTGATCGAGGCCCAGCCCGTGGACG</td><td>>hg38_21_+_8433172_8446622_RNA45SN1_45S_50ntFlanks</td><td>8174</td><td>8203</td><td>30</td><td>TCCTTCTGATCGAGGCCCAGCCCGTGGACG</td><td>149.3</td><td>-3.5</td><td>1.2</td><td>-2.8</td><td>5.0E-03</td><td>6.8E-03</td><td>MB231/MB468</td><td>MP</td></tr>
<tr><td>>MI0000447|hsa-mir-128-1&WithFlank&2|+|135665391|135665484@56.76.21;MI0000727|hsa-mir-128-2&WithFlank&3|+|35744470|35744565@58.78.21&[MIMAT0000424&hsa-miR-128-3p&offsets|0|0;m-108&2|+|135665446|135665466&offsets|0|0];[MIMAT0000424_1&hsa-miR-128-3p&offsets|0|0;m-117&3|+|35744527|35744547&offsets|0|0]|TCACAGTGAACCGGTCTCTTT</td><td>>MI0000447|hsa-mir-128-1&WithFlank&2|+|135665391|135665484</td><td>56</td><td>76</td><td>21</td><td>TCACAGTGAACCGGTCTCTTT</td><td>6031.8</td><td>2.7</td><td>0.9</td><td>2.9</td><td>3.2E-03</td><td>6.8E-03</td><td>BT20/MB468</td><td>MP</td></tr>
<tr><td>>hg38_21_+_8433172_8446622_RNA45SN1_45S_50ntFlanks@9941.9960.20|AGGACGGTGGCCATGGAAGT</td><td>>hg38_21_+_8433172_8446622_RNA45SN1_45S_50ntFlanks</td><td>9941</td><td>9960</td><td>20</td><td>AGGACGGTGGCCATGGAAGT</td><td>219.7</td><td>5.1</td><td>1.7</td><td>2.9</td><td>3.2E-03</td><td>6.8E-03</td><td>BT20/MB468</td><td>MP</td></tr>
<tr><td>>hg38_21_+_8433172_8446622_RNA45SN1_45S_50ntFlanks@11665.11692.28|TTCTGCCCAGTGCTCTGAATGTCAAAGT</td><td>>hg38_21_+_8433172_8446622_RNA45SN1_45S_50ntFlanks</td><td>11665</td><td>11692</td><td>28</td><td>TTCTGCCCAGTGCTCTGAATGTCAAAGT</td><td>266.6</td><td>2.4</td><td>0.8</td><td>3.2</td><td>1.5E-03</td><td>6.8E-03</td><td>BT20/MB231</td><td>MITO</td></tr>
<tr><td>>hg38_21_+_8433172_8446622_RNA45SN1_45S_50ntFlanks@6647.6669.23|CGTACGACTCTTAGCGGTGGATC</td><td>>hg38_21_+_8433172_8446622_RNA45SN1_45S_50ntFlanks</td><td>6647</td><td>6669</td><td>23</td><td>CGTACGACTCTTAGCGGTGGATC</td><td>17755.7</td><td>2.4</td><td>0.7</td><td>3.2</td><td>1.5E-03</td><td>6.8E-03</td><td>BT20/MB231</td><td>MITO</td></tr>
<tr><td>>hg38_MT_+_1621_3279_RNR2_16S_50ntFlanks@51.90.40|GCTAAACCTAGCCCCAAACCCACTCCACCTTACTACCAGA</td><td>>hg38_MT_+_1621_3279_RNR2_16S_50ntFlanks</td><td>51</td><td>90</td><td>40</td><td>GCTAAACCTAGCCCCAAACCCACTCCACCTTACTACCAGA</td><td>5748.5</td><td>-2.8</td><td>0.9</td><td>-3.2</td><td>1.5E-03</td><td>6.8E-03</td><td>BT20/MB231</td><td>MITO</td></tr>
<tr><td>>hg38_21_+_8433172_8446622_RNA45SN1_45S_50ntFlanks@9605.9630.26|AAGACTAATCGAACCATCTAGTAGCT</td><td>>hg38_21_+_8433172_8446622_RNA45SN1_45S_50ntFlanks</td><td>9605</td><td>9630</td><td>26</td><td>AAGACTAATCGAACCATCTAGTAGCT</td><td>304.1</td><td>4.3</td><td>1.3</td><td>3.3</td><td>1.1E-03</td><td>6.8E-03</td><td>BT20/MB231</td><td>MP</td></tr>
<tr><td>>hg38_21_+_8433172_8446622_RNA45SN1_45S_50ntFlanks@10774.10796.23|GTCTCCAAGGTGAACAGCCTCTG</td><td>>hg38_21_+_8433172_8446622_RNA45SN1_45S_50ntFlanks</td><td>10774</td><td>10796</td><td>23</td><td>GTCTCCAAGGTGAACAGCCTCTG</td><td>95.2</td><td>-4.6</td><td>1.7</td><td>-2.8</td><td>5.0E-03</td><td>6.8E-03</td><td>MB231/MB468</td><td>MP</td></tr>
<tr><td>>MI0001445|hsa-mir-423&WithFlank&17|+|30117073|30117178@59.80.22(+1G)&[MIMAT0001340&hsa-miR-423-3p&offsets|0|-1(+1G);m-100&17|+|30117131|30117153&offsets|0|-1(+1G)]|AGCTCGGTCTGAGGCCCCTCAGG</td><td>>MI0001445|hsa-mir-423&WithFlank&17|+|30117073|30117178</td><td>59</td><td>80</td><td>22(+1G)</td><td>AGCTCGGTCTGAGGCCCCTCAGG</td><td>174.7</td><td>5.3</td><td>1.8</td><td>2.9</td><td>3.2E-03</td><td>6.8E-03</td><td>BT20/MB468</td><td>MP</td></tr>
<tr><td>>MI0000650|hsa-mir-200c&WithFlank&12|+|6963693|6963772@50.72.23(+2A)&[MIMAT0000617&hsa-miR-200c-3p&offsets|0|0(+2A);m-54&12|+|6963742|6963764&offsets|0|0(+2A)]|TAATACTGCCGGGTAATGATGGAAA</td><td>>MI0000650|hsa-mir-200c&WithFlank&12|+|6963693|6963772</td><td>50</td><td>72</td><td>23(+2A)</td><td>TAATACTGCCGGGTAATGATGGAAA</td><td>113.7</td><td>3.9</td><td>1.3</td><td>2.9</td><td>3.2E-03</td><td>6.8E-03</td><td>BT20/MB468</td><td>MP</td></tr>
<tr><td>>hg38_21_+_8433172_8446622_RNA45SN1_45S_50ntFlanks@6661.6678.18|CGGTGGATCACTCGGCTC</td><td>>hg38_21_+_8433172_8446622_RNA45SN1_45S_50ntFlanks</td><td>6661</td><td>6678</td><td>18</td><td>CGGTGGATCACTCGGCTC</td><td>678.4</td><td>2.4</td><td>0.7</td><td>3.2</td><td>1.4E-03</td><td>6.8E-03</td><td>BT20/MB231</td><td>CYTO</td></tr>
<tr><td>>hg38_21_+_8433172_8446622_RNA45SN1_45S_50ntFlanks@11672.11709.38|CAGTGCTCTGAATGTCAAAGTGAAGAAATTCAATGAAG</td><td>>hg38_21_+_8433172_8446622_RNA45SN1_45S_50ntFlanks</td><td>11672</td><td>11709</td><td>38</td><td>CAGTGCTCTGAATGTCAAAGTGAAGAAATTCAATGAAG</td><td>342.4</td><td>-2.9</td><td>0.9</td><td>-3.2</td><td>1.3E-03</td><td>6.8E-03</td><td>BT20/MB231</td><td>NUC</td></tr>
<tr><td>>hg38_21_+_8433172_8446622_RNA45SN1_45S_50ntFlanks@6773.6806.34|TTCCTCCCGGGGCTACGCCTGTCTGAGCGTCGCT</td><td>>hg38_21_+_8433172_8446622_RNA45SN1_45S_50ntFlanks</td><td>6773</td><td>6806</td><td>34</td><td>TTCCTCCCGGGGCTACGCCTGTCTGAGCGTCGCT</td><td>828.7</td><td>-1.5</td><td>0.5</td><td>-2.8</td><td>5.8E-03</td><td>6.8E-03</td><td>BT20/MB468</td><td>NUC</td></tr>
<tr><td>>hg38_21_+_8433172_8446622_RNA45SN1_45S_50ntFlanks@8349.8378.30|GAAAGTTGAAAAGAACTTTGAAGAGAGAGT</td><td>>hg38_21_+_8433172_8446622_RNA45SN1_45S_50ntFlanks</td><td>8349</td><td>8378</td><td>30</td><td>GAAAGTTGAAAAGAACTTTGAAGAGAGAGT</td><td>119.1</td><td>5.8</td><td>2.0</td><td>2.9</td><td>3.2E-03</td><td>6.8E-03</td><td>BT20/MB468</td><td>MP</td></tr>
<tr><td>>hg38_21_+_8433172_8446622_RNA45SN1_45S_50ntFlanks@7975.8005.31|CGCGACCTCAGATCAGACGTGGCGACCCGCT</td><td>>hg38_21_+_8433172_8446622_RNA45SN1_45S_50ntFlanks</td><td>7975</td><td>8005</td><td>31</td><td>CGCGACCTCAGATCAGACGTGGCGACCCGCT</td><td>61770.9</td><td>4.0</td><td>1.2</td><td>3.3</td><td>1.1E-03</td><td>6.8E-03</td><td>BT20/MB231</td><td>MP</td></tr>
<tr><td>>hg38_21_+_8433172_8446622_RNA45SN1_45S_50ntFlanks@11781.11807.27|CGCATGAATGGATGAACGAGATTCCCA</td><td>>hg38_21_+_8433172_8446622_RNA45SN1_45S_50ntFlanks</td><td>11781</td><td>11807</td><td>27</td><td>CGCATGAATGGATGAACGAGATTCCCA</td><td>166.9</td><td>-5.2</td><td>1.8</td><td>-2.9</td><td>3.2E-03</td><td>6.8E-03</td><td>BT20/MB468</td><td>MP</td></tr>
<tr><td>>hg38_21_+_8433172_8446622_RNA45SN1_45S_50ntFlanks@12001.12031.31|CTGCGGGCCGCCGGTGAAATACCACTACTCT</td><td>>hg38_21_+_8433172_8446622_RNA45SN1_45S_50ntFlanks</td><td>12001</td><td>12031</td><td>31</td><td>CTGCGGGCCGCCGGTGAAATACCACTACTCT</td><td>224.7</td><td>-4.9</td><td>1.6</td><td>-2.9</td><td>3.2E-03</td><td>6.8E-03</td><td>BT20/MB468</td><td>MP</td></tr>
<tr><td>>trna10_AspGTC_12_-_125424193_125424264@36.73.38,trna12_AspGTC_12_-_125411891_125411962@36.73.38,trna144_AspGTC_6_-_27551236_27551307@36.73.38,trna38_AspGTC_17_-_8125556_8125627@36.73.38,trna45_AspGTC_6_+_27447453_27447524@36.73.38,trna48_AspGTC_6_+_27471523_27471594@36.73.38,trna4_AspGTC_12_+_96429799_96429870@36.73.38,trna69_AspGTC_1_-_161440205_161440276@36.73.38,trna72_AspGTC_1_-_161432824_161432895@36.73.38,trna75_AspGTC_1_-_161425414_161425485@36.73.38,trna78_AspGTC_1_-_161418033_161418104@36.73.38,trna81_AspGTC_1_-_161410615_161410686@36.73.38|CACGCGGGAGACCGGGGTTCGATTCCCCGACGGGGAGC</td><td>>trna10_AspGTC_12_-_125424193_125424264</td><td>36</td><td>73</td><td>38</td><td>CACGCGGGAGACCGGGGTTCGATTCCCCGACGGGGAGC</td><td>116.9</td><td>4.7</td><td>1.7</td><td>2.8</td><td>5.1E-03</td><td>6.8E-03</td><td>MB231/MB468</td><td>MP</td></tr>
<tr><td>>hg38_21_+_8433172_8446622_RNA45SN1_45S_50ntFlanks@11876.11907.32|AAGACCCTGTTGAGCTTGACTCTAGTCTGGCA</td><td>>hg38_21_+_8433172_8446622_RNA45SN1_45S_50ntFlanks</td><td>11876</td><td>11907</td><td>32</td><td>AAGACCCTGTTGAGCTTGACTCTAGTCTGGCA</td><td>87.1</td><td>-4.7</td><td>1.6</td><td>-2.9</td><td>3.2E-03</td><td>6.8E-03</td><td>BT20/MB468</td><td>MP</td></tr>
<tr><td>>hg38_21_+_8433172_8446622_RNA45SN1_45S_50ntFlanks@9862.9893.32|ATGAACCGAACGCCGGGTTAAGGCGCCCGATG</td><td>>hg38_21_+_8433172_8446622_RNA45SN1_45S_50ntFlanks</td><td>9862</td><td>9893</td><td>32</td><td>ATGAACCGAACGCCGGGTTAAGGCGCCCGATG</td><td>782.0</td><td>-3.2</td><td>1.1</td><td>-2.9</td><td>3.2E-03</td><td>6.8E-03</td><td>BT20/MB468</td><td>MP</td></tr>
<tr><td>>MI0000439|hsa-mir-23b&WithFlank&9|+|95085202|95085310@67.87.21&[MIMAT0000418&hsa-miR-23b-3p&offsets|+3|+1;m-42&9|+|95085265|95085285&offsets|+3|+3]|ACATTGCCAGGGATTACCACG</td><td>>MI0000439|hsa-mir-23b&WithFlank&9|+|95085202|95085310</td><td>67</td><td>87</td><td>21</td><td>ACATTGCCAGGGATTACCACG</td><td>29.9</td><td>8.1</td><td>2.9</td><td>2.8</td><td>5.1E-03</td><td>6.8E-03</td><td>MB231/MB468</td><td>MP</td></tr>
<tr><td>>hg38_21_+_8433172_8446622_RNA45SN1_45S_50ntFlanks@12076.12095.20|AGGGGCTCTCGCTTCTGGCG</td><td>>hg38_21_+_8433172_8446622_RNA45SN1_45S_50ntFlanks</td><td>12076</td><td>12095</td><td>20</td><td>AGGGGCTCTCGCTTCTGGCG</td><td>1407.2</td><td>-1.9</td><td>0.7</td><td>-2.8</td><td>4.7E-03</td><td>6.8E-03</td><td>BT20/MB468</td><td>MITO</td></tr>
<tr><td>>hg38_21_+_8433172_8446622_RNA45SN1_45S_50ntFlanks@7974.8003.30|ACGCGACCTCAGATCAGACGTGGCGACCCG</td><td>>hg38_21_+_8433172_8446622_RNA45SN1_45S_50ntFlanks</td><td>7974</td><td>8003</td><td>30</td><td>ACGCGACCTCAGATCAGACGTGGCGACCCG</td><td>1203.8</td><td>2.1</td><td>0.7</td><td>2.8</td><td>4.7E-03</td><td>6.8E-03</td><td>BT20/MB468</td><td>MITO</td></tr>
<tr><td>>hg38_21_+_8433172_8446622_RNA45SN1_45S_50ntFlanks@9623.9648.26|TAGTAGCTGGTTCCCTCCGAAGTTTC</td><td>>hg38_21_+_8433172_8446622_RNA45SN1_45S_50ntFlanks</td><td>9623</td><td>9648</td><td>26</td><td>TAGTAGCTGGTTCCCTCCGAAGTTTC</td><td>320.9</td><td>-5.2</td><td>1.7</td><td>-2.9</td><td>3.2E-03</td><td>6.8E-03</td><td>BT20/MB468</td><td>MP</td></tr>
<tr><td>>hg38_21_+_8433172_8446622_RNA45SN1_45S_50ntFlanks@5472.5493.22|ACGGCCCTGGCGGAGCGCTGAG</td><td>>hg38_21_+_8433172_8446622_RNA45SN1_45S_50ntFlanks</td><td>5472</td><td>5493</td><td>22</td><td>ACGGCCCTGGCGGAGCGCTGAG</td><td>82.3</td><td>-3.6</td><td>1.3</td><td>-2.8</td><td>5.1E-03</td><td>6.8E-03</td><td>MB231/MB468</td><td>MP</td></tr>
<tr><td>>hg38_21_+_8433172_8446622_RNA45SN1_45S_50ntFlanks@11672.11700.29|CAGTGCTCTGAATGTCAAAGTGAAGAAAT</td><td>>hg38_21_+_8433172_8446622_RNA45SN1_45S_50ntFlanks</td><td>11672</td><td>11700</td><td>29</td><td>CAGTGCTCTGAATGTCAAAGTGAAGAAAT</td><td>76.1</td><td>-3.5</td><td>1.3</td><td>-2.8</td><td>5.1E-03</td><td>6.8E-03</td><td>MB231/MB468</td><td>MP</td></tr>
<tr><td>>MI0005756|hsa-mir-934&WithFlank&X|+|136550872|136550966@56.77.22&[TJU_CMC_MD2.ID03472.3p-miR&offsets|-1|-1;m-3031&X|+|136550928|136550949&offsets|-1|-1]|AGAGTCTCCAGTAATGGACGGG</td><td>>MI0005756|hsa-mir-934&WithFlank&X|+|136550872|136550966</td><td>56</td><td>77</td><td>22</td><td>AGAGTCTCCAGTAATGGACGGG</td><td>55.6</td><td>9.1</td><td>2.8</td><td>3.3</td><td>1.1E-03</td><td>6.8E-03</td><td>BT20/MB231</td><td>MP</td></tr>
<tr><td>>hg38_21_+_8433172_8446622_RNA45SN1_45S_50ntFlanks@11670.11696.27|CCCAGTGCTCTGAATGTCAAAGTGAAG</td><td>>hg38_21_+_8433172_8446622_RNA45SN1_45S_50ntFlanks</td><td>11670</td><td>11696</td><td>27</td><td>CCCAGTGCTCTGAATGTCAAAGTGAAG</td><td>342.3</td><td>-5.1</td><td>1.8</td><td>-2.8</td><td>5.8E-03</td><td>6.8E-03</td><td>BT20/MB468</td><td>NUC</td></tr>
<tr><td>>hg38_21_+_8433172_8446622_RNA45SN1_45S_50ntFlanks@6771.6799.29|GGTTCCTCCCGGGGCTACGCCTGTCTGAG</td><td>>hg38_21_+_8433172_8446622_RNA45SN1_45S_50ntFlanks</td><td>6771</td><td>6799</td><td>29</td><td>GGTTCCTCCCGGGGCTACGCCTGTCTGAG</td><td>131.4</td><td>-4.0</td><td>1.4</td><td>-2.8</td><td>5.5E-03</td><td>6.8E-03</td><td>BT20/MB468</td><td>CYTO</td></tr>
<tr><td>>hg38_21_+_8433172_8446622_RNA45SN1_45S_50ntFlanks@5487.5522.36|CGCTGAGAAGACGGTCGAACTTGACTATCTAGAGGA</td><td>>hg38_21_+_8433172_8446622_RNA45SN1_45S_50ntFlanks</td><td>5487</td><td>5522</td><td>36</td><td>CGCTGAGAAGACGGTCGAACTTGACTATCTAGAGGA</td><td>198.2</td><td>-5.0</td><td>1.7</td><td>-2.9</td><td>3.2E-03</td><td>6.8E-03</td><td>BT20/MB468</td><td>MP</td></tr>
<tr><td>>MI0000803|hsa-mir-330&WithFlank&19|-|45638988|45639093@64.86.23(+1A)&[MIMAT0000751&hsa-miR-330-3p&offsets|+1|+1(+1A);m-347&19|-|45639009|45639030&offsets|0|+1(+1A)]|CAAAGCACACGGCCTGCAGAGAGA</td><td>>MI0000803|hsa-mir-330&WithFlank&19|-|45638988|45639093</td><td>64</td><td>86</td><td>23(+1A)</td><td>CAAAGCACACGGCCTGCAGAGAGA</td><td>45.6</td><td>5.3</td><td>1.9</td><td>2.8</td><td>5.1E-03</td><td>6.8E-03</td><td>MB231/MB468</td><td>MP</td></tr>
<tr><td>>hg38_MT_+_1621_3279_RNR2_16S_50ntFlanks@823.855.33|CATAAGGAAAGGTTAAAAAAAGTAAAAGGAACT</td><td>>hg38_MT_+_1621_3279_RNR2_16S_50ntFlanks</td><td>823</td><td>855</td><td>33</td><td>CATAAGGAAAGGTTAAAAAAAGTAAAAGGAACT</td><td>116.6</td><td>4.3</td><td>1.6</td><td>2.8</td><td>5.1E-03</td><td>6.8E-03</td><td>MB231/MB468</td><td>MP</td></tr>
<tr><td>>hg38_21_+_8433172_8446622_RNA45SN1_45S_50ntFlanks@3850.3871.22|GATAACTGTGGTAATTCTAGAG</td><td>>hg38_21_+_8433172_8446622_RNA45SN1_45S_50ntFlanks</td><td>3850</td><td>3871</td><td>22</td><td>GATAACTGTGGTAATTCTAGAG</td><td>201.2</td><td>-2.3</td><td>0.8</td><td>-2.8</td><td>5.5E-03</td><td>6.8E-03</td><td>BT20/MB468</td><td>CYTO</td></tr>
<tr><td>>MI0000077|hsa-mir-21&WithFlank&17|+|59841260|59841343@14.34.21(+1U)&[MIMAT0000076&hsa-miR-21-5p&offsets|0|-1(+1U);m-1&17|+|59841273|59841294&offsets|0|-1(+1U)]|TAGCTTATCAGACTGATGTTGT</td><td>>MI0000077|hsa-mir-21&WithFlank&17|+|59841260|59841343</td><td>14</td><td>34</td><td>21(+1U)</td><td>TAGCTTATCAGACTGATGTTGT</td><td>3120.4</td><td>-3.0</td><td>0.9</td><td>-3.2</td><td>1.5E-03</td><td>6.8E-03</td><td>BT20/MB231</td><td>MITO</td></tr>
<tr><td>>hg38_MT_+_1621_3279_RNR2_16S_50ntFlanks@1216.1249.34|CAACCTCCGAGCAGTACATGCTAAGACTTCACCA</td><td>>hg38_MT_+_1621_3279_RNR2_16S_50ntFlanks</td><td>1216</td><td>1249</td><td>34</td><td>CAACCTCCGAGCAGTACATGCTAAGACTTCACCA</td><td>120.1</td><td>4.3</td><td>1.6</td><td>2.8</td><td>5.9E-03</td><td>6.8E-03</td><td>MB231/MB468</td><td>MITO</td></tr>
<tr><td>>hg38_21_+_8433172_8446622_RNA45SN1_45S_50ntFlanks@12655.12673.19|ATGACTGAACGCCTCTAAG</td><td>>hg38_21_+_8433172_8446622_RNA45SN1_45S_50ntFlanks</td><td>12655</td><td>12673</td><td>19</td><td>ATGACTGAACGCCTCTAAG</td><td>354.4</td><td>-5.0</td><td>1.7</td><td>-2.9</td><td>3.2E-03</td><td>6.8E-03</td><td>BT20/MB468</td><td>MP</td></tr>
</table>

<table>
<tr><th></th><th></th><th></th><th></th><th></th><th></th><th></th><th></th><th></th><th></th><th></th><th></th><th></th><th></th></tr>
<tr><td>>hg38_21_+_8433172_8446622_RNA45SN1_45S_50ntFlanks@12635.12669.35|AAGCTACCATCTGTGGGATTATGACTGAACGCCTC</td><td>>hg38_21_+_8433172_8446622_RNA45SN1_45S_50ntFlanks</td><td>12635</td><td>12669</td><td>35</td><td>AAGCTACCATCTGTGGGATTATGACTGAACGCCTC</td><td>882.6</td><td>-2.7</td><td>1.0</td><td>-2.8</td><td>5.9E-03</td><td>6.8E-03</td><td>MB231/MB468</td><td>MITO</td></tr>
<tr><td>>hg38_21_+_8433172_8446622_RNA45SN1_45S_50ntFlanks@12615.12659.45|GCTGAGGAGCCAATGGGGCGAAGCTACCATCTGTGGGATTATGAC</td><td>>hg38_21_+_8433172_8446622_RNA45SN1_45S_50ntFlanks</td><td>12615</td><td>12659</td><td>45</td><td>GCTGAGGAGCCAATGGGGCGAAGCTACCATCTGTGGGATTATGAC</td><td>528.1</td><td>-3.7</td><td>1.3</td><td>-2.8</td><td>4.7E-03</td><td>6.8E-03</td><td>BT20/MB468</td><td>MITO</td></tr>
<tr><td>>hg38_21_+_8433172_8446622_RNA45SN1_45S_50ntFlanks@12994.13012.19|CCTCGCTGCGATCTATTGA</td><td>>hg38_21_+_8433172_8446622_RNA45SN1_45S_50ntFlanks</td><td>12994</td><td>13012</td><td>19</td><td>CCTCGCTGCGATCTATTGA</td><td>327.5</td><td>-3.2</td><td>1.1</td><td>-2.8</td><td>5.5E-03</td><td>6.8E-03</td><td>BT20/MB468</td><td>CYTO</td></tr>
<tr><td>>hg38_21_+_8433172_8446622_RNA45SN1_45S_50ntFlanks@10716.10736.21|GAAAATCCGGGGGAGAGGGTG</td><td>>hg38_21_+_8433172_8446622_RNA45SN1_45S_50ntFlanks</td><td>10716</td><td>10736</td><td>21</td><td>GAAAATCCGGGGGAGAGGGTG</td><td>271.4</td><td>-1.6</td><td>0.6</td><td>-2.8</td><td>5.5E-03</td><td>6.8E-03</td><td>BT20/MB468</td><td>CYTO</td></tr>
<tr><td>>hg38_21_+_8433172_8446622_RNA45SN1_45S_50ntFlanks@12569.12587.19|CAGTACGAGAGGAACCGCA</td><td>>hg38_21_+_8433172_8446622_RNA45SN1_45S_50ntFlanks</td><td>12569</td><td>12587</td><td>19</td><td>CAGTACGAGAGGAACCGCA</td><td>121.7</td><td>-2.5</td><td>0.9</td><td>-2.8</td><td>5.6E-03</td><td>6.8E-03</td><td>MB231/MB468</td><td>CYTO</td></tr>
<tr><td>>hg38_21_+_8433172_8446622_RNA45SN1_45S_50ntFlanks@12907.12935.29|CACCGCACGTTCGTGGGGAACCTGGCGCT</td><td>>hg38_21_+_8433172_8446622_RNA45SN1_45S_50ntFlanks</td><td>12907</td><td>12935</td><td>29</td><td>CACCGCACGTTCGTGGGGAACCTGGCGCT</td><td>278.3</td><td>-4.0</td><td>1.5</td><td>-2.8</td><td>6.0E-03</td><td>6.8E-03</td><td>MB231/MB468</td><td>NUC</td></tr>
<tr><td>>trna10_AspGTC_12_-_125424193_125424264@36.75.40,trna12_AspGTC_12_-_125411891_125411962@36.75.40,trna144_AspGTC_6_-_27551236_27551307@36.75.40,trna38_AspGTC_17_-_8125556_8125627@36.75.40,trna45_AspGTC_6_+_27447453_27447524@36.75.40,trna48_AspGTC_6_+_27471523_27471594@36.75.40,trna4_AspGTC_12_+_96429799_96429870@36.75.40,trna69_AspGTC_1_-_161440205_161440276@36.75.40,trna72_AspGTC_1_-_161432824_161432895@36.75.40,trna75_AspGTC_1_-_161425414_161425485@36.75.40,trna78_AspGTC_1_-_161418033_161418104@36.75.40,trna81_AspGTC_1_-_161410615_161410686@36.75.40|CACGCGGGAGACCGGGGTTCGATTCCCCGACGGGGAGCCA</td><td>>trna10_AspGTC_12_-_125424193_125424264</td><td>36</td><td>75</td><td>40</td><td>CACGCGGGAGACCGGGGTTCGATTCCCCGACGGGGAGCCA</td><td>22334.1</td><td>-3.6</td><td>1.1</td><td>-3.2</td><td>1.4E-03</td><td>6.8E-03</td><td>BT20/MB231</td><td>CYTO</td></tr>
<tr><td>>hg38_21_+_8433172_8446622_RNA45SN1_45S_50ntFlanks@9603.9630.28|GAAAGACTAATCGAACCATCTAGTAGCT</td><td>>hg38_21_+_8433172_8446622_RNA45SN1_45S_50ntFlanks</td><td>9603</td><td>9630</td><td>28</td><td>GAAAGACTAATCGAACCATCTAGTAGCT</td><td>128.7</td><td>4.9</td><td>1.7</td><td>2.9</td><td>3.2E-03</td><td>6.8E-03</td><td>BT20/MB468</td><td>MP</td></tr>
<tr><td>>hg38_21_+_8433172_8446622_RNA45SN1_45S_50ntFlanks@12932.12969.38|CGCTAAACCATTCGTAGACGACCTGCTTCTGGGTCGGG</td><td>>hg38_21_+_8433172_8446622_RNA45SN1_45S_50ntFlanks</td><td>12932</td><td>12969</td><td>38</td><td>CGCTAAACCATTCGTAGACGACCTGCTTCTGGGTCGGG</td><td>239.1</td><td>-4.4</td><td>1.6</td><td>-2.8</td><td>5.6E-03</td><td>6.8E-03</td><td>BT20/MB468</td><td>CYTO</td></tr>
<tr><td>>hg38_MT_+_598_1651_RNR1_12S_50ntFlanks@429.468.40|AAGTGGCTTTAACATATCTGAACACACAATAGCTAAGACC</td><td>>hg38_MT_+_598_1651_RNR1_12S_50ntFlanks</td><td>429</td><td>468</td><td>40</td><td>AAGTGGCTTTAACATATCTGAACACACAATAGCTAAGACC</td><td>58.8</td><td>5.2</td><td>1.9</td><td>2.8</td><td>5.1E-03</td><td>6.8E-03</td><td>MB231/MB468</td><td>MP</td></tr>
<tr><td>>hg38_21_+_8433172_8446622_RNA45SN1_45S_50ntFlanks@12008.12043.36|CCGCCGGTGAAATACCACTACTCTGATCGTTTTTTC</td><td>>hg38_21_+_8433172_8446622_RNA45SN1_45S_50ntFlanks</td><td>12008</td><td>12043</td><td>36</td><td>CCGCCGGTGAAATACCACTACTCTGATCGTTTTTTC</td><td>4512.8</td><td>-6.5</td><td>2.4</td><td>-2.8</td><td>5.6E-03</td><td>6.9E-03</td><td>MB231/MB468</td><td>CYTO</td></tr>
<tr><td>>hg38_21_+_8433172_8446622_RNA45SN1_45S_50ntFlanks@9928.9948.21|TGATATAGACAGCAGGACGGT</td><td>>hg38_21_+_8433172_8446622_RNA45SN1_45S_50ntFlanks</td><td>9928</td><td>9948</td><td>21</td><td>TGATATAGACAGCAGGACGGT</td><td>294.3</td><td>3.1</td><td>1.0</td><td>3.2</td><td>1.2E-03</td><td>6.9E-03</td><td>BT20/MB231</td><td>MP</td></tr>
<tr><td>>hg38_21_+_8433172_8446622_RNA45SN1_45S_50ntFlanks@12616.12653.38|CTGAGGAGCCAATGGGGCGAAGCTACCATCTGTGGGAT</td><td>>hg38_21_+_8433172_8446622_RNA45SN1_45S_50ntFlanks</td><td>12616</td><td>12653</td><td>38</td><td>CTGAGGAGCCAATGGGGCGAAGCTACCATCTGTGGGAT</td><td>259.7</td><td>-3.3</td><td>1.2</td><td>-2.8</td><td>5.6E-03</td><td>6.9E-03</td><td>BT20/MB468</td><td>CYTO</td></tr>
<tr><td>>hg38_21_+_8433172_8446622_RNA45SN1_45S_50ntFlanks@3975.4002.28|CGGGCGCCGGCGGCTTTGGTGACTCTAG</td><td>>hg38_21_+_8433172_8446622_RNA45SN1_45S_50ntFlanks</td><td>3975</td><td>4002</td><td>28</td><td>CGGGCGCCGGCGGCTTTGGTGACTCTAG</td><td>68.5</td><td>-5.0</td><td>1.7</td><td>-2.9</td><td>3.2E-03</td><td>6.9E-03</td><td>BT20/MB468</td><td>MP</td></tr>
<tr><td>>hg38_21_+_8433172_8446622_RNA45SN1_45S_50ntFlanks@9621.9656.36|TCTAGTAGCTGGTTCCCTCCGAAGTTTCCCTCAGGA</td><td>>hg38_21_+_8433172_8446622_RNA45SN1_45S_50ntFlanks</td><td>9621</td><td>9656</td><td>36</td><td>TCTAGTAGCTGGTTCCCTCCGAAGTTTCCCTCAGGA</td><td>103.1</td><td>-4.0</td><td>1.4</td><td>-2.8</td><td>4.8E-03</td><td>6.9E-03</td><td>BT20/MB468</td><td>MITO</td></tr>
<tr><td>>MI0006273|hsa-mir-1180&WithFlank&17|-|19344500|19344580@47.67.21(+1A)&[MIMAT0005825&hsa-miR-1180-3p&offsets|0|-1(+1A);m-214&17|-|19344513|19344534&offsets|0|-1(+1A)]|TTTCCGGCTCGCGTGGGTGTGA</td><td>>MI0006273|hsa-mir-1180&WithFlank&17|-|19344500|19344580</td><td>47</td><td>67</td><td>21(+1A)</td><td>TTTCCGGCTCGCGTGGGTGTGA</td><td>113.1</td><td>5.0</td><td>1.7</td><td>2.9</td><td>3.2E-03</td><td>6.9E-03</td><td>BT20/MB468</td><td>MP</td></tr>
<tr><td>>hg38_21_+_8433172_8446622_RNA45SN1_45S_50ntFlanks@10571.10591.21|GAGAAGCCGGCGGGAGCCCCG</td><td>>hg38_21_+_8433172_8446622_RNA45SN1_45S_50ntFlanks</td><td>10571</td><td>10591</td><td>21</td><td>GAGAAGCCGGCGGGAGCCCCG</td><td>522.2</td><td>-4.6</td><td>1.6</td><td>-2.8</td><td>5.6E-03</td><td>6.9E-03</td><td>BT20/MB468</td><td>CYTO</td></tr>
<tr><td>>hg38_MT_+_598_1651_RNR1_12S_50ntFlanks@428.453.26|AAAGTGGCTTTAACATATCTGAACAC</td><td>>hg38_MT_+_598_1651_RNR1_12S_50ntFlanks</td><td>428</td><td>453</td><td>26</td><td>AAAGTGGCTTTAACATATCTGAACAC</td><td>329.8</td><td>2.8</td><td>1.0</td><td>2.8</td><td>5.1E-03</td><td>6.9E-03</td><td>MB231/MB468</td><td>MP</td></tr>
<tr><td>>hg38_21_+_8433172_8446622_RNA45SN1_45S_50ntFlanks@9930.9947.18|ATATAGACAGCAGGACGG</td><td>>hg38_21_+_8433172_8446622_RNA45SN1_45S_50ntFlanks</td><td>9930</td><td>9947</td><td>18</td><td>ATATAGACAGCAGGACGG</td><td>176.2</td><td>5.9</td><td>1.8</td><td>3.2</td><td>1.2E-03</td><td>6.9E-03</td><td>BT20/MB231</td><td>MP</td></tr>
<tr><td>>hg38_21_+_8433172_8446622_RNA45SN1_45S_50ntFlanks@10872.10892.21|GCTGGGTCGGTCGGGCTGGGG</td><td>>hg38_21_+_8433172_8446622_RNA45SN1_45S_50ntFlanks</td><td>10872</td><td>10892</td><td>21</td><td>GCTGGGTCGGTCGGGCTGGGG</td><td>118.9</td><td>-2.7</td><td>1.0</td><td>-2.8</td><td>5.6E-03</td><td>6.9E-03</td><td>BT20/MB468</td><td>CYTO</td></tr>
<tr><td>>hg38_21_+_8433172_8446622_RNA45SN1_45S_50ntFlanks@10410.10431.22|TCAGTCGGTCCTGAGAGATGGG</td><td>>hg38_21_+_8433172_8446622_RNA45SN1_45S_50ntFlanks</td><td>10410</td><td>10431</td><td>22</td><td>TCAGTCGGTCCTGAGAGATGGG</td><td>839.1</td><td>-4.0</td><td>1.4</td><td>-2.9</td><td>3.2E-03</td><td>6.9E-03</td><td>BT20/MB468</td><td>MP</td></tr>
<tr><td>>hg38_21_+_8433172_8446622_RNA45SN1_45S_50ntFlanks@6733.6753.21|CATTGATCATCGACACTTCGA</td><td>>hg38_21_+_8433172_8446622_RNA45SN1_45S_50ntFlanks</td><td>6733</td><td>6753</td><td>21</td><td>CATTGATCATCGACACTTCGA</td><td>704.8</td><td>-3.0</td><td>1.0</td><td>-2.9</td><td>3.2E-03</td><td>6.9E-03</td><td>BT20/MB468</td><td>MP</td></tr>
<tr><td>>hg38_21_+_8433172_8446622_RNA45SN1_45S_50ntFlanks@11561.11603.43|GCGCCTAGCAGCCGACTTAGAACTGGTGCGGACCAGGGGAATC</td><td>>hg38_21_+_8433172_8446622_RNA45SN1_45S_50ntFlanks</td><td>11561</td><td>11603</td><td>43</td><td>GCGCCTAGCAGCCGACTTAGAACTGGTGCGGACCAGGGGAATC</td><td>811.4</td><td>-4.9</td><td>1.7</td><td>-2.9</td><td>3.2E-03</td><td>6.9E-03</td><td>BT20/MB468</td><td>MP</td></tr>
<tr><td>>hg38_21_+_8433172_8446622_RNA45SN1_45S_50ntFlanks@9604.9621.18|AAAGACTAATCGAACCAT</td><td>>hg38_21_+_8433172_8446622_RNA45SN1_45S_50ntFlanks</td><td>9604</td><td>9621</td><td>18</td><td>AAAGACTAATCGAACCAT</td><td>459.3</td><td>-5.1</td><td>1.7</td><td>-2.9</td><td>3.2E-03</td><td>6.9E-03</td><td>BT20/MB468</td><td>MP</td></tr>
<tr><td>>hg38_21_+_8433172_8446622_RNA45SN1_45S_50ntFlanks@3985.4002.18|CGGCTTTGGTGACTCTAG</td><td>>hg38_21_+_8433172_8446622_RNA45SN1_45S_50ntFlanks</td><td>3985</td><td>4002</td><td>18</td><td>CGGCTTTGGTGACTCTAG</td><td>4820.6</td><td>-5.8</td><td>2.0</td><td>-2.9</td><td>3.2E-03</td><td>6.9E-03</td><td>BT20/MB468</td><td>MP</td></tr>
<tr><td>>hg38_21_+_8433172_8446622_RNA45SN1_45S_50ntFlanks@11209.11254.46|CGGCGACTCTGGACGCGAGCCGGGCCCTTCCCGTGGATCGCCCCAG</td><td>>hg38_21_+_8433172_8446622_RNA45SN1_45S_50ntFlanks</td><td>11209</td><td>11254</td><td>46</td><td>CGGCGACTCTGGACGCGAGCCGGGCCCTTCCCGTGGATCGCCCCAG</td><td>1244.5</td><td>-3.8</td><td>1.3</td><td>-2.9</td><td>3.2E-03</td><td>6.9E-03</td><td>BT20/MB468</td><td>MP</td></tr>
<tr><td>>hg38_21_+_8433172_8446622_RNA45SN1_45S_50ntFlanks@10392.10424.33|ACAGCAGTTGAACATGGGTCAGTCGGTCCTGAG</td><td>>hg38_21_+_8433172_8446622_RNA45SN1_45S_50ntFlanks</td><td>10392</td><td>10424</td><td>33</td><td>ACAGCAGTTGAACATGGGTCAGTCGGTCCTGAG</td><td>9616.0</td><td>4.2</td><td>1.3</td><td>3.2</td><td>1.2E-03</td><td>6.9E-03</td><td>BT20/MB231</td><td>MP</td></tr>
<tr><td>>hg38_21_+_8433172_8446622_RNA45SN1_45S_50ntFlanks@8344.8373.30|TAAGGGAAAGTTGAAAAGAACTTTGAAGAG</td><td>>hg38_21_+_8433172_8446622_RNA45SN1_45S_50ntFlanks</td><td>8344</td><td>8373</td><td>30</td><td>TAAGGGAAAGTTGAAAAGAACTTTGAAGAG</td><td>1058.4</td><td>5.8</td><td>1.8</td><td>3.2</td><td>1.2E-03</td><td>6.9E-03</td><td>BT20/MB231</td><td>MP</td></tr>
<tr><td>>hg38_21_+_8433172_8446622_RNA45SN1_45S_50ntFlanks@11576.11613.38|CTTAGAACTGGTGCGGACCAGGGGAATCCGACTGTTTA</td><td>>hg38_21_+_8433172_8446622_RNA45SN1_45S_50ntFlanks</td><td>11576</td><td>11613</td><td>38</td><td>CTTAGAACTGGTGCGGACCAGGGGAATCCGACTGTTTA</td><td>201.0</td><td>-4.1</td><td>1.5</td><td>-2.8</td><td>5.1E-03</td><td>6.9E-03</td><td>MB231/MB468</td><td>MP</td></tr>
<tr><td>>trnaMT_GluTTC_MT_-_14674_14742@35.71.37,trnalookalike8_GluTTC_5_-_93905172_93905240@35.71.37|TATCATTGGTCGTGGTTGTAGTCCGTGCGAGAATACC</td><td>>trnaMT_GluTTC_MT_-_14674_14742</td><td>35</td><td>71</td><td>37</td><td>TATCATTGGTCGTGGTTGTAGTCCGTGCGAGAATACC</td><td>302.0</td><td>3.4</td><td>1.2</td><td>2.8</td><td>5.1E-03</td><td>6.9E-03</td><td>MB231/MB468</td><td>MP</td></tr>
<tr><td>>hg38_21_+_8433172_8446622_RNA45SN1_45S_50ntFlanks@11911.11943.33|TGAAGAGACATGAGAGGTGTAGAATAAGTGGGA</td><td>>hg38_21_+_8433172_8446622_RNA45SN1_45S_50ntFlanks</td><td>11911</td><td>11943</td><td>33</td><td>TGAAGAGACATGAGAGGTGTAGAATAAGTGGGA</td><td>28.2</td><td>8.6</td><td>2.9</td><td>2.9</td><td>3.2E-03</td><td>6.9E-03</td><td>BT20/MB468</td><td>MP</td></tr>
<tr><td>>hg38_21_+_8433172_8446622_RNA45SN1_45S_50ntFlanks@4740.4758.19|AGTCGGAGGTTCGAAGACG</td><td>>hg38_21_+_8433172_8446622_RNA45SN1_45S_50ntFlanks</td><td>4740</td><td>4758</td><td>19</td><td>AGTCGGAGGTTCGAAGACG</td><td>284.3</td><td>-2.1</td><td>0.7</td><td>-2.9</td><td>3.2E-03</td><td>6.9E-03</td><td>BT20/MB468</td><td>MP</td></tr>
<tr><td>>trna111_HisGTG_1_-_147774845_147774916@1.31.31,trna118_HisGTG_1_-_145396881_145396952@1.31.31,trna16_HisGTG_1_+_146544773_146544844@1.31.31,trna1_HisGTG_15_+_45493349_45493420@1.31.31,trna21_HisGTG_1_+_147753471_147753542@1.31.31,trna33_HisGTG_6_+_27125906_27125977@1.31.31,trna7_HisGTG_9_-_14433938_14434009@1.31.31,trna8_HisGTG_15_-_45492611_45492682@1.31.31,trna9_HisGTG_15_-_45490804_45490875@1.31.31|GCCGTGATCGTATAGTGGTTAGTACTCTGCG</td><td>>trna111_HisGTG_1_-_147774845_147774916</td><td>1</td><td>31</td><td>31</td><td>GCCGTGATCGTATAGTGGTTAGTACTCTGCG</td><td>10490.3</td><td>-2.1</td><td>0.7</td><td>-2.8</td><td>5.1E-03</td><td>6.9E-03</td><td>MB231/MB468</td><td>MP</td></tr>
<tr><td>>hg38_21_+_8433172_8446622_RNA45SN1_45S_50ntFlanks@485.507.23|GATGTGGTGACGTCGTGCTCTCC</td><td>>hg38_21_+_8433172_8446622_RNA45SN1_45S_50ntFlanks</td><td>485</td><td>507</td><td>23</td><td>GATGTGGTGACGTCGTGCTCTCC</td><td>29.7</td><td>8.1</td><td>2.9</td><td>2.8</td><td>5.1E-03</td><td>6.9E-03</td><td>MB231/MB468</td><td>MP</td></tr>
<tr><td>>hg38_21_+_8433172_8446622_RNA45SN1_45S_50ntFlanks@13007.13027.21|TATTGAAAGTCAGCCCTCGAC</td><td>>hg38_21_+_8433172_8446622_RNA45SN1_45S_50ntFlanks</td><td>13007</td><td>13027</td><td>21</td><td>TATTGAAAGTCAGCCCTCGAC</td><td>802.3</td><td>-2.9</td><td>1.1</td><td>-2.8</td><td>5.6E-03</td><td>6.9E-03</td><td>BT20/MB468</td><td>CYTO</td></tr>
<tr><td>>hg38_21_+_8433172_8446622_RNA45SN1_45S_50ntFlanks@9603.9626.24|GAAAGACTAATCGAACCATCTAGT</td><td>>hg38_21_+_8433172_8446622_RNA45SN1_45S_50ntFlanks</td><td>9603</td><td>9626</td><td>24</td><td>GAAAGACTAATCGAACCATCTAGT</td><td>63.6</td><td>5.0</td><td>1.7</td><td>2.9</td><td>3.3E-03</td><td>6.9E-03</td><td>BT20/MB468</td><td>MP</td></tr>
<tr><td>>hg38_21_+_8433172_8446622_RNA45SN1_45S_50ntFlanks@11668.11702.35|TGCCCAGTGCTCTGAATGTCAAAGTGAAGAAATTC</td><td>>hg38_21_+_8433172_8446622_RNA45SN1_45S_50ntFlanks</td><td>11668</td><td>11702</td><td>35</td><td>TGCCCAGTGCTCTGAATGTCAAAGTGAAGAAATTC</td><td>7354.5</td><td>-2.9</td><td>0.9</td><td>-3.2</td><td>1.5E-03</td><td>6.9E-03</td><td>BT20/MB231</td><td>CYTO</td></tr>
<tr><td>>hg38_MT_+_1621_3279_RNR2_16S_50ntFlanks@1092.1119.28|GCAAGACGAGAAGACCCTATGGAGCTTT</td><td>>hg38_MT_+_1621_3279_RNR2_16S_50ntFlanks</td><td>1092</td><td>1119</td><td>28</td><td>GCAAGACGAGAAGACCCTATGGAGCTTT</td><td>101.3</td><td>2.8</td><td>1.0</td><td>2.8</td><td>5.2E-03</td><td>6.9E-03</td><td>MB231/MB468</td><td>MP</td></tr>
<tr><td>>trnaMT_HisGTG_MT_+_12138_12206@28.60.33|ATTGTGAATCTGACAACAGAGGCTTACGACCCC</td><td>>trnaMT_HisGTG_MT_+_12138_12206</td><td>28</td><td>60</td><td>33</td><td>ATTGTGAATCTGACAACAGAGGCTTACGACCCC</td><td>225.6</td><td>3.3</td><td>1.1</td><td>2.9</td><td>3.3E-03</td><td>6.9E-03</td><td>BT20/MB468</td><td>MP</td></tr>
<tr><td>>MI0000093|hsa-mir-92a-1&WithFlank&13|+|91351308|91351397@54.75.22(+1A);MI0000094|hsa-mir-92a-2&WithFlank&X|-|134169532|134169618@54.75.22(+1A)&[MIMAT0000092&hsa-miR-92a-3p&offsets|0|0(+1A);m-9&13|+|91351361|91351382&offsets|0|0(+1A)];[MIMAT0000092_1&hsa-miR-92a-3p&offsets|0|0(+1A);m-11&X|-|134169544|134169565&offsets|0|0(+1A)]|TATTGCACTTGTCCCGGCCTGTA</td><td>>MI0000093|hsa-mir-92a-1&WithFlank&13|+|91351308|91351397</td><td>54</td><td>75</td><td>22(+1A)</td><td>TATTGCACTTGTCCCGGCCTGTA</td><td>4191.2</td><td>-1.0</td><td>0.4</td><td>-2.8</td><td>5.9E-03</td><td>6.9E-03</td><td>BT20/MB468</td><td>NUC</td></tr>
<tr><td>>hg38_21_+_8433172_8446622_RNA45SN1_45S_50ntFlanks@7983.8009.27|CAGATCAGACGTGGCGACCCGCTGAAT</td><td>>hg38_21_+_8433172_8446622_RNA45SN1_45S_50ntFlanks</td><td>7983</td><td>8009</td><td>27</td><td>CAGATCAGACGTGGCGACCCGCTGAAT</td><td>430.7</td><td>-2.2</td><td>0.8</td><td>-2.8</td><td>5.9E-03</td><td>6.9E-03</td><td>BT20/MB468</td><td>NUC</td></tr>
<tr><td>>hg38_21_+_8433172_8446622_RNA45SN1_45S_50ntFlanks@9547.9571.25|GGAGGTCCGTAGCGGTCCTGACGTG</td><td>>hg38_21_+_8433172_8446622_RNA45SN1_45S_50ntFlanks</td><td>9547</td><td>9571</td><td>25</td><td>GGAGGTCCGTAGCGGTCCTGACGTG</td><td>386.0</td><td>-2.5</td><td>0.8</td><td>-3.2</td><td>1.5E-03</td><td>6.9E-03</td><td>BT20/MB231</td><td>CYTO</td></tr>
<tr><td>>trnaMT_GluTTC_MT_-_14674_14742@39.65.27,trnalookalike8_GluTTC_5_-_93905172_93905240@39.65.27|ATTGGTCGTGGTTGTAGTCCGTGCGAG</td><td>>trnaMT_GluTTC_MT_-_14674_14742</td><td>39</td><td>65</td><td>27</td><td>ATTGGTCGTGGTTGTAGTCCGTGCGAG</td><td>182.1</td><td>2.0</td><td>0.7</td><td>2.8</td><td>5.2E-03</td><td>6.9E-03</td><td>MB231/MB468</td><td>MP</td></tr>
<tr><td>>MI0000102|hsa-mir-100&WithFlank&11|-|122152223|122152314@19.36.18&[MIMAT0000098&hsa-miR-100-5p&offsets|0|-4;m-27&11|-|122152276|122152296&offsets|0|-3]|AACCCGTAGATCCGAACT</td><td>>MI0000102|hsa-mir-100&WithFlank&11|-|122152223|122152314</td><td>19</td><td>36</td><td>18</td><td>AACCCGTAGATCCGAACT</td><td>311.8</td><td>-3.9</td><td>1.2</td><td>-3.2</td><td>1.3E-03</td><td>6.9E-03</td><td>BT20/MB231</td><td>NUC</td></tr>
<tr><td>>MI0016005|hsa-mir-3615&WithFlank&17|+|74748607|74748705@57.79.23(+1U)&[MIMAT0017994&hsa-miR-3615&offsets|0|+2(+1U);m-271&17|+|74748663|74748684&offsets|0|+1(+1U)]|TCTCTCGGCTCCTCGCGGCTCGCT</td><td>>MI0016005|hsa-mir-3615&WithFlank&17|+|74748607|74748705</td><td>57</td><td>79</td><td>23(+1U)</td><td>TCTCTCGGCTCCTCGCGGCTCGCT</td><td>125.2</td><td>3.7</td><td>1.3</td><td>2.7</td><td>6.0E-03</td><td>6.9E-03</td><td>MB231/MB468</td><td>MITO</td></tr>
</table>

<table>
<tr><td>>MI0000270|hsa-mir-181b-1&WithFlank&1|-|198858867|198858988@42.64.23(+1C);MI0000683|hsa-mir-181b-2&WithFlank&9|+|124693704|124693804@22.44.23(+1C)&[MIMAT0000257&hsa-miR-181b-5p&offsets|0|0(+1C);m-73&1|-|198858925|198858947&offsets|0|0(+1C)];[MIMAT0000257_1&hsa-miR-181b-5p&offsets|0|0(+1C);m-72&9|+|124693725|124693747&offsets|0|0(+1C)]|AACATTCATTGCTGTCGGTGGGTC</td><td>>MI0000270|hsa-mir-181b-1&WithFlank&1|-|198858867|198858988</td><td>42</td><td>64</td><td>23(+1C)</td><td>AACATTCATTGCTGTCGGTGGGTC</td><td>497.0</td><td>-2.8</td><td>0.9</td><td>-3.2</td><td>1.5E-03</td><td>6.9E-03</td><td>BT20/MB231</td><td>MITO</td></tr>
<tr><td>>hg38_21_+_8433172_8446622_RNA45SN1_45S_50ntFlanks@11601.11620.20|ATCCGACTGTTTAATTAAAA</td><td>>hg38_21_+_8433172_8446622_RNA45SN1_45S_50ntFlanks</td><td>11601</td><td>11620</td><td>20</td><td>ATCCGACTGTTTAATTAAAA</td><td>108.8</td><td>-4.5</td><td>1.6</td><td>-2.8</td><td>5.2E-03</td><td>6.9E-03</td><td>MB231/MB468</td><td>MP</td></tr>
<tr><td>>hg38_21_+_8433172_8446622_RNA45SN1_45S_50ntFlanks@10069.10090.22|AGTGGACGGGAGCGGCGGGGGC</td><td>>hg38_21_+_8433172_8446622_RNA45SN1_45S_50ntFlanks</td><td>10069</td><td>10090</td><td>22</td><td>AGTGGACGGGAGCGGCGGGGGC</td><td>73.7</td><td>-4.8</td><td>1.7</td><td>-2.8</td><td>5.2E-03</td><td>6.9E-03</td><td>MB231/MB468</td><td>MP</td></tr>
<tr><td>>hg38_21_+_8433172_8446622_RNA45SN1_45S_50ntFlanks@10785.10812.28|GAACAGCCTCTGGCATGTTGGAACAATG</td><td>>hg38_21_+_8433172_8446622_RNA45SN1_45S_50ntFlanks</td><td>10785</td><td>10812</td><td>28</td><td>GAACAGCCTCTGGCATGTTGGAACAATG</td><td>56.1</td><td>5.0</td><td>1.7</td><td>2.9</td><td>3.3E-03</td><td>6.9E-03</td><td>BT20/MB468</td><td>MP</td></tr>
<tr><td>>hg38_MT_+_1621_3279_RNR2_16S_50ntFlanks@1349.1385.37|ACAATAGGGTTTACGACCTCGATGTTGGATCAGGACA</td><td>>hg38_MT_+_1621_3279_RNR2_16S_50ntFlanks</td><td>1349</td><td>1385</td><td>37</td><td>ACAATAGGGTTTACGACCTCGATGTTGGATCAGGACA</td><td>276.0</td><td>4.4</td><td>1.5</td><td>2.9</td><td>3.3E-03</td><td>6.9E-03</td><td>BT20/MB468</td><td>MP</td></tr>
<tr><td>>hg38_21_+_8433172_8446622_RNA45SN1_45S_50ntFlanks@4976.4994.19|CCGGACACGGACAGGATTG</td><td>>hg38_21_+_8433172_8446622_RNA45SN1_45S_50ntFlanks</td><td>4976</td><td>4994</td><td>19</td><td>CCGGACACGGACAGGATTG</td><td>277.6</td><td>-4.9</td><td>1.7</td><td>-2.9</td><td>3.3E-03</td><td>6.9E-03</td><td>BT20/MB468</td><td>MP</td></tr>
<tr><td>>hg38_21_+_8433172_8446622_RNA45SN1_45S_50ntFlanks@5486.5519.34|GCGCTGAGAAGACGGTCGAACTTGACTATCTAGA</td><td>>hg38_21_+_8433172_8446622_RNA45SN1_45S_50ntFlanks</td><td>5486</td><td>5519</td><td>34</td><td>GCGCTGAGAAGACGGTCGAACTTGACTATCTAGA</td><td>120.0</td><td>2.7</td><td>1.0</td><td>2.8</td><td>6.0E-03</td><td>6.9E-03</td><td>BT20/MB468</td><td>NUC</td></tr>
<tr><td>>hg38_21_+_8433172_8446622_RNA45SN1_45S_50ntFlanks@11561.11590.30|GCGCCTAGCAGCCGACTTAGAACTGGTGCG</td><td>>hg38_21_+_8433172_8446622_RNA45SN1_45S_50ntFlanks</td><td>11561</td><td>11590</td><td>30</td><td>GCGCCTAGCAGCCGACTTAGAACTGGTGCG</td><td>323.8</td><td>-4.9</td><td>1.7</td><td>-2.9</td><td>3.3E-03</td><td>6.9E-03</td><td>BT20/MB468</td><td>MP</td></tr>
<tr><td>>hg38_21_+_8433172_8446622_RNA45SN1_45S_50ntFlanks@8177.8195.19|TTCTGATCGAGGCCCAGCC</td><td>>hg38_21_+_8433172_8446622_RNA45SN1_45S_50ntFlanks</td><td>8177</td><td>8195</td><td>19</td><td>TTCTGATCGAGGCCCAGCC</td><td>279.1</td><td>2.0</td><td>0.7</td><td>2.7</td><td>6.0E-03</td><td>6.9E-03</td><td>MB231/MB468</td><td>NUC</td></tr>
<tr><td>>hg38_21_+_8433172_8446622_RNA45SN1_45S_50ntFlanks@6648.6678.31|GTACGACTCTTAGCGGTGGATCACTCGGCTC</td><td>>hg38_21_+_8433172_8446622_RNA45SN1_45S_50ntFlanks</td><td>6648</td><td>6678</td><td>31</td><td>GTACGACTCTTAGCGGTGGATCACTCGGCTC</td><td>1036.1</td><td>4.4</td><td>1.4</td><td>3.2</td><td>1.2E-03</td><td>6.9E-03</td><td>BT20/MB231</td><td>MP</td></tr>
<tr><td>>MI0000433|hsa-let-7g&WithFlank&3|-|52268272|52268367@11.30.20&[MIMAT0000414&hsa-let-7g-5p&offsets|0|-2;m-58&3|-|52268336|52268357&offsets|0|-2]|TGAGGTAGTAGTTTGTACAG</td><td>>MI0000433|hsa-let-7g&WithFlank&3|-|52268272|52268367</td><td>11</td><td>30</td><td>20</td><td>TGAGGTAGTAGTTTGTACAG</td><td>80.6</td><td>6.0</td><td>2.1</td><td>2.9</td><td>3.3E-03</td><td>6.9E-03</td><td>BT20/MB468</td><td>MP</td></tr>
<tr><td>>hg38_21_+_8433172_8446622_RNA45SN1_45S_50ntFlanks@10458.10488.31|ATGGCCTCCGTTGCCCTCGGCCGATCGAAAG</td><td>>hg38_21_+_8433172_8446622_RNA45SN1_45S_50ntFlanks</td><td>10458</td><td>10488</td><td>31</td><td>ATGGCCTCCGTTGCCCTCGGCCGATCGAAAG</td><td>599.5</td><td>-4.9</td><td>1.8</td><td>-2.8</td><td>5.6E-03</td><td>6.9E-03</td><td>BT20/MB468</td><td>CYTO</td></tr>
<tr><td>>hg38_21_+_8433172_8446622_RNA45SN1_45S_50ntFlanks@3854.3874.21|ACTGTGGTAATTCTAGAGCTA</td><td>>hg38_21_+_8433172_8446622_RNA45SN1_45S_50ntFlanks</td><td>3854</td><td>3874</td><td>21</td><td>ACTGTGGTAATTCTAGAGCTA</td><td>218.5</td><td>-3.5</td><td>1.2</td><td>-2.8</td><td>4.8E-03</td><td>6.9E-03</td><td>BT20/MB468</td><td>MITO</td></tr>
<tr><td>>hg38_MT_+_598_1651_RNR1_12S_50ntFlanks@849.868.20|AGTGCTTAGTTGAACAGGGC</td><td>>hg38_MT_+_598_1651_RNR1_12S_50ntFlanks</td><td>849</td><td>868</td><td>20</td><td>AGTGCTTAGTTGAACAGGGC</td><td>73.9</td><td>5.2</td><td>1.8</td><td>2.9</td><td>3.3E-03</td><td>6.9E-03</td><td>BT20/MB468</td><td>MP</td></tr>
<tr><td>>MI0000466|hsa-mir-9-1&WithFlank&1|-|156420335|156420435@22.44.23;MI0000468|hsa-mir-9-3&WithFlank&15|+|89368011|89368112@22.44.23;MI0000467|hsa-mir-9-2&WithFlank&5|-|88666847|88666945@22.44.23&[MIMAT0000441&hsa-miR-9-5p&offsets|0|0;m-104&1|-|156420392|156420414&offsets|0|0];[MIMAT0000441_1&hsa-miR-9-5p&offsets|0|0;m-105&15|+|89368032|89368054&offsets|0|0];[MIMAT0000441_2&hsa-miR-9-5p&offsets|0|0;m-104&5|-|88666902|88666924&offsets|0|0]|TCTTTGGTTATCTAGCTGTATGA</td><td>>MI0000466|hsa-mir-9-1&WithFlank&1|-|156420335|156420435</td><td>22</td><td>44</td><td>23</td><td>TCTTTGGTTATCTAGCTGTATGA</td><td>260.4</td><td>2.4</td><td>0.9</td><td>2.8</td><td>5.2E-03</td><td>6.9E-03</td><td>MB231/MB468</td><td>MP</td></tr>
<tr><td>>hg38_21_+_8433172_8446622_RNA45SN1_45S_50ntFlanks@11567.11603.37|AGCAGCCGACTTAGAACTGGTGCGGACCAGGGGAATC</td><td>>hg38_21_+_8433172_8446622_RNA45SN1_45S_50ntFlanks</td><td>11567</td><td>11603</td><td>37</td><td>AGCAGCCGACTTAGAACTGGTGCGGACCAGGGGAATC</td><td>883.1</td><td>-1.8</td><td>0.6</td><td>-2.8</td><td>4.8E-03</td><td>6.9E-03</td><td>BT20/MB468</td><td>MITO</td></tr>
<tr><td>>trna143_LysTTT_6_-_27559593_27559665@14.33.20,trna149_LysTTT_6_-_27302769_27302841@14.33.20,trna14_LysTTT_11_-_59327808_59327880@14.33.20,trna2_LysTTT_17_+_8022473_8022545@14.33.20,trna54_LysTTT_1_+_204475655_204475727@14.33.20,trna5_LysTTT_11_+_59323902_59323974@14.33.20,trna62_LysTTT_1_-_204476158_204476230@14.33.20,trna76_LysTTT_6_+_28918806_28918878@14.33.20|AGTCGGTAGAGCATCAGACT</td><td>>trna143_LysTTT_6_-_27559593_27559665</td><td>14</td><td>33</td><td>20</td><td>AGTCGGTAGAGCATCAGACT</td><td>97.2</td><td>4.3</td><td>1.5</td><td>2.8</td><td>5.2E-03</td><td>6.9E-03</td><td>MB231/MB468</td><td>MP</td></tr>
<tr><td>>hg38_21_+_8433172_8446622_RNA45SN1_45S_50ntFlanks@11783.11816.34|CATGAATGGATGAACGAGATTCCCACTGTCCCTA</td><td>>hg38_21_+_8433172_8446622_RNA45SN1_45S_50ntFlanks</td><td>11783</td><td>11816</td><td>34</td><td>CATGAATGGATGAACGAGATTCCCACTGTCCCTA</td><td>260.3</td><td>-4.0</td><td>1.4</td><td>-2.9</td><td>3.3E-03</td><td>6.9E-03</td><td>BT20/MB468</td><td>MP</td></tr>
<tr><td>>hg38_21_+_8433172_8446622_RNA45SN1_45S_50ntFlanks@8402.8422.21|GAGGTAAACGGGTGGGGTCCG</td><td>>hg38_21_+_8433172_8446622_RNA45SN1_45S_50ntFlanks</td><td>8402</td><td>8422</td><td>21</td><td>GAGGTAAACGGGTGGGGTCCG</td><td>67.2</td><td>-4.7</td><td>1.6</td><td>-2.9</td><td>3.3E-03</td><td>6.9E-03</td><td>BT20/MB468</td><td>MP</td></tr>
<tr><td>>hg38_21_+_8433172_8446622_RNA45SN1_45S_50ntFlanks@12420.12457.38|ATGTCGGCTCTTCCTATCATTGTGAAGCAGAATTCACC</td><td>>hg38_21_+_8433172_8446622_RNA45SN1_45S_50ntFlanks</td><td>12420</td><td>12457</td><td>38</td><td>ATGTCGGCTCTTCCTATCATTGTGAAGCAGAATTCACC</td><td>852.9</td><td>2.1</td><td>0.8</td><td>2.7</td><td>6.1E-03</td><td>6.9E-03</td><td>MB231/MB468</td><td>NUC</td></tr>
<tr><td>>am_trna13_LysCTT_14_-_58706613_58706685@32.51.20,trna2_LysCTT_15_+_79152904_79152976@32.51.20,trna4_LysCTT_18_-_43669271_43669343@32.51.20|CTCTTAATCCCAGGGTCGTG</td><td>>am_trna13_LysCTT_14_-_58706613_58706685</td><td>32</td><td>51</td><td>20</td><td>CTCTTAATCCCAGGGTCGTG</td><td>234.0</td><td>-4.9</td><td>1.7</td><td>-2.9</td><td>3.3E-03</td><td>7.0E-03</td><td>BT20/MB468</td><td>MP</td></tr>
<tr><td>>hg38_21_+_8433172_8446622_RNA45SN1_45S_50ntFlanks@12419.12460.42|GATGTCGGCTCTTCCTATCATTGTGAAGCAGAATTCACCAAG</td><td>>hg38_21_+_8433172_8446622_RNA45SN1_45S_50ntFlanks</td><td>12419</td><td>12460</td><td>42</td><td>GATGTCGGCTCTTCCTATCATTGTGAAGCAGAATTCACCAAG</td><td>226.3</td><td>4.5</td><td>1.6</td><td>2.7</td><td>6.0E-03</td><td>7.0E-03</td><td>MB231/MB468</td><td>MITO</td></tr>
<tr><td>>am_MI0003780|hsa-mir-1296&WithFlank&10|-|63372951|63373054@22.40.19&[MIMAT0005794&hsa-miR-1296-5p&offsets|0|-3;m-304&10|-|63373012|63373033&offsets|0|-3]|TTAGGGCCCTGGCTCCATC</td><td>>am_MI0003780|hsa-mir-1296&WithFlank&10|-|63372951|63373054</td><td>22</td><td>40</td><td>19</td><td>TTAGGGCCCTGGCTCCATC</td><td>59.4</td><td>5.3</td><td>1.9</td><td>2.8</td><td>5.2E-03</td><td>7.0E-03</td><td>MB231/MB468</td><td>MP</td></tr>
<tr><td>>MI0000483|hsa-mir-186&WithFlank&1|-|71067625|71067722@21.43.23&[MIMAT0000456&hsa-miR-186-5p&offsets|0|+1;m-84&1|-|71067681|71067702&offsets|0|+1]|CAAAGAATTCTCCTTTTGGGCTT</td><td>>MI0000483|hsa-mir-186&WithFlank&1|-|71067625|71067722</td><td>21</td><td>43</td><td>23</td><td>CAAAGAATTCTCCTTTTGGGCTT</td><td>2610.2</td><td>-2.8</td><td>0.9</td><td>-3.2</td><td>1.2E-03</td><td>7.0E-03</td><td>BT20/MB231</td><td>MP</td></tr>
<tr><td>>hg38_21_+_8433172_8446622_RNA45SN1_45S_50ntFlanks@9534.9570.37|AGGAAACTCTGGTGGAGGTCCGTAGCGGTCCTGACGT</td><td>>hg38_21_+_8433172_8446622_RNA45SN1_45S_50ntFlanks</td><td>9534</td><td>9570</td><td>37</td><td>AGGAAACTCTGGTGGAGGTCCGTAGCGGTCCTGACGT</td><td>649.5</td><td>5.0</td><td>1.5</td><td>3.2</td><td>1.2E-03</td><td>7.0E-03</td><td>BT20/MB231</td><td>MP</td></tr>
<tr><td>>am_trnaMT_HisGTG_MT_+_12138_12206@18.38.21|AAAACATCAGATTGTGAATCT</td><td>>am_trnaMT_HisGTG_MT_+_12138_12206</td><td>18</td><td>38</td><td>21</td><td>AAAACATCAGATTGTGAATCT</td><td>110.2</td><td>3.4</td><td>1.2</td><td>2.8</td><td>5.2E-03</td><td>7.0E-03</td><td>MB231/MB468</td><td>MP</td></tr>
<tr><td>>trna111_HisGTG_1_-_147774845_147774916@-1T.18.19,trna118_HisGTG_1_-_145396881_145396952@-1T.18.19,trna16_HisGTG_1_+_146544773_146544844@-1T.18.19,trna1_HisGTG_15_+_45493349_45493420@-1T.18.19,trna21_HisGTG_1_+_147753471_147753542@-1T.18.19,trna33_HisGTG_6_+_27125906_27125977@-1T.18.19,trna7_HisGTG_9_-_14433938_14434009@-1T.18.19,trna8_HisGTG_15_-_45492611_45492682@-1T.18.19,trna9_HisGTG_15_-_45490804_45490875@-1T.18.19|TGCCGTGATCGTATAGTGG</td><td>>trna111_HisGTG_1_-_147774845_147774916</td><td>-1T</td><td>18</td><td>19</td><td>TGCCGTGATCGTATAGTGG</td><td>61.5</td><td>4.5</td><td>1.6</td><td>2.8</td><td>5.2E-03</td><td>7.0E-03</td><td>MB231/MB468</td><td>MP</td></tr>
<tr><td>>hg38_21_+_8433172_8446622_RNA45SN1_45S_50ntFlanks@11552.11588.37|CCTCGGCCGGCGCCTAGCAGCCGACTTAGAACTGGTG</td><td>>hg38_21_+_8433172_8446622_RNA45SN1_45S_50ntFlanks</td><td>11552</td><td>11588</td><td>37</td><td>CCTCGGCCGGCGCCTAGCAGCCGACTTAGAACTGGTG</td><td>114.8</td><td>-4.6</td><td>1.7</td><td>-2.8</td><td>5.2E-03</td><td>7.0E-03</td><td>MB231/MB468</td><td>MP</td></tr>
<tr><td>>hg38_21_+_8433172_8446622_RNA45SN1_45S_50ntFlanks@8342.8375.34|CGTAAGGGAAAGTTGAAAAGAACTTTGAAGAGAG</td><td>>hg38_21_+_8433172_8446622_RNA45SN1_45S_50ntFlanks</td><td>8342</td><td>8375</td><td>34</td><td>CGTAAGGGAAAGTTGAAAAGAACTTTGAAGAGAG</td><td>103.4</td><td>5.4</td><td>1.9</td><td>2.8</td><td>5.2E-03</td><td>7.0E-03</td><td>MB231/MB468</td><td>MP</td></tr>
<tr><td>>hg38_21_+_8433172_8446622_RNA45SN1_45S_50ntFlanks@12437.12472.36|CATTGTGAAGCAGAATTCACCAAGCGTTGGATTGTT</td><td>>hg38_21_+_8433172_8446622_RNA45SN1_45S_50ntFlanks</td><td>12437</td><td>12472</td><td>36</td><td>CATTGTGAAGCAGAATTCACCAAGCGTTGGATTGTT</td><td>383.7</td><td>-2.0</td><td>0.7</td><td>-2.8</td><td>5.7E-03</td><td>7.0E-03</td><td>BT20/MB468</td><td>CYTO</td></tr>
<tr><td>>MI0000081|hsa-mir-24-2&WithFlank&19|-|13836281|13836365@56.77.22(+1U);MI0000080|hsa-mir-24-1&WithFlank&9|+|95086015|95086094@50.71.22(+1U)&[MIMAT0000080&hsa-miR-24-3p&offsets|0|0(+1U);m-36&19|-|13836288|13836310&offsets|0|-1(+1U)];[MIMAT0000080_1&hsa-miR-24-3p&offsets|0|0(+1U);m-35&9|+|95086064|95086086&offsets|0|-1(+1U)]|TGGCTCAGTTCAGCAGGAACAGT</td><td>>MI0000081|hsa-mir-24-2&WithFlank&19|-|13836281|13836365</td><td>56</td><td>77</td><td>22(+1U)</td><td>TGGCTCAGTTCAGCAGGAACAGT</td><td>10892.5</td><td>2.7</td><td>1.0</td><td>2.8</td><td>5.2E-03</td><td>7.0E-03</td><td>MB231/MB468</td><td>MP</td></tr>
<tr><td>>trna10_LysCTT_16_+_3241501_3241573@1.35.35,trna13_LysCTT_14_-_58706613_58706685@1.35.35,trna2_LysCTT_15_+_79152904_79152976@1.35.35|GCCCGGCTAGCTCAGTCGGTAGAGCATGGGACTCT</td><td>>trna10_LysCTT_16_+_3241501_3241573</td><td>1</td><td>35</td><td>35</td><td>GCCCGGCTAGCTCAGTCGGTAGAGCATGGGACTCT</td><td>203717.1</td><td>3.7</td><td>1.2</td><td>3.2</td><td>1.5E-03</td><td>7.0E-03</td><td>BT20/MB231</td><td>CYTO</td></tr>
<tr><td>>hg38_21_+_8433172_8446622_RNA45SN1_45S_50ntFlanks@12417.12434.18|TCGATGTCGGCTCTTCCT</td><td>>hg38_21_+_8433172_8446622_RNA45SN1_45S_50ntFlanks</td><td>12417</td><td>12434</td><td>18</td><td>TCGATGTCGGCTCTTCCT</td><td>11037.6</td><td>-3.6</td><td>1.3</td><td>-2.7</td><td>6.0E-03</td><td>7.0E-03</td><td>BT20/MB468</td><td>NUC</td></tr>
<tr><td>>hg38_21_+_8433172_8446622_RNA45SN1_45S_50ntFlanks@3852.3872.21|TAACTGTGGTAATTCTAGAGC</td><td>>hg38_21_+_8433172_8446622_RNA45SN1_45S_50ntFlanks</td><td>3852</td><td>3872</td><td>21</td><td>TAACTGTGGTAATTCTAGAGC</td><td>171.5</td><td>-1.7</td><td>0.6</td><td>-2.8</td><td>5.7E-03</td><td>7.0E-03</td><td>MB231/MB468</td><td>CYTO</td></tr>
<tr><td>>hg38_21_+_8433172_8446622_RNA45SN1_45S_50ntFlanks@11573.11601.29|CGACTTAGAACTGGTGCGGACCAGGGGAA</td><td>>hg38_21_+_8433172_8446622_RNA45SN1_45S_50ntFlanks</td><td>11573</td><td>11601</td><td>29</td><td>CGACTTAGAACTGGTGCGGACCAGGGGAA</td><td>386.2</td><td>5.3</td><td>1.6</td><td>3.2</td><td>1.2E-03</td><td>7.0E-03</td><td>BT20/MB231</td><td>MP</td></tr>
<tr><td>>MI0000087|hsa-mir-29a&WithFlank&7|-|130876741|130876816@48.67.20&[MIMAT0000086&hsa-miR-29a-3p&offsets|0|-2;m-26&7|-|130876749|130876769&offsets|0|-1]|TAGCACCATCTGAAATCGGT</td><td>>MI0000087|hsa-mir-29a&WithFlank&7|-|130876741|130876816</td><td>48</td><td>67</td><td>20</td><td>TAGCACCATCTGAAATCGGT</td><td>605.3</td><td>-2.9</td><td>0.9</td><td>-3.2</td><td>1.6E-03</td><td>7.0E-03</td><td>BT20/MB231</td><td>MITO</td></tr>
<tr><td>>hg38_21_+_8433172_8446622_RNA45SN1_45S_50ntFlanks@11208.11253.46|GCGGCGACTCTGGACGCGAGCCGGGCCCTTCCCGTGGATCGCCCCA</td><td>>hg38_21_+_8433172_8446622_RNA45SN1_45S_50ntFlanks</td><td>11208</td><td>11253</td><td>46</td><td>GCGGCGACTCTGGACGCGAGCCGGGCCCTTCCCGTGGATCGCCCCA</td><td>166.9</td><td>-3.6</td><td>1.3</td><td>-2.8</td><td>5.7E-03</td><td>7.0E-03</td><td>BT20/MB468</td><td>CYTO</td></tr>
<tr><td>>MI0000067|hsa-let-7f-1&WithFlank&9|+|94176341|94176439@13.31.19(+1A);MI0000068|hsa-let-7f-2&WithFlank&X|-|53557186|53557280@14.32.19(+1A)&[MIMAT0000067&hsa-let-7f-5p&offsets|0|-3(+1A);m-20&9|+|94176353|94176374&offsets|0|-3(+1A)];[MIMAT0000067_1&hsa-let-7f-5p&offsets|0|-3(+1A);m-19&X|-|53557246|53557267&offsets|0|-3(+1A)]|TGAGGTAGTAGATTGTATAA</td><td>>MI0000067|hsa-let-7f-1&WithFlank&9|+|94176341|94176439</td><td>13</td><td>31</td><td>19(+1A)</td><td>TGAGGTAGTAGATTGTATAA</td><td>29.6</td><td>8.1</td><td>2.9</td><td>2.8</td><td>5.2E-03</td><td>7.0E-03</td><td>MB231/MB468</td><td>MP</td></tr>
<tr><td>>MI0005561|hsa-mir-877&WithFlank&6|+|30584326|30584423@7.27.21&[MIMAT0004949&hsa-miR-877-5p&offsets|0|+1;m-382&6|+|30584332|30584354&offsets|0|-2]|GTAGAGGAGATGGCGCAGGGG</td><td>>MI0005561|hsa-mir-877&WithFlank&6|+|30584326|30584423</td><td>7</td><td>27</td><td>21</td><td>GTAGAGGAGATGGCGCAGGGG</td><td>97.2</td><td>5.3</td><td>1.8</td><td>2.9</td><td>3.3E-03</td><td>7.0E-03</td><td>BT20/MB468</td><td>MP</td></tr>
<tr><td>>hg38_21_+_8433172_8446622_RNA45SN1_45S_50ntFlanks@11923.11940.18|AGAGGTGTAGAATAAGTG</td><td>>hg38_21_+_8433172_8446622_RNA45SN1_45S_50ntFlanks</td><td>11923</td><td>11940</td><td>18</td><td>AGAGGTGTAGAATAAGTG</td><td>61.7</td><td>5.0</td><td>1.7</td><td>2.9</td><td>3.3E-03</td><td>7.0E-03</td><td>BT20/MB468</td><td>MP</td></tr>
<tr><td>>hg38_21_+_8433172_8446622_RNA45SN1_45S_50ntFlanks@9563.9596.34|CCTGACGTGCAAATCGGTCGTCCGACCTGGGTAT</td><td>>hg38_21_+_8433172_8446622_RNA45SN1_45S_50ntFlanks</td><td>9563</td><td>9596</td><td>34</td><td>CCTGACGTGCAAATCGGTCGTCCGACCTGGGTAT</td><td>456.7</td><td>-5.2</td><td>1.8</td><td>-2.9</td><td>3.3E-03</td><td>7.0E-03</td><td>BT20/MB468</td><td>MP</td></tr>
<tr><td>>hg38_21_+_8433172_8446622_RNA45SN1_45S_50ntFlanks@12005.12035.31|GGGCCGCCGGTGAAATACCACTACTCTGATC</td><td>>hg38_21_+_8433172_8446622_RNA45SN1_45S_50ntFlanks</td><td>12005</td><td>12035</td><td>31</td><td>GGGCCGCCGGTGAAATACCACTACTCTGATC</td><td>164.7</td><td>4.0</td><td>1.4</td><td>2.9</td><td>3.3E-03</td><td>7.0E-03</td><td>BT20/MB468</td><td>MP</td></tr>
</table>

<table>
<tr><td>>hg38_21_+_8433172_8446622_RNA45SN1_45S_50ntFlanks@12678.12705.28|AATCCCGCCCAGGCGGAACGATACGGCA</td><td>>hg38_21_+_8433172_8446622_RNA45SN1_45S_50ntFlanks</td><td>12678</td><td>12705</td><td>28</td><td>AATCCCGCCCAGGCGGAACGATACGGCA</td><td>50.2</td><td>-4.9</td><td>1.7</td><td>-2.9</td><td>3.3E-03</td><td>7.0E-03</td><td>BT20/MB468</td><td>MP</td></tr>
<tr><td>>hg38_21_+_8433172_8446622_RNA45SN1_45S_50ntFlanks@10357.10388.32|CTTTGAAGGCCGAAGTGGAGAAGGGTTCCATG</td><td>>hg38_21_+_8433172_8446622_RNA45SN1_45S_50ntFlanks</td><td>10357</td><td>10388</td><td>32</td><td>CTTTGAAGGCCGAAGTGGAGAAGGGTTCCATG</td><td>173.4</td><td>-3.2</td><td>1.1</td><td>-2.8</td><td>4.9E-03</td><td>7.0E-03</td><td>BT20/MB468</td><td>MITO</td></tr>
<tr><td>>hg38_21_+_8433172_8446622_RNA45SN1_45S_50ntFlanks@8528.8549.22|CTTCCCCCGCCGCCCCTCCTCC</td><td>>hg38_21_+_8433172_8446622_RNA45SN1_45S_50ntFlanks</td><td>8528</td><td>8549</td><td>22</td><td>CTTCCCCCGCCGCCCCTCCTCC</td><td>380.1</td><td>-4.5</td><td>1.6</td><td>-2.8</td><td>5.7E-03</td><td>7.0E-03</td><td>MB231/MB468</td><td>CYTO</td></tr>
<tr><td>>hg38_21_+_8433172_8446622_RNA45SN1_45S_50ntFlanks@11875.11893.19|GAAGACCCTGTTGAGCTTG</td><td>>hg38_21_+_8433172_8446622_RNA45SN1_45S_50ntFlanks</td><td>11875</td><td>11893</td><td>19</td><td>GAAGACCCTGTTGAGCTTG</td><td>895.4</td><td>-2.8</td><td>1.0</td><td>-2.9</td><td>3.3E-03</td><td>7.0E-03</td><td>BT20/MB468</td><td>MP</td></tr>
<tr><td>>hg38_21_+_8433172_8446622_RNA45SN1_45S_50ntFlanks@6647.6674.28|CGTACGACTCTTAGCGGTGGATCACTCG</td><td>>hg38_21_+_8433172_8446622_RNA45SN1_45S_50ntFlanks</td><td>6647</td><td>6674</td><td>28</td><td>CGTACGACTCTTAGCGGTGGATCACTCG</td><td>1095.0</td><td>2.8</td><td>1.0</td><td>2.7</td><td>6.1E-03</td><td>7.0E-03</td><td>MB231/MB468</td><td>MITO</td></tr>
<tr><td>>hg38_MT_+_1621_3279_RNR2_16S_50ntFlanks@1356.1385.30|GGTTTACGACCTCGATGTTGGATCAGGACA</td><td>>hg38_MT_+_1621_3279_RNR2_16S_50ntFlanks</td><td>1356</td><td>1385</td><td>30</td><td>GGTTTACGACCTCGATGTTGGATCAGGACA</td><td>130.1</td><td>3.2</td><td>1.1</td><td>2.9</td><td>3.3E-03</td><td>7.0E-03</td><td>BT20/MB468</td><td>MP</td></tr>
<tr><td>>hg38_21_+_8433172_8446622_RNA45SN1_45S_50ntFlanks@11567.11601.35|AGCAGCCGACTTAGAACTGGTGCGGACCAGGGGAA</td><td>>hg38_21_+_8433172_8446622_RNA45SN1_45S_50ntFlanks</td><td>11567</td><td>11601</td><td>35</td><td>AGCAGCCGACTTAGAACTGGTGCGGACCAGGGGAA</td><td>265.5</td><td>-1.6</td><td>0.6</td><td>-2.8</td><td>5.7E-03</td><td>7.0E-03</td><td>BT20/MB468</td><td>CYTO</td></tr>
<tr><td>>hg38_MT_+_1621_3279_RNR2_16S_50ntFlanks@125.150.26|TAGGCGATAGAAATTGAAACCTGGCG</td><td>>hg38_MT_+_1621_3279_RNR2_16S_50ntFlanks</td><td>125</td><td>150</td><td>26</td><td>TAGGCGATAGAAATTGAAACCTGGCG</td><td>62.4</td><td>5.1</td><td>1.7</td><td>2.9</td><td>3.3E-03</td><td>7.0E-03</td><td>BT20/MB468</td><td>MP</td></tr>
<tr><td>>hg38_21_+_8433172_8446622_RNA45SN1_45S_50ntFlanks@8423.8442.20|CGCAGTCCGCCCGGAGGATT</td><td>>hg38_21_+_8433172_8446622_RNA45SN1_45S_50ntFlanks</td><td>8423</td><td>8442</td><td>20</td><td>CGCAGTCCGCCCGGAGGATT</td><td>1736.4</td><td>-5.4</td><td>1.8</td><td>-2.9</td><td>3.3E-03</td><td>7.0E-03</td><td>BT20/MB468</td><td>MP</td></tr>
<tr><td>>am_MI0000809|hsa-mir-151a&WithFlank&8|-|140732558|140732659@52.72.21(+1A)&[MIMAT0000757&hsa-miR-151a-3p&offsets|-1|-1(+1A);m-30&8|-|140732587|140732607&offsets|-1|-1(+1A)]|ACTAGACTGAAGCTCCTTGAGA</td><td>>am_MI0000809|hsa-mir-151a&WithFlank&8|-|140732558|140732659</td><td>52</td><td>72</td><td>21(+1A)</td><td>ACTAGACTGAAGCTCCTTGAGA</td><td>488.3</td><td>2.3</td><td>0.7</td><td>3.2</td><td>1.5E-03</td><td>7.0E-03</td><td>BT20/MB231</td><td>CYTO</td></tr>
<tr><td>>hg38_21_+_8433172_8446622_RNA45SN1_45S_50ntFlanks@4499.4519.21|AGCGTTTACTTTGAAAAAATT</td><td>>hg38_21_+_8433172_8446622_RNA45SN1_45S_50ntFlanks</td><td>4499</td><td>4519</td><td>21</td><td>AGCGTTTACTTTGAAAAAATT</td><td>370.0</td><td>2.6</td><td>0.8</td><td>3.2</td><td>1.5E-03</td><td>7.0E-03</td><td>BT20/MB231</td><td>CYTO</td></tr>
<tr><td>>hg38_21_+_8433172_8446622_RNA45SN1_45S_50ntFlanks@10099.10127.29|CGCGCGCGCGTGTGGTGTGCGTCGGAGGG</td><td>>hg38_21_+_8433172_8446622_RNA45SN1_45S_50ntFlanks</td><td>10099</td><td>10127</td><td>29</td><td>CGCGCGCGCGTGTGGTGTGCGTCGGAGGG</td><td>205.3</td><td>-4.9</td><td>1.7</td><td>-2.9</td><td>3.3E-03</td><td>7.0E-03</td><td>BT20/MB468</td><td>MP</td></tr>
<tr><td>>hg38_MT_+_1621_3279_RNR2_16S_50ntFlanks@1195.1215.21|GGGCGACCTCGGAGCAGAACC</td><td>>hg38_MT_+_1621_3279_RNR2_16S_50ntFlanks</td><td>1195</td><td>1215</td><td>21</td><td>GGGCGACCTCGGAGCAGAACC</td><td>27.7</td><td>8.0</td><td>2.9</td><td>2.8</td><td>5.2E-03</td><td>7.0E-03</td><td>MB231/MB468</td><td>MP</td></tr>
<tr><td>>MI0000803|hsa-mir-330&WithFlank&19|-|45638988|45639093@64.85.22&[MIMAT0000751&hsa-miR-330-3p&offsets|+1|0;m-347&19|-|45639009|45639030&offsets|0|0]|CAAAGCACACGGCCTGCAGAGA</td><td>>MI0000803|hsa-mir-330&WithFlank&19|-|45638988|45639093</td><td>64</td><td>85</td><td>22</td><td>CAAAGCACACGGCCTGCAGAGA</td><td>73.3</td><td>5.0</td><td>1.7</td><td>2.9</td><td>3.3E-03</td><td>7.0E-03</td><td>BT20/MB468</td><td>MP</td></tr>
<tr><td>>hg38_21_+_8433172_8446622_RNA45SN1_45S_50ntFlanks@9198.9223.26|GTTCTCTCGGGGCCACGCGCGCGTCC</td><td>>hg38_21_+_8433172_8446622_RNA45SN1_45S_50ntFlanks</td><td>9198</td><td>9223</td><td>26</td><td>GTTCTCTCGGGGCCACGCGCGCGTCC</td><td>394.5</td><td>-4.4</td><td>1.6</td><td>-2.8</td><td>4.9E-03</td><td>7.0E-03</td><td>BT20/MB468</td><td>MITO</td></tr>
<tr><td>>hg38_21_+_8433172_8446622_RNA45SN1_45S_50ntFlanks@7974.8009.36|ACGCGACCTCAGATCAGACGTGGCGACCCGCTGAAT</td><td>>hg38_21_+_8433172_8446622_RNA45SN1_45S_50ntFlanks</td><td>7974</td><td>8009</td><td>36</td><td>ACGCGACCTCAGATCAGACGTGGCGACCCGCTGAAT</td><td>20366.6</td><td>2.1</td><td>0.8</td><td>2.7</td><td>6.1E-03</td><td>7.0E-03</td><td>MB231/MB468</td><td>MITO</td></tr>
<tr><td>>hg38_21_+_8433172_8446622_RNA45SN1_45S_50ntFlanks@6655.6698.44|TCTTAGCGGTGGATCACTCGGCTCGTGCGTCGATGAAGAACGCA</td><td>>hg38_21_+_8433172_8446622_RNA45SN1_45S_50ntFlanks</td><td>6655</td><td>6698</td><td>44</td><td>TCTTAGCGGTGGATCACTCGGCTCGTGCGTCGATGAAGAACGCA</td><td>1235.3</td><td>-2.2</td><td>0.8</td><td>-2.7</td><td>6.1E-03</td><td>7.0E-03</td><td>MB231/MB468</td><td>MITO</td></tr>
<tr><td>>hg38_MT_+_598_1651_RNR1_12S_50ntFlanks@52.88.37|ATAGGTTTGGTCCTAGCCTTTCTATTAGCTCTTAGTA</td><td>>hg38_MT_+_598_1651_RNR1_12S_50ntFlanks</td><td>52</td><td>88</td><td>37</td><td>ATAGGTTTGGTCCTAGCCTTTCTATTAGCTCTTAGTA</td><td>8115.4</td><td>2.4</td><td>0.9</td><td>2.7</td><td>6.1E-03</td><td>7.0E-03</td><td>MB231/MB468</td><td>MITO</td></tr>
<tr><td>>am_trnaMT_ValTAC_MT_+_1602_1670@29.68.40|ACTTACACTTAGGAGATTTCAACTTAACTTGACCGCTCTG</td><td>>am_trnaMT_ValTAC_MT_+_1602_1670</td><td>29</td><td>68</td><td>40</td><td>ACTTACACTTAGGAGATTTCAACTTAACTTGACCGCTCTG</td><td>632.8</td><td>-5.1</td><td>1.8</td><td>-2.7</td><td>6.1E-03</td><td>7.0E-03</td><td>MB231/MB468</td><td>NUC</td></tr>
<tr><td>>MI0000085|hsa-mir-27a&WithFlank&19|-|13836434|13836523@57.77.21&[MIMAT0000084&hsa-miR-27a-3p&offsets|0|0;m-43&19|-|13836447|13836467&offsets|0|0]|TTCACAGTGGCTAAGTTCCGC</td><td>>MI0000085|hsa-mir-27a&WithFlank&19|-|13836434|13836523</td><td>57</td><td>77</td><td>21</td><td>TTCACAGTGGCTAAGTTCCGC</td><td>11896.7</td><td>2.7</td><td>1.0</td><td>2.8</td><td>5.2E-03</td><td>7.0E-03</td><td>MB231/MB468</td><td>MP</td></tr>
<tr><td>>hg38_MT_+_598_1651_RNR1_12S_50ntFlanks@634.665.32|AACCTCACCACCTCTTGCTCAGCCTATATACC</td><td>>hg38_MT_+_598_1651_RNR1_12S_50ntFlanks</td><td>634</td><td>665</td><td>32</td><td>AACCTCACCACCTCTTGCTCAGCCTATATACC</td><td>147.6</td><td>5.2</td><td>1.9</td><td>2.8</td><td>5.2E-03</td><td>7.0E-03</td><td>MB231/MB468</td><td>MP</td></tr>
<tr><td>>hg38_21_+_8433172_8446622_RNA45SN1_45S_50ntFlanks@3841.3868.28|TCCTACTTGGATAACTGTGGTAATTCTA</td><td>>hg38_21_+_8433172_8446622_RNA45SN1_45S_50ntFlanks</td><td>3841</td><td>3868</td><td>28</td><td>TCCTACTTGGATAACTGTGGTAATTCTA</td><td>370.5</td><td>-1.8</td><td>0.7</td><td>-2.7</td><td>6.0E-03</td><td>7.0E-03</td><td>BT20/MB468</td><td>NUC</td></tr>
<tr><td>>hg38_21_+_8433172_8446622_RNA45SN1_45S_50ntFlanks@8439.8470.32|GATTCAACCCGGCGGCGGGTCCGGCCGTGTCG</td><td>>hg38_21_+_8433172_8446622_RNA45SN1_45S_50ntFlanks</td><td>8439</td><td>8470</td><td>32</td><td>GATTCAACCCGGCGGCGGGTCCGGCCGTGTCG</td><td>56.0</td><td>-4.8</td><td>1.6</td><td>-2.9</td><td>3.3E-03</td><td>7.0E-03</td><td>BT20/MB468</td><td>MP</td></tr>
<tr><td>>hg38_21_+_8433172_8446622_RNA45SN1_45S_50ntFlanks@10869.10901.33|AGGGCTGGGTCGGTCGGGCTGGGGCGCGAAGCG</td><td>>hg38_21_+_8433172_8446622_RNA45SN1_45S_50ntFlanks</td><td>10869</td><td>10901</td><td>33</td><td>AGGGCTGGGTCGGTCGGGCTGGGGCGCGAAGCG</td><td>489.3</td><td>-4.5</td><td>1.6</td><td>-2.8</td><td>5.7E-03</td><td>7.0E-03</td><td>BT20/MB468</td><td>CYTO</td></tr>
<tr><td>>MI0000272|hsa-mir-182&WithFlank&7|-|129770377|129770498@29.54.26&[MIMAT0000259&hsa-miR-182-5p&offsets|0|+2;m-22&7|-|129770449|129770470&offsets|0|+4]|TTTGGCAATGGTAGAACTCACACTGG</td><td>>MI0000272|hsa-mir-182&WithFlank&7|-|129770377|129770498</td><td>29</td><td>54</td><td>26</td><td>TTTGGCAATGGTAGAACTCACACTGG</td><td>1048.4</td><td>3.2</td><td>1.1</td><td>2.9</td><td>3.3E-03</td><td>7.0E-03</td><td>BT20/MB468</td><td>MP</td></tr>
<tr><td>>hg38_21_+_8433172_8446622_RNA45SN1_45S_50ntFlanks@9564.9592.29|CTGACGTGCAAATCGGTCGTCCGACCTGG</td><td>>hg38_21_+_8433172_8446622_RNA45SN1_45S_50ntFlanks</td><td>9564</td><td>9592</td><td>29</td><td>CTGACGTGCAAATCGGTCGTCCGACCTGG</td><td>763.3</td><td>-1.7</td><td>0.6</td><td>-2.8</td><td>4.9E-03</td><td>7.0E-03</td><td>BT20/MB468</td><td>MITO</td></tr>
<tr><td>>MI0000114|hsa-mir-107&WithFlank&10|-|89592741|89592833@56.77.22&[MIMAT0000104&hsa-miR-107&offsets|0|-1;m-78&10|-|89592758|89592778&offsets|0|+1]|AGCAGCATTGTACAGGGCTATC</td><td>>MI0000114|hsa-mir-107&WithFlank&10|-|89592741|89592833</td><td>56</td><td>77</td><td>22</td><td>AGCAGCATTGTACAGGGCTATC</td><td>1029.0</td><td>-2.8</td><td>0.9</td><td>-3.2</td><td>1.6E-03</td><td>7.0E-03</td><td>BT20/MB231</td><td>MITO</td></tr>
<tr><td>>hg38_21_+_8433172_8446622_RNA45SN1_45S_50ntFlanks@7989.8010.22|AGACGTGGCGACCCGCTGAATT</td><td>>hg38_21_+_8433172_8446622_RNA45SN1_45S_50ntFlanks</td><td>7989</td><td>8010</td><td>22</td><td>AGACGTGGCGACCCGCTGAATT</td><td>1722.7</td><td>2.2</td><td>0.8</td><td>2.8</td><td>4.9E-03</td><td>7.0E-03</td><td>BT20/MB468</td><td>MITO</td></tr>
<tr><td>>MI0000102|hsa-mir-100&WithFlank&11|-|122152223|122152314@19.40.22(+1U)&[MIMAT0000098&hsa-miR-100-5p&offsets|0|0(+1U);m-27&11|-|122152276|122152296&offsets|0|+1(+1U)]|AACCCGTAGATCCGAACTTGTGT</td><td>>MI0000102|hsa-mir-100&WithFlank&11|-|122152223|122152314</td><td>19</td><td>40</td><td>22(+1U)</td><td>AACCCGTAGATCCGAACTTGTGT</td><td>48445.7</td><td>-3.8</td><td>1.2</td><td>-3.2</td><td>1.5E-03</td><td>7.0E-03</td><td>BT20/MB231</td><td>CYTO</td></tr>
<tr><td>>hg38_21_+_8433172_8446622_RNA45SN1_45S_50ntFlanks@7988.8019.32|CAGACGTGGCGACCCGCTGAATTTAAGCATAT</td><td>>hg38_21_+_8433172_8446622_RNA45SN1_45S_50ntFlanks</td><td>7988</td><td>8019</td><td>32</td><td>CAGACGTGGCGACCCGCTGAATTTAAGCATAT</td><td>403.3</td><td>-3.2</td><td>1.0</td><td>-3.2</td><td>1.5E-03</td><td>7.0E-03</td><td>BT20/MB231</td><td>CYTO</td></tr>
<tr><td>>hg38_21_+_8433172_8446622_RNA45SN1_45S_50ntFlanks@6649.6678.30|TACGACTCTTAGCGGTGGATCACTCGGCTC</td><td>>hg38_21_+_8433172_8446622_RNA45SN1_45S_50ntFlanks</td><td>6649</td><td>6678</td><td>30</td><td>TACGACTCTTAGCGGTGGATCACTCGGCTC</td><td>239.8</td><td>3.0</td><td>1.0</td><td>2.9</td><td>3.3E-03</td><td>7.0E-03</td><td>BT20/MB468</td><td>MP</td></tr>
<tr><td>>hg38_21_+_8433172_8446622_RNA45SN1_45S_50ntFlanks@10401.10422.22|GAACATGGGTCAGTCGGTCCTG</td><td>>hg38_21_+_8433172_8446622_RNA45SN1_45S_50ntFlanks</td><td>10401</td><td>10422</td><td>22</td><td>GAACATGGGTCAGTCGGTCCTG</td><td>209.4</td><td>-2.2</td><td>0.8</td><td>-2.9</td><td>3.3E-03</td><td>7.0E-03</td><td>BT20/MB468</td><td>MP</td></tr>
<tr><td>>hg38_21_+_8433172_8446622_RNA45SN1_45S_50ntFlanks@10805.10826.22|GAACAATGTAGGTAAGGGAAGT</td><td>>hg38_21_+_8433172_8446622_RNA45SN1_45S_50ntFlanks</td><td>10805</td><td>10826</td><td>22</td><td>GAACAATGTAGGTAAGGGAAGT</td><td>135.2</td><td>-4.7</td><td>1.6</td><td>-2.9</td><td>3.3E-03</td><td>7.0E-03</td><td>BT20/MB468</td><td>MP</td></tr>
<tr><td>>hg38_21_+_8433172_8446622_RNA45SN1_45S_50ntFlanks@9533.9568.36|GAGGAAACTCTGGTGGAGGTCCGTAGCGGTCCTGAC</td><td>>hg38_21_+_8433172_8446622_RNA45SN1_45S_50ntFlanks</td><td>9533</td><td>9568</td><td>36</td><td>GAGGAAACTCTGGTGGAGGTCCGTAGCGGTCCTGAC</td><td>185.8</td><td>-2.4</td><td>0.7</td><td>-3.2</td><td>1.5E-03</td><td>7.0E-03</td><td>BT20/MB231</td><td>CYTO</td></tr>
<tr><td>>hg38_21_+_8433172_8446622_RNA45SN1_45S_50ntFlanks@10789.10818.30|AGCCTCTGGCATGTTGGAACAATGTAGGTA</td><td>>hg38_21_+_8433172_8446622_RNA45SN1_45S_50ntFlanks</td><td>10789</td><td>10818</td><td>30</td><td>AGCCTCTGGCATGTTGGAACAATGTAGGTA</td><td>200.7</td><td>5.1</td><td>1.6</td><td>3.2</td><td>1.2E-03</td><td>7.0E-03</td><td>BT20/MB231</td><td>MP</td></tr>
<tr><td>>trna111_HisGTG_1_-_147774845_147774916@-1T.28.29,trna118_HisGTG_1_-_145396881_145396952@-1T.28.29,trna16_HisGTG_1_+_146544773_146544844@-1T.28.29,trna1_HisGTG_15_+_45493349_45493420@-1T.28.29,trna21_HisGTG_1_+_147753471_147753542@-1T.28.29,trna33_HisGTG_6_+_27125906_27125977@-1T.28.29,trna7_HisGTG_9_-_14433938_14434009@-1T.28.29,trna8_HisGTG_15_-_45492611_45492682@-1T.28.29,trna9_HisGTG_15_-_45490804_45490875@-1T.28.29|TGCCGTGATCGTATAGTGGTTAGTACTCT</td><td>>trna111_HisGTG_1_-_147774845_147774916</td><td>-1T</td><td>28</td><td>29</td><td>TGCCGTGATCGTATAGTGGTTAGTACTCT</td><td>617.9</td><td>2.4</td><td>0.9</td><td>2.7</td><td>6.1E-03</td><td>7.0E-03</td><td>MB231/MB468</td><td>MITO</td></tr>
<tr><td>>hg38_21_+_8433172_8446622_RNA45SN1_45S_50ntFlanks@8192.8215.24|AGCCCGTGGACGGTGTGAGGCCGG</td><td>>hg38_21_+_8433172_8446622_RNA45SN1_45S_50ntFlanks</td><td>8192</td><td>8215</td><td>24</td><td>AGCCCGTGGACGGTGTGAGGCCGG</td><td>343.5</td><td>-4.5</td><td>1.7</td><td>-2.7</td><td>6.1E-03</td><td>7.0E-03</td><td>MB231/MB468</td><td>NUC</td></tr>
<tr><td>>hg38_MT_+_1621_3279_RNR2_16S_50ntFlanks@1370.1388.19|ATGTTGGATCAGGACATCC</td><td>>hg38_MT_+_1621_3279_RNR2_16S_50ntFlanks</td><td>1370</td><td>1388</td><td>19</td><td>ATGTTGGATCAGGACATCC</td><td>220.9</td><td>3.6</td><td>1.3</td><td>2.7</td><td>6.1E-03</td><td>7.1E-03</td><td>MB231/MB468</td><td>MITO</td></tr>
<tr><td>>hg38_21_+_8433172_8446622_RNA45SN1_45S_50ntFlanks@12008.12037.30|CCGCCGGTGAAATACCACTACTCTGATCGT</td><td>>hg38_21_+_8433172_8446622_RNA45SN1_45S_50ntFlanks</td><td>12008</td><td>12037</td><td>30</td><td>CCGCCGGTGAAATACCACTACTCTGATCGT</td><td>342.8</td><td>-1.7</td><td>0.6</td><td>-2.8</td><td>4.9E-03</td><td>7.1E-03</td><td>BT20/MB468</td><td>MITO</td></tr>
<tr><td>>trna116_GluCTC_1_-_145399233_145399304@36.71.36,trna134_GluTTC_1_-_16861774_16861845@36.71.36,trna71_GluCTC_1_-_161439189_161439260@36.71.36,trna74_GluCTC_1_-_161431809_161431880@36.71.36,trna77_GluCTC_1_-_161424398_161424469@36.71.36,trna77_GluCTC_6_+_28949976_28950047@36.71.36,trna80_GluCTC_1_-_161417018_161417089@36.71.36,trna84_GluTTC_1_-_161391883_161391954@36.71.36,trna87_GluCTC_6_-_126101393_126101464@36.71.36|CACCGCCGCGGCCCGGGTTCGATTCCCGGTCAGGGA</td><td>>trna116_GluCTC_1_-_145399233_145399304</td><td>36</td><td>71</td><td>36</td><td>CACCGCCGCGGCCCGGGTTCGATTCCCGGTCAGGGA</td><td>155.3</td><td>-3.4</td><td>1.2</td><td>-2.8</td><td>5.8E-03</td><td>7.1E-03</td><td>MB231/MB468</td><td>CYTO</td></tr>
<tr><td>>hg38_21_+_8433172_8446622_RNA45SN1_45S_50ntFlanks@8189.8214.26|CCCAGCCCGTGGACGGTGTGAGGCCG</td><td>>hg38_21_+_8433172_8446622_RNA45SN1_45S_50ntFlanks</td><td>8189</td><td>8214</td><td>26</td><td>CCCAGCCCGTGGACGGTGTGAGGCCG</td><td>66.9</td><td>-4.8</td><td>1.6</td><td>-2.9</td><td>3.4E-03</td><td>7.1E-03</td><td>BT20/MB468</td><td>MP</td></tr>
<tr><td>>hg38_21_+_8433172_8446622_RNA45SN1_45S_50ntFlanks@4970.4999.30|CCCGGCCCGGACACGGACAGGATTGACAGA</td><td>>hg38_21_+_8433172_8446622_RNA45SN1_45S_50ntFlanks</td><td>4970</td><td>4999</td><td>30</td><td>CCCGGCCCGGACACGGACAGGATTGACAGA</td><td>190.2</td><td>-3.7</td><td>1.3</td><td>-2.8</td><td>4.9E-03</td><td>7.1E-03</td><td>BT20/MB468</td><td>MITO</td></tr>
<tr><td>>hg38_1_-_228634819_228635039_RNA5S12_5S_50ntFlanks@136.171.36|GGAGACCGCCTGGGAATACCGGGTGCTGTAGGCTTT</td><td>>hg38_1_-_228634819_228635039_RNA5S12_5S_50ntFlanks</td><td>136</td><td>171</td><td>36</td><td>GGAGACCGCCTGGGAATACCGGGTGCTGTAGGCTTT</td><td>28679.1</td><td>5.2</td><td>1.6</td><td>3.2</td><td>1.2E-03</td><td>7.1E-03</td><td>BT20/MB231</td><td>MP</td></tr>
<tr><td>>MI0000088|hsa-mir-30a&WithFlank&6|-|71403545|71403627@54.73.20(+1A)&[MIMAT0000088&hsa-miR-30a-3p&offsets|+1|-1(+1A);m-38&6|-|71403554|71403575&offsets|+1|-1(+1A)]|TTTCAGTCGGATGTTTGCAGA</td><td>>MI0000088|hsa-mir-30a&WithFlank&6|-|71403545|71403627</td><td>54</td><td>73</td><td>20(+1A)</td><td>TTTCAGTCGGATGTTTGCAGA</td><td>396.0</td><td>-3.5</td><td>1.1</td><td>-3.2</td><td>1.2E-03</td><td>7.1E-03</td><td>BT20/MB231</td><td>MP</td></tr>
<tr><td>>hg38_1_-_228634819_228635039_RNA5S12_5S_50ntFlanks@151.171.21|ATACCGGGTGCTGTAGGCTTT</td><td>>hg38_1_-_228634819_228635039_RNA5S12_5S_50ntFlanks</td><td>151</td><td>171</td><td>21</td><td>ATACCGGGTGCTGTAGGCTTT</td><td>9447.5</td><td>-2.8</td><td>0.9</td><td>-2.9</td><td>3.4E-03</td><td>7.1E-03</td><td>BT20/MB468</td><td>MP</td></tr>
<tr><td>>MI0000471|hsa-mir-126&WithFlank&9|+|136670596|136670692@58.79.22&[MIMAT0000445&hsa-miR-126-3p&offsets|0|0;m-40&9|+|136670653|136670674&offsets|0|0]|TCGTACCGTGAGTAATAATGCG</td><td>>MI0000471|hsa-mir-126&WithFlank&9|+|136670596|136670692</td><td>58</td><td>79</td><td>22</td><td>TCGTACCGTGAGTAATAATGCG</td><td>4954.4</td><td>3.1</td><td>1.0</td><td>3.2</td><td>1.5E-03</td><td>7.1E-03</td><td>BT20/MB231</td><td>CYTO</td></tr>
<tr><td>>am_trnaMT_ValTAC_MT_+_1602_1670@43.67.25|GATTTCAACTTAACTTGACCGCTCT</td><td>>am_trnaMT_ValTAC_MT_+_1602_1670</td><td>43</td><td>67</td><td>25</td><td>GATTTCAACTTAACTTGACCGCTCT</td><td>388.8</td><td>2.7</td><td>0.9</td><td>3.2</td><td>1.5E-03</td><td>7.1E-03</td><td>BT20/MB231</td><td>CYTO</td></tr>
<tr><td>>MI0000065|hsa-let-7d&WithFlank&9|+|94178828|94178926@14.31.18&[MIMAT0000065&hsa-let-7d-5p&offsets|0|-4;m-90&9|+|94178841|94178862&offsets|0|-4]|AGAGGTAGTAGGTTGCAT</td><td>>MI0000065|hsa-let-7d&WithFlank&9|+|94178828|94178926</td><td>14</td><td>31</td><td>18</td><td>AGAGGTAGTAGGTTGCAT</td><td>138.0</td><td>2.1</td><td>0.8</td><td>2.7</td><td>6.1E-03</td><td>7.1E-03</td><td>BT20/MB468</td><td>NUC</td></tr>
</table>

<table>
<tr><th></th><th></th><th></th><th></th><th></th><th></th><th></th><th></th><th></th><th></th><th></th><th></th><th></th><th></th></tr>
<tr><td>>trnaMT_ValTAC_MT_+_1602_1670@37.71.35|TTAGGAGATTTCAACTTAACTTGACCGCTCTGACC</td><td>>trnaMT_ValTAC_MT_+_1602_1670</td><td>37</td><td>71</td><td>35</td><td>TTAGGAGATTTCAACTTAACTTGACCGCTCTGACC</td><td>46.5</td><td>5.2</td><td>1.9</td><td>2.8</td><td>5.3E-03</td><td>7.1E-03</td><td>MB231/MB468</td><td>MP</td></tr>
<tr><td>>am_trnaMT_SerGCT_MT_+_12207_12265@14.31.18|AGAACTGCTAACTCATGC</td><td>>am_trnaMT_SerGCT_MT_+_12207_12265</td><td>14</td><td>31</td><td>18</td><td>AGAACTGCTAACTCATGC</td><td>536.8</td><td>2.4</td><td>0.7</td><td>3.2</td><td>1.5E-03</td><td>7.1E-03</td><td>BT20/MB231</td><td>CYTO</td></tr>
<tr><td>>hg38_21_+_8433172_8446622_RNA45SN1_45S_50ntFlanks@11566.11599.34|TAGCAGCCGACTTAGAACTGGTGCGGACCAGGGG</td><td>>hg38_21_+_8433172_8446622_RNA45SN1_45S_50ntFlanks</td><td>11566</td><td>11599</td><td>34</td><td>TAGCAGCCGACTTAGAACTGGTGCGGACCAGGGG</td><td>57.8</td><td>5.8</td><td>2.0</td><td>2.9</td><td>3.4E-03</td><td>7.1E-03</td><td>BT20/MB468</td><td>MP</td></tr>
<tr><td>>hg38_21_+_8433172_8446622_RNA45SN1_45S_50ntFlanks@10731.10755.25|AGGGTGTAAATCTCGCGCCGGGCCG</td><td>>hg38_21_+_8433172_8446622_RNA45SN1_45S_50ntFlanks</td><td>10731</td><td>10755</td><td>25</td><td>AGGGTGTAAATCTCGCGCCGGGCCG</td><td>341.2</td><td>-4.3</td><td>1.6</td><td>-2.8</td><td>5.8E-03</td><td>7.1E-03</td><td>BT20/MB468</td><td>CYTO</td></tr>
<tr><td>>hg38_21_+_8433172_8446622_RNA45SN1_45S_50ntFlanks@12244.12266.23|AAGGGCAAAAGCTCGCTTGATCT</td><td>>hg38_21_+_8433172_8446622_RNA45SN1_45S_50ntFlanks</td><td>12244</td><td>12266</td><td>23</td><td>AAGGGCAAAAGCTCGCTTGATCT</td><td>145.7</td><td>-4.8</td><td>1.6</td><td>-2.9</td><td>3.4E-03</td><td>7.1E-03</td><td>BT20/MB468</td><td>MP</td></tr>
<tr><td>>hg38_21_+_8433172_8446622_RNA45SN1_45S_50ntFlanks@12642.12669.28|CATCTGTGGGATTATGACTGAACGCCTC</td><td>>hg38_21_+_8433172_8446622_RNA45SN1_45S_50ntFlanks</td><td>12642</td><td>12669</td><td>28</td><td>CATCTGTGGGATTATGACTGAACGCCTC</td><td>366.4</td><td>5.7</td><td>1.8</td><td>3.2</td><td>1.2E-03</td><td>7.1E-03</td><td>BT20/MB231</td><td>MP</td></tr>
<tr><td>>hg38_21_+_8433172_8446622_RNA45SN1_45S_50ntFlanks@11569.11609.41|CAGCCGACTTAGAACTGGTGCGGACCAGGGGAATCCGACTG</td><td>>hg38_21_+_8433172_8446622_RNA45SN1_45S_50ntFlanks</td><td>11569</td><td>11609</td><td>41</td><td>CAGCCGACTTAGAACTGGTGCGGACCAGGGGAATCCGACTG</td><td>269.2</td><td>-2.5</td><td>0.9</td><td>-2.7</td><td>6.2E-03</td><td>7.1E-03</td><td>MB231/MB468</td><td>NUC</td></tr>
<tr><td>>hg38_21_+_8433172_8446622_RNA45SN1_45S_50ntFlanks@10393.10426.34|CAGCAGTTGAACATGGGTCAGTCGGTCCTGAGAG</td><td>>hg38_21_+_8433172_8446622_RNA45SN1_45S_50ntFlanks</td><td>10393</td><td>10426</td><td>34</td><td>CAGCAGTTGAACATGGGTCAGTCGGTCCTGAGAG</td><td>221.8</td><td>-1.6</td><td>0.6</td><td>-2.8</td><td>4.9E-03</td><td>7.1E-03</td><td>BT20/MB468</td><td>MITO</td></tr>
<tr><td>>hg38_21_+_8433172_8446622_RNA45SN1_45S_50ntFlanks@9738.9768.31|GAAACGATCTCAACCTATTCTCAAACTTTAA</td><td>>hg38_21_+_8433172_8446622_RNA45SN1_45S_50ntFlanks</td><td>9738</td><td>9768</td><td>31</td><td>GAAACGATCTCAACCTATTCTCAAACTTTAA</td><td>168.0</td><td>-1.8</td><td>0.6</td><td>-2.8</td><td>4.9E-03</td><td>7.1E-03</td><td>BT20/MB468</td><td>MITO</td></tr>
<tr><td>>MI0000750|hsa-mir-26a-2&WithFlank&12|-|57824603|57824698@20.42.23;MI0000083|hsa-mir-26a-1&WithFlank&3|+|37969398|37969486@16.38.23&[MIMAT0000082&hsa-miR-26a-5p&offsets|0|+1;m-33&12|-|57824658|57824679&offsets|0|+1];[MIMAT0000082_1&hsa-miR-26a-5p&offsets|0|+1;m-34&3|+|37969413|37969434&offsets|0|+1]|TTCAAGTAATCCAGGATAGGCTG</td><td>>MI0000750|hsa-mir-26a-2&WithFlank&12|-|57824603|57824698</td><td>20</td><td>42</td><td>23</td><td>TTCAAGTAATCCAGGATAGGCTG</td><td>316.8</td><td>3.4</td><td>1.2</td><td>2.9</td><td>3.4E-03</td><td>7.1E-03</td><td>BT20/MB468</td><td>MP</td></tr>
<tr><td>>hg38_21_+_8433172_8446622_RNA45SN1_45S_50ntFlanks@4724.4761.38|ATTAATCAAGAACGAAAGTCGGAGGTTCGAAGACGATC</td><td>>hg38_21_+_8433172_8446622_RNA45SN1_45S_50ntFlanks</td><td>4724</td><td>4761</td><td>38</td><td>ATTAATCAAGAACGAAAGTCGGAGGTTCGAAGACGATC</td><td>215.3</td><td>-1.9</td><td>0.7</td><td>-2.7</td><td>6.2E-03</td><td>7.1E-03</td><td>MB231/MB468</td><td>NUC</td></tr>
<tr><td>>hg38_21_+_8433172_8446622_RNA45SN1_45S_50ntFlanks@8357.8375.19|AAAAGAACTTTGAAGAGAG</td><td>>hg38_21_+_8433172_8446622_RNA45SN1_45S_50ntFlanks</td><td>8357</td><td>8375</td><td>19</td><td>AAAAGAACTTTGAAGAGAG</td><td>15864.0</td><td>4.6</td><td>1.4</td><td>3.2</td><td>1.2E-03</td><td>7.1E-03</td><td>BT20/MB231</td><td>MP</td></tr>
<tr><td>>hg38_21_+_8433172_8446622_RNA45SN1_45S_50ntFlanks@11674.11692.19|GTGCTCTGAATGTCAAAGT</td><td>>hg38_21_+_8433172_8446622_RNA45SN1_45S_50ntFlanks</td><td>11674</td><td>11692</td><td>19</td><td>GTGCTCTGAATGTCAAAGT</td><td>140.5</td><td>6.1</td><td>1.9</td><td>3.2</td><td>1.2E-03</td><td>7.1E-03</td><td>BT20/MB231</td><td>MP</td></tr>
<tr><td>>hg38_21_+_8433172_8446622_RNA45SN1_45S_50ntFlanks@8156.8181.26|GCTCGTGGGGGGCCCAAGTCCTTCTG</td><td>>hg38_21_+_8433172_8446622_RNA45SN1_45S_50ntFlanks</td><td>8156</td><td>8181</td><td>26</td><td>GCTCGTGGGGGGCCCAAGTCCTTCTG</td><td>259.2</td><td>-2.0</td><td>0.7</td><td>-2.8</td><td>5.0E-03</td><td>7.1E-03</td><td>BT20/MB468</td><td>MITO</td></tr>
<tr><td>>trnaMT_GluTTC_MT_-_14674_14742@34.66.33,trnalookalike8_GluTTC_5_-_93905172_93905240@34.66.33|ATATCATTGGTCGTGGTTGTAGTCCGTGCGAGA</td><td>>trnaMT_GluTTC_MT_-_14674_14742</td><td>34</td><td>66</td><td>33</td><td>ATATCATTGGTCGTGGTTGTAGTCCGTGCGAGA</td><td>177.4</td><td>4.0</td><td>1.4</td><td>2.9</td><td>3.4E-03</td><td>7.1E-03</td><td>BT20/MB468</td><td>MP</td></tr>
<tr><td>>MI0000086|hsa-mir-28&WithFlank&3|+|188688775|188688872@20.40.21&[MIMAT0000085&hsa-miR-28-5p&offsets|0|-1;m-133&3|+|188688794|188688815&offsets|0|-1]|AAGGAGCTCACAGTCTATTGA</td><td>>MI0000086|hsa-mir-28&WithFlank&3|+|188688775|188688872</td><td>20</td><td>40</td><td>21</td><td>AAGGAGCTCACAGTCTATTGA</td><td>314.8</td><td>3.8</td><td>1.4</td><td>2.8</td><td>5.3E-03</td><td>7.1E-03</td><td>MB231/MB468</td><td>MP</td></tr>
<tr><td>>hg38_21_+_8433172_8446622_RNA45SN1_45S_50ntFlanks@11029.11051.23|CTCTCTCTCTCTCCCCCGCTCCC</td><td>>hg38_21_+_8433172_8446622_RNA45SN1_45S_50ntFlanks</td><td>11029</td><td>11051</td><td>23</td><td>CTCTCTCTCTCTCCCCCGCTCCC</td><td>582.4</td><td>-5.5</td><td>2.0</td><td>-2.7</td><td>6.1E-03</td><td>7.1E-03</td><td>BT20/MB468</td><td>NUC</td></tr>
<tr><td>>MI0001445|hsa-mir-423&WithFlank&17|+|30117073|30117178@23.44.22(+1G)&[MIMAT0004748&hsa-miR-423-5p&offsets|0|-1(+1G);m-141&17|+|30117095|30117117&offsets|0|-1(+1G)]|TGAGGGGCAGAGAGCGAGACTTG</td><td>>MI0001445|hsa-mir-423&WithFlank&17|+|30117073|30117178</td><td>23</td><td>44</td><td>22(+1G)</td><td>TGAGGGGCAGAGAGCGAGACTTG</td><td>71.6</td><td>3.6</td><td>1.3</td><td>2.8</td><td>5.3E-03</td><td>7.1E-03</td><td>MB231/MB468</td><td>MP</td></tr>
<tr><td>>trnaMT_GlnTTG_MT_-_4329_4400@1.33.33,trnalookalike2_GlnTTG_1_-_564879_564950@1.33.33|TAGGATGGGGTGTGATAGGTGGCACGGAGAATT</td><td>>trnaMT_GlnTTG_MT_-_4329_4400</td><td>1</td><td>33</td><td>33</td><td>TAGGATGGGGTGTGATAGGTGGCACGGAGAATT</td><td>269.0</td><td>4.1</td><td>1.3</td><td>3.2</td><td>1.3E-03</td><td>7.1E-03</td><td>BT20/MB231</td><td>NUC</td></tr>
<tr><td>>hg38_21_+_8433172_8446622_RNA45SN1_45S_50ntFlanks@8378.8400.23|TTCAAGAGGGCGTGAAACCGTTA</td><td>>hg38_21_+_8433172_8446622_RNA45SN1_45S_50ntFlanks</td><td>8378</td><td>8400</td><td>23</td><td>TTCAAGAGGGCGTGAAACCGTTA</td><td>166.5</td><td>-4.9</td><td>1.7</td><td>-2.9</td><td>3.4E-03</td><td>7.1E-03</td><td>BT20/MB468</td><td>MP</td></tr>
<tr><td>>hg38_21_+_8433172_8446622_RNA45SN1_45S_50ntFlanks@9807.9830.24|GTGGAATGCGAGTGCCTAGTGGGC</td><td>>hg38_21_+_8433172_8446622_RNA45SN1_45S_50ntFlanks</td><td>9807</td><td>9830</td><td>24</td><td>GTGGAATGCGAGTGCCTAGTGGGC</td><td>177.5</td><td>5.2</td><td>1.8</td><td>2.9</td><td>3.4E-03</td><td>7.1E-03</td><td>BT20/MB468</td><td>MP</td></tr>
<tr><td>>hg38_21_+_8433172_8446622_RNA45SN1_45S_50ntFlanks@12413.12436.24|TCCTTCGATGTCGGCTCTTCCTAT</td><td>>hg38_21_+_8433172_8446622_RNA45SN1_45S_50ntFlanks</td><td>12413</td><td>12436</td><td>24</td><td>TCCTTCGATGTCGGCTCTTCCTAT</td><td>2075.3</td><td>-5.9</td><td>2.1</td><td>-2.7</td><td>6.2E-03</td><td>7.1E-03</td><td>BT20/MB468</td><td>NUC</td></tr>
<tr><td>>hg38_21_+_8433172_8446622_RNA45SN1_45S_50ntFlanks@10686.10710.25|GCGGCGTCCGGTGAGCTCTCGCTGG</td><td>>hg38_21_+_8433172_8446622_RNA45SN1_45S_50ntFlanks</td><td>10686</td><td>10710</td><td>25</td><td>GCGGCGTCCGGTGAGCTCTCGCTGG</td><td>953.6</td><td>-2.8</td><td>0.9</td><td>-2.9</td><td>3.4E-03</td><td>7.1E-03</td><td>BT20/MB468</td><td>MP</td></tr>
<tr><td>>am_trnaMT_ProTGG_MT_-_15956_16023@52.71.20|AAAGACTTTTTCTCTGACCA</td><td>>am_trnaMT_ProTGG_MT_-_15956_16023</td><td>52</td><td>71</td><td>20</td><td>AAAGACTTTTTCTCTGACCA</td><td>233.5</td><td>2.7</td><td>1.0</td><td>2.7</td><td>6.2E-03</td><td>7.1E-03</td><td>BT20/MB468</td><td>NUC</td></tr>
<tr><td>>hg38_21_+_8433172_8446622_RNA45SN1_45S_50ntFlanks@12419.12441.23|GATGTCGGCTCTTCCTATCATTG</td><td>>hg38_21_+_8433172_8446622_RNA45SN1_45S_50ntFlanks</td><td>12419</td><td>12441</td><td>23</td><td>GATGTCGGCTCTTCCTATCATTG</td><td>2668.4</td><td>-2.6</td><td>0.8</td><td>-3.2</td><td>1.5E-03</td><td>7.1E-03</td><td>BT20/MB231</td><td>CYTO</td></tr>
<tr><td>>hg38_21_+_8433172_8446622_RNA45SN1_45S_50ntFlanks@475.523.49|CTTCGTGATCGATGTGGTGACGTCGTGCTCTCCCGGGCCGGGTCCGAGC</td><td>>hg38_21_+_8433172_8446622_RNA45SN1_45S_50ntFlanks</td><td>475</td><td>523</td><td>49</td><td>CTTCGTGATCGATGTGGTGACGTCGTGCTCTCCCGGGCCGGGTCCGAGC</td><td>4610.3</td><td>3.8</td><td>1.4</td><td>2.7</td><td>6.2E-03</td><td>7.1E-03</td><td>MB231/MB468</td><td>NUC</td></tr>
<tr><td>>hg38_MT_+_1621_3279_RNR2_16S_50ntFlanks@51.81.31|GCTAAACCTAGCCCCAAACCCACTCCACCTT</td><td>>hg38_MT_+_1621_3279_RNR2_16S_50ntFlanks</td><td>51</td><td>81</td><td>31</td><td>GCTAAACCTAGCCCCAAACCCACTCCACCTT</td><td>803.6</td><td>2.4</td><td>0.9</td><td>2.7</td><td>6.2E-03</td><td>7.1E-03</td><td>MB231/MB468</td><td>MITO</td></tr>
<tr><td>>hg38_1_-_228634819_228635039_RNA5S12_5S_50ntFlanks@149.170.22|GAATACCGGGTGCTGTAGGCTT</td><td>>hg38_1_-_228634819_228635039_RNA5S12_5S_50ntFlanks</td><td>149</td><td>170</td><td>22</td><td>GAATACCGGGTGCTGTAGGCTT</td><td>1345.8</td><td>2.1</td><td>0.7</td><td>3.2</td><td>1.6E-03</td><td>7.2E-03</td><td>BT20/MB231</td><td>MITO</td></tr>
<tr><td>>MI0000289|hsa-mir-181a-1&WithFlank&1|-|198859038|198859159@30.48.19(+1A);MI0000269|hsa-mir-181a-2&WithFlank&9|+|124692436|124692557@45.63.19(+1A)&[MIMAT0000256&hsa-miR-181a-5p&offsets|0|-4(+1A);m-32&1|-|198859108|198859130&offsets|0|-4(+1A)];[MIMAT0000256_1&hsa-miR-181a-5p&offsets|0|-4(+1A);m-32&9|+|124692480|124692502&offsets|0|-4(+1A)]|AACATTCAACGCTGTCGGTA</td><td>>MI0000289|hsa-mir-181a-1&WithFlank&1|-|198859038|198859159</td><td>30</td><td>48</td><td>19(+1A)</td><td>AACATTCAACGCTGTCGGTA</td><td>165.5</td><td>3.9</td><td>1.4</td><td>2.8</td><td>5.4E-03</td><td>7.2E-03</td><td>MB231/MB468</td><td>MP</td></tr>
<tr><td>>MI0003815|hsa-mir-1301&WithFlank&2|-|25328634|25328727@54.75.22&[MIMAT0005797&hsa-miR-1301-3p&offsets|0|-2;m-256&2|-|25328650|25328674&offsets|0|-3]|TTGCAGCTGCCTGGGAGTGACT</td><td>>MI0003815|hsa-mir-1301&WithFlank&2|-|25328634|25328727</td><td>54</td><td>75</td><td>22</td><td>TTGCAGCTGCCTGGGAGTGACT</td><td>105.9</td><td>3.9</td><td>1.3</td><td>2.9</td><td>3.4E-03</td><td>7.2E-03</td><td>BT20/MB468</td><td>MP</td></tr>
<tr><td>>hg38_21_+_8433172_8446622_RNA45SN1_45S_50ntFlanks@12418.12451.34|CGATGTCGGCTCTTCCTATCATTGTGAAGCAGAA</td><td>>hg38_21_+_8433172_8446622_RNA45SN1_45S_50ntFlanks</td><td>12418</td><td>12451</td><td>34</td><td>CGATGTCGGCTCTTCCTATCATTGTGAAGCAGAA</td><td>270.9</td><td>-1.1</td><td>0.4</td><td>-2.8</td><td>5.0E-03</td><td>7.2E-03</td><td>BT20/MB468</td><td>MITO</td></tr>
<tr><td>>hg38_21_+_8433172_8446622_RNA45SN1_45S_50ntFlanks@9158.9187.30|AGTGCGCCCCGGGCGGGTCGCGCCGTCGGG</td><td>>hg38_21_+_8433172_8446622_RNA45SN1_45S_50ntFlanks</td><td>9158</td><td>9187</td><td>30</td><td>AGTGCGCCCCGGGCGGGTCGCGCCGTCGGG</td><td>161.0</td><td>-4.2</td><td>1.5</td><td>-2.7</td><td>6.2E-03</td><td>7.2E-03</td><td>MB231/MB468</td><td>NUC</td></tr>
<tr><td>>hg38_MT_+_1621_3279_RNR2_16S_50ntFlanks@120.145.26|AAGTATAGGCGATAGAAATTGAAACC</td><td>>hg38_MT_+_1621_3279_RNR2_16S_50ntFlanks</td><td>120</td><td>145</td><td>26</td><td>AAGTATAGGCGATAGAAATTGAAACC</td><td>47.6</td><td>6.0</td><td>2.0</td><td>2.9</td><td>3.4E-03</td><td>7.2E-03</td><td>BT20/MB468</td><td>MP</td></tr>
<tr><td>>hg38_21_+_8433172_8446622_RNA45SN1_45S_50ntFlanks@10514.10533.20|CGGAGTGGCGGAGATGGGCG</td><td>>hg38_21_+_8433172_8446622_RNA45SN1_45S_50ntFlanks</td><td>10514</td><td>10533</td><td>20</td><td>CGGAGTGGCGGAGATGGGCG</td><td>496.4</td><td>-4.6</td><td>1.6</td><td>-2.8</td><td>5.0E-03</td><td>7.2E-03</td><td>BT20/MB468</td><td>MITO</td></tr>
<tr><td>>hg38_21_+_8433172_8446622_RNA45SN1_45S_50ntFlanks@11567.11585.19|AGCAGCCGACTTAGAACTG</td><td>>hg38_21_+_8433172_8446622_RNA45SN1_45S_50ntFlanks</td><td>11567</td><td>11585</td><td>19</td><td>AGCAGCCGACTTAGAACTG</td><td>1371.8</td><td>-2.5</td><td>0.9</td><td>-2.8</td><td>5.4E-03</td><td>7.2E-03</td><td>MB231/MB468</td><td>MP</td></tr>
<tr><td>>hg38_21_+_8433172_8446622_RNA45SN1_45S_50ntFlanks@12407.12433.27|TTTTGATCCTTCGATGTCGGCTCTTCC</td><td>>hg38_21_+_8433172_8446622_RNA45SN1_45S_50ntFlanks</td><td>12407</td><td>12433</td><td>27</td><td>TTTTGATCCTTCGATGTCGGCTCTTCC</td><td>173.5</td><td>-3.5</td><td>1.2</td><td>-2.8</td><td>5.4E-03</td><td>7.2E-03</td><td>MB231/MB468</td><td>MP</td></tr>
<tr><td>>MI0003643|hsa-mir-629&WithFlank&15|-|70079366|70079474@28.48.21(+1C)&[MIMAT0004810&hsa-miR-629-5p&offsets|0|0(+1C);m-150&15|-|70079427|70079447&offsets|0|0(+1C)]|TGGGTTTACGTTGGGAGAACTC</td><td>>MI0003643|hsa-mir-629&WithFlank&15|-|70079366|70079474</td><td>28</td><td>48</td><td>21(+1C)</td><td>TGGGTTTACGTTGGGAGAACTC</td><td>434.2</td><td>2.6</td><td>0.8</td><td>3.2</td><td>1.5E-03</td><td>7.2E-03</td><td>BT20/MB231</td><td>CYTO</td></tr>
<tr><td>>hg38_21_+_8433172_8446622_RNA45SN1_45S_50ntFlanks@12974.12992.19|CGTACGTAGCAGAGCAGCT</td><td>>hg38_21_+_8433172_8446622_RNA45SN1_45S_50ntFlanks</td><td>12974</td><td>12992</td><td>19</td><td>CGTACGTAGCAGAGCAGCT</td><td>182.1</td><td>-3.6</td><td>1.3</td><td>-2.8</td><td>5.4E-03</td><td>7.2E-03</td><td>MB231/MB468</td><td>MP</td></tr>
<tr><td>>hg38_21_+_8433172_8446622_RNA45SN1_45S_50ntFlanks@8179.8203.25|CTGATCGAGGCCCAGCCCGTGGACG</td><td>>hg38_21_+_8433172_8446622_RNA45SN1_45S_50ntFlanks</td><td>8179</td><td>8203</td><td>25</td><td>CTGATCGAGGCCCAGCCCGTGGACG</td><td>417.1</td><td>-5.0</td><td>1.7</td><td>-2.9</td><td>3.4E-03</td><td>7.2E-03</td><td>BT20/MB468</td><td>MP</td></tr>
<tr><td>>hg38_21_+_8433172_8446622_RNA45SN1_45S_50ntFlanks@12570.12587.18|AGTACGAGAGGAACCGCA</td><td>>hg38_21_+_8433172_8446622_RNA45SN1_45S_50ntFlanks</td><td>12570</td><td>12587</td><td>18</td><td>AGTACGAGAGGAACCGCA</td><td>1410.5</td><td>-5.4</td><td>1.8</td><td>-2.9</td><td>3.4E-03</td><td>7.2E-03</td><td>BT20/MB468</td><td>MP</td></tr>
<tr><td>>hg38_21_+_8433172_8446622_RNA45SN1_45S_50ntFlanks@11569.11609.41|CAGCCGACTTAGAACTGGTGCGGACCAGGGGAATCCGACTG</td><td>>hg38_21_+_8433172_8446622_RNA45SN1_45S_50ntFlanks</td><td>11569</td><td>11609</td><td>41</td><td>CAGCCGACTTAGAACTGGTGCGGACCAGGGGAATCCGACTG</td><td>382.5</td><td>-2.3</td><td>0.8</td><td>-2.8</td><td>5.9E-03</td><td>7.2E-03</td><td>MB231/MB468</td><td>CYTO</td></tr>
<tr><td>>trna111_HisGTG_1_-_147774845_147774916@-1G.32.33,trna118_HisGTG_1_-_145396881_145396952@-1G.32.33,trna16_HisGTG_1_+_146544773_146544844@-1G.32.33,trna1_HisGTG_15_+_45493349_45493420@-1G.32.33,trna21_HisGTG_1_+_147753471_147753542@-1G.32.33,trna33_HisGTG_6_+_27125906_27125977@-1G.32.33,trna7_HisGTG_9_-_14433938_14434009@-1G.32.33,trna8_HisGTG_15_-_45492611_45492682@-1G.32.33,trna9_HisGTG_15_-_45490804_45490875@-1G.32.33|GGCCGTGATCGTATAGTGGTTAGTACTCTGCGT</td><td>>trna111_HisGTG_1_-_147774845_147774916</td><td>-1G</td><td>32</td><td>33</td><td>GGCCGTGATCGTATAGTGGTTAGTACTCTGCGT</td><td>5893.8</td><td>-4.2</td><td>1.3</td><td>-3.2</td><td>1.2E-03</td><td>7.2E-03</td><td>BT20/MB231</td><td>MP</td></tr>
<tr><td>>hg38_MT_+_598_1651_RNR1_12S_50ntFlanks@970.1004.35|ATGGTAAGTGTACTGGAAAGTGCACTTGGACGAAC</td><td>>hg38_MT_+_598_1651_RNR1_12S_50ntFlanks</td><td>970</td><td>1004</td><td>35</td><td>ATGGTAAGTGTACTGGAAAGTGCACTTGGACGAAC</td><td>406.3</td><td>-3.7</td><td>1.2</td><td>-3.2</td><td>1.2E-03</td><td>7.2E-03</td><td>BT20/MB231</td><td>MP</td></tr>
<tr><td>>hg38_21_+_8433172_8446622_RNA45SN1_45S_50ntFlanks@11569.11599.31|CAGCCGACTTAGAACTGGTGCGGACCAGGGG</td><td>>hg38_21_+_8433172_8446622_RNA45SN1_45S_50ntFlanks</td><td>11569</td><td>11599</td><td>31</td><td>CAGCCGACTTAGAACTGGTGCGGACCAGGGG</td><td>253.0</td><td>6.3</td><td>1.9</td><td>3.2</td><td>1.2E-03</td><td>7.2E-03</td><td>BT20/MB231</td><td>MP</td></tr>
<tr><td>>hg38_21_+_8433172_8446622_RNA45SN1_45S_50ntFlanks@9540.9569.30|CTCTGGTGGAGGTCCGTAGCGGTCCTGACG</td><td>>hg38_21_+_8433172_8446622_RNA45SN1_45S_50ntFlanks</td><td>9540</td><td>9569</td><td>30</td><td>CTCTGGTGGAGGTCCGTAGCGGTCCTGACG</td><td>372.8</td><td>-2.1</td><td>0.8</td><td>-2.7</td><td>6.3E-03</td><td>7.2E-03</td><td>MB231/MB468</td><td>NUC</td></tr>
<tr><td>>hg38_21_+_8433172_8446622_RNA45SN1_45S_50ntFlanks@11211.11243.33|GCGACTCTGGACGCGAGCCGGGCCCTTCCCGTG</td><td>>hg38_21_+_8433172_8446622_RNA45SN1_45S_50ntFlanks</td><td>11211</td><td>11243</td><td>33</td><td>GCGACTCTGGACGCGAGCCGGGCCCTTCCCGTG</td><td>88.0</td><td>5.0</td><td>1.7</td><td>2.9</td><td>3.4E-03</td><td>7.2E-03</td><td>BT20/MB468</td><td>MP</td></tr>
</table>

<table>
<tr><th></th><th></th><th></th><th></th><th></th><th></th><th></th><th></th><th></th><th></th><th></th><th></th><th></th><th></th></tr>
<tr><td>>trna11_SerAGA_8_-_96281885_96281966@1.23.23,trna145_SerAGA_6_-_27521192_27521273@1.23.23,trna147_SerAGA_6_-_27509554_27509635@1.23.23,trna148_SerTGA_6_-_27473607_27473688@1.23.23,trna172_SerTGA_6_-_26312824_26312905@1.23.23,trna35_SerAGA_17_-_8129928_8130009@1.23.23,trna44_SerAGA_6_+_27446591_27446672@1.23.23,trna46_SerAGA_6_+_27463593_27463674@1.23.23,trna47_SerAGA_6_+_27470818_27470899@1.23.23,trna50_SerAGA_6_+_27499987_27500068@1.23.23,trna51_SerTGA_6_+_27513468_27513549@1.23.23,trna5_SerAGA_6_+_26327817_26327898@1.23.23|GTAGTCGTGGCCGAGTGGTTAAG</td><td>>trna11_SerAGA_8_-_96281885_96281966</td><td>1</td><td>23</td><td>23</td><td>GTAGTCGTGGCCGAGTGGTTAAG</td><td>491.9</td><td>4.5</td><td>1.4</td><td>3.2</td><td>1.2E-03</td><td>7.2E-03</td><td>BT20/MB231</td><td>MP</td></tr>
<tr><td>>hg38_21_+_8433172_8446622_RNA45SN1_45S_50ntFlanks@10350.10371.22|ACGAGAACTTTGAAGGCCGAAG</td><td>>hg38_21_+_8433172_8446622_RNA45SN1_45S_50ntFlanks</td><td>10350</td><td>10371</td><td>22</td><td>ACGAGAACTTTGAAGGCCGAAG</td><td>916.9</td><td>-1.3</td><td>0.5</td><td>-2.8</td><td>5.9E-03</td><td>7.2E-03</td><td>MB231/MB468</td><td>CYTO</td></tr>
<tr><td>>hg38_21_+_8433172_8446622_RNA45SN1_45S_50ntFlanks@10348.10371.24|AAACGAGAACTTTGAAGGCCGAAG</td><td>>hg38_21_+_8433172_8446622_RNA45SN1_45S_50ntFlanks</td><td>10348</td><td>10371</td><td>24</td><td>AAACGAGAACTTTGAAGGCCGAAG</td><td>653.6</td><td>2.0</td><td>0.7</td><td>2.8</td><td>5.9E-03</td><td>7.2E-03</td><td>MB231/MB468</td><td>CYTO</td></tr>
<tr><td>>hg38_21_+_8433172_8446622_RNA45SN1_45S_50ntFlanks@12417.12437.21|TCGATGTCGGCTCTTCCTATC</td><td>>hg38_21_+_8433172_8446622_RNA45SN1_45S_50ntFlanks</td><td>12417</td><td>12437</td><td>21</td><td>TCGATGTCGGCTCTTCCTATC</td><td>3511.1</td><td>-1.6</td><td>0.6</td><td>-2.8</td><td>5.9E-03</td><td>7.2E-03</td><td>BT20/MB468</td><td>CYTO</td></tr>
<tr><td>>hg38_21_+_8433172_8446622_RNA45SN1_45S_50ntFlanks@6649.6681.33|TACGACTCTTAGCGGTGGATCACTCGGCTCGTG</td><td>>hg38_21_+_8433172_8446622_RNA45SN1_45S_50ntFlanks</td><td>6649</td><td>6681</td><td>33</td><td>TACGACTCTTAGCGGTGGATCACTCGGCTCGTG</td><td>2131.3</td><td>-1.7</td><td>0.6</td><td>-2.8</td><td>5.0E-03</td><td>7.2E-03</td><td>BT20/MB468</td><td>MITO</td></tr>
<tr><td>>hg38_21_+_8433172_8446622_RNA45SN1_45S_50ntFlanks@4685.4726.42|AGACGGACCAGAGCGAAAGCATTTGCCAAGAATGTTTTCATT</td><td>>hg38_21_+_8433172_8446622_RNA45SN1_45S_50ntFlanks</td><td>4685</td><td>4726</td><td>42</td><td>AGACGGACCAGAGCGAAAGCATTTGCCAAGAATGTTTTCATT</td><td>486.2</td><td>-3.2</td><td>1.0</td><td>-3.2</td><td>1.6E-03</td><td>7.2E-03</td><td>BT20/MB231</td><td>CYTO</td></tr>
<tr><td>>hg38_21_+_8433172_8446622_RNA45SN1_45S_50ntFlanks@12999.13027.29|CTGCGATCTATTGAAAGTCAGCCCTCGAC</td><td>>hg38_21_+_8433172_8446622_RNA45SN1_45S_50ntFlanks</td><td>12999</td><td>13027</td><td>29</td><td>CTGCGATCTATTGAAAGTCAGCCCTCGAC</td><td>630.9</td><td>-2.0</td><td>0.7</td><td>-2.8</td><td>5.9E-03</td><td>7.2E-03</td><td>BT20/MB468</td><td>CYTO</td></tr>
<tr><td>>hg38_21_+_8433172_8446622_RNA45SN1_45S_50ntFlanks@9806.9830.25|CGTGGAATGCGAGTGCCTAGTGGGC</td><td>>hg38_21_+_8433172_8446622_RNA45SN1_45S_50ntFlanks</td><td>9806</td><td>9830</td><td>25</td><td>CGTGGAATGCGAGTGCCTAGTGGGC</td><td>149.2</td><td>-3.2</td><td>1.2</td><td>-2.7</td><td>6.3E-03</td><td>7.2E-03</td><td>MB231/MB468</td><td>NUC</td></tr>
<tr><td>>MI0016590|hsa-mir-3934&WithFlank&6|+|33698122|33698240@31.52.22&[MIMAT0018349&hsa-miR-3934-5p&offsets|0|0;m-557&6|+|33698152|33698173&offsets|0|0]|TCAGGTGTGGAAACTGAGGCAG</td><td>>MI0016590|hsa-mir-3934&WithFlank&6|+|33698122|33698240</td><td>31</td><td>52</td><td>22</td><td>TCAGGTGTGGAAACTGAGGCAG</td><td>148.0</td><td>4.4</td><td>1.6</td><td>2.8</td><td>5.4E-03</td><td>7.2E-03</td><td>MB231/MB468</td><td>MP</td></tr>
<tr><td>>hg38_21_+_8433172_8446622_RNA45SN1_45S_50ntFlanks@4678.4720.43|CGGCGCAAGACGGACCAGAGCGAAAGCATTTGCCAAGAATGTT</td><td>>hg38_21_+_8433172_8446622_RNA45SN1_45S_50ntFlanks</td><td>4678</td><td>4720</td><td>43</td><td>CGGCGCAAGACGGACCAGAGCGAAAGCATTTGCCAAGAATGTT</td><td>671.6</td><td>-3.2</td><td>1.0</td><td>-3.2</td><td>1.6E-03</td><td>7.2E-03</td><td>BT20/MB231</td><td>CYTO</td></tr>
<tr><td>>MI0000085|hsa-mir-27a&WithFlank&19|-|13836434|13836523@57.75.19(+1A)&[MIMAT0000084&hsa-miR-27a-3p&offsets|0|-2(+1A);m-43&19|-|13836447|13836467&offsets|0|-2(+1A)]|TTCACAGTGGCTAAGTTCCA</td><td>>MI0000085|hsa-mir-27a&WithFlank&19|-|13836434|13836523</td><td>57</td><td>75</td><td>19(+1A)</td><td>TTCACAGTGGCTAAGTTCCA</td><td>291.2</td><td>1.8</td><td>0.7</td><td>2.7</td><td>6.2E-03</td><td>7.2E-03</td><td>BT20/MB468</td><td>NUC</td></tr>
<tr><td>>hg38_21_+_8433172_8446622_RNA45SN1_45S_50ntFlanks@12937.12966.30|AACCATTCGTAGACGACCTGCTTCTGGGTC</td><td>>hg38_21_+_8433172_8446622_RNA45SN1_45S_50ntFlanks</td><td>12937</td><td>12966</td><td>30</td><td>AACCATTCGTAGACGACCTGCTTCTGGGTC</td><td>138.8</td><td>-4.8</td><td>1.7</td><td>-2.9</td><td>3.4E-03</td><td>7.2E-03</td><td>BT20/MB468</td><td>MP</td></tr>
<tr><td>>hg38_21_+_8433172_8446622_RNA45SN1_45S_50ntFlanks@10475.10512.38|CGGCCGATCGAAAGGGAGTCGGGTTCAGATCCCCGAAT</td><td>>hg38_21_+_8433172_8446622_RNA45SN1_45S_50ntFlanks</td><td>10475</td><td>10512</td><td>38</td><td>CGGCCGATCGAAAGGGAGTCGGGTTCAGATCCCCGAAT</td><td>141.1</td><td>-2.4</td><td>0.9</td><td>-2.8</td><td>6.0E-03</td><td>7.2E-03</td><td>MB231/MB468</td><td>CYTO</td></tr>
<tr><td>>hg38_21_+_8433172_8446622_RNA45SN1_45S_50ntFlanks@12006.12035.30|GGCCGCCGGTGAAATACCACTACTCTGATC</td><td>>hg38_21_+_8433172_8446622_RNA45SN1_45S_50ntFlanks</td><td>12006</td><td>12035</td><td>30</td><td>GGCCGCCGGTGAAATACCACTACTCTGATC</td><td>261.1</td><td>4.3</td><td>1.4</td><td>3.2</td><td>1.4E-03</td><td>7.2E-03</td><td>BT20/MB231</td><td>NUC</td></tr>
<tr><td>>hg38_21_+_8433172_8446622_RNA45SN1_45S_50ntFlanks@476.506.31|TTCGTGATCGATGTGGTGACGTCGTGCTCTC</td><td>>hg38_21_+_8433172_8446622_RNA45SN1_45S_50ntFlanks</td><td>476</td><td>506</td><td>31</td><td>TTCGTGATCGATGTGGTGACGTCGTGCTCTC</td><td>88.3</td><td>3.5</td><td>1.3</td><td>2.8</td><td>5.4E-03</td><td>7.2E-03</td><td>MB231/MB468</td><td>MP</td></tr>
<tr><td>>MI0000299|hsa-mir-222&WithFlank&X|-|45747009|45747130@75.95.21(+1A)&[MIMAT0000279&hsa-miR-222-3p&offsets|0|0(+1A);m-140&X|-|45747033|45747056&offsets|0|-3(+1A)]|AGCTACATCTGGCTACTGGGTA</td><td>>MI0000299|hsa-mir-222&WithFlank&X|-|45747009|45747130</td><td>75</td><td>95</td><td>21(+1A)</td><td>AGCTACATCTGGCTACTGGGTA</td><td>194.0</td><td>3.7</td><td>1.3</td><td>2.8</td><td>5.4E-03</td><td>7.2E-03</td><td>MB231/MB468</td><td>MP</td></tr>
<tr><td>>am_trnaMT_TyrGTA_MT_-_5826_5891@13.31.19|GAGTGAAGCATTGGACTGT</td><td>>am_trnaMT_TyrGTA_MT_-_5826_5891</td><td>13</td><td>31</td><td>19</td><td>GAGTGAAGCATTGGACTGT</td><td>72.2</td><td>4.8</td><td>1.7</td><td>2.9</td><td>3.4E-03</td><td>7.2E-03</td><td>BT20/MB468</td><td>MP</td></tr>
<tr><td>>hg38_MT_+_1621_3279_RNR2_16S_50ntFlanks@1092.1120.29|GCAAGACGAGAAGACCCTATGGAGCTTTA</td><td>>hg38_MT_+_1621_3279_RNR2_16S_50ntFlanks</td><td>1092</td><td>1120</td><td>29</td><td>GCAAGACGAGAAGACCCTATGGAGCTTTA</td><td>246.6</td><td>2.6</td><td>0.9</td><td>2.8</td><td>5.4E-03</td><td>7.2E-03</td><td>MB231/MB468</td><td>MP</td></tr>
<tr><td>>trna116_GluCTC_1_-_145399233_145399304@32.75.44,trna71_GluCTC_1_-_161439189_161439260@32.75.44,trna74_GluCTC_1_-_161431809_161431880@32.75.44,trna77_GluCTC_1_-_161424398_161424469@32.75.44,trna77_GluCTC_6_+_28949976_28950047@32.75.44,trna80_GluCTC_1_-_161417018_161417089@32.75.44,trna87_GluCTC_6_-_126101393_126101464@32.75.44|CTCTCACCGCCGCGGCCCGGGTTCGATTCCCGGTCAGGGAACCA</td><td>>trna116_GluCTC_1_-_145399233_145399304</td><td>32</td><td>75</td><td>44</td><td>CTCTCACCGCCGCGGCCCGGGTTCGATTCCCGGTCAGGGAACCA</td><td>146.4</td><td>-4.5</td><td>1.6</td><td>-2.8</td><td>5.0E-03</td><td>7.2E-03</td><td>BT20/MB468</td><td>MITO</td></tr>
<tr><td>>MI0000269|hsa-mir-181a-2&WithFlank&9|+|124692436|124692557@83.103.21(+1A)&[MIMAT0004558&hsa-miR-181a-2-3p&offsets|0|-1(+1A);m-71&9|+|124692518|124692539&offsets|0|-1(+1A)]|ACCACTGACCGTTGACTGTACA</td><td>>MI0000269|hsa-mir-181a-2&WithFlank&9|+|124692436|124692557</td><td>83</td><td>103</td><td>21(+1A)</td><td>ACCACTGACCGTTGACTGTACA</td><td>75.2</td><td>5.3</td><td>1.9</td><td>2.8</td><td>5.4E-03</td><td>7.2E-03</td><td>MB231/MB468</td><td>MP</td></tr>
<tr><td>>hg38_21_+_8433172_8446622_RNA45SN1_45S_50ntFlanks@12488.12522.35|AACGTGAGCTGGGTTTAGACCGTCGTGAGACAGGT</td><td>>hg38_21_+_8433172_8446622_RNA45SN1_45S_50ntFlanks</td><td>12488</td><td>12522</td><td>35</td><td>AACGTGAGCTGGGTTTAGACCGTCGTGAGACAGGT</td><td>142.9</td><td>-2.2</td><td>0.8</td><td>-2.8</td><td>5.0E-03</td><td>7.2E-03</td><td>BT20/MB468</td><td>MITO</td></tr>
<tr><td>>hg38_21_+_8433172_8446622_RNA45SN1_45S_50ntFlanks@11580.11607.28|GAACTGGTGCGGACCAGGGGAATCCGAC</td><td>>hg38_21_+_8433172_8446622_RNA45SN1_45S_50ntFlanks</td><td>11580</td><td>11607</td><td>28</td><td>GAACTGGTGCGGACCAGGGGAATCCGAC</td><td>476.3</td><td>-3.1</td><td>1.1</td><td>-2.9</td><td>3.4E-03</td><td>7.2E-03</td><td>BT20/MB468</td><td>MP</td></tr>
<tr><td>>trnaMT_GluTTC_MT_-_14674_14742@46.72.27,trnalookalike8_GluTTC_5_-_93905172_93905240@46.72.27|GTGGTTGTAGTCCGTGCGAGAATACCA</td><td>>trnaMT_GluTTC_MT_-_14674_14742</td><td>46</td><td>72</td><td>27</td><td>GTGGTTGTAGTCCGTGCGAGAATACCA</td><td>1183.6</td><td>3.5</td><td>1.2</td><td>2.9</td><td>3.4E-03</td><td>7.2E-03</td><td>BT20/MB468</td><td>MP</td></tr>
<tr><td>>hg38_21_+_8433172_8446622_RNA45SN1_45S_50ntFlanks@10101.10124.24|CGCGCGCGTGTGGTGTGCGTCGGA</td><td>>hg38_21_+_8433172_8446622_RNA45SN1_45S_50ntFlanks</td><td>10101</td><td>10124</td><td>24</td><td>CGCGCGCGTGTGGTGTGCGTCGGA</td><td>155.5</td><td>-4.5</td><td>1.6</td><td>-2.8</td><td>5.1E-03</td><td>7.2E-03</td><td>BT20/MB468</td><td>MITO</td></tr>
<tr><td>>hg38_21_+_8433172_8446622_RNA45SN1_45S_50ntFlanks@10879.10901.23|CGGTCGGGCTGGGGCGCGAAGCG</td><td>>hg38_21_+_8433172_8446622_RNA45SN1_45S_50ntFlanks</td><td>10879</td><td>10901</td><td>23</td><td>CGGTCGGGCTGGGGCGCGAAGCG</td><td>181.2</td><td>-4.3</td><td>1.6</td><td>-2.7</td><td>6.3E-03</td><td>7.3E-03</td><td>MB231/MB468</td><td>NUC</td></tr>
<tr><td>>hg38_MT_+_598_1651_RNR1_12S_50ntFlanks@438.469.32|TAACATATCTGAACACACAATAGCTAAGACCC</td><td>>hg38_MT_+_598_1651_RNR1_12S_50ntFlanks</td><td>438</td><td>469</td><td>32</td><td>TAACATATCTGAACACACAATAGCTAAGACCC</td><td>27.5</td><td>8.0</td><td>2.9</td><td>2.8</td><td>5.4E-03</td><td>7.3E-03</td><td>MB231/MB468</td><td>MP</td></tr>
<tr><td>>hg38_21_+_8433172_8446622_RNA45SN1_45S_50ntFlanks@8179.8202.24|CTGATCGAGGCCCAGCCCGTGGAC</td><td>>hg38_21_+_8433172_8446622_RNA45SN1_45S_50ntFlanks</td><td>8179</td><td>8202</td><td>24</td><td>CTGATCGAGGCCCAGCCCGTGGAC</td><td>112.8</td><td>-4.6</td><td>1.7</td><td>-2.8</td><td>5.4E-03</td><td>7.3E-03</td><td>MB231/MB468</td><td>MP</td></tr>
<tr><td>>hg38_21_+_8433172_8446622_RNA45SN1_45S_50ntFlanks@6652.6681.30|GACTCTTAGCGGTGGATCACTCGGCTCGTG</td><td>>hg38_21_+_8433172_8446622_RNA45SN1_45S_50ntFlanks</td><td>6652</td><td>6681</td><td>30</td><td>GACTCTTAGCGGTGGATCACTCGGCTCGTG</td><td>7390.6</td><td>-1.8</td><td>0.6</td><td>-3.2</td><td>1.4E-03</td><td>7.3E-03</td><td>BT20/MB231</td><td>NUC</td></tr>
<tr><td>>hg38_21_+_8433172_8446622_RNA45SN1_45S_50ntFlanks@12416.12454.39|TTCGATGTCGGCTCTTCCTATCATTGTGAAGCAGAATTC</td><td>>hg38_21_+_8433172_8446622_RNA45SN1_45S_50ntFlanks</td><td>12416</td><td>12454</td><td>39</td><td>TTCGATGTCGGCTCTTCCTATCATTGTGAAGCAGAATTC</td><td>239.4</td><td>-2.8</td><td>0.9</td><td>-3.2</td><td>1.4E-03</td><td>7.3E-03</td><td>BT20/MB231</td><td>NUC</td></tr>
<tr><td>>hg38_21_+_8433172_8446622_RNA45SN1_45S_50ntFlanks@6654.6689.36|CTCTTAGCGGTGGATCACTCGGCTCGTGCGTCGATG</td><td>>hg38_21_+_8433172_8446622_RNA45SN1_45S_50ntFlanks</td><td>6654</td><td>6689</td><td>36</td><td>CTCTTAGCGGTGGATCACTCGGCTCGTGCGTCGATG</td><td>213.7</td><td>-2.6</td><td>0.9</td><td>-2.7</td><td>6.0E-03</td><td>7.3E-03</td><td>MB231/MB468</td><td>CYTO</td></tr>
<tr><td>>hg38_21_+_8433172_8446622_RNA45SN1_45S_50ntFlanks@480.497.18|TGATCGATGTGGTGACGT</td><td>>hg38_21_+_8433172_8446622_RNA45SN1_45S_50ntFlanks</td><td>480</td><td>497</td><td>18</td><td>TGATCGATGTGGTGACGT</td><td>97.3</td><td>4.3</td><td>1.6</td><td>2.8</td><td>5.4E-03</td><td>7.3E-03</td><td>MB231/MB468</td><td>MP</td></tr>
<tr><td>>am_trna128_GlyGCC_6_-_27870686_27870756@4.28.25,trna133_GlyCCC_1_-_16872434_16872504@4.28.25,trna18_GlyGCC_16_+_70822597_70822667@4.28.25,trna19_GlyGCC_16_+_70823410_70823480@4.28.25,trna19_GlyGCC_2_-_157257659_157257729@4.28.25,trna24_GlyGCC_16_-_70812942_70813012@4.28.25,trna25_GlyGCC_16_-_70812114_70812184@4.28.25,trna4_GlyCCC_1_+_17188416_17188486@4.28.25,trna5_GlyGCC_17_+_8029064_8029134@4.28.25,trna68_GlyGCC_1_-_161493637_161493707@4.28.25|TTGGTGGTTCAGTGGTAGAATTCTC</td><td>>am_trna128_GlyGCC_6_-_27870686_27870756</td><td>4</td><td>28</td><td>25</td><td>TTGGTGGTTCAGTGGTAGAATTCTC</td><td>227.9</td><td>-3.4</td><td>1.2</td><td>-2.7</td><td>6.3E-03</td><td>7.3E-03</td><td>MB231/MB468</td><td>NUC</td></tr>
<tr><td>>hg38_21_+_8433172_8446622_RNA45SN1_45S_50ntFlanks@3705.3745.41|TACCTGGTTGATCCTGCCAGTAGCATATGCTTGTCTCAAAG</td><td>>hg38_21_+_8433172_8446622_RNA45SN1_45S_50ntFlanks</td><td>3705</td><td>3745</td><td>41</td><td>TACCTGGTTGATCCTGCCAGTAGCATATGCTTGTCTCAAAG</td><td>822.2</td><td>2.4</td><td>0.7</td><td>3.2</td><td>1.4E-03</td><td>7.3E-03</td><td>BT20/MB231</td><td>NUC</td></tr>
<tr><td>>hg38_21_+_8433172_8446622_RNA45SN1_45S_50ntFlanks@6648.6696.49|GTACGACTCTTAGCGGTGGATCACTCGGCTCGTGCGTCGATGAAGAACG</td><td>>hg38_21_+_8433172_8446622_RNA45SN1_45S_50ntFlanks</td><td>6648</td><td>6696</td><td>49</td><td>GTACGACTCTTAGCGGTGGATCACTCGGCTCGTGCGTCGATGAAGAACG</td><td>2248.4</td><td>2.8</td><td>1.0</td><td>2.8</td><td>5.1E-03</td><td>7.3E-03</td><td>BT20/MB468</td><td>MITO</td></tr>
<tr><td>>hg38_21_+_8433172_8446622_RNA45SN1_45S_50ntFlanks@9941.9978.38|AGGACGGTGGCCATGGAAGTCGGAATCCGCTAAGGAGT</td><td>>hg38_21_+_8433172_8446622_RNA45SN1_45S_50ntFlanks</td><td>9941</td><td>9978</td><td>38</td><td>AGGACGGTGGCCATGGAAGTCGGAATCCGCTAAGGAGT</td><td>463.4</td><td>4.3</td><td>1.3</td><td>3.2</td><td>1.2E-03</td><td>7.3E-03</td><td>BT20/MB231</td><td>MP</td></tr>
<tr><td>>hg38_21_+_8433172_8446622_RNA45SN1_45S_50ntFlanks@6652.6677.26|GACTCTTAGCGGTGGATCACTCGGCT</td><td>>hg38_21_+_8433172_8446622_RNA45SN1_45S_50ntFlanks</td><td>6652</td><td>6677</td><td>26</td><td>GACTCTTAGCGGTGGATCACTCGGCT</td><td>5775.5</td><td>3.8</td><td>1.2</td><td>3.2</td><td>1.2E-03</td><td>7.3E-03</td><td>BT20/MB231</td><td>MP</td></tr>
<tr><td>>hg38_21_+_8433172_8446622_RNA45SN1_45S_50ntFlanks@11567.11588.22|AGCAGCCGACTTAGAACTGGTG</td><td>>hg38_21_+_8433172_8446622_RNA45SN1_45S_50ntFlanks</td><td>11567</td><td>11588</td><td>22</td><td>AGCAGCCGACTTAGAACTGGTG</td><td>433.8</td><td>-1.4</td><td>0.5</td><td>-2.7</td><td>6.0E-03</td><td>7.3E-03</td><td>MB231/MB468</td><td>CYTO</td></tr>
<tr><td>>hg38_21_+_8433172_8446622_RNA45SN1_45S_50ntFlanks@3854.3887.34|ACTGTGGTAATTCTAGAGCTAATACATGCCGACG</td><td>>hg38_21_+_8433172_8446622_RNA45SN1_45S_50ntFlanks</td><td>3854</td><td>3887</td><td>34</td><td>ACTGTGGTAATTCTAGAGCTAATACATGCCGACG</td><td>97.7</td><td>-5.0</td><td>1.8</td><td>-2.8</td><td>5.5E-03</td><td>7.3E-03</td><td>MB231/MB468</td><td>MP</td></tr>
<tr><td>>am_trna111_HisGTG_1_-_147774845_147774916@-1G.22.23,trna118_HisGTG_1_-_145396881_145396952@-1G.22.23,trna16_HisGTG_1_+_146544773_146544844@-1G.22.23,trna1_HisGTG_15_+_45493349_45493420@-1G.22.23,trna21_HisGTG_1_+_147753471_147753542@-1G.22.23,trna33_HisGTG_6_+_27125906_27125977@-1G.22.23,trna7_HisGTG_9_-_14433938_14434009@-1G.22.23,trna8_HisGTG_15_-_45492611_45492682@-1G.22.23,trna9_HisGTG_15_-_45490804_45490875@-1G.22.23|GGCCGTGATCGTATAGTGGTTAG</td><td>>am_trna111_HisGTG_1_-_147774845_147774916</td><td>-1G</td><td>22</td><td>23</td><td>GGCCGTGATCGTATAGTGGTTAG</td><td>253.6</td><td>-5.8</td><td>1.8</td><td>-3.2</td><td>1.2E-03</td><td>7.3E-03</td><td>BT20/MB231</td><td>MP</td></tr>
<tr><td>>hg38_21_+_8433172_8446622_RNA45SN1_45S_50ntFlanks@10712.10736.25|CCTTGAAAATCCGGGGGAGAGGGTG</td><td>>hg38_21_+_8433172_8446622_RNA45SN1_45S_50ntFlanks</td><td>10712</td><td>10736</td><td>25</td><td>CCTTGAAAATCCGGGGGAGAGGGTG</td><td>116.9</td><td>-4.7</td><td>1.6</td><td>-2.9</td><td>3.5E-03</td><td>7.3E-03</td><td>BT20/MB468</td><td>MP</td></tr>
</table>

<table>
<tr><td>>hg38_21_+_8433172_8446622_RNA45SN1_45S_50ntFlanks@11206.11232.27|CGGCGGCGACTCTGGACGCGAGCCGGG</td><td>>hg38_21_+_8433172_8446622_RNA45SN1_45S_50ntFlanks</td><td>11206</td><td>11232</td><td>27</td><td>CGGCGGCGACTCTGGACGCGAGCCGGG</td><td>316.2</td><td>-3.3</td><td>1.2</td><td>-2.7</td><td>6.4E-03</td><td>7.3E-03</td><td>MB231/MB468</td><td>NUC</td></tr>
<tr><td>>hg38_21_+_8433172_8446622_RNA45SN1_45S_50ntFlanks@9622.9639.18|CTAGTAGCTGGTTCCCTC</td><td>>hg38_21_+_8433172_8446622_RNA45SN1_45S_50ntFlanks</td><td>9622</td><td>9639</td><td>18</td><td>CTAGTAGCTGGTTCCCTC</td><td>71.9</td><td>-4.8</td><td>1.7</td><td>-2.9</td><td>3.5E-03</td><td>7.3E-03</td><td>BT20/MB468</td><td>MP</td></tr>
<tr><td>>hg38_21_+_8433172_8446622_RNA45SN1_45S_50ntFlanks@6651.6675.25|CGACTCTTAGCGGTGGATCACTCGG</td><td>>hg38_21_+_8433172_8446622_RNA45SN1_45S_50ntFlanks</td><td>6651</td><td>6675</td><td>25</td><td>CGACTCTTAGCGGTGGATCACTCGG</td><td>10226.7</td><td>2.7</td><td>1.0</td><td>2.7</td><td>6.4E-03</td><td>7.3E-03</td><td>MB231/MB468</td><td>NUC</td></tr>
<tr><td>>hg38_21_+_8433172_8446622_RNA45SN1_45S_50ntFlanks@6646.6686.41|TCGTACGACTCTTAGCGGTGGATCACTCGGCTCGTGCGTCG</td><td>>hg38_21_+_8433172_8446622_RNA45SN1_45S_50ntFlanks</td><td>6646</td><td>6686</td><td>41</td><td>TCGTACGACTCTTAGCGGTGGATCACTCGGCTCGTGCGTCG</td><td>481.5</td><td>-2.6</td><td>0.9</td><td>-2.8</td><td>5.5E-03</td><td>7.3E-03</td><td>MB231/MB468</td><td>MP</td></tr>
<tr><td>>hg38_MT_+_598_1651_RNR1_12S_50ntFlanks@908.946.39|ATACTTCAAAGGACATTTAACTAAAACCCCTACGCATTT</td><td>>hg38_MT_+_598_1651_RNR1_12S_50ntFlanks</td><td>908</td><td>946</td><td>39</td><td>ATACTTCAAAGGACATTTAACTAAAACCCCTACGCATTT</td><td>255.9</td><td>2.6</td><td>0.8</td><td>3.1</td><td>1.6E-03</td><td>7.3E-03</td><td>BT20/MB231</td><td>MITO</td></tr>
<tr><td>>MI0000114|hsa-mir-107&WithFlank&10|-|89592741|89592833@56.76.21;MI0000108|hsa-mir-103a-2&WithFlank&20|+|3917488|3917577@54.74.21;MI0000109|hsa-mir-103a-1&WithFlank&5|-|168560890|168560979@54.74.21&[MIMAT0000104&hsa-miR-107&offsets|0|-2;m-78&10|-|89592758|89592778&offsets|0|0];[MIMAT0000101&hsa-miR-103a-3p&offsets|0|-2;m-12&20|+|3917541|3917563&offsets|0|-2];[MIMAT0000101_1&hsa-miR-103a-3p&offsets|0|-2;m-13&5|-|168560904|168560926&offsets|0|-2;m-44220&5|-|168560908|168560926&offsets|0|+2]|AGCAGCATTGTACAGGGCTAT</td><td>>MI0000114|hsa-mir-107&WithFlank&10|-|89592741|89592833</td><td>56</td><td>76</td><td>21</td><td>AGCAGCATTGTACAGGGCTAT</td><td>1498.7</td><td>-2.8</td><td>0.9</td><td>-3.1</td><td>1.6E-03</td><td>7.3E-03</td><td>BT20/MB231</td><td>MITO</td></tr>
<tr><td>>hg38_21_+_8433172_8446622_RNA45SN1_45S_50ntFlanks@12005.12035.31|GGGCCGCCGGTGAAATACCACTACTCTGATC</td><td>>hg38_21_+_8433172_8446622_RNA45SN1_45S_50ntFlanks</td><td>12005</td><td>12035</td><td>31</td><td>GGGCCGCCGGTGAAATACCACTACTCTGATC</td><td>484.3</td><td>2.2</td><td>0.7</td><td>3.1</td><td>1.7E-03</td><td>7.3E-03</td><td>BT20/MB231</td><td>MITO</td></tr>
<tr><td>>hg38_21_+_8433172_8446622_RNA45SN1_45S_50ntFlanks@6661.6702.42|CGGTGGATCACTCGGCTCGTGCGTCGATGAAGAACGCAGCTA</td><td>>hg38_21_+_8433172_8446622_RNA45SN1_45S_50ntFlanks</td><td>6661</td><td>6702</td><td>42</td><td>CGGTGGATCACTCGGCTCGTGCGTCGATGAAGAACGCAGCTA</td><td>529.7</td><td>2.8</td><td>0.9</td><td>3.1</td><td>1.6E-03</td><td>7.3E-03</td><td>BT20/MB231</td><td>MITO</td></tr>
<tr><td>>MI0000298|hsa-mir-221&WithFlank&X|-|45746151|45746272@31.53.23&[MIMAT0004568&hsa-miR-221-5p&offsets|0|+1;m-405&X|-|45746220|45746242&offsets|0|0]|ACCTGGCATACAATGTAGATTTC</td><td>>MI0000298|hsa-mir-221&WithFlank&X|-|45746151|45746272</td><td>31</td><td>53</td><td>23</td><td>ACCTGGCATACAATGTAGATTTC</td><td>723.0</td><td>-2.4</td><td>0.8</td><td>-3.1</td><td>1.7E-03</td><td>7.3E-03</td><td>BT20/MB231</td><td>MITO</td></tr>
<tr><td>>hg38_21_+_8433172_8446622_RNA45SN1_45S_50ntFlanks@10350.10378.29|ACGAGAACTTTGAAGGCCGAAGTGGAGAA</td><td>>hg38_21_+_8433172_8446622_RNA45SN1_45S_50ntFlanks</td><td>10350</td><td>10378</td><td>29</td><td>ACGAGAACTTTGAAGGCCGAAGTGGAGAA</td><td>236.7</td><td>-1.7</td><td>0.6</td><td>-2.8</td><td>5.9E-03</td><td>7.3E-03</td><td>BT20/MB468</td><td>CYTO</td></tr>
<tr><td>>hg38_21_+_8433172_8446622_RNA45SN1_45S_50ntFlanks@10548.10576.29|GTGCGGTAACGCGACCGATCCCGGAGAAG</td><td>>hg38_21_+_8433172_8446622_RNA45SN1_45S_50ntFlanks</td><td>10548</td><td>10576</td><td>29</td><td>GTGCGGTAACGCGACCGATCCCGGAGAAG</td><td>84.2</td><td>2.6</td><td>0.9</td><td>2.9</td><td>3.5E-03</td><td>7.3E-03</td><td>BT20/MB468</td><td>MP</td></tr>
<tr><td>>hg38_21_+_8433172_8446622_RNA45SN1_45S_50ntFlanks@10729.10755.27|AGAGGGTGTAAATCTCGCGCCGGGCCG</td><td>>hg38_21_+_8433172_8446622_RNA45SN1_45S_50ntFlanks</td><td>10729</td><td>10755</td><td>27</td><td>AGAGGGTGTAAATCTCGCGCCGGGCCG</td><td>314.1</td><td>-4.9</td><td>1.7</td><td>-2.9</td><td>3.5E-03</td><td>7.3E-03</td><td>BT20/MB468</td><td>MP</td></tr>
<tr><td>>hg38_21_+_8433172_8446622_RNA45SN1_45S_50ntFlanks@9548.9571.24|GAGGTCCGTAGCGGTCCTGACGTG</td><td>>hg38_21_+_8433172_8446622_RNA45SN1_45S_50ntFlanks</td><td>9548</td><td>9571</td><td>24</td><td>GAGGTCCGTAGCGGTCCTGACGTG</td><td>657.9</td><td>-1.7</td><td>0.6</td><td>-2.7</td><td>6.4E-03</td><td>7.3E-03</td><td>MB231/MB468</td><td>NUC</td></tr>
<tr><td>>MI0000270|hsa-mir-181b-1&WithFlank&1|-|198858867|198858988@42.64.23(+1A);MI0000683|hsa-mir-181b-2&WithFlank&9|+|124693704|124693804@22.44.23(+1A)&[MIMAT0000257&hsa-miR-181b-5p&offsets|0|0(+1A);m-73&1|-|198858925|198858947&offsets|0|0(+1A)];[MIMAT0000257_1&hsa-miR-181b-5p&offsets|0|0(+1A);m-72&9|+|124693725|124693747&offsets|0|0(+1A)]|AACATTCATTGCTGTCGGTGGGTA</td><td>>MI0000270|hsa-mir-181b-1&WithFlank&1|-|198858867|198858988</td><td>42</td><td>64</td><td>23(+1A)</td><td>AACATTCATTGCTGTCGGTGGGTA</td><td>233.7</td><td>3.7</td><td>1.3</td><td>2.8</td><td>5.5E-03</td><td>7.3E-03</td><td>MB231/MB468</td><td>MP</td></tr>
<tr><td>>trnaMT_PheGAA_MT_+_577_647@22.53.32|AAAGCAATACACTGAAAATGTTTAGACGGGCT</td><td>>trnaMT_PheGAA_MT_+_577_647</td><td>22</td><td>53</td><td>32</td><td>AAAGCAATACACTGAAAATGTTTAGACGGGCT</td><td>176.8</td><td>5.4</td><td>1.9</td><td>2.9</td><td>3.5E-03</td><td>7.3E-03</td><td>BT20/MB468</td><td>MP</td></tr>
<tr><td>>hg38_21_+_8433172_8446622_RNA45SN1_45S_50ntFlanks@4731.4766.36|AAGAACGAAAGTCGGAGGTTCGAAGACGATCAGATA</td><td>>hg38_21_+_8433172_8446622_RNA45SN1_45S_50ntFlanks</td><td>4731</td><td>4766</td><td>36</td><td>AAGAACGAAAGTCGGAGGTTCGAAGACGATCAGATA</td><td>223.5</td><td>2.7</td><td>0.9</td><td>2.9</td><td>3.5E-03</td><td>7.3E-03</td><td>BT20/MB468</td><td>MP</td></tr>
<tr><td>>hg38_1_-_228634819_228635039_RNA5S12_5S_50ntFlanks@138.169.32|AGACCGCCTGGGAATACCGGGTGCTGTAGGCT</td><td>>hg38_1_-_228634819_228635039_RNA5S12_5S_50ntFlanks</td><td>138</td><td>169</td><td>32</td><td>AGACCGCCTGGGAATACCGGGTGCTGTAGGCT</td><td>930.3</td><td>2.4</td><td>0.8</td><td>2.9</td><td>3.5E-03</td><td>7.3E-03</td><td>BT20/MB468</td><td>MP</td></tr>
<tr><td>>hg38_21_+_8433172_8446622_RNA45SN1_45S_50ntFlanks@10808.10838.31|CAATGTAGGTAAGGGAAGTCGGCAAGCCGGA</td><td>>hg38_21_+_8433172_8446622_RNA45SN1_45S_50ntFlanks</td><td>10808</td><td>10838</td><td>31</td><td>CAATGTAGGTAAGGGAAGTCGGCAAGCCGGA</td><td>132.8</td><td>-2.7</td><td>0.9</td><td>-2.9</td><td>3.5E-03</td><td>7.3E-03</td><td>BT20/MB468</td><td>MP</td></tr>
<tr><td>>trna5_AspGTC_12_+_98897281_98897352@35.75.41|TCACGCGGGAGACCGGGGTTCAATTCCCCGACGGGGAGCCA</td><td>>trna5_AspGTC_12_+_98897281_98897352</td><td>35</td><td>75</td><td>41</td><td>TCACGCGGGAGACCGGGGTTCAATTCCCCGACGGGGAGCCA</td><td>335.8</td><td>4.5</td><td>1.7</td><td>2.7</td><td>6.4E-03</td><td>7.3E-03</td><td>MB231/MB468</td><td>NUC</td></tr>
<tr><td>>hg38_21_+_8433172_8446622_RNA45SN1_45S_50ntFlanks@4069.4097.29|CTTTCGATGGTAGTCGCCGTGCCTACCAT</td><td>>hg38_21_+_8433172_8446622_RNA45SN1_45S_50ntFlanks</td><td>4069</td><td>4097</td><td>29</td><td>CTTTCGATGGTAGTCGCCGTGCCTACCAT</td><td>48.2</td><td>-4.7</td><td>1.6</td><td>-2.9</td><td>3.5E-03</td><td>7.3E-03</td><td>BT20/MB468</td><td>MP</td></tr>
<tr><td>>MI0000234|hsa-mir-192&WithFlank&11|-|64891131|64891252@30.51.22(+1U)&[MIMAT0000222&hsa-miR-192-5p&offsets|0|+1(+1U);m-59&11|-|64891203|64891223&offsets|0|+1(+1U)]|CTGACCTATGAATTGACAGCCAT</td><td>>MI0000234|hsa-mir-192&WithFlank&11|-|64891131|64891252</td><td>30</td><td>51</td><td>22(+1U)</td><td>CTGACCTATGAATTGACAGCCAT</td><td>1657.2</td><td>-2.7</td><td>0.8</td><td>-3.2</td><td>1.3E-03</td><td>7.3E-03</td><td>BT20/MB231</td><td>MP</td></tr>
<tr><td>>MI0000088|hsa-mir-30a&WithFlank&6|-|71403545|71403627@54.74.21(+1A)&[MIMAT0000088&hsa-miR-30a-3p&offsets|+1|0(+1A);m-38&6|-|71403554|71403575&offsets|+1|0(+1A)]|TTTCAGTCGGATGTTTGCAGCA</td><td>>MI0000088|hsa-mir-30a&WithFlank&6|-|71403545|71403627</td><td>54</td><td>74</td><td>21(+1A)</td><td>TTTCAGTCGGATGTTTGCAGCA</td><td>258.0</td><td>-3.8</td><td>1.2</td><td>-3.2</td><td>1.3E-03</td><td>7.3E-03</td><td>BT20/MB231</td><td>MP</td></tr>
<tr><td>>hg38_21_+_8433172_8446622_RNA45SN1_45S_50ntFlanks@9533.9564.32|GAGGAAACTCTGGTGGAGGTCCGTAGCGGTCC</td><td>>hg38_21_+_8433172_8446622_RNA45SN1_45S_50ntFlanks</td><td>9533</td><td>9564</td><td>32</td><td>GAGGAAACTCTGGTGGAGGTCCGTAGCGGTCC</td><td>142.7</td><td>-4.7</td><td>1.7</td><td>-2.8</td><td>5.5E-03</td><td>7.3E-03</td><td>MB231/MB468</td><td>MP</td></tr>
<tr><td>>hg38_21_+_8433172_8446622_RNA45SN1_45S_50ntFlanks@12691.12711.21|CGGAACGATACGGCAGCGCCG</td><td>>hg38_21_+_8433172_8446622_RNA45SN1_45S_50ntFlanks</td><td>12691</td><td>12711</td><td>21</td><td>CGGAACGATACGGCAGCGCCG</td><td>132.1</td><td>-4.8</td><td>1.6</td><td>-2.9</td><td>3.5E-03</td><td>7.3E-03</td><td>BT20/MB468</td><td>MP</td></tr>
<tr><td>>MI0000811|hsa-mir-148b&WithFlank&12|+|54337210|54337320@69.90.22(+3U)&[MIMAT0000759&hsa-miR-148b-3p&offsets|0|0(+3U);m-77&12|+|54337278|54337299&offsets|0|0(+3U)]|TCAGTGCATCACAGAACTTTGTTTT</td><td>>MI0000811|hsa-mir-148b&WithFlank&12|+|54337210|54337320</td><td>69</td><td>90</td><td>22(+3U)</td><td>TCAGTGCATCACAGAACTTTGTTTT</td><td>326.7</td><td>-4.8</td><td>1.5</td><td>-3.2</td><td>1.3E-03</td><td>7.3E-03</td><td>BT20/MB231</td><td>MP</td></tr>
<tr><td>>hg38_21_+_8433172_8446622_RNA45SN1_45S_50ntFlanks@5490.5525.36|TGAGAAGACGGTCGAACTTGACTATCTAGAGGAAGT</td><td>>hg38_21_+_8433172_8446622_RNA45SN1_45S_50ntFlanks</td><td>5490</td><td>5525</td><td>36</td><td>TGAGAAGACGGTCGAACTTGACTATCTAGAGGAAGT</td><td>2306.9</td><td>5.1</td><td>1.6</td><td>3.2</td><td>1.3E-03</td><td>7.3E-03</td><td>BT20/MB231</td><td>MP</td></tr>
<tr><td>>hg38_21_+_8433172_8446622_RNA45SN1_45S_50ntFlanks@10792.10818.27|CTCTGGCATGTTGGAACAATGTAGGTA</td><td>>hg38_21_+_8433172_8446622_RNA45SN1_45S_50ntFlanks</td><td>10792</td><td>10818</td><td>27</td><td>CTCTGGCATGTTGGAACAATGTAGGTA</td><td>230.8</td><td>-1.7</td><td>0.6</td><td>-2.7</td><td>6.0E-03</td><td>7.3E-03</td><td>BT20/MB468</td><td>CYTO</td></tr>
<tr><td>>hg38_21_+_8433172_8446622_RNA45SN1_45S_50ntFlanks@4722.4755.34|TCATTAATCAAGAACGAAAGTCGGAGGTTCGAAG</td><td>>hg38_21_+_8433172_8446622_RNA45SN1_45S_50ntFlanks</td><td>4722</td><td>4755</td><td>34</td><td>TCATTAATCAAGAACGAAAGTCGGAGGTTCGAAG</td><td>117.5</td><td>-4.8</td><td>1.6</td><td>-2.9</td><td>3.5E-03</td><td>7.3E-03</td><td>BT20/MB468</td><td>MP</td></tr>
<tr><td>>hg38_21_+_8433172_8446622_RNA45SN1_45S_50ntFlanks@6646.6675.30|TCGTACGACTCTTAGCGGTGGATCACTCGG</td><td>>hg38_21_+_8433172_8446622_RNA45SN1_45S_50ntFlanks</td><td>6646</td><td>6675</td><td>30</td><td>TCGTACGACTCTTAGCGGTGGATCACTCGG</td><td>2003.2</td><td>2.1</td><td>0.7</td><td>2.7</td><td>6.0E-03</td><td>7.3E-03</td><td>BT20/MB468</td><td>CYTO</td></tr>
<tr><td>>hg38_21_+_8433172_8446622_RNA45SN1_45S_50ntFlanks@10446.10463.18|AAGGGACGGGCGATGGCC</td><td>>hg38_21_+_8433172_8446622_RNA45SN1_45S_50ntFlanks</td><td>10446</td><td>10463</td><td>18</td><td>AAGGGACGGGCGATGGCC</td><td>61.5</td><td>9.3</td><td>2.9</td><td>3.2</td><td>1.3E-03</td><td>7.3E-03</td><td>BT20/MB231</td><td>MP</td></tr>
<tr><td>>hg38_21_+_8433172_8446622_RNA45SN1_45S_50ntFlanks@12417.12445.29|TCGATGTCGGCTCTTCCTATCATTGTGAA</td><td>>hg38_21_+_8433172_8446622_RNA45SN1_45S_50ntFlanks</td><td>12417</td><td>12445</td><td>29</td><td>TCGATGTCGGCTCTTCCTATCATTGTGAA</td><td>95.4</td><td>-4.6</td><td>1.7</td><td>-2.8</td><td>5.5E-03</td><td>7.3E-03</td><td>MB231/MB468</td><td>MP</td></tr>
<tr><td>>hg38_21_+_8433172_8446622_RNA45SN1_45S_50ntFlanks@12927.12973.47|CCTGGCGCTAAACCATTCGTAGACGACCTGCTTCTGGGTCGGGGTTT</td><td>>hg38_21_+_8433172_8446622_RNA45SN1_45S_50ntFlanks</td><td>12927</td><td>12973</td><td>47</td><td>CCTGGCGCTAAACCATTCGTAGACGACCTGCTTCTGGGTCGGGGTTT</td><td>319.9</td><td>-3.7</td><td>1.3</td><td>-2.8</td><td>5.1E-03</td><td>7.3E-03</td><td>BT20/MB468</td><td>MITO</td></tr>
<tr><td>>MI0000433|hsa-let-7g&WithFlank&3|-|52268272|52268367@10.32.23&[MIMAT0000414&hsa-let-7g-5p&offsets|-1|0;m-58&3|-|52268336|52268357&offsets|-1|0]|CTGAGGTAGTAGTTTGTACAGTT</td><td>>MI0000433|hsa-let-7g&WithFlank&3|-|52268272|52268367</td><td>10</td><td>32</td><td>23</td><td>CTGAGGTAGTAGTTTGTACAGTT</td><td>132.8</td><td>3.7</td><td>1.3</td><td>2.8</td><td>5.5E-03</td><td>7.3E-03</td><td>MB231/MB468</td><td>MP</td></tr>
<tr><td>>hg38_21_+_8433172_8446622_RNA45SN1_45S_50ntFlanks@10787.10825.39|ACAGCCTCTGGCATGTTGGAACAATGTAGGTAAGGGAAG</td><td>>hg38_21_+_8433172_8446622_RNA45SN1_45S_50ntFlanks</td><td>10787</td><td>10825</td><td>39</td><td>ACAGCCTCTGGCATGTTGGAACAATGTAGGTAAGGGAAG</td><td>193.9</td><td>-1.9</td><td>0.7</td><td>-2.8</td><td>5.1E-03</td><td>7.4E-03</td><td>BT20/MB468</td><td>MITO</td></tr>
<tr><td>>hg38_21_+_8433172_8446622_RNA45SN1_45S_50ntFlanks@4557.4574.18|CAGCTAGGAATAATGGAA</td><td>>hg38_21_+_8433172_8446622_RNA45SN1_45S_50ntFlanks</td><td>4557</td><td>4574</td><td>18</td><td>CAGCTAGGAATAATGGAA</td><td>148.6</td><td>-2.3</td><td>0.8</td><td>-2.7</td><td>6.4E-03</td><td>7.4E-03</td><td>MB231/MB468</td><td>NUC</td></tr>
<tr><td>>hg38_21_+_8433172_8446622_RNA45SN1_45S_50ntFlanks@9540.9570.31|CTCTGGTGGAGGTCCGTAGCGGTCCTGACGT</td><td>>hg38_21_+_8433172_8446622_RNA45SN1_45S_50ntFlanks</td><td>9540</td><td>9570</td><td>31</td><td>CTCTGGTGGAGGTCCGTAGCGGTCCTGACGT</td><td>109.4</td><td>3.0</td><td>1.0</td><td>2.9</td><td>3.5E-03</td><td>7.4E-03</td><td>BT20/MB468</td><td>MP</td></tr>
<tr><td>>am_MI0000809|hsa-mir-151a&WithFlank&8|-|140732558|140732659@54.73.20&[MIMAT0000757&hsa-miR-151a-3p&offsets|+1|0;m-30&8|-|140732587|140732607&offsets|+1|0]|TAGACTGAAGCTCCTTGAGG</td><td>>am_MI0000809|hsa-mir-151a&WithFlank&8|-|140732558|140732659</td><td>54</td><td>73</td><td>20</td><td>TAGACTGAAGCTCCTTGAGG</td><td>54.6</td><td>4.4</td><td>1.6</td><td>2.8</td><td>5.5E-03</td><td>7.4E-03</td><td>MB231/MB468</td><td>MP</td></tr>
<tr><td>>hg38_21_+_8433172_8446622_RNA45SN1_45S_50ntFlanks@12454.12476.23|CACCAAGCGTTGGATTGTTCACC</td><td>>hg38_21_+_8433172_8446622_RNA45SN1_45S_50ntFlanks</td><td>12454</td><td>12476</td><td>23</td><td>CACCAAGCGTTGGATTGTTCACC</td><td>854.4</td><td>1.8</td><td>0.7</td><td>2.7</td><td>6.1E-03</td><td>7.4E-03</td><td>MB231/MB468</td><td>CYTO</td></tr>
<tr><td>>trnaMT_GluTTC_MT_-_14674_14742@34.65.32,trnalookalike8_GluTTC_5_-_93905172_93905240@34.65.32|ATATCATTGGTCGTGGTTGTAGTCCGTGCGAG</td><td>>trnaMT_GluTTC_MT_-_14674_14742</td><td>34</td><td>65</td><td>32</td><td>ATATCATTGGTCGTGGTTGTAGTCCGTGCGAG</td><td>185.4</td><td>3.7</td><td>1.3</td><td>2.9</td><td>3.5E-03</td><td>7.4E-03</td><td>BT20/MB468</td><td>MP</td></tr>
<tr><td>>hg38_21_+_8433172_8446622_RNA45SN1_45S_50ntFlanks@12625.12657.33|CAATGGGGCGAAGCTACCATCTGTGGGATTATG</td><td>>hg38_21_+_8433172_8446622_RNA45SN1_45S_50ntFlanks</td><td>12625</td><td>12657</td><td>33</td><td>CAATGGGGCGAAGCTACCATCTGTGGGATTATG</td><td>107.4</td><td>-2.9</td><td>1.1</td><td>-2.7</td><td>6.0E-03</td><td>7.4E-03</td><td>BT20/MB468</td><td>CYTO</td></tr>
<tr><td>>hg38_21_+_8433172_8446622_RNA45SN1_45S_50ntFlanks@10409.10433.25|GTCAGTCGGTCCTGAGAGATGGGCG</td><td>>hg38_21_+_8433172_8446622_RNA45SN1_45S_50ntFlanks</td><td>10409</td><td>10433</td><td>25</td><td>GTCAGTCGGTCCTGAGAGATGGGCG</td><td>221.7</td><td>6.5</td><td>2.0</td><td>3.2</td><td>1.3E-03</td><td>7.4E-03</td><td>BT20/MB231</td><td>MP</td></tr>
<tr><td>>hg38_21_+_8433172_8446622_RNA45SN1_45S_50ntFlanks@6721.6750.30|AATTGCAGGACACATTGATCATCGACACTT</td><td>>hg38_21_+_8433172_8446622_RNA45SN1_45S_50ntFlanks</td><td>6721</td><td>6750</td><td>30</td><td>AATTGCAGGACACATTGATCATCGACACTT</td><td>352.2</td><td>-1.4</td><td>0.5</td><td>-2.8</td><td>5.1E-03</td><td>7.4E-03</td><td>BT20/MB468</td><td>MITO</td></tr>
<tr><td>>hg38_21_+_8433172_8446622_RNA45SN1_45S_50ntFlanks@13022.13039.18|CTCGACACAAGGGTTTGT</td><td>>hg38_21_+_8433172_8446622_RNA45SN1_45S_50ntFlanks</td><td>13022</td><td>13039</td><td>18</td><td>CTCGACACAAGGGTTTGT</td><td>994.8</td><td>2.2</td><td>0.8</td><td>2.7</td><td>6.4E-03</td><td>7.4E-03</td><td>MB231/MB468</td><td>MITO</td></tr>
</table>

| | | | | | | | | | | | | | |
|---|---|---|---|---|---|---|---|---|---|---|---|---|---|
| >MI0000061\|hsa-let-7a-2&WithFlank&11\|-\|122146516\|122146599@11.28.18;MI0000064\|hsa-let-7c&WithFlank&21\|+\|16539822\|16539917@17.34.18;MI0000062\|hsa-let-7a-3&WithFlank&22\|+\|46112743\|46112828@10.27.18;MI0000060\|hsa-let-7a-1&WithFlank&9\|+\|94175951\|94176042@12.29.18&[MIMAT0000062&hsa-let-7a-5p&offsets\|0\|-4;m-6&11\|-\|122146568\|122146589&offsets\|0\|-4];[MIMAT0000064&hsa-let-7c-5p&offsets\|0\|-4;m-37&21\|+\|16539838\|16539859&offsets\|0\|-4];[MIMAT0000062_1&hsa-let-7a-5p&offsets\|0\|-4;m-5&22\|+\|46112752\|46112773&offsets\|0\|-4];[MIMAT0000062_2&hsa-let-7a-5p&offsets\|0\|-4;m-4&9\|+\|94175962\|94175983&offsets\|0\|-4]\|TGAGGTAGTAGGTTGTAT | >MI0000061\|hsa-let-7a-2&WithFlank&11\|-\|122146516\|122146599 | 11 | 28 | 18 | TGAGGTAGTAGGTTGTAT | 1034.4 | 2.0 | 0.7 | 2.7 | 6.4E-03 | 7.4E-03 | BT20/MB468 | NUC |
| >hg38_21_+_8433172_8446622_RNA45SN1_45S_50ntFlanks@10466.10487.22\|CGTTGCCCTCGGCCGATCGAAA | >hg38_21_+_8433172_8446622_RNA45SN1_45S_50ntFlanks | 10466 | 10487 | 22 | CGTTGCCCTCGGCCGATCGAAA | 204.0 | -3.2 | 1.2 | -2.7 | 6.0E-03 | 7.4E-03 | BT20/MB468 | CYTO |
| >hg38_21_+_8433172_8446622_RNA45SN1_45S_50ntFlanks@8197.8215.19\|GTGGACGGTGTGAGGCCGG | >hg38_21_+_8433172_8446622_RNA45SN1_45S_50ntFlanks | 8197 | 8215 | 19 | GTGGACGGTGTGAGGCCGG | 416.7 | -2.2 | 0.8 | -2.7 | 6.0E-03 | 7.4E-03 | BT20/MB468 | CYTO |
| >hg38_21_+_8433172_8446622_RNA45SN1_45S_50ntFlanks@9554.9574.21\|CGTAGCGGTCCTGACGTGCAA | >hg38_21_+_8433172_8446622_RNA45SN1_45S_50ntFlanks | 9554 | 9574 | 21 | CGTAGCGGTCCTGACGTGCAA | 3611.2 | -1.6 | 0.6 | -2.7 | 6.1E-03 | 7.4E-03 | MB231/MB468 | CYTO |
| >hg38_21_+_8433172_8446622_RNA45SN1_45S_50ntFlanks@8154.8179.26\|CCGCTCGTGGGGGGCCCAAGTCCTTC | >hg38_21_+_8433172_8446622_RNA45SN1_45S_50ntFlanks | 8154 | 8179 | 26 | CCGCTCGTGGGGGGCCCAAGTCCTTC | 63.2 | -4.7 | 1.6 | -2.9 | 3.5E-03 | 7.4E-03 | BT20/MB468 | MP |
| >hg38_21_+_8433172_8446622_RNA45SN1_45S_50ntFlanks@6652.6693.42\|GACTCTTAGCGGTGGATCACTCGGCTCGTGCGTCGATGAAGA | >hg38_21_+_8433172_8446622_RNA45SN1_45S_50ntFlanks | 6652 | 6693 | 42 | GACTCTTAGCGGTGGATCACTCGGCTCGTGCGTCGATGAAGA | 555.3 | 4.3 | 1.5 | 2.9 | 3.5E-03 | 7.4E-03 | BT20/MB468 | MP |
| >hg38_21_+_8433172_8446622_RNA45SN1_45S_50ntFlanks@5471.5514.44\|CACGGCCCTGGCGGAGCGCTGAGAAGACGGTCGAACTTGACTAT | >hg38_21_+_8433172_8446622_RNA45SN1_45S_50ntFlanks | 5471 | 5514 | 44 | CACGGCCCTGGCGGAGCGCTGAGAAGACGGTCGAACTTGACTAT | 3023.5 | -2.5 | 0.9 | -2.7 | 6.1E-03 | 7.4E-03 | MB231/MB468 | CYTO |
| >hg38_21_+_8433172_8446622_RNA45SN1_45S_50ntFlanks@9532.9563.32\|AGAGGAAACTCTGGTGGAGGTCCGTAGCGGTC | >hg38_21_+_8433172_8446622_RNA45SN1_45S_50ntFlanks | 9532 | 9563 | 32 | AGAGGAAACTCTGGTGGAGGTCCGTAGCGGTC | 305.9 | -2.6 | 0.8 | -3.2 | 1.6E-03 | 7.4E-03 | BT20/MB231 | CYTO |
| >hg38_21_+_8433172_8446622_RNA45SN1_45S_50ntFlanks@11846.11867.22\|GAACGGGCTTGGCGGAATCAGC | >hg38_21_+_8433172_8446622_RNA45SN1_45S_50ntFlanks | 11846 | 11867 | 22 | GAACGGGCTTGGCGGAATCAGC | 171.5 | 4.3 | 1.5 | 2.9 | 3.5E-03 | 7.4E-03 | BT20/MB468 | MP |
| >am_trna17_LeuCAG_16_+_57333863_57333945@1.25.25,trna26_LeuCAG_16_-_57334392_57334474@1.25.25,trna34_LeuCAG_1_+_161411323_161411405@1.25.25,trna36_LeuCAG_1_+_161418741_161418823@1.25.25,trna38_LeuCAG_1_+_161426122_161426204@1.25.25,trna40_LeuCAG_1_+_161433532_161433614@1.25.25,trna42_LeuCAG_1_+_161440913_161440995@1.25.25,trna67_LeuCAG_1_-_161500132_161500214@1.25.25,trna7_LeuCAG_6_+_26521436_26521518@1.25.25\|GTCAGGATGGCCGAGCGGTCTAAGG | >am_trna17_LeuCAG_16_+_57333863_57333945 | 1 | 25 | 25 | GTCAGGATGGCCGAGCGGTCTAAGG | 1498.5 | -3.9 | 1.2 | -3.2 | 1.6E-03 | 7.4E-03 | BT20/MB231 | CYTO |
| >hg38_1_-_228634819_228635039_RNA5S12_5S_50ntFlanks@51.86.36\|GTCTACGGCCATACCACCCTGAACGCGCCCGATCTC | >hg38_1_-_228634819_228635039_RNA5S12_5S_50ntFlanks | 51 | 86 | 36 | GTCTACGGCCATACCACCCTGAACGCGCCCGATCTC | 1738.0 | -2.9 | 0.9 | -3.1 | 1.7E-03 | 7.4E-03 | BT20/MB231 | MITO |
| >hg38_21_+_8433172_8446622_RNA45SN1_45S_50ntFlanks@10387.10410.24\|TGTGAACAGCAGTTGAACATGGGT | >hg38_21_+_8433172_8446622_RNA45SN1_45S_50ntFlanks | 10387 | 10410 | 24 | TGTGAACAGCAGTTGAACATGGGT | 331.4 | -3.0 | 1.0 | -2.9 | 3.5E-03 | 7.4E-03 | BT20/MB468 | MP |
| >hg38_21_+_8433172_8446622_RNA45SN1_45S_50ntFlanks@3900.3938.39\|CTTCGCGGGGGGGATGCGTGCATTTATCAGATCAAAACC | >hg38_21_+_8433172_8446622_RNA45SN1_45S_50ntFlanks | 3900 | 3938 | 39 | CTTCGCGGGGGGGATGCGTGCATTTATCAGATCAAAACC | 405.4 | -5.5 | 1.9 | -2.9 | 3.5E-03 | 7.4E-03 | BT20/MB468 | MP |
| >hg38_21_+_8433172_8446622_RNA45SN1_45S_50ntFlanks@12203.12221.19\|TAAGGCGAGCTCAGGGAGG | >hg38_21_+_8433172_8446622_RNA45SN1_45S_50ntFlanks | 12203 | 12221 | 19 | TAAGGCGAGCTCAGGGAGG | 164.6 | -4.5 | 1.6 | -2.8 | 5.2E-03 | 7.4E-03 | BT20/MB468 | MITO |
| >hg38_21_+_8433172_8446622_RNA45SN1_45S_50ntFlanks@9938.9968.31\|AGCAGGACGGTGGCCATGGAAGTCGGAATCC | >hg38_21_+_8433172_8446622_RNA45SN1_45S_50ntFlanks | 9938 | 9968 | 31 | AGCAGGACGGTGGCCATGGAAGTCGGAATCC | 1529.2 | 6.4 | 2.0 | 3.2 | 1.3E-03 | 7.4E-03 | BT20/MB231 | MP |
| >hg38_21_+_8433172_8446622_RNA45SN1_45S_50ntFlanks@12632.12664.33\|GCGAAGCTACCATCTGTGGGATTATGACTGAAC | >hg38_21_+_8433172_8446622_RNA45SN1_45S_50ntFlanks | 12632 | 12664 | 33 | GCGAAGCTACCATCTGTGGGATTATGACTGAAC | 131.7 | 5.1 | 1.7 | 2.9 | 3.5E-03 | 7.4E-03 | BT20/MB468 | MP |
| >hg38_21_+_8433172_8446622_RNA45SN1_45S_50ntFlanks@10466.10488.23\|CGTTGCCCTCGGCCGATCGAAAG | >hg38_21_+_8433172_8446622_RNA45SN1_45S_50ntFlanks | 10466 | 10488 | 23 | CGTTGCCCTCGGCCGATCGAAAG | 324.1 | -3.1 | 1.1 | -2.7 | 6.1E-03 | 7.4E-03 | MB231/MB468 | CYTO |
| >hg38_21_+_8433172_8446622_RNA45SN1_45S_50ntFlanks@12628.12653.26\|TGGGGCGAAGCTACCATCTGTGGGAT | >hg38_21_+_8433172_8446622_RNA45SN1_45S_50ntFlanks | 12628 | 12653 | 26 | TGGGGCGAAGCTACCATCTGTGGGAT | 820.8 | -4.6 | 1.7 | -2.7 | 6.0E-03 | 7.4E-03 | BT20/MB468 | CYTO |
| >MI0000100\|hsa-mir-98&WithFlank&X\|-\|53556217\|53556347@28.49.22&[MIMAT0000096&hsa-miR-98-5p&offsets\|0\|0;m-143&X\|-\|53556299\|53556320&offsets\|0\|0]\|TGAGGTAGTAAGTTGTATTGTT | >MI0000100\|hsa-mir-98&WithFlank&X\|-\|53556217\|53556347 | 28 | 49 | 22 | TGAGGTAGTAAGTTGTATTGTT | 6809.9 | -2.9 | 0.9 | -3.2 | 1.3E-03 | 7.4E-03 | BT20/MB231 | MP |
| >hg38_MT_+_1621_3279_RNR2_16S_50ntFlanks@837.864.28\|AAAAAAAGTAAAAGGAACTCGGCAAATC | >hg38_MT_+_1621_3279_RNR2_16S_50ntFlanks | 837 | 864 | 28 | AAAAAAAGTAAAAGGAACTCGGCAAATC | 27.2 | 8.0 | 2.9 | 2.8 | 5.6E-03 | 7.4E-03 | MB231/MB468 | MP |
| >hg38_MT_+_1621_3279_RNR2_16S_50ntFlanks@1380.1414.35\|AGGACATCCCGATGGTGCAGCCGCTATTAAAGGTT | >hg38_MT_+_1621_3279_RNR2_16S_50ntFlanks | 1380 | 1414 | 35 | AGGACATCCCGATGGTGCAGCCGCTATTAAAGGTT | 246.1 | 4.7 | 1.6 | 2.9 | 3.5E-03 | 7.4E-03 | BT20/MB468 | MP |
| >hg38_21_+_8433172_8446622_RNA45SN1_45S_50ntFlanks@10781.10817.37\|AGGTGAACAGCCTCTGGCATGTTGGAACAATGTAGGT | >hg38_21_+_8433172_8446622_RNA45SN1_45S_50ntFlanks | 10781 | 10817 | 37 | AGGTGAACAGCCTCTGGCATGTTGGAACAATGTAGGT | 326.7 | -4.0 | 1.4 | -2.9 | 3.5E-03 | 7.4E-03 | BT20/MB468 | MP |
| >hg38_MT_+_1621_3279_RNR2_16S_50ntFlanks@1507.1529.23\|GAAAGGACAAGAGAAATAAGGCC | >hg38_MT_+_1621_3279_RNR2_16S_50ntFlanks | 1507 | 1529 | 23 | GAAAGGACAAGAGAAATAAGGCC | 27.6 | 8.0 | 2.9 | 2.8 | 5.6E-03 | 7.4E-03 | MB231/MB468 | MP |
| >hg38_21_+_8433172_8446622_RNA45SN1_45S_50ntFlanks@9860.9890.31\|GGATGAACCGAACGCCGGGTTAAGGCGCCCG | >hg38_21_+_8433172_8446622_RNA45SN1_45S_50ntFlanks | 9860 | 9890 | 31 | GGATGAACCGAACGCCGGGTTAAGGCGCCCG | 57.8 | -4.6 | 1.7 | -2.8 | 5.6E-03 | 7.4E-03 | MB231/MB468 | MP |
| >hg38_1_-_228634819_228635039_RNA5S12_5S_50ntFlanks@51.83.33\|GTCTACGGCCATACCACCCTGAACGCGCCCGAT | >hg38_1_-_228634819_228635039_RNA5S12_5S_50ntFlanks | 51 | 83 | 33 | GTCTACGGCCATACCACCCTGAACGCGCCCGAT | 14019.7 | -2.2 | 0.8 | -2.7 | 6.5E-03 | 7.4E-03 | MB231/MB468 | NUC |
| >hg38_21_+_8433172_8446622_RNA45SN1_45S_50ntFlanks@12416.12456.41\|TTCGATGTCGGCTCTTCCTATCATTGTGAAGCAGAATTCAC | >hg38_21_+_8433172_8446622_RNA45SN1_45S_50ntFlanks | 12416 | 12456 | 41 | TTCGATGTCGGCTCTTCCTATCATTGTGAAGCAGAATTCAC | 292.0 | -2.7 | 0.9 | -3.1 | 1.7E-03 | 7.4E-03 | BT20/MB231 | MITO |
| >hg38_21_+_8433172_8446622_RNA45SN1_45S_50ntFlanks@9459.9494.36\|CCGGGGAGGTGGAGCACGAGCGCACGTGTTAGGACC | >hg38_21_+_8433172_8446622_RNA45SN1_45S_50ntFlanks | 9459 | 9494 | 36 | CCGGGGAGGTGGAGCACGAGCGCACGTGTTAGGACC | 107.6 | -4.1 | 1.5 | -2.8 | 5.2E-03 | 7.4E-03 | BT20/MB468 | MITO |
| >hg38_21_+_8433172_8446622_RNA45SN1_45S_50ntFlanks@12204.12221.18\|AAGGCGAGCTCAGGGAGG | >hg38_21_+_8433172_8446622_RNA45SN1_45S_50ntFlanks | 12204 | 12221 | 18 | AAGGCGAGCTCAGGGAGG | 265.0 | -4.7 | 1.7 | -2.8 | 5.2E-03 | 7.4E-03 | BT20/MB468 | MITO |
| >hg38_21_+_8433172_8446622_RNA45SN1_45S_50ntFlanks@5050.5081.32\|TTGGTGGAGCGATTTGTCTGGTTAATTCCGAT | >hg38_21_+_8433172_8446622_RNA45SN1_45S_50ntFlanks | 5050 | 5081 | 32 | TTGGTGGAGCGATTTGTCTGGTTAATTCCGAT | 56.0 | 5.9 | 2.0 | 2.9 | 3.5E-03 | 7.4E-03 | BT20/MB468 | MP |
| >trnaMT_ValTAC_MT_+_1602_1670@51.72.22\|CTTAACTTGACCGCTCTGACCA | >trnaMT_ValTAC_MT_+_1602_1670 | 51 | 72 | 22 | CTTAACTTGACCGCTCTGACCA | 786.2 | 2.5 | 0.9 | 2.7 | 6.1E-03 | 7.4E-03 | BT20/MB468 | CYTO |
| >MI0005566\|hsa-mir-374b&WithFlank&X\|-\|74218541\|74218624@17.37.21(+1A)&[MIMAT0004955&hsa-miR-374b-5p&offsets\|0\|-1(+1A);m-153&X\|-\|74218587\|74218608&offsets\|0\|-1(+1A)]\|ATATAATACAACCTGCTAAGTA | >MI0005566\|hsa-mir-374b&WithFlank&X\|-\|74218541\|74218624 | 17 | 37 | 21(+1A) | ATATAATACAACCTGCTAAGTA | 148.1 | 4.6 | 1.7 | 2.8 | 5.6E-03 | 7.4E-03 | MB231/MB468 | MP |
| >hg38_21_+_8433172_8446622_RNA45SN1_45S_50ntFlanks@9559.9584.26\|CGGTCCTGACGTGCAAATCGGTCGTC | >hg38_21_+_8433172_8446622_RNA45SN1_45S_50ntFlanks | 9559 | 9584 | 26 | CGGTCCTGACGTGCAAATCGGTCGTC | 128.8 | -4.5 | 1.6 | -2.8 | 5.6E-03 | 7.4E-03 | MB231/MB468 | MP |
| >MI0000270\|hsa-mir-181b-1&WithFlank&1\|-\|198858867\|198858988@42.64.23;MI0000683\|hsa-mir-181b-2&WithFlank&9\|+\|124693704\|124693804@22.44.23&[MIMAT0000257&hsa-miR-181b-5p&offsets\|0\|0;m-73&1\|-\|198858925\|198858947&offsets\|0\|0];[MIMAT0000257_1&hsa-miR-181b-5p&offsets\|0\|0;m-72&9\|+\|124693725\|124693747&offsets\|0\|0]\|AACATTCATTGCTGTCGGTGGGT | >MI0000270\|hsa-mir-181b-1&WithFlank&1\|-\|198858867\|198858988 | 42 | 64 | 23 | AACATTCATTGCTGTCGGTGGGT | 3716.5 | -2.9 | 0.9 | -3.2 | 1.3E-03 | 7.4E-03 | BT20/MB231 | MP |
| >hg38_21_+_8433172_8446622_RNA45SN1_45S_50ntFlanks@12324.12358.35\|TTGGGTTTTAAGCAGGAGGTGTCAGAAAAGTTACC | >hg38_21_+_8433172_8446622_RNA45SN1_45S_50ntFlanks | 12324 | 12358 | 35 | TTGGGTTTTAAGCAGGAGGTGTCAGAAAAGTTACC | 64.0 | -5.0 | 1.7 | -2.9 | 3.5E-03 | 7.4E-03 | BT20/MB468 | MP |
| >hg38_21_+_8433172_8446622_RNA45SN1_45S_50ntFlanks@6679.6696.18\|GTGCGTCGATGAAGAACG | >hg38_21_+_8433172_8446622_RNA45SN1_45S_50ntFlanks | 6679 | 6696 | 18 | GTGCGTCGATGAAGAACG | 127.2 | -3.5 | 1.2 | -2.8 | 5.6E-03 | 7.4E-03 | MB231/MB468 | MP |
| >trnaMT_ValTAC_MT_+_1602_1670@32.72.41\|TACACTTAGGAGATTTCAACTTAACTTGACCGCTCTGACCA | >trnaMT_ValTAC_MT_+_1602_1670 | 32 | 72 | 41 | TACACTTAGGAGATTTCAACTTAACTTGACCGCTCTGACCA | 506.5 | 5.1 | 1.8 | 2.9 | 3.6E-03 | 7.4E-03 | BT20/MB468 | MP |
| >hg38_21_+_8433172_8446622_RNA45SN1_45S_50ntFlanks@10392.10420.29\|ACAGCAGTTGAACATGGGTCAGTCGGTCC | >hg38_21_+_8433172_8446622_RNA45SN1_45S_50ntFlanks | 10392 | 10420 | 29 | ACAGCAGTTGAACATGGGTCAGTCGGTCC | 375.2 | -3.8 | 1.3 | -2.9 | 3.6E-03 | 7.4E-03 | BT20/MB468 | MP |
| >hg38_21_+_8433172_8446622_RNA45SN1_45S_50ntFlanks@8243.8269.27\|GTCTTCCCGGAGTCGGGTTGCTTGGGA | >hg38_21_+_8433172_8446622_RNA45SN1_45S_50ntFlanks | 8243 | 8269 | 27 | GTCTTCCCGGAGTCGGGTTGCTTGGGA | 342.9 | 4.3 | 1.3 | 3.2 | 1.3E-03 | 7.5E-03 | BT20/MB231 | MP |
| >hg38_21_+_8433172_8446622_RNA45SN1_45S_50ntFlanks@8408.8442.35\|AACGGGTGGGGTCCGCGCAGTCCGCCCGGAGGATT | >hg38_21_+_8433172_8446622_RNA45SN1_45S_50ntFlanks | 8408 | 8442 | 35 | AACGGGTGGGGTCCGCGCAGTCCGCCCGGAGGATT | 151.0 | -3.7 | 1.3 | -2.8 | 5.6E-03 | 7.5E-03 | MB231/MB468 | MP |
| >trnaMT_GluTTC_MT_-_14674_14742@40.72.33,trnalookalike8_GluTTC_5_-_93905172_93905240@40.72.33\|TTGGTCGTGGTTGTAGTCCGTGCGAGAATACCA | >trnaMT_GluTTC_MT_-_14674_14742 | 40 | 72 | 33 | TTGGTCGTGGTTGTAGTCCGTGCGAGAATACCA | 17881.2 | -2.5 | 0.8 | -3.2 | 1.4E-03 | 7.5E-03 | BT20/MB231 | NUC |
| >MI0003591\|hsa-mir-584&WithFlank&5\|-\|149062307\|149062415@22.43.22(+1A)&[MIMAT0003249&hsa-miR-584-5p&offsets\|0\|0(+1A);m-167&5\|-\|149062374\|149062394&offsets\|0\|+1(+1A)]\|TTATGGTTTGCCTGGGACTGAGA | >MI0003591\|hsa-mir-584&WithFlank&5\|-\|149062307\|149062415 | 22 | 43 | 22(+1A) | TTATGGTTTGCCTGGGACTGAGA | 168.5 | 5.0 | 1.7 | 2.9 | 3.6E-03 | 7.5E-03 | BT20/MB468 | MP |
| >hg38_MT_+_1621_3279_RNR2_16S_50ntFlanks@1427.1458.32\|GATTAAAGTCCTACGTGATCTGAGTTCAGACC | >hg38_MT_+_1621_3279_RNR2_16S_50ntFlanks | 1427 | 1458 | 32 | GATTAAAGTCCTACGTGATCTGAGTTCAGACC | 457.2 | 3.5 | 1.3 | 2.8 | 5.6E-03 | 7.5E-03 | MB231/MB468 | MP |
| >hg38_21_+_8433172_8446622_RNA45SN1_45S_50ntFlanks@4895.4917.23\|CTTAAAGGAATTGACGGAAGGGC | >hg38_21_+_8433172_8446622_RNA45SN1_45S_50ntFlanks | 4895 | 4917 | 23 | CTTAAAGGAATTGACGGAAGGGC | 133.1 | -4.9 | 1.7 | -2.9 | 3.6E-03 | 7.5E-03 | BT20/MB468 | MP |
| >trnaMT_SerGCT_MT_+_12207_12265@13.59.47\|AAGAACTGCTAACTCATGCCCCCATGTCTAACAACATGGCTTTCTCA | >trnaMT_SerGCT_MT_+_12207_12265 | 13 | 59 | 47 | AAGAACTGCTAACTCATGCCCCCATGTCTAACAACATGGCTTTCTCA | 137.8 | 6.4 | 2.3 | 2.7 | 6.5E-03 | 7.5E-03 | MB231/MB468 | MITO |
| >hg38_21_+_8433172_8446622_RNA45SN1_45S_50ntFlanks@5473.5516.44\|CGGCCCTGGCGGAGCGCTGAGAAGACGGTCGAACTTGACTATCT | >hg38_21_+_8433172_8446622_RNA45SN1_45S_50ntFlanks | 5473 | 5516 | 44 | CGGCCCTGGCGGAGCGCTGAGAAGACGGTCGAACTTGACTATCT | 190.7 | 2.4 | 0.9 | 2.7 | 6.5E-03 | 7.5E-03 | MB231/MB468 | MITO |

<table>
<tr><th></th><th></th><th></th><th></th><th></th><th></th><th></th><th></th><th></th><th></th><th></th><th></th><th></th><th></th></tr>
<tr><td>>hg38_21_+_8433172_8446622_RNA45SN1_45S_50ntFlanks@12394.12417.24|TAGCGACGTCGCTTTTTGATCCTT</td><td>>hg38_21_+_8433172_8446622_RNA45SN1_45S_50ntFlanks</td><td>12394</td><td>12417</td><td>24</td><td>TAGCGACGTCGCTTTTTGATCCTT</td><td>267.1</td><td>-3.5</td><td>1.3</td><td>-2.7</td><td>6.5E-03</td><td>7.5E-03</td><td>MB231/MB468</td><td>NUC</td></tr>
<tr><td>>hg38_21_+_8433172_8446622_RNA45SN1_45S_50ntFlanks@6660.6688.29|GCGGTGGATCACTCGGCTCGTGCGTCGAT</td><td>>hg38_21_+_8433172_8446622_RNA45SN1_45S_50ntFlanks</td><td>6660</td><td>6688</td><td>29</td><td>GCGGTGGATCACTCGGCTCGTGCGTCGAT</td><td>522.9</td><td>5.2</td><td>1.6</td><td>3.2</td><td>1.3E-03</td><td>7.5E-03</td><td>BT20/MB231</td><td>MP</td></tr>
<tr><td>>hg38_MT_+_1621_3279_RNR2_16S_50ntFlanks@285.305.21|CGAAACCAGACGAGCTACCTA</td><td>>hg38_MT_+_1621_3279_RNR2_16S_50ntFlanks</td><td>285</td><td>305</td><td>21</td><td>CGAAACCAGACGAGCTACCTA</td><td>109.6</td><td>3.4</td><td>1.2</td><td>2.8</td><td>5.6E-03</td><td>7.5E-03</td><td>MB231/MB468</td><td>MP</td></tr>
<tr><td>>hg38_21_+_8433172_8446622_RNA45SN1_45S_50ntFlanks@11662.11681.20|GATTTCTGCCCAGTGCTCTG</td><td>>hg38_21_+_8433172_8446622_RNA45SN1_45S_50ntFlanks</td><td>11662</td><td>11681</td><td>20</td><td>GATTTCTGCCCAGTGCTCTG</td><td>541.5</td><td>-3.9</td><td>1.4</td><td>-2.7</td><td>6.4E-03</td><td>7.5E-03</td><td>BT20/MB468</td><td>NUC</td></tr>
<tr><td>>MI0000750|hsa-mir-26a-2&WithFlank&12|-|57824603|57824698@20.41.22(+1U);MI0000083|hsa-mir-26a-1&WithFlank&3|+|37969398|37969486@16.37.22(+1U)&[MIMAT0000082&hsa-miR-26a-5p&offsets|0|0(+1U);m-33&12|-|57824658|57824679&offsets|0|0(+1U)];[MIMAT0000082_1&hsa-miR-26a-5p&offsets|0|0(+1U);m-34&3|+|37969413|37969434&offsets|0|0(+1U)]|TTCAAGTAATCCAGGATAGGCTT</td><td>>MI0000750|hsa-mir-26a-2&WithFlank&12|-|57824603|57824698</td><td>20</td><td>41</td><td>22(+1U)</td><td>TTCAAGTAATCCAGGATAGGCTT</td><td>6298.6</td><td>-2.6</td><td>0.8</td><td>-3.1</td><td>1.7E-03</td><td>7.5E-03</td><td>BT20/MB231</td><td>MITO</td></tr>
<tr><td>>MI0005559|hsa-mir-744&WithFlank&17|+|12081893|12082002@17.36.20(+1U)&[MIMAT0004945&hsa-miR-744-5p&offsets|0|-2(+1U);m-172&17|+|12081909|12081929&offsets|0|-1(+1U)]|TGCGGGGCTAGGGCTAACAGT</td><td>>MI0005559|hsa-mir-744&WithFlank&17|+|12081893|12082002</td><td>17</td><td>36</td><td>20(+1U)</td><td>TGCGGGGCTAGGGCTAACAGT</td><td>176.2</td><td>4.7</td><td>1.7</td><td>2.8</td><td>5.6E-03</td><td>7.5E-03</td><td>MB231/MB468</td><td>MP</td></tr>
<tr><td>>hg38_21_+_8433172_8446622_RNA45SN1_45S_50ntFlanks@8105.8142.38|CGCGGCGGGGCGCGGGACATGTGGCGTACGGAAGACCC</td><td>>hg38_21_+_8433172_8446622_RNA45SN1_45S_50ntFlanks</td><td>8105</td><td>8142</td><td>38</td><td>CGCGGCGGGGCGCGGGACATGTGGCGTACGGAAGACCC</td><td>1743.0</td><td>-3.3</td><td>1.1</td><td>-2.9</td><td>3.6E-03</td><td>7.5E-03</td><td>BT20/MB468</td><td>MP</td></tr>
<tr><td>>hg38_21_+_8433172_8446622_RNA45SN1_45S_50ntFlanks@6721.6744.24|AATTGCAGGACACATTGATCATCG</td><td>>hg38_21_+_8433172_8446622_RNA45SN1_45S_50ntFlanks</td><td>6721</td><td>6744</td><td>24</td><td>AATTGCAGGACACATTGATCATCG</td><td>4485.5</td><td>-5.6</td><td>1.9</td><td>-2.9</td><td>3.6E-03</td><td>7.5E-03</td><td>BT20/MB468</td><td>MP</td></tr>
<tr><td>>MI0000749|hsa-mir-30e&WithFlank&1|+|40754349|40754452@23.46.24&[MIMAT0000692&hsa-miR-30e-5p&offsets|0|+2;m-24&1|+|40754371|40754394&offsets|0|0]|TGTAAACATCCTTGACTGGAAGCT</td><td>>MI0000749|hsa-mir-30e&WithFlank&1|+|40754349|40754452</td><td>23</td><td>46</td><td>24</td><td>TGTAAACATCCTTGACTGGAAGCT</td><td>782.6</td><td>2.3</td><td>0.9</td><td>2.7</td><td>6.2E-03</td><td>7.5E-03</td><td>MB231/MB468</td><td>CYTO</td></tr>
<tr><td>>trnaMT_PheGAA_MT_+_577_647@35.53.19|GAAAATGTTTAGACGGGCT</td><td>>trnaMT_PheGAA_MT_+_577_647</td><td>35</td><td>53</td><td>19</td><td>GAAAATGTTTAGACGGGCT</td><td>219.2</td><td>2.8</td><td>1.0</td><td>2.7</td><td>6.5E-03</td><td>7.5E-03</td><td>MB231/MB468</td><td>MITO</td></tr>
<tr><td>>hg38_21_+_8433172_8446622_RNA45SN1_45S_50ntFlanks@7974.7995.22|ACGCGACCTCAGATCAGACGTG</td><td>>hg38_21_+_8433172_8446622_RNA45SN1_45S_50ntFlanks</td><td>7974</td><td>7995</td><td>22</td><td>ACGCGACCTCAGATCAGACGTG</td><td>2055.1</td><td>-1.9</td><td>0.7</td><td>-2.8</td><td>5.6E-03</td><td>7.5E-03</td><td>MB231/MB468</td><td>MP</td></tr>
<tr><td>>am_MI0000786|hsa-mir-378a&WithFlank&5|+|149732819|149732896@50.71.22&[MIMAT0000732&hsa-miR-378a-3p&offsets|+1|+1;m-44&5|+|149732867|149732888&offsets|+1|+1]|CTGGACTTGGAGTCAGAAGGCC</td><td>>am_MI0000786|hsa-mir-378a&WithFlank&5|+|149732819|149732896</td><td>50</td><td>71</td><td>22</td><td>CTGGACTTGGAGTCAGAAGGCC</td><td>720.8</td><td>3.2</td><td>1.1</td><td>2.9</td><td>3.6E-03</td><td>7.5E-03</td><td>BT20/MB468</td><td>MP</td></tr>
<tr><td>>trnaMT_LysTTT_MT_+_8295_8364@53.72.20,trnalookalike7_LysTTT_1_+_568844_568913@53.72.20|AACACCTCTTTACAGTGACC</td><td>>trnaMT_LysTTT_MT_+_8295_8364</td><td>53</td><td>72</td><td>20</td><td>AACACCTCTTTACAGTGACC</td><td>484.1</td><td>4.6</td><td>1.7</td><td>2.8</td><td>5.6E-03</td><td>7.5E-03</td><td>MB231/MB468</td><td>MP</td></tr>
<tr><td>>hg38_21_+_8433172_8446622_RNA45SN1_45S_50ntFlanks@6652.6701.50|GACTCTTAGCGGTGGATCACTCGGCTCGTGCGTCGATGAAGAACGCAGCT</td><td>>hg38_21_+_8433172_8446622_RNA45SN1_45S_50ntFlanks</td><td>6652</td><td>6701</td><td>50</td><td>GACTCTTAGCGGTGGATCACTCGGCTCGTGCGTCGATGAAGAACGCAGCT</td><td>1293.7</td><td>5.8</td><td>2.0</td><td>2.9</td><td>3.6E-03</td><td>7.5E-03</td><td>BT20/MB468</td><td>MP</td></tr>
<tr><td>>MI0000782|hsa-mir-374a&WithFlank&X|-|74287280|74287363@18.37.20&[MIMAT0000727&hsa-miR-374a-5p&offsets|0|-2;m-184&X|-|74287325|74287346&offsets|0|-2]|TTATAATACAACCTGATAAG</td><td>>MI0000782|hsa-mir-374a&WithFlank&X|-|74287280|74287363</td><td>18</td><td>37</td><td>20</td><td>TTATAATACAACCTGATAAG</td><td>56.5</td><td>-9.2</td><td>2.9</td><td>-3.2</td><td>1.3E-03</td><td>7.5E-03</td><td>BT20/MB231</td><td>MP</td></tr>
<tr><td>>hg38_21_+_8433172_8446622_RNA45SN1_45S_50ntFlanks@7968.7988.21|TCCGAGACGCGACCTCAGATC</td><td>>hg38_21_+_8433172_8446622_RNA45SN1_45S_50ntFlanks</td><td>7968</td><td>7988</td><td>21</td><td>TCCGAGACGCGACCTCAGATC</td><td>505.0</td><td>2.3</td><td>0.9</td><td>2.7</td><td>6.5E-03</td><td>7.5E-03</td><td>BT20/MB468</td><td>NUC</td></tr>
<tr><td>>MI0000446|hsa-mir-125b-1&WithFlank&11|-|122099751|122099850@21.41.21(+1G);MI0000470|hsa-mir-125b-2&WithFlank&21|+|16590231|16590331@23.43.21(+1G)&[MIMAT0000423&hsa-miR-125b-5p&offsets|0|-1(+1G);m-67&11|-|122099809|122099830&offsets|0|-1(+1G)];[MIMAT0000423_1&hsa-miR-125b-5p&offsets|0|-1(+1G);m-68&21|+|16590253|16590274&offsets|0|-1(+1G)]|TCCCTGAGACCCTAACTTGTGG</td><td>>MI0000446|hsa-mir-125b-1&WithFlank&11|-|122099751|122099850</td><td>21</td><td>41</td><td>21(+1G)</td><td>TCCCTGAGACCCTAACTTGTGG</td><td>263.6</td><td>4.6</td><td>1.6</td><td>2.8</td><td>5.6E-03</td><td>7.5E-03</td><td>MB231/MB468</td><td>MP</td></tr>
<tr><td>>MI0000809|hsa-mir-151a&WithFlank&8|-|140732558|140732659@51.74.24&[MIMAT0000757&hsa-miR-151a-3p&offsets|-2|+1;m-30&8|-|140732587|140732607&offsets|-2|+1]|TACTAGACTGAAGCTCCTTGAGGA</td><td>>MI0000809|hsa-mir-151a&WithFlank&8|-|140732558|140732659</td><td>51</td><td>74</td><td>24</td><td>TACTAGACTGAAGCTCCTTGAGGA</td><td>42.3</td><td>5.2</td><td>1.9</td><td>2.8</td><td>5.6E-03</td><td>7.5E-03</td><td>MB231/MB468</td><td>MP</td></tr>
<tr><td>>hg38_21_+_8433172_8446622_RNA45SN1_45S_50ntFlanks@4724.4761.38|ATTAATCAAGAACGAAAGTCGGAGGTTCGAAGACGATC</td><td>>hg38_21_+_8433172_8446622_RNA45SN1_45S_50ntFlanks</td><td>4724</td><td>4761</td><td>38</td><td>ATTAATCAAGAACGAAAGTCGGAGGTTCGAAGACGATC</td><td>149.0</td><td>-4.9</td><td>1.8</td><td>-2.8</td><td>5.6E-03</td><td>7.5E-03</td><td>MB231/MB468</td><td>MP</td></tr>
<tr><td>>hg38_21_+_8433172_8446622_RNA45SN1_45S_50ntFlanks@5431.5466.36|GTTTAGTGAGGCCCTCGGATCGGCCCCGCCGGGGTC</td><td>>hg38_21_+_8433172_8446622_RNA45SN1_45S_50ntFlanks</td><td>5431</td><td>5466</td><td>36</td><td>GTTTAGTGAGGCCCTCGGATCGGCCCCGCCGGGGTC</td><td>225.0</td><td>-3.2</td><td>1.2</td><td>-2.7</td><td>6.1E-03</td><td>7.5E-03</td><td>BT20/MB468</td><td>CYTO</td></tr>
<tr><td>>hg38_21_+_8433172_8446622_RNA45SN1_45S_50ntFlanks@11212.11251.40|CGACTCTGGACGCGAGCCGGGCCCTTCCCGTGGATCGCCC</td><td>>hg38_21_+_8433172_8446622_RNA45SN1_45S_50ntFlanks</td><td>11212</td><td>11251</td><td>40</td><td>CGACTCTGGACGCGAGCCGGGCCCTTCCCGTGGATCGCCC</td><td>355.4</td><td>-2.2</td><td>0.8</td><td>-2.7</td><td>6.2E-03</td><td>7.5E-03</td><td>MB231/MB468</td><td>CYTO</td></tr>
<tr><td>>trna116_GluCTC_1_-_145399233_145399304@13.31.19,trna25_GluCTC_2_-_71273488_71273560@13.31.19,trna59_GluCTC_1_+_249168447_249168518@13.31.19,trna71_GluCTC_1_-_161439189_161439260@13.31.19,trna74_GluCTC_1_-_161431809_161431880@13.31.19,trna77_GluCTC_1_-_161424398_161424469@13.31.19,trna77_GluCTC_6_+_28949976_28950047@13.31.19,trna80_GluCTC_1_-_161417018_161417089@13.31.19,trna87_GluCTC_6_-_126101393_126101464@13.31.19|TAGTGGTTAGGATTCGGCG</td><td>>trna116_GluCTC_1_-_145399233_145399304</td><td>13</td><td>31</td><td>19</td><td>TAGTGGTTAGGATTCGGCG</td><td>899.5</td><td>2.1</td><td>0.7</td><td>2.7</td><td>6.1E-03</td><td>7.5E-03</td><td>BT20/MB468</td><td>CYTO</td></tr>
<tr><td>>hg38_21_+_8433172_8446622_RNA45SN1_45S_50ntFlanks@9564.9601.38|CTGACGTGCAAATCGGTCGTCCGACCTGGGTATAGGGG</td><td>>hg38_21_+_8433172_8446622_RNA45SN1_45S_50ntFlanks</td><td>9564</td><td>9601</td><td>38</td><td>CTGACGTGCAAATCGGTCGTCCGACCTGGGTATAGGGG</td><td>437.1</td><td>-3.5</td><td>1.3</td><td>-2.7</td><td>6.5E-03</td><td>7.5E-03</td><td>MB231/MB468</td><td>MITO</td></tr>
<tr><td>>hg38_21_+_8433172_8446622_RNA45SN1_45S_50ntFlanks@6659.6688.30|AGCGGTGGATCACTCGGCTCGTGCGTCGAT</td><td>>hg38_21_+_8433172_8446622_RNA45SN1_45S_50ntFlanks</td><td>6659</td><td>6688</td><td>30</td><td>AGCGGTGGATCACTCGGCTCGTGCGTCGAT</td><td>228.7</td><td>5.3</td><td>1.6</td><td>3.2</td><td>1.3E-03</td><td>7.5E-03</td><td>BT20/MB231</td><td>MP</td></tr>
<tr><td>>hg38_MT_+_1621_3279_RNR2_16S_50ntFlanks@644.664.21|AATTGGACCAATCTATCACCC</td><td>>hg38_MT_+_1621_3279_RNR2_16S_50ntFlanks</td><td>644</td><td>664</td><td>21</td><td>AATTGGACCAATCTATCACCC</td><td>106.9</td><td>5.1</td><td>1.7</td><td>2.9</td><td>3.6E-03</td><td>7.5E-03</td><td>BT20/MB468</td><td>MP</td></tr>
<tr><td>>hg38_21_+_8433172_8446622_RNA45SN1_45S_50ntFlanks@5066.5098.33|TCTGGTTAATTCCGATAACGAACGAGACTCTGG</td><td>>hg38_21_+_8433172_8446622_RNA45SN1_45S_50ntFlanks</td><td>5066</td><td>5098</td><td>33</td><td>TCTGGTTAATTCCGATAACGAACGAGACTCTGG</td><td>122.7</td><td>-5.5</td><td>1.9</td><td>-2.9</td><td>3.6E-03</td><td>7.5E-03</td><td>BT20/MB468</td><td>MP</td></tr>
<tr><td>>hg38_21_+_8433172_8446622_RNA45SN1_45S_50ntFlanks@9309.9338.30|GGAGTCTAACACGTGCGCGAGTCGGGGGCT</td><td>>hg38_21_+_8433172_8446622_RNA45SN1_45S_50ntFlanks</td><td>9309</td><td>9338</td><td>30</td><td>GGAGTCTAACACGTGCGCGAGTCGGGGGCT</td><td>125.0</td><td>-4.8</td><td>1.6</td><td>-2.9</td><td>3.6E-03</td><td>7.5E-03</td><td>BT20/MB468</td><td>MP</td></tr>
<tr><td>>hg38_21_+_8433172_8446622_RNA45SN1_45S_50ntFlanks@11214.11234.21|ACTCTGGACGCGAGCCGGGCC</td><td>>hg38_21_+_8433172_8446622_RNA45SN1_45S_50ntFlanks</td><td>11214</td><td>11234</td><td>21</td><td>ACTCTGGACGCGAGCCGGGCC</td><td>64.2</td><td>-5.0</td><td>1.7</td><td>-2.9</td><td>3.6E-03</td><td>7.5E-03</td><td>BT20/MB468</td><td>MP</td></tr>
<tr><td>>trnaMT_PheGAA_MT_+_577_647@45.74.30|AGACGGGCTCACATCACCCCATAAACACCA</td><td>>trnaMT_PheGAA_MT_+_577_647</td><td>45</td><td>74</td><td>30</td><td>AGACGGGCTCACATCACCCCATAAACACCA</td><td>559.0</td><td>-1.8</td><td>0.7</td><td>-2.8</td><td>5.6E-03</td><td>7.5E-03</td><td>MB231/MB468</td><td>MP</td></tr>
<tr><td>>hg38_21_+_8433172_8446622_RNA45SN1_45S_50ntFlanks@10346.10379.34|TCAAACGAGAACTTTGAAGGCCGAAGTGGAGAAG</td><td>>hg38_21_+_8433172_8446622_RNA45SN1_45S_50ntFlanks</td><td>10346</td><td>10379</td><td>34</td><td>TCAAACGAGAACTTTGAAGGCCGAAGTGGAGAAG</td><td>74.4</td><td>-3.6</td><td>1.3</td><td>-2.8</td><td>5.6E-03</td><td>7.5E-03</td><td>MB231/MB468</td><td>MP</td></tr>
<tr><td>>hg38_21_+_8433172_8446622_RNA45SN1_45S_50ntFlanks@8171.8212.42|AAGTCCTTCTGATCGAGGCCCAGCCCGTGGACGGTGTGAGGC</td><td>>hg38_21_+_8433172_8446622_RNA45SN1_45S_50ntFlanks</td><td>8171</td><td>8212</td><td>42</td><td>AAGTCCTTCTGATCGAGGCCCAGCCCGTGGACGGTGTGAGGC</td><td>641.6</td><td>-1.9</td><td>0.7</td><td>-2.8</td><td>5.3E-03</td><td>7.5E-03</td><td>BT20/MB468</td><td>MITO</td></tr>
<tr><td>>hg38_21_+_8433172_8446622_RNA45SN1_45S_50ntFlanks@12241.12271.31|CAGAAGGGCAAAAGCTCGCTTGATCTTGATT</td><td>>hg38_21_+_8433172_8446622_RNA45SN1_45S_50ntFlanks</td><td>12241</td><td>12271</td><td>31</td><td>CAGAAGGGCAAAAGCTCGCTTGATCTTGATT</td><td>171.7</td><td>-3.7</td><td>1.3</td><td>-2.8</td><td>5.6E-03</td><td>7.5E-03</td><td>MB231/MB468</td><td>MP</td></tr>
<tr><td>>hg38_MT_+_1621_3279_RNR2_16S_50ntFlanks@1369.1404.36|GATGTTGGATCAGGACATCCCGATGGTGCAGCCGCT</td><td>>hg38_MT_+_1621_3279_RNR2_16S_50ntFlanks</td><td>1369</td><td>1404</td><td>36</td><td>GATGTTGGATCAGGACATCCCGATGGTGCAGCCGCT</td><td>109.7</td><td>5.2</td><td>1.8</td><td>2.9</td><td>3.6E-03</td><td>7.5E-03</td><td>BT20/MB468</td><td>MP</td></tr>
<tr><td>>hg38_21_+_8433172_8446622_RNA45SN1_45S_50ntFlanks@6719.6756.38|TGAATTGCAGGACACATTGATCATCGACACTTCGAACG</td><td>>hg38_21_+_8433172_8446622_RNA45SN1_45S_50ntFlanks</td><td>6719</td><td>6756</td><td>38</td><td>TGAATTGCAGGACACATTGATCATCGACACTTCGAACG</td><td>247.0</td><td>-2.3</td><td>0.8</td><td>-2.8</td><td>5.3E-03</td><td>7.5E-03</td><td>BT20/MB468</td><td>MITO</td></tr>
<tr><td>>hg38_21_+_8433172_8446622_RNA45SN1_45S_50ntFlanks@9297.9322.26|AACACGGACCAAGGAGTCTAACACGT</td><td>>hg38_21_+_8433172_8446622_RNA45SN1_45S_50ntFlanks</td><td>9297</td><td>9322</td><td>26</td><td>AACACGGACCAAGGAGTCTAACACGT</td><td>27.4</td><td>8.5</td><td>2.9</td><td>2.9</td><td>3.6E-03</td><td>7.5E-03</td><td>BT20/MB468</td><td>MP</td></tr>
<tr><td>>hg38_21_+_8433172_8446622_RNA45SN1_45S_50ntFlanks@10410.10427.18|TCAGTCGGTCCTGAGAGA</td><td>>hg38_21_+_8433172_8446622_RNA45SN1_45S_50ntFlanks</td><td>10410</td><td>10427</td><td>18</td><td>TCAGTCGGTCCTGAGAGA</td><td>160.4</td><td>-3.7</td><td>1.3</td><td>-2.8</td><td>5.7E-03</td><td>7.5E-03</td><td>MB231/MB468</td><td>MP</td></tr>
<tr><td>>hg38_21_+_8433172_8446622_RNA45SN1_45S_50ntFlanks@11540.11584.45|CGCGGCGCCTCGCCTCGGCCGGCGCCTAGCAGCCGACTTAGAACT</td><td>>hg38_21_+_8433172_8446622_RNA45SN1_45S_50ntFlanks</td><td>11540</td><td>11584</td><td>45</td><td>CGCGGCGCCTCGCCTCGGCCGGCGCCTAGCAGCCGACTTAGAACT</td><td>1860.5</td><td>-3.0</td><td>0.9</td><td>-3.2</td><td>1.4E-03</td><td>7.5E-03</td><td>BT20/MB231</td><td>NUC</td></tr>
<tr><td>>MI0000446|hsa-mir-125b-1&WithFlank&11|-|122099751|122099850@21.40.20(+1A);MI0000470|hsa-mir-125b-2&WithFlank&21|+|16590231|16590331@23.42.20(+1A)&[MIMAT0000423&hsa-miR-125b-5p&offsets|0|-2(+1A);m-67&11|-|122099809|122099830&offsets|0|-2(+1A)];[MIMAT0000423_1&hsa-miR-125b-5p&offsets|0|-2(+1A);m-68&21|+|16590253|16590274&offsets|0|-2(+1A)]|TCCCTGAGACCCTAACTTGTA</td><td>>MI0000446|hsa-mir-125b-1&WithFlank&11|-|122099751|122099850</td><td>21</td><td>40</td><td>20(+1A)</td><td>TCCCTGAGACCCTAACTTGTA</td><td>1394.6</td><td>-7.5</td><td>2.4</td><td>-3.1</td><td>1.7E-03</td><td>7.5E-03</td><td>BT20/MB231</td><td>MITO</td></tr>
<tr><td>>hg38_21_+_8433172_8446622_RNA45SN1_45S_50ntFlanks@10345.10373.29|TTCAAACGAGAACTTTGAAGGCCGAAGTG</td><td>>hg38_21_+_8433172_8446622_RNA45SN1_45S_50ntFlanks</td><td>10345</td><td>10373</td><td>29</td><td>TTCAAACGAGAACTTTGAAGGCCGAAGTG</td><td>155.0</td><td>2.4</td><td>0.8</td><td>2.9</td><td>3.6E-03</td><td>7.5E-03</td><td>BT20/MB468</td><td>MP</td></tr>
<tr><td>>MI0000063|hsa-let-7b&WithFlank&22|+|46113680|46113774@66.86.21&[MIMAT0004482&hsa-let-7b-3p&offsets|0|-1;m-171&22|+|46113745|46113766&offsets|0|-1]|CTATACAACCTACTGCCTTCC</td><td>>MI0000063|hsa-let-7b&WithFlank&22|+|46113680|46113774</td><td>66</td><td>86</td><td>21</td><td>CTATACAACCTACTGCCTTCC</td><td>39.6</td><td>5.4</td><td>2.0</td><td>2.8</td><td>5.7E-03</td><td>7.5E-03</td><td>MB231/MB468</td><td>MP</td></tr>
<tr><td>>MI0000749|hsa-mir-30e&WithFlank&1|+|40754349|40754452@23.46.24(+1C)&[MIMAT0000692&hsa-miR-30e-5p&offsets|0|+2(+1C);m-24&1|+|40754371|40754394&offsets|0|0(+1C)]|TGTAAACATCCTTGACTGGAAGCTC</td><td>>MI0000749|hsa-mir-30e&WithFlank&1|+|40754349|40754452</td><td>23</td><td>46</td><td>24(+1C)</td><td>TGTAAACATCCTTGACTGGAAGCTC</td><td>259.2</td><td>2.7</td><td>1.0</td><td>2.8</td><td>5.7E-03</td><td>7.5E-03</td><td>MB231/MB468</td><td>MP</td></tr>
<tr><td>>hg38_21_+_8433172_8446622_RNA45SN1_45S_50ntFlanks@9313.9336.24|TCTAACACGTGCGCGAGTCGGGGG</td><td>>hg38_21_+_8433172_8446622_RNA45SN1_45S_50ntFlanks</td><td>9313</td><td>9336</td><td>24</td><td>TCTAACACGTGCGCGAGTCGGGGG</td><td>374.9</td><td>-5.1</td><td>1.7</td><td>-2.9</td><td>3.6E-03</td><td>7.5E-03</td><td>BT20/MB468</td><td>MP</td></tr>
<tr><td>>hg38_21_+_8433172_8446622_RNA45SN1_45S_50ntFlanks@11580.11607.28|GAACTGGTGCGGACCAGGGGAATCCGAC</td><td>>hg38_21_+_8433172_8446622_RNA45SN1_45S_50ntFlanks</td><td>11580</td><td>11607</td><td>28</td><td>GAACTGGTGCGGACCAGGGGAATCCGAC</td><td>251.1</td><td>-1.7</td><td>0.6</td><td>-2.7</td><td>6.6E-03</td><td>7.5E-03</td><td>MB231/MB468</td><td>NUC</td></tr>
<tr><td>>hg38_21_+_8433172_8446622_RNA45SN1_45S_50ntFlanks@4732.4767.36|AGAACGAAAGTCGGAGGTTCGAAGACGATCAGATAC</td><td>>hg38_21_+_8433172_8446622_RNA45SN1_45S_50ntFlanks</td><td>4732</td><td>4767</td><td>36</td><td>AGAACGAAAGTCGGAGGTTCGAAGACGATCAGATAC</td><td>81.9</td><td>3.7</td><td>1.4</td><td>2.7</td><td>6.6E-03</td><td>7.5E-03</td><td>MB231/MB468</td><td>MITO</td></tr>
<tr><td>>hg38_21_+_8433172_8446622_RNA45SN1_45S_50ntFlanks@3854.3873.20|ACTGTGGTAATTCTAGAGCT</td><td>>hg38_21_+_8433172_8446622_RNA45SN1_45S_50ntFlanks</td><td>3854</td><td>3873</td><td>20</td><td>ACTGTGGTAATTCTAGAGCT</td><td>271.4</td><td>-3.3</td><td>1.2</td><td>-2.7</td><td>6.6E-03</td><td>7.5E-03</td><td>MB231/MB468</td><td>NUC</td></tr>
</table>

| | | | | | | | | | | | | | |
|---|---|---|---|---|---|---|---|---|---|---|---|---|---|
| >hg38_21_+_8433172_8446622_RNA45SN1_45S_50ntFlanks@7974.8012.39\|ACGCGACCTCAGATCAGACGTGGCGACCCGCTGAATTTA | >hg38_21_+_8433172_8446622_RNA45SN1_45S_50ntFlanks | 7974 | 8012 | 39 | ACGCGACCTCAGATCAGACGTGGCGACCCGCTGAATTTA | 3883.2 | -1.9 | 0.7 | -2.7 | 6.5E-03 | 7.5E-03 | BT20/MB468 | NUC |
| >MI0000299\|hsa-mir-222&WithFlank&X\|-\|45747009\|45747130@75.98.24(+1C)&[MIMAT0000279&hsa-miR-222-3p&offsets\|0\|+3(+1C);m-140&X\|-\|45747033\|45747056&offsets\|0\|0(+1C)]\|AGCTACATCTGGCTACTGGGTCTCC | >MI0000299\|hsa-mir-222&WithFlank&X\|-\|45747009\|45747130 | 75 | 98 | 24(+1C) | AGCTACATCTGGCTACTGGGTCTCC | 4114.1 | -3.0 | 0.9 | -3.2 | 1.3E-03 | 7.5E-03 | BT20/MB231 | MP |
| >hg38_21_+_8433172_8446622_RNA45SN1_45S_50ntFlanks@9035.9055.21\|CGTGGGGGGGCCGGGCCACCC | >hg38_21_+_8433172_8446622_RNA45SN1_45S_50ntFlanks | 9035 | 9055 | 21 | CGTGGGGGGGCCGGGCCACCC | 196.7 | -4.9 | 1.7 | -2.9 | 3.6E-03 | 7.6E-03 | BT20/MB468 | MP |
| >trnaMT_GluTTC_MT_-_14674_14742@24.58.35,trnalookalike8_GluTTC_5_-_93905172_93905240@24.58.35\|ATGGTTTTTCATATCATTGGTCGTGGTTGTAGTCC | >trnaMT_GluTTC_MT_-_14674_14742 | 24 | 58 | 35 | ATGGTTTTTCATATCATTGGTCGTGGTTGTAGTCC | 67.8 | 5.1 | 1.9 | 2.8 | 5.7E-03 | 7.6E-03 | MB231/MB468 | MP |
| >am_MI0016746\|hsa-mir-548o-2&WithFlank&20\|+\|38516557\|38516638@51.71.21(+1A);MI0006402\|hsa-mir-548o&WithFlank&7\|-\|102405736\|102405861@93.113.21(+1A)&[MIMAT0005919&hsa-miR-548o-3p&offsets\|0\|-1(+1A);m-431&20\|+\|38516607\|38516628&offsets\|0\|-1(+1A)];[MIMAT0005919_1&hsa-miR-548o-3p&offsets\|0\|-1(+1A);m-430&7\|-\|102405748\|102405769&offsets\|0\|-1(+1A)]\|CCAAAACTGCAGTTACTTTTGA | >am_MI0016746\|hsa-mir-548o-2&WithFlank&20\|+\|38516557\|38516638 | 51 | 71 | 21(+1A) | CCAAAACTGCAGTTACTTTTGA | 41.3 | 5.2 | 1.9 | 2.8 | 5.7E-03 | 7.6E-03 | MB231/MB468 | MP |
| >hg38_21_+_8433172_8446622_RNA45SN1_45S_50ntFlanks@12419.12439.21\|GATGTCGGCTCTTCCTATCAT | >hg38_21_+_8433172_8446622_RNA45SN1_45S_50ntFlanks | 12419 | 12439 | 21 | GATGTCGGCTCTTCCTATCAT | 405.3 | -1.5 | 0.6 | -2.7 | 6.5E-03 | 7.6E-03 | BT20/MB468 | NUC |
| >MI0000087\|hsa-mir-29a&WithFlank&7\|-\|130876741\|130876816@48.67.20&[MIMAT0000086&hsa-miR-29a-3p&offsets\|0\|-2;m-26&7\|-\|130876749\|130876769&offsets\|0\|-1]\|TAGCACCATCTGAAATCGGT | >MI0000087\|hsa-mir-29a&WithFlank&7\|-\|130876741\|130876816 | 48 | 67 | 20 | TAGCACCATCTGAAATCGGT | 517.7 | 5.7 | 1.9 | 2.9 | 3.6E-03 | 7.6E-03 | BT20/MB468 | MP |
| >hg38_MT_+_1621_3279_RNR2_16S_50ntFlanks@506.530.25\|TAGGAAAAAACCTTGTAGAGAGAGT | >hg38_MT_+_1621_3279_RNR2_16S_50ntFlanks | 506 | 530 | 25 | TAGGAAAAAACCTTGTAGAGAGAGT | 51.7 | 5.9 | 2.0 | 2.9 | 3.6E-03 | 7.6E-03 | BT20/MB468 | MP |
| >hg38_21_+_8433172_8446622_RNA45SN1_45S_50ntFlanks@4892.4910.19\|AAACTTAAAGGAATTGACG | >hg38_21_+_8433172_8446622_RNA45SN1_45S_50ntFlanks | 4892 | 4910 | 19 | AAACTTAAAGGAATTGACG | 58.5 | -4.8 | 1.7 | -2.9 | 3.6E-03 | 7.6E-03 | BT20/MB468 | MP |
| >hg38_21_+_8433172_8446622_RNA45SN1_45S_50ntFlanks@8380.8403.24\|CAAGAGGGCGTGAAACCGTTAAGA | >hg38_21_+_8433172_8446622_RNA45SN1_45S_50ntFlanks | 8380 | 8403 | 24 | CAAGAGGGCGTGAAACCGTTAAGA | 539.9 | -4.4 | 1.6 | -2.7 | 6.2E-03 | 7.6E-03 | BT20/MB468 | CYTO |
| >hg38_MT_+_598_1651_RNR1_12S_50ntFlanks@51.85.35\|AATAGGTTTGGTCCTAGCCTTTCTATTAGCTCTTA | >hg38_MT_+_598_1651_RNR1_12S_50ntFlanks | 51 | 85 | 35 | AATAGGTTTGGTCCTAGCCTTTCTATTAGCTCTTA | 914.0 | 4.1 | 1.4 | 2.9 | 3.6E-03 | 7.6E-03 | BT20/MB468 | MP |
| >hg38_21_+_8433172_8446622_RNA45SN1_45S_50ntFlanks@6652.6690.39\|GACTCTTAGCGGTGGATCACTCGGCTCGTGCGTCGATGA | >hg38_21_+_8433172_8446622_RNA45SN1_45S_50ntFlanks | 6652 | 6690 | 39 | GACTCTTAGCGGTGGATCACTCGGCTCGTGCGTCGATGA | 1573.0 | 3.1 | 1.0 | 3.2 | 1.3E-03 | 7.6E-03 | BT20/MB231 | MP |
| >trna5_AspGTC_12_+_98897281_98897352@37.75.39\|ACGCGGGAGACCGGGGTTCAATTCCCCGACGGGGAGCCA | >trna5_AspGTC_12_+_98897281_98897352 | 37 | 75 | 39 | ACGCGGGAGACCGGGGTTCAATTCCCCGACGGGGAGCCA | 2631.2 | -5.5 | 1.8 | -3.1 | 1.6E-03 | 7.6E-03 | BT20/MB231 | CYTO |
| >hg38_21_+_8433172_8446622_RNA45SN1_45S_50ntFlanks@11671.11693.23\|CCAGTGCTCTGAATGTCAAAGTG | >hg38_21_+_8433172_8446622_RNA45SN1_45S_50ntFlanks | 11671 | 11693 | 23 | CCAGTGCTCTGAATGTCAAAGTG | 496.8 | -2.9 | 1.0 | -2.9 | 3.6E-03 | 7.6E-03 | BT20/MB468 | MP |
| >MI0003205\|hsa-mir-532&WithFlank&X\|+\|50003142\|50003244@26.46.21&[MIMAT0002888&hsa-miR-532-5p&offsets\|0\|-1;m-65&X\|+\|50003167\|50003188&offsets\|0\|-1]\|CATGCCTTGAGTGTAGGACCG | >MI0003205\|hsa-mir-532&WithFlank&X\|+\|50003142\|50003244 | 26 | 46 | 21 | CATGCCTTGAGTGTAGGACCG | 143.8 | 4.3 | 1.5 | 2.9 | 3.6E-03 | 7.6E-03 | BT20/MB468 | MP |
| >hg38_1_-_228634819_228635039_RNA5S12_5S_50ntFlanks@141.171.31\|CCGCCTGGGAATACCGGGTGCTGTAGGCTTT | >hg38_1_-_228634819_228635039_RNA5S12_5S_50ntFlanks | 141 | 171 | 31 | CCGCCTGGGAATACCGGGTGCTGTAGGCTTT | 1393.9 | -2.1 | 0.7 | -3.1 | 1.7E-03 | 7.6E-03 | BT20/MB231 | MITO |
| >hg38_21_+_8433172_8446622_RNA45SN1_45S_50ntFlanks@3988.4023.36\|CTTTGGTGACTCTAGATAACCTCGGGCCGATCGCAC | >hg38_21_+_8433172_8446622_RNA45SN1_45S_50ntFlanks | 3988 | 4023 | 36 | CTTTGGTGACTCTAGATAACCTCGGGCCGATCGCAC | 761.9 | 2.4 | 0.8 | 3.1 | 1.7E-03 | 7.6E-03 | BT20/MB231 | MITO |
| >MI0000299\|hsa-mir-222&WithFlank&X\|-\|45747009\|45747130@75.98.24(+1G)&[MIMAT0000279&hsa-miR-222-3p&offsets\|0\|+3(+1G);m-140&X\|-\|45747033\|45747056&offsets\|0\|0(+1G)]\|AGCTACATCTGGCTACTGGGTCTCG | >MI0000299\|hsa-mir-222&WithFlank&X\|-\|45747009\|45747130 | 75 | 98 | 24(+1G) | AGCTACATCTGGCTACTGGGTCTCG | 921.2 | -2.4 | 0.8 | -3.1 | 1.7E-03 | 7.6E-03 | BT20/MB231 | MITO |
| >MI0000088\|hsa-mir-30a&WithFlank&6\|-\|71403545\|71403627@53.73.21(+1A)&[MIMAT0000088&hsa-miR-30a-3p&offsets\|0\|-1(+1A);m-38&6\|-\|71403554\|71403575&offsets\|0\|-1(+1A)]\|CTTTCAGTCGGATGTTTGCAGA | >MI0000088\|hsa-mir-30a&WithFlank&6\|-\|71403545\|71403627 | 53 | 73 | 21(+1A) | CTTTCAGTCGGATGTTTGCAGA | 550.5 | 3.8 | 1.4 | 2.8 | 5.7E-03 | 7.6E-03 | MB231/MB468 | MP |
| >am_trnaMT_LysTTT_MT_+_8295_8364@35.52.18,trnalookalike7_LysTTT_1_+_568844_568913@35.52.18\|TTAAAGATTAAGAGAACC | >am_trnaMT_LysTTT_MT_+_8295_8364 | 35 | 52 | 18 | TTAAAGATTAAGAGAACC | 249.4 | 2.6 | 0.8 | 3.1 | 1.6E-03 | 7.6E-03 | BT20/MB231 | CYTO |
| >trnaMT_GluTTC_MT_-_14674_14742@37.71.35,trnalookalike8_GluTTC_5_-_93905172_93905240@37.71.35\|TCATTGGTCGTGGTTGTAGTCCGTGCGAGAATACC | >trnaMT_GluTTC_MT_-_14674_14742 | 37 | 71 | 35 | TCATTGGTCGTGGTTGTAGTCCGTGCGAGAATACC | 1077.0 | -3.1 | 1.0 | -3.1 | 1.6E-03 | 7.6E-03 | BT20/MB231 | CYTO |
| >hg38_MT_+_1621_3279_RNR2_16S_50ntFlanks@51.69.19\|GCTAAACCTAGCCCCAAAC | >hg38_MT_+_1621_3279_RNR2_16S_50ntFlanks | 51 | 69 | 19 | GCTAAACCTAGCCCCAAAC | 494.5 | 2.6 | 1.0 | 2.8 | 5.7E-03 | 7.6E-03 | MB231/MB468 | MP |
| >hg38_21_+_8433172_8446622_RNA45SN1_45S_50ntFlanks@8364.8395.32\|CTTTGAAGAGAGAGTTCAAGAGGGCGTGAAAC | >hg38_21_+_8433172_8446622_RNA45SN1_45S_50ntFlanks | 8364 | 8395 | 32 | CTTTGAAGAGAGAGTTCAAGAGGGCGTGAAAC | 65.7 | 5.7 | 2.0 | 2.9 | 3.6E-03 | 7.6E-03 | BT20/MB468 | MP |
| >hg38_21_+_8433172_8446622_RNA45SN1_45S_50ntFlanks@4279.4314.36\|AATCCTTTAACGAGGATCCATTGGAGGGCAAGTCTG | >hg38_21_+_8433172_8446622_RNA45SN1_45S_50ntFlanks | 4279 | 4314 | 36 | AATCCTTTAACGAGGATCCATTGGAGGGCAAGTCTG | 192.1 | -1.7 | 0.6 | -2.7 | 6.3E-03 | 7.6E-03 | MB231/MB468 | CYTO |
| >MI0006349\|hsa-mir-1287&WithFlank&10\|-\|98395212\|98395313@22.43.22&[MIMAT0005878&hsa-miR-1287-5p&offsets\|0\|0;m-244&10\|-\|98395272\|98395292&offsets\|0\|+1]\|TGCTGGATCAGTGGTTCGAGTC | >MI0006349\|hsa-mir-1287&WithFlank&10\|-\|98395212\|98395313 | 22 | 43 | 22 | TGCTGGATCAGTGGTTCGAGTC | 549.0 | -4.5 | 1.4 | -3.2 | 1.3E-03 | 7.6E-03 | BT20/MB231 | MP |
| >am_trna111_HisGTG_1_-_147774845_147774916@-1G.29.30,trna118_HisGTG_1_-_145396881_145396952@-1G.29.30,trna16_HisGTG_1_+_146544773_146544844@-1G.29.30,trna1_HisGTG_15_+_45493349_45493420@-1G.29.30,trna21_HisGTG_1_+_147753471_147753542@-1G.29.30,trna33_HisGTG_6_+_27125906_27125977@-1G.29.30,trna7_HisGTG_9_-_14433938_14434009@-1G.29.30,trna8_HisGTG_15_-_45492611_45492682@-1G.29.30,trna9_HisGTG_15_-_45490804_45490875@-1G.29.30\|GGCCGTGATCGTATAGTGGTTAGTACTCTG | >am_trna111_HisGTG_1_-_147774845_147774916 | -1G | 29 | 30 | GGCCGTGATCGTATAGTGGTTAGTACTCTG | 194.4 | -4.0 | 1.3 | -3.2 | 1.4E-03 | 7.6E-03 | BT20/MB231 | NUC |
| >hg38_21_+_8433172_8446622_RNA45SN1_45S_50ntFlanks@4731.4761.31\|AAGAACGAAAGTCGGAGGTTCGAAGACGATC | >hg38_21_+_8433172_8446622_RNA45SN1_45S_50ntFlanks | 4731 | 4761 | 31 | AAGAACGAAAGTCGGAGGTTCGAAGACGATC | 1907.2 | 3.1 | 1.1 | 2.9 | 3.6E-03 | 7.6E-03 | BT20/MB468 | MP |
| >hg38_21_+_8433172_8446622_RNA45SN1_45S_50ntFlanks@8421.8442.22\|CGCGCAGTCCGCCCGGAGGATT | >hg38_21_+_8433172_8446622_RNA45SN1_45S_50ntFlanks | 8421 | 8442 | 22 | CGCGCAGTCCGCCCGGAGGATT | 60.9 | -4.7 | 1.6 | -2.9 | 3.6E-03 | 7.6E-03 | BT20/MB468 | MP |
| >hg38_21_+_8433172_8446622_RNA45SN1_45S_50ntFlanks@9737.9765.29\|CGAAACGATCTCAACCTATTCTCAAACTT | >hg38_21_+_8433172_8446622_RNA45SN1_45S_50ntFlanks | 9737 | 9765 | 29 | CGAAACGATCTCAACCTATTCTCAAACTT | 392.7 | -3.1 | 1.0 | -3.2 | 1.4E-03 | 7.6E-03 | BT20/MB231 | NUC |
| >hg38_21_+_8433172_8446622_RNA45SN1_45S_50ntFlanks@3843.3874.32\|CTACTTGGATAACTGTGGTAATTCTAGAGCTA | >hg38_21_+_8433172_8446622_RNA45SN1_45S_50ntFlanks | 3843 | 3874 | 32 | CTACTTGGATAACTGTGGTAATTCTAGAGCTA | 290.7 | -3.3 | 1.2 | -2.8 | 5.3E-03 | 7.6E-03 | BT20/MB468 | MITO |
| >hg38_21_+_8433172_8446622_RNA45SN1_45S_50ntFlanks@11591.11612.22\|GACCAGGGGAATCCGACTGTTT | >hg38_21_+_8433172_8446622_RNA45SN1_45S_50ntFlanks | 11591 | 11612 | 22 | GACCAGGGGAATCCGACTGTTT | 103.8 | -5.0 | 1.7 | -2.9 | 3.6E-03 | 7.6E-03 | BT20/MB468 | MP |
| >hg38_MT_+_598_1651_RNR1_12S_50ntFlanks@440.468.29\|ACATATCTGAACACACAATAGCTAAGACC | >hg38_MT_+_598_1651_RNR1_12S_50ntFlanks | 440 | 468 | 29 | ACATATCTGAACACACAATAGCTAAGACC | 129.1 | 3.7 | 1.3 | 2.8 | 5.7E-03 | 7.6E-03 | MB231/MB468 | MP |
| >hg38_21_+_8433172_8446622_RNA45SN1_45S_50ntFlanks@12927.12972.46\|CCTGGCGCTAAACCATTCGTAGACGACCTGCTTCTGGGTCGGGGTT | >hg38_21_+_8433172_8446622_RNA45SN1_45S_50ntFlanks | 12927 | 12972 | 46 | CCTGGCGCTAAACCATTCGTAGACGACCTGCTTCTGGGTCGGGGTT | 240.2 | -4.4 | 1.6 | -2.8 | 5.3E-03 | 7.6E-03 | BT20/MB468 | MITO |
| >MI0000067\|hsa-let-7f-1&WithFlank&9\|+\|94176341\|94176439@13.35.23&[MIMAT0000067&hsa-let-7f-5p&offsets\|0\|+1;m-20&9\|+\|94176353\|94176374&offsets\|0\|+1]\|TGAGGTAGTAGATTGTATAGTTG | >MI0000067\|hsa-let-7f-1&WithFlank&9\|+\|94176341\|94176439 | 13 | 35 | 23 | TGAGGTAGTAGATTGTATAGTTG | 170.1 | 3.5 | 1.3 | 2.8 | 5.7E-03 | 7.6E-03 | MB231/MB468 | MP |
| >MI0000085\|hsa-mir-27a&WithFlank&19\|-\|13836434\|13836523@57.77.21(+1A)&[MIMAT0000084&hsa-miR-27a-3p&offsets\|0\|0(+1A);m-43&19\|-\|13836447\|13836467&offsets\|0\|0(+1A)]\|TTCACAGTGGCTAAGTTCCGCA | >MI0000085\|hsa-mir-27a&WithFlank&19\|-\|13836434\|13836523 | 57 | 77 | 21(+1A) | TTCACAGTGGCTAAGTTCCGCA | 428.8 | -3.9 | 1.2 | -3.2 | 1.3E-03 | 7.6E-03 | BT20/MB231 | MP |
| >hg38_21_+_8433172_8446622_RNA45SN1_45S_50ntFlanks@5444.5466.23\|CTCGGATCGGCCCCGCCGGGGTC | >hg38_21_+_8433172_8446622_RNA45SN1_45S_50ntFlanks | 5444 | 5466 | 23 | CTCGGATCGGCCCCGCCGGGGTC | 13837.9 | -3.5 | 1.2 | -2.8 | 5.3E-03 | 7.6E-03 | BT20/MB468 | MITO |
| >hg38_21_+_8433172_8446622_RNA45SN1_45S_50ntFlanks@12420.12461.42\|ATGTCGGCTCTTCCTATCATTGTGAAGCAGAATTCACCAAGC | >hg38_21_+_8433172_8446622_RNA45SN1_45S_50ntFlanks | 12420 | 12461 | 42 | ATGTCGGCTCTTCCTATCATTGTGAAGCAGAATTCACCAAGC | 252.8 | 4.3 | 1.6 | 2.7 | 6.6E-03 | 7.6E-03 | MB231/MB468 | MITO |
| >hg38_1_-_228634819_228635039_RNA5S12_5S_50ntFlanks@152.169.18\|TACCGGGTGCTGTAGGCT | >hg38_1_-_228634819_228635039_RNA5S12_5S_50ntFlanks | 152 | 169 | 18 | TACCGGGTGCTGTAGGCT | 507.7 | 1.7 | 0.6 | 2.8 | 5.3E-03 | 7.6E-03 | BT20/MB468 | MITO |
| >MI0000299\|hsa-mir-222&WithFlank&X\|-\|45747009\|45747130@75.96.22(+1C)&[MIMAT0000279&hsa-miR-222-3p&offsets\|0\|+1(+1C);m-140&X\|-\|45747033\|45747056&offsets\|0\|-2(+1C)]\|AGCTACATCTGGCTACTGGGTCC | >MI0000299\|hsa-mir-222&WithFlank&X\|-\|45747009\|45747130 | 75 | 96 | 22(+1C) | AGCTACATCTGGCTACTGGGTCC | 323.0 | 3.3 | 1.1 | 2.9 | 3.7E-03 | 7.6E-03 | BT20/MB468 | MP |
| >hg38_21_+_8433172_8446622_RNA45SN1_45S_50ntFlanks@7979.8003.25\|ACCTCAGATCAGACGTGGCGACCCG | >hg38_21_+_8433172_8446622_RNA45SN1_45S_50ntFlanks | 7979 | 8003 | 25 | ACCTCAGATCAGACGTGGCGACCCG | 390.1 | -2.6 | 0.9 | -2.9 | 3.7E-03 | 7.6E-03 | BT20/MB468 | MP |
| >hg38_21_+_8433172_8446622_RNA45SN1_45S_50ntFlanks@5489.5514.26\|CTGAGAAGACGGTCGAACTTGACTAT | >hg38_21_+_8433172_8446622_RNA45SN1_45S_50ntFlanks | 5489 | 5514 | 26 | CTGAGAAGACGGTCGAACTTGACTAT | 869.8 | -2.0 | 0.7 | -2.7 | 6.2E-03 | 7.6E-03 | BT20/MB468 | CYTO |
| >MI0000447\|hsa-mir-128-1&WithFlank&2\|+\|135665391\|135665484@56.75.20(+1C);MI0000727\|hsa-mir-128-2&WithFlank&3\|+\|35744470\|35744565@58.77.20(+1C)&[MIMAT0000424&hsa-miR-128-3p&offsets\|0\|-1(+1C);m-108&2\|+\|135665446\|135665466&offsets\|0\|-1(+1C)];[MIMAT0000424_1&hsa-miR-128-3p&offsets\|0\|-1(+1C);m-117&3\|+\|35744527\|35744547&offsets\|0\|-1(+1C)]\|TCACAGTGAACCGGTCTCTTC | >MI0000447\|hsa-mir-128-1&WithFlank&2\|+\|135665391\|135665484 | 56 | 75 | 20(+1C) | TCACAGTGAACCGGTCTCTTC | 613.9 | 2.5 | 0.9 | 2.7 | 6.7E-03 | 7.6E-03 | MB231/MB468 | MITO |
| >MI0000734\|hsa-mir-106b&WithFlank&7\|-\|100093987\|100094080@18.39.22&[MIMAT0000680&hsa-miR-106b-5p&offsets\|0\|+1;m-103&7\|-\|100094043\|100094063&offsets\|0\|+1]\|TAAAGTGCTGACAGTGCAGATA | >MI0000734\|hsa-mir-106b&WithFlank&7\|-\|100093987\|100094080 | 18 | 39 | 22 | TAAAGTGCTGACAGTGCAGATA | 1969.9 | -2.8 | 0.9 | -3.1 | 1.8E-03 | 7.6E-03 | BT20/MB231 | MITO |
| >hg38_21_+_8433172_8446622_RNA45SN1_45S_50ntFlanks@6649.6669.21\|TACGACTCTTAGCGGTGGATC | >hg38_21_+_8433172_8446622_RNA45SN1_45S_50ntFlanks | 6649 | 6669 | 21 | TACGACTCTTAGCGGTGGATC | 4212.8 | 2.3 | 0.7 | 3.1 | 1.8E-03 | 7.7E-03 | BT20/MB231 | MITO |
| >hg38_MT_+_598_1651_RNR1_12S_50ntFlanks@401.424.24\|ACTCCAGTTGACACAAAATAGACT | >hg38_MT_+_598_1651_RNR1_12S_50ntFlanks | 401 | 424 | 24 | ACTCCAGTTGACACAAAATAGACT | 139.6 | 4.7 | 1.7 | 2.8 | 5.8E-03 | 7.7E-03 | MB231/MB468 | MP |

<table>
<tr><td>>hg38_21_+_8433172_8446622_RNA45SN1_45S_50ntFlanks@6778.6803.26|CCCGGGGCTACGCCTGTCTGAGCGTC</td><td>>hg38_21_+_8433172_8446622_RNA45SN1_45S_50ntFlanks</td><td>6778</td><td>6803</td><td>26</td><td>CCCGGGGCTACGCCTGTCTGAGCGTC</td><td>1446.4</td><td>-2.7</td><td>1.0</td><td>-2.7</td><td>6.7E-03</td><td>7.7E-03</td><td>MB231/MB468</td><td>MITO</td></tr>
<tr><td>>hg38_21_+_8433172_8446622_RNA45SN1_45S_50ntFlanks@12418.12445.28|CGATGTCGGCTCTTCCTATCATTGTGAA</td><td>>hg38_21_+_8433172_8446622_RNA45SN1_45S_50ntFlanks</td><td>12418</td><td>12445</td><td>28</td><td>CGATGTCGGCTCTTCCTATCATTGTGAA</td><td>946.8</td><td>-1.9</td><td>0.7</td><td>-2.7</td><td>6.3E-03</td><td>7.7E-03</td><td>BT20/MB468</td><td>CYTO</td></tr>
<tr><td>>hg38_21_+_8433172_8446622_RNA45SN1_45S_50ntFlanks@12408.12434.27|TTTGATCCTTCGATGTCGGCTCTTCCT</td><td>>hg38_21_+_8433172_8446622_RNA45SN1_45S_50ntFlanks</td><td>12408</td><td>12434</td><td>27</td><td>TTTGATCCTTCGATGTCGGCTCTTCCT</td><td>351.6</td><td>-1.8</td><td>0.7</td><td>-2.7</td><td>6.7E-03</td><td>7.7E-03</td><td>MB231/MB468</td><td>MITO</td></tr>
<tr><td>>MI0000471|hsa-mir-126&WithFlank&9|+|136670596|136670692@21.42.22&[MIMAT0000444&hsa-miR-126-5p&offsets|0|+1;m-114&9|+|136670616|136670636&offsets|0|+1]|CATTATTACTTTTGGTACGCGC</td><td>>MI0000471|hsa-mir-126&WithFlank&9|+|136670596|136670692</td><td>21</td><td>42</td><td>22</td><td>CATTATTACTTTTGGTACGCGC</td><td>28.4</td><td>8.1</td><td>2.9</td><td>2.8</td><td>5.8E-03</td><td>7.7E-03</td><td>MB231/MB468</td><td>MP</td></tr>
<tr><td>>hg38_21_+_8433172_8446622_RNA45SN1_45S_50ntFlanks@3765.3794.30|AGTACGCACGGCCGGTACAGTGAAACTGCG</td><td>>hg38_21_+_8433172_8446622_RNA45SN1_45S_50ntFlanks</td><td>3765</td><td>3794</td><td>30</td><td>AGTACGCACGGCCGGTACAGTGAAACTGCG</td><td>200.8</td><td>1.8</td><td>0.7</td><td>2.7</td><td>6.3E-03</td><td>7.7E-03</td><td>MB231/MB468</td><td>CYTO</td></tr>
<tr><td>>hg38_21_+_8433172_8446622_RNA45SN1_45S_50ntFlanks@9484.9520.37|GTGTTAGGACCCGAAAGATGGTGAACTATGCCTGGGC</td><td>>hg38_21_+_8433172_8446622_RNA45SN1_45S_50ntFlanks</td><td>9484</td><td>9520</td><td>37</td><td>GTGTTAGGACCCGAAAGATGGTGAACTATGCCTGGGC</td><td>60.6</td><td>9.3</td><td>2.9</td><td>3.2</td><td>1.3E-03</td><td>7.7E-03</td><td>BT20/MB231</td><td>MP</td></tr>
<tr><td>>hg38_21_+_8433172_8446622_RNA45SN1_45S_50ntFlanks@7978.8009.32|GACCTCAGATCAGACGTGGCGACCCGCTGAAT</td><td>>hg38_21_+_8433172_8446622_RNA45SN1_45S_50ntFlanks</td><td>7978</td><td>8009</td><td>32</td><td>GACCTCAGATCAGACGTGGCGACCCGCTGAAT</td><td>207.7</td><td>2.2</td><td>0.8</td><td>2.7</td><td>6.7E-03</td><td>7.7E-03</td><td>MB231/MB468</td><td>MITO</td></tr>
<tr><td>>hg38_21_+_8433172_8446622_RNA45SN1_45S_50ntFlanks@12415.12439.25|CTTCGATGTCGGCTCTTCCTATCAT</td><td>>hg38_21_+_8433172_8446622_RNA45SN1_45S_50ntFlanks</td><td>12415</td><td>12439</td><td>25</td><td>CTTCGATGTCGGCTCTTCCTATCAT</td><td>612.4</td><td>-2.7</td><td>1.0</td><td>-2.7</td><td>6.7E-03</td><td>7.7E-03</td><td>MB231/MB468</td><td>MITO</td></tr>
<tr><td>>trnaMT_GluTTC_MT_-_14674_14742@40.61.22,trnalookalike8_GluTTC_5_-_93905172_93905240@40.61.22|TTGGTCGTGGTTGTAGTCCGTG</td><td>>trnaMT_GluTTC_MT_-_14674_14742</td><td>40</td><td>61</td><td>22</td><td>TTGGTCGTGGTTGTAGTCCGTG</td><td>231.5</td><td>2.3</td><td>0.8</td><td>2.8</td><td>5.8E-03</td><td>7.7E-03</td><td>MB231/MB468</td><td>MP</td></tr>
<tr><td>>hg38_21_+_8433172_8446622_RNA45SN1_45S_50ntFlanks@11573.11609.37|CGACTTAGAACTGGTGCGGACCAGGGGAATCCGACTG</td><td>>hg38_21_+_8433172_8446622_RNA45SN1_45S_50ntFlanks</td><td>11573</td><td>11609</td><td>37</td><td>CGACTTAGAACTGGTGCGGACCAGGGGAATCCGACTG</td><td>78.4</td><td>-3.8</td><td>1.4</td><td>-2.8</td><td>5.8E-03</td><td>7.7E-03</td><td>MB231/MB468</td><td>MP</td></tr>
<tr><td>>hg38_21_+_8433172_8446622_RNA45SN1_45S_50ntFlanks@8168.8212.45|CCCAAGTCCTTCTGATCGAGGCCCAGCCCGTGGACGGTGTGAGGC</td><td>>hg38_21_+_8433172_8446622_RNA45SN1_45S_50ntFlanks</td><td>8168</td><td>8212</td><td>45</td><td>CCCAAGTCCTTCTGATCGAGGCCCAGCCCGTGGACGGTGTGAGGC</td><td>197.1</td><td>-1.8</td><td>0.7</td><td>-2.7</td><td>6.3E-03</td><td>7.7E-03</td><td>MB231/MB468</td><td>CYTO</td></tr>
<tr><td>>hg38_21_+_8433172_8446622_RNA45SN1_45S_50ntFlanks@10392.10419.28|ACAGCAGTTGAACATGGGTCAGTCGGTC</td><td>>hg38_21_+_8433172_8446622_RNA45SN1_45S_50ntFlanks</td><td>10392</td><td>10419</td><td>28</td><td>ACAGCAGTTGAACATGGGTCAGTCGGTC</td><td>440.7</td><td>-4.4</td><td>1.6</td><td>-2.7</td><td>6.3E-03</td><td>7.7E-03</td><td>BT20/MB468</td><td>CYTO</td></tr>
<tr><td>>hg38_21_+_8433172_8446622_RNA45SN1_45S_50ntFlanks@11318.11342.25|CCCCACGTCTCGTCGCGCGCGCGTC</td><td>>hg38_21_+_8433172_8446622_RNA45SN1_45S_50ntFlanks</td><td>11318</td><td>11342</td><td>25</td><td>CCCCACGTCTCGTCGCGCGCGCGTC</td><td>1469.8</td><td>-2.8</td><td>1.0</td><td>-2.8</td><td>5.4E-03</td><td>7.7E-03</td><td>BT20/MB468</td><td>MITO</td></tr>
<tr><td>>hg38_21_+_8433172_8446622_RNA45SN1_45S_50ntFlanks@12479.12500.22|CTAATAGGGAACGTGAGCTGGG</td><td>>hg38_21_+_8433172_8446622_RNA45SN1_45S_50ntFlanks</td><td>12479</td><td>12500</td><td>22</td><td>CTAATAGGGAACGTGAGCTGGG</td><td>240.7</td><td>-2.6</td><td>0.9</td><td>-2.9</td><td>3.7E-03</td><td>7.7E-03</td><td>BT20/MB468</td><td>MP</td></tr>
<tr><td>>MI0000264|hsa-mir-7-2&WithFlank&15|+|88611819|88611940@38.58.21;MI0000265|hsa-mir-7-3&WithFlank&19|+|4770664|4770785@37.57.21;MI0000263|hsa-mir-7-1&WithFlank&9|-|83969742|83969863@30.50.21&[MIMAT0000252&hsa-miR-7-5p&offsets|0|-3;m-400&15|+|88611856|88611879&offsets|0|-3];[MIMAT0000252_1&hsa-miR-7-5p&offsets|0|-3;m-402&19|+|4770700|4770723&offsets|0|-3];[MIMAT0000252_2&hsa-miR-7-5p&offsets|0|-3;m-401&9|-|83969811|83969834&offsets|0|-3]|TGGAAGACTAGTGATTTTGTT</td><td>>MI0000264|hsa-mir-7-2&WithFlank&15|+|88611819|88611940</td><td>38</td><td>58</td><td>21</td><td>TGGAAGACTAGTGATTTTGTT</td><td>543.2</td><td>2.9</td><td>1.0</td><td>2.9</td><td>3.7E-03</td><td>7.7E-03</td><td>BT20/MB468</td><td>MP</td></tr>
<tr><td>>hg38_21_+_8433172_8446622_RNA45SN1_45S_50ntFlanks@4760.4793.34|TCAGATACCGTCGTAGTTCCGACCATAAACGATG</td><td>>hg38_21_+_8433172_8446622_RNA45SN1_45S_50ntFlanks</td><td>4760</td><td>4793</td><td>34</td><td>TCAGATACCGTCGTAGTTCCGACCATAAACGATG</td><td>133.1</td><td>-4.0</td><td>1.4</td><td>-2.8</td><td>5.4E-03</td><td>7.7E-03</td><td>BT20/MB468</td><td>MITO</td></tr>
<tr><td>>am_trnaMT_PheGAA_MT_+_577_647@41.65.25|GTTTAGACGGGCTCACATCACCCCA</td><td>>am_trnaMT_PheGAA_MT_+_577_647</td><td>41</td><td>65</td><td>25</td><td>GTTTAGACGGGCTCACATCACCCCA</td><td>280.9</td><td>4.8</td><td>1.7</td><td>2.9</td><td>3.7E-03</td><td>7.7E-03</td><td>BT20/MB468</td><td>MP</td></tr>
<tr><td>>hg38_21_+_8433172_8446622_RNA45SN1_45S_50ntFlanks@12456.12499.44|CCAAGCGTTGGATTGTTCACCCACTAATAGGGAACGTGAGCTGG</td><td>>hg38_21_+_8433172_8446622_RNA45SN1_45S_50ntFlanks</td><td>12456</td><td>12499</td><td>44</td><td>CCAAGCGTTGGATTGTTCACCCACTAATAGGGAACGTGAGCTGG</td><td>330.9</td><td>-3.9</td><td>1.4</td><td>-2.8</td><td>5.4E-03</td><td>7.7E-03</td><td>BT20/MB468</td><td>MITO</td></tr>
<tr><td>>hg38_21_+_8433172_8446622_RNA45SN1_45S_50ntFlanks@7973.8017.45|GACGCGACCTCAGATCAGACGTGGCGACCCGCTGAATTTAAGCAT</td><td>>hg38_21_+_8433172_8446622_RNA45SN1_45S_50ntFlanks</td><td>7973</td><td>8017</td><td>45</td><td>GACGCGACCTCAGATCAGACGTGGCGACCCGCTGAATTTAAGCAT</td><td>177.5</td><td>3.7</td><td>1.4</td><td>2.7</td><td>6.7E-03</td><td>7.7E-03</td><td>MB231/MB468</td><td>MITO</td></tr>
<tr><td>>MI0000746|hsa-mir-99b&WithFlank&19|+|51692606|51692687@51.73.23&[MIMAT0004678&hsa-miR-99b-3p&offsets|0|+1;m-151&19|+|51692656|51692678&offsets|0|0]|CAAGCTCGTGTCTGTGGGTCCGT</td><td>>MI0000746|hsa-mir-99b&WithFlank&19|+|51692606|51692687</td><td>51</td><td>73</td><td>23</td><td>CAAGCTCGTGTCTGTGGGTCCGT</td><td>2463.9</td><td>2.5</td><td>0.8</td><td>3.1</td><td>1.7E-03</td><td>7.7E-03</td><td>BT20/MB231</td><td>CYTO</td></tr>
<tr><td>>trna2_GlyGCC_21_-_18827107_18827177@1.29.29,trna35_GlyGCC_1_+_161413094_161413164@1.29.29,trna37_GlyGCC_1_+_161420467_161420537@1.29.29,trna39_GlyGCC_1_+_161427898_161427968@1.29.29,trna41_GlyGCC_1_+_161435258_161435328@1.29.29|GCATGGGTGGTTCAGTGGTAGAATTCTCG</td><td>>trna2_GlyGCC_21_-_18827107_18827177</td><td>1</td><td>29</td><td>29</td><td>GCATGGGTGGTTCAGTGGTAGAATTCTCG</td><td>1782.7</td><td>3.2</td><td>1.0</td><td>3.1</td><td>1.7E-03</td><td>7.7E-03</td><td>BT20/MB231</td><td>CYTO</td></tr>
<tr><td>>trna16_ValTAC_11_-_59318460_59318532@1.34.34|GGTTCCATAGTGTAGCGGTTATCACGTCTGCTTT</td><td>>trna16_ValTAC_11_-_59318460_59318532</td><td>1</td><td>34</td><td>34</td><td>GGTTCCATAGTGTAGCGGTTATCACGTCTGCTTT</td><td>343.6</td><td>-3.3</td><td>1.1</td><td>-3.1</td><td>1.7E-03</td><td>7.7E-03</td><td>BT20/MB231</td><td>CYTO</td></tr>
<tr><td>>hg38_21_+_8433172_8446622_RNA45SN1_45S_50ntFlanks@12454.12477.24|CACCAAGCGTTGGATTGTTCACCC</td><td>>hg38_21_+_8433172_8446622_RNA45SN1_45S_50ntFlanks</td><td>12454</td><td>12477</td><td>24</td><td>CACCAAGCGTTGGATTGTTCACCC</td><td>403.5</td><td>-2.6</td><td>0.8</td><td>-3.1</td><td>1.7E-03</td><td>7.7E-03</td><td>BT20/MB231</td><td>CYTO</td></tr>
<tr><td>>hg38_21_+_8433172_8446622_RNA45SN1_45S_50ntFlanks@12990.13014.25|GCTCCCTCGCTGCGATCTATTGAAA</td><td>>hg38_21_+_8433172_8446622_RNA45SN1_45S_50ntFlanks</td><td>12990</td><td>13014</td><td>25</td><td>GCTCCCTCGCTGCGATCTATTGAAA</td><td>478.3</td><td>-2.8</td><td>0.9</td><td>-3.1</td><td>1.7E-03</td><td>7.7E-03</td><td>BT20/MB231</td><td>CYTO</td></tr>
<tr><td>>hg38_21_+_8433172_8446622_RNA45SN1_45S_50ntFlanks@12418.12449.32|CGATGTCGGCTCTTCCTATCATTGTGAAGCAG</td><td>>hg38_21_+_8433172_8446622_RNA45SN1_45S_50ntFlanks</td><td>12418</td><td>12449</td><td>32</td><td>CGATGTCGGCTCTTCCTATCATTGTGAAGCAG</td><td>318.8</td><td>-1.3</td><td>0.5</td><td>-2.8</td><td>5.4E-03</td><td>7.7E-03</td><td>BT20/MB468</td><td>MITO</td></tr>
<tr><td>>am_trna15_CysGCA_17_+_37023898_37023969@14.34.21,trna26_CysGCA_17_-_37310744_37310815@14.34.21,trna27_CysGCA_17_-_37309987_37310058@14.34.21,trna28_CysGCA_17_-_37025545_37025616@14.34.21,trna3_CysGCA_4_-_124430005_124430076@14.34.21,trna5_CysGCA_7_+_149007281_149007352@14.34.21,trna7_CysGCA_3_-_131947944_131948015@14.34.21|AGTGGTAGAGCATTTGACTGC</td><td>>am_trna15_CysGCA_17_+_37023898_37023969</td><td>14</td><td>34</td><td>21</td><td>AGTGGTAGAGCATTTGACTGC</td><td>125.2</td><td>-5.7</td><td>1.8</td><td>-3.2</td><td>1.3E-03</td><td>7.7E-03</td><td>BT20/MB231</td><td>MP</td></tr>
<tr><td>>hg38_21_+_8433172_8446622_RNA45SN1_45S_50ntFlanks@6721.6756.36|AATTGCAGGACACATTGATCATCGACACTTCGAACG</td><td>>hg38_21_+_8433172_8446622_RNA45SN1_45S_50ntFlanks</td><td>6721</td><td>6756</td><td>36</td><td>AATTGCAGGACACATTGATCATCGACACTTCGAACG</td><td>592.5</td><td>-1.7</td><td>0.6</td><td>-2.7</td><td>6.4E-03</td><td>7.7E-03</td><td>MB231/MB468</td><td>CYTO</td></tr>
<tr><td>>hg38_21_+_8433172_8446622_RNA45SN1_45S_50ntFlanks@8196.8214.19|CGTGGACGGTGTGAGGCCG</td><td>>hg38_21_+_8433172_8446622_RNA45SN1_45S_50ntFlanks</td><td>8196</td><td>8214</td><td>19</td><td>CGTGGACGGTGTGAGGCCG</td><td>425.1</td><td>-1.6</td><td>0.6</td><td>-2.8</td><td>5.4E-03</td><td>7.7E-03</td><td>BT20/MB468</td><td>MITO</td></tr>
<tr><td>>hg38_21_+_8433172_8446622_RNA45SN1_45S_50ntFlanks@9471.9500.30|AGCACGAGCGCACGTGTTAGGACCCGAAAG</td><td>>hg38_21_+_8433172_8446622_RNA45SN1_45S_50ntFlanks</td><td>9471</td><td>9500</td><td>30</td><td>AGCACGAGCGCACGTGTTAGGACCCGAAAG</td><td>145.9</td><td>-1.3</td><td>0.5</td><td>-2.8</td><td>5.4E-03</td><td>7.7E-03</td><td>BT20/MB468</td><td>MITO</td></tr>
<tr><td>>hg38_21_+_8433172_8446622_RNA45SN1_45S_50ntFlanks@6670.6700.31|ACTCGGCTCGTGCGTCGATGAAGAACGCAGC</td><td>>hg38_21_+_8433172_8446622_RNA45SN1_45S_50ntFlanks</td><td>6670</td><td>6700</td><td>31</td><td>ACTCGGCTCGTGCGTCGATGAAGAACGCAGC</td><td>160.3</td><td>2.5</td><td>0.9</td><td>2.7</td><td>6.7E-03</td><td>7.7E-03</td><td>MB231/MB468</td><td>MITO</td></tr>
<tr><td>>hg38_21_+_8433172_8446622_RNA45SN1_45S_50ntFlanks@10401.10429.29|GAACATGGGTCAGTCGGTCCTGAGAGATG</td><td>>hg38_21_+_8433172_8446622_RNA45SN1_45S_50ntFlanks</td><td>10401</td><td>10429</td><td>29</td><td>GAACATGGGTCAGTCGGTCCTGAGAGATG</td><td>756.8</td><td>-2.6</td><td>1.0</td><td>-2.7</td><td>6.7E-03</td><td>7.7E-03</td><td>MB231/MB468</td><td>MITO</td></tr>
<tr><td>>hg38_21_+_8433172_8446622_RNA45SN1_45S_50ntFlanks@9546.9572.27|TGGAGGTCCGTAGCGGTCCTGACGTGC</td><td>>hg38_21_+_8433172_8446622_RNA45SN1_45S_50ntFlanks</td><td>9546</td><td>9572</td><td>27</td><td>TGGAGGTCCGTAGCGGTCCTGACGTGC</td><td>292.5</td><td>3.1</td><td>1.1</td><td>2.9</td><td>3.7E-03</td><td>7.7E-03</td><td>BT20/MB468</td><td>MP</td></tr>
<tr><td>>hg38_21_+_8433172_8446622_RNA45SN1_45S_50ntFlanks@5085.5102.18|GAACGAGACTCTGGCATG</td><td>>hg38_21_+_8433172_8446622_RNA45SN1_45S_50ntFlanks</td><td>5085</td><td>5102</td><td>18</td><td>GAACGAGACTCTGGCATG</td><td>621.3</td><td>4.9</td><td>1.5</td><td>3.2</td><td>1.3E-03</td><td>7.7E-03</td><td>BT20/MB231</td><td>MP</td></tr>
<tr><td>>hg38_21_+_8433172_8446622_RNA45SN1_45S_50ntFlanks@10838.10868.31|ATCCGTAACTTCGGGATAAGGATTGGCTCTA</td><td>>hg38_21_+_8433172_8446622_RNA45SN1_45S_50ntFlanks</td><td>10838</td><td>10868</td><td>31</td><td>ATCCGTAACTTCGGGATAAGGATTGGCTCTA</td><td>167.0</td><td>-4.6</td><td>1.7</td><td>-2.8</td><td>5.8E-03</td><td>7.7E-03</td><td>MB231/MB468</td><td>MP</td></tr>
<tr><td>>hg38_MT_+_1621_3279_RNR2_16S_50ntFlanks@626.652.27|ACTGAACTCCTCACACCCAATTGGACC</td><td>>hg38_MT_+_1621_3279_RNR2_16S_50ntFlanks</td><td>626</td><td>652</td><td>27</td><td>ACTGAACTCCTCACACCCAATTGGACC</td><td>110.9</td><td>4.3</td><td>1.5</td><td>2.9</td><td>3.7E-03</td><td>7.7E-03</td><td>BT20/MB468</td><td>MP</td></tr>
<tr><td>>hg38_21_+_8433172_8446622_RNA45SN1_45S_50ntFlanks@5472.5494.23|ACGGCCCTGGCGGAGCGCTGAGA</td><td>>hg38_21_+_8433172_8446622_RNA45SN1_45S_50ntFlanks</td><td>5472</td><td>5494</td><td>23</td><td>ACGGCCCTGGCGGAGCGCTGAGA</td><td>301.4</td><td>4.3</td><td>1.5</td><td>2.9</td><td>3.7E-03</td><td>7.7E-03</td><td>BT20/MB468</td><td>MP</td></tr>
<tr><td>>hg38_21_+_8433172_8446622_RNA45SN1_45S_50ntFlanks@6652.6682.31|GACTCTTAGCGGTGGATCACTCGGCTCGTGC</td><td>>hg38_21_+_8433172_8446622_RNA45SN1_45S_50ntFlanks</td><td>6652</td><td>6682</td><td>31</td><td>GACTCTTAGCGGTGGATCACTCGGCTCGTGC</td><td>1844.6</td><td>3.0</td><td>1.1</td><td>2.7</td><td>6.7E-03</td><td>7.7E-03</td><td>MB231/MB468</td><td>MITO</td></tr>
<tr><td>>hg38_21_+_8433172_8446622_RNA45SN1_45S_50ntFlanks@3842.3881.40|CCTACTTGGATAACTGTGGTAATTCTAGAGCTAATACATG</td><td>>hg38_21_+_8433172_8446622_RNA45SN1_45S_50ntFlanks</td><td>3842</td><td>3881</td><td>40</td><td>CCTACTTGGATAACTGTGGTAATTCTAGAGCTAATACATG</td><td>188.5</td><td>-2.0</td><td>0.7</td><td>-2.7</td><td>6.8E-03</td><td>7.7E-03</td><td>MB231/MB468</td><td>NUC</td></tr>
<tr><td>>hg38_21_+_8433172_8446622_RNA45SN1_45S_50ntFlanks@12190.12211.22|AACGCAGGTGTCCTAAGGCGAG</td><td>>hg38_21_+_8433172_8446622_RNA45SN1_45S_50ntFlanks</td><td>12190</td><td>12211</td><td>22</td><td>AACGCAGGTGTCCTAAGGCGAG</td><td>109.2</td><td>-4.7</td><td>1.7</td><td>-2.8</td><td>5.8E-03</td><td>7.7E-03</td><td>MB231/MB468</td><td>MP</td></tr>
<tr><td>>hg38_21_+_8433172_8446622_RNA45SN1_45S_50ntFlanks@5487.5518.32|CGCTGAGAAGACGGTCGAACTTGACTATCTAG</td><td>>hg38_21_+_8433172_8446622_RNA45SN1_45S_50ntFlanks</td><td>5487</td><td>5518</td><td>32</td><td>CGCTGAGAAGACGGTCGAACTTGACTATCTAG</td><td>119.0</td><td>-4.0</td><td>1.4</td><td>-2.8</td><td>5.4E-03</td><td>7.7E-03</td><td>BT20/MB468</td><td>MITO</td></tr>
<tr><td>>trnaMT_SerGCT_MT_+_12207_12265@1.28.28|GAGAAAGCTCACAAGAACTGCTAACTCA</td><td>>trnaMT_SerGCT_MT_+_12207_12265</td><td>1</td><td>28</td><td>28</td><td>GAGAAAGCTCACAAGAACTGCTAACTCA</td><td>191.7</td><td>2.4</td><td>0.9</td><td>2.8</td><td>5.8E-03</td><td>7.7E-03</td><td>MB231/MB468</td><td>MP</td></tr>
<tr><td>>MI0000273|hsa-mir-183&WithFlank&7|-|129774899|129775020@33.54.22(+1A)&[MIMAT0000261&hsa-miR-183-5p&offsets|0|0(+1A);m-41&7|-|129774966|129774987&offsets|-1|-1(+1A)]|TATGGCACTGGTAGAATTCACTA</td><td>>MI0000273|hsa-mir-183&WithFlank&7|-|129774899|129775020</td><td>33</td><td>54</td><td>22(+1A)</td><td>TATGGCACTGGTAGAATTCACTA</td><td>346.6</td><td>4.2</td><td>1.5</td><td>2.9</td><td>3.7E-03</td><td>7.7E-03</td><td>BT20/MB468</td><td>MP</td></tr>
<tr><td>>hg38_MT_+_1621_3279_RNR2_16S_50ntFlanks@929.958.30|CAGTGACACATGTTTAACGGCCGCGGTACC</td><td>>hg38_MT_+_1621_3279_RNR2_16S_50ntFlanks</td><td>929</td><td>958</td><td>30</td><td>CAGTGACACATGTTTAACGGCCGCGGTACC</td><td>268.8</td><td>4.6</td><td>1.6</td><td>2.9</td><td>3.7E-03</td><td>7.7E-03</td><td>BT20/MB468</td><td>MP</td></tr>
<tr><td>>MI0000746|hsa-mir-99b&WithFlank&19|+|51692606|51692687@51.73.23(+1U)&[MIMAT0004678&hsa-miR-99b-3p&offsets|0|+1(+1U);m-151&19|+|51692656|51692678&offsets|0|0(+1U)]|CAAGCTCGTGTCTGTGGGTCCGTT</td><td>>MI0000746|hsa-mir-99b&WithFlank&19|+|51692606|51692687</td><td>51</td><td>73</td><td>23(+1U)</td><td>CAAGCTCGTGTCTGTGGGTCCGTT</td><td>127.8</td><td>5.8</td><td>2.0</td><td>2.9</td><td>3.7E-03</td><td>7.7E-03</td><td>BT20/MB468</td><td>MP</td></tr>
<tr><td>>MI0000270|hsa-mir-181b-1&WithFlank&1|-|198858867|198858988@42.61.20;MI0000683|hsa-mir-181b-2&WithFlank&9|+|124693704|124693804@22.41.20&[MIMAT0000257&hsa-miR-181b-5p&offsets|0|-3;m-73&1|-|198858925|198858947&offsets|0|-3];[MIMAT0000257_1&hsa-miR-181b-5p&offsets|0|-3;m-72&9|+|124693725|124693747&offsets|0|-3]|AACATTCATTGCTGTCGGTG</td><td>>MI0000270|hsa-mir-181b-1&WithFlank&1|-|198858867|198858988</td><td>42</td><td>61</td><td>20</td><td>AACATTCATTGCTGTCGGTG</td><td>356.8</td><td>2.4</td><td>0.9</td><td>2.8</td><td>5.8E-03</td><td>7.7E-03</td><td>MB231/MB468</td><td>MP</td></tr>
<tr><td>>hg38_MT_+_598_1651_RNR1_12S_50ntFlanks@443.465.23|TATCTGAACACACACAATAGCTAAG</td><td>>hg38_MT_+_598_1651_RNR1_12S_50ntFlanks</td><td>443</td><td>465</td><td>23</td><td>TATCTGAACACACACAATAGCTAAG</td><td>173.0</td><td>3.1</td><td>1.1</td><td>2.8</td><td>5.8E-03</td><td>7.7E-03</td><td>MB231/MB468</td><td>MP</td></tr>
<tr><td>>trnaMT_ProTGG_MT_-_15956_16023@49.70.22|GTTAAAGACTTTTTCTCTGACC</td><td>>trnaMT_ProTGG_MT_-_15956_16023</td><td>49</td><td>70</td><td>22</td><td>GTTAAAGACTTTTTCTCTGACC</td><td>87.3</td><td>2.7</td><td>1.0</td><td>2.8</td><td>5.8E-03</td><td>7.7E-03</td><td>MB231/MB468</td><td>MP</td></tr>
<tr><td>>hg38_MT_+_1621_3279_RNR2_16S_50ntFlanks@283.305.23|CCCGAAACCAGACGAGCTACCTA</td><td>>hg38_MT_+_1621_3279_RNR2_16S_50ntFlanks</td><td>283</td><td>305</td><td>23</td><td>CCCGAAACCAGACGAGCTACCTA</td><td>63.5</td><td>5.8</td><td>2.0</td><td>2.9</td><td>3.7E-03</td><td>7.7E-03</td><td>BT20/MB468</td><td>MP</td></tr>
<tr><td>>MI0000255|hsa-mir-30d&WithFlank&8|-|134804870|134804951@12.33.22&[MIMAT0000245&hsa-miR-30d-5p&offsets|0|0;m-21&8|-|134804919|134804940&offsets|0|0]|TGTAAACATCCCCGACTGGAAG</td><td>>MI0000255|hsa-mir-30d&WithFlank&8|-|134804870|134804951</td><td>12</td><td>33</td><td>22</td><td>TGTAAACATCCCCGACTGGAAG</td><td>831.6</td><td>2.2</td><td>0.8</td><td>2.8</td><td>5.8E-03</td><td>7.7E-03</td><td>MB231/MB468</td><td>MP</td></tr>
</table>

<table>
<tr><th></th><th></th><th></th><th></th><th></th><th></th><th></th><th></th><th></th><th></th><th></th><th></th><th></th><th></th></tr>
<tr><td>>hg38_21_+_8433172_8446622_RNA45SN1_45S_50ntFlanks@10107.10125.19|CGTGTGGTGTGCGTCGGAG</td><td>>hg38_21_+_8433172_8446622_RNA45SN1_45S_50ntFlanks</td><td>10107</td><td>10125</td><td>19</td><td>CGTGTGGTGTGCGTCGGAG</td><td>731.3</td><td>-3.9</td><td>1.3</td><td>-2.9</td><td>3.7E-03</td><td>7.8E-03</td><td>BT20/MB468</td><td>MP</td></tr>
<tr><td>>hg38_21_+_8433172_8446622_RNA45SN1_45S_50ntFlanks@7988.8009.22|CAGACGTGGCGACCCGCTGAAT</td><td>>hg38_21_+_8433172_8446622_RNA45SN1_45S_50ntFlanks</td><td>7988</td><td>8009</td><td>22</td><td>CAGACGTGGCGACCCGCTGAAT</td><td>713.7</td><td>3.3</td><td>1.0</td><td>3.2</td><td>1.5E-03</td><td>7.8E-03</td><td>BT20/MB231</td><td>NUC</td></tr>
<tr><td>>MI0000088|hsa-mir-30a&WithFlank&6|-|71403545|71403627@54.71.18&[MIMAT0000088&hsa-miR-30a-3p&offsets|+1|-3;m-38&6|-|71403554|71403575&offsets|+1|-3]|TTTCAGTCGGATGTTTGC</td><td>>MI0000088|hsa-mir-30a&WithFlank&6|-|71403545|71403627</td><td>54</td><td>71</td><td>18</td><td>TTTCAGTCGGATGTTTGC</td><td>1022.0</td><td>-2.0</td><td>0.6</td><td>-3.2</td><td>1.5E-03</td><td>7.8E-03</td><td>BT20/MB231</td><td>NUC</td></tr>
<tr><td>>hg38_21_+_8433172_8446622_RNA45SN1_45S_50ntFlanks@9792.9818.27|GGCGTGGAGCCGGGCGTGGAATGCGAG</td><td>>hg38_21_+_8433172_8446622_RNA45SN1_45S_50ntFlanks</td><td>9792</td><td>9818</td><td>27</td><td>GGCGTGGAGCCGGGCGTGGAATGCGAG</td><td>60.4</td><td>9.2</td><td>2.9</td><td>3.2</td><td>1.3E-03</td><td>7.8E-03</td><td>BT20/MB231</td><td>MP</td></tr>
<tr><td>>hg38_21_+_8433172_8446622_RNA45SN1_45S_50ntFlanks@12412.12431.20|ATCCTTCGATGTCGGCTCTT</td><td>>hg38_21_+_8433172_8446622_RNA45SN1_45S_50ntFlanks</td><td>12412</td><td>12431</td><td>20</td><td>ATCCTTCGATGTCGGCTCTT</td><td>634.9</td><td>-4.2</td><td>1.5</td><td>-2.7</td><td>6.4E-03</td><td>7.8E-03</td><td>MB231/MB468</td><td>CYTO</td></tr>
<tr><td>>MI0000440|hsa-mir-27b&WithFlank&9|+|95085439|95085547@67.88.22(+2U)&[MIMAT0000419&hsa-miR-27b-3p&offsets|0|+1(+2U);m-39&9|+|95085505|95085525&offsets|0|+1(+2U)]|TTCACAGTGGCTAAGTTCTGCATT</td><td>>MI0000440|hsa-mir-27b&WithFlank&9|+|95085439|95085547</td><td>67</td><td>88</td><td>22(+2U)</td><td>TTCACAGTGGCTAAGTTCTGCATT</td><td>140.8</td><td>5.2</td><td>1.9</td><td>2.7</td><td>6.8E-03</td><td>7.8E-03</td><td>MB231/MB468</td><td>MITO</td></tr>
<tr><td>>hg38_21_+_8433172_8446622_RNA45SN1_45S_50ntFlanks@11727.11744.18|AGTAACTATGACTCTCTT</td><td>>hg38_21_+_8433172_8446622_RNA45SN1_45S_50ntFlanks</td><td>11727</td><td>11744</td><td>18</td><td>AGTAACTATGACTCTCTT</td><td>863.1</td><td>-1.9</td><td>0.7</td><td>-2.7</td><td>6.8E-03</td><td>7.8E-03</td><td>MB231/MB468</td><td>NUC</td></tr>
<tr><td>>MI0000446|hsa-mir-125b-1&WithFlank&11|-|122099751|122099850@21.40.20(+1A);MI0000470|hsa-mir-125b-2&WithFlank&21|+|16590231|16590331@23.42.20(+1A)&[MIMAT0000423&hsa-miR-125b-5p&offsets|0|-2(+1A);m-67&11|-|122099809|122099830&offsets|0|-2(+1A)];[MIMAT0000423_1&hsa-miR-125b-5p&offsets|0|-2(+1A);m-68&21|+|16590253|16590274&offsets|0|-2(+1A)]|TCCCTGAGACCCTAACTTGTA</td><td>>MI0000446|hsa-mir-125b-1&WithFlank&11|-|122099751|122099850</td><td>21</td><td>40</td><td>20(+1A)</td><td>TCCCTGAGACCCTAACTTGTA</td><td>574.0</td><td>-5.2</td><td>1.6</td><td>-3.2</td><td>1.5E-03</td><td>7.8E-03</td><td>BT20/MB231</td><td>NUC</td></tr>
<tr><td>>hg38_21_+_8433172_8446622_RNA45SN1_45S_50ntFlanks@6651.6688.38|CGACTCTTAGCGGTGGATCACTCGGCTCGTGCGTCGAT</td><td>>hg38_21_+_8433172_8446622_RNA45SN1_45S_50ntFlanks</td><td>6651</td><td>6688</td><td>38</td><td>CGACTCTTAGCGGTGGATCACTCGGCTCGTGCGTCGAT</td><td>1791.3</td><td>2.8</td><td>0.9</td><td>3.1</td><td>1.7E-03</td><td>7.8E-03</td><td>BT20/MB231</td><td>CYTO</td></tr>
<tr><td>>hg38_21_+_8433172_8446622_RNA45SN1_45S_50ntFlanks@12420.12451.32|ATGTCGGCTCTTCCTATCATTGTGAAGCAGAA</td><td>>hg38_21_+_8433172_8446622_RNA45SN1_45S_50ntFlanks</td><td>12420</td><td>12451</td><td>32</td><td>ATGTCGGCTCTTCCTATCATTGTGAAGCAGAA</td><td>170.4</td><td>-2.4</td><td>0.9</td><td>-2.7</td><td>6.4E-03</td><td>7.8E-03</td><td>MB231/MB468</td><td>CYTO</td></tr>
<tr><td>>MI0000088|hsa-mir-30a&WithFlank&6|-|71403545|71403627@53.74.22(+1A)&[MIMAT0000088&hsa-miR-30a-3p&offsets|0|0(+1A);m-38&6|-|71403554|71403575&offsets|0|0(+1A)]|CTTTCAGTCGGATGTTTGCAGCA</td><td>>MI0000088|hsa-mir-30a&WithFlank&6|-|71403545|71403627</td><td>53</td><td>74</td><td>22(+1A)</td><td>CTTTCAGTCGGATGTTTGCAGCA</td><td>178.0</td><td>-3.5</td><td>1.1</td><td>-3.2</td><td>1.4E-03</td><td>7.8E-03</td><td>BT20/MB231</td><td>MP</td></tr>
<tr><td>>hg38_21_+_8433172_8446622_RNA45SN1_45S_50ntFlanks@7974.8009.36|ACGCGACCTCAGATCAGACGTGGCGACCCGCTGAAT</td><td>>hg38_21_+_8433172_8446622_RNA45SN1_45S_50ntFlanks</td><td>7974</td><td>8009</td><td>36</td><td>ACGCGACCTCAGATCAGACGTGGCGACCCGCTGAAT</td><td>47138.0</td><td>2.8</td><td>0.9</td><td>3.2</td><td>1.5E-03</td><td>7.8E-03</td><td>BT20/MB231</td><td>NUC</td></tr>
<tr><td>>hg38_21_+_8433172_8446622_RNA45SN1_45S_50ntFlanks@12241.12269.29|CAGAAGGGCAAAAGCTCGCTTGATCTTGA</td><td>>hg38_21_+_8433172_8446622_RNA45SN1_45S_50ntFlanks</td><td>12241</td><td>12269</td><td>29</td><td>CAGAAGGGCAAAAGCTCGCTTGATCTTGA</td><td>372.9</td><td>-2.1</td><td>0.8</td><td>-2.8</td><td>5.5E-03</td><td>7.8E-03</td><td>BT20/MB468</td><td>MITO</td></tr>
<tr><td>>hg38_21_+_8433172_8446622_RNA45SN1_45S_50ntFlanks@7978.8010.33|GACCTCAGATCAGACGTGGCGACCCGCTGAATT</td><td>>hg38_21_+_8433172_8446622_RNA45SN1_45S_50ntFlanks</td><td>7978</td><td>8010</td><td>33</td><td>GACCTCAGATCAGACGTGGCGACCCGCTGAATT</td><td>125.8</td><td>1.7</td><td>0.6</td><td>2.7</td><td>6.7E-03</td><td>7.8E-03</td><td>BT20/MB468</td><td>NUC</td></tr>
<tr><td>>hg38_21_+_8433172_8446622_RNA45SN1_45S_50ntFlanks@6649.6688.40|TACGACTCTTAGCGGTGGATCACTCGGCTCGTGCGTCGAT</td><td>>hg38_21_+_8433172_8446622_RNA45SN1_45S_50ntFlanks</td><td>6649</td><td>6688</td><td>40</td><td>TACGACTCTTAGCGGTGGATCACTCGGCTCGTGCGTCGAT</td><td>447.9</td><td>2.2</td><td>0.8</td><td>2.8</td><td>5.5E-03</td><td>7.8E-03</td><td>BT20/MB468</td><td>MITO</td></tr>
<tr><td>>am_trnaMT_PheGAA_MT_+_577_647@44.61.18|TAGACGGGCTCACATCAC</td><td>>am_trnaMT_PheGAA_MT_+_577_647</td><td>44</td><td>61</td><td>18</td><td>TAGACGGGCTCACATCAC</td><td>104.1</td><td>4.4</td><td>1.5</td><td>2.9</td><td>3.7E-03</td><td>7.8E-03</td><td>BT20/MB468</td><td>MP</td></tr>
<tr><td>>hg38_21_+_8433172_8446622_RNA45SN1_45S_50ntFlanks@9932.9951.20|ATAGACAGCAGGACGGTGGC</td><td>>hg38_21_+_8433172_8446622_RNA45SN1_45S_50ntFlanks</td><td>9932</td><td>9951</td><td>20</td><td>ATAGACAGCAGGACGGTGGC</td><td>399.1</td><td>5.3</td><td>1.8</td><td>2.9</td><td>3.8E-03</td><td>7.8E-03</td><td>BT20/MB468</td><td>MP</td></tr>
<tr><td>>hg38_21_+_8433172_8446622_RNA45SN1_45S_50ntFlanks@6660.6693.34|GCGGTGGATCACTCGGCTCGTGCGTCGATGAAGA</td><td>>hg38_21_+_8433172_8446622_RNA45SN1_45S_50ntFlanks</td><td>6660</td><td>6693</td><td>34</td><td>GCGGTGGATCACTCGGCTCGTGCGTCGATGAAGA</td><td>96.4</td><td>3.0</td><td>1.1</td><td>2.9</td><td>3.8E-03</td><td>7.8E-03</td><td>BT20/MB468</td><td>MP</td></tr>
<tr><td>>hg38_21_+_8433172_8446622_RNA45SN1_45S_50ntFlanks@3960.3984.25|GCCCCGGCCGGGGGGCGGGCGCCGG</td><td>>hg38_21_+_8433172_8446622_RNA45SN1_45S_50ntFlanks</td><td>3960</td><td>3984</td><td>25</td><td>GCCCCGGCCGGGGGGCGGGCGCCGG</td><td>103.5</td><td>-2.8</td><td>1.0</td><td>-2.7</td><td>6.4E-03</td><td>7.8E-03</td><td>BT20/MB468</td><td>CYTO</td></tr>
<tr><td>>hg38_21_+_8433172_8446622_RNA45SN1_45S_50ntFlanks@9545.9573.29|GTGGAGGTCCGTAGCGGTCCTGACGTGCA</td><td>>hg38_21_+_8433172_8446622_RNA45SN1_45S_50ntFlanks</td><td>9545</td><td>9573</td><td>29</td><td>GTGGAGGTCCGTAGCGGTCCTGACGTGCA</td><td>488.8</td><td>-2.6</td><td>0.9</td><td>-2.9</td><td>3.8E-03</td><td>7.8E-03</td><td>BT20/MB468</td><td>MP</td></tr>
<tr><td>>hg38_21_+_8433172_8446622_RNA45SN1_45S_50ntFlanks@5492.5510.19|AGAAGACGGTCGAACTTGA</td><td>>hg38_21_+_8433172_8446622_RNA45SN1_45S_50ntFlanks</td><td>5492</td><td>5510</td><td>19</td><td>AGAAGACGGTCGAACTTGA</td><td>388.5</td><td>-5.1</td><td>1.9</td><td>-2.7</td><td>6.8E-03</td><td>7.8E-03</td><td>BT20/MB468</td><td>NUC</td></tr>
<tr><td>>hg38_21_+_8433172_8446622_RNA45SN1_45S_50ntFlanks@9936.9960.25|ACAGCAGGACGGTGGCCATGGAAGT</td><td>>hg38_21_+_8433172_8446622_RNA45SN1_45S_50ntFlanks</td><td>9936</td><td>9960</td><td>25</td><td>ACAGCAGGACGGTGGCCATGGAAGT</td><td>61.8</td><td>4.9</td><td>1.7</td><td>2.9</td><td>3.8E-03</td><td>7.8E-03</td><td>BT20/MB468</td><td>MP</td></tr>
<tr><td>>hg38_21_+_8433172_8446622_RNA45SN1_45S_50ntFlanks@6723.6744.22|TTGCAGGACACATTGATCATCG</td><td>>hg38_21_+_8433172_8446622_RNA45SN1_45S_50ntFlanks</td><td>6723</td><td>6744</td><td>22</td><td>TTGCAGGACACATTGATCATCG</td><td>138.3</td><td>-1.8</td><td>0.6</td><td>-2.8</td><td>5.5E-03</td><td>7.8E-03</td><td>BT20/MB468</td><td>MITO</td></tr>
<tr><td>>hg38_21_+_8433172_8446622_RNA45SN1_45S_50ntFlanks@10408.10427.20|GGTCAGTCGGTCCTGAGAGA</td><td>>hg38_21_+_8433172_8446622_RNA45SN1_45S_50ntFlanks</td><td>10408</td><td>10427</td><td>20</td><td>GGTCAGTCGGTCCTGAGAGA</td><td>144.6</td><td>-4.1</td><td>1.5</td><td>-2.7</td><td>6.4E-03</td><td>7.8E-03</td><td>BT20/MB468</td><td>CYTO</td></tr>
<tr><td>>trnaMT_GluTTC_MT_-_14674_14742@40.60.21,trnalookalike8_GluTTC_5_-_93905172_93905240@40.60.21|TTGGTCGTGGTTGTAGTCCGT</td><td>>trnaMT_GluTTC_MT_-_14674_14742</td><td>40</td><td>60</td><td>21</td><td>TTGGTCGTGGTTGTAGTCCGT</td><td>66.5</td><td>4.9</td><td>1.7</td><td>2.9</td><td>3.8E-03</td><td>7.8E-03</td><td>BT20/MB468</td><td>MP</td></tr>
<tr><td>>hg38_MT_+_1621_3279_RNR2_16S_50ntFlanks@493.513.21|GCTCTTTGGACACTAGGAAAA</td><td>>hg38_MT_+_1621_3279_RNR2_16S_50ntFlanks</td><td>493</td><td>513</td><td>21</td><td>GCTCTTTGGACACTAGGAAAA</td><td>139.0</td><td>3.5</td><td>1.3</td><td>2.7</td><td>6.8E-03</td><td>7.8E-03</td><td>MB231/MB468</td><td>MITO</td></tr>
<tr><td>>hg38_21_+_8433172_8446622_RNA45SN1_45S_50ntFlanks@6662.6700.39|GGTGGATCACTCGGCTCGTGCGTCGATGAAGAACGCAGC</td><td>>hg38_21_+_8433172_8446622_RNA45SN1_45S_50ntFlanks</td><td>6662</td><td>6700</td><td>39</td><td>GGTGGATCACTCGGCTCGTGCGTCGATGAAGAACGCAGC</td><td>445.2</td><td>5.9</td><td>1.8</td><td>3.2</td><td>1.4E-03</td><td>7.8E-03</td><td>BT20/MB231</td><td>MP</td></tr>
<tr><td>>MI0000294|hsa-mir-218-1&WithFlank&4|+|20528269|20528390@31.50.20;MI0000295|hsa-mir-218-2&WithFlank&5|-|168768140|168768261@31.50.20&[MIMAT0000275&hsa-miR-218-5p&offsets|0|-1;m-185&4|+|20528299|20528319&offsets|0|-1];[MIMAT0000275_1&hsa-miR-218-5p&offsets|0|-1;m-187&5|-|168768211|168768231&offsets|0|-1]|TTGTGCTTGATCTAACCATG</td><td>>MI0000294|hsa-mir-218-1&WithFlank&4|+|20528269|20528390</td><td>31</td><td>50</td><td>20</td><td>TTGTGCTTGATCTAACCATG</td><td>28.8</td><td>8.1</td><td>2.9</td><td>2.8</td><td>5.9E-03</td><td>7.8E-03</td><td>MB231/MB468</td><td>MP</td></tr>
<tr><td>>hg38_21_+_8433172_8446622_RNA45SN1_45S_50ntFlanks@12625.12661.37|CAATGGGGCGAAGCTACCATCTGTGGGATTATGACTG</td><td>>hg38_21_+_8433172_8446622_RNA45SN1_45S_50ntFlanks</td><td>12625</td><td>12661</td><td>37</td><td>CAATGGGGCGAAGCTACCATCTGTGGGATTATGACTG</td><td>132.9</td><td>-1.0</td><td>0.4</td><td>-2.7</td><td>6.4E-03</td><td>7.9E-03</td><td>BT20/MB468</td><td>CYTO</td></tr>
<tr><td>>hg38_21_+_8433172_8446622_RNA45SN1_45S_50ntFlanks@12936.12962.27|AAACCATTCGTAGACGACCTGCTTCTG</td><td>>hg38_21_+_8433172_8446622_RNA45SN1_45S_50ntFlanks</td><td>12936</td><td>12962</td><td>27</td><td>AAACCATTCGTAGACGACCTGCTTCTG</td><td>137.6</td><td>-2.9</td><td>1.1</td><td>-2.7</td><td>6.4E-03</td><td>7.9E-03</td><td>BT20/MB468</td><td>CYTO</td></tr>
<tr><td>>hg38_21_+_8433172_8446622_RNA45SN1_45S_50ntFlanks@10794.10819.26|CTGGCATGTTGGAACAATGTAGGTAA</td><td>>hg38_21_+_8433172_8446622_RNA45SN1_45S_50ntFlanks</td><td>10794</td><td>10819</td><td>26</td><td>CTGGCATGTTGGAACAATGTAGGTAA</td><td>46.7</td><td>-4.8</td><td>1.6</td><td>-2.9</td><td>3.8E-03</td><td>7.9E-03</td><td>BT20/MB468</td><td>MP</td></tr>
<tr><td>>hg38_21_+_8433172_8446622_RNA45SN1_45S_50ntFlanks@9266.9285.20|CGACGTCGGCTACCCACCCG</td><td>>hg38_21_+_8433172_8446622_RNA45SN1_45S_50ntFlanks</td><td>9266</td><td>9285</td><td>20</td><td>CGACGTCGGCTACCCACCCG</td><td>485.6</td><td>-3.2</td><td>1.2</td><td>-2.7</td><td>6.5E-03</td><td>7.9E-03</td><td>MB231/MB468</td><td>CYTO</td></tr>
<tr><td>>hg38_21_+_8433172_8446622_RNA45SN1_45S_50ntFlanks@11571.11591.21|GCCGACTTAGAACTGGTGCGG</td><td>>hg38_21_+_8433172_8446622_RNA45SN1_45S_50ntFlanks</td><td>11571</td><td>11591</td><td>21</td><td>GCCGACTTAGAACTGGTGCGG</td><td>445.1</td><td>-3.8</td><td>1.3</td><td>-2.9</td><td>3.8E-03</td><td>7.9E-03</td><td>BT20/MB468</td><td>MP</td></tr>
<tr><td>>hg38_21_+_8433172_8446622_RNA45SN1_45S_50ntFlanks@9791.9812.22|TGGCGTGGAGCCGGGCGTGGAA</td><td>>hg38_21_+_8433172_8446622_RNA45SN1_45S_50ntFlanks</td><td>9791</td><td>9812</td><td>22</td><td>TGGCGTGGAGCCGGGCGTGGAA</td><td>133.2</td><td>6.2</td><td>1.9</td><td>3.2</td><td>1.4E-03</td><td>7.9E-03</td><td>BT20/MB231</td><td>MP</td></tr>
<tr><td>>MI0000077|hsa-mir-21&WithFlank&17|+|59841260|59841343@14.32.19&[MIMAT0000076&hsa-miR-21-5p&offsets|0|-3;m-1&17|+|59841273|59841294&offsets|0|-3]|TAGCTTATCAGACTGATGT</td><td>>MI0000077|hsa-mir-21&WithFlank&17|+|59841260|59841343</td><td>14</td><td>32</td><td>19</td><td>TAGCTTATCAGACTGATGT</td><td>1623.0</td><td>3.4</td><td>1.1</td><td>3.1</td><td>1.7E-03</td><td>7.9E-03</td><td>BT20/MB231</td><td>CYTO</td></tr>
<tr><td>>hg38_21_+_8433172_8446622_RNA45SN1_45S_50ntFlanks@6651.6696.46|CGACTCTTAGCGGTGGATCACTCGGCTCGTGCGTCGATGAAGAACG</td><td>>hg38_21_+_8433172_8446622_RNA45SN1_45S_50ntFlanks</td><td>6651</td><td>6696</td><td>46</td><td>CGACTCTTAGCGGTGGATCACTCGGCTCGTGCGTCGATGAAGAACG</td><td>6546.4</td><td>-2.8</td><td>0.9</td><td>-3.1</td><td>1.7E-03</td><td>7.9E-03</td><td>BT20/MB231</td><td>CYTO</td></tr>
<tr><td>>trnaMT_GluTTC_MT_-_14674_14742@39.71.33,trnalookalike8_GluTTC_5_-_93905172_93905240@39.71.33|ATTGGTCGTGGTTGTAGTCCGTGCGAGAATACC</td><td>>trnaMT_GluTTC_MT_-_14674_14742</td><td>39</td><td>71</td><td>33</td><td>ATTGGTCGTGGTTGTAGTCCGTGCGAGAATACC</td><td>2355.8</td><td>-3.3</td><td>1.0</td><td>-3.1</td><td>1.7E-03</td><td>7.9E-03</td><td>BT20/MB231</td><td>CYTO</td></tr>
<tr><td>>hg38_21_+_8433172_8446622_RNA45SN1_45S_50ntFlanks@4007.4042.36|CCTCGGGCCGATCGCACGCCCCCCGTGGCGGCGACG</td><td>>hg38_21_+_8433172_8446622_RNA45SN1_45S_50ntFlanks</td><td>4007</td><td>4042</td><td>36</td><td>CCTCGGGCCGATCGCACGCCCCCCGTGGCGGCGACG</td><td>285.8</td><td>-4.4</td><td>1.6</td><td>-2.7</td><td>6.9E-03</td><td>7.9E-03</td><td>MB231/MB468</td><td>NUC</td></tr>
<tr><td>>hg38_21_+_8433172_8446622_RNA45SN1_45S_50ntFlanks@9916.9947.32|AAAGGTGTTGGTTGATATAGACAGCAGGACGG</td><td>>hg38_21_+_8433172_8446622_RNA45SN1_45S_50ntFlanks</td><td>9916</td><td>9947</td><td>32</td><td>AAAGGTGTTGGTTGATATAGACAGCAGGACGG</td><td>57.2</td><td>5.8</td><td>2.0</td><td>2.9</td><td>3.8E-03</td><td>7.9E-03</td><td>BT20/MB468</td><td>MP</td></tr>
<tr><td>>hg38_21_+_8433172_8446622_RNA45SN1_45S_50ntFlanks@9822.9850.29|CTAGTGGGCCACTTTTGGTAAGCAGAACT</td><td>>hg38_21_+_8433172_8446622_RNA45SN1_45S_50ntFlanks</td><td>9822</td><td>9850</td><td>29</td><td>CTAGTGGGCCACTTTTGGTAAGCAGAACT</td><td>288.5</td><td>-3.2</td><td>1.2</td><td>-2.7</td><td>6.5E-03</td><td>7.9E-03</td><td>MB231/MB468</td><td>CYTO</td></tr>
<tr><td>>MI0000286|hsa-mir-210&WithFlank&11|-|568083|568204@72.92.21(+1U)&[MIMAT0000267&hsa-miR-210-3p&offsets|0|-1(+1U);m-102&11|-|568112|568133&offsets|0|-1(+1U)]|CTGTGCGTGTGACAGCGGCTGT</td><td>>MI0000286|hsa-mir-210&WithFlank&11|-|568083|568204</td><td>72</td><td>92</td><td>21(+1U)</td><td>CTGTGCGTGTGACAGCGGCTGT</td><td>708.8</td><td>6.6</td><td>2.3</td><td>2.9</td><td>3.8E-03</td><td>7.9E-03</td><td>BT20/MB468</td><td>MP</td></tr>
<tr><td>>MI0000095|hsa-mir-93&WithFlank&7|-|100093762|100093853@17.38.22(+1U)&[MIMAT0000093&hsa-miR-93-5p&offsets|0|-1(+1U);m-29&7|-|100093815|100093837&offsets|0|-1(+1U)]|CAAAGTGCTGTTCGTGCAGGTAT</td><td>>MI0000095|hsa-mir-93&WithFlank&7|-|100093762|100093853</td><td>17</td><td>38</td><td>22(+1U)</td><td>CAAAGTGCTGTTCGTGCAGGTAT</td><td>387.3</td><td>3.8</td><td>1.3</td><td>2.9</td><td>3.8E-03</td><td>7.9E-03</td><td>BT20/MB468</td><td>MP</td></tr>
<tr><td>>MI0000270|hsa-mir-181b-1&WithFlank&1|-|198858867|198858988@42.62.21;MI0000683|hsa-mir-181b-2&WithFlank&9|+|124693704|124693804@22.42.21&[MIMAT0000257&hsa-miR-181b-5p&offsets|0|-2;m-73&1|-|198858925|198858947&offsets|0|-2];[MIMAT0000257_1&hsa-miR-181b-5p&offsets|0|-2;m-72&9|+|124693725|124693747&offsets|0|-2]|AACATTCATTGCTGTCGGTGG</td><td>>MI0000270|hsa-mir-181b-1&WithFlank&1|-|198858867|198858988</td><td>42</td><td>62</td><td>21</td><td>AACATTCATTGCTGTCGGTGG</td><td>321.3</td><td>2.5</td><td>0.9</td><td>2.8</td><td>5.9E-03</td><td>7.9E-03</td><td>MB231/MB468</td><td>MP</td></tr>
<tr><td>>hg38_21_+_8433172_8446622_RNA45SN1_45S_50ntFlanks@12456.12493.38|CCAAGCGTTGGATTGTTCACCCACTAATAGGGAACGTG</td><td>>hg38_21_+_8433172_8446622_RNA45SN1_45S_50ntFlanks</td><td>12456</td><td>12493</td><td>38</td><td>CCAAGCGTTGGATTGTTCACCCACTAATAGGGAACGTG</td><td>488.3</td><td>-2.7</td><td>0.9</td><td>-2.9</td><td>3.8E-03</td><td>7.9E-03</td><td>BT20/MB468</td><td>MP</td></tr>
<tr><td>>hg38_21_+_8433172_8446622_RNA45SN1_45S_50ntFlanks@7992.8017.26|CGTGGCGACCCGCTGAATTTAAGCAT</td><td>>hg38_21_+_8433172_8446622_RNA45SN1_45S_50ntFlanks</td><td>7992</td><td>8017</td><td>26</td><td>CGTGGCGACCCGCTGAATTTAAGCAT</td><td>196.7</td><td>-3.6</td><td>1.2</td><td>-3.1</td><td>1.7E-03</td><td>7.9E-03</td><td>BT20/MB231</td><td>CYTO</td></tr>
<tr><td>>hg38_21_+_8433172_8446622_RNA45SN1_45S_50ntFlanks@10186.10208.23|TCCTCCCCTCCTCCCGCCCACGC</td><td>>hg38_21_+_8433172_8446622_RNA45SN1_45S_50ntFlanks</td><td>10186</td><td>10208</td><td>23</td><td>TCCTCCCCTCCTCCCGCCCACGC</td><td>96.5</td><td>-4.0</td><td>1.5</td><td>-2.8</td><td>5.5E-03</td><td>7.9E-03</td><td>BT20/MB468</td><td>MITO</td></tr>
<tr><td>>hg38_21_+_8433172_8446622_RNA45SN1_45S_50ntFlanks@8352.8378.27|AGTTGAAAAGAACTTTGAAGAGAGAGT</td><td>>hg38_21_+_8433172_8446622_RNA45SN1_45S_50ntFlanks</td><td>8352</td><td>8378</td><td>27</td><td>AGTTGAAAAGAACTTTGAAGAGAGAGT</td><td>184.8</td><td>2.5</td><td>0.9</td><td>2.9</td><td>3.8E-03</td><td>7.9E-03</td><td>BT20/MB468</td><td>MP</td></tr>
<tr><td>>trnaMT_PheGAA_MT_+_577_647@35.55.21|GAAAATGTTTAGACGGGCTCA</td><td>>trnaMT_PheGAA_MT_+_577_647</td><td>35</td><td>55</td><td>21</td><td>GAAAATGTTTAGACGGGCTCA</td><td>2081.5</td><td>2.1</td><td>0.7</td><td>3.1</td><td>1.8E-03</td><td>7.9E-03</td><td>BT20/MB231</td><td>MITO</td></tr>
</table>

<table>
<tr><th></th><th></th><th></th><th></th><th></th><th></th><th></th><th></th><th></th><th></th><th></th><th></th><th></th><th></th></tr>
<tr><td>>MI0000459|hsa-mir-143&WithFlank&5|+|149428912|149429029@67.87.21&[MIMAT0000435&hsa-miR-143-3p&offsets|0|0;m-3&5|+|149428978|149428997&offsets|0|+1]|TGAGATGAAGCACTGTAGCTC</td><td>>MI0000459|hsa-mir-143&WithFlank&5|+|149428912|149429029</td><td>67</td><td>87</td><td>21</td><td>TGAGATGAAGCACTGTAGCTC</td><td>482.0</td><td>-5.0</td><td>1.7</td><td>-2.9</td><td>3.8E-03</td><td>7.9E-03</td><td>BT20/MB468</td><td>MP</td></tr>
<tr><td>>MI0000082|hsa-mir-25&WithFlank&7|-|100093554|100093649@58.80.23&[MIMAT0000081&hsa-miR-25-3p&offsets|0|+1;m-23&7|-|100093571|100093592&offsets|0|+1]|CATTGCACTTGTCTCGGTCTGAC</td><td>>MI0000082|hsa-mir-25&WithFlank&7|-|100093554|100093649</td><td>58</td><td>80</td><td>23</td><td>CATTGCACTTGTCTCGGTCTGAC</td><td>73.8</td><td>3.7</td><td>1.3</td><td>2.7</td><td>6.0E-03</td><td>7.9E-03</td><td>MB231/MB468</td><td>MP</td></tr>
<tr><td>>hg38_21_+_8433172_8446622_RNA45SN1_45S_50ntFlanks@7975.8007.33|CGCGACCTCAGATCAGACGTGGCGACCCGCTGA</td><td>>hg38_21_+_8433172_8446622_RNA45SN1_45S_50ntFlanks</td><td>7975</td><td>8007</td><td>33</td><td>CGCGACCTCAGATCAGACGTGGCGACCCGCTGA</td><td>3501.3</td><td>-2.0</td><td>0.7</td><td>-2.7</td><td>6.0E-03</td><td>7.9E-03</td><td>MB231/MB468</td><td>MP</td></tr>
<tr><td>>hg38_21_+_8433172_8446622_RNA45SN1_45S_50ntFlanks@8381.8404.24|AAGAGGGCGTGAAACCGTTAAGAG</td><td>>hg38_21_+_8433172_8446622_RNA45SN1_45S_50ntFlanks</td><td>8381</td><td>8404</td><td>24</td><td>AAGAGGGCGTGAAACCGTTAAGAG</td><td>383.2</td><td>-4.9</td><td>1.7</td><td>-2.9</td><td>3.8E-03</td><td>7.9E-03</td><td>BT20/MB468</td><td>MP</td></tr>
<tr><td>>hg38_21_+_8433172_8446622_RNA45SN1_45S_50ntFlanks@8169.8212.44|CCAAGTCCTTCTGATCGAGGCCCAGCCCGTGGACGGTGTGAGGC</td><td>>hg38_21_+_8433172_8446622_RNA45SN1_45S_50ntFlanks</td><td>8169</td><td>8212</td><td>44</td><td>CCAAGTCCTTCTGATCGAGGCCCAGCCCGTGGACGGTGTGAGGC</td><td>139.6</td><td>-2.5</td><td>0.9</td><td>-2.7</td><td>6.6E-03</td><td>7.9E-03</td><td>MB231/MB468</td><td>CYTO</td></tr>
<tr><td>>am_MI0000786|hsa-mir-378a&WithFlank&5|+|149732819|149732896@49.70.22(+1U)&[MIMAT0000732&hsa-miR-378a-3p&offsets|0|0(+1U);m-44&5|+|149732867|149732888&offsets|0|0(+1U)]|ACTGGACTTGGAGTCAGAAGGCT</td><td>>am_MI0000786|hsa-mir-378a&WithFlank&5|+|149732819|149732896</td><td>49</td><td>70</td><td>22(+1U)</td><td>ACTGGACTTGGAGTCAGAAGGCT</td><td>771.3</td><td>2.5</td><td>0.9</td><td>2.7</td><td>6.9E-03</td><td>7.9E-03</td><td>MB231/MB468</td><td>MITO</td></tr>
<tr><td>>MI0000102|hsa-mir-100&WithFlank&11|-|122152223|122152314@22.40.19(+1U)&[MIMAT0000098&hsa-miR-100-5p&offsets|+3|0(+1U);m-27&11|-|122152276|122152296&offsets|+3|+1(+1U)]|CCGTAGATCCGAACTTGTGT</td><td>>MI0000102|hsa-mir-100&WithFlank&11|-|122152223|122152314</td><td>22</td><td>40</td><td>19(+1U)</td><td>CCGTAGATCCGAACTTGTGT</td><td>28.3</td><td>8.0</td><td>2.9</td><td>2.7</td><td>6.0E-03</td><td>7.9E-03</td><td>MB231/MB468</td><td>MP</td></tr>
<tr><td>>MI0000089|hsa-mir-31&WithFlank&9|-|21512109|21512191@14.34.21(+1A)&[MIMAT0000089&hsa-miR-31-5p&offsets|0|0(+1A);m-360&9|-|21512157|21512178&offsets|0|-1(+1A)]|AGGCAAGATGCTGGCATAGCTA</td><td>>MI0000089|hsa-mir-31&WithFlank&9|-|21512109|21512191</td><td>14</td><td>34</td><td>21(+1A)</td><td>AGGCAAGATGCTGGCATAGCTA</td><td>52.8</td><td>9.1</td><td>2.8</td><td>3.2</td><td>1.4E-03</td><td>7.9E-03</td><td>BT20/MB231</td><td>MP</td></tr>
<tr><td>>hg38_21_+_8433172_8446622_RNA45SN1_45S_50ntFlanks@8343.8365.23|GTAAGGGAAAGTTGAAAAGAACT</td><td>>hg38_21_+_8433172_8446622_RNA45SN1_45S_50ntFlanks</td><td>8343</td><td>8365</td><td>23</td><td>GTAAGGGAAAGTTGAAAAGAACT</td><td>26.6</td><td>8.0</td><td>2.9</td><td>2.7</td><td>6.0E-03</td><td>7.9E-03</td><td>MB231/MB468</td><td>MP</td></tr>
<tr><td>>hg38_21_+_8433172_8446622_RNA45SN1_45S_50ntFlanks@12365.12382.18|ATAACTGGCTTGTGGCGG</td><td>>hg38_21_+_8433172_8446622_RNA45SN1_45S_50ntFlanks</td><td>12365</td><td>12382</td><td>18</td><td>ATAACTGGCTTGTGGCGG</td><td>381.5</td><td>-4.3</td><td>1.6</td><td>-2.8</td><td>5.6E-03</td><td>7.9E-03</td><td>BT20/MB468</td><td>MITO</td></tr>
<tr><td>>hg38_21_+_8433172_8446622_RNA45SN1_45S_50ntFlanks@11575.11611.37|ACTTAGAACTGGTGCGGACCAGGGGAATCCGACTGTT</td><td>>hg38_21_+_8433172_8446622_RNA45SN1_45S_50ntFlanks</td><td>11575</td><td>11611</td><td>37</td><td>ACTTAGAACTGGTGCGGACCAGGGGAATCCGACTGTT</td><td>776.5</td><td>-3.0</td><td>1.0</td><td>-2.9</td><td>3.8E-03</td><td>7.9E-03</td><td>BT20/MB468</td><td>MP</td></tr>
<tr><td>>hg38_21_+_8433172_8446622_RNA45SN1_45S_50ntFlanks@11778.11803.26|ACGCGCATGAATGGATGAACGAGATT</td><td>>hg38_21_+_8433172_8446622_RNA45SN1_45S_50ntFlanks</td><td>11778</td><td>11803</td><td>26</td><td>ACGCGCATGAATGGATGAACGAGATT</td><td>849.6</td><td>-3.9</td><td>1.4</td><td>-2.9</td><td>3.8E-03</td><td>7.9E-03</td><td>BT20/MB468</td><td>MP</td></tr>
<tr><td>>hg38_21_+_8433172_8446622_RNA45SN1_45S_50ntFlanks@11805.11830.26|CCACTGTCCCTACCTACTATCCAGCG</td><td>>hg38_21_+_8433172_8446622_RNA45SN1_45S_50ntFlanks</td><td>11805</td><td>11830</td><td>26</td><td>CCACTGTCCCTACCTACTATCCAGCG</td><td>77.5</td><td>-4.8</td><td>1.6</td><td>-2.9</td><td>3.8E-03</td><td>7.9E-03</td><td>BT20/MB468</td><td>MP</td></tr>
<tr><td>>hg38_21_+_8433172_8446622_RNA45SN1_45S_50ntFlanks@8174.8203.30|TCCTTCTGATCGAGGCCCAGCCCGTGGACG</td><td>>hg38_21_+_8433172_8446622_RNA45SN1_45S_50ntFlanks</td><td>8174</td><td>8203</td><td>30</td><td>TCCTTCTGATCGAGGCCCAGCCCGTGGACG</td><td>365.9</td><td>1.8</td><td>0.7</td><td>2.8</td><td>5.6E-03</td><td>7.9E-03</td><td>BT20/MB468</td><td>MITO</td></tr>
<tr><td>>hg38_21_+_8433172_8446622_RNA45SN1_45S_50ntFlanks@5493.5525.33|GAAGACGGTCGAACTTGACTATCTAGAGGAAGT</td><td>>hg38_21_+_8433172_8446622_RNA45SN1_45S_50ntFlanks</td><td>5493</td><td>5525</td><td>33</td><td>GAAGACGGTCGAACTTGACTATCTAGAGGAAGT</td><td>560.9</td><td>5.7</td><td>1.8</td><td>3.2</td><td>1.4E-03</td><td>8.0E-03</td><td>BT20/MB231</td><td>MP</td></tr>
<tr><td>>hg38_21_+_8433172_8446622_RNA45SN1_45S_50ntFlanks@12407.12429.23|TTTTGATCCTTCGATGTCGGCTC</td><td>>hg38_21_+_8433172_8446622_RNA45SN1_45S_50ntFlanks</td><td>12407</td><td>12429</td><td>23</td><td>TTTTGATCCTTCGATGTCGGCTC</td><td>259.7</td><td>-1.5</td><td>0.6</td><td>-2.7</td><td>7.0E-03</td><td>8.0E-03</td><td>MB231/MB468</td><td>NUC</td></tr>
<tr><td>>hg38_21_+_8433172_8446622_RNA45SN1_45S_50ntFlanks@6663.6699.37|GTGGATCACTCGGCTCGTGCGTCGATGAAGAACGCAG</td><td>>hg38_21_+_8433172_8446622_RNA45SN1_45S_50ntFlanks</td><td>6663</td><td>6699</td><td>37</td><td>GTGGATCACTCGGCTCGTGCGTCGATGAAGAACGCAG</td><td>179.0</td><td>-1.9</td><td>0.7</td><td>-2.7</td><td>6.6E-03</td><td>8.0E-03</td><td>MB231/MB468</td><td>CYTO</td></tr>
<tr><td>>hg38_21_+_8433172_8446622_RNA45SN1_45S_50ntFlanks@12640.12659.20|ACCATCTGTGGGATTATGAC</td><td>>hg38_21_+_8433172_8446622_RNA45SN1_45S_50ntFlanks</td><td>12640</td><td>12659</td><td>20</td><td>ACCATCTGTGGGATTATGAC</td><td>1223.5</td><td>-3.4</td><td>1.2</td><td>-2.9</td><td>3.8E-03</td><td>8.0E-03</td><td>BT20/MB468</td><td>MP</td></tr>
<tr><td>>am_MI0000786|hsa-mir-378a&WithFlank&5|+|149732819|149732896@50.70.21&[MIMAT0000732&hsa-miR-378a-3p&offsets|+1|0;m-44&5|+|149732867|149732888&offsets|+1|0]|CTGGACTTGGAGTCAGAAGGC</td><td>>am_MI0000786|hsa-mir-378a&WithFlank&5|+|149732819|149732896</td><td>50</td><td>70</td><td>21</td><td>CTGGACTTGGAGTCAGAAGGC</td><td>596.1</td><td>2.3</td><td>0.9</td><td>2.7</td><td>7.0E-03</td><td>8.0E-03</td><td>MB231/MB468</td><td>MITO</td></tr>
<tr><td>>hg38_21_+_8433172_8446622_RNA45SN1_45S_50ntFlanks@7973.8005.33|GACGCGACCTCAGATCAGACGTGGCGACCCGCT</td><td>>hg38_21_+_8433172_8446622_RNA45SN1_45S_50ntFlanks</td><td>7973</td><td>8005</td><td>33</td><td>GACGCGACCTCAGATCAGACGTGGCGACCCGCT</td><td>916.3</td><td>2.0</td><td>0.7</td><td>2.7</td><td>7.0E-03</td><td>8.0E-03</td><td>MB231/MB468</td><td>MITO</td></tr>
<tr><td>>trnaMT_ThrTGT_MT_+_15888_15953@34.69.36|TAAACCGGAGATGAAAACCTTTTTCCAAGGACACCA</td><td>>trnaMT_ThrTGT_MT_+_15888_15953</td><td>34</td><td>69</td><td>36</td><td>TAAACCGGAGATGAAAACCTTTTTCCAAGGACACCA</td><td>233.1</td><td>4.6</td><td>1.6</td><td>2.9</td><td>3.8E-03</td><td>8.0E-03</td><td>BT20/MB468</td><td>MP</td></tr>
<tr><td>>trnaMT_ValTAC_MT_+_1602_1670@36.71.36|CTTAGGAGATTTCAACTTAACTTGACCGCTCTGACC</td><td>>trnaMT_ValTAC_MT_+_1602_1670</td><td>36</td><td>71</td><td>36</td><td>CTTAGGAGATTTCAACTTAACTTGACCGCTCTGACC</td><td>222.0</td><td>6.4</td><td>2.2</td><td>2.9</td><td>3.8E-03</td><td>8.0E-03</td><td>BT20/MB468</td><td>MP</td></tr>
<tr><td>>hg38_1_-_228634819_228635039_RNA5S12_5S_50ntFlanks@147.170.24|GGGAATACCGGGTGCTGTAGGCTT</td><td>>hg38_1_-_228634819_228635039_RNA5S12_5S_50ntFlanks</td><td>147</td><td>170</td><td>24</td><td>GGGAATACCGGGTGCTGTAGGCTT</td><td>52685.6</td><td>4.6</td><td>1.5</td><td>3.2</td><td>1.4E-03</td><td>8.0E-03</td><td>BT20/MB231</td><td>MP</td></tr>
<tr><td>>MI0000301|hsa-mir-224&WithFlank&X|-|151958572|151958664@14.36.23&[MIMAT0000281&hsa-miR-224-5p&offsets|+1|-1;m-238&X|-|151958630|151958651&offsets|0|+1]|CAAGTCACTAGTGGTTCCGTTTA</td><td>>MI0000301|hsa-mir-224&WithFlank&X|-|151958572|151958664</td><td>14</td><td>36</td><td>23</td><td>CAAGTCACTAGTGGTTCCGTTTA</td><td>1531.2</td><td>2.4</td><td>0.8</td><td>3.2</td><td>1.5E-03</td><td>8.0E-03</td><td>BT20/MB231</td><td>NUC</td></tr>
<tr><td>>trnaMT_GluTTC_MT_-_14674_14742@34.72.39,trnalookalike8_GluTTC_5_-_93905172_93905240@34.72.39|ATATCATTGGTCGTGGTTGTAGTCCGTGCGAGAATACCA</td><td>>trnaMT_GluTTC_MT_-_14674_14742</td><td>34</td><td>72</td><td>39</td><td>ATATCATTGGTCGTGGTTGTAGTCCGTGCGAGAATACCA</td><td>2374.4</td><td>-2.3</td><td>0.7</td><td>-3.2</td><td>1.5E-03</td><td>8.0E-03</td><td>BT20/MB231</td><td>NUC</td></tr>
<tr><td>>hg38_21_+_8433172_8446622_RNA45SN1_45S_50ntFlanks@11920.11940.21|ATGAGAGGTGTAGAATAAGTG</td><td>>hg38_21_+_8433172_8446622_RNA45SN1_45S_50ntFlanks</td><td>11920</td><td>11940</td><td>21</td><td>ATGAGAGGTGTAGAATAAGTG</td><td>102.4</td><td>5.0</td><td>1.7</td><td>2.9</td><td>3.8E-03</td><td>8.0E-03</td><td>BT20/MB468</td><td>MP</td></tr>
<tr><td>>trnaMT_SerGCT_MT_+_12207_12265@1.26.26|GAGAAAGCTCACAAGAACTGCTAACT</td><td>>trnaMT_SerGCT_MT_+_12207_12265</td><td>1</td><td>26</td><td>26</td><td>GAGAAAGCTCACAAGAACTGCTAACT</td><td>24807.5</td><td>2.4</td><td>0.9</td><td>2.7</td><td>6.0E-03</td><td>8.0E-03</td><td>MB231/MB468</td><td>MP</td></tr>
<tr><td>>hg38_21_+_8433172_8446622_RNA45SN1_45S_50ntFlanks@11845.11872.28|GGAACGGGCTTGGCGGAATCAGCGGGGA</td><td>>hg38_21_+_8433172_8446622_RNA45SN1_45S_50ntFlanks</td><td>11845</td><td>11872</td><td>28</td><td>GGAACGGGCTTGGCGGAATCAGCGGGGA</td><td>199.3</td><td>-4.1</td><td>1.5</td><td>-2.7</td><td>6.5E-03</td><td>8.0E-03</td><td>BT20/MB468</td><td>CYTO</td></tr>
<tr><td>>MI0000447|hsa-mir-128-1&WithFlank&2|+|135665391|135665484@56.76.21(+2A);MI0000727|hsa-mir-128-2&WithFlank&3|+|35744470|35744565@58.78.21(+2A)&[MIMAT0000424&hsa-miR-128-3p&offsets|0|0(+2A);m-108&2|+|135665446|135665466&offsets|0|0(+2A)];[MIMAT0000424_1&hsa-miR-128-3p&offsets|0|0(+2A);m-117&3|+|35744527|35744547&offsets|0|0(+2A)]|TCACAGTGAACCGGTCTCTTTAA</td><td>>MI0000447|hsa-mir-128-1&WithFlank&2|+|135665391|135665484</td><td>56</td><td>76</td><td>21(+2A)</td><td>TCACAGTGAACCGGTCTCTTTAA</td><td>580.8</td><td>2.0</td><td>0.7</td><td>2.7</td><td>7.0E-03</td><td>8.0E-03</td><td>MB231/MB468</td><td>NUC</td></tr>
<tr><td>>hg38_21_+_8433172_8446622_RNA45SN1_45S_50ntFlanks@10678.10711.34|CGGTTCCGGCGGCGTCCGGTGAGCTCTCGCTGGC</td><td>>hg38_21_+_8433172_8446622_RNA45SN1_45S_50ntFlanks</td><td>10678</td><td>10711</td><td>34</td><td>CGGTTCCGGCGGCGTCCGGTGAGCTCTCGCTGGC</td><td>835.7</td><td>7.1</td><td>2.2</td><td>3.2</td><td>1.4E-03</td><td>8.0E-03</td><td>BT20/MB231</td><td>MP</td></tr>
<tr><td>>am_trnaMT_ValTAC_MT_+_1602_1670@36.67.32|CTTAGGAGATTTCAACTTAACTTGACCGCTCT</td><td>>am_trnaMT_ValTAC_MT_+_1602_1670</td><td>36</td><td>67</td><td>32</td><td>CTTAGGAGATTTCAACTTAACTTGACCGCTCT</td><td>368.2</td><td>6.6</td><td>2.3</td><td>2.9</td><td>3.8E-03</td><td>8.0E-03</td><td>BT20/MB468</td><td>MP</td></tr>
<tr><td>>hg38_21_+_8433172_8446622_RNA45SN1_45S_50ntFlanks@4724.4755.32|ATTAATCAAGAACGAAAGTCGGAGGTTCGAAG</td><td>>hg38_21_+_8433172_8446622_RNA45SN1_45S_50ntFlanks</td><td>4724</td><td>4755</td><td>32</td><td>ATTAATCAAGAACGAAAGTCGGAGGTTCGAAG</td><td>162.3</td><td>-4.7</td><td>1.6</td><td>-2.9</td><td>3.8E-03</td><td>8.0E-03</td><td>BT20/MB468</td><td>MP</td></tr>
<tr><td>>hg38_21_+_8433172_8446622_RNA45SN1_45S_50ntFlanks@8345.8365.21|AAGGGAAAGTTGAAAAGAACT</td><td>>hg38_21_+_8433172_8446622_RNA45SN1_45S_50ntFlanks</td><td>8345</td><td>8365</td><td>21</td><td>AAGGGAAAGTTGAAAAGAACT</td><td>48.0</td><td>5.2</td><td>1.9</td><td>2.7</td><td>6.0E-03</td><td>8.0E-03</td><td>MB231/MB468</td><td>MP</td></tr>
<tr><td>>MI0005561|hsa-mir-877&WithFlank&6|+|30584326|30584423@7.28.22(+1U)&[MIMAT0004949&hsa-miR-877-5p&offsets|0|+2(+1U);m-382&6|+|30584332|30584354&offsets|0|-1(+1U)]|GTAGAGGAGATGGCGCAGGGGAT</td><td>>MI0005561|hsa-mir-877&WithFlank&6|+|30584326|30584423</td><td>7</td><td>28</td><td>22(+1U)</td><td>GTAGAGGAGATGGCGCAGGGGAT</td><td>44.3</td><td>4.4</td><td>1.6</td><td>2.7</td><td>6.0E-03</td><td>8.0E-03</td><td>MB231/MB468</td><td>MP</td></tr>
<tr><td>>hg38_21_+_8433172_8446622_RNA45SN1_45S_50ntFlanks@12993.13039.47|CCCTCGCTGCGATCTATTGAAAGTCAGCCCTCGACACAAGGGTTTGT</td><td>>hg38_21_+_8433172_8446622_RNA45SN1_45S_50ntFlanks</td><td>12993</td><td>13039</td><td>47</td><td>CCCTCGCTGCGATCTATTGAAAGTCAGCCCTCGACACAAGGGTTTGT</td><td>2019.4</td><td>3.3</td><td>1.2</td><td>2.7</td><td>6.5E-03</td><td>8.0E-03</td><td>BT20/MB468</td><td>CYTO</td></tr>
<tr><td>>hg38_MT_+_1621_3279_RNR2_16S_50ntFlanks@1380.1404.25|AGGACATCCCGATGGTGCAGCCGCT</td><td>>hg38_MT_+_1621_3279_RNR2_16S_50ntFlanks</td><td>1380</td><td>1404</td><td>25</td><td>AGGACATCCCGATGGTGCAGCCGCT</td><td>97.4</td><td>4.5</td><td>1.6</td><td>2.7</td><td>6.0E-03</td><td>8.0E-03</td><td>MB231/MB468</td><td>MP</td></tr>
<tr><td>>am_trna2_GlyGCC_21_-_18827107_18827177@1.24.24,trna35_GlyGCC_1_+_161413094_161413164@1.24.24,trna37_GlyGCC_1_+_161420467_161420537@1.24.24,trna39_GlyGCC_1_+_161427898_161427968@1.24.24,trna41_GlyGCC_1_+_161435258_161435328@1.24.24|GCATGGGTGGTTCAGTGGTAGAAT</td><td>>am_trna2_GlyGCC_21_-_18827107_18827177</td><td>1</td><td>24</td><td>24</td><td>GCATGGGTGGTTCAGTGGTAGAAT</td><td>182.5</td><td>3.2</td><td>1.1</td><td>2.9</td><td>3.9E-03</td><td>8.0E-03</td><td>BT20/MB468</td><td>MP</td></tr>
<tr><td>>hg38_21_+_8433172_8446622_RNA45SN1_45S_50ntFlanks@475.493.19|CTTCGTGATCGATGTGGTG</td><td>>hg38_21_+_8433172_8446622_RNA45SN1_45S_50ntFlanks</td><td>475</td><td>493</td><td>19</td><td>CTTCGTGATCGATGTGGTG</td><td>109.8</td><td>5.1</td><td>1.8</td><td>2.9</td><td>3.9E-03</td><td>8.0E-03</td><td>BT20/MB468</td><td>MP</td></tr>
<tr><td>>hg38_MT_+_598_1651_RNR1_12S_50ntFlanks@608.637.30|TGTTCTGTAATCGATAAACCCCGATCAACC</td><td>>hg38_MT_+_598_1651_RNR1_12S_50ntFlanks</td><td>608</td><td>637</td><td>30</td><td>TGTTCTGTAATCGATAAACCCCGATCAACC</td><td>55.3</td><td>5.3</td><td>1.9</td><td>2.7</td><td>6.0E-03</td><td>8.0E-03</td><td>MB231/MB468</td><td>MP</td></tr>
<tr><td>>hg38_21_+_8433172_8446622_RNA45SN1_45S_50ntFlanks@12418.12454.37|CGATGTCGGCTCTTCCTATCATTGTGAAGCAGAATTC</td><td>>hg38_21_+_8433172_8446622_RNA45SN1_45S_50ntFlanks</td><td>12418</td><td>12454</td><td>37</td><td>CGATGTCGGCTCTTCCTATCATTGTGAAGCAGAATTC</td><td>312.9</td><td>-1.8</td><td>0.7</td><td>-2.7</td><td>6.5E-03</td><td>8.0E-03</td><td>BT20/MB468</td><td>CYTO</td></tr>
<tr><td>>hg38_21_+_8433172_8446622_RNA45SN1_45S_50ntFlanks@5485.5519.35|AGCGCTGAGAAGACGGTCGAACTTGACTATCTAGA</td><td>>hg38_21_+_8433172_8446622_RNA45SN1_45S_50ntFlanks</td><td>5485</td><td>5519</td><td>35</td><td>AGCGCTGAGAAGACGGTCGAACTTGACTATCTAGA</td><td>310.0</td><td>2.9</td><td>0.9</td><td>3.1</td><td>1.8E-03</td><td>8.0E-03</td><td>BT20/MB231</td><td>CYTO</td></tr>
<tr><td>>hg38_21_+_8433172_8446622_RNA45SN1_45S_50ntFlanks@10513.10532.20|CCGGAGTGGCGGAGATGGGC</td><td>>hg38_21_+_8433172_8446622_RNA45SN1_45S_50ntFlanks</td><td>10513</td><td>10532</td><td>20</td><td>CCGGAGTGGCGGAGATGGGC</td><td>964.3</td><td>5.6</td><td>1.8</td><td>3.2</td><td>1.4E-03</td><td>8.0E-03</td><td>BT20/MB231</td><td>MP</td></tr>
<tr><td>>hg38_21_+_8433172_8446622_RNA45SN1_45S_50ntFlanks@11917.11942.26|GACATGAGAGGTGTAGAATAAGTGGG</td><td>>hg38_21_+_8433172_8446622_RNA45SN1_45S_50ntFlanks</td><td>11917</td><td>11942</td><td>26</td><td>GACATGAGAGGTGTAGAATAAGTGGG</td><td>60.0</td><td>9.2</td><td>2.9</td><td>3.2</td><td>1.4E-03</td><td>8.0E-03</td><td>BT20/MB231</td><td>MP</td></tr>
<tr><td>>hg38_21_+_8433172_8446622_RNA45SN1_45S_50ntFlanks@4727.4759.33|AATCAAGAACGAAAGTCGGAGGTTCGAAGACGA</td><td>>hg38_21_+_8433172_8446622_RNA45SN1_45S_50ntFlanks</td><td>4727</td><td>4759</td><td>33</td><td>AATCAAGAACGAAAGTCGGAGGTTCGAAGACGA</td><td>1000.1</td><td>4.7</td><td>1.5</td><td>3.2</td><td>1.4E-03</td><td>8.0E-03</td><td>BT20/MB231</td><td>MP</td></tr>
<tr><td>>hg38_21_+_8433172_8446622_RNA45SN1_45S_50ntFlanks@12420.12449.30|ATGTCGGCTCTTCCTATCATTGTGAAGCAG</td><td>>hg38_21_+_8433172_8446622_RNA45SN1_45S_50ntFlanks</td><td>12420</td><td>12449</td><td>30</td><td>ATGTCGGCTCTTCCTATCATTGTGAAGCAG</td><td>1234.5</td><td>-1.7</td><td>0.6</td><td>-2.7</td><td>7.0E-03</td><td>8.0E-03</td><td>MB231/MB468</td><td>NUC</td></tr>
<tr><td>>hg38_21_+_8433172_8446622_RNA45SN1_45S_50ntFlanks@11781.11806.26|CGCATGAATGGATGAACGAGATTCCC</td><td>>hg38_21_+_8433172_8446622_RNA45SN1_45S_50ntFlanks</td><td>11781</td><td>11806</td><td>26</td><td>CGCATGAATGGATGAACGAGATTCCC</td><td>824.5</td><td>-4.0</td><td>1.4</td><td>-2.9</td><td>3.9E-03</td><td>8.0E-03</td><td>BT20/MB468</td><td>MP</td></tr>
<tr><td>>hg38_21_+_8433172_8446622_RNA45SN1_45S_50ntFlanks@6761.6807.47|TGCGGCCCCGGGTTCCTCCCGGGGCTACGCCTGTCTGAGCGTCGCTT</td><td>>hg38_21_+_8433172_8446622_RNA45SN1_45S_50ntFlanks</td><td>6761</td><td>6807</td><td>47</td><td>TGCGGCCCCGGGTTCCTCCCGGGGCTACGCCTGTCTGAGCGTCGCTT</td><td>94.2</td><td>3.3</td><td>1.2</td><td>2.7</td><td>7.0E-03</td><td>8.0E-03</td><td>MB231/MB468</td><td>MITO</td></tr>
<tr><td>>hg38_21_+_8433172_8446622_RNA45SN1_45S_50ntFlanks@4045.4073.29|CCATTCGAACGTCTGCCCTATCAACTTTC</td><td>>hg38_21_+_8433172_8446622_RNA45SN1_45S_50ntFlanks</td><td>4045</td><td>4073</td><td>29</td><td>CCATTCGAACGTCTGCCCTATCAACTTTC</td><td>264.0</td><td>1.8</td><td>0.7</td><td>2.7</td><td>6.6E-03</td><td>8.0E-03</td><td>MB231/MB468</td><td>CYTO</td></tr>
<tr><td>>hg38_21_+_8433172_8446622_RNA45SN1_45S_50ntFlanks@12004.12026.23|CGGGCCGCCGGTGAAATACCACT</td><td>>hg38_21_+_8433172_8446622_RNA45SN1_45S_50ntFlanks</td><td>12004</td><td>12026</td><td>23</td><td>CGGGCCGCCGGTGAAATACCACT</td><td>258.4</td><td>5.0</td><td>1.7</td><td>2.9</td><td>3.9E-03</td><td>8.0E-03</td><td>BT20/MB468</td><td>MP</td></tr>
</table>

| | | | | | | | | | | | | | |
|---|---|---|---|---|---|---|---|---|---|---|---|---|---|
| >hg38_21_+_8433172_8446622_RNA45SN1_45S_50ntFlanks@11672.11702.31\|CAGTGCTCTGAATGTCAAAGTGAAGAAATTC | >hg38_21_+_8433172_8446622_RNA45SN1_45S_50ntFlanks | 11672 | 11702 | 31 | CAGTGCTCTGAATGTCAAAGTGAAGAAATTC | 326.6 | 1.3 | 0.5 | 2.8 | 5.6E-03 | 8.0E-03 | BT20/MB468 | MITO |
| >MI0000114\|hsa-mir-107&WithFlank&10\|-\|89592741\|89592833@56.76.21(+1U);MI0000108\|hsa-mir-103a-2&WithFlank&20\|+\|3917488\|3917577@54.74.21(+1U);MI0000109\|hsa-mir-103a-1&WithFlank&5\|-\|168560890\|168560979@54.74.21(+1U)&[MIMAT0000104&hsa-miR-107&offsets\|0\|-2(+1U);m-78&10\|-\|89592758\|89592778&offsets\|0\|0(+1U)];[MIMAT0000101&hsa-miR-103a-3p&offsets\|0\|-2(+1U);m-12&20\|+\|3917541\|3917563&offsets\|0\|-2(+1U)];[MIMAT0000101_1&hsa-miR-103a-3p&offsets\|0\|-2(+1U);m-13&5\|-\|168560904\|168560926&offsets\|0\|-2(+1U);m-44220&5\|-\|168560908\|168560926&offsets\|0\|+2(+1U)]\|AGCAGCATTGTACAGGGCTATT | >MI0000114\|hsa-mir-107&WithFlank&10\|-\|89592741\|89592833 | 56 | 76 | 21(+1U) | AGCAGCATTGTACAGGGCTATT | 767.8 | -4.6 | 1.4 | -3.2 | 1.4E-03 | 8.0E-03 | BT20/MB231 | MP |
| >MI0000809\|hsa-mir-151a&WithFlank&8\|-\|140732558\|140732659@51.73.23(+1U)&[MIMAT0000757&hsa-miR-151a-3p&offsets\|-2\|0(+1U);m-30&8\|-\|140732587\|140732607&offsets\|-2\|0(+1U)]\|TACTAGACTGAAGCTCCTTGAGGT | >MI0000809\|hsa-mir-151a&WithFlank&8\|-\|140732558\|140732659 | 51 | 73 | 23(+1U) | TACTAGACTGAAGCTCCTTGAGGT | 64.9 | 4.1 | 1.4 | 2.9 | 3.9E-03 | 8.0E-03 | BT20/MB468 | MP |
| >trna111_HisGTG_1_-_147774845_147774916@-1G.31.32,trna118_HisGTG_1_-_145396881_145396952@-1G.31.32,trna16_HisGTG_1_+_146544773_146544844@-1G.31.32,trna1_HisGTG_15_+_45493349_45493420@-1G.31.32,trna21_HisGTG_1_+_147753471_147753542@-1G.31.32,trna33_HisGTG_6_+_27125906_27125977@-1G.31.32,trna7_HisGTG_9_-_14433938_14434009@-1G.31.32,trna8_HisGTG_15_-_45492611_45492682@-1G.31.32,trna9_HisGTG_15_-_45490804_45490875@-1G.31.32\|GGCCGTGATCGTATAGTGGTTAGTACTCTGCG | >trna111_HisGTG_1_-_147774845_147774916 | -1G | 31 | 32 | GGCCGTGATCGTATAGTGGTTAGTACTCTGCG | 12989.1 | -3.1 | 1.1 | -2.9 | 3.9E-03 | 8.0E-03 | BT20/MB468 | MP |
| >MI0000088\|hsa-mir-30a&WithFlank&6\|-\|71403545\|71403627@12.32.21&[MIMAT0000087&hsa-miR-30a-5p&offsets\|0\|-1;m-15&6\|-\|71403595\|71403616&offsets\|0\|-1]\|TGTAAACATCCTCGACTGGAA | >MI0000088\|hsa-mir-30a&WithFlank&6\|-\|71403545\|71403627 | 12 | 32 | 21 | TGTAAACATCCTCGACTGGAA | 1115.9 | -2.4 | 0.8 | -3.1 | 1.8E-03 | 8.0E-03 | BT20/MB231 | MITO |
| >hg38_21_+_8433172_8446622_RNA45SN1_45S_50ntFlanks@6645.6669.25\|CTCGTACGACTCTTAGCGGTGGATC | >hg38_21_+_8433172_8446622_RNA45SN1_45S_50ntFlanks | 6645 | 6669 | 25 | CTCGTACGACTCTTAGCGGTGGATC | 21612.4 | 2.3 | 0.7 | 3.1 | 1.8E-03 | 8.0E-03 | BT20/MB231 | MITO |
| >hg38_21_+_8433172_8446622_RNA45SN1_45S_50ntFlanks@5490.5531.42\|TGAGAAGACGGTCGAACTTGACTATCTAGAGGAAGTAAAAGT | >hg38_21_+_8433172_8446622_RNA45SN1_45S_50ntFlanks | 5490 | 5531 | 42 | TGAGAAGACGGTCGAACTTGACTATCTAGAGGAAGTAAAAGT | 775.2 | -3.4 | 1.1 | -3.1 | 1.8E-03 | 8.0E-03 | BT20/MB231 | CYTO |
| >hg38_21_+_8433172_8446622_RNA45SN1_45S_50ntFlanks@12638.12656.19\|CTACCATCTGTGGGATTAT | >hg38_21_+_8433172_8446622_RNA45SN1_45S_50ntFlanks | 12638 | 12656 | 19 | CTACCATCTGTGGGATTAT | 70.5 | -4.6 | 1.6 | -2.9 | 3.9E-03 | 8.0E-03 | BT20/MB468 | MP |
| >hg38_21_+_8433172_8446622_RNA45SN1_45S_50ntFlanks@12644.12661.18\|TCTGTGGGATTATGACTG | >hg38_21_+_8433172_8446622_RNA45SN1_45S_50ntFlanks | 12644 | 12661 | 18 | TCTGTGGGATTATGACTG | 1677.3 | -4.2 | 1.5 | -2.9 | 3.9E-03 | 8.0E-03 | BT20/MB468 | MP |
| >hg38_21_+_8433172_8446622_RNA45SN1_45S_50ntFlanks@12974.12997.24\|CGTACGTAGCAGAGCAGCTCCCTC | >hg38_21_+_8433172_8446622_RNA45SN1_45S_50ntFlanks | 12974 | 12997 | 24 | CGTACGTAGCAGAGCAGCTCCCTC | 267.3 | -2.4 | 0.8 | -3.1 | 1.8E-03 | 8.0E-03 | BT20/MB231 | CYTO |
| >hg38_21_+_8433172_8446622_RNA45SN1_45S_50ntFlanks@4372.4398.27\|AAAAAGCTCGTAGTTGGATCTTGGGAG | >hg38_21_+_8433172_8446622_RNA45SN1_45S_50ntFlanks | 4372 | 4398 | 27 | AAAAAGCTCGTAGTTGGATCTTGGGAG | 115.8 | -4.9 | 1.7 | -2.9 | 3.9E-03 | 8.0E-03 | BT20/MB468 | MP |
| >hg38_21_+_8433172_8446622_RNA45SN1_45S_50ntFlanks@6745.6763.19\|ACACTTCGAACGCACTTGC | >hg38_21_+_8433172_8446622_RNA45SN1_45S_50ntFlanks | 6745 | 6763 | 19 | ACACTTCGAACGCACTTGC | 67.8 | -4.7 | 1.6 | -2.9 | 3.9E-03 | 8.0E-03 | BT20/MB468 | MP |
| >MI0003667\|hsa-mir-652&WithFlank&X\|+\|110055323\|110055432@67.87.21(+1A)&[MIMAT0003322&hsa-miR-652-3p&offsets\|0\|0(+1A);m-148&X\|+\|110055389\|110055410&offsets\|0\|-1(+1A)]\|AATGGCGCCACTAGGGTTGTGA | >MI0003667\|hsa-mir-652&WithFlank&X\|+\|110055323\|110055432 | 67 | 87 | 21(+1A) | AATGGCGCCACTAGGGTTGTGA | 173.4 | 4.4 | 1.5 | 2.9 | 3.9E-03 | 8.0E-03 | BT20/MB468 | MP |
| >hg38_21_+_8433172_8446622_RNA45SN1_45S_50ntFlanks@11190.11225.36\|CCCGGGGGGCCGGCGGCGGCGGCGACTCTGGACGCG | >hg38_21_+_8433172_8446622_RNA45SN1_45S_50ntFlanks | 11190 | 11225 | 36 | CCCGGGGGGCCGGCGGCGGCGGCGACTCTGGACGCG | 464.6 | -3.5 | 1.3 | -2.7 | 6.6E-03 | 8.0E-03 | BT20/MB468 | CYTO |
| >hg38_21_+_8433172_8446622_RNA45SN1_45S_50ntFlanks@5487.5515.29\|CGCTGAGAAGACGGTCGAACTTGACTATC | >hg38_21_+_8433172_8446622_RNA45SN1_45S_50ntFlanks | 5487 | 5515 | 29 | CGCTGAGAAGACGGTCGAACTTGACTATC | 492.9 | -1.8 | 0.7 | -2.8 | 5.6E-03 | 8.0E-03 | BT20/MB468 | MITO |
| >hg38_21_+_8433172_8446622_RNA45SN1_45S_50ntFlanks@4727.4758.32\|AATCAAGAACGAAAGTCGGAGGTTCGAAGACG | >hg38_21_+_8433172_8446622_RNA45SN1_45S_50ntFlanks | 4727 | 4758 | 32 | AATCAAGAACGAAAGTCGGAGGTTCGAAGACG | 1169.0 | 2.1 | 0.8 | 2.7 | 6.6E-03 | 8.0E-03 | BT20/MB468 | CYTO |
| >trna27_GlyCCC_2_-_70476123_70476193@1.33.33,trna34_GlyCCC_16_-_686736_686806@1.33.33\|GCGCCGCTGGTGTAGTGGTATCATGCAAGATTC | >trna27_GlyCCC_2_-_70476123_70476193 | 1 | 33 | 33 | GCGCCGCTGGTGTAGTGGTATCATGCAAGATTC | 285.5 | 6.6 | 2.3 | 2.9 | 3.9E-03 | 8.1E-03 | BT20/MB468 | MP |
| >hg38_MT_+_598_1651_RNR1_12S_50ntFlanks@52.98.47\|ATAGGTTTGGTCCTAGCCTTTCTATTAGCTCTTAGTAAGATTACACA | >hg38_MT_+_598_1651_RNR1_12S_50ntFlanks | 52 | 98 | 47 | ATAGGTTTGGTCCTAGCCTTTCTATTAGCTCTTAGTAAGATTACACA | 235.0 | 4.6 | 1.7 | 2.7 | 7.0E-03 | 8.1E-03 | MB231/MB468 | MITO |
| >hg38_21_+_8433172_8446622_RNA45SN1_45S_50ntFlanks@6672.6696.25\|TCGGCTCGTGCGTCGATGAAGAACG | >hg38_21_+_8433172_8446622_RNA45SN1_45S_50ntFlanks | 6672 | 6696 | 25 | TCGGCTCGTGCGTCGATGAAGAACG | 84.9 | -3.4 | 1.2 | -2.7 | 6.1E-03 | 8.1E-03 | MB231/MB468 | MP |
| >hg38_21_+_8433172_8446622_RNA45SN1_45S_50ntFlanks@10470.10497.28\|GCCCTCGGCCGATCGAAAGGGAGTCGGG | >hg38_21_+_8433172_8446622_RNA45SN1_45S_50ntFlanks | 10470 | 10497 | 28 | GCCCTCGGCCGATCGAAAGGGAGTCGGG | 54.1 | -4.6 | 1.6 | -2.9 | 3.9E-03 | 8.1E-03 | BT20/MB468 | MP |
| >MI0000815\|hsa-mir-339&WithFlank&7\|-\|1022927\|1023032@56.75.20(+1U)&[MIMAT0004702&hsa-miR-339-3p&offsets\|0\|-3(+1U);m-235&7\|-\|1022957\|1022977&offsets\|0\|-1(+1U)]\|TGAGCGCCTCGACGACAGAGT | >MI0000815\|hsa-mir-339&WithFlank&7\|-\|1022927\|1023032 | 56 | 75 | 20(+1U) | TGAGCGCCTCGACGACAGAGT | 99.6 | 5.3 | 1.9 | 2.7 | 6.1E-03 | 8.1E-03 | MB231/MB468 | MP |
| >hg38_1_-_228634819_228635039_RNA5S12_5S_50ntFlanks@85.103.19\|TCGTCTGATCTCGGAAGCT | >hg38_1_-_228634819_228635039_RNA5S12_5S_50ntFlanks | 85 | 103 | 19 | TCGTCTGATCTCGGAAGCT | 462.8 | 5.8 | 1.8 | 3.2 | 1.4E-03 | 8.1E-03 | BT20/MB231 | MP |
| >hg38_21_+_8433172_8446622_RNA45SN1_45S_50ntFlanks@6661.6693.33\|CGGTGGATCACTCGGCTCGTGCGTCGATGAAGA | >hg38_21_+_8433172_8446622_RNA45SN1_45S_50ntFlanks | 6661 | 6693 | 33 | CGGTGGATCACTCGGCTCGTGCGTCGATGAAGA | 998.7 | 3.5 | 1.1 | 3.2 | 1.4E-03 | 8.1E-03 | BT20/MB231 | MP |
| >MI0000750\|hsa-mir-26a-2&WithFlank&12\|-\|57824603\|57824698@20.39.20(+1U);MI0000083\|hsa-mir-26a-1&WithFlank&3\|+\|37969398\|37969486@16.35.20(+1U)&[MIMAT0000082&hsa-miR-26a-5p&offsets\|0\|-2(+1U);m-33&12\|-\|57824658\|57824679&offsets\|0\|-2(+1U)];[MIMAT0000082_1&hsa-miR-26a-5p&offsets\|0\|-2(+1U);m-34&3\|+\|37969413\|37969434&offsets\|0\|-2(+1U)]\|TTCAAGTAATCCAGGATAGGT | >MI0000750\|hsa-mir-26a-2&WithFlank&12\|-\|57824603\|57824698 | 20 | 39 | 20(+1U) | TTCAAGTAATCCAGGATAGGT | 823.6 | -4.9 | 1.5 | -3.2 | 1.4E-03 | 8.1E-03 | BT20/MB231 | MP |
| >MI0033423\|hsa-mir-10399&WithFlank&7\|-\|139044094\|139044163@43.63.21(+1U)&[MIMAT0041630&hsa-miR-10399-3p&offsets\|-1\|-1(+1U);TJU_CMC_MD2.ID03061.3p-miR&offsets\|+3\|+2(+1U);m-2657&7\|-\|139044103\|139044124&offsets\|+3\|+2(+1U)]\|TCTCTCGGACAAGCTGTAGGTT | >MI0033423\|hsa-mir-10399&WithFlank&7\|-\|139044094\|139044163 | 43 | 63 | 21(+1U) | TCTCTCGGACAAGCTGTAGGTT | 471.0 | -4.5 | 1.4 | -3.2 | 1.4E-03 | 8.1E-03 | BT20/MB231 | MP |
| >hg38_21_+_8433172_8446622_RNA45SN1_45S_50ntFlanks@11320.11342.23\|CCACGTCTCGTCGCGCGCGCGTC | >hg38_21_+_8433172_8446622_RNA45SN1_45S_50ntFlanks | 11320 | 11342 | 23 | CCACGTCTCGTCGCGCGCGCGTC | 309.0 | -5.0 | 1.7 | -2.9 | 3.9E-03 | 8.1E-03 | BT20/MB468 | MP |
| >hg38_MT_+_598_1651_RNR1_12S_50ntFlanks@719.743.25\|TAAAGACGTTAGGTCAAGGTGTAGC | >hg38_MT_+_598_1651_RNR1_12S_50ntFlanks | 719 | 743 | 25 | TAAAGACGTTAGGTCAAGGTGTAGC | 179.6 | -2.4 | 0.9 | -2.7 | 6.7E-03 | 8.1E-03 | MB231/MB468 | CYTO |
| >hg38_21_+_8433172_8446622_RNA45SN1_45S_50ntFlanks@5471.5512.42\|CACGGCCCTGGCGGAGCGCTGAGAAGACGGTCGAACTTGACT | >hg38_21_+_8433172_8446622_RNA45SN1_45S_50ntFlanks | 5471 | 5512 | 42 | CACGGCCCTGGCGGAGCGCTGAGAAGACGGTCGAACTTGACT | 338.1 | 2.5 | 0.9 | 2.7 | 6.6E-03 | 8.1E-03 | BT20/MB468 | CYTO |
| >hg38_21_+_8433172_8446622_RNA45SN1_45S_50ntFlanks@9252.9273.22\|CACGGGGTCGGCGGCGACGTCG | >hg38_21_+_8433172_8446622_RNA45SN1_45S_50ntFlanks | 9252 | 9273 | 22 | CACGGGGTCGGCGGCGACGTCG | 374.6 | -4.9 | 1.7 | -2.9 | 3.9E-03 | 8.1E-03 | BT20/MB468 | MP |
| >hg38_21_+_8433172_8446622_RNA45SN1_45S_50ntFlanks@10316.10340.25\|GGTGCAGATCTTGGTGGTAGTAGCA | >hg38_21_+_8433172_8446622_RNA45SN1_45S_50ntFlanks | 10316 | 10340 | 25 | GGTGCAGATCTTGGTGGTAGTAGCA | 743.6 | 4.9 | 1.7 | 2.9 | 3.9E-03 | 8.1E-03 | BT20/MB468 | MP |
| >hg38_21_+_8433172_8446622_RNA45SN1_45S_50ntFlanks@3851.3873.23\|ATAACTGTGGTAATTCTAGAGCT | >hg38_21_+_8433172_8446622_RNA45SN1_45S_50ntFlanks | 3851 | 3873 | 23 | ATAACTGTGGTAATTCTAGAGCT | 237.4 | -3.8 | 1.3 | -2.9 | 3.9E-03 | 8.1E-03 | BT20/MB468 | MP |
| >MI0000299\|hsa-mir-222&WithFlank&X\|-\|45747009\|45747130@75.95.21(+2U)&[MIMAT0000279&hsa-miR-222-3p&offsets\|0\|0(+2U);m-140&X\|-\|45747033\|45747056&offsets\|0\|-3(+2U)]\|AGCTACATCTGGCTACTGGGTTT | >MI0000299\|hsa-mir-222&WithFlank&X\|-\|45747009\|45747130 | 75 | 95 | 21(+2U) | AGCTACATCTGGCTACTGGGTTT | 43.3 | 5.3 | 1.9 | 2.7 | 6.1E-03 | 8.1E-03 | MB231/MB468 | MP |
| >hg38_21_+_8433172_8446622_RNA45SN1_45S_50ntFlanks@4894.4917.24\|ACTTAAAGGAATTGACGGAAGGGC | >hg38_21_+_8433172_8446622_RNA45SN1_45S_50ntFlanks | 4894 | 4917 | 24 | ACTTAAAGGAATTGACGGAAGGGC | 88.5 | -5.0 | 1.7 | -2.9 | 3.9E-03 | 8.1E-03 | BT20/MB468 | MP |
| >hg38_MT_+_598_1651_RNR1_12S_50ntFlanks@439.469.31\|AACATATCTGAACACACAATAGCTAAGACCC | >hg38_MT_+_598_1651_RNR1_12S_50ntFlanks | 439 | 469 | 31 | AACATATCTGAACACACAATAGCTAAGACCC | 143.9 | 3.4 | 1.2 | 2.7 | 6.1E-03 | 8.1E-03 | MB231/MB468 | MP |
| >hg38_21_+_8433172_8446622_RNA45SN1_45S_50ntFlanks@10058.10076.19\|CGGCAGTCGAGAGTGGACG | >hg38_21_+_8433172_8446622_RNA45SN1_45S_50ntFlanks | 10058 | 10076 | 19 | CGGCAGTCGAGAGTGGACG | 222.5 | 6.1 | 1.9 | 3.2 | 1.4E-03 | 8.1E-03 | BT20/MB231 | MP |
| >hg38_21_+_8433172_8446622_RNA45SN1_45S_50ntFlanks@10237.10261.25\|ACGCTACGCCGCGACGAGTAGGAGG | >hg38_21_+_8433172_8446622_RNA45SN1_45S_50ntFlanks | 10237 | 10261 | 25 | ACGCTACGCCGCGACGAGTAGGAGG | 214.0 | 5.9 | 1.9 | 3.2 | 1.4E-03 | 8.1E-03 | BT20/MB231 | MP |
| >hg38_21_+_8433172_8446622_RNA45SN1_45S_50ntFlanks@11771.11790.20\|ATTAGTGACGCGCATGAATG | >hg38_21_+_8433172_8446622_RNA45SN1_45S_50ntFlanks | 11771 | 11790 | 20 | ATTAGTGACGCGCATGAATG | 134.0 | -4.7 | 1.7 | -2.7 | 6.1E-03 | 8.1E-03 | MB231/MB468 | MP |
| >MI0016590\|hsa-mir-3934&WithFlank&6\|+\|33698122\|33698240@31.52.22&[MIMAT0018349&hsa-miR-3934-5p&offsets\|0\|0;m-557&6\|+\|33698152\|33698173&offsets\|0\|0]\|TCAGGTGTGGAAACTGAGGCAG | >MI0016590\|hsa-mir-3934&WithFlank&6\|+\|33698122\|33698240 | 31 | 52 | 22 | TCAGGTGTGGAAACTGAGGCAG | 155.7 | 5.2 | 1.8 | 2.9 | 3.9E-03 | 8.1E-03 | BT20/MB468 | MP |
| >hg38_MT_+_598_1651_RNR1_12S_50ntFlanks@424.456.33\|TACGAAAGTGGCTTTAACATATCTGAACACACA | >hg38_MT_+_598_1651_RNR1_12S_50ntFlanks | 424 | 456 | 33 | TACGAAAGTGGCTTTAACATATCTGAACACACA | 203.9 | 5.3 | 1.8 | 2.9 | 3.9E-03 | 8.1E-03 | BT20/MB468 | MP |
| >hg38_21_+_8433172_8446622_RNA45SN1_45S_50ntFlanks@4972.4994.23\|CGGCCCGGACACGGACAGGATTG | >hg38_21_+_8433172_8446622_RNA45SN1_45S_50ntFlanks | 4972 | 4994 | 23 | CGGCCCGGACACGGACAGGATTG | 143.8 | -4.6 | 1.7 | -2.8 | 5.7E-03 | 8.1E-03 | BT20/MB468 | MITO |
| >MI0000266\|hsa-mir-10a&WithFlank&17\|-\|48579832\|48579953@28.48.21(+1C)&[MIMAT0000253&hsa-miR-10a-5p&offsets\|0\|-2(+1C);m-18&17\|-\|48579905\|48579926&offsets\|0\|-1(+1C)]\|TACCCTGTAGATCCGAATTTGC | >MI0000266\|hsa-mir-10a&WithFlank&17\|-\|48579832\|48579953 | 28 | 48 | 21(+1C) | TACCCTGTAGATCCGAATTTGC | 4297.1 | 2.4 | 0.8 | 3.1 | 1.8E-03 | 8.1E-03 | BT20/MB231 | CYTO |
| >hg38_21_+_8433172_8446622_RNA45SN1_45S_50ntFlanks@11600.11627.28\|AATCCGACTGTTTAATTAAAACAAAGCA | >hg38_21_+_8433172_8446622_RNA45SN1_45S_50ntFlanks | 11600 | 11627 | 28 | AATCCGACTGTTTAATTAAAACAAAGCA | 291.8 | 6.2 | 2.0 | 3.2 | 1.4E-03 | 8.1E-03 | BT20/MB231 | MP |
| >hg38_21_+_8433172_8446622_RNA45SN1_45S_50ntFlanks@11578.11609.32\|TAGAACTGGTGCGGACCAGGGGAATCCGACTG | >hg38_21_+_8433172_8446622_RNA45SN1_45S_50ntFlanks | 11578 | 11609 | 32 | TAGAACTGGTGCGGACCAGGGGAATCCGACTG | 227.4 | 3.3 | 1.1 | 3.1 | 1.9E-03 | 8.1E-03 | BT20/MB231 | MITO |

| | | | | | | | | | | | | | |
|---|---|---|---|---|---|---|---|---|---|---|---|---|---|
| >hg38_21_+_8433172_8446622_RNA45SN1_45S_50ntFlanks@9533.9572.40\|GAGGAAACTCTGGTGGAGGTCCGTAGCGGTCCTGACGTGC | >hg38_21_+_8433172_8446622_RNA45SN1_45S_50ntFlanks | 9533 | 9572 | 40 | GAGGAAACTCTGGTGGAGGTCCGTAGCGGTCCTGACGTGC | 387.3 | 3.1 | 1.0 | 3.1 | 1.9E-03 | 8.1E-03 | BT20/MB231 | MITO |
| >hg38_21_+_8433172_8446622_RNA45SN1_45S_50ntFlanks@10348.10379.32\|AAACGAGAACTTTGAAGGCCGAAGTGGAGAAG | >hg38_21_+_8433172_8446622_RNA45SN1_45S_50ntFlanks | 10348 | 10379 | 32 | AAACGAGAACTTTGAAGGCCGAAGTGGAGAAG | 220.6 | -2.4 | 0.9 | -2.7 | 6.1E-03 | 8.1E-03 | MB231/MB468 | MP |
| >hg38_21_+_8433172_8446622_RNA45SN1_45S_50ntFlanks@12588.12616.29\|GGTTCAGACATTTGGTGTATGTGCTTGGC | >hg38_21_+_8433172_8446622_RNA45SN1_45S_50ntFlanks | 12588 | 12616 | 29 | GGTTCAGACATTTGGTGTATGTGCTTGGC | 130.4 | -4.7 | 1.6 | -2.9 | 3.9E-03 | 8.1E-03 | BT20/MB468 | MP |
| >hg38_21_+_8433172_8446622_RNA45SN1_45S_50ntFlanks@9549.9572.24\|AGGTCCGTAGCGGTCCTGACGTGC | >hg38_21_+_8433172_8446622_RNA45SN1_45S_50ntFlanks | 9549 | 9572 | 24 | AGGTCCGTAGCGGTCCTGACGTGC | 224.3 | 5.7 | 1.8 | 3.2 | 1.4E-03 | 8.1E-03 | BT20/MB231 | MP |
| >MI0000114\|hsa-mir-107&WithFlank&10\|-\|89592741\|89592833@56.78.23&[MIMAT0000104&hsa-miR-107&offsets\|0\|0;m-78&10\|-\|89592758\|89592778&offsets\|0\|+2]\|AGCAGCATTGTACAGGGCTATCA | >MI0000114\|hsa-mir-107&WithFlank&10\|-\|89592741\|89592833 | 56 | 78 | 23 | AGCAGCATTGTACAGGGCTATCA | 204.3 | -4.4 | 1.4 | -3.2 | 1.4E-03 | 8.1E-03 | BT20/MB231 | MP |
| >hg38_21_+_8433172_8446622_RNA45SN1_45S_50ntFlanks@5470.5493.24\|CCACGGCCCTGGCGGAGCGCTGAG | >hg38_21_+_8433172_8446622_RNA45SN1_45S_50ntFlanks | 5470 | 5493 | 24 | CCACGGCCCTGGCGGAGCGCTGAG | 51.9 | -4.7 | 1.6 | -2.9 | 3.9E-03 | 8.1E-03 | BT20/MB468 | MP |
| >hg38_21_+_8433172_8446622_RNA45SN1_45S_50ntFlanks@9533.9555.23\|GAGGAAACTCTGGTGGAGGTCCG | >hg38_21_+_8433172_8446622_RNA45SN1_45S_50ntFlanks | 9533 | 9555 | 23 | GAGGAAACTCTGGTGGAGGTCCG | 60.0 | -9.0 | 3.1 | -2.9 | 3.9E-03 | 8.1E-03 | BT20/MB468 | MP |
| >hg38_21_+_8433172_8446622_RNA45SN1_45S_50ntFlanks@11672.11702.31\|CAGTGCTCTGAATGTCAAAGTGAAGAAATTC | >hg38_21_+_8433172_8446622_RNA45SN1_45S_50ntFlanks | 11672 | 11702 | 31 | CAGTGCTCTGAATGTCAAAGTGAAGAAATTC | 324.3 | -1.9 | 0.7 | -2.7 | 7.1E-03 | 8.1E-03 | MB231/MB468 | MITO |
| >hg38_21_+_8433172_8446622_RNA45SN1_45S_50ntFlanks@9534.9576.43\|AGGAAACTCTGGTGGAGGTCCGTAGCGGTCCTGACGTGCAAAT | >hg38_21_+_8433172_8446622_RNA45SN1_45S_50ntFlanks | 9534 | 9576 | 43 | AGGAAACTCTGGTGGAGGTCCGTAGCGGTCCTGACGTGCAAAT | 353.9 | -3.0 | 1.0 | -3.1 | 1.8E-03 | 8.1E-03 | BT20/MB231 | CYTO |
| >trnaMT_GluTTC_MT_-_14674_14742@39.72.34,trnalookalike8_GluTTC_5_-_93905172_93905240@39.72.34\|ATTGGTCGTGGTTGTAGTCCGTGCGAGAATACCA | >trnaMT_GluTTC_MT_-_14674_14742 | 39 | 72 | 34 | ATTGGTCGTGGTTGTAGTCCGTGCGAGAATACCA | 16222.8 | 2.6 | 1.0 | 2.7 | 6.1E-03 | 8.1E-03 | MB231/MB468 | MP |
| >hg38_21_+_8433172_8446622_RNA45SN1_45S_50ntFlanks@11564.11593.30\|CCTAGCAGCCGACTTAGAACTGGTGCGGAC | >hg38_21_+_8433172_8446622_RNA45SN1_45S_50ntFlanks | 11564 | 11593 | 30 | CCTAGCAGCCGACTTAGAACTGGTGCGGAC | 124.8 | -3.9 | 1.4 | -2.8 | 5.7E-03 | 8.1E-03 | BT20/MB468 | MITO |
| >hg38_21_+_8433172_8446622_RNA45SN1_45S_50ntFlanks@10808.10839.32\|CAATGTAGGTAAGGGAAGTCGGCAAGCCGGAT | >hg38_21_+_8433172_8446622_RNA45SN1_45S_50ntFlanks | 10808 | 10839 | 32 | CAATGTAGGTAAGGGAAGTCGGCAAGCCGGAT | 149.6 | -4.7 | 1.6 | -2.9 | 3.9E-03 | 8.1E-03 | BT20/MB468 | MP |
| >am_trna116_GluCTC_1_-_145399233_145399304@1.30.30,trna59_GluCTC_1_+_249168447_249168518@1.30.30,trna71_GluCTC_1_-_161439189_161439260@1.30.30,trna74_GluCTC_1_-_161431809_161431880@1.30.30,trna77_GluCTC_1_-_161424398_161424469@1.30.30,trna77_GluCTC_6_+_28949976_28950047@1.30.30,trna80_GluCTC_1_-_161417018_161417089@1.30.30,trna87_GluCTC_6_-_126101393_126101464@1.30.30\|TCCCTGGTGGTCTAGTGGTTAGGATTCGGC | >am_trna116_GluCTC_1_-_145399233_145399304 | 1 | 30 | 30 | TCCCTGGTGGTCTAGTGGTTAGGATTCGGC | 7284.3 | -2.5 | 0.9 | -2.7 | 6.1E-03 | 8.1E-03 | MB231/MB468 | MP |
| >trna10_AspGTC_12_-_125424193_125424264@31.75.45,trna12_AspGTC_12_-_125411891_125411962@31.75.45,trna38_AspGTC_17_-_8125556_8125627@31.75.45,trna45_AspGTC_6_+_27447453_27447524@31.75.45,trna48_AspGTC_6_+_27471523_27471594@31.75.45,trna4_AspGTC_12_+_96429799_96429870@31.75.45,trna69_AspGTC_1_-_161440205_161440276@31.75.45,trna72_AspGTC_1_-_161432824_161432895@31.75.45,trna75_AspGTC_1_-_161425414_161425485@31.75.45,trna78_AspGTC_1_-_161418033_161418104@31.75.45,trna81_AspGTC_1_-_161410615_161410686@31.75.45\|CCTGTCACGCGGGAGACCGGGGTTCGATTCCCCGACGGGGAGCCA | >trna10_AspGTC_12_-_125424193_125424264 | 31 | 75 | 45 | CCTGTCACGCGGGAGACCGGGGTTCGATTCCCCGACGGGGAGCCA | 364.4 | -4.5 | 1.7 | -2.7 | 6.7E-03 | 8.1E-03 | BT20/MB468 | CYTO |
| >hg38_21_+_8433172_8446622_RNA45SN1_45S_50ntFlanks@6661.6689.29\|CGGTGGATCACTCGGCTCGTGCGTCGATG | >hg38_21_+_8433172_8446622_RNA45SN1_45S_50ntFlanks | 6661 | 6689 | 29 | CGGTGGATCACTCGGCTCGTGCGTCGATG | 30798.7 | 1.6 | 0.6 | 2.8 | 5.7E-03 | 8.1E-03 | BT20/MB468 | MITO |
| >hg38_21_+_8433172_8446622_RNA45SN1_45S_50ntFlanks@3986.4024.39\|GGCTTTGGTGACTCTAGATAACCTCGGGCCGATCGCACG | >hg38_21_+_8433172_8446622_RNA45SN1_45S_50ntFlanks | 3986 | 4024 | 39 | GGCTTTGGTGACTCTAGATAACCTCGGGCCGATCGCACG | 573.7 | -4.6 | 1.6 | -2.9 | 3.9E-03 | 8.1E-03 | BT20/MB468 | MP |
| >hg38_21_+_8433172_8446622_RNA45SN1_45S_50ntFlanks@3840.3879.40\|CTCCTACTTGGATAACTGTGGTAATTCTAGAGCTAATACA | >hg38_21_+_8433172_8446622_RNA45SN1_45S_50ntFlanks | 3840 | 3879 | 40 | CTCCTACTTGGATAACTGTGGTAATTCTAGAGCTAATACA | 487.0 | 2.7 | 0.9 | 3.1 | 1.9E-03 | 8.1E-03 | BT20/MB231 | MITO |
| >hg38_21_+_8433172_8446622_RNA45SN1_45S_50ntFlanks@7974.8005.32\|ACGCGACCTCAGATCAGACGTGGCGACCCGCT | >hg38_21_+_8433172_8446622_RNA45SN1_45S_50ntFlanks | 7974 | 8005 | 32 | ACGCGACCTCAGATCAGACGTGGCGACCCGCT | 47573.2 | 2.5 | 0.8 | 3.1 | 1.9E-03 | 8.1E-03 | BT20/MB231 | MITO |
| >MI0000078\|hsa-mir-22&WithFlank&17\|-\|1713897\|1713993@59.80.22(+1A)&[MIMAT0000077&hsa-miR-22-3p&offsets\|0\|0(+1A);m-2&17\|-\|1713914\|1713935&offsets\|0\|0(+1A)]\|AAGCTGCCAGTTGAAGAACTGTA | >MI0000078\|hsa-mir-22&WithFlank&17\|-\|1713897\|1713993 | 59 | 80 | 22(+1A) | AAGCTGCCAGTTGAAGAACTGTA | 606.2 | 3.8 | 1.4 | 2.7 | 6.1E-03 | 8.1E-03 | MB231/MB468 | MP |
| >MI0000746\|hsa-mir-99b&WithFlank&19\|+\|51692606\|51692687@13.31.19(+1A)&[MIMAT0000689&hsa-miR-99b-5p&offsets\|0\|-3(+1A);m-10&19\|+\|51692618\|51692639&offsets\|0\|-3(+1A)]\|CACCCGTAGAACCGACCTTA | >MI0000746\|hsa-mir-99b&WithFlank&19\|+\|51692606\|51692687 | 13 | 31 | 19(+1A) | CACCCGTAGAACCGACCTTA | 47.6 | 5.4 | 2.0 | 2.7 | 6.1E-03 | 8.1E-03 | MB231/MB468 | MP |
| >am_MI0000299\|hsa-mir-222&WithFlank&X\|-\|45747009\|45747130@75.92.18&[MIMAT0000279&hsa-miR-222-3p&offsets\|0\|-3;m-140&X\|-\|45747033\|45747056&offsets\|0\|-6]\|AGCTACATCTGGCTACTG | >am_MI0000299\|hsa-mir-222&WithFlank&X\|-\|45747009\|45747130 | 75 | 92 | 18 | AGCTACATCTGGCTACTG | 54.5 | 4.6 | 1.7 | 2.7 | 6.1E-03 | 8.1E-03 | MB231/MB468 | MP |
| >MI0003834\|hsa-mir-769&WithFlank&19\|+\|46018926\|46019055@36.57.22(+1U)&[MIMAT0003886&hsa-miR-769-5p&offsets\|0\|0(+1U);m-216&19\|+\|46018961\|46018982&offsets\|0\|0(+1U)]\|TGAGACCTCTGGGTTCTGAGCTT | >MI0003834\|hsa-mir-769&WithFlank&19\|+\|46018926\|46019055 | 36 | 57 | 22(+1U) | TGAGACCTCTGGGTTCTGAGCTT | 347.1 | 3.5 | 1.2 | 2.9 | 3.9E-03 | 8.2E-03 | BT20/MB468 | MP |
| >MI0000077\|hsa-mir-21&WithFlank&17\|+\|59841260\|59841343@16.36.21&[MIMAT0000076&hsa-miR-21-5p&offsets\|+2\|+1;m-1&17\|+\|59841273\|59841294&offsets\|+2\|+1]\|GCTTATCAGACTGATGTTGAC | >MI0000077\|hsa-mir-21&WithFlank&17\|+\|59841260\|59841343 | 16 | 36 | 21 | GCTTATCAGACTGATGTTGAC | 75.2 | 5.1 | 1.9 | 2.7 | 6.1E-03 | 8.2E-03 | MB231/MB468 | MP |
| >trnaMT_ProTGG_MT_-_15956_16023@38.70.33\|CTAATGGTGGAGTTAAAGACTTTTTCTCTGACC | >trnaMT_ProTGG_MT_-_15956_16023 | 38 | 70 | 33 | CTAATGGTGGAGTTAAAGACTTTTTCTCTGACC | 39.1 | 5.2 | 1.9 | 2.7 | 6.1E-03 | 8.2E-03 | MB231/MB468 | MP |
| >MI0000446\|hsa-mir-125b-1&WithFlank&11\|-\|122099751\|122099850@21.43.23&[MIMAT0000423&hsa-miR-125b-5p&offsets\|0\|+1;m-67&11\|-\|122099809\|122099830&offsets\|0\|+1]\|TCCCTGAGACCCTAACTTGTGAT | >MI0000446\|hsa-mir-125b-1&WithFlank&11\|-\|122099751\|122099850 | 21 | 43 | 23 | TCCCTGAGACCCTAACTTGTGAT | 2764.7 | -7.9 | 2.5 | -3.1 | 1.9E-03 | 8.2E-03 | BT20/MB231 | MITO |
| >trna111_HisGTG_1_-_147774845_147774916@-1G.32.33,trna118_HisGTG_1_-_145396881_145396952@-1G.32.33,trna16_HisGTG_1_+_146544773_146544844@-1G.32.33,trna1_HisGTG_15_+_45493349_45493420@-1G.32.33,trna21_HisGTG_1_+_147753471_147753542@-1G.32.33,trna33_HisGTG_6_+_27125906_27125977@-1G.32.33,trna7_HisGTG_9_-_14433938_14434009@-1G.32.33,trna8_HisGTG_15_-_45492611_45492682@-1G.32.33,trna9_HisGTG_15_-_45490804_45490875@-1G.32.33\|GGCCGTGATCGTATAGTGGTTAGTACTCTGCGT | >trna111_HisGTG_1_-_147774845_147774916 | -1G | 32 | 33 | GGCCGTGATCGTATAGTGGTTAGTACTCTGCGT | 2813.4 | -3.1 | 1.0 | -3.1 | 1.9E-03 | 8.2E-03 | BT20/MB231 | MITO |
| >hg38_21_+_8433172_8446622_RNA45SN1_45S_50ntFlanks@11573.11594.22\|CGACTTAGAACTGGTGCGGACC | >hg38_21_+_8433172_8446622_RNA45SN1_45S_50ntFlanks | 11573 | 11594 | 22 | CGACTTAGAACTGGTGCGGACC | 524.8 | 1.9 | 0.7 | 2.7 | 6.7E-03 | 8.2E-03 | MB231/MB468 | CYTO |
| >MI0003591\|hsa-mir-584&WithFlank&5\|-\|149062307\|149062415@22.42.21(+1A)&[MIMAT0003249&hsa-miR-584-5p&offsets\|0\|-1(+1A);m-167&5\|-\|149062374\|149062394&offsets\|0\|0(+1A)]\|TTATGGTTTGCCTGGGACTGAA | >MI0003591\|hsa-mir-584&WithFlank&5\|-\|149062307\|149062415 | 22 | 42 | 21(+1A) | TTATGGTTTGCCTGGGACTGAA | 137.3 | 3.5 | 1.3 | 2.7 | 6.2E-03 | 8.2E-03 | MB231/MB468 | MP |
| >hg38_21_+_8433172_8446622_RNA45SN1_45S_50ntFlanks@11208.11233.26\|GCGGCGACTCTGGACGCGAGCCGGGC | >hg38_21_+_8433172_8446622_RNA45SN1_45S_50ntFlanks | 11208 | 11233 | 26 | GCGGCGACTCTGGACGCGAGCCGGGC | 511.1 | 7.8 | 2.4 | 3.2 | 1.4E-03 | 8.2E-03 | BT20/MB231 | MP |
| >MI0000062\|hsa-let-7a-3&WithFlank&22\|+\|46112743\|46112828@58.78.21(+1U);MI0000060\|hsa-let-7a-1&WithFlank&9\|+\|94175951\|94176042@63.83.21(+1U)&[MIMAT0004481&hsa-let-7a-3p&offsets\|0\|0(+1U);m-180&22\|+\|46112800\|46112821&offsets\|0\|-1(+1U)];[MIMAT0004481_1&hsa-let-7a-3p&offsets\|0\|0(+1U);m-181&9\|+\|94176013\|94176034&offsets\|0\|-1(+1U)]\|CTATACAATCTACTGTCTTTCT | >MI0000062\|hsa-let-7a-3&WithFlank&22\|+\|46112743\|46112828 | 58 | 78 | 21(+1U) | CTATACAATCTACTGTCTTTCT | 542.5 | 3.0 | 1.1 | 2.9 | 4.0E-03 | 8.2E-03 | BT20/MB468 | MP |
| >hg38_21_+_8433172_8446622_RNA45SN1_45S_50ntFlanks@10069.10097.29\|AGTGGACGGGAGCGGCGGGGGCGGCGCGC | >hg38_21_+_8433172_8446622_RNA45SN1_45S_50ntFlanks | 10069 | 10097 | 29 | AGTGGACGGGAGCGGCGGGGGCGGCGCGC | 101.1 | 5.9 | 1.9 | 3.2 | 1.4E-03 | 8.2E-03 | BT20/MB231 | MP |
| >am_trna119_LysCTT_1_-_145395522_145395594@13.34.22,trna11_LysCTT_5_-_180648979_180649051@13.34.22,trna128_LysCTT_1_-_55423542_55423614@13.34.22,trna13_LysCTT_6_+_26556774_26556846@13.34.22,trna6_LysCTT_19_-_52425396_52425468@13.34.22,trna7_LysCTT_16_+_3225692_3225764@13.34.22,trna9_LysCTT_5_+_180634755_180634827@13.34.22\|CAGTCGGTAGAGCATGAGACTC | >am_trna119_LysCTT_1_-_145395522_145395594 | 13 | 34 | 22 | CAGTCGGTAGAGCATGAGACTC | 62.0 | 4.3 | 1.6 | 2.7 | 6.2E-03 | 8.2E-03 | MB231/MB468 | MP |
| >trnaMT_PheGAA_MT_+_577_647@36.59.24\|AAAATGTTTAGACGGGCTCACATC | >trnaMT_PheGAA_MT_+_577_647 | 36 | 59 | 24 | AAAATGTTTAGACGGGCTCACATC | 136.3 | 3.5 | 1.3 | 2.7 | 7.1E-03 | 8.2E-03 | MB231/MB468 | MITO |
| >trnaMT_SerGCT_MT_+_12207_12265@13.48.36\|AAGAACTGCTAACTCATGCCCCCATGTCTAACAACA | >trnaMT_SerGCT_MT_+_12207_12265 | 13 | 48 | 36 | AAGAACTGCTAACTCATGCCCCCATGTCTAACAACA | 264.8 | 4.6 | 1.6 | 2.9 | 4.0E-03 | 8.2E-03 | BT20/MB468 | MP |
| >hg38_MT_+_1621_3279_RNR2_16S_50ntFlanks@1370.1395.26\|ATGTTGGATCAGGACATCCCGATGGT | >hg38_MT_+_1621_3279_RNR2_16S_50ntFlanks | 1370 | 1395 | 26 | ATGTTGGATCAGGACATCCCGATGGT | 249.8 | 3.7 | 1.3 | 2.9 | 4.0E-03 | 8.2E-03 | BT20/MB468 | MP |
| >hg38_21_+_8433172_8446622_RNA45SN1_45S_50ntFlanks@9562.9592.31\|TCCTGACGTGCAAATCGGTCGTCCGACCTGG | >hg38_21_+_8433172_8446622_RNA45SN1_45S_50ntFlanks | 9562 | 9592 | 31 | TCCTGACGTGCAAATCGGTCGTCCGACCTGG | 159.4 | -3.1 | 1.2 | -2.7 | 7.2E-03 | 8.2E-03 | MB231/MB468 | NUC |
| >hg38_21_+_8433172_8446622_RNA45SN1_45S_50ntFlanks@11667.11706.40\|CTGCCCAGTGCTCTGAATGTCAAAGTGAAGAAATTCAATG | >hg38_21_+_8433172_8446622_RNA45SN1_45S_50ntFlanks | 11667 | 11706 | 40 | CTGCCCAGTGCTCTGAATGTCAAAGTGAAGAAATTCAATG | 480.5 | 3.0 | 1.0 | 3.1 | 1.9E-03 | 8.2E-03 | BT20/MB231 | MITO |
| >MI0000298\|hsa-mir-221&WithFlank&X\|-\|45746151\|45746272@71.93.23(+2U)&[MIMAT0000278&hsa-miR-221-3p&offsets\|0\|0(+2U);m-91&X\|-\|45746181\|45746202&offsets\|0\|+1(+2U)]\|AGCTACATTGTCTGCTGGGTTTCTT | >MI0000298\|hsa-mir-221&WithFlank&X\|-\|45746151\|45746272 | 71 | 93 | 23(+2U) | AGCTACATTGTCTGCTGGGTTTCTT | 165.9 | -2.2 | 0.7 | -3.2 | 1.6E-03 | 8.2E-03 | BT20/MB231 | NUC |

| | | | | | | | | | | | | | |
|---|---|---|---|---|---|---|---|---|---|---|---|---|---|
| >hg38_21_+_8433172_8446622_RNA45SN1_45S_50ntFlanks@10772.10818.47\|AGGTCTCCAAGGTGAACAGCCTCTGGCATGTTGGAACAATGTAGGTA | >hg38_21_+_8433172_8446622_RNA45SN1_45S_50ntFlanks | 10772 | 10818 | 47 | AGGTCTCCAAGGTGAACAGCCTCTGGCATGTTGGAACAATGTAGGTA | 209.7 | -2.3 | 0.8 | -2.7 | 6.8E-03 | 8.2E-03 | MB231/MB468 | CYTO |
| >hg38_MT_+_1621_3279_RNR2_16S_50ntFlanks@1432.1458.27\|AAGTCCTACGTGATCTGAGTTCAGACC | >hg38_MT_+_1621_3279_RNR2_16S_50ntFlanks | 1432 | 1458 | 27 | AAGTCCTACGTGATCTGAGTTCAGACC | 160.7 | 3.4 | 1.3 | 2.7 | 7.2E-03 | 8.2E-03 | MB231/MB468 | MITO |
| >hg38_21_+_8433172_8446622_RNA45SN1_45S_50ntFlanks@12995.13014.20\|CTCGCTGCGATCTATTGAAA | >hg38_21_+_8433172_8446622_RNA45SN1_45S_50ntFlanks | 12995 | 13014 | 20 | CTCGCTGCGATCTATTGAAA | 509.6 | 2.1 | 0.8 | 2.7 | 6.7E-03 | 8.2E-03 | BT20/MB468 | CYTO |
| >hg38_21_+_8433172_8446622_RNA45SN1_45S_50ntFlanks@12419.12440.22\|GATGTCGGCTCTTCCTATCATT | >hg38_21_+_8433172_8446622_RNA45SN1_45S_50ntFlanks | 12419 | 12440 | 22 | GATGTCGGCTCTTCCTATCATT | 168.5 | 2.2 | 0.8 | 2.7 | 7.2E-03 | 8.2E-03 | MB231/MB468 | MITO |
| >hg38_21_+_8433172_8446622_RNA45SN1_45S_50ntFlanks@5471.5516.46\|CACGGCCCTGGCGGAGCGCTGAGAAGACGGTCGAACTTGACTATCT | >hg38_21_+_8433172_8446622_RNA45SN1_45S_50ntFlanks | 5471 | 5516 | 46 | CACGGCCCTGGCGGAGCGCTGAGAAGACGGTCGAACTTGACTATCT | 395.1 | 2.5 | 0.9 | 2.7 | 6.7E-03 | 8.2E-03 | BT20/MB468 | CYTO |
| >hg38_21_+_8433172_8446622_RNA45SN1_45S_50ntFlanks@10388.10410.23\|GTGAACAGCAGTTGAACATGGGT | >hg38_21_+_8433172_8446622_RNA45SN1_45S_50ntFlanks | 10388 | 10410 | 23 | GTGAACAGCAGTTGAACATGGGT | 349.6 | -2.6 | 0.9 | -2.9 | 4.0E-03 | 8.2E-03 | BT20/MB468 | MP |
| >trna10_AspGTC_12_-_125424193_125424264@35.75.41,trna12_AspGTC_12_-_125411891_125411962@35.75.41,trna144_AspGTC_6_-_27551236_27551307@35.75.41,trna38_AspGTC_17_-_8125556_8125627@35.75.41,trna45_AspGTC_6_+_27447453_27447524@35.75.41,trna48_AspGTC_6_+_27471523_27471594@35.75.41,trna4_AspGTC_12_+_96429799_96429870@35.75.41,trna69_AspGTC_1_-_161440205_161440276@35.75.41,trna72_AspGTC_1_-_161432824_161432895@35.75.41,trna75_AspGTC_1_-_161425414_161425485@35.75.41,trna78_AspGTC_1_-_161418033_161418104@35.75.41,trna81_AspGTC_1_-_161410615_161410686@35.75.41\|TCACGCGGGAGACCGGGGTTCGATTCCCCGACGGGGAGCCA | >trna10_AspGTC_12_-_125424193_125424264 | 35 | 75 | 41 | TCACGCGGGAGACCGGGGTTCGATTCCCCGACGGGGAGCCA | 2183.7 | 4.0 | 1.5 | 2.7 | 7.2E-03 | 8.2E-03 | MB231/MB468 | NUC |
| >hg38_MT_+_1621_3279_RNR2_16S_50ntFlanks@1095.1118.24\|AGACGAGAAGACCCTATGGAGCTT | >hg38_MT_+_1621_3279_RNR2_16S_50ntFlanks | 1095 | 1118 | 24 | AGACGAGAAGACCCTATGGAGCTT | 147.9 | 2.9 | 1.1 | 2.7 | 6.2E-03 | 8.2E-03 | MB231/MB468 | MP |
| >MI0000746\|hsa-mir-99b&WithFlank&19\|+\|51692606\|51692687@13.31.19(+1A)&[MIMAT0000689&hsa-miR-99b-5p&offsets\|0\|-3(+1A);m-10&19\|+\|51692618\|51692639&offsets\|0\|-3(+1A)]\|CACCCGTAGAACCGACCTTA | >MI0000746\|hsa-mir-99b&WithFlank&19\|+\|51692606\|51692687 | 13 | 31 | 19(+1A) | CACCCGTAGAACCGACCTTA | 609.8 | 2.8 | 0.9 | 3.1 | 1.8E-03 | 8.2E-03 | BT20/MB231 | CYTO |
| >hg38_MT_+_1621_3279_RNR2_16S_50ntFlanks@931.958.28\|GTGACACATGTTTAACGGCCGCGGTACC | >hg38_MT_+_1621_3279_RNR2_16S_50ntFlanks | 931 | 958 | 28 | GTGACACATGTTTAACGGCCGCGGTACC | 87.3 | 5.1 | 1.9 | 2.7 | 6.2E-03 | 8.2E-03 | MB231/MB468 | MP |
| >MI0000749\|hsa-mir-30e&WithFlank&1\|+\|40754349\|40754452@23.45.23(+1A)&[MIMAT0000692&hsa-miR-30e-5p&offsets\|0\|+1(+1A);m-24&1\|+\|40754371\|40754394&offsets\|0\|-1(+1A)]\|TGTAAACATCCTTGACTGGAAGCA | >MI0000749\|hsa-mir-30e&WithFlank&1\|+\|40754349\|40754452 | 23 | 45 | 23(+1A) | TGTAAACATCCTTGACTGGAAGCA | 800.0 | -2.8 | 0.9 | -3.1 | 1.9E-03 | 8.2E-03 | BT20/MB231 | MITO |
| >hg38_21_+_8433172_8446622_RNA45SN1_45S_50ntFlanks@6653.6680.28\|ACTCTTAGCGGTGGATCACTCGGCTCGT | >hg38_21_+_8433172_8446622_RNA45SN1_45S_50ntFlanks | 6653 | 6680 | 28 | ACTCTTAGCGGTGGATCACTCGGCTCGT | 526.0 | 1.4 | 0.5 | 2.8 | 5.8E-03 | 8.2E-03 | BT20/MB468 | MITO |
| >MI0000298\|hsa-mir-221&WithFlank&X\|-\|45746151\|45746272@31.55.25(+1C)&[MIMAT0004568&hsa-miR-221-5p&offsets\|0\|+3(+1C);m-405&X\|-\|45746220\|45746242&offsets\|0\|+2(+1C)]\|ACCTGGCATACAATGTAGATTTCTGC | >MI0000298\|hsa-mir-221&WithFlank&X\|-\|45746151\|45746272 | 31 | 55 | 25(+1C) | ACCTGGCATACAATGTAGATTTCTGC | 183.3 | -2.2 | 0.7 | -3.2 | 1.6E-03 | 8.2E-03 | BT20/MB231 | NUC |
| >hg38_21_+_8433172_8446622_RNA45SN1_45S_50ntFlanks@5489.5509.21\|CTGAGAAGACGGTCGAACTTG | >hg38_21_+_8433172_8446622_RNA45SN1_45S_50ntFlanks | 5489 | 5509 | 21 | CTGAGAAGACGGTCGAACTTG | 154.0 | -1.4 | 0.5 | -2.8 | 5.8E-03 | 8.2E-03 | BT20/MB468 | MITO |
| >MI0000812\|hsa-mir-331&WithFlank&12\|+\|95308414\|95308519@32.52.21&[MIMAT0004700&hsa-miR-331-5p&offsets\|0\|-1;m-377&12\|+\|95308445\|95308465&offsets\|0\|0]\|CTAGGTATGGTCCCAGGGATC | >MI0000812\|hsa-mir-331&WithFlank&12\|+\|95308414\|95308519 | 32 | 52 | 21 | CTAGGTATGGTCCCAGGGATC | 119.0 | 5.3 | 1.9 | 2.7 | 6.2E-03 | 8.2E-03 | MB231/MB468 | MP |
| >hg38_21_+_8433172_8446622_RNA45SN1_45S_50ntFlanks@12420.12447.28\|ATGTCGGCTCTTCCTATCATTGTGAAGC | >hg38_21_+_8433172_8446622_RNA45SN1_45S_50ntFlanks | 12420 | 12447 | 28 | ATGTCGGCTCTTCCTATCATTGTGAAGC | 1101.2 | -2.6 | 0.8 | -3.1 | 1.8E-03 | 8.2E-03 | BT20/MB231 | CYTO |
| >trna111_HisGTG_1_-_147774845_147774916@-1G.30.31,trna118_HisGTG_1_-_145396881_145396952@-1G.30.31,trna16_HisGTG_1_+_146544773_146544844@-1G.30.31,trna1_HisGTG_15_+_45493349_45493420@-1G.30.31,trna21_HisGTG_1_+_147753471_147753542@-1G.30.31,trna33_HisGTG_6_+_27125906_27125977@-1G.30.31,trna7_HisGTG_9_-_14433938_14434009@-1G.30.31,trna8_HisGTG_15_-_45492611_45492682@-1G.30.31,trna9_HisGTG_15_-_45490804_45490875@-1G.30.31\|GGCCGTGATCGTATAGTGGTTAGTACTCTGC | >trna111_HisGTG_1_-_147774845_147774916 | -1G | 30 | 31 | GGCCGTGATCGTATAGTGGTTAGTACTCTGC | 1602.3 | 2.2 | 0.8 | 2.7 | 7.2E-03 | 8.2E-03 | MB231/MB468 | MITO |
| >hg38_21_+_8433172_8446622_RNA45SN1_45S_50ntFlanks@11920.11951.32\|ATGAGAGGTGTAGAATAAGTGGGAGGCCCCCG | >hg38_21_+_8433172_8446622_RNA45SN1_45S_50ntFlanks | 11920 | 11951 | 32 | ATGAGAGGTGTAGAATAAGTGGGAGGCCCCCG | 542.9 | -3.0 | 1.1 | -2.7 | 6.8E-03 | 8.2E-03 | MB231/MB468 | CYTO |
| >MI0000272\|hsa-mir-182&WithFlank&7\|-\|129770377\|129770498@29.53.25(+1C)&[MIMAT0000259&hsa-miR-182-5p&offsets\|0\|+1(+1C);m-22&7\|-\|129770449\|129770470&offsets\|0\|+3(+1C)]\|TTTGGCAATGGTAGAACTCACACTGC | >MI0000272\|hsa-mir-182&WithFlank&7\|-\|129770377\|129770498 | 29 | 53 | 25(+1C) | TTTGGCAATGGTAGAACTCACACTGC | 268.9 | 2.3 | 0.7 | 3.2 | 1.6E-03 | 8.2E-03 | BT20/MB231 | NUC |
| >hg38_21_+_8433172_8446622_RNA45SN1_45S_50ntFlanks@4007.4025.19\|CCTCGGGCCGATCGCACGC | >hg38_21_+_8433172_8446622_RNA45SN1_45S_50ntFlanks | 4007 | 4025 | 19 | CCTCGGGCCGATCGCACGC | 202.8 | -2.8 | 1.0 | -2.7 | 6.8E-03 | 8.3E-03 | BT20/MB468 | CYTO |
| >hg38_21_+_8433172_8446622_RNA45SN1_45S_50ntFlanks@11568.11603.36\|GCAGCCGACTTAGAACTGGTGCGGACCAGGGGAATC | >hg38_21_+_8433172_8446622_RNA45SN1_45S_50ntFlanks | 11568 | 11603 | 36 | GCAGCCGACTTAGAACTGGTGCGGACCAGGGGAATC | 916.1 | -2.5 | 0.9 | -2.9 | 4.0E-03 | 8.3E-03 | BT20/MB468 | MP |
| >am_trnaMT_PheGAA_MT_+_577_647@41.66.26\|GTTTAGACGGGCTCACATCACCCCAT | >am_trnaMT_PheGAA_MT_+_577_647 | 41 | 66 | 26 | GTTTAGACGGGCTCACATCACCCCAT | 177.3 | 4.5 | 1.6 | 2.9 | 4.0E-03 | 8.3E-03 | BT20/MB468 | MP |
| >hg38_21_+_8433172_8446622_RNA45SN1_45S_50ntFlanks@11190.11227.38\|CCCGGGGGGCCGGCGGCGGCGGCGACTCTGGACGCGAG | >hg38_21_+_8433172_8446622_RNA45SN1_45S_50ntFlanks | 11190 | 11227 | 38 | CCCGGGGGGCCGGCGGCGGCGGCGACTCTGGACGCGAG | 132.7 | -5.0 | 1.7 | -2.9 | 4.0E-03 | 8.3E-03 | BT20/MB468 | MP |
| >hg38_21_+_8433172_8446622_RNA45SN1_45S_50ntFlanks@10107.10137.31\|CGTGTGGTGTGCGTCGGAGGGCGGCGGCGGC | >hg38_21_+_8433172_8446622_RNA45SN1_45S_50ntFlanks | 10107 | 10137 | 31 | CGTGTGGTGTGCGTCGGAGGGCGGCGGCGGC | 58.9 | 9.2 | 2.9 | 3.2 | 1.5E-03 | 8.3E-03 | BT20/MB231 | MP |
| >MI0000081\|hsa-mir-24-2&WithFlank&19\|-\|13836281\|13836365@56.77.22(+2U);MI0000080\|hsa-mir-24-1&WithFlank&9\|+\|95086015\|95086094@50.71.22(+2U)&[MIMAT0000080&hsa-miR-24-3p&offsets\|0\|0(+2U);m-36&19\|-\|13836288\|13836310&offsets\|0\|-1(+2U)];[MIMAT0000080_1&hsa-miR-24-3p&offsets\|0\|0(+2U);m-35&9\|+\|95086064\|95086086&offsets\|0\|-1(+2U)]\|TGGCTCAGTTCAGCAGGAACAGTT | >MI0000081\|hsa-mir-24-2&WithFlank&19\|-\|13836281\|13836365 | 56 | 77 | 22(+2U) | TGGCTCAGTTCAGCAGGAACAGTT | 125.6 | -5.8 | 1.8 | -3.2 | 1.5E-03 | 8.3E-03 | BT20/MB231 | MP |
| >MI0000102\|hsa-mir-100&WithFlank&11\|-\|122152223\|122152314@19.36.18(+2G)&[MIMAT0000098&hsa-miR-100-5p&offsets\|0\|-4(+2G);m-27&11\|-\|122152276\|122152296&offsets\|0\|-3(+2G)]\|AACCCGTAGATCCGAACTGG | >MI0000102\|hsa-mir-100&WithFlank&11\|-\|122152223\|122152314 | 19 | 36 | 18(+2G) | AACCCGTAGATCCGAACTGG | 54.2 | -9.2 | 2.9 | -3.2 | 1.5E-03 | 8.3E-03 | BT20/MB231 | MP |
| >hg38_21_+_8433172_8446622_RNA45SN1_45S_50ntFlanks@12396.12414.19\|GCGACGTCGCTTTTTGATC | >hg38_21_+_8433172_8446622_RNA45SN1_45S_50ntFlanks | 12396 | 12414 | 19 | GCGACGTCGCTTTTTGATC | 1952.0 | -1.2 | 0.4 | -2.7 | 6.8E-03 | 8.3E-03 | BT20/MB468 | CYTO |
| >MI0000084\|hsa-mir-26b&WithFlank&2\|+\|218402640\|218402728@18.38.21&[MIMAT0000083&hsa-miR-26b-5p&offsets\|0\|0;m-53&2\|+\|218402657\|218402678&offsets\|0\|-1]\|TTCAAGTAATTCAGGATAGGT | >MI0000084\|hsa-mir-26b&WithFlank&2\|+\|218402640\|218402728 | 18 | 38 | 21 | TTCAAGTAATTCAGGATAGGT | 1589.2 | 2.8 | 1.0 | 2.7 | 6.2E-03 | 8.3E-03 | MB231/MB468 | MP |
| >hg38_MT_+_1621_3279_RNR2_16S_50ntFlanks@1216.1265.50\|CAACCTCCGAGCAGTACATGCTAAGACTTCACCAGTCAAAGCGAACTACT | >hg38_MT_+_1621_3279_RNR2_16S_50ntFlanks | 1216 | 1265 | 50 | CAACCTCCGAGCAGTACATGCTAAGACTTCACCAGTCAAAGCGAACTACT | 179.1 | 5.2 | 1.9 | 2.7 | 7.2E-03 | 8.3E-03 | MB231/MB468 | MITO |
| >hg38_21_+_8433172_8446622_RNA45SN1_45S_50ntFlanks@5473.5512.40\|CGGCCCTGGCGGAGCGCTGAGAAGACGGTCGAACTTGACT | >hg38_21_+_8433172_8446622_RNA45SN1_45S_50ntFlanks | 5473 | 5512 | 40 | CGGCCCTGGCGGAGCGCTGAGAAGACGGTCGAACTTGACT | 240.8 | 4.2 | 1.4 | 2.9 | 4.0E-03 | 8.3E-03 | BT20/MB468 | MP |
| >hg38_21_+_8433172_8446622_RNA45SN1_45S_50ntFlanks@12927.12953.27\|CCTGGCGCTAAACCATTCGTAGACGAC | >hg38_21_+_8433172_8446622_RNA45SN1_45S_50ntFlanks | 12927 | 12953 | 27 | CCTGGCGCTAAACCATTCGTAGACGAC | 187.2 | -4.8 | 1.7 | -2.9 | 4.0E-03 | 8.3E-03 | BT20/MB468 | MP |
| >hg38_21_+_8433172_8446622_RNA45SN1_45S_50ntFlanks@8638.8658.21\|GCGACCGGCCGCCGCCGGGCG | >hg38_21_+_8433172_8446622_RNA45SN1_45S_50ntFlanks | 8638 | 8658 | 21 | GCGACCGGCCGCCGCCGGGCG | 196.7 | -5.3 | 1.8 | -2.9 | 4.0E-03 | 8.3E-03 | BT20/MB468 | MP |
| >hg38_21_+_8433172_8446622_RNA45SN1_45S_50ntFlanks@9352.9371.20\|CCGTGGCGCAATGAAGGTGA | >hg38_21_+_8433172_8446622_RNA45SN1_45S_50ntFlanks | 9352 | 9371 | 20 | CCGTGGCGCAATGAAGGTGA | 280.3 | -4.9 | 1.7 | -2.9 | 4.0E-03 | 8.3E-03 | BT20/MB468 | MP |
| >hg38_21_+_8433172_8446622_RNA45SN1_45S_50ntFlanks@5487.5521.35\|CGCTGAGAAGACGGTCGAACTTGACTATCTAGAGG | >hg38_21_+_8433172_8446622_RNA45SN1_45S_50ntFlanks | 5487 | 5521 | 35 | CGCTGAGAAGACGGTCGAACTTGACTATCTAGAGG | 286.9 | -2.9 | 1.0 | -2.9 | 4.0E-03 | 8.3E-03 | BT20/MB468 | MP |
| >trnaMT_SerGCT_MT_+_12207_12265@1.18.18\|GAGAAAGCTCACAAGAAC | >trnaMT_SerGCT_MT_+_12207_12265 | 1 | 18 | 18 | GAGAAAGCTCACAAGAAC | 97215.6 | 2.3 | 0.8 | 2.7 | 6.3E-03 | 8.3E-03 | MB231/MB468 | MP |
| >hg38_21_+_8433172_8446622_RNA45SN1_45S_50ntFlanks@13009.13027.19\|TTGAAAGTCAGCCCTCGAC | >hg38_21_+_8433172_8446622_RNA45SN1_45S_50ntFlanks | 13009 | 13027 | 19 | TTGAAAGTCAGCCCTCGAC | 73.9 | -4.7 | 1.6 | -2.9 | 4.0E-03 | 8.3E-03 | BT20/MB468 | MP |
| >hg38_21_+_8433172_8446622_RNA45SN1_45S_50ntFlanks@8176.8217.42\|CTTCTGATCGAGGCCCAGCCCGTGGACGGTGTGAGGCCGGTA | >hg38_21_+_8433172_8446622_RNA45SN1_45S_50ntFlanks | 8176 | 8217 | 42 | CTTCTGATCGAGGCCCAGCCCGTGGACGGTGTGAGGCCGGTA | 229.2 | -2.4 | 0.9 | -2.7 | 7.3E-03 | 8.3E-03 | MB231/MB468 | MITO |
| >hg38_21_+_8433172_8446622_RNA45SN1_45S_50ntFlanks@11571.11603.33\|GCCGACTTAGAACTGGTGCGGACCAGGGGAATC | >hg38_21_+_8433172_8446622_RNA45SN1_45S_50ntFlanks | 11571 | 11603 | 33 | GCCGACTTAGAACTGGTGCGGACCAGGGGAATC | 447.5 | 4.3 | 1.3 | 3.2 | 1.5E-03 | 8.3E-03 | BT20/MB231 | MP |
| >hg38_21_+_8433172_8446622_RNA45SN1_45S_50ntFlanks@10393.10429.37\|CAGCAGTTGAACATGGGTCAGTCGGTCCTGAGAGATG | >hg38_21_+_8433172_8446622_RNA45SN1_45S_50ntFlanks | 10393 | 10429 | 37 | CAGCAGTTGAACATGGGTCAGTCGGTCCTGAGAGATG | 1096.2 | -3.0 | 1.1 | -2.9 | 4.0E-03 | 8.3E-03 | BT20/MB468 | MP |
| >hg38_21_+_8433172_8446622_RNA45SN1_45S_50ntFlanks@9786.9819.34\|CTCGCTGGCGTGGAGCCGGGCGTGGAATGCGAGT | >hg38_21_+_8433172_8446622_RNA45SN1_45S_50ntFlanks | 9786 | 9819 | 34 | CTCGCTGGCGTGGAGCCGGGCGTGGAATGCGAGT | 191.3 | 2.6 | 0.9 | 2.9 | 4.0E-03 | 8.3E-03 | BT20/MB468 | MP |
| >hg38_21_+_8433172_8446622_RNA45SN1_45S_50ntFlanks@12490.12518.29\|CGTGAGCTGGGTTTAGACCGTCGTGAGAC | >hg38_21_+_8433172_8446622_RNA45SN1_45S_50ntFlanks | 12490 | 12518 | 29 | CGTGAGCTGGGTTTAGACCGTCGTGAGAC | 56.7 | -4.8 | 1.7 | -2.9 | 4.0E-03 | 8.3E-03 | BT20/MB468 | MP |
| >MI0000439\|hsa-mir-23b&WithFlank&9\|+\|95085202\|95085310@67.86.20(+2U)&[MIMAT0000418&hsa-miR-23b-3p&offsets\|+3\|0(+2U);m-42&9\|+\|95085265\|95085285&offsets\|+3\|+2(+2U)]\|ACATTGCCAGGGATTACCACTT | >MI0000439\|hsa-mir-23b&WithFlank&9\|+\|95085202\|95085310 | 67 | 86 | 20(+2U) | ACATTGCCAGGGATTACCACTT | 28.1 | 8.0 | 2.9 | 2.7 | 6.3E-03 | 8.3E-03 | MB231/MB468 | MP |
| >MI0000088\|hsa-mir-30a&WithFlank&6\|-\|71403545\|71403627@54.73.20&[MIMAT0000088&hsa-miR-30a-3p&offsets\|+1\|-1;m-38&6\|-\|71403554\|71403575&offsets\|+1\|-1]\|TTTCAGTCGGATGTTTGCAG | >MI0000088\|hsa-mir-30a&WithFlank&6\|-\|71403545\|71403627 | 54 | 73 | 20 | TTTCAGTCGGATGTTTGCAG | 133.5 | 4.2 | 1.5 | 2.9 | 4.0E-03 | 8.3E-03 | BT20/MB468 | MP |

<table>
<tr><td></td><td></td><td></td><td></td><td></td><td></td><td></td><td></td><td></td><td></td><td></td><td></td><td></td><td></td></tr>
<tr><td>>MI0000750|hsa-mir-26a-2&WithFlank&12|-|57824603|57824698@20.39.20(+1U);MI0000083|hsa-mir-26a-1&WithFlank&3|+|37969398|37969486@16.35.20(+1U)&[MIMAT0000082&hsa-miR-26a-5p&offsets|0|-2(+1U);m-33&12|-|57824658|57824679&offsets|0|-2(+1U)];[MIMAT0000082_1&hsa-miR-26a-5p&offsets|0|-2(+1U);m-34&3|+|37969413|37969434&offsets|0|-2(+1U)]|TTCAAGTAATCCAGGATAGGT</td><td>>MI0000750|hsa-mir-26a-2&WithFlank&12|-|57824603|57824698</td><td>20</td><td>39</td><td>20(+1U)</td><td>TTCAAGTAATCCAGGATAGGT</td><td>493.7</td><td>3.9</td><td>1.4</td><td>2.7</td><td>6.3E-03</td><td>8.3E-03</td><td>MB231/MB468</td><td>MP</td></tr>
<tr><td>>hg38_21_+_8433172_8446622_RNA45SN1_45S_50ntFlanks@10841.10860.20|CGTAACTTCGGGATAAGGAT</td><td>>hg38_21_+_8433172_8446622_RNA45SN1_45S_50ntFlanks</td><td>10841</td><td>10860</td><td>20</td><td>CGTAACTTCGGGATAAGGAT</td><td>558.8</td><td>-2.4</td><td>0.9</td><td>-2.7</td><td>6.9E-03</td><td>8.3E-03</td><td>MB231/MB468</td><td>CYTO</td></tr>
<tr><td>>hg38_21_+_8433172_8446622_RNA45SN1_45S_50ntFlanks@6661.6690.30|CGGTGGATCACTCGGCTCGTGCGTCGATGA</td><td>>hg38_21_+_8433172_8446622_RNA45SN1_45S_50ntFlanks</td><td>6661</td><td>6690</td><td>30</td><td>CGGTGGATCACTCGGCTCGTGCGTCGATGA</td><td>152.8</td><td>-2.1</td><td>0.8</td><td>-2.7</td><td>6.3E-03</td><td>8.3E-03</td><td>MB231/MB468</td><td>MP</td></tr>
<tr><td>>hg38_21_+_8433172_8446622_RNA45SN1_45S_50ntFlanks@9567.9594.28|ACGTGCAAATCGGTCGTCCGACCTGGGT</td><td>>hg38_21_+_8433172_8446622_RNA45SN1_45S_50ntFlanks</td><td>9567</td><td>9594</td><td>28</td><td>ACGTGCAAATCGGTCGTCCGACCTGGGT</td><td>624.7</td><td>-1.7</td><td>0.6</td><td>-2.8</td><td>5.9E-03</td><td>8.3E-03</td><td>BT20/MB468</td><td>MITO</td></tr>
<tr><td>>hg38_1_-_228634819_228635039_RNA5S12_5S_50ntFlanks@137.170.34|GAGACCGCCTGGGAATACCGGGTGCTGTAGGCTT</td><td>>hg38_1_-_228634819_228635039_RNA5S12_5S_50ntFlanks</td><td>137</td><td>170</td><td>34</td><td>GAGACCGCCTGGGAATACCGGGTGCTGTAGGCTT</td><td>151.4</td><td>2.6</td><td>1.0</td><td>2.7</td><td>7.3E-03</td><td>8.3E-03</td><td>MB231/MB468</td><td>MITO</td></tr>
<tr><td>>trnaMT_ValTAC_MT_+_1602_1670@52.71.20|TTAACTTGACCGCTCTGACC</td><td>>trnaMT_ValTAC_MT_+_1602_1670</td><td>52</td><td>71</td><td>20</td><td>TTAACTTGACCGCTCTGACC</td><td>257.8</td><td>2.5</td><td>0.9</td><td>2.7</td><td>7.3E-03</td><td>8.3E-03</td><td>MB231/MB468</td><td>MITO</td></tr>
<tr><td>>trna11_GluTTC_15_-_26327381_26327452@1.29.29,trna3_GluTTC_13_-_45492062_45492133@1.29.29|TCCCACATGGTCTAGCGGTTAGGATTCCT</td><td>>trna11_GluTTC_15_-_26327381_26327452</td><td>1</td><td>29</td><td>29</td><td>TCCCACATGGTCTAGCGGTTAGGATTCCT</td><td>5230.1</td><td>-2.3</td><td>0.9</td><td>-2.7</td><td>6.9E-03</td><td>8.3E-03</td><td>MB231/MB468</td><td>CYTO</td></tr>
<tr><td>>hg38_21_+_8433172_8446622_RNA45SN1_45S_50ntFlanks@6646.6693.48|TCGTACGACTCTTAGCGGTGGATCACTCGGCTCGTGCGTCGATGAAGA</td><td>>hg38_21_+_8433172_8446622_RNA45SN1_45S_50ntFlanks</td><td>6646</td><td>6693</td><td>48</td><td>TCGTACGACTCTTAGCGGTGGATCACTCGGCTCGTGCGTCGATGAAGA</td><td>214.8</td><td>3.7</td><td>1.3</td><td>2.9</td><td>4.0E-03</td><td>8.3E-03</td><td>BT20/MB468</td><td>MP</td></tr>
<tr><td>>hg38_21_+_8433172_8446622_RNA45SN1_45S_50ntFlanks@4926.4944.19|GAGTGGAGCCTGCGGCTTA</td><td>>hg38_21_+_8433172_8446622_RNA45SN1_45S_50ntFlanks</td><td>4926</td><td>4944</td><td>19</td><td>GAGTGGAGCCTGCGGCTTA</td><td>498.1</td><td>5.2</td><td>1.6</td><td>3.2</td><td>1.5E-03</td><td>8.3E-03</td><td>BT20/MB231</td><td>MP</td></tr>
<tr><td>>hg38_21_+_8433172_8446622_RNA45SN1_45S_50ntFlanks@5490.5516.27|TGAGAAGACGGTCGAACTTGACTATCT</td><td>>hg38_21_+_8433172_8446622_RNA45SN1_45S_50ntFlanks</td><td>5490</td><td>5516</td><td>27</td><td>TGAGAAGACGGTCGAACTTGACTATCT</td><td>2269.4</td><td>3.8</td><td>1.2</td><td>3.2</td><td>1.5E-03</td><td>8.3E-03</td><td>BT20/MB231</td><td>MP</td></tr>
<tr><td>>hg38_MT_+_1621_3279_RNR2_16S_50ntFlanks@1370.1401.32|ATGTTGGATCAGGACATCCCGATGGTGCAGCC</td><td>>hg38_MT_+_1621_3279_RNR2_16S_50ntFlanks</td><td>1370</td><td>1401</td><td>32</td><td>ATGTTGGATCAGGACATCCCGATGGTGCAGCC</td><td>219.3</td><td>4.5</td><td>1.6</td><td>2.9</td><td>4.0E-03</td><td>8.3E-03</td><td>BT20/MB468</td><td>MP</td></tr>
<tr><td>>hg38_21_+_8433172_8446622_RNA45SN1_45S_50ntFlanks@10103.10125.23|CGCGCGTGTGGTGTGCGTCGGAG</td><td>>hg38_21_+_8433172_8446622_RNA45SN1_45S_50ntFlanks</td><td>10103</td><td>10125</td><td>23</td><td>CGCGCGTGTGGTGTGCGTCGGAG</td><td>171.3</td><td>-3.6</td><td>1.3</td><td>-2.9</td><td>4.0E-03</td><td>8.3E-03</td><td>BT20/MB468</td><td>MP</td></tr>
<tr><td>>hg38_21_+_8433172_8446622_RNA45SN1_45S_50ntFlanks@11199.11229.31|CCGGCGGCGGCGGCGACTCTGGACGCGAGCC</td><td>>hg38_21_+_8433172_8446622_RNA45SN1_45S_50ntFlanks</td><td>11199</td><td>11229</td><td>31</td><td>CCGGCGGCGGCGGCGACTCTGGACGCGAGCC</td><td>578.3</td><td>-5.2</td><td>1.9</td><td>-2.7</td><td>7.2E-03</td><td>8.3E-03</td><td>BT20/MB468</td><td>NUC</td></tr>
<tr><td>>MI0000065|hsa-let-7d&WithFlank&9|+|94178828|94178926@69.89.21&[MIMAT0004484&hsa-let-7d-3p&offsets|+1|0;m-98&9|+|94178895|94178916&offsets|+1|0]|TATACGACCTGCTGCCTTTCT</td><td>>MI0000065|hsa-let-7d&WithFlank&9|+|94178828|94178926</td><td>69</td><td>89</td><td>21</td><td>TATACGACCTGCTGCCTTTCT</td><td>92.1</td><td>4.2</td><td>1.5</td><td>2.9</td><td>4.0E-03</td><td>8.3E-03</td><td>BT20/MB468</td><td>MP</td></tr>
<tr><td>>hg38_21_+_8433172_8446622_RNA45SN1_45S_50ntFlanks@11568.11599.32|GCAGCCGACTTAGAACTGGTGCGGACCAGGGG</td><td>>hg38_21_+_8433172_8446622_RNA45SN1_45S_50ntFlanks</td><td>11568</td><td>11599</td><td>32</td><td>GCAGCCGACTTAGAACTGGTGCGGACCAGGGG</td><td>243.7</td><td>5.2</td><td>1.6</td><td>3.2</td><td>1.5E-03</td><td>8.4E-03</td><td>BT20/MB231</td><td>MP</td></tr>
<tr><td>>hg38_21_+_8433172_8446622_RNA45SN1_45S_50ntFlanks@8104.8137.34|CCGCGGCGGGGCGCGGGACATGTGGCGTACGGAA</td><td>>hg38_21_+_8433172_8446622_RNA45SN1_45S_50ntFlanks</td><td>8104</td><td>8137</td><td>34</td><td>CCGCGGCGGGGCGCGGGACATGTGGCGTACGGAA</td><td>446.7</td><td>6.2</td><td>1.9</td><td>3.2</td><td>1.5E-03</td><td>8.4E-03</td><td>BT20/MB231</td><td>MP</td></tr>
<tr><td>>hg38_21_+_8433172_8446622_RNA45SN1_45S_50ntFlanks@10789.10812.24|AGCCTCTGGCATGTTGGAACAATG</td><td>>hg38_21_+_8433172_8446622_RNA45SN1_45S_50ntFlanks</td><td>10789</td><td>10812</td><td>24</td><td>AGCCTCTGGCATGTTGGAACAATG</td><td>65.6</td><td>4.8</td><td>1.7</td><td>2.9</td><td>4.1E-03</td><td>8.4E-03</td><td>BT20/MB468</td><td>MP</td></tr>
<tr><td>>hg38_21_+_8433172_8446622_RNA45SN1_45S_50ntFlanks@3705.3748.44|TACCTGGTTGATCCTGCCAGTAGCATATGCTTGTCTCAAAGATT</td><td>>hg38_21_+_8433172_8446622_RNA45SN1_45S_50ntFlanks</td><td>3705</td><td>3748</td><td>44</td><td>TACCTGGTTGATCCTGCCAGTAGCATATGCTTGTCTCAAAGATT</td><td>286.8</td><td>4.6</td><td>1.6</td><td>2.9</td><td>4.1E-03</td><td>8.4E-03</td><td>BT20/MB468</td><td>MP</td></tr>
<tr><td>>am_trna111_HisGTG_1_-_147774845_147774916@-1G.28.29,trna118_HisGTG_1_-_145396881_145396952@-1G.28.29,trna16_HisGTG_1_+_146544773_146544844@-1G.28.29,trna1_HisGTG_15_+_45493349_45493420@-1G.28.29,trna21_HisGTG_1_+_147753471_147753542@-1G.28.29,trna33_HisGTG_6_+_27125906_27125977@-1G.28.29,trna7_HisGTG_9_-_14433938_14434009@-1G.28.29,trna8_HisGTG_15_-_45492611_45492682@-1G.28.29,trna9_HisGTG_15_-_45490804_45490875@-1G.28.29|GGCCGTGATCGTATAGTGGTTAGTACTCT</td><td>>am_trna111_HisGTG_1_-_147774845_147774916</td><td>-1G</td><td>28</td><td>29</td><td>GGCCGTGATCGTATAGTGGTTAGTACTCT</td><td>601.3</td><td>-2.8</td><td>0.9</td><td>-3.2</td><td>1.6E-03</td><td>8.4E-03</td><td>BT20/MB231</td><td>NUC</td></tr>
<tr><td>>hg38_21_+_8433172_8446622_RNA45SN1_45S_50ntFlanks@4002.4022.21|GATAACCTCGGGCCGATCGCA</td><td>>hg38_21_+_8433172_8446622_RNA45SN1_45S_50ntFlanks</td><td>4002</td><td>4022</td><td>21</td><td>GATAACCTCGGGCCGATCGCA</td><td>315.6</td><td>-3.3</td><td>1.2</td><td>-2.8</td><td>5.9E-03</td><td>8.4E-03</td><td>BT20/MB468</td><td>MITO</td></tr>
<tr><td>>hg38_21_+_8433172_8446622_RNA45SN1_45S_50ntFlanks@4728.4755.28|ATCAAGAACGAAAGTCGGAGGTTCGAAG</td><td>>hg38_21_+_8433172_8446622_RNA45SN1_45S_50ntFlanks</td><td>4728</td><td>4755</td><td>28</td><td>ATCAAGAACGAAAGTCGGAGGTTCGAAG</td><td>142.5</td><td>-4.0</td><td>1.5</td><td>-2.7</td><td>7.3E-03</td><td>8.4E-03</td><td>MB231/MB468</td><td>NUC</td></tr>
<tr><td>>hg38_21_+_8433172_8446622_RNA45SN1_45S_50ntFlanks@10846.10863.18|CTTCGGGATAAGGATTGG</td><td>>hg38_21_+_8433172_8446622_RNA45SN1_45S_50ntFlanks</td><td>10846</td><td>10863</td><td>18</td><td>CTTCGGGATAAGGATTGG</td><td>292.0</td><td>-4.8</td><td>1.7</td><td>-2.9</td><td>4.1E-03</td><td>8.4E-03</td><td>BT20/MB468</td><td>MP</td></tr>
<tr><td>>hg38_21_+_8433172_8446622_RNA45SN1_45S_50ntFlanks@8104.8137.34|CCGCGGCGGGGCGCGGGACATGTGGCGTACGGAA</td><td>>hg38_21_+_8433172_8446622_RNA45SN1_45S_50ntFlanks</td><td>8104</td><td>8137</td><td>34</td><td>CCGCGGCGGGGCGCGGGACATGTGGCGTACGGAA</td><td>136.8</td><td>-4.8</td><td>1.8</td><td>-2.7</td><td>6.3E-03</td><td>8.4E-03</td><td>MB231/MB468</td><td>MP</td></tr>
<tr><td>>hg38_21_+_8433172_8446622_RNA45SN1_45S_50ntFlanks@11572.11600.29|CCGACTTAGAACTGGTGCGGACCAGGGGA</td><td>>hg38_21_+_8433172_8446622_RNA45SN1_45S_50ntFlanks</td><td>11572</td><td>11600</td><td>29</td><td>CCGACTTAGAACTGGTGCGGACCAGGGGA</td><td>705.9</td><td>3.9</td><td>1.2</td><td>3.2</td><td>1.5E-03</td><td>8.4E-03</td><td>BT20/MB231</td><td>MP</td></tr>
<tr><td>>hg38_21_+_8433172_8446622_RNA45SN1_45S_50ntFlanks@10236.10261.26|GACGCTACGCCGCGACGAGTAGGAGG</td><td>>hg38_21_+_8433172_8446622_RNA45SN1_45S_50ntFlanks</td><td>10236</td><td>10261</td><td>26</td><td>GACGCTACGCCGCGACGAGTAGGAGG</td><td>169.8</td><td>3.0</td><td>1.0</td><td>2.9</td><td>4.1E-03</td><td>8.4E-03</td><td>BT20/MB468</td><td>MP</td></tr>
<tr><td>>hg38_21_+_8433172_8446622_RNA45SN1_45S_50ntFlanks@10423.10443.21|AGAGATGGGCGAGCGCCGTTC</td><td>>hg38_21_+_8433172_8446622_RNA45SN1_45S_50ntFlanks</td><td>10423</td><td>10443</td><td>21</td><td>AGAGATGGGCGAGCGCCGTTC</td><td>145.2</td><td>-6.5</td><td>2.2</td><td>-2.9</td><td>4.1E-03</td><td>8.4E-03</td><td>BT20/MB468</td><td>MP</td></tr>
<tr><td>>hg38_21_+_8433172_8446622_RNA45SN1_45S_50ntFlanks@11845.11886.42|GGAACGGGCTTGGCGGAATCAGCGGGGAAAGAAGACCCTGTT</td><td>>hg38_21_+_8433172_8446622_RNA45SN1_45S_50ntFlanks</td><td>11845</td><td>11886</td><td>42</td><td>GGAACGGGCTTGGCGGAATCAGCGGGGAAAGAAGACCCTGTT</td><td>58.4</td><td>5.7</td><td>2.0</td><td>2.9</td><td>4.1E-03</td><td>8.4E-03</td><td>BT20/MB468</td><td>MP</td></tr>
<tr><td>>hg38_21_+_8433172_8446622_RNA45SN1_45S_50ntFlanks@12416.12435.20|TTCGATGTCGGCTCTTCCTA</td><td>>hg38_21_+_8433172_8446622_RNA45SN1_45S_50ntFlanks</td><td>12416</td><td>12435</td><td>20</td><td>TTCGATGTCGGCTCTTCCTA</td><td>309.0</td><td>-1.2</td><td>0.5</td><td>-2.8</td><td>5.9E-03</td><td>8.4E-03</td><td>BT20/MB468</td><td>MITO</td></tr>
<tr><td>>hg38_MT_+_1621_3279_RNR2_16S_50ntFlanks@1089.1117.29|ACAGCAAGACGAGAAGACCCTATGGAGCT</td><td>>hg38_MT_+_1621_3279_RNR2_16S_50ntFlanks</td><td>1089</td><td>1117</td><td>29</td><td>ACAGCAAGACGAGAAGACCCTATGGAGCT</td><td>61.8</td><td>5.9</td><td>2.1</td><td>2.9</td><td>4.1E-03</td><td>8.4E-03</td><td>BT20/MB468</td><td>MP</td></tr>
<tr><td>>hg38_21_+_8433172_8446622_RNA45SN1_45S_50ntFlanks@10819.10839.21|AGGGAAGTCGGCAAGCCGGAT</td><td>>hg38_21_+_8433172_8446622_RNA45SN1_45S_50ntFlanks</td><td>10819</td><td>10839</td><td>21</td><td>AGGGAAGTCGGCAAGCCGGAT</td><td>167.7</td><td>-3.0</td><td>1.1</td><td>-2.7</td><td>6.9E-03</td><td>8.4E-03</td><td>BT20/MB468</td><td>CYTO</td></tr>
<tr><td>>MI0000253|hsa-mir-148a&WithFlank&7|-|25949913|25949992@50.71.22&[MIMAT0000243&hsa-miR-148a-3p&offsets|0|0;m-7&7|-|25949922|25949943&offsets|0|0]|TCAGTGCACTACAGAACTTTGT</td><td>>MI0000253|hsa-mir-148a&WithFlank&7|-|25949913|25949992</td><td>50</td><td>71</td><td>22</td><td>TCAGTGCACTACAGAACTTTGT</td><td>378901.1</td><td>3.5</td><td>1.1</td><td>3.2</td><td>1.5E-03</td><td>8.4E-03</td><td>BT20/MB231</td><td>MP</td></tr>
<tr><td>>hg38_21_+_8433172_8446622_RNA45SN1_45S_50ntFlanks@4588.4622.35|CTATTTTGTTGGTTTTCGGAACTGAGGCCATGATT</td><td>>hg38_21_+_8433172_8446622_RNA45SN1_45S_50ntFlanks</td><td>4588</td><td>4622</td><td>35</td><td>CTATTTTGTTGGTTTTCGGAACTGAGGCCATGATT</td><td>224.4</td><td>-2.0</td><td>0.7</td><td>-2.8</td><td>5.9E-03</td><td>8.4E-03</td><td>BT20/MB468</td><td>MITO</td></tr>
<tr><td>>hg38_MT_+_1621_3279_RNR2_16S_50ntFlanks@1181.1216.36|AAAAATTTCGGTTGGGGCGACCTCGGAGCAGAACCC</td><td>>hg38_MT_+_1621_3279_RNR2_16S_50ntFlanks</td><td>1181</td><td>1216</td><td>36</td><td>AAAAATTTCGGTTGGGGCGACCTCGGAGCAGAACCC</td><td>120.6</td><td>5.9</td><td>2.0</td><td>2.9</td><td>4.1E-03</td><td>8.4E-03</td><td>BT20/MB468</td><td>MP</td></tr>
<tr><td>>hg38_21_+_8433172_8446622_RNA45SN1_45S_50ntFlanks@9791.9813.23|TGGCGTGGAGCCGGGCGTGGAAT</td><td>>hg38_21_+_8433172_8446622_RNA45SN1_45S_50ntFlanks</td><td>9791</td><td>9813</td><td>23</td><td>TGGCGTGGAGCCGGGCGTGGAAT</td><td>182.3</td><td>6.2</td><td>1.9</td><td>3.2</td><td>1.5E-03</td><td>8.4E-03</td><td>BT20/MB231</td><td>MP</td></tr>
<tr><td>>hg38_21_+_8433172_8446622_RNA45SN1_45S_50ntFlanks@4048.4069.22|TTCGAACGTCTGCCCTATCAAC</td><td>>hg38_21_+_8433172_8446622_RNA45SN1_45S_50ntFlanks</td><td>4048</td><td>4069</td><td>22</td><td>TTCGAACGTCTGCCCTATCAAC</td><td>244.7</td><td>-2.5</td><td>0.8</td><td>-3.1</td><td>1.9E-03</td><td>8.4E-03</td><td>BT20/MB231</td><td>CYTO</td></tr>
<tr><td>>trnaMT_GluTTC_MT_-_14674_14742@45.72.28,trnalookalike8_GluTTC_5_-_93905172_93905240@45.72.28|CGTGGTTGTAGTCCGTGCGAGAATACCA</td><td>>trnaMT_GluTTC_MT_-_14674_14742</td><td>45</td><td>72</td><td>28</td><td>CGTGGTTGTAGTCCGTGCGAGAATACCA</td><td>655.0</td><td>2.9</td><td>1.0</td><td>2.9</td><td>4.1E-03</td><td>8.4E-03</td><td>BT20/MB468</td><td>MP</td></tr>
<tr><td>>hg38_21_+_8433172_8446622_RNA45SN1_45S_50ntFlanks@9931.9949.19|TATAGACAGCAGGACGGTG</td><td>>hg38_21_+_8433172_8446622_RNA45SN1_45S_50ntFlanks</td><td>9931</td><td>9949</td><td>19</td><td>TATAGACAGCAGGACGGTG</td><td>188.4</td><td>5.8</td><td>2.0</td><td>2.9</td><td>4.1E-03</td><td>8.4E-03</td><td>BT20/MB468</td><td>MP</td></tr>
<tr><td>>hg38_21_+_8433172_8446622_RNA45SN1_45S_50ntFlanks@4980.5001.22|ACACGGACAGGATTGACAGATT</td><td>>hg38_21_+_8433172_8446622_RNA45SN1_45S_50ntFlanks</td><td>4980</td><td>5001</td><td>22</td><td>ACACGGACAGGATTGACAGATT</td><td>138.4</td><td>2.4</td><td>0.8</td><td>2.9</td><td>4.1E-03</td><td>8.4E-03</td><td>BT20/MB468</td><td>MP</td></tr>
<tr><td>>hg38_21_+_8433172_8446622_RNA45SN1_45S_50ntFlanks@6661.6692.32|CGGTGGATCACTCGGCTCGTGCGTCGATGAAG</td><td>>hg38_21_+_8433172_8446622_RNA45SN1_45S_50ntFlanks</td><td>6661</td><td>6692</td><td>32</td><td>CGGTGGATCACTCGGCTCGTGCGTCGATGAAG</td><td>726.9</td><td>3.6</td><td>1.1</td><td>3.2</td><td>1.5E-03</td><td>8.4E-03</td><td>BT20/MB231</td><td>MP</td></tr>
<tr><td>>MI0000273|hsa-mir-183&WithFlank&7|-|129774899|129775020@34.54.21&[MIMAT0000261&hsa-miR-183-5p&offsets|+1|0;m-41&7|-|129774966|129774987&offsets|0|-1]|ATGGCACTGGTAGAATTCACT</td><td>>MI0000273|hsa-mir-183&WithFlank&7|-|129774899|129775020</td><td>34</td><td>54</td><td>21</td><td>ATGGCACTGGTAGAATTCACT</td><td>1374.0</td><td>2.5</td><td>0.9</td><td>2.9</td><td>4.1E-03</td><td>8.4E-03</td><td>BT20/MB468</td><td>MP</td></tr>
<tr><td>>hg38_21_+_8433172_8446622_RNA45SN1_45S_50ntFlanks@10458.10483.26|ATGGCCTCCGTTGCCCTCGGCCGATC</td><td>>hg38_21_+_8433172_8446622_RNA45SN1_45S_50ntFlanks</td><td>10458</td><td>10483</td><td>26</td><td>ATGGCCTCCGTTGCCCTCGGCCGATC</td><td>141.0</td><td>-5.0</td><td>1.8</td><td>-2.9</td><td>4.1E-03</td><td>8.4E-03</td><td>BT20/MB468</td><td>MP</td></tr>
<tr><td>>MI0000101|hsa-mir-99a&WithFlank&21|+|16539083|16539175@19.39.21&[MIMAT0000097&hsa-miR-99a-5p&offsets|0|-1;m-64&21|+|16539101|16539121&offsets|0|0]|AACCCGTAGATCCGATCTTGT</td><td>>MI0000101|hsa-mir-99a&WithFlank&21|+|16539083|16539175</td><td>19</td><td>39</td><td>21</td><td>AACCCGTAGATCCGATCTTGT</td><td>5236.1</td><td>-2.0</td><td>0.7</td><td>-2.7</td><td>6.4E-03</td><td>8.4E-03</td><td>MB231/MB468</td><td>MP</td></tr>
<tr><td>>hg38_21_+_8433172_8446622_RNA45SN1_45S_50ntFlanks@10677.10708.32|GCGGTTCCGGCGGCGTCCGGTGAGCTCTCGCT</td><td>>hg38_21_+_8433172_8446622_RNA45SN1_45S_50ntFlanks</td><td>10677</td><td>10708</td><td>32</td><td>GCGGTTCCGGCGGCGTCCGGTGAGCTCTCGCT</td><td>556.6</td><td>-1.5</td><td>0.6</td><td>-2.7</td><td>7.4E-03</td><td>8.4E-03</td><td>MB231/MB468</td><td>NUC</td></tr>
<tr><td>>hg38_21_+_8433172_8446622_RNA45SN1_45S_50ntFlanks@13005.13039.35|TCTATTGAAAGTCAGCCCTCGACACAAGGGTTTGT</td><td>>hg38_21_+_8433172_8446622_RNA45SN1_45S_50ntFlanks</td><td>13005</td><td>13039</td><td>35</td><td>TCTATTGAAAGTCAGCCCTCGACACAAGGGTTTGT</td><td>133.6</td><td>2.8</td><td>1.0</td><td>2.9</td><td>4.1E-03</td><td>8.4E-03</td><td>BT20/MB468</td><td>MP</td></tr>
<tr><td>>trnaMT_HisGTG_MT_+_12138_12206@32.69.38|TGAATCTGACAACAGAGGCTTACGACCCCTTATTTACC</td><td>>trnaMT_HisGTG_MT_+_12138_12206</td><td>32</td><td>69</td><td>38</td><td>TGAATCTGACAACAGAGGCTTACGACCCCTTATTTACC</td><td>3443.9</td><td>2.5</td><td>0.9</td><td>2.7</td><td>7.4E-03</td><td>8.4E-03</td><td>MB231/MB468</td><td>MITO</td></tr>
<tr><td>>MI0003602|hsa-mir-590&WithFlank&7|+|74191192|74191300@62.82.21&[MIMAT0004801&hsa-miR-590-3p&offsets|0|0;m-332&7|+|74191253|74191273&offsets|0|0]|TAATTTTATGTATAAGCTAGT</td><td>>MI0003602|hsa-mir-590&WithFlank&7|+|74191192|74191300</td><td>62</td><td>82</td><td>21</td><td>TAATTTTATGTATAAGCTAGT</td><td>141.2</td><td>3.5</td><td>1.3</td><td>2.7</td><td>6.4E-03</td><td>8.4E-03</td><td>MB231/MB468</td><td>MP</td></tr>
<tr><td>>hg38_21_+_8433172_8446622_RNA45SN1_45S_50ntFlanks@9352.9370.19|CCGTGGCGCAATGAAGGTG</td><td>>hg38_21_+_8433172_8446622_RNA45SN1_45S_50ntFlanks</td><td>9352</td><td>9370</td><td>19</td><td>CCGTGGCGCAATGAAGGTG</td><td>238.3</td><td>-4.3</td><td>1.6</td><td>-2.7</td><td>6.9E-03</td><td>8.4E-03</td><td>BT20/MB468</td><td>CYTO</td></tr>
<tr><td>>hg38_MT_+_598_1651_RNR1_12S_50ntFlanks@51.88.38|AATAGGTTTGGTCCTAGCCTTTCTATTAGCTCTTAGTA</td><td>>hg38_MT_+_598_1651_RNR1_12S_50ntFlanks</td><td>51</td><td>88</td><td>38</td><td>AATAGGTTTGGTCCTAGCCTTTCTATTAGCTCTTAGTA</td><td>7234.7</td><td>3.1</td><td>1.1</td><td>2.7</td><td>6.4E-03</td><td>8.4E-03</td><td>MB231/MB468</td><td>MP</td></tr>
<tr><td>>hg38_21_+_8433172_8446622_RNA45SN1_45S_50ntFlanks@9129.9151.23|CGGGTCGGGGGGCGGGGCGGACT</td><td>>hg38_21_+_8433172_8446622_RNA45SN1_45S_50ntFlanks</td><td>9129</td><td>9151</td><td>23</td><td>CGGGTCGGGGGGCGGGGCGGACT</td><td>212.1</td><td>-4.9</td><td>1.7</td><td>-2.9</td><td>4.1E-03</td><td>8.4E-03</td><td>BT20/MB468</td><td>MP</td></tr>
</table>

| | | | | | | | | | | | | | |
|---|---|---|---|---|---|---|---|---|---|---|---|---|---|
| >hg38_21_+_8433172_8446622_RNA45SN1_45S_50ntFlanks@12251.12284.34\|AAAGCTCGCTTGATCTTGATTTTCAGTACGAATA | >hg38_21_+_8433172_8446622_RNA45SN1_45S_50ntFlanks | 12251 | 12284 | 34 | AAAGCTCGCTTGATCTTGATTTTCAGTACGAATA | 204.7 | -1.5 | 0.6 | -2.7 | 7.4E-03 | 8.4E-03 | MB231/MB468 | NUC |
| >hg38_MT_+_1621_3279_RNR2_16S_50ntFlanks@1425.1453.29\|ACGATTAAAGTCCTACGTGATCTGAGTTC | >hg38_MT_+_1621_3279_RNR2_16S_50ntFlanks | 1425 | 1453 | 29 | ACGATTAAAGTCCTACGTGATCTGAGTTC | 213.4 | 3.9 | 1.4 | 2.7 | 6.4E-03 | 8.4E-03 | MB231/MB468 | MP |
| >hg38_21_+_8433172_8446622_RNA45SN1_45S_50ntFlanks@12405.12435.31\|CTTTTTGATCCTTCGATGTCGGCTCTTCCTA | >hg38_21_+_8433172_8446622_RNA45SN1_45S_50ntFlanks | 12405 | 12435 | 31 | CTTTTTGATCCTTCGATGTCGGCTCTTCCTA | 1319.1 | -5.2 | 1.8 | -2.9 | 4.1E-03 | 8.4E-03 | BT20/MB468 | MP |
| >hg38_21_+_8433172_8446622_RNA45SN1_45S_50ntFlanks@12627.12659.33\|ATGGGGCGAAGCTACCATCTGTGGGATTATGAC | >hg38_21_+_8433172_8446622_RNA45SN1_45S_50ntFlanks | 12627 | 12659 | 33 | ATGGGGCGAAGCTACCATCTGTGGGATTATGAC | 2268.5 | -4.7 | 1.6 | -2.9 | 4.1E-03 | 8.4E-03 | BT20/MB468 | MP |
| >trnaMT_ValTAC_MT_+_1602_1670@33.72.40\|ACACTTAGGAGATTTCAACTTAACTTGACCGCTCTGACCA | >trnaMT_ValTAC_MT_+_1602_1670 | 33 | 72 | 40 | ACACTTAGGAGATTTCAACTTAACTTGACCGCTCTGACCA | 368.5 | 3.9 | 1.4 | 2.7 | 6.4E-03 | 8.4E-03 | MB231/MB468 | MP |
| >hg38_21_+_8433172_8446622_RNA45SN1_45S_50ntFlanks@6742.6772.31\|TCGACACTTCGAACGCACTTGCGGCCCCGGG | >hg38_21_+_8433172_8446622_RNA45SN1_45S_50ntFlanks | 6742 | 6772 | 31 | TCGACACTTCGAACGCACTTGCGGCCCCGGG | 200.9 | -1.9 | 0.7 | -2.8 | 5.9E-03 | 8.4E-03 | BT20/MB468 | MITO |
| >hg38_21_+_8433172_8446622_RNA45SN1_45S_50ntFlanks@11200.11227.28\|CGGCGGCGGCGGCGACTCTGGACGCGAG | >hg38_21_+_8433172_8446622_RNA45SN1_45S_50ntFlanks | 11200 | 11227 | 28 | CGGCGGCGGCGGCGACTCTGGACGCGAG | 425.8 | -2.6 | 1.0 | -2.7 | 7.0E-03 | 8.5E-03 | MB231/MB468 | CYTO |
| >MI0000065\|hsa-let-7d&WithFlank&9\|+\|94178828\|94178926@14.34.21(+1A)&[MIMAT0000065&hsa-let-7d-5p&offsets\|0\|-1(+1A);m-90&9\|+\|94178841\|94178862&offsets\|0\|-1(+1A)]\|AGAGGTAGTAGGTTGCATAGTA | >MI0000065\|hsa-let-7d&WithFlank&9\|+\|94178828\|94178926 | 14 | 34 | 21(+1A) | AGAGGTAGTAGGTTGCATAGTA | 335.3 | 1.9 | 0.6 | 3.1 | 1.6E-03 | 8.5E-03 | BT20/MB231 | NUC |
| >MI0000825\|hsa-mir-345&WithFlank&14\|+\|100307853\|100307962@24.45.22&[MIMAT0000772&hsa-miR-345-5p&offsets\|0\|0;m-193&14\|+\|100307876\|100307896&offsets\|0\|+1]\|GCTGACTCCTAGTCCAGGGCTC | >MI0000825\|hsa-mir-345&WithFlank&14\|+\|100307853\|100307962 | 24 | 45 | 22 | GCTGACTCCTAGTCCAGGGCTC | 362.5 | 3.3 | 1.1 | 2.9 | 4.1E-03 | 8.5E-03 | BT20/MB468 | MP |
| >MI0000750\|hsa-mir-26a-2&WithFlank&12\|-\|57824603\|57824698@23.41.19(+2U);MI0000083\|hsa-mir-26a-1&WithFlank&3\|+\|37969398\|37969486@19.37.19(+2U)&[MIMAT0000082&hsa-miR-26a-5p&offsets\|+3\|0(+2U);m-33&12\|-\|57824658\|57824679&offsets\|+3\|0(+2U)];[MIMAT0000082_1&hsa-miR-26a-5p&offsets\|+3\|0(+2U);m-34&3\|+\|37969413\|37969434&offsets\|+3\|0(+2U)]\|AAGTAATCCAGGATAGGCTTT | >MI0000750\|hsa-mir-26a-2&WithFlank&12\|-\|57824603\|57824698 | 23 | 41 | 19(+2U) | AAGTAATCCAGGATAGGCTTT | 27.4 | 8.0 | 2.9 | 2.7 | 6.4E-03 | 8.5E-03 | MB231/MB468 | MP |
| >hg38_MT_+_1621_3279_RNR2_16S_50ntFlanks@1389.1415.27\|CGATGGTGCAGCCGCTATTAAAGGTTC | >hg38_MT_+_1621_3279_RNR2_16S_50ntFlanks | 1389 | 1415 | 27 | CGATGGTGCAGCCGCTATTAAAGGTTC | 236.7 | 4.7 | 1.7 | 2.7 | 6.4E-03 | 8.5E-03 | MB231/MB468 | MP |
| >hg38_21_+_8433172_8446622_RNA45SN1_45S_50ntFlanks@4730.4751.22\|CAAGAACGAAAGTCGGAGGTTC | >hg38_21_+_8433172_8446622_RNA45SN1_45S_50ntFlanks | 4730 | 4751 | 22 | CAAGAACGAAAGTCGGAGGTTC | 968.1 | -2.8 | 1.0 | -2.9 | 4.1E-03 | 8.5E-03 | BT20/MB468 | MP |
| >hg38_21_+_8433172_8446622_RNA45SN1_45S_50ntFlanks@10830.10851.22\|CAAGCCGGATCCGTAACTTCGG | >hg38_21_+_8433172_8446622_RNA45SN1_45S_50ntFlanks | 10830 | 10851 | 22 | CAAGCCGGATCCGTAACTTCGG | 347.9 | -2.4 | 0.8 | -2.9 | 4.1E-03 | 8.5E-03 | BT20/MB468 | MP |
| >hg38_21_+_8433172_8446622_RNA45SN1_45S_50ntFlanks@12627.12662.36\|ATGGGGCGAAGCTACCATCTGTGGGATTATGACTGA | >hg38_21_+_8433172_8446622_RNA45SN1_45S_50ntFlanks | 12627 | 12662 | 36 | ATGGGGCGAAGCTACCATCTGTGGGATTATGACTGA | 145.4 | -4.1 | 1.5 | -2.7 | 6.0E-03 | 8.5E-03 | BT20/MB468 | MITO |
| >hg38_21_+_8433172_8446622_RNA45SN1_45S_50ntFlanks@9567.9603.37\|ACGTGCAAATCGGTCGTCCGACCTGGGTATAGGGGCG | >hg38_21_+_8433172_8446622_RNA45SN1_45S_50ntFlanks | 9567 | 9603 | 37 | ACGTGCAAATCGGTCGTCCGACCTGGGTATAGGGGCG | 1119.3 | -2.2 | 0.8 | -2.8 | 6.0E-03 | 8.5E-03 | BT20/MB468 | MITO |
| >hg38_21_+_8433172_8446622_RNA45SN1_45S_50ntFlanks@11565.11592.28\|CTAGCAGCCGACTTAGAACTGGTGCGGA | >hg38_21_+_8433172_8446622_RNA45SN1_45S_50ntFlanks | 11565 | 11592 | 28 | CTAGCAGCCGACTTAGAACTGGTGCGGA | 174.5 | -4.2 | 1.5 | -2.9 | 4.1E-03 | 8.5E-03 | BT20/MB468 | MP |
| >hg38_21_+_8433172_8446622_RNA45SN1_45S_50ntFlanks@11212.11253.42\|CGACTCTGGACGCGAGCCGGGCCCTTCCCGTGGATCGCCCCA | >hg38_21_+_8433172_8446622_RNA45SN1_45S_50ntFlanks | 11212 | 11253 | 42 | CGACTCTGGACGCGAGCCGGGCCCTTCCCGTGGATCGCCCCA | 619.1 | -4.8 | 1.8 | -2.7 | 7.4E-03 | 8.5E-03 | MB231/MB468 | NUC |
| >MI0000456\|hsa-mir-140&WithFlank&16\|+\|69933075\|69933186@29.49.21(+1A)&[MIMAT0000431&hsa-miR-140-5p&offsets\|0\|-1(+1A);m-173&16\|+\|69933103\|69933124&offsets\|0\|-1(+1A)]\|CAGTGGTTTTACCCTATGGTAA | >MI0000456\|hsa-mir-140&WithFlank&16\|+\|69933075\|69933186 | 29 | 49 | 21(+1A) | CAGTGGTTTTACCCTATGGTAA | 121.7 | 6.2 | 2.1 | 2.9 | 4.1E-03 | 8.5E-03 | BT20/MB468 | MP |
| >hg38_21_+_8433172_8446622_RNA45SN1_45S_50ntFlanks@10393.10424.32\|CAGCAGTTGAACATGGGTCAGTCGGTCCTGAG | >hg38_21_+_8433172_8446622_RNA45SN1_45S_50ntFlanks | 10393 | 10424 | 32 | CAGCAGTTGAACATGGGTCAGTCGGTCCTGAG | 14608.0 | 4.5 | 1.4 | 3.2 | 1.5E-03 | 8.5E-03 | BT20/MB231 | MP |
| >MI0000108\|hsa-mir-103a-2&WithFlank&20\|+\|3917488\|3917577@54.75.22(+1C);MI0000109\|hsa-mir-103a-1&WithFlank&5\|-\|168560890\|168560979@54.75.22(+1C)&[MIMAT0000101&hsa-miR-103a-3p&offsets\|0\|-1(+1C);m-12&20\|+\|3917541\|3917563&offsets\|0\|-1(+1C)];[MIMAT0000101_1&hsa-miR-103a-3p&offsets\|0\|-1(+1C);m-13&5\|-\|168560904\|168560926&offsets\|0\|-1(+1C);m-44220&5\|-\|168560908\|168560926&offsets\|0\|+3(+1C)]\|AGCAGCATTGTACAGGGCTATGC | >MI0000108\|hsa-mir-103a-2&WithFlank&20\|+\|3917488\|3917577 | 54 | 75 | 22(+1C) | AGCAGCATTGTACAGGGCTATGC | 484.4 | -4.6 | 1.4 | -3.2 | 1.5E-03 | 8.5E-03 | BT20/MB231 | MP |
| >hg38_21_+_8433172_8446622_RNA45SN1_45S_50ntFlanks@12416.12434.19\|TTCGATGTCGGCTCTTCCT | >hg38_21_+_8433172_8446622_RNA45SN1_45S_50ntFlanks | 12416 | 12434 | 19 | TTCGATGTCGGCTCTTCCT | 3019.5 | 1.9 | 0.7 | 2.7 | 6.9E-03 | 8.5E-03 | BT20/MB468 | CYTO |
| >trnaMT_MetCAT_MT_+_4402_4469@32.71.40\|ATACCCCGAAAATGTTGGTTATACCCTTCCCGTACTACCA | >trnaMT_MetCAT_MT_+_4402_4469 | 32 | 71 | 40 | ATACCCCGAAAATGTTGGTTATACCCTTCCCGTACTACCA | 407.4 | -5.8 | 1.9 | -3.1 | 1.9E-03 | 8.5E-03 | BT20/MB231 | CYTO |
| >hg38_21_+_8433172_8446622_RNA45SN1_45S_50ntFlanks@8528.8550.23\|CTTCCCCCGCCGCCCCTCCTCCT | >hg38_21_+_8433172_8446622_RNA45SN1_45S_50ntFlanks | 8528 | 8550 | 23 | CTTCCCCCGCCGCCCCTCCTCCT | 355.0 | -4.3 | 1.6 | -2.7 | 7.0E-03 | 8.5E-03 | MB231/MB468 | CYTO |
| >MI0000085\|hsa-mir-27a&WithFlank&19\|-\|13836434\|13836523@57.75.19&[MIMAT0000084&hsa-miR-27a-3p&offsets\|0\|-2;m-43&19\|-\|13836447\|13836467&offsets\|0\|-2]\|TTCACAGTGGCTAAGTTCC | >MI0000085\|hsa-mir-27a&WithFlank&19\|-\|13836434\|13836523 | 57 | 75 | 19 | TTCACAGTGGCTAAGTTCC | 530.8 | 2.8 | 1.0 | 2.9 | 4.1E-03 | 8.5E-03 | BT20/MB468 | MP |
| >hg38_21_+_8433172_8446622_RNA45SN1_45S_50ntFlanks@6656.6686.31\|CTTAGCGGTGGATCACTCGGCTCGTGCGTCG | >hg38_21_+_8433172_8446622_RNA45SN1_45S_50ntFlanks | 6656 | 6686 | 31 | CTTAGCGGTGGATCACTCGGCTCGTGCGTCG | 192.5 | -1.8 | 0.7 | -2.7 | 7.4E-03 | 8.5E-03 | MB231/MB468 | NUC |
| >hg38_21_+_8433172_8446622_RNA45SN1_45S_50ntFlanks@12999.13029.31\|CTGCGATCTATTGAAAGTCAGCCCTCGACAC | >hg38_21_+_8433172_8446622_RNA45SN1_45S_50ntFlanks | 12999 | 13029 | 31 | CTGCGATCTATTGAAAGTCAGCCCTCGACAC | 180.4 | -3.4 | 1.1 | -3.1 | 1.9E-03 | 8.5E-03 | BT20/MB231 | CYTO |
| >hg38_21_+_8433172_8446622_RNA45SN1_45S_50ntFlanks@3705.3722.18\|TACCTGGTTGATCCTGCC | >hg38_21_+_8433172_8446622_RNA45SN1_45S_50ntFlanks | 3705 | 3722 | 18 | TACCTGGTTGATCCTGCC | 10001.2 | 2.8 | 1.0 | 2.7 | 6.0E-03 | 8.5E-03 | BT20/MB468 | MITO |
| >hg38_21_+_8433172_8446622_RNA45SN1_45S_50ntFlanks@8168.8191.24\|CCCAAGTCCTTCTGATCGAGGCCC | >hg38_21_+_8433172_8446622_RNA45SN1_45S_50ntFlanks | 8168 | 8191 | 24 | CCCAAGTCCTTCTGATCGAGGCCC | 75.5 | -4.6 | 1.6 | -2.9 | 4.1E-03 | 8.5E-03 | BT20/MB468 | MP |
| >hg38_21_+_8433172_8446622_RNA45SN1_45S_50ntFlanks@4733.4767.35\|GAACGAAAGTCGGAGGTTCGAAGACGATCAGATAC | >hg38_21_+_8433172_8446622_RNA45SN1_45S_50ntFlanks | 4733 | 4767 | 35 | GAACGAAAGTCGGAGGTTCGAAGACGATCAGATAC | 81.5 | 4.3 | 1.6 | 2.7 | 7.4E-03 | 8.5E-03 | MB231/MB468 | MITO |
| >hg38_21_+_8433172_8446622_RNA45SN1_45S_50ntFlanks@10680.10716.37\|GTTCCGGCGGCGTCCGGTGAGCTCTCGCTGGCCCTTG | >hg38_21_+_8433172_8446622_RNA45SN1_45S_50ntFlanks | 10680 | 10716 | 37 | GTTCCGGCGGCGTCCGGTGAGCTCTCGCTGGCCCTTG | 90.8 | 4.2 | 1.6 | 2.7 | 7.4E-03 | 8.5E-03 | MB231/MB468 | MITO |
| >hg38_21_+_8433172_8446622_RNA45SN1_45S_50ntFlanks@11674.11705.32\|GTGCTCTGAATGTCAAAGTGAAGAAATTCAAT | >hg38_21_+_8433172_8446622_RNA45SN1_45S_50ntFlanks | 11674 | 11705 | 32 | GTGCTCTGAATGTCAAAGTGAAGAAATTCAAT | 58.6 | 9.2 | 2.9 | 3.2 | 1.5E-03 | 8.5E-03 | BT20/MB231 | MP |
| >hg38_21_+_8433172_8446622_RNA45SN1_45S_50ntFlanks@12991.13014.24\|CTCCCTCGCTGCGATCTATTGAAA | >hg38_21_+_8433172_8446622_RNA45SN1_45S_50ntFlanks | 12991 | 13014 | 24 | CTCCCTCGCTGCGATCTATTGAAA | 1876.0 | -3.7 | 1.4 | -2.7 | 7.0E-03 | 8.5E-03 | BT20/MB468 | CYTO |
| >hg38_21_+_8433172_8446622_RNA45SN1_45S_50ntFlanks@12436.12473.38\|TCATTGTGAAGCAGAATTCACCAAGCGTTGGATTGTTC | >hg38_21_+_8433172_8446622_RNA45SN1_45S_50ntFlanks | 12436 | 12473 | 38 | TCATTGTGAAGCAGAATTCACCAAGCGTTGGATTGTTC | 110.5 | -3.4 | 1.2 | -2.7 | 6.0E-03 | 8.5E-03 | BT20/MB468 | MITO |
| >MI0000270\|hsa-mir-181b-1&WithFlank&1\|-\|198858867\|198858988@42.63.22(+1C);MI0000683\|hsa-mir-181b-2&WithFlank&9\|+\|124693704\|124693804@22.43.22(+1C)&[MIMAT0000257&hsa-miR-181b-5p&offsets\|0\|-1(+1C);m-73&1\|-\|198858925\|198858947&offsets\|0\|-1(+1C)];[MIMAT0000257_1&hsa-miR-181b-5p&offsets\|0\|-1(+1C);m-72&9\|+\|124693725\|124693747&offsets\|0\|-1(+1C)]\|AACATTCATTGCTGTCGGTGGGC | >MI0000270\|hsa-mir-181b-1&WithFlank&1\|-\|198858867\|198858988 | 42 | 63 | 22(+1C) | AACATTCATTGCTGTCGGTGGGC | 626.9 | -3.1 | 1.0 | -3.2 | 1.5E-03 | 8.5E-03 | BT20/MB231 | MP |
| >hg38_21_+_8433172_8446622_RNA45SN1_45S_50ntFlanks@10687.10707.21\|CGGCGTCCGGTGAGCTCTCGC | >hg38_21_+_8433172_8446622_RNA45SN1_45S_50ntFlanks | 10687 | 10707 | 21 | CGGCGTCCGGTGAGCTCTCGC | 172.9 | -3.1 | 1.2 | -2.7 | 7.5E-03 | 8.5E-03 | MB231/MB468 | NUC |
| >MI0000542\|hsa-mir-320a&WithFlank&8\|-\|22244960\|22245043@48.68.21(+1C)&[MIMAT0000510&hsa-miR-320a-3p&offsets\|0\|-1(+1C);m-60&8\|-\|22244975\|22244996&offsets\|0\|-1(+1C)]\|AAAAGCTGGGTTGAGAGGGCGC | >MI0000542\|hsa-mir-320a&WithFlank&8\|-\|22244960\|22245043 | 48 | 68 | 21(+1C) | AAAAGCTGGGTTGAGAGGGCGC | 598.0 | 2.2 | 0.7 | 3.1 | 1.7E-03 | 8.5E-03 | BT20/MB231 | NUC |
| >MI0000542\|hsa-mir-320a&WithFlank&8\|-\|22244960\|22245043@49.70.22&[MIMAT0000510&hsa-miR-320a-3p&offsets\|+1\|+1;m-60&8\|-\|22244975\|22244996&offsets\|+1\|+1]\|AAAGCTGGGTTGAGAGGGCGAA | >MI0000542\|hsa-mir-320a&WithFlank&8\|-\|22244960\|22245043 | 49 | 70 | 22 | AAAGCTGGGTTGAGAGGGCGAA | 452.8 | 2.0 | 0.6 | 3.1 | 1.7E-03 | 8.5E-03 | BT20/MB231 | NUC |
| >trna116_GluCTC_1_-_145399233_145399304@37.74.38,trna134_GluTTC_1_-_16861774_16861845@37.74.38,trna71_GluCTC_1_-_161439189_161439260@37.74.38,trna74_GluCTC_1_-_161431809_161431880@37.74.38,trna77_GluCTC_1_-_161424398_161424469@37.74.38,trna77_GluCTC_6_+_28949976_28950047@37.74.38,trna80_GluCTC_1_-_161417018_161417089@37.74.38,trna84_GluTTC_1_-_161391883_161391954@37.74.38,trna87_GluCTC_6_-_126101393_126101464@37.74.38\|ACCGCCGCGGCCCGGGTTCGATTCCCGGTCAGGGAACC | >trna116_GluCTC_1_-_145399233_145399304 | 37 | 74 | 38 | ACCGCCGCGGCCCGGGTTCGATTCCCGGTCAGGGAACC | 3214.6 | -4.1 | 1.3 | -3.1 | 1.7E-03 | 8.5E-03 | BT20/MB231 | NUC |
| >trnaMT_GluTTC_MT_-_14674_14742@34.71.38,trnalookalike8_GluTTC_5_-_93905172_93905240@34.71.38\|ATATCATTGGTCGTGGTTGTAGTCCGTGCGAGAATACC | >trnaMT_GluTTC_MT_-_14674_14742 | 34 | 71 | 38 | ATATCATTGGTCGTGGTTGTAGTCCGTGCGAGAATACC | 327.3 | -2.4 | 0.8 | -3.1 | 1.7E-03 | 8.5E-03 | BT20/MB231 | NUC |
| >hg38_MT_+_1621_3279_RNR2_16S_50ntFlanks@1175.1215.41\|TGCATTAAAAATTTCGGTTGGGGCGACCTCGGAGCAGAACC | >hg38_MT_+_1621_3279_RNR2_16S_50ntFlanks | 1175 | 1215 | 41 | TGCATTAAAAATTTCGGTTGGGGCGACCTCGGAGCAGAACC | 64.1 | 5.2 | 1.9 | 2.7 | 6.4E-03 | 8.5E-03 | MB231/MB468 | MP |
| >hg38_21_+_8433172_8446622_RNA45SN1_45S_50ntFlanks@9488.9505.18\|TAGGACCCGAAAGATGGT | >hg38_21_+_8433172_8446622_RNA45SN1_45S_50ntFlanks | 9488 | 9505 | 18 | TAGGACCCGAAAGATGGT | 45.9 | 4.8 | 1.7 | 2.9 | 4.2E-03 | 8.5E-03 | BT20/MB468 | MP |
| >hg38_21_+_8433172_8446622_RNA45SN1_45S_50ntFlanks@6721.6750.30\|AATTGCAGGACACATTGATCATCGACACTT | >hg38_21_+_8433172_8446622_RNA45SN1_45S_50ntFlanks | 6721 | 6750 | 30 | AATTGCAGGACACATTGATCATCGACACTT | 255.4 | -2.6 | 1.0 | -2.7 | 7.1E-03 | 8.5E-03 | MB231/MB468 | CYTO |
| >hg38_MT_+_1621_3279_RNR2_16S_50ntFlanks@499.518.20\|TGGACACTAGGAAAAAACCT | >hg38_MT_+_1621_3279_RNR2_16S_50ntFlanks | 499 | 518 | 20 | TGGACACTAGGAAAAAACCT | 59.8 | 5.7 | 2.0 | 2.9 | 4.2E-03 | 8.5E-03 | BT20/MB468 | MP |
| >hg38_MT_+_1621_3279_RNR2_16S_50ntFlanks@1254.1279.26\|AAGCGAACTACTATACTCAATTGATC | >hg38_MT_+_1621_3279_RNR2_16S_50ntFlanks | 1254 | 1279 | 26 | AAGCGAACTACTATACTCAATTGATC | 38.2 | 5.4 | 2.0 | 2.7 | 6.4E-03 | 8.5E-03 | MB231/MB468 | MP |
| >trna32_LysCTT_16_-_3207406_3207478@1.33.33\|GCCCGGCTAGCTCAGTCGGTAGAGCATGAGACC | >trna32_LysCTT_16_-_3207406_3207478 | 1 | 33 | 33 | GCCCGGCTAGCTCAGTCGGTAGAGCATGAGACC | 4991.0 | -3.1 | 1.0 | -3.1 | 1.9E-03 | 8.5E-03 | BT20/MB231 | CYTO |
| >hg38_21_+_8433172_8446622_RNA45SN1_45S_50ntFlanks@12934.12961.28\|CTAAACCATTCGTAGACGACCTGCTTCT | >hg38_21_+_8433172_8446622_RNA45SN1_45S_50ntFlanks | 12934 | 12961 | 28 | CTAAACCATTCGTAGACGACCTGCTTCT | 216.0 | -1.5 | 0.5 | -2.7 | 6.0E-03 | 8.5E-03 | BT20/MB468 | MITO |

<table>
<tr><th></th><th></th><th></th><th></th><th></th><th></th><th></th><th></th><th></th><th></th><th></th><th></th><th></th><th></th></tr>
<tr><td>>hg38_21_+_8433172_8446622_RNA45SN1_45S_50ntFlanks@4453.4471.19|TCGGCGCCCCCTCGATGCT</td><td>>hg38_21_+_8433172_8446622_RNA45SN1_45S_50ntFlanks</td><td>4453</td><td>4471</td><td>19</td><td>TCGGCGCCCCCTCGATGCT</td><td>72.3</td><td>-4.7</td><td>1.7</td><td>-2.9</td><td>4.2E-03</td><td>8.5E-03</td><td>BT20/MB468</td><td>MP</td></tr>
<tr><td>>MI0000446|hsa-mir-125b-1&WithFlank&11|-|122099751|122099850@21.39.19(+1C);MI0000470|hsa-mir-125b-2&WithFlank&21|+|16590231|16590331@23.41.19(+1C)&[MIMAT0000423&hsa-miR-125b-5p&offsets|0|-3(+1C);m-67&11|-|122099809|122099830&offsets|0|-3(+1C)];[MIMAT0000423_1&hsa-miR-125b-5p&offsets|0|-3(+1C);m-68&21|+|16590253|16590274&offsets|0|-3(+1C)]|TCCCTGAGACCCTAACTTGC</td><td>>MI0000446|hsa-mir-125b-1&WithFlank&11|-|122099751|122099850</td><td>21</td><td>39</td><td>19(+1C)</td><td>TCCCTGAGACCCTAACTTGC</td><td>170.5</td><td>-4.9</td><td>1.6</td><td>-3.1</td><td>1.7E-03</td><td>8.6E-03</td><td>BT20/MB231</td><td>NUC</td></tr>
<tr><td>>hg38_21_+_8433172_8446622_RNA45SN1_45S_50ntFlanks@10385.10424.40|CATGTGAACAGCAGTTGAACATGGGTCAGTCGGTCCTGAG</td><td>>hg38_21_+_8433172_8446622_RNA45SN1_45S_50ntFlanks</td><td>10385</td><td>10424</td><td>40</td><td>CATGTGAACAGCAGTTGAACATGGGTCAGTCGGTCCTGAG</td><td>1672.6</td><td>2.4</td><td>0.8</td><td>3.1</td><td>2.0E-03</td><td>8.6E-03</td><td>BT20/MB231</td><td>MITO</td></tr>
<tr><td>>MI0000077|hsa-mir-21&WithFlank&17|+|59841260|59841343@14.35.22(+2U)&[MIMAT0000076&hsa-miR-21-5p&offsets|0|0(+2U);m-1&17|+|59841273|59841294&offsets|0|0(+2U)]|TAGCTTATCAGACTGATGTTGATT</td><td>>MI0000077|hsa-mir-21&WithFlank&17|+|59841260|59841343</td><td>14</td><td>35</td><td>22(+2U)</td><td>TAGCTTATCAGACTGATGTTGATT</td><td>390.5</td><td>-2.5</td><td>0.8</td><td>-3.1</td><td>2.0E-03</td><td>8.6E-03</td><td>BT20/MB231</td><td>MITO</td></tr>
<tr><td>>am_MI0000786|hsa-mir-378a&WithFlank&5|+|149732819|149732896@49.70.22(+1G)&[MIMAT0000732&hsa-miR-378a-3p&offsets|0|0(+1G);m-44&5|+|149732867|149732888&offsets|0|0(+1G)]|ACTGGACTTGGAGTCAGAAGGCG</td><td>>am_MI0000786|hsa-mir-378a&WithFlank&5|+|149732819|149732896</td><td>49</td><td>70</td><td>22(+1G)</td><td>ACTGGACTTGGAGTCAGAAGGCG</td><td>1822.7</td><td>3.0</td><td>1.1</td><td>2.9</td><td>4.2E-03</td><td>8.6E-03</td><td>BT20/MB468</td><td>MP</td></tr>
<tr><td>>hg38_21_+_8433172_8446622_RNA45SN1_45S_50ntFlanks@12394.12422.29|TAGCGACGTCGCTTTTTGATCCTTCGATG</td><td>>hg38_21_+_8433172_8446622_RNA45SN1_45S_50ntFlanks</td><td>12394</td><td>12422</td><td>29</td><td>TAGCGACGTCGCTTTTTGATCCTTCGATG</td><td>67.6</td><td>-4.5</td><td>1.7</td><td>-2.7</td><td>6.5E-03</td><td>8.6E-03</td><td>MB231/MB468</td><td>MP</td></tr>
<tr><td>>hg38_21_+_8433172_8446622_RNA45SN1_45S_50ntFlanks@8358.8378.21|AAAGAACTTTGAAGAGAGAGT</td><td>>hg38_21_+_8433172_8446622_RNA45SN1_45S_50ntFlanks</td><td>8358</td><td>8378</td><td>21</td><td>AAAGAACTTTGAAGAGAGAGT</td><td>414.2</td><td>-2.3</td><td>0.9</td><td>-2.7</td><td>7.5E-03</td><td>8.6E-03</td><td>MB231/MB468</td><td>NUC</td></tr>
<tr><td>>hg38_21_+_8433172_8446622_RNA45SN1_45S_50ntFlanks@4724.4761.38|ATTAATCAAGAACGAAAGTCGGAGGTTCGAAGACGATC</td><td>>hg38_21_+_8433172_8446622_RNA45SN1_45S_50ntFlanks</td><td>4724</td><td>4761</td><td>38</td><td>ATTAATCAAGAACGAAAGTCGGAGGTTCGAAGACGATC</td><td>446.6</td><td>2.2</td><td>0.8</td><td>2.7</td><td>7.1E-03</td><td>8.6E-03</td><td>MB231/MB468</td><td>CYTO</td></tr>
<tr><td>>hg38_21_+_8433172_8446622_RNA45SN1_45S_50ntFlanks@12008.12035.28|CCGCCGGTGAAATACCACTACTCTGATC</td><td>>hg38_21_+_8433172_8446622_RNA45SN1_45S_50ntFlanks</td><td>12008</td><td>12035</td><td>28</td><td>CCGCCGGTGAAATACCACTACTCTGATC</td><td>226.3</td><td>-3.0</td><td>1.1</td><td>-2.7</td><td>7.0E-03</td><td>8.6E-03</td><td>BT20/MB468</td><td>CYTO</td></tr>
<tr><td>>hg38_21_+_8433172_8446622_RNA45SN1_45S_50ntFlanks@4099.4117.19|GTGACCACGGGTGACGGGG</td><td>>hg38_21_+_8433172_8446622_RNA45SN1_45S_50ntFlanks</td><td>4099</td><td>4117</td><td>19</td><td>GTGACCACGGGTGACGGGG</td><td>363.9</td><td>-4.9</td><td>1.7</td><td>-2.9</td><td>4.2E-03</td><td>8.6E-03</td><td>BT20/MB468</td><td>MP</td></tr>
<tr><td>>hg38_21_+_8433172_8446622_RNA45SN1_45S_50ntFlanks@9915.9938.24|AAAAGGTGTTGGTTGATATAGACA</td><td>>hg38_21_+_8433172_8446622_RNA45SN1_45S_50ntFlanks</td><td>9915</td><td>9938</td><td>24</td><td>AAAAGGTGTTGGTTGATATAGACA</td><td>106.5</td><td>5.7</td><td>2.0</td><td>2.9</td><td>4.2E-03</td><td>8.6E-03</td><td>BT20/MB468</td><td>MP</td></tr>
<tr><td>>hg38_21_+_8433172_8446622_RNA45SN1_45S_50ntFlanks@11846.11887.42|GAACGGGCTTGGCGGAATCAGCGGGGAAAGAAGACCCTGTTG</td><td>>hg38_21_+_8433172_8446622_RNA45SN1_45S_50ntFlanks</td><td>11846</td><td>11887</td><td>42</td><td>GAACGGGCTTGGCGGAATCAGCGGGGAAAGAAGACCCTGTTG</td><td>47.4</td><td>5.6</td><td>2.0</td><td>2.9</td><td>4.2E-03</td><td>8.6E-03</td><td>BT20/MB468</td><td>MP</td></tr>
<tr><td>>hg38_21_+_8433172_8446622_RNA45SN1_45S_50ntFlanks@9921.9951.31|TGTTGGTTGATATAGACAGCAGGACGGTGGC</td><td>>hg38_21_+_8433172_8446622_RNA45SN1_45S_50ntFlanks</td><td>9921</td><td>9951</td><td>31</td><td>TGTTGGTTGATATAGACAGCAGGACGGTGGC</td><td>60.5</td><td>5.6</td><td>2.0</td><td>2.9</td><td>4.2E-03</td><td>8.6E-03</td><td>BT20/MB468</td><td>MP</td></tr>
<tr><td>>hg38_21_+_8433172_8446622_RNA45SN1_45S_50ntFlanks@10737.10755.19|TAAATCTCGCGCCGGGCCG</td><td>>hg38_21_+_8433172_8446622_RNA45SN1_45S_50ntFlanks</td><td>10737</td><td>10755</td><td>19</td><td>TAAATCTCGCGCCGGGCCG</td><td>135.9</td><td>-4.6</td><td>1.6</td><td>-2.9</td><td>4.2E-03</td><td>8.6E-03</td><td>BT20/MB468</td><td>MP</td></tr>
<tr><td>>MI0000086|hsa-mir-28&WithFlank&3|+|188688775|188688872@60.82.23&[MIMAT0004502&hsa-miR-28-3p&offsets|0|+1;m-28&3|+|188688834|188688855&offsets|0|+1]|CACTAGATTGTGAGCTCCTGGAG</td><td>>MI0000086|hsa-mir-28&WithFlank&3|+|188688775|188688872</td><td>60</td><td>82</td><td>23</td><td>CACTAGATTGTGAGCTCCTGGAG</td><td>1784.8</td><td>2.7</td><td>0.9</td><td>3.1</td><td>1.9E-03</td><td>8.6E-03</td><td>BT20/MB231</td><td>CYTO</td></tr>
<tr><td>>hg38_21_+_8433172_8446622_RNA45SN1_45S_50ntFlanks@12418.12462.45|CGATGTCGGCTCTTCCTATCATTGTGAAGCAGAATTCACCAAGCG</td><td>>hg38_21_+_8433172_8446622_RNA45SN1_45S_50ntFlanks</td><td>12418</td><td>12462</td><td>45</td><td>CGATGTCGGCTCTTCCTATCATTGTGAAGCAGAATTCACCAAGCG</td><td>227.6</td><td>-5.4</td><td>1.7</td><td>-3.1</td><td>1.9E-03</td><td>8.6E-03</td><td>BT20/MB231</td><td>CYTO</td></tr>
<tr><td>>hg38_21_+_8433172_8446622_RNA45SN1_45S_50ntFlanks@9556.9581.26|TAGCGGTCCTGACGTGCAAATCGGTC</td><td>>hg38_21_+_8433172_8446622_RNA45SN1_45S_50ntFlanks</td><td>9556</td><td>9581</td><td>26</td><td>TAGCGGTCCTGACGTGCAAATCGGTC</td><td>187.0</td><td>-2.6</td><td>0.8</td><td>-3.1</td><td>1.9E-03</td><td>8.6E-03</td><td>BT20/MB231</td><td>CYTO</td></tr>
<tr><td>>hg38_21_+_8433172_8446622_RNA45SN1_45S_50ntFlanks@10909.10931.23|GCGCGCGCCGCGGCTGGACGAGG</td><td>>hg38_21_+_8433172_8446622_RNA45SN1_45S_50ntFlanks</td><td>10909</td><td>10931</td><td>23</td><td>GCGCGCGCCGCGGCTGGACGAGG</td><td>57.1</td><td>9.2</td><td>2.9</td><td>3.2</td><td>1.5E-03</td><td>8.6E-03</td><td>BT20/MB231</td><td>MP</td></tr>
<tr><td>>hg38_21_+_8433172_8446622_RNA45SN1_45S_50ntFlanks@10361.10383.23|GAAGGCCGAAGTGGAGAAGGGTT</td><td>>hg38_21_+_8433172_8446622_RNA45SN1_45S_50ntFlanks</td><td>10361</td><td>10383</td><td>23</td><td>GAAGGCCGAAGTGGAGAAGGGTT</td><td>631.0</td><td>-4.1</td><td>1.4</td><td>-2.9</td><td>4.2E-03</td><td>8.6E-03</td><td>BT20/MB468</td><td>MP</td></tr>
<tr><td>>hg38_21_+_8433172_8446622_RNA45SN1_45S_50ntFlanks@12718.12747.30|CTCGGTTGGCCTCGGATAGCCGGTCCCCCG</td><td>>hg38_21_+_8433172_8446622_RNA45SN1_45S_50ntFlanks</td><td>12718</td><td>12747</td><td>30</td><td>CTCGGTTGGCCTCGGATAGCCGGTCCCCCG</td><td>103.2</td><td>-3.7</td><td>1.3</td><td>-2.9</td><td>4.2E-03</td><td>8.6E-03</td><td>BT20/MB468</td><td>MP</td></tr>
<tr><td>>am_MI0005763|hsa-mir-941-1&WithFlank&20|+|63919443|63919526@53.73.21;MI0005764|hsa-mir-941-2&WithFlank&20|+|63919499|63919582@53.73.21;MI0005765|hsa-mir-941-3&WithFlank&20|+|63919555|63919638@53.73.21;MI0005766|hsa-mir-941-4&WithFlank&20|+|63919750|63919833@53.73.21;MI0031520|hsa-mir-941-5&WithFlank&20|+|63919862|63919945@53.73.21&[MIMAT0004984&hsa-miR-941&offsets|0|-2;m-263&20|+|63919495|63919517&offsets|0|-2];[MIMAT0004984_1&hsa-miR-941&offsets|0|-2;m-263&20|+|63919551|63919573&offsets|0|-2];[MIMAT0004984_2&hsa-miR-941&offsets|0|-2;m-263&20|+|63919607|63919629&offsets|0|-2];[MIMAT0004984_3&hsa-miR-941&offsets|0|-2;m-263&20|+|63919802|63919824&offsets|0|-2];[MIMAT0004984_4&hsa-miR-941&offsets|0|-2;m-263&20|+|63919914|63919936&offsets|0|-2]|CACCCGGCTGTGTGCACATGT</td><td>>am_MI0005763|hsa-mir-941-1&WithFlank&20|+|63919443|63919526</td><td>53</td><td>73</td><td>21</td><td>CACCCGGCTGTGTGCACATGT</td><td>55.0</td><td>4.5</td><td>1.7</td><td>2.7</td><td>6.5E-03</td><td>8.6E-03</td><td>MB231/MB468</td><td>MP</td></tr>
<tr><td>>hg38_21_+_8433172_8446622_RNA45SN1_45S_50ntFlanks@6645.6675.31|CTCGTACGACTCTTAGCGGTGGATCACTCGG</td><td>>hg38_21_+_8433172_8446622_RNA45SN1_45S_50ntFlanks</td><td>6645</td><td>6675</td><td>31</td><td>CTCGTACGACTCTTAGCGGTGGATCACTCGG</td><td>1056.4</td><td>2.3</td><td>0.9</td><td>2.7</td><td>7.5E-03</td><td>8.6E-03</td><td>MB231/MB468</td><td>NUC</td></tr>
<tr><td>>hg38_21_+_8433172_8446622_RNA45SN1_45S_50ntFlanks@6778.6806.29|CCCGGGGCTACGCCTGTCTGAGCGTCGCT</td><td>>hg38_21_+_8433172_8446622_RNA45SN1_45S_50ntFlanks</td><td>6778</td><td>6806</td><td>29</td><td>CCCGGGGCTACGCCTGTCTGAGCGTCGCT</td><td>425.2</td><td>2.0</td><td>0.8</td><td>2.7</td><td>6.5E-03</td><td>8.6E-03</td><td>MB231/MB468</td><td>MP</td></tr>
<tr><td>>am_MI0006370|hsa-mir-1303&WithFlank&5|+|154685770|154685867@58.79.22&[MIMAT0005891&hsa-miR-1303&offsets|0|0;m-516&5|+|154685826|154685848&offsets|+1|0]|TTTAGAGACGGGGTCTTGCTCT</td><td>>am_MI0006370|hsa-mir-1303&WithFlank&5|+|154685770|154685867</td><td>58</td><td>79</td><td>22</td><td>TTTAGAGACGGGGTCTTGCTCT</td><td>65.3</td><td>5.8</td><td>2.0</td><td>2.9</td><td>4.2E-03</td><td>8.6E-03</td><td>BT20/MB468</td><td>MP</td></tr>
<tr><td>>MI0000088|hsa-mir-30a&WithFlank&6|-|71403545|71403627@53.71.19&[MIMAT0000088&hsa-miR-30a-3p&offsets|0|-3;m-38&6|-|71403554|71403575&offsets|0|-3]|CTTTCAGTCGGATGTTTGC</td><td>>MI0000088|hsa-mir-30a&WithFlank&6|-|71403545|71403627</td><td>53</td><td>71</td><td>19</td><td>CTTTCAGTCGGATGTTTGC</td><td>1705.7</td><td>-2.0</td><td>0.7</td><td>-3.1</td><td>2.0E-03</td><td>8.6E-03</td><td>BT20/MB231</td><td>MITO</td></tr>
<tr><td>>hg38_21_+_8433172_8446622_RNA45SN1_45S_50ntFlanks@12001.12031.31|CTGCGGGCCGCCGGTGAAATACCACTACTCT</td><td>>hg38_21_+_8433172_8446622_RNA45SN1_45S_50ntFlanks</td><td>12001</td><td>12031</td><td>31</td><td>CTGCGGGCCGCCGGTGAAATACCACTACTCT</td><td>576.6</td><td>-2.6</td><td>1.0</td><td>-2.7</td><td>7.1E-03</td><td>8.6E-03</td><td>MB231/MB468</td><td>CYTO</td></tr>
<tr><td>>hg38_21_+_8433172_8446622_RNA45SN1_45S_50ntFlanks@5486.5515.30|GCGCTGAGAAGACGGTCGAACTTGACTATC</td><td>>hg38_21_+_8433172_8446622_RNA45SN1_45S_50ntFlanks</td><td>5486</td><td>5515</td><td>30</td><td>GCGCTGAGAAGACGGTCGAACTTGACTATC</td><td>562.8</td><td>6.5</td><td>2.1</td><td>3.2</td><td>1.5E-03</td><td>8.6E-03</td><td>BT20/MB231</td><td>MP</td></tr>
<tr><td>>hg38_21_+_8433172_8446622_RNA45SN1_45S_50ntFlanks@12655.12674.20|ATGACTGAACGCCTCTAAGT</td><td>>hg38_21_+_8433172_8446622_RNA45SN1_45S_50ntFlanks</td><td>12655</td><td>12674</td><td>20</td><td>ATGACTGAACGCCTCTAAGT</td><td>1134.3</td><td>-1.7</td><td>0.6</td><td>-2.7</td><td>6.1E-03</td><td>8.6E-03</td><td>BT20/MB468</td><td>MITO</td></tr>
<tr><td>>hg38_1_-_228634819_228635039_RNA5S12_5S_50ntFlanks@130.170.41|TGGATGGGAGACCGCCTGGGAATACCGGGTGCTGTAGGCTT</td><td>>hg38_1_-_228634819_228635039_RNA5S12_5S_50ntFlanks</td><td>130</td><td>170</td><td>41</td><td>TGGATGGGAGACCGCCTGGGAATACCGGGTGCTGTAGGCTT</td><td>486.4</td><td>2.6</td><td>0.9</td><td>2.7</td><td>6.5E-03</td><td>8.6E-03</td><td>MB231/MB468</td><td>MP</td></tr>
<tr><td>>hg38_21_+_8433172_8446622_RNA45SN1_45S_50ntFlanks@6731.6765.35|CACATTGATCATCGACACTTCGAACGCACTTGCGG</td><td>>hg38_21_+_8433172_8446622_RNA45SN1_45S_50ntFlanks</td><td>6731</td><td>6765</td><td>35</td><td>CACATTGATCATCGACACTTCGAACGCACTTGCGG</td><td>249.3</td><td>-4.6</td><td>1.7</td><td>-2.7</td><td>7.5E-03</td><td>8.6E-03</td><td>MB231/MB468</td><td>MITO</td></tr>
<tr><td>>hg38_21_+_8433172_8446622_RNA45SN1_45S_50ntFlanks@11573.11603.31|CGACTTAGAACTGGTGCGGACCAGGGGAATC</td><td>>hg38_21_+_8433172_8446622_RNA45SN1_45S_50ntFlanks</td><td>11573</td><td>11603</td><td>31</td><td>CGACTTAGAACTGGTGCGGACCAGGGGAATC</td><td>333.6</td><td>5.3</td><td>1.7</td><td>3.2</td><td>1.5E-03</td><td>8.6E-03</td><td>BT20/MB231</td><td>MP</td></tr>
<tr><td>>MI0000085|hsa-mir-27a&WithFlank&19|-|13836434|13836523@57.75.19(+3U)&[MIMAT0000084&hsa-miR-27a-3p&offsets|0|-2(+3U);m-43&19|-|13836447|13836467&offsets|0|-2(+3U)]|TTCACAGTGGCTAAGTTCCTTT</td><td>>MI0000085|hsa-mir-27a&WithFlank&19|-|13836434|13836523</td><td>57</td><td>75</td><td>19(+3U)</td><td>TTCACAGTGGCTAAGTTCCTTT</td><td>231.7</td><td>2.9</td><td>1.1</td><td>2.7</td><td>7.6E-03</td><td>8.6E-03</td><td>MB231/MB468</td><td>MITO</td></tr>
<tr><td>>MI0000086|hsa-mir-28&WithFlank&3|+|188688775|188688872@20.41.22&[MIMAT0000085&hsa-miR-28-5p&offsets|0|0;m-133&3|+|188688794|188688815&offsets|0|0]|AAGGAGCTCACAGTCTATTGAG</td><td>>MI0000086|hsa-mir-28&WithFlank&3|+|188688775|188688872</td><td>20</td><td>41</td><td>22</td><td>AAGGAGCTCACAGTCTATTGAG</td><td>838.2</td><td>1.8</td><td>0.6</td><td>3.1</td><td>1.7E-03</td><td>8.6E-03</td><td>BT20/MB231</td><td>NUC</td></tr>
<tr><td>>trna10_AspGTC_12_-_125424193_125424264@38.72.35,trna12_AspGTC_12_-_125411891_125411962@38.72.35,trna144_AspGTC_6_-_27551236_27551307@38.72.35,trna38_AspGTC_17_-_8125556_8125627@38.72.35,trna45_AspGTC_6_+_27447453_27447524@38.72.35,trna48_AspGTC_6_+_27471523_27471594@38.72.35,trna4_AspGTC_12_+_96429799_96429870@38.72.35,trna69_AspGTC_1_-_161440205_161440276@38.72.35,trna72_AspGTC_1_-_161432824_161432895@38.72.35,trna75_AspGTC_1_-_161425414_161425485@38.72.35,trna78_AspGTC_1_-_161418033_161418104@38.72.35,trna81_AspGTC_1_-_161410615_161410686@38.72.35|CGCGGGAGACCGGGGTTCGATTCCCCGACGGGGAG</td><td>>trna10_AspGTC_12_-_125424193_125424264</td><td>38</td><td>72</td><td>35</td><td>CGCGGGAGACCGGGGTTCGATTCCCCGACGGGGAG</td><td>428.5</td><td>-5.1</td><td>1.9</td><td>-2.7</td><td>7.5E-03</td><td>8.6E-03</td><td>BT20/MB468</td><td>NUC</td></tr>
<tr><td>>hg38_21_+_8433172_8446622_RNA45SN1_45S_50ntFlanks@11710.11740.31|CGCGGGTAAACGGCGGGAGTAACTATGACTC</td><td>>hg38_21_+_8433172_8446622_RNA45SN1_45S_50ntFlanks</td><td>11710</td><td>11740</td><td>31</td><td>CGCGGGTAAACGGCGGGAGTAACTATGACTC</td><td>364.1</td><td>-2.7</td><td>1.0</td><td>-2.7</td><td>7.1E-03</td><td>8.6E-03</td><td>BT20/MB468</td><td>CYTO</td></tr>
<tr><td>>hg38_21_+_8433172_8446622_RNA45SN1_45S_50ntFlanks@6661.6691.31|CGGTGGATCACTCGGCTCGTGCGTCGATGAA</td><td>>hg38_21_+_8433172_8446622_RNA45SN1_45S_50ntFlanks</td><td>6661</td><td>6691</td><td>31</td><td>CGGTGGATCACTCGGCTCGTGCGTCGATGAA</td><td>121.2</td><td>2.4</td><td>0.8</td><td>2.9</td><td>4.2E-03</td><td>8.6E-03</td><td>BT20/MB468</td><td>MP</td></tr>
<tr><td>>MI0000093|hsa-mir-92a-1&WithFlank&13|+|91351308|91351397@54.74.21;MI0000094|hsa-mir-92a-2&WithFlank&X|-|134169532|134169618@54.74.21&[MIMAT0000092&hsa-miR-92a-3p&offsets|0|-1;m-9&13|+|91351361|91351382&offsets|0|-1];[MIMAT0000092_1&hsa-miR-92a-3p&offsets|0|-1;m-11&X|-|134169544|134169565&offsets|0|-1]|TATTGCACTTGTCCCGGCCTG</td><td>>MI0000093|hsa-mir-92a-1&WithFlank&13|+|91351308|91351397</td><td>54</td><td>74</td><td>21</td><td>TATTGCACTTGTCCCGGCCTG</td><td>1010.7</td><td>-2.1</td><td>0.7</td><td>-3.1</td><td>2.0E-03</td><td>8.6E-03</td><td>BT20/MB231</td><td>MITO</td></tr>
<tr><td>>hg38_21_+_8433172_8446622_RNA45SN1_45S_50ntFlanks@8195.8212.18|CCGTGGACGGTGTGAGGC</td><td>>hg38_21_+_8433172_8446622_RNA45SN1_45S_50ntFlanks</td><td>8195</td><td>8212</td><td>18</td><td>CCGTGGACGGTGTGAGGC</td><td>364.3</td><td>-2.2</td><td>0.8</td><td>-2.7</td><td>6.5E-03</td><td>8.6E-03</td><td>MB231/MB468</td><td>MP</td></tr>
<tr><td>>hg38_21_+_8433172_8446622_RNA45SN1_45S_50ntFlanks@8189.8209.21|CCCAGCCCGTGGACGGTGTGA</td><td>>hg38_21_+_8433172_8446622_RNA45SN1_45S_50ntFlanks</td><td>8189</td><td>8209</td><td>21</td><td>CCCAGCCCGTGGACGGTGTGA</td><td>67.9</td><td>-4.6</td><td>1.6</td><td>-2.9</td><td>4.2E-03</td><td>8.6E-03</td><td>BT20/MB468</td><td>MP</td></tr>
</table>

| | | | | | | | | | | | | | |
|---|---|---|---|---|---|---|---|---|---|---|---|---|---|
| >hg38_21_+_8433172_8446622_RNA45SN1_45S_50ntFlanks@11710.11738.29\|CGCGGGTAAACGGCGGGAGTAACTATGAC | >hg38_21_+_8433172_8446622_RNA45SN1_45S_50ntFlanks | 11710 | 11738 | 29 | CGCGGGTAAACGGCGGGAGTAACTATGAC | 58.7 | -4.6 | 1.6 | -2.9 | 4.2E-03 | 8.6E-03 | BT20/MB468 | MP |
| >hg38_21_+_8433172_8446622_RNA45SN1_45S_50ntFlanks@475.503.29\|CTTCGTGATCGATGTGGTGACGTCGTGCT | >hg38_21_+_8433172_8446622_RNA45SN1_45S_50ntFlanks | 475 | 503 | 29 | CTTCGTGATCGATGTGGTGACGTCGTGCT | 441.5 | 3.8 | 1.4 | 2.7 | 6.5E-03 | 8.6E-03 | MB231/MB468 | MP |
| >hg38_21_+_8433172_8446622_RNA45SN1_45S_50ntFlanks@5084.5104.21\|CGAACGAGACTCTGGCATGCT | >hg38_21_+_8433172_8446622_RNA45SN1_45S_50ntFlanks | 5084 | 5104 | 21 | CGAACGAGACTCTGGCATGCT | 425.5 | -3.0 | 1.1 | -2.7 | 6.1E-03 | 8.6E-03 | BT20/MB468 | MITO |
| >hg38_1_-_228634819_228635039_RNA5S12_5S_50ntFlanks@129.171.43\|TTGGATGGGAGACCGCCTGGGAATACCGGGTGCTGTAGGCTTT | >hg38_1_-_228634819_228635039_RNA5S12_5S_50ntFlanks | 129 | 171 | 43 | TTGGATGGGAGACCGCCTGGGAATACCGGGTGCTGTAGGCTTT | 135.5 | 4.5 | 1.7 | 2.7 | 6.5E-03 | 8.7E-03 | MB231/MB468 | MP |
| >hg38_21_+_8433172_8446622_RNA45SN1_45S_50ntFlanks@4067.4089.23\|AACTTTCGATGGTAGTCGCCGTG | >hg38_21_+_8433172_8446622_RNA45SN1_45S_50ntFlanks | 4067 | 4089 | 23 | AACTTTCGATGGTAGTCGCCGTG | 537.6 | -4.1 | 1.5 | -2.7 | 6.1E-03 | 8.7E-03 | BT20/MB468 | MITO |
| >hg38_21_+_8433172_8446622_RNA45SN1_45S_50ntFlanks@4759.4776.18\|ATCAGATACCGTCGTAGT | >hg38_21_+_8433172_8446622_RNA45SN1_45S_50ntFlanks | 4759 | 4776 | 18 | ATCAGATACCGTCGTAGT | 169.4 | -4.5 | 1.6 | -2.7 | 6.5E-03 | 8.7E-03 | MB231/MB468 | MP |
| >hg38_21_+_8433172_8446622_RNA45SN1_45S_50ntFlanks@12444.12480.37\|AAGCAGAATTCACCAAGCGTTGGATTGTTCACCCACT | >hg38_21_+_8433172_8446622_RNA45SN1_45S_50ntFlanks | 12444 | 12480 | 37 | AAGCAGAATTCACCAAGCGTTGGATTGTTCACCCACT | 311.0 | -2.7 | 1.0 | -2.7 | 6.6E-03 | 8.7E-03 | MB231/MB468 | MP |
| >hg38_21_+_8433172_8446622_RNA45SN1_45S_50ntFlanks@7975.8009.35\|CGCGACCTCAGATCAGACGTGGCGACCCGCTGAAT | >hg38_21_+_8433172_8446622_RNA45SN1_45S_50ntFlanks | 7975 | 8009 | 35 | CGCGACCTCAGATCAGACGTGGCGACCCGCTGAAT | 681746.7 | 1.8 | 0.6 | 3.1 | 2.0E-03 | 8.7E-03 | BT20/MB231 | MITO |
| >MI0003583\|hsa-mir-576&WithFlank&4\|+\|109488692\|109488801@22.43.22&[MIMAT0003241&hsa-miR-576-5p&offsets\|0\|0;m-264&4\|+\|109488713\|109488735&offsets\|0\|-1]\|ATTCTAATTTCTCCACGTCTTT | >MI0003583\|hsa-mir-576&WithFlank&4\|+\|109488692\|109488801 | 22 | 43 | 22 | ATTCTAATTTCTCCACGTCTTT | 268.7 | 2.5 | 0.8 | 3.1 | 1.9E-03 | 8.7E-03 | BT20/MB231 | CYTO |
| >hg38_21_+_8433172_8446622_RNA45SN1_45S_50ntFlanks@4725.4768.44\|TTAATCAAGAACGAAAGTCGGAGGTTCGAAGACGATCAGATACC | >hg38_21_+_8433172_8446622_RNA45SN1_45S_50ntFlanks | 4725 | 4768 | 44 | TTAATCAAGAACGAAAGTCGGAGGTTCGAAGACGATCAGATACC | 768.0 | -3.2 | 1.0 | -3.1 | 1.9E-03 | 8.7E-03 | BT20/MB231 | CYTO |
| >hg38_21_+_8433172_8446622_RNA45SN1_45S_50ntFlanks@6661.6680.20\|CGGTGGATCACTCGGCTCGT | >hg38_21_+_8433172_8446622_RNA45SN1_45S_50ntFlanks | 6661 | 6680 | 20 | CGGTGGATCACTCGGCTCGT | 424.6 | -1.3 | 0.5 | -2.7 | 7.5E-03 | 8.7E-03 | BT20/MB468 | NUC |
| >hg38_MT_+_1621_3279_RNR2_16S_50ntFlanks@1250.1275.26\|GTCAAAGCGAACTACTATACTCAATT | >hg38_MT_+_1621_3279_RNR2_16S_50ntFlanks | 1250 | 1275 | 26 | GTCAAAGCGAACTACTATACTCAATT | 126.7 | 4.6 | 1.7 | 2.7 | 6.6E-03 | 8.7E-03 | MB231/MB468 | MP |
| >hg38_21_+_8433172_8446622_RNA45SN1_45S_50ntFlanks@4971.4999.29\|CCGGCCCGGACACGGACAGGATTGACAGA | >hg38_21_+_8433172_8446622_RNA45SN1_45S_50ntFlanks | 4971 | 4999 | 29 | CCGGCCCGGACACGGACAGGATTGACAGA | 785.1 | -5.2 | 1.8 | -2.9 | 4.2E-03 | 8.7E-03 | BT20/MB468 | MP |
| >hg38_21_+_8433172_8446622_RNA45SN1_45S_50ntFlanks@12598.12626.29\|TTTGGTGTATGTGCTTGGCTGAGGAGCCA | >hg38_21_+_8433172_8446622_RNA45SN1_45S_50ntFlanks | 12598 | 12626 | 29 | TTTGGTGTATGTGCTTGGCTGAGGAGCCA | 107.6 | -2.7 | 1.0 | -2.7 | 7.2E-03 | 8.7E-03 | MB231/MB468 | CYTO |
| >hg38_21_+_8433172_8446622_RNA45SN1_45S_50ntFlanks@12201.12221.21\|CCTAAGGCGAGCTCAGGGAGG | >hg38_21_+_8433172_8446622_RNA45SN1_45S_50ntFlanks | 12201 | 12221 | 21 | CCTAAGGCGAGCTCAGGGAGG | 84.3 | -4.6 | 1.6 | -2.9 | 4.2E-03 | 8.7E-03 | BT20/MB468 | MP |
| >trnaMT_MetCAT_MT_+_4402_4469@38.71.34\|CGAAAATGTTGGTTATACCCTTCCCGTACTACCA | >trnaMT_MetCAT_MT_+_4402_4469 | 38 | 71 | 34 | CGAAAATGTTGGTTATACCCTTCCCGTACTACCA | 263.7 | 4.4 | 1.6 | 2.7 | 6.6E-03 | 8.7E-03 | MB231/MB468 | MP |
| >trnaMT_GluTTC_MT_-_14674_14742@41.66.26,trnalookalike8_GluTTC_5_-_93905172_93905240@41.66.26\|TGGTCGTGGTTGTAGTCCGTGCGAGA | >trnaMT_GluTTC_MT_-_14674_14742 | 41 | 66 | 26 | TGGTCGTGGTTGTAGTCCGTGCGAGA | 117.0 | 5.0 | 1.8 | 2.9 | 4.2E-03 | 8.7E-03 | BT20/MB468 | MP |
| >hg38_21_+_8433172_8446622_RNA45SN1_45S_50ntFlanks@12416.12460.45\|TTCGATGTCGGCTCTTCCTATCATTGTGAAGCAGAATTCACCAAG | >hg38_21_+_8433172_8446622_RNA45SN1_45S_50ntFlanks | 12416 | 12460 | 45 | TTCGATGTCGGCTCTTCCTATCATTGTGAAGCAGAATTCACCAAG | 519.4 | -3.1 | 1.0 | -3.1 | 2.0E-03 | 8.7E-03 | BT20/MB231 | MITO |
| >hg38_21_+_8433172_8446622_RNA45SN1_45S_50ntFlanks@4499.4520.22\|AGCGTTTACTTTGAAAAAATTA | >hg38_21_+_8433172_8446622_RNA45SN1_45S_50ntFlanks | 4499 | 4520 | 22 | AGCGTTTACTTTGAAAAAATTA | 726.3 | 3.1 | 1.0 | 3.1 | 2.0E-03 | 8.7E-03 | BT20/MB231 | CYTO |
| >trnaMT_HisGTG_MT_+_12138_12206@27.61.35\|GATTGTGAATCTGACAACAGAGGCTTACGACCCCT | >trnaMT_HisGTG_MT_+_12138_12206 | 27 | 61 | 35 | GATTGTGAATCTGACAACAGAGGCTTACGACCCCT | 131.3 | 6.0 | 2.1 | 2.9 | 4.2E-03 | 8.7E-03 | BT20/MB468 | MP |
| >hg38_21_+_8433172_8446622_RNA45SN1_45S_50ntFlanks@12241.12265.25\|CAGAAGGGCAAAAGCTCGCTTGATC | >hg38_21_+_8433172_8446622_RNA45SN1_45S_50ntFlanks | 12241 | 12265 | 25 | CAGAAGGGCAAAAGCTCGCTTGATC | 270.3 | -3.7 | 1.3 | -2.9 | 4.2E-03 | 8.7E-03 | BT20/MB468 | MP |
| >trna111_HisGTG_1_-_147774845_147774916@-1A.31.32,trna118_HisGTG_1_-_145396881_145396952@-1A.31.32,trna16_HisGTG_1_+_146544773_146544844@-1A.31.32,trna1_HisGTG_15_+_45493349_45493420@-1A.31.32,trna21_HisGTG_1_+_147753471_147753542@-1A.31.32,trna33_HisGTG_6_+_27125906_27125977@-1A.31.32,trna7_HisGTG_9_-_14433938_14434009@-1A.31.32,trna8_HisGTG_15_-_45492611_45492682@-1A.31.32,trna9_HisGTG_15_-_45490804_45490875@-1A.31.32\|AGCCGTGATCGTATAGTGGTTAGTACTCTGCG | >trna111_HisGTG_1_-_147774845_147774916 | -1A | 31 | 32 | AGCCGTGATCGTATAGTGGTTAGTACTCTGCG | 873.5 | -4.3 | 1.5 | -2.9 | 4.2E-03 | 8.7E-03 | BT20/MB468 | MP |
| >hg38_21_+_8433172_8446622_RNA45SN1_45S_50ntFlanks@9929.9959.31\|GATATAGACAGCAGGACGGTGGCCATGGAAG | >hg38_21_+_8433172_8446622_RNA45SN1_45S_50ntFlanks | 9929 | 9959 | 31 | GATATAGACAGCAGGACGGTGGCCATGGAAG | 247.7 | -2.9 | 1.0 | -2.9 | 4.2E-03 | 8.7E-03 | BT20/MB468 | MP |
| >hg38_1_-_228634819_228635039_RNA5S12_5S_50ntFlanks@150.170.21\|AATACCGGGTGCTGTAGGCTT | >hg38_1_-_228634819_228635039_RNA5S12_5S_50ntFlanks | 150 | 170 | 21 | AATACCGGGTGCTGTAGGCTT | 6563.9 | 2.6 | 0.8 | 3.2 | 1.6E-03 | 8.7E-03 | BT20/MB231 | MP |
| >hg38_21_+_8433172_8446622_RNA45SN1_45S_50ntFlanks@10351.10377.27\|CGAGAACTTTGAAGGCCGAAGTGGAGA | >hg38_21_+_8433172_8446622_RNA45SN1_45S_50ntFlanks | 10351 | 10377 | 27 | CGAGAACTTTGAAGGCCGAAGTGGAGA | 972.3 | 5.6 | 1.8 | 3.2 | 1.6E-03 | 8.7E-03 | BT20/MB231 | MP |
| >hg38_21_+_8433172_8446622_RNA45SN1_45S_50ntFlanks@10508.10532.25\|CGAATCCGGAGTGGCGGAGATGGGC | >hg38_21_+_8433172_8446622_RNA45SN1_45S_50ntFlanks | 10508 | 10532 | 25 | CGAATCCGGAGTGGCGGAGATGGGC | 135.3 | -3.5 | 1.3 | -2.7 | 6.6E-03 | 8.7E-03 | MB231/MB468 | MP |
| >MI0000342\|hsa-mir-200b&WithFlank&1\|+\|1167098\|1167204@63.84.22(+1U)&[MIMAT0000318&hsa-miR-200b-3p&offsets\|0\|0(+1U);m-93&1\|+\|1167160\|1167182&offsets\|0\|-1(+1U)]\|TAATACTGCCTGGTAATGATGAT | >MI0000342\|hsa-mir-200b&WithFlank&1\|+\|1167098\|1167204 | 63 | 84 | 22(+1U) | TAATACTGCCTGGTAATGATGAT | 413.7 | 1.5 | 0.6 | 2.7 | 7.1E-03 | 8.7E-03 | BT20/MB468 | CYTO |
| >trna20_GluTTC_2_-_131094701_131094772@1.29.29,trna5_GluTTC_13_-_41634874_41634945@1.29.29\|TCCCATATGGTCTAGCGGTTAGGATTCCT | >trna20_GluTTC_2_-_131094701_131094772 | 1 | 29 | 29 | TCCCATATGGTCTAGCGGTTAGGATTCCT | 267.7 | -2.7 | 1.0 | -2.7 | 6.6E-03 | 8.7E-03 | MB231/MB468 | MP |
| >hg38_21_+_8433172_8446622_RNA45SN1_45S_50ntFlanks@6655.6699.45\|TCTTAGCGGTGGATCACTCGGCTCGTGCGTCGATGAAGAACGCAG | >hg38_21_+_8433172_8446622_RNA45SN1_45S_50ntFlanks | 6655 | 6699 | 45 | TCTTAGCGGTGGATCACTCGGCTCGTGCGTCGATGAAGAACGCAG | 211.3 | -2.6 | 1.0 | -2.7 | 7.6E-03 | 8.7E-03 | MB231/MB468 | MITO |
| >hg38_1_-_228634819_228635039_RNA5S12_5S_50ntFlanks@127.171.45\|ACTTGGATGGGAGACCGCCTGGGAATACCGGGTGCTGTAGGCTTT | >hg38_1_-_228634819_228635039_RNA5S12_5S_50ntFlanks | 127 | 171 | 45 | ACTTGGATGGGAGACCGCCTGGGAATACCGGGTGCTGTAGGCTTT | 125.9 | 4.3 | 1.6 | 2.7 | 7.6E-03 | 8.7E-03 | MB231/MB468 | MITO |
| >hg38_21_+_8433172_8446622_RNA45SN1_45S_50ntFlanks@12995.13039.45\|CTCGCTGCGATCTATTGAAAGTCAGCCCTCGACACAAGGGTTTGT | >hg38_21_+_8433172_8446622_RNA45SN1_45S_50ntFlanks | 12995 | 13039 | 45 | CTCGCTGCGATCTATTGAAAGTCAGCCCTCGACACAAGGGTTTGT | 218472.7 | -3.5 | 1.1 | -3.1 | 1.7E-03 | 8.7E-03 | BT20/MB231 | NUC |
| >am_MI0000786\|hsa-mir-378a&WithFlank&5\|+\|149732819\|149732896@50.70.21(+1U)&[MIMAT0000732&hsa-miR-378a-3p&offsets\|+1\|0(+1U);m-44&5\|+\|149732867\|149732888&offsets\|+1\|0(+1U)]\|CTGGACTTGGAGTCAGAAGGCT | >am_MI0000786\|hsa-mir-378a&WithFlank&5\|+\|149732819\|149732896 | 50 | 70 | 21(+1U) | CTGGACTTGGAGTCAGAAGGCT | 3662.3 | 3.1 | 1.1 | 2.9 | 4.3E-03 | 8.7E-03 | BT20/MB468 | MP |
| >hg38_21_+_8433172_8446622_RNA45SN1_45S_50ntFlanks@4981.4999.19\|CACGGACAGGATTGACAGA | >hg38_21_+_8433172_8446622_RNA45SN1_45S_50ntFlanks | 4981 | 4999 | 19 | CACGGACAGGATTGACAGA | 318.6 | -2.3 | 0.8 | -2.9 | 4.3E-03 | 8.7E-03 | BT20/MB468 | MP |
| >hg38_21_+_8433172_8446622_RNA45SN1_45S_50ntFlanks@6646.6675.30\|TCGTACGACTCTTAGCGGTGGATCACTCGG | >hg38_21_+_8433172_8446622_RNA45SN1_45S_50ntFlanks | 6646 | 6675 | 30 | TCGTACGACTCTTAGCGGTGGATCACTCGG | 2036.5 | 1.7 | 0.7 | 2.7 | 7.2E-03 | 8.7E-03 | MB231/MB468 | CYTO |
| >hg38_21_+_8433172_8446622_RNA45SN1_45S_50ntFlanks@10402.10431.30\|AACATGGGTCAGTCGGTCCTGAGAGATGGG | >hg38_21_+_8433172_8446622_RNA45SN1_45S_50ntFlanks | 10402 | 10431 | 30 | AACATGGGTCAGTCGGTCCTGAGAGATGGG | 726.1 | -3.9 | 1.4 | -2.9 | 4.3E-03 | 8.7E-03 | BT20/MB468 | MP |
| >hg38_21_+_8433172_8446622_RNA45SN1_45S_50ntFlanks@10444.10462.19\|CGAAGGGACGGGCGATGGC | >hg38_21_+_8433172_8446622_RNA45SN1_45S_50ntFlanks | 10444 | 10462 | 19 | CGAAGGGACGGGCGATGGC | 416.0 | -3.1 | 1.2 | -2.7 | 7.2E-03 | 8.7E-03 | MB231/MB468 | CYTO |
| >hg38_21_+_8433172_8446622_RNA45SN1_45S_50ntFlanks@3782.3810.29\|CAGTGAAACTGCGAATGGCTCATTAAATC | >hg38_21_+_8433172_8446622_RNA45SN1_45S_50ntFlanks | 3782 | 3810 | 29 | CAGTGAAACTGCGAATGGCTCATTAAATC | 238.1 | -4.3 | 1.6 | -2.7 | 6.2E-03 | 8.7E-03 | BT20/MB468 | MITO |
| >MI0000077\|hsa-mir-21&WithFlank&17\|+\|59841260\|59841343@14.34.21&[MIMAT0000076&hsa-miR-21-5p&offsets\|0\|-1;m-1&17\|+\|59841273\|59841294&offsets\|0\|-1]\|TAGCTTATCAGACTGATGTTG | >MI0000077\|hsa-mir-21&WithFlank&17\|+\|59841260\|59841343 | 14 | 34 | 21 | TAGCTTATCAGACTGATGTTG | 21391.4 | 3.3 | 1.1 | 2.9 | 4.3E-03 | 8.7E-03 | BT20/MB468 | MP |
| >hg38_21_+_8433172_8446622_RNA45SN1_45S_50ntFlanks@10678.10708.31\|CGGTTCCGGCGGCGTCCGGTGAGCTCTCGCT | >hg38_21_+_8433172_8446622_RNA45SN1_45S_50ntFlanks | 10678 | 10708 | 31 | CGGTTCCGGCGGCGTCCGGTGAGCTCTCGCT | 625.7 | 4.2 | 1.3 | 3.2 | 1.6E-03 | 8.7E-03 | BT20/MB231 | MP |
| >hg38_21_+_8433172_8446622_RNA45SN1_45S_50ntFlanks@10791.10823.33\|CCTCTGGCATGTTGGAACAATGTAGGTAAGGGA | >hg38_21_+_8433172_8446622_RNA45SN1_45S_50ntFlanks | 10791 | 10823 | 33 | CCTCTGGCATGTTGGAACAATGTAGGTAAGGGA | 472.0 | 4.9 | 1.6 | 3.2 | 1.6E-03 | 8.7E-03 | BT20/MB231 | MP |
| >hg38_21_+_8433172_8446622_RNA45SN1_45S_50ntFlanks@11648.11687.40\|TGTTGACGCGATGTGATTTCTGCCCAGTGCTCTGAATGTC | >hg38_21_+_8433172_8446622_RNA45SN1_45S_50ntFlanks | 11648 | 11687 | 40 | TGTTGACGCGATGTGATTTCTGCCCAGTGCTCTGAATGTC | 431.4 | 6.7 | 2.1 | 3.2 | 1.6E-03 | 8.7E-03 | BT20/MB231 | MP |
| >hg38_21_+_8433172_8446622_RNA45SN1_45S_50ntFlanks@6660.6681.22\|GCGGTGGATCACTCGGCTCGTG | >hg38_21_+_8433172_8446622_RNA45SN1_45S_50ntFlanks | 6660 | 6681 | 22 | GCGGTGGATCACTCGGCTCGTG | 333.1 | 5.1 | 1.6 | 3.2 | 1.6E-03 | 8.7E-03 | BT20/MB231 | MP |
| >hg38_21_+_8433172_8446622_RNA45SN1_45S_50ntFlanks@7990.8008.19\|GACGTGGCGACCCGCTGAA | >hg38_21_+_8433172_8446622_RNA45SN1_45S_50ntFlanks | 7990 | 8008 | 19 | GACGTGGCGACCCGCTGAA | 1044.7 | 4.4 | 1.4 | 3.2 | 1.6E-03 | 8.7E-03 | BT20/MB231 | MP |
| >hg38_21_+_8433172_8446622_RNA45SN1_45S_50ntFlanks@9498.9520.23\|AAGATGGTGAACTATGCCTGGGC | >hg38_21_+_8433172_8446622_RNA45SN1_45S_50ntFlanks | 9498 | 9520 | 23 | AAGATGGTGAACTATGCCTGGGC | 126.3 | 5.9 | 1.9 | 3.2 | 1.6E-03 | 8.7E-03 | BT20/MB231 | MP |
| >Antisense_hg38_21_-_8433172_8446622_RNA45SN1_45S_50ntFlanks@7674.7692.19\|CTCTCTCTCTCTCTCCCCC | >Antisense_hg38_21_-_8433172_8446622_RNA45SN1_45S_50ntFlanks | 7674 | 7692 | 19 | CTCTCTCTCTCTCTCCCCC | 365.6 | -3.5 | 1.3 | -2.7 | 7.2E-03 | 8.7E-03 | BT20/MB468 | CYTO |
| >hg38_21_+_8433172_8446622_RNA45SN1_45S_50ntFlanks@11027.11045.19\|CTCTCTCTCTCTCTCCCCC | >hg38_21_+_8433172_8446622_RNA45SN1_45S_50ntFlanks | 11027 | 11045 | 19 | CTCTCTCTCTCTCTCCCCC | 365.6 | -3.5 | 1.3 | -2.7 | 7.2E-03 | 8.7E-03 | BT20/MB468 | CYTO |
| >hg38_21_+_8433172_8446622_RNA45SN1_45S_50ntFlanks@9555.9573.19\|GTAGCGGTCCTGACGTGCA | >hg38_21_+_8433172_8446622_RNA45SN1_45S_50ntFlanks | 9555 | 9573 | 19 | GTAGCGGTCCTGACGTGCA | 75.9 | -4.6 | 1.6 | -2.9 | 4.3E-03 | 8.7E-03 | BT20/MB468 | MP |
| >am_trnaMT_MetCAT_MT_+_4402_4469@1.21.21\|AGTAAGGTCAGCTAAATAAGC | >am_trnaMT_MetCAT_MT_+_4402_4469 | 1 | 21 | 21 | AGTAAGGTCAGCTAAATAAGC | 5890.9 | -3.0 | 1.0 | -3.2 | 1.6E-03 | 8.7E-03 | BT20/MB231 | MP |
| >hg38_21_+_8433172_8446622_RNA45SN1_45S_50ntFlanks@11206.11225.20\|CGGCGGCGACTCTGGACGCG | >hg38_21_+_8433172_8446622_RNA45SN1_45S_50ntFlanks | 11206 | 11225 | 20 | CGGCGGCGACTCTGGACGCG | 390.7 | -2.3 | 0.8 | -2.9 | 4.3E-03 | 8.8E-03 | BT20/MB468 | MP |
| >hg38_21_+_8433172_8446622_RNA45SN1_45S_50ntFlanks@10357.10376.20\|CTTTGAAGGCCGAAGTGGAG | >hg38_21_+_8433172_8446622_RNA45SN1_45S_50ntFlanks | 10357 | 10376 | 20 | CTTTGAAGGCCGAAGTGGAG | 93.5 | -4.7 | 1.6 | -2.9 | 4.3E-03 | 8.8E-03 | BT20/MB468 | MP |

<table>
<tr><td>>hg38_21_+_8433172_8446622_RNA45SN1_45S_50ntFlanks@10843.10865.23|TAACTTCGGGATAAGGATTGGCT</td><td>>hg38_21_+_8433172_8446622_RNA45SN1_45S_50ntFlanks</td><td>10843</td><td>10865</td><td>23</td><td>TAACTTCGGGATAAGGATTGGCT</td><td>158.9</td><td>-1.5</td><td>0.6</td><td>-2.7</td><td>6.2E-03</td><td>8.8E-03</td><td>BT20/MB468</td><td>MITO</td></tr>
<tr><td>>hg38_21_+_8433172_8446622_RNA45SN1_45S_50ntFlanks@12406.12436.31|TTTTTGATCCTTCGATGTCGGCTCTTCCTAT</td><td>>hg38_21_+_8433172_8446622_RNA45SN1_45S_50ntFlanks</td><td>12406</td><td>12436</td><td>31</td><td>TTTTTGATCCTTCGATGTCGGCTCTTCCTAT</td><td>192.5</td><td>-4.0</td><td>1.5</td><td>-2.7</td><td>6.2E-03</td><td>8.8E-03</td><td>BT20/MB468</td><td>MITO</td></tr>
<tr><td>>hg38_21_+_8433172_8446622_RNA45SN1_45S_50ntFlanks@11562.11588.27|CGCCTAGCAGCCGACTTAGAACTGGTG</td><td>>hg38_21_+_8433172_8446622_RNA45SN1_45S_50ntFlanks</td><td>11562</td><td>11588</td><td>27</td><td>CGCCTAGCAGCCGACTTAGAACTGGTG</td><td>114.0</td><td>6.2</td><td>2.0</td><td>3.2</td><td>1.6E-03</td><td>8.8E-03</td><td>BT20/MB231</td><td>MP</td></tr>
<tr><td>>hg38_21_+_8433172_8446622_RNA45SN1_45S_50ntFlanks@12416.12459.44|TTCGATGTCGGCTCTTCCTATCATTGTGAAGCAGAATTCACCAA</td><td>>hg38_21_+_8433172_8446622_RNA45SN1_45S_50ntFlanks</td><td>12416</td><td>12459</td><td>44</td><td>TTCGATGTCGGCTCTTCCTATCATTGTGAAGCAGAATTCACCAA</td><td>515.0</td><td>-3.1</td><td>1.0</td><td>-3.1</td><td>2.1E-03</td><td>8.8E-03</td><td>BT20/MB231</td><td>MITO</td></tr>
<tr><td>>hg38_21_+_8433172_8446622_RNA45SN1_45S_50ntFlanks@12717.12739.23|CCTCGGTTGGCCTCGGATAGCCG</td><td>>hg38_21_+_8433172_8446622_RNA45SN1_45S_50ntFlanks</td><td>12717</td><td>12739</td><td>23</td><td>CCTCGGTTGGCCTCGGATAGCCG</td><td>231.8</td><td>-4.8</td><td>1.7</td><td>-2.9</td><td>4.3E-03</td><td>8.8E-03</td><td>BT20/MB468</td><td>MP</td></tr>
<tr><td>>am_trnaMT_ValTAC_MT_+_1602_1670@44.68.25|ATTTCAACTTAACTTGACCGCTCTG</td><td>>am_trnaMT_ValTAC_MT_+_1602_1670</td><td>44</td><td>68</td><td>25</td><td>ATTTCAACTTAACTTGACCGCTCTG</td><td>332.2</td><td>-3.6</td><td>1.4</td><td>-2.7</td><td>7.3E-03</td><td>8.8E-03</td><td>MB231/MB468</td><td>CYTO</td></tr>
<tr><td>>am_trnaMT_MetCAT_MT_+_4402_4469@1.22.22|AGTAAGGTCAGCTAAATAAGCT</td><td>>am_trnaMT_MetCAT_MT_+_4402_4469</td><td>1</td><td>22</td><td>22</td><td>AGTAAGGTCAGCTAAATAAGCT</td><td>851.2</td><td>2.5</td><td>0.9</td><td>2.7</td><td>6.6E-03</td><td>8.8E-03</td><td>MB231/MB468</td><td>MP</td></tr>
<tr><td>>MI0000066|hsa-let-7e&WithFlank&19|+|51692780|51692870@14.34.21(+1C)&[MIMAT0000066&hsa-let-7e-5p&offsets|0|-1(+1C);m-50&19|+|51692793|51692814&offsets|0|-1(+1C)]|TGAGGTAGGAGGTTGTATAGTC</td><td>>MI0000066|hsa-let-7e&WithFlank&19|+|51692780|51692870</td><td>14</td><td>34</td><td>21(+1C)</td><td>TGAGGTAGGAGGTTGTATAGTC</td><td>1710.7</td><td>1.6</td><td>0.5</td><td>3.1</td><td>1.7E-03</td><td>8.8E-03</td><td>BT20/MB231</td><td>NUC</td></tr>
<tr><td>>hg38_21_+_8433172_8446622_RNA45SN1_45S_50ntFlanks@10350.10380.31|ACGAGAACTTTGAAGGCCGAAGTGGAGAAGG</td><td>>hg38_21_+_8433172_8446622_RNA45SN1_45S_50ntFlanks</td><td>10350</td><td>10380</td><td>31</td><td>ACGAGAACTTTGAAGGCCGAAGTGGAGAAGG</td><td>225.2</td><td>-3.2</td><td>1.2</td><td>-2.7</td><td>7.7E-03</td><td>8.8E-03</td><td>MB231/MB468</td><td>MITO</td></tr>
<tr><td>>MI0000746|hsa-mir-99b&WithFlank&19|+|51692606|51692687@13.32.20(+1A)&[MIMAT0000689&hsa-miR-99b-5p&offsets|0|-2(+1A);m-10&19|+|51692618|51692639&offsets|0|-2(+1A)]|CACCCGTAGAACCGACCTTGA</td><td>>MI0000746|hsa-mir-99b&WithFlank&19|+|51692606|51692687</td><td>13</td><td>32</td><td>20(+1A)</td><td>CACCCGTAGAACCGACCTTGA</td><td>2963.4</td><td>2.6</td><td>0.9</td><td>3.1</td><td>2.0E-03</td><td>8.8E-03</td><td>BT20/MB231</td><td>CYTO</td></tr>
<tr><td>>trna10_AspGTC_12_-_125424193_125424264@38.74.37,trna12_AspGTC_12_-_125411891_125411962@38.74.37,trna144_AspGTC_6_-_27551236_27551307@38.74.37,trna38_AspGTC_17_-_8125556_8125627@38.74.37,trna45_AspGTC_6_+_27447453_27447524@38.74.37,trna48_AspGTC_6_+_27471523_27471594@38.74.37,trna4_AspGTC_12_+_96429799_96429870@38.74.37,trna69_AspGTC_1_-_161440205_161440276@38.74.37,trna72_AspGTC_1_-_161432824_161432895@38.74.37,trna75_AspGTC_1_-_161425414_161425485@38.74.37,trna78_AspGTC_1_-_161418033_161418104@38.74.37,trna81_AspGTC_1_-_161410615_161410686@38.74.37|CGCGGGAGACCGGGGTTCGATTCCCCGACGGGGAGCC</td><td>>trna10_AspGTC_12_-_125424193_125424264</td><td>38</td><td>74</td><td>37</td><td>CGCGGGAGACCGGGGTTCGATTCCCCGACGGGGAGCC</td><td>1588.6</td><td>-2.8</td><td>0.9</td><td>-3.1</td><td>2.0E-03</td><td>8.8E-03</td><td>BT20/MB231</td><td>CYTO</td></tr>
<tr><td>>trnaMT_PheGAA_MT_+_577_647@35.59.25|GAAAATGTTTAGACGGGCTCACATC</td><td>>trnaMT_PheGAA_MT_+_577_647</td><td>35</td><td>59</td><td>25</td><td>GAAAATGTTTAGACGGGCTCACATC</td><td>160.4</td><td>2.5</td><td>0.9</td><td>2.7</td><td>6.6E-03</td><td>8.8E-03</td><td>MB231/MB468</td><td>MP</td></tr>
<tr><td>>hg38_MT_+_598_1651_RNR1_12S_50ntFlanks@916.935.20|AAGGACATTTAACTAAAACC</td><td>>hg38_MT_+_598_1651_RNR1_12S_50ntFlanks</td><td>916</td><td>935</td><td>20</td><td>AAGGACATTTAACTAAAACC</td><td>496.2</td><td>2.9</td><td>1.1</td><td>2.7</td><td>6.6E-03</td><td>8.8E-03</td><td>MB231/MB468</td><td>MP</td></tr>
<tr><td>>hg38_21_+_8433172_8446622_RNA45SN1_45S_50ntFlanks@7973.8006.34|GACGCGACCTCAGATCAGACGTGGCGACCCGCTG</td><td>>hg38_21_+_8433172_8446622_RNA45SN1_45S_50ntFlanks</td><td>7973</td><td>8006</td><td>34</td><td>GACGCGACCTCAGATCAGACGTGGCGACCCGCTG</td><td>235.6</td><td>-1.3</td><td>0.5</td><td>-2.7</td><td>7.3E-03</td><td>8.8E-03</td><td>MB231/MB468</td><td>CYTO</td></tr>
<tr><td>>MI0000088|hsa-mir-30a&WithFlank&6|-|71403545|71403627@54.73.20(+3U)&[MIMAT0000088&hsa-miR-30a-3p&offsets|+1|-1(+3U);m-38&6|-|71403554|71403575&offsets|+1|-1(+3U)]|TTTCAGTCGGATGTTTGCAGTTT</td><td>>MI0000088|hsa-mir-30a&WithFlank&6|-|71403545|71403627</td><td>54</td><td>73</td><td>20(+3U)</td><td>TTTCAGTCGGATGTTTGCAGTTT</td><td>188.0</td><td>-5.6</td><td>1.8</td><td>-3.2</td><td>1.6E-03</td><td>8.8E-03</td><td>BT20/MB231</td><td>MP</td></tr>
<tr><td>>trnaMT_PheGAA_MT_+_577_647@47.74.28|ACGGGCTCACATCACCCCATAAACACCA</td><td>>trnaMT_PheGAA_MT_+_577_647</td><td>47</td><td>74</td><td>28</td><td>ACGGGCTCACATCACCCCATAAACACCA</td><td>85.6</td><td>1.9</td><td>0.7</td><td>2.7</td><td>6.7E-03</td><td>8.8E-03</td><td>MB231/MB468</td><td>MP</td></tr>
<tr><td>>hg38_21_+_8433172_8446622_RNA45SN1_45S_50ntFlanks@6647.6682.36|CGTACGACTCTTAGCGGTGGATCACTCGGCTCGTGC</td><td>>hg38_21_+_8433172_8446622_RNA45SN1_45S_50ntFlanks</td><td>6647</td><td>6682</td><td>36</td><td>CGTACGACTCTTAGCGGTGGATCACTCGGCTCGTGC</td><td>89.4</td><td>4.2</td><td>1.5</td><td>2.9</td><td>4.3E-03</td><td>8.8E-03</td><td>BT20/MB468</td><td>MP</td></tr>
<tr><td>>hg38_21_+_8433172_8446622_RNA45SN1_45S_50ntFlanks@9835.9854.20|TTTGGTAAGCAGAACTGGCG</td><td>>hg38_21_+_8433172_8446622_RNA45SN1_45S_50ntFlanks</td><td>9835</td><td>9854</td><td>20</td><td>TTTGGTAAGCAGAACTGGCG</td><td>86.8</td><td>7.1</td><td>2.5</td><td>2.9</td><td>4.3E-03</td><td>8.8E-03</td><td>BT20/MB468</td><td>MP</td></tr>
<tr><td>>hg38_21_+_8433172_8446622_RNA45SN1_45S_50ntFlanks@6658.6681.24|TAGCGGTGGATCACTCGGCTCGTG</td><td>>hg38_21_+_8433172_8446622_RNA45SN1_45S_50ntFlanks</td><td>6658</td><td>6681</td><td>24</td><td>TAGCGGTGGATCACTCGGCTCGTG</td><td>327.2</td><td>-3.8</td><td>1.3</td><td>-2.9</td><td>4.3E-03</td><td>8.8E-03</td><td>BT20/MB468</td><td>MP</td></tr>
<tr><td>>hg38_21_+_8433172_8446622_RNA45SN1_45S_50ntFlanks@12419.12449.31|GATGTCGGCTCTTCCTATCATTGTGAAGCAG</td><td>>hg38_21_+_8433172_8446622_RNA45SN1_45S_50ntFlanks</td><td>12419</td><td>12449</td><td>31</td><td>GATGTCGGCTCTTCCTATCATTGTGAAGCAG</td><td>483.7</td><td>-2.6</td><td>0.8</td><td>-3.1</td><td>2.1E-03</td><td>8.8E-03</td><td>BT20/MB231</td><td>MITO</td></tr>
<tr><td>>hg38_21_+_8433172_8446622_RNA45SN1_45S_50ntFlanks@10239.10267.29|GCTACGCCGCGACGAGTAGGAGGGCCGCT</td><td>>hg38_21_+_8433172_8446622_RNA45SN1_45S_50ntFlanks</td><td>10239</td><td>10267</td><td>29</td><td>GCTACGCCGCGACGAGTAGGAGGGCCGCT</td><td>96.8</td><td>2.8</td><td>1.0</td><td>2.9</td><td>4.3E-03</td><td>8.8E-03</td><td>BT20/MB468</td><td>MP</td></tr>
<tr><td>>am_trna116_GluCTC_1_-_145399233_145399304@1.26.26,trna59_GluCTC_1_+_249168447_249168518@1.26.26,trna71_GluCTC_1_-_161439189_161439260@1.26.26,trna74_GluCTC_1_-_161431809_161431880@1.26.26,trna77_GluCTC_1_-_161424398_161424469@1.26.26,trna77_GluCTC_6_+_28949976_28950047@1.26.26,trna80_GluCTC_1_-_161417018_161417089@1.26.26,trna87_GluCTC_6_-_126101393_126101464@1.26.26|TCCCTGGTGGTCTAGTGGTTAGGATT</td><td>>am_trna116_GluCTC_1_-_145399233_145399304</td><td>1</td><td>26</td><td>26</td><td>TCCCTGGTGGTCTAGTGGTTAGGATT</td><td>1232.6</td><td>3.0</td><td>1.1</td><td>2.9</td><td>4.3E-03</td><td>8.8E-03</td><td>BT20/MB468</td><td>MP</td></tr>
<tr><td>>hg38_MT_+_1621_3279_RNR2_16S_50ntFlanks@1356.1385.30|GGTTTACGACCTCGATGTTGGATCAGGACA</td><td>>hg38_MT_+_1621_3279_RNR2_16S_50ntFlanks</td><td>1356</td><td>1385</td><td>30</td><td>GGTTTACGACCTCGATGTTGGATCAGGACA</td><td>119.7</td><td>2.5</td><td>0.9</td><td>2.7</td><td>6.7E-03</td><td>8.8E-03</td><td>MB231/MB468</td><td>MP</td></tr>
<tr><td>>hg38_21_+_8433172_8446622_RNA45SN1_45S_50ntFlanks@9553.9570.18|CCGTAGCGGTCCTGACGT</td><td>>hg38_21_+_8433172_8446622_RNA45SN1_45S_50ntFlanks</td><td>9553</td><td>9570</td><td>18</td><td>CCGTAGCGGTCCTGACGT</td><td>343.0</td><td>1.8</td><td>0.7</td><td>2.7</td><td>7.2E-03</td><td>8.8E-03</td><td>BT20/MB468</td><td>CYTO</td></tr>
<tr><td>>hg38_21_+_8433172_8446622_RNA45SN1_45S_50ntFlanks@9467.9486.20|GTGGAGCACGAGCGCACGTG</td><td>>hg38_21_+_8433172_8446622_RNA45SN1_45S_50ntFlanks</td><td>9467</td><td>9486</td><td>20</td><td>GTGGAGCACGAGCGCACGTG</td><td>150.6</td><td>-5.4</td><td>1.9</td><td>-2.9</td><td>4.3E-03</td><td>8.8E-03</td><td>BT20/MB468</td><td>MP</td></tr>
<tr><td>>hg38_21_+_8433172_8446622_RNA45SN1_45S_50ntFlanks@12539.12565.27|ATGATGTGTTGTTGCCATGGTAATCCT</td><td>>hg38_21_+_8433172_8446622_RNA45SN1_45S_50ntFlanks</td><td>12539</td><td>12565</td><td>27</td><td>ATGATGTGTTGTTGCCATGGTAATCCT</td><td>221.6</td><td>-4.1</td><td>1.5</td><td>-2.7</td><td>6.2E-03</td><td>8.8E-03</td><td>BT20/MB468</td><td>MITO</td></tr>
<tr><td>>hg38_21_+_8433172_8446622_RNA45SN1_45S_50ntFlanks@4552.4572.21|TACCGCAGCTAGGAATAATGG</td><td>>hg38_21_+_8433172_8446622_RNA45SN1_45S_50ntFlanks</td><td>4552</td><td>4572</td><td>21</td><td>TACCGCAGCTAGGAATAATGG</td><td>119.2</td><td>-4.1</td><td>1.5</td><td>-2.7</td><td>7.7E-03</td><td>8.8E-03</td><td>MB231/MB468</td><td>NUC</td></tr>
<tr><td>>hg38_21_+_8433172_8446622_RNA45SN1_45S_50ntFlanks@6721.6746.26|AATTGCAGGACACATTGATCATCGAC</td><td>>hg38_21_+_8433172_8446622_RNA45SN1_45S_50ntFlanks</td><td>6721</td><td>6746</td><td>26</td><td>AATTGCAGGACACATTGATCATCGAC</td><td>2373.0</td><td>-5.4</td><td>1.9</td><td>-2.9</td><td>4.3E-03</td><td>8.8E-03</td><td>BT20/MB468</td><td>MP</td></tr>
<tr><td>>trnaMT_PheGAA_MT_+_577_647@38.56.19|AATGTTTAGACGGGCTCAC</td><td>>trnaMT_PheGAA_MT_+_577_647</td><td>38</td><td>56</td><td>19</td><td>AATGTTTAGACGGGCTCAC</td><td>91.8</td><td>3.5</td><td>1.3</td><td>2.7</td><td>6.7E-03</td><td>8.8E-03</td><td>MB231/MB468</td><td>MP</td></tr>
<tr><td>>hg38_21_+_8433172_8446622_RNA45SN1_45S_50ntFlanks@10290.10319.30|GCGCGGGCCCGGGTGGAGCCGCCGCAGGTG</td><td>>hg38_21_+_8433172_8446622_RNA45SN1_45S_50ntFlanks</td><td>10290</td><td>10319</td><td>30</td><td>GCGCGGGCCCGGGTGGAGCCGCCGCAGGTG</td><td>225.8</td><td>-4.1</td><td>1.4</td><td>-2.9</td><td>4.3E-03</td><td>8.8E-03</td><td>BT20/MB468</td><td>MP</td></tr>
<tr><td>>hg38_21_+_8433172_8446622_RNA45SN1_45S_50ntFlanks@12416.12443.28|TTCGATGTCGGCTCTTCCTATCATTGTG</td><td>>hg38_21_+_8433172_8446622_RNA45SN1_45S_50ntFlanks</td><td>12416</td><td>12443</td><td>28</td><td>TTCGATGTCGGCTCTTCCTATCATTGTG</td><td>3790.2</td><td>2.1</td><td>0.8</td><td>2.7</td><td>7.3E-03</td><td>8.8E-03</td><td>MB231/MB468</td><td>CYTO</td></tr>
<tr><td>>trna119_LysCTT_1_-_145395522_145395594@1.32.32,trna11_LysCTT_5_-_180648979_180649051@1.32.32,trna13_LysCTT_6_+_26556774_26556846@1.32.32,trna32_LysCTT_16_-_3207406_3207478@1.32.32,trna7_LysCTT_16_+_3225692_3225764@1.32.32,trna9_LysCTT_5_+_180634755_180634827@1.32.32|GCCCGGCTAGCTCAGTCGGTAGAGCATGAGAC</td><td>>trna119_LysCTT_1_-_145395522_145395594</td><td>1</td><td>32</td><td>32</td><td>GCCCGGCTAGCTCAGTCGGTAGAGCATGAGAC</td><td>1112.0</td><td>-3.3</td><td>1.1</td><td>-3.2</td><td>1.6E-03</td><td>8.8E-03</td><td>BT20/MB231</td><td>MP</td></tr>
<tr><td>>hg38_21_+_8433172_8446622_RNA45SN1_45S_50ntFlanks@6721.6760.40|AATTGCAGGACACATTGATCATCGACACTTCGAACGCACT</td><td>>hg38_21_+_8433172_8446622_RNA45SN1_45S_50ntFlanks</td><td>6721</td><td>6760</td><td>40</td><td>AATTGCAGGACACATTGATCATCGACACTTCGAACGCACT</td><td>412.8</td><td>-2.2</td><td>0.8</td><td>-2.7</td><td>7.2E-03</td><td>8.8E-03</td><td>BT20/MB468</td><td>CYTO</td></tr>
<tr><td>>hg38_21_+_8433172_8446622_RNA45SN1_45S_50ntFlanks@9568.9594.27|CGTGCAAATCGGTCGTCCGACCTGGGT</td><td>>hg38_21_+_8433172_8446622_RNA45SN1_45S_50ntFlanks</td><td>9568</td><td>9594</td><td>27</td><td>CGTGCAAATCGGTCGTCCGACCTGGGT</td><td>640.5</td><td>-3.9</td><td>1.4</td><td>-2.9</td><td>4.3E-03</td><td>8.8E-03</td><td>BT20/MB468</td><td>MP</td></tr>
<tr><td>>hg38_21_+_8433172_8446622_RNA45SN1_45S_50ntFlanks@12641.12661.21|CCATCTGTGGGATTATGACTG</td><td>>hg38_21_+_8433172_8446622_RNA45SN1_45S_50ntFlanks</td><td>12641</td><td>12661</td><td>21</td><td>CCATCTGTGGGATTATGACTG</td><td>2029.1</td><td>-3.4</td><td>1.2</td><td>-2.9</td><td>4.3E-03</td><td>8.8E-03</td><td>BT20/MB468</td><td>MP</td></tr>
<tr><td>>MI0000077|hsa-mir-21&WithFlank&17|+|59841260|59841343@14.34.21(+3C)&[MIMAT0000076&hsa-miR-21-5p&offsets|0|-1(+3C);m-1&17|+|59841273|59841294&offsets|0|-1(+3C)]|TAGCTTATCAGACTGATGTTGCCC</td><td>>MI0000077|hsa-mir-21&WithFlank&17|+|59841260|59841343</td><td>14</td><td>34</td><td>21(+3C)</td><td>TAGCTTATCAGACTGATGTTGCCC</td><td>235.3</td><td>-2.5</td><td>0.8</td><td>-3.1</td><td>1.7E-03</td><td>8.8E-03</td><td>BT20/MB231</td><td>NUC</td></tr>
<tr><td>>hg38_21_+_8433172_8446622_RNA45SN1_45S_50ntFlanks@11702.11728.27|CAATGAAGCGCGGGTAAACGGCGGGAG</td><td>>hg38_21_+_8433172_8446622_RNA45SN1_45S_50ntFlanks</td><td>11702</td><td>11728</td><td>27</td><td>CAATGAAGCGCGGGTAAACGGCGGGAG</td><td>115.1</td><td>-3.8</td><td>1.4</td><td>-2.7</td><td>7.2E-03</td><td>8.8E-03</td><td>BT20/MB468</td><td>CYTO</td></tr>
<tr><td>>hg38_21_+_8433172_8446622_RNA45SN1_45S_50ntFlanks@9494.9515.22|CCGAAAGATGGTGAACTATGCC</td><td>>hg38_21_+_8433172_8446622_RNA45SN1_45S_50ntFlanks</td><td>9494</td><td>9515</td><td>22</td><td>CCGAAAGATGGTGAACTATGCC</td><td>114.5</td><td>4.9</td><td>1.7</td><td>2.9</td><td>4.3E-03</td><td>8.8E-03</td><td>BT20/MB468</td><td>MP</td></tr>
<tr><td>>hg38_21_+_8433172_8446622_RNA45SN1_45S_50ntFlanks@4659.4678.20|AGAGGTGAAATTCTTGGACC</td><td>>hg38_21_+_8433172_8446622_RNA45SN1_45S_50ntFlanks</td><td>4659</td><td>4678</td><td>20</td><td>AGAGGTGAAATTCTTGGACC</td><td>71.4</td><td>5.2</td><td>1.9</td><td>2.7</td><td>6.7E-03</td><td>8.8E-03</td><td>MB231/MB468</td><td>MP</td></tr>
<tr><td>>hg38_MT_+_1621_3279_RNR2_16S_50ntFlanks@625.643.19|AACTGAACTCCTCACACCC</td><td>>hg38_MT_+_1621_3279_RNR2_16S_50ntFlanks</td><td>625</td><td>643</td><td>19</td><td>AACTGAACTCCTCACACCC</td><td>79.7</td><td>5.1</td><td>1.8</td><td>2.9</td><td>4.3E-03</td><td>8.8E-03</td><td>BT20/MB468</td><td>MP</td></tr>
<tr><td>>hg38_21_+_8433172_8446622_RNA45SN1_45S_50ntFlanks@10401.10424.24|GAACATGGGTCAGTCGGTCCTGAG</td><td>>hg38_21_+_8433172_8446622_RNA45SN1_45S_50ntFlanks</td><td>10401</td><td>10424</td><td>24</td><td>GAACATGGGTCAGTCGGTCCTGAG</td><td>3818.9</td><td>2.9</td><td>1.1</td><td>2.7</td><td>6.7E-03</td><td>8.8E-03</td><td>MB231/MB468</td><td>MP</td></tr>
<tr><td>>hg38_21_+_8433172_8446622_RNA45SN1_45S_50ntFlanks@6786.6806.21|TACGCCTGTCTGAGCGTCGCT</td><td>>hg38_21_+_8433172_8446622_RNA45SN1_45S_50ntFlanks</td><td>6786</td><td>6806</td><td>21</td><td>TACGCCTGTCTGAGCGTCGCT</td><td>477.9</td><td>2.3</td><td>0.8</td><td>2.7</td><td>6.7E-03</td><td>8.8E-03</td><td>MB231/MB468</td><td>MP</td></tr>
<tr><td>>hg38_21_+_8433172_8446622_RNA45SN1_45S_50ntFlanks@10770.10796.27|GCAGGTCTCCAAGGTGAACAGCCTCTG</td><td>>hg38_21_+_8433172_8446622_RNA45SN1_45S_50ntFlanks</td><td>10770</td><td>10796</td><td>27</td><td>GCAGGTCTCCAAGGTGAACAGCCTCTG</td><td>418.9</td><td>-3.6</td><td>1.3</td><td>-2.7</td><td>6.7E-03</td><td>8.8E-03</td><td>MB231/MB468</td><td>MP</td></tr>
<tr><td>>MI0000077|hsa-mir-21&WithFlank&17|+|59841260|59841343@52.71.20(+1A)&[MIMAT0004494&hsa-miR-21-3p&offsets|0|-1(+1A);m-52&17|+|59841311|59841332&offsets|0|-2(+1A)]|CAACACCAGTCGATGGGCTGA</td><td>>MI0000077|hsa-mir-21&WithFlank&17|+|59841260|59841343</td><td>52</td><td>71</td><td>20(+1A)</td><td>CAACACCAGTCGATGGGCTGA</td><td>49.8</td><td>5.2</td><td>1.9</td><td>2.7</td><td>6.7E-03</td><td>8.8E-03</td><td>MB231/MB468</td><td>MP</td></tr>
<tr><td>>hg38_21_+_8433172_8446622_RNA45SN1_45S_50ntFlanks@10402.10424.23|AACATGGGTCAGTCGGTCCTGAG</td><td>>hg38_21_+_8433172_8446622_RNA45SN1_45S_50ntFlanks</td><td>10402</td><td>10424</td><td>23</td><td>AACATGGGTCAGTCGGTCCTGAG</td><td>4379.6</td><td>-2.1</td><td>0.8</td><td>-2.7</td><td>7.7E-03</td><td>8.8E-03</td><td>MB231/MB468</td><td>MITO</td></tr>
<tr><td>>hg38_21_+_8433172_8446622_RNA45SN1_45S_50ntFlanks@12405.12446.42|CTTTTTGATCCTTCGATGTCGGCTCTTCCTATCATTGTGAAG</td><td>>hg38_21_+_8433172_8446622_RNA45SN1_45S_50ntFlanks</td><td>12405</td><td>12446</td><td>42</td><td>CTTTTTGATCCTTCGATGTCGGCTCTTCCTATCATTGTGAAG</td><td>169.4</td><td>-5.4</td><td>1.9</td><td>-2.9</td><td>4.3E-03</td><td>8.8E-03</td><td>BT20/MB468</td><td>MP</td></tr>
</table>

<table>
<tr><td>>hg38_21_+_8433172_8446622_RNA45SN1_45S_50ntFlanks@10349.10369.21|AACGAGAACTTTGAAGGCCGA</td><td>>hg38_21_+_8433172_8446622_RNA45SN1_45S_50ntFlanks</td><td>10349</td><td>10369</td><td>21</td><td>AACGAGAACTTTGAAGGCCGA</td><td>878.1</td><td>2.0</td><td>0.8</td><td>2.7</td><td>7.8E-03</td><td>8.8E-03</td><td>MB231/MB468</td><td>NUC</td></tr>
<tr><td>>hg38_21_+_8433172_8446622_RNA45SN1_45S_50ntFlanks@8175.8214.40|CCTTCTGATCGAGGCCCAGCCCGTGGACGGTGTGAGGCCG</td><td>>hg38_21_+_8433172_8446622_RNA45SN1_45S_50ntFlanks</td><td>8175</td><td>8214</td><td>40</td><td>CCTTCTGATCGAGGCCCAGCCCGTGGACGGTGTGAGGCCG</td><td>122.8</td><td>-3.0</td><td>1.1</td><td>-2.7</td><td>6.7E-03</td><td>8.8E-03</td><td>MB231/MB468</td><td>MP</td></tr>
<tr><td>>hg38_21_+_8433172_8446622_RNA45SN1_45S_50ntFlanks@4590.4616.27|ATTTTGTTGGTTTTCGGAACTGAGGCC</td><td>>hg38_21_+_8433172_8446622_RNA45SN1_45S_50ntFlanks</td><td>4590</td><td>4616</td><td>27</td><td>ATTTTGTTGGTTTTCGGAACTGAGGCC</td><td>527.1</td><td>-2.1</td><td>0.8</td><td>-2.7</td><td>7.3E-03</td><td>8.8E-03</td><td>MB231/MB468</td><td>CYTO</td></tr>
<tr><td>>hg38_21_+_8433172_8446622_RNA45SN1_45S_50ntFlanks@6650.6699.50|ACGACTCTTAGCGGTGGATCACTCGGCTCGTGCGTCGATGAAGAACGCAG</td><td>>hg38_21_+_8433172_8446622_RNA45SN1_45S_50ntFlanks</td><td>6650</td><td>6699</td><td>50</td><td>ACGACTCTTAGCGGTGGATCACTCGGCTCGTGCGTCGATGAAGAACGCAG</td><td>368.9</td><td>-5.5</td><td>2.0</td><td>-2.7</td><td>6.7E-03</td><td>8.8E-03</td><td>MB231/MB468</td><td>MP</td></tr>
<tr><td>>hg38_21_+_8433172_8446622_RNA45SN1_45S_50ntFlanks@12007.12030.24|GCCGCCGGTGAAATACCACTACTC</td><td>>hg38_21_+_8433172_8446622_RNA45SN1_45S_50ntFlanks</td><td>12007</td><td>12030</td><td>24</td><td>GCCGCCGGTGAAATACCACTACTC</td><td>942.9</td><td>-5.5</td><td>2.0</td><td>-2.7</td><td>7.7E-03</td><td>8.8E-03</td><td>BT20/MB468</td><td>NUC</td></tr>
<tr><td>>trna152_ValCAC_6_-_27248049_27248121@1.28.28|GCTTCTGTAGTGTAGTGGTTATCACGTT</td><td>>trna152_ValCAC_6_-_27248049_27248121</td><td>1</td><td>28</td><td>28</td><td>GCTTCTGTAGTGTAGTGGTTATCACGTT</td><td>264.0</td><td>-1.6</td><td>0.6</td><td>-2.7</td><td>7.3E-03</td><td>8.9E-03</td><td>BT20/MB468</td><td>CYTO</td></tr>
<tr><td>>MI0000746|hsa-mir-99b&WithFlank&19|+|51692606|51692687@51.72.22&[MIMAT0004678&hsa-miR-99b-3p&offsets|0|0;m-151&19|+|51692656|51692678&offsets|0|-1]|CAAGCTCGTGTCTGTGGGTCCG</td><td>>MI0000746|hsa-mir-99b&WithFlank&19|+|51692606|51692687</td><td>51</td><td>72</td><td>22</td><td>CAAGCTCGTGTCTGTGGGTCCG</td><td>2536.6</td><td>2.3</td><td>0.7</td><td>3.1</td><td>2.0E-03</td><td>8.9E-03</td><td>BT20/MB231</td><td>CYTO</td></tr>
<tr><td>>hg38_21_+_8433172_8446622_RNA45SN1_45S_50ntFlanks@7989.8020.32|AGACGTGGCGACCCGCTGAATTTAAGCATATT</td><td>>hg38_21_+_8433172_8446622_RNA45SN1_45S_50ntFlanks</td><td>7989</td><td>8020</td><td>32</td><td>AGACGTGGCGACCCGCTGAATTTAAGCATATT</td><td>196.8</td><td>-2.9</td><td>0.9</td><td>-3.1</td><td>2.0E-03</td><td>8.9E-03</td><td>BT20/MB231</td><td>CYTO</td></tr>
<tr><td>>hg38_21_+_8433172_8446622_RNA45SN1_45S_50ntFlanks@5473.5499.27|CGGCCCTGGCGGAGCGCTGAGAAGACG</td><td>>hg38_21_+_8433172_8446622_RNA45SN1_45S_50ntFlanks</td><td>5473</td><td>5499</td><td>27</td><td>CGGCCCTGGCGGAGCGCTGAGAAGACG</td><td>1773.1</td><td>-3.0</td><td>1.1</td><td>-2.7</td><td>7.7E-03</td><td>8.9E-03</td><td>BT20/MB468</td><td>NUC</td></tr>
<tr><td>>am_MI0003581|hsa-mir-574&WithFlank&4|+|38868026|38868133@31.52.22(+1C)&[MIMAT0004795&hsa-miR-574-5p&offsets|0|-1(+1C);m-213&4|+|38868056|38868078&offsets|0|-1(+1C)]|TGAGTGTGTGTGTGTGAGTGTGC</td><td>>am_MI0003581|hsa-mir-574&WithFlank&4|+|38868026|38868133</td><td>31</td><td>52</td><td>22(+1C)</td><td>TGAGTGTGTGTGTGTGAGTGTGC</td><td>153.0</td><td>4.2</td><td>1.5</td><td>2.9</td><td>4.3E-03</td><td>8.9E-03</td><td>BT20/MB468</td><td>MP</td></tr>
<tr><td>>hg38_21_+_8433172_8446622_RNA45SN1_45S_50ntFlanks@6651.6677.27|CGACTCTTAGCGGTGGATCACTCGGCT</td><td>>hg38_21_+_8433172_8446622_RNA45SN1_45S_50ntFlanks</td><td>6651</td><td>6677</td><td>27</td><td>CGACTCTTAGCGGTGGATCACTCGGCT</td><td>7344.2</td><td>2.7</td><td>1.0</td><td>2.9</td><td>4.3E-03</td><td>8.9E-03</td><td>BT20/MB468</td><td>MP</td></tr>
<tr><td>>hg38_21_+_8433172_8446622_RNA45SN1_45S_50ntFlanks@11666.11687.22|TCTGCCCAGTGCTCTGAATGTC</td><td>>hg38_21_+_8433172_8446622_RNA45SN1_45S_50ntFlanks</td><td>11666</td><td>11687</td><td>22</td><td>TCTGCCCAGTGCTCTGAATGTC</td><td>428.9</td><td>2.9</td><td>1.0</td><td>2.9</td><td>4.3E-03</td><td>8.9E-03</td><td>BT20/MB468</td><td>MP</td></tr>
<tr><td>>hg38_MT_+_1621_3279_RNR2_16S_50ntFlanks@1379.1401.23|CAGGACATCCCGATGGTGCAGCC</td><td>>hg38_MT_+_1621_3279_RNR2_16S_50ntFlanks</td><td>1379</td><td>1401</td><td>23</td><td>CAGGACATCCCGATGGTGCAGCC</td><td>68.6</td><td>3.8</td><td>1.4</td><td>2.7</td><td>6.7E-03</td><td>8.9E-03</td><td>MB231/MB468</td><td>MP</td></tr>
<tr><td>>trnaMT_ProTGG_MT_-_15956_16023@40.70.31|AATGGTGGAGTTAAAGACTTTTTCTCTGACC</td><td>>trnaMT_ProTGG_MT_-_15956_16023</td><td>40</td><td>70</td><td>31</td><td>AATGGTGGAGTTAAAGACTTTTTCTCTGACC</td><td>86.8</td><td>5.3</td><td>2.0</td><td>2.7</td><td>6.7E-03</td><td>8.9E-03</td><td>MB231/MB468</td><td>MP</td></tr>
<tr><td>>MI0006444|hsa-mir-1307&WithFlank&10|-|103394247|103394407@86.107.22(+1A)&[MIMAT0005951&hsa-miR-1307-3p&offsets|0|0(+1A);m-66&10|-|103394300|103394322&offsets|0|-1(+1A)]|ACTCGGCGTGGCGTCGGTCGTGA</td><td>>MI0006444|hsa-mir-1307&WithFlank&10|-|103394247|103394407</td><td>86</td><td>107</td><td>22(+1A)</td><td>ACTCGGCGTGGCGTCGGTCGTGA</td><td>213.0</td><td>2.7</td><td>1.0</td><td>2.7</td><td>6.7E-03</td><td>8.9E-03</td><td>MB231/MB468</td><td>MP</td></tr>
<tr><td>>hg38_21_+_8433172_8446622_RNA45SN1_45S_50ntFlanks@10910.10932.23|CGCGCGCCGCGGCTGGACGAGGC</td><td>>hg38_21_+_8433172_8446622_RNA45SN1_45S_50ntFlanks</td><td>10910</td><td>10932</td><td>23</td><td>CGCGCGCCGCGGCTGGACGAGGC</td><td>389.6</td><td>-2.5</td><td>0.9</td><td>-2.9</td><td>4.4E-03</td><td>8.9E-03</td><td>BT20/MB468</td><td>MP</td></tr>
<tr><td>>hg38_MT_+_1621_3279_RNR2_16S_50ntFlanks@644.677.34|AATTGGACCAATCTATCACCCTATAGAAGAACTA</td><td>>hg38_MT_+_1621_3279_RNR2_16S_50ntFlanks</td><td>644</td><td>677</td><td>34</td><td>AATTGGACCAATCTATCACCCTATAGAAGAACTA</td><td>56.2</td><td>5.1</td><td>1.9</td><td>2.7</td><td>6.7E-03</td><td>8.9E-03</td><td>MB231/MB468</td><td>MP</td></tr>
<tr><td>>MI0000077|hsa-mir-21&WithFlank&17|+|59841260|59841343@14.31.18&[MIMAT0000076&hsa-miR-21-5p&offsets|0|-4;m-1&17|+|59841273|59841294&offsets|0|-4]|TAGCTTATCAGACTGATG</td><td>>MI0000077|hsa-mir-21&WithFlank&17|+|59841260|59841343</td><td>14</td><td>31</td><td>18</td><td>TAGCTTATCAGACTGATG</td><td>747.6</td><td>1.8</td><td>0.7</td><td>2.7</td><td>6.7E-03</td><td>8.9E-03</td><td>MB231/MB468</td><td>MP</td></tr>
<tr><td>>hg38_21_+_8433172_8446622_RNA45SN1_45S_50ntFlanks@6645.6679.35|CTCGTACGACTCTTAGCGGTGGATCACTCGGCTCG</td><td>>hg38_21_+_8433172_8446622_RNA45SN1_45S_50ntFlanks</td><td>6645</td><td>6679</td><td>35</td><td>CTCGTACGACTCTTAGCGGTGGATCACTCGGCTCG</td><td>598.5</td><td>1.5</td><td>0.6</td><td>2.7</td><td>7.4E-03</td><td>8.9E-03</td><td>MB231/MB468</td><td>CYTO</td></tr>
<tr><td>>am_trnaMT_TyrGTA_MT_-_5826_5891@13.36.24|GAGTGAAGCATTGGACTGTAAATC</td><td>>am_trnaMT_TyrGTA_MT_-_5826_5891</td><td>13</td><td>36</td><td>24</td><td>GAGTGAAGCATTGGACTGTAAATC</td><td>25.7</td><td>7.9</td><td>2.9</td><td>2.7</td><td>6.7E-03</td><td>8.9E-03</td><td>MB231/MB468</td><td>MP</td></tr>
<tr><td>>hg38_21_+_8433172_8446622_RNA45SN1_45S_50ntFlanks@4659.4678.20|AGAGGTGAAATTCTTGGACC</td><td>>hg38_21_+_8433172_8446622_RNA45SN1_45S_50ntFlanks</td><td>4659</td><td>4678</td><td>20</td><td>AGAGGTGAAATTCTTGGACC</td><td>76.9</td><td>5.9</td><td>2.1</td><td>2.9</td><td>4.4E-03</td><td>8.9E-03</td><td>BT20/MB468</td><td>MP</td></tr>
<tr><td>>hg38_21_+_8433172_8446622_RNA45SN1_45S_50ntFlanks@10423.10456.34|AGAGATGGGCGAGCGCCGTTCCGAAGGGACGGGC</td><td>>hg38_21_+_8433172_8446622_RNA45SN1_45S_50ntFlanks</td><td>10423</td><td>10456</td><td>34</td><td>AGAGATGGGCGAGCGCCGTTCCGAAGGGACGGGC</td><td>1025.5</td><td>-1.5</td><td>0.5</td><td>-2.7</td><td>7.3E-03</td><td>8.9E-03</td><td>BT20/MB468</td><td>CYTO</td></tr>
<tr><td>>hg38_MT_+_598_1651_RNR1_12S_50ntFlanks@52.93.42|ATAGGTTTGGTCCTAGCCTTTCTATTAGCTCTTAGTAAGATT</td><td>>hg38_MT_+_598_1651_RNR1_12S_50ntFlanks</td><td>52</td><td>93</td><td>42</td><td>ATAGGTTTGGTCCTAGCCTTTCTATTAGCTCTTAGTAAGATT</td><td>3586.0</td><td>4.6</td><td>1.6</td><td>2.9</td><td>4.4E-03</td><td>8.9E-03</td><td>BT20/MB468</td><td>MP</td></tr>
<tr><td>>trnaMT_GluTTC_MT_-_14674_14742@32.72.41,trnalookalike8_GluTTC_5_-_93905172_93905240@32.72.41|TCATATCATTGGTCGTGGTTGTAGTCCGTGCGAGAATACCA</td><td>>trnaMT_GluTTC_MT_-_14674_14742</td><td>32</td><td>72</td><td>41</td><td>TCATATCATTGGTCGTGGTTGTAGTCCGTGCGAGAATACCA</td><td>872.9</td><td>-2.6</td><td>0.8</td><td>-3.1</td><td>2.0E-03</td><td>8.9E-03</td><td>BT20/MB231</td><td>CYTO</td></tr>
<tr><td>>hg38_21_+_8433172_8446622_RNA45SN1_45S_50ntFlanks@10841.10867.27|CGTAACTTCGGGATAAGGATTGGCTCT</td><td>>hg38_21_+_8433172_8446622_RNA45SN1_45S_50ntFlanks</td><td>10841</td><td>10867</td><td>27</td><td>CGTAACTTCGGGATAAGGATTGGCTCT</td><td>4348.4</td><td>-2.7</td><td>1.0</td><td>-2.7</td><td>7.8E-03</td><td>8.9E-03</td><td>MB231/MB468</td><td>NUC</td></tr>
<tr><td>>MI0000749|hsa-mir-30e&WithFlank&1|+|40754349|40754452@65.85.21(+2U)&[MIMAT0000693&hsa-miR-30e-3p&offsets|0|-1(+2U);m-25&1|+|40754413|40754434&offsets|0|-1(+2U)]|CTTTCAGTCGGATGTTTACAGTT</td><td>>MI0000749|hsa-mir-30e&WithFlank&1|+|40754349|40754452</td><td>65</td><td>85</td><td>21(+2U)</td><td>CTTTCAGTCGGATGTTTACAGTT</td><td>288.5</td><td>-1.0</td><td>0.4</td><td>-2.7</td><td>7.7E-03</td><td>8.9E-03</td><td>BT20/MB468</td><td>NUC</td></tr>
<tr><td>>hg38_21_+_8433172_8446622_RNA45SN1_45S_50ntFlanks@10852.10878.27|GATAAGGATTGGCTCTAAGGGCTGGGT</td><td>>hg38_21_+_8433172_8446622_RNA45SN1_45S_50ntFlanks</td><td>10852</td><td>10878</td><td>27</td><td>GATAAGGATTGGCTCTAAGGGCTGGGT</td><td>57.5</td><td>9.2</td><td>2.9</td><td>3.2</td><td>1.6E-03</td><td>8.9E-03</td><td>BT20/MB231</td><td>MP</td></tr>
<tr><td>>hg38_21_+_8433172_8446622_RNA45SN1_45S_50ntFlanks@3845.3881.37|ACTTGGATAACTGTGGTAATTCTAGAGCTAATACATG</td><td>>hg38_21_+_8433172_8446622_RNA45SN1_45S_50ntFlanks</td><td>3845</td><td>3881</td><td>37</td><td>ACTTGGATAACTGTGGTAATTCTAGAGCTAATACATG</td><td>114.5</td><td>-4.0</td><td>1.5</td><td>-2.7</td><td>6.3E-03</td><td>8.9E-03</td><td>BT20/MB468</td><td>MITO</td></tr>
<tr><td>>MI0000269|hsa-mir-181a-2&WithFlank&9|+|124692436|124692557@83.103.21(+1U)&[MIMAT0004558&hsa-miR-181a-2-3p&offsets|0|-1(+1U);m-71&9|+|124692518|124692539&offsets|0|-1(+1U)]|ACCACTGACCGTTGACTGTACT</td><td>>MI0000269|hsa-mir-181a-2&WithFlank&9|+|124692436|124692557</td><td>83</td><td>103</td><td>21(+1U)</td><td>ACCACTGACCGTTGACTGTACT</td><td>219.0</td><td>4.4</td><td>1.6</td><td>2.7</td><td>6.8E-03</td><td>8.9E-03</td><td>MB231/MB468</td><td>MP</td></tr>
<tr><td>>MI0000070|hsa-mir-16-1&WithFlank&13|-|50048967|50049067@20.40.21(+1A);MI0000115|hsa-mir-16-2&WithFlank&3|+|160404739|160404831@16.36.21(+1A)&[MIMAT0000069&hsa-miR-16-5p&offsets|0|-1(+1A);m-62&13|-|50049027|50049048&offsets|0|-1(+1A)];[MIMAT0000069_1&hsa-miR-16-5p&offsets|0|-1(+1A);m-61&3|+|160404754|160404775&offsets|0|-1(+1A)]|TAGCAGCACGTAAATATTGGCA</td><td>>MI0000070|hsa-mir-16-1&WithFlank&13|-|50048967|50049067</td><td>20</td><td>40</td><td>21(+1A)</td><td>TAGCAGCACGTAAATATTGGCA</td><td>1706.4</td><td>2.5</td><td>0.8</td><td>3.1</td><td>1.8E-03</td><td>8.9E-03</td><td>BT20/MB231</td><td>NUC</td></tr>
<tr><td>>MI0000749|hsa-mir-30e&WithFlank&1|+|40754349|40754452@65.85.21(+1U)&[MIMAT0000693&hsa-miR-30e-3p&offsets|0|-1(+1U);m-25&1|+|40754413|40754434&offsets|0|-1(+1U)]|CTTTCAGTCGGATGTTTACAGT</td><td>>MI0000749|hsa-mir-30e&WithFlank&1|+|40754349|40754452</td><td>65</td><td>85</td><td>21(+1U)</td><td>CTTTCAGTCGGATGTTTACAGT</td><td>2386.0</td><td>-2.9</td><td>0.9</td><td>-3.2</td><td>1.6E-03</td><td>8.9E-03</td><td>BT20/MB231</td><td>MP</td></tr>
<tr><td>>hg38_21_+_8433172_8446622_RNA45SN1_45S_50ntFlanks@11570.11599.30|AGCCGACTTAGAACTGGTGCGGACCAGGGG</td><td>>hg38_21_+_8433172_8446622_RNA45SN1_45S_50ntFlanks</td><td>11570</td><td>11599</td><td>30</td><td>AGCCGACTTAGAACTGGTGCGGACCAGGGG</td><td>369.5</td><td>5.3</td><td>1.7</td><td>3.2</td><td>1.6E-03</td><td>8.9E-03</td><td>BT20/MB231</td><td>MP</td></tr>
<tr><td>>trna111_HisGTG_1_-_147774845_147774916@-1T.22.23,trna118_HisGTG_1_-_145396881_145396952@-1T.22.23,trna16_HisGTG_1_+_146544773_146544844@-1T.22.23,trna1_HisGTG_15_+_45493349_45493420@-1T.22.23,trna21_HisGTG_1_+_147753471_147753542@-1T.22.23,trna33_HisGTG_6_+_27125906_27125977@-1T.22.23,trna7_HisGTG_9_-_14433938_14434009@-1T.22.23,trna8_HisGTG_15_-_45492611_45492682@-1T.22.23,trna9_HisGTG_15_-_45490804_45490875@-1T.22.23|TGCCGTGATCGTATAGTGGTTAG</td><td>>trna111_HisGTG_1_-_147774845_147774916</td><td>-1T</td><td>22</td><td>23</td><td>TGCCGTGATCGTATAGTGGTTAG</td><td>278.9</td><td>-4.1</td><td>1.3</td><td>-3.1</td><td>1.8E-03</td><td>8.9E-03</td><td>BT20/MB231</td><td>NUC</td></tr>
<tr><td>>hg38_21_+_8433172_8446622_RNA45SN1_45S_50ntFlanks@11665.11685.21|TTCTGCCCAGTGCTCTGAATG</td><td>>hg38_21_+_8433172_8446622_RNA45SN1_45S_50ntFlanks</td><td>11665</td><td>11685</td><td>21</td><td>TTCTGCCCAGTGCTCTGAATG</td><td>588.0</td><td>-2.0</td><td>0.7</td><td>-2.7</td><td>7.4E-03</td><td>8.9E-03</td><td>MB231/MB468</td><td>CYTO</td></tr>
<tr><td>>hg38_21_+_8433172_8446622_RNA45SN1_45S_50ntFlanks@12415.12456.42|CTTCGATGTCGGCTCTTCCTATCATTGTGAAGCAGAATTCAC</td><td>>hg38_21_+_8433172_8446622_RNA45SN1_45S_50ntFlanks</td><td>12415</td><td>12456</td><td>42</td><td>CTTCGATGTCGGCTCTTCCTATCATTGTGAAGCAGAATTCAC</td><td>300.9</td><td>-2.2</td><td>0.8</td><td>-2.7</td><td>7.8E-03</td><td>8.9E-03</td><td>MB231/MB468</td><td>MITO</td></tr>
<tr><td>>am_trnaMT_ValTAC_MT_+_1602_1670@36.65.30|CTTAGGAGATTTCAACTTAACTTGACCGCT</td><td>>am_trnaMT_ValTAC_MT_+_1602_1670</td><td>36</td><td>65</td><td>30</td><td>CTTAGGAGATTTCAACTTAACTTGACCGCT</td><td>288.1</td><td>7.3</td><td>2.6</td><td>2.8</td><td>4.4E-03</td><td>8.9E-03</td><td>BT20/MB468</td><td>MP</td></tr>
<tr><td>>hg38_21_+_8433172_8446622_RNA45SN1_45S_50ntFlanks@5083.5110.28|ACGAACGAGACTCTGGCATGCTAACTAG</td><td>>hg38_21_+_8433172_8446622_RNA45SN1_45S_50ntFlanks</td><td>5083</td><td>5110</td><td>28</td><td>ACGAACGAGACTCTGGCATGCTAACTAG</td><td>443.8</td><td>-2.5</td><td>1.0</td><td>-2.7</td><td>7.9E-03</td><td>8.9E-03</td><td>MB231/MB468</td><td>NUC</td></tr>
<tr><td>>hg38_21_+_8433172_8446622_RNA45SN1_45S_50ntFlanks@12160.12179.20|TTGACTGGGGCGGTACACCT</td><td>>hg38_21_+_8433172_8446622_RNA45SN1_45S_50ntFlanks</td><td>12160</td><td>12179</td><td>20</td><td>TTGACTGGGGCGGTACACCT</td><td>170.9</td><td>-2.3</td><td>0.8</td><td>-2.7</td><td>6.3E-03</td><td>9.0E-03</td><td>BT20/MB468</td><td>MITO</td></tr>
<tr><td>>hg38_21_+_8433172_8446622_RNA45SN1_45S_50ntFlanks@12006.12026.21|GGCCGCCGGTGAAATACCACT</td><td>>hg38_21_+_8433172_8446622_RNA45SN1_45S_50ntFlanks</td><td>12006</td><td>12026</td><td>21</td><td>GGCCGCCGGTGAAATACCACT</td><td>306.2</td><td>5.8</td><td>1.8</td><td>3.2</td><td>1.6E-03</td><td>9.0E-03</td><td>BT20/MB231</td><td>MP</td></tr>
<tr><td>>hg38_21_+_8433172_8446622_RNA45SN1_45S_50ntFlanks@11670.11697.28|CCCAGTGCTCTGAATGTCAAAGTGAAGA</td><td>>hg38_21_+_8433172_8446622_RNA45SN1_45S_50ntFlanks</td><td>11670</td><td>11697</td><td>28</td><td>CCCAGTGCTCTGAATGTCAAAGTGAAGA</td><td>314.6</td><td>5.4</td><td>1.7</td><td>3.2</td><td>1.6E-03</td><td>9.0E-03</td><td>BT20/MB231</td><td>MP</td></tr>
<tr><td>>hg38_21_+_8433172_8446622_RNA45SN1_45S_50ntFlanks@10291.10315.25|CGCGGGCCCGGGTGGAGCCGCCGCA</td><td>>hg38_21_+_8433172_8446622_RNA45SN1_45S_50ntFlanks</td><td>10291</td><td>10315</td><td>25</td><td>CGCGGGCCCGGGTGGAGCCGCCGCA</td><td>179.8</td><td>-5.3</td><td>1.9</td><td>-2.8</td><td>4.4E-03</td><td>9.0E-03</td><td>BT20/MB468</td><td>MP</td></tr>
<tr><td>>MI0000270|hsa-mir-181b-1&WithFlank&1|-|198858867|198858988@42.64.23(+1A);MI0000683|hsa-mir-181b-2&WithFlank&9|+|124693704|124693804@22.44.23(+1A)&[MIMAT0000257&hsa-miR-181b-5p&offsets|0|0(+1A);m-73&1|-|198858925|198858947&offsets|0|0(+1A)];[MIMAT0000257_1&hsa-miR-181b-5p&offsets|0|0(+1A);m-72&9|+|124693725|124693747&offsets|0|0(+1A)]|AACATTCATTGCTGTCGGTGGGTA</td><td>>MI0000270|hsa-mir-181b-1&WithFlank&1|-|198858867|198858988</td><td>42</td><td>64</td><td>23(+1A)</td><td>AACATTCATTGCTGTCGGTGGGTA</td><td>258.7</td><td>-2.6</td><td>0.8</td><td>-3.1</td><td>2.1E-03</td><td>9.0E-03</td><td>BT20/MB231</td><td>MITO</td></tr>
<tr><td>>MI0003684|hsa-mir-660&WithFlank&X|+|50013235|50013343@22.44.23&[MIMAT0003338&hsa-miR-660-5p&offsets|0|+1;m-157&X|+|50013256|50013278&offsets|0|0]|TACCCATTGCATATCGGAGTTGT</td><td>>MI0003684|hsa-mir-660&WithFlank&X|+|50013235|50013343</td><td>22</td><td>44</td><td>23</td><td>TACCCATTGCATATCGGAGTTGT</td><td>352.3</td><td>-3.5</td><td>1.1</td><td>-3.1</td><td>2.1E-03</td><td>9.0E-03</td><td>BT20/MB231</td><td>MITO</td></tr>
<tr><td>>hg38_21_+_8433172_8446622_RNA45SN1_45S_50ntFlanks@12990.13013.24|GCTCCCTCGCTGCGATCTATTGAA</td><td>>hg38_21_+_8433172_8446622_RNA45SN1_45S_50ntFlanks</td><td>12990</td><td>13013</td><td>24</td><td>GCTCCCTCGCTGCGATCTATTGAA</td><td>949.3</td><td>-2.8</td><td>0.9</td><td>-3.1</td><td>2.0E-03</td><td>9.0E-03</td><td>BT20/MB231</td><td>CYTO</td></tr>
<tr><td>>hg38_21_+_8433172_8446622_RNA45SN1_45S_50ntFlanks@4574.4613.40|ATAGGACCGCGGTTCTATTTTGTTGGTTTTCGGAACTGAG</td><td>>hg38_21_+_8433172_8446622_RNA45SN1_45S_50ntFlanks</td><td>4574</td><td>4613</td><td>40</td><td>ATAGGACCGCGGTTCTATTTTGTTGGTTTTCGGAACTGAG</td><td>149.1</td><td>-2.3</td><td>0.9</td><td>-2.7</td><td>7.5E-03</td><td>9.0E-03</td><td>MB231/MB468</td><td>CYTO</td></tr>
</table>

| | | | | | | | | | | | | | |
|---|---|---|---|---|---|---|---|---|---|---|---|---|---|
| >hg38_21_+_8433172_8446622_RNA45SN1_45S_50ntFlanks@6722.6746.25\|ATTGCAGGACACATTGATCATCGAC | >hg38_21_+_8433172_8446622_RNA45SN1_45S_50ntFlanks | 6722 | 6746 | 25 | ATTGCAGGACACATTGATCATCGAC | 186.6 | -3.1 | 1.2 | -2.7 | 7.4E-03 | 9.0E-03 | BT20/MB468 | CYTO |
| >hg38_21_+_8433172_8446622_RNA45SN1_45S_50ntFlanks@11316.11338.23\|CACCCCACGTCTCGTCGCGCGCG | >hg38_21_+_8433172_8446622_RNA45SN1_45S_50ntFlanks | 11316 | 11338 | 23 | CACCCCACGTCTCGTCGCGCGCG | 3276.7 | -4.0 | 1.5 | -2.7 | 6.3E-03 | 9.0E-03 | BT20/MB468 | MITO |
| >hg38_21_+_8433172_8446622_RNA45SN1_45S_50ntFlanks@11641.11667.27\|CGGCGGGTGTTGACGCGATGTGATTTC | >hg38_21_+_8433172_8446622_RNA45SN1_45S_50ntFlanks | 11641 | 11667 | 27 | CGGCGGGTGTTGACGCGATGTGATTTC | 197.9 | -4.2 | 1.6 | -2.7 | 7.4E-03 | 9.0E-03 | BT20/MB468 | CYTO |
| >hg38_21_+_8433172_8446622_RNA45SN1_45S_50ntFlanks@3880.3898.19\|TGCCGACGGGCGCTGACCC | >hg38_21_+_8433172_8446622_RNA45SN1_45S_50ntFlanks | 3880 | 3898 | 19 | TGCCGACGGGCGCTGACCC | 202.7 | -1.9 | 0.7 | -2.7 | 7.4E-03 | 9.0E-03 | BT20/MB468 | CYTO |
| >hg38_21_+_8433172_8446622_RNA45SN1_45S_50ntFlanks@11576.11593.18\|CTTAGAACTGGTGCGGAC | >hg38_21_+_8433172_8446622_RNA45SN1_45S_50ntFlanks | 11576 | 11593 | 18 | CTTAGAACTGGTGCGGAC | 286.3 | -3.2 | 1.2 | -2.7 | 6.3E-03 | 9.0E-03 | BT20/MB468 | MITO |
| >hg38_21_+_8433172_8446622_RNA45SN1_45S_50ntFlanks@12717.12757.41\|CCTCGGTTGGCCTCGGATAGCCGGTCCCCCGCCTGTCCCCG | >hg38_21_+_8433172_8446622_RNA45SN1_45S_50ntFlanks | 12717 | 12757 | 41 | CCTCGGTTGGCCTCGGATAGCCGGTCCCCCGCCTGTCCCCG | 536.5 | -4.4 | 1.6 | -2.7 | 6.3E-03 | 9.0E-03 | BT20/MB468 | MITO |
| >hg38_21_+_8433172_8446622_RNA45SN1_45S_50ntFlanks@7992.8009.18\|CGTGGCGACCCGCTGAAT | >hg38_21_+_8433172_8446622_RNA45SN1_45S_50ntFlanks | 7992 | 8009 | 18 | CGTGGCGACCCGCTGAAT | 1887.5 | -2.4 | 0.9 | -2.7 | 7.9E-03 | 9.0E-03 | MB231/MB468 | MITO |
| >am_trna128_GlyGCC_6_-_27870686_27870756@1.36.36,trna18_GlyGCC_16_+_70822597_70822667@1.36.36,trna19_GlyGCC_16_+_70823410_70823480@1.36.36,trna19_GlyGCC_2_-_157257659_157257729@1.36.36,trna24_GlyGCC_16_-_70812942_70813012@1.36.36,trna25_GlyGCC_16_-_70812114_70812184@1.36.36,trna5_GlyGCC_17_+_8029064_8029134@1.36.36,trna68_GlyGCC_1_-_161493637_161493707@1.36.36\|GCATTGGTGGTTCAGTGGTAGAATTCTCGCCTGCCA | >am_trna128_GlyGCC_6_-_27870686_27870756 | 1 | 36 | 36 | GCATTGGTGGTTCAGTGGTAGAATTCTCGCCTGCCA | 372.1 | -2.7 | 0.9 | -3.1 | 1.8E-03 | 9.0E-03 | BT20/MB231 | NUC |
| >MI0000298\|hsa-mir-221&WithFlank&X\|-\|45746151\|45746272@71.92.22(+1G)&[MIMAT0000278&hsa-miR-221-3p&offsets\|0\|-1(+1G);m-91&X\|-\|45746181\|45746202&offsets\|0\|0(+1G)]\|AGCTACATTGTCTGCTGGGTTTG | >MI0000298\|hsa-mir-221&WithFlank&X\|-\|45746151\|45746272 | 71 | 92 | 22(+1G) | AGCTACATTGTCTGCTGGGTTTG | 92.2 | 4.4 | 1.6 | 2.7 | 6.8E-03 | 9.0E-03 | MB231/MB468 | MP |
| >hg38_21_+_8433172_8446622_RNA45SN1_45S_50ntFlanks@8194.8216.23\|CCCGTGGACGGTGTGAGGCCGGT | >hg38_21_+_8433172_8446622_RNA45SN1_45S_50ntFlanks | 8194 | 8216 | 23 | CCCGTGGACGGTGTGAGGCCGGT | 195.1 | -2.9 | 1.1 | -2.7 | 7.4E-03 | 9.0E-03 | BT20/MB468 | CYTO |
| >hg38_21_+_8433172_8446622_RNA45SN1_45S_50ntFlanks@10348.10367.20\|AAACGAGAACTTTGAAGGCC | >hg38_21_+_8433172_8446622_RNA45SN1_45S_50ntFlanks | 10348 | 10367 | 20 | AAACGAGAACTTTGAAGGCC | 737.6 | 1.5 | 0.6 | 2.7 | 7.4E-03 | 9.0E-03 | BT20/MB468 | CYTO |
| >hg38_21_+_8433172_8446622_RNA45SN1_45S_50ntFlanks@12988.13013.26\|CAGCTCCCTCGCTGCGATCTATTGAA | >hg38_21_+_8433172_8446622_RNA45SN1_45S_50ntFlanks | 12988 | 13013 | 26 | CAGCTCCCTCGCTGCGATCTATTGAA | 418.1 | 1.4 | 0.5 | 2.7 | 7.4E-03 | 9.0E-03 | BT20/MB468 | CYTO |
| >MI0005566\|hsa-mir-374b&WithFlank&X\|-\|74218541\|74218624@17.39.23&[MIMAT0004955&hsa-miR-374b-5p&offsets\|0\|+1;m-153&X\|-\|74218587\|74218608&offsets\|0\|+1]\|ATATAATACAACCTGCTAAGTGT | >MI0005566\|hsa-mir-374b&WithFlank&X\|-\|74218541\|74218624 | 17 | 39 | 23 | ATATAATACAACCTGCTAAGTGT | 396.0 | 4.6 | 1.7 | 2.7 | 6.8E-03 | 9.0E-03 | MB231/MB468 | MP |
| >hg38_21_+_8433172_8446622_RNA45SN1_45S_50ntFlanks@4917.4941.25\|CACCACCAGGAGTGGAGCCTGCGGC | >hg38_21_+_8433172_8446622_RNA45SN1_45S_50ntFlanks | 4917 | 4941 | 25 | CACCACCAGGAGTGGAGCCTGCGGC | 272.4 | -2.5 | 0.9 | -2.8 | 4.4E-03 | 9.0E-03 | BT20/MB468 | MP |
| >hg38_21_+_8433172_8446622_RNA45SN1_45S_50ntFlanks@11576.11593.18\|CTTAGAACTGGTGCGGAC | >hg38_21_+_8433172_8446622_RNA45SN1_45S_50ntFlanks | 11576 | 11593 | 18 | CTTAGAACTGGTGCGGAC | 557.9 | -3.9 | 1.4 | -2.8 | 4.4E-03 | 9.0E-03 | BT20/MB468 | MP |
| >hg38_21_+_8433172_8446622_RNA45SN1_45S_50ntFlanks@10065.10082.18\|CGAGAGTGGACGGGAGCG | >hg38_21_+_8433172_8446622_RNA45SN1_45S_50ntFlanks | 10065 | 10082 | 18 | CGAGAGTGGACGGGAGCG | 66.3 | 5.7 | 2.0 | 2.8 | 4.4E-03 | 9.0E-03 | BT20/MB468 | MP |
| >hg38_MT_+_598_1651_RNR1_12S_50ntFlanks@51.88.38\|AATAGGTTTGGTCCTAGCCTTTCTATTAGCTCTTAGTA | >hg38_MT_+_598_1651_RNR1_12S_50ntFlanks | 51 | 88 | 38 | AATAGGTTTGGTCCTAGCCTTTCTATTAGCTCTTAGTA | 15171.1 | 4.6 | 1.6 | 2.8 | 4.4E-03 | 9.0E-03 | BT20/MB468 | MP |
| >hg38_21_+_8433172_8446622_RNA45SN1_45S_50ntFlanks@3705.3742.38\|TACCTGGTTGATCCTGCCAGTAGCATATGCTTGTCTCA | >hg38_21_+_8433172_8446622_RNA45SN1_45S_50ntFlanks | 3705 | 3742 | 38 | TACCTGGTTGATCCTGCCAGTAGCATATGCTTGTCTCA | 774.5 | 1.3 | 0.5 | 2.7 | 7.8E-03 | 9.0E-03 | BT20/MB468 | NUC |
| >hg38_21_+_8433172_8446622_RNA45SN1_45S_50ntFlanks@4007.4025.19\|CCTCGGGCCGATCGCACGC | >hg38_21_+_8433172_8446622_RNA45SN1_45S_50ntFlanks | 4007 | 4025 | 19 | CCTCGGGCCGATCGCACGC | 222.5 | -4.3 | 1.6 | -2.7 | 6.4E-03 | 9.0E-03 | BT20/MB468 | MITO |
| >hg38_21_+_8433172_8446622_RNA45SN1_45S_50ntFlanks@5236.5254.19\|CACTGACTGGCTCAGCGTG | >hg38_21_+_8433172_8446622_RNA45SN1_45S_50ntFlanks | 5236 | 5254 | 19 | CACTGACTGGCTCAGCGTG | 1055.0 | -5.1 | 1.9 | -2.7 | 6.4E-03 | 9.0E-03 | BT20/MB468 | MITO |
| >hg38_21_+_8433172_8446622_RNA45SN1_45S_50ntFlanks@6733.6754.22\|CATTGATCATCGACACTTCGAA | >hg38_21_+_8433172_8446622_RNA45SN1_45S_50ntFlanks | 6733 | 6754 | 22 | CATTGATCATCGACACTTCGAA | 178.7 | -4.0 | 1.4 | -2.8 | 4.4E-03 | 9.0E-03 | BT20/MB468 | MP |
| >hg38_21_+_8433172_8446622_RNA45SN1_45S_50ntFlanks@6734.6772.39\|ATTGATCATCGACACTTCGAACGCACTTGCGGCCCCGGG | >hg38_21_+_8433172_8446622_RNA45SN1_45S_50ntFlanks | 6734 | 6772 | 39 | ATTGATCATCGACACTTCGAACGCACTTGCGGCCCCGGG | 758.8 | -2.0 | 0.8 | -2.7 | 7.9E-03 | 9.0E-03 | MB231/MB468 | MITO |
| >hg38_21_+_8433172_8446622_RNA45SN1_45S_50ntFlanks@10346.10369.24\|TCAAACGAGAACTTTGAAGGCCGA | >hg38_21_+_8433172_8446622_RNA45SN1_45S_50ntFlanks | 10346 | 10369 | 24 | TCAAACGAGAACTTTGAAGGCCGA | 631.5 | 4.1 | 1.3 | 3.1 | 1.6E-03 | 9.0E-03 | BT20/MB231 | MP |
| >hg38_21_+_8433172_8446622_RNA45SN1_45S_50ntFlanks@6648.6675.28\|GTACGACTCTTAGCGGTGGATCACTCGG | >hg38_21_+_8433172_8446622_RNA45SN1_45S_50ntFlanks | 6648 | 6675 | 28 | GTACGACTCTTAGCGGTGGATCACTCGG | 106.7 | -2.5 | 0.9 | -2.7 | 6.9E-03 | 9.0E-03 | MB231/MB468 | MP |
| >hg38_MT_+_1621_3279_RNR2_16S_50ntFlanks@460.491.32\|TTGTAAATTTAACTGTTAGTCCAAAGAGGAAC | >hg38_MT_+_1621_3279_RNR2_16S_50ntFlanks | 460 | 491 | 32 | TTGTAAATTTAACTGTTAGTCCAAAGAGGAAC | 86.2 | 3.3 | 1.2 | 2.7 | 6.9E-03 | 9.0E-03 | MB231/MB468 | MP |
| >MI0000066\|hsa-let-7e&WithFlank&19\|+\|51692780\|51692870@14.34.21(+1A)&[MIMAT0000066&hsa-let-7e-5p&offsets\|0\|-1(+1A);m-50&19\|+\|51692793\|51692814&offsets\|0\|-1(+1A)]\|TGAGGTAGGAGGTTGTATAGTA | >MI0000066\|hsa-let-7e&WithFlank&19\|+\|51692780\|51692870 | 14 | 34 | 21(+1A) | TGAGGTAGGAGGTTGTATAGTA | 315.5 | 1.8 | 0.6 | 3.1 | 1.8E-03 | 9.0E-03 | BT20/MB231 | NUC |
| >hg38_21_+_8433172_8446622_RNA45SN1_45S_50ntFlanks@11875.11898.24\|GAAGACCCTGTTGAGCTTGACTCT | >hg38_21_+_8433172_8446622_RNA45SN1_45S_50ntFlanks | 11875 | 11898 | 24 | GAAGACCCTGTTGAGCTTGACTCT | 398.0 | -2.0 | 0.8 | -2.7 | 7.9E-03 | 9.1E-03 | MB231/MB468 | NUC |
| >hg38_21_+_8433172_8446622_RNA45SN1_45S_50ntFlanks@10393.10426.34\|CAGCAGTTGAACATGGGTCAGTCGGTCCTGAGAG | >hg38_21_+_8433172_8446622_RNA45SN1_45S_50ntFlanks | 10393 | 10426 | 34 | CAGCAGTTGAACATGGGTCAGTCGGTCCTGAGAG | 342.1 | -2.1 | 0.8 | -2.7 | 7.5E-03 | 9.1E-03 | MB231/MB468 | CYTO |
| >hg38_1_-_228634819_228635039_RNA5S12_5S_50ntFlanks@145.171.27\|CTGGGAATACCGGGTGCTGTAGGCTTT | >hg38_1_-_228634819_228635039_RNA5S12_5S_50ntFlanks | 145 | 171 | 27 | CTGGGAATACCGGGTGCTGTAGGCTTT | 616.2 | 2.4 | 0.9 | 2.7 | 6.4E-03 | 9.1E-03 | BT20/MB468 | MITO |
| >hg38_21_+_8433172_8446622_RNA45SN1_45S_50ntFlanks@12415.12433.19\|CTTCGATGTCGGCTCTTCC | >hg38_21_+_8433172_8446622_RNA45SN1_45S_50ntFlanks | 12415 | 12433 | 19 | CTTCGATGTCGGCTCTTCC | 3873.3 | 1.7 | 0.6 | 2.7 | 7.5E-03 | 9.1E-03 | MB231/MB468 | CYTO |
| >hg38_21_+_8433172_8446622_RNA45SN1_45S_50ntFlanks@10922.10947.26\|CTGGACGAGGCGCCGCCGCCCCCCCC | >hg38_21_+_8433172_8446622_RNA45SN1_45S_50ntFlanks | 10922 | 10947 | 26 | CTGGACGAGGCGCCGCCGCCCCCCCC | 271.9 | -4.2 | 1.6 | -2.7 | 7.5E-03 | 9.1E-03 | BT20/MB468 | CYTO |
| >MI0006444\|hsa-mir-1307&WithFlank&10\|-\|103394247\|103394407@87.107.21(+1A)&[MIMAT0005951&hsa-miR-1307-3p&offsets\|+1\|0(+1A);m-66&10\|-\|103394300\|103394322&offsets\|+1\|-1(+1A)]\|CTCGGCGTGGCGTCGGTCGTGA | >MI0006444\|hsa-mir-1307&WithFlank&10\|-\|103394247\|103394407 | 87 | 107 | 21(+1A) | CTCGGCGTGGCGTCGGTCGTGA | 132.4 | 2.6 | 1.0 | 2.7 | 6.9E-03 | 9.1E-03 | MB231/MB468 | MP |
| >MI0000483\|hsa-mir-186&WithFlank&1\|-\|71067625\|71067722@21.43.23(+1C)&[MIMAT0000456&hsa-miR-186-5p&offsets\|0\|+1(+1C);m-84&1\|-\|71067681\|71067702&offsets\|0\|+1(+1C)]\|CAAAGAATTCTCCTTTTGGGCTTC | >MI0000483\|hsa-mir-186&WithFlank&1\|-\|71067625\|71067722 | 21 | 43 | 23(+1C) | CAAAGAATTCTCCTTTTGGGCTTC | 410.3 | -4.4 | 1.4 | -3.1 | 1.6E-03 | 9.1E-03 | BT20/MB231 | MP |
| >hg38_21_+_8433172_8446622_RNA45SN1_45S_50ntFlanks@6662.6688.27\|GGTGGATCACTCGGCTCGTGCGTCGAT | >hg38_21_+_8433172_8446622_RNA45SN1_45S_50ntFlanks | 6662 | 6688 | 27 | GGTGGATCACTCGGCTCGTGCGTCGAT | 169.5 | 4.2 | 1.5 | 2.8 | 4.5E-03 | 9.1E-03 | BT20/MB468 | MP |
| >am_trnaMT_SerGCT_MT_+_12207_12265@14.31.18\|AGAACTGCTAACTCATGC | >am_trnaMT_SerGCT_MT_+_12207_12265 | 14 | 31 | 18 | AGAACTGCTAACTCATGC | 329.9 | 3.0 | 1.1 | 2.8 | 4.5E-03 | 9.1E-03 | BT20/MB468 | MP |
| >trna10_AspGTC_12_-_125424193_125424264@1.31.31,trna12_AspGTC_12_-_125411891_125411962@1.31.31,trna38_AspGTC_17_-_8125556_8125627@1.31.31,trna45_AspGTC_6_+_27447453_27447524@1.31.31,trna48_AspGTC_6_+_27471523_27471594@1.31.31,trna4_AspGTC_12_+_96429799_96429870@1.31.31,trna69_AspGTC_1_-_161440205_161440276@1.31.31,trna72_AspGTC_1_-_161432824_161432895@1.31.31,trna75_AspGTC_1_-_161425414_161425485@1.31.31,trna78_AspGTC_1_-_161418033_161418104@1.31.31,trna81_AspGTC_1_-_161410615_161410686@1.31.31\|TCCTCGTTAGTATAGTGGTGAGTATCCCCGC | >trna10_AspGTC_12_-_125424193_125424264 | 1 | 31 | 31 | TCCTCGTTAGTATAGTGGTGAGTATCCCCGC | 300.4 | -4.2 | 1.6 | -2.7 | 7.5E-03 | 9.1E-03 | BT20/MB468 | CYTO |
| >hg38_21_+_8433172_8446622_RNA45SN1_45S_50ntFlanks@11566.11603.38\|TAGCAGCCGACTTAGAACTGGTGCGGACCAGGGGAATC | >hg38_21_+_8433172_8446622_RNA45SN1_45S_50ntFlanks | 11566 | 11603 | 38 | TAGCAGCCGACTTAGAACTGGTGCGGACCAGGGGAATC | 80.5 | -3.7 | 1.4 | -2.7 | 6.9E-03 | 9.1E-03 | MB231/MB468 | MP |
| >MI0000440\|hsa-mir-27b&WithFlank&9\|+\|95085439\|95085547@67.88.22&[MIMAT0000419&hsa-miR-27b-3p&offsets\|0\|+1;m-39&9\|+\|95085505\|95085525&offsets\|0\|+1]\|TTCACAGTGGCTAAGTTCTGCA | >MI0000440\|hsa-mir-27b&WithFlank&9\|+\|95085439\|95085547 | 67 | 88 | 22 | TTCACAGTGGCTAAGTTCTGCA | 513.3 | 2.2 | 0.8 | 2.8 | 4.5E-03 | 9.1E-03 | BT20/MB468 | MP |
| >hg38_21_+_8433172_8446622_RNA45SN1_45S_50ntFlanks@3785.3802.18\|TGAAACTGCGAATGGCTC | >hg38_21_+_8433172_8446622_RNA45SN1_45S_50ntFlanks | 3785 | 3802 | 18 | TGAAACTGCGAATGGCTC | 120.4 | -3.1 | 1.1 | -2.7 | 6.4E-03 | 9.1E-03 | BT20/MB468 | MITO |
| >MI0000253\|hsa-mir-148a&WithFlank&7\|-\|25949913\|25949992@12.32.21(+1C)&[MIMAT0004549&hsa-miR-148a-5p&offsets\|0\|-1(+1C);m-189&7\|-\|25949960\|25949981&offsets\|0\|-1(+1C)]\|AAAGTTCTGAGACACTCCGACC | >MI0000253\|hsa-mir-148a&WithFlank&7\|-\|25949913\|25949992 | 12 | 32 | 21(+1C) | AAAGTTCTGAGACACTCCGACC | 65.8 | 5.0 | 1.8 | 2.8 | 4.5E-03 | 9.1E-03 | BT20/MB468 | MP |
| >hg38_21_+_8433172_8446622_RNA45SN1_45S_50ntFlanks@10420.10454.35\|CTGAGAGATGGGCGAGCGCCGTTCCGAAGGGACGG | >hg38_21_+_8433172_8446622_RNA45SN1_45S_50ntFlanks | 10420 | 10454 | 35 | CTGAGAGATGGGCGAGCGCCGTTCCGAAGGGACGG | 185.1 | -1.9 | 0.7 | -2.7 | 7.5E-03 | 9.1E-03 | MB231/MB468 | CYTO |
| >hg38_21_+_8433172_8446622_RNA45SN1_45S_50ntFlanks@10906.10930.25\|TGGGCGCGCGCCGCGGCTGGACGAG | >hg38_21_+_8433172_8446622_RNA45SN1_45S_50ntFlanks | 10906 | 10930 | 25 | TGGGCGCGCGCCGCGGCTGGACGAG | 55.6 | 9.1 | 2.9 | 3.1 | 1.6E-03 | 9.1E-03 | BT20/MB231 | MP |
| >hg38_21_+_8433172_8446622_RNA45SN1_45S_50ntFlanks@11539.11583.45\|GCGCGGCGCCTCGCCTCGGCCGGCGCCTAGCAGCCGACTTAGAAC | >hg38_21_+_8433172_8446622_RNA45SN1_45S_50ntFlanks | 11539 | 11583 | 45 | GCGCGGCGCCTCGCCTCGGCCGGCGCCTAGCAGCCGACTTAGAAC | 139.5 | 4.2 | 1.6 | 2.7 | 8.0E-03 | 9.1E-03 | MB231/MB468 | MITO |
| >hg38_21_+_8433172_8446622_RNA45SN1_45S_50ntFlanks@11205.11225.21\|GCGGCGGCGACTCTGGACGCG | >hg38_21_+_8433172_8446622_RNA45SN1_45S_50ntFlanks | 11205 | 11225 | 21 | GCGGCGGCGACTCTGGACGCG | 389.1 | -5.0 | 1.9 | -2.7 | 7.9E-03 | 9.1E-03 | BT20/MB468 | NUC |
| >hg38_MT_+_1621_3279_RNR2_16S_50ntFlanks@837.858.22\|AAAAAAAGTAAAAGGAACTCGG | >hg38_MT_+_1621_3279_RNR2_16S_50ntFlanks | 837 | 858 | 22 | AAAAAAAGTAAAAGGAACTCGG | 148.1 | -3.0 | 1.0 | -2.8 | 4.5E-03 | 9.1E-03 | BT20/MB468 | MP |
| >hg38_21_+_8433172_8446622_RNA45SN1_45S_50ntFlanks@5085.5110.26\|GAACGAGACTCTGGCATGCTAACTAG | >hg38_21_+_8433172_8446622_RNA45SN1_45S_50ntFlanks | 5085 | 5110 | 26 | GAACGAGACTCTGGCATGCTAACTAG | 336.5 | 5.5 | 1.7 | 3.1 | 1.6E-03 | 9.1E-03 | BT20/MB231 | MP |
| >hg38_21_+_8433172_8446622_RNA45SN1_45S_50ntFlanks@12455.12477.23\|ACCAAGCGTTGGATTGTTCACCC | >hg38_21_+_8433172_8446622_RNA45SN1_45S_50ntFlanks | 12455 | 12477 | 23 | ACCAAGCGTTGGATTGTTCACCC | 174.7 | -3.4 | 1.3 | -2.7 | 6.9E-03 | 9.1E-03 | MB231/MB468 | MP |

<table>
<tr><td>>hg38_21_+_8433172_8446622_RNA45SN1_45S_50ntFlanks@10677.10708.32|GCGGTTCCGGCGGCGTCCGGTGAGCTCTCGCT</td><td>>hg38_21_+_8433172_8446622_RNA45SN1_45S_50ntFlanks</td><td>10677</td><td>10708</td><td>32</td><td>GCGGTTCCGGCGGCGTCCGGTGAGCTCTCGCT</td><td>334.8</td><td>3.2</td><td>1.1</td><td>2.8</td><td>4.5E-03</td><td>9.1E-03</td><td>BT20/MB468</td><td>MP</td></tr>
<tr><td>>hg38_21_+_8433172_8446622_RNA45SN1_45S_50ntFlanks@13001.13028.28|GCGATCTATTGAAAGTCAGCCCTCGACA</td><td>>hg38_21_+_8433172_8446622_RNA45SN1_45S_50ntFlanks</td><td>13001</td><td>13028</td><td>28</td><td>GCGATCTATTGAAAGTCAGCCCTCGACA</td><td>178.0</td><td>-3.0</td><td>1.1</td><td>-2.7</td><td>8.0E-03</td><td>9.1E-03</td><td>MB231/MB468</td><td>NUC</td></tr>
<tr><td>>MI0000488|hsa-mir-194-1&WithFlank&1|-|220118151|220118247@21.42.22;MI0000732|hsa-mir-194-2&WithFlank&11|-|64891349|64891445@21.42.22&[MIMAT0000460&hsa-miR-194-5p&offsets|0|0;m-131&1|-|220118206|220118227&offsets|0|0];[MIMAT0000460_1&hsa-miR-194-5p&offsets|0|0;m-126&11|-|64891404|64891425&offsets|0|0]|TGTAACAGCAACTCCATGTGGA</td><td>>MI0000488|hsa-mir-194-1&WithFlank&1|-|220118151|220118247</td><td>21</td><td>42</td><td>22</td><td>TGTAACAGCAACTCCATGTGGA</td><td>433.0</td><td>2.8</td><td>0.9</td><td>3.1</td><td>2.1E-03</td><td>9.1E-03</td><td>BT20/MB231</td><td>CYTO</td></tr>
<tr><td>>hg38_21_+_8433172_8446622_RNA45SN1_45S_50ntFlanks@4722.4753.32|TCATTAATCAAGAACGAAAGTCGGAGGTTCGA</td><td>>hg38_21_+_8433172_8446622_RNA45SN1_45S_50ntFlanks</td><td>4722</td><td>4753</td><td>32</td><td>TCATTAATCAAGAACGAAAGTCGGAGGTTCGA</td><td>110.5</td><td>-4.7</td><td>1.6</td><td>-2.8</td><td>4.5E-03</td><td>9.1E-03</td><td>BT20/MB468</td><td>MP</td></tr>
<tr><td>>hg38_21_+_8433172_8446622_RNA45SN1_45S_50ntFlanks@6646.6664.19|TCGTACGACTCTTAGCGGT</td><td>>hg38_21_+_8433172_8446622_RNA45SN1_45S_50ntFlanks</td><td>6646</td><td>6664</td><td>19</td><td>TCGTACGACTCTTAGCGGT</td><td>28855.1</td><td>1.9</td><td>0.7</td><td>2.7</td><td>8.0E-03</td><td>9.1E-03</td><td>MB231/MB468</td><td>MITO</td></tr>
<tr><td>>hg38_21_+_8433172_8446622_RNA45SN1_45S_50ntFlanks@7991.8010.20|ACGTGGCGACCCGCTGAATT</td><td>>hg38_21_+_8433172_8446622_RNA45SN1_45S_50ntFlanks</td><td>7991</td><td>8010</td><td>20</td><td>ACGTGGCGACCCGCTGAATT</td><td>1146.8</td><td>-2.3</td><td>0.8</td><td>-2.8</td><td>4.5E-03</td><td>9.1E-03</td><td>BT20/MB468</td><td>MP</td></tr>
<tr><td>>hg38_21_+_8433172_8446622_RNA45SN1_45S_50ntFlanks@11718.11738.21|AACGGCGGGAGTAACTATGAC</td><td>>hg38_21_+_8433172_8446622_RNA45SN1_45S_50ntFlanks</td><td>11718</td><td>11738</td><td>21</td><td>AACGGCGGGAGTAACTATGAC</td><td>344.4</td><td>-4.8</td><td>1.7</td><td>-2.8</td><td>4.5E-03</td><td>9.1E-03</td><td>BT20/MB468</td><td>MP</td></tr>
<tr><td>>hg38_21_+_8433172_8446622_RNA45SN1_45S_50ntFlanks@3987.4005.19|GCTTTGGTGACTCTAGATA</td><td>>hg38_21_+_8433172_8446622_RNA45SN1_45S_50ntFlanks</td><td>3987</td><td>4005</td><td>19</td><td>GCTTTGGTGACTCTAGATA</td><td>190.6</td><td>-2.5</td><td>0.9</td><td>-2.8</td><td>4.5E-03</td><td>9.1E-03</td><td>BT20/MB468</td><td>MP</td></tr>
<tr><td>>hg38_21_+_8433172_8446622_RNA45SN1_45S_50ntFlanks@12715.12751.37|AGCCTCGGTTGGCCTCGGATAGCCGGTCCCCCGCCTG</td><td>>hg38_21_+_8433172_8446622_RNA45SN1_45S_50ntFlanks</td><td>12715</td><td>12751</td><td>37</td><td>AGCCTCGGTTGGCCTCGGATAGCCGGTCCCCCGCCTG</td><td>137.7</td><td>-5.5</td><td>1.9</td><td>-2.8</td><td>4.5E-03</td><td>9.1E-03</td><td>BT20/MB468</td><td>MP</td></tr>
<tr><td>>hg38_21_+_8433172_8446622_RNA45SN1_45S_50ntFlanks@9956.9981.26|GAAGTCGGAATCCGCTAAGGAGTGTG</td><td>>hg38_21_+_8433172_8446622_RNA45SN1_45S_50ntFlanks</td><td>9956</td><td>9981</td><td>26</td><td>GAAGTCGGAATCCGCTAAGGAGTGTG</td><td>75.7</td><td>4.8</td><td>1.7</td><td>2.8</td><td>4.5E-03</td><td>9.1E-03</td><td>BT20/MB468</td><td>MP</td></tr>
<tr><td>>hg38_21_+_8433172_8446622_RNA45SN1_45S_50ntFlanks@11539.11584.46|GCGCGGCGCCTCGCCTCGGCCGGCGCCTAGCAGCCGACTTAGAACT</td><td>>hg38_21_+_8433172_8446622_RNA45SN1_45S_50ntFlanks</td><td>11539</td><td>11584</td><td>46</td><td>GCGCGGCGCCTCGCCTCGGCCGGCGCCTAGCAGCCGACTTAGAACT</td><td>427.5</td><td>-2.9</td><td>0.9</td><td>-3.1</td><td>1.8E-03</td><td>9.1E-03</td><td>BT20/MB231</td><td>NUC</td></tr>
<tr><td>>hg38_21_+_8433172_8446622_RNA45SN1_45S_50ntFlanks@474.496.23|TCTTCGTGATCGATGTGGTGACG</td><td>>hg38_21_+_8433172_8446622_RNA45SN1_45S_50ntFlanks</td><td>474</td><td>496</td><td>23</td><td>TCTTCGTGATCGATGTGGTGACG</td><td>546.2</td><td>-2.9</td><td>0.9</td><td>-3.1</td><td>1.8E-03</td><td>9.1E-03</td><td>BT20/MB231</td><td>NUC</td></tr>
<tr><td>>hg38_21_+_8433172_8446622_RNA45SN1_45S_50ntFlanks@6643.6683.41|ACCTCGTACGACTCTTAGCGGTGGATCACTCGGCTCGTGCG</td><td>>hg38_21_+_8433172_8446622_RNA45SN1_45S_50ntFlanks</td><td>6643</td><td>6683</td><td>41</td><td>ACCTCGTACGACTCTTAGCGGTGGATCACTCGGCTCGTGCG</td><td>640.3</td><td>-2.7</td><td>0.9</td><td>-3.1</td><td>1.8E-03</td><td>9.1E-03</td><td>BT20/MB231</td><td>NUC</td></tr>
<tr><td>>MI0006349|hsa-mir-1287&WithFlank&10|-|98395212|98395313@22.42.21(+1U)&[MIMAT0005878&hsa-miR-1287-5p&offsets|0|-1(+1U);m-244&10|-|98395272|98395292&offsets|0|0(+1U)]|TGCTGGATCAGTGGTTCGAGTT</td><td>>MI0006349|hsa-mir-1287&WithFlank&10|-|98395212|98395313</td><td>22</td><td>42</td><td>21(+1U)</td><td>TGCTGGATCAGTGGTTCGAGTT</td><td>117.1</td><td>-5.8</td><td>1.8</td><td>-3.1</td><td>1.7E-03</td><td>9.1E-03</td><td>BT20/MB231</td><td>MP</td></tr>
<tr><td>>hg38_21_+_8433172_8446622_RNA45SN1_45S_50ntFlanks@4589.4616.28|TATTTTGTTGGTTTTCGGAACTGAGGCC</td><td>>hg38_21_+_8433172_8446622_RNA45SN1_45S_50ntFlanks</td><td>4589</td><td>4616</td><td>28</td><td>TATTTTGTTGGTTTTCGGAACTGAGGCC</td><td>131.6</td><td>-1.5</td><td>0.6</td><td>-2.7</td><td>7.5E-03</td><td>9.1E-03</td><td>BT20/MB468</td><td>CYTO</td></tr>
<tr><td>>hg38_21_+_8433172_8446622_RNA45SN1_45S_50ntFlanks@12640.12659.20|ACCATCTGTGGGATTATGAC</td><td>>hg38_21_+_8433172_8446622_RNA45SN1_45S_50ntFlanks</td><td>12640</td><td>12659</td><td>20</td><td>ACCATCTGTGGGATTATGAC</td><td>166.2</td><td>-1.9</td><td>0.7</td><td>-2.7</td><td>7.5E-03</td><td>9.1E-03</td><td>BT20/MB468</td><td>CYTO</td></tr>
<tr><td>>hg38_21_+_8433172_8446622_RNA45SN1_45S_50ntFlanks@10348.10366.19|AAACGAGAACTTTGAAGGC</td><td>>hg38_21_+_8433172_8446622_RNA45SN1_45S_50ntFlanks</td><td>10348</td><td>10366</td><td>19</td><td>AAACGAGAACTTTGAAGGC</td><td>990.4</td><td>-3.0</td><td>1.0</td><td>-2.8</td><td>4.5E-03</td><td>9.1E-03</td><td>BT20/MB468</td><td>MP</td></tr>
<tr><td>>hg38_21_+_8433172_8446622_RNA45SN1_45S_50ntFlanks@7973.8010.38|GACGCGACCTCAGATCAGACGTGGCGACCCGCTGAATT</td><td>>hg38_21_+_8433172_8446622_RNA45SN1_45S_50ntFlanks</td><td>7973</td><td>8010</td><td>38</td><td>GACGCGACCTCAGATCAGACGTGGCGACCCGCTGAATT</td><td>273.6</td><td>3.1</td><td>1.1</td><td>2.7</td><td>6.9E-03</td><td>9.1E-03</td><td>MB231/MB468</td><td>MP</td></tr>
<tr><td>>hg38_21_+_8433172_8446622_RNA45SN1_45S_50ntFlanks@4110.4150.41|TGACGGGGAATCAGGGTTCGATTCCGGAGAGGGAGCCTGAG</td><td>>hg38_21_+_8433172_8446622_RNA45SN1_45S_50ntFlanks</td><td>4110</td><td>4150</td><td>41</td><td>TGACGGGGAATCAGGGTTCGATTCCGGAGAGGGAGCCTGAG</td><td>76.6</td><td>7.1</td><td>2.5</td><td>2.8</td><td>4.5E-03</td><td>9.1E-03</td><td>BT20/MB468</td><td>MP</td></tr>
<tr><td>>hg38_21_+_8433172_8446622_RNA45SN1_45S_50ntFlanks@11667.11696.30|CTGCCCAGTGCTCTGAATGTCAAAGTGAAG</td><td>>hg38_21_+_8433172_8446622_RNA45SN1_45S_50ntFlanks</td><td>11667</td><td>11696</td><td>30</td><td>CTGCCCAGTGCTCTGAATGTCAAAGTGAAG</td><td>247.8</td><td>6.3</td><td>2.0</td><td>3.1</td><td>1.7E-03</td><td>9.1E-03</td><td>BT20/MB231</td><td>MP</td></tr>
<tr><td>>hg38_21_+_8433172_8446622_RNA45SN1_45S_50ntFlanks@6652.6674.23|GACTCTTAGCGGTGGATCACTCG</td><td>>hg38_21_+_8433172_8446622_RNA45SN1_45S_50ntFlanks</td><td>6652</td><td>6674</td><td>23</td><td>GACTCTTAGCGGTGGATCACTCG</td><td>840.0</td><td>3.6</td><td>1.2</td><td>3.1</td><td>1.7E-03</td><td>9.1E-03</td><td>BT20/MB231</td><td>MP</td></tr>
<tr><td>>MI0000087|hsa-mir-29a&WithFlank&7|-|130876741|130876816@10.34.25&[MIMAT0004503&hsa-miR-29a-5p&offsets|0|+3;m-315&7|-|130876785|130876807&offsets|0|+2]|ACTGATTTCTTTTGGTGTTCAGAGT</td><td>>MI0000087|hsa-mir-29a&WithFlank&7|-|130876741|130876816</td><td>10</td><td>34</td><td>25</td><td>ACTGATTTCTTTTGGTGTTCAGAGT</td><td>67.5</td><td>5.1</td><td>1.9</td><td>2.7</td><td>6.9E-03</td><td>9.1E-03</td><td>MB231/MB468</td><td>MP</td></tr>
<tr><td>>hg38_21_+_8433172_8446622_RNA45SN1_45S_50ntFlanks@12943.12972.30|TCGTAGACGACCTGCTTCTGGGTCGGGGTT</td><td>>hg38_21_+_8433172_8446622_RNA45SN1_45S_50ntFlanks</td><td>12943</td><td>12972</td><td>30</td><td>TCGTAGACGACCTGCTTCTGGGTCGGGGTT</td><td>159.0</td><td>-3.2</td><td>1.2</td><td>-2.7</td><td>7.5E-03</td><td>9.1E-03</td><td>BT20/MB468</td><td>CYTO</td></tr>
<tr><td>>hg38_21_+_8433172_8446622_RNA45SN1_45S_50ntFlanks@12973.12990.18|TCGTACGTAGCAGAGCAG</td><td>>hg38_21_+_8433172_8446622_RNA45SN1_45S_50ntFlanks</td><td>12973</td><td>12990</td><td>18</td><td>TCGTACGTAGCAGAGCAG</td><td>2627.0</td><td>-5.3</td><td>1.9</td><td>-2.8</td><td>4.5E-03</td><td>9.1E-03</td><td>BT20/MB468</td><td>MP</td></tr>
<tr><td>>hg38_1_-_228634819_228635039_RNA5S12_5S_50ntFlanks@142.171.30|CGCCTGGGAATACCGGGTGCTGTAGGCTTT</td><td>>hg38_1_-_228634819_228635039_RNA5S12_5S_50ntFlanks</td><td>142</td><td>171</td><td>30</td><td>CGCCTGGGAATACCGGGTGCTGTAGGCTTT</td><td>1187.1</td><td>-2.1</td><td>0.7</td><td>-3.1</td><td>1.8E-03</td><td>9.2E-03</td><td>BT20/MB231</td><td>NUC</td></tr>
<tr><td>>hg38_21_+_8433172_8446622_RNA45SN1_45S_50ntFlanks@4367.4407.41|CAGTTAAAAAGCTCGTAGTTGGATCTTGGGAGCGGGCGGGC</td><td>>hg38_21_+_8433172_8446622_RNA45SN1_45S_50ntFlanks</td><td>4367</td><td>4407</td><td>41</td><td>CAGTTAAAAAGCTCGTAGTTGGATCTTGGGAGCGGGCGGGC</td><td>112.9</td><td>-3.3</td><td>1.3</td><td>-2.7</td><td>7.5E-03</td><td>9.2E-03</td><td>BT20/MB468</td><td>CYTO</td></tr>
<tr><td>>hg38_MT_+_1621_3279_RNR2_16S_50ntFlanks@1095.1116.22|AGACGAGAAGACCCTATGGAGC</td><td>>hg38_MT_+_1621_3279_RNR2_16S_50ntFlanks</td><td>1095</td><td>1116</td><td>22</td><td>AGACGAGAAGACCCTATGGAGC</td><td>89.5</td><td>4.9</td><td>1.7</td><td>2.8</td><td>4.5E-03</td><td>9.2E-03</td><td>BT20/MB468</td><td>MP</td></tr>
<tr><td>>hg38_21_+_8433172_8446622_RNA45SN1_45S_50ntFlanks@12416.12453.38|TTCGATGTCGGCTCTTCCTATCATTGTGAAGCAGAATT</td><td>>hg38_21_+_8433172_8446622_RNA45SN1_45S_50ntFlanks</td><td>12416</td><td>12453</td><td>38</td><td>TTCGATGTCGGCTCTTCCTATCATTGTGAAGCAGAATT</td><td>162.0</td><td>4.1</td><td>1.6</td><td>2.7</td><td>8.0E-03</td><td>9.2E-03</td><td>MB231/MB468</td><td>MITO</td></tr>
<tr><td>>hg38_21_+_8433172_8446622_RNA45SN1_45S_50ntFlanks@12453.12476.24|TCACCAAGCGTTGGATTGTTCACC</td><td>>hg38_21_+_8433172_8446622_RNA45SN1_45S_50ntFlanks</td><td>12453</td><td>12476</td><td>24</td><td>TCACCAAGCGTTGGATTGTTCACC</td><td>669.0</td><td>-2.5</td><td>0.8</td><td>-3.1</td><td>2.1E-03</td><td>9.2E-03</td><td>BT20/MB231</td><td>CYTO</td></tr>
<tr><td>>trna27_GlyCCC_2_-_70476123_70476193@1.32.32,trna34_GlyCCC_16_-_686736_686806@1.32.32|GCGCCGCTGGTGTAGTGGTATCATGCAAGATT</td><td>>trna27_GlyCCC_2_-_70476123_70476193</td><td>1</td><td>32</td><td>32</td><td>GCGCCGCTGGTGTAGTGGTATCATGCAAGATT</td><td>780.0</td><td>9.4</td><td>3.5</td><td>2.7</td><td>7.6E-03</td><td>9.2E-03</td><td>MB231/MB468</td><td>CYTO</td></tr>
<tr><td>>trna6_TrpCCA_12_+_98898030_98898101@30.52.23|ACTCCAGATCAGAAGGCTGCGTG</td><td>>trna6_TrpCCA_12_+_98898030_98898101</td><td>30</td><td>52</td><td>23</td><td>ACTCCAGATCAGAAGGCTGCGTG</td><td>128.7</td><td>-4.6</td><td>1.6</td><td>-2.8</td><td>4.5E-03</td><td>9.2E-03</td><td>BT20/MB468</td><td>MP</td></tr>
<tr><td>>trna14_LysTTT_11_-_59327808_59327880@3.33.31,trna2_LysTTT_17_+_8022473_8022545@3.33.31,trna54_LysTTT_1_+_204475655_204475727@3.33.31,trna5_LysTTT_11_+_59323902_59323974@3.33.31,trna62_LysTTT_1_-_204476158_204476230@3.33.31,trna76_LysTTT_6_+_28918806_28918878@3.33.31|CCGGATAGCTCAGTCGGTAGAGCATCAGACT</td><td>>trna14_LysTTT_11_-_59327808_59327880</td><td>3</td><td>33</td><td>31</td><td>CCGGATAGCTCAGTCGGTAGAGCATCAGACT</td><td>228.6</td><td>-4.4</td><td>1.4</td><td>-3.1</td><td>1.7E-03</td><td>9.2E-03</td><td>BT20/MB231</td><td>MP</td></tr>
<tr><td>>am_trna117_GlyTCC_1_-_145397864_145397935@1.28.28,trna45_GlyTCC_1_+_161500903_161500974@1.28.28,trna70_GlyTCC_1_-_161439547_161439618@1.28.28,trna73_GlyTCC_1_-_161432166_161432237@1.28.28,trna76_GlyTCC_1_-_161424756_161424827@1.28.28,trna79_GlyTCC_1_-_161417375_161417446@1.28.28|GCGTTGGTGGTATAGTGGTGAGCATAGC</td><td>>am_trna117_GlyTCC_1_-_145397864_145397935</td><td>1</td><td>28</td><td>28</td><td>GCGTTGGTGGTATAGTGGTGAGCATAGC</td><td>174.3</td><td>-1.8</td><td>0.7</td><td>-2.7</td><td>7.5E-03</td><td>9.2E-03</td><td>BT20/MB468</td><td>CYTO</td></tr>
<tr><td>>hg38_21_+_8433172_8446622_RNA45SN1_45S_50ntFlanks@6646.6667.22|TCGTACGACTCTTAGCGGTGGA</td><td>>hg38_21_+_8433172_8446622_RNA45SN1_45S_50ntFlanks</td><td>6646</td><td>6667</td><td>22</td><td>TCGTACGACTCTTAGCGGTGGA</td><td>11304.3</td><td>2.2</td><td>0.7</td><td>3.1</td><td>1.8E-03</td><td>9.2E-03</td><td>BT20/MB231</td><td>NUC</td></tr>
<tr><td>>hg38_21_+_8433172_8446622_RNA45SN1_45S_50ntFlanks@11571.11590.20|GCCGACTTAGAACTGGTGCG</td><td>>hg38_21_+_8433172_8446622_RNA45SN1_45S_50ntFlanks</td><td>11571</td><td>11590</td><td>20</td><td>GCCGACTTAGAACTGGTGCG</td><td>187.9</td><td>-2.3</td><td>0.8</td><td>-3.1</td><td>2.1E-03</td><td>9.2E-03</td><td>BT20/MB231</td><td>CYTO</td></tr>
<tr><td>>hg38_21_+_8433172_8446622_RNA45SN1_45S_50ntFlanks@6699.6721.23|GCTAGCTGCGAGAATTAATGTGA</td><td>>hg38_21_+_8433172_8446622_RNA45SN1_45S_50ntFlanks</td><td>6699</td><td>6721</td><td>23</td><td>GCTAGCTGCGAGAATTAATGTGA</td><td>684.7</td><td>-2.0</td><td>0.8</td><td>-2.6</td><td>8.1E-03</td><td>9.2E-03</td><td>MB231/MB468</td><td>NUC</td></tr>
<tr><td>>trna14_LysTTT_11_-_59327808_59327880@4.32.29,trna2_LysTTT_17_+_8022473_8022545@4.32.29,trna54_LysTTT_1_+_204475655_204475727@4.32.29,trna5_LysTTT_11_+_59323902_59323974@4.32.29,trna62_LysTTT_1_-_204476158_204476230@4.32.29,trna76_LysTTT_6_+_28918806_28918878@4.32.29|CGGATAGCTCAGTCGGTAGAGCATCAGAC</td><td>>trna14_LysTTT_11_-_59327808_59327880</td><td>4</td><td>32</td><td>29</td><td>CGGATAGCTCAGTCGGTAGAGCATCAGAC</td><td>78.9</td><td>3.8</td><td>1.4</td><td>2.7</td><td>7.0E-03</td><td>9.2E-03</td><td>MB231/MB468</td><td>MP</td></tr>
<tr><td>>hg38_21_+_8433172_8446622_RNA45SN1_45S_50ntFlanks@11666.11684.19|TCTGCCCAGTGCTCTGAAT</td><td>>hg38_21_+_8433172_8446622_RNA45SN1_45S_50ntFlanks</td><td>11666</td><td>11684</td><td>19</td><td>TCTGCCCAGTGCTCTGAAT</td><td>140.9</td><td>-2.6</td><td>1.0</td><td>-2.7</td><td>7.0E-03</td><td>9.2E-03</td><td>MB231/MB468</td><td>MP</td></tr>
<tr><td>>hg38_1_-_228634819_228635039_RNA5S12_5S_50ntFlanks@140.169.30|ACCGCCTGGGAATACCGGGTGCTGTAGGCT</td><td>>hg38_1_-_228634819_228635039_RNA5S12_5S_50ntFlanks</td><td>140</td><td>169</td><td>30</td><td>ACCGCCTGGGAATACCGGGTGCTGTAGGCT</td><td>3786.2</td><td>1.8</td><td>0.6</td><td>2.7</td><td>6.5E-03</td><td>9.2E-03</td><td>BT20/MB468</td><td>MITO</td></tr>
<tr><td>>am_trnaMT_ProTGG_MT_-_15956_16023@53.70.18|AAGACTTTTTCTCTGACC</td><td>>am_trnaMT_ProTGG_MT_-_15956_16023</td><td>53</td><td>70</td><td>18</td><td>AAGACTTTTTCTCTGACC</td><td>579.8</td><td>3.7</td><td>1.4</td><td>2.7</td><td>7.6E-03</td><td>9.2E-03</td><td>MB231/MB468</td><td>CYTO</td></tr>
<tr><td>>hg38_21_+_8433172_8446622_RNA45SN1_45S_50ntFlanks@4000.4024.25|TAGATAACCTCGGGCCGATCGCACG</td><td>>hg38_21_+_8433172_8446622_RNA45SN1_45S_50ntFlanks</td><td>4000</td><td>4024</td><td>25</td><td>TAGATAACCTCGGGCCGATCGCACG</td><td>213.8</td><td>-4.8</td><td>1.7</td><td>-2.8</td><td>4.5E-03</td><td>9.2E-03</td><td>BT20/MB468</td><td>MP</td></tr>
<tr><td>>hg38_21_+_8433172_8446622_RNA45SN1_45S_50ntFlanks@7980.8004.25|CCTCAGATCAGACGTGGCGACCCGC</td><td>>hg38_21_+_8433172_8446622_RNA45SN1_45S_50ntFlanks</td><td>7980</td><td>8004</td><td>25</td><td>CCTCAGATCAGACGTGGCGACCCGC</td><td>304.0</td><td>-1.8</td><td>0.7</td><td>-2.7</td><td>6.5E-03</td><td>9.2E-03</td><td>BT20/MB468</td><td>MITO</td></tr>
<tr><td>>hg38_1_-_228634819_228635039_RNA5S12_5S_50ntFlanks@51.86.36|GTCTACGGCCATACCACCCTGAACGCGCCCGATCTC</td><td>>hg38_1_-_228634819_228635039_RNA5S12_5S_50ntFlanks</td><td>51</td><td>86</td><td>36</td><td>GTCTACGGCCATACCACCCTGAACGCGCCCGATCTC</td><td>950.1</td><td>3.5</td><td>1.3</td><td>2.6</td><td>8.1E-03</td><td>9.2E-03</td><td>MB231/MB468</td><td>MITO</td></tr>
<tr><td>>MI0000298|hsa-mir-221&WithFlank&X|-|45746151|45746272@71.93.23(+1G)&[MIMAT0000278&hsa-miR-221-3p&offsets|0|0(+1G);m-91&X|-|45746181|45746202&offsets|0|+1(+1G)]|AGCTACATTGTCTGCTGGGTTTCG</td><td>>MI0000298|hsa-mir-221&WithFlank&X|-|45746151|45746272</td><td>71</td><td>93</td><td>23(+1G)</td><td>AGCTACATTGTCTGCTGGGTTTCG</td><td>364.4</td><td>-3.2</td><td>1.0</td><td>-3.1</td><td>1.7E-03</td><td>9.2E-03</td><td>BT20/MB231</td><td>MP</td></tr>
<tr><td>>hg38_21_+_8433172_8446622_RNA45SN1_45S_50ntFlanks@11707.11728.22|AAGCGCGGGTAAACGGCGGGAG</td><td>>hg38_21_+_8433172_8446622_RNA45SN1_45S_50ntFlanks</td><td>11707</td><td>11728</td><td>22</td><td>AAGCGCGGGTAAACGGCGGGAG</td><td>263.3</td><td>-4.1</td><td>1.5</td><td>-2.7</td><td>7.6E-03</td><td>9.2E-03</td><td>BT20/MB468</td><td>CYTO</td></tr>
<tr><td>>MI0000469|hsa-mir-125a&WithFlank&19|+|51693248|51693345@21.41.21&[MIMAT0000443&hsa-miR-125a-5p&offsets|0|-3;m-63&19|+|51693268|51693289&offsets|0|-1]|TCCCTGAGACCCTTTAACCTG</td><td>>MI0000469|hsa-mir-125a&WithFlank&19|+|51693248|51693345</td><td>21</td><td>41</td><td>21</td><td>TCCCTGAGACCCTTTAACCTG</td><td>99.0</td><td>2.1</td><td>0.8</td><td>2.7</td><td>8.0E-03</td><td>9.2E-03</td><td>BT20/MB468</td><td>NUC</td></tr>
</table>

| | | | | | | | | | | | | | |
|---|---|---|---|---|---|---|---|---|---|---|---|---|---|
| >MI0000289\|hsa-mir-181a-1&WithFlank&1\|-\|198859038\|198859159@30.50.21(+1A);MI0000269\|hsa-mir-181a-2&WithFlank&9\|+\|124692436\|124692557@45.65.21(+1A)&[MIMAT0000256&hsa-miR-181a-5p&offsets\|0\|-2(+1A);m-32&1\|-\|198859108\|198859130&offsets\|0\|-2(+1A)];[MIMAT0000256_1&hsa-miR-181a-5p&offsets\|0\|-2(+1A);m-32&9\|+\|124692480\|124692502&offsets\|0\|-2(+1A)]\|AACATTCAACGCTGTCGGTGAA | >MI0000289\|hsa-mir-181a-1&WithFlank&1\|-\|198859038\|198859159 | 30 | 50 | 21(+1A) | AACATTCAACGCTGTCGGTGAA | 96.6 | 3.7 | 1.4 | 2.7 | 7.0E-03 | 9.2E-03 | MB231/MB468 | MP |
| >hg38_21_+_8433172_8446622_RNA45SN1_45S_50ntFlanks@12418.12454.37\|CGATGTCGGCTCTTCCTATCATTGTGAAGCAGAATTC | >hg38_21_+_8433172_8446622_RNA45SN1_45S_50ntFlanks | 12418 | 12454 | 37 | CGATGTCGGCTCTTCCTATCATTGTGAAGCAGAATTC | 158.0 | -3.6 | 1.4 | -2.7 | 7.0E-03 | 9.2E-03 | MB231/MB468 | MP |
| >hg38_21_+_8433172_8446622_RNA45SN1_45S_50ntFlanks@9498.9525.28\|AAGATGGTGAACTATGCCTGGGCAGGGC | >hg38_21_+_8433172_8446622_RNA45SN1_45S_50ntFlanks | 9498 | 9525 | 28 | AAGATGGTGAACTATGCCTGGGCAGGGC | 195.7 | 6.3 | 2.0 | 3.1 | 1.7E-03 | 9.2E-03 | BT20/MB231 | MP |
| >am_trnaMT_ValTAC_MT_+_1602_1670@48.68.21\|CAACTTAACTTGACCGCTCTG | >am_trnaMT_ValTAC_MT_+_1602_1670 | 48 | 68 | 21 | CAACTTAACTTGACCGCTCTG | 778.6 | -2.7 | 1.0 | -2.6 | 8.1E-03 | 9.2E-03 | MB231/MB468 | NUC |
| >hg38_21_+_8433172_8446622_RNA45SN1_45S_50ntFlanks@9504.9522.19\|GTGAACTATGCCTGGGCAG | >hg38_21_+_8433172_8446622_RNA45SN1_45S_50ntFlanks | 9504 | 9522 | 19 | GTGAACTATGCCTGGGCAG | 817.8 | -5.1 | 1.8 | -2.8 | 4.5E-03 | 9.2E-03 | BT20/MB468 | MP |
| >hg38_1_-_228634819_228635039_RNA5S12_5S_50ntFlanks@51.81.31\|GTCTACGGCCATACCACCCTGAACGCGCCCG | >hg38_1_-_228634819_228635039_RNA5S12_5S_50ntFlanks | 51 | 81 | 31 | GTCTACGGCCATACCACCCTGAACGCGCCCG | 1264.0 | -2.9 | 1.0 | -3.1 | 2.2E-03 | 9.2E-03 | BT20/MB231 | MITO |
| >hg38_21_+_8433172_8446622_RNA45SN1_45S_50ntFlanks@10241.10273.33\|TACGCCGCGACGAGTAGGAGGGCCGCTGCGGTG | >hg38_21_+_8433172_8446622_RNA45SN1_45S_50ntFlanks | 10241 | 10273 | 33 | TACGCCGCGACGAGTAGGAGGGCCGCTGCGGTG | 69.0 | 6.0 | 2.1 | 2.8 | 4.5E-03 | 9.2E-03 | BT20/MB468 | MP |
| >hg38_21_+_8433172_8446622_RNA45SN1_45S_50ntFlanks@4660.4682.23\|GAGGTGAAATTCTTGGACCGGCG | >hg38_21_+_8433172_8446622_RNA45SN1_45S_50ntFlanks | 4660 | 4682 | 23 | GAGGTGAAATTCTTGGACCGGCG | 136.3 | -4.7 | 1.6 | -2.8 | 4.5E-03 | 9.2E-03 | BT20/MB468 | MP |
| >trnaMT_MetCAT_MT_+_4402_4469@39.71.33\|GAAAATGTTGGTTATACCCTTCCCGTACTACCA | >trnaMT_MetCAT_MT_+_4402_4469 | 39 | 71 | 33 | GAAAATGTTGGTTATACCCTTCCCGTACTACCA | 228.8 | 5.2 | 1.8 | 2.8 | 4.5E-03 | 9.2E-03 | BT20/MB468 | MP |
| >MI0000095\|hsa-mir-93&WithFlank&7\|-\|100093762\|100093853@17.40.24&[MIMAT0000093&hsa-miR-93-5p&offsets\|0\|+1;m-29&7\|-\|100093815\|100093837&offsets\|0\|+1]\|CAAAGTGCTGTTCGTGCAGGTAGT | >MI0000095\|hsa-mir-93&WithFlank&7\|-\|100093762\|100093853 | 17 | 40 | 24 | CAAAGTGCTGTTCGTGCAGGTAGT | 3645.2 | -2.6 | 0.9 | -3.1 | 2.2E-03 | 9.2E-03 | BT20/MB231 | MITO |
| >hg38_21_+_8433172_8446622_RNA45SN1_45S_50ntFlanks@6661.6695.35\|CGGTGGATCACTCGGCTCGTGCGTCGATGAAGAAC | >hg38_21_+_8433172_8446622_RNA45SN1_45S_50ntFlanks | 6661 | 6695 | 35 | CGGTGGATCACTCGGCTCGTGCGTCGATGAAGAAC | 3728.4 | 2.4 | 0.8 | 3.1 | 2.2E-03 | 9.2E-03 | BT20/MB231 | MITO |
| >hg38_21_+_8433172_8446622_RNA45SN1_45S_50ntFlanks@12399.12433.35\|ACGTCGCTTTTTGATCCTTCGATGTCGGCTCTTCC | >hg38_21_+_8433172_8446622_RNA45SN1_45S_50ntFlanks | 12399 | 12433 | 35 | ACGTCGCTTTTTGATCCTTCGATGTCGGCTCTTCC | 378.1 | 2.0 | 0.8 | 2.6 | 8.1E-03 | 9.2E-03 | MB231/MB468 | NUC |
| >hg38_21_+_8433172_8446622_RNA45SN1_45S_50ntFlanks@10868.10890.23\|AAGGGCTGGGTCGGTCGGGCTGG | >hg38_21_+_8433172_8446622_RNA45SN1_45S_50ntFlanks | 10868 | 10890 | 23 | AAGGGCTGGGTCGGTCGGGCTGG | 103.8 | -3.6 | 1.3 | -2.7 | 7.0E-03 | 9.2E-03 | MB231/MB468 | MP |
| >trnaMT_ValTAC_MT_+_1602_1670@36.71.36\|CTTAGGAGATTTCAACTTAACTTGACCGCTCTGACC | >trnaMT_ValTAC_MT_+_1602_1670 | 36 | 71 | 36 | CTTAGGAGATTTCAACTTAACTTGACCGCTCTGACC | 78.4 | 4.2 | 1.6 | 2.7 | 7.0E-03 | 9.2E-03 | MB231/MB468 | MP |
| >hg38_21_+_8433172_8446622_RNA45SN1_45S_50ntFlanks@12420.12448.29\|ATGTCGGCTCTTCCTATCATTGTGAAGCA | >hg38_21_+_8433172_8446622_RNA45SN1_45S_50ntFlanks | 12420 | 12448 | 29 | ATGTCGGCTCTTCCTATCATTGTGAAGCA | 991.6 | 1.8 | 0.7 | 2.7 | 7.6E-03 | 9.2E-03 | BT20/MB468 | CYTO |
| >MI0000239\|hsa-mir-197&WithFlank&1\|+\|109598887\|109598973@54.75.22&[MIMAT0000227&hsa-miR-197-3p&offsets\|0\|0;m-99&1\|+\|109598940\|109598961&offsets\|0\|0]\|TTCACCACCTTCTCCACCCAGC | >MI0000239\|hsa-mir-197&WithFlank&1\|+\|109598887\|109598973 | 54 | 75 | 22 | TTCACCACCTTCTCCACCCAGC | 1131.6 | 2.8 | 1.0 | 2.8 | 4.6E-03 | 9.2E-03 | BT20/MB468 | MP |
| >hg38_21_+_8433172_8446622_RNA45SN1_45S_50ntFlanks@475.507.33\|CTTCGTGATCGATGTGGTGACGTCGTGCTCTCC | >hg38_21_+_8433172_8446622_RNA45SN1_45S_50ntFlanks | 475 | 507 | 33 | CTTCGTGATCGATGTGGTGACGTCGTGCTCTCC | 157.6 | 5.1 | 1.8 | 2.8 | 4.6E-03 | 9.2E-03 | BT20/MB468 | MP |
| >MI0000065\|hsa-let-7d&WithFlank&9\|+\|94178828\|94178926@14.33.20(+1C)&[MIMAT0000065&hsa-let-7d-5p&offsets\|0\|-2(+1C);m-90&9\|+\|94178841\|94178862&offsets\|0\|-2(+1C)]\|AGAGGTAGTAGGTTGCATAGC | >MI0000065\|hsa-let-7d&WithFlank&9\|+\|94178828\|94178926 | 14 | 33 | 20(+1C) | AGAGGTAGTAGGTTGCATAGC | 130.5 | 5.3 | 2.0 | 2.7 | 7.0E-03 | 9.2E-03 | MB231/MB468 | MP |
| >MI0000438\|hsa-mir-15b&WithFlank&3\|+\|160404582\|160404691@64.83.20(+1C)&[MIMAT0004586&hsa-miR-15b-3p&offsets\|0\|-2(+1C);m-204&3\|+\|160404645\|160404665&offsets\|0\|-1(+1C)]\|CGAATCATTATTTGCTGCTCC | >MI0000438\|hsa-mir-15b&WithFlank&3\|+\|160404582\|160404691 | 64 | 83 | 20(+1C) | CGAATCATTATTTGCTGCTCC | 80.7 | 5.4 | 2.0 | 2.7 | 7.0E-03 | 9.2E-03 | MB231/MB468 | MP |
| >hg38_21_+_8433172_8446622_RNA45SN1_45S_50ntFlanks@6734.6771.38\|ATTGATCATCGACACTTCGAACGCACTTGCGGCCCCGG | >hg38_21_+_8433172_8446622_RNA45SN1_45S_50ntFlanks | 6734 | 6771 | 38 | ATTGATCATCGACACTTCGAACGCACTTGCGGCCCCGG | 10147.6 | 5.2 | 1.8 | 2.8 | 4.6E-03 | 9.3E-03 | BT20/MB468 | MP |
| >hg38_21_+_8433172_8446622_RNA45SN1_45S_50ntFlanks@6647.6676.30\|CGTACGACTCTTAGCGGTGGATCACTCGGC | >hg38_21_+_8433172_8446622_RNA45SN1_45S_50ntFlanks | 6647 | 6676 | 30 | CGTACGACTCTTAGCGGTGGATCACTCGGC | 272.4 | 1.6 | 0.6 | 2.7 | 8.0E-03 | 9.3E-03 | BT20/MB468 | NUC |
| >hg38_21_+_8433172_8446622_RNA45SN1_45S_50ntFlanks@3705.3742.38\|TACCTGGTTGATCCTGCCAGTAGCATATGCTTGTCTCA | >hg38_21_+_8433172_8446622_RNA45SN1_45S_50ntFlanks | 3705 | 3742 | 38 | TACCTGGTTGATCCTGCCAGTAGCATATGCTTGTCTCA | 129.6 | 3.1 | 1.1 | 2.8 | 4.6E-03 | 9.3E-03 | BT20/MB468 | MP |
| >hg38_21_+_8433172_8446622_RNA45SN1_45S_50ntFlanks@11321.11342.22\|CACGTCTCGTCGCGCGCGCGTC | >hg38_21_+_8433172_8446622_RNA45SN1_45S_50ntFlanks | 11321 | 11342 | 22 | CACGTCTCGTCGCGCGCGCGTC | 56.4 | 9.2 | 2.9 | 3.1 | 1.7E-03 | 9.3E-03 | BT20/MB231 | MP |
| >hg38_21_+_8433172_8446622_RNA45SN1_45S_50ntFlanks@7976.8009.34\|GCGACCTCAGATCAGACGTGGCGACCCGCTGAAT | >hg38_21_+_8433172_8446622_RNA45SN1_45S_50ntFlanks | 7976 | 8009 | 34 | GCGACCTCAGATCAGACGTGGCGACCCGCTGAAT | 264.8 | 2.6 | 1.0 | 2.7 | 7.0E-03 | 9.3E-03 | MB231/MB468 | MP |
| >trna111_HisGTG_1_-_147774845_147774916@-1T.31.32,trna118_HisGTG_1_-_145396881_145396952@-1T.31.32,trna16_HisGTG_1_+_146544773_146544844@-1T.31.32,trna1_HisGTG_15_+_45493349_45493420@-1T.31.32,trna21_HisGTG_1_+_147753471_147753542@-1T.31.32,trna33_HisGTG_6_+_27125906_27125977@-1T.31.32,trna7_HisGTG_9_-_14433938_14434009@-1T.31.32,trna8_HisGTG_15_-_45492611_45492682@-1T.31.32,trna9_HisGTG_15_-_45490804_45490875@-1T.31.32\|TGCCGTGATCGTATAGTGGTTAGTACTCTGCG | >trna111_HisGTG_1_-_147774845_147774916 | -1T | 31 | 32 | TGCCGTGATCGTATAGTGGTTAGTACTCTGCG | 545.8 | -3.4 | 1.1 | -3.1 | 2.2E-03 | 9.3E-03 | BT20/MB231 | MITO |
| >hg38_21_+_8433172_8446622_RNA45SN1_45S_50ntFlanks@9572.9604.33\|CAAATCGGTCGTCCGACCTGGGTATAGGGGCGA | >hg38_21_+_8433172_8446622_RNA45SN1_45S_50ntFlanks | 9572 | 9604 | 33 | CAAATCGGTCGTCCGACCTGGGTATAGGGGCGA | 185.0 | -4.0 | 1.4 | -2.8 | 4.6E-03 | 9.3E-03 | BT20/MB468 | MP |
| >hg38_MT_+_1621_3279_RNR2_16S_50ntFlanks@827.856.30\|AGGAAAGGTTAAAAAAAGTAAAAGGAACTC | >hg38_MT_+_1621_3279_RNR2_16S_50ntFlanks | 827 | 856 | 30 | AGGAAAGGTTAAAAAAAGTAAAAGGAACTC | 44.1 | 5.2 | 1.9 | 2.7 | 7.1E-03 | 9.3E-03 | MB231/MB468 | MP |
| >MI0000299\|hsa-mir-222&WithFlank&X\|-\|45747009\|45747130@75.93.19(+1A)&[MIMAT0000279&hsa-miR-222-3p&offsets\|0\|-2(+1A);m-140&X\|-\|45747033\|45747056&offsets\|0\|-5(+1A)]\|AGCTACATCTGGCTACTGGA | >MI0000299\|hsa-mir-222&WithFlank&X\|-\|45747009\|45747130 | 75 | 93 | 19(+1A) | AGCTACATCTGGCTACTGGA | 99.0 | 5.3 | 2.0 | 2.7 | 7.1E-03 | 9.3E-03 | MB231/MB468 | MP |
| >hg38_21_+_8433172_8446622_RNA45SN1_45S_50ntFlanks@6646.6666.21\|TCGTACGACTCTTAGCGGTGG | >hg38_21_+_8433172_8446622_RNA45SN1_45S_50ntFlanks | 6646 | 6666 | 21 | TCGTACGACTCTTAGCGGTGG | 16120.6 | 3.3 | 1.0 | 3.1 | 1.7E-03 | 9.3E-03 | BT20/MB231 | MP |
| >hg38_21_+_8433172_8446622_RNA45SN1_45S_50ntFlanks@12418.12451.34\|CGATGTCGGCTCTTCCTATCATTGTGAAGCAGAA | >hg38_21_+_8433172_8446622_RNA45SN1_45S_50ntFlanks | 12418 | 12451 | 34 | CGATGTCGGCTCTTCCTATCATTGTGAAGCAGAA | 245.6 | -1.4 | 0.5 | -2.6 | 8.2E-03 | 9.3E-03 | MB231/MB468 | NUC |
| >hg38_21_+_8433172_8446622_RNA45SN1_45S_50ntFlanks@4131.4150.20\|TTCCGGAGAGGGAGCCTGAG | >hg38_21_+_8433172_8446622_RNA45SN1_45S_50ntFlanks | 4131 | 4150 | 20 | TTCCGGAGAGGGAGCCTGAG | 75.3 | 5.0 | 1.8 | 2.8 | 4.6E-03 | 9.3E-03 | BT20/MB468 | MP |
| >hg38_21_+_8433172_8446622_RNA45SN1_45S_50ntFlanks@8114.8141.28\|GCGCGGGACATGTGGCGTACGGAAGACC | >hg38_21_+_8433172_8446622_RNA45SN1_45S_50ntFlanks | 8114 | 8141 | 28 | GCGCGGGACATGTGGCGTACGGAAGACC | 990.0 | -2.0 | 0.7 | -2.7 | 6.6E-03 | 9.3E-03 | BT20/MB468 | MITO |
| >hg38_21_+_8433172_8446622_RNA45SN1_45S_50ntFlanks@12996.13036.41\|TCGCTGCGATCTATTGAAAGTCAGCCCTCGACACAAGGGTT | >hg38_21_+_8433172_8446622_RNA45SN1_45S_50ntFlanks | 12996 | 13036 | 41 | TCGCTGCGATCTATTGAAAGTCAGCCCTCGACACAAGGGTT | 764.8 | 3.7 | 1.4 | 2.6 | 8.2E-03 | 9.3E-03 | MB231/MB468 | MITO |
| >hg38_21_+_8433172_8446622_RNA45SN1_45S_50ntFlanks@10841.10866.26\|CGTAACTTCGGGATAAGGATTGGCTC | >hg38_21_+_8433172_8446622_RNA45SN1_45S_50ntFlanks | 10841 | 10866 | 26 | CGTAACTTCGGGATAAGGATTGGCTC | 1865.6 | -2.1 | 0.8 | -2.7 | 7.7E-03 | 9.3E-03 | MB231/MB468 | CYTO |
| >hg38_21_+_8433172_8446622_RNA45SN1_45S_50ntFlanks@12632.12667.36\|GCGAAGCTACCATCTGTGGGATTATGACTGAACGCC | >hg38_21_+_8433172_8446622_RNA45SN1_45S_50ntFlanks | 12632 | 12667 | 36 | GCGAAGCTACCATCTGTGGGATTATGACTGAACGCC | 104.0 | 4.2 | 1.6 | 2.6 | 8.2E-03 | 9.3E-03 | MB231/MB468 | MITO |
| >hg38_21_+_8433172_8446622_RNA45SN1_45S_50ntFlanks@11581.11607.27\|AACTGGTGCGGACCAGGGGAATCCGAC | >hg38_21_+_8433172_8446622_RNA45SN1_45S_50ntFlanks | 11581 | 11607 | 27 | AACTGGTGCGGACCAGGGGAATCCGAC | 401.3 | -1.6 | 0.6 | -2.7 | 7.7E-03 | 9.3E-03 | BT20/MB468 | CYTO |
| >hg38_21_+_8433172_8446622_RNA45SN1_45S_50ntFlanks@4738.4758.21\|AAAGTCGGAGGTTCGAAGACG | >hg38_21_+_8433172_8446622_RNA45SN1_45S_50ntFlanks | 4738 | 4758 | 21 | AAAGTCGGAGGTTCGAAGACG | 4758.2 | -2.3 | 0.9 | -2.6 | 8.2E-03 | 9.3E-03 | MB231/MB468 | MITO |
| >hg38_21_+_8433172_8446622_RNA45SN1_45S_50ntFlanks@6772.6806.35\|GTTCCTCCCGGGGCTACGCCTGTCTGAGCGTCGCT | >hg38_21_+_8433172_8446622_RNA45SN1_45S_50ntFlanks | 6772 | 6806 | 35 | GTTCCTCCCGGGGCTACGCCTGTCTGAGCGTCGCT | 1855.8 | -2.4 | 0.9 | -2.8 | 4.6E-03 | 9.3E-03 | BT20/MB468 | MP |
| >MI0000285\|hsa-mir-205&WithFlank&1\|+\|209432127\|209432248@40.58.19(+1U)&[MIMAT0000266&hsa-miR-205-5p&offsets\|0\|-3(+1U);m-351&1\|+\|209432166\|209432187&offsets\|0\|-3(+1U)]\|TCCTTCATTCCACCGGAGTT | >MI0000285\|hsa-mir-205&WithFlank&1\|+\|209432127\|209432248 | 40 | 58 | 19(+1U) | TCCTTCATTCCACCGGAGTT | 50.0 | 9.0 | 2.9 | 3.1 | 1.7E-03 | 9.3E-03 | BT20/MB231 | MP |
| >hg38_21_+_8433172_8446622_RNA45SN1_45S_50ntFlanks@7990.8008.19\|GACGTGGCGACCCGCTGAA | >hg38_21_+_8433172_8446622_RNA45SN1_45S_50ntFlanks | 7990 | 8008 | 19 | GACGTGGCGACCCGCTGAA | 4517.0 | -2.8 | 1.0 | -2.8 | 4.6E-03 | 9.3E-03 | BT20/MB468 | MP |
| >hg38_21_+_8433172_8446622_RNA45SN1_45S_50ntFlanks@12996.13017.22\|TCGCTGCGATCTATTGAAAGTC | >hg38_21_+_8433172_8446622_RNA45SN1_45S_50ntFlanks | 12996 | 13017 | 22 | TCGCTGCGATCTATTGAAAGTC | 73.6 | 4.4 | 1.6 | 2.7 | 7.1E-03 | 9.3E-03 | MB231/MB468 | MP |
| >MI0000071\|hsa-mir-17&WithFlank&13\|+\|91350599\|91350694@20.43.24&[MIMAT0000070&hsa-miR-17-5p&offsets\|0\|+1;m-88&13\|+\|91350618\|91350640&offsets\|0\|+1]\|CAAAGTGCTTACAGTGCAGGTAGT | >MI0000071\|hsa-mir-17&WithFlank&13\|+\|91350599\|91350694 | 20 | 43 | 24 | CAAAGTGCTTACAGTGCAGGTAGT | 1065.6 | 2.7 | 1.0 | 2.7 | 7.1E-03 | 9.3E-03 | MB231/MB468 | MP |
| >MI0005566\|hsa-mir-374b&WithFlank&X\|-\|74218541\|74218624@17.38.22&[MIMAT0004955&hsa-miR-374b-5p&offsets\|0\|0;m-153&X\|-\|74218587\|74218608&offsets\|0\|0]\|ATATAATACAACCTGCTAAGTG | >MI0005566\|hsa-mir-374b&WithFlank&X\|-\|74218541\|74218624 | 17 | 38 | 22 | ATATAATACAACCTGCTAAGTG | 2492.1 | -2.5 | 0.8 | -3.1 | 2.2E-03 | 9.3E-03 | BT20/MB231 | MITO |
| >hg38_21_+_8433172_8446622_RNA45SN1_45S_50ntFlanks@4770.4793.24\|TCGTAGTTCCGACCATAAACGATG | >hg38_21_+_8433172_8446622_RNA45SN1_45S_50ntFlanks | 4770 | 4793 | 24 | TCGTAGTTCCGACCATAAACGATG | 642.9 | -1.6 | 0.6 | -2.7 | 7.8E-03 | 9.3E-03 | MB231/MB468 | CYTO |
| >MI0000095\|hsa-mir-93&WithFlank&7\|-\|100093762\|100093853@17.38.22&[MIMAT0000093&hsa-miR-93-5p&offsets\|0\|-1;m-29&7\|-\|100093815\|100093837&offsets\|0\|-1]\|CAAAGTGCTGTTCGTGCAGGTA | >MI0000095\|hsa-mir-93&WithFlank&7\|-\|100093762\|100093853 | 17 | 38 | 22 | CAAAGTGCTGTTCGTGCAGGTA | 1008.0 | -2.6 | 0.8 | -3.1 | 2.2E-03 | 9.3E-03 | BT20/MB231 | MITO |
| >MI0000063\|hsa-let-7b&WithFlank&22\|+\|46113680\|46113774@12.31.20&[MIMAT0000063&hsa-let-7b-5p&offsets\|0\|-2;m-14&22\|+\|46113691\|46113712&offsets\|0\|-2]\|TGAGGTAGTAGGTTGTGTGG | >MI0000063\|hsa-let-7b&WithFlank&22\|+\|46113680\|46113774 | 12 | 31 | 20 | TGAGGTAGTAGGTTGTGTGG | 352.8 | 2.1 | 0.7 | 3.1 | 2.2E-03 | 9.3E-03 | BT20/MB231 | MITO |

| | | | | | | | | | | | | | |
|---|---|---|---|---|---|---|---|---|---|---|---|---|---|
| >hg38_MT_+_1621_3279_RNR2_16S_50ntFlanks@51.93.43\|GCTAAACCTAGCCCCAAACCCACTCCACCTTACTACCAGACAA | >hg38_MT_+_1621_3279_RNR2_16S_50ntFlanks | 51 | 93 | 43 | GCTAAACCTAGCCCCAAACCCACTCCACCTTACTACCAGACAA | 2168.6 | -2.8 | 0.9 | -3.1 | 2.2E-03 | 9.3E-03 | BT20/MB231 | MITO |
| >MI0000062\|hsa-let-7a-3&WithFlank&22\|+\|46112743\|46112828@58.79.22;MI0000060\|hsa-let-7a-1&WithFlank&9\|+\|94175951\|94176042@63.84.22&[MIMAT0004481&hsa-let-7a-3p&offsets\|0\|+1;m-180&22\|+\|46112800\|46112821&offsets\|0\|0];[MIMAT0004481_1&hsa-let-7a-3p&offsets\|0\|+1;m-181&9\|+\|94176013\|94176034&offsets\|0\|0]\|CTATACAATCTACTGTCTTTCC | >MI0000062\|hsa-let-7a-3&WithFlank&22\|+\|46112743\|46112828 | 58 | 79 | 22 | CTATACAATCTACTGTCTTTCC | 200.9 | 3.3 | 1.2 | 2.8 | 4.6E-03 | 9.3E-03 | BT20/MB468 | MP |
| >hg38_21_+_8433172_8446622_RNA45SN1_45S_50ntFlanks@9307.9330.24\|AAGGAGTCTAACACGTGCGCGAGT | >hg38_21_+_8433172_8446622_RNA45SN1_45S_50ntFlanks | 9307 | 9330 | 24 | AAGGAGTCTAACACGTGCGCGAGT | 384.9 | -3.0 | 1.0 | -3.1 | 2.1E-03 | 9.3E-03 | BT20/MB231 | CYTO |
| >hg38_21_+_8433172_8446622_RNA45SN1_45S_50ntFlanks@9532.9572.41\|AGAGGAAACTCTGGTGGAGGTCCGTAGCGGTCCTGACGTGC | >hg38_21_+_8433172_8446622_RNA45SN1_45S_50ntFlanks | 9532 | 9572 | 41 | AGAGGAAACTCTGGTGGAGGTCCGTAGCGGTCCTGACGTGC | 457.8 | -2.4 | 0.8 | -3.1 | 2.1E-03 | 9.3E-03 | BT20/MB231 | CYTO |
| >hg38_21_+_8433172_8446622_RNA45SN1_45S_50ntFlanks@6646.6678.33\|TCGTACGACTCTTAGCGGTGGATCACTCGGCTC | >hg38_21_+_8433172_8446622_RNA45SN1_45S_50ntFlanks | 6646 | 6678 | 33 | TCGTACGACTCTTAGCGGTGGATCACTCGGCTC | 2103.5 | 2.3 | 0.9 | 2.6 | 8.2E-03 | 9.3E-03 | MB231/MB468 | NUC |
| >MI0000281\|hsa-mir-199a-2&WithFlank&1\|-\|172144529\|172144650@76.96.21;MI0000242\|hsa-mir-199a-1&WithFlank&19\|-\|10817420\|10817502@53.73.21;MI0000282\|hsa-mir-199b&WithFlank&9\|-\|128244715\|128244836@71.91.21&[MIMAT0000232&hsa-miR-199a-3p&offsets\|0\|-1;m-46&1\|-\|172144555\|172144575&offsets\|0\|0];[MIMAT0000232_1&hsa-miR-199a-3p&offsets\|0\|-1;m-47&19\|-\|10817430\|10817450&offsets\|0\|0];[MIMAT0004563&hsa-miR-199b-3p&offsets\|0\|-1;m-47&9\|-\|128244746\|128244766&offsets\|0\|0]\|ACAGTAGTCTGCACATTGGTT | >MI0000281\|hsa-mir-199a-2&WithFlank&1\|-\|172144529\|172144650 | 76 | 96 | 21 | ACAGTAGTCTGCACATTGGTT | 141.3 | 6.1 | 2.2 | 2.8 | 4.6E-03 | 9.3E-03 | BT20/MB468 | MP |
| >MI0000093\|hsa-mir-92a-1&WithFlank&13\|+\|91351308\|91351397@54.72.19;MI0000094\|hsa-mir-92a-2&WithFlank&X\|-\|134169532\|134169618@54.72.19&[MIMAT0000092&hsa-miR-92a-3p&offsets\|0\|-3;m-9&13\|+\|91351361\|91351382&offsets\|0\|-3];[MIMAT0000092_1&hsa-miR-92a-3p&offsets\|0\|-3;m-11&X\|-\|134169544\|134169565&offsets\|0\|-3]\|TATTGCACTTGTCCCGGCC | >MI0000093\|hsa-mir-92a-1&WithFlank&13\|+\|91351308\|91351397 | 54 | 72 | 19 | TATTGCACTTGTCCCGGCC | 35.6 | 5.2 | 1.9 | 2.7 | 7.1E-03 | 9.3E-03 | MB231/MB468 | MP |
| >hg38_21_+_8433172_8446622_RNA45SN1_45S_50ntFlanks@5488.5512.25\|GCTGAGAAGACGGTCGAACTTGACT | >hg38_21_+_8433172_8446622_RNA45SN1_45S_50ntFlanks | 5488 | 5512 | 25 | GCTGAGAAGACGGTCGAACTTGACT | 950.9 | 2.8 | 0.9 | 3.1 | 2.1E-03 | 9.3E-03 | BT20/MB231 | CYTO |
| >MI0000434\|hsa-let-7i&WithFlank&12\|+\|62603680\|62603775@13.33.21&[MIMAT0000415&hsa-let-7i-5p&offsets\|+1\|0;m-49&12\|+\|62603691\|62603712&offsets\|+1\|0]\|GAGGTAGTAGTTTGTGCTGTT | >MI0000434\|hsa-let-7i&WithFlank&12\|+\|62603680\|62603775 | 13 | 33 | 21 | GAGGTAGTAGTTTGTGCTGTT | 39.8 | 5.3 | 2.0 | 2.7 | 7.1E-03 | 9.4E-03 | MB231/MB468 | MP |
| >hg38_21_+_8433172_8446622_RNA45SN1_45S_50ntFlanks@5496.5525.30\|GACGGTCGAACTTGACTATCTAGAGGAAGT | >hg38_21_+_8433172_8446622_RNA45SN1_45S_50ntFlanks | 5496 | 5525 | 30 | GACGGTCGAACTTGACTATCTAGAGGAAGT | 311.4 | 5.6 | 1.8 | 3.1 | 1.7E-03 | 9.4E-03 | BT20/MB231 | MP |
| >hg38_21_+_8433172_8446622_RNA45SN1_45S_50ntFlanks@11568.11588.21\|GCAGCCGACTTAGAACTGGTG | >hg38_21_+_8433172_8446622_RNA45SN1_45S_50ntFlanks | 11568 | 11588 | 21 | GCAGCCGACTTAGAACTGGTG | 475.2 | -2.0 | 0.8 | -2.6 | 8.2E-03 | 9.4E-03 | MB231/MB468 | MITO |
| >hg38_21_+_8433172_8446622_RNA45SN1_45S_50ntFlanks@7975.8001.27\|CGCGACCTCAGATCAGACGTGGCGACC | >hg38_21_+_8433172_8446622_RNA45SN1_45S_50ntFlanks | 7975 | 8001 | 27 | CGCGACCTCAGATCAGACGTGGCGACC | 865.8 | 2.4 | 0.9 | 2.6 | 8.2E-03 | 9.4E-03 | MB231/MB468 | MITO |
| >hg38_21_+_8433172_8446622_RNA45SN1_45S_50ntFlanks@9536.9571.36\|GAAACTCTGGTGGAGGTCCGTAGCGGTCCTGACGTG | >hg38_21_+_8433172_8446622_RNA45SN1_45S_50ntFlanks | 9536 | 9571 | 36 | GAAACTCTGGTGGAGGTCCGTAGCGGTCCTGACGTG | 449.2 | -1.7 | 0.6 | -2.7 | 7.7E-03 | 9.4E-03 | BT20/MB468 | CYTO |
| >hg38_21_+_8433172_8446622_RNA45SN1_45S_50ntFlanks@12909.12933.25\|CCGCACGTTCGTGGGGAACCTGGCG | >hg38_21_+_8433172_8446622_RNA45SN1_45S_50ntFlanks | 12909 | 12933 | 25 | CCGCACGTTCGTGGGGAACCTGGCG | 247.6 | -3.4 | 1.3 | -2.7 | 7.8E-03 | 9.4E-03 | MB231/MB468 | CYTO |
| >trnaMT_ProTGG_MT_-_15956_16023@37.70.34\|GCTAATGGTGGAGTTAAAGACTTTTTCTCTGACC | >trnaMT_ProTGG_MT_-_15956_16023 | 37 | 70 | 34 | GCTAATGGTGGAGTTAAAGACTTTTTCTCTGACC | 47.5 | 6.0 | 2.1 | 2.8 | 4.6E-03 | 9.4E-03 | BT20/MB468 | MP |
| >hg38_21_+_8433172_8446622_RNA45SN1_45S_50ntFlanks@11911.11943.33\|TGAAGAGACATGAGAGGTGTAGAATAAGTGGGA | >hg38_21_+_8433172_8446622_RNA45SN1_45S_50ntFlanks | 11911 | 11943 | 33 | TGAAGAGACATGAGAGGTGTAGAATAAGTGGGA | 56.4 | 9.1 | 2.9 | 3.1 | 1.7E-03 | 9.4E-03 | BT20/MB231 | MP |
| >hg38_21_+_8433172_8446622_RNA45SN1_45S_50ntFlanks@10392.10430.39\|ACAGCAGTTGAACATGGGTCAGTCGGTCCTGAGAGATGG | >hg38_21_+_8433172_8446622_RNA45SN1_45S_50ntFlanks | 10392 | 10430 | 39 | ACAGCAGTTGAACATGGGTCAGTCGGTCCTGAGAGATGG | 480.8 | -5.1 | 1.8 | -2.8 | 4.6E-03 | 9.4E-03 | BT20/MB468 | MP |
| >hg38_21_+_8433172_8446622_RNA45SN1_45S_50ntFlanks@9313.9335.23\|TCTAACACGTGCGCGAGTCGGGG | >hg38_21_+_8433172_8446622_RNA45SN1_45S_50ntFlanks | 9313 | 9335 | 23 | TCTAACACGTGCGCGAGTCGGGG | 353.9 | -5.0 | 1.8 | -2.8 | 4.6E-03 | 9.4E-03 | BT20/MB468 | MP |
| >MI0000447\|hsa-mir-128-1&WithFlank&2\|+\|135665391\|135665484@56.76.21;MI0000727\|hsa-mir-128-2&WithFlank&3\|+\|35744470\|35744565@58.78.21&[MIMAT0000424&hsa-miR-128-3p&offsets\|0\|0;m-108&2\|+\|135665446\|135665466&offsets\|0\|0];[MIMAT0000424_1&hsa-miR-128-3p&offsets\|0\|0;m-117&3\|+\|35744527\|35744547&offsets\|0\|0]\|TCACAGTGAACCGGTCTCTTT | >MI0000447\|hsa-mir-128-1&WithFlank&2\|+\|135665391\|135665484 | 56 | 76 | 21 | TCACAGTGAACCGGTCTCTTT | 5049.2 | 2.3 | 0.9 | 2.6 | 8.2E-03 | 9.4E-03 | MB231/MB468 | MITO |
| >hg38_1_-_228634819_228635039_RNA5S12_5S_50ntFlanks@132.170.39\|GATGGGAGACCGCCTGGGAATACCGGGTGCTGTAGGCTT | >hg38_1_-_228634819_228635039_RNA5S12_5S_50ntFlanks | 132 | 170 | 39 | GATGGGAGACCGCCTGGGAATACCGGGTGCTGTAGGCTT | 58896.9 | 2.6 | 1.0 | 2.7 | 7.7E-03 | 9.4E-03 | BT20/MB468 | CYTO |
| >hg38_21_+_8433172_8446622_RNA45SN1_45S_50ntFlanks@12419.12449.31\|GATGTCGGCTCTTCCTATCATTGTGAAGCAG | >hg38_21_+_8433172_8446622_RNA45SN1_45S_50ntFlanks | 12419 | 12449 | 31 | GATGTCGGCTCTTCCTATCATTGTGAAGCAG | 287.4 | 2.3 | 0.9 | 2.6 | 8.2E-03 | 9.4E-03 | MB231/MB468 | MITO |
| >hg38_21_+_8433172_8446622_RNA45SN1_45S_50ntFlanks@12940.12972.33\|CATTCGTAGACGACCTGCTTCTGGGTCGGGGTT | >hg38_21_+_8433172_8446622_RNA45SN1_45S_50ntFlanks | 12940 | 12972 | 33 | CATTCGTAGACGACCTGCTTCTGGGTCGGGGTT | 100.7 | -3.0 | 1.1 | -2.7 | 6.6E-03 | 9.4E-03 | BT20/MB468 | MITO |
| >MI0000077\|hsa-mir-21&WithFlank&17\|+\|59841260\|59841343@52.74.23&[MIMAT0004494&hsa-miR-21-3p&offsets\|0\|+2;m-52&17\|+\|59841311\|59841332&offsets\|0\|+1]\|CAACACCAGTCGATGGGCTGTCT | >MI0000077\|hsa-mir-21&WithFlank&17\|+\|59841260\|59841343 | 52 | 74 | 23 | CAACACCAGTCGATGGGCTGTCT | 383.3 | 4.5 | 1.6 | 2.8 | 4.6E-03 | 9.4E-03 | BT20/MB468 | MP |
| >hg38_21_+_8433172_8446622_RNA45SN1_45S_50ntFlanks@6647.6668.22\|CGTACGACTCTTAGCGGTGGAT | >hg38_21_+_8433172_8446622_RNA45SN1_45S_50ntFlanks | 6647 | 6668 | 22 | CGTACGACTCTTAGCGGTGGAT | 706.9 | 1.8 | 0.7 | 2.6 | 8.1E-03 | 9.4E-03 | BT20/MB468 | NUC |
| >hg38_21_+_8433172_8446622_RNA45SN1_45S_50ntFlanks@4727.4753.27\|AATCAAGAACGAAAGTCGGAGGTTCGA | >hg38_21_+_8433172_8446622_RNA45SN1_45S_50ntFlanks | 4727 | 4753 | 27 | AATCAAGAACGAAAGTCGGAGGTTCGA | 80.1 | -3.4 | 1.3 | -2.7 | 7.1E-03 | 9.4E-03 | MB231/MB468 | MP |
| >MI0000289\|hsa-mir-181a-1&WithFlank&1\|-\|198859038\|198859159@30.54.25(+1U);MI0000269\|hsa-mir-181a-2&WithFlank&9\|+\|124692436\|124692557@45.69.25(+1U)&[MIMAT0000256&hsa-miR-181a-5p&offsets\|0\|+2(+1U);m-32&1\|-\|198859108\|198859130&offsets\|0\|+2(+1U)];[MIMAT0000256_1&hsa-miR-181a-5p&offsets\|0\|+2(+1U);m-32&9\|+\|124692480\|124692502&offsets\|0\|+2(+1U)]\|AACATTCAACGCTGTCGGTGAGTTTT | >MI0000289\|hsa-mir-181a-1&WithFlank&1\|-\|198859038\|198859159 | 30 | 54 | 25(+1U) | AACATTCAACGCTGTCGGTGAGTTTT | 304.3 | -5.1 | 1.6 | -3.1 | 1.9E-03 | 9.4E-03 | BT20/MB231 | NUC |
| >hg38_21_+_8433172_8446622_RNA45SN1_45S_50ntFlanks@12631.12659.29\|GGCGAAGCTACCATCTGTGGGATTATGAC | >hg38_21_+_8433172_8446622_RNA45SN1_45S_50ntFlanks | 12631 | 12659 | 29 | GGCGAAGCTACCATCTGTGGGATTATGAC | 510.7 | -2.4 | 0.9 | -2.7 | 6.6E-03 | 9.4E-03 | BT20/MB468 | MITO |
| >hg38_21_+_8433172_8446622_RNA45SN1_45S_50ntFlanks@12645.12669.25\|CTGTGGGATTATGACTGAACGCCTC | >hg38_21_+_8433172_8446622_RNA45SN1_45S_50ntFlanks | 12645 | 12669 | 25 | CTGTGGGATTATGACTGAACGCCTC | 231.3 | 3.6 | 1.3 | 2.8 | 4.6E-03 | 9.4E-03 | BT20/MB468 | MP |
| >MI0000065\|hsa-let-7d&WithFlank&9\|+\|94178828\|94178926@14.33.20&[MIMAT0000065&hsa-let-7d-5p&offsets\|0\|-2;m-90&9\|+\|94178841\|94178862&offsets\|0\|-2]\|AGAGGTAGTAGGTTGCATAG | >MI0000065\|hsa-let-7d&WithFlank&9\|+\|94178828\|94178926 | 14 | 33 | 20 | AGAGGTAGTAGGTTGCATAG | 48.9 | 4.9 | 1.7 | 2.8 | 4.6E-03 | 9.4E-03 | BT20/MB468 | MP |
| >hg38_21_+_8433172_8446622_RNA45SN1_45S_50ntFlanks@11890.11909.20\|CTTGACTCTAGTCTGGCACG | >hg38_21_+_8433172_8446622_RNA45SN1_45S_50ntFlanks | 11890 | 11909 | 20 | CTTGACTCTAGTCTGGCACG | 186.5 | -4.8 | 1.7 | -2.8 | 4.6E-03 | 9.4E-03 | BT20/MB468 | MP |
| >hg38_21_+_8433172_8446622_RNA45SN1_45S_50ntFlanks@12399.12419.21\|ACGTCGCTTTTTGATCCTTCG | >hg38_21_+_8433172_8446622_RNA45SN1_45S_50ntFlanks | 12399 | 12419 | 21 | ACGTCGCTTTTTGATCCTTCG | 358.3 | -2.8 | 1.0 | -2.7 | 7.8E-03 | 9.4E-03 | MB231/MB468 | CYTO |
| >hg38_21_+_8433172_8446622_RNA45SN1_45S_50ntFlanks@12258.12284.27\|GCTTGATCTTGATTTTCAGTACGAATA | >hg38_21_+_8433172_8446622_RNA45SN1_45S_50ntFlanks | 12258 | 12284 | 27 | GCTTGATCTTGATTTTCAGTACGAATA | 135.1 | -2.0 | 0.8 | -2.7 | 7.7E-03 | 9.4E-03 | BT20/MB468 | CYTO |
| >hg38_21_+_8433172_8446622_RNA45SN1_45S_50ntFlanks@10767.10785.19\|GCAGCAGGTCTCCAAGGTG | >hg38_21_+_8433172_8446622_RNA45SN1_45S_50ntFlanks | 10767 | 10785 | 19 | GCAGCAGGTCTCCAAGGTG | 208.9 | -4.8 | 1.7 | -2.8 | 4.6E-03 | 9.4E-03 | BT20/MB468 | MP |
| >hg38_1_-_228634819_228635039_RNA5S12_5S_50ntFlanks@131.171.41\|GGATGGGAGACCGCCTGGGAATACCGGGTGCTGTAGGCTTT | >hg38_1_-_228634819_228635039_RNA5S12_5S_50ntFlanks | 131 | 171 | 41 | GGATGGGAGACCGCCTGGGAATACCGGGTGCTGTAGGCTTT | 871.9 | 2.7 | 0.9 | 3.1 | 2.3E-03 | 9.4E-03 | BT20/MB231 | MITO |
| >hg38_21_+_8433172_8446622_RNA45SN1_45S_50ntFlanks@12960.12982.23\|CTGGGTCGGGGTTTCGTACGTAG | >hg38_21_+_8433172_8446622_RNA45SN1_45S_50ntFlanks | 12960 | 12982 | 23 | CTGGGTCGGGGTTTCGTACGTAG | 213.0 | -3.0 | 1.1 | -2.7 | 7.8E-03 | 9.4E-03 | BT20/MB468 | CYTO |
| >hg38_21_+_8433172_8446622_RNA45SN1_45S_50ntFlanks@9554.9571.18\|CGTAGCGGTCCTGACGTG | >hg38_21_+_8433172_8446622_RNA45SN1_45S_50ntFlanks | 9554 | 9571 | 18 | CGTAGCGGTCCTGACGTG | 10379.5 | -2.7 | 0.9 | -3.1 | 2.1E-03 | 9.4E-03 | BT20/MB231 | CYTO |
| >MI0000077\|hsa-mir-21&WithFlank&17\|+\|59841260\|59841343@14.33.20(+1A)&[MIMAT0000076&hsa-miR-21-5p&offsets\|0\|-2(+1A);m-1&17\|+\|59841273\|59841294&offsets\|0\|-2(+1A)]\|TAGCTTATCAGACTGATGTTA | >MI0000077\|hsa-mir-21&WithFlank&17\|+\|59841260\|59841343 | 14 | 33 | 20(+1A) | TAGCTTATCAGACTGATGTTA | 2626.7 | 3.4 | 1.2 | 2.8 | 4.6E-03 | 9.4E-03 | BT20/MB468 | MP |
| >am_trnaMT_GluTTC_MT_-_14674_14742@24.48.25,trnalookalike8_GluTTC_5_-_93905172_93905240@24.48.25\|ATGGTTTTTCATATCATTGGTCGTG | >am_trnaMT_GluTTC_MT_-_14674_14742 | 24 | 48 | 25 | ATGGTTTTTCATATCATTGGTCGTG | 148.8 | 5.9 | 2.1 | 2.8 | 4.6E-03 | 9.4E-03 | BT20/MB468 | MP |
| >MI0016887\|hsa-mir-4521&WithFlank&17\|+\|8186939\|8187010@13.34.22&[MIMAT0019058&hsa-miR-4521&offsets\|+3\|+3;m-765&17\|+\|8186948\|8186969&offsets\|+3\|+3]\|AAGGAAGTCCTGTGCTCAGTTT | >MI0016887\|hsa-mir-4521&WithFlank&17\|+\|8186939\|8187010 | 13 | 34 | 22 | AAGGAAGTCCTGTGCTCAGTTT | 428.8 | 3.2 | 1.0 | 3.1 | 1.9E-03 | 9.4E-03 | BT20/MB231 | NUC |
| >trnaMT_MetCAT_MT_+_4402_4469@41.71.31\|AAATGTTGGTTATACCCTTCCCGTACTACCA | >trnaMT_MetCAT_MT_+_4402_4469 | 41 | 71 | 31 | AAATGTTGGTTATACCCTTCCCGTACTACCA | 350.1 | -2.4 | 0.9 | -2.6 | 8.3E-03 | 9.4E-03 | MB231/MB468 | MITO |
| >hg38_21_+_8433172_8446622_RNA45SN1_45S_50ntFlanks@10678.10696.19\|CGGTTCCGGCGGCGTCCGG | >hg38_21_+_8433172_8446622_RNA45SN1_45S_50ntFlanks | 10678 | 10696 | 19 | CGGTTCCGGCGGCGTCCGG | 209.5 | -4.1 | 1.5 | -2.7 | 7.8E-03 | 9.4E-03 | BT20/MB468 | CYTO |
| >hg38_21_+_8433172_8446622_RNA45SN1_45S_50ntFlanks@4367.4402.36\|CAGTTAAAAAGCTCGTAGTTGGATCTTGGGAGCGGG | >hg38_21_+_8433172_8446622_RNA45SN1_45S_50ntFlanks | 4367 | 4402 | 36 | CAGTTAAAAAGCTCGTAGTTGGATCTTGGGAGCGGG | 114.9 | -3.2 | 1.2 | -2.7 | 7.8E-03 | 9.4E-03 | BT20/MB468 | CYTO |
| >trnaMT_ValTAC_MT_+_1602_1670@41.71.31\|GAGATTTCAACTTAACTTGACCGCTCTGACC | >trnaMT_ValTAC_MT_+_1602_1670 | 41 | 71 | 31 | GAGATTTCAACTTAACTTGACCGCTCTGACC | 1092.7 | 2.7 | 1.0 | 2.7 | 6.7E-03 | 9.4E-03 | BT20/MB468 | MITO |
| >hg38_21_+_8433172_8446622_RNA45SN1_45S_50ntFlanks@12416.12446.31\|TTCGATGTCGGCTCTTCCTATCATTGTGAAG | >hg38_21_+_8433172_8446622_RNA45SN1_45S_50ntFlanks | 12416 | 12446 | 31 | TTCGATGTCGGCTCTTCCTATCATTGTGAAG | 513.7 | -2.5 | 0.8 | -3.1 | 2.3E-03 | 9.4E-03 | BT20/MB231 | MITO |

<table>
<tr><td>>MI0001448|hsa-mir-425&WithFlank&3|-|49020142|49020240@20.43.24(+1C)&[MIMAT0003393&hsa-miR-425-5p&offsets|0|+1(+1C);m-95&3|-|49020200|49020221&offsets|0|+2(+1C)]|AATGACACGATCACTCCCGTTGAGC</td><td>>MI0001448|hsa-mir-425&WithFlank&3|-|49020142|49020240</td><td>20</td><td>43</td><td>24(+1C)</td><td>AATGACACGATCACTCCCGTTGAGC</td><td>885.3</td><td>-2.7</td><td>0.9</td><td>-3.1</td><td>2.3E-03</td><td>9.4E-03</td><td>BT20/MB231</td><td>MITO</td></tr>
<tr><td>>hg38_21_+_8433172_8446622_RNA45SN1_45S_50ntFlanks@9533.9562.30|GAGGAAACTCTGGTGGAGGTCCGTAGCGGT</td><td>>hg38_21_+_8433172_8446622_RNA45SN1_45S_50ntFlanks</td><td>9533</td><td>9562</td><td>30</td><td>GAGGAAACTCTGGTGGAGGTCCGTAGCGGT</td><td>342.4</td><td>-2.0</td><td>0.8</td><td>-2.7</td><td>7.8E-03</td><td>9.4E-03</td><td>BT20/MB468</td><td>CYTO</td></tr>
<tr><td>>MI0000066|hsa-let-7e&WithFlank&19|+|51692780|51692870@14.31.18&[MIMAT0000066&hsa-let-7e-5p&offsets|0|-4;m-50&19|+|51692793|51692814&offsets|0|-4]|TGAGGTAGGAGGTTGTAT</td><td>>MI0000066|hsa-let-7e&WithFlank&19|+|51692780|51692870</td><td>14</td><td>31</td><td>18</td><td>TGAGGTAGGAGGTTGTAT</td><td>125.6</td><td>1.5</td><td>0.6</td><td>2.6</td><td>8.2E-03</td><td>9.4E-03</td><td>BT20/MB468</td><td>NUC</td></tr>
<tr><td>>hg38_21_+_8433172_8446622_RNA45SN1_45S_50ntFlanks@10247.10276.30|GCGACGAGTAGGAGGGCCGCTGCGGTGAGC</td><td>>hg38_21_+_8433172_8446622_RNA45SN1_45S_50ntFlanks</td><td>10247</td><td>10276</td><td>30</td><td>GCGACGAGTAGGAGGGCCGCTGCGGTGAGC</td><td>174.8</td><td>5.0</td><td>1.6</td><td>3.1</td><td>1.7E-03</td><td>9.4E-03</td><td>BT20/MB231</td><td>MP</td></tr>
<tr><td>>hg38_21_+_8433172_8446622_RNA45SN1_45S_50ntFlanks@10852.10875.24|GATAAGGATTGGCTCTAAGGGCTG</td><td>>hg38_21_+_8433172_8446622_RNA45SN1_45S_50ntFlanks</td><td>10852</td><td>10875</td><td>24</td><td>GATAAGGATTGGCTCTAAGGGCTG</td><td>56.5</td><td>9.2</td><td>2.9</td><td>3.1</td><td>1.7E-03</td><td>9.4E-03</td><td>BT20/MB231</td><td>MP</td></tr>
<tr><td>>hg38_21_+_8433172_8446622_RNA45SN1_45S_50ntFlanks@6651.6675.25|CGACTCTTAGCGGTGGATCACTCGG</td><td>>hg38_21_+_8433172_8446622_RNA45SN1_45S_50ntFlanks</td><td>6651</td><td>6675</td><td>25</td><td>CGACTCTTAGCGGTGGATCACTCGG</td><td>6121.1</td><td>4.4</td><td>1.4</td><td>3.1</td><td>1.7E-03</td><td>9.4E-03</td><td>BT20/MB231</td><td>MP</td></tr>
<tr><td>>hg38_21_+_8433172_8446622_RNA45SN1_45S_50ntFlanks@8346.8375.30|AGGGAAAGTTGAAAAGAACTTTGAAGAGAG</td><td>>hg38_21_+_8433172_8446622_RNA45SN1_45S_50ntFlanks</td><td>8346</td><td>8375</td><td>30</td><td>AGGGAAAGTTGAAAAGAACTTTGAAGAGAG</td><td>3366.7</td><td>5.5</td><td>1.8</td><td>3.1</td><td>1.7E-03</td><td>9.4E-03</td><td>BT20/MB231</td><td>MP</td></tr>
<tr><td>>hg38_21_+_8433172_8446622_RNA45SN1_45S_50ntFlanks@9494.9531.38|CCGAAAGATGGTGAACTATGCCTGGGCAGGGCGAAGCC</td><td>>hg38_21_+_8433172_8446622_RNA45SN1_45S_50ntFlanks</td><td>9494</td><td>9531</td><td>38</td><td>CCGAAAGATGGTGAACTATGCCTGGGCAGGGCGAAGCC</td><td>260.8</td><td>4.2</td><td>1.3</td><td>3.1</td><td>1.7E-03</td><td>9.4E-03</td><td>BT20/MB231</td><td>MP</td></tr>
<tr><td>>MI0000448|hsa-mir-130a&WithFlank&11|+|57641192|57641292@21.42.22&[MIMAT0004593&hsa-miR-130a-5p&offsets|0|0;m-622&11|+|57641212|57641233&offsets|0|0]|GCTCTTTTCACATTGTGCTACT</td><td>>MI0000448|hsa-mir-130a&WithFlank&11|+|57641192|57641292</td><td>21</td><td>42</td><td>22</td><td>GCTCTTTTCACATTGTGCTACT</td><td>188.7</td><td>-5.6</td><td>1.8</td><td>-3.1</td><td>1.7E-03</td><td>9.4E-03</td><td>BT20/MB231</td><td>MP</td></tr>
<tr><td>>MI0003180|hsa-mir-516a-1&WithFlank&19|+|53756735|53756836@22.44.23;MI0003181|hsa-mir-516a-2&WithFlank&19|+|53761127|53761228@22.44.23&[MIMAT0004770&hsa-miR-516a-5p&offsets|0|0;m-852&19|+|53756756|53756777&offsets|0|+1];[MIMAT0004770_1&hsa-miR-516a-5p&offsets|0|0;m-852&19|+|53761148|53761169&offsets|0|+1]|TTCTCGAGGAAAGAAGCACTTTC</td><td>>MI0003180|hsa-mir-516a-1&WithFlank&19|+|53756735|53756836</td><td>22</td><td>44</td><td>23</td><td>TTCTCGAGGAAAGAAGCACTTTC</td><td>123.9</td><td>-4.4</td><td>1.4</td><td>-3.1</td><td>1.7E-03</td><td>9.4E-03</td><td>BT20/MB231</td><td>MP</td></tr>
<tr><td>>MI0000289|hsa-mir-181a-1&WithFlank&1|-|198859038|198859159@30.48.19;MI0000269|hsa-mir-181a-2&WithFlank&9|+|124692436|124692557@45.63.19&[MIMAT0000256&hsa-miR-181a-5p&offsets|0|-4;m-32&1|-|198859108|198859130&offsets|0|-4];[MIMAT0000256_1&hsa-miR-181a-5p&offsets|0|-4;m-32&9|+|124692480|124692502&offsets|0|-4]|AACATTCAACGCTGTCGGT</td><td>>MI0000289|hsa-mir-181a-1&WithFlank&1|-|198859038|198859159</td><td>30</td><td>48</td><td>19</td><td>AACATTCAACGCTGTCGGT</td><td>111.5</td><td>2.4</td><td>0.9</td><td>2.7</td><td>7.2E-03</td><td>9.4E-03</td><td>MB231/MB468</td><td>MP</td></tr>
<tr><td>>MI0000086|hsa-mir-28&WithFlank&3|+|188688775|188688872@61.80.20&[MIMAT0004502&hsa-miR-28-3p&offsets|+1|-1;m-28&3|+|188688834|188688855&offsets|+1|-1]|ACTAGATTGTGAGCTCCTGG</td><td>>MI0000086|hsa-mir-28&WithFlank&3|+|188688775|188688872</td><td>61</td><td>80</td><td>20</td><td>ACTAGATTGTGAGCTCCTGG</td><td>54.4</td><td>5.1</td><td>1.9</td><td>2.7</td><td>7.2E-03</td><td>9.4E-03</td><td>MB231/MB468</td><td>MP</td></tr>
<tr><td>>hg38_21_+_8433172_8446622_RNA45SN1_45S_50ntFlanks@11665.11687.23|TTCTGCCCAGTGCTCTGAATGTC</td><td>>hg38_21_+_8433172_8446622_RNA45SN1_45S_50ntFlanks</td><td>11665</td><td>11687</td><td>23</td><td>TTCTGCCCAGTGCTCTGAATGTC</td><td>93.0</td><td>3.8</td><td>1.3</td><td>2.8</td><td>4.7E-03</td><td>9.4E-03</td><td>BT20/MB468</td><td>MP</td></tr>
<tr><td>>hg38_21_+_8433172_8446622_RNA45SN1_45S_50ntFlanks@6731.6758.28|CACATTGATCATCGACACTTCGAACGCA</td><td>>hg38_21_+_8433172_8446622_RNA45SN1_45S_50ntFlanks</td><td>6731</td><td>6758</td><td>28</td><td>CACATTGATCATCGACACTTCGAACGCA</td><td>147.5</td><td>-4.7</td><td>1.6</td><td>-2.8</td><td>4.7E-03</td><td>9.4E-03</td><td>BT20/MB468</td><td>MP</td></tr>
<tr><td>>am_MI0000815|hsa-mir-339&WithFlank&7|-|1022927|1023032@21.39.19(+1U)&[MIMAT0000764&hsa-miR-339-5p&offsets|0|-4(+1U);m-163&7|-|1022990|1023012&offsets|0|-4(+1U)]|TCCCTGTCCTCCAGGAGCTT</td><td>>am_MI0000815|hsa-mir-339&WithFlank&7|-|1022927|1023032</td><td>21</td><td>39</td><td>19(+1U)</td><td>TCCCTGTCCTCCAGGAGCTT</td><td>37.2</td><td>5.3</td><td>2.0</td><td>2.7</td><td>7.2E-03</td><td>9.4E-03</td><td>MB231/MB468</td><td>MP</td></tr>
<tr><td>>trna10_GlyTCC_17_+_8124866_8124937@31.53.23,trna117_GlyTCC_1_-_145397864_145397935@31.53.23,trna2_GlyTCC_19_+_4724082_4724153@31.53.23,trna45_GlyTCC_1_+_161500903_161500974@31.53.23,trna70_GlyTCC_1_-_161439547_161439618@31.53.23,trna73_GlyTCC_1_-_161432166_161432237@31.53.23,trna76_GlyTCC_1_-_161424756_161424827@31.53.23,trna79_GlyTCC_1_-_161417375_161417446@31.53.23|CCTTCCAAGCAGTTGACCCGGGT</td><td>>trna10_GlyTCC_17_+_8124866_8124937</td><td>31</td><td>53</td><td>23</td><td>CCTTCCAAGCAGTTGACCCGGGT</td><td>83.8</td><td>-3.5</td><td>1.3</td><td>-2.7</td><td>7.2E-03</td><td>9.4E-03</td><td>MB231/MB468</td><td>MP</td></tr>
<tr><td>>hg38_21_+_8433172_8446622_RNA45SN1_45S_50ntFlanks@10391.10433.43|AACAGCAGTTGAACATGGGTCAGTCGGTCCTGAGAGATGGGCG</td><td>>hg38_21_+_8433172_8446622_RNA45SN1_45S_50ntFlanks</td><td>10391</td><td>10433</td><td>43</td><td>AACAGCAGTTGAACATGGGTCAGTCGGTCCTGAGAGATGGGCG</td><td>347.0</td><td>-2.8</td><td>1.0</td><td>-2.7</td><td>6.7E-03</td><td>9.4E-03</td><td>BT20/MB468</td><td>MITO</td></tr>
<tr><td>>trna111_HisGTG_1_-_147774845_147774916@-1T.33.34,trna118_HisGTG_1_-_145396881_145396952@-1T.33.34,trna16_HisGTG_1_+_146544773_146544844@-1T.33.34,trna1_HisGTG_15_+_45493349_45493420@-1T.33.34,trna21_HisGTG_1_+_147753471_147753542@-1T.33.34,trna33_HisGTG_6_+_27125906_27125977@-1T.33.34,trna7_HisGTG_9_-_14433938_14434009@-1T.33.34,trna8_HisGTG_15_-_45492611_45492682@-1T.33.34,trna9_HisGTG_15_-_45490804_45490875@-1T.33.34|TGCCGTGATCGTATAGTGGTTAGTACTCTGCGTT</td><td>>trna111_HisGTG_1_-_147774845_147774916</td><td>-1T</td><td>33</td><td>34</td><td>TGCCGTGATCGTATAGTGGTTAGTACTCTGCGTT</td><td>130.0</td><td>2.0</td><td>0.7</td><td>2.6</td><td>8.2E-03</td><td>9.4E-03</td><td>BT20/MB468</td><td>NUC</td></tr>
<tr><td>>hg38_21_+_8433172_8446622_RNA45SN1_45S_50ntFlanks@11322.11347.26|ACGTCTCGTCGCGCGCGCGTCCGCTG</td><td>>hg38_21_+_8433172_8446622_RNA45SN1_45S_50ntFlanks</td><td>11322</td><td>11347</td><td>26</td><td>ACGTCTCGTCGCGCGCGCGTCCGCTG</td><td>2432.1</td><td>8.6</td><td>2.8</td><td>3.1</td><td>2.3E-03</td><td>9.4E-03</td><td>BT20/MB231</td><td>MITO</td></tr>
<tr><td>>hg38_21_+_8433172_8446622_RNA45SN1_45S_50ntFlanks@9301.9323.23|CGGACCAAGGAGTCTAACACGTG</td><td>>hg38_21_+_8433172_8446622_RNA45SN1_45S_50ntFlanks</td><td>9301</td><td>9323</td><td>23</td><td>CGGACCAAGGAGTCTAACACGTG</td><td>55.8</td><td>9.1</td><td>2.9</td><td>3.1</td><td>1.7E-03</td><td>9.5E-03</td><td>BT20/MB231</td><td>MP</td></tr>
<tr><td>>MI0000081|hsa-mir-24-2&WithFlank&19|-|13836281|13836365@18.41.24&[MIMAT0004497&hsa-miR-24-2-5p&offsets|-1|+1;m-226&19|-|13836326|13836348&offsets|0|+1]|GTGCCTACTGAGCTGAAACACAGT</td><td>>MI0000081|hsa-mir-24-2&WithFlank&19|-|13836281|13836365</td><td>18</td><td>41</td><td>24</td><td>GTGCCTACTGAGCTGAAACACAGT</td><td>294.7</td><td>-3.3</td><td>1.1</td><td>-3.1</td><td>1.7E-03</td><td>9.5E-03</td><td>BT20/MB231</td><td>MP</td></tr>
<tr><td>>trnaMT_GluTTC_MT_-_14674_14742@41.65.25,trnalookalike8_GluTTC_5_-_93905172_93905240@41.65.25|TGGTCGTGGTTGTAGTCCGTGCGAG</td><td>>trnaMT_GluTTC_MT_-_14674_14742</td><td>41</td><td>65</td><td>25</td><td>TGGTCGTGGTTGTAGTCCGTGCGAG</td><td>103.3</td><td>4.3</td><td>1.5</td><td>2.8</td><td>4.7E-03</td><td>9.5E-03</td><td>BT20/MB468</td><td>MP</td></tr>
<tr><td>>MI0000101|hsa-mir-99a&WithFlank&21|+|16539083|16539175@19.40.22&[MIMAT0000097&hsa-miR-99a-5p&offsets|0|0;m-64&21|+|16539101|16539121&offsets|0|+1]|AACCCGTAGATCCGATCTTGTG</td><td>>MI0000101|hsa-mir-99a&WithFlank&21|+|16539083|16539175</td><td>19</td><td>40</td><td>22</td><td>AACCCGTAGATCCGATCTTGTG</td><td>921.7</td><td>-2.7</td><td>0.9</td><td>-3.1</td><td>2.2E-03</td><td>9.5E-03</td><td>BT20/MB231</td><td>CYTO</td></tr>
<tr><td>>MI0000087|hsa-mir-29a&WithFlank&7|-|130876741|130876816@48.67.20(+1C)&[MIMAT0000086&hsa-miR-29a-3p&offsets|0|-2(+1C);m-26&7|-|130876749|130876769&offsets|0|-1(+1C)]|TAGCACCATCTGAAATCGGTC</td><td>>MI0000087|hsa-mir-29a&WithFlank&7|-|130876741|130876816</td><td>48</td><td>67</td><td>20(+1C)</td><td>TAGCACCATCTGAAATCGGTC</td><td>2189.4</td><td>-2.8</td><td>0.9</td><td>-3.1</td><td>2.3E-03</td><td>9.5E-03</td><td>BT20/MB231</td><td>MITO</td></tr>
<tr><td>>MI0000086|hsa-mir-28&WithFlank&3|+|188688775|188688872@60.81.22(+1A)&[MIMAT0004502&hsa-miR-28-3p&offsets|0|0(+1A);m-28&3|+|188688834|188688855&offsets|0|0(+1A)]|CACTAGATTGTGAGCTCCTGGAA</td><td>>MI0000086|hsa-mir-28&WithFlank&3|+|188688775|188688872</td><td>60</td><td>81</td><td>22(+1A)</td><td>CACTAGATTGTGAGCTCCTGGAA</td><td>4532.8</td><td>2.6</td><td>0.9</td><td>3.1</td><td>2.2E-03</td><td>9.5E-03</td><td>BT20/MB231</td><td>CYTO</td></tr>
<tr><td>>hg38_21_+_8433172_8446622_RNA45SN1_45S_50ntFlanks@6646.6688.43|TCGTACGACTCTTAGCGGTGGATCACTCGGCTCGTGCGTCGAT</td><td>>hg38_21_+_8433172_8446622_RNA45SN1_45S_50ntFlanks</td><td>6646</td><td>6688</td><td>43</td><td>TCGTACGACTCTTAGCGGTGGATCACTCGGCTCGTGCGTCGAT</td><td>1931.0</td><td>2.4</td><td>0.9</td><td>2.7</td><td>6.7E-03</td><td>9.5E-03</td><td>BT20/MB468</td><td>MITO</td></tr>
<tr><td>>hg38_21_+_8433172_8446622_RNA45SN1_45S_50ntFlanks@11452.11469.18|CTCCTCGGCGCGCGGCGG</td><td>>hg38_21_+_8433172_8446622_RNA45SN1_45S_50ntFlanks</td><td>11452</td><td>11469</td><td>18</td><td>CTCCTCGGCGCGCGGCGG</td><td>584.1</td><td>-5.4</td><td>2.0</td><td>-2.6</td><td>8.2E-03</td><td>9.5E-03</td><td>BT20/MB468</td><td>NUC</td></tr>
<tr><td>>hg38_21_+_8433172_8446622_RNA45SN1_45S_50ntFlanks@5000.5024.25|TTGATAGCTCTTTCTCGATTCCGTG</td><td>>hg38_21_+_8433172_8446622_RNA45SN1_45S_50ntFlanks</td><td>5000</td><td>5024</td><td>25</td><td>TTGATAGCTCTTTCTCGATTCCGTG</td><td>57.0</td><td>-4.6</td><td>1.6</td><td>-2.8</td><td>4.7E-03</td><td>9.5E-03</td><td>BT20/MB468</td><td>MP</td></tr>
<tr><td>>hg38_21_+_8433172_8446622_RNA45SN1_45S_50ntFlanks@8171.8215.45|AAGTCCTTCTGATCGAGGCCCAGCCCGTGGACGGTGTGAGGCCGG</td><td>>hg38_21_+_8433172_8446622_RNA45SN1_45S_50ntFlanks</td><td>8171</td><td>8215</td><td>45</td><td>AAGTCCTTCTGATCGAGGCCCAGCCCGTGGACGGTGTGAGGCCGG</td><td>336.5</td><td>-3.7</td><td>1.4</td><td>-2.7</td><td>7.8E-03</td><td>9.5E-03</td><td>BT20/MB468</td><td>CYTO</td></tr>
<tr><td>>MI0000270|hsa-mir-181b-1&WithFlank&1|-|198858867|198858988@42.65.24(+1A);MI0000683|hsa-mir-181b-2&WithFlank&9|+|124693704|124693804@22.45.24(+1A)&[MIMAT0000257&hsa-miR-181b-5p&offsets|0|+1(+1A);m-73&1|-|198858925|198858947&offsets|0|+1(+1A)];[MIMAT0000257_1&hsa-miR-181b-5p&offsets|0|+1(+1A);m-72&9|+|124693725|124693747&offsets|0|+1(+1A)]|AACATTCATTGCTGTCGGTGGGTTA</td><td>>MI0000270|hsa-mir-181b-1&WithFlank&1|-|198858867|198858988</td><td>42</td><td>65</td><td>24(+1A)</td><td>AACATTCATTGCTGTCGGTGGGTTA</td><td>64.7</td><td>4.4</td><td>1.6</td><td>2.7</td><td>7.2E-03</td><td>9.5E-03</td><td>MB231/MB468</td><td>MP</td></tr>
<tr><td>>hg38_21_+_8433172_8446622_RNA45SN1_45S_50ntFlanks@8176.8218.43|CTTCTGATCGAGGCCCAGCCCGTGGACGGTGTGAGGCCGGTAG</td><td>>hg38_21_+_8433172_8446622_RNA45SN1_45S_50ntFlanks</td><td>8176</td><td>8218</td><td>43</td><td>CTTCTGATCGAGGCCCAGCCCGTGGACGGTGTGAGGCCGGTAG</td><td>166.6</td><td>-4.2</td><td>1.5</td><td>-2.8</td><td>4.7E-03</td><td>9.5E-03</td><td>BT20/MB468</td><td>MP</td></tr>
<tr><td>>MI0005566|hsa-mir-374b&WithFlank&X|-|74218541|74218624@17.38.22(+1C)&[MIMAT0004955&hsa-miR-374b-5p&offsets|0|0(+1C);m-153&X|-|74218587|74218608&offsets|0|0(+1C)]|ATATAATACAACCTGCTAAGTGC</td><td>>MI0005566|hsa-mir-374b&WithFlank&X|-|74218541|74218624</td><td>17</td><td>38</td><td>22(+1C)</td><td>ATATAATACAACCTGCTAAGTGC</td><td>73.1</td><td>5.2</td><td>2.0</td><td>2.7</td><td>7.2E-03</td><td>9.5E-03</td><td>MB231/MB468</td><td>MP</td></tr>
<tr><td>>hg38_21_+_8433172_8446622_RNA45SN1_45S_50ntFlanks@6646.6681.36|TCGTACGACTCTTAGCGGTGGATCACTCGGCTCGTG</td><td>>hg38_21_+_8433172_8446622_RNA45SN1_45S_50ntFlanks</td><td>6646</td><td>6681</td><td>36</td><td>TCGTACGACTCTTAGCGGTGGATCACTCGGCTCGTG</td><td>21795.7</td><td>-1.9</td><td>0.6</td><td>-3.1</td><td>1.9E-03</td><td>9.5E-03</td><td>BT20/MB231</td><td>NUC</td></tr>
<tr><td>>hg38_21_+_8433172_8446622_RNA45SN1_45S_50ntFlanks@10352.10377.26|GAGAACTTTGAAGGCCGAAGTGGAGA</td><td>>hg38_21_+_8433172_8446622_RNA45SN1_45S_50ntFlanks</td><td>10352</td><td>10377</td><td>26</td><td>GAGAACTTTGAAGGCCGAAGTGGAGA</td><td>124.7</td><td>-3.6</td><td>1.3</td><td>-2.7</td><td>7.2E-03</td><td>9.5E-03</td><td>MB231/MB468</td><td>MP</td></tr>
<tr><td>>hg38_21_+_8433172_8446622_RNA45SN1_45S_50ntFlanks@12997.13035.39|CGCTGCGATCTATTGAAAGTCAGCCCTCGACACAAGGGT</td><td>>hg38_21_+_8433172_8446622_RNA45SN1_45S_50ntFlanks</td><td>12997</td><td>13035</td><td>39</td><td>CGCTGCGATCTATTGAAAGTCAGCCCTCGACACAAGGGT</td><td>178.1</td><td>4.2</td><td>1.6</td><td>2.6</td><td>8.4E-03</td><td>9.5E-03</td><td>MB231/MB468</td><td>MITO</td></tr>
<tr><td>>trnaMT_GlnTTG_MT_-_4329_4400@1.34.34,trnalookalike2_GlnTTG_1_-_564879_564950@1.34.34|TAGGATGGGGTGTGATAGGTGGCACGGAGAATTT</td><td>>trnaMT_GlnTTG_MT_-_4329_4400</td><td>1</td><td>34</td><td>34</td><td>TAGGATGGGGTGTGATAGGTGGCACGGAGAATTT</td><td>203.5</td><td>3.3</td><td>1.2</td><td>2.7</td><td>7.2E-03</td><td>9.5E-03</td><td>MB231/MB468</td><td>MP</td></tr>
<tr><td>>hg38_21_+_8433172_8446622_RNA45SN1_45S_50ntFlanks@10792.10817.26|CTCTGGCATGTTGGAACAATGTAGGT</td><td>>hg38_21_+_8433172_8446622_RNA45SN1_45S_50ntFlanks</td><td>10792</td><td>10817</td><td>26</td><td>CTCTGGCATGTTGGAACAATGTAGGT</td><td>123.7</td><td>-2.9</td><td>1.0</td><td>-2.8</td><td>4.7E-03</td><td>9.5E-03</td><td>BT20/MB468</td><td>MP</td></tr>
<tr><td>>hg38_1_-_228634819_228635039_RNA5S12_5S_50ntFlanks@146.170.25|TGGGAATACCGGGTGCTGTAGGCTT</td><td>>hg38_1_-_228634819_228635039_RNA5S12_5S_50ntFlanks</td><td>146</td><td>170</td><td>25</td><td>TGGGAATACCGGGTGCTGTAGGCTT</td><td>9946.9</td><td>4.1</td><td>1.3</td><td>3.1</td><td>1.8E-03</td><td>9.5E-03</td><td>BT20/MB231</td><td>MP</td></tr>
<tr><td>>am_trnaMT_PheGAA_MT_+_577_647@43.62.20|TTAGACGGGCTCACATCACC</td><td>>am_trnaMT_PheGAA_MT_+_577_647</td><td>43</td><td>62</td><td>20</td><td>TTAGACGGGCTCACATCACC</td><td>60.5</td><td>2.1</td><td>0.8</td><td>2.7</td><td>7.2E-03</td><td>9.5E-03</td><td>MB231/MB468</td><td>MP</td></tr>
<tr><td>>hg38_21_+_8433172_8446622_RNA45SN1_45S_50ntFlanks@6678.6696.19|CGTGCGTCGATGAAGAACG</td><td>>hg38_21_+_8433172_8446622_RNA45SN1_45S_50ntFlanks</td><td>6678</td><td>6696</td><td>19</td><td>CGTGCGTCGATGAAGAACG</td><td>428.1</td><td>4.8</td><td>1.5</td><td>3.1</td><td>1.8E-03</td><td>9.5E-03</td><td>BT20/MB231</td><td>MP</td></tr>
<tr><td>>hg38_21_+_8433172_8446622_RNA45SN1_45S_50ntFlanks@12606.12626.21|ATGTGCTTGGCTGAGGAGCCA</td><td>>hg38_21_+_8433172_8446622_RNA45SN1_45S_50ntFlanks</td><td>12606</td><td>12626</td><td>21</td><td>ATGTGCTTGGCTGAGGAGCCA</td><td>322.6</td><td>-3.0</td><td>1.1</td><td>-2.7</td><td>7.9E-03</td><td>9.5E-03</td><td>BT20/MB468</td><td>CYTO</td></tr>
</table>

<table>
<tr><td>>hg38_21_+_8433172_8446622_RNA45SN1_45S_50ntFlanks@4006.4023.18|ACCTCGGGCCGATCGCAC</td><td>>hg38_21_+_8433172_8446622_RNA45SN1_45S_50ntFlanks</td><td>4006</td><td>4023</td><td>18</td><td>ACCTCGGGCCGATCGCAC</td><td>465.0</td><td>-3.3</td><td>1.3</td><td>-2.6</td><td>8.4E-03</td><td>9.5E-03</td><td>MB231/MB468</td><td>MITO</td></tr>
<tr><td>>hg38_21_+_8433172_8446622_RNA45SN1_45S_50ntFlanks@7982.8005.24|TCAGATCAGACGTGGCGACCCGCT</td><td>>hg38_21_+_8433172_8446622_RNA45SN1_45S_50ntFlanks</td><td>7982</td><td>8005</td><td>24</td><td>TCAGATCAGACGTGGCGACCCGCT</td><td>112.6</td><td>5.8</td><td>1.8</td><td>3.1</td><td>1.8E-03</td><td>9.5E-03</td><td>BT20/MB231</td><td>MP</td></tr>
<tr><td>>hg38_21_+_8433172_8446622_RNA45SN1_45S_50ntFlanks@9593.9610.18|GTATAGGGGCGAAAGACT</td><td>>hg38_21_+_8433172_8446622_RNA45SN1_45S_50ntFlanks</td><td>9593</td><td>9610</td><td>18</td><td>GTATAGGGGCGAAAGACT</td><td>81.2</td><td>5.7</td><td>2.0</td><td>2.8</td><td>4.7E-03</td><td>9.5E-03</td><td>BT20/MB468</td><td>MP</td></tr>
<tr><td>>hg38_21_+_8433172_8446622_RNA45SN1_45S_50ntFlanks@10866.10891.26|CTAAGGGCTGGGTCGGTCGGGCTGGG</td><td>>hg38_21_+_8433172_8446622_RNA45SN1_45S_50ntFlanks</td><td>10866</td><td>10891</td><td>26</td><td>CTAAGGGCTGGGTCGGTCGGGCTGGG</td><td>576.2</td><td>-4.4</td><td>1.6</td><td>-2.7</td><td>7.9E-03</td><td>9.5E-03</td><td>BT20/MB468</td><td>CYTO</td></tr>
<tr><td>>MI0000299|hsa-mir-222&WithFlank&X|-|45747009|45747130@75.94.20(+1A)&[MIMAT0000279&hsa-miR-222-3p&offsets|0|-1(+1A);m-140&X|-|45747033|45747056&offsets|0|-4(+1A)]|AGCTACATCTGGCTACTGGGA</td><td>>MI0000299|hsa-mir-222&WithFlank&X|-|45747009|45747130</td><td>75</td><td>94</td><td>20(+1A)</td><td>AGCTACATCTGGCTACTGGGA</td><td>277.3</td><td>4.2</td><td>1.5</td><td>2.8</td><td>4.7E-03</td><td>9.5E-03</td><td>BT20/MB468</td><td>MP</td></tr>
<tr><td>>hg38_21_+_8433172_8446622_RNA45SN1_45S_50ntFlanks@9493.9515.23|CCCGAAAGATGGTGAACTATGCC</td><td>>hg38_21_+_8433172_8446622_RNA45SN1_45S_50ntFlanks</td><td>9493</td><td>9515</td><td>23</td><td>CCCGAAAGATGGTGAACTATGCC</td><td>138.0</td><td>5.7</td><td>2.0</td><td>2.8</td><td>4.7E-03</td><td>9.5E-03</td><td>BT20/MB468</td><td>MP</td></tr>
<tr><td>>hg38_21_+_8433172_8446622_RNA45SN1_45S_50ntFlanks@12394.12419.26|TAGCGACGTCGCTTTTTGATCCTTCG</td><td>>hg38_21_+_8433172_8446622_RNA45SN1_45S_50ntFlanks</td><td>12394</td><td>12419</td><td>26</td><td>TAGCGACGTCGCTTTTTGATCCTTCG</td><td>776.7</td><td>-1.9</td><td>0.7</td><td>-2.7</td><td>7.9E-03</td><td>9.6E-03</td><td>MB231/MB468</td><td>CYTO</td></tr>
<tr><td>>hg38_21_+_8433172_8446622_RNA45SN1_45S_50ntFlanks@3785.3810.26|TGAAACTGCGAATGGCTCATTAAATC</td><td>>hg38_21_+_8433172_8446622_RNA45SN1_45S_50ntFlanks</td><td>3785</td><td>3810</td><td>26</td><td>TGAAACTGCGAATGGCTCATTAAATC</td><td>298.4</td><td>-1.6</td><td>0.6</td><td>-2.7</td><td>8.0E-03</td><td>9.6E-03</td><td>MB231/MB468</td><td>CYTO</td></tr>
<tr><td>>hg38_21_+_8433172_8446622_RNA45SN1_45S_50ntFlanks@9027.9064.38|CGTCTCCTCGTGGGGGGGCCGGGCCACCCCTCCCACGG</td><td>>hg38_21_+_8433172_8446622_RNA45SN1_45S_50ntFlanks</td><td>9027</td><td>9064</td><td>38</td><td>CGTCTCCTCGTGGGGGGGCCGGGCCACCCCTCCCACGG</td><td>286.3</td><td>-4.8</td><td>1.7</td><td>-2.8</td><td>4.7E-03</td><td>9.6E-03</td><td>BT20/MB468</td><td>MP</td></tr>
<tr><td>>hg38_21_+_8433172_8446622_RNA45SN1_45S_50ntFlanks@10318.10340.23|TGCAGATCTTGGTGGTAGTAGCA</td><td>>hg38_21_+_8433172_8446622_RNA45SN1_45S_50ntFlanks</td><td>10318</td><td>10340</td><td>23</td><td>TGCAGATCTTGGTGGTAGTAGCA</td><td>313.1</td><td>4.3</td><td>1.4</td><td>3.1</td><td>1.8E-03</td><td>9.6E-03</td><td>BT20/MB231</td><td>MP</td></tr>
<tr><td>>MI0000811|hsa-mir-148b&WithFlank&12|+|54337210|54337320@30.52.23&[MIMAT0004699&hsa-miR-148b-5p&offsets|-1|0;m-371&12|+|54337239|54337262&offsets|0|-1]|GAAGTTCTGTTATACACTCAGGC</td><td>>MI0000811|hsa-mir-148b&WithFlank&12|+|54337210|54337320</td><td>30</td><td>52</td><td>23</td><td>GAAGTTCTGTTATACACTCAGGC</td><td>180.2</td><td>-3.2</td><td>1.0</td><td>-3.1</td><td>1.8E-03</td><td>9.6E-03</td><td>BT20/MB231</td><td>MP</td></tr>
<tr><td>>am_MI0003639|hsa-mir-625&WithFlank&14|+|65471096|65471192@58.78.21(+1U)&[MIMAT0004808&hsa-miR-625-3p&offsets|0|-1(+1U);m-137&14|+|65471153|65471174&offsets|0|-1(+1U)]|GACTATAGAACTTTCCCCCTCT</td><td>>am_MI0003639|hsa-mir-625&WithFlank&14|+|65471096|65471192</td><td>58</td><td>78</td><td>21(+1U)</td><td>GACTATAGAACTTTCCCCCTCT</td><td>98.1</td><td>5.1</td><td>1.8</td><td>2.8</td><td>4.7E-03</td><td>9.6E-03</td><td>BT20/MB468</td><td>MP</td></tr>
<tr><td>>MI0000253|hsa-mir-148a&WithFlank&7|-|25949913|25949992@12.31.20&[MIMAT0004549&hsa-miR-148a-5p&offsets|0|-2;m-189&7|-|25949960|25949981&offsets|0|-2]|AAAGTTCTGAGACACTCCGA</td><td>>MI0000253|hsa-mir-148a&WithFlank&7|-|25949913|25949992</td><td>12</td><td>31</td><td>20</td><td>AAAGTTCTGAGACACTCCGA</td><td>81.4</td><td>5.7</td><td>1.8</td><td>3.1</td><td>1.8E-03</td><td>9.6E-03</td><td>BT20/MB231</td><td>MP</td></tr>
<tr><td>>hg38_21_+_8433172_8446622_RNA45SN1_45S_50ntFlanks@12419.12448.30|GATGTCGGCTCTTCCTATCATTGTGAAGCA</td><td>>hg38_21_+_8433172_8446622_RNA45SN1_45S_50ntFlanks</td><td>12419</td><td>12448</td><td>30</td><td>GATGTCGGCTCTTCCTATCATTGTGAAGCA</td><td>158.8</td><td>4.0</td><td>1.4</td><td>2.8</td><td>4.7E-03</td><td>9.6E-03</td><td>BT20/MB468</td><td>MP</td></tr>
<tr><td>>am_MI0005763|hsa-mir-941-1&WithFlank&20|+|63919443|63919526@53.75.23(+1U);MI0005764|hsa-mir-941-2&WithFlank&20|+|63919499|63919582@53.75.23(+1U);MI0005765|hsa-mir-941-3&WithFlank&20|+|63919555|63919638@53.75.23(+1U);MI0005766|hsa-mir-941-4&WithFlank&20|+|63919750|63919833@53.75.23(+1U);MI0031520|hsa-mir-941-5&WithFlank&20|+|63919862|63919945@53.75.23(+1U)&[MIMAT0004984&hsa-miR-941&offsets|0|0(+1U);m-263&20|+|63919495|63919517&offsets|0|0(+1U)];[MIMAT0004984_1&hsa-miR-941&offsets|0|0(+1U);m-263&20|+|63919551|63919573&offsets|0|0(+1U)];[MIMAT0004984_2&hsa-miR-941&offsets|0|0(+1U);m-263&20|+|63919607|63919629&offsets|0|0(+1U)];[MIMAT0004984_3&hsa-miR-941&offsets|0|0(+1U);m-263&20|+|63919802|63919824&offsets|0|0(+1U)];[MIMAT0004984_4&hsa-miR-941&offsets|0|0(+1U);m-263&20|+|63919914|63919936&offsets|0|0(+1U)]|CACCCGGCTGTGTGCACATGTGCT</td><td>>am_MI0005763|hsa-mir-941-1&WithFlank&20|+|63919443|63919526</td><td>53</td><td>75</td><td>23(+1U)</td><td>CACCCGGCTGTGTGCACATGTGCT</td><td>78.6</td><td>5.1</td><td>1.9</td><td>2.7</td><td>7.3E-03</td><td>9.6E-03</td><td>MB231/MB468</td><td>MP</td></tr>
<tr><td>>hg38_21_+_8433172_8446622_RNA45SN1_45S_50ntFlanks@12219.12241.23|AGGACAGAAACCTCCCGTGGAGC</td><td>>hg38_21_+_8433172_8446622_RNA45SN1_45S_50ntFlanks</td><td>12219</td><td>12241</td><td>23</td><td>AGGACAGAAACCTCCCGTGGAGC</td><td>170.0</td><td>3.0</td><td>1.0</td><td>2.8</td><td>4.7E-03</td><td>9.6E-03</td><td>BT20/MB468</td><td>MP</td></tr>
<tr><td>>hg38_21_+_8433172_8446622_RNA45SN1_45S_50ntFlanks@11321.11344.24|CACGTCTCGTCGCGCGCGCGTCCG</td><td>>hg38_21_+_8433172_8446622_RNA45SN1_45S_50ntFlanks</td><td>11321</td><td>11344</td><td>24</td><td>CACGTCTCGTCGCGCGCGCGTCCG</td><td>1270.5</td><td>-5.2</td><td>1.8</td><td>-2.8</td><td>4.7E-03</td><td>9.6E-03</td><td>BT20/MB468</td><td>MP</td></tr>
<tr><td>>hg38_21_+_8433172_8446622_RNA45SN1_45S_50ntFlanks@12436.12454.19|TCATTGTGAAGCAGAATTC</td><td>>hg38_21_+_8433172_8446622_RNA45SN1_45S_50ntFlanks</td><td>12436</td><td>12454</td><td>19</td><td>TCATTGTGAAGCAGAATTC</td><td>171.9</td><td>-4.0</td><td>1.5</td><td>-2.7</td><td>6.8E-03</td><td>9.6E-03</td><td>BT20/MB468</td><td>MITO</td></tr>
<tr><td>>hg38_MT_+_598_1651_RNR1_12S_50ntFlanks@443.469.27|TATCTGAACACACAATAGCTAAGACCC</td><td>>hg38_MT_+_598_1651_RNR1_12S_50ntFlanks</td><td>443</td><td>469</td><td>27</td><td>TATCTGAACACACAATAGCTAAGACCC</td><td>179.8</td><td>4.8</td><td>1.7</td><td>2.8</td><td>4.7E-03</td><td>9.6E-03</td><td>BT20/MB468</td><td>MP</td></tr>
<tr><td>>hg38_21_+_8433172_8446622_RNA45SN1_45S_50ntFlanks@12081.12109.29|CTCTCGCTTCTGGCGCCAAGCGCCCGGCC</td><td>>hg38_21_+_8433172_8446622_RNA45SN1_45S_50ntFlanks</td><td>12081</td><td>12109</td><td>29</td><td>CTCTCGCTTCTGGCGCCAAGCGCCCGGCC</td><td>81.1</td><td>-4.6</td><td>1.6</td><td>-2.8</td><td>4.7E-03</td><td>9.6E-03</td><td>BT20/MB468</td><td>MP</td></tr>
<tr><td>>MI0000088|hsa-mir-30a&WithFlank&6|-|71403545|71403627@13.34.22(+1A)&[MIMAT0000087&hsa-miR-30a-5p&offsets|+1|+1(+1A);m-15&6|-|71403595|71403616&offsets|+1|+1(+1A)]|GTAAACATCCTCGACTGGAAGCA</td><td>>MI0000088|hsa-mir-30a&WithFlank&6|-|71403545|71403627</td><td>13</td><td>34</td><td>22(+1A)</td><td>GTAAACATCCTCGACTGGAAGCA</td><td>249.2</td><td>4.5</td><td>1.6</td><td>2.8</td><td>4.8E-03</td><td>9.6E-03</td><td>BT20/MB468</td><td>MP</td></tr>
<tr><td>>hg38_21_+_8433172_8446622_RNA45SN1_45S_50ntFlanks@6650.6676.27|ACGACTCTTAGCGGTGGATCACTCGGC</td><td>>hg38_21_+_8433172_8446622_RNA45SN1_45S_50ntFlanks</td><td>6650</td><td>6676</td><td>27</td><td>ACGACTCTTAGCGGTGGATCACTCGGC</td><td>2533.0</td><td>3.5</td><td>1.2</td><td>3.1</td><td>2.2E-03</td><td>9.6E-03</td><td>BT20/MB231</td><td>CYTO</td></tr>
<tr><td>>hg38_21_+_8433172_8446622_RNA45SN1_45S_50ntFlanks@11569.11592.24|CAGCCGACTTAGAACTGGTGCGGA</td><td>>hg38_21_+_8433172_8446622_RNA45SN1_45S_50ntFlanks</td><td>11569</td><td>11592</td><td>24</td><td>CAGCCGACTTAGAACTGGTGCGGA</td><td>308.1</td><td>-2.0</td><td>0.7</td><td>-2.7</td><td>6.8E-03</td><td>9.6E-03</td><td>BT20/MB468</td><td>MITO</td></tr>
<tr><td>>hg38_21_+_8433172_8446622_RNA45SN1_45S_50ntFlanks@7988.8010.23|CAGACGTGGCGACCCGCTGAATT</td><td>>hg38_21_+_8433172_8446622_RNA45SN1_45S_50ntFlanks</td><td>7988</td><td>8010</td><td>23</td><td>CAGACGTGGCGACCCGCTGAATT</td><td>520.8</td><td>2.7</td><td>0.9</td><td>3.1</td><td>1.9E-03</td><td>9.6E-03</td><td>BT20/MB231</td><td>NUC</td></tr>
<tr><td>>hg38_21_+_8433172_8446622_RNA45SN1_45S_50ntFlanks@12935.12955.21|TAAACCATTCGTAGACGACCT</td><td>>hg38_21_+_8433172_8446622_RNA45SN1_45S_50ntFlanks</td><td>12935</td><td>12955</td><td>21</td><td>TAAACCATTCGTAGACGACCT</td><td>178.0</td><td>-3.9</td><td>1.5</td><td>-2.7</td><td>7.9E-03</td><td>9.6E-03</td><td>BT20/MB468</td><td>CYTO</td></tr>
<tr><td>>hg38_21_+_8433172_8446622_RNA45SN1_45S_50ntFlanks@3705.3747.43|TACCTGGTTGATCCTGCCAGTAGCATATGCTTGTCTCAAAGAT</td><td>>hg38_21_+_8433172_8446622_RNA45SN1_45S_50ntFlanks</td><td>3705</td><td>3747</td><td>43</td><td>TACCTGGTTGATCCTGCCAGTAGCATATGCTTGTCTCAAAGAT</td><td>233.7</td><td>3.5</td><td>1.3</td><td>2.6</td><td>8.4E-03</td><td>9.6E-03</td><td>MB231/MB468</td><td>MITO</td></tr>
<tr><td>>MI0006444|hsa-mir-1307&WithFlank&10|-|103394247|103394407@86.107.22&[MIMAT0005951&hsa-miR-1307-3p&offsets|0|0;m-66&10|-|103394300|103394322&offsets|0|-1]|ACTCGGCGTGGCGTCGGTCGTG</td><td>>MI0006444|hsa-mir-1307&WithFlank&10|-|103394247|103394407</td><td>86</td><td>107</td><td>22</td><td>ACTCGGCGTGGCGTCGGTCGTG</td><td>351.0</td><td>2.4</td><td>0.9</td><td>2.7</td><td>7.3E-03</td><td>9.6E-03</td><td>MB231/MB468</td><td>MP</td></tr>
<tr><td>>hg38_21_+_8433172_8446622_RNA45SN1_45S_50ntFlanks@12926.12953.28|ACCTGGCGCTAAACCATTCGTAGACGAC</td><td>>hg38_21_+_8433172_8446622_RNA45SN1_45S_50ntFlanks</td><td>12926</td><td>12953</td><td>28</td><td>ACCTGGCGCTAAACCATTCGTAGACGAC</td><td>187.7</td><td>-1.5</td><td>0.6</td><td>-2.7</td><td>6.8E-03</td><td>9.6E-03</td><td>BT20/MB468</td><td>MITO</td></tr>
<tr><td>>am_MI0003581|hsa-mir-574&WithFlank&4|+|38868026|38868133@31.54.24(+1C)&[MIMAT0004795&hsa-miR-574-5p&offsets|0|+1(+1C);m-213&4|+|38868056|38868078&offsets|0|+1(+1C)]|TGAGTGTGTGTGTGTGAGTGTGTGC</td><td>>am_MI0003581|hsa-mir-574&WithFlank&4|+|38868026|38868133</td><td>31</td><td>54</td><td>24(+1C)</td><td>TGAGTGTGTGTGTGTGAGTGTGTGC</td><td>115.9</td><td>4.3</td><td>1.6</td><td>2.7</td><td>7.3E-03</td><td>9.6E-03</td><td>MB231/MB468</td><td>MP</td></tr>
<tr><td>>MI0000077|hsa-mir-21&WithFlank&17|+|59841260|59841343@14.35.22(+2A)&[MIMAT0000076&hsa-miR-21-5p&offsets|0|0(+2A);m-1&17|+|59841273|59841294&offsets|0|0(+2A)]|TAGCTTATCAGACTGATGTTGAAA</td><td>>MI0000077|hsa-mir-21&WithFlank&17|+|59841260|59841343</td><td>14</td><td>35</td><td>22(+2A)</td><td>TAGCTTATCAGACTGATGTTGAAA</td><td>1414.5</td><td>2.9</td><td>0.9</td><td>3.1</td><td>2.2E-03</td><td>9.6E-03</td><td>BT20/MB231</td><td>CYTO</td></tr>
<tr><td>>MI0000078|hsa-mir-22&WithFlank&17|-|1713897|1713993@59.80.22(+1A)&[MIMAT0000077&hsa-miR-22-3p&offsets|0|0(+1A);m-2&17|-|1713914|1713935&offsets|0|0(+1A)]|AAGCTGCCAGTTGAAGAACTGTA</td><td>>MI0000078|hsa-mir-22&WithFlank&17|-|1713897|1713993</td><td>59</td><td>80</td><td>22(+1A)</td><td>AAGCTGCCAGTTGAAGAACTGTA</td><td>458.3</td><td>2.7</td><td>0.9</td><td>3.1</td><td>2.2E-03</td><td>9.6E-03</td><td>BT20/MB231</td><td>CYTO</td></tr>
<tr><td>>hg38_21_+_8433172_8446622_RNA45SN1_45S_50ntFlanks@11669.11702.34|GCCCAGTGCTCTGAATGTCAAAGTGAAGAAATTC</td><td>>hg38_21_+_8433172_8446622_RNA45SN1_45S_50ntFlanks</td><td>11669</td><td>11702</td><td>34</td><td>GCCCAGTGCTCTGAATGTCAAAGTGAAGAAATTC</td><td>5858.6</td><td>-2.9</td><td>0.9</td><td>-3.1</td><td>2.2E-03</td><td>9.6E-03</td><td>BT20/MB231</td><td>CYTO</td></tr>
<tr><td>>hg38_21_+_8433172_8446622_RNA45SN1_45S_50ntFlanks@10410.10427.18|TCAGTCGGTCCTGAGAGA</td><td>>hg38_21_+_8433172_8446622_RNA45SN1_45S_50ntFlanks</td><td>10410</td><td>10427</td><td>18</td><td>TCAGTCGGTCCTGAGAGA</td><td>111.6</td><td>-3.6</td><td>1.3</td><td>-2.8</td><td>4.8E-03</td><td>9.6E-03</td><td>BT20/MB468</td><td>MP</td></tr>
<tr><td>>hg38_21_+_8433172_8446622_RNA45SN1_45S_50ntFlanks@11674.11701.28|GTGCTCTGAATGTCAAAGTGAAGAAATT</td><td>>hg38_21_+_8433172_8446622_RNA45SN1_45S_50ntFlanks</td><td>11674</td><td>11701</td><td>28</td><td>GTGCTCTGAATGTCAAAGTGAAGAAATT</td><td>80.8</td><td>4.9</td><td>1.7</td><td>2.8</td><td>4.8E-03</td><td>9.6E-03</td><td>BT20/MB468</td><td>MP</td></tr>
<tr><td>>hg38_1_-_228634819_228635039_RNA5S12_5S_50ntFlanks@154.171.18|CCGGGTGCTGTAGGCTTT</td><td>>hg38_1_-_228634819_228635039_RNA5S12_5S_50ntFlanks</td><td>154</td><td>171</td><td>18</td><td>CCGGGTGCTGTAGGCTTT</td><td>1748.8</td><td>2.3</td><td>0.9</td><td>2.6</td><td>8.5E-03</td><td>9.6E-03</td><td>MB231/MB468</td><td>NUC</td></tr>
<tr><td>>hg38_21_+_8433172_8446622_RNA45SN1_45S_50ntFlanks@12974.13000.27|CGTACGTAGCAGAGCAGCTCCCTCGCT</td><td>>hg38_21_+_8433172_8446622_RNA45SN1_45S_50ntFlanks</td><td>12974</td><td>13000</td><td>27</td><td>CGTACGTAGCAGAGCAGCTCCCTCGCT</td><td>111.6</td><td>-4.4</td><td>1.6</td><td>-2.7</td><td>7.3E-03</td><td>9.6E-03</td><td>MB231/MB468</td><td>MP</td></tr>
<tr><td>>hg38_21_+_8433172_8446622_RNA45SN1_45S_50ntFlanks@6699.6722.24|GCTAGCTGCGAGAATTAATGTGAA</td><td>>hg38_21_+_8433172_8446622_RNA45SN1_45S_50ntFlanks</td><td>6699</td><td>6722</td><td>24</td><td>GCTAGCTGCGAGAATTAATGTGAA</td><td>100.6</td><td>-3.4</td><td>1.2</td><td>-2.8</td><td>4.8E-03</td><td>9.6E-03</td><td>BT20/MB468</td><td>MP</td></tr>
<tr><td>>hg38_21_+_8433172_8446622_RNA45SN1_45S_50ntFlanks@5491.5516.26|GAGAAGACGGTCGAACTTGACTATCT</td><td>>hg38_21_+_8433172_8446622_RNA45SN1_45S_50ntFlanks</td><td>5491</td><td>5516</td><td>26</td><td>GAGAAGACGGTCGAACTTGACTATCT</td><td>359.8</td><td>4.0</td><td>1.5</td><td>2.7</td><td>7.3E-03</td><td>9.6E-03</td><td>MB231/MB468</td><td>MP</td></tr>
<tr><td>>hg38_21_+_8433172_8446622_RNA45SN1_45S_50ntFlanks@6649.6675.27|TACGACTCTTAGCGGTGGATCACTCGG</td><td>>hg38_21_+_8433172_8446622_RNA45SN1_45S_50ntFlanks</td><td>6649</td><td>6675</td><td>27</td><td>TACGACTCTTAGCGGTGGATCACTCGG</td><td>646.9</td><td>2.1</td><td>0.8</td><td>2.6</td><td>8.5E-03</td><td>9.6E-03</td><td>MB231/MB468</td><td>NUC</td></tr>
<tr><td>>MI0001445|hsa-mir-423&WithFlank&17|+|30117073|30117178@23.42.20&[MIMAT0004748&hsa-miR-423-5p&offsets|0|-3;m-141&17|+|30117095|30117117&offsets|0|-3]|TGAGGGGCAGAGAGCGAGAC</td><td>>MI0001445|hsa-mir-423&WithFlank&17|+|30117073|30117178</td><td>23</td><td>42</td><td>20</td><td>TGAGGGGCAGAGAGCGAGAC</td><td>71.7</td><td>4.4</td><td>1.6</td><td>2.7</td><td>7.3E-03</td><td>9.6E-03</td><td>MB231/MB468</td><td>MP</td></tr>
<tr><td>>hg38_21_+_8433172_8446622_RNA45SN1_45S_50ntFlanks@9792.9830.39|GGCGTGGAGCCGGGCGTGGAATGCGAGTGCCTAGTGGGC</td><td>>hg38_21_+_8433172_8446622_RNA45SN1_45S_50ntFlanks</td><td>9792</td><td>9830</td><td>39</td><td>GGCGTGGAGCCGGGCGTGGAATGCGAGTGCCTAGTGGGC</td><td>94.1</td><td>5.7</td><td>2.0</td><td>2.8</td><td>4.8E-03</td><td>9.7E-03</td><td>BT20/MB468</td><td>MP</td></tr>
<tr><td>>hg38_21_+_8433172_8446622_RNA45SN1_45S_50ntFlanks@8371.8391.21|GAGAGAGTTCAAGAGGGCGTG</td><td>>hg38_21_+_8433172_8446622_RNA45SN1_45S_50ntFlanks</td><td>8371</td><td>8391</td><td>21</td><td>GAGAGAGTTCAAGAGGGCGTG</td><td>37.3</td><td>5.6</td><td>2.0</td><td>2.8</td><td>4.8E-03</td><td>9.7E-03</td><td>BT20/MB468</td><td>MP</td></tr>
<tr><td>>MI0000071|hsa-mir-17&WithFlank&13|+|91350599|91350694@20.42.23(+1A)&[MIMAT0000070&hsa-miR-17-5p&offsets|0|0(+1A);m-88&13|+|91350618|91350640&offsets|0|0(+1A)]|CAAAGTGCTTACAGTGCAGGTAGA</td><td>>MI0000071|hsa-mir-17&WithFlank&13|+|91350599|91350694</td><td>20</td><td>42</td><td>23(+1A)</td><td>CAAAGTGCTTACAGTGCAGGTAGA</td><td>1831.6</td><td>2.3</td><td>0.9</td><td>2.7</td><td>7.4E-03</td><td>9.7E-03</td><td>MB231/MB468</td><td>MP</td></tr>
</table>

<table>
<tr><td>>hg38_21_+_8433172_8446622_RNA45SN1_45S_50ntFlanks@7993.8016.24|GTGGCGACCCGCTGAATTTAAGCA</td><td>>hg38_21_+_8433172_8446622_RNA45SN1_45S_50ntFlanks</td><td>7993</td><td>8016</td><td>24</td><td>GTGGCGACCCGCTGAATTTAAGCA</td><td>178.9</td><td>6.6</td><td>2.5</td><td>2.7</td><td>8.0E-03</td><td>9.7E-03</td><td>MB231/MB468</td><td>CYTO</td></tr>
<tr><td>>hg38_21_+_8433172_8446622_RNA45SN1_45S_50ntFlanks@10866.10886.21|CTAAGGGCTGGGTCGGTCGGG</td><td>>hg38_21_+_8433172_8446622_RNA45SN1_45S_50ntFlanks</td><td>10866</td><td>10886</td><td>21</td><td>CTAAGGGCTGGGTCGGTCGGG</td><td>384.2</td><td>-3.8</td><td>1.3</td><td>-2.8</td><td>4.8E-03</td><td>9.7E-03</td><td>BT20/MB468</td><td>MP</td></tr>
<tr><td>>hg38_21_+_8433172_8446622_RNA45SN1_45S_50ntFlanks@12602.12625.24|GTGTATGTGCTTGGCTGAGGAGCC</td><td>>hg38_21_+_8433172_8446622_RNA45SN1_45S_50ntFlanks</td><td>12602</td><td>12625</td><td>24</td><td>GTGTATGTGCTTGGCTGAGGAGCC</td><td>245.4</td><td>-1.8</td><td>0.7</td><td>-2.7</td><td>8.0E-03</td><td>9.7E-03</td><td>BT20/MB468</td><td>CYTO</td></tr>
<tr><td>>hg38_MT_+_1621_3279_RNR2_16S_50ntFlanks@1095.1122.28|AGACGAGAAGACCCTATGGAGCTTTAAT</td><td>>hg38_MT_+_1621_3279_RNR2_16S_50ntFlanks</td><td>1095</td><td>1122</td><td>28</td><td>AGACGAGAAGACCCTATGGAGCTTTAAT</td><td>60.9</td><td>5.7</td><td>2.0</td><td>2.8</td><td>4.8E-03</td><td>9.7E-03</td><td>BT20/MB468</td><td>MP</td></tr>
<tr><td>>hg38_21_+_8433172_8446622_RNA45SN1_45S_50ntFlanks@8114.8141.28|GCGCGGGACATGTGGCGTACGGAAGACC</td><td>>hg38_21_+_8433172_8446622_RNA45SN1_45S_50ntFlanks</td><td>8114</td><td>8141</td><td>28</td><td>GCGCGGGACATGTGGCGTACGGAAGACC</td><td>464.3</td><td>-5.1</td><td>1.9</td><td>-2.6</td><td>8.4E-03</td><td>9.7E-03</td><td>BT20/MB468</td><td>NUC</td></tr>
<tr><td>>hg38_21_+_8433172_8446622_RNA45SN1_45S_50ntFlanks@10391.10424.34|AACAGCAGTTGAACATGGGTCAGTCGGTCCTGAG</td><td>>hg38_21_+_8433172_8446622_RNA45SN1_45S_50ntFlanks</td><td>10391</td><td>10424</td><td>34</td><td>AACAGCAGTTGAACATGGGTCAGTCGGTCCTGAG</td><td>4025.3</td><td>-2.3</td><td>0.9</td><td>-2.6</td><td>8.5E-03</td><td>9.7E-03</td><td>MB231/MB468</td><td>MITO</td></tr>
<tr><td>>am_trna128_GlyGCC_6_-_27870686_27870756@3.30.28,trna133_GlyCCC_1_-_16872434_16872504@3.30.28,trna18_GlyGCC_16_+_70822597_70822667@3.30.28,trna19_GlyGCC_16_+_70823410_70823480@3.30.28,trna19_GlyGCC_2_-_157257659_157257729@3.30.28,trna24_GlyGCC_16_-_70812942_70813012@3.30.28,trna25_GlyGCC_16_-_70812114_70812184@3.30.28,trna4_GlyCCC_1_+_17188416_17188486@3.30.28,trna5_GlyGCC_17_+_8029064_8029134@3.30.28,trna68_GlyGCC_1_-_161493637_161493707@3.30.28|ATTGGTGGTTCAGTGGTAGAATTCTCGC</td><td>>am_trna128_GlyGCC_6_-_27870686_27870756</td><td>3</td><td>30</td><td>28</td><td>ATTGGTGGTTCAGTGGTAGAATTCTCGC</td><td>120.4</td><td>3.6</td><td>1.3</td><td>2.7</td><td>7.4E-03</td><td>9.7E-03</td><td>MB231/MB468</td><td>MP</td></tr>
<tr><td>>hg38_21_+_8433172_8446622_RNA45SN1_45S_50ntFlanks@8170.8216.47|CAAGTCCTTCTGATCGAGGCCCAGCCCGTGGACGGTGTGAGGCCGGT</td><td>>hg38_21_+_8433172_8446622_RNA45SN1_45S_50ntFlanks</td><td>8170</td><td>8216</td><td>47</td><td>CAAGTCCTTCTGATCGAGGCCCAGCCCGTGGACGGTGTGAGGCCGGT</td><td>509.8</td><td>-3.6</td><td>1.4</td><td>-2.6</td><td>8.5E-03</td><td>9.7E-03</td><td>MB231/MB468</td><td>NUC</td></tr>
<tr><td>>hg38_21_+_8433172_8446622_RNA45SN1_45S_50ntFlanks@12412.12437.26|ATCCTTCGATGTCGGCTCTTCCTATC</td><td>>hg38_21_+_8433172_8446622_RNA45SN1_45S_50ntFlanks</td><td>12412</td><td>12437</td><td>26</td><td>ATCCTTCGATGTCGGCTCTTCCTATC</td><td>489.4</td><td>-4.5</td><td>1.7</td><td>-2.6</td><td>8.5E-03</td><td>9.7E-03</td><td>MB231/MB468</td><td>MITO</td></tr>
<tr><td>>hg38_21_+_8433172_8446622_RNA45SN1_45S_50ntFlanks@3988.4023.36|CTTTGGTGACTCTAGATAACCTCGGGCCGATCGCAC</td><td>>hg38_21_+_8433172_8446622_RNA45SN1_45S_50ntFlanks</td><td>3988</td><td>4023</td><td>36</td><td>CTTTGGTGACTCTAGATAACCTCGGGCCGATCGCAC</td><td>447.8</td><td>2.1</td><td>0.8</td><td>2.6</td><td>8.5E-03</td><td>9.7E-03</td><td>MB231/MB468</td><td>NUC</td></tr>
<tr><td>>hg38_21_+_8433172_8446622_RNA45SN1_45S_50ntFlanks@11569.11612.44|CAGCCGACTTAGAACTGGTGCGGACCAGGGGAATCCGACTGTTT</td><td>>hg38_21_+_8433172_8446622_RNA45SN1_45S_50ntFlanks</td><td>11569</td><td>11612</td><td>44</td><td>CAGCCGACTTAGAACTGGTGCGGACCAGGGGAATCCGACTGTTT</td><td>856.2</td><td>-2.2</td><td>0.8</td><td>-2.6</td><td>8.1E-03</td><td>9.7E-03</td><td>MB231/MB468</td><td>CYTO</td></tr>
<tr><td>>hg38_21_+_8433172_8446622_RNA45SN1_45S_50ntFlanks@4047.4067.21|ATTCGAACGTCTGCCCTATCA</td><td>>hg38_21_+_8433172_8446622_RNA45SN1_45S_50ntFlanks</td><td>4047</td><td>4067</td><td>21</td><td>ATTCGAACGTCTGCCCTATCA</td><td>409.3</td><td>-1.7</td><td>0.6</td><td>-2.6</td><td>8.1E-03</td><td>9.7E-03</td><td>MB231/MB468</td><td>CYTO</td></tr>
<tr><td>>hg38_21_+_8433172_8446622_RNA45SN1_45S_50ntFlanks@11771.11790.20|ATTAGTGACGCGCATGAATG</td><td>>hg38_21_+_8433172_8446622_RNA45SN1_45S_50ntFlanks</td><td>11771</td><td>11790</td><td>20</td><td>ATTAGTGACGCGCATGAATG</td><td>232.2</td><td>-1.5</td><td>0.6</td><td>-2.6</td><td>8.5E-03</td><td>9.7E-03</td><td>MB231/MB468</td><td>NUC</td></tr>
<tr><td>>hg38_21_+_8433172_8446622_RNA45SN1_45S_50ntFlanks@10402.10435.34|AACATGGGTCAGTCGGTCCTGAGAGATGGGCGAG</td><td>>hg38_21_+_8433172_8446622_RNA45SN1_45S_50ntFlanks</td><td>10402</td><td>10435</td><td>34</td><td>AACATGGGTCAGTCGGTCCTGAGAGATGGGCGAG</td><td>811.5</td><td>-5.1</td><td>1.8</td><td>-2.8</td><td>4.8E-03</td><td>9.7E-03</td><td>BT20/MB468</td><td>MP</td></tr>
<tr><td>>hg38_21_+_8433172_8446622_RNA45SN1_45S_50ntFlanks@11665.11685.21|TTCTGCCCAGTGCTCTGAATG</td><td>>hg38_21_+_8433172_8446622_RNA45SN1_45S_50ntFlanks</td><td>11665</td><td>11685</td><td>21</td><td>TTCTGCCCAGTGCTCTGAATG</td><td>506.0</td><td>-1.8</td><td>0.7</td><td>-2.7</td><td>8.0E-03</td><td>9.7E-03</td><td>BT20/MB468</td><td>CYTO</td></tr>
<tr><td>>hg38_1_-_228634819_228635039_RNA5S12_5S_50ntFlanks@51.82.32|GTCTACGGCCATACCACCCTGAACGCGCCCGA</td><td>>hg38_1_-_228634819_228635039_RNA5S12_5S_50ntFlanks</td><td>51</td><td>82</td><td>32</td><td>GTCTACGGCCATACCACCCTGAACGCGCCCGA</td><td>725.1</td><td>2.5</td><td>1.0</td><td>2.6</td><td>8.5E-03</td><td>9.7E-03</td><td>MB231/MB468</td><td>MITO</td></tr>
<tr><td>>hg38_21_+_8433172_8446622_RNA45SN1_45S_50ntFlanks@11769.11806.38|TAATTAGTGACGCGCATGAATGGATGAACGAGATTCCC</td><td>>hg38_21_+_8433172_8446622_RNA45SN1_45S_50ntFlanks</td><td>11769</td><td>11806</td><td>38</td><td>TAATTAGTGACGCGCATGAATGGATGAACGAGATTCCC</td><td>67.5</td><td>-4.6</td><td>1.7</td><td>-2.7</td><td>7.4E-03</td><td>9.7E-03</td><td>MB231/MB468</td><td>MP</td></tr>
<tr><td>>trnaMT_SerGCT_MT_+_12207_12265@14.41.28|AGAACTGCTAACTCATGCCCCCATGTCT</td><td>>trnaMT_SerGCT_MT_+_12207_12265</td><td>14</td><td>41</td><td>28</td><td>AGAACTGCTAACTCATGCCCCCATGTCT</td><td>131.2</td><td>3.4</td><td>1.3</td><td>2.6</td><td>8.5E-03</td><td>9.7E-03</td><td>MB231/MB468</td><td>MITO</td></tr>
<tr><td>>hg38_21_+_8433172_8446622_RNA45SN1_45S_50ntFlanks@10401.10439.39|GAACATGGGTCAGTCGGTCCTGAGAGATGGGCGAGCGCC</td><td>>hg38_21_+_8433172_8446622_RNA45SN1_45S_50ntFlanks</td><td>10401</td><td>10439</td><td>39</td><td>GAACATGGGTCAGTCGGTCCTGAGAGATGGGCGAGCGCC</td><td>142.8</td><td>4.8</td><td>1.7</td><td>2.8</td><td>4.8E-03</td><td>9.7E-03</td><td>BT20/MB468</td><td>MP</td></tr>
<tr><td>>hg38_21_+_8433172_8446622_RNA45SN1_45S_50ntFlanks@10458.10495.38|ATGGCCTCCGTTGCCCTCGGCCGATCGAAAGGGAGTCG</td><td>>hg38_21_+_8433172_8446622_RNA45SN1_45S_50ntFlanks</td><td>10458</td><td>10495</td><td>38</td><td>ATGGCCTCCGTTGCCCTCGGCCGATCGAAAGGGAGTCG</td><td>391.2</td><td>-4.9</td><td>1.7</td><td>-2.8</td><td>4.8E-03</td><td>9.7E-03</td><td>BT20/MB468</td><td>MP</td></tr>
<tr><td>>hg38_21_+_8433172_8446622_RNA45SN1_45S_50ntFlanks@4969.4999.31|ACCCGGCCCGGACACGGACAGGATTGACAGA</td><td>>hg38_21_+_8433172_8446622_RNA45SN1_45S_50ntFlanks</td><td>4969</td><td>4999</td><td>31</td><td>ACCCGGCCCGGACACGGACAGGATTGACAGA</td><td>241.0</td><td>-4.0</td><td>1.5</td><td>-2.7</td><td>6.9E-03</td><td>9.7E-03</td><td>BT20/MB468</td><td>MITO</td></tr>
<tr><td>>hg38_21_+_8433172_8446622_RNA45SN1_45S_50ntFlanks@5085.5104.20|GAACGAGACTCTGGCATGCT</td><td>>hg38_21_+_8433172_8446622_RNA45SN1_45S_50ntFlanks</td><td>5085</td><td>5104</td><td>20</td><td>GAACGAGACTCTGGCATGCT</td><td>1011.2</td><td>4.8</td><td>1.5</td><td>3.1</td><td>1.8E-03</td><td>9.7E-03</td><td>BT20/MB231</td><td>MP</td></tr>
<tr><td>>MI0000736|hsa-mir-30c-1&WithFlank&1|+|40757278|40757378@23.48.26;MI0000254|hsa-mir-30c-2&WithFlank&6|-|71376954|71377037@13.38.26&[MIMAT0000244&hsa-miR-30c-5p&offsets|0|+3;m-57&1|+|40757300|40757323&offsets|0|+2];[MIMAT0000244_1&hsa-miR-30c-5p&offsets|0|+3;m-56&6|-|71377002|71377025&offsets|0|+2]|TGTAAACATCCTACACTCTCAGCTGT</td><td>>MI0000736|hsa-mir-30c-1&WithFlank&1|+|40757278|40757378</td><td>23</td><td>48</td><td>26</td><td>TGTAAACATCCTACACTCTCAGCTGT</td><td>173.2</td><td>-5.8</td><td>1.9</td><td>-3.1</td><td>1.8E-03</td><td>9.7E-03</td><td>BT20/MB231</td><td>MP</td></tr>
<tr><td>>hg38_21_+_8433172_8446622_RNA45SN1_45S_50ntFlanks@10095.10124.30|CGCGCGCGCGCGCGTGTGGTGTGCGTCGGA</td><td>>hg38_21_+_8433172_8446622_RNA45SN1_45S_50ntFlanks</td><td>10095</td><td>10124</td><td>30</td><td>CGCGCGCGCGCGCGTGTGGTGTGCGTCGGA</td><td>54.9</td><td>9.1</td><td>2.9</td><td>3.1</td><td>1.8E-03</td><td>9.7E-03</td><td>BT20/MB231</td><td>MP</td></tr>
<tr><td>>MI0000102|hsa-mir-100&WithFlank&11|-|122152223|122152314@19.38.20(+1C)&[MIMAT0000098&hsa-miR-100-5p&offsets|0|-2(+1C);m-27&11|-|122152276|122152296&offsets|0|-1(+1C)]|AACCCGTAGATCCGAACTTGC</td><td>>MI0000102|hsa-mir-100&WithFlank&11|-|122152223|122152314</td><td>19</td><td>38</td><td>20(+1C)</td><td>AACCCGTAGATCCGAACTTGC</td><td>19100.7</td><td>-3.7</td><td>1.2</td><td>-3.1</td><td>2.2E-03</td><td>9.7E-03</td><td>BT20/MB231</td><td>CYTO</td></tr>
<tr><td>>hg38_21_+_8433172_8446622_RNA45SN1_45S_50ntFlanks@12396.12414.19|GCGACGTCGCTTTTTGATC</td><td>>hg38_21_+_8433172_8446622_RNA45SN1_45S_50ntFlanks</td><td>12396</td><td>12414</td><td>19</td><td>GCGACGTCGCTTTTTGATC</td><td>2067.6</td><td>-2.6</td><td>0.8</td><td>-3.1</td><td>2.2E-03</td><td>9.7E-03</td><td>BT20/MB231</td><td>CYTO</td></tr>
<tr><td>>MI0003667|hsa-mir-652&WithFlank&X|+|110055323|110055432@67.88.22&[MIMAT0003322&hsa-miR-652-3p&offsets|0|+1;m-148&X|+|110055389|110055410&offsets|0|0]|AATGGCGCCACTAGGGTTGTGC</td><td>>MI0003667|hsa-mir-652&WithFlank&X|+|110055323|110055432</td><td>67</td><td>88</td><td>22</td><td>AATGGCGCCACTAGGGTTGTGC</td><td>237.2</td><td>3.4</td><td>1.1</td><td>3.1</td><td>2.2E-03</td><td>9.7E-03</td><td>BT20/MB231</td><td>CYTO</td></tr>
<tr><td>>am_MI0000809|hsa-mir-151a&WithFlank&8|-|140732558|140732659@53.72.20&[MIMAT0000757&hsa-miR-151a-3p&offsets|0|-1;m-30&8|-|140732587|140732607&offsets|0|-1]|CTAGACTGAAGCTCCTTGAG</td><td>>am_MI0000809|hsa-mir-151a&WithFlank&8|-|140732558|140732659</td><td>53</td><td>72</td><td>20</td><td>CTAGACTGAAGCTCCTTGAG</td><td>3982.6</td><td>2.3</td><td>0.8</td><td>3.1</td><td>2.2E-03</td><td>9.7E-03</td><td>BT20/MB231</td><td>CYTO</td></tr>
<tr><td>>hg38_21_+_8433172_8446622_RNA45SN1_45S_50ntFlanks@6651.6676.26|CGACTCTTAGCGGTGGATCACTCGGC</td><td>>hg38_21_+_8433172_8446622_RNA45SN1_45S_50ntFlanks</td><td>6651</td><td>6676</td><td>26</td><td>CGACTCTTAGCGGTGGATCACTCGGC</td><td>39161.3</td><td>3.5</td><td>1.1</td><td>3.1</td><td>2.2E-03</td><td>9.7E-03</td><td>BT20/MB231</td><td>CYTO</td></tr>
<tr><td>>trnaMT_HisGTG_MT_+_12138_12206@32.72.41|TGAATCTGACAACAGAGGCTTACGACCCCTTATTTACCCCA</td><td>>trnaMT_HisGTG_MT_+_12138_12206</td><td>32</td><td>72</td><td>41</td><td>TGAATCTGACAACAGAGGCTTACGACCCCTTATTTACCCCA</td><td>594.0</td><td>-3.2</td><td>1.0</td><td>-3.1</td><td>2.2E-03</td><td>9.7E-03</td><td>BT20/MB231</td><td>CYTO</td></tr>
<tr><td>>hg38_21_+_8433172_8446622_RNA45SN1_45S_50ntFlanks@11574.11603.30|GACTTAGAACTGGTGCGGACCAGGGGAATC</td><td>>hg38_21_+_8433172_8446622_RNA45SN1_45S_50ntFlanks</td><td>11574</td><td>11603</td><td>30</td><td>GACTTAGAACTGGTGCGGACCAGGGGAATC</td><td>207.5</td><td>-2.4</td><td>0.9</td><td>-2.7</td><td>8.0E-03</td><td>9.7E-03</td><td>BT20/MB468</td><td>CYTO</td></tr>
<tr><td>>hg38_21_+_8433172_8446622_RNA45SN1_45S_50ntFlanks@8156.8181.26|GCTCGTGGGGGGCCCAAGTCCTTCTG</td><td>>hg38_21_+_8433172_8446622_RNA45SN1_45S_50ntFlanks</td><td>8156</td><td>8181</td><td>26</td><td>GCTCGTGGGGGGCCCAAGTCCTTCTG</td><td>142.8</td><td>-4.7</td><td>1.7</td><td>-2.8</td><td>4.8E-03</td><td>9.7E-03</td><td>BT20/MB468</td><td>MP</td></tr>
<tr><td>>hg38_21_+_8433172_8446622_RNA45SN1_45S_50ntFlanks@3841.3861.21|TCCTACTTGGATAACTGTGGT</td><td>>hg38_21_+_8433172_8446622_RNA45SN1_45S_50ntFlanks</td><td>3841</td><td>3861</td><td>21</td><td>TCCTACTTGGATAACTGTGGT</td><td>268.4</td><td>2.2</td><td>0.7</td><td>3.1</td><td>2.3E-03</td><td>9.7E-03</td><td>BT20/MB231</td><td>CYTO</td></tr>
<tr><td>>hg38_21_+_8433172_8446622_RNA45SN1_45S_50ntFlanks@8168.8215.48|CCCAAGTCCTTCTGATCGAGGCCCAGCCCGTGGACGGTGTGAGGCCGG</td><td>>hg38_21_+_8433172_8446622_RNA45SN1_45S_50ntFlanks</td><td>8168</td><td>8215</td><td>48</td><td>CCCAAGTCCTTCTGATCGAGGCCCAGCCCGTGGACGGTGTGAGGCCGG</td><td>213.2</td><td>-3.9</td><td>1.5</td><td>-2.7</td><td>8.0E-03</td><td>9.7E-03</td><td>BT20/MB468</td><td>CYTO</td></tr>
<tr><td>>hg38_21_+_8433172_8446622_RNA45SN1_45S_50ntFlanks@12990.13017.28|GCTCCCTCGCTGCGATCTATTGAAAGTC</td><td>>hg38_21_+_8433172_8446622_RNA45SN1_45S_50ntFlanks</td><td>12990</td><td>13017</td><td>28</td><td>GCTCCCTCGCTGCGATCTATTGAAAGTC</td><td>569.7</td><td>-2.9</td><td>1.0</td><td>-3.1</td><td>2.3E-03</td><td>9.7E-03</td><td>BT20/MB231</td><td>CYTO</td></tr>
<tr><td>>hg38_21_+_8433172_8446622_RNA45SN1_45S_50ntFlanks@11540.11583.44|CGCGGCGCCTCGCCTCGGCCGGCGCCTAGCAGCCGACTTAGAAC</td><td>>hg38_21_+_8433172_8446622_RNA45SN1_45S_50ntFlanks</td><td>11540</td><td>11583</td><td>44</td><td>CGCGGCGCCTCGCCTCGGCCGGCGCCTAGCAGCCGACTTAGAAC</td><td>892.3</td><td>3.1</td><td>1.2</td><td>2.6</td><td>8.6E-03</td><td>9.7E-03</td><td>MB231/MB468</td><td>NUC</td></tr>
<tr><td>>hg38_21_+_8433172_8446622_RNA45SN1_45S_50ntFlanks@11572.11592.21|CCGACTTAGAACTGGTGCGGA</td><td>>hg38_21_+_8433172_8446622_RNA45SN1_45S_50ntFlanks</td><td>11572</td><td>11592</td><td>21</td><td>CCGACTTAGAACTGGTGCGGA</td><td>305.9</td><td>-1.7</td><td>0.6</td><td>-2.7</td><td>6.9E-03</td><td>9.7E-03</td><td>BT20/MB468</td><td>MITO</td></tr>
<tr><td>>hg38_MT_+_1621_3279_RNR2_16S_50ntFlanks@1404.1421.18|TATTAAAGGTTCGTTTGT</td><td>>hg38_MT_+_1621_3279_RNR2_16S_50ntFlanks</td><td>1404</td><td>1421</td><td>18</td><td>TATTAAAGGTTCGTTTGT</td><td>1267.9</td><td>-4.7</td><td>1.7</td><td>-2.8</td><td>4.8E-03</td><td>9.7E-03</td><td>BT20/MB468</td><td>MP</td></tr>
<tr><td>>hg38_21_+_8433172_8446622_RNA45SN1_45S_50ntFlanks@10097.10117.21|CGCGCGCGCGCGTGTGGTGTG</td><td>>hg38_21_+_8433172_8446622_RNA45SN1_45S_50ntFlanks</td><td>10097</td><td>10117</td><td>21</td><td>CGCGCGCGCGCGTGTGGTGTG</td><td>299.2</td><td>-3.7</td><td>1.3</td><td>-2.8</td><td>4.8E-03</td><td>9.7E-03</td><td>BT20/MB468</td><td>MP</td></tr>
<tr><td>>MI0000095|hsa-mir-93&WithFlank&7|-|100093762|100093853@17.36.20(+1A)&[MIMAT0000093&hsa-miR-93-5p&offsets|0|-3(+1A);m-29&7|-|100093815|100093837&offsets|0|-3(+1A)]|CAAAGTGCTGTTCGTGCAGGA</td><td>>MI0000095|hsa-mir-93&WithFlank&7|-|100093762|100093853</td><td>17</td><td>36</td><td>20(+1A)</td><td>CAAAGTGCTGTTCGTGCAGGA</td><td>199.8</td><td>3.6</td><td>1.3</td><td>2.7</td><td>7.4E-03</td><td>9.7E-03</td><td>MB231/MB468</td><td>MP</td></tr>
<tr><td>>hg38_21_+_8433172_8446622_RNA45SN1_45S_50ntFlanks@12404.12441.38|GCTTTTTGATCCTTCGATGTCGGCTCTTCCTATCATTG</td><td>>hg38_21_+_8433172_8446622_RNA45SN1_45S_50ntFlanks</td><td>12404</td><td>12441</td><td>38</td><td>GCTTTTTGATCCTTCGATGTCGGCTCTTCCTATCATTG</td><td>453.1</td><td>-2.4</td><td>0.9</td><td>-2.6</td><td>8.6E-03</td><td>9.7E-03</td><td>MB231/MB468</td><td>NUC</td></tr>
<tr><td>>trnaMT_SerGCT_MT_+_12207_12265@8.31.24|CTCACAAGAACTGCTAACTCATGC</td><td>>trnaMT_SerGCT_MT_+_12207_12265</td><td>8</td><td>31</td><td>24</td><td>CTCACAAGAACTGCTAACTCATGC</td><td>218.0</td><td>-3.3</td><td>1.3</td><td>-2.6</td><td>8.1E-03</td><td>9.7E-03</td><td>MB231/MB468</td><td>CYTO</td></tr>
<tr><td>>MI0000077|hsa-mir-21&WithFlank&17|+|59841260|59841343@13.37.25&[MIMAT0000076&hsa-miR-21-5p&offsets|-1|+2;m-1&17|+|59841273|59841294&offsets|-1|+2]|GTAGCTTATCAGACTGATGTTGACT</td><td>>MI0000077|hsa-mir-21&WithFlank&17|+|59841260|59841343</td><td>13</td><td>37</td><td>25</td><td>GTAGCTTATCAGACTGATGTTGACT</td><td>48.5</td><td>4.3</td><td>1.6</td><td>2.7</td><td>7.4E-03</td><td>9.7E-03</td><td>MB231/MB468</td><td>MP</td></tr>
<tr><td>>hg38_21_+_8433172_8446622_RNA45SN1_45S_50ntFlanks@10869.10903.35|AGGGCTGGGTCGGTCGGGCTGGGGCGCGAAGCGGG</td><td>>hg38_21_+_8433172_8446622_RNA45SN1_45S_50ntFlanks</td><td>10869</td><td>10903</td><td>35</td><td>AGGGCTGGGTCGGTCGGGCTGGGGCGCGAAGCGGG</td><td>160.3</td><td>-3.6</td><td>1.3</td><td>-2.8</td><td>4.8E-03</td><td>9.7E-03</td><td>BT20/MB468</td><td>MP</td></tr>
<tr><td>>hg38_21_+_8433172_8446622_RNA45SN1_45S_50ntFlanks@6726.6743.18|CAGGACACATTGATCATC</td><td>>hg38_21_+_8433172_8446622_RNA45SN1_45S_50ntFlanks</td><td>6726</td><td>6743</td><td>18</td><td>CAGGACACATTGATCATC</td><td>154.6</td><td>-3.4</td><td>1.3</td><td>-2.6</td><td>8.6E-03</td><td>9.7E-03</td><td>MB231/MB468</td><td>NUC</td></tr>
<tr><td>>hg38_21_+_8433172_8446622_RNA45SN1_45S_50ntFlanks@11602.11619.18|TCCGACTGTTTAATTAAA</td><td>>hg38_21_+_8433172_8446622_RNA45SN1_45S_50ntFlanks</td><td>11602</td><td>11619</td><td>18</td><td>TCCGACTGTTTAATTAAA</td><td>1720.2</td><td>-2.3</td><td>0.9</td><td>-2.6</td><td>8.1E-03</td><td>9.8E-03</td><td>MB231/MB468</td><td>CYTO</td></tr>
<tr><td>>MI0000088|hsa-mir-30a&WithFlank&6|-|71403545|71403627@53.72.20&[MIMAT0000088&hsa-miR-30a-3p&offsets|0|-2;m-38&6|-|71403554|71403575&offsets|0|-2]|CTTTCAGTCGGATGTTTGCA</td><td>>MI0000088|hsa-mir-30a&WithFlank&6|-|71403545|71403627</td><td>53</td><td>72</td><td>20</td><td>CTTTCAGTCGGATGTTTGCA</td><td>210.9</td><td>-2.2</td><td>0.7</td><td>-3.0</td><td>2.3E-03</td><td>9.8E-03</td><td>BT20/MB231</td><td>MITO</td></tr>
<tr><td>>MI0000736|hsa-mir-30c-1&WithFlank&1|+|40757278|40757378@23.46.24(+1U);MI0000254|hsa-mir-30c-2&WithFlank&6|-|71376954|71377037@13.36.24(+1U)&[MIMAT0000244&hsa-miR-30c-5p&offsets|0|+1(+1U);m-57&1|+|40757300|40757323&offsets|0|0(+1U)];[MIMAT0000244_1&hsa-miR-30c-5p&offsets|0|+1(+1U);m-56&6|-|71377002|71377025&offsets|0|0(+1U)]|TGTAAACATCCTACACTCTCAGCTT</td><td>>MI0000736|hsa-mir-30c-1&WithFlank&1|+|40757278|40757378</td><td>23</td><td>46</td><td>24(+1U)</td><td>TGTAAACATCCTACACTCTCAGCTT</td><td>860.6</td><td>-3.5</td><td>1.1</td><td>-3.1</td><td>1.8E-03</td><td>9.8E-03</td><td>BT20/MB231</td><td>MP</td></tr>
<tr><td>>MI0006444|hsa-mir-1307&WithFlank&10|-|103394247|103394407@86.106.21&[MIMAT0005951&hsa-miR-1307-3p&offsets|0|-1;m-66&10|-|103394300|103394322&offsets|0|-2]|ACTCGGCGTGGCGTCGGTCGT</td><td>>MI0006444|hsa-mir-1307&WithFlank&10|-|103394247|103394407</td><td>86</td><td>106</td><td>21</td><td>ACTCGGCGTGGCGTCGGTCGT</td><td>211.1</td><td>4.7</td><td>1.7</td><td>2.7</td><td>7.4E-03</td><td>9.8E-03</td><td>MB231/MB468</td><td>MP</td></tr>
</table>

| | | | | | | | | | | | | | |
|---|---|---|---|---|---|---|---|---|---|---|---|---|---|
| >hg38_21_+_8433172_8446622_RNA45SN1_45S_50ntFlanks@9928.9949.22\|TGATATAGACAGCAGGACGGTG | >hg38_21_+_8433172_8446622_RNA45SN1_45S_50ntFlanks | 9928 | 9949 | 22 | TGATATAGACAGCAGGACGGTG | 238.3 | 4.5 | 1.4 | 3.1 | 1.8E-03 | 9.8E-03 | BT20/MB231 | MP |
| >MI0000085\|hsa-mir-27a&WithFlank&19\|-\|13836434\|13836523@57.78.22&[MIMAT0000084&hsa-miR-27a-3p&offsets\|0\|+1;m-43&19\|-\|13836447\|13836467&offsets\|0\|+1]\|TTCACAGTGGCTAAGTTCCGCC | >MI0000085\|hsa-mir-27a&WithFlank&19\|-\|13836434\|13836523 | 57 | 78 | 22 | TTCACAGTGGCTAAGTTCCGCC | 220.2 | -5.8 | 1.9 | -3.1 | 1.8E-03 | 9.8E-03 | BT20/MB231 | MP |
| >hg38_21_+_8433172_8446622_RNA45SN1_45S_50ntFlanks@9539.9571.33\|ACTCTGGTGGAGGTCCGTAGCGGTCCTGACGTG | >hg38_21_+_8433172_8446622_RNA45SN1_45S_50ntFlanks | 9539 | 9571 | 33 | ACTCTGGTGGAGGTCCGTAGCGGTCCTGACGTG | 1124.8 | -2.1 | 0.8 | -2.6 | 8.1E-03 | 9.8E-03 | BT20/MB468 | CYTO |
| >trnaMT_MetCAT_MT_+_4402_4469@39.71.33\|GAAAATGTTGGTTATACCCTTCCCGTACTACCA | >trnaMT_MetCAT_MT_+_4402_4469 | 39 | 71 | 33 | GAAAATGTTGGTTATACCCTTCCCGTACTACCA | 139.4 | 4.2 | 1.6 | 2.7 | 7.4E-03 | 9.8E-03 | MB231/MB468 | MP |
| >hg38_21_+_8433172_8446622_RNA45SN1_45S_50ntFlanks@3999.4023.25\|CTAGATAACCTCGGGCCGATCGCAC | >hg38_21_+_8433172_8446622_RNA45SN1_45S_50ntFlanks | 3999 | 4023 | 25 | CTAGATAACCTCGGGCCGATCGCAC | 216.8 | -1.7 | 0.6 | -2.6 | 8.6E-03 | 9.8E-03 | MB231/MB468 | NUC |
| >hg38_21_+_8433172_8446622_RNA45SN1_45S_50ntFlanks@6646.6674.29\|TCGTACGACTCTTAGCGGTGGATCACTCG | >hg38_21_+_8433172_8446622_RNA45SN1_45S_50ntFlanks | 6646 | 6674 | 29 | TCGTACGACTCTTAGCGGTGGATCACTCG | 5908.6 | 2.6 | 1.0 | 2.6 | 8.6E-03 | 9.8E-03 | MB231/MB468 | MITO |
| >hg38_MT_+_1621_3279_RNR2_16S_50ntFlanks@1369.1401.33\|GATGTTGGATCAGGACATCCCGATGGTGCAGCC | >hg38_MT_+_1621_3279_RNR2_16S_50ntFlanks | 1369 | 1401 | 33 | GATGTTGGATCAGGACATCCCGATGGTGCAGCC | 252.3 | 5.1 | 1.8 | 2.8 | 4.9E-03 | 9.8E-03 | BT20/MB468 | MP |
| >hg38_21_+_8433172_8446622_RNA45SN1_45S_50ntFlanks@10404.10426.23\|CATGGGTCAGTCGGTCCTGAGAG | >hg38_21_+_8433172_8446622_RNA45SN1_45S_50ntFlanks | 10404 | 10426 | 23 | CATGGGTCAGTCGGTCCTGAGAG | 132.7 | -4.6 | 1.6 | -2.8 | 4.9E-03 | 9.8E-03 | BT20/MB468 | MP |
| >hg38_21_+_8433172_8446622_RNA45SN1_45S_50ntFlanks@11570.11604.35\|AGCCGACTTAGAACTGGTGCGGACCAGGGGAATCC | >hg38_21_+_8433172_8446622_RNA45SN1_45S_50ntFlanks | 11570 | 11604 | 35 | AGCCGACTTAGAACTGGTGCGGACCAGGGGAATCC | 116.8 | 3.1 | 1.1 | 2.8 | 4.9E-03 | 9.8E-03 | BT20/MB468 | MP |
| >hg38_21_+_8433172_8446622_RNA45SN1_45S_50ntFlanks@11207.11232.26\|GGCGGCGACTCTGGACGCGAGCCGGG | >hg38_21_+_8433172_8446622_RNA45SN1_45S_50ntFlanks | 11207 | 11232 | 26 | GGCGGCGACTCTGGACGCGAGCCGGG | 513.9 | -3.8 | 1.4 | -2.8 | 4.9E-03 | 9.8E-03 | BT20/MB468 | MP |
| >trna116_GluCTC_1_-_145399233_145399304@38.75.38,trna134_GluTTC_1_-_16861774_16861845@38.75.38,trna71_GluCTC_1_-_161439189_161439260@38.75.38,trna74_GluCTC_1_-_161431809_161431880@38.75.38,trna77_GluCTC_1_-_161424398_161424469@38.75.38,trna77_GluCTC_6_+_28949976_28950047@38.75.38,trna80_GluCTC_1_-_161417018_161417089@38.75.38,trna84_GluTTC_1_-_161391883_161391954@38.75.38,trna87_GluCTC_6_-_126101393_126101464@38.75.38\|CCGCCGCGGCCCGGGTTCGATTCCCGGTCAGGGAACCA | >trna116_GluCTC_1_-_145399233_145399304 | 38 | 75 | 38 | CCGCCGCGGCCCGGGTTCGATTCCCGGTCAGGGAACCA | 669.2 | 2.8 | 1.1 | 2.6 | 8.2E-03 | 9.8E-03 | MB231/MB468 | CYTO |
| >hg38_21_+_8433172_8446622_RNA45SN1_45S_50ntFlanks@10468.10495.28\|TTGCCCTCGGCCGATCGAAAGGGAGTCG | >hg38_21_+_8433172_8446622_RNA45SN1_45S_50ntFlanks | 10468 | 10495 | 28 | TTGCCCTCGGCCGATCGAAAGGGAGTCG | 276.2 | -2.1 | 0.8 | -2.6 | 8.2E-03 | 9.8E-03 | MB231/MB468 | CYTO |
| >hg38_21_+_8433172_8446622_RNA45SN1_45S_50ntFlanks@11662.11681.20\|GATTTCTGCCCAGTGCTCTG | >hg38_21_+_8433172_8446622_RNA45SN1_45S_50ntFlanks | 11662 | 11681 | 20 | GATTTCTGCCCAGTGCTCTG | 210.3 | -1.6 | 0.6 | -2.6 | 8.2E-03 | 9.8E-03 | MB231/MB468 | CYTO |
| >hg38_21_+_8433172_8446622_RNA45SN1_45S_50ntFlanks@5489.5516.28\|CTGAGAAGACGGTCGAACTTGACTATCT | >hg38_21_+_8433172_8446622_RNA45SN1_45S_50ntFlanks | 5489 | 5516 | 28 | CTGAGAAGACGGTCGAACTTGACTATCT | 826.5 | 1.7 | 0.7 | 2.6 | 8.6E-03 | 9.8E-03 | MB231/MB468 | NUC |
| >MI0000299\|hsa-mir-222&WithFlank&X\|-\|45747009\|45747130@75.96.22(+1A)&[MIMAT0000279&hsa-miR-222-3p&offsets\|0\|+1(+1A);m-140&X\|-\|45747033\|45747056&offsets\|0\|-2(+1A)]\|AGCTACATCTGGCTACTGGGTCA | >MI0000299\|hsa-mir-222&WithFlank&X\|-\|45747009\|45747130 | 75 | 96 | 22(+1A) | AGCTACATCTGGCTACTGGGTCA | 1928.6 | -3.4 | 1.1 | -3.1 | 1.8E-03 | 9.8E-03 | BT20/MB231 | MP |
| >hg38_21_+_8433172_8446622_RNA45SN1_45S_50ntFlanks@11569.11602.34\|CAGCCGACTTAGAACTGGTGCGGACCAGGGGAAT | >hg38_21_+_8433172_8446622_RNA45SN1_45S_50ntFlanks | 11569 | 11602 | 34 | CAGCCGACTTAGAACTGGTGCGGACCAGGGGAAT | 264.6 | 2.5 | 0.8 | 3.0 | 2.4E-03 | 9.8E-03 | BT20/MB231 | MITO |
| >hg38_21_+_8433172_8446622_RNA45SN1_45S_50ntFlanks@11208.11227.20\|GCGGCGACTCTGGACGCGAG | >hg38_21_+_8433172_8446622_RNA45SN1_45S_50ntFlanks | 11208 | 11227 | 20 | GCGGCGACTCTGGACGCGAG | 1536.9 | -5.5 | 2.1 | -2.6 | 8.5E-03 | 9.8E-03 | BT20/MB468 | NUC |
| >trnaMT_ValTAC_MT_+_1602_1670@53.72.20\|TAACTTGACCGCTCTGACCA | >trnaMT_ValTAC_MT_+_1602_1670 | 53 | 72 | 20 | TAACTTGACCGCTCTGACCA | 1685.8 | 2.7 | 0.9 | 3.1 | 2.3E-03 | 9.8E-03 | BT20/MB231 | CYTO |
| >hg38_21_+_8433172_8446622_RNA45SN1_45S_50ntFlanks@11036.11060.25\|TCTCTCCCCCGCTCCCCGTCCTCCC | >hg38_21_+_8433172_8446622_RNA45SN1_45S_50ntFlanks | 11036 | 11060 | 25 | TCTCTCCCCCGCTCCCCGTCCTCCC | 693.0 | -5.4 | 2.1 | -2.6 | 8.5E-03 | 9.8E-03 | BT20/MB468 | NUC |
| >hg38_21_+_8433172_8446622_RNA45SN1_45S_50ntFlanks@8196.8225.30\|CGTGGACGGTGTGAGGCCGGTAGCGGCCCC | >hg38_21_+_8433172_8446622_RNA45SN1_45S_50ntFlanks | 8196 | 8225 | 30 | CGTGGACGGTGTGAGGCCGGTAGCGGCCCC | 553.7 | -3.4 | 1.1 | -3.1 | 2.3E-03 | 9.8E-03 | BT20/MB231 | CYTO |
| >hg38_21_+_8433172_8446622_RNA45SN1_45S_50ntFlanks@10781.10822.42\|AGGTGAACAGCCTCTGGCATGTTGGAACAATGTAGGTAAGGG | >hg38_21_+_8433172_8446622_RNA45SN1_45S_50ntFlanks | 10781 | 10822 | 42 | AGGTGAACAGCCTCTGGCATGTTGGAACAATGTAGGTAAGGG | 138.0 | -3.3 | 1.2 | -2.6 | 8.1E-03 | 9.8E-03 | BT20/MB468 | CYTO |
| >hg38_MT_+_598_1651_RNR1_12S_50ntFlanks@447.480.34\|TGAACACACAATAGCTAAGACCCAAACTGGGATT | >hg38_MT_+_598_1651_RNR1_12S_50ntFlanks | 447 | 480 | 34 | TGAACACACAATAGCTAAGACCCAAACTGGGATT | 253.6 | 6.2 | 2.2 | 2.8 | 4.9E-03 | 9.8E-03 | BT20/MB468 | MP |
| >hg38_21_+_8433172_8446622_RNA45SN1_45S_50ntFlanks@8170.8212.43\|CAAGTCCTTCTGATCGAGGCCCAGCCCGTGGACGGTGTGAGGC | >hg38_21_+_8433172_8446622_RNA45SN1_45S_50ntFlanks | 8170 | 8212 | 43 | CAAGTCCTTCTGATCGAGGCCCAGCCCGTGGACGGTGTGAGGC | 1704.0 | -4.7 | 1.7 | -2.8 | 4.9E-03 | 9.8E-03 | BT20/MB468 | MP |
| >hg38_21_+_8433172_8446622_RNA45SN1_45S_50ntFlanks@11037.11060.24\|CTCTCCCCCGCTCCCCGTCCTCCC | >hg38_21_+_8433172_8446622_RNA45SN1_45S_50ntFlanks | 11037 | 11060 | 24 | CTCTCCCCCGCTCCCCGTCCTCCC | 3519.5 | -5.9 | 2.3 | -2.6 | 8.5E-03 | 9.8E-03 | BT20/MB468 | NUC |
| >hg38_21_+_8433172_8446622_RNA45SN1_45S_50ntFlanks@11672.11705.34\|CAGTGCTCTGAATGTCAAAGTGAAGAAATTCAAT | >hg38_21_+_8433172_8446622_RNA45SN1_45S_50ntFlanks | 11672 | 11705 | 34 | CAGTGCTCTGAATGTCAAAGTGAAGAAATTCAAT | 44.7 | 5.7 | 2.0 | 2.8 | 4.9E-03 | 9.8E-03 | BT20/MB468 | MP |
| >hg38_21_+_8433172_8446622_RNA45SN1_45S_50ntFlanks@4760.4777.18\|TCAGATACCGTCGTAGTT | >hg38_21_+_8433172_8446622_RNA45SN1_45S_50ntFlanks | 4760 | 4777 | 18 | TCAGATACCGTCGTAGTT | 165.9 | -4.7 | 1.7 | -2.8 | 4.9E-03 | 9.8E-03 | BT20/MB468 | MP |
| >hg38_21_+_8433172_8446622_RNA45SN1_45S_50ntFlanks@12639.12659.21\|TACCATCTGTGGGATTATGAC | >hg38_21_+_8433172_8446622_RNA45SN1_45S_50ntFlanks | 12639 | 12659 | 21 | TACCATCTGTGGGATTATGAC | 268.9 | -2.3 | 0.9 | -2.7 | 7.0E-03 | 9.8E-03 | BT20/MB468 | MITO |
| >hg38_21_+_8433172_8446622_RNA45SN1_45S_50ntFlanks@11821.11854.34\|CTATCCAGCGAAACCACAGCCAAGGGAACGGGCT | >hg38_21_+_8433172_8446622_RNA45SN1_45S_50ntFlanks | 11821 | 11854 | 34 | CTATCCAGCGAAACCACAGCCAAGGGAACGGGCT | 74.7 | -4.6 | 1.6 | -2.8 | 4.9E-03 | 9.8E-03 | BT20/MB468 | MP |
| >hg38_21_+_8433172_8446622_RNA45SN1_45S_50ntFlanks@9926.9949.24\|GTTGATATAGACAGCAGGACGGTG | >hg38_21_+_8433172_8446622_RNA45SN1_45S_50ntFlanks | 9926 | 9949 | 24 | GTTGATATAGACAGCAGGACGGTG | 132.9 | 5.7 | 1.8 | 3.1 | 1.8E-03 | 9.8E-03 | BT20/MB231 | MP |
| >am_trna116_GluCTC_1_-_145399233_145399304@1.26.26,trna59_GluCTC_1_+_249168447_249168518@1.26.26,trna71_GluCTC_1_-_161439189_161439260@1.26.26,trna74_GluCTC_1_-_161431809_161431880@1.26.26,trna77_GluCTC_1_-_161424398_161424469@1.26.26,trna77_GluCTC_6_+_28949976_28950047@1.26.26,trna80_GluCTC_1_-_161417018_161417089@1.26.26,trna87_GluCTC_6_-_126101393_126101464@1.26.26\|TCCCTGGTGGTCTAGTGGTTAGGATT | >am_trna116_GluCTC_1_-_145399233_145399304 | 1 | 26 | 26 | TCCCTGGTGGTCTAGTGGTTAGGATT | 810.8 | -2.1 | 0.8 | -2.6 | 8.7E-03 | 9.8E-03 | MB231/MB468 | NUC |
| >hg38_21_+_8433172_8446622_RNA45SN1_45S_50ntFlanks@5490.5512.23\|TGAGAAGACGGTCGAACTTGACT | >hg38_21_+_8433172_8446622_RNA45SN1_45S_50ntFlanks | 5490 | 5512 | 23 | TGAGAAGACGGTCGAACTTGACT | 123.3 | 4.8 | 1.7 | 2.8 | 4.9E-03 | 9.8E-03 | BT20/MB468 | MP |
| >hg38_21_+_8433172_8446622_RNA45SN1_45S_50ntFlanks@7977.8018.42\|CGACCTCAGATCAGACGTGGCGACCCGCTGAATTTAAGCATA | >hg38_21_+_8433172_8446622_RNA45SN1_45S_50ntFlanks | 7977 | 8018 | 42 | CGACCTCAGATCAGACGTGGCGACCCGCTGAATTTAAGCATA | 269.0 | 2.7 | 1.0 | 2.6 | 8.7E-03 | 9.8E-03 | MB231/MB468 | MITO |
| >MI0000811\|hsa-mir-148b&WithFlank&12\|+\|54337210\|54337320@30.52.23&[MIMAT0004699&hsa-miR-148b-5p&offsets\|-1\|0;m-371&12\|+\|54337239\|54337262&offsets\|0\|-1]\|GAAGTTCTGTTATACACTCAGGC | >MI0000811\|hsa-mir-148b&WithFlank&12\|+\|54337210\|54337320 | 30 | 52 | 23 | GAAGTTCTGTTATACACTCAGGC | 97.9 | 3.5 | 1.3 | 2.7 | 7.5E-03 | 9.8E-03 | MB231/MB468 | MP |
| >MI0000087\|hsa-mir-29a&WithFlank&7\|-\|130876741\|130876816@48.68.21&[MIMAT0000086&hsa-miR-29a-3p&offsets\|0\|-1;m-26&7\|-\|130876749\|130876769&offsets\|0\|0]\|TAGCACCATCTGAAATCGGTT | >MI0000087\|hsa-mir-29a&WithFlank&7\|-\|130876741\|130876816 | 48 | 68 | 21 | TAGCACCATCTGAAATCGGTT | 15960.9 | -2.7 | 0.9 | -3.0 | 2.4E-03 | 9.8E-03 | BT20/MB231 | MITO |
| >hg38_21_+_8433172_8446622_RNA45SN1_45S_50ntFlanks@6646.6674.29\|TCGTACGACTCTTAGCGGTGGATCACTCG | >hg38_21_+_8433172_8446622_RNA45SN1_45S_50ntFlanks | 6646 | 6674 | 29 | TCGTACGACTCTTAGCGGTGGATCACTCG | 4129.9 | 4.1 | 1.3 | 3.1 | 1.8E-03 | 9.8E-03 | BT20/MB231 | MP |
| >hg38_21_+_8433172_8446622_RNA45SN1_45S_50ntFlanks@8356.8375.20\|GAAAAGAACTTTGAAGAGAG | >hg38_21_+_8433172_8446622_RNA45SN1_45S_50ntFlanks | 8356 | 8375 | 20 | GAAAAGAACTTTGAAGAGAG | 3077.6 | -2.8 | 1.0 | -2.7 | 7.5E-03 | 9.8E-03 | MB231/MB468 | MP |
| >hg38_21_+_8433172_8446622_RNA45SN1_45S_50ntFlanks@6646.6665.20\|TCGTACGACTCTTAGCGGTG | >hg38_21_+_8433172_8446622_RNA45SN1_45S_50ntFlanks | 6646 | 6665 | 20 | TCGTACGACTCTTAGCGGTG | 21885.6 | 2.3 | 0.9 | 2.6 | 8.7E-03 | 9.8E-03 | MB231/MB468 | MITO |
| >trna20_GluTTC_2_-_131094701_131094772@1.29.29,trna5_GluTTC_13_-_41634874_41634945@1.29.29\|TCCCATATGGTCTAGCGGTTAGGATTCCT | >trna20_GluTTC_2_-_131094701_131094772 | 1 | 29 | 29 | TCCCATATGGTCTAGCGGTTAGGATTCCT | 342.1 | -2.0 | 0.8 | -2.6 | 8.7E-03 | 9.8E-03 | MB231/MB468 | MITO |
| >hg38_21_+_8433172_8446622_RNA45SN1_45S_50ntFlanks@4738.4767.30\|AAAGTCGGAGGTTCGAAGACGATCAGATAC | >hg38_21_+_8433172_8446622_RNA45SN1_45S_50ntFlanks | 4738 | 4767 | 30 | AAAGTCGGAGGTTCGAAGACGATCAGATAC | 189.4 | 2.5 | 0.9 | 2.7 | 7.5E-03 | 9.8E-03 | MB231/MB468 | MP |
| >hg38_MT_+_1621_3279_RNR2_16S_50ntFlanks@1428.1453.26\|ATTAAAGTCCTACGTGATCTGAGTTC | >hg38_MT_+_1621_3279_RNR2_16S_50ntFlanks | 1428 | 1453 | 26 | ATTAAAGTCCTACGTGATCTGAGTTC | 416.5 | 2.3 | 0.9 | 2.7 | 7.5E-03 | 9.8E-03 | MB231/MB468 | MP |
| >hg38_21_+_8433172_8446622_RNA45SN1_45S_50ntFlanks@7988.8016.29\|CAGACGTGGCGACCCGCTGAATTTAAGCA | >hg38_21_+_8433172_8446622_RNA45SN1_45S_50ntFlanks | 7988 | 8016 | 29 | CAGACGTGGCGACCCGCTGAATTTAAGCA | 772.2 | -1.7 | 0.6 | -2.7 | 7.0E-03 | 9.8E-03 | BT20/MB468 | MITO |
| >hg38_21_+_8433172_8446622_RNA45SN1_45S_50ntFlanks@6723.6740.18\|TTGCAGGACACATTGATC | >hg38_21_+_8433172_8446622_RNA45SN1_45S_50ntFlanks | 6723 | 6740 | 18 | TTGCAGGACACATTGATC | 357.6 | -2.5 | 0.9 | -2.6 | 8.1E-03 | 9.8E-03 | BT20/MB468 | CYTO |
| >hg38_21_+_8433172_8446622_RNA45SN1_45S_50ntFlanks@9547.9571.25\|GGAGGTCCGTAGCGGTCCTGACGTG | >hg38_21_+_8433172_8446622_RNA45SN1_45S_50ntFlanks | 9547 | 9571 | 25 | GGAGGTCCGTAGCGGTCCTGACGTG | 490.5 | 1.8 | 0.7 | 2.6 | 8.2E-03 | 9.8E-03 | MB231/MB468 | CYTO |
| >trnaMT_PheGAA_MT_+_577_647@42.73.32\|TTTAGACGGGCTCACATCACCCCATAAACACC | >trnaMT_PheGAA_MT_+_577_647 | 42 | 73 | 32 | TTTAGACGGGCTCACATCACCCCATAAACACC | 198.5 | 3.8 | 1.3 | 2.8 | 4.9E-03 | 9.8E-03 | BT20/MB468 | MP |
| >hg38_21_+_8433172_8446622_RNA45SN1_45S_50ntFlanks@4895.4916.22\|CTTAAAGGAATTGACGGAAGGG | >hg38_21_+_8433172_8446622_RNA45SN1_45S_50ntFlanks | 4895 | 4916 | 22 | CTTAAAGGAATTGACGGAAGGG | 88.5 | -4.6 | 1.7 | -2.8 | 4.9E-03 | 9.8E-03 | BT20/MB468 | MP |
| >hg38_21_+_8433172_8446622_RNA45SN1_45S_50ntFlanks@9913.9938.26\|AGAAAAGGTGTTGGTTGATATAGACA | >hg38_21_+_8433172_8446622_RNA45SN1_45S_50ntFlanks | 9913 | 9938 | 26 | AGAAAAGGTGTTGGTTGATATAGACA | 158.0 | 4.6 | 1.5 | 3.1 | 1.8E-03 | 9.8E-03 | BT20/MB231 | MP |
| >MI0000077\|hsa-mir-21&WithFlank&17\|+\|59841260\|59841343@14.33.20(+1C)&[MIMAT0000076&hsa-miR-21-5p&offsets\|0\|-2(+1C);m-1&17\|+\|59841273\|59841294&offsets\|0\|-2(+1C)]\|TAGCTTATCAGACTGATGTTC | >MI0000077\|hsa-mir-21&WithFlank&17\|+\|59841260\|59841343 | 14 | 33 | 20(+1C) | TAGCTTATCAGACTGATGTTC | 150.0 | 3.8 | 1.4 | 2.7 | 7.5E-03 | 9.8E-03 | MB231/MB468 | MP |
| >am_trnaMT_TyrGTA_MT_-_5826_5891@1.32.32\|GGTAAAATGGCTGAGTGAAGCATTGGACTGTA | >am_trnaMT_TyrGTA_MT_-_5826_5891 | 1 | 32 | 32 | GGTAAAATGGCTGAGTGAAGCATTGGACTGTA | 2515.3 | -3.5 | 1.2 | -3.0 | 2.4E-03 | 9.8E-03 | BT20/MB231 | MITO |

<table>
<tr><td>>hg38_21_+_8433172_8446622_RNA45SN1_45S_50ntFlanks@11907.11929.23|ACGGTGAAGAGACATGAGAGGTG</td><td>>hg38_21_+_8433172_8446622_RNA45SN1_45S_50ntFlanks</td><td>11907</td><td>11929</td><td>23</td><td>ACGGTGAAGAGACATGAGAGGTG</td><td>103.5</td><td>-3.3</td><td>1.3</td><td>-2.6</td><td>8.2E-03</td><td>9.9E-03</td><td>MB231/MB468</td><td>CYTO</td></tr>
<tr><td>>hg38_21_+_8433172_8446622_RNA45SN1_45S_50ntFlanks@6644.6681.38|CCTCGTACGACTCTTAGCGGTGGATCACTCGGCTCGTG</td><td>>hg38_21_+_8433172_8446622_RNA45SN1_45S_50ntFlanks</td><td>6644</td><td>6681</td><td>38</td><td>CCTCGTACGACTCTTAGCGGTGGATCACTCGGCTCGTG</td><td>526.1</td><td>-1.4</td><td>0.5</td><td>-2.7</td><td>7.0E-03</td><td>9.9E-03</td><td>BT20/MB468</td><td>MITO</td></tr>
<tr><td>>hg38_21_+_8433172_8446622_RNA45SN1_45S_50ntFlanks@7974.8002.29|ACGCGACCTCAGATCAGACGTGGCGACCC</td><td>>hg38_21_+_8433172_8446622_RNA45SN1_45S_50ntFlanks</td><td>7974</td><td>8002</td><td>29</td><td>ACGCGACCTCAGATCAGACGTGGCGACCC</td><td>354.0</td><td>2.2</td><td>0.8</td><td>2.6</td><td>8.7E-03</td><td>9.9E-03</td><td>MB231/MB468</td><td>MITO</td></tr>
<tr><td>>MI0000465|hsa-mir-191&WithFlank&3|-|49020612|49020715@22.42.21&[MIMAT0000440&hsa-miR-191-5p&offsets|0|-2;m-48&3|-|49020672|49020694&offsets|0|-2]|CAACGGAATCCCAAAAGCAGC</td><td>>MI0000465|hsa-mir-191&WithFlank&3|-|49020612|49020715</td><td>22</td><td>42</td><td>21</td><td>CAACGGAATCCCAAAAGCAGC</td><td>581.8</td><td>2.5</td><td>0.9</td><td>2.7</td><td>7.5E-03</td><td>9.9E-03</td><td>MB231/MB468</td><td>MP</td></tr>
<tr><td>>hg38_1_-_228634819_228635039_RNA5S12_5S_50ntFlanks@51.71.21|GTCTACGGCCATACCACCCTG</td><td>>hg38_1_-_228634819_228635039_RNA5S12_5S_50ntFlanks</td><td>51</td><td>71</td><td>21</td><td>GTCTACGGCCATACCACCCTG</td><td>208.0</td><td>-3.7</td><td>1.4</td><td>-2.7</td><td>7.5E-03</td><td>9.9E-03</td><td>MB231/MB468</td><td>MP</td></tr>
<tr><td>>trnaMT_ValTAC_MT_+_1602_1670@34.71.38|CACTTAGGAGATTTCAACTTAACTTGACCGCTCTGACC</td><td>>trnaMT_ValTAC_MT_+_1602_1670</td><td>34</td><td>71</td><td>38</td><td>CACTTAGGAGATTTCAACTTAACTTGACCGCTCTGACC</td><td>41.5</td><td>5.1</td><td>1.9</td><td>2.7</td><td>7.5E-03</td><td>9.9E-03</td><td>MB231/MB468</td><td>MP</td></tr>
<tr><td>>hg38_21_+_8433172_8446622_RNA45SN1_45S_50ntFlanks@4735.4761.27|ACGAAAGTCGGAGGTTCGAAGACGATC</td><td>>hg38_21_+_8433172_8446622_RNA45SN1_45S_50ntFlanks</td><td>4735</td><td>4761</td><td>27</td><td>ACGAAAGTCGGAGGTTCGAAGACGATC</td><td>199.8</td><td>2.1</td><td>0.8</td><td>2.8</td><td>4.9E-03</td><td>9.9E-03</td><td>BT20/MB468</td><td>MP</td></tr>
<tr><td>>MI0000077|hsa-mir-21&WithFlank&17|+|59841260|59841343@14.34.21(+2U)&[MIMAT0000076&hsa-miR-21-5p&offsets|0|-1(+2U);m-1&17|+|59841273|59841294&offsets|0|-1(+2U)]|TAGCTTATCAGACTGATGTTGTT</td><td>>MI0000077|hsa-mir-21&WithFlank&17|+|59841260|59841343</td><td>14</td><td>34</td><td>21(+2U)</td><td>TAGCTTATCAGACTGATGTTGTT</td><td>266.8</td><td>3.2</td><td>1.0</td><td>3.0</td><td>2.3E-03</td><td>9.9E-03</td><td>BT20/MB231</td><td>CYTO</td></tr>
<tr><td>>trnaMT_ValTAC_MT_+_1602_1670@49.70.22|AACTTAACTTGACCGCTCTGAC</td><td>>trnaMT_ValTAC_MT_+_1602_1670</td><td>49</td><td>70</td><td>22</td><td>AACTTAACTTGACCGCTCTGAC</td><td>234.0</td><td>2.3</td><td>0.8</td><td>3.0</td><td>2.3E-03</td><td>9.9E-03</td><td>BT20/MB231</td><td>CYTO</td></tr>
<tr><td>>hg38_21_+_8433172_8446622_RNA45SN1_45S_50ntFlanks@10420.10453.34|CTGAGAGATGGGCGAGCGCCGTTCCGAAGGGACG</td><td>>hg38_21_+_8433172_8446622_RNA45SN1_45S_50ntFlanks</td><td>10420</td><td>10453</td><td>34</td><td>CTGAGAGATGGGCGAGCGCCGTTCCGAAGGGACG</td><td>216.3</td><td>-2.6</td><td>0.9</td><td>-3.0</td><td>2.3E-03</td><td>9.9E-03</td><td>BT20/MB231</td><td>CYTO</td></tr>
<tr><td>>hg38_21_+_8433172_8446622_RNA45SN1_45S_50ntFlanks@12424.12457.34|CGGCTCTTCCTATCATTGTGAAGCAGAATTCACC</td><td>>hg38_21_+_8433172_8446622_RNA45SN1_45S_50ntFlanks</td><td>12424</td><td>12457</td><td>34</td><td>CGGCTCTTCCTATCATTGTGAAGCAGAATTCACC</td><td>300.6</td><td>-2.5</td><td>0.8</td><td>-3.0</td><td>2.3E-03</td><td>9.9E-03</td><td>BT20/MB231</td><td>CYTO</td></tr>
<tr><td>>hg38_21_+_8433172_8446622_RNA45SN1_45S_50ntFlanks@12001.12043.43|CTGCGGGCCGCCGGTGAAATACCACTACTCTGATCGTTTTTTC</td><td>>hg38_21_+_8433172_8446622_RNA45SN1_45S_50ntFlanks</td><td>12001</td><td>12043</td><td>43</td><td>CTGCGGGCCGCCGGTGAAATACCACTACTCTGATCGTTTTTTC</td><td>402.6</td><td>-4.4</td><td>1.6</td><td>-2.7</td><td>7.0E-03</td><td>9.9E-03</td><td>BT20/MB468</td><td>MITO</td></tr>
<tr><td>>hg38_21_+_8433172_8446622_RNA45SN1_45S_50ntFlanks@6734.6756.23|ATTGATCATCGACACTTCGAACG</td><td>>hg38_21_+_8433172_8446622_RNA45SN1_45S_50ntFlanks</td><td>6734</td><td>6756</td><td>23</td><td>ATTGATCATCGACACTTCGAACG</td><td>1396.9</td><td>5.1</td><td>1.6</td><td>3.1</td><td>1.8E-03</td><td>9.9E-03</td><td>BT20/MB231</td><td>MP</td></tr>
<tr><td>>hg38_21_+_8433172_8446622_RNA45SN1_45S_50ntFlanks@4297.4315.19|CATTGGAGGGCAAGTCTGG</td><td>>hg38_21_+_8433172_8446622_RNA45SN1_45S_50ntFlanks</td><td>4297</td><td>4315</td><td>19</td><td>CATTGGAGGGCAAGTCTGG</td><td>125.8</td><td>-3.5</td><td>1.3</td><td>-2.7</td><td>7.5E-03</td><td>9.9E-03</td><td>MB231/MB468</td><td>MP</td></tr>
<tr><td>>trnaMT_SerGCT_MT_+_12207_12265@15.62.48|GAACTGCTAACTCATGCCCCCATGTCTAACAACATGGCTTTCTCACCA</td><td>>trnaMT_SerGCT_MT_+_12207_12265</td><td>15</td><td>62</td><td>48</td><td>GAACTGCTAACTCATGCCCCCATGTCTAACAACATGGCTTTCTCACCA</td><td>738.7</td><td>4.7</td><td>1.8</td><td>2.6</td><td>8.7E-03</td><td>9.9E-03</td><td>MB231/MB468</td><td>MITO</td></tr>
<tr><td>>hg38_1_-_228634819_228635039_RNA5S12_5S_50ntFlanks@51.88.38|GTCTACGGCCATACCACCCTGAACGCGCCCGATCTCGT</td><td>>hg38_1_-_228634819_228635039_RNA5S12_5S_50ntFlanks</td><td>51</td><td>88</td><td>38</td><td>GTCTACGGCCATACCACCCTGAACGCGCCCGATCTCGT</td><td>502.0</td><td>2.8</td><td>1.0</td><td>2.7</td><td>7.5E-03</td><td>9.9E-03</td><td>MB231/MB468</td><td>MP</td></tr>
<tr><td>>hg38_21_+_8433172_8446622_RNA45SN1_45S_50ntFlanks@6648.6686.39|GTACGACTCTTAGCGGTGGATCACTCGGCTCGTGCGTCG</td><td>>hg38_21_+_8433172_8446622_RNA45SN1_45S_50ntFlanks</td><td>6648</td><td>6686</td><td>39</td><td>GTACGACTCTTAGCGGTGGATCACTCGGCTCGTGCGTCG</td><td>114.3</td><td>-2.4</td><td>0.9</td><td>-2.7</td><td>7.5E-03</td><td>9.9E-03</td><td>MB231/MB468</td><td>MP</td></tr>
<tr><td>>trna10_AspGTC_12_-_125424193_125424264@39.73.35,trna12_AspGTC_12_-_125411891_125411962@39.73.35,trna144_AspGTC_6_-_27551236_27551307@39.73.35,trna38_AspGTC_17_-_8125556_8125627@39.73.35,trna45_AspGTC_6_+_27447453_27447524@39.73.35,trna48_AspGTC_6_+_27471523_27471594@39.73.35,trna4_AspGTC_12_+_96429799_96429870@39.73.35,trna69_AspGTC_1_-_161440205_161440276@39.73.35,trna72_AspGTC_1_-_161432824_161432895@39.73.35,trna75_AspGTC_1_-_161425414_161425485@39.73.35,trna78_AspGTC_1_-_161418033_161418104@39.73.35,trna81_AspGTC_1_-_161410615_161410686@39.73.35|GCGGGAGACCGGGGTTCGATTCCCCGACGGGGAGC</td><td>>trna10_AspGTC_12_-_125424193_125424264</td><td>39</td><td>73</td><td>35</td><td>GCGGGAGACCGGGGTTCGATTCCCCGACGGGGAGC</td><td>103.0</td><td>2.2</td><td>0.8</td><td>2.7</td><td>7.5E-03</td><td>9.9E-03</td><td>MB231/MB468</td><td>MP</td></tr>
<tr><td>>MI0000456|hsa-mir-140&WithFlank&16|+|69933075|69933186@68.87.20&[MIMAT0004597&hsa-miR-140-3p&offsets|0|-1;m-45&16|+|69933143|69933164&offsets|-1|-3]|TACCACAGGGTAGAACCACG</td><td>>MI0000456|hsa-mir-140&WithFlank&16|+|69933075|69933186</td><td>68</td><td>87</td><td>20</td><td>TACCACAGGGTAGAACCACG</td><td>39.7</td><td>5.1</td><td>1.9</td><td>2.7</td><td>7.5E-03</td><td>9.9E-03</td><td>MB231/MB468</td><td>MP</td></tr>
<tr><td>>hg38_21_+_8433172_8446622_RNA45SN1_45S_50ntFlanks@483.505.23|TCGATGTGGTGACGTCGTGCTCT</td><td>>hg38_21_+_8433172_8446622_RNA45SN1_45S_50ntFlanks</td><td>483</td><td>505</td><td>23</td><td>TCGATGTGGTGACGTCGTGCTCT</td><td>121.5</td><td>2.3</td><td>0.9</td><td>2.7</td><td>7.6E-03</td><td>9.9E-03</td><td>MB231/MB468</td><td>MP</td></tr>
<tr><td>>hg38_21_+_8433172_8446622_RNA45SN1_45S_50ntFlanks@9790.9820.31|CTGGCGTGGAGCCGGGCGTGGAATGCGAGTG</td><td>>hg38_21_+_8433172_8446622_RNA45SN1_45S_50ntFlanks</td><td>9790</td><td>9820</td><td>31</td><td>CTGGCGTGGAGCCGGGCGTGGAATGCGAGTG</td><td>354.8</td><td>-5.2</td><td>2.0</td><td>-2.6</td><td>8.6E-03</td><td>9.9E-03</td><td>BT20/MB468</td><td>NUC</td></tr>
<tr><td>>hg38_MT_+_1621_3279_RNR2_16S_50ntFlanks@1369.1397.29|GATGTTGGATCAGGACATCCCGATGGTGC</td><td>>hg38_MT_+_1621_3279_RNR2_16S_50ntFlanks</td><td>1369</td><td>1397</td><td>29</td><td>GATGTTGGATCAGGACATCCCGATGGTGC</td><td>81.0</td><td>3.5</td><td>1.3</td><td>2.7</td><td>7.6E-03</td><td>9.9E-03</td><td>MB231/MB468</td><td>MP</td></tr>
<tr><td>>MI0000803|hsa-mir-330&WithFlank&19|-|45638988|45639093@64.86.23(+1A)&[MIMAT0000751&hsa-miR-330-3p&offsets|+1|+1(+1A);m-347&19|-|45639009|45639030&offsets|0|+1(+1A)]|CAAAGCACACGGCCTGCAGAGAGA</td><td>>MI0000803|hsa-mir-330&WithFlank&19|-|45638988|45639093</td><td>64</td><td>86</td><td>23(+1A)</td><td>CAAAGCACACGGCCTGCAGAGAGA</td><td>47.0</td><td>5.9</td><td>2.1</td><td>2.8</td><td>4.9E-03</td><td>9.9E-03</td><td>BT20/MB468</td><td>MP</td></tr>
<tr><td>>hg38_21_+_8433172_8446622_RNA45SN1_45S_50ntFlanks@8198.8224.27|TGGACGGTGTGAGGCCGGTAGCGGCCC</td><td>>hg38_21_+_8433172_8446622_RNA45SN1_45S_50ntFlanks</td><td>8198</td><td>8224</td><td>27</td><td>TGGACGGTGTGAGGCCGGTAGCGGCCC</td><td>291.8</td><td>-2.2</td><td>0.8</td><td>-2.8</td><td>4.9E-03</td><td>9.9E-03</td><td>BT20/MB468</td><td>MP</td></tr>
<tr><td>>MI0000095|hsa-mir-93&WithFlank&7|-|100093762|100093853@17.39.23(+1A)&[MIMAT0000093&hsa-miR-93-5p&offsets|0|0(+1A);m-29&7|-|100093815|100093837&offsets|0|0(+1A)]|CAAAGTGCTGTTCGTGCAGGTAGA</td><td>>MI0000095|hsa-mir-93&WithFlank&7|-|100093762|100093853</td><td>17</td><td>39</td><td>23(+1A)</td><td>CAAAGTGCTGTTCGTGCAGGTAGA</td><td>636.9</td><td>1.3</td><td>0.5</td><td>2.6</td><td>8.6E-03</td><td>9.9E-03</td><td>BT20/MB468</td><td>NUC</td></tr>
<tr><td>>hg38_21_+_8433172_8446622_RNA45SN1_45S_50ntFlanks@10441.10460.20|TTCCGAAGGGACGGGCGATG</td><td>>hg38_21_+_8433172_8446622_RNA45SN1_45S_50ntFlanks</td><td>10441</td><td>10460</td><td>20</td><td>TTCCGAAGGGACGGGCGATG</td><td>269.3</td><td>-2.0</td><td>0.8</td><td>-2.6</td><td>8.2E-03</td><td>9.9E-03</td><td>BT20/MB468</td><td>CYTO</td></tr>
<tr><td>>hg38_21_+_8433172_8446622_RNA45SN1_45S_50ntFlanks@10391.10424.34|AACAGCAGTTGAACATGGGTCAGTCGGTCCTGAG</td><td>>hg38_21_+_8433172_8446622_RNA45SN1_45S_50ntFlanks</td><td>10391</td><td>10424</td><td>34</td><td>AACAGCAGTTGAACATGGGTCAGTCGGTCCTGAG</td><td>3974.2</td><td>1.9</td><td>0.7</td><td>2.6</td><td>8.2E-03</td><td>9.9E-03</td><td>BT20/MB468</td><td>CYTO</td></tr>
<tr><td>>hg38_21_+_8433172_8446622_RNA45SN1_45S_50ntFlanks@5237.5255.19|ACTGACTGGCTCAGCGTGT</td><td>>hg38_21_+_8433172_8446622_RNA45SN1_45S_50ntFlanks</td><td>5237</td><td>5255</td><td>19</td><td>ACTGACTGGCTCAGCGTGT</td><td>133.0</td><td>2.2</td><td>0.8</td><td>2.7</td><td>7.6E-03</td><td>9.9E-03</td><td>MB231/MB468</td><td>MP</td></tr>
<tr><td>>MI0003834|hsa-mir-769&WithFlank&19|+|46018926|46019055@36.57.22(+3U)&[MIMAT0003886&hsa-miR-769-5p&offsets|0|0(+3U);m-216&19|+|46018961|46018982&offsets|0|0(+3U)]|TGAGACCTCTGGGTTCTGAGCTTTT</td><td>>MI0003834|hsa-mir-769&WithFlank&19|+|46018926|46019055</td><td>36</td><td>57</td><td>22(+3U)</td><td>TGAGACCTCTGGGTTCTGAGCTTTT</td><td>61.3</td><td>3.4</td><td>1.3</td><td>2.7</td><td>7.6E-03</td><td>9.9E-03</td><td>MB231/MB468</td><td>MP</td></tr>
<tr><td>>trna10_AspGTC_12_-_125424193_125424264@38.69.32,trna12_AspGTC_12_-_125411891_125411962@38.69.32,trna144_AspGTC_6_-_27551236_27551307@38.69.32,trna38_AspGTC_17_-_8125556_8125627@38.69.32,trna45_AspGTC_6_+_27447453_27447524@38.69.32,trna48_AspGTC_6_+_27471523_27471594@38.69.32,trna4_AspGTC_12_+_96429799_96429870@38.69.32,trna69_AspGTC_1_-_161440205_161440276@38.69.32,trna72_AspGTC_1_-_161432824_161432895@38.69.32,trna75_AspGTC_1_-_161425414_161425485@38.69.32,trna78_AspGTC_1_-_161418033_161418104@38.69.32,trna81_AspGTC_1_-_161410615_161410686@38.69.32|CGCGGGAGACCGGGGTTCGATTCCCCGACGGG</td><td>>trna10_AspGTC_12_-_125424193_125424264</td><td>38</td><td>69</td><td>32</td><td>CGCGGGAGACCGGGGTTCGATTCCCCGACGGG</td><td>702.5</td><td>3.5</td><td>1.1</td><td>3.0</td><td>2.3E-03</td><td>9.9E-03</td><td>BT20/MB231</td><td>CYTO</td></tr>
<tr><td>>trna116_GluCTC_1_-_145399233_145399304@37.75.39,trna134_GluTTC_1_-_16861774_16861845@37.75.39,trna71_GluCTC_1_-_161439189_161439260@37.75.39,trna74_GluCTC_1_-_161431809_161431880@37.75.39,trna77_GluCTC_1_-_161424398_161424469@37.75.39,trna77_GluCTC_6_+_28949976_28950047@37.75.39,trna80_GluCTC_1_-_161417018_161417089@37.75.39,trna84_GluTTC_1_-_161391883_161391954@37.75.39,trna87_GluCTC_6_-_126101393_126101464@37.75.39|ACCGCCGCGGCCCGGGTTCGATTCCCGGTCAGGGAACCA</td><td>>trna116_GluCTC_1_-_145399233_145399304</td><td>37</td><td>75</td><td>39</td><td>ACCGCCGCGGCCCGGGTTCGATTCCCGGTCAGGGAACCA</td><td>514.9</td><td>4.6</td><td>1.7</td><td>2.6</td><td>8.8E-03</td><td>9.9E-03</td><td>MB231/MB468</td><td>MITO</td></tr>
<tr><td>>hg38_21_+_8433172_8446622_RNA45SN1_45S_50ntFlanks@10353.10388.36|AGAACTTTGAAGGCCGAAGTGGAGAAGGGTTCCATG</td><td>>hg38_21_+_8433172_8446622_RNA45SN1_45S_50ntFlanks</td><td>10353</td><td>10388</td><td>36</td><td>AGAACTTTGAAGGCCGAAGTGGAGAAGGGTTCCATG</td><td>376.7</td><td>-2.5</td><td>1.0</td><td>-2.6</td><td>8.2E-03</td><td>9.9E-03</td><td>BT20/MB468</td><td>CYTO</td></tr>
<tr><td>>hg38_21_+_8433172_8446622_RNA45SN1_45S_50ntFlanks@3837.3861.25|CCTCTCCTACTTGGATAACTGTGGT</td><td>>hg38_21_+_8433172_8446622_RNA45SN1_45S_50ntFlanks</td><td>3837</td><td>3861</td><td>25</td><td>CCTCTCCTACTTGGATAACTGTGGT</td><td>588.5</td><td>-2.6</td><td>1.0</td><td>-2.6</td><td>8.8E-03</td><td>9.9E-03</td><td>MB231/MB468</td><td>MITO</td></tr>
<tr><td>>hg38_1_-_228634819_228635039_RNA5S12_5S_50ntFlanks@135.169.35|GGGAGACCGCCTGGGAATACCGGGTGCTGTAGGCT</td><td>>hg38_1_-_228634819_228635039_RNA5S12_5S_50ntFlanks</td><td>135</td><td>169</td><td>35</td><td>GGGAGACCGCCTGGGAATACCGGGTGCTGTAGGCT</td><td>1141.5</td><td>2.6</td><td>0.9</td><td>2.8</td><td>4.9E-03</td><td>9.9E-03</td><td>BT20/MB468</td><td>MP</td></tr>
<tr><td>>MI0000081|hsa-mir-24-2&WithFlank&19|-|13836281|13836365@56.75.20(+1G);MI0000080|hsa-mir-24-1&WithFlank&9|+|95086015|95086094@50.69.20(+1G)&[MIMAT0000080&hsa-miR-24-3p&offsets|0|-2(+1G);m-36&19|-|13836288|13836310&offsets|0|-3(+1G)];[MIMAT0000080_1&hsa-miR-24-3p&offsets|0|-2(+1G);m-35&9|+|95086064|95086086&offsets|0|-3(+1G)]|TGGCTCAGTTCAGCAGGAACG</td><td>>MI0000081|hsa-mir-24-2&WithFlank&19|-|13836281|13836365</td><td>56</td><td>75</td><td>20(+1G)</td><td>TGGCTCAGTTCAGCAGGAACG</td><td>140.4</td><td>2.4</td><td>0.9</td><td>2.7</td><td>7.6E-03</td><td>1.0E-02</td><td>MB231/MB468</td><td>MP</td></tr>
<tr><td>>hg38_21_+_8433172_8446622_RNA45SN1_45S_50ntFlanks@5073.5115.43|AATTCCGATAACGAACGAGACTCTGGCATGCTAACTAGTTACG</td><td>>hg38_21_+_8433172_8446622_RNA45SN1_45S_50ntFlanks</td><td>5073</td><td>5115</td><td>43</td><td>AATTCCGATAACGAACGAGACTCTGGCATGCTAACTAGTTACG</td><td>186.9</td><td>-2.1</td><td>0.8</td><td>-2.6</td><td>8.8E-03</td><td>1.0E-02</td><td>MB231/MB468</td><td>NUC</td></tr>
<tr><td>>MI0000434|hsa-let-7i&WithFlank&12|+|62603680|62603775@12.34.23&[MIMAT0000415&hsa-let-7i-5p&offsets|0|+1;m-49&12|+|62603691|62603712&offsets|0|+1]|TGAGGTAGTAGTTTGTGCTGTTG</td><td>>MI0000434|hsa-let-7i&WithFlank&12|+|62603680|62603775</td><td>12</td><td>34</td><td>23</td><td>TGAGGTAGTAGTTTGTGCTGTTG</td><td>347.0</td><td>3.5</td><td>1.3</td><td>2.7</td><td>7.6E-03</td><td>1.0E-02</td><td>MB231/MB468</td><td>MP</td></tr>
<tr><td>>hg38_21_+_8433172_8446622_RNA45SN1_45S_50ntFlanks@12990.13012.23|GCTCCCTCGCTGCGATCTATTGA</td><td>>hg38_21_+_8433172_8446622_RNA45SN1_45S_50ntFlanks</td><td>12990</td><td>13012</td><td>23</td><td>GCTCCCTCGCTGCGATCTATTGA</td><td>365.6</td><td>-2.7</td><td>0.9</td><td>-3.0</td><td>2.3E-03</td><td>1.0E-02</td><td>BT20/MB231</td><td>CYTO</td></tr>
</table>

<table>
<tr><td>>hg38_21_+_8433172_8446622_RNA45SN1_45S_50ntFlanks@7981.8009.29|CTCAGATCAGACGTGGCGACCCGCTGAAT</td><td>>hg38_21_+_8433172_8446622_RNA45SN1_45S_50ntFlanks</td><td>7981</td><td>8009</td><td>29</td><td>CTCAGATCAGACGTGGCGACCCGCTGAAT</td><td>373.6</td><td>2.1</td><td>0.7</td><td>3.0</td><td>2.4E-03</td><td>1.0E-02</td><td>BT20/MB231</td><td>MITO</td></tr>
<tr><td>>hg38_MT_+_1621_3279_RNR2_16S_50ntFlanks@1367.1405.39|TCGATGTTGGATCAGGACATCCCGATGGTGCAGCCGCTA</td><td>>hg38_MT_+_1621_3279_RNR2_16S_50ntFlanks</td><td>1367</td><td>1405</td><td>39</td><td>TCGATGTTGGATCAGGACATCCCGATGGTGCAGCCGCTA</td><td>24.5</td><td>7.8</td><td>2.9</td><td>2.7</td><td>7.6E-03</td><td>1.0E-02</td><td>MB231/MB468</td><td>MP</td></tr>
<tr><td>>hg38_21_+_8433172_8446622_RNA45SN1_45S_50ntFlanks@10248.10279.32|CGACGAGTAGGAGGGCCGCTGCGGTGAGCCTT</td><td>>hg38_21_+_8433172_8446622_RNA45SN1_45S_50ntFlanks</td><td>10248</td><td>10279</td><td>32</td><td>CGACGAGTAGGAGGGCCGCTGCGGTGAGCCTT</td><td>487.4</td><td>3.9</td><td>1.2</td><td>3.1</td><td>1.9E-03</td><td>1.0E-02</td><td>BT20/MB231</td><td>MP</td></tr>
<tr><td>>hg38_21_+_8433172_8446622_RNA45SN1_45S_50ntFlanks@9297.9322.26|AACACGGACCAAGGAGTCTAACACGT</td><td>>hg38_21_+_8433172_8446622_RNA45SN1_45S_50ntFlanks</td><td>9297</td><td>9322</td><td>26</td><td>AACACGGACCAAGGAGTCTAACACGT</td><td>55.1</td><td>9.1</td><td>2.9</td><td>3.1</td><td>1.9E-03</td><td>1.0E-02</td><td>BT20/MB231</td><td>MP</td></tr>
<tr><td>>trnaMT_GluTTC_MT_-_14674_14742@38.72.35,trnalookalike8_GluTTC_5_-_93905172_93905240@38.72.35|CATTGGTCGTGGTTGTAGTCCGTGCGAGAATACCA</td><td>>trnaMT_GluTTC_MT_-_14674_14742</td><td>38</td><td>72</td><td>35</td><td>CATTGGTCGTGGTTGTAGTCCGTGCGAGAATACCA</td><td>9087.0</td><td>3.7</td><td>1.3</td><td>2.8</td><td>5.0E-03</td><td>1.0E-02</td><td>BT20/MB468</td><td>MP</td></tr>
<tr><td>>hg38_21_+_8433172_8446622_RNA45SN1_45S_50ntFlanks@6781.6799.19|GGGGCTACGCCTGTCTGAG</td><td>>hg38_21_+_8433172_8446622_RNA45SN1_45S_50ntFlanks</td><td>6781</td><td>6799</td><td>19</td><td>GGGGCTACGCCTGTCTGAG</td><td>102.4</td><td>-3.5</td><td>1.2</td><td>-2.8</td><td>5.0E-03</td><td>1.0E-02</td><td>BT20/MB468</td><td>MP</td></tr>
<tr><td>>hg38_21_+_8433172_8446622_RNA45SN1_45S_50ntFlanks@8198.8225.28|TGGACGGTGTGAGGCCGGTAGCGGCCCC</td><td>>hg38_21_+_8433172_8446622_RNA45SN1_45S_50ntFlanks</td><td>8198</td><td>8225</td><td>28</td><td>TGGACGGTGTGAGGCCGGTAGCGGCCCC</td><td>304.8</td><td>-2.4</td><td>0.9</td><td>-2.8</td><td>5.0E-03</td><td>1.0E-02</td><td>BT20/MB468</td><td>MP</td></tr>
<tr><td>>hg38_21_+_8433172_8446622_RNA45SN1_45S_50ntFlanks@4663.4695.33|GTGAAATTCTTGGACCGGCGCAAGACGGACCAG</td><td>>hg38_21_+_8433172_8446622_RNA45SN1_45S_50ntFlanks</td><td>4663</td><td>4695</td><td>33</td><td>GTGAAATTCTTGGACCGGCGCAAGACGGACCAG</td><td>54.8</td><td>-4.5</td><td>1.6</td><td>-2.8</td><td>5.0E-03</td><td>1.0E-02</td><td>BT20/MB468</td><td>MP</td></tr>
<tr><td>>hg38_21_+_8433172_8446622_RNA45SN1_45S_50ntFlanks@5474.5506.33|GGCCCTGGCGGAGCGCTGAGAAGACGGTCGAAC</td><td>>hg38_21_+_8433172_8446622_RNA45SN1_45S_50ntFlanks</td><td>5474</td><td>5506</td><td>33</td><td>GGCCCTGGCGGAGCGCTGAGAAGACGGTCGAAC</td><td>46.3</td><td>5.6</td><td>2.0</td><td>2.8</td><td>5.0E-03</td><td>1.0E-02</td><td>BT20/MB468</td><td>MP</td></tr>
<tr><td>>MI0000071|hsa-mir-17&WithFlank&13|+|91350599|91350694@20.39.20(+1A)&[MIMAT0000070&hsa-miR-17-5p&offsets|0|-3(+1A);m-88&13|+|91350618|91350640&offsets|0|-3(+1A)]|CAAAGTGCTTACAGTGCAGGA</td><td>>MI0000071|hsa-mir-17&WithFlank&13|+|91350599|91350694</td><td>20</td><td>39</td><td>20(+1A)</td><td>CAAAGTGCTTACAGTGCAGGA</td><td>91.5</td><td>3.6</td><td>1.4</td><td>2.7</td><td>7.6E-03</td><td>1.0E-02</td><td>MB231/MB468</td><td>MP</td></tr>
<tr><td>>MI0000077|hsa-mir-21&WithFlank&17|+|59841260|59841343@14.32.19(+1A)&[MIMAT0000076&hsa-miR-21-5p&offsets|0|-3(+1A);m-1&17|+|59841273|59841294&offsets|0|-3(+1A)]|TAGCTTATCAGACTGATGTA</td><td>>MI0000077|hsa-mir-21&WithFlank&17|+|59841260|59841343</td><td>14</td><td>32</td><td>19(+1A)</td><td>TAGCTTATCAGACTGATGTA</td><td>92.2</td><td>3.6</td><td>1.4</td><td>2.7</td><td>7.6E-03</td><td>1.0E-02</td><td>MB231/MB468</td><td>MP</td></tr>
<tr><td>>hg38_21_+_8433172_8446622_RNA45SN1_45S_50ntFlanks@10866.10889.24|CTAAGGGCTGGGTCGGTCGGGCTG</td><td>>hg38_21_+_8433172_8446622_RNA45SN1_45S_50ntFlanks</td><td>10866</td><td>10889</td><td>24</td><td>CTAAGGGCTGGGTCGGTCGGGCTG</td><td>446.1</td><td>-2.9</td><td>1.0</td><td>-2.8</td><td>5.0E-03</td><td>1.0E-02</td><td>BT20/MB468</td><td>MP</td></tr>
<tr><td>>hg38_21_+_8433172_8446622_RNA45SN1_45S_50ntFlanks@3863.3898.36|ATTCTAGAGCTAATACATGCCGACGGGCGCTGACCC</td><td>>hg38_21_+_8433172_8446622_RNA45SN1_45S_50ntFlanks</td><td>3863</td><td>3898</td><td>36</td><td>ATTCTAGAGCTAATACATGCCGACGGGCGCTGACCC</td><td>214.4</td><td>-3.0</td><td>1.1</td><td>-2.6</td><td>8.3E-03</td><td>1.0E-02</td><td>BT20/MB468</td><td>CYTO</td></tr>
<tr><td>>hg38_21_+_8433172_8446622_RNA45SN1_45S_50ntFlanks@11572.11604.33|CCGACTTAGAACTGGTGCGGACCAGGGGAATCC</td><td>>hg38_21_+_8433172_8446622_RNA45SN1_45S_50ntFlanks</td><td>11572</td><td>11604</td><td>33</td><td>CCGACTTAGAACTGGTGCGGACCAGGGGAATCC</td><td>177.9</td><td>-1.0</td><td>0.4</td><td>-2.6</td><td>8.3E-03</td><td>1.0E-02</td><td>BT20/MB468</td><td>CYTO</td></tr>
<tr><td>>hg38_21_+_8433172_8446622_RNA45SN1_45S_50ntFlanks@6646.6663.18|TCGTACGACTCTTAGCGG</td><td>>hg38_21_+_8433172_8446622_RNA45SN1_45S_50ntFlanks</td><td>6646</td><td>6663</td><td>18</td><td>TCGTACGACTCTTAGCGG</td><td>20457.6</td><td>2.0</td><td>0.7</td><td>2.8</td><td>5.0E-03</td><td>1.0E-02</td><td>BT20/MB468</td><td>MP</td></tr>
<tr><td>>hg38_21_+_8433172_8446622_RNA45SN1_45S_50ntFlanks@6730.6771.42|ACACATTGATCATCGACACTTCGAACGCACTTGCGGCCCCGG</td><td>>hg38_21_+_8433172_8446622_RNA45SN1_45S_50ntFlanks</td><td>6730</td><td>6771</td><td>42</td><td>ACACATTGATCATCGACACTTCGAACGCACTTGCGGCCCCGG</td><td>118.3</td><td>-5.2</td><td>1.9</td><td>-2.8</td><td>5.0E-03</td><td>1.0E-02</td><td>BT20/MB468</td><td>MP</td></tr>
<tr><td>>hg38_21_+_8433172_8446622_RNA45SN1_45S_50ntFlanks@11567.11611.45|AGCAGCCGACTTAGAACTGGTGCGGACCAGGGGAATCCGACTGTT</td><td>>hg38_21_+_8433172_8446622_RNA45SN1_45S_50ntFlanks</td><td>11567</td><td>11611</td><td>45</td><td>AGCAGCCGACTTAGAACTGGTGCGGACCAGGGGAATCCGACTGTT</td><td>507.3</td><td>3.9</td><td>1.4</td><td>2.8</td><td>5.0E-03</td><td>1.0E-02</td><td>BT20/MB468</td><td>MP</td></tr>
<tr><td>>hg38_21_+_8433172_8446622_RNA45SN1_45S_50ntFlanks@12936.12962.27|AAACCATTCGTAGACGACCTGCTTCTG</td><td>>hg38_21_+_8433172_8446622_RNA45SN1_45S_50ntFlanks</td><td>12936</td><td>12962</td><td>27</td><td>AAACCATTCGTAGACGACCTGCTTCTG</td><td>69.3</td><td>-4.5</td><td>1.6</td><td>-2.8</td><td>5.0E-03</td><td>1.0E-02</td><td>BT20/MB468</td><td>MP</td></tr>
<tr><td>>trna11_GluTTC_15_-_26327381_26327452@1.30.30,trna3_GluTTC_13_-_45492062_45492133@1.30.30|TCCCACATGGTCTAGCGGTTAGGATTCCTG</td><td>>trna11_GluTTC_15_-_26327381_26327452</td><td>1</td><td>30</td><td>30</td><td>TCCCACATGGTCTAGCGGTTAGGATTCCTG</td><td>1052.8</td><td>-2.7</td><td>1.0</td><td>-2.6</td><td>8.8E-03</td><td>1.0E-02</td><td>MB231/MB468</td><td>NUC</td></tr>
<tr><td>>hg38_21_+_8433172_8446622_RNA45SN1_45S_50ntFlanks@12407.12447.41|TTTTGATCCTTCGATGTCGGCTCTTCCTATCATTGTGAAGC</td><td>>hg38_21_+_8433172_8446622_RNA45SN1_45S_50ntFlanks</td><td>12407</td><td>12447</td><td>41</td><td>TTTTGATCCTTCGATGTCGGCTCTTCCTATCATTGTGAAGC</td><td>142.4</td><td>5.3</td><td>2.0</td><td>2.6</td><td>8.3E-03</td><td>1.0E-02</td><td>MB231/MB468</td><td>CYTO</td></tr>
<tr><td>>hg38_21_+_8433172_8446622_RNA45SN1_45S_50ntFlanks@10240.10259.20|CTACGCCGCGACGAGTAGGA</td><td>>hg38_21_+_8433172_8446622_RNA45SN1_45S_50ntFlanks</td><td>10240</td><td>10259</td><td>20</td><td>CTACGCCGCGACGAGTAGGA</td><td>476.2</td><td>2.1</td><td>0.7</td><td>3.1</td><td>2.0E-03</td><td>1.0E-02</td><td>BT20/MB231</td><td>NUC</td></tr>
<tr><td>>hg38_MT_+_1621_3279_RNR2_16S_50ntFlanks@1357.1388.32|GTTTACGACCTCGATGTTGGATCAGGACATCC</td><td>>hg38_MT_+_1621_3279_RNR2_16S_50ntFlanks</td><td>1357</td><td>1388</td><td>32</td><td>GTTTACGACCTCGATGTTGGATCAGGACATCC</td><td>143.7</td><td>3.4</td><td>1.3</td><td>2.6</td><td>8.8E-03</td><td>1.0E-02</td><td>MB231/MB468</td><td>MITO</td></tr>
<tr><td>>hg38_21_+_8433172_8446622_RNA45SN1_45S_50ntFlanks@6656.6680.25|CTTAGCGGTGGATCACTCGGCTCGT</td><td>>hg38_21_+_8433172_8446622_RNA45SN1_45S_50ntFlanks</td><td>6656</td><td>6680</td><td>25</td><td>CTTAGCGGTGGATCACTCGGCTCGT</td><td>114.8</td><td>-3.8</td><td>1.4</td><td>-2.6</td><td>8.3E-03</td><td>1.0E-02</td><td>BT20/MB468</td><td>CYTO</td></tr>
<tr><td>>hg38_21_+_8433172_8446622_RNA45SN1_45S_50ntFlanks@4243.4263.21|CTTTCGAGGCCCTGTAATTGG</td><td>>hg38_21_+_8433172_8446622_RNA45SN1_45S_50ntFlanks</td><td>4243</td><td>4263</td><td>21</td><td>CTTTCGAGGCCCTGTAATTGG</td><td>411.8</td><td>-1.6</td><td>0.6</td><td>-2.6</td><td>8.3E-03</td><td>1.0E-02</td><td>MB231/MB468</td><td>CYTO</td></tr>
<tr><td>>hg38_21_+_8433172_8446622_RNA45SN1_45S_50ntFlanks@4568.4589.22|AATGGAATAGGACCGCGGTTCT</td><td>>hg38_21_+_8433172_8446622_RNA45SN1_45S_50ntFlanks</td><td>4568</td><td>4589</td><td>22</td><td>AATGGAATAGGACCGCGGTTCT</td><td>155.1</td><td>2.5</td><td>0.9</td><td>2.8</td><td>5.0E-03</td><td>1.0E-02</td><td>BT20/MB468</td><td>MP</td></tr>
<tr><td>>hg38_21_+_8433172_8446622_RNA45SN1_45S_50ntFlanks@12154.12179.26|GGGAGTTTGACTGGGGCGGTACACCT</td><td>>hg38_21_+_8433172_8446622_RNA45SN1_45S_50ntFlanks</td><td>12154</td><td>12179</td><td>26</td><td>GGGAGTTTGACTGGGGCGGTACACCT</td><td>404.4</td><td>-4.8</td><td>1.7</td><td>-2.8</td><td>5.0E-03</td><td>1.0E-02</td><td>BT20/MB468</td><td>MP</td></tr>
<tr><td>>hg38_21_+_8433172_8446622_RNA45SN1_45S_50ntFlanks@6721.6754.34|AATTGCAGGACACATTGATCATCGACACTTCGAA</td><td>>hg38_21_+_8433172_8446622_RNA45SN1_45S_50ntFlanks</td><td>6721</td><td>6754</td><td>34</td><td>AATTGCAGGACACATTGATCATCGACACTTCGAA</td><td>141.7</td><td>-1.8</td><td>0.7</td><td>-2.7</td><td>7.1E-03</td><td>1.0E-02</td><td>BT20/MB468</td><td>MITO</td></tr>
<tr><td>>hg38_21_+_8433172_8446622_RNA45SN1_45S_50ntFlanks@4975.4993.19|CCCGGACACGGACAGGATT</td><td>>hg38_21_+_8433172_8446622_RNA45SN1_45S_50ntFlanks</td><td>4975</td><td>4993</td><td>19</td><td>CCCGGACACGGACAGGATT</td><td>148.9</td><td>1.8</td><td>0.7</td><td>2.6</td><td>8.7E-03</td><td>1.0E-02</td><td>BT20/MB468</td><td>NUC</td></tr>
<tr><td>>trnaMT_PheGAA_MT_+_577_647@35.65.31|GAAAATGTTTAGACGGGCTCACATCACCCCA</td><td>>trnaMT_PheGAA_MT_+_577_647</td><td>35</td><td>65</td><td>31</td><td>GAAAATGTTTAGACGGGCTCACATCACCCCA</td><td>179.4</td><td>4.2</td><td>1.6</td><td>2.7</td><td>7.7E-03</td><td>1.0E-02</td><td>MB231/MB468</td><td>MP</td></tr>
<tr><td>>am_trnaMT_ValTAC_MT_+_1602_1670@49.67.19|AACTTAACTTGACCGCTCT</td><td>>am_trnaMT_ValTAC_MT_+_1602_1670</td><td>49</td><td>67</td><td>19</td><td>AACTTAACTTGACCGCTCT</td><td>604.5</td><td>3.6</td><td>1.4</td><td>2.6</td><td>8.4E-03</td><td>1.0E-02</td><td>MB231/MB468</td><td>CYTO</td></tr>
<tr><td>>hg38_21_+_8433172_8446622_RNA45SN1_45S_50ntFlanks@11235.11259.25|CTTCCCGTGGATCGCCCCAGCTGCG</td><td>>hg38_21_+_8433172_8446622_RNA45SN1_45S_50ntFlanks</td><td>11235</td><td>11259</td><td>25</td><td>CTTCCCGTGGATCGCCCCAGCTGCG</td><td>346.9</td><td>-4.9</td><td>1.9</td><td>-2.6</td><td>8.7E-03</td><td>1.0E-02</td><td>BT20/MB468</td><td>NUC</td></tr>
<tr><td>>hg38_21_+_8433172_8446622_RNA45SN1_45S_50ntFlanks@11786.11803.18|GAATGGATGAACGAGATT</td><td>>hg38_21_+_8433172_8446622_RNA45SN1_45S_50ntFlanks</td><td>11786</td><td>11803</td><td>18</td><td>GAATGGATGAACGAGATT</td><td>581.6</td><td>-2.5</td><td>0.9</td><td>-2.6</td><td>8.4E-03</td><td>1.0E-02</td><td>MB231/MB468</td><td>CYTO</td></tr>
<tr><td>>trnaMT_GluTTC_MT_-_14674_14742@47.71.25,trnalookalike8_GluTTC_5_-_93905172_93905240@47.71.25|TGGTTGTAGTCCGTGCGAGAATACC</td><td>>trnaMT_GluTTC_MT_-_14674_14742</td><td>47</td><td>71</td><td>25</td><td>TGGTTGTAGTCCGTGCGAGAATACC</td><td>136.0</td><td>3.3</td><td>1.2</td><td>2.8</td><td>5.0E-03</td><td>1.0E-02</td><td>BT20/MB468</td><td>MP</td></tr>
<tr><td>>hg38_21_+_8433172_8446622_RNA45SN1_45S_50ntFlanks@5491.5512.22|GAGAAGACGGTCGAACTTGACT</td><td>>hg38_21_+_8433172_8446622_RNA45SN1_45S_50ntFlanks</td><td>5491</td><td>5512</td><td>22</td><td>GAGAAGACGGTCGAACTTGACT</td><td>311.7</td><td>4.2</td><td>1.5</td><td>2.8</td><td>5.0E-03</td><td>1.0E-02</td><td>BT20/MB468</td><td>MP</td></tr>
<tr><td>>trna16_LeuAAG_16_+_22308461_22308542@1.20.20,trna16_LeuAAG_5_-_180601044_180601125@1.20.20,trna19_LeuAAG_5_-_180524474_180524555@1.20.20,trna1_LeuAAG_14_+_21078291_21078372@1.20.20,trna3_LeuAAG_5_+_180528840_180528921@1.20.20,trna42_LeuTAG_17_-_8023632_8023713@1.20.20,trna78_LeuAAG_6_+_28956779_28956860@1.20.20,trna7_LeuAAG_5_+_180614701_180614782@1.20.20,trna98_LeuAAG_6_-_28911399_28911480@1.20.20|GGTAGCGTGGCCGAGCGGTC</td><td>>trna16_LeuAAG_16_+_22308461_22308542</td><td>1</td><td>20</td><td>20</td><td>GGTAGCGTGGCCGAGCGGTC</td><td>157.1</td><td>5.4</td><td>1.9</td><td>2.8</td><td>5.0E-03</td><td>1.0E-02</td><td>BT20/MB468</td><td>MP</td></tr>
<tr><td>>hg38_21_+_8433172_8446622_RNA45SN1_45S_50ntFlanks@4726.4755.30|TAATCAAGAACGAAAGTCGGAGGTTCGAAG</td><td>>hg38_21_+_8433172_8446622_RNA45SN1_45S_50ntFlanks</td><td>4726</td><td>4755</td><td>30</td><td>TAATCAAGAACGAAAGTCGGAGGTTCGAAG</td><td>72.3</td><td>-4.6</td><td>1.6</td><td>-2.8</td><td>5.0E-03</td><td>1.0E-02</td><td>BT20/MB468</td><td>MP</td></tr>
<tr><td>>hg38_21_+_8433172_8446622_RNA45SN1_45S_50ntFlanks@9930.9959.30|ATATAGACAGCAGGACGGTGGCCATGGAAG</td><td>>hg38_21_+_8433172_8446622_RNA45SN1_45S_50ntFlanks</td><td>9930</td><td>9959</td><td>30</td><td>ATATAGACAGCAGGACGGTGGCCATGGAAG</td><td>230.0</td><td>-4.1</td><td>1.5</td><td>-2.8</td><td>5.0E-03</td><td>1.0E-02</td><td>BT20/MB468</td><td>MP</td></tr>
<tr><td>>hg38_21_+_8433172_8446622_RNA45SN1_45S_50ntFlanks@10866.10891.26|CTAAGGGCTGGGTCGGTCGGGCTGGG</td><td>>hg38_21_+_8433172_8446622_RNA45SN1_45S_50ntFlanks</td><td>10866</td><td>10891</td><td>26</td><td>CTAAGGGCTGGGTCGGTCGGGCTGGG</td><td>497.6</td><td>-3.3</td><td>1.2</td><td>-2.7</td><td>7.1E-03</td><td>1.0E-02</td><td>BT20/MB468</td><td>MITO</td></tr>
<tr><td>>hg38_21_+_8433172_8446622_RNA45SN1_45S_50ntFlanks@11667.11702.36|CTGCCCAGTGCTCTGAATGTCAAAGTGAAGAAATTC</td><td>>hg38_21_+_8433172_8446622_RNA45SN1_45S_50ntFlanks</td><td>11667</td><td>11702</td><td>36</td><td>CTGCCCAGTGCTCTGAATGTCAAAGTGAAGAAATTC</td><td>2371.1</td><td>-2.8</td><td>0.9</td><td>-3.0</td><td>2.4E-03</td><td>1.0E-02</td><td>BT20/MB231</td><td>CYTO</td></tr>
<tr><td>>hg38_21_+_8433172_8446622_RNA45SN1_45S_50ntFlanks@6648.6681.34|GTACGACTCTTAGCGGTGGATCACTCGGCTCGTG</td><td>>hg38_21_+_8433172_8446622_RNA45SN1_45S_50ntFlanks</td><td>6648</td><td>6681</td><td>34</td><td>GTACGACTCTTAGCGGTGGATCACTCGGCTCGTG</td><td>3880.6</td><td>-2.8</td><td>1.0</td><td>-2.8</td><td>5.0E-03</td><td>1.0E-02</td><td>BT20/MB468</td><td>MP</td></tr>
<tr><td>>hg38_21_+_8433172_8446622_RNA45SN1_45S_50ntFlanks@3876.3899.24|TACATGCCGACGGGCGCTGACCCC</td><td>>hg38_21_+_8433172_8446622_RNA45SN1_45S_50ntFlanks</td><td>3876</td><td>3899</td><td>24</td><td>TACATGCCGACGGGCGCTGACCCC</td><td>315.5</td><td>-1.4</td><td>0.5</td><td>-2.6</td><td>8.3E-03</td><td>1.0E-02</td><td>BT20/MB468</td><td>CYTO</td></tr>
<tr><td>>hg38_21_+_8433172_8446622_RNA45SN1_45S_50ntFlanks@11663.11692.30|ATTTCTGCCCAGTGCTCTGAATGTCAAAGT</td><td>>hg38_21_+_8433172_8446622_RNA45SN1_45S_50ntFlanks</td><td>11663</td><td>11692</td><td>30</td><td>ATTTCTGCCCAGTGCTCTGAATGTCAAAGT</td><td>186.7</td><td>5.1</td><td>1.7</td><td>3.1</td><td>1.9E-03</td><td>1.0E-02</td><td>BT20/MB231</td><td>MP</td></tr>
<tr><td>>MI0000299|hsa-mir-222&WithFlank&X|-|45747009|45747130@75.99.25(+2U)&[MIMAT0000279&hsa-miR-222-3p&offsets|0|+4(+2U);m-140&X|-|45747033|45747056&offsets|0|+1(+2U)]|AGCTACATCTGGCTACTGGGTCTCTTT</td><td>>MI0000299|hsa-mir-222&WithFlank&X|-|45747009|45747130</td><td>75</td><td>99</td><td>25(+2U)</td><td>AGCTACATCTGGCTACTGGGTCTCTTT</td><td>605.0</td><td>-3.3</td><td>1.0</td><td>-3.1</td><td>1.9E-03</td><td>1.0E-02</td><td>BT20/MB231</td><td>MP</td></tr>
<tr><td>>hg38_21_+_8433172_8446622_RNA45SN1_45S_50ntFlanks@9554.9580.27|CGTAGCGGTCCTGACGTGCAAATCGGT</td><td>>hg38_21_+_8433172_8446622_RNA45SN1_45S_50ntFlanks</td><td>9554</td><td>9580</td><td>27</td><td>CGTAGCGGTCCTGACGTGCAAATCGGT</td><td>258.1</td><td>-3.3</td><td>1.3</td><td>-2.6</td><td>8.9E-03</td><td>1.0E-02</td><td>MB231/MB468</td><td>MITO</td></tr>
<tr><td>>hg38_21_+_8433172_8446622_RNA45SN1_45S_50ntFlanks@12957.12987.31|CTTCTGGGTCGGGGTTTCGTACGTAGCAGAG</td><td>>hg38_21_+_8433172_8446622_RNA45SN1_45S_50ntFlanks</td><td>12957</td><td>12987</td><td>31</td><td>CTTCTGGGTCGGGGTTTCGTACGTAGCAGAG</td><td>105.1</td><td>-4.6</td><td>1.6</td><td>-2.8</td><td>5.0E-03</td><td>1.0E-02</td><td>BT20/MB468</td><td>MP</td></tr>
<tr><td>>hg38_21_+_8433172_8446622_RNA45SN1_45S_50ntFlanks@10574.10592.19|AAGCCGGCGGGAGCCCCGG</td><td>>hg38_21_+_8433172_8446622_RNA45SN1_45S_50ntFlanks</td><td>10574</td><td>10592</td><td>19</td><td>AAGCCGGCGGGAGCCCCGG</td><td>193.8</td><td>-4.6</td><td>1.6</td><td>-2.8</td><td>5.0E-03</td><td>1.0E-02</td><td>BT20/MB468</td><td>MP</td></tr>
<tr><td>>hg38_21_+_8433172_8446622_RNA45SN1_45S_50ntFlanks@8344.8365.22|TAAGGGAAAGTTGAAAAGAACT</td><td>>hg38_21_+_8433172_8446622_RNA45SN1_45S_50ntFlanks</td><td>8344</td><td>8365</td><td>22</td><td>TAAGGGAAAGTTGAAAAGAACT</td><td>46.7</td><td>4.8</td><td>1.7</td><td>2.8</td><td>5.0E-03</td><td>1.0E-02</td><td>BT20/MB468</td><td>MP</td></tr>
<tr><td>>hg38_MT_+_1621_3279_RNR2_16S_50ntFlanks@505.531.27|CTAGGAAAAAACCTTGTAGAGAGAGTA</td><td>>hg38_MT_+_1621_3279_RNR2_16S_50ntFlanks</td><td>505</td><td>531</td><td>27</td><td>CTAGGAAAAAACCTTGTAGAGAGAGTA</td><td>176.4</td><td>3.8</td><td>1.4</td><td>2.8</td><td>5.0E-03</td><td>1.0E-02</td><td>BT20/MB468</td><td>MP</td></tr>
<tr><td>>hg38_21_+_8433172_8446622_RNA45SN1_45S_50ntFlanks@12996.13014.19|TCGCTGCGATCTATTGAAA</td><td>>hg38_21_+_8433172_8446622_RNA45SN1_45S_50ntFlanks</td><td>12996</td><td>13014</td><td>19</td><td>TCGCTGCGATCTATTGAAA</td><td>374.7</td><td>2.0</td><td>0.8</td><td>2.6</td><td>8.9E-03</td><td>1.0E-02</td><td>MB231/MB468</td><td>MITO</td></tr>
<tr><td>>MI0000478|hsa-mir-149&WithFlank&2|+|240455995|240456095@21.42.22&[MIMAT0000450&hsa-miR-149-5p&offsets|0|-1;m-195&2|+|240456015|240456037&offsets|0|-1]|TCTGGCTCCGTGTCTTCACTCC</td><td>>MI0000478|hsa-mir-149&WithFlank&2|+|240455995|240456095</td><td>21</td><td>42</td><td>22</td><td>TCTGGCTCCGTGTCTTCACTCC</td><td>102.0</td><td>4.3</td><td>1.6</td><td>2.7</td><td>7.7E-03</td><td>1.0E-02</td><td>MB231/MB468</td><td>MP</td></tr>
<tr><td>>hg38_21_+_8433172_8446622_RNA45SN1_45S_50ntFlanks@9167.9197.31|CGGGCGGGTCGCGCCGTCGGGCCCGGGGGAG</td><td>>hg38_21_+_8433172_8446622_RNA45SN1_45S_50ntFlanks</td><td>9167</td><td>9197</td><td>31</td><td>CGGGCGGGTCGCGCCGTCGGGCCCGGGGGAG</td><td>822.3</td><td>-4.8</td><td>1.8</td><td>-2.7</td><td>7.2E-03</td><td>1.0E-02</td><td>BT20/MB468</td><td>MITO</td></tr>
</table>

<table>
<tr><th></th><th></th><th></th><th></th><th></th><th></th><th></th><th></th><th></th><th></th><th></th><th></th><th></th><th></th></tr>
<tr><td>>hg38_21_+_8433172_8446622_RNA45SN1_45S_50ntFlanks@8105.8139.35|CGCGGCGGGGCGCGGGACATGTGGCGTACGGAAGA</td><td>>hg38_21_+_8433172_8446622_RNA45SN1_45S_50ntFlanks</td><td>8105</td><td>8139</td><td>35</td><td>CGCGGCGGGGCGCGGGACATGTGGCGTACGGAAGA</td><td>1476.1</td><td>5.5</td><td>1.8</td><td>3.1</td><td>1.9E-03</td><td>1.0E-02</td><td>BT20/MB231</td><td>MP</td></tr>
<tr><td>>hg38_21_+_8433172_8446622_RNA45SN1_45S_50ntFlanks@8112.8139.28|GGGCGCGGGACATGTGGCGTACGGAAGA</td><td>>hg38_21_+_8433172_8446622_RNA45SN1_45S_50ntFlanks</td><td>8112</td><td>8139</td><td>28</td><td>GGGCGCGGGACATGTGGCGTACGGAAGA</td><td>533.5</td><td>6.6</td><td>2.1</td><td>3.1</td><td>1.9E-03</td><td>1.0E-02</td><td>BT20/MB231</td><td>MP</td></tr>
<tr><td>>MI0000095|hsa-mir-93&WithFlank&7|-|100093762|100093853@17.39.23(+1A)&[MIMAT0000093&hsa-miR-93-5p&offsets|0|0(+1A);m-29&7|-|100093815|100093837&offsets|0|0(+1A)]|CAAAGTGCTGTTCGTGCAGGTAGA</td><td>>MI0000095|hsa-mir-93&WithFlank&7|-|100093762|100093853</td><td>17</td><td>39</td><td>23(+1A)</td><td>CAAAGTGCTGTTCGTGCAGGTAGA</td><td>827.9</td><td>2.0</td><td>0.8</td><td>2.7</td><td>7.7E-03</td><td>1.0E-02</td><td>MB231/MB468</td><td>MP</td></tr>
<tr><td>>hg38_21_+_8433172_8446622_RNA45SN1_45S_50ntFlanks@6655.6681.27|TCTTAGCGGTGGATCACTCGGCTCGTG</td><td>>hg38_21_+_8433172_8446622_RNA45SN1_45S_50ntFlanks</td><td>6655</td><td>6681</td><td>27</td><td>TCTTAGCGGTGGATCACTCGGCTCGTG</td><td>487.3</td><td>5.1</td><td>1.7</td><td>3.1</td><td>1.9E-03</td><td>1.0E-02</td><td>BT20/MB231</td><td>MP</td></tr>
<tr><td>>hg38_21_+_8433172_8446622_RNA45SN1_45S_50ntFlanks@11921.11951.31|TGAGAGGTGTAGAATAAGTGGGAGGCCCCCG</td><td>>hg38_21_+_8433172_8446622_RNA45SN1_45S_50ntFlanks</td><td>11921</td><td>11951</td><td>31</td><td>TGAGAGGTGTAGAATAAGTGGGAGGCCCCCG</td><td>413.2</td><td>-3.8</td><td>1.3</td><td>-2.8</td><td>5.0E-03</td><td>1.0E-02</td><td>BT20/MB468</td><td>MP</td></tr>
<tr><td>>MI0000078|hsa-mir-22&WithFlank&17|-|1713897|1713993@59.81.23&[MIMAT0000077&hsa-miR-22-3p&offsets|0|+1;m-2&17|-|1713914|1713935&offsets|0|+1]|AAGCTGCCAGTTGAAGAACTGTT</td><td>>MI0000078|hsa-mir-22&WithFlank&17|-|1713897|1713993</td><td>59</td><td>81</td><td>23</td><td>AAGCTGCCAGTTGAAGAACTGTT</td><td>1646.5</td><td>-3.4</td><td>1.1</td><td>-3.1</td><td>1.9E-03</td><td>1.0E-02</td><td>BT20/MB231</td><td>MP</td></tr>
<tr><td>>hg38_21_+_8433172_8446622_RNA45SN1_45S_50ntFlanks@11209.11252.44|CGGCGACTCTGGACGCGAGCCGGGCCCTTCCCGTGGATCGCCCC</td><td>>hg38_21_+_8433172_8446622_RNA45SN1_45S_50ntFlanks</td><td>11209</td><td>11252</td><td>44</td><td>CGGCGACTCTGGACGCGAGCCGGGCCCTTCCCGTGGATCGCCCC</td><td>553.4</td><td>5.8</td><td>1.9</td><td>3.0</td><td>2.5E-03</td><td>1.0E-02</td><td>BT20/MB231</td><td>MITO</td></tr>
<tr><td>>MI0000089|hsa-mir-31&WithFlank&9|-|21512109|21512191@14.36.23&[MIMAT0000089&hsa-miR-31-5p&offsets|0|+2;m-360&9|-|21512157|21512178&offsets|0|+1]|AGGCAAGATGCTGGCATAGCTGT</td><td>>MI0000089|hsa-mir-31&WithFlank&9|-|21512109|21512191</td><td>14</td><td>36</td><td>23</td><td>AGGCAAGATGCTGGCATAGCTGT</td><td>1826.6</td><td>3.7</td><td>1.3</td><td>2.8</td><td>5.0E-03</td><td>1.0E-02</td><td>BT20/MB468</td><td>MP</td></tr>
<tr><td>>hg38_21_+_8433172_8446622_RNA45SN1_45S_50ntFlanks@5238.5283.46|CTGACTGGCTCAGCGTGTGCCTACCCTACGCCGGCAGGCGCGGGTA</td><td>>hg38_21_+_8433172_8446622_RNA45SN1_45S_50ntFlanks</td><td>5238</td><td>5283</td><td>46</td><td>CTGACTGGCTCAGCGTGTGCCTACCCTACGCCGGCAGGCGCGGGTA</td><td>73.3</td><td>5.9</td><td>2.1</td><td>2.8</td><td>5.1E-03</td><td>1.0E-02</td><td>BT20/MB468</td><td>MP</td></tr>
<tr><td>>MI0000063|hsa-let-7b&WithFlank&22|+|46113680|46113774@12.31.20&[MIMAT0000063&hsa-let-7b-5p&offsets|0|-2;m-14&22|+|46113691|46113712&offsets|0|-2]|TGAGGTAGTAGGTTGTGTGG</td><td>>MI0000063|hsa-let-7b&WithFlank&22|+|46113680|46113774</td><td>12</td><td>31</td><td>20</td><td>TGAGGTAGTAGGTTGTGTGG</td><td>370.4</td><td>2.3</td><td>0.9</td><td>2.7</td><td>7.7E-03</td><td>1.0E-02</td><td>MB231/MB468</td><td>MP</td></tr>
<tr><td>>hg38_21_+_8433172_8446622_RNA45SN1_45S_50ntFlanks@9716.9735.20|AATGATTAGAGGTCTTGGGG</td><td>>hg38_21_+_8433172_8446622_RNA45SN1_45S_50ntFlanks</td><td>9716</td><td>9735</td><td>20</td><td>AATGATTAGAGGTCTTGGGG</td><td>205.2</td><td>-2.4</td><td>0.8</td><td>-2.8</td><td>5.1E-03</td><td>1.0E-02</td><td>BT20/MB468</td><td>MP</td></tr>
<tr><td>>hg38_21_+_8433172_8446622_RNA45SN1_45S_50ntFlanks@12397.12418.22|CGACGTCGCTTTTTGATCCTTC</td><td>>hg38_21_+_8433172_8446622_RNA45SN1_45S_50ntFlanks</td><td>12397</td><td>12418</td><td>22</td><td>CGACGTCGCTTTTTGATCCTTC</td><td>2569.0</td><td>-2.8</td><td>0.9</td><td>-3.0</td><td>2.4E-03</td><td>1.0E-02</td><td>BT20/MB231</td><td>CYTO</td></tr>
<tr><td>>MI0000767|hsa-mir-365a&WithFlank&16|+|14309279|14309377@62.83.22;MI0000769|hsa-mir-365b&WithFlank&17|+|31575405|31575527@74.95.22&[MIMAT0000710&hsa-miR-365a-3p&offsets|0|0;m-176&16|+|14309340|14309361&offsets|0|0];[MIMAT0022834&hsa-miR-365b-3p&offsets|0|0;m-175&17|+|31575478|31575499&offsets|0|0]|TAATGCCCCTAAAAATCCTTAT</td><td>>MI0000767|hsa-mir-365a&WithFlank&16|+|14309279|14309377</td><td>62</td><td>83</td><td>22</td><td>TAATGCCCCTAAAAATCCTTAT</td><td>344.4</td><td>4.3</td><td>1.5</td><td>2.8</td><td>5.1E-03</td><td>1.0E-02</td><td>BT20/MB468</td><td>MP</td></tr>
<tr><td>>hg38_21_+_8433172_8446622_RNA45SN1_45S_50ntFlanks@11663.11687.25|ATTTCTGCCCAGTGCTCTGAATGTC</td><td>>hg38_21_+_8433172_8446622_RNA45SN1_45S_50ntFlanks</td><td>11663</td><td>11687</td><td>25</td><td>ATTTCTGCCCAGTGCTCTGAATGTC</td><td>219.9</td><td>2.1</td><td>0.7</td><td>3.0</td><td>2.5E-03</td><td>1.0E-02</td><td>BT20/MB231</td><td>MITO</td></tr>
<tr><td>>hg38_21_+_8433172_8446622_RNA45SN1_45S_50ntFlanks@6751.6772.22|CGAACGCACTTGCGGCCCCGGG</td><td>>hg38_21_+_8433172_8446622_RNA45SN1_45S_50ntFlanks</td><td>6751</td><td>6772</td><td>22</td><td>CGAACGCACTTGCGGCCCCGGG</td><td>77.4</td><td>-4.6</td><td>1.6</td><td>-2.8</td><td>5.1E-03</td><td>1.0E-02</td><td>BT20/MB468</td><td>MP</td></tr>
<tr><td>>trna119_LysCTT_1_-_145395522_145395594@4.33.30,trna11_LysCTT_5_-_180648979_180649051@4.33.30,trna13_LysCTT_6_+_26556774_26556846@4.33.30,trna7_LysCTT_16_+_3225692_3225764@4.33.30,trna9_LysCTT_5_+_180634755_180634827@4.33.30|CGGCTAGCTCAGTCGGTAGAGCATGAGACT</td><td>>trna119_LysCTT_1_-_145395522_145395594</td><td>4</td><td>33</td><td>30</td><td>CGGCTAGCTCAGTCGGTAGAGCATGAGACT</td><td>931.5</td><td>-4.6</td><td>1.5</td><td>-3.1</td><td>1.9E-03</td><td>1.0E-02</td><td>BT20/MB231</td><td>MP</td></tr>
<tr><td>>trnaMT_LysTTT_MT_+_8295_8364@54.72.19,trnalookalike7_LysTTT_1_+_568844_568913@54.72.19|ACACCTCTTTACAGTGACC</td><td>>trnaMT_LysTTT_MT_+_8295_8364</td><td>54</td><td>72</td><td>19</td><td>ACACCTCTTTACAGTGACC</td><td>397.8</td><td>-4.5</td><td>1.4</td><td>-3.1</td><td>1.9E-03</td><td>1.0E-02</td><td>BT20/MB231</td><td>MP</td></tr>
<tr><td>>hg38_MT_+_1621_3279_RNR2_16S_50ntFlanks@622.652.31|TATAACTGAACTCCTCACACCCAATTGGACC</td><td>>hg38_MT_+_1621_3279_RNR2_16S_50ntFlanks</td><td>622</td><td>652</td><td>31</td><td>TATAACTGAACTCCTCACACCCAATTGGACC</td><td>138.5</td><td>3.3</td><td>1.2</td><td>2.7</td><td>7.7E-03</td><td>1.0E-02</td><td>MB231/MB468</td><td>MP</td></tr>
<tr><td>>hg38_21_+_8433172_8446622_RNA45SN1_45S_50ntFlanks@10513.10533.21|CCGGAGTGGCGGAGATGGGCG</td><td>>hg38_21_+_8433172_8446622_RNA45SN1_45S_50ntFlanks</td><td>10513</td><td>10533</td><td>21</td><td>CCGGAGTGGCGGAGATGGGCG</td><td>318.8</td><td>6.3</td><td>2.0</td><td>3.1</td><td>1.9E-03</td><td>1.0E-02</td><td>BT20/MB231</td><td>MP</td></tr>
<tr><td>>hg38_21_+_8433172_8446622_RNA45SN1_45S_50ntFlanks@4733.4765.33|GAACGAAAGTCGGAGGTTCGAAGACGATCAGAT</td><td>>hg38_21_+_8433172_8446622_RNA45SN1_45S_50ntFlanks</td><td>4733</td><td>4765</td><td>33</td><td>GAACGAAAGTCGGAGGTTCGAAGACGATCAGAT</td><td>117.4</td><td>2.0</td><td>0.8</td><td>2.7</td><td>7.8E-03</td><td>1.0E-02</td><td>MB231/MB468</td><td>MP</td></tr>
<tr><td>>hg38_21_+_8433172_8446622_RNA45SN1_45S_50ntFlanks@12405.12442.38|CTTTTTGATCCTTCGATGTCGGCTCTTCCTATCATTGT</td><td>>hg38_21_+_8433172_8446622_RNA45SN1_45S_50ntFlanks</td><td>12405</td><td>12442</td><td>38</td><td>CTTTTTGATCCTTCGATGTCGGCTCTTCCTATCATTGT</td><td>183.0</td><td>-2.0</td><td>0.7</td><td>-2.7</td><td>7.2E-03</td><td>1.0E-02</td><td>BT20/MB468</td><td>MITO</td></tr>
<tr><td>>hg38_21_+_8433172_8446622_RNA45SN1_45S_50ntFlanks@5424.5466.43|TTGGATGGTTTAGTGAGGCCCTCGGATCGGCCCCGCCGGGGTC</td><td>>hg38_21_+_8433172_8446622_RNA45SN1_45S_50ntFlanks</td><td>5424</td><td>5466</td><td>43</td><td>TTGGATGGTTTAGTGAGGCCCTCGGATCGGCCCCGCCGGGGTC</td><td>208.5</td><td>-2.4</td><td>0.9</td><td>-2.6</td><td>8.4E-03</td><td>1.0E-02</td><td>BT20/MB468</td><td>CYTO</td></tr>
<tr><td>>MI0000750|hsa-mir-26a-2&WithFlank&12|-|57824603|57824698@20.41.22(+2U);MI0000083|hsa-mir-26a-1&WithFlank&3|+|37969398|37969486@16.37.22(+2U)&[MIMAT0000082&hsa-miR-26a-5p&offsets|0|0(+2U);m-33&12|-|57824658|57824679&offsets|0|0(+2U)];[MIMAT0000082_1&hsa-miR-26a-5p&offsets|0|0(+2U);m-34&3|+|37969413|37969434&offsets|0|0(+2U)]|TTCAAGTAATCCAGGATAGGCTTT</td><td>>MI0000750|hsa-mir-26a-2&WithFlank&12|-|57824603|57824698</td><td>20</td><td>41</td><td>22(+2U)</td><td>TTCAAGTAATCCAGGATAGGCTTT</td><td>4918.0</td><td>-3.3</td><td>1.1</td><td>-3.1</td><td>1.9E-03</td><td>1.0E-02</td><td>BT20/MB231</td><td>MP</td></tr>
<tr><td>>hg38_21_+_8433172_8446622_RNA45SN1_45S_50ntFlanks@6662.6698.37|GGTGGATCACTCGGCTCGTGCGTCGATGAAGAACGCA</td><td>>hg38_21_+_8433172_8446622_RNA45SN1_45S_50ntFlanks</td><td>6662</td><td>6698</td><td>37</td><td>GGTGGATCACTCGGCTCGTGCGTCGATGAAGAACGCA</td><td>239.5</td><td>5.9</td><td>1.9</td><td>3.1</td><td>1.9E-03</td><td>1.0E-02</td><td>BT20/MB231</td><td>MP</td></tr>
<tr><td>>hg38_21_+_8433172_8446622_RNA45SN1_45S_50ntFlanks@7975.8004.30|CGCGACCTCAGATCAGACGTGGCGACCCGC</td><td>>hg38_21_+_8433172_8446622_RNA45SN1_45S_50ntFlanks</td><td>7975</td><td>8004</td><td>30</td><td>CGCGACCTCAGATCAGACGTGGCGACCCGC</td><td>15439.6</td><td>3.7</td><td>1.2</td><td>3.1</td><td>1.9E-03</td><td>1.0E-02</td><td>BT20/MB231</td><td>MP</td></tr>
<tr><td>>MI0000087|hsa-mir-29a&WithFlank&7|-|130876741|130876816@48.68.21(+1U)&[MIMAT0000086&hsa-miR-29a-3p&offsets|0|-1(+1U);m-26&7|-|130876749|130876769&offsets|0|0(+1U)]|TAGCACCATCTGAAATCGGTTT</td><td>>MI0000087|hsa-mir-29a&WithFlank&7|-|130876741|130876816</td><td>48</td><td>68</td><td>21(+1U)</td><td>TAGCACCATCTGAAATCGGTTT</td><td>794.6</td><td>-2.9</td><td>0.9</td><td>-3.0</td><td>2.5E-03</td><td>1.0E-02</td><td>BT20/MB231</td><td>MITO</td></tr>
<tr><td>>hg38_21_+_8433172_8446622_RNA45SN1_45S_50ntFlanks@10386.10408.23|ATGTGAACAGCAGTTGAACATGG</td><td>>hg38_21_+_8433172_8446622_RNA45SN1_45S_50ntFlanks</td><td>10386</td><td>10408</td><td>23</td><td>ATGTGAACAGCAGTTGAACATGG</td><td>171.4</td><td>-7.0</td><td>2.6</td><td>-2.7</td><td>7.8E-03</td><td>1.0E-02</td><td>MB231/MB468</td><td>MP</td></tr>
<tr><td>>hg38_21_+_8433172_8446622_RNA45SN1_45S_50ntFlanks@6647.6681.35|CGTACGACTCTTAGCGGTGGATCACTCGGCTCGTG</td><td>>hg38_21_+_8433172_8446622_RNA45SN1_45S_50ntFlanks</td><td>6647</td><td>6681</td><td>35</td><td>CGTACGACTCTTAGCGGTGGATCACTCGGCTCGTG</td><td>7335.1</td><td>-1.9</td><td>0.6</td><td>-3.1</td><td>2.1E-03</td><td>1.0E-02</td><td>BT20/MB231</td><td>NUC</td></tr>
<tr><td>>trna119_LysCTT_1_-_145395522_145395594@4.36.33,trna11_LysCTT_5_-_180648979_180649051@4.36.33,trna13_LysCTT_6_+_26556774_26556846@4.36.33,trna7_LysCTT_16_+_3225692_3225764@4.36.33,trna9_LysCTT_5_+_180634755_180634827@4.36.33|CGGCTAGCTCAGTCGGTAGAGCATGAGACTCTT</td><td>>trna119_LysCTT_1_-_145395522_145395594</td><td>4</td><td>36</td><td>33</td><td>CGGCTAGCTCAGTCGGTAGAGCATGAGACTCTT</td><td>394.1</td><td>-4.2</td><td>1.4</td><td>-3.1</td><td>2.1E-03</td><td>1.0E-02</td><td>BT20/MB231</td><td>NUC</td></tr>
<tr><td>>hg38_21_+_8433172_8446622_RNA45SN1_45S_50ntFlanks@12911.12932.22|GCACGTTCGTGGGGAACCTGGC</td><td>>hg38_21_+_8433172_8446622_RNA45SN1_45S_50ntFlanks</td><td>12911</td><td>12932</td><td>22</td><td>GCACGTTCGTGGGGAACCTGGC</td><td>746.9</td><td>5.6</td><td>1.8</td><td>3.1</td><td>1.9E-03</td><td>1.0E-02</td><td>BT20/MB231</td><td>MP</td></tr>
<tr><td>>trnaMT_ProTGG_MT_-_15956_16023@40.62.23|AATGGTGGAGTTAAAGACTTTTT</td><td>>trnaMT_ProTGG_MT_-_15956_16023</td><td>40</td><td>62</td><td>23</td><td>AATGGTGGAGTTAAAGACTTTTT</td><td>38.8</td><td>5.2</td><td>2.0</td><td>2.7</td><td>7.8E-03</td><td>1.0E-02</td><td>MB231/MB468</td><td>MP</td></tr>
<tr><td>>hg38_21_+_8433172_8446622_RNA45SN1_45S_50ntFlanks@10905.10928.24|CTGGGCGCGCGCCGCGGCTGGACG</td><td>>hg38_21_+_8433172_8446622_RNA45SN1_45S_50ntFlanks</td><td>10905</td><td>10928</td><td>24</td><td>CTGGGCGCGCGCCGCGGCTGGACG</td><td>123.3</td><td>-3.6</td><td>1.3</td><td>-2.8</td><td>5.1E-03</td><td>1.0E-02</td><td>BT20/MB468</td><td>MP</td></tr>
<tr><td>>hg38_21_+_8433172_8446622_RNA45SN1_45S_50ntFlanks@10403.10420.18|ACATGGGTCAGTCGGTCC</td><td>>hg38_21_+_8433172_8446622_RNA45SN1_45S_50ntFlanks</td><td>10403</td><td>10420</td><td>18</td><td>ACATGGGTCAGTCGGTCC</td><td>189.7</td><td>-3.7</td><td>1.3</td><td>-2.8</td><td>5.1E-03</td><td>1.0E-02</td><td>BT20/MB468</td><td>MP</td></tr>
<tr><td>>hg38_21_+_8433172_8446622_RNA45SN1_45S_50ntFlanks@10770.10796.27|GCAGGTCTCCAAGGTGAACAGCCTCTG</td><td>>hg38_21_+_8433172_8446622_RNA45SN1_45S_50ntFlanks</td><td>10770</td><td>10796</td><td>27</td><td>GCAGGTCTCCAAGGTGAACAGCCTCTG</td><td>568.3</td><td>-1.5</td><td>0.6</td><td>-2.6</td><td>8.4E-03</td><td>1.0E-02</td><td>BT20/MB468</td><td>CYTO</td></tr>
<tr><td>>hg38_21_+_8433172_8446622_RNA45SN1_45S_50ntFlanks@10873.10902.30|CTGGGTCGGTCGGGCTGGGGCGCGAAGCGG</td><td>>hg38_21_+_8433172_8446622_RNA45SN1_45S_50ntFlanks</td><td>10873</td><td>10902</td><td>30</td><td>CTGGGTCGGTCGGGCTGGGGCGCGAAGCGG</td><td>664.3</td><td>-4.4</td><td>1.7</td><td>-2.6</td><td>8.4E-03</td><td>1.0E-02</td><td>BT20/MB468</td><td>CYTO</td></tr>
<tr><td>>trna10_AspGTC_12_-_125424193_125424264@37.72.36,trna12_AspGTC_12_-_125411891_125411962@37.72.36,trna144_AspGTC_6_-_27551236_27551307@37.72.36,trna38_AspGTC_17_-_8125556_8125627@37.72.36,trna45_AspGTC_6_+_27447453_27447524@37.72.36,trna48_AspGTC_6_+_27471523_27471594@37.72.36,trna4_AspGTC_12_+_96429799_96429870@37.72.36,trna69_AspGTC_1_-_161440205_161440276@37.72.36,trna72_AspGTC_1_-_161432824_161432895@37.72.36,trna75_AspGTC_1_-_161425414_161425485@37.72.36,trna78_AspGTC_1_-_161418033_161418104@37.72.36,trna81_AspGTC_1_-_161410615_161410686@37.72.36|ACGCGGGAGACCGGGGTTCGATTCCCCGACGGGGAG</td><td>>trna10_AspGTC_12_-_125424193_125424264</td><td>37</td><td>72</td><td>36</td><td>ACGCGGGAGACCGGGGTTCGATTCCCCGACGGGGAG</td><td>594.5</td><td>-5.2</td><td>1.9</td><td>-2.8</td><td>5.1E-03</td><td>1.0E-02</td><td>BT20/MB468</td><td>MP</td></tr>
<tr><td>>trna111_HisGTG_1_-_147774845_147774916@1.30.30,trna118_HisGTG_1_-_145396881_145396952@1.30.30,trna16_HisGTG_1_+_146544773_146544844@1.30.30,trna1_HisGTG_15_+_45493349_45493420@1.30.30,trna21_HisGTG_1_+_147753471_147753542@1.30.30,trna33_HisGTG_6_+_27125906_27125977@1.30.30,trna7_HisGTG_9_-_14433938_14434009@1.30.30,trna8_HisGTG_15_-_45492611_45492682@1.30.30,trna9_HisGTG_15_-_45490804_45490875@1.30.30|GCCGTGATCGTATAGTGGTTAGTACTCTGC</td><td>>trna111_HisGTG_1_-_147774845_147774916</td><td>1</td><td>30</td><td>30</td><td>GCCGTGATCGTATAGTGGTTAGTACTCTGC</td><td>959.1</td><td>2.6</td><td>1.0</td><td>2.6</td><td>9.0E-03</td><td>1.0E-02</td><td>MB231/MB468</td><td>MITO</td></tr>
<tr><td>>hg38_21_+_8433172_8446622_RNA45SN1_45S_50ntFlanks@6661.6706.46|CGGTGGATCACTCGGCTCGTGCGTCGATGAAGAACGCAGCTAGCTG</td><td>>hg38_21_+_8433172_8446622_RNA45SN1_45S_50ntFlanks</td><td>6661</td><td>6706</td><td>46</td><td>CGGTGGATCACTCGGCTCGTGCGTCGATGAAGAACGCAGCTAGCTG</td><td>220.8</td><td>-3.3</td><td>1.2</td><td>-2.7</td><td>7.8E-03</td><td>1.0E-02</td><td>MB231/MB468</td><td>MP</td></tr>
<tr><td>>hg38_21_+_8433172_8446622_RNA45SN1_45S_50ntFlanks@9790.9820.31|CTGGCGTGGAGCCGGGCGTGGAATGCGAGTG</td><td>>hg38_21_+_8433172_8446622_RNA45SN1_45S_50ntFlanks</td><td>9790</td><td>9820</td><td>31</td><td>CTGGCGTGGAGCCGGGCGTGGAATGCGAGTG</td><td>121.0</td><td>-2.5</td><td>0.9</td><td>-2.6</td><td>8.5E-03</td><td>1.0E-02</td><td>MB231/MB468</td><td>CYTO</td></tr>
<tr><td>>trna11_GluTTC_15_-_26327381_26327452@1.30.30,trna3_GluTTC_13_-_45492062_45492133@1.30.30|TCCCACATGGTCTAGCGGTTAGGATTCCTG</td><td>>trna11_GluTTC_15_-_26327381_26327452</td><td>1</td><td>30</td><td>30</td><td>TCCCACATGGTCTAGCGGTTAGGATTCCTG</td><td>1368.7</td><td>-4.2</td><td>1.5</td><td>-2.8</td><td>5.1E-03</td><td>1.0E-02</td><td>BT20/MB468</td><td>MP</td></tr>
<tr><td>>hg38_21_+_8433172_8446622_RNA45SN1_45S_50ntFlanks@6733.6772.40|CATTGATCATCGACACTTCGAACGCACTTGCGGCCCCGGG</td><td>>hg38_21_+_8433172_8446622_RNA45SN1_45S_50ntFlanks</td><td>6733</td><td>6772</td><td>40</td><td>CATTGATCATCGACACTTCGAACGCACTTGCGGCCCCGGG</td><td>798.3</td><td>-1.5</td><td>0.6</td><td>-2.7</td><td>7.2E-03</td><td>1.0E-02</td><td>BT20/MB468</td><td>MITO</td></tr>
</table>

| | | | | | | | | | | | | | |
|---|---|---|---|---|---|---|---|---|---|---|---|---|---|
| >trna116_GluCTC_1_-_145399233_145399304@32.75.44,trna71_GluCTC_1_-_161439189_161439260@32.75.44,trna74_GluCTC_1_-_161431809_161431880@32.75.44,trna77_GluCTC_1_-_161424398_161424469@32.75.44,trna77_GluCTC_6_+_28949976_28950047@32.75.44,trna80_GluCTC_1_-_161417018_161417089@32.75.44,trna87_GluCTC_6_-_126101393_126101464@32.75.44\|CTCTCACCGCCGCGGCCCGGGTTCGATTCCCGGTCAGGGAACCA | >trna116_GluCTC_1_-_145399233_145399304 | 32 | 75 | 44 | CTCTCACCGCCGCGGCCCGGGTTCGATTCCCGGTCAGGGAACCA | 921.6 | -4.6 | 1.7 | -2.6 | 8.4E-03 | 1.0E-02 | BT20/MB468 | CYTO |
| >MI0003760\|hsa-mir-671&WithFlank&7\|+\|151238415\|151238544@35.59.25(+2U)&[MIMAT0003880&hsa-miR-671-5p&offsets\|0\|+2(+2U);m-316&7\|+\|151238449\|151238472&offsets\|0\|+1(+2U)]\|AGGAAGCCCTGGAGGGGCTGGAGGTTT | >MI0003760\|hsa-mir-671&WithFlank&7\|+\|151238415\|151238544 | 35 | 59 | 25(+2U) | AGGAAGCCCTGGAGGGGCTGGAGGTTT | 32.5 | 5.1 | 1.9 | 2.7 | 7.8E-03 | 1.0E-02 | MB231/MB468 | MP |
| >trnaMT_SerGCT_MT_+_12207_12265@13.61.49\|AAGAACTGCTAACTCATGCCCCCATGTCTAACAACATGGCTTTCTCACC | >trnaMT_SerGCT_MT_+_12207_12265 | 13 | 61 | 49 | AAGAACTGCTAACTCATGCCCCCATGTCTAACAACATGGCTTTCTCACC | 190.5 | 4.6 | 1.8 | 2.6 | 9.0E-03 | 1.0E-02 | MB231/MB468 | MITO |
| >hg38_21_+_8433172_8446622_RNA45SN1_45S_50ntFlanks@12490.12520.31\|CGTGAGCTGGGTTTAGACCGTCGTGAGACAG | >hg38_21_+_8433172_8446622_RNA45SN1_45S_50ntFlanks | 12490 | 12520 | 31 | CGTGAGCTGGGTTTAGACCGTCGTGAGACAG | 114.2 | -4.6 | 1.6 | -2.8 | 5.1E-03 | 1.0E-02 | BT20/MB468 | MP |
| >hg38_21_+_8433172_8446622_RNA45SN1_45S_50ntFlanks@12405.12448.44\|CTTTTTGATCCTTCGATGTCGGCTCTTCCTATCATTGTGAAGCA | >hg38_21_+_8433172_8446622_RNA45SN1_45S_50ntFlanks | 12405 | 12448 | 44 | CTTTTTGATCCTTCGATGTCGGCTCTTCCTATCATTGTGAAGCA | 359.1 | -2.1 | 0.8 | -2.6 | 9.0E-03 | 1.0E-02 | MB231/MB468 | MITO |
| >MI0000084\|hsa-mir-26b&WithFlank&2\|+\|218402640\|218402728@18.38.21&[MIMAT0000083&hsa-miR-26b-5p&offsets\|0\|0;m-53&2\|+\|218402657\|218402678&offsets\|0\|-1]\|TTCAAGTAATTCAGGATAGGT | >MI0000084\|hsa-mir-26b&WithFlank&2\|+\|218402640\|218402728 | 18 | 38 | 21 | TTCAAGTAATTCAGGATAGGT | 1166.9 | 3.0 | 1.0 | 3.0 | 2.4E-03 | 1.0E-02 | BT20/MB231 | CYTO |
| >hg38_21_+_8433172_8446622_RNA45SN1_45S_50ntFlanks@8919.8936.18\|TCCCGGGGCCGAGGGAGC | >hg38_21_+_8433172_8446622_RNA45SN1_45S_50ntFlanks | 8919 | 8936 | 18 | TCCCGGGGCCGAGGGAGC | 364.9 | -2.3 | 0.8 | -3.0 | 2.4E-03 | 1.0E-02 | BT20/MB231 | CYTO |
| >MI0000447\|hsa-mir-128-1&WithFlank&2\|+\|135665391\|135665484@56.75.20(+1A);MI0000727\|hsa-mir-128-2&WithFlank&3\|+\|35744470\|35744565@58.77.20(+1A)&[MIMAT0000424&hsa-miR-128-3p&offsets\|0\|-1(+1A);m-108&2\|+\|135665446\|135665466&offsets\|0\|-1(+1A)];[MIMAT0000424_1&hsa-miR-128-3p&offsets\|0\|-1(+1A);m-117&3\|+\|35744527\|35744547&offsets\|0\|-1(+1A)]\|TCACAGTGAACCGGTCTCTTA | >MI0000447\|hsa-mir-128-1&WithFlank&2\|+\|135665391\|135665484 | 56 | 75 | 20(+1A) | TCACAGTGAACCGGTCTCTTA | 359.9 | 2.4 | 0.8 | 3.0 | 2.4E-03 | 1.0E-02 | BT20/MB231 | CYTO |
| >hg38_21_+_8433172_8446622_RNA45SN1_45S_50ntFlanks@12027.12055.29\|ACTCTGATCGTTTTTTCACTGACCCGGTG | >hg38_21_+_8433172_8446622_RNA45SN1_45S_50ntFlanks | 12027 | 12055 | 29 | ACTCTGATCGTTTTTTCACTGACCCGGTG | 190.4 | 6.0 | 1.9 | 3.1 | 1.9E-03 | 1.0E-02 | BT20/MB231 | MP |
| >hg38_21_+_8433172_8446622_RNA45SN1_45S_50ntFlanks@6652.6672.21\|GACTCTTAGCGGTGGATCACT | >hg38_21_+_8433172_8446622_RNA45SN1_45S_50ntFlanks | 6652 | 6672 | 21 | GACTCTTAGCGGTGGATCACT | 328.0 | 2.7 | 1.0 | 2.6 | 9.0E-03 | 1.0E-02 | MB231/MB468 | MITO |
| >hg38_21_+_8433172_8446622_RNA45SN1_45S_50ntFlanks@10401.10429.29\|GAACATGGGTCAGTCGGTCCTGAGAGATG | >hg38_21_+_8433172_8446622_RNA45SN1_45S_50ntFlanks | 10401 | 10429 | 29 | GAACATGGGTCAGTCGGTCCTGAGAGATG | 1141.1 | -2.2 | 0.8 | -2.6 | 9.0E-03 | 1.0E-02 | MB231/MB468 | NUC |
| >MI0000070\|hsa-mir-16-1&WithFlank&13\|-\|50048967\|50049067@20.38.19(+1A);MI0000115\|hsa-mir-16-2&WithFlank&3\|+\|160404739\|160404831@16.34.19(+1A)&[MIMAT0000069&hsa-miR-16-5p&offsets\|0\|-3(+1A);m-62&13\|-\|50049027\|50049048&offsets\|0\|-3(+1A)];[MIMAT0000069_1&hsa-miR-16-5p&offsets\|0\|-3(+1A);m-61&3\|+\|160404754\|160404775&offsets\|0\|-3(+1A)]\|TAGCAGCACGTAAATATTGA | >MI0000070\|hsa-mir-16-1&WithFlank&13\|-\|50048967\|50049067 | 20 | 38 | 19(+1A) | TAGCAGCACGTAAATATTGA | 101.1 | 4.7 | 1.8 | 2.7 | 7.8E-03 | 1.0E-02 | MB231/MB468 | MP |
| >hg38_21_+_8433172_8446622_RNA45SN1_45S_50ntFlanks@11027.11050.24\|CTCTCTCTCTCTCTCCCCCGCTCC | >hg38_21_+_8433172_8446622_RNA45SN1_45S_50ntFlanks | 11027 | 11050 | 24 | CTCTCTCTCTCTCTCCCCCGCTCC | 66.8 | -4.7 | 1.7 | -2.8 | 5.1E-03 | 1.0E-02 | BT20/MB468 | MP |
| >hg38_21_+_8433172_8446622_RNA45SN1_45S_50ntFlanks@4555.4573.19\|CGCAGCTAGGAATAATGGA | >hg38_21_+_8433172_8446622_RNA45SN1_45S_50ntFlanks | 4555 | 4573 | 19 | CGCAGCTAGGAATAATGGA | 435.8 | -4.8 | 1.7 | -2.8 | 5.1E-03 | 1.0E-02 | BT20/MB468 | MP |
| >hg38_21_+_8433172_8446622_RNA45SN1_45S_50ntFlanks@12481.12499.19\|AATAGGGAACGTGAGCTGG | >hg38_21_+_8433172_8446622_RNA45SN1_45S_50ntFlanks | 12481 | 12499 | 19 | AATAGGGAACGTGAGCTGG | 611.4 | 2.3 | 0.9 | 2.6 | 8.5E-03 | 1.0E-02 | BT20/MB468 | CYTO |
| >hg38_21_+_8433172_8446622_RNA45SN1_45S_50ntFlanks@10399.10424.26\|TTGAACATGGGTCAGTCGGTCCTGAG | >hg38_21_+_8433172_8446622_RNA45SN1_45S_50ntFlanks | 10399 | 10424 | 26 | TTGAACATGGGTCAGTCGGTCCTGAG | 1949.3 | 3.9 | 1.3 | 3.1 | 1.9E-03 | 1.0E-02 | BT20/MB231 | MP |
| >hg38_21_+_8433172_8446622_RNA45SN1_45S_50ntFlanks@11569.11602.34\|CAGCCGACTTAGAACTGGTGCGGACCAGGGGAAT | >hg38_21_+_8433172_8446622_RNA45SN1_45S_50ntFlanks | 11569 | 11602 | 34 | CAGCCGACTTAGAACTGGTGCGGACCAGGGGAAT | 364.6 | 3.8 | 1.2 | 3.1 | 1.9E-03 | 1.0E-02 | BT20/MB231 | MP |
| >hg38_21_+_8433172_8446622_RNA45SN1_45S_50ntFlanks@9554.9577.24\|CGTAGCGGTCCTGACGTGCAAATC | >hg38_21_+_8433172_8446622_RNA45SN1_45S_50ntFlanks | 9554 | 9577 | 24 | CGTAGCGGTCCTGACGTGCAAATC | 542.3 | 1.2 | 0.4 | 2.7 | 7.3E-03 | 1.0E-02 | BT20/MB468 | MITO |
| >hg38_21_+_8433172_8446622_RNA45SN1_45S_50ntFlanks@4729.4755.27\|TCAAGAACGAAAGTCGGAGGTTCGAAG | >hg38_21_+_8433172_8446622_RNA45SN1_45S_50ntFlanks | 4729 | 4755 | 27 | TCAAGAACGAAAGTCGGAGGTTCGAAG | 73.0 | -4.5 | 1.6 | -2.8 | 5.1E-03 | 1.0E-02 | BT20/MB468 | MP |
| >hg38_MT_+_1621_3279_RNR2_16S_50ntFlanks@1368.1404.37\|CGATGTTGGATCAGGACATCCCGATGGTGCAGCCGCT | >hg38_MT_+_1621_3279_RNR2_16S_50ntFlanks | 1368 | 1404 | 37 | CGATGTTGGATCAGGACATCCCGATGGTGCAGCCGCT | 75.4 | 4.9 | 1.8 | 2.8 | 5.1E-03 | 1.0E-02 | BT20/MB468 | MP |
| >hg38_21_+_8433172_8446622_RNA45SN1_45S_50ntFlanks@12633.12651.19\|CGAAGCTACCATCTGTGGG | >hg38_21_+_8433172_8446622_RNA45SN1_45S_50ntFlanks | 12633 | 12651 | 19 | CGAAGCTACCATCTGTGGG | 1670.9 | -2.5 | 0.9 | -2.8 | 5.1E-03 | 1.0E-02 | BT20/MB468 | MP |
| >hg38_21_+_8433172_8446622_RNA45SN1_45S_50ntFlanks@11564.11582.19\|CCTAGCAGCCGACTTAGAA | >hg38_21_+_8433172_8446622_RNA45SN1_45S_50ntFlanks | 11564 | 11582 | 19 | CCTAGCAGCCGACTTAGAA | 460.0 | -4.8 | 1.7 | -2.8 | 5.1E-03 | 1.0E-02 | BT20/MB468 | MP |
| >hg38_21_+_8433172_8446622_RNA45SN1_45S_50ntFlanks@6645.6675.31\|CTCGTACGACTCTTAGCGGTGGATCACTCGG | >hg38_21_+_8433172_8446622_RNA45SN1_45S_50ntFlanks | 6645 | 6675 | 31 | CTCGTACGACTCTTAGCGGTGGATCACTCGG | 210.4 | -2.6 | 1.0 | -2.7 | 7.8E-03 | 1.0E-02 | MB231/MB468 | MP |
| >hg38_21_+_8433172_8446622_RNA45SN1_45S_50ntFlanks@10830.10859.30\|CAAGCCGGATCCGTAACTTCGGGATAAGGA | >hg38_21_+_8433172_8446622_RNA45SN1_45S_50ntFlanks | 10830 | 10859 | 30 | CAAGCCGGATCCGTAACTTCGGGATAAGGA | 245.7 | -4.6 | 1.6 | -2.8 | 5.1E-03 | 1.0E-02 | BT20/MB468 | MP |
| >hg38_21_+_8433172_8446622_RNA45SN1_45S_50ntFlanks@7971.8010.40\|GAGACGCGACCTCAGATCAGACGTGGCGACCCGCTGAATT | >hg38_21_+_8433172_8446622_RNA45SN1_45S_50ntFlanks | 7971 | 8010 | 40 | GAGACGCGACCTCAGATCAGACGTGGCGACCCGCTGAATT | 1329.8 | 3.2 | 1.0 | 3.1 | 2.1E-03 | 1.0E-02 | BT20/MB231 | NUC |
| >hg38_21_+_8433172_8446622_RNA45SN1_45S_50ntFlanks@7988.8011.24\|CAGACGTGGCGACCCGCTGAATTT | >hg38_21_+_8433172_8446622_RNA45SN1_45S_50ntFlanks | 7988 | 8011 | 24 | CAGACGTGGCGACCCGCTGAATTT | 1179.3 | 2.5 | 0.8 | 3.1 | 2.1E-03 | 1.0E-02 | BT20/MB231 | NUC |
| >hg38_21_+_8433172_8446622_RNA45SN1_45S_50ntFlanks@12997.13038.42\|CGCTGCGATCTATTGAAAGTCAGCCCTCGACACAAGGGTTTG | >hg38_21_+_8433172_8446622_RNA45SN1_45S_50ntFlanks | 12997 | 13038 | 42 | CGCTGCGATCTATTGAAAGTCAGCCCTCGACACAAGGGTTTG | 1084.4 | -2.3 | 0.7 | -3.1 | 2.1E-03 | 1.0E-02 | BT20/MB231 | NUC |
| >hg38_21_+_8433172_8446622_RNA45SN1_45S_50ntFlanks@9495.9523.29\|CGAAAGATGGTGAACTATGCCTGGGCAGG | >hg38_21_+_8433172_8446622_RNA45SN1_45S_50ntFlanks | 9495 | 9523 | 29 | CGAAAGATGGTGAACTATGCCTGGGCAGG | 404.9 | 7.6 | 2.4 | 3.1 | 1.9E-03 | 1.0E-02 | BT20/MB231 | MP |
| >hg38_21_+_8433172_8446622_RNA45SN1_45S_50ntFlanks@6778.6802.25\|CCCGGGGCTACGCCTGTCTGAGCGT | >hg38_21_+_8433172_8446622_RNA45SN1_45S_50ntFlanks | 6778 | 6802 | 25 | CCCGGGGCTACGCCTGTCTGAGCGT | 203.6 | 4.2 | 1.5 | 2.8 | 5.1E-03 | 1.0E-02 | BT20/MB468 | MP |
| >MI0000078\|hsa-mir-22&WithFlank&17\|-\|1713897\|1713993@59.81.23&[MIMAT0000077&hsa-miR-22-3p&offsets\|0\|+1;m-2&17\|-\|1713914\|1713935&offsets\|0\|+1]\|AAGCTGCCAGTTGAAGAACTGTT | >MI0000078\|hsa-mir-22&WithFlank&17\|-\|1713897\|1713993 | 59 | 81 | 23 | AAGCTGCCAGTTGAAGAACTGTT | 203.8 | 4.2 | 1.5 | 2.8 | 5.1E-03 | 1.0E-02 | BT20/MB468 | MP |
| >hg38_21_+_8433172_8446622_RNA45SN1_45S_50ntFlanks@8197.8222.26\|GTGGACGGTGTGAGGCCGGTAGCGGC | >hg38_21_+_8433172_8446622_RNA45SN1_45S_50ntFlanks | 8197 | 8222 | 26 | GTGGACGGTGTGAGGCCGGTAGCGGC | 83.1 | -4.5 | 1.7 | -2.7 | 7.8E-03 | 1.0E-02 | MB231/MB468 | MP |
| >hg38_21_+_8433172_8446622_RNA45SN1_45S_50ntFlanks@11671.11701.31\|CCAGTGCTCTGAATGTCAAAGTGAAGAAATT | >hg38_21_+_8433172_8446622_RNA45SN1_45S_50ntFlanks | 11671 | 11701 | 31 | CCAGTGCTCTGAATGTCAAAGTGAAGAAATT | 435.2 | -1.6 | 0.6 | -2.6 | 8.6E-03 | 1.0E-02 | MB231/MB468 | CYTO |
| >hg38_21_+_8433172_8446622_RNA45SN1_45S_50ntFlanks@6751.6771.21\|CGAACGCACTTGCGGCCCCGG | >hg38_21_+_8433172_8446622_RNA45SN1_45S_50ntFlanks | 6751 | 6771 | 21 | CGAACGCACTTGCGGCCCCGG | 236.5 | -4.7 | 1.7 | -2.8 | 5.1E-03 | 1.0E-02 | BT20/MB468 | MP |
| >hg38_21_+_8433172_8446622_RNA45SN1_45S_50ntFlanks@10389.10424.36\|TGAACAGCAGTTGAACATGGGTCAGTCGGTCCTGAG | >hg38_21_+_8433172_8446622_RNA45SN1_45S_50ntFlanks | 10389 | 10424 | 36 | TGAACAGCAGTTGAACATGGGTCAGTCGGTCCTGAG | 12988.6 | 3.8 | 1.2 | 3.1 | 1.9E-03 | 1.0E-02 | BT20/MB231 | MP |
| >MI0000078\|hsa-mir-22&WithFlank&17\|-\|1713897\|1713993@21.41.21&[MIMAT0004495&hsa-miR-22-5p&offsets\|0\|-1;m-217&17\|-\|1713953\|1713973&offsets\|0\|0]\|AGTTCTTCAGTGGCAAGCTTT | >MI0000078\|hsa-mir-22&WithFlank&17\|-\|1713897\|1713993 | 21 | 41 | 21 | AGTTCTTCAGTGGCAAGCTTT | 5336.1 | -2.9 | 0.9 | -3.1 | 1.9E-03 | 1.0E-02 | BT20/MB231 | MP |
| >hg38_21_+_8433172_8446622_RNA45SN1_45S_50ntFlanks@6719.6742.24\|TGAATTGCAGGACACATTGATCAT | >hg38_21_+_8433172_8446622_RNA45SN1_45S_50ntFlanks | 6719 | 6742 | 24 | TGAATTGCAGGACACATTGATCAT | 138.6 | -4.6 | 1.6 | -2.8 | 5.1E-03 | 1.0E-02 | BT20/MB468 | MP |
| >MI0000804\|hsa-mir-328&WithFlank&16\|-\|67202315\|67202401@54.75.22&[MIMAT0000752&hsa-miR-328-3p&offsets\|0\|0;m-160&16\|-\|67202327\|67202348&offsets\|0\|0]\|CTGGCCCTCTCTGCCCTTCCGT | >MI0000804\|hsa-mir-328&WithFlank&16\|-\|67202315\|67202401 | 54 | 75 | 22 | CTGGCCCTCTCTGCCCTTCCGT | 772.8 | 2.3 | 0.8 | 3.0 | 2.5E-03 | 1.0E-02 | BT20/MB231 | MITO |
| >hg38_21_+_8433172_8446622_RNA45SN1_45S_50ntFlanks@7990.8008.19\|GACGTGGCGACCCGCTGAA | >hg38_21_+_8433172_8446622_RNA45SN1_45S_50ntFlanks | 7990 | 8008 | 19 | GACGTGGCGACCCGCTGAA | 381.6 | 2.1 | 0.8 | 2.6 | 8.6E-03 | 1.0E-02 | MB231/MB468 | CYTO |
| >hg38_21_+_8433172_8446622_RNA45SN1_45S_50ntFlanks@10401.10430.30\|GAACATGGGTCAGTCGGTCCTGAGAGATGG | >hg38_21_+_8433172_8446622_RNA45SN1_45S_50ntFlanks | 10401 | 10430 | 30 | GAACATGGGTCAGTCGGTCCTGAGAGATGG | 254.4 | -2.4 | 0.9 | -2.8 | 5.1E-03 | 1.0E-02 | BT20/MB468 | MP |
| >hg38_21_+_8433172_8446622_RNA45SN1_45S_50ntFlanks@10781.10800.20\|AGGTGAACAGCCTCTGGCAT | >hg38_21_+_8433172_8446622_RNA45SN1_45S_50ntFlanks | 10781 | 10800 | 20 | AGGTGAACAGCCTCTGGCAT | 144.2 | -3.5 | 1.3 | -2.8 | 5.1E-03 | 1.0E-02 | BT20/MB468 | MP |
| >hg38_21_+_8433172_8446622_RNA45SN1_45S_50ntFlanks@10407.10430.24\|GGGTCAGTCGGTCCTGAGAGATGG | >hg38_21_+_8433172_8446622_RNA45SN1_45S_50ntFlanks | 10407 | 10430 | 24 | GGGTCAGTCGGTCCTGAGAGATGG | 99.6 | -4.6 | 1.7 | -2.8 | 5.1E-03 | 1.0E-02 | BT20/MB468 | MP |
| >MI0000101\|hsa-mir-99a&WithFlank&21\|+\|16539083\|16539175@19.39.21&[MIMAT0000097&hsa-miR-99a-5p&offsets\|0\|-1;m-64&21\|+\|16539101\|16539121&offsets\|0\|0]\|AACCCGTAGATCCGATCTTGT | >MI0000101\|hsa-mir-99a&WithFlank&21\|+\|16539083\|16539175 | 19 | 39 | 21 | AACCCGTAGATCCGATCTTGT | 703.1 | -2.6 | 0.8 | -3.0 | 2.4E-03 | 1.0E-02 | BT20/MB231 | CYTO |
| >hg38_MT_+_598_1651_RNR1_12S_50ntFlanks@608.633.26\|TGTTCTGTAATCGATAAACCCCGATC | >hg38_MT_+_598_1651_RNR1_12S_50ntFlanks | 608 | 633 | 26 | TGTTCTGTAATCGATAAACCCCGATC | 107.2 | 3.4 | 1.3 | 2.6 | 9.1E-03 | 1.0E-02 | MB231/MB468 | MITO |
| >hg38_21_+_8433172_8446622_RNA45SN1_45S_50ntFlanks@9929.9950.22\|GATATAGACAGCAGGACGGTGG | >hg38_21_+_8433172_8446622_RNA45SN1_45S_50ntFlanks | 9929 | 9950 | 22 | GATATAGACAGCAGGACGGTGG | 154.0 | 5.7 | 1.8 | 3.1 | 1.9E-03 | 1.0E-02 | BT20/MB231 | MP |
| >MI0000078\|hsa-mir-22&WithFlank&17\|-\|1713897\|1713993@21.40.20(+1C)&[MIMAT0004495&hsa-miR-22-5p&offsets\|0\|-2(+1C);m-217&17\|-\|1713953\|1713973&offsets\|0\|-1(+1C)]\|AGTTCTTCAGTGGCAAGCTTC | >MI0000078\|hsa-mir-22&WithFlank&17\|-\|1713897\|1713993 | 21 | 40 | 20(+1C) | AGTTCTTCAGTGGCAAGCTTC | 720.4 | -3.2 | 1.0 | -3.1 | 1.9E-03 | 1.0E-02 | BT20/MB231 | MP |
| >MI0000234\|hsa-mir-192&WithFlank&11\|-\|64891131\|64891252@31.53.23&[MIMAT0000222&hsa-miR-192-5p&offsets\|+1\|+3;m-59&11\|-\|64891203\|64891223&offsets\|+1\|+3]\|TGACCTATGAATTGACAGCCAGT | >MI0000234\|hsa-mir-192&WithFlank&11\|-\|64891131\|64891252 | 31 | 53 | 23 | TGACCTATGAATTGACAGCCAGT | 1419.9 | -3.0 | 1.0 | -3.1 | 1.9E-03 | 1.0E-02 | BT20/MB231 | MP |

<table>
<tr><th></th><th></th><th></th><th></th><th></th><th></th><th></th><th></th><th></th><th></th><th></th><th></th><th></th><th></th><th></th></tr>
<tr><td>>trnaMT_GluTTC_MT_-_14674_14742@37.71.35,trnalookalike8_GluTTC_5_-_93905172_93905240@37.71.35|TCATTGGTCGTGGTTGTAGTCCGTGCGAGAATACC</td><td>>trnaMT_GluTTC_MT_-_14674_14742</td><td>37</td><td>71</td><td>35</td><td>TCATTGGTCGTGGTTGTAGTCCGTGCGAGAATACC</td><td>148.8</td><td>2.7</td><td>1.0</td><td>2.6</td><td>9.1E-03</td><td>1.0E-02</td><td>MB231/MB468</td><td>MITO</td></tr>
<tr><td>>hg38_21_+_8433172_8446622_RNA45SN1_45S_50ntFlanks@10821.10839.19|GGAAGTCGGCAAGCCGGAT</td><td>>hg38_21_+_8433172_8446622_RNA45SN1_45S_50ntFlanks</td><td>10821</td><td>10839</td><td>19</td><td>GGAAGTCGGCAAGCCGGAT</td><td>556.4</td><td>-2.9</td><td>1.0</td><td>-2.8</td><td>5.2E-03</td><td>1.0E-02</td><td>BT20/MB468</td><td>MP</td></tr>
<tr><td>>hg38_21_+_8433172_8446622_RNA45SN1_45S_50ntFlanks@12932.12953.22|CGCTAAACCATTCGTAGACGAC</td><td>>hg38_21_+_8433172_8446622_RNA45SN1_45S_50ntFlanks</td><td>12932</td><td>12953</td><td>22</td><td>CGCTAAACCATTCGTAGACGAC</td><td>146.6</td><td>-4.6</td><td>1.6</td><td>-2.8</td><td>5.2E-03</td><td>1.0E-02</td><td>BT20/MB468</td><td>MP</td></tr>
<tr><td>>hg38_21_+_8433172_8446622_RNA45SN1_45S_50ntFlanks@10869.10901.33|AGGGCTGGGTCGGTCGGGCTGGGGCGCGAAGCG</td><td>>hg38_21_+_8433172_8446622_RNA45SN1_45S_50ntFlanks</td><td>10869</td><td>10901</td><td>33</td><td>AGGGCTGGGTCGGTCGGGCTGGGGCGCGAAGCG</td><td>470.2</td><td>-4.3</td><td>1.6</td><td>-2.7</td><td>7.3E-03</td><td>1.0E-02</td><td>BT20/MB468</td><td>MITO</td></tr>
<tr><td>>hg38_21_+_8433172_8446622_RNA45SN1_45S_50ntFlanks@10240.10273.34|CTACGCCGCGACGAGTAGGAGGGCCGCTGCGGTG</td><td>>hg38_21_+_8433172_8446622_RNA45SN1_45S_50ntFlanks</td><td>10240</td><td>10273</td><td>34</td><td>CTACGCCGCGACGAGTAGGAGGGCCGCTGCGGTG</td><td>474.8</td><td>3.6</td><td>1.3</td><td>2.8</td><td>5.2E-03</td><td>1.0E-02</td><td>BT20/MB468</td><td>MP</td></tr>
<tr><td>>hg38_21_+_8433172_8446622_RNA45SN1_45S_50ntFlanks@12474.12497.24|ACCCACTAATAGGGAACGTGAGCT</td><td>>hg38_21_+_8433172_8446622_RNA45SN1_45S_50ntFlanks</td><td>12474</td><td>12497</td><td>24</td><td>ACCCACTAATAGGGAACGTGAGCT</td><td>393.8</td><td>2.0</td><td>0.8</td><td>2.6</td><td>9.1E-03</td><td>1.0E-02</td><td>MB231/MB468</td><td>NUC</td></tr>
<tr><td>>hg38_MT_+_1621_3279_RNR2_16S_50ntFlanks@1349.1382.34|ACAATAGGGTTTACGACCTCGATGTTGGATCAGG</td><td>>hg38_MT_+_1621_3279_RNR2_16S_50ntFlanks</td><td>1349</td><td>1382</td><td>34</td><td>ACAATAGGGTTTACGACCTCGATGTTGGATCAGG</td><td>207.1</td><td>3.3</td><td>1.2</td><td>2.7</td><td>7.9E-03</td><td>1.0E-02</td><td>MB231/MB468</td><td>MP</td></tr>
<tr><td>>hg38_MT_+_1621_3279_RNR2_16S_50ntFlanks@51.84.34|GCTAAACCTAGCCCCAAACCCACTCCACCTTACT</td><td>>hg38_MT_+_1621_3279_RNR2_16S_50ntFlanks</td><td>51</td><td>84</td><td>34</td><td>GCTAAACCTAGCCCCAAACCCACTCCACCTTACT</td><td>259.6</td><td>1.9</td><td>0.7</td><td>2.6</td><td>9.0E-03</td><td>1.0E-02</td><td>BT20/MB468</td><td>NUC</td></tr>
<tr><td>>hg38_21_+_8433172_8446622_RNA45SN1_45S_50ntFlanks@11207.11225.19|GGCGGCGACTCTGGACGCG</td><td>>hg38_21_+_8433172_8446622_RNA45SN1_45S_50ntFlanks</td><td>11207</td><td>11225</td><td>19</td><td>GGCGGCGACTCTGGACGCG</td><td>933.0</td><td>-3.8</td><td>1.4</td><td>-2.8</td><td>5.2E-03</td><td>1.0E-02</td><td>BT20/MB468</td><td>MP</td></tr>
<tr><td>>hg38_MT_+_1621_3279_RNR2_16S_50ntFlanks@929.959.31|CAGTGACACATGTTTAACGGCCGCGGTACCC</td><td>>hg38_MT_+_1621_3279_RNR2_16S_50ntFlanks</td><td>929</td><td>959</td><td>31</td><td>CAGTGACACATGTTTAACGGCCGCGGTACCC</td><td>178.3</td><td>5.3</td><td>1.9</td><td>2.8</td><td>5.2E-03</td><td>1.0E-02</td><td>BT20/MB468</td><td>MP</td></tr>
<tr><td>>hg38_21_+_8433172_8446622_RNA45SN1_45S_50ntFlanks@11587.11611.25|TGCGGACCAGGGGAATCCGACTGTT</td><td>>hg38_21_+_8433172_8446622_RNA45SN1_45S_50ntFlanks</td><td>11587</td><td>11611</td><td>25</td><td>TGCGGACCAGGGGAATCCGACTGTT</td><td>490.4</td><td>-1.6</td><td>0.6</td><td>-2.6</td><td>9.1E-03</td><td>1.0E-02</td><td>MB231/MB468</td><td>NUC</td></tr>
<tr><td>>hg38_21_+_8433172_8446622_RNA45SN1_45S_50ntFlanks@9496.9524.29|GAAAGATGGTGAACTATGCCTGGGCAGGG</td><td>>hg38_21_+_8433172_8446622_RNA45SN1_45S_50ntFlanks</td><td>9496</td><td>9524</td><td>29</td><td>GAAAGATGGTGAACTATGCCTGGGCAGGG</td><td>55.9</td><td>5.6</td><td>2.0</td><td>2.8</td><td>5.2E-03</td><td>1.0E-02</td><td>BT20/MB468</td><td>MP</td></tr>
<tr><td>>trnaMT_SerGCT_MT_+_12207_12265@13.62.50|AAGAACTGCTAACTCATGCCCCCATGTCTAACAACATGGCTTTCTCACCA</td><td>>trnaMT_SerGCT_MT_+_12207_12265</td><td>13</td><td>62</td><td>50</td><td>AAGAACTGCTAACTCATGCCCCCATGTCTAACAACATGGCTTTCTCACCA</td><td>1004.8</td><td>7.6</td><td>2.7</td><td>2.8</td><td>5.2E-03</td><td>1.0E-02</td><td>BT20/MB468</td><td>MP</td></tr>
<tr><td>>hg38_21_+_8433172_8446622_RNA45SN1_45S_50ntFlanks@10677.10711.35|GCGGTTCCGGCGGCGTCCGGTGAGCTCTCGCTGGC</td><td>>hg38_21_+_8433172_8446622_RNA45SN1_45S_50ntFlanks</td><td>10677</td><td>10711</td><td>35</td><td>GCGGTTCCGGCGGCGTCCGGTGAGCTCTCGCTGGC</td><td>1057.5</td><td>3.2</td><td>1.1</td><td>3.0</td><td>2.5E-03</td><td>1.0E-02</td><td>BT20/MB231</td><td>MITO</td></tr>
<tr><td>>trna143_LysTTT_6_-_27559593_27559665@14.31.18,trna149_LysTTT_6_-_27302769_27302841@14.31.18,trna14_LysTTT_11_-_59327808_59327880@14.31.18,trna2_LysTTT_17_+_8022473_8022545@14.31.18,trna54_LysTTT_1_+_204475655_204475727@14.31.18,trna5_LysTTT_11_+_59323902_59323974@14.31.18,trna62_LysTTT_1_-_204476158_204476230@14.31.18,trna76_LysTTT_6_+_28918806_28918878@14.31.18|AGTCGGTAGAGCATCAGA</td><td>>trna143_LysTTT_6_-_27559593_27559665</td><td>14</td><td>31</td><td>18</td><td>AGTCGGTAGAGCATCAGA</td><td>1432.3</td><td>2.4</td><td>0.9</td><td>2.6</td><td>8.6E-03</td><td>1.0E-02</td><td>BT20/MB468</td><td>CYTO</td></tr>
<tr><td>>hg38_21_+_8433172_8446622_RNA45SN1_45S_50ntFlanks@7975.8013.39|CGCGACCTCAGATCAGACGTGGCGACCCGCTGAATTTAA</td><td>>hg38_21_+_8433172_8446622_RNA45SN1_45S_50ntFlanks</td><td>7975</td><td>8013</td><td>39</td><td>CGCGACCTCAGATCAGACGTGGCGACCCGCTGAATTTAA</td><td>10668.8</td><td>2.3</td><td>0.9</td><td>2.6</td><td>9.1E-03</td><td>1.0E-02</td><td>MB231/MB468</td><td>MITO</td></tr>
<tr><td>>hg38_21_+_8433172_8446622_RNA45SN1_45S_50ntFlanks@9555.9584.30|GTAGCGGTCCTGACGTGCAAATCGGTCGTC</td><td>>hg38_21_+_8433172_8446622_RNA45SN1_45S_50ntFlanks</td><td>9555</td><td>9584</td><td>30</td><td>GTAGCGGTCCTGACGTGCAAATCGGTCGTC</td><td>341.6</td><td>-2.0</td><td>0.8</td><td>-2.6</td><td>9.1E-03</td><td>1.0E-02</td><td>MB231/MB468</td><td>NUC</td></tr>
<tr><td>>am_trnaMT_ValTAC_MT_+_1602_1670@28.58.31|AACTTACACTTAGGAGATTTCAACTTAACTT</td><td>>am_trnaMT_ValTAC_MT_+_1602_1670</td><td>28</td><td>58</td><td>31</td><td>AACTTACACTTAGGAGATTTCAACTTAACTT</td><td>247.5</td><td>5.5</td><td>2.0</td><td>2.8</td><td>5.2E-03</td><td>1.0E-02</td><td>BT20/MB468</td><td>MP</td></tr>
<tr><td>>MI0000082|hsa-mir-25&WithFlank&7|-|100093554|100093649@58.76.19&[MIMAT0000081&hsa-miR-25-3p&offsets|0|-3;m-23&7|-|100093571|100093592&offsets|0|-3]|CATTGCACTTGTCTCGGTC</td><td>>MI0000082|hsa-mir-25&WithFlank&7|-|100093554|100093649</td><td>58</td><td>76</td><td>19</td><td>CATTGCACTTGTCTCGGTC</td><td>179.1</td><td>6.3</td><td>2.2</td><td>2.8</td><td>5.2E-03</td><td>1.0E-02</td><td>BT20/MB468</td><td>MP</td></tr>
<tr><td>>am_trna116_GluCTC_1_-_145399233_145399304@1.26.26,trna59_GluCTC_1_+_249168447_249168518@1.26.26,trna71_GluCTC_1_-_161439189_161439260@1.26.26,trna74_GluCTC_1_-_161431809_161431880@1.26.26,trna77_GluCTC_1_-_161424398_161424469@1.26.26,trna77_GluCTC_6_+_28949976_28950047@1.26.26,trna80_GluCTC_1_-_161417018_161417089@1.26.26,trna87_GluCTC_6_-_126101393_126101464@1.26.26|TCCCTGGTGGTCTAGTGGTTAGGATT</td><td>>am_trna116_GluCTC_1_-_145399233_145399304</td><td>1</td><td>26</td><td>26</td><td>TCCCTGGTGGTCTAGTGGTTAGGATT</td><td>2307.5</td><td>1.4</td><td>0.5</td><td>2.6</td><td>8.7E-03</td><td>1.0E-02</td><td>MB231/MB468</td><td>CYTO</td></tr>
<tr><td>>trna152_ValCAC_6_-_27248049_27248121@1.24.24|GCTTCTGTAGTGTAGTGGTTATCA</td><td>>trna152_ValCAC_6_-_27248049_27248121</td><td>1</td><td>24</td><td>24</td><td>GCTTCTGTAGTGTAGTGGTTATCA</td><td>273.8</td><td>-4.8</td><td>1.8</td><td>-2.6</td><td>9.1E-03</td><td>1.0E-02</td><td>MB231/MB468</td><td>NUC</td></tr>
<tr><td>>MI0000070|hsa-mir-16-1&WithFlank&13|-|50048967|50049067@20.42.23;MI0000115|hsa-mir-16-2&WithFlank&3|+|160404739|160404831@16.38.23&[MIMAT0000069&hsa-miR-16-5p&offsets|0|+1;m-62&13|-|50049027|50049048&offsets|0|+1];[MIMAT0000069_1&hsa-miR-16-5p&offsets|0|+1;m-61&3|+|160404754|160404775&offsets|0|+1]|TAGCAGCACGTAAATATTGGCGT</td><td>>MI0000070|hsa-mir-16-1&WithFlank&13|-|50048967|50049067</td><td>20</td><td>42</td><td>23</td><td>TAGCAGCACGTAAATATTGGCGT</td><td>2538.5</td><td>-2.8</td><td>0.9</td><td>-3.0</td><td>2.5E-03</td><td>1.0E-02</td><td>BT20/MB231</td><td>MITO</td></tr>
<tr><td>>MI0000085|hsa-mir-27a&WithFlank&19|-|13836434|13836523@57.76.20(+1A)&[MIMAT0000084&hsa-miR-27a-3p&offsets|0|-1(+1A);m-43&19|-|13836447|13836467&offsets|0|-1(+1A)]|TTCACAGTGGCTAAGTTCCGA</td><td>>MI0000085|hsa-mir-27a&WithFlank&19|-|13836434|13836523</td><td>57</td><td>76</td><td>20(+1A)</td><td>TTCACAGTGGCTAAGTTCCGA</td><td>3713.2</td><td>-2.2</td><td>0.7</td><td>-3.0</td><td>2.5E-03</td><td>1.0E-02</td><td>BT20/MB231</td><td>MITO</td></tr>
<tr><td>>hg38_21_+_8433172_8446622_RNA45SN1_45S_50ntFlanks@4733.4767.35|GAACGAAAGTCGGAGGTTCGAAGACGATCAGATAC</td><td>>hg38_21_+_8433172_8446622_RNA45SN1_45S_50ntFlanks</td><td>4733</td><td>4767</td><td>35</td><td>GAACGAAAGTCGGAGGTTCGAAGACGATCAGATAC</td><td>105.5</td><td>4.3</td><td>1.6</td><td>2.7</td><td>7.9E-03</td><td>1.0E-02</td><td>MB231/MB468</td><td>MP</td></tr>
<tr><td>>hg38_21_+_8433172_8446622_RNA45SN1_45S_50ntFlanks@12907.12930.24|CACCGCACGTTCGTGGGGAACCTG</td><td>>hg38_21_+_8433172_8446622_RNA45SN1_45S_50ntFlanks</td><td>12907</td><td>12930</td><td>24</td><td>CACCGCACGTTCGTGGGGAACCTG</td><td>274.1</td><td>-4.9</td><td>1.8</td><td>-2.8</td><td>5.2E-03</td><td>1.0E-02</td><td>BT20/MB468</td><td>MP</td></tr>
<tr><td>>hg38_21_+_8433172_8446622_RNA45SN1_45S_50ntFlanks@4731.4765.35|AAGAACGAAAGTCGGAGGTTCGAAGACGATCAGAT</td><td>>hg38_21_+_8433172_8446622_RNA45SN1_45S_50ntFlanks</td><td>4731</td><td>4765</td><td>35</td><td>AAGAACGAAAGTCGGAGGTTCGAAGACGATCAGAT</td><td>276.5</td><td>1.3</td><td>0.5</td><td>2.6</td><td>9.0E-03</td><td>1.0E-02</td><td>BT20/MB468</td><td>NUC</td></tr>
<tr><td>>trna116_GluCTC_1_-_145399233_145399304@2.31.30,trna59_GluCTC_1_+_249168447_249168518@2.31.30,trna71_GluCTC_1_-_161439189_161439260@2.31.30,trna74_GluCTC_1_-_161431809_161431880@2.31.30,trna77_GluCTC_1_-_161424398_161424469@2.31.30,trna77_GluCTC_6_+_28949976_28950047@2.31.30,trna80_GluCTC_1_-_161417018_161417089@2.31.30,trna87_GluCTC_6_-_126101393_126101464@2.31.30|CCCTGGTGGTCTAGTGGTTAGGATTCGGCG</td><td>>trna116_GluCTC_1_-_145399233_145399304</td><td>2</td><td>31</td><td>30</td><td>CCCTGGTGGTCTAGTGGTTAGGATTCGGCG</td><td>406.1</td><td>-2.4</td><td>0.9</td><td>-2.6</td><td>9.2E-03</td><td>1.0E-02</td><td>MB231/MB468</td><td>NUC</td></tr>
<tr><td>>hg38_21_+_8433172_8446622_RNA45SN1_45S_50ntFlanks@12419.12439.21|GATGTCGGCTCTTCCTATCAT</td><td>>hg38_21_+_8433172_8446622_RNA45SN1_45S_50ntFlanks</td><td>12419</td><td>12439</td><td>21</td><td>GATGTCGGCTCTTCCTATCAT</td><td>297.2</td><td>2.1</td><td>0.8</td><td>2.6</td><td>9.2E-03</td><td>1.0E-02</td><td>MB231/MB468</td><td>MITO</td></tr>
<tr><td>>hg38_21_+_8433172_8446622_RNA45SN1_45S_50ntFlanks@3993.4024.32|GTGACTCTAGATAACCTCGGGCCGATCGCACG</td><td>>hg38_21_+_8433172_8446622_RNA45SN1_45S_50ntFlanks</td><td>3993</td><td>4024</td><td>32</td><td>GTGACTCTAGATAACCTCGGGCCGATCGCACG</td><td>916.5</td><td>2.3</td><td>0.9</td><td>2.6</td><td>8.7E-03</td><td>1.0E-02</td><td>MB231/MB468</td><td>CYTO</td></tr>
<tr><td>>MI0000299|hsa-mir-222&WithFlank&X|-|45747009|45747130@75.96.22(+1C)&[MIMAT0000279&hsa-miR-222-3p&offsets|0|+1(+1C);m-140&X|-|45747033|45747056&offsets|0|-2(+1C)]|AGCTACATCTGGCTACTGGGTCC</td><td>>MI0000299|hsa-mir-222&WithFlank&X|-|45747009|45747130</td><td>75</td><td>96</td><td>22(+1C)</td><td>AGCTACATCTGGCTACTGGGTCC</td><td>2291.7</td><td>-3.4</td><td>1.1</td><td>-3.1</td><td>2.0E-03</td><td>1.0E-02</td><td>BT20/MB231</td><td>MP</td></tr>
<tr><td>>hg38_21_+_8433172_8446622_RNA45SN1_45S_50ntFlanks@12418.12436.19|CGATGTCGGCTCTTCCTAT</td><td>>hg38_21_+_8433172_8446622_RNA45SN1_45S_50ntFlanks</td><td>12418</td><td>12436</td><td>19</td><td>CGATGTCGGCTCTTCCTAT</td><td>1938.3</td><td>-5.2</td><td>1.9</td><td>-2.8</td><td>5.2E-03</td><td>1.0E-02</td><td>BT20/MB468</td><td>MP</td></tr>
<tr><td>>hg38_21_+_8433172_8446622_RNA45SN1_45S_50ntFlanks@4353.4371.19|TATTAAAGTTGCTGCAGTT</td><td>>hg38_21_+_8433172_8446622_RNA45SN1_45S_50ntFlanks</td><td>4353</td><td>4371</td><td>19</td><td>TATTAAAGTTGCTGCAGTT</td><td>195.1</td><td>-2.3</td><td>0.8</td><td>-2.7</td><td>8.0E-03</td><td>1.0E-02</td><td>MB231/MB468</td><td>MP</td></tr>
<tr><td>>hg38_21_+_8433172_8446622_RNA45SN1_45S_50ntFlanks@12417.12438.22|TCGATGTCGGCTCTTCCTATCA</td><td>>hg38_21_+_8433172_8446622_RNA45SN1_45S_50ntFlanks</td><td>12417</td><td>12438</td><td>22</td><td>TCGATGTCGGCTCTTCCTATCA</td><td>1045.4</td><td>-2.6</td><td>1.0</td><td>-2.6</td><td>9.0E-03</td><td>1.0E-02</td><td>BT20/MB468</td><td>NUC</td></tr>
<tr><td>>hg38_21_+_8433172_8446622_RNA45SN1_45S_50ntFlanks@12927.12951.25|CCTGGCGCTAAACCATTCGTAGACG</td><td>>hg38_21_+_8433172_8446622_RNA45SN1_45S_50ntFlanks</td><td>12927</td><td>12951</td><td>25</td><td>CCTGGCGCTAAACCATTCGTAGACG</td><td>172.3</td><td>-4.7</td><td>1.7</td><td>-2.8</td><td>5.2E-03</td><td>1.0E-02</td><td>BT20/MB468</td><td>MP</td></tr>
<tr><td>>hg38_21_+_8433172_8446622_RNA45SN1_45S_50ntFlanks@10355.10382.28|AACTTTGAAGGCCGAAGTGGAGAAGGGT</td><td>>hg38_21_+_8433172_8446622_RNA45SN1_45S_50ntFlanks</td><td>10355</td><td>10382</td><td>28</td><td>AACTTTGAAGGCCGAAGTGGAGAAGGGT</td><td>126.5</td><td>-3.4</td><td>1.3</td><td>-2.7</td><td>8.0E-03</td><td>1.0E-02</td><td>MB231/MB468</td><td>MP</td></tr>
<tr><td>>hg38_21_+_8433172_8446622_RNA45SN1_45S_50ntFlanks@4003.4026.24|ATAACCTCGGGCCGATCGCACGCC</td><td>>hg38_21_+_8433172_8446622_RNA45SN1_45S_50ntFlanks</td><td>4003</td><td>4026</td><td>24</td><td>ATAACCTCGGGCCGATCGCACGCC</td><td>390.2</td><td>-4.8</td><td>1.7</td><td>-2.8</td><td>5.2E-03</td><td>1.0E-02</td><td>BT20/MB468</td><td>MP</td></tr>
<tr><td>>hg38_21_+_8433172_8446622_RNA45SN1_45S_50ntFlanks@9576.9594.19|TCGGTCGTCCGACCTGGGT</td><td>>hg38_21_+_8433172_8446622_RNA45SN1_45S_50ntFlanks</td><td>9576</td><td>9594</td><td>19</td><td>TCGGTCGTCCGACCTGGGT</td><td>178.9</td><td>-3.8</td><td>1.4</td><td>-2.8</td><td>5.2E-03</td><td>1.0E-02</td><td>BT20/MB468</td><td>MP</td></tr>
<tr><td>>hg38_21_+_8433172_8446622_RNA45SN1_45S_50ntFlanks@12932.12966.35|CGCTAAACCATTCGTAGACGACCTGCTTCTGGGTC</td><td>>hg38_21_+_8433172_8446622_RNA45SN1_45S_50ntFlanks</td><td>12932</td><td>12966</td><td>35</td><td>CGCTAAACCATTCGTAGACGACCTGCTTCTGGGTC</td><td>136.2</td><td>-3.9</td><td>1.5</td><td>-2.6</td><td>8.6E-03</td><td>1.0E-02</td><td>BT20/MB468</td><td>CYTO</td></tr>
<tr><td>>trnaMT_TrpTCA_MT_+_5512_5579@17.34.18,trnalookalike3_TrpTCA_1_+_566062_566129@17.34.18|TACAGACCAAGAGCCTTC</td><td>>trnaMT_TrpTCA_MT_+_5512_5579</td><td>17</td><td>34</td><td>18</td><td>TACAGACCAAGAGCCTTC</td><td>40.5</td><td>5.0</td><td>1.9</td><td>2.7</td><td>8.0E-03</td><td>1.0E-02</td><td>MB231/MB468</td><td>MP</td></tr>
<tr><td>>trnaMT_ValTAC_MT_+_1602_1670@36.72.37|CTTAGGAGATTTCAACTTAACTTGACCGCTCTGACCA</td><td>>trnaMT_ValTAC_MT_+_1602_1670</td><td>36</td><td>72</td><td>37</td><td>CTTAGGAGATTTCAACTTAACTTGACCGCTCTGACCA</td><td>119.5</td><td>3.4</td><td>1.3</td><td>2.7</td><td>8.0E-03</td><td>1.0E-02</td><td>MB231/MB468</td><td>MP</td></tr>
<tr><td>>hg38_21_+_8433172_8446622_RNA45SN1_45S_50ntFlanks@3835.3861.27|CTCCTCTCCTACTTGGATAACTGTGGT</td><td>>hg38_21_+_8433172_8446622_RNA45SN1_45S_50ntFlanks</td><td>3835</td><td>3861</td><td>27</td><td>CTCCTCTCCTACTTGGATAACTGTGGT</td><td>336.2</td><td>-1.2</td><td>0.5</td><td>-2.6</td><td>8.7E-03</td><td>1.0E-02</td><td>BT20/MB468</td><td>CYTO</td></tr>
<tr><td>>MI0000253|hsa-mir-148a&WithFlank&7|-|25949913|25949992@50.69.20&[MIMAT0000243&hsa-miR-148a-3p&offsets|0|-2;m-7&7|-|25949922|25949943&offsets|0|-2]|TCAGTGCACTACAGAACTTT</td><td>>MI0000253|hsa-mir-148a&WithFlank&7|-|25949913|25949992</td><td>50</td><td>69</td><td>20</td><td>TCAGTGCACTACAGAACTTT</td><td>87.6</td><td>6.2</td><td>2.2</td><td>2.8</td><td>5.2E-03</td><td>1.0E-02</td><td>BT20/MB468</td><td>MP</td></tr>
<tr><td>>hg38_21_+_8433172_8446622_RNA45SN1_45S_50ntFlanks@9533.9572.40|GAGGAAACTCTGGTGGAGGTCCGTAGCGGTCCTGACGTGC</td><td>>hg38_21_+_8433172_8446622_RNA45SN1_45S_50ntFlanks</td><td>9533</td><td>9572</td><td>40</td><td>GAGGAAACTCTGGTGGAGGTCCGTAGCGGTCCTGACGTGC</td><td>180.6</td><td>-2.8</td><td>1.1</td><td>-2.7</td><td>8.0E-03</td><td>1.0E-02</td><td>MB231/MB468</td><td>MP</td></tr>
<tr><td>>hg38_21_+_8433172_8446622_RNA45SN1_45S_50ntFlanks@7973.7994.22|GACGCGACCTCAGATCAGACGT</td><td>>hg38_21_+_8433172_8446622_RNA45SN1_45S_50ntFlanks</td><td>7973</td><td>7994</td><td>22</td><td>GACGCGACCTCAGATCAGACGT</td><td>918.4</td><td>2.6</td><td>0.9</td><td>2.8</td><td>5.2E-03</td><td>1.0E-02</td><td>BT20/MB468</td><td>MP</td></tr>
<tr><td>>hg38_MT_+_598_1651_RNR1_12S_50ntFlanks@425.453.29|ACGAAAGTGGCTTTAACATATCTGAACAC</td><td>>hg38_MT_+_598_1651_RNR1_12S_50ntFlanks</td><td>425</td><td>453</td><td>29</td><td>ACGAAAGTGGCTTTAACATATCTGAACAC</td><td>304.1</td><td>4.3</td><td>1.5</td><td>2.8</td><td>5.2E-03</td><td>1.0E-02</td><td>BT20/MB468</td><td>MP</td></tr>
<tr><td>>hg38_21_+_8433172_8446622_RNA45SN1_45S_50ntFlanks@7991.8017.27|ACGTGGCGACCCGCTGAATTTAAGCAT</td><td>>hg38_21_+_8433172_8446622_RNA45SN1_45S_50ntFlanks</td><td>7991</td><td>8017</td><td>27</td><td>ACGTGGCGACCCGCTGAATTTAAGCAT</td><td>831.3</td><td>-2.1</td><td>0.8</td><td>-2.7</td><td>7.4E-03</td><td>1.0E-02</td><td>BT20/MB468</td><td>MITO</td></tr>
</table>

<table>
<tr><th></th><th></th><th></th><th></th><th></th><th></th><th></th><th></th><th></th><th></th><th></th><th></th><th></th></tr>
<tr><td>>hg38_21_+_8433172_8446622_RNA45SN1_45S_50ntFlanks@9305.9335.31|CCAAGGAGTCTAACACGTGCGCGAGTCGGGG</td><td>>hg38_21_+_8433172_8446622_RNA45SN1_45S_50ntFlanks</td><td>9305</td><td>9335</td><td>31</td><td>CCAAGGAGTCTAACACGTGCGCGAGTCGGGG</td><td>728.4</td><td>-3.9</td><td>1.4</td><td>-2.8</td><td>5.3E-03</td><td>1.0E-02</td><td>BT20/MB468</td><td>MP</td></tr>
<tr><td>>hg38_21_+_8433172_8446622_RNA45SN1_45S_50ntFlanks@3854.3880.27|ACTGTGGTAATTCTAGAGCTAATACAT</td><td>>hg38_21_+_8433172_8446622_RNA45SN1_45S_50ntFlanks</td><td>3854</td><td>3880</td><td>27</td><td>ACTGTGGTAATTCTAGAGCTAATACAT</td><td>222.0</td><td>-2.0</td><td>0.8</td><td>-2.6</td><td>8.7E-03</td><td>1.0E-02</td><td>BT20/MB468</td><td>CYTO</td></tr>
<tr><td>>hg38_21_+_8433172_8446622_RNA45SN1_45S_50ntFlanks@8176.8196.21|CTTCTGATCGAGGCCCAGCCC</td><td>>hg38_21_+_8433172_8446622_RNA45SN1_45S_50ntFlanks</td><td>8176</td><td>8196</td><td>21</td><td>CTTCTGATCGAGGCCCAGCCC</td><td>938.5</td><td>1.8</td><td>0.7</td><td>2.6</td><td>8.8E-03</td><td>1.0E-02</td><td>MB231/MB468</td><td>CYTO</td></tr>
<tr><td>>hg38_21_+_8433172_8446622_RNA45SN1_45S_50ntFlanks@8342.8379.38|CGTAAGGGAAAGTTGAAAAGAACTTTGAAGAGAGAGTT</td><td>>hg38_21_+_8433172_8446622_RNA45SN1_45S_50ntFlanks</td><td>8342</td><td>8379</td><td>38</td><td>CGTAAGGGAAAGTTGAAAAGAACTTTGAAGAGAGAGTT</td><td>121.8</td><td>3.4</td><td>1.3</td><td>2.7</td><td>8.0E-03</td><td>1.0E-02</td><td>MB231/MB468</td><td>MP</td></tr>
<tr><td>>hg38_21_+_8433172_8446622_RNA45SN1_45S_50ntFlanks@8186.8211.26|AGGCCCAGCCCGTGGACGGTGTGAGG</td><td>>hg38_21_+_8433172_8446622_RNA45SN1_45S_50ntFlanks</td><td>8186</td><td>8211</td><td>26</td><td>AGGCCCAGCCCGTGGACGGTGTGAGG</td><td>255.7</td><td>-3.8</td><td>1.4</td><td>-2.7</td><td>8.0E-03</td><td>1.0E-02</td><td>MB231/MB468</td><td>MP</td></tr>
<tr><td>>hg38_21_+_8433172_8446622_RNA45SN1_45S_50ntFlanks@7974.8006.33|ACGCGACCTCAGATCAGACGTGGCGACCCGCTG</td><td>>hg38_21_+_8433172_8446622_RNA45SN1_45S_50ntFlanks</td><td>7974</td><td>8006</td><td>33</td><td>ACGCGACCTCAGATCAGACGTGGCGACCCGCTG</td><td>16872.2</td><td>2.5</td><td>0.8</td><td>3.0</td><td>2.6E-03</td><td>1.0E-02</td><td>BT20/MB231</td><td>MITO</td></tr>
<tr><td>>am_trna128_GlyGCC_6_-_27870686_27870756@7.34.28,trna18_GlyGCC_16_+_70822597_70822667@7.34.28,trna19_GlyGCC_16_+_70823410_70823480@7.34.28,trna19_GlyGCC_2_-_157257659_157257729@7.34.28,trna24_GlyGCC_16_-_70812942_70813012@7.34.28,trna25_GlyGCC_16_-_70812114_70812184@7.34.28,trna2_GlyGCC_21_-_18827107_18827177@7.34.28,trna35_GlyGCC_1_+_161413094_161413164@7.34.28,trna37_GlyGCC_1_+_161420467_161420537@7.34.28,trna39_GlyGCC_1_+_161427898_161427968@7.34.28,trna41_GlyGCC_1_+_161435258_161435328@7.34.28,trna5_GlyGCC_17_+_8029064_8029134@7.34.28,trna68_GlyGCC_1_-_161493637_161493707@7.34.28|GTGGTTCAGTGGTAGAATTCTCGCCTGC</td><td>>am_trna128_GlyGCC_6_-_27870686_27870756</td><td>7</td><td>34</td><td>28</td><td>GTGGTTCAGTGGTAGAATTCTCGCCTGC</td><td>212.5</td><td>-3.1</td><td>1.0</td><td>-3.0</td><td>2.6E-03</td><td>1.0E-02</td><td>BT20/MB231</td><td>MITO</td></tr>
<tr><td>>am_trnaMT_MetCAT_MT_+_4402_4469@1.21.21|AGTAAGGTCAGCTAAATAAGC</td><td>>am_trnaMT_MetCAT_MT_+_4402_4469</td><td>1</td><td>21</td><td>21</td><td>AGTAAGGTCAGCTAAATAAGC</td><td>4121.8</td><td>-2.8</td><td>0.9</td><td>-3.0</td><td>2.6E-03</td><td>1.0E-02</td><td>BT20/MB231</td><td>MITO</td></tr>
<tr><td>>hg38_21_+_8433172_8446622_RNA45SN1_45S_50ntFlanks@3858.3898.41|TGGTAATTCTAGAGCTAATACATGCCGACGGGCGCTGACCC</td><td>>hg38_21_+_8433172_8446622_RNA45SN1_45S_50ntFlanks</td><td>3858</td><td>3898</td><td>41</td><td>TGGTAATTCTAGAGCTAATACATGCCGACGGGCGCTGACCC</td><td>170.3</td><td>-1.9</td><td>0.7</td><td>-2.6</td><td>8.8E-03</td><td>1.0E-02</td><td>MB231/MB468</td><td>CYTO</td></tr>
<tr><td>>hg38_21_+_8433172_8446622_RNA45SN1_45S_50ntFlanks@11027.11052.26|CTCTCTCTCTCTCTCCCCCGCTCCCC</td><td>>hg38_21_+_8433172_8446622_RNA45SN1_45S_50ntFlanks</td><td>11027</td><td>11052</td><td>26</td><td>CTCTCTCTCTCTCTCCCCCGCTCCCC</td><td>148.9</td><td>-2.7</td><td>1.0</td><td>-2.6</td><td>8.7E-03</td><td>1.0E-02</td><td>BT20/MB468</td><td>CYTO</td></tr>
<tr><td>>hg38_21_+_8433172_8446622_RNA45SN1_45S_50ntFlanks@10237.10258.22|ACGCTACGCCGCGACGAGTAGG</td><td>>hg38_21_+_8433172_8446622_RNA45SN1_45S_50ntFlanks</td><td>10237</td><td>10258</td><td>22</td><td>ACGCTACGCCGCGACGAGTAGG</td><td>174.6</td><td>2.9</td><td>1.0</td><td>2.8</td><td>5.3E-03</td><td>1.1E-02</td><td>BT20/MB468</td><td>MP</td></tr>
<tr><td>>hg38_21_+_8433172_8446622_RNA45SN1_45S_50ntFlanks@9532.9570.39|AGAGGAAACTCTGGTGGAGGTCCGTAGCGGTCCTGACGT</td><td>>hg38_21_+_8433172_8446622_RNA45SN1_45S_50ntFlanks</td><td>9532</td><td>9570</td><td>39</td><td>AGAGGAAACTCTGGTGGAGGTCCGTAGCGGTCCTGACGT</td><td>307.5</td><td>-2.4</td><td>0.8</td><td>-3.0</td><td>2.5E-03</td><td>1.1E-02</td><td>BT20/MB231</td><td>CYTO</td></tr>
<tr><td>>hg38_21_+_8433172_8446622_RNA45SN1_45S_50ntFlanks@10770.10805.36|GCAGGTCTCCAAGGTGAACAGCCTCTGGCATGTTGG</td><td>>hg38_21_+_8433172_8446622_RNA45SN1_45S_50ntFlanks</td><td>10770</td><td>10805</td><td>36</td><td>GCAGGTCTCCAAGGTGAACAGCCTCTGGCATGTTGG</td><td>156.2</td><td>-3.1</td><td>1.2</td><td>-2.7</td><td>7.5E-03</td><td>1.1E-02</td><td>BT20/MB468</td><td>MITO</td></tr>
<tr><td>>hg38_21_+_8433172_8446622_RNA45SN1_45S_50ntFlanks@12997.13036.40|CGCTGCGATCTATTGAAAGTCAGCCCTCGACACAAGGGTT</td><td>>hg38_21_+_8433172_8446622_RNA45SN1_45S_50ntFlanks</td><td>12997</td><td>13036</td><td>40</td><td>CGCTGCGATCTATTGAAAGTCAGCCCTCGACACAAGGGTT</td><td>366.6</td><td>2.0</td><td>0.8</td><td>2.6</td><td>9.1E-03</td><td>1.1E-02</td><td>BT20/MB468</td><td>NUC</td></tr>
<tr><td>>hg38_21_+_8433172_8446622_RNA45SN1_45S_50ntFlanks@11666.11692.27|TCTGCCCAGTGCTCTGAATGTCAAAGT</td><td>>hg38_21_+_8433172_8446622_RNA45SN1_45S_50ntFlanks</td><td>11666</td><td>11692</td><td>27</td><td>TCTGCCCAGTGCTCTGAATGTCAAAGT</td><td>304.3</td><td>3.9</td><td>1.2</td><td>3.1</td><td>2.0E-03</td><td>1.1E-02</td><td>BT20/MB231</td><td>MP</td></tr>
<tr><td>>hg38_21_+_8433172_8446622_RNA45SN1_45S_50ntFlanks@6732.6756.25|ACATTGATCATCGACACTTCGAACG</td><td>>hg38_21_+_8433172_8446622_RNA45SN1_45S_50ntFlanks</td><td>6732</td><td>6756</td><td>25</td><td>ACATTGATCATCGACACTTCGAACG</td><td>782.3</td><td>-2.2</td><td>0.8</td><td>-2.6</td><td>9.3E-03</td><td>1.1E-02</td><td>MB231/MB468</td><td>MITO</td></tr>
<tr><td>>hg38_21_+_8433172_8446622_RNA45SN1_45S_50ntFlanks@12029.12050.22|TCTGATCGTTTTTTCACTGACC</td><td>>hg38_21_+_8433172_8446622_RNA45SN1_45S_50ntFlanks</td><td>12029</td><td>12050</td><td>22</td><td>TCTGATCGTTTTTTCACTGACC</td><td>263.2</td><td>6.3</td><td>2.0</td><td>3.1</td><td>2.0E-03</td><td>1.1E-02</td><td>BT20/MB231</td><td>MP</td></tr>
<tr><td>>hg38_21_+_8433172_8446622_RNA45SN1_45S_50ntFlanks@12423.12456.34|TCGGCTCTTCCTATCATTGTGAAGCAGAATTCAC</td><td>>hg38_21_+_8433172_8446622_RNA45SN1_45S_50ntFlanks</td><td>12423</td><td>12456</td><td>34</td><td>TCGGCTCTTCCTATCATTGTGAAGCAGAATTCAC</td><td>346.4</td><td>-1.7</td><td>0.7</td><td>-2.6</td><td>9.3E-03</td><td>1.1E-02</td><td>MB231/MB468</td><td>NUC</td></tr>
<tr><td>>MI0000767|hsa-mir-365a&WithFlank&16|+|14309279|14309377@62.82.21;MI0000769|hsa-mir-365b&WithFlank&17|+|31575405|31575527@74.94.21&[MIMAT0000710&hsa-miR-365a-3p&offsets|0|-1;m-176&16|+|14309340|14309361&offsets|0|-1];[MIMAT0022834&hsa-miR-365b-3p&offsets|0|-1;m-175&17|+|31575478|31575499&offsets|0|-1]|TAATGCCCCTAAAAATCCTTA</td><td>>MI0000767|hsa-mir-365a&WithFlank&16|+|14309279|14309377</td><td>62</td><td>82</td><td>21</td><td>TAATGCCCCTAAAAATCCTTA</td><td>448.9</td><td>2.5</td><td>0.8</td><td>3.0</td><td>2.6E-03</td><td>1.1E-02</td><td>BT20/MB231</td><td>MITO</td></tr>
<tr><td>>hg38_21_+_8433172_8446622_RNA45SN1_45S_50ntFlanks@12073.12096.24|CCGAGGGGCTCTCGCTTCTGGCGC</td><td>>hg38_21_+_8433172_8446622_RNA45SN1_45S_50ntFlanks</td><td>12073</td><td>12096</td><td>24</td><td>CCGAGGGGCTCTCGCTTCTGGCGC</td><td>520.2</td><td>-4.9</td><td>1.7</td><td>-2.8</td><td>5.3E-03</td><td>1.1E-02</td><td>BT20/MB468</td><td>MP</td></tr>
<tr><td>>hg38_21_+_8433172_8446622_RNA45SN1_45S_50ntFlanks@3996.4026.31|ACTCTAGATAACCTCGGGCCGATCGCACGCC</td><td>>hg38_21_+_8433172_8446622_RNA45SN1_45S_50ntFlanks</td><td>3996</td><td>4026</td><td>31</td><td>ACTCTAGATAACCTCGGGCCGATCGCACGCC</td><td>68.5</td><td>-4.5</td><td>1.6</td><td>-2.8</td><td>5.3E-03</td><td>1.1E-02</td><td>BT20/MB468</td><td>MP</td></tr>
<tr><td>>hg38_21_+_8433172_8446622_RNA45SN1_45S_50ntFlanks@10841.10866.26|CGTAACTTCGGGATAAGGATTGGCTC</td><td>>hg38_21_+_8433172_8446622_RNA45SN1_45S_50ntFlanks</td><td>10841</td><td>10866</td><td>26</td><td>CGTAACTTCGGGATAAGGATTGGCTC</td><td>545.2</td><td>-2.5</td><td>0.9</td><td>-2.8</td><td>5.3E-03</td><td>1.1E-02</td><td>BT20/MB468</td><td>MP</td></tr>
<tr><td>>MI0000465|hsa-mir-191&WithFlank&3|-|49020612|49020715@23.43.21&[MIMAT0000440&hsa-miR-191-5p&offsets|+1|-1;m-48&3|-|49020672|49020694&offsets|+1|-1]|AACGGAATCCCAAAAGCAGCT</td><td>>MI0000465|hsa-mir-191&WithFlank&3|-|49020612|49020715</td><td>23</td><td>43</td><td>21</td><td>AACGGAATCCCAAAAGCAGCT</td><td>124.6</td><td>3.4</td><td>1.3</td><td>2.6</td><td>8.1E-03</td><td>1.1E-02</td><td>MB231/MB468</td><td>MP</td></tr>
<tr><td>>MI0006444|hsa-mir-1307&WithFlank&10|-|103394247|103394407@86.106.21(+1A)&[MIMAT0005951&hsa-miR-1307-3p&offsets|0|-1(+1A);m-66&10|-|103394300|103394322&offsets|0|-2(+1A)]|ACTCGGCGTGGCGTCGGTCGTA</td><td>>MI0006444|hsa-mir-1307&WithFlank&10|-|103394247|103394407</td><td>86</td><td>106</td><td>21(+1A)</td><td>ACTCGGCGTGGCGTCGGTCGTA</td><td>55.7</td><td>4.4</td><td>1.7</td><td>2.6</td><td>8.1E-03</td><td>1.1E-02</td><td>MB231/MB468</td><td>MP</td></tr>
<tr><td>>hg38_21_+_8433172_8446622_RNA45SN1_45S_50ntFlanks@7977.8005.29|CGACCTCAGATCAGACGTGGCGACCCGCT</td><td>>hg38_21_+_8433172_8446622_RNA45SN1_45S_50ntFlanks</td><td>7977</td><td>8005</td><td>29</td><td>CGACCTCAGATCAGACGTGGCGACCCGCT</td><td>83.0</td><td>-4.3</td><td>1.6</td><td>-2.6</td><td>8.1E-03</td><td>1.1E-02</td><td>MB231/MB468</td><td>MP</td></tr>
<tr><td>>hg38_MT_+_1621_3279_RNR2_16S_50ntFlanks@1182.1215.34|AAAATTTCGGTTGGGGCGACCTCGGAGCAGAACC</td><td>>hg38_MT_+_1621_3279_RNR2_16S_50ntFlanks</td><td>1182</td><td>1215</td><td>34</td><td>AAAATTTCGGTTGGGGCGACCTCGGAGCAGAACC</td><td>122.2</td><td>3.2</td><td>1.2</td><td>2.6</td><td>8.1E-03</td><td>1.1E-02</td><td>MB231/MB468</td><td>MP</td></tr>
<tr><td>>hg38_21_+_8433172_8446622_RNA45SN1_45S_50ntFlanks@8354.8375.22|TTGAAAAGAACTTTGAAGAGAG</td><td>>hg38_21_+_8433172_8446622_RNA45SN1_45S_50ntFlanks</td><td>8354</td><td>8375</td><td>22</td><td>TTGAAAAGAACTTTGAAGAGAG</td><td>2718.4</td><td>4.3</td><td>1.4</td><td>3.1</td><td>2.0E-03</td><td>1.1E-02</td><td>BT20/MB231</td><td>MP</td></tr>
<tr><td>>MI0000102|hsa-mir-100&WithFlank&11|-|122152223|122152314@19.40.22(+1U)&[MIMAT0000098&hsa-miR-100-5p&offsets|0|0(+1U);m-27&11|-|122152276|122152296&offsets|0|+1(+1U)]|AACCCGTAGATCCGAACTTGTGT</td><td>>MI0000102|hsa-mir-100&WithFlank&11|-|122152223|122152314</td><td>19</td><td>40</td><td>22(+1U)</td><td>AACCCGTAGATCCGAACTTGTGT</td><td>203.2</td><td>4.3</td><td>1.5</td><td>2.8</td><td>5.3E-03</td><td>1.1E-02</td><td>BT20/MB468</td><td>MP</td></tr>
<tr><td>>hg38_21_+_8433172_8446622_RNA45SN1_45S_50ntFlanks@10679.10708.30|GGTTCCGGCGGCGTCCGGTGAGCTCTCGCT</td><td>>hg38_21_+_8433172_8446622_RNA45SN1_45S_50ntFlanks</td><td>10679</td><td>10708</td><td>30</td><td>GGTTCCGGCGGCGTCCGGTGAGCTCTCGCT</td><td>3570.2</td><td>-3.7</td><td>1.4</td><td>-2.6</td><td>8.8E-03</td><td>1.1E-02</td><td>BT20/MB468</td><td>CYTO</td></tr>
<tr><td>>MI0000272|hsa-mir-182&WithFlank&7|-|129770377|129770498@29.51.23(+1A)&[MIMAT0000259&hsa-miR-182-5p&offsets|0|-1(+1A);m-22&7|-|129770449|129770470&offsets|0|+1(+1A)]|TTTGGCAATGGTAGAACTCACACA</td><td>>MI0000272|hsa-mir-182&WithFlank&7|-|129770377|129770498</td><td>29</td><td>51</td><td>23(+1A)</td><td>TTTGGCAATGGTAGAACTCACACA</td><td>583.0</td><td>3.2</td><td>1.2</td><td>2.8</td><td>5.3E-03</td><td>1.1E-02</td><td>BT20/MB468</td><td>MP</td></tr>
<tr><td>>trnaMT_SerGCT_MT_+_12207_12265@14.47.34|AGAACTGCTAACTCATGCCCCCATGTCTAACAAC</td><td>>trnaMT_SerGCT_MT_+_12207_12265</td><td>14</td><td>47</td><td>34</td><td>AGAACTGCTAACTCATGCCCCCATGTCTAACAAC</td><td>179.7</td><td>4.4</td><td>1.6</td><td>2.8</td><td>5.3E-03</td><td>1.1E-02</td><td>BT20/MB468</td><td>MP</td></tr>
<tr><td>>am_MI0003780|hsa-mir-1296&WithFlank&10|-|63372951|63373054@22.40.19&[MIMAT0005794&hsa-miR-1296-5p&offsets|0|-3;m-304&10|-|63373012|63373033&offsets|0|-3]|TTAGGGCCCTGGCTCCATC</td><td>>am_MI0003780|hsa-mir-1296&WithFlank&10|-|63372951|63373054</td><td>22</td><td>40</td><td>19</td><td>TTAGGGCCCTGGCTCCATC</td><td>57.1</td><td>5.8</td><td>2.1</td><td>2.8</td><td>5.3E-03</td><td>1.1E-02</td><td>BT20/MB468</td><td>MP</td></tr>
<tr><td>>hg38_21_+_8433172_8446622_RNA45SN1_45S_50ntFlanks@8159.8188.30|CGTGGGGGGCCCAAGTCCTTCTGATCGAGG</td><td>>hg38_21_+_8433172_8446622_RNA45SN1_45S_50ntFlanks</td><td>8159</td><td>8188</td><td>30</td><td>CGTGGGGGGCCCAAGTCCTTCTGATCGAGG</td><td>120.9</td><td>-3.5</td><td>1.3</td><td>-2.6</td><td>8.1E-03</td><td>1.1E-02</td><td>MB231/MB468</td><td>MP</td></tr>
<tr><td>>hg38_21_+_8433172_8446622_RNA45SN1_45S_50ntFlanks@10351.10385.35|CGAGAACTTTGAAGGCCGAAGTGGAGAAGGGTTCC</td><td>>hg38_21_+_8433172_8446622_RNA45SN1_45S_50ntFlanks</td><td>10351</td><td>10385</td><td>35</td><td>CGAGAACTTTGAAGGCCGAAGTGGAGAAGGGTTCC</td><td>406.3</td><td>-4.0</td><td>1.4</td><td>-2.8</td><td>5.3E-03</td><td>1.1E-02</td><td>BT20/MB468</td><td>MP</td></tr>
<tr><td>>hg38_21_+_8433172_8446622_RNA45SN1_45S_50ntFlanks@6648.6669.22|GTACGACTCTTAGCGGTGGATC</td><td>>hg38_21_+_8433172_8446622_RNA45SN1_45S_50ntFlanks</td><td>6648</td><td>6669</td><td>22</td><td>GTACGACTCTTAGCGGTGGATC</td><td>691.8</td><td>-2.6</td><td>1.0</td><td>-2.6</td><td>8.1E-03</td><td>1.1E-02</td><td>MB231/MB468</td><td>MP</td></tr>
<tr><td>>hg38_MT_+_1621_3279_RNR2_16S_50ntFlanks@1094.1120.27|AAGACGAGAAGACCCTATGGAGCTTTA</td><td>>hg38_MT_+_1621_3279_RNR2_16S_50ntFlanks</td><td>1094</td><td>1120</td><td>27</td><td>AAGACGAGAAGACCCTATGGAGCTTTA</td><td>153.6</td><td>2.2</td><td>0.8</td><td>2.6</td><td>8.1E-03</td><td>1.1E-02</td><td>MB231/MB468</td><td>MP</td></tr>
<tr><td>>hg38_21_+_8433172_8446622_RNA45SN1_45S_50ntFlanks@5475.5506.32|GCCCTGGCGGAGCGCTGAGAAGACGGTCGAAC</td><td>>hg38_21_+_8433172_8446622_RNA45SN1_45S_50ntFlanks</td><td>5475</td><td>5506</td><td>32</td><td>GCCCTGGCGGAGCGCTGAGAAGACGGTCGAAC</td><td>122.3</td><td>6.2</td><td>2.0</td><td>3.1</td><td>2.0E-03</td><td>1.1E-02</td><td>BT20/MB231</td><td>MP</td></tr>
<tr><td>>trna111_HisGTG_1_-_147774845_147774916@1.35.35,trna118_HisGTG_1_-_145396881_145396952@1.35.35,trna16_HisGTG_1_+_146544773_146544844@1.35.35,trna1_HisGTG_15_+_45493349_45493420@1.35.35,trna21_HisGTG_1_+_147753471_147753542@1.35.35,trna33_HisGTG_6_+_27125906_27125977@1.35.35,trna7_HisGTG_9_-_14433938_14434009@1.35.35,trna8_HisGTG_15_-_45492611_45492682@1.35.35,trna9_HisGTG_15_-_45490804_45490875@1.35.35|GCCGTGATCGTATAGTGGTTAGTACTCTGCGTTGT</td><td>>trna111_HisGTG_1_-_147774845_147774916</td><td>1</td><td>35</td><td>35</td><td>GCCGTGATCGTATAGTGGTTAGTACTCTGCGTTGT</td><td>1075.6</td><td>3.7</td><td>1.2</td><td>3.1</td><td>2.2E-03</td><td>1.1E-02</td><td>BT20/MB231</td><td>NUC</td></tr>
<tr><td>>hg38_21_+_8433172_8446622_RNA45SN1_45S_50ntFlanks@475.495.21|CTTCGTGATCGATGTGGTGAC</td><td>>hg38_21_+_8433172_8446622_RNA45SN1_45S_50ntFlanks</td><td>475</td><td>495</td><td>21</td><td>CTTCGTGATCGATGTGGTGAC</td><td>5715.7</td><td>-1.8</td><td>0.6</td><td>-3.1</td><td>2.2E-03</td><td>1.1E-02</td><td>BT20/MB231</td><td>NUC</td></tr>
<tr><td>>hg38_21_+_8433172_8446622_RNA45SN1_45S_50ntFlanks@11540.11586.47|CGCGGCGCCTCGCCTCGGCCGGCGCCTAGCAGCCGACTTAGAACTGG</td><td>>hg38_21_+_8433172_8446622_RNA45SN1_45S_50ntFlanks</td><td>11540</td><td>11586</td><td>47</td><td>CGCGGCGCCTCGCCTCGGCCGGCGCCTAGCAGCCGACTTAGAACTGG</td><td>238.7</td><td>-3.2</td><td>1.1</td><td>-3.1</td><td>2.2E-03</td><td>1.1E-02</td><td>BT20/MB231</td><td>NUC</td></tr>
<tr><td>>hg38_21_+_8433172_8446622_RNA45SN1_45S_50ntFlanks@11580.11611.32|GAACTGGTGCGGACCAGGGGAATCCGACTGTT</td><td>>hg38_21_+_8433172_8446622_RNA45SN1_45S_50ntFlanks</td><td>11580</td><td>11611</td><td>32</td><td>GAACTGGTGCGGACCAGGGGAATCCGACTGTT</td><td>1562.5</td><td>-1.7</td><td>0.7</td><td>-2.6</td><td>8.8E-03</td><td>1.1E-02</td><td>MB231/MB468</td><td>CYTO</td></tr>
<tr><td>>trnaMT_ValTAC_MT_+_1602_1670@50.72.23|ACTTAACTTGACCGCTCTGACCA</td><td>>trnaMT_ValTAC_MT_+_1602_1670</td><td>50</td><td>72</td><td>23</td><td>ACTTAACTTGACCGCTCTGACCA</td><td>2491.1</td><td>2.5</td><td>0.8</td><td>3.0</td><td>2.5E-03</td><td>1.1E-02</td><td>BT20/MB231</td><td>CYTO</td></tr>
<tr><td>>MI0000255|hsa-mir-30d&WithFlank&8|-|134804870|134804951@12.34.23&[MIMAT0000245&hsa-miR-30d-5p&offsets|0|+1;m-21&8|-|134804919|134804940&offsets|0|+1]|TGTAAACATCCCCGACTGGAAGC</td><td>>MI0000255|hsa-mir-30d&WithFlank&8|-|134804870|134804951</td><td>12</td><td>34</td><td>23</td><td>TGTAAACATCCCCGACTGGAAGC</td><td>39919.5</td><td>2.5</td><td>0.8</td><td>3.0</td><td>2.5E-03</td><td>1.1E-02</td><td>BT20/MB231</td><td>CYTO</td></tr>
<tr><td>>trnaMT_ValTAC_MT_+_1602_1670@50.70.21|ACTTAACTTGACCGCTCTGAC</td><td>>trnaMT_ValTAC_MT_+_1602_1670</td><td>50</td><td>70</td><td>21</td><td>ACTTAACTTGACCGCTCTGAC</td><td>397.2</td><td>2.3</td><td>0.8</td><td>3.0</td><td>2.5E-03</td><td>1.1E-02</td><td>BT20/MB231</td><td>CYTO</td></tr>
<tr><td>>hg38_21_+_8433172_8446622_RNA45SN1_45S_50ntFlanks@7978.8017.40|GACCTCAGATCAGACGTGGCGACCCGCTGAATTTAAGCAT</td><td>>hg38_21_+_8433172_8446622_RNA45SN1_45S_50ntFlanks</td><td>7978</td><td>8017</td><td>40</td><td>GACCTCAGATCAGACGTGGCGACCCGCTGAATTTAAGCAT</td><td>440.3</td><td>-5.2</td><td>1.7</td><td>-3.0</td><td>2.5E-03</td><td>1.1E-02</td><td>BT20/MB231</td><td>CYTO</td></tr>
<tr><td>>hg38_21_+_8433172_8446622_RNA45SN1_45S_50ntFlanks@10061.10078.18|CAGTCGAGAGTGGACGGG</td><td>>hg38_21_+_8433172_8446622_RNA45SN1_45S_50ntFlanks</td><td>10061</td><td>10078</td><td>18</td><td>CAGTCGAGAGTGGACGGG</td><td>200.3</td><td>5.1</td><td>1.6</td><td>3.1</td><td>2.0E-03</td><td>1.1E-02</td><td>BT20/MB231</td><td>MP</td></tr>
</table>

| | | | | | | | | | | | | | |
|---|---|---|---|---|---|---|---|---|---|---|---|---|---|
| >MI0003137\|hsa-mir-193b&WithFlank&16\|+\|14303961\|14304055@57.78.22&[MIMAT0002819&hsa-miR-193b-3p&offsets\|0\|0;m-166&16\|+\|14304017\|14304038&offsets\|0\|0]\|AACTGGCCCTCAAAGTCCCGCT | >MI0003137\|hsa-mir-193b&WithFlank&16\|+\|14303961\|14304055 | 57 | 78 | 22 | AACTGGCCCTCAAAGTCCCGCT | 2251.7 | 2.5 | 0.8 | 3.0 | 2.6E-03 | 1.1E-02 | BT20/MB231 | MITO |
| >hg38_MT_+_598_1651_RNR1_12S_50ntFlanks@439.468.30\|AACATATCTGAACACACAATAGCTAAGACC | >hg38_MT_+_598_1651_RNR1_12S_50ntFlanks | 439 | 468 | 30 | AACATATCTGAACACACAATAGCTAAGACC | 139.7 | 3.4 | 1.3 | 2.6 | 8.1E-03 | 1.1E-02 | MB231/MB468 | MP |
| >hg38_21_+_8433172_8446622_RNA45SN1_45S_50ntFlanks@10403.10424.22\|ACATGGGTCAGTCGGTCCTGAG | >hg38_21_+_8433172_8446622_RNA45SN1_45S_50ntFlanks | 10403 | 10424 | 22 | ACATGGGTCAGTCGGTCCTGAG | 10576.4 | 3.3 | 1.1 | 3.1 | 2.0E-03 | 1.1E-02 | BT20/MB231 | MP |
| >hg38_21_+_8433172_8446622_RNA45SN1_45S_50ntFlanks@10409.10432.24\|GTCAGTCGGTCCTGAGAGATGGGC | >hg38_21_+_8433172_8446622_RNA45SN1_45S_50ntFlanks | 10409 | 10432 | 24 | GTCAGTCGGTCCTGAGAGATGGGC | 747.6 | 3.6 | 1.3 | 2.8 | 5.3E-03 | 1.1E-02 | BT20/MB468 | MP |
| >trnaMT_ValTAC_MT_+_1602_1670@45.71.27\|TTTCAACTTAACTTGACCGCTCTGACC | >trnaMT_ValTAC_MT_+_1602_1670 | 45 | 71 | 27 | TTTCAACTTAACTTGACCGCTCTGACC | 360.0 | -3.1 | 1.0 | -3.0 | 2.5E-03 | 1.1E-02 | BT20/MB231 | CYTO |
| >am_trna10_ValCAC_5_-_180649395_180649467@1.23.23,trna12_ValAAC_5_-_180645270_180645342@1.23.23,trna132_ValAAC_6_-_27721179_27721251@1.23.23,trna136_ValAAC_6_-_27648885_27648957@1.23.23,trna139_ValAAC_6_-_27618707_27618779@1.23.23,trna18_ValCAC_5_-_180529253_180529325@1.23.23,trna2_ValAAC_3_+_169490018_169490090@1.23.23,trna2_ValCAC_5_+_180524070_180524142@1.23.23,trna37_ValAAC_6_+_27203288_27203360@1.23.23,trna4_ValAAC_5_+_180591154_180591226@1.23.23,trna5_ValAAC_5_+_180596610_180596682@1.23.23,trna6_ValCAC_5_+_180600650_180600722@1.23.23,trna85_ValCAC_1_-_161369490_161369562@1.23.23,trna90_ValCAC_1_-_149684088_149684161@1.23.23,trna98_ValCAC_1_-_149298555_149298627@1.23.23,trna9_ValCAC_6_+_26538282_26538354@1.23.23\|GTTTCCGTAGTGTAGTGGTTATC | >am_trna10_ValCAC_5_-_180649395_180649467 | 1 | 23 | 23 | GTTTCCGTAGTGTAGTGGTTATC | 3932.9 | 3.5 | 1.1 | 3.1 | 2.0E-03 | 1.1E-02 | BT20/MB231 | MP |
| >trnaMT_PheGAA_MT_+_577_647@1.18.18\|GTTTATGTAGCTTACCTC | >trnaMT_PheGAA_MT_+_577_647 | 1 | 18 | 18 | GTTTATGTAGCTTACCTC | 363.2 | -2.7 | 0.9 | -3.1 | 2.2E-03 | 1.1E-02 | BT20/MB231 | NUC |
| >hg38_21_+_8433172_8446622_RNA45SN1_45S_50ntFlanks@12996.13040.45\|TCGCTGCGATCTATTGAAAGTCAGCCCTCGACACAAGGGTTTGTC | >hg38_21_+_8433172_8446622_RNA45SN1_45S_50ntFlanks | 12996 | 13040 | 45 | TCGCTGCGATCTATTGAAAGTCAGCCCTCGACACAAGGGTTTGTC | 636.6 | -2.6 | 0.9 | -3.0 | 2.6E-03 | 1.1E-02 | BT20/MB231 | MITO |
| >hg38_MT_+_1621_3279_RNR2_16S_50ntFlanks@617.634.18\|AAACATATAACTGAACTC | >hg38_MT_+_1621_3279_RNR2_16S_50ntFlanks | 617 | 634 | 18 | AAACATATAACTGAACTC | 48.9 | -9.0 | 2.9 | -3.1 | 2.0E-03 | 1.1E-02 | BT20/MB231 | MP |
| >hg38_21_+_8433172_8446622_RNA45SN1_45S_50ntFlanks@10014.10038.25\|AAAATGGATGGCGCTGGAGCGTCGG | >hg38_21_+_8433172_8446622_RNA45SN1_45S_50ntFlanks | 10014 | 10038 | 25 | AAAATGGATGGCGCTGGAGCGTCGG | 52.1 | -8.8 | 3.2 | -2.8 | 5.3E-03 | 1.1E-02 | BT20/MB468 | MP |
| >hg38_21_+_8433172_8446622_RNA45SN1_45S_50ntFlanks@10498.10521.24\|TTCAGATCCCCGAATCCGGAGTGG | >hg38_21_+_8433172_8446622_RNA45SN1_45S_50ntFlanks | 10498 | 10521 | 24 | TTCAGATCCCCGAATCCGGAGTGG | 176.8 | -4.8 | 1.7 | -2.8 | 5.3E-03 | 1.1E-02 | BT20/MB468 | MP |
| >hg38_21_+_8433172_8446622_RNA45SN1_45S_50ntFlanks@8189.8216.28\|CCCAGCCCGTGGACGGTGTGAGGCCGGT | >hg38_21_+_8433172_8446622_RNA45SN1_45S_50ntFlanks | 8189 | 8216 | 28 | CCCAGCCCGTGGACGGTGTGAGGCCGGT | 78.1 | -4.3 | 1.6 | -2.6 | 8.1E-03 | 1.1E-02 | MB231/MB468 | MP |
| >trnaMT_HisGTG_MT_+_12138_12206@28.61.34\|ATTGTGAATCTGACAACAGAGGCTTACGACCCCT | >trnaMT_HisGTG_MT_+_12138_12206 | 28 | 61 | 34 | ATTGTGAATCTGACAACAGAGGCTTACGACCCCT | 123.1 | -3.1 | 1.2 | -2.6 | 8.9E-03 | 1.1E-02 | MB231/MB468 | CYTO |
| >hg38_21_+_8433172_8446622_RNA45SN1_45S_50ntFlanks@9481.9513.33\|CACGTGTTAGGACCCGAAAGATGGTGAACTATG | >hg38_21_+_8433172_8446622_RNA45SN1_45S_50ntFlanks | 9481 | 9513 | 33 | CACGTGTTAGGACCCGAAAGATGGTGAACTATG | 195.8 | 6.1 | 2.0 | 3.1 | 2.0E-03 | 1.1E-02 | BT20/MB231 | MP |
| >hg38_MT_+_1621_3279_RNR2_16S_50ntFlanks@1099.1117.19\|GAGAAGACCCTATGGAGCT | >hg38_MT_+_1621_3279_RNR2_16S_50ntFlanks | 1099 | 1117 | 19 | GAGAAGACCCTATGGAGCT | 89.1 | 3.0 | 1.1 | 2.8 | 5.4E-03 | 1.1E-02 | BT20/MB468 | MP |
| >hg38_21_+_8433172_8446622_RNA45SN1_45S_50ntFlanks@10686.10707.22\|GCGGCGTCCGGTGAGCTCTCGC | >hg38_21_+_8433172_8446622_RNA45SN1_45S_50ntFlanks | 10686 | 10707 | 22 | GCGGCGTCCGGTGAGCTCTCGC | 938.1 | 2.9 | 1.0 | 3.0 | 2.6E-03 | 1.1E-02 | BT20/MB231 | MITO |
| >hg38_21_+_8433172_8446622_RNA45SN1_45S_50ntFlanks@6641.6681.41\|CGACCTCGTACGACTCTTAGCGGTGGATCACTCGGCTCGTG | >hg38_21_+_8433172_8446622_RNA45SN1_45S_50ntFlanks | 6641 | 6681 | 41 | CGACCTCGTACGACTCTTAGCGGTGGATCACTCGGCTCGTG | 150.8 | -3.8 | 1.2 | -3.1 | 2.2E-03 | 1.1E-02 | BT20/MB231 | NUC |
| >hg38_21_+_8433172_8446622_RNA45SN1_45S_50ntFlanks@10420.10437.18\|CTGAGAGATGGGCGAGCG | >hg38_21_+_8433172_8446622_RNA45SN1_45S_50ntFlanks | 10420 | 10437 | 18 | CTGAGAGATGGGCGAGCG | 102.7 | -4.6 | 1.7 | -2.8 | 5.4E-03 | 1.1E-02 | BT20/MB468 | MP |
| >hg38_21_+_8433172_8446622_RNA45SN1_45S_50ntFlanks@4006.4026.21\|ACCTCGGGCCGATCGCACGCC | >hg38_21_+_8433172_8446622_RNA45SN1_45S_50ntFlanks | 4006 | 4026 | 21 | ACCTCGGGCCGATCGCACGCC | 155.5 | -4.6 | 1.6 | -2.8 | 5.4E-03 | 1.1E-02 | BT20/MB468 | MP |
| >hg38_21_+_8433172_8446622_RNA45SN1_45S_50ntFlanks@7986.8020.35\|ATCAGACGTGGCGACCCGCTGAATTTAAGCATATT | >hg38_21_+_8433172_8446622_RNA45SN1_45S_50ntFlanks | 7986 | 8020 | 35 | ATCAGACGTGGCGACCCGCTGAATTTAAGCATATT | 209.2 | -4.5 | 1.7 | -2.6 | 9.4E-03 | 1.1E-02 | MB231/MB468 | MITO |
| >hg38_21_+_8433172_8446622_RNA45SN1_45S_50ntFlanks@11596.11621.26\|GGGGAATCCGACTGTTTAATTAAAAC | >hg38_21_+_8433172_8446622_RNA45SN1_45S_50ntFlanks | 11596 | 11621 | 26 | GGGGAATCCGACTGTTTAATTAAAAC | 61.2 | 5.6 | 2.0 | 2.8 | 5.4E-03 | 1.1E-02 | BT20/MB468 | MP |
| >hg38_21_+_8433172_8446622_RNA45SN1_45S_50ntFlanks@10402.10425.24\|AACATGGGTCAGTCGGTCCTGAGA | >hg38_21_+_8433172_8446622_RNA45SN1_45S_50ntFlanks | 10402 | 10425 | 24 | AACATGGGTCAGTCGGTCCTGAGA | 167.9 | -3.5 | 1.3 | -2.8 | 5.4E-03 | 1.1E-02 | BT20/MB468 | MP |
| >hg38_21_+_8433172_8446622_RNA45SN1_45S_50ntFlanks@7974.7992.19\|ACGCGACCTCAGATCAGAC | >hg38_21_+_8433172_8446622_RNA45SN1_45S_50ntFlanks | 7974 | 7992 | 19 | ACGCGACCTCAGATCAGAC | 6054.7 | 4.1 | 1.3 | 3.1 | 2.0E-03 | 1.1E-02 | BT20/MB231 | MP |
| >MI0000107\|hsa-mir-29b-2&WithFlank&1\|-\|207802437\|207802529@58.80.23;MI0000105\|hsa-mir-29b-1&WithFlank&7\|-\|130877453\|130877545@57.79.23&[MIMAT0000100&hsa-miR-29b-3p&offsets\|0\|0;m-81&1\|-\|207802450\|207802472&offsets\|0\|0];[MIMAT0000100_1&hsa-miR-29b-3p&offsets\|0\|0;m-82&7\|-\|130877467\|130877489&offsets\|0\|0]\|TAGCACCATTTGAAATCAGTGTT | >MI0000107\|hsa-mir-29b-2&WithFlank&1\|-\|207802437\|207802529 | 58 | 80 | 23 | TAGCACCATTTGAAATCAGTGTT | 745.1 | -2.7 | 0.9 | -3.0 | 2.6E-03 | 1.1E-02 | BT20/MB231 | MITO |
| >trnaMT_LysTTT_MT_+_8295_8364@37.73.37,trnalookalike7_LysTTT_1_+_568844_568913@37.73.37\|AAAGATTAAGAGAACCAACACCTCTTTACAGTGACCA | >trnaMT_LysTTT_MT_+_8295_8364 | 37 | 73 | 37 | AAAGATTAAGAGAACCAACACCTCTTTACAGTGACCA | 359.6 | 3.6 | 1.4 | 2.6 | 8.2E-03 | 1.1E-02 | MB231/MB468 | MP |
| >trna10_AspGTC_12_-_125424193_125424264@39.69.31,trna12_AspGTC_12_-_125411891_125411962@39.69.31,trna144_AspGTC_6_-_27551236_27551307@39.69.31,trna38_AspGTC_17_-_8125556_8125627@39.69.31,trna45_AspGTC_6_+_27447453_27447524@39.69.31,trna48_AspGTC_6_+_27471523_27471594@39.69.31,trna4_AspGTC_12_+_96429799_96429870@39.69.31,trna69_AspGTC_1_-_161440205_161440276@39.69.31,trna72_AspGTC_1_-_161432824_161432895@39.69.31,trna75_AspGTC_1_-_161425414_161425485@39.69.31,trna78_AspGTC_1_-_161418033_161418104@39.69.31,trna81_AspGTC_1_-_161410615_161410686@39.69.31\|GCGGGAGACCGGGGTTCGATTCCCCGACGGG | >trna10_AspGTC_12_-_125424193_125424264 | 39 | 69 | 31 | GCGGGAGACCGGGGTTCGATTCCCCGACGGG | 177.2 | -2.1 | 0.8 | -2.6 | 8.2E-03 | 1.1E-02 | MB231/MB468 | MP |
| >hg38_21_+_8433172_8446622_RNA45SN1_45S_50ntFlanks@10349.10383.35\|AACGAGAACTTTGAAGGCCGAAGTGGAGAAGGGTT | >hg38_21_+_8433172_8446622_RNA45SN1_45S_50ntFlanks | 10349 | 10383 | 35 | AACGAGAACTTTGAAGGCCGAAGTGGAGAAGGGTT | 5144.3 | -2.8 | 1.0 | -2.8 | 5.4E-03 | 1.1E-02 | BT20/MB468 | MP |
| >MI0000289\|hsa-mir-181a-1&WithFlank&1\|-\|198859038\|198859159@30.48.19(+1U);MI0000269\|hsa-mir-181a-2&WithFlank&9\|+\|124692436\|124692557@45.63.19(+1U)&[MIMAT0000256&hsa-miR-181a-5p&offsets\|0\|-4(+1U);m-32&1\|-\|198859108\|198859130&offsets\|0\|-4(+1U)];[MIMAT0000256_1&hsa-miR-181a-5p&offsets\|0\|-4(+1U);m-32&9\|+\|124692480\|124692502&offsets\|0\|-4(+1U)]\|AACATTCAACGCTGTCGGTT | >MI0000289\|hsa-mir-181a-1&WithFlank&1\|-\|198859038\|198859159 | 30 | 48 | 19(+1U) | AACATTCAACGCTGTCGGTT | 37.1 | 5.3 | 2.0 | 2.6 | 8.2E-03 | 1.1E-02 | MB231/MB468 | MP |
| >hg38_21_+_8433172_8446622_RNA45SN1_45S_50ntFlanks@6652.6675.24\|GACTCTTAGCGGTGGATCACTCGG | >hg38_21_+_8433172_8446622_RNA45SN1_45S_50ntFlanks | 6652 | 6675 | 24 | GACTCTTAGCGGTGGATCACTCGG | 666.3 | 2.7 | 1.0 | 2.6 | 9.4E-03 | 1.1E-02 | MB231/MB468 | NUC |
| >hg38_21_+_8433172_8446622_RNA45SN1_45S_50ntFlanks@12636.12661.26\|AGCTACCATCTGTGGGATTATGACTG | >hg38_21_+_8433172_8446622_RNA45SN1_45S_50ntFlanks | 12636 | 12661 | 26 | AGCTACCATCTGTGGGATTATGACTG | 244.9 | -2.8 | 1.1 | -2.6 | 8.9E-03 | 1.1E-02 | BT20/MB468 | CYTO |
| >MI0000088\|hsa-mir-30a&WithFlank&6\|-\|71403545\|71403627@12.34.23(+1C)&[MIMAT0000087&hsa-miR-30a-5p&offsets\|0\|+1(+1C);m-15&6\|-\|71403595\|71403616&offsets\|0\|+1(+1C)]\|TGTAAACATCCTCGACTGGAAGCC | >MI0000088\|hsa-mir-30a&WithFlank&6\|-\|71403545\|71403627 | 12 | 34 | 23(+1C) | TGTAAACATCCTCGACTGGAAGCC | 75910.1 | -3.2 | 1.0 | -3.1 | 2.0E-03 | 1.1E-02 | BT20/MB231 | MP |
| >hg38_21_+_8433172_8446622_RNA45SN1_45S_50ntFlanks@10266.10284.19\|CTGCGGTGAGCCTTGAAGC | >hg38_21_+_8433172_8446622_RNA45SN1_45S_50ntFlanks | 10266 | 10284 | 19 | CTGCGGTGAGCCTTGAAGC | 147.7 | 5.0 | 1.6 | 3.1 | 2.0E-03 | 1.1E-02 | BT20/MB231 | MP |
| >MI0000816\|hsa-mir-335&WithFlank&7\|+\|130496105\|130496210@58.79.22&[MIMAT0004703&hsa-miR-335-3p&offsets\|0\|0;m-159&7\|+\|130496162\|130496183&offsets\|0\|0]\|TTTTTCATTATTGCTCCTGACC | >MI0000816\|hsa-mir-335&WithFlank&7\|+\|130496105\|130496210 | 58 | 79 | 22 | TTTTTCATTATTGCTCCTGACC | 372.9 | 3.2 | 1.1 | 2.8 | 5.4E-03 | 1.1E-02 | BT20/MB468 | MP |
| >hg38_21_+_8433172_8446622_RNA45SN1_45S_50ntFlanks@8349.8378.30\|GAAAGTTGAAAAGAACTTTGAAGAGAGAGT | >hg38_21_+_8433172_8446622_RNA45SN1_45S_50ntFlanks | 8349 | 8378 | 30 | GAAAGTTGAAAAGAACTTTGAAGAGAGAGT | 170.1 | 4.7 | 1.5 | 3.1 | 2.0E-03 | 1.1E-02 | BT20/MB231 | MP |
| >MI0000253\|hsa-mir-148a&WithFlank&7\|-\|25949913\|25949992@50.70.21(+1C)&[MIMAT0000243&hsa-miR-148a-3p&offsets\|0\|-1(+1C);m-7&7\|-\|25949922\|25949943&offsets\|0\|-1(+1C)]\|TCAGTGCACTACAGAACTTTGC | >MI0000253\|hsa-mir-148a&WithFlank&7\|-\|25949913\|25949992 | 50 | 70 | 21(+1C) | TCAGTGCACTACAGAACTTTGC | 55514.6 | 3.6 | 1.2 | 3.1 | 2.0E-03 | 1.1E-02 | BT20/MB231 | MP |
| >MI0033423\|hsa-mir-10399&WithFlank&7\|-\|139044094\|139044163@7.27.21&[MIMAT0041629&hsa-miR-10399-5p&offsets\|0\|0;TJU_CMC_MD2.ID03061.5p-miR&offsets\|0\|0;m-2644&7\|-\|139044137\|139044157&offsets\|0\|0]\|AATTACAGATTGTCTCAGAGA | >MI0033423\|hsa-mir-10399&WithFlank&7\|-\|139044094\|139044163 | 7 | 27 | 21 | AATTACAGATTGTCTCAGAGA | 158.3 | -5.6 | 1.8 | -3.1 | 2.0E-03 | 1.1E-02 | BT20/MB231 | MP |
| >MI0000067\|hsa-let-7f-1&WithFlank&9\|+\|94176341\|94176439@13.32.20;MI0000068\|hsa-let-7f-2&WithFlank&X\|-\|53557186\|53557280@14.33.20&[MIMAT0000067&hsa-let-7f-5p&offsets\|0\|-2;m-20&9\|+\|94176353\|94176374&offsets\|0\|-2];[MIMAT0000067_1&hsa-let-7f-5p&offsets\|0\|-2;m-19&X\|-\|53557246\|53557267&offsets\|0\|-2]\|TGAGGTAGTAGATTGTATAG | >MI0000067\|hsa-let-7f-1&WithFlank&9\|+\|94176341\|94176439 | 13 | 32 | 20 | TGAGGTAGTAGATTGTATAG | 229.9 | 4.5 | 1.7 | 2.6 | 8.2E-03 | 1.1E-02 | MB231/MB468 | MP |
| >MI0000440\|hsa-mir-27b&WithFlank&9\|+\|95085439\|95085547@67.88.22(+1U)&[MIMAT0000419&hsa-miR-27b-3p&offsets\|0\|+1(+1U);m-39&9\|+\|95085505\|95085525&offsets\|0\|+1(+1U)]\|TTCACAGTGGCTAAGTTCTGCAT | >MI0000440\|hsa-mir-27b&WithFlank&9\|+\|95085439\|95085547 | 67 | 88 | 22(+1U) | TTCACAGTGGCTAAGTTCTGCAT | 62.2 | 3.9 | 1.4 | 2.8 | 5.4E-03 | 1.1E-02 | BT20/MB468 | MP |
| >trna119_LysCTT_1_-_145395522_145395594@4.35.32,trna11_LysCTT_5_-_180648979_180649051@4.35.32,trna13_LysCTT_6_+_26556774_26556846@4.35.32,trna7_LysCTT_16_+_3225692_3225764@4.35.32,trna9_LysCTT_5_+_180634755_180634827@4.35.32\|CGGCTAGCTCAGTCGGTAGAGCATGAGACTCT | >trna119_LysCTT_1_-_145395522_145395594 | 4 | 35 | 32 | CGGCTAGCTCAGTCGGTAGAGCATGAGACTCT | 148.0 | 3.3 | 1.2 | 2.8 | 5.4E-03 | 1.1E-02 | BT20/MB468 | MP |

<table>
<tr><td>>trnaMT_GlnTTG_MT_-_4329_4400@1.32.32,trnalookalike2_GlnTTG_1_-_564879_564950@1.32.32|TAGGATGGGGTGTGATAGGTGGCACGGAGAAT</td><td>>trnaMT_GlnTTG_MT_-_4329_4400</td><td>1</td><td>32</td><td>32</td><td>TAGGATGGGGTGTGATAGGTGGCACGGAGAAT</td><td>692.3</td><td>3.9</td><td>1.4</td><td>2.8</td><td>5.4E-03</td><td>1.1E-02</td><td>BT20/MB468</td><td>MP</td></tr>
<tr><td>>hg38_21_+_8433172_8446622_RNA45SN1_45S_50ntFlanks@7969.7986.18|CCGAGACGCGACCTCAGA</td><td>>hg38_21_+_8433172_8446622_RNA45SN1_45S_50ntFlanks</td><td>7969</td><td>7986</td><td>18</td><td>CCGAGACGCGACCTCAGA</td><td>64.7</td><td>1.9</td><td>0.7</td><td>2.6</td><td>9.3E-03</td><td>1.1E-02</td><td>BT20/MB468</td><td>NUC</td></tr>
<tr><td>>MI0000746|hsa-mir-99b&WithFlank&19|+|51692606|51692687@14.34.21&[MIMAT0000689&hsa-miR-99b-5p&offsets|+1|0;m-10&19|+|51692618|51692639&offsets|+1|0]|ACCCGTAGAACCGACCTTGCG</td><td>>MI0000746|hsa-mir-99b&WithFlank&19|+|51692606|51692687</td><td>14</td><td>34</td><td>21</td><td>ACCCGTAGAACCGACCTTGCG</td><td>72.0</td><td>3.8</td><td>1.4</td><td>2.8</td><td>5.4E-03</td><td>1.1E-02</td><td>BT20/MB468</td><td>MP</td></tr>
<tr><td>>hg38_21_+_8433172_8446622_RNA45SN1_45S_50ntFlanks@6790.6807.18|CCTGTCTGAGCGTCGCTT</td><td>>hg38_21_+_8433172_8446622_RNA45SN1_45S_50ntFlanks</td><td>6790</td><td>6807</td><td>18</td><td>CCTGTCTGAGCGTCGCTT</td><td>2884.6</td><td>4.9</td><td>1.6</td><td>3.1</td><td>2.0E-03</td><td>1.1E-02</td><td>BT20/MB231</td><td>MP</td></tr>
<tr><td>>hg38_21_+_8433172_8446622_RNA45SN1_45S_50ntFlanks@12141.12166.26|ACAGTGCCAGGTGGGGAGTTTGACTG</td><td>>hg38_21_+_8433172_8446622_RNA45SN1_45S_50ntFlanks</td><td>12141</td><td>12166</td><td>26</td><td>ACAGTGCCAGGTGGGGAGTTTGACTG</td><td>58.3</td><td>-4.6</td><td>1.7</td><td>-2.8</td><td>5.4E-03</td><td>1.1E-02</td><td>BT20/MB468</td><td>MP</td></tr>
<tr><td>>MI0000087|hsa-mir-29a&WithFlank&7|-|130876741|130876816@47.67.21(+1G)&[MIMAT0000086&hsa-miR-29a-3p&offsets|-1|-2(+1G);m-26&7|-|130876749|130876769&offsets|-1|-1(+1G)]|CTAGCACCATCTGAAATCGGTG</td><td>>MI0000087|hsa-mir-29a&WithFlank&7|-|130876741|130876816</td><td>47</td><td>67</td><td>21(+1G)</td><td>CTAGCACCATCTGAAATCGGTG</td><td>182.6</td><td>5.2</td><td>1.9</td><td>2.8</td><td>5.4E-03</td><td>1.1E-02</td><td>BT20/MB468</td><td>MP</td></tr>
<tr><td>>hg38_21_+_8433172_8446622_RNA45SN1_45S_50ntFlanks@11933.11968.36|AATAAGTGGGAGGCCCCCGGCGCCCCCCCGGTGTCC</td><td>>hg38_21_+_8433172_8446622_RNA45SN1_45S_50ntFlanks</td><td>11933</td><td>11968</td><td>36</td><td>AATAAGTGGGAGGCCCCCGGCGCCCCCCCGGTGTCC</td><td>115.4</td><td>-4.4</td><td>1.7</td><td>-2.7</td><td>7.6E-03</td><td>1.1E-02</td><td>BT20/MB468</td><td>MITO</td></tr>
<tr><td>>hg38_21_+_8433172_8446622_RNA45SN1_45S_50ntFlanks@6653.6681.29|ACTCTTAGCGGTGGATCACTCGGCTCGTG</td><td>>hg38_21_+_8433172_8446622_RNA45SN1_45S_50ntFlanks</td><td>6653</td><td>6681</td><td>29</td><td>ACTCTTAGCGGTGGATCACTCGGCTCGTG</td><td>3169.0</td><td>-1.3</td><td>0.5</td><td>-2.7</td><td>7.6E-03</td><td>1.1E-02</td><td>BT20/MB468</td><td>MITO</td></tr>
<tr><td>>hg38_21_+_8433172_8446622_RNA45SN1_45S_50ntFlanks@9622.9658.37|CTAGTAGCTGGTTCCCTCCGAAGTTTCCCTCAGGATA</td><td>>hg38_21_+_8433172_8446622_RNA45SN1_45S_50ntFlanks</td><td>9622</td><td>9658</td><td>37</td><td>CTAGTAGCTGGTTCCCTCCGAAGTTTCCCTCAGGATA</td><td>196.2</td><td>-4.7</td><td>1.7</td><td>-2.8</td><td>5.4E-03</td><td>1.1E-02</td><td>BT20/MB468</td><td>MP</td></tr>
<tr><td>>am_MI0000786|hsa-mir-378a&WithFlank&5|+|149732819|149732896@50.70.21(+1U)&[MIMAT0000732&hsa-miR-378a-3p&offsets|+1|0(+1U);m-44&5|+|149732867|149732888&offsets|+1|0(+1U)]|CTGGACTTGGAGTCAGAAGGCT</td><td>>am_MI0000786|hsa-mir-378a&WithFlank&5|+|149732819|149732896</td><td>50</td><td>70</td><td>21(+1U)</td><td>CTGGACTTGGAGTCAGAAGGCT</td><td>320.3</td><td>-1.0</td><td>0.4</td><td>-2.6</td><td>9.3E-03</td><td>1.1E-02</td><td>BT20/MB468</td><td>NUC</td></tr>
<tr><td>>hg38_21_+_8433172_8446622_RNA45SN1_45S_50ntFlanks@10807.10829.23|ACAATGTAGGTAAGGGAAGTCGG</td><td>>hg38_21_+_8433172_8446622_RNA45SN1_45S_50ntFlanks</td><td>10807</td><td>10829</td><td>23</td><td>ACAATGTAGGTAAGGGAAGTCGG</td><td>116.5</td><td>-3.7</td><td>1.4</td><td>-2.6</td><td>8.9E-03</td><td>1.1E-02</td><td>BT20/MB468</td><td>CYTO</td></tr>
<tr><td>>hg38_21_+_8433172_8446622_RNA45SN1_45S_50ntFlanks@10344.10369.26|ATTCAAACGAGAACTTTGAAGGCCGA</td><td>>hg38_21_+_8433172_8446622_RNA45SN1_45S_50ntFlanks</td><td>10344</td><td>10369</td><td>26</td><td>ATTCAAACGAGAACTTTGAAGGCCGA</td><td>331.4</td><td>3.8</td><td>1.2</td><td>3.1</td><td>2.1E-03</td><td>1.1E-02</td><td>BT20/MB231</td><td>MP</td></tr>
<tr><td>>hg38_21_+_8433172_8446622_RNA45SN1_45S_50ntFlanks@9198.9222.25|GTTCTCTCGGGGCCACGCGCGCGTC</td><td>>hg38_21_+_8433172_8446622_RNA45SN1_45S_50ntFlanks</td><td>9198</td><td>9222</td><td>25</td><td>GTTCTCTCGGGGCCACGCGCGCGTC</td><td>2541.6</td><td>-3.9</td><td>1.5</td><td>-2.7</td><td>7.7E-03</td><td>1.1E-02</td><td>BT20/MB468</td><td>MITO</td></tr>
<tr><td>>hg38_MT_+_1621_3279_RNR2_16S_50ntFlanks@1369.1395.27|GATGTTGGATCAGGACATCCCGATGGT</td><td>>hg38_MT_+_1621_3279_RNR2_16S_50ntFlanks</td><td>1369</td><td>1395</td><td>27</td><td>GATGTTGGATCAGGACATCCCGATGGT</td><td>166.4</td><td>3.9</td><td>1.4</td><td>2.8</td><td>5.4E-03</td><td>1.1E-02</td><td>BT20/MB468</td><td>MP</td></tr>
<tr><td>>MI0000272|hsa-mir-182&WithFlank&7|-|129770377|129770498@29.48.20&[MIMAT0000259&hsa-miR-182-5p&offsets|0|-4;m-22&7|-|129770449|129770470&offsets|0|-2]|TTTGGCAATGGTAGAACTCA</td><td>>MI0000272|hsa-mir-182&WithFlank&7|-|129770377|129770498</td><td>29</td><td>48</td><td>20</td><td>TTTGGCAATGGTAGAACTCA</td><td>174.4</td><td>5.0</td><td>1.8</td><td>2.8</td><td>5.4E-03</td><td>1.1E-02</td><td>BT20/MB468</td><td>MP</td></tr>
<tr><td>>hg38_21_+_8433172_8446622_RNA45SN1_45S_50ntFlanks@9937.9957.21|CAGCAGGACGGTGGCCATGGA</td><td>>hg38_21_+_8433172_8446622_RNA45SN1_45S_50ntFlanks</td><td>9937</td><td>9957</td><td>21</td><td>CAGCAGGACGGTGGCCATGGA</td><td>125.5</td><td>4.8</td><td>1.7</td><td>2.8</td><td>5.4E-03</td><td>1.1E-02</td><td>BT20/MB468</td><td>MP</td></tr>
<tr><td>>MI0000071|hsa-mir-17&WithFlank&13|+|91350599|91350694@20.40.21&[MIMAT0000070&hsa-miR-17-5p&offsets|0|-2;m-88&13|+|91350618|91350640&offsets|0|-2]|CAAAGTGCTTACAGTGCAGGT</td><td>>MI0000071|hsa-mir-17&WithFlank&13|+|91350599|91350694</td><td>20</td><td>40</td><td>21</td><td>CAAAGTGCTTACAGTGCAGGT</td><td>3099.7</td><td>-4.8</td><td>1.5</td><td>-3.1</td><td>2.1E-03</td><td>1.1E-02</td><td>BT20/MB231</td><td>MP</td></tr>
<tr><td>>MI0000071|hsa-mir-17&WithFlank&13|+|91350599|91350694@20.42.23&[MIMAT0000070&hsa-miR-17-5p&offsets|0|0;m-88&13|+|91350618|91350640&offsets|0|0]|CAAAGTGCTTACAGTGCAGGTAG</td><td>>MI0000071|hsa-mir-17&WithFlank&13|+|91350599|91350694</td><td>20</td><td>42</td><td>23</td><td>CAAAGTGCTTACAGTGCAGGTAG</td><td>22445.7</td><td>-3.5</td><td>1.1</td><td>-3.1</td><td>2.1E-03</td><td>1.1E-02</td><td>BT20/MB231</td><td>MP</td></tr>
<tr><td>>MI0000650|hsa-mir-200c&WithFlank&12|+|6963693|6963772@49.71.23&[MIMAT0000617&hsa-miR-200c-3p&offsets|-1|-1;m-54&12|+|6963742|6963764&offsets|-1|-1]|CTAATACTGCCGGGTAATGATGG</td><td>>MI0000650|hsa-mir-200c&WithFlank&12|+|6963693|6963772</td><td>49</td><td>71</td><td>23</td><td>CTAATACTGCCGGGTAATGATGG</td><td>211.4</td><td>4.2</td><td>1.5</td><td>2.8</td><td>5.4E-03</td><td>1.1E-02</td><td>BT20/MB468</td><td>MP</td></tr>
<tr><td>>MI0000095|hsa-mir-93&WithFlank&7|-|100093762|100093853@17.39.23(+1C)&[MIMAT0000093&hsa-miR-93-5p&offsets|0|0(+1C);m-29&7|-|100093815|100093837&offsets|0|0(+1C)]|CAAAGTGCTGTTCGTGCAGGTAGC</td><td>>MI0000095|hsa-mir-93&WithFlank&7|-|100093762|100093853</td><td>17</td><td>39</td><td>23(+1C)</td><td>CAAAGTGCTGTTCGTGCAGGTAGC</td><td>176.0</td><td>3.7</td><td>1.4</td><td>2.6</td><td>8.2E-03</td><td>1.1E-02</td><td>MB231/MB468</td><td>MP</td></tr>
<tr><td>>hg38_21_+_8433172_8446622_RNA45SN1_45S_50ntFlanks@6774.6807.34|TCCTCCCGGGGCTACGCCTGTCTGAGCGTCGCTT</td><td>>hg38_21_+_8433172_8446622_RNA45SN1_45S_50ntFlanks</td><td>6774</td><td>6807</td><td>34</td><td>TCCTCCCGGGGCTACGCCTGTCTGAGCGTCGCTT</td><td>173.6</td><td>2.3</td><td>0.9</td><td>2.6</td><td>9.5E-03</td><td>1.1E-02</td><td>MB231/MB468</td><td>MITO</td></tr>
<tr><td>>hg38_21_+_8433172_8446622_RNA45SN1_45S_50ntFlanks@4742.4759.18|TCGGAGGTTCGAAGACGA</td><td>>hg38_21_+_8433172_8446622_RNA45SN1_45S_50ntFlanks</td><td>4742</td><td>4759</td><td>18</td><td>TCGGAGGTTCGAAGACGA</td><td>477.9</td><td>-2.1</td><td>0.7</td><td>-3.1</td><td>2.2E-03</td><td>1.1E-02</td><td>BT20/MB231</td><td>NUC</td></tr>
<tr><td>>hg38_MT_+_598_1651_RNR1_12S_50ntFlanks@971.1004.34|TGGTAAGTGTACTGGAAAGTGCACTTGGACGAAC</td><td>>hg38_MT_+_598_1651_RNR1_12S_50ntFlanks</td><td>971</td><td>1004</td><td>34</td><td>TGGTAAGTGTACTGGAAAGTGCACTTGGACGAAC</td><td>162.2</td><td>-4.0</td><td>1.3</td><td>-3.1</td><td>2.2E-03</td><td>1.1E-02</td><td>BT20/MB231</td><td>NUC</td></tr>
<tr><td>>hg38_21_+_8433172_8446622_RNA45SN1_45S_50ntFlanks@11564.11602.39|CCTAGCAGCCGACTTAGAACTGGTGCGGACCAGGGGAAT</td><td>>hg38_21_+_8433172_8446622_RNA45SN1_45S_50ntFlanks</td><td>11564</td><td>11602</td><td>39</td><td>CCTAGCAGCCGACTTAGAACTGGTGCGGACCAGGGGAAT</td><td>1650.5</td><td>-3.0</td><td>1.1</td><td>-2.8</td><td>5.4E-03</td><td>1.1E-02</td><td>BT20/MB468</td><td>MP</td></tr>
<tr><td>>hg38_21_+_8433172_8446622_RNA45SN1_45S_50ntFlanks@3985.4025.41|CGGCTTTGGTGACTCTAGATAACCTCGGGCCGATCGCACGC</td><td>>hg38_21_+_8433172_8446622_RNA45SN1_45S_50ntFlanks</td><td>3985</td><td>4025</td><td>41</td><td>CGGCTTTGGTGACTCTAGATAACCTCGGGCCGATCGCACGC</td><td>121.4</td><td>-5.1</td><td>1.8</td><td>-2.8</td><td>5.4E-03</td><td>1.1E-02</td><td>BT20/MB468</td><td>MP</td></tr>
<tr><td>>hg38_21_+_8433172_8446622_RNA45SN1_45S_50ntFlanks@4729.4751.23|TCAAGAACGAAAGTCGGAGGTTC</td><td>>hg38_21_+_8433172_8446622_RNA45SN1_45S_50ntFlanks</td><td>4729</td><td>4751</td><td>23</td><td>TCAAGAACGAAAGTCGGAGGTTC</td><td>287.6</td><td>-1.5</td><td>0.6</td><td>-2.6</td><td>8.9E-03</td><td>1.1E-02</td><td>BT20/MB468</td><td>CYTO</td></tr>
<tr><td>>hg38_21_+_8433172_8446622_RNA45SN1_45S_50ntFlanks@11572.11613.42|CCGACTTAGAACTGGTGCGGACCAGGGGAATCCGACTGTTTA</td><td>>hg38_21_+_8433172_8446622_RNA45SN1_45S_50ntFlanks</td><td>11572</td><td>11613</td><td>42</td><td>CCGACTTAGAACTGGTGCGGACCAGGGGAATCCGACTGTTTA</td><td>1047.0</td><td>3.0</td><td>1.0</td><td>3.0</td><td>2.7E-03</td><td>1.1E-02</td><td>BT20/MB231</td><td>MITO</td></tr>
<tr><td>>hg38_21_+_8433172_8446622_RNA45SN1_45S_50ntFlanks@12472.12496.25|TCACCCACTAATAGGGAACGTGAGC</td><td>>hg38_21_+_8433172_8446622_RNA45SN1_45S_50ntFlanks</td><td>12472</td><td>12496</td><td>25</td><td>TCACCCACTAATAGGGAACGTGAGC</td><td>280.0</td><td>2.3</td><td>0.8</td><td>3.0</td><td>2.7E-03</td><td>1.1E-02</td><td>BT20/MB231</td><td>MITO</td></tr>
<tr><td>>hg38_21_+_8433172_8446622_RNA45SN1_45S_50ntFlanks@6671.6720.50|CTCGGCTCGTGCGTCGATGAAGAACGCAGCTAGCTGCGAGAATTAATGTG</td><td>>hg38_21_+_8433172_8446622_RNA45SN1_45S_50ntFlanks</td><td>6671</td><td>6720</td><td>50</td><td>CTCGGCTCGTGCGTCGATGAAGAACGCAGCTAGCTGCGAGAATTAATGTG</td><td>138.6</td><td>-3.1</td><td>1.2</td><td>-2.7</td><td>7.7E-03</td><td>1.1E-02</td><td>BT20/MB468</td><td>MITO</td></tr>
<tr><td>>hg38_21_+_8433172_8446622_RNA45SN1_45S_50ntFlanks@3837.3871.35|CCTCTCCTACTTGGATAACTGTGGTAATTCTAGAG</td><td>>hg38_21_+_8433172_8446622_RNA45SN1_45S_50ntFlanks</td><td>3837</td><td>3871</td><td>35</td><td>CCTCTCCTACTTGGATAACTGTGGTAATTCTAGAG</td><td>311.7</td><td>-2.1</td><td>0.8</td><td>-2.6</td><td>9.0E-03</td><td>1.1E-02</td><td>BT20/MB468</td><td>CYTO</td></tr>
<tr><td>>hg38_21_+_8433172_8446622_RNA45SN1_45S_50ntFlanks@4102.4124.23|ACCACGGGTGACGGGGAATCAGG</td><td>>hg38_21_+_8433172_8446622_RNA45SN1_45S_50ntFlanks</td><td>4102</td><td>4124</td><td>23</td><td>ACCACGGGTGACGGGGAATCAGG</td><td>247.4</td><td>2.1</td><td>0.8</td><td>2.6</td><td>8.3E-03</td><td>1.1E-02</td><td>MB231/MB468</td><td>MP</td></tr>
<tr><td>>hg38_21_+_8433172_8446622_RNA45SN1_45S_50ntFlanks@8915.8941.27|CGAATCCCGGGGCCGAGGGAGCGAGAC</td><td>>hg38_21_+_8433172_8446622_RNA45SN1_45S_50ntFlanks</td><td>8915</td><td>8941</td><td>27</td><td>CGAATCCCGGGGCCGAGGGAGCGAGAC</td><td>96.2</td><td>5.0</td><td>1.8</td><td>2.8</td><td>5.4E-03</td><td>1.1E-02</td><td>BT20/MB468</td><td>MP</td></tr>
<tr><td>>am_trnaMT_SerGCT_MT_+_12207_12265@14.32.19|AGAACTGCTAACTCATGCC</td><td>>am_trnaMT_SerGCT_MT_+_12207_12265</td><td>14</td><td>32</td><td>19</td><td>AGAACTGCTAACTCATGCC</td><td>118.4</td><td>4.1</td><td>1.5</td><td>2.8</td><td>5.4E-03</td><td>1.1E-02</td><td>BT20/MB468</td><td>MP</td></tr>
<tr><td>>hg38_21_+_8433172_8446622_RNA45SN1_45S_50ntFlanks@10869.10891.23|AGGGCTGGGTCGGTCGGGCTGGG</td><td>>hg38_21_+_8433172_8446622_RNA45SN1_45S_50ntFlanks</td><td>10869</td><td>10891</td><td>23</td><td>AGGGCTGGGTCGGTCGGGCTGGG</td><td>1015.4</td><td>-2.7</td><td>1.0</td><td>-2.6</td><td>9.0E-03</td><td>1.1E-02</td><td>BT20/MB468</td><td>CYTO</td></tr>
<tr><td>>trnaMT_LeuTAA_MT_+_3230_3304@18.42.25|CGGTAATCGCATAAAACTTAAAACT</td><td>>trnaMT_LeuTAA_MT_+_3230_3304</td><td>18</td><td>42</td><td>25</td><td>CGGTAATCGCATAAAACTTAAAACT</td><td>65.6</td><td>4.4</td><td>1.6</td><td>2.8</td><td>5.5E-03</td><td>1.1E-02</td><td>BT20/MB468</td><td>MP</td></tr>
<tr><td>>hg38_21_+_8433172_8446622_RNA45SN1_45S_50ntFlanks@12713.12742.30|GGAGCCTCGGTTGGCCTCGGATAGCCGGTC</td><td>>hg38_21_+_8433172_8446622_RNA45SN1_45S_50ntFlanks</td><td>12713</td><td>12742</td><td>30</td><td>GGAGCCTCGGTTGGCCTCGGATAGCCGGTC</td><td>90.6</td><td>-4.4</td><td>1.7</td><td>-2.6</td><td>8.3E-03</td><td>1.1E-02</td><td>MB231/MB468</td><td>MP</td></tr>
<tr><td>>hg38_21_+_8433172_8446622_RNA45SN1_45S_50ntFlanks@10777.10818.42|TCCAAGGTGAACAGCCTCTGGCATGTTGGAACAATGTAGGTA</td><td>>hg38_21_+_8433172_8446622_RNA45SN1_45S_50ntFlanks</td><td>10777</td><td>10818</td><td>42</td><td>TCCAAGGTGAACAGCCTCTGGCATGTTGGAACAATGTAGGTA</td><td>217.5</td><td>-2.2</td><td>0.8</td><td>-2.7</td><td>7.7E-03</td><td>1.1E-02</td><td>BT20/MB468</td><td>MITO</td></tr>
<tr><td>>MI0000542|hsa-mir-320a&WithFlank&8|-|22244960|22245043@48.68.21&[MIMAT0000510&hsa-miR-320a-3p&offsets|0|-1;m-60&8|-|22244975|22244996&offsets|0|-1]|AAAAGCTGGGTTGAGAGGGCG</td><td>>MI0000542|hsa-mir-320a&WithFlank&8|-|22244960|22245043</td><td>48</td><td>68</td><td>21</td><td>AAAAGCTGGGTTGAGAGGGCG</td><td>20237.5</td><td>2.8</td><td>0.9</td><td>3.1</td><td>2.1E-03</td><td>1.1E-02</td><td>BT20/MB231</td><td>MP</td></tr>
<tr><td>>hg38_21_+_8433172_8446622_RNA45SN1_45S_50ntFlanks@11670.11690.21|CCCAGTGCTCTGAATGTCAAA</td><td>>hg38_21_+_8433172_8446622_RNA45SN1_45S_50ntFlanks</td><td>11670</td><td>11690</td><td>21</td><td>CCCAGTGCTCTGAATGTCAAA</td><td>171.9</td><td>-1.5</td><td>0.6</td><td>-2.6</td><td>9.0E-03</td><td>1.1E-02</td><td>MB231/MB468</td><td>CYTO</td></tr>
<tr><td>>hg38_21_+_8433172_8446622_RNA45SN1_45S_50ntFlanks@3987.4023.37|GCTTTGGTGACTCTAGATAACCTCGGGCCGATCGCAC</td><td>>hg38_21_+_8433172_8446622_RNA45SN1_45S_50ntFlanks</td><td>3987</td><td>4023</td><td>37</td><td>GCTTTGGTGACTCTAGATAACCTCGGGCCGATCGCAC</td><td>392.3</td><td>2.9</td><td>1.0</td><td>3.1</td><td>2.3E-03</td><td>1.1E-02</td><td>BT20/MB231</td><td>NUC</td></tr>
<tr><td>>hg38_21_+_8433172_8446622_RNA45SN1_45S_50ntFlanks@5482.5501.20|CGGAGCGCTGAGAAGACGGT</td><td>>hg38_21_+_8433172_8446622_RNA45SN1_45S_50ntFlanks</td><td>5482</td><td>5501</td><td>20</td><td>CGGAGCGCTGAGAAGACGGT</td><td>170.1</td><td>6.0</td><td>1.9</td><td>3.1</td><td>2.1E-03</td><td>1.1E-02</td><td>BT20/MB231</td><td>MP</td></tr>
<tr><td>>hg38_21_+_8433172_8446622_RNA45SN1_45S_50ntFlanks@12995.13015.21|CTCGCTGCGATCTATTGAAAG</td><td>>hg38_21_+_8433172_8446622_RNA45SN1_45S_50ntFlanks</td><td>12995</td><td>13015</td><td>21</td><td>CTCGCTGCGATCTATTGAAAG</td><td>938.3</td><td>-2.6</td><td>1.0</td><td>-2.6</td><td>9.4E-03</td><td>1.1E-02</td><td>BT20/MB468</td><td>NUC</td></tr>
<tr><td>>am_trna18_AlaTGC_11_-_50233879_50233950@58.75.18,trna67_AlaAGC_6_+_28678366_28678437@58.75.18,trna87_AlaAGC_1_-_150017382_150017452@57.74.18|TCCCCAGCACCTCCACCA</td><td>>am_trna18_AlaTGC_11_-_50233879_50233950</td><td>58</td><td>75</td><td>18</td><td>TCCCCAGCACCTCCACCA</td><td>286.3</td><td>2.5</td><td>0.8</td><td>3.0</td><td>2.6E-03</td><td>1.1E-02</td><td>BT20/MB231</td><td>CYTO</td></tr>
<tr><td>>MI0000289|hsa-mir-181a-1&WithFlank&1|-|198859038|198859159@30.53.24;MI0000269|hsa-mir-181a-2&WithFlank&9|+|124692436|124692557@45.68.24&[MIMAT0000256&hsa-miR-181a-5p&offsets|0|+1;m-32&1|-|198859108|198859130&offsets|0|+1];[MIMAT0000256_1&hsa-miR-181a-5p&offsets|0|+1;m-32&9|+|124692480|124692502&offsets|0|+1]|AACATTCAACGCTGTCGGTGAGTT</td><td>>MI0000289|hsa-mir-181a-1&WithFlank&1|-|198859038|198859159</td><td>30</td><td>53</td><td>24</td><td>AACATTCAACGCTGTCGGTGAGTT</td><td>9621.7</td><td>-5.0</td><td>1.6</td><td>-3.1</td><td>2.1E-03</td><td>1.1E-02</td><td>BT20/MB231</td><td>MP</td></tr>
<tr><td>>MI0000471|hsa-mir-126&WithFlank&9|+|136670596|136670692@21.42.22&[MIMAT0000444&hsa-miR-126-5p&offsets|0|+1;m-114&9|+|136670616|136670636&offsets|0|+1]|CATTATTACTTTTGGTACGCGC</td><td>>MI0000471|hsa-mir-126&WithFlank&9|+|136670596|136670692</td><td>21</td><td>42</td><td>22</td><td>CATTATTACTTTTGGTACGCGC</td><td>48.7</td><td>-9.0</td><td>2.9</td><td>-3.1</td><td>2.1E-03</td><td>1.1E-02</td><td>BT20/MB231</td><td>MP</td></tr>
<tr><td>>hg38_21_+_8433172_8446622_RNA45SN1_45S_50ntFlanks@9544.9571.28|GGTGGAGGTCCGTAGCGGTCCTGACGTG</td><td>>hg38_21_+_8433172_8446622_RNA45SN1_45S_50ntFlanks</td><td>9544</td><td>9571</td><td>28</td><td>GGTGGAGGTCCGTAGCGGTCCTGACGTG</td><td>1912.0</td><td>-3.0</td><td>1.0</td><td>-3.0</td><td>2.6E-03</td><td>1.1E-02</td><td>BT20/MB231</td><td>CYTO</td></tr>
<tr><td>>hg38_21_+_8433172_8446622_RNA45SN1_45S_50ntFlanks@10786.10815.30|AACAGCCTCTGGCATGTTGGAACAATGTAG</td><td>>hg38_21_+_8433172_8446622_RNA45SN1_45S_50ntFlanks</td><td>10786</td><td>10815</td><td>30</td><td>AACAGCCTCTGGCATGTTGGAACAATGTAG</td><td>255.1</td><td>-3.8</td><td>1.4</td><td>-2.8</td><td>5.5E-03</td><td>1.1E-02</td><td>BT20/MB468</td><td>MP</td></tr>
<tr><td>>hg38_21_+_8433172_8446622_RNA45SN1_45S_50ntFlanks@12421.12443.23|TGTCGGCTCTTCCTATCATTGTG</td><td>>hg38_21_+_8433172_8446622_RNA45SN1_45S_50ntFlanks</td><td>12421</td><td>12443</td><td>23</td><td>TGTCGGCTCTTCCTATCATTGTG</td><td>181.0</td><td>-3.2</td><td>1.2</td><td>-2.6</td><td>9.6E-03</td><td>1.1E-02</td><td>MB231/MB468</td><td>NUC</td></tr>
</table>

| | | | | | | | | | | | | | |
|---|---|---|---|---|---|---|---|---|---|---|---|---|---|
| >MI0000093\|hsa-mir-92a-1&WithFlank&13\|+\|91351308\|91351397@54.74.21(+1G);MI0000094\|hsa-mir-92a-2&WithFlank&X\|-\|134169532\|134169618@54.74.21(+1G)&[MIMAT0000092&hsa-miR-92a-3p&offsets\|0\|-1(+1G);m-9&13\|+\|91351361\|91351382&offsets\|0\|-1(+1G)];[MIMAT0000092_1&hsa-miR-92a-3p&offsets\|0\|-1(+1G);m-11&X\|-\|134169544\|134169565&offsets\|0\|-1(+1G)]\|TATTGCACTTGTCCCGGCCTGG | >MI0000093\|hsa-mir-92a-1&WithFlank&13\|+\|91351308\|91351397 | 54 | 74 | 21(+1G) | TATTGCACTTGTCCCGGCCTGG | 1206.5 | -2.4 | 0.8 | -3.0 | 2.7E-03 | 1.1E-02 | BT20/MB231 | MITO |
| >hg38_21_+_8433172_8446622_RNA45SN1_45S_50ntFlanks@12419.12442.24\|GATGTCGGCTCTTCCTATCATTGT | >hg38_21_+_8433172_8446622_RNA45SN1_45S_50ntFlanks | 12419 | 12442 | 24 | GATGTCGGCTCTTCCTATCATTGT | 4460.6 | -2.5 | 0.8 | -3.0 | 2.6E-03 | 1.1E-02 | BT20/MB231 | CYTO |
| >hg38_21_+_8433172_8446622_RNA45SN1_45S_50ntFlanks@12405.12439.35\|CTTTTTGATCCTTCGATGTCGGCTCTTCCTATCAT | >hg38_21_+_8433172_8446622_RNA45SN1_45S_50ntFlanks | 12405 | 12439 | 35 | CTTTTTGATCCTTCGATGTCGGCTCTTCCTATCAT | 247.0 | -1.8 | 0.7 | -2.6 | 9.1E-03 | 1.1E-02 | MB231/MB468 | CYTO |
| >hg38_21_+_8433172_8446622_RNA45SN1_45S_50ntFlanks@8176.8199.24\|CTTCTGATCGAGGCCCAGCCCGTG | >hg38_21_+_8433172_8446622_RNA45SN1_45S_50ntFlanks | 8176 | 8199 | 24 | CTTCTGATCGAGGCCCAGCCCGTG | 265.4 | -1.4 | 0.5 | -2.6 | 9.0E-03 | 1.1E-02 | BT20/MB468 | CYTO |
| >MI0000067\|hsa-let-7f-1&WithFlank&9\|+\|94176341\|94176439@13.33.21(+1G);MI0000068\|hsa-let-7f-2&WithFlank&X\|-\|53557186\|53557280@14.34.21(+1G)&[MIMAT0000067&hsa-let-7f-5p&offsets\|0\|-1(+1G);m-20&9\|+\|94176353\|94176374&offsets\|0\|-1(+1G)];[MIMAT0000067_1&hsa-let-7f-5p&offsets\|0\|-1(+1G);m-19&X\|-\|53557246\|53557267&offsets\|0\|-1(+1G)]\|TGAGGTAGTAGATTGTATAGTG | >MI0000067\|hsa-let-7f-1&WithFlank&9\|+\|94176341\|94176439 | 13 | 33 | 21(+1G) | TGAGGTAGTAGATTGTATAGTG | 550.1 | 2.5 | 1.0 | 2.6 | 9.1E-03 | 1.1E-02 | MB231/MB468 | CYTO |
| >hg38_21_+_8433172_8446622_RNA45SN1_45S_50ntFlanks@11875.11898.24\|GAAGACCCTGTTGAGCTTGACTCT | >hg38_21_+_8433172_8446622_RNA45SN1_45S_50ntFlanks | 11875 | 11898 | 24 | GAAGACCCTGTTGAGCTTGACTCT | 313.0 | -1.4 | 0.5 | -2.6 | 9.0E-03 | 1.1E-02 | BT20/MB468 | CYTO |
| >hg38_MT_+_1621_3279_RNR2_16S_50ntFlanks@999.1030.32\|AGGGACCTGTATGAATGGCTCCACGAGGGTTC | >hg38_MT_+_1621_3279_RNR2_16S_50ntFlanks | 999 | 1030 | 32 | AGGGACCTGTATGAATGGCTCCACGAGGGTTC | 82.0 | 3.1 | 1.2 | 2.6 | 8.3E-03 | 1.1E-02 | MB231/MB468 | MP |
| >hg38_21_+_8433172_8446622_RNA45SN1_45S_50ntFlanks@4733.4761.29\|GAACGAAAGTCGGAGGTTCGAAGACGATC | >hg38_21_+_8433172_8446622_RNA45SN1_45S_50ntFlanks | 4733 | 4761 | 29 | GAACGAAAGTCGGAGGTTCGAAGACGATC | 5878.0 | 5.0 | 1.6 | 3.1 | 2.1E-03 | 1.1E-02 | BT20/MB231 | MP |
| >hg38_21_+_8433172_8446622_RNA45SN1_45S_50ntFlanks@11210.11234.25\|GGCGACTCTGGACGCGAGCCGGGCC | >hg38_21_+_8433172_8446622_RNA45SN1_45S_50ntFlanks | 11210 | 11234 | 25 | GGCGACTCTGGACGCGAGCCGGGCC | 145.4 | -4.6 | 1.7 | -2.8 | 5.5E-03 | 1.1E-02 | BT20/MB468 | MP |
| >MI0000071\|hsa-mir-17&WithFlank&13\|+\|91350599\|91350694@20.41.22(+1A)&[MIMAT0000070&hsa-miR-17-5p&offsets\|0\|-1(+1A);m-88&13\|+\|91350618\|91350640&offsets\|0\|-1(+1A)]\|CAAAGTGCTTACAGTGCAGGTAA | >MI0000071\|hsa-mir-17&WithFlank&13\|+\|91350599\|91350694 | 20 | 41 | 22(+1A) | CAAAGTGCTTACAGTGCAGGTAA | 3058.3 | -3.8 | 1.2 | -3.1 | 2.1E-03 | 1.1E-02 | BT20/MB231 | MP |
| >MI0000088\|hsa-mir-30a&WithFlank&6\|-\|71403545\|71403627@12.35.24&[MIMAT0000087&hsa-miR-30a-5p&offsets\|0\|+2;m-15&6\|-\|71403595\|71403616&offsets\|0\|+2]\|TGTAAACATCCTCGACTGGAAGCT | >MI0000088\|hsa-mir-30a&WithFlank&6\|-\|71403545\|71403627 | 12 | 35 | 24 | TGTAAACATCCTCGACTGGAAGCT | 84008.8 | 3.2 | 1.2 | 2.8 | 5.5E-03 | 1.1E-02 | BT20/MB468 | MP |
| >hg38_MT_+_1621_3279_RNR2_16S_50ntFlanks@1426.1458.33\|CGATTAAAGTCCTACGTGATCTGAGTTCAGACC | >hg38_MT_+_1621_3279_RNR2_16S_50ntFlanks | 1426 | 1458 | 33 | CGATTAAAGTCCTACGTGATCTGAGTTCAGACC | 83.1 | 7.0 | 2.5 | 2.8 | 5.5E-03 | 1.1E-02 | BT20/MB468 | MP |
| >hg38_21_+_8433172_8446622_RNA45SN1_45S_50ntFlanks@10247.10289.43\|GCGACGAGTAGGAGGGCCGCTGCGGTGAGCCTTGAAGCCTAGG | >hg38_21_+_8433172_8446622_RNA45SN1_45S_50ntFlanks | 10247 | 10289 | 43 | GCGACGAGTAGGAGGGCCGCTGCGGTGAGCCTTGAAGCCTAGG | 179.9 | 6.0 | 2.0 | 3.1 | 2.1E-03 | 1.1E-02 | BT20/MB231 | MP |
| >trnaMT_HisGTG_MT_+_12138_12206@32.63.32\|TGAATCTGACAACAGAGGCTTACGACCCCTTA | >trnaMT_HisGTG_MT_+_12138_12206 | 32 | 63 | 32 | TGAATCTGACAACAGAGGCTTACGACCCCTTA | 1593.2 | 2.6 | 1.0 | 2.6 | 9.6E-03 | 1.1E-02 | MB231/MB468 | MITO |
| >hg38_21_+_8433172_8446622_RNA45SN1_45S_50ntFlanks@12073.12106.34\|CCGAGGGGCTCTCGCTTCTGGCGCCAAGCGCCCG | >hg38_21_+_8433172_8446622_RNA45SN1_45S_50ntFlanks | 12073 | 12106 | 34 | CCGAGGGGCTCTCGCTTCTGGCGCCAAGCGCCCG | 498.6 | -3.2 | 1.2 | -2.7 | 7.7E-03 | 1.1E-02 | BT20/MB468 | MITO |
| >hg38_21_+_8433172_8446622_RNA45SN1_45S_50ntFlanks@11669.11691.23\|GCCCAGTGCTCTGAATGTCAAAG | >hg38_21_+_8433172_8446622_RNA45SN1_45S_50ntFlanks | 11669 | 11691 | 23 | GCCCAGTGCTCTGAATGTCAAAG | 124.0 | 2.7 | 1.0 | 2.8 | 5.5E-03 | 1.1E-02 | BT20/MB468 | MP |
| >hg38_21_+_8433172_8446622_RNA45SN1_45S_50ntFlanks@6650.6685.36\|ACGACTCTTAGCGGTGGATCACTCGGCTCGTGCGTC | >hg38_21_+_8433172_8446622_RNA45SN1_45S_50ntFlanks | 6650 | 6685 | 36 | ACGACTCTTAGCGGTGGATCACTCGGCTCGTGCGTC | 263.7 | 2.1 | 0.7 | 3.1 | 2.3E-03 | 1.1E-02 | BT20/MB231 | NUC |
| >MI0001448\|hsa-mir-425&WithFlank&3\|-\|49020142\|49020240@20.44.25&[MIMAT0003393&hsa-miR-425-5p&offsets\|0\|+2;m-95&3\|-\|49020200\|49020221&offsets\|0\|+3]\|AATGACACGATCACTCCCGTTGAGT | >MI0001448\|hsa-mir-425&WithFlank&3\|-\|49020142\|49020240 | 20 | 44 | 25 | AATGACACGATCACTCCCGTTGAGT | 1299.1 | 2.8 | 1.1 | 2.6 | 8.3E-03 | 1.1E-02 | MB231/MB468 | MP |
| >trnaMT_LeuTAA_MT_+_3230_3304@16.45.30\|CCCGGTAATCGCATAAAACTTAAAACTTTA | >trnaMT_LeuTAA_MT_+_3230_3304 | 16 | 45 | 30 | CCCGGTAATCGCATAAAACTTAAAACTTTA | 203.4 | -4.1 | 1.5 | -2.7 | 7.7E-03 | 1.1E-02 | BT20/MB468 | MITO |
| >hg38_21_+_8433172_8446622_RNA45SN1_45S_50ntFlanks@12993.13039.47\|CCCTCGCTGCGATCTATTGAAAGTCAGCCCTCGACACAAGGGTTTGT | >hg38_21_+_8433172_8446622_RNA45SN1_45S_50ntFlanks | 12993 | 13039 | 47 | CCCTCGCTGCGATCTATTGAAAGTCAGCCCTCGACACAAGGGTTTGT | 3862.6 | -4.2 | 1.4 | -3.1 | 2.3E-03 | 1.1E-02 | BT20/MB231 | NUC |
| >hg38_21_+_8433172_8446622_RNA45SN1_45S_50ntFlanks@10392.10432.41\|ACAGCAGTTGAACATGGGTCAGTCGGTCCTGAGAGATGGGC | >hg38_21_+_8433172_8446622_RNA45SN1_45S_50ntFlanks | 10392 | 10432 | 41 | ACAGCAGTTGAACATGGGTCAGTCGGTCCTGAGAGATGGGC | 204.0 | -3.4 | 1.3 | -2.6 | 9.0E-03 | 1.1E-02 | BT20/MB468 | CYTO |
| >trna152_ValCAC_6_-_27248049_27248121@1.24.24\|GCTTCTGTAGTGTAGTGGTTATCA | >trna152_ValCAC_6_-_27248049_27248121 | 1 | 24 | 24 | GCTTCTGTAGTGTAGTGGTTATCA | 357.0 | -2.5 | 1.0 | -2.6 | 9.1E-03 | 1.1E-02 | MB231/MB468 | CYTO |
| >hg38_21_+_8433172_8446622_RNA45SN1_45S_50ntFlanks@4760.4783.24\|TCAGATACCGTCGTAGTTCCGACC | >hg38_21_+_8433172_8446622_RNA45SN1_45S_50ntFlanks | 4760 | 4783 | 24 | TCAGATACCGTCGTAGTTCCGACC | 628.0 | -3.9 | 1.4 | -2.8 | 5.5E-03 | 1.1E-02 | BT20/MB468 | MP |
| >hg38_21_+_8433172_8446622_RNA45SN1_45S_50ntFlanks@11587.11613.27\|TGCGGACCAGGGGAATCCGACTGTTTA | >hg38_21_+_8433172_8446622_RNA45SN1_45S_50ntFlanks | 11587 | 11613 | 27 | TGCGGACCAGGGGAATCCGACTGTTTA | 92.7 | -3.4 | 1.3 | -2.6 | 8.4E-03 | 1.1E-02 | MB231/MB468 | MP |
| >MI0000061\|hsa-let-7a-2&WithFlank&11\|-\|122146516\|122146599@11.31.21(+1C);MI0000062\|hsa-let-7a-3&WithFlank&22\|+\|46112743\|46112828@10.30.21(+1C);MI0000060\|hsa-let-7a-1&WithFlank&9\|+\|94175951\|94176042@12.32.21(+1C)&[MIMAT0000062&hsa-let-7a-5p&offsets\|0\|-1(+1C);m-6&11\|-\|122146568\|122146589&offsets\|0\|-1(+1C)];[MIMAT0000062_1&hsa-let-7a-5p&offsets\|0\|-1(+1C);m-5&22\|+\|46112752\|46112773&offsets\|0\|-1(+1C)];[MIMAT0000062_2&hsa-let-7a-5p&offsets\|0\|-1(+1C);m-4&9\|+\|94175962\|94175983&offsets\|0\|-1(+1C)]\|TGAGGTAGTAGGTTGTATAGTC | >MI0000061\|hsa-let-7a-2&WithFlank&11\|-\|122146516\|122146599 | 11 | 31 | 21(+1C) | TGAGGTAGTAGGTTGTATAGTC | 2008.0 | 2.7 | 1.0 | 2.8 | 5.5E-03 | 1.1E-02 | BT20/MB468 | MP |
| >hg38_21_+_8433172_8446622_RNA45SN1_45S_50ntFlanks@12615.12653.39\|GCTGAGGAGCCAATGGGGCGAAGCTACCATCTGTGGGAT | >hg38_21_+_8433172_8446622_RNA45SN1_45S_50ntFlanks | 12615 | 12653 | 39 | GCTGAGGAGCCAATGGGGCGAAGCTACCATCTGTGGGAT | 958.9 | -3.7 | 1.4 | -2.6 | 9.1E-03 | 1.1E-02 | BT20/MB468 | CYTO |
| >hg38_21_+_8433172_8446622_RNA45SN1_45S_50ntFlanks@10871.10892.22\|GGCTGGGTCGGTCGGGCTGGGG | >hg38_21_+_8433172_8446622_RNA45SN1_45S_50ntFlanks | 10871 | 10892 | 22 | GGCTGGGTCGGTCGGGCTGGGG | 275.9 | -2.5 | 0.9 | -2.8 | 5.5E-03 | 1.1E-02 | BT20/MB468 | MP |
| >hg38_21_+_8433172_8446622_RNA45SN1_45S_50ntFlanks@8355.8373.19\|TGAAAAGAACTTTGAAGAG | >hg38_21_+_8433172_8446622_RNA45SN1_45S_50ntFlanks | 8355 | 8373 | 19 | TGAAAAGAACTTTGAAGAG | 3930.8 | 5.3 | 1.7 | 3.1 | 2.1E-03 | 1.1E-02 | BT20/MB231 | MP |
| >MI0000086\|hsa-mir-28&WithFlank&3\|+\|188688775\|188688872@61.81.21&[MIMAT0004502&hsa-miR-28-3p&offsets\|+1\|0;m-28&3\|+\|188688834\|188688855&offsets\|+1\|0]\|ACTAGATTGTGAGCTCCTGGA | >MI0000086\|hsa-mir-28&WithFlank&3\|+\|188688775\|188688872 | 61 | 81 | 21 | ACTAGATTGTGAGCTCCTGGA | 319.9 | 2.5 | 0.8 | 3.0 | 2.6E-03 | 1.1E-02 | BT20/MB231 | CYTO |
| >hg38_21_+_8433172_8446622_RNA45SN1_45S_50ntFlanks@12415.12432.18\|CTTCGATGTCGGCTCTTC | >hg38_21_+_8433172_8446622_RNA45SN1_45S_50ntFlanks | 12415 | 12432 | 18 | CTTCGATGTCGGCTCTTC | 1764.1 | -2.5 | 0.8 | -3.0 | 2.6E-03 | 1.1E-02 | BT20/MB231 | CYTO |
| >MI0000101\|hsa-mir-99a&WithFlank&21\|+\|16539083\|16539175@19.39.21&[MIMAT0000097&hsa-miR-99a-5p&offsets\|0\|-1;m-64&21\|+\|16539101\|16539121&offsets\|0\|0]\|AACCCGTAGATCCGATCTTGT | >MI0000101\|hsa-mir-99a&WithFlank&21\|+\|16539083\|16539175 | 19 | 39 | 21 | AACCCGTAGATCCGATCTTGT | 645.3 | -3.2 | 1.2 | -2.7 | 7.8E-03 | 1.1E-02 | BT20/MB468 | MITO |
| >hg38_21_+_8433172_8446622_RNA45SN1_45S_50ntFlanks@5073.5098.26\|AATTCCGATAACGAACGAGACTCTGG | >hg38_21_+_8433172_8446622_RNA45SN1_45S_50ntFlanks | 5073 | 5098 | 26 | AATTCCGATAACGAACGAGACTCTGG | 366.5 | -1.9 | 0.7 | -2.6 | 9.1E-03 | 1.1E-02 | MB231/MB468 | CYTO |
| >MI0000294\|hsa-mir-218-1&WithFlank&4\|+\|20528269\|20528390@31.50.20;MI0000295\|hsa-mir-218-2&WithFlank&5\|-\|168768140\|168768261@31.50.20&[MIMAT0000275&hsa-miR-218-5p&offsets\|0\|-1;m-185&4\|+\|20528299\|20528319&offsets\|0\|-1];[MIMAT0000275_1&hsa-miR-218-5p&offsets\|0\|-1;m-187&5\|-\|168768211\|168768231&offsets\|0\|-1]\|TTGTGCTTGATCTAACCATG | >MI0000294\|hsa-mir-218-1&WithFlank&4\|+\|20528269\|20528390 | 31 | 50 | 20 | TTGTGCTTGATCTAACCATG | 49.5 | -9.0 | 2.9 | -3.1 | 2.1E-03 | 1.1E-02 | BT20/MB231 | MP |
| >hg38_21_+_8433172_8446622_RNA45SN1_45S_50ntFlanks@12071.12092.22\|CCCCGAGGGGCTCTCGCTTCTG | >hg38_21_+_8433172_8446622_RNA45SN1_45S_50ntFlanks | 12071 | 12092 | 22 | CCCCGAGGGGCTCTCGCTTCTG | 379.0 | -1.8 | 0.7 | -2.6 | 9.1E-03 | 1.1E-02 | BT20/MB468 | CYTO |
| >MI0000273\|hsa-mir-183&WithFlank&7\|-\|129774899\|129775020@34.56.23&[MIMAT0000261&hsa-miR-183-5p&offsets\|+1\|+2;m-41&7\|-\|129774966\|129774987&offsets\|0\|+1]\|ATGGCACTGGTAGAATTCACTGT | >MI0000273\|hsa-mir-183&WithFlank&7\|-\|129774899\|129775020 | 34 | 56 | 23 | ATGGCACTGGTAGAATTCACTGT | 477.1 | 2.9 | 1.1 | 2.8 | 5.5E-03 | 1.1E-02 | BT20/MB468 | MP |
| >hg38_21_+_8433172_8446622_RNA45SN1_45S_50ntFlanks@3813.3834.22\|TTATGGTTCCTTTGGTCGCTCG | >hg38_21_+_8433172_8446622_RNA45SN1_45S_50ntFlanks | 3813 | 3834 | 22 | TTATGGTTCCTTTGGTCGCTCG | 350.7 | -1.9 | 0.7 | -2.6 | 9.1E-03 | 1.1E-02 | BT20/MB468 | CYTO |
| >hg38_21_+_8433172_8446622_RNA45SN1_45S_50ntFlanks@6651.6697.47\|CGACTCTTAGCGGTGGATCACTCGGCTCGTGCGTCGATGAAGAACGC | >hg38_21_+_8433172_8446622_RNA45SN1_45S_50ntFlanks | 6651 | 6697 | 47 | CGACTCTTAGCGGTGGATCACTCGGCTCGTGCGTCGATGAAGAACGC | 5314.5 | -2.8 | 1.1 | -2.6 | 9.5E-03 | 1.1E-02 | BT20/MB468 | NUC |
| >MI0000286\|hsa-mir-210&WithFlank&11\|-\|568083\|568204@72.91.20&[MIMAT0000267&hsa-miR-210-3p&offsets\|0\|-2;m-102&11\|-\|568112\|568133&offsets\|0\|-2]\|CTGTGCGTGTGACAGCGGCT | >MI0000286\|hsa-mir-210&WithFlank&11\|-\|568083\|568204 | 72 | 91 | 20 | CTGTGCGTGTGACAGCGGCT | 48.7 | 5.2 | 2.0 | 2.6 | 8.4E-03 | 1.1E-02 | MB231/MB468 | MP |
| >hg38_21_+_8433172_8446622_RNA45SN1_45S_50ntFlanks@6650.6698.49\|ACGACTCTTAGCGGTGGATCACTCGGCTCGTGCGTCGATGAAGAACGCA | >hg38_21_+_8433172_8446622_RNA45SN1_45S_50ntFlanks | 6650 | 6698 | 49 | ACGACTCTTAGCGGTGGATCACTCGGCTCGTGCGTCGATGAAGAACGCA | 720.8 | 2.3 | 0.9 | 2.6 | 9.2E-03 | 1.1E-02 | MB231/MB468 | CYTO |
| >hg38_21_+_8433172_8446622_RNA45SN1_45S_50ntFlanks@9574.9609.36\|AATCGGTCGTCCGACCTGGGTATAGGGGCGAAAGAC | >hg38_21_+_8433172_8446622_RNA45SN1_45S_50ntFlanks | 9574 | 9609 | 36 | AATCGGTCGTCCGACCTGGGTATAGGGGCGAAAGAC | 188.8 | 3.3 | 1.3 | 2.6 | 9.7E-03 | 1.1E-02 | MB231/MB468 | MITO |
| >MI0003834\|hsa-mir-769&WithFlank&19\|+\|46018926\|46019055@36.57.22(+2U)&[MIMAT0003886&hsa-miR-769-5p&offsets\|0\|0(+2U);m-216&19\|+\|46018961\|46018982&offsets\|0\|0(+2U)]\|TGAGACCTCTGGGTTCTGAGCTTT | >MI0003834\|hsa-mir-769&WithFlank&19\|+\|46018926\|46019055 | 36 | 57 | 22(+2U) | TGAGACCTCTGGGTTCTGAGCTTT | 156.1 | 3.3 | 1.3 | 2.6 | 8.4E-03 | 1.1E-02 | MB231/MB468 | MP |
| >hg38_21_+_8433172_8446622_RNA45SN1_45S_50ntFlanks@12407.12444.38\|TTTTGATCCTTCGATGTCGGCTCTTCCTATCATTGTGA | >hg38_21_+_8433172_8446622_RNA45SN1_45S_50ntFlanks | 12407 | 12444 | 38 | TTTTGATCCTTCGATGTCGGCTCTTCCTATCATTGTGA | 409.8 | -2.2 | 0.9 | -2.6 | 9.7E-03 | 1.1E-02 | MB231/MB468 | MITO |
| >hg38_21_+_8433172_8446622_RNA45SN1_45S_50ntFlanks@4574.4611.38\|ATAGGACCGCGGTTCTATTTTGTTGGTTTTCGGAACTG | >hg38_21_+_8433172_8446622_RNA45SN1_45S_50ntFlanks | 4574 | 4611 | 38 | ATAGGACCGCGGTTCTATTTTGTTGGTTTTCGGAACTG | 222.4 | -2.3 | 0.9 | -2.6 | 9.2E-03 | 1.1E-02 | MB231/MB468 | CYTO |
| >hg38_21_+_8433172_8446622_RNA45SN1_45S_50ntFlanks@11999.12036.38\|CCCTGCGGGCCGCCGGTGAAATACCACTACTCTGATCG | >hg38_21_+_8433172_8446622_RNA45SN1_45S_50ntFlanks | 11999 | 12036 | 38 | CCCTGCGGGCCGCCGGTGAAATACCACTACTCTGATCG | 193.4 | -3.0 | 1.1 | -2.7 | 7.8E-03 | 1.1E-02 | BT20/MB468 | MITO |
| >MI0000299\|hsa-mir-222&WithFlank&X\|-\|45747009\|45747130@75.94.20&[MIMAT0000279&hsa-miR-222-3p&offsets\|0\|-1;m-140&X\|-\|45747033\|45747056&offsets\|0\|-4]\|AGCTACATCTGGCTACTGGG | >MI0000299\|hsa-mir-222&WithFlank&X\|-\|45747009\|45747130 | 75 | 94 | 20 | AGCTACATCTGGCTACTGGG | 357.6 | 4.4 | 1.6 | 2.8 | 5.5E-03 | 1.1E-02 | BT20/MB468 | MP |

| | | | | | | | | | | | | | |
|---|---|---|---|---|---|---|---|---|---|---|---|---|---|
| >hg38_21_+_8433172_8446622_RNA45SN1_45S_50ntFlanks@12488.12518.31\|AACGTGAGCTGGGTTTAGACCGTCGTGAGAC | >hg38_21_+_8433172_8446622_RNA45SN1_45S_50ntFlanks | 12488 | 12518 | 31 | AACGTGAGCTGGGTTTAGACCGTCGTGAGAC | 213.8 | -4.0 | 1.5 | -2.6 | 9.1E-03 | 1.1E-02 | BT20/MB468 | CYTO |
| >hg38_21_+_8433172_8446622_RNA45SN1_45S_50ntFlanks@3705.3722.18\|TACCTGGTTGATCCTGCC | >hg38_21_+_8433172_8446622_RNA45SN1_45S_50ntFlanks | 3705 | 3722 | 18 | TACCTGGTTGATCCTGCC | 1343.9 | -2.2 | 0.8 | -2.6 | 8.4E-03 | 1.1E-02 | MB231/MB468 | MP |
| >MI0000299\|hsa-mir-222&WithFlank&X\|-\|45747009\|45747130@75.97.23&[MIMAT0000279&hsa-miR-222-3p&offsets\|0\|+2;m-140&X\|-\|45747033\|45747056&offsets\|0\|-1]\|AGCTACATCTGGCTACTGGGTCT | >MI0000299\|hsa-mir-222&WithFlank&X\|-\|45747009\|45747130 | 75 | 97 | 23 | AGCTACATCTGGCTACTGGGTCT | 12629.3 | -3.1 | 1.0 | -3.1 | 2.1E-03 | 1.1E-02 | BT20/MB231 | MP |
| >MI0000062\|hsa-let-7a-3&WithFlank&22\|+\|46112743\|46112828@58.78.21(+1U);MI0000060\|hsa-let-7a-1&WithFlank&9\|+\|94175951\|94176042@63.83.21(+1U)&[MIMAT0004481&hsa-let-7a-3p&offsets\|0\|0(+1U);m-180&22\|+\|46112800\|46112821&offsets\|0\|-1(+1U)];[MIMAT0004481_1&hsa-let-7a-3p&offsets\|0\|0(+1U);m-181&9\|+\|94176013\|94176034&offsets\|0\|-1(+1U)]\|CTATACAATCTACTGTCTTTCT | >MI0000062\|hsa-let-7a-3&WithFlank&22\|+\|46112743\|46112828 | 58 | 78 | 21(+1U) | CTATACAATCTACTGTCTTTCT | 2468.6 | -2.6 | 0.9 | -3.0 | 2.7E-03 | 1.1E-02 | BT20/MB231 | MITO |
| >hg38_21_+_8433172_8446622_RNA45SN1_45S_50ntFlanks@3878.3898.21\|CATGCCGACGGGCGCTGACCC | >hg38_21_+_8433172_8446622_RNA45SN1_45S_50ntFlanks | 3878 | 3898 | 21 | CATGCCGACGGGCGCTGACCC | 509.5 | -1.4 | 0.6 | -2.6 | 9.2E-03 | 1.1E-02 | MB231/MB468 | CYTO |
| >hg38_MT_+_598_1651_RNR1_12S_50ntFlanks@970.1004.35\|ATGGTAAGTGTACTGGAAAGTGCACTTGGACGAAC | >hg38_MT_+_598_1651_RNR1_12S_50ntFlanks | 970 | 1004 | 35 | ATGGTAAGTGTACTGGAAAGTGCACTTGGACGAAC | 123.9 | -3.9 | 1.3 | -3.0 | 2.3E-03 | 1.1E-02 | BT20/MB231 | NUC |
| >hg38_21_+_8433172_8446622_RNA45SN1_45S_50ntFlanks@4588.4619.32\|CTATTTTGTTGGTTTTCGGAACTGAGGCCATG | >hg38_21_+_8433172_8446622_RNA45SN1_45S_50ntFlanks | 4588 | 4619 | 32 | CTATTTTGTTGGTTTTCGGAACTGAGGCCATG | 388.4 | -2.2 | 0.9 | -2.6 | 9.2E-03 | 1.1E-02 | MB231/MB468 | CYTO |
| >hg38_21_+_8433172_8446622_RNA45SN1_45S_50ntFlanks@8194.8212.19\|CCCGTGGACGGTGTGAGGC | >hg38_21_+_8433172_8446622_RNA45SN1_45S_50ntFlanks | 8194 | 8212 | 19 | CCCGTGGACGGTGTGAGGC | 2372.6 | -3.1 | 1.1 | -2.8 | 5.6E-03 | 1.1E-02 | BT20/MB468 | MP |
| >am_trnaMT_ValTAC_MT_+_1602_1670@27.57.31\|CAACTTACACTTAGGAGATTTCAACTTAACT | >am_trnaMT_ValTAC_MT_+_1602_1670 | 27 | 57 | 31 | CAACTTACACTTAGGAGATTTCAACTTAACT | 212.4 | 7.1 | 2.6 | 2.8 | 5.6E-03 | 1.1E-02 | BT20/MB468 | MP |
| >hg38_21_+_8433172_8446622_RNA45SN1_45S_50ntFlanks@5082.5104.23\|AACGAACGAGACTCTGGCATGCT | >hg38_21_+_8433172_8446622_RNA45SN1_45S_50ntFlanks | 5082 | 5104 | 23 | AACGAACGAGACTCTGGCATGCT | 532.3 | 2.9 | 1.0 | 2.8 | 5.6E-03 | 1.1E-02 | BT20/MB468 | MP |
| >MI0000272\|hsa-mir-182&WithFlank&7\|-\|129770377\|129770498@29.50.22(+1U)&[MIMAT0000259&hsa-miR-182-5p&offsets\|0\|-2(+1U);m-22&7\|-\|129770449\|129770470&offsets\|0\|0(+1U)]\|TTTGGCAATGGTAGAACTCACAT | >MI0000272\|hsa-mir-182&WithFlank&7\|-\|129770377\|129770498 | 29 | 50 | 22(+1U) | TTTGGCAATGGTAGAACTCACAT | 182.6 | 4.1 | 1.5 | 2.8 | 5.6E-03 | 1.1E-02 | BT20/MB468 | MP |
| >trnaMT_TyrGTA_MT_-_5826_5891@37.69.33\|TAAAGACAGGGGTTAGGCCTCTTTTTACCACCA | >trnaMT_TyrGTA_MT_-_5826_5891 | 37 | 69 | 33 | TAAAGACAGGGGTTAGGCCTCTTTTTACCACCA | 194.2 | 5.4 | 1.9 | 2.8 | 5.6E-03 | 1.1E-02 | BT20/MB468 | MP |
| >MI0000062\|hsa-let-7a-3&WithFlank&22\|+\|46112743\|46112828@9.31.23&[MIMAT0000062_1&hsa-let-7a-5p&offsets\|-1\|0;m-5&22\|+\|46112752\|46112773&offsets\|-1\|0]\|GTGAGGTAGTAGGTTGTATAGTT | >MI0000062\|hsa-let-7a-3&WithFlank&22\|+\|46112743\|46112828 | 9 | 31 | 23 | GTGAGGTAGTAGGTTGTATAGTT | 293.9 | 2.3 | 0.8 | 3.0 | 2.6E-03 | 1.1E-02 | BT20/MB231 | CYTO |
| >hg38_21_+_8433172_8446622_RNA45SN1_45S_50ntFlanks@12416.12441.26\|TTCGATGTCGGCTCTTCCTATCATTG | >hg38_21_+_8433172_8446622_RNA45SN1_45S_50ntFlanks | 12416 | 12441 | 26 | TTCGATGTCGGCTCTTCCTATCATTG | 2641.2 | -2.7 | 0.9 | -3.0 | 2.6E-03 | 1.1E-02 | BT20/MB231 | CYTO |
| >hg38_21_+_8433172_8446622_RNA45SN1_45S_50ntFlanks@9545.9571.27\|GTGGAGGTCCGTAGCGGTCCTGACGTG | >hg38_21_+_8433172_8446622_RNA45SN1_45S_50ntFlanks | 9545 | 9571 | 27 | GTGGAGGTCCGTAGCGGTCCTGACGTG | 1944.6 | -2.7 | 0.9 | -3.0 | 2.6E-03 | 1.1E-02 | BT20/MB231 | CYTO |
| >trna10_AspGTC_12_-_125424193_125424264@36.72.37,trna12_AspGTC_12_-_125411891_125411962@36.72.37,trna144_AspGTC_6_-_27551236_27551307@36.72.37,trna38_AspGTC_17_-_8125556_8125627@36.72.37,trna45_AspGTC_6_+_27447453_27447524@36.72.37,trna48_AspGTC_6_+_27471523_27471594@36.72.37,trna4_AspGTC_12_+_96429799_96429870@36.72.37,trna69_AspGTC_1_-_161440205_161440276@36.72.37,trna72_AspGTC_1_-_161432824_161432895@36.72.37,trna75_AspGTC_1_-_161425414_161425485@36.72.37,trna78_AspGTC_1_-_161418033_161418104@36.72.37,trna81_AspGTC_1_-_161410615_161410686@36.72.37\|CACGCGGGAGACCGGGGTTCGATTCCCCGACGGGGAG | >trna10_AspGTC_12_-_125424193_125424264 | 36 | 72 | 37 | CACGCGGGAGACCGGGGTTCGATTCCCCGACGGGGAG | 596.8 | -2.9 | 1.0 | -3.0 | 2.6E-03 | 1.1E-02 | BT20/MB231 | CYTO |
| >hg38_21_+_8433172_8446622_RNA45SN1_45S_50ntFlanks@6787.6806.20\|ACGCCTGTCTGAGCGTCGCT | >hg38_21_+_8433172_8446622_RNA45SN1_45S_50ntFlanks | 6787 | 6806 | 20 | ACGCCTGTCTGAGCGTCGCT | 657.7 | 1.9 | 0.7 | 2.6 | 8.4E-03 | 1.1E-02 | MB231/MB468 | MP |
| >hg38_MT_+_598_1651_RNR1_12S_50ntFlanks@373.404.32\|AAGCTAAAACTCACCTGAGTTGTAAAAAACTC | >hg38_MT_+_598_1651_RNR1_12S_50ntFlanks | 373 | 404 | 32 | AAGCTAAAACTCACCTGAGTTGTAAAAAACTC | 292.1 | 2.3 | 0.9 | 2.6 | 9.7E-03 | 1.1E-02 | MB231/MB468 | MITO |
| >hg38_21_+_8433172_8446622_RNA45SN1_45S_50ntFlanks@10830.10848.19\|CAAGCCGGATCCGTAACTT | >hg38_21_+_8433172_8446622_RNA45SN1_45S_50ntFlanks | 10830 | 10848 | 19 | CAAGCCGGATCCGTAACTT | 92.2 | -4.5 | 1.6 | -2.8 | 5.6E-03 | 1.1E-02 | BT20/MB468 | MP |
| >hg38_21_+_8433172_8446622_RNA45SN1_45S_50ntFlanks@8384.8405.22\|AGGGCGTGAAACCGTTAAGAGG | >hg38_21_+_8433172_8446622_RNA45SN1_45S_50ntFlanks | 8384 | 8405 | 22 | AGGGCGTGAAACCGTTAAGAGG | 359.7 | -2.3 | 0.8 | -2.8 | 5.6E-03 | 1.1E-02 | BT20/MB468 | MP |
| >hg38_21_+_8433172_8446622_RNA45SN1_45S_50ntFlanks@11317.11342.26\|ACCCCACGTCTCGTCGCGCGCGCGTC | >hg38_21_+_8433172_8446622_RNA45SN1_45S_50ntFlanks | 11317 | 11342 | 26 | ACCCCACGTCTCGTCGCGCGCGCGTC | 2070.9 | -2.9 | 1.1 | -2.7 | 7.9E-03 | 1.1E-02 | BT20/MB468 | MITO |
| >hg38_21_+_8433172_8446622_RNA45SN1_45S_50ntFlanks@10382.10424.43\|TTCCATGTGAACAGCAGTTGAACATGGGTCAGTCGGTCCTGAG | >hg38_21_+_8433172_8446622_RNA45SN1_45S_50ntFlanks | 10382 | 10424 | 43 | TTCCATGTGAACAGCAGTTGAACATGGGTCAGTCGGTCCTGAG | 1352.4 | 3.6 | 1.2 | 3.1 | 2.1E-03 | 1.1E-02 | BT20/MB231 | MP |
| >hg38_21_+_8433172_8446622_RNA45SN1_45S_50ntFlanks@11539.11583.45\|GCGCGGCGCCTCGCCTCGGCCGGCGCCTAGCAGCCGACTTAGAAC | >hg38_21_+_8433172_8446622_RNA45SN1_45S_50ntFlanks | 11539 | 11583 | 45 | GCGCGGCGCCTCGCCTCGGCCGGCGCCTAGCAGCCGACTTAGAAC | 222.1 | -2.7 | 0.9 | -3.0 | 2.3E-03 | 1.1E-02 | BT20/MB231 | NUC |
| >hg38_21_+_8433172_8446622_RNA45SN1_45S_50ntFlanks@5486.5519.34\|GCGCTGAGAAGACGGTCGAACTTGACTATCTAGA | >hg38_21_+_8433172_8446622_RNA45SN1_45S_50ntFlanks | 5486 | 5519 | 34 | GCGCTGAGAAGACGGTCGAACTTGACTATCTAGA | 284.8 | 2.5 | 0.8 | 3.0 | 2.6E-03 | 1.1E-02 | BT20/MB231 | CYTO |
| >hg38_21_+_8433172_8446622_RNA45SN1_45S_50ntFlanks@12003.12032.30\|GCGGGCCGCCGGTGAAATACCACTACTCTG | >hg38_21_+_8433172_8446622_RNA45SN1_45S_50ntFlanks | 12003 | 12032 | 30 | GCGGGCCGCCGGTGAAATACCACTACTCTG | 584.1 | -3.0 | 1.0 | -3.0 | 2.6E-03 | 1.1E-02 | BT20/MB231 | CYTO |
| >hg38_21_+_8433172_8446622_RNA45SN1_45S_50ntFlanks@7978.8019.42\|GACCTCAGATCAGACGTGGCGACCCGCTGAATTTAAGCATAT | >hg38_21_+_8433172_8446622_RNA45SN1_45S_50ntFlanks | 7978 | 8019 | 42 | GACCTCAGATCAGACGTGGCGACCCGCTGAATTTAAGCATAT | 486.0 | -7.2 | 2.4 | -3.0 | 2.7E-03 | 1.1E-02 | BT20/MB231 | CYTO |
| >hg38_21_+_8433172_8446622_RNA45SN1_45S_50ntFlanks@10468.10487.20\|TTGCCCTCGGCCGATCGAAA | >hg38_21_+_8433172_8446622_RNA45SN1_45S_50ntFlanks | 10468 | 10487 | 20 | TTGCCCTCGGCCGATCGAAA | 338.7 | -2.2 | 0.7 | -3.0 | 2.7E-03 | 1.1E-02 | BT20/MB231 | CYTO |
| >hg38_21_+_8433172_8446622_RNA45SN1_45S_50ntFlanks@3876.3899.24\|TACATGCCGACGGGCGCTGACCCC | >hg38_21_+_8433172_8446622_RNA45SN1_45S_50ntFlanks | 3876 | 3899 | 24 | TACATGCCGACGGGCGCTGACCCC | 357.8 | -1.7 | 0.7 | -2.6 | 9.2E-03 | 1.1E-02 | MB231/MB468 | CYTO |
| >hg38_21_+_8433172_8446622_RNA45SN1_45S_50ntFlanks@9485.9503.19\|TGTTAGGACCCGAAAGATG | >hg38_21_+_8433172_8446622_RNA45SN1_45S_50ntFlanks | 9485 | 9503 | 19 | TGTTAGGACCCGAAAGATG | 76.7 | -4.3 | 1.6 | -2.6 | 8.5E-03 | 1.1E-02 | MB231/MB468 | MP |
| >hg38_21_+_8433172_8446622_RNA45SN1_45S_50ntFlanks@11573.11612.40\|CGACTTAGAACTGGTGCGGACCAGGGGAATCCGACTGTTT | >hg38_21_+_8433172_8446622_RNA45SN1_45S_50ntFlanks | 11573 | 11612 | 40 | CGACTTAGAACTGGTGCGGACCAGGGGAATCCGACTGTTT | 632.5 | -2.3 | 0.9 | -2.6 | 9.8E-03 | 1.1E-02 | MB231/MB468 | MITO |
| >am_trnaMT_ValTAC_MT_+_1602_1670@39.65.27\|AGGAGATTTCAACTTAACTTGACCGCT | >am_trnaMT_ValTAC_MT_+_1602_1670 | 39 | 65 | 27 | AGGAGATTTCAACTTAACTTGACCGCT | 102.3 | 7.0 | 2.5 | 2.8 | 5.6E-03 | 1.1E-02 | BT20/MB468 | MP |
| >hg38_21_+_8433172_8446622_RNA45SN1_45S_50ntFlanks@9554.9580.27\|CGTAGCGGTCCTGACGTGCAAATCGGT | >hg38_21_+_8433172_8446622_RNA45SN1_45S_50ntFlanks | 9554 | 9580 | 27 | CGTAGCGGTCCTGACGTGCAAATCGGT | 115.6 | -4.3 | 1.6 | -2.6 | 8.5E-03 | 1.1E-02 | MB231/MB468 | MP |
| >MI0000077\|hsa-mir-21&WithFlank&17\|+\|59841260\|59841343@52.72.21(+1U)&[MIMAT0004494&hsa-miR-21-3p&offsets\|0\|0(+1U);m-52&17\|+\|59841311\|59841332&offsets\|0\|-1(+1U)]\|CAACACCAGTCGATGGGCTGTT | >MI0000077\|hsa-mir-21&WithFlank&17\|+\|59841260\|59841343 | 52 | 72 | 21(+1U) | CAACACCAGTCGATGGGCTGTT | 351.4 | 4.4 | 1.7 | 2.6 | 8.5E-03 | 1.1E-02 | MB231/MB468 | MP |
| >hg38_MT_+_1621_3279_RNR2_16S_50ntFlanks@51.86.36\|GCTAAACCTAGCCCCAAACCCACTCCACCTTACTAC | >hg38_MT_+_1621_3279_RNR2_16S_50ntFlanks | 51 | 86 | 36 | GCTAAACCTAGCCCCAAACCCACTCCACCTTACTAC | 487.3 | 4.8 | 1.7 | 2.8 | 5.6E-03 | 1.1E-02 | BT20/MB468 | MP |
| >hg38_21_+_8433172_8446622_RNA45SN1_45S_50ntFlanks@3771.3788.18\|CACGGCCGGTACAGTGAA | >hg38_21_+_8433172_8446622_RNA45SN1_45S_50ntFlanks | 3771 | 3788 | 18 | CACGGCCGGTACAGTGAA | 57.6 | -4.5 | 1.6 | -2.8 | 5.6E-03 | 1.1E-02 | BT20/MB468 | MP |
| >hg38_21_+_8433172_8446622_RNA45SN1_45S_50ntFlanks@9533.9569.37\|GAGGAAACTCTGGTGGAGGTCCGTAGCGGTCCTGACG | >hg38_21_+_8433172_8446622_RNA45SN1_45S_50ntFlanks | 9533 | 9569 | 37 | GAGGAAACTCTGGTGGAGGTCCGTAGCGGTCCTGACG | 314.0 | -3.7 | 1.4 | -2.6 | 8.5E-03 | 1.1E-02 | MB231/MB468 | MP |
| >trnaMT_GluTTC_MT_-_14674_14742@31.71.41,trnalookalike8_GluTTC_5_-_93905172_93905240@31.71.41\|TTCATATCATTGGTCGTGGTTGTAGTCCGTGCGAGAATACC | >trnaMT_GluTTC_MT_-_14674_14742 | 31 | 71 | 41 | TTCATATCATTGGTCGTGGTTGTAGTCCGTGCGAGAATACC | 294.0 | -2.3 | 0.8 | -3.0 | 2.7E-03 | 1.1E-02 | BT20/MB231 | MITO |
| >trnaMT_PheGAA_MT_+_577_647@35.64.30\|GAAAATGTTTAGACGGGCTCACATCACCCC | >trnaMT_PheGAA_MT_+_577_647 | 35 | 64 | 30 | GAAAATGTTTAGACGGGCTCACATCACCCC | 133.4 | 4.2 | 1.6 | 2.6 | 8.5E-03 | 1.1E-02 | MB231/MB468 | MP |
| >hg38_21_+_8433172_8446622_RNA45SN1_45S_50ntFlanks@7973.7990.18\|GACGCGACCTCAGATCAG | >hg38_21_+_8433172_8446622_RNA45SN1_45S_50ntFlanks | 7973 | 7990 | 18 | GACGCGACCTCAGATCAG | 431.8 | 2.1 | 0.8 | 2.7 | 7.9E-03 | 1.1E-02 | BT20/MB468 | MITO |
| >hg38_21_+_8433172_8446622_RNA45SN1_45S_50ntFlanks@12997.13017.21\|CGCTGCGATCTATTGAAAGTC | >hg38_21_+_8433172_8446622_RNA45SN1_45S_50ntFlanks | 12997 | 13017 | 21 | CGCTGCGATCTATTGAAAGTC | 158.1 | 3.5 | 1.3 | 2.6 | 8.5E-03 | 1.1E-02 | MB231/MB468 | MP |
| >hg38_21_+_8433172_8446622_RNA45SN1_45S_50ntFlanks@12625.12667.43\|CAATGGGGCGAAGCTACCATCTGTGGGATTATGACTGAACGCC | >hg38_21_+_8433172_8446622_RNA45SN1_45S_50ntFlanks | 12625 | 12667 | 43 | CAATGGGGCGAAGCTACCATCTGTGGGATTATGACTGAACGCC | 175.1 | 3.4 | 1.3 | 2.6 | 9.8E-03 | 1.1E-02 | MB231/MB468 | MITO |
| >hg38_21_+_8433172_8446622_RNA45SN1_45S_50ntFlanks@11553.11588.36\|CTCGGCCGGCGCCTAGCAGCCGACTTAGAACTGGTG | >hg38_21_+_8433172_8446622_RNA45SN1_45S_50ntFlanks | 11553 | 11588 | 36 | CTCGGCCGGCGCCTAGCAGCCGACTTAGAACTGGTG | 289.8 | 1.5 | 0.6 | 2.7 | 7.9E-03 | 1.1E-02 | BT20/MB468 | MITO |
| >hg38_21_+_8433172_8446622_RNA45SN1_45S_50ntFlanks@8170.8199.30\|CAAGTCCTTCTGATCGAGGCCCAGCCCGTG | >hg38_21_+_8433172_8446622_RNA45SN1_45S_50ntFlanks | 8170 | 8199 | 30 | CAAGTCCTTCTGATCGAGGCCCAGCCCGTG | 229.5 | -4.6 | 1.7 | -2.8 | 5.6E-03 | 1.1E-02 | BT20/MB468 | MP |
| >hg38_21_+_8433172_8446622_RNA45SN1_45S_50ntFlanks@12306.12326.21\|CACGATCCTTCTGACCTTTTG | >hg38_21_+_8433172_8446622_RNA45SN1_45S_50ntFlanks | 12306 | 12326 | 21 | CACGATCCTTCTGACCTTTTG | 865.7 | -2.5 | 0.9 | -2.7 | 7.9E-03 | 1.1E-02 | BT20/MB468 | MITO |
| >hg38_21_+_8433172_8446622_RNA45SN1_45S_50ntFlanks@6652.6699.48\|GACTCTTAGCGGTGGATCACTCGGCTCGTGCGTCGATGAAGAACGCAG | >hg38_21_+_8433172_8446622_RNA45SN1_45S_50ntFlanks | 6652 | 6699 | 48 | GACTCTTAGCGGTGGATCACTCGGCTCGTGCGTCGATGAAGAACGCAG | 1472.1 | -3.9 | 1.5 | -2.6 | 8.5E-03 | 1.1E-02 | MB231/MB468 | MP |
| >trna11_GluTTC_15_-_26327381_26327452@39.73.35,trna3_GluTTC_13_-_45492062_45492133@39.73.35\|CCAGGCGGCCCGGGTTCGACTCCCGGTGTGGGAAC | >trna11_GluTTC_15_-_26327381_26327452 | 39 | 73 | 35 | CCAGGCGGCCCGGGTTCGACTCCCGGTGTGGGAAC | 201.8 | -3.4 | 1.1 | -3.0 | 2.7E-03 | 1.1E-02 | BT20/MB231 | CYTO |
| >hg38_21_+_8433172_8446622_RNA45SN1_45S_50ntFlanks@12159.12178.20\|TTTGACTGGGGCGGTACACC | >hg38_21_+_8433172_8446622_RNA45SN1_45S_50ntFlanks | 12159 | 12178 | 20 | TTTGACTGGGGCGGTACACC | 1114.0 | -4.0 | 1.4 | -2.8 | 5.6E-03 | 1.1E-02 | BT20/MB468 | MP |
| >hg38_21_+_8433172_8446622_RNA45SN1_45S_50ntFlanks@5525.5549.25\|TAAAAGTCGTAACAAGGTTTCCGTA | >hg38_21_+_8433172_8446622_RNA45SN1_45S_50ntFlanks | 5525 | 5549 | 25 | TAAAAGTCGTAACAAGGTTTCCGTA | 112.0 | -4.5 | 1.6 | -2.8 | 5.6E-03 | 1.1E-02 | BT20/MB468 | MP |
| >hg38_21_+_8433172_8446622_RNA45SN1_45S_50ntFlanks@11668.11693.26\|TGCCCAGTGCTCTGAATGTCAAAGTG | >hg38_21_+_8433172_8446622_RNA45SN1_45S_50ntFlanks | 11668 | 11693 | 26 | TGCCCAGTGCTCTGAATGTCAAAGTG | 74.3 | 3.7 | 1.3 | 2.8 | 5.6E-03 | 1.1E-02 | BT20/MB468 | MP |
| >hg38_21_+_8433172_8446622_RNA45SN1_45S_50ntFlanks@3705.3743.39\|TACCTGGTTGATCCTGCCAGTAGCATATGCTTGTCTCAA | >hg38_21_+_8433172_8446622_RNA45SN1_45S_50ntFlanks | 3705 | 3743 | 39 | TACCTGGTTGATCCTGCCAGTAGCATATGCTTGTCTCAA | 1841.3 | 2.1 | 0.7 | 3.0 | 2.3E-03 | 1.1E-02 | BT20/MB231 | NUC |

| | | | | | | | | | | | | | |
|---|---|---|---|---|---|---|---|---|---|---|---|---|---|
| >trna10_AspGTC_12_-_125424193_125424264@37.74.38,trna12_AspGTC_12_-_125411891_125411962@37.74.38,trna144_AspGTC_6_-_27551236_27551307@37.74.38,trna38_AspGTC_17_-_8125556_8125627@37.74.38,trna45_AspGTC_6_+_27447453_27447524@37.74.38,trna48_AspGTC_6_+_27471523_27471594@37.74.38,trna4_AspGTC_12_+_96429799_96429870@37.74.38,trna69_AspGTC_1_-_161440205_161440276@37.74.38,trna72_AspGTC_1_-_161432824_161432895@37.74.38,trna75_AspGTC_1_-_161425414_161425485@37.74.38,trna78_AspGTC_1_-_161418033_161418104@37.74.38,trna81_AspGTC_1_-_161410615_161410686@37.74.38\|ACGCGGGAGACCGGGGTTCGATTCCCCGACGGGGAGCC | >trna10_AspGTC_12_-_125424193_125424264 | 37 | 74 | 38 | ACGCGGGAGACCGGGGTTCGATTCCCCGACGGGGAGCC | 794.6 | -4.9 | 1.6 | -3.0 | 2.3E-03 | 1.1E-02 | BT20/MB231 | NUC |
| >hg38_21_+_8433172_8446622_RNA45SN1_45S_50ntFlanks@12954.12972.19\|CTGCTTCTGGGTCGGGGTT | >hg38_21_+_8433172_8446622_RNA45SN1_45S_50ntFlanks | 12954 | 12972 | 19 | CTGCTTCTGGGTCGGGGTT | 464.4 | -4.3 | 1.6 | -2.6 | 9.2E-03 | 1.1E-02 | BT20/MB468 | CYTO |
| >MI0003834\|hsa-mir-769&WithFlank&19\|+\|46018926\|46019055@36.57.22&[MIMAT0003886&hsa-miR-769-5p&offsets\|0\|0;m-216&19\|+\|46018961\|46018982&offsets\|0\|0]\|TGAGACCTCTGGGTTCTGAGCT | >MI0003834\|hsa-mir-769&WithFlank&19\|+\|46018926\|46019055 | 36 | 57 | 22 | TGAGACCTCTGGGTTCTGAGCT | 1468.6 | 3.0 | 1.1 | 2.8 | 5.6E-03 | 1.1E-02 | BT20/MB468 | MP |
| >MI0000266\|hsa-mir-10a&WithFlank&17\|-\|48579832\|48579953@29.50.22(+1A)&[MIMAT0000253&hsa-miR-10a-5p&offsets\|+1\|0(+1A);m-18&17\|-\|48579905\|48579926&offsets\|+1\|+1(+1A)]\|ACCCTGTAGATCCGAATTTGTGA | >MI0000266\|hsa-mir-10a&WithFlank&17\|-\|48579832\|48579953 | 29 | 50 | 22(+1A) | ACCCTGTAGATCCGAATTTGTGA | 1514.9 | 2.3 | 0.8 | 3.0 | 2.7E-03 | 1.1E-02 | BT20/MB231 | CYTO |
| >MI0000440\|hsa-mir-27b&WithFlank&9\|+\|95085439\|95085547@67.87.21(+1G)&[MIMAT0000419&hsa-miR-27b-3p&offsets\|0\|0(+1G);m-39&9\|+\|95085505\|95085525&offsets\|0\|0(+1G)]\|TTCACAGTGGCTAAGTTCTGCG | >MI0000440\|hsa-mir-27b&WithFlank&9\|+\|95085439\|95085547 | 67 | 87 | 21(+1G) | TTCACAGTGGCTAAGTTCTGCG | 297.6 | 2.7 | 0.9 | 3.0 | 2.7E-03 | 1.1E-02 | BT20/MB231 | CYTO |
| >MI0006383\|hsa-mir-1248&WithFlank&3\|+\|186786666\|186786783@10.32.23&[MIMAT0005900&hsa-miR-1248&offsets\|0\|-4;m-827&3\|+\|186786675\|186786701&offsets\|0\|-4]\|ACCTTCTTGTATAAGCACTGTGC | >MI0006383\|hsa-mir-1248&WithFlank&3\|+\|186786666\|186786783 | 10 | 32 | 23 | ACCTTCTTGTATAAGCACTGTGC | 367.2 | 2.6 | 0.9 | 3.0 | 2.7E-03 | 1.1E-02 | BT20/MB231 | CYTO |
| >hg38_21_+_8433172_8446622_RNA45SN1_45S_50ntFlanks@9534.9571.38\|AGGAAACTCTGGTGGAGGTCCGTAGCGGTCCTGACGTG | >hg38_21_+_8433172_8446622_RNA45SN1_45S_50ntFlanks | 9534 | 9571 | 38 | AGGAAACTCTGGTGGAGGTCCGTAGCGGTCCTGACGTG | 1981.0 | -2.8 | 0.9 | -3.0 | 2.7E-03 | 1.1E-02 | BT20/MB231 | CYTO |
| >hg38_21_+_8433172_8446622_RNA45SN1_45S_50ntFlanks@6721.6760.40\|AATTGCAGGACACATTGATCATCGACACTTCGAACGCACT | >hg38_21_+_8433172_8446622_RNA45SN1_45S_50ntFlanks | 6721 | 6760 | 40 | AATTGCAGGACACATTGATCATCGACACTTCGAACGCACT | 231.3 | -2.6 | 0.9 | -3.0 | 2.7E-03 | 1.1E-02 | BT20/MB231 | CYTO |
| >hg38_21_+_8433172_8446622_RNA45SN1_45S_50ntFlanks@10351.10387.37\|CGAGAACTTTGAAGGCCGAAGTGGAGAAGGGTTCCAT | >hg38_21_+_8433172_8446622_RNA45SN1_45S_50ntFlanks | 10351 | 10387 | 37 | CGAGAACTTTGAAGGCCGAAGTGGAGAAGGGTTCCAT | 140.7 | -4.7 | 1.7 | -2.8 | 5.6E-03 | 1.1E-02 | BT20/MB468 | MP |
| >am_MI0000786\|hsa-mir-378a&WithFlank&5\|+\|149732819\|149732896@50.70.21(+1G)&[MIMAT0000732&hsa-miR-378a-3p&offsets\|+1\|0(+1G);m-44&5\|+\|149732867\|149732888&offsets\|+1\|0(+1G)]\|CTGGACTTGGAGTCAGAAGGCG | >am_MI0000786\|hsa-mir-378a&WithFlank&5\|+\|149732819\|149732896 | 50 | 70 | 21(+1G) | CTGGACTTGGAGTCAGAAGGCG | 294.0 | 3.1 | 1.1 | 2.8 | 5.6E-03 | 1.1E-02 | BT20/MB468 | MP |
| >hg38_21_+_8433172_8446622_RNA45SN1_45S_50ntFlanks@10350.10373.24\|ACGAGAACTTTGAAGGCCGAAGTG | >hg38_21_+_8433172_8446622_RNA45SN1_45S_50ntFlanks | 10350 | 10373 | 24 | ACGAGAACTTTGAAGGCCGAAGTG | 616.9 | -1.6 | 0.6 | -2.6 | 9.3E-03 | 1.1E-02 | MB231/MB468 | CYTO |
| >hg38_MT_+_598_1651_RNR1_12S_50ntFlanks@438.468.31\|TAACATATCTGAACACACAATAGCTAAGACC | >hg38_MT_+_598_1651_RNR1_12S_50ntFlanks | 438 | 468 | 31 | TAACATATCTGAACACACAATAGCTAAGACC | 23.5 | 7.8 | 3.0 | 2.6 | 8.5E-03 | 1.1E-02 | MB231/MB468 | MP |
| >hg38_21_+_8433172_8446622_RNA45SN1_45S_50ntFlanks@8528.8551.24\|CTTCCCCCGCCGCCCCTCCTCCTC | >hg38_21_+_8433172_8446622_RNA45SN1_45S_50ntFlanks | 8528 | 8551 | 24 | CTTCCCCCGCCGCCCCTCCTCCTC | 535.4 | -4.3 | 1.7 | -2.6 | 9.3E-03 | 1.1E-02 | MB231/MB468 | CYTO |
| >hg38_21_+_8433172_8446622_RNA45SN1_45S_50ntFlanks@6785.6807.23\|CTACGCCTGTCTGAGCGTCGCTT | >hg38_21_+_8433172_8446622_RNA45SN1_45S_50ntFlanks | 6785 | 6807 | 23 | CTACGCCTGTCTGAGCGTCGCTT | 235.6 | -1.9 | 0.7 | -2.6 | 8.5E-03 | 1.1E-02 | MB231/MB468 | MP |
| >hg38_21_+_8433172_8446622_RNA45SN1_45S_50ntFlanks@5515.5532.18\|CTAGAGGAAGTAAAAGTC | >hg38_21_+_8433172_8446622_RNA45SN1_45S_50ntFlanks | 5515 | 5532 | 18 | CTAGAGGAAGTAAAAGTC | 140.5 | -4.3 | 1.7 | -2.6 | 8.5E-03 | 1.1E-02 | MB231/MB468 | MP |
| >MI0000102\|hsa-mir-100&WithFlank&11\|-\|122152223\|122152314@22.40.19(+1U)&[MIMAT0000098&hsa-miR-100-5p&offsets\|+3\|0(+1U);m-27&11\|-\|122152276\|122152296&offsets\|+3\|+1(+1U)]\|CCGTAGATCCGAACTTGTGT | >MI0000102\|hsa-mir-100&WithFlank&11\|-\|122152223\|122152314 | 22 | 40 | 19(+1U) | CCGTAGATCCGAACTTGTGT | 48.7 | -9.0 | 2.9 | -3.1 | 2.1E-03 | 1.1E-02 | BT20/MB231 | MP |
| >hg38_21_+_8433172_8446622_RNA45SN1_45S_50ntFlanks@10476.10493.18\|GGCCGATCGAAAGGGAGT | >hg38_21_+_8433172_8446622_RNA45SN1_45S_50ntFlanks | 10476 | 10493 | 18 | GGCCGATCGAAAGGGAGT | 91.7 | -4.3 | 1.6 | -2.6 | 8.6E-03 | 1.1E-02 | MB231/MB468 | MP |
| >hg38_21_+_8433172_8446622_RNA45SN1_45S_50ntFlanks@9929.9946.18\|GATATAGACAGCAGGACG | >hg38_21_+_8433172_8446622_RNA45SN1_45S_50ntFlanks | 9929 | 9946 | 18 | GATATAGACAGCAGGACG | 137.0 | 4.5 | 1.5 | 3.1 | 2.1E-03 | 1.1E-02 | BT20/MB231 | MP |
| >hg38_21_+_8433172_8446622_RNA45SN1_45S_50ntFlanks@6664.6686.23\|TGGATCACTCGGCTCGTGCGTCG | >hg38_21_+_8433172_8446622_RNA45SN1_45S_50ntFlanks | 6664 | 6686 | 23 | TGGATCACTCGGCTCGTGCGTCG | 443.0 | -3.8 | 1.4 | -2.8 | 5.6E-03 | 1.1E-02 | BT20/MB468 | MP |
| >MI0003137\|hsa-mir-193b&WithFlank&16\|+\|14303961\|14304055@57.77.21(+1G)&[MIMAT0002819&hsa-miR-193b-3p&offsets\|0\|-1(+1G);m-166&16\|+\|14304017\|14304038&offsets\|0\|-1(+1G)]\|AACTGGCCCTCAAAGTCCCGCG | >MI0003137\|hsa-mir-193b&WithFlank&16\|+\|14303961\|14304055 | 57 | 77 | 21(+1G) | AACTGGCCCTCAAAGTCCCGCG | 178.2 | 5.2 | 1.9 | 2.8 | 5.7E-03 | 1.1E-02 | BT20/MB468 | MP |
| >trnaMT_HisGTG_MT_+_12138_12206@32.61.30\|TGAATCTGACAACAGAGGCTTACGACCCCT | >trnaMT_HisGTG_MT_+_12138_12206 | 32 | 61 | 30 | TGAATCTGACAACAGAGGCTTACGACCCCT | 401.7 | 2.5 | 1.0 | 2.6 | 9.9E-03 | 1.1E-02 | MB231/MB468 | MITO |
| >hg38_21_+_8433172_8446622_RNA45SN1_45S_50ntFlanks@12686.12708.23\|CCAGGCGGAACGATACGGCAGCG | >hg38_21_+_8433172_8446622_RNA45SN1_45S_50ntFlanks | 12686 | 12708 | 23 | CCAGGCGGAACGATACGGCAGCG | 132.2 | -4.5 | 1.6 | -2.8 | 5.7E-03 | 1.1E-02 | BT20/MB468 | MP |
| >MI0000085\|hsa-mir-27a&WithFlank&19\|-\|13836434\|13836523@57.76.20(+1U)&[MIMAT0000084&hsa-miR-27a-3p&offsets\|0\|-1(+1U);m-43&19\|-\|13836447\|13836467&offsets\|0\|-1(+1U)]\|TTCACAGTGGCTAAGTTCCGT | >MI0000085\|hsa-mir-27a&WithFlank&19\|-\|13836434\|13836523 | 57 | 76 | 20(+1U) | TTCACAGTGGCTAAGTTCCGT | 20605.0 | -3.5 | 1.1 | -3.1 | 2.2E-03 | 1.1E-02 | BT20/MB231 | MP |
| >hg38_21_+_8433172_8446622_RNA45SN1_45S_50ntFlanks@7989.8013.25\|AGACGTGGCGACCCGCTGAATTTAA | >hg38_21_+_8433172_8446622_RNA45SN1_45S_50ntFlanks | 7989 | 8013 | 25 | AGACGTGGCGACCCGCTGAATTTAA | 257.8 | 3.7 | 1.2 | 3.1 | 2.2E-03 | 1.1E-02 | BT20/MB231 | MP |
| >trna10_AlaCGC_6_+_26553731_26553802@1.19.19,trna119_AlaCGC_6_-_28641613_28641684@1.19.19,trna120_AlaAGC_6_-_28626014_28626085@1.19.19,trna13_AlaCGC_2_+_157257281_157257352@1.19.19,trna13_AlaTGC_12_-_125406301_125406372@1.19.19,trna66_AlaTGC_6_+_28611222_28611293@1.19.19,trna8_AlaTGC_12_+_125424512_125424583@1.19.19,trna8_AlaTGC_5_+_180633868_180633939@1.19.19\|GGGGATGTAGCTCAGTGGT | >trna10_AlaCGC_6_+_26553731_26553802 | 1 | 19 | 19 | GGGGATGTAGCTCAGTGGT | 418.9 | 3.0 | 1.1 | 2.8 | 5.7E-03 | 1.1E-02 | BT20/MB468 | MP |
| >MI0000073\|hsa-mir-19a&WithFlank&13\|+\|91350885\|91350978@55.76.22(+1U)&[MIMAT0000073&hsa-miR-19a-3p&offsets\|0\|-1(+1U);m-201&13\|+\|91350939\|91350961&offsets\|0\|-1(+1U)]\|TGTGCAAATCTATGCAAAACTGT | >MI0000073\|hsa-mir-19a&WithFlank&13\|+\|91350885\|91350978 | 55 | 76 | 22(+1U) | TGTGCAAATCTATGCAAAACTGT | 821.4 | 2.4 | 0.9 | 2.6 | 8.6E-03 | 1.1E-02 | MB231/MB468 | MP |
| >hg38_21_+_8433172_8446622_RNA45SN1_45S_50ntFlanks@10058.10076.19\|CGGCAGTCGAGAGTGGACG | >hg38_21_+_8433172_8446622_RNA45SN1_45S_50ntFlanks | 10058 | 10076 | 19 | CGGCAGTCGAGAGTGGACG | 136.9 | 3.8 | 1.4 | 2.8 | 5.7E-03 | 1.1E-02 | BT20/MB468 | MP |
| >MI0003763\|hsa-mir-767&WithFlank&X\|-\|152393415\|152393535@33.54.22&[MIMAT0003882&hsa-miR-767-5p&offsets\|0\|-1;m-788&X\|-\|152393481\|152393503&offsets\|0\|-1]\|TGCACCATGGTTGTCTGAGCAT | >MI0003763\|hsa-mir-767&WithFlank&X\|-\|152393415\|152393535 | 33 | 54 | 22 | TGCACCATGGTTGTCTGAGCAT | 157.4 | 3.1 | 1.1 | 2.8 | 5.7E-03 | 1.1E-02 | BT20/MB468 | MP |
| >hg38_21_+_8433172_8446622_RNA45SN1_45S_50ntFlanks@6663.6689.27\|GTGGATCACTCGGCTCGTGCGTCGATG | >hg38_21_+_8433172_8446622_RNA45SN1_45S_50ntFlanks | 6663 | 6689 | 27 | GTGGATCACTCGGCTCGTGCGTCGATG | 520.2 | 5.1 | 1.7 | 3.1 | 2.2E-03 | 1.1E-02 | BT20/MB231 | MP |
| >hg38_21_+_8433172_8446622_RNA45SN1_45S_50ntFlanks@12455.12487.33\|ACCAAGCGTTGGATTGTTCACCCACTAATAGGG | >hg38_21_+_8433172_8446622_RNA45SN1_45S_50ntFlanks | 12455 | 12487 | 33 | ACCAAGCGTTGGATTGTTCACCCACTAATAGGG | 206.1 | -4.1 | 1.6 | -2.6 | 9.3E-03 | 1.1E-02 | BT20/MB468 | CYTO |
| >hg38_21_+_8433172_8446622_RNA45SN1_45S_50ntFlanks@11319.11343.25\|CCCACGTCTCGTCGCGCGCGCGTCC | >hg38_21_+_8433172_8446622_RNA45SN1_45S_50ntFlanks | 11319 | 11343 | 25 | CCCACGTCTCGTCGCGCGCGCGTCC | 281.2 | -2.5 | 1.0 | -2.6 | 9.4E-03 | 1.1E-02 | MB231/MB468 | CYTO |
| >hg38_21_+_8433172_8446622_RNA45SN1_45S_50ntFlanks@12628.12661.34\|TGGGGCGAAGCTACCATCTGTGGGATTATGACTG | >hg38_21_+_8433172_8446622_RNA45SN1_45S_50ntFlanks | 12628 | 12661 | 34 | TGGGGCGAAGCTACCATCTGTGGGATTATGACTG | 1536.4 | -4.4 | 1.6 | -2.8 | 5.7E-03 | 1.1E-02 | BT20/MB468 | MP |
| >hg38_21_+_8433172_8446622_RNA45SN1_45S_50ntFlanks@4726.4767.42\|TAATCAAGAACGAAAGTCGGAGGTTCGAAGACGATCAGATAC | >hg38_21_+_8433172_8446622_RNA45SN1_45S_50ntFlanks | 4726 | 4767 | 42 | TAATCAAGAACGAAAGTCGGAGGTTCGAAGACGATCAGATAC | 887.1 | -2.8 | 0.9 | -3.0 | 2.7E-03 | 1.1E-02 | BT20/MB231 | CYTO |
| >MI0000108\|hsa-mir-103a-2&WithFlank&20\|+\|3917488\|3917577@54.76.23(+1C);MI0000109\|hsa-mir-103a-1&WithFlank&5\|-\|168560890\|168560979@54.76.23(+1C)&[MIMAT0000101&hsa-miR-103a-3p&offsets\|0\|0(+1C);m-12&20\|+\|3917541\|3917563&offsets\|0\|0(+1C)];[MIMAT0000101_1&hsa-miR-103a-3p&offsets\|0\|0(+1C);m-13&5\|-\|168560904\|168560926&offsets\|0\|0(+1C);m-44220&5\|-\|168560908\|168560926&offsets\|0\|+4(+1C)]\|AGCAGCATTGTACAGGGCTATGAC | >MI0000108\|hsa-mir-103a-2&WithFlank&20\|+\|3917488\|3917577 | 54 | 76 | 23(+1C) | AGCAGCATTGTACAGGGCTATGAC | 774.6 | -2.6 | 0.9 | -3.0 | 2.8E-03 | 1.1E-02 | BT20/MB231 | MITO |
| >hg38_21_+_8433172_8446622_RNA45SN1_45S_50ntFlanks@12419.12448.30\|GATGTCGGCTCTTCCTATCATTGTGAAGCA | >hg38_21_+_8433172_8446622_RNA45SN1_45S_50ntFlanks | 12419 | 12448 | 30 | GATGTCGGCTCTTCCTATCATTGTGAAGCA | 8482.4 | -2.7 | 0.9 | -3.0 | 2.7E-03 | 1.1E-02 | BT20/MB231 | CYTO |
| >hg38_1_-_228634819_228635039_RNA5S12_5S_50ntFlanks@85.103.19\|TCGTCTGATCTCGGAAGCT | >hg38_1_-_228634819_228635039_RNA5S12_5S_50ntFlanks | 85 | 103 | 19 | TCGTCTGATCTCGGAAGCT | 197.5 | 4.2 | 1.5 | 2.8 | 5.7E-03 | 1.1E-02 | BT20/MB468 | MP |
| >hg38_MT_+_1621_3279_RNR2_16S_50ntFlanks@719.740.22\|GCCTGCGTCAGATTAAAACACT | >hg38_MT_+_1621_3279_RNR2_16S_50ntFlanks | 719 | 740 | 22 | GCCTGCGTCAGATTAAAACACT | 23.4 | 7.8 | 3.0 | 2.6 | 8.6E-03 | 1.1E-02 | MB231/MB468 | MP |
| >hg38_21_+_8433172_8446622_RNA45SN1_45S_50ntFlanks@6649.6698.50\|TACGACTCTTAGCGGTGGATCACTCGGCTCGTGCGTCGATGAAGAACGCA | >hg38_21_+_8433172_8446622_RNA45SN1_45S_50ntFlanks | 6649 | 6698 | 50 | TACGACTCTTAGCGGTGGATCACTCGGCTCGTGCGTCGATGAAGAACGCA | 228.2 | -5.9 | 2.2 | -2.6 | 8.6E-03 | 1.1E-02 | MB231/MB468 | MP |
| >hg38_21_+_8433172_8446622_RNA45SN1_45S_50ntFlanks@12463.12496.34\|TTGGATTGTTCACCCACTAATAGGGAACGTGAGC | >hg38_21_+_8433172_8446622_RNA45SN1_45S_50ntFlanks | 12463 | 12496 | 34 | TTGGATTGTTCACCCACTAATAGGGAACGTGAGC | 55.6 | 4.6 | 1.7 | 2.8 | 5.7E-03 | 1.1E-02 | BT20/MB468 | MP |
| >trna119_LysCTT_1_-_145395522_145395594@2.33.32,trna11_LysCTT_5_-_180648979_180649051@2.33.32,trna13_LysCTT_6_+_26556774_26556846@2.33.32,trna7_LysCTT_16_+_3225692_3225764@2.33.32,trna9_LysCTT_5_+_180634755_180634827@2.33.32\|CCCGGCTAGCTCAGTCGGTAGAGCATGAGACT | >trna119_LysCTT_1_-_145395522_145395594 | 2 | 33 | 32 | CCCGGCTAGCTCAGTCGGTAGAGCATGAGACT | 337.7 | -5.4 | 1.8 | -3.0 | 2.7E-03 | 1.1E-02 | BT20/MB231 | CYTO |

<table>
<tr><th></th><th></th><th></th><th></th><th></th><th></th><th></th><th></th><th></th><th></th><th></th><th></th><th></th><th></th></tr>
<tr><td>>hg38_21_+_8433172_8446622_RNA45SN1_45S_50ntFlanks@12455.12475.21|ACCAAGCGTTGGATTGTTCAC</td><td>>hg38_21_+_8433172_8446622_RNA45SN1_45S_50ntFlanks</td><td>12455</td><td>12475</td><td>21</td><td>ACCAAGCGTTGGATTGTTCAC</td><td>77.7</td><td>-4.6</td><td>1.7</td><td>-2.8</td><td>5.7E-03</td><td>1.1E-02</td><td>BT20/MB468</td><td>MP</td></tr>
<tr><td>>am_trna128_GlyGCC_6_-_27870686_27870756@1.34.34,trna18_GlyGCC_16_+_70822597_70822667@1.34.34,trna19_GlyGCC_16_+_70823410_70823480@1.34.34,trna19_GlyGCC_2_-_157257659_157257729@1.34.34,trna24_GlyGCC_16_-_70812942_70813012@1.34.34,trna25_GlyGCC_16_-_70812114_70812184@1.34.34,trna5_GlyGCC_17_+_8029064_8029134@1.34.34,trna68_GlyGCC_1_-_161493637_161493707@1.34.34|GCATTGGTGGTTCAGTGGTAGAATTCTCGCCTGC</td><td>>am_trna128_GlyGCC_6_-_27870686_27870756</td><td>1</td><td>34</td><td>34</td><td>GCATTGGTGGTTCAGTGGTAGAATTCTCGCCTGC</td><td>290.3</td><td>3.5</td><td>1.3</td><td>2.8</td><td>5.7E-03</td><td>1.1E-02</td><td>BT20/MB468</td><td>MP</td></tr>
<tr><td>>am_trna2_GlyGCC_21_-_18827107_18827177@1.18.18,trna35_GlyGCC_1_+_161413094_161413164@1.18.18,trna37_GlyGCC_1_+_161420467_161420537@1.18.18,trna39_GlyGCC_1_+_161427898_161427968@1.18.18,trna41_GlyGCC_1_+_161435258_161435328@1.18.18|GCATGGGTGGTTCAGTGG</td><td>>am_trna2_GlyGCC_21_-_18827107_18827177</td><td>1</td><td>18</td><td>18</td><td>GCATGGGTGGTTCAGTGG</td><td>207.9</td><td>4.3</td><td>1.6</td><td>2.8</td><td>5.7E-03</td><td>1.1E-02</td><td>BT20/MB468</td><td>MP</td></tr>
<tr><td>>hg38_21_+_8433172_8446622_RNA45SN1_45S_50ntFlanks@6725.6744.20|GCAGGACACATTGATCATCG</td><td>>hg38_21_+_8433172_8446622_RNA45SN1_45S_50ntFlanks</td><td>6725</td><td>6744</td><td>20</td><td>GCAGGACACATTGATCATCG</td><td>479.9</td><td>-2.3</td><td>0.9</td><td>-2.6</td><td>9.9E-03</td><td>1.1E-02</td><td>MB231/MB468</td><td>NUC</td></tr>
<tr><td>>hg38_MT_+_598_1651_RNR1_12S_50ntFlanks@599.616.18|AGAGGAGCCTGTTCTGTA</td><td>>hg38_MT_+_598_1651_RNR1_12S_50ntFlanks</td><td>599</td><td>616</td><td>18</td><td>AGAGGAGCCTGTTCTGTA</td><td>170.2</td><td>3.0</td><td>1.1</td><td>2.8</td><td>5.7E-03</td><td>1.1E-02</td><td>BT20/MB468</td><td>MP</td></tr>
<tr><td>>MI0000812|hsa-mir-331&WithFlank&12|+|95308414|95308519@67.86.20&[MIMAT0000760&hsa-miR-331-3p&offsets|0|-1;m-192&12|+|95308480|95308501&offsets|0|-2]|GCCCCTGGGCCTATCCTAGA</td><td>>MI0000812|hsa-mir-331&WithFlank&12|+|95308414|95308519</td><td>67</td><td>86</td><td>20</td><td>GCCCCTGGGCCTATCCTAGA</td><td>288.4</td><td>2.6</td><td>0.8</td><td>3.0</td><td>2.4E-03</td><td>1.1E-02</td><td>BT20/MB231</td><td>NUC</td></tr>
<tr><td>>trnaMT_GluTTC_MT_-_14674_14742@46.72.27,trnalookalike8_GluTTC_5_-_93905172_93905240@46.72.27|GTGGTTGTAGTCCGTGCGAGAATACCA</td><td>>trnaMT_GluTTC_MT_-_14674_14742</td><td>46</td><td>72</td><td>27</td><td>GTGGTTGTAGTCCGTGCGAGAATACCA</td><td>560.9</td><td>-1.7</td><td>0.5</td><td>-3.0</td><td>2.4E-03</td><td>1.1E-02</td><td>BT20/MB231</td><td>NUC</td></tr>
<tr><td>>hg38_21_+_8433172_8446622_RNA45SN1_45S_50ntFlanks@10348.10390.43|AAACGAGAACTTTGAAGGCCGAAGTGGAGAAGGGTTCCATGTG</td><td>>hg38_21_+_8433172_8446622_RNA45SN1_45S_50ntFlanks</td><td>10348</td><td>10390</td><td>43</td><td>AAACGAGAACTTTGAAGGCCGAAGTGGAGAAGGGTTCCATGTG</td><td>91.0</td><td>-2.4</td><td>0.9</td><td>-2.6</td><td>8.7E-03</td><td>1.1E-02</td><td>MB231/MB468</td><td>MP</td></tr>
<tr><td>>hg38_21_+_8433172_8446622_RNA45SN1_45S_50ntFlanks@11565.11602.38|CTAGCAGCCGACTTAGAACTGGTGCGGACCAGGGGAAT</td><td>>hg38_21_+_8433172_8446622_RNA45SN1_45S_50ntFlanks</td><td>11565</td><td>11602</td><td>38</td><td>CTAGCAGCCGACTTAGAACTGGTGCGGACCAGGGGAAT</td><td>117.3</td><td>-4.7</td><td>1.8</td><td>-2.6</td><td>8.7E-03</td><td>1.1E-02</td><td>MB231/MB468</td><td>MP</td></tr>
<tr><td>>MI0000078|hsa-mir-22&WithFlank&17|-|1713897|1713993@59.78.20(+1A)&[MIMAT0000077&hsa-miR-22-3p&offsets|0|-2(+1A);m-2&17|-|1713914|1713935&offsets|0|-2(+1A)]|AAGCTGCCAGTTGAAGAACTA</td><td>>MI0000078|hsa-mir-22&WithFlank&17|-|1713897|1713993</td><td>59</td><td>78</td><td>20(+1A)</td><td>AAGCTGCCAGTTGAAGAACTA</td><td>24.7</td><td>7.8</td><td>3.0</td><td>2.6</td><td>8.7E-03</td><td>1.1E-02</td><td>MB231/MB468</td><td>MP</td></tr>
<tr><td>>hg38_21_+_8433172_8446622_RNA45SN1_45S_50ntFlanks@3705.3737.33|TACCTGGTTGATCCTGCCAGTAGCATATGCTTG</td><td>>hg38_21_+_8433172_8446622_RNA45SN1_45S_50ntFlanks</td><td>3705</td><td>3737</td><td>33</td><td>TACCTGGTTGATCCTGCCAGTAGCATATGCTTG</td><td>690.1</td><td>2.3</td><td>0.9</td><td>2.6</td><td>9.5E-03</td><td>1.1E-02</td><td>MB231/MB468</td><td>CYTO</td></tr>
<tr><td>>hg38_21_+_8433172_8446622_RNA45SN1_45S_50ntFlanks@12419.12451.33|GATGTCGGCTCTTCCTATCATTGTGAAGCAGAA</td><td>>hg38_21_+_8433172_8446622_RNA45SN1_45S_50ntFlanks</td><td>12419</td><td>12451</td><td>33</td><td>GATGTCGGCTCTTCCTATCATTGTGAAGCAGAA</td><td>250.2</td><td>-2.4</td><td>0.8</td><td>-3.0</td><td>2.7E-03</td><td>1.1E-02</td><td>BT20/MB231</td><td>CYTO</td></tr>
<tr><td>>MI0000088|hsa-mir-30a&WithFlank&6|-|71403545|71403627@12.34.23(+1A)&[MIMAT0000087&hsa-miR-30a-5p&offsets|0|+1(+1A);m-15&6|-|71403595|71403616&offsets|0|+1(+1A)]|TGTAAACATCCTCGACTGGAAGCA</td><td>>MI0000088|hsa-mir-30a&WithFlank&6|-|71403545|71403627</td><td>12</td><td>34</td><td>23(+1A)</td><td>TGTAAACATCCTCGACTGGAAGCA</td><td>51249.8</td><td>-3.3</td><td>1.1</td><td>-3.1</td><td>2.2E-03</td><td>1.1E-02</td><td>BT20/MB231</td><td>MP</td></tr>
<tr><td>>hg38_21_+_8433172_8446622_RNA45SN1_45S_50ntFlanks@6650.6673.24|ACGACTCTTAGCGGTGGATCACTC</td><td>>hg38_21_+_8433172_8446622_RNA45SN1_45S_50ntFlanks</td><td>6650</td><td>6673</td><td>24</td><td>ACGACTCTTAGCGGTGGATCACTC</td><td>434.7</td><td>3.9</td><td>1.3</td><td>3.1</td><td>2.2E-03</td><td>1.1E-02</td><td>BT20/MB231</td><td>MP</td></tr>
<tr><td>>MI0000749|hsa-mir-30e&WithFlank&1|+|40754349|40754452@65.86.22&[MIMAT0000693&hsa-miR-30e-3p&offsets|0|0;m-25&1|+|40754413|40754434&offsets|0|0]|CTTTCAGTCGGATGTTTACAGC</td><td>>MI0000749|hsa-mir-30e&WithFlank&1|+|40754349|40754452</td><td>65</td><td>86</td><td>22</td><td>CTTTCAGTCGGATGTTTACAGC</td><td>1780.5</td><td>-2.8</td><td>0.9</td><td>-3.1</td><td>2.2E-03</td><td>1.1E-02</td><td>BT20/MB231</td><td>MP</td></tr>
<tr><td>>hg38_21_+_8433172_8446622_RNA45SN1_45S_50ntFlanks@10800.10818.19|TGTTGGAACAATGTAGGTA</td><td>>hg38_21_+_8433172_8446622_RNA45SN1_45S_50ntFlanks</td><td>10800</td><td>10818</td><td>19</td><td>TGTTGGAACAATGTAGGTA</td><td>91.2</td><td>-4.6</td><td>1.7</td><td>-2.8</td><td>5.7E-03</td><td>1.1E-02</td><td>BT20/MB468</td><td>MP</td></tr>
<tr><td>>hg38_21_+_8433172_8446622_RNA45SN1_45S_50ntFlanks@12941.12958.18|ATTCGTAGACGACCTGCT</td><td>>hg38_21_+_8433172_8446622_RNA45SN1_45S_50ntFlanks</td><td>12941</td><td>12958</td><td>18</td><td>ATTCGTAGACGACCTGCT</td><td>96.1</td><td>-3.3</td><td>1.3</td><td>-2.6</td><td>8.7E-03</td><td>1.1E-02</td><td>MB231/MB468</td><td>MP</td></tr>
<tr><td>>hg38_21_+_8433172_8446622_RNA45SN1_45S_50ntFlanks@9738.9765.28|GAAACGATCTCAACCTATTCTCAAACTT</td><td>>hg38_21_+_8433172_8446622_RNA45SN1_45S_50ntFlanks</td><td>9738</td><td>9765</td><td>28</td><td>GAAACGATCTCAACCTATTCTCAAACTT</td><td>632.7</td><td>-3.5</td><td>1.3</td><td>-2.6</td><td>9.4E-03</td><td>1.1E-02</td><td>BT20/MB468</td><td>CYTO</td></tr>
<tr><td>>MI0000289|hsa-mir-181a-1&WithFlank&1|-|198859038|198859159@30.51.22(+1C);MI0000269|hsa-mir-181a-2&WithFlank&9|+|124692436|124692557@45.66.22(+1C)&[MIMAT0000256&hsa-miR-181a-5p&offsets|0|-1(+1C);m-32&1|-|198859108|198859130&offsets|0|-1(+1C)];[MIMAT0000256_1&hsa-miR-181a-5p&offsets|0|-1(+1C);m-32&9|+|124692480|124692502&offsets|0|-1(+1C)]|AACATTCAACGCTGTCGGTGAGC</td><td>>MI0000289|hsa-mir-181a-1&WithFlank&1|-|198859038|198859159</td><td>30</td><td>51</td><td>22(+1C)</td><td>AACATTCAACGCTGTCGGTGAGC</td><td>69.8</td><td>1.7</td><td>0.6</td><td>2.6</td><td>9.8E-03</td><td>1.1E-02</td><td>BT20/MB468</td><td>NUC</td></tr>
<tr><td>>hg38_21_+_8433172_8446622_RNA45SN1_45S_50ntFlanks@8359.8378.20|AAGAACTTTGAAGAGAGAGT</td><td>>hg38_21_+_8433172_8446622_RNA45SN1_45S_50ntFlanks</td><td>8359</td><td>8378</td><td>20</td><td>AAGAACTTTGAAGAGAGAGT</td><td>334.5</td><td>-2.3</td><td>0.9</td><td>-2.6</td><td>1.0E-02</td><td>1.1E-02</td><td>MB231/MB468</td><td>NUC</td></tr>
<tr><td>>hg38_21_+_8433172_8446622_RNA45SN1_45S_50ntFlanks@12071.12092.22|CCCCGAGGGGCTCTCGCTTCTG</td><td>>hg38_21_+_8433172_8446622_RNA45SN1_45S_50ntFlanks</td><td>12071</td><td>12092</td><td>22</td><td>CCCCGAGGGGCTCTCGCTTCTG</td><td>458.0</td><td>-1.6</td><td>0.6</td><td>-2.6</td><td>9.5E-03</td><td>1.1E-02</td><td>MB231/MB468</td><td>CYTO</td></tr>
<tr><td>>trna119_LysCTT_1_-_145395522_145395594@1.34.34,trna11_LysCTT_5_-_180648979_180649051@1.34.34,trna13_LysCTT_6_+_26556774_26556846@1.34.34,trna7_LysCTT_16_+_3225692_3225764@1.34.34,trna9_LysCTT_5_+_180634755_180634827@1.34.34|GCCCGGCTAGCTCAGTCGGTAGAGCATGAGACTC</td><td>>trna119_LysCTT_1_-_145395522_145395594</td><td>1</td><td>34</td><td>34</td><td>GCCCGGCTAGCTCAGTCGGTAGAGCATGAGACTC</td><td>886.3</td><td>4.0</td><td>1.4</td><td>2.8</td><td>5.7E-03</td><td>1.1E-02</td><td>BT20/MB468</td><td>MP</td></tr>
<tr><td>>hg38_21_+_8433172_8446622_RNA45SN1_45S_50ntFlanks@6658.6683.26|TAGCGGTGGATCACTCGGCTCGTGCG</td><td>>hg38_21_+_8433172_8446622_RNA45SN1_45S_50ntFlanks</td><td>6658</td><td>6683</td><td>26</td><td>TAGCGGTGGATCACTCGGCTCGTGCG</td><td>256.5</td><td>-2.3</td><td>0.8</td><td>-3.0</td><td>2.4E-03</td><td>1.1E-02</td><td>BT20/MB231</td><td>NUC</td></tr>
<tr><td>>hg38_MT_+_598_1651_RNR1_12S_50ntFlanks@428.452.25|AAAGTGGCTTTAACATATCTGAACA</td><td>>hg38_MT_+_598_1651_RNR1_12S_50ntFlanks</td><td>428</td><td>452</td><td>25</td><td>AAAGTGGCTTTAACATATCTGAACA</td><td>397.0</td><td>3.0</td><td>1.1</td><td>2.6</td><td>8.7E-03</td><td>1.1E-02</td><td>MB231/MB468</td><td>MP</td></tr>
<tr><td>>trna16_LeuAAG_16_+_22308461_22308542@1.18.18,trna16_LeuAAG_5_-_180601044_180601125@1.18.18,trna19_LeuAAG_5_-_180524474_180524555@1.18.18,trna1_LeuAAG_14_+_21078291_21078372@1.18.18,trna3_LeuAAG_5_+_180528840_180528921@1.18.18,trna42_LeuTAG_17_-_8023632_8023713@1.18.18,trna78_LeuAAG_6_+_28956779_28956860@1.18.18,trna7_LeuAAG_5_+_180614701_180614782@1.18.18,trna98_LeuAAG_6_-_28911399_28911480@1.18.18|GGTAGCGTGGCCGAGCGG</td><td>>trna16_LeuAAG_16_+_22308461_22308542</td><td>1</td><td>18</td><td>18</td><td>GGTAGCGTGGCCGAGCGG</td><td>193.1</td><td>4.9</td><td>1.8</td><td>2.8</td><td>5.7E-03</td><td>1.1E-02</td><td>BT20/MB468</td><td>MP</td></tr>
<tr><td>>hg38_21_+_8433172_8446622_RNA45SN1_45S_50ntFlanks@5312.5330.19|TCGGGGATTGCAATTATTC</td><td>>hg38_21_+_8433172_8446622_RNA45SN1_45S_50ntFlanks</td><td>5312</td><td>5330</td><td>19</td><td>TCGGGGATTGCAATTATTC</td><td>82.4</td><td>-4.8</td><td>1.7</td><td>-2.8</td><td>5.7E-03</td><td>1.1E-02</td><td>BT20/MB468</td><td>MP</td></tr>
<tr><td>>MI0000469|hsa-mir-125a&WithFlank&19|+|51693248|51693345@21.42.22(+1U)&[MIMAT0000443&hsa-miR-125a-5p&offsets|0|-2(+1U);m-63&19|+|51693268|51693289&offsets|0|0(+1U)]|TCCCTGAGACCCTTTAACCTGTT</td><td>>MI0000469|hsa-mir-125a&WithFlank&19|+|51693248|51693345</td><td>21</td><td>42</td><td>22(+1U)</td><td>TCCCTGAGACCCTTTAACCTGTT</td><td>456.6</td><td>2.7</td><td>0.9</td><td>3.0</td><td>2.8E-03</td><td>1.1E-02</td><td>BT20/MB231</td><td>CYTO</td></tr>
<tr><td>>am_trnaMT_PheGAA_MT_+_577_647@45.62.18|AGACGGGCTCACATCACC</td><td>>am_trnaMT_PheGAA_MT_+_577_647</td><td>45</td><td>62</td><td>18</td><td>AGACGGGCTCACATCACC</td><td>662.7</td><td>2.3</td><td>0.8</td><td>3.0</td><td>2.8E-03</td><td>1.1E-02</td><td>BT20/MB231</td><td>CYTO</td></tr>
<tr><td>>hg38_21_+_8433172_8446622_RNA45SN1_45S_50ntFlanks@7974.8013.40|ACGCGACCTCAGATCAGACGTGGCGACCCGCTGAATTTAA</td><td>>hg38_21_+_8433172_8446622_RNA45SN1_45S_50ntFlanks</td><td>7974</td><td>8013</td><td>40</td><td>ACGCGACCTCAGATCAGACGTGGCGACCCGCTGAATTTAA</td><td>1773.7</td><td>-2.7</td><td>0.9</td><td>-3.0</td><td>2.8E-03</td><td>1.1E-02</td><td>BT20/MB231</td><td>CYTO</td></tr>
<tr><td>>hg38_MT_+_1621_3279_RNR2_16S_50ntFlanks@1387.1414.28|CCCGATGGTGCAGCCGCTATTAAAGGTT</td><td>>hg38_MT_+_1621_3279_RNR2_16S_50ntFlanks</td><td>1387</td><td>1414</td><td>28</td><td>CCCGATGGTGCAGCCGCTATTAAAGGTT</td><td>23.3</td><td>7.8</td><td>3.0</td><td>2.6</td><td>8.7E-03</td><td>1.1E-02</td><td>MB231/MB468</td><td>MP</td></tr>
<tr><td>>hg38_21_+_8433172_8446622_RNA45SN1_45S_50ntFlanks@3854.3884.31|ACTGTGGTAATTCTAGAGCTAATACATGCCG</td><td>>hg38_21_+_8433172_8446622_RNA45SN1_45S_50ntFlanks</td><td>3854</td><td>3884</td><td>31</td><td>ACTGTGGTAATTCTAGAGCTAATACATGCCG</td><td>233.8</td><td>-1.5</td><td>0.6</td><td>-2.6</td><td>9.4E-03</td><td>1.1E-02</td><td>BT20/MB468</td><td>CYTO</td></tr>
<tr><td>>am_trnaMT_ArgTCG_MT_+_10405_10469@29.48.20|TTTCGACTCATTAAATTATG</td><td>>am_trnaMT_ArgTCG_MT_+_10405_10469</td><td>29</td><td>48</td><td>20</td><td>TTTCGACTCATTAAATTATG</td><td>206.5</td><td>-3.5</td><td>1.4</td><td>-2.6</td><td>9.4E-03</td><td>1.1E-02</td><td>BT20/MB468</td><td>CYTO</td></tr>
<tr><td>>hg38_21_+_8433172_8446622_RNA45SN1_45S_50ntFlanks@4452.4471.20|CTCGGCGCCCCCTCGATGCT</td><td>>hg38_21_+_8433172_8446622_RNA45SN1_45S_50ntFlanks</td><td>4452</td><td>4471</td><td>20</td><td>CTCGGCGCCCCCTCGATGCT</td><td>115.2</td><td>-3.9</td><td>1.5</td><td>-2.6</td><td>8.1E-03</td><td>1.1E-02</td><td>BT20/MB468</td><td>MITO</td></tr>
<tr><td>>hg38_21_+_8433172_8446622_RNA45SN1_45S_50ntFlanks@10681.10712.32|TTCCGGCGGCGTCCGGTGAGCTCTCGCTGGCC</td><td>>hg38_21_+_8433172_8446622_RNA45SN1_45S_50ntFlanks</td><td>10681</td><td>10712</td><td>32</td><td>TTCCGGCGGCGTCCGGTGAGCTCTCGCTGGCC</td><td>317.7</td><td>-4.7</td><td>1.7</td><td>-2.8</td><td>5.8E-03</td><td>1.1E-02</td><td>BT20/MB468</td><td>MP</td></tr>
<tr><td>>MI0000813|hsa-mir-324&WithFlank&17|-|7223291|7223385@59.78.20(+2U)&[MIMAT0000762&hsa-miR-324-3p&offsets|+3|0(+2U);m-203&17|-|7223308|7223329&offsets|+2|0(+2U)]|ACTGCCCCAGGTGCTGCTGGTT</td><td>>MI0000813|hsa-mir-324&WithFlank&17|-|7223291|7223385</td><td>59</td><td>78</td><td>20(+2U)</td><td>ACTGCCCCAGGTGCTGCTGGTT</td><td>97.7</td><td>4.3</td><td>1.6</td><td>2.6</td><td>8.7E-03</td><td>1.1E-02</td><td>MB231/MB468</td><td>MP</td></tr>
<tr><td>>hg38_21_+_8433172_8446622_RNA45SN1_45S_50ntFlanks@8380.8403.24|CAAGAGGGCGTGAAACCGTTAAGA</td><td>>hg38_21_+_8433172_8446622_RNA45SN1_45S_50ntFlanks</td><td>8380</td><td>8403</td><td>24</td><td>CAAGAGGGCGTGAAACCGTTAAGA</td><td>1694.8</td><td>-2.9</td><td>1.1</td><td>-2.8</td><td>5.8E-03</td><td>1.1E-02</td><td>BT20/MB468</td><td>MP</td></tr>
<tr><td>>hg38_1_-_228634819_228635039_RNA5S12_5S_50ntFlanks@132.171.40|GATGGGAGACCGCCTGGGAATACCGGGTGCTGTAGGCTTT</td><td>>hg38_1_-_228634819_228635039_RNA5S12_5S_50ntFlanks</td><td>132</td><td>171</td><td>40</td><td>GATGGGAGACCGCCTGGGAATACCGGGTGCTGTAGGCTTT</td><td>14090.5</td><td>3.1</td><td>1.2</td><td>2.6</td><td>8.1E-03</td><td>1.1E-02</td><td>BT20/MB468</td><td>MITO</td></tr>
<tr><td>>hg38_21_+_8433172_8446622_RNA45SN1_45S_50ntFlanks@8177.8211.35|TTCTGATCGAGGCCCAGCCCGTGGACGGTGTGAGG</td><td>>hg38_21_+_8433172_8446622_RNA45SN1_45S_50ntFlanks</td><td>8177</td><td>8211</td><td>35</td><td>TTCTGATCGAGGCCCAGCCCGTGGACGGTGTGAGG</td><td>538.2</td><td>-5.1</td><td>2.0</td><td>-2.6</td><td>9.9E-03</td><td>1.1E-02</td><td>BT20/MB468</td><td>NUC</td></tr>
<tr><td>>hg38_21_+_8433172_8446622_RNA45SN1_45S_50ntFlanks@9940.9975.36|CAGGACGGTGGCCATGGAAGTCGGAATCCGCTAAGG</td><td>>hg38_21_+_8433172_8446622_RNA45SN1_45S_50ntFlanks</td><td>9940</td><td>9975</td><td>36</td><td>CAGGACGGTGGCCATGGAAGTCGGAATCCGCTAAGG</td><td>210.5</td><td>3.5</td><td>1.1</td><td>3.1</td><td>2.2E-03</td><td>1.1E-02</td><td>BT20/MB231</td><td>MP</td></tr>
<tr><td>>Antisense_hg38_21_-_8433172_8446622_RNA45SN1_45S_50ntFlanks@7674.7691.18|CTCTCTCTCTCTCTCCCC</td><td>>Antisense_hg38_21_-_8433172_8446622_RNA45SN1_45S_50ntFlanks</td><td>7674</td><td>7691</td><td>18</td><td>CTCTCTCTCTCTCTCCCC</td><td>575.6</td><td>-5.2</td><td>2.0</td><td>-2.6</td><td>9.9E-03</td><td>1.1E-02</td><td>BT20/MB468</td><td>NUC</td></tr>
<tr><td>>hg38_21_+_8433172_8446622_RNA45SN1_45S_50ntFlanks@11027.11044.18|CTCTCTCTCTCTCTCCCC</td><td>>hg38_21_+_8433172_8446622_RNA45SN1_45S_50ntFlanks</td><td>11027</td><td>11044</td><td>18</td><td>CTCTCTCTCTCTCTCCCC</td><td>575.6</td><td>-5.2</td><td>2.0</td><td>-2.6</td><td>9.9E-03</td><td>1.1E-02</td><td>BT20/MB468</td><td>NUC</td></tr>
<tr><td>>am_trna128_GlyGCC_6_-_27870686_27870756@3.35.33,trna18_GlyGCC_16_+_70822597_70822667@3.35.33,trna19_GlyGCC_16_+_70823410_70823480@3.35.33,trna19_GlyGCC_2_-_157257659_157257729@3.35.33,trna24_GlyGCC_16_-_70812942_70813012@3.35.33,trna25_GlyGCC_16_-_70812114_70812184@3.35.33,trna5_GlyGCC_17_+_8029064_8029134@3.35.33,trna68_GlyGCC_1_-_161493637_161493707@3.35.33|ATTGGTGGTTCAGTGGTAGAATTCTCGCCTGCC</td><td>>am_trna128_GlyGCC_6_-_27870686_27870756</td><td>3</td><td>35</td><td>33</td><td>ATTGGTGGTTCAGTGGTAGAATTCTCGCCTGCC</td><td>531.9</td><td>-3.3</td><td>1.1</td><td>-3.1</td><td>2.2E-03</td><td>1.1E-02</td><td>BT20/MB231</td><td>MP</td></tr>
</table>

| | | | | | | | | | | | | | |
|---|---|---|---|---|---|---|---|---|---|---|---|---|---|
| >hg38_21_+_8433172_8446622_RNA45SN1_45S_50ntFlanks@6653.6689.37\|ACTCTTAGCGGTGGATCACTCGGCTCGTGCGTCGATG | >hg38_21_+_8433172_8446622_RNA45SN1_45S_50ntFlanks | 6653 | 6689 | 37 | ACTCTTAGCGGTGGATCACTCGGCTCGTGCGTCGATG | 783.3 | 3.7 | 1.2 | 3.1 | 2.2E-03 | 1.1E-02 | BT20/MB231 | MP |
| >hg38_1_-_228634819_228635039_RNA5S12_5S_50ntFlanks@51.88.38\|GTCTACGGCCATACCACCCTGAACGCGCCCGATCTCGT | >hg38_1_-_228634819_228635039_RNA5S12_5S_50ntFlanks | 51 | 88 | 38 | GTCTACGGCCATACCACCCTGAACGCGCCCGATCTCGT | 1264.0 | 2.3 | 0.9 | 2.6 | 9.9E-03 | 1.1E-02 | BT20/MB468 | NUC |
| >hg38_21_+_8433172_8446622_RNA45SN1_45S_50ntFlanks@10349.10375.27\|AACGAGAACTTTGAAGGCCGAAGTGGA | >hg38_21_+_8433172_8446622_RNA45SN1_45S_50ntFlanks | 10349 | 10375 | 27 | AACGAGAACTTTGAAGGCCGAAGTGGA | 92.0 | -4.9 | 1.8 | -2.8 | 5.8E-03 | 1.1E-02 | BT20/MB468 | MP |
| >hg38_21_+_8433172_8446622_RNA45SN1_45S_50ntFlanks@12458.12476.19\|AAGCGTTGGATTGTTCACC | >hg38_21_+_8433172_8446622_RNA45SN1_45S_50ntFlanks | 12458 | 12476 | 19 | AAGCGTTGGATTGTTCACC | 4219.5 | -1.8 | 0.7 | -2.6 | 9.5E-03 | 1.1E-02 | BT20/MB468 | CYTO |
| >hg38_MT_+_1621_3279_RNR2_16S_50ntFlanks@841.864.24\|AAAGTAAAAGGAACTCGGCAAATC | >hg38_MT_+_1621_3279_RNR2_16S_50ntFlanks | 841 | 864 | 24 | AAAGTAAAAGGAACTCGGCAAATC | 71.9 | 3.6 | 1.4 | 2.6 | 8.7E-03 | 1.1E-02 | MB231/MB468 | MP |
| >hg38_MT_+_598_1651_RNR1_12S_50ntFlanks@454.480.27\|ACAATAGCTAAGACCCAAACTGGGATT | >hg38_MT_+_598_1651_RNR1_12S_50ntFlanks | 454 | 480 | 27 | ACAATAGCTAAGACCCAAACTGGGATT | 169.8 | 2.1 | 0.8 | 2.6 | 8.7E-03 | 1.1E-02 | MB231/MB468 | MP |
| >MI0000266\|hsa-mir-10a&WithFlank&17\|-\|48579832\|48579953@29.49.21(+1A)&[MIMAT0000253&hsa-miR-10a-5p&offsets\|+1\|-1(+1A);m-18&17\|-\|48579905\|48579926&offsets\|+1\|0(+1A)]\|ACCCTGTAGATCCGAATTTGTA | >MI0000266\|hsa-mir-10a&WithFlank&17\|-\|48579832\|48579953 | 29 | 49 | 21(+1A) | ACCCTGTAGATCCGAATTTGTA | 892.9 | 1.5 | 0.5 | 3.0 | 2.4E-03 | 1.1E-02 | BT20/MB231 | NUC |
| >hg38_21_+_8433172_8446622_RNA45SN1_45S_50ntFlanks@11675.11707.33\|TGCTCTGAATGTCAAAGTGAAGAAATTCAATGA | >hg38_21_+_8433172_8446622_RNA45SN1_45S_50ntFlanks | 11675 | 11707 | 33 | TGCTCTGAATGTCAAAGTGAAGAAATTCAATGA | 416.9 | -2.5 | 0.8 | -3.0 | 2.4E-03 | 1.1E-02 | BT20/MB231 | NUC |
| >hg38_21_+_8433172_8446622_RNA45SN1_45S_50ntFlanks@5054.5081.28\|TGGAGCGATTTGTCTGGTTAATTCCGAT | >hg38_21_+_8433172_8446622_RNA45SN1_45S_50ntFlanks | 5054 | 5081 | 28 | TGGAGCGATTTGTCTGGTTAATTCCGAT | 182.4 | 5.1 | 1.7 | 3.1 | 2.2E-03 | 1.1E-02 | BT20/MB231 | MP |
| >hg38_21_+_8433172_8446622_RNA45SN1_45S_50ntFlanks@6788.6807.20\|CGCCTGTCTGAGCGTCGCTT | >hg38_21_+_8433172_8446622_RNA45SN1_45S_50ntFlanks | 6788 | 6807 | 20 | CGCCTGTCTGAGCGTCGCTT | 1857.7 | 4.6 | 1.5 | 3.1 | 2.2E-03 | 1.1E-02 | BT20/MB231 | MP |
| >hg38_21_+_8433172_8446622_RNA45SN1_45S_50ntFlanks@3854.3884.31\|ACTGTGGTAATTCTAGAGCTAATACATGCCG | >hg38_21_+_8433172_8446622_RNA45SN1_45S_50ntFlanks | 3854 | 3884 | 31 | ACTGTGGTAATTCTAGAGCTAATACATGCCG | 261.9 | -2.0 | 0.8 | -2.6 | 9.5E-03 | 1.1E-02 | MB231/MB468 | CYTO |
| >hg38_21_+_8433172_8446622_RNA45SN1_45S_50ntFlanks@6733.6770.38\|CATTGATCATCGACACTTCGAACGCACTTGCGGCCCCG | >hg38_21_+_8433172_8446622_RNA45SN1_45S_50ntFlanks | 6733 | 6770 | 38 | CATTGATCATCGACACTTCGAACGCACTTGCGGCCCCG | 1494.8 | -5.4 | 1.9 | -2.8 | 5.8E-03 | 1.1E-02 | BT20/MB468 | MP |
| >MI0000299\|hsa-mir-222&WithFlank&X\|-\|45747009\|45747130@75.97.23(+1A)&[MIMAT0000279&hsa-miR-222-3p&offsets\|0\|+2(+1A);m-140&X\|-\|45747033\|45747056&offsets\|0\|-1(+1A)]\|AGCTACATCTGGCTACTGGGTCTA | >MI0000299\|hsa-mir-222&WithFlank&X\|-\|45747009\|45747130 | 75 | 97 | 23(+1A) | AGCTACATCTGGCTACTGGGTCTA | 217.7 | 4.0 | 1.5 | 2.8 | 5.8E-03 | 1.1E-02 | BT20/MB468 | MP |
| >hg38_21_+_8433172_8446622_RNA45SN1_45S_50ntFlanks@4884.4918.35\|CAAAGCTGAAACTTAAAGGAATTGACGGAAGGGCA | >hg38_21_+_8433172_8446622_RNA45SN1_45S_50ntFlanks | 4884 | 4918 | 35 | CAAAGCTGAAACTTAAAGGAATTGACGGAAGGGCA | 191.3 | -4.7 | 1.7 | -2.8 | 5.8E-03 | 1.1E-02 | BT20/MB468 | MP |
| >hg38_21_+_8433172_8446622_RNA45SN1_45S_50ntFlanks@11208.11228.21\|GCGGCGACTCTGGACGCGAGC | >hg38_21_+_8433172_8446622_RNA45SN1_45S_50ntFlanks | 11208 | 11228 | 21 | GCGGCGACTCTGGACGCGAGC | 351.8 | -3.2 | 1.2 | -2.6 | 1.0E-02 | 1.1E-02 | MB231/MB468 | NUC |
| >hg38_21_+_8433172_8446622_RNA45SN1_45S_50ntFlanks@11571.11604.34\|GCCGACTTAGAACTGGTGCGGACCAGGGGAATCC | >hg38_21_+_8433172_8446622_RNA45SN1_45S_50ntFlanks | 11571 | 11604 | 34 | GCCGACTTAGAACTGGTGCGGACCAGGGGAATCC | 164.5 | 4.9 | 1.8 | 2.8 | 5.8E-03 | 1.1E-02 | BT20/MB468 | MP |
| >MI0000299\|hsa-mir-222&WithFlank&X\|-\|45747009\|45747130@75.97.23(+1U)&[MIMAT0000279&hsa-miR-222-3p&offsets\|0\|+2(+1U);m-140&X\|-\|45747033\|45747056&offsets\|0\|-1(+1U)]\|AGCTACATCTGGCTACTGGGTCTT | >MI0000299\|hsa-mir-222&WithFlank&X\|-\|45747009\|45747130 | 75 | 97 | 23(+1U) | AGCTACATCTGGCTACTGGGTCTT | 5593.5 | -2.3 | 0.8 | -3.0 | 2.8E-03 | 1.1E-02 | BT20/MB231 | MITO |
| >hg38_21_+_8433172_8446622_RNA45SN1_45S_50ntFlanks@10797.10819.23\|GCATGTTGGAACAATGTAGGTAA | >hg38_21_+_8433172_8446622_RNA45SN1_45S_50ntFlanks | 10797 | 10819 | 23 | GCATGTTGGAACAATGTAGGTAA | 83.3 | -4.3 | 1.6 | -2.6 | 8.8E-03 | 1.1E-02 | MB231/MB468 | MP |
| >hg38_21_+_8433172_8446622_RNA45SN1_45S_50ntFlanks@6652.6669.18\|GACTCTTAGCGGTGGATC | >hg38_21_+_8433172_8446622_RNA45SN1_45S_50ntFlanks | 6652 | 6669 | 18 | GACTCTTAGCGGTGGATC | 440.4 | 2.1 | 0.7 | 3.0 | 2.8E-03 | 1.1E-02 | BT20/MB231 | CYTO |
| >am_trnaMT_PheGAA_MT_+_577_647@41.64.24\|GTTTAGACGGGCTCACATCACCCC | >am_trnaMT_PheGAA_MT_+_577_647 | 41 | 64 | 24 | GTTTAGACGGGCTCACATCACCCC | 318.5 | 5.6 | 2.0 | 2.8 | 5.8E-03 | 1.1E-02 | BT20/MB468 | MP |
| >hg38_21_+_8433172_8446622_RNA45SN1_45S_50ntFlanks@7991.8016.26\|ACGTGGCGACCCGCTGAATTTAAGCA | >hg38_21_+_8433172_8446622_RNA45SN1_45S_50ntFlanks | 7991 | 8016 | 26 | ACGTGGCGACCCGCTGAATTTAAGCA | 1004.3 | -1.8 | 0.7 | -2.6 | 8.1E-03 | 1.1E-02 | BT20/MB468 | MITO |
| >hg38_21_+_8433172_8446622_RNA45SN1_45S_50ntFlanks@12937.12957.21\|AACCATTCGTAGACGACCTGC | >hg38_21_+_8433172_8446622_RNA45SN1_45S_50ntFlanks | 12937 | 12957 | 21 | AACCATTCGTAGACGACCTGC | 647.1 | 1.8 | 0.7 | 2.6 | 9.6E-03 | 1.1E-02 | MB231/MB468 | CYTO |
| >hg38_21_+_8433172_8446622_RNA45SN1_45S_50ntFlanks@12008.12035.28\|CCGCCGGTGAAATACCACTACTCTGATC | >hg38_21_+_8433172_8446622_RNA45SN1_45S_50ntFlanks | 12008 | 12035 | 28 | CCGCCGGTGAAATACCACTACTCTGATC | 369.4 | -1.6 | 0.6 | -2.6 | 8.1E-03 | 1.1E-02 | BT20/MB468 | MITO |
| >hg38_21_+_8433172_8446622_RNA45SN1_45S_50ntFlanks@10386.10410.25\|ATGTGAACAGCAGTTGAACATGGGT | >hg38_21_+_8433172_8446622_RNA45SN1_45S_50ntFlanks | 10386 | 10410 | 25 | ATGTGAACAGCAGTTGAACATGGGT | 222.4 | -2.7 | 1.0 | -2.6 | 8.2E-03 | 1.1E-02 | BT20/MB468 | MITO |
| >MI0000082\|hsa-mir-25&WithFlank&7\|-\|100093554\|100093649@58.76.19(+1C)&[MIMAT0000081&hsa-miR-25-3p&offsets\|0\|-3(+1C);m-23&7\|-\|100093571\|100093592&offsets\|0\|-3(+1C)]\|CATTGCACTTGTCTCGGTCC | >MI0000082\|hsa-mir-25&WithFlank&7\|-\|100093554\|100093649 | 58 | 76 | 19(+1C) | CATTGCACTTGTCTCGGTCC | 45.7 | 5.3 | 2.0 | 2.6 | 8.8E-03 | 1.1E-02 | MB231/MB468 | MP |
| >hg38_21_+_8433172_8446622_RNA45SN1_45S_50ntFlanks@10874.10894.21\|TGGGTCGGTCGGGCTGGGGCG | >hg38_21_+_8433172_8446622_RNA45SN1_45S_50ntFlanks | 10874 | 10894 | 21 | TGGGTCGGTCGGGCTGGGGCG | 71.0 | -4.5 | 1.6 | -2.8 | 5.8E-03 | 1.1E-02 | BT20/MB468 | MP |
| >hg38_21_+_8433172_8446622_RNA45SN1_45S_50ntFlanks@474.505.32\|TCTTCGTGATCGATGTGGTGACGTCGTGCTCT | >hg38_21_+_8433172_8446622_RNA45SN1_45S_50ntFlanks | 474 | 505 | 32 | TCTTCGTGATCGATGTGGTGACGTCGTGCTCT | 32.6 | 5.1 | 2.0 | 2.6 | 8.8E-03 | 1.1E-02 | MB231/MB468 | MP |
| >hg38_21_+_8433172_8446622_RNA45SN1_45S_50ntFlanks@4903.4941.39\|AATTGACGGAAGGGCACCACCAGGAGTGGAGCCTGCGGC | >hg38_21_+_8433172_8446622_RNA45SN1_45S_50ntFlanks | 4903 | 4941 | 39 | AATTGACGGAAGGGCACCACCAGGAGTGGAGCCTGCGGC | 111.0 | 3.1 | 1.1 | 2.8 | 5.8E-03 | 1.1E-02 | BT20/MB468 | MP |
| >hg38_21_+_8433172_8446622_RNA45SN1_45S_50ntFlanks@10852.10879.28\|GATAAGGATTGGCTCTAAGGGCTGGGTC | >hg38_21_+_8433172_8446622_RNA45SN1_45S_50ntFlanks | 10852 | 10879 | 28 | GATAAGGATTGGCTCTAAGGGCTGGGTC | 284.5 | 5.1 | 1.7 | 3.1 | 2.2E-03 | 1.1E-02 | BT20/MB231 | MP |
| >hg38_MT_+_1621_3279_RNR2_16S_50ntFlanks@1416.1452.37\|GTTTGTTCAACGATTAAAGTCCTACGTGATCTGAGTT | >hg38_MT_+_1621_3279_RNR2_16S_50ntFlanks | 1416 | 1452 | 37 | GTTTGTTCAACGATTAAAGTCCTACGTGATCTGAGTT | 47.2 | 5.1 | 1.9 | 2.6 | 8.8E-03 | 1.1E-02 | MB231/MB468 | MP |
| >hg38_21_+_8433172_8446622_RNA45SN1_45S_50ntFlanks@10351.10388.38\|CGAGAACTTTGAAGGCCGAAGTGGAGAAGGGTTCCATG | >hg38_21_+_8433172_8446622_RNA45SN1_45S_50ntFlanks | 10351 | 10388 | 38 | CGAGAACTTTGAAGGCCGAAGTGGAGAAGGGTTCCATG | 274.3 | 6.1 | 2.0 | 3.1 | 2.2E-03 | 1.1E-02 | BT20/MB231 | MP |
| >hg38_21_+_8433172_8446622_RNA45SN1_45S_50ntFlanks@4899.4916.18\|AAGGAATTGACGGAAGGG | >hg38_21_+_8433172_8446622_RNA45SN1_45S_50ntFlanks | 4899 | 4916 | 18 | AAGGAATTGACGGAAGGG | 191.3 | -4.6 | 1.7 | -2.8 | 5.8E-03 | 1.1E-02 | BT20/MB468 | MP |
| >trna116_GluCTC_1_-_145399233_145399304@32.74.43,trna71_GluCTC_1_-_161439189_161439260@32.74.43,trna74_GluCTC_1_-_161431809_161431880@32.74.43,trna77_GluCTC_1_-_161424398_161424469@32.74.43,trna77_GluCTC_6_+_28949976_28950047@32.74.43,trna80_GluCTC_1_-_161417018_161417089@32.74.43,trna87_GluCTC_6_-_126101393_126101464@32.74.43\|CTCTCACCGCCGCGGCCCGGGTTCGATTCCCGGTCAGGGAACC | >trna116_GluCTC_1_-_145399233_145399304 | 32 | 74 | 43 | CTCTCACCGCCGCGGCCCGGGTTCGATTCCCGGTCAGGGAACC | 180.5 | -4.0 | 1.5 | -2.6 | 9.5E-03 | 1.1E-02 | BT20/MB468 | CYTO |
| >hg38_21_+_8433172_8446622_RNA45SN1_45S_50ntFlanks@10346.10367.22\|TCAAACGAGAACTTTGAAGGCC | >hg38_21_+_8433172_8446622_RNA45SN1_45S_50ntFlanks | 10346 | 10367 | 22 | TCAAACGAGAACTTTGAAGGCC | 270.3 | -2.8 | 0.9 | -3.0 | 2.4E-03 | 1.1E-02 | BT20/MB231 | NUC |
| >hg38_21_+_8433172_8446622_RNA45SN1_45S_50ntFlanks@12156.12181.26\|GAGTTTGACTGGGGCGGTACACCTGT | >hg38_21_+_8433172_8446622_RNA45SN1_45S_50ntFlanks | 12156 | 12181 | 26 | GAGTTTGACTGGGGCGGTACACCTGT | 210.4 | -4.6 | 1.7 | -2.8 | 5.8E-03 | 1.1E-02 | BT20/MB468 | MP |
| >MI0000483\|hsa-mir-186&WithFlank&1\|-\|71067625\|71067722@21.41.21&[MIMAT0000456&hsa-miR-186-5p&offsets\|0\|-1;m-84&1\|-\|71067681\|71067702&offsets\|0\|-1]\|CAAAGAATTCTCCTTTTGGGC | >MI0000483\|hsa-mir-186&WithFlank&1\|-\|71067625\|71067722 | 21 | 41 | 21 | CAAAGAATTCTCCTTTTGGGC | 516.0 | 2.7 | 0.9 | 3.0 | 2.8E-03 | 1.1E-02 | BT20/MB231 | CYTO |
| >trnaMT_ValTAC_MT_+_1602_1670@37.71.35\|TTAGGAGATTTCAACTTAACTTGACCGCTCTGACC | >trnaMT_ValTAC_MT_+_1602_1670 | 37 | 71 | 35 | TTAGGAGATTTCAACTTAACTTGACCGCTCTGACC | 78.7 | 6.9 | 2.5 | 2.8 | 5.8E-03 | 1.1E-02 | BT20/MB468 | MP |
| >hg38_21_+_8433172_8446622_RNA45SN1_45S_50ntFlanks@9564.9603.40\|CTGACGTGCAAATCGGTCGTCCGACCTGGGTATAGGGGCG | >hg38_21_+_8433172_8446622_RNA45SN1_45S_50ntFlanks | 9564 | 9603 | 40 | CTGACGTGCAAATCGGTCGTCCGACCTGGGTATAGGGGCG | 1121.8 | -2.5 | 0.9 | -2.6 | 8.2E-03 | 1.1E-02 | BT20/MB468 | MITO |
| >hg38_21_+_8433172_8446622_RNA45SN1_45S_50ntFlanks@6661.6708.48\|CGGTGGATCACTCGGCTCGTGCGTCGATGAAGAACGCAGCTAGCTGCG | >hg38_21_+_8433172_8446622_RNA45SN1_45S_50ntFlanks | 6661 | 6708 | 48 | CGGTGGATCACTCGGCTCGTGCGTCGATGAAGAACGCAGCTAGCTGCG | 1336.6 | -2.4 | 0.9 | -2.6 | 1.0E-02 | 1.1E-02 | MB231/MB468 | MITO |
| >hg38_21_+_8433172_8446622_RNA45SN1_45S_50ntFlanks@11563.11584.22\|GCCTAGCAGCCGACTTAGAACT | >hg38_21_+_8433172_8446622_RNA45SN1_45S_50ntFlanks | 11563 | 11584 | 22 | GCCTAGCAGCCGACTTAGAACT | 37.1 | 5.1 | 2.0 | 2.6 | 8.8E-03 | 1.1E-02 | MB231/MB468 | MP |
| >hg38_21_+_8433172_8446622_RNA45SN1_45S_50ntFlanks@51.72.22\|GCTGACACGCTGTCCTCTGGCG | >hg38_21_+_8433172_8446622_RNA45SN1_45S_50ntFlanks | 51 | 72 | 22 | GCTGACACGCTGTCCTCTGGCG | 133.2 | -3.8 | 1.5 | -2.6 | 9.6E-03 | 1.1E-02 | BT20/MB468 | CYTO |
| >hg38_21_+_8433172_8446622_RNA45SN1_45S_50ntFlanks@10341.10362.22\|AATATTCAAACGAGAACTTTGA | >hg38_21_+_8433172_8446622_RNA45SN1_45S_50ntFlanks | 10341 | 10362 | 22 | AATATTCAAACGAGAACTTTGA | 187.5 | 4.8 | 1.6 | 3.1 | 2.2E-03 | 1.1E-02 | BT20/MB231 | MP |
| >hg38_21_+_8433172_8446622_RNA45SN1_45S_50ntFlanks@12999.13028.30\|CTGCGATCTATTGAAAGTCAGCCCTCGACA | >hg38_21_+_8433172_8446622_RNA45SN1_45S_50ntFlanks | 12999 | 13028 | 30 | CTGCGATCTATTGAAAGTCAGCCCTCGACA | 700.5 | -1.9 | 0.7 | -2.6 | 1.0E-02 | 1.1E-02 | MB231/MB468 | NUC |
| >hg38_21_+_8433172_8446622_RNA45SN1_45S_50ntFlanks@4620.4638.19\|ATTAAGAGGGACGGCCGGG | >hg38_21_+_8433172_8446622_RNA45SN1_45S_50ntFlanks | 4620 | 4638 | 19 | ATTAAGAGGGACGGCCGGG | 138.9 | -2.6 | 1.0 | -2.6 | 8.8E-03 | 1.1E-02 | MB231/MB468 | MP |
| >hg38_21_+_8433172_8446622_RNA45SN1_45S_50ntFlanks@10341.10361.21\|AATATTCAAACGAGAACTTTG | >hg38_21_+_8433172_8446622_RNA45SN1_45S_50ntFlanks | 10341 | 10361 | 21 | AATATTCAAACGAGAACTTTG | 51.1 | 9.0 | 2.9 | 3.1 | 2.2E-03 | 1.1E-02 | BT20/MB231 | MP |
| >hg38_21_+_8433172_8446622_RNA45SN1_45S_50ntFlanks@11568.11611.44\|GCAGCCGACTTAGAACTGGTGCGGACCAGGGGAATCCGACTGTT | >hg38_21_+_8433172_8446622_RNA45SN1_45S_50ntFlanks | 11568 | 11611 | 44 | GCAGCCGACTTAGAACTGGTGCGGACCAGGGGAATCCGACTGTT | 13287.6 | 3.4 | 1.1 | 3.0 | 2.9E-03 | 1.1E-02 | BT20/MB231 | MITO |
| >hg38_21_+_8433172_8446622_RNA45SN1_45S_50ntFlanks@10393.10419.27\|CAGCAGTTGAACATGGGTCAGTCGGTC | >hg38_21_+_8433172_8446622_RNA45SN1_45S_50ntFlanks | 10393 | 10419 | 27 | CAGCAGTTGAACATGGGTCAGTCGGTC | 255.6 | -4.0 | 1.5 | -2.6 | 9.6E-03 | 1.2E-02 | BT20/MB468 | CYTO |
| >hg38_21_+_8433172_8446622_RNA45SN1_45S_50ntFlanks@10348.10384.37\|AAACGAGAACTTTGAAGGCCGAAGTGGAGAAGGGTTC | >hg38_21_+_8433172_8446622_RNA45SN1_45S_50ntFlanks | 10348 | 10384 | 37 | AAACGAGAACTTTGAAGGCCGAAGTGGAGAAGGGTTC | 256.4 | -3.6 | 1.4 | -2.6 | 8.9E-03 | 1.2E-02 | MB231/MB468 | MP |
| >hg38_21_+_8433172_8446622_RNA45SN1_45S_50ntFlanks@10768.10792.25\|CAGCAGGTCTCCAAGGTGAACAGCC | >hg38_21_+_8433172_8446622_RNA45SN1_45S_50ntFlanks | 10768 | 10792 | 25 | CAGCAGGTCTCCAAGGTGAACAGCC | 59.8 | -4.6 | 1.7 | -2.6 | 8.9E-03 | 1.2E-02 | MB231/MB468 | MP |
| >hg38_MT_+_598_1651_RNR1_12S_50ntFlanks@978.1004.27\|TGTACTGGAAAGTGCACTTGGACGAAC | >hg38_MT_+_598_1651_RNR1_12S_50ntFlanks | 978 | 1004 | 27 | TGTACTGGAAAGTGCACTTGGACGAAC | 90.4 | 2.5 | 1.0 | 2.6 | 8.9E-03 | 1.2E-02 | MB231/MB468 | MP |
| >hg38_21_+_8433172_8446622_RNA45SN1_45S_50ntFlanks@9739.9760.22\|AAACGATCTCAACCTATTCTCA | >hg38_21_+_8433172_8446622_RNA45SN1_45S_50ntFlanks | 9739 | 9760 | 22 | AAACGATCTCAACCTATTCTCA | 211.2 | -1.5 | 0.6 | -2.6 | 1.0E-02 | 1.2E-02 | MB231/MB468 | NUC |
| >hg38_MT_+_598_1651_RNR1_12S_50ntFlanks@718.744.27\|GTAAAGACGTTAGGTCAAGGTGTAGCC | >hg38_MT_+_598_1651_RNR1_12S_50ntFlanks | 718 | 744 | 27 | GTAAAGACGTTAGGTCAAGGTGTAGCC | 297.7 | 3.5 | 1.3 | 2.6 | 8.9E-03 | 1.2E-02 | MB231/MB468 | MP |

| | | | | | | | | | | | | | |
|---|---|---|---|---|---|---|---|---|---|---|---|---|---|
| >MI0000067\|hsa-let-7f-1&WithFlank&9\|+\|94176341\|94176439@13.33.21(+1A);MI0000068\|hsa-let-7f-2&WithFlank&X\|-\|53557186\|53557280@14.34.21(+1A)&[MIMAT0000067&hsa-let-7f-5p&offsets\|0\|-1(+1A);m-20&9\|+\|94176353\|94176374&offsets\|0\|-1(+1A)];[MIMAT0000067_1&hsa-let-7f-5p&offsets\|0\|-1(+1A);m-19&X\|-\|53557246\|53557267&offsets\|0\|-1(+1A)]\|TGAGGTAGTAGATTGTATAGTA | >MI0000067\|hsa-let-7f-1&WithFlank&9\|+\|94176341\|94176439 | 13 | 33 | 21(+1A) | TGAGGTAGTAGATTGTATAGTA | 3265.3 | -3.4 | 1.1 | -3.1 | 2.2E-03 | 1.2E-02 | BT20/MB231 | MP |
| >hg38_21_+_8433172_8446622_RNA45SN1_45S_50ntFlanks@5073.5108.36\|AATTCCGATAACGAACGAGACTCTGGCATGCTAACT | >hg38_21_+_8433172_8446622_RNA45SN1_45S_50ntFlanks | 5073 | 5108 | 36 | AATTCCGATAACGAACGAGACTCTGGCATGCTAACT | 379.6 | -2.8 | 1.1 | -2.6 | 8.2E-03 | 1.2E-02 | BT20/MB468 | MITO |
| >hg38_21_+_8433172_8446622_RNA45SN1_45S_50ntFlanks@7975.8015.41\|CGCGACCTCAGATCAGACGTGGCGACCCGCTGAATTTAAGC | >hg38_21_+_8433172_8446622_RNA45SN1_45S_50ntFlanks | 7975 | 8015 | 41 | CGCGACCTCAGATCAGACGTGGCGACCCGCTGAATTTAAGC | 1151.3 | 3.2 | 1.2 | 2.6 | 8.9E-03 | 1.2E-02 | MB231/MB468 | MP |
| >hg38_21_+_8433172_8446622_RNA45SN1_45S_50ntFlanks@12457.12475.19\|CAAGCGTTGGATTGTTCAC | >hg38_21_+_8433172_8446622_RNA45SN1_45S_50ntFlanks | 12457 | 12475 | 19 | CAAGCGTTGGATTGTTCAC | 321.6 | -4.7 | 1.7 | -2.8 | 5.9E-03 | 1.2E-02 | BT20/MB468 | MP |
| >hg38_21_+_8433172_8446622_RNA45SN1_45S_50ntFlanks@12625.12661.37\|CAATGGGGCGAAGCTACCATCTGTGGGATTATGACTG | >hg38_21_+_8433172_8446622_RNA45SN1_45S_50ntFlanks | 12625 | 12661 | 37 | CAATGGGGCGAAGCTACCATCTGTGGGATTATGACTG | 540.3 | -4.1 | 1.5 | -2.8 | 5.9E-03 | 1.2E-02 | BT20/MB468 | MP |
| >am_trna128_GlyGCC_6_-_27870686_27870756@1.29.29,trna133_GlyCCC_1_-_16872434_16872504@1.29.29,trna18_GlyGCC_16_+_70822597_70822667@1.29.29,trna19_GlyGCC_16_+_70823410_70823480@1.29.29,trna19_GlyGCC_2_-_157257659_157257729@1.29.29,trna24_GlyGCC_16_-_70812942_70813012@1.29.29,trna25_GlyGCC_16_-_70812114_70812184@1.29.29,trna4_GlyCCC_1_+_17188416_17188486@1.29.29,trna5_GlyGCC_17_+_8029064_8029134@1.29.29,trna68_GlyGCC_1_-_161493637_161493707@1.29.29\|GCATTGGTGGTTCAGTGGTAGAATTCTCG | >am_trna128_GlyGCC_6_-_27870686_27870756 | 1 | 29 | 29 | GCATTGGTGGTTCAGTGGTAGAATTCTCG | 529.0 | -2.3 | 0.8 | -3.0 | 2.9E-03 | 1.2E-02 | BT20/MB231 | MITO |
| >hg38_21_+_8433172_8446622_RNA45SN1_45S_50ntFlanks@5487.5526.40\|CGCTGAGAAGACGGTCGAACTTGACTATCTAGAGGAAGTA | >hg38_21_+_8433172_8446622_RNA45SN1_45S_50ntFlanks | 5487 | 5526 | 40 | CGCTGAGAAGACGGTCGAACTTGACTATCTAGAGGAAGTA | 654.6 | -2.3 | 0.9 | -2.6 | 9.6E-03 | 1.2E-02 | BT20/MB468 | CYTO |
| >hg38_MT_+_1621_3279_RNR2_16S_50ntFlanks@1383.1414.32\|ACATCCCGATGGTGCAGCCGCTATTAAAGGTT | >hg38_MT_+_1621_3279_RNR2_16S_50ntFlanks | 1383 | 1414 | 32 | ACATCCCGATGGTGCAGCCGCTATTAAAGGTT | 61.2 | 4.2 | 1.6 | 2.6 | 8.9E-03 | 1.2E-02 | MB231/MB468 | MP |
| >hg38_21_+_8433172_8446622_RNA45SN1_45S_50ntFlanks@5473.5505.33\|CGGCCCTGGCGGAGCGCTGAGAAGACGGTCGAA | >hg38_21_+_8433172_8446622_RNA45SN1_45S_50ntFlanks | 5473 | 5505 | 33 | CGGCCCTGGCGGAGCGCTGAGAAGACGGTCGAA | 1779.4 | 3.0 | 1.0 | 3.0 | 2.8E-03 | 1.2E-02 | BT20/MB231 | CYTO |
| >hg38_21_+_8433172_8446622_RNA45SN1_45S_50ntFlanks@7991.8013.23\|ACGTGGCGACCCGCTGAATTTAA | >hg38_21_+_8433172_8446622_RNA45SN1_45S_50ntFlanks | 7991 | 8013 | 23 | ACGTGGCGACCCGCTGAATTTAA | 501.0 | -2.5 | 0.9 | -2.8 | 5.9E-03 | 1.2E-02 | BT20/MB468 | MP |
| >hg38_1_-_228634819_228635039_RNA5S12_5S_50ntFlanks@124.147.24\|AGTACTTGGATGGGAGACCGCCTG | >hg38_1_-_228634819_228635039_RNA5S12_5S_50ntFlanks | 124 | 147 | 24 | AGTACTTGGATGGGAGACCGCCTG | 158.7 | -1.9 | 0.7 | -2.6 | 8.3E-03 | 1.2E-02 | BT20/MB468 | MITO |
| >hg38_21_+_8433172_8446622_RNA45SN1_45S_50ntFlanks@6646.6668.23\|TCGTACGACTCTTAGCGGTGGAT | >hg38_21_+_8433172_8446622_RNA45SN1_45S_50ntFlanks | 6646 | 6668 | 23 | TCGTACGACTCTTAGCGGTGGAT | 6800.6 | 1.7 | 0.7 | 2.6 | 1.0E-02 | 1.2E-02 | MB231/MB468 | MITO |
| >hg38_21_+_8433172_8446622_RNA45SN1_45S_50ntFlanks@5472.5514.43\|ACGGCCCTGGCGGAGCGCTGAGAAGACGGTCGAACTTGACTAT | >hg38_21_+_8433172_8446622_RNA45SN1_45S_50ntFlanks | 5472 | 5514 | 43 | ACGGCCCTGGCGGAGCGCTGAGAAGACGGTCGAACTTGACTAT | 490.6 | -3.4 | 1.3 | -2.6 | 8.9E-03 | 1.2E-02 | MB231/MB468 | MP |
| >hg38_MT_+_598_1651_RNR1_12S_50ntFlanks@410.429.20\|GACACAAAATAGACTACGAA | >hg38_MT_+_598_1651_RNR1_12S_50ntFlanks | 410 | 429 | 20 | GACACAAAATAGACTACGAA | 46.4 | -8.9 | 2.9 | -3.1 | 2.3E-03 | 1.2E-02 | BT20/MB231 | MP |
| >hg38_21_+_8433172_8446622_RNA45SN1_45S_50ntFlanks@4050.4082.33\|CGAACGTCTGCCCTATCAACTTTCGATGGTAGT | >hg38_21_+_8433172_8446622_RNA45SN1_45S_50ntFlanks | 4050 | 4082 | 33 | CGAACGTCTGCCCTATCAACTTTCGATGGTAGT | 48.7 | 4.9 | 1.9 | 2.6 | 8.9E-03 | 1.2E-02 | MB231/MB468 | MP |
| >hg38_21_+_8433172_8446622_RNA45SN1_45S_50ntFlanks@4590.4616.27\|ATTTTGTTGGTTTTCGGAACTGAGGCC | >hg38_21_+_8433172_8446622_RNA45SN1_45S_50ntFlanks | 4590 | 4616 | 27 | ATTTTGTTGGTTTTCGGAACTGAGGCC | 236.7 | -1.7 | 0.7 | -2.6 | 8.3E-03 | 1.2E-02 | BT20/MB468 | MITO |
| >MI0000234\|hsa-mir-192&WithFlank&11\|-\|64891131\|64891252@31.53.23&[MIMAT0000222&hsa-miR-192-5p&offsets\|+1\|+3;m-59&11\|-\|64891203\|64891223&offsets\|+1\|+3]\|TGACCTATGAATTGACAGCCAGT | >MI0000234\|hsa-mir-192&WithFlank&11\|-\|64891131\|64891252 | 31 | 53 | 23 | TGACCTATGAATTGACAGCCAGT | 941.5 | 1.9 | 0.7 | 2.6 | 8.9E-03 | 1.2E-02 | MB231/MB468 | MP |
| >hg38_21_+_8433172_8446622_RNA45SN1_45S_50ntFlanks@12007.12033.27\|GCCGCCGGTGAAATACCACTACTCTGA | >hg38_21_+_8433172_8446622_RNA45SN1_45S_50ntFlanks | 12007 | 12033 | 27 | GCCGCCGGTGAAATACCACTACTCTGA | 631.3 | -2.2 | 0.9 | -2.6 | 1.0E-02 | 1.2E-02 | MB231/MB468 | MITO |
| >hg38_21_+_8433172_8446622_RNA45SN1_45S_50ntFlanks@8355.8375.21\|TGAAAAGAACTTTGAAGAGAG | >hg38_21_+_8433172_8446622_RNA45SN1_45S_50ntFlanks | 8355 | 8375 | 21 | TGAAAAGAACTTTGAAGAGAG | 164.9 | 3.8 | 1.4 | 2.6 | 8.9E-03 | 1.2E-02 | MB231/MB468 | MP |
| >hg38_21_+_8433172_8446622_RNA45SN1_45S_50ntFlanks@8184.8204.21\|CGAGGCCCAGCCCGTGGACGG | >hg38_21_+_8433172_8446622_RNA45SN1_45S_50ntFlanks | 8184 | 8204 | 21 | CGAGGCCCAGCCCGTGGACGG | 153.9 | -3.1 | 1.2 | -2.6 | 8.3E-03 | 1.2E-02 | BT20/MB468 | MITO |
| >hg38_21_+_8433172_8446622_RNA45SN1_45S_50ntFlanks@4753.4783.31\|AAGACGATCAGATACCGTCGTAGTTCCGACC | >hg38_21_+_8433172_8446622_RNA45SN1_45S_50ntFlanks | 4753 | 4783 | 31 | AAGACGATCAGATACCGTCGTAGTTCCGACC | 767.8 | -2.2 | 0.9 | -2.6 | 1.0E-02 | 1.2E-02 | MB231/MB468 | NUC |
| >hg38_21_+_8433172_8446622_RNA45SN1_45S_50ntFlanks@4732.4759.28\|AGAACGAAAGTCGGAGGTTCGAAGACGA | >hg38_21_+_8433172_8446622_RNA45SN1_45S_50ntFlanks | 4732 | 4759 | 28 | AGAACGAAAGTCGGAGGTTCGAAGACGA | 2140.6 | 4.9 | 1.6 | 3.1 | 2.3E-03 | 1.2E-02 | BT20/MB231 | MP |
| >hg38_21_+_8433172_8446622_RNA45SN1_45S_50ntFlanks@12605.12634.30\|TATGTGCTTGGCTGAGGAGCCAATGGGGCG | >hg38_21_+_8433172_8446622_RNA45SN1_45S_50ntFlanks | 12605 | 12634 | 30 | TATGTGCTTGGCTGAGGAGCCAATGGGGCG | 101.8 | -4.5 | 1.7 | -2.8 | 5.9E-03 | 1.2E-02 | BT20/MB468 | MP |
| >hg38_21_+_8433172_8446622_RNA45SN1_45S_50ntFlanks@8104.8140.37\|CCGCGGCGGGGCGCGGGACATGTGGCGTACGGAAGAC | >hg38_21_+_8433172_8446622_RNA45SN1_45S_50ntFlanks | 8104 | 8140 | 37 | CCGCGGCGGGGCGCGGGACATGTGGCGTACGGAAGAC | 1433.3 | -4.3 | 1.7 | -2.6 | 1.0E-02 | 1.2E-02 | BT20/MB468 | NUC |
| >hg38_21_+_8433172_8446622_RNA45SN1_45S_50ntFlanks@3985.4005.21\|CGGCTTTGGTGACTCTAGATA | >hg38_21_+_8433172_8446622_RNA45SN1_45S_50ntFlanks | 3985 | 4005 | 21 | CGGCTTTGGTGACTCTAGATA | 927.1 | -5.0 | 1.8 | -2.8 | 5.9E-03 | 1.2E-02 | BT20/MB468 | MP |
| >hg38_21_+_8433172_8446622_RNA45SN1_45S_50ntFlanks@11672.11692.21\|CAGTGCTCTGAATGTCAAAGT | >hg38_21_+_8433172_8446622_RNA45SN1_45S_50ntFlanks | 11672 | 11692 | 21 | CAGTGCTCTGAATGTCAAAGT | 507.8 | 2.4 | 0.8 | 3.0 | 2.9E-03 | 1.2E-02 | BT20/MB231 | MITO |
| >hg38_21_+_8433172_8446622_RNA45SN1_45S_50ntFlanks@6650.6698.49\|ACGACTCTTAGCGGTGGATCACTCGGCTCGTGCGTCGATGAAGAACGCA | >hg38_21_+_8433172_8446622_RNA45SN1_45S_50ntFlanks | 6650 | 6698 | 49 | ACGACTCTTAGCGGTGGATCACTCGGCTCGTGCGTCGATGAAGAACGCA | 7536.3 | -2.5 | 1.0 | -2.6 | 1.0E-02 | 1.2E-02 | MB231/MB468 | MITO |
| >am_MI0039740\|hsa-mir-12136&WithFlank&1\|-\|632609\|632691@7.31.25&[MIMAT0049032&hsa-miR-12136&offsets\|0\|+7]\|GAAAAAGTCATGGAGGCCATGGGGT | >am_MI0039740\|hsa-mir-12136&WithFlank&1\|-\|632609\|632691 | 7 | 31 | 25 | GAAAAAGTCATGGAGGCCATGGGGT | 3555.5 | 4.2 | 1.5 | 2.8 | 5.9E-03 | 1.2E-02 | BT20/MB468 | MP |
| >trnaMT_SerTGA_MT_-_7446_7514@1.25.25,trnalookalike6_SerTGA_1_-_567997_568065@1.25.25\|GAAAAAGTCATGGAGGCCATGGGGT | >trnaMT_SerTGA_MT_-_7446_7514 | 1 | 25 | 25 | GAAAAAGTCATGGAGGCCATGGGGT | 3555.5 | 4.2 | 1.5 | 2.8 | 5.9E-03 | 1.2E-02 | BT20/MB468 | MP |
| >hg38_21_+_8433172_8446622_RNA45SN1_45S_50ntFlanks@3984.4014.31\|GCGGCTTTGGTGACTCTAGATAACCTCGGGC | >hg38_21_+_8433172_8446622_RNA45SN1_45S_50ntFlanks | 3984 | 4014 | 31 | GCGGCTTTGGTGACTCTAGATAACCTCGGGC | 76.1 | -4.6 | 1.7 | -2.8 | 5.9E-03 | 1.2E-02 | BT20/MB468 | MP |
| >MI0000061\|hsa-let-7a-2&WithFlank&11\|-\|122146516\|122146599@11.32.22;MI0000062\|hsa-let-7a-3&WithFlank&22\|+\|46112743\|46112828@10.31.22;MI0000060\|hsa-let-7a-1&WithFlank&9\|+\|94175951\|94176042@12.33.22&[MIMAT0000062&hsa-let-7a-5p&offsets\|0\|0;m-6&11\|-\|122146568\|122146589&offsets\|0\|0];[MIMAT0000062_1&hsa-let-7a-5p&offsets\|0\|0;m-5&22\|+\|46112752\|46112773&offsets\|0\|0];[MIMAT0000062_2&hsa-let-7a-5p&offsets\|0\|0;m-4&9\|+\|94175962\|94175983&offsets\|0\|0]\|TGAGGTAGTAGGTTGTATAGTT | >MI0000061\|hsa-let-7a-2&WithFlank&11\|-\|122146516\|122146599 | 11 | 32 | 22 | TGAGGTAGTAGGTTGTATAGTT | 14556.7 | 2.5 | 0.9 | 2.8 | 5.9E-03 | 1.2E-02 | BT20/MB468 | MP |
| >hg38_21_+_8433172_8446622_RNA45SN1_45S_50ntFlanks@11920.11948.29\|ATGAGAGGTGTAGAATAAGTGGGAGGCCC | >hg38_21_+_8433172_8446622_RNA45SN1_45S_50ntFlanks | 11920 | 11948 | 29 | ATGAGAGGTGTAGAATAAGTGGGAGGCCC | 1261.4 | -3.9 | 1.4 | -2.8 | 5.9E-03 | 1.2E-02 | BT20/MB468 | MP |
| >hg38_21_+_8433172_8446622_RNA45SN1_45S_50ntFlanks@4005.4039.35\|AACCTCGGGCCGATCGCACGCCCCCCGTGGCGGCG | >hg38_21_+_8433172_8446622_RNA45SN1_45S_50ntFlanks | 4005 | 4039 | 35 | AACCTCGGGCCGATCGCACGCCCCCCGTGGCGGCG | 94.3 | -2.7 | 1.1 | -2.6 | 9.7E-03 | 1.2E-02 | BT20/MB468 | CYTO |
| >hg38_MT_+_1621_3279_RNR2_16S_50ntFlanks@1249.1274.26\|AGTCAAAGCGAACTACTATACTCAAT | >hg38_MT_+_1621_3279_RNR2_16S_50ntFlanks | 1249 | 1274 | 26 | AGTCAAAGCGAACTACTATACTCAAT | 142.0 | 3.5 | 1.3 | 2.6 | 9.0E-03 | 1.2E-02 | MB231/MB468 | MP |
| >hg38_21_+_8433172_8446622_RNA45SN1_45S_50ntFlanks@11565.11599.35\|CTAGCAGCCGACTTAGAACTGGTGCGGACCAGGGG | >hg38_21_+_8433172_8446622_RNA45SN1_45S_50ntFlanks | 11565 | 11599 | 35 | CTAGCAGCCGACTTAGAACTGGTGCGGACCAGGGG | 101.3 | 3.1 | 1.1 | 2.8 | 5.9E-03 | 1.2E-02 | BT20/MB468 | MP |
| >hg38_21_+_8433172_8446622_RNA45SN1_45S_50ntFlanks@7970.7988.19\|CGAGACGCGACCTCAGATC | >hg38_21_+_8433172_8446622_RNA45SN1_45S_50ntFlanks | 7970 | 7988 | 19 | CGAGACGCGACCTCAGATC | 79.9 | 3.9 | 1.4 | 2.8 | 5.9E-03 | 1.2E-02 | BT20/MB468 | MP |
| >hg38_21_+_8433172_8446622_RNA45SN1_45S_50ntFlanks@6700.6721.22\|CTAGCTGCGAGAATTAATGTGA | >hg38_21_+_8433172_8446622_RNA45SN1_45S_50ntFlanks | 6700 | 6721 | 22 | CTAGCTGCGAGAATTAATGTGA | 140.2 | -4.5 | 1.6 | -2.8 | 5.9E-03 | 1.2E-02 | BT20/MB468 | MP |
| >hg38_21_+_8433172_8446622_RNA45SN1_45S_50ntFlanks@6650.6668.19\|ACGACTCTTAGCGGTGGAT | >hg38_21_+_8433172_8446622_RNA45SN1_45S_50ntFlanks | 6650 | 6668 | 19 | ACGACTCTTAGCGGTGGAT | 1148.4 | 1.9 | 0.8 | 2.6 | 1.0E-02 | 1.2E-02 | MB231/MB468 | MITO |
| >MI0000270\|hsa-mir-181b-1&WithFlank&1\|-\|198858867\|198858988@42.64.23(+1C);MI0000683\|hsa-mir-181b-2&WithFlank&9\|+\|124693704\|124693804@22.44.23(+1C)&[MIMAT0000257&hsa-miR-181b-5p&offsets\|0\|0(+1C);m-73&1\|-\|198858925\|198858947&offsets\|0\|0(+1C)];[MIMAT0000257_1&hsa-miR-181b-5p&offsets\|0\|0(+1C);m-72&9\|+\|124693725\|124693747&offsets\|0\|0(+1C)]\|AACATTCATTGCTGTCGGTGGGTC | >MI0000270\|hsa-mir-181b-1&WithFlank&1\|-\|198858867\|198858988 | 42 | 64 | 23(+1C) | AACATTCATTGCTGTCGGTGGGTC | 1020.1 | -3.4 | 1.1 | -3.1 | 2.3E-03 | 1.2E-02 | BT20/MB231 | MP |
| >hg38_21_+_8433172_8446622_RNA45SN1_45S_50ntFlanks@10387.10424.38\|TGTGAACAGCAGTTGAACATGGGTCAGTCGGTCCTGAG | >hg38_21_+_8433172_8446622_RNA45SN1_45S_50ntFlanks | 10387 | 10424 | 38 | TGTGAACAGCAGTTGAACATGGGTCAGTCGGTCCTGAG | 6725.3 | 3.9 | 1.3 | 3.1 | 2.3E-03 | 1.2E-02 | BT20/MB231 | MP |
| >MI0000088\|hsa-mir-30a&WithFlank&6\|-\|71403545\|71403627@12.34.23&[MIMAT0000087&hsa-miR-30a-5p&offsets\|0\|+1;m-15&6\|-\|71403595\|71403616&offsets\|0\|+1]\|TGTAAACATCCTCGACTGGAAGC | >MI0000088\|hsa-mir-30a&WithFlank&6\|-\|71403545\|71403627 | 12 | 34 | 23 | TGTAAACATCCTCGACTGGAAGC | 29465.4 | 3.2 | 1.2 | 2.8 | 5.9E-03 | 1.2E-02 | BT20/MB468 | MP |
| >MI0003581\|hsa-mir-574&WithFlank&4\|+\|38868026\|38868133@67.86.20(+1U)&[MIMAT0003239&hsa-miR-574-3p&offsets\|0\|-2(+1U);m-149&4\|+\|38868092\|38868113&offsets\|0\|-2(+1U)]\|CACGCTCATGCACACACCCAT | >MI0003581\|hsa-mir-574&WithFlank&4\|+\|38868026\|38868133 | 67 | 86 | 20(+1U) | CACGCTCATGCACACACCCAT | 86.0 | 5.1 | 1.9 | 2.6 | 9.0E-03 | 1.2E-02 | MB231/MB468 | MP |
| >hg38_21_+_8433172_8446622_RNA45SN1_45S_50ntFlanks@6733.6753.21\|CATTGATCATCGACACTTCGA | >hg38_21_+_8433172_8446622_RNA45SN1_45S_50ntFlanks | 6733 | 6753 | 21 | CATTGATCATCGACACTTCGA | 567.3 | -2.1 | 0.8 | -2.6 | 1.0E-02 | 1.2E-02 | MB231/MB468 | MITO |
| >hg38_21_+_8433172_8446622_RNA45SN1_45S_50ntFlanks@4687.4720.34\|ACGGACCAGAGCGAAAGCATTTGCCAAGAATGTT | >hg38_21_+_8433172_8446622_RNA45SN1_45S_50ntFlanks | 4687 | 4720 | 34 | ACGGACCAGAGCGAAAGCATTTGCCAAGAATGTT | 383.2 | 3.5 | 1.3 | 2.6 | 9.7E-03 | 1.2E-02 | BT20/MB468 | CYTO |
| >hg38_21_+_8433172_8446622_RNA45SN1_45S_50ntFlanks@11672.11703.32\|CAGTGCTCTGAATGTCAAAGTGAAGAAATTCA | >hg38_21_+_8433172_8446622_RNA45SN1_45S_50ntFlanks | 11672 | 11703 | 32 | CAGTGCTCTGAATGTCAAAGTGAAGAAATTCA | 220.4 | -3.5 | 1.3 | -2.6 | 9.0E-03 | 1.2E-02 | MB231/MB468 | MP |

<table>
<tr><th></th><th></th><th></th><th></th><th></th><th></th><th></th><th></th><th></th><th></th><th></th><th></th><th></th><th></th></tr>
<tr><td>>hg38_21_+_8433172_8446622_RNA45SN1_45S_50ntFlanks@483.529.47|TCGATGTGGTGACGTCGTGCTCTCCCGGGCCGGGTCCGAGCCGCGAC</td><td>>hg38_21_+_8433172_8446622_RNA45SN1_45S_50ntFlanks</td><td>483</td><td>529</td><td>47</td><td>TCGATGTGGTGACGTCGTGCTCTCCCGGGCCGGGTCCGAGCCGCGAC</td><td>1249.2</td><td>3.2</td><td>1.3</td><td>2.6</td><td>1.0E-02</td><td>1.2E-02</td><td>MB231/MB468</td><td>NUC</td></tr>
<tr><td>>hg38_21_+_8433172_8446622_RNA45SN1_45S_50ntFlanks@11592.11613.22|ACCAGGGGAATCCGACTGTTTA</td><td>>hg38_21_+_8433172_8446622_RNA45SN1_45S_50ntFlanks</td><td>11592</td><td>11613</td><td>22</td><td>ACCAGGGGAATCCGACTGTTTA</td><td>188.0</td><td>-3.7</td><td>1.4</td><td>-2.8</td><td>5.9E-03</td><td>1.2E-02</td><td>BT20/MB468</td><td>MP</td></tr>
<tr><td>>hg38_21_+_8433172_8446622_RNA45SN1_45S_50ntFlanks@4557.4579.23|CAGCTAGGAATAATGGAATAGGA</td><td>>hg38_21_+_8433172_8446622_RNA45SN1_45S_50ntFlanks</td><td>4557</td><td>4579</td><td>23</td><td>CAGCTAGGAATAATGGAATAGGA</td><td>67.0</td><td>-4.5</td><td>1.6</td><td>-2.8</td><td>5.9E-03</td><td>1.2E-02</td><td>BT20/MB468</td><td>MP</td></tr>
<tr><td>>hg38_21_+_8433172_8446622_RNA45SN1_45S_50ntFlanks@12081.12109.29|CTCTCGCTTCTGGCGCCAAGCGCCCGGCC</td><td>>hg38_21_+_8433172_8446622_RNA45SN1_45S_50ntFlanks</td><td>12081</td><td>12109</td><td>29</td><td>CTCTCGCTTCTGGCGCCAAGCGCCCGGCC</td><td>130.4</td><td>-3.9</td><td>1.5</td><td>-2.6</td><td>8.3E-03</td><td>1.2E-02</td><td>BT20/MB468</td><td>MITO</td></tr>
<tr><td>>hg38_21_+_8433172_8446622_RNA45SN1_45S_50ntFlanks@6652.6673.22|GACTCTTAGCGGTGGATCACTC</td><td>>hg38_21_+_8433172_8446622_RNA45SN1_45S_50ntFlanks</td><td>6652</td><td>6673</td><td>22</td><td>GACTCTTAGCGGTGGATCACTC</td><td>644.3</td><td>2.7</td><td>1.1</td><td>2.6</td><td>1.0E-02</td><td>1.2E-02</td><td>MB231/MB468</td><td>MITO</td></tr>
<tr><td>>MI0000462|hsa-mir-152&WithFlank&17|-|48037155|48037253@60.81.22(+1A)&[MIMAT0000438&hsa-miR-152-3p&offsets|0|+1(+1A);m-97&17|-|48037174|48037194&offsets|0|+1(+1A)]|TCAGTGCATGACAGAACTTGGGA</td><td>>MI0000462|hsa-mir-152&WithFlank&17|-|48037155|48037253</td><td>60</td><td>81</td><td>22(+1A)</td><td>TCAGTGCATGACAGAACTTGGGA</td><td>508.8</td><td>2.5</td><td>0.8</td><td>3.0</td><td>2.9E-03</td><td>1.2E-02</td><td>BT20/MB231</td><td>CYTO</td></tr>
<tr><td>>hg38_21_+_8433172_8446622_RNA45SN1_45S_50ntFlanks@7975.7994.20|CGCGACCTCAGATCAGACGT</td><td>>hg38_21_+_8433172_8446622_RNA45SN1_45S_50ntFlanks</td><td>7975</td><td>7994</td><td>20</td><td>CGCGACCTCAGATCAGACGT</td><td>194886.8</td><td>2.3</td><td>0.8</td><td>3.0</td><td>2.5E-03</td><td>1.2E-02</td><td>BT20/MB231</td><td>NUC</td></tr>
<tr><td>>MI0000070|hsa-mir-16-1&WithFlank&13|-|50048967|50049067@20.39.20(+1U);MI0000115|hsa-mir-16-2&WithFlank&3|+|160404739|160404831@16.35.20(+1U)&[MIMAT0000069&hsa-miR-16-5p&offsets|0|-2(+1U);m-62&13|-|50049027|50049048&offsets|0|-2(+1U)];[MIMAT0000069_1&hsa-miR-16-5p&offsets|0|-2(+1U);m-61&3|+|160404754|160404775&offsets|0|-2(+1U)]|TAGCAGCACGTAAATATTGGT</td><td>>MI0000070|hsa-mir-16-1&WithFlank&13|-|50048967|50049067</td><td>20</td><td>39</td><td>20(+1U)</td><td>TAGCAGCACGTAAATATTGGT</td><td>523.7</td><td>3.4</td><td>1.2</td><td>2.8</td><td>5.9E-03</td><td>1.2E-02</td><td>BT20/MB468</td><td>MP</td></tr>
<tr><td>>hg38_21_+_8433172_8446622_RNA45SN1_45S_50ntFlanks@7988.8005.18|CAGACGTGGCGACCCGCT</td><td>>hg38_21_+_8433172_8446622_RNA45SN1_45S_50ntFlanks</td><td>7988</td><td>8005</td><td>18</td><td>CAGACGTGGCGACCCGCT</td><td>143.9</td><td>-4.0</td><td>1.5</td><td>-2.6</td><td>8.3E-03</td><td>1.2E-02</td><td>BT20/MB468</td><td>MITO</td></tr>
<tr><td>>hg38_21_+_8433172_8446622_RNA45SN1_45S_50ntFlanks@12603.12624.22|TGTATGTGCTTGGCTGAGGAGC</td><td>>hg38_21_+_8433172_8446622_RNA45SN1_45S_50ntFlanks</td><td>12603</td><td>12624</td><td>22</td><td>TGTATGTGCTTGGCTGAGGAGC</td><td>366.8</td><td>5.5</td><td>1.8</td><td>3.1</td><td>2.3E-03</td><td>1.2E-02</td><td>BT20/MB231</td><td>MP</td></tr>
<tr><td>>hg38_21_+_8433172_8446622_RNA45SN1_45S_50ntFlanks@9739.9766.28|AAACGATCTCAACCTATTCTCAAACTTT</td><td>>hg38_21_+_8433172_8446622_RNA45SN1_45S_50ntFlanks</td><td>9739</td><td>9766</td><td>28</td><td>AAACGATCTCAACCTATTCTCAAACTTT</td><td>274.6</td><td>-1.6</td><td>0.6</td><td>-2.6</td><td>1.0E-02</td><td>1.2E-02</td><td>MB231/MB468</td><td>NUC</td></tr>
<tr><td>>hg38_21_+_8433172_8446622_RNA45SN1_45S_50ntFlanks@4857.4876.20|CTTTGGGTTCCGGGGGGAGT</td><td>>hg38_21_+_8433172_8446622_RNA45SN1_45S_50ntFlanks</td><td>4857</td><td>4876</td><td>20</td><td>CTTTGGGTTCCGGGGGGAGT</td><td>63.8</td><td>4.5</td><td>1.7</td><td>2.6</td><td>9.0E-03</td><td>1.2E-02</td><td>MB231/MB468</td><td>MP</td></tr>
<tr><td>>hg38_21_+_8433172_8446622_RNA45SN1_45S_50ntFlanks@6647.6670.24|CGTACGACTCTTAGCGGTGGATCA</td><td>>hg38_21_+_8433172_8446622_RNA45SN1_45S_50ntFlanks</td><td>6647</td><td>6670</td><td>24</td><td>CGTACGACTCTTAGCGGTGGATCA</td><td>233.0</td><td>-1.5</td><td>0.6</td><td>-2.6</td><td>9.8E-03</td><td>1.2E-02</td><td>MB231/MB468</td><td>CYTO</td></tr>
<tr><td>>hg38_21_+_8433172_8446622_RNA45SN1_45S_50ntFlanks@4103.4128.26|CCACGGGTGACGGGGAATCAGGGTTC</td><td>>hg38_21_+_8433172_8446622_RNA45SN1_45S_50ntFlanks</td><td>4103</td><td>4128</td><td>26</td><td>CCACGGGTGACGGGGAATCAGGGTTC</td><td>47.1</td><td>-4.5</td><td>1.6</td><td>-2.7</td><td>6.0E-03</td><td>1.2E-02</td><td>BT20/MB468</td><td>MP</td></tr>
<tr><td>>hg38_21_+_8433172_8446622_RNA45SN1_45S_50ntFlanks@7973.8010.38|GACGCGACCTCAGATCAGACGTGGCGACCCGCTGAATT</td><td>>hg38_21_+_8433172_8446622_RNA45SN1_45S_50ntFlanks</td><td>7973</td><td>8010</td><td>38</td><td>GACGCGACCTCAGATCAGACGTGGCGACCCGCTGAATT</td><td>14966.7</td><td>3.0</td><td>1.0</td><td>3.0</td><td>2.5E-03</td><td>1.2E-02</td><td>BT20/MB231</td><td>NUC</td></tr>
<tr><td>>trnaMT_SerGCT_MT_+_12207_12265@10.38.29|CACAAGAACTGCTAACTCATGCCCCCATG</td><td>>trnaMT_SerGCT_MT_+_12207_12265</td><td>10</td><td>38</td><td>29</td><td>CACAAGAACTGCTAACTCATGCCCCCATG</td><td>514.7</td><td>-2.7</td><td>1.0</td><td>-2.6</td><td>8.4E-03</td><td>1.2E-02</td><td>BT20/MB468</td><td>MITO</td></tr>
<tr><td>>MI0000084|hsa-mir-26b&WithFlank&2|+|218402640|218402728@18.38.21(+1C)&[MIMAT0000083&hsa-miR-26b-5p&offsets|0|0(+1C);m-53&2|+|218402657|218402678&offsets|0|-1(+1C)]|TTCAAGTAATTCAGGATAGGTC</td><td>>MI0000084|hsa-mir-26b&WithFlank&2|+|218402640|218402728</td><td>18</td><td>38</td><td>21(+1C)</td><td>TTCAAGTAATTCAGGATAGGTC</td><td>1465.8</td><td>-3.6</td><td>1.2</td><td>-3.1</td><td>2.3E-03</td><td>1.2E-02</td><td>BT20/MB231</td><td>MP</td></tr>
<tr><td>>hg38_21_+_8433172_8446622_RNA45SN1_45S_50ntFlanks@4499.4517.19|AGCGTTTACTTTGAAAAAA</td><td>>hg38_21_+_8433172_8446622_RNA45SN1_45S_50ntFlanks</td><td>4499</td><td>4517</td><td>19</td><td>AGCGTTTACTTTGAAAAAA</td><td>481.8</td><td>2.6</td><td>0.9</td><td>3.0</td><td>2.9E-03</td><td>1.2E-02</td><td>BT20/MB231</td><td>CYTO</td></tr>
<tr><td>>hg38_21_+_8433172_8446622_RNA45SN1_45S_50ntFlanks@6650.6676.27|ACGACTCTTAGCGGTGGATCACTCGGC</td><td>>hg38_21_+_8433172_8446622_RNA45SN1_45S_50ntFlanks</td><td>6650</td><td>6676</td><td>27</td><td>ACGACTCTTAGCGGTGGATCACTCGGC</td><td>2745.5</td><td>4.1</td><td>1.3</td><td>3.0</td><td>2.3E-03</td><td>1.2E-02</td><td>BT20/MB231</td><td>MP</td></tr>
<tr><td>>hg38_21_+_8433172_8446622_RNA45SN1_45S_50ntFlanks@475.492.18|CTTCGTGATCGATGTGGT</td><td>>hg38_21_+_8433172_8446622_RNA45SN1_45S_50ntFlanks</td><td>475</td><td>492</td><td>18</td><td>CTTCGTGATCGATGTGGT</td><td>192.0</td><td>4.2</td><td>1.5</td><td>2.7</td><td>6.0E-03</td><td>1.2E-02</td><td>BT20/MB468</td><td>MP</td></tr>
<tr><td>>trna119_LysCTT_1_-_145395522_145395594@1.32.32,trna11_LysCTT_5_-_180648979_180649051@1.32.32,trna13_LysCTT_6_+_26556774_26556846@1.32.32,trna32_LysCTT_16_-_3207406_3207478@1.32.32,trna7_LysCTT_16_+_3225692_3225764@1.32.32,trna9_LysCTT_5_+_180634755_180634827@1.32.32|GCCCGGCTAGCTCAGTCGGTAGAGCATGAGAC</td><td>>trna119_LysCTT_1_-_145395522_145395594</td><td>1</td><td>32</td><td>32</td><td>GCCCGGCTAGCTCAGTCGGTAGAGCATGAGAC</td><td>1760.0</td><td>2.9</td><td>1.1</td><td>2.6</td><td>1.0E-02</td><td>1.2E-02</td><td>MB231/MB468</td><td>NUC</td></tr>
<tr><td>>hg38_1_-_228634819_228635039_RNA5S12_5S_50ntFlanks@51.79.29|GTCTACGGCCATACCACCCTGAACGCGCC</td><td>>hg38_1_-_228634819_228635039_RNA5S12_5S_50ntFlanks</td><td>51</td><td>79</td><td>29</td><td>GTCTACGGCCATACCACCCTGAACGCGCC</td><td>1872.9</td><td>2.6</td><td>1.0</td><td>2.6</td><td>1.0E-02</td><td>1.2E-02</td><td>MB231/MB468</td><td>NUC</td></tr>
<tr><td>>hg38_21_+_8433172_8446622_RNA45SN1_45S_50ntFlanks@8238.8266.29|CCCGGGTCTTCCCGGAGTCGGGTTGCTTG</td><td>>hg38_21_+_8433172_8446622_RNA45SN1_45S_50ntFlanks</td><td>8238</td><td>8266</td><td>29</td><td>CCCGGGTCTTCCCGGAGTCGGGTTGCTTG</td><td>195.1</td><td>-1.7</td><td>0.6</td><td>-2.6</td><td>8.4E-03</td><td>1.2E-02</td><td>BT20/MB468</td><td>MITO</td></tr>
<tr><td>>hg38_21_+_8433172_8446622_RNA45SN1_45S_50ntFlanks@9304.9336.33|ACCAAGGAGTCTAACACGTGCGCGAGTCGGGGG</td><td>>hg38_21_+_8433172_8446622_RNA45SN1_45S_50ntFlanks</td><td>9304</td><td>9336</td><td>33</td><td>ACCAAGGAGTCTAACACGTGCGCGAGTCGGGGG</td><td>165.8</td><td>-3.0</td><td>1.1</td><td>-2.6</td><td>8.4E-03</td><td>1.2E-02</td><td>BT20/MB468</td><td>MITO</td></tr>
<tr><td>>hg38_21_+_8433172_8446622_RNA45SN1_45S_50ntFlanks@11576.11609.34|CTTAGAACTGGTGCGGACCAGGGGAATCCGACTG</td><td>>hg38_21_+_8433172_8446622_RNA45SN1_45S_50ntFlanks</td><td>11576</td><td>11609</td><td>34</td><td>CTTAGAACTGGTGCGGACCAGGGGAATCCGACTG</td><td>826.2</td><td>-3.0</td><td>1.1</td><td>-2.7</td><td>6.0E-03</td><td>1.2E-02</td><td>BT20/MB468</td><td>MP</td></tr>
<tr><td>>hg38_21_+_8433172_8446622_RNA45SN1_45S_50ntFlanks@12073.12110.38|CCGAGGGGCTCTCGCTTCTGGCGCCAAGCGCCCGGCCG</td><td>>hg38_21_+_8433172_8446622_RNA45SN1_45S_50ntFlanks</td><td>12073</td><td>12110</td><td>38</td><td>CCGAGGGGCTCTCGCTTCTGGCGCCAAGCGCCCGGCCG</td><td>302.8</td><td>-4.8</td><td>1.7</td><td>-2.7</td><td>6.0E-03</td><td>1.2E-02</td><td>BT20/MB468</td><td>MP</td></tr>
<tr><td>>hg38_21_+_8433172_8446622_RNA45SN1_45S_50ntFlanks@8183.8211.29|TCGAGGCCCAGCCCGTGGACGGTGTGAGG</td><td>>hg38_21_+_8433172_8446622_RNA45SN1_45S_50ntFlanks</td><td>8183</td><td>8211</td><td>29</td><td>TCGAGGCCCAGCCCGTGGACGGTGTGAGG</td><td>103.7</td><td>-4.6</td><td>1.7</td><td>-2.7</td><td>6.0E-03</td><td>1.2E-02</td><td>BT20/MB468</td><td>MP</td></tr>
<tr><td>>hg38_21_+_8433172_8446622_RNA45SN1_45S_50ntFlanks@11575.11611.37|ACTTAGAACTGGTGCGGACCAGGGGAATCCGACTGTT</td><td>>hg38_21_+_8433172_8446622_RNA45SN1_45S_50ntFlanks</td><td>11575</td><td>11611</td><td>37</td><td>ACTTAGAACTGGTGCGGACCAGGGGAATCCGACTGTT</td><td>306.7</td><td>-2.3</td><td>0.9</td><td>-2.6</td><td>9.8E-03</td><td>1.2E-02</td><td>BT20/MB468</td><td>CYTO</td></tr>
<tr><td>>hg38_21_+_8433172_8446622_RNA45SN1_45S_50ntFlanks@5471.5499.29|CACGGCCCTGGCGGAGCGCTGAGAAGACG</td><td>>hg38_21_+_8433172_8446622_RNA45SN1_45S_50ntFlanks</td><td>5471</td><td>5499</td><td>29</td><td>CACGGCCCTGGCGGAGCGCTGAGAAGACG</td><td>949.5</td><td>-2.3</td><td>0.9</td><td>-2.6</td><td>8.4E-03</td><td>1.2E-02</td><td>BT20/MB468</td><td>MITO</td></tr>
<tr><td>>MI0005531|hsa-mir-450b&WithFlank&X|-|134540179|134540268@17.37.21&[MIMAT0004909&hsa-miR-450b-5p&offsets|0|-1;m-266&X|-|134540232|134540252&offsets|0|0]|TTTTGCAATATGTTCCTGAAT</td><td>>MI0005531|hsa-mir-450b&WithFlank&X|-|134540179|134540268</td><td>17</td><td>37</td><td>21</td><td>TTTTGCAATATGTTCCTGAAT</td><td>501.1</td><td>1.7</td><td>0.6</td><td>3.0</td><td>2.5E-03</td><td>1.2E-02</td><td>BT20/MB231</td><td>NUC</td></tr>
<tr><td>>am_MI0039740|hsa-mir-12136&WithFlank&1|-|632609|632691@7.31.25&[MIMAT0049032&hsa-miR-12136&offsets|0|+7]|GAAAAAGTCATGGAGGCCATGGGGT</td><td>>am_MI0039740|hsa-mir-12136&WithFlank&1|-|632609|632691</td><td>7</td><td>31</td><td>25</td><td>GAAAAAGTCATGGAGGCCATGGGGT</td><td>364.0</td><td>3.9</td><td>1.3</td><td>3.0</td><td>2.5E-03</td><td>1.2E-02</td><td>BT20/MB231</td><td>NUC</td></tr>
<tr><td>>trnaMT_SerTGA_MT_-_7446_7514@1.25.25,trnalookalike6_SerTGA_1_-_567997_568065@1.25.25|GAAAAAGTCATGGAGGCCATGGGGT</td><td>>trnaMT_SerTGA_MT_-_7446_7514</td><td>1</td><td>25</td><td>25</td><td>GAAAAAGTCATGGAGGCCATGGGGT</td><td>364.0</td><td>3.9</td><td>1.3</td><td>3.0</td><td>2.5E-03</td><td>1.2E-02</td><td>BT20/MB231</td><td>NUC</td></tr>
<tr><td>>MI0000088|hsa-mir-30a&WithFlank&6|-|71403545|71403627@53.73.21(+2U)&[MIMAT0000088&hsa-miR-30a-3p&offsets|0|-1(+2U);m-38&6|-|71403554|71403575&offsets|0|-1(+2U)]|CTTTCAGTCGGATGTTTGCAGTT</td><td>>MI0000088|hsa-mir-30a&WithFlank&6|-|71403545|71403627</td><td>53</td><td>73</td><td>21(+2U)</td><td>CTTTCAGTCGGATGTTTGCAGTT</td><td>698.7</td><td>-1.9</td><td>0.6</td><td>-3.0</td><td>2.5E-03</td><td>1.2E-02</td><td>BT20/MB231</td><td>NUC</td></tr>
<tr><td>>hg38_21_+_8433172_8446622_RNA45SN1_45S_50ntFlanks@12928.12953.26|CTGGCGCTAAACCATTCGTAGACGAC</td><td>>hg38_21_+_8433172_8446622_RNA45SN1_45S_50ntFlanks</td><td>12928</td><td>12953</td><td>26</td><td>CTGGCGCTAAACCATTCGTAGACGAC</td><td>107.5</td><td>-4.5</td><td>1.6</td><td>-2.7</td><td>6.0E-03</td><td>1.2E-02</td><td>BT20/MB468</td><td>MP</td></tr>
<tr><td>>MI0000299|hsa-mir-222&WithFlank&X|-|45747009|45747130@75.97.23(+1A)&[MIMAT0000279&hsa-miR-222-3p&offsets|0|+2(+1A);m-140&X|-|45747033|45747056&offsets|0|-1(+1A)]|AGCTACATCTGGCTACTGGGTCTA</td><td>>MI0000299|hsa-mir-222&WithFlank&X|-|45747009|45747130</td><td>75</td><td>97</td><td>23(+1A)</td><td>AGCTACATCTGGCTACTGGGTCTA</td><td>1591.2</td><td>-2.2</td><td>0.7</td><td>-3.0</td><td>2.9E-03</td><td>1.2E-02</td><td>BT20/MB231</td><td>MITO</td></tr>
<tr><td>>MI0000433|hsa-let-7g&WithFlank&3|-|52268272|52268367@11.29.19(+1A)&[MIMAT0000414&hsa-let-7g-5p&offsets|0|-3(+1A);m-58&3|-|52268336|52268357&offsets|0|-3(+1A)]|TGAGGTAGTAGTTTGTACAA</td><td>>MI0000433|hsa-let-7g&WithFlank&3|-|52268272|52268367</td><td>11</td><td>29</td><td>19(+1A)</td><td>TGAGGTAGTAGTTTGTACAA</td><td>24.4</td><td>7.8</td><td>3.0</td><td>2.6</td><td>9.0E-03</td><td>1.2E-02</td><td>MB231/MB468</td><td>MP</td></tr>
<tr><td>>MI0000088|hsa-mir-30a&WithFlank&6|-|71403545|71403627@12.35.24&[MIMAT0000087&hsa-miR-30a-5p&offsets|0|+2;m-15&6|-|71403595|71403616&offsets|0|+2]|TGTAAACATCCTCGACTGGAAGCT</td><td>>MI0000088|hsa-mir-30a&WithFlank&6|-|71403545|71403627</td><td>12</td><td>35</td><td>24</td><td>TGTAAACATCCTCGACTGGAAGCT</td><td>456646.3</td><td>-3.1</td><td>1.0</td><td>-3.0</td><td>2.3E-03</td><td>1.2E-02</td><td>BT20/MB231</td><td>MP</td></tr>
<tr><td>>hg38_21_+_8433172_8446622_RNA45SN1_45S_50ntFlanks@3939.3959.21|AACCCGGTCAGCCCCTCTCCG</td><td>>hg38_21_+_8433172_8446622_RNA45SN1_45S_50ntFlanks</td><td>3939</td><td>3959</td><td>21</td><td>AACCCGGTCAGCCCCTCTCCG</td><td>286.7</td><td>-4.7</td><td>1.7</td><td>-2.7</td><td>6.0E-03</td><td>1.2E-02</td><td>BT20/MB468</td><td>MP</td></tr>
<tr><td>>trna119_LysCTT_1_-_145395522_145395594@3.33.31,trna11_LysCTT_5_-_180648979_180649051@3.33.31,trna13_LysCTT_6_+_26556774_26556846@3.33.31,trna7_LysCTT_16_+_3225692_3225764@3.33.31,trna9_LysCTT_5_+_180634755_180634827@3.33.31|CCGGCTAGCTCAGTCGGTAGAGCATGAGACT</td><td>>trna119_LysCTT_1_-_145395522_145395594</td><td>3</td><td>33</td><td>31</td><td>CCGGCTAGCTCAGTCGGTAGAGCATGAGACT</td><td>1019.3</td><td>-4.4</td><td>1.5</td><td>-3.0</td><td>2.5E-03</td><td>1.2E-02</td><td>BT20/MB231</td><td>NUC</td></tr>
<tr><td>>trna14_LysTTT_11_-_59327808_59327880@3.36.34,trna2_LysTTT_17_+_8022473_8022545@3.36.34,trna54_LysTTT_1_+_204475655_204475727@3.36.34,trna5_LysTTT_11_+_59323902_59323974@3.36.34,trna62_LysTTT_1_-_204476158_204476230@3.36.34,trna76_LysTTT_6_+_28918806_28918878@3.36.34|CCGGATAGCTCAGTCGGTAGAGCATCAGACTTTT</td><td>>trna14_LysTTT_11_-_59327808_59327880</td><td>3</td><td>36</td><td>34</td><td>CCGGATAGCTCAGTCGGTAGAGCATCAGACTTTT</td><td>133.9</td><td>4.3</td><td>1.6</td><td>2.6</td><td>9.0E-03</td><td>1.2E-02</td><td>MB231/MB468</td><td>MP</td></tr>
<tr><td>>MI0000063|hsa-let-7b&WithFlank&22|+|46113680|46113774@12.33.22(+1C)&[MIMAT0000063&hsa-let-7b-5p&offsets|0|0(+1C);m-14&22|+|46113691|46113712&offsets|0|0(+1C)]|TGAGGTAGTAGGTTGTGTGGTTC</td><td>>MI0000063|hsa-let-7b&WithFlank&22|+|46113680|46113774</td><td>12</td><td>33</td><td>22(+1C)</td><td>TGAGGTAGTAGGTTGTGTGGTTC</td><td>1215.4</td><td>2.4</td><td>0.8</td><td>3.0</td><td>2.9E-03</td><td>1.2E-02</td><td>BT20/MB231</td><td>CYTO</td></tr>
<tr><td>>trna111_HisGTG_1_-_147774845_147774916@-1G.35.36,trna118_HisGTG_1_-_145396881_145396952@-1G.35.36,trna16_HisGTG_1_+_146544773_146544844@-1G.35.36,trna1_HisGTG_15_+_45493349_45493420@-1G.35.36,trna21_HisGTG_1_+_147753471_147753542@-1G.35.36,trna33_HisGTG_6_+_27125906_27125977@-1G.35.36,trna7_HisGTG_9_-_14433938_14434009@-1G.35.36,trna8_HisGTG_15_-_45492611_45492682@-1G.35.36,trna9_HisGTG_15_-_45490804_45490875@-1G.35.36|GGCCGTGATCGTATAGTGGTTAGTACTCTGCGTTGT</td><td>>trna111_HisGTG_1_-_147774845_147774916</td><td>-1G</td><td>35</td><td>36</td><td>GGCCGTGATCGTATAGTGGTTAGTACTCTGCGTTGT</td><td>4595.5</td><td>-3.2</td><td>1.1</td><td>-3.0</td><td>2.9E-03</td><td>1.2E-02</td><td>BT20/MB231</td><td>CYTO</td></tr>
</table>

<table>
<tr><td>>MI0000299|hsa-mir-222&WithFlank&X|-|45747009|45747130@77.97.21&[MIMAT0000279&hsa-miR-222-3p&offsets|+2|+2;m-140&X|-|45747033|45747056&offsets|+2|-1]|CTACATCTGGCTACTGGGTCT</td><td>>MI0000299|hsa-mir-222&WithFlank&X|-|45747009|45747130</td><td>77</td><td>97</td><td>21</td><td>CTACATCTGGCTACTGGGTCT</td><td>34.1</td><td>5.2</td><td>2.0</td><td>2.6</td><td>9.1E-03</td><td>1.2E-02</td><td>MB231/MB468</td><td>MP</td></tr>
<tr><td>>hg38_21_+_8433172_8446622_RNA45SN1_45S_50ntFlanks@10841.10873.33|CGTAACTTCGGGATAAGGATTGGCTCTAAGGGC</td><td>>hg38_21_+_8433172_8446622_RNA45SN1_45S_50ntFlanks</td><td>10841</td><td>10873</td><td>33</td><td>CGTAACTTCGGGATAAGGATTGGCTCTAAGGGC</td><td>562.6</td><td>-1.5</td><td>0.6</td><td>-2.6</td><td>9.8E-03</td><td>1.2E-02</td><td>BT20/MB468</td><td>CYTO</td></tr>
<tr><td>>hg38_21_+_8433172_8446622_RNA45SN1_45S_50ntFlanks@6648.6668.21|GTACGACTCTTAGCGGTGGAT</td><td>>hg38_21_+_8433172_8446622_RNA45SN1_45S_50ntFlanks</td><td>6648</td><td>6668</td><td>21</td><td>GTACGACTCTTAGCGGTGGAT</td><td>1306.9</td><td>1.9</td><td>0.8</td><td>2.6</td><td>1.0E-02</td><td>1.2E-02</td><td>MB231/MB468</td><td>MITO</td></tr>
<tr><td>>hg38_21_+_8433172_8446622_RNA45SN1_45S_50ntFlanks@10375.10424.50|AGAAGGGTTCCATGTGAACAGCAGTTGAACATGGGTCAGTCGGTCCTGAG</td><td>>hg38_21_+_8433172_8446622_RNA45SN1_45S_50ntFlanks</td><td>10375</td><td>10424</td><td>50</td><td>AGAAGGGTTCCATGTGAACAGCAGTTGAACATGGGTCAGTCGGTCCTGAG</td><td>182.1</td><td>4.9</td><td>1.8</td><td>2.7</td><td>6.0E-03</td><td>1.2E-02</td><td>BT20/MB468</td><td>MP</td></tr>
<tr><td>>trna10_ValCAC_5_-_180649395_180649467@36.54.19,trna12_ValAAC_5_-_180645270_180645342@36.54.19,trna132_ValAAC_6_-_27721179_27721251@36.54.19,trna13_ValCAC_19_-_4724647_4724719@36.54.19,trna152_ValCAC_6_-_27248049_27248121@36.54.19,trna157_ValCAC_6_-_27173867_27173939@36.54.19,trna15_ValAAC_5_-_180615416_180615488@36.54.19,trna18_ValCAC_5_-_180529253_180529325@36.54.19,trna2_ValAAC_3_+_169490018_169490090@36.54.19,trna2_ValCAC_5_+_180524070_180524142@36.54.19,trna37_ValAAC_6_+_27203288_27203360@36.54.19,trna4_ValAAC_5_+_180591154_180591226@36.54.19,trna5_ValAAC_5_+_180596610_180596682@36.54.19,trna6_ValCAC_5_+_180600650_180600722@36.54.19,trna85_ValCAC_1_-_161369490_161369562@36.54.19,trna98_ValCAC_1_-_149298555_149298627@36.54.19,trna9_ValCAC_6_+_26538282_26538354@36.54.19|CACGCGAAAGGTCCCCGGT</td><td>>trna10_ValCAC_5_-_180649395_180649467</td><td>36</td><td>54</td><td>19</td><td>CACGCGAAAGGTCCCCGGT</td><td>263.7</td><td>-5.1</td><td>1.9</td><td>-2.7</td><td>6.0E-03</td><td>1.2E-02</td><td>BT20/MB468</td><td>MP</td></tr>
<tr><td>>am_trnaMT_SerGCT_MT_+_12207_12265@15.32.18|GAACTGCTAACTCATGCC</td><td>>am_trnaMT_SerGCT_MT_+_12207_12265</td><td>15</td><td>32</td><td>18</td><td>GAACTGCTAACTCATGCC</td><td>174.9</td><td>2.4</td><td>0.8</td><td>3.0</td><td>2.9E-03</td><td>1.2E-02</td><td>BT20/MB231</td><td>CYTO</td></tr>
<tr><td>>hg38_21_+_8433172_8446622_RNA45SN1_45S_50ntFlanks@12445.12473.29|AGCAGAATTCACCAAGCGTTGGATTGTTC</td><td>>hg38_21_+_8433172_8446622_RNA45SN1_45S_50ntFlanks</td><td>12445</td><td>12473</td><td>29</td><td>AGCAGAATTCACCAAGCGTTGGATTGTTC</td><td>200.6</td><td>-4.8</td><td>1.7</td><td>-2.7</td><td>6.0E-03</td><td>1.2E-02</td><td>BT20/MB468</td><td>MP</td></tr>
<tr><td>>hg38_21_+_8433172_8446622_RNA45SN1_45S_50ntFlanks@7975.7997.23|CGCGACCTCAGATCAGACGTGGC</td><td>>hg38_21_+_8433172_8446622_RNA45SN1_45S_50ntFlanks</td><td>7975</td><td>7997</td><td>23</td><td>CGCGACCTCAGATCAGACGTGGC</td><td>16708.0</td><td>2.1</td><td>0.8</td><td>2.6</td><td>1.0E-02</td><td>1.2E-02</td><td>MB231/MB468</td><td>MITO</td></tr>
<tr><td>>hg38_21_+_8433172_8446622_RNA45SN1_45S_50ntFlanks@7973.7994.22|GACGCGACCTCAGATCAGACGT</td><td>>hg38_21_+_8433172_8446622_RNA45SN1_45S_50ntFlanks</td><td>7973</td><td>7994</td><td>22</td><td>GACGCGACCTCAGATCAGACGT</td><td>2253.6</td><td>3.0</td><td>1.0</td><td>3.0</td><td>2.5E-03</td><td>1.2E-02</td><td>BT20/MB231</td><td>NUC</td></tr>
<tr><td>>trna10_AspGTC_12_-_125424193_125424264@35.73.39,trna12_AspGTC_12_-_125411891_125411962@35.73.39,trna144_AspGTC_6_-_27551236_27551307@35.73.39,trna38_AspGTC_17_-_8125556_8125627@35.73.39,trna45_AspGTC_6_+_27447453_27447524@35.73.39,trna48_AspGTC_6_+_27471523_27471594@35.73.39,trna4_AspGTC_12_+_96429799_96429870@35.73.39,trna69_AspGTC_1_-_161440205_161440276@35.73.39,trna72_AspGTC_1_-_161432824_161432895@35.73.39,trna75_AspGTC_1_-_161425414_161425485@35.73.39,trna78_AspGTC_1_-_161418033_161418104@35.73.39,trna81_AspGTC_1_-_161410615_161410686@35.73.39|TCACGCGGGAGACCGGGGTTCGATTCCCCGACGGGGAGC</td><td>>trna10_AspGTC_12_-_125424193_125424264</td><td>35</td><td>73</td><td>39</td><td>TCACGCGGGAGACCGGGGTTCGATTCCCCGACGGGGAGC</td><td>335.5</td><td>2.8</td><td>1.1</td><td>2.6</td><td>1.0E-02</td><td>1.2E-02</td><td>MB231/MB468</td><td>NUC</td></tr>
<tr><td>>hg38_21_+_8433172_8446622_RNA45SN1_45S_50ntFlanks@12415.12455.41|CTTCGATGTCGGCTCTTCCTATCATTGTGAAGCAGAATTCA</td><td>>hg38_21_+_8433172_8446622_RNA45SN1_45S_50ntFlanks</td><td>12415</td><td>12455</td><td>41</td><td>CTTCGATGTCGGCTCTTCCTATCATTGTGAAGCAGAATTCA</td><td>372.4</td><td>-2.4</td><td>0.9</td><td>-2.6</td><td>1.0E-02</td><td>1.2E-02</td><td>MB231/MB468</td><td>NUC</td></tr>
<tr><td>>hg38_21_+_8433172_8446622_RNA45SN1_45S_50ntFlanks@11668.11703.36|TGCCCAGTGCTCTGAATGTCAAAGTGAAGAAATTCA</td><td>>hg38_21_+_8433172_8446622_RNA45SN1_45S_50ntFlanks</td><td>11668</td><td>11703</td><td>36</td><td>TGCCCAGTGCTCTGAATGTCAAAGTGAAGAAATTCA</td><td>2370.0</td><td>-2.5</td><td>0.8</td><td>-3.0</td><td>2.9E-03</td><td>1.2E-02</td><td>BT20/MB231</td><td>CYTO</td></tr>
<tr><td>>MI0006273|hsa-mir-1180&WithFlank&17|-|19344500|19344580@47.65.19&[MIMAT0005825&hsa-miR-1180-3p&offsets|0|-3;m-214&17|-|19344513|19344534&offsets|0|-3]|TTTCCGGCTCGCGTGGGTG</td><td>>MI0006273|hsa-mir-1180&WithFlank&17|-|19344500|19344580</td><td>47</td><td>65</td><td>19</td><td>TTTCCGGCTCGCGTGGGTG</td><td>72.9</td><td>5.6</td><td>2.1</td><td>2.7</td><td>6.0E-03</td><td>1.2E-02</td><td>BT20/MB468</td><td>MP</td></tr>
<tr><td>>trna119_LysCTT_1_-_145395522_145395594@5.35.31,trna11_LysCTT_5_-_180648979_180649051@5.35.31,trna13_LysCTT_6_+_26556774_26556846@5.35.31,trna7_LysCTT_16_+_3225692_3225764@5.35.31,trna9_LysCTT_5_+_180634755_180634827@5.35.31|GGCTAGCTCAGTCGGTAGAGCATGAGACTCT</td><td>>trna119_LysCTT_1_-_145395522_145395594</td><td>5</td><td>35</td><td>31</td><td>GGCTAGCTCAGTCGGTAGAGCATGAGACTCT</td><td>400.6</td><td>-3.4</td><td>1.1</td><td>-3.0</td><td>2.3E-03</td><td>1.2E-02</td><td>BT20/MB231</td><td>MP</td></tr>
<tr><td>>hg38_1_-_228634819_228635039_RNA5S12_5S_50ntFlanks@126.169.44|TACTTGGATGGGAGACCGCCTGGGAATACCGGGTGCTGTAGGCT</td><td>>hg38_1_-_228634819_228635039_RNA5S12_5S_50ntFlanks</td><td>126</td><td>169</td><td>44</td><td>TACTTGGATGGGAGACCGCCTGGGAATACCGGGTGCTGTAGGCT</td><td>154.8</td><td>5.7</td><td>2.1</td><td>2.7</td><td>6.0E-03</td><td>1.2E-02</td><td>BT20/MB468</td><td>MP</td></tr>
<tr><td>>am_MI0000786|hsa-mir-378a&WithFlank&5|+|149732819|149732896@49.71.23&[MIMAT0000732&hsa-miR-378a-3p&offsets|0|+1;m-44&5|+|149732867|149732888&offsets|0|+1]|ACTGGACTTGGAGTCAGAAGGCC</td><td>>am_MI0000786|hsa-mir-378a&WithFlank&5|+|149732819|149732896</td><td>49</td><td>71</td><td>23</td><td>ACTGGACTTGGAGTCAGAAGGCC</td><td>1195.1</td><td>2.9</td><td>1.0</td><td>2.7</td><td>6.0E-03</td><td>1.2E-02</td><td>BT20/MB468</td><td>MP</td></tr>
<tr><td>>hg38_21_+_8433172_8446622_RNA45SN1_45S_50ntFlanks@10349.10390.42|AACGAGAACTTTGAAGGCCGAAGTGGAGAAGGGTTCCATGTG</td><td>>hg38_21_+_8433172_8446622_RNA45SN1_45S_50ntFlanks</td><td>10349</td><td>10390</td><td>42</td><td>AACGAGAACTTTGAAGGCCGAAGTGGAGAAGGGTTCCATGTG</td><td>469.8</td><td>-2.6</td><td>0.9</td><td>-3.0</td><td>2.9E-03</td><td>1.2E-02</td><td>BT20/MB231</td><td>CYTO</td></tr>
<tr><td>>hg38_21_+_8433172_8446622_RNA45SN1_45S_50ntFlanks@5489.5516.28|CTGAGAAGACGGTCGAACTTGACTATCT</td><td>>hg38_21_+_8433172_8446622_RNA45SN1_45S_50ntFlanks</td><td>5489</td><td>5516</td><td>28</td><td>CTGAGAAGACGGTCGAACTTGACTATCT</td><td>1317.6</td><td>-2.8</td><td>1.1</td><td>-2.6</td><td>9.1E-03</td><td>1.2E-02</td><td>MB231/MB468</td><td>MP</td></tr>
<tr><td>>MI0000239|hsa-mir-197&WithFlank&1|+|109598887|109598973@54.75.22&[MIMAT0000227&hsa-miR-197-3p&offsets|0|0;m-99&1|+|109598940|109598961&offsets|0|0]|TTCACCACCTTCTCCACCCAGC</td><td>>MI0000239|hsa-mir-197&WithFlank&1|+|109598887|109598973</td><td>54</td><td>75</td><td>22</td><td>TTCACCACCTTCTCCACCCAGC</td><td>9053.8</td><td>2.3</td><td>0.8</td><td>3.0</td><td>3.0E-03</td><td>1.2E-02</td><td>BT20/MB231</td><td>MITO</td></tr>
<tr><td>>hg38_21_+_8433172_8446622_RNA45SN1_45S_50ntFlanks@4685.4723.39|AGACGGACCAGAGCGAAAGCATTTGCCAAGAATGTTTTC</td><td>>hg38_21_+_8433172_8446622_RNA45SN1_45S_50ntFlanks</td><td>4685</td><td>4723</td><td>39</td><td>AGACGGACCAGAGCGAAAGCATTTGCCAAGAATGTTTTC</td><td>419.6</td><td>-2.2</td><td>0.8</td><td>-2.6</td><td>8.5E-03</td><td>1.2E-02</td><td>BT20/MB468</td><td>MITO</td></tr>
<tr><td>>hg38_21_+_8433172_8446622_RNA45SN1_45S_50ntFlanks@6650.6688.39|ACGACTCTTAGCGGTGGATCACTCGGCTCGTGCGTCGAT</td><td>>hg38_21_+_8433172_8446622_RNA45SN1_45S_50ntFlanks</td><td>6650</td><td>6688</td><td>39</td><td>ACGACTCTTAGCGGTGGATCACTCGGCTCGTGCGTCGAT</td><td>261.6</td><td>2.2</td><td>0.7</td><td>3.0</td><td>2.6E-03</td><td>1.2E-02</td><td>BT20/MB231</td><td>NUC</td></tr>
<tr><td>>hg38_21_+_8433172_8446622_RNA45SN1_45S_50ntFlanks@12456.12477.22|CCAAGCGTTGGATTGTTCACCC</td><td>>hg38_21_+_8433172_8446622_RNA45SN1_45S_50ntFlanks</td><td>12456</td><td>12477</td><td>22</td><td>CCAAGCGTTGGATTGTTCACCC</td><td>6759.4</td><td>-4.7</td><td>1.7</td><td>-2.7</td><td>6.0E-03</td><td>1.2E-02</td><td>BT20/MB468</td><td>MP</td></tr>
<tr><td>>MI0000746|hsa-mir-99b&WithFlank&19|+|51692606|51692687@51.72.22&[MIMAT0004678&hsa-miR-99b-3p&offsets|0|0;m-151&19|+|51692656|51692678&offsets|0|-1]|CAAGCTCGTGTCTGTGGGTCCG</td><td>>MI0000746|hsa-mir-99b&WithFlank&19|+|51692606|51692687</td><td>51</td><td>72</td><td>22</td><td>CAAGCTCGTGTCTGTGGGTCCG</td><td>1962.8</td><td>2.2</td><td>0.7</td><td>3.0</td><td>3.0E-03</td><td>1.2E-02</td><td>BT20/MB231</td><td>MITO</td></tr>
<tr><td>>hg38_21_+_8433172_8446622_RNA45SN1_45S_50ntFlanks@5488.5515.28|GCTGAGAAGACGGTCGAACTTGACTATC</td><td>>hg38_21_+_8433172_8446622_RNA45SN1_45S_50ntFlanks</td><td>5488</td><td>5515</td><td>28</td><td>GCTGAGAAGACGGTCGAACTTGACTATC</td><td>655.3</td><td>5.6</td><td>1.8</td><td>3.0</td><td>2.3E-03</td><td>1.2E-02</td><td>BT20/MB231</td><td>MP</td></tr>
<tr><td>>hg38_21_+_8433172_8446622_RNA45SN1_45S_50ntFlanks@11318.11344.27|CCCCACGTCTCGTCGCGCGCGCGTCCG</td><td>>hg38_21_+_8433172_8446622_RNA45SN1_45S_50ntFlanks</td><td>11318</td><td>11344</td><td>27</td><td>CCCCACGTCTCGTCGCGCGCGCGTCCG</td><td>223.2</td><td>-4.7</td><td>1.7</td><td>-2.7</td><td>6.0E-03</td><td>1.2E-02</td><td>BT20/MB468</td><td>MP</td></tr>
<tr><td>>trnaMT_ValTAC_MT_+_1602_1670@29.71.43|ACTTACACTTAGGAGATTTCAACTTAACTTGACCGCTCTGACC</td><td>>trnaMT_ValTAC_MT_+_1602_1670</td><td>29</td><td>71</td><td>43</td><td>ACTTACACTTAGGAGATTTCAACTTAACTTGACCGCTCTGACC</td><td>278.1</td><td>3.7</td><td>1.4</td><td>2.6</td><td>1.0E-02</td><td>1.2E-02</td><td>MB231/MB468</td><td>MITO</td></tr>
<tr><td>>MI0000067|hsa-let-7f-1&WithFlank&9|+|94176341|94176439@13.32.20(+1A);MI0000068|hsa-let-7f-2&WithFlank&X|-|53557186|53557280@14.33.20(+1A)&[MIMAT0000067&hsa-let-7f-5p&offsets|0|-2(+1A);m-20&9|+|94176353|94176374&offsets|0|-2(+1A)];[MIMAT0000067_1&hsa-let-7f-5p&offsets|0|-2(+1A);m-19&X|-|53557246|53557267&offsets|0|-2(+1A)]|TGAGGTAGTAGATTGTATAGA</td><td>>MI0000067|hsa-let-7f-1&WithFlank&9|+|94176341|94176439</td><td>13</td><td>32</td><td>20(+1A)</td><td>TGAGGTAGTAGATTGTATAGA</td><td>310.9</td><td>2.5</td><td>0.8</td><td>3.0</td><td>2.9E-03</td><td>1.2E-02</td><td>BT20/MB231</td><td>CYTO</td></tr>
<tr><td>>trnaMT_GluTTC_MT_-_14674_14742@25.72.48,trnalookalike8_GluTTC_5_-_93905172_93905240@25.72.48|TGGTTTTTCATATCATTGGTCGTGGTTGTAGTCCGTGCGAGAATACCA</td><td>>trnaMT_GluTTC_MT_-_14674_14742</td><td>25</td><td>72</td><td>48</td><td>TGGTTTTTCATATCATTGGTCGTGGTTGTAGTCCGTGCGAGAATACCA</td><td>358.4</td><td>-6.8</td><td>2.3</td><td>-3.0</td><td>2.9E-03</td><td>1.2E-02</td><td>BT20/MB231</td><td>CYTO</td></tr>
<tr><td>>trnaMT_HisGTG_MT_+_12138_12206@32.70.39|TGAATCTGACAACAGAGGCTTACGACCCCTTATTTACCC</td><td>>trnaMT_HisGTG_MT_+_12138_12206</td><td>32</td><td>70</td><td>39</td><td>TGAATCTGACAACAGAGGCTTACGACCCCTTATTTACCC</td><td>610.0</td><td>-3.2</td><td>1.1</td><td>-3.0</td><td>2.9E-03</td><td>1.2E-02</td><td>BT20/MB231</td><td>CYTO</td></tr>
<tr><td>>hg38_21_+_8433172_8446622_RNA45SN1_45S_50ntFlanks@11670.11702.33|CCCAGTGCTCTGAATGTCAAAGTGAAGAAATTC</td><td>>hg38_21_+_8433172_8446622_RNA45SN1_45S_50ntFlanks</td><td>11670</td><td>11702</td><td>33</td><td>CCCAGTGCTCTGAATGTCAAAGTGAAGAAATTC</td><td>448.0</td><td>-2.9</td><td>1.0</td><td>-3.0</td><td>2.9E-03</td><td>1.2E-02</td><td>BT20/MB231</td><td>CYTO</td></tr>
<tr><td>>hg38_21_+_8433172_8446622_RNA45SN1_45S_50ntFlanks@12423.12455.33|TCGGCTCTTCCTATCATTGTGAAGCAGAATTCA</td><td>>hg38_21_+_8433172_8446622_RNA45SN1_45S_50ntFlanks</td><td>12423</td><td>12455</td><td>33</td><td>TCGGCTCTTCCTATCATTGTGAAGCAGAATTCA</td><td>148.2</td><td>-3.1</td><td>1.2</td><td>-2.6</td><td>8.5E-03</td><td>1.2E-02</td><td>BT20/MB468</td><td>MITO</td></tr>
<tr><td>>hg38_21_+_8433172_8446622_RNA45SN1_45S_50ntFlanks@6719.6751.33|TGAATTGCAGGACACATTGATCATCGACACTTC</td><td>>hg38_21_+_8433172_8446622_RNA45SN1_45S_50ntFlanks</td><td>6719</td><td>6751</td><td>33</td><td>TGAATTGCAGGACACATTGATCATCGACACTTC</td><td>232.7</td><td>-2.5</td><td>1.0</td><td>-2.6</td><td>9.9E-03</td><td>1.2E-02</td><td>MB231/MB468</td><td>CYTO</td></tr>
<tr><td>>hg38_21_+_8433172_8446622_RNA45SN1_45S_50ntFlanks@9597.9617.21|AGGGGCGAAAGACTAATCGAA</td><td>>hg38_21_+_8433172_8446622_RNA45SN1_45S_50ntFlanks</td><td>9597</td><td>9617</td><td>21</td><td>AGGGGCGAAAGACTAATCGAA</td><td>146.2</td><td>-3.7</td><td>1.4</td><td>-2.7</td><td>6.1E-03</td><td>1.2E-02</td><td>BT20/MB468</td><td>MP</td></tr>
<tr><td>>MI0000239|hsa-mir-197&WithFlank&1|+|109598887|109598973@54.74.21&[MIMAT0000227&hsa-miR-197-3p&offsets|0|-1;m-99&1|+|109598940|109598961&offsets|0|-1]|TTCACCACCTTCTCCACCCAG</td><td>>MI0000239|hsa-mir-197&WithFlank&1|+|109598887|109598973</td><td>54</td><td>74</td><td>21</td><td>TTCACCACCTTCTCCACCCAG</td><td>139.0</td><td>4.9</td><td>1.8</td><td>2.7</td><td>6.1E-03</td><td>1.2E-02</td><td>BT20/MB468</td><td>MP</td></tr>
<tr><td>>hg38_21_+_8433172_8446622_RNA45SN1_45S_50ntFlanks@11582.11600.19|ACTGGTGCGGACCAGGGGA</td><td>>hg38_21_+_8433172_8446622_RNA45SN1_45S_50ntFlanks</td><td>11582</td><td>11600</td><td>19</td><td>ACTGGTGCGGACCAGGGGA</td><td>352.3</td><td>-2.0</td><td>0.8</td><td>-2.6</td><td>9.9E-03</td><td>1.2E-02</td><td>MB231/MB468</td><td>CYTO</td></tr>
<tr><td>>hg38_21_+_8433172_8446622_RNA45SN1_45S_50ntFlanks@9821.9850.30|CCTAGTGGGCCACTTTTGGTAAGCAGAACT</td><td>>hg38_21_+_8433172_8446622_RNA45SN1_45S_50ntFlanks</td><td>9821</td><td>9850</td><td>30</td><td>CCTAGTGGGCCACTTTTGGTAAGCAGAACT</td><td>1508.5</td><td>-2.1</td><td>0.8</td><td>-2.6</td><td>8.5E-03</td><td>1.2E-02</td><td>BT20/MB468</td><td>MITO</td></tr>
<tr><td>>MI0003581|hsa-mir-574&WithFlank&4|+|38868026|38868133@67.87.21(+1U)&[MIMAT0003239&hsa-miR-574-3p&offsets|0|-1(+1U);m-149&4|+|38868092|38868113&offsets|0|-1(+1U)]|CACGCTCATGCACACACCCACT</td><td>>MI0003581|hsa-mir-574&WithFlank&4|+|38868026|38868133</td><td>67</td><td>87</td><td>21(+1U)</td><td>CACGCTCATGCACACACCCACT</td><td>110.2</td><td>4.3</td><td>1.6</td><td>2.6</td><td>9.1E-03</td><td>1.2E-02</td><td>MB231/MB468</td><td>MP</td></tr>
<tr><td>>am_trnaMT_SerGCT_MT_+_12207_12265@11.32.22|ACAAGAACTGCTAACTCATGCC</td><td>>am_trnaMT_SerGCT_MT_+_12207_12265</td><td>11</td><td>32</td><td>22</td><td>ACAAGAACTGCTAACTCATGCC</td><td>72.0</td><td>2.1</td><td>0.8</td><td>2.6</td><td>9.1E-03</td><td>1.2E-02</td><td>MB231/MB468</td><td>MP</td></tr>
<tr><td>>trnaMT_GluTTC_MT_-_14674_14742@37.72.36,trnalookalike8_GluTTC_5_-_93905172_93905240@37.72.36|TCATTGGTCGTGGTTGTAGTCCGTGCGAGAATACCA</td><td>>trnaMT_GluTTC_MT_-_14674_14742</td><td>37</td><td>72</td><td>36</td><td>TCATTGGTCGTGGTTGTAGTCCGTGCGAGAATACCA</td><td>1693.7</td><td>3.4</td><td>1.2</td><td>2.7</td><td>6.1E-03</td><td>1.2E-02</td><td>BT20/MB468</td><td>MP</td></tr>
<tr><td>>hg38_21_+_8433172_8446622_RNA45SN1_45S_50ntFlanks@9860.9885.26|GGATGAACCGAACGCCGGGTTAAGGC</td><td>>hg38_21_+_8433172_8446622_RNA45SN1_45S_50ntFlanks</td><td>9860</td><td>9885</td><td>26</td><td>GGATGAACCGAACGCCGGGTTAAGGC</td><td>61.9</td><td>5.6</td><td>2.0</td><td>2.7</td><td>6.1E-03</td><td>1.2E-02</td><td>BT20/MB468</td><td>MP</td></tr>
</table>

| | | | | | | | | | | | | | |
|---|---|---|---|---|---|---|---|---|---|---|---|---|---|
| >hg38_21_+_8433172_8446622_RNA45SN1_45S_50ntFlanks@12936.12953.18\|AAACCATTCGTAGACGAC | >hg38_21_+_8433172_8446622_RNA45SN1_45S_50ntFlanks | 12936 | 12953 | 18 | AAACCATTCGTAGACGAC | 448.8 | -4.7 | 1.7 | -2.7 | 6.1E-03 | 1.2E-02 | BT20/MB468 | MP |
| >hg38_21_+_8433172_8446622_RNA45SN1_45S_50ntFlanks@11767.11803.37\|TCTAATTAGTGACGCGCATGAATGGATGAACGAGATT | >hg38_21_+_8433172_8446622_RNA45SN1_45S_50ntFlanks | 11767 | 11803 | 37 | TCTAATTAGTGACGCGCATGAATGGATGAACGAGATT | 383.6 | 1.8 | 0.7 | 2.6 | 8.5E-03 | 1.2E-02 | BT20/MB468 | MITO |
| >trnaMT_MetCAT_MT_+_4402_4469@33.71.39\|TACCCCGAAAATGTTGGTTATACCCTTCCCGTACTACCA | >trnaMT_MetCAT_MT_+_4402_4469 | 33 | 71 | 39 | TACCCCGAAAATGTTGGTTATACCCTTCCCGTACTACCA | 326.6 | -4.5 | 1.5 | -3.0 | 2.6E-03 | 1.2E-02 | BT20/MB231 | NUC |
| >hg38_21_+_8433172_8446622_RNA45SN1_45S_50ntFlanks@7969.7988.20\|CCGAGACGCGACCTCAGATC | >hg38_21_+_8433172_8446622_RNA45SN1_45S_50ntFlanks | 7969 | 7988 | 20 | CCGAGACGCGACCTCAGATC | 167.5 | 2.2 | 0.9 | 2.6 | 1.0E-02 | 1.2E-02 | BT20/MB468 | NUC |
| >MI0001641\|hsa-mir-429&WithFlank&1\|+\|1168999\|1169093@57.78.22&[MIMAT0001536&hsa-miR-429&offsets\|0\|0;m-207&1\|+\|1169055\|1169076&offsets\|0\|0]\|TAATACTGTCTGGTAAAACCGT | >MI0001641\|hsa-mir-429&WithFlank&1\|+\|1168999\|1169093 | 57 | 78 | 22 | TAATACTGTCTGGTAAAACCGT | 257.9 | -3.0 | 1.2 | -2.6 | 1.1E-02 | 1.2E-02 | MB231/MB468 | NUC |
| >MI0000253\|hsa-mir-148a&WithFlank&7\|-\|25949913\|25949992@12.33.22&[MIMAT0004549&hsa-miR-148a-5p&offsets\|0\|0;m-189&7\|-\|25949960\|25949981&offsets\|0\|0]\|AAAGTTCTGAGACACTCCGACT | >MI0000253\|hsa-mir-148a&WithFlank&7\|-\|25949913\|25949992 | 12 | 33 | 22 | AAAGTTCTGAGACACTCCGACT | 351.1 | 3.0 | 1.0 | 3.0 | 2.3E-03 | 1.2E-02 | BT20/MB231 | MP |
| >hg38_21_+_8433172_8446622_RNA45SN1_45S_50ntFlanks@12942.12959.18\|TTCGTAGACGACCTGCTT | >hg38_21_+_8433172_8446622_RNA45SN1_45S_50ntFlanks | 12942 | 12959 | 18 | TTCGTAGACGACCTGCTT | 1102.9 | -1.5 | 0.6 | -2.6 | 1.0E-02 | 1.2E-02 | MB231/MB468 | CYTO |
| >hg38_21_+_8433172_8446622_RNA45SN1_45S_50ntFlanks@6652.6676.25\|GACTCTTAGCGGTGGATCACTCGGC | >hg38_21_+_8433172_8446622_RNA45SN1_45S_50ntFlanks | 6652 | 6676 | 25 | GACTCTTAGCGGTGGATCACTCGGC | 3139.3 | 2.6 | 1.0 | 2.6 | 1.1E-02 | 1.2E-02 | MB231/MB468 | MITO |
| >MI0000746\|hsa-mir-99b&WithFlank&19\|+\|51692606\|51692687@13.32.20(+1A)&[MIMAT0000689&hsa-miR-99b-5p&offsets\|0\|-2(+1A);m-10&19\|+\|51692618\|51692639&offsets\|0\|-2(+1A)]\|CACCCGTAGAACCGACCTTGA | >MI0000746\|hsa-mir-99b&WithFlank&19\|+\|51692606\|51692687 | 13 | 32 | 20(+1A) | CACCCGTAGAACCGACCTTGA | 234.5 | 3.4 | 1.3 | 2.6 | 9.2E-03 | 1.2E-02 | MB231/MB468 | MP |
| >hg38_21_+_8433172_8446622_RNA45SN1_45S_50ntFlanks@9789.9819.31\|GCTGGCGTGGAGCCGGGCGTGGAATGCGAGT | >hg38_21_+_8433172_8446622_RNA45SN1_45S_50ntFlanks | 9789 | 9819 | 31 | GCTGGCGTGGAGCCGGGCGTGGAATGCGAGT | 185.5 | 4.2 | 1.5 | 2.7 | 6.1E-03 | 1.2E-02 | BT20/MB468 | MP |
| >trnaMT_GluTTC_MT_-_14674_14742@34.63.30,trnalookalike8_GluTTC_5_-_93905172_93905240@34.63.30\|ATATCATTGGTCGTGGTTGTAGTCCGTGCG | >trnaMT_GluTTC_MT_-_14674_14742 | 34 | 63 | 30 | ATATCATTGGTCGTGGTTGTAGTCCGTGCG | 101.5 | 2.3 | 0.9 | 2.6 | 9.2E-03 | 1.2E-02 | MB231/MB468 | MP |
| >hg38_MT_+_1621_3279_RNR2_16S_50ntFlanks@622.651.30\|TATAACTGAACTCCTCACACCCAATTGGAC | >hg38_MT_+_1621_3279_RNR2_16S_50ntFlanks | 622 | 651 | 30 | TATAACTGAACTCCTCACACCCAATTGGAC | 65.7 | 5.7 | 2.1 | 2.7 | 6.1E-03 | 1.2E-02 | BT20/MB468 | MP |
| >hg38_21_+_8433172_8446622_RNA45SN1_45S_50ntFlanks@7975.8011.37\|CGCGACCTCAGATCAGACGTGGCGACCCGCTGAATTT | >hg38_21_+_8433172_8446622_RNA45SN1_45S_50ntFlanks | 7975 | 8011 | 37 | CGCGACCTCAGATCAGACGTGGCGACCCGCTGAATTT | 113331.6 | 1.8 | 0.7 | 2.6 | 1.1E-02 | 1.2E-02 | MB231/MB468 | MITO |
| >trna2_GlyGCC_21_-_18827107_18827177@1.28.28,trna35_GlyGCC_1_+_161413094_161413164@1.28.28,trna37_GlyGCC_1_+_161420467_161420537@1.28.28,trna39_GlyGCC_1_+_161427898_161427968@1.28.28,trna41_GlyGCC_1_+_161435258_161435328@1.28.28\|GCATGGGTGGTTCAGTGGTAGAATTCTC | >trna2_GlyGCC_21_-_18827107_18827177 | 1 | 28 | 28 | GCATGGGTGGTTCAGTGGTAGAATTCTC | 47142.3 | -1.8 | 0.7 | -2.6 | 1.0E-02 | 1.2E-02 | BT20/MB468 | CYTO |
| >hg38_21_+_8433172_8446622_RNA45SN1_45S_50ntFlanks@10060.10081.22\|GCAGTCGAGAGTGGACGGGAGC | >hg38_21_+_8433172_8446622_RNA45SN1_45S_50ntFlanks | 10060 | 10081 | 22 | GCAGTCGAGAGTGGACGGGAGC | 119.6 | 4.7 | 1.7 | 2.7 | 6.1E-03 | 1.2E-02 | BT20/MB468 | MP |
| >hg38_21_+_8433172_8446622_RNA45SN1_45S_50ntFlanks@4049.4066.18\|TCGAACGTCTGCCCTATC | >hg38_21_+_8433172_8446622_RNA45SN1_45S_50ntFlanks | 4049 | 4066 | 18 | TCGAACGTCTGCCCTATC | 210.7 | -3.5 | 1.3 | -2.6 | 8.6E-03 | 1.2E-02 | BT20/MB468 | MITO |
| >MI0001448\|hsa-mir-425&WithFlank&3\|-\|49020142\|49020240@20.43.24(+1C)&[MIMAT0003393&hsa-miR-425-5p&offsets\|0\|+1(+1C);m-95&3\|-\|49020200\|49020221&offsets\|0\|+2(+1C)]\|AATGACACGATCACTCCCGTTGAGC | >MI0001448\|hsa-mir-425&WithFlank&3\|-\|49020142\|49020240 | 20 | 43 | 24(+1C) | AATGACACGATCACTCCCGTTGAGC | 329.5 | -2.9 | 1.0 | -3.0 | 2.3E-03 | 1.2E-02 | BT20/MB231 | MP |
| >MI0000289\|hsa-mir-181a-1&WithFlank&1\|-\|198859038\|198859159@30.52.23(+1C);MI0000269\|hsa-mir-181a-2&WithFlank&9\|+\|124692436\|124692557@45.67.23(+1C)&[MIMAT0000256&hsa-miR-181a-5p&offsets\|0\|0(+1C);m-32&1\|-\|198859108\|198859130&offsets\|0\|0(+1C)];[MIMAT0000256_1&hsa-miR-181a-5p&offsets\|0\|0(+1C);m-32&9\|+\|124692480\|124692502&offsets\|0\|0(+1C)]\|AACATTCAACGCTGTCGGTGAGTC | >MI0000289\|hsa-mir-181a-1&WithFlank&1\|-\|198859038\|198859159 | 30 | 52 | 23(+1C) | AACATTCAACGCTGTCGGTGAGTC | 946.2 | 3.9 | 1.5 | 2.6 | 9.2E-03 | 1.2E-02 | MB231/MB468 | MP |
| >MI0000736\|hsa-mir-30c-1&WithFlank&1\|+\|40757278\|40757378@23.44.22(+1U);MI0000254\|hsa-mir-30c-2&WithFlank&6\|-\|71376954\|71377037@13.34.22(+1U)&[MIMAT0000244&hsa-miR-30c-5p&offsets\|0\|-1(+1U);m-57&1\|+\|40757300\|40757323&offsets\|0\|-2(+1U)];[MIMAT0000244_1&hsa-miR-30c-5p&offsets\|0\|-1(+1U);m-56&6\|-\|71377002\|71377025&offsets\|0\|-2(+1U)]\|TGTAAACATCCTACACTCTCAGT | >MI0000736\|hsa-mir-30c-1&WithFlank&1\|+\|40757278\|40757378 | 23 | 44 | 22(+1U) | TGTAAACATCCTACACTCTCAGT | 3691.9 | -3.5 | 1.1 | -3.0 | 2.3E-03 | 1.2E-02 | BT20/MB231 | MP |
| >hg38_21_+_8433172_8446622_RNA45SN1_45S_50ntFlanks@11563.11600.38\|GCCTAGCAGCCGACTTAGAACTGGTGCGGACCAGGGGA | >hg38_21_+_8433172_8446622_RNA45SN1_45S_50ntFlanks | 11563 | 11600 | 38 | GCCTAGCAGCCGACTTAGAACTGGTGCGGACCAGGGGA | 114.6 | 3.8 | 1.4 | 2.7 | 6.1E-03 | 1.2E-02 | BT20/MB468 | MP |
| >MI0003184\|hsa-mir-500a&WithFlank&X\|+\|50008425\|50008520@58.77.20(+1C);MI0003186\|hsa-mir-502&WithFlank&X\|+\|50014592\|50014689@59.78.20(+1C)&[MIMAT0002871&hsa-miR-500a-3p&offsets\|0\|-2(+1C);m-111&X\|+\|50008482\|50008504&offsets\|0\|-3(+1C)];[MIMAT0004775&hsa-miR-502-3p&offsets\|+1\|-1(+1C);m-236&X\|+\|50014649\|50014669&offsets\|+1\|0(+1C)]\|ATGCACCTGGGCAAGGATTCC | >MI0003184\|hsa-mir-500a&WithFlank&X\|+\|50008425\|50008520 | 58 | 77 | 20(+1C) | ATGCACCTGGGCAAGGATTCC | 50.1 | 5.0 | 1.8 | 2.7 | 6.1E-03 | 1.2E-02 | BT20/MB468 | MP |
| >hg38_21_+_8433172_8446622_RNA45SN1_45S_50ntFlanks@4007.4042.36\|CCTCGGGCCGATCGCACGCCCCCCGTGGCGGCGACG | >hg38_21_+_8433172_8446622_RNA45SN1_45S_50ntFlanks | 4007 | 4042 | 36 | CCTCGGGCCGATCGCACGCCCCCCGTGGCGGCGACG | 190.3 | -1.3 | 0.5 | -2.6 | 8.6E-03 | 1.2E-02 | BT20/MB468 | MITO |
| >hg38_21_+_8433172_8446622_RNA45SN1_45S_50ntFlanks@10872.10893.22\|GCTGGGTCGGTCGGGCTGGGGC | >hg38_21_+_8433172_8446622_RNA45SN1_45S_50ntFlanks | 10872 | 10893 | 22 | GCTGGGTCGGTCGGGCTGGGGC | 148.0 | 5.9 | 2.0 | 3.0 | 2.4E-03 | 1.2E-02 | BT20/MB231 | MP |
| >hg38_21_+_8433172_8446622_RNA45SN1_45S_50ntFlanks@8381.8403.23\|AAGAGGGCGTGAAACCGTTAAGA | >hg38_21_+_8433172_8446622_RNA45SN1_45S_50ntFlanks | 8381 | 8403 | 23 | AAGAGGGCGTGAAACCGTTAAGA | 223.7 | -3.9 | 1.5 | -2.6 | 1.0E-02 | 1.2E-02 | BT20/MB468 | CYTO |
| >hg38_21_+_8433172_8446622_RNA45SN1_45S_50ntFlanks@3850.3881.32\|GATAACTGTGGTAATTCTAGAGCTAATACATG | >hg38_21_+_8433172_8446622_RNA45SN1_45S_50ntFlanks | 3850 | 3881 | 32 | GATAACTGTGGTAATTCTAGAGCTAATACATG | 160.0 | -4.7 | 1.7 | -2.7 | 6.1E-03 | 1.2E-02 | BT20/MB468 | MP |
| >hg38_21_+_8433172_8446622_RNA45SN1_45S_50ntFlanks@6657.6689.33\|TTAGCGGTGGATCACTCGGCTCGTGCGTCGATG | >hg38_21_+_8433172_8446622_RNA45SN1_45S_50ntFlanks | 6657 | 6689 | 33 | TTAGCGGTGGATCACTCGGCTCGTGCGTCGATG | 267.5 | -2.5 | 0.9 | -2.7 | 6.1E-03 | 1.2E-02 | BT20/MB468 | MP |
| >hg38_21_+_8433172_8446622_RNA45SN1_45S_50ntFlanks@7981.8002.22\|CTCAGATCAGACGTGGCGACCC | >hg38_21_+_8433172_8446622_RNA45SN1_45S_50ntFlanks | 7981 | 8002 | 22 | CTCAGATCAGACGTGGCGACCC | 79.8 | 2.6 | 0.9 | 2.7 | 6.1E-03 | 1.2E-02 | BT20/MB468 | MP |
| >trna10_AlaCGC_6_+_26553731_26553802@54.75.22,trna119_AlaCGC_6_-_28641613_28641684@54.75.22,trna13_AlaTGC_12_-_125406301_125406372@54.75.22,trna66_AlaTGC_6_+_28611222_28611293@54.75.22,trna8_AlaTGC_5_+_180633868_180633939@54.75.22\|TCGATCCCCGGCATCTCCACCA | >trna10_AlaCGC_6_+_26553731_26553802 | 54 | 75 | 22 | TCGATCCCCGGCATCTCCACCA | 494.7 | 2.6 | 0.9 | 3.0 | 3.0E-03 | 1.2E-02 | BT20/MB231 | CYTO |
| >hg38_21_+_8433172_8446622_RNA45SN1_45S_50ntFlanks@4003.4023.21\|ATAACCTCGGGCCGATCGCAC | >hg38_21_+_8433172_8446622_RNA45SN1_45S_50ntFlanks | 4003 | 4023 | 21 | ATAACCTCGGGCCGATCGCAC | 523.5 | -2.0 | 0.8 | -2.6 | 8.6E-03 | 1.2E-02 | BT20/MB468 | MITO |
| >trnaMT_SerGCT_MT_+_12207_12265@13.61.49\|AAGAACTGCTAACTCATGCCCCCATGTCTAACAACATGGCTTTCTCACC | >trnaMT_SerGCT_MT_+_12207_12265 | 13 | 61 | 49 | AAGAACTGCTAACTCATGCCCCCATGTCTAACAACATGGCTTTCTCACC | 495.5 | 7.2 | 2.6 | 2.7 | 6.1E-03 | 1.2E-02 | BT20/MB468 | MP |
| >MI0003643\|hsa-mir-629&WithFlank&15\|-\|70079366\|70079474@28.49.22&[MIMAT0004810&hsa-miR-629-5p&offsets\|0\|+1;m-150&15\|-\|70079427\|70079447&offsets\|0\|+1]\|TGGGTTTACGTTGGGAGAACTT | >MI0003643\|hsa-mir-629&WithFlank&15\|-\|70079366\|70079474 | 28 | 49 | 22 | TGGGTTTACGTTGGGAGAACTT | 3325.7 | 2.3 | 0.8 | 3.0 | 3.0E-03 | 1.2E-02 | BT20/MB231 | CYTO |
| >hg38_21_+_8433172_8446622_RNA45SN1_45S_50ntFlanks@9568.9605.38\|CGTGCAAATCGGTCGTCCGACCTGGGTATAGGGGCGAA | >hg38_21_+_8433172_8446622_RNA45SN1_45S_50ntFlanks | 9568 | 9605 | 38 | CGTGCAAATCGGTCGTCCGACCTGGGTATAGGGGCGAA | 58.5 | -5.6 | 2.2 | -2.6 | 9.3E-03 | 1.2E-02 | MB231/MB468 | MP |
| >trnaMT_HisGTG_MT_+_12138_12206@28.62.35\|ATTGTGAATCTGACAACAGAGGCTTACGACCCCTT | >trnaMT_HisGTG_MT_+_12138_12206 | 28 | 62 | 35 | ATTGTGAATCTGACAACAGAGGCTTACGACCCCTT | 102.2 | -4.3 | 1.7 | -2.6 | 1.0E-02 | 1.2E-02 | BT20/MB468 | CYTO |
| >hg38_21_+_8433172_8446622_RNA45SN1_45S_50ntFlanks@7989.8016.28\|AGACGTGGCGACCCGCTGAATTTAAGCA | >hg38_21_+_8433172_8446622_RNA45SN1_45S_50ntFlanks | 7989 | 8016 | 28 | AGACGTGGCGACCCGCTGAATTTAAGCA | 848.7 | 2.0 | 0.8 | 2.6 | 8.6E-03 | 1.2E-02 | BT20/MB468 | MITO |
| >hg38_21_+_8433172_8446622_RNA45SN1_45S_50ntFlanks@11914.11940.27\|AGAGACATGAGAGGTGTAGAATAAGTG | >hg38_21_+_8433172_8446622_RNA45SN1_45S_50ntFlanks | 11914 | 11940 | 27 | AGAGACATGAGAGGTGTAGAATAAGTG | 317.7 | 5.2 | 1.7 | 3.0 | 2.4E-03 | 1.2E-02 | BT20/MB231 | MP |
| >hg38_21_+_8433172_8446622_RNA45SN1_45S_50ntFlanks@11660.11693.34\|GTGATTTCTGCCCAGTGCTCTGAATGTCAAAGTG | >hg38_21_+_8433172_8446622_RNA45SN1_45S_50ntFlanks | 11660 | 11693 | 34 | GTGATTTCTGCCCAGTGCTCTGAATGTCAAAGTG | 183.2 | 4.3 | 1.6 | 2.7 | 6.2E-03 | 1.2E-02 | BT20/MB468 | MP |
| >hg38_21_+_8433172_8446622_RNA45SN1_45S_50ntFlanks@7984.8009.26\|AGATCAGACGTGGCGACCCGCTGAAT | >hg38_21_+_8433172_8446622_RNA45SN1_45S_50ntFlanks | 7984 | 8009 | 26 | AGATCAGACGTGGCGACCCGCTGAAT | 983.9 | -2.1 | 0.8 | -2.7 | 6.2E-03 | 1.2E-02 | BT20/MB468 | MP |
| >MI0000749\|hsa-mir-30e&WithFlank&1\|+\|40754349\|40754452@65.85.21&[MIMAT0000693&hsa-miR-30e-3p&offsets\|0\|-1;m-25&1\|+\|40754413\|40754434&offsets\|0\|-1]\|CTTTCAGTCGGATGTTTACAG | >MI0000749\|hsa-mir-30e&WithFlank&1\|+\|40754349\|40754452 | 65 | 85 | 21 | CTTTCAGTCGGATGTTTACAG | 341.0 | -3.1 | 1.0 | -3.0 | 2.4E-03 | 1.2E-02 | BT20/MB231 | MP |
| >hg38_21_+_8433172_8446622_RNA45SN1_45S_50ntFlanks@9940.9961.22\|CAGGACGGTGGCCATGGAAGTC | >hg38_21_+_8433172_8446622_RNA45SN1_45S_50ntFlanks | 9940 | 9961 | 22 | CAGGACGGTGGCCATGGAAGTC | 445.2 | 4.8 | 1.6 | 3.0 | 2.4E-03 | 1.2E-02 | BT20/MB231 | MP |
| >MI0000286\|hsa-mir-210&WithFlank&11\|-\|568083\|568204@72.94.23&[MIMAT0000267&hsa-miR-210-3p&offsets\|0\|+1;m-102&11\|-\|568112\|568133&offsets\|0\|+1]\|CTGTGCGTGTGACAGCGGCTGAT | >MI0000286\|hsa-mir-210&WithFlank&11\|-\|568083\|568204 | 72 | 94 | 23 | CTGTGCGTGTGACAGCGGCTGAT | 343.8 | 3.1 | 1.0 | 3.0 | 3.0E-03 | 1.2E-02 | BT20/MB231 | CYTO |
| >hg38_21_+_8433172_8446622_RNA45SN1_45S_50ntFlanks@5086.5108.23\|AACGAGACTCTGGCATGCTAACT | >hg38_21_+_8433172_8446622_RNA45SN1_45S_50ntFlanks | 5086 | 5108 | 23 | AACGAGACTCTGGCATGCTAACT | 842.6 | 4.6 | 1.5 | 3.0 | 2.4E-03 | 1.2E-02 | BT20/MB231 | MP |
| >MI0000076\|hsa-mir-20a&WithFlank&13\|+\|91351059\|91351141@14.35.22(+1U)&[MIMAT0000075&hsa-miR-20a-5p&offsets\|0\|-1(+1U);m-96&13\|+\|91351072\|91351094&offsets\|0\|-1(+1U)]\|TAAAGTGCTTATAGTGCAGGTAT | >MI0000076\|hsa-mir-20a&WithFlank&13\|+\|91351059\|91351141 | 14 | 35 | 22(+1U) | TAAAGTGCTTATAGTGCAGGTAT | 459.5 | -4.4 | 1.4 | -3.0 | 2.4E-03 | 1.2E-02 | BT20/MB231 | MP |
| >MI0000088\|hsa-mir-30a&WithFlank&6\|-\|71403545\|71403627@53.73.21(+1A)&[MIMAT0000088&hsa-miR-30a-3p&offsets\|0\|-1(+1A);m-38&6\|-\|71403554\|71403575&offsets\|0\|-1(+1A)]\|CTTTCAGTCGGATGTTTGCAGA | >MI0000088\|hsa-mir-30a&WithFlank&6\|-\|71403545\|71403627 | 53 | 73 | 21(+1A) | CTTTCAGTCGGATGTTTGCAGA | 107.3 | 1.4 | 0.6 | 2.6 | 1.0E-02 | 1.2E-02 | BT20/MB468 | NUC |
| >MI0000825\|hsa-mir-345&WithFlank&14\|+\|100307853\|100307962@24.44.21&[MIMAT0000772&hsa-miR-345-5p&offsets\|0\|-1;m-193&14\|+\|100307876\|100307896&offsets\|0\|0]\|GCTGACTCCTAGTCCAGGGCT | >MI0000825\|hsa-mir-345&WithFlank&14\|+\|100307853\|100307962 | 24 | 44 | 21 | GCTGACTCCTAGTCCAGGGCT | 241.9 | 3.5 | 1.3 | 2.6 | 9.3E-03 | 1.2E-02 | MB231/MB468 | MP |
| >hg38_21_+_8433172_8446622_RNA45SN1_45S_50ntFlanks@4762.4782.21\|AGATACCGTCGTAGTTCCGAC | >hg38_21_+_8433172_8446622_RNA45SN1_45S_50ntFlanks | 4762 | 4782 | 21 | AGATACCGTCGTAGTTCCGAC | 205.3 | -3.4 | 1.3 | -2.6 | 9.3E-03 | 1.2E-02 | MB231/MB468 | MP |
| >hg38_21_+_8433172_8446622_RNA45SN1_45S_50ntFlanks@12006.12043.38\|GGCCGCCGGTGAAATACCACTACTCTGATCGTTTTTTC | >hg38_21_+_8433172_8446622_RNA45SN1_45S_50ntFlanks | 12006 | 12043 | 38 | GGCCGCCGGTGAAATACCACTACTCTGATCGTTTTTTC | 291.7 | -3.2 | 1.2 | -2.6 | 8.6E-03 | 1.2E-02 | BT20/MB468 | MITO |

| | | | | | | | | | | | | | |
|---|---|---|---|---|---|---|---|---|---|---|---|---|---|
| >hg38_21_+_8433172_8446622_RNA45SN1_45S_50ntFlanks@7989.8011.23\|AGACGTGGCGACCCGCTGAATTT | >hg38_21_+_8433172_8446622_RNA45SN1_45S_50ntFlanks | 7989 | 8011 | 23 | AGACGTGGCGACCCGCTGAATTT | 3682.1 | 2.1 | 0.8 | 2.6 | 8.7E-03 | 1.2E-02 | BT20/MB468 | MITO |
| >hg38_21_+_8433172_8446622_RNA45SN1_45S_50ntFlanks@5484.5512.29\|GAGCGCTGAGAAGACGGTCGAACTTGACT | >hg38_21_+_8433172_8446622_RNA45SN1_45S_50ntFlanks | 5484 | 5512 | 29 | GAGCGCTGAGAAGACGGTCGAACTTGACT | 402.1 | 5.5 | 1.8 | 3.0 | 2.4E-03 | 1.2E-02 | BT20/MB231 | MP |
| >hg38_21_+_8433172_8446622_RNA45SN1_45S_50ntFlanks@6661.6687.27\|CGGTGGATCACTCGGCTCGTGCGTCGA | >hg38_21_+_8433172_8446622_RNA45SN1_45S_50ntFlanks | 6661 | 6687 | 27 | CGGTGGATCACTCGGCTCGTGCGTCGA | 800.2 | 2.7 | 1.0 | 2.7 | 6.2E-03 | 1.2E-02 | BT20/MB468 | MP |
| >MI0000482\|hsa-mir-185&WithFlank&22\|+\|20033133\|20033226@21.42.22&[MIMAT0000455&hsa-miR-185-5p&offsets\|0\|0;m-145&22\|+\|20033153\|20033174&offsets\|0\|0]\|TGGAGAGAAAGGCAGTTCCTGA | >MI0000482\|hsa-mir-185&WithFlank&22\|+\|20033133\|20033226 | 21 | 42 | 22 | TGGAGAGAAAGGCAGTTCCTGA | 1850.0 | 1.6 | 0.5 | 3.0 | 2.6E-03 | 1.2E-02 | BT20/MB231 | NUC |
| >hg38_21_+_8433172_8446622_RNA45SN1_45S_50ntFlanks@8105.8145.41\|CGCGGCGGGGCGCGGGACATGTGGCGTACGGAAGACCCGCT | >hg38_21_+_8433172_8446622_RNA45SN1_45S_50ntFlanks | 8105 | 8145 | 41 | CGCGGCGGGGCGCGGGACATGTGGCGTACGGAAGACCCGCT | 610.7 | 2.8 | 1.1 | 2.6 | 8.7E-03 | 1.2E-02 | BT20/MB468 | MITO |
| >hg38_21_+_8433172_8446622_RNA45SN1_45S_50ntFlanks@7982.8007.26\|TCAGATCAGACGTGGCGACCCGCTGA | >hg38_21_+_8433172_8446622_RNA45SN1_45S_50ntFlanks | 7982 | 8007 | 26 | TCAGATCAGACGTGGCGACCCGCTGA | 369.7 | -1.8 | 0.7 | -2.6 | 8.7E-03 | 1.2E-02 | BT20/MB468 | MITO |
| >hg38_21_+_8433172_8446622_RNA45SN1_45S_50ntFlanks@11566.11583.18\|TAGCAGCCGACTTAGAAC | >hg38_21_+_8433172_8446622_RNA45SN1_45S_50ntFlanks | 11566 | 11583 | 18 | TAGCAGCCGACTTAGAAC | 756.0 | -2.2 | 0.7 | -3.0 | 3.0E-03 | 1.2E-02 | BT20/MB231 | CYTO |
| >hg38_21_+_8433172_8446622_RNA45SN1_45S_50ntFlanks@4367.4407.41\|CAGTTAAAAAGCTCGTAGTTGGATCTTGGGAGCGGGCGGGC | >hg38_21_+_8433172_8446622_RNA45SN1_45S_50ntFlanks | 4367 | 4407 | 41 | CAGTTAAAAAGCTCGTAGTTGGATCTTGGGAGCGGGCGGGC | 146.4 | -2.1 | 0.8 | -2.6 | 1.0E-02 | 1.2E-02 | MB231/MB468 | CYTO |
| >hg38_1_-_228634819_228635039_RNA5S12_5S_50ntFlanks@132.171.40\|GATGGGAGACCGCCTGGGAATACCGGGTGCTGTAGGCTTT | >hg38_1_-_228634819_228635039_RNA5S12_5S_50ntFlanks | 132 | 171 | 40 | GATGGGAGACCGCCTGGGAATACCGGGTGCTGTAGGCTTT | 17268.0 | 2.5 | 1.0 | 2.6 | 1.0E-02 | 1.2E-02 | MB231/MB468 | CYTO |
| >hg38_21_+_8433172_8446622_RNA45SN1_45S_50ntFlanks@11318.11340.23\|CCCCACGTCTCGTCGCGCGCGCG | >hg38_21_+_8433172_8446622_RNA45SN1_45S_50ntFlanks | 11318 | 11340 | 23 | CCCCACGTCTCGTCGCGCGCGCG | 366.2 | -4.8 | 1.9 | -2.6 | 1.1E-02 | 1.2E-02 | BT20/MB468 | NUC |
| >hg38_21_+_8433172_8446622_RNA45SN1_45S_50ntFlanks@11541.11583.43\|GCGGCGCCTCGCCTCGGCCGGCGCCTAGCAGCCGACTTAGAAC | >hg38_21_+_8433172_8446622_RNA45SN1_45S_50ntFlanks | 11541 | 11583 | 43 | GCGGCGCCTCGCCTCGGCCGGCGCCTAGCAGCCGACTTAGAAC | 77.3 | 4.1 | 1.6 | 2.6 | 1.1E-02 | 1.2E-02 | MB231/MB468 | MITO |
| >trnaMT_SerGCT_MT_+_12207_12265@13.40.28\|AAGAACTGCTAACTCATGCCCCCATGTC | >trnaMT_SerGCT_MT_+_12207_12265 | 13 | 40 | 28 | AAGAACTGCTAACTCATGCCCCCATGTC | 326.7 | 3.3 | 1.2 | 2.7 | 6.2E-03 | 1.2E-02 | BT20/MB468 | MP |
| >hg38_MT_+_1621_3279_RNR2_16S_50ntFlanks@1367.1401.35\|TCGATGTTGGATCAGGACATCCCGATGGTGCAGCC | >hg38_MT_+_1621_3279_RNR2_16S_50ntFlanks | 1367 | 1401 | 35 | TCGATGTTGGATCAGGACATCCCGATGGTGCAGCC | 156.4 | 3.2 | 1.2 | 2.6 | 9.3E-03 | 1.2E-02 | MB231/MB468 | MP |
| >hg38_21_+_8433172_8446622_RNA45SN1_45S_50ntFlanks@6661.6698.38\|CGGTGGATCACTCGGCTCGTGCGTCGATGAAGAACGCA | >hg38_21_+_8433172_8446622_RNA45SN1_45S_50ntFlanks | 6661 | 6698 | 38 | CGGTGGATCACTCGGCTCGTGCGTCGATGAAGAACGCA | 874.3 | 2.3 | 0.8 | 3.0 | 3.0E-03 | 1.2E-02 | BT20/MB231 | MITO |
| >hg38_21_+_8433172_8446622_RNA45SN1_45S_50ntFlanks@9545.9562.18\|GTGGAGGTCCGTAGCGGT | >hg38_21_+_8433172_8446622_RNA45SN1_45S_50ntFlanks | 9545 | 9562 | 18 | GTGGAGGTCCGTAGCGGT | 344.1 | -2.7 | 1.0 | -2.6 | 1.0E-02 | 1.2E-02 | MB231/MB468 | CYTO |
| >hg38_21_+_8433172_8446622_RNA45SN1_45S_50ntFlanks@4051.4076.26\|GAACGTCTGCCCTATCAACTTTCGAT | >hg38_21_+_8433172_8446622_RNA45SN1_45S_50ntFlanks | 4051 | 4076 | 26 | GAACGTCTGCCCTATCAACTTTCGAT | 222.0 | -2.3 | 0.8 | -3.0 | 3.0E-03 | 1.2E-02 | BT20/MB231 | CYTO |
| >hg38_21_+_8433172_8446622_RNA45SN1_45S_50ntFlanks@8381.8406.26\|AAGAGGGCGTGAAACCGTTAAGAGGT | >hg38_21_+_8433172_8446622_RNA45SN1_45S_50ntFlanks | 8381 | 8406 | 26 | AAGAGGGCGTGAAACCGTTAAGAGGT | 580.6 | -2.7 | 0.9 | -3.0 | 3.0E-03 | 1.2E-02 | BT20/MB231 | CYTO |
| >hg38_1_-_228634819_228635039_RNA5S12_5S_50ntFlanks@151.169.19\|ATACCGGGTGCTGTAGGCT | >hg38_1_-_228634819_228635039_RNA5S12_5S_50ntFlanks | 151 | 169 | 19 | ATACCGGGTGCTGTAGGCT | 273.4 | 2.1 | 0.7 | 3.0 | 3.0E-03 | 1.2E-02 | BT20/MB231 | MITO |
| >hg38_MT_+_1621_3279_RNR2_16S_50ntFlanks@378.411.34\|TACCGAGCCTGGTGATAGCTGGTTGTCCAAGATA | >hg38_MT_+_1621_3279_RNR2_16S_50ntFlanks | 378 | 411 | 34 | TACCGAGCCTGGTGATAGCTGGTTGTCCAAGATA | 57.3 | 5.0 | 1.9 | 2.6 | 9.3E-03 | 1.2E-02 | MB231/MB468 | MP |
| >am_MI0006370\|hsa-mir-1303&WithFlank&5\|+\|154685770\|154685867@58.79.22&[MIMAT0005891&hsa-miR-1303&offsets\|0\|0;m-516&5\|+\|154685826\|154685848&offsets\|+1\|0]\|TTTAGAGACGGGGTCTTGCTCT | >am_MI0006370\|hsa-mir-1303&WithFlank&5\|+\|154685770\|154685867 | 58 | 79 | 22 | TTTAGAGACGGGGTCTTGCTCT | 69.4 | 5.0 | 1.9 | 2.6 | 9.3E-03 | 1.2E-02 | MB231/MB468 | MP |
| >MI0000061\|hsa-let-7a-2&WithFlank&11\|-\|122146516\|122146599@11.33.23;MI0000062\|hsa-let-7a-3&WithFlank&22\|+\|46112743\|46112828@10.32.23;MI0000060\|hsa-let-7a-1&WithFlank&9\|+\|94175951\|94176042@12.34.23&[MIMAT0000062&hsa-let-7a-5p&offsets\|0\|+1;m-6&11\|-\|122146568\|122146589&offsets\|0\|+1];[MIMAT0000062_1&hsa-let-7a-5p&offsets\|0\|+1;m-5&22\|+\|46112752\|46112773&offsets\|0\|+1];[MIMAT0000062_2&hsa-let-7a-5p&offsets\|0\|+1;m-4&9\|+\|94175962\|94175983&offsets\|0\|+1]\|TGAGGTAGTAGGTTGTATAGTTT | >MI0000061\|hsa-let-7a-2&WithFlank&11\|-\|122146516\|122146599 | 11 | 33 | 23 | TGAGGTAGTAGGTTGTATAGTTT | 8560.8 | -2.8 | 0.9 | -3.0 | 2.4E-03 | 1.2E-02 | BT20/MB231 | MP |
| >hg38_21_+_8433172_8446622_RNA45SN1_45S_50ntFlanks@6650.6681.32\|ACGACTCTTAGCGGTGGATCACTCGGCTCGTG | >hg38_21_+_8433172_8446622_RNA45SN1_45S_50ntFlanks | 6650 | 6681 | 32 | ACGACTCTTAGCGGTGGATCACTCGGCTCGTG | 1342.2 | -1.5 | 0.6 | -2.6 | 1.0E-02 | 1.2E-02 | MB231/MB468 | CYTO |
| >hg38_21_+_8433172_8446622_RNA45SN1_45S_50ntFlanks@10014.10036.23\|AAAATGGATGGCGCTGGAGCGTC | >hg38_21_+_8433172_8446622_RNA45SN1_45S_50ntFlanks | 10014 | 10036 | 23 | AAAATGGATGGCGCTGGAGCGTC | 180.9 | -3.5 | 1.3 | -2.7 | 6.2E-03 | 1.2E-02 | BT20/MB468 | MP |
| >MI0000114\|hsa-mir-107&WithFlank&10\|-\|89592741\|89592833@56.77.22(+1U)&[MIMAT0000104&hsa-miR-107&offsets\|0\|-1(+1U);m-78&10\|-\|89592758\|89592778&offsets\|0\|+1(+1U)]\|AGCAGCATTGTACAGGGCTATCT | >MI0000114\|hsa-mir-107&WithFlank&10\|-\|89592741\|89592833 | 56 | 77 | 22(+1U) | AGCAGCATTGTACAGGGCTATCT | 116.8 | -5.6 | 1.8 | -3.0 | 2.4E-03 | 1.2E-02 | BT20/MB231 | MP |
| >hg38_1_-_228634819_228635039_RNA5S12_5S_50ntFlanks@144.171.28\|CCTGGGAATACCGGGTGCTGTAGGCTTT | >hg38_1_-_228634819_228635039_RNA5S12_5S_50ntFlanks | 144 | 171 | 28 | CCTGGGAATACCGGGTGCTGTAGGCTTT | 2836.5 | -2.1 | 0.7 | -3.0 | 3.1E-03 | 1.2E-02 | BT20/MB231 | MITO |
| >hg38_21_+_8433172_8446622_RNA45SN1_45S_50ntFlanks@11933.11952.20\|AATAAGTGGGAGGCCCCCGG | >hg38_21_+_8433172_8446622_RNA45SN1_45S_50ntFlanks | 11933 | 11952 | 20 | AATAAGTGGGAGGCCCCCGG | 180.9 | -3.8 | 1.4 | -2.7 | 6.2E-03 | 1.2E-02 | BT20/MB468 | MP |
| >hg38_21_+_8433172_8446622_RNA45SN1_45S_50ntFlanks@4917.4941.25\|CACCACCAGGAGTGGAGCCTGCGGC | >hg38_21_+_8433172_8446622_RNA45SN1_45S_50ntFlanks | 4917 | 4941 | 25 | CACCACCAGGAGTGGAGCCTGCGGC | 294.6 | -3.6 | 1.4 | -2.6 | 8.7E-03 | 1.2E-02 | BT20/MB468 | MITO |
| >hg38_21_+_8433172_8446622_RNA45SN1_45S_50ntFlanks@12243.12268.26\|GAAGGGCAAAAGCTCGCTTGATCTTG | >hg38_21_+_8433172_8446622_RNA45SN1_45S_50ntFlanks | 12243 | 12268 | 26 | GAAGGGCAAAAGCTCGCTTGATCTTG | 181.8 | -3.1 | 1.2 | -2.6 | 1.1E-02 | 1.2E-02 | MB231/MB468 | NUC |
| >hg38_MT_+_1621_3279_RNR2_16S_50ntFlanks@1389.1410.22\|CGATGGTGCAGCCGCTATTAAA | >hg38_MT_+_1621_3279_RNR2_16S_50ntFlanks | 1389 | 1410 | 22 | CGATGGTGCAGCCGCTATTAAA | 22.8 | 7.7 | 3.0 | 2.6 | 9.4E-03 | 1.2E-02 | MB231/MB468 | MP |
| >MI0003776\|hsa-mir-320b-1&WithFlank&1\|+\|116671740\|116671823@48.66.19(+1A);MI0003839\|hsa-mir-320b-2&WithFlank&1\|-\|224257034\|224257116@47.65.19(+1A);MI0003778\|hsa-mir-320c-1&WithFlank&18\|+\|21683512\|21683595@48.66.19(+1A);MI0008191\|hsa-mir-320c-2&WithFlank&18\|+\|24321669\|24321752@48.66.19(+1A);MI0000542\|hsa-mir-320a&WithFlank&8\|-\|22244960\|22245043@48.66.19(+1A)&[MIMAT0005792&hsa-miR-320b&offsets\|0\|-3(+1A);m-183&1\|+\|116671787\|116671806&offsets\|0\|-1(+1A)];[MIMAT0005792_1&hsa-miR-320b&offsets\|0\|-3(+1A);m-183&1\|-\|224257051\|224257070&offsets\|0\|-1(+1A)];[MIMAT0005793&hsa-miR-320c&offsets\|0\|-1(+1A);m-222&18\|+\|21683559\|21683578&offsets\|0\|-1(+1A)];[MIMAT0005793_1&hsa-miR-320c&offsets\|0\|-1(+1A);m-219&18\|+\|24321716\|24321735&offsets\|0\|-1(+1A)];[MIMAT0000510&hsa-miR-320a-3p&offsets\|0\|-3(+1A);m-60&8\|-\|22244975\|22244996&offsets\|0\|-3(+1A)]\|AAAAGCTGGGTTGAGAGGGA | >MI0003776\|hsa-mir-320b-1&WithFlank&1\|+\|116671740\|116671823 | 48 | 66 | 19(+1A) | AAAAGCTGGGTTGAGAGGGA | 1440.0 | 2.9 | 1.0 | 3.0 | 2.4E-03 | 1.2E-02 | BT20/MB231 | MP |
| >trna111_HisGTG_1_-_147774845_147774916@1.30.30,trna118_HisGTG_1_-_145396881_145396952@1.30.30,trna16_HisGTG_1_+_146544773_146544844@1.30.30,trna1_HisGTG_15_+_45493349_45493420@1.30.30,trna21_HisGTG_1_+_147753471_147753542@1.30.30,trna33_HisGTG_6_+_27125906_27125977@1.30.30,trna7_HisGTG_9_-_14433938_14434009@1.30.30,trna8_HisGTG_15_-_45492611_45492682@1.30.30,trna9_HisGTG_15_-_45490804_45490875@1.30.30\|GCCGTGATCGTATAGTGGTTAGTACTCTGC | >trna111_HisGTG_1_-_147774845_147774916 | 1 | 30 | 30 | GCCGTGATCGTATAGTGGTTAGTACTCTGC | 1699.3 | -3.4 | 1.1 | -3.0 | 2.4E-03 | 1.2E-02 | BT20/MB231 | MP |
| >hg38_21_+_8433172_8446622_RNA45SN1_45S_50ntFlanks@7985.8004.20\|GATCAGACGTGGCGACCCGC | >hg38_21_+_8433172_8446622_RNA45SN1_45S_50ntFlanks | 7985 | 8004 | 20 | GATCAGACGTGGCGACCCGC | 181.7 | -4.0 | 1.6 | -2.6 | 1.1E-02 | 1.2E-02 | MB231/MB468 | NUC |
| >hg38_21_+_8433172_8446622_RNA45SN1_45S_50ntFlanks@12603.12624.22\|TGTATGTGCTTGGCTGAGGAGC | >hg38_21_+_8433172_8446622_RNA45SN1_45S_50ntFlanks | 12603 | 12624 | 22 | TGTATGTGCTTGGCTGAGGAGC | 142.8 | 4.2 | 1.5 | 2.7 | 6.2E-03 | 1.2E-02 | BT20/MB468 | MP |
| >hg38_21_+_8433172_8446622_RNA45SN1_45S_50ntFlanks@11581.11600.20\|AACTGGTGCGGACCAGGGGA | >hg38_21_+_8433172_8446622_RNA45SN1_45S_50ntFlanks | 11581 | 11600 | 20 | AACTGGTGCGGACCAGGGGA | 137.6 | -1.7 | 0.7 | -2.6 | 8.7E-03 | 1.2E-02 | BT20/MB468 | MITO |
| >hg38_21_+_8433172_8446622_RNA45SN1_45S_50ntFlanks@4927.4947.21\|AGTGGAGCCTGCGGCTTAATT | >hg38_21_+_8433172_8446622_RNA45SN1_45S_50ntFlanks | 4927 | 4947 | 21 | AGTGGAGCCTGCGGCTTAATT | 57.8 | 4.7 | 1.7 | 2.7 | 6.2E-03 | 1.2E-02 | BT20/MB468 | MP |
| >hg38_21_+_8433172_8446622_RNA45SN1_45S_50ntFlanks@6661.6687.27\|CGGTGGATCACTCGGCTCGTGCGTCGA | >hg38_21_+_8433172_8446622_RNA45SN1_45S_50ntFlanks | 6661 | 6687 | 27 | CGGTGGATCACTCGGCTCGTGCGTCGA | 242.8 | -2.3 | 0.9 | -2.6 | 9.4E-03 | 1.2E-02 | MB231/MB468 | MP |
| >hg38_21_+_8433172_8446622_RNA45SN1_45S_50ntFlanks@12948.12989.42\|GACGACCTGCTTCTGGGTCGGGGTTTCGTACGTAGCAGAGCA | >hg38_21_+_8433172_8446622_RNA45SN1_45S_50ntFlanks | 12948 | 12989 | 42 | GACGACCTGCTTCTGGGTCGGGGTTTCGTACGTAGCAGAGCA | 236.1 | -2.4 | 0.9 | -2.5 | 1.1E-02 | 1.2E-02 | MB231/MB468 | NUC |
| >MI0000299\|hsa-mir-222&WithFlank&X\|-\|45747009\|45747130@75.97.23(+1U)&[MIMAT0000279&hsa-miR-222-3p&offsets\|0\|+2(+1U);m-140&X\|-\|45747033\|45747056&offsets\|0\|-1(+1U)]\|AGCTACATCTGGCTACTGGGTCTT | >MI0000299\|hsa-mir-222&WithFlank&X\|-\|45747009\|45747130 | 75 | 97 | 23(+1U) | AGCTACATCTGGCTACTGGGTCTT | 3984.1 | -2.9 | 1.0 | -3.0 | 2.4E-03 | 1.2E-02 | BT20/MB231 | MP |
| >hg38_21_+_8433172_8446622_RNA45SN1_45S_50ntFlanks@6646.6678.33\|TCGTACGACTCTTAGCGGTGGATCACTCGGCTC | >hg38_21_+_8433172_8446622_RNA45SN1_45S_50ntFlanks | 6646 | 6678 | 33 | TCGTACGACTCTTAGCGGTGGATCACTCGGCTC | 4405.1 | 3.6 | 1.2 | 3.0 | 2.4E-03 | 1.2E-02 | BT20/MB231 | MP |
| >hg38_21_+_8433172_8446622_RNA45SN1_45S_50ntFlanks@12569.12587.19\|CAGTACGAGAGGAACCGCA | >hg38_21_+_8433172_8446622_RNA45SN1_45S_50ntFlanks | 12569 | 12587 | 19 | CAGTACGAGAGGAACCGCA | 94.4 | -3.8 | 1.5 | -2.6 | 1.0E-02 | 1.2E-02 | BT20/MB468 | CYTO |
| >hg38_21_+_8433172_8446622_RNA45SN1_45S_50ntFlanks@12946.12982.37\|TAGACGACCTGCTTCTGGGTCGGGGTTTCGTACGTAG | >hg38_21_+_8433172_8446622_RNA45SN1_45S_50ntFlanks | 12946 | 12982 | 37 | TAGACGACCTGCTTCTGGGTCGGGGTTTCGTACGTAG | 250.1 | -2.6 | 1.0 | -2.6 | 1.0E-02 | 1.2E-02 | MB231/MB468 | CYTO |
| >hg38_21_+_8433172_8446622_RNA45SN1_45S_50ntFlanks@9304.9338.35\|ACCAAGGAGTCTAACACGTGCGCGAGTCGGGGGCT | >hg38_21_+_8433172_8446622_RNA45SN1_45S_50ntFlanks | 9304 | 9338 | 35 | ACCAAGGAGTCTAACACGTGCGCGAGTCGGGGGCT | 1082.2 | -5.0 | 1.8 | -2.7 | 6.3E-03 | 1.2E-02 | BT20/MB468 | MP |

<table>
<tr><td>>MI0000446|hsa-mir-125b-1&WithFlank&11|-|122099751|122099850@21.41.21(+1G);MI0000470|hsa-mir-125b-2&WithFlank&21|+|16590231|16590331@23.43.21(+1G)&[MIMAT0000423&hsa-miR-125b-5p&offsets|0|-1(+1G);m-67&11|-|122099809|122099830&offsets|0|-1(+1G)];[MIMAT0000423_1&hsa-miR-125b-5p&offsets|0|-1(+1G);m-68&21|+|16590253|16590274&offsets|0|-1(+1G)]|TCCCTGAGACCCTAACTTGTGG</td><td>>MI0000446|hsa-mir-125b-1&WithFlank&11|-|122099751|122099850</td><td>21</td><td>41</td><td>21(+1G)</td><td>TCCCTGAGACCCTAACTTGTGG</td><td>280.1</td><td>-4.9</td><td>1.6</td><td>-3.0</td><td>2.7E-03</td><td>1.2E-02</td><td>BT20/MB231</td><td>NUC</td></tr>
<tr><td>>hg38_MT_+_598_1651_RNR1_12S_50ntFlanks@441.468.28|CATATCTGAACACACAATAGCTAAGACC</td><td>>hg38_MT_+_598_1651_RNR1_12S_50ntFlanks</td><td>441</td><td>468</td><td>28</td><td>CATATCTGAACACACAATAGCTAAGACC</td><td>48.4</td><td>5.1</td><td>2.0</td><td>2.6</td><td>9.4E-03</td><td>1.2E-02</td><td>MB231/MB468</td><td>MP</td></tr>
<tr><td>>MI0000825|hsa-mir-345&WithFlank&14|+|100307853|100307962@24.43.20(+1C)&[MIMAT0000772&hsa-miR-345-5p&offsets|0|-2(+1C);m-193&14|+|100307876|100307896&offsets|0|-1(+1C)]|GCTGACTCCTAGTCCAGGGCC</td><td>>MI0000825|hsa-mir-345&WithFlank&14|+|100307853|100307962</td><td>24</td><td>43</td><td>20(+1C)</td><td>GCTGACTCCTAGTCCAGGGCC</td><td>35.3</td><td>5.2</td><td>2.0</td><td>2.6</td><td>9.4E-03</td><td>1.2E-02</td><td>MB231/MB468</td><td>MP</td></tr>
<tr><td>>hg38_21_+_8433172_8446622_RNA45SN1_45S_50ntFlanks@8081.8099.19|GAGCCCAGCGCCGAATCCC</td><td>>hg38_21_+_8433172_8446622_RNA45SN1_45S_50ntFlanks</td><td>8081</td><td>8099</td><td>19</td><td>GAGCCCAGCGCCGAATCCC</td><td>93.2</td><td>6.9</td><td>2.5</td><td>2.7</td><td>6.3E-03</td><td>1.2E-02</td><td>BT20/MB468</td><td>MP</td></tr>
<tr><td>>hg38_21_+_8433172_8446622_RNA45SN1_45S_50ntFlanks@8381.8401.21|AAGAGGGCGTGAAACCGTTAA</td><td>>hg38_21_+_8433172_8446622_RNA45SN1_45S_50ntFlanks</td><td>8381</td><td>8401</td><td>21</td><td>AAGAGGGCGTGAAACCGTTAA</td><td>264.4</td><td>-4.7</td><td>1.7</td><td>-2.7</td><td>6.3E-03</td><td>1.2E-02</td><td>BT20/MB468</td><td>MP</td></tr>
<tr><td>>trna11_GluTTC_15_-_26327381_26327452@1.31.31,trna3_GluTTC_13_-_45492062_45492133@1.31.31|TCCCACATGGTCTAGCGGTTAGGATTCCTGG</td><td>>trna11_GluTTC_15_-_26327381_26327452</td><td>1</td><td>31</td><td>31</td><td>TCCCACATGGTCTAGCGGTTAGGATTCCTGG</td><td>2405.1</td><td>-1.7</td><td>0.7</td><td>-2.6</td><td>1.0E-02</td><td>1.2E-02</td><td>BT20/MB468</td><td>CYTO</td></tr>
<tr><td>>hg38_21_+_8433172_8446622_RNA45SN1_45S_50ntFlanks@12488.12525.38|AACGTGAGCTGGGTTTAGACCGTCGTGAGACAGGTTAG</td><td>>hg38_21_+_8433172_8446622_RNA45SN1_45S_50ntFlanks</td><td>12488</td><td>12525</td><td>38</td><td>AACGTGAGCTGGGTTTAGACCGTCGTGAGACAGGTTAG</td><td>100.3</td><td>-4.0</td><td>1.5</td><td>-2.6</td><td>8.8E-03</td><td>1.2E-02</td><td>BT20/MB468</td><td>MITO</td></tr>
<tr><td>>hg38_21_+_8433172_8446622_RNA45SN1_45S_50ntFlanks@4995.5023.29|ACAGATTGATAGCTCTTTCTCGATTCCGT</td><td>>hg38_21_+_8433172_8446622_RNA45SN1_45S_50ntFlanks</td><td>4995</td><td>5023</td><td>29</td><td>ACAGATTGATAGCTCTTTCTCGATTCCGT</td><td>168.8</td><td>-4.3</td><td>1.7</td><td>-2.6</td><td>8.8E-03</td><td>1.2E-02</td><td>BT20/MB468</td><td>MITO</td></tr>
<tr><td>>MI0000061|hsa-let-7a-2&WithFlank&11|-|122146516|122146599@11.31.21(+1A);MI0000062|hsa-let-7a-3&WithFlank&22|+|46112743|46112828@10.30.21(+1A);MI0000060|hsa-let-7a-1&WithFlank&9|+|94175951|94176042@12.32.21(+1A)&[MIMAT0000062&hsa-let-7a-5p&offsets|0|-1(+1A);m-6&11|-|122146568|122146589&offsets|0|-1(+1A)];[MIMAT0000062_1&hsa-let-7a-5p&offsets|0|-1(+1A);m-5&22|+|46112752|46112773&offsets|0|-1(+1A)];[MIMAT0000062_2&hsa-let-7a-5p&offsets|0|-1(+1A);m-4&9|+|94175962|94175983&offsets|0|-1(+1A)]|TGAGGTAGTAGGTTGTATAGTA</td><td>>MI0000061|hsa-let-7a-2&WithFlank&11|-|122146516|122146599</td><td>11</td><td>31</td><td>21(+1A)</td><td>TGAGGTAGTAGGTTGTATAGTA</td><td>3009.8</td><td>-3.1</td><td>1.0</td><td>-3.0</td><td>2.4E-03</td><td>1.2E-02</td><td>BT20/MB231</td><td>MP</td></tr>
<tr><td>>hg38_21_+_8433172_8446622_RNA45SN1_45S_50ntFlanks@10788.10823.36|CAGCCTCTGGCATGTTGGAACAATGTAGGTAAGGGA</td><td>>hg38_21_+_8433172_8446622_RNA45SN1_45S_50ntFlanks</td><td>10788</td><td>10823</td><td>36</td><td>CAGCCTCTGGCATGTTGGAACAATGTAGGTAAGGGA</td><td>213.4</td><td>-1.3</td><td>0.5</td><td>-2.6</td><td>8.8E-03</td><td>1.2E-02</td><td>BT20/MB468</td><td>MITO</td></tr>
<tr><td>>hg38_21_+_8433172_8446622_RNA45SN1_45S_50ntFlanks@10116.10138.23|TGCGTCGGAGGGCGGCGGCGGCG</td><td>>hg38_21_+_8433172_8446622_RNA45SN1_45S_50ntFlanks</td><td>10116</td><td>10138</td><td>23</td><td>TGCGTCGGAGGGCGGCGGCGGCG</td><td>214.4</td><td>-4.8</td><td>1.8</td><td>-2.7</td><td>6.3E-03</td><td>1.2E-02</td><td>BT20/MB468</td><td>MP</td></tr>
<tr><td>>MI0000298|hsa-mir-221&WithFlank&X|-|45746151|45746272@31.53.23(+1C)&[MIMAT0004568&hsa-miR-221-5p&offsets|0|+1(+1C);m-405&X|-|45746220|45746242&offsets|0|0(+1C)]|ACCTGGCATACAATGTAGATTTCC</td><td>>MI0000298|hsa-mir-221&WithFlank&X|-|45746151|45746272</td><td>31</td><td>53</td><td>23(+1C)</td><td>ACCTGGCATACAATGTAGATTTCC</td><td>321.3</td><td>-3.4</td><td>1.1</td><td>-3.0</td><td>3.1E-03</td><td>1.2E-02</td><td>BT20/MB231</td><td>MITO</td></tr>
<tr><td>>hg38_21_+_8433172_8446622_RNA45SN1_45S_50ntFlanks@11568.11593.26|GCAGCCGACTTAGAACTGGTGCGGAC</td><td>>hg38_21_+_8433172_8446622_RNA45SN1_45S_50ntFlanks</td><td>11568</td><td>11593</td><td>26</td><td>GCAGCCGACTTAGAACTGGTGCGGAC</td><td>246.9</td><td>-4.1</td><td>1.5</td><td>-2.7</td><td>6.3E-03</td><td>1.2E-02</td><td>BT20/MB468</td><td>MP</td></tr>
<tr><td>>hg38_21_+_8433172_8446622_RNA45SN1_45S_50ntFlanks@9934.9952.19|AGACAGCAGGACGGTGGCC</td><td>>hg38_21_+_8433172_8446622_RNA45SN1_45S_50ntFlanks</td><td>9934</td><td>9952</td><td>19</td><td>AGACAGCAGGACGGTGGCC</td><td>181.2</td><td>5.9</td><td>1.9</td><td>3.0</td><td>2.4E-03</td><td>1.2E-02</td><td>BT20/MB231</td><td>MP</td></tr>
<tr><td>>hg38_21_+_8433172_8446622_RNA45SN1_45S_50ntFlanks@3705.3725.21|TACCTGGTTGATCCTGCCAGT</td><td>>hg38_21_+_8433172_8446622_RNA45SN1_45S_50ntFlanks</td><td>3705</td><td>3725</td><td>21</td><td>TACCTGGTTGATCCTGCCAGT</td><td>1519.8</td><td>2.6</td><td>1.0</td><td>2.5</td><td>1.1E-02</td><td>1.2E-02</td><td>MB231/MB468</td><td>MITO</td></tr>
<tr><td>>hg38_21_+_8433172_8446622_RNA45SN1_45S_50ntFlanks@10574.10599.26|AAGCCGGCGGGAGCCCCGGGGAGAGT</td><td>>hg38_21_+_8433172_8446622_RNA45SN1_45S_50ntFlanks</td><td>10574</td><td>10599</td><td>26</td><td>AAGCCGGCGGGAGCCCCGGGGAGAGT</td><td>94.2</td><td>-3.5</td><td>1.3</td><td>-2.6</td><td>9.5E-03</td><td>1.2E-02</td><td>MB231/MB468</td><td>MP</td></tr>
<tr><td>>MI0000067|hsa-let-7f-1&WithFlank&9|+|94176341|94176439@13.34.22;MI0000068|hsa-let-7f-2&WithFlank&X|-|53557186|53557280@14.35.22&[MIMAT0000067&hsa-let-7f-5p&offsets|0|0;m-20&9|+|94176353|94176374&offsets|0|0];[MIMAT0000067_1&hsa-let-7f-5p&offsets|0|0;m-19&X|-|53557246|53557267&offsets|0|0]|TGAGGTAGTAGATTGTATAGTT</td><td>>MI0000067|hsa-let-7f-1&WithFlank&9|+|94176341|94176439</td><td>13</td><td>34</td><td>22</td><td>TGAGGTAGTAGATTGTATAGTT</td><td>109420.5</td><td>-3.0</td><td>1.0</td><td>-3.0</td><td>2.4E-03</td><td>1.2E-02</td><td>BT20/MB231</td><td>MP</td></tr>
<tr><td>>hg38_21_+_8433172_8446622_RNA45SN1_45S_50ntFlanks@6661.6684.24|CGGTGGATCACTCGGCTCGTGCGT</td><td>>hg38_21_+_8433172_8446622_RNA45SN1_45S_50ntFlanks</td><td>6661</td><td>6684</td><td>24</td><td>CGGTGGATCACTCGGCTCGTGCGT</td><td>501.1</td><td>2.6</td><td>0.9</td><td>3.0</td><td>2.4E-03</td><td>1.2E-02</td><td>BT20/MB231</td><td>MP</td></tr>
<tr><td>>hg38_21_+_8433172_8446622_RNA45SN1_45S_50ntFlanks@10404.10435.32|CATGGGTCAGTCGGTCCTGAGAGATGGGCGAG</td><td>>hg38_21_+_8433172_8446622_RNA45SN1_45S_50ntFlanks</td><td>10404</td><td>10435</td><td>32</td><td>CATGGGTCAGTCGGTCCTGAGAGATGGGCGAG</td><td>194.9</td><td>-2.4</td><td>0.9</td><td>-2.7</td><td>6.3E-03</td><td>1.2E-02</td><td>BT20/MB468</td><td>MP</td></tr>
<tr><td>>hg38_MT_+_1621_3279_RNR2_16S_50ntFlanks@620.652.33|CATATAACTGAACTCCTCACACCCAATTGGACC</td><td>>hg38_MT_+_1621_3279_RNR2_16S_50ntFlanks</td><td>620</td><td>652</td><td>33</td><td>CATATAACTGAACTCCTCACACCCAATTGGACC</td><td>104.2</td><td>5.9</td><td>2.1</td><td>2.7</td><td>6.3E-03</td><td>1.2E-02</td><td>BT20/MB468</td><td>MP</td></tr>
<tr><td>>hg38_21_+_8433172_8446622_RNA45SN1_45S_50ntFlanks@8552.8577.26|CTCCCCGGAGGGGGCGGGCTCCGGCG</td><td>>hg38_21_+_8433172_8446622_RNA45SN1_45S_50ntFlanks</td><td>8552</td><td>8577</td><td>26</td><td>CTCCCCGGAGGGGGCGGGCTCCGGCG</td><td>486.5</td><td>-4.2</td><td>1.6</td><td>-2.6</td><td>1.0E-02</td><td>1.2E-02</td><td>BT20/MB468</td><td>CYTO</td></tr>
<tr><td>>hg38_21_+_8433172_8446622_RNA45SN1_45S_50ntFlanks@10240.10259.20|CTACGCCGCGACGAGTAGGA</td><td>>hg38_21_+_8433172_8446622_RNA45SN1_45S_50ntFlanks</td><td>10240</td><td>10259</td><td>20</td><td>CTACGCCGCGACGAGTAGGA</td><td>4931.0</td><td>3.8</td><td>1.3</td><td>3.0</td><td>2.4E-03</td><td>1.2E-02</td><td>BT20/MB231</td><td>MP</td></tr>
<tr><td>>hg38_21_+_8433172_8446622_RNA45SN1_45S_50ntFlanks@12990.13014.25|GCTCCCTCGCTGCGATCTATTGAAA</td><td>>hg38_21_+_8433172_8446622_RNA45SN1_45S_50ntFlanks</td><td>12990</td><td>13014</td><td>25</td><td>GCTCCCTCGCTGCGATCTATTGAAA</td><td>275.2</td><td>-1.7</td><td>0.7</td><td>-2.6</td><td>8.8E-03</td><td>1.2E-02</td><td>BT20/MB468</td><td>MITO</td></tr>
<tr><td>>hg38_21_+_8433172_8446622_RNA45SN1_45S_50ntFlanks@3785.3802.18|TGAAACTGCGAATGGCTC</td><td>>hg38_21_+_8433172_8446622_RNA45SN1_45S_50ntFlanks</td><td>3785</td><td>3802</td><td>18</td><td>TGAAACTGCGAATGGCTC</td><td>167.3</td><td>-1.3</td><td>0.5</td><td>-2.6</td><td>1.0E-02</td><td>1.2E-02</td><td>MB231/MB468</td><td>CYTO</td></tr>
<tr><td>>MI0000088|hsa-mir-30a&WithFlank&6|-|71403545|71403627@12.33.22(+1U)&[MIMAT0000087&hsa-miR-30a-5p&offsets|0|0(+1U);m-15&6|-|71403595|71403616&offsets|0|0(+1U)]|TGTAAACATCCTCGACTGGAAGT</td><td>>MI0000088|hsa-mir-30a&WithFlank&6|-|71403545|71403627</td><td>12</td><td>33</td><td>22(+1U)</td><td>TGTAAACATCCTCGACTGGAAGT</td><td>8436.3</td><td>3.4</td><td>1.3</td><td>2.7</td><td>6.3E-03</td><td>1.2E-02</td><td>BT20/MB468</td><td>MP</td></tr>
<tr><td>>hg38_21_+_8433172_8446622_RNA45SN1_45S_50ntFlanks@11874.11900.27|AGAAGACCCTGTTGAGCTTGACTCTAG</td><td>>hg38_21_+_8433172_8446622_RNA45SN1_45S_50ntFlanks</td><td>11874</td><td>11900</td><td>27</td><td>AGAAGACCCTGTTGAGCTTGACTCTAG</td><td>564.4</td><td>-4.8</td><td>1.7</td><td>-2.7</td><td>6.3E-03</td><td>1.2E-02</td><td>BT20/MB468</td><td>MP</td></tr>
<tr><td>>hg38_21_+_8433172_8446622_RNA45SN1_45S_50ntFlanks@12423.12445.23|TCGGCTCTTCCTATCATTGTGAA</td><td>>hg38_21_+_8433172_8446622_RNA45SN1_45S_50ntFlanks</td><td>12423</td><td>12445</td><td>23</td><td>TCGGCTCTTCCTATCATTGTGAA</td><td>181.5</td><td>-1.4</td><td>0.5</td><td>-2.6</td><td>1.0E-02</td><td>1.2E-02</td><td>MB231/MB468</td><td>CYTO</td></tr>
<tr><td>>MI0000434|hsa-let-7i&WithFlank&12|+|62603680|62603775@12.31.20(+2C)&[MIMAT0000415&hsa-let-7i-5p&offsets|0|-2(+2C);m-49&12|+|62603691|62603712&offsets|0|-2(+2C)]|TGAGGTAGTAGTTTGTGCTGCC</td><td>>MI0000434|hsa-let-7i&WithFlank&12|+|62603680|62603775</td><td>12</td><td>31</td><td>20(+2C)</td><td>TGAGGTAGTAGTTTGTGCTGCC</td><td>61.9</td><td>5.1</td><td>2.0</td><td>2.6</td><td>9.5E-03</td><td>1.2E-02</td><td>MB231/MB468</td><td>MP</td></tr>
<tr><td>>hg38_21_+_8433172_8446622_RNA45SN1_45S_50ntFlanks@11161.11179.19|CACGGTCCCCCGCGAGGGG</td><td>>hg38_21_+_8433172_8446622_RNA45SN1_45S_50ntFlanks</td><td>11161</td><td>11179</td><td>19</td><td>CACGGTCCCCCGCGAGGGG</td><td>1390.7</td><td>-4.6</td><td>1.8</td><td>-2.6</td><td>1.0E-02</td><td>1.2E-02</td><td>BT20/MB468</td><td>CYTO</td></tr>
<tr><td>>hg38_21_+_8433172_8446622_RNA45SN1_45S_50ntFlanks@9549.9573.25|AGGTCCGTAGCGGTCCTGACGTGCA</td><td>>hg38_21_+_8433172_8446622_RNA45SN1_45S_50ntFlanks</td><td>9549</td><td>9573</td><td>25</td><td>AGGTCCGTAGCGGTCCTGACGTGCA</td><td>408.3</td><td>2.1</td><td>0.8</td><td>2.6</td><td>1.0E-02</td><td>1.2E-02</td><td>BT20/MB468</td><td>CYTO</td></tr>
<tr><td>>hg38_21_+_8433172_8446622_RNA45SN1_45S_50ntFlanks@9573.9605.33|AAATCGGTCGTCCGACCTGGGTATAGGGGCGAA</td><td>>hg38_21_+_8433172_8446622_RNA45SN1_45S_50ntFlanks</td><td>9573</td><td>9605</td><td>33</td><td>AAATCGGTCGTCCGACCTGGGTATAGGGGCGAA</td><td>854.9</td><td>3.5</td><td>1.1</td><td>3.0</td><td>2.4E-03</td><td>1.2E-02</td><td>BT20/MB231</td><td>MP</td></tr>
<tr><td>>hg38_21_+_8433172_8446622_RNA45SN1_45S_50ntFlanks@10856.10873.18|AGGATTGGCTCTAAGGGC</td><td>>hg38_21_+_8433172_8446622_RNA45SN1_45S_50ntFlanks</td><td>10856</td><td>10873</td><td>18</td><td>AGGATTGGCTCTAAGGGC</td><td>179.4</td><td>-1.3</td><td>0.5</td><td>-2.6</td><td>1.0E-02</td><td>1.2E-02</td><td>BT20/MB468</td><td>CYTO</td></tr>
<tr><td>>hg38_21_+_8433172_8446622_RNA45SN1_45S_50ntFlanks@10239.10263.25|GCTACGCCGCGACGAGTAGGAGGGC</td><td>>hg38_21_+_8433172_8446622_RNA45SN1_45S_50ntFlanks</td><td>10239</td><td>10263</td><td>25</td><td>GCTACGCCGCGACGAGTAGGAGGGC</td><td>174.7</td><td>4.0</td><td>1.4</td><td>2.7</td><td>6.3E-03</td><td>1.2E-02</td><td>BT20/MB468</td><td>MP</td></tr>
<tr><td>>hg38_21_+_8433172_8446622_RNA45SN1_45S_50ntFlanks@5190.5225.36|AACAGGTCTGTGATGCCCTTAGATGTCCGGGGCTGC</td><td>>hg38_21_+_8433172_8446622_RNA45SN1_45S_50ntFlanks</td><td>5190</td><td>5225</td><td>36</td><td>AACAGGTCTGTGATGCCCTTAGATGTCCGGGGCTGC</td><td>183.7</td><td>5.1</td><td>1.9</td><td>2.7</td><td>6.3E-03</td><td>1.2E-02</td><td>BT20/MB468</td><td>MP</td></tr>
<tr><td>>hg38_21_+_8433172_8446622_RNA45SN1_45S_50ntFlanks@10795.10826.32|TGGCATGTTGGAACAATGTAGGTAAGGGAAGT</td><td>>hg38_21_+_8433172_8446622_RNA45SN1_45S_50ntFlanks</td><td>10795</td><td>10826</td><td>32</td><td>TGGCATGTTGGAACAATGTAGGTAAGGGAAGT</td><td>141.7</td><td>-2.3</td><td>0.8</td><td>-2.7</td><td>6.3E-03</td><td>1.2E-02</td><td>BT20/MB468</td><td>MP</td></tr>
<tr><td>>hg38_21_+_8433172_8446622_RNA45SN1_45S_50ntFlanks@6731.6754.24|CACATTGATCATCGACACTTCGAA</td><td>>hg38_21_+_8433172_8446622_RNA45SN1_45S_50ntFlanks</td><td>6731</td><td>6754</td><td>24</td><td>CACATTGATCATCGACACTTCGAA</td><td>74.3</td><td>-4.5</td><td>1.7</td><td>-2.7</td><td>6.3E-03</td><td>1.2E-02</td><td>BT20/MB468</td><td>MP</td></tr>
<tr><td>>MI0000289|hsa-mir-181a-1&WithFlank&1|-|198859038|198859159@30.54.25(+1C);MI0000269|hsa-mir-181a-2&WithFlank&9|+|124692436|124692557@45.69.25(+1C)&[MIMAT0000256&hsa-miR-181a-5p&offsets|0|+2(+1C);m-32&1|-|198859108|198859130&offsets|0|+2(+1C)];[MIMAT0000256_1&hsa-miR-181a-5p&offsets|0|+2(+1C);m-32&9|+|124692480|124692502&offsets|0|+2(+1C)]|AACATTCAACGCTGTCGGTGAGTTTC</td><td>>MI0000289|hsa-mir-181a-1&WithFlank&1|-|198859038|198859159</td><td>30</td><td>54</td><td>25(+1C)</td><td>AACATTCAACGCTGTCGGTGAGTTTC</td><td>108.6</td><td>5.0</td><td>1.9</td><td>2.6</td><td>9.5E-03</td><td>1.2E-02</td><td>MB231/MB468</td><td>MP</td></tr>
<tr><td>>hg38_21_+_8433172_8446622_RNA45SN1_45S_50ntFlanks@12416.12462.47|TTCGATGTCGGCTCTTCCTATCATTGTGAAGCAGAATTCACCAAGCG</td><td>>hg38_21_+_8433172_8446622_RNA45SN1_45S_50ntFlanks</td><td>12416</td><td>12462</td><td>47</td><td>TTCGATGTCGGCTCTTCCTATCATTGTGAAGCAGAATTCACCAAGCG</td><td>584.7</td><td>-5.5</td><td>1.9</td><td>-3.0</td><td>3.1E-03</td><td>1.2E-02</td><td>BT20/MB231</td><td>CYTO</td></tr>
<tr><td>>hg38_21_+_8433172_8446622_RNA45SN1_45S_50ntFlanks@12627.12652.26|ATGGGGCGAAGCTACCATCTGTGGGA</td><td>>hg38_21_+_8433172_8446622_RNA45SN1_45S_50ntFlanks</td><td>12627</td><td>12652</td><td>26</td><td>ATGGGGCGAAGCTACCATCTGTGGGA</td><td>314.6</td><td>-3.0</td><td>1.2</td><td>-2.6</td><td>1.0E-02</td><td>1.2E-02</td><td>BT20/MB468</td><td>CYTO</td></tr>
<tr><td>>hg38_21_+_8433172_8446622_RNA45SN1_45S_50ntFlanks@8345.8379.35|AAGGGAAAGTTGAAAAGAACTTTGAAGAGAGAGTT</td><td>>hg38_21_+_8433172_8446622_RNA45SN1_45S_50ntFlanks</td><td>8345</td><td>8379</td><td>35</td><td>AAGGGAAAGTTGAAAAGAACTTTGAAGAGAGAGTT</td><td>77.3</td><td>3.8</td><td>1.4</td><td>2.7</td><td>6.3E-03</td><td>1.2E-02</td><td>BT20/MB468</td><td>MP</td></tr>
<tr><td>>hg38_21_+_8433172_8446622_RNA45SN1_45S_50ntFlanks@9459.9506.48|CCGGGGAGGTGGAGCACGAGCGCACGTGTTAGGACCCGAAAGATGGTG</td><td>>hg38_21_+_8433172_8446622_RNA45SN1_45S_50ntFlanks</td><td>9459</td><td>9506</td><td>48</td><td>CCGGGGAGGTGGAGCACGAGCGCACGTGTTAGGACCCGAAAGATGGTG</td><td>46.4</td><td>-8.6</td><td>3.2</td><td>-2.7</td><td>6.3E-03</td><td>1.2E-02</td><td>BT20/MB468</td><td>MP</td></tr>
<tr><td>>trnaMT_ValTAC_MT_+_1602_1670@33.72.40|ACACTTAGGAGATTTCAACTTAACTTGACCGCTCTGACCA</td><td>>trnaMT_ValTAC_MT_+_1602_1670</td><td>33</td><td>72</td><td>40</td><td>ACACTTAGGAGATTTCAACTTAACTTGACCGCTCTGACCA</td><td>520.0</td><td>4.8</td><td>1.7</td><td>2.7</td><td>6.3E-03</td><td>1.2E-02</td><td>BT20/MB468</td><td>MP</td></tr>
<tr><td>>MI0000088|hsa-mir-30a&WithFlank&6|-|71403545|71403627@12.33.22(+2U)&[MIMAT0000087&hsa-miR-30a-5p&offsets|0|0(+2U);m-15&6|-|71403595|71403616&offsets|0|0(+2U)]|TGTAAACATCCTCGACTGGAAGTT</td><td>>MI0000088|hsa-mir-30a&WithFlank&6|-|71403545|71403627</td><td>12</td><td>33</td><td>22(+2U)</td><td>TGTAAACATCCTCGACTGGAAGTT</td><td>336.0</td><td>3.4</td><td>1.3</td><td>2.6</td><td>9.5E-03</td><td>1.2E-02</td><td>MB231/MB468</td><td>MP</td></tr>
<tr><td>>hg38_21_+_8433172_8446622_RNA45SN1_45S_50ntFlanks@9915.9938.24|AAAAGGTGTTGGTTGATATAGACA</td><td>>hg38_21_+_8433172_8446622_RNA45SN1_45S_50ntFlanks</td><td>9915</td><td>9938</td><td>24</td><td>AAAAGGTGTTGGTTGATATAGACA</td><td>123.8</td><td>4.5</td><td>1.5</td><td>3.0</td><td>2.5E-03</td><td>1.2E-02</td><td>BT20/MB231</td><td>MP</td></tr>
<tr><td>>MI0000088|hsa-mir-30a&WithFlank&6|-|71403545|71403627@54.73.20&[MIMAT0000088&hsa-miR-30a-3p&offsets|+1|-1;m-38&6|-|71403554|71403575&offsets|+1|-1]|TTTCAGTCGGATGTTTGCAG</td><td>>MI0000088|hsa-mir-30a&WithFlank&6|-|71403545|71403627</td><td>54</td><td>73</td><td>20</td><td>TTTCAGTCGGATGTTTGCAG</td><td>846.8</td><td>-3.0</td><td>1.0</td><td>-3.0</td><td>2.5E-03</td><td>1.2E-02</td><td>BT20/MB231</td><td>MP</td></tr>
</table>

<table>
<tr><td>>hg38_21_+_8433172_8446622_RNA45SN1_45S_50ntFlanks@11569.11613.45|CAGCCGACTTAGAACTGGTGCGGACCAGGGGAATCCGACTGTTTA</td><td>>hg38_21_+_8433172_8446622_RNA45SN1_45S_50ntFlanks</td><td>11569</td><td>11613</td><td>45</td><td>CAGCCGACTTAGAACTGGTGCGGACCAGGGGAATCCGACTGTTTA</td><td>118.5</td><td>-5.0</td><td>1.9</td><td>-2.6</td><td>9.5E-03</td><td>1.2E-02</td><td>MB231/MB468</td><td>MP</td></tr>
<tr><td>>hg38_21_+_8433172_8446622_RNA45SN1_45S_50ntFlanks@12931.12951.21|GCGCTAAACCATTCGTAGACG</td><td>>hg38_21_+_8433172_8446622_RNA45SN1_45S_50ntFlanks</td><td>12931</td><td>12951</td><td>21</td><td>GCGCTAAACCATTCGTAGACG</td><td>51.7</td><td>-4.4</td><td>1.6</td><td>-2.7</td><td>6.4E-03</td><td>1.2E-02</td><td>BT20/MB468</td><td>MP</td></tr>
<tr><td>>trnaMT_LeuTAG_MT_+_12266_12336@40.74.35|CAAAAATTTTGGTGCAACTCCAAATAAAAGTACCA</td><td>>trnaMT_LeuTAG_MT_+_12266_12336</td><td>40</td><td>74</td><td>35</td><td>CAAAAATTTTGGTGCAACTCCAAATAAAAGTACCA</td><td>134.0</td><td>3.2</td><td>1.2</td><td>2.6</td><td>9.5E-03</td><td>1.2E-02</td><td>MB231/MB468</td><td>MP</td></tr>
<tr><td>>hg38_21_+_8433172_8446622_RNA45SN1_45S_50ntFlanks@12933.12965.33|GCTAAACCATTCGTAGACGACCTGCTTCTGGGT</td><td>>hg38_21_+_8433172_8446622_RNA45SN1_45S_50ntFlanks</td><td>12933</td><td>12965</td><td>33</td><td>GCTAAACCATTCGTAGACGACCTGCTTCTGGGT</td><td>243.5</td><td>-2.2</td><td>0.9</td><td>-2.6</td><td>1.0E-02</td><td>1.2E-02</td><td>BT20/MB468</td><td>CYTO</td></tr>
<tr><td>>hg38_21_+_8433172_8446622_RNA45SN1_45S_50ntFlanks@11570.11597.28|AGCCGACTTAGAACTGGTGCGGACCAGG</td><td>>hg38_21_+_8433172_8446622_RNA45SN1_45S_50ntFlanks</td><td>11570</td><td>11597</td><td>28</td><td>AGCCGACTTAGAACTGGTGCGGACCAGG</td><td>178.3</td><td>2.3</td><td>0.8</td><td>2.7</td><td>6.4E-03</td><td>1.2E-02</td><td>BT20/MB468</td><td>MP</td></tr>
<tr><td>>MI0000082|hsa-mir-25&WithFlank&7|-|100093554|100093649@20.42.23&[MIMAT0004498&hsa-miR-25-5p&offsets|0|+2;m-572&7|-|100093608|100093630&offsets|0|0]|AGGCGGAGACTTGGGCAATTGCT</td><td>>MI0000082|hsa-mir-25&WithFlank&7|-|100093554|100093649</td><td>20</td><td>42</td><td>23</td><td>AGGCGGAGACTTGGGCAATTGCT</td><td>131.0</td><td>3.9</td><td>1.4</td><td>2.7</td><td>6.4E-03</td><td>1.2E-02</td><td>BT20/MB468</td><td>MP</td></tr>
<tr><td>>hg38_21_+_8433172_8446622_RNA45SN1_45S_50ntFlanks@8692.8714.23|CCGGGACGGCTGGGAAGGCCCGG</td><td>>hg38_21_+_8433172_8446622_RNA45SN1_45S_50ntFlanks</td><td>8692</td><td>8714</td><td>23</td><td>CCGGGACGGCTGGGAAGGCCCGG</td><td>109.2</td><td>6.1</td><td>2.2</td><td>2.7</td><td>6.4E-03</td><td>1.2E-02</td><td>BT20/MB468</td><td>MP</td></tr>
<tr><td>>trna111_HisGTG_1_-_147774845_147774916@1.35.35,trna118_HisGTG_1_-_145396881_145396952@1.35.35,trna16_HisGTG_1_+_146544773_146544844@1.35.35,trna1_HisGTG_15_+_45493349_45493420@1.35.35,trna21_HisGTG_1_+_147753471_147753542@1.35.35,trna33_HisGTG_6_+_27125906_27125977@1.35.35,trna7_HisGTG_9_-_14433938_14434009@1.35.35,trna8_HisGTG_15_-_45492611_45492682@1.35.35,trna9_HisGTG_15_-_45490804_45490875@1.35.35|GCCGTGATCGTATAGTGGTTAGTACTCTGCGTTGT</td><td>>trna111_HisGTG_1_-_147774845_147774916</td><td>1</td><td>35</td><td>35</td><td>GCCGTGATCGTATAGTGGTTAGTACTCTGCGTTGT</td><td>47.7</td><td>5.1</td><td>2.0</td><td>2.6</td><td>9.6E-03</td><td>1.2E-02</td><td>MB231/MB468</td><td>MP</td></tr>
<tr><td>>hg38_21_+_8433172_8446622_RNA45SN1_45S_50ntFlanks@11208.11229.22|GCGGCGACTCTGGACGCGAGCC</td><td>>hg38_21_+_8433172_8446622_RNA45SN1_45S_50ntFlanks</td><td>11208</td><td>11229</td><td>22</td><td>GCGGCGACTCTGGACGCGAGCC</td><td>1707.1</td><td>-4.3</td><td>1.7</td><td>-2.5</td><td>1.1E-02</td><td>1.2E-02</td><td>BT20/MB468</td><td>NUC</td></tr>
<tr><td>>MI0000088|hsa-mir-30a&WithFlank&6|-|71403545|71403627@12.34.23(+1C)&[MIMAT0000087&hsa-miR-30a-5p&offsets|0|+1(+1C);m-15&6|-|71403595|71403616&offsets|0|+1(+1C)]|TGTAAACATCCTCGACTGGAAGCC</td><td>>MI0000088|hsa-mir-30a&WithFlank&6|-|71403545|71403627</td><td>12</td><td>34</td><td>23(+1C)</td><td>TGTAAACATCCTCGACTGGAAGCC</td><td>12620.7</td><td>3.2</td><td>1.2</td><td>2.7</td><td>6.4E-03</td><td>1.2E-02</td><td>BT20/MB468</td><td>MP</td></tr>
<tr><td>>hg38_21_+_8433172_8446622_RNA45SN1_45S_50ntFlanks@12615.12652.38|GCTGAGGAGCCAATGGGGCGAAGCTACCATCTGTGGGA</td><td>>hg38_21_+_8433172_8446622_RNA45SN1_45S_50ntFlanks</td><td>12615</td><td>12652</td><td>38</td><td>GCTGAGGAGCCAATGGGGCGAAGCTACCATCTGTGGGA</td><td>75.9</td><td>-4.6</td><td>1.7</td><td>-2.7</td><td>6.4E-03</td><td>1.2E-02</td><td>BT20/MB468</td><td>MP</td></tr>
<tr><td>>hg38_21_+_8433172_8446622_RNA45SN1_45S_50ntFlanks@12913.12933.21|ACGTTCGTGGGGAACCTGGCG</td><td>>hg38_21_+_8433172_8446622_RNA45SN1_45S_50ntFlanks</td><td>12913</td><td>12933</td><td>21</td><td>ACGTTCGTGGGGAACCTGGCG</td><td>492.0</td><td>-4.1</td><td>1.5</td><td>-2.7</td><td>6.4E-03</td><td>1.2E-02</td><td>BT20/MB468</td><td>MP</td></tr>
<tr><td>>hg38_21_+_8433172_8446622_RNA45SN1_45S_50ntFlanks@4623.4640.18|AAGAGGGACGGCCGGGGG</td><td>>hg38_21_+_8433172_8446622_RNA45SN1_45S_50ntFlanks</td><td>4623</td><td>4640</td><td>18</td><td>AAGAGGGACGGCCGGGGG</td><td>156.0</td><td>-3.6</td><td>1.4</td><td>-2.6</td><td>9.6E-03</td><td>1.2E-02</td><td>MB231/MB468</td><td>MP</td></tr>
<tr><td>>am_trnaMT_MetCAT_MT_+_4402_4469@1.32.32|AGTAAGGTCAGCTAAATAAGCTATCGGGCCCA</td><td>>am_trnaMT_MetCAT_MT_+_4402_4469</td><td>1</td><td>32</td><td>32</td><td>AGTAAGGTCAGCTAAATAAGCTATCGGGCCCA</td><td>223.5</td><td>2.5</td><td>1.0</td><td>2.6</td><td>9.6E-03</td><td>1.2E-02</td><td>MB231/MB468</td><td>MP</td></tr>
<tr><td>>MI0003205|hsa-mir-532&WithFlank&X|+|50003142|50003244@26.46.21(+1C)&[MIMAT0002888&hsa-miR-532-5p&offsets|0|-1(+1C);m-65&X|+|50003167|50003188&offsets|0|-1(+1C)]|CATGCCTTGAGTGTAGGACCGC</td><td>>MI0003205|hsa-mir-532&WithFlank&X|+|50003142|50003244</td><td>26</td><td>46</td><td>21(+1C)</td><td>CATGCCTTGAGTGTAGGACCGC</td><td>545.9</td><td>2.0</td><td>0.8</td><td>2.6</td><td>9.6E-03</td><td>1.2E-02</td><td>MB231/MB468</td><td>MP</td></tr>
<tr><td>>MI0006444|hsa-mir-1307&WithFlank&10|-|103394247|103394407@86.108.23&[MIMAT0005951&hsa-miR-1307-3p&offsets|0|+1;m-66&10|-|103394300|103394322&offsets|0|0]|ACTCGGCGTGGCGTCGGTCGTGG</td><td>>MI0006444|hsa-mir-1307&WithFlank&10|-|103394247|103394407</td><td>86</td><td>108</td><td>23</td><td>ACTCGGCGTGGCGTCGGTCGTGG</td><td>998.3</td><td>2.3</td><td>0.9</td><td>2.6</td><td>9.6E-03</td><td>1.2E-02</td><td>MB231/MB468</td><td>MP</td></tr>
<tr><td>>hg38_21_+_8433172_8446622_RNA45SN1_45S_50ntFlanks@8182.8206.25|ATCGAGGCCCAGCCCGTGGACGGTG</td><td>>hg38_21_+_8433172_8446622_RNA45SN1_45S_50ntFlanks</td><td>8182</td><td>8206</td><td>25</td><td>ATCGAGGCCCAGCCCGTGGACGGTG</td><td>240.5</td><td>-3.5</td><td>1.4</td><td>-2.6</td><td>1.0E-02</td><td>1.2E-02</td><td>MB231/MB468</td><td>CYTO</td></tr>
<tr><td>>hg38_1_-_228634819_228635039_RNA5S12_5S_50ntFlanks@131.170.40|GGATGGGAGACCGCCTGGGAATACCGGGTGCTGTAGGCTT</td><td>>hg38_1_-_228634819_228635039_RNA5S12_5S_50ntFlanks</td><td>131</td><td>170</td><td>40</td><td>GGATGGGAGACCGCCTGGGAATACCGGGTGCTGTAGGCTT</td><td>6570.3</td><td>3.6</td><td>1.2</td><td>3.0</td><td>2.5E-03</td><td>1.2E-02</td><td>BT20/MB231</td><td>MP</td></tr>
<tr><td>>hg38_21_+_8433172_8446622_RNA45SN1_45S_50ntFlanks@8244.8269.26|TCTTCCCGGAGTCGGGTTGCTTGGGA</td><td>>hg38_21_+_8433172_8446622_RNA45SN1_45S_50ntFlanks</td><td>8244</td><td>8269</td><td>26</td><td>TCTTCCCGGAGTCGGGTTGCTTGGGA</td><td>304.9</td><td>5.0</td><td>1.7</td><td>3.0</td><td>2.5E-03</td><td>1.2E-02</td><td>BT20/MB231</td><td>MP</td></tr>
<tr><td>>hg38_21_+_8433172_8446622_RNA45SN1_45S_50ntFlanks@9556.9582.27|TAGCGGTCCTGACGTGCAAATCGGTCG</td><td>>hg38_21_+_8433172_8446622_RNA45SN1_45S_50ntFlanks</td><td>9556</td><td>9582</td><td>27</td><td>TAGCGGTCCTGACGTGCAAATCGGTCG</td><td>104.8</td><td>-4.5</td><td>1.7</td><td>-2.7</td><td>6.4E-03</td><td>1.2E-02</td><td>BT20/MB468</td><td>MP</td></tr>
<tr><td>>hg38_21_+_8433172_8446622_RNA45SN1_45S_50ntFlanks@9201.9225.25|CTCTCGGGGCCACGCGCGCGTCCCC</td><td>>hg38_21_+_8433172_8446622_RNA45SN1_45S_50ntFlanks</td><td>9201</td><td>9225</td><td>25</td><td>CTCTCGGGGCCACGCGCGCGTCCCC</td><td>191.0</td><td>-4.1</td><td>1.6</td><td>-2.6</td><td>8.9E-03</td><td>1.2E-02</td><td>BT20/MB468</td><td>MITO</td></tr>
<tr><td>>am_trnaMT_ValTAC_MT_+_1602_1670@49.68.20|AACTTAACTTGACCGCTCTG</td><td>>am_trnaMT_ValTAC_MT_+_1602_1670</td><td>49</td><td>68</td><td>20</td><td>AACTTAACTTGACCGCTCTG</td><td>542.3</td><td>-2.8</td><td>1.1</td><td>-2.6</td><td>1.0E-02</td><td>1.2E-02</td><td>MB231/MB468</td><td>CYTO</td></tr>
<tr><td>>hg38_1_-_228634819_228635039_RNA5S12_5S_50ntFlanks@145.170.26|CTGGGAATACCGGGTGCTGTAGGCTT</td><td>>hg38_1_-_228634819_228635039_RNA5S12_5S_50ntFlanks</td><td>145</td><td>170</td><td>26</td><td>CTGGGAATACCGGGTGCTGTAGGCTT</td><td>1162.6</td><td>-2.0</td><td>0.8</td><td>-2.6</td><td>9.6E-03</td><td>1.2E-02</td><td>MB231/MB468</td><td>MP</td></tr>
<tr><td>>hg38_MT_+_1621_3279_RNR2_16S_50ntFlanks@1372.1396.25|GTTGGATCAGGACATCCCGATGGTG</td><td>>hg38_MT_+_1621_3279_RNR2_16S_50ntFlanks</td><td>1372</td><td>1396</td><td>25</td><td>GTTGGATCAGGACATCCCGATGGTG</td><td>126.0</td><td>3.0</td><td>1.1</td><td>2.7</td><td>6.4E-03</td><td>1.2E-02</td><td>BT20/MB468</td><td>MP</td></tr>
<tr><td>>hg38_21_+_8433172_8446622_RNA45SN1_45S_50ntFlanks@11564.11600.37|CCTAGCAGCCGACTTAGAACTGGTGCGGACCAGGGGA</td><td>>hg38_21_+_8433172_8446622_RNA45SN1_45S_50ntFlanks</td><td>11564</td><td>11600</td><td>37</td><td>CCTAGCAGCCGACTTAGAACTGGTGCGGACCAGGGGA</td><td>1420.3</td><td>-2.8</td><td>1.0</td><td>-2.7</td><td>6.4E-03</td><td>1.2E-02</td><td>BT20/MB468</td><td>MP</td></tr>
<tr><td>>hg38_MT_+_1621_3279_RNR2_16S_50ntFlanks@1366.1396.31|CTCGATGTTGGATCAGGACATCCCGATGGTG</td><td>>hg38_MT_+_1621_3279_RNR2_16S_50ntFlanks</td><td>1366</td><td>1396</td><td>31</td><td>CTCGATGTTGGATCAGGACATCCCGATGGTG</td><td>86.7</td><td>3.1</td><td>1.1</td><td>2.7</td><td>6.4E-03</td><td>1.2E-02</td><td>BT20/MB468</td><td>MP</td></tr>
<tr><td>>hg38_21_+_8433172_8446622_RNA45SN1_45S_50ntFlanks@12407.12445.39|TTTTGATCCTTCGATGTCGGCTCTTCCTATCATTGTGAA</td><td>>hg38_21_+_8433172_8446622_RNA45SN1_45S_50ntFlanks</td><td>12407</td><td>12445</td><td>39</td><td>TTTTGATCCTTCGATGTCGGCTCTTCCTATCATTGTGAA</td><td>296.3</td><td>-1.8</td><td>0.7</td><td>-2.5</td><td>1.1E-02</td><td>1.2E-02</td><td>MB231/MB468</td><td>NUC</td></tr>
<tr><td>>trnaMT_HisGTG_MT_+_12138_12206@20.53.34|AACATCAGATTGTGAATCTGACAACAGAGGCTTA</td><td>>trnaMT_HisGTG_MT_+_12138_12206</td><td>20</td><td>53</td><td>34</td><td>AACATCAGATTGTGAATCTGACAACAGAGGCTTA</td><td>178.3</td><td>5.2</td><td>1.9</td><td>2.7</td><td>6.4E-03</td><td>1.2E-02</td><td>BT20/MB468</td><td>MP</td></tr>
<tr><td>>MI0000063|hsa-let-7b&WithFlank&22|+|46113680|46113774@12.34.23(+2U)&[MIMAT0000063&hsa-let-7b-5p&offsets|0|+1(+2U);m-14&22|+|46113691|46113712&offsets|0|+1(+2U)]|TGAGGTAGTAGGTTGTGTGGTTTTT</td><td>>MI0000063|hsa-let-7b&WithFlank&22|+|46113680|46113774</td><td>12</td><td>34</td><td>23(+2U)</td><td>TGAGGTAGTAGGTTGTGTGGTTTTT</td><td>40.5</td><td>4.3</td><td>1.7</td><td>2.6</td><td>9.6E-03</td><td>1.2E-02</td><td>MB231/MB468</td><td>MP</td></tr>
<tr><td>>hg38_21_+_8433172_8446622_RNA45SN1_45S_50ntFlanks@13021.13039.19|CCTCGACACAAGGGTTTGT</td><td>>hg38_21_+_8433172_8446622_RNA45SN1_45S_50ntFlanks</td><td>13021</td><td>13039</td><td>19</td><td>CCTCGACACAAGGGTTTGT</td><td>135.3</td><td>2.5</td><td>1.0</td><td>2.5</td><td>1.1E-02</td><td>1.2E-02</td><td>MB231/MB468</td><td>MITO</td></tr>
<tr><td>>hg38_21_+_8433172_8446622_RNA45SN1_45S_50ntFlanks@9310.9330.21|GAGTCTAACACGTGCGCGAGT</td><td>>hg38_21_+_8433172_8446622_RNA45SN1_45S_50ntFlanks</td><td>9310</td><td>9330</td><td>21</td><td>GAGTCTAACACGTGCGCGAGT</td><td>82.5</td><td>4.9</td><td>1.8</td><td>2.7</td><td>6.4E-03</td><td>1.2E-02</td><td>BT20/MB468</td><td>MP</td></tr>
<tr><td>>hg38_MT_+_598_1651_RNR1_12S_50ntFlanks@916.946.31|AAGGACATTTAACTAAAACCCCTACGCATTT</td><td>>hg38_MT_+_598_1651_RNR1_12S_50ntFlanks</td><td>916</td><td>946</td><td>31</td><td>AAGGACATTTAACTAAAACCCCTACGCATTT</td><td>370.2</td><td>3.4</td><td>1.3</td><td>2.6</td><td>9.6E-03</td><td>1.2E-02</td><td>MB231/MB468</td><td>MP</td></tr>
<tr><td>>hg38_21_+_8433172_8446622_RNA45SN1_45S_50ntFlanks@11668.11691.24|TGCCCAGTGCTCTGAATGTCAAAG</td><td>>hg38_21_+_8433172_8446622_RNA45SN1_45S_50ntFlanks</td><td>11668</td><td>11691</td><td>24</td><td>TGCCCAGTGCTCTGAATGTCAAAG</td><td>117.1</td><td>2.7</td><td>1.0</td><td>2.7</td><td>6.4E-03</td><td>1.2E-02</td><td>BT20/MB468</td><td>MP</td></tr>
<tr><td>>hg38_21_+_8433172_8446622_RNA45SN1_45S_50ntFlanks@9306.9336.31|CAAGGAGTCTAACACGTGCGCGAGTCGGGGG</td><td>>hg38_21_+_8433172_8446622_RNA45SN1_45S_50ntFlanks</td><td>9306</td><td>9336</td><td>31</td><td>CAAGGAGTCTAACACGTGCGCGAGTCGGGGG</td><td>921.2</td><td>-4.9</td><td>1.8</td><td>-2.7</td><td>6.4E-03</td><td>1.2E-02</td><td>BT20/MB468</td><td>MP</td></tr>
<tr><td>>MI0000269|hsa-mir-181a-2&WithFlank&9|+|124692436|124692557@83.104.22&[MIMAT0004558&hsa-miR-181a-2-3p&offsets|0|0;m-71&9|+|124692518|124692539&offsets|0|0]|ACCACTGACCGTTGACTGTACC</td><td>>MI0000269|hsa-mir-181a-2&WithFlank&9|+|124692436|124692557</td><td>83</td><td>104</td><td>22</td><td>ACCACTGACCGTTGACTGTACC</td><td>447.8</td><td>3.5</td><td>1.4</td><td>2.6</td><td>9.6E-03</td><td>1.2E-02</td><td>MB231/MB468</td><td>MP</td></tr>
<tr><td>>MI0003834|hsa-mir-769&WithFlank&19|+|46018926|46019055@36.55.20(+1U)&[MIMAT0003886&hsa-miR-769-5p&offsets|0|-2(+1U);m-216&19|+|46018961|46018982&offsets|0|-2(+1U)]|TGAGACCTCTGGGTTCTGAGT</td><td>>MI0003834|hsa-mir-769&WithFlank&19|+|46018926|46019055</td><td>36</td><td>55</td><td>20(+1U)</td><td>TGAGACCTCTGGGTTCTGAGT</td><td>133.1</td><td>3.4</td><td>1.3</td><td>2.6</td><td>9.6E-03</td><td>1.2E-02</td><td>MB231/MB468</td><td>MP</td></tr>
<tr><td>>hg38_21_+_8433172_8446622_RNA45SN1_45S_50ntFlanks@8240.8260.21|CGGGTCTTCCCGGAGTCGGGT</td><td>>hg38_21_+_8433172_8446622_RNA45SN1_45S_50ntFlanks</td><td>8240</td><td>8260</td><td>21</td><td>CGGGTCTTCCCGGAGTCGGGT</td><td>308.9</td><td>-2.6</td><td>0.9</td><td>-3.0</td><td>3.1E-03</td><td>1.2E-02</td><td>BT20/MB231</td><td>CYTO</td></tr>
<tr><td>>hg38_21_+_8433172_8446622_RNA45SN1_45S_50ntFlanks@7976.8008.33|GCGACCTCAGATCAGACGTGGCGACCCGCTGAA</td><td>>hg38_21_+_8433172_8446622_RNA45SN1_45S_50ntFlanks</td><td>7976</td><td>8008</td><td>33</td><td>GCGACCTCAGATCAGACGTGGCGACCCGCTGAA</td><td>1890.5</td><td>4.8</td><td>1.6</td><td>3.0</td><td>2.5E-03</td><td>1.2E-02</td><td>BT20/MB231</td><td>MP</td></tr>
<tr><td>>hg38_21_+_8433172_8446622_RNA45SN1_45S_50ntFlanks@9839.9864.26|GTAAGCAGAACTGGCGCTGCGGGATG</td><td>>hg38_21_+_8433172_8446622_RNA45SN1_45S_50ntFlanks</td><td>9839</td><td>9864</td><td>26</td><td>GTAAGCAGAACTGGCGCTGCGGGATG</td><td>244.4</td><td>-3.0</td><td>1.1</td><td>-2.7</td><td>6.4E-03</td><td>1.2E-02</td><td>BT20/MB468</td><td>MP</td></tr>
<tr><td>>hg38_MT_+_598_1651_RNR1_12S_50ntFlanks@431.468.38|GTGGCTTTAACATATCTGAACACACAATAGCTAAGACC</td><td>>hg38_MT_+_598_1651_RNR1_12S_50ntFlanks</td><td>431</td><td>468</td><td>38</td><td>GTGGCTTTAACATATCTGAACACACAATAGCTAAGACC</td><td>50.6</td><td>5.0</td><td>1.9</td><td>2.6</td><td>9.6E-03</td><td>1.2E-02</td><td>MB231/MB468</td><td>MP</td></tr>
<tr><td>>hg38_21_+_8433172_8446622_RNA45SN1_45S_50ntFlanks@12235.12265.31|GTGGAGCAGAAGGGCAAAAGCTCGCTTGATC</td><td>>hg38_21_+_8433172_8446622_RNA45SN1_45S_50ntFlanks</td><td>12235</td><td>12265</td><td>31</td><td>GTGGAGCAGAAGGGCAAAAGCTCGCTTGATC</td><td>47.3</td><td>5.5</td><td>2.0</td><td>2.7</td><td>6.4E-03</td><td>1.2E-02</td><td>BT20/MB468</td><td>MP</td></tr>
<tr><td>>hg38_MT_+_1621_3279_RNR2_16S_50ntFlanks@51.70.20|GCTAAACCTAGCCCCAAACC</td><td>>hg38_MT_+_1621_3279_RNR2_16S_50ntFlanks</td><td>51</td><td>70</td><td>20</td><td>GCTAAACCTAGCCCCAAACC</td><td>2366.8</td><td>-2.9</td><td>1.0</td><td>-3.0</td><td>3.2E-03</td><td>1.2E-02</td><td>BT20/MB231</td><td>MITO</td></tr>
<tr><td>>hg38_MT_+_1621_3279_RNR2_16S_50ntFlanks@1372.1395.24|GTTGGATCAGGACATCCCGATGGT</td><td>>hg38_MT_+_1621_3279_RNR2_16S_50ntFlanks</td><td>1372</td><td>1395</td><td>24</td><td>GTTGGATCAGGACATCCCGATGGT</td><td>22.6</td><td>7.7</td><td>3.0</td><td>2.6</td><td>9.6E-03</td><td>1.2E-02</td><td>MB231/MB468</td><td>MP</td></tr>
<tr><td>>hg38_21_+_8433172_8446622_RNA45SN1_45S_50ntFlanks@11567.11587.21|AGCAGCCGACTTAGAACTGGT</td><td>>hg38_21_+_8433172_8446622_RNA45SN1_45S_50ntFlanks</td><td>11567</td><td>11587</td><td>21</td><td>AGCAGCCGACTTAGAACTGGT</td><td>306.1</td><td>-2.4</td><td>0.8</td><td>-3.0</td><td>2.7E-03</td><td>1.2E-02</td><td>BT20/MB231</td><td>NUC</td></tr>
<tr><td>>hg38_21_+_8433172_8446622_RNA45SN1_45S_50ntFlanks@8170.8213.44|CAAGTCCTTCTGATCGAGGCCCAGCCCGTGGACGGTGTGAGGCC</td><td>>hg38_21_+_8433172_8446622_RNA45SN1_45S_50ntFlanks</td><td>8170</td><td>8213</td><td>44</td><td>CAAGTCCTTCTGATCGAGGCCCAGCCCGTGGACGGTGTGAGGCC</td><td>844.1</td><td>-2.0</td><td>0.8</td><td>-2.6</td><td>9.0E-03</td><td>1.2E-02</td><td>BT20/MB468</td><td>MITO</td></tr>
<tr><td>>hg38_21_+_8433172_8446622_RNA45SN1_45S_50ntFlanks@6645.6663.19|CTCGTACGACTCTTAGCGG</td><td>>hg38_21_+_8433172_8446622_RNA45SN1_45S_50ntFlanks</td><td>6645</td><td>6663</td><td>19</td><td>CTCGTACGACTCTTAGCGG</td><td>9250.8</td><td>2.7</td><td>1.1</td><td>2.5</td><td>1.1E-02</td><td>1.2E-02</td><td>MB231/MB468</td><td>MITO</td></tr>
<tr><td>>hg38_21_+_8433172_8446622_RNA45SN1_45S_50ntFlanks@11885.11903.19|TTGAGCTTGACTCTAGTCT</td><td>>hg38_21_+_8433172_8446622_RNA45SN1_45S_50ntFlanks</td><td>11885</td><td>11903</td><td>19</td><td>TTGAGCTTGACTCTAGTCT</td><td>342.7</td><td>-4.6</td><td>1.7</td><td>-2.7</td><td>6.4E-03</td><td>1.2E-02</td><td>BT20/MB468</td><td>MP</td></tr>
<tr><td>>trnaMT_SerGCT_MT_+_12207_12265@14.45.32|AGAACTGCTAACTCATGCCCCCATGTCTAACA</td><td>>trnaMT_SerGCT_MT_+_12207_12265</td><td>14</td><td>45</td><td>32</td><td>AGAACTGCTAACTCATGCCCCCATGTCTAACA</td><td>233.8</td><td>2.0</td><td>0.8</td><td>2.5</td><td>1.1E-02</td><td>1.2E-02</td><td>MB231/MB468</td><td>MITO</td></tr>
<tr><td>>hg38_21_+_8433172_8446622_RNA45SN1_45S_50ntFlanks@5471.5514.44|CACGGCCCTGGCGGAGCGCTGAGAAGACGGTCGAACTTGACTAT</td><td>>hg38_21_+_8433172_8446622_RNA45SN1_45S_50ntFlanks</td><td>5471</td><td>5514</td><td>44</td><td>CACGGCCCTGGCGGAGCGCTGAGAAGACGGTCGAACTTGACTAT</td><td>703.1</td><td>-3.7</td><td>1.4</td><td>-2.6</td><td>9.0E-03</td><td>1.2E-02</td><td>BT20/MB468</td><td>MITO</td></tr>
<tr><td>>hg38_21_+_8433172_8446622_RNA45SN1_45S_50ntFlanks@10342.10369.28|ATATTCAAACGAGAACTTTGAAGGCCGA</td><td>>hg38_21_+_8433172_8446622_RNA45SN1_45S_50ntFlanks</td><td>10342</td><td>10369</td><td>28</td><td>ATATTCAAACGAGAACTTTGAAGGCCGA</td><td>234.7</td><td>4.9</td><td>1.6</td><td>3.0</td><td>2.5E-03</td><td>1.3E-02</td><td>BT20/MB231</td><td>MP</td></tr>
<tr><td>>hg38_1_-_228634819_228635039_RNA5S12_5S_50ntFlanks@84.112.29|CTCGTCTGATCTCGGAAGCTAAGCAGGGT</td><td>>hg38_1_-_228634819_228635039_RNA5S12_5S_50ntFlanks</td><td>84</td><td>112</td><td>29</td><td>CTCGTCTGATCTCGGAAGCTAAGCAGGGT</td><td>274.9</td><td>-3.0</td><td>1.2</td><td>-2.6</td><td>1.0E-02</td><td>1.3E-02</td><td>BT20/MB468</td><td>CYTO</td></tr>
<tr><td>>hg38_21_+_8433172_8446622_RNA45SN1_45S_50ntFlanks@5486.5513.28|GCGCTGAGAAGACGGTCGAACTTGACTA</td><td>>hg38_21_+_8433172_8446622_RNA45SN1_45S_50ntFlanks</td><td>5486</td><td>5513</td><td>28</td><td>GCGCTGAGAAGACGGTCGAACTTGACTA</td><td>305.7</td><td>-2.4</td><td>0.9</td><td>-2.7</td><td>6.4E-03</td><td>1.3E-02</td><td>BT20/MB468</td><td>MP</td></tr>
</table>

<table>
<tr><td>>hg38_21_+_8433172_8446622_RNA45SN1_45S_50ntFlanks@12396.12417.22|GCGACGTCGCTTTTTGATCCTT</td><td>>hg38_21_+_8433172_8446622_RNA45SN1_45S_50ntFlanks</td><td>12396</td><td>12417</td><td>22</td><td>GCGACGTCGCTTTTTGATCCTT</td><td>320.9</td><td>-3.0</td><td>1.2</td><td>-2.6</td><td>1.0E-02</td><td>1.3E-02</td><td>BT20/MB468</td><td>CYTO</td></tr>
<tr><td>>hg38_21_+_8433172_8446622_RNA45SN1_45S_50ntFlanks@9545.9572.28|GTGGAGGTCCGTAGCGGTCCTGACGTGC</td><td>>hg38_21_+_8433172_8446622_RNA45SN1_45S_50ntFlanks</td><td>9545</td><td>9572</td><td>28</td><td>GTGGAGGTCCGTAGCGGTCCTGACGTGC</td><td>729.8</td><td>4.5</td><td>1.5</td><td>3.0</td><td>2.5E-03</td><td>1.3E-02</td><td>BT20/MB231</td><td>MP</td></tr>
<tr><td>>trnaMT_ValTAC_MT_+_1602_1670@55.72.18|ACTTGACCGCTCTGACCA</td><td>>trnaMT_ValTAC_MT_+_1602_1670</td><td>55</td><td>72</td><td>18</td><td>ACTTGACCGCTCTGACCA</td><td>2622.4</td><td>2.7</td><td>0.9</td><td>3.0</td><td>3.1E-03</td><td>1.3E-02</td><td>BT20/MB231</td><td>CYTO</td></tr>
<tr><td>>trna10_AspGTC_12_-_125424193_125424264@1.35.35,trna12_AspGTC_12_-_125411891_125411962@1.35.35,trna38_AspGTC_17_-_8125556_8125627@1.35.35,trna45_AspGTC_6_+_27447453_27447524@1.35.35,trna48_AspGTC_6_+_27471523_27471594@1.35.35,trna4_AspGTC_12_+_96429799_96429870@1.35.35,trna69_AspGTC_1_-_161440205_161440276@1.35.35,trna72_AspGTC_1_-_161432824_161432895@1.35.35,trna75_AspGTC_1_-_161425414_161425485@1.35.35,trna78_AspGTC_1_-_161418033_161418104@1.35.35,trna81_AspGTC_1_-_161410615_161410686@1.35.35|TCCTCGTTAGTATAGTGGTGAGTATCCCCGCCTGT</td><td>>trna10_AspGTC_12_-_125424193_125424264</td><td>1</td><td>35</td><td>35</td><td>TCCTCGTTAGTATAGTGGTGAGTATCCCCGCCTGT</td><td>254.1</td><td>-3.1</td><td>1.0</td><td>-3.0</td><td>3.1E-03</td><td>1.3E-02</td><td>BT20/MB231</td><td>CYTO</td></tr>
<tr><td>>hg38_21_+_8433172_8446622_RNA45SN1_45S_50ntFlanks@12418.12452.35|CGATGTCGGCTCTTCCTATCATTGTGAAGCAGAAT</td><td>>hg38_21_+_8433172_8446622_RNA45SN1_45S_50ntFlanks</td><td>12418</td><td>12452</td><td>35</td><td>CGATGTCGGCTCTTCCTATCATTGTGAAGCAGAAT</td><td>362.2</td><td>-2.3</td><td>0.8</td><td>-3.0</td><td>3.1E-03</td><td>1.3E-02</td><td>BT20/MB231</td><td>CYTO</td></tr>
<tr><td>>hg38_MT_+_1621_3279_RNR2_16S_50ntFlanks@1355.1385.31|GGGTTTACGACCTCGATGTTGGATCAGGACA</td><td>>hg38_MT_+_1621_3279_RNR2_16S_50ntFlanks</td><td>1355</td><td>1385</td><td>31</td><td>GGGTTTACGACCTCGATGTTGGATCAGGACA</td><td>98.4</td><td>4.6</td><td>1.7</td><td>2.7</td><td>6.5E-03</td><td>1.3E-02</td><td>BT20/MB468</td><td>MP</td></tr>
<tr><td>>hg38_21_+_8433172_8446622_RNA45SN1_45S_50ntFlanks@9569.9609.41|GTGCAAATCGGTCGTCCGACCTGGGTATAGGGGCGAAAGAC</td><td>>hg38_21_+_8433172_8446622_RNA45SN1_45S_50ntFlanks</td><td>9569</td><td>9609</td><td>41</td><td>GTGCAAATCGGTCGTCCGACCTGGGTATAGGGGCGAAAGAC</td><td>36.3</td><td>5.1</td><td>2.0</td><td>2.6</td><td>9.7E-03</td><td>1.3E-02</td><td>MB231/MB468</td><td>MP</td></tr>
<tr><td>>hg38_21_+_8433172_8446622_RNA45SN1_45S_50ntFlanks@11779.11803.25|CGCGCATGAATGGATGAACGAGATT</td><td>>hg38_21_+_8433172_8446622_RNA45SN1_45S_50ntFlanks</td><td>11779</td><td>11803</td><td>25</td><td>CGCGCATGAATGGATGAACGAGATT</td><td>172.2</td><td>-3.6</td><td>1.4</td><td>-2.6</td><td>9.7E-03</td><td>1.3E-02</td><td>MB231/MB468</td><td>MP</td></tr>
<tr><td>>MI0000446|hsa-mir-125b-1&WithFlank&11|-|122099751|122099850@21.39.19(+1A);MI0000470|hsa-mir-125b-2&WithFlank&21|+|16590231|16590331@23.41.19(+1A)&[MIMAT0000423&hsa-miR-125b-5p&offsets|0|-3(+1A);m-67&11|-|122099809|122099830&offsets|0|-3(+1A)];[MIMAT0000423_1&hsa-miR-125b-5p&offsets|0|-3(+1A);m-68&21|+|16590253|16590274&offsets|0|-3(+1A)]|TCCCTGAGACCCTAACTTGA</td><td>>MI0000446|hsa-mir-125b-1&WithFlank&11|-|122099751|122099850</td><td>21</td><td>39</td><td>19(+1A)</td><td>TCCCTGAGACCCTAACTTGA</td><td>23.8</td><td>7.8</td><td>3.0</td><td>2.6</td><td>9.7E-03</td><td>1.3E-02</td><td>MB231/MB468</td><td>MP</td></tr>
<tr><td>>trna10_AspGTC_12_-_125424193_125424264@31.71.41,trna12_AspGTC_12_-_125411891_125411962@31.71.41,trna38_AspGTC_17_-_8125556_8125627@31.71.41,trna45_AspGTC_6_+_27447453_27447524@31.71.41,trna48_AspGTC_6_+_27471523_27471594@31.71.41,trna4_AspGTC_12_+_96429799_96429870@31.71.41,trna69_AspGTC_1_-_161440205_161440276@31.71.41,trna72_AspGTC_1_-_161432824_161432895@31.71.41,trna75_AspGTC_1_-_161425414_161425485@31.71.41,trna78_AspGTC_1_-_161418033_161418104@31.71.41,trna81_AspGTC_1_-_161410615_161410686@31.71.41|CCTGTCACGCGGGAGACCGGGGTTCGATTCCCCGACGGGGA</td><td>>trna10_AspGTC_12_-_125424193_125424264</td><td>31</td><td>71</td><td>41</td><td>CCTGTCACGCGGGAGACCGGGGTTCGATTCCCCGACGGGGA</td><td>111.1</td><td>-5.0</td><td>1.9</td><td>-2.6</td><td>9.0E-03</td><td>1.3E-02</td><td>BT20/MB468</td><td>MITO</td></tr>
<tr><td>>hg38_21_+_8433172_8446622_RNA45SN1_45S_50ntFlanks@11314.11344.31|CCCACCCCACGTCTCGTCGCGCGCGCGTCCG</td><td>>hg38_21_+_8433172_8446622_RNA45SN1_45S_50ntFlanks</td><td>11314</td><td>11344</td><td>31</td><td>CCCACCCCACGTCTCGTCGCGCGCGCGTCCG</td><td>767.6</td><td>-4.3</td><td>1.7</td><td>-2.6</td><td>9.0E-03</td><td>1.3E-02</td><td>BT20/MB468</td><td>MITO</td></tr>
<tr><td>>MI0000447|hsa-mir-128-1&WithFlank&2|+|135665391|135665484@56.75.20(+1C);MI0000727|hsa-mir-128-2&WithFlank&3|+|35744470|35744565@58.77.20(+1C)&[MIMAT0000424&hsa-miR-128-3p&offsets|0|-1(+1C);m-108&2|+|135665446|135665466&offsets|0|-1(+1C)];[MIMAT0000424_1&hsa-miR-128-3p&offsets|0|-1(+1C);m-117&3|+|35744527|35744547&offsets|0|-1(+1C)]|TCACAGTGAACCGGTCTCTTC</td><td>>MI0000447|hsa-mir-128-1&WithFlank&2|+|135665391|135665484</td><td>56</td><td>75</td><td>20(+1C)</td><td>TCACAGTGAACCGGTCTCTTC</td><td>1066.6</td><td>2.3</td><td>0.8</td><td>3.0</td><td>3.2E-03</td><td>1.3E-02</td><td>BT20/MB231</td><td>CYTO</td></tr>
<tr><td>>am_trna116_GluCTC_1_-_145399233_145399304@13.30.18,trna25_GluCTC_2_-_71273488_71273560@13.30.18,trna59_GluCTC_1_+_249168447_249168518@13.30.18,trna71_GluCTC_1_-_161439189_161439260@13.30.18,trna74_GluCTC_1_-_161431809_161431880@13.30.18,trna77_GluCTC_1_-_161424398_161424469@13.30.18,trna77_GluCTC_6_+_28949976_28950047@13.30.18,trna80_GluCTC_1_-_161417018_161417089@13.30.18,trna87_GluCTC_6_-_126101393_126101464@13.30.18|TAGTGGTTAGGATTCGGC</td><td>>am_trna116_GluCTC_1_-_145399233_145399304</td><td>13</td><td>30</td><td>18</td><td>TAGTGGTTAGGATTCGGC</td><td>1663.7</td><td>2.7</td><td>0.9</td><td>3.0</td><td>3.2E-03</td><td>1.3E-02</td><td>BT20/MB231</td><td>CYTO</td></tr>
<tr><td>>am_trnaMT_LysTTT_MT_+_8295_8364@33.52.20,trnalookalike7_LysTTT_1_+_568844_568913@33.52.20|AGTTAAAGATTAAGAGAACC</td><td>>am_trnaMT_LysTTT_MT_+_8295_8364</td><td>33</td><td>52</td><td>20</td><td>AGTTAAAGATTAAGAGAACC</td><td>381.2</td><td>2.7</td><td>0.9</td><td>3.0</td><td>3.2E-03</td><td>1.3E-02</td><td>BT20/MB231</td><td>CYTO</td></tr>
<tr><td>>hg38_21_+_8433172_8446622_RNA45SN1_45S_50ntFlanks@9720.9749.30|ATTAGAGGTCTTGGGGCCGAAACGATCTCA</td><td>>hg38_21_+_8433172_8446622_RNA45SN1_45S_50ntFlanks</td><td>9720</td><td>9749</td><td>30</td><td>ATTAGAGGTCTTGGGGCCGAAACGATCTCA</td><td>46.3</td><td>-4.5</td><td>1.6</td><td>-2.7</td><td>6.5E-03</td><td>1.3E-02</td><td>BT20/MB468</td><td>MP</td></tr>
<tr><td>>hg38_21_+_8433172_8446622_RNA45SN1_45S_50ntFlanks@3990.4024.35|TTGGTGACTCTAGATAACCTCGGGCCGATCGCACG</td><td>>hg38_21_+_8433172_8446622_RNA45SN1_45S_50ntFlanks</td><td>3990</td><td>4024</td><td>35</td><td>TTGGTGACTCTAGATAACCTCGGGCCGATCGCACG</td><td>717.5</td><td>-1.4</td><td>0.6</td><td>-2.5</td><td>1.1E-02</td><td>1.3E-02</td><td>MB231/MB468</td><td>NUC</td></tr>
<tr><td>>hg38_MT_+_1621_3279_RNR2_16S_50ntFlanks@1095.1113.19|AGACGAGAAGACCCTATGG</td><td>>hg38_MT_+_1621_3279_RNR2_16S_50ntFlanks</td><td>1095</td><td>1113</td><td>19</td><td>AGACGAGAAGACCCTATGG</td><td>109.9</td><td>2.9</td><td>1.1</td><td>2.7</td><td>6.5E-03</td><td>1.3E-02</td><td>BT20/MB468</td><td>MP</td></tr>
<tr><td>>hg38_21_+_8433172_8446622_RNA45SN1_45S_50ntFlanks@7988.8012.25|CAGACGTGGCGACCCGCTGAATTTA</td><td>>hg38_21_+_8433172_8446622_RNA45SN1_45S_50ntFlanks</td><td>7988</td><td>8012</td><td>25</td><td>CAGACGTGGCGACCCGCTGAATTTA</td><td>405.2</td><td>-1.6</td><td>0.6</td><td>-2.6</td><td>1.1E-02</td><td>1.3E-02</td><td>MB231/MB468</td><td>CYTO</td></tr>
<tr><td>>hg38_21_+_8433172_8446622_RNA45SN1_45S_50ntFlanks@6652.6698.47|GACTCTTAGCGGTGGATCACTCGGCTCGTGCGTCGATGAAGAACGCA</td><td>>hg38_21_+_8433172_8446622_RNA45SN1_45S_50ntFlanks</td><td>6652</td><td>6698</td><td>47</td><td>GACTCTTAGCGGTGGATCACTCGGCTCGTGCGTCGATGAAGAACGCA</td><td>1972.5</td><td>-2.8</td><td>0.9</td><td>-3.0</td><td>3.2E-03</td><td>1.3E-02</td><td>BT20/MB231</td><td>CYTO</td></tr>
<tr><td>>hg38_MT_+_1621_3279_RNR2_16S_50ntFlanks@1187.1215.29|TTCGGTTGGGGCGACCTCGGAGCAGAACC</td><td>>hg38_MT_+_1621_3279_RNR2_16S_50ntFlanks</td><td>1187</td><td>1215</td><td>29</td><td>TTCGGTTGGGGCGACCTCGGAGCAGAACC</td><td>157.3</td><td>4.8</td><td>1.8</td><td>2.7</td><td>6.5E-03</td><td>1.3E-02</td><td>BT20/MB468</td><td>MP</td></tr>
<tr><td>>hg38_21_+_8433172_8446622_RNA45SN1_45S_50ntFlanks@9553.9577.25|CCGTAGCGGTCCTGACGTGCAAATC</td><td>>hg38_21_+_8433172_8446622_RNA45SN1_45S_50ntFlanks</td><td>9553</td><td>9577</td><td>25</td><td>CCGTAGCGGTCCTGACGTGCAAATC</td><td>285.8</td><td>-2.3</td><td>0.8</td><td>-2.9</td><td>3.2E-03</td><td>1.3E-02</td><td>BT20/MB231</td><td>CYTO</td></tr>
<tr><td>>hg38_21_+_8433172_8446622_RNA45SN1_45S_50ntFlanks@12627.12662.36|ATGGGGCGAAGCTACCATCTGTGGGATTATGACTGA</td><td>>hg38_21_+_8433172_8446622_RNA45SN1_45S_50ntFlanks</td><td>12627</td><td>12662</td><td>36</td><td>ATGGGGCGAAGCTACCATCTGTGGGATTATGACTGA</td><td>75.8</td><td>-4.6</td><td>1.7</td><td>-2.7</td><td>6.5E-03</td><td>1.3E-02</td><td>BT20/MB468</td><td>MP</td></tr>
<tr><td>>hg38_21_+_8433172_8446622_RNA45SN1_45S_50ntFlanks@11568.11611.44|GCAGCCGACTTAGAACTGGTGCGGACCAGGGGAATCCGACTGTT</td><td>>hg38_21_+_8433172_8446622_RNA45SN1_45S_50ntFlanks</td><td>11568</td><td>11611</td><td>44</td><td>GCAGCCGACTTAGAACTGGTGCGGACCAGGGGAATCCGACTGTT</td><td>1050.8</td><td>2.8</td><td>1.1</td><td>2.6</td><td>1.1E-02</td><td>1.3E-02</td><td>MB231/MB468</td><td>CYTO</td></tr>
<tr><td>>hg38_21_+_8433172_8446622_RNA45SN1_45S_50ntFlanks@8198.8220.23|TGGACGGTGTGAGGCCGGTAGCG</td><td>>hg38_21_+_8433172_8446622_RNA45SN1_45S_50ntFlanks</td><td>8198</td><td>8220</td><td>23</td><td>TGGACGGTGTGAGGCCGGTAGCG</td><td>126.8</td><td>-4.7</td><td>1.7</td><td>-2.7</td><td>6.5E-03</td><td>1.3E-02</td><td>BT20/MB468</td><td>MP</td></tr>
<tr><td>>hg38_21_+_8433172_8446622_RNA45SN1_45S_50ntFlanks@6645.6679.35|CTCGTACGACTCTTAGCGGTGGATCACTCGGCTCG</td><td>>hg38_21_+_8433172_8446622_RNA45SN1_45S_50ntFlanks</td><td>6645</td><td>6679</td><td>35</td><td>CTCGTACGACTCTTAGCGGTGGATCACTCGGCTCG</td><td>1749.9</td><td>4.5</td><td>1.5</td><td>3.0</td><td>2.5E-03</td><td>1.3E-02</td><td>BT20/MB231</td><td>MP</td></tr>
<tr><td>>hg38_21_+_8433172_8446622_RNA45SN1_45S_50ntFlanks@6645.6666.22|CTCGTACGACTCTTAGCGGTGG</td><td>>hg38_21_+_8433172_8446622_RNA45SN1_45S_50ntFlanks</td><td>6645</td><td>6666</td><td>22</td><td>CTCGTACGACTCTTAGCGGTGG</td><td>3951.8</td><td>3.3</td><td>1.1</td><td>3.0</td><td>2.5E-03</td><td>1.3E-02</td><td>BT20/MB231</td><td>MP</td></tr>
<tr><td>>hg38_21_+_8433172_8446622_RNA45SN1_45S_50ntFlanks@9943.9978.36|GACGGTGGCCATGGAAGTCGGAATCCGCTAAGGAGT</td><td>>hg38_21_+_8433172_8446622_RNA45SN1_45S_50ntFlanks</td><td>9943</td><td>9978</td><td>36</td><td>GACGGTGGCCATGGAAGTCGGAATCCGCTAAGGAGT</td><td>72.9</td><td>4.6</td><td>1.7</td><td>2.7</td><td>6.5E-03</td><td>1.3E-02</td><td>BT20/MB468</td><td>MP</td></tr>
<tr><td>>hg38_21_+_8433172_8446622_RNA45SN1_45S_50ntFlanks@11208.11228.21|GCGGCGACTCTGGACGCGAGC</td><td>>hg38_21_+_8433172_8446622_RNA45SN1_45S_50ntFlanks</td><td>11208</td><td>11228</td><td>21</td><td>GCGGCGACTCTGGACGCGAGC</td><td>1038.9</td><td>2.9</td><td>1.1</td><td>2.6</td><td>9.1E-03</td><td>1.3E-02</td><td>BT20/MB468</td><td>MITO</td></tr>
<tr><td>>hg38_21_+_8433172_8446622_RNA45SN1_45S_50ntFlanks@12995.13037.43|CTCGCTGCGATCTATTGAAAGTCAGCCCTCGACACAAGGGTTT</td><td>>hg38_21_+_8433172_8446622_RNA45SN1_45S_50ntFlanks</td><td>12995</td><td>13037</td><td>43</td><td>CTCGCTGCGATCTATTGAAAGTCAGCCCTCGACACAAGGGTTT</td><td>602.2</td><td>-2.2</td><td>0.8</td><td>-2.9</td><td>3.2E-03</td><td>1.3E-02</td><td>BT20/MB231</td><td>MITO</td></tr>
<tr><td>>hg38_21_+_8433172_8446622_RNA45SN1_45S_50ntFlanks@9737.9765.29|CGAAACGATCTCAACCTATTCTCAAACTT</td><td>>hg38_21_+_8433172_8446622_RNA45SN1_45S_50ntFlanks</td><td>9737</td><td>9765</td><td>29</td><td>CGAAACGATCTCAACCTATTCTCAAACTT</td><td>296.9</td><td>-2.3</td><td>0.8</td><td>-2.9</td><td>3.2E-03</td><td>1.3E-02</td><td>BT20/MB231</td><td>MITO</td></tr>
<tr><td>>hg38_1_-_228634819_228635039_RNA5S12_5S_50ntFlanks@132.168.37|GATGGGAGACCGCCTGGGAATACCGGGTGCTGTAGGC</td><td>>hg38_1_-_228634819_228635039_RNA5S12_5S_50ntFlanks</td><td>132</td><td>168</td><td>37</td><td>GATGGGAGACCGCCTGGGAATACCGGGTGCTGTAGGC</td><td>258.6</td><td>5.0</td><td>1.7</td><td>3.0</td><td>2.5E-03</td><td>1.3E-02</td><td>BT20/MB231</td><td>MP</td></tr>
<tr><td>>hg38_21_+_8433172_8446622_RNA45SN1_45S_50ntFlanks@11876.11893.18|AAGACCCTGTTGAGCTTG</td><td>>hg38_21_+_8433172_8446622_RNA45SN1_45S_50ntFlanks</td><td>11876</td><td>11893</td><td>18</td><td>AAGACCCTGTTGAGCTTG</td><td>171.5</td><td>4.8</td><td>1.6</td><td>3.0</td><td>2.5E-03</td><td>1.3E-02</td><td>BT20/MB231</td><td>MP</td></tr>
<tr><td>>MI0000077|hsa-mir-21&WithFlank&17|+|59841260|59841343@14.33.20(+1U)&[MIMAT0000076&hsa-miR-21-5p&offsets|0|-2(+1U);m-1&17|+|59841273|59841294&offsets|0|-2(+1U)]|TAGCTTATCAGACTGATGTTT</td><td>>MI0000077|hsa-mir-21&WithFlank&17|+|59841260|59841343</td><td>14</td><td>33</td><td>20(+1U)</td><td>TAGCTTATCAGACTGATGTTT</td><td>409.0</td><td>3.6</td><td>1.3</td><td>2.7</td><td>6.5E-03</td><td>1.3E-02</td><td>BT20/MB468</td><td>MP</td></tr>
<tr><td>>hg38_21_+_8433172_8446622_RNA45SN1_45S_50ntFlanks@6674.6696.23|GGCTCGTGCGTCGATGAAGAACG</td><td>>hg38_21_+_8433172_8446622_RNA45SN1_45S_50ntFlanks</td><td>6674</td><td>6696</td><td>23</td><td>GGCTCGTGCGTCGATGAAGAACG</td><td>162.5</td><td>2.5</td><td>0.9</td><td>2.7</td><td>6.5E-03</td><td>1.3E-02</td><td>BT20/MB468</td><td>MP</td></tr>
<tr><td>>hg38_21_+_8433172_8446622_RNA45SN1_45S_50ntFlanks@11569.11601.33|CAGCCGACTTAGAACTGGTGCGGACCAGGGGAA</td><td>>hg38_21_+_8433172_8446622_RNA45SN1_45S_50ntFlanks</td><td>11569</td><td>11601</td><td>33</td><td>CAGCCGACTTAGAACTGGTGCGGACCAGGGGAA</td><td>303.1</td><td>3.9</td><td>1.3</td><td>3.0</td><td>2.5E-03</td><td>1.3E-02</td><td>BT20/MB231</td><td>MP</td></tr>
<tr><td>>hg38_21_+_8433172_8446622_RNA45SN1_45S_50ntFlanks@10107.10125.19|CGTGTGGTGTGCGTCGGAG</td><td>>hg38_21_+_8433172_8446622_RNA45SN1_45S_50ntFlanks</td><td>10107</td><td>10125</td><td>19</td><td>CGTGTGGTGTGCGTCGGAG</td><td>930.5</td><td>-5.2</td><td>2.1</td><td>-2.5</td><td>1.1E-02</td><td>1.3E-02</td><td>BT20/MB468</td><td>NUC</td></tr>
<tr><td>>hg38_21_+_8433172_8446622_RNA45SN1_45S_50ntFlanks@12423.12443.21|TCGGCTCTTCCTATCATTGTG</td><td>>hg38_21_+_8433172_8446622_RNA45SN1_45S_50ntFlanks</td><td>12423</td><td>12443</td><td>21</td><td>TCGGCTCTTCCTATCATTGTG</td><td>174.4</td><td>-4.0</td><td>1.5</td><td>-2.6</td><td>9.1E-03</td><td>1.3E-02</td><td>BT20/MB468</td><td>MITO</td></tr>
<tr><td>>MI0000076|hsa-mir-20a&WithFlank&13|+|91351059|91351141@14.36.23(+1A)&[MIMAT0000075&hsa-miR-20a-5p&offsets|0|0(+1A);m-96&13|+|91351072|91351094&offsets|0|0(+1A)]|TAAAGTGCTTATAGTGCAGGTAGA</td><td>>MI0000076|hsa-mir-20a&WithFlank&13|+|91351059|91351141</td><td>14</td><td>36</td><td>23(+1A)</td><td>TAAAGTGCTTATAGTGCAGGTAGA</td><td>1111.1</td><td>2.1</td><td>0.8</td><td>2.6</td><td>9.8E-03</td><td>1.3E-02</td><td>MB231/MB468</td><td>MP</td></tr>
</table>

<table>
<tr><td>>hg38_21_+_8433172_8446622_RNA45SN1_45S_50ntFlanks@6646.6680.35|TCGTACGACTCTTAGCGGTGGATCACTCGGCTCGT</td><td>>hg38_21_+_8433172_8446622_RNA45SN1_45S_50ntFlanks</td><td>6646</td><td>6680</td><td>35</td><td>TCGTACGACTCTTAGCGGTGGATCACTCGGCTCGT</td><td>4904.7</td><td>1.8</td><td>0.7</td><td>2.5</td><td>1.1E-02</td><td>1.3E-02</td><td>MB231/MB468</td><td>NUC</td></tr>
<tr><td>>am_trna119_LysCTT_1_-_145395522_145395594@14.31.18,trna11_LysCTT_5_-_180648979_180649051@14.31.18,trna128_LysCTT_1_-_55423542_55423614@14.31.18,trna13_LysCTT_6_+_26556774_26556846@14.31.18,trna32_LysCTT_16_-_3207406_3207478@14.31.18,trna6_LysCTT_19_-_52425396_52425468@14.31.18,trna7_LysCTT_16_+_3225692_3225764@14.31.18,trna9_LysCTT_5_+_180634755_180634827@14.31.18|AGTCGGTAGAGCATGAGA</td><td>>am_trna119_LysCTT_1_-_145395522_145395594</td><td>14</td><td>31</td><td>18</td><td>AGTCGGTAGAGCATGAGA</td><td>197.7</td><td>-3.5</td><td>1.3</td><td>-2.7</td><td>6.5E-03</td><td>1.3E-02</td><td>BT20/MB468</td><td>MP</td></tr>
<tr><td>>hg38_21_+_8433172_8446622_RNA45SN1_45S_50ntFlanks@6722.6743.22|ATTGCAGGACACATTGATCATC</td><td>>hg38_21_+_8433172_8446622_RNA45SN1_45S_50ntFlanks</td><td>6722</td><td>6743</td><td>22</td><td>ATTGCAGGACACATTGATCATC</td><td>399.5</td><td>-2.7</td><td>1.0</td><td>-2.7</td><td>6.5E-03</td><td>1.3E-02</td><td>BT20/MB468</td><td>MP</td></tr>
<tr><td>>hg38_21_+_8433172_8446622_RNA45SN1_45S_50ntFlanks@10393.10422.30|CAGCAGTTGAACATGGGTCAGTCGGTCCTG</td><td>>hg38_21_+_8433172_8446622_RNA45SN1_45S_50ntFlanks</td><td>10393</td><td>10422</td><td>30</td><td>CAGCAGTTGAACATGGGTCAGTCGGTCCTG</td><td>357.9</td><td>-2.6</td><td>0.9</td><td>-2.7</td><td>6.5E-03</td><td>1.3E-02</td><td>BT20/MB468</td><td>MP</td></tr>
<tr><td>>MI0006444|hsa-mir-1307&WithFlank&10|-|103394247|103394407@86.106.21&[MIMAT0005951&hsa-miR-1307-3p&offsets|0|-1;m-66&10|-|103394300|103394322&offsets|0|-2]|ACTCGGCGTGGCGTCGGTCGT</td><td>>MI0006444|hsa-mir-1307&WithFlank&10|-|103394247|103394407</td><td>86</td><td>106</td><td>21</td><td>ACTCGGCGTGGCGTCGGTCGT</td><td>144.9</td><td>4.9</td><td>1.8</td><td>2.7</td><td>6.5E-03</td><td>1.3E-02</td><td>BT20/MB468</td><td>MP</td></tr>
<tr><td>>hg38_21_+_8433172_8446622_RNA45SN1_45S_50ntFlanks@5473.5494.22|CGGCCCTGGCGGAGCGCTGAGA</td><td>>hg38_21_+_8433172_8446622_RNA45SN1_45S_50ntFlanks</td><td>5473</td><td>5494</td><td>22</td><td>CGGCCCTGGCGGAGCGCTGAGA</td><td>2414.8</td><td>3.2</td><td>1.1</td><td>2.9</td><td>3.2E-03</td><td>1.3E-02</td><td>BT20/MB231</td><td>CYTO</td></tr>
<tr><td>>hg38_21_+_8433172_8446622_RNA45SN1_45S_50ntFlanks@12418.12453.36|CGATGTCGGCTCTTCCTATCATTGTGAAGCAGAATT</td><td>>hg38_21_+_8433172_8446622_RNA45SN1_45S_50ntFlanks</td><td>12418</td><td>12453</td><td>36</td><td>CGATGTCGGCTCTTCCTATCATTGTGAAGCAGAATT</td><td>271.4</td><td>-2.2</td><td>0.8</td><td>-2.9</td><td>3.2E-03</td><td>1.3E-02</td><td>BT20/MB231</td><td>CYTO</td></tr>
<tr><td>>trna20_GluTTC_2_-_131094701_131094772@1.33.33,trna5_GluTTC_13_-_41634874_41634945@1.33.33|TCCCATATGGTCTAGCGGTTAGGATTCCTGGTT</td><td>>trna20_GluTTC_2_-_131094701_131094772</td><td>1</td><td>33</td><td>33</td><td>TCCCATATGGTCTAGCGGTTAGGATTCCTGGTT</td><td>566.7</td><td>-2.5</td><td>0.9</td><td>-3.0</td><td>2.8E-03</td><td>1.3E-02</td><td>BT20/MB231</td><td>NUC</td></tr>
<tr><td>>MI0000264|hsa-mir-7-2&WithFlank&15|+|88611819|88611940@38.58.21;MI0000265|hsa-mir-7-3&WithFlank&19|+|4770664|4770785@37.57.21;MI0000263|hsa-mir-7-1&WithFlank&9|-|83969742|83969863@30.50.21&[MIMAT0000252&hsa-miR-7-5p&offsets|0|-3;m-400&15|+|88611856|88611879&offsets|0|-3];[MIMAT0000252_1&hsa-miR-7-5p&offsets|0|-3;m-402&19|+|4770700|4770723&offsets|0|-3];[MIMAT0000252_2&hsa-miR-7-5p&offsets|0|-3;m-401&9|-|83969811|83969834&offsets|0|-3]|TGGAAGACTAGTGATTTTGTT</td><td>>MI0000264|hsa-mir-7-2&WithFlank&15|+|88611819|88611940</td><td>38</td><td>58</td><td>21</td><td>TGGAAGACTAGTGATTTTGTT</td><td>3658.2</td><td>-3.0</td><td>1.0</td><td>-3.0</td><td>2.5E-03</td><td>1.3E-02</td><td>BT20/MB231</td><td>MP</td></tr>
<tr><td>>am_MI0003581|hsa-mir-574&WithFlank&4|+|38868026|38868133@31.54.24&[MIMAT0004795&hsa-miR-574-5p&offsets|0|+1;m-213&4|+|38868056|38868078&offsets|0|+1]|TGAGTGTGTGTGTGTGAGTGTGTG</td><td>>am_MI0003581|hsa-mir-574&WithFlank&4|+|38868026|38868133</td><td>31</td><td>54</td><td>24</td><td>TGAGTGTGTGTGTGTGAGTGTGTG</td><td>926.1</td><td>-3.0</td><td>1.0</td><td>-3.0</td><td>2.5E-03</td><td>1.3E-02</td><td>BT20/MB231</td><td>MP</td></tr>
<tr><td>>trnaMT_ValTAC_MT_+_1602_1670@42.71.30|AGATTTCAACTTAACTTGACCGCTCTGACC</td><td>>trnaMT_ValTAC_MT_+_1602_1670</td><td>42</td><td>71</td><td>30</td><td>AGATTTCAACTTAACTTGACCGCTCTGACC</td><td>7099.6</td><td>-2.8</td><td>0.9</td><td>-3.0</td><td>2.5E-03</td><td>1.3E-02</td><td>BT20/MB231</td><td>MP</td></tr>
<tr><td>>hg38_21_+_8433172_8446622_RNA45SN1_45S_50ntFlanks@9927.9949.23|TTGATATAGACAGCAGGACGGTG</td><td>>hg38_21_+_8433172_8446622_RNA45SN1_45S_50ntFlanks</td><td>9927</td><td>9949</td><td>23</td><td>TTGATATAGACAGCAGGACGGTG</td><td>242.7</td><td>4.5</td><td>1.5</td><td>3.0</td><td>2.5E-03</td><td>1.3E-02</td><td>BT20/MB231</td><td>MP</td></tr>
<tr><td>>hg38_MT_+_1621_3279_RNR2_16S_50ntFlanks@51.92.42|GCTAAACCTAGCCCCAAACCCACTCCACCTTACTACCAGACA</td><td>>hg38_MT_+_1621_3279_RNR2_16S_50ntFlanks</td><td>51</td><td>92</td><td>42</td><td>GCTAAACCTAGCCCCAAACCCACTCCACCTTACTACCAGACA</td><td>8213.3</td><td>-2.8</td><td>0.9</td><td>-2.9</td><td>3.2E-03</td><td>1.3E-02</td><td>BT20/MB231</td><td>MITO</td></tr>
<tr><td>>hg38_1_-_228634819_228635039_RNA5S12_5S_50ntFlanks@131.169.39|GGATGGGAGACCGCCTGGGAATACCGGGTGCTGTAGGCT</td><td>>hg38_1_-_228634819_228635039_RNA5S12_5S_50ntFlanks</td><td>131</td><td>169</td><td>39</td><td>GGATGGGAGACCGCCTGGGAATACCGGGTGCTGTAGGCT</td><td>1094.1</td><td>3.5</td><td>1.2</td><td>3.0</td><td>2.5E-03</td><td>1.3E-02</td><td>BT20/MB231</td><td>MP</td></tr>
<tr><td>>hg38_21_+_8433172_8446622_RNA45SN1_45S_50ntFlanks@4567.4587.21|TAATGGAATAGGACCGCGGTT</td><td>>hg38_21_+_8433172_8446622_RNA45SN1_45S_50ntFlanks</td><td>4567</td><td>4587</td><td>21</td><td>TAATGGAATAGGACCGCGGTT</td><td>193.6</td><td>-3.1</td><td>1.2</td><td>-2.6</td><td>9.1E-03</td><td>1.3E-02</td><td>BT20/MB468</td><td>MITO</td></tr>
<tr><td>>hg38_21_+_8433172_8446622_RNA45SN1_45S_50ntFlanks@11843.11873.31|AGGGAACGGGCTTGGCGGAATCAGCGGGGAA</td><td>>hg38_21_+_8433172_8446622_RNA45SN1_45S_50ntFlanks</td><td>11843</td><td>11873</td><td>31</td><td>AGGGAACGGGCTTGGCGGAATCAGCGGGGAA</td><td>318.9</td><td>-4.7</td><td>1.7</td><td>-2.7</td><td>6.6E-03</td><td>1.3E-02</td><td>BT20/MB468</td><td>MP</td></tr>
<tr><td>>MI0000255|hsa-mir-30d&WithFlank&8|-|134804870|134804951@12.32.21(+1A)&[MIMAT0000245&hsa-miR-30d-5p&offsets|0|-1(+1A);m-21&8|-|134804919|134804940&offsets|0|-1(+1A)]|TGTAAACATCCCCGACTGGAAA</td><td>>MI0000255|hsa-mir-30d&WithFlank&8|-|134804870|134804951</td><td>12</td><td>32</td><td>21(+1A)</td><td>TGTAAACATCCCCGACTGGAAA</td><td>148.9</td><td>3.8</td><td>1.4</td><td>2.7</td><td>6.6E-03</td><td>1.3E-02</td><td>BT20/MB468</td><td>MP</td></tr>
<tr><td>>hg38_21_+_8433172_8446622_RNA45SN1_45S_50ntFlanks@10099.10117.19|CGCGCGCGCGTGTGGTGTG</td><td>>hg38_21_+_8433172_8446622_RNA45SN1_45S_50ntFlanks</td><td>10099</td><td>10117</td><td>19</td><td>CGCGCGCGCGTGTGGTGTG</td><td>582.0</td><td>-5.0</td><td>2.0</td><td>-2.5</td><td>1.1E-02</td><td>1.3E-02</td><td>BT20/MB468</td><td>NUC</td></tr>
<tr><td>>hg38_MT_+_598_1651_RNR1_12S_50ntFlanks@442.468.27|ATATCTGAACACACAATAGCTAAGACC</td><td>>hg38_MT_+_598_1651_RNR1_12S_50ntFlanks</td><td>442</td><td>468</td><td>27</td><td>ATATCTGAACACACAATAGCTAAGACC</td><td>77.2</td><td>4.2</td><td>1.6</td><td>2.7</td><td>6.6E-03</td><td>1.3E-02</td><td>BT20/MB468</td><td>MP</td></tr>
<tr><td>>hg38_21_+_8433172_8446622_RNA45SN1_45S_50ntFlanks@11575.11600.26|ACTTAGAACTGGTGCGGACCAGGGGA</td><td>>hg38_21_+_8433172_8446622_RNA45SN1_45S_50ntFlanks</td><td>11575</td><td>11600</td><td>26</td><td>ACTTAGAACTGGTGCGGACCAGGGGA</td><td>181.2</td><td>-1.6</td><td>0.6</td><td>-2.6</td><td>9.2E-03</td><td>1.3E-02</td><td>BT20/MB468</td><td>MITO</td></tr>
<tr><td>>hg38_21_+_8433172_8446622_RNA45SN1_45S_50ntFlanks@10097.10115.19|CGCGCGCGCGCGTGTGGTG</td><td>>hg38_21_+_8433172_8446622_RNA45SN1_45S_50ntFlanks</td><td>10097</td><td>10115</td><td>19</td><td>CGCGCGCGCGCGTGTGGTG</td><td>171.7</td><td>-4.5</td><td>1.7</td><td>-2.7</td><td>6.6E-03</td><td>1.3E-02</td><td>BT20/MB468</td><td>MP</td></tr>
<tr><td>>hg38_21_+_8433172_8446622_RNA45SN1_45S_50ntFlanks@11784.11804.21|ATGAATGGATGAACGAGATTC</td><td>>hg38_21_+_8433172_8446622_RNA45SN1_45S_50ntFlanks</td><td>11784</td><td>11804</td><td>21</td><td>ATGAATGGATGAACGAGATTC</td><td>1171.6</td><td>-1.5</td><td>0.6</td><td>-2.6</td><td>9.2E-03</td><td>1.3E-02</td><td>BT20/MB468</td><td>MITO</td></tr>
<tr><td>>MI0000067|hsa-let-7f-1&WithFlank&9|+|94176341|94176439@69.90.22&[MIMAT0004486&hsa-let-7f-1-3p&offsets|0|0;m-275&9|+|94176409|94176430&offsets|0|0]|CTATACAATCTATTGCCTTCCC</td><td>>MI0000067|hsa-let-7f-1&WithFlank&9|+|94176341|94176439</td><td>69</td><td>90</td><td>22</td><td>CTATACAATCTATTGCCTTCCC</td><td>65.0</td><td>6.9</td><td>2.5</td><td>2.7</td><td>6.6E-03</td><td>1.3E-02</td><td>BT20/MB468</td><td>MP</td></tr>
<tr><td>>hg38_21_+_8433172_8446622_RNA45SN1_45S_50ntFlanks@13002.13039.38|CGATCTATTGAAAGTCAGCCCTCGACACAAGGGTTTGT</td><td>>hg38_21_+_8433172_8446622_RNA45SN1_45S_50ntFlanks</td><td>13002</td><td>13039</td><td>38</td><td>CGATCTATTGAAAGTCAGCCCTCGACACAAGGGTTTGT</td><td>3016.3</td><td>-4.0</td><td>1.6</td><td>-2.6</td><td>1.1E-02</td><td>1.3E-02</td><td>MB231/MB468</td><td>CYTO</td></tr>
<tr><td>>MI0000734|hsa-mir-106b&WithFlank&7|-|100093987|100094080@58.79.22&[MIMAT0004672&hsa-miR-106b-3p&offsets|0|0;m-87&7|-|100094003|100094023&offsets|0|+1]|CCGCACTGTGGGTACTTGCTGC</td><td>>MI0000734|hsa-mir-106b&WithFlank&7|-|100093987|100094080</td><td>58</td><td>79</td><td>22</td><td>CCGCACTGTGGGTACTTGCTGC</td><td>2066.7</td><td>2.5</td><td>1.0</td><td>2.6</td><td>9.8E-03</td><td>1.3E-02</td><td>MB231/MB468</td><td>MP</td></tr>
<tr><td>>hg38_MT_+_1621_3279_RNR2_16S_50ntFlanks@1370.1388.19|ATGTTGGATCAGGACATCC</td><td>>hg38_MT_+_1621_3279_RNR2_16S_50ntFlanks</td><td>1370</td><td>1388</td><td>19</td><td>ATGTTGGATCAGGACATCC</td><td>636.8</td><td>2.5</td><td>1.0</td><td>2.6</td><td>9.8E-03</td><td>1.3E-02</td><td>MB231/MB468</td><td>MP</td></tr>
<tr><td>>MI0000071|hsa-mir-17&WithFlank&13|+|91350599|91350694@20.39.20(+1C)&[MIMAT0000070&hsa-miR-17-5p&offsets|0|-3(+1C);m-88&13|+|91350618|91350640&offsets|0|-3(+1C)]|CAAAGTGCTTACAGTGCAGGC</td><td>>MI0000071|hsa-mir-17&WithFlank&13|+|91350599|91350694</td><td>20</td><td>39</td><td>20(+1C)</td><td>CAAAGTGCTTACAGTGCAGGC</td><td>340.2</td><td>3.6</td><td>1.4</td><td>2.6</td><td>9.9E-03</td><td>1.3E-02</td><td>MB231/MB468</td><td>MP</td></tr>
<tr><td>>hg38_21_+_8433172_8446622_RNA45SN1_45S_50ntFlanks@11604.11623.20|CGACTGTTTAATTAAAACAA</td><td>>hg38_21_+_8433172_8446622_RNA45SN1_45S_50ntFlanks</td><td>11604</td><td>11623</td><td>20</td><td>CGACTGTTTAATTAAAACAA</td><td>450.0</td><td>2.3</td><td>0.9</td><td>2.5</td><td>1.1E-02</td><td>1.3E-02</td><td>MB231/MB468</td><td>NUC</td></tr>
<tr><td>>hg38_21_+_8433172_8446622_RNA45SN1_45S_50ntFlanks@10781.10809.29|AGGTGAACAGCCTCTGGCATGTTGGAACA</td><td>>hg38_21_+_8433172_8446622_RNA45SN1_45S_50ntFlanks</td><td>10781</td><td>10809</td><td>29</td><td>AGGTGAACAGCCTCTGGCATGTTGGAACA</td><td>208.4</td><td>-2.1</td><td>0.8</td><td>-2.6</td><td>1.1E-02</td><td>1.3E-02</td><td>BT20/MB468</td><td>CYTO</td></tr>
<tr><td>>hg38_21_+_8433172_8446622_RNA45SN1_45S_50ntFlanks@10686.10709.24|GCGGCGTCCGGTGAGCTCTCGCTG</td><td>>hg38_21_+_8433172_8446622_RNA45SN1_45S_50ntFlanks</td><td>10686</td><td>10709</td><td>24</td><td>GCGGCGTCCGGTGAGCTCTCGCTG</td><td>2017.7</td><td>-3.3</td><td>1.3</td><td>-2.5</td><td>1.1E-02</td><td>1.3E-02</td><td>BT20/MB468</td><td>NUC</td></tr>
<tr><td>>hg38_21_+_8433172_8446622_RNA45SN1_45S_50ntFlanks@6698.6720.23|AGCTAGCTGCGAGAATTAATGTG</td><td>>hg38_21_+_8433172_8446622_RNA45SN1_45S_50ntFlanks</td><td>6698</td><td>6720</td><td>23</td><td>AGCTAGCTGCGAGAATTAATGTG</td><td>22608.4</td><td>-3.1</td><td>1.2</td><td>-2.7</td><td>6.6E-03</td><td>1.3E-02</td><td>BT20/MB468</td><td>MP</td></tr>
<tr><td>>MI0000063|hsa-let-7b&WithFlank&22|+|46113680|46113774@66.87.22&[MIMAT0004482&hsa-let-7b-3p&offsets|0|0;m-171&22|+|46113745|46113766&offsets|0|0]|CTATACAACCTACTGCCTTCCC</td><td>>MI0000063|hsa-let-7b&WithFlank&22|+|46113680|46113774</td><td>66</td><td>87</td><td>22</td><td>CTATACAACCTACTGCCTTCCC</td><td>74.6</td><td>6.8</td><td>2.5</td><td>2.7</td><td>6.6E-03</td><td>1.3E-02</td><td>BT20/MB468</td><td>MP</td></tr>
<tr><td>>trna119_LysCTT_1_-_145395522_145395594@2.35.34,trna11_LysCTT_5_-_180648979_180649051@2.35.34,trna13_LysCTT_6_+_26556774_26556846@2.35.34,trna7_LysCTT_16_+_3225692_3225764@2.35.34,trna9_LysCTT_5_+_180634755_180634827@2.35.34|CCCGGCTAGCTCAGTCGGTAGAGCATGAGACTCT</td><td>>trna119_LysCTT_1_-_145395522_145395594</td><td>2</td><td>35</td><td>34</td><td>CCCGGCTAGCTCAGTCGGTAGAGCATGAGACTCT</td><td>198.1</td><td>-5.9</td><td>2.0</td><td>-2.9</td><td>3.2E-03</td><td>1.3E-02</td><td>BT20/MB231</td><td>MITO</td></tr>
<tr><td>>trna152_ValCAC_6_-_27248049_27248121@1.33.33|GCTTCTGTAGTGTAGTGGTTATCACGTTCGCCT</td><td>>trna152_ValCAC_6_-_27248049_27248121</td><td>1</td><td>33</td><td>33</td><td>GCTTCTGTAGTGTAGTGGTTATCACGTTCGCCT</td><td>758.7</td><td>-3.6</td><td>1.2</td><td>-3.0</td><td>2.8E-03</td><td>1.3E-02</td><td>BT20/MB231</td><td>NUC</td></tr>
<tr><td>>hg38_21_+_8433172_8446622_RNA45SN1_45S_50ntFlanks@6773.6807.35|TTCCTCCCGGGGCTACGCCTGTCTGAGCGTCGCTT</td><td>>hg38_21_+_8433172_8446622_RNA45SN1_45S_50ntFlanks</td><td>6773</td><td>6807</td><td>35</td><td>TTCCTCCCGGGGCTACGCCTGTCTGAGCGTCGCTT</td><td>438.2</td><td>-1.9</td><td>0.7</td><td>-2.5</td><td>1.1E-02</td><td>1.3E-02</td><td>MB231/MB468</td><td>NUC</td></tr>
<tr><td>>trnaMT_GluTTC_MT_-_14674_14742@40.71.32,trnalookalike8_GluTTC_5_-_93905172_93905240@40.71.32|TTGGTCGTGGTTGTAGTCCGTGCGAGAATACC</td><td>>trnaMT_GluTTC_MT_-_14674_14742</td><td>40</td><td>71</td><td>32</td><td>TTGGTCGTGGTTGTAGTCCGTGCGAGAATACC</td><td>1271.2</td><td>-2.4</td><td>0.8</td><td>-3.0</td><td>2.8E-03</td><td>1.3E-02</td><td>BT20/MB231</td><td>NUC</td></tr>
<tr><td>>trnaMT_PheGAA_MT_+_577_647@36.53.18|AAAATGTTTAGACGGGCT</td><td>>trnaMT_PheGAA_MT_+_577_647</td><td>36</td><td>53</td><td>18</td><td>AAAATGTTTAGACGGGCT</td><td>334.7</td><td>3.4</td><td>1.2</td><td>2.7</td><td>6.6E-03</td><td>1.3E-02</td><td>BT20/MB468</td><td>MP</td></tr>
<tr><td>>hg38_21_+_8433172_8446622_RNA45SN1_45S_50ntFlanks@9546.9570.25|TGGAGGTCCGTAGCGGTCCTGACGT</td><td>>hg38_21_+_8433172_8446622_RNA45SN1_45S_50ntFlanks</td><td>9546</td><td>9570</td><td>25</td><td>TGGAGGTCCGTAGCGGTCCTGACGT</td><td>154.7</td><td>2.5</td><td>0.9</td><td>2.7</td><td>6.6E-03</td><td>1.3E-02</td><td>BT20/MB468</td><td>MP</td></tr>
<tr><td>>hg38_21_+_8433172_8446622_RNA45SN1_45S_50ntFlanks@9567.9588.22|ACGTGCAAATCGGTCGTCCGAC</td><td>>hg38_21_+_8433172_8446622_RNA45SN1_45S_50ntFlanks</td><td>9567</td><td>9588</td><td>22</td><td>ACGTGCAAATCGGTCGTCCGAC</td><td>183.8</td><td>-4.5</td><td>1.7</td><td>-2.7</td><td>6.6E-03</td><td>1.3E-02</td><td>BT20/MB468</td><td>MP</td></tr>
<tr><td>>hg38_21_+_8433172_8446622_RNA45SN1_45S_50ntFlanks@11579.11609.31|AGAACTGGTGCGGACCAGGGGAATCCGACTG</td><td>>hg38_21_+_8433172_8446622_RNA45SN1_45S_50ntFlanks</td><td>11579</td><td>11609</td><td>31</td><td>AGAACTGGTGCGGACCAGGGGAATCCGACTG</td><td>353.1</td><td>-1.9</td><td>0.8</td><td>-2.5</td><td>1.1E-02</td><td>1.3E-02</td><td>MB231/MB468</td><td>NUC</td></tr>
<tr><td>>hg38_21_+_8433172_8446622_RNA45SN1_45S_50ntFlanks@4739.4759.21|AAGTCGGAGGTTCGAAGACGA</td><td>>hg38_21_+_8433172_8446622_RNA45SN1_45S_50ntFlanks</td><td>4739</td><td>4759</td><td>21</td><td>AAGTCGGAGGTTCGAAGACGA</td><td>260.7</td><td>-3.7</td><td>1.4</td><td>-2.7</td><td>6.6E-03</td><td>1.3E-02</td><td>BT20/MB468</td><td>MP</td></tr>
<tr><td>>am_trnaMT_PheGAA_MT_+_577_647@45.62.18|AGACGGGCTCACATCACC</td><td>>am_trnaMT_PheGAA_MT_+_577_647</td><td>45</td><td>62</td><td>18</td><td>AGACGGGCTCACATCACC</td><td>452.2</td><td>2.9</td><td>1.1</td><td>2.7</td><td>6.6E-03</td><td>1.3E-02</td><td>BT20/MB468</td><td>MP</td></tr>
<tr><td>>hg38_21_+_8433172_8446622_RNA45SN1_45S_50ntFlanks@11572.11612.41|CCGACTTAGAACTGGTGCGGACCAGGGGAATCCGACTGTTT</td><td>>hg38_21_+_8433172_8446622_RNA45SN1_45S_50ntFlanks</td><td>11572</td><td>11612</td><td>41</td><td>CCGACTTAGAACTGGTGCGGACCAGGGGAATCCGACTGTTT</td><td>4804.0</td><td>-2.5</td><td>0.9</td><td>-2.6</td><td>9.2E-03</td><td>1.3E-02</td><td>BT20/MB468</td><td>MITO</td></tr>
<tr><td>>hg38_21_+_8433172_8446622_RNA45SN1_45S_50ntFlanks@11602.11619.18|TCCGACTGTTTAATTAAA</td><td>>hg38_21_+_8433172_8446622_RNA45SN1_45S_50ntFlanks</td><td>11602</td><td>11619</td><td>18</td><td>TCCGACTGTTTAATTAAA</td><td>474.9</td><td>-1.5</td><td>0.6</td><td>-2.6</td><td>9.2E-03</td><td>1.3E-02</td><td>BT20/MB468</td><td>MITO</td></tr>
<tr><td>>MI0000082|hsa-mir-25&WithFlank&7|-|100093554|100093649@58.79.22(+1A)&[MIMAT0000081&hsa-miR-25-3p&offsets|0|0(+1A);m-23&7|-|100093571|100093592&offsets|0|0(+1A)]|CATTGCACTTGTCTCGGTCTGAA</td><td>>MI0000082|hsa-mir-25&WithFlank&7|-|100093554|100093649</td><td>58</td><td>79</td><td>22(+1A)</td><td>CATTGCACTTGTCTCGGTCTGAA</td><td>345.1</td><td>3.8</td><td>1.4</td><td>2.7</td><td>6.6E-03</td><td>1.3E-02</td><td>BT20/MB468</td><td>MP</td></tr>
<tr><td>>hg38_21_+_8433172_8446622_RNA45SN1_45S_50ntFlanks@10348.10365.18|AAACGAGAACTTTGAAGG</td><td>>hg38_21_+_8433172_8446622_RNA45SN1_45S_50ntFlanks</td><td>10348</td><td>10365</td><td>18</td><td>AAACGAGAACTTTGAAGG</td><td>392.5</td><td>-2.7</td><td>1.0</td><td>-2.7</td><td>6.6E-03</td><td>1.3E-02</td><td>BT20/MB468</td><td>MP</td></tr>
<tr><td>>hg38_MT_+_1621_3279_RNR2_16S_50ntFlanks@1000.1018.19|GGGACCTGTATGAATGGCT</td><td>>hg38_MT_+_1621_3279_RNR2_16S_50ntFlanks</td><td>1000</td><td>1018</td><td>19</td><td>GGGACCTGTATGAATGGCT</td><td>170.3</td><td>3.3</td><td>1.3</td><td>2.6</td><td>9.9E-03</td><td>1.3E-02</td><td>MB231/MB468</td><td>MP</td></tr>
</table>

| | | | | | | | | | | | | | |
|---|---|---|---|---|---|---|---|---|---|---|---|---|---|
| >MI0000760\|hsa-mir-361&WithFlank&X\|-\|85903630\|85903713@12.31.20&[MIMAT0000703&hsa-miR-361-5p&offsets\|0\|-2;m-89&X\|-\|85903681\|85903702&offsets\|0\|-2]\|TTATCAGAATCTCCAGGGGT | >MI0000760\|hsa-mir-361&WithFlank&X\|-\|85903630\|85903713 | 12 | 31 | 20 | TTATCAGAATCTCCAGGGGT | 91.0 | 3.4 | 1.3 | 2.6 | 9.9E-03 | 1.3E-02 | MB231/MB468 | MP |
| >hg38_21_+_8433172_8446622_RNA45SN1_45S_50ntFlanks@10388.10424.37\|GTGAACAGCAGTTGAACATGGGTCAGTCGGTCCTGAG | >hg38_21_+_8433172_8446622_RNA45SN1_45S_50ntFlanks | 10388 | 10424 | 37 | GTGAACAGCAGTTGAACATGGGTCAGTCGGTCCTGAG | 11773.8 | 3.8 | 1.3 | 3.0 | 2.6E-03 | 1.3E-02 | BT20/MB231 | MP |
| >hg38_21_+_8433172_8446622_RNA45SN1_45S_50ntFlanks@10677.10712.36\|GCGGTTCCGGCGGCGTCCGGTGAGCTCTCGCTGGCC | >hg38_21_+_8433172_8446622_RNA45SN1_45S_50ntFlanks | 10677 | 10712 | 36 | GCGGTTCCGGCGGCGTCCGGTGAGCTCTCGCTGGCC | 2253.5 | -3.2 | 1.3 | -2.5 | 1.1E-02 | 1.3E-02 | BT20/MB468 | NUC |
| >hg38_MT_+_1621_3279_RNR2_16S_50ntFlanks@1389.1414.26\|CGATGGTGCAGCCGCTATTAAAGGTT | >hg38_MT_+_1621_3279_RNR2_16S_50ntFlanks | 1389 | 1414 | 26 | CGATGGTGCAGCCGCTATTAAAGGTT | 791.5 | -3.5 | 1.2 | -3.0 | 2.6E-03 | 1.3E-02 | BT20/MB231 | MP |
| >hg38_21_+_8433172_8446622_RNA45SN1_45S_50ntFlanks@11211.11252.42\|GCGACTCTGGACGCGAGCCGGGCCCTTCCCGTGGATCGCCCC | >hg38_21_+_8433172_8446622_RNA45SN1_45S_50ntFlanks | 11211 | 11252 | 42 | GCGACTCTGGACGCGAGCCGGGCCCTTCCCGTGGATCGCCCC | 682.2 | -4.0 | 1.6 | -2.5 | 1.1E-02 | 1.3E-02 | BT20/MB468 | NUC |
| >hg38_21_+_8433172_8446622_RNA45SN1_45S_50ntFlanks@12258.12284.27\|GCTTGATCTTGATTTTCAGTACGAATA | >hg38_21_+_8433172_8446622_RNA45SN1_45S_50ntFlanks | 12258 | 12284 | 27 | GCTTGATCTTGATTTTCAGTACGAATA | 155.3 | -2.6 | 1.0 | -2.6 | 1.1E-02 | 1.3E-02 | MB231/MB468 | CYTO |
| >MI0000082\|hsa-mir-25&WithFlank&7\|-\|100093554\|100093649@58.75.18(+1U)&[MIMAT0000081&hsa-miR-25-3p&offsets\|0\|-4(+1U);m-23&7\|-\|100093571\|100093592&offsets\|0\|-4(+1U)]\|CATTGCACTTGTCTCGGTT | >MI0000082\|hsa-mir-25&WithFlank&7\|-\|100093554\|100093649 | 58 | 75 | 18(+1U) | CATTGCACTTGTCTCGGTT | 23.5 | 7.8 | 3.0 | 2.6 | 9.9E-03 | 1.3E-02 | MB231/MB468 | MP |
| >hg38_21_+_8433172_8446622_RNA45SN1_45S_50ntFlanks@12598.12626.29\|TTTGGTGTATGTGCTTGGCTGAGGAGCCA | >hg38_21_+_8433172_8446622_RNA45SN1_45S_50ntFlanks | 12598 | 12626 | 29 | TTTGGTGTATGTGCTTGGCTGAGGAGCCA | 83.1 | -3.8 | 1.5 | -2.6 | 1.1E-02 | 1.3E-02 | BT20/MB468 | CYTO |
| >hg38_21_+_8433172_8446622_RNA45SN1_45S_50ntFlanks@9574.9594.21\|AATCGGTCGTCCGACCTGGGT | >hg38_21_+_8433172_8446622_RNA45SN1_45S_50ntFlanks | 9574 | 9594 | 21 | AATCGGTCGTCCGACCTGGGT | 542.7 | -2.3 | 0.9 | -2.6 | 9.2E-03 | 1.3E-02 | BT20/MB468 | MITO |
| >hg38_21_+_8433172_8446622_RNA45SN1_45S_50ntFlanks@9307.9324.18\|AAGGAGTCTAACACGTGC | >hg38_21_+_8433172_8446622_RNA45SN1_45S_50ntFlanks | 9307 | 9324 | 18 | AAGGAGTCTAACACGTGC | 111.2 | 5.9 | 2.0 | 3.0 | 2.6E-03 | 1.3E-02 | BT20/MB231 | MP |
| >hg38_21_+_8433172_8446622_RNA45SN1_45S_50ntFlanks@11542.11584.43\|CGGCGCCTCGCCTCGGCCGGCGCCTAGCAGCCGACTTAGAACT | >hg38_21_+_8433172_8446622_RNA45SN1_45S_50ntFlanks | 11542 | 11584 | 43 | CGGCGCCTCGCCTCGGCCGGCGCCTAGCAGCCGACTTAGAACT | 145.0 | 4.0 | 1.6 | 2.5 | 1.1E-02 | 1.3E-02 | MB231/MB468 | MITO |
| >hg38_1_-_228634819_228635039_RNA5S12_5S_50ntFlanks@51.86.36\|GTCTACGGCCATACCACCCTGAACGCGCCCGATCTC | >hg38_1_-_228634819_228635039_RNA5S12_5S_50ntFlanks | 51 | 86 | 36 | GTCTACGGCCATACCACCCTGAACGCGCCCGATCTC | 3463.7 | 5.5 | 2.2 | 2.6 | 1.1E-02 | 1.3E-02 | BT20/MB468 | CYTO |
| >MI0000813\|hsa-mir-324&WithFlank&17\|-\|7223291\|7223385@59.78.20(+1U)&[MIMAT0000762&hsa-miR-324-3p&offsets\|+3\|0(+1U);m-203&17\|-\|7223308\|7223329&offsets\|+2\|0(+1U)]\|ACTGCCCCAGGTGCTGCTGGT | >MI0000813\|hsa-mir-324&WithFlank&17\|-\|7223291\|7223385 | 59 | 78 | 20(+1U) | ACTGCCCCAGGTGCTGCTGGT | 262.6 | 4.3 | 1.6 | 2.7 | 6.6E-03 | 1.3E-02 | BT20/MB468 | MP |
| >hg38_MT_+_1621_3279_RNR2_16S_50ntFlanks@51.78.28\|GCTAAACCTAGCCCCAAACCCACTCCAC | >hg38_MT_+_1621_3279_RNR2_16S_50ntFlanks | 51 | 78 | 28 | GCTAAACCTAGCCCCAAACCCACTCCAC | 114.6 | 2.5 | 1.0 | 2.5 | 1.1E-02 | 1.3E-02 | MB231/MB468 | MITO |
| >MI0000746\|hsa-mir-99b&WithFlank&19\|+\|51692606\|51692687@13.34.22(+1C)&[MIMAT0000689&hsa-miR-99b-5p&offsets\|0\|0(+1C);m-10&19\|+\|51692618\|51692639&offsets\|0\|0(+1C)]\|CACCCGTAGAACCGACCTTGCGC | >MI0000746\|hsa-mir-99b&WithFlank&19\|+\|51692606\|51692687 | 13 | 34 | 22(+1C) | CACCCGTAGAACCGACCTTGCGC | 127.4 | 3.4 | 1.3 | 2.6 | 9.9E-03 | 1.3E-02 | MB231/MB468 | MP |
| >am_MI0006370\|hsa-mir-1303&WithFlank&5\|+\|154685770\|154685867@57.79.23&[MIMAT0005891&hsa-miR-1303&offsets\|-1\|0;m-516&5\|+\|154685826\|154685848&offsets\|0\|0]\|TTTTAGAGACGGGGTCTTGCTCT | >am_MI0006370\|hsa-mir-1303&WithFlank&5\|+\|154685770\|154685867 | 57 | 79 | 23 | TTTTAGAGACGGGGTCTTGCTCT | 141.8 | 3.4 | 1.3 | 2.6 | 9.9E-03 | 1.3E-02 | MB231/MB468 | MP |
| >hg38_21_+_8433172_8446622_RNA45SN1_45S_50ntFlanks@5467.5497.31\|GGCCCACGGCCCTGGCGGAGCGCTGAGAAGA | >hg38_21_+_8433172_8446622_RNA45SN1_45S_50ntFlanks | 5467 | 5497 | 31 | GGCCCACGGCCCTGGCGGAGCGCTGAGAAGA | 436.4 | -4.9 | 1.9 | -2.5 | 1.1E-02 | 1.3E-02 | BT20/MB468 | NUC |
| >hg38_21_+_8433172_8446622_RNA45SN1_45S_50ntFlanks@9551.9571.21\|GTCCGTAGCGGTCCTGACGTG | >hg38_21_+_8433172_8446622_RNA45SN1_45S_50ntFlanks | 9551 | 9571 | 21 | GTCCGTAGCGGTCCTGACGTG | 102.1 | -3.5 | 1.4 | -2.6 | 9.9E-03 | 1.3E-02 | MB231/MB468 | MP |
| >hg38_21_+_8433172_8446622_RNA45SN1_45S_50ntFlanks@10797.10823.27\|GCATGTTGGAACAATGTAGGTAAGGGA | >hg38_21_+_8433172_8446622_RNA45SN1_45S_50ntFlanks | 10797 | 10823 | 27 | GCATGTTGGAACAATGTAGGTAAGGGA | 179.8 | -1.1 | 0.4 | -2.6 | 9.3E-03 | 1.3E-02 | BT20/MB468 | MITO |
| >hg38_1_-_228634819_228635039_RNA5S12_5S_50ntFlanks@132.171.40\|GATGGGAGACCGCCTGGGAATACCGGGTGCTGTAGGCTTT | >hg38_1_-_228634819_228635039_RNA5S12_5S_50ntFlanks | 132 | 171 | 40 | GATGGGAGACCGCCTGGGAATACCGGGTGCTGTAGGCTTT | 37935.5 | 4.2 | 1.4 | 3.0 | 2.6E-03 | 1.3E-02 | BT20/MB231 | MP |
| >hg38_21_+_8433172_8446622_RNA45SN1_45S_50ntFlanks@6661.6700.40\|CGGTGGATCACTCGGCTCGTGCGTCGATGAAGAACGCAGC | >hg38_21_+_8433172_8446622_RNA45SN1_45S_50ntFlanks | 6661 | 6700 | 40 | CGGTGGATCACTCGGCTCGTGCGTCGATGAAGAACGCAGC | 5653.9 | 3.7 | 1.2 | 3.0 | 2.6E-03 | 1.3E-02 | BT20/MB231 | MP |
| >hg38_21_+_8433172_8446622_RNA45SN1_45S_50ntFlanks@8106.8143.38\|GCGGCGGGGCGCGGGACATGTGGCGTACGGAAGACCCG | >hg38_21_+_8433172_8446622_RNA45SN1_45S_50ntFlanks | 8106 | 8143 | 38 | GCGGCGGGGCGCGGGACATGTGGCGTACGGAAGACCCG | 609.0 | -4.3 | 1.6 | -2.6 | 9.3E-03 | 1.3E-02 | BT20/MB468 | MITO |
| >MI0003815\|hsa-mir-1301&WithFlank&2\|-\|25328634\|25328727@54.77.24&[MIMAT0005797&hsa-miR-1301-3p&offsets\|0\|0;m-256&2\|-\|25328650\|25328674&offsets\|0\|-1]\|TTGCAGCTGCCTGGGAGTGACTTC | >MI0003815\|hsa-mir-1301&WithFlank&2\|-\|25328634\|25328727 | 54 | 77 | 24 | TTGCAGCTGCCTGGGAGTGACTTC | 188.0 | 2.4 | 0.8 | 2.9 | 3.3E-03 | 1.3E-02 | BT20/MB231 | CYTO |
| >MI0000440\|hsa-mir-27b&WithFlank&9\|+\|95085439\|95085547@67.88.22(+1A)&[MIMAT0000419&hsa-miR-27b-3p&offsets\|0\|+1(+1A);m-39&9\|+\|95085505\|95085525&offsets\|0\|+1(+1A)]\|TTCACAGTGGCTAAGTTCTGCAA | >MI0000440\|hsa-mir-27b&WithFlank&9\|+\|95085439\|95085547 | 67 | 88 | 22(+1A) | TTCACAGTGGCTAAGTTCTGCAA | 87.5 | 2.3 | 0.8 | 2.7 | 6.7E-03 | 1.3E-02 | BT20/MB468 | MP |
| >hg38_1_-_228634819_228635039_RNA5S12_5S_50ntFlanks@51.82.32\|GTCTACGGCCATACCACCCTGAACGCGCCCGA | >hg38_1_-_228634819_228635039_RNA5S12_5S_50ntFlanks | 51 | 82 | 32 | GTCTACGGCCATACCACCCTGAACGCGCCCGA | 2006.5 | 2.1 | 0.8 | 2.5 | 1.1E-02 | 1.3E-02 | MB231/MB468 | NUC |
| >am_trna128_GlyGCC_6_-_27870686_27870756@1.30.30,trna133_GlyCCC_1_-_16872434_16872504@1.30.30,trna18_GlyGCC_16_+_70822597_70822667@1.30.30,trna19_GlyGCC_16_+_70823410_70823480@1.30.30,trna19_GlyGCC_2_-_157257659_157257729@1.30.30,trna24_GlyGCC_16_-_70812942_70813012@1.30.30,trna25_GlyGCC_16_-_70812114_70812184@1.30.30,trna4_GlyCCC_1_+_17188416_17188486@1.30.30,trna5_GlyGCC_17_+_8029064_8029134@1.30.30,trna68_GlyGCC_1_-_161493637_161493707@1.30.30\|GCATTGGTGGTTCAGTGGTAGAATTCTCGC | >am_trna128_GlyGCC_6_-_27870686_27870756 | 1 | 30 | 30 | GCATTGGTGGTTCAGTGGTAGAATTCTCGC | 35695.1 | -3.6 | 1.2 | -3.0 | 2.6E-03 | 1.3E-02 | BT20/MB231 | MP |
| >trnaMT_ValTAC_MT_+_1602_1670@51.71.21\|CTTAACTTGACCGCTCTGACC | >trnaMT_ValTAC_MT_+_1602_1670 | 51 | 71 | 21 | CTTAACTTGACCGCTCTGACC | 535.4 | -2.5 | 0.9 | -2.7 | 6.7E-03 | 1.3E-02 | BT20/MB468 | MP |
| >hg38_21_+_8433172_8446622_RNA45SN1_45S_50ntFlanks@7974.8010.37\|ACGCGACCTCAGATCAGACGTGGCGACCCGCTGAATT | >hg38_21_+_8433172_8446622_RNA45SN1_45S_50ntFlanks | 7974 | 8010 | 37 | ACGCGACCTCAGATCAGACGTGGCGACCCGCTGAATT | 1539.2 | 2.7 | 1.0 | 2.6 | 1.0E-02 | 1.3E-02 | MB231/MB468 | MP |
| >hg38_21_+_8433172_8446622_RNA45SN1_45S_50ntFlanks@12474.12496.23\|ACCCACTAATAGGGAACGTGAGC | >hg38_21_+_8433172_8446622_RNA45SN1_45S_50ntFlanks | 12474 | 12496 | 23 | ACCCACTAATAGGGAACGTGAGC | 768.5 | 1.4 | 0.6 | 2.6 | 1.1E-02 | 1.3E-02 | BT20/MB468 | CYTO |
| >hg38_1_-_228634819_228635039_RNA5S12_5S_50ntFlanks@51.83.33\|GTCTACGGCCATACCACCCTGAACGCGCCCGAT | >hg38_1_-_228634819_228635039_RNA5S12_5S_50ntFlanks | 51 | 83 | 33 | GTCTACGGCCATACCACCCTGAACGCGCCCGAT | 3407.4 | -2.0 | 0.8 | -2.5 | 1.1E-02 | 1.3E-02 | MB231/MB468 | MITO |
| >hg38_21_+_8433172_8446622_RNA45SN1_45S_50ntFlanks@4564.4587.24\|GAATAATGGAATAGGACCGCGGTT | >hg38_21_+_8433172_8446622_RNA45SN1_45S_50ntFlanks | 4564 | 4587 | 24 | GAATAATGGAATAGGACCGCGGTT | 168.8 | -3.8 | 1.5 | -2.5 | 1.1E-02 | 1.3E-02 | BT20/MB468 | CYTO |
| >hg38_21_+_8433172_8446622_RNA45SN1_45S_50ntFlanks@6660.6681.22\|GCGGTGGATCACTCGGCTCGTG | >hg38_21_+_8433172_8446622_RNA45SN1_45S_50ntFlanks | 6660 | 6681 | 22 | GCGGTGGATCACTCGGCTCGTG | 308.7 | -1.8 | 0.7 | -2.5 | 1.1E-02 | 1.3E-02 | BT20/MB468 | CYTO |
| >hg38_21_+_8433172_8446622_RNA45SN1_45S_50ntFlanks@11027.11053.27\|CTCTCTCTCTCTCTCCCCCGCTCCCCG | >hg38_21_+_8433172_8446622_RNA45SN1_45S_50ntFlanks | 11027 | 11053 | 27 | CTCTCTCTCTCTCTCCCCCGCTCCCCG | 1217.4 | -3.5 | 1.4 | -2.5 | 1.1E-02 | 1.3E-02 | BT20/MB468 | CYTO |
| >hg38_21_+_8433172_8446622_RNA45SN1_45S_50ntFlanks@10760.10785.26\|CATATCCGCAGCAGGTCTCCAAGGTG | >hg38_21_+_8433172_8446622_RNA45SN1_45S_50ntFlanks | 10760 | 10785 | 26 | CATATCCGCAGCAGGTCTCCAAGGTG | 198.8 | -4.6 | 1.7 | -2.7 | 6.7E-03 | 1.3E-02 | BT20/MB468 | MP |
| >hg38_MT_+_1621_3279_RNR2_16S_50ntFlanks@843.864.22\|AGTAAAAGGAACTCGGCAAATC | >hg38_MT_+_1621_3279_RNR2_16S_50ntFlanks | 843 | 864 | 22 | AGTAAAAGGAACTCGGCAAATC | 22.3 | 7.7 | 3.0 | 2.6 | 1.0E-02 | 1.3E-02 | MB231/MB468 | MP |
| >hg38_MT_+_1621_3279_RNR2_16S_50ntFlanks@616.652.37\|CAAACATATAACTGAACTCCTCACACCCAATTGGACC | >hg38_MT_+_1621_3279_RNR2_16S_50ntFlanks | 616 | 652 | 37 | CAAACATATAACTGAACTCCTCACACCCAATTGGACC | 45.8 | 5.7 | 2.1 | 2.7 | 6.7E-03 | 1.3E-02 | BT20/MB468 | MP |
| >hg38_21_+_8433172_8446622_RNA45SN1_45S_50ntFlanks@3996.4023.28\|ACTCTAGATAACCTCGGGCCGATCGCAC | >hg38_21_+_8433172_8446622_RNA45SN1_45S_50ntFlanks | 3996 | 4023 | 28 | ACTCTAGATAACCTCGGGCCGATCGCAC | 421.1 | -1.6 | 0.6 | -2.5 | 1.1E-02 | 1.3E-02 | MB231/MB468 | NUC |
| >hg38_21_+_8433172_8446622_RNA45SN1_45S_50ntFlanks@10352.10369.18\|GAGAACTTTGAAGGCCGA | >hg38_21_+_8433172_8446622_RNA45SN1_45S_50ntFlanks | 10352 | 10369 | 18 | GAGAACTTTGAAGGCCGA | 241.1 | 2.3 | 0.8 | 2.7 | 6.7E-03 | 1.3E-02 | BT20/MB468 | MP |
| >hg38_21_+_8433172_8446622_RNA45SN1_45S_50ntFlanks@10547.10570.24\|AGTGCGGTAACGCGACCGATCCCG | >hg38_21_+_8433172_8446622_RNA45SN1_45S_50ntFlanks | 10547 | 10570 | 24 | AGTGCGGTAACGCGACCGATCCCG | 175.3 | -2.5 | 0.9 | -2.7 | 6.7E-03 | 1.3E-02 | BT20/MB468 | MP |
| >hg38_MT_+_1621_3279_RNR2_16S_50ntFlanks@51.87.37\|GCTAAACCTAGCCCCAAACCCACTCCACCTTACTACC | >hg38_MT_+_1621_3279_RNR2_16S_50ntFlanks | 51 | 87 | 37 | GCTAAACCTAGCCCCAAACCCACTCCACCTTACTACC | 138.7 | 1.4 | 0.6 | 2.5 | 1.1E-02 | 1.3E-02 | BT20/MB468 | NUC |
| >hg38_21_+_8433172_8446622_RNA45SN1_45S_50ntFlanks@10248.10269.22\|CGACGAGTAGGAGGGCCGCTGC | >hg38_21_+_8433172_8446622_RNA45SN1_45S_50ntFlanks | 10248 | 10269 | 22 | CGACGAGTAGGAGGGCCGCTGC | 52.7 | 5.5 | 2.0 | 2.7 | 6.7E-03 | 1.3E-02 | BT20/MB468 | MP |
| >hg38_21_+_8433172_8446622_RNA45SN1_45S_50ntFlanks@4565.4589.25\|AATAATGGAATAGGACCGCGGTTCT | >hg38_21_+_8433172_8446622_RNA45SN1_45S_50ntFlanks | 4565 | 4589 | 25 | AATAATGGAATAGGACCGCGGTTCT | 592.1 | 3.0 | 1.1 | 2.7 | 6.7E-03 | 1.3E-02 | BT20/MB468 | MP |
| >hg38_21_+_8433172_8446622_RNA45SN1_45S_50ntFlanks@8178.8207.30\|TCTGATCGAGGCCCAGCCCGTGGACGGTGT | >hg38_21_+_8433172_8446622_RNA45SN1_45S_50ntFlanks | 8178 | 8207 | 30 | TCTGATCGAGGCCCAGCCCGTGGACGGTGT | 207.0 | 7.1 | 2.4 | 3.0 | 2.6E-03 | 1.3E-02 | BT20/MB231 | MP |
| >hg38_MT_+_1621_3279_RNR2_16S_50ntFlanks@753.777.25\|AACAGCCCAATATCTACAATCAACC | >hg38_MT_+_1621_3279_RNR2_16S_50ntFlanks | 753 | 777 | 25 | AACAGCCCAATATCTACAATCAACC | 57.4 | 4.9 | 1.9 | 2.6 | 1.0E-02 | 1.3E-02 | MB231/MB468 | MP |
| >MI0000283\|hsa-mir-203a&WithFlank&14\|+\|104117399\|104117520@71.93.23&[MIMAT0000264&hsa-miR-203a-3p&offsets\|0\|+1;m-79&14\|+\|104117469\|104117490&offsets\|0\|+1]\|GTGAAATGTTTAGGACCACTAGA | >MI0000283\|hsa-mir-203a&WithFlank&14\|+\|104117399\|104117520 | 71 | 93 | 23 | GTGAAATGTTTAGGACCACTAGA | 76.5 | 4.7 | 1.7 | 2.7 | 6.7E-03 | 1.3E-02 | BT20/MB468 | MP |
| >hg38_21_+_8433172_8446622_RNA45SN1_45S_50ntFlanks@10058.10078.21\|CGGCAGTCGAGAGTGGACGGG | >hg38_21_+_8433172_8446622_RNA45SN1_45S_50ntFlanks | 10058 | 10078 | 21 | CGGCAGTCGAGAGTGGACGGG | 137.2 | 3.8 | 1.4 | 2.7 | 6.7E-03 | 1.3E-02 | BT20/MB468 | MP |
| >MI0000065\|hsa-let-7d&WithFlank&9\|+\|94178828\|94178926@68.87.20&[MIMAT0004484&hsa-let-7d-3p&offsets\|0\|-2;m-98&9\|+\|94178895\|94178916&offsets\|0\|-2]\|CTATACGACCTGCTGCCTTT | >MI0000065\|hsa-let-7d&WithFlank&9\|+\|94178828\|94178926 | 68 | 87 | 20 | CTATACGACCTGCTGCCTTT | 65.0 | 6.9 | 2.5 | 2.7 | 6.7E-03 | 1.3E-02 | BT20/MB468 | MP |

<table>
<tr><td>>trna10_AspGTC_12_-_125424193_125424264@37.70.34,trna12_AspGTC_12_-_125411891_125411962@37.70.34,trna144_AspGTC_6_-_27551236_27551307@37.70.34,trna38_AspGTC_17_-_8125556_8125627@37.70.34,trna45_AspGTC_6_+_27447453_27447524@37.70.34,trna48_AspGTC_6_+_27471523_27471594@37.70.34,trna4_AspGTC_12_+_96429799_96429870@37.70.34,trna69_AspGTC_1_-_161440205_161440276@37.70.34,trna72_AspGTC_1_-_161432824_161432895@37.70.34,trna75_AspGTC_1_-_161425414_161425485@37.70.34,trna78_AspGTC_1_-_161418033_161418104@37.70.34,trna81_AspGTC_1_-_161410615_161410686@37.70.34|ACGCGGGAGACCGGGGTTCGATTCCCCGACGGGG</td><td>>trna10_AspGTC_12_-_125424193_125424264</td><td>37</td><td>70</td><td>34</td><td>ACGCGGGAGACCGGGGTTCGATTCCCCGACGGGG</td><td>2273.1</td><td>2.6</td><td>1.0</td><td>2.5</td><td>1.1E-02</td><td>1.3E-02</td><td>MB231/MB468</td><td>CYTO</td></tr>
<tr><td>>hg38_21_+_8433172_8446622_RNA45SN1_45S_50ntFlanks@3853.3898.46|AACTGTGGTAATTCTAGAGCTAATACATGCCGACGGGCGCTGACCC</td><td>>hg38_21_+_8433172_8446622_RNA45SN1_45S_50ntFlanks</td><td>3853</td><td>3898</td><td>46</td><td>AACTGTGGTAATTCTAGAGCTAATACATGCCGACGGGCGCTGACCC</td><td>262.3</td><td>-2.2</td><td>0.9</td><td>-2.5</td><td>1.1E-02</td><td>1.3E-02</td><td>MB231/MB468</td><td>CYTO</td></tr>
<tr><td>>am_trnaMT_LysTTT_MT_+_8295_8364@34.56.23,trnalookalike7_LysTTT_1_+_568844_568913@34.56.23|GTTAAAGATTAAGAGAACCAACA</td><td>>am_trnaMT_LysTTT_MT_+_8295_8364</td><td>34</td><td>56</td><td>23</td><td>GTTAAAGATTAAGAGAACCAACA</td><td>22.3</td><td>7.7</td><td>3.0</td><td>2.6</td><td>1.0E-02</td><td>1.3E-02</td><td>MB231/MB468</td><td>MP</td></tr>
<tr><td>>hg38_21_+_8433172_8446622_RNA45SN1_45S_50ntFlanks@12994.13039.46|CCTCGCTGCGATCTATTGAAAGTCAGCCCTCGACACAAGGGTTTGT</td><td>>hg38_21_+_8433172_8446622_RNA45SN1_45S_50ntFlanks</td><td>12994</td><td>13039</td><td>46</td><td>CCTCGCTGCGATCTATTGAAAGTCAGCCCTCGACACAAGGGTTTGT</td><td>35430.2</td><td>-3.1</td><td>1.1</td><td>-2.9</td><td>3.3E-03</td><td>1.3E-02</td><td>BT20/MB231</td><td>CYTO</td></tr>
<tr><td>>hg38_MT_+_598_1651_RNR1_12S_50ntFlanks@52.84.33|ATAGGTTTGGTCCTAGCCTTTCTATTAGCTCTT</td><td>>hg38_MT_+_598_1651_RNR1_12S_50ntFlanks</td><td>52</td><td>84</td><td>33</td><td>ATAGGTTTGGTCCTAGCCTTTCTATTAGCTCTT</td><td>22072.7</td><td>3.6</td><td>1.3</td><td>2.7</td><td>6.7E-03</td><td>1.3E-02</td><td>BT20/MB468</td><td>MP</td></tr>
<tr><td>>hg38_21_+_8433172_8446622_RNA45SN1_45S_50ntFlanks@3843.3872.30|CTACTTGGATAACTGTGGTAATTCTAGAGC</td><td>>hg38_21_+_8433172_8446622_RNA45SN1_45S_50ntFlanks</td><td>3843</td><td>3872</td><td>30</td><td>CTACTTGGATAACTGTGGTAATTCTAGAGC</td><td>205.3</td><td>-4.6</td><td>1.7</td><td>-2.7</td><td>6.7E-03</td><td>1.3E-02</td><td>BT20/MB468</td><td>MP</td></tr>
<tr><td>>hg38_21_+_8433172_8446622_RNA45SN1_45S_50ntFlanks@5085.5102.18|GAACGAGACTCTGGCATG</td><td>>hg38_21_+_8433172_8446622_RNA45SN1_45S_50ntFlanks</td><td>5085</td><td>5102</td><td>18</td><td>GAACGAGACTCTGGCATG</td><td>248.5</td><td>3.4</td><td>1.2</td><td>2.7</td><td>6.7E-03</td><td>1.3E-02</td><td>BT20/MB468</td><td>MP</td></tr>
<tr><td>>trna37_ValAAC_6_+_27203288_27203360@1.29.29|GTTTCCGTAGTGTAGTGGTTATCACGTTT</td><td>>trna37_ValAAC_6_+_27203288_27203360</td><td>1</td><td>29</td><td>29</td><td>GTTTCCGTAGTGTAGTGGTTATCACGTTT</td><td>1103.6</td><td>-2.7</td><td>1.0</td><td>-2.7</td><td>6.7E-03</td><td>1.3E-02</td><td>BT20/MB468</td><td>MP</td></tr>
<tr><td>>hg38_MT_+_598_1651_RNR1_12S_50ntFlanks@52.88.37|ATAGGTTTGGTCCTAGCCTTTCTATTAGCTCTTAGTA</td><td>>hg38_MT_+_598_1651_RNR1_12S_50ntFlanks</td><td>52</td><td>88</td><td>37</td><td>ATAGGTTTGGTCCTAGCCTTTCTATTAGCTCTTAGTA</td><td>70256.8</td><td>2.8</td><td>1.1</td><td>2.6</td><td>1.0E-02</td><td>1.3E-02</td><td>MB231/MB468</td><td>MP</td></tr>
<tr><td>>hg38_MT_+_598_1651_RNR1_12S_50ntFlanks@52.84.33|ATAGGTTTGGTCCTAGCCTTTCTATTAGCTCTT</td><td>>hg38_MT_+_598_1651_RNR1_12S_50ntFlanks</td><td>52</td><td>84</td><td>33</td><td>ATAGGTTTGGTCCTAGCCTTTCTATTAGCTCTT</td><td>418.8</td><td>1.8</td><td>0.6</td><td>3.0</td><td>2.8E-03</td><td>1.3E-02</td><td>BT20/MB231</td><td>NUC</td></tr>
<tr><td>>trnaMT_HisGTG_MT_+_12138_12206@34.72.39|AATCTGACAACAGAGGCTTACGACCCCTTATTTACCCCA</td><td>>trnaMT_HisGTG_MT_+_12138_12206</td><td>34</td><td>72</td><td>39</td><td>AATCTGACAACAGAGGCTTACGACCCCTTATTTACCCCA</td><td>264.5</td><td>6.2</td><td>2.3</td><td>2.7</td><td>6.7E-03</td><td>1.3E-02</td><td>BT20/MB468</td><td>MP</td></tr>
<tr><td>>hg38_21_+_8433172_8446622_RNA45SN1_45S_50ntFlanks@10391.10426.36|AACAGCAGTTGAACATGGGTCAGTCGGTCCTGAGAG</td><td>>hg38_21_+_8433172_8446622_RNA45SN1_45S_50ntFlanks</td><td>10391</td><td>10426</td><td>36</td><td>AACAGCAGTTGAACATGGGTCAGTCGGTCCTGAGAG</td><td>217.6</td><td>-4.7</td><td>1.7</td><td>-2.7</td><td>6.7E-03</td><td>1.3E-02</td><td>BT20/MB468</td><td>MP</td></tr>
<tr><td>>hg38_21_+_8433172_8446622_RNA45SN1_45S_50ntFlanks@13000.13039.40|TGCGATCTATTGAAAGTCAGCCCTCGACACAAGGGTTTGT</td><td>>hg38_21_+_8433172_8446622_RNA45SN1_45S_50ntFlanks</td><td>13000</td><td>13039</td><td>40</td><td>TGCGATCTATTGAAAGTCAGCCCTCGACACAAGGGTTTGT</td><td>854.1</td><td>2.9</td><td>1.1</td><td>2.6</td><td>9.4E-03</td><td>1.3E-02</td><td>BT20/MB468</td><td>MITO</td></tr>
<tr><td>>MI0000487|hsa-mir-193a&WithFlank&17|+|31559990|31560089@27.46.20(+1U)&[MIMAT0004614&hsa-miR-193a-5p&offsets|0|-2(+1U);m-123&17|+|31560016|31560037&offsets|0|-2(+1U)]|TGGGTCTTTGCGGGCGAGATT</td><td>>MI0000487|hsa-mir-193a&WithFlank&17|+|31559990|31560089</td><td>27</td><td>46</td><td>20(+1U)</td><td>TGGGTCTTTGCGGGCGAGATT</td><td>87.1</td><td>5.7</td><td>1.9</td><td>3.0</td><td>2.6E-03</td><td>1.3E-02</td><td>BT20/MB231</td><td>MP</td></tr>
<tr><td>>hg38_21_+_8433172_8446622_RNA45SN1_45S_50ntFlanks@7976.8011.36|GCGACCTCAGATCAGACGTGGCGACCCGCTGAATTT</td><td>>hg38_21_+_8433172_8446622_RNA45SN1_45S_50ntFlanks</td><td>7976</td><td>8011</td><td>36</td><td>GCGACCTCAGATCAGACGTGGCGACCCGCTGAATTT</td><td>135.1</td><td>3.5</td><td>1.4</td><td>2.6</td><td>1.0E-02</td><td>1.3E-02</td><td>MB231/MB468</td><td>MP</td></tr>
<tr><td>>hg38_21_+_8433172_8446622_RNA45SN1_45S_50ntFlanks@4473.4497.25|TTAGCTGAGTGTCCCGCGGGGCCCG</td><td>>hg38_21_+_8433172_8446622_RNA45SN1_45S_50ntFlanks</td><td>4473</td><td>4497</td><td>25</td><td>TTAGCTGAGTGTCCCGCGGGGCCCG</td><td>71.9</td><td>-4.5</td><td>1.7</td><td>-2.7</td><td>6.7E-03</td><td>1.3E-02</td><td>BT20/MB468</td><td>MP</td></tr>
<tr><td>>hg38_21_+_8433172_8446622_RNA45SN1_45S_50ntFlanks@3782.3810.29|CAGTGAAACTGCGAATGGCTCATTAAATC</td><td>>hg38_21_+_8433172_8446622_RNA45SN1_45S_50ntFlanks</td><td>3782</td><td>3810</td><td>29</td><td>CAGTGAAACTGCGAATGGCTCATTAAATC</td><td>206.3</td><td>-3.8</td><td>1.4</td><td>-2.7</td><td>6.7E-03</td><td>1.3E-02</td><td>BT20/MB468</td><td>MP</td></tr>
<tr><td>>hg38_1_-_228634819_228635039_RNA5S12_5S_50ntFlanks@134.172.39|TGGGAGACCGCCTGGGAATACCGGGTGCTGTAGGCTTTT</td><td>>hg38_1_-_228634819_228635039_RNA5S12_5S_50ntFlanks</td><td>134</td><td>172</td><td>39</td><td>TGGGAGACCGCCTGGGAATACCGGGTGCTGTAGGCTTTT</td><td>659.4</td><td>-4.7</td><td>1.6</td><td>-3.0</td><td>2.9E-03</td><td>1.3E-02</td><td>BT20/MB231</td><td>NUC</td></tr>
<tr><td>>hg38_21_+_8433172_8446622_RNA45SN1_45S_50ntFlanks@12475.12496.22|CCCACTAATAGGGAACGTGAGC</td><td>>hg38_21_+_8433172_8446622_RNA45SN1_45S_50ntFlanks</td><td>12475</td><td>12496</td><td>22</td><td>CCCACTAATAGGGAACGTGAGC</td><td>465.0</td><td>-2.8</td><td>0.9</td><td>-3.0</td><td>2.9E-03</td><td>1.3E-02</td><td>BT20/MB231</td><td>NUC</td></tr>
<tr><td>>hg38_21_+_8433172_8446622_RNA45SN1_45S_50ntFlanks@8673.8703.31|CGGTGCGCCGCGACCGGCTCCGGGACGGCTG</td><td>>hg38_21_+_8433172_8446622_RNA45SN1_45S_50ntFlanks</td><td>8673</td><td>8703</td><td>31</td><td>CGGTGCGCCGCGACCGGCTCCGGGACGGCTG</td><td>45.6</td><td>4.6</td><td>1.7</td><td>2.7</td><td>6.7E-03</td><td>1.3E-02</td><td>BT20/MB468</td><td>MP</td></tr>
<tr><td>>hg38_21_+_8433172_8446622_RNA45SN1_45S_50ntFlanks@11857.11875.19|GCGGAATCAGCGGGGAAAG</td><td>>hg38_21_+_8433172_8446622_RNA45SN1_45S_50ntFlanks</td><td>11857</td><td>11875</td><td>19</td><td>GCGGAATCAGCGGGGAAAG</td><td>91.9</td><td>-4.5</td><td>1.6</td><td>-2.7</td><td>6.7E-03</td><td>1.3E-02</td><td>BT20/MB468</td><td>MP</td></tr>
<tr><td>>hg38_21_+_8433172_8446622_RNA45SN1_45S_50ntFlanks@7976.8010.35|GCGACCTCAGATCAGACGTGGCGACCCGCTGAATT</td><td>>hg38_21_+_8433172_8446622_RNA45SN1_45S_50ntFlanks</td><td>7976</td><td>8010</td><td>35</td><td>GCGACCTCAGATCAGACGTGGCGACCCGCTGAATT</td><td>2096.8</td><td>2.1</td><td>0.7</td><td>2.9</td><td>3.3E-03</td><td>1.3E-02</td><td>BT20/MB231</td><td>MITO</td></tr>
<tr><td>>hg38_21_+_8433172_8446622_RNA45SN1_45S_50ntFlanks@6646.6673.28|TCGTACGACTCTTAGCGGTGGATCACTC</td><td>>hg38_21_+_8433172_8446622_RNA45SN1_45S_50ntFlanks</td><td>6646</td><td>6673</td><td>28</td><td>TCGTACGACTCTTAGCGGTGGATCACTC</td><td>2131.4</td><td>2.3</td><td>0.9</td><td>2.5</td><td>1.2E-02</td><td>1.3E-02</td><td>MB231/MB468</td><td>MITO</td></tr>
<tr><td>>hg38_21_+_8433172_8446622_RNA45SN1_45S_50ntFlanks@6646.6695.50|TCGTACGACTCTTAGCGGTGGATCACTCGGCTCGTGCGTCGATGAAGAAC</td><td>>hg38_21_+_8433172_8446622_RNA45SN1_45S_50ntFlanks</td><td>6646</td><td>6695</td><td>50</td><td>TCGTACGACTCTTAGCGGTGGATCACTCGGCTCGTGCGTCGATGAAGAAC</td><td>1232.3</td><td>5.4</td><td>2.0</td><td>2.7</td><td>6.7E-03</td><td>1.3E-02</td><td>BT20/MB468</td><td>MP</td></tr>
<tr><td>>MI0001445|hsa-mir-423&WithFlank&17|+|30117073|30117178@58.79.22&[MIMAT0001340&hsa-miR-423-3p&offsets|-1|-2;m-100&17|+|30117131|30117153&offsets|-1|-2]|AAGCTCGGTCTGAGGCCCCTCA</td><td>>MI0001445|hsa-mir-423&WithFlank&17|+|30117073|30117178</td><td>58</td><td>79</td><td>22</td><td>AAGCTCGGTCTGAGGCCCCTCA</td><td>576.4</td><td>2.2</td><td>0.7</td><td>2.9</td><td>3.3E-03</td><td>1.3E-02</td><td>BT20/MB231</td><td>CYTO</td></tr>
<tr><td>>hg38_21_+_8433172_8446622_RNA45SN1_45S_50ntFlanks@12892.12932.41|CCCTCGCCCGTCACGCACCGCACGTTCGTGGGGAACCTGGC</td><td>>hg38_21_+_8433172_8446622_RNA45SN1_45S_50ntFlanks</td><td>12892</td><td>12932</td><td>41</td><td>CCCTCGCCCGTCACGCACCGCACGTTCGTGGGGAACCTGGC</td><td>371.3</td><td>-5.3</td><td>2.0</td><td>-2.7</td><td>6.8E-03</td><td>1.3E-02</td><td>BT20/MB468</td><td>MP</td></tr>
<tr><td>>MI0000114|hsa-mir-107&WithFlank&10|-|89592741|89592833@56.74.19;MI0000108|hsa-mir-103a-2&WithFlank&20|+|3917488|3917577@54.72.19;MI0000109|hsa-mir-103a-1&WithFlank&5|-|168560890|168560979@54.72.19&[MIMAT0000104&hsa-miR-107&offsets|0|-4;m-78&10|-|89592758|89592778&offsets|0|-2];[MIMAT0000101&hsa-miR-103a-3p&offsets|0|-4;m-12&20|+|3917541|3917563&offsets|0|-4];[MIMAT0000101_1&hsa-miR-103a-3p&offsets|0|-4;m-13&5|-|168560904|168560926&offsets|0|-4;m-44220&5|-|168560908|168560926&offsets|0|0]|AGCAGCATTGTACAGGGCT</td><td>>MI0000114|hsa-mir-107&WithFlank&10|-|89592741|89592833</td><td>56</td><td>74</td><td>19</td><td>AGCAGCATTGTACAGGGCT</td><td>112.5</td><td>3.4</td><td>1.3</td><td>2.6</td><td>1.0E-02</td><td>1.3E-02</td><td>MB231/MB468</td><td>MP</td></tr>
<tr><td>>hg38_21_+_8433172_8446622_RNA45SN1_45S_50ntFlanks@11848.11871.24|ACGGGCTTGGCGGAATCAGCGGGG</td><td>>hg38_21_+_8433172_8446622_RNA45SN1_45S_50ntFlanks</td><td>11848</td><td>11871</td><td>24</td><td>ACGGGCTTGGCGGAATCAGCGGGG</td><td>271.6</td><td>-2.2</td><td>0.8</td><td>-2.7</td><td>6.8E-03</td><td>1.3E-02</td><td>BT20/MB468</td><td>MP</td></tr>
<tr><td>>MI0000465|hsa-mir-191&WithFlank&3|-|49020612|49020715@22.43.22(+1C)&[MIMAT0000440&hsa-miR-191-5p&offsets|0|-1(+1C);m-48&3|-|49020672|49020694&offsets|0|-1(+1C)]|CAACGGAATCCCAAAAGCAGCTC</td><td>>MI0000465|hsa-mir-191&WithFlank&3|-|49020612|49020715</td><td>22</td><td>43</td><td>22(+1C)</td><td>CAACGGAATCCCAAAAGCAGCTC</td><td>107.7</td><td>3.5</td><td>1.4</td><td>2.6</td><td>1.0E-02</td><td>1.3E-02</td><td>MB231/MB468</td><td>MP</td></tr>
<tr><td>>hg38_21_+_8433172_8446622_RNA45SN1_45S_50ntFlanks@11694.11728.35|AAGAAATTCAATGAAGCGCGGGTAAACGGCGGGAG</td><td>>hg38_21_+_8433172_8446622_RNA45SN1_45S_50ntFlanks</td><td>11694</td><td>11728</td><td>35</td><td>AAGAAATTCAATGAAGCGCGGGTAAACGGCGGGAG</td><td>233.0</td><td>-4.2</td><td>1.6</td><td>-2.6</td><td>9.4E-03</td><td>1.3E-02</td><td>BT20/MB468</td><td>MITO</td></tr>
<tr><td>>MI0003667|hsa-mir-652&WithFlank&X|+|110055323|110055432@67.86.20(+1U)&[MIMAT0003322&hsa-miR-652-3p&offsets|0|-1(+1U);m-148&X|+|110055389|110055410&offsets|0|-2(+1U)]|AATGGCGCCACTAGGGTTGTT</td><td>>MI0003667|hsa-mir-652&WithFlank&X|+|110055323|110055432</td><td>67</td><td>86</td><td>20(+1U)</td><td>AATGGCGCCACTAGGGTTGTT</td><td>86.8</td><td>5.8</td><td>2.1</td><td>2.7</td><td>6.8E-03</td><td>1.3E-02</td><td>BT20/MB468</td><td>MP</td></tr>
<tr><td>>hg38_21_+_8433172_8446622_RNA45SN1_45S_50ntFlanks@9568.9586.19|CGTGCAAATCGGTCGTCCG</td><td>>hg38_21_+_8433172_8446622_RNA45SN1_45S_50ntFlanks</td><td>9568</td><td>9586</td><td>19</td><td>CGTGCAAATCGGTCGTCCG</td><td>124.9</td><td>-4.5</td><td>1.6</td><td>-2.7</td><td>6.8E-03</td><td>1.3E-02</td><td>BT20/MB468</td><td>MP</td></tr>
<tr><td>>hg38_21_+_8433172_8446622_RNA45SN1_45S_50ntFlanks@5083.5120.38|ACGAACGAGACTCTGGCATGCTAACTAGTTACGCGACC</td><td>>hg38_21_+_8433172_8446622_RNA45SN1_45S_50ntFlanks</td><td>5083</td><td>5120</td><td>38</td><td>ACGAACGAGACTCTGGCATGCTAACTAGTTACGCGACC</td><td>185.6</td><td>-1.7</td><td>0.7</td><td>-2.6</td><td>9.4E-03</td><td>1.3E-02</td><td>BT20/MB468</td><td>MITO</td></tr>
<tr><td>>hg38_21_+_8433172_8446622_RNA45SN1_45S_50ntFlanks@11668.11706.39|TGCCCAGTGCTCTGAATGTCAAAGTGAAGAAATTCAATG</td><td>>hg38_21_+_8433172_8446622_RNA45SN1_45S_50ntFlanks</td><td>11668</td><td>11706</td><td>39</td><td>TGCCCAGTGCTCTGAATGTCAAAGTGAAGAAATTCAATG</td><td>659.3</td><td>-3.0</td><td>1.0</td><td>-3.0</td><td>2.9E-03</td><td>1.3E-02</td><td>BT20/MB231</td><td>NUC</td></tr>
<tr><td>>hg38_21_+_8433172_8446622_RNA45SN1_45S_50ntFlanks@10425.10453.29|AGATGGGCGAGCGCCGTTCCGAAGGGACG</td><td>>hg38_21_+_8433172_8446622_RNA45SN1_45S_50ntFlanks</td><td>10425</td><td>10453</td><td>29</td><td>AGATGGGCGAGCGCCGTTCCGAAGGGACG</td><td>79.7</td><td>-4.4</td><td>1.6</td><td>-2.7</td><td>6.8E-03</td><td>1.3E-02</td><td>BT20/MB468</td><td>MP</td></tr>
<tr><td>>hg38_21_+_8433172_8446622_RNA45SN1_45S_50ntFlanks@11672.11702.31|CAGTGCTCTGAATGTCAAAGTGAAGAAATTC</td><td>>hg38_21_+_8433172_8446622_RNA45SN1_45S_50ntFlanks</td><td>11672</td><td>11702</td><td>31</td><td>CAGTGCTCTGAATGTCAAAGTGAAGAAATTC</td><td>577.8</td><td>-1.4</td><td>0.6</td><td>-2.5</td><td>1.1E-02</td><td>1.3E-02</td><td>BT20/MB468</td><td>CYTO</td></tr>
<tr><td>>trnaMT_ValTAC_MT_+_1602_1670@28.72.45|AACTTACACTTAGGAGATTTCAACTTAACTTGACCGCTCTGACCA</td><td>>trnaMT_ValTAC_MT_+_1602_1670</td><td>28</td><td>72</td><td>45</td><td>AACTTACACTTAGGAGATTTCAACTTAACTTGACCGCTCTGACCA</td><td>205.4</td><td>-2.7</td><td>0.9</td><td>-2.9</td><td>3.3E-03</td><td>1.3E-02</td><td>BT20/MB231</td><td>CYTO</td></tr>
<tr><td>>trnaMT_ProTGG_MT_-_15956_16023@46.70.25|GGAGTTAAAGACTTTTTCTCTGACC</td><td>>trnaMT_ProTGG_MT_-_15956_16023</td><td>46</td><td>70</td><td>25</td><td>GGAGTTAAAGACTTTTTCTCTGACC</td><td>158.8</td><td>2.7</td><td>1.0</td><td>2.6</td><td>1.0E-02</td><td>1.3E-02</td><td>MB231/MB468</td><td>MP</td></tr>
<tr><td>>hg38_21_+_8433172_8446622_RNA45SN1_45S_50ntFlanks@7973.8015.43|GACGCGACCTCAGATCAGACGTGGCGACCCGCTGAATTTAAGC</td><td>>hg38_21_+_8433172_8446622_RNA45SN1_45S_50ntFlanks</td><td>7973</td><td>8015</td><td>43</td><td>GACGCGACCTCAGATCAGACGTGGCGACCCGCTGAATTTAAGC</td><td>344.4</td><td>4.1</td><td>1.6</td><td>2.5</td><td>1.2E-02</td><td>1.3E-02</td><td>MB231/MB468</td><td>MITO</td></tr>
<tr><td>>trna119_LysCTT_1_-_145395522_145395594@1.40.40,trna11_LysCTT_5_-_180648979_180649051@1.40.40,trna13_LysCTT_6_+_26556774_26556846@1.40.40,trna7_LysCTT_16_+_3225692_3225764@1.40.40,trna9_LysCTT_5_+_180634755_180634827@1.40.40|GCCCGGCTAGCTCAGTCGGTAGAGCATGAGACTCTTAATC</td><td>>trna119_LysCTT_1_-_145395522_145395594</td><td>1</td><td>40</td><td>40</td><td>GCCCGGCTAGCTCAGTCGGTAGAGCATGAGACTCTTAATC</td><td>491.7</td><td>5.0</td><td>2.0</td><td>2.5</td><td>1.1E-02</td><td>1.3E-02</td><td>BT20/MB468</td><td>CYTO</td></tr>
<tr><td>>hg38_21_+_8433172_8446622_RNA45SN1_45S_50ntFlanks@9304.9335.32|ACCAAGGAGTCTAACACGTGCGCGAGTCGGGG</td><td>>hg38_21_+_8433172_8446622_RNA45SN1_45S_50ntFlanks</td><td>9304</td><td>9335</td><td>32</td><td>ACCAAGGAGTCTAACACGTGCGCGAGTCGGGG</td><td>220.0</td><td>-3.6</td><td>1.3</td><td>-2.7</td><td>6.8E-03</td><td>1.3E-02</td><td>BT20/MB468</td><td>MP</td></tr>
<tr><td>>hg38_MT_+_598_1651_RNR1_12S_50ntFlanks@274.294.21|AGGGTTGGTCAATTTCGTGCC</td><td>>hg38_MT_+_598_1651_RNR1_12S_50ntFlanks</td><td>274</td><td>294</td><td>21</td><td>AGGGTTGGTCAATTTCGTGCC</td><td>71.5</td><td>5.6</td><td>2.1</td><td>2.7</td><td>6.8E-03</td><td>1.3E-02</td><td>BT20/MB468</td><td>MP</td></tr>
<tr><td>>hg38_21_+_8433172_8446622_RNA45SN1_45S_50ntFlanks@6660.6681.22|GCGGTGGATCACTCGGCTCGTG</td><td>>hg38_21_+_8433172_8446622_RNA45SN1_45S_50ntFlanks</td><td>6660</td><td>6681</td><td>22</td><td>GCGGTGGATCACTCGGCTCGTG</td><td>1024.1</td><td>-1.6</td><td>0.6</td><td>-2.6</td><td>9.5E-03</td><td>1.3E-02</td><td>BT20/MB468</td><td>MITO</td></tr>
<tr><td>>hg38_21_+_8433172_8446622_RNA45SN1_45S_50ntFlanks@4727.4758.32|AATCAAGAACGAAAGTCGGAGGTTCGAAGACG</td><td>>hg38_21_+_8433172_8446622_RNA45SN1_45S_50ntFlanks</td><td>4727</td><td>4758</td><td>32</td><td>AATCAAGAACGAAAGTCGGAGGTTCGAAGACG</td><td>1568.7</td><td>-1.6</td><td>0.6</td><td>-2.6</td><td>1.0E-02</td><td>1.3E-02</td><td>MB231/MB468</td><td>MP</td></tr>
</table>

<table>
<tr><td>>hg38_21_+_8433172_8446622_RNA45SN1_45S_50ntFlanks@11037.11061.25|CTCTCCCCCGCTCCCCGTCCTCCCC</td><td>>hg38_21_+_8433172_8446622_RNA45SN1_45S_50ntFlanks</td><td>11037</td><td>11061</td><td>25</td><td>CTCTCCCCCGCTCCCCGTCCTCCCC</td><td>721.4</td><td>-5.2</td><td>2.1</td><td>-2.5</td><td>1.1E-02</td><td>1.3E-02</td><td>BT20/MB468</td><td>NUC</td></tr>
<tr><td>>hg38_21_+_8433172_8446622_RNA45SN1_45S_50ntFlanks@11200.11233.34|CGGCGGCGGCGGCGACTCTGGACGCGAGCCGGGC</td><td>>hg38_21_+_8433172_8446622_RNA45SN1_45S_50ntFlanks</td><td>11200</td><td>11233</td><td>34</td><td>CGGCGGCGGCGGCGACTCTGGACGCGAGCCGGGC</td><td>144.2</td><td>-2.6</td><td>1.0</td><td>-2.7</td><td>6.8E-03</td><td>1.3E-02</td><td>BT20/MB468</td><td>MP</td></tr>
<tr><td>>hg38_MT_+_1621_3279_RNR2_16S_50ntFlanks@1369.1395.27|GATGTTGGATCAGGACATCCCGATGGT</td><td>>hg38_MT_+_1621_3279_RNR2_16S_50ntFlanks</td><td>1369</td><td>1395</td><td>27</td><td>GATGTTGGATCAGGACATCCCGATGGT</td><td>149.2</td><td>3.2</td><td>1.2</td><td>2.6</td><td>1.0E-02</td><td>1.3E-02</td><td>MB231/MB468</td><td>MP</td></tr>
<tr><td>>hg38_21_+_8433172_8446622_RNA45SN1_45S_50ntFlanks@4753.4786.34|AAGACGATCAGATACCGTCGTAGTTCCGACCATA</td><td>>hg38_21_+_8433172_8446622_RNA45SN1_45S_50ntFlanks</td><td>4753</td><td>4786</td><td>34</td><td>AAGACGATCAGATACCGTCGTAGTTCCGACCATA</td><td>467.0</td><td>-3.6</td><td>1.4</td><td>-2.6</td><td>9.5E-03</td><td>1.3E-02</td><td>BT20/MB468</td><td>MITO</td></tr>
<tr><td>>MI0000825|hsa-mir-345&WithFlank&14|+|100307853|100307962@24.45.22&[MIMAT0000772&hsa-miR-345-5p&offsets|0|0;m-193&14|+|100307876|100307896&offsets|0|+1]|GCTGACTCCTAGTCCAGGGCTC</td><td>>MI0000825|hsa-mir-345&WithFlank&14|+|100307853|100307962</td><td>24</td><td>45</td><td>22</td><td>GCTGACTCCTAGTCCAGGGCTC</td><td>349.1</td><td>-2.1</td><td>0.7</td><td>-2.9</td><td>3.3E-03</td><td>1.3E-02</td><td>BT20/MB231</td><td>MITO</td></tr>
<tr><td>>trna14_LysTTT_11_-_59327808_59327880@1.36.36,trna2_LysTTT_17_+_8022473_8022545@1.36.36,trna54_LysTTT_1_+_204475655_204475727@1.36.36,trna5_LysTTT_11_+_59323902_59323974@1.36.36,trna62_LysTTT_1_-_204476158_204476230@1.36.36,trna76_LysTTT_6_+_28918806_28918878@1.36.36|GCCCGGATAGCTCAGTCGGTAGAGCATCAGACTTTT</td><td>>trna14_LysTTT_11_-_59327808_59327880</td><td>1</td><td>36</td><td>36</td><td>GCCCGGATAGCTCAGTCGGTAGAGCATCAGACTTTT</td><td>325.7</td><td>-2.8</td><td>0.9</td><td>-2.9</td><td>3.3E-03</td><td>1.3E-02</td><td>BT20/MB231</td><td>MITO</td></tr>
<tr><td>>hg38_21_+_8433172_8446622_RNA45SN1_45S_50ntFlanks@11595.11621.27|AGGGGAATCCGACTGTTTAATTAAAAC</td><td>>hg38_21_+_8433172_8446622_RNA45SN1_45S_50ntFlanks</td><td>11595</td><td>11621</td><td>27</td><td>AGGGGAATCCGACTGTTTAATTAAAAC</td><td>144.0</td><td>5.1</td><td>1.7</td><td>3.0</td><td>2.6E-03</td><td>1.3E-02</td><td>BT20/MB231</td><td>MP</td></tr>
<tr><td>>hg38_21_+_8433172_8446622_RNA45SN1_45S_50ntFlanks@9940.9958.19|CAGGACGGTGGCCATGGAA</td><td>>hg38_21_+_8433172_8446622_RNA45SN1_45S_50ntFlanks</td><td>9940</td><td>9958</td><td>19</td><td>CAGGACGGTGGCCATGGAA</td><td>68.2</td><td>5.4</td><td>2.0</td><td>2.7</td><td>6.8E-03</td><td>1.3E-02</td><td>BT20/MB468</td><td>MP</td></tr>
<tr><td>>MI0000072|hsa-mir-18a&WithFlank&13|+|91350745|91350827@53.74.22(+1U)&[MIMAT0002891&hsa-miR-18a-3p&offsets|0|-1(+1U);m-389&13|+|91350797|91350819&offsets|0|-1(+1U)]|ACTGCCCTAAGTGCTCCTTCTGT</td><td>>MI0000072|hsa-mir-18a&WithFlank&13|+|91350745|91350827</td><td>53</td><td>74</td><td>22(+1U)</td><td>ACTGCCCTAAGTGCTCCTTCTGT</td><td>82.9</td><td>5.0</td><td>2.0</td><td>2.6</td><td>1.0E-02</td><td>1.3E-02</td><td>MB231/MB468</td><td>MP</td></tr>
<tr><td>>hg38_21_+_8433172_8446622_RNA45SN1_45S_50ntFlanks@6744.6772.29|GACACTTCGAACGCACTTGCGGCCCCGGG</td><td>>hg38_21_+_8433172_8446622_RNA45SN1_45S_50ntFlanks</td><td>6744</td><td>6772</td><td>29</td><td>GACACTTCGAACGCACTTGCGGCCCCGGG</td><td>277.4</td><td>-2.0</td><td>0.8</td><td>-2.6</td><td>9.5E-03</td><td>1.3E-02</td><td>BT20/MB468</td><td>MITO</td></tr>
<tr><td>>hg38_MT_+_598_1651_RNR1_12S_50ntFlanks@906.929.24|GTATACTTCAAAGGACATTTAACT</td><td>>hg38_MT_+_598_1651_RNR1_12S_50ntFlanks</td><td>906</td><td>929</td><td>24</td><td>GTATACTTCAAAGGACATTTAACT</td><td>76.8</td><td>5.0</td><td>2.0</td><td>2.6</td><td>1.0E-02</td><td>1.3E-02</td><td>MB231/MB468</td><td>MP</td></tr>
<tr><td>>am_MI0000786|hsa-mir-378a&WithFlank&5|+|149732819|149732896@49.70.22(+1A)&[MIMAT0000732&hsa-miR-378a-3p&offsets|0|0(+1A);m-44&5|+|149732867|149732888&offsets|0|0(+1A)]|ACTGGACTTGGAGTCAGAAGGCA</td><td>>am_MI0000786|hsa-mir-378a&WithFlank&5|+|149732819|149732896</td><td>49</td><td>70</td><td>22(+1A)</td><td>ACTGGACTTGGAGTCAGAAGGCA</td><td>505.9</td><td>2.2</td><td>0.9</td><td>2.5</td><td>1.2E-02</td><td>1.3E-02</td><td>MB231/MB468</td><td>MITO</td></tr>
<tr><td>>hg38_21_+_8433172_8446622_RNA45SN1_45S_50ntFlanks@6652.6674.23|GACTCTTAGCGGTGGATCACTCG</td><td>>hg38_21_+_8433172_8446622_RNA45SN1_45S_50ntFlanks</td><td>6652</td><td>6674</td><td>23</td><td>GACTCTTAGCGGTGGATCACTCG</td><td>967.5</td><td>2.8</td><td>1.1</td><td>2.5</td><td>1.2E-02</td><td>1.3E-02</td><td>MB231/MB468</td><td>MITO</td></tr>
<tr><td>>MI0000088|hsa-mir-30a&WithFlank&6|-|71403545|71403627@53.74.22(+1C)&[MIMAT0000088&hsa-miR-30a-3p&offsets|0|0(+1C);m-38&6|-|71403554|71403575&offsets|0|0(+1C)]|CTTTCAGTCGGATGTTTGCAGCC</td><td>>MI0000088|hsa-mir-30a&WithFlank&6|-|71403545|71403627</td><td>53</td><td>74</td><td>22(+1C)</td><td>CTTTCAGTCGGATGTTTGCAGCC</td><td>38.1</td><td>5.1</td><td>2.0</td><td>2.6</td><td>1.0E-02</td><td>1.3E-02</td><td>MB231/MB468</td><td>MP</td></tr>
<tr><td>>hg38_21_+_8433172_8446622_RNA45SN1_45S_50ntFlanks@12416.12441.26|TTCGATGTCGGCTCTTCCTATCATTG</td><td>>hg38_21_+_8433172_8446622_RNA45SN1_45S_50ntFlanks</td><td>12416</td><td>12441</td><td>26</td><td>TTCGATGTCGGCTCTTCCTATCATTG</td><td>487.7</td><td>1.4</td><td>0.5</td><td>2.6</td><td>9.5E-03</td><td>1.3E-02</td><td>BT20/MB468</td><td>MITO</td></tr>
<tr><td>>hg38_21_+_8433172_8446622_RNA45SN1_45S_50ntFlanks@6739.6756.18|TCATCGACACTTCGAACG</td><td>>hg38_21_+_8433172_8446622_RNA45SN1_45S_50ntFlanks</td><td>6739</td><td>6756</td><td>18</td><td>TCATCGACACTTCGAACG</td><td>85.7</td><td>-4.6</td><td>1.7</td><td>-2.7</td><td>6.8E-03</td><td>1.3E-02</td><td>BT20/MB468</td><td>MP</td></tr>
<tr><td>>trna119_LysCTT_1_-_145395522_145395594@4.34.31,trna11_LysCTT_5_-_180648979_180649051@4.34.31,trna13_LysCTT_6_+_26556774_26556846@4.34.31,trna7_LysCTT_16_+_3225692_3225764@4.34.31,trna9_LysCTT_5_+_180634755_180634827@4.34.31|CGGCTAGCTCAGTCGGTAGAGCATGAGACTC</td><td>>trna119_LysCTT_1_-_145395522_145395594</td><td>4</td><td>34</td><td>31</td><td>CGGCTAGCTCAGTCGGTAGAGCATGAGACTC</td><td>287.5</td><td>-3.3</td><td>1.1</td><td>-3.0</td><td>2.7E-03</td><td>1.3E-02</td><td>BT20/MB231</td><td>MP</td></tr>
<tr><td>>hg38_21_+_8433172_8446622_RNA45SN1_45S_50ntFlanks@4047.4065.19|ATTCGAACGTCTGCCCTAT</td><td>>hg38_21_+_8433172_8446622_RNA45SN1_45S_50ntFlanks</td><td>4047</td><td>4065</td><td>19</td><td>ATTCGAACGTCTGCCCTAT</td><td>103.6</td><td>-4.5</td><td>1.7</td><td>-2.7</td><td>6.8E-03</td><td>1.3E-02</td><td>BT20/MB468</td><td>MP</td></tr>
<tr><td>>hg38_21_+_8433172_8446622_RNA45SN1_45S_50ntFlanks@7974.8006.33|ACGCGACCTCAGATCAGACGTGGCGACCCGCTG</td><td>>hg38_21_+_8433172_8446622_RNA45SN1_45S_50ntFlanks</td><td>7974</td><td>8006</td><td>33</td><td>ACGCGACCTCAGATCAGACGTGGCGACCCGCTG</td><td>10287.7</td><td>3.2</td><td>1.3</td><td>2.5</td><td>1.1E-02</td><td>1.3E-02</td><td>BT20/MB468</td><td>CYTO</td></tr>
<tr><td>>hg38_21_+_8433172_8446622_RNA45SN1_45S_50ntFlanks@11212.11253.42|CGACTCTGGACGCGAGCCGGGCCCTTCCCGTGGATCGCCCCA</td><td>>hg38_21_+_8433172_8446622_RNA45SN1_45S_50ntFlanks</td><td>11212</td><td>11253</td><td>42</td><td>CGACTCTGGACGCGAGCCGGGCCCTTCCCGTGGATCGCCCCA</td><td>304.7</td><td>-2.4</td><td>0.9</td><td>-2.5</td><td>1.1E-02</td><td>1.3E-02</td><td>BT20/MB468</td><td>CYTO</td></tr>
<tr><td>>hg38_21_+_8433172_8446622_RNA45SN1_45S_50ntFlanks@4569.4587.19|ATGGAATAGGACCGCGGTT</td><td>>hg38_21_+_8433172_8446622_RNA45SN1_45S_50ntFlanks</td><td>4569</td><td>4587</td><td>19</td><td>ATGGAATAGGACCGCGGTT</td><td>217.7</td><td>-2.2</td><td>0.8</td><td>-2.7</td><td>6.8E-03</td><td>1.3E-02</td><td>BT20/MB468</td><td>MP</td></tr>
<tr><td>>hg38_21_+_8433172_8446622_RNA45SN1_45S_50ntFlanks@4728.4748.21|ATCAAGAACGAAAGTCGGAGG</td><td>>hg38_21_+_8433172_8446622_RNA45SN1_45S_50ntFlanks</td><td>4728</td><td>4748</td><td>21</td><td>ATCAAGAACGAAAGTCGGAGG</td><td>246.4</td><td>-3.6</td><td>1.4</td><td>-2.5</td><td>1.1E-02</td><td>1.3E-02</td><td>MB231/MB468</td><td>CYTO</td></tr>
<tr><td>>trnaMT_GluTTC_MT_-_14674_14742@35.65.31,trnalookalike8_GluTTC_5_-_93905172_93905240@35.65.31|TATCATTGGTCGTGGTTGTAGTCCGTGCGAG</td><td>>trnaMT_GluTTC_MT_-_14674_14742</td><td>35</td><td>65</td><td>31</td><td>TATCATTGGTCGTGGTTGTAGTCCGTGCGAG</td><td>117.2</td><td>4.8</td><td>1.8</td><td>2.7</td><td>6.8E-03</td><td>1.3E-02</td><td>BT20/MB468</td><td>MP</td></tr>
<tr><td>>hg38_21_+_8433172_8446622_RNA45SN1_45S_50ntFlanks@9927.9948.22|TTGATATAGACAGCAGGACGGT</td><td>>hg38_21_+_8433172_8446622_RNA45SN1_45S_50ntFlanks</td><td>9927</td><td>9948</td><td>22</td><td>TTGATATAGACAGCAGGACGGT</td><td>306.4</td><td>4.8</td><td>1.8</td><td>2.7</td><td>6.8E-03</td><td>1.3E-02</td><td>BT20/MB468</td><td>MP</td></tr>
<tr><td>>hg38_21_+_8433172_8446622_RNA45SN1_45S_50ntFlanks@6646.6690.45|TCGTACGACTCTTAGCGGTGGATCACTCGGCTCGTGCGTCGATGA</td><td>>hg38_21_+_8433172_8446622_RNA45SN1_45S_50ntFlanks</td><td>6646</td><td>6690</td><td>45</td><td>TCGTACGACTCTTAGCGGTGGATCACTCGGCTCGTGCGTCGATGA</td><td>1320.3</td><td>2.1</td><td>0.7</td><td>2.9</td><td>3.4E-03</td><td>1.3E-02</td><td>BT20/MB231</td><td>MITO</td></tr>
<tr><td>>hg38_21_+_8433172_8446622_RNA45SN1_45S_50ntFlanks@11853.11873.21|CTTGGCGGAATCAGCGGGGAA</td><td>>hg38_21_+_8433172_8446622_RNA45SN1_45S_50ntFlanks</td><td>11853</td><td>11873</td><td>21</td><td>CTTGGCGGAATCAGCGGGGAA</td><td>70.7</td><td>-4.4</td><td>1.6</td><td>-2.7</td><td>6.8E-03</td><td>1.3E-02</td><td>BT20/MB468</td><td>MP</td></tr>
<tr><td>>hg38_21_+_8433172_8446622_RNA45SN1_45S_50ntFlanks@10391.10419.29|AACAGCAGTTGAACATGGGTCAGTCGGTC</td><td>>hg38_21_+_8433172_8446622_RNA45SN1_45S_50ntFlanks</td><td>10391</td><td>10419</td><td>29</td><td>AACAGCAGTTGAACATGGGTCAGTCGGTC</td><td>138.6</td><td>-3.9</td><td>1.5</td><td>-2.6</td><td>9.5E-03</td><td>1.3E-02</td><td>BT20/MB468</td><td>MITO</td></tr>
<tr><td>>trnaMT_HisGTG_MT_+_12138_12206@32.66.35|TGAATCTGACAACAGAGGCTTACGACCCCTTATTT</td><td>>trnaMT_HisGTG_MT_+_12138_12206</td><td>32</td><td>66</td><td>35</td><td>TGAATCTGACAACAGAGGCTTACGACCCCTTATTT</td><td>339.0</td><td>2.5</td><td>1.0</td><td>2.5</td><td>1.2E-02</td><td>1.3E-02</td><td>MB231/MB468</td><td>MITO</td></tr>
<tr><td>>MI0000285|hsa-mir-205&WithFlank&1|+|209432127|209432248@40.60.21(+1U)&[MIMAT0000266&hsa-miR-205-5p&offsets|0|-1(+1U);m-351&1|+|209432166|209432187&offsets|0|-1(+1U)]|TCCTTCATTCCACCGGAGTCTT</td><td>>MI0000285|hsa-mir-205&WithFlank&1|+|209432127|209432248</td><td>40</td><td>60</td><td>21(+1U)</td><td>TCCTTCATTCCACCGGAGTCTT</td><td>1154.5</td><td>3.3</td><td>1.2</td><td>2.7</td><td>6.9E-03</td><td>1.3E-02</td><td>BT20/MB468</td><td>MP</td></tr>
<tr><td>>MI0000298|hsa-mir-221&WithFlank&X|-|45746151|45746272@71.92.22&[MIMAT0000278&hsa-miR-221-3p&offsets|0|-1;m-91&X|-|45746181|45746202&offsets|0|0]|AGCTACATTGTCTGCTGGGTTT</td><td>>MI0000298|hsa-mir-221&WithFlank&X|-|45746151|45746272</td><td>71</td><td>92</td><td>22</td><td>AGCTACATTGTCTGCTGGGTTT</td><td>17811.6</td><td>-1.8</td><td>0.6</td><td>-3.0</td><td>2.9E-03</td><td>1.3E-02</td><td>BT20/MB231</td><td>NUC</td></tr>
<tr><td>>hg38_21_+_8433172_8446622_RNA45SN1_45S_50ntFlanks@12419.12456.38|GATGTCGGCTCTTCCTATCATTGTGAAGCAGAATTCAC</td><td>>hg38_21_+_8433172_8446622_RNA45SN1_45S_50ntFlanks</td><td>12419</td><td>12456</td><td>38</td><td>GATGTCGGCTCTTCCTATCATTGTGAAGCAGAATTCAC</td><td>204.4</td><td>-2.6</td><td>0.9</td><td>-3.0</td><td>2.9E-03</td><td>1.3E-02</td><td>BT20/MB231</td><td>NUC</td></tr>
<tr><td>>hg38_21_+_8433172_8446622_RNA45SN1_45S_50ntFlanks@4738.4758.21|AAAGTCGGAGGTTCGAAGACG</td><td>>hg38_21_+_8433172_8446622_RNA45SN1_45S_50ntFlanks</td><td>4738</td><td>4758</td><td>21</td><td>AAAGTCGGAGGTTCGAAGACG</td><td>9513.9</td><td>-1.9</td><td>0.7</td><td>-2.7</td><td>6.9E-03</td><td>1.3E-02</td><td>BT20/MB468</td><td>MP</td></tr>
<tr><td>>hg38_MT_+_598_1651_RNR1_12S_50ntFlanks@428.465.38|AAAGTGGCTTTAACATATCTGAACACACAATAGCTAAG</td><td>>hg38_MT_+_598_1651_RNR1_12S_50ntFlanks</td><td>428</td><td>465</td><td>38</td><td>AAAGTGGCTTTAACATATCTGAACACACAATAGCTAAG</td><td>252.7</td><td>4.4</td><td>1.6</td><td>2.7</td><td>6.9E-03</td><td>1.3E-02</td><td>BT20/MB468</td><td>MP</td></tr>
<tr><td>>hg38_21_+_8433172_8446622_RNA45SN1_45S_50ntFlanks@13002.13039.38|CGATCTATTGAAAGTCAGCCCTCGACACAAGGGTTTGT</td><td>>hg38_21_+_8433172_8446622_RNA45SN1_45S_50ntFlanks</td><td>13002</td><td>13039</td><td>38</td><td>CGATCTATTGAAAGTCAGCCCTCGACACAAGGGTTTGT</td><td>8630.4</td><td>-4.9</td><td>1.9</td><td>-2.5</td><td>1.2E-02</td><td>1.3E-02</td><td>MB231/MB468</td><td>MITO</td></tr>
<tr><td>>trna111_HisGTG_1_-_147774845_147774916@-1A.23.24,trna118_HisGTG_1_-_145396881_145396952@-1A.23.24,trna16_HisGTG_1_+_146544773_146544844@-1A.23.24,trna1_HisGTG_15_+_45493349_45493420@-1A.23.24,trna21_HisGTG_1_+_147753471_147753542@-1A.23.24,trna33_HisGTG_6_+_27125906_27125977@-1A.23.24,trna7_HisGTG_9_-_14433938_14434009@-1A.23.24,trna8_HisGTG_15_-_45492611_45492682@-1A.23.24,trna9_HisGTG_15_-_45490804_45490875@-1A.23.24|AGCCGTGATCGTATAGTGGTTAGT</td><td>>trna111_HisGTG_1_-_147774845_147774916</td><td>-1A</td><td>23</td><td>24</td><td>AGCCGTGATCGTATAGTGGTTAGT</td><td>214.9</td><td>-5.8</td><td>1.9</td><td>-3.0</td><td>2.7E-03</td><td>1.3E-02</td><td>BT20/MB231</td><td>MP</td></tr>
<tr><td>>hg38_21_+_8433172_8446622_RNA45SN1_45S_50ntFlanks@5059.5080.22|CGATTTGTCTGGTTAATTCCGA</td><td>>hg38_21_+_8433172_8446622_RNA45SN1_45S_50ntFlanks</td><td>5059</td><td>5080</td><td>22</td><td>CGATTTGTCTGGTTAATTCCGA</td><td>584.1</td><td>-3.7</td><td>1.4</td><td>-2.6</td><td>9.5E-03</td><td>1.3E-02</td><td>BT20/MB468</td><td>MITO</td></tr>
<tr><td>>MI0000084|hsa-mir-26b&WithFlank&2|+|218402640|218402728@18.38.21(+1A)&[MIMAT0000083&hsa-miR-26b-5p&offsets|0|0(+1A);m-53&2|+|218402657|218402678&offsets|0|-1(+1A)]|TTCAAGTAATTCAGGATAGGTA</td><td>>MI0000084|hsa-mir-26b&WithFlank&2|+|218402640|218402728</td><td>18</td><td>38</td><td>21(+1A)</td><td>TTCAAGTAATTCAGGATAGGTA</td><td>467.9</td><td>2.9</td><td>1.0</td><td>2.9</td><td>3.4E-03</td><td>1.3E-02</td><td>BT20/MB231</td><td>CYTO</td></tr>
<tr><td>>am_MI0000786|hsa-mir-378a&WithFlank&5|+|149732819|149732896@49.70.22(+1G)&[MIMAT0000732&hsa-miR-378a-3p&offsets|0|0(+1G);m-44&5|+|149732867|149732888&offsets|0|0(+1G)]|ACTGGACTTGGAGTCAGAAGGCG</td><td>>am_MI0000786|hsa-mir-378a&WithFlank&5|+|149732819|149732896</td><td>49</td><td>70</td><td>22(+1G)</td><td>ACTGGACTTGGAGTCAGAAGGCG</td><td>541.0</td><td>2.2</td><td>0.8</td><td>2.9</td><td>3.4E-03</td><td>1.3E-02</td><td>BT20/MB231</td><td>CYTO</td></tr>
<tr><td>>hg38_21_+_8433172_8446622_RNA45SN1_45S_50ntFlanks@6661.6680.20|CGGTGGATCACTCGGCTCGT</td><td>>hg38_21_+_8433172_8446622_RNA45SN1_45S_50ntFlanks</td><td>6661</td><td>6680</td><td>20</td><td>CGGTGGATCACTCGGCTCGT</td><td>317.1</td><td>2.0</td><td>0.7</td><td>2.9</td><td>3.4E-03</td><td>1.3E-02</td><td>BT20/MB231</td><td>CYTO</td></tr>
<tr><td>>hg38_21_+_8433172_8446622_RNA45SN1_45S_50ntFlanks@12446.12476.31|GCAGAATTCACCAAGCGTTGGATTGTTCACC</td><td>>hg38_21_+_8433172_8446622_RNA45SN1_45S_50ntFlanks</td><td>12446</td><td>12476</td><td>31</td><td>GCAGAATTCACCAAGCGTTGGATTGTTCACC</td><td>362.5</td><td>-2.5</td><td>0.8</td><td>-2.9</td><td>3.4E-03</td><td>1.3E-02</td><td>BT20/MB231</td><td>CYTO</td></tr>
<tr><td>>hg38_21_+_8433172_8446622_RNA45SN1_45S_50ntFlanks@12637.12661.25|GCTACCATCTGTGGGATTATGACTG</td><td>>hg38_21_+_8433172_8446622_RNA45SN1_45S_50ntFlanks</td><td>12637</td><td>12661</td><td>25</td><td>GCTACCATCTGTGGGATTATGACTG</td><td>2276.2</td><td>-3.5</td><td>1.3</td><td>-2.7</td><td>6.9E-03</td><td>1.3E-02</td><td>BT20/MB468</td><td>MP</td></tr>
<tr><td>>MI0000811|hsa-mir-148b&WithFlank&12|+|54337210|54337320@69.90.22(+1U)&[MIMAT0000759&hsa-miR-148b-3p&offsets|0|0(+1U);m-77&12|+|54337278|54337299&offsets|0|0(+1U)]|TCAGTGCATCACAGAACTTTGTT</td><td>>MI0000811|hsa-mir-148b&WithFlank&12|+|54337210|54337320</td><td>69</td><td>90</td><td>22(+1U)</td><td>TCAGTGCATCACAGAACTTTGTT</td><td>4414.0</td><td>-3.3</td><td>1.1</td><td>-3.0</td><td>2.7E-03</td><td>1.3E-02</td><td>BT20/MB231</td><td>MP</td></tr>
<tr><td>>hg38_21_+_8433172_8446622_RNA45SN1_45S_50ntFlanks@3863.3898.36|ATTCTAGAGCTAATACATGCCGACGGGCGCTGACCC</td><td>>hg38_21_+_8433172_8446622_RNA45SN1_45S_50ntFlanks</td><td>3863</td><td>3898</td><td>36</td><td>ATTCTAGAGCTAATACATGCCGACGGGCGCTGACCC</td><td>262.3</td><td>-3.9</td><td>1.5</td><td>-2.6</td><td>9.5E-03</td><td>1.3E-02</td><td>BT20/MB468</td><td>MITO</td></tr>
</table>

| | | | | | | | | | | | | | |
|---|---|---|---|---|---|---|---|---|---|---|---|---|---|
| >hg38_21_+_8433172_8446622_RNA45SN1_45S_50ntFlanks@9932.9952.21\|ATAGACAGCAGGACGGTGGCC | >hg38_21_+_8433172_8446622_RNA45SN1_45S_50ntFlanks | 9932 | 9952 | 21 | ATAGACAGCAGGACGGTGGCC | 165.7 | 4.7 | 1.8 | 2.7 | 6.9E-03 | 1.3E-02 | BT20/MB468 | MP |
| >hg38_MT_+_1621_3279_RNR2_16S_50ntFlanks@1374.1400.27\|TGGATCAGGACATCCCGATGGTGCAGC | >hg38_MT_+_1621_3279_RNR2_16S_50ntFlanks | 1374 | 1400 | 27 | TGGATCAGGACATCCCGATGGTGCAGC | 22.1 | 7.7 | 3.0 | 2.6 | 1.0E-02 | 1.3E-02 | MB231/MB468 | MP |
| >hg38_21_+_8433172_8446622_RNA45SN1_45S_50ntFlanks@11872.11904.33\|AAAGAAGACCCTGTTGAGCTTGACTCTAGTCTG | >hg38_21_+_8433172_8446622_RNA45SN1_45S_50ntFlanks | 11872 | 11904 | 33 | AAAGAAGACCCTGTTGAGCTTGACTCTAGTCTG | 102.1 | -3.5 | 1.3 | -2.7 | 6.9E-03 | 1.3E-02 | BT20/MB468 | MP |
| >trnaMT_HisGTG_MT_+_12138_12206@46.71.26\|GAGGCTTACGACCCCTTATTTACCCC | >trnaMT_HisGTG_MT_+_12138_12206 | 46 | 71 | 26 | GAGGCTTACGACCCCTTATTTACCCC | 82.6 | 4.1 | 1.6 | 2.6 | 1.0E-02 | 1.3E-02 | MB231/MB468 | MP |
| >trna111_HisGTG_1_-_147774845_147774916@-1A.31.32,trna118_HisGTG_1_-_145396881_145396952@-1A.31.32,trna16_HisGTG_1_+_146544773_146544844@-1A.31.32,trna1_HisGTG_15_+_45493349_45493420@-1A.31.32,trna21_HisGTG_1_+_147753471_147753542@-1A.31.32,trna33_HisGTG_6_+_27125906_27125977@-1A.31.32,trna7_HisGTG_9_-_14433938_14434009@-1A.31.32,trna8_HisGTG_15_-_45492611_45492682@-1A.31.32,trna9_HisGTG_15_-_45490804_45490875@-1A.31.32\|AGCCGTGATCGTATAGTGGTTAGTACTCTGCG | >trna111_HisGTG_1_-_147774845_147774916 | -1A | 31 | 32 | AGCCGTGATCGTATAGTGGTTAGTACTCTGCG | 3041.0 | -2.8 | 1.1 | -2.5 | 1.1E-02 | 1.3E-02 | BT20/MB468 | CYTO |
| >hg38_MT_+_1621_3279_RNR2_16S_50ntFlanks@51.83.33\|GCTAAACCTAGCCCCAAACCCACTCCACCTTAC | >hg38_MT_+_1621_3279_RNR2_16S_50ntFlanks | 51 | 83 | 33 | GCTAAACCTAGCCCCAAACCCACTCCACCTTAC | 220.9 | 3.7 | 1.4 | 2.7 | 6.9E-03 | 1.3E-02 | BT20/MB468 | MP |
| >hg38_21_+_8433172_8446622_RNA45SN1_45S_50ntFlanks@11781.11802.22\|CGCATGAATGGATGAACGAGAT | >hg38_21_+_8433172_8446622_RNA45SN1_45S_50ntFlanks | 11781 | 11802 | 22 | CGCATGAATGGATGAACGAGAT | 85.2 | -4.5 | 1.7 | -2.7 | 6.9E-03 | 1.3E-02 | BT20/MB468 | MP |
| >hg38_21_+_8433172_8446622_RNA45SN1_45S_50ntFlanks@8112.8140.29\|GGGCGCGGGACATGTGGCGTACGGAAGAC | >hg38_21_+_8433172_8446622_RNA45SN1_45S_50ntFlanks | 8112 | 8140 | 29 | GGGCGCGGGACATGTGGCGTACGGAAGAC | 1320.5 | -3.0 | 1.1 | -2.7 | 6.9E-03 | 1.3E-02 | BT20/MB468 | MP |
| >hg38_21_+_8433172_8446622_RNA45SN1_45S_50ntFlanks@8176.8218.43\|CTTCTGATCGAGGCCCAGCCCGTGGACGGTGTGAGGCCGGTAG | >hg38_21_+_8433172_8446622_RNA45SN1_45S_50ntFlanks | 8176 | 8218 | 43 | CTTCTGATCGAGGCCCAGCCCGTGGACGGTGTGAGGCCGGTAG | 366.2 | -3.0 | 1.2 | -2.5 | 1.1E-02 | 1.3E-02 | MB231/MB468 | CYTO |
| >hg38_21_+_8433172_8446622_RNA45SN1_45S_50ntFlanks@9915.9948.34\|AAAAGGTGTTGGTTGATATAGACAGCAGGACGGT | >hg38_21_+_8433172_8446622_RNA45SN1_45S_50ntFlanks | 9915 | 9948 | 34 | AAAAGGTGTTGGTTGATATAGACAGCAGGACGGT | 164.1 | 3.7 | 1.2 | 3.0 | 2.7E-03 | 1.3E-02 | BT20/MB231 | MP |
| >hg38_21_+_8433172_8446622_RNA45SN1_45S_50ntFlanks@5444.5469.26\|CTCGGATCGGCCCCGCCGGGGTCGGC | >hg38_21_+_8433172_8446622_RNA45SN1_45S_50ntFlanks | 5444 | 5469 | 26 | CTCGGATCGGCCCCGCCGGGGTCGGC | 45.1 | -8.9 | 3.0 | -3.0 | 2.7E-03 | 1.3E-02 | BT20/MB231 | MP |
| >MI0000088\|hsa-mir-30a&WithFlank&6\|-\|71403545\|71403627@13.35.23&[MIMAT0000087&hsa-miR-30a-5p&offsets\|+1\|+2;m-15&6\|-\|71403595\|71403616&offsets\|+1\|+2]\|GTAAACATCCTCGACTGGAAGCT | >MI0000088\|hsa-mir-30a&WithFlank&6\|-\|71403545\|71403627 | 13 | 35 | 23 | GTAAACATCCTCGACTGGAAGCT | 1915.6 | -2.4 | 0.8 | -2.9 | 3.4E-03 | 1.3E-02 | BT20/MB231 | MITO |
| >trna10_AspGTC_12_-_125424193_125424264@1.31.31,trna12_AspGTC_12_-_125411891_125411962@1.31.31,trna38_AspGTC_17_-_8125556_8125627@1.31.31,trna45_AspGTC_6_+_27447453_27447524@1.31.31,trna48_AspGTC_6_+_27471523_27471594@1.31.31,trna4_AspGTC_12_+_96429799_96429870@1.31.31,trna69_AspGTC_1_-_161440205_161440276@1.31.31,trna72_AspGTC_1_-_161432824_161432895@1.31.31,trna75_AspGTC_1_-_161425414_161425485@1.31.31,trna78_AspGTC_1_-_161418033_161418104@1.31.31,trna81_AspGTC_1_-_161410615_161410686@1.31.31\|TCCTCGTTAGTATAGTGGTGAGTATCCCCGC | >trna10_AspGTC_12_-_125424193_125424264 | 1 | 31 | 31 | TCCTCGTTAGTATAGTGGTGAGTATCCCCGC | 122.0 | -2.7 | 1.1 | -2.6 | 1.0E-02 | 1.3E-02 | MB231/MB468 | MP |
| >MI0000103\|hsa-mir-101-1&WithFlank&1\|-\|65058428\|65058514@53.72.20;MI0000739\|hsa-mir-101-2&WithFlank&9\|+\|4850291\|4850381@55.74.20&[MIMAT0000099&hsa-miR-101-3p&offsets\|0\|-1;m-17&1\|-\|65058443\|65058462&offsets\|0\|0];[MIMAT0000099_1&hsa-miR-101-3p&offsets\|0\|-1;m-16&9\|+\|4850345\|4850364&offsets\|0\|0]\|TACAGTACTGTGATAACTGA | >MI0000103\|hsa-mir-101-1&WithFlank&1\|-\|65058428\|65058514 | 53 | 72 | 20 | TACAGTACTGTGATAACTGA | 594.6 | 2.3 | 0.8 | 2.9 | 3.4E-03 | 1.3E-02 | BT20/MB231 | CYTO |
| >MI0000469\|hsa-mir-125a&WithFlank&19\|+\|51693248\|51693345@21.41.21(+1A)&[MIMAT0000443&hsa-miR-125a-5p&offsets\|0\|-3(+1A);m-63&19\|+\|51693268\|51693289&offsets\|0\|-1(+1A)]\|TCCCTGAGACCCTTTAACCTGA | >MI0000469\|hsa-mir-125a&WithFlank&19\|+\|51693248\|51693345 | 21 | 41 | 21(+1A) | TCCCTGAGACCCTTTAACCTGA | 304.0 | 2.5 | 0.9 | 2.9 | 3.4E-03 | 1.3E-02 | BT20/MB231 | CYTO |
| >hg38_21_+_8433172_8446622_RNA45SN1_45S_50ntFlanks@12408.12448.41\|TTTGATCCTTCGATGTCGGCTCTTCCTATCATTGTGAAGCA | >hg38_21_+_8433172_8446622_RNA45SN1_45S_50ntFlanks | 12408 | 12448 | 41 | TTTGATCCTTCGATGTCGGCTCTTCCTATCATTGTGAAGCA | 388.4 | -3.0 | 1.0 | -2.9 | 3.4E-03 | 1.3E-02 | BT20/MB231 | CYTO |
| >hg38_21_+_8433172_8446622_RNA45SN1_45S_50ntFlanks@9533.9572.40\|GAGGAAACTCTGGTGGAGGTCCGTAGCGGTCCTGACGTGC | >hg38_21_+_8433172_8446622_RNA45SN1_45S_50ntFlanks | 9533 | 9572 | 40 | GAGGAAACTCTGGTGGAGGTCCGTAGCGGTCCTGACGTGC | 529.9 | -2.5 | 0.8 | -2.9 | 3.4E-03 | 1.3E-02 | BT20/MB231 | CYTO |
| >hg38_21_+_8433172_8446622_RNA45SN1_45S_50ntFlanks@8673.8706.34\|CGGTGCGCCGCGACCGGCTCCGGGACGGCTGGGA | >hg38_21_+_8433172_8446622_RNA45SN1_45S_50ntFlanks | 8673 | 8706 | 34 | CGGTGCGCCGCGACCGGCTCCGGGACGGCTGGGA | 221.2 | 5.1 | 1.9 | 2.7 | 6.9E-03 | 1.3E-02 | BT20/MB468 | MP |
| >hg38_21_+_8433172_8446622_RNA45SN1_45S_50ntFlanks@10393.10410.18\|CAGCAGTTGAACATGGGT | >hg38_21_+_8433172_8446622_RNA45SN1_45S_50ntFlanks | 10393 | 10410 | 18 | CAGCAGTTGAACATGGGT | 394.5 | -3.2 | 1.2 | -2.6 | 9.6E-03 | 1.3E-02 | BT20/MB468 | MITO |
| >hg38_21_+_8433172_8446622_RNA45SN1_45S_50ntFlanks@9562.9597.36\|TCCTGACGTGCAAATCGGTCGTCCGACCTGGGTATA | >hg38_21_+_8433172_8446622_RNA45SN1_45S_50ntFlanks | 9562 | 9597 | 36 | TCCTGACGTGCAAATCGGTCGTCCGACCTGGGTATA | 179.8 | -4.8 | 1.8 | -2.7 | 6.9E-03 | 1.3E-02 | BT20/MB468 | MP |
| >hg38_21_+_8433172_8446622_RNA45SN1_45S_50ntFlanks@5103.5136.34\|CTAACTAGTTACGCGACCCCCGAGCGGTCGGCGT | >hg38_21_+_8433172_8446622_RNA45SN1_45S_50ntFlanks | 5103 | 5136 | 34 | CTAACTAGTTACGCGACCCCCGAGCGGTCGGCGT | 44.6 | -8.6 | 3.2 | -2.7 | 6.9E-03 | 1.3E-02 | BT20/MB468 | MP |
| >hg38_21_+_8433172_8446622_RNA45SN1_45S_50ntFlanks@6732.6758.27\|ACATTGATCATCGACACTTCGAACGCA | >hg38_21_+_8433172_8446622_RNA45SN1_45S_50ntFlanks | 6732 | 6758 | 27 | ACATTGATCATCGACACTTCGAACGCA | 401.3 | -2.1 | 0.8 | -2.5 | 1.2E-02 | 1.3E-02 | MB231/MB468 | MITO |
| >hg38_MT_+_598_1651_RNR1_12S_50ntFlanks@428.454.27\|AAAGTGGCTTTAACATATCTGAACACA | >hg38_MT_+_598_1651_RNR1_12S_50ntFlanks | 428 | 454 | 27 | AAAGTGGCTTTAACATATCTGAACACA | 180.4 | 3.4 | 1.3 | 2.6 | 1.0E-02 | 1.3E-02 | MB231/MB468 | MP |
| >MI0000082\|hsa-mir-25&WithFlank&7\|-\|100093554\|100093649@58.76.19&[MIMAT0000081&hsa-miR-25-3p&offsets\|0\|-3;m-23&7\|-\|100093571\|100093592&offsets\|0\|-3]\|CATTGCACTTGTCTCGGTC | >MI0000082\|hsa-mir-25&WithFlank&7\|-\|100093554\|100093649 | 58 | 76 | 19 | CATTGCACTTGTCTCGGTC | 87.9 | 4.5 | 1.7 | 2.6 | 1.0E-02 | 1.3E-02 | MB231/MB468 | MP |
| >hg38_MT_+_1621_3279_RNR2_16S_50ntFlanks@1431.1450.20\|AAAGTCCTACGTGATCTGAG | >hg38_MT_+_1621_3279_RNR2_16S_50ntFlanks | 1431 | 1450 | 20 | AAAGTCCTACGTGATCTGAG | 85.8 | 2.5 | 1.0 | 2.6 | 1.0E-02 | 1.3E-02 | MB231/MB468 | MP |
| >hg38_21_+_8433172_8446622_RNA45SN1_45S_50ntFlanks@7973.7996.24\|GACGCGACCTCAGATCAGACGTGG | >hg38_21_+_8433172_8446622_RNA45SN1_45S_50ntFlanks | 7973 | 7996 | 24 | GACGCGACCTCAGATCAGACGTGG | 102.0 | -1.8 | 0.7 | -2.6 | 1.0E-02 | 1.3E-02 | MB231/MB468 | MP |
| >hg38_21_+_8433172_8446622_RNA45SN1_45S_50ntFlanks@7991.8009.19\|ACGTGGCGACCCGCTGAAT | >hg38_21_+_8433172_8446622_RNA45SN1_45S_50ntFlanks | 7991 | 8009 | 19 | ACGTGGCGACCCGCTGAAT | 1507.6 | -2.2 | 0.9 | -2.5 | 1.2E-02 | 1.3E-02 | BT20/MB468 | NUC |
| >hg38_21_+_8433172_8446622_RNA45SN1_45S_50ntFlanks@5442.5466.25\|CCCTCGGATCGGCCCCGCCGGGGTC | >hg38_21_+_8433172_8446622_RNA45SN1_45S_50ntFlanks | 5442 | 5466 | 25 | CCCTCGGATCGGCCCCGCCGGGGTC | 1045.6 | -5.2 | 2.1 | -2.5 | 1.2E-02 | 1.3E-02 | BT20/MB468 | NUC |
| >hg38_21_+_8433172_8446622_RNA45SN1_45S_50ntFlanks@10868.10889.22\|AAGGGCTGGGTCGGTCGGGCTG | >hg38_21_+_8433172_8446622_RNA45SN1_45S_50ntFlanks | 10868 | 10889 | 22 | AAGGGCTGGGTCGGTCGGGCTG | 80.9 | -2.3 | 0.9 | -2.6 | 1.0E-02 | 1.3E-02 | MB231/MB468 | MP |
| >hg38_21_+_8433172_8446622_RNA45SN1_45S_50ntFlanks@6700.6718.19\|CTAGCTGCGAGAATTAATG | >hg38_21_+_8433172_8446622_RNA45SN1_45S_50ntFlanks | 6700 | 6718 | 19 | CTAGCTGCGAGAATTAATG | 254.6 | -2.6 | 0.9 | -2.9 | 3.4E-03 | 1.3E-02 | BT20/MB231 | CYTO |
| >MI0000114\|hsa-mir-107&WithFlank&10\|-\|89592741\|89592833@56.75.20(+1A);MI0000108\|hsa-mir-103a-2&WithFlank&20\|+\|3917488\|3917577@54.73.20(+1A);MI0000109\|hsa-mir-103a-1&WithFlank&5\|-\|168560890\|168560979@54.73.20(+1A)&[MIMAT0000104&hsa-miR-107&offsets\|0\|-3(+1A);m-78&10\|-\|89592758\|89592778&offsets\|0\|-1(+1A)];[MIMAT0000101&hsa-miR-103a-3p&offsets\|0\|-3(+1A);m-12&20\|+\|3917541\|3917563&offsets\|0\|-3(+1A)];[MIMAT0000101_1&hsa-miR-103a-3p&offsets\|0\|-3(+1A);m-13&5\|-\|168560904\|168560926&offsets\|0\|-3(+1A);m-44220&5\|-\|168560908\|168560926&offsets\|0\|+1(+1A)]\|AGCAGCATTGTACAGGGCTAA | >MI0000114\|hsa-mir-107&WithFlank&10\|-\|89592741\|89592833 | 56 | 75 | 20(+1A) | AGCAGCATTGTACAGGGCTAA | 85.2 | 4.2 | 1.6 | 2.6 | 1.0E-02 | 1.3E-02 | MB231/MB468 | MP |
| >trnaMT_PheGAA_MT_+_577_647@36.62.27\|AAAATGTTTAGACGGGCTCACATCACC | >trnaMT_PheGAA_MT_+_577_647 | 36 | 62 | 27 | AAAATGTTTAGACGGGCTCACATCACC | 154.0 | 4.2 | 1.7 | 2.6 | 1.0E-02 | 1.3E-02 | MB231/MB468 | MP |
| >hg38_21_+_8433172_8446622_RNA45SN1_45S_50ntFlanks@6732.6752.21\|ACATTGATCATCGACACTTCG | >hg38_21_+_8433172_8446622_RNA45SN1_45S_50ntFlanks | 6732 | 6752 | 21 | ACATTGATCATCGACACTTCG | 708.8 | 1.7 | 0.7 | 2.6 | 9.6E-03 | 1.3E-02 | BT20/MB468 | MITO |
| >hg38_21_+_8433172_8446622_RNA45SN1_45S_50ntFlanks@9623.9652.30\|TAGTAGCTGGTTCCCTCCGAAGTTTCCCTC | >hg38_21_+_8433172_8446622_RNA45SN1_45S_50ntFlanks | 9623 | 9652 | 30 | TAGTAGCTGGTTCCCTCCGAAGTTTCCCTC | 253.2 | -4.6 | 1.7 | -2.7 | 6.9E-03 | 1.3E-02 | BT20/MB468 | MP |
| >hg38_21_+_8433172_8446622_RNA45SN1_45S_50ntFlanks@11853.11872.20\|CTTGGCGGAATCAGCGGGGA | >hg38_21_+_8433172_8446622_RNA45SN1_45S_50ntFlanks | 11853 | 11872 | 20 | CTTGGCGGAATCAGCGGGGA | 154.3 | -2.9 | 1.1 | -2.6 | 9.6E-03 | 1.3E-02 | BT20/MB468 | MITO |
| >hg38_21_+_8433172_8446622_RNA45SN1_45S_50ntFlanks@9533.9570.38\|GAGGAAACTCTGGTGGAGGTCCGTAGCGGTCCTGACGT | >hg38_21_+_8433172_8446622_RNA45SN1_45S_50ntFlanks | 9533 | 9570 | 38 | GAGGAAACTCTGGTGGAGGTCCGTAGCGGTCCTGACGT | 616.9 | -2.3 | 0.8 | -2.9 | 3.4E-03 | 1.3E-02 | BT20/MB231 | CYTO |
| >hg38_21_+_8433172_8446622_RNA45SN1_45S_50ntFlanks@11666.11703.38\|TCTGCCCAGTGCTCTGAATGTCAAAGTGAAGAAATTCA | >hg38_21_+_8433172_8446622_RNA45SN1_45S_50ntFlanks | 11666 | 11703 | 38 | TCTGCCCAGTGCTCTGAATGTCAAAGTGAAGAAATTCA | 667.6 | -2.4 | 0.8 | -2.9 | 3.4E-03 | 1.3E-02 | BT20/MB231 | CYTO |
| >hg38_21_+_8433172_8446622_RNA45SN1_45S_50ntFlanks@6734.6777.44\|ATTGATCATCGACACTTCGAACGCACTTGCGGCCCCGGGTTCCT | >hg38_21_+_8433172_8446622_RNA45SN1_45S_50ntFlanks | 6734 | 6777 | 44 | ATTGATCATCGACACTTCGAACGCACTTGCGGCCCCGGGTTCCT | 168.6 | -6.5 | 2.2 | -2.9 | 3.4E-03 | 1.3E-02 | BT20/MB231 | CYTO |
| >hg38_21_+_8433172_8446622_RNA45SN1_45S_50ntFlanks@3993.4022.30\|GTGACTCTAGATAACCTCGGGCCGATCGCA | >hg38_21_+_8433172_8446622_RNA45SN1_45S_50ntFlanks | 3993 | 4022 | 30 | GTGACTCTAGATAACCTCGGGCCGATCGCA | 158.7 | -3.2 | 1.2 | -2.6 | 1.0E-02 | 1.3E-02 | MB231/MB468 | MP |
| >hg38_MT_+_1621_3279_RNR2_16S_50ntFlanks@930.958.29\|AGTGACACATGTTTAACGGCCGCGGTACC | >hg38_MT_+_1621_3279_RNR2_16S_50ntFlanks | 930 | 958 | 29 | AGTGACACATGTTTAACGGCCGCGGTACC | 325.4 | 4.4 | 1.6 | 2.7 | 7.0E-03 | 1.3E-02 | BT20/MB468 | MP |
| >am_trna13_LysCTT_14_-_58706613_58706685@31.51.21,trna2_LysCTT_15_+_79152904_79152976@31.51.21,trna4_LysCTT_18_-_43669271_43669343@31.51.21\|ACTCTTAATCCCAGGGTCGTG | >am_trna13_LysCTT_14_-_58706613_58706685 | 31 | 51 | 21 | ACTCTTAATCCCAGGGTCGTG | 266.7 | -4.6 | 1.7 | -2.7 | 7.0E-03 | 1.3E-02 | BT20/MB468 | MP |
| >hg38_21_+_8433172_8446622_RNA45SN1_45S_50ntFlanks@10349.10371.23\|AACGAGAACTTTGAAGGCCGAAG | >hg38_21_+_8433172_8446622_RNA45SN1_45S_50ntFlanks | 10349 | 10371 | 23 | AACGAGAACTTTGAAGGCCGAAG | 1915.3 | -2.1 | 0.8 | -2.6 | 1.0E-02 | 1.3E-02 | MB231/MB468 | MP |
| >MI0000063\|hsa-let-7b&WithFlank&22\|+\|46113680\|46113774@12.34.23&[MIMAT0000063&hsa-let-7b-5p&offsets\|0\|+1;m-14&22\|+\|46113691\|46113712&offsets\|0\|+1]\|TGAGGTAGTAGGTTGTGTGGTTT | >MI0000063\|hsa-let-7b&WithFlank&22\|+\|46113680\|46113774 | 12 | 34 | 23 | TGAGGTAGTAGGTTGTGTGGTTT | 9833.3 | 2.3 | 0.8 | 2.9 | 3.4E-03 | 1.3E-02 | BT20/MB231 | CYTO |
| >hg38_21_+_8433172_8446622_RNA45SN1_45S_50ntFlanks@11768.11798.31\|CTAATTAGTGACGCGCATGAATGGATGAACG | >hg38_21_+_8433172_8446622_RNA45SN1_45S_50ntFlanks | 11768 | 11798 | 31 | CTAATTAGTGACGCGCATGAATGGATGAACG | 70.8 | -4.4 | 1.6 | -2.7 | 7.0E-03 | 1.3E-02 | BT20/MB468 | MP |

| | | | | | | | | | | | | | |
|---|---|---|---|---|---|---|---|---|---|---|---|---|---|
| >hg38_21_+_8433172_8446622_RNA45SN1_45S_50ntFlanks@10845.10865.21\|ACTTCGGGATAAGGATTGGCT | >hg38_21_+_8433172_8446622_RNA45SN1_45S_50ntFlanks | 10845 | 10865 | 21 | ACTTCGGGATAAGGATTGGCT | 2543.6 | -5.2 | 1.9 | -2.7 | 7.0E-03 | 1.3E-02 | BT20/MB468 | MP |
| >MI0000465\|hsa-mir-191&WithFlank&3\|-\|49020612\|49020715@22.41.20(+1U)&[MIMAT0000440&hsa-miR-191-5p&offsets\|0\|-3(+1U);m-48&3\|-\|49020672\|49020694&offsets\|0\|-3(+1U)]\|CAACGGAATCCCAAAAGCAGT | >MI0000465\|hsa-mir-191&WithFlank&3\|-\|49020612\|49020715 | 22 | 41 | 20(+1U) | CAACGGAATCCCAAAAGCAGT | 162.2 | 3.6 | 1.4 | 2.6 | 1.0E-02 | 1.3E-02 | MB231/MB468 | MP |
| >hg38_21_+_8433172_8446622_RNA45SN1_45S_50ntFlanks@12488.12522.35\|AACGTGAGCTGGGTTTAGACCGTCGTGAGACAGGT | >hg38_21_+_8433172_8446622_RNA45SN1_45S_50ntFlanks | 12488 | 12522 | 35 | AACGTGAGCTGGGTTTAGACCGTCGTGAGACAGGT | 184.0 | -3.8 | 1.5 | -2.5 | 1.1E-02 | 1.3E-02 | BT20/MB468 | CYTO |
| >am_trnaMT_SerGCT_MT_+_12207_12265@13.31.19\|AAGAACTGCTAACTCATGC | >am_trnaMT_SerGCT_MT_+_12207_12265 | 13 | 31 | 19 | AAGAACTGCTAACTCATGC | 242.3 | 2.1 | 0.7 | 2.9 | 3.4E-03 | 1.3E-02 | BT20/MB231 | CYTO |
| >am_MI0016746\|hsa-mir-548o-2&WithFlank&20\|+\|38516557\|38516638@51.71.21(+1U);MI0006402\|hsa-mir-548o&WithFlank&7\|-\|102405736\|102405861@93.113.21(+1U)&[MIMAT0005919&hsa-miR-548o-3p&offsets\|0\|-1(+1U);m-431&20\|+\|38516607\|38516628&offsets\|0\|-1(+1U)];[MIMAT0005919_1&hsa-miR-548o-3p&offsets\|0\|-1(+1U);m-430&7\|-\|102405748\|102405769&offsets\|0\|-1(+1U)]\|CCAAAACTGCAGTTACTTTTGT | >am_MI0016746\|hsa-mir-548o-2&WithFlank&20\|+\|38516557\|38516638 | 51 | 71 | 21(+1U) | CCAAAACTGCAGTTACTTTTGT | 432.2 | 2.3 | 0.8 | 2.9 | 3.5E-03 | 1.3E-02 | BT20/MB231 | CYTO |
| >hg38_21_+_8433172_8446622_RNA45SN1_45S_50ntFlanks@10383.10424.42\|TCCATGTGAACAGCAGTTGAACATGGGTCAGTCGGTCCTGAG | >hg38_21_+_8433172_8446622_RNA45SN1_45S_50ntFlanks | 10383 | 10424 | 42 | TCCATGTGAACAGCAGTTGAACATGGGTCAGTCGGTCCTGAG | 105.9 | 4.1 | 1.6 | 2.6 | 1.0E-02 | 1.3E-02 | MB231/MB468 | MP |
| >MI0000082\|hsa-mir-25&WithFlank&7\|-\|100093554\|100093649@58.79.22(+1U)&[MIMAT0000081&hsa-miR-25-3p&offsets\|0\|0(+1U);m-23&7\|-\|100093571\|100093592&offsets\|0\|0(+1U)]\|CATTGCACTTGTCTCGGTCTGAT | >MI0000082\|hsa-mir-25&WithFlank&7\|-\|100093554\|100093649 | 58 | 79 | 22(+1U) | CATTGCACTTGTCTCGGTCTGAT | 148.4 | 2.3 | 0.9 | 2.7 | 7.0E-03 | 1.3E-02 | BT20/MB468 | MP |
| >hg38_21_+_8433172_8446622_RNA45SN1_45S_50ntFlanks@9930.9952.23\|ATATAGACAGCAGGACGGTGGCC | >hg38_21_+_8433172_8446622_RNA45SN1_45S_50ntFlanks | 9930 | 9952 | 23 | ATATAGACAGCAGGACGGTGGCC | 211.1 | 5.8 | 1.9 | 3.0 | 2.7E-03 | 1.3E-02 | BT20/MB231 | MP |
| >hg38_21_+_8433172_8446622_RNA45SN1_45S_50ntFlanks@11670.11687.18\|CCCAGTGCTCTGAATGTC | >hg38_21_+_8433172_8446622_RNA45SN1_45S_50ntFlanks | 11670 | 11687 | 18 | CCCAGTGCTCTGAATGTC | 1515.4 | -1.9 | 0.8 | -2.5 | 1.1E-02 | 1.3E-02 | MB231/MB468 | CYTO |
| >MI0000088\|hsa-mir-30a&WithFlank&6\|-\|71403545\|71403627@55.74.20&[MIMAT0000088&hsa-miR-30a-3p&offsets\|+2\|0;m-38&6\|-\|71403554\|71403575&offsets\|+2\|0]\|TTCAGTCGGATGTTTGCAGC | >MI0000088\|hsa-mir-30a&WithFlank&6\|-\|71403545\|71403627 | 55 | 74 | 20 | TTCAGTCGGATGTTTGCAGC | 53.6 | 3.2 | 1.3 | 2.6 | 1.0E-02 | 1.3E-02 | MB231/MB468 | MP |
| >hg38_21_+_8433172_8446622_RNA45SN1_45S_50ntFlanks@10781.10826.46\|AGGTGAACAGCCTCTGGCATGTTGGAACAATGTAGGTAAGGGAAGT | >hg38_21_+_8433172_8446622_RNA45SN1_45S_50ntFlanks | 10781 | 10826 | 46 | AGGTGAACAGCCTCTGGCATGTTGGAACAATGTAGGTAAGGGAAGT | 142.5 | -4.2 | 1.6 | -2.6 | 9.7E-03 | 1.3E-02 | BT20/MB468 | MITO |
| >hg38_21_+_8433172_8446622_RNA45SN1_45S_50ntFlanks@11570.11602.33\|AGCCGACTTAGAACTGGTGCGGACCAGGGGAAT | >hg38_21_+_8433172_8446622_RNA45SN1_45S_50ntFlanks | 11570 | 11602 | 33 | AGCCGACTTAGAACTGGTGCGGACCAGGGGAAT | 529.9 | 3.9 | 1.3 | 3.0 | 2.7E-03 | 1.3E-02 | BT20/MB231 | MP |
| >trnaMT_ProTGG_MT_-_15956_16023@1.32.32\|CAGAGAATAGTTTAAATTAGAATCTTAGCTTT | >trnaMT_ProTGG_MT_-_15956_16023 | 1 | 32 | 32 | CAGAGAATAGTTTAAATTAGAATCTTAGCTTT | 254.1 | 2.6 | 1.0 | 2.6 | 1.0E-02 | 1.3E-02 | MB231/MB468 | MP |
| >hg38_21_+_8433172_8446622_RNA45SN1_45S_50ntFlanks@12607.12624.18\|TGTGCTTGGCTGAGGAGC | >hg38_21_+_8433172_8446622_RNA45SN1_45S_50ntFlanks | 12607 | 12624 | 18 | TGTGCTTGGCTGAGGAGC | 281.1 | 2.9 | 1.1 | 2.7 | 7.0E-03 | 1.3E-02 | BT20/MB468 | MP |
| >trna133_ValCAC_6_-_27696327_27696399@1.23.23\|GTTTCCGTAGTGTAGTGGTTATT | >trna133_ValCAC_6_-_27696327_27696399 | 1 | 23 | 23 | GTTTCCGTAGTGTAGTGGTTATT | 215.9 | 5.8 | 2.2 | 2.7 | 7.0E-03 | 1.3E-02 | BT20/MB468 | MP |
| >hg38_21_+_8433172_8446622_RNA45SN1_45S_50ntFlanks@5471.5491.21\|CACGGCCCTGGCGGAGCGCTG | >hg38_21_+_8433172_8446622_RNA45SN1_45S_50ntFlanks | 5471 | 5491 | 21 | CACGGCCCTGGCGGAGCGCTG | 134.5 | -3.6 | 1.3 | -2.7 | 7.0E-03 | 1.3E-02 | BT20/MB468 | MP |
| >hg38_21_+_8433172_8446622_RNA45SN1_45S_50ntFlanks@4367.4396.30\|CAGTTAAAAAGCTCGTAGTTGGATCTTGGG | >hg38_21_+_8433172_8446622_RNA45SN1_45S_50ntFlanks | 4367 | 4396 | 30 | CAGTTAAAAAGCTCGTAGTTGGATCTTGGG | 114.0 | -4.5 | 1.7 | -2.7 | 7.0E-03 | 1.3E-02 | BT20/MB468 | MP |
| >hg38_21_+_8433172_8446622_RNA45SN1_45S_50ntFlanks@5472.5499.28\|ACGGCCCTGGCGGAGCGCTGAGAAGACG | >hg38_21_+_8433172_8446622_RNA45SN1_45S_50ntFlanks | 5472 | 5499 | 28 | ACGGCCCTGGCGGAGCGCTGAGAAGACG | 2255.7 | -2.5 | 1.0 | -2.5 | 1.2E-02 | 1.3E-02 | MB231/MB468 | MITO |
| >am_trnaMT_SerGCT_MT_+_12207_12265@12.32.21\|CAAGAACTGCTAACTCATGCC | >am_trnaMT_SerGCT_MT_+_12207_12265 | 12 | 32 | 21 | CAAGAACTGCTAACTCATGCC | 142.1 | 3.3 | 1.2 | 2.7 | 7.0E-03 | 1.3E-02 | BT20/MB468 | MP |
| >hg38_21_+_8433172_8446622_RNA45SN1_45S_50ntFlanks@12078.12095.18\|GGGCTCTCGCTTCTGGCG | >hg38_21_+_8433172_8446622_RNA45SN1_45S_50ntFlanks | 12078 | 12095 | 18 | GGGCTCTCGCTTCTGGCG | 83.1 | -4.5 | 1.7 | -2.7 | 7.0E-03 | 1.3E-02 | BT20/MB468 | MP |
| >hg38_21_+_8433172_8446622_RNA45SN1_45S_50ntFlanks@8401.8422.22\|AGAGGTAAACGGGTGGGGTCCG | >hg38_21_+_8433172_8446622_RNA45SN1_45S_50ntFlanks | 8401 | 8422 | 22 | AGAGGTAAACGGGTGGGGTCCG | 87.4 | -4.4 | 1.6 | -2.7 | 7.0E-03 | 1.3E-02 | BT20/MB468 | MP |
| >hg38_21_+_8433172_8446622_RNA45SN1_45S_50ntFlanks@3787.3805.19\|AAACTGCGAATGGCTCATT | >hg38_21_+_8433172_8446622_RNA45SN1_45S_50ntFlanks | 3787 | 3805 | 19 | AAACTGCGAATGGCTCATT | 348.2 | -2.4 | 1.0 | -2.5 | 1.2E-02 | 1.3E-02 | BT20/MB468 | NUC |
| >hg38_1_-_228634819_228635039_RNA5S12_5S_50ntFlanks@135.171.37\|GGGAGACCGCCTGGGAATACCGGGTGCTGTAGGCTTT | >hg38_1_-_228634819_228635039_RNA5S12_5S_50ntFlanks | 135 | 171 | 37 | GGGAGACCGCCTGGGAATACCGGGTGCTGTAGGCTTT | 2076.0 | 1.5 | 0.6 | 2.5 | 1.2E-02 | 1.3E-02 | BT20/MB468 | NUC |
| >hg38_MT_+_1621_3279_RNR2_16S_50ntFlanks@829.859.31\|GAAAGGTTAAAAAAAGTAAAAGGAACTCGGC | >hg38_MT_+_1621_3279_RNR2_16S_50ntFlanks | 829 | 859 | 31 | GAAAGGTTAAAAAAAGTAAAAGGAACTCGGC | 34.2 | 5.0 | 2.0 | 2.6 | 1.0E-02 | 1.3E-02 | MB231/MB468 | MP |
| >hg38_21_+_8433172_8446622_RNA45SN1_45S_50ntFlanks@12419.12455.37\|GATGTCGGCTCTTCCTATCATTGTGAAGCAGAATTCA | >hg38_21_+_8433172_8446622_RNA45SN1_45S_50ntFlanks | 12419 | 12455 | 37 | GATGTCGGCTCTTCCTATCATTGTGAAGCAGAATTCA | 769.7 | -2.6 | 1.1 | -2.5 | 1.2E-02 | 1.3E-02 | MB231/MB468 | MITO |
| >hg38_21_+_8433172_8446622_RNA45SN1_45S_50ntFlanks@11539.11586.48\|GCGCGGCGCCTCGCCTCGGCCGGCGCCTAGCAGCCGACTTAGAACTGG | >hg38_21_+_8433172_8446622_RNA45SN1_45S_50ntFlanks | 11539 | 11586 | 48 | GCGCGGCGCCTCGCCTCGGCCGGCGCCTAGCAGCCGACTTAGAACTGG | 106.0 | 4.8 | 1.9 | 2.5 | 1.2E-02 | 1.3E-02 | MB231/MB468 | MITO |
| >hg38_21_+_8433172_8446622_RNA45SN1_45S_50ntFlanks@12411.12436.26\|GATCCTTCGATGTCGGCTCTTCCTAT | >hg38_21_+_8433172_8446622_RNA45SN1_45S_50ntFlanks | 12411 | 12436 | 26 | GATCCTTCGATGTCGGCTCTTCCTAT | 436.2 | -4.3 | 1.7 | -2.5 | 1.2E-02 | 1.3E-02 | MB231/MB468 | NUC |
| >hg38_21_+_8433172_8446622_RNA45SN1_45S_50ntFlanks@9790.9820.31\|CTGGCGTGGAGCCGGGCGTGGAATGCGAGTG | >hg38_21_+_8433172_8446622_RNA45SN1_45S_50ntFlanks | 9790 | 9820 | 31 | CTGGCGTGGAGCCGGGCGTGGAATGCGAGTG | 167.2 | -3.3 | 1.3 | -2.5 | 1.2E-02 | 1.3E-02 | MB231/MB468 | NUC |
| >trna10_AspGTC_12_-_125424193_125424264@32.71.40,trna12_AspGTC_12_-_125411891_125411962@32.71.40,trna144_AspGTC_6_-_27551236_27551307@32.71.40,trna38_AspGTC_17_-_8125556_8125627@32.71.40,trna45_AspGTC_6_+_27447453_27447524@32.71.40,trna48_AspGTC_6_+_27471523_27471594@32.71.40,trna4_AspGTC_12_+_96429799_96429870@32.71.40,trna69_AspGTC_1_-_161440205_161440276@32.71.40,trna72_AspGTC_1_-_161432824_161432895@32.71.40,trna75_AspGTC_1_-_161425414_161425485@32.71.40,trna78_AspGTC_1_-_161418033_161418104@32.71.40,trna81_AspGTC_1_-_161410615_161410686@32.71.40\|CTGTCACGCGGGAGACCGGGGTTCGATTCCCCGACGGGGA | >trna10_AspGTC_12_-_125424193_125424264 | 32 | 71 | 40 | CTGTCACGCGGGAGACCGGGGTTCGATTCCCCGACGGGGA | 106.8 | -4.7 | 1.7 | -2.7 | 7.0E-03 | 1.3E-02 | BT20/MB468 | MP |
| >hg38_21_+_8433172_8446622_RNA45SN1_45S_50ntFlanks@10343.10361.19\|TATTCAAACGAGAACTTTG | >hg38_21_+_8433172_8446622_RNA45SN1_45S_50ntFlanks | 10343 | 10361 | 19 | TATTCAAACGAGAACTTTG | 256.1 | -3.4 | 1.3 | -2.7 | 7.0E-03 | 1.3E-02 | BT20/MB468 | MP |
| >trnaMT_ValTAC_MT_+_1602_1670@27.71.45\|CAACTTACACTTAGGAGATTTCAACTTAACTTGACCGCTCTGACC | >trnaMT_ValTAC_MT_+_1602_1670 | 27 | 71 | 45 | CAACTTACACTTAGGAGATTTCAACTTAACTTGACCGCTCTGACC | 188.1 | 5.0 | 2.0 | 2.5 | 1.2E-02 | 1.3E-02 | MB231/MB468 | MITO |
| >hg38_21_+_8433172_8446622_RNA45SN1_45S_50ntFlanks@12007.12031.25\|GCCGCCGGTGAAATACCACTACTCT | >hg38_21_+_8433172_8446622_RNA45SN1_45S_50ntFlanks | 12007 | 12031 | 25 | GCCGCCGGTGAAATACCACTACTCT | 3660.3 | -2.2 | 0.9 | -2.6 | 9.8E-03 | 1.3E-02 | BT20/MB468 | MITO |
| >MI0000101\|hsa-mir-99a&WithFlank&21\|+\|16539083\|16539175@19.38.20(+1C)&[MIMAT0000097&hsa-miR-99a-5p&offsets\|0\|-2(+1C);m-64&21\|+\|16539101\|16539121&offsets\|0\|-1(+1C)]\|AACCCGTAGATCCGATCTTGC | >MI0000101\|hsa-mir-99a&WithFlank&21\|+\|16539083\|16539175 | 19 | 38 | 20(+1C) | AACCCGTAGATCCGATCTTGC | 128.5 | -1.6 | 0.6 | -2.5 | 1.1E-02 | 1.3E-02 | BT20/MB468 | CYTO |
| >MI0000088\|hsa-mir-30a&WithFlank&6\|-\|71403545\|71403627@54.73.20(+1U)&[MIMAT0000088&hsa-miR-30a-3p&offsets\|+1\|-1(+1U);m-38&6\|-\|71403554\|71403575&offsets\|+1\|-1(+1U)]\|TTTCAGTCGGATGTTTGCAGT | >MI0000088\|hsa-mir-30a&WithFlank&6\|-\|71403545\|71403627 | 54 | 73 | 20(+1U) | TTTCAGTCGGATGTTTGCAGT | 512.1 | 2.6 | 1.0 | 2.7 | 7.0E-03 | 1.3E-02 | BT20/MB468 | MP |
| >hg38_1_-_228634819_228635039_RNA5S12_5S_50ntFlanks@150.172.23\|AATACCGGGTGCTGTAGGCTTTT | >hg38_1_-_228634819_228635039_RNA5S12_5S_50ntFlanks | 150 | 172 | 23 | AATACCGGGTGCTGTAGGCTTTT | 201.9 | -5.0 | 1.9 | -2.6 | 1.0E-02 | 1.3E-02 | MB231/MB468 | MP |
| >hg38_21_+_8433172_8446622_RNA45SN1_45S_50ntFlanks@7974.8012.39\|ACGCGACCTCAGATCAGACGTGGCGACCCGCTGAATTTA | >hg38_21_+_8433172_8446622_RNA45SN1_45S_50ntFlanks | 7974 | 8012 | 39 | ACGCGACCTCAGATCAGACGTGGCGACCCGCTGAATTTA | 7511.4 | 1.3 | 0.5 | 2.6 | 9.8E-03 | 1.3E-02 | BT20/MB468 | MITO |
| >hg38_21_+_8433172_8446622_RNA45SN1_45S_50ntFlanks@9567.9586.20\|ACGTGCAAATCGGTCGTCCG | >hg38_21_+_8433172_8446622_RNA45SN1_45S_50ntFlanks | 9567 | 9586 | 20 | ACGTGCAAATCGGTCGTCCG | 154.7 | -1.4 | 0.6 | -2.6 | 9.8E-03 | 1.3E-02 | BT20/MB468 | MITO |
| >hg38_21_+_8433172_8446622_RNA45SN1_45S_50ntFlanks@4734.4751.18\|AACGAAAGTCGGAGGTTC | >hg38_21_+_8433172_8446622_RNA45SN1_45S_50ntFlanks | 4734 | 4751 | 18 | AACGAAAGTCGGAGGTTC | 1802.4 | 4.9 | 1.6 | 3.0 | 2.7E-03 | 1.3E-02 | BT20/MB231 | MP |
| >hg38_21_+_8433172_8446622_RNA45SN1_45S_50ntFlanks@12324.12351.28\|TTGGGTTTTAAGCAGGAGGTGTCAGAAA | >hg38_21_+_8433172_8446622_RNA45SN1_45S_50ntFlanks | 12324 | 12351 | 28 | TTGGGTTTTAAGCAGGAGGTGTCAGAAA | 58.3 | -4.4 | 1.6 | -2.7 | 7.0E-03 | 1.3E-02 | BT20/MB468 | MP |
| >hg38_21_+_8433172_8446622_RNA45SN1_45S_50ntFlanks@10236.10273.38\|GACGCTACGCCGCGACGAGTAGGAGGGCCGCTGCGGTG | >hg38_21_+_8433172_8446622_RNA45SN1_45S_50ntFlanks | 10236 | 10273 | 38 | GACGCTACGCCGCGACGAGTAGGAGGGCCGCTGCGGTG | 341.3 | 2.9 | 1.1 | 2.7 | 7.0E-03 | 1.3E-02 | BT20/MB468 | MP |
| >hg38_21_+_8433172_8446622_RNA45SN1_45S_50ntFlanks@10802.10828.27\|TTGGAACAATGTAGGTAAGGGAAGTCG | >hg38_21_+_8433172_8446622_RNA45SN1_45S_50ntFlanks | 10802 | 10828 | 27 | TTGGAACAATGTAGGTAAGGGAAGTCG | 134.6 | -4.4 | 1.6 | -2.7 | 7.0E-03 | 1.3E-02 | BT20/MB468 | MP |
| >hg38_21_+_8433172_8446622_RNA45SN1_45S_50ntFlanks@12997.13016.20\|CGCTGCGATCTATTGAAAGT | >hg38_21_+_8433172_8446622_RNA45SN1_45S_50ntFlanks | 12997 | 13016 | 20 | CGCTGCGATCTATTGAAAGT | 116.7 | 4.4 | 1.7 | 2.6 | 1.0E-02 | 1.3E-02 | MB231/MB468 | MP |
| >hg38_21_+_8433172_8446622_RNA45SN1_45S_50ntFlanks@12628.12661.34\|TGGGGCGAAGCTACCATCTGTGGGATTATGACTG | >hg38_21_+_8433172_8446622_RNA45SN1_45S_50ntFlanks | 12628 | 12661 | 34 | TGGGGCGAAGCTACCATCTGTGGGATTATGACTG | 190.5 | -2.8 | 1.1 | -2.5 | 1.1E-02 | 1.3E-02 | BT20/MB468 | CYTO |
| >hg38_21_+_8433172_8446622_RNA45SN1_45S_50ntFlanks@9940.9979.40\|CAGGACGGTGGCCATGGAAGTCGGAATCCGCTAAGGAGTG | >hg38_21_+_8433172_8446622_RNA45SN1_45S_50ntFlanks | 9940 | 9979 | 40 | CAGGACGGTGGCCATGGAAGTCGGAATCCGCTAAGGAGTG | 109.6 | 5.4 | 2.0 | 2.7 | 7.1E-03 | 1.4E-02 | BT20/MB468 | MP |
| >hg38_21_+_8433172_8446622_RNA45SN1_45S_50ntFlanks@12244.12272.29\|AAGGGCAAAAGCTCGCTTGATCTTGATTT | >hg38_21_+_8433172_8446622_RNA45SN1_45S_50ntFlanks | 12244 | 12272 | 29 | AAGGGCAAAAGCTCGCTTGATCTTGATTT | 314.1 | -1.7 | 0.7 | -2.5 | 1.2E-02 | 1.4E-02 | MB231/MB468 | NUC |
| >hg38_21_+_8433172_8446622_RNA45SN1_45S_50ntFlanks@12444.12484.41\|AAGCAGAATTCACCAAGCGTTGGATTGTTCACCCACTAATA | >hg38_21_+_8433172_8446622_RNA45SN1_45S_50ntFlanks | 12444 | 12484 | 41 | AAGCAGAATTCACCAAGCGTTGGATTGTTCACCCACTAATA | 317.4 | -2.2 | 0.8 | -2.5 | 1.1E-02 | 1.4E-02 | MB231/MB468 | CYTO |
| >hg38_21_+_8433172_8446622_RNA45SN1_45S_50ntFlanks@11669.11687.19\|GCCCAGTGCTCTGAATGTC | >hg38_21_+_8433172_8446622_RNA45SN1_45S_50ntFlanks | 11669 | 11687 | 19 | GCCCAGTGCTCTGAATGTC | 323.4 | 4.0 | 1.5 | 2.7 | 7.1E-03 | 1.4E-02 | BT20/MB468 | MP |
| >hg38_21_+_8433172_8446622_RNA45SN1_45S_50ntFlanks@11595.11612.18\|AGGGGAATCCGACTGTTT | >hg38_21_+_8433172_8446622_RNA45SN1_45S_50ntFlanks | 11595 | 11612 | 18 | AGGGGAATCCGACTGTTT | 594.8 | -2.6 | 0.9 | -2.9 | 3.5E-03 | 1.4E-02 | BT20/MB231 | CYTO |

<table>
<tr><td>>am_trna119_LysCTT_1_-_145395522_145395594@14.33.20,trna11_LysCTT_5_-_180648979_180649051@14.33.20,trna128_LysCTT_1_-_55423542_55423614@14.33.20,trna13_LysCTT_6_+_26556774_26556846@14.33.20,trna6_LysCTT_19_-_52425396_52425468@14.33.20,trna7_LysCTT_16_+_3225692_3225764@14.33.20,trna9_LysCTT_5_+_180634755_180634827@14.33.20|AGTCGGTAGAGCATGAGACT</td><td>>am_trna119_LysCTT_1_-_145395522_145395594</td><td>14</td><td>33</td><td>20</td><td>AGTCGGTAGAGCATGAGACT</td><td>194.9</td><td>-2.9</td><td>1.0</td><td>-3.0</td><td>2.7E-03</td><td>1.4E-02</td><td>BT20/MB231</td><td>MP</td></tr>
<tr><td>>hg38_21_+_8433172_8446622_RNA45SN1_45S_50ntFlanks@11667.11692.26|CTGCCCAGTGCTCTGAATGTCAAAGT</td><td>>hg38_21_+_8433172_8446622_RNA45SN1_45S_50ntFlanks</td><td>11667</td><td>11692</td><td>26</td><td>CTGCCCAGTGCTCTGAATGTCAAAGT</td><td>386.1</td><td>4.3</td><td>1.5</td><td>3.0</td><td>2.7E-03</td><td>1.4E-02</td><td>BT20/MB231</td><td>MP</td></tr>
<tr><td>>hg38_21_+_8433172_8446622_RNA45SN1_45S_50ntFlanks@10789.10816.28|AGCCTCTGGCATGTTGGAACAATGTAGG</td><td>>hg38_21_+_8433172_8446622_RNA45SN1_45S_50ntFlanks</td><td>10789</td><td>10816</td><td>28</td><td>AGCCTCTGGCATGTTGGAACAATGTAGG</td><td>167.6</td><td>-1.8</td><td>0.7</td><td>-2.5</td><td>1.2E-02</td><td>1.4E-02</td><td>MB231/MB468</td><td>NUC</td></tr>
<tr><td>>trnaMT_SerGCT_MT_+_12207_12265@1.26.26|GAGAAAGCTCACAAGAACTGCTAACT</td><td>>trnaMT_SerGCT_MT_+_12207_12265</td><td>1</td><td>26</td><td>26</td><td>GAGAAAGCTCACAAGAACTGCTAACT</td><td>569.2</td><td>2.6</td><td>1.0</td><td>2.5</td><td>1.1E-02</td><td>1.4E-02</td><td>MB231/MB468</td><td>CYTO</td></tr>
<tr><td>>hg38_21_+_8433172_8446622_RNA45SN1_45S_50ntFlanks@12002.12035.34|TGCGGGCCGCCGGTGAAATACCACTACTCTGATC</td><td>>hg38_21_+_8433172_8446622_RNA45SN1_45S_50ntFlanks</td><td>12002</td><td>12035</td><td>34</td><td>TGCGGGCCGCCGGTGAAATACCACTACTCTGATC</td><td>100.6</td><td>-2.5</td><td>1.0</td><td>-2.6</td><td>1.1E-02</td><td>1.4E-02</td><td>MB231/MB468</td><td>MP</td></tr>
<tr><td>>trna11_GluTTC_15_-_26327381_26327452@1.31.31,trna3_GluTTC_13_-_45492062_45492133@1.31.31|TCCCACATGGTCTAGCGGTTAGGATTCCTGG</td><td>>trna11_GluTTC_15_-_26327381_26327452</td><td>1</td><td>31</td><td>31</td><td>TCCCACATGGTCTAGCGGTTAGGATTCCTGG</td><td>612.8</td><td>-3.3</td><td>1.1</td><td>-2.9</td><td>3.5E-03</td><td>1.4E-02</td><td>BT20/MB231</td><td>MITO</td></tr>
<tr><td>>hg38_21_+_8433172_8446622_RNA45SN1_45S_50ntFlanks@6661.6683.23|CGGTGGATCACTCGGCTCGTGCG</td><td>>hg38_21_+_8433172_8446622_RNA45SN1_45S_50ntFlanks</td><td>6661</td><td>6683</td><td>23</td><td>CGGTGGATCACTCGGCTCGTGCG</td><td>195.3</td><td>-1.6</td><td>0.6</td><td>-2.6</td><td>1.1E-02</td><td>1.4E-02</td><td>MB231/MB468</td><td>MP</td></tr>
<tr><td>>hg38_21_+_8433172_8446622_RNA45SN1_45S_50ntFlanks@10392.10424.33|ACAGCAGTTGAACATGGGTCAGTCGGTCCTGAG</td><td>>hg38_21_+_8433172_8446622_RNA45SN1_45S_50ntFlanks</td><td>10392</td><td>10424</td><td>33</td><td>ACAGCAGTTGAACATGGGTCAGTCGGTCCTGAG</td><td>4143.6</td><td>-1.8</td><td>0.7</td><td>-2.5</td><td>1.2E-02</td><td>1.4E-02</td><td>MB231/MB468</td><td>NUC</td></tr>
<tr><td>>hg38_21_+_8433172_8446622_RNA45SN1_45S_50ntFlanks@7990.8009.20|GACGTGGCGACCCGCTGAAT</td><td>>hg38_21_+_8433172_8446622_RNA45SN1_45S_50ntFlanks</td><td>7990</td><td>8009</td><td>20</td><td>GACGTGGCGACCCGCTGAAT</td><td>916.5</td><td>1.8</td><td>0.7</td><td>2.6</td><td>9.8E-03</td><td>1.4E-02</td><td>BT20/MB468</td><td>MITO</td></tr>
<tr><td>>MI0000102|hsa-mir-100&WithFlank&11|-|122152223|122152314@20.39.20&[MIMAT0000098&hsa-miR-100-5p&offsets|+1|-1;m-27&11|-|122152276|122152296&offsets|+1|0]|ACCCGTAGATCCGAACTTGT</td><td>>MI0000102|hsa-mir-100&WithFlank&11|-|122152223|122152314</td><td>20</td><td>39</td><td>20</td><td>ACCCGTAGATCCGAACTTGT</td><td>324.7</td><td>-5.2</td><td>1.8</td><td>-2.9</td><td>3.5E-03</td><td>1.4E-02</td><td>BT20/MB231</td><td>CYTO</td></tr>
<tr><td>>am_MI0015825|hsa-mir-378c&WithFlank&10|-|130962582|130962674@17.34.18;MI0016749|hsa-mir-378d-1&WithFlank&4|-|5923269|5923334@37.54.18;MI0000786|hsa-mir-378a&WithFlank&5|+|149732819|149732896@49.66.18;MI0003840|hsa-mir-378d-2&WithFlank&8|-|93916016|93916125@28.45.18&[MIMAT0016847&hsa-miR-378c&offsets|0|-7;m-130&10|-|130962639|130962658&offsets|0|-2];[MIMAT0018926&hsa-miR-378d&offsets|0|-2;m-228&4|-|5923281|5923298&offsets|0|0];[MIMAT0000732&hsa-miR-378a-3p&offsets|0|-4;m-44&5|+|149732867|149732888&offsets|0|-4];[MIMAT0018926_1&hsa-miR-378d&offsets|0|-2;m-227&8|-|93916081|93916098&offsets|0|0]|ACTGGACTTGGAGTCAGA</td><td>>am_MI0015825|hsa-mir-378c&WithFlank&10|-|130962582|130962674</td><td>17</td><td>34</td><td>18</td><td>ACTGGACTTGGAGTCAGA</td><td>1423.7</td><td>3.3</td><td>1.2</td><td>2.7</td><td>7.1E-03</td><td>1.4E-02</td><td>BT20/MB468</td><td>MP</td></tr>
<tr><td>>hg38_21_+_8433172_8446622_RNA45SN1_45S_50ntFlanks@11188.11206.19|CACCCGGGGGGCCGGCGGC</td><td>>hg38_21_+_8433172_8446622_RNA45SN1_45S_50ntFlanks</td><td>11188</td><td>11206</td><td>19</td><td>CACCCGGGGGGCCGGCGGC</td><td>81.2</td><td>-4.7</td><td>1.7</td><td>-2.7</td><td>7.1E-03</td><td>1.4E-02</td><td>BT20/MB468</td><td>MP</td></tr>
<tr><td>>hg38_MT_+_598_1651_RNR1_12S_50ntFlanks@905.929.25|AGTATACTTCAAAGGACATTTAACT</td><td>>hg38_MT_+_598_1651_RNR1_12S_50ntFlanks</td><td>905</td><td>929</td><td>25</td><td>AGTATACTTCAAAGGACATTTAACT</td><td>5382.1</td><td>-3.7</td><td>1.2</td><td>-3.0</td><td>2.8E-03</td><td>1.4E-02</td><td>BT20/MB231</td><td>MP</td></tr>
<tr><td>>hg38_21_+_8433172_8446622_RNA45SN1_45S_50ntFlanks@13000.13039.40|TGCGATCTATTGAAAGTCAGCCCTCGACACAAGGGTTTGT</td><td>>hg38_21_+_8433172_8446622_RNA45SN1_45S_50ntFlanks</td><td>13000</td><td>13039</td><td>40</td><td>TGCGATCTATTGAAAGTCAGCCCTCGACACAAGGGTTTGT</td><td>1768.2</td><td>2.8</td><td>1.1</td><td>2.5</td><td>1.2E-02</td><td>1.4E-02</td><td>MB231/MB468</td><td>MITO</td></tr>
<tr><td>>trnaMT_LeuTAG_MT_+_12266_12336@54.74.21|CAACTCCAAATAAAAGTACCA</td><td>>trnaMT_LeuTAG_MT_+_12266_12336</td><td>54</td><td>74</td><td>21</td><td>CAACTCCAAATAAAAGTACCA</td><td>139.3</td><td>-4.1</td><td>1.6</td><td>-2.6</td><td>9.8E-03</td><td>1.4E-02</td><td>BT20/MB468</td><td>MITO</td></tr>
<tr><td>>am_MI0000786|hsa-mir-378a&WithFlank&5|+|149732819|149732896@49.70.22(+1A)&[MIMAT0000732&hsa-miR-378a-3p&offsets|0|0(+1A);m-44&5|+|149732867|149732888&offsets|0|0(+1A)]|ACTGGACTTGGAGTCAGAAGGCA</td><td>>am_MI0000786|hsa-mir-378a&WithFlank&5|+|149732819|149732896</td><td>49</td><td>70</td><td>22(+1A)</td><td>ACTGGACTTGGAGTCAGAAGGCA</td><td>3309.1</td><td>2.3</td><td>0.8</td><td>2.9</td><td>3.5E-03</td><td>1.4E-02</td><td>BT20/MB231</td><td>CYTO</td></tr>
<tr><td>>trna10_AspGTC_12_-_125424193_125424264@39.73.35,trna12_AspGTC_12_-_125411891_125411962@39.73.35,trna144_AspGTC_6_-_27551236_27551307@39.73.35,trna38_AspGTC_17_-_8125556_8125627@39.73.35,trna45_AspGTC_6_+_27447453_27447524@39.73.35,trna48_AspGTC_6_+_27471523_27471594@39.73.35,trna4_AspGTC_12_+_96429799_96429870@39.73.35,trna69_AspGTC_1_-_161440205_161440276@39.73.35,trna72_AspGTC_1_-_161432824_161432895@39.73.35,trna75_AspGTC_1_-_161425414_161425485@39.73.35,trna78_AspGTC_1_-_161418033_161418104@39.73.35,trna81_AspGTC_1_-_161410615_161410686@39.73.35|GCGGGAGACCGGGGTTCGATTCCCCGACGGGGAGC</td><td>>trna10_AspGTC_12_-_125424193_125424264</td><td>39</td><td>73</td><td>35</td><td>GCGGGAGACCGGGGTTCGATTCCCCGACGGGGAGC</td><td>2798.7</td><td>-2.7</td><td>0.9</td><td>-2.9</td><td>3.5E-03</td><td>1.4E-02</td><td>BT20/MB231</td><td>CYTO</td></tr>
<tr><td>>trna10_AspGTC_12_-_125424193_125424264@38.72.35,trna12_AspGTC_12_-_125411891_125411962@38.72.35,trna144_AspGTC_6_-_27551236_27551307@38.72.35,trna38_AspGTC_17_-_8125556_8125627@38.72.35,trna45_AspGTC_6_+_27447453_27447524@38.72.35,trna48_AspGTC_6_+_27471523_27471594@38.72.35,trna4_AspGTC_12_+_96429799_96429870@38.72.35,trna69_AspGTC_1_-_161440205_161440276@38.72.35,trna72_AspGTC_1_-_161432824_161432895@38.72.35,trna75_AspGTC_1_-_161425414_161425485@38.72.35,trna78_AspGTC_1_-_161418033_161418104@38.72.35,trna81_AspGTC_1_-_161410615_161410686@38.72.35|CGCGGGAGACCGGGGTTCGATTCCCCGACGGGGAG</td><td>>trna10_AspGTC_12_-_125424193_125424264</td><td>38</td><td>72</td><td>35</td><td>CGCGGGAGACCGGGGTTCGATTCCCCGACGGGGAG</td><td>412.2</td><td>-2.6</td><td>0.9</td><td>-2.9</td><td>3.5E-03</td><td>1.4E-02</td><td>BT20/MB231</td><td>CYTO</td></tr>
<tr><td>>trnaMT_ValTAC_MT_+_1602_1670@53.70.18|TAACTTGACCGCTCTGAC</td><td>>trnaMT_ValTAC_MT_+_1602_1670</td><td>53</td><td>70</td><td>18</td><td>TAACTTGACCGCTCTGAC</td><td>238.8</td><td>2.4</td><td>0.8</td><td>2.9</td><td>3.5E-03</td><td>1.4E-02</td><td>BT20/MB231</td><td>CYTO</td></tr>
<tr><td>>MI0000066|hsa-let-7e&WithFlank&19|+|51692780|51692870@14.35.22(+1U)&[MIMAT0000066&hsa-let-7e-5p&offsets|0|0(+1U);m-50&19|+|51692793|51692814&offsets|0|0(+1U)]|TGAGGTAGGAGGTTGTATAGTTT</td><td>>MI0000066|hsa-let-7e&WithFlank&19|+|51692780|51692870</td><td>14</td><td>35</td><td>22(+1U)</td><td>TGAGGTAGGAGGTTGTATAGTTT</td><td>336.7</td><td>2.7</td><td>1.0</td><td>2.7</td><td>7.1E-03</td><td>1.4E-02</td><td>BT20/MB468</td><td>MP</td></tr>
<tr><td>>hg38_21_+_8433172_8446622_RNA45SN1_45S_50ntFlanks@11567.11599.33|AGCAGCCGACTTAGAACTGGTGCGGACCAGGGG</td><td>>hg38_21_+_8433172_8446622_RNA45SN1_45S_50ntFlanks</td><td>11567</td><td>11599</td><td>33</td><td>AGCAGCCGACTTAGAACTGGTGCGGACCAGGGG</td><td>215.4</td><td>5.1</td><td>1.7</td><td>3.0</td><td>2.8E-03</td><td>1.4E-02</td><td>BT20/MB231</td><td>MP</td></tr>
<tr><td>>hg38_21_+_8433172_8446622_RNA45SN1_45S_50ntFlanks@10107.10127.21|CGTGTGGTGTGCGTCGGAGGG</td><td>>hg38_21_+_8433172_8446622_RNA45SN1_45S_50ntFlanks</td><td>10107</td><td>10127</td><td>21</td><td>CGTGTGGTGTGCGTCGGAGGG</td><td>1030.7</td><td>-5.2</td><td>2.1</td><td>-2.5</td><td>1.2E-02</td><td>1.4E-02</td><td>BT20/MB468</td><td>NUC</td></tr>
<tr><td>>MI0001446|hsa-mir-424&WithFlank&X|-|134546608|134546717@17.37.21&[MIMAT0001341&hsa-miR-424-5p&offsets|0|-1;m-152&X|-|134546681|134546701&offsets|0|0]|CAGCAGCAATTCATGTTTTGA</td><td>>MI0001446|hsa-mir-424&WithFlank&X|-|134546608|134546717</td><td>17</td><td>37</td><td>21</td><td>CAGCAGCAATTCATGTTTTGA</td><td>246.0</td><td>6.1</td><td>2.3</td><td>2.7</td><td>7.1E-03</td><td>1.4E-02</td><td>BT20/MB468</td><td>MP</td></tr>
<tr><td>>hg38_21_+_8433172_8446622_RNA45SN1_45S_50ntFlanks@11568.11601.34|GCAGCCGACTTAGAACTGGTGCGGACCAGGGGAA</td><td>>hg38_21_+_8433172_8446622_RNA45SN1_45S_50ntFlanks</td><td>11568</td><td>11601</td><td>34</td><td>GCAGCCGACTTAGAACTGGTGCGGACCAGGGGAA</td><td>1111.2</td><td>-2.3</td><td>0.9</td><td>-2.7</td><td>7.1E-03</td><td>1.4E-02</td><td>BT20/MB468</td><td>MP</td></tr>
<tr><td>>hg38_21_+_8433172_8446622_RNA45SN1_45S_50ntFlanks@4555.4577.23|CGCAGCTAGGAATAATGGAATAG</td><td>>hg38_21_+_8433172_8446622_RNA45SN1_45S_50ntFlanks</td><td>4555</td><td>4577</td><td>23</td><td>CGCAGCTAGGAATAATGGAATAG</td><td>62.7</td><td>-4.4</td><td>1.7</td><td>-2.7</td><td>7.1E-03</td><td>1.4E-02</td><td>BT20/MB468</td><td>MP</td></tr>
<tr><td>>hg38_21_+_8433172_8446622_RNA45SN1_45S_50ntFlanks@6684.6706.23|TCGATGAAGAACGCAGCTAGCTG</td><td>>hg38_21_+_8433172_8446622_RNA45SN1_45S_50ntFlanks</td><td>6684</td><td>6706</td><td>23</td><td>TCGATGAAGAACGCAGCTAGCTG</td><td>128.4</td><td>-4.4</td><td>1.6</td><td>-2.7</td><td>7.1E-03</td><td>1.4E-02</td><td>BT20/MB468</td><td>MP</td></tr>
<tr><td>>hg38_21_+_8433172_8446622_RNA45SN1_45S_50ntFlanks@7973.7997.25|GACGCGACCTCAGATCAGACGTGGC</td><td>>hg38_21_+_8433172_8446622_RNA45SN1_45S_50ntFlanks</td><td>7973</td><td>7997</td><td>25</td><td>GACGCGACCTCAGATCAGACGTGGC</td><td>1668.0</td><td>2.9</td><td>1.0</td><td>3.0</td><td>3.0E-03</td><td>1.4E-02</td><td>BT20/MB231</td><td>NUC</td></tr>
<tr><td>>hg38_21_+_8433172_8446622_RNA45SN1_45S_50ntFlanks@5082.5099.18|AACGAACGAGACTCTGGC</td><td>>hg38_21_+_8433172_8446622_RNA45SN1_45S_50ntFlanks</td><td>5082</td><td>5099</td><td>18</td><td>AACGAACGAGACTCTGGC</td><td>336.5</td><td>4.4</td><td>1.5</td><td>3.0</td><td>2.8E-03</td><td>1.4E-02</td><td>BT20/MB231</td><td>MP</td></tr>
<tr><td>>hg38_21_+_8433172_8446622_RNA45SN1_45S_50ntFlanks@7972.8009.38|AGACGCGACCTCAGATCAGACGTGGCGACCCGCTGAAT</td><td>>hg38_21_+_8433172_8446622_RNA45SN1_45S_50ntFlanks</td><td>7972</td><td>8009</td><td>38</td><td>AGACGCGACCTCAGATCAGACGTGGCGACCCGCTGAAT</td><td>65.9</td><td>4.9</td><td>1.9</td><td>2.6</td><td>1.1E-02</td><td>1.4E-02</td><td>MB231/MB468</td><td>MP</td></tr>
<tr><td>>hg38_MT_+_1621_3279_RNR2_16S_50ntFlanks@643.676.34|CAATTGGACCAATCTATCACCCTATAGAAGAACT</td><td>>hg38_MT_+_1621_3279_RNR2_16S_50ntFlanks</td><td>643</td><td>676</td><td>34</td><td>CAATTGGACCAATCTATCACCCTATAGAAGAACT</td><td>84.1</td><td>4.0</td><td>1.6</td><td>2.6</td><td>1.1E-02</td><td>1.4E-02</td><td>MB231/MB468</td><td>MP</td></tr>
<tr><td>>hg38_21_+_8433172_8446622_RNA45SN1_45S_50ntFlanks@12456.12496.41|CCAAGCGTTGGATTGTTCACCCACTAATAGGGAACGTGAGC</td><td>>hg38_21_+_8433172_8446622_RNA45SN1_45S_50ntFlanks</td><td>12456</td><td>12496</td><td>41</td><td>CCAAGCGTTGGATTGTTCACCCACTAATAGGGAACGTGAGC</td><td>668.8</td><td>-1.9</td><td>0.8</td><td>-2.5</td><td>1.1E-02</td><td>1.4E-02</td><td>BT20/MB468</td><td>CYTO</td></tr>
<tr><td>>MI0000750|hsa-mir-26a-2&WithFlank&12|-|57824603|57824698@20.40.21(+1G);MI0000083|hsa-mir-26a-1&WithFlank&3|+|37969398|37969486@16.36.21(+1G)&[MIMAT0000082&hsa-miR-26a-5p&offsets|0|-1(+1G);m-33&12|-|57824658|57824679&offsets|0|-1(+1G)];[MIMAT0000082_1&hsa-miR-26a-5p&offsets|0|-1(+1G);m-34&3|+|37969413|37969434&offsets|0|-1(+1G)]|TTCAAGTAATCCAGGATAGGCG</td><td>>MI0000750|hsa-mir-26a-2&WithFlank&12|-|57824603|57824698</td><td>20</td><td>40</td><td>21(+1G)</td><td>TTCAAGTAATCCAGGATAGGCG</td><td>2854.1</td><td>-3.4</td><td>1.1</td><td>-3.0</td><td>2.8E-03</td><td>1.4E-02</td><td>BT20/MB231</td><td>MP</td></tr>
<tr><td>>hg38_21_+_8433172_8446622_RNA45SN1_45S_50ntFlanks@11671.11692.22|CCAGTGCTCTGAATGTCAAAGT</td><td>>hg38_21_+_8433172_8446622_RNA45SN1_45S_50ntFlanks</td><td>11671</td><td>11692</td><td>22</td><td>CCAGTGCTCTGAATGTCAAAGT</td><td>446.6</td><td>4.2</td><td>1.4</td><td>3.0</td><td>2.8E-03</td><td>1.4E-02</td><td>BT20/MB231</td><td>MP</td></tr>
<tr><td>>hg38_21_+_8433172_8446622_RNA45SN1_45S_50ntFlanks@8527.8550.24|CCTTCCCCCGCCGCCCCTCCTCCT</td><td>>hg38_21_+_8433172_8446622_RNA45SN1_45S_50ntFlanks</td><td>8527</td><td>8550</td><td>24</td><td>CCTTCCCCCGCCGCCCCTCCTCCT</td><td>265.6</td><td>-3.9</td><td>1.6</td><td>-2.5</td><td>1.1E-02</td><td>1.4E-02</td><td>MB231/MB468</td><td>CYTO</td></tr>
<tr><td>>hg38_21_+_8433172_8446622_RNA45SN1_45S_50ntFlanks@4724.4764.41|ATTAATCAAGAACGAAAGTCGGAGGTTCGAAGACGATCAGA</td><td>>hg38_21_+_8433172_8446622_RNA45SN1_45S_50ntFlanks</td><td>4724</td><td>4764</td><td>41</td><td>ATTAATCAAGAACGAAAGTCGGAGGTTCGAAGACGATCAGA</td><td>371.5</td><td>-3.3</td><td>1.3</td><td>-2.6</td><td>9.9E-03</td><td>1.4E-02</td><td>BT20/MB468</td><td>MITO</td></tr>
<tr><td>>hg38_21_+_8433172_8446622_RNA45SN1_45S_50ntFlanks@11602.11619.18|TCCGACTGTTTAATTAAA</td><td>>hg38_21_+_8433172_8446622_RNA45SN1_45S_50ntFlanks</td><td>11602</td><td>11619</td><td>18</td><td>TCCGACTGTTTAATTAAA</td><td>453.5</td><td>-3.5</td><td>1.3</td><td>-2.7</td><td>7.1E-03</td><td>1.4E-02</td><td>BT20/MB468</td><td>MP</td></tr>
<tr><td>>trnaMT_LeuTAG_MT_+_12266_12336@39.74.36|CCAAAAATTTTGGTGCAACTCCAAATAAAAGTACCA</td><td>>trnaMT_LeuTAG_MT_+_12266_12336</td><td>39</td><td>74</td><td>36</td><td>CCAAAAATTTTGGTGCAACTCCAAATAAAAGTACCA</td><td>91.9</td><td>4.1</td><td>1.6</td><td>2.6</td><td>1.1E-02</td><td>1.4E-02</td><td>MB231/MB468</td><td>MP</td></tr>
<tr><td>>hg38_21_+_8433172_8446622_RNA45SN1_45S_50ntFlanks@4738.4755.18|AAAGTCGGAGGTTCGAAG</td><td>>hg38_21_+_8433172_8446622_RNA45SN1_45S_50ntFlanks</td><td>4738</td><td>4755</td><td>18</td><td>AAAGTCGGAGGTTCGAAG</td><td>238.1</td><td>-1.4</td><td>0.6</td><td>-2.5</td><td>1.2E-02</td><td>1.4E-02</td><td>MB231/MB468</td><td>NUC</td></tr>
<tr><td>>hg38_21_+_8433172_8446622_RNA45SN1_45S_50ntFlanks@10822.10842.21|GAAGTCGGCAAGCCGGATCCG</td><td>>hg38_21_+_8433172_8446622_RNA45SN1_45S_50ntFlanks</td><td>10822</td><td>10842</td><td>21</td><td>GAAGTCGGCAAGCCGGATCCG</td><td>91.2</td><td>-3.5</td><td>1.3</td><td>-2.7</td><td>7.1E-03</td><td>1.4E-02</td><td>BT20/MB468</td><td>MP</td></tr>
</table>

| | | | | | | | | | | | | | |
|---|---|---|---|---|---|---|---|---|---|---|---|---|---|
| >hg38_21_+_8433172_8446622_RNA45SN1_45S_50ntFlanks@12415.12445.31\|CTTCGATGTCGGCTCTTCCTATCATTGTGAA | >hg38_21_+_8433172_8446622_RNA45SN1_45S_50ntFlanks | 12415 | 12445 | 31 | CTTCGATGTCGGCTCTTCCTATCATTGTGAA | 55.0 | -4.4 | 1.7 | -2.6 | 1.1E-02 | 1.4E-02 | MB231/MB468 | MP |
| >hg38_21_+_8433172_8446622_RNA45SN1_45S_50ntFlanks@6700.6720.21\|CTAGCTGCGAGAATTAATGTG | >hg38_21_+_8433172_8446622_RNA45SN1_45S_50ntFlanks | 6700 | 6720 | 21 | CTAGCTGCGAGAATTAATGTG | 457.5 | -3.2 | 1.1 | -3.0 | 3.0E-03 | 1.4E-02 | BT20/MB231 | NUC |
| >hg38_21_+_8433172_8446622_RNA45SN1_45S_50ntFlanks@9747.9769.23\|TCAACCTATTCTCAAACTTTAAA | >hg38_21_+_8433172_8446622_RNA45SN1_45S_50ntFlanks | 9747 | 9769 | 23 | TCAACCTATTCTCAAACTTTAAA | 263.1 | 3.3 | 1.1 | 2.9 | 3.5E-03 | 1.4E-02 | BT20/MB231 | MITO |
| >hg38_21_+_8433172_8446622_RNA45SN1_45S_50ntFlanks@8380.8398.19\|CAAGAGGGCGTGAAACCGT | >hg38_21_+_8433172_8446622_RNA45SN1_45S_50ntFlanks | 8380 | 8398 | 19 | CAAGAGGGCGTGAAACCGT | 220.0 | -3.7 | 1.4 | -2.7 | 7.2E-03 | 1.4E-02 | BT20/MB468 | MP |
| >trna13_ValCAC_19_-_4724647_4724719@1.32.32\|GTTTCCGTAGTGTAGCGGTTATCACATTCGCC | >trna13_ValCAC_19_-_4724647_4724719 | 1 | 32 | 32 | GTTTCCGTAGTGTAGCGGTTATCACATTCGCC | 446.0 | -4.5 | 1.5 | -3.0 | 2.8E-03 | 1.4E-02 | BT20/MB231 | MP |
| >hg38_21_+_8433172_8446622_RNA45SN1_45S_50ntFlanks@8351.8375.25\|AAGTTGAAAAGAACTTTGAAGAGAG | >hg38_21_+_8433172_8446622_RNA45SN1_45S_50ntFlanks | 8351 | 8375 | 25 | AAGTTGAAAAGAACTTTGAAGAGAG | 174.9 | 3.9 | 1.5 | 2.6 | 1.1E-02 | 1.4E-02 | MB231/MB468 | MP |
| >hg38_21_+_8433172_8446622_RNA45SN1_45S_50ntFlanks@9033.9075.43\|CTCGTGGGGGGGCCGGGCCACCCCTCCCACGGCGCGACCGCTC | >hg38_21_+_8433172_8446622_RNA45SN1_45S_50ntFlanks | 9033 | 9075 | 43 | CTCGTGGGGGGGCCGGGCCACCCCTCCCACGGCGCGACCGCTC | 43.9 | -8.5 | 3.2 | -2.7 | 7.2E-03 | 1.4E-02 | BT20/MB468 | MP |
| >hg38_21_+_8433172_8446622_RNA45SN1_45S_50ntFlanks@10058.10077.20\|CGGCAGTCGAGAGTGGACGG | >hg38_21_+_8433172_8446622_RNA45SN1_45S_50ntFlanks | 10058 | 10077 | 20 | CGGCAGTCGAGAGTGGACGG | 141.7 | 3.7 | 1.4 | 2.7 | 7.2E-03 | 1.4E-02 | BT20/MB468 | MP |
| >hg38_21_+_8433172_8446622_RNA45SN1_45S_50ntFlanks@10775.10818.44\|TCTCCAAGGTGAACAGCCTCTGGCATGTTGGAACAATGTAGGTA | >hg38_21_+_8433172_8446622_RNA45SN1_45S_50ntFlanks | 10775 | 10818 | 44 | TCTCCAAGGTGAACAGCCTCTGGCATGTTGGAACAATGTAGGTA | 303.4 | -2.3 | 0.9 | -2.5 | 1.2E-02 | 1.4E-02 | MB231/MB468 | CYTO |
| >hg38_21_+_8433172_8446622_RNA45SN1_45S_50ntFlanks@10420.10457.38\|CTGAGAGATGGGCGAGCGCCGTTCCGAAGGGACGGGCG | >hg38_21_+_8433172_8446622_RNA45SN1_45S_50ntFlanks | 10420 | 10457 | 38 | CTGAGAGATGGGCGAGCGCCGTTCCGAAGGGACGGGCG | 90.3 | -4.5 | 1.7 | -2.7 | 7.2E-03 | 1.4E-02 | BT20/MB468 | MP |
| >hg38_21_+_8433172_8446622_RNA45SN1_45S_50ntFlanks@7989.8014.26\|AGACGTGGCGACCCGCTGAATTTAAG | >hg38_21_+_8433172_8446622_RNA45SN1_45S_50ntFlanks | 7989 | 8014 | 26 | AGACGTGGCGACCCGCTGAATTTAAG | 242.8 | 3.8 | 1.3 | 3.0 | 2.8E-03 | 1.4E-02 | BT20/MB231 | MP |
| >hg38_21_+_8433172_8446622_RNA45SN1_45S_50ntFlanks@6663.6695.33\|GTGGATCACTCGGCTCGTGCGTCGATGAAGAAC | >hg38_21_+_8433172_8446622_RNA45SN1_45S_50ntFlanks | 6663 | 6695 | 33 | GTGGATCACTCGGCTCGTGCGTCGATGAAGAAC | 117.7 | -3.3 | 1.3 | -2.6 | 1.1E-02 | 1.4E-02 | MB231/MB468 | MP |
| >hg38_21_+_8433172_8446622_RNA45SN1_45S_50ntFlanks@10788.10822.35\|CAGCCTCTGGCATGTTGGAACAATGTAGGTAAGGG | >hg38_21_+_8433172_8446622_RNA45SN1_45S_50ntFlanks | 10788 | 10822 | 35 | CAGCCTCTGGCATGTTGGAACAATGTAGGTAAGGG | 228.1 | -2.2 | 0.8 | -2.7 | 7.2E-03 | 1.4E-02 | BT20/MB468 | MP |
| >hg38_21_+_8433172_8446622_RNA45SN1_45S_50ntFlanks@12475.12497.23\|CCCACTAATAGGGAACGTGAGCT | >hg38_21_+_8433172_8446622_RNA45SN1_45S_50ntFlanks | 12475 | 12497 | 23 | CCCACTAATAGGGAACGTGAGCT | 669.5 | 1.5 | 0.6 | 2.5 | 1.2E-02 | 1.4E-02 | MB231/MB468 | CYTO |
| >hg38_21_+_8433172_8446622_RNA45SN1_45S_50ntFlanks@11641.11682.42\|CGGCGGGTGTTGACGCGATGTGATTTCTGCCCAGTGCTCTGA | >hg38_21_+_8433172_8446622_RNA45SN1_45S_50ntFlanks | 11641 | 11682 | 42 | CGGCGGGTGTTGACGCGATGTGATTTCTGCCCAGTGCTCTGA | 380.8 | -3.4 | 1.3 | -2.6 | 1.0E-02 | 1.4E-02 | BT20/MB468 | MITO |
| >hg38_21_+_8433172_8446622_RNA45SN1_45S_50ntFlanks@10229.10252.24\|CCCCGCGGACGCTACGCCGCGACG | >hg38_21_+_8433172_8446622_RNA45SN1_45S_50ntFlanks | 10229 | 10252 | 24 | CCCCGCGGACGCTACGCCGCGACG | 76.6 | 6.4 | 2.5 | 2.6 | 1.1E-02 | 1.4E-02 | MB231/MB468 | MP |
| >MI0000434\|hsa-let-7i&WithFlank&12\|+\|62603680\|62603775@12.31.20(+1G)&[MIMAT0000415&hsa-let-7i-5p&offsets\|0\|-2(+1G);m-49&12\|+\|62603691\|62603712&offsets\|0\|-2(+1G)]\|TGAGGTAGTAGTTTGTGCTGG | >MI0000434\|hsa-let-7i&WithFlank&12\|+\|62603680\|62603775 | 12 | 31 | 20(+1G) | TGAGGTAGTAGTTTGTGCTGG | 107.9 | 3.6 | 1.4 | 2.6 | 1.1E-02 | 1.4E-02 | MB231/MB468 | MP |
| >MI0000272\|hsa-mir-182&WithFlank&7\|-\|129770377\|129770498@29.54.26(+1U)&[MIMAT0000259&hsa-miR-182-5p&offsets\|0\|+2(+1U);m-22&7\|-\|129770449\|129770470&offsets\|0\|+4(+1U)]\|TTTGGCAATGGTAGAACTCACACTGGT | >MI0000272\|hsa-mir-182&WithFlank&7\|-\|129770377\|129770498 | 29 | 54 | 26(+1U) | TTTGGCAATGGTAGAACTCACACTGGT | 199.4 | 4.0 | 1.5 | 2.7 | 7.2E-03 | 1.4E-02 | BT20/MB468 | MP |
| >MI0000289\|hsa-mir-181a-1&WithFlank&1\|-\|198859038\|198859159@30.52.23(+1A);MI0000269\|hsa-mir-181a-2&WithFlank&9\|+\|124692436\|124692557@45.67.23(+1A)&[MIMAT0000256&hsa-miR-181a-5p&offsets\|0\|0(+1A);m-32&1\|-\|198859108\|198859130&offsets\|0\|0(+1A)];[MIMAT0000256_1&hsa-miR-181a-5p&offsets\|0\|0(+1A);m-32&9\|+\|124692480\|124692502&offsets\|0\|0(+1A)]\|AACATTCAACGCTGTCGGTGAGTA | >MI0000289\|hsa-mir-181a-1&WithFlank&1\|-\|198859038\|198859159 | 30 | 52 | 23(+1A) | AACATTCAACGCTGTCGGTGAGTA | 269.3 | 3.6 | 1.4 | 2.6 | 1.1E-02 | 1.4E-02 | MB231/MB468 | MP |
| >hg38_21_+_8433172_8446622_RNA45SN1_45S_50ntFlanks@11783.11809.27\|CATGAATGGATGAACGAGATTCCCACT | >hg38_21_+_8433172_8446622_RNA45SN1_45S_50ntFlanks | 11783 | 11809 | 27 | CATGAATGGATGAACGAGATTCCCACT | 94.5 | -5.6 | 2.2 | -2.6 | 1.1E-02 | 1.4E-02 | MB231/MB468 | MP |
| >hg38_21_+_8433172_8446622_RNA45SN1_45S_50ntFlanks@11776.11794.19\|TGACGCGCATGAATGGATG | >hg38_21_+_8433172_8446622_RNA45SN1_45S_50ntFlanks | 11776 | 11794 | 19 | TGACGCGCATGAATGGATG | 196.2 | -2.3 | 0.9 | -2.6 | 1.1E-02 | 1.4E-02 | MB231/MB468 | MP |
| >hg38_21_+_8433172_8446622_RNA45SN1_45S_50ntFlanks@7978.8019.42\|GACCTCAGATCAGACGTGGCGACCCGCTGAATTTAAGCATAT | >hg38_21_+_8433172_8446622_RNA45SN1_45S_50ntFlanks | 7978 | 8019 | 42 | GACCTCAGATCAGACGTGGCGACCCGCTGAATTTAAGCATAT | 167.0 | 5.8 | 2.3 | 2.5 | 1.2E-02 | 1.4E-02 | MB231/MB468 | MITO |
| >hg38_21_+_8433172_8446622_RNA45SN1_45S_50ntFlanks@10680.10710.31\|GTTCCGGCGGCGTCCGGTGAGCTCTCGCTGG | >hg38_21_+_8433172_8446622_RNA45SN1_45S_50ntFlanks | 10680 | 10710 | 31 | GTTCCGGCGGCGTCCGGTGAGCTCTCGCTGG | 381.7 | 5.5 | 1.8 | 3.0 | 2.8E-03 | 1.4E-02 | BT20/MB231 | MP |
| >hg38_21_+_8433172_8446622_RNA45SN1_45S_50ntFlanks@9738.9764.27\|GAAACGATCTCAACCTATTCTCAAACT | >hg38_21_+_8433172_8446622_RNA45SN1_45S_50ntFlanks | 9738 | 9764 | 27 | GAAACGATCTCAACCTATTCTCAAACT | 396.9 | -2.0 | 0.8 | -2.6 | 1.0E-02 | 1.4E-02 | BT20/MB468 | MITO |
| >MI0000811\|hsa-mir-148b&WithFlank&12\|+\|54337210\|54337320@69.90.22(+2U)&[MIMAT0000759&hsa-miR-148b-3p&offsets\|0\|0(+2U);m-77&12\|+\|54337278\|54337299&offsets\|0\|0(+2U)]\|TCAGTGCATCACAGAACTTTGTTT | >MI0000811\|hsa-mir-148b&WithFlank&12\|+\|54337210\|54337320 | 69 | 90 | 22(+2U) | TCAGTGCATCACAGAACTTTGTTT | 411.5 | 3.5 | 1.4 | 2.6 | 1.1E-02 | 1.4E-02 | MB231/MB468 | MP |
| >MI0000085\|hsa-mir-27a&WithFlank&19\|-\|13836434\|13836523@16.36.21&[MIMAT0004501&hsa-miR-27a-5p&offsets\|0\|-1;m-270&19\|-\|13836487\|13836508&offsets\|0\|-1]\|AGGGCTTAGCTGCTTGTGAGC | >MI0000085\|hsa-mir-27a&WithFlank&19\|-\|13836434\|13836523 | 16 | 36 | 21 | AGGGCTTAGCTGCTTGTGAGC | 246.9 | 4.3 | 1.7 | 2.6 | 1.1E-02 | 1.4E-02 | MB231/MB468 | MP |
| >MI0000103\|hsa-mir-101-1&WithFlank&1\|-\|65058428\|65058514@53.73.21(+1C);MI0000739\|hsa-mir-101-2&WithFlank&9\|+\|4850291\|4850381@55.75.21(+1C)&[MIMAT0000099&hsa-miR-101-3p&offsets\|0\|0(+1C);m-17&1\|-\|65058443\|65058462&offsets\|0\|+1(+1C)];[MIMAT0000099_1&hsa-miR-101-3p&offsets\|0\|0(+1C);m-16&9\|+\|4850345\|4850364&offsets\|0\|+1(+1C)]\|TACAGTACTGTGATAACTGAAC | >MI0000103\|hsa-mir-101-1&WithFlank&1\|-\|65058428\|65058514 | 53 | 73 | 21(+1C) | TACAGTACTGTGATAACTGAAC | 71.4 | 4.9 | 1.9 | 2.6 | 1.1E-02 | 1.4E-02 | MB231/MB468 | MP |
| >hg38_MT_+_1621_3279_RNR2_16S_50ntFlanks@1372.1396.25\|GTTGGATCAGGACATCCCGATGGTG | >hg38_MT_+_1621_3279_RNR2_16S_50ntFlanks | 1372 | 1396 | 25 | GTTGGATCAGGACATCCCGATGGTG | 139.4 | 2.5 | 1.0 | 2.6 | 1.1E-02 | 1.4E-02 | MB231/MB468 | MP |
| >MI0000088\|hsa-mir-30a&WithFlank&6\|-\|71403545\|71403627@53.74.22(+1A)&[MIMAT0000088&hsa-miR-30a-3p&offsets\|0\|0(+1A);m-38&6\|-\|71403554\|71403575&offsets\|0\|0(+1A)]\|CTTTCAGTCGGATGTTTGCAGCA | >MI0000088\|hsa-mir-30a&WithFlank&6\|-\|71403545\|71403627 | 53 | 74 | 22(+1A) | CTTTCAGTCGGATGTTTGCAGCA | 222.6 | -2.1 | 0.7 | -2.9 | 3.5E-03 | 1.4E-02 | BT20/MB231 | MITO |
| >hg38_21_+_8433172_8446622_RNA45SN1_45S_50ntFlanks@11666.11702.37\|TCTGCCCAGTGCTCTGAATGTCAAAGTGAAGAAATTC | >hg38_21_+_8433172_8446622_RNA45SN1_45S_50ntFlanks | 11666 | 11702 | 37 | TCTGCCCAGTGCTCTGAATGTCAAAGTGAAGAAATTC | 296.2 | 1.2 | 0.5 | 2.6 | 1.0E-02 | 1.4E-02 | BT20/MB468 | MITO |
| >hg38_MT_+_598_1651_RNR1_12S_50ntFlanks@439.469.31\|AACATATCTGAACACACAATAGCTAAGACCC | >hg38_MT_+_598_1651_RNR1_12S_50ntFlanks | 439 | 469 | 31 | AACATATCTGAACACACAATAGCTAAGACCC | 186.5 | 4.3 | 1.6 | 2.7 | 7.2E-03 | 1.4E-02 | BT20/MB468 | MP |
| >hg38_21_+_8433172_8446622_RNA45SN1_45S_50ntFlanks@10680.10711.32\|GTTCCGGCGGCGTCCGGTGAGCTCTCGCTGGC | >hg38_21_+_8433172_8446622_RNA45SN1_45S_50ntFlanks | 10680 | 10711 | 32 | GTTCCGGCGGCGTCCGGTGAGCTCTCGCTGGC | 2556.2 | -5.1 | 1.9 | -2.7 | 7.2E-03 | 1.4E-02 | BT20/MB468 | MP |
| >hg38_21_+_8433172_8446622_RNA45SN1_45S_50ntFlanks@10099.10117.19\|CGCGCGCGCGTGTGGTGTG | >hg38_21_+_8433172_8446622_RNA45SN1_45S_50ntFlanks | 10099 | 10117 | 19 | CGCGCGCGCGTGTGGTGTG | 916.7 | -4.4 | 1.7 | -2.6 | 1.0E-02 | 1.4E-02 | BT20/MB468 | MITO |
| >hg38_21_+_8433172_8446622_RNA45SN1_45S_50ntFlanks@10781.10804.24\|AGGTGAACAGCCTCTGGCATGTTG | >hg38_21_+_8433172_8446622_RNA45SN1_45S_50ntFlanks | 10781 | 10804 | 24 | AGGTGAACAGCCTCTGGCATGTTG | 116.6 | -3.8 | 1.5 | -2.6 | 1.0E-02 | 1.4E-02 | BT20/MB468 | MITO |
| >hg38_21_+_8433172_8446622_RNA45SN1_45S_50ntFlanks@9556.9581.26\|TAGCGGTCCTGACGTGCAAATCGGTC | >hg38_21_+_8433172_8446622_RNA45SN1_45S_50ntFlanks | 9556 | 9581 | 26 | TAGCGGTCCTGACGTGCAAATCGGTC | 327.7 | -1.9 | 0.7 | -2.5 | 1.2E-02 | 1.4E-02 | MB231/MB468 | NUC |
| >MI0000077\|hsa-mir-21&WithFlank&17\|+\|59841260\|59841343@14.35.22(+1A)&[MIMAT0000076&hsa-miR-21-5p&offsets\|0\|0(+1A);m-1&17\|+\|59841273\|59841294&offsets\|0\|0(+1A)]\|TAGCTTATCAGACTGATGTTGAA | >MI0000077\|hsa-mir-21&WithFlank&17\|+\|59841260\|59841343 | 14 | 35 | 22(+1A) | TAGCTTATCAGACTGATGTTGAA | 23610.0 | -2.3 | 0.8 | -2.9 | 3.5E-03 | 1.4E-02 | BT20/MB231 | MITO |
| >hg38_21_+_8433172_8446622_RNA45SN1_45S_50ntFlanks@4924.4943.20\|AGGAGTGGAGCCTGCGGCTT | >hg38_21_+_8433172_8446622_RNA45SN1_45S_50ntFlanks | 4924 | 4943 | 20 | AGGAGTGGAGCCTGCGGCTT | 358.0 | -2.5 | 0.9 | -2.9 | 3.6E-03 | 1.4E-02 | BT20/MB231 | CYTO |
| >hg38_21_+_8433172_8446622_RNA45SN1_45S_50ntFlanks@7975.8003.29\|CGCGACCTCAGATCAGACGTGGCGACCCG | >hg38_21_+_8433172_8446622_RNA45SN1_45S_50ntFlanks | 7975 | 8003 | 29 | CGCGACCTCAGATCAGACGTGGCGACCCG | 4489.3 | 2.3 | 0.8 | 3.0 | 3.1E-03 | 1.4E-02 | BT20/MB231 | NUC |
| >hg38_21_+_8433172_8446622_RNA45SN1_45S_50ntFlanks@6646.6664.19\|TCGTACGACTCTTAGCGGT | >hg38_21_+_8433172_8446622_RNA45SN1_45S_50ntFlanks | 6646 | 6664 | 19 | TCGTACGACTCTTAGCGGT | 13150.0 | 1.4 | 0.6 | 2.5 | 1.2E-02 | 1.4E-02 | BT20/MB468 | NUC |
| >hg38_21_+_8433172_8446622_RNA45SN1_45S_50ntFlanks@4760.4778.19\|TCAGATACCGTCGTAGTTC | >hg38_21_+_8433172_8446622_RNA45SN1_45S_50ntFlanks | 4760 | 4778 | 19 | TCAGATACCGTCGTAGTTC | 476.6 | -4.7 | 1.7 | -2.7 | 7.2E-03 | 1.4E-02 | BT20/MB468 | MP |
| >hg38_21_+_8433172_8446622_RNA45SN1_45S_50ntFlanks@4728.4759.32\|ATCAAGAACGAAAGTCGGAGGTTCGAAGACGA | >hg38_21_+_8433172_8446622_RNA45SN1_45S_50ntFlanks | 4728 | 4759 | 32 | ATCAAGAACGAAAGTCGGAGGTTCGAAGACGA | 395.4 | -3.7 | 1.5 | -2.5 | 1.2E-02 | 1.4E-02 | MB231/MB468 | CYTO |
| >hg38_21_+_8433172_8446622_RNA45SN1_45S_50ntFlanks@3841.3873.33\|TCCTACTTGGATAACTGTGGTAATTCTAGAGCT | >hg38_21_+_8433172_8446622_RNA45SN1_45S_50ntFlanks | 3841 | 3873 | 33 | TCCTACTTGGATAACTGTGGTAATTCTAGAGCT | 70.4 | -5.4 | 2.1 | -2.5 | 1.1E-02 | 1.4E-02 | MB231/MB468 | MP |
| >trnaMT_GluTTC_MT_-_14674_14742@33.61.29,trnalookalike8_GluTTC_5_-_93905172_93905240@33.61.29\|CATATCATTGGTCGTGGTTGTAGTCCGTG | >trnaMT_GluTTC_MT_-_14674_14742 | 33 | 61 | 29 | CATATCATTGGTCGTGGTTGTAGTCCGTG | 87.1 | 2.8 | 1.0 | 2.7 | 7.3E-03 | 1.4E-02 | BT20/MB468 | MP |
| >hg38_21_+_8433172_8446622_RNA45SN1_45S_50ntFlanks@10103.10127.25\|CGCGCGTGTGGTGTGCGTCGGAGGG | >hg38_21_+_8433172_8446622_RNA45SN1_45S_50ntFlanks | 10103 | 10127 | 25 | CGCGCGTGTGGTGTGCGTCGGAGGG | 118.3 | -4.5 | 1.7 | -2.7 | 7.3E-03 | 1.4E-02 | BT20/MB468 | MP |
| >trnaMT_HisGTG_MT_+_12138_12206@32.62.31\|TGAATCTGACAACAGAGGCTTACGACCCCTT | >trnaMT_HisGTG_MT_+_12138_12206 | 32 | 62 | 31 | TGAATCTGACAACAGAGGCTTACGACCCCTT | 288.2 | 2.4 | 0.9 | 2.5 | 1.2E-02 | 1.4E-02 | MB231/MB468 | MITO |
| >hg38_1_-_228634819_228635039_RNA5S12_5S_50ntFlanks@150.171.22\|AATACCGGGTGCTGTAGGCTTT | >hg38_1_-_228634819_228635039_RNA5S12_5S_50ntFlanks | 150 | 171 | 22 | AATACCGGGTGCTGTAGGCTTT | 17635.0 | -2.9 | 1.1 | -2.7 | 7.3E-03 | 1.4E-02 | BT20/MB468 | MP |
| >hg38_21_+_8433172_8446622_RNA45SN1_45S_50ntFlanks@11567.11588.22\|AGCAGCCGACTTAGAACTGGTG | >hg38_21_+_8433172_8446622_RNA45SN1_45S_50ntFlanks | 11567 | 11588 | 22 | AGCAGCCGACTTAGAACTGGTG | 719.0 | -2.2 | 0.8 | -2.7 | 7.3E-03 | 1.4E-02 | BT20/MB468 | MP |
| >hg38_21_+_8433172_8446622_RNA45SN1_45S_50ntFlanks@11326.11345.20\|CTCGTCGCGCGCGCGTCCGC | >hg38_21_+_8433172_8446622_RNA45SN1_45S_50ntFlanks | 11326 | 11345 | 20 | CTCGTCGCGCGCGCGTCCGC | 1266.1 | -3.5 | 1.2 | -3.0 | 3.1E-03 | 1.4E-02 | BT20/MB231 | NUC |
| >trnaMT_GluTTC_MT_-_14674_14742@39.71.33,trnalookalike8_GluTTC_5_-_93905172_93905240@39.71.33\|ATTGGTCGTGGTTGTAGTCCGTGCGAGAATACC | >trnaMT_GluTTC_MT_-_14674_14742 | 39 | 71 | 33 | ATTGGTCGTGGTTGTAGTCCGTGCGAGAATACC | 1959.9 | -2.6 | 0.9 | -3.0 | 3.1E-03 | 1.4E-02 | BT20/MB231 | NUC |

| | | | | | | | | | | | | | |
|---|---|---|---|---|---|---|---|---|---|---|---|---|---|
| >hg38_21_+_8433172_8446622_RNA45SN1_45S_50ntFlanks@11207.11225.19\|GGCGGCGACTCTGGACGCG | >hg38_21_+_8433172_8446622_RNA45SN1_45S_50ntFlanks | 11207 | 11225 | 19 | GGCGGCGACTCTGGACGCG | 429.4 | -4.8 | 1.9 | -2.5 | 1.2E-02 | 1.4E-02 | BT20/MB468 | NUC |
| >MI0000086\|hsa-mir-28&WithFlank&3\|+\|188688775\|188688872@61.82.22&[MIMAT0004502&hsa-miR-28-3p&offsets\|+1\|+1;m-28&3\|+\|188688834\|188688855&offsets\|+1\|+1]\|ACTAGATTGTGAGCTCCTGGAG | >MI0000086\|hsa-mir-28&WithFlank&3\|+\|188688775\|188688872 | 61 | 82 | 22 | ACTAGATTGTGAGCTCCTGGAG | 173.3 | 4.2 | 1.6 | 2.5 | 1.1E-02 | 1.4E-02 | MB231/MB468 | MP |
| >hg38_21_+_8433172_8446622_RNA45SN1_45S_50ntFlanks@11875.11906.32\|GAAGACCCTGTTGAGCTTGACTCTAGTCTGGC | >hg38_21_+_8433172_8446622_RNA45SN1_45S_50ntFlanks | 11875 | 11906 | 32 | GAAGACCCTGTTGAGCTTGACTCTAGTCTGGC | 184.1 | -1.8 | 0.7 | -2.6 | 1.0E-02 | 1.4E-02 | BT20/MB468 | MITO |
| >MI0000441\|hsa-mir-30b&WithFlank&8\|-\|134800514\|134800613@23.44.22&[MIMAT0000420&hsa-miR-30b-5p&offsets\|0\|0;m-76&8\|-\|134800570\|134800591&offsets\|0\|0]\|TGTAAACATCCTACACTCAGCT | >MI0000441\|hsa-mir-30b&WithFlank&8\|-\|134800514\|134800613 | 23 | 44 | 22 | TGTAAACATCCTACACTCAGCT | 2831.3 | -3.0 | 1.0 | -3.0 | 2.8E-03 | 1.4E-02 | BT20/MB231 | MP |
| >hg38_21_+_8433172_8446622_RNA45SN1_45S_50ntFlanks@8917.8940.24\|AATCCCGGGGCCGAGGGAGCGAGA | >hg38_21_+_8433172_8446622_RNA45SN1_45S_50ntFlanks | 8917 | 8940 | 24 | AATCCCGGGGCCGAGGGAGCGAGA | 100.0 | 2.4 | 0.9 | 2.7 | 7.3E-03 | 1.4E-02 | BT20/MB468 | MP |
| >hg38_21_+_8433172_8446622_RNA45SN1_45S_50ntFlanks@6771.6807.37\|GGTTCCTCCCGGGGCTACGCCTGTCTGAGCGTCGCTT | >hg38_21_+_8433172_8446622_RNA45SN1_45S_50ntFlanks | 6771 | 6807 | 37 | GGTTCCTCCCGGGGCTACGCCTGTCTGAGCGTCGCTT | 243.4 | -2.3 | 0.9 | -2.7 | 7.3E-03 | 1.4E-02 | BT20/MB468 | MP |
| >hg38_MT_+_1621_3279_RNR2_16S_50ntFlanks@1424.1448.25\|AACGATTAAAGTCCTACGTGATCTG | >hg38_MT_+_1621_3279_RNR2_16S_50ntFlanks | 1424 | 1448 | 25 | AACGATTAAAGTCCTACGTGATCTG | 130.1 | 2.9 | 1.1 | 2.5 | 1.1E-02 | 1.4E-02 | MB231/MB468 | MP |
| >MI0000809\|hsa-mir-151a&WithFlank&8\|-\|140732558\|140732659@51.72.22(+1A)&[MIMAT0000757&hsa-miR-151a-3p&offsets\|-2\|-1(+1A);m-30&8\|-\|140732587\|140732607&offsets\|-2\|-1(+1A)]\|TACTAGACTGAAGCTCCTTGAGA | >MI0000809\|hsa-mir-151a&WithFlank&8\|-\|140732558\|140732659 | 51 | 72 | 22(+1A) | TACTAGACTGAAGCTCCTTGAGA | 1464.5 | 2.1 | 0.7 | 2.9 | 3.6E-03 | 1.4E-02 | BT20/MB231 | CYTO |
| >hg38_21_+_8433172_8446622_RNA45SN1_45S_50ntFlanks@9308.9330.23\|AGGAGTCTAACACGTGCGCGAGT | >hg38_21_+_8433172_8446622_RNA45SN1_45S_50ntFlanks | 9308 | 9330 | 23 | AGGAGTCTAACACGTGCGCGAGT | 129.8 | -2.8 | 1.1 | -2.5 | 1.1E-02 | 1.4E-02 | MB231/MB468 | MP |
| >hg38_21_+_8433172_8446622_RNA45SN1_45S_50ntFlanks@12934.12968.35\|CTAAACCATTCGTAGACGACCTGCTTCTGGGTCGG | >hg38_21_+_8433172_8446622_RNA45SN1_45S_50ntFlanks | 12934 | 12968 | 35 | CTAAACCATTCGTAGACGACCTGCTTCTGGGTCGG | 60.3 | -4.9 | 1.8 | -2.7 | 7.3E-03 | 1.4E-02 | BT20/MB468 | MP |
| >MI0000239\|hsa-mir-197&WithFlank&1\|+\|109598887\|109598973@54.72.19&[MIMAT0000227&hsa-miR-197-3p&offsets\|0\|-3;m-99&1\|+\|109598940\|109598961&offsets\|0\|-3]\|TTCACCACCTTCTCCACCC | >MI0000239\|hsa-mir-197&WithFlank&1\|+\|109598887\|109598973 | 54 | 72 | 19 | TTCACCACCTTCTCCACCC | 373.2 | 2.1 | 0.7 | 2.9 | 3.6E-03 | 1.4E-02 | BT20/MB231 | MITO |
| >MI0000084\|hsa-mir-26b&WithFlank&2\|+\|218402640\|218402728@18.38.21(+1C)&[MIMAT0000083&hsa-miR-26b-5p&offsets\|0\|0(+1C);m-53&2\|+\|218402657\|218402678&offsets\|0\|-1(+1C)]\|TTCAAGTAATTCAGGATAGGTC | >MI0000084\|hsa-mir-26b&WithFlank&2\|+\|218402640\|218402728 | 18 | 38 | 21(+1C) | TTCAAGTAATTCAGGATAGGTC | 1040.8 | -2.4 | 0.8 | -2.9 | 3.6E-03 | 1.4E-02 | BT20/MB231 | MITO |
| >MI0000446\|hsa-mir-125b-1&WithFlank&11\|-\|122099751\|122099850@21.41.21(+1U);MI0000470\|hsa-mir-125b-2&WithFlank&21\|+\|16590231\|16590331@23.43.21(+1U)&[MIMAT0000423&hsa-miR-125b-5p&offsets\|0\|-1(+1U);m-67&11\|-\|122099809\|122099830&offsets\|0\|-1(+1U)];[MIMAT0000423_1&hsa-miR-125b-5p&offsets\|0\|-1(+1U);m-68&21\|+\|16590253\|16590274&offsets\|0\|-1(+1U)]\|TCCCTGAGACCCTAACTTGTGT | >MI0000446\|hsa-mir-125b-1&WithFlank&11\|-\|122099751\|122099850 | 21 | 41 | 21(+1U) | TCCCTGAGACCCTAACTTGTGT | 1459.3 | -7.2 | 2.5 | -2.9 | 3.6E-03 | 1.4E-02 | BT20/MB231 | MITO |
| >trna10_AspGTC_12_-_125424193_125424264@37.74.38,trna12_AspGTC_12_-_125411891_125411962@37.74.38,trna144_AspGTC_6_-_27551236_27551307@37.74.38,trna38_AspGTC_17_-_8125556_8125627@37.74.38,trna45_AspGTC_6_+_27447453_27447524@37.74.38,trna48_AspGTC_6_+_27471523_27471594@37.74.38,trna4_AspGTC_12_+_96429799_96429870@37.74.38,trna69_AspGTC_1_-_161440205_161440276@37.74.38,trna72_AspGTC_1_-_161432824_161432895@37.74.38,trna75_AspGTC_1_-_161425414_161425485@37.74.38,trna78_AspGTC_1_-_161418033_161418104@37.74.38,trna81_AspGTC_1_-_161410615_161410686@37.74.38\|ACGCGGGAGACCGGGGTTCGATTCCCCGACGGGGAGCC | >trna10_AspGTC_12_-_125424193_125424264 | 37 | 74 | 38 | ACGCGGGAGACCGGGGTTCGATTCCCCGACGGGGAGCC | 182.0 | -3.3 | 1.1 | -2.9 | 3.6E-03 | 1.4E-02 | BT20/MB231 | MITO |
| >hg38_21_+_8433172_8446622_RNA45SN1_45S_50ntFlanks@12420.12453.34\|ATGTCGGCTCTTCCTATCATTGTGAAGCAGAATT | >hg38_21_+_8433172_8446622_RNA45SN1_45S_50ntFlanks | 12420 | 12453 | 34 | ATGTCGGCTCTTCCTATCATTGTGAAGCAGAATT | 494.3 | -2.3 | 0.8 | -2.9 | 3.6E-03 | 1.4E-02 | BT20/MB231 | MITO |
| >hg38_21_+_8433172_8446622_RNA45SN1_45S_50ntFlanks@11583.11612.30\|CTGGTGCGGACCAGGGGAATCCGACTGTTT | >hg38_21_+_8433172_8446622_RNA45SN1_45S_50ntFlanks | 11583 | 11612 | 30 | CTGGTGCGGACCAGGGGAATCCGACTGTTT | 429.8 | 2.1 | 0.7 | 2.9 | 3.6E-03 | 1.4E-02 | BT20/MB231 | MITO |
| >hg38_21_+_8433172_8446622_RNA45SN1_45S_50ntFlanks@11582.11616.35\|ACTGGTGCGGACCAGGGGAATCCGACTGTTTAATT | >hg38_21_+_8433172_8446622_RNA45SN1_45S_50ntFlanks | 11582 | 11616 | 35 | ACTGGTGCGGACCAGGGGAATCCGACTGTTTAATT | 81.5 | 2.5 | 1.0 | 2.5 | 1.1E-02 | 1.4E-02 | MB231/MB468 | MP |
| >hg38_21_+_8433172_8446622_RNA45SN1_45S_50ntFlanks@12479.12497.19\|CTAATAGGGAACGTGAGCT | >hg38_21_+_8433172_8446622_RNA45SN1_45S_50ntFlanks | 12479 | 12497 | 19 | CTAATAGGGAACGTGAGCT | 515.8 | -2.0 | 0.8 | -2.5 | 1.1E-02 | 1.4E-02 | MB231/MB468 | MP |
| >hg38_21_+_8433172_8446622_RNA45SN1_45S_50ntFlanks@4731.4751.21\|AAGAACGAAAGTCGGAGGTTC | >hg38_21_+_8433172_8446622_RNA45SN1_45S_50ntFlanks | 4731 | 4751 | 21 | AAGAACGAAAGTCGGAGGTTC | 2467.1 | -2.5 | 0.9 | -2.7 | 7.3E-03 | 1.4E-02 | BT20/MB468 | MP |
| >hg38_21_+_8433172_8446622_RNA45SN1_45S_50ntFlanks@4585.4619.35\|GTTCTATTTTGTTGGTTTTCGGAACTGAGGCCATG | >hg38_21_+_8433172_8446622_RNA45SN1_45S_50ntFlanks | 4585 | 4619 | 35 | GTTCTATTTTGTTGGTTTTCGGAACTGAGGCCATG | 209.2 | -2.3 | 0.8 | -2.9 | 3.6E-03 | 1.4E-02 | BT20/MB231 | CYTO |
| >hg38_21_+_8433172_8446622_RNA45SN1_45S_50ntFlanks@12633.12656.24\|CGAAGCTACCATCTGTGGGATTAT | >hg38_21_+_8433172_8446622_RNA45SN1_45S_50ntFlanks | 12633 | 12656 | 24 | CGAAGCTACCATCTGTGGGATTAT | 323.9 | -2.2 | 0.9 | -2.6 | 1.0E-02 | 1.4E-02 | BT20/MB468 | MITO |
| >hg38_21_+_8433172_8446622_RNA45SN1_45S_50ntFlanks@11582.11607.26\|ACTGGTGCGGACCAGGGGAATCCGAC | >hg38_21_+_8433172_8446622_RNA45SN1_45S_50ntFlanks | 11582 | 11607 | 26 | ACTGGTGCGGACCAGGGGAATCCGAC | 96.1 | -2.6 | 1.0 | -2.5 | 1.1E-02 | 1.4E-02 | MB231/MB468 | MP |
| >hg38_21_+_8433172_8446622_RNA45SN1_45S_50ntFlanks@5066.5093.28\|TCTGGTTAATTCCGATAACGAACGAGAC | >hg38_21_+_8433172_8446622_RNA45SN1_45S_50ntFlanks | 5066 | 5093 | 28 | TCTGGTTAATTCCGATAACGAACGAGAC | 43.3 | -8.5 | 3.2 | -2.7 | 7.3E-03 | 1.4E-02 | BT20/MB468 | MP |
| >trnaMT_GluTTC_MT_-_14674_14742@46.65.20,trnalookalike8_GluTTC_5_-_93905172_93905240@46.65.20\|GTGGTTGTAGTCCGTGCGAG | >trnaMT_GluTTC_MT_-_14674_14742 | 46 | 65 | 20 | GTGGTTGTAGTCCGTGCGAG | 56.7 | 5.5 | 2.0 | 2.7 | 7.3E-03 | 1.4E-02 | BT20/MB468 | MP |
| >hg38_21_+_8433172_8446622_RNA45SN1_45S_50ntFlanks@8196.8216.21\|CGTGGACGGTGTGAGGCCGGT | >hg38_21_+_8433172_8446622_RNA45SN1_45S_50ntFlanks | 8196 | 8216 | 21 | CGTGGACGGTGTGAGGCCGGT | 748.8 | -2.3 | 0.9 | -2.7 | 7.3E-03 | 1.4E-02 | BT20/MB468 | MP |
| >hg38_21_+_8433172_8446622_RNA45SN1_45S_50ntFlanks@4378.4396.19\|CTCGTAGTTGGATCTTGGG | >hg38_21_+_8433172_8446622_RNA45SN1_45S_50ntFlanks | 4378 | 4396 | 19 | CTCGTAGTTGGATCTTGGG | 88.3 | -4.4 | 1.6 | -2.7 | 7.3E-03 | 1.4E-02 | BT20/MB468 | MP |
| >hg38_21_+_8433172_8446622_RNA45SN1_45S_50ntFlanks@6734.6772.39\|ATTGATCATCGACACTTCGAACGCACTTGCGGCCCCGGG | >hg38_21_+_8433172_8446622_RNA45SN1_45S_50ntFlanks | 6734 | 6772 | 39 | ATTGATCATCGACACTTCGAACGCACTTGCGGCCCCGGG | 3510.9 | 5.8 | 2.1 | 2.7 | 7.3E-03 | 1.4E-02 | BT20/MB468 | MP |
| >am_MI0000809\|hsa-mir-151a&WithFlank&8\|-\|140732558\|140732659@53.72.20(+1C)&[MIMAT0000757&hsa-miR-151a-3p&offsets\|0\|-1(+1C);m-30&8\|-\|140732587\|140732607&offsets\|0\|-1(+1C)]\|CTAGACTGAAGCTCCTTGAGC | >am_MI0000809\|hsa-mir-151a&WithFlank&8\|-\|140732558\|140732659 | 53 | 72 | 20(+1C) | CTAGACTGAAGCTCCTTGAGC | 1157.2 | 2.1 | 0.7 | 2.9 | 3.6E-03 | 1.4E-02 | BT20/MB231 | CYTO |
| >hg38_MT_+_598_1651_RNR1_12S_50ntFlanks@434.468.35\|GCTTTAACATATCTGAACACACAATAGCTAAGACC | >hg38_MT_+_598_1651_RNR1_12S_50ntFlanks | 434 | 468 | 35 | GCTTTAACATATCTGAACACACAATAGCTAAGACC | 21.6 | 7.7 | 3.0 | 2.5 | 1.1E-02 | 1.4E-02 | MB231/MB468 | MP |
| >hg38_21_+_8433172_8446622_RNA45SN1_45S_50ntFlanks@6660.6692.33\|GCGGTGGATCACTCGGCTCGTGCGTCGATGAAG | >hg38_21_+_8433172_8446622_RNA45SN1_45S_50ntFlanks | 6660 | 6692 | 33 | GCGGTGGATCACTCGGCTCGTGCGTCGATGAAG | 78.9 | -4.4 | 1.7 | -2.5 | 1.1E-02 | 1.4E-02 | MB231/MB468 | MP |
| >am_trna2_GlyTCC_19_+_4724082_4724153@1.32.32\|GCGTTGGTGGTATAGTGGTTAGCATAGCTGCC | >am_trna2_GlyTCC_19_+_4724082_4724153 | 1 | 32 | 32 | GCGTTGGTGGTATAGTGGTTAGCATAGCTGCC | 58.3 | 4.9 | 1.9 | 2.5 | 1.1E-02 | 1.4E-02 | MB231/MB468 | MP |
| >hg38_21_+_8433172_8446622_RNA45SN1_45S_50ntFlanks@11562.11584.23\|CGCCTAGCAGCCGACTTAGAACT | >hg38_21_+_8433172_8446622_RNA45SN1_45S_50ntFlanks | 11562 | 11584 | 23 | CGCCTAGCAGCCGACTTAGAACT | 123.5 | 5.8 | 2.0 | 3.0 | 2.9E-03 | 1.4E-02 | BT20/MB231 | MP |
| >hg38_21_+_8433172_8446622_RNA45SN1_45S_50ntFlanks@4661.4682.22\|AGGTGAAATTCTTGGACCGGCG | >hg38_21_+_8433172_8446622_RNA45SN1_45S_50ntFlanks | 4661 | 4682 | 22 | AGGTGAAATTCTTGGACCGGCG | 84.4 | -2.8 | 1.1 | -2.5 | 1.2E-02 | 1.4E-02 | BT20/MB468 | CYTO |
| >hg38_21_+_8433172_8446622_RNA45SN1_45S_50ntFlanks@10350.10391.42\|ACGAGAACTTTGAAGGCCGAAGTGGAGAAGGGTTCCATGTGA | >hg38_21_+_8433172_8446622_RNA45SN1_45S_50ntFlanks | 10350 | 10391 | 42 | ACGAGAACTTTGAAGGCCGAAGTGGAGAAGGGTTCCATGTGA | 167.0 | -2.7 | 1.0 | -2.7 | 7.4E-03 | 1.4E-02 | BT20/MB468 | MP |
| >hg38_21_+_8433172_8446622_RNA45SN1_45S_50ntFlanks@11570.11603.34\|AGCCGACTTAGAACTGGTGCGGACCAGGGGAATC | >hg38_21_+_8433172_8446622_RNA45SN1_45S_50ntFlanks | 11570 | 11603 | 34 | AGCCGACTTAGAACTGGTGCGGACCAGGGGAATC | 368.6 | 5.1 | 1.7 | 3.0 | 2.9E-03 | 1.4E-02 | BT20/MB231 | MP |
| >hg38_21_+_8433172_8446622_RNA45SN1_45S_50ntFlanks@7993.8012.20\|GTGGCGACCCGCTGAATTTA | >hg38_21_+_8433172_8446622_RNA45SN1_45S_50ntFlanks | 7993 | 8012 | 20 | GTGGCGACCCGCTGAATTTA | 246.8 | -3.1 | 1.2 | -2.6 | 1.0E-02 | 1.4E-02 | BT20/MB468 | MITO |
| >hg38_21_+_8433172_8446622_RNA45SN1_45S_50ntFlanks@4565.4587.23\|AATAATGGAATAGGACCGCGGTT | >hg38_21_+_8433172_8446622_RNA45SN1_45S_50ntFlanks | 4565 | 4587 | 23 | AATAATGGAATAGGACCGCGGTT | 587.1 | -1.2 | 0.5 | -2.5 | 1.2E-02 | 1.4E-02 | BT20/MB468 | CYTO |
| >trnaMT_HisGTG_MT_+_12138_12206@32.71.40\|TGAATCTGACAACAGAGGCTTACGACCCCTTATTTACCCC | >trnaMT_HisGTG_MT_+_12138_12206 | 32 | 71 | 40 | TGAATCTGACAACAGAGGCTTACGACCCCTTATTTACCCC | 533.5 | -3.8 | 1.5 | -2.5 | 1.2E-02 | 1.4E-02 | BT20/MB468 | NUC |
| >hg38_21_+_8433172_8446622_RNA45SN1_45S_50ntFlanks@9623.9659.37\|TAGTAGCTGGTTCCCTCCGAAGTTTCCCTCAGGATAG | >hg38_21_+_8433172_8446622_RNA45SN1_45S_50ntFlanks | 9623 | 9659 | 37 | TAGTAGCTGGTTCCCTCCGAAGTTTCCCTCAGGATAG | 99.9 | -3.8 | 1.5 | -2.6 | 1.0E-02 | 1.4E-02 | BT20/MB468 | MITO |
| >hg38_21_+_8433172_8446622_RNA45SN1_45S_50ntFlanks@5071.5093.23\|TTAATTCCGATAACGAACGAGAC | >hg38_21_+_8433172_8446622_RNA45SN1_45S_50ntFlanks | 5071 | 5093 | 23 | TTAATTCCGATAACGAACGAGAC | 118.5 | -5.0 | 1.8 | -2.7 | 7.4E-03 | 1.4E-02 | BT20/MB468 | MP |
| >hg38_21_+_8433172_8446622_RNA45SN1_45S_50ntFlanks@6723.6740.18\|TTGCAGGACACATTGATC | >hg38_21_+_8433172_8446622_RNA45SN1_45S_50ntFlanks | 6723 | 6740 | 18 | TTGCAGGACACATTGATC | 401.1 | -3.1 | 1.2 | -2.5 | 1.2E-02 | 1.4E-02 | MB231/MB468 | CYTO |
| >trna27_GlyCCC_2_-_70476123_70476193@1.29.29,trna34_GlyCCC_16_-_686736_686806@1.29.29\|GCGCCGCTGGTGTAGTGGTATCATGCAAG | >trna27_GlyCCC_2_-_70476123_70476193 | 1 | 29 | 29 | GCGCCGCTGGTGTAGTGGTATCATGCAAG | 203.2 | -3.3 | 1.3 | -2.5 | 1.2E-02 | 1.4E-02 | MB231/MB468 | MITO |
| >MI0000441\|hsa-mir-30b&WithFlank&8\|-\|134800514\|134800613@23.43.21&[MIMAT0000420&hsa-miR-30b-5p&offsets\|0\|-1;m-76&8\|-\|134800570\|134800591&offsets\|0\|-1]\|TGTAAACATCCTACACTCAGC | >MI0000441\|hsa-mir-30b&WithFlank&8\|-\|134800514\|134800613 | 23 | 43 | 21 | TGTAAACATCCTACACTCAGC | 295.6 | 3.5 | 1.4 | 2.5 | 1.1E-02 | 1.4E-02 | MB231/MB468 | MP |
| >MI0000447\|hsa-mir-128-1&WithFlank&2\|+\|135665391\|135665484@56.75.20(+2A);MI0000727\|hsa-mir-128-2&WithFlank&3\|+\|35744470\|35744565@58.77.20(+2A)&[MIMAT0000424&hsa-miR-128-3p&offsets\|0\|-1(+2A);m-108&2\|+\|135665446\|135665466&offsets\|0\|-1(+2A)];[MIMAT0000424_1&hsa-miR-128-3p&offsets\|0\|-1(+2A);m-117&3\|+\|35744527\|35744547&offsets\|0\|-1(+2A)]\|TCACAGTGAACCGGTCTCTTAA | >MI0000447\|hsa-mir-128-1&WithFlank&2\|+\|135665391\|135665484 | 56 | 75 | 20(+2A) | TCACAGTGAACCGGTCTCTTAA | 132.5 | 2.8 | 1.0 | 2.7 | 7.4E-03 | 1.4E-02 | BT20/MB468 | MP |

<table>
<tr><td>>hg38_21_+_8433172_8446622_RNA45SN1_45S_50ntFlanks@6646.6689.44|TCGTACGACTCTTAGCGGTGGATCACTCGGCTCGTGCGTCGATG</td><td>>hg38_21_+_8433172_8446622_RNA45SN1_45S_50ntFlanks</td><td>6646</td><td>6689</td><td>44</td><td>TCGTACGACTCTTAGCGGTGGATCACTCGGCTCGTGCGTCGATG</td><td>6772.8</td><td>-2.9</td><td>1.1</td><td>-2.5</td><td>1.1E-02</td><td>1.4E-02</td><td>MB231/MB468</td><td>MP</td></tr>
<tr><td>>hg38_21_+_8433172_8446622_RNA45SN1_45S_50ntFlanks@4297.4314.18|CATTGGAGGGCAAGTCTG</td><td>>hg38_21_+_8433172_8446622_RNA45SN1_45S_50ntFlanks</td><td>4297</td><td>4314</td><td>18</td><td>CATTGGAGGGCAAGTCTG</td><td>409.5</td><td>-1.7</td><td>0.7</td><td>-2.5</td><td>1.2E-02</td><td>1.4E-02</td><td>BT20/MB468</td><td>CYTO</td></tr>
<tr><td>>MI0000061|hsa-let-7a-2&WithFlank&11|-|122146516|122146599@11.30.20(+1C);MI0000062|hsa-let-7a-3&WithFlank&22|+|46112743|46112828@10.29.20(+1C);MI0000060|hsa-let-7a-1&WithFlank&9|+|94175951|94176042@12.31.20(+1C)&[MIMAT0000062&hsa-let-7a-5p&offsets|0|-2(+1C);m-6&11|-|122146568|122146589&offsets|0|-2(+1C)];[MIMAT0000062_1&hsa-let-7a-5p&offsets|0|-2(+1C);m-5&22|+|46112752|46112773&offsets|0|-2(+1C)];[MIMAT0000062_2&hsa-let-7a-5p&offsets|0|-2(+1C);m-4&9|+|94175962|94175983&offsets|0|-2(+1C)]|TGAGGTAGTAGGTTGTATAGC</td><td>>MI0000061|hsa-let-7a-2&WithFlank&11|-|122146516|122146599</td><td>11</td><td>30</td><td>20(+1C)</td><td>TGAGGTAGTAGGTTGTATAGC</td><td>223.2</td><td>3.7</td><td>1.4</td><td>2.7</td><td>7.4E-03</td><td>1.4E-02</td><td>BT20/MB468</td><td>MP</td></tr>
<tr><td>>hg38_21_+_8433172_8446622_RNA45SN1_45S_50ntFlanks@8380.8418.39|CAAGAGGGCGTGAAACCGTTAAGAGGTAAACGGGTGGGG</td><td>>hg38_21_+_8433172_8446622_RNA45SN1_45S_50ntFlanks</td><td>8380</td><td>8418</td><td>39</td><td>CAAGAGGGCGTGAAACCGTTAAGAGGTAAACGGGTGGGG</td><td>71.7</td><td>-4.8</td><td>1.8</td><td>-2.7</td><td>7.4E-03</td><td>1.4E-02</td><td>BT20/MB468</td><td>MP</td></tr>
<tr><td>>MI0000077|hsa-mir-21&WithFlank&17|+|59841260|59841343@14.33.20(+1A)&[MIMAT0000076&hsa-miR-21-5p&offsets|0|-2(+1A);m-1&17|+|59841273|59841294&offsets|0|-2(+1A)]|TAGCTTATCAGACTGATGTTA</td><td>>MI0000077|hsa-mir-21&WithFlank&17|+|59841260|59841343</td><td>14</td><td>33</td><td>20(+1A)</td><td>TAGCTTATCAGACTGATGTTA</td><td>3074.0</td><td>3.2</td><td>1.1</td><td>2.9</td><td>3.7E-03</td><td>1.4E-02</td><td>BT20/MB231</td><td>CYTO</td></tr>
<tr><td>>hg38_21_+_8433172_8446622_RNA45SN1_45S_50ntFlanks@7979.7999.21|ACCTCAGATCAGACGTGGCGA</td><td>>hg38_21_+_8433172_8446622_RNA45SN1_45S_50ntFlanks</td><td>7979</td><td>7999</td><td>21</td><td>ACCTCAGATCAGACGTGGCGA</td><td>150.6</td><td>-3.4</td><td>1.3</td><td>-2.7</td><td>7.4E-03</td><td>1.4E-02</td><td>BT20/MB468</td><td>MP</td></tr>
<tr><td>>hg38_1_-_228634819_228635039_RNA5S12_5S_50ntFlanks@133.169.37|ATGGGAGACCGCCTGGGAATACCGGGTGCTGTAGGCT</td><td>>hg38_1_-_228634819_228635039_RNA5S12_5S_50ntFlanks</td><td>133</td><td>169</td><td>37</td><td>ATGGGAGACCGCCTGGGAATACCGGGTGCTGTAGGCT</td><td>651.2</td><td>1.8</td><td>0.7</td><td>2.5</td><td>1.2E-02</td><td>1.4E-02</td><td>MB231/MB468</td><td>NUC</td></tr>
<tr><td>>hg38_21_+_8433172_8446622_RNA45SN1_45S_50ntFlanks@9544.9569.26|GGTGGAGGTCCGTAGCGGTCCTGACG</td><td>>hg38_21_+_8433172_8446622_RNA45SN1_45S_50ntFlanks</td><td>9544</td><td>9569</td><td>26</td><td>GGTGGAGGTCCGTAGCGGTCCTGACG</td><td>172.6</td><td>-3.4</td><td>1.3</td><td>-2.5</td><td>1.1E-02</td><td>1.4E-02</td><td>MB231/MB468</td><td>MP</td></tr>
<tr><td>>MI0000086|hsa-mir-28&WithFlank&3|+|188688775|188688872@61.82.22&[MIMAT0004502&hsa-miR-28-3p&offsets|+1|+1;m-28&3|+|188688834|188688855&offsets|+1|+1]|ACTAGATTGTGAGCTCCTGGAG</td><td>>MI0000086|hsa-mir-28&WithFlank&3|+|188688775|188688872</td><td>61</td><td>82</td><td>22</td><td>ACTAGATTGTGAGCTCCTGGAG</td><td>1282.7</td><td>2.4</td><td>0.8</td><td>2.9</td><td>3.7E-03</td><td>1.4E-02</td><td>BT20/MB231</td><td>CYTO</td></tr>
<tr><td>>hg38_21_+_8433172_8446622_RNA45SN1_45S_50ntFlanks@8176.8201.26|CTTCTGATCGAGGCCCAGCCCGTGGA</td><td>>hg38_21_+_8433172_8446622_RNA45SN1_45S_50ntFlanks</td><td>8176</td><td>8201</td><td>26</td><td>CTTCTGATCGAGGCCCAGCCCGTGGA</td><td>575.6</td><td>-1.8</td><td>0.7</td><td>-2.5</td><td>1.2E-02</td><td>1.4E-02</td><td>MB231/MB468</td><td>CYTO</td></tr>
<tr><td>>trnaMT_ThrTGT_MT_+_15888_15953@38.69.32|CCGGAGATGAAAACCTTTTTCCAAGGACACCA</td><td>>trnaMT_ThrTGT_MT_+_15888_15953</td><td>38</td><td>69</td><td>32</td><td>CCGGAGATGAAAACCTTTTTCCAAGGACACCA</td><td>188.4</td><td>2.5</td><td>1.0</td><td>2.5</td><td>1.1E-02</td><td>1.4E-02</td><td>MB231/MB468</td><td>MP</td></tr>
<tr><td>>hg38_21_+_8433172_8446622_RNA45SN1_45S_50ntFlanks@5082.5112.31|AACGAACGAGACTCTGGCATGCTAACTAGTT</td><td>>hg38_21_+_8433172_8446622_RNA45SN1_45S_50ntFlanks</td><td>5082</td><td>5112</td><td>31</td><td>AACGAACGAGACTCTGGCATGCTAACTAGTT</td><td>196.9</td><td>-2.6</td><td>1.0</td><td>-2.5</td><td>1.1E-02</td><td>1.4E-02</td><td>MB231/MB468</td><td>MP</td></tr>
<tr><td>>MI0000272|hsa-mir-182&WithFlank&7|-|129770377|129770498@29.50.22(+1U)&[MIMAT0000259&hsa-miR-182-5p&offsets|0|-2(+1U);m-22&7|-|129770449|129770470&offsets|0|0(+1U)]|TTTGGCAATGGTAGAACTCACAT</td><td>>MI0000272|hsa-mir-182&WithFlank&7|-|129770377|129770498</td><td>29</td><td>50</td><td>22(+1U)</td><td>TTTGGCAATGGTAGAACTCACAT</td><td>223.2</td><td>2.3</td><td>0.8</td><td>2.9</td><td>3.7E-03</td><td>1.4E-02</td><td>BT20/MB231</td><td>MITO</td></tr>
<tr><td>>am_trnaMT_ValTAC_MT_+_1602_1670@28.57.30|AACTTACACTTAGGAGATTTCAACTTAACT</td><td>>am_trnaMT_ValTAC_MT_+_1602_1670</td><td>28</td><td>57</td><td>30</td><td>AACTTACACTTAGGAGATTTCAACTTAACT</td><td>169.6</td><td>4.3</td><td>1.7</td><td>2.5</td><td>1.1E-02</td><td>1.4E-02</td><td>MB231/MB468</td><td>MP</td></tr>
<tr><td>>hg38_21_+_8433172_8446622_RNA45SN1_45S_50ntFlanks@12657.12674.18|GACTGAACGCCTCTAAGT</td><td>>hg38_21_+_8433172_8446622_RNA45SN1_45S_50ntFlanks</td><td>12657</td><td>12674</td><td>18</td><td>GACTGAACGCCTCTAAGT</td><td>169.7</td><td>-4.6</td><td>1.7</td><td>-2.7</td><td>7.4E-03</td><td>1.4E-02</td><td>BT20/MB468</td><td>MP</td></tr>
<tr><td>>am_trnaMT_HisGTG_MT_+_12138_12206@19.38.20|AAACATCAGATTGTGAATCT</td><td>>am_trnaMT_HisGTG_MT_+_12138_12206</td><td>19</td><td>38</td><td>20</td><td>AAACATCAGATTGTGAATCT</td><td>88.6</td><td>2.3</td><td>0.9</td><td>2.5</td><td>1.1E-02</td><td>1.4E-02</td><td>MB231/MB468</td><td>MP</td></tr>
<tr><td>>MI0000070|hsa-mir-16-1&WithFlank&13|-|50048967|50049067@20.40.21(+1A);MI0000115|hsa-mir-16-2&WithFlank&3|+|160404739|160404831@16.36.21(+1A)&[MIMAT0000069&hsa-miR-16-5p&offsets|0|-1(+1A);m-62&13|-|50049027|50049048&offsets|0|-1(+1A)];[MIMAT0000069_1&hsa-miR-16-5p&offsets|0|-1(+1A);m-61&3|+|160404754|160404775&offsets|0|-1(+1A)]|TAGCAGCACGTAAATATTGGCA</td><td>>MI0000070|hsa-mir-16-1&WithFlank&13|-|50048967|50049067</td><td>20</td><td>40</td><td>21(+1A)</td><td>TAGCAGCACGTAAATATTGGCA</td><td>1578.2</td><td>2.9</td><td>1.0</td><td>2.9</td><td>3.7E-03</td><td>1.4E-02</td><td>BT20/MB231</td><td>CYTO</td></tr>
<tr><td>>trnaMT_ValTAC_MT_+_1602_1670@47.71.25|TCAACTTAACTTGACCGCTCTGACC</td><td>>trnaMT_ValTAC_MT_+_1602_1670</td><td>47</td><td>71</td><td>25</td><td>TCAACTTAACTTGACCGCTCTGACC</td><td>638.9</td><td>2.4</td><td>0.9</td><td>2.6</td><td>1.0E-02</td><td>1.4E-02</td><td>BT20/MB468</td><td>MITO</td></tr>
<tr><td>>MI0000101|hsa-mir-99a&WithFlank&21|+|16539083|16539175@18.39.22&[MIMAT0000097&hsa-miR-99a-5p&offsets|-1|-1;m-64&21|+|16539101|16539121&offsets|-1|0]|AAACCCGTAGATCCGATCTTGT</td><td>>MI0000101|hsa-mir-99a&WithFlank&21|+|16539083|16539175</td><td>18</td><td>39</td><td>22</td><td>AAACCCGTAGATCCGATCTTGT</td><td>395.4</td><td>-2.0</td><td>0.8</td><td>-2.5</td><td>1.1E-02</td><td>1.4E-02</td><td>MB231/MB468</td><td>MP</td></tr>
<tr><td>>MI0000809|hsa-mir-151a&WithFlank&8|-|140732558|140732659@53.75.23&[MIMAT0000757&hsa-miR-151a-3p&offsets|0|+2;m-30&8|-|140732587|140732607&offsets|0|+2]|CTAGACTGAAGCTCCTTGAGGAC</td><td>>MI0000809|hsa-mir-151a&WithFlank&8|-|140732558|140732659</td><td>53</td><td>75</td><td>23</td><td>CTAGACTGAAGCTCCTTGAGGAC</td><td>320.4</td><td>2.0</td><td>0.7</td><td>2.9</td><td>3.7E-03</td><td>1.4E-02</td><td>BT20/MB231</td><td>CYTO</td></tr>
<tr><td>>MI0000115|hsa-mir-16-2&WithFlank&3|+|160404739|160404831@58.78.21(+1C)&[MIMAT0004518&hsa-miR-16-2-3p&offsets|-1|-2(+1C);m-252&3|+|160404796|160404816&offsets|0|0(+1C)]|ACCAATATTACTGTGCTGCTTC</td><td>>MI0000115|hsa-mir-16-2&WithFlank&3|+|160404739|160404831</td><td>58</td><td>78</td><td>21(+1C)</td><td>ACCAATATTACTGTGCTGCTTC</td><td>142.7</td><td>4.4</td><td>1.7</td><td>2.5</td><td>1.1E-02</td><td>1.4E-02</td><td>MB231/MB468</td><td>MP</td></tr>
<tr><td>>hg38_21_+_8433172_8446622_RNA45SN1_45S_50ntFlanks@6726.6743.18|CAGGACACATTGATCATC</td><td>>hg38_21_+_8433172_8446622_RNA45SN1_45S_50ntFlanks</td><td>6726</td><td>6743</td><td>18</td><td>CAGGACACATTGATCATC</td><td>125.0</td><td>-4.5</td><td>1.7</td><td>-2.7</td><td>7.4E-03</td><td>1.4E-02</td><td>BT20/MB468</td><td>MP</td></tr>
<tr><td>>hg38_1_-_228634819_228635039_RNA5S12_5S_50ntFlanks@152.171.20|TACCGGGTGCTGTAGGCTTT</td><td>>hg38_1_-_228634819_228635039_RNA5S12_5S_50ntFlanks</td><td>152</td><td>171</td><td>20</td><td>TACCGGGTGCTGTAGGCTTT</td><td>5611.6</td><td>-2.5</td><td>0.9</td><td>-2.7</td><td>7.4E-03</td><td>1.4E-02</td><td>BT20/MB468</td><td>MP</td></tr>
<tr><td>>MI0000095|hsa-mir-93&WithFlank&7|-|100093762|100093853@17.37.21&[MIMAT0000093&hsa-miR-93-5p&offsets|0|-2;m-29&7|-|100093815|100093837&offsets|0|-2]|CAAAGTGCTGTTCGTGCAGGT</td><td>>MI0000095|hsa-mir-93&WithFlank&7|-|100093762|100093853</td><td>17</td><td>37</td><td>21</td><td>CAAAGTGCTGTTCGTGCAGGT</td><td>1724.9</td><td>-2.3</td><td>0.8</td><td>-2.9</td><td>3.7E-03</td><td>1.4E-02</td><td>BT20/MB231</td><td>MITO</td></tr>
<tr><td>>am_trnaMT_MetCAT_MT_+_4402_4469@1.19.19|AGTAAGGTCAGCTAAATAA</td><td>>am_trnaMT_MetCAT_MT_+_4402_4469</td><td>1</td><td>19</td><td>19</td><td>AGTAAGGTCAGCTAAATAA</td><td>263.6</td><td>-5.5</td><td>1.8</td><td>-3.0</td><td>2.9E-03</td><td>1.4E-02</td><td>BT20/MB231</td><td>MP</td></tr>
<tr><td>>hg38_21_+_8433172_8446622_RNA45SN1_45S_50ntFlanks@6651.6673.23|CGACTCTTAGCGGTGGATCACTC</td><td>>hg38_21_+_8433172_8446622_RNA45SN1_45S_50ntFlanks</td><td>6651</td><td>6673</td><td>23</td><td>CGACTCTTAGCGGTGGATCACTC</td><td>2601.6</td><td>3.6</td><td>1.2</td><td>3.0</td><td>2.9E-03</td><td>1.4E-02</td><td>BT20/MB231</td><td>MP</td></tr>
<tr><td>>am_MI0000815|hsa-mir-339&WithFlank&7|-|1022927|1023032@21.40.20&[MIMAT0000764&hsa-miR-339-5p&offsets|0|-3;m-163&7|-|1022990|1023012&offsets|0|-3]|TCCCTGTCCTCCAGGAGCTC</td><td>>am_MI0000815|hsa-mir-339&WithFlank&7|-|1022927|1023032</td><td>21</td><td>40</td><td>20</td><td>TCCCTGTCCTCCAGGAGCTC</td><td>169.4</td><td>2.2</td><td>0.9</td><td>2.5</td><td>1.1E-02</td><td>1.4E-02</td><td>MB231/MB468</td><td>MP</td></tr>
<tr><td>>hg38_21_+_8433172_8446622_RNA45SN1_45S_50ntFlanks@10854.10873.20|TAAGGATTGGCTCTAAGGGC</td><td>>hg38_21_+_8433172_8446622_RNA45SN1_45S_50ntFlanks</td><td>10854</td><td>10873</td><td>20</td><td>TAAGGATTGGCTCTAAGGGC</td><td>143.7</td><td>5.8</td><td>2.0</td><td>3.0</td><td>2.9E-03</td><td>1.4E-02</td><td>BT20/MB231</td><td>MP</td></tr>
<tr><td>>am_trnaMT_GluTTC_MT_-_14674_14742@41.58.18,trnalookalike8_GluTTC_5_-_93905172_93905240@41.58.18|TGGTCGTGGTTGTAGTCC</td><td>>am_trnaMT_GluTTC_MT_-_14674_14742</td><td>41</td><td>58</td><td>18</td><td>TGGTCGTGGTTGTAGTCC</td><td>137.3</td><td>3.4</td><td>1.3</td><td>2.5</td><td>1.1E-02</td><td>1.4E-02</td><td>MB231/MB468</td><td>MP</td></tr>
<tr><td>>hg38_MT_+_1621_3279_RNR2_16S_50ntFlanks@459.492.34|CTTGTAAATTTAACTGTTAGTCCAAAGAGGAACA</td><td>>hg38_MT_+_1621_3279_RNR2_16S_50ntFlanks</td><td>459</td><td>492</td><td>34</td><td>CTTGTAAATTTAACTGTTAGTCCAAAGAGGAACA</td><td>285.1</td><td>3.1</td><td>1.2</td><td>2.5</td><td>1.1E-02</td><td>1.4E-02</td><td>MB231/MB468</td><td>MP</td></tr>
<tr><td>>MI0000448|hsa-mir-130a&WithFlank&11|+|57641192|57641292@61.83.23&[MIMAT0000425&hsa-miR-130a-3p&offsets|0|+1;m-139&11|+|57641252|57641273&offsets|0|+1]|CAGTGCAATGTTAAAAGGGCATT</td><td>>MI0000448|hsa-mir-130a&WithFlank&11|+|57641192|57641292</td><td>61</td><td>83</td><td>23</td><td>CAGTGCAATGTTAAAAGGGCATT</td><td>289.4</td><td>-4.4</td><td>1.5</td><td>-3.0</td><td>2.9E-03</td><td>1.4E-02</td><td>BT20/MB231</td><td>MP</td></tr>
<tr><td>>hg38_21_+_8433172_8446622_RNA45SN1_45S_50ntFlanks@12416.12441.26|TTCGATGTCGGCTCTTCCTATCATTG</td><td>>hg38_21_+_8433172_8446622_RNA45SN1_45S_50ntFlanks</td><td>12416</td><td>12441</td><td>26</td><td>TTCGATGTCGGCTCTTCCTATCATTG</td><td>958.8</td><td>1.0</td><td>0.4</td><td>2.5</td><td>1.2E-02</td><td>1.4E-02</td><td>BT20/MB468</td><td>CYTO</td></tr>
<tr><td>>hg38_21_+_8433172_8446622_RNA45SN1_45S_50ntFlanks@11921.11948.28|TGAGAGGTGTAGAATAAGTGGGAGGCCC</td><td>>hg38_21_+_8433172_8446622_RNA45SN1_45S_50ntFlanks</td><td>11921</td><td>11948</td><td>28</td><td>TGAGAGGTGTAGAATAAGTGGGAGGCCC</td><td>1001.6</td><td>-3.7</td><td>1.4</td><td>-2.7</td><td>7.5E-03</td><td>1.4E-02</td><td>BT20/MB468</td><td>MP</td></tr>
<tr><td>>hg38_MT_+_1621_3279_RNR2_16S_50ntFlanks@116.145.30|AATAAAGTATAGGCGATAGAAATTGAAACC</td><td>>hg38_MT_+_1621_3279_RNR2_16S_50ntFlanks</td><td>116</td><td>145</td><td>30</td><td>AATAAAGTATAGGCGATAGAAATTGAAACC</td><td>66.6</td><td>4.9</td><td>1.8</td><td>2.7</td><td>7.5E-03</td><td>1.4E-02</td><td>BT20/MB468</td><td>MP</td></tr>
<tr><td>>hg38_21_+_8433172_8446622_RNA45SN1_45S_50ntFlanks@6655.6676.22|TCTTAGCGGTGGATCACTCGGC</td><td>>hg38_21_+_8433172_8446622_RNA45SN1_45S_50ntFlanks</td><td>6655</td><td>6676</td><td>22</td><td>TCTTAGCGGTGGATCACTCGGC</td><td>103.5</td><td>4.6</td><td>1.7</td><td>2.7</td><td>7.5E-03</td><td>1.4E-02</td><td>BT20/MB468</td><td>MP</td></tr>
<tr><td>>trna119_LysCTT_1_-_145395522_145395594@3.36.34,trna11_LysCTT_5_-_180648979_180649051@3.36.34,trna13_LysCTT_6_+_26556774_26556846@3.36.34,trna7_LysCTT_16_+_3225692_3225764@3.36.34,trna9_LysCTT_5_+_180634755_180634827@3.36.34|CCGGCTAGCTCAGTCGGTAGAGCATGAGACTCTT</td><td>>trna119_LysCTT_1_-_145395522_145395594</td><td>3</td><td>36</td><td>34</td><td>CCGGCTAGCTCAGTCGGTAGAGCATGAGACTCTT</td><td>132.9</td><td>4.2</td><td>1.6</td><td>2.5</td><td>1.1E-02</td><td>1.4E-02</td><td>MB231/MB468</td><td>MP</td></tr>
<tr><td>>hg38_21_+_8433172_8446622_RNA45SN1_45S_50ntFlanks@9956.9978.23|GAAGTCGGAATCCGCTAAGGAGT</td><td>>hg38_21_+_8433172_8446622_RNA45SN1_45S_50ntFlanks</td><td>9956</td><td>9978</td><td>23</td><td>GAAGTCGGAATCCGCTAAGGAGT</td><td>93.9</td><td>4.5</td><td>1.7</td><td>2.7</td><td>7.5E-03</td><td>1.4E-02</td><td>BT20/MB468</td><td>MP</td></tr>
<tr><td>>hg38_21_+_8433172_8446622_RNA45SN1_45S_50ntFlanks@6657.6681.25|TTAGCGGTGGATCACTCGGCTCGTG</td><td>>hg38_21_+_8433172_8446622_RNA45SN1_45S_50ntFlanks</td><td>6657</td><td>6681</td><td>25</td><td>TTAGCGGTGGATCACTCGGCTCGTG</td><td>529.2</td><td>-1.5</td><td>0.6</td><td>-2.5</td><td>1.3E-02</td><td>1.4E-02</td><td>MB231/MB468</td><td>NUC</td></tr>
<tr><td>>MI0000809|hsa-mir-151a&WithFlank&8|-|140732558|140732659@51.72.22&[MIMAT0000757&hsa-miR-151a-3p&offsets|-2|-1;m-30&8|-|140732587|140732607&offsets|-2|-1]|TACTAGACTGAAGCTCCTTGAG</td><td>>MI0000809|hsa-mir-151a&WithFlank&8|-|140732558|140732659</td><td>51</td><td>72</td><td>22</td><td>TACTAGACTGAAGCTCCTTGAG</td><td>1044.3</td><td>2.0</td><td>0.7</td><td>2.9</td><td>3.7E-03</td><td>1.4E-02</td><td>BT20/MB231</td><td>CYTO</td></tr>
<tr><td>>am_trnaMT_ValTAC_MT_+_1602_1670@49.68.20|AACTTAACTTGACCGCTCTG</td><td>>am_trnaMT_ValTAC_MT_+_1602_1670</td><td>49</td><td>68</td><td>20</td><td>AACTTAACTTGACCGCTCTG</td><td>286.1</td><td>2.4</td><td>0.8</td><td>2.9</td><td>3.7E-03</td><td>1.4E-02</td><td>BT20/MB231</td><td>CYTO</td></tr>
<tr><td>>hg38_21_+_8433172_8446622_RNA45SN1_45S_50ntFlanks@475.494.20|CTTCGTGATCGATGTGGTGA</td><td>>hg38_21_+_8433172_8446622_RNA45SN1_45S_50ntFlanks</td><td>475</td><td>494</td><td>20</td><td>CTTCGTGATCGATGTGGTGA</td><td>68.6</td><td>5.2</td><td>2.0</td><td>2.5</td><td>1.1E-02</td><td>1.4E-02</td><td>MB231/MB468</td><td>MP</td></tr>
<tr><td>>hg38_21_+_8433172_8446622_RNA45SN1_45S_50ntFlanks@7979.8007.29|ACCTCAGATCAGACGTGGCGACCCGCTGA</td><td>>hg38_21_+_8433172_8446622_RNA45SN1_45S_50ntFlanks</td><td>7979</td><td>8007</td><td>29</td><td>ACCTCAGATCAGACGTGGCGACCCGCTGA</td><td>506.6</td><td>-1.6</td><td>0.6</td><td>-2.6</td><td>1.0E-02</td><td>1.4E-02</td><td>BT20/MB468</td><td>MITO</td></tr>
<tr><td>>trnaMT_SerGCT_MT_+_12207_12265@1.26.26|GAGAAAGCTCACAAGAACTGCTAACT</td><td>>trnaMT_SerGCT_MT_+_12207_12265</td><td>1</td><td>26</td><td>26</td><td>GAGAAAGCTCACAAGAACTGCTAACT</td><td>7599.4</td><td>2.1</td><td>0.9</td><td>2.5</td><td>1.3E-02</td><td>1.4E-02</td><td>MB231/MB468</td><td>MITO</td></tr>
<tr><td>>trnaMT_AspGTC_MT_+_7518_7585@47.71.25|AGGCTAAATCCTATATATCTTACCA</td><td>>trnaMT_AspGTC_MT_+_7518_7585</td><td>47</td><td>71</td><td>25</td><td>AGGCTAAATCCTATATATCTTACCA</td><td>69.0</td><td>4.8</td><td>1.9</td><td>2.5</td><td>1.1E-02</td><td>1.4E-02</td><td>MB231/MB468</td><td>MP</td></tr>
<tr><td>>hg38_21_+_8433172_8446622_RNA45SN1_45S_50ntFlanks@11918.11939.22|ACATGAGAGGTGTAGAATAAGT</td><td>>hg38_21_+_8433172_8446622_RNA45SN1_45S_50ntFlanks</td><td>11918</td><td>11939</td><td>22</td><td>ACATGAGAGGTGTAGAATAAGT</td><td>269.4</td><td>3.7</td><td>1.3</td><td>3.0</td><td>2.9E-03</td><td>1.4E-02</td><td>BT20/MB231</td><td>MP</td></tr>
<tr><td>>hg38_21_+_8433172_8446622_RNA45SN1_45S_50ntFlanks@12646.12666.21|TGTGGGATTATGACTGAACGC</td><td>>hg38_21_+_8433172_8446622_RNA45SN1_45S_50ntFlanks</td><td>12646</td><td>12666</td><td>21</td><td>TGTGGGATTATGACTGAACGC</td><td>572.7</td><td>5.2</td><td>1.8</td><td>3.0</td><td>2.9E-03</td><td>1.4E-02</td><td>BT20/MB231</td><td>MP</td></tr>
<tr><td>>hg38_21_+_8433172_8446622_RNA45SN1_45S_50ntFlanks@11583.11607.25|CTGGTGCGGACCAGGGGAATCCGAC</td><td>>hg38_21_+_8433172_8446622_RNA45SN1_45S_50ntFlanks</td><td>11583</td><td>11607</td><td>25</td><td>CTGGTGCGGACCAGGGGAATCCGAC</td><td>571.0</td><td>1.6</td><td>0.6</td><td>2.6</td><td>1.0E-02</td><td>1.4E-02</td><td>BT20/MB468</td><td>MITO</td></tr>
<tr><td>>hg38_MT_+_1621_3279_RNR2_16S_50ntFlanks@1428.1458.31|ATTAAAGTCCTACGTGATCTGAGTTCAGACC</td><td>>hg38_MT_+_1621_3279_RNR2_16S_50ntFlanks</td><td>1428</td><td>1458</td><td>31</td><td>ATTAAAGTCCTACGTGATCTGAGTTCAGACC</td><td>281.5</td><td>-2.6</td><td>0.9</td><td>-2.9</td><td>3.2E-03</td><td>1.4E-02</td><td>BT20/MB231</td><td>NUC</td></tr>
<tr><td>>MI0000298|hsa-mir-221&WithFlank&X|-|45746151|45746272@71.91.21(+1C)&[MIMAT0000278&hsa-miR-221-3p&offsets|0|-2(+1C);m-91&X|-|45746181|45746202&offsets|0|-1(+1C)]|AGCTACATTGTCTGCTGGGTTC</td><td>>MI0000298|hsa-mir-221&WithFlank&X|-|45746151|45746272</td><td>71</td><td>91</td><td>21(+1C)</td><td>AGCTACATTGTCTGCTGGGTTC</td><td>12613.6</td><td>-3.2</td><td>1.1</td><td>-3.0</td><td>2.9E-03</td><td>1.4E-02</td><td>BT20/MB231</td><td>MP</td></tr>
</table>

<table>
<tr><td>>hg38_MT_+_1621_3279_RNR2_16S_50ntFlanks@618.652.35|AACATATAACTGAACTCCTCACACCCAATTGGACC</td><td>>hg38_MT_+_1621_3279_RNR2_16S_50ntFlanks</td><td>618</td><td>652</td><td>35</td><td>AACATATAACTGAACTCCTCACACCCAATTGGACC</td><td>97.4</td><td>2.9</td><td>1.2</td><td>2.5</td><td>1.1E-02</td><td>1.4E-02</td><td>MB231/MB468</td><td>MP</td></tr>
<tr><td>>trna10_AspGTC_12_-_125424193_125424264@38.71.34,trna12_AspGTC_12_-_125411891_125411962@38.71.34,trna144_AspGTC_6_-_27551236_27551307@38.71.34,trna38_AspGTC_17_-_8125556_8125627@38.71.34,trna45_AspGTC_6_+_27447453_27447524@38.71.34,trna48_AspGTC_6_+_27471523_27471594@38.71.34,trna4_AspGTC_12_+_96429799_96429870@38.71.34,trna69_AspGTC_1_-_161440205_161440276@38.71.34,trna72_AspGTC_1_-_161432824_161432895@38.71.34,trna75_AspGTC_1_-_161425414_161425485@38.71.34,trna78_AspGTC_1_-_161418033_161418104@38.71.34,trna81_AspGTC_1_-_161410615_161410686@38.71.34|CGCGGGAGACCGGGGTTCGATTCCCCGACGGGGA</td><td>>trna10_AspGTC_12_-_125424193_125424264</td><td>38</td><td>71</td><td>34</td><td>CGCGGGAGACCGGGGTTCGATTCCCCGACGGGGA</td><td>297.9</td><td>-2.6</td><td>0.9</td><td>-2.9</td><td>3.8E-03</td><td>1.4E-02</td><td>BT20/MB231</td><td>CYTO</td></tr>
<tr><td>>hg38_MT_+_598_1651_RNR1_12S_50ntFlanks@428.453.26|AAAGTGGCTTTAACATATCTGAACAC</td><td>>hg38_MT_+_598_1651_RNR1_12S_50ntFlanks</td><td>428</td><td>453</td><td>26</td><td>AAAGTGGCTTTAACATATCTGAACAC</td><td>276.9</td><td>3.0</td><td>1.1</td><td>2.7</td><td>7.5E-03</td><td>1.4E-02</td><td>BT20/MB468</td><td>MP</td></tr>
<tr><td>>hg38_21_+_8433172_8446622_RNA45SN1_45S_50ntFlanks@12035.12053.19|CGTTTTTTCACTGACCCGG</td><td>>hg38_21_+_8433172_8446622_RNA45SN1_45S_50ntFlanks</td><td>12035</td><td>12053</td><td>19</td><td>CGTTTTTTCACTGACCCGG</td><td>54.2</td><td>-4.6</td><td>1.7</td><td>-2.7</td><td>7.5E-03</td><td>1.4E-02</td><td>BT20/MB468</td><td>MP</td></tr>
<tr><td>>hg38_21_+_8433172_8446622_RNA45SN1_45S_50ntFlanks@8359.8395.37|AAGAACTTTGAAGAGAGAGTTCAAGAGGGCGTGAAAC</td><td>>hg38_21_+_8433172_8446622_RNA45SN1_45S_50ntFlanks</td><td>8359</td><td>8395</td><td>37</td><td>AAGAACTTTGAAGAGAGAGTTCAAGAGGGCGTGAAAC</td><td>155.1</td><td>5.9</td><td>2.0</td><td>3.0</td><td>2.9E-03</td><td>1.4E-02</td><td>BT20/MB231</td><td>MP</td></tr>
<tr><td>>hg38_21_+_8433172_8446622_RNA45SN1_45S_50ntFlanks@7980.8007.28|CCTCAGATCAGACGTGGCGACCCGCTGA</td><td>>hg38_21_+_8433172_8446622_RNA45SN1_45S_50ntFlanks</td><td>7980</td><td>8007</td><td>28</td><td>CCTCAGATCAGACGTGGCGACCCGCTGA</td><td>162.0</td><td>-3.9</td><td>1.6</td><td>-2.5</td><td>1.3E-02</td><td>1.4E-02</td><td>MB231/MB468</td><td>NUC</td></tr>
<tr><td>>hg38_21_+_8433172_8446622_RNA45SN1_45S_50ntFlanks@6648.6677.30|GTACGACTCTTAGCGGTGGATCACTCGGCT</td><td>>hg38_21_+_8433172_8446622_RNA45SN1_45S_50ntFlanks</td><td>6648</td><td>6677</td><td>30</td><td>GTACGACTCTTAGCGGTGGATCACTCGGCT</td><td>1006.8</td><td>3.9</td><td>1.3</td><td>3.0</td><td>2.9E-03</td><td>1.4E-02</td><td>BT20/MB231</td><td>MP</td></tr>
<tr><td>>MI0000746|hsa-mir-99b&WithFlank&19|+|51692606|51692687@13.32.20&[MIMAT0000689&hsa-miR-99b-5p&offsets|0|-2;m-10&19|+|51692618|51692639&offsets|0|-2]|CACCCGTAGAACCGACCTTG</td><td>>MI0000746|hsa-mir-99b&WithFlank&19|+|51692606|51692687</td><td>13</td><td>32</td><td>20</td><td>CACCCGTAGAACCGACCTTG</td><td>1654.9</td><td>1.9</td><td>0.7</td><td>2.9</td><td>3.7E-03</td><td>1.4E-02</td><td>BT20/MB231</td><td>MITO</td></tr>
<tr><td>>hg38_21_+_8433172_8446622_RNA45SN1_45S_50ntFlanks@11785.11806.22|TGAATGGATGAACGAGATTCCC</td><td>>hg38_21_+_8433172_8446622_RNA45SN1_45S_50ntFlanks</td><td>11785</td><td>11806</td><td>22</td><td>TGAATGGATGAACGAGATTCCC</td><td>262.4</td><td>-3.1</td><td>1.2</td><td>-2.6</td><td>1.0E-02</td><td>1.4E-02</td><td>BT20/MB468</td><td>MITO</td></tr>
<tr><td>>hg38_21_+_8433172_8446622_RNA45SN1_45S_50ntFlanks@10664.10706.43|CTTGGAAAGCGTCGCGGTTCCGGCGGCGTCCGGTGAGCTCTCG</td><td>>hg38_21_+_8433172_8446622_RNA45SN1_45S_50ntFlanks</td><td>10664</td><td>10706</td><td>43</td><td>CTTGGAAAGCGTCGCGGTTCCGGCGGCGTCCGGTGAGCTCTCG</td><td>683.4</td><td>-3.3</td><td>1.3</td><td>-2.6</td><td>1.0E-02</td><td>1.4E-02</td><td>BT20/MB468</td><td>MITO</td></tr>
<tr><td>>MI0000270|hsa-mir-181b-1&WithFlank&1|-|198858867|198858988@42.65.24;MI0000683|hsa-mir-181b-2&WithFlank&9|+|124693704|124693804@22.45.24&[MIMAT0000257&hsa-miR-181b-5p&offsets|0|+1;m-73&1|-|198858925|198858947&offsets|0|+1];[MIMAT0000257_1&hsa-miR-181b-5p&offsets|0|+1;m-72&9|+|124693725|124693747&offsets|0|+1]|AACATTCATTGCTGTCGGTGGGTT</td><td>>MI0000270|hsa-mir-181b-1&WithFlank&1|-|198858867|198858988</td><td>42</td><td>65</td><td>24</td><td>AACATTCATTGCTGTCGGTGGGTT</td><td>6226.2</td><td>-3.2</td><td>1.1</td><td>-3.0</td><td>2.9E-03</td><td>1.4E-02</td><td>BT20/MB231</td><td>MP</td></tr>
<tr><td>>hg38_21_+_8433172_8446622_RNA45SN1_45S_50ntFlanks@10345.10362.18|TTCAAACGAGAACTTTGA</td><td>>hg38_21_+_8433172_8446622_RNA45SN1_45S_50ntFlanks</td><td>10345</td><td>10362</td><td>18</td><td>TTCAAACGAGAACTTTGA</td><td>104.4</td><td>2.4</td><td>0.9</td><td>2.7</td><td>7.5E-03</td><td>1.4E-02</td><td>BT20/MB468</td><td>MP</td></tr>
<tr><td>>hg38_21_+_8433172_8446622_RNA45SN1_45S_50ntFlanks@12936.12961.26|AAACCATTCGTAGACGACCTGCTTCT</td><td>>hg38_21_+_8433172_8446622_RNA45SN1_45S_50ntFlanks</td><td>12936</td><td>12961</td><td>26</td><td>AAACCATTCGTAGACGACCTGCTTCT</td><td>231.5</td><td>-1.6</td><td>0.7</td><td>-2.5</td><td>1.2E-02</td><td>1.4E-02</td><td>MB231/MB468</td><td>CYTO</td></tr>
<tr><td>>am_trnaMT_PheGAA_MT_+_577_647@41.63.23|GTTTAGACGGGCTCACATCACCC</td><td>>am_trnaMT_PheGAA_MT_+_577_647</td><td>41</td><td>63</td><td>23</td><td>GTTTAGACGGGCTCACATCACCC</td><td>61.9</td><td>2.4</td><td>0.9</td><td>2.5</td><td>1.1E-02</td><td>1.4E-02</td><td>MB231/MB468</td><td>MP</td></tr>
<tr><td>>hg38_21_+_8433172_8446622_RNA45SN1_45S_50ntFlanks@10401.10424.24|GAACATGGGTCAGTCGGTCCTGAG</td><td>>hg38_21_+_8433172_8446622_RNA45SN1_45S_50ntFlanks</td><td>10401</td><td>10424</td><td>24</td><td>GAACATGGGTCAGTCGGTCCTGAG</td><td>15920.9</td><td>2.7</td><td>0.9</td><td>2.9</td><td>3.7E-03</td><td>1.4E-02</td><td>BT20/MB231</td><td>MITO</td></tr>
<tr><td>>hg38_21_+_8433172_8446622_RNA45SN1_45S_50ntFlanks@9939.9968.30|GCAGGACGGTGGCCATGGAAGTCGGAATCC</td><td>>hg38_21_+_8433172_8446622_RNA45SN1_45S_50ntFlanks</td><td>9939</td><td>9968</td><td>30</td><td>GCAGGACGGTGGCCATGGAAGTCGGAATCC</td><td>192.6</td><td>6.0</td><td>2.1</td><td>2.9</td><td>3.7E-03</td><td>1.4E-02</td><td>BT20/MB231</td><td>MITO</td></tr>
<tr><td>>MI0000253|hsa-mir-148a&WithFlank&7|-|25949913|25949992@50.70.21(+1G)&[MIMAT0000243&hsa-miR-148a-3p&offsets|0|-1(+1G);m-7&7|-|25949922|25949943&offsets|0|-1(+1G)]|TCAGTGCACTACAGAACTTTGG</td><td>>MI0000253|hsa-mir-148a&WithFlank&7|-|25949913|25949992</td><td>50</td><td>70</td><td>21(+1G)</td><td>TCAGTGCACTACAGAACTTTGG</td><td>1215.8</td><td>-2.0</td><td>0.8</td><td>-2.5</td><td>1.1E-02</td><td>1.4E-02</td><td>MB231/MB468</td><td>MP</td></tr>
<tr><td>>hg38_21_+_8433172_8446622_RNA45SN1_45S_50ntFlanks@7974.8006.33|ACGCGACCTCAGATCAGACGTGGCGACCCGCTG</td><td>>hg38_21_+_8433172_8446622_RNA45SN1_45S_50ntFlanks</td><td>7974</td><td>8006</td><td>33</td><td>ACGCGACCTCAGATCAGACGTGGCGACCCGCTG</td><td>5806.7</td><td>3.4</td><td>1.2</td><td>2.9</td><td>3.8E-03</td><td>1.4E-02</td><td>BT20/MB231</td><td>CYTO</td></tr>
<tr><td>>MI0000081|hsa-mir-24-2&WithFlank&19|-|13836281|13836365@56.77.22(+1A);MI0000080|hsa-mir-24-1&WithFlank&9|+|95086015|95086094@50.71.22(+1A)&[MIMAT0000080&hsa-miR-24-3p&offsets|0|0(+1A);m-36&19|-|13836288|13836310&offsets|0|-1(+1A)];[MIMAT0000080_1&hsa-miR-24-3p&offsets|0|0(+1A);m-35&9|+|95086064|95086086&offsets|0|-1(+1A)]|TGGCTCAGTTCAGCAGGAACAGA</td><td>>MI0000081|hsa-mir-24-2&WithFlank&19|-|13836281|13836365</td><td>56</td><td>77</td><td>22(+1A)</td><td>TGGCTCAGTTCAGCAGGAACAGA</td><td>857.8</td><td>1.8</td><td>0.6</td><td>2.9</td><td>3.2E-03</td><td>1.4E-02</td><td>BT20/MB231</td><td>NUC</td></tr>
<tr><td>>hg38_21_+_8433172_8446622_RNA45SN1_45S_50ntFlanks@10854.10873.20|TAAGGATTGGCTCTAAGGGC</td><td>>hg38_21_+_8433172_8446622_RNA45SN1_45S_50ntFlanks</td><td>10854</td><td>10873</td><td>20</td><td>TAAGGATTGGCTCTAAGGGC</td><td>208.3</td><td>-1.7</td><td>0.7</td><td>-2.5</td><td>1.2E-02</td><td>1.4E-02</td><td>MB231/MB468</td><td>CYTO</td></tr>
<tr><td>>hg38_21_+_8433172_8446622_RNA45SN1_45S_50ntFlanks@6719.6741.23|TGAATTGCAGGACACATTGATCA</td><td>>hg38_21_+_8433172_8446622_RNA45SN1_45S_50ntFlanks</td><td>6719</td><td>6741</td><td>23</td><td>TGAATTGCAGGACACATTGATCA</td><td>193.7</td><td>-4.5</td><td>1.7</td><td>-2.7</td><td>7.5E-03</td><td>1.4E-02</td><td>BT20/MB468</td><td>MP</td></tr>
<tr><td>>hg38_21_+_8433172_8446622_RNA45SN1_45S_50ntFlanks@12413.12433.21|TCCTTCGATGTCGGCTCTTCC</td><td>>hg38_21_+_8433172_8446622_RNA45SN1_45S_50ntFlanks</td><td>12413</td><td>12433</td><td>21</td><td>TCCTTCGATGTCGGCTCTTCC</td><td>325.0</td><td>-1.3</td><td>0.5</td><td>-2.5</td><td>1.3E-02</td><td>1.4E-02</td><td>MB231/MB468</td><td>NUC</td></tr>
<tr><td>>hg38_21_+_8433172_8446622_RNA45SN1_45S_50ntFlanks@11325.11344.20|TCTCGTCGCGCGCGCGTCCG</td><td>>hg38_21_+_8433172_8446622_RNA45SN1_45S_50ntFlanks</td><td>11325</td><td>11344</td><td>20</td><td>TCTCGTCGCGCGCGCGTCCG</td><td>736.3</td><td>-3.3</td><td>1.3</td><td>-2.5</td><td>1.2E-02</td><td>1.4E-02</td><td>BT20/MB468</td><td>CYTO</td></tr>
<tr><td>>am_trnaMT_GluTTC_MT_-_14674_14742@27.53.27,trnalookalike8_GluTTC_5_-_93905172_93905240@27.53.27|GTTTTTCATATCATTGGTCGTGGTTGT</td><td>>am_trnaMT_GluTTC_MT_-_14674_14742</td><td>27</td><td>53</td><td>27</td><td>GTTTTTCATATCATTGGTCGTGGTTGT</td><td>162.2</td><td>-2.5</td><td>0.9</td><td>-2.7</td><td>7.5E-03</td><td>1.4E-02</td><td>BT20/MB468</td><td>MP</td></tr>
<tr><td>>MI0000108|hsa-mir-103a-2&WithFlank&20|+|3917488|3917577@54.76.23(+1C);MI0000109|hsa-mir-103a-1&WithFlank&5|-|168560890|168560979@54.76.23(+1C)&[MIMAT0000101&hsa-miR-103a-3p&offsets|0|0(+1C);m-12&20|+|3917541|3917563&offsets|0|0(+1C)];[MIMAT0000101_1&hsa-miR-103a-3p&offsets|0|0(+1C);m-13&5|-|168560904|168560926&offsets|0|0(+1C);m-44220&5|-|168560908|168560926&offsets|0|+4(+1C)]|AGCAGCATTGTACAGGGCTATGAC</td><td>>MI0000108|hsa-mir-103a-2&WithFlank&20|+|3917488|3917577</td><td>54</td><td>76</td><td>23(+1C)</td><td>AGCAGCATTGTACAGGGCTATGAC</td><td>194.9</td><td>-3.1</td><td>1.1</td><td>-3.0</td><td>3.0E-03</td><td>1.4E-02</td><td>BT20/MB231</td><td>MP</td></tr>
<tr><td>>hg38_21_+_8433172_8446622_RNA45SN1_45S_50ntFlanks@12973.13000.28|TCGTACGTAGCAGAGCAGCTCCCTCGCT</td><td>>hg38_21_+_8433172_8446622_RNA45SN1_45S_50ntFlanks</td><td>12973</td><td>13000</td><td>28</td><td>TCGTACGTAGCAGAGCAGCTCCCTCGCT</td><td>192.4</td><td>-2.3</td><td>0.9</td><td>-2.5</td><td>1.2E-02</td><td>1.4E-02</td><td>MB231/MB468</td><td>CYTO</td></tr>
<tr><td>>MI0000736|hsa-mir-30c-1&WithFlank&1|+|40757278|40757378@23.44.22(+1G);MI0000254|hsa-mir-30c-2&WithFlank&6|-|71376954|71377037@13.34.22(+1G)&[MIMAT0000244&hsa-miR-30c-5p&offsets|0|-1(+1G);m-57&1|+|40757300|40757323&offsets|0|-2(+1G)];[MIMAT0000244_1&hsa-miR-30c-5p&offsets|0|-1(+1G);m-56&6|-|71377002|71377025&offsets|0|-2(+1G)]|TGTAAACATCCTACACTCTCAGG</td><td>>MI0000736|hsa-mir-30c-1&WithFlank&1|+|40757278|40757378</td><td>23</td><td>44</td><td>22(+1G)</td><td>TGTAAACATCCTACACTCTCAGG</td><td>22.7</td><td>7.7</td><td>3.0</td><td>2.5</td><td>1.1E-02</td><td>1.4E-02</td><td>MB231/MB468</td><td>MP</td></tr>
<tr><td>>hg38_21_+_8433172_8446622_RNA45SN1_45S_50ntFlanks@8191.8214.24|CAGCCCGTGGACGGTGTGAGGCCG</td><td>>hg38_21_+_8433172_8446622_RNA45SN1_45S_50ntFlanks</td><td>8191</td><td>8214</td><td>24</td><td>CAGCCCGTGGACGGTGTGAGGCCG</td><td>431.3</td><td>-3.1</td><td>1.2</td><td>-2.5</td><td>1.2E-02</td><td>1.4E-02</td><td>BT20/MB468</td><td>CYTO</td></tr>
<tr><td>>hg38_MT_+_1621_3279_RNR2_16S_50ntFlanks@1352.1385.34|ATAGGGTTTACGACCTCGATGTTGGATCAGGACA</td><td>>hg38_MT_+_1621_3279_RNR2_16S_50ntFlanks</td><td>1352</td><td>1385</td><td>34</td><td>ATAGGGTTTACGACCTCGATGTTGGATCAGGACA</td><td>160.4</td><td>4.1</td><td>1.5</td><td>2.7</td><td>7.6E-03</td><td>1.4E-02</td><td>BT20/MB468</td><td>MP</td></tr>
<tr><td>>hg38_21_+_8433172_8446622_RNA45SN1_45S_50ntFlanks@3910.3932.23|GGGATGCGTGCATTTATCAGATC</td><td>>hg38_21_+_8433172_8446622_RNA45SN1_45S_50ntFlanks</td><td>3910</td><td>3932</td><td>23</td><td>GGGATGCGTGCATTTATCAGATC</td><td>260.9</td><td>-3.0</td><td>1.1</td><td>-2.7</td><td>7.6E-03</td><td>1.4E-02</td><td>BT20/MB468</td><td>MP</td></tr>
<tr><td>>hg38_21_+_8433172_8446622_RNA45SN1_45S_50ntFlanks@9532.9576.45|AGAGGAAACTCTGGTGGAGGTCCGTAGCGGTCCTGACGTGCAAAT</td><td>>hg38_21_+_8433172_8446622_RNA45SN1_45S_50ntFlanks</td><td>9532</td><td>9576</td><td>45</td><td>AGAGGAAACTCTGGTGGAGGTCCGTAGCGGTCCTGACGTGCAAAT</td><td>164.4</td><td>-3.4</td><td>1.2</td><td>-2.9</td><td>3.8E-03</td><td>1.4E-02</td><td>BT20/MB231</td><td>CYTO</td></tr>
<tr><td>>hg38_21_+_8433172_8446622_RNA45SN1_45S_50ntFlanks@7979.8010.32|ACCTCAGATCAGACGTGGCGACCCGCTGAATT</td><td>>hg38_21_+_8433172_8446622_RNA45SN1_45S_50ntFlanks</td><td>7979</td><td>8010</td><td>32</td><td>ACCTCAGATCAGACGTGGCGACCCGCTGAATT</td><td>448.5</td><td>2.0</td><td>0.7</td><td>2.9</td><td>3.2E-03</td><td>1.4E-02</td><td>BT20/MB231</td><td>NUC</td></tr>
<tr><td>>hg38_21_+_8433172_8446622_RNA45SN1_45S_50ntFlanks@4732.4759.28|AGAACGAAAGTCGGAGGTTCGAAGACGA</td><td>>hg38_21_+_8433172_8446622_RNA45SN1_45S_50ntFlanks</td><td>4732</td><td>4759</td><td>28</td><td>AGAACGAAAGTCGGAGGTTCGAAGACGA</td><td>302.6</td><td>-1.8</td><td>0.7</td><td>-2.6</td><td>1.0E-02</td><td>1.4E-02</td><td>BT20/MB468</td><td>MITO</td></tr>
<tr><td>>hg38_21_+_8433172_8446622_RNA45SN1_45S_50ntFlanks@10677.10710.34|GCGGTTCCGGCGGCGTCCGGTGAGCTCTCGCTGG</td><td>>hg38_21_+_8433172_8446622_RNA45SN1_45S_50ntFlanks</td><td>10677</td><td>10710</td><td>34</td><td>GCGGTTCCGGCGGCGTCCGGTGAGCTCTCGCTGG</td><td>2922.5</td><td>3.4</td><td>1.2</td><td>2.9</td><td>3.8E-03</td><td>1.4E-02</td><td>BT20/MB231</td><td>MITO</td></tr>
<tr><td>>hg38_21_+_8433172_8446622_RNA45SN1_45S_50ntFlanks@3840.3868.29|CTCCTACTTGGATAACTGTGGTAATTCTA</td><td>>hg38_21_+_8433172_8446622_RNA45SN1_45S_50ntFlanks</td><td>3840</td><td>3868</td><td>29</td><td>CTCCTACTTGGATAACTGTGGTAATTCTA</td><td>607.8</td><td>2.6</td><td>0.9</td><td>2.9</td><td>3.8E-03</td><td>1.4E-02</td><td>BT20/MB231</td><td>MITO</td></tr>
<tr><td>>MI0000108|hsa-mir-103a-2&WithFlank&20|+|3917488|3917577@56.76.21;MI0000109|hsa-mir-103a-1&WithFlank&5|-|168560890|168560979@56.76.21&[MIMAT0000101&hsa-miR-103a-3p&offsets|+2|0;m-12&20|+|3917541|3917563&offsets|+2|0];[MIMAT0000101_1&hsa-miR-103a-3p&offsets|+2|0;m-13&5|-|168560904|168560926&offsets|+2|0;m-44220&5|-|168560908|168560926&offsets|+2|+4]|CAGCATTGTACAGGGCTATGA</td><td>>MI0000108|hsa-mir-103a-2&WithFlank&20|+|3917488|3917577</td><td>56</td><td>76</td><td>21</td><td>CAGCATTGTACAGGGCTATGA</td><td>379.4</td><td>-2.4</td><td>0.8</td><td>-2.9</td><td>3.8E-03</td><td>1.4E-02</td><td>BT20/MB231</td><td>MITO</td></tr>
<tr><td>>hg38_MT_+_1621_3279_RNR2_16S_50ntFlanks@1428.1458.31|ATTAAAGTCCTACGTGATCTGAGTTCAGACC</td><td>>hg38_MT_+_1621_3279_RNR2_16S_50ntFlanks</td><td>1428</td><td>1458</td><td>31</td><td>ATTAAAGTCCTACGTGATCTGAGTTCAGACC</td><td>454.0</td><td>2.6</td><td>1.0</td><td>2.5</td><td>1.1E-02</td><td>1.4E-02</td><td>MB231/MB468</td><td>MP</td></tr>
<tr><td>>MI0000062|hsa-let-7a-3&WithFlank&22|+|46112743|46112828@58.78.21;MI0000060|hsa-let-7a-1&WithFlank&9|+|94175951|94176042@63.83.21&[MIMAT0004481&hsa-let-7a-3p&offsets|0|0;m-180&22|+|46112800|46112821&offsets|0|-1];[MIMAT0004481_1&hsa-let-7a-3p&offsets|0|0;m-181&9|+|94176013|94176034&offsets|0|-1]|CTATACAATCTACTGTCTTTC</td><td>>MI0000062|hsa-let-7a-3&WithFlank&22|+|46112743|46112828</td><td>58</td><td>78</td><td>21</td><td>CTATACAATCTACTGTCTTTC</td><td>71.3</td><td>4.1</td><td>1.5</td><td>2.7</td><td>7.6E-03</td><td>1.4E-02</td><td>BT20/MB468</td><td>MP</td></tr>
<tr><td>>hg38_21_+_8433172_8446622_RNA45SN1_45S_50ntFlanks@12418.12449.32|CGATGTCGGCTCTTCCTATCATTGTGAAGCAG</td><td>>hg38_21_+_8433172_8446622_RNA45SN1_45S_50ntFlanks</td><td>12418</td><td>12449</td><td>32</td><td>CGATGTCGGCTCTTCCTATCATTGTGAAGCAG</td><td>177.3</td><td>-2.2</td><td>0.8</td><td>-2.7</td><td>7.6E-03</td><td>1.4E-02</td><td>BT20/MB468</td><td>MP</td></tr>
<tr><td>>hg38_21_+_8433172_8446622_RNA45SN1_45S_50ntFlanks@9482.9513.32|ACGTGTTAGGACCCGAAAGATGGTGAACTATG</td><td>>hg38_21_+_8433172_8446622_RNA45SN1_45S_50ntFlanks</td><td>9482</td><td>9513</td><td>32</td><td>ACGTGTTAGGACCCGAAAGATGGTGAACTATG</td><td>254.4</td><td>6.0</td><td>2.0</td><td>3.0</td><td>3.0E-03</td><td>1.4E-02</td><td>BT20/MB231</td><td>MP</td></tr>
</table>

| | | | | | | | | | | | | | |
|---|---|---|---|---|---|---|---|---|---|---|---|---|---|
| >hg38_21_+_8433172_8446622_RNA45SN1_45S_50ntFlanks@6652.6698.47\|GACTCTTAGCGGTGGATCACTCGGCTCGTGCGTCGATGAAGAACGCA | >hg38_21_+_8433172_8446622_RNA45SN1_45S_50ntFlanks | 6652 | 6698 | 47 | GACTCTTAGCGGTGGATCACTCGGCTCGTGCGTCGATGAAGAACGCA | 2510.4 | 1.9 | 0.8 | 2.5 | 1.2E-02 | 1.4E-02 | MB231/MB468 | CYTO |
| >hg38_21_+_8433172_8446622_RNA45SN1_45S_50ntFlanks@12404.12436.33\|GCTTTTTGATCCTTCGATGTCGGCTCTTCCTAT | >hg38_21_+_8433172_8446622_RNA45SN1_45S_50ntFlanks | 12404 | 12436 | 33 | GCTTTTTGATCCTTCGATGTCGGCTCTTCCTAT | 125.7 | -3.0 | 1.2 | -2.6 | 1.0E-02 | 1.4E-02 | BT20/MB468 | MITO |
| >MI0000285\|hsa-mir-205&WithFlank&1\|+\|209432127\|209432248@40.61.22&[MIMAT0000266&hsa-miR-205-5p&offsets\|0\|0;m-351&1\|+\|209432166\|209432187&offsets\|0\|0]\|TCCTTCATTCCACCGGAGTCTG | >MI0000285\|hsa-mir-205&WithFlank&1\|+\|209432127\|209432248 | 40 | 61 | 22 | TCCTTCATTCCACCGGAGTCTG | 43227.0 | 3.1 | 1.2 | 2.7 | 7.6E-03 | 1.4E-02 | BT20/MB468 | MP |
| >hg38_21_+_8433172_8446622_RNA45SN1_45S_50ntFlanks@11205.11232.28\|GCGGCGGCGACTCTGGACGCGAGCCGGG | >hg38_21_+_8433172_8446622_RNA45SN1_45S_50ntFlanks | 11205 | 11232 | 28 | GCGGCGGCGACTCTGGACGCGAGCCGGG | 625.6 | -3.7 | 1.4 | -2.7 | 7.6E-03 | 1.4E-02 | BT20/MB468 | MP |
| >hg38_21_+_8433172_8446622_RNA45SN1_45S_50ntFlanks@12626.12667.42\|AATGGGGCGAAGCTACCATCTGTGGGATTATGACTGAACGCC | >hg38_21_+_8433172_8446622_RNA45SN1_45S_50ntFlanks | 12626 | 12667 | 42 | AATGGGGCGAAGCTACCATCTGTGGGATTATGACTGAACGCC | 90.2 | 3.1 | 1.2 | 2.5 | 1.1E-02 | 1.4E-02 | MB231/MB468 | MP |
| >am_trnaMT_SerGCT_MT_+_12207_12265@13.31.19\|AAGAACTGCTAACTCATGC | >am_trnaMT_SerGCT_MT_+_12207_12265 | 13 | 31 | 19 | AAGAACTGCTAACTCATGC | 363.2 | 3.3 | 1.3 | 2.5 | 1.2E-02 | 1.4E-02 | BT20/MB468 | CYTO |
| >MI0000077\|hsa-mir-21&WithFlank&17\|+\|59841260\|59841343@15.37.23&[MIMAT0000076&hsa-miR-21-5p&offsets\|+1\|+2;m-1&17\|+\|59841273\|59841294&offsets\|+1\|+2]\|AGCTTATCAGACTGATGTTGACT | >MI0000077\|hsa-mir-21&WithFlank&17\|+\|59841260\|59841343 | 15 | 37 | 23 | AGCTTATCAGACTGATGTTGACT | 293.5 | -3.0 | 1.0 | -3.0 | 3.0E-03 | 1.4E-02 | BT20/MB231 | MP |
| >trnaMT_SerGCT_MT_+_12207_12265@20.62.43\|GCTAACTCATGCCCCCATGTCTAACAACATGGCTTTCTCACCA | >trnaMT_SerGCT_MT_+_12207_12265 | 20 | 62 | 43 | GCTAACTCATGCCCCCATGTCTAACAACATGGCTTTCTCACCA | 233.0 | 4.1 | 1.6 | 2.5 | 1.3E-02 | 1.4E-02 | MB231/MB468 | MITO |
| >hg38_21_+_8433172_8446622_RNA45SN1_45S_50ntFlanks@3838.3859.22\|CTCTCCTACTTGGATAACTGTG | >hg38_21_+_8433172_8446622_RNA45SN1_45S_50ntFlanks | 3838 | 3859 | 22 | CTCTCCTACTTGGATAACTGTG | 159.2 | -1.5 | 0.6 | -2.5 | 1.2E-02 | 1.4E-02 | BT20/MB468 | CYTO |
| >hg38_21_+_8433172_8446622_RNA45SN1_45S_50ntFlanks@12419.12455.37\|GATGTCGGCTCTTCCTATCATTGTGAAGCAGAATTCA | >hg38_21_+_8433172_8446622_RNA45SN1_45S_50ntFlanks | 12419 | 12455 | 37 | GATGTCGGCTCTTCCTATCATTGTGAAGCAGAATTCA | 223.4 | -2.4 | 0.8 | -2.9 | 3.2E-03 | 1.4E-02 | BT20/MB231 | NUC |
| >hg38_21_+_8433172_8446622_RNA45SN1_45S_50ntFlanks@13000.13039.40\|TGCGATCTATTGAAAGTCAGCCCTCGACACAAGGGTTTGT | >hg38_21_+_8433172_8446622_RNA45SN1_45S_50ntFlanks | 13000 | 13039 | 40 | TGCGATCTATTGAAAGTCAGCCCTCGACACAAGGGTTTGT | 623.2 | -2.1 | 0.7 | -2.9 | 3.2E-03 | 1.4E-02 | BT20/MB231 | NUC |
| >hg38_21_+_8433172_8446622_RNA45SN1_45S_50ntFlanks@6728.6758.31\|GGACACATTGATCATCGACACTTCGAACGCA | >hg38_21_+_8433172_8446622_RNA45SN1_45S_50ntFlanks | 6728 | 6758 | 31 | GGACACATTGATCATCGACACTTCGAACGCA | 147.6 | -3.2 | 1.1 | -2.9 | 3.8E-03 | 1.4E-02 | BT20/MB231 | CYTO |
| >MI0000465\|hsa-mir-191&WithFlank&3\|-\|49020612\|49020715@22.39.18&[MIMAT0000440&hsa-miR-191-5p&offsets\|0\|-5;m-48&3\|-\|49020672\|49020694&offsets\|0\|-5]\|CAACGGAATCCCAAAAGC | >MI0000465\|hsa-mir-191&WithFlank&3\|-\|49020612\|49020715 | 22 | 39 | 18 | CAACGGAATCCCAAAAGC | 123.0 | 3.4 | 1.4 | 2.5 | 1.3E-02 | 1.4E-02 | MB231/MB468 | MITO |
| >hg38_21_+_8433172_8446622_RNA45SN1_45S_50ntFlanks@4048.4066.19\|TTCGAACGTCTGCCCTATC | >hg38_21_+_8433172_8446622_RNA45SN1_45S_50ntFlanks | 4048 | 4066 | 19 | TTCGAACGTCTGCCCTATC | 603.4 | -1.8 | 0.7 | -2.5 | 1.3E-02 | 1.4E-02 | MB231/MB468 | NUC |
| >hg38_21_+_8433172_8446622_RNA45SN1_45S_50ntFlanks@6730.6752.23\|ACACATTGATCATCGACACTTCG | >hg38_21_+_8433172_8446622_RNA45SN1_45S_50ntFlanks | 6730 | 6752 | 23 | ACACATTGATCATCGACACTTCG | 239.6 | -2.4 | 0.9 | -2.5 | 1.2E-02 | 1.4E-02 | MB231/MB468 | CYTO |
| >hg38_21_+_8433172_8446622_RNA45SN1_45S_50ntFlanks@4725.4752.28\|TTAATCAAGAACGAAAGTCGGAGGTTCG | >hg38_21_+_8433172_8446622_RNA45SN1_45S_50ntFlanks | 4725 | 4752 | 28 | TTAATCAAGAACGAAAGTCGGAGGTTCG | 106.9 | -4.4 | 1.6 | -2.7 | 7.6E-03 | 1.4E-02 | BT20/MB468 | MP |
| >hg38_21_+_8433172_8446622_RNA45SN1_45S_50ntFlanks@12474.12498.25\|ACCCACTAATAGGGAACGTGAGCTG | >hg38_21_+_8433172_8446622_RNA45SN1_45S_50ntFlanks | 12474 | 12498 | 25 | ACCCACTAATAGGGAACGTGAGCTG | 174.2 | -1.7 | 0.7 | -2.6 | 1.1E-02 | 1.4E-02 | BT20/MB468 | MITO |
| >hg38_21_+_8433172_8446622_RNA45SN1_45S_50ntFlanks@9484.9503.20\|GTGTTAGGACCCGAAAGATG | >hg38_21_+_8433172_8446622_RNA45SN1_45S_50ntFlanks | 9484 | 9503 | 20 | GTGTTAGGACCCGAAAGATG | 137.2 | 2.5 | 0.9 | 2.7 | 7.6E-03 | 1.4E-02 | BT20/MB468 | MP |
| >hg38_21_+_8433172_8446622_RNA45SN1_45S_50ntFlanks@10797.10820.24\|GCATGTTGGAACAATGTAGGTAAG | >hg38_21_+_8433172_8446622_RNA45SN1_45S_50ntFlanks | 10797 | 10820 | 24 | GCATGTTGGAACAATGTAGGTAAG | 184.5 | -3.6 | 1.3 | -2.7 | 7.6E-03 | 1.4E-02 | BT20/MB468 | MP |
| >hg38_MT_+_1621_3279_RNR2_16S_50ntFlanks@284.305.22\|CCGAAACCAGACGAGCTACCTA | >hg38_MT_+_1621_3279_RNR2_16S_50ntFlanks | 284 | 305 | 22 | CCGAAACCAGACGAGCTACCTA | 38.3 | 5.0 | 2.0 | 2.5 | 1.1E-02 | 1.4E-02 | MB231/MB468 | MP |
| >trna134_GluTTC_1_-_16861774_16861845@1.33.33,trna5_GluTTC_1_+_17199078_17199149@1.33.33,trna84_GluTTC_1_-_161391883_161391954@1.33.33,trna94_GluTTC_1_-_149664355_149664427@1.33.33\|TCCCTGGTGGTCTAGTGGCTAGGATTCGGCGCT | >trna134_GluTTC_1_-_16861774_16861845 | 1 | 33 | 33 | TCCCTGGTGGTCTAGTGGCTAGGATTCGGCGCT | 223.8 | 1.6 | 0.6 | 2.5 | 1.3E-02 | 1.4E-02 | BT20/MB468 | NUC |
| >hg38_21_+_8433172_8446622_RNA45SN1_45S_50ntFlanks@9554.9573.20\|CGTAGCGGTCCTGACGTGCA | >hg38_21_+_8433172_8446622_RNA45SN1_45S_50ntFlanks | 9554 | 9573 | 20 | CGTAGCGGTCCTGACGTGCA | 15410.6 | -1.4 | 0.6 | -2.5 | 1.2E-02 | 1.4E-02 | MB231/MB468 | CYTO |
| >MI0000285\|hsa-mir-205&WithFlank&1\|+\|209432127\|209432248@40.60.21(+1A)&[MIMAT0000266&hsa-miR-205-5p&offsets\|0\|-1(+1A);m-351&1\|+\|209432166\|209432187&offsets\|0\|-1(+1A)]\|TCCTTCATTCCACCGGAGTCTA | >MI0000285\|hsa-mir-205&WithFlank&1\|+\|209432127\|209432248 | 40 | 60 | 21(+1A) | TCCTTCATTCCACCGGAGTCTA | 5551.7 | 3.2 | 1.2 | 2.7 | 7.6E-03 | 1.4E-02 | BT20/MB468 | MP |
| >hg38_21_+_8433172_8446622_RNA45SN1_45S_50ntFlanks@10347.10369.23\|CAAACGAGAACTTTGAAGGCCGA | >hg38_21_+_8433172_8446622_RNA45SN1_45S_50ntFlanks | 10347 | 10369 | 23 | CAAACGAGAACTTTGAAGGCCGA | 506.1 | 3.9 | 1.3 | 3.0 | 3.0E-03 | 1.4E-02 | BT20/MB231 | MP |
| >am_trna119_LysCTT_1_-_145395522_145395594@14.32.19,trna11_LysCTT_5_-_180648979_180649051@14.32.19,trna128_LysCTT_1_-_55423542_55423614@14.32.19,trna13_LysCTT_6_+_26556774_26556846@14.32.19,trna32_LysCTT_16_-_3207406_3207478@14.32.19,trna6_LysCTT_19_-_52425396_52425468@14.32.19,trna7_LysCTT_16_+_3225692_3225764@14.32.19,trna9_LysCTT_5_+_180634755_180634827@14.32.19\|AGTCGGTAGAGCATGAGAC | >am_trna119_LysCTT_1_-_145395522_145395594 | 14 | 32 | 19 | AGTCGGTAGAGCATGAGAC | 200.8 | -3.0 | 1.0 | -3.0 | 3.0E-03 | 1.4E-02 | BT20/MB231 | MP |
| >MI0000750\|hsa-mir-26a-2&WithFlank&12\|-\|57824603\|57824698@20.41.22;MI0000083\|hsa-mir-26a-1&WithFlank&3\|+\|37969398\|37969486@16.37.22&[MIMAT0000082&hsa-miR-26a-5p&offsets\|0\|0;m-33&12\|-\|57824658\|57824679&offsets\|0\|0];[MIMAT0000082_1&hsa-miR-26a-5p&offsets\|0\|0;m-34&3\|+\|37969413\|37969434&offsets\|0\|0]\|TTCAAGTAATCCAGGATAGGCT | >MI0000750\|hsa-mir-26a-2&WithFlank&12\|-\|57824603\|57824698 | 20 | 41 | 22 | TTCAAGTAATCCAGGATAGGCT | 19406.5 | 1.1 | 0.5 | 2.5 | 1.3E-02 | 1.4E-02 | BT20/MB468 | NUC |
| >hg38_21_+_8433172_8446622_RNA45SN1_45S_50ntFlanks@10788.10825.38\|CAGCCTCTGGCATGTTGGAACAATGTAGGTAAGGGAAG | >hg38_21_+_8433172_8446622_RNA45SN1_45S_50ntFlanks | 10788 | 10825 | 38 | CAGCCTCTGGCATGTTGGAACAATGTAGGTAAGGGAAG | 221.3 | -1.7 | 0.7 | -2.6 | 1.1E-02 | 1.4E-02 | BT20/MB468 | MITO |
| >hg38_21_+_8433172_8446622_RNA45SN1_45S_50ntFlanks@11601.11619.19\|ATCCGACTGTTTAATTAAA | >hg38_21_+_8433172_8446622_RNA45SN1_45S_50ntFlanks | 11601 | 11619 | 19 | ATCCGACTGTTTAATTAAA | 125.2 | -4.2 | 1.7 | -2.5 | 1.1E-02 | 1.4E-02 | MB231/MB468 | MP |
| >hg38_21_+_8433172_8446622_RNA45SN1_45S_50ntFlanks@12907.12931.25\|CACCGCACGTTCGTGGGGAACCTGG | >hg38_21_+_8433172_8446622_RNA45SN1_45S_50ntFlanks | 12907 | 12931 | 25 | CACCGCACGTTCGTGGGGAACCTGG | 189.4 | -4.6 | 1.7 | -2.7 | 7.6E-03 | 1.5E-02 | BT20/MB468 | MP |
| >hg38_21_+_8433172_8446622_RNA45SN1_45S_50ntFlanks@6678.6700.23\|CGTGCGTCGATGAAGAACGCAGC | >hg38_21_+_8433172_8446622_RNA45SN1_45S_50ntFlanks | 6678 | 6700 | 23 | CGTGCGTCGATGAAGAACGCAGC | 118.7 | 5.6 | 2.1 | 2.7 | 7.6E-03 | 1.5E-02 | BT20/MB468 | MP |
| >hg38_21_+_8433172_8446622_RNA45SN1_45S_50ntFlanks@4731.4761.31\|AAGAACGAAAGTCGGAGGTTCGAAGACGATC | >hg38_21_+_8433172_8446622_RNA45SN1_45S_50ntFlanks | 4731 | 4761 | 31 | AAGAACGAAAGTCGGAGGTTCGAAGACGATC | 676.2 | -1.2 | 0.5 | -2.5 | 1.3E-02 | 1.5E-02 | BT20/MB468 | NUC |
| >hg38_21_+_8433172_8446622_RNA45SN1_45S_50ntFlanks@10239.10258.20\|GCTACGCCGCGACGAGTAGG | >hg38_21_+_8433172_8446622_RNA45SN1_45S_50ntFlanks | 10239 | 10258 | 20 | GCTACGCCGCGACGAGTAGG | 134.9 | 3.9 | 1.4 | 2.7 | 7.6E-03 | 1.5E-02 | BT20/MB468 | MP |
| >hg38_21_+_8433172_8446622_RNA45SN1_45S_50ntFlanks@12475.12496.22\|CCCACTAATAGGGAACGTGAGC | >hg38_21_+_8433172_8446622_RNA45SN1_45S_50ntFlanks | 12475 | 12496 | 22 | CCCACTAATAGGGAACGTGAGC | 1081.8 | 1.6 | 0.6 | 2.5 | 1.2E-02 | 1.5E-02 | MB231/MB468 | CYTO |
| >hg38_21_+_8433172_8446622_RNA45SN1_45S_50ntFlanks@11104.11123.20\|AAGGGTCGGGGCGGCAGGGG | >hg38_21_+_8433172_8446622_RNA45SN1_45S_50ntFlanks | 11104 | 11123 | 20 | AAGGGTCGGGGCGGCAGGGG | 493.3 | -3.2 | 1.3 | -2.5 | 1.2E-02 | 1.5E-02 | BT20/MB468 | CYTO |
| >MI0000750\|hsa-mir-26a-2&WithFlank&12\|-\|57824603\|57824698@20.41.22(+1C);MI0000083\|hsa-mir-26a-1&WithFlank&3\|+\|37969398\|37969486@16.37.22(+1C)&[MIMAT0000082&hsa-miR-26a-5p&offsets\|0\|0(+1C);m-33&12\|-\|57824658\|57824679&offsets\|0\|0(+1C)];[MIMAT0000082_1&hsa-miR-26a-5p&offsets\|0\|0(+1C);m-34&3\|+\|37969413\|37969434&offsets\|0\|0(+1C)]\|TTCAAGTAATCCAGGATAGGCTC | >MI0000750\|hsa-mir-26a-2&WithFlank&12\|-\|57824603\|57824698 | 20 | 41 | 22(+1C) | TTCAAGTAATCCAGGATAGGCTC | 220.4 | 4.1 | 1.5 | 2.7 | 7.7E-03 | 1.5E-02 | BT20/MB468 | MP |
| >hg38_21_+_8433172_8446622_RNA45SN1_45S_50ntFlanks@475.511.37\|CTTCGTGATCGATGTGGTGACGTCGTGCTCTCCCGGG | >hg38_21_+_8433172_8446622_RNA45SN1_45S_50ntFlanks | 475 | 511 | 37 | CTTCGTGATCGATGTGGTGACGTCGTGCTCTCCCGGG | 57.8 | 4.9 | 1.9 | 2.5 | 1.1E-02 | 1.5E-02 | MB231/MB468 | MP |
| >hg38_1_-_228634819_228635039_RNA5S12_5S_50ntFlanks@51.86.36\|GTCTACGGCCATACCACCCTGAACGCGCCCGATCTC | >hg38_1_-_228634819_228635039_RNA5S12_5S_50ntFlanks | 51 | 86 | 36 | GTCTACGGCCATACCACCCTGAACGCGCCCGATCTC | 3559.4 | 3.8 | 1.5 | 2.5 | 1.3E-02 | 1.5E-02 | MB231/MB468 | NUC |
| >trnaMT_LysTTT_MT_+_8295_8364@53.72.20,trnalookalike7_LysTTT_1_+_568844_568913@53.72.20\|AACACCTCTTTACAGTGACC | >trnaMT_LysTTT_MT_+_8295_8364 | 53 | 72 | 20 | AACACCTCTTTACAGTGACC | 973.4 | -4.4 | 1.5 | -3.0 | 3.0E-03 | 1.5E-02 | BT20/MB231 | MP |
| >hg38_21_+_8433172_8446622_RNA45SN1_45S_50ntFlanks@9561.9597.37\|GTCCTGACGTGCAAATCGGTCGTCCGACCTGGGTATA | >hg38_21_+_8433172_8446622_RNA45SN1_45S_50ntFlanks | 9561 | 9597 | 37 | GTCCTGACGTGCAAATCGGTCGTCCGACCTGGGTATA | 211.8 | -2.6 | 1.0 | -2.5 | 1.2E-02 | 1.5E-02 | MB231/MB468 | CYTO |
| >hg38_21_+_8433172_8446622_RNA45SN1_45S_50ntFlanks@9553.9571.19\|CCGTAGCGGTCCTGACGTG | >hg38_21_+_8433172_8446622_RNA45SN1_45S_50ntFlanks | 9553 | 9571 | 19 | CCGTAGCGGTCCTGACGTG | 231.9 | -2.7 | 1.1 | -2.5 | 1.2E-02 | 1.5E-02 | BT20/MB468 | CYTO |
| >hg38_21_+_8433172_8446622_RNA45SN1_45S_50ntFlanks@6646.6665.20\|TCGTACGACTCTTAGCGGTG | >hg38_21_+_8433172_8446622_RNA45SN1_45S_50ntFlanks | 6646 | 6665 | 20 | TCGTACGACTCTTAGCGGTG | 51725.8 | 2.2 | 0.7 | 2.9 | 3.3E-03 | 1.5E-02 | BT20/MB231 | NUC |
| >hg38_21_+_8433172_8446622_RNA45SN1_45S_50ntFlanks@12418.12457.40\|CGATGTCGGCTCTTCCTATCATTGTGAAGCAGAATTCACC | >hg38_21_+_8433172_8446622_RNA45SN1_45S_50ntFlanks | 12418 | 12457 | 40 | CGATGTCGGCTCTTCCTATCATTGTGAAGCAGAATTCACC | 232.0 | -4.0 | 1.4 | -2.9 | 3.3E-03 | 1.5E-02 | BT20/MB231 | NUC |
| >hg38_21_+_8433172_8446622_RNA45SN1_45S_50ntFlanks@11567.11584.18\|AGCAGCCGACTTAGAACT | >hg38_21_+_8433172_8446622_RNA45SN1_45S_50ntFlanks | 11567 | 11584 | 18 | AGCAGCCGACTTAGAACT | 379.5 | 2.1 | 0.8 | 2.5 | 1.1E-02 | 1.5E-02 | MB231/MB468 | MP |
| >hg38_21_+_8433172_8446622_RNA45SN1_45S_50ntFlanks@12479.12509.31\|CTAATAGGGAACGTGAGCTGGGTTTAGACCG | >hg38_21_+_8433172_8446622_RNA45SN1_45S_50ntFlanks | 12479 | 12509 | 31 | CTAATAGGGAACGTGAGCTGGGTTTAGACCG | 176.8 | -3.8 | 1.5 | -2.5 | 1.2E-02 | 1.5E-02 | BT20/MB468 | CYTO |
| >MI0000272\|hsa-mir-182&WithFlank&7\|-\|129770377\|129770498@29.53.25(+1U)&[MIMAT0000259&hsa-miR-182-5p&offsets\|0\|+1(+1U);m-22&7\|-\|129770449\|129770470&offsets\|0\|+3(+1U)]\|TTTGGCAATGGTAGAACTCACACTGT | >MI0000272\|hsa-mir-182&WithFlank&7\|-\|129770377\|129770498 | 29 | 53 | 25(+1U) | TTTGGCAATGGTAGAACTCACACTGT | 1664.1 | 1.8 | 0.6 | 2.9 | 3.3E-03 | 1.5E-02 | BT20/MB231 | NUC |
| >MI0016005\|hsa-mir-3615&WithFlank&17\|+\|74748607\|74748705@57.78.22&[MIMAT0017994&hsa-miR-3615&offsets\|0\|+1;m-271&17\|+\|74748663\|74748684&offsets\|0\|0]\|TCTCTCGGCTCCTCGCGGCTCG | >MI0016005\|hsa-mir-3615&WithFlank&17\|+\|74748607\|74748705 | 57 | 78 | 22 | TCTCTCGGCTCCTCGCGGCTCG | 754.3 | -2.0 | 0.7 | -2.9 | 3.3E-03 | 1.5E-02 | BT20/MB231 | NUC |
| >hg38_21_+_8433172_8446622_RNA45SN1_45S_50ntFlanks@11665.11702.38\|TTCTGCCCAGTGCTCTGAATGTCAAAGTGAAGAAATTC | >hg38_21_+_8433172_8446622_RNA45SN1_45S_50ntFlanks | 11665 | 11702 | 38 | TTCTGCCCAGTGCTCTGAATGTCAAAGTGAAGAAATTC | 361.4 | -2.9 | 1.0 | -2.9 | 3.3E-03 | 1.5E-02 | BT20/MB231 | NUC |
| >hg38_21_+_8433172_8446622_RNA45SN1_45S_50ntFlanks@7981.8005.25\|CTCAGATCAGACGTGGCGACCCGCT | >hg38_21_+_8433172_8446622_RNA45SN1_45S_50ntFlanks | 7981 | 8005 | 25 | CTCAGATCAGACGTGGCGACCCGCT | 156.9 | -1.5 | 0.6 | -2.5 | 1.2E-02 | 1.5E-02 | MB231/MB468 | CYTO |

| | | | | | | | | | | | | | |
|---|---|---|---|---|---|---|---|---|---|---|---|---|---|
| >hg38_21_+_8433172_8446622_RNA45SN1_45S_50ntFlanks@4283.4317.35\|CTTTAACGAGGATCCATTGGAGGGCAAGTCTGGTG | >hg38_21_+_8433172_8446622_RNA45SN1_45S_50ntFlanks | 4283 | 4317 | 35 | CTTTAACGAGGATCCATTGGAGGGCAAGTCTGGTG | 3148.4 | -3.8 | 1.4 | -2.7 | 7.7E-03 | 1.5E-02 | BT20/MB468 | MP |
| >hg38_21_+_8433172_8446622_RNA45SN1_45S_50ntFlanks@4067.4101.35\|AACTTTCGATGGTAGTCGCCGTGCCTACCATGGTG | >hg38_21_+_8433172_8446622_RNA45SN1_45S_50ntFlanks | 4067 | 4101 | 35 | AACTTTCGATGGTAGTCGCCGTGCCTACCATGGTG | 394.4 | 1.9 | 0.8 | 2.5 | 1.2E-02 | 1.5E-02 | BT20/MB468 | CYTO |
| >hg38_21_+_8433172_8446622_RNA45SN1_45S_50ntFlanks@11578.11613.36\|TAGAACTGGTGCGGACCAGGGGAATCCGACTGTTTA | >hg38_21_+_8433172_8446622_RNA45SN1_45S_50ntFlanks | 11578 | 11613 | 36 | TAGAACTGGTGCGGACCAGGGGAATCCGACTGTTTA | 98.7 | 3.5 | 1.3 | 2.7 | 7.7E-03 | 1.5E-02 | BT20/MB468 | MP |
| >hg38_21_+_8433172_8446622_RNA45SN1_45S_50ntFlanks@11587.11609.23\|TGCGGACCAGGGGAATCCGACTG | >hg38_21_+_8433172_8446622_RNA45SN1_45S_50ntFlanks | 11587 | 11609 | 23 | TGCGGACCAGGGGAATCCGACTG | 228.9 | -2.6 | 1.0 | -2.7 | 7.7E-03 | 1.5E-02 | BT20/MB468 | MP |
| >hg38_21_+_8433172_8446622_RNA45SN1_45S_50ntFlanks@12627.12658.32\|ATGGGGCGAAGCTACCATCTGTGGGATTATGA | >hg38_21_+_8433172_8446622_RNA45SN1_45S_50ntFlanks | 12627 | 12658 | 32 | ATGGGGCGAAGCTACCATCTGTGGGATTATGA | 39.0 | -4.5 | 1.7 | -2.7 | 7.7E-03 | 1.5E-02 | BT20/MB468 | MP |
| >am_MI0000786\|hsa-mir-378a&WithFlank&5\|+\|149732819\|149732896@49.70.22(+1U)&[MIMAT0000732&hsa-miR-378a-3p&offsets\|0\|0(+1U);m-44&5\|+\|149732867\|149732888&offsets\|0\|0(+1U)]\|ACTGGACTTGGAGTCAGAAGGCT | >am_MI0000786\|hsa-mir-378a&WithFlank&5\|+\|149732819\|149732896 | 49 | 70 | 22(+1U) | ACTGGACTTGGAGTCAGAAGGCT | 7466.8 | 2.7 | 1.0 | 2.7 | 7.7E-03 | 1.5E-02 | BT20/MB468 | MP |
| >MI0000066\|hsa-let-7e&WithFlank&19\|+\|51692780\|51692870@14.35.22(+1A)&[MIMAT0000066&hsa-let-7e-5p&offsets\|0\|0(+1A);m-50&19\|+\|51692793\|51692814&offsets\|0\|0(+1A)]\|TGAGGTAGGAGGTTGTATAGTTA | >MI0000066\|hsa-let-7e&WithFlank&19\|+\|51692780\|51692870 | 14 | 35 | 22(+1A) | TGAGGTAGGAGGTTGTATAGTTA | 378.2 | 1.8 | 0.6 | 2.9 | 3.3E-03 | 1.5E-02 | BT20/MB231 | NUC |
| >MI0003686\|hsa-mir-542&WithFlank&X\|-\|134541335\|134541443@59.80.22&[MIMAT0003389&hsa-miR-542-3p&offsets\|0\|0;m-136&X\|-\|134541364\|134541385&offsets\|0\|0]\|TGTGACAGATTGATAACTGAAA | >MI0003686\|hsa-mir-542&WithFlank&X\|-\|134541335\|134541443 | 59 | 80 | 22 | TGTGACAGATTGATAACTGAAA | 269.0 | 2.0 | 0.7 | 2.9 | 3.3E-03 | 1.5E-02 | BT20/MB231 | NUC |
| >hg38_MT_+_1621_3279_RNR2_16S_50ntFlanks@1425.1457.33\|ACGATTAAAGTCCTACGTGATCTGAGTTCAGAC | >hg38_MT_+_1621_3279_RNR2_16S_50ntFlanks | 1425 | 1457 | 33 | ACGATTAAAGTCCTACGTGATCTGAGTTCAGAC | 58.4 | 4.0 | 1.6 | 2.5 | 1.1E-02 | 1.5E-02 | MB231/MB468 | MP |
| >hg38_21_+_8433172_8446622_RNA45SN1_45S_50ntFlanks@10786.10827.42\|AACAGCCTCTGGCATGTTGGAACAATGTAGGTAAGGGAAGTC | >hg38_21_+_8433172_8446622_RNA45SN1_45S_50ntFlanks | 10786 | 10827 | 42 | AACAGCCTCTGGCATGTTGGAACAATGTAGGTAAGGGAAGTC | 129.5 | -3.0 | 1.2 | -2.5 | 1.2E-02 | 1.5E-02 | BT20/MB468 | CYTO |
| >hg38_21_+_8433172_8446622_RNA45SN1_45S_50ntFlanks@9930.9949.20\|ATATAGACAGCAGGACGGTG | >hg38_21_+_8433172_8446622_RNA45SN1_45S_50ntFlanks | 9930 | 9949 | 20 | ATATAGACAGCAGGACGGTG | 453.7 | 4.4 | 1.5 | 3.0 | 3.0E-03 | 1.5E-02 | BT20/MB231 | MP |
| >trna14_LysTTT_11_-_59327808_59327880@4.32.29,trna2_LysTTT_17_+_8022473_8022545@4.32.29,trna54_LysTTT_1_+_204475655_204475727@4.32.29,trna5_LysTTT_11_+_59323902_59323974@4.32.29,trna62_LysTTT_1_-_204476158_204476230@4.32.29,trna76_LysTTT_6_+_28918806_28918878@4.32.29\|CGGATAGCTCAGTCGGTAGAGCATCAGAC | >trna14_LysTTT_11_-_59327808_59327880 | 4 | 32 | 29 | CGGATAGCTCAGTCGGTAGAGCATCAGAC | 132.1 | -4.6 | 1.5 | -3.0 | 3.0E-03 | 1.5E-02 | BT20/MB231 | MP |
| >hg38_21_+_8433172_8446622_RNA45SN1_45S_50ntFlanks@12001.12031.31\|CTGCGGGCCGCCGGTGAAATACCACTACTCT | >hg38_21_+_8433172_8446622_RNA45SN1_45S_50ntFlanks | 12001 | 12031 | 31 | CTGCGGGCCGCCGGTGAAATACCACTACTCT | 448.8 | -4.1 | 1.6 | -2.5 | 1.2E-02 | 1.5E-02 | BT20/MB468 | CYTO |
| >hg38_21_+_8433172_8446622_RNA45SN1_45S_50ntFlanks@12635.12665.31\|AAGCTACCATCTGTGGGATTATGACTGAACG | >hg38_21_+_8433172_8446622_RNA45SN1_45S_50ntFlanks | 12635 | 12665 | 31 | AAGCTACCATCTGTGGGATTATGACTGAACG | 104.5 | -4.3 | 1.7 | -2.5 | 1.1E-02 | 1.5E-02 | MB231/MB468 | MP |
| >hg38_21_+_8433172_8446622_RNA45SN1_45S_50ntFlanks@11572.11603.32\|CCGACTTAGAACTGGTGCGGACCAGGGGAATC | >hg38_21_+_8433172_8446622_RNA45SN1_45S_50ntFlanks | 11572 | 11603 | 32 | CCGACTTAGAACTGGTGCGGACCAGGGGAATC | 404.8 | 4.0 | 1.3 | 3.0 | 3.0E-03 | 1.5E-02 | BT20/MB231 | MP |
| >hg38_MT_+_1621_3279_RNR2_16S_50ntFlanks@1263.1284.22\|ACTATACTCAATTGATCCAATA | >hg38_MT_+_1621_3279_RNR2_16S_50ntFlanks | 1263 | 1284 | 22 | ACTATACTCAATTGATCCAATA | 114.2 | -5.8 | 1.9 | -3.0 | 3.0E-03 | 1.5E-02 | BT20/MB231 | MP |
| >trna2_GlyGCC_21_-_18827107_18827177@3.35.33,trna35_GlyGCC_1_+_161413094_161413164@3.35.33,trna37_GlyGCC_1_+_161420467_161420537@3.35.33,trna39_GlyGCC_1_+_161427898_161427968@3.35.33,trna41_GlyGCC_1_+_161435258_161435328@3.35.33\|ATGGGTGGTTCAGTGGTAGAATTCTCGCCTGCC | >trna2_GlyGCC_21_-_18827107_18827177 | 3 | 35 | 33 | ATGGGTGGTTCAGTGGTAGAATTCTCGCCTGCC | 115.3 | 5.9 | 2.2 | 2.7 | 7.7E-03 | 1.5E-02 | BT20/MB468 | MP |
| >hg38_21_+_8433172_8446622_RNA45SN1_45S_50ntFlanks@8198.8222.25\|TGGACGGTGTGAGGCCGGTAGCGGC | >hg38_21_+_8433172_8446622_RNA45SN1_45S_50ntFlanks | 8198 | 8222 | 25 | TGGACGGTGTGAGGCCGGTAGCGGC | 105.1 | 6.1 | 2.1 | 3.0 | 3.0E-03 | 1.5E-02 | BT20/MB231 | MP |
| >MI0000087\|hsa-mir-29a&WithFlank&7\|-\|130876741\|130876816@47.69.23&[MIMAT0000086&hsa-miR-29a-3p&offsets\|-1\|0;m-26&7\|-\|130876749\|130876769&offsets\|-1\|+1]\|CTAGCACCATCTGAAATCGGTTA | >MI0000087\|hsa-mir-29a&WithFlank&7\|-\|130876741\|130876816 | 47 | 69 | 23 | CTAGCACCATCTGAAATCGGTTA | 2957.3 | -2.7 | 0.9 | -2.9 | 3.8E-03 | 1.5E-02 | BT20/MB231 | MITO |
| >hg38_1_-_228634819_228635039_RNA5S12_5S_50ntFlanks@138.171.34\|AGACCGCCTGGGAATACCGGGTGCTGTAGGCTTT | >hg38_1_-_228634819_228635039_RNA5S12_5S_50ntFlanks | 138 | 171 | 34 | AGACCGCCTGGGAATACCGGGTGCTGTAGGCTTT | 1768.1 | -2.3 | 0.8 | -2.9 | 3.8E-03 | 1.5E-02 | BT20/MB231 | MITO |
| >hg38_21_+_8433172_8446622_RNA45SN1_45S_50ntFlanks@12417.12460.44\|TCGATGTCGGCTCTTCCTATCATTGTGAAGCAGAATTCACCAAG | >hg38_21_+_8433172_8446622_RNA45SN1_45S_50ntFlanks | 12417 | 12460 | 44 | TCGATGTCGGCTCTTCCTATCATTGTGAAGCAGAATTCACCAAG | 415.8 | -2.9 | 1.0 | -2.9 | 3.8E-03 | 1.5E-02 | BT20/MB231 | MITO |
| >hg38_MT_+_1621_3279_RNR2_16S_50ntFlanks@51.89.39\|GCTAAACCTAGCCCCAAACCCACTCCACCTTACTACCAG | >hg38_MT_+_1621_3279_RNR2_16S_50ntFlanks | 51 | 89 | 39 | GCTAAACCTAGCCCCAAACCCACTCCACCTTACTACCAG | 951.1 | -2.3 | 0.8 | -2.9 | 3.8E-03 | 1.5E-02 | BT20/MB231 | MITO |
| >hg38_21_+_8433172_8446622_RNA45SN1_45S_50ntFlanks@10408.10430.23\|GGTCAGTCGGTCCTGAGAGATGG | >hg38_21_+_8433172_8446622_RNA45SN1_45S_50ntFlanks | 10408 | 10430 | 23 | GGTCAGTCGGTCCTGAGAGATGG | 750.4 | -4.9 | 1.8 | -2.7 | 7.7E-03 | 1.5E-02 | BT20/MB468 | MP |
| >hg38_21_+_8433172_8446622_RNA45SN1_45S_50ntFlanks@11576.11600.25\|CTTAGAACTGGTGCGGACCAGGGGA | >hg38_21_+_8433172_8446622_RNA45SN1_45S_50ntFlanks | 11576 | 11600 | 25 | CTTAGAACTGGTGCGGACCAGGGGA | 129.7 | -1.6 | 0.6 | -2.6 | 1.1E-02 | 1.5E-02 | BT20/MB468 | MITO |
| >hg38_21_+_8433172_8446622_RNA45SN1_45S_50ntFlanks@9551.9571.21\|GTCCGTAGCGGTCCTGACGTG | >hg38_21_+_8433172_8446622_RNA45SN1_45S_50ntFlanks | 9551 | 9571 | 21 | GTCCGTAGCGGTCCTGACGTG | 626.6 | -2.4 | 0.8 | -2.9 | 3.9E-03 | 1.5E-02 | BT20/MB231 | CYTO |
| >trnaMT_PheGAA_MT_+_577_647@43.74.32\|TTAGACGGGCTCACATCACCCCATAAACACCA | >trnaMT_PheGAA_MT_+_577_647 | 43 | 74 | 32 | TTAGACGGGCTCACATCACCCCATAAACACCA | 838.4 | 4.5 | 1.7 | 2.7 | 7.7E-03 | 1.5E-02 | BT20/MB468 | MP |
| >hg38_21_+_8433172_8446622_RNA45SN1_45S_50ntFlanks@10793.10818.26\|TCTGGCATGTTGGAACAATGTAGGTA | >hg38_21_+_8433172_8446622_RNA45SN1_45S_50ntFlanks | 10793 | 10818 | 26 | TCTGGCATGTTGGAACAATGTAGGTA | 244.9 | -1.7 | 0.7 | -2.6 | 1.1E-02 | 1.5E-02 | BT20/MB468 | MITO |
| >MI0000066\|hsa-let-7e&WithFlank&19\|+\|51692780\|51692870@14.35.22(+1U)&[MIMAT0000066&hsa-let-7e-5p&offsets\|0\|0(+1U);m-50&19\|+\|51692793\|51692814&offsets\|0\|0(+1U)]\|TGAGGTAGGAGGTTGTATAGTTT | >MI0000066\|hsa-let-7e&WithFlank&19\|+\|51692780\|51692870 | 14 | 35 | 22(+1U) | TGAGGTAGGAGGTTGTATAGTTT | 268.5 | 1.0 | 0.4 | 2.5 | 1.3E-02 | 1.5E-02 | BT20/MB468 | NUC |
| >hg38_21_+_8433172_8446622_RNA45SN1_45S_50ntFlanks@4287.4316.30\|AACGAGGATCCATTGGAGGGCAAGTCTGGT | >hg38_21_+_8433172_8446622_RNA45SN1_45S_50ntFlanks | 4287 | 4316 | 30 | AACGAGGATCCATTGGAGGGCAAGTCTGGT | 231.2 | 4.2 | 1.6 | 2.7 | 7.7E-03 | 1.5E-02 | BT20/MB468 | MP |
| >hg38_21_+_8433172_8446622_RNA45SN1_45S_50ntFlanks@11587.11612.26\|TGCGGACCAGGGGAATCCGACTGTTT | >hg38_21_+_8433172_8446622_RNA45SN1_45S_50ntFlanks | 11587 | 11612 | 26 | TGCGGACCAGGGGAATCCGACTGTTT | 417.8 | -2.6 | 1.0 | -2.7 | 7.7E-03 | 1.5E-02 | BT20/MB468 | MP |
| >MI0000542\|hsa-mir-320a&WithFlank&8\|-\|22244960\|22245043@47.68.22&[MIMAT0000510&hsa-miR-320a-3p&offsets\|-1\|-1;m-60&8\|-\|22244975\|22244996&offsets\|-1\|-1]\|GAAAAGCTGGGTTGAGAGGGCG | >MI0000542\|hsa-mir-320a&WithFlank&8\|-\|22244960\|22245043 | 47 | 68 | 22 | GAAAAGCTGGGTTGAGAGGGCG | 282.4 | 3.1 | 1.0 | 3.0 | 3.0E-03 | 1.5E-02 | BT20/MB231 | MP |
| >hg38_21_+_8433172_8446622_RNA45SN1_45S_50ntFlanks@10403.10429.27\|ACATGGGTCAGTCGGTCCTGAGAGATG | >hg38_21_+_8433172_8446622_RNA45SN1_45S_50ntFlanks | 10403 | 10429 | 27 | ACATGGGTCAGTCGGTCCTGAGAGATG | 349.9 | 4.9 | 1.7 | 3.0 | 3.0E-03 | 1.5E-02 | BT20/MB231 | MP |
| >hg38_21_+_8433172_8446622_RNA45SN1_45S_50ntFlanks@11562.11592.31\|CGCCTAGCAGCCGACTTAGAACTGGTGCGGA | >hg38_21_+_8433172_8446622_RNA45SN1_45S_50ntFlanks | 11562 | 11592 | 31 | CGCCTAGCAGCCGACTTAGAACTGGTGCGGA | 149.9 | 5.9 | 2.0 | 3.0 | 3.0E-03 | 1.5E-02 | BT20/MB231 | MP |
| >MI0000749\|hsa-mir-30e&WithFlank&1\|+\|40754349\|40754452@23.46.24(+2U)&[MIMAT0000692&hsa-miR-30e-5p&offsets\|0\|+2(+2U);m-24&1\|+\|40754371\|40754394&offsets\|0\|0(+2U)]\|TGTAAACATCCTTGACTGGAAGCTTT | >MI0000749\|hsa-mir-30e&WithFlank&1\|+\|40754349\|40754452 | 23 | 46 | 24(+2U) | TGTAAACATCCTTGACTGGAAGCTTT | 308.6 | -6.4 | 2.2 | -2.9 | 3.9E-03 | 1.5E-02 | BT20/MB231 | MITO |
| >MI0003820\|hsa-mir-454&WithFlank&17\|-\|59137752\|59137878@70.92.23(+1A)&[MIMAT0003885&hsa-miR-454-3p&offsets\|0\|0(+1A);m-272&17\|-\|59137787\|59137809&offsets\|0\|0(+1A)]\|TAGTGCAATATTGCTTATAGGGTA | >MI0003820\|hsa-mir-454&WithFlank&17\|-\|59137752\|59137878 | 70 | 92 | 23(+1A) | TAGTGCAATATTGCTTATAGGGTA | 34.8 | 5.0 | 2.0 | 2.5 | 1.1E-02 | 1.5E-02 | MB231/MB468 | MP |
| >hg38_21_+_8433172_8446622_RNA45SN1_45S_50ntFlanks@5190.5227.38\|AACAGGTCTGTGATGCCCTTAGATGTCCGGGGCTGCAC | >hg38_21_+_8433172_8446622_RNA45SN1_45S_50ntFlanks | 5190 | 5227 | 38 | AACAGGTCTGTGATGCCCTTAGATGTCCGGGGCTGCAC | 149.4 | 5.7 | 2.1 | 2.7 | 7.7E-03 | 1.5E-02 | BT20/MB468 | MP |
| >hg38_21_+_8433172_8446622_RNA45SN1_45S_50ntFlanks@12395.12414.20\|AGCGACGTCGCTTTTTGATC | >hg38_21_+_8433172_8446622_RNA45SN1_45S_50ntFlanks | 12395 | 12414 | 20 | AGCGACGTCGCTTTTTGATC | 1053.0 | -2.2 | 0.8 | -2.9 | 3.9E-03 | 1.5E-02 | BT20/MB231 | CYTO |
| >MI0003685\|hsa-mir-421&WithFlank&X\|-\|74218371\|74218467@57.75.19(+1A)&[MIMAT0003339&hsa-miR-421&offsets\|+3\|-1(+1A);m-309&X\|-\|74218392\|74218414&offsets\|+3\|-1(+1A)]\|AACAGACATTAATTGGGCGA | >MI0003685\|hsa-mir-421&WithFlank&X\|-\|74218371\|74218467 | 57 | 75 | 19(+1A) | AACAGACATTAATTGGGCGA | 22.5 | 7.7 | 3.0 | 2.5 | 1.1E-02 | 1.5E-02 | MB231/MB468 | MP |
| >MI0003139\|hsa-mir-181d&WithFlank&19\|+\|13874869\|13875017@42.64.23&[MIMAT0002821&hsa-miR-181d-5p&offsets\|0\|0;m-194&19\|+\|13874910\|13874932&offsets\|0\|0]\|AACATTCATTGTTGTCGGTGGGT | >MI0003139\|hsa-mir-181d&WithFlank&19\|+\|13874869\|13875017 | 42 | 64 | 23 | AACATTCATTGTTGTCGGTGGGT | 137.8 | 2.9 | 1.1 | 2.7 | 7.7E-03 | 1.5E-02 | BT20/MB468 | MP |
| >hg38_21_+_8433172_8446622_RNA45SN1_45S_50ntFlanks@9623.9658.36\|TAGTAGCTGGTTCCCTCCGAAGTTTCCCTCAGGATA | >hg38_21_+_8433172_8446622_RNA45SN1_45S_50ntFlanks | 9623 | 9658 | 36 | TAGTAGCTGGTTCCCTCCGAAGTTTCCCTCAGGATA | 252.9 | -3.9 | 1.5 | -2.7 | 7.7E-03 | 1.5E-02 | BT20/MB468 | MP |
| >hg38_21_+_8433172_8446622_RNA45SN1_45S_50ntFlanks@9938.9978.41\|AGCAGGACGGTGGCCATGGAAGTCGGAATCCGCTAAGGAGT | >hg38_21_+_8433172_8446622_RNA45SN1_45S_50ntFlanks | 9938 | 9978 | 41 | AGCAGGACGGTGGCCATGGAAGTCGGAATCCGCTAAGGAGT | 44.2 | 5.0 | 2.0 | 2.5 | 1.1E-02 | 1.5E-02 | MB231/MB468 | MP |
| >hg38_21_+_8433172_8446622_RNA45SN1_45S_50ntFlanks@4099.4121.23\|GTGACCACGGGTGACGGGGAATC | >hg38_21_+_8433172_8446622_RNA45SN1_45S_50ntFlanks | 4099 | 4121 | 23 | GTGACCACGGGTGACGGGGAATC | 346.6 | -4.6 | 1.7 | -2.7 | 7.7E-03 | 1.5E-02 | BT20/MB468 | MP |
| >hg38_21_+_8433172_8446622_RNA45SN1_45S_50ntFlanks@11575.11594.20\|ACTTAGAACTGGTGCGGACC | >hg38_21_+_8433172_8446622_RNA45SN1_45S_50ntFlanks | 11575 | 11594 | 20 | ACTTAGAACTGGTGCGGACC | 387.3 | -2.2 | 0.8 | -2.9 | 3.9E-03 | 1.5E-02 | BT20/MB231 | CYTO |
| >MI0000764\|hsa-mir-363&WithFlank&X\|-\|134169372\|134169458@56.77.22&[MIMAT0000707&hsa-miR-363-3p&offsets\|0\|0;m-178&X\|-\|134169381\|134169403&offsets\|0\|-1]\|AATTGCACGGTATCCATCTGTA | >MI0000764\|hsa-mir-363&WithFlank&X\|-\|134169372\|134169458 | 56 | 77 | 22 | AATTGCACGGTATCCATCTGTA | 1884.8 | 2.4 | 0.9 | 2.7 | 7.8E-03 | 1.5E-02 | BT20/MB468 | MP |
| >hg38_21_+_8433172_8446622_RNA45SN1_45S_50ntFlanks@475.509.35\|CTTCGTGATCGATGTGGTGACGTCGTGCTCTCCCG | >hg38_21_+_8433172_8446622_RNA45SN1_45S_50ntFlanks | 475 | 509 | 35 | CTTCGTGATCGATGTGGTGACGTCGTGCTCTCCCG | 109.9 | 4.9 | 1.8 | 2.7 | 7.8E-03 | 1.5E-02 | BT20/MB468 | MP |
| >MI0003656\|hsa-mir-641&WithFlank&19\|-\|40282537\|40282647@23.46.24&[MIMAT0003311&hsa-miR-641&offsets\|+1\|+1;m-786&19\|-\|40282603\|40282626&offsets\|+1\|+1]\|AAGACATAGGATAGAGTCACCTCT | >MI0003656\|hsa-mir-641&WithFlank&19\|-\|40282537\|40282647 | 23 | 46 | 24 | AAGACATAGGATAGAGTCACCTCT | 31.4 | 5.1 | 2.0 | 2.5 | 1.1E-02 | 1.5E-02 | MB231/MB468 | MP |

| | | | | | | | | | | | | | |
|---|---|---|---|---|---|---|---|---|---|---|---|---|---|
| >trna111_HisGTG_1_-_147774845_147774916@16.33.18,trna118_HisGTG_1_-_145396881_145396952@16.33.18,trna16_HisGTG_1_+_146544773_146544844@16.33.18,trna1_HisGTG_15_+_45493349_45493420@16.33.18,trna21_HisGTG_1_+_147753471_147753542@16.33.18,trna33_HisGTG_6_+_27125906_27125977@16.33.18,trna7_HisGTG_9_-_14433938_14434009@16.33.18,trna8_HisGTG_15_-_45492611_45492682@16.33.18,trna9_HisGTG_15_-_45490804_45490875@16.33.18\|TGGTTAGTACTCTGCGTT | >trna111_HisGTG_1_-_147774845_147774916 | 16 | 33 | 18 | TGGTTAGTACTCTGCGTT | 567.2 | 2.3 | 0.8 | 2.9 | 3.9E-03 | 1.5E-02 | BT20/MB231 | CYTO |
| >hg38_21_+_8433172_8446622_RNA45SN1_45S_50ntFlanks@11583.11609.27\|CTGGTGCGGACCAGGGGAATCCGACTG | >hg38_21_+_8433172_8446622_RNA45SN1_45S_50ntFlanks | 11583 | 11609 | 27 | CTGGTGCGGACCAGGGGAATCCGACTG | 826.5 | -1.6 | 0.7 | -2.5 | 1.2E-02 | 1.5E-02 | MB231/MB468 | CYTO |
| >hg38_21_+_8433172_8446622_RNA45SN1_45S_50ntFlanks@11874.11907.34\|AGAAGACCCTGTTGAGCTTGACTCTAGTCTGGCA | >hg38_21_+_8433172_8446622_RNA45SN1_45S_50ntFlanks | 11874 | 11907 | 34 | AGAAGACCCTGTTGAGCTTGACTCTAGTCTGGCA | 271.7 | -4.6 | 1.7 | -2.7 | 7.8E-03 | 1.5E-02 | BT20/MB468 | MP |
| >hg38_21_+_8433172_8446622_RNA45SN1_45S_50ntFlanks@8184.8204.21\|CGAGGCCCAGCCCGTGGACGG | >hg38_21_+_8433172_8446622_RNA45SN1_45S_50ntFlanks | 8184 | 8204 | 21 | CGAGGCCCAGCCCGTGGACGG | 220.3 | -4.5 | 1.7 | -2.7 | 7.8E-03 | 1.5E-02 | BT20/MB468 | MP |
| >hg38_21_+_8433172_8446622_RNA45SN1_45S_50ntFlanks@9954.9981.28\|TGGAAGTCGGAATCCGCTAAGGAGTGTG | >hg38_21_+_8433172_8446622_RNA45SN1_45S_50ntFlanks | 9954 | 9981 | 28 | TGGAAGTCGGAATCCGCTAAGGAGTGTG | 68.3 | 4.6 | 1.7 | 2.7 | 7.8E-03 | 1.5E-02 | BT20/MB468 | MP |
| >hg38_21_+_8433172_8446622_RNA45SN1_45S_50ntFlanks@12001.12030.30\|CTGCGGGCCGCCGGTGAAATACCACTACTC | >hg38_21_+_8433172_8446622_RNA45SN1_45S_50ntFlanks | 12001 | 12030 | 30 | CTGCGGGCCGCCGGTGAAATACCACTACTC | 143.0 | -2.8 | 1.1 | -2.6 | 1.1E-02 | 1.5E-02 | BT20/MB468 | MITO |
| >hg38_21_+_8433172_8446622_RNA45SN1_45S_50ntFlanks@7976.8010.35\|GCGACCTCAGATCAGACGTGGCGACCCGCTGAATT | >hg38_21_+_8433172_8446622_RNA45SN1_45S_50ntFlanks | 7976 | 8010 | 35 | GCGACCTCAGATCAGACGTGGCGACCCGCTGAATT | 3233.6 | 5.0 | 1.7 | 3.0 | 3.0E-03 | 1.5E-02 | BT20/MB231 | MP |
| >trna10_AspGTC_12_-_125424193_125424264@37.74.38,trna12_AspGTC_12_-_125411891_125411962@37.74.38,trna144_AspGTC_6_-_27551236_27551307@37.74.38,trna38_AspGTC_17_-_8125556_8125627@37.74.38,trna45_AspGTC_6_+_27447453_27447524@37.74.38,trna48_AspGTC_6_+_27471523_27471594@37.74.38,trna4_AspGTC_12_+_96429799_96429870@37.74.38,trna69_AspGTC_1_-_161440205_161440276@37.74.38,trna72_AspGTC_1_-_161432824_161432895@37.74.38,trna75_AspGTC_1_-_161425414_161425485@37.74.38,trna78_AspGTC_1_-_161418033_161418104@37.74.38,trna81_AspGTC_1_-_161410615_161410686@37.74.38\|ACGCGGGAGACCGGGGTTCGATTCCCCGACGGGGAGCC | >trna10_AspGTC_12_-_125424193_125424264 | 37 | 74 | 38 | ACGCGGGAGACCGGGGTTCGATTCCCCGACGGGGAGCC | 494.4 | -3.8 | 1.3 | -3.0 | 3.0E-03 | 1.5E-02 | BT20/MB231 | MP |
| >am_trna111_HisGTG_1_-_147774845_147774916@3.28.26,trna118_HisGTG_1_-_145396881_145396952@3.28.26,trna16_HisGTG_1_+_146544773_146544844@3.28.26,trna1_HisGTG_15_+_45493349_45493420@3.28.26,trna21_HisGTG_1_+_147753471_147753542@3.28.26,trna33_HisGTG_6_+_27125906_27125977@3.28.26,trna7_HisGTG_9_-_14433938_14434009@3.28.26,trna8_HisGTG_15_-_45492611_45492682@3.28.26,trna9_HisGTG_15_-_45490804_45490875@3.28.26\|CGTGATCGTATAGTGGTTAGTACTCT | >am_trna111_HisGTG_1_-_147774845_147774916 | 3 | 28 | 26 | CGTGATCGTATAGTGGTTAGTACTCT | 847.8 | 4.5 | 1.5 | 3.0 | 3.1E-03 | 1.5E-02 | BT20/MB231 | MP |
| >hg38_21_+_8433172_8446622_RNA45SN1_45S_50ntFlanks@5485.5526.42\|AGCGCTGAGAAGACGGTCGAACTTGACTATCTAGAGGAAGTA | >hg38_21_+_8433172_8446622_RNA45SN1_45S_50ntFlanks | 5485 | 5526 | 42 | AGCGCTGAGAAGACGGTCGAACTTGACTATCTAGAGGAAGTA | 451.4 | -2.5 | 1.0 | -2.5 | 1.2E-02 | 1.5E-02 | BT20/MB468 | CYTO |
| >trnaMT_SerGCT_MT_+_12207_12265@10.31.22\|CACAAGAACTGCTAACTCATGC | >trnaMT_SerGCT_MT_+_12207_12265 | 10 | 31 | 22 | CACAAGAACTGCTAACTCATGC | 265.0 | -3.3 | 1.3 | -2.5 | 1.1E-02 | 1.5E-02 | MB231/MB468 | MP |
| >hg38_1_-_228634819_228635039_RNA5S12_5S_50ntFlanks@139.169.31\|GACCGCCTGGGAATACCGGGTGCTGTAGGCT | >hg38_1_-_228634819_228635039_RNA5S12_5S_50ntFlanks | 139 | 169 | 31 | GACCGCCTGGGAATACCGGGTGCTGTAGGCT | 3443.4 | 3.9 | 1.3 | 3.0 | 3.1E-03 | 1.5E-02 | BT20/MB231 | MP |
| >hg38_21_+_8433172_8446622_RNA45SN1_45S_50ntFlanks@4727.4766.40\|AATCAAGAACGAAAGTCGGAGGTTCGAAGACGATCAGATA | >hg38_21_+_8433172_8446622_RNA45SN1_45S_50ntFlanks | 4727 | 4766 | 40 | AATCAAGAACGAAAGTCGGAGGTTCGAAGACGATCAGATA | 79.0 | 5.4 | 2.0 | 2.7 | 7.8E-03 | 1.5E-02 | BT20/MB468 | MP |
| >hg38_21_+_8433172_8446622_RNA45SN1_45S_50ntFlanks@6659.6695.37\|AGCGGTGGATCACTCGGCTCGTGCGTCGATGAAGAAC | >hg38_21_+_8433172_8446622_RNA45SN1_45S_50ntFlanks | 6659 | 6695 | 37 | AGCGGTGGATCACTCGGCTCGTGCGTCGATGAAGAAC | 124.3 | 4.6 | 1.7 | 2.7 | 7.8E-03 | 1.5E-02 | BT20/MB468 | MP |
| >MI0000077\|hsa-mir-21&WithFlank&17\|+\|59841260\|59841343@14.35.22&[MIMAT0000076&hsa-miR-21-5p&offsets\|0\|0;m-1&17\|+\|59841273\|59841294&offsets\|0\|0]\|TAGCTTATCAGACTGATGTTGA | >MI0000077\|hsa-mir-21&WithFlank&17\|+\|59841260\|59841343 | 14 | 35 | 22 | TAGCTTATCAGACTGATGTTGA | 132003.2 | 2.9 | 1.0 | 2.9 | 3.9E-03 | 1.5E-02 | BT20/MB231 | CYTO |
| >trnaMT_GluTTC_MT_-_14674_14742@42.72.31,trnalookalike8_GluTTC_5_-_93905172_93905240@42.72.31\|GGTCGTGGTTGTAGTCCGTGCGAGAATACCA | >trnaMT_GluTTC_MT_-_14674_14742 | 42 | 72 | 31 | GGTCGTGGTTGTAGTCCGTGCGAGAATACCA | 456.7 | 3.0 | 1.1 | 2.7 | 7.8E-03 | 1.5E-02 | BT20/MB468 | MP |
| >hg38_21_+_8433172_8446622_RNA45SN1_45S_50ntFlanks@12394.12414.21\|TAGCGACGTCGCTTTTTGATC | >hg38_21_+_8433172_8446622_RNA45SN1_45S_50ntFlanks | 12394 | 12414 | 21 | TAGCGACGTCGCTTTTTGATC | 550.3 | -2.7 | 1.0 | -2.7 | 7.8E-03 | 1.5E-02 | BT20/MB468 | MP |
| >hg38_21_+_8433172_8446622_RNA45SN1_45S_50ntFlanks@3839.3869.31\|TCTCCTACTTGGATAACTGTGGTAATTCTAG | >hg38_21_+_8433172_8446622_RNA45SN1_45S_50ntFlanks | 3839 | 3869 | 31 | TCTCCTACTTGGATAACTGTGGTAATTCTAG | 311.7 | -2.9 | 1.0 | -2.9 | 3.9E-03 | 1.5E-02 | BT20/MB231 | CYTO |
| >hg38_21_+_8433172_8446622_RNA45SN1_45S_50ntFlanks@7973.7995.23\|GACGCGACCTCAGATCAGACGTG | >hg38_21_+_8433172_8446622_RNA45SN1_45S_50ntFlanks | 7973 | 7995 | 23 | GACGCGACCTCAGATCAGACGTG | 537.2 | 1.7 | 0.7 | 2.5 | 1.2E-02 | 1.5E-02 | MB231/MB468 | CYTO |
| >hg38_21_+_8433172_8446622_RNA45SN1_45S_50ntFlanks@3838.3873.36\|CTCTCCTACTTGGATAACTGTGGTAATTCTAGAGCT | >hg38_21_+_8433172_8446622_RNA45SN1_45S_50ntFlanks | 3838 | 3873 | 36 | CTCTCCTACTTGGATAACTGTGGTAATTCTAGAGCT | 100.4 | -5.0 | 1.9 | -2.7 | 7.8E-03 | 1.5E-02 | BT20/MB468 | MP |
| >MI0000813\|hsa-mir-324&WithFlank&17\|-\|7223291\|7223385@59.78.20(+1C)&[MIMAT0000762&hsa-miR-324-3p&offsets\|+3\|0(+1C);m-203&17\|-\|7223308\|7223329&offsets\|+2\|0(+1C)]\|ACTGCCCCAGGTGCTGCTGGC | >MI0000813\|hsa-mir-324&WithFlank&17\|-\|7223291\|7223385 | 59 | 78 | 20(+1C) | ACTGCCCCAGGTGCTGCTGGC | 57.7 | 5.1 | 2.0 | 2.5 | 1.2E-02 | 1.5E-02 | MB231/MB468 | MP |
| >hg38_21_+_8433172_8446622_RNA45SN1_45S_50ntFlanks@11570.11588.19\|AGCCGACTTAGAACTGGTG | >hg38_21_+_8433172_8446622_RNA45SN1_45S_50ntFlanks | 11570 | 11588 | 19 | AGCCGACTTAGAACTGGTG | 347.6 | -2.8 | 1.1 | -2.7 | 7.8E-03 | 1.5E-02 | BT20/MB468 | MP |
| >hg38_21_+_8433172_8446622_RNA45SN1_45S_50ntFlanks@10793.10825.33\|TCTGGCATGTTGGAACAATGTAGGTAAGGGAAG | >hg38_21_+_8433172_8446622_RNA45SN1_45S_50ntFlanks | 10793 | 10825 | 33 | TCTGGCATGTTGGAACAATGTAGGTAAGGGAAG | 66.0 | -4.4 | 1.6 | -2.7 | 7.8E-03 | 1.5E-02 | BT20/MB468 | MP |
| >hg38_21_+_8433172_8446622_RNA45SN1_45S_50ntFlanks@11774.11794.21\|AGTGACGCGCATGAATGGATG | >hg38_21_+_8433172_8446622_RNA45SN1_45S_50ntFlanks | 11774 | 11794 | 21 | AGTGACGCGCATGAATGGATG | 428.3 | -2.3 | 0.9 | -2.5 | 1.2E-02 | 1.5E-02 | MB231/MB468 | MP |
| >hg38_21_+_8433172_8446622_RNA45SN1_45S_50ntFlanks@4996.5013.18\|CAGATTGATAGCTCTTTC | >hg38_21_+_8433172_8446622_RNA45SN1_45S_50ntFlanks | 4996 | 5013 | 18 | CAGATTGATAGCTCTTTC | 177.7 | -4.6 | 1.7 | -2.7 | 7.8E-03 | 1.5E-02 | BT20/MB468 | MP |
| >trnaMT_PheGAA_MT_+_577_647@52.74.23\|CTCACATCACCCCATAAACACCA | >trnaMT_PheGAA_MT_+_577_647 | 52 | 74 | 23 | CTCACATCACCCCATAAACACCA | 313.1 | -4.9 | 2.0 | -2.5 | 1.3E-02 | 1.5E-02 | BT20/MB468 | NUC |
| >hg38_MT_+_1621_3279_RNR2_16S_50ntFlanks@1369.1398.30\|GATGTTGGATCAGGACATCCCGATGGTGCA | >hg38_MT_+_1621_3279_RNR2_16S_50ntFlanks | 1369 | 1398 | 30 | GATGTTGGATCAGGACATCCCGATGGTGCA | 122.9 | 4.6 | 1.7 | 2.7 | 7.8E-03 | 1.5E-02 | BT20/MB468 | MP |
| >trnaMT_TyrGTA_MT_-_5826_5891@37.69.33\|TAAAGACAGGGGTTAGGCCTCTTTTTACCACCA | >trnaMT_TyrGTA_MT_-_5826_5891 | 37 | 69 | 33 | TAAAGACAGGGGTTAGGCCTCTTTTTACCACCA | 112.1 | 4.2 | 1.6 | 2.5 | 1.2E-02 | 1.5E-02 | MB231/MB468 | MP |
| >hg38_21_+_8433172_8446622_RNA45SN1_45S_50ntFlanks@11666.11692.27\|TCTGCCCAGTGCTCTGAATGTCAAAGT | >hg38_21_+_8433172_8446622_RNA45SN1_45S_50ntFlanks | 11666 | 11692 | 27 | TCTGCCCAGTGCTCTGAATGTCAAAGT | 312.1 | 2.3 | 0.8 | 2.9 | 3.9E-03 | 1.5E-02 | BT20/MB231 | MITO |
| >hg38_21_+_8433172_8446622_RNA45SN1_45S_50ntFlanks@4981.5002.22\|CACGGACAGGATTGACAGATTG | >hg38_21_+_8433172_8446622_RNA45SN1_45S_50ntFlanks | 4981 | 5002 | 22 | CACGGACAGGATTGACAGATTG | 360.0 | 4.5 | 1.5 | 3.0 | 3.1E-03 | 1.5E-02 | BT20/MB231 | MP |
| >MI0000266\|hsa-mir-10a&WithFlank&17\|-\|48579832\|48579953@28.50.23(+1A)&[MIMAT0000253&hsa-miR-10a-5p&offsets\|0\|0(+1A);m-18&17\|-\|48579905\|48579926&offsets\|0\|+1(+1A)]\|TACCCTGTAGATCCGAATTTGTGA | >MI0000266\|hsa-mir-10a&WithFlank&17\|-\|48579832\|48579953 | 28 | 50 | 23(+1A) | TACCCTGTAGATCCGAATTTGTGA | 1658.7 | 2.3 | 0.8 | 2.9 | 3.9E-03 | 1.5E-02 | BT20/MB231 | CYTO |
| >hg38_MT_+_1621_3279_RNR2_16S_50ntFlanks@1293.1320.28\|AACGGAACAAGTTACCCTAGGGATAACA | >hg38_MT_+_1621_3279_RNR2_16S_50ntFlanks | 1293 | 1320 | 28 | AACGGAACAAGTTACCCTAGGGATAACA | 169.5 | 3.2 | 1.2 | 2.7 | 7.8E-03 | 1.5E-02 | BT20/MB468 | MP |
| >hg38_21_+_8433172_8446622_RNA45SN1_45S_50ntFlanks@10571.10588.18\|GAGAAGCCGGCGGGAGCC | >hg38_21_+_8433172_8446622_RNA45SN1_45S_50ntFlanks | 10571 | 10588 | 18 | GAGAAGCCGGCGGGAGCC | 77.9 | -4.3 | 1.6 | -2.7 | 7.8E-03 | 1.5E-02 | BT20/MB468 | MP |
| >hg38_MT_+_1621_3279_RNR2_16S_50ntFlanks@459.492.34\|CTTGTAAATTTAACTGTTAGTCCAAAGAGGAACA | >hg38_MT_+_1621_3279_RNR2_16S_50ntFlanks | 459 | 492 | 34 | CTTGTAAATTTAACTGTTAGTCCAAAGAGGAACA | 424.5 | 4.2 | 1.6 | 2.7 | 7.8E-03 | 1.5E-02 | BT20/MB468 | MP |
| >trna10_LysCTT_16_+_3241501_3241573@3.35.33,trna13_LysCTT_14_-_58706613_58706685@3.35.33,trna2_LysCTT_15_+_79152904_79152976@3.35.33\|CCGGCTAGCTCAGTCGGTAGAGCATGGGACTCT | >trna10_LysCTT_16_+_3241501_3241573 | 3 | 35 | 33 | CCGGCTAGCTCAGTCGGTAGAGCATGGGACTCT | 509.3 | -2.4 | 0.8 | -2.9 | 3.9E-03 | 1.5E-02 | BT20/MB231 | CYTO |
| >hg38_21_+_8433172_8446622_RNA45SN1_45S_50ntFlanks@12408.12441.34\|TTTGATCCTTCGATGTCGGCTCTTCCTATCATTG | >hg38_21_+_8433172_8446622_RNA45SN1_45S_50ntFlanks | 12408 | 12441 | 34 | TTTGATCCTTCGATGTCGGCTCTTCCTATCATTG | 302.4 | -1.4 | 0.5 | -2.5 | 1.3E-02 | 1.5E-02 | MB231/MB468 | NUC |
| >MI0000088\|hsa-mir-30a&WithFlank&6\|-\|71403545\|71403627@12.32.21&[MIMAT0000087&hsa-miR-30a-5p&offsets\|0\|-1;m-15&6\|-\|71403595\|71403616&offsets\|0\|-1]\|TGTAAACATCCTCGACTGGAA | >MI0000088\|hsa-mir-30a&WithFlank&6\|-\|71403545\|71403627 | 12 | 32 | 21 | TGTAAACATCCTCGACTGGAA | 3115.9 | -2.8 | 1.0 | -3.0 | 3.1E-03 | 1.5E-02 | BT20/MB231 | MP |
| >MI0003513\|hsa-mir-455&WithFlank&9\|+\|114209428\|114209535@22.43.22(+1C)&[MIMAT0003150&hsa-miR-455-5p&offsets\|0\|0(+1C);m-232&9\|+\|114209449\|114209470&offsets\|0\|0(+1C)]\|TATGTGCCTTTGGACTACATCGC | >MI0003513\|hsa-mir-455&WithFlank&9\|+\|114209428\|114209535 | 22 | 43 | 22(+1C) | TATGTGCCTTTGGACTACATCGC | 283.3 | -4.5 | 1.5 | -3.0 | 3.1E-03 | 1.5E-02 | BT20/MB231 | MP |
| >hg38_21_+_8433172_8446622_RNA45SN1_45S_50ntFlanks@12418.12443.26\|CGATGTCGGCTCTTCCTATCATTGTG | >hg38_21_+_8433172_8446622_RNA45SN1_45S_50ntFlanks | 12418 | 12443 | 26 | CGATGTCGGCTCTTCCTATCATTGTG | 946.0 | -2.4 | 0.8 | -2.9 | 4.0E-03 | 1.5E-02 | BT20/MB231 | CYTO |
| >MI0000749\|hsa-mir-30e&WithFlank&1\|+\|40754349\|40754452@24.46.23(+2U)&[MIMAT0000692&hsa-miR-30e-5p&offsets\|+1\|+2(+2U);m-24&1\|+\|40754371\|40754394&offsets\|+1\|0(+2U)]\|GTAAACATCCTTGACTGGAAGCTTT | >MI0000749\|hsa-mir-30e&WithFlank&1\|+\|40754349\|40754452 | 24 | 46 | 23(+2U) | GTAAACATCCTTGACTGGAAGCTTT | 47.4 | 4.9 | 2.0 | 2.5 | 1.2E-02 | 1.5E-02 | MB231/MB468 | MP |
| >hg38_21_+_8433172_8446622_RNA45SN1_45S_50ntFlanks@11582.11601.20\|ACTGGTGCGGACCAGGGGAA | >hg38_21_+_8433172_8446622_RNA45SN1_45S_50ntFlanks | 11582 | 11601 | 20 | ACTGGTGCGGACCAGGGGAA | 480.9 | -2.0 | 0.8 | -2.5 | 1.2E-02 | 1.5E-02 | BT20/MB468 | CYTO |

<table>
<tr><td>>MI0000469|hsa-mir-125a&WithFlank&19|+|51693248|51693345@21.42.22(+1A)&[MIMAT0000443&hsa-miR-125a-5p&offsets|0|-2(+1A);m-63&19|+|51693268|51693289&offsets|0|0(+1A)]|TCCCTGAGACCCTTTAACCTGTA</td><td>>MI0000469|hsa-mir-125a&WithFlank&19|+|51693248|51693345</td><td>21</td><td>42</td><td>22(+1A)</td><td>TCCCTGAGACCCTTTAACCTGTA</td><td>5091.3</td><td>2.6</td><td>0.9</td><td>2.9</td><td>4.0E-03</td><td>1.5E-02</td><td>BT20/MB231</td><td>CYTO</td></tr>
<tr><td>>hg38_21_+_8433172_8446622_RNA45SN1_45S_50ntFlanks@12420.12449.30|ATGTCGGCTCTTCCTATCATTGTGAAGCAG</td><td>>hg38_21_+_8433172_8446622_RNA45SN1_45S_50ntFlanks</td><td>12420</td><td>12449</td><td>30</td><td>ATGTCGGCTCTTCCTATCATTGTGAAGCAG</td><td>803.2</td><td>-2.4</td><td>0.8</td><td>-2.9</td><td>4.0E-03</td><td>1.5E-02</td><td>BT20/MB231</td><td>CYTO</td></tr>
<tr><td>>hg38_1_-_228634819_228635039_RNA5S12_5S_50ntFlanks@151.172.22|ATACCGGGTGCTGTAGGCTTTT</td><td>>hg38_1_-_228634819_228635039_RNA5S12_5S_50ntFlanks</td><td>151</td><td>172</td><td>22</td><td>ATACCGGGTGCTGTAGGCTTTT</td><td>104.2</td><td>-5.6</td><td>2.2</td><td>-2.5</td><td>1.2E-02</td><td>1.5E-02</td><td>MB231/MB468</td><td>MP</td></tr>
<tr><td>>MI0000542|hsa-mir-320a&WithFlank&8|-|22244960|22245043@48.72.25&[MIMAT0000510&hsa-miR-320a-3p&offsets|0|+3;m-60&8|-|22244975|22244996&offsets|0|+3]|AAAAGCTGGGTTGAGAGGGCGAAAA</td><td>>MI0000542|hsa-mir-320a&WithFlank&8|-|22244960|22245043</td><td>48</td><td>72</td><td>25</td><td>AAAAGCTGGGTTGAGAGGGCGAAAA</td><td>45.3</td><td>5.0</td><td>2.0</td><td>2.5</td><td>1.2E-02</td><td>1.5E-02</td><td>MB231/MB468</td><td>MP</td></tr>
<tr><td>>trnaMT_HisGTG_MT_+_12138_12206@32.69.38|TGAATCTGACAACAGAGGCTTACGACCCCTTATTTACC</td><td>>trnaMT_HisGTG_MT_+_12138_12206</td><td>32</td><td>69</td><td>38</td><td>TGAATCTGACAACAGAGGCTTACGACCCCTTATTTACC</td><td>4915.0</td><td>4.5</td><td>1.8</td><td>2.5</td><td>1.1E-02</td><td>1.5E-02</td><td>BT20/MB468</td><td>MITO</td></tr>
<tr><td>>MI0000298|hsa-mir-221&WithFlank&X|-|45746151|45746272@31.56.26&[MIMAT0004568&hsa-miR-221-5p&offsets|0|+4;m-405&X|-|45746220|45746242&offsets|0|+3]|ACCTGGCATACAATGTAGATTTCTGT</td><td>>MI0000298|hsa-mir-221&WithFlank&X|-|45746151|45746272</td><td>31</td><td>56</td><td>26</td><td>ACCTGGCATACAATGTAGATTTCTGT</td><td>98.4</td><td>4.9</td><td>1.9</td><td>2.5</td><td>1.2E-02</td><td>1.5E-02</td><td>MB231/MB468</td><td>MP</td></tr>
<tr><td>>MI0000289|hsa-mir-181a-1&WithFlank&1|-|198859038|198859159@30.48.19(+1A);MI0000269|hsa-mir-181a-2&WithFlank&9|+|124692436|124692557@45.63.19(+1A)&[MIMAT0000256&hsa-miR-181a-5p&offsets|0|-4(+1A);m-32&1|-|198859108|198859130&offsets|0|-4(+1A)];[MIMAT0000256_1&hsa-miR-181a-5p&offsets|0|-4(+1A);m-32&9|+|124692480|124692502&offsets|0|-4(+1A)]|AACATTCAACGCTGTCGGTA</td><td>>MI0000289|hsa-mir-181a-1&WithFlank&1|-|198859038|198859159</td><td>30</td><td>48</td><td>19(+1A)</td><td>AACATTCAACGCTGTCGGTA</td><td>278.2</td><td>-4.4</td><td>1.5</td><td>-3.0</td><td>3.1E-03</td><td>1.5E-02</td><td>BT20/MB231</td><td>MP</td></tr>
<tr><td>>hg38_21_+_8433172_8446622_RNA45SN1_45S_50ntFlanks@12448.12473.26|AGAATTCACCAAGCGTTGGATTGTTC</td><td>>hg38_21_+_8433172_8446622_RNA45SN1_45S_50ntFlanks</td><td>12448</td><td>12473</td><td>26</td><td>AGAATTCACCAAGCGTTGGATTGTTC</td><td>210.3</td><td>-3.3</td><td>1.1</td><td>-3.0</td><td>3.1E-03</td><td>1.5E-02</td><td>BT20/MB231</td><td>MP</td></tr>
<tr><td>>hg38_21_+_8433172_8446622_RNA45SN1_45S_50ntFlanks@10791.10818.28|CCTCTGGCATGTTGGAACAATGTAGGTA</td><td>>hg38_21_+_8433172_8446622_RNA45SN1_45S_50ntFlanks</td><td>10791</td><td>10818</td><td>28</td><td>CCTCTGGCATGTTGGAACAATGTAGGTA</td><td>185.4</td><td>-1.1</td><td>0.4</td><td>-2.5</td><td>1.1E-02</td><td>1.5E-02</td><td>BT20/MB468</td><td>MITO</td></tr>
<tr><td>>hg38_21_+_8433172_8446622_RNA45SN1_45S_50ntFlanks@4554.4574.21|CCGCAGCTAGGAATAATGGAA</td><td>>hg38_21_+_8433172_8446622_RNA45SN1_45S_50ntFlanks</td><td>4554</td><td>4574</td><td>21</td><td>CCGCAGCTAGGAATAATGGAA</td><td>148.3</td><td>-1.8</td><td>0.7</td><td>-2.5</td><td>1.3E-02</td><td>1.5E-02</td><td>MB231/MB468</td><td>NUC</td></tr>
<tr><td>>hg38_21_+_8433172_8446622_RNA45SN1_45S_50ntFlanks@11587.11611.25|TGCGGACCAGGGGAATCCGACTGTT</td><td>>hg38_21_+_8433172_8446622_RNA45SN1_45S_50ntFlanks</td><td>11587</td><td>11611</td><td>25</td><td>TGCGGACCAGGGGAATCCGACTGTT</td><td>708.6</td><td>-2.3</td><td>0.9</td><td>-2.7</td><td>7.9E-03</td><td>1.5E-02</td><td>BT20/MB468</td><td>MP</td></tr>
<tr><td>>am_trnaMT_ProTGG_MT_-_15956_16023@53.70.18|AAGACTTTTTCTCTGACC</td><td>>am_trnaMT_ProTGG_MT_-_15956_16023</td><td>53</td><td>70</td><td>18</td><td>AAGACTTTTTCTCTGACC</td><td>1230.5</td><td>2.0</td><td>0.8</td><td>2.5</td><td>1.2E-02</td><td>1.5E-02</td><td>MB231/MB468</td><td>MP</td></tr>
<tr><td>>hg38_21_+_8433172_8446622_RNA45SN1_45S_50ntFlanks@5472.5490.19|ACGGCCCTGGCGGAGCGCT</td><td>>hg38_21_+_8433172_8446622_RNA45SN1_45S_50ntFlanks</td><td>5472</td><td>5490</td><td>19</td><td>ACGGCCCTGGCGGAGCGCT</td><td>129.7</td><td>3.1</td><td>1.3</td><td>2.5</td><td>1.3E-02</td><td>1.5E-02</td><td>BT20/MB468</td><td>NUC</td></tr>
<tr><td>>MI0000067|hsa-let-7f-1&WithFlank&9|+|94176341|94176439@13.33.21(+1C);MI0000068|hsa-let-7f-2&WithFlank&X|-|53557186|53557280@14.34.21(+1C)&[MIMAT0000067&hsa-let-7f-5p&offsets|0|-1(+1C);m-20&9|+|94176353|94176374&offsets|0|-1(+1C)];[MIMAT0000067_1&hsa-let-7f-5p&offsets|0|-1(+1C);m-19&X|-|53557246|53557267&offsets|0|-1(+1C)]|TGAGGTAGTAGATTGTATAGTC</td><td>>MI0000067|hsa-let-7f-1&WithFlank&9|+|94176341|94176439</td><td>13</td><td>33</td><td>21(+1C)</td><td>TGAGGTAGTAGATTGTATAGTC</td><td>17830.8</td><td>-3.2</td><td>1.1</td><td>-3.0</td><td>3.1E-03</td><td>1.5E-02</td><td>BT20/MB231</td><td>MP</td></tr>
<tr><td>>hg38_21_+_8433172_8446622_RNA45SN1_45S_50ntFlanks@10347.10366.20|CAAACGAGAACTTTGAAGGC</td><td>>hg38_21_+_8433172_8446622_RNA45SN1_45S_50ntFlanks</td><td>10347</td><td>10366</td><td>20</td><td>CAAACGAGAACTTTGAAGGC</td><td>194.5</td><td>-3.4</td><td>1.3</td><td>-2.5</td><td>1.2E-02</td><td>1.5E-02</td><td>MB231/MB468</td><td>MP</td></tr>
<tr><td>>hg38_1_-_228634819_228635039_RNA5S12_5S_50ntFlanks@134.169.36|TGGGAGACCGCCTGGGAATACCGGGTGCTGTAGGCT</td><td>>hg38_1_-_228634819_228635039_RNA5S12_5S_50ntFlanks</td><td>134</td><td>169</td><td>36</td><td>TGGGAGACCGCCTGGGAATACCGGGTGCTGTAGGCT</td><td>1215.7</td><td>2.2</td><td>0.9</td><td>2.5</td><td>1.2E-02</td><td>1.5E-02</td><td>MB231/MB468</td><td>MP</td></tr>
<tr><td>>hg38_1_-_228634819_228635039_RNA5S12_5S_50ntFlanks@129.171.43|TTGGATGGGAGACCGCCTGGGAATACCGGGTGCTGTAGGCTTT</td><td>>hg38_1_-_228634819_228635039_RNA5S12_5S_50ntFlanks</td><td>129</td><td>171</td><td>43</td><td>TTGGATGGGAGACCGCCTGGGAATACCGGGTGCTGTAGGCTTT</td><td>182.8</td><td>4.0</td><td>1.6</td><td>2.5</td><td>1.3E-02</td><td>1.5E-02</td><td>MB231/MB468</td><td>MITO</td></tr>
<tr><td>>MI0000812|hsa-mir-331&WithFlank&12|+|95308414|95308519@31.52.22&[MIMAT0004700&hsa-miR-331-5p&offsets|-1|-1;TJU_CMC_MD2.ID00734.5p-miR&offsets|+19|+21;m-2747&12|+|95308425|95308444&offsets|+19|+21;m-377&12|+|95308445|95308465&offsets|-1|0]|TCTAGGTATGGTCCCAGGGATC</td><td>>MI0000812|hsa-mir-331&WithFlank&12|+|95308414|95308519</td><td>31</td><td>52</td><td>22</td><td>TCTAGGTATGGTCCCAGGGATC</td><td>62.2</td><td>5.0</td><td>2.0</td><td>2.5</td><td>1.2E-02</td><td>1.5E-02</td><td>MB231/MB468</td><td>MP</td></tr>
<tr><td>>hg38_21_+_8433172_8446622_RNA45SN1_45S_50ntFlanks@12412.12429.18|ATCCTTCGATGTCGGCTC</td><td>>hg38_21_+_8433172_8446622_RNA45SN1_45S_50ntFlanks</td><td>12412</td><td>12429</td><td>18</td><td>ATCCTTCGATGTCGGCTC</td><td>324.2</td><td>-3.6</td><td>1.5</td><td>-2.5</td><td>1.3E-02</td><td>1.5E-02</td><td>MB231/MB468</td><td>CYTO</td></tr>
<tr><td>>hg38_1_-_228634819_228635039_RNA5S12_5S_50ntFlanks@126.170.45|TACTTGGATGGGAGACCGCCTGGGAATACCGGGTGCTGTAGGCTT</td><td>>hg38_1_-_228634819_228635039_RNA5S12_5S_50ntFlanks</td><td>126</td><td>170</td><td>45</td><td>TACTTGGATGGGAGACCGCCTGGGAATACCGGGTGCTGTAGGCTT</td><td>1144.1</td><td>2.9</td><td>1.0</td><td>2.9</td><td>3.9E-03</td><td>1.5E-02</td><td>BT20/MB231</td><td>MITO</td></tr>
<tr><td>>hg38_21_+_8433172_8446622_RNA45SN1_45S_50ntFlanks@7970.7997.28|CGAGACGCGACCTCAGATCAGACGTGGC</td><td>>hg38_21_+_8433172_8446622_RNA45SN1_45S_50ntFlanks</td><td>7970</td><td>7997</td><td>28</td><td>CGAGACGCGACCTCAGATCAGACGTGGC</td><td>90.8</td><td>1.9</td><td>0.7</td><td>2.5</td><td>1.3E-02</td><td>1.5E-02</td><td>BT20/MB468</td><td>NUC</td></tr>
<tr><td>>hg38_21_+_8433172_8446622_RNA45SN1_45S_50ntFlanks@12418.12456.39|CGATGTCGGCTCTTCCTATCATTGTGAAGCAGAATTCAC</td><td>>hg38_21_+_8433172_8446622_RNA45SN1_45S_50ntFlanks</td><td>12418</td><td>12456</td><td>39</td><td>CGATGTCGGCTCTTCCTATCATTGTGAAGCAGAATTCAC</td><td>1423.8</td><td>-1.9</td><td>0.7</td><td>-2.5</td><td>1.3E-02</td><td>1.5E-02</td><td>MB231/MB468</td><td>CYTO</td></tr>
<tr><td>>hg38_21_+_8433172_8446622_RNA45SN1_45S_50ntFlanks@6645.6686.42|CTCGTACGACTCTTAGCGGTGGATCACTCGGCTCGTGCGTCG</td><td>>hg38_21_+_8433172_8446622_RNA45SN1_45S_50ntFlanks</td><td>6645</td><td>6686</td><td>42</td><td>CTCGTACGACTCTTAGCGGTGGATCACTCGGCTCGTGCGTCG</td><td>400.6</td><td>-1.9</td><td>0.8</td><td>-2.5</td><td>1.3E-02</td><td>1.5E-02</td><td>MB231/MB468</td><td>MITO</td></tr>
<tr><td>>hg38_21_+_8433172_8446622_RNA45SN1_45S_50ntFlanks@11705.11728.24|TGAAGCGCGGGTAAACGGCGGGAG</td><td>>hg38_21_+_8433172_8446622_RNA45SN1_45S_50ntFlanks</td><td>11705</td><td>11728</td><td>24</td><td>TGAAGCGCGGGTAAACGGCGGGAG</td><td>533.9</td><td>-4.1</td><td>1.7</td><td>-2.5</td><td>1.3E-02</td><td>1.5E-02</td><td>BT20/MB468</td><td>CYTO</td></tr>
<tr><td>>hg38_21_+_8433172_8446622_RNA45SN1_45S_50ntFlanks@476.522.47|TTCGTGATCGATGTGGTGACGTCGTGCTCTCCCGGGCCGGGTCCGAG</td><td>>hg38_21_+_8433172_8446622_RNA45SN1_45S_50ntFlanks</td><td>476</td><td>522</td><td>47</td><td>TTCGTGATCGATGTGGTGACGTCGTGCTCTCCCGGGCCGGGTCCGAG</td><td>137.3</td><td>4.9</td><td>2.0</td><td>2.5</td><td>1.3E-02</td><td>1.5E-02</td><td>MB231/MB468</td><td>MITO</td></tr>
<tr><td>>hg38_21_+_8433172_8446622_RNA45SN1_45S_50ntFlanks@10386.10410.25|ATGTGAACAGCAGTTGAACATGGGT</td><td>>hg38_21_+_8433172_8446622_RNA45SN1_45S_50ntFlanks</td><td>10386</td><td>10410</td><td>25</td><td>ATGTGAACAGCAGTTGAACATGGGT</td><td>481.1</td><td>-3.0</td><td>1.2</td><td>-2.5</td><td>1.3E-02</td><td>1.5E-02</td><td>MB231/MB468</td><td>MITO</td></tr>
<tr><td>>hg38_21_+_8433172_8446622_RNA45SN1_45S_50ntFlanks@10841.10867.27|CGTAACTTCGGGATAAGGATTGGCTCT</td><td>>hg38_21_+_8433172_8446622_RNA45SN1_45S_50ntFlanks</td><td>10841</td><td>10867</td><td>27</td><td>CGTAACTTCGGGATAAGGATTGGCTCT</td><td>684.4</td><td>-2.3</td><td>0.9</td><td>-2.5</td><td>1.3E-02</td><td>1.5E-02</td><td>MB231/MB468</td><td>MITO</td></tr>
<tr><td>>MI0000447|hsa-mir-128-1&WithFlank&2|+|135665391|135665484@56.75.20(+2A);MI0000727|hsa-mir-128-2&WithFlank&3|+|35744470|35744565@58.77.20(+2A)&[MIMAT0000424&hsa-miR-128-3p&offsets|0|-1(+2A);m-108&2|+|135665446|135665466&offsets|0|-1(+2A)];[MIMAT0000424_1&hsa-miR-128-3p&offsets|0|-1(+2A);m-117&3|+|35744527|35744547&offsets|0|-1(+2A)]|TCACAGTGAACCGGTCTCTTAA</td><td>>MI0000447|hsa-mir-128-1&WithFlank&2|+|135665391|135665484</td><td>56</td><td>75</td><td>20(+2A)</td><td>TCACAGTGAACCGGTCTCTTAA</td><td>242.8</td><td>2.1</td><td>0.7</td><td>2.9</td><td>4.0E-03</td><td>1.5E-02</td><td>BT20/MB231</td><td>CYTO</td></tr>
<tr><td>>hg38_21_+_8433172_8446622_RNA45SN1_45S_50ntFlanks@10786.10812.27|AACAGCCTCTGGCATGTTGGAACAATG</td><td>>hg38_21_+_8433172_8446622_RNA45SN1_45S_50ntFlanks</td><td>10786</td><td>10812</td><td>27</td><td>AACAGCCTCTGGCATGTTGGAACAATG</td><td>128.1</td><td>5.6</td><td>1.9</td><td>3.0</td><td>3.1E-03</td><td>1.5E-02</td><td>BT20/MB231</td><td>MP</td></tr>
<tr><td>>hg38_21_+_8433172_8446622_RNA45SN1_45S_50ntFlanks@11868.11893.26|GGGGAAAGAAGACCCTGTTGAGCTTG</td><td>>hg38_21_+_8433172_8446622_RNA45SN1_45S_50ntFlanks</td><td>11868</td><td>11893</td><td>26</td><td>GGGGAAAGAAGACCCTGTTGAGCTTG</td><td>96.6</td><td>5.8</td><td>2.0</td><td>3.0</td><td>3.1E-03</td><td>1.5E-02</td><td>BT20/MB231</td><td>MP</td></tr>
<tr><td>>hg38_21_+_8433172_8446622_RNA45SN1_45S_50ntFlanks@8191.8215.25|CAGCCCGTGGACGGTGTGAGGCCGG</td><td>>hg38_21_+_8433172_8446622_RNA45SN1_45S_50ntFlanks</td><td>8191</td><td>8215</td><td>25</td><td>CAGCCCGTGGACGGTGTGAGGCCGG</td><td>1030.7</td><td>-3.5</td><td>1.4</td><td>-2.5</td><td>1.3E-02</td><td>1.5E-02</td><td>BT20/MB468</td><td>CYTO</td></tr>
<tr><td>>hg38_21_+_8433172_8446622_RNA45SN1_45S_50ntFlanks@10873.10894.22|CTGGGTCGGTCGGGCTGGGGCG</td><td>>hg38_21_+_8433172_8446622_RNA45SN1_45S_50ntFlanks</td><td>10873</td><td>10894</td><td>22</td><td>CTGGGTCGGTCGGGCTGGGGCG</td><td>208.8</td><td>-2.9</td><td>1.1</td><td>-2.5</td><td>1.1E-02</td><td>1.5E-02</td><td>BT20/MB468</td><td>MITO</td></tr>
<tr><td>>hg38_21_+_8433172_8446622_RNA45SN1_45S_50ntFlanks@12417.12439.23|TCGATGTCGGCTCTTCCTATCAT</td><td>>hg38_21_+_8433172_8446622_RNA45SN1_45S_50ntFlanks</td><td>12417</td><td>12439</td><td>23</td><td>TCGATGTCGGCTCTTCCTATCAT</td><td>731.2</td><td>-2.6</td><td>1.1</td><td>-2.5</td><td>1.3E-02</td><td>1.5E-02</td><td>BT20/MB468</td><td>NUC</td></tr>
<tr><td>>hg38_MT_+_598_1651_RNR1_12S_50ntFlanks@905.935.31|AGTATACTTCAAAGGACATTTAACTAAAACC</td><td>>hg38_MT_+_598_1651_RNR1_12S_50ntFlanks</td><td>905</td><td>935</td><td>31</td><td>AGTATACTTCAAAGGACATTTAACTAAAACC</td><td>220.1</td><td>3.4</td><td>1.3</td><td>2.5</td><td>1.2E-02</td><td>1.5E-02</td><td>MB231/MB468</td><td>MP</td></tr>
<tr><td>>am_trnaMT_ValTAC_MT_+_1602_1670@51.69.19|CTTAACTTGACCGCTCTGA</td><td>>am_trnaMT_ValTAC_MT_+_1602_1670</td><td>51</td><td>69</td><td>19</td><td>CTTAACTTGACCGCTCTGA</td><td>258.2</td><td>-3.5</td><td>1.4</td><td>-2.5</td><td>1.3E-02</td><td>1.5E-02</td><td>MB231/MB468</td><td>MITO</td></tr>
<tr><td>>trnaMT_SerGCT_MT_+_12207_12265@13.48.36|AAGAACTGCTAACTCATGCCCCCATGTCTAACAACA</td><td>>trnaMT_SerGCT_MT_+_12207_12265</td><td>13</td><td>48</td><td>36</td><td>AAGAACTGCTAACTCATGCCCCCATGTCTAACAACA</td><td>150.7</td><td>3.4</td><td>1.3</td><td>2.5</td><td>1.2E-02</td><td>1.5E-02</td><td>MB231/MB468</td><td>MP</td></tr>
<tr><td>>hg38_MT_+_1621_3279_RNR2_16S_50ntFlanks@1389.1422.34|CGATGGTGCAGCCGCTATTAAAGGTTCGTTTGTT</td><td>>hg38_MT_+_1621_3279_RNR2_16S_50ntFlanks</td><td>1389</td><td>1422</td><td>34</td><td>CGATGGTGCAGCCGCTATTAAAGGTTCGTTTGTT</td><td>82.0</td><td>3.2</td><td>1.3</td><td>2.5</td><td>1.2E-02</td><td>1.5E-02</td><td>MB231/MB468</td><td>MP</td></tr>
<tr><td>>MI0003184|hsa-mir-500a&WithFlank&X|+|50008425|50008520@57.77.21;MI0003186|hsa-mir-502&WithFlank&X|+|50014592|50014689@58.78.21&[MIMAT0002871&hsa-miR-500a-3p&offsets|-1|-2;m-111&X|+|50008482|50008504&offsets|-1|-3];[MIMAT0004775&hsa-miR-502-3p&offsets|0|-1;m-236&X|+|50014649|50014669&offsets|0|0]|AATGCACCTGGGCAAGGATTC</td><td>>MI0003184|hsa-mir-500a&WithFlank&X|+|50008425|50008520</td><td>57</td><td>77</td><td>21</td><td>AATGCACCTGGGCAAGGATTC</td><td>248.2</td><td>4.2</td><td>1.6</td><td>2.7</td><td>7.9E-03</td><td>1.5E-02</td><td>BT20/MB468</td><td>MP</td></tr>
<tr><td>>hg38_21_+_8433172_8446622_RNA45SN1_45S_50ntFlanks@9454.9502.49|CCGCGCCGGGGAGGTGGAGCACGAGCGCACGTGTTAGGACCCGAAAGAT</td><td>>hg38_21_+_8433172_8446622_RNA45SN1_45S_50ntFlanks</td><td>9454</td><td>9502</td><td>49</td><td>CCGCGCCGGGGAGGTGGAGCACGAGCGCACGTGTTAGGACCCGAAAGAT</td><td>42.0</td><td>-8.5</td><td>3.2</td><td>-2.7</td><td>7.9E-03</td><td>1.5E-02</td><td>BT20/MB468</td><td>MP</td></tr>
<tr><td>>MI0000082|hsa-mir-25&WithFlank&7|-|100093554|100093649@20.41.22&[MIMAT0004498&hsa-miR-25-5p&offsets|0|+1;m-572&7|-|100093608|100093630&offsets|0|-1]|AGGCGGAGACTTGGGCAATTGC</td><td>>MI0000082|hsa-mir-25&WithFlank&7|-|100093554|100093649</td><td>20</td><td>41</td><td>22</td><td>AGGCGGAGACTTGGGCAATTGC</td><td>54.9</td><td>5.5</td><td>2.1</td><td>2.7</td><td>7.9E-03</td><td>1.5E-02</td><td>BT20/MB468</td><td>MP</td></tr>
<tr><td>>hg38_21_+_8433172_8446622_RNA45SN1_45S_50ntFlanks@6652.6681.30|GACTCTTAGCGGTGGATCACTCGGCTCGTG</td><td>>hg38_21_+_8433172_8446622_RNA45SN1_45S_50ntFlanks</td><td>6652</td><td>6681</td><td>30</td><td>GACTCTTAGCGGTGGATCACTCGGCTCGTG</td><td>3809.6</td><td>3.4</td><td>1.1</td><td>3.0</td><td>3.1E-03</td><td>1.5E-02</td><td>BT20/MB231</td><td>MP</td></tr>
<tr><td>>hg38_21_+_8433172_8446622_RNA45SN1_45S_50ntFlanks@11728.11751.24|GTAACTATGACTCTCTTAAGGTAG</td><td>>hg38_21_+_8433172_8446622_RNA45SN1_45S_50ntFlanks</td><td>11728</td><td>11751</td><td>24</td><td>GTAACTATGACTCTCTTAAGGTAG</td><td>72.8</td><td>-4.5</td><td>1.7</td><td>-2.7</td><td>8.0E-03</td><td>1.5E-02</td><td>BT20/MB468</td><td>MP</td></tr>
<tr><td>>trnaMT_ProTGG_MT_-_15956_16023@50.70.21|TTAAAGACTTTTTCTCTGACC</td><td>>trnaMT_ProTGG_MT_-_15956_16023</td><td>50</td><td>70</td><td>21</td><td>TTAAAGACTTTTTCTCTGACC</td><td>156.2</td><td>4.1</td><td>1.7</td><td>2.5</td><td>1.3E-02</td><td>1.5E-02</td><td>MB231/MB468</td><td>MITO</td></tr>
<tr><td>>hg38_21_+_8433172_8446622_RNA45SN1_45S_50ntFlanks@8175.8213.39|CCTTCTGATCGAGGCCCAGCCCGTGGACGGTGTGAGGCC</td><td>>hg38_21_+_8433172_8446622_RNA45SN1_45S_50ntFlanks</td><td>8175</td><td>8213</td><td>39</td><td>CCTTCTGATCGAGGCCCAGCCCGTGGACGGTGTGAGGCC</td><td>172.8</td><td>3.8</td><td>1.4</td><td>2.7</td><td>8.0E-03</td><td>1.5E-02</td><td>BT20/MB468</td><td>MP</td></tr>
<tr><td>>hg38_21_+_8433172_8446622_RNA45SN1_45S_50ntFlanks@3782.3802.21|CAGTGAAACTGCGAATGGCTC</td><td>>hg38_21_+_8433172_8446622_RNA45SN1_45S_50ntFlanks</td><td>3782</td><td>3802</td><td>21</td><td>CAGTGAAACTGCGAATGGCTC</td><td>393.7</td><td>-3.9</td><td>1.5</td><td>-2.7</td><td>8.0E-03</td><td>1.5E-02</td><td>BT20/MB468</td><td>MP</td></tr>
</table>

<table>
<tr><th></th><th></th><th></th><th></th><th></th><th></th><th></th><th></th><th></th><th></th><th></th><th></th><th></th><th></th></tr>
<tr><td>>hg38_21_+_8433172_8446622_RNA45SN1_45S_50ntFlanks@12893.12932.40|CCTCGCCCGTCACGCACCGCACGTTCGTGGGGAACCTGGC</td><td>>hg38_21_+_8433172_8446622_RNA45SN1_45S_50ntFlanks</td><td>12893</td><td>12932</td><td>40</td><td>CCTCGCCCGTCACGCACCGCACGTTCGTGGGGAACCTGGC</td><td>1121.3</td><td>-5.3</td><td>2.0</td><td>-2.7</td><td>8.0E-03</td><td>1.5E-02</td><td>BT20/MB468</td><td>MP</td></tr>
<tr><td>>hg38_21_+_8433172_8446622_RNA45SN1_45S_50ntFlanks@3853.3873.21|AACTGTGGTAATTCTAGAGCT</td><td>>hg38_21_+_8433172_8446622_RNA45SN1_45S_50ntFlanks</td><td>3853</td><td>3873</td><td>21</td><td>AACTGTGGTAATTCTAGAGCT</td><td>206.0</td><td>-2.5</td><td>1.0</td><td>-2.5</td><td>1.1E-02</td><td>1.5E-02</td><td>BT20/MB468</td><td>MITO</td></tr>
<tr><td>>trnaMT_GluTTC_MT_-_14674_14742@26.58.33,trnalookalike8_GluTTC_5_-_93905172_93905240@26.58.33|GGTTTTTCATATCATTGGTCGTGGTTGTAGTCC</td><td>>trnaMT_GluTTC_MT_-_14674_14742</td><td>26</td><td>58</td><td>33</td><td>GGTTTTTCATATCATTGGTCGTGGTTGTAGTCC</td><td>45.7</td><td>4.8</td><td>1.9</td><td>2.5</td><td>1.2E-02</td><td>1.5E-02</td><td>MB231/MB468</td><td>MP</td></tr>
<tr><td>>MI0010629|hsa-mir-2110&WithFlank&10|-|114174099|114174185@14.34.21&[MIMAT0010133&hsa-miR-2110&offsets|0|-1;m-424&10|-|114174152|114174172&offsets|0|0]|TTGGGGAAACGGCCGCTGAGT</td><td>>MI0010629|hsa-mir-2110&WithFlank&10|-|114174099|114174185</td><td>14</td><td>34</td><td>21</td><td>TTGGGGAAACGGCCGCTGAGT</td><td>70.8</td><td>4.8</td><td>1.8</td><td>2.7</td><td>8.0E-03</td><td>1.5E-02</td><td>BT20/MB468</td><td>MP</td></tr>
<tr><td>>trna116_GluCTC_1_-_145399233_145399304@37.74.38,trna134_GluTTC_1_-_16861774_16861845@37.74.38,trna71_GluCTC_1_-_161439189_161439260@37.74.38,trna74_GluCTC_1_-_161431809_161431880@37.74.38,trna77_GluCTC_1_-_161424398_161424469@37.74.38,trna77_GluCTC_6_+_28949976_28950047@37.74.38,trna80_GluCTC_1_-_161417018_161417089@37.74.38,trna84_GluTTC_1_-_161391883_161391954@37.74.38,trna87_GluCTC_6_-_126101393_126101464@37.74.38|ACCGCCGCGGCCCGGGTTCGATTCCCGGTCAGGGAACC</td><td>>trna116_GluCTC_1_-_145399233_145399304</td><td>37</td><td>74</td><td>38</td><td>ACCGCCGCGGCCCGGGTTCGATTCCCGGTCAGGGAACC</td><td>699.5</td><td>-2.7</td><td>0.9</td><td>-2.9</td><td>4.0E-03</td><td>1.5E-02</td><td>BT20/MB231</td><td>MITO</td></tr>
<tr><td>>trnaMT_MetCAT_MT_+_4402_4469@52.71.20|ATACCCTTCCCGTACTACCA</td><td>>trnaMT_MetCAT_MT_+_4402_4469</td><td>52</td><td>71</td><td>20</td><td>ATACCCTTCCCGTACTACCA</td><td>304.1</td><td>2.8</td><td>1.1</td><td>2.5</td><td>1.3E-02</td><td>1.5E-02</td><td>MB231/MB468</td><td>MITO</td></tr>
<tr><td>>hg38_21_+_8433172_8446622_RNA45SN1_45S_50ntFlanks@12006.12032.27|GGCCGCCGGTGAAATACCACTACTCTG</td><td>>hg38_21_+_8433172_8446622_RNA45SN1_45S_50ntFlanks</td><td>12006</td><td>12032</td><td>27</td><td>GGCCGCCGGTGAAATACCACTACTCTG</td><td>699.6</td><td>-1.6</td><td>0.6</td><td>-2.5</td><td>1.1E-02</td><td>1.5E-02</td><td>BT20/MB468</td><td>MITO</td></tr>
<tr><td>>hg38_21_+_8433172_8446622_RNA45SN1_45S_50ntFlanks@12934.12958.25|CTAAACCATTCGTAGACGACCTGCT</td><td>>hg38_21_+_8433172_8446622_RNA45SN1_45S_50ntFlanks</td><td>12934</td><td>12958</td><td>25</td><td>CTAAACCATTCGTAGACGACCTGCT</td><td>336.1</td><td>-1.5</td><td>0.6</td><td>-2.5</td><td>1.3E-02</td><td>1.5E-02</td><td>MB231/MB468</td><td>CYTO</td></tr>
<tr><td>>MI0000446|hsa-mir-125b-1&WithFlank&11|-|122099751|122099850@61.80.20(+2U)&[MIMAT0004592&hsa-miR-125b-1-3p&offsets|0|-2(+2U);m-485&11|-|122099769|122099790&offsets|0|-2(+2U)]|ACGGGTTAGGCTCTTGGGAGTT</td><td>>MI0000446|hsa-mir-125b-1&WithFlank&11|-|122099751|122099850</td><td>61</td><td>80</td><td>20(+2U)</td><td>ACGGGTTAGGCTCTTGGGAGTT</td><td>188.8</td><td>-5.6</td><td>1.9</td><td>-3.0</td><td>3.1E-03</td><td>1.5E-02</td><td>BT20/MB231</td><td>MP</td></tr>
<tr><td>>hg38_21_+_8433172_8446622_RNA45SN1_45S_50ntFlanks@10717.10736.20|AAAATCCGGGGGAGAGGGTG</td><td>>hg38_21_+_8433172_8446622_RNA45SN1_45S_50ntFlanks</td><td>10717</td><td>10736</td><td>20</td><td>AAAATCCGGGGGAGAGGGTG</td><td>293.4</td><td>-2.4</td><td>0.9</td><td>-2.7</td><td>8.0E-03</td><td>1.5E-02</td><td>BT20/MB468</td><td>MP</td></tr>
<tr><td>>hg38_21_+_8433172_8446622_RNA45SN1_45S_50ntFlanks@11572.11594.23|CCGACTTAGAACTGGTGCGGACC</td><td>>hg38_21_+_8433172_8446622_RNA45SN1_45S_50ntFlanks</td><td>11572</td><td>11594</td><td>23</td><td>CCGACTTAGAACTGGTGCGGACC</td><td>678.2</td><td>-2.2</td><td>0.9</td><td>-2.5</td><td>1.3E-02</td><td>1.5E-02</td><td>BT20/MB468</td><td>CYTO</td></tr>
<tr><td>>hg38_21_+_8433172_8446622_RNA45SN1_45S_50ntFlanks@10871.10890.20|GGCTGGGTCGGTCGGGCTGG</td><td>>hg38_21_+_8433172_8446622_RNA45SN1_45S_50ntFlanks</td><td>10871</td><td>10890</td><td>20</td><td>GGCTGGGTCGGTCGGGCTGG</td><td>481.5</td><td>6.5</td><td>2.2</td><td>3.0</td><td>3.2E-03</td><td>1.5E-02</td><td>BT20/MB231</td><td>MP</td></tr>
<tr><td>>MI0000108|hsa-mir-103a-2&WithFlank&20|+|3917488|3917577@54.77.24;MI0000109|hsa-mir-103a-1&WithFlank&5|-|168560890|168560979@54.77.24&[MIMAT0000101&hsa-miR-103a-3p&offsets|0|+1;m-12&20|+|3917541|3917563&offsets|0|+1];[MIMAT0000101_1&hsa-miR-103a-3p&offsets|0|+1;m-13&5|-|168560904|168560926&offsets|0|+1;m-44220&5|-|168560908|168560926&offsets|0|+5]|AGCAGCATTGTACAGGGCTATGAA</td><td>>MI0000108|hsa-mir-103a-2&WithFlank&20|+|3917488|3917577</td><td>54</td><td>77</td><td>24</td><td>AGCAGCATTGTACAGGGCTATGAA</td><td>428.7</td><td>-3.0</td><td>1.0</td><td>-3.0</td><td>3.2E-03</td><td>1.5E-02</td><td>BT20/MB231</td><td>MP</td></tr>
<tr><td>>MI0000750|hsa-mir-26a-2&WithFlank&12|-|57824603|57824698@20.39.20(+1A);MI0000083|hsa-mir-26a-1&WithFlank&3|+|37969398|37969486@16.35.20(+1A)&[MIMAT0000082&hsa-miR-26a-5p&offsets|0|-2(+1A);m-33&12|-|57824658|57824679&offsets|0|-2(+1A)];[MIMAT0000082_1&hsa-miR-26a-5p&offsets|0|-2(+1A);m-34&3|+|37969413|37969434&offsets|0|-2(+1A)]|TTCAAGTAATCCAGGATAGGA</td><td>>MI0000750|hsa-mir-26a-2&WithFlank&12|-|57824603|57824698</td><td>20</td><td>39</td><td>20(+1A)</td><td>TTCAAGTAATCCAGGATAGGA</td><td>196.9</td><td>-4.5</td><td>1.5</td><td>-3.0</td><td>3.2E-03</td><td>1.5E-02</td><td>BT20/MB231</td><td>MP</td></tr>
<tr><td>>hg38_21_+_8433172_8446622_RNA45SN1_45S_50ntFlanks@8357.8395.39|AAAAGAACTTTGAAGAGAGAGTTCAAGAGGGCGTGAAAC</td><td>>hg38_21_+_8433172_8446622_RNA45SN1_45S_50ntFlanks</td><td>8357</td><td>8395</td><td>39</td><td>AAAAGAACTTTGAAGAGAGAGTTCAAGAGGGCGTGAAAC</td><td>121.8</td><td>2.9</td><td>1.1</td><td>2.7</td><td>8.0E-03</td><td>1.5E-02</td><td>BT20/MB468</td><td>MP</td></tr>
<tr><td>>hg38_21_+_8433172_8446622_RNA45SN1_45S_50ntFlanks@12397.12417.21|CGACGTCGCTTTTTGATCCTT</td><td>>hg38_21_+_8433172_8446622_RNA45SN1_45S_50ntFlanks</td><td>12397</td><td>12417</td><td>21</td><td>CGACGTCGCTTTTTGATCCTT</td><td>796.4</td><td>-3.0</td><td>1.2</td><td>-2.5</td><td>1.3E-02</td><td>1.5E-02</td><td>BT20/MB468</td><td>CYTO</td></tr>
<tr><td>>hg38_21_+_8433172_8446622_RNA45SN1_45S_50ntFlanks@6653.6688.36|ACTCTTAGCGGTGGATCACTCGGCTCGTGCGTCGAT</td><td>>hg38_21_+_8433172_8446622_RNA45SN1_45S_50ntFlanks</td><td>6653</td><td>6688</td><td>36</td><td>ACTCTTAGCGGTGGATCACTCGGCTCGTGCGTCGAT</td><td>1242.4</td><td>1.7</td><td>0.7</td><td>2.5</td><td>1.3E-02</td><td>1.5E-02</td><td>MB231/MB468</td><td>NUC</td></tr>
<tr><td>>trna16_LeuAAG_16_+_22308461_22308542@1.24.24,trna16_LeuAAG_5_-_180601044_180601125@1.24.24,trna19_LeuAAG_5_-_180524474_180524555@1.24.24,trna1_LeuAAG_14_+_21078291_21078372@1.24.24,trna3_LeuAAG_5_+_180528840_180528921@1.24.24,trna42_LeuTAG_17_-_8023632_8023713@1.24.24,trna78_LeuAAG_6_+_28956779_28956860@1.24.24,trna7_LeuAAG_5_+_180614701_180614782@1.24.24,trna98_LeuAAG_6_-_28911399_28911480@1.24.24|GGTAGCGTGGCCGAGCGGTCTAAG</td><td>>trna16_LeuAAG_16_+_22308461_22308542</td><td>1</td><td>24</td><td>24</td><td>GGTAGCGTGGCCGAGCGGTCTAAG</td><td>60.8</td><td>5.5</td><td>2.1</td><td>2.7</td><td>8.0E-03</td><td>1.5E-02</td><td>BT20/MB468</td><td>MP</td></tr>
<tr><td>>hg38_21_+_8433172_8446622_RNA45SN1_45S_50ntFlanks@10236.10263.28|GACGCTACGCCGCGACGAGTAGGAGGGC</td><td>>hg38_21_+_8433172_8446622_RNA45SN1_45S_50ntFlanks</td><td>10236</td><td>10263</td><td>28</td><td>GACGCTACGCCGCGACGAGTAGGAGGGC</td><td>218.8</td><td>2.1</td><td>0.8</td><td>2.7</td><td>8.0E-03</td><td>1.5E-02</td><td>BT20/MB468</td><td>MP</td></tr>
<tr><td>>hg38_21_+_8433172_8446622_RNA45SN1_45S_50ntFlanks@11666.11693.28|TCTGCCCAGTGCTCTGAATGTCAAAGTG</td><td>>hg38_21_+_8433172_8446622_RNA45SN1_45S_50ntFlanks</td><td>11666</td><td>11693</td><td>28</td><td>TCTGCCCAGTGCTCTGAATGTCAAAGTG</td><td>112.9</td><td>2.3</td><td>0.9</td><td>2.7</td><td>8.0E-03</td><td>1.5E-02</td><td>BT20/MB468</td><td>MP</td></tr>
<tr><td>>hg38_21_+_8433172_8446622_RNA45SN1_45S_50ntFlanks@10421.10443.23|TGAGAGATGGGCGAGCGCCGTTC</td><td>>hg38_21_+_8433172_8446622_RNA45SN1_45S_50ntFlanks</td><td>10421</td><td>10443</td><td>23</td><td>TGAGAGATGGGCGAGCGCCGTTC</td><td>248.7</td><td>-5.0</td><td>1.9</td><td>-2.7</td><td>8.0E-03</td><td>1.5E-02</td><td>BT20/MB468</td><td>MP</td></tr>
<tr><td>>hg38_21_+_8433172_8446622_RNA45SN1_45S_50ntFlanks@10342.10369.28|ATATTCAAACGAGAACTTTGAAGGCCGA</td><td>>hg38_21_+_8433172_8446622_RNA45SN1_45S_50ntFlanks</td><td>10342</td><td>10369</td><td>28</td><td>ATATTCAAACGAGAACTTTGAAGGCCGA</td><td>126.1</td><td>3.7</td><td>1.4</td><td>2.7</td><td>8.0E-03</td><td>1.5E-02</td><td>BT20/MB468</td><td>MP</td></tr>
<tr><td>>hg38_MT_+_1621_3279_RNR2_16S_50ntFlanks@1363.1393.31|GACCTCGATGTTGGATCAGGACATCCCGATG</td><td>>hg38_MT_+_1621_3279_RNR2_16S_50ntFlanks</td><td>1363</td><td>1393</td><td>31</td><td>GACCTCGATGTTGGATCAGGACATCCCGATG</td><td>87.0</td><td>3.1</td><td>1.2</td><td>2.5</td><td>1.2E-02</td><td>1.5E-02</td><td>MB231/MB468</td><td>MP</td></tr>
<tr><td>>am_trna10_ValCAC_5_-_180649395_180649467@1.29.29,trna12_ValAAC_5_-_180645270_180645342@1.29.29,trna132_ValAAC_6_-_27721179_27721251@1.29.29,trna136_ValAAC_6_-_27648885_27648957@1.29.29,trna139_ValAAC_6_-_27618707_27618779@1.29.29,trna18_ValCAC_5_-_180529253_180529325@1.29.29,trna2_ValAAC_3_+_169490018_169490090@1.29.29,trna2_ValCAC_5_+_180524070_180524142@1.29.29,trna4_ValAAC_5_+_180591154_180591226@1.29.29,trna5_ValAAC_5_+_180596610_180596682@1.29.29,trna6_ValCAC_5_+_180600650_180600722@1.29.29,trna85_ValCAC_1_-_161369490_161369562@1.29.29,trna90_ValCAC_1_-_149684088_149684161@1.29.29,trna98_ValCAC_1_-_149298555_149298627@1.29.29,trna9_ValCAC_6_+_26538282_26538354@1.29.29|GTTTCCGTAGTGTAGTGGTTATCACGTTC</td><td>>am_trna10_ValCAC_5_-_180649395_180649467</td><td>1</td><td>29</td><td>29</td><td>GTTTCCGTAGTGTAGTGGTTATCACGTTC</td><td>18907.1</td><td>-3.0</td><td>1.1</td><td>-2.7</td><td>8.0E-03</td><td>1.5E-02</td><td>BT20/MB468</td><td>MP</td></tr>
<tr><td>>hg38_21_+_8433172_8446622_RNA45SN1_45S_50ntFlanks@5073.5108.36|AATTCCGATAACGAACGAGACTCTGGCATGCTAACT</td><td>>hg38_21_+_8433172_8446622_RNA45SN1_45S_50ntFlanks</td><td>5073</td><td>5108</td><td>36</td><td>AATTCCGATAACGAACGAGACTCTGGCATGCTAACT</td><td>409.1</td><td>-2.4</td><td>1.0</td><td>-2.5</td><td>1.3E-02</td><td>1.5E-02</td><td>MB231/MB468</td><td>NUC</td></tr>
<tr><td>>trna111_HisGTG_1_-_147774845_147774916@15.33.19,trna118_HisGTG_1_-_145396881_145396952@15.33.19,trna16_HisGTG_1_+_146544773_146544844@15.33.19,trna1_HisGTG_15_+_45493349_45493420@15.33.19,trna21_HisGTG_1_+_147753471_147753542@15.33.19,trna33_HisGTG_6_+_27125906_27125977@15.33.19,trna7_HisGTG_9_-_14433938_14434009@15.33.19,trna8_HisGTG_15_-_45492611_45492682@15.33.19,trna9_HisGTG_15_-_45490804_45490875@15.33.19|GTGGTTAGTACTCTGCGTT</td><td>>trna111_HisGTG_1_-_147774845_147774916</td><td>15</td><td>33</td><td>19</td><td>GTGGTTAGTACTCTGCGTT</td><td>214.4</td><td>2.5</td><td>0.9</td><td>2.9</td><td>4.0E-03</td><td>1.5E-02</td><td>BT20/MB231</td><td>CYTO</td></tr>
<tr><td>>MI0001445|hsa-mir-423&WithFlank&17|+|30117073|30117178@59.77.19&[MIMAT0001340&hsa-miR-423-3p&offsets|0|-4;m-100&17|+|30117131|30117153&offsets|0|-4]|AGCTCGGTCTGAGGCCCCT</td><td>>MI0001445|hsa-mir-423&WithFlank&17|+|30117073|30117178</td><td>59</td><td>77</td><td>19</td><td>AGCTCGGTCTGAGGCCCCT</td><td>75.3</td><td>4.2</td><td>1.7</td><td>2.5</td><td>1.2E-02</td><td>1.5E-02</td><td>MB231/MB468</td><td>MP</td></tr>
<tr><td>>hg38_21_+_8433172_8446622_RNA45SN1_45S_50ntFlanks@9307.9330.24|AAGGAGTCTAACACGTGCGCGAGT</td><td>>hg38_21_+_8433172_8446622_RNA45SN1_45S_50ntFlanks</td><td>9307</td><td>9330</td><td>24</td><td>AAGGAGTCTAACACGTGCGCGAGT</td><td>218.9</td><td>2.3</td><td>0.9</td><td>2.5</td><td>1.2E-02</td><td>1.5E-02</td><td>MB231/MB468</td><td>MP</td></tr>
<tr><td>>trna16_GlnTTG_17_+_47269890_47269961@1.29.29,trna64_GlnTTG_6_+_28557156_28557227@1.29.29,trna84_GlnTTG_6_+_145503859_145503930@1.29.29|GGTCCCATGGTGTAATGGTTAGCACTCTG</td><td>>trna16_GlnTTG_17_+_47269890_47269961</td><td>1</td><td>29</td><td>29</td><td>GGTCCCATGGTGTAATGGTTAGCACTCTG</td><td>441.6</td><td>-4.3</td><td>1.7</td><td>-2.5</td><td>1.1E-02</td><td>1.5E-02</td><td>BT20/MB468</td><td>MITO</td></tr>
<tr><td>>hg38_21_+_8433172_8446622_RNA45SN1_45S_50ntFlanks@6789.6806.18|GCCTGTCTGAGCGTCGCT</td><td>>hg38_21_+_8433172_8446622_RNA45SN1_45S_50ntFlanks</td><td>6789</td><td>6806</td><td>18</td><td>GCCTGTCTGAGCGTCGCT</td><td>272.2</td><td>2.1</td><td>0.8</td><td>2.5</td><td>1.2E-02</td><td>1.5E-02</td><td>MB231/MB468</td><td>MP</td></tr>
<tr><td>>trnaMT_PheGAA_MT_+_577_647@36.55.20|AAAATGTTTAGACGGGCTCA</td><td>>trnaMT_PheGAA_MT_+_577_647</td><td>36</td><td>55</td><td>20</td><td>AAAATGTTTAGACGGGCTCA</td><td>852.1</td><td>2.0</td><td>0.7</td><td>2.9</td><td>4.0E-03</td><td>1.5E-02</td><td>BT20/MB231</td><td>MITO</td></tr>
<tr><td>>hg38_21_+_8433172_8446622_RNA45SN1_45S_50ntFlanks@4560.4579.20|CTAGGAATAATGGAATAGGA</td><td>>hg38_21_+_8433172_8446622_RNA45SN1_45S_50ntFlanks</td><td>4560</td><td>4579</td><td>20</td><td>CTAGGAATAATGGAATAGGA</td><td>232.0</td><td>-4.5</td><td>1.7</td><td>-2.7</td><td>8.0E-03</td><td>1.5E-02</td><td>BT20/MB468</td><td>MP</td></tr>
<tr><td>>MI0000073|hsa-mir-19a&WithFlank&13|+|91350885|91350978@55.78.24&[MIMAT0000073&hsa-miR-19a-3p&offsets|0|+1;m-201&13|+|91350939|91350961&offsets|0|+1]|TGTGCAAATCTATGCAAAACTGAT</td><td>>MI0000073|hsa-mir-19a&WithFlank&13|+|91350885|91350978</td><td>55</td><td>78</td><td>24</td><td>TGTGCAAATCTATGCAAAACTGAT</td><td>564.5</td><td>2.7</td><td>1.1</td><td>2.5</td><td>1.2E-02</td><td>1.5E-02</td><td>MB231/MB468</td><td>MP</td></tr>
<tr><td>>MI0000486|hsa-mir-190a&WithFlank&15|+|62823951|62824047@21.44.24&[MIMAT0000458&hsa-miR-190a-5p&offsets|0|+2;m-349&15|+|62823971|62823993&offsets|0|+1]|TGATATGTTTGATATATTAGGTTG</td><td>>MI0000486|hsa-mir-190a&WithFlank&15|+|62823951|62824047</td><td>21</td><td>44</td><td>24</td><td>TGATATGTTTGATATATTAGGTTG</td><td>82.4</td><td>4.0</td><td>1.6</td><td>2.5</td><td>1.2E-02</td><td>1.5E-02</td><td>MB231/MB468</td><td>MP</td></tr>
</table>

<table>
<tr><td></td><td></td><td></td><td></td><td></td><td></td><td></td><td></td><td></td><td></td><td></td><td></td><td></td><td></td></tr>
<tr><td>>MI0000062|hsa-let-7a-3&WithFlank&22|+|46112743|46112828@58.78.21(+1A);MI0000060|hsa-let-7a-1&WithFlank&9|+|94175951|94176042@63.83.21(+1A)&[MIMAT0004481&hsa-let-7a-3p&offsets|0|0(+1A);m-180&22|+|46112800|46112821&offsets|0|-1(+1A)];[MIMAT0004481_1&hsa-let-7a-3p&offsets|0|0(+1A);m-181&9|+|94176013|94176034&offsets|0|-1(+1A)]|CTATACAATCTACTGTCTTTCA</td><td>>MI0000062|hsa-let-7a-3&WithFlank&22|+|46112743|46112828</td><td>58</td><td>78</td><td>21(+1A)</td><td>CTATACAATCTACTGTCTTTCA</td><td>82.0</td><td>3.5</td><td>1.4</td><td>2.5</td><td>1.2E-02</td><td>1.5E-02</td><td>MB231/MB468</td><td>MP</td></tr>
<tr><td>>MI0000270|hsa-mir-181b-1&WithFlank&1|-|198858867|198858988@42.62.21;MI0000683|hsa-mir-181b-2&WithFlank&9|+|124693704|124693804@22.42.21&[MIMAT0000257&hsa-miR-181b-5p&offsets|0|-2;m-73&1|-|198858925|198858947&offsets|0|-2];[MIMAT0000257_1&hsa-miR-181b-5p&offsets|0|-2;m-72&9|+|124693725|124693747&offsets|0|-2]|AACATTCATTGCTGTCGGTGG</td><td>>MI0000270|hsa-mir-181b-1&WithFlank&1|-|198858867|198858988</td><td>42</td><td>62</td><td>21</td><td>AACATTCATTGCTGTCGGTGG</td><td>509.4</td><td>-2.1</td><td>0.7</td><td>-2.9</td><td>4.0E-03</td><td>1.5E-02</td><td>BT20/MB231</td><td>MITO</td></tr>
<tr><td>>MI0000750|hsa-mir-26a-2&WithFlank&12|-|57824603|57824698@20.41.22(+1C);MI0000083|hsa-mir-26a-1&WithFlank&3|+|37969398|37969486@16.37.22(+1C)&[MIMAT0000082&hsa-miR-26a-5p&offsets|0|0(+1C);m-33&12|-|57824658|57824679&offsets|0|0(+1C)];[MIMAT0000082_1&hsa-miR-26a-5p&offsets|0|0(+1C);m-34&3|+|37969413|37969434&offsets|0|0(+1C)]|TTCAAGTAATCCAGGATAGGCTC</td><td>>MI0000750|hsa-mir-26a-2&WithFlank&12|-|57824603|57824698</td><td>20</td><td>41</td><td>22(+1C)</td><td>TTCAAGTAATCCAGGATAGGCTC</td><td>931.6</td><td>-2.5</td><td>0.9</td><td>-2.9</td><td>4.0E-03</td><td>1.5E-02</td><td>BT20/MB231</td><td>MITO</td></tr>
<tr><td>>hg38_MT_+_598_1651_RNR1_12S_50ntFlanks@965.1004.40|GTAACATGGTAAGTGTACTGGAAAGTGCACTTGGACGAAC</td><td>>hg38_MT_+_598_1651_RNR1_12S_50ntFlanks</td><td>965</td><td>1004</td><td>40</td><td>GTAACATGGTAAGTGTACTGGAAAGTGCACTTGGACGAAC</td><td>256.6</td><td>-2.5</td><td>0.9</td><td>-2.9</td><td>4.0E-03</td><td>1.5E-02</td><td>BT20/MB231</td><td>MITO</td></tr>
<tr><td>>MI0000465|hsa-mir-191&WithFlank&3|-|49020612|49020715@22.45.24&[MIMAT0000440&hsa-miR-191-5p&offsets|0|+1;m-48&3|-|49020672|49020694&offsets|0|+1]|CAACGGAATCCCAAAAGCAGCTGT</td><td>>MI0000465|hsa-mir-191&WithFlank&3|-|49020612|49020715</td><td>22</td><td>45</td><td>24</td><td>CAACGGAATCCCAAAAGCAGCTGT</td><td>2845.7</td><td>3.5</td><td>1.3</td><td>2.6</td><td>8.1E-03</td><td>1.5E-02</td><td>BT20/MB468</td><td>MP</td></tr>
<tr><td>>MI0000269|hsa-mir-181a-2&WithFlank&9|+|124692436|124692557@83.103.21&[MIMAT0004558&hsa-miR-181a-2-3p&offsets|0|-1;m-71&9|+|124692518|124692539&offsets|0|-1]|ACCACTGACCGTTGACTGTAC</td><td>>MI0000269|hsa-mir-181a-2&WithFlank&9|+|124692436|124692557</td><td>83</td><td>103</td><td>21</td><td>ACCACTGACCGTTGACTGTAC</td><td>103.4</td><td>4.0</td><td>1.5</td><td>2.6</td><td>8.1E-03</td><td>1.5E-02</td><td>BT20/MB468</td><td>MP</td></tr>
<tr><td>>am_trnaMT_ValTAC_MT_+_1602_1670@46.67.22|TTCAACTTAACTTGACCGCTCT</td><td>>am_trnaMT_ValTAC_MT_+_1602_1670</td><td>46</td><td>67</td><td>22</td><td>TTCAACTTAACTTGACCGCTCT</td><td>563.9</td><td>2.6</td><td>0.9</td><td>2.9</td><td>4.1E-03</td><td>1.5E-02</td><td>BT20/MB231</td><td>CYTO</td></tr>
<tr><td>>am_MI0016800|hsa-mir-4454&WithFlank&4|-|163093568|163093634@12.30.19&[MIMAT0018976&hsa-miR-4454&offsets|+3|+2;m-444&4|-|163093607|163093623&offsets|0|+2]|TCCGAGTCACGGCACCAAA</td><td>>am_MI0016800|hsa-mir-4454&WithFlank&4|-|163093568|163093634</td><td>12</td><td>30</td><td>19</td><td>TCCGAGTCACGGCACCAAA</td><td>233.5</td><td>6.9</td><td>2.6</td><td>2.6</td><td>8.1E-03</td><td>1.5E-02</td><td>BT20/MB468</td><td>MP</td></tr>
<tr><td>>hg38_21_+_8433172_8446622_RNA45SN1_45S_50ntFlanks@4068.4095.28|ACTTTCGATGGTAGTCGCCGTGCCTACC</td><td>>hg38_21_+_8433172_8446622_RNA45SN1_45S_50ntFlanks</td><td>4068</td><td>4095</td><td>28</td><td>ACTTTCGATGGTAGTCGCCGTGCCTACC</td><td>162.7</td><td>-4.5</td><td>1.7</td><td>-2.6</td><td>8.1E-03</td><td>1.5E-02</td><td>BT20/MB468</td><td>MP</td></tr>
<tr><td>>hg38_21_+_8433172_8446622_RNA45SN1_45S_50ntFlanks@8105.8137.33|CGCGGCGGGGCGCGGGACATGTGGCGTACGGAA</td><td>>hg38_21_+_8433172_8446622_RNA45SN1_45S_50ntFlanks</td><td>8105</td><td>8137</td><td>33</td><td>CGCGGCGGGGCGCGGGACATGTGGCGTACGGAA</td><td>481.4</td><td>5.9</td><td>2.0</td><td>2.9</td><td>3.2E-03</td><td>1.5E-02</td><td>BT20/MB231</td><td>MP</td></tr>
<tr><td>>hg38_21_+_8433172_8446622_RNA45SN1_45S_50ntFlanks@9533.9552.20|GAGGAAACTCTGGTGGAGGT</td><td>>hg38_21_+_8433172_8446622_RNA45SN1_45S_50ntFlanks</td><td>9533</td><td>9552</td><td>20</td><td>GAGGAAACTCTGGTGGAGGT</td><td>367.4</td><td>-1.5</td><td>0.6</td><td>-2.5</td><td>1.3E-02</td><td>1.5E-02</td><td>BT20/MB468</td><td>CYTO</td></tr>
<tr><td>>hg38_21_+_8433172_8446622_RNA45SN1_45S_50ntFlanks@10458.10498.41|ATGGCCTCCGTTGCCCTCGGCCGATCGAAAGGGAGTCGGGT</td><td>>hg38_21_+_8433172_8446622_RNA45SN1_45S_50ntFlanks</td><td>10458</td><td>10498</td><td>41</td><td>ATGGCCTCCGTTGCCCTCGGCCGATCGAAAGGGAGTCGGGT</td><td>82.3</td><td>-4.5</td><td>1.7</td><td>-2.6</td><td>8.1E-03</td><td>1.5E-02</td><td>BT20/MB468</td><td>MP</td></tr>
<tr><td>>hg38_21_+_8433172_8446622_RNA45SN1_45S_50ntFlanks@7988.8019.32|CAGACGTGGCGACCCGCTGAATTTAAGCATAT</td><td>>hg38_21_+_8433172_8446622_RNA45SN1_45S_50ntFlanks</td><td>7988</td><td>8019</td><td>32</td><td>CAGACGTGGCGACCCGCTGAATTTAAGCATAT</td><td>47.7</td><td>-4.7</td><td>1.8</td><td>-2.6</td><td>8.1E-03</td><td>1.5E-02</td><td>BT20/MB468</td><td>MP</td></tr>
<tr><td>>hg38_21_+_8433172_8446622_RNA45SN1_45S_50ntFlanks@3846.3874.29|CTTGGATAACTGTGGTAATTCTAGAGCTA</td><td>>hg38_21_+_8433172_8446622_RNA45SN1_45S_50ntFlanks</td><td>3846</td><td>3874</td><td>29</td><td>CTTGGATAACTGTGGTAATTCTAGAGCTA</td><td>152.1</td><td>-3.9</td><td>1.5</td><td>-2.5</td><td>1.1E-02</td><td>1.5E-02</td><td>BT20/MB468</td><td>MITO</td></tr>
<tr><td>>MI0000065|hsa-let-7d&WithFlank&9|+|94178828|94178926@14.34.21(+1C)&[MIMAT0000065&hsa-let-7d-5p&offsets|0|-1(+1C);m-90&9|+|94178841|94178862&offsets|0|-1(+1C)]|AGAGGTAGTAGGTTGCATAGTC</td><td>>MI0000065|hsa-let-7d&WithFlank&9|+|94178828|94178926</td><td>14</td><td>34</td><td>21(+1C)</td><td>AGAGGTAGTAGGTTGCATAGTC</td><td>2071.8</td><td>1.6</td><td>0.5</td><td>2.9</td><td>3.5E-03</td><td>1.5E-02</td><td>BT20/MB231</td><td>NUC</td></tr>
<tr><td>>hg38_21_+_8433172_8446622_RNA45SN1_45S_50ntFlanks@12332.12349.18|TAAGCAGGAGGTGTCAGA</td><td>>hg38_21_+_8433172_8446622_RNA45SN1_45S_50ntFlanks</td><td>12332</td><td>12349</td><td>18</td><td>TAAGCAGGAGGTGTCAGA</td><td>73.9</td><td>-4.3</td><td>1.6</td><td>-2.6</td><td>8.1E-03</td><td>1.5E-02</td><td>BT20/MB468</td><td>MP</td></tr>
<tr><td>>hg38_21_+_8433172_8446622_RNA45SN1_45S_50ntFlanks@8381.8406.26|AAGAGGGCGTGAAACCGTTAAGAGGT</td><td>>hg38_21_+_8433172_8446622_RNA45SN1_45S_50ntFlanks</td><td>8381</td><td>8406</td><td>26</td><td>AAGAGGGCGTGAAACCGTTAAGAGGT</td><td>672.5</td><td>-2.6</td><td>1.0</td><td>-2.5</td><td>1.4E-02</td><td>1.5E-02</td><td>MB231/MB468</td><td>MITO</td></tr>
<tr><td>>hg38_1_-_228634819_228635039_RNA5S12_5S_50ntFlanks@51.85.35|GTCTACGGCCATACCACCCTGAACGCGCCCGATCT</td><td>>hg38_1_-_228634819_228635039_RNA5S12_5S_50ntFlanks</td><td>51</td><td>85</td><td>35</td><td>GTCTACGGCCATACCACCCTGAACGCGCCCGATCT</td><td>423.8</td><td>2.0</td><td>0.8</td><td>2.5</td><td>1.4E-02</td><td>1.5E-02</td><td>MB231/MB468</td><td>MITO</td></tr>
<tr><td>>hg38_21_+_8433172_8446622_RNA45SN1_45S_50ntFlanks@10400.10424.25|TGAACATGGGTCAGTCGGTCCTGAG</td><td>>hg38_21_+_8433172_8446622_RNA45SN1_45S_50ntFlanks</td><td>10400</td><td>10424</td><td>25</td><td>TGAACATGGGTCAGTCGGTCCTGAG</td><td>9364.9</td><td>3.7</td><td>1.2</td><td>2.9</td><td>3.2E-03</td><td>1.5E-02</td><td>BT20/MB231</td><td>MP</td></tr>
<tr><td>>hg38_21_+_8433172_8446622_RNA45SN1_45S_50ntFlanks@9483.9513.31|CGTGTTAGGACCCGAAAGATGGTGAACTATG</td><td>>hg38_21_+_8433172_8446622_RNA45SN1_45S_50ntFlanks</td><td>9483</td><td>9513</td><td>31</td><td>CGTGTTAGGACCCGAAAGATGGTGAACTATG</td><td>147.3</td><td>5.9</td><td>2.0</td><td>2.9</td><td>3.2E-03</td><td>1.5E-02</td><td>BT20/MB231</td><td>MP</td></tr>
<tr><td>>hg38_21_+_8433172_8446622_RNA45SN1_45S_50ntFlanks@10234.10255.22|CGGACGCTACGCCGCGACGAGT</td><td>>hg38_21_+_8433172_8446622_RNA45SN1_45S_50ntFlanks</td><td>10234</td><td>10255</td><td>22</td><td>CGGACGCTACGCCGCGACGAGT</td><td>94.4</td><td>2.5</td><td>0.9</td><td>2.6</td><td>8.1E-03</td><td>1.5E-02</td><td>BT20/MB468</td><td>MP</td></tr>
<tr><td>>hg38_21_+_8433172_8446622_RNA45SN1_45S_50ntFlanks@10731.10756.26|AGGGTGTAAATCTCGCGCCGGGCCGT</td><td>>hg38_21_+_8433172_8446622_RNA45SN1_45S_50ntFlanks</td><td>10731</td><td>10756</td><td>26</td><td>AGGGTGTAAATCTCGCGCCGGGCCGT</td><td>84.1</td><td>-2.1</td><td>0.8</td><td>-2.6</td><td>8.1E-03</td><td>1.5E-02</td><td>BT20/MB468</td><td>MP</td></tr>
<tr><td>>hg38_21_+_8433172_8446622_RNA45SN1_45S_50ntFlanks@4047.4081.35|ATTCGAACGTCTGCCCTATCAACTTTCGATGGTAG</td><td>>hg38_21_+_8433172_8446622_RNA45SN1_45S_50ntFlanks</td><td>4047</td><td>4081</td><td>35</td><td>ATTCGAACGTCTGCCCTATCAACTTTCGATGGTAG</td><td>185.4</td><td>-4.7</td><td>1.8</td><td>-2.6</td><td>8.1E-03</td><td>1.5E-02</td><td>BT20/MB468</td><td>MP</td></tr>
<tr><td>>hg38_1_-_228634819_228635039_RNA5S12_5S_50ntFlanks@141.171.31|CCGCCTGGGAATACCGGGTGCTGTAGGCTTT</td><td>>hg38_1_-_228634819_228635039_RNA5S12_5S_50ntFlanks</td><td>141</td><td>171</td><td>31</td><td>CCGCCTGGGAATACCGGGTGCTGTAGGCTTT</td><td>3666.6</td><td>-2.2</td><td>0.9</td><td>-2.5</td><td>1.2E-02</td><td>1.5E-02</td><td>MB231/MB468</td><td>MP</td></tr>
<tr><td>>hg38_21_+_8433172_8446622_RNA45SN1_45S_50ntFlanks@12419.12441.23|GATGTCGGCTCTTCCTATCATTG</td><td>>hg38_21_+_8433172_8446622_RNA45SN1_45S_50ntFlanks</td><td>12419</td><td>12441</td><td>23</td><td>GATGTCGGCTCTTCCTATCATTG</td><td>694.3</td><td>-2.5</td><td>0.8</td><td>-2.9</td><td>3.5E-03</td><td>1.5E-02</td><td>BT20/MB231</td><td>NUC</td></tr>
<tr><td>>MI0001448|hsa-mir-425&WithFlank&3|-|49020142|49020240@20.43.24&[MIMAT0003393&hsa-miR-425-5p&offsets|0|+1;m-95&3|-|49020200|49020221&offsets|0|+2]|AATGACACGATCACTCCCGTTGAG</td><td>>MI0001448|hsa-mir-425&WithFlank&3|-|49020142|49020240</td><td>20</td><td>43</td><td>24</td><td>AATGACACGATCACTCCCGTTGAG</td><td>531.9</td><td>2.4</td><td>1.0</td><td>2.5</td><td>1.2E-02</td><td>1.5E-02</td><td>MB231/MB468</td><td>MP</td></tr>
<tr><td>>am_MI0015825|hsa-mir-378c&WithFlank&10|-|130962582|130962674@17.34.18;MI0016749|hsa-mir-378d-1&WithFlank&4|-|5923269|5923334@37.54.18;MI0000786|hsa-mir-378a&WithFlank&5|+|149732819|149732896@49.66.18;MI0003840|hsa-mir-378d-2&WithFlank&8|-|93916016|93916125@28.45.18&[MIMAT0016847&hsa-miR-378c&offsets|0|-7;m-130&10|-|130962639|130962658&offsets|0|-2];[MIMAT0018926&hsa-miR-378d&offsets|0|-2;m-228&4|-|5923281|5923298&offsets|0|0];[MIMAT0000732&hsa-miR-378a-3p&offsets|0|-4;m-44&5|+|149732867|149732888&offsets|0|-4];[MIMAT0018926_1&hsa-miR-378d&offsets|0|-2;m-227&8|-|93916081|93916098&offsets|0|0]|ACTGGACTTGGAGTCAGA</td><td>>am_MI0015825|hsa-mir-378c&WithFlank&10|-|130962582|130962674</td><td>17</td><td>34</td><td>18</td><td>ACTGGACTTGGAGTCAGA</td><td>1049.5</td><td>2.1</td><td>0.8</td><td>2.5</td><td>1.2E-02</td><td>1.5E-02</td><td>MB231/MB468</td><td>MP</td></tr>
<tr><td>>hg38_21_+_8433172_8446622_RNA45SN1_45S_50ntFlanks@11666.11702.37|TCTGCCCAGTGCTCTGAATGTCAAAGTGAAGAAATTC</td><td>>hg38_21_+_8433172_8446622_RNA45SN1_45S_50ntFlanks</td><td>11666</td><td>11702</td><td>37</td><td>TCTGCCCAGTGCTCTGAATGTCAAAGTGAAGAAATTC</td><td>2212.3</td><td>-2.3</td><td>0.8</td><td>-2.9</td><td>4.1E-03</td><td>1.5E-02</td><td>BT20/MB231</td><td>CYTO</td></tr>
<tr><td>>hg38_21_+_8433172_8446622_RNA45SN1_45S_50ntFlanks@9569.9603.35|GTGCAAATCGGTCGTCCGACCTGGGTATAGGGGCG</td><td>>hg38_21_+_8433172_8446622_RNA45SN1_45S_50ntFlanks</td><td>9569</td><td>9603</td><td>35</td><td>GTGCAAATCGGTCGTCCGACCTGGGTATAGGGGCG</td><td>438.6</td><td>-2.9</td><td>1.0</td><td>-2.9</td><td>4.1E-03</td><td>1.5E-02</td><td>BT20/MB231</td><td>CYTO</td></tr>
<tr><td>>MI0003776|hsa-mir-320b-1&WithFlank&1|+|116671740|116671823@48.67.20(+2U);MI0003839|hsa-mir-320b-2&WithFlank&1|-|224257034|224257116@47.66.20(+2U);MI0000542|hsa-mir-320a&WithFlank&8|-|22244960|22245043@48.67.20(+2U)&[MIMAT0005792&hsa-miR-320b&offsets|0|-2(+2U);m-183&1|+|116671787|116671806&offsets|0|0(+2U)];[MIMAT0005792_1&hsa-miR-320b&offsets|0|-2(+2U);m-183&1|-|224257051|224257070&offsets|0|0(+2U)];[MIMAT0000510&hsa-miR-320a-3p&offsets|0|-2(+2U);m-60&8|-|22244975|22244996&offsets|0|-2(+2U)]|AAAAGCTGGGTTGAGAGGGCTT</td><td>>MI0003776|hsa-mir-320b-1&WithFlank&1|+|116671740|116671823</td><td>48</td><td>67</td><td>20(+2U)</td><td>AAAAGCTGGGTTGAGAGGGCTT</td><td>37.8</td><td>5.0</td><td>2.0</td><td>2.5</td><td>1.2E-02</td><td>1.5E-02</td><td>MB231/MB468</td><td>MP</td></tr>
<tr><td>>hg38_21_+_8433172_8446622_RNA45SN1_45S_50ntFlanks@6785.6803.19|CTACGCCTGTCTGAGCGTC</td><td>>hg38_21_+_8433172_8446622_RNA45SN1_45S_50ntFlanks</td><td>6785</td><td>6803</td><td>19</td><td>CTACGCCTGTCTGAGCGTC</td><td>623.1</td><td>-5.0</td><td>1.9</td><td>-2.6</td><td>8.1E-03</td><td>1.5E-02</td><td>BT20/MB468</td><td>MP</td></tr>
<tr><td>>hg38_21_+_8433172_8446622_RNA45SN1_45S_50ntFlanks@12934.12956.23|CTAAACCATTCGTAGACGACCTG</td><td>>hg38_21_+_8433172_8446622_RNA45SN1_45S_50ntFlanks</td><td>12934</td><td>12956</td><td>23</td><td>CTAAACCATTCGTAGACGACCTG</td><td>3135.4</td><td>-5.1</td><td>1.9</td><td>-2.6</td><td>8.1E-03</td><td>1.5E-02</td><td>BT20/MB468</td><td>MP</td></tr>
<tr><td>>hg38_21_+_8433172_8446622_RNA45SN1_45S_50ntFlanks@11885.11905.21|TTGAGCTTGACTCTAGTCTGG</td><td>>hg38_21_+_8433172_8446622_RNA45SN1_45S_50ntFlanks</td><td>11885</td><td>11905</td><td>21</td><td>TTGAGCTTGACTCTAGTCTGG</td><td>332.5</td><td>-3.2</td><td>1.2</td><td>-2.5</td><td>1.1E-02</td><td>1.5E-02</td><td>BT20/MB468</td><td>MITO</td></tr>
<tr><td>>hg38_21_+_8433172_8446622_RNA45SN1_45S_50ntFlanks@11217.11250.34|CTGGACGCGAGCCGGGCCCTTCCCGTGGATCGCC</td><td>>hg38_21_+_8433172_8446622_RNA45SN1_45S_50ntFlanks</td><td>11217</td><td>11250</td><td>34</td><td>CTGGACGCGAGCCGGGCCCTTCCCGTGGATCGCC</td><td>175.0</td><td>-3.3</td><td>1.3</td><td>-2.6</td><td>8.1E-03</td><td>1.5E-02</td><td>BT20/MB468</td><td>MP</td></tr>
<tr><td>>am_trnaMT_HisGTG_MT_+_12138_12206@20.51.32|AACATCAGATTGTGAATCTGACAACAGAGGCT</td><td>>am_trnaMT_HisGTG_MT_+_12138_12206</td><td>20</td><td>51</td><td>32</td><td>AACATCAGATTGTGAATCTGACAACAGAGGCT</td><td>207.1</td><td>3.1</td><td>1.2</td><td>2.5</td><td>1.2E-02</td><td>1.5E-02</td><td>MB231/MB468</td><td>MP</td></tr>
<tr><td>>hg38_21_+_8433172_8446622_RNA45SN1_45S_50ntFlanks@9198.9226.29|GTTCTCTCGGGGCCACGCGCGCGTCCCCC</td><td>>hg38_21_+_8433172_8446622_RNA45SN1_45S_50ntFlanks</td><td>9198</td><td>9226</td><td>29</td><td>GTTCTCTCGGGGCCACGCGCGCGTCCCCC</td><td>182.6</td><td>-4.4</td><td>1.7</td><td>-2.6</td><td>8.1E-03</td><td>1.5E-02</td><td>BT20/MB468</td><td>MP</td></tr>
<tr><td>>hg38_21_+_8433172_8446622_RNA45SN1_45S_50ntFlanks@7980.8003.24|CCTCAGATCAGACGTGGCGACCCG</td><td>>hg38_21_+_8433172_8446622_RNA45SN1_45S_50ntFlanks</td><td>7980</td><td>8003</td><td>24</td><td>CCTCAGATCAGACGTGGCGACCCG</td><td>308.9</td><td>-1.7</td><td>0.7</td><td>-2.5</td><td>1.1E-02</td><td>1.5E-02</td><td>BT20/MB468</td><td>MITO</td></tr>
<tr><td>>hg38_MT_+_1621_3279_RNR2_16S_50ntFlanks@619.652.34|ACATATAACTGAACTCCTCACACCCAATTGGACC</td><td>>hg38_MT_+_1621_3279_RNR2_16S_50ntFlanks</td><td>619</td><td>652</td><td>34</td><td>ACATATAACTGAACTCCTCACACCCAATTGGACC</td><td>95.7</td><td>6.7</td><td>2.5</td><td>2.6</td><td>8.1E-03</td><td>1.5E-02</td><td>BT20/MB468</td><td>MP</td></tr>
<tr><td>>hg38_21_+_8433172_8446622_RNA45SN1_45S_50ntFlanks@12408.12434.27|TTTGATCCTTCGATGTCGGCTCTTCCT</td><td>>hg38_21_+_8433172_8446622_RNA45SN1_45S_50ntFlanks</td><td>12408</td><td>12434</td><td>27</td><td>TTTGATCCTTCGATGTCGGCTCTTCCT</td><td>1412.8</td><td>-2.5</td><td>1.0</td><td>-2.5</td><td>1.3E-02</td><td>1.5E-02</td><td>BT20/MB468</td><td>NUC</td></tr>
<tr><td>>hg38_MT_+_1621_3279_RNR2_16S_50ntFlanks@489.518.30|AACAGCTCTTTGGACACTAGGAAAAAACCT</td><td>>hg38_MT_+_1621_3279_RNR2_16S_50ntFlanks</td><td>489</td><td>518</td><td>30</td><td>AACAGCTCTTTGGACACTAGGAAAAAACCT</td><td>37.0</td><td>4.9</td><td>2.0</td><td>2.5</td><td>1.2E-02</td><td>1.5E-02</td><td>MB231/MB468</td><td>MP</td></tr>
<tr><td>>trnaMT_ValTAC_MT_+_1602_1670@48.72.25|CAACTTAACTTGACCGCTCTGACCA</td><td>>trnaMT_ValTAC_MT_+_1602_1670</td><td>48</td><td>72</td><td>25</td><td>CAACTTAACTTGACCGCTCTGACCA</td><td>9108.0</td><td>2.9</td><td>1.2</td><td>2.5</td><td>1.1E-02</td><td>1.5E-02</td><td>BT20/MB468</td><td>MITO</td></tr>
<tr><td>>hg38_21_+_8433172_8446622_RNA45SN1_45S_50ntFlanks@3854.3871.18|ACTGTGGTAATTCTAGAG</td><td>>hg38_21_+_8433172_8446622_RNA45SN1_45S_50ntFlanks</td><td>3854</td><td>3871</td><td>18</td><td>ACTGTGGTAATTCTAGAG</td><td>255.1</td><td>-2.4</td><td>1.0</td><td>-2.5</td><td>1.3E-02</td><td>1.5E-02</td><td>BT20/MB468</td><td>CYTO</td></tr>
<tr><td>>am_trnaMT_ValTAC_MT_+_1602_1670@28.58.31|AACTTACACTTAGGAGATTTCAACTTAACTT</td><td>>am_trnaMT_ValTAC_MT_+_1602_1670</td><td>28</td><td>58</td><td>31</td><td>AACTTACACTTAGGAGATTTCAACTTAACTT</td><td>89.5</td><td>3.4</td><td>1.3</td><td>2.5</td><td>1.2E-02</td><td>1.5E-02</td><td>MB231/MB468</td><td>MP</td></tr>
</table>

<table>
<tr><th></th><th></th><th></th><th></th><th></th><th></th><th></th><th></th><th></th><th></th><th></th><th></th><th></th><th></th></tr>
<tr><td>>hg38_MT_+_1621_3279_RNR2_16S_50ntFlanks@622.643.22|TATAACTGAACTCCTCACACCC</td><td>>hg38_MT_+_1621_3279_RNR2_16S_50ntFlanks</td><td>622</td><td>643</td><td>22</td><td>TATAACTGAACTCCTCACACCC</td><td>110.8</td><td>4.9</td><td>1.9</td><td>2.6</td><td>8.2E-03</td><td>1.5E-02</td><td>BT20/MB468</td><td>MP</td></tr>
<tr><td>>MI0001448|hsa-mir-425&WithFlank&3|-|49020142|49020240@20.40.21&[MIMAT0003393&hsa-miR-425-5p&offsets|0|-2;m-95&3|-|49020200|49020221&offsets|0|-1]|AATGACACGATCACTCCCGTT</td><td>>MI0001448|hsa-mir-425&WithFlank&3|-|49020142|49020240</td><td>20</td><td>40</td><td>21</td><td>AATGACACGATCACTCCCGTT</td><td>82.1</td><td>4.1</td><td>1.6</td><td>2.5</td><td>1.2E-02</td><td>1.5E-02</td><td>MB231/MB468</td><td>MP</td></tr>
<tr><td>>hg38_21_+_8433172_8446622_RNA45SN1_45S_50ntFlanks@6652.6675.24|GACTCTTAGCGGTGGATCACTCGG</td><td>>hg38_21_+_8433172_8446622_RNA45SN1_45S_50ntFlanks</td><td>6652</td><td>6675</td><td>24</td><td>GACTCTTAGCGGTGGATCACTCGG</td><td>745.6</td><td>3.5</td><td>1.2</td><td>2.9</td><td>3.2E-03</td><td>1.5E-02</td><td>BT20/MB231</td><td>MP</td></tr>
<tr><td>>hg38_21_+_8433172_8446622_RNA45SN1_45S_50ntFlanks@12414.12447.34|CCTTCGATGTCGGCTCTTCCTATCATTGTGAAGC</td><td>>hg38_21_+_8433172_8446622_RNA45SN1_45S_50ntFlanks</td><td>12414</td><td>12447</td><td>34</td><td>CCTTCGATGTCGGCTCTTCCTATCATTGTGAAGC</td><td>401.2</td><td>-2.1</td><td>0.7</td><td>-2.9</td><td>4.1E-03</td><td>1.5E-02</td><td>BT20/MB231</td><td>CYTO</td></tr>
<tr><td>>hg38_21_+_8433172_8446622_RNA45SN1_45S_50ntFlanks@10347.10369.23|CAAACGAGAACTTTGAAGGCCGA</td><td>>hg38_21_+_8433172_8446622_RNA45SN1_45S_50ntFlanks</td><td>10347</td><td>10369</td><td>23</td><td>CAAACGAGAACTTTGAAGGCCGA</td><td>104.7</td><td>-2.0</td><td>0.8</td><td>-2.5</td><td>1.2E-02</td><td>1.5E-02</td><td>MB231/MB468</td><td>MP</td></tr>
<tr><td>>MI0006273|hsa-mir-1180&WithFlank&17|-|19344500|19344580@47.68.22&[MIMAT0005825&hsa-miR-1180-3p&offsets|0|0;m-214&17|-|19344513|19344534&offsets|0|0]|TTTCCGGCTCGCGTGGGTGTGT</td><td>>MI0006273|hsa-mir-1180&WithFlank&17|-|19344500|19344580</td><td>47</td><td>68</td><td>22</td><td>TTTCCGGCTCGCGTGGGTGTGT</td><td>472.0</td><td>1.8</td><td>0.7</td><td>2.5</td><td>1.3E-02</td><td>1.5E-02</td><td>BT20/MB468</td><td>NUC</td></tr>
<tr><td>>hg38_MT_+_1621_3279_RNR2_16S_50ntFlanks@1217.1262.46|AACCTCCGAGCAGTACATGCTAAGACTTCACCAGTCAAAGCGAACT</td><td>>hg38_MT_+_1621_3279_RNR2_16S_50ntFlanks</td><td>1217</td><td>1262</td><td>46</td><td>AACCTCCGAGCAGTACATGCTAAGACTTCACCAGTCAAAGCGAACT</td><td>353.2</td><td>4.2</td><td>1.7</td><td>2.5</td><td>1.4E-02</td><td>1.5E-02</td><td>MB231/MB468</td><td>MITO</td></tr>
<tr><td>>hg38_21_+_8433172_8446622_RNA45SN1_45S_50ntFlanks@7975.8000.26|CGCGACCTCAGATCAGACGTGGCGAC</td><td>>hg38_21_+_8433172_8446622_RNA45SN1_45S_50ntFlanks</td><td>7975</td><td>8000</td><td>26</td><td>CGCGACCTCAGATCAGACGTGGCGAC</td><td>593.1</td><td>2.3</td><td>0.9</td><td>2.5</td><td>1.4E-02</td><td>1.5E-02</td><td>MB231/MB468</td><td>MITO</td></tr>
<tr><td>>hg38_21_+_8433172_8446622_RNA45SN1_45S_50ntFlanks@10797.10827.31|GCATGTTGGAACAATGTAGGTAAGGGAAGTC</td><td>>hg38_21_+_8433172_8446622_RNA45SN1_45S_50ntFlanks</td><td>10797</td><td>10827</td><td>31</td><td>GCATGTTGGAACAATGTAGGTAAGGGAAGTC</td><td>291.5</td><td>-2.2</td><td>0.8</td><td>-2.6</td><td>8.2E-03</td><td>1.5E-02</td><td>BT20/MB468</td><td>MP</td></tr>
<tr><td>>hg38_21_+_8433172_8446622_RNA45SN1_45S_50ntFlanks@12004.12038.35|CGGGCCGCCGGTGAAATACCACTACTCTGATCGTT</td><td>>hg38_21_+_8433172_8446622_RNA45SN1_45S_50ntFlanks</td><td>12004</td><td>12038</td><td>35</td><td>CGGGCCGCCGGTGAAATACCACTACTCTGATCGTT</td><td>99.2</td><td>2.6</td><td>1.0</td><td>2.6</td><td>8.2E-03</td><td>1.5E-02</td><td>BT20/MB468</td><td>MP</td></tr>
<tr><td>>hg38_21_+_8433172_8446622_RNA45SN1_45S_50ntFlanks@6652.6676.25|GACTCTTAGCGGTGGATCACTCGGC</td><td>>hg38_21_+_8433172_8446622_RNA45SN1_45S_50ntFlanks</td><td>6652</td><td>6676</td><td>25</td><td>GACTCTTAGCGGTGGATCACTCGGC</td><td>5026.4</td><td>3.4</td><td>1.2</td><td>2.9</td><td>3.2E-03</td><td>1.5E-02</td><td>BT20/MB231</td><td>MP</td></tr>
<tr><td>>MI0003591|hsa-mir-584&WithFlank&5|-|149062307|149062415@21.42.22&[MIMAT0003249&hsa-miR-584-5p&offsets|-1|-1;m-167&5|-|149062374|149062394&offsets|-1|0]|ATTATGGTTTGCCTGGGACTGA</td><td>>MI0003591|hsa-mir-584&WithFlank&5|-|149062307|149062415</td><td>21</td><td>42</td><td>22</td><td>ATTATGGTTTGCCTGGGACTGA</td><td>85.9</td><td>6.7</td><td>2.5</td><td>2.6</td><td>8.2E-03</td><td>1.5E-02</td><td>BT20/MB468</td><td>MP</td></tr>
<tr><td>>hg38_21_+_8433172_8446622_RNA45SN1_45S_50ntFlanks@8183.8208.26|TCGAGGCCCAGCCCGTGGACGGTGTG</td><td>>hg38_21_+_8433172_8446622_RNA45SN1_45S_50ntFlanks</td><td>8183</td><td>8208</td><td>26</td><td>TCGAGGCCCAGCCCGTGGACGGTGTG</td><td>408.4</td><td>-4.8</td><td>1.9</td><td>-2.5</td><td>1.3E-02</td><td>1.5E-02</td><td>BT20/MB468</td><td>NUC</td></tr>
<tr><td>>MI0000811|hsa-mir-148b&WithFlank&12|+|54337210|54337320@69.90.22(+1U)&[MIMAT0000759&hsa-miR-148b-3p&offsets|0|0(+1U);m-77&12|+|54337278|54337299&offsets|0|0(+1U)]|TCAGTGCATCACAGAACTTTGTT</td><td>>MI0000811|hsa-mir-148b&WithFlank&12|+|54337210|54337320</td><td>69</td><td>90</td><td>22(+1U)</td><td>TCAGTGCATCACAGAACTTTGTT</td><td>5946.5</td><td>-2.5</td><td>0.9</td><td>-2.9</td><td>4.1E-03</td><td>1.5E-02</td><td>BT20/MB231</td><td>MITO</td></tr>
<tr><td>>hg38_21_+_8433172_8446622_RNA45SN1_45S_50ntFlanks@7976.8009.34|GCGACCTCAGATCAGACGTGGCGACCCGCTGAAT</td><td>>hg38_21_+_8433172_8446622_RNA45SN1_45S_50ntFlanks</td><td>7976</td><td>8009</td><td>34</td><td>GCGACCTCAGATCAGACGTGGCGACCCGCTGAAT</td><td>2216.4</td><td>2.1</td><td>0.7</td><td>2.9</td><td>4.1E-03</td><td>1.5E-02</td><td>BT20/MB231</td><td>MITO</td></tr>
<tr><td>>MI0000088|hsa-mir-30a&WithFlank&6|-|71403545|71403627@54.73.20(+2U)&[MIMAT0000088&hsa-miR-30a-3p&offsets|+1|-1(+2U);m-38&6|-|71403554|71403575&offsets|+1|-1(+2U)]|TTTCAGTCGGATGTTTGCAGTT</td><td>>MI0000088|hsa-mir-30a&WithFlank&6|-|71403545|71403627</td><td>54</td><td>73</td><td>20(+2U)</td><td>TTTCAGTCGGATGTTTGCAGTT</td><td>414.5</td><td>-2.1</td><td>0.7</td><td>-2.9</td><td>4.1E-03</td><td>1.5E-02</td><td>BT20/MB231</td><td>MITO</td></tr>
<tr><td>>hg38_MT_+_1621_3279_RNR2_16S_50ntFlanks@1347.1389.43|CAACAATAGGGTTTACGACCTCGATGTTGGATCAGGACATCCC</td><td>>hg38_MT_+_1621_3279_RNR2_16S_50ntFlanks</td><td>1347</td><td>1389</td><td>43</td><td>CAACAATAGGGTTTACGACCTCGATGTTGGATCAGGACATCCC</td><td>119.0</td><td>5.0</td><td>1.9</td><td>2.6</td><td>8.2E-03</td><td>1.5E-02</td><td>BT20/MB468</td><td>MP</td></tr>
<tr><td>>MI0000298|hsa-mir-221&WithFlank&X|-|45746151|45746272@71.92.22(+1U)&[MIMAT0000278&hsa-miR-221-3p&offsets|0|-1(+1U);m-91&X|-|45746181|45746202&offsets|0|0(+1U)]|AGCTACATTGTCTGCTGGGTTTT</td><td>>MI0000298|hsa-mir-221&WithFlank&X|-|45746151|45746272</td><td>71</td><td>92</td><td>22(+1U)</td><td>AGCTACATTGTCTGCTGGGTTTT</td><td>3376.6</td><td>-1.8</td><td>0.6</td><td>-2.9</td><td>3.5E-03</td><td>1.5E-02</td><td>BT20/MB231</td><td>NUC</td></tr>
<tr><td>>hg38_21_+_8433172_8446622_RNA45SN1_45S_50ntFlanks@12474.12497.24|ACCCACTAATAGGGAACGTGAGCT</td><td>>hg38_21_+_8433172_8446622_RNA45SN1_45S_50ntFlanks</td><td>12474</td><td>12497</td><td>24</td><td>ACCCACTAATAGGGAACGTGAGCT</td><td>302.9</td><td>-2.9</td><td>1.0</td><td>-2.9</td><td>3.5E-03</td><td>1.5E-02</td><td>BT20/MB231</td><td>NUC</td></tr>
<tr><td>>hg38_21_+_8433172_8446622_RNA45SN1_45S_50ntFlanks@4590.4616.27|ATTTTGTTGGTTTTCGGAACTGAGGCC</td><td>>hg38_21_+_8433172_8446622_RNA45SN1_45S_50ntFlanks</td><td>4590</td><td>4616</td><td>27</td><td>ATTTTGTTGGTTTTCGGAACTGAGGCC</td><td>461.2</td><td>-1.7</td><td>0.7</td><td>-2.5</td><td>1.3E-02</td><td>1.5E-02</td><td>BT20/MB468</td><td>CYTO</td></tr>
<tr><td>>hg38_21_+_8433172_8446622_RNA45SN1_45S_50ntFlanks@10394.10424.31|AGCAGTTGAACATGGGTCAGTCGGTCCTGAG</td><td>>hg38_21_+_8433172_8446622_RNA45SN1_45S_50ntFlanks</td><td>10394</td><td>10424</td><td>31</td><td>AGCAGTTGAACATGGGTCAGTCGGTCCTGAG</td><td>934.6</td><td>-1.5</td><td>0.6</td><td>-2.5</td><td>1.4E-02</td><td>1.5E-02</td><td>BT20/MB468</td><td>NUC</td></tr>
<tr><td>>hg38_21_+_8433172_8446622_RNA45SN1_45S_50ntFlanks@12073.12108.36|CCGAGGGGCTCTCGCTTCTGGCGCCAAGCGCCCGGC</td><td>>hg38_21_+_8433172_8446622_RNA45SN1_45S_50ntFlanks</td><td>12073</td><td>12108</td><td>36</td><td>CCGAGGGGCTCTCGCTTCTGGCGCCAAGCGCCCGGC</td><td>278.2</td><td>-3.9</td><td>1.5</td><td>-2.5</td><td>1.1E-02</td><td>1.5E-02</td><td>BT20/MB468</td><td>MITO</td></tr>
<tr><td>>am_trnaMT_LysTTT_MT_+_8295_8364@45.69.25,trnalookalike7_LysTTT_1_+_568844_568913@45.69.25|AGAGAACCAACACCTCTTTACAGTG</td><td>>am_trnaMT_LysTTT_MT_+_8295_8364</td><td>45</td><td>69</td><td>25</td><td>AGAGAACCAACACCTCTTTACAGTG</td><td>80.5</td><td>-4.2</td><td>1.7</td><td>-2.5</td><td>1.2E-02</td><td>1.5E-02</td><td>MB231/MB468</td><td>MP</td></tr>
<tr><td>>am_trna10_ValCAC_5_-_180649395_180649467@1.24.24,trna12_ValAAC_5_-_180645270_180645342@1.24.24,trna132_ValAAC_6_-_27721179_27721251@1.24.24,trna136_ValAAC_6_-_27648885_27648957@1.24.24,trna139_ValAAC_6_-_27618707_27618779@1.24.24,trna18_ValCAC_5_-_180529253_180529325@1.24.24,trna2_ValAAC_3_+_169490018_169490090@1.24.24,trna2_ValCAC_5_+_180524070_180524142@1.24.24,trna37_ValAAC_6_+_27203288_27203360@1.24.24,trna4_ValAAC_5_+_180591154_180591226@1.24.24,trna5_ValAAC_5_+_180596610_180596682@1.24.24,trna6_ValCAC_5_+_180600650_180600722@1.24.24,trna85_ValCAC_1_-_161369490_161369562@1.24.24,trna90_ValCAC_1_-_149684088_149684161@1.24.24,trna98_ValCAC_1_-_149298555_149298627@1.24.24,trna9_ValCAC_6_+_26538282_26538354@1.24.24|GTTTCCGTAGTGTAGTGGTTATCA</td><td>>am_trna10_ValCAC_5_-_180649395_180649467</td><td>1</td><td>24</td><td>24</td><td>GTTTCCGTAGTGTAGTGGTTATCA</td><td>106.3</td><td>-4.7</td><td>1.9</td><td>-2.5</td><td>1.2E-02</td><td>1.5E-02</td><td>MB231/MB468</td><td>MP</td></tr>
<tr><td>>MI0000270|hsa-mir-181b-1&WithFlank&1|-|198858867|198858988@42.64.23(+1A);MI0000683|hsa-mir-181b-2&WithFlank&9|+|124693704|124693804@22.44.23(+1A)&[MIMAT0000257&hsa-miR-181b-5p&offsets|0|0(+1A);m-73&1|-|198858925|198858947&offsets|0|0(+1A)];[MIMAT0000257_1&hsa-miR-181b-5p&offsets|0|0(+1A);m-72&9|+|124693725|124693747&offsets|0|0(+1A)]|AACATTCATTGCTGTCGGTGGGTA</td><td>>MI0000270|hsa-mir-181b-1&WithFlank&1|-|198858867|198858988</td><td>42</td><td>64</td><td>23(+1A)</td><td>AACATTCATTGCTGTCGGTGGGTA</td><td>420.2</td><td>-3.1</td><td>1.1</td><td>-2.9</td><td>3.2E-03</td><td>1.5E-02</td><td>BT20/MB231</td><td>MP</td></tr>
<tr><td>>hg38_21_+_8433172_8446622_RNA45SN1_45S_50ntFlanks@6648.6687.40|GTACGACTCTTAGCGGTGGATCACTCGGCTCGTGCGTCGA</td><td>>hg38_21_+_8433172_8446622_RNA45SN1_45S_50ntFlanks</td><td>6648</td><td>6687</td><td>40</td><td>GTACGACTCTTAGCGGTGGATCACTCGGCTCGTGCGTCGA</td><td>346.5</td><td>2.4</td><td>1.0</td><td>2.5</td><td>1.4E-02</td><td>1.5E-02</td><td>MB231/MB468</td><td>MITO</td></tr>
<tr><td>>hg38_21_+_8433172_8446622_RNA45SN1_45S_50ntFlanks@12994.13039.46|CCTCGCTGCGATCTATTGAAAGTCAGCCCTCGACACAAGGGTTTGT</td><td>>hg38_21_+_8433172_8446622_RNA45SN1_45S_50ntFlanks</td><td>12994</td><td>13039</td><td>46</td><td>CCTCGCTGCGATCTATTGAAAGTCAGCCCTCGACACAAGGGTTTGT</td><td>14846.1</td><td>3.1</td><td>1.3</td><td>2.5</td><td>1.4E-02</td><td>1.5E-02</td><td>MB231/MB468</td><td>NUC</td></tr>
<tr><td>>hg38_21_+_8433172_8446622_RNA45SN1_45S_50ntFlanks@5444.5463.20|CTCGGATCGGCCCCGCCGGG</td><td>>hg38_21_+_8433172_8446622_RNA45SN1_45S_50ntFlanks</td><td>5444</td><td>5463</td><td>20</td><td>CTCGGATCGGCCCCGCCGGG</td><td>62.6</td><td>-4.7</td><td>1.8</td><td>-2.6</td><td>8.2E-03</td><td>1.5E-02</td><td>BT20/MB468</td><td>MP</td></tr>
<tr><td>>hg38_21_+_8433172_8446622_RNA45SN1_45S_50ntFlanks@6646.6680.35|TCGTACGACTCTTAGCGGTGGATCACTCGGCTCGT</td><td>>hg38_21_+_8433172_8446622_RNA45SN1_45S_50ntFlanks</td><td>6646</td><td>6680</td><td>35</td><td>TCGTACGACTCTTAGCGGTGGATCACTCGGCTCGT</td><td>5090.4</td><td>3.5</td><td>1.2</td><td>2.9</td><td>3.3E-03</td><td>1.5E-02</td><td>BT20/MB231</td><td>MP</td></tr>
<tr><td>>hg38_21_+_8433172_8446622_RNA45SN1_45S_50ntFlanks@6651.6681.31|CGACTCTTAGCGGTGGATCACTCGGCTCGTG</td><td>>hg38_21_+_8433172_8446622_RNA45SN1_45S_50ntFlanks</td><td>6651</td><td>6681</td><td>31</td><td>CGACTCTTAGCGGTGGATCACTCGGCTCGTG</td><td>17537.9</td><td>4.0</td><td>1.4</td><td>2.9</td><td>3.3E-03</td><td>1.5E-02</td><td>BT20/MB231</td><td>MP</td></tr>
<tr><td>>MI0000103|hsa-mir-101-1&WithFlank&1|-|65058428|65058514@52.73.22;MI0000739|hsa-mir-101-2&WithFlank&9|+|4850291|4850381@54.75.22&[MIMAT0000099&hsa-miR-101-3p&offsets|-1|0;m-17&1|-|65058443|65058462&offsets|-1|+1];[MIMAT0000099_1&hsa-miR-101-3p&offsets|-1|0;m-16&9|+|4850345|4850364&offsets|-1|+1]|GTACAGTACTGTGATAACTGAA</td><td>>MI0000103|hsa-mir-101-1&WithFlank&1|-|65058428|65058514</td><td>52</td><td>73</td><td>22</td><td>GTACAGTACTGTGATAACTGAA</td><td>453.9</td><td>2.5</td><td>0.9</td><td>2.9</td><td>4.2E-03</td><td>1.5E-02</td><td>BT20/MB231</td><td>CYTO</td></tr>
<tr><td>>hg38_21_+_8433172_8446622_RNA45SN1_45S_50ntFlanks@12945.12988.44|GTAGACGACCTGCTTCTGGGTCGGGGTTTCGTACGTAGCAGAGC</td><td>>hg38_21_+_8433172_8446622_RNA45SN1_45S_50ntFlanks</td><td>12945</td><td>12988</td><td>44</td><td>GTAGACGACCTGCTTCTGGGTCGGGGTTTCGTACGTAGCAGAGC</td><td>835.2</td><td>-5.1</td><td>1.8</td><td>-2.9</td><td>4.2E-03</td><td>1.5E-02</td><td>BT20/MB231</td><td>CYTO</td></tr>
<tr><td>>hg38_MT_+_1621_3279_RNR2_16S_50ntFlanks@1426.1458.33|CGATTAAAGTCCTACGTGATCTGAGTTCAGACC</td><td>>hg38_MT_+_1621_3279_RNR2_16S_50ntFlanks</td><td>1426</td><td>1458</td><td>33</td><td>CGATTAAAGTCCTACGTGATCTGAGTTCAGACC</td><td>37.2</td><td>4.9</td><td>2.0</td><td>2.5</td><td>1.2E-02</td><td>1.5E-02</td><td>MB231/MB468</td><td>MP</td></tr>
<tr><td>>hg38_MT_+_1621_3279_RNR2_16S_50ntFlanks@1431.1451.21|AAAGTCCTACGTGATCTGAGT</td><td>>hg38_MT_+_1621_3279_RNR2_16S_50ntFlanks</td><td>1431</td><td>1451</td><td>21</td><td>AAAGTCCTACGTGATCTGAGT</td><td>446.4</td><td>-2.7</td><td>0.9</td><td>-2.9</td><td>3.3E-03</td><td>1.5E-02</td><td>BT20/MB231</td><td>MP</td></tr>
<tr><td>>hg38_21_+_8433172_8446622_RNA45SN1_45S_50ntFlanks@8356.8376.21|GAAAAGAACTTTGAAGAGAGA</td><td>>hg38_21_+_8433172_8446622_RNA45SN1_45S_50ntFlanks</td><td>8356</td><td>8376</td><td>21</td><td>GAAAAGAACTTTGAAGAGAGA</td><td>384.1</td><td>-2.6</td><td>1.0</td><td>-2.6</td><td>8.2E-03</td><td>1.5E-02</td><td>BT20/MB468</td><td>MP</td></tr>
<tr><td>>hg38_21_+_8433172_8446622_RNA45SN1_45S_50ntFlanks@9343.9369.27|CGAAAGCCGCCGTGGCGCAATGAAGGT</td><td>>hg38_21_+_8433172_8446622_RNA45SN1_45S_50ntFlanks</td><td>9343</td><td>9369</td><td>27</td><td>CGAAAGCCGCCGTGGCGCAATGAAGGT</td><td>67.3</td><td>-4.4</td><td>1.7</td><td>-2.6</td><td>8.2E-03</td><td>1.5E-02</td><td>BT20/MB468</td><td>MP</td></tr>
<tr><td>>hg38_21_+_8433172_8446622_RNA45SN1_45S_50ntFlanks@4565.4588.24|AATAATGGAATAGGACCGCGGTTC</td><td>>hg38_21_+_8433172_8446622_RNA45SN1_45S_50ntFlanks</td><td>4565</td><td>4588</td><td>24</td><td>AATAATGGAATAGGACCGCGGTTC</td><td>167.0</td><td>3.3</td><td>1.2</td><td>2.6</td><td>8.3E-03</td><td>1.5E-02</td><td>BT20/MB468</td><td>MP</td></tr>
<tr><td>>hg38_21_+_8433172_8446622_RNA45SN1_45S_50ntFlanks@12892.12930.39|CCCTCGCCCGTCACGCACCGCACGTTCGTGGGGAACCTG</td><td>>hg38_21_+_8433172_8446622_RNA45SN1_45S_50ntFlanks</td><td>12892</td><td>12930</td><td>39</td><td>CCCTCGCCCGTCACGCACCGCACGTTCGTGGGGAACCTG</td><td>289.4</td><td>-4.9</td><td>1.9</td><td>-2.6</td><td>8.3E-03</td><td>1.5E-02</td><td>BT20/MB468</td><td>MP</td></tr>
<tr><td>>hg38_21_+_8433172_8446622_RNA45SN1_45S_50ntFlanks@7974.8012.39|ACGCGACCTCAGATCAGACGTGGCGACCCGCTGAATTTA</td><td>>hg38_21_+_8433172_8446622_RNA45SN1_45S_50ntFlanks</td><td>7974</td><td>8012</td><td>39</td><td>ACGCGACCTCAGATCAGACGTGGCGACCCGCTGAATTTA</td><td>5368.6</td><td>1.7</td><td>0.7</td><td>2.5</td><td>1.4E-02</td><td>1.6E-02</td><td>MB231/MB468</td><td>NUC</td></tr>
<tr><td>>hg38_1_-_228634819_228635039_RNA5S12_5S_50ntFlanks@139.171.33|GACCGCCTGGGAATACCGGGTGCTGTAGGCTTT</td><td>>hg38_1_-_228634819_228635039_RNA5S12_5S_50ntFlanks</td><td>139</td><td>171</td><td>33</td><td>GACCGCCTGGGAATACCGGGTGCTGTAGGCTTT</td><td>907.0</td><td>2.6</td><td>1.0</td><td>2.6</td><td>8.3E-03</td><td>1.6E-02</td><td>BT20/MB468</td><td>MP</td></tr>
<tr><td>>MI0000299|hsa-mir-222&WithFlank&X|-|45747009|45747130@75.95.21(+1U)&[MIMAT0000279&hsa-miR-222-3p&offsets|0|0(+1U);m-140&X|-|45747033|45747056&offsets|0|-3(+1U)]|AGCTACATCTGGCTACTGGGTT</td><td>>MI0000299|hsa-mir-222&WithFlank&X|-|45747009|45747130</td><td>75</td><td>95</td><td>21(+1U)</td><td>AGCTACATCTGGCTACTGGGTT</td><td>197.7</td><td>3.9</td><td>1.5</td><td>2.6</td><td>8.3E-03</td><td>1.6E-02</td><td>BT20/MB468</td><td>MP</td></tr>
<tr><td>>hg38_21_+_8433172_8446622_RNA45SN1_45S_50ntFlanks@6651.6674.24|CGACTCTTAGCGGTGGATCACTCG</td><td>>hg38_21_+_8433172_8446622_RNA45SN1_45S_50ntFlanks</td><td>6651</td><td>6674</td><td>24</td><td>CGACTCTTAGCGGTGGATCACTCG</td><td>4450.1</td><td>4.0</td><td>1.3</td><td>2.9</td><td>3.3E-03</td><td>1.6E-02</td><td>BT20/MB231</td><td>MP</td></tr>
</table>

| | | | | | | | | | | | | | |
|---|---|---|---|---|---|---|---|---|---|---|---|---|---|
| >hg38_21_+_8433172_8446622_RNA45SN1_45S_50ntFlanks@10069.10087.19\|AGTGGACGGGAGCGGCGGG | >hg38_21_+_8433172_8446622_RNA45SN1_45S_50ntFlanks | 10069 | 10087 | 19 | AGTGGACGGGAGCGGCGGG | 113.0 | -4.4 | 1.7 | -2.5 | 1.2E-02 | 1.6E-02 | MB231/MB468 | MP |
| >trnaMT_ProTGG_MT_-_15956_16023@40.70.31\|AATGGTGGAGTTAAAGACTTTTTCTCTGACC | >trnaMT_ProTGG_MT_-_15956_16023 | 40 | 70 | 31 | AATGGTGGAGTTAAAGACTTTTTCTCTGACC | 71.3 | 5.5 | 2.1 | 2.6 | 8.3E-03 | 1.6E-02 | BT20/MB468 | MP |
| >trnaMT_PheGAA_MT_+_577_647@38.57.20\|AATGTTTAGACGGGCTCACA | >trnaMT_PheGAA_MT_+_577_647 | 38 | 57 | 20 | AATGTTTAGACGGGCTCACA | 75.7 | 2.0 | 0.8 | 2.5 | 1.2E-02 | 1.6E-02 | MB231/MB468 | MP |
| >hg38_21_+_8433172_8446622_RNA45SN1_45S_50ntFlanks@10064.10081.18\|TCGAGAGTGGACGGGAGC | >hg38_21_+_8433172_8446622_RNA45SN1_45S_50ntFlanks | 10064 | 10081 | 18 | TCGAGAGTGGACGGGAGC | 123.8 | 4.6 | 1.6 | 2.9 | 3.3E-03 | 1.6E-02 | BT20/MB231 | MP |
| >hg38_MT_+_1621_3279_RNR2_16S_50ntFlanks@695.727.33\|ATGAAAACATTCTCCTCCGCATAAGCCTGCGTC | >hg38_MT_+_1621_3279_RNR2_16S_50ntFlanks | 695 | 727 | 33 | ATGAAAACATTCTCCTCCGCATAAGCCTGCGTC | 41.7 | 8.7 | 3.0 | 2.9 | 3.3E-03 | 1.6E-02 | BT20/MB231 | MP |
| >MI0000465\|hsa-mir-191&WithFlank&3\|-\|49020612\|49020715@22.42.21&[MIMAT0000440&hsa-miR-191-5p&offsets\|0\|-2;m-48&3\|-\|49020672\|49020694&offsets\|0\|-2]\|CAACGGAATCCCAAAAGCAGC | >MI0000465\|hsa-mir-191&WithFlank&3\|-\|49020612\|49020715 | 22 | 42 | 21 | CAACGGAATCCCAAAAGCAGC | 614.6 | 3.3 | 1.2 | 2.6 | 8.3E-03 | 1.6E-02 | BT20/MB468 | MP |
| >hg38_21_+_8433172_8446622_RNA45SN1_45S_50ntFlanks@6733.6752.20\|CATTGATCATCGACACTTCG | >hg38_21_+_8433172_8446622_RNA45SN1_45S_50ntFlanks | 6733 | 6752 | 20 | CATTGATCATCGACACTTCG | 378.4 | -1.5 | 0.6 | -2.5 | 1.4E-02 | 1.6E-02 | MB231/MB468 | NUC |
| >hg38_21_+_8433172_8446622_RNA45SN1_45S_50ntFlanks@9564.9593.30\|CTGACGTGCAAATCGGTCGTCCGACCTGGG | >hg38_21_+_8433172_8446622_RNA45SN1_45S_50ntFlanks | 9564 | 9593 | 30 | CTGACGTGCAAATCGGTCGTCCGACCTGGG | 141.9 | -2.2 | 0.9 | -2.5 | 1.3E-02 | 1.6E-02 | MB231/MB468 | CYTO |
| >trnaMT_ValTAC_MT_+_1602_1670@51.72.22\|CTTAACTTGACCGCTCTGACCA | >trnaMT_ValTAC_MT_+_1602_1670 | 51 | 72 | 22 | CTTAACTTGACCGCTCTGACCA | 1537.1 | -2.5 | 1.0 | -2.6 | 8.3E-03 | 1.6E-02 | BT20/MB468 | MP |
| >hg38_21_+_8433172_8446622_RNA45SN1_45S_50ntFlanks@9546.9571.26\|TGGAGGTCCGTAGCGGTCCTGACGTG | >hg38_21_+_8433172_8446622_RNA45SN1_45S_50ntFlanks | 9546 | 9571 | 26 | TGGAGGTCCGTAGCGGTCCTGACGTG | 479.4 | 1.1 | 0.4 | 2.5 | 1.1E-02 | 1.6E-02 | BT20/MB468 | MITO |
| >hg38_MT_+_1621_3279_RNR2_16S_50ntFlanks@470.491.22\|AACTGTTAGTCCAAAGAGGAAC | >hg38_MT_+_1621_3279_RNR2_16S_50ntFlanks | 470 | 491 | 22 | AACTGTTAGTCCAAAGAGGAAC | 84.2 | 2.9 | 1.1 | 2.6 | 8.3E-03 | 1.6E-02 | BT20/MB468 | MP |
| >trnaMT_ProTGG_MT_-_15956_16023@40.69.30\|AATGGTGGAGTTAAAGACTTTTTCTCTGAC | >trnaMT_ProTGG_MT_-_15956_16023 | 40 | 69 | 30 | AATGGTGGAGTTAAAGACTTTTTCTCTGAC | 30.9 | 4.9 | 1.9 | 2.5 | 1.2E-02 | 1.6E-02 | MB231/MB468 | MP |
| >MI0000077\|hsa-mir-21&WithFlank&17\|+\|59841260\|59841343@14.35.22(+2U)&[MIMAT0000076&hsa-miR-21-5p&offsets\|0\|0(+2U);m-1&17\|+\|59841273\|59841294&offsets\|0\|0(+2U)]\|TAGCTTATCAGACTGATGTTGATT | >MI0000077\|hsa-mir-21&WithFlank&17\|+\|59841260\|59841343 | 14 | 35 | 22(+2U) | TAGCTTATCAGACTGATGTTGATT | 50.3 | 4.0 | 1.6 | 2.5 | 1.2E-02 | 1.6E-02 | MB231/MB468 | MP |
| >hg38_1_-_228634819_228635039_RNA5S12_5S_50ntFlanks@51.96.46\|GTCTACGGCCATACCACCCTGAACGCGCCCGATCTCGTCTGATCTC | >hg38_1_-_228634819_228635039_RNA5S12_5S_50ntFlanks | 51 | 96 | 46 | GTCTACGGCCATACCACCCTGAACGCGCCCGATCTCGTCTGATCTC | 201.6 | 5.8 | 2.2 | 2.6 | 8.3E-03 | 1.6E-02 | BT20/MB468 | MP |
| >hg38_MT_+_1621_3279_RNR2_16S_50ntFlanks@1221.1262.42\|TCCGAGCAGTACATGCTAAGACTTCACCAGTCAAAGCGAACT | >hg38_MT_+_1621_3279_RNR2_16S_50ntFlanks | 1221 | 1262 | 42 | TCCGAGCAGTACATGCTAAGACTTCACCAGTCAAAGCGAACT | 35.8 | 4.9 | 1.9 | 2.5 | 1.2E-02 | 1.6E-02 | MB231/MB468 | MP |
| >hg38_21_+_8433172_8446622_RNA45SN1_45S_50ntFlanks@9465.9503.39\|AGGTGGAGCACGAGCGCACGTGTTAGGACCCGAAAGATG | >hg38_21_+_8433172_8446622_RNA45SN1_45S_50ntFlanks | 9465 | 9503 | 39 | AGGTGGAGCACGAGCGCACGTGTTAGGACCCGAAAGATG | 464.6 | -4.8 | 1.8 | -2.6 | 8.3E-03 | 1.6E-02 | BT20/MB468 | MP |
| >trna2_GlyGCC_21_-_18827107_18827177@1.29.29,trna35_GlyGCC_1_+_161413094_161413164@1.29.29,trna37_GlyGCC_1_+_161420467_161420537@1.29.29,trna39_GlyGCC_1_+_161427898_161427968@1.29.29,trna41_GlyGCC_1_+_161435258_161435328@1.29.29\|GCATGGGTGGTTCAGTGGTAGAATTCTCG | >trna2_GlyGCC_21_-_18827107_18827177 | 1 | 29 | 29 | GCATGGGTGGTTCAGTGGTAGAATTCTCG | 319.5 | 6.7 | 2.4 | 2.9 | 4.2E-03 | 1.6E-02 | BT20/MB231 | MITO |
| >hg38_21_+_8433172_8446622_RNA45SN1_45S_50ntFlanks@12997.13038.42\|CGCTGCGATCTATTGAAAGTCAGCCCTCGACACAAGGGTTTG | >hg38_21_+_8433172_8446622_RNA45SN1_45S_50ntFlanks | 12997 | 13038 | 42 | CGCTGCGATCTATTGAAAGTCAGCCCTCGACACAAGGGTTTG | 2387.5 | -2.0 | 0.7 | -2.9 | 4.2E-03 | 1.6E-02 | BT20/MB231 | MITO |
| >MI0000809\|hsa-mir-151a&WithFlank&8\|-\|140732558\|140732659@55.75.21(+1U)&[MIMAT0000757&hsa-miR-151a-3p&offsets\|+2\|+2(+1U);m-30&8\|-\|140732587\|140732607&offsets\|+2\|+2(+1U)]\|AGACTGAAGCTCCTTGAGGACT | >MI0000809\|hsa-mir-151a&WithFlank&8\|-\|140732558\|140732659 | 55 | 75 | 21(+1U) | AGACTGAAGCTCCTTGAGGACT | 50.3 | 4.8 | 1.9 | 2.5 | 1.2E-02 | 1.6E-02 | MB231/MB468 | MP |
| >hg38_1_-_228634819_228635039_RNA5S12_5S_50ntFlanks@139.170.32\|GACCGCCTGGGAATACCGGGTGCTGTAGGCTT | >hg38_1_-_228634819_228635039_RNA5S12_5S_50ntFlanks | 139 | 170 | 32 | GACCGCCTGGGAATACCGGGTGCTGTAGGCTT | 437.3 | 2.1 | 0.8 | 2.5 | 1.4E-02 | 1.6E-02 | MB231/MB468 | MITO |
| >hg38_21_+_8433172_8446622_RNA45SN1_45S_50ntFlanks@7971.8011.41\|GAGACGCGACCTCAGATCAGACGTGGCGACCCGCTGAATTT | >hg38_21_+_8433172_8446622_RNA45SN1_45S_50ntFlanks | 7971 | 8011 | 41 | GAGACGCGACCTCAGATCAGACGTGGCGACCCGCTGAATTT | 113.0 | 3.9 | 1.6 | 2.5 | 1.4E-02 | 1.6E-02 | BT20/MB468 | NUC |
| >hg38_21_+_8433172_8446622_RNA45SN1_45S_50ntFlanks@12454.12474.21\|CACCAAGCGTTGGATTGTTCA | >hg38_21_+_8433172_8446622_RNA45SN1_45S_50ntFlanks | 12454 | 12474 | 21 | CACCAAGCGTTGGATTGTTCA | 78.5 | -4.4 | 1.8 | -2.5 | 1.2E-02 | 1.6E-02 | MB231/MB468 | MP |
| >hg38_1_-_228634819_228635039_RNA5S12_5S_50ntFlanks@51.69.19\|GTCTACGGCCATACCACCC | >hg38_1_-_228634819_228635039_RNA5S12_5S_50ntFlanks | 51 | 69 | 19 | GTCTACGGCCATACCACCC | 161.2 | -4.5 | 1.8 | -2.5 | 1.2E-02 | 1.6E-02 | MB231/MB468 | MP |
| >trnaMT_SerGCT_MT_+_12207_12265@14.45.32\|AGAACTGCTAACTCATGCCCCCATGTCTAACA | >trnaMT_SerGCT_MT_+_12207_12265 | 14 | 45 | 32 | AGAACTGCTAACTCATGCCCCCATGTCTAACA | 369.6 | 4.1 | 1.5 | 2.6 | 8.3E-03 | 1.6E-02 | BT20/MB468 | MP |
| >trnaMT_SerGCT_MT_+_12207_12265@15.62.48\|GAACTGCTAACTCATGCCCCCATGTCTAACAACATGGCTTTCTCACCA | >trnaMT_SerGCT_MT_+_12207_12265 | 15 | 62 | 48 | GAACTGCTAACTCATGCCCCCATGTCTAACAACATGGCTTTCTCACCA | 334.7 | 6.9 | 2.6 | 2.6 | 8.3E-03 | 1.6E-02 | BT20/MB468 | MP |
| >hg38_MT_+_1621_3279_RNR2_16S_50ntFlanks@1374.1414.41\|TGGATCAGGACATCCCGATGGTGCAGCCGCTATTAAAGGTT | >hg38_MT_+_1621_3279_RNR2_16S_50ntFlanks | 1374 | 1414 | 41 | TGGATCAGGACATCCCGATGGTGCAGCCGCTATTAAAGGTT | 104.7 | 4.0 | 1.6 | 2.5 | 1.2E-02 | 1.6E-02 | MB231/MB468 | MP |
| >hg38_21_+_8433172_8446622_RNA45SN1_45S_50ntFlanks@12242.12265.24\|AGAAGGGCAAAAGCTCGCTTGATC | >hg38_21_+_8433172_8446622_RNA45SN1_45S_50ntFlanks | 12242 | 12265 | 24 | AGAAGGGCAAAAGCTCGCTTGATC | 324.0 | -2.0 | 0.8 | -2.6 | 8.3E-03 | 1.6E-02 | BT20/MB468 | MP |
| >hg38_21_+_8433172_8446622_RNA45SN1_45S_50ntFlanks@12075.12095.21\|GAGGGGCTCTCGCTTCTGGCG | >hg38_21_+_8433172_8446622_RNA45SN1_45S_50ntFlanks | 12075 | 12095 | 21 | GAGGGGCTCTCGCTTCTGGCG | 390.6 | -1.8 | 0.7 | -2.5 | 1.3E-02 | 1.6E-02 | BT20/MB468 | CYTO |
| >hg38_21_+_8433172_8446622_RNA45SN1_45S_50ntFlanks@10792.10816.25\|CTCTGGCATGTTGGAACAATGTAGG | >hg38_21_+_8433172_8446622_RNA45SN1_45S_50ntFlanks | 10792 | 10816 | 25 | CTCTGGCATGTTGGAACAATGTAGG | 87.3 | 5.8 | 2.0 | 2.9 | 3.3E-03 | 1.6E-02 | BT20/MB231 | MP |
| >hg38_21_+_8433172_8446622_RNA45SN1_45S_50ntFlanks@10841.10861.21\|CGTAACTTCGGGATAAGGATT | >hg38_21_+_8433172_8446622_RNA45SN1_45S_50ntFlanks | 10841 | 10861 | 21 | CGTAACTTCGGGATAAGGATT | 706.0 | 4.3 | 1.5 | 2.9 | 3.3E-03 | 1.6E-02 | BT20/MB231 | MP |
| >hg38_21_+_8433172_8446622_RNA45SN1_45S_50ntFlanks@9300.9334.35\|ACGGACCAAGGAGTCTAACACGTGCGCGAGTCGGG | >hg38_21_+_8433172_8446622_RNA45SN1_45S_50ntFlanks | 9300 | 9334 | 35 | ACGGACCAAGGAGTCTAACACGTGCGCGAGTCGGG | 107.2 | 5.5 | 1.9 | 2.9 | 3.3E-03 | 1.6E-02 | BT20/MB231 | MP |
| >MI0000234\|hsa-mir-192&WithFlank&11\|-\|64891131\|64891252@31.51.21(+1U)&[MIMAT0000222&hsa-miR-192-5p&offsets\|+1\|+1(+1U);m-59&11\|-\|64891203\|64891223&offsets\|+1\|+1(+1U)]\|TGACCTATGAATTGACAGCCAT | >MI0000234\|hsa-mir-192&WithFlank&11\|-\|64891131\|64891252 | 31 | 51 | 21(+1U) | TGACCTATGAATTGACAGCCAT | 441.9 | 1.6 | 0.6 | 2.5 | 1.2E-02 | 1.6E-02 | MB231/MB468 | MP |
| >hg38_21_+_8433172_8446622_RNA45SN1_45S_50ntFlanks@7968.7997.30\|TCCGAGACGCGACCTCAGATCAGACGTGGC | >hg38_21_+_8433172_8446622_RNA45SN1_45S_50ntFlanks | 7968 | 7997 | 30 | TCCGAGACGCGACCTCAGATCAGACGTGGC | 108.3 | 2.0 | 0.8 | 2.5 | 1.4E-02 | 1.6E-02 | BT20/MB468 | NUC |
| >hg38_21_+_8433172_8446622_RNA45SN1_45S_50ntFlanks@11104.11121.18\|AAGGGTCGGGGCGGCAGG | >hg38_21_+_8433172_8446622_RNA45SN1_45S_50ntFlanks | 11104 | 11121 | 18 | AAGGGTCGGGGCGGCAGG | 767.8 | -5.0 | 2.0 | -2.5 | 1.4E-02 | 1.6E-02 | BT20/MB468 | NUC |
| >hg38_21_+_8433172_8446622_RNA45SN1_45S_50ntFlanks@12397.12414.18\|CGACGTCGCTTTTTGATC | >hg38_21_+_8433172_8446622_RNA45SN1_45S_50ntFlanks | 12397 | 12414 | 18 | CGACGTCGCTTTTTGATC | 1184.1 | -2.8 | 1.0 | -2.6 | 8.4E-03 | 1.6E-02 | BT20/MB468 | MP |
| >hg38_21_+_8433172_8446622_RNA45SN1_45S_50ntFlanks@5472.5491.20\|ACGGCCCTGGCGGAGCGCTG | >hg38_21_+_8433172_8446622_RNA45SN1_45S_50ntFlanks | 5472 | 5491 | 20 | ACGGCCCTGGCGGAGCGCTG | 144.7 | -2.3 | 0.9 | -2.5 | 1.2E-02 | 1.6E-02 | MB231/MB468 | MP |
| >am_trnaMT_PheGAA_MT_+_577_647@42.65.24\|TTTAGACGGGCTCACATCACCCCA | >am_trnaMT_PheGAA_MT_+_577_647 | 42 | 65 | 24 | TTTAGACGGGCTCACATCACCCCA | 58.8 | 6.7 | 2.5 | 2.6 | 8.4E-03 | 1.6E-02 | BT20/MB468 | MP |
| >trnaMT_PheGAA_MT_+_577_647@1.29.29\|GTTTATGTAGCTTACCTCCTCAAAGCAAT | >trnaMT_PheGAA_MT_+_577_647 | 1 | 29 | 29 | GTTTATGTAGCTTACCTCCTCAAAGCAAT | 2492.9 | 4.3 | 1.6 | 2.6 | 8.4E-03 | 1.6E-02 | BT20/MB468 | MP |
| >hg38_21_+_8433172_8446622_RNA45SN1_45S_50ntFlanks@12414.12432.19\|CCTTCGATGTCGGCTCTTC | >hg38_21_+_8433172_8446622_RNA45SN1_45S_50ntFlanks | 12414 | 12432 | 19 | CCTTCGATGTCGGCTCTTC | 424.7 | -2.4 | 0.8 | -2.9 | 4.2E-03 | 1.6E-02 | BT20/MB231 | CYTO |
| >hg38_21_+_8433172_8446622_RNA45SN1_45S_50ntFlanks@12409.12431.23\|TTGATCCTTCGATGTCGGCTCTT | >hg38_21_+_8433172_8446622_RNA45SN1_45S_50ntFlanks | 12409 | 12431 | 23 | TTGATCCTTCGATGTCGGCTCTT | 297.4 | -2.5 | 1.0 | -2.5 | 1.3E-02 | 1.6E-02 | BT20/MB468 | CYTO |
| >MI0006444\|hsa-mir-1307&WithFlank&10\|-\|103394247\|103394407@87.107.21(+1U)&[MIMAT0005951&hsa-miR-1307-3p&offsets\|+1\|0(+1U);m-66&10\|-\|103394300\|103394322&offsets\|+1\|-1(+1U)]\|CTCGGCGTGGCGTCGGTCGTGT | >MI0006444\|hsa-mir-1307&WithFlank&10\|-\|103394247\|103394407 | 87 | 107 | 21(+1U) | CTCGGCGTGGCGTCGGTCGTGT | 179.3 | 2.3 | 0.9 | 2.5 | 1.2E-02 | 1.6E-02 | MB231/MB468 | MP |
| >hg38_21_+_8433172_8446622_RNA45SN1_45S_50ntFlanks@10346.10390.45\|TCAAACGAGAACTTTGAAGGCCGAAGTGGAGAAGGGTTCCATGTG | >hg38_21_+_8433172_8446622_RNA45SN1_45S_50ntFlanks | 10346 | 10390 | 45 | TCAAACGAGAACTTTGAAGGCCGAAGTGGAGAAGGGTTCCATGTG | 94.8 | -4.4 | 1.8 | -2.5 | 1.2E-02 | 1.6E-02 | MB231/MB468 | MP |
| >hg38_21_+_8433172_8446622_RNA45SN1_45S_50ntFlanks@9860.9879.20\|GGATGAACCGAACGCCGGGT | >hg38_21_+_8433172_8446622_RNA45SN1_45S_50ntFlanks | 9860 | 9879 | 20 | GGATGAACCGAACGCCGGGT | 89.6 | -5.6 | 2.2 | -2.5 | 1.2E-02 | 1.6E-02 | MB231/MB468 | MP |
| >hg38_21_+_8433172_8446622_RNA45SN1_45S_50ntFlanks@10781.10822.42\|AGGTGAACAGCCTCTGGCATGTTGGAACAATGTAGGTAAGGG | >hg38_21_+_8433172_8446622_RNA45SN1_45S_50ntFlanks | 10781 | 10822 | 42 | AGGTGAACAGCCTCTGGCATGTTGGAACAATGTAGGTAAGGG | 124.6 | -3.7 | 1.4 | -2.6 | 8.4E-03 | 1.6E-02 | BT20/MB468 | MP |
| >MI0000298\|hsa-mir-221&WithFlank&X\|-\|45746151\|45746272@71.90.20(+1C)&[MIMAT0000278&hsa-miR-221-3p&offsets\|0\|-3(+1C);m-91&X\|-\|45746181\|45746202&offsets\|0\|-2(+1C)]\|AGCTACATTGTCTGCTGGGTC | >MI0000298\|hsa-mir-221&WithFlank&X\|-\|45746151\|45746272 | 71 | 90 | 20(+1C) | AGCTACATTGTCTGCTGGGTC | 273.6 | 4.2 | 1.6 | 2.6 | 8.4E-03 | 1.6E-02 | BT20/MB468 | MP |
| >hg38_21_+_8433172_8446622_RNA45SN1_45S_50ntFlanks@6772.6806.35\|GTTCCTCCCGGGGCTACGCCTGTCTGAGCGTCGCT | >hg38_21_+_8433172_8446622_RNA45SN1_45S_50ntFlanks | 6772 | 6806 | 35 | GTTCCTCCCGGGGCTACGCCTGTCTGAGCGTCGCT | 878.2 | -1.5 | 0.6 | -2.5 | 1.4E-02 | 1.6E-02 | MB231/MB468 | NUC |
| >hg38_21_+_8433172_8446622_RNA45SN1_45S_50ntFlanks@3885.3915.31\|ACGGGCGCTGACCCCCTTCGCGGGGGGGATG | >hg38_21_+_8433172_8446622_RNA45SN1_45S_50ntFlanks | 3885 | 3915 | 31 | ACGGGCGCTGACCCCCTTCGCGGGGGGGATG | 129.4 | -3.7 | 1.5 | -2.5 | 1.1E-02 | 1.6E-02 | BT20/MB468 | MITO |
| >hg38_21_+_8433172_8446622_RNA45SN1_45S_50ntFlanks@11029.11050.22\|CTCTCTCTCTCTCCCCCGCTCC | >hg38_21_+_8433172_8446622_RNA45SN1_45S_50ntFlanks | 11029 | 11050 | 22 | CTCTCTCTCTCTCCCCCGCTCC | 619.6 | -5.1 | 2.1 | -2.5 | 1.4E-02 | 1.6E-02 | BT20/MB468 | NUC |
| >hg38_21_+_8433172_8446622_RNA45SN1_45S_50ntFlanks@11571.11613.43\|GCCGACTTAGAACTGGTGCGGACCAGGGGAATCCGACTGTTTA | >hg38_21_+_8433172_8446622_RNA45SN1_45S_50ntFlanks | 11571 | 11613 | 43 | GCCGACTTAGAACTGGTGCGGACCAGGGGAATCCGACTGTTTA | 275.5 | -2.9 | 1.0 | -2.9 | 4.2E-03 | 1.6E-02 | BT20/MB231 | CYTO |
| >hg38_21_+_8433172_8446622_RNA45SN1_45S_50ntFlanks@11211.11232.22\|GCGACTCTGGACGCGAGCCGGG | >hg38_21_+_8433172_8446622_RNA45SN1_45S_50ntFlanks | 11211 | 11232 | 22 | GCGACTCTGGACGCGAGCCGGG | 365.4 | -1.5 | 0.6 | -2.5 | 1.1E-02 | 1.6E-02 | BT20/MB468 | MITO |
| >hg38_21_+_8433172_8446622_RNA45SN1_45S_50ntFlanks@475.510.36\|CTTCGTGATCGATGTGGTGACGTCGTGCTCTCCCGG | >hg38_21_+_8433172_8446622_RNA45SN1_45S_50ntFlanks | 475 | 510 | 36 | CTTCGTGATCGATGTGGTGACGTCGTGCTCTCCCGG | 5564.3 | -1.1 | 0.4 | -2.5 | 1.4E-02 | 1.6E-02 | BT20/MB468 | NUC |
| >MI0000102\|hsa-mir-100&WithFlank&11\|-\|122152223\|122152314@19.40.22&[MIMAT0000098&hsa-miR-100-5p&offsets\|0\|0;m-27&11\|-\|122152276\|122152296&offsets\|0\|+1]\|AACCCGTAGATCCGAACTTGTG | >MI0000102\|hsa-mir-100&WithFlank&11\|-\|122152223\|122152314 | 19 | 40 | 22 | AACCCGTAGATCCGAACTTGTG | 5133.4 | 4.7 | 1.8 | 2.6 | 8.4E-03 | 1.6E-02 | BT20/MB468 | MP |

| | | | | | | | | | | | | | |
|---|---|---|---|---|---|---|---|---|---|---|---|---|---|
| >MI0000085\|hsa-mir-27a&WithFlank&19\|-\|13836434\|13836523@57.74.18(+1A);MI0000440\|hsa-mir-27b&WithFlank&9\|+\|95085439\|95085547@67.84.18(+1A)&[MIMAT0000084&hsa-miR-27a-3p&offsets\|0\|-3(+1A);m-43&19\|-\|13836447\|13836467&offsets\|0\|-3(+1A)];[MIMAT0000419&hsa-miR-27b-3p&offsets\|0\|-3(+1A);m-39&9\|+\|95085505\|95085525&offsets\|0\|-3(+1A)]\|TTCACAGTGGCTAAGTTCA | >MI0000085\|hsa-mir-27a&WithFlank&19\|-\|13836434\|13836523 | 57 | 74 | 18(+1A) | TTCACAGTGGCTAAGTTCA | 136.0 | 3.8 | 1.4 | 2.6 | 8.4E-03 | 1.6E-02 | BT20/MB468 | MP |
| >MI0000081\|hsa-mir-24-2&WithFlank&19\|-\|13836281\|13836365@56.76.21;MI0000080\|hsa-mir-24-1&WithFlank&9\|+\|95086015\|95086094@50.70.21&[MIMAT0000080&hsa-miR-24-3p&offsets\|0\|-1;m-36&19\|-\|13836288\|13836310&offsets\|0\|-2];[MIMAT0000080_1&hsa-miR-24-3p&offsets\|0\|-1;m-35&9\|+\|95086064\|95086086&offsets\|0\|-2]\|TGGCTCAGTTCAGCAGGAACA | >MI0000081\|hsa-mir-24-2&WithFlank&19\|-\|13836281\|13836365 | 56 | 76 | 21 | TGGCTCAGTTCAGCAGGAACA | 976.1 | 1.0 | 0.4 | 2.5 | 1.4E-02 | 1.6E-02 | BT20/MB468 | NUC |
| >hg38_21_+_8433172_8446622_RNA45SN1_45S_50ntFlanks@11570.11603.34\|AGCCGACTTAGAACTGGTGCGGACCAGGGGAATC | >hg38_21_+_8433172_8446622_RNA45SN1_45S_50ntFlanks | 11570 | 11603 | 34 | AGCCGACTTAGAACTGGTGCGGACCAGGGGAATC | 1229.0 | -2.3 | 0.9 | -2.6 | 8.4E-03 | 1.6E-02 | BT20/MB468 | MP |
| >hg38_21_+_8433172_8446622_RNA45SN1_45S_50ntFlanks@12408.12426.19\|TTTGATCCTTCGATGTCGG | >hg38_21_+_8433172_8446622_RNA45SN1_45S_50ntFlanks | 12408 | 12426 | 19 | TTTGATCCTTCGATGTCGG | 83.2 | -4.4 | 1.7 | -2.6 | 8.4E-03 | 1.6E-02 | BT20/MB468 | MP |
| >hg38_21_+_8433172_8446622_RNA45SN1_45S_50ntFlanks@10796.10816.21\|GGCATGTTGGAACAATGTAGG | >hg38_21_+_8433172_8446622_RNA45SN1_45S_50ntFlanks | 10796 | 10816 | 21 | GGCATGTTGGAACAATGTAGG | 57.1 | -4.4 | 1.7 | -2.6 | 8.4E-03 | 1.6E-02 | BT20/MB468 | MP |
| >hg38_21_+_8433172_8446622_RNA45SN1_45S_50ntFlanks@11670.11696.27\|CCCAGTGCTCTGAATGTCAAAGTGAAG | >hg38_21_+_8433172_8446622_RNA45SN1_45S_50ntFlanks | 11670 | 11696 | 27 | CCCAGTGCTCTGAATGTCAAAGTGAAG | 173.9 | -2.9 | 1.2 | -2.5 | 1.4E-02 | 1.6E-02 | MB231/MB468 | NUC |
| >hg38_21_+_8433172_8446622_RNA45SN1_45S_50ntFlanks@11602.11621.20\|TCCGACTGTTTAATTAAAAC | >hg38_21_+_8433172_8446622_RNA45SN1_45S_50ntFlanks | 11602 | 11621 | 20 | TCCGACTGTTTAATTAAAAC | 424.6 | -1.7 | 0.7 | -2.5 | 1.3E-02 | 1.6E-02 | BT20/MB468 | CYTO |
| >hg38_21_+_8433172_8446622_RNA45SN1_45S_50ntFlanks@12163.12182.20\|ACTGGGGCGGTACACCTGTC | >hg38_21_+_8433172_8446622_RNA45SN1_45S_50ntFlanks | 12163 | 12182 | 20 | ACTGGGGCGGTACACCTGTC | 160.3 | 4.9 | 1.7 | 2.9 | 3.3E-03 | 1.6E-02 | BT20/MB231 | MP |
| >hg38_21_+_8433172_8446622_RNA45SN1_45S_50ntFlanks@9592.9610.19\|GGTATAGGGGCGAAAGACT | >hg38_21_+_8433172_8446622_RNA45SN1_45S_50ntFlanks | 9592 | 9610 | 19 | GGTATAGGGGCGAAAGACT | 178.1 | 5.0 | 1.7 | 2.9 | 3.3E-03 | 1.6E-02 | BT20/MB231 | MP |
| >hg38_21_+_8433172_8446622_RNA45SN1_45S_50ntFlanks@12419.12438.20\|GATGTCGGCTCTTCCTATCA | >hg38_21_+_8433172_8446622_RNA45SN1_45S_50ntFlanks | 12419 | 12438 | 20 | GATGTCGGCTCTTCCTATCA | 4397.0 | -2.4 | 0.9 | -2.9 | 4.2E-03 | 1.6E-02 | BT20/MB231 | CYTO |
| >hg38_21_+_8433172_8446622_RNA45SN1_45S_50ntFlanks@5481.5516.36\|GCGGAGCGCTGAGAAGACGGTCGAACTTGACTATCT | >hg38_21_+_8433172_8446622_RNA45SN1_45S_50ntFlanks | 5481 | 5516 | 36 | GCGGAGCGCTGAGAAGACGGTCGAACTTGACTATCT | 308.8 | 5.2 | 1.8 | 2.9 | 3.3E-03 | 1.6E-02 | BT20/MB231 | MP |
| >hg38_21_+_8433172_8446622_RNA45SN1_45S_50ntFlanks@4006.4042.37\|ACCTCGGGCCGATCGCACGCCCCCCGTGGCGGCGACG | >hg38_21_+_8433172_8446622_RNA45SN1_45S_50ntFlanks | 4006 | 4042 | 37 | ACCTCGGGCCGATCGCACGCCCCCCGTGGCGGCGACG | 198.7 | -2.9 | 1.1 | -2.5 | 1.2E-02 | 1.6E-02 | BT20/MB468 | MITO |
| >hg38_21_+_8433172_8446622_RNA45SN1_45S_50ntFlanks@12413.12435.23\|TCCTTCGATGTCGGCTCTTCCTA | >hg38_21_+_8433172_8446622_RNA45SN1_45S_50ntFlanks | 12413 | 12435 | 23 | TCCTTCGATGTCGGCTCTTCCTA | 1074.0 | -5.2 | 2.1 | -2.5 | 1.4E-02 | 1.6E-02 | BT20/MB468 | NUC |
| >MI0000299\|hsa-mir-222&WithFlank&X\|-\|45747009\|45747130@75.98.24(+1G)&[MIMAT0000279&hsa-miR-222-3p&offsets\|0\|+3(+1G);m-140&X\|-\|45747033\|45747056&offsets\|0\|0(+1G)]\|AGCTACATCTGGCTACTGGGTCTCG | >MI0000299\|hsa-mir-222&WithFlank&X\|-\|45747009\|45747130 | 75 | 98 | 24(+1G) | AGCTACATCTGGCTACTGGGTCTCG | 491.4 | -2.8 | 1.0 | -2.9 | 3.3E-03 | 1.6E-02 | BT20/MB231 | MP |
| >hg38_1_-_228634819_228635039_RNA5S12_5S_50ntFlanks@131.171.41\|GGATGGGAGACCGCCTGGGAATACCGGGTGCTGTAGGCTTT | >hg38_1_-_228634819_228635039_RNA5S12_5S_50ntFlanks | 131 | 171 | 41 | GGATGGGAGACCGCCTGGGAATACCGGGTGCTGTAGGCTTT | 124.0 | 1.8 | 0.7 | 2.5 | 1.4E-02 | 1.6E-02 | BT20/MB468 | NUC |
| >hg38_21_+_8433172_8446622_RNA45SN1_45S_50ntFlanks@11663.11681.19\|ATTTCTGCCCAGTGCTCTG | >hg38_21_+_8433172_8446622_RNA45SN1_45S_50ntFlanks | 11663 | 11681 | 19 | ATTTCTGCCCAGTGCTCTG | 225.3 | -2.7 | 1.1 | -2.5 | 1.3E-02 | 1.6E-02 | BT20/MB468 | CYTO |
| >hg38_21_+_8433172_8446622_RNA45SN1_45S_50ntFlanks@9458.9503.46\|GCCGGGGAGGTGGAGCACGAGCGCACGTGTTAGGACCCGAAAGATG | >hg38_21_+_8433172_8446622_RNA45SN1_45S_50ntFlanks | 9458 | 9503 | 46 | GCCGGGGAGGTGGAGCACGAGCGCACGTGTTAGGACCCGAAAGATG | 40.5 | -8.4 | 3.2 | -2.6 | 8.4E-03 | 1.6E-02 | BT20/MB468 | MP |
| >MI0000440\|hsa-mir-27b&WithFlank&9\|+\|95085439\|95085547@67.88.22(+1A)&[MIMAT0000419&hsa-miR-27b-3p&offsets\|0\|+1(+1A);m-39&9\|+\|95085505\|95085525&offsets\|0\|+1(+1A)]\|TTCACAGTGGCTAAGTTCTGCAA | >MI0000440\|hsa-mir-27b&WithFlank&9\|+\|95085439\|95085547 | 67 | 88 | 22(+1A) | TTCACAGTGGCTAAGTTCTGCAA | 628.7 | 2.4 | 0.8 | 2.9 | 4.3E-03 | 1.6E-02 | BT20/MB231 | CYTO |
| >hg38_21_+_8433172_8446622_RNA45SN1_45S_50ntFlanks@10835.10858.24\|CGGATCCGTAACTTCGGGATAAGG | >hg38_21_+_8433172_8446622_RNA45SN1_45S_50ntFlanks | 10835 | 10858 | 24 | CGGATCCGTAACTTCGGGATAAGG | 209.1 | -2.3 | 0.8 | -2.9 | 4.3E-03 | 1.6E-02 | BT20/MB231 | CYTO |
| >MI0003667\|hsa-mir-652&WithFlank&X\|+\|110055323\|110055432@67.88.22&[MIMAT0003322&hsa-miR-652-3p&offsets\|0\|+1;m-148&X\|+\|110055389\|110055410&offsets\|0\|0]\|AATGGCGCCACTAGGGTTGTGC | >MI0003667\|hsa-mir-652&WithFlank&X\|+\|110055323\|110055432 | 67 | 88 | 22 | AATGGCGCCACTAGGGTTGTGC | 480.6 | 4.5 | 1.7 | 2.6 | 8.4E-03 | 1.6E-02 | BT20/MB468 | MP |
| >hg38_21_+_8433172_8446622_RNA45SN1_45S_50ntFlanks@8527.8551.25\|CCTTCCCCCGCCGCCCCTCCTCCTC | >hg38_21_+_8433172_8446622_RNA45SN1_45S_50ntFlanks | 8527 | 8551 | 25 | CCTTCCCCCGCCGCCCCTCCTCCTC | 416.9 | -4.0 | 1.6 | -2.5 | 1.3E-02 | 1.6E-02 | MB231/MB468 | CYTO |
| >trna11_SerAGA_8_-_96281885_96281966@1.18.18,trna145_SerAGA_6_-_27521192_27521273@1.18.18,trna147_SerAGA_6_-_27509554_27509635@1.18.18,trna148_SerTGA_6_-_27473607_27473688@1.18.18,trna172_SerTGA_6_-_26312824_26312905@1.18.18,trna35_SerAGA_17_-_8129928_8130009@1.18.18,trna44_SerAGA_6_+_27446591_27446672@1.18.18,trna46_SerAGA_6_+_27463593_27463674@1.18.18,trna47_SerAGA_6_+_27470818_27470899@1.18.18,trna50_SerAGA_6_+_27499987_27500068@1.18.18,trna51_SerTGA_6_+_27513468_27513549@1.18.18,trna5_SerAGA_6_+_26327817_26327898@1.18.18\|GTAGTCGTGGCCGAGTGG | >trna11_SerAGA_8_-_96281885_96281966 | 1 | 18 | 18 | GTAGTCGTGGCCGAGTGG | 94.8 | 2.0 | 0.8 | 2.5 | 1.2E-02 | 1.6E-02 | MB231/MB468 | MP |
| >hg38_MT_+_1621_3279_RNR2_16S_50ntFlanks@1369.1404.36\|GATGTTGGATCAGGACATCCCGATGGTGCAGCCGCT | >hg38_MT_+_1621_3279_RNR2_16S_50ntFlanks | 1369 | 1404 | 36 | GATGTTGGATCAGGACATCCCGATGGTGCAGCCGCT | 75.2 | 4.0 | 1.6 | 2.5 | 1.2E-02 | 1.6E-02 | MB231/MB468 | MP |
| >hg38_21_+_8433172_8446622_RNA45SN1_45S_50ntFlanks@6733.6771.39\|CATTGATCATCGACACTTCGAACGCACTTGCGGCCCCGG | >hg38_21_+_8433172_8446622_RNA45SN1_45S_50ntFlanks | 6733 | 6771 | 39 | CATTGATCATCGACACTTCGAACGCACTTGCGGCCCCGG | 589.3 | -2.8 | 1.1 | -2.5 | 1.3E-02 | 1.6E-02 | MB231/MB468 | CYTO |
| >hg38_21_+_8433172_8446622_RNA45SN1_45S_50ntFlanks@12072.12096.25\|CCCGAGGGGCTCTCGCTTCTGGCGC | >hg38_21_+_8433172_8446622_RNA45SN1_45S_50ntFlanks | 12072 | 12096 | 25 | CCCGAGGGGCTCTCGCTTCTGGCGC | 123.9 | -4.6 | 1.7 | -2.6 | 8.5E-03 | 1.6E-02 | BT20/MB468 | MP |
| >hg38_21_+_8433172_8446622_RNA45SN1_45S_50ntFlanks@3999.4024.26\|CTAGATAACCTCGGGCCGATCGCACG | >hg38_21_+_8433172_8446622_RNA45SN1_45S_50ntFlanks | 3999 | 4024 | 26 | CTAGATAACCTCGGGCCGATCGCACG | 96.7 | -4.4 | 1.7 | -2.6 | 8.5E-03 | 1.6E-02 | BT20/MB468 | MP |
| >hg38_21_+_8433172_8446622_RNA45SN1_45S_50ntFlanks@5420.5445.26\|CCGATTGGATGGTTTAGTGAGGCCCT | >hg38_21_+_8433172_8446622_RNA45SN1_45S_50ntFlanks | 5420 | 5445 | 26 | CCGATTGGATGGTTTAGTGAGGCCCT | 706.2 | -4.8 | 1.8 | -2.6 | 8.5E-03 | 1.6E-02 | BT20/MB468 | MP |
| >hg38_MT_+_1621_3279_RNR2_16S_50ntFlanks@750.777.28\|ATTAACAGCCCAATATCTACAATCAACC | >hg38_MT_+_1621_3279_RNR2_16S_50ntFlanks | 750 | 777 | 28 | ATTAACAGCCCAATATCTACAATCAACC | 60.7 | 4.8 | 1.9 | 2.5 | 1.2E-02 | 1.6E-02 | MB231/MB468 | MP |
| >hg38_21_+_8433172_8446622_RNA45SN1_45S_50ntFlanks@6772.6804.33\|GTTCCTCCCGGGGCTACGCCTGTCTGAGCGTCG | >hg38_21_+_8433172_8446622_RNA45SN1_45S_50ntFlanks | 6772 | 6804 | 33 | GTTCCTCCCGGGGCTACGCCTGTCTGAGCGTCG | 64.3 | -4.3 | 1.6 | -2.6 | 8.5E-03 | 1.6E-02 | BT20/MB468 | MP |
| >hg38_21_+_8433172_8446622_RNA45SN1_45S_50ntFlanks@9305.9323.19\|CCAAGGAGTCTAACACGTG | >hg38_21_+_8433172_8446622_RNA45SN1_45S_50ntFlanks | 9305 | 9323 | 19 | CCAAGGAGTCTAACACGTG | 564.2 | -2.6 | 1.0 | -2.6 | 8.5E-03 | 1.6E-02 | BT20/MB468 | MP |
| >hg38_21_+_8433172_8446622_RNA45SN1_45S_50ntFlanks@12645.12669.25\|CTGTGGGATTATGACTGAACGCCTC | >hg38_21_+_8433172_8446622_RNA45SN1_45S_50ntFlanks | 12645 | 12669 | 25 | CTGTGGGATTATGACTGAACGCCTC | 446.8 | 2.5 | 0.9 | 2.9 | 4.2E-03 | 1.6E-02 | BT20/MB231 | MITO |
| >am_trnaMT_PheGAA_MT_+_577_647@41.60.20\|GTTTAGACGGGCTCACATCA | >am_trnaMT_PheGAA_MT_+_577_647 | 41 | 60 | 20 | GTTTAGACGGGCTCACATCA | 93.1 | 3.2 | 1.3 | 2.5 | 1.2E-02 | 1.6E-02 | MB231/MB468 | MP |
| >hg38_21_+_8433172_8446622_RNA45SN1_45S_50ntFlanks@7991.8011.21\|ACGTGGCGACCCGCTGAATTT | >hg38_21_+_8433172_8446622_RNA45SN1_45S_50ntFlanks | 7991 | 8011 | 21 | ACGTGGCGACCCGCTGAATTT | 658.6 | 1.5 | 0.6 | 2.5 | 1.3E-02 | 1.6E-02 | BT20/MB468 | CYTO |
| >hg38_21_+_8433172_8446622_RNA45SN1_45S_50ntFlanks@3899.3927.29\|CCTTCGCGGGGGGGATGCGTGCATTTATC | >hg38_21_+_8433172_8446622_RNA45SN1_45S_50ntFlanks | 3899 | 3927 | 29 | CCTTCGCGGGGGGGATGCGTGCATTTATC | 456.9 | -2.2 | 0.9 | -2.5 | 1.3E-02 | 1.6E-02 | BT20/MB468 | CYTO |
| >hg38_21_+_8433172_8446622_RNA45SN1_45S_50ntFlanks@7975.8007.33\|CGCGACCTCAGATCAGACGTGGCGACCCGCTGA | >hg38_21_+_8433172_8446622_RNA45SN1_45S_50ntFlanks | 7975 | 8007 | 33 | CGCGACCTCAGATCAGACGTGGCGACCCGCTGA | 72504.6 | 2.2 | 0.8 | 2.9 | 4.2E-03 | 1.6E-02 | BT20/MB231 | MITO |
| >hg38_21_+_8433172_8446622_RNA45SN1_45S_50ntFlanks@10683.10705.23\|CCGGCGGCGTCCGGTGAGCTCTC | >hg38_21_+_8433172_8446622_RNA45SN1_45S_50ntFlanks | 10683 | 10705 | 23 | CCGGCGGCGTCCGGTGAGCTCTC | 614.5 | -1.6 | 0.6 | -2.5 | 1.2E-02 | 1.6E-02 | BT20/MB468 | MITO |
| >hg38_21_+_8433172_8446622_RNA45SN1_45S_50ntFlanks@12418.12458.41\|CGATGTCGGCTCTTCCTATCATTGTGAAGCAGAATTCACCA | >hg38_21_+_8433172_8446622_RNA45SN1_45S_50ntFlanks | 12418 | 12458 | 41 | CGATGTCGGCTCTTCCTATCATTGTGAAGCAGAATTCACCA | 566.4 | 2.1 | 0.8 | 2.5 | 1.3E-02 | 1.6E-02 | MB231/MB468 | CYTO |
| >MI0001448\|hsa-mir-425&WithFlank&3\|-\|49020142\|49020240@20.44.25&[MIMAT0003393&hsa-miR-425-5p&offsets\|0\|+2;m-95&3\|-\|49020200\|49020221&offsets\|0\|+3]\|AATGACACGATCACTCCCGTTGAGT | >MI0001448\|hsa-mir-425&WithFlank&3\|-\|49020142\|49020240 | 20 | 44 | 25 | AATGACACGATCACTCCCGTTGAGT | 2458.9 | -2.7 | 0.9 | -2.9 | 3.4E-03 | 1.6E-02 | BT20/MB231 | MP |
| >MI0000087\|hsa-mir-29a&WithFlank&7\|-\|130876741\|130876816@48.66.19(+1C)&[MIMAT0000086&hsa-miR-29a-3p&offsets\|0\|-3(+1C);m-26&7\|-\|130876749\|130876769&offsets\|0\|-2(+1C)]\|TAGCACCATCTGAAATCGGC | >MI0000087\|hsa-mir-29a&WithFlank&7\|-\|130876741\|130876816 | 48 | 66 | 19(+1C) | TAGCACCATCTGAAATCGGC | 67.8 | 6.7 | 2.6 | 2.6 | 8.5E-03 | 1.6E-02 | BT20/MB468 | MP |
| >MI0003591\|hsa-mir-584&WithFlank&5\|-\|149062307\|149062415@21.42.22&[MIMAT0003249&hsa-miR-584-5p&offsets\|-1\|-1;m-167&5\|-\|149062374\|149062394&offsets\|-1\|0]\|ATTATGGTTTGCCTGGGACTGA | >MI0003591\|hsa-mir-584&WithFlank&5\|-\|149062307\|149062415 | 21 | 42 | 22 | ATTATGGTTTGCCTGGGACTGA | 45.9 | 4.8 | 1.9 | 2.5 | 1.2E-02 | 1.6E-02 | MB231/MB468 | MP |
| >MI0000093\|hsa-mir-92a-1&WithFlank&13\|+\|91351308\|91351397@54.75.22;MI0000094\|hsa-mir-92a-2&WithFlank&X\|-\|134169532\|134169618@54.75.22&[MIMAT0000092&hsa-miR-92a-3p&offsets\|0\|0;m-9&13\|+\|91351361\|91351382&offsets\|0\|0];[MIMAT0000092_1&hsa-miR-92a-3p&offsets\|0\|0;m-11&X\|-\|134169544\|134169565&offsets\|0\|0]\|TATTGCACTTGTCCCGGCCTGT | >MI0000093\|hsa-mir-92a-1&WithFlank&13\|+\|91351308\|91351397 | 54 | 75 | 22 | TATTGCACTTGTCCCGGCCTGT | 95131.7 | -2.0 | 0.7 | -2.9 | 4.3E-03 | 1.6E-02 | BT20/MB231 | MITO |
| >hg38_21_+_8433172_8446622_RNA45SN1_45S_50ntFlanks@11206.11225.20\|CGGCGGCGACTCTGGACGCG | >hg38_21_+_8433172_8446622_RNA45SN1_45S_50ntFlanks | 11206 | 11225 | 20 | CGGCGGCGACTCTGGACGCG | 87.7 | 5.8 | 2.0 | 2.9 | 3.4E-03 | 1.6E-02 | BT20/MB231 | MP |
| >hg38_1_-_228634819_228635039_RNA5S12_5S_50ntFlanks@135.171.37\|GGGAGACCGCCTGGGAATACCGGGTGCTGTAGGCTTT | >hg38_1_-_228634819_228635039_RNA5S12_5S_50ntFlanks | 135 | 171 | 37 | GGGAGACCGCCTGGGAATACCGGGTGCTGTAGGCTTT | 3796.6 | 2.1 | 0.9 | 2.5 | 1.4E-02 | 1.6E-02 | MB231/MB468 | MITO |
| >hg38_21_+_8433172_8446622_RNA45SN1_45S_50ntFlanks@8176.8199.24\|CTTCTGATCGAGGCCCAGCCCGTG | >hg38_21_+_8433172_8446622_RNA45SN1_45S_50ntFlanks | 8176 | 8199 | 24 | CTTCTGATCGAGGCCCAGCCCGTG | 313.4 | -1.5 | 0.6 | -2.5 | 1.3E-02 | 1.6E-02 | MB231/MB468 | CYTO |
| >MI0003763\|hsa-mir-767&WithFlank&X\|-\|152393415\|152393535@33.52.20&[MIMAT0003882&hsa-miR-767-5p&offsets\|0\|-3;m-788&X\|-\|152393481\|152393503&offsets\|0\|-3]\|TGCACCATGGTTGTCTGAGC | >MI0003763\|hsa-mir-767&WithFlank&X\|-\|152393415\|152393535 | 33 | 52 | 20 | TGCACCATGGTTGTCTGAGC | 52.7 | 5.5 | 2.1 | 2.6 | 8.5E-03 | 1.6E-02 | BT20/MB468 | MP |
| >trnaMT_SerGCT_MT_+_12207_12265@8.38.31\|CTCACAAGAACTGCTAACTCATGCCCCCATG | >trnaMT_SerGCT_MT_+_12207_12265 | 8 | 38 | 31 | CTCACAAGAACTGCTAACTCATGCCCCCATG | 602.1 | -4.3 | 1.7 | -2.5 | 1.2E-02 | 1.6E-02 | BT20/MB468 | MITO |

<table>
<tr><td>>hg38_21_+_8433172_8446622_RNA45SN1_45S_50ntFlanks@11602.11620.19|TCCGACTGTTTAATTAAAA</td><td>>hg38_21_+_8433172_8446622_RNA45SN1_45S_50ntFlanks</td><td>11602</td><td>11620</td><td>19</td><td>TCCGACTGTTTAATTAAAA</td><td>382.2</td><td>-1.5</td><td>0.6</td><td>-2.5</td><td>1.2E-02</td><td>1.6E-02</td><td>BT20/MB468</td><td>MITO</td></tr>
<tr><td>>hg38_21_+_8433172_8446622_RNA45SN1_45S_50ntFlanks@9471.9503.33|AGCACGAGCGCACGTGTTAGGACCCGAAAGATG</td><td>>hg38_21_+_8433172_8446622_RNA45SN1_45S_50ntFlanks</td><td>9471</td><td>9503</td><td>33</td><td>AGCACGAGCGCACGTGTTAGGACCCGAAAGATG</td><td>1218.0</td><td>-3.0</td><td>1.1</td><td>-2.6</td><td>8.5E-03</td><td>1.6E-02</td><td>BT20/MB468</td><td>MP</td></tr>
<tr><td>>hg38_MT_+_598_1651_RNR1_12S_50ntFlanks@720.744.25|AAAGACGTTAGGTCAAGGTGTAGCC</td><td>>hg38_MT_+_598_1651_RNR1_12S_50ntFlanks</td><td>720</td><td>744</td><td>25</td><td>AAAGACGTTAGGTCAAGGTGTAGCC</td><td>122.4</td><td>3.3</td><td>1.3</td><td>2.5</td><td>1.2E-02</td><td>1.6E-02</td><td>MB231/MB468</td><td>MP</td></tr>
<tr><td>>hg38_21_+_8433172_8446622_RNA45SN1_45S_50ntFlanks@10805.10823.19|GAACAATGTAGGTAAGGGA</td><td>>hg38_21_+_8433172_8446622_RNA45SN1_45S_50ntFlanks</td><td>10805</td><td>10823</td><td>19</td><td>GAACAATGTAGGTAAGGGA</td><td>234.8</td><td>-3.4</td><td>1.3</td><td>-2.6</td><td>8.5E-03</td><td>1.6E-02</td><td>BT20/MB468</td><td>MP</td></tr>
<tr><td>>MI0003126|hsa-mir-491&WithFlank&9|+|20716099|20716194@22.44.23&[MIMAT0002807&hsa-miR-491-5p&offsets|0|+1;m-451&9|+|20716120|20716141&offsets|0|+1]|AGTGGGGAACCCTTCCATGAGGA</td><td>>MI0003126|hsa-mir-491&WithFlank&9|+|20716099|20716194</td><td>22</td><td>44</td><td>23</td><td>AGTGGGGAACCCTTCCATGAGGA</td><td>85.1</td><td>4.9</td><td>1.9</td><td>2.5</td><td>1.3E-02</td><td>1.6E-02</td><td>MB231/MB468</td><td>MP</td></tr>
<tr><td>>trnaMT_LysTTT_MT_+_8295_8364@41.72.32,trnalookalike7_LysTTT_1_+_568844_568913@41.72.32|ATTAAGAGAACCAACACCTCTTTACAGTGACC</td><td>>trnaMT_LysTTT_MT_+_8295_8364</td><td>41</td><td>72</td><td>32</td><td>ATTAAGAGAACCAACACCTCTTTACAGTGACC</td><td>63.8</td><td>4.0</td><td>1.6</td><td>2.5</td><td>1.3E-02</td><td>1.6E-02</td><td>MB231/MB468</td><td>MP</td></tr>
<tr><td>>hg38_MT_+_1621_3279_RNR2_16S_50ntFlanks@999.1018.20|AGGGACCTGTATGAATGGCT</td><td>>hg38_MT_+_1621_3279_RNR2_16S_50ntFlanks</td><td>999</td><td>1018</td><td>20</td><td>AGGGACCTGTATGAATGGCT</td><td>167.3</td><td>3.3</td><td>1.2</td><td>2.6</td><td>8.5E-03</td><td>1.6E-02</td><td>BT20/MB468</td><td>MP</td></tr>
<tr><td>>hg38_21_+_8433172_8446622_RNA45SN1_45S_50ntFlanks@11718.11744.27|AACGGCGGGAGTAACTATGACTCTCTT</td><td>>hg38_21_+_8433172_8446622_RNA45SN1_45S_50ntFlanks</td><td>11718</td><td>11744</td><td>27</td><td>AACGGCGGGAGTAACTATGACTCTCTT</td><td>169.0</td><td>-3.8</td><td>1.5</td><td>-2.5</td><td>1.2E-02</td><td>1.6E-02</td><td>BT20/MB468</td><td>MITO</td></tr>
<tr><td>>MI0000264|hsa-mir-7-2&WithFlank&15|+|88611819|88611940@38.60.23(+1C);MI0000265|hsa-mir-7-3&WithFlank&19|+|4770664|4770785@37.59.23(+1C);MI0000263|hsa-mir-7-1&WithFlank&9|-|83969742|83969863@30.52.23(+1C)&[MIMAT0000252&hsa-miR-7-5p&offsets|0|-1(+1C);m-400&15|+|88611856|88611879&offsets|0|-1(+1C)];[MIMAT0000252_1&hsa-miR-7-5p&offsets|0|-1(+1C);m-402&19|+|4770700|4770723&offsets|0|-1(+1C)];[MIMAT0000252_2&hsa-miR-7-5p&offsets|0|-1(+1C);m-401&9|-|83969811|83969834&offsets|0|-1(+1C)]|TGGAAGACTAGTGATTTTGTTGTC</td><td>>MI0000264|hsa-mir-7-2&WithFlank&15|+|88611819|88611940</td><td>38</td><td>60</td><td>23(+1C)</td><td>TGGAAGACTAGTGATTTTGTTGTC</td><td>2010.8</td><td>2.6</td><td>1.0</td><td>2.6</td><td>8.5E-03</td><td>1.6E-02</td><td>BT20/MB468</td><td>MP</td></tr>
<tr><td>>hg38_21_+_8433172_8446622_RNA45SN1_45S_50ntFlanks@10922.10946.25|CTGGACGAGGCGCCGCCGCCCCCCC</td><td>>hg38_21_+_8433172_8446622_RNA45SN1_45S_50ntFlanks</td><td>10922</td><td>10946</td><td>25</td><td>CTGGACGAGGCGCCGCCGCCCCCCC</td><td>592.5</td><td>-3.3</td><td>1.3</td><td>-2.5</td><td>1.2E-02</td><td>1.6E-02</td><td>BT20/MB468</td><td>MITO</td></tr>
<tr><td>>hg38_21_+_8433172_8446622_RNA45SN1_45S_50ntFlanks@4730.4751.22|CAAGAACGAAAGTCGGAGGTTC</td><td>>hg38_21_+_8433172_8446622_RNA45SN1_45S_50ntFlanks</td><td>4730</td><td>4751</td><td>22</td><td>CAAGAACGAAAGTCGGAGGTTC</td><td>184.4</td><td>-2.8</td><td>1.1</td><td>-2.5</td><td>1.2E-02</td><td>1.6E-02</td><td>BT20/MB468</td><td>MITO</td></tr>
<tr><td>>hg38_1_-_228634819_228635039_RNA5S12_5S_50ntFlanks@144.171.28|CCTGGGAATACCGGGTGCTGTAGGCTTT</td><td>>hg38_1_-_228634819_228635039_RNA5S12_5S_50ntFlanks</td><td>144</td><td>171</td><td>28</td><td>CCTGGGAATACCGGGTGCTGTAGGCTTT</td><td>3445.2</td><td>-2.0</td><td>0.8</td><td>-2.6</td><td>8.5E-03</td><td>1.6E-02</td><td>BT20/MB468</td><td>MP</td></tr>
<tr><td>>hg38_21_+_8433172_8446622_RNA45SN1_45S_50ntFlanks@6655.6688.34|TCTTAGCGGTGGATCACTCGGCTCGTGCGTCGAT</td><td>>hg38_21_+_8433172_8446622_RNA45SN1_45S_50ntFlanks</td><td>6655</td><td>6688</td><td>34</td><td>TCTTAGCGGTGGATCACTCGGCTCGTGCGTCGAT</td><td>252.5</td><td>5.1</td><td>1.7</td><td>2.9</td><td>3.4E-03</td><td>1.6E-02</td><td>BT20/MB231</td><td>MP</td></tr>
<tr><td>>hg38_21_+_8433172_8446622_RNA45SN1_45S_50ntFlanks@11572.11597.26|CCGACTTAGAACTGGTGCGGACCAGG</td><td>>hg38_21_+_8433172_8446622_RNA45SN1_45S_50ntFlanks</td><td>11572</td><td>11597</td><td>26</td><td>CCGACTTAGAACTGGTGCGGACCAGG</td><td>478.7</td><td>1.6</td><td>0.6</td><td>2.5</td><td>1.2E-02</td><td>1.6E-02</td><td>BT20/MB468</td><td>MITO</td></tr>
<tr><td>>hg38_21_+_8433172_8446622_RNA45SN1_45S_50ntFlanks@4045.4070.26|CCATTCGAACGTCTGCCCTATCAACT</td><td>>hg38_21_+_8433172_8446622_RNA45SN1_45S_50ntFlanks</td><td>4045</td><td>4070</td><td>26</td><td>CCATTCGAACGTCTGCCCTATCAACT</td><td>162.3</td><td>-2.4</td><td>0.9</td><td>-2.9</td><td>4.3E-03</td><td>1.6E-02</td><td>BT20/MB231</td><td>CYTO</td></tr>
<tr><td>>hg38_21_+_8433172_8446622_RNA45SN1_45S_50ntFlanks@3705.3737.33|TACCTGGTTGATCCTGCCAGTAGCATATGCTTG</td><td>>hg38_21_+_8433172_8446622_RNA45SN1_45S_50ntFlanks</td><td>3705</td><td>3737</td><td>33</td><td>TACCTGGTTGATCCTGCCAGTAGCATATGCTTG</td><td>559.5</td><td>-2.3</td><td>0.8</td><td>-2.9</td><td>4.3E-03</td><td>1.6E-02</td><td>BT20/MB231</td><td>MITO</td></tr>
<tr><td>>hg38_21_+_8433172_8446622_RNA45SN1_45S_50ntFlanks@7977.8012.36|CGACCTCAGATCAGACGTGGCGACCCGCTGAATTTA</td><td>>hg38_21_+_8433172_8446622_RNA45SN1_45S_50ntFlanks</td><td>7977</td><td>8012</td><td>36</td><td>CGACCTCAGATCAGACGTGGCGACCCGCTGAATTTA</td><td>200.4</td><td>-3.6</td><td>1.4</td><td>-2.5</td><td>1.3E-02</td><td>1.6E-02</td><td>MB231/MB468</td><td>MP</td></tr>
<tr><td>>hg38_21_+_8433172_8446622_RNA45SN1_45S_50ntFlanks@7983.8017.35|CAGATCAGACGTGGCGACCCGCTGAATTTAAGCAT</td><td>>hg38_21_+_8433172_8446622_RNA45SN1_45S_50ntFlanks</td><td>7983</td><td>8017</td><td>35</td><td>CAGATCAGACGTGGCGACCCGCTGAATTTAAGCAT</td><td>145.3</td><td>-4.4</td><td>1.8</td><td>-2.5</td><td>1.3E-02</td><td>1.6E-02</td><td>MB231/MB468</td><td>MP</td></tr>
<tr><td>>hg38_21_+_8433172_8446622_RNA45SN1_45S_50ntFlanks@10060.10078.19|GCAGTCGAGAGTGGACGGG</td><td>>hg38_21_+_8433172_8446622_RNA45SN1_45S_50ntFlanks</td><td>10060</td><td>10078</td><td>19</td><td>GCAGTCGAGAGTGGACGGG</td><td>224.8</td><td>4.6</td><td>1.6</td><td>2.9</td><td>3.4E-03</td><td>1.6E-02</td><td>BT20/MB231</td><td>MP</td></tr>
<tr><td>>hg38_21_+_8433172_8446622_RNA45SN1_45S_50ntFlanks@3852.3872.21|TAACTGTGGTAATTCTAGAGC</td><td>>hg38_21_+_8433172_8446622_RNA45SN1_45S_50ntFlanks</td><td>3852</td><td>3872</td><td>21</td><td>TAACTGTGGTAATTCTAGAGC</td><td>148.9</td><td>-1.4</td><td>0.6</td><td>-2.5</td><td>1.3E-02</td><td>1.6E-02</td><td>BT20/MB468</td><td>CYTO</td></tr>
<tr><td>>hg38_21_+_8433172_8446622_RNA45SN1_45S_50ntFlanks@6645.6667.23|CTCGTACGACTCTTAGCGGTGGA</td><td>>hg38_21_+_8433172_8446622_RNA45SN1_45S_50ntFlanks</td><td>6645</td><td>6667</td><td>23</td><td>CTCGTACGACTCTTAGCGGTGGA</td><td>805.4</td><td>1.7</td><td>0.7</td><td>2.5</td><td>1.4E-02</td><td>1.6E-02</td><td>BT20/MB468</td><td>NUC</td></tr>
<tr><td>>MI0000272|hsa-mir-182&WithFlank&7|-|129770377|129770498@29.53.25&[MIMAT0000259&hsa-miR-182-5p&offsets|0|+1;m-22&7|-|129770449|129770470&offsets|0|+3]|TTTGGCAATGGTAGAACTCACACTG</td><td>>MI0000272|hsa-mir-182&WithFlank&7|-|129770377|129770498</td><td>29</td><td>53</td><td>25</td><td>TTTGGCAATGGTAGAACTCACACTG</td><td>1544.7</td><td>2.8</td><td>1.1</td><td>2.6</td><td>8.6E-03</td><td>1.6E-02</td><td>BT20/MB468</td><td>MP</td></tr>
<tr><td>>hg38_21_+_8433172_8446622_RNA45SN1_45S_50ntFlanks@7970.8011.42|CGAGACGCGACCTCAGATCAGACGTGGCGACCCGCTGAATTT</td><td>>hg38_21_+_8433172_8446622_RNA45SN1_45S_50ntFlanks</td><td>7970</td><td>8011</td><td>42</td><td>CGAGACGCGACCTCAGATCAGACGTGGCGACCCGCTGAATTT</td><td>150.3</td><td>2.4</td><td>1.0</td><td>2.5</td><td>1.4E-02</td><td>1.6E-02</td><td>BT20/MB468</td><td>NUC</td></tr>
<tr><td>>hg38_21_+_8433172_8446622_RNA45SN1_45S_50ntFlanks@11672.11692.21|CAGTGCTCTGAATGTCAAAGT</td><td>>hg38_21_+_8433172_8446622_RNA45SN1_45S_50ntFlanks</td><td>11672</td><td>11692</td><td>21</td><td>CAGTGCTCTGAATGTCAAAGT</td><td>307.8</td><td>1.7</td><td>0.7</td><td>2.5</td><td>1.4E-02</td><td>1.6E-02</td><td>MB231/MB468</td><td>CYTO</td></tr>
<tr><td>>am_MI0003772|hsa-mir-151b&WithFlank&14|-|100109413|100109520@66.85.20(+1C);MI0000809|hsa-mir-151a&WithFlank&8|-|140732558|140732659@17.36.20(+1C)&[MIMAT0010214&hsa-miR-151b&offsets|0|+2(+1C);m-179&14|-|100109436|100109455&offsets|0|0(+1C)];[MIMAT0004697&hsa-miR-151a-5p&offsets|0|-1(+1C);m-75&8|-|140732623|140732643&offsets|0|-1(+1C)]|TCGAGGAGCTCACAGTCTAGC</td><td>>am_MI0003772|hsa-mir-151b&WithFlank&14|-|100109413|100109520</td><td>66</td><td>85</td><td>20(+1C)</td><td>TCGAGGAGCTCACAGTCTAGC</td><td>467.2</td><td>2.2</td><td>0.8</td><td>2.9</td><td>3.7E-03</td><td>1.6E-02</td><td>BT20/MB231</td><td>NUC</td></tr>
<tr><td>>hg38_21_+_8433172_8446622_RNA45SN1_45S_50ntFlanks@10781.10809.29|AGGTGAACAGCCTCTGGCATGTTGGAACA</td><td>>hg38_21_+_8433172_8446622_RNA45SN1_45S_50ntFlanks</td><td>10781</td><td>10809</td><td>29</td><td>AGGTGAACAGCCTCTGGCATGTTGGAACA</td><td>171.6</td><td>-2.1</td><td>0.8</td><td>-2.6</td><td>8.6E-03</td><td>1.6E-02</td><td>BT20/MB468</td><td>MP</td></tr>
<tr><td>>hg38_21_+_8433172_8446622_RNA45SN1_45S_50ntFlanks@12393.12414.22|ATAGCGACGTCGCTTTTTGATC</td><td>>hg38_21_+_8433172_8446622_RNA45SN1_45S_50ntFlanks</td><td>12393</td><td>12414</td><td>22</td><td>ATAGCGACGTCGCTTTTTGATC</td><td>227.6</td><td>-2.3</td><td>0.9</td><td>-2.6</td><td>8.6E-03</td><td>1.6E-02</td><td>BT20/MB468</td><td>MP</td></tr>
<tr><td>>hg38_MT_+_1621_3279_RNR2_16S_50ntFlanks@51.88.38|GCTAAACCTAGCCCCAAACCCACTCCACCTTACTACCA</td><td>>hg38_MT_+_1621_3279_RNR2_16S_50ntFlanks</td><td>51</td><td>88</td><td>38</td><td>GCTAAACCTAGCCCCAAACCCACTCCACCTTACTACCA</td><td>363.1</td><td>-2.4</td><td>0.9</td><td>-2.9</td><td>4.3E-03</td><td>1.6E-02</td><td>BT20/MB231</td><td>CYTO</td></tr>
<tr><td>>hg38_21_+_8433172_8446622_RNA45SN1_45S_50ntFlanks@10248.10283.36|CGACGAGTAGGAGGGCCGCTGCGGTGAGCCTTGAAG</td><td>>hg38_21_+_8433172_8446622_RNA45SN1_45S_50ntFlanks</td><td>10248</td><td>10283</td><td>36</td><td>CGACGAGTAGGAGGGCCGCTGCGGTGAGCCTTGAAG</td><td>66.3</td><td>5.4</td><td>2.0</td><td>2.6</td><td>8.6E-03</td><td>1.6E-02</td><td>BT20/MB468</td><td>MP</td></tr>
<tr><td>>trnaMT_GluTTC_MT_-_14674_14742@35.61.27,trnalookalike8_GluTTC_5_-_93905172_93905240@35.61.27|TATCATTGGTCGTGGTTGTAGTCCGTG</td><td>>trnaMT_GluTTC_MT_-_14674_14742</td><td>35</td><td>61</td><td>27</td><td>TATCATTGGTCGTGGTTGTAGTCCGTG</td><td>228.8</td><td>3.0</td><td>1.1</td><td>2.6</td><td>8.6E-03</td><td>1.6E-02</td><td>BT20/MB468</td><td>MP</td></tr>
<tr><td>>hg38_21_+_8433172_8446622_RNA45SN1_45S_50ntFlanks@12907.12933.27|CACCGCACGTTCGTGGGGAACCTGGCG</td><td>>hg38_21_+_8433172_8446622_RNA45SN1_45S_50ntFlanks</td><td>12907</td><td>12933</td><td>27</td><td>CACCGCACGTTCGTGGGGAACCTGGCG</td><td>1101.9</td><td>-3.7</td><td>1.5</td><td>-2.5</td><td>1.4E-02</td><td>1.6E-02</td><td>MB231/MB468</td><td>CYTO</td></tr>
<tr><td>>hg38_MT_+_598_1651_RNR1_12S_50ntFlanks@428.463.36|AAAGTGGCTTTAACATATCTGAACACACAATAGCTA</td><td>>hg38_MT_+_598_1651_RNR1_12S_50ntFlanks</td><td>428</td><td>463</td><td>36</td><td>AAAGTGGCTTTAACATATCTGAACACACAATAGCTA</td><td>164.4</td><td>4.1</td><td>1.6</td><td>2.6</td><td>8.6E-03</td><td>1.6E-02</td><td>BT20/MB468</td><td>MP</td></tr>
<tr><td>>hg38_21_+_8433172_8446622_RNA45SN1_45S_50ntFlanks@10777.10818.42|TCCAAGGTGAACAGCCTCTGGCATGTTGGAACAATGTAGGTA</td><td>>hg38_21_+_8433172_8446622_RNA45SN1_45S_50ntFlanks</td><td>10777</td><td>10818</td><td>42</td><td>TCCAAGGTGAACAGCCTCTGGCATGTTGGAACAATGTAGGTA</td><td>383.1</td><td>-3.5</td><td>1.4</td><td>-2.5</td><td>1.4E-02</td><td>1.6E-02</td><td>BT20/MB468</td><td>CYTO</td></tr>
<tr><td>>hg38_21_+_8433172_8446622_RNA45SN1_45S_50ntFlanks@9310.9338.29|GAGTCTAACACGTGCGCGAGTCGGGGGCT</td><td>>hg38_21_+_8433172_8446622_RNA45SN1_45S_50ntFlanks</td><td>9310</td><td>9338</td><td>29</td><td>GAGTCTAACACGTGCGCGAGTCGGGGGCT</td><td>257.6</td><td>-3.6</td><td>1.4</td><td>-2.6</td><td>8.6E-03</td><td>1.6E-02</td><td>BT20/MB468</td><td>MP</td></tr>
<tr><td>>MI0000736|hsa-mir-30c-1&WithFlank&1|+|40757278|40757378@23.45.23;MI0000254|hsa-mir-30c-2&WithFlank&6|-|71376954|71377037@13.35.23&[MIMAT0000244&hsa-miR-30c-5p&offsets|0|0;m-57&1|+|40757300|40757323&offsets|0|-1];[MIMAT0000244_1&hsa-miR-30c-5p&offsets|0|0;m-56&6|-|71377002|71377025&offsets|0|-1]|TGTAAACATCCTACACTCTCAGC</td><td>>MI0000736|hsa-mir-30c-1&WithFlank&1|+|40757278|40757378</td><td>23</td><td>45</td><td>23</td><td>TGTAAACATCCTACACTCTCAGC</td><td>9572.5</td><td>-2.9</td><td>1.0</td><td>-2.9</td><td>3.4E-03</td><td>1.6E-02</td><td>BT20/MB231</td><td>MP</td></tr>
<tr><td>>hg38_21_+_8433172_8446622_RNA45SN1_45S_50ntFlanks@11576.11594.19|CTTAGAACTGGTGCGGACC</td><td>>hg38_21_+_8433172_8446622_RNA45SN1_45S_50ntFlanks</td><td>11576</td><td>11594</td><td>19</td><td>CTTAGAACTGGTGCGGACC</td><td>332.9</td><td>-2.1</td><td>0.7</td><td>-2.9</td><td>4.3E-03</td><td>1.6E-02</td><td>BT20/MB231</td><td>CYTO</td></tr>
<tr><td>>hg38_21_+_8433172_8446622_RNA45SN1_45S_50ntFlanks@11199.11223.25|CCGGCGGCGGCGGCGACTCTGGACG</td><td>>hg38_21_+_8433172_8446622_RNA45SN1_45S_50ntFlanks</td><td>11199</td><td>11223</td><td>25</td><td>CCGGCGGCGGCGGCGACTCTGGACG</td><td>353.8</td><td>-4.6</td><td>1.7</td><td>-2.6</td><td>8.6E-03</td><td>1.6E-02</td><td>BT20/MB468</td><td>MP</td></tr>
<tr><td>>hg38_21_+_8433172_8446622_RNA45SN1_45S_50ntFlanks@12296.12323.28|AGCGGGGCCTCACGATCCTTCTGACCTT</td><td>>hg38_21_+_8433172_8446622_RNA45SN1_45S_50ntFlanks</td><td>12296</td><td>12323</td><td>28</td><td>AGCGGGGCCTCACGATCCTTCTGACCTT</td><td>146.6</td><td>-4.2</td><td>1.6</td><td>-2.5</td><td>1.2E-02</td><td>1.6E-02</td><td>BT20/MB468</td><td>MITO</td></tr>
<tr><td>>trnaMT_SerGCT_MT_+_12207_12265@11.40.30|ACAAGAACTGCTAACTCATGCCCCCATGTC</td><td>>trnaMT_SerGCT_MT_+_12207_12265</td><td>11</td><td>40</td><td>30</td><td>ACAAGAACTGCTAACTCATGCCCCCATGTC</td><td>102.4</td><td>4.1</td><td>1.5</td><td>2.6</td><td>8.6E-03</td><td>1.6E-02</td><td>BT20/MB468</td><td>MP</td></tr>
<tr><td>>hg38_21_+_8433172_8446622_RNA45SN1_45S_50ntFlanks@10402.10432.31|AACATGGGTCAGTCGGTCCTGAGAGATGGGC</td><td>>hg38_21_+_8433172_8446622_RNA45SN1_45S_50ntFlanks</td><td>10402</td><td>10432</td><td>31</td><td>AACATGGGTCAGTCGGTCCTGAGAGATGGGC</td><td>158.5</td><td>6.0</td><td>2.0</td><td>2.9</td><td>3.4E-03</td><td>1.6E-02</td><td>BT20/MB231</td><td>MP</td></tr>
<tr><td>>hg38_21_+_8433172_8446622_RNA45SN1_45S_50ntFlanks@10788.10826.39|CAGCCTCTGGCATGTTGGAACAATGTAGGTAAGGGAAGT</td><td>>hg38_21_+_8433172_8446622_RNA45SN1_45S_50ntFlanks</td><td>10788</td><td>10826</td><td>39</td><td>CAGCCTCTGGCATGTTGGAACAATGTAGGTAAGGGAAGT</td><td>242.6</td><td>-1.6</td><td>0.7</td><td>-2.5</td><td>1.2E-02</td><td>1.6E-02</td><td>BT20/MB468</td><td>MITO</td></tr>
<tr><td>>hg38_21_+_8433172_8446622_RNA45SN1_45S_50ntFlanks@8340.8373.34|ACCGTAAGGGAAAGTTGAAAAGAACTTTGAAGAG</td><td>>hg38_21_+_8433172_8446622_RNA45SN1_45S_50ntFlanks</td><td>8340</td><td>8373</td><td>34</td><td>ACCGTAAGGGAAAGTTGAAAAGAACTTTGAAGAG</td><td>554.5</td><td>5.2</td><td>1.8</td><td>2.9</td><td>3.4E-03</td><td>1.6E-02</td><td>BT20/MB231</td><td>MP</td></tr>
<tr><td>>hg38_21_+_8433172_8446622_RNA45SN1_45S_50ntFlanks@9496.9520.25|GAAAGATGGTGAACTATGCCTGGGC</td><td>>hg38_21_+_8433172_8446622_RNA45SN1_45S_50ntFlanks</td><td>9496</td><td>9520</td><td>25</td><td>GAAAGATGGTGAACTATGCCTGGGC</td><td>136.9</td><td>4.6</td><td>1.7</td><td>2.6</td><td>8.6E-03</td><td>1.6E-02</td><td>BT20/MB468</td><td>MP</td></tr>
<tr><td>>trnaMT_MetCAT_MT_+_4402_4469@34.71.38|ACCCCGAAAATGTTGGTTATACCCTTCCCGTACTACCA</td><td>>trnaMT_MetCAT_MT_+_4402_4469</td><td>34</td><td>71</td><td>38</td><td>ACCCCGAAAATGTTGGTTATACCCTTCCCGTACTACCA</td><td>373.4</td><td>-4.2</td><td>1.5</td><td>-2.9</td><td>3.7E-03</td><td>1.6E-02</td><td>BT20/MB231</td><td>NUC</td></tr>
<tr><td>>MI0000266|hsa-mir-10a&WithFlank&17|-|48579832|48579953@28.49.22&[MIMAT0000253&hsa-miR-10a-5p&offsets|0|-1;m-18&17|-|48579905|48579926&offsets|0|0]|TACCCTGTAGATCCGAATTTGT</td><td>>MI0000266|hsa-mir-10a&WithFlank&17|-|48579832|48579953</td><td>28</td><td>49</td><td>22</td><td>TACCCTGTAGATCCGAATTTGT</td><td>33007.7</td><td>2.2</td><td>0.8</td><td>2.9</td><td>4.4E-03</td><td>1.6E-02</td><td>BT20/MB231</td><td>CYTO</td></tr>
<tr><td>>hg38_21_+_8433172_8446622_RNA45SN1_45S_50ntFlanks@11209.11229.21|CGGCGACTCTGGACGCGAGCC</td><td>>hg38_21_+_8433172_8446622_RNA45SN1_45S_50ntFlanks</td><td>11209</td><td>11229</td><td>21</td><td>CGGCGACTCTGGACGCGAGCC</td><td>921.9</td><td>2.1</td><td>0.8</td><td>2.5</td><td>1.2E-02</td><td>1.6E-02</td><td>BT20/MB468</td><td>MITO</td></tr>
<tr><td>>hg38_1_-_228634819_228635039_RNA5S12_5S_50ntFlanks@153.170.18|ACCGGGTGCTGTAGGCTT</td><td>>hg38_1_-_228634819_228635039_RNA5S12_5S_50ntFlanks</td><td>153</td><td>170</td><td>18</td><td>ACCGGGTGCTGTAGGCTT</td><td>6728.0</td><td>3.2</td><td>1.3</td><td>2.5</td><td>1.4E-02</td><td>1.6E-02</td><td>BT20/MB468</td><td>NUC</td></tr>
<tr><td>>hg38_21_+_8433172_8446622_RNA45SN1_45S_50ntFlanks@7983.8004.22|CAGATCAGACGTGGCGACCCGC</td><td>>hg38_21_+_8433172_8446622_RNA45SN1_45S_50ntFlanks</td><td>7983</td><td>8004</td><td>22</td><td>CAGATCAGACGTGGCGACCCGC</td><td>234.2</td><td>-2.9</td><td>1.2</td><td>-2.5</td><td>1.4E-02</td><td>1.6E-02</td><td>BT20/MB468</td><td>CYTO</td></tr>
</table>

<table>
<tr><th></th><th></th><th></th><th></th><th></th><th></th><th></th><th></th><th></th><th></th><th></th><th></th><th></th><th></th></tr>
<tr><td>>hg38_21_+_8433172_8446622_RNA45SN1_45S_50ntFlanks@11999.12038.40|CCCTGCGGGCCGCCGGTGAAATACCACTACTCTGATCGTT</td><td>>hg38_21_+_8433172_8446622_RNA45SN1_45S_50ntFlanks</td><td>11999</td><td>12038</td><td>40</td><td>CCCTGCGGGCCGCCGGTGAAATACCACTACTCTGATCGTT</td><td>215.0</td><td>-3.1</td><td>1.2</td><td>-2.5</td><td>1.2E-02</td><td>1.6E-02</td><td>BT20/MB468</td><td>MITO</td></tr>
<tr><td>>hg38_21_+_8433172_8446622_RNA45SN1_45S_50ntFlanks@9457.9486.30|CGCCGGGGAGGTGGAGCACGAGCGCACGTG</td><td>>hg38_21_+_8433172_8446622_RNA45SN1_45S_50ntFlanks</td><td>9457</td><td>9486</td><td>30</td><td>CGCCGGGGAGGTGGAGCACGAGCGCACGTG</td><td>99.7</td><td>-4.4</td><td>1.7</td><td>-2.6</td><td>8.6E-03</td><td>1.6E-02</td><td>BT20/MB468</td><td>MP</td></tr>
<tr><td>>hg38_21_+_8433172_8446622_RNA45SN1_45S_50ntFlanks@6649.6680.32|TACGACTCTTAGCGGTGGATCACTCGGCTCGT</td><td>>hg38_21_+_8433172_8446622_RNA45SN1_45S_50ntFlanks</td><td>6649</td><td>6680</td><td>32</td><td>TACGACTCTTAGCGGTGGATCACTCGGCTCGT</td><td>1079.8</td><td>1.3</td><td>0.5</td><td>2.5</td><td>1.2E-02</td><td>1.6E-02</td><td>BT20/MB468</td><td>MITO</td></tr>
<tr><td>>hg38_21_+_8433172_8446622_RNA45SN1_45S_50ntFlanks@4787.4812.26|AACGATGCCGACCGGCGATGCGGCGG</td><td>>hg38_21_+_8433172_8446622_RNA45SN1_45S_50ntFlanks</td><td>4787</td><td>4812</td><td>26</td><td>AACGATGCCGACCGGCGATGCGGCGG</td><td>124.0</td><td>-4.6</td><td>1.7</td><td>-2.6</td><td>8.6E-03</td><td>1.6E-02</td><td>BT20/MB468</td><td>MP</td></tr>
<tr><td>>hg38_21_+_8433172_8446622_RNA45SN1_45S_50ntFlanks@7975.8006.32|CGCGACCTCAGATCAGACGTGGCGACCCGCTG</td><td>>hg38_21_+_8433172_8446622_RNA45SN1_45S_50ntFlanks</td><td>7975</td><td>8006</td><td>32</td><td>CGCGACCTCAGATCAGACGTGGCGACCCGCTG</td><td>99932.0</td><td>-1.6</td><td>0.6</td><td>-2.5</td><td>1.4E-02</td><td>1.6E-02</td><td>BT20/MB468</td><td>NUC</td></tr>
<tr><td>>hg38_21_+_8433172_8446622_RNA45SN1_45S_50ntFlanks@5472.5515.44|ACGGCCCTGGCGGAGCGCTGAGAAGACGGTCGAACTTGACTATC</td><td>>hg38_21_+_8433172_8446622_RNA45SN1_45S_50ntFlanks</td><td>5472</td><td>5515</td><td>44</td><td>ACGGCCCTGGCGGAGCGCTGAGAAGACGGTCGAACTTGACTATC</td><td>106.8</td><td>5.5</td><td>2.1</td><td>2.6</td><td>8.6E-03</td><td>1.6E-02</td><td>BT20/MB468</td><td>MP</td></tr>
<tr><td>>trnaMT_LeuTAA_MT_+_3230_3304@1.35.35|GTTAAGATGGCAGAGCCCGGTAATCGCATAAAACT</td><td>>trnaMT_LeuTAA_MT_+_3230_3304</td><td>1</td><td>35</td><td>35</td><td>GTTAAGATGGCAGAGCCCGGTAATCGCATAAAACT</td><td>214.9</td><td>3.8</td><td>1.3</td><td>2.9</td><td>3.7E-03</td><td>1.6E-02</td><td>BT20/MB231</td><td>NUC</td></tr>
<tr><td>>hg38_21_+_8433172_8446622_RNA45SN1_45S_50ntFlanks@12418.12445.28|CGATGTCGGCTCTTCCTATCATTGTGAA</td><td>>hg38_21_+_8433172_8446622_RNA45SN1_45S_50ntFlanks</td><td>12418</td><td>12445</td><td>28</td><td>CGATGTCGGCTCTTCCTATCATTGTGAA</td><td>656.3</td><td>-1.2</td><td>0.5</td><td>-2.5</td><td>1.2E-02</td><td>1.6E-02</td><td>BT20/MB468</td><td>MITO</td></tr>
<tr><td>>hg38_21_+_8433172_8446622_RNA45SN1_45S_50ntFlanks@5078.5097.20|CGATAACGAACGAGACTCTG</td><td>>hg38_21_+_8433172_8446622_RNA45SN1_45S_50ntFlanks</td><td>5078</td><td>5097</td><td>20</td><td>CGATAACGAACGAGACTCTG</td><td>334.6</td><td>-2.0</td><td>0.8</td><td>-2.5</td><td>1.2E-02</td><td>1.6E-02</td><td>BT20/MB468</td><td>MITO</td></tr>
<tr><td>>hg38_21_+_8433172_8446622_RNA45SN1_45S_50ntFlanks@12416.12433.18|TTCGATGTCGGCTCTTCC</td><td>>hg38_21_+_8433172_8446622_RNA45SN1_45S_50ntFlanks</td><td>12416</td><td>12433</td><td>18</td><td>TTCGATGTCGGCTCTTCC</td><td>226.9</td><td>1.7</td><td>0.7</td><td>2.4</td><td>1.4E-02</td><td>1.6E-02</td><td>MB231/MB468</td><td>MITO</td></tr>
<tr><td>>hg38_21_+_8433172_8446622_RNA45SN1_45S_50ntFlanks@12624.12669.46|CCAATGGGGCGAAGCTACCATCTGTGGGATTATGACTGAACGCCTC</td><td>>hg38_21_+_8433172_8446622_RNA45SN1_45S_50ntFlanks</td><td>12624</td><td>12669</td><td>46</td><td>CCAATGGGGCGAAGCTACCATCTGTGGGATTATGACTGAACGCCTC</td><td>489.0</td><td>-2.6</td><td>1.1</td><td>-2.4</td><td>1.4E-02</td><td>1.6E-02</td><td>MB231/MB468</td><td>MITO</td></tr>
<tr><td>>MI0001445|hsa-mir-423&WithFlank&17|+|30117073|30117178@59.77.19(+2U)&[MIMAT0001340&hsa-miR-423-3p&offsets|0|-4(+2U);m-100&17|+|30117131|30117153&offsets|0|-4(+2U)]|AGCTCGGTCTGAGGCCCCTTT</td><td>>MI0001445|hsa-mir-423&WithFlank&17|+|30117073|30117178</td><td>59</td><td>77</td><td>19(+2U)</td><td>AGCTCGGTCTGAGGCCCCTTT</td><td>32.7</td><td>5.0</td><td>2.0</td><td>2.5</td><td>1.3E-02</td><td>1.6E-02</td><td>MB231/MB468</td><td>MP</td></tr>
<tr><td>>hg38_MT_+_1621_3279_RNR2_16S_50ntFlanks@1216.1248.33|CAACCTCCGAGCAGTACATGCTAAGACTTCACC</td><td>>hg38_MT_+_1621_3279_RNR2_16S_50ntFlanks</td><td>1216</td><td>1248</td><td>33</td><td>CAACCTCCGAGCAGTACATGCTAAGACTTCACC</td><td>327.4</td><td>2.5</td><td>1.0</td><td>2.5</td><td>1.3E-02</td><td>1.6E-02</td><td>MB231/MB468</td><td>MP</td></tr>
<tr><td>>hg38_21_+_8433172_8446622_RNA45SN1_45S_50ntFlanks@10771.10790.20|CAGGTCTCCAAGGTGAACAG</td><td>>hg38_21_+_8433172_8446622_RNA45SN1_45S_50ntFlanks</td><td>10771</td><td>10790</td><td>20</td><td>CAGGTCTCCAAGGTGAACAG</td><td>92.8</td><td>-4.3</td><td>1.6</td><td>-2.6</td><td>8.7E-03</td><td>1.6E-02</td><td>BT20/MB468</td><td>MP</td></tr>
<tr><td>>MI0000542|hsa-mir-320a&WithFlank&8|-|22244960|22245043@48.68.21&[MIMAT0000510&hsa-miR-320a-3p&offsets|0|-1;m-60&8|-|22244975|22244996&offsets|0|-1]|AAAAGCTGGGTTGAGAGGGCG</td><td>>MI0000542|hsa-mir-320a&WithFlank&8|-|22244960|22245043</td><td>48</td><td>68</td><td>21</td><td>AAAAGCTGGGTTGAGAGGGCG</td><td>3121.0</td><td>1.9</td><td>0.7</td><td>2.9</td><td>4.3E-03</td><td>1.6E-02</td><td>BT20/MB231</td><td>MITO</td></tr>
<tr><td>>trna116_GluCTC_1_-_145399233_145399304@1.37.37,trna59_GluCTC_1_+_249168447_249168518@1.37.37,trna71_GluCTC_1_-_161439189_161439260@1.37.37,trna74_GluCTC_1_-_161431809_161431880@1.37.37,trna77_GluCTC_1_-_161424398_161424469@1.37.37,trna77_GluCTC_6_+_28949976_28950047@1.37.37,trna80_GluCTC_1_-_161417018_161417089@1.37.37,trna87_GluCTC_6_-_126101393_126101464@1.37.37|TCCCTGGTGGTCTAGTGGTTAGGATTCGGCGCTCTCA</td><td>>trna116_GluCTC_1_-_145399233_145399304</td><td>1</td><td>37</td><td>37</td><td>TCCCTGGTGGTCTAGTGGTTAGGATTCGGCGCTCTCA</td><td>284.2</td><td>2.3</td><td>0.8</td><td>2.9</td><td>4.4E-03</td><td>1.6E-02</td><td>BT20/MB231</td><td>MITO</td></tr>
<tr><td>>hg38_21_+_8433172_8446622_RNA45SN1_45S_50ntFlanks@11565.11611.47|CTAGCAGCCGACTTAGAACTGGTGCGGACCAGGGGAATCCGACTGTT</td><td>>hg38_21_+_8433172_8446622_RNA45SN1_45S_50ntFlanks</td><td>11565</td><td>11611</td><td>47</td><td>CTAGCAGCCGACTTAGAACTGGTGCGGACCAGGGGAATCCGACTGTT</td><td>2351.6</td><td>3.4</td><td>1.2</td><td>2.9</td><td>4.4E-03</td><td>1.6E-02</td><td>BT20/MB231</td><td>MITO</td></tr>
<tr><td>>MI0000074|hsa-mir-19b-1&WithFlank&13|+|91351186|91351284@60.81.22;MI0000075|hsa-mir-19b-2&WithFlank&X|-|134169665|134169772@68.89.22&[MIMAT0000074&hsa-miR-19b-3p&offsets|0|-1;m-122&13|+|91351245|91351267&offsets|0|-1];[MIMAT0000074_1&hsa-miR-19b-3p&offsets|0|-1;m-121&X|-|134169683|134169705&offsets|0|-1]|TGTGCAAATCCATGCAAAACTG</td><td>>MI0000074|hsa-mir-19b-1&WithFlank&13|+|91351186|91351284</td><td>60</td><td>81</td><td>22</td><td>TGTGCAAATCCATGCAAAACTG</td><td>621.2</td><td>-2.6</td><td>0.9</td><td>-2.9</td><td>4.3E-03</td><td>1.6E-02</td><td>BT20/MB231</td><td>MITO</td></tr>
<tr><td>>MI0000085|hsa-mir-27a&WithFlank&19|-|13836434|13836523@57.75.19&[MIMAT0000084&hsa-miR-27a-3p&offsets|0|-2;m-43&19|-|13836447|13836467&offsets|0|-2]|TTCACAGTGGCTAAGTTCC</td><td>>MI0000085|hsa-mir-27a&WithFlank&19|-|13836434|13836523</td><td>57</td><td>75</td><td>19</td><td>TTCACAGTGGCTAAGTTCC</td><td>866.0</td><td>-2.1</td><td>0.7</td><td>-2.9</td><td>4.4E-03</td><td>1.6E-02</td><td>BT20/MB231</td><td>MITO</td></tr>
<tr><td>>hg38_21_+_8433172_8446622_RNA45SN1_45S_50ntFlanks@6647.6675.29|CGTACGACTCTTAGCGGTGGATCACTCGG</td><td>>hg38_21_+_8433172_8446622_RNA45SN1_45S_50ntFlanks</td><td>6647</td><td>6675</td><td>29</td><td>CGTACGACTCTTAGCGGTGGATCACTCGG</td><td>573.8</td><td>4.0</td><td>1.4</td><td>2.9</td><td>3.4E-03</td><td>1.6E-02</td><td>BT20/MB231</td><td>MP</td></tr>
<tr><td>>am_trna119_LysCTT_1_-_145395522_145395594@6.36.31,trna11_LysCTT_5_-_180648979_180649051@6.36.31,trna128_LysCTT_1_-_55423542_55423614@6.36.31,trna13_LysCTT_6_+_26556774_26556846@6.36.31,trna6_LysCTT_19_-_52425396_52425468@6.36.31,trna7_LysCTT_16_+_3225692_3225764@6.36.31,trna9_LysCTT_5_+_180634755_180634827@6.36.31|GCTAGCTCAGTCGGTAGAGCATGAGACTCTT</td><td>>am_trna119_LysCTT_1_-_145395522_145395594</td><td>6</td><td>36</td><td>31</td><td>GCTAGCTCAGTCGGTAGAGCATGAGACTCTT</td><td>52.1</td><td>4.8</td><td>1.9</td><td>2.5</td><td>1.3E-02</td><td>1.6E-02</td><td>MB231/MB468</td><td>MP</td></tr>
<tr><td>>hg38_21_+_8433172_8446622_RNA45SN1_45S_50ntFlanks@12539.12580.42|ATGATGTGTTGTTGCCATGGTAATCCTGCTCAGTACGAGAGG</td><td>>hg38_21_+_8433172_8446622_RNA45SN1_45S_50ntFlanks</td><td>12539</td><td>12580</td><td>42</td><td>ATGATGTGTTGTTGCCATGGTAATCCTGCTCAGTACGAGAGG</td><td>172.5</td><td>-4.4</td><td>1.7</td><td>-2.5</td><td>1.2E-02</td><td>1.6E-02</td><td>BT20/MB468</td><td>MITO</td></tr>
<tr><td>>hg38_21_+_8433172_8446622_RNA45SN1_45S_50ntFlanks@10349.10388.40|AACGAGAACTTTGAAGGCCGAAGTGGAGAAGGGTTCCATG</td><td>>hg38_21_+_8433172_8446622_RNA45SN1_45S_50ntFlanks</td><td>10349</td><td>10388</td><td>40</td><td>AACGAGAACTTTGAAGGCCGAAGTGGAGAAGGGTTCCATG</td><td>738.8</td><td>-2.4</td><td>0.9</td><td>-2.6</td><td>8.7E-03</td><td>1.6E-02</td><td>BT20/MB468</td><td>MP</td></tr>
<tr><td>>hg38_21_+_8433172_8446622_RNA45SN1_45S_50ntFlanks@10684.10705.22|CGGCGGCGTCCGGTGAGCTCTC</td><td>>hg38_21_+_8433172_8446622_RNA45SN1_45S_50ntFlanks</td><td>10684</td><td>10705</td><td>22</td><td>CGGCGGCGTCCGGTGAGCTCTC</td><td>236.4</td><td>-2.6</td><td>1.1</td><td>-2.5</td><td>1.4E-02</td><td>1.6E-02</td><td>BT20/MB468</td><td>CYTO</td></tr>
<tr><td>>hg38_21_+_8433172_8446622_RNA45SN1_45S_50ntFlanks@3854.3873.20|ACTGTGGTAATTCTAGAGCT</td><td>>hg38_21_+_8433172_8446622_RNA45SN1_45S_50ntFlanks</td><td>3854</td><td>3873</td><td>20</td><td>ACTGTGGTAATTCTAGAGCT</td><td>302.8</td><td>-3.2</td><td>1.3</td><td>-2.5</td><td>1.2E-02</td><td>1.6E-02</td><td>BT20/MB468</td><td>MITO</td></tr>
<tr><td>>hg38_21_+_8433172_8446622_RNA45SN1_45S_50ntFlanks@8120.8141.22|GACATGTGGCGTACGGAAGACC</td><td>>hg38_21_+_8433172_8446622_RNA45SN1_45S_50ntFlanks</td><td>8120</td><td>8141</td><td>22</td><td>GACATGTGGCGTACGGAAGACC</td><td>91.8</td><td>4.6</td><td>1.8</td><td>2.6</td><td>8.7E-03</td><td>1.6E-02</td><td>BT20/MB468</td><td>MP</td></tr>
<tr><td>>hg38_21_+_8433172_8446622_RNA45SN1_45S_50ntFlanks@10878.10904.27|TCGGTCGGGCTGGGGCGCGAAGCGGGG</td><td>>hg38_21_+_8433172_8446622_RNA45SN1_45S_50ntFlanks</td><td>10878</td><td>10904</td><td>27</td><td>TCGGTCGGGCTGGGGCGCGAAGCGGGG</td><td>211.9</td><td>-4.0</td><td>1.6</td><td>-2.4</td><td>1.4E-02</td><td>1.6E-02</td><td>MB231/MB468</td><td>NUC</td></tr>
<tr><td>>hg38_21_+_8433172_8446622_RNA45SN1_45S_50ntFlanks@11315.11340.26|CCACCCCACGTCTCGTCGCGCGCGCG</td><td>>hg38_21_+_8433172_8446622_RNA45SN1_45S_50ntFlanks</td><td>11315</td><td>11340</td><td>26</td><td>CCACCCCACGTCTCGTCGCGCGCGCG</td><td>536.3</td><td>-4.9</td><td>2.0</td><td>-2.5</td><td>1.4E-02</td><td>1.6E-02</td><td>BT20/MB468</td><td>NUC</td></tr>
<tr><td>>hg38_21_+_8433172_8446622_RNA45SN1_45S_50ntFlanks@11873.11907.35|AAGAAGACCCTGTTGAGCTTGACTCTAGTCTGGCA</td><td>>hg38_21_+_8433172_8446622_RNA45SN1_45S_50ntFlanks</td><td>11873</td><td>11907</td><td>35</td><td>AAGAAGACCCTGTTGAGCTTGACTCTAGTCTGGCA</td><td>256.6</td><td>-4.5</td><td>1.7</td><td>-2.6</td><td>8.7E-03</td><td>1.6E-02</td><td>BT20/MB468</td><td>MP</td></tr>
<tr><td>>hg38_21_+_8433172_8446622_RNA45SN1_45S_50ntFlanks@12416.12436.21|TTCGATGTCGGCTCTTCCTAT</td><td>>hg38_21_+_8433172_8446622_RNA45SN1_45S_50ntFlanks</td><td>12416</td><td>12436</td><td>21</td><td>TTCGATGTCGGCTCTTCCTAT</td><td>1280.5</td><td>-1.5</td><td>0.6</td><td>-2.5</td><td>1.4E-02</td><td>1.6E-02</td><td>BT20/MB468</td><td>CYTO</td></tr>
<tr><td>>hg38_21_+_8433172_8446622_RNA45SN1_45S_50ntFlanks@8436.8467.32|GAGGATTCAACCCGGCGGCGGGTCCGGCCGTG</td><td>>hg38_21_+_8433172_8446622_RNA45SN1_45S_50ntFlanks</td><td>8436</td><td>8467</td><td>32</td><td>GAGGATTCAACCCGGCGGCGGGTCCGGCCGTG</td><td>60.7</td><td>-4.5</td><td>1.7</td><td>-2.6</td><td>8.7E-03</td><td>1.6E-02</td><td>BT20/MB468</td><td>MP</td></tr>
<tr><td>>hg38_21_+_8433172_8446622_RNA45SN1_45S_50ntFlanks@6650.6673.24|ACGACTCTTAGCGGTGGATCACTC</td><td>>hg38_21_+_8433172_8446622_RNA45SN1_45S_50ntFlanks</td><td>6650</td><td>6673</td><td>24</td><td>ACGACTCTTAGCGGTGGATCACTC</td><td>120.7</td><td>2.0</td><td>0.8</td><td>2.5</td><td>1.4E-02</td><td>1.6E-02</td><td>BT20/MB468</td><td>NUC</td></tr>
<tr><td>>hg38_21_+_8433172_8446622_RNA45SN1_45S_50ntFlanks@8157.8185.29|CTCGTGGGGGGCCCAAGTCCTTCTGATCG</td><td>>hg38_21_+_8433172_8446622_RNA45SN1_45S_50ntFlanks</td><td>8157</td><td>8185</td><td>29</td><td>CTCGTGGGGGGCCCAAGTCCTTCTGATCG</td><td>250.6</td><td>-3.5</td><td>1.3</td><td>-2.6</td><td>8.7E-03</td><td>1.6E-02</td><td>BT20/MB468</td><td>MP</td></tr>
<tr><td>>MI0000272|hsa-mir-182&WithFlank&7|-|129770377|129770498@29.50.22&[MIMAT0000259&hsa-miR-182-5p&offsets|0|-2;m-22&7|-|129770449|129770470&offsets|0|0]|TTTGGCAATGGTAGAACTCACA</td><td>>MI0000272|hsa-mir-182&WithFlank&7|-|129770377|129770498</td><td>29</td><td>50</td><td>22</td><td>TTTGGCAATGGTAGAACTCACA</td><td>1496.4</td><td>2.0</td><td>0.7</td><td>2.8</td><td>4.4E-03</td><td>1.6E-02</td><td>BT20/MB231</td><td>MITO</td></tr>
<tr><td>>hg38_21_+_8433172_8446622_RNA45SN1_45S_50ntFlanks@10685.10708.24|GGCGGCGTCCGGTGAGCTCTCGCT</td><td>>hg38_21_+_8433172_8446622_RNA45SN1_45S_50ntFlanks</td><td>10685</td><td>10708</td><td>24</td><td>GGCGGCGTCCGGTGAGCTCTCGCT</td><td>221.5</td><td>2.6</td><td>1.0</td><td>2.6</td><td>8.7E-03</td><td>1.6E-02</td><td>BT20/MB468</td><td>MP</td></tr>
<tr><td>>hg38_21_+_8433172_8446622_RNA45SN1_45S_50ntFlanks@4602.4622.21|TTCGGAACTGAGGCCATGATT</td><td>>hg38_21_+_8433172_8446622_RNA45SN1_45S_50ntFlanks</td><td>4602</td><td>4622</td><td>21</td><td>TTCGGAACTGAGGCCATGATT</td><td>94.8</td><td>-4.3</td><td>1.6</td><td>-2.6</td><td>8.7E-03</td><td>1.6E-02</td><td>BT20/MB468</td><td>MP</td></tr>
<tr><td>>hg38_21_+_8433172_8446622_RNA45SN1_45S_50ntFlanks@12275.12293.19|AGTACGAATACAGACCGTG</td><td>>hg38_21_+_8433172_8446622_RNA45SN1_45S_50ntFlanks</td><td>12275</td><td>12293</td><td>19</td><td>AGTACGAATACAGACCGTG</td><td>84.0</td><td>-4.7</td><td>1.8</td><td>-2.6</td><td>8.7E-03</td><td>1.6E-02</td><td>BT20/MB468</td><td>MP</td></tr>
<tr><td>>hg38_21_+_8433172_8446622_RNA45SN1_45S_50ntFlanks@11315.11334.20|CCACCCCACGTCTCGTCGCG</td><td>>hg38_21_+_8433172_8446622_RNA45SN1_45S_50ntFlanks</td><td>11315</td><td>11334</td><td>20</td><td>CCACCCCACGTCTCGTCGCG</td><td>1006.6</td><td>-5.3</td><td>2.2</td><td>-2.5</td><td>1.4E-02</td><td>1.6E-02</td><td>BT20/MB468</td><td>NUC</td></tr>
<tr><td>>hg38_21_+_8433172_8446622_RNA45SN1_45S_50ntFlanks@12626.12657.32|AATGGGGCGAAGCTACCATCTGTGGGATTATG</td><td>>hg38_21_+_8433172_8446622_RNA45SN1_45S_50ntFlanks</td><td>12626</td><td>12657</td><td>32</td><td>AATGGGGCGAAGCTACCATCTGTGGGATTATG</td><td>319.1</td><td>-2.6</td><td>1.1</td><td>-2.5</td><td>1.4E-02</td><td>1.6E-02</td><td>MB231/MB468</td><td>CYTO</td></tr>
<tr><td>>hg38_MT_+_1621_3279_RNR2_16S_50ntFlanks@1209.1231.23|CAGAACCCAACCTCCGAGCAGTA</td><td>>hg38_MT_+_1621_3279_RNR2_16S_50ntFlanks</td><td>1209</td><td>1231</td><td>23</td><td>CAGAACCCAACCTCCGAGCAGTA</td><td>204.5</td><td>-4.2</td><td>1.7</td><td>-2.5</td><td>1.2E-02</td><td>1.6E-02</td><td>BT20/MB468</td><td>MITO</td></tr>
<tr><td>>hg38_21_+_8433172_8446622_RNA45SN1_45S_50ntFlanks@8114.8134.21|GCGCGGGACATGTGGCGTACG</td><td>>hg38_21_+_8433172_8446622_RNA45SN1_45S_50ntFlanks</td><td>8114</td><td>8134</td><td>21</td><td>GCGCGGGACATGTGGCGTACG</td><td>351.1</td><td>-2.1</td><td>0.8</td><td>-2.6</td><td>8.7E-03</td><td>1.6E-02</td><td>BT20/MB468</td><td>MP</td></tr>
<tr><td>>hg38_21_+_8433172_8446622_RNA45SN1_45S_50ntFlanks@11573.11600.28|CGACTTAGAACTGGTGCGGACCAGGGGA</td><td>>hg38_21_+_8433172_8446622_RNA45SN1_45S_50ntFlanks</td><td>11573</td><td>11600</td><td>28</td><td>CGACTTAGAACTGGTGCGGACCAGGGGA</td><td>513.4</td><td>4.8</td><td>1.6</td><td>2.9</td><td>3.5E-03</td><td>1.6E-02</td><td>BT20/MB231</td><td>MP</td></tr>
<tr><td>>hg38_21_+_8433172_8446622_RNA45SN1_45S_50ntFlanks@5156.5185.30|ACAAGTGGCGTTCAGCCACCCGAGATTGAG</td><td>>hg38_21_+_8433172_8446622_RNA45SN1_45S_50ntFlanks</td><td>5156</td><td>5185</td><td>30</td><td>ACAAGTGGCGTTCAGCCACCCGAGATTGAG</td><td>88.8</td><td>-4.6</td><td>1.7</td><td>-2.6</td><td>8.7E-03</td><td>1.6E-02</td><td>BT20/MB468</td><td>MP</td></tr>
</table>

<table>
<tr><td>>trna10_AspGTC_12_-_125424193_125424264@37.72.36,trna12_AspGTC_12_-_125411891_125411962@37.72.36,trna144_AspGTC_6_-_27551236_27551307@37.72.36,trna38_AspGTC_17_-_8125556_8125627@37.72.36,trna45_AspGTC_6_+_27447453_27447524@37.72.36,trna48_AspGTC_6_+_27471523_27471594@37.72.36,trna4_AspGTC_12_+_96429799_96429870@37.72.36,trna69_AspGTC_1_-_161440205_161440276@37.72.36,trna72_AspGTC_1_-_161432824_161432895@37.72.36,trna75_AspGTC_1_-_161425414_161425485@37.72.36,trna78_AspGTC_1_-_161418033_161418104@37.72.36,trna81_AspGTC_1_-_161410615_161410686@37.72.36|ACGCGGGAGACCGGGGTTCGATTCCCCGACGGGGAG</td><td>>trna10_AspGTC_12_-_125424193_125424264</td><td>37</td><td>72</td><td>36</td><td>ACGCGGGAGACCGGGGTTCGATTCCCCGACGGGGAG</td><td>4027.2</td><td>2.6</td><td>1.1</td><td>2.5</td><td>1.4E-02</td><td>1.6E-02</td><td>MB231/MB468</td><td>CYTO</td></tr>
<tr><td>>hg38_21_+_8433172_8446622_RNA45SN1_45S_50ntFlanks@6647.6695.49|CGTACGACTCTTAGCGGTGGATCACTCGGCTCGTGCGTCGATGAAGAAC</td><td>>hg38_21_+_8433172_8446622_RNA45SN1_45S_50ntFlanks</td><td>6647</td><td>6695</td><td>49</td><td>CGTACGACTCTTAGCGGTGGATCACTCGGCTCGTGCGTCGATGAAGAAC</td><td>277.5</td><td>4.9</td><td>1.9</td><td>2.6</td><td>8.7E-03</td><td>1.6E-02</td><td>BT20/MB468</td><td>MP</td></tr>
<tr><td>>hg38_21_+_8433172_8446622_RNA45SN1_45S_50ntFlanks@11876.11903.28|AAGACCCTGTTGAGCTTGACTCTAGTCT</td><td>>hg38_21_+_8433172_8446622_RNA45SN1_45S_50ntFlanks</td><td>11876</td><td>11903</td><td>28</td><td>AAGACCCTGTTGAGCTTGACTCTAGTCT</td><td>117.8</td><td>-4.4</td><td>1.7</td><td>-2.6</td><td>8.7E-03</td><td>1.6E-02</td><td>BT20/MB468</td><td>MP</td></tr>
<tr><td>>hg38_21_+_8433172_8446622_RNA45SN1_45S_50ntFlanks@10794.10827.34|CTGGCATGTTGGAACAATGTAGGTAAGGGAAGTC</td><td>>hg38_21_+_8433172_8446622_RNA45SN1_45S_50ntFlanks</td><td>10794</td><td>10827</td><td>34</td><td>CTGGCATGTTGGAACAATGTAGGTAAGGGAAGTC</td><td>171.4</td><td>-3.3</td><td>1.3</td><td>-2.6</td><td>8.7E-03</td><td>1.6E-02</td><td>BT20/MB468</td><td>MP</td></tr>
<tr><td>>hg38_21_+_8433172_8446622_RNA45SN1_45S_50ntFlanks@4892.4911.20|AAACTTAAAGGAATTGACGG</td><td>>hg38_21_+_8433172_8446622_RNA45SN1_45S_50ntFlanks</td><td>4892</td><td>4911</td><td>20</td><td>AAACTTAAAGGAATTGACGG</td><td>222.3</td><td>-2.3</td><td>0.9</td><td>-2.6</td><td>8.7E-03</td><td>1.6E-02</td><td>BT20/MB468</td><td>MP</td></tr>
<tr><td>>hg38_21_+_8433172_8446622_RNA45SN1_45S_50ntFlanks@5056.5082.27|GAGCGATTTGTCTGGTTAATTCCGATA</td><td>>hg38_21_+_8433172_8446622_RNA45SN1_45S_50ntFlanks</td><td>5056</td><td>5082</td><td>27</td><td>GAGCGATTTGTCTGGTTAATTCCGATA</td><td>323.4</td><td>-2.4</td><td>1.0</td><td>-2.4</td><td>1.4E-02</td><td>1.6E-02</td><td>MB231/MB468</td><td>NUC</td></tr>
<tr><td>>hg38_21_+_8433172_8446622_RNA45SN1_45S_50ntFlanks@3775.3798.24|GCCGGTACAGTGAAACTGCGAATG</td><td>>hg38_21_+_8433172_8446622_RNA45SN1_45S_50ntFlanks</td><td>3775</td><td>3798</td><td>24</td><td>GCCGGTACAGTGAAACTGCGAATG</td><td>133.8</td><td>-3.9</td><td>1.5</td><td>-2.5</td><td>1.2E-02</td><td>1.6E-02</td><td>BT20/MB468</td><td>MITO</td></tr>
<tr><td>>hg38_MT_+_1621_3279_RNR2_16S_50ntFlanks@1430.1458.29|TAAAGTCCTACGTGATCTGAGTTCAGACC</td><td>>hg38_MT_+_1621_3279_RNR2_16S_50ntFlanks</td><td>1430</td><td>1458</td><td>29</td><td>TAAAGTCCTACGTGATCTGAGTTCAGACC</td><td>93.2</td><td>3.3</td><td>1.3</td><td>2.4</td><td>1.5E-02</td><td>1.6E-02</td><td>MB231/MB468</td><td>MITO</td></tr>
<tr><td>>MI0000448|hsa-mir-130a&WithFlank&11|+|57641192|57641292@61.80.20&[MIMAT0000425&hsa-miR-130a-3p&offsets|0|-2;m-139&11|+|57641252|57641273&offsets|0|-2]|CAGTGCAATGTTAAAAGGGC</td><td>>MI0000448|hsa-mir-130a&WithFlank&11|+|57641192|57641292</td><td>61</td><td>80</td><td>20</td><td>CAGTGCAATGTTAAAAGGGC</td><td>338.9</td><td>-3.3</td><td>1.1</td><td>-2.9</td><td>3.5E-03</td><td>1.6E-02</td><td>BT20/MB231</td><td>MP</td></tr>
<tr><td>>hg38_21_+_8433172_8446622_RNA45SN1_45S_50ntFlanks@5020.5044.25|CCGTGGGTGGTGGTGCATGGCCGTT</td><td>>hg38_21_+_8433172_8446622_RNA45SN1_45S_50ntFlanks</td><td>5020</td><td>5044</td><td>25</td><td>CCGTGGGTGGTGGTGCATGGCCGTT</td><td>108.2</td><td>4.8</td><td>1.8</td><td>2.6</td><td>8.7E-03</td><td>1.6E-02</td><td>BT20/MB468</td><td>MP</td></tr>
<tr><td>>hg38_21_+_8433172_8446622_RNA45SN1_45S_50ntFlanks@11769.11791.23|TAATTAGTGACGCGCATGAATGG</td><td>>hg38_21_+_8433172_8446622_RNA45SN1_45S_50ntFlanks</td><td>11769</td><td>11791</td><td>23</td><td>TAATTAGTGACGCGCATGAATGG</td><td>228.6</td><td>-4.5</td><td>1.7</td><td>-2.6</td><td>8.8E-03</td><td>1.6E-02</td><td>BT20/MB468</td><td>MP</td></tr>
<tr><td>>hg38_21_+_8433172_8446622_RNA45SN1_45S_50ntFlanks@4972.5001.30|CGGCCCGGACACGGACAGGATTGACAGATT</td><td>>hg38_21_+_8433172_8446622_RNA45SN1_45S_50ntFlanks</td><td>4972</td><td>5001</td><td>30</td><td>CGGCCCGGACACGGACAGGATTGACAGATT</td><td>58.5</td><td>-3.3</td><td>1.3</td><td>-2.5</td><td>1.3E-02</td><td>1.6E-02</td><td>MB231/MB468</td><td>MP</td></tr>
<tr><td>>hg38_21_+_8433172_8446622_RNA45SN1_45S_50ntFlanks@12627.12660.34|ATGGGGCGAAGCTACCATCTGTGGGATTATGACT</td><td>>hg38_21_+_8433172_8446622_RNA45SN1_45S_50ntFlanks</td><td>12627</td><td>12660</td><td>34</td><td>ATGGGGCGAAGCTACCATCTGTGGGATTATGACT</td><td>190.0</td><td>-2.9</td><td>1.2</td><td>-2.5</td><td>1.2E-02</td><td>1.6E-02</td><td>BT20/MB468</td><td>MITO</td></tr>
<tr><td>>hg38_21_+_8433172_8446622_RNA45SN1_45S_50ntFlanks@11563.11600.38|GCCTAGCAGCCGACTTAGAACTGGTGCGGACCAGGGGA</td><td>>hg38_21_+_8433172_8446622_RNA45SN1_45S_50ntFlanks</td><td>11563</td><td>11600</td><td>38</td><td>GCCTAGCAGCCGACTTAGAACTGGTGCGGACCAGGGGA</td><td>246.7</td><td>4.0</td><td>1.4</td><td>2.9</td><td>3.5E-03</td><td>1.6E-02</td><td>BT20/MB231</td><td>MP</td></tr>
<tr><td>>hg38_21_+_8433172_8446622_RNA45SN1_45S_50ntFlanks@10345.10381.37|TTCAAACGAGAACTTTGAAGGCCGAAGTGGAGAAGGG</td><td>>hg38_21_+_8433172_8446622_RNA45SN1_45S_50ntFlanks</td><td>10345</td><td>10381</td><td>37</td><td>TTCAAACGAGAACTTTGAAGGCCGAAGTGGAGAAGGG</td><td>77.4</td><td>5.3</td><td>2.0</td><td>2.6</td><td>8.8E-03</td><td>1.6E-02</td><td>BT20/MB468</td><td>MP</td></tr>
<tr><td>>hg38_21_+_8433172_8446622_RNA45SN1_45S_50ntFlanks@9503.9526.24|GGTGAACTATGCCTGGGCAGGGCG</td><td>>hg38_21_+_8433172_8446622_RNA45SN1_45S_50ntFlanks</td><td>9503</td><td>9526</td><td>24</td><td>GGTGAACTATGCCTGGGCAGGGCG</td><td>691.1</td><td>-4.9</td><td>1.9</td><td>-2.6</td><td>8.8E-03</td><td>1.6E-02</td><td>BT20/MB468</td><td>MP</td></tr>
<tr><td>>hg38_21_+_8433172_8446622_RNA45SN1_45S_50ntFlanks@4732.4762.31|AGAACGAAAGTCGGAGGTTCGAAGACGATCA</td><td>>hg38_21_+_8433172_8446622_RNA45SN1_45S_50ntFlanks</td><td>4732</td><td>4762</td><td>31</td><td>AGAACGAAAGTCGGAGGTTCGAAGACGATCA</td><td>60.4</td><td>3.8</td><td>1.4</td><td>2.6</td><td>8.8E-03</td><td>1.6E-02</td><td>BT20/MB468</td><td>MP</td></tr>
<tr><td>>hg38_21_+_8433172_8446622_RNA45SN1_45S_50ntFlanks@4738.4768.31|AAAGTCGGAGGTTCGAAGACGATCAGATACC</td><td>>hg38_21_+_8433172_8446622_RNA45SN1_45S_50ntFlanks</td><td>4738</td><td>4768</td><td>31</td><td>AAAGTCGGAGGTTCGAAGACGATCAGATACC</td><td>249.4</td><td>3.8</td><td>1.5</td><td>2.6</td><td>8.8E-03</td><td>1.6E-02</td><td>BT20/MB468</td><td>MP</td></tr>
<tr><td>>trnaMT_GluTTC_MT_-_14674_14742@34.61.28,trnalookalike8_GluTTC_5_-_93905172_93905240@34.61.28|ATATCATTGGTCGTGGTTGTAGTCCGTG</td><td>>trnaMT_GluTTC_MT_-_14674_14742</td><td>34</td><td>61</td><td>28</td><td>ATATCATTGGTCGTGGTTGTAGTCCGTG</td><td>323.9</td><td>2.8</td><td>1.1</td><td>2.6</td><td>8.8E-03</td><td>1.6E-02</td><td>BT20/MB468</td><td>MP</td></tr>
<tr><td>>hg38_21_+_8433172_8446622_RNA45SN1_45S_50ntFlanks@10771.10818.48|CAGGTCTCCAAGGTGAACAGCCTCTGGCATGTTGGAACAATGTAGGTA</td><td>>hg38_21_+_8433172_8446622_RNA45SN1_45S_50ntFlanks</td><td>10771</td><td>10818</td><td>48</td><td>CAGGTCTCCAAGGTGAACAGCCTCTGGCATGTTGGAACAATGTAGGTA</td><td>822.5</td><td>-4.7</td><td>1.9</td><td>-2.5</td><td>1.4E-02</td><td>1.6E-02</td><td>BT20/MB468</td><td>CYTO</td></tr>
<tr><td>>hg38_MT_+_1621_3279_RNR2_16S_50ntFlanks@1370.1407.38|ATGTTGGATCAGGACATCCCGATGGTGCAGCCGCTATT</td><td>>hg38_MT_+_1621_3279_RNR2_16S_50ntFlanks</td><td>1370</td><td>1407</td><td>38</td><td>ATGTTGGATCAGGACATCCCGATGGTGCAGCCGCTATT</td><td>88.6</td><td>3.1</td><td>1.2</td><td>2.5</td><td>1.3E-02</td><td>1.6E-02</td><td>MB231/MB468</td><td>MP</td></tr>
<tr><td>>MI0000469|hsa-mir-125a&WithFlank&19|+|51693248|51693345@21.42.22&[MIMAT0000443&hsa-miR-125a-5p&offsets|0|-2;m-63&19|+|51693268|51693289&offsets|0|0]|TCCCTGAGACCCTTTAACCTGT</td><td>>MI0000469|hsa-mir-125a&WithFlank&19|+|51693248|51693345</td><td>21</td><td>42</td><td>22</td><td>TCCCTGAGACCCTTTAACCTGT</td><td>5318.9</td><td>1.6</td><td>0.6</td><td>2.9</td><td>3.8E-03</td><td>1.6E-02</td><td>BT20/MB231</td><td>NUC</td></tr>
<tr><td>>trnaMT_ProTGG_MT_-_15956_16023@50.70.21|TTAAAGACTTTTTCTCTGACC</td><td>>trnaMT_ProTGG_MT_-_15956_16023</td><td>50</td><td>70</td><td>21</td><td>TTAAAGACTTTTTCTCTGACC</td><td>195.7</td><td>-3.8</td><td>1.3</td><td>-2.9</td><td>3.8E-03</td><td>1.6E-02</td><td>BT20/MB231</td><td>NUC</td></tr>
<tr><td>>hg38_21_+_8433172_8446622_RNA45SN1_45S_50ntFlanks@10341.10372.32|AATATTCAAACGAGAACTTTGAAGGCCGAAGT</td><td>>hg38_21_+_8433172_8446622_RNA45SN1_45S_50ntFlanks</td><td>10341</td><td>10372</td><td>32</td><td>AATATTCAAACGAGAACTTTGAAGGCCGAAGT</td><td>63.7</td><td>4.5</td><td>1.7</td><td>2.6</td><td>8.8E-03</td><td>1.6E-02</td><td>BT20/MB468</td><td>MP</td></tr>
<tr><td>>trnaMT_PheGAA_MT_+_577_647@35.61.27|GAAAATGTTTAGACGGGCTCACATCAC</td><td>>trnaMT_PheGAA_MT_+_577_647</td><td>35</td><td>61</td><td>27</td><td>GAAAATGTTTAGACGGGCTCACATCAC</td><td>107.8</td><td>4.7</td><td>1.9</td><td>2.4</td><td>1.5E-02</td><td>1.6E-02</td><td>MB231/MB468</td><td>MITO</td></tr>
<tr><td>>trnaMT_GluTTC_MT_-_14674_14742@40.72.33,trnalookalike8_GluTTC_5_-_93905172_93905240@40.72.33|TTGGTCGTGGTTGTAGTCCGTGCGAGAATACCA</td><td>>trnaMT_GluTTC_MT_-_14674_14742</td><td>40</td><td>72</td><td>33</td><td>TTGGTCGTGGTTGTAGTCCGTGCGAGAATACCA</td><td>4705.9</td><td>-1.9</td><td>0.8</td><td>-2.4</td><td>1.4E-02</td><td>1.6E-02</td><td>BT20/MB468</td><td>NUC</td></tr>
<tr><td>>hg38_21_+_8433172_8446622_RNA45SN1_45S_50ntFlanks@8121.8141.21|ACATGTGGCGTACGGAAGACC</td><td>>hg38_21_+_8433172_8446622_RNA45SN1_45S_50ntFlanks</td><td>8121</td><td>8141</td><td>21</td><td>ACATGTGGCGTACGGAAGACC</td><td>179.5</td><td>4.6</td><td>1.8</td><td>2.6</td><td>8.8E-03</td><td>1.6E-02</td><td>BT20/MB468</td><td>MP</td></tr>
<tr><td>>MI0000088|hsa-mir-30a&WithFlank&6|-|71403545|71403627@13.34.22&[MIMAT0000087&hsa-miR-30a-5p&offsets|+1|+1;m-15&6|-|71403595|71403616&offsets|+1|+1]|GTAAACATCCTCGACTGGAAGC</td><td>>MI0000088|hsa-mir-30a&WithFlank&6|-|71403545|71403627</td><td>13</td><td>34</td><td>22</td><td>GTAAACATCCTCGACTGGAAGC</td><td>254.4</td><td>4.3</td><td>1.6</td><td>2.6</td><td>8.8E-03</td><td>1.6E-02</td><td>BT20/MB468</td><td>MP</td></tr>
<tr><td>>hg38_21_+_8433172_8446622_RNA45SN1_45S_50ntFlanks@9312.9330.19|GTCTAACACGTGCGCGAGT</td><td>>hg38_21_+_8433172_8446622_RNA45SN1_45S_50ntFlanks</td><td>9312</td><td>9330</td><td>19</td><td>GTCTAACACGTGCGCGAGT</td><td>55.1</td><td>4.6</td><td>1.7</td><td>2.6</td><td>8.8E-03</td><td>1.6E-02</td><td>BT20/MB468</td><td>MP</td></tr>
<tr><td>>MI0000273|hsa-mir-183&WithFlank&7|-|129774899|129775020@34.54.21(+1A)&[MIMAT0000261&hsa-miR-183-5p&offsets|+1|0(+1A);m-41&7|-|129774966|129774987&offsets|0|-1(+1A)]|ATGGCACTGGTAGAATTCACTA</td><td>>MI0000273|hsa-mir-183&WithFlank&7|-|129774899|129775020</td><td>34</td><td>54</td><td>21(+1A)</td><td>ATGGCACTGGTAGAATTCACTA</td><td>347.2</td><td>3.9</td><td>1.5</td><td>2.6</td><td>8.8E-03</td><td>1.6E-02</td><td>BT20/MB468</td><td>MP</td></tr>
<tr><td>>MI0000082|hsa-mir-25&WithFlank&7|-|100093554|100093649@58.79.22(+1G)&[MIMAT0000081&hsa-miR-25-3p&offsets|0|0(+1G);m-23&7|-|100093571|100093592&offsets|0|0(+1G)]|CATTGCACTTGTCTCGGTCTGAG</td><td>>MI0000082|hsa-mir-25&WithFlank&7|-|100093554|100093649</td><td>58</td><td>79</td><td>22(+1G)</td><td>CATTGCACTTGTCTCGGTCTGAG</td><td>40.7</td><td>4.9</td><td>2.0</td><td>2.5</td><td>1.3E-02</td><td>1.6E-02</td><td>MB231/MB468</td><td>MP</td></tr>
<tr><td>>hg38_21_+_8433172_8446622_RNA45SN1_45S_50ntFlanks@11670.11698.29|CCCAGTGCTCTGAATGTCAAAGTGAAGAA</td><td>>hg38_21_+_8433172_8446622_RNA45SN1_45S_50ntFlanks</td><td>11670</td><td>11698</td><td>29</td><td>CCCAGTGCTCTGAATGTCAAAGTGAAGAA</td><td>263.5</td><td>-2.6</td><td>1.1</td><td>-2.5</td><td>1.4E-02</td><td>1.6E-02</td><td>MB231/MB468</td><td>CYTO</td></tr>
<tr><td>>trnaMT_GluTTC_MT_-_14674_14742@33.66.34,trnalookalike8_GluTTC_5_-_93905172_93905240@33.66.34|CATATCATTGGTCGTGGTTGTAGTCCGTGCGAGA</td><td>>trnaMT_GluTTC_MT_-_14674_14742</td><td>33</td><td>66</td><td>34</td><td>CATATCATTGGTCGTGGTTGTAGTCCGTGCGAGA</td><td>62.4</td><td>5.4</td><td>2.1</td><td>2.6</td><td>8.8E-03</td><td>1.6E-02</td><td>BT20/MB468</td><td>MP</td></tr>
<tr><td>>hg38_21_+_8433172_8446622_RNA45SN1_45S_50ntFlanks@10412.10431.20|AGTCGGTCCTGAGAGATGGG</td><td>>hg38_21_+_8433172_8446622_RNA45SN1_45S_50ntFlanks</td><td>10412</td><td>10431</td><td>20</td><td>AGTCGGTCCTGAGAGATGGG</td><td>127.8</td><td>-4.3</td><td>1.7</td><td>-2.6</td><td>8.8E-03</td><td>1.6E-02</td><td>BT20/MB468</td><td>MP</td></tr>
<tr><td>>hg38_21_+_8433172_8446622_RNA45SN1_45S_50ntFlanks@7978.8009.32|GACCTCAGATCAGACGTGGCGACCCGCTGAAT</td><td>>hg38_21_+_8433172_8446622_RNA45SN1_45S_50ntFlanks</td><td>7978</td><td>8009</td><td>32</td><td>GACCTCAGATCAGACGTGGCGACCCGCTGAAT</td><td>238.1</td><td>2.4</td><td>1.0</td><td>2.5</td><td>1.3E-02</td><td>1.6E-02</td><td>MB231/MB468</td><td>MP</td></tr>
<tr><td>>hg38_MT_+_1621_3279_RNR2_16S_50ntFlanks@1250.1271.22|GTCAAAGCGAACTACTATACTC</td><td>>hg38_MT_+_1621_3279_RNR2_16S_50ntFlanks</td><td>1250</td><td>1271</td><td>22</td><td>GTCAAAGCGAACTACTATACTC</td><td>83.8</td><td>4.1</td><td>1.7</td><td>2.5</td><td>1.3E-02</td><td>1.6E-02</td><td>MB231/MB468</td><td>MP</td></tr>
<tr><td>>hg38_21_+_8433172_8446622_RNA45SN1_45S_50ntFlanks@4353.4383.31|TATTAAAGTTGCTGCAGTTAAAAAGCTCGTA</td><td>>hg38_21_+_8433172_8446622_RNA45SN1_45S_50ntFlanks</td><td>4353</td><td>4383</td><td>31</td><td>TATTAAAGTTGCTGCAGTTAAAAAGCTCGTA</td><td>91.4</td><td>-2.2</td><td>0.9</td><td>-2.5</td><td>1.3E-02</td><td>1.6E-02</td><td>MB231/MB468</td><td>MP</td></tr>
<tr><td>>hg38_21_+_8433172_8446622_RNA45SN1_45S_50ntFlanks@9736.9753.18|CCGAAACGATCTCAACCT</td><td>>hg38_21_+_8433172_8446622_RNA45SN1_45S_50ntFlanks</td><td>9736</td><td>9753</td><td>18</td><td>CCGAAACGATCTCAACCT</td><td>137.2</td><td>-4.4</td><td>1.7</td><td>-2.6</td><td>8.8E-03</td><td>1.6E-02</td><td>BT20/MB468</td><td>MP</td></tr>
<tr><td>>hg38_21_+_8433172_8446622_RNA45SN1_45S_50ntFlanks@12405.12445.41|CTTTTTGATCCTTCGATGTCGGCTCTTCCTATCATTGTGAA</td><td>>hg38_21_+_8433172_8446622_RNA45SN1_45S_50ntFlanks</td><td>12405</td><td>12445</td><td>41</td><td>CTTTTTGATCCTTCGATGTCGGCTCTTCCTATCATTGTGAA</td><td>352.6</td><td>-1.9</td><td>0.8</td><td>-2.5</td><td>1.2E-02</td><td>1.6E-02</td><td>BT20/MB468</td><td>MITO</td></tr>
<tr><td>>hg38_21_+_8433172_8446622_RNA45SN1_45S_50ntFlanks@10128.10148.21|CGGCGGCGGCGGCGGCGGCGG</td><td>>hg38_21_+_8433172_8446622_RNA45SN1_45S_50ntFlanks</td><td>10128</td><td>10148</td><td>21</td><td>CGGCGGCGGCGGCGGCGGCGG</td><td>345.0</td><td>-4.1</td><td>1.6</td><td>-2.5</td><td>1.2E-02</td><td>1.6E-02</td><td>BT20/MB468</td><td>MITO</td></tr>
<tr><td>>hg38_21_+_8433172_8446622_RNA45SN1_45S_50ntFlanks@8845.8865.21|CGGCGGCGGCGGCGGCGGCGG</td><td>>hg38_21_+_8433172_8446622_RNA45SN1_45S_50ntFlanks</td><td>8845</td><td>8865</td><td>21</td><td>CGGCGGCGGCGGCGGCGGCGG</td><td>345.0</td><td>-4.1</td><td>1.6</td><td>-2.5</td><td>1.2E-02</td><td>1.6E-02</td><td>BT20/MB468</td><td>MITO</td></tr>
<tr><td>>hg38_MT_+_1621_3279_RNR2_16S_50ntFlanks@1354.1385.32|AGGGTTTACGACCTCGATGTTGGATCAGGACA</td><td>>hg38_MT_+_1621_3279_RNR2_16S_50ntFlanks</td><td>1354</td><td>1385</td><td>32</td><td>AGGGTTTACGACCTCGATGTTGGATCAGGACA</td><td>74.8</td><td>5.5</td><td>2.1</td><td>2.6</td><td>8.8E-03</td><td>1.6E-02</td><td>BT20/MB468</td><td>MP</td></tr>
<tr><td>>hg38_21_+_8433172_8446622_RNA45SN1_45S_50ntFlanks@9574.9598.25|AATCGGTCGTCCGACCTGGGTATAG</td><td>>hg38_21_+_8433172_8446622_RNA45SN1_45S_50ntFlanks</td><td>9574</td><td>9598</td><td>25</td><td>AATCGGTCGTCCGACCTGGGTATAG</td><td>203.5</td><td>-2.8</td><td>1.1</td><td>-2.5</td><td>1.2E-02</td><td>1.6E-02</td><td>BT20/MB468</td><td>MITO</td></tr>
<tr><td>>hg38_21_+_8433172_8446622_RNA45SN1_45S_50ntFlanks@6721.6757.37|AATTGCAGGACACATTGATCATCGACACTTCGAACGC</td><td>>hg38_21_+_8433172_8446622_RNA45SN1_45S_50ntFlanks</td><td>6721</td><td>6757</td><td>37</td><td>AATTGCAGGACACATTGATCATCGACACTTCGAACGC</td><td>299.9</td><td>-2.1</td><td>0.8</td><td>-2.5</td><td>1.2E-02</td><td>1.6E-02</td><td>BT20/MB468</td><td>MITO</td></tr>
<tr><td>>hg38_21_+_8433172_8446622_RNA45SN1_45S_50ntFlanks@3938.3959.22|CAACCCGGTCAGCCCCTCTCCG</td><td>>hg38_21_+_8433172_8446622_RNA45SN1_45S_50ntFlanks</td><td>3938</td><td>3959</td><td>22</td><td>CAACCCGGTCAGCCCCTCTCCG</td><td>160.5</td><td>-3.8</td><td>1.5</td><td>-2.5</td><td>1.2E-02</td><td>1.6E-02</td><td>BT20/MB468</td><td>MITO</td></tr>
<tr><td>>hg38_21_+_8433172_8446622_RNA45SN1_45S_50ntFlanks@4585.4621.37|GTTCTATTTTGTTGGTTTTCGGAACTGAGGCCATGAT</td><td>>hg38_21_+_8433172_8446622_RNA45SN1_45S_50ntFlanks</td><td>4585</td><td>4621</td><td>37</td><td>GTTCTATTTTGTTGGTTTTCGGAACTGAGGCCATGAT</td><td>556.3</td><td>-2.4</td><td>0.8</td><td>-2.8</td><td>4.5E-03</td><td>1.6E-02</td><td>BT20/MB231</td><td>CYTO</td></tr>
<tr><td>>hg38_MT_+_1621_3279_RNR2_16S_50ntFlanks@623.652.30|ATAACTGAACTCCTCACACCCAATTGGACC</td><td>>hg38_MT_+_1621_3279_RNR2_16S_50ntFlanks</td><td>623</td><td>652</td><td>30</td><td>ATAACTGAACTCCTCACACCCAATTGGACC</td><td>50.2</td><td>4.8</td><td>1.9</td><td>2.5</td><td>1.3E-02</td><td>1.6E-02</td><td>MB231/MB468</td><td>MP</td></tr>
<tr><td>>hg38_21_+_8433172_8446622_RNA45SN1_45S_50ntFlanks@9570.9598.29|TGCAAATCGGTCGTCCGACCTGGGTATAG</td><td>>hg38_21_+_8433172_8446622_RNA45SN1_45S_50ntFlanks</td><td>9570</td><td>9598</td><td>29</td><td>TGCAAATCGGTCGTCCGACCTGGGTATAG</td><td>49.3</td><td>-4.4</td><td>1.7</td><td>-2.6</td><td>8.9E-03</td><td>1.6E-02</td><td>BT20/MB468</td><td>MP</td></tr>
<tr><td>>hg38_21_+_8433172_8446622_RNA45SN1_45S_50ntFlanks@11561.11600.40|GCGCCTAGCAGCCGACTTAGAACTGGTGCGGACCAGGGGA</td><td>>hg38_21_+_8433172_8446622_RNA45SN1_45S_50ntFlanks</td><td>11561</td><td>11600</td><td>40</td><td>GCGCCTAGCAGCCGACTTAGAACTGGTGCGGACCAGGGGA</td><td>860.3</td><td>-1.7</td><td>0.7</td><td>-2.5</td><td>1.2E-02</td><td>1.6E-02</td><td>BT20/MB468</td><td>MITO</td></tr>
</table>

| | | | | | | | | | | | | | |
|---|---|---|---|---|---|---|---|---|---|---|---|---|---|
| >hg38_21_+_8433172_8446622_RNA45SN1_45S_50ntFlanks@10116.10135.20\|TGCGTCGGAGGGCGGCGGCG | >hg38_21_+_8433172_8446622_RNA45SN1_45S_50ntFlanks | 10116 | 10135 | 20 | TGCGTCGGAGGGCGGCGGCG | 279.3 | -3.9 | 1.5 | -2.5 | 1.2E-02 | 1.6E-02 | BT20/MB468 | MITO |
| >hg38_21_+_8433172_8446622_RNA45SN1_45S_50ntFlanks@12406.12432.27\|TTTTTGATCCTTCGATGTCGGCTCTTC | >hg38_21_+_8433172_8446622_RNA45SN1_45S_50ntFlanks | 12406 | 12432 | 27 | TTTTTGATCCTTCGATGTCGGCTCTTC | 289.1 | -3.0 | 1.2 | -2.4 | 1.5E-02 | 1.6E-02 | MB231/MB468 | NUC |
| >trnaMT_GluTTC_MT_-_14674_14742@46.71.26,trnalookalike8_GluTTC_5_-_93905172_93905240@46.71.26\|GTGGTTGTAGTCCGTGCGAGAATACC | >trnaMT_GluTTC_MT_-_14674_14742 | 46 | 71 | 26 | GTGGTTGTAGTCCGTGCGAGAATACC | 171.8 | -2.1 | 0.7 | -2.9 | 3.8E-03 | 1.6E-02 | BT20/MB231 | NUC |
| >MI0000440\|hsa-mir-27b&WithFlank&9\|+\|95085439\|95085547@67.87.21(+1U)&[MIMAT0000419&hsa-miR-27b-3p&offsets\|0\|0(+1U);m-39&9\|+\|95085505\|95085525&offsets\|0\|0(+1U)]\|TTCACAGTGGCTAAGTTCTGCT | >MI0000440\|hsa-mir-27b&WithFlank&9\|+\|95085439\|95085547 | 67 | 87 | 21(+1U) | TTCACAGTGGCTAAGTTCTGCT | 2589.5 | -3.2 | 1.1 | -2.9 | 3.5E-03 | 1.6E-02 | BT20/MB231 | MP |
| >hg38_21_+_8433172_8446622_RNA45SN1_45S_50ntFlanks@10515.10532.18\|GGAGTGGCGGAGATGGGC | >hg38_21_+_8433172_8446622_RNA45SN1_45S_50ntFlanks | 10515 | 10532 | 18 | GGAGTGGCGGAGATGGGC | 361.1 | 6.0 | 2.1 | 2.9 | 3.5E-03 | 1.6E-02 | BT20/MB231 | MP |
| >hg38_MT_+_1621_3279_RNR2_16S_50ntFlanks@1405.1426.22\|ATTAAAGGTTCGTTTGTTCAAC | >hg38_MT_+_1621_3279_RNR2_16S_50ntFlanks | 1405 | 1426 | 22 | ATTAAAGGTTCGTTTGTTCAAC | 123.9 | 2.5 | 1.0 | 2.5 | 1.3E-02 | 1.6E-02 | MB231/MB468 | MP |
| >MI0000088\|hsa-mir-30a&WithFlank&6\|-\|71403545\|71403627@53.75.23&[MIMAT0000088&hsa-miR-30a-3p&offsets\|0\|+1;m-38&6\|-\|71403554\|71403575&offsets\|0\|+1]\|CTTTCAGTCGGATGTTTGCAGCT | >MI0000088\|hsa-mir-30a&WithFlank&6\|-\|71403545\|71403627 | 53 | 75 | 23 | CTTTCAGTCGGATGTTTGCAGCT | 325.2 | -2.0 | 0.7 | -2.9 | 3.8E-03 | 1.6E-02 | BT20/MB231 | NUC |
| >hg38_21_+_8433172_8446622_RNA45SN1_45S_50ntFlanks@12433.12452.20\|CTATCATTGTGAAGCAGAAT | >hg38_21_+_8433172_8446622_RNA45SN1_45S_50ntFlanks | 12433 | 12452 | 20 | CTATCATTGTGAAGCAGAAT | 65.8 | -4.3 | 1.6 | -2.6 | 8.9E-03 | 1.6E-02 | BT20/MB468 | MP |
| >hg38_21_+_8433172_8446622_RNA45SN1_45S_50ntFlanks@4570.4589.20\|TGGAATAGGACCGCGGTTCT | >hg38_21_+_8433172_8446622_RNA45SN1_45S_50ntFlanks | 4570 | 4589 | 20 | TGGAATAGGACCGCGGTTCT | 180.3 | -1.4 | 0.6 | -2.5 | 1.4E-02 | 1.6E-02 | MB231/MB468 | CYTO |
| >trnaMT_GluTTC_MT_-_14674_14742@29.72.44,trnalookalike8_GluTTC_5_-_93905172_93905240@29.72.44\|TTTTCATATCATTGGTCGTGGTTGTAGTCCGTGCGAGAATACCA | >trnaMT_GluTTC_MT_-_14674_14742 | 29 | 72 | 44 | TTTTCATATCATTGGTCGTGGTTGTAGTCCGTGCGAGAATACCA | 165.3 | 4.9 | 1.9 | 2.6 | 8.9E-03 | 1.6E-02 | BT20/MB468 | MP |
| >trnaMT_SerGCT_MT_+_12207_12265@1.26.26\|GAGAAAGCTCACAAGAACTGCTAACT | >trnaMT_SerGCT_MT_+_12207_12265 | 1 | 26 | 26 | GAGAAAGCTCACAAGAACTGCTAACT | 35537.4 | 3.5 | 1.3 | 2.6 | 8.9E-03 | 1.6E-02 | BT20/MB468 | MP |
| >hg38_21_+_8433172_8446622_RNA45SN1_45S_50ntFlanks@5487.5516.30\|CGCTGAGAAGACGGTCGAACTTGACTATCT | >hg38_21_+_8433172_8446622_RNA45SN1_45S_50ntFlanks | 5487 | 5516 | 30 | CGCTGAGAAGACGGTCGAACTTGACTATCT | 1184.7 | 3.2 | 1.1 | 2.9 | 3.8E-03 | 1.6E-02 | BT20/MB231 | NUC |
| >hg38_21_+_8433172_8446622_RNA45SN1_45S_50ntFlanks@9931.9948.18\|TATAGACAGCAGGACGGT | >hg38_21_+_8433172_8446622_RNA45SN1_45S_50ntFlanks | 9931 | 9948 | 18 | TATAGACAGCAGGACGGT | 424.3 | 3.1 | 1.1 | 2.9 | 3.5E-03 | 1.6E-02 | BT20/MB231 | MP |
| >hg38_MT_+_1621_3279_RNR2_16S_50ntFlanks@716.741.26\|TAAGCCTGCGTCAGATTAAAACACTG | >hg38_MT_+_1621_3279_RNR2_16S_50ntFlanks | 716 | 741 | 26 | TAAGCCTGCGTCAGATTAAAACACTG | 20.3 | 7.6 | 3.0 | 2.5 | 1.3E-02 | 1.6E-02 | MB231/MB468 | MP |
| >hg38_21_+_8433172_8446622_RNA45SN1_45S_50ntFlanks@12717.12753.37\|CCTCGGTTGGCCTCGGATAGCCGGTCCCCCGCCTGTC | >hg38_21_+_8433172_8446622_RNA45SN1_45S_50ntFlanks | 12717 | 12753 | 37 | CCTCGGTTGGCCTCGGATAGCCGGTCCCCCGCCTGTC | 78.1 | -5.0 | 1.9 | -2.6 | 8.9E-03 | 1.6E-02 | BT20/MB468 | MP |
| >MI0000734\|hsa-mir-106b&WithFlank&7\|-\|100093987\|100094080@58.77.20(+1A)&[MIMAT0004672&hsa-miR-106b-3p&offsets\|0\|-2(+1A);m-87&7\|-\|100094003\|100094023&offsets\|0\|-1(+1A)]\|CCGCACTGTGGGTACTTGCTA | >MI0000734\|hsa-mir-106b&WithFlank&7\|-\|100093987\|100094080 | 58 | 77 | 20(+1A) | CCGCACTGTGGGTACTTGCTA | 149.9 | 5.0 | 2.0 | 2.5 | 1.3E-02 | 1.7E-02 | MB231/MB468 | MP |
| >hg38_21_+_8433172_8446622_RNA45SN1_45S_50ntFlanks@12283.12318.36\|TACAGACCGTGAAAGCGGGGCCTCACGATCCTTCTG | >hg38_21_+_8433172_8446622_RNA45SN1_45S_50ntFlanks | 12283 | 12318 | 36 | TACAGACCGTGAAAGCGGGGCCTCACGATCCTTCTG | 79.5 | -4.4 | 1.7 | -2.6 | 8.9E-03 | 1.7E-02 | BT20/MB468 | MP |
| >trna116_GluCTC_1_-_145399233_145399304@1.34.34,trna59_GluCTC_1_+_249168447_249168518@1.34.34,trna71_GluCTC_1_-_161439189_161439260@1.34.34,trna74_GluCTC_1_-_161431809_161431880@1.34.34,trna77_GluCTC_1_-_161424398_161424469@1.34.34,trna77_GluCTC_6_+_28949976_28950047@1.34.34,trna80_GluCTC_1_-_161417018_161417089@1.34.34,trna87_GluCTC_6_-_126101393_126101464@1.34.34\|TCCCTGGTGGTCTAGTGGTTAGGATTCGGCGCTC | >trna116_GluCTC_1_-_145399233_145399304 | 1 | 34 | 34 | TCCCTGGTGGTCTAGTGGTTAGGATTCGGCGCTC | 1106.1 | 1.9 | 0.8 | 2.4 | 1.5E-02 | 1.7E-02 | BT20/MB468 | NUC |
| >MI0000077\|hsa-mir-21&WithFlank&17\|+\|59841260\|59841343@15.36.22(+1A)&[MIMAT0000076&hsa-miR-21-5p&offsets\|+1\|+1(+1A);m-1&17\|+\|59841273\|59841294&offsets\|+1\|+1(+1A)]\|AGCTTATCAGACTGATGTTGACA | >MI0000077\|hsa-mir-21&WithFlank&17\|+\|59841260\|59841343 | 15 | 36 | 22(+1A) | AGCTTATCAGACTGATGTTGACA | 392.5 | 1.8 | 0.6 | 2.9 | 3.8E-03 | 1.7E-02 | BT20/MB231 | NUC |
| >hg38_21_+_8433172_8446622_RNA45SN1_45S_50ntFlanks@12418.12439.22\|CGATGTCGGCTCTTCCTATCAT | >hg38_21_+_8433172_8446622_RNA45SN1_45S_50ntFlanks | 12418 | 12439 | 22 | CGATGTCGGCTCTTCCTATCAT | 855.8 | -1.4 | 0.6 | -2.4 | 1.5E-02 | 1.7E-02 | MB231/MB468 | NUC |
| >hg38_21_+_8433172_8446622_RNA45SN1_45S_50ntFlanks@12937.12955.19\|AACCATTCGTAGACGACCT | >hg38_21_+_8433172_8446622_RNA45SN1_45S_50ntFlanks | 12937 | 12955 | 19 | AACCATTCGTAGACGACCT | 290.1 | -3.9 | 1.6 | -2.5 | 1.4E-02 | 1.7E-02 | BT20/MB468 | CYTO |
| >hg38_21_+_8433172_8446622_RNA45SN1_45S_50ntFlanks@10410.10453.44\|TCAGTCGGTCCTGAGAGATGGGCGAGCGCCGTTCCGAAGGGACG | >hg38_21_+_8433172_8446622_RNA45SN1_45S_50ntFlanks | 10410 | 10453 | 44 | TCAGTCGGTCCTGAGAGATGGGCGAGCGCCGTTCCGAAGGGACG | 162.3 | -3.0 | 1.1 | -2.6 | 8.9E-03 | 1.7E-02 | BT20/MB468 | MP |
| >MI0000469\|hsa-mir-125a&WithFlank&19\|+\|51693248\|51693345@59.78.20(+1U)&[MIMAT0004602&hsa-miR-125a-3p&offsets\|0\|-2(+1U);m-333&19\|+\|51693306\|51693326&offsets\|0\|-1(+1U)]\|ACAGGTGAGGTTCTTGGGAGT | >MI0000469\|hsa-mir-125a&WithFlank&19\|+\|51693248\|51693345 | 59 | 78 | 20(+1U) | ACAGGTGAGGTTCTTGGGAGT | 270.9 | 2.2 | 0.8 | 2.8 | 4.5E-03 | 1.7E-02 | BT20/MB231 | CYTO |
| >trna116_GluCTC_1_-_145399233_145399304@36.73.38,trna134_GluTTC_1_-_16861774_16861845@36.73.38,trna71_GluCTC_1_-_161439189_161439260@36.73.38,trna74_GluCTC_1_-_161431809_161431880@36.73.38,trna77_GluCTC_1_-_161424398_161424469@36.73.38,trna77_GluCTC_6_+_28949976_28950047@36.73.38,trna80_GluCTC_1_-_161417018_161417089@36.73.38,trna84_GluTTC_1_-_161391883_161391954@36.73.38,trna87_GluCTC_6_-_126101393_126101464@36.73.38\|CACCGCCGCGGCCCGGGTTCGATTCCCGGTCAGGGAAC | >trna116_GluCTC_1_-_145399233_145399304 | 36 | 73 | 38 | CACCGCCGCGGCCCGGGTTCGATTCCCGGTCAGGGAAC | 834.5 | 3.1 | 1.3 | 2.5 | 1.4E-02 | 1.7E-02 | MB231/MB468 | CYTO |
| >trnaMT_ThrTGT_MT_+_15888_15953@33.69.37\|GTAAACCGGAGATGAAAACCTTTTTCCAAGGACACCA | >trnaMT_ThrTGT_MT_+_15888_15953 | 33 | 69 | 37 | GTAAACCGGAGATGAAAACCTTTTTCCAAGGACACCA | 182.3 | 5.9 | 2.3 | 2.6 | 8.9E-03 | 1.7E-02 | BT20/MB468 | MP |
| >am_trnaMT_PheGAA_MT_+_577_647@41.61.21\|GTTTAGACGGGCTCACATCAC | >am_trnaMT_PheGAA_MT_+_577_647 | 41 | 61 | 21 | GTTTAGACGGGCTCACATCAC | 130.8 | 4.9 | 1.9 | 2.6 | 8.9E-03 | 1.7E-02 | BT20/MB468 | MP |
| >hg38_21_+_8433172_8446622_RNA45SN1_45S_50ntFlanks@12458.12477.20\|AAGCGTTGGATTGTTCACCC | >hg38_21_+_8433172_8446622_RNA45SN1_45S_50ntFlanks | 12458 | 12477 | 20 | AAGCGTTGGATTGTTCACCC | 584.5 | -2.0 | 0.8 | -2.4 | 1.5E-02 | 1.7E-02 | MB231/MB468 | MITO |
| >MI0000483\|hsa-mir-186&WithFlank&1\|-\|71067625\|71067722@21.42.22&[MIMAT0000456&hsa-miR-186-5p&offsets\|0\|0;m-84&1\|-\|71067681\|71067702&offsets\|0\|0]\|CAAAGAATTCTCCTTTTGGGCT | >MI0000483\|hsa-mir-186&WithFlank&1\|-\|71067625\|71067722 | 21 | 42 | 22 | CAAAGAATTCTCCTTTTGGGCT | 1580.1 | 2.6 | 0.9 | 2.8 | 4.5E-03 | 1.7E-02 | BT20/MB231 | CYTO |
| >MI0001445\|hsa-mir-423&WithFlank&17\|+\|30117073\|30117178@60.81.22&[MIMAT0001340&hsa-miR-423-3p&offsets\|+1\|0;m-100&17\|+\|30117131\|30117153&offsets\|+1\|0]\|GCTCGGTCTGAGGCCCCTCAGT | >MI0001445\|hsa-mir-423&WithFlank&17\|+\|30117073\|30117178 | 60 | 81 | 22 | GCTCGGTCTGAGGCCCCTCAGT | 55.1 | 4.7 | 1.8 | 2.6 | 8.9E-03 | 1.7E-02 | BT20/MB468 | MP |
| >hg38_21_+_8433172_8446622_RNA45SN1_45S_50ntFlanks@5482.5503.22\|CGGAGCGCTGAGAAGACGGTCG | >hg38_21_+_8433172_8446622_RNA45SN1_45S_50ntFlanks | 5482 | 5503 | 22 | CGGAGCGCTGAGAAGACGGTCG | 88.7 | -3.7 | 1.4 | -2.6 | 8.9E-03 | 1.7E-02 | BT20/MB468 | MP |
| >hg38_21_+_8433172_8446622_RNA45SN1_45S_50ntFlanks@11703.11740.38\|AATGAAGCGCGGGTAAACGGCGGGAGTAACTATGACTC | >hg38_21_+_8433172_8446622_RNA45SN1_45S_50ntFlanks | 11703 | 11740 | 38 | AATGAAGCGCGGGTAAACGGCGGGAGTAACTATGACTC | 181.7 | -1.9 | 0.7 | -2.5 | 1.2E-02 | 1.7E-02 | BT20/MB468 | MITO |
| >hg38_21_+_8433172_8446622_RNA45SN1_45S_50ntFlanks@8174.8214.41\|TCCTTCTGATCGAGGCCCAGCCCGTGGACGGTGTGAGGCCG | >hg38_21_+_8433172_8446622_RNA45SN1_45S_50ntFlanks | 8174 | 8214 | 41 | TCCTTCTGATCGAGGCCCAGCCCGTGGACGGTGTGAGGCCG | 122.0 | -3.6 | 1.4 | -2.5 | 1.3E-02 | 1.7E-02 | MB231/MB468 | MP |
| >hg38_21_+_8433172_8446622_RNA45SN1_45S_50ntFlanks@12420.12447.28\|ATGTCGGCTCTTCCTATCATTGTGAAGC | >hg38_21_+_8433172_8446622_RNA45SN1_45S_50ntFlanks | 12420 | 12447 | 28 | ATGTCGGCTCTTCCTATCATTGTGAAGC | 269.8 | -2.5 | 0.9 | -2.9 | 3.9E-03 | 1.7E-02 | BT20/MB231 | NUC |
| >hg38_21_+_8433172_8446622_RNA45SN1_45S_50ntFlanks@11672.11702.31\|CAGTGCTCTGAATGTCAAAGTGAAGAAATTC | >hg38_21_+_8433172_8446622_RNA45SN1_45S_50ntFlanks | 11672 | 11702 | 31 | CAGTGCTCTGAATGTCAAAGTGAAGAAATTC | 561.4 | -2.6 | 0.9 | -2.8 | 4.5E-03 | 1.7E-02 | BT20/MB231 | CYTO |
| >hg38_21_+_8433172_8446622_RNA45SN1_45S_50ntFlanks@8196.8217.22\|CGTGGACGGTGTGAGGCCGGTA | >hg38_21_+_8433172_8446622_RNA45SN1_45S_50ntFlanks | 8196 | 8217 | 22 | CGTGGACGGTGTGAGGCCGGTA | 345.7 | 1.4 | 0.6 | 2.5 | 1.4E-02 | 1.7E-02 | BT20/MB468 | CYTO |
| >hg38_21_+_8433172_8446622_RNA45SN1_45S_50ntFlanks@9313.9338.26\|TCTAACACGTGCGCGAGTCGGGGGCT | >hg38_21_+_8433172_8446622_RNA45SN1_45S_50ntFlanks | 9313 | 9338 | 26 | TCTAACACGTGCGCGAGTCGGGGGCT | 679.8 | -4.8 | 1.8 | -2.6 | 9.0E-03 | 1.7E-02 | BT20/MB468 | MP |
| >hg38_21_+_8433172_8446622_RNA45SN1_45S_50ntFlanks@12420.12438.19\|ATGTCGGCTCTTCCTATCA | >hg38_21_+_8433172_8446622_RNA45SN1_45S_50ntFlanks | 12420 | 12438 | 19 | ATGTCGGCTCTTCCTATCA | 1314.7 | 1.8 | 0.7 | 2.5 | 1.4E-02 | 1.7E-02 | MB231/MB468 | CYTO |
| >hg38_21_+_8433172_8446622_RNA45SN1_45S_50ntFlanks@3850.3869.20\|GATAACTGTGGTAATTCTAG | >hg38_21_+_8433172_8446622_RNA45SN1_45S_50ntFlanks | 3850 | 3869 | 20 | GATAACTGTGGTAATTCTAG | 162.1 | -4.3 | 1.7 | -2.6 | 9.0E-03 | 1.7E-02 | BT20/MB468 | MP |
| >hg38_21_+_8433172_8446622_RNA45SN1_45S_50ntFlanks@5487.5506.20\|CGCTGAGAAGACGGTCGAAC | >hg38_21_+_8433172_8446622_RNA45SN1_45S_50ntFlanks | 5487 | 5506 | 20 | CGCTGAGAAGACGGTCGAAC | 239.6 | -4.4 | 1.7 | -2.6 | 9.0E-03 | 1.7E-02 | BT20/MB468 | MP |
| >hg38_21_+_8433172_8446622_RNA45SN1_45S_50ntFlanks@4006.4023.18\|ACCTCGGGCCGATCGCAC | >hg38_21_+_8433172_8446622_RNA45SN1_45S_50ntFlanks | 4006 | 4023 | 18 | ACCTCGGGCCGATCGCAC | 1230.5 | 2.0 | 0.8 | 2.5 | 1.4E-02 | 1.7E-02 | MB231/MB468 | CYTO |
| >hg38_21_+_8433172_8446622_RNA45SN1_45S_50ntFlanks@12893.12933.41\|CCTCGCCCGTCACGCACCGCACGTTCGTGGGGAACCTGGCG | >hg38_21_+_8433172_8446622_RNA45SN1_45S_50ntFlanks | 12893 | 12933 | 41 | CCTCGCCCGTCACGCACCGCACGTTCGTGGGGAACCTGGCG | 160.0 | -3.7 | 1.5 | -2.5 | 1.4E-02 | 1.7E-02 | BT20/MB468 | CYTO |
| >trna134_GluTTC_1_-_16861774_16861845@1.33.33,trna5_GluTTC_1_+_17199078_17199149@1.33.33,trna84_GluTTC_1_-_161391883_161391954@1.33.33,trna94_GluTTC_1_-_149664355_149664427@1.33.33\|TCCCTGGTGGTCTAGTGGCTAGGATTCGGCGCT | >trna134_GluTTC_1_-_16861774_16861845 | 1 | 33 | 33 | TCCCTGGTGGTCTAGTGGCTAGGATTCGGCGCT | 330.4 | 3.5 | 1.2 | 2.9 | 3.5E-03 | 1.7E-02 | BT20/MB231 | MP |
| >hg38_21_+_8433172_8446622_RNA45SN1_45S_50ntFlanks@10240.10264.25\|CTACGCCGCGACGAGTAGGAGGGCC | >hg38_21_+_8433172_8446622_RNA45SN1_45S_50ntFlanks | 10240 | 10264 | 25 | CTACGCCGCGACGAGTAGGAGGGCC | 130.1 | 3.8 | 1.4 | 2.6 | 9.0E-03 | 1.7E-02 | BT20/MB468 | MP |
| >hg38_21_+_8433172_8446622_RNA45SN1_45S_50ntFlanks@12419.12460.42\|GATGTCGGCTCTTCCTATCATTGTGAAGCAGAATTCACCAAG | >hg38_21_+_8433172_8446622_RNA45SN1_45S_50ntFlanks | 12419 | 12460 | 42 | GATGTCGGCTCTTCCTATCATTGTGAAGCAGAATTCACCAAG | 453.9 | -2.7 | 0.9 | -2.8 | 4.5E-03 | 1.7E-02 | BT20/MB231 | MITO |
| >hg38_21_+_8433172_8446622_RNA45SN1_45S_50ntFlanks@11745.11768.24\|AAGGTAGCCAAATGCCTCGTCATC | >hg38_21_+_8433172_8446622_RNA45SN1_45S_50ntFlanks | 11745 | 11768 | 24 | AAGGTAGCCAAATGCCTCGTCATC | 71.1 | -5.4 | 2.2 | -2.5 | 1.3E-02 | 1.7E-02 | MB231/MB468 | MP |
| >hg38_21_+_8433172_8446622_RNA45SN1_45S_50ntFlanks@12053.12072.20\|GTGAGGCGGGGGGGCGAGCC | >hg38_21_+_8433172_8446622_RNA45SN1_45S_50ntFlanks | 12053 | 12072 | 20 | GTGAGGCGGGGGGGCGAGCC | 74.4 | -4.4 | 1.7 | -2.6 | 9.0E-03 | 1.7E-02 | BT20/MB468 | MP |
| >hg38_21_+_8433172_8446622_RNA45SN1_45S_50ntFlanks@9940.9980.41\|CAGGACGGTGGCCATGGAAGTCGGAATCCGCTAAGGAGTGT | >hg38_21_+_8433172_8446622_RNA45SN1_45S_50ntFlanks | 9940 | 9980 | 41 | CAGGACGGTGGCCATGGAAGTCGGAATCCGCTAAGGAGTGT | 259.9 | 3.7 | 1.3 | 2.9 | 3.6E-03 | 1.7E-02 | BT20/MB231 | MP |

| | | | | | | | | | | | | | |
|---|---|---|---|---|---|---|---|---|---|---|---|---|---|
| >MI0000102\|hsa-mir-100&WithFlank&11\|-\|122152223\|122152314@19.40.22(+1C)&[MIMAT0000098&hsa-miR-100-5p&offsets\|0\|0(+1C);m-27&11\|-\|122152276\|122152296&offsets\|0\|+1(+1C)]\|AACCCGTAGATCCGAACTTGTGC | >MI0000102\|hsa-mir-100&WithFlank&11\|-\|122152223\|122152314 | 19 | 40 | 22(+1C) | AACCCGTAGATCCGAACTTGTGC | 5694.0 | -3.4 | 1.2 | -2.8 | 4.5E-03 | 1.7E-02 | BT20/MB231 | CYTO |
| >hg38_21_+_8433172_8446622_RNA45SN1_45S_50ntFlanks@12072.12092.21\|CCCGAGGGGCTCTCGCTTCTG | >hg38_21_+_8433172_8446622_RNA45SN1_45S_50ntFlanks | 12072 | 12092 | 21 | CCCGAGGGGCTCTCGCTTCTG | 67.4 | -4.3 | 1.7 | -2.6 | 9.0E-03 | 1.7E-02 | BT20/MB468 | MP |
| >hg38_21_+_8433172_8446622_RNA45SN1_45S_50ntFlanks@4724.4768.45\|ATTAATCAAGAACGAAAGTCGGAGGTTCGAAGACGATCAGATACC | >hg38_21_+_8433172_8446622_RNA45SN1_45S_50ntFlanks | 4724 | 4768 | 45 | ATTAATCAAGAACGAAAGTCGGAGGTTCGAAGACGATCAGATACC | 359.8 | -3.0 | 1.0 | -2.8 | 4.6E-03 | 1.7E-02 | BT20/MB231 | CYTO |
| >trna15_ValAAC_5_-_180615416_180615488@1.32.32\|GTTTCCGTAGTGTAGTGGTCATCACGTTCGCC | >trna15_ValAAC_5_-_180615416_180615488 | 1 | 32 | 32 | GTTTCCGTAGTGTAGTGGTCATCACGTTCGCC | 229.7 | 6.9 | 2.6 | 2.6 | 9.0E-03 | 1.7E-02 | BT20/MB468 | MP |
| >MI0000457\|hsa-mir-141&WithFlank&12\|+\|6964091\|6964197@65.84.20&[MIMAT0000432&hsa-miR-141-3p&offsets\|0\|-2;m-134&12\|+\|6964155\|6964175&offsets\|0\|-1]\|TAACACTGTCTGGTAAAGAT | >MI0000457\|hsa-mir-141&WithFlank&12\|+\|6964091\|6964197 | 65 | 84 | 20 | TAACACTGTCTGGTAAAGAT | 166.5 | 3.9 | 1.5 | 2.6 | 9.0E-03 | 1.7E-02 | BT20/MB468 | MP |
| >hg38_21_+_8433172_8446622_RNA45SN1_45S_50ntFlanks@7975.8003.29\|CGCGACCTCAGATCAGACGTGGCGACCCG | >hg38_21_+_8433172_8446622_RNA45SN1_45S_50ntFlanks | 7975 | 8003 | 29 | CGCGACCTCAGATCAGACGTGGCGACCCG | 4273.1 | 3.6 | 1.2 | 2.9 | 3.6E-03 | 1.7E-02 | BT20/MB231 | MP |
| >hg38_21_+_8433172_8446622_RNA45SN1_45S_50ntFlanks@12251.12269.19\|AAAGCTCGCTTGATCTTGA | >hg38_21_+_8433172_8446622_RNA45SN1_45S_50ntFlanks | 12251 | 12269 | 19 | AAAGCTCGCTTGATCTTGA | 237.5 | -2.7 | 1.1 | -2.5 | 1.2E-02 | 1.7E-02 | BT20/MB468 | MITO |
| >hg38_21_+_8433172_8446622_RNA45SN1_45S_50ntFlanks@12446.12477.32\|GCAGAATTCACCAAGCGTTGGATTGTTCACCC | >hg38_21_+_8433172_8446622_RNA45SN1_45S_50ntFlanks | 12446 | 12477 | 32 | GCAGAATTCACCAAGCGTTGGATTGTTCACCC | 315.2 | -2.5 | 0.9 | -2.8 | 4.6E-03 | 1.7E-02 | BT20/MB231 | CYTO |
| >hg38_21_+_8433172_8446622_RNA45SN1_45S_50ntFlanks@12241.12269.29\|CAGAAGGGCAAAAGCTCGCTTGATCTTGA | >hg38_21_+_8433172_8446622_RNA45SN1_45S_50ntFlanks | 12241 | 12269 | 29 | CAGAAGGGCAAAAGCTCGCTTGATCTTGA | 215.7 | -4.4 | 1.7 | -2.6 | 9.0E-03 | 1.7E-02 | BT20/MB468 | MP |
| >trna10_LysCTT_16_+_3241501_3241573@8.35.28,trna13_LysCTT_14_-_58706613_58706685@8.35.28,trna2_LysCTT_15_+_79152904_79152976@8.35.28,trna4_LysCTT_18_-_43669271_43669343@8.35.28\|TAGCTCAGTCGGTAGAGCATGGGACTCT | >trna10_LysCTT_16_+_3241501_3241573 | 8 | 35 | 28 | TAGCTCAGTCGGTAGAGCATGGGACTCT | 280.5 | -5.9 | 2.1 | -2.8 | 4.5E-03 | 1.7E-02 | BT20/MB231 | MITO |
| >hg38_21_+_8433172_8446622_RNA45SN1_45S_50ntFlanks@12748.12774.27\|CCTGTCCCCGCCGGCGGGCCGCCCCCC | >hg38_21_+_8433172_8446622_RNA45SN1_45S_50ntFlanks | 12748 | 12774 | 27 | CCTGTCCCCGCCGGCGGGCCGCCCCCC | 265.6 | -3.0 | 1.2 | -2.5 | 1.2E-02 | 1.7E-02 | BT20/MB468 | MITO |
| >hg38_21_+_8433172_8446622_RNA45SN1_45S_50ntFlanks@7973.7994.22\|GACGCGACCTCAGATCAGACGT | >hg38_21_+_8433172_8446622_RNA45SN1_45S_50ntFlanks | 7973 | 7994 | 22 | GACGCGACCTCAGATCAGACGT | 797.8 | 2.1 | 0.9 | 2.4 | 1.5E-02 | 1.7E-02 | BT20/MB468 | NUC |
| >hg38_21_+_8433172_8446622_RNA45SN1_45S_50ntFlanks@3711.3729.19\|GTTGATCCTGCCAGTAGCA | >hg38_21_+_8433172_8446622_RNA45SN1_45S_50ntFlanks | 3711 | 3729 | 19 | GTTGATCCTGCCAGTAGCA | 285.6 | 1.3 | 0.5 | 2.5 | 1.4E-02 | 1.7E-02 | BT20/MB468 | CYTO |
| >MI0000289\|hsa-mir-181a-1&WithFlank&1\|-\|198859038\|198859159@30.51.22(+1A);MI0000269\|hsa-mir-181a-2&WithFlank&9\|+\|124692436\|124692557@45.66.22(+1A)&[MIMAT0000256&hsa-miR-181a-5p&offsets\|0\|-1(+1A);m-32&1\|-\|198859108\|198859130&offsets\|0\|-1(+1A)];[MIMAT0000256_1&hsa-miR-181a-5p&offsets\|0\|-1(+1A);m-32&9\|+\|124692480\|124692502&offsets\|0\|-1(+1A)]\|AACATTCAACGCTGTCGGTGAGA | >MI0000289\|hsa-mir-181a-1&WithFlank&1\|-\|198859038\|198859159 | 30 | 51 | 22(+1A) | AACATTCAACGCTGTCGGTGAGA | 1004.0 | -3.0 | 1.0 | -2.9 | 3.6E-03 | 1.7E-02 | BT20/MB231 | MP |
| >MI0000103\|hsa-mir-101-1&WithFlank&1\|-\|65058428\|65058514@53.73.21;MI0000739\|hsa-mir-101-2&WithFlank&9\|+\|4850291\|4850381@55.75.21&[MIMAT0000099&hsa-miR-101-3p&offsets\|0\|0;m-17&1\|-\|65058443\|65058462&offsets\|0\|+1];[MIMAT0000099_1&hsa-miR-101-3p&offsets\|0\|0;m-16&9\|+\|4850345\|4850364&offsets\|0\|+1]\|TACAGTACTGTGATAACTGAA | >MI0000103\|hsa-mir-101-1&WithFlank&1\|-\|65058428\|65058514 | 53 | 73 | 21 | TACAGTACTGTGATAACTGAA | 459.2 | -2.5 | 0.9 | -2.8 | 4.5E-03 | 1.7E-02 | BT20/MB231 | MITO |
| >hg38_21_+_8433172_8446622_RNA45SN1_45S_50ntFlanks@10841.10858.18\|CGTAACTTCGGGATAAGG | >hg38_21_+_8433172_8446622_RNA45SN1_45S_50ntFlanks | 10841 | 10858 | 18 | CGTAACTTCGGGATAAGG | 463.4 | -3.7 | 1.4 | -2.6 | 9.0E-03 | 1.7E-02 | BT20/MB468 | MP |
| >hg38_MT_+_1621_3279_RNR2_16S_50ntFlanks@493.518.26\|GCTCTTTGGACACTAGGAAAAAACCT | >hg38_MT_+_1621_3279_RNR2_16S_50ntFlanks | 493 | 518 | 26 | GCTCTTTGGACACTAGGAAAAAACCT | 299.1 | 2.9 | 1.1 | 2.6 | 9.1E-03 | 1.7E-02 | BT20/MB468 | MP |
| >MI0000734\|hsa-mir-106b&WithFlank&7\|-\|100093987\|100094080@18.37.20(+1C)&[MIMAT0000680&hsa-miR-106b-5p&offsets\|0\|-1(+1C);m-103&7\|-\|100094043\|100094063&offsets\|0\|-1(+1C)]\|TAAAGTGCTGACAGTGCAGAC | >MI0000734\|hsa-mir-106b&WithFlank&7\|-\|100093987\|100094080 | 18 | 37 | 20(+1C) | TAAAGTGCTGACAGTGCAGAC | 206.5 | -4.3 | 1.5 | -2.9 | 3.6E-03 | 1.7E-02 | BT20/MB231 | MP |
| >hg38_21_+_8433172_8446622_RNA45SN1_45S_50ntFlanks@4069.4095.27\|CTTTCGATGGTAGTCGCCGTGCCTACC | >hg38_21_+_8433172_8446622_RNA45SN1_45S_50ntFlanks | 4069 | 4095 | 27 | CTTTCGATGGTAGTCGCCGTGCCTACC | 655.3 | -4.7 | 1.8 | -2.6 | 9.1E-03 | 1.7E-02 | BT20/MB468 | MP |
| >MI0000087\|hsa-mir-29a&WithFlank&7\|-\|130876741\|130876816@47.70.24&[MIMAT0000086&hsa-miR-29a-3p&offsets\|-1\|+1;m-26&7\|-\|130876749\|130876769&offsets\|-1\|+2]\|CTAGCACCATCTGAAATCGGTTAT | >MI0000087\|hsa-mir-29a&WithFlank&7\|-\|130876741\|130876816 | 47 | 70 | 24 | CTAGCACCATCTGAAATCGGTTAT | 1087.5 | -3.3 | 1.1 | -2.9 | 3.6E-03 | 1.7E-02 | BT20/MB231 | MP |
| >hg38_21_+_8433172_8446622_RNA45SN1_45S_50ntFlanks@11671.11702.32\|CCAGTGCTCTGAATGTCAAAGTGAAGAAATTC | >hg38_21_+_8433172_8446622_RNA45SN1_45S_50ntFlanks | 11671 | 11702 | 32 | CCAGTGCTCTGAATGTCAAAGTGAAGAAATTC | 368.2 | -2.6 | 0.9 | -2.8 | 4.6E-03 | 1.7E-02 | BT20/MB231 | CYTO |
| >hg38_21_+_8433172_8446622_RNA45SN1_45S_50ntFlanks@8176.8214.39\|CTTCTGATCGAGGCCCAGCCCGTGGACGGTGTGAGGCCG | >hg38_21_+_8433172_8446622_RNA45SN1_45S_50ntFlanks | 8176 | 8214 | 39 | CTTCTGATCGAGGCCCAGCCCGTGGACGGTGTGAGGCCG | 905.6 | -3.3 | 1.4 | -2.5 | 1.4E-02 | 1.7E-02 | MB231/MB468 | CYTO |
| >hg38_21_+_8433172_8446622_RNA45SN1_45S_50ntFlanks@11207.11233.27\|GGCGGCGACTCTGGACGCGAGCCGGGC | >hg38_21_+_8433172_8446622_RNA45SN1_45S_50ntFlanks | 11207 | 11233 | 27 | GGCGGCGACTCTGGACGCGAGCCGGGC | 418.6 | -4.7 | 1.9 | -2.4 | 1.5E-02 | 1.7E-02 | BT20/MB468 | NUC |
| >trnaMT_GluTTC_MT_-_14674_14742@38.71.34,trnalookalike8_GluTTC_5_-_93905172_93905240@38.71.34\|CATTGGTCGTGGTTGTAGTCCGTGCGAGAATACC | >trnaMT_GluTTC_MT_-_14674_14742 | 38 | 71 | 34 | CATTGGTCGTGGTTGTAGTCCGTGCGAGAATACC | 582.1 | 3.9 | 1.5 | 2.6 | 9.1E-03 | 1.7E-02 | BT20/MB468 | MP |
| >MI0000067\|hsa-let-7f-1&WithFlank&9\|+\|94176341\|94176439@13.34.22(+1A);MI0000068\|hsa-let-7f-2&WithFlank&X\|-\|53557186\|53557280@14.35.22(+1A)&[MIMAT0000067&hsa-let-7f-5p&offsets\|0\|0(+1A);m-20&9\|+\|94176353\|94176374&offsets\|0\|0(+1A)];[MIMAT0000067_1&hsa-let-7f-5p&offsets\|0\|0(+1A);m-19&X\|-\|53557246\|53557267&offsets\|0\|0(+1A)]\|TGAGGTAGTAGATTGTATAGTTA | >MI0000067\|hsa-let-7f-1&WithFlank&9\|+\|94176341\|94176439 | 13 | 34 | 22(+1A) | TGAGGTAGTAGATTGTATAGTTA | 2524.5 | 2.3 | 0.8 | 2.8 | 4.6E-03 | 1.7E-02 | BT20/MB231 | CYTO |
| >am_trnaMT_HisGTG_MT_+_12138_12206@26.51.26\|AGATTGTGAATCTGACAACAGAGGCT | >am_trnaMT_HisGTG_MT_+_12138_12206 | 26 | 51 | 26 | AGATTGTGAATCTGACAACAGAGGCT | 57.1 | 4.1 | 1.7 | 2.5 | 1.3E-02 | 1.7E-02 | MB231/MB468 | MP |
| >hg38_21_+_8433172_8446622_RNA45SN1_45S_50ntFlanks@7987.8009.23\|TCAGACGTGGCGACCCGCTGAAT | >hg38_21_+_8433172_8446622_RNA45SN1_45S_50ntFlanks | 7987 | 8009 | 23 | TCAGACGTGGCGACCCGCTGAAT | 159.5 | -2.4 | 0.9 | -2.6 | 9.1E-03 | 1.7E-02 | BT20/MB468 | MP |
| >hg38_21_+_8433172_8446622_RNA45SN1_45S_50ntFlanks@12999.13039.41\|CTGCGATCTATTGAAAGTCAGCCCTCGACACAAGGGTTTGT | >hg38_21_+_8433172_8446622_RNA45SN1_45S_50ntFlanks | 12999 | 13039 | 41 | CTGCGATCTATTGAAAGTCAGCCCTCGACACAAGGGTTTGT | 5520.9 | 2.4 | 1.0 | 2.4 | 1.5E-02 | 1.7E-02 | MB231/MB468 | MITO |
| >hg38_21_+_8433172_8446622_RNA45SN1_45S_50ntFlanks@12394.12418.25\|TAGCGACGTCGCTTTTTGATCCTTC | >hg38_21_+_8433172_8446622_RNA45SN1_45S_50ntFlanks | 12394 | 12418 | 25 | TAGCGACGTCGCTTTTTGATCCTTC | 284.7 | -2.2 | 0.9 | -2.4 | 1.5E-02 | 1.7E-02 | MB231/MB468 | MITO |
| >trna111_HisGTG_1_-_147774845_147774916@-1C.31.32,trna118_HisGTG_1_-_145396881_145396952@-1C.31.32,trna16_HisGTG_1_+_146544773_146544844@-1C.31.32,trna1_HisGTG_15_+_45493349_45493420@-1C.31.32,trna21_HisGTG_1_+_147753471_147753542@-1C.31.32,trna33_HisGTG_6_+_27125906_27125977@-1C.31.32,trna7_HisGTG_9_-_14433938_14434009@-1C.31.32,trna8_HisGTG_15_-_45492611_45492682@-1C.31.32,trna9_HisGTG_15_-_45490804_45490875@-1C.31.32\|CGCCGTGATCGTATAGTGGTTAGTACTCTGCG | >trna111_HisGTG_1_-_147774845_147774916 | -1C | 31 | 32 | CGCCGTGATCGTATAGTGGTTAGTACTCTGCG | 216.2 | -1.9 | 0.8 | -2.5 | 1.4E-02 | 1.7E-02 | BT20/MB468 | CYTO |
| >hg38_21_+_8433172_8446622_RNA45SN1_45S_50ntFlanks@12455.12476.22\|ACCAAGCGTTGGATTGTTCACC | >hg38_21_+_8433172_8446622_RNA45SN1_45S_50ntFlanks | 12455 | 12476 | 22 | ACCAAGCGTTGGATTGTTCACC | 509.1 | -2.2 | 0.8 | -2.8 | 4.6E-03 | 1.7E-02 | BT20/MB231 | CYTO |
| >hg38_21_+_8433172_8446622_RNA45SN1_45S_50ntFlanks@9251.9271.21\|GCACGGGGTCGGCGGCGACGT | >hg38_21_+_8433172_8446622_RNA45SN1_45S_50ntFlanks | 9251 | 9271 | 21 | GCACGGGGTCGGCGGCGACGT | 94.0 | 3.7 | 1.4 | 2.6 | 9.1E-03 | 1.7E-02 | BT20/MB468 | MP |
| >MI0005561\|hsa-mir-877&WithFlank&6\|+\|30584326\|30584423@7.25.19&[MIMAT0004949&hsa-miR-877-5p&offsets\|0\|-1;m-382&6\|+\|30584332\|30584354&offsets\|0\|-4]\|GTAGAGGAGATGGCGCAGG | >MI0005561\|hsa-mir-877&WithFlank&6\|+\|30584326\|30584423 | 7 | 25 | 19 | GTAGAGGAGATGGCGCAGG | 52.9 | 4.1 | 1.7 | 2.5 | 1.3E-02 | 1.7E-02 | MB231/MB468 | MP |
| >hg38_MT_+_1621_3279_RNR2_16S_50ntFlanks@292.313.22\|AGACGAGCTACCTAAGAACAGC | >hg38_MT_+_1621_3279_RNR2_16S_50ntFlanks | 292 | 313 | 22 | AGACGAGCTACCTAAGAACAGC | 64.4 | 3.4 | 1.4 | 2.5 | 1.3E-02 | 1.7E-02 | MB231/MB468 | MP |
| >trnaMT_AspGTC_MT_+_7518_7585@46.71.26\|TAGGCTAAATCCTATATATCTTACCA | >trnaMT_AspGTC_MT_+_7518_7585 | 46 | 71 | 26 | TAGGCTAAATCCTATATATCTTACCA | 56.7 | 4.1 | 1.6 | 2.5 | 1.3E-02 | 1.7E-02 | MB231/MB468 | MP |
| >trna10_LysCTT_16_+_3241501_3241573@4.36.33,trna13_LysCTT_14_-_58706613_58706685@4.36.33,trna2_LysCTT_15_+_79152904_79152976@4.36.33\|CGGCTAGCTCAGTCGGTAGAGCATGGGACTCTT | >trna10_LysCTT_16_+_3241501_3241573 | 4 | 36 | 33 | CGGCTAGCTCAGTCGGTAGAGCATGGGACTCTT | 140.4 | 5.1 | 1.9 | 2.6 | 9.1E-03 | 1.7E-02 | BT20/MB468 | MP |
| >hg38_MT_+_1621_3279_RNR2_16S_50ntFlanks@482.504.23\|AAAGAGGAACAGCTCTTTGGACA | >hg38_MT_+_1621_3279_RNR2_16S_50ntFlanks | 482 | 504 | 23 | AAAGAGGAACAGCTCTTTGGACA | 154.5 | 2.6 | 1.0 | 2.6 | 9.1E-03 | 1.7E-02 | BT20/MB468 | MP |
| >hg38_21_+_8433172_8446622_RNA45SN1_45S_50ntFlanks@9821.9850.30\|CCTAGTGGGCCACTTTTGGTAAGCAGAACT | >hg38_21_+_8433172_8446622_RNA45SN1_45S_50ntFlanks | 9821 | 9850 | 30 | CCTAGTGGGCCACTTTTGGTAAGCAGAACT | 320.6 | -2.9 | 1.2 | -2.4 | 1.4E-02 | 1.7E-02 | MB231/MB468 | CYTO |
| >hg38_21_+_8433172_8446622_RNA45SN1_45S_50ntFlanks@5059.5079.21\|CGATTTGTCTGGTTAATTCCG | >hg38_21_+_8433172_8446622_RNA45SN1_45S_50ntFlanks | 5059 | 5079 | 21 | CGATTTGTCTGGTTAATTCCG | 64.8 | -4.7 | 1.8 | -2.6 | 9.1E-03 | 1.7E-02 | BT20/MB468 | MP |
| >hg38_21_+_8433172_8446622_RNA45SN1_45S_50ntFlanks@10401.10420.20\|GAACATGGGTCAGTCGGTCC | >hg38_21_+_8433172_8446622_RNA45SN1_45S_50ntFlanks | 10401 | 10420 | 20 | GAACATGGGTCAGTCGGTCC | 147.8 | -3.7 | 1.5 | -2.5 | 1.4E-02 | 1.7E-02 | BT20/MB468 | CYTO |
| >hg38_21_+_8433172_8446622_RNA45SN1_45S_50ntFlanks@12417.12447.31\|TCGATGTCGGCTCTTCCTATCATTGTGAAGC | >hg38_21_+_8433172_8446622_RNA45SN1_45S_50ntFlanks | 12417 | 12447 | 31 | TCGATGTCGGCTCTTCCTATCATTGTGAAGC | 1002.3 | -2.4 | 0.8 | -2.8 | 4.6E-03 | 1.7E-02 | BT20/MB231 | CYTO |
| >hg38_MT_+_1621_3279_RNR2_16S_50ntFlanks@493.517.25\|GCTCTTTGGACACTAGGAAAAAACC | >hg38_MT_+_1621_3279_RNR2_16S_50ntFlanks | 493 | 517 | 25 | GCTCTTTGGACACTAGGAAAAAACC | 319.7 | -2.6 | 0.9 | -2.9 | 3.9E-03 | 1.7E-02 | BT20/MB231 | NUC |
| >hg38_21_+_8433172_8446622_RNA45SN1_45S_50ntFlanks@8175.8202.28\|CCTTCTGATCGAGGCCCAGCCCGTGGAC | >hg38_21_+_8433172_8446622_RNA45SN1_45S_50ntFlanks | 8175 | 8202 | 28 | CCTTCTGATCGAGGCCCAGCCCGTGGAC | 85.8 | 2.6 | 1.1 | 2.4 | 1.5E-02 | 1.7E-02 | MB231/MB468 | MITO |
| >am_trnaMT_SerGCT_MT_+_12207_12265@14.36.23\|AGAACTGCTAACTCATGCCCCCA | >am_trnaMT_SerGCT_MT_+_12207_12265 | 14 | 36 | 23 | AGAACTGCTAACTCATGCCCCCA | 60.5 | 5.4 | 2.1 | 2.6 | 9.1E-03 | 1.7E-02 | BT20/MB468 | MP |
| >hg38_MT_+_1621_3279_RNR2_16S_50ntFlanks@1370.1389.20\|ATGTTGGATCAGGACATCCC | >hg38_MT_+_1621_3279_RNR2_16S_50ntFlanks | 1370 | 1389 | 20 | ATGTTGGATCAGGACATCCC | 183.1 | 2.7 | 1.1 | 2.5 | 1.3E-02 | 1.7E-02 | MB231/MB468 | MP |
| >hg38_21_+_8433172_8446622_RNA45SN1_45S_50ntFlanks@6662.6696.35\|GGTGGATCACTCGGCTCGTGCGTCGATGAAGAACG | >hg38_21_+_8433172_8446622_RNA45SN1_45S_50ntFlanks | 6662 | 6696 | 35 | GGTGGATCACTCGGCTCGTGCGTCGATGAAGAACG | 180.9 | 4.6 | 1.6 | 2.9 | 3.6E-03 | 1.7E-02 | BT20/MB231 | MP |
| >MI0000816\|hsa-mir-335&WithFlank&7\|+\|130496105\|130496210@22.44.23&[MIMAT0000765&hsa-miR-335-5p&offsets\|0\|0;m-249&7\|+\|130496126\|130496147&offsets\|0\|+1]\|TCAAGAGCAATAACGAAAAATGT | >MI0000816\|hsa-mir-335&WithFlank&7\|+\|130496105\|130496210 | 22 | 44 | 23 | TCAAGAGCAATAACGAAAAATGT | 488.4 | -4.2 | 1.4 | -2.9 | 3.6E-03 | 1.7E-02 | BT20/MB231 | MP |
| >MI0000077\|hsa-mir-21&WithFlank&17\|+\|59841260\|59841343@15.36.22&[MIMAT0000076&hsa-miR-21-5p&offsets\|+1\|+1;m-1&17\|+\|59841273\|59841294&offsets\|+1\|+1]\|AGCTTATCAGACTGATGTTGAC | >MI0000077\|hsa-mir-21&WithFlank&17\|+\|59841260\|59841343 | 15 | 36 | 22 | AGCTTATCAGACTGATGTTGAC | 1625.5 | -2.8 | 1.0 | -2.9 | 3.6E-03 | 1.7E-02 | BT20/MB231 | MP |

| | | | | | | | | | | | | | |
|---|---|---|---|---|---|---|---|---|---|---|---|---|---|
| >MI0000108\|hsa-mir-103a-2&WithFlank&20\|+\|3917488\|3917577@54.75.22(+1G);MI0000109\|hsa-mir-103a-1&WithFlank&5\|-\|168560890\|168560979@54.75.22(+1G)&[MIMAT0000101&hsa-miR-103a-3p&offsets\|0\|-1(+1G);m-12&20\|+\|3917541\|3917563&offsets\|0\|-1(+1G)];[MIMAT0000101_1&hsa-miR-103a-3p&offsets\|0\|-1(+1G);m-13&5\|-\|168560904\|168560926&offsets\|0\|-1(+1G);m-44220&5\|-\|168560908\|168560926&offsets\|0\|+3(+1G)]\|AGCAGCATTGTACAGGGCTATGG | >MI0000108\|hsa-mir-103a-2&WithFlank&20\|+\|3917488\|3917577 | 54 | 75 | 22(+1G) | AGCAGCATTGTACAGGGCTATGG | 761.1 | -3.1 | 1.1 | -2.9 | 3.6E-03 | 1.7E-02 | BT20/MB231 | MP |
| >trnaMT_MetCAT_MT_+_4402_4469@40.71.32\|AAAATGTTGGTTATACCCTTCCCGTACTACCA | >trnaMT_MetCAT_MT_+_4402_4469 | 40 | 71 | 32 | AAAATGTTGGTTATACCCTTCCCGTACTACCA | 475.7 | 3.0 | 1.2 | 2.6 | 9.2E-03 | 1.7E-02 | BT20/MB468 | MP |
| >hg38_21_+_8433172_8446622_RNA45SN1_45S_50ntFlanks@12453.12477.25\|TCACCAAGCGTTGGATTGTTCACCC | >hg38_21_+_8433172_8446622_RNA45SN1_45S_50ntFlanks | 12453 | 12477 | 25 | TCACCAAGCGTTGGATTGTTCACCC | 489.7 | -2.4 | 0.8 | -2.8 | 4.7E-03 | 1.7E-02 | BT20/MB231 | CYTO |
| >hg38_21_+_8433172_8446622_RNA45SN1_45S_50ntFlanks@8176.8220.45\|CTTCTGATCGAGGCCCAGCCCGTGGACGGTGTGAGGCCGGTAGCG | >hg38_21_+_8433172_8446622_RNA45SN1_45S_50ntFlanks | 8176 | 8220 | 45 | CTTCTGATCGAGGCCCAGCCCGTGGACGGTGTGAGGCCGGTAGCG | 481.4 | -2.6 | 1.0 | -2.5 | 1.2E-02 | 1.7E-02 | BT20/MB468 | MITO |
| >hg38_21_+_8433172_8446622_RNA45SN1_45S_50ntFlanks@9157.9187.31\|CAGTGCGCCCCGGGCGGGTCGCGCCGTCGGG | >hg38_21_+_8433172_8446622_RNA45SN1_45S_50ntFlanks | 9157 | 9187 | 31 | CAGTGCGCCCCGGGCGGGTCGCGCCGTCGGG | 420.8 | -4.8 | 2.0 | -2.4 | 1.5E-02 | 1.7E-02 | BT20/MB468 | NUC |
| >hg38_21_+_8433172_8446622_RNA45SN1_45S_50ntFlanks@12423.12443.21\|TCGGCTCTTCCTATCATTGTG | >hg38_21_+_8433172_8446622_RNA45SN1_45S_50ntFlanks | 12423 | 12443 | 21 | TCGGCTCTTCCTATCATTGTG | 520.9 | -2.7 | 1.1 | -2.4 | 1.4E-02 | 1.7E-02 | MB231/MB468 | CYTO |
| >hg38_21_+_8433172_8446622_RNA45SN1_45S_50ntFlanks@12907.12934.28\|CACCGCACGTTCGTGGGGAACCTGGCGC | >hg38_21_+_8433172_8446622_RNA45SN1_45S_50ntFlanks | 12907 | 12934 | 28 | CACCGCACGTTCGTGGGGAACCTGGCGC | 110.6 | -1.7 | 0.7 | -2.5 | 1.2E-02 | 1.7E-02 | BT20/MB468 | MITO |
| >hg38_MT_+_598_1651_RNR1_12S_50ntFlanks@855.872.18\|TAGTTGAACAGGGCCCTG | >hg38_MT_+_598_1651_RNR1_12S_50ntFlanks | 855 | 872 | 18 | TAGTTGAACAGGGCCCTG | 138.3 | 4.1 | 1.6 | 2.6 | 9.2E-03 | 1.7E-02 | BT20/MB468 | MP |
| >MI0000809\|hsa-mir-151a&WithFlank&8\|-\|140732558\|140732659@53.74.22&[MIMAT0000757&hsa-miR-151a-3p&offsets\|0\|+1;m-30&8\|-\|140732587\|140732607&offsets\|0\|+1]\|CTAGACTGAAGCTCCTTGAGGA | >MI0000809\|hsa-mir-151a&WithFlank&8\|-\|140732558\|140732659 | 53 | 74 | 22 | CTAGACTGAAGCTCCTTGAGGA | 25366.3 | 2.0 | 0.7 | 2.8 | 4.7E-03 | 1.7E-02 | BT20/MB231 | CYTO |
| >hg38_21_+_8433172_8446622_RNA45SN1_45S_50ntFlanks@5236.5255.20\|CACTGACTGGCTCAGCGTGT | >hg38_21_+_8433172_8446622_RNA45SN1_45S_50ntFlanks | 5236 | 5255 | 20 | CACTGACTGGCTCAGCGTGT | 159.1 | -1.6 | 0.7 | -2.5 | 1.3E-02 | 1.7E-02 | MB231/MB468 | MP |
| >hg38_21_+_8433172_8446622_RNA45SN1_45S_50ntFlanks@4049.4066.18\|TCGAACGTCTGCCCTATC | >hg38_21_+_8433172_8446622_RNA45SN1_45S_50ntFlanks | 4049 | 4066 | 18 | TCGAACGTCTGCCCTATC | 829.6 | -2.5 | 0.9 | -2.8 | 4.7E-03 | 1.7E-02 | BT20/MB231 | CYTO |
| >MI0000077\|hsa-mir-21&WithFlank&17\|+\|59841260\|59841343@14.36.23(+1G)&[MIMAT0000076&hsa-miR-21-5p&offsets\|0\|+1(+1G);m-1&17\|+\|59841273\|59841294&offsets\|0\|+1(+1G)]\|TAGCTTATCAGACTGATGTTGACG | >MI0000077\|hsa-mir-21&WithFlank&17\|+\|59841260\|59841343 | 14 | 36 | 23(+1G) | TAGCTTATCAGACTGATGTTGACG | 1877.3 | 2.3 | 0.9 | 2.5 | 1.3E-02 | 1.7E-02 | MB231/MB468 | MP |
| >hg38_21_+_8433172_8446622_RNA45SN1_45S_50ntFlanks@12992.13017.26\|TCCCTCGCTGCGATCTATTGAAAGTC | >hg38_21_+_8433172_8446622_RNA45SN1_45S_50ntFlanks | 12992 | 13017 | 26 | TCCCTCGCTGCGATCTATTGAAAGTC | 493.0 | -2.6 | 0.9 | -2.8 | 4.7E-03 | 1.7E-02 | BT20/MB231 | CYTO |
| >MI0000087\|hsa-mir-29a&WithFlank&7\|-\|130876741\|130876816@48.67.20(+1A)&[MIMAT0000086&hsa-miR-29a-3p&offsets\|0\|-2(+1A);m-26&7\|-\|130876749\|130876769&offsets\|0\|-1(+1A)]\|TAGCACCATCTGAAATCGGTA | >MI0000087\|hsa-mir-29a&WithFlank&7\|-\|130876741\|130876816 | 48 | 67 | 20(+1A) | TAGCACCATCTGAAATCGGTA | 560.9 | 6.1 | 2.3 | 2.6 | 9.2E-03 | 1.7E-02 | BT20/MB468 | MP |
| >hg38_21_+_8433172_8446622_RNA45SN1_45S_50ntFlanks@12415.12433.19\|CTTCGATGTCGGCTCTTCC | >hg38_21_+_8433172_8446622_RNA45SN1_45S_50ntFlanks | 12415 | 12433 | 19 | CTTCGATGTCGGCTCTTCC | 777.0 | -1.9 | 0.8 | -2.4 | 1.5E-02 | 1.7E-02 | MB231/MB468 | MITO |
| >hg38_21_+_8433172_8446622_RNA45SN1_45S_50ntFlanks@8185.8202.18\|GAGGCCCAGCCCGTGGAC | >hg38_21_+_8433172_8446622_RNA45SN1_45S_50ntFlanks | 8185 | 8202 | 18 | GAGGCCCAGCCCGTGGAC | 89.4 | 5.4 | 2.1 | 2.6 | 9.2E-03 | 1.7E-02 | BT20/MB468 | MP |
| >hg38_21_+_8433172_8446622_RNA45SN1_45S_50ntFlanks@4733.4758.26\|GAACGAAAGTCGGAGGTTCGAAGACG | >hg38_21_+_8433172_8446622_RNA45SN1_45S_50ntFlanks | 4733 | 4758 | 26 | GAACGAAAGTCGGAGGTTCGAAGACG | 1237.8 | 1.8 | 0.7 | 2.4 | 1.4E-02 | 1.7E-02 | BT20/MB468 | CYTO |
| >hg38_21_+_8433172_8446622_RNA45SN1_45S_50ntFlanks@9546.9563.18\|TGGAGGTCCGTAGCGGTC | >hg38_21_+_8433172_8446622_RNA45SN1_45S_50ntFlanks | 9546 | 9563 | 18 | TGGAGGTCCGTAGCGGTC | 187.2 | -1.4 | 0.6 | -2.4 | 1.4E-02 | 1.7E-02 | BT20/MB468 | CYTO |
| >MI0000286\|hsa-mir-210&WithFlank&11\|-\|568083\|568204@72.94.23(+1U)&[MIMAT0000267&hsa-miR-210-3p&offsets\|0\|+1(+1U);m-102&11\|-\|568112\|568133&offsets\|0\|+1(+1U)]\|CTGTGCGTGTGACAGCGGCTGATT | >MI0000286\|hsa-mir-210&WithFlank&11\|-\|568083\|568204 | 72 | 94 | 23(+1U) | CTGTGCGTGTGACAGCGGCTGATT | 139.8 | 4.9 | 2.0 | 2.5 | 1.3E-02 | 1.7E-02 | MB231/MB468 | MP |
| >hg38_1_-_228634819_228635039_RNA5S12_5S_50ntFlanks@137.171.35\|GAGACCGCCTGGGAATACCGGGTGCTGTAGGCTTT | >hg38_1_-_228634819_228635039_RNA5S12_5S_50ntFlanks | 137 | 171 | 35 | GAGACCGCCTGGGAATACCGGGTGCTGTAGGCTTT | 1490.2 | -1.9 | 0.7 | -2.9 | 4.0E-03 | 1.7E-02 | BT20/MB231 | NUC |
| >hg38_21_+_8433172_8446622_RNA45SN1_45S_50ntFlanks@7974.8006.33\|ACGCGACCTCAGATCAGACGTGGCGACCCGCTG | >hg38_21_+_8433172_8446622_RNA45SN1_45S_50ntFlanks | 7974 | 8006 | 33 | ACGCGACCTCAGATCAGACGTGGCGACCCGCTG | 38940.6 | 1.8 | 0.7 | 2.5 | 1.3E-02 | 1.7E-02 | BT20/MB468 | MITO |
| >MI0000447\|hsa-mir-128-1&WithFlank&2\|+\|135665391\|135665484@56.76.21(+1A);MI0000727\|hsa-mir-128-2&WithFlank&3\|+\|35744470\|35744565@58.78.21(+1A)&[MIMAT0000424&hsa-miR-128-3p&offsets\|0\|0(+1A);m-108&2\|+\|135665446\|135665466&offsets\|0\|0(+1A)];[MIMAT0000424_1&hsa-miR-128-3p&offsets\|0\|0(+1A);m-117&3\|+\|35744527\|35744547&offsets\|0\|0(+1A)]\|TCACAGTGAACCGGTCTCTTTA | >MI0000447\|hsa-mir-128-1&WithFlank&2\|+\|135665391\|135665484 | 56 | 76 | 21(+1A) | TCACAGTGAACCGGTCTCTTTA | 925.3 | 1.8 | 0.8 | 2.4 | 1.5E-02 | 1.7E-02 | MB231/MB468 | NUC |
| >MI0000077\|hsa-mir-21&WithFlank&17\|+\|59841260\|59841343@14.36.23(+1A)&[MIMAT0000076&hsa-miR-21-5p&offsets\|0\|+1(+1A);m-1&17\|+\|59841273\|59841294&offsets\|0\|+1(+1A)]\|TAGCTTATCAGACTGATGTTGACA | >MI0000077\|hsa-mir-21&WithFlank&17\|+\|59841260\|59841343 | 14 | 36 | 23(+1A) | TAGCTTATCAGACTGATGTTGACA | 12667.1 | 2.5 | 0.9 | 2.6 | 9.2E-03 | 1.7E-02 | BT20/MB468 | MP |
| >hg38_21_+_8433172_8446622_RNA45SN1_45S_50ntFlanks@5487.5513.27\|CGCTGAGAAGACGGTCGAACTTGACTA | >hg38_21_+_8433172_8446622_RNA45SN1_45S_50ntFlanks | 5487 | 5513 | 27 | CGCTGAGAAGACGGTCGAACTTGACTA | 177.0 | -2.8 | 1.1 | -2.5 | 1.3E-02 | 1.7E-02 | BT20/MB468 | MITO |
| >hg38_21_+_8433172_8446622_RNA45SN1_45S_50ntFlanks@6660.6686.27\|GCGGTGGATCACTCGGCTCGTGCGTCG | >hg38_21_+_8433172_8446622_RNA45SN1_45S_50ntFlanks | 6660 | 6686 | 27 | GCGGTGGATCACTCGGCTCGTGCGTCG | 155.9 | 4.8 | 1.6 | 2.9 | 3.7E-03 | 1.7E-02 | BT20/MB231 | MP |
| >trnaMT_TrpTCA_MT_+_5512_5579@13.34.22,trnalookalike3_TrpTCA_1_+_566062_566129@13.34.22\|TAAATACAGACCAAGAGCCTTC | >trnaMT_TrpTCA_MT_+_5512_5579 | 13 | 34 | 22 | TAAATACAGACCAAGAGCCTTC | 146.3 | 3.4 | 1.4 | 2.4 | 1.5E-02 | 1.7E-02 | MB231/MB468 | MITO |
| >hg38_21_+_8433172_8446622_RNA45SN1_45S_50ntFlanks@11671.11691.21\|CCAGTGCTCTGAATGTCAAAG | >hg38_21_+_8433172_8446622_RNA45SN1_45S_50ntFlanks | 11671 | 11691 | 21 | CCAGTGCTCTGAATGTCAAAG | 391.3 | -2.5 | 1.0 | -2.4 | 1.5E-02 | 1.7E-02 | MB231/MB468 | MITO |
| >hg38_MT_+_1621_3279_RNR2_16S_50ntFlanks@51.81.31\|GCTAAACCTAGCCCCAAACCCACTCCACCTT | >hg38_MT_+_1621_3279_RNR2_16S_50ntFlanks | 51 | 81 | 31 | GCTAAACCTAGCCCCAAACCCACTCCACCTT | 790.2 | 2.6 | 0.9 | 2.9 | 4.0E-03 | 1.7E-02 | BT20/MB231 | NUC |
| >hg38_21_+_8433172_8446622_RNA45SN1_45S_50ntFlanks@6649.6667.19\|TACGACTCTTAGCGGTGGA | >hg38_21_+_8433172_8446622_RNA45SN1_45S_50ntFlanks | 6649 | 6667 | 19 | TACGACTCTTAGCGGTGGA | 258.9 | 2.2 | 0.9 | 2.4 | 1.5E-02 | 1.7E-02 | MB231/MB468 | MITO |
| >hg38_21_+_8433172_8446622_RNA45SN1_45S_50ntFlanks@10687.10711.25\|CGGCGTCCGGTGAGCTCTCGCTGGC | >hg38_21_+_8433172_8446622_RNA45SN1_45S_50ntFlanks | 10687 | 10711 | 25 | CGGCGTCCGGTGAGCTCTCGCTGGC | 1958.6 | -4.1 | 1.6 | -2.6 | 9.2E-03 | 1.7E-02 | BT20/MB468 | MP |
| >am_trna119_LysCTT_1_-_145395522_145395594@15.33.19,trna11_LysCTT_5_-_180648979_180649051@15.33.19,trna128_LysCTT_1_-_55423542_55423614@15.33.19,trna13_LysCTT_6_+_26556774_26556846@15.33.19,trna6_LysCTT_19_-_52425396_52425468@15.33.19,trna7_LysCTT_16_+_3225692_3225764@15.33.19,trna9_LysCTT_5_+_180634755_180634827@15.33.19\|GTCGGTAGAGCATGAGACT | >am_trna119_LysCTT_1_-_145395522_145395594 | 15 | 33 | 19 | GTCGGTAGAGCATGAGACT | 29.6 | 5.0 | 2.0 | 2.5 | 1.3E-02 | 1.7E-02 | MB231/MB468 | MP |
| >hg38_21_+_8433172_8446622_RNA45SN1_45S_50ntFlanks@12444.12478.35\|AAGCAGAATTCACCAAGCGTTGGATTGTTCACCCA | >hg38_21_+_8433172_8446622_RNA45SN1_45S_50ntFlanks | 12444 | 12478 | 35 | AAGCAGAATTCACCAAGCGTTGGATTGTTCACCCA | 182.7 | -3.9 | 1.5 | -2.6 | 9.2E-03 | 1.7E-02 | BT20/MB468 | MP |
| >MI0000482\|hsa-mir-185&WithFlank&22\|+\|20033133\|20033226@21.43.23&[MIMAT0000455&hsa-miR-185-5p&offsets\|0\|+1;m-145&22\|+\|20033153\|20033174&offsets\|0\|+1]\|TGGAGAGAAAGGCAGTTCCTGAT | >MI0000482\|hsa-mir-185&WithFlank&22\|+\|20033133\|20033226 | 21 | 43 | 23 | TGGAGAGAAAGGCAGTTCCTGAT | 402.6 | 1.6 | 0.6 | 2.9 | 4.0E-03 | 1.7E-02 | BT20/MB231 | NUC |
| >trnaMT_HisGTG_MT_+_12138_12206@27.61.35\|GATTGTGAATCTGACAACAGAGGCTTACGACCCCT | >trnaMT_HisGTG_MT_+_12138_12206 | 27 | 61 | 35 | GATTGTGAATCTGACAACAGAGGCTTACGACCCCT | 77.9 | 4.8 | 1.9 | 2.5 | 1.3E-02 | 1.7E-02 | MB231/MB468 | MP |
| >MI0000482\|hsa-mir-185&WithFlank&22\|+\|20033133\|20033226@21.41.21&[MIMAT0000455&hsa-miR-185-5p&offsets\|0\|-1;m-145&22\|+\|20033153\|20033174&offsets\|0\|-1]\|TGGAGAGAAAGGCAGTTCCTG | >MI0000482\|hsa-mir-185&WithFlank&22\|+\|20033133\|20033226 | 21 | 41 | 21 | TGGAGAGAAAGGCAGTTCCTG | 384.2 | 3.0 | 1.1 | 2.6 | 9.2E-03 | 1.7E-02 | BT20/MB468 | MP |
| >hg38_21_+_8433172_8446622_RNA45SN1_45S_50ntFlanks@9788.9819.32\|CGCTGGCGTGGAGCCGGGCGTGGAATGCGAGT | >hg38_21_+_8433172_8446622_RNA45SN1_45S_50ntFlanks | 9788 | 9819 | 32 | CGCTGGCGTGGAGCCGGGCGTGGAATGCGAGT | 356.9 | 3.0 | 1.2 | 2.6 | 9.2E-03 | 1.7E-02 | BT20/MB468 | MP |
| >trnaMT_GluTTC_MT_-_14674_14742@46.63.18,trnalookalike8_GluTTC_5_-_93905172_93905240@46.63.18\|GTGGTTGTAGTCCGTGCG | >trnaMT_GluTTC_MT_-_14674_14742 | 46 | 63 | 18 | GTGGTTGTAGTCCGTGCG | 60.3 | 4.1 | 1.7 | 2.5 | 1.3E-02 | 1.7E-02 | MB231/MB468 | MP |
| >hg38_21_+_8433172_8446622_RNA45SN1_45S_50ntFlanks@5074.5098.25\|ATTCCGATAACGAACGAGACTCTGG | >hg38_21_+_8433172_8446622_RNA45SN1_45S_50ntFlanks | 5074 | 5098 | 25 | ATTCCGATAACGAACGAGACTCTGG | 361.0 | -3.6 | 1.4 | -2.5 | 1.3E-02 | 1.7E-02 | BT20/MB468 | MITO |
| >hg38_MT_+_1621_3279_RNR2_16S_50ntFlanks@1351.1388.38\|AATAGGGTTTACGACCTCGATGTTGGATCAGGACATCC | >hg38_MT_+_1621_3279_RNR2_16S_50ntFlanks | 1351 | 1388 | 38 | AATAGGGTTTACGACCTCGATGTTGGATCAGGACATCC | 73.1 | 3.1 | 1.3 | 2.5 | 1.3E-02 | 1.7E-02 | MB231/MB468 | MP |
| >MI0000736\|hsa-mir-30c-1&WithFlank&1\|+\|40757278\|40757378@23.43.21(+1A);MI0000254\|hsa-mir-30c-2&WithFlank&6\|-\|71376954\|71377037@13.33.21(+1A)&[MIMAT0000244&hsa-miR-30c-5p&offsets\|0\|-2(+1A);m-57&1\|+\|40757300\|40757323&offsets\|0\|-3(+1A)];[MIMAT0000244_1&hsa-miR-30c-5p&offsets\|0\|-2(+1A);m-56&6\|-\|71377002\|71377025&offsets\|0\|-3(+1A)]\|TGTAAACATCCTACACTCTCAA | >MI0000736\|hsa-mir-30c-1&WithFlank&1\|+\|40757278\|40757378 | 23 | 43 | 21(+1A) | TGTAAACATCCTACACTCTCAA | 58.5 | 4.9 | 2.0 | 2.5 | 1.3E-02 | 1.7E-02 | MB231/MB468 | MP |
| >hg38_21_+_8433172_8446622_RNA45SN1_45S_50ntFlanks@12003.12032.30\|GCGGGCCGCCGGTGAAATACCACTACTCTG | >hg38_21_+_8433172_8446622_RNA45SN1_45S_50ntFlanks | 12003 | 12032 | 30 | GCGGGCCGCCGGTGAAATACCACTACTCTG | 3188.4 | -2.3 | 1.0 | -2.4 | 1.5E-02 | 1.7E-02 | MB231/MB468 | NUC |
| >hg38_21_+_8433172_8446622_RNA45SN1_45S_50ntFlanks@12576.12604.29\|AGAGGAACCGCAGGTTCAGACATTTGGTG | >hg38_21_+_8433172_8446622_RNA45SN1_45S_50ntFlanks | 12576 | 12604 | 29 | AGAGGAACCGCAGGTTCAGACATTTGGTG | 134.5 | -4.5 | 1.7 | -2.6 | 9.2E-03 | 1.7E-02 | BT20/MB468 | MP |
| >hg38_21_+_8433172_8446622_RNA45SN1_45S_50ntFlanks@11999.12039.41\|CCCTGCGGGCCGCCGGTGAAATACCACTACTCTGATCGTTT | >hg38_21_+_8433172_8446622_RNA45SN1_45S_50ntFlanks | 11999 | 12039 | 41 | CCCTGCGGGCCGCCGGTGAAATACCACTACTCTGATCGTTT | 158.1 | -3.9 | 1.6 | -2.4 | 1.4E-02 | 1.7E-02 | BT20/MB468 | CYTO |
| >hg38_21_+_8433172_8446622_RNA45SN1_45S_50ntFlanks@10680.10708.29\|GTTCCGGCGGCGTCCGGTGAGCTCTCGCT | >hg38_21_+_8433172_8446622_RNA45SN1_45S_50ntFlanks | 10680 | 10708 | 29 | GTTCCGGCGGCGTCCGGTGAGCTCTCGCT | 779.5 | 4.3 | 1.5 | 2.9 | 3.7E-03 | 1.7E-02 | BT20/MB231 | MP |
| >trnaMT_GluTTC_MT_-_14674_14742@31.72.42,trnalookalike8_GluTTC_5_-_93905172_93905240@31.72.42\|TTCATATCATTGGTCGTGGTTGTAGTCCGTGCGAGAATACCA | >trnaMT_GluTTC_MT_-_14674_14742 | 31 | 72 | 42 | TTCATATCATTGGTCGTGGTTGTAGTCCGTGCGAGAATACCA | 569.3 | 3.7 | 1.5 | 2.5 | 1.3E-02 | 1.7E-02 | MB231/MB468 | MP |
| >hg38_21_+_8433172_8446622_RNA45SN1_45S_50ntFlanks@10735.10755.21\|TGTAAATCTCGCGCCGGGCCG | >hg38_21_+_8433172_8446622_RNA45SN1_45S_50ntFlanks | 10735 | 10755 | 21 | TGTAAATCTCGCGCCGGGCCG | 235.1 | -4.4 | 1.7 | -2.6 | 9.3E-03 | 1.7E-02 | BT20/MB468 | MP |

<table>
<tr><td>>hg38_21_+_8433172_8446622_RNA45SN1_45S_50ntFlanks@6718.6744.27|GTGAATTGCAGGACACATTGATCATCG</td><td>>hg38_21_+_8433172_8446622_RNA45SN1_45S_50ntFlanks</td><td>6718</td><td>6744</td><td>27</td><td>GTGAATTGCAGGACACATTGATCATCG</td><td>169.0</td><td>-1.7</td><td>0.7</td><td>-2.4</td><td>1.4E-02</td><td>1.7E-02</td><td>BT20/MB468</td><td>CYTO</td></tr>
<tr><td>>MI0003780|hsa-mir-1296&WithFlank&10|-|63372951|63373054@22.42.21&[MIMAT0005794&hsa-miR-1296-5p&offsets|0|-1;m-304&10|-|63373012|63373033&offsets|0|-1]|TTAGGGCCCTGGCTCCATCTC</td><td>>MI0003780|hsa-mir-1296&WithFlank&10|-|63372951|63373054</td><td>22</td><td>42</td><td>21</td><td>TTAGGGCCCTGGCTCCATCTC</td><td>99.3</td><td>4.1</td><td>1.7</td><td>2.5</td><td>1.3E-02</td><td>1.7E-02</td><td>MB231/MB468</td><td>MP</td></tr>
<tr><td>>hg38_MT_+_1621_3279_RNR2_16S_50ntFlanks@1424.1453.30|AACGATTAAAGTCCTACGTGATCTGAGTTC</td><td>>hg38_MT_+_1621_3279_RNR2_16S_50ntFlanks</td><td>1424</td><td>1453</td><td>30</td><td>AACGATTAAAGTCCTACGTGATCTGAGTTC</td><td>55.7</td><td>4.0</td><td>1.6</td><td>2.5</td><td>1.4E-02</td><td>1.7E-02</td><td>MB231/MB468</td><td>MP</td></tr>
<tr><td>>hg38_21_+_8433172_8446622_RNA45SN1_45S_50ntFlanks@11581.11616.36|AACTGGTGCGGACCAGGGGAATCCGACTGTTTAATT</td><td>>hg38_21_+_8433172_8446622_RNA45SN1_45S_50ntFlanks</td><td>11581</td><td>11616</td><td>36</td><td>AACTGGTGCGGACCAGGGGAATCCGACTGTTTAATT</td><td>303.4</td><td>3.0</td><td>1.1</td><td>2.6</td><td>9.3E-03</td><td>1.7E-02</td><td>BT20/MB468</td><td>MP</td></tr>
<tr><td>>hg38_21_+_8433172_8446622_RNA45SN1_45S_50ntFlanks@12511.12546.36|CGTGAGACAGGTTAGTTTTACCCTACTGATGATGTG</td><td>>hg38_21_+_8433172_8446622_RNA45SN1_45S_50ntFlanks</td><td>12511</td><td>12546</td><td>36</td><td>CGTGAGACAGGTTAGTTTTACCCTACTGATGATGTG</td><td>38.7</td><td>-4.5</td><td>1.7</td><td>-2.6</td><td>9.3E-03</td><td>1.7E-02</td><td>BT20/MB468</td><td>MP</td></tr>
<tr><td>>MI0000270|hsa-mir-181b-1&WithFlank&1|-|198858867|198858988@42.63.22(+1C);MI0000683|hsa-mir-181b-2&WithFlank&9|+|124693704|124693804@22.43.22(+1C)&[MIMAT0000257&hsa-miR-181b-5p&offsets|0|-1(+1C);m-73&1|-|198858925|198858947&offsets|0|-1(+1C)];[MIMAT0000257_1&hsa-miR-181b-5p&offsets|0|-1(+1C);m-72&9|+|124693725|124693747&offsets|0|-1(+1C)]|AACATTCATTGCTGTCGGTGGGC</td><td>>MI0000270|hsa-mir-181b-1&WithFlank&1|-|198858867|198858988</td><td>42</td><td>63</td><td>22(+1C)</td><td>AACATTCATTGCTGTCGGTGGGC</td><td>356.3</td><td>3.4</td><td>1.4</td><td>2.5</td><td>1.4E-02</td><td>1.7E-02</td><td>MB231/MB468</td><td>MP</td></tr>
<tr><td>>hg38_21_+_8433172_8446622_RNA45SN1_45S_50ntFlanks@7974.8011.38|ACGCGACCTCAGATCAGACGTGGCGACCCGCTGAATTT</td><td>>hg38_21_+_8433172_8446622_RNA45SN1_45S_50ntFlanks</td><td>7974</td><td>8011</td><td>38</td><td>ACGCGACCTCAGATCAGACGTGGCGACCCGCTGAATTT</td><td>2576.7</td><td>3.2</td><td>1.2</td><td>2.6</td><td>9.3E-03</td><td>1.7E-02</td><td>BT20/MB468</td><td>MP</td></tr>
<tr><td>>trnaMT_HisGTG_MT_+_12138_12206@27.60.34|GATTGTGAATCTGACAACAGAGGCTTACGACCCC</td><td>>trnaMT_HisGTG_MT_+_12138_12206</td><td>27</td><td>60</td><td>34</td><td>GATTGTGAATCTGACAACAGAGGCTTACGACCCC</td><td>170.9</td><td>6.6</td><td>2.6</td><td>2.6</td><td>9.3E-03</td><td>1.7E-02</td><td>BT20/MB468</td><td>MP</td></tr>
<tr><td>>hg38_MT_+_598_1651_RNR1_12S_50ntFlanks@327.352.26|ATAGAAGCCGGCGTAAAGAGTGTTTT</td><td>>hg38_MT_+_598_1651_RNR1_12S_50ntFlanks</td><td>327</td><td>352</td><td>26</td><td>ATAGAAGCCGGCGTAAAGAGTGTTTT</td><td>171.7</td><td>4.1</td><td>1.6</td><td>2.6</td><td>9.3E-03</td><td>1.7E-02</td><td>BT20/MB468</td><td>MP</td></tr>
<tr><td>>trna10_AspGTC_12_-_125424193_125424264@1.30.30,trna12_AspGTC_12_-_125411891_125411962@1.30.30,trna38_AspGTC_17_-_8125556_8125627@1.30.30,trna45_AspGTC_6_+_27447453_27447524@1.30.30,trna48_AspGTC_6_+_27471523_27471594@1.30.30,trna4_AspGTC_12_+_96429799_96429870@1.30.30,trna69_AspGTC_1_-_161440205_161440276@1.30.30,trna72_AspGTC_1_-_161432824_161432895@1.30.30,trna75_AspGTC_1_-_161425414_161425485@1.30.30,trna78_AspGTC_1_-_161418033_161418104@1.30.30,trna81_AspGTC_1_-_161410615_161410686@1.30.30|TCCTCGTTAGTATAGTGGTGAGTATCCCCG</td><td>>trna10_AspGTC_12_-_125424193_125424264</td><td>1</td><td>30</td><td>30</td><td>TCCTCGTTAGTATAGTGGTGAGTATCCCCG</td><td>152.3</td><td>-3.8</td><td>1.5</td><td>-2.5</td><td>1.3E-02</td><td>1.7E-02</td><td>BT20/MB468</td><td>MITO</td></tr>
<tr><td>>hg38_21_+_8433172_8446622_RNA45SN1_45S_50ntFlanks@10781.10809.29|AGGTGAACAGCCTCTGGCATGTTGGAACA</td><td>>hg38_21_+_8433172_8446622_RNA45SN1_45S_50ntFlanks</td><td>10781</td><td>10809</td><td>29</td><td>AGGTGAACAGCCTCTGGCATGTTGGAACA</td><td>176.8</td><td>-1.4</td><td>0.6</td><td>-2.5</td><td>1.3E-02</td><td>1.7E-02</td><td>BT20/MB468</td><td>MITO</td></tr>
<tr><td>>hg38_21_+_8433172_8446622_RNA45SN1_45S_50ntFlanks@3843.3869.27|CTACTTGGATAACTGTGGTAATTCTAG</td><td>>hg38_21_+_8433172_8446622_RNA45SN1_45S_50ntFlanks</td><td>3843</td><td>3869</td><td>27</td><td>CTACTTGGATAACTGTGGTAATTCTAG</td><td>252.0</td><td>-1.7</td><td>0.7</td><td>-2.4</td><td>1.5E-02</td><td>1.7E-02</td><td>BT20/MB468</td><td>CYTO</td></tr>
<tr><td>>hg38_MT_+_598_1651_RNR1_12S_50ntFlanks@52.72.21|ATAGGTTTGGTCCTAGCCTTT</td><td>>hg38_MT_+_598_1651_RNR1_12S_50ntFlanks</td><td>52</td><td>72</td><td>21</td><td>ATAGGTTTGGTCCTAGCCTTT</td><td>1704.9</td><td>-3.2</td><td>1.3</td><td>-2.5</td><td>1.4E-02</td><td>1.7E-02</td><td>MB231/MB468</td><td>MP</td></tr>
<tr><td>>hg38_21_+_8433172_8446622_RNA45SN1_45S_50ntFlanks@12431.12455.25|TCCTATCATTGTGAAGCAGAATTCA</td><td>>hg38_21_+_8433172_8446622_RNA45SN1_45S_50ntFlanks</td><td>12431</td><td>12455</td><td>25</td><td>TCCTATCATTGTGAAGCAGAATTCA</td><td>69.9</td><td>-4.3</td><td>1.7</td><td>-2.6</td><td>9.3E-03</td><td>1.7E-02</td><td>BT20/MB468</td><td>MP</td></tr>
<tr><td>>hg38_21_+_8433172_8446622_RNA45SN1_45S_50ntFlanks@7991.8017.27|ACGTGGCGACCCGCTGAATTTAAGCAT</td><td>>hg38_21_+_8433172_8446622_RNA45SN1_45S_50ntFlanks</td><td>7991</td><td>8017</td><td>27</td><td>ACGTGGCGACCCGCTGAATTTAAGCAT</td><td>194.3</td><td>-2.6</td><td>0.9</td><td>-2.8</td><td>4.7E-03</td><td>1.7E-02</td><td>BT20/MB231</td><td>CYTO</td></tr>
<tr><td>>MI0000749|hsa-mir-30e&WithFlank&1|+|40754349|40754452@23.45.23(+1C)&[MIMAT0000692&hsa-miR-30e-5p&offsets|0|+1(+1C);m-24&1|+|40754371|40754394&offsets|0|-1(+1C)]|TGTAAACATCCTTGACTGGAAGCC</td><td>>MI0000749|hsa-mir-30e&WithFlank&1|+|40754349|40754452</td><td>23</td><td>45</td><td>23(+1C)</td><td>TGTAAACATCCTTGACTGGAAGCC</td><td>1399.4</td><td>-2.6</td><td>0.9</td><td>-2.8</td><td>4.7E-03</td><td>1.7E-02</td><td>BT20/MB231</td><td>MITO</td></tr>
<tr><td>>hg38_MT_+_1621_3279_RNR2_16S_50ntFlanks@1216.1261.46|CAACCTCCGAGCAGTACATGCTAAGACTTCACCAGTCAAAGCGAAC</td><td>>hg38_MT_+_1621_3279_RNR2_16S_50ntFlanks</td><td>1216</td><td>1261</td><td>46</td><td>CAACCTCCGAGCAGTACATGCTAAGACTTCACCAGTCAAAGCGAAC</td><td>638.2</td><td>-2.5</td><td>0.9</td><td>-2.8</td><td>4.7E-03</td><td>1.7E-02</td><td>BT20/MB231</td><td>MITO</td></tr>
<tr><td>>hg38_MT_+_1621_3279_RNR2_16S_50ntFlanks@1293.1320.28|AACGGAACAAGTTACCCTAGGGATAACA</td><td>>hg38_MT_+_1621_3279_RNR2_16S_50ntFlanks</td><td>1293</td><td>1320</td><td>28</td><td>AACGGAACAAGTTACCCTAGGGATAACA</td><td>154.6</td><td>2.4</td><td>1.0</td><td>2.5</td><td>1.4E-02</td><td>1.7E-02</td><td>MB231/MB468</td><td>MP</td></tr>
<tr><td>>hg38_21_+_8433172_8446622_RNA45SN1_45S_50ntFlanks@12420.12451.32|ATGTCGGCTCTTCCTATCATTGTGAAGCAGAA</td><td>>hg38_21_+_8433172_8446622_RNA45SN1_45S_50ntFlanks</td><td>12420</td><td>12451</td><td>32</td><td>ATGTCGGCTCTTCCTATCATTGTGAAGCAGAA</td><td>90.9</td><td>-4.1</td><td>1.7</td><td>-2.5</td><td>1.4E-02</td><td>1.7E-02</td><td>MB231/MB468</td><td>MP</td></tr>
<tr><td>>MI0000254|hsa-mir-30c-2&WithFlank&6|-|71376954|71377037@53.75.23&[MIMAT0004550&hsa-miR-30c-2-3p&offsets|0|+1;m-206&6|-|71376964|71376985&offsets|0|+1]|CTGGGAGAAGGCTGTTTACTCTT</td><td>>MI0000254|hsa-mir-30c-2&WithFlank&6|-|71376954|71377037</td><td>53</td><td>75</td><td>23</td><td>CTGGGAGAAGGCTGTTTACTCTT</td><td>78.8</td><td>4.0</td><td>1.6</td><td>2.5</td><td>1.4E-02</td><td>1.7E-02</td><td>MB231/MB468</td><td>MP</td></tr>
<tr><td>>hg38_21_+_8433172_8446622_RNA45SN1_45S_50ntFlanks@12640.12661.22|ACCATCTGTGGGATTATGACTG</td><td>>hg38_21_+_8433172_8446622_RNA45SN1_45S_50ntFlanks</td><td>12640</td><td>12661</td><td>22</td><td>ACCATCTGTGGGATTATGACTG</td><td>1178.9</td><td>-2.0</td><td>0.8</td><td>-2.5</td><td>1.3E-02</td><td>1.7E-02</td><td>BT20/MB468</td><td>MITO</td></tr>
<tr><td>>hg38_21_+_8433172_8446622_RNA45SN1_45S_50ntFlanks@4732.4751.20|AGAACGAAAGTCGGAGGTTC</td><td>>hg38_21_+_8433172_8446622_RNA45SN1_45S_50ntFlanks</td><td>4732</td><td>4751</td><td>20</td><td>AGAACGAAAGTCGGAGGTTC</td><td>487.5</td><td>1.3</td><td>0.5</td><td>2.4</td><td>1.5E-02</td><td>1.7E-02</td><td>BT20/MB468</td><td>CYTO</td></tr>
<tr><td>>MI0003610|hsa-mir-598&WithFlank&8|-|11035200|11035308@67.87.21&[MIMAT0003266&hsa-miR-598-3p&offsets|0|-1;m-231&8|-|11035221|11035242&offsets|0|-1]|TACGTCATCGTTGTCATCGTC</td><td>>MI0003610|hsa-mir-598&WithFlank&8|-|11035200|11035308</td><td>67</td><td>87</td><td>21</td><td>TACGTCATCGTTGTCATCGTC</td><td>105.1</td><td>-4.2</td><td>1.5</td><td>-2.9</td><td>3.7E-03</td><td>1.7E-02</td><td>BT20/MB231</td><td>MP</td></tr>
<tr><td>>hg38_21_+_8433172_8446622_RNA45SN1_45S_50ntFlanks@6730.6752.23|ACACATTGATCATCGACACTTCG</td><td>>hg38_21_+_8433172_8446622_RNA45SN1_45S_50ntFlanks</td><td>6730</td><td>6752</td><td>23</td><td>ACACATTGATCATCGACACTTCG</td><td>186.2</td><td>-3.7</td><td>1.5</td><td>-2.4</td><td>1.5E-02</td><td>1.7E-02</td><td>BT20/MB468</td><td>CYTO</td></tr>
<tr><td>>am_MI0003581|hsa-mir-574&WithFlank&4|+|38868026|38868133@31.51.21&[MIMAT0004795&hsa-miR-574-5p&offsets|0|-2;m-213&4|+|38868056|38868078&offsets|0|-2]|TGAGTGTGTGTGTGTGAGTGT</td><td>>am_MI0003581|hsa-mir-574&WithFlank&4|+|38868026|38868133</td><td>31</td><td>51</td><td>21</td><td>TGAGTGTGTGTGTGTGAGTGT</td><td>268.6</td><td>3.9</td><td>1.5</td><td>2.6</td><td>9.3E-03</td><td>1.7E-02</td><td>BT20/MB468</td><td>MP</td></tr>
<tr><td>>hg38_21_+_8433172_8446622_RNA45SN1_45S_50ntFlanks@11672.11697.26|CAGTGCTCTGAATGTCAAAGTGAAGA</td><td>>hg38_21_+_8433172_8446622_RNA45SN1_45S_50ntFlanks</td><td>11672</td><td>11697</td><td>26</td><td>CAGTGCTCTGAATGTCAAAGTGAAGA</td><td>132.2</td><td>2.5</td><td>1.0</td><td>2.6</td><td>9.3E-03</td><td>1.7E-02</td><td>BT20/MB468</td><td>MP</td></tr>
<tr><td>>hg38_21_+_8433172_8446622_RNA45SN1_45S_50ntFlanks@10401.10426.26|GAACATGGGTCAGTCGGTCCTGAGAG</td><td>>hg38_21_+_8433172_8446622_RNA45SN1_45S_50ntFlanks</td><td>10401</td><td>10426</td><td>26</td><td>GAACATGGGTCAGTCGGTCCTGAGAG</td><td>148.2</td><td>-3.3</td><td>1.3</td><td>-2.6</td><td>9.3E-03</td><td>1.7E-02</td><td>BT20/MB468</td><td>MP</td></tr>
<tr><td>>hg38_1_-_228634819_228635039_RNA5S12_5S_50ntFlanks@136.170.35|GGAGACCGCCTGGGAATACCGGGTGCTGTAGGCTT</td><td>>hg38_1_-_228634819_228635039_RNA5S12_5S_50ntFlanks</td><td>136</td><td>170</td><td>35</td><td>GGAGACCGCCTGGGAATACCGGGTGCTGTAGGCTT</td><td>3321.0</td><td>2.4</td><td>0.8</td><td>2.8</td><td>4.7E-03</td><td>1.7E-02</td><td>BT20/MB231</td><td>MITO</td></tr>
<tr><td>>hg38_21_+_8433172_8446622_RNA45SN1_45S_50ntFlanks@9573.9594.22|AAATCGGTCGTCCGACCTGGGT</td><td>>hg38_21_+_8433172_8446622_RNA45SN1_45S_50ntFlanks</td><td>9573</td><td>9594</td><td>22</td><td>AAATCGGTCGTCCGACCTGGGT</td><td>778.2</td><td>-2.8</td><td>1.2</td><td>-2.4</td><td>1.5E-02</td><td>1.7E-02</td><td>MB231/MB468</td><td>MITO</td></tr>
<tr><td>>MI0000100|hsa-mir-98&WithFlank&X|-|53556217|53556347@28.48.21(+1C)&[MIMAT0000096&hsa-miR-98-5p&offsets|0|-1(+1C);m-143&X|-|53556299|53556320&offsets|0|-1(+1C)]|TGAGGTAGTAAGTTGTATTGTC</td><td>>MI0000100|hsa-mir-98&WithFlank&X|-|53556217|53556347</td><td>28</td><td>48</td><td>21(+1C)</td><td>TGAGGTAGTAAGTTGTATTGTC</td><td>1046.0</td><td>-3.0</td><td>1.0</td><td>-2.9</td><td>3.7E-03</td><td>1.7E-02</td><td>BT20/MB231</td><td>MP</td></tr>
<tr><td>>hg38_21_+_8433172_8446622_RNA45SN1_45S_50ntFlanks@10361.10384.24|GAAGGCCGAAGTGGAGAAGGGTTC</td><td>>hg38_21_+_8433172_8446622_RNA45SN1_45S_50ntFlanks</td><td>10361</td><td>10384</td><td>24</td><td>GAAGGCCGAAGTGGAGAAGGGTTC</td><td>165.1</td><td>-4.4</td><td>1.7</td><td>-2.6</td><td>9.4E-03</td><td>1.7E-02</td><td>BT20/MB468</td><td>MP</td></tr>
<tr><td>>trnaMT_HisGTG_MT_+_12138_12206@32.67.36|TGAATCTGACAACAGAGGCTTACGACCCCTTATTTA</td><td>>trnaMT_HisGTG_MT_+_12138_12206</td><td>32</td><td>67</td><td>36</td><td>TGAATCTGACAACAGAGGCTTACGACCCCTTATTTA</td><td>815.6</td><td>2.5</td><td>1.0</td><td>2.4</td><td>1.5E-02</td><td>1.7E-02</td><td>MB231/MB468</td><td>MITO</td></tr>
<tr><td>>trnaMT_GluTTC_MT_-_14674_14742@34.57.24,trnalookalike8_GluTTC_5_-_93905172_93905240@34.57.24|ATATCATTGGTCGTGGTTGTAGTC</td><td>>trnaMT_GluTTC_MT_-_14674_14742</td><td>34</td><td>57</td><td>24</td><td>ATATCATTGGTCGTGGTTGTAGTC</td><td>155.0</td><td>2.1</td><td>0.9</td><td>2.5</td><td>1.4E-02</td><td>1.7E-02</td><td>MB231/MB468</td><td>MP</td></tr>
<tr><td>>hg38_21_+_8433172_8446622_RNA45SN1_45S_50ntFlanks@10686.10707.22|GCGGCGTCCGGTGAGCTCTCGC</td><td>>hg38_21_+_8433172_8446622_RNA45SN1_45S_50ntFlanks</td><td>10686</td><td>10707</td><td>22</td><td>GCGGCGTCCGGTGAGCTCTCGC</td><td>902.4</td><td>4.2</td><td>1.5</td><td>2.9</td><td>3.7E-03</td><td>1.7E-02</td><td>BT20/MB231</td><td>MP</td></tr>
<tr><td>>hg38_21_+_8433172_8446622_RNA45SN1_45S_50ntFlanks@11999.12034.36|CCCTGCGGGCCGCCGGTGAAATACCACTACTCTGAT</td><td>>hg38_21_+_8433172_8446622_RNA45SN1_45S_50ntFlanks</td><td>11999</td><td>12034</td><td>36</td><td>CCCTGCGGGCCGCCGGTGAAATACCACTACTCTGAT</td><td>185.9</td><td>-3.9</td><td>1.6</td><td>-2.5</td><td>1.3E-02</td><td>1.7E-02</td><td>BT20/MB468</td><td>MITO</td></tr>
<tr><td>>hg38_21_+_8433172_8446622_RNA45SN1_45S_50ntFlanks@11785.11802.18|TGAATGGATGAACGAGAT</td><td>>hg38_21_+_8433172_8446622_RNA45SN1_45S_50ntFlanks</td><td>11785</td><td>11802</td><td>18</td><td>TGAATGGATGAACGAGAT</td><td>320.9</td><td>-4.8</td><td>1.8</td><td>-2.6</td><td>9.4E-03</td><td>1.7E-02</td><td>BT20/MB468</td><td>MP</td></tr>
<tr><td>>hg38_21_+_8433172_8446622_RNA45SN1_45S_50ntFlanks@12438.12473.36|ATTGTGAAGCAGAATTCACCAAGCGTTGGATTGTTC</td><td>>hg38_21_+_8433172_8446622_RNA45SN1_45S_50ntFlanks</td><td>12438</td><td>12473</td><td>36</td><td>ATTGTGAAGCAGAATTCACCAAGCGTTGGATTGTTC</td><td>364.1</td><td>-1.7</td><td>0.7</td><td>-2.5</td><td>1.4E-02</td><td>1.7E-02</td><td>MB231/MB468</td><td>MP</td></tr>
<tr><td>>hg38_21_+_8433172_8446622_RNA45SN1_45S_50ntFlanks@5085.5112.28|GAACGAGACTCTGGCATGCTAACTAGTT</td><td>>hg38_21_+_8433172_8446622_RNA45SN1_45S_50ntFlanks</td><td>5085</td><td>5112</td><td>28</td><td>GAACGAGACTCTGGCATGCTAACTAGTT</td><td>300.5</td><td>4.9</td><td>1.9</td><td>2.6</td><td>9.4E-03</td><td>1.7E-02</td><td>BT20/MB468</td><td>MP</td></tr>
<tr><td>>hg38_21_+_8433172_8446622_RNA45SN1_45S_50ntFlanks@5497.5526.30|ACGGTCGAACTTGACTATCTAGAGGAAGTA</td><td>>hg38_21_+_8433172_8446622_RNA45SN1_45S_50ntFlanks</td><td>5497</td><td>5526</td><td>30</td><td>ACGGTCGAACTTGACTATCTAGAGGAAGTA</td><td>357.0</td><td>-2.5</td><td>1.0</td><td>-2.6</td><td>9.4E-03</td><td>1.7E-02</td><td>BT20/MB468</td><td>MP</td></tr>
<tr><td>>hg38_21_+_8433172_8446622_RNA45SN1_45S_50ntFlanks@6645.6685.41|CTCGTACGACTCTTAGCGGTGGATCACTCGGCTCGTGCGTC</td><td>>hg38_21_+_8433172_8446622_RNA45SN1_45S_50ntFlanks</td><td>6645</td><td>6685</td><td>41</td><td>CTCGTACGACTCTTAGCGGTGGATCACTCGGCTCGTGCGTC</td><td>385.6</td><td>3.1</td><td>1.2</td><td>2.5</td><td>1.4E-02</td><td>1.7E-02</td><td>MB231/MB468</td><td>MP</td></tr>
<tr><td>>hg38_21_+_8433172_8446622_RNA45SN1_45S_50ntFlanks@9546.9571.26|TGGAGGTCCGTAGCGGTCCTGACGTG</td><td>>hg38_21_+_8433172_8446622_RNA45SN1_45S_50ntFlanks</td><td>9546</td><td>9571</td><td>26</td><td>TGGAGGTCCGTAGCGGTCCTGACGTG</td><td>2261.9</td><td>1.6</td><td>0.6</td><td>2.4</td><td>1.5E-02</td><td>1.7E-02</td><td>MB231/MB468</td><td>CYTO</td></tr>
<tr><td>>hg38_21_+_8433172_8446622_RNA45SN1_45S_50ntFlanks@3705.3735.31|TACCTGGTTGATCCTGCCAGTAGCATATGCT</td><td>>hg38_21_+_8433172_8446622_RNA45SN1_45S_50ntFlanks</td><td>3705</td><td>3735</td><td>31</td><td>TACCTGGTTGATCCTGCCAGTAGCATATGCT</td><td>2069.2</td><td>2.3</td><td>0.8</td><td>2.8</td><td>4.8E-03</td><td>1.7E-02</td><td>BT20/MB231</td><td>CYTO</td></tr>
<tr><td>>hg38_21_+_8433172_8446622_RNA45SN1_45S_50ntFlanks@11571.11604.34|GCCGACTTAGAACTGGTGCGGACCAGGGGAATCC</td><td>>hg38_21_+_8433172_8446622_RNA45SN1_45S_50ntFlanks</td><td>11571</td><td>11604</td><td>34</td><td>GCCGACTTAGAACTGGTGCGGACCAGGGGAATCC</td><td>414.1</td><td>4.1</td><td>1.4</td><td>2.9</td><td>3.7E-03</td><td>1.7E-02</td><td>BT20/MB231</td><td>MP</td></tr>
<tr><td>>hg38_21_+_8433172_8446622_RNA45SN1_45S_50ntFlanks@12912.12931.20|CACGTTCGTGGGGAACCTGG</td><td>>hg38_21_+_8433172_8446622_RNA45SN1_45S_50ntFlanks</td><td>12912</td><td>12931</td><td>20</td><td>CACGTTCGTGGGGAACCTGG</td><td>146.8</td><td>-3.6</td><td>1.5</td><td>-2.4</td><td>1.5E-02</td><td>1.7E-02</td><td>BT20/MB468</td><td>CYTO</td></tr>
<tr><td>>MI0000063|hsa-let-7b&WithFlank&22|+|46113680|46113774@12.31.20(+1A)&[MIMAT0000063&hsa-let-7b-5p&offsets|0|-2(+1A);m-14&22|+|46113691|46113712&offsets|0|-2(+1A)]|TGAGGTAGTAGGTTGTGTGGA</td><td>>MI0000063|hsa-let-7b&WithFlank&22|+|46113680|46113774</td><td>12</td><td>31</td><td>20(+1A)</td><td>TGAGGTAGTAGGTTGTGTGGA</td><td>161.4</td><td>3.3</td><td>1.3</td><td>2.5</td><td>1.4E-02</td><td>1.7E-02</td><td>MB231/MB468</td><td>MP</td></tr>
<tr><td>>hg38_21_+_8433172_8446622_RNA45SN1_45S_50ntFlanks@5473.5499.27|CGGCCCTGGCGGAGCGCTGAGAAGACG</td><td>>hg38_21_+_8433172_8446622_RNA45SN1_45S_50ntFlanks</td><td>5473</td><td>5499</td><td>27</td><td>CGGCCCTGGCGGAGCGCTGAGAAGACG</td><td>2478.8</td><td>-2.2</td><td>0.9</td><td>-2.6</td><td>9.4E-03</td><td>1.7E-02</td><td>BT20/MB468</td><td>MP</td></tr>
<tr><td>>hg38_MT_+_1621_3279_RNR2_16S_50ntFlanks@1090.1117.28|CAGCAAGACGAGAAGACCCTATGGAGCT</td><td>>hg38_MT_+_1621_3279_RNR2_16S_50ntFlanks</td><td>1090</td><td>1117</td><td>28</td><td>CAGCAAGACGAGAAGACCCTATGGAGCT</td><td>103.9</td><td>3.5</td><td>1.4</td><td>2.5</td><td>1.4E-02</td><td>1.7E-02</td><td>MB231/MB468</td><td>MP</td></tr>
</table>

| | | | | | | | | | | | | | |
|---|---|---|---|---|---|---|---|---|---|---|---|---|---|
| >hg38_21_+_8433172_8446622_RNA45SN1_45S_50ntFlanks@3993.4031.39\|GTGACTCTAGATAACCTCGGGCCGATCGCACGCCCCCCG | >hg38_21_+_8433172_8446622_RNA45SN1_45S_50ntFlanks | 3993 | 4031 | 39 | GTGACTCTAGATAACCTCGGGCCGATCGCACGCCCCCCG | 711.1 | -3.2 | 1.1 | -2.8 | 4.8E-03 | 1.7E-02 | BT20/MB231 | CYTO |
| >hg38_21_+_8433172_8446622_RNA45SN1_45S_50ntFlanks@10391.10430.40\|AACAGCAGTTGAACATGGGTCAGTCGGTCCTGAGAGATGG | >hg38_21_+_8433172_8446622_RNA45SN1_45S_50ntFlanks | 10391 | 10430 | 40 | AACAGCAGTTGAACATGGGTCAGTCGGTCCTGAGAGATGG | 258.1 | -3.7 | 1.4 | -2.6 | 9.4E-03 | 1.7E-02 | BT20/MB468 | MP |
| >trna10_LysCTT_16_+_3241501_3241573@3.36.34,trna13_LysCTT_14_-_58706613_58706685@3.36.34,trna2_LysCTT_15_+_79152904_79152976@3.36.34\|CCGGCTAGCTCAGTCGGTAGAGCATGGGACTCTT | >trna10_LysCTT_16_+_3241501_3241573 | 3 | 36 | 34 | CCGGCTAGCTCAGTCGGTAGAGCATGGGACTCTT | 282.0 | 5.2 | 2.0 | 2.6 | 9.4E-03 | 1.7E-02 | BT20/MB468 | MP |
| >hg38_21_+_8433172_8446622_RNA45SN1_45S_50ntFlanks@10783.10813.31\|GTGAACAGCCTCTGGCATGTTGGAACAATGT | >hg38_21_+_8433172_8446622_RNA45SN1_45S_50ntFlanks | 10783 | 10813 | 31 | GTGAACAGCCTCTGGCATGTTGGAACAATGT | 89.1 | -4.5 | 1.7 | -2.6 | 9.4E-03 | 1.7E-02 | BT20/MB468 | MP |
| >hg38_21_+_8433172_8446622_RNA45SN1_45S_50ntFlanks@6742.6772.31\|TCGACACTTCGAACGCACTTGCGGCCCCGGG | >hg38_21_+_8433172_8446622_RNA45SN1_45S_50ntFlanks | 6742 | 6772 | 31 | TCGACACTTCGAACGCACTTGCGGCCCCGGG | 86.1 | -4.5 | 1.7 | -2.6 | 9.4E-03 | 1.7E-02 | BT20/MB468 | MP |
| >hg38_21_+_8433172_8446622_RNA45SN1_45S_50ntFlanks@8197.8222.26\|GTGGACGGTGTGAGGCCGGTAGCGGC | >hg38_21_+_8433172_8446622_RNA45SN1_45S_50ntFlanks | 8197 | 8222 | 26 | GTGGACGGTGTGAGGCCGGTAGCGGC | 289.8 | 6.0 | 2.1 | 2.9 | 3.7E-03 | 1.7E-02 | BT20/MB231 | MP |
| >trnaMT_ProTGG_MT_-_15956_16023@40.60.21\|AATGGTGGAGTTAAAGACTTT | >trnaMT_ProTGG_MT_-_15956_16023 | 40 | 60 | 21 | AATGGTGGAGTTAAAGACTTT | 83.7 | 3.8 | 1.4 | 2.6 | 9.4E-03 | 1.7E-02 | BT20/MB468 | MP |
| >hg38_21_+_8433172_8446622_RNA45SN1_45S_50ntFlanks@6649.6698.50\|TACGACTCTTAGCGGTGGATCACTCGGCTCGTGCGTCGATGAAGAACGCA | >hg38_21_+_8433172_8446622_RNA45SN1_45S_50ntFlanks | 6649 | 6698 | 50 | TACGACTCTTAGCGGTGGATCACTCGGCTCGTGCGTCGATGAAGAACGCA | 2895.7 | -2.6 | 1.1 | -2.4 | 1.6E-02 | 1.7E-02 | MB231/MB468 | MITO |
| >hg38_21_+_8433172_8446622_RNA45SN1_45S_50ntFlanks@5494.5514.21\|AAGACGGTCGAACTTGACTAT | >hg38_21_+_8433172_8446622_RNA45SN1_45S_50ntFlanks | 5494 | 5514 | 21 | AAGACGGTCGAACTTGACTAT | 301.2 | -2.3 | 1.0 | -2.4 | 1.5E-02 | 1.7E-02 | MB231/MB468 | CYTO |
| >MI0000067\|hsa-let-7f-1&WithFlank&9\|+\|94176341\|94176439@13.33.21;MI0000068\|hsa-let-7f-2&WithFlank&X\|-\|53557186\|53557280@14.34.21&[MIMAT0000067&hsa-let-7f-5p&offsets\|0\|-1;m-20&9\|+\|94176353\|94176374&offsets\|0\|-1];[MIMAT0000067_1&hsa-let-7f-5p&offsets\|0\|-1;m-19&X\|-\|53557246\|53557267&offsets\|0\|-1]\|TGAGGTAGTAGATTGTATAGT | >MI0000067\|hsa-let-7f-1&WithFlank&9\|+\|94176341\|94176439 | 13 | 33 | 21 | TGAGGTAGTAGATTGTATAGT | 10187.7 | -2.9 | 1.0 | -2.9 | 3.8E-03 | 1.7E-02 | BT20/MB231 | MP |
| >hg38_21_+_8433172_8446622_RNA45SN1_45S_50ntFlanks@5073.5094.22\|AATTCCGATAACGAACGAGACT | >hg38_21_+_8433172_8446622_RNA45SN1_45S_50ntFlanks | 5073 | 5094 | 22 | AATTCCGATAACGAACGAGACT | 837.8 | -4.3 | 1.6 | -2.6 | 9.5E-03 | 1.7E-02 | BT20/MB468 | MP |
| >hg38_21_+_8433172_8446622_RNA45SN1_45S_50ntFlanks@12989.13013.25\|AGCTCCCTCGCTGCGATCTATTGAA | >hg38_21_+_8433172_8446622_RNA45SN1_45S_50ntFlanks | 12989 | 13013 | 25 | AGCTCCCTCGCTGCGATCTATTGAA | 266.2 | 1.8 | 0.7 | 2.4 | 1.6E-02 | 1.7E-02 | MB231/MB468 | NUC |
| >hg38_21_+_8433172_8446622_RNA45SN1_45S_50ntFlanks@11597.11621.25\|GGGAATCCGACTGTTTAATTAAAAC | >hg38_21_+_8433172_8446622_RNA45SN1_45S_50ntFlanks | 11597 | 11621 | 25 | GGGAATCCGACTGTTTAATTAAAAC | 89.8 | 3.7 | 1.4 | 2.6 | 9.5E-03 | 1.7E-02 | BT20/MB468 | MP |
| >hg38_21_+_8433172_8446622_RNA45SN1_45S_50ntFlanks@11843.11871.29\|AGGGAACGGGCTTGGCGGAATCAGCGGGG | >hg38_21_+_8433172_8446622_RNA45SN1_45S_50ntFlanks | 11843 | 11871 | 29 | AGGGAACGGGCTTGGCGGAATCAGCGGGG | 297.9 | -3.4 | 1.3 | -2.6 | 9.5E-03 | 1.7E-02 | BT20/MB468 | MP |
| >hg38_21_+_8433172_8446622_RNA45SN1_45S_50ntFlanks@12479.12500.22\|CTAATAGGGAACGTGAGCTGGG | >hg38_21_+_8433172_8446622_RNA45SN1_45S_50ntFlanks | 12479 | 12500 | 22 | CTAATAGGGAACGTGAGCTGGG | 572.9 | -1.4 | 0.6 | -2.4 | 1.5E-02 | 1.7E-02 | MB231/MB468 | CYTO |
| >hg38_21_+_8433172_8446622_RNA45SN1_45S_50ntFlanks@10240.10261.22\|CTACGCCGCGACGAGTAGGAGG | >hg38_21_+_8433172_8446622_RNA45SN1_45S_50ntFlanks | 10240 | 10261 | 22 | CTACGCCGCGACGAGTAGGAGG | 222.2 | 2.4 | 0.9 | 2.6 | 9.5E-03 | 1.7E-02 | BT20/MB468 | MP |
| >MI0000102\|hsa-mir-100&WithFlank&11\|-\|122152223\|122152314@19.39.21(+1A)&[MIMAT0000098&hsa-miR-100-5p&offsets\|0\|-1(+1A);m-27&11\|-\|122152276\|122152296&offsets\|0\|0(+1A)]\|AACCCGTAGATCCGAACTTGTA | >MI0000102\|hsa-mir-100&WithFlank&11\|-\|122152223\|122152314 | 19 | 39 | 21(+1A) | AACCCGTAGATCCGAACTTGTA | 495.8 | 4.4 | 1.7 | 2.6 | 9.5E-03 | 1.7E-02 | BT20/MB468 | MP |
| >hg38_21_+_8433172_8446622_RNA45SN1_45S_50ntFlanks@10806.10844.39\|AACAATGTAGGTAAGGGAAGTCGGCAAGCCGGATCCGTA | >hg38_21_+_8433172_8446622_RNA45SN1_45S_50ntFlanks | 10806 | 10844 | 39 | AACAATGTAGGTAAGGGAAGTCGGCAAGCCGGATCCGTA | 142.6 | -3.0 | 1.2 | -2.5 | 1.3E-02 | 1.7E-02 | BT20/MB468 | MITO |
| >hg38_21_+_8433172_8446622_RNA45SN1_45S_50ntFlanks@4977.4999.23\|CGGACACGGACAGGATTGACAGA | >hg38_21_+_8433172_8446622_RNA45SN1_45S_50ntFlanks | 4977 | 4999 | 23 | CGGACACGGACAGGATTGACAGA | 357.9 | -4.5 | 1.7 | -2.6 | 9.5E-03 | 1.8E-02 | BT20/MB468 | MP |
| >hg38_21_+_8433172_8446622_RNA45SN1_45S_50ntFlanks@6648.6665.18\|GTACGACTCTTAGCGGTG | >hg38_21_+_8433172_8446622_RNA45SN1_45S_50ntFlanks | 6648 | 6665 | 18 | GTACGACTCTTAGCGGTG | 925.8 | 2.2 | 0.9 | 2.4 | 1.6E-02 | 1.8E-02 | MB231/MB468 | MITO |
| >trnaMT_PheGAA_MT_+_577_647@39.65.27\|ATGTTTAGACGGGCTCACATCACCCCA | >trnaMT_PheGAA_MT_+_577_647 | 39 | 65 | 27 | ATGTTTAGACGGGCTCACATCACCCCA | 92.4 | 2.3 | 0.9 | 2.5 | 1.4E-02 | 1.8E-02 | MB231/MB468 | MP |
| >hg38_21_+_8433172_8446622_RNA45SN1_45S_50ntFlanks@11920.11949.30\|ATGAGAGGTGTAGAATAAGTGGGAGGCCCC | >hg38_21_+_8433172_8446622_RNA45SN1_45S_50ntFlanks | 11920 | 11949 | 30 | ATGAGAGGTGTAGAATAAGTGGGAGGCCCC | 195.2 | -2.3 | 1.0 | -2.4 | 1.5E-02 | 1.8E-02 | MB231/MB468 | CYTO |
| >hg38_21_+_8433172_8446622_RNA45SN1_45S_50ntFlanks@10689.10708.20\|GCGTCCGGTGAGCTCTCGCT | >hg38_21_+_8433172_8446622_RNA45SN1_45S_50ntFlanks | 10689 | 10708 | 20 | GCGTCCGGTGAGCTCTCGCT | 633.5 | -1.7 | 0.7 | -2.5 | 1.3E-02 | 1.8E-02 | BT20/MB468 | MITO |
| >hg38_21_+_8433172_8446622_RNA45SN1_45S_50ntFlanks@12419.12437.19\|GATGTCGGCTCTTCCTATC | >hg38_21_+_8433172_8446622_RNA45SN1_45S_50ntFlanks | 12419 | 12437 | 19 | GATGTCGGCTCTTCCTATC | 8948.0 | -2.2 | 0.8 | -2.8 | 4.9E-03 | 1.8E-02 | BT20/MB231 | CYTO |
| >hg38_1_-_228634819_228635039_RNA5S12_5S_50ntFlanks@129.170.42\|TTGGATGGGAGACCGCCTGGGAATACCGGGTGCTGTAGGCTT | >hg38_1_-_228634819_228635039_RNA5S12_5S_50ntFlanks | 129 | 170 | 42 | TTGGATGGGAGACCGCCTGGGAATACCGGGTGCTGTAGGCTT | 193.7 | 3.1 | 1.3 | 2.4 | 1.6E-02 | 1.8E-02 | MB231/MB468 | MITO |
| >hg38_21_+_8433172_8446622_RNA45SN1_45S_50ntFlanks@11315.11344.30\|CCACCCCACGTCTCGTCGCGCGCGCGTCCG | >hg38_21_+_8433172_8446622_RNA45SN1_45S_50ntFlanks | 11315 | 11344 | 30 | CCACCCCACGTCTCGTCGCGCGCGCGTCCG | 3374.3 | -4.6 | 1.8 | -2.5 | 1.3E-02 | 1.8E-02 | BT20/MB468 | MITO |
| >hg38_21_+_8433172_8446622_RNA45SN1_45S_50ntFlanks@10855.10879.25\|AAGGATTGGCTCTAAGGGCTGGGTC | >hg38_21_+_8433172_8446622_RNA45SN1_45S_50ntFlanks | 10855 | 10879 | 25 | AAGGATTGGCTCTAAGGGCTGGGTC | 179.3 | 3.8 | 1.3 | 2.9 | 3.8E-03 | 1.8E-02 | BT20/MB231 | MP |
| >MI0000446\|hsa-mir-125b-1&WithFlank&11\|-\|122099751\|122099850@21.39.19(+1A);MI0000470\|hsa-mir-125b-2&WithFlank&21\|+\|16590231\|16590331@23.41.19(+1A)&[MIMAT0000423&hsa-miR-125b-5p&offsets\|0\|-3(+1A);m-67&11\|-\|122099809\|122099830&offsets\|0\|-3(+1A)];[MIMAT0000423_1&hsa-miR-125b-5p&offsets\|0\|-3(+1A);m-68&21\|+\|16590253\|16590274&offsets\|0\|-3(+1A)]\|TCCCTGAGACCCTAACTTGA | >MI0000446\|hsa-mir-125b-1&WithFlank&11\|-\|122099751\|122099850 | 21 | 39 | 19(+1A) | TCCCTGAGACCCTAACTTGA | 40.7 | -8.7 | 3.0 | -2.9 | 3.8E-03 | 1.8E-02 | BT20/MB231 | MP |
| >hg38_21_+_8433172_8446622_RNA45SN1_45S_50ntFlanks@4130.4151.22\|ATTCCGGAGAGGGAGCCTGAGA | >hg38_21_+_8433172_8446622_RNA45SN1_45S_50ntFlanks | 4130 | 4151 | 22 | ATTCCGGAGAGGGAGCCTGAGA | 294.1 | -3.5 | 1.5 | -2.4 | 1.5E-02 | 1.8E-02 | BT20/MB468 | NUC |
| >hg38_21_+_8433172_8446622_RNA45SN1_45S_50ntFlanks@11580.11604.25\|GAACTGGTGCGGACCAGGGGAATCC | >hg38_21_+_8433172_8446622_RNA45SN1_45S_50ntFlanks | 11580 | 11604 | 25 | GAACTGGTGCGGACCAGGGGAATCC | 136.6 | -3.5 | 1.4 | -2.6 | 9.5E-03 | 1.8E-02 | BT20/MB468 | MP |
| >hg38_21_+_8433172_8446622_RNA45SN1_45S_50ntFlanks@11846.11867.22\|GAACGGGCTTGGCGGAATCAGC | >hg38_21_+_8433172_8446622_RNA45SN1_45S_50ntFlanks | 11846 | 11867 | 22 | GAACGGGCTTGGCGGAATCAGC | 433.9 | 5.2 | 1.8 | 2.9 | 3.8E-03 | 1.8E-02 | BT20/MB231 | MP |
| >hg38_MT_+_1621_3279_RNR2_16S_50ntFlanks@1216.1233.18\|CAACCTCCGAGCAGTACA | >hg38_MT_+_1621_3279_RNR2_16S_50ntFlanks | 1216 | 1233 | 18 | CAACCTCCGAGCAGTACA | 145.3 | 3.2 | 1.3 | 2.4 | 1.6E-02 | 1.8E-02 | MB231/MB468 | MITO |
| >hg38_21_+_8433172_8446622_RNA45SN1_45S_50ntFlanks@11570.11593.24\|AGCCGACTTAGAACTGGTGCGGAC | >hg38_21_+_8433172_8446622_RNA45SN1_45S_50ntFlanks | 11570 | 11593 | 24 | AGCCGACTTAGAACTGGTGCGGAC | 120.5 | -3.6 | 1.4 | -2.6 | 9.5E-03 | 1.8E-02 | BT20/MB468 | MP |
| >hg38_21_+_8433172_8446622_RNA45SN1_45S_50ntFlanks@10809.10826.18\|AATGTAGGTAAGGGAAGT | >hg38_21_+_8433172_8446622_RNA45SN1_45S_50ntFlanks | 10809 | 10826 | 18 | AATGTAGGTAAGGGAAGT | 756.3 | 2.3 | 1.0 | 2.4 | 1.6E-02 | 1.8E-02 | MB231/MB468 | NUC |
| >hg38_21_+_8433172_8446622_RNA45SN1_45S_50ntFlanks@12006.12035.30\|GGCCGCCGGTGAAATACCACTACTCTGATC | >hg38_21_+_8433172_8446622_RNA45SN1_45S_50ntFlanks | 12006 | 12035 | 30 | GGCCGCCGGTGAAATACCACTACTCTGATC | 292.6 | 3.8 | 1.3 | 2.9 | 3.8E-03 | 1.8E-02 | BT20/MB231 | MP |
| >hg38_MT_+_1621_3279_RNR2_16S_50ntFlanks@1428.1451.24\|ATTAAAGTCCTACGTGATCTGAGT | >hg38_MT_+_1621_3279_RNR2_16S_50ntFlanks | 1428 | 1451 | 24 | ATTAAAGTCCTACGTGATCTGAGT | 206.2 | 2.2 | 0.9 | 2.5 | 1.4E-02 | 1.8E-02 | MB231/MB468 | MP |
| >hg38_21_+_8433172_8446622_RNA45SN1_45S_50ntFlanks@6661.6689.29\|CGGTGGATCACTCGGCTCGTGCGTCGATG | >hg38_21_+_8433172_8446622_RNA45SN1_45S_50ntFlanks | 6661 | 6689 | 29 | CGGTGGATCACTCGGCTCGTGCGTCGATG | 2802.2 | 1.9 | 0.8 | 2.4 | 1.5E-02 | 1.8E-02 | BT20/MB468 | CYTO |
| >hg38_21_+_8433172_8446622_RNA45SN1_45S_50ntFlanks@9555.9584.30\|GTAGCGGTCCTGACGTGCAAATCGGTCGTC | >hg38_21_+_8433172_8446622_RNA45SN1_45S_50ntFlanks | 9555 | 9584 | 30 | GTAGCGGTCCTGACGTGCAAATCGGTCGTC | 244.7 | -1.9 | 0.8 | -2.4 | 1.5E-02 | 1.8E-02 | BT20/MB468 | CYTO |
| >hg38_21_+_8433172_8446622_RNA45SN1_45S_50ntFlanks@4725.4767.43\|TTAATCAAGAACGAAAGTCGGAGGTTCGAAGACGATCAGATAC | >hg38_21_+_8433172_8446622_RNA45SN1_45S_50ntFlanks | 4725 | 4767 | 43 | TTAATCAAGAACGAAAGTCGGAGGTTCGAAGACGATCAGATAC | 1191.7 | -2.8 | 1.0 | -2.8 | 4.9E-03 | 1.8E-02 | BT20/MB231 | CYTO |
| >trna2_GlyGCC_21_-_18827107_18827177@1.32.32,trna35_GlyGCC_1_+_161413094_161413164@1.32.32,trna37_GlyGCC_1_+_161420467_161420537@1.32.32,trna39_GlyGCC_1_+_161427898_161427968@1.32.32,trna41_GlyGCC_1_+_161435258_161435328@1.32.32\|GCATGGGTGGTTCAGTGGTAGAATTCTCGCCT | >trna2_GlyGCC_21_-_18827107_18827177 | 1 | 32 | 32 | GCATGGGTGGTTCAGTGGTAGAATTCTCGCCT | 851.0 | 2.5 | 1.0 | 2.4 | 1.5E-02 | 1.8E-02 | MB231/MB468 | CYTO |
| >hg38_21_+_8433172_8446622_RNA45SN1_45S_50ntFlanks@8176.8213.38\|CTTCTGATCGAGGCCCAGCCCGTGGACGGTGTGAGGCC | >hg38_21_+_8433172_8446622_RNA45SN1_45S_50ntFlanks | 8176 | 8213 | 38 | CTTCTGATCGAGGCCCAGCCCGTGGACGGTGTGAGGCC | 727.1 | -2.7 | 1.1 | -2.6 | 9.6E-03 | 1.8E-02 | BT20/MB468 | MP |
| >hg38_1_-_228634819_228635039_RNA5S12_5S_50ntFlanks@146.169.24\|TGGGAATACCGGGTGCTGTAGGCT | >hg38_1_-_228634819_228635039_RNA5S12_5S_50ntFlanks | 146 | 169 | 24 | TGGGAATACCGGGTGCTGTAGGCT | 277.8 | -1.9 | 0.8 | -2.5 | 1.4E-02 | 1.8E-02 | MB231/MB468 | MP |
| >hg38_MT_+_1621_3279_RNR2_16S_50ntFlanks@1401.1423.23\|CGCTATTAAAGGTTCGTTTGTTC | >hg38_MT_+_1621_3279_RNR2_16S_50ntFlanks | 1401 | 1423 | 23 | CGCTATTAAAGGTTCGTTTGTTC | 19.8 | 7.5 | 3.1 | 2.5 | 1.4E-02 | 1.8E-02 | MB231/MB468 | MP |
| >hg38_21_+_8433172_8446622_RNA45SN1_45S_50ntFlanks@6648.6665.18\|GTACGACTCTTAGCGGTG | >hg38_21_+_8433172_8446622_RNA45SN1_45S_50ntFlanks | 6648 | 6665 | 18 | GTACGACTCTTAGCGGTG | 521.1 | 2.4 | 0.8 | 2.8 | 4.9E-03 | 1.8E-02 | BT20/MB231 | CYTO |
| >hg38_21_+_8433172_8446622_RNA45SN1_45S_50ntFlanks@9553.9570.18\|CCGTAGCGGTCCTGACGT | >hg38_21_+_8433172_8446622_RNA45SN1_45S_50ntFlanks | 9553 | 9570 | 18 | CCGTAGCGGTCCTGACGT | 77.3 | -2.3 | 0.9 | -2.5 | 1.4E-02 | 1.8E-02 | MB231/MB468 | MP |
| >hg38_21_+_8433172_8446622_RNA45SN1_45S_50ntFlanks@10427.10456.30\|ATGGGCGAGCGCCGTTCCGAAGGGACGGGC | >hg38_21_+_8433172_8446622_RNA45SN1_45S_50ntFlanks | 10427 | 10456 | 30 | ATGGGCGAGCGCCGTTCCGAAGGGACGGGC | 650.6 | -2.3 | 0.9 | -2.6 | 9.6E-03 | 1.8E-02 | BT20/MB468 | MP |
| >MI0000088\|hsa-mir-30a&WithFlank&6\|-\|71403545\|71403627@54.71.18&[MIMAT0000088&hsa-miR-30a-3p&offsets\|+1\|-3;m-38&6\|-\|71403554\|71403575&offsets\|+1\|-3]\|TTTCAGTCGGATGTTTGC | >MI0000088\|hsa-mir-30a&WithFlank&6\|-\|71403545\|71403627 | 54 | 71 | 18 | TTTCAGTCGGATGTTTGC | 1733.1 | -2.9 | 1.0 | -2.9 | 3.8E-03 | 1.8E-02 | BT20/MB231 | MP |
| >hg38_21_+_8433172_8446622_RNA45SN1_45S_50ntFlanks@4603.4622.20\|TCGGAACTGAGGCCATGATT | >hg38_21_+_8433172_8446622_RNA45SN1_45S_50ntFlanks | 4603 | 4622 | 20 | TCGGAACTGAGGCCATGATT | 60.9 | -4.2 | 1.6 | -2.6 | 9.6E-03 | 1.8E-02 | BT20/MB468 | MP |
| >MI0000111\|hsa-mir-105-1&WithFlank&X\|-\|152392213\|152392305@21.42.22;MI0000112\|hsa-mir-105-2&WithFlank&X\|-\|152394406\|152394498@21.42.22&[MIMAT0000102&hsa-miR-105-5p&offsets\|+2\|+1;m-735&X\|-\|152392266\|152392287&offsets\|+2\|+2];[MIMAT0000102_1&hsa-miR-105-5p&offsets\|+2\|+1;m-735&X\|-\|152394459\|152394480&offsets\|+2\|+2]\|AAATGCTCAGACTCCTGTGGTG | >MI0000111\|hsa-mir-105-1&WithFlank&X\|-\|152392213\|152392305 | 21 | 42 | 22 | AAATGCTCAGACTCCTGTGGTG | 492.0 | 3.1 | 1.2 | 2.6 | 9.6E-03 | 1.8E-02 | BT20/MB468 | MP |
| >hg38_21_+_8433172_8446622_RNA45SN1_45S_50ntFlanks@5479.5499.21\|TGGCGGAGCGCTGAGAAGACG | >hg38_21_+_8433172_8446622_RNA45SN1_45S_50ntFlanks | 5479 | 5499 | 21 | TGGCGGAGCGCTGAGAAGACG | 103.2 | 2.3 | 0.9 | 2.6 | 9.6E-03 | 1.8E-02 | BT20/MB468 | MP |

| | | | | | | | | | | | | | |
|---|---|---|---|---|---|---|---|---|---|---|---|---|---|
| >hg38_21_+_8433172_8446622_RNA45SN1_45S_50ntFlanks@11913.11939.27\|AAGAGACATGAGAGGTGTAGAATAAGT | >hg38_21_+_8433172_8446622_RNA45SN1_45S_50ntFlanks | 11913 | 11939 | 27 | AAGAGACATGAGAGGTGTAGAATAAGT | 41.6 | 4.5 | 1.7 | 2.6 | 9.6E-03 | 1.8E-02 | BT20/MB468 | MP |
| >hg38_21_+_8433172_8446622_RNA45SN1_45S_50ntFlanks@7975.8006.32\|CGCGACCTCAGATCAGACGTGGCGACCCGCTG | >hg38_21_+_8433172_8446622_RNA45SN1_45S_50ntFlanks | 7975 | 8006 | 32 | CGCGACCTCAGATCAGACGTGGCGACCCGCTG | 68593.4 | 3.0 | 1.2 | 2.4 | 1.5E-02 | 1.8E-02 | BT20/MB468 | CYTO |
| >hg38_21_+_8433172_8446622_RNA45SN1_45S_50ntFlanks@10386.10424.39\|ATGTGAACAGCAGTTGAACATGGGTCAGTCGGTCCTGAG | >hg38_21_+_8433172_8446622_RNA45SN1_45S_50ntFlanks | 10386 | 10424 | 39 | ATGTGAACAGCAGTTGAACATGGGTCAGTCGGTCCTGAG | 6781.5 | 3.6 | 1.2 | 2.9 | 3.8E-03 | 1.8E-02 | BT20/MB231 | MP |
| >hg38_21_+_8433172_8446622_RNA45SN1_45S_50ntFlanks@9603.9630.28\|GAAAGACTAATCGAACCATCTAGTAGCT | >hg38_21_+_8433172_8446622_RNA45SN1_45S_50ntFlanks | 9603 | 9630 | 28 | GAAAGACTAATCGAACCATCTAGTAGCT | 237.6 | 3.6 | 1.2 | 2.9 | 3.8E-03 | 1.8E-02 | BT20/MB231 | MP |
| >trna111_HisGTG_1_-_147774845_147774916@-1G.31.32,trna118_HisGTG_1_-_145396881_145396952@-1G.31.32,trna16_HisGTG_1_+_146544773_146544844@-1G.31.32,trna1_HisGTG_15_+_45493349_45493420@-1G.31.32,trna21_HisGTG_1_+_147753471_147753542@-1G.31.32,trna33_HisGTG_6_+_27125906_27125977@-1G.31.32,trna7_HisGTG_9_-_14433938_14434009@-1G.31.32,trna8_HisGTG_15_-_45492611_45492682@-1G.31.32,trna9_HisGTG_15_-_45490804_45490875@-1G.31.32\|GGCCGTGATCGTATAGTGGTTAGTACTCTGCG | >trna111_HisGTG_1_-_147774845_147774916 | -1G | 31 | 32 | GGCCGTGATCGTATAGTGGTTAGTACTCTGCG | 24989.1 | -2.8 | 1.1 | -2.4 | 1.5E-02 | 1.8E-02 | BT20/MB468 | CYTO |
| >MI0000084\|hsa-mir-26b&WithFlank&2\|+\|218402640\|218402728@18.39.22&[MIMAT0000083&hsa-miR-26b-5p&offsets\|0\|+1;m-53&2\|+\|218402657\|218402678&offsets\|0\|0]\|TTCAAGTAATTCAGGATAGGTT | >MI0000084\|hsa-mir-26b&WithFlank&2\|+\|218402640\|218402728 | 18 | 39 | 22 | TTCAAGTAATTCAGGATAGGTT | 7658.9 | -2.3 | 0.8 | -2.8 | 4.8E-03 | 1.8E-02 | BT20/MB231 | MITO |
| >hg38_21_+_8433172_8446622_RNA45SN1_45S_50ntFlanks@10408.10453.46\|GGTCAGTCGGTCCTGAGAGATGGGCGAGCGCCGTTCCGAAGGGACG | >hg38_21_+_8433172_8446622_RNA45SN1_45S_50ntFlanks | 10408 | 10453 | 46 | GGTCAGTCGGTCCTGAGAGATGGGCGAGCGCCGTTCCGAAGGGACG | 397.6 | -3.3 | 1.4 | -2.4 | 1.6E-02 | 1.8E-02 | MB231/MB468 | NUC |
| >hg38_21_+_8433172_8446622_RNA45SN1_45S_50ntFlanks@12606.12624.19\|ATGTGCTTGGCTGAGGAGC | >hg38_21_+_8433172_8446622_RNA45SN1_45S_50ntFlanks | 12606 | 12624 | 19 | ATGTGCTTGGCTGAGGAGC | 457.5 | 5.3 | 1.8 | 2.9 | 3.8E-03 | 1.8E-02 | BT20/MB231 | MP |
| >hg38_21_+_8433172_8446622_RNA45SN1_45S_50ntFlanks@5489.5526.38\|CTGAGAAGACGGTCGAACTTGACTATCTAGAGGAAGTA | >hg38_21_+_8433172_8446622_RNA45SN1_45S_50ntFlanks | 5489 | 5526 | 38 | CTGAGAAGACGGTCGAACTTGACTATCTAGAGGAAGTA | 111.2 | -3.7 | 1.5 | -2.5 | 1.3E-02 | 1.8E-02 | BT20/MB468 | MITO |
| >trnaMT_ValTAC_MT_+_1602_1670@28.71.44\|AACTTACACTTAGGAGATTTCAACTTAACTTGACCGCTCTGACC | >trnaMT_ValTAC_MT_+_1602_1670 | 28 | 71 | 44 | AACTTACACTTAGGAGATTTCAACTTAACTTGACCGCTCTGACC | 289.4 | 3.5 | 1.5 | 2.4 | 1.6E-02 | 1.8E-02 | MB231/MB468 | MITO |
| >trnaMT_ValTAC_MT_+_1602_1670@39.71.33\|AGGAGATTTCAACTTAACTTGACCGCTCTGACC | >trnaMT_ValTAC_MT_+_1602_1670 | 39 | 71 | 33 | AGGAGATTTCAACTTAACTTGACCGCTCTGACC | 169.0 | 2.6 | 1.0 | 2.5 | 1.4E-02 | 1.8E-02 | MB231/MB468 | MP |
| >hg38_21_+_8433172_8446622_RNA45SN1_45S_50ntFlanks@4657.4682.26\|CTAGAGGTGAAATTCTTGGACCGGCG | >hg38_21_+_8433172_8446622_RNA45SN1_45S_50ntFlanks | 4657 | 4682 | 26 | CTAGAGGTGAAATTCTTGGACCGGCG | 205.6 | -4.4 | 1.7 | -2.6 | 9.6E-03 | 1.8E-02 | BT20/MB468 | MP |
| >am_trnaMT_ProTGG_MT_-_15956_16023@54.71.18\|AGACTTTTTCTCTGACCA | >am_trnaMT_ProTGG_MT_-_15956_16023 | 54 | 71 | 18 | AGACTTTTTCTCTGACCA | 4832.6 | 2.8 | 1.1 | 2.5 | 1.3E-02 | 1.8E-02 | BT20/MB468 | MITO |
| >hg38_21_+_8433172_8446622_RNA45SN1_45S_50ntFlanks@3993.4013.21\|GTGACTCTAGATAACCTCGGG | >hg38_21_+_8433172_8446622_RNA45SN1_45S_50ntFlanks | 3993 | 4013 | 21 | GTGACTCTAGATAACCTCGGG | 432.5 | -2.1 | 0.8 | -2.5 | 1.3E-02 | 1.8E-02 | BT20/MB468 | MITO |
| >hg38_21_+_8433172_8446622_RNA45SN1_45S_50ntFlanks@8357.8391.35\|AAAAGAACTTTGAAGAGAGAGTTCAAGAGGGCGTG | >hg38_21_+_8433172_8446622_RNA45SN1_45S_50ntFlanks | 8357 | 8391 | 35 | AAAAGAACTTTGAAGAGAGAGTTCAAGAGGGCGTG | 552.0 | -1.7 | 0.7 | -2.4 | 1.5E-02 | 1.8E-02 | BT20/MB468 | CYTO |
| >hg38_21_+_8433172_8446622_RNA45SN1_45S_50ntFlanks@7965.8005.41\|CCCTCCGAGACGCGACCTCAGATCAGACGTGGCGACCCGCT | >hg38_21_+_8433172_8446622_RNA45SN1_45S_50ntFlanks | 7965 | 8005 | 41 | CCCTCCGAGACGCGACCTCAGATCAGACGTGGCGACCCGCT | 402.9 | 3.6 | 1.3 | 2.9 | 4.2E-03 | 1.8E-02 | BT20/MB231 | NUC |
| >hg38_21_+_8433172_8446622_RNA45SN1_45S_50ntFlanks@8194.8212.19\|CCCGTGGACGGTGTGAGGC | >hg38_21_+_8433172_8446622_RNA45SN1_45S_50ntFlanks | 8194 | 8212 | 19 | CCCGTGGACGGTGTGAGGC | 524.6 | 5.3 | 1.8 | 2.9 | 3.8E-03 | 1.8E-02 | BT20/MB231 | MP |
| >hg38_21_+_8433172_8446622_RNA45SN1_45S_50ntFlanks@11321.11342.22\|CACGTCTCGTCGCGCGCGCGTC | >hg38_21_+_8433172_8446622_RNA45SN1_45S_50ntFlanks | 11321 | 11342 | 22 | CACGTCTCGTCGCGCGCGCGTC | 441.6 | -3.4 | 1.3 | -2.6 | 9.7E-03 | 1.8E-02 | BT20/MB468 | MP |
| >hg38_21_+_8433172_8446622_RNA45SN1_45S_50ntFlanks@10685.10711.27\|GGCGGCGTCCGGTGAGCTCTCGCTGGC | >hg38_21_+_8433172_8446622_RNA45SN1_45S_50ntFlanks | 10685 | 10711 | 27 | GGCGGCGTCCGGTGAGCTCTCGCTGGC | 541.3 | -4.8 | 2.0 | -2.4 | 1.6E-02 | 1.8E-02 | BT20/MB468 | NUC |
| >trna116_GluCTC_1_-_145399233_145399304@37.73.37,trna134_GluTTC_1_-_16861774_16861845@37.73.37,trna71_GluCTC_1_-_161439189_161439260@37.73.37,trna74_GluCTC_1_-_161431809_161431880@37.73.37,trna77_GluCTC_1_-_161424398_161424469@37.73.37,trna77_GluCTC_6_+_28949976_28950047@37.73.37,trna80_GluCTC_1_-_161417018_161417089@37.73.37,trna84_GluTTC_1_-_161391883_161391954@37.73.37,trna87_GluCTC_6_-_126101393_126101464@37.73.37\|ACCGCCGCGGCCCGGGTTCGATTCCCGGTCAGGGAAC | >trna116_GluCTC_1_-_145399233_145399304 | 37 | 73 | 37 | ACCGCCGCGGCCCGGGTTCGATTCCCGGTCAGGGAAC | 424.7 | 1.8 | 0.7 | 2.4 | 1.5E-02 | 1.8E-02 | BT20/MB468 | CYTO |
| >hg38_21_+_8433172_8446622_RNA45SN1_45S_50ntFlanks@4727.4765.39\|AATCAAGAACGAAAGTCGGAGGTTCGAAGACGATCAGAT | >hg38_21_+_8433172_8446622_RNA45SN1_45S_50ntFlanks | 4727 | 4765 | 39 | AATCAAGAACGAAAGTCGGAGGTTCGAAGACGATCAGAT | 230.7 | 1.2 | 0.5 | 2.4 | 1.6E-02 | 1.8E-02 | BT20/MB468 | NUC |
| >hg38_21_+_8433172_8446622_RNA45SN1_45S_50ntFlanks@11570.11602.33\|AGCCGACTTAGAACTGGTGCGGACCAGGGGAAT | >hg38_21_+_8433172_8446622_RNA45SN1_45S_50ntFlanks | 11570 | 11602 | 33 | AGCCGACTTAGAACTGGTGCGGACCAGGGGAAT | 387.1 | -1.4 | 0.6 | -2.4 | 1.5E-02 | 1.8E-02 | BT20/MB468 | CYTO |
| >MI0000299\|hsa-mir-222&WithFlank&X\|-\|45747009\|45747130@75.96.22(+1G)&[MIMAT0000279&hsa-miR-222-3p&offsets\|0\|+1(+1G);m-140&X\|-\|45747033\|45747056&offsets\|0\|-2(+1G)]\|AGCTACATCTGGCTACTGGGTCG | >MI0000299\|hsa-mir-222&WithFlank&X\|-\|45747009\|45747130 | 75 | 96 | 22(+1G) | AGCTACATCTGGCTACTGGGTCG | 170.9 | 4.2 | 1.7 | 2.5 | 1.4E-02 | 1.8E-02 | MB231/MB468 | MP |
| >trna10_LysCTT_16_+_3241501_3241573@14.31.18,trna13_LysCTT_14_-_58706613_58706685@14.31.18,trna2_LysCTT_15_+_79152904_79152976@14.31.18,trna4_LysCTT_18_-_43669271_43669343@14.31.18\|AGTCGGTAGAGCATGGGA | >trna10_LysCTT_16_+_3241501_3241573 | 14 | 31 | 18 | AGTCGGTAGAGCATGGGA | 1304.1 | -1.8 | 0.8 | -2.4 | 1.5E-02 | 1.8E-02 | MB231/MB468 | CYTO |
| >hg38_21_+_8433172_8446622_RNA45SN1_45S_50ntFlanks@11582.11613.32\|ACTGGTGCGGACCAGGGGAATCCGACTGTTTA | >hg38_21_+_8433172_8446622_RNA45SN1_45S_50ntFlanks | 11582 | 11613 | 32 | ACTGGTGCGGACCAGGGGAATCCGACTGTTTA | 347.6 | 1.4 | 0.6 | 2.5 | 1.3E-02 | 1.8E-02 | BT20/MB468 | MITO |
| >MI0000478\|hsa-mir-149&WithFlank&2\|+\|240455995\|240456095@21.43.23&[MIMAT0000450&hsa-miR-149-5p&offsets\|0\|0;m-195&2\|+\|240456015\|240456037&offsets\|0\|0]\|TCTGGCTCCGTGTCTTCACTCCC | >MI0000478\|hsa-mir-149&WithFlank&2\|+\|240455995\|240456095 | 21 | 43 | 23 | TCTGGCTCCGTGTCTTCACTCCC | 1082.5 | 3.2 | 1.1 | 2.9 | 3.9E-03 | 1.8E-02 | BT20/MB231 | MP |
| >trnaMT_GluTTC_MT_-_14674_14742@35.71.37,trnalookalike8_GluTTC_5_-_93905172_93905240@35.71.37\|TATCATTGGTCGTGGTTGTAGTCCGTGCGAGAATACC | >trnaMT_GluTTC_MT_-_14674_14742 | 35 | 71 | 37 | TATCATTGGTCGTGGTTGTAGTCCGTGCGAGAATACC | 321.8 | 4.0 | 1.5 | 2.6 | 9.7E-03 | 1.8E-02 | BT20/MB468 | MP |
| >hg38_21_+_8433172_8446622_RNA45SN1_45S_50ntFlanks@12447.12486.40\|CAGAATTCACCAAGCGTTGGATTGTTCACCCACTAATAGG | >hg38_21_+_8433172_8446622_RNA45SN1_45S_50ntFlanks | 12447 | 12486 | 40 | CAGAATTCACCAAGCGTTGGATTGTTCACCCACTAATAGG | 93.2 | -4.2 | 1.7 | -2.5 | 1.3E-02 | 1.8E-02 | BT20/MB468 | MITO |
| >hg38_21_+_8433172_8446622_RNA45SN1_45S_50ntFlanks@12473.12496.24\|CACCCACTAATAGGGAACGTGAGC | >hg38_21_+_8433172_8446622_RNA45SN1_45S_50ntFlanks | 12473 | 12496 | 24 | CACCCACTAATAGGGAACGTGAGC | 256.2 | 1.9 | 0.8 | 2.4 | 1.6E-02 | 1.8E-02 | MB231/MB468 | NUC |
| >hg38_21_+_8433172_8446622_RNA45SN1_45S_50ntFlanks@11568.11609.42\|GCAGCCGACTTAGAACTGGTGCGGACCAGGGGAATCCGACTG | >hg38_21_+_8433172_8446622_RNA45SN1_45S_50ntFlanks | 11568 | 11609 | 42 | GCAGCCGACTTAGAACTGGTGCGGACCAGGGGAATCCGACTG | 416.9 | -2.4 | 1.0 | -2.4 | 1.6E-02 | 1.8E-02 | MB231/MB468 | MITO |
| >MI0000064\|hsa-let-7c&WithFlank&21\|+\|16539822\|16539917@17.38.22&[MIMAT0000064&hsa-let-7c-5p&offsets\|0\|0;m-37&21\|+\|16539838\|16539859&offsets\|0\|0]\|TGAGGTAGTAGGTTGTATGGTT | >MI0000064\|hsa-let-7c&WithFlank&21\|+\|16539822\|16539917 | 17 | 38 | 22 | TGAGGTAGTAGGTTGTATGGTT | 772.1 | -3.1 | 1.1 | -2.9 | 3.9E-03 | 1.8E-02 | BT20/MB231 | MP |
| >hg38_21_+_8433172_8446622_RNA45SN1_45S_50ntFlanks@6734.6771.38\|ATTGATCATCGACACTTCGAACGCACTTGCGGCCCCGG | >hg38_21_+_8433172_8446622_RNA45SN1_45S_50ntFlanks | 6734 | 6771 | 38 | ATTGATCATCGACACTTCGAACGCACTTGCGGCCCCGG | 1874.1 | -2.9 | 1.2 | -2.4 | 1.6E-02 | 1.8E-02 | BT20/MB468 | NUC |
| >hg38_21_+_8433172_8446622_RNA45SN1_45S_50ntFlanks@6647.6669.23\|CGTACGACTCTTAGCGGTGGATC | >hg38_21_+_8433172_8446622_RNA45SN1_45S_50ntFlanks | 6647 | 6669 | 23 | CGTACGACTCTTAGCGGTGGATC | 3285.9 | 2.3 | 0.8 | 2.8 | 4.9E-03 | 1.8E-02 | BT20/MB231 | CYTO |
| >MI0000095\|hsa-mir-93&WithFlank&7\|-\|100093762\|100093853@17.39.23(+1C)&[MIMAT0000093&hsa-miR-93-5p&offsets\|0\|0(+1C);m-29&7\|-\|100093815\|100093837&offsets\|0\|0(+1C)]\|CAAAGTGCTGTTCGTGCAGGTAGC | >MI0000095\|hsa-mir-93&WithFlank&7\|-\|100093762\|100093853 | 17 | 39 | 23(+1C) | CAAAGTGCTGTTCGTGCAGGTAGC | 501.3 | -2.6 | 0.9 | -2.8 | 4.9E-03 | 1.8E-02 | BT20/MB231 | MITO |
| >hg38_21_+_8433172_8446622_RNA45SN1_45S_50ntFlanks@13000.13039.40\|TGCGATCTATTGAAAGTCAGCCCTCGACACAAGGGTTTGT | >hg38_21_+_8433172_8446622_RNA45SN1_45S_50ntFlanks | 13000 | 13039 | 40 | TGCGATCTATTGAAAGTCAGCCCTCGACACAAGGGTTTGT | 126.5 | 3.7 | 1.4 | 2.6 | 9.7E-03 | 1.8E-02 | BT20/MB468 | MP |
| >hg38_21_+_8433172_8446622_RNA45SN1_45S_50ntFlanks@5442.5466.25\|CCCTCGGATCGGCCCCGCCGGGGTC | >hg38_21_+_8433172_8446622_RNA45SN1_45S_50ntFlanks | 5442 | 5466 | 25 | CCCTCGGATCGGCCCCGCCGGGGTC | 84.4 | -4.4 | 1.7 | -2.6 | 9.7E-03 | 1.8E-02 | BT20/MB468 | MP |
| >hg38_21_+_8433172_8446622_RNA45SN1_45S_50ntFlanks@11783.11814.32\|CATGAATGGATGAACGAGATTCCCACTGTCCC | >hg38_21_+_8433172_8446622_RNA45SN1_45S_50ntFlanks | 11783 | 11814 | 32 | CATGAATGGATGAACGAGATTCCCACTGTCCC | 145.1 | -3.4 | 1.3 | -2.6 | 9.7E-03 | 1.8E-02 | BT20/MB468 | MP |
| >hg38_21_+_8433172_8446622_RNA45SN1_45S_50ntFlanks@10465.10495.31\|CCGTTGCCCTCGGCCGATCGAAAGGGAGTCG | >hg38_21_+_8433172_8446622_RNA45SN1_45S_50ntFlanks | 10465 | 10495 | 31 | CCGTTGCCCTCGGCCGATCGAAAGGGAGTCG | 77.7 | -4.3 | 1.6 | -2.6 | 9.7E-03 | 1.8E-02 | BT20/MB468 | MP |
| >hg38_21_+_8433172_8446622_RNA45SN1_45S_50ntFlanks@12424.12456.33\|CGGCTCTTCCTATCATTGTGAAGCAGAATTCAC | >hg38_21_+_8433172_8446622_RNA45SN1_45S_50ntFlanks | 12424 | 12456 | 33 | CGGCTCTTCCTATCATTGTGAAGCAGAATTCAC | 226.3 | -1.5 | 0.6 | -2.4 | 1.6E-02 | 1.8E-02 | MB231/MB468 | NUC |
| >trnaMT_PheGAA_MT_+_577_647@39.64.26\|ATGTTTAGACGGGCTCACATCACCCC | >trnaMT_PheGAA_MT_+_577_647 | 39 | 64 | 26 | ATGTTTAGACGGGCTCACATCACCCC | 306.5 | 4.5 | 1.7 | 2.6 | 9.7E-03 | 1.8E-02 | BT20/MB468 | MP |
| >hg38_21_+_8433172_8446622_RNA45SN1_45S_50ntFlanks@6648.6666.19\|GTACGACTCTTAGCGGTGG | >hg38_21_+_8433172_8446622_RNA45SN1_45S_50ntFlanks | 6648 | 6666 | 19 | GTACGACTCTTAGCGGTGG | 211.7 | -1.9 | 0.8 | -2.5 | 1.4E-02 | 1.8E-02 | MB231/MB468 | MP |
| >hg38_21_+_8433172_8446622_RNA45SN1_45S_50ntFlanks@12911.12932.22\|GCACGTTCGTGGGGAACCTGGC | >hg38_21_+_8433172_8446622_RNA45SN1_45S_50ntFlanks | 12911 | 12932 | 22 | GCACGTTCGTGGGGAACCTGGC | 301.4 | 3.9 | 1.5 | 2.6 | 9.7E-03 | 1.8E-02 | BT20/MB468 | MP |
| >hg38_MT_+_598_1651_RNR1_12S_50ntFlanks@905.922.18\|AGTATACTTCAAAGGACA | >hg38_MT_+_598_1651_RNR1_12S_50ntFlanks | 905 | 922 | 18 | AGTATACTTCAAAGGACA | 142.6 | 3.3 | 1.4 | 2.5 | 1.4E-02 | 1.8E-02 | MB231/MB468 | MP |
| >hg38_21_+_8433172_8446622_RNA45SN1_45S_50ntFlanks@10395.10424.30\|GCAGTTGAACATGGGTCAGTCGGTCCTGAG | >hg38_21_+_8433172_8446622_RNA45SN1_45S_50ntFlanks | 10395 | 10424 | 30 | GCAGTTGAACATGGGTCAGTCGGTCCTGAG | 10409.7 | 3.6 | 1.2 | 2.9 | 3.9E-03 | 1.8E-02 | BT20/MB231 | MP |
| >hg38_21_+_8433172_8446622_RNA45SN1_45S_50ntFlanks@11604.11626.23\|CGACTGTTTAATTAAAACAAAGC | >hg38_21_+_8433172_8446622_RNA45SN1_45S_50ntFlanks | 11604 | 11626 | 23 | CGACTGTTTAATTAAAACAAAGC | 296.1 | 5.7 | 2.0 | 2.9 | 3.9E-03 | 1.8E-02 | BT20/MB231 | MP |
| >hg38_21_+_8433172_8446622_RNA45SN1_45S_50ntFlanks@10787.10822.36\|ACAGCCTCTGGCATGTTGGAACAATGTAGGTAAGGG | >hg38_21_+_8433172_8446622_RNA45SN1_45S_50ntFlanks | 10787 | 10822 | 36 | ACAGCCTCTGGCATGTTGGAACAATGTAGGTAAGGG | 207.4 | -2.2 | 0.8 | -2.6 | 9.7E-03 | 1.8E-02 | BT20/MB468 | MP |

| | | | | | | | | | | | | | |
|---|---|---|---|---|---|---|---|---|---|---|---|---|---|
| >MI0000088\|hsa-mir-30a&WithFlank&6\|-\|71403545\|71403627@12.35.24(+1U)&[MIMAT0000087&hsa-miR-30a-5p&offsets\|0\|+2(+1U);m-15&6\|-\|71403595\|71403616&offsets\|0\|+2(+1U)]\|TGTAAACATCCTCGACTGGAAGCTT | >MI0000088\|hsa-mir-30a&WithFlank&6\|-\|71403545\|71403627 | 12 | 35 | 24(+1U) | TGTAAACATCCTCGACTGGAAGCTT | 3404.3 | 3.2 | 1.3 | 2.6 | 9.8E-03 | 1.8E-02 | BT20/MB468 | MP |
| >hg38_21_+_8433172_8446622_RNA45SN1_45S_50ntFlanks@7975.8012.38\|CGCGACCTCAGATCAGACGTGGCGACCCGCTGAATTTA | >hg38_21_+_8433172_8446622_RNA45SN1_45S_50ntFlanks | 7975 | 8012 | 38 | CGCGACCTCAGATCAGACGTGGCGACCCGCTGAATTTA | 76586.6 | 1.5 | 0.6 | 2.5 | 1.3E-02 | 1.8E-02 | BT20/MB468 | MITO |
| >hg38_21_+_8433172_8446622_RNA45SN1_45S_50ntFlanks@12479.12498.20\|CTAATAGGGAACGTGAGCTG | >hg38_21_+_8433172_8446622_RNA45SN1_45S_50ntFlanks | 12479 | 12498 | 20 | CTAATAGGGAACGTGAGCTG | 165.2 | -1.5 | 0.6 | -2.5 | 1.3E-02 | 1.8E-02 | BT20/MB468 | MITO |
| >hg38_21_+_8433172_8446622_RNA45SN1_45S_50ntFlanks@4726.4761.36\|TAATCAAGAACGAAAGTCGGAGGTTCGAAGACGATC | >hg38_21_+_8433172_8446622_RNA45SN1_45S_50ntFlanks | 4726 | 4761 | 36 | TAATCAAGAACGAAAGTCGGAGGTTCGAAGACGATC | 270.8 | 2.7 | 1.1 | 2.4 | 1.5E-02 | 1.8E-02 | MB231/MB468 | CYTO |
| >hg38_21_+_8433172_8446622_RNA45SN1_45S_50ntFlanks@10350.10385.36\|ACGAGAACTTTGAAGGCCGAAGTGGAGAAGGGTTCC | >hg38_21_+_8433172_8446622_RNA45SN1_45S_50ntFlanks | 10350 | 10385 | 36 | ACGAGAACTTTGAAGGCCGAAGTGGAGAAGGGTTCC | 145.9 | -2.7 | 1.0 | -2.6 | 9.8E-03 | 1.8E-02 | BT20/MB468 | MP |
| >hg38_21_+_8433172_8446622_RNA45SN1_45S_50ntFlanks@12416.12438.23\|TTCGATGTCGGCTCTTCCTATCA | >hg38_21_+_8433172_8446622_RNA45SN1_45S_50ntFlanks | 12416 | 12438 | 23 | TTCGATGTCGGCTCTTCCTATCA | 4512.7 | -2.5 | 0.9 | -2.8 | 5.0E-03 | 1.8E-02 | BT20/MB231 | CYTO |
| >hg38_21_+_8433172_8446622_RNA45SN1_45S_50ntFlanks@6647.6665.19\|CGTACGACTCTTAGCGGTG | >hg38_21_+_8433172_8446622_RNA45SN1_45S_50ntFlanks | 6647 | 6665 | 19 | CGTACGACTCTTAGCGGTG | 5295.1 | 2.1 | 0.7 | 2.8 | 5.0E-03 | 1.8E-02 | BT20/MB231 | CYTO |
| >trnaMT_HisGTG_MT_+_12138_12206@34.72.39\|AATCTGACAACAGAGGCTTACGACCCCTTATTTACCCCA | >trnaMT_HisGTG_MT_+_12138_12206 | 34 | 72 | 39 | AATCTGACAACAGAGGCTTACGACCCCTTATTTACCCCA | 174.5 | 4.1 | 1.7 | 2.4 | 1.6E-02 | 1.8E-02 | MB231/MB468 | MITO |
| >am_trna119_LysCTT_1_-_145395522_145395594@16.33.18,trna11_LysCTT_5_-_180648979_180649051@16.33.18,trna128_LysCTT_1_-_55423542_55423614@16.33.18,trna13_LysCTT_6_+_26556774_26556846@16.33.18,trna6_LysCTT_19_-_52425396_52425468@16.33.18,trna7_LysCTT_16_+_3225692_3225764@16.33.18,trna9_LysCTT_5_+_180634755_180634827@16.33.18\|TCGGTAGAGCATGAGACT | >am_trna119_LysCTT_1_-_145395522_145395594 | 16 | 33 | 18 | TCGGTAGAGCATGAGACT | 44.0 | 4.8 | 2.0 | 2.5 | 1.4E-02 | 1.8E-02 | MB231/MB468 | MP |
| >hg38_21_+_8433172_8446622_RNA45SN1_45S_50ntFlanks@12641.12664.24\|CCATCTGTGGGATTATGACTGAAC | >hg38_21_+_8433172_8446622_RNA45SN1_45S_50ntFlanks | 12641 | 12664 | 24 | CCATCTGTGGGATTATGACTGAAC | 86.2 | 4.6 | 1.8 | 2.6 | 9.8E-03 | 1.8E-02 | BT20/MB468 | MP |
| >MI0000077\|hsa-mir-21&WithFlank&17\|+\|59841260\|59841343@13.35.23(+1U)&[MIMAT0000076&hsa-miR-21-5p&offsets\|-1\|0(+1U);m-1&17\|+\|59841273\|59841294&offsets\|-1\|0(+1U)]\|GTAGCTTATCAGACTGATGTTGAT | >MI0000077\|hsa-mir-21&WithFlank&17\|+\|59841260\|59841343 | 13 | 35 | 23(+1U) | GTAGCTTATCAGACTGATGTTGAT | 40.7 | 4.9 | 2.0 | 2.5 | 1.4E-02 | 1.8E-02 | MB231/MB468 | MP |
| >hg38_MT_+_598_1651_RNR1_12S_50ntFlanks@428.465.38\|AAAGTGGCTTTAACATATCTGAACACACAATAGCTAAG | >hg38_MT_+_598_1651_RNR1_12S_50ntFlanks | 428 | 465 | 38 | AAAGTGGCTTTAACATATCTGAACACACAATAGCTAAG | 146.0 | 3.1 | 1.2 | 2.5 | 1.4E-02 | 1.8E-02 | MB231/MB468 | MP |
| >hg38_21_+_8433172_8446622_RNA45SN1_45S_50ntFlanks@11573.11594.22\|CGACTTAGAACTGGTGCGGACC | >hg38_21_+_8433172_8446622_RNA45SN1_45S_50ntFlanks | 11573 | 11594 | 22 | CGACTTAGAACTGGTGCGGACC | 213.0 | -3.1 | 1.3 | -2.5 | 1.4E-02 | 1.8E-02 | MB231/MB468 | MP |
| >hg38_21_+_8433172_8446622_RNA45SN1_45S_50ntFlanks@5056.5097.42\|GAGCGATTTGTCTGGTTAATTCCGATAACGAACGAGACTCTG | >hg38_21_+_8433172_8446622_RNA45SN1_45S_50ntFlanks | 5056 | 5097 | 42 | GAGCGATTTGTCTGGTTAATTCCGATAACGAACGAGACTCTG | 37.6 | -8.3 | 3.2 | -2.6 | 9.8E-03 | 1.8E-02 | BT20/MB468 | MP |
| >hg38_21_+_8433172_8446622_RNA45SN1_45S_50ntFlanks@10342.10362.21\|ATATTCAAACGAGAACTTTGA | >hg38_21_+_8433172_8446622_RNA45SN1_45S_50ntFlanks | 10342 | 10362 | 21 | ATATTCAAACGAGAACTTTGA | 127.0 | 2.1 | 0.8 | 2.6 | 9.8E-03 | 1.8E-02 | BT20/MB468 | MP |
| >hg38_21_+_8433172_8446622_RNA45SN1_45S_50ntFlanks@8325.8354.30\|CGATAGTCAACAAGTACCGTAAGGGAAAGT | >hg38_21_+_8433172_8446622_RNA45SN1_45S_50ntFlanks | 8325 | 8354 | 30 | CGATAGTCAACAAGTACCGTAAGGGAAAGT | 44.0 | -4.5 | 1.8 | -2.6 | 9.8E-03 | 1.8E-02 | BT20/MB468 | MP |
| >hg38_21_+_8433172_8446622_RNA45SN1_45S_50ntFlanks@8196.8224.29\|CGTGGACGGTGTGAGGCCGGTAGCGGCCC | >hg38_21_+_8433172_8446622_RNA45SN1_45S_50ntFlanks | 8196 | 8224 | 29 | CGTGGACGGTGTGAGGCCGGTAGCGGCCC | 289.1 | -1.4 | 0.6 | -2.5 | 1.3E-02 | 1.8E-02 | BT20/MB468 | MITO |
| >hg38_21_+_8433172_8446622_RNA45SN1_45S_50ntFlanks@11603.11620.18\|CCGACTGTTTAATTAAAA | >hg38_21_+_8433172_8446622_RNA45SN1_45S_50ntFlanks | 11603 | 11620 | 18 | CCGACTGTTTAATTAAAA | 785.6 | -2.1 | 0.8 | -2.4 | 1.5E-02 | 1.8E-02 | MB231/MB468 | CYTO |
| >hg38_21_+_8433172_8446622_RNA45SN1_45S_50ntFlanks@11873.11909.37\|AAGAAGACCCTGTTGAGCTTGACTCTAGTCTGGCACG | >hg38_21_+_8433172_8446622_RNA45SN1_45S_50ntFlanks | 11873 | 11909 | 37 | AAGAAGACCCTGTTGAGCTTGACTCTAGTCTGGCACG | 134.7 | -3.6 | 1.4 | -2.6 | 9.8E-03 | 1.8E-02 | BT20/MB468 | MP |
| >am_trna128_GlyGCC_6_-_27870686_27870756@1.26.26,trna133_GlyCCC_1_-_16872434_16872504@1.26.26,trna18_GlyGCC_16_+_70822597_70822667@1.26.26,trna19_GlyGCC_16_+_70823410_70823480@1.26.26,trna19_GlyGCC_2_-_157257659_157257729@1.26.26,trna24_GlyGCC_16_-_70812942_70813012@1.26.26,trna25_GlyGCC_16_-_70812114_70812184@1.26.26,trna4_GlyCCC_1_+_17188416_17188486@1.26.26,trna5_GlyGCC_17_+_8029064_8029134@1.26.26,trna68_GlyGCC_1_-_161493637_161493707@1.26.26\|GCATTGGTGGTTCAGTGGTAGAATTC | >am_trna128_GlyGCC_6_-_27870686_27870756 | 1 | 26 | 26 | GCATTGGTGGTTCAGTGGTAGAATTC | 544.7 | -2.1 | 0.9 | -2.4 | 1.6E-02 | 1.8E-02 | MB231/MB468 | NUC |
| >trnaMT_PheGAA_MT_+_577_647@36.63.28\|AAAATGTTTAGACGGGCTCACATCACCC | >trnaMT_PheGAA_MT_+_577_647 | 36 | 63 | 28 | AAAATGTTTAGACGGGCTCACATCACCC | 161.9 | 5.7 | 2.2 | 2.6 | 9.8E-03 | 1.8E-02 | BT20/MB468 | MP |
| >MI0005559\|hsa-mir-744&WithFlank&17\|+\|12081893\|12082002@17.36.20(+1A)&[MIMAT0004945&hsa-miR-744-5p&offsets\|0\|-2(+1A);m-172&17\|+\|12081909\|12081929&offsets\|0\|-1(+1A)]\|TGCGGGGCTAGGGCTAACAGA | >MI0005559\|hsa-mir-744&WithFlank&17\|+\|12081893\|12082002 | 17 | 36 | 20(+1A) | TGCGGGGCTAGGGCTAACAGA | 230.5 | 2.1 | 0.8 | 2.8 | 5.0E-03 | 1.8E-02 | BT20/MB231 | CYTO |
| >hg38_1_-_228634819_228635039_RNA5S12_5S_50ntFlanks@153.171.19\|ACCGGGTGCTGTAGGCTTT | >hg38_1_-_228634819_228635039_RNA5S12_5S_50ntFlanks | 153 | 171 | 19 | ACCGGGTGCTGTAGGCTTT | 3965.2 | 3.0 | 1.3 | 2.4 | 1.6E-02 | 1.8E-02 | BT20/MB468 | NUC |
| >MI0000103\|hsa-mir-101-1&WithFlank&1\|-\|65058428\|65058514@53.72.20(+2U);MI0000739\|hsa-mir-101-2&WithFlank&9\|+\|4850291\|4850381@55.74.20(+2U)&[MIMAT0000099&hsa-miR-101-3p&offsets\|0\|-1(+2U);m-17&1\|-\|65058443\|65058462&offsets\|0\|0(+2U)];[MIMAT0000099_1&hsa-miR-101-3p&offsets\|0\|-1(+2U);m-16&9\|+\|4850345\|4850364&offsets\|0\|0(+2U)]\|TACAGTACTGTGATAACTGATT | >MI0000103\|hsa-mir-101-1&WithFlank&1\|-\|65058428\|65058514 | 53 | 72 | 20(+2U) | TACAGTACTGTGATAACTGATT | 95.6 | 3.2 | 1.3 | 2.5 | 1.4E-02 | 1.8E-02 | MB231/MB468 | MP |
| >hg38_21_+_8433172_8446622_RNA45SN1_45S_50ntFlanks@3992.4026.35\|GGTGACTCTAGATAACCTCGGGCCGATCGCACGCC | >hg38_21_+_8433172_8446622_RNA45SN1_45S_50ntFlanks | 3992 | 4026 | 35 | GGTGACTCTAGATAACCTCGGGCCGATCGCACGCC | 294.7 | -1.9 | 0.8 | -2.4 | 1.6E-02 | 1.8E-02 | MB231/MB468 | NUC |
| >hg38_21_+_8433172_8446622_RNA45SN1_45S_50ntFlanks@12241.12272.32\|CAGAAGGGCAAAAGCTCGCTTGATCTTGATTT | >hg38_21_+_8433172_8446622_RNA45SN1_45S_50ntFlanks | 12241 | 12272 | 32 | CAGAAGGGCAAAAGCTCGCTTGATCTTGATTT | 369.7 | -2.7 | 1.1 | -2.6 | 9.8E-03 | 1.8E-02 | BT20/MB468 | MP |
| >hg38_21_+_8433172_8446622_RNA45SN1_45S_50ntFlanks@9300.9322.23\|ACGGACCAAGGAGTCTAACACGT | >hg38_21_+_8433172_8446622_RNA45SN1_45S_50ntFlanks | 9300 | 9322 | 23 | ACGGACCAAGGAGTCTAACACGT | 67.8 | 4.5 | 1.7 | 2.6 | 9.8E-03 | 1.8E-02 | BT20/MB468 | MP |
| >hg38_21_+_8433172_8446622_RNA45SN1_45S_50ntFlanks@6654.6685.32\|CTCTTAGCGGTGGATCACTCGGCTCGTGCGTC | >hg38_21_+_8433172_8446622_RNA45SN1_45S_50ntFlanks | 6654 | 6685 | 32 | CTCTTAGCGGTGGATCACTCGGCTCGTGCGTC | 101.7 | 3.6 | 1.4 | 2.6 | 9.8E-03 | 1.8E-02 | BT20/MB468 | MP |
| >hg38_21_+_8433172_8446622_RNA45SN1_45S_50ntFlanks@9940.9957.18\|CAGGACGGTGGCCATGGA | >hg38_21_+_8433172_8446622_RNA45SN1_45S_50ntFlanks | 9940 | 9957 | 18 | CAGGACGGTGGCCATGGA | 559.8 | 3.1 | 1.1 | 2.9 | 3.9E-03 | 1.8E-02 | BT20/MB231 | MP |
| >hg38_21_+_8433172_8446622_RNA45SN1_45S_50ntFlanks@9956.9983.28\|GAAGTCGGAATCCGCTAAGGAGTGTGTA | >hg38_21_+_8433172_8446622_RNA45SN1_45S_50ntFlanks | 9956 | 9983 | 28 | GAAGTCGGAATCCGCTAAGGAGTGTGTA | 143.2 | 5.7 | 2.0 | 2.9 | 3.9E-03 | 1.8E-02 | BT20/MB231 | MP |
| >hg38_21_+_8433172_8446622_RNA45SN1_45S_50ntFlanks@12006.12035.30\|GGCCGCCGGTGAAATACCACTACTCTGATC | >hg38_21_+_8433172_8446622_RNA45SN1_45S_50ntFlanks | 12006 | 12035 | 30 | GGCCGCCGGTGAAATACCACTACTCTGATC | 102.9 | 1.6 | 0.7 | 2.4 | 1.6E-02 | 1.8E-02 | BT20/MB468 | NUC |
| >hg38_21_+_8433172_8446622_RNA45SN1_45S_50ntFlanks@13006.13039.34\|CTATTGAAAGTCAGCCCTCGACACAAGGGTTTGT | >hg38_21_+_8433172_8446622_RNA45SN1_45S_50ntFlanks | 13006 | 13039 | 34 | CTATTGAAAGTCAGCCCTCGACACAAGGGTTTGT | 702.2 | 2.6 | 1.0 | 2.6 | 9.8E-03 | 1.8E-02 | BT20/MB468 | MP |
| >hg38_MT_+_598_1651_RNR1_12S_50ntFlanks@638.665.28\|TCACCACCTCTTGCTCAGCCTATATACC | >hg38_MT_+_598_1651_RNR1_12S_50ntFlanks | 638 | 665 | 28 | TCACCACCTCTTGCTCAGCCTATATACC | 95.5 | 5.6 | 2.2 | 2.6 | 9.8E-03 | 1.8E-02 | BT20/MB468 | MP |
| >hg38_21_+_8433172_8446622_RNA45SN1_45S_50ntFlanks@4722.4757.36\|TCATTAATCAAGAACGAAAGTCGGAGGTTCGAAGAC | >hg38_21_+_8433172_8446622_RNA45SN1_45S_50ntFlanks | 4722 | 4757 | 36 | TCATTAATCAAGAACGAAAGTCGGAGGTTCGAAGAC | 142.1 | -1.8 | 0.8 | -2.4 | 1.5E-02 | 1.8E-02 | MB231/MB468 | CYTO |
| >trna119_LysCTT_1_-_145395522_145395594@4.34.31,trna11_LysCTT_5_-_180648979_180649051@4.34.31,trna13_LysCTT_6_+_26556774_26556846@4.34.31,trna7_LysCTT_16_+_3225692_3225764@4.34.31,trna9_LysCTT_5_+_180634755_180634827@4.34.31\|CGGCTAGCTCAGTCGGTAGAGCATGAGACTC | >trna119_LysCTT_1_-_145395522_145395594 | 4 | 34 | 31 | CGGCTAGCTCAGTCGGTAGAGCATGAGACTC | 285.8 | -3.9 | 1.3 | -2.9 | 4.2E-03 | 1.8E-02 | BT20/MB231 | NUC |
| >hg38_21_+_8433172_8446622_RNA45SN1_45S_50ntFlanks@12419.12450.32\|GATGTCGGCTCTTCCTATCATTGTGAAGCAGA | >hg38_21_+_8433172_8446622_RNA45SN1_45S_50ntFlanks | 12419 | 12450 | 32 | GATGTCGGCTCTTCCTATCATTGTGAAGCAGA | 780.9 | -2.2 | 0.8 | -2.8 | 5.0E-03 | 1.8E-02 | BT20/MB231 | CYTO |
| >hg38_21_+_8433172_8446622_RNA45SN1_45S_50ntFlanks@8104.8146.43\|CCGCGGCGGGGCGCGGGACATGTGGCGTACGGAAGACCCGCTC | >hg38_21_+_8433172_8446622_RNA45SN1_45S_50ntFlanks | 8104 | 8146 | 43 | CCGCGGCGGGGCGCGGGACATGTGGCGTACGGAAGACCCGCTC | 37.5 | -8.3 | 3.2 | -2.6 | 9.9E-03 | 1.8E-02 | BT20/MB468 | MP |
| >hg38_21_+_8433172_8446622_RNA45SN1_45S_50ntFlanks@9201.9226.26\|CTCTCGGGGCCACGCGCGCGTCCCCC | >hg38_21_+_8433172_8446622_RNA45SN1_45S_50ntFlanks | 9201 | 9226 | 26 | CTCTCGGGGCCACGCGCGCGTCCCCC | 118.9 | -4.3 | 1.7 | -2.6 | 9.9E-03 | 1.8E-02 | BT20/MB468 | MP |
| >hg38_21_+_8433172_8446622_RNA45SN1_45S_50ntFlanks@6661.6686.26\|CGGTGGATCACTCGGCTCGTGCGTCG | >hg38_21_+_8433172_8446622_RNA45SN1_45S_50ntFlanks | 6661 | 6686 | 26 | CGGTGGATCACTCGGCTCGTGCGTCG | 4442.2 | 1.6 | 0.6 | 2.5 | 1.3E-02 | 1.8E-02 | BT20/MB468 | MITO |
| >hg38_21_+_8433172_8446622_RNA45SN1_45S_50ntFlanks@11874.11910.37\|AGAAGACCCTGTTGAGCTTGACTCTAGTCTGGCACGG | >hg38_21_+_8433172_8446622_RNA45SN1_45S_50ntFlanks | 11874 | 11910 | 37 | AGAAGACCCTGTTGAGCTTGACTCTAGTCTGGCACGG | 213.4 | -4.4 | 1.7 | -2.6 | 9.9E-03 | 1.8E-02 | BT20/MB468 | MP |
| >hg38_21_+_8433172_8446622_RNA45SN1_45S_50ntFlanks@12997.13017.21\|CGCTGCGATCTATTGAAAGTC | >hg38_21_+_8433172_8446622_RNA45SN1_45S_50ntFlanks | 12997 | 13017 | 21 | CGCTGCGATCTATTGAAAGTC | 1568.5 | -2.2 | 0.8 | -2.9 | 4.3E-03 | 1.8E-02 | BT20/MB231 | NUC |
| >hg38_21_+_8433172_8446622_RNA45SN1_45S_50ntFlanks@7977.8016.40\|CGACCTCAGATCAGACGTGGCGACCCGCTGAATTTAAGCA | >hg38_21_+_8433172_8446622_RNA45SN1_45S_50ntFlanks | 7977 | 8016 | 40 | CGACCTCAGATCAGACGTGGCGACCCGCTGAATTTAAGCA | 1032.7 | -3.3 | 1.2 | -2.9 | 4.3E-03 | 1.8E-02 | BT20/MB231 | NUC |
| >hg38_21_+_8433172_8446622_RNA45SN1_45S_50ntFlanks@10350.10371.22\|ACGAGAACTTTGAAGGCCGAAG | >hg38_21_+_8433172_8446622_RNA45SN1_45S_50ntFlanks | 10350 | 10371 | 22 | ACGAGAACTTTGAAGGCCGAAG | 2517.5 | -2.3 | 0.9 | -2.6 | 9.9E-03 | 1.8E-02 | BT20/MB468 | MP |
| >hg38_MT_+_1621_3279_RNR2_16S_50ntFlanks@999.1017.19\|AGGGACCTGTATGAATGGC | >hg38_MT_+_1621_3279_RNR2_16S_50ntFlanks | 999 | 1017 | 19 | AGGGACCTGTATGAATGGC | 128.9 | 3.2 | 1.3 | 2.4 | 1.4E-02 | 1.8E-02 | MB231/MB468 | MP |
| >hg38_21_+_8433172_8446622_RNA45SN1_45S_50ntFlanks@6646.6689.44\|TCGTACGACTCTTAGCGGTGGATCACTCGGCTCGTGCGTCGATG | >hg38_21_+_8433172_8446622_RNA45SN1_45S_50ntFlanks | 6646 | 6689 | 44 | TCGTACGACTCTTAGCGGTGGATCACTCGGCTCGTGCGTCGATG | 7766.3 | 2.7 | 1.0 | 2.9 | 4.3E-03 | 1.8E-02 | BT20/MB231 | NUC |
| >hg38_21_+_8433172_8446622_RNA45SN1_45S_50ntFlanks@6773.6806.34\|TTCCTCCCGGGGCTACGCCTGTCTGAGCGTCGCT | >hg38_21_+_8433172_8446622_RNA45SN1_45S_50ntFlanks | 6773 | 6806 | 34 | TTCCTCCCGGGGCTACGCCTGTCTGAGCGTCGCT | 781.0 | 1.6 | 0.6 | 2.9 | 4.3E-03 | 1.8E-02 | BT20/MB231 | NUC |

<table>
<tr><td>>trna10_LysCTT_16_+_3241501_3241573@1.28.28,trna119_LysCTT_1_-_145395522_145395594@1.28.28,trna11_LysCTT_5_-_180648979_180649051@1.28.28,trna13_LysCTT_14_-_58706613_58706685@1.28.28,trna13_LysCTT_6_+_26556774_26556846@1.28.28,trna2_LysCTT_15_+_79152904_79152976@1.28.28,trna32_LysCTT_16_-_3207406_3207478@1.28.28,trna7_LysCTT_16_+_3225692_3225764@1.28.28,trna9_LysCTT_5_+_180634755_180634827@1.28.28|GCCCGGCTAGCTCAGTCGGTAGAGCATG</td><td>>trna10_LysCTT_16_+_3241501_3241573</td><td>1</td><td>28</td><td>28</td><td>GCCCGGCTAGCTCAGTCGGTAGAGCATG</td><td>1556.8</td><td>-3.6</td><td>1.4</td><td>-2.5</td><td>1.3E-02</td><td>1.8E-02</td><td>BT20/MB468</td><td>MITO</td></tr>
<tr><td>>hg38_21_+_8433172_8446622_RNA45SN1_45S_50ntFlanks@11835.11872.38|CACAGCCAAGGGAACGGGCTTGGCGGAATCAGCGGGGA</td><td>>hg38_21_+_8433172_8446622_RNA45SN1_45S_50ntFlanks</td><td>11835</td><td>11872</td><td>38</td><td>CACAGCCAAGGGAACGGGCTTGGCGGAATCAGCGGGGA</td><td>67.7</td><td>-4.5</td><td>1.7</td><td>-2.6</td><td>9.9E-03</td><td>1.8E-02</td><td>BT20/MB468</td><td>MP</td></tr>
<tr><td>>hg38_21_+_8433172_8446622_RNA45SN1_45S_50ntFlanks@11891.11912.22|TTGACTCTAGTCTGGCACGGTG</td><td>>hg38_21_+_8433172_8446622_RNA45SN1_45S_50ntFlanks</td><td>11891</td><td>11912</td><td>22</td><td>TTGACTCTAGTCTGGCACGGTG</td><td>782.3</td><td>-2.8</td><td>1.1</td><td>-2.6</td><td>9.9E-03</td><td>1.8E-02</td><td>BT20/MB468</td><td>MP</td></tr>
<tr><td>>hg38_21_+_8433172_8446622_RNA45SN1_45S_50ntFlanks@12420.12450.31|ATGTCGGCTCTTCCTATCATTGTGAAGCAGA</td><td>>hg38_21_+_8433172_8446622_RNA45SN1_45S_50ntFlanks</td><td>12420</td><td>12450</td><td>31</td><td>ATGTCGGCTCTTCCTATCATTGTGAAGCAGA</td><td>181.6</td><td>-3.2</td><td>1.3</td><td>-2.4</td><td>1.4E-02</td><td>1.8E-02</td><td>MB231/MB468</td><td>MP</td></tr>
<tr><td>>hg38_21_+_8433172_8446622_RNA45SN1_45S_50ntFlanks@10794.10813.20|CTGGCATGTTGGAACAATGT</td><td>>hg38_21_+_8433172_8446622_RNA45SN1_45S_50ntFlanks</td><td>10794</td><td>10813</td><td>20</td><td>CTGGCATGTTGGAACAATGT</td><td>141.0</td><td>-3.4</td><td>1.3</td><td>-2.6</td><td>9.9E-03</td><td>1.8E-02</td><td>BT20/MB468</td><td>MP</td></tr>
<tr><td>>hg38_21_+_8433172_8446622_RNA45SN1_45S_50ntFlanks@12413.12434.22|TCCTTCGATGTCGGCTCTTCCT</td><td>>hg38_21_+_8433172_8446622_RNA45SN1_45S_50ntFlanks</td><td>12413</td><td>12434</td><td>22</td><td>TCCTTCGATGTCGGCTCTTCCT</td><td>296.7</td><td>1.2</td><td>0.5</td><td>2.5</td><td>1.3E-02</td><td>1.8E-02</td><td>BT20/MB468</td><td>MITO</td></tr>
<tr><td>>hg38_21_+_8433172_8446622_RNA45SN1_45S_50ntFlanks@8349.8373.25|GAAAGTTGAAAAGAACTTTGAAGAG</td><td>>hg38_21_+_8433172_8446622_RNA45SN1_45S_50ntFlanks</td><td>8349</td><td>8373</td><td>25</td><td>GAAAGTTGAAAAGAACTTTGAAGAG</td><td>2160.7</td><td>5.0</td><td>1.7</td><td>2.9</td><td>3.9E-03</td><td>1.8E-02</td><td>BT20/MB231</td><td>MP</td></tr>
<tr><td>>hg38_21_+_8433172_8446622_RNA45SN1_45S_50ntFlanks@6661.6701.41|CGGTGGATCACTCGGCTCGTGCGTCGATGAAGAACGCAGCT</td><td>>hg38_21_+_8433172_8446622_RNA45SN1_45S_50ntFlanks</td><td>6661</td><td>6701</td><td>41</td><td>CGGTGGATCACTCGGCTCGTGCGTCGATGAAGAACGCAGCT</td><td>301.3</td><td>2.1</td><td>0.9</td><td>2.4</td><td>1.4E-02</td><td>1.8E-02</td><td>MB231/MB468</td><td>MP</td></tr>
<tr><td>>hg38_21_+_8433172_8446622_RNA45SN1_45S_50ntFlanks@3841.3871.31|TCCTACTTGGATAACTGTGGTAATTCTAGAG</td><td>>hg38_21_+_8433172_8446622_RNA45SN1_45S_50ntFlanks</td><td>3841</td><td>3871</td><td>31</td><td>TCCTACTTGGATAACTGTGGTAATTCTAGAG</td><td>222.3</td><td>2.2</td><td>0.8</td><td>2.8</td><td>5.1E-03</td><td>1.8E-02</td><td>BT20/MB231</td><td>CYTO</td></tr>
<tr><td>>MI0000239|hsa-mir-197&WithFlank&1|+|109598887|109598973@54.74.21(+1U)&[MIMAT0000227&hsa-miR-197-3p&offsets|0|-1(+1U);m-99&1|+|109598940|109598961&offsets|0|-1(+1U)]|TTCACCACCTTCTCCACCCAGT</td><td>>MI0000239|hsa-mir-197&WithFlank&1|+|109598887|109598973</td><td>54</td><td>74</td><td>21(+1U)</td><td>TTCACCACCTTCTCCACCCAGT</td><td>267.1</td><td>3.7</td><td>1.4</td><td>2.6</td><td>9.9E-03</td><td>1.8E-02</td><td>BT20/MB468</td><td>MP</td></tr>
<tr><td>>hg38_21_+_8433172_8446622_RNA45SN1_45S_50ntFlanks@12016.12035.20|GAAATACCACTACTCTGATC</td><td>>hg38_21_+_8433172_8446622_RNA45SN1_45S_50ntFlanks</td><td>12016</td><td>12035</td><td>20</td><td>GAAATACCACTACTCTGATC</td><td>37.9</td><td>5.4</td><td>2.1</td><td>2.6</td><td>9.9E-03</td><td>1.8E-02</td><td>BT20/MB468</td><td>MP</td></tr>
<tr><td>>am_trnaMT_PheGAA_MT_+_577_647@43.65.23|TTAGACGGGCTCACATCACCCCA</td><td>>am_trnaMT_PheGAA_MT_+_577_647</td><td>43</td><td>65</td><td>23</td><td>TTAGACGGGCTCACATCACCCCA</td><td>55.7</td><td>6.6</td><td>2.6</td><td>2.6</td><td>9.9E-03</td><td>1.8E-02</td><td>BT20/MB468</td><td>MP</td></tr>
<tr><td>>hg38_21_+_8433172_8446622_RNA45SN1_45S_50ntFlanks@4657.4683.27|CTAGAGGTGAAATTCTTGGACCGGCGC</td><td>>hg38_21_+_8433172_8446622_RNA45SN1_45S_50ntFlanks</td><td>4657</td><td>4683</td><td>27</td><td>CTAGAGGTGAAATTCTTGGACCGGCGC</td><td>86.3</td><td>-4.3</td><td>1.7</td><td>-2.6</td><td>9.9E-03</td><td>1.8E-02</td><td>BT20/MB468</td><td>MP</td></tr>
<tr><td>>am_trnaMT_GlyTCC_MT_+_9991_10058@28.48.21|ACTTCCAATTAACTAGTTTTG</td><td>>am_trnaMT_GlyTCC_MT_+_9991_10058</td><td>28</td><td>48</td><td>21</td><td>ACTTCCAATTAACTAGTTTTG</td><td>110.0</td><td>-6.3</td><td>2.6</td><td>-2.4</td><td>1.5E-02</td><td>1.8E-02</td><td>MB231/MB468</td><td>CYTO</td></tr>
<tr><td>>hg38_21_+_8433172_8446622_RNA45SN1_45S_50ntFlanks@3987.4026.40|GCTTTGGTGACTCTAGATAACCTCGGGCCGATCGCACGCC</td><td>>hg38_21_+_8433172_8446622_RNA45SN1_45S_50ntFlanks</td><td>3987</td><td>4026</td><td>40</td><td>GCTTTGGTGACTCTAGATAACCTCGGGCCGATCGCACGCC</td><td>373.6</td><td>-3.4</td><td>1.4</td><td>-2.4</td><td>1.6E-02</td><td>1.8E-02</td><td>BT20/MB468</td><td>NUC</td></tr>
<tr><td>>hg38_21_+_8433172_8446622_RNA45SN1_45S_50ntFlanks@11570.11612.43|AGCCGACTTAGAACTGGTGCGGACCAGGGGAATCCGACTGTTT</td><td>>hg38_21_+_8433172_8446622_RNA45SN1_45S_50ntFlanks</td><td>11570</td><td>11612</td><td>43</td><td>AGCCGACTTAGAACTGGTGCGGACCAGGGGAATCCGACTGTTT</td><td>188.8</td><td>-4.6</td><td>1.9</td><td>-2.4</td><td>1.4E-02</td><td>1.8E-02</td><td>MB231/MB468</td><td>MP</td></tr>
<tr><td>>hg38_21_+_8433172_8446622_RNA45SN1_45S_50ntFlanks@10798.10822.25|CATGTTGGAACAATGTAGGTAAGGG</td><td>>hg38_21_+_8433172_8446622_RNA45SN1_45S_50ntFlanks</td><td>10798</td><td>10822</td><td>25</td><td>CATGTTGGAACAATGTAGGTAAGGG</td><td>252.7</td><td>-1.5</td><td>0.6</td><td>-2.5</td><td>1.3E-02</td><td>1.8E-02</td><td>BT20/MB468</td><td>MITO</td></tr>
<tr><td>>hg38_21_+_8433172_8446622_RNA45SN1_45S_50ntFlanks@7973.8011.39|GACGCGACCTCAGATCAGACGTGGCGACCCGCTGAATTT</td><td>>hg38_21_+_8433172_8446622_RNA45SN1_45S_50ntFlanks</td><td>7973</td><td>8011</td><td>39</td><td>GACGCGACCTCAGATCAGACGTGGCGACCCGCTGAATTT</td><td>2365.0</td><td>-2.4</td><td>0.9</td><td>-2.8</td><td>5.1E-03</td><td>1.8E-02</td><td>BT20/MB231</td><td>CYTO</td></tr>
<tr><td>>hg38_21_+_8433172_8446622_RNA45SN1_45S_50ntFlanks@12989.13013.25|AGCTCCCTCGCTGCGATCTATTGAA</td><td>>hg38_21_+_8433172_8446622_RNA45SN1_45S_50ntFlanks</td><td>12989</td><td>13013</td><td>25</td><td>AGCTCCCTCGCTGCGATCTATTGAA</td><td>458.6</td><td>-2.3</td><td>0.8</td><td>-2.8</td><td>5.1E-03</td><td>1.8E-02</td><td>BT20/MB231</td><td>CYTO</td></tr>
<tr><td>>am_MI0000809|hsa-mir-151a&WithFlank&8|-|140732558|140732659@53.70.18(+3G)&[MIMAT0000757&hsa-miR-151a-3p&offsets|0|-3(+3G);m-30&8|-|140732587|140732607&offsets|0|-3(+3G)]|CTAGACTGAAGCTCCTTGGGG</td><td>>am_MI0000809|hsa-mir-151a&WithFlank&8|-|140732558|140732659</td><td>53</td><td>70</td><td>18(+3G)</td><td>CTAGACTGAAGCTCCTTGGGG</td><td>139.3</td><td>3.1</td><td>1.3</td><td>2.4</td><td>1.4E-02</td><td>1.8E-02</td><td>MB231/MB468</td><td>MP</td></tr>
<tr><td>>hg38_21_+_8433172_8446622_RNA45SN1_45S_50ntFlanks@7990.8010.21|GACGTGGCGACCCGCTGAATT</td><td>>hg38_21_+_8433172_8446622_RNA45SN1_45S_50ntFlanks</td><td>7990</td><td>8010</td><td>21</td><td>GACGTGGCGACCCGCTGAATT</td><td>269.5</td><td>-1.9</td><td>0.8</td><td>-2.4</td><td>1.4E-02</td><td>1.8E-02</td><td>MB231/MB468</td><td>MP</td></tr>
<tr><td>>hg38_21_+_8433172_8446622_RNA45SN1_45S_50ntFlanks@11671.11702.32|CCAGTGCTCTGAATGTCAAAGTGAAGAAATTC</td><td>>hg38_21_+_8433172_8446622_RNA45SN1_45S_50ntFlanks</td><td>11671</td><td>11702</td><td>32</td><td>CCAGTGCTCTGAATGTCAAAGTGAAGAAATTC</td><td>782.8</td><td>-2.4</td><td>0.9</td><td>-2.6</td><td>1.0E-02</td><td>1.8E-02</td><td>BT20/MB468</td><td>MP</td></tr>
<tr><td>>MI0000736|hsa-mir-30c-1&WithFlank&1|+|40757278|40757378@23.43.21;MI0000254|hsa-mir-30c-2&WithFlank&6|-|71376954|71377037@13.33.21&[MIMAT0000244&hsa-miR-30c-5p&offsets|0|-2;m-57&1|+|40757300|40757323&offsets|0|-3];[MIMAT0000244_1&hsa-miR-30c-5p&offsets|0|-2;m-56&6|-|71377002|71377025&offsets|0|-3]|TGTAAACATCCTACACTCTCA</td><td>>MI0000736|hsa-mir-30c-1&WithFlank&1|+|40757278|40757378</td><td>23</td><td>43</td><td>21</td><td>TGTAAACATCCTACACTCTCA</td><td>111.6</td><td>4.1</td><td>1.6</td><td>2.6</td><td>1.0E-02</td><td>1.8E-02</td><td>BT20/MB468</td><td>MP</td></tr>
<tr><td>>hg38_21_+_8433172_8446622_RNA45SN1_45S_50ntFlanks@7973.8009.37|GACGCGACCTCAGATCAGACGTGGCGACCCGCTGAAT</td><td>>hg38_21_+_8433172_8446622_RNA45SN1_45S_50ntFlanks</td><td>7973</td><td>8009</td><td>37</td><td>GACGCGACCTCAGATCAGACGTGGCGACCCGCTGAAT</td><td>8210.8</td><td>2.3</td><td>0.8</td><td>2.9</td><td>4.3E-03</td><td>1.8E-02</td><td>BT20/MB231</td><td>NUC</td></tr>
<tr><td>>MI0006371|hsa-mir-1304&WithFlank&11|-|93733668|93733770@23.44.22&[MIMAT0005892&hsa-miR-1304-5p&offsets|-3|-3;m-632&11|-|93733727|93733748&offsets|0|0]|CGGTTTGAGGCTACAGTGAGAT</td><td>>MI0006371|hsa-mir-1304&WithFlank&11|-|93733668|93733770</td><td>23</td><td>44</td><td>22</td><td>CGGTTTGAGGCTACAGTGAGAT</td><td>204.1</td><td>2.3</td><td>0.8</td><td>2.8</td><td>5.1E-03</td><td>1.8E-02</td><td>BT20/MB231</td><td>CYTO</td></tr>
<tr><td>>hg38_21_+_8433172_8446622_RNA45SN1_45S_50ntFlanks@3843.3862.20|CTACTTGGATAACTGTGGTA</td><td>>hg38_21_+_8433172_8446622_RNA45SN1_45S_50ntFlanks</td><td>3843</td><td>3862</td><td>20</td><td>CTACTTGGATAACTGTGGTA</td><td>236.3</td><td>2.3</td><td>0.8</td><td>2.8</td><td>5.1E-03</td><td>1.8E-02</td><td>BT20/MB231</td><td>CYTO</td></tr>
<tr><td>>hg38_21_+_8433172_8446622_RNA45SN1_45S_50ntFlanks@9938.9959.22|AGCAGGACGGTGGCCATGGAAG</td><td>>hg38_21_+_8433172_8446622_RNA45SN1_45S_50ntFlanks</td><td>9938</td><td>9959</td><td>22</td><td>AGCAGGACGGTGGCCATGGAAG</td><td>183.3</td><td>3.6</td><td>1.4</td><td>2.6</td><td>1.0E-02</td><td>1.8E-02</td><td>BT20/MB468</td><td>MP</td></tr>
<tr><td>>hg38_MT_+_1621_3279_RNR2_16S_50ntFlanks@1374.1401.28|TGGATCAGGACATCCCGATGGTGCAGCC</td><td>>hg38_MT_+_1621_3279_RNR2_16S_50ntFlanks</td><td>1374</td><td>1401</td><td>28</td><td>TGGATCAGGACATCCCGATGGTGCAGCC</td><td>111.8</td><td>3.9</td><td>1.5</td><td>2.6</td><td>1.0E-02</td><td>1.8E-02</td><td>BT20/MB468</td><td>MP</td></tr>
<tr><td>>MI0003776|hsa-mir-320b-1&WithFlank&1|+|116671740|116671823@48.67.20(+1U);MI0003839|hsa-mir-320b-2&WithFlank&1|-|224257034|224257116@47.66.20(+1U);MI0000542|hsa-mir-320a&WithFlank&8|-|22244960|22245043@48.67.20(+1U)&[MIMAT0005792&hsa-miR-320b&offsets|0|-2(+1U);m-183&1|+|116671787|116671806&offsets|0|0(+1U)];[MIMAT0005792_1&hsa-miR-320b&offsets|0|-2(+1U);m-183&1|-|224257051|224257070&offsets|0|0(+1U)];[MIMAT0000510&hsa-miR-320a-3p&offsets|0|-2(+1U);m-60&8|-|22244975|22244996&offsets|0|-2(+1U)]|AAAAGCTGGGTTGAGAGGGCT</td><td>>MI0003776|hsa-mir-320b-1&WithFlank&1|+|116671740|116671823</td><td>48</td><td>67</td><td>20(+1U)</td><td>AAAAGCTGGGTTGAGAGGGCT</td><td>239.0</td><td>3.5</td><td>1.4</td><td>2.4</td><td>1.4E-02</td><td>1.8E-02</td><td>MB231/MB468</td><td>MP</td></tr>
<tr><td>>hg38_21_+_8433172_8446622_RNA45SN1_45S_50ntFlanks@4760.4793.34|TCAGATACCGTCGTAGTTCCGACCATAAACGATG</td><td>>hg38_21_+_8433172_8446622_RNA45SN1_45S_50ntFlanks</td><td>4760</td><td>4793</td><td>34</td><td>TCAGATACCGTCGTAGTTCCGACCATAAACGATG</td><td>168.3</td><td>-3.6</td><td>1.4</td><td>-2.6</td><td>1.0E-02</td><td>1.8E-02</td><td>BT20/MB468</td><td>MP</td></tr>
<tr><td>>hg38_21_+_8433172_8446622_RNA45SN1_45S_50ntFlanks@11209.11231.23|CGGCGACTCTGGACGCGAGCCGG</td><td>>hg38_21_+_8433172_8446622_RNA45SN1_45S_50ntFlanks</td><td>11209</td><td>11231</td><td>23</td><td>CGGCGACTCTGGACGCGAGCCGG</td><td>852.9</td><td>-2.9</td><td>1.1</td><td>-2.6</td><td>1.0E-02</td><td>1.8E-02</td><td>BT20/MB468</td><td>MP</td></tr>
<tr><td>>MI0000469|hsa-mir-125a&WithFlank&19|+|51693248|51693345@21.42.22(+1U)&[MIMAT0000443&hsa-miR-125a-5p&offsets|0|-2(+1U);m-63&19|+|51693268|51693289&offsets|0|0(+1U)]|TCCCTGAGACCCTTTAACCTGTT</td><td>>MI0000469|hsa-mir-125a&WithFlank&19|+|51693248|51693345</td><td>21</td><td>42</td><td>22(+1U)</td><td>TCCCTGAGACCCTTTAACCTGTT</td><td>284.2</td><td>1.9</td><td>0.7</td><td>2.9</td><td>4.4E-03</td><td>1.8E-02</td><td>BT20/MB231</td><td>NUC</td></tr>
<tr><td>>hg38_21_+_8433172_8446622_RNA45SN1_45S_50ntFlanks@9549.9572.24|AGGTCCGTAGCGGTCCTGACGTGC</td><td>>hg38_21_+_8433172_8446622_RNA45SN1_45S_50ntFlanks</td><td>9549</td><td>9572</td><td>24</td><td>AGGTCCGTAGCGGTCCTGACGTGC</td><td>280.3</td><td>1.2</td><td>0.5</td><td>2.4</td><td>1.6E-02</td><td>1.8E-02</td><td>MB231/MB468</td><td>CYTO</td></tr>
<tr><td>>hg38_21_+_8433172_8446622_RNA45SN1_45S_50ntFlanks@8383.8405.23|GAGGGCGTGAAACCGTTAAGAGG</td><td>>hg38_21_+_8433172_8446622_RNA45SN1_45S_50ntFlanks</td><td>8383</td><td>8405</td><td>23</td><td>GAGGGCGTGAAACCGTTAAGAGG</td><td>110.4</td><td>-3.5</td><td>1.4</td><td>-2.4</td><td>1.4E-02</td><td>1.8E-02</td><td>MB231/MB468</td><td>MP</td></tr>
<tr><td>>hg38_21_+_8433172_8446622_RNA45SN1_45S_50ntFlanks@12416.12435.20|TTCGATGTCGGCTCTTCCTA</td><td>>hg38_21_+_8433172_8446622_RNA45SN1_45S_50ntFlanks</td><td>12416</td><td>12435</td><td>20</td><td>TTCGATGTCGGCTCTTCCTA</td><td>129.4</td><td>-4.3</td><td>1.7</td><td>-2.6</td><td>1.0E-02</td><td>1.8E-02</td><td>BT20/MB468</td><td>MP</td></tr>
<tr><td>>hg38_MT_+_598_1651_RNR1_12S_50ntFlanks@903.929.27|CAAGTATACTTCAAAGGACATTTAACT</td><td>>hg38_MT_+_598_1651_RNR1_12S_50ntFlanks</td><td>903</td><td>929</td><td>27</td><td>CAAGTATACTTCAAAGGACATTTAACT</td><td>136.4</td><td>4.2</td><td>1.7</td><td>2.4</td><td>1.4E-02</td><td>1.8E-02</td><td>MB231/MB468</td><td>MP</td></tr>
<tr><td>>hg38_21_+_8433172_8446622_RNA45SN1_45S_50ntFlanks@8236.8266.31|GGCCCGGGTCTTCCCGGAGTCGGGTTGCTTG</td><td>>hg38_21_+_8433172_8446622_RNA45SN1_45S_50ntFlanks</td><td>8236</td><td>8266</td><td>31</td><td>GGCCCGGGTCTTCCCGGAGTCGGGTTGCTTG</td><td>73.7</td><td>-4.3</td><td>1.7</td><td>-2.6</td><td>1.0E-02</td><td>1.8E-02</td><td>BT20/MB468</td><td>MP</td></tr>
<tr><td>>hg38_1_-_228634819_228635039_RNA5S12_5S_50ntFlanks@151.170.20|ATACCGGGTGCTGTAGGCTT</td><td>>hg38_1_-_228634819_228635039_RNA5S12_5S_50ntFlanks</td><td>151</td><td>170</td><td>20</td><td>ATACCGGGTGCTGTAGGCTT</td><td>6937.8</td><td>2.2</td><td>0.8</td><td>2.9</td><td>4.0E-03</td><td>1.8E-02</td><td>BT20/MB231</td><td>MP</td></tr>
<tr><td>>MI0003776|hsa-mir-320b-1&WithFlank&1|+|116671740|116671823@49.67.19;MI0003839|hsa-mir-320b-2&WithFlank&1|-|224257034|224257116@48.66.19;MI0000542|hsa-mir-320a&WithFlank&8|-|22244960|22245043@49.67.19&[MIMAT0005792&hsa-miR-320b&offsets|+1|-2;m-183&1|+|116671787|116671806&offsets|+1|0];[MIMAT0005792_1&hsa-miR-320b&offsets|+1|-2;m-183&1|-|224257051|224257070&offsets|+1|0];[MIMAT0000510&hsa-miR-320a-3p&offsets|+1|-2;m-60&8|-|22244975|22244996&offsets|+1|-2]|AAAGCTGGGTTGAGAGGGC</td><td>>MI0003776|hsa-mir-320b-1&WithFlank&1|+|116671740|116671823</td><td>49</td><td>67</td><td>19</td><td>AAAGCTGGGTTGAGAGGGC</td><td>424.9</td><td>2.9</td><td>1.0</td><td>2.9</td><td>4.0E-03</td><td>1.8E-02</td><td>BT20/MB231</td><td>MP</td></tr>
<tr><td>>hg38_21_+_8433172_8446622_RNA45SN1_45S_50ntFlanks@10797.10819.23|GCATGTTGGAACAATGTAGGTAA</td><td>>hg38_21_+_8433172_8446622_RNA45SN1_45S_50ntFlanks</td><td>10797</td><td>10819</td><td>23</td><td>GCATGTTGGAACAATGTAGGTAA</td><td>57.9</td><td>-4.2</td><td>1.6</td><td>-2.6</td><td>1.0E-02</td><td>1.8E-02</td><td>BT20/MB468</td><td>MP</td></tr>
<tr><td>>hg38_21_+_8433172_8446622_RNA45SN1_45S_50ntFlanks@12003.12030.28|GCGGGCCGCCGGTGAAATACCACTACTC</td><td>>hg38_21_+_8433172_8446622_RNA45SN1_45S_50ntFlanks</td><td>12003</td><td>12030</td><td>28</td><td>GCGGGCCGCCGGTGAAATACCACTACTC</td><td>217.5</td><td>-2.4</td><td>0.8</td><td>-2.8</td><td>5.1E-03</td><td>1.8E-02</td><td>BT20/MB231</td><td>CYTO</td></tr>
<tr><td>>hg38_21_+_8433172_8446622_RNA45SN1_45S_50ntFlanks@12396.12417.22|GCGACGTCGCTTTTTGATCCTT</td><td>>hg38_21_+_8433172_8446622_RNA45SN1_45S_50ntFlanks</td><td>12396</td><td>12417</td><td>22</td><td>GCGACGTCGCTTTTTGATCCTT</td><td>395.9</td><td>-2.5</td><td>1.0</td><td>-2.4</td><td>1.6E-02</td><td>1.8E-02</td><td>MB231/MB468</td><td>CYTO</td></tr>
<tr><td>>hg38_21_+_8433172_8446622_RNA45SN1_45S_50ntFlanks@10468.10486.19|TTGCCCTCGGCCGATCGAA</td><td>>hg38_21_+_8433172_8446622_RNA45SN1_45S_50ntFlanks</td><td>10468</td><td>10486</td><td>19</td><td>TTGCCCTCGGCCGATCGAA</td><td>362.9</td><td>-1.8</td><td>0.8</td><td>-2.4</td><td>1.6E-02</td><td>1.8E-02</td><td>MB231/MB468</td><td>NUC</td></tr>
<tr><td>>hg38_21_+_8433172_8446622_RNA45SN1_45S_50ntFlanks@9202.9220.19|TCTCGGGGCCACGCGCGCG</td><td>>hg38_21_+_8433172_8446622_RNA45SN1_45S_50ntFlanks</td><td>9202</td><td>9220</td><td>19</td><td>TCTCGGGGCCACGCGCGCG</td><td>371.8</td><td>-4.0</td><td>1.6</td><td>-2.5</td><td>1.4E-02</td><td>1.8E-02</td><td>BT20/MB468</td><td>MITO</td></tr>
</table>

<table>
<tr><th></th><th></th><th></th><th></th><th></th><th></th><th></th><th></th><th></th><th></th><th></th><th></th><th></th><th></th></tr>
<tr><td>>MI0000266|hsa-mir-10a&WithFlank&17|-|48579832|48579953@29.50.22&[MIMAT0000253&hsa-miR-10a-5p&offsets|+1|0;m-18&17|-|48579905|48579926&offsets|+1|+1]|ACCCTGTAGATCCGAATTTGTG</td><td>>MI0000266|hsa-mir-10a&WithFlank&17|-|48579832|48579953</td><td>29</td><td>50</td><td>22</td><td>ACCCTGTAGATCCGAATTTGTG</td><td>13085.6</td><td>2.1</td><td>0.8</td><td>2.8</td><td>5.2E-03</td><td>1.8E-02</td><td>BT20/MB231</td><td>CYTO</td></tr>
<tr><td>>hg38_21_+_8433172_8446622_RNA45SN1_45S_50ntFlanks@11255.11281.27|CTGCGGCGGGCGTCGCGGCCGCCCCCG</td><td>>hg38_21_+_8433172_8446622_RNA45SN1_45S_50ntFlanks</td><td>11255</td><td>11281</td><td>27</td><td>CTGCGGCGGGCGTCGCGGCCGCCCCCG</td><td>91.9</td><td>-4.3</td><td>1.7</td><td>-2.6</td><td>1.0E-02</td><td>1.8E-02</td><td>BT20/MB468</td><td>MP</td></tr>
<tr><td>>hg38_21_+_8433172_8446622_RNA45SN1_45S_50ntFlanks@9957.9981.25|AAGTCGGAATCCGCTAAGGAGTGTG</td><td>>hg38_21_+_8433172_8446622_RNA45SN1_45S_50ntFlanks</td><td>9957</td><td>9981</td><td>25</td><td>AAGTCGGAATCCGCTAAGGAGTGTG</td><td>41.5</td><td>4.5</td><td>1.8</td><td>2.6</td><td>1.0E-02</td><td>1.8E-02</td><td>BT20/MB468</td><td>MP</td></tr>
<tr><td>>hg38_21_+_8433172_8446622_RNA45SN1_45S_50ntFlanks@12539.12570.32|ATGATGTGTTGTTGCCATGGTAATCCTGCTCA</td><td>>hg38_21_+_8433172_8446622_RNA45SN1_45S_50ntFlanks</td><td>12539</td><td>12570</td><td>32</td><td>ATGATGTGTTGTTGCCATGGTAATCCTGCTCA</td><td>55.2</td><td>-4.3</td><td>1.7</td><td>-2.6</td><td>1.0E-02</td><td>1.8E-02</td><td>BT20/MB468</td><td>MP</td></tr>
<tr><td>>hg38_21_+_8433172_8446622_RNA45SN1_45S_50ntFlanks@9033.9076.44|CTCGTGGGGGGGCCGGGCCACCCCTCCCACGGCGCGACCGCTCT</td><td>>hg38_21_+_8433172_8446622_RNA45SN1_45S_50ntFlanks</td><td>9033</td><td>9076</td><td>44</td><td>CTCGTGGGGGGGCCGGGCCACCCCTCCCACGGCGCGACCGCTCT</td><td>375.4</td><td>-4.7</td><td>2.0</td><td>-2.4</td><td>1.6E-02</td><td>1.8E-02</td><td>BT20/MB468</td><td>NUC</td></tr>
<tr><td>>hg38_21_+_8433172_8446622_RNA45SN1_45S_50ntFlanks@10922.10939.18|CTGGACGAGGCGCCGCCG</td><td>>hg38_21_+_8433172_8446622_RNA45SN1_45S_50ntFlanks</td><td>10922</td><td>10939</td><td>18</td><td>CTGGACGAGGCGCCGCCG</td><td>1460.0</td><td>-2.6</td><td>1.1</td><td>-2.5</td><td>1.4E-02</td><td>1.8E-02</td><td>BT20/MB468</td><td>MITO</td></tr>
<tr><td>>hg38_21_+_8433172_8446622_RNA45SN1_45S_50ntFlanks@11586.11607.22|GTGCGGACCAGGGGAATCCGAC</td><td>>hg38_21_+_8433172_8446622_RNA45SN1_45S_50ntFlanks</td><td>11586</td><td>11607</td><td>22</td><td>GTGCGGACCAGGGGAATCCGAC</td><td>164.8</td><td>-4.6</td><td>1.8</td><td>-2.6</td><td>1.0E-02</td><td>1.8E-02</td><td>BT20/MB468</td><td>MP</td></tr>
<tr><td>>hg38_21_+_8433172_8446622_RNA45SN1_45S_50ntFlanks@11584.11623.40|TGGTGCGGACCAGGGGAATCCGACTGTTTAATTAAAACAA</td><td>>hg38_21_+_8433172_8446622_RNA45SN1_45S_50ntFlanks</td><td>11584</td><td>11623</td><td>40</td><td>TGGTGCGGACCAGGGGAATCCGACTGTTTAATTAAAACAA</td><td>132.3</td><td>4.1</td><td>1.7</td><td>2.4</td><td>1.6E-02</td><td>1.8E-02</td><td>MB231/MB468</td><td>MITO</td></tr>
<tr><td>>am_trna116_GluCTC_1_-_145399233_145399304@1.30.30,trna59_GluCTC_1_+_249168447_249168518@1.30.30,trna71_GluCTC_1_-_161439189_161439260@1.30.30,trna74_GluCTC_1_-_161431809_161431880@1.30.30,trna77_GluCTC_1_-_161424398_161424469@1.30.30,trna77_GluCTC_6_+_28949976_28950047@1.30.30,trna80_GluCTC_1_-_161417018_161417089@1.30.30,trna87_GluCTC_6_-_126101393_126101464@1.30.30|TCCCTGGTGGTCTAGTGGTTAGGATTCGGC</td><td>>am_trna116_GluCTC_1_-_145399233_145399304</td><td>1</td><td>30</td><td>30</td><td>TCCCTGGTGGTCTAGTGGTTAGGATTCGGC</td><td>6807.7</td><td>2.3</td><td>0.9</td><td>2.5</td><td>1.4E-02</td><td>1.8E-02</td><td>BT20/MB468</td><td>MITO</td></tr>
<tr><td>>hg38_21_+_8433172_8446622_RNA45SN1_45S_50ntFlanks@9033.9056.24|CTCGTGGGGGGGCCGGGCCACCCC</td><td>>hg38_21_+_8433172_8446622_RNA45SN1_45S_50ntFlanks</td><td>9033</td><td>9056</td><td>24</td><td>CTCGTGGGGGGGCCGGGCCACCCC</td><td>407.2</td><td>-4.2</td><td>1.7</td><td>-2.5</td><td>1.4E-02</td><td>1.8E-02</td><td>BT20/MB468</td><td>MITO</td></tr>
<tr><td>>hg38_21_+_8433172_8446622_RNA45SN1_45S_50ntFlanks@10908.10930.23|GGCGCGCGCCGCGGCTGGACGAG</td><td>>hg38_21_+_8433172_8446622_RNA45SN1_45S_50ntFlanks</td><td>10908</td><td>10930</td><td>23</td><td>GGCGCGCGCCGCGGCTGGACGAG</td><td>35.6</td><td>5.4</td><td>2.1</td><td>2.6</td><td>1.0E-02</td><td>1.8E-02</td><td>BT20/MB468</td><td>MP</td></tr>
<tr><td>>MI0000077|hsa-mir-21&WithFlank&17|+|59841260|59841343@14.35.22(+2A)&[MIMAT0000076&hsa-miR-21-5p&offsets|0|0(+2A);m-1&17|+|59841273|59841294&offsets|0|0(+2A)]|TAGCTTATCAGACTGATGTTGAAA</td><td>>MI0000077|hsa-mir-21&WithFlank&17|+|59841260|59841343</td><td>14</td><td>35</td><td>22(+2A)</td><td>TAGCTTATCAGACTGATGTTGAAA</td><td>172.6</td><td>2.3</td><td>0.9</td><td>2.4</td><td>1.5E-02</td><td>1.8E-02</td><td>MB231/MB468</td><td>MP</td></tr>
<tr><td>>hg38_21_+_8433172_8446622_RNA45SN1_45S_50ntFlanks@475.496.22|CTTCGTGATCGATGTGGTGACG</td><td>>hg38_21_+_8433172_8446622_RNA45SN1_45S_50ntFlanks</td><td>475</td><td>496</td><td>22</td><td>CTTCGTGATCGATGTGGTGACG</td><td>1448.5</td><td>3.8</td><td>1.5</td><td>2.4</td><td>1.5E-02</td><td>1.8E-02</td><td>MB231/MB468</td><td>MP</td></tr>
<tr><td>>hg38_21_+_8433172_8446622_RNA45SN1_45S_50ntFlanks@12628.12649.22|TGGGGCGAAGCTACCATCTGTG</td><td>>hg38_21_+_8433172_8446622_RNA45SN1_45S_50ntFlanks</td><td>12628</td><td>12649</td><td>22</td><td>TGGGGCGAAGCTACCATCTGTG</td><td>229.6</td><td>-4.4</td><td>1.7</td><td>-2.6</td><td>1.0E-02</td><td>1.8E-02</td><td>BT20/MB468</td><td>MP</td></tr>
<tr><td>>hg38_21_+_8433172_8446622_RNA45SN1_45S_50ntFlanks@5494.5516.23|AAGACGGTCGAACTTGACTATCT</td><td>>hg38_21_+_8433172_8446622_RNA45SN1_45S_50ntFlanks</td><td>5494</td><td>5516</td><td>23</td><td>AAGACGGTCGAACTTGACTATCT</td><td>1967.8</td><td>4.0</td><td>1.4</td><td>2.9</td><td>4.0E-03</td><td>1.8E-02</td><td>BT20/MB231</td><td>MP</td></tr>
<tr><td>>MI0000255|hsa-mir-30d&WithFlank&8|-|134804870|134804951@12.32.21&[MIMAT0000245&hsa-miR-30d-5p&offsets|0|-1;m-21&8|-|134804919|134804940&offsets|0|-1]|TGTAAACATCCCCGACTGGAA</td><td>>MI0000255|hsa-mir-30d&WithFlank&8|-|134804870|134804951</td><td>12</td><td>32</td><td>21</td><td>TGTAAACATCCCCGACTGGAA</td><td>1136.7</td><td>1.8</td><td>0.6</td><td>2.8</td><td>4.4E-03</td><td>1.8E-02</td><td>BT20/MB231</td><td>NUC</td></tr>
<tr><td>>hg38_21_+_8433172_8446622_RNA45SN1_45S_50ntFlanks@11714.11741.28|GGTAAACGGCGGGAGTAACTATGACTCT</td><td>>hg38_21_+_8433172_8446622_RNA45SN1_45S_50ntFlanks</td><td>11714</td><td>11741</td><td>28</td><td>GGTAAACGGCGGGAGTAACTATGACTCT</td><td>277.0</td><td>-2.6</td><td>0.9</td><td>-2.8</td><td>5.2E-03</td><td>1.8E-02</td><td>BT20/MB231</td><td>CYTO</td></tr>
<tr><td>>hg38_21_+_8433172_8446622_RNA45SN1_45S_50ntFlanks@9918.9948.31|AGGTGTTGGTTGATATAGACAGCAGGACGGT</td><td>>hg38_21_+_8433172_8446622_RNA45SN1_45S_50ntFlanks</td><td>9918</td><td>9948</td><td>31</td><td>AGGTGTTGGTTGATATAGACAGCAGGACGGT</td><td>64.8</td><td>5.2</td><td>2.0</td><td>2.6</td><td>1.0E-02</td><td>1.8E-02</td><td>BT20/MB468</td><td>MP</td></tr>
<tr><td>>hg38_21_+_8433172_8446622_RNA45SN1_45S_50ntFlanks@5192.5209.18|CAGGTCTGTGATGCCCTT</td><td>>hg38_21_+_8433172_8446622_RNA45SN1_45S_50ntFlanks</td><td>5192</td><td>5209</td><td>18</td><td>CAGGTCTGTGATGCCCTT</td><td>73.6</td><td>3.5</td><td>1.3</td><td>2.6</td><td>1.0E-02</td><td>1.8E-02</td><td>BT20/MB468</td><td>MP</td></tr>
<tr><td>>hg38_21_+_8433172_8446622_RNA45SN1_45S_50ntFlanks@10348.10373.26|AAACGAGAACTTTGAAGGCCGAAGTG</td><td>>hg38_21_+_8433172_8446622_RNA45SN1_45S_50ntFlanks</td><td>10348</td><td>10373</td><td>26</td><td>AAACGAGAACTTTGAAGGCCGAAGTG</td><td>556.6</td><td>-3.5</td><td>1.4</td><td>-2.4</td><td>1.5E-02</td><td>1.8E-02</td><td>MB231/MB468</td><td>MP</td></tr>
<tr><td>>hg38_21_+_8433172_8446622_RNA45SN1_45S_50ntFlanks@7992.8020.29|CGTGGCGACCCGCTGAATTTAAGCATATT</td><td>>hg38_21_+_8433172_8446622_RNA45SN1_45S_50ntFlanks</td><td>7992</td><td>8020</td><td>29</td><td>CGTGGCGACCCGCTGAATTTAAGCATATT</td><td>156.3</td><td>-3.2</td><td>1.2</td><td>-2.8</td><td>5.2E-03</td><td>1.8E-02</td><td>BT20/MB231</td><td>CYTO</td></tr>
<tr><td>>hg38_MT_+_1621_3279_RNR2_16S_50ntFlanks@625.663.39|AACTGAACTCCTCACACCCAATTGGACCAATCTATCACC</td><td>>hg38_MT_+_1621_3279_RNR2_16S_50ntFlanks</td><td>625</td><td>663</td><td>39</td><td>AACTGAACTCCTCACACCCAATTGGACCAATCTATCACC</td><td>43.8</td><td>4.8</td><td>2.0</td><td>2.4</td><td>1.5E-02</td><td>1.8E-02</td><td>MB231/MB468</td><td>MP</td></tr>
<tr><td>>hg38_21_+_8433172_8446622_RNA45SN1_45S_50ntFlanks@6656.6689.34|CTTAGCGGTGGATCACTCGGCTCGTGCGTCGATG</td><td>>hg38_21_+_8433172_8446622_RNA45SN1_45S_50ntFlanks</td><td>6656</td><td>6689</td><td>34</td><td>CTTAGCGGTGGATCACTCGGCTCGTGCGTCGATG</td><td>688.8</td><td>-2.5</td><td>1.0</td><td>-2.6</td><td>1.0E-02</td><td>1.8E-02</td><td>BT20/MB468</td><td>MP</td></tr>
<tr><td>>MI0000072|hsa-mir-18a&WithFlank&13|+|91350745|91350827@53.75.23&[MIMAT0002891&hsa-miR-18a-3p&offsets|0|0;m-389&13|+|91350797|91350819&offsets|0|0]|ACTGCCCTAAGTGCTCCTTCTGG</td><td>>MI0000072|hsa-mir-18a&WithFlank&13|+|91350745|91350827</td><td>53</td><td>75</td><td>23</td><td>ACTGCCCTAAGTGCTCCTTCTGG</td><td>89.4</td><td>3.3</td><td>1.4</td><td>2.4</td><td>1.5E-02</td><td>1.8E-02</td><td>MB231/MB468</td><td>MP</td></tr>
<tr><td>>hg38_21_+_8433172_8446622_RNA45SN1_45S_50ntFlanks@6650.6695.46|ACGACTCTTAGCGGTGGATCACTCGGCTCGTGCGTCGATGAAGAAC</td><td>>hg38_21_+_8433172_8446622_RNA45SN1_45S_50ntFlanks</td><td>6650</td><td>6695</td><td>46</td><td>ACGACTCTTAGCGGTGGATCACTCGGCTCGTGCGTCGATGAAGAAC</td><td>278.7</td><td>4.6</td><td>1.8</td><td>2.6</td><td>1.0E-02</td><td>1.8E-02</td><td>BT20/MB468</td><td>MP</td></tr>
<tr><td>>hg38_21_+_8433172_8446622_RNA45SN1_45S_50ntFlanks@9564.9597.34|CTGACGTGCAAATCGGTCGTCCGACCTGGGTATA</td><td>>hg38_21_+_8433172_8446622_RNA45SN1_45S_50ntFlanks</td><td>9564</td><td>9597</td><td>34</td><td>CTGACGTGCAAATCGGTCGTCCGACCTGGGTATA</td><td>2389.7</td><td>-5.0</td><td>2.0</td><td>-2.6</td><td>1.0E-02</td><td>1.8E-02</td><td>BT20/MB468</td><td>MP</td></tr>
<tr><td>>hg38_21_+_8433172_8446622_RNA45SN1_45S_50ntFlanks@4588.4612.25|CTATTTTGTTGGTTTTCGGAACTGA</td><td>>hg38_21_+_8433172_8446622_RNA45SN1_45S_50ntFlanks</td><td>4588</td><td>4612</td><td>25</td><td>CTATTTTGTTGGTTTTCGGAACTGA</td><td>116.1</td><td>-4.3</td><td>1.7</td><td>-2.6</td><td>1.0E-02</td><td>1.8E-02</td><td>BT20/MB468</td><td>MP</td></tr>
<tr><td>>MI0000804|hsa-mir-328&WithFlank&16|-|67202315|67202401@54.75.22(+1U)&[MIMAT0000752&hsa-miR-328-3p&offsets|0|0(+1U);m-160&16|-|67202327|67202348&offsets|0|0(+1U)]|CTGGCCCTCTCTGCCCTTCCGTT</td><td>>MI0000804|hsa-mir-328&WithFlank&16|-|67202315|67202401</td><td>54</td><td>75</td><td>22(+1U)</td><td>CTGGCCCTCTCTGCCCTTCCGTT</td><td>195.4</td><td>2.4</td><td>0.9</td><td>2.8</td><td>5.2E-03</td><td>1.9E-02</td><td>BT20/MB231</td><td>CYTO</td></tr>
<tr><td>>MI0003666|hsa-mir-651&WithFlank&X|+|8126959|8127067@22.43.22&[MIMAT0003321&hsa-miR-651-5p&offsets|0|0;m-399&X|+|8126980|8127001&offsets|0|0]|TTTAGGATAAGCTTGACTTTTG</td><td>>MI0003666|hsa-mir-651&WithFlank&X|+|8126959|8127067</td><td>22</td><td>43</td><td>22</td><td>TTTAGGATAAGCTTGACTTTTG</td><td>237.5</td><td>2.5</td><td>0.9</td><td>2.8</td><td>5.2E-03</td><td>1.9E-02</td><td>BT20/MB231</td><td>CYTO</td></tr>
<tr><td>>MI0000734|hsa-mir-106b&WithFlank&7|-|100093987|100094080@18.38.21&[MIMAT0000680&hsa-miR-106b-5p&offsets|0|0;m-103&7|-|100094043|100094063&offsets|0|0]|TAAAGTGCTGACAGTGCAGAT</td><td>>MI0000734|hsa-mir-106b&WithFlank&7|-|100093987|100094080</td><td>18</td><td>38</td><td>21</td><td>TAAAGTGCTGACAGTGCAGAT</td><td>1294.0</td><td>-4.3</td><td>1.5</td><td>-2.9</td><td>4.0E-03</td><td>1.9E-02</td><td>BT20/MB231</td><td>MP</td></tr>
<tr><td>>hg38_21_+_8433172_8446622_RNA45SN1_45S_50ntFlanks@10237.10261.25|ACGCTACGCCGCGACGAGTAGGAGG</td><td>>hg38_21_+_8433172_8446622_RNA45SN1_45S_50ntFlanks</td><td>10237</td><td>10261</td><td>25</td><td>ACGCTACGCCGCGACGAGTAGGAGG</td><td>215.9</td><td>2.6</td><td>1.0</td><td>2.6</td><td>1.0E-02</td><td>1.9E-02</td><td>BT20/MB468</td><td>MP</td></tr>
<tr><td>>hg38_21_+_8433172_8446622_RNA45SN1_45S_50ntFlanks@12941.12961.21|ATTCGTAGACGACCTGCTTCT</td><td>>hg38_21_+_8433172_8446622_RNA45SN1_45S_50ntFlanks</td><td>12941</td><td>12961</td><td>21</td><td>ATTCGTAGACGACCTGCTTCT</td><td>357.9</td><td>1.3</td><td>0.5</td><td>2.4</td><td>1.7E-02</td><td>1.9E-02</td><td>MB231/MB468</td><td>NUC</td></tr>
<tr><td>>hg38_21_+_8433172_8446622_RNA45SN1_45S_50ntFlanks@9158.9187.30|AGTGCGCCCCGGGCGGGTCGCGCCGTCGGG</td><td>>hg38_21_+_8433172_8446622_RNA45SN1_45S_50ntFlanks</td><td>9158</td><td>9187</td><td>30</td><td>AGTGCGCCCCGGGCGGGTCGCGCCGTCGGG</td><td>337.5</td><td>-4.6</td><td>1.9</td><td>-2.4</td><td>1.6E-02</td><td>1.9E-02</td><td>BT20/MB468</td><td>NUC</td></tr>
<tr><td>>MI0000440|hsa-mir-27b&WithFlank&9|+|95085439|95085547@67.86.20(+1U)&[MIMAT0000419&hsa-miR-27b-3p&offsets|0|-1(+1U);m-39&9|+|95085505|95085525&offsets|0|-1(+1U)]|TTCACAGTGGCTAAGTTCTGT</td><td>>MI0000440|hsa-mir-27b&WithFlank&9|+|95085439|95085547</td><td>67</td><td>86</td><td>20(+1U)</td><td>TTCACAGTGGCTAAGTTCTGT</td><td>13472.5</td><td>2.5</td><td>0.9</td><td>2.8</td><td>5.2E-03</td><td>1.9E-02</td><td>BT20/MB231</td><td>CYTO</td></tr>
<tr><td>>hg38_21_+_8433172_8446622_RNA45SN1_45S_50ntFlanks@6648.6673.26|GTACGACTCTTAGCGGTGGATCACTC</td><td>>hg38_21_+_8433172_8446622_RNA45SN1_45S_50ntFlanks</td><td>6648</td><td>6673</td><td>26</td><td>GTACGACTCTTAGCGGTGGATCACTC</td><td>88.6</td><td>1.5</td><td>0.6</td><td>2.4</td><td>1.6E-02</td><td>1.9E-02</td><td>BT20/MB468</td><td>NUC</td></tr>
<tr><td>>trnaMT_GluTTC_MT_-_14674_14742@40.65.26,trnalookalike8_GluTTC_5_-_93905172_93905240@40.65.26|TTGGTCGTGGTTGTAGTCCGTGCGAG</td><td>>trnaMT_GluTTC_MT_-_14674_14742</td><td>40</td><td>65</td><td>26</td><td>TTGGTCGTGGTTGTAGTCCGTGCGAG</td><td>146.9</td><td>2.0</td><td>0.8</td><td>2.4</td><td>1.5E-02</td><td>1.9E-02</td><td>MB231/MB468</td><td>MP</td></tr>
<tr><td>>hg38_21_+_8433172_8446622_RNA45SN1_45S_50ntFlanks@10234.10273.40|CGGACGCTACGCCGCGACGAGTAGGAGGGCCGCTGCGGTG</td><td>>hg38_21_+_8433172_8446622_RNA45SN1_45S_50ntFlanks</td><td>10234</td><td>10273</td><td>40</td><td>CGGACGCTACGCCGCGACGAGTAGGAGGGCCGCTGCGGTG</td><td>748.0</td><td>-4.4</td><td>1.8</td><td>-2.4</td><td>1.5E-02</td><td>1.9E-02</td><td>MB231/MB468</td><td>MP</td></tr>
<tr><td>>hg38_21_+_8433172_8446622_RNA45SN1_45S_50ntFlanks@476.524.49|TTCGTGATCGATGTGGTGACGTCGTGCTCTCCCGGGCCGGGTCCGAGCC</td><td>>hg38_21_+_8433172_8446622_RNA45SN1_45S_50ntFlanks</td><td>476</td><td>524</td><td>49</td><td>TTCGTGATCGATGTGGTGACGTCGTGCTCTCCCGGGCCGGGTCCGAGCC</td><td>7631.2</td><td>-4.1</td><td>1.4</td><td>-2.8</td><td>4.4E-03</td><td>1.9E-02</td><td>BT20/MB231</td><td>NUC</td></tr>
<tr><td>>hg38_21_+_8433172_8446622_RNA45SN1_45S_50ntFlanks@5443.5466.24|CCTCGGATCGGCCCCGCCGGGGTC</td><td>>hg38_21_+_8433172_8446622_RNA45SN1_45S_50ntFlanks</td><td>5443</td><td>5466</td><td>24</td><td>CCTCGGATCGGCCCCGCCGGGGTC</td><td>2194.6</td><td>-5.2</td><td>2.2</td><td>-2.4</td><td>1.6E-02</td><td>1.9E-02</td><td>BT20/MB468</td><td>NUC</td></tr>
<tr><td>>hg38_21_+_8433172_8446622_RNA45SN1_45S_50ntFlanks@5472.5509.38|ACGGCCCTGGCGGAGCGCTGAGAAGACGGTCGAACTTG</td><td>>hg38_21_+_8433172_8446622_RNA45SN1_45S_50ntFlanks</td><td>5472</td><td>5509</td><td>38</td><td>ACGGCCCTGGCGGAGCGCTGAGAAGACGGTCGAACTTG</td><td>383.4</td><td>2.3</td><td>1.0</td><td>2.4</td><td>1.7E-02</td><td>1.9E-02</td><td>MB231/MB468</td><td>MITO</td></tr>
<tr><td>>hg38_21_+_8433172_8446622_RNA45SN1_45S_50ntFlanks@12907.12930.24|CACCGCACGTTCGTGGGGAACCTG</td><td>>hg38_21_+_8433172_8446622_RNA45SN1_45S_50ntFlanks</td><td>12907</td><td>12930</td><td>24</td><td>CACCGCACGTTCGTGGGGAACCTG</td><td>106.3</td><td>-2.9</td><td>1.2</td><td>-2.4</td><td>1.6E-02</td><td>1.9E-02</td><td>MB231/MB468</td><td>CYTO</td></tr>
<tr><td>>hg38_21_+_8433172_8446622_RNA45SN1_45S_50ntFlanks@476.497.22|TTCGTGATCGATGTGGTGACGT</td><td>>hg38_21_+_8433172_8446622_RNA45SN1_45S_50ntFlanks</td><td>476</td><td>497</td><td>22</td><td>TTCGTGATCGATGTGGTGACGT</td><td>215.5</td><td>4.0</td><td>1.6</td><td>2.4</td><td>1.5E-02</td><td>1.9E-02</td><td>MB231/MB468</td><td>MP</td></tr>
<tr><td>>hg38_21_+_8433172_8446622_RNA45SN1_45S_50ntFlanks@6702.6726.25|AGCTGCGAGAATTAATGTGAATTGC</td><td>>hg38_21_+_8433172_8446622_RNA45SN1_45S_50ntFlanks</td><td>6702</td><td>6726</td><td>25</td><td>AGCTGCGAGAATTAATGTGAATTGC</td><td>52.8</td><td>4.1</td><td>1.7</td><td>2.4</td><td>1.5E-02</td><td>1.9E-02</td><td>MB231/MB468</td><td>MP</td></tr>
<tr><td>>hg38_MT_+_598_1651_RNR1_12S_50ntFlanks@638.665.28|TCACCACCTCTTGCTCAGCCTATATACC</td><td>>hg38_MT_+_598_1651_RNR1_12S_50ntFlanks</td><td>638</td><td>665</td><td>28</td><td>TCACCACCTCTTGCTCAGCCTATATACC</td><td>71.6</td><td>4.7</td><td>1.9</td><td>2.4</td><td>1.5E-02</td><td>1.9E-02</td><td>MB231/MB468</td><td>MP</td></tr>
<tr><td>>MI0000542|hsa-mir-320a&WithFlank&8|-|22244960|22245043@47.69.23&[MIMAT0000510&hsa-miR-320a-3p&offsets|-1|0;m-60&8|-|22244975|22244996&offsets|-1|0]|GAAAAGCTGGGTTGAGAGGGCGA</td><td>>MI0000542|hsa-mir-320a&WithFlank&8|-|22244960|22245043</td><td>47</td><td>69</td><td>23</td><td>GAAAAGCTGGGTTGAGAGGGCGA</td><td>60.2</td><td>4.1</td><td>1.7</td><td>2.4</td><td>1.5E-02</td><td>1.9E-02</td><td>MB231/MB468</td><td>MP</td></tr>
<tr><td>>hg38_21_+_8433172_8446622_RNA45SN1_45S_50ntFlanks@7981.8004.24|CTCAGATCAGACGTGGCGACCCGC</td><td>>hg38_21_+_8433172_8446622_RNA45SN1_45S_50ntFlanks</td><td>7981</td><td>8004</td><td>24</td><td>CTCAGATCAGACGTGGCGACCCGC</td><td>228.8</td><td>-2.2</td><td>0.9</td><td>-2.4</td><td>1.7E-02</td><td>1.9E-02</td><td>MB231/MB468</td><td>NUC</td></tr>
<tr><td>>hg38_21_+_8433172_8446622_RNA45SN1_45S_50ntFlanks@11567.11602.36|AGCAGCCGACTTAGAACTGGTGCGGACCAGGGGAAT</td><td>>hg38_21_+_8433172_8446622_RNA45SN1_45S_50ntFlanks</td><td>11567</td><td>11602</td><td>36</td><td>AGCAGCCGACTTAGAACTGGTGCGGACCAGGGGAAT</td><td>209.4</td><td>-2.4</td><td>0.8</td><td>-2.8</td><td>5.2E-03</td><td>1.9E-02</td><td>BT20/MB231</td><td>CYTO</td></tr>
<tr><td>>hg38_1_-_228634819_228635039_RNA5S12_5S_50ntFlanks@132.171.40|GATGGGAGACCGCCTGGGAATACCGGGTGCTGTAGGCTTT</td><td>>hg38_1_-_228634819_228635039_RNA5S12_5S_50ntFlanks</td><td>132</td><td>171</td><td>40</td><td>GATGGGAGACCGCCTGGGAATACCGGGTGCTGTAGGCTTT</td><td>17142.0</td><td>2.8</td><td>1.2</td><td>2.4</td><td>1.6E-02</td><td>1.9E-02</td><td>BT20/MB468</td><td>CYTO</td></tr>
</table>

| | | | | | | | | | | | | | |
|---|---|---|---|---|---|---|---|---|---|---|---|---|---|
| >hg38_21_+_8433172_8446622_RNA45SN1_45S_50ntFlanks@10910.10928.19\|CGCGCGCCGCGGCTGGACG | >hg38_21_+_8433172_8446622_RNA45SN1_45S_50ntFlanks | 10910 | 10928 | 19 | CGCGCGCCGCGGCTGGACG | 261.3 | -2.1 | 0.9 | -2.4 | 1.6E-02 | 1.9E-02 | MB231/MB468 | CYTO |
| >hg38_21_+_8433172_8446622_RNA45SN1_45S_50ntFlanks@3854.3880.27\|ACTGTGGTAATTCTAGAGCTAAT ACAT | >hg38_21_+_8433172_8446622_RNA45SN1_45S_50ntFlanks | 3854 | 3880 | 27 | ACTGTGGTAATTCTAGAGCTAATACAT | 271.0 | -1.7 | 0.7 | -2.4 | 1.6E-02 | 1.9E-02 | MB231/MB468 | CYTO |
| >hg38_21_+_8433172_8446622_RNA45SN1_45S_50ntFlanks@12417.12446.30\|TCGATGTCGGCTCTTCCTATCA TTGTGAAG | >hg38_21_+_8433172_8446622_RNA45SN1_45S_50ntFlanks | 12417 | 12446 | 30 | TCGATGTCGGCTCTTCCTATCATTGTGAAG | 367.2 | -4.5 | 1.7 | -2.6 | 1.0E-02 | 1.9E-02 | BT20/MB468 | MP |
| >hg38_21_+_8433172_8446622_RNA45SN1_45S_50ntFlanks@10853.10879.27\|ATAAGGATTGGCTCTAAGGGC TGGGTC | >hg38_21_+_8433172_8446622_RNA45SN1_45S_50ntFlanks | 10853 | 10879 | 27 | ATAAGGATTGGCTCTAAGGGCTGGGTC | 144.3 | 5.6 | 2.0 | 2.9 | 4.1E-03 | 1.9E-02 | BT20/MB231 | MP |
| >hg38_21_+_8433172_8446622_RNA45SN1_45S_50ntFlanks@4666.4689.24\|AAATTCTTGGACCGGCGCAAGAC G | >hg38_21_+_8433172_8446622_RNA45SN1_45S_50ntFlanks | 4666 | 4689 | 24 | AAATTCTTGGACCGGCGCAAGACG | 68.3 | -4.4 | 1.7 | -2.6 | 1.0E-02 | 1.9E-02 | BT20/MB468 | MP |
| >trnaMT_GlnTTG_MT_-_4329_4400@1.19.19,trnalookalike2_GlnTTG_1_-_564879_564950@1.19.19\|TAGGATGGGGTGTGATAGG | >trnaMT_GlnTTG_MT_-_4329_4400 | 1 | 19 | 19 | TAGGATGGGGTGTGATAGG | 516.6 | 3.2 | 1.2 | 2.6 | 1.0E-02 | 1.9E-02 | BT20/MB468 | MP |
| >hg38_21_+_8433172_8446622_RNA45SN1_45S_50ntFlanks@8180.8210.31\|TGATCGAGGCCCAGCCCGTGGAC GGTGTGAG | >hg38_21_+_8433172_8446622_RNA45SN1_45S_50ntFlanks | 8180 | 8210 | 31 | TGATCGAGGCCCAGCCCGTGGACGGTGTGAG | 65.9 | -4.1 | 1.7 | -2.4 | 1.5E-02 | 1.9E-02 | MB231/MB468 | MP |
| >hg38_21_+_8433172_8446622_RNA45SN1_45S_50ntFlanks@12203.12240.38\|TAAGGCGAGCTCAGGGAGGA CAGAAACCTCCCGTGGAG | >hg38_21_+_8433172_8446622_RNA45SN1_45S_50ntFlanks | 12203 | 12240 | 38 | TAAGGCGAGCTCAGGGAGGACAGAAACCTCCCGTGGAG | 36.8 | -8.3 | 3.2 | -2.6 | 1.0E-02 | 1.9E-02 | BT20/MB468 | MP |
| >hg38_21_+_8433172_8446622_RNA45SN1_45S_50ntFlanks@8384.8402.19\|AGGGCGTGAAACCGTTAAG | >hg38_21_+_8433172_8446622_RNA45SN1_45S_50ntFlanks | 8384 | 8402 | 19 | AGGGCGTGAAACCGTTAAG | 129.7 | -4.0 | 1.6 | -2.5 | 1.4E-02 | 1.9E-02 | BT20/MB468 | MITO |
| >hg38_21_+_8433172_8446622_RNA45SN1_45S_50ntFlanks@6652.6695.44\|GACTCTTAGCGGTGGATCACTCG GCTCGTGCGTCGATGAAGAAC | >hg38_21_+_8433172_8446622_RNA45SN1_45S_50ntFlanks | 6652 | 6695 | 44 | GACTCTTAGCGGTGGATCACTCGGCTCGTGCGTCGATGAAGAAC | 613.1 | 2.1 | 0.9 | 2.4 | 1.6E-02 | 1.9E-02 | BT20/MB468 | NUC |
| >hg38_21_+_8433172_8446622_RNA45SN1_45S_50ntFlanks@6646.6672.27\|TCGTACGACTCTTAGCGGTGGAT CACT | >hg38_21_+_8433172_8446622_RNA45SN1_45S_50ntFlanks | 6646 | 6672 | 27 | TCGTACGACTCTTAGCGGTGGATCACT | 10333.7 | 1.8 | 0.6 | 2.8 | 4.5E-03 | 1.9E-02 | BT20/MB231 | NUC |
| >hg38_21_+_8433172_8446622_RNA45SN1_45S_50ntFlanks@9955.9980.26\|GGAAGTCGGAATCCGCTAAGGA GTGT | >hg38_21_+_8433172_8446622_RNA45SN1_45S_50ntFlanks | 9955 | 9980 | 26 | GGAAGTCGGAATCCGCTAAGGAGTGT | 163.2 | 5.6 | 2.0 | 2.9 | 4.1E-03 | 1.9E-02 | BT20/MB231 | MP |
| >hg38_21_+_8433172_8446622_RNA45SN1_45S_50ntFlanks@7975.8002.28\|CGCGACCTCAGATCAGACGTGGC GACCC | >hg38_21_+_8433172_8446622_RNA45SN1_45S_50ntFlanks | 7975 | 8002 | 28 | CGCGACCTCAGATCAGACGTGGCGACCC | 2089.7 | 1.9 | 0.8 | 2.4 | 1.7E-02 | 1.9E-02 | MB231/MB468 | MITO |
| >hg38_MT_+_598_1651_RNR1_12S_50ntFlanks@915.935.21\|AAAGGACATTTAACTAAAACC | >hg38_MT_+_598_1651_RNR1_12S_50ntFlanks | 915 | 935 | 21 | AAAGGACATTTAACTAAAACC | 189.4 | 2.6 | 1.1 | 2.4 | 1.5E-02 | 1.9E-02 | MB231/MB468 | MP |
| >trnaMT_PheGAA_MT_+_577_647@35.57.23\|GAAAATGTTTAGACGGGCTCACA | >trnaMT_PheGAA_MT_+_577_647 | 35 | 57 | 23 | GAAAATGTTTAGACGGGCTCACA | 640.5 | 2.6 | 0.9 | 2.8 | 5.3E-03 | 1.9E-02 | BT20/MB231 | CYTO |
| >hg38_21_+_8433172_8446622_RNA45SN1_45S_50ntFlanks@6648.6672.25\|GTACGACTCTTAGCGGTGGATCA CT | >hg38_21_+_8433172_8446622_RNA45SN1_45S_50ntFlanks | 6648 | 6672 | 25 | GTACGACTCTTAGCGGTGGATCACT | 126.8 | 3.7 | 1.4 | 2.6 | 1.0E-02 | 1.9E-02 | BT20/MB468 | MP |
| >MI0000298\|hsa-mir-221&WithFlank&X\|-\|45746151\|45746272@71.89.19(+1C)&[MIMAT0000278&hsa-miR-221-3p&offsets\|0\|-4(+1C);m-91&X\|-\|45746181\|45746202&offsets\|0\|-3(+1C)]\|AGCTACATTGTCTGCTGGGC | >MI0000298\|hsa-mir-221&WithFlank&X\|-\|45746151\|45746272 | 71 | 89 | 19(+1C) | AGCTACATTGTCTGCTGGGC | 84.3 | 4.8 | 1.9 | 2.6 | 1.0E-02 | 1.9E-02 | BT20/MB468 | MP |
| >hg38_21_+_8433172_8446622_RNA45SN1_45S_50ntFlanks@10869.10902.34\|AGGGCTGGGTCGGTCGGGCT GGGGCGCGAAGCGG | >hg38_21_+_8433172_8446622_RNA45SN1_45S_50ntFlanks | 10869 | 10902 | 34 | AGGGCTGGGTCGGTCGGGCTGGGGCGCGAAGCGG | 900.4 | -4.2 | 1.8 | -2.4 | 1.6E-02 | 1.9E-02 | BT20/MB468 | CYTO |
| >hg38_21_+_8433172_8446622_RNA45SN1_45S_50ntFlanks@11577.11600.24\|TTAGAACTGGTGCGGACCAG GGGA | >hg38_21_+_8433172_8446622_RNA45SN1_45S_50ntFlanks | 11577 | 11600 | 24 | TTAGAACTGGTGCGGACCAGGGGA | 181.0 | -3.1 | 1.3 | -2.4 | 1.6E-02 | 1.9E-02 | BT20/MB468 | CYTO |
| >hg38_21_+_8433172_8446622_RNA45SN1_45S_50ntFlanks@9545.9570.26\|GTGGAGGTCCGTAGCGGTCCTG ACGT | >hg38_21_+_8433172_8446622_RNA45SN1_45S_50ntFlanks | 9545 | 9570 | 26 | GTGGAGGTCCGTAGCGGTCCTGACGT | 643.7 | -2.3 | 0.9 | -2.6 | 1.0E-02 | 1.9E-02 | BT20/MB468 | MP |
| >hg38_MT_+_598_1651_RNR1_12S_50ntFlanks@416.456.41\|AAATAGACTACGAAAGTGGCTTTAACATATCTGA ACACACA | >hg38_MT_+_598_1651_RNR1_12S_50ntFlanks | 416 | 456 | 41 | AAATAGACTACGAAAGTGGCTTTAACATATCTGAACACACA | 153.6 | 4.6 | 1.9 | 2.4 | 1.7E-02 | 1.9E-02 | MB231/MB468 | MITO |
| >hg38_21_+_8433172_8446622_RNA45SN1_45S_50ntFlanks@12155.12183.29\|GGAGTTTGACTGGGGCGGTA CACCTGTCA | >hg38_21_+_8433172_8446622_RNA45SN1_45S_50ntFlanks | 12155 | 12183 | 29 | GGAGTTTGACTGGGGCGGTACACCTGTCA | 100.0 | -3.6 | 1.4 | -2.6 | 1.0E-02 | 1.9E-02 | BT20/MB468 | MP |
| >hg38_21_+_8433172_8446622_RNA45SN1_45S_50ntFlanks@5078.5104.27\|CGATAACGAACGAGACTCTGGCA TGCT | >hg38_21_+_8433172_8446622_RNA45SN1_45S_50ntFlanks | 5078 | 5104 | 27 | CGATAACGAACGAGACTCTGGCATGCT | 147.1 | -3.3 | 1.4 | -2.5 | 1.4E-02 | 1.9E-02 | BT20/MB468 | MITO |
| >hg38_21_+_8433172_8446622_RNA45SN1_45S_50ntFlanks@7990.8012.23\|GACGTGGCGACCCGCTGAATTTA | >hg38_21_+_8433172_8446622_RNA45SN1_45S_50ntFlanks | 7990 | 8012 | 23 | GACGTGGCGACCCGCTGAATTTA | 642.5 | 4.1 | 1.4 | 2.9 | 4.1E-03 | 1.9E-02 | BT20/MB231 | MP |
| >hg38_21_+_8433172_8446622_RNA45SN1_45S_50ntFlanks@7977.8010.34\|CGACCTCAGATCAGACGTGGCGA CCCGCTGAATT | >hg38_21_+_8433172_8446622_RNA45SN1_45S_50ntFlanks | 7977 | 8010 | 34 | CGACCTCAGATCAGACGTGGCGACCCGCTGAATT | 1458.3 | -2.6 | 0.9 | -2.8 | 5.3E-03 | 1.9E-02 | BT20/MB231 | CYTO |
| >hg38_21_+_8433172_8446622_RNA45SN1_45S_50ntFlanks@11561.11583.23\|GCGCCTAGCAGCCGACTTAGA AC | >hg38_21_+_8433172_8446622_RNA45SN1_45S_50ntFlanks | 11561 | 11583 | 23 | GCGCCTAGCAGCCGACTTAGAAC | 384.2 | -1.8 | 0.7 | -2.5 | 1.4E-02 | 1.9E-02 | BT20/MB468 | MITO |
| >hg38_21_+_8433172_8446622_RNA45SN1_45S_50ntFlanks@6678.6695.18\|CGTGCGTCGATGAAGAAC | >hg38_21_+_8433172_8446622_RNA45SN1_45S_50ntFlanks | 6678 | 6695 | 18 | CGTGCGTCGATGAAGAAC | 185.1 | 5.8 | 2.0 | 2.9 | 4.1E-03 | 1.9E-02 | BT20/MB231 | MP |
| >MI0000076\|hsa-mir-20a&WithFlank&13\|+\|91351059\|91351141@14.35.22(+1C)&[MIMAT0000075&hsa-miR-20a-5p&offsets\|0\|-1(+1C);m-96&13\|+\|91351072\|91351094&offsets\|0\|-1(+1C)]\|TAAAGTGCTTATAGTGCAGGTAC | >MI0000076\|hsa-mir-20a&WithFlank&13\|+\|91351059\|91351141 | 14 | 35 | 22(+1C) | TAAAGTGCTTATAGTGCAGGTAC | 146.5 | -4.2 | 1.5 | -2.9 | 4.1E-03 | 1.9E-02 | BT20/MB231 | MP |
| >hg38_21_+_8433172_8446622_RNA45SN1_45S_50ntFlanks@8116.8141.26\|GCGGGACATGTGGCGTACGGAA GACC | >hg38_21_+_8433172_8446622_RNA45SN1_45S_50ntFlanks | 8116 | 8141 | 26 | GCGGGACATGTGGCGTACGGAAGACC | 572.7 | 2.5 | 1.0 | 2.6 | 1.0E-02 | 1.9E-02 | BT20/MB468 | MP |
| >hg38_21_+_8433172_8446622_RNA45SN1_45S_50ntFlanks@8115.8132.18\|CGCGGGACATGTGGCGTA | >hg38_21_+_8433172_8446622_RNA45SN1_45S_50ntFlanks | 8115 | 8132 | 18 | CGCGGGACATGTGGCGTA | 124.9 | 5.6 | 2.0 | 2.9 | 4.1E-03 | 1.9E-02 | BT20/MB231 | MP |
| >hg38_MT_+_1621_3279_RNR2_16S_50ntFlanks@841.864.24\|AAAGTAAAAGGAACTCGGCAAATC | >hg38_MT_+_1621_3279_RNR2_16S_50ntFlanks | 841 | 864 | 24 | AAAGTAAAAGGAACTCGGCAAATC | 61.8 | 4.1 | 1.6 | 2.6 | 1.0E-02 | 1.9E-02 | BT20/MB468 | MP |
| >hg38_21_+_8433172_8446622_RNA45SN1_45S_50ntFlanks@12932.12972.41\|CGCTAAACCATTCGTAGACGA CCTGCTTCTGGGTCGGGGTT | >hg38_21_+_8433172_8446622_RNA45SN1_45S_50ntFlanks | 12932 | 12972 | 41 | CGCTAAACCATTCGTAGACGACCTGCTTCTGGGTCGGGGTT | 1005.3 | -3.5 | 1.5 | -2.4 | 1.6E-02 | 1.9E-02 | BT20/MB468 | CYTO |
| >hg38_1_-_228634819_228635039_RNA5S12_5S_50ntFlanks@51.80.30\|GTCTACGGCCATACCACCCTGAACGCGCCC | >hg38_1_-_228634819_228635039_RNA5S12_5S_50ntFlanks | 51 | 80 | 30 | GTCTACGGCCATACCACCCTGAACGCGCCC | 1748.9 | 3.2 | 1.1 | 2.9 | 4.1E-03 | 1.9E-02 | BT20/MB231 | MP |
| >hg38_21_+_8433172_8446622_RNA45SN1_45S_50ntFlanks@9737.9765.29\|CGAAACGATCTCAACCTATTCTCA AACTT | >hg38_21_+_8433172_8446622_RNA45SN1_45S_50ntFlanks | 9737 | 9765 | 29 | CGAAACGATCTCAACCTATTCTCAAACTT | 1889.7 | -2.4 | 1.0 | -2.4 | 1.7E-02 | 1.9E-02 | MB231/MB468 | MITO |
| >hg38_21_+_8433172_8446622_RNA45SN1_45S_50ntFlanks@6661.6706.46\|CGGTGGATCACTCGGCTCGTGCG TCGATGAAGAACGCAGCTAGCTG | >hg38_21_+_8433172_8446622_RNA45SN1_45S_50ntFlanks | 6661 | 6706 | 46 | CGGTGGATCACTCGGCTCGTGCGTCGATGAAGAACGCAGCTAGCTG | 2264.4 | 3.1 | 1.1 | 2.8 | 4.5E-03 | 1.9E-02 | BT20/MB231 | NUC |
| >hg38_21_+_8433172_8446622_RNA45SN1_45S_50ntFlanks@9554.9576.23\|CGTAGCGGTCCTGACGTGCAAAT | >hg38_21_+_8433172_8446622_RNA45SN1_45S_50ntFlanks | 9554 | 9576 | 23 | CGTAGCGGTCCTGACGTGCAAAT | 764.5 | 1.5 | 0.6 | 2.4 | 1.7E-02 | 1.9E-02 | MB231/MB468 | NUC |
| >hg38_21_+_8433172_8446622_RNA45SN1_45S_50ntFlanks@9549.9571.23\|AGGTCCGTAGCGGTCCTGACGT G | >hg38_21_+_8433172_8446622_RNA45SN1_45S_50ntFlanks | 9549 | 9571 | 23 | AGGTCCGTAGCGGTCCTGACGTG | 307.2 | -4.6 | 1.9 | -2.4 | 1.7E-02 | 1.9E-02 | BT20/MB468 | NUC |
| >hg38_21_+_8433172_8446622_RNA45SN1_45S_50ntFlanks@4742.4765.24\|TCGGAGGTTCGAAGACGATCAG AT | >hg38_21_+_8433172_8446622_RNA45SN1_45S_50ntFlanks | 4742 | 4765 | 24 | TCGGAGGTTCGAAGACGATCAGAT | 53.2 | 5.2 | 2.0 | 2.6 | 1.0E-02 | 1.9E-02 | BT20/MB468 | MP |
| >hg38_21_+_8433172_8446622_RNA45SN1_45S_50ntFlanks@9934.9957.24\|AGACAGCAGGACGGTGGCCATG GA | >hg38_21_+_8433172_8446622_RNA45SN1_45S_50ntFlanks | 9934 | 9957 | 24 | AGACAGCAGGACGGTGGCCATGGA | 72.1 | 5.2 | 2.0 | 2.6 | 1.0E-02 | 1.9E-02 | BT20/MB468 | MP |
| >hg38_21_+_8433172_8446622_RNA45SN1_45S_50ntFlanks@9027.9065.39\|CGTCTCCTCGTGGGGGGGCCGG GCCACCCCTCCCACGGC | >hg38_21_+_8433172_8446622_RNA45SN1_45S_50ntFlanks | 9027 | 9065 | 39 | CGTCTCCTCGTGGGGGGGCCGGGCCACCCCTCCCACGGC | 269.7 | -4.5 | 1.8 | -2.6 | 1.0E-02 | 1.9E-02 | BT20/MB468 | MP |
| >trna11_GluTTC_15_-_26327381_26327452@41.74.34,trna3_GluTTC_13_-_45492062_45492133@41.74.34\|AGGCGGCCCGGGTTCGACTCCCGGTGTGGGAACC | >trna11_GluTTC_15_-_26327381_26327452 | 41 | 74 | 34 | AGGCGGCCCGGGTTCGACTCCCGGTGTGGGAACC | 98.3 | 5.0 | 2.1 | 2.4 | 1.5E-02 | 1.9E-02 | MB231/MB468 | MP |
| >hg38_21_+_8433172_8446622_RNA45SN1_45S_50ntFlanks@10476.10512.37\|GGCCGATCGAAAGGGAGTCG GGTTCAGATCCCCGAAT | >hg38_21_+_8433172_8446622_RNA45SN1_45S_50ntFlanks | 10476 | 10512 | 37 | GGCCGATCGAAAGGGAGTCGGGTTCAGATCCCCGAAT | 226.9 | -2.4 | 1.0 | -2.4 | 1.6E-02 | 1.9E-02 | MB231/MB468 | CYTO |
| >hg38_21_+_8433172_8446622_RNA45SN1_45S_50ntFlanks@6651.6674.24\|CGACTCTTAGCGGTGGATCACTC G | >hg38_21_+_8433172_8446622_RNA45SN1_45S_50ntFlanks | 6651 | 6674 | 24 | CGACTCTTAGCGGTGGATCACTCG | 11409.8 | 2.5 | 1.0 | 2.4 | 1.7E-02 | 1.9E-02 | MB231/MB468 | MITO |
| >hg38_21_+_8433172_8446622_RNA45SN1_45S_50ntFlanks@7992.8010.19\|CGTGGCGACCCGCTGAATT | >hg38_21_+_8433172_8446622_RNA45SN1_45S_50ntFlanks | 7992 | 8010 | 19 | CGTGGCGACCCGCTGAATT | 196.6 | -2.3 | 1.0 | -2.4 | 1.5E-02 | 1.9E-02 | MB231/MB468 | MP |
| >hg38_21_+_8433172_8446622_RNA45SN1_45S_50ntFlanks@9790.9813.24\|CTGGCGTGGAGCCGGGCGTGGA AT | >hg38_21_+_8433172_8446622_RNA45SN1_45S_50ntFlanks | 9790 | 9813 | 24 | CTGGCGTGGAGCCGGGCGTGGAAT | 61.0 | 2.7 | 1.1 | 2.6 | 1.0E-02 | 1.9E-02 | BT20/MB468 | MP |
| >hg38_21_+_8433172_8446622_RNA45SN1_45S_50ntFlanks@9931.9948.18\|TATAGACAGCAGGACGGT | >hg38_21_+_8433172_8446622_RNA45SN1_45S_50ntFlanks | 9931 | 9948 | 18 | TATAGACAGCAGGACGGT | 358.1 | 4.0 | 1.5 | 2.6 | 1.0E-02 | 1.9E-02 | BT20/MB468 | MP |
| >trnaMT_ProTGG_MT_-_15956_16023@38.58.21\|CTAATGGTGGAGTTAAAGACT | >trnaMT_ProTGG_MT_-_15956_16023 | 38 | 58 | 21 | CTAATGGTGGAGTTAAAGACT | 105.8 | 3.0 | 1.3 | 2.4 | 1.5E-02 | 1.9E-02 | MB231/MB468 | MP |
| >hg38_21_+_8433172_8446622_RNA45SN1_45S_50ntFlanks@12838.12856.19\|GTGCGGAGTGCCCTTCGTC | >hg38_21_+_8433172_8446622_RNA45SN1_45S_50ntFlanks | 12838 | 12856 | 19 | GTGCGGAGTGCCCTTCGTC | 87.2 | -4.6 | 1.8 | -2.6 | 1.0E-02 | 1.9E-02 | BT20/MB468 | MP |
| >hg38_21_+_8433172_8446622_RNA45SN1_45S_50ntFlanks@10291.10317.27\|CGCGGGCCCGGGTGGAGCCG CCGCAGG | >hg38_21_+_8433172_8446622_RNA45SN1_45S_50ntFlanks | 10291 | 10317 | 27 | CGCGGGCCCGGGTGGAGCCGCCGCAGG | 189.0 | -4.8 | 1.9 | -2.6 | 1.0E-02 | 1.9E-02 | BT20/MB468 | MP |
| >hg38_21_+_8433172_8446622_RNA45SN1_45S_50ntFlanks@7989.8012.24\|AGACGTGGCGACCCGCTGAATTT A | >hg38_21_+_8433172_8446622_RNA45SN1_45S_50ntFlanks | 7989 | 8012 | 24 | AGACGTGGCGACCCGCTGAATTTA | 4418.3 | -2.0 | 0.8 | -2.5 | 1.4E-02 | 1.9E-02 | BT20/MB468 | MITO |
| >hg38_21_+_8433172_8446622_RNA45SN1_45S_50ntFlanks@4731.4765.35\|AAGAACGAAAGTCGGAGGTTCG AAGACGATCAGAT | >hg38_21_+_8433172_8446622_RNA45SN1_45S_50ntFlanks | 4731 | 4765 | 35 | AAGAACGAAAGTCGGAGGTTCGAAGACGATCAGAT | 223.1 | 1.9 | 0.8 | 2.4 | 1.5E-02 | 1.9E-02 | MB231/MB468 | MP |
| >hg38_21_+_8433172_8446622_RNA45SN1_45S_50ntFlanks@11873.11898.26\|AAGAAGACCCTGTTGAGCTTG ACTCT | >hg38_21_+_8433172_8446622_RNA45SN1_45S_50ntFlanks | 11873 | 11898 | 26 | AAGAAGACCCTGTTGAGCTTGACTCT | 801.4 | -2.6 | 1.0 | -2.6 | 1.0E-02 | 1.9E-02 | BT20/MB468 | MP |
| >trnaMT_PheGAA_MT_+_577_647@38.64.27\|AATGTTTAGACGGGCTCACATCACCCC | >trnaMT_PheGAA_MT_+_577_647 | 38 | 64 | 27 | AATGTTTAGACGGGCTCACATCACCCC | 29.1 | 4.8 | 2.0 | 2.4 | 1.5E-02 | 1.9E-02 | MB231/MB468 | MP |
| >MI0003628\|hsa-mir-615&WithFlank&12\|+\|54033944\|54034051@67.86.20(+1C)&[MIMAT0003283&hsa-miR-615-3p&offsets\|0\|-2(+1C);m-495&12\|+\|54034010\|54034030&offsets\|0\|-1(+1C)]\|TCCGAGCCTGGGTCTCCCTCC | >MI0003628\|hsa-mir-615&WithFlank&12\|+\|54033944\|54034051 | 67 | 86 | 20(+1C) | TCCGAGCCTGGGTCTCCCTCC | 111.6 | 1.5 | 0.6 | 2.4 | 1.7E-02 | 1.9E-02 | BT20/MB468 | NUC |

| | | | | | | | | | | | | | |
|---|---|---|---|---|---|---|---|---|---|---|---|---|---|
| >MI0000803\|hsa-mir-330&WithFlank&19\|-\|45638988\|45639093@63.85.23&[MIMAT0000751&hsa-miR-330-3p&offsets\|0\|0;m-347&19\|-\|45639009\|45639030&offsets\|-1\|0]\|GCAAAGCACACGGCCTGCAGAGA | >MI0000803\|hsa-mir-330&WithFlank&19\|-\|45638988\|45639093 | 63 | 85 | 23 | GCAAAGCACACGGCCTGCAGAGA | 172.0 | 3.8 | 1.5 | 2.6 | 1.0E-02 | 1.9E-02 | BT20/MB468 | MP |
| >hg38_21_+_8433172_8446622_RNA45SN1_45S_50ntFlanks@9299.9322.24\|CACGGACCAAGGAGTCTAACACGT | >hg38_21_+_8433172_8446622_RNA45SN1_45S_50ntFlanks | 9299 | 9322 | 24 | CACGGACCAAGGAGTCTAACACGT | 41.3 | 5.3 | 2.1 | 2.6 | 1.0E-02 | 1.9E-02 | BT20/MB468 | MP |
| >MI0000234\|hsa-mir-192&WithFlank&11\|-\|64891131\|64891252@31.53.23&[MIMAT0000222&hsa-miR-192-5p&offsets\|+1\|+3;m-59&11\|-\|64891203\|64891223&offsets\|+1\|+3]\|TGACCTATGAATTGACAGCCAGT | >MI0000234\|hsa-mir-192&WithFlank&11\|-\|64891131\|64891252 | 31 | 53 | 23 | TGACCTATGAATTGACAGCCAGT | 522.2 | -2.2 | 0.8 | -2.8 | 5.1E-03 | 1.9E-02 | BT20/MB231 | MITO |
| >hg38_21_+_8433172_8446622_RNA45SN1_45S_50ntFlanks@10798.10828.31\|CATGTTGGAACAATGTAGGTAAGGGAAGTCG | >hg38_21_+_8433172_8446622_RNA45SN1_45S_50ntFlanks | 10798 | 10828 | 31 | CATGTTGGAACAATGTAGGTAAGGGAAGTCG | 119.1 | -3.2 | 1.2 | -2.6 | 1.0E-02 | 1.9E-02 | BT20/MB468 | MP |
| >hg38_MT_+_1621_3279_RNR2_16S_50ntFlanks@1431.1453.23\|AAAGTCCTACGTGATCTGAGTTC | >hg38_MT_+_1621_3279_RNR2_16S_50ntFlanks | 1431 | 1453 | 23 | AAAGTCCTACGTGATCTGAGTTC | 220.8 | 3.1 | 1.2 | 2.6 | 1.0E-02 | 1.9E-02 | BT20/MB468 | MP |
| >hg38_MT_+_1621_3279_RNR2_16S_50ntFlanks@929.959.31\|CAGTGACACATGTTTAACGGCCGCGGTACCC | >hg38_MT_+_1621_3279_RNR2_16S_50ntFlanks | 929 | 959 | 31 | CAGTGACACATGTTTAACGGCCGCGGTACCC | 112.2 | 4.0 | 1.6 | 2.4 | 1.5E-02 | 1.9E-02 | MB231/MB468 | MP |
| >trnaMT_ProTGG_MT_-_15956_16023@29.70.42\|CTTTGGGTGCTAATGGTGGAGTTAAAGACTTTTTCTCTGACC | >trnaMT_ProTGG_MT_-_15956_16023 | 29 | 70 | 42 | CTTTGGGTGCTAATGGTGGAGTTAAAGACTTTTTCTCTGACC | 70.4 | 5.5 | 2.2 | 2.6 | 1.0E-02 | 1.9E-02 | BT20/MB468 | MP |
| >hg38_21_+_8433172_8446622_RNA45SN1_45S_50ntFlanks@11589.11611.23\|CGGACCAGGGGAATCCGACTGTT | >hg38_21_+_8433172_8446622_RNA45SN1_45S_50ntFlanks | 11589 | 11611 | 23 | CGGACCAGGGGAATCCGACTGTT | 929.8 | -2.2 | 0.9 | -2.6 | 1.0E-02 | 1.9E-02 | BT20/MB468 | MP |
| >hg38_21_+_8433172_8446622_RNA45SN1_45S_50ntFlanks@8241.8261.21\|GGGTCTTCCCGGAGTCGGGTT | >hg38_21_+_8433172_8446622_RNA45SN1_45S_50ntFlanks | 8241 | 8261 | 21 | GGGTCTTCCCGGAGTCGGGTT | 86.5 | 4.5 | 1.7 | 2.6 | 1.0E-02 | 1.9E-02 | BT20/MB468 | MP |
| >hg38_21_+_8433172_8446622_RNA45SN1_45S_50ntFlanks@9494.9525.32\|CCGAAAGATGGTGAACTATGCCTGGGCAGGGC | >hg38_21_+_8433172_8446622_RNA45SN1_45S_50ntFlanks | 9494 | 9525 | 32 | CCGAAAGATGGTGAACTATGCCTGGGCAGGGC | 130.3 | 3.8 | 1.5 | 2.6 | 1.0E-02 | 1.9E-02 | BT20/MB468 | MP |
| >MI0016887\|hsa-mir-4521&WithFlank&17\|+\|8186939\|8187010@10.34.25&[MIMAT0019058&hsa-miR-4521&offsets\|0\|+3;m-765&17\|+\|8186948\|8186969&offsets\|0\|+3]\|GCTAAGGAAGTCCTGTGCTCAGTTT | >MI0016887\|hsa-mir-4521&WithFlank&17\|+\|8186939\|8187010 | 10 | 34 | 25 | GCTAAGGAAGTCCTGTGCTCAGTTT | 48.9 | 4.6 | 1.9 | 2.4 | 1.5E-02 | 1.9E-02 | MB231/MB468 | MP |
| >hg38_21_+_8433172_8446622_RNA45SN1_45S_50ntFlanks@12415.12435.21\|CTTCGATGTCGGCTCTTCCTA | >hg38_21_+_8433172_8446622_RNA45SN1_45S_50ntFlanks | 12415 | 12435 | 21 | CTTCGATGTCGGCTCTTCCTA | 483.7 | -4.5 | 1.8 | -2.6 | 1.0E-02 | 1.9E-02 | BT20/MB468 | MP |
| >hg38_21_+_8433172_8446622_RNA45SN1_45S_50ntFlanks@11208.11230.23\|GCGGCGACTCTGGACGCGAGCCG | >hg38_21_+_8433172_8446622_RNA45SN1_45S_50ntFlanks | 11208 | 11230 | 23 | GCGGCGACTCTGGACGCGAGCCG | 1036.9 | -3.8 | 1.5 | -2.6 | 1.0E-02 | 1.9E-02 | BT20/MB468 | MP |
| >MI0000088\|hsa-mir-30a&WithFlank&6\|-\|71403545\|71403627@12.30.19(+1G)&[MIMAT0000087&hsa-miR-30a-5p&offsets\|0\|-3(+1G);m-15&6\|-\|71403595\|71403616&offsets\|0\|-3(+1G)]\|TGTAAACATCCTCGACTGGG | >MI0000088\|hsa-mir-30a&WithFlank&6\|-\|71403545\|71403627 | 12 | 30 | 19(+1G) | TGTAAACATCCTCGACTGGG | 180.6 | 4.0 | 1.7 | 2.4 | 1.5E-02 | 1.9E-02 | MB231/MB468 | MP |
| >hg38_21_+_8433172_8446622_RNA45SN1_45S_50ntFlanks@6651.6675.25\|CGACTCTTAGCGGTGGATCACTCGG | >hg38_21_+_8433172_8446622_RNA45SN1_45S_50ntFlanks | 6651 | 6675 | 25 | CGACTCTTAGCGGTGGATCACTCGG | 16451.4 | 2.0 | 0.8 | 2.5 | 1.4E-02 | 1.9E-02 | BT20/MB468 | MITO |
| >hg38_21_+_8433172_8446622_RNA45SN1_45S_50ntFlanks@11671.11700.30\|CCAGTGCTCTGAATGTCAAAGTGAAGAAAT | >hg38_21_+_8433172_8446622_RNA45SN1_45S_50ntFlanks | 11671 | 11700 | 30 | CCAGTGCTCTGAATGTCAAAGTGAAGAAAT | 207.9 | -2.3 | 1.0 | -2.4 | 1.6E-02 | 1.9E-02 | MB231/MB468 | CYTO |
| >hg38_21_+_8433172_8446622_RNA45SN1_45S_50ntFlanks@11573.11613.41\|CGACTTAGAACTGGTGCGGACCAGGGGAATCCGACTGTTTA | >hg38_21_+_8433172_8446622_RNA45SN1_45S_50ntFlanks | 11573 | 11613 | 41 | CGACTTAGAACTGGTGCGGACCAGGGGAATCCGACTGTTTA | 1648.4 | 2.6 | 0.9 | 2.8 | 5.2E-03 | 1.9E-02 | BT20/MB231 | MITO |
| >hg38_21_+_8433172_8446622_RNA45SN1_45S_50ntFlanks@3705.3742.38\|TACCTGGTTGATCCTGCCAGTAGCATATGCTTGTCTCA | >hg38_21_+_8433172_8446622_RNA45SN1_45S_50ntFlanks | 3705 | 3742 | 38 | TACCTGGTTGATCCTGCCAGTAGCATATGCTTGTCTCA | 7062.2 | 2.3 | 0.8 | 2.8 | 5.4E-03 | 1.9E-02 | BT20/MB231 | CYTO |
| >hg38_21_+_8433172_8446622_RNA45SN1_45S_50ntFlanks@12451.12476.26\|ATTCACCAAGCGTTGGATTGTTCACC | >hg38_21_+_8433172_8446622_RNA45SN1_45S_50ntFlanks | 12451 | 12476 | 26 | ATTCACCAAGCGTTGGATTGTTCACC | 409.6 | -2.2 | 0.8 | -2.8 | 5.3E-03 | 1.9E-02 | BT20/MB231 | CYTO |
| >hg38_21_+_8433172_8446622_RNA45SN1_45S_50ntFlanks@11666.11702.37\|TCTGCCCAGTGCTCTGAATGTCAAAGTGAAGAAATTC | >hg38_21_+_8433172_8446622_RNA45SN1_45S_50ntFlanks | 11666 | 11702 | 37 | TCTGCCCAGTGCTCTGAATGTCAAAGTGAAGAAATTC | 396.3 | -2.7 | 1.0 | -2.8 | 4.5E-03 | 1.9E-02 | BT20/MB231 | NUC |
| >hg38_21_+_8433172_8446622_RNA45SN1_45S_50ntFlanks@4732.4758.27\|AGAACGAAAGTCGGAGGTTCGAAGACG | >hg38_21_+_8433172_8446622_RNA45SN1_45S_50ntFlanks | 4732 | 4758 | 27 | AGAACGAAAGTCGGAGGTTCGAAGACG | 2208.7 | 1.8 | 0.8 | 2.4 | 1.6E-02 | 1.9E-02 | BT20/MB468 | CYTO |
| >hg38_21_+_8433172_8446622_RNA45SN1_45S_50ntFlanks@11576.11609.34\|CTTAGAACTGGTGCGGACCAGGGGAATCCGACTG | >hg38_21_+_8433172_8446622_RNA45SN1_45S_50ntFlanks | 11576 | 11609 | 34 | CTTAGAACTGGTGCGGACCAGGGGAATCCGACTG | 168.6 | 5.0 | 1.7 | 2.9 | 4.1E-03 | 1.9E-02 | BT20/MB231 | MP |
| >MI0000107\|hsa-mir-29b-2&WithFlank&1\|-\|207802437\|207802529@58.77.20;MI0000105\|hsa-mir-29b-1&WithFlank&7\|-\|130877453\|130877545@57.76.20&[MIMAT0000100&hsa-miR-29b-3p&offsets\|0\|-3;m-81&1\|-\|207802450\|207802472&offsets\|0\|-3];[MIMAT0000100_1&hsa-miR-29b-3p&offsets\|0\|-3;m-82&7\|-\|130877467\|130877489&offsets\|0\|-3]\|TAGCACCATTTGAAATCAGT | >MI0000107\|hsa-mir-29b-2&WithFlank&1\|-\|207802437\|207802529 | 58 | 77 | 20 | TAGCACCATTTGAAATCAGT | 220.3 | -3.0 | 1.1 | -2.9 | 4.1E-03 | 1.9E-02 | BT20/MB231 | MP |
| >hg38_21_+_8433172_8446622_RNA45SN1_45S_50ntFlanks@9547.9572.26\|GGAGGTCCGTAGCGGTCCTGACGTGC | >hg38_21_+_8433172_8446622_RNA45SN1_45S_50ntFlanks | 9547 | 9572 | 26 | GGAGGTCCGTAGCGGTCCTGACGTGC | 84.3 | 4.5 | 1.7 | 2.6 | 1.0E-02 | 1.9E-02 | BT20/MB468 | MP |
| >hg38_21_+_8433172_8446622_RNA45SN1_45S_50ntFlanks@7994.8012.19\|TGGCGACCCGCTGAATTTA | >hg38_21_+_8433172_8446622_RNA45SN1_45S_50ntFlanks | 7994 | 8012 | 19 | TGGCGACCCGCTGAATTTA | 52.0 | -4.3 | 1.7 | -2.6 | 1.0E-02 | 1.9E-02 | BT20/MB468 | MP |
| >hg38_21_+_8433172_8446622_RNA45SN1_45S_50ntFlanks@9862.9880.19\|ATGAACCGAACGCCGGGTT | >hg38_21_+_8433172_8446622_RNA45SN1_45S_50ntFlanks | 9862 | 9880 | 19 | ATGAACCGAACGCCGGGTT | 327.3 | -4.4 | 1.7 | -2.6 | 1.0E-02 | 1.9E-02 | BT20/MB468 | MP |
| >hg38_21_+_8433172_8446622_RNA45SN1_45S_50ntFlanks@10905.10936.32\|CTGGGCGCGCGCCGCGGCTGGACGAGGCGCCG | >hg38_21_+_8433172_8446622_RNA45SN1_45S_50ntFlanks | 10905 | 10936 | 32 | CTGGGCGCGCGCCGCGGCTGGACGAGGCGCCG | 218.4 | -3.8 | 1.5 | -2.5 | 1.4E-02 | 1.9E-02 | BT20/MB468 | MITO |
| >hg38_21_+_8433172_8446622_RNA45SN1_45S_50ntFlanks@12416.12446.31\|TTCGATGTCGGCTCTTCCTATCATTGTGAAG | >hg38_21_+_8433172_8446622_RNA45SN1_45S_50ntFlanks | 12416 | 12446 | 31 | TTCGATGTCGGCTCTTCCTATCATTGTGAAG | 3134.2 | -2.7 | 1.0 | -2.8 | 5.4E-03 | 1.9E-02 | BT20/MB231 | CYTO |
| >hg38_MT_+_1621_3279_RNR2_16S_50ntFlanks@617.642.26\|AAACATATAACTGAACTCCTCACACC | >hg38_MT_+_1621_3279_RNR2_16S_50ntFlanks | 617 | 642 | 26 | AAACATATAACTGAACTCCTCACACC | 151.0 | 3.4 | 1.3 | 2.6 | 1.0E-02 | 1.9E-02 | BT20/MB468 | MP |
| >hg38_21_+_8433172_8446622_RNA45SN1_45S_50ntFlanks@9914.9938.25\|GAAAAGGTGTTGGTTGATATAGACA | >hg38_21_+_8433172_8446622_RNA45SN1_45S_50ntFlanks | 9914 | 9938 | 25 | GAAAAGGTGTTGGTTGATATAGACA | 166.7 | 4.5 | 1.6 | 2.9 | 4.2E-03 | 1.9E-02 | BT20/MB231 | MP |
| >hg38_21_+_8433172_8446622_RNA45SN1_45S_50ntFlanks@5494.5526.33\|AAGACGGTCGAACTTGACTATCTAGAGGAAGTA | >hg38_21_+_8433172_8446622_RNA45SN1_45S_50ntFlanks | 5494 | 5526 | 33 | AAGACGGTCGAACTTGACTATCTAGAGGAAGTA | 756.8 | -2.7 | 1.1 | -2.6 | 1.0E-02 | 1.9E-02 | BT20/MB468 | MP |
| >MI0003776\|hsa-mir-320b-1&WithFlank&1\|+\|116671740\|116671823@48.68.21;MI0003839\|hsa-mir-320b-2&WithFlank&1\|-\|224257034\|224257116@47.67.21&[MIMAT0005792&hsa-miR-320b&offsets\|0\|-1;m-183&1\|+\|116671787\|116671806&offsets\|0\|+1];[MIMAT0005792_1&hsa-miR-320b&offsets\|0\|-1;m-183&1\|-\|224257051\|224257070&offsets\|0\|+1]\|AAAAGCTGGGTTGAGAGGGCA | >MI0003776\|hsa-mir-320b-1&WithFlank&1\|+\|116671740\|116671823 | 48 | 68 | 21 | AAAAGCTGGGTTGAGAGGGCA | 452.2 | 3.5 | 1.4 | 2.4 | 1.5E-02 | 1.9E-02 | MB231/MB468 | MP |
| >hg38_21_+_8433172_8446622_RNA45SN1_45S_50ntFlanks@11876.11893.18\|AAGACCCTGTTGAGCTTG | >hg38_21_+_8433172_8446622_RNA45SN1_45S_50ntFlanks | 11876 | 11893 | 18 | AAGACCCTGTTGAGCTTG | 3339.1 | -5.4 | 2.3 | -2.4 | 1.7E-02 | 1.9E-02 | BT20/MB468 | NUC |
| >trnaMT_ValTAC_MT_+_1602_1670@53.71.19\|TAACTTGACCGCTCTGACC | >trnaMT_ValTAC_MT_+_1602_1670 | 53 | 71 | 19 | TAACTTGACCGCTCTGACC | 278.1 | 2.7 | 1.1 | 2.6 | 1.0E-02 | 1.9E-02 | BT20/MB468 | MP |
| >hg38_21_+_8433172_8446622_RNA45SN1_45S_50ntFlanks@11668.11704.37\|TGCCCAGTGCTCTGAATGTCAAAGTGAAGAAATTCAA | >hg38_21_+_8433172_8446622_RNA45SN1_45S_50ntFlanks | 11668 | 11704 | 37 | TGCCCAGTGCTCTGAATGTCAAAGTGAAGAAATTCAA | 697.0 | -2.2 | 0.8 | -2.8 | 5.4E-03 | 1.9E-02 | BT20/MB231 | CYTO |
| >hg38_21_+_8433172_8446622_RNA45SN1_45S_50ntFlanks@10247.10289.43\|GCGACGAGTAGGAGGGCCGCTGCGGTGAGCCTTGAAGCCTAGG | >hg38_21_+_8433172_8446622_RNA45SN1_45S_50ntFlanks | 10247 | 10289 | 43 | GCGACGAGTAGGAGGGCCGCTGCGGTGAGCCTTGAAGCCTAGG | 90.7 | 5.3 | 2.1 | 2.6 | 1.0E-02 | 1.9E-02 | BT20/MB468 | MP |
| >hg38_21_+_8433172_8446622_RNA45SN1_45S_50ntFlanks@10387.10408.22\|TGTGAACAGCAGTTGAACATGG | >hg38_21_+_8433172_8446622_RNA45SN1_45S_50ntFlanks | 10387 | 10408 | 22 | TGTGAACAGCAGTTGAACATGG | 37.3 | -8.3 | 3.2 | -2.6 | 1.0E-02 | 1.9E-02 | BT20/MB468 | MP |
| >hg38_21_+_8433172_8446622_RNA45SN1_45S_50ntFlanks@6742.6771.30\|TCGACACTTCGAACGCACTTGCGGCCCCGG | >hg38_21_+_8433172_8446622_RNA45SN1_45S_50ntFlanks | 6742 | 6771 | 30 | TCGACACTTCGAACGCACTTGCGGCCCCGG | 308.0 | -4.4 | 1.7 | -2.6 | 1.0E-02 | 1.9E-02 | BT20/MB468 | MP |
| >hg38_21_+_8433172_8446622_RNA45SN1_45S_50ntFlanks@7975.7996.22\|CGCGACCTCAGATCAGACGTGG | >hg38_21_+_8433172_8446622_RNA45SN1_45S_50ntFlanks | 7975 | 7996 | 22 | CGCGACCTCAGATCAGACGTGG | 6411.4 | 1.6 | 0.7 | 2.4 | 1.6E-02 | 1.9E-02 | MB231/MB468 | CYTO |
| >hg38_21_+_8433172_8446622_RNA45SN1_45S_50ntFlanks@11667.11691.25\|CTGCCCAGTGCTCTGAATGTCAAAG | >hg38_21_+_8433172_8446622_RNA45SN1_45S_50ntFlanks | 11667 | 11691 | 25 | CTGCCCAGTGCTCTGAATGTCAAAG | 458.6 | 4.0 | 1.4 | 2.9 | 4.2E-03 | 1.9E-02 | BT20/MB231 | MP |
| >hg38_21_+_8433172_8446622_RNA45SN1_45S_50ntFlanks@12406.12443.38\|TTTTTGATCCTTCGATGTCGGCTCTTCCTATCATTGTG | >hg38_21_+_8433172_8446622_RNA45SN1_45S_50ntFlanks | 12406 | 12443 | 38 | TTTTTGATCCTTCGATGTCGGCTCTTCCTATCATTGTG | 147.7 | -2.5 | 1.0 | -2.4 | 1.6E-02 | 1.9E-02 | BT20/MB468 | CYTO |
| >MI0000108\|hsa-mir-103a-2&WithFlank&20\|+\|3917488\|3917577@54.76.23(+1U);MI0000109\|hsa-mir-103a-1&WithFlank&5\|-\|168560890\|168560979@54.76.23(+1U)&[MIMAT0000101&hsa-miR-103a-3p&offsets\|0\|0(+1U);m-12&20\|+\|3917541\|3917563&offsets\|0\|0(+1U)];[MIMAT0000101_1&hsa-miR-103a-3p&offsets\|0\|0(+1U);m-13&5\|-\|168560904\|168560926&offsets\|0\|0(+1U);m-44220&5\|-\|168560908\|168560926&offsets\|0\|+4(+1U)]\|AGCAGCATTGTACAGGGCTATGAT | >MI0000108\|hsa-mir-103a-2&WithFlank&20\|+\|3917488\|3917577 | 54 | 76 | 23(+1U) | AGCAGCATTGTACAGGGCTATGAT | 738.3 | 2.4 | 1.0 | 2.4 | 1.5E-02 | 1.9E-02 | MB231/MB468 | MP |
| >hg38_21_+_8433172_8446622_RNA45SN1_45S_50ntFlanks@10391.10426.36\|AACAGCAGTTGAACATGGGTCAGTCGGTCCTGAGAG | >hg38_21_+_8433172_8446622_RNA45SN1_45S_50ntFlanks | 10391 | 10426 | 36 | AACAGCAGTTGAACATGGGTCAGTCGGTCCTGAGAG | 238.0 | -1.9 | 0.8 | -2.5 | 1.4E-02 | 1.9E-02 | BT20/MB468 | MITO |
| >MI0000434\|hsa-let-7i&WithFlank&12\|+\|62603680\|62603775@11.33.23&[MIMAT0000415&hsa-let-7i-5p&offsets\|-1\|0;m-49&12\|+\|62603691\|62603712&offsets\|-1\|0]\|CTGAGGTAGTAGTTTGTGCTGTT | >MI0000434\|hsa-let-7i&WithFlank&12\|+\|62603680\|62603775 | 11 | 33 | 23 | CTGAGGTAGTAGTTTGTGCTGTT | 73.6 | 4.1 | 1.7 | 2.4 | 1.5E-02 | 1.9E-02 | MB231/MB468 | MP |
| >hg38_21_+_8433172_8446622_RNA45SN1_45S_50ntFlanks@6661.6686.26\|CGGTGGATCACTCGGCTCGTGCGTCG | >hg38_21_+_8433172_8446622_RNA45SN1_45S_50ntFlanks | 6661 | 6686 | 26 | CGGTGGATCACTCGGCTCGTGCGTCG | 1540.9 | 1.4 | 0.6 | 2.4 | 1.7E-02 | 1.9E-02 | MB231/MB468 | NUC |
| >hg38_21_+_8433172_8446622_RNA45SN1_45S_50ntFlanks@11212.11230.19\|CGACTCTGGACGCGAGCCG | >hg38_21_+_8433172_8446622_RNA45SN1_45S_50ntFlanks | 11212 | 11230 | 19 | CGACTCTGGACGCGAGCCG | 237.4 | -4.4 | 1.7 | -2.6 | 1.0E-02 | 1.9E-02 | BT20/MB468 | MP |
| >hg38_21_+_8433172_8446622_RNA45SN1_45S_50ntFlanks@12417.12443.27\|TCGATGTCGGCTCTTCCTATCATTGTG | >hg38_21_+_8433172_8446622_RNA45SN1_45S_50ntFlanks | 12417 | 12443 | 27 | TCGATGTCGGCTCTTCCTATCATTGTG | 1360.3 | -2.5 | 0.9 | -2.8 | 5.4E-03 | 1.9E-02 | BT20/MB231 | CYTO |
| >hg38_21_+_8433172_8446622_RNA45SN1_45S_50ntFlanks@12896.12935.40\|CGCCCGTCACGCACCGCACGTTCGTGGGGAACCTGGCGCT | >hg38_21_+_8433172_8446622_RNA45SN1_45S_50ntFlanks | 12896 | 12935 | 40 | CGCCCGTCACGCACCGCACGTTCGTGGGGAACCTGGCGCT | 190.0 | -4.2 | 1.8 | -2.4 | 1.7E-02 | 1.9E-02 | MB231/MB468 | MITO |
| >hg38_21_+_8433172_8446622_RNA45SN1_45S_50ntFlanks@10822.10847.26\|GAAGTCGGCAAGCCGGATCCGTAACT | >hg38_21_+_8433172_8446622_RNA45SN1_45S_50ntFlanks | 10822 | 10847 | 26 | GAAGTCGGCAAGCCGGATCCGTAACT | 197.3 | -2.0 | 0.7 | -2.8 | 5.4E-03 | 1.9E-02 | BT20/MB231 | CYTO |
| >hg38_21_+_8433172_8446622_RNA45SN1_45S_50ntFlanks@8169.8215.47\|CCAAGTCCTTCTGATCGAGGCCCAGCCCGTGGACGGTGTGAGGCCGG | >hg38_21_+_8433172_8446622_RNA45SN1_45S_50ntFlanks | 8169 | 8215 | 47 | CCAAGTCCTTCTGATCGAGGCCCAGCCCGTGGACGGTGTGAGGCCGG | 163.0 | -3.5 | 1.5 | -2.4 | 1.6E-02 | 1.9E-02 | BT20/MB468 | CYTO |

<table>
<tr><td>>hg38_21_+_8433172_8446622_RNA45SN1_45S_50ntFlanks@9554.9572.19|CGTAGCGGTCCTGACGTGC</td><td>>hg38_21_+_8433172_8446622_RNA45SN1_45S_50ntFlanks</td><td>9554</td><td>9572</td><td>19</td><td>CGTAGCGGTCCTGACGTGC</td><td>5787.4</td><td>-3.0</td><td>1.2</td><td>-2.6</td><td>1.0E-02</td><td>1.9E-02</td><td>BT20/MB468</td><td>MP</td></tr>
<tr><td>>hg38_21_+_8433172_8446622_RNA45SN1_45S_50ntFlanks@12957.12977.21|CTTCTGGGTCGGGGTTTCGTA</td><td>>hg38_21_+_8433172_8446622_RNA45SN1_45S_50ntFlanks</td><td>12957</td><td>12977</td><td>21</td><td>CTTCTGGGTCGGGGTTTCGTA</td><td>108.6</td><td>-4.3</td><td>1.7</td><td>-2.6</td><td>1.0E-02</td><td>1.9E-02</td><td>BT20/MB468</td><td>MP</td></tr>
<tr><td>>hg38_21_+_8433172_8446622_RNA45SN1_45S_50ntFlanks@5485.5510.26|AGCGCTGAGAAGACGGTCGAACTTGA</td><td>>hg38_21_+_8433172_8446622_RNA45SN1_45S_50ntFlanks</td><td>5485</td><td>5510</td><td>26</td><td>AGCGCTGAGAAGACGGTCGAACTTGA</td><td>305.1</td><td>-1.9</td><td>0.8</td><td>-2.5</td><td>1.4E-02</td><td>1.9E-02</td><td>BT20/MB468</td><td>MITO</td></tr>
<tr><td>>MI0000299|hsa-mir-222&WithFlank&X|-|45747009|45747130@75.95.21&[MIMAT0000279&hsa-miR-222-3p&offsets|0|0;m-140&X|-|45747033|45747056&offsets|0|-3]|AGCTACATCTGGCTACTGGGT</td><td>>MI0000299|hsa-mir-222&WithFlank&X|-|45747009|45747130</td><td>75</td><td>95</td><td>21</td><td>AGCTACATCTGGCTACTGGGT</td><td>4002.2</td><td>-2.0</td><td>0.7</td><td>-2.8</td><td>5.2E-03</td><td>1.9E-02</td><td>BT20/MB231</td><td>MITO</td></tr>
<tr><td>>MI0000065|hsa-let-7d&WithFlank&9|+|94178828|94178926@14.35.22(+1A)&[MIMAT0000065&hsa-let-7d-5p&offsets|0|0(+1A);m-90&9|+|94178841|94178862&offsets|0|0(+1A)]|AGAGGTAGTAGGTTGCATAGTTA</td><td>>MI0000065|hsa-let-7d&WithFlank&9|+|94178828|94178926</td><td>14</td><td>35</td><td>22(+1A)</td><td>AGAGGTAGTAGGTTGCATAGTTA</td><td>333.3</td><td>2.4</td><td>0.9</td><td>2.8</td><td>5.4E-03</td><td>1.9E-02</td><td>BT20/MB231</td><td>CYTO</td></tr>
<tr><td>>MI0003667|hsa-mir-652&WithFlank&X|+|110055323|110055432@67.87.21(+1U)&[MIMAT0003322&hsa-miR-652-3p&offsets|0|0(+1U);m-148&X|+|110055389|110055410&offsets|0|-1(+1U)]|AATGGCGCCACTAGGGTTGTGT</td><td>>MI0003667|hsa-mir-652&WithFlank&X|+|110055323|110055432</td><td>67</td><td>87</td><td>21(+1U)</td><td>AATGGCGCCACTAGGGTTGTGT</td><td>1307.6</td><td>3.0</td><td>1.1</td><td>2.8</td><td>5.5E-03</td><td>1.9E-02</td><td>BT20/MB231</td><td>CYTO</td></tr>
<tr><td>>hg38_1_-_228634819_228635039_RNA5S12_5S_50ntFlanks@134.171.38|TGGGAGACCGCCTGGGAATACCGGGTGCTGTAGGCTTT</td><td>>hg38_1_-_228634819_228635039_RNA5S12_5S_50ntFlanks</td><td>134</td><td>171</td><td>38</td><td>TGGGAGACCGCCTGGGAATACCGGGTGCTGTAGGCTTT</td><td>54191.2</td><td>-2.6</td><td>0.9</td><td>-2.8</td><td>5.4E-03</td><td>1.9E-02</td><td>BT20/MB231</td><td>CYTO</td></tr>
<tr><td>>hg38_21_+_8433172_8446622_RNA45SN1_45S_50ntFlanks@12416.12443.28|TTCGATGTCGGCTCTTCCTATCATTGTG</td><td>>hg38_21_+_8433172_8446622_RNA45SN1_45S_50ntFlanks</td><td>12416</td><td>12443</td><td>28</td><td>TTCGATGTCGGCTCTTCCTATCATTGTG</td><td>3283.9</td><td>-2.7</td><td>1.0</td><td>-2.8</td><td>5.4E-03</td><td>1.9E-02</td><td>BT20/MB231</td><td>CYTO</td></tr>
<tr><td>>hg38_21_+_8433172_8446622_RNA45SN1_45S_50ntFlanks@5487.5512.26|CGCTGAGAAGACGGTCGAACTTGACT</td><td>>hg38_21_+_8433172_8446622_RNA45SN1_45S_50ntFlanks</td><td>5487</td><td>5512</td><td>26</td><td>CGCTGAGAAGACGGTCGAACTTGACT</td><td>544.0</td><td>2.9</td><td>1.1</td><td>2.8</td><td>5.4E-03</td><td>1.9E-02</td><td>BT20/MB231</td><td>CYTO</td></tr>
<tr><td>>hg38_21_+_8433172_8446622_RNA45SN1_45S_50ntFlanks@5491.5516.26|GAGAAGACGGTCGAACTTGACTATCT</td><td>>hg38_21_+_8433172_8446622_RNA45SN1_45S_50ntFlanks</td><td>5491</td><td>5516</td><td>26</td><td>GAGAAGACGGTCGAACTTGACTATCT</td><td>55038.3</td><td>2.8</td><td>1.0</td><td>2.8</td><td>5.5E-03</td><td>1.9E-02</td><td>BT20/MB231</td><td>CYTO</td></tr>
<tr><td>>MI0003513|hsa-mir-455&WithFlank&9|+|114209428|114209535@22.44.23&[MIMAT0003150&hsa-miR-455-5p&offsets|0|+1;m-232&9|+|114209449|114209470&offsets|0|+1]|TATGTGCCTTTGGACTACATCGT</td><td>>MI0003513|hsa-mir-455&WithFlank&9|+|114209428|114209535</td><td>22</td><td>44</td><td>23</td><td>TATGTGCCTTTGGACTACATCGT</td><td>269.0</td><td>-2.2</td><td>0.8</td><td>-2.8</td><td>5.5E-03</td><td>1.9E-02</td><td>BT20/MB231</td><td>CYTO</td></tr>
<tr><td>>hg38_21_+_8433172_8446622_RNA45SN1_45S_50ntFlanks@4680.4720.41|GCGCAAGACGGACCAGAGCGAAAGCATTTGCCAAGAATGTT</td><td>>hg38_21_+_8433172_8446622_RNA45SN1_45S_50ntFlanks</td><td>4680</td><td>4720</td><td>41</td><td>GCGCAAGACGGACCAGAGCGAAAGCATTTGCCAAGAATGTT</td><td>383.3</td><td>-2.5</td><td>0.9</td><td>-2.8</td><td>5.5E-03</td><td>1.9E-02</td><td>BT20/MB231</td><td>CYTO</td></tr>
<tr><td>>hg38_21_+_8433172_8446622_RNA45SN1_45S_50ntFlanks@7992.8018.27|CGTGGCGACCCGCTGAATTTAAGCATA</td><td>>hg38_21_+_8433172_8446622_RNA45SN1_45S_50ntFlanks</td><td>7992</td><td>8018</td><td>27</td><td>CGTGGCGACCCGCTGAATTTAAGCATA</td><td>167.2</td><td>-3.3</td><td>1.2</td><td>-2.8</td><td>5.5E-03</td><td>1.9E-02</td><td>BT20/MB231</td><td>CYTO</td></tr>
<tr><td>>hg38_21_+_8433172_8446622_RNA45SN1_45S_50ntFlanks@12409.12436.28|TTGATCCTTCGATGTCGGCTCTTCCTAT</td><td>>hg38_21_+_8433172_8446622_RNA45SN1_45S_50ntFlanks</td><td>12409</td><td>12436</td><td>28</td><td>TTGATCCTTCGATGTCGGCTCTTCCTAT</td><td>571.7</td><td>-4.8</td><td>2.0</td><td>-2.4</td><td>1.7E-02</td><td>1.9E-02</td><td>BT20/MB468</td><td>NUC</td></tr>
<tr><td>>hg38_21_+_8433172_8446622_RNA45SN1_45S_50ntFlanks@6651.6668.18|CGACTCTTAGCGGTGGAT</td><td>>hg38_21_+_8433172_8446622_RNA45SN1_45S_50ntFlanks</td><td>6651</td><td>6668</td><td>18</td><td>CGACTCTTAGCGGTGGAT</td><td>61232.4</td><td>1.9</td><td>0.7</td><td>2.8</td><td>5.2E-03</td><td>1.9E-02</td><td>BT20/MB231</td><td>MITO</td></tr>
<tr><td>>hg38_21_+_8433172_8446622_RNA45SN1_45S_50ntFlanks@6663.6686.24|GTGGATCACTCGGCTCGTGCGTCG</td><td>>hg38_21_+_8433172_8446622_RNA45SN1_45S_50ntFlanks</td><td>6663</td><td>6686</td><td>24</td><td>GTGGATCACTCGGCTCGTGCGTCG</td><td>967.5</td><td>-2.9</td><td>1.1</td><td>-2.6</td><td>1.1E-02</td><td>1.9E-02</td><td>BT20/MB468</td><td>MP</td></tr>
<tr><td>>hg38_MT_+_1621_3279_RNR2_16S_50ntFlanks@754.777.24|ACAGCCCAATATCTACAATCAACC</td><td>>hg38_MT_+_1621_3279_RNR2_16S_50ntFlanks</td><td>754</td><td>777</td><td>24</td><td>ACAGCCCAATATCTACAATCAACC</td><td>54.2</td><td>4.7</td><td>1.9</td><td>2.4</td><td>1.5E-02</td><td>1.9E-02</td><td>MB231/MB468</td><td>MP</td></tr>
<tr><td>>hg38_21_+_8433172_8446622_RNA45SN1_45S_50ntFlanks@12995.13012.18|CTCGCTGCGATCTATTGA</td><td>>hg38_21_+_8433172_8446622_RNA45SN1_45S_50ntFlanks</td><td>12995</td><td>13012</td><td>18</td><td>CTCGCTGCGATCTATTGA</td><td>571.2</td><td>1.3</td><td>0.6</td><td>2.4</td><td>1.6E-02</td><td>1.9E-02</td><td>BT20/MB468</td><td>CYTO</td></tr>
<tr><td>>hg38_21_+_8433172_8446622_RNA45SN1_45S_50ntFlanks@11188.11225.38|CACCCGGGGGGCCGGCGGCGGCGGCGACTCTGGACGCG</td><td>>hg38_21_+_8433172_8446622_RNA45SN1_45S_50ntFlanks</td><td>11188</td><td>11225</td><td>38</td><td>CACCCGGGGGGCCGGCGGCGGCGGCGACTCTGGACGCG</td><td>280.3</td><td>-3.9</td><td>1.6</td><td>-2.4</td><td>1.6E-02</td><td>1.9E-02</td><td>BT20/MB468</td><td>CYTO</td></tr>
<tr><td>>hg38_21_+_8433172_8446622_RNA45SN1_45S_50ntFlanks@10346.10369.24|TCAAACGAGAACTTTGAAGGCCGA</td><td>>hg38_21_+_8433172_8446622_RNA45SN1_45S_50ntFlanks</td><td>10346</td><td>10369</td><td>24</td><td>TCAAACGAGAACTTTGAAGGCCGA</td><td>468.5</td><td>2.5</td><td>0.9</td><td>2.8</td><td>5.2E-03</td><td>1.9E-02</td><td>BT20/MB231</td><td>MITO</td></tr>
<tr><td>>hg38_21_+_8433172_8446622_RNA45SN1_45S_50ntFlanks@12488.12519.32|AACGTGAGCTGGGTTTAGACCGTCGTGAGACA</td><td>>hg38_21_+_8433172_8446622_RNA45SN1_45S_50ntFlanks</td><td>12488</td><td>12519</td><td>32</td><td>AACGTGAGCTGGGTTTAGACCGTCGTGAGACA</td><td>116.4</td><td>-3.5</td><td>1.5</td><td>-2.4</td><td>1.6E-02</td><td>1.9E-02</td><td>BT20/MB468</td><td>CYTO</td></tr>
<tr><td>>hg38_21_+_8433172_8446622_RNA45SN1_45S_50ntFlanks@10843.10865.23|TAACTTCGGGATAAGGATTGGCT</td><td>>hg38_21_+_8433172_8446622_RNA45SN1_45S_50ntFlanks</td><td>10843</td><td>10865</td><td>23</td><td>TAACTTCGGGATAAGGATTGGCT</td><td>292.5</td><td>-4.4</td><td>1.7</td><td>-2.6</td><td>1.1E-02</td><td>1.9E-02</td><td>BT20/MB468</td><td>MP</td></tr>
<tr><td>>hg38_21_+_8433172_8446622_RNA45SN1_45S_50ntFlanks@4049.4066.18|TCGAACGTCTGCCCTATC</td><td>>hg38_21_+_8433172_8446622_RNA45SN1_45S_50ntFlanks</td><td>4049</td><td>4066</td><td>18</td><td>TCGAACGTCTGCCCTATC</td><td>328.6</td><td>1.1</td><td>0.5</td><td>2.4</td><td>1.6E-02</td><td>1.9E-02</td><td>BT20/MB468</td><td>CYTO</td></tr>
<tr><td>>hg38_21_+_8433172_8446622_RNA45SN1_45S_50ntFlanks@8383.8406.24|GAGGGCGTGAAACCGTTAAGAGGT</td><td>>hg38_21_+_8433172_8446622_RNA45SN1_45S_50ntFlanks</td><td>8383</td><td>8406</td><td>24</td><td>GAGGGCGTGAAACCGTTAAGAGGT</td><td>123.7</td><td>2.1</td><td>0.8</td><td>2.6</td><td>1.1E-02</td><td>1.9E-02</td><td>BT20/MB468</td><td>MP</td></tr>
<tr><td>>hg38_21_+_8433172_8446622_RNA45SN1_45S_50ntFlanks@4128.4148.21|CGATTCCGGAGAGGGAGCCTG</td><td>>hg38_21_+_8433172_8446622_RNA45SN1_45S_50ntFlanks</td><td>4128</td><td>4148</td><td>21</td><td>CGATTCCGGAGAGGGAGCCTG</td><td>218.8</td><td>-2.1</td><td>0.8</td><td>-2.6</td><td>1.1E-02</td><td>1.9E-02</td><td>BT20/MB468</td><td>MP</td></tr>
<tr><td>>hg38_21_+_8433172_8446622_RNA45SN1_45S_50ntFlanks@4614.4633.20|GCCATGATTAAGAGGGACGG</td><td>>hg38_21_+_8433172_8446622_RNA45SN1_45S_50ntFlanks</td><td>4614</td><td>4633</td><td>20</td><td>GCCATGATTAAGAGGGACGG</td><td>81.1</td><td>-4.4</td><td>1.7</td><td>-2.6</td><td>1.1E-02</td><td>1.9E-02</td><td>BT20/MB468</td><td>MP</td></tr>
<tr><td>>trna37_ValAAC_6_+_27203288_27203360@1.29.29|GTTTCCGTAGTGTAGTGGTTATCACGTTT</td><td>>trna37_ValAAC_6_+_27203288_27203360</td><td>1</td><td>29</td><td>29</td><td>GTTTCCGTAGTGTAGTGGTTATCACGTTT</td><td>270.8</td><td>-3.3</td><td>1.3</td><td>-2.5</td><td>1.4E-02</td><td>1.9E-02</td><td>BT20/MB468</td><td>MITO</td></tr>
<tr><td>>hg38_21_+_8433172_8446622_RNA45SN1_45S_50ntFlanks@6661.6678.18|CGGTGGATCACTCGGCTC</td><td>>hg38_21_+_8433172_8446622_RNA45SN1_45S_50ntFlanks</td><td>6661</td><td>6678</td><td>18</td><td>CGGTGGATCACTCGGCTC</td><td>185.0</td><td>2.1</td><td>0.9</td><td>2.4</td><td>1.7E-02</td><td>1.9E-02</td><td>MB231/MB468</td><td>MITO</td></tr>
<tr><td>>hg38_21_+_8433172_8446622_RNA45SN1_45S_50ntFlanks@6651.6688.38|CGACTCTTAGCGGTGGATCACTCGGCTCGTGCGTCGAT</td><td>>hg38_21_+_8433172_8446622_RNA45SN1_45S_50ntFlanks</td><td>6651</td><td>6688</td><td>38</td><td>CGACTCTTAGCGGTGGATCACTCGGCTCGTGCGTCGAT</td><td>10739.0</td><td>-1.7</td><td>0.7</td><td>-2.4</td><td>1.7E-02</td><td>1.9E-02</td><td>MB231/MB468</td><td>MITO</td></tr>
<tr><td>>MI0003656|hsa-mir-641&WithFlank&19|-|40282537|40282647@22.46.25&[MIMAT0003311&hsa-miR-641&offsets|0|+1;m-786&19|-|40282603|40282626&offsets|0|+1]|AAAGACATAGGATAGAGTCACCTCT</td><td>>MI0003656|hsa-mir-641&WithFlank&19|-|40282537|40282647</td><td>22</td><td>46</td><td>25</td><td>AAAGACATAGGATAGAGTCACCTCT</td><td>82.5</td><td>-5.5</td><td>1.9</td><td>-2.9</td><td>4.2E-03</td><td>1.9E-02</td><td>BT20/MB231</td><td>MP</td></tr>
<tr><td>>MI0000734|hsa-mir-106b&WithFlank&7|-|100093987|100094080@18.38.21(+1U)&[MIMAT0000680&hsa-miR-106b-5p&offsets|0|0(+1U);m-103&7|-|100094043|100094063&offsets|0|0(+1U)]|TAAAGTGCTGACAGTGCAGATT</td><td>>MI0000734|hsa-mir-106b&WithFlank&7|-|100093987|100094080</td><td>18</td><td>38</td><td>21(+1U)</td><td>TAAAGTGCTGACAGTGCAGATT</td><td>149.0</td><td>3.2</td><td>1.3</td><td>2.4</td><td>1.5E-02</td><td>1.9E-02</td><td>MB231/MB468</td><td>MP</td></tr>
<tr><td>>hg38_21_+_8433172_8446622_RNA45SN1_45S_50ntFlanks@3775.3797.23|GCCGGTACAGTGAAACTGCGAAT</td><td>>hg38_21_+_8433172_8446622_RNA45SN1_45S_50ntFlanks</td><td>3775</td><td>3797</td><td>23</td><td>GCCGGTACAGTGAAACTGCGAAT</td><td>147.3</td><td>-4.8</td><td>1.9</td><td>-2.6</td><td>1.1E-02</td><td>1.9E-02</td><td>BT20/MB468</td><td>MP</td></tr>
<tr><td>>hg38_21_+_8433172_8446622_RNA45SN1_45S_50ntFlanks@6648.6686.39|GTACGACTCTTAGCGGTGGATCACTCGGCTCGTGCGTCG</td><td>>hg38_21_+_8433172_8446622_RNA45SN1_45S_50ntFlanks</td><td>6648</td><td>6686</td><td>39</td><td>GTACGACTCTTAGCGGTGGATCACTCGGCTCGTGCGTCG</td><td>386.3</td><td>-1.9</td><td>0.8</td><td>-2.4</td><td>1.7E-02</td><td>1.9E-02</td><td>MB231/MB468</td><td>MITO</td></tr>
<tr><td>>hg38_21_+_8433172_8446622_RNA45SN1_45S_50ntFlanks@7989.8006.18|AGACGTGGCGACCCGCTG</td><td>>hg38_21_+_8433172_8446622_RNA45SN1_45S_50ntFlanks</td><td>7989</td><td>8006</td><td>18</td><td>AGACGTGGCGACCCGCTG</td><td>416.4</td><td>-1.8</td><td>0.8</td><td>-2.4</td><td>1.6E-02</td><td>1.9E-02</td><td>MB231/MB468</td><td>CYTO</td></tr>
<tr><td>>hg38_21_+_8433172_8446622_RNA45SN1_45S_50ntFlanks@11919.11940.22|CATGAGAGGTGTAGAATAAGTG</td><td>>hg38_21_+_8433172_8446622_RNA45SN1_45S_50ntFlanks</td><td>11919</td><td>11940</td><td>22</td><td>CATGAGAGGTGTAGAATAAGTG</td><td>130.0</td><td>4.6</td><td>1.8</td><td>2.6</td><td>1.1E-02</td><td>1.9E-02</td><td>BT20/MB468</td><td>MP</td></tr>
<tr><td>>trnaMT_ValTAC_MT_+_1602_1670@48.72.25|CAACTTAACTTGACCGCTCTGACCA</td><td>>trnaMT_ValTAC_MT_+_1602_1670</td><td>48</td><td>72</td><td>25</td><td>CAACTTAACTTGACCGCTCTGACCA</td><td>980.7</td><td>2.8</td><td>1.1</td><td>2.6</td><td>1.1E-02</td><td>1.9E-02</td><td>BT20/MB468</td><td>MP</td></tr>
<tr><td>>hg38_21_+_8433172_8446622_RNA45SN1_45S_50ntFlanks@9739.9757.19|AAACGATCTCAACCTATTC</td><td>>hg38_21_+_8433172_8446622_RNA45SN1_45S_50ntFlanks</td><td>9739</td><td>9757</td><td>19</td><td>AAACGATCTCAACCTATTC</td><td>635.5</td><td>-3.2</td><td>1.3</td><td>-2.4</td><td>1.6E-02</td><td>1.9E-02</td><td>BT20/MB468</td><td>CYTO</td></tr>
<tr><td>>hg38_21_+_8433172_8446622_RNA45SN1_45S_50ntFlanks@10346.10368.23|TCAAACGAGAACTTTGAAGGCCG</td><td>>hg38_21_+_8433172_8446622_RNA45SN1_45S_50ntFlanks</td><td>10346</td><td>10368</td><td>23</td><td>TCAAACGAGAACTTTGAAGGCCG</td><td>418.3</td><td>-2.7</td><td>1.0</td><td>-2.8</td><td>4.6E-03</td><td>1.9E-02</td><td>BT20/MB231</td><td>NUC</td></tr>
<tr><td>>MI0000114|hsa-mir-107&WithFlank&10|-|89592741|89592833@56.76.21(+1U);MI0000108|hsa-mir-103a-2&WithFlank&20|+|3917488|3917577@54.74.21(+1U);MI0000109|hsa-mir-103a-1&WithFlank&5|-|168560890|168560979@54.74.21(+1U)&[MIMAT0000104&hsa-miR-107&offsets|0|-2(+1U);m-78&10|-|89592758|89592778&offsets|0|0(+1U)];[MIMAT0000101&hsa-miR-103a-3p&offsets|0|-2(+1U);m-12&20|+|3917541|3917563&offsets|0|-2(+1U)];[MIMAT0000101_1&hsa-miR-103a-3p&offsets|0|-2(+1U);m-13&5|-|168560904|168560926&offsets|0|-2(+1U);m-44220&5|-|168560908|168560926&offsets|0|+2(+1U)]|AGCAGCATTGTACAGGGCTATT</td><td>>MI0000114|hsa-mir-107&WithFlank&10|-|89592741|89592833</td><td>56</td><td>76</td><td>21(+1U)</td><td>AGCAGCATTGTACAGGGCTATT</td><td>454.8</td><td>3.4</td><td>1.4</td><td>2.4</td><td>1.5E-02</td><td>1.9E-02</td><td>MB231/MB468</td><td>MP</td></tr>
<tr><td>>MI0003610|hsa-mir-598&WithFlank&8|-|11035200|11035308@67.88.22&[MIMAT0003266&hsa-miR-598-3p&offsets|0|0;m-231&8|-|11035221|11035242&offsets|0|0]|TACGTCATCGTTGTCATCGTCA</td><td>>MI0003610|hsa-mir-598&WithFlank&8|-|11035200|11035308</td><td>67</td><td>88</td><td>22</td><td>TACGTCATCGTTGTCATCGTCA</td><td>124.5</td><td>-5.3</td><td>1.8</td><td>-2.9</td><td>4.2E-03</td><td>1.9E-02</td><td>BT20/MB231</td><td>MP</td></tr>
<tr><td>>hg38_21_+_8433172_8446622_RNA45SN1_45S_50ntFlanks@9924.9948.25|TGGTTGATATAGACAGCAGGACGGT</td><td>>hg38_21_+_8433172_8446622_RNA45SN1_45S_50ntFlanks</td><td>9924</td><td>9948</td><td>25</td><td>TGGTTGATATAGACAGCAGGACGGT</td><td>66.2</td><td>5.2</td><td>2.0</td><td>2.6</td><td>1.1E-02</td><td>1.9E-02</td><td>BT20/MB468</td><td>MP</td></tr>
<tr><td>>MI0000068|hsa-let-7f-2&WithFlank&X|-|53557186|53557280@64.84.21(+1U)&[MIMAT0004487&hsa-let-7f-2-3p&offsets|0|-1(+1U);m-453&X|-|53557196|53557217&offsets|0|-1(+1U)]|CTATACAGTCTACTGTCTTTCT</td><td>>MI0000068|hsa-let-7f-2&WithFlank&X|-|53557186|53557280</td><td>64</td><td>84</td><td>21(+1U)</td><td>CTATACAGTCTACTGTCTTTCT</td><td>220.1</td><td>2.1</td><td>0.9</td><td>2.4</td><td>1.5E-02</td><td>1.9E-02</td><td>MB231/MB468</td><td>MP</td></tr>
<tr><td>>MI0000085|hsa-mir-27a&WithFlank&19|-|13836434|13836523@57.76.20(+2A)&[MIMAT0000084&hsa-miR-27a-3p&offsets|0|-1(+2A);m-43&19|-|13836447|13836467&offsets|0|-1(+2A)]|TTCACAGTGGCTAAGTTCCGAA</td><td>>MI0000085|hsa-mir-27a&WithFlank&19|-|13836434|13836523</td><td>57</td><td>76</td><td>20(+2A)</td><td>TTCACAGTGGCTAAGTTCCGAA</td><td>99.1</td><td>3.3</td><td>1.4</td><td>2.4</td><td>1.5E-02</td><td>1.9E-02</td><td>MB231/MB468</td><td>MP</td></tr>
<tr><td>>hg38_21_+_8433172_8446622_RNA45SN1_45S_50ntFlanks@11572.11602.31|CCGACTTAGAACTGGTGCGGACCAGGGGAAT</td><td>>hg38_21_+_8433172_8446622_RNA45SN1_45S_50ntFlanks</td><td>11572</td><td>11602</td><td>31</td><td>CCGACTTAGAACTGGTGCGGACCAGGGGAAT</td><td>579.9</td><td>3.5</td><td>1.2</td><td>2.9</td><td>4.2E-03</td><td>1.9E-02</td><td>BT20/MB231</td><td>MP</td></tr>
<tr><td>>MI0000440|hsa-mir-27b&WithFlank&9|+|95085439|95085547@66.87.22&[MIMAT0000419&hsa-miR-27b-3p&offsets|-1|0;m-39&9|+|95085505|95085525&offsets|-1|0]|GTTCACAGTGGCTAAGTTCTGC</td><td>>MI0000440|hsa-mir-27b&WithFlank&9|+|95085439|95085547</td><td>66</td><td>87</td><td>22</td><td>GTTCACAGTGGCTAAGTTCTGC</td><td>852.6</td><td>2.3</td><td>0.8</td><td>2.8</td><td>5.5E-03</td><td>1.9E-02</td><td>BT20/MB231</td><td>CYTO</td></tr>
<tr><td>>hg38_21_+_8433172_8446622_RNA45SN1_45S_50ntFlanks@12420.12452.33|ATGTCGGCTCTTCCTATCATTGTGAAGCAGAAT</td><td>>hg38_21_+_8433172_8446622_RNA45SN1_45S_50ntFlanks</td><td>12420</td><td>12452</td><td>33</td><td>ATGTCGGCTCTTCCTATCATTGTGAAGCAGAAT</td><td>243.8</td><td>-2.1</td><td>0.8</td><td>-2.8</td><td>5.5E-03</td><td>1.9E-02</td><td>BT20/MB231</td><td>CYTO</td></tr>
<tr><td>>hg38_21_+_8433172_8446622_RNA45SN1_45S_50ntFlanks@4047.4069.23|ATTCGAACGTCTGCCCTATCAAC</td><td>>hg38_21_+_8433172_8446622_RNA45SN1_45S_50ntFlanks</td><td>4047</td><td>4069</td><td>23</td><td>ATTCGAACGTCTGCCCTATCAAC</td><td>238.2</td><td>-2.3</td><td>0.8</td><td>-2.8</td><td>5.5E-03</td><td>1.9E-02</td><td>BT20/MB231</td><td>CYTO</td></tr>
<tr><td>>hg38_MT_+_1621_3279_RNR2_16S_50ntFlanks@51.92.42|GCTAAACCTAGCCCCAAACCCACTCCACCTTACTACCAGACA</td><td>>hg38_MT_+_1621_3279_RNR2_16S_50ntFlanks</td><td>51</td><td>92</td><td>42</td><td>GCTAAACCTAGCCCCAAACCCACTCCACCTTACTACCAGACA</td><td>597.6</td><td>-2.9</td><td>1.1</td><td>-2.8</td><td>5.5E-03</td><td>1.9E-02</td><td>BT20/MB231</td><td>CYTO</td></tr>
</table>

<table>
<tr><th></th><th></th><th></th><th></th><th></th><th></th><th></th><th></th><th></th><th></th><th></th><th></th><th></th><th></th></tr>
<tr><td>>MI0000114|hsa-mir-107&WithFlank&10|-|89592741|89592833@56.76.21(+1U);MI0000108|hsa-mir-103a-2&WithFlank&20|+|3917488|3917577@54.74.21(+1U);MI0000109|hsa-mir-103a-1&WithFlank&5|-|168560890|168560979@54.74.21(+1U)&[MIMAT0000104&hsa-miR-107&offsets|0|-2(+1U);m-78&10|-|89592758|89592778&offsets|0|0(+1U)];[MIMAT0000101&hsa-miR-103a-3p&offsets|0|-2(+1U);m-12&20|+|3917541|3917563&offsets|0|-2(+1U)];[MIMAT0000101_1&hsa-miR-103a-3p&offsets|0|-2(+1U);m-13&5|-|168560904|168560926&offsets|0|-2(+1U);m-44220&5|-|168560908|168560926&offsets|0|+2(+1U)]|AGCAGCATTGTACAGGGCTATT</td><td>>MI0000114|hsa-mir-107&WithFlank&10|-|89592741|89592833</td><td>56</td><td>76</td><td>21(+1U)</td><td>AGCAGCATTGTACAGGGCTATT</td><td>405.0</td><td>-2.5</td><td>0.9</td><td>-2.8</td><td>5.3E-03</td><td>1.9E-02</td><td>BT20/MB231</td><td>MITO</td></tr>
<tr><td>>am_MI0003583|hsa-mir-576&WithFlank&4|+|109488692|109488801@61.82.22&[MIMAT0004796&hsa-miR-576-3p&offsets|0|0;m-524&4|+|109488752|109488772&offsets|0|+1]|AAGATGTGGAAAAATTGGAATC</td><td>>am_MI0003583|hsa-mir-576&WithFlank&4|+|109488692|109488801</td><td>61</td><td>82</td><td>22</td><td>AAGATGTGGAAAAATTGGAATC</td><td>166.8</td><td>5.0</td><td>1.9</td><td>2.6</td><td>1.1E-02</td><td>1.9E-02</td><td>BT20/MB468</td><td>MP</td></tr>
<tr><td>>hg38_21_+_8433172_8446622_RNA45SN1_45S_50ntFlanks@9862.9900.39|ATGAACCGAACGCCGGGTTAAGGCGCCCGATGCCGACGC</td><td>>hg38_21_+_8433172_8446622_RNA45SN1_45S_50ntFlanks</td><td>9862</td><td>9900</td><td>39</td><td>ATGAACCGAACGCCGGGTTAAGGCGCCCGATGCCGACGC</td><td>73.6</td><td>-4.6</td><td>1.8</td><td>-2.6</td><td>1.1E-02</td><td>1.9E-02</td><td>BT20/MB468</td><td>MP</td></tr>
<tr><td>>hg38_21_+_8433172_8446622_RNA45SN1_45S_50ntFlanks@11849.11873.25|CGGGCTTGGCGGAATCAGCGGGGAA</td><td>>hg38_21_+_8433172_8446622_RNA45SN1_45S_50ntFlanks</td><td>11849</td><td>11873</td><td>25</td><td>CGGGCTTGGCGGAATCAGCGGGGAA</td><td>162.8</td><td>-3.7</td><td>1.5</td><td>-2.4</td><td>1.4E-02</td><td>1.9E-02</td><td>BT20/MB468</td><td>MITO</td></tr>
<tr><td>>hg38_21_+_8433172_8446622_RNA45SN1_45S_50ntFlanks@12605.12624.20|TATGTGCTTGGCTGAGGAGC</td><td>>hg38_21_+_8433172_8446622_RNA45SN1_45S_50ntFlanks</td><td>12605</td><td>12624</td><td>20</td><td>TATGTGCTTGGCTGAGGAGC</td><td>371.1</td><td>-1.2</td><td>0.5</td><td>-2.4</td><td>1.4E-02</td><td>1.9E-02</td><td>BT20/MB468</td><td>MITO</td></tr>
<tr><td>>hg38_21_+_8433172_8446622_RNA45SN1_45S_50ntFlanks@483.524.42|TCGATGTGGTGACGTCGTGCTCTCCCGGGCCGGGTCCGAGCC</td><td>>hg38_21_+_8433172_8446622_RNA45SN1_45S_50ntFlanks</td><td>483</td><td>524</td><td>42</td><td>TCGATGTGGTGACGTCGTGCTCTCCCGGGCCGGGTCCGAGCC</td><td>1098.5</td><td>-2.4</td><td>0.9</td><td>-2.8</td><td>4.6E-03</td><td>1.9E-02</td><td>BT20/MB231</td><td>NUC</td></tr>
<tr><td>>hg38_21_+_8433172_8446622_RNA45SN1_45S_50ntFlanks@6661.6683.23|CGGTGGATCACTCGGCTCGTGCG</td><td>>hg38_21_+_8433172_8446622_RNA45SN1_45S_50ntFlanks</td><td>6661</td><td>6683</td><td>23</td><td>CGGTGGATCACTCGGCTCGTGCG</td><td>7057.8</td><td>-1.7</td><td>0.6</td><td>-2.8</td><td>4.6E-03</td><td>1.9E-02</td><td>BT20/MB231</td><td>NUC</td></tr>
<tr><td>>hg38_21_+_8433172_8446622_RNA45SN1_45S_50ntFlanks@8178.8203.26|TCTGATCGAGGCCCAGCCCGTGGACG</td><td>>hg38_21_+_8433172_8446622_RNA45SN1_45S_50ntFlanks</td><td>8178</td><td>8203</td><td>26</td><td>TCTGATCGAGGCCCAGCCCGTGGACG</td><td>340.5</td><td>-1.9</td><td>0.8</td><td>-2.4</td><td>1.7E-02</td><td>1.9E-02</td><td>MB231/MB468</td><td>MITO</td></tr>
<tr><td>>MI0000078|hsa-mir-22&WithFlank&17|-|1713897|1713993@59.79.21&[MIMAT0000077&hsa-miR-22-3p&offsets|0|-1;m-2&17|-|1713914|1713935&offsets|0|-1]|AAGCTGCCAGTTGAAGAACTG</td><td>>MI0000078|hsa-mir-22&WithFlank&17|-|1713897|1713993</td><td>59</td><td>79</td><td>21</td><td>AAGCTGCCAGTTGAAGAACTG</td><td>99.7</td><td>5.5</td><td>2.2</td><td>2.6</td><td>1.1E-02</td><td>1.9E-02</td><td>BT20/MB468</td><td>MP</td></tr>
<tr><td>>MI0000102|hsa-mir-100&WithFlank&11|-|122152223|122152314@19.40.22&[MIMAT0000098&hsa-miR-100-5p&offsets|0|0;m-27&11|-|122152276|122152296&offsets|0|+1]|AACCCGTAGATCCGAACTTGTG</td><td>>MI0000102|hsa-mir-100&WithFlank&11|-|122152223|122152314</td><td>19</td><td>40</td><td>22</td><td>AACCCGTAGATCCGAACTTGTG</td><td>578433.4</td><td>-3.4</td><td>1.2</td><td>-2.8</td><td>5.6E-03</td><td>1.9E-02</td><td>BT20/MB231</td><td>CYTO</td></tr>
<tr><td>>hg38_21_+_8433172_8446622_RNA45SN1_45S_50ntFlanks@10802.10829.28|TTGGAACAATGTAGGTAAGGGAAGTCGG</td><td>>hg38_21_+_8433172_8446622_RNA45SN1_45S_50ntFlanks</td><td>10802</td><td>10829</td><td>28</td><td>TTGGAACAATGTAGGTAAGGGAAGTCGG</td><td>74.2</td><td>-4.3</td><td>1.7</td><td>-2.6</td><td>1.1E-02</td><td>1.9E-02</td><td>BT20/MB468</td><td>MP</td></tr>
<tr><td>>hg38_21_+_8433172_8446622_RNA45SN1_45S_50ntFlanks@10684.10712.29|CGGCGGCGTCCGGTGAGCTCTCGCTGGCC</td><td>>hg38_21_+_8433172_8446622_RNA45SN1_45S_50ntFlanks</td><td>10684</td><td>10712</td><td>29</td><td>CGGCGGCGTCCGGTGAGCTCTCGCTGGCC</td><td>393.7</td><td>-2.9</td><td>1.2</td><td>-2.4</td><td>1.4E-02</td><td>1.9E-02</td><td>BT20/MB468</td><td>MITO</td></tr>
<tr><td>>MI0000767|hsa-mir-365a&WithFlank&16|+|14309279|14309377@62.82.21;MI0000769|hsa-mir-365b&WithFlank&17|+|31575405|31575527@74.94.21&[MIMAT0000710&hsa-miR-365a-3p&offsets|0|-1;m-176&16|+|14309340|14309361&offsets|0|-1];[MIMAT0022834&hsa-miR-365b-3p&offsets|0|-1;m-175&17|+|31575478|31575499&offsets|0|-1]|TAATGCCCCTAAAAATCCTTA</td><td>>MI0000767|hsa-mir-365a&WithFlank&16|+|14309279|14309377</td><td>62</td><td>82</td><td>21</td><td>TAATGCCCCTAAAAATCCTTA</td><td>77.5</td><td>4.6</td><td>1.8</td><td>2.6</td><td>1.1E-02</td><td>1.9E-02</td><td>BT20/MB468</td><td>MP</td></tr>
<tr><td>>hg38_21_+_8433172_8446622_RNA45SN1_45S_50ntFlanks@11781.11800.20|CGCATGAATGGATGAACGAG</td><td>>hg38_21_+_8433172_8446622_RNA45SN1_45S_50ntFlanks</td><td>11781</td><td>11800</td><td>20</td><td>CGCATGAATGGATGAACGAG</td><td>125.0</td><td>-4.3</td><td>1.7</td><td>-2.6</td><td>1.1E-02</td><td>1.9E-02</td><td>BT20/MB468</td><td>MP</td></tr>
<tr><td>>hg38_21_+_8433172_8446622_RNA45SN1_45S_50ntFlanks@10819.10836.18|AGGGAAGTCGGCAAGCCG</td><td>>hg38_21_+_8433172_8446622_RNA45SN1_45S_50ntFlanks</td><td>10819</td><td>10836</td><td>18</td><td>AGGGAAGTCGGCAAGCCG</td><td>158.2</td><td>-3.8</td><td>1.6</td><td>-2.4</td><td>1.6E-02</td><td>1.9E-02</td><td>BT20/MB468</td><td>CYTO</td></tr>
<tr><td>>hg38_21_+_8433172_8446622_RNA45SN1_45S_50ntFlanks@3876.3898.23|TACATGCCGACGGGCGCTGACCC</td><td>>hg38_21_+_8433172_8446622_RNA45SN1_45S_50ntFlanks</td><td>3876</td><td>3898</td><td>23</td><td>TACATGCCGACGGGCGCTGACCC</td><td>419.7</td><td>-4.5</td><td>1.8</td><td>-2.6</td><td>1.1E-02</td><td>1.9E-02</td><td>BT20/MB468</td><td>MP</td></tr>
<tr><td>>hg38_21_+_8433172_8446622_RNA45SN1_45S_50ntFlanks@10685.10711.27|GGCGGCGTCCGGTGAGCTCTCGCTGGC</td><td>>hg38_21_+_8433172_8446622_RNA45SN1_45S_50ntFlanks</td><td>10685</td><td>10711</td><td>27</td><td>GGCGGCGTCCGGTGAGCTCTCGCTGGC</td><td>131.3</td><td>-3.1</td><td>1.3</td><td>-2.4</td><td>1.5E-02</td><td>1.9E-02</td><td>MB231/MB468</td><td>MP</td></tr>
<tr><td>>MI0000750|hsa-mir-26a-2&WithFlank&12|-|57824603|57824698@20.42.23(+1C);MI0000083|hsa-mir-26a-1&WithFlank&3|+|37969398|37969486@16.38.23(+1C)&[MIMAT0000082&hsa-miR-26a-5p&offsets|0|+1(+1C);m-33&12|-|57824658|57824679&offsets|0|+1(+1C)];[MIMAT0000082_1&hsa-miR-26a-5p&offsets|0|+1(+1C);m-34&3|+|37969413|37969434&offsets|0|+1(+1C)]|TTCAAGTAATCCAGGATAGGCTGC</td><td>>MI0000750|hsa-mir-26a-2&WithFlank&12|-|57824603|57824698</td><td>20</td><td>42</td><td>23(+1C)</td><td>TTCAAGTAATCCAGGATAGGCTGC</td><td>34.2</td><td>4.9</td><td>2.0</td><td>2.4</td><td>1.5E-02</td><td>1.9E-02</td><td>MB231/MB468</td><td>MP</td></tr>
<tr><td>>MI0000253|hsa-mir-148a&WithFlank&7|-|25949913|25949992@50.70.21(+1A)&[MIMAT0000243&hsa-miR-148a-3p&offsets|0|-1(+1A);m-7&7|-|25949922|25949943&offsets|0|-1(+1A)]|TCAGTGCACTACAGAACTTTGA</td><td>>MI0000253|hsa-mir-148a&WithFlank&7|-|25949913|25949992</td><td>50</td><td>70</td><td>21(+1A)</td><td>TCAGTGCACTACAGAACTTTGA</td><td>12274.3</td><td>3.5</td><td>1.2</td><td>2.9</td><td>4.3E-03</td><td>1.9E-02</td><td>BT20/MB231</td><td>MP</td></tr>
<tr><td>>hg38_21_+_8433172_8446622_RNA45SN1_45S_50ntFlanks@4733.4758.26|GAACGAAAGTCGGAGGTTCGAAGACG</td><td>>hg38_21_+_8433172_8446622_RNA45SN1_45S_50ntFlanks</td><td>4733</td><td>4758</td><td>26</td><td>GAACGAAAGTCGGAGGTTCGAAGACG</td><td>1389.6</td><td>1.6</td><td>0.7</td><td>2.4</td><td>1.6E-02</td><td>1.9E-02</td><td>MB231/MB468</td><td>CYTO</td></tr>
<tr><td>>MI0000447|hsa-mir-128-1&WithFlank&2|+|135665391|135665484@56.76.21;MI0000727|hsa-mir-128-2&WithFlank&3|+|35744470|35744565@58.78.21&[MIMAT0000424&hsa-miR-128-3p&offsets|0|0;m-108&2|+|135665446|135665466&offsets|0|0];[MIMAT0000424_1&hsa-miR-128-3p&offsets|0|0;m-117&3|+|35744527|35744547&offsets|0|0]|TCACAGTGAACCGGTCTCTTT</td><td>>MI0000447|hsa-mir-128-1&WithFlank&2|+|135665391|135665484</td><td>56</td><td>76</td><td>21</td><td>TCACAGTGAACCGGTCTCTTT</td><td>9646.5</td><td>2.1</td><td>0.8</td><td>2.8</td><td>5.6E-03</td><td>1.9E-02</td><td>BT20/MB231</td><td>CYTO</td></tr>
<tr><td>>hg38_21_+_8433172_8446622_RNA45SN1_45S_50ntFlanks@5487.5525.39|CGCTGAGAAGACGGTCGAACTTGACTATCTAGAGGAAGT</td><td>>hg38_21_+_8433172_8446622_RNA45SN1_45S_50ntFlanks</td><td>5487</td><td>5525</td><td>39</td><td>CGCTGAGAAGACGGTCGAACTTGACTATCTAGAGGAAGT</td><td>170.9</td><td>-2.8</td><td>1.1</td><td>-2.4</td><td>1.4E-02</td><td>1.9E-02</td><td>BT20/MB468</td><td>MITO</td></tr>
<tr><td>>MI0000447|hsa-mir-128-1&WithFlank&2|+|135665391|135665484@56.76.21(+2A);MI0000727|hsa-mir-128-2&WithFlank&3|+|35744470|35744565@58.78.21(+2A)&[MIMAT0000424&hsa-miR-128-3p&offsets|0|0(+2A);m-108&2|+|135665446|135665466&offsets|0|0(+2A)];[MIMAT0000424_1&hsa-miR-128-3p&offsets|0|0(+2A);m-117&3|+|35744527|35744547&offsets|0|0(+2A)]|TCACAGTGAACCGGTCTCTTTAA</td><td>>MI0000447|hsa-mir-128-1&WithFlank&2|+|135665391|135665484</td><td>56</td><td>76</td><td>21(+2A)</td><td>TCACAGTGAACCGGTCTCTTTAA</td><td>840.6</td><td>2.1</td><td>0.8</td><td>2.8</td><td>5.6E-03</td><td>1.9E-02</td><td>BT20/MB231</td><td>CYTO</td></tr>
<tr><td>>hg38_21_+_8433172_8446622_RNA45SN1_45S_50ntFlanks@6647.6668.22|CGTACGACTCTTAGCGGTGGAT</td><td>>hg38_21_+_8433172_8446622_RNA45SN1_45S_50ntFlanks</td><td>6647</td><td>6668</td><td>22</td><td>CGTACGACTCTTAGCGGTGGAT</td><td>1474.0</td><td>1.9</td><td>0.7</td><td>2.8</td><td>5.6E-03</td><td>1.9E-02</td><td>BT20/MB231</td><td>CYTO</td></tr>
<tr><td>>hg38_21_+_8433172_8446622_RNA45SN1_45S_50ntFlanks@12991.13017.27|CTCCCTCGCTGCGATCTATTGAAAGTC</td><td>>hg38_21_+_8433172_8446622_RNA45SN1_45S_50ntFlanks</td><td>12991</td><td>13017</td><td>27</td><td>CTCCCTCGCTGCGATCTATTGAAAGTC</td><td>524.8</td><td>-2.5</td><td>0.9</td><td>-2.8</td><td>5.6E-03</td><td>1.9E-02</td><td>BT20/MB231</td><td>CYTO</td></tr>
<tr><td>>hg38_21_+_8433172_8446622_RNA45SN1_45S_50ntFlanks@12997.13037.41|CGCTGCGATCTATTGAAAGTCAGCCCTCGACACAAGGGTTT</td><td>>hg38_21_+_8433172_8446622_RNA45SN1_45S_50ntFlanks</td><td>12997</td><td>13037</td><td>41</td><td>CGCTGCGATCTATTGAAAGTCAGCCCTCGACACAAGGGTTT</td><td>1529.5</td><td>-3.1</td><td>1.1</td><td>-2.8</td><td>5.6E-03</td><td>1.9E-02</td><td>BT20/MB231</td><td>CYTO</td></tr>
<tr><td>>hg38_21_+_8433172_8446622_RNA45SN1_45S_50ntFlanks@11579.11609.31|AGAACTGGTGCGGACCAGGGGAATCCGACTG</td><td>>hg38_21_+_8433172_8446622_RNA45SN1_45S_50ntFlanks</td><td>11579</td><td>11609</td><td>31</td><td>AGAACTGGTGCGGACCAGGGGAATCCGACTG</td><td>353.0</td><td>-1.7</td><td>0.7</td><td>-2.4</td><td>1.6E-02</td><td>1.9E-02</td><td>MB231/MB468</td><td>CYTO</td></tr>
<tr><td>>hg38_21_+_8433172_8446622_RNA45SN1_45S_50ntFlanks@9473.9503.31|CACGAGCGCACGTGTTAGGACCCGAAAGATG</td><td>>hg38_21_+_8433172_8446622_RNA45SN1_45S_50ntFlanks</td><td>9473</td><td>9503</td><td>31</td><td>CACGAGCGCACGTGTTAGGACCCGAAAGATG</td><td>666.8</td><td>-2.3</td><td>0.9</td><td>-2.4</td><td>1.4E-02</td><td>1.9E-02</td><td>BT20/MB468</td><td>MITO</td></tr>
<tr><td>>hg38_21_+_8433172_8446622_RNA45SN1_45S_50ntFlanks@6772.6803.32|GTTCCTCCCGGGGCTACGCCTGTCTGAGCGTC</td><td>>hg38_21_+_8433172_8446622_RNA45SN1_45S_50ntFlanks</td><td>6772</td><td>6803</td><td>32</td><td>GTTCCTCCCGGGGCTACGCCTGTCTGAGCGTC</td><td>198.3</td><td>-3.7</td><td>1.6</td><td>-2.4</td><td>1.6E-02</td><td>1.9E-02</td><td>BT20/MB468</td><td>CYTO</td></tr>
<tr><td>>hg38_MT_+_598_1651_RNR1_12S_50ntFlanks@966.1004.39|TAACATGGTAAGTGTACTGGAAAGTGCACTTGGACGAAC</td><td>>hg38_MT_+_598_1651_RNR1_12S_50ntFlanks</td><td>966</td><td>1004</td><td>39</td><td>TAACATGGTAAGTGTACTGGAAAGTGCACTTGGACGAAC</td><td>485.0</td><td>-2.2</td><td>0.8</td><td>-2.8</td><td>5.3E-03</td><td>1.9E-02</td><td>BT20/MB231</td><td>MITO</td></tr>
<tr><td>>hg38_1_-_228634819_228635039_RNA5S12_5S_50ntFlanks@150.171.22|AATACCGGGTGCTGTAGGCTTT</td><td>>hg38_1_-_228634819_228635039_RNA5S12_5S_50ntFlanks</td><td>150</td><td>171</td><td>22</td><td>AATACCGGGTGCTGTAGGCTTT</td><td>2220.8</td><td>2.6</td><td>1.1</td><td>2.4</td><td>1.7E-02</td><td>1.9E-02</td><td>MB231/MB468</td><td>NUC</td></tr>
<tr><td>>am_trna116_GluCTC_1_-_145399233_145399304@7.28.22,trna59_GluCTC_1_+_249168447_249168518@7.28.22,trna71_GluCTC_1_-_161439189_161439260@7.28.22,trna74_GluCTC_1_-_161431809_161431880@7.28.22,trna77_GluCTC_1_-_161424398_161424469@7.28.22,trna77_GluCTC_6_+_28949976_28950047@7.28.22,trna80_GluCTC_1_-_161417018_161417089@7.28.22,trna87_GluCTC_6_-_126101393_126101464@7.28.22|GTGGTCTAGTGGTTAGGATTCG</td><td>>am_trna116_GluCTC_1_-_145399233_145399304</td><td>7</td><td>28</td><td>22</td><td>GTGGTCTAGTGGTTAGGATTCG</td><td>153.4</td><td>-3.8</td><td>1.5</td><td>-2.6</td><td>1.1E-02</td><td>1.9E-02</td><td>BT20/MB468</td><td>MP</td></tr>
<tr><td>>MI0000465|hsa-mir-191&WithFlank&3|-|49020612|49020715@22.41.20(+1A)&[MIMAT0000440&hsa-miR-191-5p&offsets|0|-3(+1A);m-48&3|-|49020672|49020694&offsets|0|-3(+1A)]|CAACGGAATCCCAAAAGCAGA</td><td>>MI0000465|hsa-mir-191&WithFlank&3|-|49020612|49020715</td><td>22</td><td>41</td><td>20(+1A)</td><td>CAACGGAATCCCAAAAGCAGA</td><td>35.9</td><td>5.0</td><td>2.0</td><td>2.4</td><td>1.5E-02</td><td>1.9E-02</td><td>MB231/MB468</td><td>MP</td></tr>
<tr><td>>hg38_21_+_8433172_8446622_RNA45SN1_45S_50ntFlanks@8324.8355.32|CCGATAGTCAACAAGTACCGTAAGGGAAAGTT</td><td>>hg38_21_+_8433172_8446622_RNA45SN1_45S_50ntFlanks</td><td>8324</td><td>8355</td><td>32</td><td>CCGATAGTCAACAAGTACCGTAAGGGAAAGTT</td><td>46.1</td><td>-4.3</td><td>1.7</td><td>-2.6</td><td>1.1E-02</td><td>1.9E-02</td><td>BT20/MB468</td><td>MP</td></tr>
<tr><td>>hg38_21_+_8433172_8446622_RNA45SN1_45S_50ntFlanks@10349.10367.19|AACGAGAACTTTGAAGGCC</td><td>>hg38_21_+_8433172_8446622_RNA45SN1_45S_50ntFlanks</td><td>10349</td><td>10367</td><td>19</td><td>AACGAGAACTTTGAAGGCC</td><td>896.6</td><td>-2.5</td><td>1.0</td><td>-2.4</td><td>1.5E-02</td><td>1.9E-02</td><td>MB231/MB468</td><td>MP</td></tr>
<tr><td>>trna111_HisGTG_1_-_147774845_147774916@-1T.31.32,trna118_HisGTG_1_-_145396881_145396952@-1T.31.32,trna16_HisGTG_1_+_146544773_146544844@-1T.31.32,trna1_HisGTG_15_+_45493349_45493420@-1T.31.32,trna21_HisGTG_1_+_147753471_147753542@-1T.31.32,trna33_HisGTG_6_+_27125906_27125977@-1T.31.32,trna7_HisGTG_9_-_14433938_14434009@-1T.31.32,trna8_HisGTG_15_-_45492611_45492682@-1T.31.32,trna9_HisGTG_15_-_45490804_45490875@-1T.31.32|TGCCGTGATCGTATAGTGGTTAGTACTCTGCG</td><td>>trna111_HisGTG_1_-_147774845_147774916</td><td>-1T</td><td>31</td><td>32</td><td>TGCCGTGATCGTATAGTGGTTAGTACTCTGCG</td><td>1673.7</td><td>-3.4</td><td>1.4</td><td>-2.4</td><td>1.4E-02</td><td>1.9E-02</td><td>BT20/MB468</td><td>MITO</td></tr>
<tr><td>>hg38_MT_+_1621_3279_RNR2_16S_50ntFlanks@999.1026.28|AGGGACCTGTATGAATGGCTCCACGAGG</td><td>>hg38_MT_+_1621_3279_RNR2_16S_50ntFlanks</td><td>999</td><td>1026</td><td>28</td><td>AGGGACCTGTATGAATGGCTCCACGAGG</td><td>70.9</td><td>4.7</td><td>1.9</td><td>2.6</td><td>1.1E-02</td><td>1.9E-02</td><td>BT20/MB468</td><td>MP</td></tr>
<tr><td>>hg38_21_+_8433172_8446622_RNA45SN1_45S_50ntFlanks@10922.10947.26|CTGGACGAGGCGCCGCCGCCCCCCCC</td><td>>hg38_21_+_8433172_8446622_RNA45SN1_45S_50ntFlanks</td><td>10922</td><td>10947</td><td>26</td><td>CTGGACGAGGCGCCGCCGCCCCCCCC</td><td>233.4</td><td>-4.5</td><td>1.8</td><td>-2.6</td><td>1.1E-02</td><td>1.9E-02</td><td>BT20/MB468</td><td>MP</td></tr>
</table>

<table>
<tr><td>>hg38_21_+_8433172_8446622_RNA45SN1_45S_50ntFlanks@9565.9595.31|TGACGTGCAAATCGGTCGTCCGACCTGGGTA</td><td>>hg38_21_+_8433172_8446622_RNA45SN1_45S_50ntFlanks</td><td>9565</td><td>9595</td><td>31</td><td>TGACGTGCAAATCGGTCGTCCGACCTGGGTA</td><td>887.9</td><td>-1.9</td><td>0.8</td><td>-2.4</td><td>1.4E-02</td><td>1.9E-02</td><td>BT20/MB468</td><td>MITO</td></tr>
<tr><td>>hg38_21_+_8433172_8446622_RNA45SN1_45S_50ntFlanks@10922.10944.23|CTGGACGAGGCGCCGCCGCCCCC</td><td>>hg38_21_+_8433172_8446622_RNA45SN1_45S_50ntFlanks</td><td>10922</td><td>10944</td><td>23</td><td>CTGGACGAGGCGCCGCCGCCCCC</td><td>335.6</td><td>-4.5</td><td>1.9</td><td>-2.4</td><td>1.7E-02</td><td>1.9E-02</td><td>BT20/MB468</td><td>NUC</td></tr>
<tr><td>>hg38_21_+_8433172_8446622_RNA45SN1_45S_50ntFlanks@6645.6670.26|CTCGTACGACTCTTAGCGGTGGATCA</td><td>>hg38_21_+_8433172_8446622_RNA45SN1_45S_50ntFlanks</td><td>6645</td><td>6670</td><td>26</td><td>CTCGTACGACTCTTAGCGGTGGATCA</td><td>538.4</td><td>-1.1</td><td>0.5</td><td>-2.4</td><td>1.7E-02</td><td>1.9E-02</td><td>MB231/MB468</td><td>NUC</td></tr>
<tr><td>>hg38_MT_+_1621_3279_RNR2_16S_50ntFlanks@1428.1458.31|ATTAAAGTCCTACGTGATCTGAGTTCAGACC</td><td>>hg38_MT_+_1621_3279_RNR2_16S_50ntFlanks</td><td>1428</td><td>1458</td><td>31</td><td>ATTAAAGTCCTACGTGATCTGAGTTCAGACC</td><td>558.0</td><td>3.4</td><td>1.3</td><td>2.6</td><td>1.1E-02</td><td>1.9E-02</td><td>BT20/MB468</td><td>MP</td></tr>
<tr><td>>hg38_21_+_8433172_8446622_RNA45SN1_45S_50ntFlanks@10839.10868.30|TCCGTAACTTCGGGATAAGGATTGGCTCTA</td><td>>hg38_21_+_8433172_8446622_RNA45SN1_45S_50ntFlanks</td><td>10839</td><td>10868</td><td>30</td><td>TCCGTAACTTCGGGATAAGGATTGGCTCTA</td><td>100.3</td><td>-3.3</td><td>1.3</td><td>-2.6</td><td>1.1E-02</td><td>1.9E-02</td><td>BT20/MB468</td><td>MP</td></tr>
<tr><td>>hg38_21_+_8433172_8446622_RNA45SN1_45S_50ntFlanks@11603.11626.24|CCGACTGTTTAATTAAAACAAAGC</td><td>>hg38_21_+_8433172_8446622_RNA45SN1_45S_50ntFlanks</td><td>11603</td><td>11626</td><td>24</td><td>CCGACTGTTTAATTAAAACAAAGC</td><td>201.8</td><td>5.7</td><td>2.0</td><td>2.9</td><td>4.3E-03</td><td>1.9E-02</td><td>BT20/MB231</td><td>MP</td></tr>
<tr><td>>hg38_21_+_8433172_8446622_RNA45SN1_45S_50ntFlanks@10682.10706.25|TCCGGCGGCGTCCGGTGAGCTCTCG</td><td>>hg38_21_+_8433172_8446622_RNA45SN1_45S_50ntFlanks</td><td>10682</td><td>10706</td><td>25</td><td>TCCGGCGGCGTCCGGTGAGCTCTCG</td><td>793.4</td><td>-4.7</td><td>1.8</td><td>-2.6</td><td>1.1E-02</td><td>1.9E-02</td><td>BT20/MB468</td><td>MP</td></tr>
<tr><td>>hg38_21_+_8433172_8446622_RNA45SN1_45S_50ntFlanks@11570.11601.32|AGCCGACTTAGAACTGGTGCGGACCAGGGGAA</td><td>>hg38_21_+_8433172_8446622_RNA45SN1_45S_50ntFlanks</td><td>11570</td><td>11601</td><td>32</td><td>AGCCGACTTAGAACTGGTGCGGACCAGGGGAA</td><td>967.3</td><td>-1.7</td><td>0.7</td><td>-2.4</td><td>1.4E-02</td><td>1.9E-02</td><td>BT20/MB468</td><td>MITO</td></tr>
<tr><td>>hg38_MT_+_1621_3279_RNR2_16S_50ntFlanks@1183.1215.33|AAATTTCGGTTGGGGCGACCTCGGAGCAGAACC</td><td>>hg38_MT_+_1621_3279_RNR2_16S_50ntFlanks</td><td>1183</td><td>1215</td><td>33</td><td>AAATTTCGGTTGGGGCGACCTCGGAGCAGAACC</td><td>96.1</td><td>2.9</td><td>1.2</td><td>2.4</td><td>1.5E-02</td><td>1.9E-02</td><td>MB231/MB468</td><td>MP</td></tr>
<tr><td>>hg38_21_+_8433172_8446622_RNA45SN1_45S_50ntFlanks@11703.11728.26|AATGAAGCGCGGGTAAACGGCGGGAG</td><td>>hg38_21_+_8433172_8446622_RNA45SN1_45S_50ntFlanks</td><td>11703</td><td>11728</td><td>26</td><td>AATGAAGCGCGGGTAAACGGCGGGAG</td><td>1567.6</td><td>-4.4</td><td>1.8</td><td>-2.4</td><td>1.7E-02</td><td>1.9E-02</td><td>BT20/MB468</td><td>CYTO</td></tr>
<tr><td>>MI0000483|hsa-mir-186&WithFlank&1|-|71067625|71067722@21.42.22(+1C)&[MIMAT0000456&hsa-miR-186-5p&offsets|0|0(+1C);m-84&1|-|71067681|71067702&offsets|0|0(+1C)]|CAAAGAATTCTCCTTTTGGGCTC</td><td>>MI0000483|hsa-mir-186&WithFlank&1|-|71067625|71067722</td><td>21</td><td>42</td><td>22(+1C)</td><td>CAAAGAATTCTCCTTTTGGGCTC</td><td>380.8</td><td>-2.9</td><td>1.0</td><td>-2.9</td><td>4.3E-03</td><td>1.9E-02</td><td>BT20/MB231</td><td>MP</td></tr>
<tr><td>>trnaMT_PheGAA_MT_+_577_647@44.74.31|TAGACGGGCTCACATCACCCCATAAACACCA</td><td>>trnaMT_PheGAA_MT_+_577_647</td><td>44</td><td>74</td><td>31</td><td>TAGACGGGCTCACATCACCCCATAAACACCA</td><td>977.0</td><td>2.8</td><td>1.1</td><td>2.4</td><td>1.4E-02</td><td>1.9E-02</td><td>BT20/MB468</td><td>MITO</td></tr>
<tr><td>>hg38_21_+_8433172_8446622_RNA45SN1_45S_50ntFlanks@9201.9218.18|CTCTCGGGGCCACGCGCG</td><td>>hg38_21_+_8433172_8446622_RNA45SN1_45S_50ntFlanks</td><td>9201</td><td>9218</td><td>18</td><td>CTCTCGGGGCCACGCGCG</td><td>555.5</td><td>-4.1</td><td>1.7</td><td>-2.4</td><td>1.4E-02</td><td>1.9E-02</td><td>BT20/MB468</td><td>MITO</td></tr>
<tr><td>>hg38_21_+_8433172_8446622_RNA45SN1_45S_50ntFlanks@9534.9566.33|AGGAAACTCTGGTGGAGGTCCGTAGCGGTCCTG</td><td>>hg38_21_+_8433172_8446622_RNA45SN1_45S_50ntFlanks</td><td>9534</td><td>9566</td><td>33</td><td>AGGAAACTCTGGTGGAGGTCCGTAGCGGTCCTG</td><td>1435.2</td><td>-4.8</td><td>1.9</td><td>-2.6</td><td>1.1E-02</td><td>1.9E-02</td><td>BT20/MB468</td><td>MP</td></tr>
<tr><td>>hg38_21_+_8433172_8446622_RNA45SN1_45S_50ntFlanks@12439.12473.35|TTGTGAAGCAGAATTCACCAAGCGTTGGATTGTTC</td><td>>hg38_21_+_8433172_8446622_RNA45SN1_45S_50ntFlanks</td><td>12439</td><td>12473</td><td>35</td><td>TTGTGAAGCAGAATTCACCAAGCGTTGGATTGTTC</td><td>632.7</td><td>1.8</td><td>0.7</td><td>2.4</td><td>1.7E-02</td><td>1.9E-02</td><td>MB231/MB468</td><td>CYTO</td></tr>
<tr><td>>hg38_21_+_8433172_8446622_RNA45SN1_45S_50ntFlanks@12999.13029.31|CTGCGATCTATTGAAAGTCAGCCCTCGACAC</td><td>>hg38_21_+_8433172_8446622_RNA45SN1_45S_50ntFlanks</td><td>12999</td><td>13029</td><td>31</td><td>CTGCGATCTATTGAAAGTCAGCCCTCGACAC</td><td>261.2</td><td>1.4</td><td>0.6</td><td>2.4</td><td>1.7E-02</td><td>1.9E-02</td><td>MB231/MB468</td><td>CYTO</td></tr>
<tr><td>>hg38_21_+_8433172_8446622_RNA45SN1_45S_50ntFlanks@11027.11062.36|CTCTCTCTCTCTCTCCCCCGCTCCCCGTCCTCCCCC</td><td>>hg38_21_+_8433172_8446622_RNA45SN1_45S_50ntFlanks</td><td>11027</td><td>11062</td><td>36</td><td>CTCTCTCTCTCTCTCCCCCGCTCCCCGTCCTCCCCC</td><td>186.6</td><td>-3.9</td><td>1.6</td><td>-2.4</td><td>1.7E-02</td><td>1.9E-02</td><td>BT20/MB468</td><td>CYTO</td></tr>
<tr><td>>hg38_21_+_8433172_8446622_RNA45SN1_45S_50ntFlanks@4723.4747.25|CATTAATCAAGAACGAAAGTCGGAG</td><td>>hg38_21_+_8433172_8446622_RNA45SN1_45S_50ntFlanks</td><td>4723</td><td>4747</td><td>25</td><td>CATTAATCAAGAACGAAAGTCGGAG</td><td>111.1</td><td>-4.3</td><td>1.7</td><td>-2.6</td><td>1.1E-02</td><td>1.9E-02</td><td>BT20/MB468</td><td>MP</td></tr>
<tr><td>>MI0000115|hsa-mir-16-2&WithFlank&3|+|160404739|160404831@58.78.21&[MIMAT0004518&hsa-miR-16-2-3p&offsets|-1|-2;m-252&3|+|160404796|160404816&offsets|0|0]|ACCAATATTACTGTGCTGCTT</td><td>>MI0000115|hsa-mir-16-2&WithFlank&3|+|160404739|160404831</td><td>58</td><td>78</td><td>21</td><td>ACCAATATTACTGTGCTGCTT</td><td>76.1</td><td>4.3</td><td>1.8</td><td>2.4</td><td>1.5E-02</td><td>1.9E-02</td><td>MB231/MB468</td><td>MP</td></tr>
<tr><td>>MI0000750|hsa-mir-26a-2&WithFlank&12|-|57824603|57824698@20.38.19(+1A);MI0000083|hsa-mir-26a-1&WithFlank&3|+|37969398|37969486@16.34.19(+1A)&[MIMAT0000082&hsa-miR-26a-5p&offsets|0|-3(+1A);m-33&12|-|57824658|57824679&offsets|0|-3(+1A)];[MIMAT0000082_1&hsa-miR-26a-5p&offsets|0|-3(+1A);m-34&3|+|37969413|37969434&offsets|0|-3(+1A)]|TTCAAGTAATCCAGGATAGA</td><td>>MI0000750|hsa-mir-26a-2&WithFlank&12|-|57824603|57824698</td><td>20</td><td>38</td><td>19(+1A)</td><td>TTCAAGTAATCCAGGATAGA</td><td>51.3</td><td>4.9</td><td>2.0</td><td>2.4</td><td>1.5E-02</td><td>1.9E-02</td><td>MB231/MB468</td><td>MP</td></tr>
<tr><td>>hg38_21_+_8433172_8446622_RNA45SN1_45S_50ntFlanks@11552.11569.18|CCTCGGCCGGCGCCTAGC</td><td>>hg38_21_+_8433172_8446622_RNA45SN1_45S_50ntFlanks</td><td>11552</td><td>11569</td><td>18</td><td>CCTCGGCCGGCGCCTAGC</td><td>595.5</td><td>-1.7</td><td>0.7</td><td>-2.4</td><td>1.4E-02</td><td>1.9E-02</td><td>BT20/MB468</td><td>MITO</td></tr>
<tr><td>>MI0000088|hsa-mir-30a&WithFlank&6|-|71403545|71403627@54.74.21(+1C)&[MIMAT0000088&hsa-miR-30a-3p&offsets|+1|0(+1C);m-38&6|-|71403554|71403575&offsets|+1|0(+1C)]|TTTCAGTCGGATGTTTGCAGCC</td><td>>MI0000088|hsa-mir-30a&WithFlank&6|-|71403545|71403627</td><td>54</td><td>74</td><td>21(+1C)</td><td>TTTCAGTCGGATGTTTGCAGCC</td><td>201.4</td><td>-5.3</td><td>1.9</td><td>-2.9</td><td>4.3E-03</td><td>1.9E-02</td><td>BT20/MB231</td><td>MP</td></tr>
<tr><td>>MI0000815|hsa-mir-339&WithFlank&7|-|1022927|1023032@56.78.23&[MIMAT0004702&hsa-miR-339-3p&offsets|0|0;m-235&7|-|1022957|1022977&offsets|0|+2]|TGAGCGCCTCGACGACAGAGCCG</td><td>>MI0000815|hsa-mir-339&WithFlank&7|-|1022927|1023032</td><td>56</td><td>78</td><td>23</td><td>TGAGCGCCTCGACGACAGAGCCG</td><td>189.6</td><td>3.1</td><td>1.3</td><td>2.4</td><td>1.5E-02</td><td>1.9E-02</td><td>MB231/MB468</td><td>MP</td></tr>
<tr><td>>hg38_21_+_8433172_8446622_RNA45SN1_45S_50ntFlanks@6646.6663.18|TCGTACGACTCTTAGCGG</td><td>>hg38_21_+_8433172_8446622_RNA45SN1_45S_50ntFlanks</td><td>6646</td><td>6663</td><td>18</td><td>TCGTACGACTCTTAGCGG</td><td>39676.7</td><td>2.4</td><td>1.0</td><td>2.4</td><td>1.7E-02</td><td>1.9E-02</td><td>MB231/MB468</td><td>MITO</td></tr>
<tr><td>>hg38_21_+_8433172_8446622_RNA45SN1_45S_50ntFlanks@4762.4783.22|AGATACCGTCGTAGTTCCGACC</td><td>>hg38_21_+_8433172_8446622_RNA45SN1_45S_50ntFlanks</td><td>4762</td><td>4783</td><td>22</td><td>AGATACCGTCGTAGTTCCGACC</td><td>211.3</td><td>2.6</td><td>1.0</td><td>2.5</td><td>1.1E-02</td><td>2.0E-02</td><td>BT20/MB468</td><td>MP</td></tr>
<tr><td>>hg38_21_+_8433172_8446622_RNA45SN1_45S_50ntFlanks@9554.9579.26|CGTAGCGGTCCTGACGTGCAAATCGG</td><td>>hg38_21_+_8433172_8446622_RNA45SN1_45S_50ntFlanks</td><td>9554</td><td>9579</td><td>26</td><td>CGTAGCGGTCCTGACGTGCAAATCGG</td><td>62.4</td><td>-4.2</td><td>1.6</td><td>-2.5</td><td>1.1E-02</td><td>2.0E-02</td><td>BT20/MB468</td><td>MP</td></tr>
<tr><td>>hg38_21_+_8433172_8446622_RNA45SN1_45S_50ntFlanks@6663.6700.38|GTGGATCACTCGGCTCGTGCGTCGATGAAGAACGCAGC</td><td>>hg38_21_+_8433172_8446622_RNA45SN1_45S_50ntFlanks</td><td>6663</td><td>6700</td><td>38</td><td>GTGGATCACTCGGCTCGTGCGTCGATGAAGAACGCAGC</td><td>110.0</td><td>3.5</td><td>1.4</td><td>2.5</td><td>1.1E-02</td><td>2.0E-02</td><td>BT20/MB468</td><td>MP</td></tr>
<tr><td>>hg38_21_+_8433172_8446622_RNA45SN1_45S_50ntFlanks@12893.12930.38|CCTCGCCCGTCACGCACCGCACGTTCGTGGGGAACCTG</td><td>>hg38_21_+_8433172_8446622_RNA45SN1_45S_50ntFlanks</td><td>12893</td><td>12930</td><td>38</td><td>CCTCGCCCGTCACGCACCGCACGTTCGTGGGGAACCTG</td><td>404.5</td><td>-3.8</td><td>1.6</td><td>-2.4</td><td>1.7E-02</td><td>2.0E-02</td><td>BT20/MB468</td><td>NUC</td></tr>
<tr><td>>MI0000234|hsa-mir-192&WithFlank&11|-|64891131|64891252@30.49.20(+1U)&[MIMAT0000222&hsa-miR-192-5p&offsets|0|-1(+1U);m-59&11|-|64891203|64891223&offsets|0|-1(+1U)]|CTGACCTATGAATTGACAGCT</td><td>>MI0000234|hsa-mir-192&WithFlank&11|-|64891131|64891252</td><td>30</td><td>49</td><td>20(+1U)</td><td>CTGACCTATGAATTGACAGCT</td><td>368.6</td><td>2.3</td><td>0.8</td><td>2.8</td><td>5.7E-03</td><td>2.0E-02</td><td>BT20/MB231</td><td>CYTO</td></tr>
<tr><td>>am_MI0000786|hsa-mir-378a&WithFlank&5|+|149732819|149732896@50.70.21(+1A)&[MIMAT0000732&hsa-miR-378a-3p&offsets|+1|0(+1A);m-44&5|+|149732867|149732888&offsets|+1|0(+1A)]|CTGGACTTGGAGTCAGAAGGCA</td><td>>am_MI0000786|hsa-mir-378a&WithFlank&5|+|149732819|149732896</td><td>50</td><td>70</td><td>21(+1A)</td><td>CTGGACTTGGAGTCAGAAGGCA</td><td>490.2</td><td>2.1</td><td>0.8</td><td>2.8</td><td>5.7E-03</td><td>2.0E-02</td><td>BT20/MB231</td><td>CYTO</td></tr>
<tr><td>>trnaMT_ValTAC_MT_+_1602_1670@51.71.21|CTTAACTTGACCGCTCTGACC</td><td>>trnaMT_ValTAC_MT_+_1602_1670</td><td>51</td><td>71</td><td>21</td><td>CTTAACTTGACCGCTCTGACC</td><td>886.6</td><td>-2.6</td><td>1.1</td><td>-2.4</td><td>1.7E-02</td><td>2.0E-02</td><td>MB231/MB468</td><td>NUC</td></tr>
<tr><td>>MI0000299|hsa-mir-222&WithFlank&X|-|45747009|45747130@75.99.25(+1C)&[MIMAT0000279&hsa-miR-222-3p&offsets|0|+4(+1C);m-140&X|-|45747033|45747056&offsets|0|+1(+1C)]|AGCTACATCTGGCTACTGGGTCTCTC</td><td>>MI0000299|hsa-mir-222&WithFlank&X|-|45747009|45747130</td><td>75</td><td>99</td><td>25(+1C)</td><td>AGCTACATCTGGCTACTGGGTCTCTC</td><td>295.2</td><td>-2.2</td><td>0.8</td><td>-2.8</td><td>4.7E-03</td><td>2.0E-02</td><td>BT20/MB231</td><td>NUC</td></tr>
<tr><td>>hg38_21_+_8433172_8446622_RNA45SN1_45S_50ntFlanks@10349.10380.32|AACGAGAACTTTGAAGGCCGAAGTGGAGAAGG</td><td>>hg38_21_+_8433172_8446622_RNA45SN1_45S_50ntFlanks</td><td>10349</td><td>10380</td><td>32</td><td>AACGAGAACTTTGAAGGCCGAAGTGGAGAAGG</td><td>336.8</td><td>1.6</td><td>0.7</td><td>2.4</td><td>1.7E-02</td><td>2.0E-02</td><td>MB231/MB468</td><td>CYTO</td></tr>
<tr><td>>trnaMT_ProTGG_MT_-_15956_16023@40.57.18|AATGGTGGAGTTAAAGAC</td><td>>trnaMT_ProTGG_MT_-_15956_16023</td><td>40</td><td>57</td><td>18</td><td>AATGGTGGAGTTAAAGAC</td><td>76.0</td><td>3.9</td><td>1.6</td><td>2.4</td><td>1.6E-02</td><td>2.0E-02</td><td>MB231/MB468</td><td>MP</td></tr>
<tr><td>>hg38_21_+_8433172_8446622_RNA45SN1_45S_50ntFlanks@5191.5226.36|ACAGGTCTGTGATGCCCTTAGATGTCCGGGGCTGCA</td><td>>hg38_21_+_8433172_8446622_RNA45SN1_45S_50ntFlanks</td><td>5191</td><td>5226</td><td>36</td><td>ACAGGTCTGTGATGCCCTTAGATGTCCGGGGCTGCA</td><td>66.7</td><td>-5.3</td><td>2.2</td><td>-2.4</td><td>1.6E-02</td><td>2.0E-02</td><td>MB231/MB468</td><td>MP</td></tr>
<tr><td>>hg38_21_+_8433172_8446622_RNA45SN1_45S_50ntFlanks@9604.9626.23|AAAGACTAATCGAACCATCTAGT</td><td>>hg38_21_+_8433172_8446622_RNA45SN1_45S_50ntFlanks</td><td>9604</td><td>9626</td><td>23</td><td>AAAGACTAATCGAACCATCTAGT</td><td>142.6</td><td>-1.9</td><td>0.8</td><td>-2.4</td><td>1.6E-02</td><td>2.0E-02</td><td>MB231/MB468</td><td>MP</td></tr>
<tr><td>>hg38_21_+_8433172_8446622_RNA45SN1_45S_50ntFlanks@11212.11233.22|CGACTCTGGACGCGAGCCGGGC</td><td>>hg38_21_+_8433172_8446622_RNA45SN1_45S_50ntFlanks</td><td>11212</td><td>11233</td><td>22</td><td>CGACTCTGGACGCGAGCCGGGC</td><td>64.6</td><td>-4.2</td><td>1.6</td><td>-2.5</td><td>1.1E-02</td><td>2.0E-02</td><td>BT20/MB468</td><td>MP</td></tr>
<tr><td>>hg38_MT_+_598_1651_RNR1_12S_50ntFlanks@905.924.20|AGTATACTTCAAAGGACATT</td><td>>hg38_MT_+_598_1651_RNR1_12S_50ntFlanks</td><td>905</td><td>924</td><td>20</td><td>AGTATACTTCAAAGGACATT</td><td>138.4</td><td>4.2</td><td>1.7</td><td>2.4</td><td>1.6E-02</td><td>2.0E-02</td><td>MB231/MB468</td><td>MP</td></tr>
<tr><td>>MI0000065|hsa-let-7d&WithFlank&9|+|94178828|94178926@14.34.21(+1A)&[MIMAT0000065&hsa-let-7d-5p&offsets|0|-1(+1A);m-90&9|+|94178841|94178862&offsets|0|-1(+1A)]|AGAGGTAGTAGGTTGCATAGTA</td><td>>MI0000065|hsa-let-7d&WithFlank&9|+|94178828|94178926</td><td>14</td><td>34</td><td>21(+1A)</td><td>AGAGGTAGTAGGTTGCATAGTA</td><td>608.8</td><td>2.5</td><td>0.9</td><td>2.8</td><td>5.7E-03</td><td>2.0E-02</td><td>BT20/MB231</td><td>CYTO</td></tr>
<tr><td>>MI0005559|hsa-mir-744&WithFlank&17|+|12081893|12082002@17.36.20(+1U)&[MIMAT0004945&hsa-miR-744-5p&offsets|0|-2(+1U);m-172&17|+|12081909|12081929&offsets|0|-1(+1U)]|TGCGGGGCTAGGGCTAACAGT</td><td>>MI0005559|hsa-mir-744&WithFlank&17|+|12081893|12082002</td><td>17</td><td>36</td><td>20(+1U)</td><td>TGCGGGGCTAGGGCTAACAGT</td><td>816.7</td><td>2.1</td><td>0.7</td><td>2.8</td><td>5.7E-03</td><td>2.0E-02</td><td>BT20/MB231</td><td>CYTO</td></tr>
<tr><td>>hg38_21_+_8433172_8446622_RNA45SN1_45S_50ntFlanks@3882.3913.32|CCGACGGGCGCTGACCCCCTTCGCGGGGGGGA</td><td>>hg38_21_+_8433172_8446622_RNA45SN1_45S_50ntFlanks</td><td>3882</td><td>3913</td><td>32</td><td>CCGACGGGCGCTGACCCCCTTCGCGGGGGGGA</td><td>281.6</td><td>-2.5</td><td>0.9</td><td>-2.8</td><td>5.7E-03</td><td>2.0E-02</td><td>BT20/MB231</td><td>CYTO</td></tr>
<tr><td>>hg38_21_+_8433172_8446622_RNA45SN1_45S_50ntFlanks@10727.10757.31|GGAGAGGGTGTAAATCTCGCGCCGGGCCGTA</td><td>>hg38_21_+_8433172_8446622_RNA45SN1_45S_50ntFlanks</td><td>10727</td><td>10757</td><td>31</td><td>GGAGAGGGTGTAAATCTCGCGCCGGGCCGTA</td><td>193.3</td><td>-3.7</td><td>1.5</td><td>-2.4</td><td>1.5E-02</td><td>2.0E-02</td><td>BT20/MB468</td><td>MITO</td></tr>
<tr><td>>MI0000734|hsa-mir-106b&WithFlank&7|-|100093987|100094080@18.38.21&[MIMAT0000680&hsa-miR-106b-5p&offsets|0|0;m-103&7|-|100094043|100094063&offsets|0|0]|TAAAGTGCTGACAGTGCAGAT</td><td>>MI0000734|hsa-mir-106b&WithFlank&7|-|100093987|100094080</td><td>18</td><td>38</td><td>21</td><td>TAAAGTGCTGACAGTGCAGAT</td><td>817.0</td><td>2.6</td><td>1.1</td><td>2.4</td><td>1.6E-02</td><td>2.0E-02</td><td>MB231/MB468</td><td>MP</td></tr>
<tr><td>>hg38_21_+_8433172_8446622_RNA45SN1_45S_50ntFlanks@6734.6772.39|ATTGATCATCGACACTTCGAACGCACTTGCGGCCCCGGG</td><td>>hg38_21_+_8433172_8446622_RNA45SN1_45S_50ntFlanks</td><td>6734</td><td>6772</td><td>39</td><td>ATTGATCATCGACACTTCGAACGCACTTGCGGCCCCGGG</td><td>426.6</td><td>-2.2</td><td>0.9</td><td>-2.4</td><td>1.7E-02</td><td>2.0E-02</td><td>BT20/MB468</td><td>NUC</td></tr>
<tr><td>>hg38_21_+_8433172_8446622_RNA45SN1_45S_50ntFlanks@12419.12444.26|GATGTCGGCTCTTCCTATCATTGTGA</td><td>>hg38_21_+_8433172_8446622_RNA45SN1_45S_50ntFlanks</td><td>12419</td><td>12444</td><td>26</td><td>GATGTCGGCTCTTCCTATCATTGTGA</td><td>2726.3</td><td>-2.3</td><td>0.8</td><td>-2.8</td><td>5.7E-03</td><td>2.0E-02</td><td>BT20/MB231</td><td>CYTO</td></tr>
<tr><td>>hg38_MT_+_1621_3279_RNR2_16S_50ntFlanks@125.150.26|TAGGCGATAGAAATTGAAACCTGGCG</td><td>>hg38_MT_+_1621_3279_RNR2_16S_50ntFlanks</td><td>125</td><td>150</td><td>26</td><td>TAGGCGATAGAAATTGAAACCTGGCG</td><td>54.5</td><td>4.1</td><td>1.7</td><td>2.4</td><td>1.6E-02</td><td>2.0E-02</td><td>MB231/MB468</td><td>MP</td></tr>
<tr><td>>hg38_21_+_8433172_8446622_RNA45SN1_45S_50ntFlanks@9459.9494.36|CCGGGGAGGTGGAGCACGAGCGCACGTGTTAGGACC</td><td>>hg38_21_+_8433172_8446622_RNA45SN1_45S_50ntFlanks</td><td>9459</td><td>9494</td><td>36</td><td>CCGGGGAGGTGGAGCACGAGCGCACGTGTTAGGACC</td><td>252.3</td><td>-4.5</td><td>1.8</td><td>-2.5</td><td>1.1E-02</td><td>2.0E-02</td><td>BT20/MB468</td><td>MP</td></tr>
<tr><td>>hg38_21_+_8433172_8446622_RNA45SN1_45S_50ntFlanks@9157.9187.31|CAGTGCGCCCCGGGCGGGTCGCGCCGTCGGG</td><td>>hg38_21_+_8433172_8446622_RNA45SN1_45S_50ntFlanks</td><td>9157</td><td>9187</td><td>31</td><td>CAGTGCGCCCCGGGCGGGTCGCGCCGTCGGG</td><td>208.1</td><td>-3.0</td><td>1.3</td><td>-2.4</td><td>1.8E-02</td><td>2.0E-02</td><td>MB231/MB468</td><td>NUC</td></tr>
</table>

| | | | | | | | | | | | | | |
|---|---|---|---|---|---|---|---|---|---|---|---|---|---|
| >hg38_21_+_8433172_8446622_RNA45SN1_45S_50ntFlanks@11667.11704.38\|CTGCCCAGTGCTCTGAATGTCAAAGTGAAGAAATTCAA | >hg38_21_+_8433172_8446622_RNA45SN1_45S_50ntFlanks | 11667 | 11704 | 38 | CTGCCCAGTGCTCTGAATGTCAAAGTGAAGAAATTCAA | 225.2 | -2.0 | 0.7 | -2.8 | 5.7E-03 | 2.0E-02 | BT20/MB231 | CYTO |
| >hg38_21_+_8433172_8446622_RNA45SN1_45S_50ntFlanks@4773.4793.21\|TAGTTCCGACCATAAACGATG | >hg38_21_+_8433172_8446622_RNA45SN1_45S_50ntFlanks | 4773 | 4793 | 21 | TAGTTCCGACCATAAACGATG | 799.1 | -4.6 | 1.8 | -2.5 | 1.1E-02 | 2.0E-02 | BT20/MB468 | MP |
| >MI0000764\|hsa-mir-363&WithFlank&X\|-\|134169372\|134169458@56.76.21&[MIMAT0000707&hsa-miR-363-3p&offsets\|0\|-1;m-178&X\|-\|134169381\|134169403&offsets\|0\|-2]\|AATTGCACGGTATCCATCTGT | >MI0000764\|hsa-mir-363&WithFlank&X\|-\|134169372\|134169458 | 56 | 76 | 21 | AATTGCACGGTATCCATCTGT | 233.7 | 2.9 | 1.2 | 2.5 | 1.1E-02 | 2.0E-02 | BT20/MB468 | MP |
| >hg38_21_+_8433172_8446622_RNA45SN1_45S_50ntFlanks@4739.4765.27\|AAGTCGGAGGTTCGAAGACGATCAGAT | >hg38_21_+_8433172_8446622_RNA45SN1_45S_50ntFlanks | 4739 | 4765 | 27 | AAGTCGGAGGTTCGAAGACGATCAGAT | 110.3 | 3.6 | 1.4 | 2.5 | 1.1E-02 | 2.0E-02 | BT20/MB468 | MP |
| >MI0000070\|hsa-mir-16-1&WithFlank&13\|-\|50048967\|50049067@20.40.21;MI0000115\|hsa-mir-16-2&WithFlank&3\|+\|160404739\|160404831@16.36.21&[MIMAT0000069&hsa-miR-16-5p&offsets\|0\|-1;m-62&13\|-\|50049027\|50049048&offsets\|0\|-1];[MIMAT0000069_1&hsa-miR-16-5p&offsets\|0\|-1;m-61&3\|+\|160404754\|160404775&offsets\|0\|-1]\|TAGCAGCACGTAAATATTGGC | >MI0000070\|hsa-mir-16-1&WithFlank&13\|-\|50048967\|50049067 | 20 | 40 | 21 | TAGCAGCACGTAAATATTGGC | 1848.5 | 3.0 | 1.2 | 2.5 | 1.1E-02 | 2.0E-02 | BT20/MB468 | MP |
| >hg38_21_+_8433172_8446622_RNA45SN1_45S_50ntFlanks@8552.8577.26\|CTCCCCGGAGGGGGCGGGCTCCGGCG | >hg38_21_+_8433172_8446622_RNA45SN1_45S_50ntFlanks | 8552 | 8577 | 26 | CTCCCCGGAGGGGGCGGGCTCCGGCG | 353.6 | -3.9 | 1.6 | -2.4 | 1.5E-02 | 2.0E-02 | BT20/MB468 | MITO |
| >MI0000736\|hsa-mir-30c-1&WithFlank&1\|+\|40757278\|40757378@23.45.23(+1A);MI0000254\|hsa-mir-30c-2&WithFlank&6\|-\|71376954\|71377037@13.35.23(+1A)&[MIMAT0000244&hsa-miR-30c-5p&offsets\|0\|0(+1A);m-57&1\|+\|40757300\|40757323&offsets\|0\|-1(+1A)];[MIMAT0000244_1&hsa-miR-30c-5p&offsets\|0\|0(+1A);m-56&6\|-\|71377002\|71377025&offsets\|0\|-1(+1A)]\|TGTAAACATCCTACACTCTCAGCA | >MI0000736\|hsa-mir-30c-1&WithFlank&1\|+\|40757278\|40757378 | 23 | 45 | 23(+1A) | TGTAAACATCCTACACTCTCAGCA | 1972.1 | -3.4 | 1.2 | -2.9 | 4.3E-03 | 2.0E-02 | BT20/MB231 | MP |
| >hg38_MT_+_598_1651_RNR1_12S_50ntFlanks@428.462.35\|AAAGTGGCTTTAACATATCTGAACACACAATAGCT | >hg38_MT_+_598_1651_RNR1_12S_50ntFlanks | 428 | 462 | 35 | AAAGTGGCTTTAACATATCTGAACACACAATAGCT | 147.2 | 3.9 | 1.5 | 2.5 | 1.1E-02 | 2.0E-02 | BT20/MB468 | MP |
| >hg38_21_+_8433172_8446622_RNA45SN1_45S_50ntFlanks@6646.6666.21\|TCGTACGACTCTTAGCGGTGG | >hg38_21_+_8433172_8446622_RNA45SN1_45S_50ntFlanks | 6646 | 6666 | 21 | TCGTACGACTCTTAGCGGTGG | 3788.3 | -1.7 | 0.7 | -2.4 | 1.6E-02 | 2.0E-02 | MB231/MB468 | MP |
| >hg38_MT_+_1621_3279_RNR2_16S_50ntFlanks@827.855.29\|AGGAAAGGTTAAAAAAAGTAAAAGGAACT | >hg38_MT_+_1621_3279_RNR2_16S_50ntFlanks | 827 | 855 | 29 | AGGAAAGGTTAAAAAAAGTAAAAGGAACT | 117.2 | 4.5 | 1.8 | 2.5 | 1.1E-02 | 2.0E-02 | BT20/MB468 | MP |
| >MI0000067\|hsa-let-7f-1&WithFlank&9\|+\|94176341\|94176439@69.89.21(+1A)&[MIMAT0004486&hsa-let-7f-1-3p&offsets\|0\|-1(+1A);m-275&9\|+\|94176409\|94176430&offsets\|0\|-1(+1A)]\|CTATACAATCTATTGCCTTCCA | >MI0000067\|hsa-let-7f-1&WithFlank&9\|+\|94176341\|94176439 | 69 | 89 | 21(+1A) | CTATACAATCTATTGCCTTCCA | 50.6 | 4.9 | 2.0 | 2.4 | 1.6E-02 | 2.0E-02 | MB231/MB468 | MP |
| >hg38_21_+_8433172_8446622_RNA45SN1_45S_50ntFlanks@6758.6806.49\|ACTTGCGGCCCCGGGTTCCTCCCGGGGCTACGCCTGTCTGAGCGTCGCT | >hg38_21_+_8433172_8446622_RNA45SN1_45S_50ntFlanks | 6758 | 6806 | 49 | ACTTGCGGCCCCGGGTTCCTCCCGGGGCTACGCCTGTCTGAGCGTCGCT | 308.6 | 3.4 | 1.4 | 2.4 | 1.8E-02 | 2.0E-02 | MB231/MB468 | MITO |
| >MI0000273\|hsa-mir-183&WithFlank&7\|-\|129774899\|129775020@34.56.23&[MIMAT0000261&hsa-miR-183-5p&offsets\|+1\|+2;m-41&7\|-\|129774966\|129774987&offsets\|0\|+1]\|ATGGCACTGGTAGAATTCACTGT | >MI0000273\|hsa-mir-183&WithFlank&7\|-\|129774899\|129775020 | 34 | 56 | 23 | ATGGCACTGGTAGAATTCACTGT | 669.1 | 1.9 | 0.7 | 2.8 | 5.4E-03 | 2.0E-02 | BT20/MB231 | MITO |
| >MI0000070\|hsa-mir-16-1&WithFlank&13\|-\|50048967\|50049067@20.40.21(+1A);MI0000115\|hsa-mir-16-2&WithFlank&3\|+\|160404739\|160404831@16.36.21(+1A)&[MIMAT0000069&hsa-miR-16-5p&offsets\|0\|-1(+1A);m-62&13\|-\|50049027\|50049048&offsets\|0\|-1(+1A)];[MIMAT0000069_1&hsa-miR-16-5p&offsets\|0\|-1(+1A);m-61&3\|+\|160404754\|160404775&offsets\|0\|-1(+1A)]\|TAGCAGCACGTAAATATTGGCA | >MI0000070\|hsa-mir-16-1&WithFlank&13\|-\|50048967\|50049067 | 20 | 40 | 21(+1A) | TAGCAGCACGTAAATATTGGCA | 699.7 | -2.7 | 1.0 | -2.8 | 5.4E-03 | 2.0E-02 | BT20/MB231 | MITO |
| >MI0000298\|hsa-mir-221&WithFlank&X\|-\|45746151\|45746272@71.93.23(+1G)&[MIMAT0000278&hsa-miR-221-3p&offsets\|0\|0(+1G);m-91&X\|-\|45746181\|45746202&offsets\|0\|+1(+1G)]\|AGCTACATTGTCTGCTGGGTTTCG | >MI0000298\|hsa-mir-221&WithFlank&X\|-\|45746151\|45746272 | 71 | 93 | 23(+1G) | AGCTACATTGTCTGCTGGGTTTCG | 346.3 | -2.0 | 0.7 | -2.8 | 5.4E-03 | 2.0E-02 | BT20/MB231 | MITO |
| >trnaMT_SerGCT_MT_+_12207_12265@1.47.47\|GAGAAAGCTCACAAGAACTGCTAACTCATGCCCCCATGTCTAACAAC | >trnaMT_SerGCT_MT_+_12207_12265 | 1 | 47 | 47 | GAGAAAGCTCACAAGAACTGCTAACTCATGCCCCCATGTCTAACAAC | 446.4 | -3.0 | 1.1 | -2.8 | 5.4E-03 | 2.0E-02 | BT20/MB231 | MITO |
| >hg38_21_+_8433172_8446622_RNA45SN1_45S_50ntFlanks@9352.9369.18\|CCGTGGCGCAATGAAGGT | >hg38_21_+_8433172_8446622_RNA45SN1_45S_50ntFlanks | 9352 | 9369 | 18 | CCGTGGCGCAATGAAGGT | 205.5 | -4.4 | 1.7 | -2.5 | 1.1E-02 | 2.0E-02 | BT20/MB468 | MP |
| >hg38_21_+_8433172_8446622_RNA45SN1_45S_50ntFlanks@4731.4768.38\|AAGAACGAAAGTCGGAGGTTCGAAGACGATCAGATACC | >hg38_21_+_8433172_8446622_RNA45SN1_45S_50ntFlanks | 4731 | 4768 | 38 | AAGAACGAAAGTCGGAGGTTCGAAGACGATCAGATACC | 77.0 | 3.9 | 1.6 | 2.4 | 1.6E-02 | 2.0E-02 | MB231/MB468 | MP |
| >hg38_MT_+_1621_3279_RNR2_16S_50ntFlanks@293.310.18\|GACGAGCTACCTAAGAAC | >hg38_MT_+_1621_3279_RNR2_16S_50ntFlanks | 293 | 310 | 18 | GACGAGCTACCTAAGAAC | 630.2 | 3.8 | 1.5 | 2.5 | 1.1E-02 | 2.0E-02 | BT20/MB468 | MP |
| >MI0003834\|hsa-mir-769&WithFlank&19\|+\|46018926\|46019055@36.57.22(+1C)&[MIMAT0003886&hsa-miR-769-5p&offsets\|0\|0(+1C);m-216&19\|+\|46018961\|46018982&offsets\|0\|0(+1C)]\|TGAGACCTCTGGGTTCTGAGCTC | >MI0003834\|hsa-mir-769&WithFlank&19\|+\|46018926\|46019055 | 36 | 57 | 22(+1C) | TGAGACCTCTGGGTTCTGAGCTC | 56.4 | 3.2 | 1.3 | 2.4 | 1.6E-02 | 2.0E-02 | MB231/MB468 | MP |
| >am_trnaMT_SerGCT_MT_+_12207_12265@14.31.18\|AGAACTGCTAACTCATGC | >am_trnaMT_SerGCT_MT_+_12207_12265 | 14 | 31 | 18 | AGAACTGCTAACTCATGC | 273.1 | 2.1 | 0.9 | 2.4 | 1.6E-02 | 2.0E-02 | MB231/MB468 | MP |
| >hg38_21_+_8433172_8446622_RNA45SN1_45S_50ntFlanks@10839.10865.27\|TCCGTAACTTCGGGATAAGGATTGGCT | >hg38_21_+_8433172_8446622_RNA45SN1_45S_50ntFlanks | 10839 | 10865 | 27 | TCCGTAACTTCGGGATAAGGATTGGCT | 182.0 | -2.7 | 1.1 | -2.4 | 1.5E-02 | 2.0E-02 | BT20/MB468 | MITO |
| >MI0000457\|hsa-mir-141&WithFlank&12\|+\|6964091\|6964197@64.85.22&[MIMAT0000432&hsa-miR-141-3p&offsets\|-1\|-1;m-134&12\|+\|6964155\|6964175&offsets\|-1\|0]\|CTAACACTGTCTGGTAAAGATG | >MI0000457\|hsa-mir-141&WithFlank&12\|+\|6964091\|6964197 | 64 | 85 | 22 | CTAACACTGTCTGGTAAAGATG | 117.7 | 5.5 | 2.1 | 2.5 | 1.1E-02 | 2.0E-02 | BT20/MB468 | MP |
| >hg38_21_+_8433172_8446622_RNA45SN1_45S_50ntFlanks@12151.12181.31\|GTGGGGAGTTTGACTGGGGCGGTACACCTGT | >hg38_21_+_8433172_8446622_RNA45SN1_45S_50ntFlanks | 12151 | 12181 | 31 | GTGGGGAGTTTGACTGGGGCGGTACACCTGT | 364.4 | -4.4 | 1.7 | -2.5 | 1.1E-02 | 2.0E-02 | BT20/MB468 | MP |
| >trnaMT_PheGAA_MT_+_577_647@36.57.22\|AAAATGTTTAGACGGGCTCACA | >trnaMT_PheGAA_MT_+_577_647 | 36 | 57 | 22 | AAAATGTTTAGACGGGCTCACA | 702.8 | 2.8 | 1.0 | 2.8 | 4.8E-03 | 2.0E-02 | BT20/MB231 | NUC |
| >trnaMT_ThrTGT_MT_+_15888_15953@37.68.32\|ACCGGAGATGAAAACCTTTTTCCAAGGACACC | >trnaMT_ThrTGT_MT_+_15888_15953 | 37 | 68 | 32 | ACCGGAGATGAAAACCTTTTTCCAAGGACACC | 72.2 | 3.9 | 1.6 | 2.4 | 1.6E-02 | 2.0E-02 | MB231/MB468 | MP |
| >MI0005566\|hsa-mir-374b&WithFlank&X\|-\|74218541\|74218624@17.37.21&[MIMAT0004955&hsa-miR-374b-5p&offsets\|0\|-1;m-153&X\|-\|74218587\|74218608&offsets\|0\|-1]\|ATATAATACAACCTGCTAAGT | >MI0005566\|hsa-mir-374b&WithFlank&X\|-\|74218541\|74218624 | 17 | 37 | 21 | ATATAATACAACCTGCTAAGT | 764.0 | -3.3 | 1.2 | -2.9 | 4.3E-03 | 2.0E-02 | BT20/MB231 | MP |
| >hg38_1_-_228634819_228635039_RNA5S12_5S_50ntFlanks@132.170.39\|GATGGGAGACCGCCTGGGAATACCGGGTGCTGTAGGCTT | >hg38_1_-_228634819_228635039_RNA5S12_5S_50ntFlanks | 132 | 170 | 39 | GATGGGAGACCGCCTGGGAATACCGGGTGCTGTAGGCTT | 15325.5 | -2.3 | 1.0 | -2.4 | 1.8E-02 | 2.0E-02 | MB231/MB468 | MITO |
| >hg38_21_+_8433172_8446622_RNA45SN1_45S_50ntFlanks@11580.11612.33\|GAACTGGTGCGGACCAGGGGAATCCGACTGTTT | >hg38_21_+_8433172_8446622_RNA45SN1_45S_50ntFlanks | 11580 | 11612 | 33 | GAACTGGTGCGGACCAGGGGAATCCGACTGTTT | 72.0 | -4.4 | 1.7 | -2.5 | 1.1E-02 | 2.0E-02 | BT20/MB468 | MP |
| >trnaMT_ValTAC_MT_+_1602_1670@31.71.41\|TTACACTTAGGAGATTTCAACTTAACTTGACCGCTCTGACC | >trnaMT_ValTAC_MT_+_1602_1670 | 31 | 71 | 41 | TTACACTTAGGAGATTTCAACTTAACTTGACCGCTCTGACC | 183.4 | 4.2 | 1.7 | 2.4 | 1.6E-02 | 2.0E-02 | MB231/MB468 | MP |
| >hg38_21_+_8433172_8446622_RNA45SN1_45S_50ntFlanks@12412.12430.19\|ATCCTTCGATGTCGGCTCT | >hg38_21_+_8433172_8446622_RNA45SN1_45S_50ntFlanks | 12412 | 12430 | 19 | ATCCTTCGATGTCGGCTCT | 333.9 | -3.5 | 1.5 | -2.4 | 1.7E-02 | 2.0E-02 | MB231/MB468 | CYTO |
| >hg38_21_+_8433172_8446622_RNA45SN1_45S_50ntFlanks@6772.6799.28\|GTTCCTCCCGGGGCTACGCCTGTCTGAG | >hg38_21_+_8433172_8446622_RNA45SN1_45S_50ntFlanks | 6772 | 6799 | 28 | GTTCCTCCCGGGGCTACGCCTGTCTGAG | 577.0 | -4.5 | 1.8 | -2.5 | 1.1E-02 | 2.0E-02 | BT20/MB468 | MP |
| >hg38_21_+_8433172_8446622_RNA45SN1_45S_50ntFlanks@10351.10381.31\|CGAGAACTTTGAAGGCCGAAGTGGAGAAGGG | >hg38_21_+_8433172_8446622_RNA45SN1_45S_50ntFlanks | 10351 | 10381 | 31 | CGAGAACTTTGAAGGCCGAAGTGGAGAAGGG | 232.9 | 4.8 | 1.7 | 2.9 | 4.4E-03 | 2.0E-02 | BT20/MB231 | MP |
| >MI0000114\|hsa-mir-107&WithFlank&10\|-\|89592741\|89592833@56.76.21;MI0000108\|hsa-mir-103a-2&WithFlank&20\|+\|3917488\|3917577@54.74.21;MI0000109\|hsa-mir-103a-1&WithFlank&5\|-\|168560890\|168560979@54.74.21&[MIMAT0000104&hsa-miR-107&offsets\|0\|-2;m-78&10\|-\|89592758\|89592778&offsets\|0\|0];[MIMAT0000101&hsa-miR-103a-3p&offsets\|0\|-2;m-12&20\|+\|3917541\|3917563&offsets\|0\|-2];[MIMAT0000101_1&hsa-miR-103a-3p&offsets\|0\|-2;m-13&5\|-\|168560904\|168560926&offsets\|0\|-2;m-44220&5\|-\|168560908\|168560926&offsets\|0\|+2]\|AGCAGCATTGTACAGGGCTAT | >MI0000114\|hsa-mir-107&WithFlank&10\|-\|89592741\|89592833 | 56 | 76 | 21 | AGCAGCATTGTACAGGGCTAT | 1332.8 | 3.4 | 1.3 | 2.5 | 1.1E-02 | 2.0E-02 | BT20/MB468 | MP |
| >hg38_21_+_8433172_8446622_RNA45SN1_45S_50ntFlanks@12420.12460.41\|ATGTCGGCTCTTCCTATCATTGTGAAGCAGAATTCACCAAG | >hg38_21_+_8433172_8446622_RNA45SN1_45S_50ntFlanks | 12420 | 12460 | 41 | ATGTCGGCTCTTCCTATCATTGTGAAGCAGAATTCACCAAG | 1451.5 | 2.4 | 1.0 | 2.4 | 1.8E-02 | 2.0E-02 | MB231/MB468 | MITO |
| >trnaMT_SerGCT_MT_+_12207_12265@12.45.34\|CAAGAACTGCTAACTCATGCCCCCATGTCTAACA | >trnaMT_SerGCT_MT_+_12207_12265 | 12 | 45 | 34 | CAAGAACTGCTAACTCATGCCCCCATGTCTAACA | 243.0 | 4.1 | 1.6 | 2.5 | 1.1E-02 | 2.0E-02 | BT20/MB468 | MP |
| >hg38_MT_+_1621_3279_RNR2_16S_50ntFlanks@618.643.26\|AACATATAACTGAACTCCTCACACCC | >hg38_MT_+_1621_3279_RNR2_16S_50ntFlanks | 618 | 643 | 26 | AACATATAACTGAACTCCTCACACCC | 143.3 | 4.9 | 1.9 | 2.5 | 1.1E-02 | 2.0E-02 | BT20/MB468 | MP |
| >hg38_21_+_8433172_8446622_RNA45SN1_45S_50ntFlanks@12072.12090.19\|CCCGAGGGGCTCTCGCTTC | >hg38_21_+_8433172_8446622_RNA45SN1_45S_50ntFlanks | 12072 | 12090 | 19 | CCCGAGGGGCTCTCGCTTC | 90.6 | -4.2 | 1.7 | -2.5 | 1.1E-02 | 2.0E-02 | BT20/MB468 | MP |
| >MI0000478\|hsa-mir-149&WithFlank&2\|+\|240455995\|240456095@21.41.21&[MIMAT0000450&hsa-miR-149-5p&offsets\|0\|-2;m-195&2\|+\|240456015\|240456037&offsets\|0\|-2]\|TCTGGCTCCGTGTCTTCACTC | >MI0000478\|hsa-mir-149&WithFlank&2\|+\|240455995\|240456095 | 21 | 41 | 21 | TCTGGCTCCGTGTCTTCACTC | 396.3 | 2.1 | 0.7 | 2.8 | 4.8E-03 | 2.0E-02 | BT20/MB231 | NUC |
| >hg38_21_+_8433172_8446622_RNA45SN1_45S_50ntFlanks@9940.9959.20\|CAGGACGGTGGCCATGGAAG | >hg38_21_+_8433172_8446622_RNA45SN1_45S_50ntFlanks | 9940 | 9959 | 20 | CAGGACGGTGGCCATGGAAG | 266.9 | 3.1 | 1.1 | 2.9 | 4.4E-03 | 2.0E-02 | BT20/MB231 | MP |
| >am_MI0003583\|hsa-mir-576&WithFlank&4\|+\|109488692\|109488801@61.82.22(+1A)&[MIMAT0004796&hsa-miR-576-3p&offsets\|0\|0(+1A);m-524&4\|+\|109488752\|109488772&offsets\|0\|+1(+1A)]\|AAGATGTGGAAAAATTGGAATCA | >am_MI0003583\|hsa-mir-576&WithFlank&4\|+\|109488692\|109488801 | 61 | 82 | 22(+1A) | AAGATGTGGAAAAATTGGAATCA | 134.6 | 5.5 | 2.2 | 2.5 | 1.1E-02 | 2.0E-02 | BT20/MB468 | MP |
| >hg38_21_+_8433172_8446622_RNA45SN1_45S_50ntFlanks@4976.4993.18\|CCGGACACGGACAGGATT | >hg38_21_+_8433172_8446622_RNA45SN1_45S_50ntFlanks | 4976 | 4993 | 18 | CCGGACACGGACAGGATT | 157.7 | 2.7 | 1.0 | 2.5 | 1.1E-02 | 2.0E-02 | BT20/MB468 | MP |
| >hg38_21_+_8433172_8446622_RNA45SN1_45S_50ntFlanks@9533.9569.37\|GAGGAAACTCTGGTGGAGGTCCGTAGCGGTCCTGACG | >hg38_21_+_8433172_8446622_RNA45SN1_45S_50ntFlanks | 9533 | 9569 | 37 | GAGGAAACTCTGGTGGAGGTCCGTAGCGGTCCTGACG | 227.5 | -2.5 | 1.0 | -2.5 | 1.1E-02 | 2.0E-02 | BT20/MB468 | MP |
| >am_MI0000815\|hsa-mir-339&WithFlank&7\|-\|1022927\|1023032@21.41.21&[MIMAT0000764&hsa-miR-339-5p&offsets\|0\|-2;m-163&7\|-\|1022990\|1023012&offsets\|0\|-2]\|TCCCTGTCCTCCAGGAGCTCA | >am_MI0000815\|hsa-mir-339&WithFlank&7\|-\|1022927\|1023032 | 21 | 41 | 21 | TCCCTGTCCTCCAGGAGCTCA | 221.0 | 3.9 | 1.5 | 2.5 | 1.1E-02 | 2.0E-02 | BT20/MB468 | MP |
| >hg38_21_+_8433172_8446622_RNA45SN1_45S_50ntFlanks@3988.4005.18\|CTTTGGTGACTCTAGATA | >hg38_21_+_8433172_8446622_RNA45SN1_45S_50ntFlanks | 3988 | 4005 | 18 | CTTTGGTGACTCTAGATA | 310.6 | -3.3 | 1.3 | -2.5 | 1.1E-02 | 2.0E-02 | BT20/MB468 | MP |
| >hg38_21_+_8433172_8446622_RNA45SN1_45S_50ntFlanks@8527.8549.23\|CCTTCCCCCGCCGCCCCTCCTCC | >hg38_21_+_8433172_8446622_RNA45SN1_45S_50ntFlanks | 8527 | 8549 | 23 | CCTTCCCCCGCCGCCCCTCCTCC | 211.4 | -3.3 | 1.4 | -2.4 | 1.7E-02 | 2.0E-02 | BT20/MB468 | CYTO |
| >trna5_AspGTC_12_+_98897281_98897352@38.70.33\|CGCGGGAGACCGGGGTTCAATTCCCCGACGGGG | >trna5_AspGTC_12_+_98897281_98897352 | 38 | 70 | 33 | CGCGGGAGACCGGGGTTCAATTCCCCGACGGGG | 100.1 | -4.1 | 1.7 | -2.4 | 1.6E-02 | 2.0E-02 | MB231/MB468 | MP |

<table>
<tr><td>>am_trna128_GlyGCC_6_-_27870686_27870756@3.35.33,trna18_GlyGCC_16_+_70822597_70822667@3.35.33,trna19_GlyGCC_16_+_70823410_70823480@3.35.33,trna19_GlyGCC_2_-_157257659_157257729@3.35.33,trna24_GlyGCC_16_-_70812942_70813012@3.35.33,trna25_GlyGCC_16_-_70812114_70812184@3.35.33,trna5_GlyGCC_17_+_8029064_8029134@3.35.33,trna68_GlyGCC_1_-_161493637_161493707@3.35.33|ATTGGTGGTTCAGTGGTAGAATTCTCGCCTGCC</td><td>>am_trna128_GlyGCC_6_-_27870686_27870756</td><td>3</td><td>35</td><td>33</td><td>ATTGGTGGTTCAGTGGTAGAATTCTCGCCTGCC</td><td>363.5</td><td>-4.9</td><td>1.7</td><td>-2.8</td><td>4.8E-03</td><td>2.0E-02</td><td>BT20/MB231</td><td>NUC</td></tr>
<tr><td>>am_MI0000809|hsa-mir-151a&WithFlank&8|-|140732558|140732659@53.72.20(+1A)&[MIMAT0000757&hsa-miR-151a-3p&offsets|0|-1(+1A);m-30&8|-|140732587|140732607&offsets|0|-1(+1A)]|CTAGACTGAAGCTCCTTGAGA</td><td>>am_MI0000809|hsa-mir-151a&WithFlank&8|-|140732558|140732659</td><td>53</td><td>72</td><td>20(+1A)</td><td>CTAGACTGAAGCTCCTTGAGA</td><td>11539.5</td><td>1.6</td><td>0.6</td><td>2.8</td><td>4.8E-03</td><td>2.0E-02</td><td>BT20/MB231</td><td>NUC</td></tr>
<tr><td>>hg38_21_+_8433172_8446622_RNA45SN1_45S_50ntFlanks@11675.11709.35|TGCTCTGAATGTCAAAGTGAAGAAATTCAATGAAG</td><td>>hg38_21_+_8433172_8446622_RNA45SN1_45S_50ntFlanks</td><td>11675</td><td>11709</td><td>35</td><td>TGCTCTGAATGTCAAAGTGAAGAAATTCAATGAAG</td><td>404.7</td><td>-2.5</td><td>0.9</td><td>-2.8</td><td>4.8E-03</td><td>2.0E-02</td><td>BT20/MB231</td><td>NUC</td></tr>
<tr><td>>hg38_21_+_8433172_8446622_RNA45SN1_45S_50ntFlanks@12644.12675.32|TCTGTGGGATTATGACTGAACGCCTCTAAGTC</td><td>>hg38_21_+_8433172_8446622_RNA45SN1_45S_50ntFlanks</td><td>12644</td><td>12675</td><td>32</td><td>TCTGTGGGATTATGACTGAACGCCTCTAAGTC</td><td>109.7</td><td>-2.2</td><td>0.9</td><td>-2.4</td><td>1.7E-02</td><td>2.0E-02</td><td>MB231/MB468</td><td>CYTO</td></tr>
<tr><td>>hg38_21_+_8433172_8446622_RNA45SN1_45S_50ntFlanks@10684.10709.26|CGGCGGCGTCCGGTGAGCTCTCGCTG</td><td>>hg38_21_+_8433172_8446622_RNA45SN1_45S_50ntFlanks</td><td>10684</td><td>10709</td><td>26</td><td>CGGCGGCGTCCGGTGAGCTCTCGCTG</td><td>1272.2</td><td>-3.7</td><td>1.5</td><td>-2.5</td><td>1.1E-02</td><td>2.0E-02</td><td>BT20/MB468</td><td>MP</td></tr>
<tr><td>>hg38_21_+_8433172_8446622_RNA45SN1_45S_50ntFlanks@10806.10827.22|AACAATGTAGGTAAGGGAAGTC</td><td>>hg38_21_+_8433172_8446622_RNA45SN1_45S_50ntFlanks</td><td>10806</td><td>10827</td><td>22</td><td>AACAATGTAGGTAAGGGAAGTC</td><td>156.0</td><td>-3.4</td><td>1.4</td><td>-2.5</td><td>1.1E-02</td><td>2.0E-02</td><td>BT20/MB468</td><td>MP</td></tr>
<tr><td>>hg38_21_+_8433172_8446622_RNA45SN1_45S_50ntFlanks@11210.11232.23|GGCGACTCTGGACGCGAGCCGGG</td><td>>hg38_21_+_8433172_8446622_RNA45SN1_45S_50ntFlanks</td><td>11210</td><td>11232</td><td>23</td><td>GGCGACTCTGGACGCGAGCCGGG</td><td>581.6</td><td>-2.2</td><td>0.9</td><td>-2.4</td><td>1.5E-02</td><td>2.0E-02</td><td>BT20/MB468</td><td>MITO</td></tr>
<tr><td>>hg38_21_+_8433172_8446622_RNA45SN1_45S_50ntFlanks@10914.10931.18|CGCCGCGGCTGGACGAGG</td><td>>hg38_21_+_8433172_8446622_RNA45SN1_45S_50ntFlanks</td><td>10914</td><td>10931</td><td>18</td><td>CGCCGCGGCTGGACGAGG</td><td>459.9</td><td>-2.1</td><td>0.9</td><td>-2.4</td><td>1.5E-02</td><td>2.0E-02</td><td>BT20/MB468</td><td>MITO</td></tr>
<tr><td>>hg38_21_+_8433172_8446622_RNA45SN1_45S_50ntFlanks@6719.6751.33|TGAATTGCAGGACACATTGATCATCGACACTTC</td><td>>hg38_21_+_8433172_8446622_RNA45SN1_45S_50ntFlanks</td><td>6719</td><td>6751</td><td>33</td><td>TGAATTGCAGGACACATTGATCATCGACACTTC</td><td>347.1</td><td>-1.4</td><td>0.6</td><td>-2.4</td><td>1.5E-02</td><td>2.0E-02</td><td>BT20/MB468</td><td>MITO</td></tr>
<tr><td>>am_MI0003581|hsa-mir-574&WithFlank&4|+|38868026|38868133@31.54.24(+1A)&[MIMAT0004795&hsa-miR-574-5p&offsets|0|+1(+1A);m-213&4|+|38868056|38868078&offsets|0|+1(+1A)]|TGAGTGTGTGTGTGTGAGTGTGTGA</td><td>>am_MI0003581|hsa-mir-574&WithFlank&4|+|38868026|38868133</td><td>31</td><td>54</td><td>24(+1A)</td><td>TGAGTGTGTGTGTGTGAGTGTGTGA</td><td>122.1</td><td>3.2</td><td>1.3</td><td>2.4</td><td>1.6E-02</td><td>2.0E-02</td><td>MB231/MB468</td><td>MP</td></tr>
<tr><td>>hg38_21_+_8433172_8446622_RNA45SN1_45S_50ntFlanks@10405.10432.28|ATGGGTCAGTCGGTCCTGAGAGATGGGC</td><td>>hg38_21_+_8433172_8446622_RNA45SN1_45S_50ntFlanks</td><td>10405</td><td>10432</td><td>28</td><td>ATGGGTCAGTCGGTCCTGAGAGATGGGC</td><td>340.5</td><td>3.7</td><td>1.3</td><td>2.8</td><td>4.4E-03</td><td>2.0E-02</td><td>BT20/MB231</td><td>MP</td></tr>
<tr><td>>hg38_21_+_8433172_8446622_RNA45SN1_45S_50ntFlanks@8355.8373.19|TGAAAAGAACTTTGAAGAG</td><td>>hg38_21_+_8433172_8446622_RNA45SN1_45S_50ntFlanks</td><td>8355</td><td>8373</td><td>19</td><td>TGAAAAGAACTTTGAAGAG</td><td>530.3</td><td>6.8</td><td>2.5</td><td>2.8</td><td>5.5E-03</td><td>2.0E-02</td><td>BT20/MB231</td><td>MITO</td></tr>
<tr><td>>hg38_21_+_8433172_8446622_RNA45SN1_45S_50ntFlanks@10829.10851.23|GCAAGCCGGATCCGTAACTTCGG</td><td>>hg38_21_+_8433172_8446622_RNA45SN1_45S_50ntFlanks</td><td>10829</td><td>10851</td><td>23</td><td>GCAAGCCGGATCCGTAACTTCGG</td><td>205.2</td><td>-1.6</td><td>0.7</td><td>-2.4</td><td>1.7E-02</td><td>2.0E-02</td><td>MB231/MB468</td><td>CYTO</td></tr>
<tr><td>>hg38_21_+_8433172_8446622_RNA45SN1_45S_50ntFlanks@7991.8008.18|ACGTGGCGACCCGCTGAA</td><td>>hg38_21_+_8433172_8446622_RNA45SN1_45S_50ntFlanks</td><td>7991</td><td>8008</td><td>18</td><td>ACGTGGCGACCCGCTGAA</td><td>2326.6</td><td>-2.1</td><td>0.9</td><td>-2.4</td><td>1.8E-02</td><td>2.0E-02</td><td>MB231/MB468</td><td>NUC</td></tr>
<tr><td>>hg38_21_+_8433172_8446622_RNA45SN1_45S_50ntFlanks@10057.10079.23|CCGGCAGTCGAGAGTGGACGGGA</td><td>>hg38_21_+_8433172_8446622_RNA45SN1_45S_50ntFlanks</td><td>10057</td><td>10079</td><td>23</td><td>CCGGCAGTCGAGAGTGGACGGGA</td><td>79.2</td><td>-4.5</td><td>1.9</td><td>-2.4</td><td>1.6E-02</td><td>2.0E-02</td><td>MB231/MB468</td><td>MP</td></tr>
<tr><td>>hg38_21_+_8433172_8446622_RNA45SN1_45S_50ntFlanks@7992.8011.20|CGTGGCGACCCGCTGAATTT</td><td>>hg38_21_+_8433172_8446622_RNA45SN1_45S_50ntFlanks</td><td>7992</td><td>8011</td><td>20</td><td>CGTGGCGACCCGCTGAATTT</td><td>339.3</td><td>1.3</td><td>0.5</td><td>2.4</td><td>1.8E-02</td><td>2.0E-02</td><td>BT20/MB468</td><td>NUC</td></tr>
<tr><td>>hg38_21_+_8433172_8446622_RNA45SN1_45S_50ntFlanks@6707.6736.30|CGAGAATTAATGTGAATTGCAGGACACATT</td><td>>hg38_21_+_8433172_8446622_RNA45SN1_45S_50ntFlanks</td><td>6707</td><td>6736</td><td>30</td><td>CGAGAATTAATGTGAATTGCAGGACACATT</td><td>92.7</td><td>-2.1</td><td>0.8</td><td>-2.5</td><td>1.1E-02</td><td>2.0E-02</td><td>BT20/MB468</td><td>MP</td></tr>
<tr><td>>hg38_21_+_8433172_8446622_RNA45SN1_45S_50ntFlanks@6653.6697.45|ACTCTTAGCGGTGGATCACTCGGCTCGTGCGTCGATGAAGAACGC</td><td>>hg38_21_+_8433172_8446622_RNA45SN1_45S_50ntFlanks</td><td>6653</td><td>6697</td><td>45</td><td>ACTCTTAGCGGTGGATCACTCGGCTCGTGCGTCGATGAAGAACGC</td><td>322.4</td><td>1.9</td><td>0.8</td><td>2.4</td><td>1.5E-02</td><td>2.0E-02</td><td>BT20/MB468</td><td>MITO</td></tr>
<tr><td>>MI0000542|hsa-mir-320a&WithFlank&8|-|22244960|22245043@49.72.24&[MIMAT0000510&hsa-miR-320a-3p&offsets|+1|+3;m-60&8|-|22244975|22244996&offsets|+1|+3]|AAAGCTGGGTTGAGAGGGCGAAAA</td><td>>MI0000542|hsa-mir-320a&WithFlank&8|-|22244960|22245043</td><td>49</td><td>72</td><td>24</td><td>AAAGCTGGGTTGAGAGGGCGAAAA</td><td>71.4</td><td>5.4</td><td>1.9</td><td>2.8</td><td>4.4E-03</td><td>2.0E-02</td><td>BT20/MB231</td><td>MP</td></tr>
<tr><td>>hg38_21_+_8433172_8446622_RNA45SN1_45S_50ntFlanks@6648.6669.22|GTACGACTCTTAGCGGTGGATC</td><td>>hg38_21_+_8433172_8446622_RNA45SN1_45S_50ntFlanks</td><td>6648</td><td>6669</td><td>22</td><td>GTACGACTCTTAGCGGTGGATC</td><td>5623.3</td><td>2.0</td><td>0.7</td><td>2.8</td><td>5.5E-03</td><td>2.0E-02</td><td>BT20/MB231</td><td>MITO</td></tr>
<tr><td>>hg38_MT_+_1621_3279_RNR2_16S_50ntFlanks@732.759.28|TAAAACACTGAACTGACAATTAACAGCC</td><td>>hg38_MT_+_1621_3279_RNR2_16S_50ntFlanks</td><td>732</td><td>759</td><td>28</td><td>TAAAACACTGAACTGACAATTAACAGCC</td><td>383.8</td><td>-3.3</td><td>1.2</td><td>-2.8</td><td>4.4E-03</td><td>2.0E-02</td><td>BT20/MB231</td><td>MP</td></tr>
<tr><td>>trna111_HisGTG_1_-_147774845_147774916@-1T.34.35,trna118_HisGTG_1_-_145396881_145396952@-1T.34.35,trna16_HisGTG_1_+_146544773_146544844@-1T.34.35,trna1_HisGTG_15_+_45493349_45493420@-1T.34.35,trna21_HisGTG_1_+_147753471_147753542@-1T.34.35,trna33_HisGTG_6_+_27125906_27125977@-1T.34.35,trna7_HisGTG_9_-_14433938_14434009@-1T.34.35,trna8_HisGTG_15_-_45492611_45492682@-1T.34.35,trna9_HisGTG_15_-_45490804_45490875@-1T.34.35|TGCCGTGATCGTATAGTGGTTAGTACTCTGCGTTG</td><td>>trna111_HisGTG_1_-_147774845_147774916</td><td>-1T</td><td>34</td><td>35</td><td>TGCCGTGATCGTATAGTGGTTAGTACTCTGCGTTG</td><td>744.4</td><td>-2.7</td><td>1.0</td><td>-2.8</td><td>4.9E-03</td><td>2.0E-02</td><td>BT20/MB231</td><td>NUC</td></tr>
<tr><td>>hg38_21_+_8433172_8446622_RNA45SN1_45S_50ntFlanks@11586.11611.26|GTGCGGACCAGGGGAATCCGACTGTT</td><td>>hg38_21_+_8433172_8446622_RNA45SN1_45S_50ntFlanks</td><td>11586</td><td>11611</td><td>26</td><td>GTGCGGACCAGGGGAATCCGACTGTT</td><td>1854.0</td><td>-1.2</td><td>0.5</td><td>-2.4</td><td>1.5E-02</td><td>2.0E-02</td><td>BT20/MB468</td><td>MITO</td></tr>
<tr><td>>hg38_21_+_8433172_8446622_RNA45SN1_45S_50ntFlanks@12415.12446.32|CTTCGATGTCGGCTCTTCCTATCATTGTGAAG</td><td>>hg38_21_+_8433172_8446622_RNA45SN1_45S_50ntFlanks</td><td>12415</td><td>12446</td><td>32</td><td>CTTCGATGTCGGCTCTTCCTATCATTGTGAAG</td><td>1257.7</td><td>-1.4</td><td>0.6</td><td>-2.4</td><td>1.5E-02</td><td>2.0E-02</td><td>BT20/MB468</td><td>MITO</td></tr>
<tr><td>>MI0000061|hsa-let-7a-2&WithFlank&11|-|122146516|122146599@11.29.19;MI0000062|hsa-let-7a-3&WithFlank&22|+|46112743|46112828@10.28.19;MI0000060|hsa-let-7a-1&WithFlank&9|+|94175951|94176042@12.30.19&[MIMAT0000062&hsa-let-7a-5p&offsets|0|-3;m-6&11|-|122146568|122146589&offsets|0|-3];[MIMAT0000062_1&hsa-let-7a-5p&offsets|0|-3;m-5&22|+|46112752|46112773&offsets|0|-3];[MIMAT0000062_2&hsa-let-7a-5p&offsets|0|-3;m-4&9|+|94175962|94175983&offsets|0|-3]|TGAGGTAGTAGGTTGTATA</td><td>>MI0000061|hsa-let-7a-2&WithFlank&11|-|122146516|122146599</td><td>11</td><td>29</td><td>19</td><td>TGAGGTAGTAGGTTGTATA</td><td>3372.4</td><td>2.7</td><td>1.0</td><td>2.8</td><td>5.8E-03</td><td>2.0E-02</td><td>BT20/MB231</td><td>CYTO</td></tr>
<tr><td>>hg38_21_+_8433172_8446622_RNA45SN1_45S_50ntFlanks@11769.11794.26|TAATTAGTGACGCGCATGAATGGATG</td><td>>hg38_21_+_8433172_8446622_RNA45SN1_45S_50ntFlanks</td><td>11769</td><td>11794</td><td>26</td><td>TAATTAGTGACGCGCATGAATGGATG</td><td>630.1</td><td>-2.3</td><td>0.8</td><td>-2.8</td><td>5.8E-03</td><td>2.0E-02</td><td>BT20/MB231</td><td>CYTO</td></tr>
<tr><td>>hg38_21_+_8433172_8446622_RNA45SN1_45S_50ntFlanks@11670.11687.18|CCCAGTGCTCTGAATGTC</td><td>>hg38_21_+_8433172_8446622_RNA45SN1_45S_50ntFlanks</td><td>11670</td><td>11687</td><td>18</td><td>CCCAGTGCTCTGAATGTC</td><td>787.1</td><td>4.4</td><td>1.6</td><td>2.8</td><td>4.4E-03</td><td>2.0E-02</td><td>BT20/MB231</td><td>MP</td></tr>
<tr><td>>hg38_21_+_8433172_8446622_RNA45SN1_45S_50ntFlanks@9547.9570.24|GGAGGTCCGTAGCGGTCCTGACGT</td><td>>hg38_21_+_8433172_8446622_RNA45SN1_45S_50ntFlanks</td><td>9547</td><td>9570</td><td>24</td><td>GGAGGTCCGTAGCGGTCCTGACGT</td><td>65.7</td><td>4.4</td><td>1.7</td><td>2.5</td><td>1.1E-02</td><td>2.0E-02</td><td>BT20/MB468</td><td>MP</td></tr>
<tr><td>>hg38_21_+_8433172_8446622_RNA45SN1_45S_50ntFlanks@10241.10261.21|TACGCCGCGACGAGTAGGAGG</td><td>>hg38_21_+_8433172_8446622_RNA45SN1_45S_50ntFlanks</td><td>10241</td><td>10261</td><td>21</td><td>TACGCCGCGACGAGTAGGAGG</td><td>90.6</td><td>5.3</td><td>2.1</td><td>2.5</td><td>1.1E-02</td><td>2.0E-02</td><td>BT20/MB468</td><td>MP</td></tr>
<tr><td>>hg38_21_+_8433172_8446622_RNA45SN1_45S_50ntFlanks@11705.11726.22|TGAAGCGCGGGTAAACGGCGGG</td><td>>hg38_21_+_8433172_8446622_RNA45SN1_45S_50ntFlanks</td><td>11705</td><td>11726</td><td>22</td><td>TGAAGCGCGGGTAAACGGCGGG</td><td>90.7</td><td>-4.3</td><td>1.7</td><td>-2.5</td><td>1.1E-02</td><td>2.0E-02</td><td>BT20/MB468</td><td>MP</td></tr>
<tr><td>>MI0000299|hsa-mir-222&WithFlank&X|-|45747009|45747130@75.97.23(+1A)&[MIMAT0000279&hsa-miR-222-3p&offsets|0|+2(+1A);m-140&X|-|45747033|45747056&offsets|0|-1(+1A)]|AGCTACATCTGGCTACTGGGTCTA</td><td>>MI0000299|hsa-mir-222&WithFlank&X|-|45747009|45747130</td><td>75</td><td>97</td><td>23(+1A)</td><td>AGCTACATCTGGCTACTGGGTCTA</td><td>1194.1</td><td>-2.9</td><td>1.0</td><td>-2.8</td><td>4.4E-03</td><td>2.0E-02</td><td>BT20/MB231</td><td>MP</td></tr>
<tr><td>>hg38_21_+_8433172_8446622_RNA45SN1_45S_50ntFlanks@5485.5516.32|AGCGCTGAGAAGACGGTCGAACTTGACTATCT</td><td>>hg38_21_+_8433172_8446622_RNA45SN1_45S_50ntFlanks</td><td>5485</td><td>5516</td><td>32</td><td>AGCGCTGAGAAGACGGTCGAACTTGACTATCT</td><td>2701.7</td><td>3.7</td><td>1.3</td><td>2.8</td><td>4.4E-03</td><td>2.0E-02</td><td>BT20/MB231</td><td>MP</td></tr>
<tr><td>>hg38_21_+_8433172_8446622_RNA45SN1_45S_50ntFlanks@6650.6672.23|ACGACTCTTAGCGGTGGATCACT</td><td>>hg38_21_+_8433172_8446622_RNA45SN1_45S_50ntFlanks</td><td>6650</td><td>6672</td><td>23</td><td>ACGACTCTTAGCGGTGGATCACT</td><td>569.8</td><td>3.8</td><td>1.3</td><td>2.8</td><td>4.4E-03</td><td>2.0E-02</td><td>BT20/MB231</td><td>MP</td></tr>
<tr><td>>hg38_21_+_8433172_8446622_RNA45SN1_45S_50ntFlanks@9796.9819.24|TGGAGCCGGGCGTGGAATGCGAGT</td><td>>hg38_21_+_8433172_8446622_RNA45SN1_45S_50ntFlanks</td><td>9796</td><td>9819</td><td>24</td><td>TGGAGCCGGGCGTGGAATGCGAGT</td><td>106.4</td><td>5.5</td><td>1.9</td><td>2.8</td><td>4.4E-03</td><td>2.0E-02</td><td>BT20/MB231</td><td>MP</td></tr>
<tr><td>>trna12_ArgCCT_16_+_3243918_3243990@35.54.20,trna18_ArgCCT_17_+_73030001_73030073@35.54.20,trna21_ArgCCT_17_-_73030526_73030598@35.54.20,trna2_ArgCCT_16_+_3202901_3202973@35.54.20,trna3_ArgCCT_7_+_139025446_139025518@35.54.20|CTAAGCCAGGGATTGTGGGT</td><td>>trna12_ArgCCT_16_+_3243918_3243990</td><td>35</td><td>54</td><td>20</td><td>CTAAGCCAGGGATTGTGGGT</td><td>1626.3</td><td>2.1</td><td>0.9</td><td>2.4</td><td>1.7E-02</td><td>2.0E-02</td><td>MB231/MB468</td><td>CYTO</td></tr>
<tr><td>>MI0000088|hsa-mir-30a&WithFlank&6|-|71403545|71403627@12.35.24(+1C)&[MIMAT0000087&hsa-miR-30a-5p&offsets|0|+2(+1C);m-15&6|-|71403595|71403616&offsets|0|+2(+1C)]|TGTAAACATCCTCGACTGGAAGCTC</td><td>>MI0000088|hsa-mir-30a&WithFlank&6|-|71403545|71403627</td><td>12</td><td>35</td><td>24(+1C)</td><td>TGTAAACATCCTCGACTGGAAGCTC</td><td>424.9</td><td>3.2</td><td>1.3</td><td>2.5</td><td>1.1E-02</td><td>2.0E-02</td><td>BT20/MB468</td><td>MP</td></tr>
<tr><td>>trnaMT_PheGAA_MT_+_577_647@1.19.19|GTTTATGTAGCTTACCTCC</td><td>>trnaMT_PheGAA_MT_+_577_647</td><td>1</td><td>19</td><td>19</td><td>GTTTATGTAGCTTACCTCC</td><td>612.6</td><td>-2.1</td><td>0.7</td><td>-2.8</td><td>4.9E-03</td><td>2.0E-02</td><td>BT20/MB231</td><td>NUC</td></tr>
<tr><td>>hg38_21_+_8433172_8446622_RNA45SN1_45S_50ntFlanks@6650.6675.26|ACGACTCTTAGCGGTGGATCACTCGG</td><td>>hg38_21_+_8433172_8446622_RNA45SN1_45S_50ntFlanks</td><td>6650</td><td>6675</td><td>26</td><td>ACGACTCTTAGCGGTGGATCACTCGG</td><td>612.0</td><td>-1.3</td><td>0.5</td><td>-2.4</td><td>1.8E-02</td><td>2.0E-02</td><td>BT20/MB468</td><td>NUC</td></tr>
<tr><td>>am_trna10_ValCAC_5_-_180649395_180649467@1.32.32,trna12_ValAAC_5_-_180645270_180645342@1.32.32,trna132_ValAAC_6_-_27721179_27721251@1.32.32,trna136_ValAAC_6_-_27648885_27648957@1.32.32,trna139_ValAAC_6_-_27618707_27618779@1.32.32,trna18_ValCAC_5_-_180529253_180529325@1.32.32,trna2_ValAAC_3_+_169490018_169490090@1.32.32,trna2_ValCAC_5_+_180524070_180524142@1.32.32,trna4_ValAAC_5_+_180591154_180591226@1.32.32,trna5_ValAAC_5_+_180596610_180596682@1.32.32,trna6_ValCAC_5_+_180600650_180600722@1.32.32,trna85_ValCAC_1_-_161369490_161369562@1.32.32,trna90_ValCAC_1_-_149684088_149684161@1.32.32,trna98_ValCAC_1_-_149298555_149298627@1.32.32,trna9_ValCAC_6_+_26538282_26538354@1.32.32|GTTTCCGTAGTGTAGTGGTTATCACGTTCGCC</td><td>>am_trna10_ValCAC_5_-_180649395_180649467</td><td>1</td><td>32</td><td>32</td><td>GTTTCCGTAGTGTAGTGGTTATCACGTTCGCC</td><td>3042.8</td><td>-2.7</td><td>1.0</td><td>-2.8</td><td>4.9E-03</td><td>2.0E-02</td><td>BT20/MB231</td><td>NUC</td></tr>
</table>

| | | | | | | | | | | | | | |
|---|---|---|---|---|---|---|---|---|---|---|---|---|---|
| >hg38_21_+_8433172_8446622_RNA45SN1_45S_50ntFlanks@4735.4758.24\|ACGAAAGTCGGAGGTTCGAAGACG | >hg38_21_+_8433172_8446622_RNA45SN1_45S_50ntFlanks | 4735 | 4758 | 24 | ACGAAAGTCGGAGGTTCGAAGACG | 1695.1 | -1.6 | 0.7 | -2.4 | 1.8E-02 | 2.0E-02 | MB231/MB468 | NUC |
| >MI0003780\|hsa-mir-1296&WithFlank&10\|-\|63372951\|63373054@22.43.22&[MIMAT0005794&hsa-miR-1296-5p&offsets\|0\|0;m-304&10\|-\|63373012\|63373033&offsets\|0\|0]\|TTAGGGCCCTGGCTCCATCTCC | >MI0003780\|hsa-mir-1296&WithFlank&10\|-\|63372951\|63373054 | 22 | 43 | 22 | TTAGGGCCCTGGCTCCATCTCC | 257.3 | 3.2 | 1.3 | 2.4 | 1.6E-02 | 2.0E-02 | MB231/MB468 | MP |
| >hg38_MT_+_1621_3279_RNR2_16S_50ntFlanks@1425.1458.34\|ACGATTAAAGTCCTACGTGATCTGAGTTCAGACC | >hg38_MT_+_1621_3279_RNR2_16S_50ntFlanks | 1425 | 1458 | 34 | ACGATTAAAGTCCTACGTGATCTGAGTTCAGACC | 84.5 | 3.1 | 1.3 | 2.4 | 1.6E-02 | 2.0E-02 | MB231/MB468 | MP |
| >hg38_21_+_8433172_8446622_RNA45SN1_45S_50ntFlanks@4046.4065.20\|CATTCGAACGTCTGCCCTAT | >hg38_21_+_8433172_8446622_RNA45SN1_45S_50ntFlanks | 4046 | 4065 | 20 | CATTCGAACGTCTGCCCTAT | 134.6 | -3.9 | 1.6 | -2.4 | 1.5E-02 | 2.0E-02 | BT20/MB468 | MITO |
| >hg38_MT_+_1621_3279_RNR2_16S_50ntFlanks@1349.1379.31\|ACAATAGGGTTTACGACCTCGATGTTGGATC | >hg38_MT_+_1621_3279_RNR2_16S_50ntFlanks | 1349 | 1379 | 31 | ACAATAGGGTTTACGACCTCGATGTTGGATC | 52.5 | 3.9 | 1.6 | 2.4 | 1.6E-02 | 2.0E-02 | MB231/MB468 | MP |
| >hg38_21_+_8433172_8446622_RNA45SN1_45S_50ntFlanks@12716.12747.32\|GCCTCGGTTGGCCTCGGATAGCCGGTCCCCCG | >hg38_21_+_8433172_8446622_RNA45SN1_45S_50ntFlanks | 12716 | 12747 | 32 | GCCTCGGTTGGCCTCGGATAGCCGGTCCCCCG | 127.0 | -4.4 | 1.7 | -2.5 | 1.1E-02 | 2.0E-02 | BT20/MB468 | MP |
| >hg38_21_+_8433172_8446622_RNA45SN1_45S_50ntFlanks@12631.12654.24\|GGCGAAGCTACCATCTGTGGGATT | >hg38_21_+_8433172_8446622_RNA45SN1_45S_50ntFlanks | 12631 | 12654 | 24 | GGCGAAGCTACCATCTGTGGGATT | 432.9 | -4.5 | 1.8 | -2.5 | 1.1E-02 | 2.0E-02 | BT20/MB468 | MP |
| >hg38_MT_+_1621_3279_RNR2_16S_50ntFlanks@1369.1407.39\|GATGTTGGATCAGGACATCCCGATGGTGCAGCCGCTATT | >hg38_MT_+_1621_3279_RNR2_16S_50ntFlanks | 1369 | 1407 | 39 | GATGTTGGATCAGGACATCCCGATGGTGCAGCCGCTATT | 38.1 | 4.7 | 2.0 | 2.4 | 1.6E-02 | 2.0E-02 | MB231/MB468 | MP |
| >hg38_21_+_8433172_8446622_RNA45SN1_45S_50ntFlanks@12603.12626.24\|TGTATGTGCTTGGCTGAGGAGCCA | >hg38_21_+_8433172_8446622_RNA45SN1_45S_50ntFlanks | 12603 | 12626 | 24 | TGTATGTGCTTGGCTGAGGAGCCA | 104.0 | -2.6 | 1.1 | -2.4 | 1.7E-02 | 2.0E-02 | BT20/MB468 | CYTO |
| >hg38_21_+_8433172_8446622_RNA45SN1_45S_50ntFlanks@11718.11743.26\|AACGGCGGGAGTAACTATGACTCTCT | >hg38_21_+_8433172_8446622_RNA45SN1_45S_50ntFlanks | 11718 | 11743 | 26 | AACGGCGGGAGTAACTATGACTCTCT | 804.6 | -2.8 | 1.2 | -2.4 | 1.7E-02 | 2.0E-02 | BT20/MB468 | CYTO |
| >hg38_21_+_8433172_8446622_RNA45SN1_45S_50ntFlanks@12988.13013.26\|CAGCTCCCTCGCTGCGATCTATTGAA | >hg38_21_+_8433172_8446622_RNA45SN1_45S_50ntFlanks | 12988 | 13013 | 26 | CAGCTCCCTCGCTGCGATCTATTGAA | 944.9 | -2.2 | 0.8 | -2.8 | 5.9E-03 | 2.0E-02 | BT20/MB231 | CYTO |
| >hg38_21_+_8433172_8446622_RNA45SN1_45S_50ntFlanks@12691.12711.21\|CGGAACGATACGGCAGCGCCG | >hg38_21_+_8433172_8446622_RNA45SN1_45S_50ntFlanks | 12691 | 12711 | 21 | CGGAACGATACGGCAGCGCCG | 153.6 | -3.9 | 1.6 | -2.4 | 1.5E-02 | 2.0E-02 | BT20/MB468 | MITO |
| >hg38_21_+_8433172_8446622_RNA45SN1_45S_50ntFlanks@10466.10487.22\|CGTTGCCCTCGGCCGATCGAAA | >hg38_21_+_8433172_8446622_RNA45SN1_45S_50ntFlanks | 10466 | 10487 | 22 | CGTTGCCCTCGGCCGATCGAAA | 193.0 | -1.9 | 0.8 | -2.4 | 1.5E-02 | 2.0E-02 | BT20/MB468 | MITO |
| >hg38_21_+_8433172_8446622_RNA45SN1_45S_50ntFlanks@11876.11899.24\|AAGACCCTGTTGAGCTTGACTCTA | >hg38_21_+_8433172_8446622_RNA45SN1_45S_50ntFlanks | 11876 | 11899 | 24 | AAGACCCTGTTGAGCTTGACTCTA | 1079.9 | -5.0 | 2.1 | -2.4 | 1.8E-02 | 2.0E-02 | BT20/MB468 | NUC |
| >hg38_21_+_8433172_8446622_RNA45SN1_45S_50ntFlanks@4884.4911.28\|CAAAGCTGAAACTTAAAGGAATTGACGG | >hg38_21_+_8433172_8446622_RNA45SN1_45S_50ntFlanks | 4884 | 4911 | 28 | CAAAGCTGAAACTTAAAGGAATTGACGG | 215.8 | -2.0 | 0.8 | -2.5 | 1.1E-02 | 2.0E-02 | BT20/MB468 | MP |
| >hg38_21_+_8433172_8446622_RNA45SN1_45S_50ntFlanks@3984.4002.19\|GCGGCTTTGGTGACTCTAG | >hg38_21_+_8433172_8446622_RNA45SN1_45S_50ntFlanks | 3984 | 4002 | 19 | GCGGCTTTGGTGACTCTAG | 1880.7 | -5.0 | 2.0 | -2.5 | 1.1E-02 | 2.0E-02 | BT20/MB468 | MP |
| >MI0000736\|hsa-mir-30c-1&WithFlank&1\|+\|40757278\|40757378@23.44.22(+1G);MI0000254\|hsa-mir-30c-2&WithFlank&6\|-\|71376954\|71377037@13.34.22(+1G)&[MIMAT0000244&hsa-miR-30c-5p&offsets\|0\|-1(+1G);m-57&1\|+\|40757300\|40757323&offsets\|0\|-2(+1G)];[MIMAT0000244_1&hsa-miR-30c-5p&offsets\|0\|-1(+1G);m-56&6\|-\|71377002\|71377025&offsets\|0\|-2(+1G)]\|TGTAAACATCCTACACTCTCAGG | >MI0000736\|hsa-mir-30c-1&WithFlank&1\|+\|40757278\|40757378 | 23 | 44 | 22(+1G) | TGTAAACATCCTACACTCTCAGG | 38.9 | -8.7 | 3.1 | -2.8 | 4.5E-03 | 2.0E-02 | BT20/MB231 | MP |
| >hg38_21_+_8433172_8446622_RNA45SN1_45S_50ntFlanks@4976.4993.18\|CCGGACACGGACAGGATT | >hg38_21_+_8433172_8446622_RNA45SN1_45S_50ntFlanks | 4976 | 4993 | 18 | CCGGACACGGACAGGATT | 320.9 | 2.4 | 0.9 | 2.8 | 4.9E-03 | 2.0E-02 | BT20/MB231 | NUC |
| >hg38_MT_+_1621_3279_RNR2_16S_50ntFlanks@837.858.22\|AAAAAAAGTAAAAGGAACTCGG | >hg38_MT_+_1621_3279_RNR2_16S_50ntFlanks | 837 | 858 | 22 | AAAAAAAGTAAAAGGAACTCGG | 186.8 | -3.2 | 1.3 | -2.4 | 1.6E-02 | 2.0E-02 | MB231/MB468 | MP |
| >hg38_MT_+_598_1651_RNR1_12S_50ntFlanks@428.453.26\|AAAGTGGCTTTAACATATCTGAACAC | >hg38_MT_+_598_1651_RNR1_12S_50ntFlanks | 428 | 453 | 26 | AAAGTGGCTTTAACATATCTGAACAC | 150.4 | 2.4 | 1.0 | 2.4 | 1.8E-02 | 2.0E-02 | MB231/MB468 | MITO |
| >hg38_21_+_8433172_8446622_RNA45SN1_45S_50ntFlanks@12203.12220.18\|TAAGGCGAGCTCAGGGAG | >hg38_21_+_8433172_8446622_RNA45SN1_45S_50ntFlanks | 12203 | 12220 | 18 | TAAGGCGAGCTCAGGGAG | 224.9 | -4.3 | 1.8 | -2.4 | 1.5E-02 | 2.0E-02 | BT20/MB468 | MITO |
| >hg38_MT_+_598_1651_RNR1_12S_50ntFlanks@51.87.37\|AATAGGTTTGGTCCTAGCCTTTCTATTAGCTCTTAGT | >hg38_MT_+_598_1651_RNR1_12S_50ntFlanks | 51 | 87 | 37 | AATAGGTTTGGTCCTAGCCTTTCTATTAGCTCTTAGT | 837.7 | 3.6 | 1.4 | 2.5 | 1.1E-02 | 2.0E-02 | BT20/MB468 | MP |
| >MI0000478\|hsa-mir-149&WithFlank&2\|+\|240455995\|240456095@21.43.23&[MIMAT0000450&hsa-miR-149-5p&offsets\|0\|0;m-195&2\|+\|240456015\|240456037&offsets\|0\|0]\|TCTGGCTCCGTGTCTTCACTCCC | >MI0000478\|hsa-mir-149&WithFlank&2\|+\|240455995\|240456095 | 21 | 43 | 23 | TCTGGCTCCGTGTCTTCACTCCC | 63.7 | 4.8 | 2.0 | 2.4 | 1.6E-02 | 2.0E-02 | MB231/MB468 | MP |
| >hg38_21_+_8433172_8446622_RNA45SN1_45S_50ntFlanks@4732.4752.21\|AGAACGAAAGTCGGAGGTTCG | >hg38_21_+_8433172_8446622_RNA45SN1_45S_50ntFlanks | 4732 | 4752 | 21 | AGAACGAAAGTCGGAGGTTCG | 362.8 | -3.4 | 1.3 | -2.5 | 1.1E-02 | 2.0E-02 | BT20/MB468 | MP |
| >hg38_21_+_8433172_8446622_RNA45SN1_45S_50ntFlanks@9029.9065.37\|TCTCCTCGTGGGGGGGCCGGGCCACCCCTCCCACGGC | >hg38_21_+_8433172_8446622_RNA45SN1_45S_50ntFlanks | 9029 | 9065 | 37 | TCTCCTCGTGGGGGGGCCGGGCCACCCCTCCCACGGC | 140.4 | -3.8 | 1.6 | -2.4 | 1.5E-02 | 2.0E-02 | BT20/MB468 | MITO |
| >hg38_21_+_8433172_8446622_RNA45SN1_45S_50ntFlanks@10407.10424.18\|GGGTCAGTCGGTCCTGAG | >hg38_21_+_8433172_8446622_RNA45SN1_45S_50ntFlanks | 10407 | 10424 | 18 | GGGTCAGTCGGTCCTGAG | 859.1 | 2.3 | 0.9 | 2.5 | 1.1E-02 | 2.0E-02 | BT20/MB468 | MP |
| >hg38_21_+_8433172_8446622_RNA45SN1_45S_50ntFlanks@7973.7991.19\|GACGCGACCTCAGATCAGA | >hg38_21_+_8433172_8446622_RNA45SN1_45S_50ntFlanks | 7973 | 7991 | 19 | GACGCGACCTCAGATCAGA | 1217.8 | -2.8 | 1.0 | -2.8 | 5.9E-03 | 2.0E-02 | BT20/MB231 | CYTO |
| >am_trnaMT_SerGCT_MT_+_12207_12265@15.41.27\|GAACTGCTAACTCATGCCCCCATGTCT | >am_trnaMT_SerGCT_MT_+_12207_12265 | 15 | 41 | 27 | GAACTGCTAACTCATGCCCCCATGTCT | 84.0 | 4.7 | 1.9 | 2.5 | 1.1E-02 | 2.0E-02 | BT20/MB468 | MP |
| >hg38_21_+_8433172_8446622_RNA45SN1_45S_50ntFlanks@6676.6696.21\|CTCGTGCGTCGATGAAGAACG | >hg38_21_+_8433172_8446622_RNA45SN1_45S_50ntFlanks | 6676 | 6696 | 21 | CTCGTGCGTCGATGAAGAACG | 352.3 | 4.5 | 1.6 | 2.8 | 4.5E-03 | 2.0E-02 | BT20/MB231 | MP |
| >hg38_21_+_8433172_8446622_RNA45SN1_45S_50ntFlanks@3842.3868.27\|CCTACTTGGATAACTGTGGTAATTCTA | >hg38_21_+_8433172_8446622_RNA45SN1_45S_50ntFlanks | 3842 | 3868 | 27 | CCTACTTGGATAACTGTGGTAATTCTA | 947.1 | 1.0 | 0.4 | 2.4 | 1.7E-02 | 2.0E-02 | BT20/MB468 | CYTO |
| >hg38_21_+_8433172_8446622_RNA45SN1_45S_50ntFlanks@7991.8016.26\|ACGTGGCGACCCGCTGAATTTAAGCA | >hg38_21_+_8433172_8446622_RNA45SN1_45S_50ntFlanks | 7991 | 8016 | 26 | ACGTGGCGACCCGCTGAATTTAAGCA | 542.5 | -2.6 | 1.0 | -2.8 | 5.9E-03 | 2.0E-02 | BT20/MB231 | CYTO |
| >hg38_21_+_8433172_8446622_RNA45SN1_45S_50ntFlanks@9534.9573.40\|AGGAAACTCTGGTGGAGGTCCGTAGCGGTCCTGACGTGCA | >hg38_21_+_8433172_8446622_RNA45SN1_45S_50ntFlanks | 9534 | 9573 | 40 | AGGAAACTCTGGTGGAGGTCCGTAGCGGTCCTGACGTGCA | 705.9 | -1.8 | 0.7 | -2.4 | 1.5E-02 | 2.0E-02 | BT20/MB468 | MITO |
| >hg38_21_+_8433172_8446622_RNA45SN1_45S_50ntFlanks@5071.5094.24\|TTAATTCCGATAACGAACGAGACT | >hg38_21_+_8433172_8446622_RNA45SN1_45S_50ntFlanks | 5071 | 5094 | 24 | TTAATTCCGATAACGAACGAGACT | 252.2 | -4.1 | 1.6 | -2.5 | 1.1E-02 | 2.0E-02 | BT20/MB468 | MP |
| >hg38_21_+_8433172_8446622_RNA45SN1_45S_50ntFlanks@8115.8142.28\|CGCGGGACATGTGGCGTACGGAAGACCC | >hg38_21_+_8433172_8446622_RNA45SN1_45S_50ntFlanks | 8115 | 8142 | 28 | CGCGGGACATGTGGCGTACGGAAGACCC | 1169.3 | -2.8 | 1.1 | -2.5 | 1.1E-02 | 2.0E-02 | BT20/MB468 | MP |
| >trnaMT_ValTAC_MT_+_1602_1670@44.71.28\|ATTTCAACTTAACTTGACCGCTCTGACC | >trnaMT_ValTAC_MT_+_1602_1670 | 44 | 71 | 28 | ATTTCAACTTAACTTGACCGCTCTGACC | 1488.3 | -2.6 | 0.9 | -2.8 | 4.5E-03 | 2.0E-02 | BT20/MB231 | MP |
| >hg38_21_+_8433172_8446622_RNA45SN1_45S_50ntFlanks@9534.9574.41\|AGGAAACTCTGGTGGAGGTCCGTAGCGGTCCTGACGTGCAA | >hg38_21_+_8433172_8446622_RNA45SN1_45S_50ntFlanks | 9534 | 9574 | 41 | AGGAAACTCTGGTGGAGGTCCGTAGCGGTCCTGACGTGCAA | 610.1 | -1.5 | 0.6 | -2.4 | 1.7E-02 | 2.0E-02 | MB231/MB468 | CYTO |
| >MI0000746\|hsa-mir-99b&WithFlank&19\|+\|51692606\|51692687@13.34.22(+1C)&[MIMAT0000689&hsa-miR-99b-5p&offsets\|0\|0(+1C);m-10&19\|+\|51692618\|51692639&offsets\|0\|0(+1C)]\|CACCCGTAGAACCGACCTTGCGC | >MI0000746\|hsa-mir-99b&WithFlank&19\|+\|51692606\|51692687 | 13 | 34 | 22(+1C) | CACCCGTAGAACCGACCTTGCGC | 87.4 | 3.5 | 1.4 | 2.5 | 1.1E-02 | 2.0E-02 | BT20/MB468 | MP |
| >trnaMT_ThrTGT_MT_+_15888_15953@39.69.31\|CGGAGATGAAAACCTTTTTCCAAGGACACCA | >trnaMT_ThrTGT_MT_+_15888_15953 | 39 | 69 | 31 | CGGAGATGAAAACCTTTTTCCAAGGACACCA | 212.4 | 3.7 | 1.5 | 2.5 | 1.1E-02 | 2.0E-02 | BT20/MB468 | MP |
| >hg38_21_+_8433172_8446622_RNA45SN1_45S_50ntFlanks@4733.4766.34\|GAACGAAAGTCGGAGGTTCGAAGACGATCAGATA | >hg38_21_+_8433172_8446622_RNA45SN1_45S_50ntFlanks | 4733 | 4766 | 34 | GAACGAAAGTCGGAGGTTCGAAGACGATCAGATA | 118.7 | 4.4 | 1.7 | 2.5 | 1.1E-02 | 2.0E-02 | BT20/MB468 | MP |
| >trna116_GluCTC_1_-_145399233_145399304@4.33.30,trna59_GluCTC_1_+_249168447_249168518@4.33.30,trna71_GluCTC_1_-_161439189_161439260@4.33.30,trna74_GluCTC_1_-_161431809_161431880@4.33.30,trna77_GluCTC_1_-_161424398_161424469@4.33.30,trna77_GluCTC_6_+_28949976_28950047@4.33.30,trna80_GluCTC_1_-_161417018_161417089@4.33.30,trna87_GluCTC_6_-_126101393_126101464@4.33.30\|CTGGTGGTCTAGTGGTTAGGATTCGGCGCT | >trna116_GluCTC_1_-_145399233_145399304 | 4 | 33 | 30 | CTGGTGGTCTAGTGGTTAGGATTCGGCGCT | 109.2 | 5.3 | 2.1 | 2.5 | 1.1E-02 | 2.0E-02 | BT20/MB468 | MP |
| >hg38_21_+_8433172_8446622_RNA45SN1_45S_50ntFlanks@3838.3859.22\|CTCTCCTACTTGGATAACTGTG | >hg38_21_+_8433172_8446622_RNA45SN1_45S_50ntFlanks | 3838 | 3859 | 22 | CTCTCCTACTTGGATAACTGTG | 107.2 | -4.5 | 1.8 | -2.5 | 1.1E-02 | 2.0E-02 | BT20/MB468 | MP |
| >hg38_21_+_8433172_8446622_RNA45SN1_45S_50ntFlanks@9533.9554.22\|GAGGAAACTCTGGTGGAGGTCC | >hg38_21_+_8433172_8446622_RNA45SN1_45S_50ntFlanks | 9533 | 9554 | 22 | GAGGAAACTCTGGTGGAGGTCC | 217.0 | -3.7 | 1.4 | -2.5 | 1.1E-02 | 2.0E-02 | BT20/MB468 | MP |
| >hg38_21_+_8433172_8446622_RNA45SN1_45S_50ntFlanks@8113.8138.26\|GGCGCGGGACATGTGGCGTACGGAAG | >hg38_21_+_8433172_8446622_RNA45SN1_45S_50ntFlanks | 8113 | 8138 | 26 | GGCGCGGGACATGTGGCGTACGGAAG | 510.2 | -3.2 | 1.3 | -2.4 | 1.5E-02 | 2.0E-02 | BT20/MB468 | MITO |
| >hg38_MT_+_1621_3279_RNR2_16S_50ntFlanks@1189.1216.28\|CGGTTGGGGCGACCTCGGAGCAGAACCC | >hg38_MT_+_1621_3279_RNR2_16S_50ntFlanks | 1189 | 1216 | 28 | CGGTTGGGGCGACCTCGGAGCAGAACCC | 91.3 | 4.5 | 1.8 | 2.5 | 1.1E-02 | 2.0E-02 | BT20/MB468 | MP |
| >hg38_21_+_8433172_8446622_RNA45SN1_45S_50ntFlanks@12420.12456.37\|ATGTCGGCTCTTCCTATCATTGTGAAGCAGAATTCAC | >hg38_21_+_8433172_8446622_RNA45SN1_45S_50ntFlanks | 12420 | 12456 | 37 | ATGTCGGCTCTTCCTATCATTGTGAAGCAGAATTCAC | 118.5 | -3.4 | 1.4 | -2.4 | 1.6E-02 | 2.0E-02 | MB231/MB468 | MP |
| >hg38_21_+_8433172_8446622_RNA45SN1_45S_50ntFlanks@11581.11604.24\|AACTGGTGCGGACCAGGGGAATCC | >hg38_21_+_8433172_8446622_RNA45SN1_45S_50ntFlanks | 11581 | 11604 | 24 | AACTGGTGCGGACCAGGGGAATCC | 242.1 | -1.1 | 0.5 | -2.4 | 1.7E-02 | 2.0E-02 | BT20/MB468 | CYTO |
| >hg38_21_+_8433172_8446622_RNA45SN1_45S_50ntFlanks@6660.6678.19\|GCGGTGGATCACTCGGCTC | >hg38_21_+_8433172_8446622_RNA45SN1_45S_50ntFlanks | 6660 | 6678 | 19 | GCGGTGGATCACTCGGCTC | 163.7 | 4.5 | 1.8 | 2.5 | 1.1E-02 | 2.0E-02 | BT20/MB468 | MP |
| >hg38_21_+_8433172_8446622_RNA45SN1_45S_50ntFlanks@10688.10706.19\|GGCGTCCGGTGAGCTCTCG | >hg38_21_+_8433172_8446622_RNA45SN1_45S_50ntFlanks | 10688 | 10706 | 19 | GGCGTCCGGTGAGCTCTCG | 108.8 | -4.3 | 1.7 | -2.5 | 1.1E-02 | 2.0E-02 | BT20/MB468 | MP |
| >MI0000067\|hsa-let-7f-1&WithFlank&9\|+\|94176341\|94176439@13.32.20(+1C);MI0000068\|hsa-let-7f-2&WithFlank&X\|-\|53557186\|53557280@14.33.20(+1C)&[MIMAT0000067&hsa-let-7f-5p&offsets\|0\|-2(+1C);m-20&9\|+\|94176353\|94176374&offsets\|0\|-2(+1C)];[MIMAT0000067_1&hsa-let-7f-5p&offsets\|0\|-2(+1C);m-19&X\|-\|53557246\|53557267&offsets\|0\|-2(+1C)]\|TGAGGTAGTAGATTGTATAGC | >MI0000067\|hsa-let-7f-1&WithFlank&9\|+\|94176341\|94176439 | 13 | 32 | 20(+1C) | TGAGGTAGTAGATTGTATAGC | 1446.4 | -3.1 | 1.1 | -2.8 | 4.5E-03 | 2.0E-02 | BT20/MB231 | MP |
| >am_MI0000786\|hsa-mir-378a&WithFlank&5\|+\|149732819\|149732896@49.69.21(+1A)&[MIMAT0000732&hsa-miR-378a-3p&offsets\|0\|-1(+1A);m-44&5\|+\|149732867\|149732888&offsets\|0\|-1(+1A)]\|ACTGGACTTGGAGTCAGAAGGA | >am_MI0000786\|hsa-mir-378a&WithFlank&5\|+\|149732819\|149732896 | 49 | 69 | 21(+1A) | ACTGGACTTGGAGTCAGAAGGA | 2404.0 | 2.4 | 0.9 | 2.8 | 5.9E-03 | 2.0E-02 | BT20/MB231 | CYTO |

| | | | | | | | | | | | | | |
|---|---|---|---|---|---|---|---|---|---|---|---|---|---|
| >am_trna10_ValCAC_5_-_180649395_180649467@1.36.36,trna18_ValCAC_5_-_180529253_180529325@1.36.36,trna2_ValCAC_5_+_180524070_180524142@1.36.36,trna6_ValCAC_5_+_180600650_180600722@1.36.36,trna85_ValCAC_1_-_161369490_161369562@1.36.36,trna90_ValCAC_1_-_149684088_149684161@1.36.36,trna98_ValCAC_1_-_149298555_149298627@1.36.36,trna9_ValCAC_6_+_26538282_26538354@1.36.36\|GTTTCCGTAGTGTAGTGGTTATCACGTTCGCCTCAC | >am_trna10_ValCAC_5_-_180649395_180649467 | 1 | 36 | 36 | GTTTCCGTAGTGTAGTGGTTATCACGTTCGCCTCAC | 1212.5 | -3.1 | 1.1 | -2.8 | 5.9E-03 | 2.0E-02 | BT20/MB231 | CYTO |
| >hg38_MT_+_598_1651_RNR1_12S_50ntFlanks@443.480.38\|TATCTGAACACACAATAGCTAAGACCCAAACTGGGATT | >hg38_MT_+_598_1651_RNR1_12S_50ntFlanks | 443 | 480 | 38 | TATCTGAACACACAATAGCTAAGACCCAAACTGGGATT | 1120.6 | -2.4 | 0.9 | -2.8 | 5.6E-03 | 2.0E-02 | BT20/MB231 | MITO |
| >hg38_21_+_8433172_8446622_RNA45SN1_45S_50ntFlanks@5472.5490.19\|ACGGCCCTGGCGGAGCGCT | >hg38_21_+_8433172_8446622_RNA45SN1_45S_50ntFlanks | 5472 | 5490 | 19 | ACGGCCCTGGCGGAGCGCT | 65.2 | 4.6 | 1.8 | 2.5 | 1.1E-02 | 2.0E-02 | BT20/MB468 | MP |
| >hg38_21_+_8433172_8446622_RNA45SN1_45S_50ntFlanks@9622.9655.34\|CTAGTAGCTGGTTCCCTCCGAAGTTTCCCTCAGG | >hg38_21_+_8433172_8446622_RNA45SN1_45S_50ntFlanks | 9622 | 9655 | 34 | CTAGTAGCTGGTTCCCTCCGAAGTTTCCCTCAGG | 104.1 | -4.2 | 1.7 | -2.5 | 1.1E-02 | 2.0E-02 | BT20/MB468 | MP |
| >hg38_21_+_8433172_8446622_RNA45SN1_45S_50ntFlanks@10351.10385.35\|CGAGAACTTTGAAGGCCGAAGTGGAGAAGGGTTCC | >hg38_21_+_8433172_8446622_RNA45SN1_45S_50ntFlanks | 10351 | 10385 | 35 | CGAGAACTTTGAAGGCCGAAGTGGAGAAGGGTTCC | 176.5 | -1.6 | 0.7 | -2.4 | 1.5E-02 | 2.0E-02 | BT20/MB468 | MITO |
| >hg38_21_+_8433172_8446622_RNA45SN1_45S_50ntFlanks@11768.11806.39\|CTAATTAGTGACGCGCATGAATGGATGAACGAGATTCCC | >hg38_21_+_8433172_8446622_RNA45SN1_45S_50ntFlanks | 11768 | 11806 | 39 | CTAATTAGTGACGCGCATGAATGGATGAACGAGATTCCC | 216.4 | -2.8 | 1.1 | -2.4 | 1.5E-02 | 2.0E-02 | BT20/MB468 | MITO |
| >hg38_MT_+_1621_3279_RNR2_16S_50ntFlanks@1380.1407.28\|AGGACATCCCGATGGTGCAGCCGCTATT | >hg38_MT_+_1621_3279_RNR2_16S_50ntFlanks | 1380 | 1407 | 28 | AGGACATCCCGATGGTGCAGCCGCTATT | 58.3 | 3.3 | 1.4 | 2.4 | 1.6E-02 | 2.0E-02 | MB231/MB468 | MP |
| >MI0000289\|hsa-mir-181a-1&WithFlank&1\|-\|198859038\|198859159@70.89.20&[MIMAT0000270&hsa-miR-181a-3p&offsets\|0\|-2;m-125&1\|-\|198859069\|198859090&offsets\|0\|-2]\|ACCATCGACCGTTGATTGTA | >MI0000289\|hsa-mir-181a-1&WithFlank&1\|-\|198859038\|198859159 | 70 | 89 | 20 | ACCATCGACCGTTGATTGTA | 36.0 | 4.0 | 1.7 | 2.4 | 1.6E-02 | 2.0E-02 | MB231/MB468 | MP |
| >hg38_21_+_8433172_8446622_RNA45SN1_45S_50ntFlanks@10868.10905.38\|AAGGGCTGGGTCGGTCGGGCTGGGGCGCGAAGCGGGGC | >hg38_21_+_8433172_8446622_RNA45SN1_45S_50ntFlanks | 10868 | 10905 | 38 | AAGGGCTGGGTCGGTCGGGCTGGGGCGCGAAGCGGGGC | 62.1 | 4.3 | 1.8 | 2.4 | 1.6E-02 | 2.0E-02 | MB231/MB468 | MP |
| >hg38_21_+_8433172_8446622_RNA45SN1_45S_50ntFlanks@10916.10936.21\|CCGCGGCTGGACGAGGCGCCG | >hg38_21_+_8433172_8446622_RNA45SN1_45S_50ntFlanks | 10916 | 10936 | 21 | CCGCGGCTGGACGAGGCGCCG | 246.4 | -3.9 | 1.6 | -2.4 | 1.5E-02 | 2.0E-02 | BT20/MB468 | MITO |
| >hg38_21_+_8433172_8446622_RNA45SN1_45S_50ntFlanks@12626.12661.36\|AATGGGGCGAAGCTACCATCTGTGGGATTATGACTG | >hg38_21_+_8433172_8446622_RNA45SN1_45S_50ntFlanks | 12626 | 12661 | 36 | AATGGGGCGAAGCTACCATCTGTGGGATTATGACTG | 622.0 | -1.4 | 0.6 | -2.4 | 1.7E-02 | 2.0E-02 | MB231/MB468 | CYTO |
| >hg38_21_+_8433172_8446622_RNA45SN1_45S_50ntFlanks@10786.10827.42\|AACAGCCTCTGGCATGTTGGAACAATGTAGGTAAGGGAAGTC | >hg38_21_+_8433172_8446622_RNA45SN1_45S_50ntFlanks | 10786 | 10827 | 42 | AACAGCCTCTGGCATGTTGGAACAATGTAGGTAAGGGAAGTC | 176.9 | -2.9 | 1.2 | -2.5 | 1.1E-02 | 2.0E-02 | BT20/MB468 | MP |
| >hg38_21_+_8433172_8446622_RNA45SN1_45S_50ntFlanks@4451.4479.29\|TCTCGGCGCCCCCTCGATGCTCTTAGCTG | >hg38_21_+_8433172_8446622_RNA45SN1_45S_50ntFlanks | 4451 | 4479 | 29 | TCTCGGCGCCCCCTCGATGCTCTTAGCTG | 163.8 | -2.7 | 1.0 | -2.7 | 6.0E-03 | 2.0E-02 | BT20/MB231 | CYTO |
| >hg38_21_+_8433172_8446622_RNA45SN1_45S_50ntFlanks@11875.11897.23\|GAAGACCCTGTTGAGCTTGACTC | >hg38_21_+_8433172_8446622_RNA45SN1_45S_50ntFlanks | 11875 | 11897 | 23 | GAAGACCCTGTTGAGCTTGACTC | 276.5 | -1.1 | 0.5 | -2.4 | 1.7E-02 | 2.0E-02 | BT20/MB468 | CYTO |
| >trnaMT_ThrTGT_MT_+_15888_15953@37.66.30\|ACCGGAGATGAAAACCTTTTTCCAAGGACA | >trnaMT_ThrTGT_MT_+_15888_15953 | 37 | 66 | 30 | ACCGGAGATGAAAACCTTTTTCCAAGGACA | 71.2 | 6.5 | 2.6 | 2.5 | 1.1E-02 | 2.0E-02 | BT20/MB468 | MP |
| >hg38_21_+_8433172_8446622_RNA45SN1_45S_50ntFlanks@9527.9571.45\|AAGCCAGAGGAAACTCTGGTGGAGGTCCGTAGCGGTCCTGACGTG | >hg38_21_+_8433172_8446622_RNA45SN1_45S_50ntFlanks | 9527 | 9571 | 45 | AAGCCAGAGGAAACTCTGGTGGAGGTCCGTAGCGGTCCTGACGTG | 250.4 | -2.7 | 1.0 | -2.7 | 6.0E-03 | 2.0E-02 | BT20/MB231 | CYTO |
| >hg38_21_+_8433172_8446622_RNA45SN1_45S_50ntFlanks@9544.9573.30\|GGTGGAGGTCCGTAGCGGTCCTGACGTGCA | >hg38_21_+_8433172_8446622_RNA45SN1_45S_50ntFlanks | 9544 | 9573 | 30 | GGTGGAGGTCCGTAGCGGTCCTGACGTGCA | 323.6 | 1.8 | 0.7 | 2.4 | 1.7E-02 | 2.0E-02 | BT20/MB468 | CYTO |
| >hg38_21_+_8433172_8446622_RNA45SN1_45S_50ntFlanks@6671.6698.28\|CTCGGCTCGTGCGTCGATGAAGAACGCA | >hg38_21_+_8433172_8446622_RNA45SN1_45S_50ntFlanks | 6671 | 6698 | 28 | CTCGGCTCGTGCGTCGATGAAGAACGCA | 1045.3 | -4.7 | 1.9 | -2.5 | 1.1E-02 | 2.0E-02 | BT20/MB468 | MP |
| >hg38_MT_+_1621_3279_RNR2_16S_50ntFlanks@1416.1446.31\|GTTTGTTCAACGATTAAAGTCCTACGTGATC | >hg38_MT_+_1621_3279_RNR2_16S_50ntFlanks | 1416 | 1446 | 31 | GTTTGTTCAACGATTAAAGTCCTACGTGATC | 190.4 | 3.0 | 1.2 | 2.4 | 1.6E-02 | 2.0E-02 | MB231/MB468 | MP |
| >hg38_21_+_8433172_8446622_RNA45SN1_45S_50ntFlanks@8176.8193.18\|CTTCTGATCGAGGCCCAG | >hg38_21_+_8433172_8446622_RNA45SN1_45S_50ntFlanks | 8176 | 8193 | 18 | CTTCTGATCGAGGCCCAG | 444.4 | -1.9 | 0.8 | -2.4 | 1.7E-02 | 2.0E-02 | MB231/MB468 | CYTO |
| >hg38_21_+_8433172_8446622_RNA45SN1_45S_50ntFlanks@9497.9525.29\|AAAGATGGTGAACTATGCCTGGGCAGGGC | >hg38_21_+_8433172_8446622_RNA45SN1_45S_50ntFlanks | 9497 | 9525 | 29 | AAAGATGGTGAACTATGCCTGGGCAGGGC | 67.8 | 5.2 | 2.1 | 2.5 | 1.1E-02 | 2.0E-02 | BT20/MB468 | MP |
| >hg38_21_+_8433172_8446622_RNA45SN1_45S_50ntFlanks@6650.6669.20\|ACGACTCTTAGCGGTGGATC | >hg38_21_+_8433172_8446622_RNA45SN1_45S_50ntFlanks | 6650 | 6669 | 20 | ACGACTCTTAGCGGTGGATC | 632.1 | -2.4 | 1.0 | -2.4 | 1.6E-02 | 2.0E-02 | MB231/MB468 | MP |
| >hg38_21_+_8433172_8446622_RNA45SN1_45S_50ntFlanks@12420.12455.36\|ATGTCGGCTCTTCCTATCATTGTGAAGCAGAATTCA | >hg38_21_+_8433172_8446622_RNA45SN1_45S_50ntFlanks | 12420 | 12455 | 36 | ATGTCGGCTCTTCCTATCATTGTGAAGCAGAATTCA | 164.5 | -2.4 | 0.9 | -2.7 | 6.0E-03 | 2.0E-02 | BT20/MB231 | CYTO |
| >hg38_21_+_8433172_8446622_RNA45SN1_45S_50ntFlanks@10404.10429.26\|CATGGGTCAGTCGGTCCTGAGAGATG | >hg38_21_+_8433172_8446622_RNA45SN1_45S_50ntFlanks | 10404 | 10429 | 26 | CATGGGTCAGTCGGTCCTGAGAGATG | 305.4 | 2.3 | 0.8 | 2.8 | 5.7E-03 | 2.0E-02 | BT20/MB231 | MITO |
| >MI0000270\|hsa-mir-181b-1&WithFlank&1\|-\|198858867\|198858988@42.63.22(+1C);MI0000683\|hsa-mir-181b-2&WithFlank&9\|+\|124693704\|124693804@22.43.22(+1C)&[MIMAT0000257&hsa-miR-181b-5p&offsets\|0\|-1(+1C);m-73&1\|-\|198858925\|198858947&offsets\|0\|-1(+1C)];[MIMAT0000257_1&hsa-miR-181b-5p&offsets\|0\|-1(+1C);m-72&9\|+\|124693725\|124693747&offsets\|0\|-1(+1C)]\|AACATTCATTGCTGTCGGTGGGC | >MI0000270\|hsa-mir-181b-1&WithFlank&1\|-\|198858867\|198858988 | 42 | 63 | 22(+1C) | AACATTCATTGCTGTCGGTGGGC | 450.7 | -2.2 | 0.8 | -2.8 | 5.7E-03 | 2.0E-02 | BT20/MB231 | MITO |
| >hg38_21_+_8433172_8446622_RNA45SN1_45S_50ntFlanks@11203.11232.30\|CGGCGGCGGCGACTCTGGACGCGAGCCGGG | >hg38_21_+_8433172_8446622_RNA45SN1_45S_50ntFlanks | 11203 | 11232 | 30 | CGGCGGCGGCGACTCTGGACGCGAGCCGGG | 539.1 | -1.3 | 0.6 | -2.4 | 1.7E-02 | 2.0E-02 | BT20/MB468 | CYTO |
| >hg38_21_+_8433172_8446622_RNA45SN1_45S_50ntFlanks@12151.12182.32\|GTGGGGAGTTTGACTGGGGCGGTACACCTGTC | >hg38_21_+_8433172_8446622_RNA45SN1_45S_50ntFlanks | 12151 | 12182 | 32 | GTGGGGAGTTTGACTGGGGCGGTACACCTGTC | 214.4 | -3.7 | 1.6 | -2.4 | 1.7E-02 | 2.0E-02 | BT20/MB468 | CYTO |
| >hg38_21_+_8433172_8446622_RNA45SN1_45S_50ntFlanks@6651.6695.45\|CGACTCTTAGCGGTGGATCACTCGGCTCGTGCGTCGATGAAGAAC | >hg38_21_+_8433172_8446622_RNA45SN1_45S_50ntFlanks | 6651 | 6695 | 45 | CGACTCTTAGCGGTGGATCACTCGGCTCGTGCGTCGATGAAGAAC | 2212.8 | 2.5 | 1.1 | 2.4 | 1.8E-02 | 2.0E-02 | BT20/MB468 | NUC |
| >hg38_21_+_8433172_8446622_RNA45SN1_45S_50ntFlanks@4983.5001.19\|CGGACAGGATTGACAGATT | >hg38_21_+_8433172_8446622_RNA45SN1_45S_50ntFlanks | 4983 | 5001 | 19 | CGGACAGGATTGACAGATT | 238.7 | 2.2 | 0.8 | 2.7 | 6.0E-03 | 2.0E-02 | BT20/MB231 | CYTO |
| >hg38_21_+_8433172_8446622_RNA45SN1_45S_50ntFlanks@11844.11871.28\|GGGAACGGGCTTGGCGGAATCAGCGGGG | >hg38_21_+_8433172_8446622_RNA45SN1_45S_50ntFlanks | 11844 | 11871 | 28 | GGGAACGGGCTTGGCGGAATCAGCGGGG | 231.4 | -4.3 | 1.7 | -2.5 | 1.1E-02 | 2.0E-02 | BT20/MB468 | MP |
| >hg38_21_+_8433172_8446622_RNA45SN1_45S_50ntFlanks@10395.10432.38\|GCAGTTGAACATGGGTCAGTCGGTCCTGAGAGATGGGC | >hg38_21_+_8433172_8446622_RNA45SN1_45S_50ntFlanks | 10395 | 10432 | 38 | GCAGTTGAACATGGGTCAGTCGGTCCTGAGAGATGGGC | 313.3 | 3.9 | 1.4 | 2.8 | 4.6E-03 | 2.0E-02 | BT20/MB231 | MP |
| >hg38_21_+_8433172_8446622_RNA45SN1_45S_50ntFlanks@7976.7993.18\|GCGACCTCAGATCAGACG | >hg38_21_+_8433172_8446622_RNA45SN1_45S_50ntFlanks | 7976 | 7993 | 18 | GCGACCTCAGATCAGACG | 240.7 | 3.6 | 1.3 | 2.8 | 4.5E-03 | 2.0E-02 | BT20/MB231 | MP |
| >MI0000270\|hsa-mir-181b-1&WithFlank&1\|-\|198858867\|198858988@42.60.19;MI0000683\|hsa-mir-181b-2&WithFlank&9\|+\|124693704\|124693804@22.40.19&[MIMAT0000257&hsa-miR-181b-5p&offsets\|0\|-4;m-73&1\|-\|198858925\|198858947&offsets\|0\|-4];[MIMAT0000257_1&hsa-miR-181b-5p&offsets\|0\|-4;m-72&9\|+\|124693725\|124693747&offsets\|0\|-4]\|AACATTCATTGCTGTCGGT | >MI0000270\|hsa-mir-181b-1&WithFlank&1\|-\|198858867\|198858988 | 42 | 60 | 19 | AACATTCATTGCTGTCGGT | 336.3 | -3.3 | 1.1 | -2.8 | 4.6E-03 | 2.0E-02 | BT20/MB231 | MP |
| >hg38_21_+_8433172_8446622_RNA45SN1_45S_50ntFlanks@12406.12429.24\|TTTTTGATCCTTCGATGTCGGCTC | >hg38_21_+_8433172_8446622_RNA45SN1_45S_50ntFlanks | 12406 | 12429 | 24 | TTTTTGATCCTTCGATGTCGGCTC | 159.7 | -2.8 | 1.2 | -2.4 | 1.8E-02 | 2.0E-02 | MB231/MB468 | NUC |
| >hg38_21_+_8433172_8446622_RNA45SN1_45S_50ntFlanks@11208.11239.32\|GCGGCGACTCTGGACGCGAGCCGGGCCCTTCC | >hg38_21_+_8433172_8446622_RNA45SN1_45S_50ntFlanks | 11208 | 11239 | 32 | GCGGCGACTCTGGACGCGAGCCGGGCCCTTCC | 474.1 | 2.1 | 0.8 | 2.4 | 1.5E-02 | 2.0E-02 | BT20/MB468 | MITO |
| >trnaMT_GluTTC_MT_-_14674_14742@33.72.40,trnalookalike8_GluTTC_5_-_93905172_93905240@33.72.40\|CATATCATTGGTCGTGGTTGTAGTCCGTGCGAGAATACCA | >trnaMT_GluTTC_MT_-_14674_14742 | 33 | 72 | 40 | CATATCATTGGTCGTGGTTGTAGTCCGTGCGAGAATACCA | 776.7 | 4.0 | 1.6 | 2.5 | 1.1E-02 | 2.0E-02 | BT20/MB468 | MP |
| >am_trnaMT_ValTAC_MT_+_1602_1670@38.62.25\|TAGGAGATTTCAACTTAACTTGACC | >am_trnaMT_ValTAC_MT_+_1602_1670 | 38 | 62 | 25 | TAGGAGATTTCAACTTAACTTGACC | 122.3 | 2.2 | 0.9 | 2.4 | 1.6E-02 | 2.0E-02 | MB231/MB468 | MP |
| >hg38_21_+_8433172_8446622_RNA45SN1_45S_50ntFlanks@7981.8003.23\|CTCAGATCAGACGTGGCGACCCG | >hg38_21_+_8433172_8446622_RNA45SN1_45S_50ntFlanks | 7981 | 8003 | 23 | CTCAGATCAGACGTGGCGACCCG | 186.7 | -2.3 | 1.0 | -2.4 | 1.8E-02 | 2.1E-02 | MB231/MB468 | CYTO |
| >hg38_21_+_8433172_8446622_RNA45SN1_45S_50ntFlanks@476.499.24\|TTCGTGATCGATGTGGTGACGTCG | >hg38_21_+_8433172_8446622_RNA45SN1_45S_50ntFlanks | 476 | 499 | 24 | TTCGTGATCGATGTGGTGACGTCG | 534.6 | 3.3 | 1.4 | 2.4 | 1.6E-02 | 2.1E-02 | MB231/MB468 | MP |
| >MI0000264\|hsa-mir-7-2&WithFlank&15\|+\|88611819\|88611940@38.61.24;MI0000265\|hsa-mir-7-3&WithFlank&19\|+\|4770664\|4770785@37.60.24;MI0000263\|hsa-mir-7-1&WithFlank&9\|-\|83969742\|83969863@30.53.24&[MIMAT0000252&hsa-miR-7-5p&offsets\|0\|0;m-400&15\|+\|88611856\|88611879&offsets\|0\|0];[MIMAT0000252_1&hsa-miR-7-5p&offsets\|0\|0;m-402&19\|+\|4770700\|4770723&offsets\|0\|0];[MIMAT0000252_2&hsa-miR-7-5p&offsets\|0\|0;m-401&9\|-\|83969811\|83969834&offsets\|0\|0]\|TGGAAGACTAGTGATTTTGTTGTT | >MI0000264\|hsa-mir-7-2&WithFlank&15\|+\|88611819\|88611940 | 38 | 61 | 24 | TGGAAGACTAGTGATTTTGTTGTT | 14318.2 | 2.4 | 0.9 | 2.5 | 1.1E-02 | 2.1E-02 | BT20/MB468 | MP |
| >hg38_21_+_8433172_8446622_RNA45SN1_45S_50ntFlanks@11529.11546.18\|CCGCGGTTCCGCGCGGCG | >hg38_21_+_8433172_8446622_RNA45SN1_45S_50ntFlanks | 11529 | 11546 | 18 | CCGCGGTTCCGCGCGGCG | 35.7 | -8.2 | 3.3 | -2.5 | 1.1E-02 | 2.1E-02 | BT20/MB468 | MP |
| >MI0000090\|hsa-mir-32&WithFlank&9\|-\|109046223\|109046304@53.73.21&[MIMAT0004505&hsa-miR-32-3p&offsets\|0\|-1;m-518&9\|-\|109046232\|109046252&offsets\|0\|0]\|CAATTTAGTGTGTGTGATATT | >MI0000090\|hsa-mir-32&WithFlank&9\|-\|109046223\|109046304 | 53 | 73 | 21 | CAATTTAGTGTGTGTGATATT | 92.2 | 4.1 | 1.7 | 2.4 | 1.6E-02 | 2.1E-02 | MB231/MB468 | MP |
| >hg38_21_+_8433172_8446622_RNA45SN1_45S_50ntFlanks@12414.12443.30\|CCTTCGATGTCGGCTCTTCCTATCATTGTG | >hg38_21_+_8433172_8446622_RNA45SN1_45S_50ntFlanks | 12414 | 12443 | 30 | CCTTCGATGTCGGCTCTTCCTATCATTGTG | 572.3 | -2.3 | 0.8 | -2.7 | 6.0E-03 | 2.1E-02 | BT20/MB231 | CYTO |
| >hg38_21_+_8433172_8446622_RNA45SN1_45S_50ntFlanks@5492.5509.18\|AGAAGACGGTCGAACTTG | >hg38_21_+_8433172_8446622_RNA45SN1_45S_50ntFlanks | 5492 | 5509 | 18 | AGAAGACGGTCGAACTTG | 540.4 | -2.1 | 0.9 | -2.4 | 1.8E-02 | 2.1E-02 | MB231/MB468 | CYTO |
| >hg38_21_+_8433172_8446622_RNA45SN1_45S_50ntFlanks@11586.11612.27\|GTGCGGACCAGGGGAATCCGACTGTTT | >hg38_21_+_8433172_8446622_RNA45SN1_45S_50ntFlanks | 11586 | 11612 | 27 | GTGCGGACCAGGGGAATCCGACTGTTT | 1008.5 | -1.5 | 0.6 | -2.4 | 1.5E-02 | 2.1E-02 | BT20/MB468 | MITO |

<table>
<tr><th></th><th></th><th></th><th></th><th></th><th></th><th></th><th></th><th></th><th></th><th></th><th></th><th></th><th></th></tr>
<tr><td>>trna10_AspGTC_12_-_125424193_125424264@38.72.35,trna12_AspGTC_12_-_125411891_125411962@38.72.35,trna144_AspGTC_6_-_27551236_27551307@38.72.35,trna38_AspGTC_17_-_8125556_8125627@38.72.35,trna45_AspGTC_6_+_27447453_27447524@38.72.35,trna48_AspGTC_6_+_27471523_27471594@38.72.35,trna4_AspGTC_12_+_96429799_96429870@38.72.35,trna69_AspGTC_1_-_161440205_161440276@38.72.35,trna72_AspGTC_1_-_161432824_161432895@38.72.35,trna75_AspGTC_1_-_161425414_161425485@38.72.35,trna78_AspGTC_1_-_161418033_161418104@38.72.35,trna81_AspGTC_1_-_161410615_161410686@38.72.35|CGCGGGAGACCGGGGTTCGATTCCCCGACGGGGAG</td><td>>trna10_AspGTC_12_-_125424193_125424264</td><td>38</td><td>72</td><td>35</td><td>CGCGGGAGACCGGGGTTCGATTCCCCGACGGGGAG</td><td>224.6</td><td>-3.0</td><td>1.3</td><td>-2.4</td><td>1.8E-02</td><td>2.1E-02</td><td>MB231/MB468</td><td>NUC</td></tr>
<tr><td>>hg38_21_+_8433172_8446622_RNA45SN1_45S_50ntFlanks@5468.5514.47|GCCCACGGCCCTGGCGGAGCGCTGAGAAGACGGTCGAACTTGACTAT</td><td>>hg38_21_+_8433172_8446622_RNA45SN1_45S_50ntFlanks</td><td>5468</td><td>5514</td><td>47</td><td>GCCCACGGCCCTGGCGGAGCGCTGAGAAGACGGTCGAACTTGACTAT</td><td>128.9</td><td>-2.8</td><td>1.2</td><td>-2.4</td><td>1.8E-02</td><td>2.1E-02</td><td>MB231/MB468</td><td>CYTO</td></tr>
<tr><td>>hg38_21_+_8433172_8446622_RNA45SN1_45S_50ntFlanks@10350.10373.24|ACGAGAACTTTGAAGGCCGAAGTG</td><td>>hg38_21_+_8433172_8446622_RNA45SN1_45S_50ntFlanks</td><td>10350</td><td>10373</td><td>24</td><td>ACGAGAACTTTGAAGGCCGAAGTG</td><td>263.5</td><td>-1.4</td><td>0.6</td><td>-2.4</td><td>1.5E-02</td><td>2.1E-02</td><td>BT20/MB468</td><td>MITO</td></tr>
<tr><td>>hg38_21_+_8433172_8446622_RNA45SN1_45S_50ntFlanks@8358.8395.38|AAAGAACTTTGAAGAGAGAGTTCAAGAGGGCGTGAAAC</td><td>>hg38_21_+_8433172_8446622_RNA45SN1_45S_50ntFlanks</td><td>8358</td><td>8395</td><td>38</td><td>AAAGAACTTTGAAGAGAGAGTTCAAGAGGGCGTGAAAC</td><td>135.4</td><td>5.5</td><td>1.9</td><td>2.8</td><td>4.6E-03</td><td>2.1E-02</td><td>BT20/MB231</td><td>MP</td></tr>
<tr><td>>hg38_MT_+_1621_3279_RNR2_16S_50ntFlanks@1368.1401.34|CGATGTTGGATCAGGACATCCCGATGGTGCAGCC</td><td>>hg38_MT_+_1621_3279_RNR2_16S_50ntFlanks</td><td>1368</td><td>1401</td><td>34</td><td>CGATGTTGGATCAGGACATCCCGATGGTGCAGCC</td><td>87.0</td><td>4.0</td><td>1.6</td><td>2.4</td><td>1.6E-02</td><td>2.1E-02</td><td>MB231/MB468</td><td>MP</td></tr>
<tr><td>>MI0000749|hsa-mir-30e&WithFlank&1|+|40754349|40754452@23.46.24(+2U)&[MIMAT0000692&hsa-miR-30e-5p&offsets|0|+2(+2U);m-24&1|+|40754371|40754394&offsets|0|0(+2U)]|TGTAAACATCCTTGACTGGAAGCTTT</td><td>>MI0000749|hsa-mir-30e&WithFlank&1|+|40754349|40754452</td><td>23</td><td>46</td><td>24(+2U)</td><td>TGTAAACATCCTTGACTGGAAGCTTT</td><td>386.9</td><td>2.1</td><td>0.9</td><td>2.4</td><td>1.6E-02</td><td>2.1E-02</td><td>MB231/MB468</td><td>MP</td></tr>
<tr><td>>hg38_21_+_8433172_8446622_RNA45SN1_45S_50ntFlanks@12008.12039.32|CCGCCGGTGAAATACCACTACTCTGATCGTTT</td><td>>hg38_21_+_8433172_8446622_RNA45SN1_45S_50ntFlanks</td><td>12008</td><td>12039</td><td>32</td><td>CCGCCGGTGAAATACCACTACTCTGATCGTTT</td><td>362.8</td><td>-4.7</td><td>1.9</td><td>-2.5</td><td>1.1E-02</td><td>2.1E-02</td><td>BT20/MB468</td><td>MP</td></tr>
<tr><td>>hg38_21_+_8433172_8446622_RNA45SN1_45S_50ntFlanks@12155.12181.27|GGAGTTTGACTGGGGCGGTACACCTGT</td><td>>hg38_21_+_8433172_8446622_RNA45SN1_45S_50ntFlanks</td><td>12155</td><td>12181</td><td>27</td><td>GGAGTTTGACTGGGGCGGTACACCTGT</td><td>116.6</td><td>-4.2</td><td>1.7</td><td>-2.5</td><td>1.1E-02</td><td>2.1E-02</td><td>BT20/MB468</td><td>MP</td></tr>
<tr><td>>MI0000750|hsa-mir-26a-2&WithFlank&12|-|57824603|57824698@20.39.20(+1A);MI0000083|hsa-mir-26a-1&WithFlank&3|+|37969398|37969486@16.35.20(+1A)&[MIMAT0000082&hsa-miR-26a-5p&offsets|0|-2(+1A);m-33&12|-|57824658|57824679&offsets|0|-2(+1A)];[MIMAT0000082_1&hsa-miR-26a-5p&offsets|0|-2(+1A);m-34&3|+|37969413|37969434&offsets|0|-2(+1A)]|TTCAAGTAATCCAGGATAGGA</td><td>>MI0000750|hsa-mir-26a-2&WithFlank&12|-|57824603|57824698</td><td>20</td><td>39</td><td>20(+1A)</td><td>TTCAAGTAATCCAGGATAGGA</td><td>116.3</td><td>4.1</td><td>1.7</td><td>2.4</td><td>1.6E-02</td><td>2.1E-02</td><td>MB231/MB468</td><td>MP</td></tr>
<tr><td>>hg38_21_+_8433172_8446622_RNA45SN1_45S_50ntFlanks@8528.8549.22|CTTCCCCCGCCGCCCCTCCTCC</td><td>>hg38_21_+_8433172_8446622_RNA45SN1_45S_50ntFlanks</td><td>8528</td><td>8549</td><td>22</td><td>CTTCCCCCGCCGCCCCTCCTCC</td><td>317.7</td><td>-3.5</td><td>1.5</td><td>-2.4</td><td>1.8E-02</td><td>2.1E-02</td><td>BT20/MB468</td><td>CYTO</td></tr>
<tr><td>>hg38_21_+_8433172_8446622_RNA45SN1_45S_50ntFlanks@9548.9573.26|GAGGTCCGTAGCGGTCCTGACGTGCA</td><td>>hg38_21_+_8433172_8446622_RNA45SN1_45S_50ntFlanks</td><td>9548</td><td>9573</td><td>26</td><td>GAGGTCCGTAGCGGTCCTGACGTGCA</td><td>666.1</td><td>1.6</td><td>0.7</td><td>2.4</td><td>1.8E-02</td><td>2.1E-02</td><td>MB231/MB468</td><td>CYTO</td></tr>
<tr><td>>hg38_21_+_8433172_8446622_RNA45SN1_45S_50ntFlanks@11774.11791.18|AGTGACGCGCATGAATGG</td><td>>hg38_21_+_8433172_8446622_RNA45SN1_45S_50ntFlanks</td><td>11774</td><td>11791</td><td>18</td><td>AGTGACGCGCATGAATGG</td><td>113.8</td><td>-4.1</td><td>1.7</td><td>-2.4</td><td>1.6E-02</td><td>2.1E-02</td><td>MB231/MB468</td><td>MP</td></tr>
<tr><td>>hg38_21_+_8433172_8446622_RNA45SN1_45S_50ntFlanks@8171.8213.43|AAGTCCTTCTGATCGAGGCCCAGCCCGTGGACGGTGTGAGGCC</td><td>>hg38_21_+_8433172_8446622_RNA45SN1_45S_50ntFlanks</td><td>8171</td><td>8213</td><td>43</td><td>AAGTCCTTCTGATCGAGGCCCAGCCCGTGGACGGTGTGAGGCC</td><td>510.0</td><td>-1.8</td><td>0.8</td><td>-2.4</td><td>1.8E-02</td><td>2.1E-02</td><td>MB231/MB468</td><td>NUC</td></tr>
<tr><td>>hg38_21_+_8433172_8446622_RNA45SN1_45S_50ntFlanks@12948.12988.41|GACGACCTGCTTCTGGGTCGGGGTTTCGTACGTAGCAGAGC</td><td>>hg38_21_+_8433172_8446622_RNA45SN1_45S_50ntFlanks</td><td>12948</td><td>12988</td><td>41</td><td>GACGACCTGCTTCTGGGTCGGGGTTTCGTACGTAGCAGAGC</td><td>682.8</td><td>-1.9</td><td>0.8</td><td>-2.4</td><td>1.8E-02</td><td>2.1E-02</td><td>MB231/MB468</td><td>NUC</td></tr>
<tr><td>>hg38_21_+_8433172_8446622_RNA45SN1_45S_50ntFlanks@10878.10902.25|TCGGTCGGGCTGGGGCGCGAAGCGG</td><td>>hg38_21_+_8433172_8446622_RNA45SN1_45S_50ntFlanks</td><td>10878</td><td>10902</td><td>25</td><td>TCGGTCGGGCTGGGGCGCGAAGCGG</td><td>297.5</td><td>-4.4</td><td>1.7</td><td>-2.5</td><td>1.2E-02</td><td>2.1E-02</td><td>BT20/MB468</td><td>MP</td></tr>
<tr><td>>MI0000264|hsa-mir-7-2&WithFlank&15|+|88611819|88611940@38.61.24(+2A);MI0000265|hsa-mir-7-3&WithFlank&19|+|4770664|4770785@37.60.24(+2A);MI0000263|hsa-mir-7-1&WithFlank&9|-|83969742|83969863@30.53.24(+2A)&[MIMAT0000252&hsa-miR-7-5p&offsets|0|0(+2A);m-400&15|+|88611856|88611879&offsets|0|0(+2A)];[MIMAT0000252_1&hsa-miR-7-5p&offsets|0|0(+2A);m-402&19|+|4770700|4770723&offsets|0|0(+2A)];[MIMAT0000252_2&hsa-miR-7-5p&offsets|0|0(+2A);m-401&9|-|83969811|83969834&offsets|0|0(+2A)]|TGGAAGACTAGTGATTTTGTTGTTAA</td><td>>MI0000264|hsa-mir-7-2&WithFlank&15|+|88611819|88611940</td><td>38</td><td>61</td><td>24(+2A)</td><td>TGGAAGACTAGTGATTTTGTTGTTAA</td><td>32.5</td><td>4.8</td><td>2.0</td><td>2.4</td><td>1.6E-02</td><td>2.1E-02</td><td>MB231/MB468</td><td>MP</td></tr>
<tr><td>>hg38_21_+_8433172_8446622_RNA45SN1_45S_50ntFlanks@3992.4023.32|GGTGACTCTAGATAACCTCGGGCCGATCGCAC</td><td>>hg38_21_+_8433172_8446622_RNA45SN1_45S_50ntFlanks</td><td>3992</td><td>4023</td><td>32</td><td>GGTGACTCTAGATAACCTCGGGCCGATCGCAC</td><td>168.6</td><td>2.3</td><td>0.9</td><td>2.5</td><td>1.2E-02</td><td>2.1E-02</td><td>BT20/MB468</td><td>MP</td></tr>
<tr><td>>MI0000750|hsa-mir-26a-2&WithFlank&12|-|57824603|57824698@21.41.21;MI0000083|hsa-mir-26a-1&WithFlank&3|+|37969398|37969486@17.37.21&[MIMAT0000082&hsa-miR-26a-5p&offsets|+1|0;m-33&12|-|57824658|57824679&offsets|+1|0];[MIMAT0000082_1&hsa-miR-26a-5p&offsets|+1|0;m-34&3|+|37969413|37969434&offsets|+1|0]|TCAAGTAATCCAGGATAGGCT</td><td>>MI0000750|hsa-mir-26a-2&WithFlank&12|-|57824603|57824698</td><td>21</td><td>41</td><td>21</td><td>TCAAGTAATCCAGGATAGGCT</td><td>201.1</td><td>-4.2</td><td>1.5</td><td>-2.8</td><td>4.6E-03</td><td>2.1E-02</td><td>BT20/MB231</td><td>MP</td></tr>
<tr><td>>hg38_21_+_8433172_8446622_RNA45SN1_45S_50ntFlanks@8341.8375.35|CCGTAAGGGAAAGTTGAAAAGAACTTTGAAGAGAG</td><td>>hg38_21_+_8433172_8446622_RNA45SN1_45S_50ntFlanks</td><td>8341</td><td>8375</td><td>35</td><td>CCGTAAGGGAAAGTTGAAAAGAACTTTGAAGAGAG</td><td>2538.5</td><td>5.2</td><td>1.8</td><td>2.8</td><td>4.6E-03</td><td>2.1E-02</td><td>BT20/MB231</td><td>MP</td></tr>
<tr><td>>hg38_21_+_8433172_8446622_RNA45SN1_45S_50ntFlanks@11197.11227.31|GGCCGGCGGCGGCGGCGACTCTGGACGCGAG</td><td>>hg38_21_+_8433172_8446622_RNA45SN1_45S_50ntFlanks</td><td>11197</td><td>11227</td><td>31</td><td>GGCCGGCGGCGGCGGCGACTCTGGACGCGAG</td><td>61.1</td><td>-4.2</td><td>1.7</td><td>-2.5</td><td>1.2E-02</td><td>2.1E-02</td><td>BT20/MB468</td><td>MP</td></tr>
<tr><td>>hg38_21_+_8433172_8446622_RNA45SN1_45S_50ntFlanks@12912.12930.19|CACGTTCGTGGGGAACCTG</td><td>>hg38_21_+_8433172_8446622_RNA45SN1_45S_50ntFlanks</td><td>12912</td><td>12930</td><td>19</td><td>CACGTTCGTGGGGAACCTG</td><td>148.7</td><td>-2.8</td><td>1.2</td><td>-2.4</td><td>1.8E-02</td><td>2.1E-02</td><td>MB231/MB468</td><td>CYTO</td></tr>
<tr><td>>hg38_21_+_8433172_8446622_RNA45SN1_45S_50ntFlanks@7968.7988.21|TCCGAGACGCGACCTCAGATC</td><td>>hg38_21_+_8433172_8446622_RNA45SN1_45S_50ntFlanks</td><td>7968</td><td>7988</td><td>21</td><td>TCCGAGACGCGACCTCAGATC</td><td>238.6</td><td>2.1</td><td>0.8</td><td>2.8</td><td>5.8E-03</td><td>2.1E-02</td><td>BT20/MB231</td><td>MITO</td></tr>
<tr><td>>hg38_21_+_8433172_8446622_RNA45SN1_45S_50ntFlanks@12071.12097.27|CCCCGAGGGGCTCTCGCTTCTGGCGCC</td><td>>hg38_21_+_8433172_8446622_RNA45SN1_45S_50ntFlanks</td><td>12071</td><td>12097</td><td>27</td><td>CCCCGAGGGGCTCTCGCTTCTGGCGCC</td><td>300.2</td><td>-3.1</td><td>1.3</td><td>-2.4</td><td>1.5E-02</td><td>2.1E-02</td><td>BT20/MB468</td><td>MITO</td></tr>
<tr><td>>hg38_21_+_8433172_8446622_RNA45SN1_45S_50ntFlanks@11582.11599.18|ACTGGTGCGGACCAGGGG</td><td>>hg38_21_+_8433172_8446622_RNA45SN1_45S_50ntFlanks</td><td>11582</td><td>11599</td><td>18</td><td>ACTGGTGCGGACCAGGGG</td><td>88.6</td><td>-4.2</td><td>1.8</td><td>-2.4</td><td>1.6E-02</td><td>2.1E-02</td><td>MB231/MB468</td><td>MP</td></tr>
<tr><td>>hg38_21_+_8433172_8446622_RNA45SN1_45S_50ntFlanks@11603.11627.25|CCGACTGTTTAATTAAAACAAAGCA</td><td>>hg38_21_+_8433172_8446622_RNA45SN1_45S_50ntFlanks</td><td>11603</td><td>11627</td><td>25</td><td>CCGACTGTTTAATTAAAACAAAGCA</td><td>171.4</td><td>5.9</td><td>2.1</td><td>2.8</td><td>4.6E-03</td><td>2.1E-02</td><td>BT20/MB231</td><td>MP</td></tr>
<tr><td>>MI0001445|hsa-mir-423&WithFlank&17|+|30117073|30117178@59.80.22&[MIMAT0001340&hsa-miR-423-3p&offsets|0|-1;m-100&17|+|30117131|30117153&offsets|0|-1]|AGCTCGGTCTGAGGCCCCTCAG</td><td>>MI0001445|hsa-mir-423&WithFlank&17|+|30117073|30117178</td><td>59</td><td>80</td><td>22</td><td>AGCTCGGTCTGAGGCCCCTCAG</td><td>292.8</td><td>1.5</td><td>0.6</td><td>2.4</td><td>1.8E-02</td><td>2.1E-02</td><td>BT20/MB468</td><td>NUC</td></tr>
<tr><td>>hg38_21_+_8433172_8446622_RNA45SN1_45S_50ntFlanks@6650.6669.20|ACGACTCTTAGCGGTGGATC</td><td>>hg38_21_+_8433172_8446622_RNA45SN1_45S_50ntFlanks</td><td>6650</td><td>6669</td><td>20</td><td>ACGACTCTTAGCGGTGGATC</td><td>1868.5</td><td>2.4</td><td>0.9</td><td>2.5</td><td>1.2E-02</td><td>2.1E-02</td><td>BT20/MB468</td><td>MP</td></tr>
<tr><td>>hg38_21_+_8433172_8446622_RNA45SN1_45S_50ntFlanks@9492.9515.24|ACCCGAAAGATGGTGAACTATGCC</td><td>>hg38_21_+_8433172_8446622_RNA45SN1_45S_50ntFlanks</td><td>9492</td><td>9515</td><td>24</td><td>ACCCGAAAGATGGTGAACTATGCC</td><td>46.0</td><td>5.2</td><td>2.1</td><td>2.5</td><td>1.2E-02</td><td>2.1E-02</td><td>BT20/MB468</td><td>MP</td></tr>
<tr><td>>hg38_21_+_8433172_8446622_RNA45SN1_45S_50ntFlanks@12841.12871.31|CGGAGTGCCCTTCGTCCTGGGAAACGGGGCG</td><td>>hg38_21_+_8433172_8446622_RNA45SN1_45S_50ntFlanks</td><td>12841</td><td>12871</td><td>31</td><td>CGGAGTGCCCTTCGTCCTGGGAAACGGGGCG</td><td>106.8</td><td>-4.3</td><td>1.7</td><td>-2.5</td><td>1.2E-02</td><td>2.1E-02</td><td>BT20/MB468</td><td>MP</td></tr>
<tr><td>>am_trna111_HisGTG_1_-_147774845_147774916@-1G.29.30,trna118_HisGTG_1_-_145396881_145396952@-1G.29.30,trna16_HisGTG_1_+_146544773_146544844@-1G.29.30,trna1_HisGTG_15_+_45493349_45493420@-1G.29.30,trna21_HisGTG_1_+_147753471_147753542@-1G.29.30,trna33_HisGTG_6_+_27125906_27125977@-1G.29.30,trna7_HisGTG_9_-_14433938_14434009@-1G.29.30,trna8_HisGTG_15_-_45492611_45492682@-1G.29.30,trna9_HisGTG_15_-_45490804_45490875@-1G.29.30|GGCCGTGATCGTATAGTGGTTAGTACTCTG</td><td>>am_trna111_HisGTG_1_-_147774845_147774916</td><td>-1G</td><td>29</td><td>30</td><td>GGCCGTGATCGTATAGTGGTTAGTACTCTG</td><td>682.0</td><td>-2.7</td><td>1.0</td><td>-2.7</td><td>6.1E-03</td><td>2.1E-02</td><td>BT20/MB231</td><td>CYTO</td></tr>
<tr><td>>MI0000081|hsa-mir-24-2&WithFlank&19|-|13836281|13836365@18.41.24&[MIMAT0004497&hsa-miR-24-2-5p&offsets|-1|+1;m-226&19|-|13836326|13836348&offsets|0|+1]|GTGCCTACTGAGCTGAAACACAGT</td><td>>MI0000081|hsa-mir-24-2&WithFlank&19|-|13836281|13836365</td><td>18</td><td>41</td><td>24</td><td>GTGCCTACTGAGCTGAAACACAGT</td><td>472.6</td><td>-2.2</td><td>0.8</td><td>-2.8</td><td>5.8E-03</td><td>2.1E-02</td><td>BT20/MB231</td><td>MITO</td></tr>
<tr><td>>hg38_21_+_8433172_8446622_RNA45SN1_45S_50ntFlanks@12416.12436.21|TTCGATGTCGGCTCTTCCTAT</td><td>>hg38_21_+_8433172_8446622_RNA45SN1_45S_50ntFlanks</td><td>12416</td><td>12436</td><td>21</td><td>TTCGATGTCGGCTCTTCCTAT</td><td>106.5</td><td>-4.2</td><td>1.7</td><td>-2.5</td><td>1.2E-02</td><td>2.1E-02</td><td>BT20/MB468</td><td>MP</td></tr>
<tr><td>>hg38_21_+_8433172_8446622_RNA45SN1_45S_50ntFlanks@12456.12486.31|CCAAGCGTTGGATTGTTCACCCACTAATAGG</td><td>>hg38_21_+_8433172_8446622_RNA45SN1_45S_50ntFlanks</td><td>12456</td><td>12486</td><td>31</td><td>CCAAGCGTTGGATTGTTCACCCACTAATAGG</td><td>5065.5</td><td>-5.1</td><td>2.0</td><td>-2.5</td><td>1.2E-02</td><td>2.1E-02</td><td>BT20/MB468</td><td>MP</td></tr>
<tr><td>>hg38_21_+_8433172_8446622_RNA45SN1_45S_50ntFlanks@4449.4466.18|CCTCTCGGCGCCCCCTCG</td><td>>hg38_21_+_8433172_8446622_RNA45SN1_45S_50ntFlanks</td><td>4449</td><td>4466</td><td>18</td><td>CCTCTCGGCGCCCCCTCG</td><td>376.3</td><td>-1.9</td><td>0.8</td><td>-2.4</td><td>1.6E-02</td><td>2.1E-02</td><td>BT20/MB468</td><td>MITO</td></tr>
<tr><td>>MI0000067|hsa-let-7f-1&WithFlank&9|+|94176341|94176439@13.33.21(+1A);MI0000068|hsa-let-7f-2&WithFlank&X|-|53557186|53557280@14.34.21(+1A)&[MIMAT0000067&hsa-let-7f-5p&offsets|0|-1(+1A);m-20&9|+|94176353|94176374&offsets|0|-1(+1A)];[MIMAT0000067_1&hsa-let-7f-5p&offsets|0|-1(+1A);m-19&X|-|53557246|53557267&offsets|0|-1(+1A)]|TGAGGTAGTAGATTGTATAGTA</td><td>>MI0000067|hsa-let-7f-1&WithFlank&9|+|94176341|94176439</td><td>13</td><td>33</td><td>21(+1A)</td><td>TGAGGTAGTAGATTGTATAGTA</td><td>5087.9</td><td>2.3</td><td>0.8</td><td>2.7</td><td>6.1E-03</td><td>2.1E-02</td><td>BT20/MB231</td><td>CYTO</td></tr>
<tr><td>>MI0000076|hsa-mir-20a&WithFlank&13|+|91351059|91351141@14.35.22(+1A)&[MIMAT0000075&hsa-miR-20a-5p&offsets|0|-1(+1A);m-96&13|+|91351072|91351094&offsets|0|-1(+1A)]|TAAAGTGCTTATAGTGCAGGTAA</td><td>>MI0000076|hsa-mir-20a&WithFlank&13|+|91351059|91351141</td><td>14</td><td>35</td><td>22(+1A)</td><td>TAAAGTGCTTATAGTGCAGGTAA</td><td>4462.0</td><td>-4.5</td><td>1.6</td><td>-2.8</td><td>4.6E-03</td><td>2.1E-02</td><td>BT20/MB231</td><td>MP</td></tr>
<tr><td>>MI0000088|hsa-mir-30a&WithFlank&6|-|71403545|71403627@12.30.19&[MIMAT0000087&hsa-miR-30a-5p&offsets|0|-3;m-15&6|-|71403595|71403616&offsets|0|-3]|TGTAAACATCCTCGACTGG</td><td>>MI0000088|hsa-mir-30a&WithFlank&6|-|71403545|71403627</td><td>12</td><td>30</td><td>19</td><td>TGTAAACATCCTCGACTGG</td><td>208.7</td><td>3.1</td><td>1.3</td><td>2.4</td><td>1.7E-02</td><td>2.1E-02</td><td>MB231/MB468</td><td>MP</td></tr>
<tr><td>>hg38_21_+_8433172_8446622_RNA45SN1_45S_50ntFlanks@5472.5517.46|ACGGCCCTGGCGGAGCGCTGAGAAGACGGTCGAACTTGACTATCTA</td><td>>hg38_21_+_8433172_8446622_RNA45SN1_45S_50ntFlanks</td><td>5472</td><td>5517</td><td>46</td><td>ACGGCCCTGGCGGAGCGCTGAGAAGACGGTCGAACTTGACTATCTA</td><td>237.2</td><td>-2.7</td><td>1.0</td><td>-2.7</td><td>6.1E-03</td><td>2.1E-02</td><td>BT20/MB231</td><td>CYTO</td></tr>
<tr><td>>MI0000542|hsa-mir-320a&WithFlank&8|-|22244960|22245043@48.70.23(+1C)&[MIMAT0000510&hsa-miR-320a-3p&offsets|0|+1(+1C);m-60&8|-|22244975|22244996&offsets|0|+1(+1C)]|AAAAGCTGGGTTGAGAGGGCGAAC</td><td>>MI0000542|hsa-mir-320a&WithFlank&8|-|22244960|22245043</td><td>48</td><td>70</td><td>23(+1C)</td><td>AAAAGCTGGGTTGAGAGGGCGAAC</td><td>446.6</td><td>2.1</td><td>0.8</td><td>2.8</td><td>5.8E-03</td><td>2.1E-02</td><td>BT20/MB231</td><td>MITO</td></tr>
</table>

<table>
<tr><th></th><th></th><th></th><th></th><th></th><th></th><th></th><th></th><th></th><th></th><th></th><th></th><th></th><th></th></tr>
<tr><td>>hg38_21_+_8433172_8446622_RNA45SN1_45S_50ntFlanks@12072.12110.39|CCCGAGGGGCTCTCGCTTCTGGCGCCAAGCGCCCGGCCG</td><td>>hg38_21_+_8433172_8446622_RNA45SN1_45S_50ntFlanks</td><td>12072</td><td>12110</td><td>39</td><td>CCCGAGGGGCTCTCGCTTCTGGCGCCAAGCGCCCGGCCG</td><td>185.8</td><td>-3.8</td><td>1.6</td><td>-2.4</td><td>1.6E-02</td><td>2.1E-02</td><td>BT20/MB468</td><td>MITO</td></tr>
<tr><td>>hg38_21_+_8433172_8446622_RNA45SN1_45S_50ntFlanks@7978.8018.41|GACCTCAGATCAGACGTGGCGACCCGCTGAATTTAAGCATA</td><td>>hg38_21_+_8433172_8446622_RNA45SN1_45S_50ntFlanks</td><td>7978</td><td>8018</td><td>41</td><td>GACCTCAGATCAGACGTGGCGACCCGCTGAATTTAAGCATA</td><td>274.7</td><td>-6.6</td><td>2.4</td><td>-2.7</td><td>6.1E-03</td><td>2.1E-02</td><td>BT20/MB231</td><td>CYTO</td></tr>
<tr><td>>MI0000736|hsa-mir-30c-1&WithFlank&1|+|40757278|40757378@23.46.24;MI0000254|hsa-mir-30c-2&WithFlank&6|-|71376954|71377037@13.36.24&[MIMAT0000244&hsa-miR-30c-5p&offsets|0|+1;m-57&1|+|40757300|40757323&offsets|0|0];[MIMAT0000244_1&hsa-miR-30c-5p&offsets|0|+1;m-56&6|-|71377002|71377025&offsets|0|0]|TGTAAACATCCTACACTCTCAGCT</td><td>>MI0000736|hsa-mir-30c-1&WithFlank&1|+|40757278|40757378</td><td>23</td><td>46</td><td>24</td><td>TGTAAACATCCTACACTCTCAGCT</td><td>15539.8</td><td>-3.0</td><td>1.1</td><td>-2.8</td><td>4.6E-03</td><td>2.1E-02</td><td>BT20/MB231</td><td>MP</td></tr>
<tr><td>>MI0003760|hsa-mir-671&WithFlank&7|+|151238415|151238544@35.59.25&[MIMAT0003880&hsa-miR-671-5p&offsets|0|+2;m-316&7|+|151238449|151238472&offsets|0|+1]|AGGAAGCCCTGGAGGGGCTGGAGGT</td><td>>MI0003760|hsa-mir-671&WithFlank&7|+|151238415|151238544</td><td>35</td><td>59</td><td>25</td><td>AGGAAGCCCTGGAGGGGCTGGAGGT</td><td>250.7</td><td>4.1</td><td>1.6</td><td>2.5</td><td>1.2E-02</td><td>2.1E-02</td><td>BT20/MB468</td><td>MP</td></tr>
<tr><td>>MI0000076|hsa-mir-20a&WithFlank&13|+|91351059|91351141@14.35.22(+1A)&[MIMAT0000075&hsa-miR-20a-5p&offsets|0|-1(+1A);m-96&13|+|91351072|91351094&offsets|0|-1(+1A)]|TAAAGTGCTTATAGTGCAGGTAA</td><td>>MI0000076|hsa-mir-20a&WithFlank&13|+|91351059|91351141</td><td>14</td><td>35</td><td>22(+1A)</td><td>TAAAGTGCTTATAGTGCAGGTAA</td><td>226.9</td><td>2.2</td><td>0.9</td><td>2.4</td><td>1.8E-02</td><td>2.1E-02</td><td>BT20/MB468</td><td>NUC</td></tr>
<tr><td>>hg38_21_+_8433172_8446622_RNA45SN1_45S_50ntFlanks@11834.11851.18|CCACAGCCAAGGGAACGG</td><td>>hg38_21_+_8433172_8446622_RNA45SN1_45S_50ntFlanks</td><td>11834</td><td>11851</td><td>18</td><td>CCACAGCCAAGGGAACGG</td><td>55.8</td><td>-4.2</td><td>1.7</td><td>-2.5</td><td>1.2E-02</td><td>2.1E-02</td><td>BT20/MB468</td><td>MP</td></tr>
<tr><td>>hg38_MT_+_1621_3279_RNR2_16S_50ntFlanks@644.664.21|AATTGGACCAATCTATCACCC</td><td>>hg38_MT_+_1621_3279_RNR2_16S_50ntFlanks</td><td>644</td><td>664</td><td>21</td><td>AATTGGACCAATCTATCACCC</td><td>81.5</td><td>3.9</td><td>1.6</td><td>2.4</td><td>1.7E-02</td><td>2.1E-02</td><td>MB231/MB468</td><td>MP</td></tr>
<tr><td>>hg38_21_+_8433172_8446622_RNA45SN1_45S_50ntFlanks@4555.4589.35|CGCAGCTAGGAATAATGGAATAGGACCGCGGTTCT</td><td>>hg38_21_+_8433172_8446622_RNA45SN1_45S_50ntFlanks</td><td>4555</td><td>4589</td><td>35</td><td>CGCAGCTAGGAATAATGGAATAGGACCGCGGTTCT</td><td>53.5</td><td>-5.2</td><td>2.2</td><td>-2.4</td><td>1.7E-02</td><td>2.1E-02</td><td>MB231/MB468</td><td>MP</td></tr>
<tr><td>>hg38_21_+_8433172_8446622_RNA45SN1_45S_50ntFlanks@6699.6720.22|GCTAGCTGCGAGAATTAATGTG</td><td>>hg38_21_+_8433172_8446622_RNA45SN1_45S_50ntFlanks</td><td>6699</td><td>6720</td><td>22</td><td>GCTAGCTGCGAGAATTAATGTG</td><td>47965.6</td><td>-3.3</td><td>1.3</td><td>-2.5</td><td>1.2E-02</td><td>2.1E-02</td><td>BT20/MB468</td><td>MP</td></tr>
<tr><td>>hg38_21_+_8433172_8446622_RNA45SN1_45S_50ntFlanks@10855.10888.34|AAGGATTGGCTCTAAGGGCTGGGTCGGTCGGGCT</td><td>>hg38_21_+_8433172_8446622_RNA45SN1_45S_50ntFlanks</td><td>10855</td><td>10888</td><td>34</td><td>AAGGATTGGCTCTAAGGGCTGGGTCGGTCGGGCT</td><td>36.4</td><td>5.2</td><td>2.1</td><td>2.5</td><td>1.2E-02</td><td>2.1E-02</td><td>BT20/MB468</td><td>MP</td></tr>
<tr><td>>hg38_21_+_8433172_8446622_RNA45SN1_45S_50ntFlanks@10344.10362.19|ATTCAAACGAGAACTTTGA</td><td>>hg38_21_+_8433172_8446622_RNA45SN1_45S_50ntFlanks</td><td>10344</td><td>10362</td><td>19</td><td>ATTCAAACGAGAACTTTGA</td><td>274.2</td><td>3.5</td><td>1.2</td><td>2.8</td><td>4.7E-03</td><td>2.1E-02</td><td>BT20/MB231</td><td>MP</td></tr>
<tr><td>>hg38_21_+_8433172_8446622_RNA45SN1_45S_50ntFlanks@4444.4472.29|CCTTGCCTCTCGGCGCCCCCTCGATGCTC</td><td>>hg38_21_+_8433172_8446622_RNA45SN1_45S_50ntFlanks</td><td>4444</td><td>4472</td><td>29</td><td>CCTTGCCTCTCGGCGCCCCCTCGATGCTC</td><td>232.6</td><td>-4.3</td><td>1.7</td><td>-2.5</td><td>1.2E-02</td><td>2.1E-02</td><td>BT20/MB468</td><td>MP</td></tr>
<tr><td>>hg38_21_+_8433172_8446622_RNA45SN1_45S_50ntFlanks@12072.12108.37|CCCGAGGGGCTCTCGCTTCTGGCGCCAAGCGCCCGGC</td><td>>hg38_21_+_8433172_8446622_RNA45SN1_45S_50ntFlanks</td><td>12072</td><td>12108</td><td>37</td><td>CCCGAGGGGCTCTCGCTTCTGGCGCCAAGCGCCCGGC</td><td>60.2</td><td>-4.3</td><td>1.7</td><td>-2.5</td><td>1.2E-02</td><td>2.1E-02</td><td>BT20/MB468</td><td>MP</td></tr>
<tr><td>>hg38_21_+_8433172_8446622_RNA45SN1_45S_50ntFlanks@9307.9335.29|AAGGAGTCTAACACGTGCGCGAGTCGGGG</td><td>>hg38_21_+_8433172_8446622_RNA45SN1_45S_50ntFlanks</td><td>9307</td><td>9335</td><td>29</td><td>AAGGAGTCTAACACGTGCGCGAGTCGGGG</td><td>327.0</td><td>5.7</td><td>2.0</td><td>2.8</td><td>4.7E-03</td><td>2.1E-02</td><td>BT20/MB231</td><td>MP</td></tr>
<tr><td>>hg38_21_+_8433172_8446622_RNA45SN1_45S_50ntFlanks@11668.11687.20|TGCCCAGTGCTCTGAATGTC</td><td>>hg38_21_+_8433172_8446622_RNA45SN1_45S_50ntFlanks</td><td>11668</td><td>11687</td><td>20</td><td>TGCCCAGTGCTCTGAATGTC</td><td>163.2</td><td>2.0</td><td>0.8</td><td>2.4</td><td>1.7E-02</td><td>2.1E-02</td><td>MB231/MB468</td><td>MP</td></tr>
<tr><td>>MI0000542|hsa-mir-320a&WithFlank&8|-|22244960|22245043@49.69.21(+1U)&[MIMAT0000510&hsa-miR-320a-3p&offsets|+1|0(+1U);m-60&8|-|22244975|22244996&offsets|+1|0(+1U)]|AAAGCTGGGTTGAGAGGGCGAT</td><td>>MI0000542|hsa-mir-320a&WithFlank&8|-|22244960|22245043</td><td>49</td><td>69</td><td>21(+1U)</td><td>AAAGCTGGGTTGAGAGGGCGAT</td><td>153.1</td><td>3.3</td><td>1.4</td><td>2.4</td><td>1.7E-02</td><td>2.1E-02</td><td>MB231/MB468</td><td>MP</td></tr>
<tr><td>>hg38_21_+_8433172_8446622_RNA45SN1_45S_50ntFlanks@6698.6721.24|AGCTAGCTGCGAGAATTAATGTGA</td><td>>hg38_21_+_8433172_8446622_RNA45SN1_45S_50ntFlanks</td><td>6698</td><td>6721</td><td>24</td><td>AGCTAGCTGCGAGAATTAATGTGA</td><td>214.3</td><td>-2.1</td><td>0.8</td><td>-2.5</td><td>1.2E-02</td><td>2.1E-02</td><td>BT20/MB468</td><td>MP</td></tr>
<tr><td>>hg38_21_+_8433172_8446622_RNA45SN1_45S_50ntFlanks@7978.8018.41|GACCTCAGATCAGACGTGGCGACCCGCTGAATTTAAGCATA</td><td>>hg38_21_+_8433172_8446622_RNA45SN1_45S_50ntFlanks</td><td>7978</td><td>8018</td><td>41</td><td>GACCTCAGATCAGACGTGGCGACCCGCTGAATTTAAGCATA</td><td>142.8</td><td>3.9</td><td>1.7</td><td>2.4</td><td>1.9E-02</td><td>2.1E-02</td><td>MB231/MB468</td><td>MITO</td></tr>
<tr><td>>MI0000111|hsa-mir-105-1&WithFlank&X|-|152392213|152392305@21.42.22(+1U);MI0000112|hsa-mir-105-2&WithFlank&X|-|152394406|152394498@21.42.22(+1U)&[MIMAT0000102&hsa-miR-105-5p&offsets|+2|+1(+1U);m-735&X|-|152392266|152392287&offsets|+2|+2(+1U)];[MIMAT0000102_1&hsa-miR-105-5p&offsets|+2|+1(+1U);m-735&X|-|152394459|152394480&offsets|+2|+2(+1U)]|AAATGCTCAGACTCCTGTGGTGT</td><td>>MI0000111|hsa-mir-105-1&WithFlank&X|-|152392213|152392305</td><td>21</td><td>42</td><td>22(+1U)</td><td>AAATGCTCAGACTCCTGTGGTGT</td><td>37.0</td><td>8.5</td><td>3.0</td><td>2.8</td><td>4.7E-03</td><td>2.1E-02</td><td>BT20/MB231</td><td>MP</td></tr>
<tr><td>>hg38_21_+_8433172_8446622_RNA45SN1_45S_50ntFlanks@2959.2979.21|CGACGTGTGGCGTGGGTCGAC</td><td>>hg38_21_+_8433172_8446622_RNA45SN1_45S_50ntFlanks</td><td>2959</td><td>2979</td><td>21</td><td>CGACGTGTGGCGTGGGTCGAC</td><td>75.8</td><td>-4.4</td><td>1.7</td><td>-2.5</td><td>1.2E-02</td><td>2.1E-02</td><td>BT20/MB468</td><td>MP</td></tr>
<tr><td>>hg38_21_+_8433172_8446622_RNA45SN1_45S_50ntFlanks@5471.5493.23|CACGGCCCTGGCGGAGCGCTGAG</td><td>>hg38_21_+_8433172_8446622_RNA45SN1_45S_50ntFlanks</td><td>5471</td><td>5493</td><td>23</td><td>CACGGCCCTGGCGGAGCGCTGAG</td><td>139.5</td><td>-3.3</td><td>1.3</td><td>-2.5</td><td>1.2E-02</td><td>2.1E-02</td><td>BT20/MB468</td><td>MP</td></tr>
<tr><td>>hg38_21_+_8433172_8446622_RNA45SN1_45S_50ntFlanks@11848.11872.25|ACGGGCTTGGCGGAATCAGCGGGGA</td><td>>hg38_21_+_8433172_8446622_RNA45SN1_45S_50ntFlanks</td><td>11848</td><td>11872</td><td>25</td><td>ACGGGCTTGGCGGAATCAGCGGGGA</td><td>120.6</td><td>-3.5</td><td>1.5</td><td>-2.4</td><td>1.8E-02</td><td>2.1E-02</td><td>BT20/MB468</td><td>CYTO</td></tr>
<tr><td>>hg38_21_+_8433172_8446622_RNA45SN1_45S_50ntFlanks@12409.12431.23|TTGATCCTTCGATGTCGGCTCTT</td><td>>hg38_21_+_8433172_8446622_RNA45SN1_45S_50ntFlanks</td><td>12409</td><td>12431</td><td>23</td><td>TTGATCCTTCGATGTCGGCTCTT</td><td>351.2</td><td>-2.4</td><td>1.0</td><td>-2.4</td><td>1.8E-02</td><td>2.1E-02</td><td>MB231/MB468</td><td>CYTO</td></tr>
<tr><td>>trnaMT_ValTAC_MT_+_1602_1670@44.71.28|ATTTCAACTTAACTTGACCGCTCTGACC</td><td>>trnaMT_ValTAC_MT_+_1602_1670</td><td>44</td><td>71</td><td>28</td><td>ATTTCAACTTAACTTGACCGCTCTGACC</td><td>837.6</td><td>2.1</td><td>0.9</td><td>2.4</td><td>1.7E-02</td><td>2.1E-02</td><td>MB231/MB468</td><td>MP</td></tr>
<tr><td>>MI0000081|hsa-mir-24-2&WithFlank&19|-|13836281|13836365@56.77.22(+1A);MI0000080|hsa-mir-24-1&WithFlank&9|+|95086015|95086094@50.71.22(+1A)&[MIMAT0000080&hsa-miR-24-3p&offsets|0|0(+1A);m-36&19|-|13836288|13836310&offsets|0|-1(+1A)];[MIMAT0000080_1&hsa-miR-24-3p&offsets|0|0(+1A);m-35&9|+|95086064|95086086&offsets|0|-1(+1A)]|TGGCTCAGTTCAGCAGGAACAGA</td><td>>MI0000081|hsa-mir-24-2&WithFlank&19|-|13836281|13836365</td><td>56</td><td>77</td><td>22(+1A)</td><td>TGGCTCAGTTCAGCAGGAACAGA</td><td>425.2</td><td>2.6</td><td>1.0</td><td>2.7</td><td>6.2E-03</td><td>2.1E-02</td><td>BT20/MB231</td><td>CYTO</td></tr>
<tr><td>>hg38_21_+_8433172_8446622_RNA45SN1_45S_50ntFlanks@10689.10707.19|GCGTCCGGTGAGCTCTCGC</td><td>>hg38_21_+_8433172_8446622_RNA45SN1_45S_50ntFlanks</td><td>10689</td><td>10707</td><td>19</td><td>GCGTCCGGTGAGCTCTCGC</td><td>126.4</td><td>-1.7</td><td>0.7</td><td>-2.4</td><td>1.8E-02</td><td>2.1E-02</td><td>BT20/MB468</td><td>CYTO</td></tr>
<tr><td>>trna116_GluCTC_1_-_145399233_145399304@36.70.35,trna134_GluTTC_1_-_16861774_16861845@36.70.35,trna71_GluCTC_1_-_161439189_161439260@36.70.35,trna74_GluCTC_1_-_161431809_161431880@36.70.35,trna77_GluCTC_1_-_161424398_161424469@36.70.35,trna77_GluCTC_6_+_28949976_28950047@36.70.35,trna80_GluCTC_1_-_161417018_161417089@36.70.35,trna84_GluTTC_1_-_161391883_161391954@36.70.35,trna87_GluCTC_6_-_126101393_126101464@36.70.35|CACCGCCGCGGCCCGGGTTCGATTCCCGGTCAGGG</td><td>>trna116_GluCTC_1_-_145399233_145399304</td><td>36</td><td>70</td><td>35</td><td>CACCGCCGCGGCCCGGGTTCGATTCCCGGTCAGGG</td><td>426.9</td><td>-3.2</td><td>1.3</td><td>-2.4</td><td>1.8E-02</td><td>2.1E-02</td><td>MB231/MB468</td><td>CYTO</td></tr>
<tr><td>>hg38_MT_+_598_1651_RNR1_12S_50ntFlanks@52.92.41|ATAGGTTTGGTCCTAGCCTTTCTATTAGCTCTTAGTAAGAT</td><td>>hg38_MT_+_598_1651_RNR1_12S_50ntFlanks</td><td>52</td><td>92</td><td>41</td><td>ATAGGTTTGGTCCTAGCCTTTCTATTAGCTCTTAGTAAGAT</td><td>370.6</td><td>3.2</td><td>1.3</td><td>2.3</td><td>1.9E-02</td><td>2.1E-02</td><td>MB231/MB468</td><td>MITO</td></tr>
<tr><td>>hg38_21_+_8433172_8446622_RNA45SN1_45S_50ntFlanks@10392.10428.37|ACAGCAGTTGAACATGGGTCAGTCGGTCCTGAGAGAT</td><td>>hg38_21_+_8433172_8446622_RNA45SN1_45S_50ntFlanks</td><td>10392</td><td>10428</td><td>37</td><td>ACAGCAGTTGAACATGGGTCAGTCGGTCCTGAGAGAT</td><td>142.5</td><td>-4.5</td><td>1.8</td><td>-2.5</td><td>1.2E-02</td><td>2.1E-02</td><td>BT20/MB468</td><td>MP</td></tr>
<tr><td>>hg38_21_+_8433172_8446622_RNA45SN1_45S_50ntFlanks@9496.9531.36|GAAAGATGGTGAACTATGCCTGGGCAGGGCGAAGCC</td><td>>hg38_21_+_8433172_8446622_RNA45SN1_45S_50ntFlanks</td><td>9496</td><td>9531</td><td>36</td><td>GAAAGATGGTGAACTATGCCTGGGCAGGGCGAAGCC</td><td>182.9</td><td>4.7</td><td>1.7</td><td>2.8</td><td>4.7E-03</td><td>2.1E-02</td><td>BT20/MB231</td><td>MP</td></tr>
<tr><td>>MI0001448|hsa-mir-425&WithFlank&3|-|49020142|49020240@60.80.21(+1A)&[MIMAT0001343&hsa-miR-425-3p&offsets|-1|-2(+1A);m-258&3|-|49020160|49020181&offsets|0|-1(+1A)]|CATCGGGAATGTCGTGTCCGCA</td><td>>MI0001448|hsa-mir-425&WithFlank&3|-|49020142|49020240</td><td>60</td><td>80</td><td>21(+1A)</td><td>CATCGGGAATGTCGTGTCCGCA</td><td>84.1</td><td>5.6</td><td>2.2</td><td>2.5</td><td>1.2E-02</td><td>2.1E-02</td><td>BT20/MB468</td><td>MP</td></tr>
<tr><td>>hg38_21_+_8433172_8446622_RNA45SN1_45S_50ntFlanks@3769.3791.23|CGCACGGCCGGTACAGTGAAACT</td><td>>hg38_21_+_8433172_8446622_RNA45SN1_45S_50ntFlanks</td><td>3769</td><td>3791</td><td>23</td><td>CGCACGGCCGGTACAGTGAAACT</td><td>360.3</td><td>-1.7</td><td>0.7</td><td>-2.4</td><td>1.9E-02</td><td>2.1E-02</td><td>MB231/MB468</td><td>NUC</td></tr>
<tr><td>>MI0000071|hsa-mir-17&WithFlank&13|+|91350599|91350694@20.41.22(+1U)&[MIMAT0000070&hsa-miR-17-5p&offsets|0|-1(+1U);m-88&13|+|91350618|91350640&offsets|0|-1(+1U)]|CAAAGTGCTTACAGTGCAGGTAT</td><td>>MI0000071|hsa-mir-17&WithFlank&13|+|91350599|91350694</td><td>20</td><td>41</td><td>22(+1U)</td><td>CAAAGTGCTTACAGTGCAGGTAT</td><td>337.9</td><td>-4.1</td><td>1.4</td><td>-2.8</td><td>4.7E-03</td><td>2.1E-02</td><td>BT20/MB231</td><td>MP</td></tr>
<tr><td>>hg38_21_+_8433172_8446622_RNA45SN1_45S_50ntFlanks@4283.4306.24|CTTTAACGAGGATCCATTGGAGGG</td><td>>hg38_21_+_8433172_8446622_RNA45SN1_45S_50ntFlanks</td><td>4283</td><td>4306</td><td>24</td><td>CTTTAACGAGGATCCATTGGAGGG</td><td>82.7</td><td>-4.2</td><td>1.7</td><td>-2.5</td><td>1.2E-02</td><td>2.1E-02</td><td>BT20/MB468</td><td>MP</td></tr>
<tr><td>>hg38_21_+_8433172_8446622_RNA45SN1_45S_50ntFlanks@7974.7992.19|ACGCGACCTCAGATCAGAC</td><td>>hg38_21_+_8433172_8446622_RNA45SN1_45S_50ntFlanks</td><td>7974</td><td>7992</td><td>19</td><td>ACGCGACCTCAGATCAGAC</td><td>2076.2</td><td>1.6</td><td>0.7</td><td>2.3</td><td>1.9E-02</td><td>2.1E-02</td><td>MB231/MB468</td><td>NUC</td></tr>
<tr><td>>MI0000299|hsa-mir-222&WithFlank&X|-|45747009|45747130@75.99.25(+1U)&[MIMAT0000279&hsa-miR-222-3p&offsets|0|+4(+1U);m-140&X|-|45747033|45747056&offsets|0|+1(+1U)]|AGCTACATCTGGCTACTGGGTCTCTT</td><td>>MI0000299|hsa-mir-222&WithFlank&X|-|45747009|45747130</td><td>75</td><td>99</td><td>25(+1U)</td><td>AGCTACATCTGGCTACTGGGTCTCTT</td><td>1155.5</td><td>-2.8</td><td>1.0</td><td>-2.8</td><td>4.7E-03</td><td>2.1E-02</td><td>BT20/MB231</td><td>MP</td></tr>
<tr><td>>MI0000269|hsa-mir-181a-2&WithFlank&9|+|124692436|124692557@83.103.21(+1U)&[MIMAT0004558&hsa-miR-181a-2-3p&offsets|0|-1(+1U);m-71&9|+|124692518|124692539&offsets|0|-1(+1U)]|ACCACTGACCGTTGACTGTACT</td><td>>MI0000269|hsa-mir-181a-2&WithFlank&9|+|124692436|124692557</td><td>83</td><td>103</td><td>21(+1U)</td><td>ACCACTGACCGTTGACTGTACT</td><td>424.6</td><td>-2.9</td><td>1.0</td><td>-2.8</td><td>4.7E-03</td><td>2.1E-02</td><td>BT20/MB231</td><td>MP</td></tr>
<tr><td>>hg38_21_+_8433172_8446622_RNA45SN1_45S_50ntFlanks@4724.4758.35|ATTAATCAAGAACGAAAGTCGGAGGTTCGAAGACG</td><td>>hg38_21_+_8433172_8446622_RNA45SN1_45S_50ntFlanks</td><td>4724</td><td>4758</td><td>35</td><td>ATTAATCAAGAACGAAAGTCGGAGGTTCGAAGACG</td><td>12051.5</td><td>-3.5</td><td>1.4</td><td>-2.5</td><td>1.2E-02</td><td>2.1E-02</td><td>BT20/MB468</td><td>MP</td></tr>
<tr><td>>MI0000734|hsa-mir-106b&WithFlank&7|-|100093987|100094080@18.38.21(+1C)&[MIMAT0000680&hsa-miR-106b-5p&offsets|0|0(+1C);m-103&7|-|100094043|100094063&offsets|0|0(+1C)]|TAAAGTGCTGACAGTGCAGATC</td><td>>MI0000734|hsa-mir-106b&WithFlank&7|-|100093987|100094080</td><td>18</td><td>38</td><td>21(+1C)</td><td>TAAAGTGCTGACAGTGCAGATC</td><td>39.0</td><td>4.9</td><td>2.0</td><td>2.4</td><td>1.7E-02</td><td>2.1E-02</td><td>MB231/MB468</td><td>MP</td></tr>
<tr><td>>MI0000070|hsa-mir-16-1&WithFlank&13|-|50048967|50049067@20.41.22;MI0000115|hsa-mir-16-2&WithFlank&3|+|160404739|160404831@16.37.22&[MIMAT0000069&hsa-miR-16-5p&offsets|0|0;m-62&13|-|50049027|50049048&offsets|0|0];[MIMAT0000069_1&hsa-miR-16-5p&offsets|0|0;m-61&3|+|160404754|160404775&offsets|0|0]|TAGCAGCACGTAAATATTGGCG</td><td>>MI0000070|hsa-mir-16-1&WithFlank&13|-|50048967|50049067</td><td>20</td><td>41</td><td>22</td><td>TAGCAGCACGTAAATATTGGCG</td><td>2605.4</td><td>-2.4</td><td>0.9</td><td>-2.8</td><td>5.9E-03</td><td>2.1E-02</td><td>BT20/MB231</td><td>MITO</td></tr>
<tr><td>>MI0000272|hsa-mir-182&WithFlank&7|-|129770377|129770498@29.50.22&[MIMAT0000259&hsa-miR-182-5p&offsets|0|-2;m-22&7|-|129770449|129770470&offsets|0|0]|TTTGGCAATGGTAGAACTCACA</td><td>>MI0000272|hsa-mir-182&WithFlank&7|-|129770377|129770498</td><td>29</td><td>50</td><td>22</td><td>TTTGGCAATGGTAGAACTCACA</td><td>1338.8</td><td>2.6</td><td>1.1</td><td>2.5</td><td>1.2E-02</td><td>2.1E-02</td><td>BT20/MB468</td><td>MP</td></tr>
</table>

| | | | | | | | | | | | | | |
|---|---|---|---|---|---|---|---|---|---|---|---|---|---|
| >hg38_MT_+_598_1651_RNR1_12S_50ntFlanks@970.1004.35\|ATGGTAAGTGTACTGGAAAGTGCACTTGGACGAAC | >hg38_MT_+_598_1651_RNR1_12S_50ntFlanks | 970 | 1004 | 35 | ATGGTAAGTGTACTGGAAAGTGCACTTGGACGAAC | 578.8 | -2.3 | 0.8 | -2.8 | 5.9E-03 | 2.1E-02 | BT20/MB231 | MITO |
| >hg38_21_+_8433172_8446622_RNA45SN1_45S_50ntFlanks@6647.6669.23\|CGTACGACTCTTAGCGGTGGATC | >hg38_21_+_8433172_8446622_RNA45SN1_45S_50ntFlanks | 6647 | 6669 | 23 | CGTACGACTCTTAGCGGTGGATC | 1839.0 | -2.2 | 0.9 | -2.4 | 1.7E-02 | 2.1E-02 | MB231/MB468 | MP |
| >trna10_AspGTC_12_-_125424193_125424264@40.68.29,trna12_AspGTC_12_-_125411891_125411962@40.68.29,trna144_AspGTC_6_-_27551236_27551307@40.68.29,trna38_AspGTC_17_-_8125556_8125627@40.68.29,trna45_AspGTC_6_+_27447453_27447524@40.68.29,trna48_AspGTC_6_+_27471523_27471594@40.68.29,trna4_AspGTC_12_+_96429799_96429870@40.68.29,trna69_AspGTC_1_-_161440205_161440276@40.68.29,trna72_AspGTC_1_-_161432824_161432895@40.68.29,trna75_AspGTC_1_-_161425414_161425485@40.68.29,trna78_AspGTC_1_-_161418033_161418104@40.68.29,trna81_AspGTC_1_-_161410615_161410686@40.68.29\|CGGGAGACCGGGGTTCGATTCCCCGACGG | >trna10_AspGTC_12_-_125424193_125424264 | 40 | 68 | 29 | CGGGAGACCGGGGTTCGATTCCCCGACGG | 95.2 | -3.2 | 1.3 | -2.4 | 1.7E-02 | 2.1E-02 | MB231/MB468 | MP |
| >hg38_21_+_8433172_8446622_RNA45SN1_45S_50ntFlanks@10786.10829.44\|AACAGCCTCTGGCATGTTGGAACAATGTAGGTAAGGGAAGTCGG | >hg38_21_+_8433172_8446622_RNA45SN1_45S_50ntFlanks | 10786 | 10829 | 44 | AACAGCCTCTGGCATGTTGGAACAATGTAGGTAAGGGAAGTCGG | 101.1 | -4.5 | 1.8 | -2.5 | 1.2E-02 | 2.1E-02 | BT20/MB468 | MP |
| >hg38_21_+_8433172_8446622_RNA45SN1_45S_50ntFlanks@7975.8006.32\|CGCGACCTCAGATCAGACGTGGCGACCCGCTG | >hg38_21_+_8433172_8446622_RNA45SN1_45S_50ntFlanks | 7975 | 8006 | 32 | CGCGACCTCAGATCAGACGTGGCGACCCGCTG | 38640.0 | 3.1 | 1.1 | 2.7 | 6.2E-03 | 2.1E-02 | BT20/MB231 | CYTO |
| >hg38_21_+_8433172_8446622_RNA45SN1_45S_50ntFlanks@6653.6698.46\|ACTCTTAGCGGTGGATCACTCGGCTCGTGCGTCGATGAAGAACGCA | >hg38_21_+_8433172_8446622_RNA45SN1_45S_50ntFlanks | 6653 | 6698 | 46 | ACTCTTAGCGGTGGATCACTCGGCTCGTGCGTCGATGAAGAACGCA | 221.1 | -2.3 | 0.8 | -2.7 | 6.2E-03 | 2.1E-02 | BT20/MB231 | CYTO |
| >trna10_AspGTC_12_-_125424193_125424264@37.73.37,trna12_AspGTC_12_-_125411891_125411962@37.73.37,trna144_AspGTC_6_-_27551236_27551307@37.73.37,trna38_AspGTC_17_-_8125556_8125627@37.73.37,trna45_AspGTC_6_+_27447453_27447524@37.73.37,trna48_AspGTC_6_+_27471523_27471594@37.73.37,trna4_AspGTC_12_+_96429799_96429870@37.73.37,trna69_AspGTC_1_-_161440205_161440276@37.73.37,trna72_AspGTC_1_-_161432824_161432895@37.73.37,trna75_AspGTC_1_-_161425414_161425485@37.73.37,trna78_AspGTC_1_-_161418033_161418104@37.73.37,trna81_AspGTC_1_-_161410615_161410686@37.73.37\|ACGCGGGAGACCGGGGTTCGATTCCCCGACGGGGAGC | >trna10_AspGTC_12_-_125424193_125424264 | 37 | 73 | 37 | ACGCGGGAGACCGGGGTTCGATTCCCCGACGGGGAGC | 386.9 | 2.5 | 1.0 | 2.4 | 1.7E-02 | 2.1E-02 | MB231/MB468 | MP |
| >hg38_21_+_8433172_8446622_RNA45SN1_45S_50ntFlanks@12008.12036.29\|CCGCCGGTGAAATACCACTACTCTGATCG | >hg38_21_+_8433172_8446622_RNA45SN1_45S_50ntFlanks | 12008 | 12036 | 29 | CCGCCGGTGAAATACCACTACTCTGATCG | 356.5 | -4.4 | 1.8 | -2.5 | 1.2E-02 | 2.1E-02 | BT20/MB468 | MP |
| >hg38_21_+_8433172_8446622_RNA45SN1_45S_50ntFlanks@12625.12651.27\|CAATGGGGCGAAGCTACCATCTGTGGG | >hg38_21_+_8433172_8446622_RNA45SN1_45S_50ntFlanks | 12625 | 12651 | 27 | CAATGGGGCGAAGCTACCATCTGTGGG | 107.1 | -4.2 | 1.7 | -2.5 | 1.2E-02 | 2.1E-02 | BT20/MB468 | MP |
| >hg38_21_+_8433172_8446622_RNA45SN1_45S_50ntFlanks@12999.13039.41\|CTGCGATCTATTGAAAGTCAGCCCTCGACACAAGGGTTTGT | >hg38_21_+_8433172_8446622_RNA45SN1_45S_50ntFlanks | 12999 | 13039 | 41 | CTGCGATCTATTGAAAGTCAGCCCTCGACACAAGGGTTTGT | 7534.7 | -2.4 | 0.9 | -2.8 | 5.2E-03 | 2.1E-02 | BT20/MB231 | NUC |
| >hg38_21_+_8433172_8446622_RNA45SN1_45S_50ntFlanks@12996.13013.18\|TCGCTGCGATCTATTGAA | >hg38_21_+_8433172_8446622_RNA45SN1_45S_50ntFlanks | 12996 | 13013 | 18 | TCGCTGCGATCTATTGAA | 836.5 | -2.1 | 0.7 | -2.8 | 5.9E-03 | 2.1E-02 | BT20/MB231 | MITO |
| >MI0000078\|hsa-mir-22&WithFlank&17\|-\|1713897\|1713993@59.79.21(+1G)&[MIMAT0000077&hsa-miR-22-3p&offsets\|0\|-1(+1G);m-2&17\|-\|1713914\|1713935&offsets\|0\|-1(+1G)]\|AAGCTGCCAGTTGAAGAACTGG | >MI0000078\|hsa-mir-22&WithFlank&17\|-\|1713897\|1713993 | 59 | 79 | 21(+1G) | AAGCTGCCAGTTGAAGAACTGG | 353.4 | 2.8 | 1.0 | 2.7 | 6.3E-03 | 2.1E-02 | BT20/MB231 | CYTO |
| >trnaMT_SerGCT_MT_+_12207_12265@14.44.31\|AGAACTGCTAACTCATGCCCCCATGTCTAAC | >trnaMT_SerGCT_MT_+_12207_12265 | 14 | 44 | 31 | AGAACTGCTAACTCATGCCCCCATGTCTAAC | 172.4 | 2.2 | 1.0 | 2.3 | 1.9E-02 | 2.1E-02 | MB231/MB468 | MITO |
| >hg38_21_+_8433172_8446622_RNA45SN1_45S_50ntFlanks@6649.6677.29\|TACGACTCTTAGCGGTGGATCACTCGGCT | >hg38_21_+_8433172_8446622_RNA45SN1_45S_50ntFlanks | 6649 | 6677 | 29 | TACGACTCTTAGCGGTGGATCACTCGGCT | 92.5 | -2.2 | 0.9 | -2.4 | 1.7E-02 | 2.1E-02 | MB231/MB468 | MP |
| >hg38_21_+_8433172_8446622_RNA45SN1_45S_50ntFlanks@11210.11232.23\|GGCGACTCTGGACGCGAGCCGGG | >hg38_21_+_8433172_8446622_RNA45SN1_45S_50ntFlanks | 11210 | 11232 | 23 | GGCGACTCTGGACGCGAGCCGGG | 1489.3 | -3.9 | 1.5 | -2.5 | 1.2E-02 | 2.1E-02 | BT20/MB468 | MP |
| >hg38_21_+_8433172_8446622_RNA45SN1_45S_50ntFlanks@9916.9938.23\|AAAGGTGTTGGTTGATATAGACA | >hg38_21_+_8433172_8446622_RNA45SN1_45S_50ntFlanks | 9916 | 9938 | 23 | AAAGGTGTTGGTTGATATAGACA | 66.6 | 4.5 | 1.8 | 2.5 | 1.2E-02 | 2.1E-02 | BT20/MB468 | MP |
| >hg38_21_+_8433172_8446622_RNA45SN1_45S_50ntFlanks@6723.6743.21\|TTGCAGGACACATTGATCATC | >hg38_21_+_8433172_8446622_RNA45SN1_45S_50ntFlanks | 6723 | 6743 | 21 | TTGCAGGACACATTGATCATC | 161.3 | -3.2 | 1.3 | -2.5 | 1.2E-02 | 2.1E-02 | BT20/MB468 | MP |
| >hg38_21_+_8433172_8446622_RNA45SN1_45S_50ntFlanks@6730.6772.43\|ACACATTGATCATCGACACTTCGAACGCACTTGCGGCCCCGGG | >hg38_21_+_8433172_8446622_RNA45SN1_45S_50ntFlanks | 6730 | 6772 | 43 | ACACATTGATCATCGACACTTCGAACGCACTTGCGGCCCCGGG | 34.5 | -8.2 | 3.3 | -2.5 | 1.2E-02 | 2.1E-02 | BT20/MB468 | MP |
| >hg38_21_+_8433172_8446622_RNA45SN1_45S_50ntFlanks@4723.4748.26\|CATTAATCAAGAACGAAAGTCGGAGG | >hg38_21_+_8433172_8446622_RNA45SN1_45S_50ntFlanks | 4723 | 4748 | 26 | CATTAATCAAGAACGAAAGTCGGAGG | 552.5 | -1.9 | 0.8 | -2.4 | 1.8E-02 | 2.1E-02 | BT20/MB468 | CYTO |
| >MI0000736\|hsa-mir-30c-1&WithFlank&1\|+\|40757278\|40757378@23.40.18;MI0000254\|hsa-mir-30c-2&WithFlank&6\|-\|71376954\|71377037@13.30.18;MI0000441\|hsa-mir-30b&WithFlank&8\|-\|134800514\|134800613@23.40.18&[MIMAT0000244&hsa-miR-30c-5p&offsets\|0\|-5;m-57&1\|+\|40757300\|40757323&offsets\|0\|-6];[MIMAT0000244_1&hsa-miR-30c-5p&offsets\|0\|-5;m-56&6\|-\|71377002\|71377025&offsets\|0\|-6];[MIMAT0000420&hsa-miR-30b-5p&offsets\|0\|-4;m-76&8\|-\|134800570\|134800591&offsets\|0\|-4]\|TGTAAACATCCTACACTC | >MI0000736\|hsa-mir-30c-1&WithFlank&1\|+\|40757278\|40757378 | 23 | 40 | 18 | TGTAAACATCCTACACTC | 57.4 | 4.9 | 2.0 | 2.4 | 1.7E-02 | 2.1E-02 | MB231/MB468 | MP |
| >hg38_21_+_8433172_8446622_RNA45SN1_45S_50ntFlanks@11205.11232.28\|GCGGCGGCGACTCTGGACGCGAGCCGGG | >hg38_21_+_8433172_8446622_RNA45SN1_45S_50ntFlanks | 11205 | 11232 | 28 | GCGGCGGCGACTCTGGACGCGAGCCGGG | 1016.8 | -4.9 | 2.1 | -2.4 | 1.9E-02 | 2.1E-02 | BT20/MB468 | NUC |
| >hg38_21_+_8433172_8446622_RNA45SN1_45S_50ntFlanks@10396.10430.35\|CAGTTGAACATGGGTCAGTCGGTCCTGAGAGATGG | >hg38_21_+_8433172_8446622_RNA45SN1_45S_50ntFlanks | 10396 | 10430 | 35 | CAGTTGAACATGGGTCAGTCGGTCCTGAGAGATGG | 119.1 | -4.3 | 1.7 | -2.5 | 1.2E-02 | 2.1E-02 | BT20/MB468 | MP |
| >am_trna10_SerGCT_15_-_40886023_40886104@1.19.19,trna123_SerGCT_6_-_28565117_28565198@1.19.19,trna31_SerGCT_6_+_27065085_27065166@1.19.19,trna43_SerGCT_6_+_27265775_27265856@1.19.19,trna62_SerGCT_6_+_28180815_28180896@1.19.19,trna7_SerGCT_17_+_8090184_8090265@1.19.19,trna8_SerGCT_11_+_66115591_66115672@1.19.19\|GACGAGGTGGCCGAGTGGT | >am_trna10_SerGCT_15_-_40886023_40886104 | 1 | 19 | 19 | GACGAGGTGGCCGAGTGGT | 28.9 | 4.9 | 2.1 | 2.4 | 1.7E-02 | 2.1E-02 | MB231/MB468 | MP |
| >hg38_21_+_8433172_8446622_RNA45SN1_45S_50ntFlanks@12303.12323.21\|CCTCACGATCCTTCTGACCTT | >hg38_21_+_8433172_8446622_RNA45SN1_45S_50ntFlanks | 12303 | 12323 | 21 | CCTCACGATCCTTCTGACCTT | 244.8 | -3.5 | 1.5 | -2.4 | 1.6E-02 | 2.1E-02 | BT20/MB468 | MITO |
| >hg38_21_+_8433172_8446622_RNA45SN1_45S_50ntFlanks@11671.11707.37\|CCAGTGCTCTGAATGTCAAAGTGAAGAAATTCAATGA | >hg38_21_+_8433172_8446622_RNA45SN1_45S_50ntFlanks | 11671 | 11707 | 37 | CCAGTGCTCTGAATGTCAAAGTGAAGAAATTCAATGA | 309.4 | -2.1 | 0.8 | -2.7 | 6.3E-03 | 2.1E-02 | BT20/MB231 | CYTO |
| >hg38_21_+_8433172_8446622_RNA45SN1_45S_50ntFlanks@10392.10422.31\|ACAGCAGTTGAACATGGGTCAGTCGGTCCTG | >hg38_21_+_8433172_8446622_RNA45SN1_45S_50ntFlanks | 10392 | 10422 | 31 | ACAGCAGTTGAACATGGGTCAGTCGGTCCTG | 441.7 | -3.6 | 1.4 | -2.5 | 1.2E-02 | 2.1E-02 | BT20/MB468 | MP |
| >MI0000084\|hsa-mir-26b&WithFlank&2\|+\|218402640\|218402728@18.38.21(+1A)&[MIMAT0000083&hsa-miR-26b-5p&offsets\|0\|0(+1A);m-53&2\|+\|218402657\|218402678&offsets\|0\|-1(+1A)]\|TTCAAGTAATTCAGGATAGGTA | >MI0000084\|hsa-mir-26b&WithFlank&2\|+\|218402640\|218402728 | 18 | 38 | 21(+1A) | TTCAAGTAATTCAGGATAGGTA | 313.4 | 1.8 | 0.7 | 2.8 | 5.2E-03 | 2.1E-02 | BT20/MB231 | NUC |
| >hg38_21_+_8433172_8446622_RNA45SN1_45S_50ntFlanks@10350.10373.24\|ACGAGAACTTTGAAGGCCGAAGTG | >hg38_21_+_8433172_8446622_RNA45SN1_45S_50ntFlanks | 10350 | 10373 | 24 | ACGAGAACTTTGAAGGCCGAAGTG | 1300.5 | -3.8 | 1.5 | -2.5 | 1.2E-02 | 2.1E-02 | BT20/MB468 | MP |
| >hg38_21_+_8433172_8446622_RNA45SN1_45S_50ntFlanks@11924.11941.18\|GAGGTGTAGAATAAGTGG | >hg38_21_+_8433172_8446622_RNA45SN1_45S_50ntFlanks | 11924 | 11941 | 18 | GAGGTGTAGAATAAGTGG | 68.1 | 5.2 | 2.1 | 2.5 | 1.2E-02 | 2.1E-02 | BT20/MB468 | MP |
| >hg38_21_+_8433172_8446622_RNA45SN1_45S_50ntFlanks@7977.7994.18\|CGACCTCAGATCAGACGT | >hg38_21_+_8433172_8446622_RNA45SN1_45S_50ntFlanks | 7977 | 7994 | 18 | CGACCTCAGATCAGACGT | 314.6 | -1.6 | 0.7 | -2.4 | 1.8E-02 | 2.1E-02 | BT20/MB468 | CYTO |
| >hg38_21_+_8433172_8446622_RNA45SN1_45S_50ntFlanks@10684.10712.29\|CGGCGGCGTCCGGTGAGCTCTCGCTGGCC | >hg38_21_+_8433172_8446622_RNA45SN1_45S_50ntFlanks | 10684 | 10712 | 29 | CGGCGGCGTCCGGTGAGCTCTCGCTGGCC | 361.0 | -4.4 | 1.7 | -2.5 | 1.2E-02 | 2.1E-02 | BT20/MB468 | MP |
| >MI0000446\|hsa-mir-125b-1&WithFlank&11\|-\|122099751\|122099850@21.40.20(+1A);MI0000470\|hsa-mir-125b-2&WithFlank&21\|+\|16590231\|16590331@23.42.20(+1A)&[MIMAT0000423&hsa-miR-125b-5p&offsets\|0\|-2(+1A);m-67&11\|-\|122099809\|122099830&offsets\|0\|-2(+1A)];[MIMAT0000423_1&hsa-miR-125b-5p&offsets\|0\|-2(+1A);m-68&21\|+\|16590253\|16590274&offsets\|0\|-2(+1A)]\|TCCCTGAGACCCTAACTTGTA | >MI0000446\|hsa-mir-125b-1&WithFlank&11\|-\|122099751\|122099850 | 21 | 40 | 20(+1A) | TCCCTGAGACCCTAACTTGTA | 1319.6 | -3.0 | 1.1 | -2.7 | 6.3E-03 | 2.1E-02 | BT20/MB231 | CYTO |
| >am_trna128_GlyGCC_6_-_27870686_27870756@1.29.29,trna133_GlyCCC_1_-_16872434_16872504@1.29.29,trna18_GlyGCC_16_+_70822597_70822667@1.29.29,trna19_GlyGCC_16_+_70823410_70823480@1.29.29,trna19_GlyGCC_2_-_157257659_157257729@1.29.29,trna24_GlyGCC_16_-_70812942_70813012@1.29.29,trna25_GlyGCC_16_-_70812114_70812184@1.29.29,trna4_GlyCCC_1_+_17188416_17188486@1.29.29,trna5_GlyGCC_17_+_8029064_8029134@1.29.29,trna68_GlyGCC_1_-_161493637_161493707@1.29.29\|GCATTGGTGGTTCAGTGGTAGAATTCTCG | >am_trna128_GlyGCC_6_-_27870686_27870756 | 1 | 29 | 29 | GCATTGGTGGTTCAGTGGTAGAATTCTCG | 2821.4 | -2.3 | 0.8 | -2.7 | 6.3E-03 | 2.1E-02 | BT20/MB231 | CYTO |
| >hg38_21_+_8433172_8446622_RNA45SN1_45S_50ntFlanks@12410.12434.25\|TGATCCTTCGATGTCGGCTCTTCCT | >hg38_21_+_8433172_8446622_RNA45SN1_45S_50ntFlanks | 12410 | 12434 | 25 | TGATCCTTCGATGTCGGCTCTTCCT | 369.0 | 2.2 | 0.9 | 2.4 | 1.8E-02 | 2.1E-02 | MB231/MB468 | CYTO |
| >hg38_21_+_8433172_8446622_RNA45SN1_45S_50ntFlanks@11576.11611.36\|CTTAGAACTGGTGCGGACCAGGGGAATCCGACTGTT | >hg38_21_+_8433172_8446622_RNA45SN1_45S_50ntFlanks | 11576 | 11611 | 36 | CTTAGAACTGGTGCGGACCAGGGGAATCCGACTGTT | 839.0 | -1.3 | 0.6 | -2.4 | 1.6E-02 | 2.1E-02 | BT20/MB468 | MITO |

| | | | | | | | | | | | | | |
|---|---|---|---|---|---|---|---|---|---|---|---|---|---|
| >hg38_21_+_8433172_8446622_RNA45SN1_45S_50ntFlanks@4728.4751.24\|ATCAAGAACGAAAGTCGGAGGTTC | >hg38_21_+_8433172_8446622_RNA45SN1_45S_50ntFlanks | 4728 | 4751 | 24 | ATCAAGAACGAAAGTCGGAGGTTC | 602.4 | -3.6 | 1.5 | -2.4 | 1.8E-02 | 2.1E-02 | MB231/MB468 | CYTO |
| >MI0000727\|hsa-mir-128-2&WithFlank&3\|+\|35744470\|35744565@58.79.22&[MIMAT0000424_1&hsa-miR-128-3p&offsets\|0\|+1;m-117&3\|+\|35744527\|35744547&offsets\|0\|+1]\|TCACAGTGAACCGGTCTCTTTC | >MI0000727\|hsa-mir-128-2&WithFlank&3\|+\|35744470\|35744565 | 58 | 79 | 22 | TCACAGTGAACCGGTCTCTTTC | 311.6 | 1.9 | 0.7 | 2.7 | 6.3E-03 | 2.1E-02 | BT20/MB231 | CYTO |
| >hg38_MT_+_598_1651_RNR1_12S_50ntFlanks@52.83.32\|ATAGGTTTGGTCCTAGCCTTTCTATTAGCTCT | >hg38_MT_+_598_1651_RNR1_12S_50ntFlanks | 52 | 83 | 32 | ATAGGTTTGGTCCTAGCCTTTCTATTAGCTCT | 3074.2 | 2.1 | 0.9 | 2.3 | 1.9E-02 | 2.1E-02 | MB231/MB468 | MITO |
| >hg38_21_+_8433172_8446622_RNA45SN1_45S_50ntFlanks@11203.11232.30\|CGGCGGCGGCGACTCTGGACGCGAGCCGGG | >hg38_21_+_8433172_8446622_RNA45SN1_45S_50ntFlanks | 11203 | 11232 | 30 | CGGCGGCGGCGACTCTGGACGCGAGCCGGG | 535.4 | -3.0 | 1.3 | -2.4 | 1.8E-02 | 2.1E-02 | MB231/MB468 | CYTO |
| >hg38_21_+_8433172_8446622_RNA45SN1_45S_50ntFlanks@6652.6691.40\|GACTCTTAGCGGTGGATCACTCGGCTCGTGCGTCGATGAA | >hg38_21_+_8433172_8446622_RNA45SN1_45S_50ntFlanks | 6652 | 6691 | 40 | GACTCTTAGCGGTGGATCACTCGGCTCGTGCGTCGATGAA | 188.4 | 4.5 | 1.8 | 2.5 | 1.2E-02 | 2.1E-02 | BT20/MB468 | MP |
| >hg38_MT_+_598_1651_RNR1_12S_50ntFlanks@52.82.31\|ATAGGTTTGGTCCTAGCCTTTCTATTAGCTC | >hg38_MT_+_598_1651_RNR1_12S_50ntFlanks | 52 | 82 | 31 | ATAGGTTTGGTCCTAGCCTTTCTATTAGCTC | 1913.3 | 3.6 | 1.4 | 2.5 | 1.2E-02 | 2.1E-02 | BT20/MB468 | MP |
| >hg38_21_+_8433172_8446622_RNA45SN1_45S_50ntFlanks@7976.7994.19\|GCGACCTCAGATCAGACGT | >hg38_21_+_8433172_8446622_RNA45SN1_45S_50ntFlanks | 7976 | 7994 | 19 | GCGACCTCAGATCAGACGT | 534.7 | 2.1 | 0.8 | 2.8 | 5.9E-03 | 2.1E-02 | BT20/MB231 | MITO |
| >hg38_21_+_8433172_8446622_RNA45SN1_45S_50ntFlanks@5445.5466.22\|TCGGATCGGCCCCGCCGGGGTC | >hg38_21_+_8433172_8446622_RNA45SN1_45S_50ntFlanks | 5445 | 5466 | 22 | TCGGATCGGCCCCGCCGGGGTC | 3358.8 | -3.6 | 1.5 | -2.4 | 1.6E-02 | 2.1E-02 | BT20/MB468 | MITO |
| >hg38_21_+_8433172_8446622_RNA45SN1_45S_50ntFlanks@11783.11816.34\|CATGAATGGATGAACGAGATTCCCACTGTCCCTA | >hg38_21_+_8433172_8446622_RNA45SN1_45S_50ntFlanks | 11783 | 11816 | 34 | CATGAATGGATGAACGAGATTCCCACTGTCCCTA | 111.1 | -3.1 | 1.3 | -2.4 | 1.6E-02 | 2.1E-02 | BT20/MB468 | MITO |
| >hg38_21_+_8433172_8446622_RNA45SN1_45S_50ntFlanks@10849.10867.19\|CGGGATAAGGATTGGCTCT | >hg38_21_+_8433172_8446622_RNA45SN1_45S_50ntFlanks | 10849 | 10867 | 19 | CGGGATAAGGATTGGCTCT | 381.3 | -2.0 | 0.8 | -2.4 | 1.8E-02 | 2.1E-02 | BT20/MB468 | CYTO |
| >hg38_21_+_8433172_8446622_RNA45SN1_45S_50ntFlanks@12405.12439.35\|CTTTTTGATCCTTCGATGTCGGCTCTTCCTATCAT | >hg38_21_+_8433172_8446622_RNA45SN1_45S_50ntFlanks | 12405 | 12439 | 35 | CTTTTTGATCCTTCGATGTCGGCTCTTCCTATCAT | 191.4 | -2.9 | 1.2 | -2.4 | 1.8E-02 | 2.1E-02 | BT20/MB468 | CYTO |
| >hg38_21_+_8433172_8446622_RNA45SN1_45S_50ntFlanks@9565.9594.30\|TGACGTGCAAATCGGTCGTCCGACCTGGGT | >hg38_21_+_8433172_8446622_RNA45SN1_45S_50ntFlanks | 9565 | 9594 | 30 | TGACGTGCAAATCGGTCGTCCGACCTGGGT | 246.5 | -1.5 | 0.6 | -2.4 | 1.8E-02 | 2.1E-02 | BT20/MB468 | CYTO |
| >hg38_21_+_8433172_8446622_RNA45SN1_45S_50ntFlanks@12415.12449.35\|CTTCGATGTCGGCTCTTCCTATCATTGTGAAGCAG | >hg38_21_+_8433172_8446622_RNA45SN1_45S_50ntFlanks | 12415 | 12449 | 35 | CTTCGATGTCGGCTCTTCCTATCATTGTGAAGCAG | 387.5 | 1.7 | 0.7 | 2.3 | 1.9E-02 | 2.1E-02 | MB231/MB468 | NUC |
| >hg38_21_+_8433172_8446622_RNA45SN1_45S_50ntFlanks@10390.10424.35\|GAACAGCAGTTGAACATGGGTCAGTCGGTCCTGAG | >hg38_21_+_8433172_8446622_RNA45SN1_45S_50ntFlanks | 10390 | 10424 | 35 | GAACAGCAGTTGAACATGGGTCAGTCGGTCCTGAG | 217.9 | 4.1 | 1.7 | 2.4 | 1.7E-02 | 2.1E-02 | MB231/MB468 | MP |
| >hg38_21_+_8433172_8446622_RNA45SN1_45S_50ntFlanks@4731.4751.21\|AAGAACGAAAGTCGGAGGTTC | >hg38_21_+_8433172_8446622_RNA45SN1_45S_50ntFlanks | 4731 | 4751 | 21 | AAGAACGAAAGTCGGAGGTTC | 382.1 | -1.3 | 0.5 | -2.4 | 1.6E-02 | 2.1E-02 | BT20/MB468 | MITO |
| >MI0000542\|hsa-mir-320a&WithFlank&8\|-\|22244960\|22245043@48.69.22(+1C)&[MIMAT0000510&hsa-miR-320a-3p&offsets\|0\|0(+1C);m-60&8\|-\|22244975\|22244996&offsets\|0\|0(+1C)]\|AAAAGCTGGGTTGAGAGGGCGAC | >MI0000542\|hsa-mir-320a&WithFlank&8\|-\|22244960\|22245043 | 48 | 69 | 22(+1C) | AAAAGCTGGGTTGAGAGGGCGAC | 1113.3 | 2.3 | 0.8 | 2.8 | 5.2E-03 | 2.1E-02 | BT20/MB231 | NUC |
| >hg38_1_-_228634819_228635039_RNA5S12_5S_50ntFlanks@132.170.39\|GATGGGAGACCGCCTGGGAATACCGGGTGCTGTAGGCTT | >hg38_1_-_228634819_228635039_RNA5S12_5S_50ntFlanks | 132 | 170 | 39 | GATGGGAGACCGCCTGGGAATACCGGGTGCTGTAGGCTT | 17146.6 | -1.7 | 0.7 | -2.4 | 1.7E-02 | 2.1E-02 | MB231/MB468 | MP |
| >hg38_21_+_8433172_8446622_RNA45SN1_45S_50ntFlanks@5489.5513.25\|CTGAGAAGACGGTCGAACTTGACTA | >hg38_21_+_8433172_8446622_RNA45SN1_45S_50ntFlanks | 5489 | 5513 | 25 | CTGAGAAGACGGTCGAACTTGACTA | 602.6 | -3.7 | 1.5 | -2.5 | 1.2E-02 | 2.1E-02 | BT20/MB468 | MP |
| >hg38_21_+_8433172_8446622_RNA45SN1_45S_50ntFlanks@6698.6721.24\|AGCTAGCTGCGAGAATTAATGTGA | >hg38_21_+_8433172_8446622_RNA45SN1_45S_50ntFlanks | 6698 | 6721 | 24 | AGCTAGCTGCGAGAATTAATGTGA | 410.8 | -1.8 | 0.8 | -2.3 | 1.9E-02 | 2.1E-02 | MB231/MB468 | NUC |
| >hg38_21_+_8433172_8446622_RNA45SN1_45S_50ntFlanks@8350.8373.24\|AAAGTTGAAAAGAACTTTGAAGAG | >hg38_21_+_8433172_8446622_RNA45SN1_45S_50ntFlanks | 8350 | 8373 | 24 | AAAGTTGAAAAGAACTTTGAAGAG | 1959.0 | 4.9 | 1.7 | 2.8 | 4.8E-03 | 2.1E-02 | BT20/MB231 | MP |
| >MI0000093\|hsa-mir-92a-1&WithFlank&13\|+\|91351308\|91351397@54.74.21;MI0000094\|hsa-mir-92a-2&WithFlank&X\|-\|134169532\|134169618@54.74.21&[MIMAT0000092&hsa-miR-92a-3p&offsets\|0\|-1;m-9&13\|+\|91351361\|91351382&offsets\|0\|-1];[MIMAT0000092_1&hsa-miR-92a-3p&offsets\|0\|-1;m-11&X\|-\|134169544\|134169565&offsets\|0\|-1]\|TATTGCACTTGTCCCGGCCTG | >MI0000093\|hsa-mir-92a-1&WithFlank&13\|+\|91351308\|91351397 | 54 | 74 | 21 | TATTGCACTTGTCCCGGCCTG | 1013.2 | -1.5 | 0.6 | -2.8 | 5.2E-03 | 2.1E-02 | BT20/MB231 | NUC |
| >trna10_AspGTC_12_-_125424193_125424264@39.72.34,trna12_AspGTC_12_-_125411891_125411962@39.72.34,trna144_AspGTC_6_-_27551236_27551307@39.72.34,trna38_AspGTC_17_-_8125556_8125627@39.72.34,trna45_AspGTC_6_+_27447453_27447524@39.72.34,trna48_AspGTC_6_+_27471523_27471594@39.72.34,trna4_AspGTC_12_+_96429799_96429870@39.72.34,trna69_AspGTC_1_-_161440205_161440276@39.72.34,trna72_AspGTC_1_-_161432824_161432895@39.72.34,trna75_AspGTC_1_-_161425414_161425485@39.72.34,trna78_AspGTC_1_-_161418033_161418104@39.72.34,trna81_AspGTC_1_-_161410615_161410686@39.72.34\|GCGGGAGACCGGGGTTCGATTCCCCGACGGGGAG | >trna10_AspGTC_12_-_125424193_125424264 | 39 | 72 | 34 | GCGGGAGACCGGGGTTCGATTCCCCGACGGGGAG | 543.6 | -2.7 | 1.0 | -2.7 | 6.3E-03 | 2.1E-02 | BT20/MB231 | CYTO |
| >hg38_21_+_8433172_8446622_RNA45SN1_45S_50ntFlanks@5491.5525.35\|GAGAAGACGGTCGAACTTGACTATCTAGAGGAAGT | >hg38_21_+_8433172_8446622_RNA45SN1_45S_50ntFlanks | 5491 | 5525 | 35 | GAGAAGACGGTCGAACTTGACTATCTAGAGGAAGT | 1346.0 | 2.8 | 1.0 | 2.7 | 6.4E-03 | 2.1E-02 | BT20/MB231 | CYTO |
| >hg38_21_+_8433172_8446622_RNA45SN1_45S_50ntFlanks@6646.6666.21\|TCGTACGACTCTTAGCGGTGG | >hg38_21_+_8433172_8446622_RNA45SN1_45S_50ntFlanks | 6646 | 6666 | 21 | TCGTACGACTCTTAGCGGTGG | 12203.8 | 1.9 | 0.7 | 2.7 | 6.4E-03 | 2.1E-02 | BT20/MB231 | CYTO |
| >hg38_21_+_8433172_8446622_RNA45SN1_45S_50ntFlanks@11542.11584.43\|CGGCGCCTCGCCTCGGCCGGCGCCTAGCAGCCGACTTAGAACT | >hg38_21_+_8433172_8446622_RNA45SN1_45S_50ntFlanks | 11542 | 11584 | 43 | CGGCGCCTCGCCTCGGCCGGCGCCTAGCAGCCGACTTAGAACT | 278.6 | 4.0 | 1.7 | 2.3 | 1.9E-02 | 2.1E-02 | MB231/MB468 | NUC |
| >hg38_21_+_8433172_8446622_RNA45SN1_45S_50ntFlanks@6646.6677.32\|TCGTACGACTCTTAGCGGTGGATCACTCGGCT | >hg38_21_+_8433172_8446622_RNA45SN1_45S_50ntFlanks | 6646 | 6677 | 32 | TCGTACGACTCTTAGCGGTGGATCACTCGGCT | 6379.0 | 3.5 | 1.2 | 2.8 | 4.8E-03 | 2.1E-02 | BT20/MB231 | MP |
| >hg38_21_+_8433172_8446622_RNA45SN1_45S_50ntFlanks@11591.11613.23\|GACCAGGGGAATCCGACTGTTTA | >hg38_21_+_8433172_8446622_RNA45SN1_45S_50ntFlanks | 11591 | 11613 | 23 | GACCAGGGGAATCCGACTGTTTA | 102.6 | -3.2 | 1.3 | -2.4 | 1.7E-02 | 2.1E-02 | MB231/MB468 | MP |
| >hg38_21_+_8433172_8446622_RNA45SN1_45S_50ntFlanks@11666.11702.37\|TCTGCCCAGTGCTCTGAATGTCAAAGTGAAGAAATTC | >hg38_21_+_8433172_8446622_RNA45SN1_45S_50ntFlanks | 11666 | 11702 | 37 | TCTGCCCAGTGCTCTGAATGTCAAAGTGAAGAAATTC | 313.4 | -1.7 | 0.7 | -2.3 | 1.9E-02 | 2.1E-02 | MB231/MB468 | MITO |
| >hg38_21_+_8433172_8446622_RNA45SN1_45S_50ntFlanks@12644.12669.26\|TCTGTGGGATTATGACTGAACGCCTC | >hg38_21_+_8433172_8446622_RNA45SN1_45S_50ntFlanks | 12644 | 12669 | 26 | TCTGTGGGATTATGACTGAACGCCTC | 579.3 | 1.9 | 0.8 | 2.4 | 1.6E-02 | 2.1E-02 | BT20/MB468 | MITO |
| >am_trnaMT_ProTGG_MT_-_15956_16023@53.71.19\|AAGACTTTTTCTCTGACCA | >am_trnaMT_ProTGG_MT_-_15956_16023 | 53 | 71 | 19 | AAGACTTTTTCTCTGACCA | 1845.0 | 2.3 | 1.0 | 2.3 | 1.9E-02 | 2.1E-02 | MB231/MB468 | NUC |
| >hg38_21_+_8433172_8446622_RNA45SN1_45S_50ntFlanks@7989.8017.29\|AGACGTGGCGACCCGCTGAATTTAAGCAT | >hg38_21_+_8433172_8446622_RNA45SN1_45S_50ntFlanks | 7989 | 8017 | 29 | AGACGTGGCGACCCGCTGAATTTAAGCAT | 373.0 | -1.7 | 0.7 | -2.5 | 1.2E-02 | 2.1E-02 | BT20/MB468 | MP |
| >hg38_21_+_8433172_8446622_RNA45SN1_45S_50ntFlanks@9482.9503.22\|ACGTGTTAGGACCCGAAAGATG | >hg38_21_+_8433172_8446622_RNA45SN1_45S_50ntFlanks | 9482 | 9503 | 22 | ACGTGTTAGGACCCGAAAGATG | 122.5 | 2.5 | 1.0 | 2.5 | 1.2E-02 | 2.1E-02 | BT20/MB468 | MP |
| >hg38_21_+_8433172_8446622_RNA45SN1_45S_50ntFlanks@9602.9630.29\|CGAAAGACTAATCGAACCATCTAGTAGCT | >hg38_21_+_8433172_8446622_RNA45SN1_45S_50ntFlanks | 9602 | 9630 | 29 | CGAAAGACTAATCGAACCATCTAGTAGCT | 218.6 | 3.8 | 1.3 | 2.8 | 4.8E-03 | 2.1E-02 | BT20/MB231 | MP |
| >hg38_MT_+_598_1651_RNR1_12S_50ntFlanks@726.744.19\|GTTAGGTCAAGGTGTAGCC | >hg38_MT_+_598_1651_RNR1_12S_50ntFlanks | 726 | 744 | 19 | GTTAGGTCAAGGTGTAGCC | 91.4 | 4.0 | 1.7 | 2.4 | 1.7E-02 | 2.1E-02 | MB231/MB468 | MP |
| >hg38_21_+_8433172_8446622_RNA45SN1_45S_50ntFlanks@12417.12449.33\|TCGATGTCGGCTCTTCCTATCATTGTGAAGCAG | >hg38_21_+_8433172_8446622_RNA45SN1_45S_50ntFlanks | 12417 | 12449 | 33 | TCGATGTCGGCTCTTCCTATCATTGTGAAGCAG | 291.5 | -2.2 | 0.8 | -2.7 | 6.4E-03 | 2.1E-02 | BT20/MB231 | CYTO |
| >hg38_21_+_8433172_8446622_RNA45SN1_45S_50ntFlanks@12825.12871.47\|GGACCGGGGTCCGGTGCGGAGTGCCCTTCGTCCTGGGAAACGGGGCG | >hg38_21_+_8433172_8446622_RNA45SN1_45S_50ntFlanks | 12825 | 12871 | 47 | GGACCGGGGTCCGGTGCGGAGTGCCCTTCGTCCTGGGAAACGGGGCG | 34.2 | -8.2 | 3.3 | -2.5 | 1.2E-02 | 2.1E-02 | BT20/MB468 | MP |
| >hg38_1_-_228634819_228635039_RNA5S12_5S_50ntFlanks@51.83.33\|GTCTACGGCCATACCACCCTGAACGCGCCCGAT | >hg38_1_-_228634819_228635039_RNA5S12_5S_50ntFlanks | 51 | 83 | 33 | GTCTACGGCCATACCACCCTGAACGCGCCCGAT | 1036.3 | -2.3 | 1.0 | -2.4 | 1.7E-02 | 2.1E-02 | MB231/MB468 | MP |
| >hg38_MT_+_1621_3279_RNR2_16S_50ntFlanks@825.855.31\|TAAGGAAAGGTTAAAAAAAGTAAAAGGAACT | >hg38_MT_+_1621_3279_RNR2_16S_50ntFlanks | 825 | 855 | 31 | TAAGGAAAGGTTAAAAAAAGTAAAAGGAACT | 108.1 | 4.5 | 1.8 | 2.5 | 1.2E-02 | 2.1E-02 | BT20/MB468 | MP |
| >hg38_21_+_8433172_8446622_RNA45SN1_45S_50ntFlanks@5074.5115.42\|ATTCCGATAACGAACGAGACTCTGGCATGCTAACTAGTTACG | >hg38_21_+_8433172_8446622_RNA45SN1_45S_50ntFlanks | 5074 | 5115 | 42 | ATTCCGATAACGAACGAGACTCTGGCATGCTAACTAGTTACG | 56.6 | -5.2 | 2.2 | -2.4 | 1.7E-02 | 2.1E-02 | MB231/MB468 | MP |
| >MI0000298\|hsa-mir-221&WithFlank&X\|-\|45746151\|45746272@31.53.23(+1C)&[MIMAT0004568&hsa-miR-221-5p&offsets\|0\|+1(+1C);m-405&X\|-\|45746220\|45746242&offsets\|0\|0(+1C)]\|ACCTGGCATACAATGTAGATTTCC | >MI0000298\|hsa-mir-221&WithFlank&X\|-\|45746151\|45746272 | 31 | 53 | 23(+1C) | ACCTGGCATACAATGTAGATTTCC | 84.3 | -5.3 | 1.9 | -2.8 | 4.8E-03 | 2.1E-02 | BT20/MB231 | MP |
| >MI0000650\|hsa-mir-200c&WithFlank&12\|+\|6963693\|6963772@49.72.24&[MIMAT0000617&hsa-miR-200c-3p&offsets\|-1\|0;m-54&12\|+\|6963742\|6963764&offsets\|-1\|0]\|CTAATACTGCCGGGTAATGATGGA | >MI0000650\|hsa-mir-200c&WithFlank&12\|+\|6963693\|6963772 | 49 | 72 | 24 | CTAATACTGCCGGGTAATGATGGA | 117.0 | 3.5 | 1.4 | 2.5 | 1.2E-02 | 2.1E-02 | BT20/MB468 | MP |
| >hg38_21_+_8433172_8446622_RNA45SN1_45S_50ntFlanks@7976.8006.31\|GCGACCTCAGATCAGACGTGGCGACCCGCTG | >hg38_21_+_8433172_8446622_RNA45SN1_45S_50ntFlanks | 7976 | 8006 | 31 | GCGACCTCAGATCAGACGTGGCGACCCGCTG | 277.7 | -1.9 | 0.8 | -2.3 | 1.9E-02 | 2.1E-02 | MB231/MB468 | NUC |
| >hg38_21_+_8433172_8446622_RNA45SN1_45S_50ntFlanks@11561.11590.30\|GCGCCTAGCAGCCGACTTAGAACTGGTGCG | >hg38_21_+_8433172_8446622_RNA45SN1_45S_50ntFlanks | 11561 | 11590 | 30 | GCGCCTAGCAGCCGACTTAGAACTGGTGCG | 239.6 | -3.7 | 1.6 | -2.4 | 1.8E-02 | 2.1E-02 | BT20/MB468 | CYTO |
| >hg38_21_+_8433172_8446622_RNA45SN1_45S_50ntFlanks@10398.10424.27\|GTTGAACATGGGTCAGTCGGTCCTGAG | >hg38_21_+_8433172_8446622_RNA45SN1_45S_50ntFlanks | 10398 | 10424 | 27 | GTTGAACATGGGTCAGTCGGTCCTGAG | 620.6 | 3.6 | 1.5 | 2.4 | 1.7E-02 | 2.1E-02 | MB231/MB468 | MP |

| | | | | | | | | | | | | | |
|---|---|---|---|---|---|---|---|---|---|---|---|---|---|
| >MI0000736\|hsa-mir-30c-1&WithFlank&1\|+\|40757278\|40757378@23.44.22(+1U);MI0000254\|hsa-mir-30c-2&WithFlank&6\|-\|71376954\|71377037@13.34.22(+1U)&[MIMAT0000244&hsa-miR-30c-5p&offsets\|0\|-1(+1U);m-57&1\|+\|40757300\|40757323&offsets\|0\|-2(+1U)];[MIMAT0000244_1&hsa-miR-30c-5p&offsets\|0\|-1(+1U);m-56&6\|-\|71377002\|71377025&offsets\|0\|-2(+1U)]\|TGTAAACATCCTACACTCTCAGT | >MI0000736\|hsa-mir-30c-1&WithFlank&1\|+\|40757278\|40757378 | 23 | 44 | 22(+1U) | TGTAAACATCCTACACTCTCAGT | 457.0 | 2.5 | 1.0 | 2.5 | 1.2E-02 | 2.1E-02 | BT20/MB468 | MP |
| >hg38_21_+_8433172_8446622_RNA45SN1_45S_50ntFlanks@11664.11691.28\|TTTCTGCCCAGTGCTCTGAATGTCAAAG | >hg38_21_+_8433172_8446622_RNA45SN1_45S_50ntFlanks | 11664 | 11691 | 28 | TTTCTGCCCAGTGCTCTGAATGTCAAAG | 71.2 | 5.3 | 2.1 | 2.5 | 1.2E-02 | 2.1E-02 | BT20/MB468 | MP |
| >MI0000077\|hsa-mir-21&WithFlank&17\|+\|59841260\|59841343@52.72.21(+1A)&[MIMAT0004494&hsa-miR-21-3p&offsets\|0\|0(+1A);m-52&17\|+\|59841311\|59841332&offsets\|0\|-1(+1A)]\|CAACACCAGTCGATGGGCTGTA | >MI0000077\|hsa-mir-21&WithFlank&17\|+\|59841260\|59841343 | 52 | 72 | 21(+1A) | CAACACCAGTCGATGGGCTGTA | 199.2 | 3.9 | 1.6 | 2.4 | 1.7E-02 | 2.1E-02 | MB231/MB468 | MP |
| >hg38_21_+_8433172_8446622_RNA45SN1_45S_50ntFlanks@7975.8010.36\|CGCGACCTCAGATCAGACGTGGCGACCCGCTGAATT | >hg38_21_+_8433172_8446622_RNA45SN1_45S_50ntFlanks | 7975 | 8010 | 36 | CGCGACCTCAGATCAGACGTGGCGACCCGCTGAATT | 9752.3 | 2.5 | 1.0 | 2.4 | 1.7E-02 | 2.1E-02 | MB231/MB468 | MP |
| >hg38_21_+_8433172_8446622_RNA45SN1_45S_50ntFlanks@11315.11334.20\|CCACCCCACGTCTCGTCGCG | >hg38_21_+_8433172_8446622_RNA45SN1_45S_50ntFlanks | 11315 | 11334 | 20 | CCACCCCACGTCTCGTCGCG | 57.9 | -4.2 | 1.7 | -2.5 | 1.2E-02 | 2.1E-02 | BT20/MB468 | MP |
| >MI0005566\|hsa-mir-374b&WithFlank&X\|-\|74218541\|74218624@17.34.18&[MIMAT0004955&hsa-miR-374b-5p&offsets\|0\|-4;m-153&X\|-\|74218587\|74218608&offsets\|0\|-4]\|ATATAATACAACCTGCTA | >MI0005566\|hsa-mir-374b&WithFlank&X\|-\|74218541\|74218624 | 17 | 34 | 18 | ATATAATACAACCTGCTA | 294.9 | -2.2 | 0.8 | -2.7 | 6.0E-03 | 2.1E-02 | BT20/MB231 | MITO |
| >hg38_21_+_8433172_8446622_RNA45SN1_45S_50ntFlanks@12995.13037.43\|CTCGCTGCGATCTATTGAAAGTCAGCCCTCGACACAAGGGTTT | >hg38_21_+_8433172_8446622_RNA45SN1_45S_50ntFlanks | 12995 | 13037 | 43 | CTCGCTGCGATCTATTGAAAGTCAGCCCTCGACACAAGGGTTT | 301.1 | -3.8 | 1.4 | -2.8 | 5.3E-03 | 2.1E-02 | BT20/MB231 | NUC |
| >am_trnaMT_LysTTT_MT_+_8295_8364@28.52.25,trnalookalike7_LysTTT_1_+_568844_568913@28.52.25\|TTTTAAGTTAAAGATTAAGAGAACC | >am_trnaMT_LysTTT_MT_+_8295_8364 | 28 | 52 | 25 | TTTTAAGTTAAAGATTAAGAGAACC | 141.6 | 3.5 | 1.4 | 2.4 | 1.7E-02 | 2.1E-02 | MB231/MB468 | MP |
| >hg38_21_+_8433172_8446622_RNA45SN1_45S_50ntFlanks@3987.4023.37\|GCTTTGGTGACTCTAGATAACCTCGGGCCGATCGCAC | >hg38_21_+_8433172_8446622_RNA45SN1_45S_50ntFlanks | 3987 | 4023 | 37 | GCTTTGGTGACTCTAGATAACCTCGGGCCGATCGCAC | 436.0 | 3.6 | 1.4 | 2.5 | 1.2E-02 | 2.1E-02 | BT20/MB468 | MP |
| >hg38_21_+_8433172_8446622_RNA45SN1_45S_50ntFlanks@7989.8015.27\|AGACGTGGCGACCCGCTGAATTTAAGC | >hg38_21_+_8433172_8446622_RNA45SN1_45S_50ntFlanks | 7989 | 8015 | 27 | AGACGTGGCGACCCGCTGAATTTAAGC | 198.7 | 2.7 | 1.2 | 2.3 | 1.9E-02 | 2.1E-02 | BT20/MB468 | NUC |
| >hg38_21_+_8433172_8446622_RNA45SN1_45S_50ntFlanks@8348.8373.26\|GGAAAGTTGAAAAGAACTTTGAAGAG | >hg38_21_+_8433172_8446622_RNA45SN1_45S_50ntFlanks | 8348 | 8373 | 26 | GGAAAGTTGAAAAGAACTTTGAAGAG | 2028.2 | 4.7 | 1.7 | 2.8 | 4.8E-03 | 2.1E-02 | BT20/MB231 | MP |
| >hg38_21_+_8433172_8446622_RNA45SN1_45S_50ntFlanks@6785.6806.22\|CTACGCCTGTCTGAGCGTCGCT | >hg38_21_+_8433172_8446622_RNA45SN1_45S_50ntFlanks | 6785 | 6806 | 22 | CTACGCCTGTCTGAGCGTCGCT | 2087.3 | -1.4 | 0.6 | -2.4 | 1.7E-02 | 2.1E-02 | MB231/MB468 | MP |
| >hg38_21_+_8433172_8446622_RNA45SN1_45S_50ntFlanks@6649.6677.29\|TACGACTCTTAGCGGTGGATCACTCGGCT | >hg38_21_+_8433172_8446622_RNA45SN1_45S_50ntFlanks | 6649 | 6677 | 29 | TACGACTCTTAGCGGTGGATCACTCGGCT | 323.6 | 2.7 | 1.1 | 2.5 | 1.2E-02 | 2.1E-02 | BT20/MB468 | MP |
| >hg38_21_+_8433172_8446622_RNA45SN1_45S_50ntFlanks@8358.8386.29\|AAAGAACTTTGAAGAGAGAGTTCAAGAGG | >hg38_21_+_8433172_8446622_RNA45SN1_45S_50ntFlanks | 8358 | 8386 | 29 | AAAGAACTTTGAAGAGAGAGTTCAAGAGG | 80.8 | 3.5 | 1.4 | 2.5 | 1.2E-02 | 2.1E-02 | BT20/MB468 | MP |
| >hg38_21_+_8433172_8446622_RNA45SN1_45S_50ntFlanks@12417.12454.38\|TCGATGTCGGCTCTTCCTATCATTGTGAAGCAGAATTC | >hg38_21_+_8433172_8446622_RNA45SN1_45S_50ntFlanks | 12417 | 12454 | 38 | TCGATGTCGGCTCTTCCTATCATTGTGAAGCAGAATTC | 428.6 | 1.6 | 0.7 | 2.4 | 1.8E-02 | 2.1E-02 | MB231/MB468 | CYTO |
| >hg38_21_+_8433172_8446622_RNA45SN1_45S_50ntFlanks@6648.6675.28\|GTACGACTCTTAGCGGTGGATCACTCGG | >hg38_21_+_8433172_8446622_RNA45SN1_45S_50ntFlanks | 6648 | 6675 | 28 | GTACGACTCTTAGCGGTGGATCACTCGG | 860.5 | 1.9 | 0.8 | 2.4 | 1.6E-02 | 2.1E-02 | BT20/MB468 | MITO |
| >hg38_21_+_8433172_8446622_RNA45SN1_45S_50ntFlanks@11672.11710.39\|CAGTGCTCTGAATGTCAAAGTGAAGAAATTCAATGAAGC | >hg38_21_+_8433172_8446622_RNA45SN1_45S_50ntFlanks | 11672 | 11710 | 39 | CAGTGCTCTGAATGTCAAAGTGAAGAAATTCAATGAAGC | 223.4 | -2.5 | 0.9 | -2.7 | 6.4E-03 | 2.1E-02 | BT20/MB231 | CYTO |
| >hg38_1_-_228634819_228635039_RNA5S12_5S_50ntFlanks@133.171.39\|ATGGGAGACCGCCTGGGAATACCGGGTGCTGTAGGCTTT | >hg38_1_-_228634819_228635039_RNA5S12_5S_50ntFlanks | 133 | 171 | 39 | ATGGGAGACCGCCTGGGAATACCGGGTGCTGTAGGCTTT | 5457.2 | 2.8 | 1.1 | 2.5 | 1.2E-02 | 2.1E-02 | BT20/MB468 | MP |
| >hg38_21_+_8433172_8446622_RNA45SN1_45S_50ntFlanks@10351.10382.32\|CGAGAACTTTGAAGGCCGAAGTGGAGAAGGGT | >hg38_21_+_8433172_8446622_RNA45SN1_45S_50ntFlanks | 10351 | 10382 | 32 | CGAGAACTTTGAAGGCCGAAGTGGAGAAGGGT | 1488.1 | -1.5 | 0.7 | -2.4 | 1.8E-02 | 2.1E-02 | MB231/MB468 | CYTO |
| >hg38_21_+_8433172_8446622_RNA45SN1_45S_50ntFlanks@11568.11600.33\|GCAGCCGACTTAGAACTGGTGCGGACCAGGGGA | >hg38_21_+_8433172_8446622_RNA45SN1_45S_50ntFlanks | 11568 | 11600 | 33 | GCAGCCGACTTAGAACTGGTGCGGACCAGGGGA | 441.7 | 3.8 | 1.4 | 2.8 | 4.8E-03 | 2.1E-02 | BT20/MB231 | MP |
| >hg38_21_+_8433172_8446622_RNA45SN1_45S_50ntFlanks@7974.8006.33\|ACGCGACCTCAGATCAGACGTGGCGACCCGCTG | >hg38_21_+_8433172_8446622_RNA45SN1_45S_50ntFlanks | 7974 | 8006 | 33 | ACGCGACCTCAGATCAGACGTGGCGACCCGCTG | 3143.4 | 3.6 | 1.3 | 2.8 | 4.8E-03 | 2.1E-02 | BT20/MB231 | MP |
| >hg38_21_+_8433172_8446622_RNA45SN1_45S_50ntFlanks@9844.9864.21\|CAGAACTGGCGCTGCGGGATG | >hg38_21_+_8433172_8446622_RNA45SN1_45S_50ntFlanks | 9844 | 9864 | 21 | CAGAACTGGCGCTGCGGGATG | 187.4 | 3.7 | 1.3 | 2.8 | 4.8E-03 | 2.1E-02 | BT20/MB231 | MP |
| >hg38_21_+_8433172_8446622_RNA45SN1_45S_50ntFlanks@9916.9938.23\|AAAGGTGTTGGTTGATATAGACA | >hg38_21_+_8433172_8446622_RNA45SN1_45S_50ntFlanks | 9916 | 9938 | 23 | AAAGGTGTTGGTTGATATAGACA | 72.0 | 5.4 | 1.9 | 2.8 | 4.8E-03 | 2.1E-02 | BT20/MB231 | MP |
| >hg38_21_+_8433172_8446622_RNA45SN1_45S_50ntFlanks@10060.10077.18\|GCAGTCGAGAGTGGACGG | >hg38_21_+_8433172_8446622_RNA45SN1_45S_50ntFlanks | 10060 | 10077 | 18 | GCAGTCGAGAGTGGACGG | 75.5 | -4.1 | 1.7 | -2.4 | 1.7E-02 | 2.1E-02 | MB231/MB468 | MP |
| >hg38_21_+_8433172_8446622_RNA45SN1_45S_50ntFlanks@9917.9947.31\|AAGGTGTTGGTTGATATAGACAGCAGGACGG | >hg38_21_+_8433172_8446622_RNA45SN1_45S_50ntFlanks | 9917 | 9947 | 31 | AAGGTGTTGGTTGATATAGACAGCAGGACGG | 63.2 | 5.1 | 2.0 | 2.5 | 1.2E-02 | 2.1E-02 | BT20/MB468 | MP |
| >MI0000825\|hsa-mir-345&WithFlank&14\|+\|100307853\|100307962@24.43.20(+1A)&[MIMAT0000772&hsa-miR-345-5p&offsets\|0\|-2(+1A);m-193&14\|+\|100307876\|100307896&offsets\|0\|-1(+1A)]\|GCTGACTCCTAGTCCAGGGCA | >MI0000825\|hsa-mir-345&WithFlank&14\|+\|100307853\|100307962 | 24 | 43 | 20(+1A) | GCTGACTCCTAGTCCAGGGCA | 28.9 | 4.9 | 2.1 | 2.4 | 1.7E-02 | 2.1E-02 | MB231/MB468 | MP |
| >hg38_21_+_8433172_8446622_RNA45SN1_45S_50ntFlanks@51.74.24\|GCTGACACGCTGTCCTCTGGCGAC | >hg38_21_+_8433172_8446622_RNA45SN1_45S_50ntFlanks | 51 | 74 | 24 | GCTGACACGCTGTCCTCTGGCGAC | 423.8 | -2.0 | 0.9 | -2.3 | 1.9E-02 | 2.1E-02 | BT20/MB468 | NUC |
| >hg38_MT_+_598_1651_RNR1_12S_50ntFlanks@52.84.33\|ATAGGTTTGGTCCTAGCCTTTCTATTAGCTCTT | >hg38_MT_+_598_1651_RNR1_12S_50ntFlanks | 52 | 84 | 33 | ATAGGTTTGGTCCTAGCCTTTCTATTAGCTCTT | 1226.7 | 2.2 | 0.9 | 2.3 | 1.9E-02 | 2.1E-02 | MB231/MB468 | MITO |
| >hg38_21_+_8433172_8446622_RNA45SN1_45S_50ntFlanks@4735.4758.24\|ACGAAAGTCGGAGGTTCGAAGACG | >hg38_21_+_8433172_8446622_RNA45SN1_45S_50ntFlanks | 4735 | 4758 | 24 | ACGAAAGTCGGAGGTTCGAAGACG | 984.7 | 1.8 | 0.8 | 2.4 | 1.8E-02 | 2.1E-02 | BT20/MB468 | CYTO |
| >MI0000734\|hsa-mir-106b&WithFlank&7\|-\|100093987\|100094080@58.76.19(+1A)&[MIMAT0004672&hsa-miR-106b-3p&offsets\|0\|-3(+1A);m-87&7\|-\|100094003\|100094023&offsets\|0\|-2(+1A)]\|CCGCACTGTGGGTACTTGCA | >MI0000734\|hsa-mir-106b&WithFlank&7\|-\|100093987\|100094080 | 58 | 76 | 19(+1A) | CCGCACTGTGGGTACTTGCA | 64.3 | 4.8 | 2.0 | 2.4 | 1.7E-02 | 2.1E-02 | MB231/MB468 | MP |
| >hg38_21_+_8433172_8446622_RNA45SN1_45S_50ntFlanks@12995.13038.44\|CTCGCTGCGATCTATTGAAAGTCAGCCCTCGACACAAGGGTTTG | >hg38_21_+_8433172_8446622_RNA45SN1_45S_50ntFlanks | 12995 | 13038 | 44 | CTCGCTGCGATCTATTGAAAGTCAGCCCTCGACACAAGGGTTTG | 8414.4 | -3.0 | 1.1 | -2.7 | 6.4E-03 | 2.1E-02 | BT20/MB231 | CYTO |
| >MI0033423\|hsa-mir-10399&WithFlank&7\|-\|139044094\|139044163@43.63.21(+1U)&[MIMAT0041630&hsa-miR-10399-3p&offsets\|-1\|-1(+1U);TJU_CMC_MD2.ID03061.3p-miR&offsets\|+3\|+2(+1U);m-2657&7\|-\|139044103\|139044124&offsets\|+3\|+2(+1U)]\|TCTCTCGGACAAGCTGTAGGTT | >MI0033423\|hsa-mir-10399&WithFlank&7\|-\|139044094\|139044163 | 43 | 63 | 21(+1U) | TCTCTCGGACAAGCTGTAGGTT | 315.5 | -2.1 | 0.8 | -2.7 | 6.4E-03 | 2.1E-02 | BT20/MB231 | CYTO |
| >MI0000093\|hsa-mir-92a-1&WithFlank&13\|+\|91351308\|91351397@54.73.20(+2U);MI0000094\|hsa-mir-92a-2&WithFlank&X\|-\|134169532\|134169618@54.73.20(+2U)&[MIMAT0000092&hsa-miR-92a-3p&offsets\|0\|-2(+2U);m-9&13\|+\|91351361\|91351382&offsets\|0\|-2(+2U)];[MIMAT0000092_1&hsa-miR-92a-3p&offsets\|0\|-2(+2U);m-11&X\|-\|134169544\|134169565&offsets\|0\|-2(+2U)]\|TATTGCACTTGTCCCGGCCTTT | >MI0000093\|hsa-mir-92a-1&WithFlank&13\|+\|91351308\|91351397 | 54 | 73 | 20(+2U) | TATTGCACTTGTCCCGGCCTTT | 54.7 | 4.0 | 1.7 | 2.4 | 1.7E-02 | 2.2E-02 | MB231/MB468 | MP |
| >hg38_21_+_8433172_8446622_RNA45SN1_45S_50ntFlanks@6647.6695.49\|CGTACGACTCTTAGCGGTGGATCACTCGGCTCGTGCGTCGATGAAGAAC | >hg38_21_+_8433172_8446622_RNA45SN1_45S_50ntFlanks | 6647 | 6695 | 49 | CGTACGACTCTTAGCGGTGGATCACTCGGCTCGTGCGTCGATGAAGAAC | 569.4 | 3.3 | 1.4 | 2.3 | 1.9E-02 | 2.2E-02 | MB231/MB468 | MITO |
| >hg38_21_+_8433172_8446622_RNA45SN1_45S_50ntFlanks@4046.4074.29\|CATTCGAACGTCTGCCCTATCAACTTTCG | >hg38_21_+_8433172_8446622_RNA45SN1_45S_50ntFlanks | 4046 | 4074 | 29 | CATTCGAACGTCTGCCCTATCAACTTTCG | 59.7 | -4.3 | 1.7 | -2.5 | 1.2E-02 | 2.2E-02 | BT20/MB468 | MP |
| >am_trnaMT_ProTGG_MT_-_15956_16023@52.71.20\|AAAGACTTTTTCTCTGACCA | >am_trnaMT_ProTGG_MT_-_15956_16023 | 52 | 71 | 20 | AAAGACTTTTTCTCTGACCA | 2462.9 | 2.6 | 1.1 | 2.4 | 1.7E-02 | 2.2E-02 | MB231/MB468 | MP |
| >hg38_21_+_8433172_8446622_RNA45SN1_45S_50ntFlanks@4132.4151.20\|TCCGGAGAGGGAGCCTGAGA | >hg38_21_+_8433172_8446622_RNA45SN1_45S_50ntFlanks | 4132 | 4151 | 20 | TCCGGAGAGGGAGCCTGAGA | 69.8 | 3.1 | 1.3 | 2.4 | 1.7E-02 | 2.2E-02 | MB231/MB468 | MP |
| >hg38_21_+_8433172_8446622_RNA45SN1_45S_50ntFlanks@10137.10157.21\|CGGCGGCGGCGGGGGTGTGGG | >hg38_21_+_8433172_8446622_RNA45SN1_45S_50ntFlanks | 10137 | 10157 | 21 | CGGCGGCGGCGGGGGTGTGGG | 116.3 | -4.3 | 1.7 | -2.5 | 1.2E-02 | 2.2E-02 | BT20/MB468 | MP |
| >hg38_1_-_228634819_228635039_RNA5S12_5S_50ntFlanks@141.169.29\|CCGCCTGGGAATACCGGGTGCTGTAGGCT | >hg38_1_-_228634819_228635039_RNA5S12_5S_50ntFlanks | 141 | 169 | 29 | CCGCCTGGGAATACCGGGTGCTGTAGGCT | 961.4 | 1.7 | 0.7 | 2.4 | 1.6E-02 | 2.2E-02 | BT20/MB468 | MITO |
| >hg38_MT_+_1621_3279_RNR2_16S_50ntFlanks@1366.1396.31\|CTCGATGTTGGATCAGGACATCCCGATGGTG | >hg38_MT_+_1621_3279_RNR2_16S_50ntFlanks | 1366 | 1396 | 31 | CTCGATGTTGGATCAGGACATCCCGATGGTG | 82.4 | 2.4 | 1.0 | 2.4 | 1.7E-02 | 2.2E-02 | MB231/MB468 | MP |
| >hg38_21_+_8433172_8446622_RNA45SN1_45S_50ntFlanks@9862.9879.18\|ATGAACCGAACGCCGGGT | >hg38_21_+_8433172_8446622_RNA45SN1_45S_50ntFlanks | 9862 | 9879 | 18 | ATGAACCGAACGCCGGGT | 213.6 | -3.4 | 1.4 | -2.5 | 1.2E-02 | 2.2E-02 | BT20/MB468 | MP |
| >hg38_21_+_8433172_8446622_RNA45SN1_45S_50ntFlanks@12003.12035.33\|GCGGGCCGCCGGTGAAATACCACTACTCTGATC | >hg38_21_+_8433172_8446622_RNA45SN1_45S_50ntFlanks | 12003 | 12035 | 33 | GCGGGCCGCCGGTGAAATACCACTACTCTGATC | 692.1 | 4.2 | 1.5 | 2.8 | 4.9E-03 | 2.2E-02 | BT20/MB231 | MP |
| >hg38_MT_+_598_1651_RNR1_12S_50ntFlanks@427.461.35\|GAAAGTGGCTTTAACATATCTGAACACACAATAGC | >hg38_MT_+_598_1651_RNR1_12S_50ntFlanks | 427 | 461 | 35 | GAAAGTGGCTTTAACATATCTGAACACACAATAGC | 18.3 | 7.4 | 3.1 | 2.4 | 1.7E-02 | 2.2E-02 | MB231/MB468 | MP |
| >MI0000061\|hsa-let-7a-2&WithFlank&11\|-\|122146516\|122146599@11.31.21(+1C);MI0000062\|hsa-let-7a-3&WithFlank&22\|+\|46112743\|46112828@10.30.21(+1C);MI0000060\|hsa-let-7a-1&WithFlank&9\|+\|94175951\|94176042@12.32.21(+1C)&[MIMAT0000062&hsa-let-7a-5p&offsets\|0\|-1(+1C);m-6&11\|-\|122146568\|122146589&offsets\|0\|-1(+1C)];[MIMAT0000062_1&hsa-let-7a-5p&offsets\|0\|-1(+1C);m-5&22\|+\|46112752\|46112773&offsets\|0\|-1(+1C)];[MIMAT0000062_2&hsa-let-7a-5p&offsets\|0\|-1(+1C);m-4&9\|+\|94175962\|94175983&offsets\|0\|-1(+1C)]\|TGAGGTAGTAGGTTGTATAGTC | >MI0000061\|hsa-let-7a-2&WithFlank&11\|-\|122146516\|122146599 | 11 | 31 | 21(+1C) | TGAGGTAGTAGGTTGTATAGTC | 15661.0 | -2.9 | 1.0 | -2.8 | 4.9E-03 | 2.2E-02 | BT20/MB231 | MP |

<table>
<tr><th></th><th></th><th></th><th></th><th></th><th></th><th></th><th></th><th></th><th></th><th></th><th></th><th></th><th></th></tr>
<tr><td>>hg38_21_+_8433172_8446622_RNA45SN1_45S_50ntFlanks@11603.11622.20|CCGACTGTTTAATTAAAACA</td><td>>hg38_21_+_8433172_8446622_RNA45SN1_45S_50ntFlanks</td><td>11603</td><td>11622</td><td>20</td><td>CCGACTGTTTAATTAAAACA</td><td>146.2</td><td>5.6</td><td>2.0</td><td>2.8</td><td>4.9E-03</td><td>2.2E-02</td><td>BT20/MB231</td><td>MP</td></tr>
<tr><td>>MI0000070|hsa-mir-16-1&WithFlank&13|-|50048967|50049067@20.42.23;MI0000115|hsa-mir-16-2&WithFlank&3|+|160404739|160404831@16.38.23&[MIMAT0000069&hsa-miR-16-5p&offsets|0|+1;m-62&13|-|50049027|50049048&offsets|0|+1];[MIMAT0000069_1&hsa-miR-16-5p&offsets|0|+1;m-61&3|+|160404754|160404775&offsets|0|+1]|TAGCAGCACGTAAATATTGGCGT</td><td>>MI0000070|hsa-mir-16-1&WithFlank&13|-|50048967|50049067</td><td>20</td><td>42</td><td>23</td><td>TAGCAGCACGTAAATATTGGCGT</td><td>3761.1</td><td>3.1</td><td>1.2</td><td>2.5</td><td>1.2E-02</td><td>2.2E-02</td><td>BT20/MB468</td><td>MP</td></tr>
<tr><td>>MI0000734|hsa-mir-106b&WithFlank&7|-|100093987|100094080@58.80.23&[MIMAT0004672&hsa-miR-106b-3p&offsets|0|+1;m-87&7|-|100094003|100094023&offsets|0|+2]|CCGCACTGTGGGTACTTGCTGCT</td><td>>MI0000734|hsa-mir-106b&WithFlank&7|-|100093987|100094080</td><td>58</td><td>80</td><td>23</td><td>CCGCACTGTGGGTACTTGCTGCT</td><td>88.5</td><td>3.7</td><td>1.5</td><td>2.5</td><td>1.2E-02</td><td>2.2E-02</td><td>BT20/MB468</td><td>MP</td></tr>
<tr><td>>hg38_21_+_8433172_8446622_RNA45SN1_45S_50ntFlanks@10344.10371.28|ATTCAAACGAGAACTTTGAAGGCCGAAG</td><td>>hg38_21_+_8433172_8446622_RNA45SN1_45S_50ntFlanks</td><td>10344</td><td>10371</td><td>28</td><td>ATTCAAACGAGAACTTTGAAGGCCGAAG</td><td>83.6</td><td>4.5</td><td>1.8</td><td>2.5</td><td>1.2E-02</td><td>2.2E-02</td><td>BT20/MB468</td><td>MP</td></tr>
<tr><td>>hg38_21_+_8433172_8446622_RNA45SN1_45S_50ntFlanks@10797.10817.21|GCATGTTGGAACAATGTAGGT</td><td>>hg38_21_+_8433172_8446622_RNA45SN1_45S_50ntFlanks</td><td>10797</td><td>10817</td><td>21</td><td>GCATGTTGGAACAATGTAGGT</td><td>436.2</td><td>-2.0</td><td>0.8</td><td>-2.4</td><td>1.8E-02</td><td>2.2E-02</td><td>BT20/MB468</td><td>CYTO</td></tr>
<tr><td>>hg38_MT_+_598_1651_RNR1_12S_50ntFlanks@440.469.30|ACATATCTGAACACACAATAGCTAAGACCC</td><td>>hg38_MT_+_598_1651_RNR1_12S_50ntFlanks</td><td>440</td><td>469</td><td>30</td><td>ACATATCTGAACACACAATAGCTAAGACCC</td><td>126.3</td><td>3.9</td><td>1.7</td><td>2.4</td><td>1.7E-02</td><td>2.2E-02</td><td>MB231/MB468</td><td>MP</td></tr>
<tr><td>>hg38_21_+_8433172_8446622_RNA45SN1_45S_50ntFlanks@4050.4069.20|CGAACGTCTGCCCTATCAAC</td><td>>hg38_21_+_8433172_8446622_RNA45SN1_45S_50ntFlanks</td><td>4050</td><td>4069</td><td>20</td><td>CGAACGTCTGCCCTATCAAC</td><td>424.7</td><td>-2.1</td><td>0.8</td><td>-2.7</td><td>6.5E-03</td><td>2.2E-02</td><td>BT20/MB231</td><td>CYTO</td></tr>
<tr><td>>hg38_21_+_8433172_8446622_RNA45SN1_45S_50ntFlanks@8168.8195.28|CCCAAGTCCTTCTGATCGAGGCCCAGCC</td><td>>hg38_21_+_8433172_8446622_RNA45SN1_45S_50ntFlanks</td><td>8168</td><td>8195</td><td>28</td><td>CCCAAGTCCTTCTGATCGAGGCCCAGCC</td><td>221.1</td><td>-2.3</td><td>1.0</td><td>-2.4</td><td>1.8E-02</td><td>2.2E-02</td><td>MB231/MB468</td><td>CYTO</td></tr>
<tr><td>>MI0000749|hsa-mir-30e&WithFlank&1|+|40754349|40754452@23.42.20&[MIMAT0000692&hsa-miR-30e-5p&offsets|0|-2;m-24&1|+|40754371|40754394&offsets|0|-4]|TGTAAACATCCTTGACTGGA</td><td>>MI0000749|hsa-mir-30e&WithFlank&1|+|40754349|40754452</td><td>23</td><td>42</td><td>20</td><td>TGTAAACATCCTTGACTGGA</td><td>373.5</td><td>2.2</td><td>0.9</td><td>2.4</td><td>1.7E-02</td><td>2.2E-02</td><td>MB231/MB468</td><td>MP</td></tr>
<tr><td>>hg38_21_+_8433172_8446622_RNA45SN1_45S_50ntFlanks@12907.12935.29|CACCGCACGTTCGTGGGGAACCTGGCGCT</td><td>>hg38_21_+_8433172_8446622_RNA45SN1_45S_50ntFlanks</td><td>12907</td><td>12935</td><td>29</td><td>CACCGCACGTTCGTGGGGAACCTGGCGCT</td><td>556.5</td><td>-1.4</td><td>0.6</td><td>-2.4</td><td>1.6E-02</td><td>2.2E-02</td><td>BT20/MB468</td><td>MITO</td></tr>
<tr><td>>hg38_21_+_8433172_8446622_RNA45SN1_45S_50ntFlanks@11704.11725.22|ATGAAGCGCGGGTAAACGGCGG</td><td>>hg38_21_+_8433172_8446622_RNA45SN1_45S_50ntFlanks</td><td>11704</td><td>11725</td><td>22</td><td>ATGAAGCGCGGGTAAACGGCGG</td><td>70.8</td><td>-4.3</td><td>1.7</td><td>-2.5</td><td>1.2E-02</td><td>2.2E-02</td><td>BT20/MB468</td><td>MP</td></tr>
<tr><td>>hg38_21_+_8433172_8446622_RNA45SN1_45S_50ntFlanks@8352.8373.22|AGTTGAAAAGAACTTTGAAGAG</td><td>>hg38_21_+_8433172_8446622_RNA45SN1_45S_50ntFlanks</td><td>8352</td><td>8373</td><td>22</td><td>AGTTGAAAAGAACTTTGAAGAG</td><td>3228.4</td><td>4.9</td><td>1.7</td><td>2.8</td><td>4.9E-03</td><td>2.2E-02</td><td>BT20/MB231</td><td>MP</td></tr>
<tr><td>>hg38_21_+_8433172_8446622_RNA45SN1_45S_50ntFlanks@6661.6685.25|CGGTGGATCACTCGGCTCGTGCGTC</td><td>>hg38_21_+_8433172_8446622_RNA45SN1_45S_50ntFlanks</td><td>6661</td><td>6685</td><td>25</td><td>CGGTGGATCACTCGGCTCGTGCGTC</td><td>7993.4</td><td>3.7</td><td>1.3</td><td>2.8</td><td>4.9E-03</td><td>2.2E-02</td><td>BT20/MB231</td><td>MP</td></tr>
<tr><td>>hg38_21_+_8433172_8446622_RNA45SN1_45S_50ntFlanks@4735.4766.32|ACGAAAGTCGGAGGTTCGAAGACGATCAGATA</td><td>>hg38_21_+_8433172_8446622_RNA45SN1_45S_50ntFlanks</td><td>4735</td><td>4766</td><td>32</td><td>ACGAAAGTCGGAGGTTCGAAGACGATCAGATA</td><td>49.7</td><td>3.2</td><td>1.3</td><td>2.4</td><td>1.7E-02</td><td>2.2E-02</td><td>MB231/MB468</td><td>MP</td></tr>
<tr><td>>MI0003205|hsa-mir-532&WithFlank&X|+|50003142|50003244@26.46.21(+1A)&[MIMAT0002888&hsa-miR-532-5p&offsets|0|-1(+1A);m-65&X|+|50003167|50003188&offsets|0|-1(+1A)]|CATGCCTTGAGTGTAGGACCGA</td><td>>MI0003205|hsa-mir-532&WithFlank&X|+|50003142|50003244</td><td>26</td><td>46</td><td>21(+1A)</td><td>CATGCCTTGAGTGTAGGACCGA</td><td>241.1</td><td>3.8</td><td>1.5</td><td>2.5</td><td>1.2E-02</td><td>2.2E-02</td><td>BT20/MB468</td><td>MP</td></tr>
<tr><td>>hg38_MT_+_1621_3279_RNR2_16S_50ntFlanks@1186.1216.31|TTTCGGTTGGGGCGACCTCGGAGCAGAACCC</td><td>>hg38_MT_+_1621_3279_RNR2_16S_50ntFlanks</td><td>1186</td><td>1216</td><td>31</td><td>TTTCGGTTGGGGCGACCTCGGAGCAGAACCC</td><td>120.0</td><td>4.4</td><td>1.8</td><td>2.5</td><td>1.2E-02</td><td>2.2E-02</td><td>BT20/MB468</td><td>MP</td></tr>
<tr><td>>hg38_21_+_8433172_8446622_RNA45SN1_45S_50ntFlanks@8381.8415.35|AAGAGGGCGTGAAACCGTTAAGAGGTAAACGGGTG</td><td>>hg38_21_+_8433172_8446622_RNA45SN1_45S_50ntFlanks</td><td>8381</td><td>8415</td><td>35</td><td>AAGAGGGCGTGAAACCGTTAAGAGGTAAACGGGTG</td><td>77.4</td><td>-4.2</td><td>1.7</td><td>-2.5</td><td>1.2E-02</td><td>2.2E-02</td><td>BT20/MB468</td><td>MP</td></tr>
<tr><td>>hg38_21_+_8433172_8446622_RNA45SN1_45S_50ntFlanks@4661.4689.29|AGGTGAAATTCTTGGACCGGCGCAAGACG</td><td>>hg38_21_+_8433172_8446622_RNA45SN1_45S_50ntFlanks</td><td>4661</td><td>4689</td><td>29</td><td>AGGTGAAATTCTTGGACCGGCGCAAGACG</td><td>268.7</td><td>-2.2</td><td>0.9</td><td>-2.4</td><td>1.9E-02</td><td>2.2E-02</td><td>MB231/MB468</td><td>CYTO</td></tr>
<tr><td>>am_MI0003583|hsa-mir-576&WithFlank&4|+|109488692|109488801@61.83.23&[MIMAT0004796&hsa-miR-576-3p&offsets|0|+1;m-524&4|+|109488752|109488772&offsets|0|+2]|AAGATGTGGAAAAATTGGAATCC</td><td>>am_MI0003583|hsa-mir-576&WithFlank&4|+|109488692|109488801</td><td>61</td><td>83</td><td>23</td><td>AAGATGTGGAAAAATTGGAATCC</td><td>536.6</td><td>3.3</td><td>1.4</td><td>2.4</td><td>1.7E-02</td><td>2.2E-02</td><td>MB231/MB468</td><td>MP</td></tr>
<tr><td>>hg38_21_+_8433172_8446622_RNA45SN1_45S_50ntFlanks@10397.10424.28|AGTTGAACATGGGTCAGTCGGTCCTGAG</td><td>>hg38_21_+_8433172_8446622_RNA45SN1_45S_50ntFlanks</td><td>10397</td><td>10424</td><td>28</td><td>AGTTGAACATGGGTCAGTCGGTCCTGAG</td><td>2799.8</td><td>3.7</td><td>1.3</td><td>2.8</td><td>4.9E-03</td><td>2.2E-02</td><td>BT20/MB231</td><td>MP</td></tr>
<tr><td>>hg38_21_+_8433172_8446622_RNA45SN1_45S_50ntFlanks@11570.11601.32|AGCCGACTTAGAACTGGTGCGGACCAGGGGAA</td><td>>hg38_21_+_8433172_8446622_RNA45SN1_45S_50ntFlanks</td><td>11570</td><td>11601</td><td>32</td><td>AGCCGACTTAGAACTGGTGCGGACCAGGGGAA</td><td>384.2</td><td>4.7</td><td>1.7</td><td>2.8</td><td>4.9E-03</td><td>2.2E-02</td><td>BT20/MB231</td><td>MP</td></tr>
<tr><td>>hg38_21_+_8433172_8446622_RNA45SN1_45S_50ntFlanks@12234.12265.32|CGTGGAGCAGAAGGGCAAAAGCTCGCTTGATC</td><td>>hg38_21_+_8433172_8446622_RNA45SN1_45S_50ntFlanks</td><td>12234</td><td>12265</td><td>32</td><td>CGTGGAGCAGAAGGGCAAAAGCTCGCTTGATC</td><td>287.3</td><td>4.8</td><td>1.7</td><td>2.8</td><td>4.9E-03</td><td>2.2E-02</td><td>BT20/MB231</td><td>MP</td></tr>
<tr><td>>hg38_21_+_8433172_8446622_RNA45SN1_45S_50ntFlanks@12934.12960.27|CTAAACCATTCGTAGACGACCTGCTTC</td><td>>hg38_21_+_8433172_8446622_RNA45SN1_45S_50ntFlanks</td><td>12934</td><td>12960</td><td>27</td><td>CTAAACCATTCGTAGACGACCTGCTTC</td><td>397.8</td><td>-1.6</td><td>0.7</td><td>-2.4</td><td>1.9E-02</td><td>2.2E-02</td><td>MB231/MB468</td><td>CYTO</td></tr>
<tr><td>>hg38_21_+_8433172_8446622_RNA45SN1_45S_50ntFlanks@10726.10753.28|GGGAGAGGGTGTAAATCTCGCGCCGGGC</td><td>>hg38_21_+_8433172_8446622_RNA45SN1_45S_50ntFlanks</td><td>10726</td><td>10753</td><td>28</td><td>GGGAGAGGGTGTAAATCTCGCGCCGGGC</td><td>60.9</td><td>-4.2</td><td>1.7</td><td>-2.5</td><td>1.2E-02</td><td>2.2E-02</td><td>BT20/MB468</td><td>MP</td></tr>
<tr><td>>trna14_LysTTT_11_-_59327808_59327880@3.36.34,trna2_LysTTT_17_+_8022473_8022545@3.36.34,trna54_LysTTT_1_+_204475655_204475727@3.36.34,trna5_LysTTT_11_+_59323902_59323974@3.36.34,trna62_LysTTT_1_-_204476158_204476230@3.36.34,trna76_LysTTT_6_+_28918806_28918878@3.36.34|CCGGATAGCTCAGTCGGTAGAGCATCAGACTTTT</td><td>>trna14_LysTTT_11_-_59327808_59327880</td><td>3</td><td>36</td><td>34</td><td>CCGGATAGCTCAGTCGGTAGAGCATCAGACTTTT</td><td>248.5</td><td>-4.1</td><td>1.4</td><td>-2.8</td><td>4.9E-03</td><td>2.2E-02</td><td>BT20/MB231</td><td>MP</td></tr>
<tr><td>>hg38_21_+_8433172_8446622_RNA45SN1_45S_50ntFlanks@6657.6683.27|TTAGCGGTGGATCACTCGGCTCGTGCG</td><td>>hg38_21_+_8433172_8446622_RNA45SN1_45S_50ntFlanks</td><td>6657</td><td>6683</td><td>27</td><td>TTAGCGGTGGATCACTCGGCTCGTGCG</td><td>167.9</td><td>-2.2</td><td>0.8</td><td>-2.8</td><td>5.4E-03</td><td>2.2E-02</td><td>BT20/MB231</td><td>NUC</td></tr>
<tr><td>>hg38_21_+_8433172_8446622_RNA45SN1_45S_50ntFlanks@12418.12459.42|CGATGTCGGCTCTTCCTATCATTGTGAAGCAGAATTCACCAA</td><td>>hg38_21_+_8433172_8446622_RNA45SN1_45S_50ntFlanks</td><td>12418</td><td>12459</td><td>42</td><td>CGATGTCGGCTCTTCCTATCATTGTGAAGCAGAATTCACCAA</td><td>228.8</td><td>3.2</td><td>1.4</td><td>2.3</td><td>1.9E-02</td><td>2.2E-02</td><td>MB231/MB468</td><td>MITO</td></tr>
<tr><td>>hg38_21_+_8433172_8446622_RNA45SN1_45S_50ntFlanks@4243.4275.33|CTTTCGAGGCCCTGTAATTGGAATGAGTCCACT</td><td>>hg38_21_+_8433172_8446622_RNA45SN1_45S_50ntFlanks</td><td>4243</td><td>4275</td><td>33</td><td>CTTTCGAGGCCCTGTAATTGGAATGAGTCCACT</td><td>1374.1</td><td>4.1</td><td>1.6</td><td>2.5</td><td>1.2E-02</td><td>2.2E-02</td><td>BT20/MB468</td><td>MP</td></tr>
<tr><td>>MI0002468|hsa-mir-484&WithFlank&16|+|15643288|15643378@14.33.20(+2U)&[MIMAT0002174&hsa-miR-484&offsets|0|-2(+2U);m-127&16|+|15643301|15643321&offsets|0|-1(+2U)]|TCAGGCTCAGTCCCCTCCCGTT</td><td>>MI0002468|hsa-mir-484&WithFlank&16|+|15643288|15643378</td><td>14</td><td>33</td><td>20(+2U)</td><td>TCAGGCTCAGTCCCCTCCCGTT</td><td>240.3</td><td>2.1</td><td>0.8</td><td>2.7</td><td>6.5E-03</td><td>2.2E-02</td><td>BT20/MB231</td><td>CYTO</td></tr>
<tr><td>>hg38_21_+_8433172_8446622_RNA45SN1_45S_50ntFlanks@11670.11706.37|CCCAGTGCTCTGAATGTCAAAGTGAAGAAATTCAATG</td><td>>hg38_21_+_8433172_8446622_RNA45SN1_45S_50ntFlanks</td><td>11670</td><td>11706</td><td>37</td><td>CCCAGTGCTCTGAATGTCAAAGTGAAGAAATTCAATG</td><td>282.7</td><td>-2.3</td><td>0.8</td><td>-2.7</td><td>6.5E-03</td><td>2.2E-02</td><td>BT20/MB231</td><td>CYTO</td></tr>
<tr><td>>MI0001448|hsa-mir-425&WithFlank&3|-|49020142|49020240@21.44.24&[MIMAT0003393&hsa-miR-425-5p&offsets|+1|+2;m-95&3|-|49020200|49020221&offsets|+1|+3]|ATGACACGATCACTCCCGTTGAGT</td><td>>MI0001448|hsa-mir-425&WithFlank&3|-|49020142|49020240</td><td>21</td><td>44</td><td>24</td><td>ATGACACGATCACTCCCGTTGAGT</td><td>774.5</td><td>2.4</td><td>0.9</td><td>2.7</td><td>6.5E-03</td><td>2.2E-02</td><td>BT20/MB231</td><td>CYTO</td></tr>
<tr><td>>hg38_21_+_8433172_8446622_RNA45SN1_45S_50ntFlanks@8342.8373.32|CGTAAGGGAAAGTTGAAAAGAACTTTGAAGAG</td><td>>hg38_21_+_8433172_8446622_RNA45SN1_45S_50ntFlanks</td><td>8342</td><td>8373</td><td>32</td><td>CGTAAGGGAAAGTTGAAAAGAACTTTGAAGAG</td><td>1487.8</td><td>4.9</td><td>1.7</td><td>2.8</td><td>4.9E-03</td><td>2.2E-02</td><td>BT20/MB231</td><td>MP</td></tr>
<tr><td>>hg38_21_+_8433172_8446622_RNA45SN1_45S_50ntFlanks@11729.11761.33|TAACTATGACTCTCTTAAGGTAGCCAAATGCCT</td><td>>hg38_21_+_8433172_8446622_RNA45SN1_45S_50ntFlanks</td><td>11729</td><td>11761</td><td>33</td><td>TAACTATGACTCTCTTAAGGTAGCCAAATGCCT</td><td>117.0</td><td>-4.3</td><td>1.7</td><td>-2.5</td><td>1.2E-02</td><td>2.2E-02</td><td>BT20/MB468</td><td>MP</td></tr>
<tr><td>>hg38_21_+_8433172_8446622_RNA45SN1_45S_50ntFlanks@5492.5532.41|AGAAGACGGTCGAACTTGACTATCTAGAGGAAGTAAAAGTC</td><td>>hg38_21_+_8433172_8446622_RNA45SN1_45S_50ntFlanks</td><td>5492</td><td>5532</td><td>41</td><td>AGAAGACGGTCGAACTTGACTATCTAGAGGAAGTAAAAGTC</td><td>205.1</td><td>3.9</td><td>1.5</td><td>2.5</td><td>1.2E-02</td><td>2.2E-02</td><td>BT20/MB468</td><td>MP</td></tr>
<tr><td>>hg38_21_+_8433172_8446622_RNA45SN1_45S_50ntFlanks@6781.6803.23|GGGGCTACGCCTGTCTGAGCGTC</td><td>>hg38_21_+_8433172_8446622_RNA45SN1_45S_50ntFlanks</td><td>6781</td><td>6803</td><td>23</td><td>GGGGCTACGCCTGTCTGAGCGTC</td><td>127.7</td><td>-2.1</td><td>0.8</td><td>-2.5</td><td>1.2E-02</td><td>2.2E-02</td><td>BT20/MB468</td><td>MP</td></tr>
<tr><td>>trna143_LysTTT_6_-_27559593_27559665@14.32.19,trna149_LysTTT_6_-_27302769_27302841@14.32.19,trna14_LysTTT_11_-_59327808_59327880@14.32.19,trna2_LysTTT_17_+_8022473_8022545@14.32.19,trna54_LysTTT_1_+_204475655_204475727@14.32.19,trna5_LysTTT_11_+_59323902_59323974@14.32.19,trna62_LysTTT_1_-_204476158_204476230@14.32.19,trna76_LysTTT_6_+_28918806_28918878@14.32.19|AGTCGGTAGAGCATCAGAC</td><td>>trna143_LysTTT_6_-_27559593_27559665</td><td>14</td><td>32</td><td>19</td><td>AGTCGGTAGAGCATCAGAC</td><td>172.9</td><td>-2.8</td><td>1.0</td><td>-2.8</td><td>4.9E-03</td><td>2.2E-02</td><td>BT20/MB231</td><td>MP</td></tr>
<tr><td>>MI0000255|hsa-mir-30d&WithFlank&8|-|134804870|134804951@12.34.23(+1G)&[MIMAT0000245&hsa-miR-30d-5p&offsets|0|+1(+1G);m-21&8|-|134804919|134804940&offsets|0|+1(+1G)]|TGTAAACATCCCCGACTGGAAGCG</td><td>>MI0000255|hsa-mir-30d&WithFlank&8|-|134804870|134804951</td><td>12</td><td>34</td><td>23(+1G)</td><td>TGTAAACATCCCCGACTGGAAGCG</td><td>525.3</td><td>3.9</td><td>1.6</td><td>2.5</td><td>1.2E-02</td><td>2.2E-02</td><td>BT20/MB468</td><td>MP</td></tr>
<tr><td>>hg38_21_+_8433172_8446622_RNA45SN1_45S_50ntFlanks@12395.12414.20|AGCGACGTCGCTTTTTGATC</td><td>>hg38_21_+_8433172_8446622_RNA45SN1_45S_50ntFlanks</td><td>12395</td><td>12414</td><td>20</td><td>AGCGACGTCGCTTTTTGATC</td><td>159.1</td><td>2.6</td><td>1.0</td><td>2.5</td><td>1.2E-02</td><td>2.2E-02</td><td>BT20/MB468</td><td>MP</td></tr>
<tr><td>>hg38_1_-_228634819_228635039_RNA5S12_5S_50ntFlanks@149.171.23|GAATACCGGGTGCTGTAGGCTTT</td><td>>hg38_1_-_228634819_228635039_RNA5S12_5S_50ntFlanks</td><td>149</td><td>171</td><td>23</td><td>GAATACCGGGTGCTGTAGGCTTT</td><td>1156.4</td><td>-1.6</td><td>0.7</td><td>-2.3</td><td>2.0E-02</td><td>2.2E-02</td><td>MB231/MB468</td><td>MITO</td></tr>
<tr><td>>hg38_21_+_8433172_8446622_RNA45SN1_45S_50ntFlanks@10922.10942.21|CTGGACGAGGCGCCGCCGCCC</td><td>>hg38_21_+_8433172_8446622_RNA45SN1_45S_50ntFlanks</td><td>10922</td><td>10942</td><td>21</td><td>CTGGACGAGGCGCCGCCGCCC</td><td>120.2</td><td>-2.5</td><td>1.1</td><td>-2.4</td><td>1.9E-02</td><td>2.2E-02</td><td>BT20/MB468</td><td>CYTO</td></tr>
<tr><td>>hg38_21_+_8433172_8446622_RNA45SN1_45S_50ntFlanks@8114.8142.29|GCGCGGGACATGTGGCGTACGGAAGACCC</td><td>>hg38_21_+_8433172_8446622_RNA45SN1_45S_50ntFlanks</td><td>8114</td><td>8142</td><td>29</td><td>GCGCGGGACATGTGGCGTACGGAAGACCC</td><td>1040.3</td><td>-3.8</td><td>1.5</td><td>-2.5</td><td>1.2E-02</td><td>2.2E-02</td><td>BT20/MB468</td><td>MP</td></tr>
<tr><td>>MI0000067|hsa-let-7f-1&WithFlank&9|+|94176341|94176439@13.33.21(+1G);MI0000068|hsa-let-7f-2&WithFlank&X|-|53557186|53557280@14.34.21(+1G)&[MIMAT0000067&hsa-let-7f-5p&offsets|0|-1(+1G);m-20&9|+|94176353|94176374&offsets|0|-1(+1G)];[MIMAT0000067_1&hsa-let-7f-5p&offsets|0|-1(+1G);m-19&X|-|53557246|53557267&offsets|0|-1(+1G)]|TGAGGTAGTAGATTGTATAGTG</td><td>>MI0000067|hsa-let-7f-1&WithFlank&9|+|94176341|94176439</td><td>13</td><td>33</td><td>21(+1G)</td><td>TGAGGTAGTAGATTGTATAGTG</td><td>662.9</td><td>1.8</td><td>0.8</td><td>2.3</td><td>1.9E-02</td><td>2.2E-02</td><td>BT20/MB468</td><td>NUC</td></tr>
<tr><td>>MI0000299|hsa-mir-222&WithFlank&X|-|45747009|45747130@75.97.23(+3U)&[MIMAT0000279&hsa-miR-222-3p&offsets|0|+2(+3U);m-140&X|-|45747033|45747056&offsets|0|-1(+3U)]|AGCTACATCTGGCTACTGGGTCTTTT</td><td>>MI0000299|hsa-mir-222&WithFlank&X|-|45747009|45747130</td><td>75</td><td>97</td><td>23(+3U)</td><td>AGCTACATCTGGCTACTGGGTCTTTT</td><td>157.5</td><td>-5.2</td><td>1.9</td><td>-2.8</td><td>4.9E-03</td><td>2.2E-02</td><td>BT20/MB231</td><td>MP</td></tr>
</table>

<table>
<tr><td>>hg38_21_+_8433172_8446622_RNA45SN1_45S_50ntFlanks@10912.10931.20|CGCGCCGCGGCTGGACGAGG</td><td>>hg38_21_+_8433172_8446622_RNA45SN1_45S_50ntFlanks</td><td>10912</td><td>10931</td><td>20</td><td>CGCGCCGCGGCTGGACGAGG</td><td>306.9</td><td>-2.3</td><td>0.9</td><td>-2.5</td><td>1.2E-02</td><td>2.2E-02</td><td>BT20/MB468</td><td>MP</td></tr>
<tr><td>>hg38_21_+_8433172_8446622_RNA45SN1_45S_50ntFlanks@9739.9769.31|AAACGATCTCAACCTATTCTCAAACTTTAAA</td><td>>hg38_21_+_8433172_8446622_RNA45SN1_45S_50ntFlanks</td><td>9739</td><td>9769</td><td>31</td><td>AAACGATCTCAACCTATTCTCAAACTTTAAA</td><td>518.1</td><td>-4.8</td><td>1.9</td><td>-2.5</td><td>1.2E-02</td><td>2.2E-02</td><td>BT20/MB468</td><td>MP</td></tr>
<tr><td>>hg38_MT_+_598_1651_RNR1_12S_50ntFlanks@429.455.27|AAGTGGCTTTAACATATCTGAACACAC</td><td>>hg38_MT_+_598_1651_RNR1_12S_50ntFlanks</td><td>429</td><td>455</td><td>27</td><td>AAGTGGCTTTAACATATCTGAACACAC</td><td>94.5</td><td>3.5</td><td>1.4</td><td>2.5</td><td>1.2E-02</td><td>2.2E-02</td><td>BT20/MB468</td><td>MP</td></tr>
<tr><td>>hg38_21_+_8433172_8446622_RNA45SN1_45S_50ntFlanks@11669.11703.35|GCCCAGTGCTCTGAATGTCAAAGTGAAGAAATTCA</td><td>>hg38_21_+_8433172_8446622_RNA45SN1_45S_50ntFlanks</td><td>11669</td><td>11703</td><td>35</td><td>GCCCAGTGCTCTGAATGTCAAAGTGAAGAAATTCA</td><td>1701.8</td><td>-2.4</td><td>0.9</td><td>-2.7</td><td>6.6E-03</td><td>2.2E-02</td><td>BT20/MB231</td><td>CYTO</td></tr>
<tr><td>>hg38_21_+_8433172_8446622_RNA45SN1_45S_50ntFlanks@8171.8216.46|AAGTCCTTCTGATCGAGGCCCAGCCCGTGGACGGTGTGAGGCCGGT</td><td>>hg38_21_+_8433172_8446622_RNA45SN1_45S_50ntFlanks</td><td>8171</td><td>8216</td><td>46</td><td>AAGTCCTTCTGATCGAGGCCCAGCCCGTGGACGGTGTGAGGCCGGT</td><td>229.1</td><td>-2.6</td><td>1.0</td><td>-2.7</td><td>6.6E-03</td><td>2.2E-02</td><td>BT20/MB231</td><td>CYTO</td></tr>
<tr><td>>hg38_21_+_8433172_8446622_RNA45SN1_45S_50ntFlanks@9539.9569.31|ACTCTGGTGGAGGTCCGTAGCGGTCCTGACG</td><td>>hg38_21_+_8433172_8446622_RNA45SN1_45S_50ntFlanks</td><td>9539</td><td>9569</td><td>31</td><td>ACTCTGGTGGAGGTCCGTAGCGGTCCTGACG</td><td>256.9</td><td>-2.3</td><td>1.0</td><td>-2.4</td><td>1.9E-02</td><td>2.2E-02</td><td>BT20/MB468</td><td>CYTO</td></tr>
<tr><td>>MI0000803|hsa-mir-330&WithFlank&19|-|45638988|45639093@64.86.23&[MIMAT0000751&hsa-miR-330-3p&offsets|+1|+1;m-347&19|-|45639009|45639030&offsets|0|+1]|CAAAGCACACGGCCTGCAGAGAG</td><td>>MI0000803|hsa-mir-330&WithFlank&19|-|45638988|45639093</td><td>64</td><td>86</td><td>23</td><td>CAAAGCACACGGCCTGCAGAGAG</td><td>51.6</td><td>5.3</td><td>2.1</td><td>2.5</td><td>1.2E-02</td><td>2.2E-02</td><td>BT20/MB468</td><td>MP</td></tr>
<tr><td>>MI0003137|hsa-mir-193b&WithFlank&16|+|14303961|14304055@57.77.21(+1C)&[MIMAT0002819&hsa-miR-193b-3p&offsets|0|-1(+1C);m-166&16|+|14304017|14304038&offsets|0|-1(+1C)]|AACTGGCCCTCAAAGTCCCGCC</td><td>>MI0003137|hsa-mir-193b&WithFlank&16|+|14303961|14304055</td><td>57</td><td>77</td><td>21(+1C)</td><td>AACTGGCCCTCAAAGTCCCGCC</td><td>369.3</td><td>2.4</td><td>0.9</td><td>2.7</td><td>6.1E-03</td><td>2.2E-02</td><td>BT20/MB231</td><td>MITO</td></tr>
<tr><td>>hg38_21_+_8433172_8446622_RNA45SN1_45S_50ntFlanks@11582.11612.31|ACTGGTGCGGACCAGGGGAATCCGACTGTTT</td><td>>hg38_21_+_8433172_8446622_RNA45SN1_45S_50ntFlanks</td><td>11582</td><td>11612</td><td>31</td><td>ACTGGTGCGGACCAGGGGAATCCGACTGTTT</td><td>561.0</td><td>-2.0</td><td>0.9</td><td>-2.3</td><td>1.9E-02</td><td>2.2E-02</td><td>BT20/MB468</td><td>NUC</td></tr>
<tr><td>>hg38_21_+_8433172_8446622_RNA45SN1_45S_50ntFlanks@11594.11612.19|CAGGGGAATCCGACTGTTT</td><td>>hg38_21_+_8433172_8446622_RNA45SN1_45S_50ntFlanks</td><td>11594</td><td>11612</td><td>19</td><td>CAGGGGAATCCGACTGTTT</td><td>584.3</td><td>-4.5</td><td>1.8</td><td>-2.5</td><td>1.2E-02</td><td>2.2E-02</td><td>BT20/MB468</td><td>MP</td></tr>
<tr><td>>hg38_21_+_8433172_8446622_RNA45SN1_45S_50ntFlanks@12008.12026.19|CCGCCGGTGAAATACCACT</td><td>>hg38_21_+_8433172_8446622_RNA45SN1_45S_50ntFlanks</td><td>12008</td><td>12026</td><td>19</td><td>CCGCCGGTGAAATACCACT</td><td>87.7</td><td>-3.4</td><td>1.4</td><td>-2.5</td><td>1.2E-02</td><td>2.2E-02</td><td>BT20/MB468</td><td>MP</td></tr>
<tr><td>>am_trna128_GlyGCC_6_-_27870686_27870756@1.26.26,trna133_GlyCCC_1_-_16872434_16872504@1.26.26,trna18_GlyGCC_16_+_70822597_70822667@1.26.26,trna19_GlyGCC_16_+_70823410_70823480@1.26.26,trna19_GlyGCC_2_-_157257659_157257729@1.26.26,trna24_GlyGCC_16_-_70812942_70813012@1.26.26,trna25_GlyGCC_16_-_70812114_70812184@1.26.26,trna4_GlyCCC_1_+_17188416_17188486@1.26.26,trna5_GlyGCC_17_+_8029064_8029134@1.26.26,trna68_GlyGCC_1_-_161493637_161493707@1.26.26|GCATTGGTGGTTCAGTGGTAGAATTC</td><td>>am_trna128_GlyGCC_6_-_27870686_27870756</td><td>1</td><td>26</td><td>26</td><td>GCATTGGTGGTTCAGTGGTAGAATTC</td><td>835.4</td><td>-2.0</td><td>0.7</td><td>-2.7</td><td>6.6E-03</td><td>2.2E-02</td><td>BT20/MB231</td><td>CYTO</td></tr>
<tr><td>>hg38_21_+_8433172_8446622_RNA45SN1_45S_50ntFlanks@4976.4993.18|CCGGACACGGACAGGATT</td><td>>hg38_21_+_8433172_8446622_RNA45SN1_45S_50ntFlanks</td><td>4976</td><td>4993</td><td>18</td><td>CCGGACACGGACAGGATT</td><td>466.7</td><td>2.0</td><td>0.7</td><td>2.7</td><td>6.6E-03</td><td>2.2E-02</td><td>BT20/MB231</td><td>CYTO</td></tr>
<tr><td>>MI0000266|hsa-mir-10a&WithFlank&17|-|48579832|48579953@29.48.20(+1C)&[MIMAT0000253&hsa-miR-10a-5p&offsets|+1|-2(+1C);m-18&17|-|48579905|48579926&offsets|+1|-1(+1C)]|ACCCTGTAGATCCGAATTTGC</td><td>>MI0000266|hsa-mir-10a&WithFlank&17|-|48579832|48579953</td><td>29</td><td>48</td><td>20(+1C)</td><td>ACCCTGTAGATCCGAATTTGC</td><td>592.4</td><td>2.2</td><td>0.8</td><td>2.7</td><td>6.6E-03</td><td>2.2E-02</td><td>BT20/MB231</td><td>CYTO</td></tr>
<tr><td>>hg38_21_+_8433172_8446622_RNA45SN1_45S_50ntFlanks@12414.12441.28|CCTTCGATGTCGGCTCTTCCTATCATTG</td><td>>hg38_21_+_8433172_8446622_RNA45SN1_45S_50ntFlanks</td><td>12414</td><td>12441</td><td>28</td><td>CCTTCGATGTCGGCTCTTCCTATCATTG</td><td>311.8</td><td>-2.4</td><td>0.9</td><td>-2.7</td><td>6.6E-03</td><td>2.2E-02</td><td>BT20/MB231</td><td>CYTO</td></tr>
<tr><td>>hg38_21_+_8433172_8446622_RNA45SN1_45S_50ntFlanks@10786.10827.42|AACAGCCTCTGGCATGTTGGAACAATGTAGGTAAGGGAAGTC</td><td>>hg38_21_+_8433172_8446622_RNA45SN1_45S_50ntFlanks</td><td>10786</td><td>10827</td><td>42</td><td>AACAGCCTCTGGCATGTTGGAACAATGTAGGTAAGGGAAGTC</td><td>160.5</td><td>-2.3</td><td>1.0</td><td>-2.4</td><td>1.9E-02</td><td>2.2E-02</td><td>MB231/MB468</td><td>CYTO</td></tr>
<tr><td>>MI0000264|hsa-mir-7-2&WithFlank&15|+|88611819|88611940@38.58.21(+1A);MI0000265|hsa-mir-7-3&WithFlank&19|+|4770664|4770785@37.57.21(+1A);MI0000263|hsa-mir-7-1&WithFlank&9|-|83969742|83969863@30.50.21(+1A)&[MIMAT0000252&hsa-miR-7-5p&offsets|0|-3(+1A);m-400&15|+|88611856|88611879&offsets|0|-3(+1A)];[MIMAT0000252_1&hsa-miR-7-5p&offsets|0|-3(+1A);m-402&19|+|4770700|4770723&offsets|0|-3(+1A)];[MIMAT0000252_2&hsa-miR-7-5p&offsets|0|-3(+1A);m-401&9|-|83969811|83969834&offsets|0|-3(+1A)]|TGGAAGACTAGTGATTTTGTTA</td><td>>MI0000264|hsa-mir-7-2&WithFlank&15|+|88611819|88611940</td><td>38</td><td>58</td><td>21(+1A)</td><td>TGGAAGACTAGTGATTTTGTTA</td><td>98.5</td><td>3.9</td><td>1.5</td><td>2.5</td><td>1.2E-02</td><td>2.2E-02</td><td>BT20/MB468</td><td>MP</td></tr>
<tr><td>>hg38_21_+_8433172_8446622_RNA45SN1_45S_50ntFlanks@4570.4587.18|TGGAATAGGACCGCGGTT</td><td>>hg38_21_+_8433172_8446622_RNA45SN1_45S_50ntFlanks</td><td>4570</td><td>4587</td><td>18</td><td>TGGAATAGGACCGCGGTT</td><td>135.8</td><td>-2.8</td><td>1.1</td><td>-2.4</td><td>1.6E-02</td><td>2.2E-02</td><td>BT20/MB468</td><td>MITO</td></tr>
<tr><td>>hg38_21_+_8433172_8446622_RNA45SN1_45S_50ntFlanks@8169.8212.44|CCAAGTCCTTCTGATCGAGGCCCAGCCCGTGGACGGTGTGAGGC</td><td>>hg38_21_+_8433172_8446622_RNA45SN1_45S_50ntFlanks</td><td>8169</td><td>8212</td><td>44</td><td>CCAAGTCCTTCTGATCGAGGCCCAGCCCGTGGACGGTGTGAGGC</td><td>386.6</td><td>-2.0</td><td>0.8</td><td>-2.4</td><td>1.6E-02</td><td>2.2E-02</td><td>BT20/MB468</td><td>MITO</td></tr>
<tr><td>>hg38_21_+_8433172_8446622_RNA45SN1_45S_50ntFlanks@10024.10072.49|GCGCTGGAGCGTCGGGCCCATACCCGGCCGTCGCCGGCAGTCGAGAGTG</td><td>>hg38_21_+_8433172_8446622_RNA45SN1_45S_50ntFlanks</td><td>10024</td><td>10072</td><td>49</td><td>GCGCTGGAGCGTCGGGCCCATACCCGGCCGTCGCCGGCAGTCGAGAGTG</td><td>105.7</td><td>-4.7</td><td>1.9</td><td>-2.5</td><td>1.2E-02</td><td>2.2E-02</td><td>BT20/MB468</td><td>MP</td></tr>
<tr><td>>hg38_21_+_8433172_8446622_RNA45SN1_45S_50ntFlanks@12615.12632.18|GCTGAGGAGCCAATGGGG</td><td>>hg38_21_+_8433172_8446622_RNA45SN1_45S_50ntFlanks</td><td>12615</td><td>12632</td><td>18</td><td>GCTGAGGAGCCAATGGGG</td><td>75.0</td><td>-4.1</td><td>1.7</td><td>-2.5</td><td>1.2E-02</td><td>2.2E-02</td><td>BT20/MB468</td><td>MP</td></tr>
<tr><td>>hg38_21_+_8433172_8446622_RNA45SN1_45S_50ntFlanks@11027.11055.29|CTCTCTCTCTCTCTCCCCCGCTCCCCGTC</td><td>>hg38_21_+_8433172_8446622_RNA45SN1_45S_50ntFlanks</td><td>11027</td><td>11055</td><td>29</td><td>CTCTCTCTCTCTCTCCCCCGCTCCCCGTC</td><td>324.3</td><td>-3.4</td><td>1.5</td><td>-2.3</td><td>2.0E-02</td><td>2.2E-02</td><td>MB231/MB468</td><td>MITO</td></tr>
<tr><td>>MI0000750|hsa-mir-26a-2&WithFlank&12|-|57824603|57824698@20.41.22(+1U);MI0000083|hsa-mir-26a-1&WithFlank&3|+|37969398|37969486@16.37.22(+1U)&[MIMAT0000082&hsa-miR-26a-5p&offsets|0|0(+1U);m-33&12|-|57824658|57824679&offsets|0|0(+1U)];[MIMAT0000082_1&hsa-miR-26a-5p&offsets|0|0(+1U);m-34&3|+|37969413|37969434&offsets|0|0(+1U)]|TTCAAGTAATCCAGGATAGGCTT</td><td>>MI0000750|hsa-mir-26a-2&WithFlank&12|-|57824603|57824698</td><td>20</td><td>41</td><td>22(+1U)</td><td>TTCAAGTAATCCAGGATAGGCTT</td><td>8742.1</td><td>-3.0</td><td>1.1</td><td>-2.8</td><td>5.0E-03</td><td>2.2E-02</td><td>BT20/MB231</td><td>MP</td></tr>
<tr><td>>hg38_21_+_8433172_8446622_RNA45SN1_45S_50ntFlanks@12992.13014.23|TCCCTCGCTGCGATCTATTGAAA</td><td>>hg38_21_+_8433172_8446622_RNA45SN1_45S_50ntFlanks</td><td>12992</td><td>13014</td><td>23</td><td>TCCCTCGCTGCGATCTATTGAAA</td><td>148.4</td><td>-2.8</td><td>1.2</td><td>-2.4</td><td>1.6E-02</td><td>2.2E-02</td><td>BT20/MB468</td><td>MITO</td></tr>
<tr><td>>hg38_21_+_8433172_8446622_RNA45SN1_45S_50ntFlanks@10817.10840.24|TAAGGGAAGTCGGCAAGCCGGATC</td><td>>hg38_21_+_8433172_8446622_RNA45SN1_45S_50ntFlanks</td><td>10817</td><td>10840</td><td>24</td><td>TAAGGGAAGTCGGCAAGCCGGATC</td><td>179.1</td><td>-1.6</td><td>0.6</td><td>-2.4</td><td>1.6E-02</td><td>2.2E-02</td><td>BT20/MB468</td><td>MITO</td></tr>
<tr><td>>MI0000782|hsa-mir-374a&WithFlank&X|-|74287280|74287363@18.38.21&[MIMAT0000727&hsa-miR-374a-5p&offsets|0|-1;m-184&X|-|74287325|74287346&offsets|0|-1]|TTATAATACAACCTGATAAGT</td><td>>MI0000782|hsa-mir-374a&WithFlank&X|-|74287280|74287363</td><td>18</td><td>38</td><td>21</td><td>TTATAATACAACCTGATAAGT</td><td>694.8</td><td>-4.4</td><td>1.6</td><td>-2.8</td><td>5.0E-03</td><td>2.2E-02</td><td>BT20/MB231</td><td>MP</td></tr>
<tr><td>>hg38_21_+_8433172_8446622_RNA45SN1_45S_50ntFlanks@7974.8005.32|ACGCGACCTCAGATCAGACGTGGCGACCCGCT</td><td>>hg38_21_+_8433172_8446622_RNA45SN1_45S_50ntFlanks</td><td>7974</td><td>8005</td><td>32</td><td>ACGCGACCTCAGATCAGACGTGGCGACCCGCT</td><td>5599.1</td><td>1.9</td><td>0.8</td><td>2.3</td><td>2.0E-02</td><td>2.2E-02</td><td>MB231/MB468</td><td>MITO</td></tr>
<tr><td>>hg38_21_+_8433172_8446622_RNA45SN1_45S_50ntFlanks@6667.6699.33|ATCACTCGGCTCGTGCGTCGATGAAGAACGCAG</td><td>>hg38_21_+_8433172_8446622_RNA45SN1_45S_50ntFlanks</td><td>6667</td><td>6699</td><td>33</td><td>ATCACTCGGCTCGTGCGTCGATGAAGAACGCAG</td><td>166.5</td><td>-3.7</td><td>1.6</td><td>-2.3</td><td>2.0E-02</td><td>2.2E-02</td><td>MB231/MB468</td><td>NUC</td></tr>
<tr><td>>hg38_21_+_8433172_8446622_RNA45SN1_45S_50ntFlanks@7992.8016.25|CGTGGCGACCCGCTGAATTTAAGCA</td><td>>hg38_21_+_8433172_8446622_RNA45SN1_45S_50ntFlanks</td><td>7992</td><td>8016</td><td>25</td><td>CGTGGCGACCCGCTGAATTTAAGCA</td><td>157.7</td><td>-2.1</td><td>0.9</td><td>-2.5</td><td>1.2E-02</td><td>2.2E-02</td><td>BT20/MB468</td><td>MP</td></tr>
<tr><td>>hg38_21_+_8433172_8446622_RNA45SN1_45S_50ntFlanks@11326.11345.20|CTCGTCGCGCGCGCGTCCGC</td><td>>hg38_21_+_8433172_8446622_RNA45SN1_45S_50ntFlanks</td><td>11326</td><td>11345</td><td>20</td><td>CTCGTCGCGCGCGCGTCCGC</td><td>1457.7</td><td>-5.8</td><td>2.1</td><td>-2.7</td><td>6.6E-03</td><td>2.2E-02</td><td>BT20/MB231</td><td>CYTO</td></tr>
<tr><td>>MI0000266|hsa-mir-10a&WithFlank&17|-|48579832|48579953@27.49.23&[MIMAT0000253&hsa-miR-10a-5p&offsets|-1|-1;m-18&17|-|48579905|48579926&offsets|-1|0]|ATACCCTGTAGATCCGAATTTGT</td><td>>MI0000266|hsa-mir-10a&WithFlank&17|-|48579832|48579953</td><td>27</td><td>49</td><td>23</td><td>ATACCCTGTAGATCCGAATTTGT</td><td>263.0</td><td>2.0</td><td>0.7</td><td>2.7</td><td>6.7E-03</td><td>2.2E-02</td><td>BT20/MB231</td><td>CYTO</td></tr>
<tr><td>>hg38_21_+_8433172_8446622_RNA45SN1_45S_50ntFlanks@12416.12452.37|TTCGATGTCGGCTCTTCCTATCATTGTGAAGCAGAAT</td><td>>hg38_21_+_8433172_8446622_RNA45SN1_45S_50ntFlanks</td><td>12416</td><td>12452</td><td>37</td><td>TTCGATGTCGGCTCTTCCTATCATTGTGAAGCAGAAT</td><td>18.1</td><td>7.4</td><td>3.1</td><td>2.4</td><td>1.8E-02</td><td>2.2E-02</td><td>MB231/MB468</td><td>MP</td></tr>
<tr><td>>hg38_21_+_8433172_8446622_RNA45SN1_45S_50ntFlanks@8159.8178.20|CGTGGGGGGCCCAAGTCCTT</td><td>>hg38_21_+_8433172_8446622_RNA45SN1_45S_50ntFlanks</td><td>8159</td><td>8178</td><td>20</td><td>CGTGGGGGGCCCAAGTCCTT</td><td>105.0</td><td>-3.5</td><td>1.4</td><td>-2.5</td><td>1.2E-02</td><td>2.2E-02</td><td>BT20/MB468</td><td>MP</td></tr>
<tr><td>>hg38_21_+_8433172_8446622_RNA45SN1_45S_50ntFlanks@11212.11252.41|CGACTCTGGACGCGAGCCGGGCCCTTCCCGTGGATCGCCCC</td><td>>hg38_21_+_8433172_8446622_RNA45SN1_45S_50ntFlanks</td><td>11212</td><td>11252</td><td>41</td><td>CGACTCTGGACGCGAGCCGGGCCCTTCCCGTGGATCGCCCC</td><td>598.7</td><td>-2.1</td><td>0.9</td><td>-2.3</td><td>1.9E-02</td><td>2.2E-02</td><td>BT20/MB468</td><td>CYTO</td></tr>
<tr><td>>MI0006444|hsa-mir-1307&WithFlank&10|-|103394247|103394407@87.107.21&[MIMAT0005951&hsa-miR-1307-3p&offsets|+1|0;m-66&10|-|103394300|103394322&offsets|+1|-1]|CTCGGCGTGGCGTCGGTCGTG</td><td>>MI0006444|hsa-mir-1307&WithFlank&10|-|103394247|103394407</td><td>87</td><td>107</td><td>21</td><td>CTCGGCGTGGCGTCGGTCGTG</td><td>201.6</td><td>2.7</td><td>1.1</td><td>2.5</td><td>1.2E-02</td><td>2.2E-02</td><td>BT20/MB468</td><td>MP</td></tr>
<tr><td>>hg38_21_+_8433172_8446622_RNA45SN1_45S_50ntFlanks@6706.6732.27|GCGAGAATTAATGTGAATTGCAGGACA</td><td>>hg38_21_+_8433172_8446622_RNA45SN1_45S_50ntFlanks</td><td>6706</td><td>6732</td><td>27</td><td>GCGAGAATTAATGTGAATTGCAGGACA</td><td>515.1</td><td>3.8</td><td>1.4</td><td>2.8</td><td>5.0E-03</td><td>2.2E-02</td><td>BT20/MB231</td><td>MP</td></tr>
<tr><td>>hg38_21_+_8433172_8446622_RNA45SN1_45S_50ntFlanks@4761.4780.20|CAGATACCGTCGTAGTTCCG</td><td>>hg38_21_+_8433172_8446622_RNA45SN1_45S_50ntFlanks</td><td>4761</td><td>4780</td><td>20</td><td>CAGATACCGTCGTAGTTCCG</td><td>184.8</td><td>-4.3</td><td>1.7</td><td>-2.5</td><td>1.2E-02</td><td>2.2E-02</td><td>BT20/MB468</td><td>MP</td></tr>
<tr><td>>hg38_21_+_8433172_8446622_RNA45SN1_45S_50ntFlanks@9551.9571.21|GTCCGTAGCGGTCCTGACGTG</td><td>>hg38_21_+_8433172_8446622_RNA45SN1_45S_50ntFlanks</td><td>9551</td><td>9571</td><td>21</td><td>GTCCGTAGCGGTCCTGACGTG</td><td>806.3</td><td>1.5</td><td>0.6</td><td>2.3</td><td>1.9E-02</td><td>2.2E-02</td><td>MB231/MB468</td><td>CYTO</td></tr>
<tr><td>>hg38_21_+_8433172_8446622_RNA45SN1_45S_50ntFlanks@12396.12418.23|GCGACGTCGCTTTTTGATCCTTC</td><td>>hg38_21_+_8433172_8446622_RNA45SN1_45S_50ntFlanks</td><td>12396</td><td>12418</td><td>23</td><td>GCGACGTCGCTTTTTGATCCTTC</td><td>257.1</td><td>-2.8</td><td>1.0</td><td>-2.8</td><td>5.4E-03</td><td>2.2E-02</td><td>BT20/MB231</td><td>NUC</td></tr>
<tr><td>>hg38_21_+_8433172_8446622_RNA45SN1_45S_50ntFlanks@4207.4242.36|CGGGGAGGTAGTGACGAAAAATAACAATACAGGACT</td><td>>hg38_21_+_8433172_8446622_RNA45SN1_45S_50ntFlanks</td><td>4207</td><td>4242</td><td>36</td><td>CGGGGAGGTAGTGACGAAAAATAACAATACAGGACT</td><td>372.8</td><td>3.9</td><td>1.4</td><td>2.8</td><td>5.0E-03</td><td>2.2E-02</td><td>BT20/MB231</td><td>MP</td></tr>
<tr><td>>hg38_21_+_8433172_8446622_RNA45SN1_45S_50ntFlanks@3776.3805.30|CCGGTACAGTGAAACTGCGAATGGCTCATT</td><td>>hg38_21_+_8433172_8446622_RNA45SN1_45S_50ntFlanks</td><td>3776</td><td>3805</td><td>30</td><td>CCGGTACAGTGAAACTGCGAATGGCTCATT</td><td>305.8</td><td>-1.5</td><td>0.6</td><td>-2.4</td><td>1.7E-02</td><td>2.2E-02</td><td>BT20/MB468</td><td>MITO</td></tr>
<tr><td>>hg38_21_+_8433172_8446622_RNA45SN1_45S_50ntFlanks@6645.6674.30|CTCGTACGACTCTTAGCGGTGGATCACTCG</td><td>>hg38_21_+_8433172_8446622_RNA45SN1_45S_50ntFlanks</td><td>6645</td><td>6674</td><td>30</td><td>CTCGTACGACTCTTAGCGGTGGATCACTCG</td><td>1966.2</td><td>3.8</td><td>1.4</td><td>2.8</td><td>5.0E-03</td><td>2.2E-02</td><td>BT20/MB231</td><td>MP</td></tr>
<tr><td>>hg38_21_+_8433172_8446622_RNA45SN1_45S_50ntFlanks@9489.9515.27|AGGACCCGAAAGATGGTGAACTATGCC</td><td>>hg38_21_+_8433172_8446622_RNA45SN1_45S_50ntFlanks</td><td>9489</td><td>9515</td><td>27</td><td>AGGACCCGAAAGATGGTGAACTATGCC</td><td>123.3</td><td>5.6</td><td>2.0</td><td>2.8</td><td>5.0E-03</td><td>2.2E-02</td><td>BT20/MB231</td><td>MP</td></tr>
<tr><td>>hg38_21_+_8433172_8446622_RNA45SN1_45S_50ntFlanks@12417.12442.26|TCGATGTCGGCTCTTCCTATCATTGT</td><td>>hg38_21_+_8433172_8446622_RNA45SN1_45S_50ntFlanks</td><td>12417</td><td>12442</td><td>26</td><td>TCGATGTCGGCTCTTCCTATCATTGT</td><td>927.0</td><td>-2.3</td><td>0.9</td><td>-2.7</td><td>6.7E-03</td><td>2.2E-02</td><td>BT20/MB231</td><td>CYTO</td></tr>
</table>

<table>
<tr><td>>am_MI0000786|hsa-mir-378a&WithFlank&5|+|149732819|149732896@50.70.21(+1U)&[MIMAT0000732&hsa-miR-378a-3p&offsets|+1|0(+1U);m-44&5|+|149732867|149732888&offsets|+1|0(+1U)]|CTGGACTTGGAGTCAGAAGGCT</td><td>>am_MI0000786|hsa-mir-378a&WithFlank&5|+|149732819|149732896</td><td>50</td><td>70</td><td>21(+1U)</td><td>CTGGACTTGGAGTCAGAAGGCT</td><td>269.7</td><td>2.3</td><td>1.0</td><td>2.3</td><td>2.0E-02</td><td>2.2E-02</td><td>MB231/MB468</td><td>MITO</td></tr>
<tr><td>>hg38_21_+_8433172_8446622_RNA45SN1_45S_50ntFlanks@12416.12452.37|TTCGATGTCGGCTCTTCCTATCATTGTGAAGCAGAAT</td><td>>hg38_21_+_8433172_8446622_RNA45SN1_45S_50ntFlanks</td><td>12416</td><td>12452</td><td>37</td><td>TTCGATGTCGGCTCTTCCTATCATTGTGAAGCAGAAT</td><td>270.2</td><td>-2.5</td><td>0.9</td><td>-2.8</td><td>5.5E-03</td><td>2.2E-02</td><td>BT20/MB231</td><td>NUC</td></tr>
<tr><td>>hg38_MT_+_598_1651_RNR1_12S_50ntFlanks@442.480.39|ATATCTGAACACACAATAGCTAAGACCCAAACTGGGATT</td><td>>hg38_MT_+_598_1651_RNR1_12S_50ntFlanks</td><td>442</td><td>480</td><td>39</td><td>ATATCTGAACACACAATAGCTAAGACCCAAACTGGGATT</td><td>329.0</td><td>3.6</td><td>1.5</td><td>2.4</td><td>1.8E-02</td><td>2.2E-02</td><td>MB231/MB468</td><td>MP</td></tr>
<tr><td>>MI0001448|hsa-mir-425&WithFlank&3|-|49020142|49020240@20.43.24&[MIMAT0003393&hsa-miR-425-5p&offsets|0|+1;m-95&3|-|49020200|49020221&offsets|0|+2]|AATGACACGATCACTCCCGTTGAG</td><td>>MI0001448|hsa-mir-425&WithFlank&3|-|49020142|49020240</td><td>20</td><td>43</td><td>24</td><td>AATGACACGATCACTCCCGTTGAG</td><td>939.8</td><td>-2.6</td><td>0.9</td><td>-2.8</td><td>5.0E-03</td><td>2.2E-02</td><td>BT20/MB231</td><td>MP</td></tr>
<tr><td>>hg38_21_+_8433172_8446622_RNA45SN1_45S_50ntFlanks@7979.8004.26|ACCTCAGATCAGACGTGGCGACCCGC</td><td>>hg38_21_+_8433172_8446622_RNA45SN1_45S_50ntFlanks</td><td>7979</td><td>8004</td><td>26</td><td>ACCTCAGATCAGACGTGGCGACCCGC</td><td>141.4</td><td>-3.3</td><td>1.4</td><td>-2.4</td><td>1.8E-02</td><td>2.2E-02</td><td>MB231/MB468</td><td>MP</td></tr>
<tr><td>>hg38_21_+_8433172_8446622_RNA45SN1_45S_50ntFlanks@6658.6685.28|TAGCGGTGGATCACTCGGCTCGTGCGTC</td><td>>hg38_21_+_8433172_8446622_RNA45SN1_45S_50ntFlanks</td><td>6658</td><td>6685</td><td>28</td><td>TAGCGGTGGATCACTCGGCTCGTGCGTC</td><td>67.0</td><td>4.3</td><td>1.7</td><td>2.5</td><td>1.2E-02</td><td>2.2E-02</td><td>BT20/MB468</td><td>MP</td></tr>
<tr><td>>hg38_21_+_8433172_8446622_RNA45SN1_45S_50ntFlanks@12044.12068.25|ACTGACCCGGTGAGGCGGGGGGGCG</td><td>>hg38_21_+_8433172_8446622_RNA45SN1_45S_50ntFlanks</td><td>12044</td><td>12068</td><td>25</td><td>ACTGACCCGGTGAGGCGGGGGGGCG</td><td>228.5</td><td>-3.7</td><td>1.6</td><td>-2.3</td><td>1.9E-02</td><td>2.2E-02</td><td>BT20/MB468</td><td>CYTO</td></tr>
<tr><td>>MI0000239|hsa-mir-197&WithFlank&1|+|109598887|109598973@54.74.21(+1A)&[MIMAT0000227&hsa-miR-197-3p&offsets|0|-1(+1A);m-99&1|+|109598940|109598961&offsets|0|-1(+1A)]|TTCACCACCTTCTCCACCCAGA</td><td>>MI0000239|hsa-mir-197&WithFlank&1|+|109598887|109598973</td><td>54</td><td>74</td><td>21(+1A)</td><td>TTCACCACCTTCTCCACCCAGA</td><td>72.4</td><td>3.5</td><td>1.4</td><td>2.5</td><td>1.2E-02</td><td>2.2E-02</td><td>BT20/MB468</td><td>MP</td></tr>
<tr><td>>MI0000070|hsa-mir-16-1&WithFlank&13|-|50048967|50049067@20.39.20(+1A);MI0000115|hsa-mir-16-2&WithFlank&3|+|160404739|160404831@16.35.20(+1A)&[MIMAT0000069&hsa-miR-16-5p&offsets|0|-2(+1A);m-62&13|-|50049027|50049048&offsets|0|-2(+1A)];[MIMAT0000069_1&hsa-miR-16-5p&offsets|0|-2(+1A);m-61&3|+|160404754|160404775&offsets|0|-2(+1A)]|TAGCAGCACGTAAATATTGGA</td><td>>MI0000070|hsa-mir-16-1&WithFlank&13|-|50048967|50049067</td><td>20</td><td>39</td><td>20(+1A)</td><td>TAGCAGCACGTAAATATTGGA</td><td>235.3</td><td>3.3</td><td>1.4</td><td>2.4</td><td>1.8E-02</td><td>2.2E-02</td><td>MB231/MB468</td><td>MP</td></tr>
<tr><td>>hg38_21_+_8433172_8446622_RNA45SN1_45S_50ntFlanks@6671.6695.25|CTCGGCTCGTGCGTCGATGAAGAAC</td><td>>hg38_21_+_8433172_8446622_RNA45SN1_45S_50ntFlanks</td><td>6671</td><td>6695</td><td>25</td><td>CTCGGCTCGTGCGTCGATGAAGAAC</td><td>182.4</td><td>-2.9</td><td>1.2</td><td>-2.4</td><td>1.8E-02</td><td>2.2E-02</td><td>MB231/MB468</td><td>MP</td></tr>
<tr><td>>hg38_21_+_8433172_8446622_RNA45SN1_45S_50ntFlanks@11566.11585.20|TAGCAGCCGACTTAGAACTG</td><td>>hg38_21_+_8433172_8446622_RNA45SN1_45S_50ntFlanks</td><td>11566</td><td>11585</td><td>20</td><td>TAGCAGCCGACTTAGAACTG</td><td>309.8</td><td>-2.7</td><td>1.0</td><td>-2.8</td><td>5.5E-03</td><td>2.2E-02</td><td>BT20/MB231</td><td>NUC</td></tr>
<tr><td>>hg38_21_+_8433172_8446622_RNA45SN1_45S_50ntFlanks@12606.12625.20|ATGTGCTTGGCTGAGGAGCC</td><td>>hg38_21_+_8433172_8446622_RNA45SN1_45S_50ntFlanks</td><td>12606</td><td>12625</td><td>20</td><td>ATGTGCTTGGCTGAGGAGCC</td><td>374.8</td><td>-2.3</td><td>0.9</td><td>-2.5</td><td>1.2E-02</td><td>2.2E-02</td><td>BT20/MB468</td><td>MP</td></tr>
<tr><td>>hg38_21_+_8433172_8446622_RNA45SN1_45S_50ntFlanks@5478.5500.23|CTGGCGGAGCGCTGAGAAGACGG</td><td>>hg38_21_+_8433172_8446622_RNA45SN1_45S_50ntFlanks</td><td>5478</td><td>5500</td><td>23</td><td>CTGGCGGAGCGCTGAGAAGACGG</td><td>239.4</td><td>2.6</td><td>1.0</td><td>2.5</td><td>1.2E-02</td><td>2.2E-02</td><td>BT20/MB468</td><td>MP</td></tr>
<tr><td>>trnaMT_SerGCT_MT_+_12207_12265@1.22.22|GAGAAAGCTCACAAGAACTGCT</td><td>>trnaMT_SerGCT_MT_+_12207_12265</td><td>1</td><td>22</td><td>22</td><td>GAGAAAGCTCACAAGAACTGCT</td><td>18500.5</td><td>1.9</td><td>0.8</td><td>2.4</td><td>1.8E-02</td><td>2.2E-02</td><td>MB231/MB468</td><td>MP</td></tr>
<tr><td>>hg38_1_-_228634819_228635039_RNA5S12_5S_50ntFlanks@137.169.33|GAGACCGCCTGGGAATACCGGGTGCTGTAGGCT</td><td>>hg38_1_-_228634819_228635039_RNA5S12_5S_50ntFlanks</td><td>137</td><td>169</td><td>33</td><td>GAGACCGCCTGGGAATACCGGGTGCTGTAGGCT</td><td>1119.3</td><td>3.6</td><td>1.3</td><td>2.8</td><td>5.0E-03</td><td>2.2E-02</td><td>BT20/MB231</td><td>MP</td></tr>
<tr><td>>trnaMT_GluTTC_MT_-_14674_14742@38.72.35,trnalookalike8_GluTTC_5_-_93905172_93905240@38.72.35|CATTGGTCGTGGTTGTAGTCCGTGCGAGAATACCA</td><td>>trnaMT_GluTTC_MT_-_14674_14742</td><td>38</td><td>72</td><td>35</td><td>CATTGGTCGTGGTTGTAGTCCGTGCGAGAATACCA</td><td>4553.7</td><td>-2.1</td><td>0.7</td><td>-2.8</td><td>5.5E-03</td><td>2.2E-02</td><td>BT20/MB231</td><td>NUC</td></tr>
<tr><td>>hg38_21_+_8433172_8446622_RNA45SN1_45S_50ntFlanks@12419.12453.35|GATGTCGGCTCTTCCTATCATTGTGAAGCAGAATT</td><td>>hg38_21_+_8433172_8446622_RNA45SN1_45S_50ntFlanks</td><td>12419</td><td>12453</td><td>35</td><td>GATGTCGGCTCTTCCTATCATTGTGAAGCAGAATT</td><td>384.7</td><td>-2.4</td><td>0.9</td><td>-2.8</td><td>5.5E-03</td><td>2.2E-02</td><td>BT20/MB231</td><td>NUC</td></tr>
<tr><td>>hg38_21_+_8433172_8446622_RNA45SN1_45S_50ntFlanks@9913.9948.36|AGAAAAGGTGTTGGTTGATATAGACAGCAGGACGGT</td><td>>hg38_21_+_8433172_8446622_RNA45SN1_45S_50ntFlanks</td><td>9913</td><td>9948</td><td>36</td><td>AGAAAAGGTGTTGGTTGATATAGACAGCAGGACGGT</td><td>127.6</td><td>5.4</td><td>1.9</td><td>2.8</td><td>5.0E-03</td><td>2.2E-02</td><td>BT20/MB231</td><td>MP</td></tr>
<tr><td>>hg38_MT_+_1621_3279_RNR2_16S_50ntFlanks@1198.1215.18|CGACCTCGGAGCAGAACC</td><td>>hg38_MT_+_1621_3279_RNR2_16S_50ntFlanks</td><td>1198</td><td>1215</td><td>18</td><td>CGACCTCGGAGCAGAACC</td><td>18.1</td><td>7.4</td><td>3.1</td><td>2.4</td><td>1.8E-02</td><td>2.2E-02</td><td>MB231/MB468</td><td>MP</td></tr>
<tr><td>>hg38_21_+_8433172_8446622_RNA45SN1_45S_50ntFlanks@11672.11703.32|CAGTGCTCTGAATGTCAAAGTGAAGAAATTCA</td><td>>hg38_21_+_8433172_8446622_RNA45SN1_45S_50ntFlanks</td><td>11672</td><td>11703</td><td>32</td><td>CAGTGCTCTGAATGTCAAAGTGAAGAAATTCA</td><td>299.5</td><td>-2.5</td><td>0.9</td><td>-2.7</td><td>6.7E-03</td><td>2.2E-02</td><td>BT20/MB231</td><td>CYTO</td></tr>
<tr><td>>hg38_21_+_8433172_8446622_RNA45SN1_45S_50ntFlanks@10409.10435.27|GTCAGTCGGTCCTGAGAGATGGGCGAG</td><td>>hg38_21_+_8433172_8446622_RNA45SN1_45S_50ntFlanks</td><td>10409</td><td>10435</td><td>27</td><td>GTCAGTCGGTCCTGAGAGATGGGCGAG</td><td>257.8</td><td>-3.3</td><td>1.3</td><td>-2.5</td><td>1.2E-02</td><td>2.2E-02</td><td>BT20/MB468</td><td>MP</td></tr>
<tr><td>>hg38_21_+_8433172_8446622_RNA45SN1_45S_50ntFlanks@8171.8214.44|AAGTCCTTCTGATCGAGGCCCAGCCCGTGGACGGTGTGAGGCCG</td><td>>hg38_21_+_8433172_8446622_RNA45SN1_45S_50ntFlanks</td><td>8171</td><td>8214</td><td>44</td><td>AAGTCCTTCTGATCGAGGCCCAGCCCGTGGACGGTGTGAGGCCG</td><td>308.8</td><td>-3.0</td><td>1.2</td><td>-2.5</td><td>1.2E-02</td><td>2.2E-02</td><td>BT20/MB468</td><td>MP</td></tr>
<tr><td>>hg38_21_+_8433172_8446622_RNA45SN1_45S_50ntFlanks@9956.9979.24|GAAGTCGGAATCCGCTAAGGAGTG</td><td>>hg38_21_+_8433172_8446622_RNA45SN1_45S_50ntFlanks</td><td>9956</td><td>9979</td><td>24</td><td>GAAGTCGGAATCCGCTAAGGAGTG</td><td>158.5</td><td>5.6</td><td>2.0</td><td>2.8</td><td>5.1E-03</td><td>2.2E-02</td><td>BT20/MB231</td><td>MP</td></tr>
<tr><td>>hg38_21_+_8433172_8446622_RNA45SN1_45S_50ntFlanks@10797.10817.21|GCATGTTGGAACAATGTAGGT</td><td>>hg38_21_+_8433172_8446622_RNA45SN1_45S_50ntFlanks</td><td>10797</td><td>10817</td><td>21</td><td>GCATGTTGGAACAATGTAGGT</td><td>154.2</td><td>-1.9</td><td>0.8</td><td>-2.3</td><td>2.0E-02</td><td>2.2E-02</td><td>MB231/MB468</td><td>NUC</td></tr>
<tr><td>>hg38_21_+_8433172_8446622_RNA45SN1_45S_50ntFlanks@9952.9979.28|CATGGAAGTCGGAATCCGCTAAGGAGTG</td><td>>hg38_21_+_8433172_8446622_RNA45SN1_45S_50ntFlanks</td><td>9952</td><td>9979</td><td>28</td><td>CATGGAAGTCGGAATCCGCTAAGGAGTG</td><td>227.3</td><td>2.7</td><td>1.1</td><td>2.5</td><td>1.3E-02</td><td>2.2E-02</td><td>BT20/MB468</td><td>MP</td></tr>
<tr><td>>trna146_GlnCTG_6_-_27515531_27515602@17.54.38,trna1_GlnCTG_6_+_18836402_18836473@17.54.38,trna3_GlnCTG_17_+_8023070_8023141@17.54.38,trna49_GlnCTG_6_+_27487308_27487379@17.54.38,trna7_GlnCTG_15_-_66161400_66161471@17.54.38,trna99_GlnCTG_6_-_28909378_28909449@17.54.38|GGTTAGCACTCTGGACTCTGAATCCAGCGATCCGAGTT</td><td>>trna146_GlnCTG_6_-_27515531_27515602</td><td>17</td><td>54</td><td>38</td><td>GGTTAGCACTCTGGACTCTGAATCCAGCGATCCGAGTT</td><td>711.9</td><td>2.5</td><td>0.9</td><td>2.7</td><td>6.8E-03</td><td>2.2E-02</td><td>BT20/MB231</td><td>CYTO</td></tr>
<tr><td>>hg38_21_+_8433172_8446622_RNA45SN1_45S_50ntFlanks@11321.11338.18|CACGTCTCGTCGCGCGCG</td><td>>hg38_21_+_8433172_8446622_RNA45SN1_45S_50ntFlanks</td><td>11321</td><td>11338</td><td>18</td><td>CACGTCTCGTCGCGCGCG</td><td>1399.4</td><td>-5.0</td><td>2.1</td><td>-2.3</td><td>2.0E-02</td><td>2.2E-02</td><td>BT20/MB468</td><td>NUC</td></tr>
<tr><td>>hg38_21_+_8433172_8446622_RNA45SN1_45S_50ntFlanks@12396.12415.20|GCGACGTCGCTTTTTGATCC</td><td>>hg38_21_+_8433172_8446622_RNA45SN1_45S_50ntFlanks</td><td>12396</td><td>12415</td><td>20</td><td>GCGACGTCGCTTTTTGATCC</td><td>148.9</td><td>2.5</td><td>1.1</td><td>2.3</td><td>1.9E-02</td><td>2.2E-02</td><td>MB231/MB468</td><td>CYTO</td></tr>
<tr><td>>trnaMT_PheGAA_MT_+_577_647@39.58.20|ATGTTTAGACGGGCTCACAT</td><td>>trnaMT_PheGAA_MT_+_577_647</td><td>39</td><td>58</td><td>20</td><td>ATGTTTAGACGGGCTCACAT</td><td>61.8</td><td>4.7</td><td>2.0</td><td>2.4</td><td>1.8E-02</td><td>2.2E-02</td><td>MB231/MB468</td><td>MP</td></tr>
<tr><td>>hg38_21_+_8433172_8446622_RNA45SN1_45S_50ntFlanks@6661.6697.37|CGGTGGATCACTCGGCTCGTGCGTCGATGAAGAACGC</td><td>>hg38_21_+_8433172_8446622_RNA45SN1_45S_50ntFlanks</td><td>6661</td><td>6697</td><td>37</td><td>CGGTGGATCACTCGGCTCGTGCGTCGATGAAGAACGC</td><td>913.0</td><td>1.5</td><td>0.5</td><td>2.8</td><td>5.5E-03</td><td>2.2E-02</td><td>BT20/MB231</td><td>NUC</td></tr>
<tr><td>>hg38_21_+_8433172_8446622_RNA45SN1_45S_50ntFlanks@10771.10816.46|CAGGTCTCCAAGGTGAACAGCCTCTGGCATGTTGGAACAATGTAGG</td><td>>hg38_21_+_8433172_8446622_RNA45SN1_45S_50ntFlanks</td><td>10771</td><td>10816</td><td>46</td><td>CAGGTCTCCAAGGTGAACAGCCTCTGGCATGTTGGAACAATGTAGG</td><td>371.8</td><td>-3.7</td><td>1.6</td><td>-2.3</td><td>1.9E-02</td><td>2.2E-02</td><td>BT20/MB468</td><td>CYTO</td></tr>
<tr><td>>hg38_21_+_8433172_8446622_RNA45SN1_45S_50ntFlanks@12458.12496.39|AAGCGTTGGATTGTTCACCCACTAATAGGGAACGTGAGC</td><td>>hg38_21_+_8433172_8446622_RNA45SN1_45S_50ntFlanks</td><td>12458</td><td>12496</td><td>39</td><td>AAGCGTTGGATTGTTCACCCACTAATAGGGAACGTGAGC</td><td>55.6</td><td>5.1</td><td>2.0</td><td>2.5</td><td>1.3E-02</td><td>2.2E-02</td><td>BT20/MB468</td><td>MP</td></tr>
<tr><td>>hg38_21_+_8433172_8446622_RNA45SN1_45S_50ntFlanks@4964.4994.31|ACCTCACCCGGCCCGGACACGGACAGGATTG</td><td>>hg38_21_+_8433172_8446622_RNA45SN1_45S_50ntFlanks</td><td>4964</td><td>4994</td><td>31</td><td>ACCTCACCCGGCCCGGACACGGACAGGATTG</td><td>45.0</td><td>-4.5</td><td>1.8</td><td>-2.5</td><td>1.3E-02</td><td>2.2E-02</td><td>BT20/MB468</td><td>MP</td></tr>
<tr><td>>MI0000077|hsa-mir-21&WithFlank&17|+|59841260|59841343@14.35.22(+1A)&[MIMAT0000076&hsa-miR-21-5p&offsets|0|0(+1A);m-1&17|+|59841273|59841294&offsets|0|0(+1A)]|TAGCTTATCAGACTGATGTTGAA</td><td>>MI0000077|hsa-mir-21&WithFlank&17|+|59841260|59841343</td><td>14</td><td>35</td><td>22(+1A)</td><td>TAGCTTATCAGACTGATGTTGAA</td><td>39683.7</td><td>2.7</td><td>1.0</td><td>2.7</td><td>6.8E-03</td><td>2.2E-02</td><td>BT20/MB231</td><td>CYTO</td></tr>
<tr><td>>hg38_21_+_8433172_8446622_RNA45SN1_45S_50ntFlanks@7992.8016.25|CGTGGCGACCCGCTGAATTTAAGCA</td><td>>hg38_21_+_8433172_8446622_RNA45SN1_45S_50ntFlanks</td><td>7992</td><td>8016</td><td>25</td><td>CGTGGCGACCCGCTGAATTTAAGCA</td><td>815.5</td><td>-2.7</td><td>1.0</td><td>-2.7</td><td>6.8E-03</td><td>2.2E-02</td><td>BT20/MB231</td><td>CYTO</td></tr>
<tr><td>>trna10_LysCTT_16_+_3241501_3241573@1.28.28,trna119_LysCTT_1_-_145395522_145395594@1.28.28,trna11_LysCTT_5_-_180648979_180649051@1.28.28,trna13_LysCTT_14_-_58706613_58706685@1.28.28,trna13_LysCTT_6_+_26556774_26556846@1.28.28,trna2_LysCTT_15_+_79152904_79152976@1.28.28,trna32_LysCTT_16_-_3207406_3207478@1.28.28,trna7_LysCTT_16_+_3225692_3225764@1.28.28,trna9_LysCTT_5_+_180634755_180634827@1.28.28|GCCCGGCTAGCTCAGTCGGTAGAGCATG</td><td>>trna10_LysCTT_16_+_3241501_3241573</td><td>1</td><td>28</td><td>28</td><td>GCCCGGCTAGCTCAGTCGGTAGAGCATG</td><td>972.8</td><td>-3.7</td><td>1.5</td><td>-2.5</td><td>1.3E-02</td><td>2.2E-02</td><td>BT20/MB468</td><td>MP</td></tr>
<tr><td>>hg38_21_+_8433172_8446622_RNA45SN1_45S_50ntFlanks@6674.6700.27|GGCTCGTGCGTCGATGAAGAACGCAGC</td><td>>hg38_21_+_8433172_8446622_RNA45SN1_45S_50ntFlanks</td><td>6674</td><td>6700</td><td>27</td><td>GGCTCGTGCGTCGATGAAGAACGCAGC</td><td>98.3</td><td>4.4</td><td>1.8</td><td>2.5</td><td>1.3E-02</td><td>2.2E-02</td><td>BT20/MB468</td><td>MP</td></tr>
<tr><td>>hg38_MT_+_1621_3279_RNR2_16S_50ntFlanks@1249.1271.23|AGTCAAAGCGAACTACTATACTC</td><td>>hg38_MT_+_1621_3279_RNR2_16S_50ntFlanks</td><td>1249</td><td>1271</td><td>23</td><td>AGTCAAAGCGAACTACTATACTC</td><td>283.9</td><td>4.2</td><td>1.8</td><td>2.4</td><td>1.8E-02</td><td>2.2E-02</td><td>MB231/MB468</td><td>MP</td></tr>
<tr><td>>hg38_MT_+_598_1651_RNR1_12S_50ntFlanks@427.452.26|GAAAGTGGCTTTAACATATCTGAACA</td><td>>hg38_MT_+_598_1651_RNR1_12S_50ntFlanks</td><td>427</td><td>452</td><td>26</td><td>GAAAGTGGCTTTAACATATCTGAACA</td><td>143.6</td><td>4.5</td><td>1.8</td><td>2.5</td><td>1.3E-02</td><td>2.2E-02</td><td>BT20/MB468</td><td>MP</td></tr>
<tr><td>>hg38_21_+_8433172_8446622_RNA45SN1_45S_50ntFlanks@10348.10376.29|AAACGAGAACTTTGAAGGCCGAAGTGGAG</td><td>>hg38_21_+_8433172_8446622_RNA45SN1_45S_50ntFlanks</td><td>10348</td><td>10376</td><td>29</td><td>AAACGAGAACTTTGAAGGCCGAAGTGGAG</td><td>386.9</td><td>-2.9</td><td>1.2</td><td>-2.5</td><td>1.3E-02</td><td>2.2E-02</td><td>BT20/MB468</td><td>MP</td></tr>
<tr><td>>hg38_21_+_8433172_8446622_RNA45SN1_45S_50ntFlanks@11321.11344.24|CACGTCTCGTCGCGCGCGCGTCCG</td><td>>hg38_21_+_8433172_8446622_RNA45SN1_45S_50ntFlanks</td><td>11321</td><td>11344</td><td>24</td><td>CACGTCTCGTCGCGCGCGCGTCCG</td><td>7156.3</td><td>-2.8</td><td>1.2</td><td>-2.4</td><td>1.7E-02</td><td>2.2E-02</td><td>BT20/MB468</td><td>MITO</td></tr>
<tr><td>>hg38_21_+_8433172_8446622_RNA45SN1_45S_50ntFlanks@11873.11899.27|AAGAAGACCCTGTTGAGCTTGACTCTA</td><td>>hg38_21_+_8433172_8446622_RNA45SN1_45S_50ntFlanks</td><td>11873</td><td>11899</td><td>27</td><td>AAGAAGACCCTGTTGAGCTTGACTCTA</td><td>297.4</td><td>-1.3</td><td>0.5</td><td>-2.4</td><td>1.7E-02</td><td>2.2E-02</td><td>BT20/MB468</td><td>MITO</td></tr>
<tr><td>>MI0000746|hsa-mir-99b&WithFlank&19|+|51692606|51692687@51.71.21(+1A)&[MIMAT0004678&hsa-miR-99b-3p&offsets|0|-1(+1A);m-151&19|+|51692656|51692678&offsets|0|-2(+1A)]|CAAGCTCGTGTCTGTGGGTCCA</td><td>>MI0000746|hsa-mir-99b&WithFlank&19|+|51692606|51692687</td><td>51</td><td>71</td><td>21(+1A)</td><td>CAAGCTCGTGTCTGTGGGTCCA</td><td>420.4</td><td>2.2</td><td>0.8</td><td>2.8</td><td>5.6E-03</td><td>2.2E-02</td><td>BT20/MB231</td><td>NUC</td></tr>
<tr><td>>trnaMT_GluTTC_MT_-_14674_14742@39.72.34,trnalookalike8_GluTTC_5_-_93905172_93905240@39.72.34|ATTGGTCGTGGTTGTAGTCCGTGCGAGAATACCA</td><td>>trnaMT_GluTTC_MT_-_14674_14742</td><td>39</td><td>72</td><td>34</td><td>ATTGGTCGTGGTTGTAGTCCGTGCGAGAATACCA</td><td>20284.9</td><td>-2.2</td><td>0.8</td><td>-2.8</td><td>5.6E-03</td><td>2.2E-02</td><td>BT20/MB231</td><td>NUC</td></tr>
</table>

<table>
<tr><td></td><td></td><td></td><td></td><td></td><td></td><td></td><td></td><td></td><td></td><td></td><td></td><td></td><td></td></tr>
<tr><td>>MI0000299|hsa-mir-222&WithFlank&X|-|45747009|45747130@75.99.25(+1U)&[MIMAT0000279&hsa-miR-222-3p&offsets|0|+4(+1U);m-140&X|-|45747033|45747056&offsets|0|+1(+1U)]|AGCTACATCTGGCTACTGGGTCTCTT</td><td>>MI0000299|hsa-mir-222&WithFlank&X|-|45747009|45747130</td><td>75</td><td>99</td><td>25(+1U)</td><td>AGCTACATCTGGCTACTGGGTCTCTT</td><td>195.7</td><td>3.7</td><td>1.5</td><td>2.5</td><td>1.3E-02</td><td>2.2E-02</td><td>BT20/MB468</td><td>MP</td></tr>
<tr><td>>hg38_21_+_8433172_8446622_RNA45SN1_45S_50ntFlanks@6660.6681.22|GCGGTGGATCACTCGGCTCGTG</td><td>>hg38_21_+_8433172_8446622_RNA45SN1_45S_50ntFlanks</td><td>6660</td><td>6681</td><td>22</td><td>GCGGTGGATCACTCGGCTCGTG</td><td>1382.0</td><td>-2.4</td><td>1.0</td><td>-2.5</td><td>1.3E-02</td><td>2.2E-02</td><td>BT20/MB468</td><td>MP</td></tr>
<tr><td>>hg38_21_+_8433172_8446622_RNA45SN1_45S_50ntFlanks@10397.10424.28|AGTTGAACATGGGTCAGTCGGTCCTGAG</td><td>>hg38_21_+_8433172_8446622_RNA45SN1_45S_50ntFlanks</td><td>10397</td><td>10424</td><td>28</td><td>AGTTGAACATGGGTCAGTCGGTCCTGAG</td><td>106.4</td><td>1.3</td><td>0.6</td><td>2.3</td><td>2.0E-02</td><td>2.2E-02</td><td>BT20/MB468</td><td>NUC</td></tr>
<tr><td>>hg38_MT_+_1621_3279_RNR2_16S_50ntFlanks@825.856.32|TAAGGAAAGGTTAAAAAAAGTAAAAGGAACTC</td><td>>hg38_MT_+_1621_3279_RNR2_16S_50ntFlanks</td><td>825</td><td>856</td><td>32</td><td>TAAGGAAAGGTTAAAAAAAGTAAAAGGAACTC</td><td>32.8</td><td>4.7</td><td>2.0</td><td>2.4</td><td>1.8E-02</td><td>2.2E-02</td><td>MB231/MB468</td><td>MP</td></tr>
<tr><td>>hg38_21_+_8433172_8446622_RNA45SN1_45S_50ntFlanks@11029.11053.25|CTCTCTCTCTCTCCCCCGCTCCCCG</td><td>>hg38_21_+_8433172_8446622_RNA45SN1_45S_50ntFlanks</td><td>11029</td><td>11053</td><td>25</td><td>CTCTCTCTCTCTCCCCCGCTCCCCG</td><td>815.7</td><td>-5.1</td><td>2.2</td><td>-2.3</td><td>2.0E-02</td><td>2.2E-02</td><td>BT20/MB468</td><td>NUC</td></tr>
<tr><td>>MI0000734|hsa-mir-106b&WithFlank&7|-|100093987|100094080@18.39.22&[MIMAT0000680&hsa-miR-106b-5p&offsets|0|+1;m-103&7|-|100094043|100094063&offsets|0|+1]|TAAAGTGCTGACAGTGCAGATA</td><td>>MI0000734|hsa-mir-106b&WithFlank&7|-|100093987|100094080</td><td>18</td><td>39</td><td>22</td><td>TAAAGTGCTGACAGTGCAGATA</td><td>2868.2</td><td>-3.3</td><td>1.2</td><td>-2.8</td><td>5.1E-03</td><td>2.2E-02</td><td>BT20/MB231</td><td>MP</td></tr>
<tr><td>>hg38_21_+_8433172_8446622_RNA45SN1_45S_50ntFlanks@9792.9821.30|GGCGTGGAGCCGGGCGTGGAATGCGAGTGC</td><td>>hg38_21_+_8433172_8446622_RNA45SN1_45S_50ntFlanks</td><td>9792</td><td>9821</td><td>30</td><td>GGCGTGGAGCCGGGCGTGGAATGCGAGTGC</td><td>94.3</td><td>4.5</td><td>1.8</td><td>2.5</td><td>1.3E-02</td><td>2.2E-02</td><td>BT20/MB468</td><td>MP</td></tr>
<tr><td>>hg38_21_+_8433172_8446622_RNA45SN1_45S_50ntFlanks@12644.12666.23|TCTGTGGGATTATGACTGAACGC</td><td>>hg38_21_+_8433172_8446622_RNA45SN1_45S_50ntFlanks</td><td>12644</td><td>12666</td><td>23</td><td>TCTGTGGGATTATGACTGAACGC</td><td>229.2</td><td>5.7</td><td>2.1</td><td>2.8</td><td>5.1E-03</td><td>2.2E-02</td><td>BT20/MB231</td><td>MP</td></tr>
<tr><td>>hg38_1_-_228634819_228635039_RNA5S12_5S_50ntFlanks@86.103.18|CGTCTGATCTCGGAAGCT</td><td>>hg38_1_-_228634819_228635039_RNA5S12_5S_50ntFlanks</td><td>86</td><td>103</td><td>18</td><td>CGTCTGATCTCGGAAGCT</td><td>249.3</td><td>-1.7</td><td>0.7</td><td>-2.3</td><td>1.9E-02</td><td>2.2E-02</td><td>BT20/MB468</td><td>CYTO</td></tr>
<tr><td>>am_trna162_MetCAT_6_-_26758550_26758622@31.54.24,trna164_MetCAT_6_-_26735574_26735646@31.54.24,trna20_MetCAT_16_+_71460396_71460468@31.54.24,trna21_MetCAT_6_+_26701712_26701784@31.54.24,trna27_MetCAT_6_+_26766444_26766516@31.54.24,trna75_MetCAT_6_+_28912352_28912424@31.54.24,trna97_MetCAT_6_-_28921042_28921114@31.54.24|TCTCATAATCTGAAGGTCCTGAGT</td><td>>am_trna162_MetCAT_6_-_26758550_26758622</td><td>31</td><td>54</td><td>24</td><td>TCTCATAATCTGAAGGTCCTGAGT</td><td>466.2</td><td>-3.1</td><td>1.3</td><td>-2.3</td><td>1.9E-02</td><td>2.2E-02</td><td>BT20/MB468</td><td>CYTO</td></tr>
<tr><td>>hg38_MT_+_1621_3279_RNR2_16S_50ntFlanks@622.653.32|TATAACTGAACTCCTCACACCCAATTGGACCA</td><td>>hg38_MT_+_1621_3279_RNR2_16S_50ntFlanks</td><td>622</td><td>653</td><td>32</td><td>TATAACTGAACTCCTCACACCCAATTGGACCA</td><td>63.1</td><td>6.4</td><td>2.6</td><td>2.5</td><td>1.3E-02</td><td>2.2E-02</td><td>BT20/MB468</td><td>MP</td></tr>
<tr><td>>hg38_MT_+_598_1651_RNR1_12S_50ntFlanks@428.456.29|AAAGTGGCTTTAACATATCTGAACACACA</td><td>>hg38_MT_+_598_1651_RNR1_12S_50ntFlanks</td><td>428</td><td>456</td><td>29</td><td>AAAGTGGCTTTAACATATCTGAACACACA</td><td>315.7</td><td>3.2</td><td>1.3</td><td>2.4</td><td>1.8E-02</td><td>2.2E-02</td><td>MB231/MB468</td><td>MP</td></tr>
<tr><td>>hg38_21_+_8433172_8446622_RNA45SN1_45S_50ntFlanks@10877.10894.18|GTCGGTCGGGCTGGGGCG</td><td>>hg38_21_+_8433172_8446622_RNA45SN1_45S_50ntFlanks</td><td>10877</td><td>10894</td><td>18</td><td>GTCGGTCGGGCTGGGGCG</td><td>82.9</td><td>-3.1</td><td>1.3</td><td>-2.5</td><td>1.3E-02</td><td>2.2E-02</td><td>BT20/MB468</td><td>MP</td></tr>
<tr><td>>hg38_MT_+_1621_3279_RNR2_16S_50ntFlanks@1390.1414.25|GATGGTGCAGCCGCTATTAAAGGTT</td><td>>hg38_MT_+_1621_3279_RNR2_16S_50ntFlanks</td><td>1390</td><td>1414</td><td>25</td><td>GATGGTGCAGCCGCTATTAAAGGTT</td><td>193.5</td><td>4.0</td><td>1.7</td><td>2.4</td><td>1.8E-02</td><td>2.2E-02</td><td>MB231/MB468</td><td>MP</td></tr>
<tr><td>>hg38_21_+_8433172_8446622_RNA45SN1_45S_50ntFlanks@10384.10424.41|CCATGTGAACAGCAGTTGAACATGGGTCAGTCGGTCCTGAG</td><td>>hg38_21_+_8433172_8446622_RNA45SN1_45S_50ntFlanks</td><td>10384</td><td>10424</td><td>41</td><td>CCATGTGAACAGCAGTTGAACATGGGTCAGTCGGTCCTGAG</td><td>1848.4</td><td>2.3</td><td>0.8</td><td>2.7</td><td>6.3E-03</td><td>2.2E-02</td><td>BT20/MB231</td><td>MITO</td></tr>
<tr><td>>MI0000088|hsa-mir-30a&WithFlank&6|-|71403545|71403627@12.36.25&[MIMAT0000087&hsa-miR-30a-5p&offsets|0|+3;m-15&6|-|71403595|71403616&offsets|0|+3]|TGTAAACATCCTCGACTGGAAGCTG</td><td>>MI0000088|hsa-mir-30a&WithFlank&6|-|71403545|71403627</td><td>12</td><td>36</td><td>25</td><td>TGTAAACATCCTCGACTGGAAGCTG</td><td>586.2</td><td>-4.0</td><td>1.4</td><td>-2.8</td><td>5.1E-03</td><td>2.2E-02</td><td>BT20/MB231</td><td>MP</td></tr>
<tr><td>>MI0003600|hsa-mir-550a-1&WithFlank&7|+|30289788|30289896@28.48.21;MI0003601|hsa-mir-550a-2&WithFlank&7|+|32732975|32733083@28.48.21&[MIMAT0004800&hsa-miR-550a-5p&offsets|0|-2;m-340&7|+|30289815|30289835&offsets|0|0];[MIMAT0004800_1&hsa-miR-550a-5p&offsets|0|-2;m-340&7|+|32733002|32733022&offsets|0|0]|AGTGCCTGAGGGAGTAAGAGC</td><td>>MI0003600|hsa-mir-550a-1&WithFlank&7|+|30289788|30289896</td><td>28</td><td>48</td><td>21</td><td>AGTGCCTGAGGGAGTAAGAGC</td><td>183.9</td><td>2.9</td><td>1.0</td><td>2.8</td><td>5.1E-03</td><td>2.2E-02</td><td>BT20/MB231</td><td>MP</td></tr>
<tr><td>>hg38_21_+_8433172_8446622_RNA45SN1_45S_50ntFlanks@12912.12930.19|CACGTTCGTGGGGAACCTG</td><td>>hg38_21_+_8433172_8446622_RNA45SN1_45S_50ntFlanks</td><td>12912</td><td>12930</td><td>19</td><td>CACGTTCGTGGGGAACCTG</td><td>240.1</td><td>-2.6</td><td>1.0</td><td>-2.5</td><td>1.3E-02</td><td>2.2E-02</td><td>BT20/MB468</td><td>MP</td></tr>
<tr><td>>hg38_21_+_8433172_8446622_RNA45SN1_45S_50ntFlanks@12147.12169.23|CCAGGTGGGGAGTTTGACTGGGG</td><td>>hg38_21_+_8433172_8446622_RNA45SN1_45S_50ntFlanks</td><td>12147</td><td>12169</td><td>23</td><td>CCAGGTGGGGAGTTTGACTGGGG</td><td>254.2</td><td>-3.2</td><td>1.3</td><td>-2.5</td><td>1.3E-02</td><td>2.2E-02</td><td>BT20/MB468</td><td>MP</td></tr>
<tr><td>>hg38_21_+_8433172_8446622_RNA45SN1_45S_50ntFlanks@9938.9957.20|AGCAGGACGGTGGCCATGGA</td><td>>hg38_21_+_8433172_8446622_RNA45SN1_45S_50ntFlanks</td><td>9938</td><td>9957</td><td>20</td><td>AGCAGGACGGTGGCCATGGA</td><td>703.2</td><td>2.6</td><td>0.9</td><td>2.8</td><td>5.1E-03</td><td>2.2E-02</td><td>BT20/MB231</td><td>MP</td></tr>
<tr><td>>hg38_21_+_8433172_8446622_RNA45SN1_45S_50ntFlanks@10717.10735.19|AAAATCCGGGGGAGAGGGT</td><td>>hg38_21_+_8433172_8446622_RNA45SN1_45S_50ntFlanks</td><td>10717</td><td>10735</td><td>19</td><td>AAAATCCGGGGGAGAGGGT</td><td>149.4</td><td>-3.1</td><td>1.3</td><td>-2.4</td><td>1.8E-02</td><td>2.2E-02</td><td>MB231/MB468</td><td>MP</td></tr>
<tr><td>>MI0000746|hsa-mir-99b&WithFlank&19|+|51692606|51692687@51.72.22&[MIMAT0004678&hsa-miR-99b-3p&offsets|0|0;m-151&19|+|51692656|51692678&offsets|0|-1]|CAAGCTCGTGTCTGTGGGTCCG</td><td>>MI0000746|hsa-mir-99b&WithFlank&19|+|51692606|51692687</td><td>51</td><td>72</td><td>22</td><td>CAAGCTCGTGTCTGTGGGTCCG</td><td>572.3</td><td>1.6</td><td>0.7</td><td>2.3</td><td>2.0E-02</td><td>2.2E-02</td><td>BT20/MB468</td><td>NUC</td></tr>
<tr><td>>hg38_21_+_8433172_8446622_RNA45SN1_45S_50ntFlanks@11359.11383.25|CGGTCGGGCGGCGGCGGTCGGCGGG</td><td>>hg38_21_+_8433172_8446622_RNA45SN1_45S_50ntFlanks</td><td>11359</td><td>11383</td><td>25</td><td>CGGTCGGGCGGCGGCGGTCGGCGGG</td><td>62.2</td><td>-4.4</td><td>1.7</td><td>-2.5</td><td>1.3E-02</td><td>2.2E-02</td><td>BT20/MB468</td><td>MP</td></tr>
<tr><td>>hg38_MT_+_598_1651_RNR1_12S_50ntFlanks@907.946.40|TATACTTCAAAGGACATTTAACTAAAACCCCTACGCATTT</td><td>>hg38_MT_+_598_1651_RNR1_12S_50ntFlanks</td><td>907</td><td>946</td><td>40</td><td>TATACTTCAAAGGACATTTAACTAAAACCCCTACGCATTT</td><td>35.0</td><td>4.8</td><td>2.0</td><td>2.4</td><td>1.8E-02</td><td>2.2E-02</td><td>MB231/MB468</td><td>MP</td></tr>
<tr><td>>am_trna12_ArgCCT_16_+_3243918_3243990@36.53.18,trna18_ArgCCT_17_+_73030001_73030073@36.53.18,trna21_ArgCCT_17_-_73030526_73030598@36.53.18,trna2_ArgCCT_16_+_3202901_3202973@36.53.18,trna3_ArgCCT_7_+_139025446_139025518@36.53.18|TAAGCCAGGGATTGTGGG</td><td>>am_trna12_ArgCCT_16_+_3243918_3243990</td><td>36</td><td>53</td><td>18</td><td>TAAGCCAGGGATTGTGGG</td><td>176.0</td><td>-2.8</td><td>1.2</td><td>-2.3</td><td>1.9E-02</td><td>2.2E-02</td><td>BT20/MB468</td><td>CYTO</td></tr>
<tr><td>>hg38_21_+_8433172_8446622_RNA45SN1_45S_50ntFlanks@10405.10429.25|ATGGGTCAGTCGGTCCTGAGAGATG</td><td>>hg38_21_+_8433172_8446622_RNA45SN1_45S_50ntFlanks</td><td>10405</td><td>10429</td><td>25</td><td>ATGGGTCAGTCGGTCCTGAGAGATG</td><td>213.7</td><td>-2.3</td><td>1.0</td><td>-2.4</td><td>1.8E-02</td><td>2.2E-02</td><td>MB231/MB468</td><td>MP</td></tr>
<tr><td>>hg38_21_+_8433172_8446622_RNA45SN1_45S_50ntFlanks@474.496.23|TCTTCGTGATCGATGTGGTGACG</td><td>>hg38_21_+_8433172_8446622_RNA45SN1_45S_50ntFlanks</td><td>474</td><td>496</td><td>23</td><td>TCTTCGTGATCGATGTGGTGACG</td><td>47.2</td><td>3.9</td><td>1.6</td><td>2.4</td><td>1.8E-02</td><td>2.2E-02</td><td>MB231/MB468</td><td>MP</td></tr>
<tr><td>>hg38_MT_+_1621_3279_RNR2_16S_50ntFlanks@643.663.21|CAATTGGACCAATCTATCACC</td><td>>hg38_MT_+_1621_3279_RNR2_16S_50ntFlanks</td><td>643</td><td>663</td><td>21</td><td>CAATTGGACCAATCTATCACC</td><td>103.3</td><td>2.9</td><td>1.2</td><td>2.4</td><td>1.8E-02</td><td>2.2E-02</td><td>MB231/MB468</td><td>MP</td></tr>
<tr><td>>hg38_MT_+_1621_3279_RNR2_16S_50ntFlanks@51.91.41|GCTAAACCTAGCCCCAAACCCACTCCACCTTACTACCAGAC</td><td>>hg38_MT_+_1621_3279_RNR2_16S_50ntFlanks</td><td>51</td><td>91</td><td>41</td><td>GCTAAACCTAGCCCCAAACCCACTCCACCTTACTACCAGAC</td><td>866.5</td><td>4.5</td><td>1.8</td><td>2.5</td><td>1.3E-02</td><td>2.2E-02</td><td>BT20/MB468</td><td>MP</td></tr>
<tr><td>>hg38_21_+_8433172_8446622_RNA45SN1_45S_50ntFlanks@6653.6673.21|ACTCTTAGCGGTGGATCACTC</td><td>>hg38_21_+_8433172_8446622_RNA45SN1_45S_50ntFlanks</td><td>6653</td><td>6673</td><td>21</td><td>ACTCTTAGCGGTGGATCACTC</td><td>57.6</td><td>6.5</td><td>2.6</td><td>2.5</td><td>1.3E-02</td><td>2.2E-02</td><td>BT20/MB468</td><td>MP</td></tr>
<tr><td>>hg38_MT_+_1621_3279_RNR2_16S_50ntFlanks@1355.1396.42|GGGTTTACGACCTCGATGTTGGATCAGGACATCCCGATGGTG</td><td>>hg38_MT_+_1621_3279_RNR2_16S_50ntFlanks</td><td>1355</td><td>1396</td><td>42</td><td>GGGTTTACGACCTCGATGTTGGATCAGGACATCCCGATGGTG</td><td>55.3</td><td>5.4</td><td>2.2</td><td>2.5</td><td>1.3E-02</td><td>2.2E-02</td><td>BT20/MB468</td><td>MP</td></tr>
<tr><td>>MI0000465|hsa-mir-191&WithFlank&3|-|49020612|49020715@22.46.25&[MIMAT0000440&hsa-miR-191-5p&offsets|0|+2;m-48&3|-|49020672|49020694&offsets|0|+2]|CAACGGAATCCCAAAAGCAGCTGTT</td><td>>MI0000465|hsa-mir-191&WithFlank&3|-|49020612|49020715</td><td>22</td><td>46</td><td>25</td><td>CAACGGAATCCCAAAAGCAGCTGTT</td><td>174.9</td><td>-4.4</td><td>1.6</td><td>-2.8</td><td>5.1E-03</td><td>2.2E-02</td><td>BT20/MB231</td><td>MP</td></tr>
<tr><td>>hg38_21_+_8433172_8446622_RNA45SN1_45S_50ntFlanks@6789.6807.19|GCCTGTCTGAGCGTCGCTT</td><td>>hg38_21_+_8433172_8446622_RNA45SN1_45S_50ntFlanks</td><td>6789</td><td>6807</td><td>19</td><td>GCCTGTCTGAGCGTCGCTT</td><td>826.4</td><td>4.4</td><td>1.6</td><td>2.8</td><td>5.1E-03</td><td>2.3E-02</td><td>BT20/MB231</td><td>MP</td></tr>
<tr><td>>am_MI0003583|hsa-mir-576&WithFlank&4|+|109488692|109488801@61.82.22&[MIMAT0004796&hsa-miR-576-3p&offsets|0|0;m-524&4|+|109488752|109488772&offsets|0|+1]|AAGATGTGGAAAAATTGGAATC</td><td>>am_MI0003583|hsa-mir-576&WithFlank&4|+|109488692|109488801</td><td>61</td><td>82</td><td>22</td><td>AAGATGTGGAAAAATTGGAATC</td><td>131.6</td><td>4.0</td><td>1.7</td><td>2.4</td><td>1.8E-02</td><td>2.3E-02</td><td>MB231/MB468</td><td>MP</td></tr>
<tr><td>>hg38_21_+_8433172_8446622_RNA45SN1_45S_50ntFlanks@10907.10928.22|GGGCGCGCGCCGCGGCTGGACG</td><td>>hg38_21_+_8433172_8446622_RNA45SN1_45S_50ntFlanks</td><td>10907</td><td>10928</td><td>22</td><td>GGGCGCGCGCCGCGGCTGGACG</td><td>112.6</td><td>5.5</td><td>2.0</td><td>2.8</td><td>5.2E-03</td><td>2.3E-02</td><td>BT20/MB231</td><td>MP</td></tr>
<tr><td>>hg38_21_+_8433172_8446622_RNA45SN1_45S_50ntFlanks@11570.11605.36|AGCCGACTTAGAACTGGTGCGGACCAGGGGAATCCG</td><td>>hg38_21_+_8433172_8446622_RNA45SN1_45S_50ntFlanks</td><td>11570</td><td>11605</td><td>36</td><td>AGCCGACTTAGAACTGGTGCGGACCAGGGGAATCCG</td><td>400.3</td><td>-1.9</td><td>0.8</td><td>-2.4</td><td>1.7E-02</td><td>2.3E-02</td><td>BT20/MB468</td><td>MITO</td></tr>
<tr><td>>hg38_21_+_8433172_8446622_RNA45SN1_45S_50ntFlanks@9572.9601.30|CAAATCGGTCGTCCGACCTGGGTATAGGGG</td><td>>hg38_21_+_8433172_8446622_RNA45SN1_45S_50ntFlanks</td><td>9572</td><td>9601</td><td>30</td><td>CAAATCGGTCGTCCGACCTGGGTATAGGGG</td><td>152.9</td><td>-2.6</td><td>1.1</td><td>-2.3</td><td>1.9E-02</td><td>2.3E-02</td><td>BT20/MB468</td><td>CYTO</td></tr>
<tr><td>>hg38_MT_+_1621_3279_RNR2_16S_50ntFlanks@469.491.23|TAACTGTTAGTCCAAAGAGGAAC</td><td>>hg38_MT_+_1621_3279_RNR2_16S_50ntFlanks</td><td>469</td><td>491</td><td>23</td><td>TAACTGTTAGTCCAAAGAGGAAC</td><td>36.1</td><td>4.7</td><td>2.0</td><td>2.4</td><td>1.8E-02</td><td>2.3E-02</td><td>MB231/MB468</td><td>MP</td></tr>
<tr><td>>hg38_21_+_8433172_8446622_RNA45SN1_45S_50ntFlanks@8382.8400.19|AGAGGGCGTGAAACCGTTA</td><td>>hg38_21_+_8433172_8446622_RNA45SN1_45S_50ntFlanks</td><td>8382</td><td>8400</td><td>19</td><td>AGAGGGCGTGAAACCGTTA</td><td>207.2</td><td>-3.7</td><td>1.5</td><td>-2.5</td><td>1.3E-02</td><td>2.3E-02</td><td>BT20/MB468</td><td>MP</td></tr>
<tr><td>>hg38_21_+_8433172_8446622_RNA45SN1_45S_50ntFlanks@11714.11736.23|GGTAAACGGCGGGAGTAACTATG</td><td>>hg38_21_+_8433172_8446622_RNA45SN1_45S_50ntFlanks</td><td>11714</td><td>11736</td><td>23</td><td>GGTAAACGGCGGGAGTAACTATG</td><td>372.1</td><td>2.0</td><td>0.9</td><td>2.3</td><td>1.9E-02</td><td>2.3E-02</td><td>MB231/MB468</td><td>CYTO</td></tr>
<tr><td>>trna14_LysTTT_11_-_59327808_59327880@1.32.32,trna2_LysTTT_17_+_8022473_8022545@1.32.32,trna54_LysTTT_1_+_204475655_204475727@1.32.32,trna5_LysTTT_11_+_59323902_59323974@1.32.32,trna62_LysTTT_1_-_204476158_204476230@1.32.32,trna76_LysTTT_6_+_28918806_28918878@1.32.32|GCCCGGATAGCTCAGTCGGTAGAGCATCAGAC</td><td>>trna14_LysTTT_11_-_59327808_59327880</td><td>1</td><td>32</td><td>32</td><td>GCCCGGATAGCTCAGTCGGTAGAGCATCAGAC</td><td>464.4</td><td>2.1</td><td>0.9</td><td>2.4</td><td>1.8E-02</td><td>2.3E-02</td><td>MB231/MB468</td><td>MP</td></tr>
<tr><td>>hg38_21_+_8433172_8446622_RNA45SN1_45S_50ntFlanks@11578.11602.25|TAGAACTGGTGCGGACCAGGGGAAT</td><td>>hg38_21_+_8433172_8446622_RNA45SN1_45S_50ntFlanks</td><td>11578</td><td>11602</td><td>25</td><td>TAGAACTGGTGCGGACCAGGGGAAT</td><td>231.8</td><td>-1.7</td><td>0.7</td><td>-2.3</td><td>1.9E-02</td><td>2.3E-02</td><td>BT20/MB468</td><td>CYTO</td></tr>
<tr><td>>hg38_21_+_8433172_8446622_RNA45SN1_45S_50ntFlanks@8388.8405.18|CGTGAAACCGTTAAGAGG</td><td>>hg38_21_+_8433172_8446622_RNA45SN1_45S_50ntFlanks</td><td>8388</td><td>8405</td><td>18</td><td>CGTGAAACCGTTAAGAGG</td><td>117.7</td><td>-3.6</td><td>1.5</td><td>-2.4</td><td>1.7E-02</td><td>2.3E-02</td><td>BT20/MB468</td><td>MITO</td></tr>
<tr><td>>hg38_21_+_8433172_8446622_RNA45SN1_45S_50ntFlanks@10922.10946.25|CTGGACGAGGCGCCGCCGCCCCCCC</td><td>>hg38_21_+_8433172_8446622_RNA45SN1_45S_50ntFlanks</td><td>10922</td><td>10946</td><td>25</td><td>CTGGACGAGGCGCCGCCGCCCCCCC</td><td>1476.8</td><td>-4.8</td><td>1.9</td><td>-2.5</td><td>1.3E-02</td><td>2.3E-02</td><td>BT20/MB468</td><td>MP</td></tr>
<tr><td>>hg38_21_+_8433172_8446622_RNA45SN1_45S_50ntFlanks@7990.8008.19|GACGTGGCGACCCGCTGAA</td><td>>hg38_21_+_8433172_8446622_RNA45SN1_45S_50ntFlanks</td><td>7990</td><td>8008</td><td>19</td><td>GACGTGGCGACCCGCTGAA</td><td>1792.9</td><td>-1.9</td><td>0.8</td><td>-2.4</td><td>1.7E-02</td><td>2.3E-02</td><td>BT20/MB468</td><td>MITO</td></tr>
<tr><td>>MI0000286|hsa-mir-210&WithFlank&11|-|568083|568204@72.92.21(+1C)&[MIMAT0000267&hsa-miR-210-3p&offsets|0|-1(+1C);m-102&11|-|568112|568133&offsets|0|-1(+1C)]|CTGTGCGTGTGACAGCGGCTGC</td><td>>MI0000286|hsa-mir-210&WithFlank&11|-|568083|568204</td><td>72</td><td>92</td><td>21(+1C)</td><td>CTGTGCGTGTGACAGCGGCTGC</td><td>39.8</td><td>4.9</td><td>2.1</td><td>2.4</td><td>1.8E-02</td><td>2.3E-02</td><td>MB231/MB468</td><td>MP</td></tr>
<tr><td>>hg38_21_+_8433172_8446622_RNA45SN1_45S_50ntFlanks@10679.10711.33|GGTTCCGGCGGCGTCCGGTGAGCTCTCGCTGGC</td><td>>hg38_21_+_8433172_8446622_RNA45SN1_45S_50ntFlanks</td><td>10679</td><td>10711</td><td>33</td><td>GGTTCCGGCGGCGTCCGGTGAGCTCTCGCTGGC</td><td>2741.8</td><td>-4.0</td><td>1.6</td><td>-2.5</td><td>1.3E-02</td><td>2.3E-02</td><td>BT20/MB468</td><td>MP</td></tr>
<tr><td>>hg38_21_+_8433172_8446622_RNA45SN1_45S_50ntFlanks@11600.11622.23|AATCCGACTGTTTAATTAAAACA</td><td>>hg38_21_+_8433172_8446622_RNA45SN1_45S_50ntFlanks</td><td>11600</td><td>11622</td><td>23</td><td>AATCCGACTGTTTAATTAAAACA</td><td>219.8</td><td>5.6</td><td>2.0</td><td>2.8</td><td>5.2E-03</td><td>2.3E-02</td><td>BT20/MB231</td><td>MP</td></tr>
<tr><td>>MI0008329|hsa-mir-1908&WithFlank&11|-|61815155|61815246@18.38.21&[MIMAT0007881&hsa-miR-1908-5p&offsets|0|0;m-915&11|-|61815209|61815229&offsets|0|0]|CGGCGGGGACGGCGATTGGTC</td><td>>MI0008329|hsa-mir-1908&WithFlank&11|-|61815155|61815246</td><td>18</td><td>38</td><td>21</td><td>CGGCGGGGACGGCGATTGGTC</td><td>118.3</td><td>-5.3</td><td>1.9</td><td>-2.8</td><td>5.2E-03</td><td>2.3E-02</td><td>BT20/MB231</td><td>MP</td></tr>
</table>

<table>
<tr><th></th><th></th><th></th><th></th><th></th><th></th><th></th><th></th><th></th><th></th><th></th><th></th><th></th><th></th></tr>
<tr><td>>hg38_21_+_8433172_8446622_RNA45SN1_45S_50ntFlanks@8349.8379.31|GAAAGTTGAAAAGAACTTTGAAGAGAGAGTT</td><td>>hg38_21_+_8433172_8446622_RNA45SN1_45S_50ntFlanks</td><td>8349</td><td>8379</td><td>31</td><td>GAAAGTTGAAAAGAACTTTGAAGAGAGAGTT</td><td>99.9</td><td>4.3</td><td>1.5</td><td>2.8</td><td>5.2E-03</td><td>2.3E-02</td><td>BT20/MB231</td><td>MP</td></tr>
<tr><td>>hg38_21_+_8433172_8446622_RNA45SN1_45S_50ntFlanks@12992.13015.24|TCCCTCGCTGCGATCTATTGAAAG</td><td>>hg38_21_+_8433172_8446622_RNA45SN1_45S_50ntFlanks</td><td>12992</td><td>13015</td><td>24</td><td>TCCCTCGCTGCGATCTATTGAAAG</td><td>501.5</td><td>-1.6</td><td>0.7</td><td>-2.3</td><td>1.9E-02</td><td>2.3E-02</td><td>MB231/MB468</td><td>CYTO</td></tr>
<tr><td>>hg38_21_+_8433172_8446622_RNA45SN1_45S_50ntFlanks@10797.10816.20|GCATGTTGGAACAATGTAGG</td><td>>hg38_21_+_8433172_8446622_RNA45SN1_45S_50ntFlanks</td><td>10797</td><td>10816</td><td>20</td><td>GCATGTTGGAACAATGTAGG</td><td>233.3</td><td>-2.3</td><td>1.0</td><td>-2.3</td><td>1.9E-02</td><td>2.3E-02</td><td>BT20/MB468</td><td>CYTO</td></tr>
<tr><td>>hg38_1_-_228634819_228635039_RNA5S12_5S_50ntFlanks@127.170.44|ACTTGGATGGGAGACCGCCTGGGAATACCGGGTGCTGTAGGCTT</td><td>>hg38_1_-_228634819_228635039_RNA5S12_5S_50ntFlanks</td><td>127</td><td>170</td><td>44</td><td>ACTTGGATGGGAGACCGCCTGGGAATACCGGGTGCTGTAGGCTT</td><td>135.4</td><td>3.8</td><td>1.6</td><td>2.3</td><td>2.0E-02</td><td>2.3E-02</td><td>MB231/MB468</td><td>MITO</td></tr>
<tr><td>>hg38_21_+_8433172_8446622_RNA45SN1_45S_50ntFlanks@11255.11281.27|CTGCGGCGGGCGTCGCGGCCGCCCCCG</td><td>>hg38_21_+_8433172_8446622_RNA45SN1_45S_50ntFlanks</td><td>11255</td><td>11281</td><td>27</td><td>CTGCGGCGGGCGTCGCGGCCGCCCCCG</td><td>170.0</td><td>-3.7</td><td>1.6</td><td>-2.3</td><td>1.9E-02</td><td>2.3E-02</td><td>BT20/MB468</td><td>CYTO</td></tr>
<tr><td>>MI0000255|hsa-mir-30d&WithFlank&8|-|134804870|134804951@12.33.22&[MIMAT0000245&hsa-miR-30d-5p&offsets|0|0;m-21&8|-|134804919|134804940&offsets|0|0]|TGTAAACATCCCCGACTGGAAG</td><td>>MI0000255|hsa-mir-30d&WithFlank&8|-|134804870|134804951</td><td>12</td><td>33</td><td>22</td><td>TGTAAACATCCCCGACTGGAAG</td><td>6048.3</td><td>1.8</td><td>0.6</td><td>2.8</td><td>5.7E-03</td><td>2.3E-02</td><td>BT20/MB231</td><td>NUC</td></tr>
<tr><td>>hg38_21_+_8433172_8446622_RNA45SN1_45S_50ntFlanks@4975.4993.19|CCCGGACACGGACAGGATT</td><td>>hg38_21_+_8433172_8446622_RNA45SN1_45S_50ntFlanks</td><td>4975</td><td>4993</td><td>19</td><td>CCCGGACACGGACAGGATT</td><td>423.9</td><td>2.8</td><td>1.0</td><td>2.8</td><td>5.7E-03</td><td>2.3E-02</td><td>BT20/MB231</td><td>NUC</td></tr>
<tr><td>>hg38_21_+_8433172_8446622_RNA45SN1_45S_50ntFlanks@10830.10856.27|CAAGCCGGATCCGTAACTTCGGGATAA</td><td>>hg38_21_+_8433172_8446622_RNA45SN1_45S_50ntFlanks</td><td>10830</td><td>10856</td><td>27</td><td>CAAGCCGGATCCGTAACTTCGGGATAA</td><td>126.3</td><td>-4.4</td><td>1.8</td><td>-2.5</td><td>1.3E-02</td><td>2.3E-02</td><td>BT20/MB468</td><td>MP</td></tr>
<tr><td>>hg38_21_+_8433172_8446622_RNA45SN1_45S_50ntFlanks@11728.11765.38|GTAACTATGACTCTCTTAAGGTAGCCAAATGCCTCGTC</td><td>>hg38_21_+_8433172_8446622_RNA45SN1_45S_50ntFlanks</td><td>11728</td><td>11765</td><td>38</td><td>GTAACTATGACTCTCTTAAGGTAGCCAAATGCCTCGTC</td><td>237.4</td><td>-4.0</td><td>1.7</td><td>-2.3</td><td>1.9E-02</td><td>2.3E-02</td><td>BT20/MB468</td><td>CYTO</td></tr>
<tr><td>>MI0000115|hsa-mir-16-2&WithFlank&3|+|160404739|160404831@58.79.22&[MIMAT0004518&hsa-miR-16-2-3p&offsets|-1|-1;m-252&3|+|160404796|160404816&offsets|0|+1]|ACCAATATTACTGTGCTGCTTT</td><td>>MI0000115|hsa-mir-16-2&WithFlank&3|+|160404739|160404831</td><td>58</td><td>79</td><td>22</td><td>ACCAATATTACTGTGCTGCTTT</td><td>288.3</td><td>1.6</td><td>0.7</td><td>2.3</td><td>2.0E-02</td><td>2.3E-02</td><td>BT20/MB468</td><td>NUC</td></tr>
<tr><td>>hg38_21_+_8433172_8446622_RNA45SN1_45S_50ntFlanks@11569.11589.21|CAGCCGACTTAGAACTGGTGC</td><td>>hg38_21_+_8433172_8446622_RNA45SN1_45S_50ntFlanks</td><td>11569</td><td>11589</td><td>21</td><td>CAGCCGACTTAGAACTGGTGC</td><td>117.9</td><td>-3.7</td><td>1.5</td><td>-2.5</td><td>1.3E-02</td><td>2.3E-02</td><td>BT20/MB468</td><td>MP</td></tr>
<tr><td>>hg38_21_+_8433172_8446622_RNA45SN1_45S_50ntFlanks@3782.3801.20|CAGTGAAACTGCGAATGGCT</td><td>>hg38_21_+_8433172_8446622_RNA45SN1_45S_50ntFlanks</td><td>3782</td><td>3801</td><td>20</td><td>CAGTGAAACTGCGAATGGCT</td><td>2181.7</td><td>-3.3</td><td>1.3</td><td>-2.5</td><td>1.3E-02</td><td>2.3E-02</td><td>BT20/MB468</td><td>MP</td></tr>
<tr><td>>hg38_21_+_8433172_8446622_RNA45SN1_45S_50ntFlanks@12151.12183.33|GTGGGGAGTTTGACTGGGGCGGTACACCTGTCA</td><td>>hg38_21_+_8433172_8446622_RNA45SN1_45S_50ntFlanks</td><td>12151</td><td>12183</td><td>33</td><td>GTGGGGAGTTTGACTGGGGCGGTACACCTGTCA</td><td>293.7</td><td>-3.4</td><td>1.4</td><td>-2.5</td><td>1.3E-02</td><td>2.3E-02</td><td>BT20/MB468</td><td>MP</td></tr>
<tr><td>>hg38_21_+_8433172_8446622_RNA45SN1_45S_50ntFlanks@11569.11586.18|CAGCCGACTTAGAACTGG</td><td>>hg38_21_+_8433172_8446622_RNA45SN1_45S_50ntFlanks</td><td>11569</td><td>11586</td><td>18</td><td>CAGCCGACTTAGAACTGG</td><td>667.7</td><td>-1.9</td><td>0.8</td><td>-2.4</td><td>1.7E-02</td><td>2.3E-02</td><td>BT20/MB468</td><td>MITO</td></tr>
<tr><td>>hg38_21_+_8433172_8446622_RNA45SN1_45S_50ntFlanks@6651.6697.47|CGACTCTTAGCGGTGGATCACTCGGCTCGTGCGTCGATGAAGAACGC</td><td>>hg38_21_+_8433172_8446622_RNA45SN1_45S_50ntFlanks</td><td>6651</td><td>6697</td><td>47</td><td>CGACTCTTAGCGGTGGATCACTCGGCTCGTGCGTCGATGAAGAACGC</td><td>3387.5</td><td>-2.5</td><td>0.9</td><td>-2.7</td><td>6.9E-03</td><td>2.3E-02</td><td>BT20/MB231</td><td>CYTO</td></tr>
<tr><td>>hg38_21_+_8433172_8446622_RNA45SN1_45S_50ntFlanks@11666.11704.39|TCTGCCCAGTGCTCTGAATGTCAAAGTGAAGAAATTCAA</td><td>>hg38_21_+_8433172_8446622_RNA45SN1_45S_50ntFlanks</td><td>11666</td><td>11704</td><td>39</td><td>TCTGCCCAGTGCTCTGAATGTCAAAGTGAAGAAATTCAA</td><td>226.5</td><td>-2.1</td><td>0.8</td><td>-2.7</td><td>6.9E-03</td><td>2.3E-02</td><td>BT20/MB231</td><td>CYTO</td></tr>
<tr><td>>MI0000434|hsa-let-7i&WithFlank&12|+|62603680|62603775@12.30.19(+3U)&[MIMAT0000415&hsa-let-7i-5p&offsets|0|-3(+3U);m-49&12|+|62603691|62603712&offsets|0|-3(+3U)]|TGAGGTAGTAGTTTGTGCTTTT</td><td>>MI0000434|hsa-let-7i&WithFlank&12|+|62603680|62603775</td><td>12</td><td>30</td><td>19(+3U)</td><td>TGAGGTAGTAGTTTGTGCTTTT</td><td>74.3</td><td>4.8</td><td>2.0</td><td>2.4</td><td>1.8E-02</td><td>2.3E-02</td><td>MB231/MB468</td><td>MP</td></tr>
<tr><td>>MI0000542|hsa-mir-320a&WithFlank&8|-|22244960|22245043@50.69.20&[MIMAT0000510&hsa-miR-320a-3p&offsets|+2|0;m-60&8|-|22244975|22244996&offsets|+2|0]|AAGCTGGGTTGAGAGGGCGA</td><td>>MI0000542|hsa-mir-320a&WithFlank&8|-|22244960|22245043</td><td>50</td><td>69</td><td>20</td><td>AAGCTGGGTTGAGAGGGCGA</td><td>90.8</td><td>4.0</td><td>1.7</td><td>2.4</td><td>1.8E-02</td><td>2.3E-02</td><td>MB231/MB468</td><td>MP</td></tr>
<tr><td>>hg38_21_+_8433172_8446622_RNA45SN1_45S_50ntFlanks@6665.6689.25|GGATCACTCGGCTCGTGCGTCGATG</td><td>>hg38_21_+_8433172_8446622_RNA45SN1_45S_50ntFlanks</td><td>6665</td><td>6689</td><td>25</td><td>GGATCACTCGGCTCGTGCGTCGATG</td><td>47.4</td><td>-4.2</td><td>1.7</td><td>-2.5</td><td>1.3E-02</td><td>2.3E-02</td><td>BT20/MB468</td><td>MP</td></tr>
<tr><td>>hg38_21_+_8433172_8446622_RNA45SN1_45S_50ntFlanks@6651.6682.32|CGACTCTTAGCGGTGGATCACTCGGCTCGTGC</td><td>>hg38_21_+_8433172_8446622_RNA45SN1_45S_50ntFlanks</td><td>6651</td><td>6682</td><td>32</td><td>CGACTCTTAGCGGTGGATCACTCGGCTCGTGC</td><td>3174.9</td><td>3.4</td><td>1.2</td><td>2.8</td><td>5.2E-03</td><td>2.3E-02</td><td>BT20/MB231</td><td>MP</td></tr>
<tr><td>>hg38_MT_+_1621_3279_RNR2_16S_50ntFlanks@1432.1458.27|AAGTCCTACGTGATCTGAGTTCAGACC</td><td>>hg38_MT_+_1621_3279_RNR2_16S_50ntFlanks</td><td>1432</td><td>1458</td><td>27</td><td>AAGTCCTACGTGATCTGAGTTCAGACC</td><td>154.0</td><td>3.1</td><td>1.2</td><td>2.5</td><td>1.3E-02</td><td>2.3E-02</td><td>BT20/MB468</td><td>MP</td></tr>
<tr><td>>hg38_21_+_8433172_8446622_RNA45SN1_45S_50ntFlanks@10686.10709.24|GCGGCGTCCGGTGAGCTCTCGCTG</td><td>>hg38_21_+_8433172_8446622_RNA45SN1_45S_50ntFlanks</td><td>10686</td><td>10709</td><td>24</td><td>GCGGCGTCCGGTGAGCTCTCGCTG</td><td>264.2</td><td>5.6</td><td>2.0</td><td>2.8</td><td>5.2E-03</td><td>2.3E-02</td><td>BT20/MB231</td><td>MP</td></tr>
<tr><td>>MI0000077|hsa-mir-21&WithFlank&17|+|59841260|59841343@14.37.24&[MIMAT0000076&hsa-miR-21-5p&offsets|0|+2;m-1&17|+|59841273|59841294&offsets|0|+2]|TAGCTTATCAGACTGATGTTGACT</td><td>>MI0000077|hsa-mir-21&WithFlank&17|+|59841260|59841343</td><td>14</td><td>37</td><td>24</td><td>TAGCTTATCAGACTGATGTTGACT</td><td>248337.0</td><td>-1.8</td><td>0.7</td><td>-2.7</td><td>6.4E-03</td><td>2.3E-02</td><td>BT20/MB231</td><td>MITO</td></tr>
<tr><td>>hg38_21_+_8433172_8446622_RNA45SN1_45S_50ntFlanks@8381.8405.25|AAGAGGGCGTGAAACCGTTAAGAGG</td><td>>hg38_21_+_8433172_8446622_RNA45SN1_45S_50ntFlanks</td><td>8381</td><td>8405</td><td>25</td><td>AAGAGGGCGTGAAACCGTTAAGAGG</td><td>712.1</td><td>-2.1</td><td>0.9</td><td>-2.3</td><td>1.9E-02</td><td>2.3E-02</td><td>MB231/MB468</td><td>CYTO</td></tr>
<tr><td>>hg38_21_+_8433172_8446622_RNA45SN1_45S_50ntFlanks@12603.12626.24|TGTATGTGCTTGGCTGAGGAGCCA</td><td>>hg38_21_+_8433172_8446622_RNA45SN1_45S_50ntFlanks</td><td>12603</td><td>12626</td><td>24</td><td>TGTATGTGCTTGGCTGAGGAGCCA</td><td>129.9</td><td>-2.2</td><td>1.0</td><td>-2.3</td><td>1.9E-02</td><td>2.3E-02</td><td>MB231/MB468</td><td>CYTO</td></tr>
<tr><td>>hg38_21_+_8433172_8446622_RNA45SN1_45S_50ntFlanks@4899.4917.19|AAGGAATTGACGGAAGGGC</td><td>>hg38_21_+_8433172_8446622_RNA45SN1_45S_50ntFlanks</td><td>4899</td><td>4917</td><td>19</td><td>AAGGAATTGACGGAAGGGC</td><td>194.0</td><td>-3.5</td><td>1.4</td><td>-2.5</td><td>1.3E-02</td><td>2.3E-02</td><td>BT20/MB468</td><td>MP</td></tr>
<tr><td>>hg38_MT_+_1621_3279_RNR2_16S_50ntFlanks@1096.1120.25|GACGAGAAGACCCTATGGAGCTTTA</td><td>>hg38_MT_+_1621_3279_RNR2_16S_50ntFlanks</td><td>1096</td><td>1120</td><td>25</td><td>GACGAGAAGACCCTATGGAGCTTTA</td><td>107.1</td><td>2.1</td><td>0.9</td><td>2.4</td><td>1.8E-02</td><td>2.3E-02</td><td>MB231/MB468</td><td>MP</td></tr>
<tr><td>>hg38_21_+_8433172_8446622_RNA45SN1_45S_50ntFlanks@9787.9819.33|TCGCTGGCGTGGAGCCGGGCGTGGAATGCGAGT</td><td>>hg38_21_+_8433172_8446622_RNA45SN1_45S_50ntFlanks</td><td>9787</td><td>9819</td><td>33</td><td>TCGCTGGCGTGGAGCCGGGCGTGGAATGCGAGT</td><td>300.5</td><td>2.3</td><td>0.9</td><td>2.5</td><td>1.3E-02</td><td>2.3E-02</td><td>BT20/MB468</td><td>MP</td></tr>
<tr><td>>hg38_21_+_8433172_8446622_RNA45SN1_45S_50ntFlanks@10866.10902.37|CTAAGGGCTGGGTCGGTCGGGCTGGGGCGCGAAGCGG</td><td>>hg38_21_+_8433172_8446622_RNA45SN1_45S_50ntFlanks</td><td>10866</td><td>10902</td><td>37</td><td>CTAAGGGCTGGGTCGGTCGGGCTGGGGCGCGAAGCGG</td><td>872.7</td><td>-4.1</td><td>1.8</td><td>-2.3</td><td>1.9E-02</td><td>2.3E-02</td><td>BT20/MB468</td><td>CYTO</td></tr>
<tr><td>>hg38_21_+_8433172_8446622_RNA45SN1_45S_50ntFlanks@3783.3800.18|AGTGAAACTGCGAATGGC</td><td>>hg38_21_+_8433172_8446622_RNA45SN1_45S_50ntFlanks</td><td>3783</td><td>3800</td><td>18</td><td>AGTGAAACTGCGAATGGC</td><td>293.7</td><td>-1.9</td><td>0.8</td><td>-2.3</td><td>2.0E-02</td><td>2.3E-02</td><td>MB231/MB468</td><td>NUC</td></tr>
<tr><td>>hg38_21_+_8433172_8446622_RNA45SN1_45S_50ntFlanks@11669.11705.37|GCCCAGTGCTCTGAATGTCAAAGTGAAGAAATTCAAT</td><td>>hg38_21_+_8433172_8446622_RNA45SN1_45S_50ntFlanks</td><td>11669</td><td>11705</td><td>37</td><td>GCCCAGTGCTCTGAATGTCAAAGTGAAGAAATTCAAT</td><td>454.8</td><td>-2.2</td><td>0.8</td><td>-2.7</td><td>7.0E-03</td><td>2.3E-02</td><td>BT20/MB231</td><td>CYTO</td></tr>
<tr><td>>MI0000077|hsa-mir-21&WithFlank&17|+|59841260|59841343@15.35.21(+1U)&[MIMAT0000076&hsa-miR-21-5p&offsets|+1|0(+1U);m-1&17|+|59841273|59841294&offsets|+1|0(+1U)]|AGCTTATCAGACTGATGTTGAT</td><td>>MI0000077|hsa-mir-21&WithFlank&17|+|59841260|59841343</td><td>15</td><td>35</td><td>21(+1U)</td><td>AGCTTATCAGACTGATGTTGAT</td><td>281.1</td><td>-3.1</td><td>1.1</td><td>-2.8</td><td>5.2E-03</td><td>2.3E-02</td><td>BT20/MB231</td><td>MP</td></tr>
<tr><td>>hg38_21_+_8433172_8446622_RNA45SN1_45S_50ntFlanks@10241.10258.18|TACGCCGCGACGAGTAGG</td><td>>hg38_21_+_8433172_8446622_RNA45SN1_45S_50ntFlanks</td><td>10241</td><td>10258</td><td>18</td><td>TACGCCGCGACGAGTAGG</td><td>110.8</td><td>4.4</td><td>1.6</td><td>2.8</td><td>5.2E-03</td><td>2.3E-02</td><td>BT20/MB231</td><td>MP</td></tr>
<tr><td>>hg38_21_+_8433172_8446622_RNA45SN1_45S_50ntFlanks@10347.10374.28|CAAACGAGAACTTTGAAGGCCGAAGTGG</td><td>>hg38_21_+_8433172_8446622_RNA45SN1_45S_50ntFlanks</td><td>10347</td><td>10374</td><td>28</td><td>CAAACGAGAACTTTGAAGGCCGAAGTGG</td><td>914.7</td><td>-3.8</td><td>1.5</td><td>-2.5</td><td>1.3E-02</td><td>2.3E-02</td><td>BT20/MB468</td><td>MP</td></tr>
<tr><td>>hg38_21_+_8433172_8446622_RNA45SN1_45S_50ntFlanks@3815.3834.20|ATGGTTCCTTTGGTCGCTCG</td><td>>hg38_21_+_8433172_8446622_RNA45SN1_45S_50ntFlanks</td><td>3815</td><td>3834</td><td>20</td><td>ATGGTTCCTTTGGTCGCTCG</td><td>61.4</td><td>-4.2</td><td>1.7</td><td>-2.5</td><td>1.3E-02</td><td>2.3E-02</td><td>BT20/MB468</td><td>MP</td></tr>
<tr><td>>hg38_21_+_8433172_8446622_RNA45SN1_45S_50ntFlanks@6729.6770.42|GACACATTGATCATCGACACTTCGAACGCACTTGCGGCCCCG</td><td>>hg38_21_+_8433172_8446622_RNA45SN1_45S_50ntFlanks</td><td>6729</td><td>6770</td><td>42</td><td>GACACATTGATCATCGACACTTCGAACGCACTTGCGGCCCCG</td><td>184.7</td><td>-5.6</td><td>2.2</td><td>-2.5</td><td>1.3E-02</td><td>2.3E-02</td><td>BT20/MB468</td><td>MP</td></tr>
<tr><td>>hg38_21_+_8433172_8446622_RNA45SN1_45S_50ntFlanks@10827.10853.27|CGGCAAGCCGGATCCGTAACTTCGGGA</td><td>>hg38_21_+_8433172_8446622_RNA45SN1_45S_50ntFlanks</td><td>10827</td><td>10853</td><td>27</td><td>CGGCAAGCCGGATCCGTAACTTCGGGA</td><td>226.8</td><td>-3.5</td><td>1.4</td><td>-2.5</td><td>1.3E-02</td><td>2.3E-02</td><td>BT20/MB468</td><td>MP</td></tr>
<tr><td>>trnaMT_MetCAT_MT_+_4402_4469@40.67.28|AAAATGTTGGTTATACCCTTCCCGTACT</td><td>>trnaMT_MetCAT_MT_+_4402_4469</td><td>40</td><td>67</td><td>28</td><td>AAAATGTTGGTTATACCCTTCCCGTACT</td><td>91.4</td><td>-4.0</td><td>1.7</td><td>-2.4</td><td>1.8E-02</td><td>2.3E-02</td><td>MB231/MB468</td><td>MP</td></tr>
<tr><td>>MI0002468|hsa-mir-484&WithFlank&16|+|15643288|15643378@14.35.22&[MIMAT0002174&hsa-miR-484&offsets|0|0;m-127&16|+|15643301|15643321&offsets|0|+1]|TCAGGCTCAGTCCCCTCCCGAT</td><td>>MI0002468|hsa-mir-484&WithFlank&16|+|15643288|15643378</td><td>14</td><td>35</td><td>22</td><td>TCAGGCTCAGTCCCCTCCCGAT</td><td>168.5</td><td>2.0</td><td>0.8</td><td>2.4</td><td>1.8E-02</td><td>2.3E-02</td><td>MB231/MB468</td><td>MP</td></tr>
<tr><td>>hg38_21_+_8433172_8446622_RNA45SN1_45S_50ntFlanks@6661.6694.34|CGGTGGATCACTCGGCTCGTGCGTCGATGAAGAA</td><td>>hg38_21_+_8433172_8446622_RNA45SN1_45S_50ntFlanks</td><td>6661</td><td>6694</td><td>34</td><td>CGGTGGATCACTCGGCTCGTGCGTCGATGAAGAA</td><td>394.0</td><td>1.7</td><td>0.6</td><td>2.8</td><td>5.7E-03</td><td>2.3E-02</td><td>BT20/MB231</td><td>NUC</td></tr>
<tr><td>>hg38_21_+_8433172_8446622_RNA45SN1_45S_50ntFlanks@12332.12356.25|TAAGCAGGAGGTGTCAGAAAAGTTA</td><td>>hg38_21_+_8433172_8446622_RNA45SN1_45S_50ntFlanks</td><td>12332</td><td>12356</td><td>25</td><td>TAAGCAGGAGGTGTCAGAAAAGTTA</td><td>67.0</td><td>-4.1</td><td>1.6</td><td>-2.5</td><td>1.3E-02</td><td>2.3E-02</td><td>BT20/MB468</td><td>MP</td></tr>
<tr><td>>hg38_MT_+_598_1651_RNR1_12S_50ntFlanks@719.744.26|TAAAGACGTTAGGTCAAGGTGTAGCC</td><td>>hg38_MT_+_598_1651_RNR1_12S_50ntFlanks</td><td>719</td><td>744</td><td>26</td><td>TAAAGACGTTAGGTCAAGGTGTAGCC</td><td>588.7</td><td>-2.1</td><td>0.8</td><td>-2.7</td><td>6.5E-03</td><td>2.3E-02</td><td>BT20/MB231</td><td>MITO</td></tr>
<tr><td>>hg38_21_+_8433172_8446622_RNA45SN1_45S_50ntFlanks@6649.6669.21|TACGACTCTTAGCGGTGGATC</td><td>>hg38_21_+_8433172_8446622_RNA45SN1_45S_50ntFlanks</td><td>6649</td><td>6669</td><td>21</td><td>TACGACTCTTAGCGGTGGATC</td><td>729.6</td><td>3.6</td><td>1.3</td><td>2.8</td><td>5.3E-03</td><td>2.3E-02</td><td>BT20/MB231</td><td>MP</td></tr>
<tr><td>>hg38_MT_+_1621_3279_RNR2_16S_50ntFlanks@1351.1385.35|AATAGGGTTTACGACCTCGATGTTGGATCAGGACA</td><td>>hg38_MT_+_1621_3279_RNR2_16S_50ntFlanks</td><td>1351</td><td>1385</td><td>35</td><td>AATAGGGTTTACGACCTCGATGTTGGATCAGGACA</td><td>137.3</td><td>4.5</td><td>1.8</td><td>2.5</td><td>1.3E-02</td><td>2.3E-02</td><td>BT20/MB468</td><td>MP</td></tr>
<tr><td>>am_trnaMT_ProTGG_MT_-_15956_16023@52.70.19|AAAGACTTTTTCTCTGACC</td><td>>am_trnaMT_ProTGG_MT_-_15956_16023</td><td>52</td><td>70</td><td>19</td><td>AAAGACTTTTTCTCTGACC</td><td>666.8</td><td>2.7</td><td>1.1</td><td>2.5</td><td>1.3E-02</td><td>2.3E-02</td><td>BT20/MB468</td><td>MP</td></tr>
<tr><td>>hg38_21_+_8433172_8446622_RNA45SN1_45S_50ntFlanks@4102.4127.26|ACCACGGGTGACGGGGAATCAGGGTT</td><td>>hg38_21_+_8433172_8446622_RNA45SN1_45S_50ntFlanks</td><td>4102</td><td>4127</td><td>26</td><td>ACCACGGGTGACGGGGAATCAGGGTT</td><td>176.0</td><td>-3.1</td><td>1.3</td><td>-2.4</td><td>1.7E-02</td><td>2.3E-02</td><td>BT20/MB468</td><td>MITO</td></tr>
<tr><td>>am_MI0003581|hsa-mir-574&WithFlank&4|+|38868026|38868133@31.52.22&[MIMAT0004795&hsa-miR-574-5p&offsets|0|-1;m-213&4|+|38868056|38868078&offsets|0|-1]|TGAGTGTGTGTGTGTGAGTGTG</td><td>>am_MI0003581|hsa-mir-574&WithFlank&4|+|38868026|38868133</td><td>31</td><td>52</td><td>22</td><td>TGAGTGTGTGTGTGTGAGTGTG</td><td>160.0</td><td>3.6</td><td>1.5</td><td>2.5</td><td>1.3E-02</td><td>2.3E-02</td><td>BT20/MB468</td><td>MP</td></tr>
<tr><td>>hg38_21_+_8433172_8446622_RNA45SN1_45S_50ntFlanks@7975.7993.19|CGCGACCTCAGATCAGACG</td><td>>hg38_21_+_8433172_8446622_RNA45SN1_45S_50ntFlanks</td><td>7975</td><td>7993</td><td>19</td><td>CGCGACCTCAGATCAGACG</td><td>34331.2</td><td>2.0</td><td>0.7</td><td>2.8</td><td>5.8E-03</td><td>2.3E-02</td><td>BT20/MB231</td><td>NUC</td></tr>
<tr><td>>hg38_21_+_8433172_8446622_RNA45SN1_45S_50ntFlanks@10347.10369.23|CAAACGAGAACTTTGAAGGCCGA</td><td>>hg38_21_+_8433172_8446622_RNA45SN1_45S_50ntFlanks</td><td>10347</td><td>10369</td><td>23</td><td>CAAACGAGAACTTTGAAGGCCGA</td><td>284.8</td><td>2.3</td><td>0.9</td><td>2.5</td><td>1.3E-02</td><td>2.3E-02</td><td>BT20/MB468</td><td>MP</td></tr>
<tr><td>>hg38_21_+_8433172_8446622_RNA45SN1_45S_50ntFlanks@12836.12862.27|CGGTGCGGAGTGCCCTTCGTCCTGGGA</td><td>>hg38_21_+_8433172_8446622_RNA45SN1_45S_50ntFlanks</td><td>12836</td><td>12862</td><td>27</td><td>CGGTGCGGAGTGCCCTTCGTCCTGGGA</td><td>188.8</td><td>-4.6</td><td>1.8</td><td>-2.5</td><td>1.3E-02</td><td>2.3E-02</td><td>BT20/MB468</td><td>MP</td></tr>
<tr><td>>hg38_MT_+_1621_3279_RNR2_16S_50ntFlanks@518.543.26|TTGTAGAGAGAGTAAAAAATTTAACA</td><td>>hg38_MT_+_1621_3279_RNR2_16S_50ntFlanks</td><td>518</td><td>543</td><td>26</td><td>TTGTAGAGAGAGTAAAAAATTTAACA</td><td>211.8</td><td>-4.3</td><td>1.6</td><td>-2.8</td><td>5.3E-03</td><td>2.3E-02</td><td>BT20/MB231</td><td>MP</td></tr>
<tr><td>>hg38_21_+_8433172_8446622_RNA45SN1_45S_50ntFlanks@4723.4758.36|CATTAATCAAGAACGAAAGTCGGAGGTTCGAAGACG</td><td>>hg38_21_+_8433172_8446622_RNA45SN1_45S_50ntFlanks</td><td>4723</td><td>4758</td><td>36</td><td>CATTAATCAAGAACGAAAGTCGGAGGTTCGAAGACG</td><td>7199.4</td><td>-3.5</td><td>1.4</td><td>-2.5</td><td>1.3E-02</td><td>2.3E-02</td><td>BT20/MB468</td><td>MP</td></tr>
<tr><td>>hg38_21_+_8433172_8446622_RNA45SN1_45S_50ntFlanks@8168.8185.18|CCCAAGTCCTTCTGATCG</td><td>>hg38_21_+_8433172_8446622_RNA45SN1_45S_50ntFlanks</td><td>8168</td><td>8185</td><td>18</td><td>CCCAAGTCCTTCTGATCG</td><td>438.7</td><td>-3.0</td><td>1.3</td><td>-2.3</td><td>2.0E-02</td><td>2.3E-02</td><td>BT20/MB468</td><td>CYTO</td></tr>
<tr><td>>hg38_MT_+_598_1651_RNR1_12S_50ntFlanks@428.458.31|AAAGTGGCTTTAACATATCTGAACACACAAT</td><td>>hg38_MT_+_598_1651_RNR1_12S_50ntFlanks</td><td>428</td><td>458</td><td>31</td><td>AAAGTGGCTTTAACATATCTGAACACACAAT</td><td>86.2</td><td>4.7</td><td>2.0</td><td>2.4</td><td>1.8E-02</td><td>2.3E-02</td><td>MB231/MB468</td><td>MP</td></tr>
<tr><td>>hg38_MT_+_1621_3279_RNR2_16S_50ntFlanks@194.214.21|AAGCATAATATAGCAAGGACT</td><td>>hg38_MT_+_1621_3279_RNR2_16S_50ntFlanks</td><td>194</td><td>214</td><td>21</td><td>AAGCATAATATAGCAAGGACT</td><td>175.6</td><td>2.3</td><td>1.0</td><td>2.4</td><td>1.8E-02</td><td>2.3E-02</td><td>MB231/MB468</td><td>MP</td></tr>
<tr><td>>hg38_21_+_8433172_8446622_RNA45SN1_45S_50ntFlanks@12944.12961.18|CGTAGACGACCTGCTTCT</td><td>>hg38_21_+_8433172_8446622_RNA45SN1_45S_50ntFlanks</td><td>12944</td><td>12961</td><td>18</td><td>CGTAGACGACCTGCTTCT</td><td>149.2</td><td>-4.1</td><td>1.7</td><td>-2.4</td><td>1.8E-02</td><td>2.3E-02</td><td>MB231/MB468</td><td>MP</td></tr>
<tr><td>>hg38_1_-_228634819_228635039_RNA5S12_5S_50ntFlanks@51.68.18|GTCTACGGCCATACCACC</td><td>>hg38_1_-_228634819_228635039_RNA5S12_5S_50ntFlanks</td><td>51</td><td>68</td><td>18</td><td>GTCTACGGCCATACCACC</td><td>429.0</td><td>1.8</td><td>0.8</td><td>2.3</td><td>2.0E-02</td><td>2.3E-02</td><td>MB231/MB468</td><td>CYTO</td></tr>
</table>

| | | | | | | | | | | | | | |
|---|---|---|---|---|---|---|---|---|---|---|---|---|---|
| >hg38_21_+_8433172_8446622_RNA45SN1_45S_50ntFlanks@4451.4472.22\|TCTCGGCGCCCCCTCGATGCTC | >hg38_21_+_8433172_8446622_RNA45SN1_45S_50ntFlanks | 4451 | 4472 | 22 | TCTCGGCGCCCCCTCGATGCTC | 62.6 | -4.1 | 1.7 | -2.5 | 1.3E-02 | 2.3E-02 | BT20/MB468 | MP |
| >hg38_21_+_8433172_8446622_RNA45SN1_45S_50ntFlanks@7983.8007.25\|CAGATCAGACGTGGCGACCCGCTGA | >hg38_21_+_8433172_8446622_RNA45SN1_45S_50ntFlanks | 7983 | 8007 | 25 | CAGATCAGACGTGGCGACCCGCTGA | 796.6 | -3.7 | 1.5 | -2.5 | 1.3E-02 | 2.3E-02 | BT20/MB468 | MP |
| >MI0000066\|hsa-let-7e&WithFlank&19\|+\|51692780\|51692870@14.31.18&[MIMAT0000066&hsa-let-7e-5p&offsets\|0\|-4;m-50&19\|+\|51692793\|51692814&offsets\|0\|-4]\|TGAGGTAGGAGGTTGTAT | >MI0000066\|hsa-let-7e&WithFlank&19\|+\|51692780\|51692870 | 14 | 31 | 18 | TGAGGTAGGAGGTTGTAT | 110.9 | 4.1 | 1.7 | 2.4 | 1.8E-02 | 2.3E-02 | MB231/MB468 | MP |
| >hg38_21_+_8433172_8446622_RNA45SN1_45S_50ntFlanks@6649.6688.40\|TACGACTCTTAGCGGTGGATCACTCGGCTCGTGCGTCGAT | >hg38_21_+_8433172_8446622_RNA45SN1_45S_50ntFlanks | 6649 | 6688 | 40 | TACGACTCTTAGCGGTGGATCACTCGGCTCGTGCGTCGAT | 52.8 | -3.4 | 1.4 | -2.4 | 1.8E-02 | 2.3E-02 | MB231/MB468 | MP |
| >MI0010629\|hsa-mir-2110&WithFlank&10\|-\|114174099\|114174185@14.36.23&[MIMAT0010133&hsa-miR-2110&offsets\|0\|+1;m-424&10\|-\|114174152\|114174172&offsets\|0\|+2]\|TTGGGGAAACGGCCGCTGAGTGA | >MI0010629\|hsa-mir-2110&WithFlank&10\|-\|114174099\|114174185 | 14 | 36 | 23 | TTGGGGAAACGGCCGCTGAGTGA | 73.7 | 3.7 | 1.5 | 2.5 | 1.3E-02 | 2.3E-02 | BT20/MB468 | MP |
| >hg38_21_+_8433172_8446622_RNA45SN1_45S_50ntFlanks@11320.11343.24\|CCACGTCTCGTCGCGCGCGCGTCC | >hg38_21_+_8433172_8446622_RNA45SN1_45S_50ntFlanks | 11320 | 11343 | 24 | CCACGTCTCGTCGCGCGCGCGTCC | 1947.5 | -5.1 | 2.2 | -2.3 | 2.0E-02 | 2.3E-02 | BT20/MB468 | NUC |
| >MI0000734\|hsa-mir-106b&WithFlank&7\|-\|100093987\|100094080@58.76.19(+1C)&[MIMAT0004672&hsa-miR-106b-3p&offsets\|0\|-3(+1C);m-87&7\|-\|100094003\|100094023&offsets\|0\|-2(+1C)]\|CCGCACTGTGGGTACTTGCC | >MI0000734\|hsa-mir-106b&WithFlank&7\|-\|100093987\|100094080 | 58 | 76 | 19(+1C) | CCGCACTGTGGGTACTTGCC | 87.2 | 4.8 | 2.0 | 2.4 | 1.8E-02 | 2.3E-02 | MB231/MB468 | MP |
| >MI0000100\|hsa-mir-98&WithFlank&X\|-\|53556217\|53556347@28.48.21&[MIMAT0000096&hsa-miR-98-5p&offsets\|0\|-1;m-143&X\|-\|53556299\|53556320&offsets\|0\|-1]\|TGAGGTAGTAAGTTGTATTGT | >MI0000100\|hsa-mir-98&WithFlank&X\|-\|53556217\|53556347 | 28 | 48 | 21 | TGAGGTAGTAAGTTGTATTGT | 431.5 | 1.9 | 0.8 | 2.4 | 1.8E-02 | 2.3E-02 | MB231/MB468 | MP |
| >hg38_21_+_8433172_8446622_RNA45SN1_45S_50ntFlanks@3987.4019.33\|GCTTTGGTGACTCTAGATAACCTCGGGCCGATC | >hg38_21_+_8433172_8446622_RNA45SN1_45S_50ntFlanks | 3987 | 4019 | 33 | GCTTTGGTGACTCTAGATAACCTCGGGCCGATC | 229.1 | 3.4 | 1.4 | 2.5 | 1.3E-02 | 2.3E-02 | BT20/MB468 | MP |
| >hg38_21_+_8433172_8446622_RNA45SN1_45S_50ntFlanks@10351.10391.41\|CGAGAACTTTGAAGGCCGAAGTGGAGAAGGGTTCCATGTGA | >hg38_21_+_8433172_8446622_RNA45SN1_45S_50ntFlanks | 10351 | 10391 | 41 | CGAGAACTTTGAAGGCCGAAGTGGAGAAGGGTTCCATGTGA | 477.1 | -3.9 | 1.6 | -2.5 | 1.3E-02 | 2.3E-02 | BT20/MB468 | MP |
| >hg38_21_+_8433172_8446622_RNA45SN1_45S_50ntFlanks@10869.10888.20\|AGGGCTGGGTCGGTCGGGCT | >hg38_21_+_8433172_8446622_RNA45SN1_45S_50ntFlanks | 10869 | 10888 | 20 | AGGGCTGGGTCGGTCGGGCT | 121.0 | 5.6 | 2.0 | 2.8 | 5.3E-03 | 2.3E-02 | BT20/MB231 | MP |
| >hg38_MT_+_1621_3279_RNR2_16S_50ntFlanks@1091.1120.30\|AGCAAGACGAGAAGACCCTATGGAGCTTTA | >hg38_MT_+_1621_3279_RNR2_16S_50ntFlanks | 1091 | 1120 | 30 | AGCAAGACGAGAAGACCCTATGGAGCTTTA | 108.7 | 2.3 | 1.0 | 2.4 | 1.8E-02 | 2.3E-02 | MB231/MB468 | MP |
| >hg38_21_+_8433172_8446622_RNA45SN1_45S_50ntFlanks@11572.11613.42\|CCGACTTAGAACTGGTGCGGACCAGGGGAATCCGACTGTTTA | >hg38_21_+_8433172_8446622_RNA45SN1_45S_50ntFlanks | 11572 | 11613 | 42 | CCGACTTAGAACTGGTGCGGACCAGGGGAATCCGACTGTTTA | 257.7 | -4.5 | 1.9 | -2.4 | 1.8E-02 | 2.3E-02 | MB231/MB468 | MP |
| >hg38_21_+_8433172_8446622_RNA45SN1_45S_50ntFlanks@5157.5185.29\|CAAGTGGCGTTCAGCCACCCGAGATTGAG | >hg38_21_+_8433172_8446622_RNA45SN1_45S_50ntFlanks | 5157 | 5185 | 29 | CAAGTGGCGTTCAGCCACCCGAGATTGAG | 77.8 | -4.5 | 1.8 | -2.5 | 1.3E-02 | 2.3E-02 | BT20/MB468 | MP |
| >hg38_21_+_8433172_8446622_RNA45SN1_45S_50ntFlanks@4927.4944.18\|AGTGGAGCCTGCGGCTTA | >hg38_21_+_8433172_8446622_RNA45SN1_45S_50ntFlanks | 4927 | 4944 | 18 | AGTGGAGCCTGCGGCTTA | 187.3 | 2.6 | 1.0 | 2.5 | 1.3E-02 | 2.3E-02 | BT20/MB468 | MP |
| >hg38_21_+_8433172_8446622_RNA45SN1_45S_50ntFlanks@6654.6689.36\|CTCTTAGCGGTGGATCACTCGGCTCGTGCGTCGATG | >hg38_21_+_8433172_8446622_RNA45SN1_45S_50ntFlanks | 6654 | 6689 | 36 | CTCTTAGCGGTGGATCACTCGGCTCGTGCGTCGATG | 297.8 | 2.1 | 0.8 | 2.7 | 6.5E-03 | 2.3E-02 | BT20/MB231 | MITO |
| >hg38_MT_+_1621_3279_RNR2_16S_50ntFlanks@51.92.42\|GCTAAACCTAGCCCCAAACCCACTCCACCTTACTACCAGACA | >hg38_MT_+_1621_3279_RNR2_16S_50ntFlanks | 51 | 92 | 42 | GCTAAACCTAGCCCCAAACCCACTCCACCTTACTACCAGACA | 1376.3 | 3.9 | 1.6 | 2.5 | 1.3E-02 | 2.3E-02 | BT20/MB468 | MP |
| >hg38_MT_+_1621_3279_RNR2_16S_50ntFlanks@1095.1120.26\|AGACGAGAAGACCCTATGGAGCTTTA | >hg38_MT_+_1621_3279_RNR2_16S_50ntFlanks | 1095 | 1120 | 26 | AGACGAGAAGACCCTATGGAGCTTTA | 200.1 | 3.1 | 1.3 | 2.3 | 2.1E-02 | 2.3E-02 | MB231/MB468 | MITO |
| >hg38_21_+_8433172_8446622_RNA45SN1_45S_50ntFlanks@9740.9769.30\|AACGATCTCAACCTATTCTCAAACTTTAAA | >hg38_21_+_8433172_8446622_RNA45SN1_45S_50ntFlanks | 9740 | 9769 | 30 | AACGATCTCAACCTATTCTCAAACTTTAAA | 460.2 | 2.0 | 0.8 | 2.4 | 1.7E-02 | 2.3E-02 | BT20/MB468 | MITO |
| >trnaMT_LeuTAA_MT_+_3230_3304@1.30.30\|GTTAAGATGGCAGAGCCCGGTAATCGCATA | >trnaMT_LeuTAA_MT_+_3230_3304 | 1 | 30 | 30 | GTTAAGATGGCAGAGCCCGGTAATCGCATA | 466.0 | 1.6 | 0.7 | 2.4 | 1.8E-02 | 2.3E-02 | MB231/MB468 | MP |
| >trna10_AspGTC_12_-_125424193_125424264@41.69.29,trna12_AspGTC_12_-_125411891_125411962@41.69.29,trna144_AspGTC_6_-_27551236_27551307@41.69.29,trna38_AspGTC_17_-_8125556_8125627@41.69.29,trna45_AspGTC_6_+_27447453_27447524@41.69.29,trna48_AspGTC_6_+_27471523_27471594@41.69.29,trna4_AspGTC_12_+_96429799_96429870@41.69.29,trna69_AspGTC_1_-_161440205_161440276@41.69.29,trna72_AspGTC_1_-_161432824_161432895@41.69.29,trna75_AspGTC_1_-_161425414_161425485@41.69.29,trna78_AspGTC_1_-_161418033_161418104@41.69.29,trna81_AspGTC_1_-_161410615_161410686@41.69.29\|GGGAGACCGGGGTTCGATTCCCCGACGGG | >trna10_AspGTC_12_-_125424193_125424264 | 41 | 69 | 29 | GGGAGACCGGGGTTCGATTCCCCGACGGG | 162.9 | 4.6 | 1.6 | 2.8 | 5.3E-03 | 2.3E-02 | BT20/MB231 | MP |
| >hg38_21_+_8433172_8446622_RNA45SN1_45S_50ntFlanks@12073.12098.26\|CCGAGGGGCTCTCGCTTCTGGCGCCA | >hg38_21_+_8433172_8446622_RNA45SN1_45S_50ntFlanks | 12073 | 12098 | 26 | CCGAGGGGCTCTCGCTTCTGGCGCCA | 215.2 | -2.8 | 1.2 | -2.4 | 1.7E-02 | 2.3E-02 | BT20/MB468 | MITO |
| >hg38_21_+_8433172_8446622_RNA45SN1_45S_50ntFlanks@6787.6807.21\|ACGCCTGTCTGAGCGTCGCTT | >hg38_21_+_8433172_8446622_RNA45SN1_45S_50ntFlanks | 6787 | 6807 | 21 | ACGCCTGTCTGAGCGTCGCTT | 309.5 | 2.0 | 0.8 | 2.3 | 2.1E-02 | 2.3E-02 | MB231/MB468 | MITO |
| >hg38_21_+_8433172_8446622_RNA45SN1_45S_50ntFlanks@3788.3810.23\|AACTGCGAATGGCTCATTAAATC | >hg38_21_+_8433172_8446622_RNA45SN1_45S_50ntFlanks | 3788 | 3810 | 23 | AACTGCGAATGGCTCATTAAATC | 165.8 | -2.9 | 1.2 | -2.4 | 1.7E-02 | 2.3E-02 | BT20/MB468 | MITO |
| >hg38_21_+_8433172_8446622_RNA45SN1_45S_50ntFlanks@8347.8375.29\|GGGAAAGTTGAAAAGAACTTTGAAGAGAG | >hg38_21_+_8433172_8446622_RNA45SN1_45S_50ntFlanks | 8347 | 8375 | 29 | GGGAAAGTTGAAAAGAACTTTGAAGAGAG | 62.5 | 4.3 | 1.8 | 2.4 | 1.8E-02 | 2.3E-02 | MB231/MB468 | MP |
| >hg38_21_+_8433172_8446622_RNA45SN1_45S_50ntFlanks@7975.7992.18\|CGCGACCTCAGATCAGAC | >hg38_21_+_8433172_8446622_RNA45SN1_45S_50ntFlanks | 7975 | 7992 | 18 | CGCGACCTCAGATCAGAC | 10644.2 | 1.7 | 0.7 | 2.3 | 2.1E-02 | 2.3E-02 | MB231/MB468 | NUC |
| >hg38_21_+_8433172_8446622_RNA45SN1_45S_50ntFlanks@12324.12355.32\|TTGGGTTTTAAGCAGGAGGTGTCAGAAAAGTT | >hg38_21_+_8433172_8446622_RNA45SN1_45S_50ntFlanks | 12324 | 12355 | 32 | TTGGGTTTTAAGCAGGAGGTGTCAGAAAAGTT | 106.8 | -3.6 | 1.5 | -2.4 | 1.7E-02 | 2.3E-02 | BT20/MB468 | MITO |
| >hg38_21_+_8433172_8446622_RNA45SN1_45S_50ntFlanks@6773.6803.31\|TTCCTCCCGGGGCTACGCCTGTCTGAGCGTC | >hg38_21_+_8433172_8446622_RNA45SN1_45S_50ntFlanks | 6773 | 6803 | 31 | TTCCTCCCGGGGCTACGCCTGTCTGAGCGTC | 906.9 | -4.5 | 1.8 | -2.5 | 1.3E-02 | 2.3E-02 | BT20/MB468 | MP |
| >hg38_21_+_8433172_8446622_RNA45SN1_45S_50ntFlanks@11200.11227.28\|CGGCGGCGGCGGCGACTCTGGACGCGAG | >hg38_21_+_8433172_8446622_RNA45SN1_45S_50ntFlanks | 11200 | 11227 | 28 | CGGCGGCGGCGGCGACTCTGGACGCGAG | 180.9 | -3.2 | 1.3 | -2.5 | 1.3E-02 | 2.3E-02 | BT20/MB468 | MP |
| >MI0000542\|hsa-mir-320a&WithFlank&8\|-\|22244960\|22245043@49.71.23&[MIMAT0000510&hsa-miR-320a-3p&offsets\|+1\|+2;m-60&8\|-\|22244975\|22244996&offsets\|+1\|+2]\|AAAGCTGGGTTGAGAGGGCGAAA | >MI0000542\|hsa-mir-320a&WithFlank&8\|-\|22244960\|22245043 | 49 | 71 | 23 | AAAGCTGGGTTGAGAGGGCGAAA | 406.0 | 2.8 | 1.0 | 2.8 | 5.3E-03 | 2.3E-02 | BT20/MB231 | MP |
| >MI0003581\|hsa-mir-574&WithFlank&4\|+\|38868026\|38868133@67.88.22&[MIMAT0003239&hsa-miR-574-3p&offsets\|0\|0;m-149&4\|+\|38868092\|38868113&offsets\|0\|0]\|CACGCTCATGCACACACCCACA | >MI0003581\|hsa-mir-574&WithFlank&4\|+\|38868026\|38868133 | 67 | 88 | 22 | CACGCTCATGCACACACCCACA | 1366.7 | -2.9 | 1.0 | -2.8 | 5.3E-03 | 2.3E-02 | BT20/MB231 | MP |
| >hg38_21_+_8433172_8446622_RNA45SN1_45S_50ntFlanks@8339.8373.35\|TACCGTAAGGGAAAGTTGAAAAGAACTTTGAAGAG | >hg38_21_+_8433172_8446622_RNA45SN1_45S_50ntFlanks | 8339 | 8373 | 35 | TACCGTAAGGGAAAGTTGAAAAGAACTTTGAAGAG | 1237.2 | 5.0 | 1.8 | 2.8 | 5.3E-03 | 2.3E-02 | BT20/MB231 | MP |
| >hg38_21_+_8433172_8446622_RNA45SN1_45S_50ntFlanks@6651.6698.48\|CGACTCTTAGCGGTGGATCACTCGGCTCGTGCGTCGATGAAGAACGCA | >hg38_21_+_8433172_8446622_RNA45SN1_45S_50ntFlanks | 6651 | 6698 | 48 | CGACTCTTAGCGGTGGATCACTCGGCTCGTGCGTCGATGAAGAACGCA | 3462.7 | 1.7 | 0.7 | 2.3 | 2.0E-02 | 2.3E-02 | MB231/MB468 | CYTO |
| >hg38_21_+_8433172_8446622_RNA45SN1_45S_50ntFlanks@3705.3744.40\|TACCTGGTTGATCCTGCCAGTAGCATATGCTTGTCTCAAA | >hg38_21_+_8433172_8446622_RNA45SN1_45S_50ntFlanks | 3705 | 3744 | 40 | TACCTGGTTGATCCTGCCAGTAGCATATGCTTGTCTCAAA | 344.7 | 1.9 | 0.8 | 2.3 | 2.1E-02 | 2.3E-02 | MB231/MB468 | MITO |
| >hg38_21_+_8433172_8446622_RNA45SN1_45S_50ntFlanks@9459.9493.35\|CCGGGGAGGTGGAGCACGAGCGCACGTGTTAGGAC | >hg38_21_+_8433172_8446622_RNA45SN1_45S_50ntFlanks | 9459 | 9493 | 35 | CCGGGGAGGTGGAGCACGAGCGCACGTGTTAGGAC | 104.1 | -4.1 | 1.7 | -2.5 | 1.3E-02 | 2.3E-02 | BT20/MB468 | MP |
| >hg38_21_+_8433172_8446622_RNA45SN1_45S_50ntFlanks@12281.12321.41\|AATACAGACCGTGAAAGCGGGGCCTCACGATCCTTCTGACC | >hg38_21_+_8433172_8446622_RNA45SN1_45S_50ntFlanks | 12281 | 12321 | 41 | AATACAGACCGTGAAAGCGGGGCCTCACGATCCTTCTGACC | 83.5 | 3.1 | 1.4 | 2.3 | 2.1E-02 | 2.3E-02 | MB231/MB468 | MITO |
| >hg38_21_+_8433172_8446622_RNA45SN1_45S_50ntFlanks@12457.12486.30\|CAAGCGTTGGATTGTTCACCCACTAATAGG | >hg38_21_+_8433172_8446622_RNA45SN1_45S_50ntFlanks | 12457 | 12486 | 30 | CAAGCGTTGGATTGTTCACCCACTAATAGG | 177.3 | -2.9 | 1.2 | -2.4 | 1.7E-02 | 2.3E-02 | BT20/MB468 | MITO |
| >MI0000087\|hsa-mir-29a&WithFlank&7\|-\|130876741\|130876816@47.68.22(+2U)&[MIMAT0000086&hsa-miR-29a-3p&offsets\|-1\|-1(+2U);m-26&7\|-\|130876749\|130876769&offsets\|-1\|0(+2U)]\|CTAGCACCATCTGAAATCGGTTTT | >MI0000087\|hsa-mir-29a&WithFlank&7\|-\|130876741\|130876816 | 47 | 68 | 22(+2U) | CTAGCACCATCTGAAATCGGTTTT | 389.1 | -2.8 | 1.0 | -2.8 | 5.3E-03 | 2.3E-02 | BT20/MB231 | MP |
| >hg38_21_+_8433172_8446622_RNA45SN1_45S_50ntFlanks@10405.10433.29\|ATGGGTCAGTCGGTCCTGAGAGATGGGCG | >hg38_21_+_8433172_8446622_RNA45SN1_45S_50ntFlanks | 10405 | 10433 | 29 | ATGGGTCAGTCGGTCCTGAGAGATGGGCG | 131.2 | 5.0 | 1.8 | 2.8 | 5.3E-03 | 2.3E-02 | BT20/MB231 | MP |
| >hg38_21_+_8433172_8446622_RNA45SN1_45S_50ntFlanks@9953.9980.28\|ATGGAAGTCGGAATCCGCTAAGGAGTGT | >hg38_21_+_8433172_8446622_RNA45SN1_45S_50ntFlanks | 9953 | 9980 | 28 | ATGGAAGTCGGAATCCGCTAAGGAGTGT | 217.7 | 5.7 | 2.0 | 2.8 | 5.3E-03 | 2.3E-02 | BT20/MB231 | MP |
| >trnaMT_ProTGG_MT_-_15956_16023@44.71.28\|GTGGAGTTAAAGACTTTTTCTCTGACCA | >trnaMT_ProTGG_MT_-_15956_16023 | 44 | 71 | 28 | GTGGAGTTAAAGACTTTTTCTCTGACCA | 162.4 | 2.7 | 1.2 | 2.3 | 2.1E-02 | 2.3E-02 | MB231/MB468 | MITO |
| >hg38_21_+_8433172_8446622_RNA45SN1_45S_50ntFlanks@6647.6688.42\|CGTACGACTCTTAGCGGTGGATCACTCGGCTCGTGCGTCGAT | >hg38_21_+_8433172_8446622_RNA45SN1_45S_50ntFlanks | 6647 | 6688 | 42 | CGTACGACTCTTAGCGGTGGATCACTCGGCTCGTGCGTCGAT | 385.8 | 2.2 | 0.9 | 2.4 | 1.7E-02 | 2.3E-02 | BT20/MB468 | MITO |
| >trna10_AspGTC_12_-_125424193_125424264@38.73.36,trna12_AspGTC_12_-_125411891_125411962@38.73.36,trna144_AspGTC_6_-_27551236_27551307@38.73.36,trna38_AspGTC_17_-_8125556_8125627@38.73.36,trna45_AspGTC_6_+_27447453_27447524@38.73.36,trna48_AspGTC_6_+_27471523_27471594@38.73.36,trna4_AspGTC_12_+_96429799_96429870@38.73.36,trna69_AspGTC_1_-_161440205_161440276@38.73.36,trna72_AspGTC_1_-_161432824_161432895@38.73.36,trna75_AspGTC_1_-_161425414_161425485@38.73.36,trna78_AspGTC_1_-_161418033_161418104@38.73.36,trna81_AspGTC_1_-_161410615_161410686@38.73.36\|CGCGGGAGACCGGGGTTCGATTCCCCGACGGGGAGC | >trna10_AspGTC_12_-_125424193_125424264 | 38 | 73 | 36 | CGCGGGAGACCGGGGTTCGATTCCCCGACGGGGAGC | 3678.3 | -2.8 | 1.0 | -2.7 | 7.1E-03 | 2.3E-02 | BT20/MB231 | CYTO |
| >trnaMT_LeuTAA_MT_+_3230_3304@1.25.25\|GTTAAGATGGCAGAGCCCGGTAATC | >trnaMT_LeuTAA_MT_+_3230_3304 | 1 | 25 | 25 | GTTAAGATGGCAGAGCCCGGTAATC | 159.5 | 2.5 | 1.1 | 2.4 | 1.9E-02 | 2.3E-02 | MB231/MB468 | MP |

<table>
<tr><th></th><th></th><th></th><th></th><th></th><th></th><th></th><th></th><th></th><th></th><th></th><th></th><th></th><th></th></tr>
<tr><td>>trna116_GluCTC_1_-_145399233_145399304@2.31.30,trna59_GluCTC_1_+_249168447_249168518@2.31.30,trna71_GluCTC_1_-_161439189_161439260@2.31.30,trna74_GluCTC_1_-_161431809_161431880@2.31.30,trna77_GluCTC_1_-_161424398_161424469@2.31.30,trna77_GluCTC_6_+_28949976_28950047@2.31.30,trna80_GluCTC_1_-_161417018_161417089@2.31.30,trna87_GluCTC_6_-_126101393_126101464@2.31.30|CCCTGGTGGTCTAGTGGTTAGGATTCGGCG</td><td>>trna116_GluCTC_1_-_145399233_145399304</td><td>2</td><td>31</td><td>30</td><td>CCCTGGTGGTCTAGTGGTTAGGATTCGGCG</td><td>481.3</td><td>-1.6</td><td>0.7</td><td>-2.3</td><td>2.0E-02</td><td>2.3E-02</td><td>BT20/MB468</td><td>CYTO</td></tr>
<tr><td>>hg38_21_+_8433172_8446622_RNA45SN1_45S_50ntFlanks@6652.6688.37|GACTCTTAGCGGTGGATCACTCGGCTCGTGCGTCGAT</td><td>>hg38_21_+_8433172_8446622_RNA45SN1_45S_50ntFlanks</td><td>6652</td><td>6688</td><td>37</td><td>GACTCTTAGCGGTGGATCACTCGGCTCGTGCGTCGAT</td><td>7979.7</td><td>2.0</td><td>0.7</td><td>2.7</td><td>6.5E-03</td><td>2.3E-02</td><td>BT20/MB231</td><td>MITO</td></tr>
<tr><td>>hg38_21_+_8433172_8446622_RNA45SN1_45S_50ntFlanks@10351.10382.32|CGAGAACTTTGAAGGCCGAAGTGGAGAAGGGT</td><td>>hg38_21_+_8433172_8446622_RNA45SN1_45S_50ntFlanks</td><td>10351</td><td>10382</td><td>32</td><td>CGAGAACTTTGAAGGCCGAAGTGGAGAAGGGT</td><td>972.6</td><td>4.1</td><td>1.5</td><td>2.8</td><td>5.3E-03</td><td>2.3E-02</td><td>BT20/MB231</td><td>MP</td></tr>
<tr><td>>hg38_21_+_8433172_8446622_RNA45SN1_45S_50ntFlanks@11820.11856.37|ACTATCCAGCGAAACCACAGCCAAGGGAACGGGCTTG</td><td>>hg38_21_+_8433172_8446622_RNA45SN1_45S_50ntFlanks</td><td>11820</td><td>11856</td><td>37</td><td>ACTATCCAGCGAAACCACAGCCAAGGGAACGGGCTTG</td><td>38.2</td><td>-4.3</td><td>1.7</td><td>-2.5</td><td>1.3E-02</td><td>2.3E-02</td><td>BT20/MB468</td><td>MP</td></tr>
<tr><td>>MI0000263|hsa-mir-7-1&WithFlank&9|-|83969742|83969863@72.92.21(+1U)&[MIMAT0004553&hsa-miR-7-1-3p&offsets|0|-1(+1U);m-237&9|-|83969771|83969792&offsets|0|-1(+1U)]|CAACAAATCACAGTCTGCCATT</td><td>>MI0000263|hsa-mir-7-1&WithFlank&9|-|83969742|83969863</td><td>72</td><td>92</td><td>21(+1U)</td><td>CAACAAATCACAGTCTGCCATT</td><td>32.5</td><td>4.8</td><td>2.0</td><td>2.4</td><td>1.9E-02</td><td>2.3E-02</td><td>MB231/MB468</td><td>MP</td></tr>
<tr><td>>hg38_21_+_8433172_8446622_RNA45SN1_45S_50ntFlanks@12996.13017.22|TCGCTGCGATCTATTGAAAGTC</td><td>>hg38_21_+_8433172_8446622_RNA45SN1_45S_50ntFlanks</td><td>12996</td><td>13017</td><td>22</td><td>TCGCTGCGATCTATTGAAAGTC</td><td>1108.6</td><td>-2.3</td><td>0.8</td><td>-2.8</td><td>5.8E-03</td><td>2.3E-02</td><td>BT20/MB231</td><td>NUC</td></tr>
<tr><td>>hg38_21_+_8433172_8446622_RNA45SN1_45S_50ntFlanks@10346.10385.40|TCAAACGAGAACTTTGAAGGCCGAAGTGGAGAAGGGTTCC</td><td>>hg38_21_+_8433172_8446622_RNA45SN1_45S_50ntFlanks</td><td>10346</td><td>10385</td><td>40</td><td>TCAAACGAGAACTTTGAAGGCCGAAGTGGAGAAGGGTTCC</td><td>70.2</td><td>4.2</td><td>1.7</td><td>2.5</td><td>1.3E-02</td><td>2.3E-02</td><td>BT20/MB468</td><td>MP</td></tr>
<tr><td>>hg38_21_+_8433172_8446622_RNA45SN1_45S_50ntFlanks@6652.6684.33|GACTCTTAGCGGTGGATCACTCGGCTCGTGCGT</td><td>>hg38_21_+_8433172_8446622_RNA45SN1_45S_50ntFlanks</td><td>6652</td><td>6684</td><td>33</td><td>GACTCTTAGCGGTGGATCACTCGGCTCGTGCGT</td><td>523.1</td><td>1.1</td><td>0.5</td><td>2.3</td><td>2.1E-02</td><td>2.3E-02</td><td>BT20/MB468</td><td>NUC</td></tr>
<tr><td>>hg38_21_+_8433172_8446622_RNA45SN1_45S_50ntFlanks@11669.11687.19|GCCCAGTGCTCTGAATGTC</td><td>>hg38_21_+_8433172_8446622_RNA45SN1_45S_50ntFlanks</td><td>11669</td><td>11687</td><td>19</td><td>GCCCAGTGCTCTGAATGTC</td><td>3274.0</td><td>-2.3</td><td>0.9</td><td>-2.7</td><td>7.1E-03</td><td>2.3E-02</td><td>BT20/MB231</td><td>CYTO</td></tr>
<tr><td>>MI0000085|hsa-mir-27a&WithFlank&19|-|13836434|13836523@57.75.19(+1A)&[MIMAT0000084&hsa-miR-27a-3p&offsets|0|-2(+1A);m-43&19|-|13836447|13836467&offsets|0|-2(+1A)]|TTCACAGTGGCTAAGTTCCA</td><td>>MI0000085|hsa-mir-27a&WithFlank&19|-|13836434|13836523</td><td>57</td><td>75</td><td>19(+1A)</td><td>TTCACAGTGGCTAAGTTCCA</td><td>6527.3</td><td>-3.2</td><td>1.2</td><td>-2.8</td><td>5.4E-03</td><td>2.3E-02</td><td>BT20/MB231</td><td>MP</td></tr>
<tr><td>>hg38_21_+_8433172_8446622_RNA45SN1_45S_50ntFlanks@10014.10039.26|AAAATGGATGGCGCTGGAGCGTCGGG</td><td>>hg38_21_+_8433172_8446622_RNA45SN1_45S_50ntFlanks</td><td>10014</td><td>10039</td><td>26</td><td>AAAATGGATGGCGCTGGAGCGTCGGG</td><td>139.3</td><td>-4.7</td><td>1.9</td><td>-2.5</td><td>1.3E-02</td><td>2.3E-02</td><td>BT20/MB468</td><td>MP</td></tr>
<tr><td>>hg38_21_+_8433172_8446622_RNA45SN1_45S_50ntFlanks@3771.3799.29|CACGGCCGGTACAGTGAAACTGCGAATGG</td><td>>hg38_21_+_8433172_8446622_RNA45SN1_45S_50ntFlanks</td><td>3771</td><td>3799</td><td>29</td><td>CACGGCCGGTACAGTGAAACTGCGAATGG</td><td>111.8</td><td>-3.6</td><td>1.5</td><td>-2.4</td><td>1.7E-02</td><td>2.3E-02</td><td>BT20/MB468</td><td>MITO</td></tr>
<tr><td>>trna16_LeuAAG_16_+_22308461_22308542@1.20.20,trna16_LeuAAG_5_-_180601044_180601125@1.20.20,trna19_LeuAAG_5_-_180524474_180524555@1.20.20,trna1_LeuAAG_14_+_21078291_21078372@1.20.20,trna3_LeuAAG_5_+_180528840_180528921@1.20.20,trna42_LeuTAG_17_-_8023632_8023713@1.20.20,trna78_LeuAAG_6_+_28956779_28956860@1.20.20,trna7_LeuAAG_5_+_180614701_180614782@1.20.20,trna98_LeuAAG_6_-_28911399_28911480@1.20.20|GGTAGCGTGGCCGAGCGGTC</td><td>>trna16_LeuAAG_16_+_22308461_22308542</td><td>1</td><td>20</td><td>20</td><td>GGTAGCGTGGCCGAGCGGTC</td><td>239.2</td><td>2.4</td><td>0.9</td><td>2.7</td><td>7.1E-03</td><td>2.3E-02</td><td>BT20/MB231</td><td>CYTO</td></tr>
<tr><td>>hg38_21_+_8433172_8446622_RNA45SN1_45S_50ntFlanks@3993.4011.19|GTGACTCTAGATAACCTCG</td><td>>hg38_21_+_8433172_8446622_RNA45SN1_45S_50ntFlanks</td><td>3993</td><td>4011</td><td>19</td><td>GTGACTCTAGATAACCTCG</td><td>302.1</td><td>1.6</td><td>0.7</td><td>2.3</td><td>2.0E-02</td><td>2.3E-02</td><td>MB231/MB468</td><td>CYTO</td></tr>
<tr><td>>hg38_21_+_8433172_8446622_RNA45SN1_45S_50ntFlanks@11600.11623.24|AATCCGACTGTTTAATTAAAACAA</td><td>>hg38_21_+_8433172_8446622_RNA45SN1_45S_50ntFlanks</td><td>11600</td><td>11623</td><td>24</td><td>AATCCGACTGTTTAATTAAAACAA</td><td>433.4</td><td>4.8</td><td>1.7</td><td>2.8</td><td>5.4E-03</td><td>2.3E-02</td><td>BT20/MB231</td><td>MP</td></tr>
<tr><td>>hg38_21_+_8433172_8446622_RNA45SN1_45S_50ntFlanks@8346.8378.33|AGGGAAAGTTGAAAAGAACTTTGAAGAGAGAGT</td><td>>hg38_21_+_8433172_8446622_RNA45SN1_45S_50ntFlanks</td><td>8346</td><td>8378</td><td>33</td><td>AGGGAAAGTTGAAAAGAACTTTGAAGAGAGAGT</td><td>312.4</td><td>3.3</td><td>1.2</td><td>2.8</td><td>5.4E-03</td><td>2.3E-02</td><td>BT20/MB231</td><td>MP</td></tr>
<tr><td>>hg38_21_+_8433172_8446622_RNA45SN1_45S_50ntFlanks@6651.6668.18|CGACTCTTAGCGGTGGAT</td><td>>hg38_21_+_8433172_8446622_RNA45SN1_45S_50ntFlanks</td><td>6651</td><td>6668</td><td>18</td><td>CGACTCTTAGCGGTGGAT</td><td>11339.7</td><td>1.7</td><td>0.7</td><td>2.3</td><td>2.1E-02</td><td>2.3E-02</td><td>MB231/MB468</td><td>MITO</td></tr>
<tr><td>>hg38_21_+_8433172_8446622_RNA45SN1_45S_50ntFlanks@9527.9571.45|AAGCCAGAGGAAACTCTGGTGGAGGTCCGTAGCGGTCCTGACGTG</td><td>>hg38_21_+_8433172_8446622_RNA45SN1_45S_50ntFlanks</td><td>9527</td><td>9571</td><td>45</td><td>AAGCCAGAGGAAACTCTGGTGGAGGTCCGTAGCGGTCCTGACGTG</td><td>441.7</td><td>-2.4</td><td>1.0</td><td>-2.3</td><td>2.1E-02</td><td>2.3E-02</td><td>MB231/MB468</td><td>NUC</td></tr>
<tr><td>>MI0000440|hsa-mir-27b&WithFlank&9|+|95085439|95085547@67.86.20(+1A)&[MIMAT0000419&hsa-miR-27b-3p&offsets|0|-1(+1A);m-39&9|+|95085505|95085525&offsets|0|-1(+1A)]|TTCACAGTGGCTAAGTTCTGA</td><td>>MI0000440|hsa-mir-27b&WithFlank&9|+|95085439|95085547</td><td>67</td><td>86</td><td>20(+1A)</td><td>TTCACAGTGGCTAAGTTCTGA</td><td>5233.5</td><td>2.5</td><td>0.9</td><td>2.7</td><td>7.1E-03</td><td>2.3E-02</td><td>BT20/MB231</td><td>CYTO</td></tr>
<tr><td>>am_trna116_GluCTC_1_-_145399233_145399304@1.27.27,trna59_GluCTC_1_+_249168447_249168518@1.27.27,trna71_GluCTC_1_-_161439189_161439260@1.27.27,trna74_GluCTC_1_-_161431809_161431880@1.27.27,trna77_GluCTC_1_-_161424398_161424469@1.27.27,trna77_GluCTC_6_+_28949976_28950047@1.27.27,trna80_GluCTC_1_-_161417018_161417089@1.27.27,trna87_GluCTC_6_-_126101393_126101464@1.27.27|TCCCTGGTGGTCTAGTGGTTAGGATTC</td><td>>am_trna116_GluCTC_1_-_145399233_145399304</td><td>1</td><td>27</td><td>27</td><td>TCCCTGGTGGTCTAGTGGTTAGGATTC</td><td>886.8</td><td>-1.6</td><td>0.7</td><td>-2.3</td><td>2.1E-02</td><td>2.3E-02</td><td>MB231/MB468</td><td>MITO</td></tr>
<tr><td>>hg38_MT_+_598_1651_RNR1_12S_50ntFlanks@909.935.27|TACTTCAAAGGACATTTAACTAAAACC</td><td>>hg38_MT_+_598_1651_RNR1_12S_50ntFlanks</td><td>909</td><td>935</td><td>27</td><td>TACTTCAAAGGACATTTAACTAAAACC</td><td>111.2</td><td>3.2</td><td>1.4</td><td>2.4</td><td>1.9E-02</td><td>2.3E-02</td><td>MB231/MB468</td><td>MP</td></tr>
<tr><td>>hg38_21_+_8433172_8446622_RNA45SN1_45S_50ntFlanks@3989.4023.35|TTTGGTGACTCTAGATAACCTCGGGCCGATCGCAC</td><td>>hg38_21_+_8433172_8446622_RNA45SN1_45S_50ntFlanks</td><td>3989</td><td>4023</td><td>35</td><td>TTTGGTGACTCTAGATAACCTCGGGCCGATCGCAC</td><td>341.2</td><td>3.8</td><td>1.4</td><td>2.8</td><td>5.4E-03</td><td>2.3E-02</td><td>BT20/MB231</td><td>MP</td></tr>
<tr><td>>hg38_21_+_8433172_8446622_RNA45SN1_45S_50ntFlanks@6646.6672.27|TCGTACGACTCTTAGCGGTGGATCACT</td><td>>hg38_21_+_8433172_8446622_RNA45SN1_45S_50ntFlanks</td><td>6646</td><td>6672</td><td>27</td><td>TCGTACGACTCTTAGCGGTGGATCACT</td><td>3659.8</td><td>3.5</td><td>1.3</td><td>2.8</td><td>5.4E-03</td><td>2.3E-02</td><td>BT20/MB231</td><td>MP</td></tr>
<tr><td>>hg38_21_+_8433172_8446622_RNA45SN1_45S_50ntFlanks@9307.9331.25|AAGGAGTCTAACACGTGCGCGAGTC</td><td>>hg38_21_+_8433172_8446622_RNA45SN1_45S_50ntFlanks</td><td>9307</td><td>9331</td><td>25</td><td>AAGGAGTCTAACACGTGCGCGAGTC</td><td>109.2</td><td>2.3</td><td>0.9</td><td>2.5</td><td>1.3E-02</td><td>2.3E-02</td><td>BT20/MB468</td><td>MP</td></tr>
<tr><td>>hg38_21_+_8433172_8446622_RNA45SN1_45S_50ntFlanks@12323.12344.22|TTTGGGTTTTAAGCAGGAGGTG</td><td>>hg38_21_+_8433172_8446622_RNA45SN1_45S_50ntFlanks</td><td>12323</td><td>12344</td><td>22</td><td>TTTGGGTTTTAAGCAGGAGGTG</td><td>86.5</td><td>-4.2</td><td>1.7</td><td>-2.5</td><td>1.3E-02</td><td>2.3E-02</td><td>BT20/MB468</td><td>MP</td></tr>
<tr><td>>trnaMT_HisGTG_MT_+_12138_12206@32.71.40|TGAATCTGACAACAGAGGCTTACGACCCCTTATTTACCCC</td><td>>trnaMT_HisGTG_MT_+_12138_12206</td><td>32</td><td>71</td><td>40</td><td>TGAATCTGACAACAGAGGCTTACGACCCCTTATTTACCCC</td><td>1756.4</td><td>-4.3</td><td>1.5</td><td>-2.8</td><td>5.9E-03</td><td>2.3E-02</td><td>BT20/MB231</td><td>NUC</td></tr>
<tr><td>>hg38_21_+_8433172_8446622_RNA45SN1_45S_50ntFlanks@7972.8009.38|AGACGCGACCTCAGATCAGACGTGGCGACCCGCTGAAT</td><td>>hg38_21_+_8433172_8446622_RNA45SN1_45S_50ntFlanks</td><td>7972</td><td>8009</td><td>38</td><td>AGACGCGACCTCAGATCAGACGTGGCGACCCGCTGAAT</td><td>690.8</td><td>2.3</td><td>0.8</td><td>2.8</td><td>5.9E-03</td><td>2.3E-02</td><td>BT20/MB231</td><td>NUC</td></tr>
<tr><td>>hg38_21_+_8433172_8446622_RNA45SN1_45S_50ntFlanks@10384.10424.41|CCATGTGAACAGCAGTTGAACATGGGTCAGTCGGTCCTGAG</td><td>>hg38_21_+_8433172_8446622_RNA45SN1_45S_50ntFlanks</td><td>10384</td><td>10424</td><td>41</td><td>CCATGTGAACAGCAGTTGAACATGGGTCAGTCGGTCCTGAG</td><td>585.0</td><td>-1.6</td><td>0.6</td><td>-2.8</td><td>5.9E-03</td><td>2.3E-02</td><td>BT20/MB231</td><td>NUC</td></tr>
<tr><td>>hg38_21_+_8433172_8446622_RNA45SN1_45S_50ntFlanks@11920.11939.20|ATGAGAGGTGTAGAATAAGT</td><td>>hg38_21_+_8433172_8446622_RNA45SN1_45S_50ntFlanks</td><td>11920</td><td>11939</td><td>20</td><td>ATGAGAGGTGTAGAATAAGT</td><td>58.4</td><td>4.4</td><td>1.8</td><td>2.5</td><td>1.3E-02</td><td>2.3E-02</td><td>BT20/MB468</td><td>MP</td></tr>
<tr><td>>hg38_21_+_8433172_8446622_RNA45SN1_45S_50ntFlanks@10346.10372.27|TCAAACGAGAACTTTGAAGGCCGAAGT</td><td>>hg38_21_+_8433172_8446622_RNA45SN1_45S_50ntFlanks</td><td>10346</td><td>10372</td><td>27</td><td>TCAAACGAGAACTTTGAAGGCCGAAGT</td><td>671.9</td><td>-2.5</td><td>1.0</td><td>-2.5</td><td>1.3E-02</td><td>2.3E-02</td><td>BT20/MB468</td><td>MP</td></tr>
<tr><td>>hg38_21_+_8433172_8446622_RNA45SN1_45S_50ntFlanks@12416.12440.25|TTCGATGTCGGCTCTTCCTATCATT</td><td>>hg38_21_+_8433172_8446622_RNA45SN1_45S_50ntFlanks</td><td>12416</td><td>12440</td><td>25</td><td>TTCGATGTCGGCTCTTCCTATCATT</td><td>4514.9</td><td>-2.4</td><td>0.9</td><td>-2.7</td><td>7.1E-03</td><td>2.3E-02</td><td>BT20/MB231</td><td>CYTO</td></tr>
<tr><td>>hg38_MT_+_1621_3279_RNR2_16S_50ntFlanks@1352.1379.28|ATAGGGTTTACGACCTCGATGTTGGATC</td><td>>hg38_MT_+_1621_3279_RNR2_16S_50ntFlanks</td><td>1352</td><td>1379</td><td>28</td><td>ATAGGGTTTACGACCTCGATGTTGGATC</td><td>109.1</td><td>2.5</td><td>1.1</td><td>2.4</td><td>1.9E-02</td><td>2.3E-02</td><td>MB231/MB468</td><td>MP</td></tr>
<tr><td>>hg38_21_+_8433172_8446622_RNA45SN1_45S_50ntFlanks@5236.5267.32|CACTGACTGGCTCAGCGTGTGCCTACCCTACG</td><td>>hg38_21_+_8433172_8446622_RNA45SN1_45S_50ntFlanks</td><td>5236</td><td>5267</td><td>32</td><td>CACTGACTGGCTCAGCGTGTGCCTACCCTACG</td><td>141.0</td><td>-1.4</td><td>0.6</td><td>-2.3</td><td>2.0E-02</td><td>2.3E-02</td><td>BT20/MB468</td><td>CYTO</td></tr>
<tr><td>>hg38_21_+_8433172_8446622_RNA45SN1_45S_50ntFlanks@8352.8375.24|AGTTGAAAAGAACTTTGAAGAGAG</td><td>>hg38_21_+_8433172_8446622_RNA45SN1_45S_50ntFlanks</td><td>8352</td><td>8375</td><td>24</td><td>AGTTGAAAAGAACTTTGAAGAGAG</td><td>982.7</td><td>7.1</td><td>2.6</td><td>2.7</td><td>6.6E-03</td><td>2.3E-02</td><td>BT20/MB231</td><td>MITO</td></tr>
<tr><td>>MI0000255|hsa-mir-30d&WithFlank&8|-|134804870|134804951@12.33.22(+2U)&[MIMAT0000245&hsa-miR-30d-5p&offsets|0|0(+2U);m-21&8|-|134804919|134804940&offsets|0|0(+2U)]|TGTAAACATCCCCGACTGGAAGTT</td><td>>MI0000255|hsa-mir-30d&WithFlank&8|-|134804870|134804951</td><td>12</td><td>33</td><td>22(+2U)</td><td>TGTAAACATCCCCGACTGGAAGTT</td><td>312.4</td><td>2.4</td><td>0.9</td><td>2.7</td><td>7.1E-03</td><td>2.3E-02</td><td>BT20/MB231</td><td>CYTO</td></tr>
<tr><td>>hg38_MT_+_1621_3279_RNR2_16S_50ntFlanks@1184.1215.32|AATTTCGGTTGGGGCGACCTCGGAGCAGAACC</td><td>>hg38_MT_+_1621_3279_RNR2_16S_50ntFlanks</td><td>1184</td><td>1215</td><td>32</td><td>AATTTCGGTTGGGGCGACCTCGGAGCAGAACC</td><td>195.6</td><td>2.9</td><td>1.2</td><td>2.4</td><td>1.9E-02</td><td>2.3E-02</td><td>MB231/MB468</td><td>MP</td></tr>
<tr><td>>hg38_21_+_8433172_8446622_RNA45SN1_45S_50ntFlanks@7974.8007.34|ACGCGACCTCAGATCAGACGTGGCGACCCGCTGA</td><td>>hg38_21_+_8433172_8446622_RNA45SN1_45S_50ntFlanks</td><td>7974</td><td>8007</td><td>34</td><td>ACGCGACCTCAGATCAGACGTGGCGACCCGCTGA</td><td>604.3</td><td>-1.6</td><td>0.7</td><td>-2.4</td><td>1.9E-02</td><td>2.3E-02</td><td>MB231/MB468</td><td>MP</td></tr>
<tr><td>>MI0000087|hsa-mir-29a&WithFlank&7|-|130876741|130876816@48.69.22&[MIMAT0000086&hsa-miR-29a-3p&offsets|0|0;m-26&7|-|130876749|130876769&offsets|0|+1]|TAGCACCATCTGAAATCGGTTA</td><td>>MI0000087|hsa-mir-29a&WithFlank&7|-|130876741|130876816</td><td>48</td><td>69</td><td>22</td><td>TAGCACCATCTGAAATCGGTTA</td><td>18965.6</td><td>-2.6</td><td>1.0</td><td>-2.7</td><td>6.6E-03</td><td>2.3E-02</td><td>BT20/MB231</td><td>MITO</td></tr>
<tr><td>>hg38_21_+_8433172_8446622_RNA45SN1_45S_50ntFlanks@4927.4944.18|AGTGGAGCCTGCGGCTTA</td><td>>hg38_21_+_8433172_8446622_RNA45SN1_45S_50ntFlanks</td><td>4927</td><td>4944</td><td>18</td><td>AGTGGAGCCTGCGGCTTA</td><td>281.8</td><td>1.9</td><td>0.7</td><td>2.7</td><td>6.6E-03</td><td>2.3E-02</td><td>BT20/MB231</td><td>MITO</td></tr>
<tr><td>>hg38_21_+_8433172_8446622_RNA45SN1_45S_50ntFlanks@12714.12747.34|GAGCCTCGGTTGGCCTCGGATAGCCGGTCCCCCG</td><td>>hg38_21_+_8433172_8446622_RNA45SN1_45S_50ntFlanks</td><td>12714</td><td>12747</td><td>34</td><td>GAGCCTCGGTTGGCCTCGGATAGCCGGTCCCCCG</td><td>345.9</td><td>-1.7</td><td>0.7</td><td>-2.4</td><td>1.8E-02</td><td>2.3E-02</td><td>BT20/MB468</td><td>MITO</td></tr>
<tr><td>>hg38_21_+_8433172_8446622_RNA45SN1_45S_50ntFlanks@6651.6676.26|CGACTCTTAGCGGTGGATCACTCGGC</td><td>>hg38_21_+_8433172_8446622_RNA45SN1_45S_50ntFlanks</td><td>6651</td><td>6676</td><td>26</td><td>CGACTCTTAGCGGTGGATCACTCGGC</td><td>20816.1</td><td>3.9</td><td>1.4</td><td>2.8</td><td>5.4E-03</td><td>2.3E-02</td><td>BT20/MB231</td><td>MP</td></tr>
<tr><td>>MI0001652|hsa-mir-450a-1&WithFlank&X|-|134540335|134540437@24.45.22;MI0003187|hsa-mir-450a-2&WithFlank&X|-|134540502|134540613@29.50.22&[MIMAT0001545&hsa-miR-450a-5p&offsets|0|0;m-288&X|-|134540393|134540414&offsets|0|0];[MIMAT0001545_1&hsa-miR-450a-5p&offsets|0|0;m-287&X|-|134540564|134540585&offsets|0|0]|TTTTGCGATGTGTTCCTAATAT</td><td>>MI0001652|hsa-mir-450a-1&WithFlank&X|-|134540335|134540437</td><td>24</td><td>45</td><td>22</td><td>TTTTGCGATGTGTTCCTAATAT</td><td>44.7</td><td>4.6</td><td>2.0</td><td>2.4</td><td>1.9E-02</td><td>2.3E-02</td><td>MB231/MB468</td><td>MP</td></tr>
</table>

| | | | | | | | | | | | | | |
|---|---|---|---|---|---|---|---|---|---|---|---|---|---|
| >trna111_HisGTG_1_-_147774845_147774916@-1T.30.31,trna118_HisGTG_1_-_145396881_145396952@-1T.30.31,trna16_HisGTG_1_+_146544773_146544844@-1T.30.31,trna1_HisGTG_15_+_45493349_45493420@-1T.30.31,trna21_HisGTG_1_+_147753471_147753542@-1T.30.31,trna33_HisGTG_6_+_27125906_27125977@-1T.30.31,trna7_HisGTG_9_-_14433938_14434009@-1T.30.31,trna8_HisGTG_15_-_45492611_45492682@-1T.30.31,trna9_HisGTG_15_-_45490804_45490875@-1T.30.31\|TGCCGTGATCGTATAGTGGTTAGTACTCTGC | >trna111_HisGTG_1_-_147774845_147774916 | -1T | 30 | 31 | TGCCGTGATCGTATAGTGGTTAGTACTCTGC | 216.9 | -2.8 | 1.0 | -2.7 | 6.6E-03 | 2.3E-02 | BT20/MB231 | MITO |
| >MI0000108\|hsa-mir-103a-2&WithFlank&20\|+\|3917488\|3917577@54.75.22(+1U);MI0000109\|hsa-mir-103a-1&WithFlank&5\|-\|168560890\|168560979@54.75.22(+1U)&[MIMAT0000101&hsa-miR-103a-3p&offsets\|0\|-1(+1U);m-12&20\|+\|3917541\|3917563&offsets\|0\|-1(+1U)];[MIMAT0000101_1&hsa-miR-103a-3p&offsets\|0\|-1(+1U);m-13&5\|-\|168560904\|168560926&offsets\|0\|-1(+1U);m-44220&5\|-\|168560908\|168560926&offsets\|0\|+3(+1U)]\|AGCAGCATTGTACAGGGCTATGT | >MI0000108\|hsa-mir-103a-2&WithFlank&20\|+\|3917488\|3917577 | 54 | 75 | 22(+1U) | AGCAGCATTGTACAGGGCTATGT | 2335.9 | -2.4 | 0.9 | -2.7 | 6.6E-03 | 2.3E-02 | BT20/MB231 | MITO |
| >hg38_21_+_8433172_8446622_RNA45SN1_45S_50ntFlanks@12997.13015.19\|CGCTGCGATCTATTGAAAG | >hg38_21_+_8433172_8446622_RNA45SN1_45S_50ntFlanks | 12997 | 13015 | 19 | CGCTGCGATCTATTGAAAG | 476.1 | -1.9 | 0.7 | -2.7 | 6.6E-03 | 2.3E-02 | BT20/MB231 | MITO |
| >hg38_21_+_8433172_8446622_RNA45SN1_45S_50ntFlanks@5487.5520.34\|CGCTGAGAAGACGGTCGAACTTGACTATCTAGAG | >hg38_21_+_8433172_8446622_RNA45SN1_45S_50ntFlanks | 5487 | 5520 | 34 | CGCTGAGAAGACGGTCGAACTTGACTATCTAGAG | 84.6 | -4.3 | 1.7 | -2.5 | 1.3E-02 | 2.3E-02 | BT20/MB468 | MP |
| >am_trna2_ArgCCT_16_+_3202901_3202973@14.31.18,trna3_ArgCCT_7_+_139025446_139025518@14.31.18\|AATGGATAAGGCATTGGC | >am_trna2_ArgCCT_16_+_3202901_3202973 | 14 | 31 | 18 | AATGGATAAGGCATTGGC | 152.6 | -3.5 | 1.5 | -2.3 | 2.0E-02 | 2.3E-02 | BT20/MB468 | CYTO |
| >hg38_21_+_8433172_8446622_RNA45SN1_45S_50ntFlanks@3771.3799.29\|CACGGCCGGTACAGTGAAACTGCGAATGG | >hg38_21_+_8433172_8446622_RNA45SN1_45S_50ntFlanks | 3771 | 3799 | 29 | CACGGCCGGTACAGTGAAACTGCGAATGG | 180.9 | -4.2 | 1.7 | -2.5 | 1.3E-02 | 2.3E-02 | BT20/MB468 | MP |
| >hg38_21_+_8433172_8446622_RNA45SN1_45S_50ntFlanks@12397.12414.18\|CGACGTCGCTTTTTGATC | >hg38_21_+_8433172_8446622_RNA45SN1_45S_50ntFlanks | 12397 | 12414 | 18 | CGACGTCGCTTTTTGATC | 2723.6 | -1.2 | 0.5 | -2.3 | 2.0E-02 | 2.3E-02 | MB231/MB468 | CYTO |
| >hg38_21_+_8433172_8446622_RNA45SN1_45S_50ntFlanks@6648.6681.34\|GTACGACTCTTAGCGGTGGATCACTCGGCTCGTG | >hg38_21_+_8433172_8446622_RNA45SN1_45S_50ntFlanks | 6648 | 6681 | 34 | GTACGACTCTTAGCGGTGGATCACTCGGCTCGTG | 11364.2 | -1.3 | 0.6 | -2.4 | 1.8E-02 | 2.3E-02 | BT20/MB468 | MITO |
| >hg38_21_+_8433172_8446622_RNA45SN1_45S_50ntFlanks@3846.3872.27\|CTTGGATAACTGTGGTAATTCTAGAGC | >hg38_21_+_8433172_8446622_RNA45SN1_45S_50ntFlanks | 3846 | 3872 | 27 | CTTGGATAACTGTGGTAATTCTAGAGC | 158.2 | -4.2 | 1.7 | -2.5 | 1.3E-02 | 2.3E-02 | BT20/MB468 | MP |
| >hg38_21_+_8433172_8446622_RNA45SN1_45S_50ntFlanks@4048.4066.19\|TTCGAACGTCTGCCCTATC | >hg38_21_+_8433172_8446622_RNA45SN1_45S_50ntFlanks | 4048 | 4066 | 19 | TTCGAACGTCTGCCCTATC | 780.5 | -2.5 | 0.9 | -2.7 | 7.2E-03 | 2.3E-02 | BT20/MB231 | CYTO |
| >hg38_21_+_8433172_8446622_RNA45SN1_45S_50ntFlanks@10685.10707.23\|GGCGGCGTCCGGTGAGCTCTCGC | >hg38_21_+_8433172_8446622_RNA45SN1_45S_50ntFlanks | 10685 | 10707 | 23 | GGCGGCGTCCGGTGAGCTCTCGC | 752.8 | 3.0 | 1.1 | 2.7 | 6.6E-03 | 2.3E-02 | BT20/MB231 | MITO |
| >hg38_21_+_8433172_8446622_RNA45SN1_45S_50ntFlanks@10820.10837.18\|GGGAAGTCGGCAAGCCGG | >hg38_21_+_8433172_8446622_RNA45SN1_45S_50ntFlanks | 10820 | 10837 | 18 | GGGAAGTCGGCAAGCCGG | 175.1 | -3.2 | 1.3 | -2.5 | 1.3E-02 | 2.3E-02 | BT20/MB468 | MP |
| >hg38_21_+_8433172_8446622_RNA45SN1_45S_50ntFlanks@12054.12072.19\|TGAGGCGGGGGGGCGAGCC | >hg38_21_+_8433172_8446622_RNA45SN1_45S_50ntFlanks | 12054 | 12072 | 19 | TGAGGCGGGGGGGCGAGCC | 242.7 | -3.9 | 1.7 | -2.3 | 2.0E-02 | 2.3E-02 | BT20/MB468 | CYTO |
| >MI0003188\|hsa-mir-503&WithFlank&X\|-\|134546322\|134546404@12.31.20(+3A)&[MIMAT0002874&hsa-miR-503-5p&offsets\|0\|-3(+3A);m-297&X\|-\|134546371\|134546393&offsets\|0\|-3(+3A)]\|TAGCAGCGGGAACAGTTCTGAAA | >MI0003188\|hsa-mir-503&WithFlank&X\|-\|134546322\|134546404 | 12 | 31 | 20(+3A) | TAGCAGCGGGAACAGTTCTGAAA | 236.4 | 2.4 | 0.9 | 2.7 | 7.2E-03 | 2.3E-02 | BT20/MB231 | CYTO |
| >hg38_21_+_8433172_8446622_RNA45SN1_45S_50ntFlanks@9549.9571.23\|AGGTCCGTAGCGGTCCTGACGTG | >hg38_21_+_8433172_8446622_RNA45SN1_45S_50ntFlanks | 9549 | 9571 | 23 | AGGTCCGTAGCGGTCCTGACGTG | 161.1 | 3.0 | 1.1 | 2.8 | 5.4E-03 | 2.3E-02 | BT20/MB231 | MP |
| >MI0000070\|hsa-mir-16-1&WithFlank&13\|-\|50048967\|50049067@20.39.20(+1U);MI0000115\|hsa-mir-16-2&WithFlank&3\|+\|160404739\|160404831@16.35.20(+1U)&[MIMAT0000069&hsa-miR-16-5p&offsets\|0\|-2(+1U);m-62&13\|-\|50049027\|50049048&offsets\|0\|-2(+1U)];[MIMAT0000069_1&hsa-miR-16-5p&offsets\|0\|-2(+1U);m-61&3\|+\|160404754\|160404775&offsets\|0\|-2(+1U)]\|TAGCAGCACGTAAATATTGGT | >MI0000070\|hsa-mir-16-1&WithFlank&13\|-\|50048967\|50049067 | 20 | 39 | 20(+1U) | TAGCAGCACGTAAATATTGGT | 691.3 | 2.5 | 0.9 | 2.7 | 7.2E-03 | 2.3E-02 | BT20/MB231 | CYTO |
| >hg38_21_+_8433172_8446622_RNA45SN1_45S_50ntFlanks@5489.5532.44\|CTGAGAAGACGGTCGAACTTGACTATCTAGAGGAAGTAAAAGTC | >hg38_21_+_8433172_8446622_RNA45SN1_45S_50ntFlanks | 5489 | 5532 | 44 | CTGAGAAGACGGTCGAACTTGACTATCTAGAGGAAGTAAAAGTC | 713.7 | -2.7 | 1.0 | -2.7 | 7.2E-03 | 2.3E-02 | BT20/MB231 | CYTO |
| >hg38_21_+_8433172_8446622_RNA45SN1_45S_50ntFlanks@6648.6673.26\|GTACGACTCTTAGCGGTGGATCACTC | >hg38_21_+_8433172_8446622_RNA45SN1_45S_50ntFlanks | 6648 | 6673 | 26 | GTACGACTCTTAGCGGTGGATCACTC | 117.8 | 4.3 | 1.7 | 2.5 | 1.3E-02 | 2.3E-02 | BT20/MB468 | MP |
| >hg38_21_+_8433172_8446622_RNA45SN1_45S_50ntFlanks@11540.11583.44\|CGCGGCGCCTCGCCTCGGCCGGCGCCTAGCAGCCGACTTAGAAC | >hg38_21_+_8433172_8446622_RNA45SN1_45S_50ntFlanks | 11540 | 11583 | 44 | CGCGGCGCCTCGCCTCGGCCGGCGCCTAGCAGCCGACTTAGAAC | 800.5 | -2.5 | 0.9 | -2.8 | 5.9E-03 | 2.3E-02 | BT20/MB231 | NUC |
| >MI0000760\|hsa-mir-361&WithFlank&X\|-\|85903630\|85903713@52.75.24&[MIMAT0004682&hsa-miR-361-3p&offsets\|+1\|+2;m-124&X\|-\|85903642\|85903663&offsets\|+1\|+3]\|CCCCCAGGTGTGATTCTGATTTGC | >MI0000760\|hsa-mir-361&WithFlank&X\|-\|85903630\|85903713 | 52 | 75 | 24 | CCCCCAGGTGTGATTCTGATTTGC | 420.8 | -1.8 | 0.6 | -2.8 | 5.9E-03 | 2.3E-02 | BT20/MB231 | NUC |
| >MI0000298\|hsa-mir-221&WithFlank&X\|-\|45746151\|45746272@71.91.21(+1A)&[MIMAT0000278&hsa-miR-221-3p&offsets\|0\|-2(+1A);m-91&X\|-\|45746181\|45746202&offsets\|0\|-1(+1A)]\|AGCTACATTGTCTGCTGGGTTA | >MI0000298\|hsa-mir-221&WithFlank&X\|-\|45746151\|45746272 | 71 | 91 | 21(+1A) | AGCTACATTGTCTGCTGGGTTA | 3148.8 | -3.2 | 1.2 | -2.8 | 5.4E-03 | 2.3E-02 | BT20/MB231 | MP |
| >hg38_21_+_8433172_8446622_RNA45SN1_45S_50ntFlanks@8353.8373.21\|GTTGAAAAGAACTTTGAAGAG | >hg38_21_+_8433172_8446622_RNA45SN1_45S_50ntFlanks | 8353 | 8373 | 21 | GTTGAAAAGAACTTTGAAGAG | 2895.5 | 4.7 | 1.7 | 2.8 | 5.4E-03 | 2.3E-02 | BT20/MB231 | MP |
| >hg38_21_+_8433172_8446622_RNA45SN1_45S_50ntFlanks@11656.11685.30\|CGATGTGATTTCTGCCCAGTGCTCTGAATG | >hg38_21_+_8433172_8446622_RNA45SN1_45S_50ntFlanks | 11656 | 11685 | 30 | CGATGTGATTTCTGCCCAGTGCTCTGAATG | 158.4 | -1.5 | 0.6 | -2.3 | 2.1E-02 | 2.3E-02 | MB231/MB468 | NUC |
| >hg38_21_+_8433172_8446622_RNA45SN1_45S_50ntFlanks@12242.12268.27\|AGAAGGGCAAAAGCTCGCTTGATCTTG | >hg38_21_+_8433172_8446622_RNA45SN1_45S_50ntFlanks | 12242 | 12268 | 27 | AGAAGGGCAAAAGCTCGCTTGATCTTG | 419.2 | -2.5 | 1.0 | -2.5 | 1.3E-02 | 2.3E-02 | BT20/MB468 | MP |
| >hg38_21_+_8433172_8446622_RNA45SN1_45S_50ntFlanks@4069.4101.33\|CTTTCGATGGTAGTCGCCGTGCCTACCATGGTG | >hg38_21_+_8433172_8446622_RNA45SN1_45S_50ntFlanks | 4069 | 4101 | 33 | CTTTCGATGGTAGTCGCCGTGCCTACCATGGTG | 646.4 | -1.6 | 0.7 | -2.3 | 2.0E-02 | 2.3E-02 | BT20/MB468 | CYTO |
| >hg38_21_+_8433172_8446622_RNA45SN1_45S_50ntFlanks@10344.10361.18\|ATTCAAACGAGAACTTTG | >hg38_21_+_8433172_8446622_RNA45SN1_45S_50ntFlanks | 10344 | 10361 | 18 | ATTCAAACGAGAACTTTG | 465.0 | -2.2 | 0.9 | -2.5 | 1.3E-02 | 2.3E-02 | BT20/MB468 | MP |
| >hg38_21_+_8433172_8446622_RNA45SN1_45S_50ntFlanks@12417.12434.18\|TCGATGTCGGCTCTTCCT | >hg38_21_+_8433172_8446622_RNA45SN1_45S_50ntFlanks | 12417 | 12434 | 18 | TCGATGTCGGCTCTTCCT | 997.5 | -2.5 | 1.0 | -2.5 | 1.3E-02 | 2.3E-02 | BT20/MB468 | MP |
| >MI0000811\|hsa-mir-148b&WithFlank&12\|+\|54337210\|54337320@69.91.23&[MIMAT0000759&hsa-miR-148b-3p&offsets\|0\|+1;m-77&12\|+\|54337278\|54337299&offsets\|0\|+1]\|TCAGTGCATCACAGAACTTTGTC | >MI0000811\|hsa-mir-148b&WithFlank&12\|+\|54337210\|54337320 | 69 | 91 | 23 | TCAGTGCATCACAGAACTTTGTC | 857.0 | -2.4 | 0.9 | -2.7 | 6.7E-03 | 2.3E-02 | BT20/MB231 | MITO |
| >hg38_21_+_8433172_8446622_RNA45SN1_45S_50ntFlanks@8380.8407.28\|CAAGAGGGCGTGAAACCGTTAAGAGGTA | >hg38_21_+_8433172_8446622_RNA45SN1_45S_50ntFlanks | 8380 | 8407 | 28 | CAAGAGGGCGTGAAACCGTTAAGAGGTA | 183.1 | -1.2 | 0.5 | -2.3 | 2.0E-02 | 2.4E-02 | MB231/MB468 | CYTO |
| >MI0000088\|hsa-mir-30a&WithFlank&6\|-\|71403545\|71403627@54.74.21&[MIMAT0000088&hsa-miR-30a-3p&offsets\|+1\|0;m-38&6\|-\|71403554\|71403575&offsets\|+1\|0]\|TTTCAGTCGGATGTTTGCAGC | >MI0000088\|hsa-mir-30a&WithFlank&6\|-\|71403545\|71403627 | 54 | 74 | 21 | TTTCAGTCGGATGTTTGCAGC | 596.1 | 2.4 | 1.0 | 2.5 | 1.3E-02 | 2.4E-02 | BT20/MB468 | MP |
| >hg38_21_+_8433172_8446622_RNA45SN1_45S_50ntFlanks@7973.8015.43\|GACGCGACCTCAGATCAGACGTGGCGACCCGCTGAATTTAAGC | >hg38_21_+_8433172_8446622_RNA45SN1_45S_50ntFlanks | 7973 | 8015 | 43 | GACGCGACCTCAGATCAGACGTGGCGACCCGCTGAATTTAAGC | 139.2 | 4.0 | 1.7 | 2.3 | 2.1E-02 | 2.4E-02 | BT20/MB468 | NUC |
| >hg38_21_+_8433172_8446622_RNA45SN1_45S_50ntFlanks@11703.11724.22\|AATGAAGCGCGGGTAAACGGCG | >hg38_21_+_8433172_8446622_RNA45SN1_45S_50ntFlanks | 11703 | 11724 | 22 | AATGAAGCGCGGGTAAACGGCG | 172.6 | -4.5 | 1.8 | -2.5 | 1.3E-02 | 2.4E-02 | BT20/MB468 | MP |
| >MI0000465\|hsa-mir-191&WithFlank&3\|-\|49020612\|49020715@22.41.20&[MIMAT0000440&hsa-miR-191-5p&offsets\|0\|-3;m-48&3\|-\|49020672\|49020694&offsets\|0\|-3]\|CAACGGAATCCCAAAAGCAG | >MI0000465\|hsa-mir-191&WithFlank&3\|-\|49020612\|49020715 | 22 | 41 | 20 | CAACGGAATCCCAAAAGCAG | 63.7 | 4.5 | 1.8 | 2.5 | 1.3E-02 | 2.4E-02 | BT20/MB468 | MP |
| >hg38_21_+_8433172_8446622_RNA45SN1_45S_50ntFlanks@12006.12035.30\|GGCCGCCGGTGAAATACCACTACTCTGATC | >hg38_21_+_8433172_8446622_RNA45SN1_45S_50ntFlanks | 12006 | 12035 | 30 | GGCCGCCGGTGAAATACCACTACTCTGATC | 164.9 | 3.4 | 1.4 | 2.5 | 1.3E-02 | 2.4E-02 | BT20/MB468 | MP |
| >hg38_21_+_8433172_8446622_RNA45SN1_45S_50ntFlanks@6771.6806.36\|GGTTCCTCCCGGGGCTACGCCTGTCTGAGCGTCGCT | >hg38_21_+_8433172_8446622_RNA45SN1_45S_50ntFlanks | 6771 | 6806 | 36 | GGTTCCTCCCGGGGCTACGCCTGTCTGAGCGTCGCT | 1347.3 | -2.1 | 0.9 | -2.5 | 1.3E-02 | 2.4E-02 | BT20/MB468 | MP |
| >trnaMT_ProTGG_MT_-_15956_16023@51.70.20\|TAAAGACTTTTTCTCTGACC | >trnaMT_ProTGG_MT_-_15956_16023 | 51 | 70 | 20 | TAAAGACTTTTTCTCTGACC | 144.3 | 2.5 | 1.1 | 2.3 | 2.1E-02 | 2.4E-02 | MB231/MB468 | MITO |
| >hg38_21_+_8433172_8446622_RNA45SN1_45S_50ntFlanks@11772.11794.23\|TTAGTGACGCGCATGAATGGATG | >hg38_21_+_8433172_8446622_RNA45SN1_45S_50ntFlanks | 11772 | 11794 | 23 | TTAGTGACGCGCATGAATGGATG | 563.5 | -2.4 | 0.9 | -2.7 | 7.3E-03 | 2.4E-02 | BT20/MB231 | CYTO |
| >hg38_21_+_8433172_8446622_RNA45SN1_45S_50ntFlanks@12447.12476.30\|CAGAATTCACCAAGCGTTGGATTGTTCACC | >hg38_21_+_8433172_8446622_RNA45SN1_45S_50ntFlanks | 12447 | 12476 | 30 | CAGAATTCACCAAGCGTTGGATTGTTCACC | 104.5 | -2.7 | 1.2 | -2.4 | 1.8E-02 | 2.4E-02 | BT20/MB468 | MITO |
| >hg38_21_+_8433172_8446622_RNA45SN1_45S_50ntFlanks@12942.12959.18\|TTCGTAGACGACCTGCTT | >hg38_21_+_8433172_8446622_RNA45SN1_45S_50ntFlanks | 12942 | 12959 | 18 | TTCGTAGACGACCTGCTT | 154.8 | -1.5 | 0.6 | -2.4 | 1.8E-02 | 2.4E-02 | BT20/MB468 | MITO |
| >trnaMT_ProTGG_MT_-_15956_16023@1.30.30\|CAGAGAATAGTTTAAATTAGAATCTTAGCT | >trnaMT_ProTGG_MT_-_15956_16023 | 1 | 30 | 30 | CAGAGAATAGTTTAAATTAGAATCTTAGCT | 267.3 | 2.4 | 1.0 | 2.3 | 1.9E-02 | 2.4E-02 | MB231/MB468 | MP |
| >MI0000077\|hsa-mir-21&WithFlank&17\|+\|59841260\|59841343@52.71.20(+1A)&[MIMAT0004494&hsa-miR-21-3p&offsets\|0\|-1(+1A);m-52&17\|+\|59841311\|59841332&offsets\|0\|-2(+1A)]\|CAACACCAGTCGATGGGCTGA | >MI0000077\|hsa-mir-21&WithFlank&17\|+\|59841260\|59841343 | 52 | 71 | 20(+1A) | CAACACCAGTCGATGGGCTGA | 76.0 | 6.4 | 2.6 | 2.5 | 1.3E-02 | 2.4E-02 | BT20/MB468 | MP |
| >hg38_21_+_8433172_8446622_RNA45SN1_45S_50ntFlanks@8380.8407.28\|CAAGAGGGCGTGAAACCGTTAAGAGGTA | >hg38_21_+_8433172_8446622_RNA45SN1_45S_50ntFlanks | 8380 | 8407 | 28 | CAAGAGGGCGTGAAACCGTTAAGAGGTA | 133.8 | -2.3 | 1.0 | -2.3 | 2.0E-02 | 2.4E-02 | BT20/MB468 | CYTO |
| >hg38_21_+_8433172_8446622_RNA45SN1_45S_50ntFlanks@10240.10258.19\|CTACGCCGCGACGAGTAGG | >hg38_21_+_8433172_8446622_RNA45SN1_45S_50ntFlanks | 10240 | 10258 | 19 | CTACGCCGCGACGAGTAGG | 255.1 | 2.4 | 1.0 | 2.5 | 1.3E-02 | 2.4E-02 | BT20/MB468 | MP |
| >trna20_GluTTC_2_-_131094701_131094772@1.33.33,trna5_GluTTC_13_-_41634874_41634945@1.33.33\|TCCCATATGGTCTAGCGGTTAGGATTCCTGGTT | >trna20_GluTTC_2_-_131094701_131094772 | 1 | 33 | 33 | TCCCATATGGTCTAGCGGTTAGGATTCCTGGTT | 2921.6 | -3.4 | 1.2 | -2.8 | 5.5E-03 | 2.4E-02 | BT20/MB231 | MP |
| >MI0000082\|hsa-mir-25&WithFlank&7\|-\|100093554\|100093649@59.79.21&[MIMAT0000081&hsa-miR-25-3p&offsets\|+1\|0;m-23&7\|-\|100093571\|100093592&offsets\|+1\|0]\|ATTGCACTTGTCTCGGTCTGA | >MI0000082\|hsa-mir-25&WithFlank&7\|-\|100093554\|100093649 | 59 | 79 | 21 | ATTGCACTTGTCTCGGTCTGA | 50.9 | 4.0 | 1.7 | 2.3 | 1.9E-02 | 2.4E-02 | MB231/MB468 | MP |
| >MI0000441\|hsa-mir-30b&WithFlank&8\|-\|134800514\|134800613@23.44.22&[MIMAT0000420&hsa-miR-30b-5p&offsets\|0\|0;m-76&8\|-\|134800570\|134800591&offsets\|0\|0]\|TGTAAACATCCTACACTCAGCT | >MI0000441\|hsa-mir-30b&WithFlank&8\|-\|134800514\|134800613 | 23 | 44 | 22 | TGTAAACATCCTACACTCAGCT | 1678.4 | 2.6 | 1.1 | 2.3 | 1.9E-02 | 2.4E-02 | MB231/MB468 | MP |
| >hg38_21_+_8433172_8446622_RNA45SN1_45S_50ntFlanks@9952.9978.27\|CATGGAAGTCGGAATCCGCTAAGGAGT | >hg38_21_+_8433172_8446622_RNA45SN1_45S_50ntFlanks | 9952 | 9978 | 27 | CATGGAAGTCGGAATCCGCTAAGGAGT | 279.6 | 3.9 | 1.4 | 2.8 | 5.5E-03 | 2.4E-02 | BT20/MB231 | MP |

<table>
<tr><th></th><th></th><th></th><th></th><th></th><th></th><th></th><th></th><th></th><th></th><th></th><th></th><th></th><th></th></tr>
<tr><td>>hg38_21_+_8433172_8446622_RNA45SN1_45S_50ntFlanks@10409.10436.28|GTCAGTCGGTCCTGAGAGATGGGCGAGC</td><td>>hg38_21_+_8433172_8446622_RNA45SN1_45S_50ntFlanks</td><td>10409</td><td>10436</td><td>28</td><td>GTCAGTCGGTCCTGAGAGATGGGCGAGC</td><td>145.6</td><td>4.9</td><td>1.8</td><td>2.8</td><td>5.5E-03</td><td>2.4E-02</td><td>BT20/MB231</td><td>MP</td></tr>
<tr><td>>hg38_21_+_8433172_8446622_RNA45SN1_45S_50ntFlanks@8353.8375.23|GTTGAAAAGAACTTTGAAGAGAG</td><td>>hg38_21_+_8433172_8446622_RNA45SN1_45S_50ntFlanks</td><td>8353</td><td>8375</td><td>23</td><td>GTTGAAAAGAACTTTGAAGAGAG</td><td>182.9</td><td>4.4</td><td>1.9</td><td>2.3</td><td>1.9E-02</td><td>2.4E-02</td><td>MB231/MB468</td><td>MP</td></tr>
<tr><td>>hg38_21_+_8433172_8446622_RNA45SN1_45S_50ntFlanks@10423.10453.31|AGAGATGGGCGAGCGCCGTTCCGAAGGGACG</td><td>>hg38_21_+_8433172_8446622_RNA45SN1_45S_50ntFlanks</td><td>10423</td><td>10453</td><td>31</td><td>AGAGATGGGCGAGCGCCGTTCCGAAGGGACG</td><td>68.4</td><td>-4.1</td><td>1.7</td><td>-2.5</td><td>1.3E-02</td><td>2.4E-02</td><td>BT20/MB468</td><td>MP</td></tr>
<tr><td>>hg38_21_+_8433172_8446622_RNA45SN1_45S_50ntFlanks@5470.5499.30|CCACGGCCCTGGCGGAGCGCTGAGAAGACG</td><td>>hg38_21_+_8433172_8446622_RNA45SN1_45S_50ntFlanks</td><td>5470</td><td>5499</td><td>30</td><td>CCACGGCCCTGGCGGAGCGCTGAGAAGACG</td><td>760.0</td><td>-4.5</td><td>1.8</td><td>-2.5</td><td>1.3E-02</td><td>2.4E-02</td><td>BT20/MB468</td><td>MP</td></tr>
<tr><td>>hg38_21_+_8433172_8446622_RNA45SN1_45S_50ntFlanks@11669.11702.34|GCCCAGTGCTCTGAATGTCAAAGTGAAGAAATTC</td><td>>hg38_21_+_8433172_8446622_RNA45SN1_45S_50ntFlanks</td><td>11669</td><td>11702</td><td>34</td><td>GCCCAGTGCTCTGAATGTCAAAGTGAAGAAATTC</td><td>1482.9</td><td>1.9</td><td>0.8</td><td>2.3</td><td>2.1E-02</td><td>2.4E-02</td><td>MB231/MB468</td><td>NUC</td></tr>
<tr><td>>hg38_21_+_8433172_8446622_RNA45SN1_45S_50ntFlanks@12488.12512.25|AACGTGAGCTGGGTTTAGACCGTCG</td><td>>hg38_21_+_8433172_8446622_RNA45SN1_45S_50ntFlanks</td><td>12488</td><td>12512</td><td>25</td><td>AACGTGAGCTGGGTTTAGACCGTCG</td><td>106.7</td><td>-4.4</td><td>1.8</td><td>-2.5</td><td>1.3E-02</td><td>2.4E-02</td><td>BT20/MB468</td><td>MP</td></tr>
<tr><td>>hg38_21_+_8433172_8446622_RNA45SN1_45S_50ntFlanks@6652.6681.30|GACTCTTAGCGGTGGATCACTCGGCTCGTG</td><td>>hg38_21_+_8433172_8446622_RNA45SN1_45S_50ntFlanks</td><td>6652</td><td>6681</td><td>30</td><td>GACTCTTAGCGGTGGATCACTCGGCTCGTG</td><td>20464.6</td><td>-1.4</td><td>0.6</td><td>-2.4</td><td>1.8E-02</td><td>2.4E-02</td><td>BT20/MB468</td><td>MITO</td></tr>
<tr><td>>hg38_21_+_8433172_8446622_RNA45SN1_45S_50ntFlanks@7990.8012.23|GACGTGGCGACCCGCTGAATTTA</td><td>>hg38_21_+_8433172_8446622_RNA45SN1_45S_50ntFlanks</td><td>7990</td><td>8012</td><td>23</td><td>GACGTGGCGACCCGCTGAATTTA</td><td>354.9</td><td>-2.3</td><td>1.0</td><td>-2.3</td><td>2.1E-02</td><td>2.4E-02</td><td>BT20/MB468</td><td>NUC</td></tr>
<tr><td>>hg38_21_+_8433172_8446622_RNA45SN1_45S_50ntFlanks@3987.4024.38|GCTTTGGTGACTCTAGATAACCTCGGGCCGATCGCACG</td><td>>hg38_21_+_8433172_8446622_RNA45SN1_45S_50ntFlanks</td><td>3987</td><td>4024</td><td>38</td><td>GCTTTGGTGACTCTAGATAACCTCGGGCCGATCGCACG</td><td>2058.2</td><td>-3.5</td><td>1.4</td><td>-2.5</td><td>1.3E-02</td><td>2.4E-02</td><td>BT20/MB468</td><td>MP</td></tr>
<tr><td>>hg38_21_+_8433172_8446622_RNA45SN1_45S_50ntFlanks@6645.6669.25|CTCGTACGACTCTTAGCGGTGGATC</td><td>>hg38_21_+_8433172_8446622_RNA45SN1_45S_50ntFlanks</td><td>6645</td><td>6669</td><td>25</td><td>CTCGTACGACTCTTAGCGGTGGATC</td><td>2936.8</td><td>1.8</td><td>0.7</td><td>2.5</td><td>1.3E-02</td><td>2.4E-02</td><td>BT20/MB468</td><td>MP</td></tr>
<tr><td>>hg38_21_+_8433172_8446622_RNA45SN1_45S_50ntFlanks@8354.8375.22|TTGAAAAGAACTTTGAAGAGAG</td><td>>hg38_21_+_8433172_8446622_RNA45SN1_45S_50ntFlanks</td><td>8354</td><td>8375</td><td>22</td><td>TTGAAAAGAACTTTGAAGAGAG</td><td>639.6</td><td>6.9</td><td>2.5</td><td>2.7</td><td>6.8E-03</td><td>2.4E-02</td><td>BT20/MB231</td><td>MITO</td></tr>
<tr><td>>hg38_21_+_8433172_8446622_RNA45SN1_45S_50ntFlanks@12418.12456.39|CGATGTCGGCTCTTCCTATCATTGTGAAGCAGAATTCAC</td><td>>hg38_21_+_8433172_8446622_RNA45SN1_45S_50ntFlanks</td><td>12418</td><td>12456</td><td>39</td><td>CGATGTCGGCTCTTCCTATCATTGTGAAGCAGAATTCAC</td><td>309.3</td><td>-2.3</td><td>0.9</td><td>-2.7</td><td>6.8E-03</td><td>2.4E-02</td><td>BT20/MB231</td><td>MITO</td></tr>
<tr><td>>hg38_21_+_8433172_8446622_RNA45SN1_45S_50ntFlanks@9564.9594.31|CTGACGTGCAAATCGGTCGTCCGACCTGGGT</td><td>>hg38_21_+_8433172_8446622_RNA45SN1_45S_50ntFlanks</td><td>9564</td><td>9594</td><td>31</td><td>CTGACGTGCAAATCGGTCGTCCGACCTGGGT</td><td>315.5</td><td>-2.2</td><td>0.9</td><td>-2.5</td><td>1.3E-02</td><td>2.4E-02</td><td>BT20/MB468</td><td>MP</td></tr>
<tr><td>>hg38_21_+_8433172_8446622_RNA45SN1_45S_50ntFlanks@12259.12287.29|CTTGATCTTGATTTTCAGTACGAATACAG</td><td>>hg38_21_+_8433172_8446622_RNA45SN1_45S_50ntFlanks</td><td>12259</td><td>12287</td><td>29</td><td>CTTGATCTTGATTTTCAGTACGAATACAG</td><td>115.9</td><td>-3.2</td><td>1.3</td><td>-2.5</td><td>1.3E-02</td><td>2.4E-02</td><td>BT20/MB468</td><td>MP</td></tr>
<tr><td>>trnaMT_PheGAA_MT_+_577_647@38.65.28|AATGTTTAGACGGGCTCACATCACCCCA</td><td>>trnaMT_PheGAA_MT_+_577_647</td><td>38</td><td>65</td><td>28</td><td>AATGTTTAGACGGGCTCACATCACCCCA</td><td>232.5</td><td>4.9</td><td>2.0</td><td>2.5</td><td>1.3E-02</td><td>2.4E-02</td><td>BT20/MB468</td><td>MP</td></tr>
<tr><td>>MI0000298|hsa-mir-221&WithFlank&X|-|45746151|45746272@71.92.22&[MIMAT0000278&hsa-miR-221-3p&offsets|0|-1;m-91&X|-|45746181|45746202&offsets|0|0]|AGCTACATTGTCTGCTGGGTTT</td><td>>MI0000298|hsa-mir-221&WithFlank&X|-|45746151|45746272</td><td>71</td><td>92</td><td>22</td><td>AGCTACATTGTCTGCTGGGTTT</td><td>75842.1</td><td>-2.9</td><td>1.1</td><td>-2.8</td><td>5.5E-03</td><td>2.4E-02</td><td>BT20/MB231</td><td>MP</td></tr>
<tr><td>>hg38_MT_+_1621_3279_RNR2_16S_50ntFlanks@1367.1402.36|TCGATGTTGGATCAGGACATCCCGATGGTGCAGCCG</td><td>>hg38_MT_+_1621_3279_RNR2_16S_50ntFlanks</td><td>1367</td><td>1402</td><td>36</td><td>TCGATGTTGGATCAGGACATCCCGATGGTGCAGCCG</td><td>66.6</td><td>5.3</td><td>2.2</td><td>2.5</td><td>1.3E-02</td><td>2.4E-02</td><td>BT20/MB468</td><td>MP</td></tr>
<tr><td>>MI0003205|hsa-mir-532&WithFlank&X|+|50003142|50003244@26.47.22&[MIMAT0002888&hsa-miR-532-5p&offsets|0|0;m-65&X|+|50003167|50003188&offsets|0|0]|CATGCCTTGAGTGTAGGACCGT</td><td>>MI0003205|hsa-mir-532&WithFlank&X|+|50003142|50003244</td><td>26</td><td>47</td><td>22</td><td>CATGCCTTGAGTGTAGGACCGT</td><td>4766.7</td><td>2.0</td><td>0.8</td><td>2.7</td><td>7.3E-03</td><td>2.4E-02</td><td>BT20/MB231</td><td>CYTO</td></tr>
<tr><td>>MI0001445|hsa-mir-423&WithFlank&17|+|30117073|30117178@23.42.20(+1G)&[MIMAT0004748&hsa-miR-423-5p&offsets|0|-3(+1G);m-141&17|+|30117095|30117117&offsets|0|-3(+1G)]|TGAGGGGCAGAGAGCGAGACG</td><td>>MI0001445|hsa-mir-423&WithFlank&17|+|30117073|30117178</td><td>23</td><td>42</td><td>20(+1G)</td><td>TGAGGGGCAGAGAGCGAGACG</td><td>44.7</td><td>4.7</td><td>2.0</td><td>2.3</td><td>1.9E-02</td><td>2.4E-02</td><td>MB231/MB468</td><td>MP</td></tr>
<tr><td>>hg38_21_+_8433172_8446622_RNA45SN1_45S_50ntFlanks@10241.10258.18|TACGCCGCGACGAGTAGG</td><td>>hg38_21_+_8433172_8446622_RNA45SN1_45S_50ntFlanks</td><td>10241</td><td>10258</td><td>18</td><td>TACGCCGCGACGAGTAGG</td><td>88.1</td><td>4.3</td><td>1.7</td><td>2.5</td><td>1.3E-02</td><td>2.4E-02</td><td>BT20/MB468</td><td>MP</td></tr>
<tr><td>>hg38_21_+_8433172_8446622_RNA45SN1_45S_50ntFlanks@11920.11956.37|ATGAGAGGTGTAGAATAAGTGGGAGGCCCCCGGCGCC</td><td>>hg38_21_+_8433172_8446622_RNA45SN1_45S_50ntFlanks</td><td>11920</td><td>11956</td><td>37</td><td>ATGAGAGGTGTAGAATAAGTGGGAGGCCCCCGGCGCC</td><td>194.1</td><td>-3.8</td><td>1.7</td><td>-2.3</td><td>2.1E-02</td><td>2.4E-02</td><td>MB231/MB468</td><td>NUC</td></tr>
<tr><td>>trnaMT_GlnTTG_MT_-_4329_4400@57.74.18,trnalookalike2_GlnTTG_1_-_564879_564950@57.74.18|ATTCTCATAGTCCTAGCC</td><td>>trnaMT_GlnTTG_MT_-_4329_4400</td><td>57</td><td>74</td><td>18</td><td>ATTCTCATAGTCCTAGCC</td><td>84.4</td><td>3.2</td><td>1.4</td><td>2.3</td><td>1.9E-02</td><td>2.4E-02</td><td>MB231/MB468</td><td>MP</td></tr>
<tr><td>>hg38_21_+_8433172_8446622_RNA45SN1_45S_50ntFlanks@12633.12659.27|CGAAGCTACCATCTGTGGGATTATGAC</td><td>>hg38_21_+_8433172_8446622_RNA45SN1_45S_50ntFlanks</td><td>12633</td><td>12659</td><td>27</td><td>CGAAGCTACCATCTGTGGGATTATGAC</td><td>2534.4</td><td>-3.1</td><td>1.2</td><td>-2.5</td><td>1.4E-02</td><td>2.4E-02</td><td>BT20/MB468</td><td>MP</td></tr>
<tr><td>>hg38_21_+_8433172_8446622_RNA45SN1_45S_50ntFlanks@9306.9334.29|CAAGGAGTCTAACACGTGCGCGAGTCGGG</td><td>>hg38_21_+_8433172_8446622_RNA45SN1_45S_50ntFlanks</td><td>9306</td><td>9334</td><td>29</td><td>CAAGGAGTCTAACACGTGCGCGAGTCGGG</td><td>590.9</td><td>-2.4</td><td>1.0</td><td>-2.5</td><td>1.4E-02</td><td>2.4E-02</td><td>BT20/MB468</td><td>MP</td></tr>
<tr><td>>hg38_21_+_8433172_8446622_RNA45SN1_45S_50ntFlanks@11918.11941.24|ACATGAGAGGTGTAGAATAAGTGG</td><td>>hg38_21_+_8433172_8446622_RNA45SN1_45S_50ntFlanks</td><td>11918</td><td>11941</td><td>24</td><td>ACATGAGAGGTGTAGAATAAGTGG</td><td>71.9</td><td>5.1</td><td>2.1</td><td>2.5</td><td>1.4E-02</td><td>2.4E-02</td><td>BT20/MB468</td><td>MP</td></tr>
<tr><td>>trnaMT_ValTAC_MT_+_1602_1670@52.72.21|TTAACTTGACCGCTCTGACCA</td><td>>trnaMT_ValTAC_MT_+_1602_1670</td><td>52</td><td>72</td><td>21</td><td>TTAACTTGACCGCTCTGACCA</td><td>426.4</td><td>2.5</td><td>1.0</td><td>2.5</td><td>1.4E-02</td><td>2.4E-02</td><td>BT20/MB468</td><td>MP</td></tr>
<tr><td>>hg38_21_+_8433172_8446622_RNA45SN1_45S_50ntFlanks@6652.6675.24|GACTCTTAGCGGTGGATCACTCGG</td><td>>hg38_21_+_8433172_8446622_RNA45SN1_45S_50ntFlanks</td><td>6652</td><td>6675</td><td>24</td><td>GACTCTTAGCGGTGGATCACTCGG</td><td>507.6</td><td>2.8</td><td>1.1</td><td>2.5</td><td>1.4E-02</td><td>2.4E-02</td><td>BT20/MB468</td><td>MP</td></tr>
<tr><td>>hg38_21_+_8433172_8446622_RNA45SN1_45S_50ntFlanks@6661.6697.37|CGGTGGATCACTCGGCTCGTGCGTCGATGAAGAACGC</td><td>>hg38_21_+_8433172_8446622_RNA45SN1_45S_50ntFlanks</td><td>6661</td><td>6697</td><td>37</td><td>CGGTGGATCACTCGGCTCGTGCGTCGATGAAGAACGC</td><td>2321.0</td><td>2.3</td><td>0.9</td><td>2.5</td><td>1.4E-02</td><td>2.4E-02</td><td>BT20/MB468</td><td>MP</td></tr>
<tr><td>>hg38_21_+_8433172_8446622_RNA45SN1_45S_50ntFlanks@3838.3859.22|CTCTCCTACTTGGATAACTGTG</td><td>>hg38_21_+_8433172_8446622_RNA45SN1_45S_50ntFlanks</td><td>3838</td><td>3859</td><td>22</td><td>CTCTCCTACTTGGATAACTGTG</td><td>186.7</td><td>-1.7</td><td>0.7</td><td>-2.3</td><td>2.0E-02</td><td>2.4E-02</td><td>MB231/MB468</td><td>CYTO</td></tr>
<tr><td>>hg38_21_+_8433172_8446622_RNA45SN1_45S_50ntFlanks@11729.11765.37|TAACTATGACTCTCTTAAGGTAGCCAAATGCCTCGTC</td><td>>hg38_21_+_8433172_8446622_RNA45SN1_45S_50ntFlanks</td><td>11729</td><td>11765</td><td>37</td><td>TAACTATGACTCTCTTAAGGTAGCCAAATGCCTCGTC</td><td>623.9</td><td>-4.6</td><td>1.9</td><td>-2.5</td><td>1.4E-02</td><td>2.4E-02</td><td>BT20/MB468</td><td>MP</td></tr>
<tr><td>>hg38_21_+_8433172_8446622_RNA45SN1_45S_50ntFlanks@12907.12931.25|CACCGCACGTTCGTGGGGAACCTGG</td><td>>hg38_21_+_8433172_8446622_RNA45SN1_45S_50ntFlanks</td><td>12907</td><td>12931</td><td>25</td><td>CACCGCACGTTCGTGGGGAACCTGG</td><td>160.6</td><td>-2.0</td><td>0.8</td><td>-2.4</td><td>1.8E-02</td><td>2.4E-02</td><td>BT20/MB468</td><td>MITO</td></tr>
<tr><td>>MI0000736|hsa-mir-30c-1&WithFlank&1|+|40757278|40757378@23.45.23(+1C);MI0000254|hsa-mir-30c-2&WithFlank&6|-|71376954|71377037@13.35.23(+1C)&[MIMAT0000244&hsa-miR-30c-5p&offsets|0|0(+1C);m-57&1|+|40757300|40757323&offsets|0|-1(+1C)];[MIMAT0000244_1&hsa-miR-30c-5p&offsets|0|0(+1C);m-56&6|-|71377002|71377025&offsets|0|-1(+1C)]|TGTAAACATCCTACACTCTCAGCC</td><td>>MI0000736|hsa-mir-30c-1&WithFlank&1|+|40757278|40757378</td><td>23</td><td>45</td><td>23(+1C)</td><td>TGTAAACATCCTACACTCTCAGCC</td><td>2688.4</td><td>-3.2</td><td>1.2</td><td>-2.8</td><td>5.5E-03</td><td>2.4E-02</td><td>BT20/MB231</td><td>MP</td></tr>
<tr><td>>hg38_21_+_8433172_8446622_RNA45SN1_45S_50ntFlanks@5482.5501.20|CGGAGCGCTGAGAAGACGGT</td><td>>hg38_21_+_8433172_8446622_RNA45SN1_45S_50ntFlanks</td><td>5482</td><td>5501</td><td>20</td><td>CGGAGCGCTGAGAAGACGGT</td><td>117.7</td><td>2.1</td><td>0.9</td><td>2.5</td><td>1.4E-02</td><td>2.4E-02</td><td>BT20/MB468</td><td>MP</td></tr>
<tr><td>>MI0000115|hsa-mir-16-2&WithFlank&3|+|160404739|160404831@58.80.23&[MIMAT0004518&hsa-miR-16-2-3p&offsets|-1|0;m-252&3|+|160404796|160404816&offsets|0|+2]|ACCAATATTACTGTGCTGCTTTA</td><td>>MI0000115|hsa-mir-16-2&WithFlank&3|+|160404739|160404831</td><td>58</td><td>80</td><td>23</td><td>ACCAATATTACTGTGCTGCTTTA</td><td>93.1</td><td>4.1</td><td>1.8</td><td>2.3</td><td>1.9E-02</td><td>2.4E-02</td><td>MB231/MB468</td><td>MP</td></tr>
<tr><td>>hg38_21_+_8433172_8446622_RNA45SN1_45S_50ntFlanks@4722.4759.38|TCATTAATCAAGAACGAAAGTCGGAGGTTCGAAGACGA</td><td>>hg38_21_+_8433172_8446622_RNA45SN1_45S_50ntFlanks</td><td>4722</td><td>4759</td><td>38</td><td>TCATTAATCAAGAACGAAAGTCGGAGGTTCGAAGACGA</td><td>4042.0</td><td>-2.3</td><td>1.0</td><td>-2.3</td><td>2.0E-02</td><td>2.4E-02</td><td>MB231/MB468</td><td>CYTO</td></tr>
<tr><td>>hg38_MT_+_1621_3279_RNR2_16S_50ntFlanks@1363.1388.26|GACCTCGATGTTGGATCAGGACATCC</td><td>>hg38_MT_+_1621_3279_RNR2_16S_50ntFlanks</td><td>1363</td><td>1388</td><td>26</td><td>GACCTCGATGTTGGATCAGGACATCC</td><td>284.2</td><td>3.1</td><td>1.2</td><td>2.5</td><td>1.4E-02</td><td>2.4E-02</td><td>BT20/MB468</td><td>MP</td></tr>
<tr><td>>MI0000070|hsa-mir-16-1&WithFlank&13|-|50048967|50049067@20.41.22(+1C);MI0000115|hsa-mir-16-2&WithFlank&3|+|160404739|160404831@16.37.22(+1C)&[MIMAT0000069&hsa-miR-16-5p&offsets|0|0(+1C);m-62&13|-|50049027|50049048&offsets|0|0(+1C)];[MIMAT0000069_1&hsa-miR-16-5p&offsets|0|0(+1C);m-61&3|+|160404754|160404775&offsets|0|0(+1C)]|TAGCAGCACGTAAATATTGGCGC</td><td>>MI0000070|hsa-mir-16-1&WithFlank&13|-|50048967|50049067</td><td>20</td><td>41</td><td>22(+1C)</td><td>TAGCAGCACGTAAATATTGGCGC</td><td>308.4</td><td>2.6</td><td>1.0</td><td>2.7</td><td>7.3E-03</td><td>2.4E-02</td><td>BT20/MB231</td><td>CYTO</td></tr>
<tr><td>>hg38_21_+_8433172_8446622_RNA45SN1_45S_50ntFlanks@6698.6719.22|AGCTAGCTGCGAGAATTAATGT</td><td>>hg38_21_+_8433172_8446622_RNA45SN1_45S_50ntFlanks</td><td>6698</td><td>6719</td><td>22</td><td>AGCTAGCTGCGAGAATTAATGT</td><td>964.8</td><td>-3.8</td><td>1.5</td><td>-2.5</td><td>1.4E-02</td><td>2.4E-02</td><td>BT20/MB468</td><td>MP</td></tr>
<tr><td>>hg38_21_+_8433172_8446622_RNA45SN1_45S_50ntFlanks@8194.8211.18|CCCGTGGACGGTGTGAGG</td><td>>hg38_21_+_8433172_8446622_RNA45SN1_45S_50ntFlanks</td><td>8194</td><td>8211</td><td>18</td><td>CCCGTGGACGGTGTGAGG</td><td>241.7</td><td>5.7</td><td>2.1</td><td>2.8</td><td>5.5E-03</td><td>2.4E-02</td><td>BT20/MB231</td><td>MP</td></tr>
<tr><td>>hg38_21_+_8433172_8446622_RNA45SN1_45S_50ntFlanks@3843.3872.30|CTACTTGGATAACTGTGGTAATTCTAGAGC</td><td>>hg38_21_+_8433172_8446622_RNA45SN1_45S_50ntFlanks</td><td>3843</td><td>3872</td><td>30</td><td>CTACTTGGATAACTGTGGTAATTCTAGAGC</td><td>204.0</td><td>-2.1</td><td>0.9</td><td>-2.4</td><td>1.8E-02</td><td>2.4E-02</td><td>BT20/MB468</td><td>MITO</td></tr>
<tr><td>>trna152_ValCAC_6_-_27248049_27248121@1.23.23|GCTTCTGTAGTGTAGTGGTTATC</td><td>>trna152_ValCAC_6_-_27248049_27248121</td><td>1</td><td>23</td><td>23</td><td>GCTTCTGTAGTGTAGTGGTTATC</td><td>814.5</td><td>3.3</td><td>1.4</td><td>2.5</td><td>1.4E-02</td><td>2.4E-02</td><td>BT20/MB468</td><td>MP</td></tr>
<tr><td>>MI0000093|hsa-mir-92a-1&WithFlank&13|+|91351308|91351397@54.75.22;MI0000094|hsa-mir-92a-2&WithFlank&X|-|134169532|134169618@54.75.22&[MIMAT0000092&hsa-miR-92a-3p&offsets|0|0;m-9&13|+|91351361|91351382&offsets|0|0];[MIMAT0000092_1&hsa-miR-92a-3p&offsets|0|0;m-11&X|-|134169544|134169565&offsets|0|0]|TATTGCACTTGTCCCGGCCTGT</td><td>>MI0000093|hsa-mir-92a-1&WithFlank&13|+|91351308|91351397</td><td>54</td><td>75</td><td>22</td><td>TATTGCACTTGTCCCGGCCTGT</td><td>23479.5</td><td>2.6</td><td>1.0</td><td>2.5</td><td>1.4E-02</td><td>2.4E-02</td><td>BT20/MB468</td><td>MP</td></tr>
<tr><td>>hg38_21_+_8433172_8446622_RNA45SN1_45S_50ntFlanks@6745.6769.25|ACACTTCGAACGCACTTGCGGCCCC</td><td>>hg38_21_+_8433172_8446622_RNA45SN1_45S_50ntFlanks</td><td>6745</td><td>6769</td><td>25</td><td>ACACTTCGAACGCACTTGCGGCCCC</td><td>251.7</td><td>-4.3</td><td>1.7</td><td>-2.5</td><td>1.4E-02</td><td>2.4E-02</td><td>BT20/MB468</td><td>MP</td></tr>
<tr><td>>trnaMT_GluTTC_MT_-_14674_14742@44.66.23,trnalookalike8_GluTTC_5_-_93905172_93905240@44.66.23|TCGTGGTTGTAGTCCGTGCGAGA</td><td>>trnaMT_GluTTC_MT_-_14674_14742</td><td>44</td><td>66</td><td>23</td><td>TCGTGGTTGTAGTCCGTGCGAGA</td><td>143.3</td><td>2.7</td><td>1.1</td><td>2.5</td><td>1.4E-02</td><td>2.4E-02</td><td>BT20/MB468</td><td>MP</td></tr>
<tr><td>>hg38_21_+_8433172_8446622_RNA45SN1_45S_50ntFlanks@12896.12932.37|CGCCCGTCACGCACCGCACGTTCGTGGGGAACCTGGC</td><td>>hg38_21_+_8433172_8446622_RNA45SN1_45S_50ntFlanks</td><td>12896</td><td>12932</td><td>37</td><td>CGCCCGTCACGCACCGCACGTTCGTGGGGAACCTGGC</td><td>241.5</td><td>-1.8</td><td>0.8</td><td>-2.4</td><td>1.8E-02</td><td>2.4E-02</td><td>BT20/MB468</td><td>MITO</td></tr>
<tr><td>>hg38_21_+_8433172_8446622_RNA45SN1_45S_50ntFlanks@9598.9616.19|GGGGCGAAAGACTAATCGA</td><td>>hg38_21_+_8433172_8446622_RNA45SN1_45S_50ntFlanks</td><td>9598</td><td>9616</td><td>19</td><td>GGGGCGAAAGACTAATCGA</td><td>130.5</td><td>-3.2</td><td>1.3</td><td>-2.5</td><td>1.4E-02</td><td>2.4E-02</td><td>BT20/MB468</td><td>MP</td></tr>
<tr><td>>MI0000683|hsa-mir-181b-2&WithFlank&9|+|124693704|124693804@22.46.25(+2U)&[MIMAT0000257_1&hsa-miR-181b-5p&offsets|0|+2(+2U);m-72&9|+|124693725|124693747&offsets|0|+2(+2U)]|AACATTCATTGCTGTCGGTGGGTTTTT</td><td>>MI0000683|hsa-mir-181b-2&WithFlank&9|+|124693704|124693804</td><td>22</td><td>46</td><td>25(+2U)</td><td>AACATTCATTGCTGTCGGTGGGTTTTT</td><td>116.8</td><td>-5.3</td><td>1.9</td><td>-2.8</td><td>5.6E-03</td><td>2.4E-02</td><td>BT20/MB231</td><td>MP</td></tr>
</table>

<table>
<tr><td>>trna119_LysCTT_1_-_145395522_145395594@2.33.32,trna11_LysCTT_5_-_180648979_180649051@2.33.32,trna13_LysCTT_6_+_26556774_26556846@2.33.32,trna7_LysCTT_16_+_3225692_3225764@2.33.32,trna9_LysCTT_5_+_180634755_180634827@2.33.32|CCCGGCTAGCTCAGTCGGTAGAGCATGAGACT</td><td>>trna119_LysCTT_1_-_145395522_145395594</td><td>2</td><td>33</td><td>32</td><td>CCCGGCTAGCTCAGTCGGTAGAGCATGAGACT</td><td>113.3</td><td>-5.2</td><td>1.9</td><td>-2.8</td><td>5.6E-03</td><td>2.4E-02</td><td>BT20/MB231</td><td>MP</td></tr>
<tr><td>>trnaMT_GluTTC_MT_-_14674_14742@46.70.25,trnalookalike8_GluTTC_5_-_93905172_93905240@46.70.25|GTGGTTGTAGTCCGTGCGAGAATAC</td><td>>trnaMT_GluTTC_MT_-_14674_14742</td><td>46</td><td>70</td><td>25</td><td>GTGGTTGTAGTCCGTGCGAGAATAC</td><td>77.5</td><td>4.6</td><td>2.0</td><td>2.3</td><td>1.9E-02</td><td>2.4E-02</td><td>MB231/MB468</td><td>MP</td></tr>
<tr><td>>hg38_21_+_8433172_8446622_RNA45SN1_45S_50ntFlanks@10404.10429.26|CATGGGTCAGTCGGTCCTGAGAGATG</td><td>>hg38_21_+_8433172_8446622_RNA45SN1_45S_50ntFlanks</td><td>10404</td><td>10429</td><td>26</td><td>CATGGGTCAGTCGGTCCTGAGAGATG</td><td>560.3</td><td>3.5</td><td>1.3</td><td>2.8</td><td>5.6E-03</td><td>2.4E-02</td><td>BT20/MB231</td><td>MP</td></tr>
<tr><td>>hg38_21_+_8433172_8446622_RNA45SN1_45S_50ntFlanks@10855.10873.19|AAGGATTGGCTCTAAGGGC</td><td>>hg38_21_+_8433172_8446622_RNA45SN1_45S_50ntFlanks</td><td>10855</td><td>10873</td><td>19</td><td>AAGGATTGGCTCTAAGGGC</td><td>275.0</td><td>3.6</td><td>1.3</td><td>2.8</td><td>5.6E-03</td><td>2.4E-02</td><td>BT20/MB231</td><td>MP</td></tr>
<tr><td>>hg38_21_+_8433172_8446622_RNA45SN1_45S_50ntFlanks@4734.4765.32|AACGAAAGTCGGAGGTTCGAAGACGATCAGAT</td><td>>hg38_21_+_8433172_8446622_RNA45SN1_45S_50ntFlanks</td><td>4734</td><td>4765</td><td>32</td><td>AACGAAAGTCGGAGGTTCGAAGACGATCAGAT</td><td>170.8</td><td>3.0</td><td>1.3</td><td>2.3</td><td>1.9E-02</td><td>2.4E-02</td><td>MB231/MB468</td><td>MP</td></tr>
<tr><td>>trnaMT_HisGTG_MT_+_12138_12206@20.53.34|AACATCAGATTGTGAATCTGACAACAGAGGCTTA</td><td>>trnaMT_HisGTG_MT_+_12138_12206</td><td>20</td><td>53</td><td>34</td><td>AACATCAGATTGTGAATCTGACAACAGAGGCTTA</td><td>94.5</td><td>3.9</td><td>1.7</td><td>2.3</td><td>1.9E-02</td><td>2.4E-02</td><td>MB231/MB468</td><td>MP</td></tr>
<tr><td>>hg38_21_+_8433172_8446622_RNA45SN1_45S_50ntFlanks@10248.10284.37|CGACGAGTAGGAGGGCCGCTGCGGTGAGCCTTGAAGC</td><td>>hg38_21_+_8433172_8446622_RNA45SN1_45S_50ntFlanks</td><td>10248</td><td>10284</td><td>37</td><td>CGACGAGTAGGAGGGCCGCTGCGGTGAGCCTTGAAGC</td><td>199.1</td><td>5.5</td><td>2.0</td><td>2.8</td><td>5.6E-03</td><td>2.4E-02</td><td>BT20/MB231</td><td>MP</td></tr>
<tr><td>>MI0000298|hsa-mir-221&WithFlank&X|-|45746151|45746272@71.90.20(+1C)&[MIMAT0000278&hsa-miR-221-3p&offsets|0|-3(+1C);m-91&X|-|45746181|45746202&offsets|0|-2(+1C)]|AGCTACATTGTCTGCTGGGTC</td><td>>MI0000298|hsa-mir-221&WithFlank&X|-|45746151|45746272</td><td>71</td><td>90</td><td>20(+1C)</td><td>AGCTACATTGTCTGCTGGGTC</td><td>1613.6</td><td>-3.2</td><td>1.2</td><td>-2.8</td><td>5.6E-03</td><td>2.4E-02</td><td>BT20/MB231</td><td>MP</td></tr>
<tr><td>>hg38_21_+_8433172_8446622_RNA45SN1_45S_50ntFlanks@11579.11606.28|AGAACTGGTGCGGACCAGGGGAATCCGA</td><td>>hg38_21_+_8433172_8446622_RNA45SN1_45S_50ntFlanks</td><td>11579</td><td>11606</td><td>28</td><td>AGAACTGGTGCGGACCAGGGGAATCCGA</td><td>269.2</td><td>-2.3</td><td>0.9</td><td>-2.5</td><td>1.4E-02</td><td>2.4E-02</td><td>BT20/MB468</td><td>MP</td></tr>
<tr><td>>hg38_21_+_8433172_8446622_RNA45SN1_45S_50ntFlanks@12420.12444.25|ATGTCGGCTCTTCCTATCATTGTGA</td><td>>hg38_21_+_8433172_8446622_RNA45SN1_45S_50ntFlanks</td><td>12420</td><td>12444</td><td>25</td><td>ATGTCGGCTCTTCCTATCATTGTGA</td><td>1250.5</td><td>-2.0</td><td>0.9</td><td>-2.3</td><td>2.2E-02</td><td>2.4E-02</td><td>MB231/MB468</td><td>MITO</td></tr>
<tr><td>>hg38_21_+_8433172_8446622_RNA45SN1_45S_50ntFlanks@7975.8010.36|CGCGACCTCAGATCAGACGTGGCGACCCGCTGAATT</td><td>>hg38_21_+_8433172_8446622_RNA45SN1_45S_50ntFlanks</td><td>7975</td><td>8010</td><td>36</td><td>CGCGACCTCAGATCAGACGTGGCGACCCGCTGAATT</td><td>616058.6</td><td>1.6</td><td>0.6</td><td>2.7</td><td>6.8E-03</td><td>2.4E-02</td><td>BT20/MB231</td><td>MITO</td></tr>
<tr><td>>MI0000078|hsa-mir-22&WithFlank&17|-|1713897|1713993@59.79.21(+1C)&[MIMAT0000077&hsa-miR-22-3p&offsets|0|-1(+1C);m-2&17|-|1713914|1713935&offsets|0|-1(+1C)]|AAGCTGCCAGTTGAAGAACTGC</td><td>>MI0000078|hsa-mir-22&WithFlank&17|-|1713897|1713993</td><td>59</td><td>79</td><td>21(+1C)</td><td>AAGCTGCCAGTTGAAGAACTGC</td><td>1931.7</td><td>2.6</td><td>1.0</td><td>2.7</td><td>7.4E-03</td><td>2.4E-02</td><td>BT20/MB231</td><td>CYTO</td></tr>
<tr><td>>hg38_21_+_8433172_8446622_RNA45SN1_45S_50ntFlanks@8171.8216.46|AAGTCCTTCTGATCGAGGCCCAGCCCGTGGACGGTGTGAGGCCGGT</td><td>>hg38_21_+_8433172_8446622_RNA45SN1_45S_50ntFlanks</td><td>8171</td><td>8216</td><td>46</td><td>AAGTCCTTCTGATCGAGGCCCAGCCCGTGGACGGTGTGAGGCCGGT</td><td>351.2</td><td>1.9</td><td>0.8</td><td>2.3</td><td>2.1E-02</td><td>2.4E-02</td><td>MB231/MB468</td><td>CYTO</td></tr>
<tr><td>>hg38_21_+_8433172_8446622_RNA45SN1_45S_50ntFlanks@10349.10366.18|AACGAGAACTTTGAAGGC</td><td>>hg38_21_+_8433172_8446622_RNA45SN1_45S_50ntFlanks</td><td>10349</td><td>10366</td><td>18</td><td>AACGAGAACTTTGAAGGC</td><td>1325.7</td><td>-3.7</td><td>1.5</td><td>-2.5</td><td>1.4E-02</td><td>2.4E-02</td><td>BT20/MB468</td><td>MP</td></tr>
<tr><td>>hg38_21_+_8433172_8446622_RNA45SN1_45S_50ntFlanks@10350.10369.20|ACGAGAACTTTGAAGGCCGA</td><td>>hg38_21_+_8433172_8446622_RNA45SN1_45S_50ntFlanks</td><td>10350</td><td>10369</td><td>20</td><td>ACGAGAACTTTGAAGGCCGA</td><td>605.8</td><td>-1.8</td><td>0.8</td><td>-2.3</td><td>2.2E-02</td><td>2.4E-02</td><td>MB231/MB468</td><td>MITO</td></tr>
<tr><td>>MI0000085|hsa-mir-27a&WithFlank&19|-|13836434|13836523@16.37.22&[MIMAT0004501&hsa-miR-27a-5p&offsets|0|0;m-270&19|-|13836487|13836508&offsets|0|0]|AGGGCTTAGCTGCTTGTGAGCA</td><td>>MI0000085|hsa-mir-27a&WithFlank&19|-|13836434|13836523</td><td>16</td><td>37</td><td>22</td><td>AGGGCTTAGCTGCTTGTGAGCA</td><td>56.8</td><td>4.4</td><td>1.8</td><td>2.5</td><td>1.4E-02</td><td>2.4E-02</td><td>BT20/MB468</td><td>MP</td></tr>
<tr><td>>hg38_21_+_8433172_8446622_RNA45SN1_45S_50ntFlanks@8358.8377.20|AAAGAACTTTGAAGAGAGAG</td><td>>hg38_21_+_8433172_8446622_RNA45SN1_45S_50ntFlanks</td><td>8358</td><td>8377</td><td>20</td><td>AAAGAACTTTGAAGAGAGAG</td><td>249.1</td><td>-3.9</td><td>1.6</td><td>-2.5</td><td>1.4E-02</td><td>2.4E-02</td><td>BT20/MB468</td><td>MP</td></tr>
<tr><td>>hg38_21_+_8433172_8446622_RNA45SN1_45S_50ntFlanks@5472.5509.38|ACGGCCCTGGCGGAGCGCTGAGAAGACGGTCGAACTTG</td><td>>hg38_21_+_8433172_8446622_RNA45SN1_45S_50ntFlanks</td><td>5472</td><td>5509</td><td>38</td><td>ACGGCCCTGGCGGAGCGCTGAGAAGACGGTCGAACTTG</td><td>310.1</td><td>2.5</td><td>1.0</td><td>2.5</td><td>1.4E-02</td><td>2.4E-02</td><td>BT20/MB468</td><td>MP</td></tr>
<tr><td>>MI0000298|hsa-mir-221&WithFlank&X|-|45746151|45746272@71.89.19&[MIMAT0000278&hsa-miR-221-3p&offsets|0|-4;m-91&X|-|45746181|45746202&offsets|0|-3]|AGCTACATTGTCTGCTGGG</td><td>>MI0000298|hsa-mir-221&WithFlank&X|-|45746151|45746272</td><td>71</td><td>89</td><td>19</td><td>AGCTACATTGTCTGCTGGG</td><td>124.3</td><td>3.1</td><td>1.3</td><td>2.3</td><td>1.9E-02</td><td>2.4E-02</td><td>MB231/MB468</td><td>MP</td></tr>
<tr><td>>hg38_21_+_8433172_8446622_RNA45SN1_45S_50ntFlanks@476.503.28|TTCGTGATCGATGTGGTGACGTCGTGCT</td><td>>hg38_21_+_8433172_8446622_RNA45SN1_45S_50ntFlanks</td><td>476</td><td>503</td><td>28</td><td>TTCGTGATCGATGTGGTGACGTCGTGCT</td><td>136.7</td><td>3.8</td><td>1.6</td><td>2.3</td><td>1.9E-02</td><td>2.4E-02</td><td>MB231/MB468</td><td>MP</td></tr>
<tr><td>>MI0000483|hsa-mir-186&WithFlank&1|-|71067625|71067722@21.40.20&[MIMAT0000456&hsa-miR-186-5p&offsets|0|-2;m-84&1|-|71067681|71067702&offsets|0|-2]|CAAAGAATTCTCCTTTTGGG</td><td>>MI0000483|hsa-mir-186&WithFlank&1|-|71067625|71067722</td><td>21</td><td>40</td><td>20</td><td>CAAAGAATTCTCCTTTTGGG</td><td>341.4</td><td>1.7</td><td>0.7</td><td>2.3</td><td>1.9E-02</td><td>2.4E-02</td><td>MB231/MB468</td><td>MP</td></tr>
<tr><td>>hg38_21_+_8433172_8446622_RNA45SN1_45S_50ntFlanks@12481.12500.20|AATAGGGAACGTGAGCTGGG</td><td>>hg38_21_+_8433172_8446622_RNA45SN1_45S_50ntFlanks</td><td>12481</td><td>12500</td><td>20</td><td>AATAGGGAACGTGAGCTGGG</td><td>323.9</td><td>1.5</td><td>0.7</td><td>2.3</td><td>2.1E-02</td><td>2.4E-02</td><td>BT20/MB468</td><td>CYTO</td></tr>
<tr><td>>hg38_21_+_8433172_8446622_RNA45SN1_45S_50ntFlanks@3987.4023.37|GCTTTGGTGACTCTAGATAACCTCGGGCCGATCGCAC</td><td>>hg38_21_+_8433172_8446622_RNA45SN1_45S_50ntFlanks</td><td>3987</td><td>4023</td><td>37</td><td>GCTTTGGTGACTCTAGATAACCTCGGGCCGATCGCAC</td><td>568.6</td><td>3.2</td><td>1.2</td><td>2.8</td><td>5.6E-03</td><td>2.4E-02</td><td>BT20/MB231</td><td>MP</td></tr>
<tr><td>>hg38_21_+_8433172_8446622_RNA45SN1_45S_50ntFlanks@6646.6663.18|TCGTACGACTCTTAGCGG</td><td>>hg38_21_+_8433172_8446622_RNA45SN1_45S_50ntFlanks</td><td>6646</td><td>6663</td><td>18</td><td>TCGTACGACTCTTAGCGG</td><td>24477.6</td><td>3.0</td><td>1.1</td><td>2.8</td><td>5.6E-03</td><td>2.4E-02</td><td>BT20/MB231</td><td>MP</td></tr>
<tr><td>>hg38_21_+_8433172_8446622_RNA45SN1_45S_50ntFlanks@12415.12447.33|CTTCGATGTCGGCTCTTCCTATCATTGTGAAGC</td><td>>hg38_21_+_8433172_8446622_RNA45SN1_45S_50ntFlanks</td><td>12415</td><td>12447</td><td>33</td><td>CTTCGATGTCGGCTCTTCCTATCATTGTGAAGC</td><td>1107.2</td><td>-2.1</td><td>0.8</td><td>-2.7</td><td>7.4E-03</td><td>2.4E-02</td><td>BT20/MB231</td><td>CYTO</td></tr>
<tr><td>>MI0000762|hsa-mir-362&WithFlank&X|+|50008958|50009034@11.34.24&[MIMAT0000705&hsa-miR-362-5p&offsets|0|0;m-267&X|+|50008968|50008991&offsets|0|0]|AATCCTTGGAACCTAGGTGTGAGT</td><td>>MI0000762|hsa-mir-362&WithFlank&X|+|50008958|50009034</td><td>11</td><td>34</td><td>24</td><td>AATCCTTGGAACCTAGGTGTGAGT</td><td>350.7</td><td>2.6</td><td>1.0</td><td>2.7</td><td>7.4E-03</td><td>2.4E-02</td><td>BT20/MB231</td><td>CYTO</td></tr>
<tr><td>>hg38_21_+_8433172_8446622_RNA45SN1_45S_50ntFlanks@12456.12476.21|CCAAGCGTTGGATTGTTCACC</td><td>>hg38_21_+_8433172_8446622_RNA45SN1_45S_50ntFlanks</td><td>12456</td><td>12476</td><td>21</td><td>CCAAGCGTTGGATTGTTCACC</td><td>690.3</td><td>-2.6</td><td>1.0</td><td>-2.7</td><td>7.4E-03</td><td>2.4E-02</td><td>BT20/MB231</td><td>CYTO</td></tr>
<tr><td>>hg38_21_+_8433172_8446622_RNA45SN1_45S_50ntFlanks@6645.6686.42|CTCGTACGACTCTTAGCGGTGGATCACTCGGCTCGTGCGTCG</td><td>>hg38_21_+_8433172_8446622_RNA45SN1_45S_50ntFlanks</td><td>6645</td><td>6686</td><td>42</td><td>CTCGTACGACTCTTAGCGGTGGATCACTCGGCTCGTGCGTCG</td><td>187.2</td><td>-2.3</td><td>1.0</td><td>-2.3</td><td>1.9E-02</td><td>2.4E-02</td><td>MB231/MB468</td><td>MP</td></tr>
<tr><td>>trna119_LysCTT_1_-_145395522_145395594@1.31.31,trna11_LysCTT_5_-_180648979_180649051@1.31.31,trna13_LysCTT_6_+_26556774_26556846@1.31.31,trna32_LysCTT_16_-_3207406_3207478@1.31.31,trna7_LysCTT_16_+_3225692_3225764@1.31.31,trna9_LysCTT_5_+_180634755_180634827@1.31.31|GCCCGGCTAGCTCAGTCGGTAGAGCATGAGA</td><td>>trna119_LysCTT_1_-_145395522_145395594</td><td>1</td><td>31</td><td>31</td><td>GCCCGGCTAGCTCAGTCGGTAGAGCATGAGA</td><td>2814.9</td><td>-2.0</td><td>0.9</td><td>-2.3</td><td>2.2E-02</td><td>2.4E-02</td><td>MB231/MB468</td><td>MITO</td></tr>
<tr><td>>hg38_21_+_8433172_8446622_RNA45SN1_45S_50ntFlanks@11572.11594.23|CCGACTTAGAACTGGTGCGGACC</td><td>>hg38_21_+_8433172_8446622_RNA45SN1_45S_50ntFlanks</td><td>11572</td><td>11594</td><td>23</td><td>CCGACTTAGAACTGGTGCGGACC</td><td>387.3</td><td>1.5</td><td>0.6</td><td>2.4</td><td>1.8E-02</td><td>2.4E-02</td><td>BT20/MB468</td><td>MITO</td></tr>
<tr><td>>hg38_21_+_8433172_8446622_RNA45SN1_45S_50ntFlanks@6703.6720.18|GCTGCGAGAATTAATGTG</td><td>>hg38_21_+_8433172_8446622_RNA45SN1_45S_50ntFlanks</td><td>6703</td><td>6720</td><td>18</td><td>GCTGCGAGAATTAATGTG</td><td>154.8</td><td>-3.6</td><td>1.5</td><td>-2.3</td><td>2.1E-02</td><td>2.4E-02</td><td>BT20/MB468</td><td>CYTO</td></tr>
<tr><td>>hg38_21_+_8433172_8446622_RNA45SN1_45S_50ntFlanks@6646.6686.41|TCGTACGACTCTTAGCGGTGGATCACTCGGCTCGTGCGTCG</td><td>>hg38_21_+_8433172_8446622_RNA45SN1_45S_50ntFlanks</td><td>6646</td><td>6686</td><td>41</td><td>TCGTACGACTCTTAGCGGTGGATCACTCGGCTCGTGCGTCG</td><td>1000.6</td><td>2.0</td><td>0.8</td><td>2.4</td><td>1.8E-02</td><td>2.4E-02</td><td>BT20/MB468</td><td>MITO</td></tr>
<tr><td>>hg38_21_+_8433172_8446622_RNA45SN1_45S_50ntFlanks@7978.8010.33|GACCTCAGATCAGACGTGGCGACCCGCTGAATT</td><td>>hg38_21_+_8433172_8446622_RNA45SN1_45S_50ntFlanks</td><td>7978</td><td>8010</td><td>33</td><td>GACCTCAGATCAGACGTGGCGACCCGCTGAATT</td><td>780.1</td><td>-2.5</td><td>0.9</td><td>-2.7</td><td>7.5E-03</td><td>2.4E-02</td><td>BT20/MB231</td><td>CYTO</td></tr>
<tr><td>>hg38_21_+_8433172_8446622_RNA45SN1_45S_50ntFlanks@9544.9577.34|GGTGGAGGTCCGTAGCGGTCCTGACGTGCAAATC</td><td>>hg38_21_+_8433172_8446622_RNA45SN1_45S_50ntFlanks</td><td>9544</td><td>9577</td><td>34</td><td>GGTGGAGGTCCGTAGCGGTCCTGACGTGCAAATC</td><td>234.7</td><td>-2.5</td><td>0.9</td><td>-2.7</td><td>7.5E-03</td><td>2.4E-02</td><td>BT20/MB231</td><td>CYTO</td></tr>
<tr><td>>hg38_21_+_8433172_8446622_RNA45SN1_45S_50ntFlanks@9555.9584.30|GTAGCGGTCCTGACGTGCAAATCGGTCGTC</td><td>>hg38_21_+_8433172_8446622_RNA45SN1_45S_50ntFlanks</td><td>9555</td><td>9584</td><td>30</td><td>GTAGCGGTCCTGACGTGCAAATCGGTCGTC</td><td>300.5</td><td>-1.5</td><td>0.7</td><td>-2.3</td><td>2.1E-02</td><td>2.4E-02</td><td>MB231/MB468</td><td>CYTO</td></tr>
<tr><td>>hg38_21_+_8433172_8446622_RNA45SN1_45S_50ntFlanks@12044.12070.27|ACTGACCCGGTGAGGCGGGGGGGCGAG</td><td>>hg38_21_+_8433172_8446622_RNA45SN1_45S_50ntFlanks</td><td>12044</td><td>12070</td><td>27</td><td>ACTGACCCGGTGAGGCGGGGGGGCGAG</td><td>278.5</td><td>-3.7</td><td>1.6</td><td>-2.3</td><td>2.1E-02</td><td>2.4E-02</td><td>BT20/MB468</td><td>CYTO</td></tr>
<tr><td>>hg38_21_+_8433172_8446622_RNA45SN1_45S_50ntFlanks@11572.11611.40|CCGACTTAGAACTGGTGCGGACCAGGGGAATCCGACTGTT</td><td>>hg38_21_+_8433172_8446622_RNA45SN1_45S_50ntFlanks</td><td>11572</td><td>11611</td><td>40</td><td>CCGACTTAGAACTGGTGCGGACCAGGGGAATCCGACTGTT</td><td>582.4</td><td>-2.4</td><td>1.0</td><td>-2.3</td><td>2.1E-02</td><td>2.4E-02</td><td>BT20/MB468</td><td>CYTO</td></tr>
<tr><td>>hg38_21_+_8433172_8446622_RNA45SN1_45S_50ntFlanks@11561.11583.23|GCGCCTAGCAGCCGACTTAGAAC</td><td>>hg38_21_+_8433172_8446622_RNA45SN1_45S_50ntFlanks</td><td>11561</td><td>11583</td><td>23</td><td>GCGCCTAGCAGCCGACTTAGAAC</td><td>205.6</td><td>-1.7</td><td>0.7</td><td>-2.3</td><td>2.2E-02</td><td>2.4E-02</td><td>MB231/MB468</td><td>NUC</td></tr>
<tr><td>>hg38_21_+_8433172_8446622_RNA45SN1_45S_50ntFlanks@11569.11603.35|CAGCCGACTTAGAACTGGTGCGGACCAGGGGAATC</td><td>>hg38_21_+_8433172_8446622_RNA45SN1_45S_50ntFlanks</td><td>11569</td><td>11603</td><td>35</td><td>CAGCCGACTTAGAACTGGTGCGGACCAGGGGAATC</td><td>252.1</td><td>5.5</td><td>2.0</td><td>2.8</td><td>5.6E-03</td><td>2.4E-02</td><td>BT20/MB231</td><td>MP</td></tr>
<tr><td>>hg38_21_+_8433172_8446622_RNA45SN1_45S_50ntFlanks@9939.9957.19|GCAGGACGGTGGCCATGGA</td><td>>hg38_21_+_8433172_8446622_RNA45SN1_45S_50ntFlanks</td><td>9939</td><td>9957</td><td>19</td><td>GCAGGACGGTGGCCATGGA</td><td>475.0</td><td>3.3</td><td>1.2</td><td>2.8</td><td>5.6E-03</td><td>2.4E-02</td><td>BT20/MB231</td><td>MP</td></tr>
<tr><td>>hg38_21_+_8433172_8446622_RNA45SN1_45S_50ntFlanks@9952.9980.29|CATGGAAGTCGGAATCCGCTAAGGAGTGT</td><td>>hg38_21_+_8433172_8446622_RNA45SN1_45S_50ntFlanks</td><td>9952</td><td>9980</td><td>29</td><td>CATGGAAGTCGGAATCCGCTAAGGAGTGT</td><td>166.8</td><td>4.8</td><td>1.7</td><td>2.8</td><td>5.6E-03</td><td>2.4E-02</td><td>BT20/MB231</td><td>MP</td></tr>
<tr><td>>MI0000100|hsa-mir-98&WithFlank&X|-|53556217|53556347@28.48.21(+1A)&[MIMAT0000096&hsa-miR-98-5p&offsets|0|-1(+1A);m-143&X|-|53556299|53556320&offsets|0|-1(+1A)]|TGAGGTAGTAAGTTGTATTGTA</td><td>>MI0000100|hsa-mir-98&WithFlank&X|-|53556217|53556347</td><td>28</td><td>48</td><td>21(+1A)</td><td>TGAGGTAGTAAGTTGTATTGTA</td><td>251.1</td><td>2.6</td><td>1.0</td><td>2.7</td><td>7.5E-03</td><td>2.4E-02</td><td>BT20/MB231</td><td>CYTO</td></tr>
<tr><td>>hg38_21_+_8433172_8446622_RNA45SN1_45S_50ntFlanks@12416.12437.22|TTCGATGTCGGCTCTTCCTATC</td><td>>hg38_21_+_8433172_8446622_RNA45SN1_45S_50ntFlanks</td><td>12416</td><td>12437</td><td>22</td><td>TTCGATGTCGGCTCTTCCTATC</td><td>7418.3</td><td>-2.4</td><td>0.9</td><td>-2.7</td><td>7.5E-03</td><td>2.4E-02</td><td>BT20/MB231</td><td>CYTO</td></tr>
<tr><td>>hg38_21_+_8433172_8446622_RNA45SN1_45S_50ntFlanks@9573.9606.34|AAATCGGTCGTCCGACCTGGGTATAGGGGCGAAA</td><td>>hg38_21_+_8433172_8446622_RNA45SN1_45S_50ntFlanks</td><td>9573</td><td>9606</td><td>34</td><td>AAATCGGTCGTCCGACCTGGGTATAGGGGCGAAA</td><td>52.5</td><td>5.1</td><td>2.1</td><td>2.5</td><td>1.4E-02</td><td>2.4E-02</td><td>BT20/MB468</td><td>MP</td></tr>
<tr><td>>MI0000746|hsa-mir-99b&WithFlank&19|+|51692606|51692687@13.33.21(+1U)&[MIMAT0000689&hsa-miR-99b-5p&offsets|0|-1(+1U);m-10&19|+|51692618|51692639&offsets|0|-1(+1U)]|CACCCGTAGAACCGACCTTGCT</td><td>>MI0000746|hsa-mir-99b&WithFlank&19|+|51692606|51692687</td><td>13</td><td>33</td><td>21(+1U)</td><td>CACCCGTAGAACCGACCTTGCT</td><td>2628.2</td><td>2.4</td><td>0.9</td><td>2.7</td><td>7.5E-03</td><td>2.4E-02</td><td>BT20/MB231</td><td>CYTO</td></tr>
<tr><td>>MI0000477|hsa-mir-146a&WithFlank&5|+|160485346|160485456@27.48.22&[MIMAT0000449&hsa-miR-146a-5p&offsets|0|0;m-129&5|+|160485372|160485393&offsets|0|0]|TGAGAACTGAATTCCATGGGTT</td><td>>MI0000477|hsa-mir-146a&WithFlank&5|+|160485346|160485456</td><td>27</td><td>48</td><td>22</td><td>TGAGAACTGAATTCCATGGGTT</td><td>9439.6</td><td>-5.1</td><td>1.9</td><td>-2.7</td><td>7.5E-03</td><td>2.4E-02</td><td>BT20/MB231</td><td>CYTO</td></tr>
<tr><td>>hg38_21_+_8433172_8446622_RNA45SN1_45S_50ntFlanks@7988.8017.30|CAGACGTGGCGACCCGCTGAATTTAAGCAT</td><td>>hg38_21_+_8433172_8446622_RNA45SN1_45S_50ntFlanks</td><td>7988</td><td>8017</td><td>30</td><td>CAGACGTGGCGACCCGCTGAATTTAAGCAT</td><td>326.7</td><td>-2.3</td><td>0.9</td><td>-2.7</td><td>7.5E-03</td><td>2.4E-02</td><td>BT20/MB231</td><td>CYTO</td></tr>
<tr><td>>hg38_21_+_8433172_8446622_RNA45SN1_45S_50ntFlanks@3771.3791.21|CACGGCCGGTACAGTGAAACT</td><td>>hg38_21_+_8433172_8446622_RNA45SN1_45S_50ntFlanks</td><td>3771</td><td>3791</td><td>21</td><td>CACGGCCGGTACAGTGAAACT</td><td>410.3</td><td>-1.8</td><td>0.8</td><td>-2.3</td><td>2.2E-02</td><td>2.4E-02</td><td>MB231/MB468</td><td>NUC</td></tr>
<tr><td>>hg38_21_+_8433172_8446622_RNA45SN1_45S_50ntFlanks@9568.9603.36|CGTGCAAATCGGTCGTCCGACCTGGGTATAGGGGCG</td><td>>hg38_21_+_8433172_8446622_RNA45SN1_45S_50ntFlanks</td><td>9568</td><td>9603</td><td>36</td><td>CGTGCAAATCGGTCGTCCGACCTGGGTATAGGGGCG</td><td>391.7</td><td>-2.4</td><td>1.1</td><td>-2.3</td><td>2.1E-02</td><td>2.4E-02</td><td>MB231/MB468</td><td>CYTO</td></tr>
</table>

<table>
<tr><th></th><th></th><th></th><th></th><th></th><th></th><th></th><th></th><th></th><th></th><th></th><th></th><th></th><th></th></tr>
<tr><td>>hg38_21_+_8433172_8446622_RNA45SN1_45S_50ntFlanks@9505.9523.19|TGAACTATGCCTGGGCAGG</td><td>>hg38_21_+_8433172_8446622_RNA45SN1_45S_50ntFlanks</td><td>9505</td><td>9523</td><td>19</td><td>TGAACTATGCCTGGGCAGG</td><td>206.2</td><td>-3.6</td><td>1.5</td><td>-2.5</td><td>1.4E-02</td><td>2.4E-02</td><td>BT20/MB468</td><td>MP</td></tr>
<tr><td>>MI0000255|hsa-mir-30d&WithFlank&8|-|134804870|134804951@12.34.23(+1A)&[MIMAT0000245&hsa-miR-30d-5p&offsets|0|+1(+1A);m-21&8|-|134804919|134804940&offsets|0|+1(+1A)]|TGTAAACATCCCCGACTGGAAGCA</td><td>>MI0000255|hsa-mir-30d&WithFlank&8|-|134804870|134804951</td><td>12</td><td>34</td><td>23(+1A)</td><td>TGTAAACATCCCCGACTGGAAGCA</td><td>9091.3</td><td>2.3</td><td>0.9</td><td>2.7</td><td>7.5E-03</td><td>2.4E-02</td><td>BT20/MB231</td><td>CYTO</td></tr>
<tr><td>>trnaMT_GluTTC_MT_-_14674_14742@45.72.28,trnalookalike8_GluTTC_5_-_93905172_93905240@45.72.28|CGTGGTTGTAGTCCGTGCGAGAATACCA</td><td>>trnaMT_GluTTC_MT_-_14674_14742</td><td>45</td><td>72</td><td>28</td><td>CGTGGTTGTAGTCCGTGCGAGAATACCA</td><td>664.7</td><td>-2.1</td><td>0.7</td><td>-2.7</td><td>6.1E-03</td><td>2.4E-02</td><td>BT20/MB231</td><td>NUC</td></tr>
<tr><td>>hg38_21_+_8433172_8446622_RNA45SN1_45S_50ntFlanks@4399.4423.25|CGGGCGGGCGGTCCGCCGCGAGGCG</td><td>>hg38_21_+_8433172_8446622_RNA45SN1_45S_50ntFlanks</td><td>4399</td><td>4423</td><td>25</td><td>CGGGCGGGCGGTCCGCCGCGAGGCG</td><td>149.0</td><td>-4.2</td><td>1.7</td><td>-2.5</td><td>1.4E-02</td><td>2.4E-02</td><td>BT20/MB468</td><td>MP</td></tr>
<tr><td>>hg38_21_+_8433172_8446622_RNA45SN1_45S_50ntFlanks@9858.9890.33|CGGGATGAACCGAACGCCGGGTTAAGGCGCCCG</td><td>>hg38_21_+_8433172_8446622_RNA45SN1_45S_50ntFlanks</td><td>9858</td><td>9890</td><td>33</td><td>CGGGATGAACCGAACGCCGGGTTAAGGCGCCCG</td><td>57.1</td><td>-4.4</td><td>1.8</td><td>-2.5</td><td>1.4E-02</td><td>2.4E-02</td><td>BT20/MB468</td><td>MP</td></tr>
<tr><td>>trnaMT_IleGAT_MT_+_4263_4331@43.67.25|ATAGGAGCTTAAACCCCCTTATTTC</td><td>>trnaMT_IleGAT_MT_+_4263_4331</td><td>43</td><td>67</td><td>25</td><td>ATAGGAGCTTAAACCCCCTTATTTC</td><td>107.7</td><td>6.3</td><td>2.6</td><td>2.5</td><td>1.4E-02</td><td>2.4E-02</td><td>BT20/MB468</td><td>MP</td></tr>
<tr><td>>MI0000299|hsa-mir-222&WithFlank&X|-|45747009|45747130@75.98.24&[MIMAT0000279&hsa-miR-222-3p&offsets|0|+3;m-140&X|-|45747033|45747056&offsets|0|0]|AGCTACATCTGGCTACTGGGTCTC</td><td>>MI0000299|hsa-mir-222&WithFlank&X|-|45747009|45747130</td><td>75</td><td>98</td><td>24</td><td>AGCTACATCTGGCTACTGGGTCTC</td><td>24826.3</td><td>-2.0</td><td>0.7</td><td>-2.7</td><td>6.9E-03</td><td>2.4E-02</td><td>BT20/MB231</td><td>MITO</td></tr>
<tr><td>>hg38_21_+_8433172_8446622_RNA45SN1_45S_50ntFlanks@13011.13039.29|GAAAGTCAGCCCTCGACACAAGGGTTTGT</td><td>>hg38_21_+_8433172_8446622_RNA45SN1_45S_50ntFlanks</td><td>13011</td><td>13039</td><td>29</td><td>GAAAGTCAGCCCTCGACACAAGGGTTTGT</td><td>341.5</td><td>2.4</td><td>0.9</td><td>2.7</td><td>6.9E-03</td><td>2.4E-02</td><td>BT20/MB231</td><td>MITO</td></tr>
<tr><td>>hg38_21_+_8433172_8446622_RNA45SN1_45S_50ntFlanks@10787.10826.40|ACAGCCTCTGGCATGTTGGAACAATGTAGGTAAGGGAAGT</td><td>>hg38_21_+_8433172_8446622_RNA45SN1_45S_50ntFlanks</td><td>10787</td><td>10826</td><td>40</td><td>ACAGCCTCTGGCATGTTGGAACAATGTAGGTAAGGGAAGT</td><td>171.2</td><td>-2.7</td><td>1.1</td><td>-2.5</td><td>1.4E-02</td><td>2.4E-02</td><td>BT20/MB468</td><td>MP</td></tr>
<tr><td>>hg38_21_+_8433172_8446622_RNA45SN1_45S_50ntFlanks@10114.10131.18|TGTGCGTCGGAGGGCGGC</td><td>>hg38_21_+_8433172_8446622_RNA45SN1_45S_50ntFlanks</td><td>10114</td><td>10131</td><td>18</td><td>TGTGCGTCGGAGGGCGGC</td><td>50.8</td><td>6.3</td><td>2.6</td><td>2.5</td><td>1.4E-02</td><td>2.4E-02</td><td>BT20/MB468</td><td>MP</td></tr>
<tr><td>>am_trna111_HisGTG_1_-_147774845_147774916@2.29.28,trna118_HisGTG_1_-_145396881_145396952@2.29.28,trna16_HisGTG_1_+_146544773_146544844@2.29.28,trna1_HisGTG_15_+_45493349_45493420@2.29.28,trna21_HisGTG_1_+_147753471_147753542@2.29.28,trna33_HisGTG_6_+_27125906_27125977@2.29.28,trna7_HisGTG_9_-_14433938_14434009@2.29.28,trna8_HisGTG_15_-_45492611_45492682@2.29.28,trna9_HisGTG_15_-_45490804_45490875@2.29.28|CCGTGATCGTATAGTGGTTAGTACTCTG</td><td>>am_trna111_HisGTG_1_-_147774845_147774916</td><td>2</td><td>29</td><td>28</td><td>CCGTGATCGTATAGTGGTTAGTACTCTG</td><td>427.1</td><td>-4.3</td><td>1.8</td><td>-2.5</td><td>1.4E-02</td><td>2.4E-02</td><td>BT20/MB468</td><td>MP</td></tr>
<tr><td>>trna129_MetCAT_6_-_27870271_27870342@12.31.20,trna142_MetCAT_6_-_27560600_27560671@12.31.20,trna150_MetCAT_6_-_27300764_27300835@12.31.20,trna151_ThrCGT_6_-_27271568_27271639@12.31.20,trna169_MetCAT_6_-_26330529_26330600@12.31.20,trna171_MetCAT_6_-_26313352_26313423@12.31.20,trna20_MetCAT_17_-_80452597_80452668@12.31.20,trna2_MetCAT_6_+_26286754_26286825@12.31.20,trna32_MetCAT_1_+_153643726_153643797@12.31.20,trna61_MetCAT_6_+_27745664_27745735@12.31.20|GCAGCGGAAGCGTGCTGGGC</td><td>>trna129_MetCAT_6_-_27870271_27870342</td><td>12</td><td>31</td><td>20</td><td>GCAGCGGAAGCGTGCTGGGC</td><td>40.8</td><td>4.9</td><td>2.1</td><td>2.3</td><td>1.9E-02</td><td>2.4E-02</td><td>MB231/MB468</td><td>MP</td></tr>
<tr><td>>hg38_21_+_8433172_8446622_RNA45SN1_45S_50ntFlanks@9855.9874.20|CTGCGGGATGAACCGAACGC</td><td>>hg38_21_+_8433172_8446622_RNA45SN1_45S_50ntFlanks</td><td>9855</td><td>9874</td><td>20</td><td>CTGCGGGATGAACCGAACGC</td><td>461.6</td><td>4.6</td><td>1.7</td><td>2.8</td><td>5.6E-03</td><td>2.4E-02</td><td>BT20/MB231</td><td>MP</td></tr>
<tr><td>>MI0000088|hsa-mir-30a&WithFlank&6|-|71403545|71403627@12.34.23(+1A)&[MIMAT0000087&hsa-miR-30a-5p&offsets|0|+1(+1A);m-15&6|-|71403595|71403616&offsets|0|+1(+1A)]|TGTAAACATCCTCGACTGGAAGCA</td><td>>MI0000088|hsa-mir-30a&WithFlank&6|-|71403545|71403627</td><td>12</td><td>34</td><td>23(+1A)</td><td>TGTAAACATCCTCGACTGGAAGCA</td><td>8445.5</td><td>3.0</td><td>1.2</td><td>2.5</td><td>1.4E-02</td><td>2.4E-02</td><td>BT20/MB468</td><td>MP</td></tr>
<tr><td>>hg38_21_+_8433172_8446622_RNA45SN1_45S_50ntFlanks@8111.8136.26|GGGGCGCGGGACATGTGGCGTACGGA</td><td>>hg38_21_+_8433172_8446622_RNA45SN1_45S_50ntFlanks</td><td>8111</td><td>8136</td><td>26</td><td>GGGGCGCGGGACATGTGGCGTACGGA</td><td>626.8</td><td>5.1</td><td>2.1</td><td>2.5</td><td>1.4E-02</td><td>2.4E-02</td><td>BT20/MB468</td><td>MP</td></tr>
<tr><td>>hg38_21_+_8433172_8446622_RNA45SN1_45S_50ntFlanks@5492.5525.34|AGAAGACGGTCGAACTTGACTATCTAGAGGAAGT</td><td>>hg38_21_+_8433172_8446622_RNA45SN1_45S_50ntFlanks</td><td>5492</td><td>5525</td><td>34</td><td>AGAAGACGGTCGAACTTGACTATCTAGAGGAAGT</td><td>640.5</td><td>3.6</td><td>1.3</td><td>2.8</td><td>5.7E-03</td><td>2.4E-02</td><td>BT20/MB231</td><td>MP</td></tr>
<tr><td>>hg38_21_+_8433172_8446622_RNA45SN1_45S_50ntFlanks@10806.10828.23|AACAATGTAGGTAAGGGAAGTCG</td><td>>hg38_21_+_8433172_8446622_RNA45SN1_45S_50ntFlanks</td><td>10806</td><td>10828</td><td>23</td><td>AACAATGTAGGTAAGGGAAGTCG</td><td>208.9</td><td>-4.2</td><td>1.7</td><td>-2.5</td><td>1.4E-02</td><td>2.4E-02</td><td>BT20/MB468</td><td>MP</td></tr>
<tr><td>>hg38_21_+_8433172_8446622_RNA45SN1_45S_50ntFlanks@12997.13015.19|CGCTGCGATCTATTGAAAG</td><td>>hg38_21_+_8433172_8446622_RNA45SN1_45S_50ntFlanks</td><td>12997</td><td>13015</td><td>19</td><td>CGCTGCGATCTATTGAAAG</td><td>478.2</td><td>-1.7</td><td>0.7</td><td>-2.3</td><td>2.1E-02</td><td>2.4E-02</td><td>BT20/MB468</td><td>NUC</td></tr>
<tr><td>>trnaMT_ValTAC_MT_+_1602_1670@33.71.39|ACACTTAGGAGATTTCAACTTAACTTGACCGCTCTGACC</td><td>>trnaMT_ValTAC_MT_+_1602_1670</td><td>33</td><td>71</td><td>39</td><td>ACACTTAGGAGATTTCAACTTAACTTGACCGCTCTGACC</td><td>175.8</td><td>3.9</td><td>1.6</td><td>2.5</td><td>1.4E-02</td><td>2.4E-02</td><td>BT20/MB468</td><td>MP</td></tr>
<tr><td>>hg38_21_+_8433172_8446622_RNA45SN1_45S_50ntFlanks@11675.11702.28|TGCTCTGAATGTCAAAGTGAAGAAATTC</td><td>>hg38_21_+_8433172_8446622_RNA45SN1_45S_50ntFlanks</td><td>11675</td><td>11702</td><td>28</td><td>TGCTCTGAATGTCAAAGTGAAGAAATTC</td><td>659.9</td><td>-2.0</td><td>0.9</td><td>-2.3</td><td>2.2E-02</td><td>2.4E-02</td><td>MB231/MB468</td><td>MITO</td></tr>
<tr><td>>hg38_21_+_8433172_8446622_RNA45SN1_45S_50ntFlanks@12644.12666.23|TCTGTGGGATTATGACTGAACGC</td><td>>hg38_21_+_8433172_8446622_RNA45SN1_45S_50ntFlanks</td><td>12644</td><td>12666</td><td>23</td><td>TCTGTGGGATTATGACTGAACGC</td><td>102.0</td><td>3.6</td><td>1.5</td><td>2.5</td><td>1.4E-02</td><td>2.4E-02</td><td>BT20/MB468</td><td>MP</td></tr>
<tr><td>>hg38_21_+_8433172_8446622_RNA45SN1_45S_50ntFlanks@6661.6688.28|CGGTGGATCACTCGGCTCGTGCGTCGAT</td><td>>hg38_21_+_8433172_8446622_RNA45SN1_45S_50ntFlanks</td><td>6661</td><td>6688</td><td>28</td><td>CGGTGGATCACTCGGCTCGTGCGTCGAT</td><td>787.0</td><td>2.4</td><td>0.9</td><td>2.7</td><td>7.5E-03</td><td>2.4E-02</td><td>BT20/MB231</td><td>CYTO</td></tr>
<tr><td>>hg38_21_+_8433172_8446622_RNA45SN1_45S_50ntFlanks@6773.6802.30|TTCCTCCCGGGGCTACGCCTGTCTGAGCGT</td><td>>hg38_21_+_8433172_8446622_RNA45SN1_45S_50ntFlanks</td><td>6773</td><td>6802</td><td>30</td><td>TTCCTCCCGGGGCTACGCCTGTCTGAGCGT</td><td>290.6</td><td>-1.3</td><td>0.6</td><td>-2.3</td><td>2.2E-02</td><td>2.4E-02</td><td>MB231/MB468</td><td>NUC</td></tr>
<tr><td>>hg38_21_+_8433172_8446622_RNA45SN1_45S_50ntFlanks@8358.8375.18|AAAGAACTTTGAAGAGAG</td><td>>hg38_21_+_8433172_8446622_RNA45SN1_45S_50ntFlanks</td><td>8358</td><td>8375</td><td>18</td><td>AAAGAACTTTGAAGAGAG</td><td>8020.8</td><td>3.7</td><td>1.3</td><td>2.8</td><td>5.7E-03</td><td>2.4E-02</td><td>BT20/MB231</td><td>MP</td></tr>
<tr><td>>am_trnaMT_ValTAC_MT_+_1602_1670@29.67.39|ACTTACACTTAGGAGATTTCAACTTAACTTGACCGCTCT</td><td>>am_trnaMT_ValTAC_MT_+_1602_1670</td><td>29</td><td>67</td><td>39</td><td>ACTTACACTTAGGAGATTTCAACTTAACTTGACCGCTCT</td><td>84.2</td><td>3.7</td><td>1.6</td><td>2.3</td><td>1.9E-02</td><td>2.4E-02</td><td>MB231/MB468</td><td>MP</td></tr>
<tr><td>>hg38_21_+_8433172_8446622_RNA45SN1_45S_50ntFlanks@8115.8140.26|CGCGGGACATGTGGCGTACGGAAGAC</td><td>>hg38_21_+_8433172_8446622_RNA45SN1_45S_50ntFlanks</td><td>8115</td><td>8140</td><td>26</td><td>CGCGGGACATGTGGCGTACGGAAGAC</td><td>482.1</td><td>-3.5</td><td>1.5</td><td>-2.3</td><td>2.1E-02</td><td>2.4E-02</td><td>BT20/MB468</td><td>NUC</td></tr>
<tr><td>>am_MI0000809|hsa-mir-151a&WithFlank&8|-|140732558|140732659@17.38.22&[MIMAT0004697&hsa-miR-151a-5p&offsets|0|+1;m-75&8|-|140732623|140732643&offsets|0|+1]|TCGAGGAGCTCACAGTCTAGTA</td><td>>am_MI0000809|hsa-mir-151a&WithFlank&8|-|140732558|140732659</td><td>17</td><td>38</td><td>22</td><td>TCGAGGAGCTCACAGTCTAGTA</td><td>1047.8</td><td>2.4</td><td>0.9</td><td>2.7</td><td>7.6E-03</td><td>2.4E-02</td><td>BT20/MB231</td><td>CYTO</td></tr>
<tr><td>>hg38_21_+_8433172_8446622_RNA45SN1_45S_50ntFlanks@10819.10840.22|AGGGAAGTCGGCAAGCCGGATC</td><td>>hg38_21_+_8433172_8446622_RNA45SN1_45S_50ntFlanks</td><td>10819</td><td>10840</td><td>22</td><td>AGGGAAGTCGGCAAGCCGGATC</td><td>761.9</td><td>5.2</td><td>1.9</td><td>2.8</td><td>5.7E-03</td><td>2.4E-02</td><td>BT20/MB231</td><td>MP</td></tr>
<tr><td>>hg38_21_+_8433172_8446622_RNA45SN1_45S_50ntFlanks@12475.12499.25|CCCACTAATAGGGAACGTGAGCTGG</td><td>>hg38_21_+_8433172_8446622_RNA45SN1_45S_50ntFlanks</td><td>12475</td><td>12499</td><td>25</td><td>CCCACTAATAGGGAACGTGAGCTGG</td><td>183.7</td><td>-2.7</td><td>1.1</td><td>-2.3</td><td>2.1E-02</td><td>2.4E-02</td><td>BT20/MB468</td><td>CYTO</td></tr>
<tr><td>>am_trnaMT_GlyTCC_MT_+_9991_10058@28.48.21|ACTTCCAATTAACTAGTTTTG</td><td>>am_trnaMT_GlyTCC_MT_+_9991_10058</td><td>28</td><td>48</td><td>21</td><td>ACTTCCAATTAACTAGTTTTG</td><td>128.4</td><td>-4.4</td><td>1.8</td><td>-2.5</td><td>1.4E-02</td><td>2.4E-02</td><td>BT20/MB468</td><td>MP</td></tr>
<tr><td>>hg38_21_+_8433172_8446622_RNA45SN1_45S_50ntFlanks@13004.13039.36|ATCTATTGAAAGTCAGCCCTCGACACAAGGGTTTGT</td><td>>hg38_21_+_8433172_8446622_RNA45SN1_45S_50ntFlanks</td><td>13004</td><td>13039</td><td>36</td><td>ATCTATTGAAAGTCAGCCCTCGACACAAGGGTTTGT</td><td>139.5</td><td>-4.0</td><td>1.7</td><td>-2.3</td><td>1.9E-02</td><td>2.4E-02</td><td>MB231/MB468</td><td>MP</td></tr>
<tr><td>>hg38_21_+_8433172_8446622_RNA45SN1_45S_50ntFlanks@11199.11220.22|CCGGCGGCGGCGGCGACTCTGG</td><td>>hg38_21_+_8433172_8446622_RNA45SN1_45S_50ntFlanks</td><td>11199</td><td>11220</td><td>22</td><td>CCGGCGGCGGCGGCGACTCTGG</td><td>197.7</td><td>-4.2</td><td>1.7</td><td>-2.5</td><td>1.4E-02</td><td>2.4E-02</td><td>BT20/MB468</td><td>MP</td></tr>
<tr><td>>hg38_21_+_8433172_8446622_RNA45SN1_45S_50ntFlanks@12003.12031.29|GCGGGCCGCCGGTGAAATACCACTACTCT</td><td>>hg38_21_+_8433172_8446622_RNA45SN1_45S_50ntFlanks</td><td>12003</td><td>12031</td><td>29</td><td>GCGGGCCGCCGGTGAAATACCACTACTCT</td><td>402.5</td><td>-2.4</td><td>0.9</td><td>-2.7</td><td>7.6E-03</td><td>2.4E-02</td><td>BT20/MB231</td><td>CYTO</td></tr>
<tr><td>>hg38_21_+_8433172_8446622_RNA45SN1_45S_50ntFlanks@12676.12704.29|AGAATCCCGCCCAGGCGGAACGATACGGC</td><td>>hg38_21_+_8433172_8446622_RNA45SN1_45S_50ntFlanks</td><td>12676</td><td>12704</td><td>29</td><td>AGAATCCCGCCCAGGCGGAACGATACGGC</td><td>151.3</td><td>-2.7</td><td>1.2</td><td>-2.3</td><td>2.1E-02</td><td>2.4E-02</td><td>MB231/MB468</td><td>CYTO</td></tr>
<tr><td>>MI0000108|hsa-mir-103a-2&WithFlank&20|+|3917488|3917577@54.77.24(+1U);MI0000109|hsa-mir-103a-1&WithFlank&5|-|168560890|168560979@54.77.24(+1U)&[MIMAT0000101&hsa-miR-103a-3p&offsets|0|+1(+1U);m-12&20|+|3917541|3917563&offsets|0|+1(+1U)];[MIMAT0000101_1&hsa-miR-103a-3p&offsets|0|+1(+1U);m-13&5|-|168560904|168560926&offsets|0|+1(+1U);m-44220&5|-|168560908|168560926&offsets|0|+5(+1U)]|AGCAGCATTGTACAGGGCTATGAAT</td><td>>MI0000108|hsa-mir-103a-2&WithFlank&20|+|3917488|3917577</td><td>54</td><td>77</td><td>24(+1U)</td><td>AGCAGCATTGTACAGGGCTATGAAT</td><td>135.7</td><td>-5.2</td><td>1.9</td><td>-2.8</td><td>5.7E-03</td><td>2.4E-02</td><td>BT20/MB231</td><td>MP</td></tr>
<tr><td>>hg38_MT_+_598_1651_RNR1_12S_50ntFlanks@913.946.34|TCAAAGGACATTTAACTAAAACCCCTACGCATTT</td><td>>hg38_MT_+_598_1651_RNR1_12S_50ntFlanks</td><td>913</td><td>946</td><td>34</td><td>TCAAAGGACATTTAACTAAAACCCCTACGCATTT</td><td>412.9</td><td>4.6</td><td>1.9</td><td>2.5</td><td>1.4E-02</td><td>2.4E-02</td><td>BT20/MB468</td><td>MP</td></tr>
<tr><td>>hg38_MT_+_598_1651_RNR1_12S_50ntFlanks@905.950.46|AGTATACTTCAAAGGACATTTAACTAAAACCCCTACGCATTTATAT</td><td>>hg38_MT_+_598_1651_RNR1_12S_50ntFlanks</td><td>905</td><td>950</td><td>46</td><td>AGTATACTTCAAAGGACATTTAACTAAAACCCCTACGCATTTATAT</td><td>34.8</td><td>8.5</td><td>3.1</td><td>2.8</td><td>5.7E-03</td><td>2.4E-02</td><td>BT20/MB231</td><td>MP</td></tr>
<tr><td>>hg38_21_+_8433172_8446622_RNA45SN1_45S_50ntFlanks@4385.4423.39|TTGGATCTTGGGAGCGGGCGGGCGGTCCGCCGCGAGGCG</td><td>>hg38_21_+_8433172_8446622_RNA45SN1_45S_50ntFlanks</td><td>4385</td><td>4423</td><td>39</td><td>TTGGATCTTGGGAGCGGGCGGGCGGTCCGCCGCGAGGCG</td><td>177.8</td><td>-3.7</td><td>1.6</td><td>-2.3</td><td>2.1E-02</td><td>2.4E-02</td><td>BT20/MB468</td><td>CYTO</td></tr>
<tr><td>>trna10_AspGTC_12_-_125424193_125424264@38.64.27,trna12_AspGTC_12_-_125411891_125411962@38.64.27,trna144_AspGTC_6_-_27551236_27551307@38.64.27,trna38_AspGTC_17_-_8125556_8125627@38.64.27,trna45_AspGTC_6_+_27447453_27447524@38.64.27,trna48_AspGTC_6_+_27471523_27471594@38.64.27,trna4_AspGTC_12_+_96429799_96429870@38.64.27,trna69_AspGTC_1_-_161440205_161440276@38.64.27,trna72_AspGTC_1_-_161432824_161432895@38.64.27,trna75_AspGTC_1_-_161425414_161425485@38.64.27,trna78_AspGTC_1_-_161418033_161418104@38.64.27,trna81_AspGTC_1_-_161410615_161410686@38.64.27|CGCGGGAGACCGGGGTTCGATTCCCCG</td><td>>trna10_AspGTC_12_-_125424193_125424264</td><td>38</td><td>64</td><td>27</td><td>CGCGGGAGACCGGGGTTCGATTCCCCG</td><td>233.9</td><td>-4.5</td><td>1.8</td><td>-2.5</td><td>1.4E-02</td><td>2.4E-02</td><td>BT20/MB468</td><td>MP</td></tr>
<tr><td>>hg38_21_+_8433172_8446622_RNA45SN1_45S_50ntFlanks@10772.10796.25|AGGTCTCCAAGGTGAACAGCCTCTG</td><td>>hg38_21_+_8433172_8446622_RNA45SN1_45S_50ntFlanks</td><td>10772</td><td>10796</td><td>25</td><td>AGGTCTCCAAGGTGAACAGCCTCTG</td><td>327.0</td><td>-1.7</td><td>0.7</td><td>-2.3</td><td>2.1E-02</td><td>2.4E-02</td><td>BT20/MB468</td><td>CYTO</td></tr>
<tr><td>>hg38_21_+_8433172_8446622_RNA45SN1_45S_50ntFlanks@12071.12110.40|CCCCGAGGGGCTCTCGCTTCTGGCGCCAAGCGCCCGGCCG</td><td>>hg38_21_+_8433172_8446622_RNA45SN1_45S_50ntFlanks</td><td>12071</td><td>12110</td><td>40</td><td>CCCCGAGGGGCTCTCGCTTCTGGCGCCAAGCGCCCGGCCG</td><td>303.5</td><td>-3.8</td><td>1.6</td><td>-2.4</td><td>1.8E-02</td><td>2.4E-02</td><td>BT20/MB468</td><td>MITO</td></tr>
<tr><td>>hg38_21_+_8433172_8446622_RNA45SN1_45S_50ntFlanks@10384.10424.41|CCATGTGAACAGCAGTTGAACATGGGTCAGTCGGTCCTGAG</td><td>>hg38_21_+_8433172_8446622_RNA45SN1_45S_50ntFlanks</td><td>10384</td><td>10424</td><td>41</td><td>CCATGTGAACAGCAGTTGAACATGGGTCAGTCGGTCCTGAG</td><td>989.8</td><td>-2.3</td><td>0.9</td><td>-2.7</td><td>7.6E-03</td><td>2.4E-02</td><td>BT20/MB231</td><td>CYTO</td></tr>
<tr><td>>hg38_21_+_8433172_8446622_RNA45SN1_45S_50ntFlanks@12419.12463.45|GATGTCGGCTCTTCCTATCATTGTGAAGCAGAATTCACCAAGCGT</td><td>>hg38_21_+_8433172_8446622_RNA45SN1_45S_50ntFlanks</td><td>12419</td><td>12463</td><td>45</td><td>GATGTCGGCTCTTCCTATCATTGTGAAGCAGAATTCACCAAGCGT</td><td>469.4</td><td>-4.8</td><td>1.8</td><td>-2.7</td><td>7.6E-03</td><td>2.4E-02</td><td>BT20/MB231</td><td>CYTO</td></tr>
</table>

| | | | | | | | | | | | | | |
|---|---|---|---|---|---|---|---|---|---|---|---|---|---|
| >MI0000750\|hsa-mir-26a-2&WithFlank&12\|-\|57824603\|57824698@20.39.20;MI0000083\|hsa-mir-26a-1&WithFlank&3\|+\|37969398\|37969486@16.35.20&[MIMAT0000082&hsa-miR-26a-5p&offsets\|0\|-2;m-33&12\|-\|57824658\|57824679&offsets\|0\|-2];[MIMAT0000082_1&hsa-miR-26a-5p&offsets\|0\|-2;m-34&3\|+\|37969413\|37969434&offsets\|0\|-2]\|TTCAAGTAATCCAGGATAGG | >MI0000750\|hsa-mir-26a-2&WithFlank&12\|-\|57824603\|57824698 | 20 | 39 | 20 | TTCAAGTAATCCAGGATAGG | 557.1 | -4.1 | 1.5 | -2.8 | 5.7E-03 | 2.4E-02 | BT20/MB231 | MP |
| >trnaMT_SerGCT_MT_+_12207_12265@8.38.31\|CTCACAAGAACTGCTAACTCATGCCCCCATG | >trnaMT_SerGCT_MT_+_12207_12265 | 8 | 38 | 31 | CTCACAAGAACTGCTAACTCATGCCCCCATG | 357.6 | -4.3 | 1.9 | -2.3 | 2.1E-02 | 2.4E-02 | BT20/MB468 | CYTO |
| >hg38_21_+_8433172_8446622_RNA45SN1_45S_50ntFlanks@12685.12703.19\|CCCAGGCGGAACGATACGG | >hg38_21_+_8433172_8446622_RNA45SN1_45S_50ntFlanks | 12685 | 12703 | 19 | CCCAGGCGGAACGATACGG | 180.2 | -1.9 | 0.8 | -2.4 | 1.8E-02 | 2.4E-02 | BT20/MB468 | MITO |
| >trna10_AspGTC_12_-_125424193_125424264@37.75.39,trna12_AspGTC_12_-_125411891_125411962@37.75.39,trna144_AspGTC_6_-_27551236_27551307@37.75.39,trna38_AspGTC_17_-_8125556_8125627@37.75.39,trna45_AspGTC_6_+_27447453_27447524@37.75.39,trna48_AspGTC_6_+_27471523_27471594@37.75.39,trna4_AspGTC_12_+_96429799_96429870@37.75.39,trna69_AspGTC_1_-_161440205_161440276@37.75.39,trna72_AspGTC_1_-_161432824_161432895@37.75.39,trna75_AspGTC_1_-_161425414_161425485@37.75.39,trna78_AspGTC_1_-_161418033_161418104@37.75.39,trna81_AspGTC_1_-_161410615_161410686@37.75.39\|ACGCGGGAGACCGGGGTTCGATTCCCCGACGGGGAGCCA | >trna10_AspGTC_12_-_125424193_125424264 | 37 | 75 | 39 | ACGCGGGAGACCGGGGTTCGATTCCCCGACGGGGAGCCA | 764.6 | -3.4 | 1.2 | -2.7 | 6.2E-03 | 2.4E-02 | BT20/MB231 | NUC |
| >hg38_21_+_8433172_8446622_RNA45SN1_45S_50ntFlanks@5236.5263.28\|CACTGACTGGCTCAGCGTGTGCCTACCC | >hg38_21_+_8433172_8446622_RNA45SN1_45S_50ntFlanks | 5236 | 5263 | 28 | CACTGACTGGCTCAGCGTGTGCCTACCC | 141.5 | -3.4 | 1.5 | -2.4 | 1.8E-02 | 2.4E-02 | BT20/MB468 | MITO |
| >hg38_21_+_8433172_8446622_RNA45SN1_45S_50ntFlanks@3782.3810.29\|CAGTGAAACTGCGAATGGCTCATTAAATC | >hg38_21_+_8433172_8446622_RNA45SN1_45S_50ntFlanks | 3782 | 3810 | 29 | CAGTGAAACTGCGAATGGCTCATTAAATC | 280.0 | -1.1 | 0.5 | -2.3 | 2.1E-02 | 2.4E-02 | BT20/MB468 | CYTO |
| >hg38_21_+_8433172_8446622_RNA45SN1_45S_50ntFlanks@12202.12220.19\|CTAAGGCGAGCTCAGGGAG | >hg38_21_+_8433172_8446622_RNA45SN1_45S_50ntFlanks | 12202 | 12220 | 19 | CTAAGGCGAGCTCAGGGAG | 140.3 | -2.8 | 1.2 | -2.3 | 2.1E-02 | 2.4E-02 | BT20/MB468 | CYTO |
| >hg38_21_+_8433172_8446622_RNA45SN1_45S_50ntFlanks@11188.11225.38\|CACCCGGGGGGCCGGCGGCGGCGGCGACTCTGGACGCG | >hg38_21_+_8433172_8446622_RNA45SN1_45S_50ntFlanks | 11188 | 11225 | 38 | CACCCGGGGGGCCGGCGGCGGCGGCGACTCTGGACGCG | 109.6 | -4.3 | 1.8 | -2.5 | 1.4E-02 | 2.4E-02 | BT20/MB468 | MP |
| >hg38_21_+_8433172_8446622_RNA45SN1_45S_50ntFlanks@10353.10390.38\|AGAACTTTGAAGGCCGAAGTGGAGAAGGGTTCCATGTG | >hg38_21_+_8433172_8446622_RNA45SN1_45S_50ntFlanks | 10353 | 10390 | 38 | AGAACTTTGAAGGCCGAAGTGGAGAAGGGTTCCATGTG | 512.1 | -2.3 | 1.0 | -2.3 | 2.2E-02 | 2.4E-02 | MB231/MB468 | MITO |
| >hg38_MT_+_598_1651_RNR1_12S_50ntFlanks@910.935.26\|ACTTCAAAGGACATTTAACTAAAACC | >hg38_MT_+_598_1651_RNR1_12S_50ntFlanks | 910 | 935 | 26 | ACTTCAAAGGACATTTAACTAAAACC | 109.3 | 3.2 | 1.4 | 2.3 | 2.0E-02 | 2.4E-02 | MB231/MB468 | MP |
| >hg38_21_+_8433172_8446622_RNA45SN1_45S_50ntFlanks@11906.11929.24\|CACGGTGAAGAGACATGAGAGGTG | >hg38_21_+_8433172_8446622_RNA45SN1_45S_50ntFlanks | 11906 | 11929 | 24 | CACGGTGAAGAGACATGAGAGGTG | 130.9 | -1.8 | 0.8 | -2.3 | 2.1E-02 | 2.4E-02 | BT20/MB468 | CYTO |
| >hg38_21_+_8433172_8446622_RNA45SN1_45S_50ntFlanks@9918.9938.21\|AGGTGTTGGTTGATATAGACA | >hg38_21_+_8433172_8446622_RNA45SN1_45S_50ntFlanks | 9918 | 9938 | 21 | AGGTGTTGGTTGATATAGACA | 153.6 | 5.2 | 2.1 | 2.5 | 1.4E-02 | 2.4E-02 | BT20/MB468 | MP |
| >hg38_21_+_8433172_8446622_RNA45SN1_45S_50ntFlanks@10259.10276.18\|AGGGCCGCTGCGGTGAGC | >hg38_21_+_8433172_8446622_RNA45SN1_45S_50ntFlanks | 10259 | 10276 | 18 | AGGGCCGCTGCGGTGAGC | 60.3 | 4.4 | 1.8 | 2.5 | 1.4E-02 | 2.4E-02 | BT20/MB468 | MP |
| >hg38_21_+_8433172_8446622_RNA45SN1_45S_50ntFlanks@10342.10373.32\|ATATTCAAACGAGAACTTTGAAGGCCGAAGTG | >hg38_21_+_8433172_8446622_RNA45SN1_45S_50ntFlanks | 10342 | 10373 | 32 | ATATTCAAACGAGAACTTTGAAGGCCGAAGTG | 193.5 | 2.1 | 0.9 | 2.5 | 1.4E-02 | 2.4E-02 | BT20/MB468 | MP |
| >MI0006444\|hsa-mir-1307&WithFlank&10\|-\|103394247\|103394407@86.107.22(+1C)&[MIMAT0005951&hsa-miR-1307-3p&offsets\|0\|0(+1C);m-66&10\|-\|103394300\|103394322&offsets\|0\|-1(+1C)]\|ACTCGGCGTGGCGTCGGTCGTGC | >MI0006444\|hsa-mir-1307&WithFlank&10\|-\|103394247\|103394407 | 86 | 107 | 22(+1C) | ACTCGGCGTGGCGTCGGTCGTGC | 67.0 | 4.0 | 1.7 | 2.3 | 2.0E-02 | 2.4E-02 | MB231/MB468 | MP |
| >hg38_MT_+_1621_3279_RNR2_16S_50ntFlanks@1390.1415.26\|GATGGTGCAGCCGCTATTAAAGGTTC | >hg38_MT_+_1621_3279_RNR2_16S_50ntFlanks | 1390 | 1415 | 26 | GATGGTGCAGCCGCTATTAAAGGTTC | 142.6 | 4.0 | 1.7 | 2.3 | 2.0E-02 | 2.4E-02 | MB231/MB468 | MP |
| >hg38_21_+_8433172_8446622_RNA45SN1_45S_50ntFlanks@4661.4696.36\|AGGTGAAATTCTTGGACCGGCGCAAGACGGACCAGA | >hg38_21_+_8433172_8446622_RNA45SN1_45S_50ntFlanks | 4661 | 4696 | 36 | AGGTGAAATTCTTGGACCGGCGCAAGACGGACCAGA | 320.2 | 3.0 | 1.1 | 2.7 | 7.7E-03 | 2.4E-02 | BT20/MB231 | CYTO |
| >hg38_21_+_8433172_8446622_RNA45SN1_45S_50ntFlanks@6646.6669.24\|TCGTACGACTCTTAGCGGTGGATC | >hg38_21_+_8433172_8446622_RNA45SN1_45S_50ntFlanks | 6646 | 6669 | 24 | TCGTACGACTCTTAGCGGTGGATC | 26255.1 | 2.3 | 0.9 | 2.7 | 7.7E-03 | 2.4E-02 | BT20/MB231 | CYTO |
| >hg38_21_+_8433172_8446622_RNA45SN1_45S_50ntFlanks@6645.6669.25\|CTCGTACGACTCTTAGCGGTGGATC | >hg38_21_+_8433172_8446622_RNA45SN1_45S_50ntFlanks | 6645 | 6669 | 25 | CTCGTACGACTCTTAGCGGTGGATC | 6007.3 | 2.4 | 0.9 | 2.7 | 6.2E-03 | 2.4E-02 | BT20/MB231 | NUC |
| >hg38_21_+_8433172_8446622_RNA45SN1_45S_50ntFlanks@6661.6701.41\|CGGTGGATCACTCGGCTCGTGCGTCGATGAAGAACGCAGCT | >hg38_21_+_8433172_8446622_RNA45SN1_45S_50ntFlanks | 6661 | 6701 | 41 | CGGTGGATCACTCGGCTCGTGCGTCGATGAAGAACGCAGCT | 1720.6 | 1.9 | 0.7 | 2.7 | 6.2E-03 | 2.4E-02 | BT20/MB231 | NUC |
| >hg38_21_+_8433172_8446622_RNA45SN1_45S_50ntFlanks@4005.4023.19\|AACCTCGGGCCGATCGCAC | >hg38_21_+_8433172_8446622_RNA45SN1_45S_50ntFlanks | 4005 | 4023 | 19 | AACCTCGGGCCGATCGCAC | 359.4 | -1.9 | 0.8 | -2.3 | 2.2E-02 | 2.4E-02 | MB231/MB468 | MITO |
| >hg38_21_+_8433172_8446622_RNA45SN1_45S_50ntFlanks@7977.8009.33\|CGACCTCAGATCAGACGTGGCGACCCGCTGAAT | >hg38_21_+_8433172_8446622_RNA45SN1_45S_50ntFlanks | 7977 | 8009 | 33 | CGACCTCAGATCAGACGTGGCGACCCGCTGAAT | 754.5 | 2.9 | 1.2 | 2.5 | 1.4E-02 | 2.4E-02 | BT20/MB468 | MP |
| >hg38_21_+_8433172_8446622_RNA45SN1_45S_50ntFlanks@10355.10384.30\|AACTTTGAAGGCCGAAGTGGAGAAGGGTTC | >hg38_21_+_8433172_8446622_RNA45SN1_45S_50ntFlanks | 10355 | 10384 | 30 | AACTTTGAAGGCCGAAGTGGAGAAGGGTTC | 107.6 | -3.6 | 1.5 | -2.5 | 1.4E-02 | 2.4E-02 | BT20/MB468 | MP |
| >hg38_21_+_8433172_8446622_RNA45SN1_45S_50ntFlanks@12836.12869.34\|CGGTGCGGAGTGCCCTTCGTCCTGGGAAACGGGG | >hg38_21_+_8433172_8446622_RNA45SN1_45S_50ntFlanks | 12836 | 12869 | 34 | CGGTGCGGAGTGCCCTTCGTCCTGGGAAACGGGG | 278.9 | -4.2 | 1.7 | -2.5 | 1.4E-02 | 2.4E-02 | BT20/MB468 | MP |
| >hg38_21_+_8433172_8446622_RNA45SN1_45S_50ntFlanks@6653.6688.36\|ACTCTTAGCGGTGGATCACTCGGCTCGTGCGTCGAT | >hg38_21_+_8433172_8446622_RNA45SN1_45S_50ntFlanks | 6653 | 6688 | 36 | ACTCTTAGCGGTGGATCACTCGGCTCGTGCGTCGAT | 132.8 | -2.7 | 1.1 | -2.3 | 2.0E-02 | 2.4E-02 | MB231/MB468 | MP |
| >hg38_21_+_8433172_8446622_RNA45SN1_45S_50ntFlanks@12934.12958.25\|CTAAACCATTCGTAGACGACCTGCT | >hg38_21_+_8433172_8446622_RNA45SN1_45S_50ntFlanks | 12934 | 12958 | 25 | CTAAACCATTCGTAGACGACCTGCT | 128.2 | -4.1 | 1.7 | -2.5 | 1.4E-02 | 2.4E-02 | BT20/MB468 | MP |
| >hg38_21_+_8433172_8446622_RNA45SN1_45S_50ntFlanks@10427.10453.27\|ATGGGCGAGCGCCGTTCCGAAGGGACG | >hg38_21_+_8433172_8446622_RNA45SN1_45S_50ntFlanks | 10427 | 10453 | 27 | ATGGGCGAGCGCCGTTCCGAAGGGACG | 269.0 | -3.4 | 1.4 | -2.5 | 1.4E-02 | 2.4E-02 | BT20/MB468 | MP |
| >MI0000088\|hsa-mir-30a&WithFlank&6\|-\|71403545\|71403627@12.35.24(+2U)&[MIMAT0000087&hsa-miR-30a-5p&offsets\|0\|+2(+2U);m-15&6\|-\|71403595\|71403616&offsets\|0\|+2(+2U)]\|TGTAAACATCCTCGACTGGAAGCTTT | >MI0000088\|hsa-mir-30a&WithFlank&6\|-\|71403545\|71403627 | 12 | 35 | 24(+2U) | TGTAAACATCCTCGACTGGAAGCTTT | 397.2 | -2.2 | 0.8 | -2.7 | 7.7E-03 | 2.4E-02 | BT20/MB231 | CYTO |
| >hg38_21_+_8433172_8446622_RNA45SN1_45S_50ntFlanks@10468.10488.21\|TTGCCCTCGGCCGATCGAAAG | >hg38_21_+_8433172_8446622_RNA45SN1_45S_50ntFlanks | 10468 | 10488 | 21 | TTGCCCTCGGCCGATCGAAAG | 184.9 | -2.4 | 0.9 | -2.7 | 7.7E-03 | 2.4E-02 | BT20/MB231 | CYTO |
| >MI0000108\|hsa-mir-103a-2&WithFlank&20\|+\|3917488\|3917577@54.75.22(+2U);MI0000109\|hsa-mir-103a-1&WithFlank&5\|-\|168560890\|168560979@54.75.22(+2U)&[MIMAT0000101&hsa-miR-103a-3p&offsets\|0\|-1(+2U);m-12&20\|+\|3917541\|3917563&offsets\|0\|-1(+2U)];[MIMAT0000101_1&hsa-miR-103a-3p&offsets\|0\|-1(+2U);m-13&5\|-\|168560904\|168560926&offsets\|0\|-1(+2U);m-44220&5\|-\|168560908\|168560926&offsets\|0\|+3(+2U)]\|AGCAGCATTGTACAGGGCTATGTT | >MI0000108\|hsa-mir-103a-2&WithFlank&20\|+\|3917488\|3917577 | 54 | 75 | 22(+2U) | AGCAGCATTGTACAGGGCTATGTT | 91.2 | 4.7 | 2.0 | 2.3 | 2.0E-02 | 2.5E-02 | MB231/MB468 | MP |
| >MI0000269\|hsa-mir-181a-2&WithFlank&9\|+\|124692436\|124692557@83.104.22&[MIMAT0004558&hsa-miR-181a-2-3p&offsets\|0\|0;m-71&9\|+\|124692518\|124692539&offsets\|0\|0]\|ACCACTGACCGTTGACTGTACC | >MI0000269\|hsa-mir-181a-2&WithFlank&9\|+\|124692436\|124692557 | 83 | 104 | 22 | ACCACTGACCGTTGACTGTACC | 1298.7 | -1.6 | 0.6 | -2.7 | 6.3E-03 | 2.5E-02 | BT20/MB231 | NUC |
| >hg38_21_+_8433172_8446622_RNA45SN1_45S_50ntFlanks@10347.10374.28\|CAAACGAGAACTTTGAAGGCCGAAGTGG | >hg38_21_+_8433172_8446622_RNA45SN1_45S_50ntFlanks | 10347 | 10374 | 28 | CAAACGAGAACTTTGAAGGCCGAAGTGG | 192.9 | -1.5 | 0.6 | -2.4 | 1.9E-02 | 2.5E-02 | BT20/MB468 | MITO |
| >trnaMT_HisGTG_MT_+_12138_12206@43.71.29\|ACAGAGGCTTACGACCCCTTATTTACCCC | >trnaMT_HisGTG_MT_+_12138_12206 | 43 | 71 | 29 | ACAGAGGCTTACGACCCCTTATTTACCCC | 42.7 | 4.6 | 2.0 | 2.3 | 2.0E-02 | 2.5E-02 | MB231/MB468 | MP |
| >MI0033423\|hsa-mir-10399&WithFlank&7\|-\|139044094\|139044163@43.63.21&[MIMAT0041630&hsa-miR-10399-3p&offsets\|-1\|-1;TJU_CMC_MD2.ID03061.3p-miR&offsets\|+3\|+2;m-2657&7\|-\|139044103\|139044124&offsets\|+3\|+2]\|TCTCTCGGACAAGCTGTAGGT | >MI0033423\|hsa-mir-10399&WithFlank&7\|-\|139044094\|139044163 | 43 | 63 | 21 | TCTCTCGGACAAGCTGTAGGT | 29.8 | 4.7 | 2.0 | 2.3 | 2.0E-02 | 2.5E-02 | MB231/MB468 | MP |
| >hg38_21_+_8433172_8446622_RNA45SN1_45S_50ntFlanks@5515.5532.18\|CTAGAGGAAGTAAAAGTC | >hg38_21_+_8433172_8446622_RNA45SN1_45S_50ntFlanks | 5515 | 5532 | 18 | CTAGAGGAAGTAAAAGTC | 94.9 | -4.1 | 1.7 | -2.5 | 1.4E-02 | 2.5E-02 | BT20/MB468 | MP |
| >trnaMT_MetCAT_MT_+_4402_4469@42.71.30\|AATGTTGGTTATACCCTTCCCGTACTACCA | >trnaMT_MetCAT_MT_+_4402_4469 | 42 | 71 | 30 | AATGTTGGTTATACCCTTCCCGTACTACCA | 599.2 | 1.8 | 0.8 | 2.3 | 2.0E-02 | 2.5E-02 | MB231/MB468 | MP |
| >hg38_21_+_8433172_8446622_RNA45SN1_45S_50ntFlanks@9533.9571.39\|GAGGAAACTCTGGTGGAGGTCCGTAGCGGTCCTGACGTG | >hg38_21_+_8433172_8446622_RNA45SN1_45S_50ntFlanks | 9533 | 9571 | 39 | GAGGAAACTCTGGTGGAGGTCCGTAGCGGTCCTGACGTG | 1443.0 | -2.8 | 1.1 | -2.5 | 1.4E-02 | 2.5E-02 | BT20/MB468 | MP |
| >hg38_MT_+_598_1651_RNR1_12S_50ntFlanks@904.935.32\|AAGTATACTTCAAAGGACATTTAACTAAAACC | >hg38_MT_+_598_1651_RNR1_12S_50ntFlanks | 904 | 935 | 32 | AAGTATACTTCAAAGGACATTTAACTAAAACC | 77.1 | 2.9 | 1.3 | 2.3 | 2.0E-02 | 2.5E-02 | MB231/MB468 | MP |
| >MI0000542\|hsa-mir-320a&WithFlank&8\|-\|22244960\|22245043@48.68.21(+1G)&[MIMAT0000510&hsa-miR-320a-3p&offsets\|0\|-1(+1G);m-60&8\|-\|22244975\|22244996&offsets\|0\|-1(+1G)]\|AAAAGCTGGGTTGAGAGGGCGG | >MI0000542\|hsa-mir-320a&WithFlank&8\|-\|22244960\|22245043 | 48 | 68 | 21(+1G) | AAAAGCTGGGTTGAGAGGGCGG | 186.5 | 3.2 | 1.4 | 2.3 | 2.0E-02 | 2.5E-02 | MB231/MB468 | MP |
| >hg38_21_+_8433172_8446622_RNA45SN1_45S_50ntFlanks@7975.8012.38\|CGCGACCTCAGATCAGACGTGGCGACCCGCTGAATTTA | >hg38_21_+_8433172_8446622_RNA45SN1_45S_50ntFlanks | 7975 | 8012 | 38 | CGCGACCTCAGATCAGACGTGGCGACCCGCTGAATTTA | 5525.9 | -2.5 | 1.1 | -2.3 | 2.0E-02 | 2.5E-02 | MB231/MB468 | MP |
| >trnaMT_ValTAC_MT_+_1602_1670@34.72.39\|CACTTAGGAGATTTCAACTTAACTTGACCGCTCTGACCA | >trnaMT_ValTAC_MT_+_1602_1670 | 34 | 72 | 39 | CACTTAGGAGATTTCAACTTAACTTGACCGCTCTGACCA | 171.9 | 2.4 | 1.1 | 2.3 | 2.2E-02 | 2.5E-02 | MB231/MB468 | MITO |
| >MI0000065\|hsa-let-7d&WithFlank&9\|+\|94178828\|94178926@14.34.21(+1G)&[MIMAT0000065&hsa-let-7d-5p&offsets\|0\|-1(+1G);m-90&9\|+\|94178841\|94178862&offsets\|0\|-1(+1G)]\|AGAGGTAGTAGGTTGCATAGTG | >MI0000065\|hsa-let-7d&WithFlank&9\|+\|94178828\|94178926 | 14 | 34 | 21(+1G) | AGAGGTAGTAGGTTGCATAGTG | 227.4 | 2.4 | 0.9 | 2.7 | 7.8E-03 | 2.5E-02 | BT20/MB231 | CYTO |
| >MI0000483\|hsa-mir-186&WithFlank&1\|-\|71067625\|71067722@21.43.23&[MIMAT0000456&hsa-miR-186-5p&offsets\|0\|+1;m-84&1\|-\|71067681\|71067702&offsets\|0\|+1]\|CAAAGAATTCTCCTTTTGGGCTT | >MI0000483\|hsa-mir-186&WithFlank&1\|-\|71067625\|71067722 | 21 | 43 | 23 | CAAAGAATTCTCCTTTTGGGCTT | 546.9 | 2.4 | 0.9 | 2.7 | 7.8E-03 | 2.5E-02 | BT20/MB231 | CYTO |
| >hg38_21_+_8433172_8446622_RNA45SN1_45S_50ntFlanks@7991.8018.28\|ACGTGGCGACCCGCTGAATTTAAGCATA | >hg38_21_+_8433172_8446622_RNA45SN1_45S_50ntFlanks | 7991 | 8018 | 28 | ACGTGGCGACCCGCTGAATTTAAGCATA | 153.7 | -3.2 | 1.2 | -2.7 | 7.8E-03 | 2.5E-02 | BT20/MB231 | CYTO |
| >hg38_21_+_8433172_8446622_RNA45SN1_45S_50ntFlanks@10351.10369.19\|CGAGAACTTTGAAGGCCGA | >hg38_21_+_8433172_8446622_RNA45SN1_45S_50ntFlanks | 10351 | 10369 | 19 | CGAGAACTTTGAAGGCCGA | 671.6 | -1.4 | 0.6 | -2.3 | 2.2E-02 | 2.5E-02 | MB231/MB468 | NUC |
| >hg38_21_+_8433172_8446622_RNA45SN1_45S_50ntFlanks@10350.10369.20\|ACGAGAACTTTGAAGGCCGA | >hg38_21_+_8433172_8446622_RNA45SN1_45S_50ntFlanks | 10350 | 10369 | 20 | ACGAGAACTTTGAAGGCCGA | 279.1 | -2.3 | 0.8 | -2.7 | 6.3E-03 | 2.5E-02 | BT20/MB231 | NUC |

<table>
<tr><td>>MI0000076|hsa-mir-20a&WithFlank&13|+|91351059|91351141@14.33.20(+1A)&[MIMAT0000075&hsa-miR-20a-5p&offsets|0|-3(+1A);m-96&13|+|91351072|91351094&offsets|0|-3(+1A)]|TAAAGTGCTTATAGTGCAGGA</td><td>>MI0000076|hsa-mir-20a&WithFlank&13|+|91351059|91351141</td><td>14</td><td>33</td><td>20(+1A)</td><td>TAAAGTGCTTATAGTGCAGGA</td><td>89.2</td><td>3.9</td><td>1.7</td><td>2.3</td><td>2.0E-02</td><td>2.5E-02</td><td>MB231/MB468</td><td>MP</td></tr>
<tr><td>>am_trnaMT_SerGCT_MT_+_12207_12265@13.32.20|AAGAACTGCTAACTCATGCC</td><td>>am_trnaMT_SerGCT_MT_+_12207_12265</td><td>13</td><td>32</td><td>20</td><td>AAGAACTGCTAACTCATGCC</td><td>134.8</td><td>3.0</td><td>1.2</td><td>2.5</td><td>1.4E-02</td><td>2.5E-02</td><td>BT20/MB468</td><td>MP</td></tr>
<tr><td>>hg38_21_+_8433172_8446622_RNA45SN1_45S_50ntFlanks@10064.10081.18|TCGAGAGTGGACGGGAGC</td><td>>hg38_21_+_8433172_8446622_RNA45SN1_45S_50ntFlanks</td><td>10064</td><td>10081</td><td>18</td><td>TCGAGAGTGGACGGGAGC</td><td>82.1</td><td>5.1</td><td>2.1</td><td>2.5</td><td>1.4E-02</td><td>2.5E-02</td><td>BT20/MB468</td><td>MP</td></tr>
<tr><td>>hg38_21_+_8433172_8446622_RNA45SN1_45S_50ntFlanks@12418.12442.25|CGATGTCGGCTCTTCCTATCATTGT</td><td>>hg38_21_+_8433172_8446622_RNA45SN1_45S_50ntFlanks</td><td>12418</td><td>12442</td><td>25</td><td>CGATGTCGGCTCTTCCTATCATTGT</td><td>713.2</td><td>-2.1</td><td>0.8</td><td>-2.7</td><td>7.8E-03</td><td>2.5E-02</td><td>BT20/MB231</td><td>CYTO</td></tr>
<tr><td>>hg38_21_+_8433172_8446622_RNA45SN1_45S_50ntFlanks@11669.11704.36|GCCCAGTGCTCTGAATGTCAAAGTGAAGAAATTCAA</td><td>>hg38_21_+_8433172_8446622_RNA45SN1_45S_50ntFlanks</td><td>11669</td><td>11704</td><td>36</td><td>GCCCAGTGCTCTGAATGTCAAAGTGAAGAAATTCAA</td><td>554.5</td><td>-2.2</td><td>0.8</td><td>-2.7</td><td>7.8E-03</td><td>2.5E-02</td><td>BT20/MB231</td><td>CYTO</td></tr>
<tr><td>>MI0000749|hsa-mir-30e&WithFlank&1|+|40754349|40754452@24.46.23&[MIMAT0000692&hsa-miR-30e-5p&offsets|+1|+2;m-24&1|+|40754371|40754394&offsets|+1|0]|GTAAACATCCTTGACTGGAAGCT</td><td>>MI0000749|hsa-mir-30e&WithFlank&1|+|40754349|40754452</td><td>24</td><td>46</td><td>23</td><td>GTAAACATCCTTGACTGGAAGCT</td><td>1448.9</td><td>3.1</td><td>1.2</td><td>2.5</td><td>1.4E-02</td><td>2.5E-02</td><td>BT20/MB468</td><td>MP</td></tr>
<tr><td>>MI0000746|hsa-mir-99b&WithFlank&19|+|51692606|51692687@13.32.20&[MIMAT0000689&hsa-miR-99b-5p&offsets|0|-2;m-10&19|+|51692618|51692639&offsets|0|-2]|CACCCGTAGAACCGACCTTG</td><td>>MI0000746|hsa-mir-99b&WithFlank&19|+|51692606|51692687</td><td>13</td><td>32</td><td>20</td><td>CACCCGTAGAACCGACCTTG</td><td>2085.2</td><td>2.0</td><td>0.7</td><td>2.7</td><td>6.3E-03</td><td>2.5E-02</td><td>BT20/MB231</td><td>NUC</td></tr>
<tr><td>>hg38_21_+_8433172_8446622_RNA45SN1_45S_50ntFlanks@5474.5516.43|GGCCCTGGCGGAGCGCTGAGAAGACGGTCGAACTTGACTATCT</td><td>>hg38_21_+_8433172_8446622_RNA45SN1_45S_50ntFlanks</td><td>5474</td><td>5516</td><td>43</td><td>GGCCCTGGCGGAGCGCTGAGAAGACGGTCGAACTTGACTATCT</td><td>295.6</td><td>-1.9</td><td>0.7</td><td>-2.7</td><td>7.8E-03</td><td>2.5E-02</td><td>BT20/MB231</td><td>CYTO</td></tr>
<tr><td>>trnaMT_GluTTC_MT_-_14674_14742@35.72.38,trnalookalike8_GluTTC_5_-_93905172_93905240@35.72.38|TATCATTGGTCGTGGTTGTAGTCCGTGCGAGAATACCA</td><td>>trnaMT_GluTTC_MT_-_14674_14742</td><td>35</td><td>72</td><td>38</td><td>TATCATTGGTCGTGGTTGTAGTCCGTGCGAGAATACCA</td><td>3536.1</td><td>4.0</td><td>1.6</td><td>2.5</td><td>1.4E-02</td><td>2.5E-02</td><td>BT20/MB468</td><td>MP</td></tr>
<tr><td>>MI0000088|hsa-mir-30a&WithFlank&6|-|71403545|71403627@53.74.22&[MIMAT0000088&hsa-miR-30a-3p&offsets|0|0;m-38&6|-|71403554|71403575&offsets|0|0]|CTTTCAGTCGGATGTTTGCAGC</td><td>>MI0000088|hsa-mir-30a&WithFlank&6|-|71403545|71403627</td><td>53</td><td>74</td><td>22</td><td>CTTTCAGTCGGATGTTTGCAGC</td><td>14590.0</td><td>-1.6</td><td>0.6</td><td>-2.7</td><td>6.4E-03</td><td>2.5E-02</td><td>BT20/MB231</td><td>NUC</td></tr>
<tr><td>>hg38_21_+_8433172_8446622_RNA45SN1_45S_50ntFlanks@6654.6683.30|CTCTTAGCGGTGGATCACTCGGCTCGTGCG</td><td>>hg38_21_+_8433172_8446622_RNA45SN1_45S_50ntFlanks</td><td>6654</td><td>6683</td><td>30</td><td>CTCTTAGCGGTGGATCACTCGGCTCGTGCG</td><td>1363.1</td><td>-1.9</td><td>0.7</td><td>-2.7</td><td>6.4E-03</td><td>2.5E-02</td><td>BT20/MB231</td><td>NUC</td></tr>
<tr><td>>hg38_21_+_8433172_8446622_RNA45SN1_45S_50ntFlanks@4919.4938.20|CCACCAGGAGTGGAGCCTGC</td><td>>hg38_21_+_8433172_8446622_RNA45SN1_45S_50ntFlanks</td><td>4919</td><td>4938</td><td>20</td><td>CCACCAGGAGTGGAGCCTGC</td><td>254.8</td><td>-3.0</td><td>1.3</td><td>-2.4</td><td>1.9E-02</td><td>2.5E-02</td><td>BT20/MB468</td><td>MITO</td></tr>
<tr><td>>hg38_21_+_8433172_8446622_RNA45SN1_45S_50ntFlanks@6661.6706.46|CGGTGGATCACTCGGCTCGTGCGTCGATGAAGAACGCAGCTAGCTG</td><td>>hg38_21_+_8433172_8446622_RNA45SN1_45S_50ntFlanks</td><td>6661</td><td>6706</td><td>46</td><td>CGGTGGATCACTCGGCTCGTGCGTCGATGAAGAACGCAGCTAGCTG</td><td>406.9</td><td>3.4</td><td>1.2</td><td>2.8</td><td>5.8E-03</td><td>2.5E-02</td><td>BT20/MB231</td><td>MP</td></tr>
<tr><td>>hg38_21_+_8433172_8446622_RNA45SN1_45S_50ntFlanks@9605.9630.26|AAGACTAATCGAACCATCTAGTAGCT</td><td>>hg38_21_+_8433172_8446622_RNA45SN1_45S_50ntFlanks</td><td>9605</td><td>9630</td><td>26</td><td>AAGACTAATCGAACCATCTAGTAGCT</td><td>190.4</td><td>2.2</td><td>0.9</td><td>2.5</td><td>1.4E-02</td><td>2.5E-02</td><td>BT20/MB468</td><td>MP</td></tr>
<tr><td>>hg38_21_+_8433172_8446622_RNA45SN1_45S_50ntFlanks@9951.9980.30|CCATGGAAGTCGGAATCCGCTAAGGAGTGT</td><td>>hg38_21_+_8433172_8446622_RNA45SN1_45S_50ntFlanks</td><td>9951</td><td>9980</td><td>30</td><td>CCATGGAAGTCGGAATCCGCTAAGGAGTGT</td><td>38.7</td><td>5.1</td><td>2.1</td><td>2.5</td><td>1.4E-02</td><td>2.5E-02</td><td>BT20/MB468</td><td>MP</td></tr>
<tr><td>>hg38_21_+_8433172_8446622_RNA45SN1_45S_50ntFlanks@8355.8390.36|TGAAAAGAACTTTGAAGAGAGAGTTCAAGAGGGCGT</td><td>>hg38_21_+_8433172_8446622_RNA45SN1_45S_50ntFlanks</td><td>8355</td><td>8390</td><td>36</td><td>TGAAAAGAACTTTGAAGAGAGAGTTCAAGAGGGCGT</td><td>176.5</td><td>5.3</td><td>1.9</td><td>2.8</td><td>5.8E-03</td><td>2.5E-02</td><td>BT20/MB231</td><td>MP</td></tr>
<tr><td>>hg38_21_+_8433172_8446622_RNA45SN1_45S_50ntFlanks@12218.12236.19|GAGGACAGAAACCTCCCGT</td><td>>hg38_21_+_8433172_8446622_RNA45SN1_45S_50ntFlanks</td><td>12218</td><td>12236</td><td>19</td><td>GAGGACAGAAACCTCCCGT</td><td>410.3</td><td>6.2</td><td>2.3</td><td>2.7</td><td>7.1E-03</td><td>2.5E-02</td><td>BT20/MB231</td><td>MITO</td></tr>
<tr><td>>hg38_21_+_8433172_8446622_RNA45SN1_45S_50ntFlanks@12934.12960.27|CTAAACCATTCGTAGACGACCTGCTTC</td><td>>hg38_21_+_8433172_8446622_RNA45SN1_45S_50ntFlanks</td><td>12934</td><td>12960</td><td>27</td><td>CTAAACCATTCGTAGACGACCTGCTTC</td><td>243.1</td><td>-4.2</td><td>1.7</td><td>-2.5</td><td>1.4E-02</td><td>2.5E-02</td><td>BT20/MB468</td><td>MP</td></tr>
<tr><td>>am_MI0000786|hsa-mir-378a&WithFlank&5|+|149732819|149732896@49.69.21(+1U)&[MIMAT0000732&hsa-miR-378a-3p&offsets|0|-1(+1U);m-44&5|+|149732867|149732888&offsets|0|-1(+1U)]|ACTGGACTTGGAGTCAGAAGGT</td><td>>am_MI0000786|hsa-mir-378a&WithFlank&5|+|149732819|149732896</td><td>49</td><td>69</td><td>21(+1U)</td><td>ACTGGACTTGGAGTCAGAAGGT</td><td>6430.3</td><td>2.2</td><td>0.8</td><td>2.7</td><td>7.9E-03</td><td>2.5E-02</td><td>BT20/MB231</td><td>CYTO</td></tr>
<tr><td>>trnaMT_ValTAC_MT_+_1602_1670@54.72.19|AACTTGACCGCTCTGACCA</td><td>>trnaMT_ValTAC_MT_+_1602_1670</td><td>54</td><td>72</td><td>19</td><td>AACTTGACCGCTCTGACCA</td><td>2362.1</td><td>2.5</td><td>0.9</td><td>2.7</td><td>7.8E-03</td><td>2.5E-02</td><td>BT20/MB231</td><td>CYTO</td></tr>
<tr><td>>hg38_21_+_8433172_8446622_RNA45SN1_45S_50ntFlanks@10427.10456.30|ATGGGCGAGCGCCGTTCCGAAGGGACGGGC</td><td>>hg38_21_+_8433172_8446622_RNA45SN1_45S_50ntFlanks</td><td>10427</td><td>10456</td><td>30</td><td>ATGGGCGAGCGCCGTTCCGAAGGGACGGGC</td><td>335.8</td><td>-2.1</td><td>0.8</td><td>-2.7</td><td>7.9E-03</td><td>2.5E-02</td><td>BT20/MB231</td><td>CYTO</td></tr>
<tr><td>>hg38_21_+_8433172_8446622_RNA45SN1_45S_50ntFlanks@4585.4622.38|GTTCTATTTTGTTGGTTTTCGGAACTGAGGCCATGATT</td><td>>hg38_21_+_8433172_8446622_RNA45SN1_45S_50ntFlanks</td><td>4585</td><td>4622</td><td>38</td><td>GTTCTATTTTGTTGGTTTTCGGAACTGAGGCCATGATT</td><td>532.5</td><td>-2.6</td><td>1.0</td><td>-2.7</td><td>7.9E-03</td><td>2.5E-02</td><td>BT20/MB231</td><td>CYTO</td></tr>
<tr><td>>hg38_MT_+_1621_3279_RNR2_16S_50ntFlanks@1347.1385.39|CAACAATAGGGTTTACGACCTCGATGTTGGATCAGGACA</td><td>>hg38_MT_+_1621_3279_RNR2_16S_50ntFlanks</td><td>1347</td><td>1385</td><td>39</td><td>CAACAATAGGGTTTACGACCTCGATGTTGGATCAGGACA</td><td>53.3</td><td>6.3</td><td>2.6</td><td>2.5</td><td>1.4E-02</td><td>2.5E-02</td><td>BT20/MB468</td><td>MP</td></tr>
<tr><td>>hg38_MT_+_1621_3279_RNR2_16S_50ntFlanks@826.855.30|AAGGAAAGGTTAAAAAAAGTAAAAGGAACT</td><td>>hg38_MT_+_1621_3279_RNR2_16S_50ntFlanks</td><td>826</td><td>855</td><td>30</td><td>AAGGAAAGGTTAAAAAAAGTAAAAGGAACT</td><td>141.0</td><td>3.6</td><td>1.5</td><td>2.5</td><td>1.4E-02</td><td>2.5E-02</td><td>BT20/MB468</td><td>MP</td></tr>
<tr><td>>hg38_21_+_8433172_8446622_RNA45SN1_45S_50ntFlanks@11540.11584.45|CGCGGCGCCTCGCCTCGGCCGGCGCCTAGCAGCCGACTTAGAACT</td><td>>hg38_21_+_8433172_8446622_RNA45SN1_45S_50ntFlanks</td><td>11540</td><td>11584</td><td>45</td><td>CGCGGCGCCTCGCCTCGGCCGGCGCCTAGCAGCCGACTTAGAACT</td><td>2073.0</td><td>3.2</td><td>1.4</td><td>2.3</td><td>2.2E-02</td><td>2.5E-02</td><td>MB231/MB468</td><td>NUC</td></tr>
<tr><td>>hg38_21_+_8433172_8446622_RNA45SN1_45S_50ntFlanks@11565.11603.39|CTAGCAGCCGACTTAGAACTGGTGCGGACCAGGGGAATC</td><td>>hg38_21_+_8433172_8446622_RNA45SN1_45S_50ntFlanks</td><td>11565</td><td>11603</td><td>39</td><td>CTAGCAGCCGACTTAGAACTGGTGCGGACCAGGGGAATC</td><td>374.1</td><td>-1.8</td><td>0.8</td><td>-2.4</td><td>1.9E-02</td><td>2.5E-02</td><td>BT20/MB468</td><td>MITO</td></tr>
<tr><td>>hg38_21_+_8433172_8446622_RNA45SN1_45S_50ntFlanks@9739.9762.24|AAACGATCTCAACCTATTCTCAAA</td><td>>hg38_21_+_8433172_8446622_RNA45SN1_45S_50ntFlanks</td><td>9739</td><td>9762</td><td>24</td><td>AAACGATCTCAACCTATTCTCAAA</td><td>354.0</td><td>-2.2</td><td>0.9</td><td>-2.4</td><td>1.9E-02</td><td>2.5E-02</td><td>BT20/MB468</td><td>MITO</td></tr>
<tr><td>>hg38_21_+_8433172_8446622_RNA45SN1_45S_50ntFlanks@10738.10755.18|AAATCTCGCGCCGGGCCG</td><td>>hg38_21_+_8433172_8446622_RNA45SN1_45S_50ntFlanks</td><td>10738</td><td>10755</td><td>18</td><td>AAATCTCGCGCCGGGCCG</td><td>331.5</td><td>-1.7</td><td>0.7</td><td>-2.4</td><td>1.9E-02</td><td>2.5E-02</td><td>BT20/MB468</td><td>MITO</td></tr>
<tr><td>>hg38_21_+_8433172_8446622_RNA45SN1_45S_50ntFlanks@10350.10371.22|ACGAGAACTTTGAAGGCCGAAG</td><td>>hg38_21_+_8433172_8446622_RNA45SN1_45S_50ntFlanks</td><td>10350</td><td>10371</td><td>22</td><td>ACGAGAACTTTGAAGGCCGAAG</td><td>726.4</td><td>-1.7</td><td>0.7</td><td>-2.3</td><td>2.1E-02</td><td>2.5E-02</td><td>BT20/MB468</td><td>CYTO</td></tr>
<tr><td>>trnaMT_LeuTAA_MT_+_3230_3304@1.30.30|GTTAAGATGGCAGAGCCCGGTAATCGCATA</td><td>>trnaMT_LeuTAA_MT_+_3230_3304</td><td>1</td><td>30</td><td>30</td><td>GTTAAGATGGCAGAGCCCGGTAATCGCATA</td><td>155.6</td><td>1.9</td><td>0.8</td><td>2.3</td><td>2.2E-02</td><td>2.5E-02</td><td>BT20/MB468</td><td>NUC</td></tr>
<tr><td>>hg38_21_+_8433172_8446622_RNA45SN1_45S_50ntFlanks@10408.10456.49|GGTCAGTCGGTCCTGAGAGATGGGCGAGCGCCGTTCCGAAGGGACGGGC</td><td>>hg38_21_+_8433172_8446622_RNA45SN1_45S_50ntFlanks</td><td>10408</td><td>10456</td><td>49</td><td>GGTCAGTCGGTCCTGAGAGATGGGCGAGCGCCGTTCCGAAGGGACGGGC</td><td>458.8</td><td>-3.6</td><td>1.6</td><td>-2.3</td><td>2.2E-02</td><td>2.5E-02</td><td>MB231/MB468</td><td>NUC</td></tr>
<tr><td>>MI0000285|hsa-mir-205&WithFlank&1|+|209432127|209432248@40.62.23(+1A)&[MIMAT0000266&hsa-miR-205-5p&offsets|0|+1(+1A);m-351&1|+|209432166|209432187&offsets|0|+1(+1A)]|TCCTTCATTCCACCGGAGTCTGTA</td><td>>MI0000285|hsa-mir-205&WithFlank&1|+|209432127|209432248</td><td>40</td><td>62</td><td>23(+1A)</td><td>TCCTTCATTCCACCGGAGTCTGTA</td><td>590.5</td><td>1.6</td><td>0.7</td><td>2.3</td><td>2.2E-02</td><td>2.5E-02</td><td>BT20/MB468</td><td>NUC</td></tr>
<tr><td>>trnaMT_ValTAC_MT_+_1602_1670@50.72.23|ACTTAACTTGACCGCTCTGACCA</td><td>>trnaMT_ValTAC_MT_+_1602_1670</td><td>50</td><td>72</td><td>23</td><td>ACTTAACTTGACCGCTCTGACCA</td><td>1871.3</td><td>2.7</td><td>1.1</td><td>2.5</td><td>1.4E-02</td><td>2.5E-02</td><td>BT20/MB468</td><td>MP</td></tr>
<tr><td>>hg38_21_+_8433172_8446622_RNA45SN1_45S_50ntFlanks@10466.10486.21|CGTTGCCCTCGGCCGATCGAA</td><td>>hg38_21_+_8433172_8446622_RNA45SN1_45S_50ntFlanks</td><td>10466</td><td>10486</td><td>21</td><td>CGTTGCCCTCGGCCGATCGAA</td><td>161.7</td><td>-1.7</td><td>0.7</td><td>-2.3</td><td>1.9E-02</td><td>2.5E-02</td><td>BT20/MB468</td><td>MITO</td></tr>
<tr><td>>hg38_MT_+_1621_3279_RNR2_16S_50ntFlanks@1388.1414.27|CCGATGGTGCAGCCGCTATTAAAGGTT</td><td>>hg38_MT_+_1621_3279_RNR2_16S_50ntFlanks</td><td>1388</td><td>1414</td><td>27</td><td>CCGATGGTGCAGCCGCTATTAAAGGTT</td><td>72.1</td><td>4.0</td><td>1.7</td><td>2.3</td><td>2.0E-02</td><td>2.5E-02</td><td>MB231/MB468</td><td>MP</td></tr>
<tr><td>>hg38_21_+_8433172_8446622_RNA45SN1_45S_50ntFlanks@4003.4039.37|ATAACCTCGGGCCGATCGCACGCCCCCCGTGGCGGCG</td><td>>hg38_21_+_8433172_8446622_RNA45SN1_45S_50ntFlanks</td><td>4003</td><td>4039</td><td>37</td><td>ATAACCTCGGGCCGATCGCACGCCCCCCGTGGCGGCG</td><td>115.8</td><td>-2.5</td><td>1.1</td><td>-2.3</td><td>1.9E-02</td><td>2.5E-02</td><td>BT20/MB468</td><td>MITO</td></tr>
<tr><td>>hg38_21_+_8433172_8446622_RNA45SN1_45S_50ntFlanks@8114.8136.23|GCGCGGGACATGTGGCGTACGGA</td><td>>hg38_21_+_8433172_8446622_RNA45SN1_45S_50ntFlanks</td><td>8114</td><td>8136</td><td>23</td><td>GCGCGGGACATGTGGCGTACGGA</td><td>332.5</td><td>-1.9</td><td>0.8</td><td>-2.3</td><td>1.9E-02</td><td>2.5E-02</td><td>BT20/MB468</td><td>MITO</td></tr>
<tr><td>>MI0000093|hsa-mir-92a-1&WithFlank&13|+|91351308|91351397@54.74.21(+1A);MI0000094|hsa-mir-92a-2&WithFlank&X|-|134169532|134169618@54.74.21(+1A)&[MIMAT0000092&hsa-miR-92a-3p&offsets|0|-1(+1A);m-9&13|+|91351361|91351382&offsets|0|-1(+1A)];[MIMAT0000092_1&hsa-miR-92a-3p&offsets|0|-1(+1A);m-11&X|-|134169544|134169565&offsets|0|-1(+1A)]|TATTGCACTTGTCCCGGCCTGA</td><td>>MI0000093|hsa-mir-92a-1&WithFlank&13|+|91351308|91351397</td><td>54</td><td>74</td><td>21(+1A)</td><td>TATTGCACTTGTCCCGGCCTGA</td><td>3282.3</td><td>-2.0</td><td>0.8</td><td>-2.7</td><td>7.2E-03</td><td>2.5E-02</td><td>BT20/MB231</td><td>MITO</td></tr>
<tr><td>>hg38_21_+_8433172_8446622_RNA45SN1_45S_50ntFlanks@11663.11691.29|ATTTCTGCCCAGTGCTCTGAATGTCAAAG</td><td>>hg38_21_+_8433172_8446622_RNA45SN1_45S_50ntFlanks</td><td>11663</td><td>11691</td><td>29</td><td>ATTTCTGCCCAGTGCTCTGAATGTCAAAG</td><td>714.1</td><td>2.2</td><td>0.8</td><td>2.7</td><td>7.2E-03</td><td>2.5E-02</td><td>BT20/MB231</td><td>MITO</td></tr>
<tr><td>>MI0000234|hsa-mir-192&WithFlank&11|-|64891131|64891252@30.51.22(+1C)&[MIMAT0000222&hsa-miR-192-5p&offsets|0|+1(+1C);m-59&11|-|64891203|64891223&offsets|0|+1(+1C)]|CTGACCTATGAATTGACAGCCAC</td><td>>MI0000234|hsa-mir-192&WithFlank&11|-|64891131|64891252</td><td>30</td><td>51</td><td>22(+1C)</td><td>CTGACCTATGAATTGACAGCCAC</td><td>244.9</td><td>-2.8</td><td>1.0</td><td>-2.8</td><td>5.9E-03</td><td>2.5E-02</td><td>BT20/MB231</td><td>MP</td></tr>
<tr><td>>hg38_21_+_8433172_8446622_RNA45SN1_45S_50ntFlanks@12444.12477.34|AAGCAGAATTCACCAAGCGTTGGATTGTTCACCC</td><td>>hg38_21_+_8433172_8446622_RNA45SN1_45S_50ntFlanks</td><td>12444</td><td>12477</td><td>34</td><td>AAGCAGAATTCACCAAGCGTTGGATTGTTCACCC</td><td>257.1</td><td>-2.2</td><td>0.9</td><td>-2.3</td><td>2.0E-02</td><td>2.5E-02</td><td>MB231/MB468</td><td>MP</td></tr>
<tr><td>>MI0000736|hsa-mir-30c-1&WithFlank&1|+|40757278|40757378@23.44.22(+1A);MI0000254|hsa-mir-30c-2&WithFlank&6|-|71376954|71377037@13.34.22(+1A)&[MIMAT0000244&hsa-miR-30c-5p&offsets|0|-1(+1A);m-57&1|+|40757300|40757323&offsets|0|-2(+1A)];[MIMAT0000244_1&hsa-miR-30c-5p&offsets|0|-1(+1A);m-56&6|-|71377002|71377025&offsets|0|-2(+1A)]|TGTAAACATCCTACACTCTCAGA</td><td>>MI0000736|hsa-mir-30c-1&WithFlank&1|+|40757278|40757378</td><td>23</td><td>44</td><td>22(+1A)</td><td>TGTAAACATCCTACACTCTCAGA</td><td>847.4</td><td>-3.3</td><td>1.2</td><td>-2.8</td><td>5.9E-03</td><td>2.5E-02</td><td>BT20/MB231</td><td>MP</td></tr>
<tr><td>>MI0000063|hsa-let-7b&WithFlank&22|+|46113680|46113774@12.34.23(+1U)&[MIMAT0000063&hsa-let-7b-5p&offsets|0|+1(+1U);m-14&22|+|46113691|46113712&offsets|0|+1(+1U)]|TGAGGTAGTAGGTTGTGTGGTTTT</td><td>>MI0000063|hsa-let-7b&WithFlank&22|+|46113680|46113774</td><td>12</td><td>34</td><td>23(+1U)</td><td>TGAGGTAGTAGGTTGTGTGGTTTT</td><td>604.5</td><td>1.6</td><td>0.6</td><td>2.7</td><td>6.4E-03</td><td>2.5E-02</td><td>BT20/MB231</td><td>NUC</td></tr>
<tr><td>>hg38_21_+_8433172_8446622_RNA45SN1_45S_50ntFlanks@10394.10424.31|AGCAGTTGAACATGGGTCAGTCGGTCCTGAG</td><td>>hg38_21_+_8433172_8446622_RNA45SN1_45S_50ntFlanks</td><td>10394</td><td>10424</td><td>31</td><td>AGCAGTTGAACATGGGTCAGTCGGTCCTGAG</td><td>1191.3</td><td>1.6</td><td>0.7</td><td>2.3</td><td>2.2E-02</td><td>2.5E-02</td><td>MB231/MB468</td><td>NUC</td></tr>
<tr><td>>hg38_21_+_8433172_8446622_RNA45SN1_45S_50ntFlanks@11874.11901.28|AGAAGACCCTGTTGAGCTTGACTCTAGT</td><td>>hg38_21_+_8433172_8446622_RNA45SN1_45S_50ntFlanks</td><td>11874</td><td>11901</td><td>28</td><td>AGAAGACCCTGTTGAGCTTGACTCTAGT</td><td>167.6</td><td>-2.7</td><td>1.2</td><td>-2.3</td><td>2.2E-02</td><td>2.5E-02</td><td>MB231/MB468</td><td>NUC</td></tr>
<tr><td>>MI0000070|hsa-mir-16-1&WithFlank&13|-|50048967|50049067@20.41.22(+1C);MI0000115|hsa-mir-16-2&WithFlank&3|+|160404739|160404831@16.37.22(+1C)&[MIMAT0000069&hsa-miR-16-5p&offsets|0|0(+1C);m-62&13|-|50049027|50049048&offsets|0|0(+1C)];[MIMAT0000069_1&hsa-miR-16-5p&offsets|0|0(+1C);m-61&3|+|160404754|160404775&offsets|0|0(+1C)]|TAGCAGCACGTAAATATTGGCGC</td><td>>MI0000070|hsa-mir-16-1&WithFlank&13|-|50048967|50049067</td><td>20</td><td>41</td><td>22(+1C)</td><td>TAGCAGCACGTAAATATTGGCGC</td><td>501.9</td><td>3.1</td><td>1.3</td><td>2.4</td><td>1.4E-02</td><td>2.5E-02</td><td>BT20/MB468</td><td>MP</td></tr>
</table>

<table>
<tr><th></th><th></th><th></th><th></th><th></th><th></th><th></th><th></th><th></th><th></th><th></th><th></th><th></th><th></th></tr>
<tr><td>>hg38_21_+_8433172_8446622_RNA45SN1_45S_50ntFlanks@12999.13027.29|CTGCGATCTATTGAAAGTCAGCCCTCGAC</td><td>>hg38_21_+_8433172_8446622_RNA45SN1_45S_50ntFlanks</td><td>12999</td><td>13027</td><td>29</td><td>CTGCGATCTATTGAAAGTCAGCCCTCGAC</td><td>155.6</td><td>-4.2</td><td>1.7</td><td>-2.4</td><td>1.4E-02</td><td>2.5E-02</td><td>BT20/MB468</td><td>MP</td></tr>
<tr><td>>hg38_21_+_8433172_8446622_RNA45SN1_45S_50ntFlanks@12603.12624.22|TGTATGTGCTTGGCTGAGGAGC</td><td>>hg38_21_+_8433172_8446622_RNA45SN1_45S_50ntFlanks</td><td>12603</td><td>12624</td><td>22</td><td>TGTATGTGCTTGGCTGAGGAGC</td><td>436.5</td><td>-2.3</td><td>0.9</td><td>-2.7</td><td>7.9E-03</td><td>2.5E-02</td><td>BT20/MB231</td><td>CYTO</td></tr>
<tr><td>>trna17_ValTAC_11_-_59318102_59318174@1.30.30,trna4_ValTAC_X_-_18693029_18693101@1.30.30|GGTTCCATAGTGTAGTGGTTATCACGTCTG</td><td>>trna17_ValTAC_11_-_59318102_59318174</td><td>1</td><td>30</td><td>30</td><td>GGTTCCATAGTGTAGTGGTTATCACGTCTG</td><td>90.8</td><td>-4.2</td><td>1.7</td><td>-2.4</td><td>1.4E-02</td><td>2.5E-02</td><td>BT20/MB468</td><td>MP</td></tr>
<tr><td>>hg38_21_+_8433172_8446622_RNA45SN1_45S_50ntFlanks@5152.5185.34|AGGGACAAGTGGCGTTCAGCCACCCGAGATTGAG</td><td>>hg38_21_+_8433172_8446622_RNA45SN1_45S_50ntFlanks</td><td>5152</td><td>5185</td><td>34</td><td>AGGGACAAGTGGCGTTCAGCCACCCGAGATTGAG</td><td>565.2</td><td>-4.9</td><td>2.0</td><td>-2.4</td><td>1.4E-02</td><td>2.5E-02</td><td>BT20/MB468</td><td>MP</td></tr>
<tr><td>>hg38_21_+_8433172_8446622_RNA45SN1_45S_50ntFlanks@12640.12659.20|ACCATCTGTGGGATTATGAC</td><td>>hg38_21_+_8433172_8446622_RNA45SN1_45S_50ntFlanks</td><td>12640</td><td>12659</td><td>20</td><td>ACCATCTGTGGGATTATGAC</td><td>212.4</td><td>-1.5</td><td>0.7</td><td>-2.3</td><td>2.1E-02</td><td>2.5E-02</td><td>MB231/MB468</td><td>CYTO</td></tr>
<tr><td>>hg38_MT_+_1621_3279_RNR2_16S_50ntFlanks@1427.1452.26|GATTAAAGTCCTACGTGATCTGAGTT</td><td>>hg38_MT_+_1621_3279_RNR2_16S_50ntFlanks</td><td>1427</td><td>1452</td><td>26</td><td>GATTAAAGTCCTACGTGATCTGAGTT</td><td>172.4</td><td>4.2</td><td>1.8</td><td>2.3</td><td>2.0E-02</td><td>2.5E-02</td><td>MB231/MB468</td><td>MP</td></tr>
<tr><td>>hg38_1_-_228634819_228635039_RNA5S12_5S_50ntFlanks@137.171.35|GAGACCGCCTGGGAATACCGGGTGCTGTAGGCTTT</td><td>>hg38_1_-_228634819_228635039_RNA5S12_5S_50ntFlanks</td><td>137</td><td>171</td><td>35</td><td>GAGACCGCCTGGGAATACCGGGTGCTGTAGGCTTT</td><td>494.1</td><td>1.9</td><td>0.8</td><td>2.3</td><td>2.0E-02</td><td>2.5E-02</td><td>MB231/MB468</td><td>MP</td></tr>
<tr><td>>hg38_21_+_8433172_8446622_RNA45SN1_45S_50ntFlanks@6649.6680.32|TACGACTCTTAGCGGTGGATCACTCGGCTCGT</td><td>>hg38_21_+_8433172_8446622_RNA45SN1_45S_50ntFlanks</td><td>6649</td><td>6680</td><td>32</td><td>TACGACTCTTAGCGGTGGATCACTCGGCTCGT</td><td>321.0</td><td>1.8</td><td>0.8</td><td>2.3</td><td>2.2E-02</td><td>2.5E-02</td><td>MB231/MB468</td><td>NUC</td></tr>
<tr><td>>hg38_21_+_8433172_8446622_RNA45SN1_45S_50ntFlanks@9950.9982.33|GCCATGGAAGTCGGAATCCGCTAAGGAGTGTGT</td><td>>hg38_21_+_8433172_8446622_RNA45SN1_45S_50ntFlanks</td><td>9950</td><td>9982</td><td>33</td><td>GCCATGGAAGTCGGAATCCGCTAAGGAGTGTGT</td><td>94.2</td><td>5.6</td><td>2.0</td><td>2.8</td><td>5.9E-03</td><td>2.5E-02</td><td>BT20/MB231</td><td>MP</td></tr>
<tr><td>>hg38_MT_+_1621_3279_RNR2_16S_50ntFlanks@1390.1422.33|GATGGTGCAGCCGCTATTAAAGGTTCGTTTGTT</td><td>>hg38_MT_+_1621_3279_RNR2_16S_50ntFlanks</td><td>1390</td><td>1422</td><td>33</td><td>GATGGTGCAGCCGCTATTAAAGGTTCGTTTGTT</td><td>58.4</td><td>4.6</td><td>2.0</td><td>2.3</td><td>2.0E-02</td><td>2.5E-02</td><td>MB231/MB468</td><td>MP</td></tr>
<tr><td>>hg38_MT_+_598_1651_RNR1_12S_50ntFlanks@994.1016.23|CTTGGACGAACCAGAGTGTAGCT</td><td>>hg38_MT_+_598_1651_RNR1_12S_50ntFlanks</td><td>994</td><td>1016</td><td>23</td><td>CTTGGACGAACCAGAGTGTAGCT</td><td>93.3</td><td>-5.2</td><td>1.9</td><td>-2.8</td><td>5.9E-03</td><td>2.5E-02</td><td>BT20/MB231</td><td>MP</td></tr>
<tr><td>>hg38_MT_+_1621_3279_RNR2_16S_50ntFlanks@643.663.21|CAATTGGACCAATCTATCACC</td><td>>hg38_MT_+_1621_3279_RNR2_16S_50ntFlanks</td><td>643</td><td>663</td><td>21</td><td>CAATTGGACCAATCTATCACC</td><td>130.2</td><td>3.9</td><td>1.6</td><td>2.4</td><td>1.4E-02</td><td>2.5E-02</td><td>BT20/MB468</td><td>MP</td></tr>
<tr><td>>hg38_21_+_8433172_8446622_RNA45SN1_45S_50ntFlanks@6674.6696.23|GGCTCGTGCGTCGATGAAGAACG</td><td>>hg38_21_+_8433172_8446622_RNA45SN1_45S_50ntFlanks</td><td>6674</td><td>6696</td><td>23</td><td>GGCTCGTGCGTCGATGAAGAACG</td><td>285.7</td><td>4.6</td><td>1.7</td><td>2.8</td><td>5.9E-03</td><td>2.5E-02</td><td>BT20/MB231</td><td>MP</td></tr>
<tr><td>>hg38_21_+_8433172_8446622_RNA45SN1_45S_50ntFlanks@8196.8213.18|CGTGGACGGTGTGAGGCC</td><td>>hg38_21_+_8433172_8446622_RNA45SN1_45S_50ntFlanks</td><td>8196</td><td>8213</td><td>18</td><td>CGTGGACGGTGTGAGGCC</td><td>6914.4</td><td>4.1</td><td>1.5</td><td>2.8</td><td>5.9E-03</td><td>2.5E-02</td><td>BT20/MB231</td><td>MP</td></tr>
<tr><td>>hg38_21_+_8433172_8446622_RNA45SN1_45S_50ntFlanks@7985.8017.33|GATCAGACGTGGCGACCCGCTGAATTTAAGCAT</td><td>>hg38_21_+_8433172_8446622_RNA45SN1_45S_50ntFlanks</td><td>7985</td><td>8017</td><td>33</td><td>GATCAGACGTGGCGACCCGCTGAATTTAAGCAT</td><td>479.4</td><td>-3.9</td><td>1.7</td><td>-2.3</td><td>2.3E-02</td><td>2.5E-02</td><td>MB231/MB468</td><td>MITO</td></tr>
<tr><td>>hg38_21_+_8433172_8446622_RNA45SN1_45S_50ntFlanks@12394.12414.21|TAGCGACGTCGCTTTTTGATC</td><td>>hg38_21_+_8433172_8446622_RNA45SN1_45S_50ntFlanks</td><td>12394</td><td>12414</td><td>21</td><td>TAGCGACGTCGCTTTTTGATC</td><td>933.3</td><td>-2.3</td><td>0.9</td><td>-2.7</td><td>8.0E-03</td><td>2.5E-02</td><td>BT20/MB231</td><td>CYTO</td></tr>
<tr><td>>hg38_21_+_8433172_8446622_RNA45SN1_45S_50ntFlanks@6652.6687.36|GACTCTTAGCGGTGGATCACTCGGCTCGTGCGTCGA</td><td>>hg38_21_+_8433172_8446622_RNA45SN1_45S_50ntFlanks</td><td>6652</td><td>6687</td><td>36</td><td>GACTCTTAGCGGTGGATCACTCGGCTCGTGCGTCGA</td><td>522.7</td><td>2.5</td><td>1.0</td><td>2.4</td><td>1.4E-02</td><td>2.5E-02</td><td>BT20/MB468</td><td>MP</td></tr>
<tr><td>>hg38_21_+_8433172_8446622_RNA45SN1_45S_50ntFlanks@11564.11587.24|CCTAGCAGCCGACTTAGAACTGGT</td><td>>hg38_21_+_8433172_8446622_RNA45SN1_45S_50ntFlanks</td><td>11564</td><td>11587</td><td>24</td><td>CCTAGCAGCCGACTTAGAACTGGT</td><td>154.5</td><td>4.6</td><td>1.7</td><td>2.8</td><td>5.9E-03</td><td>2.5E-02</td><td>BT20/MB231</td><td>MP</td></tr>
<tr><td>>hg38_21_+_8433172_8446622_RNA45SN1_45S_50ntFlanks@7977.8011.35|CGACCTCAGATCAGACGTGGCGACCCGCTGAATTT</td><td>>hg38_21_+_8433172_8446622_RNA45SN1_45S_50ntFlanks</td><td>7977</td><td>8011</td><td>35</td><td>CGACCTCAGATCAGACGTGGCGACCCGCTGAATTT</td><td>3295.5</td><td>4.5</td><td>1.6</td><td>2.8</td><td>5.9E-03</td><td>2.5E-02</td><td>BT20/MB231</td><td>MP</td></tr>
<tr><td>>hg38_MT_+_598_1651_RNR1_12S_50ntFlanks@428.461.34|AAAGTGGCTTTAACATATCTGAACACACAATAGC</td><td>>hg38_MT_+_598_1651_RNR1_12S_50ntFlanks</td><td>428</td><td>461</td><td>34</td><td>AAAGTGGCTTTAACATATCTGAACACACAATAGC</td><td>122.5</td><td>3.8</td><td>1.6</td><td>2.4</td><td>1.4E-02</td><td>2.5E-02</td><td>BT20/MB468</td><td>MP</td></tr>
<tr><td>>hg38_21_+_8433172_8446622_RNA45SN1_45S_50ntFlanks@9177.9197.21|GCGCCGTCGGGCCCGGGGGAG</td><td>>hg38_21_+_8433172_8446622_RNA45SN1_45S_50ntFlanks</td><td>9177</td><td>9197</td><td>21</td><td>GCGCCGTCGGGCCCGGGGGAG</td><td>154.6</td><td>-4.4</td><td>1.8</td><td>-2.4</td><td>1.4E-02</td><td>2.5E-02</td><td>BT20/MB468</td><td>MP</td></tr>
<tr><td>>MI0000234|hsa-mir-192&WithFlank&11|-|64891131|64891252@30.51.22(+1U)&[MIMAT0000222&hsa-miR-192-5p&offsets|0|+1(+1U);m-59&11|-|64891203|64891223&offsets|0|+1(+1U)]|CTGACCTATGAATTGACAGCCAT</td><td>>MI0000234|hsa-mir-192&WithFlank&11|-|64891131|64891252</td><td>30</td><td>51</td><td>22(+1U)</td><td>CTGACCTATGAATTGACAGCCAT</td><td>1106.5</td><td>-1.9</td><td>0.7</td><td>-2.7</td><td>7.2E-03</td><td>2.5E-02</td><td>BT20/MB231</td><td>MITO</td></tr>
<tr><td>>MI0000749|hsa-mir-30e&WithFlank&1|+|40754349|40754452@24.45.22(+1C)&[MIMAT0000692&hsa-miR-30e-5p&offsets|+1|+1(+1C);m-24&1|+|40754371|40754394&offsets|+1|-1(+1C)]|GTAAACATCCTTGACTGGAAGCC</td><td>>MI0000749|hsa-mir-30e&WithFlank&1|+|40754349|40754452</td><td>24</td><td>45</td><td>22(+1C)</td><td>GTAAACATCCTTGACTGGAAGCC</td><td>174.1</td><td>2.1</td><td>0.9</td><td>2.3</td><td>2.0E-02</td><td>2.5E-02</td><td>MB231/MB468</td><td>MP</td></tr>
<tr><td>>hg38_21_+_8433172_8446622_RNA45SN1_45S_50ntFlanks@9533.9566.34|GAGGAAACTCTGGTGGAGGTCCGTAGCGGTCCTG</td><td>>hg38_21_+_8433172_8446622_RNA45SN1_45S_50ntFlanks</td><td>9533</td><td>9566</td><td>34</td><td>GAGGAAACTCTGGTGGAGGTCCGTAGCGGTCCTG</td><td>130.5</td><td>-3.4</td><td>1.4</td><td>-2.4</td><td>1.4E-02</td><td>2.5E-02</td><td>BT20/MB468</td><td>MP</td></tr>
<tr><td>>hg38_MT_+_1621_3279_RNR2_16S_50ntFlanks@485.517.33|GAGGAACAGCTCTTTGGACACTAGGAAAAAACC</td><td>>hg38_MT_+_1621_3279_RNR2_16S_50ntFlanks</td><td>485</td><td>517</td><td>33</td><td>GAGGAACAGCTCTTTGGACACTAGGAAAAAACC</td><td>64.0</td><td>3.8</td><td>1.6</td><td>2.3</td><td>2.0E-02</td><td>2.5E-02</td><td>MB231/MB468</td><td>MP</td></tr>
<tr><td>>hg38_1_-_228634819_228635039_RNA5S12_5S_50ntFlanks@152.169.18|TACCGGGTGCTGTAGGCT</td><td>>hg38_1_-_228634819_228635039_RNA5S12_5S_50ntFlanks</td><td>152</td><td>169</td><td>18</td><td>TACCGGGTGCTGTAGGCT</td><td>1010.5</td><td>2.4</td><td>0.9</td><td>2.8</td><td>5.9E-03</td><td>2.5E-02</td><td>BT20/MB231</td><td>MP</td></tr>
<tr><td>>MI0000270|hsa-mir-181b-1&WithFlank&1|-|198858867|198858988@42.64.23;MI0000683|hsa-mir-181b-2&WithFlank&9|+|124693704|124693804@22.44.23&[MIMAT0000257&hsa-miR-181b-5p&offsets|0|0;m-73&1|-|198858925|198858947&offsets|0|0];[MIMAT0000257_1&hsa-miR-181b-5p&offsets|0|0;m-72&9|+|124693725|124693747&offsets|0|0]|AACATTCATTGCTGTCGGTGGGT</td><td>>MI0000270|hsa-mir-181b-1&WithFlank&1|-|198858867|198858988</td><td>42</td><td>64</td><td>23</td><td>AACATTCATTGCTGTCGGTGGGT</td><td>3159.4</td><td>-2.0</td><td>0.7</td><td>-2.7</td><td>7.3E-03</td><td>2.5E-02</td><td>BT20/MB231</td><td>MITO</td></tr>
<tr><td>>MI0000749|hsa-mir-30e&WithFlank&1|+|40754349|40754452@23.46.24&[MIMAT0000692&hsa-miR-30e-5p&offsets|0|+2;m-24&1|+|40754371|40754394&offsets|0|0]|TGTAAACATCCTTGACTGGAAGCT</td><td>>MI0000749|hsa-mir-30e&WithFlank&1|+|40754349|40754452</td><td>23</td><td>46</td><td>24</td><td>TGTAAACATCCTTGACTGGAAGCT</td><td>8202.9</td><td>-2.4</td><td>0.9</td><td>-2.7</td><td>7.3E-03</td><td>2.5E-02</td><td>BT20/MB231</td><td>MITO</td></tr>
<tr><td>>hg38_21_+_8433172_8446622_RNA45SN1_45S_50ntFlanks@11208.11252.45|GCGGCGACTCTGGACGCGAGCCGGGCCCTTCCCGTGGATCGCCCC</td><td>>hg38_21_+_8433172_8446622_RNA45SN1_45S_50ntFlanks</td><td>11208</td><td>11252</td><td>45</td><td>GCGGCGACTCTGGACGCGAGCCGGGCCCTTCCCGTGGATCGCCCC</td><td>528.9</td><td>7.1</td><td>2.7</td><td>2.7</td><td>7.3E-03</td><td>2.5E-02</td><td>BT20/MB231</td><td>MITO</td></tr>
<tr><td>>MI0000750|hsa-mir-26a-2&WithFlank&12|-|57824603|57824698@20.43.24;MI0000083|hsa-mir-26a-1&WithFlank&3|+|37969398|37969486@16.39.24&[MIMAT0000082&hsa-miR-26a-5p&offsets|0|+2;m-33&12|-|57824658|57824679&offsets|0|+2];[MIMAT0000082_1&hsa-miR-26a-5p&offsets|0|+2;m-34&3|+|37969413|37969434&offsets|0|+2]|TTCAAGTAATCCAGGATAGGCTGT</td><td>>MI0000750|hsa-mir-26a-2&WithFlank&12|-|57824603|57824698</td><td>20</td><td>43</td><td>24</td><td>TTCAAGTAATCCAGGATAGGCTGT</td><td>307.6</td><td>-2.3</td><td>0.9</td><td>-2.7</td><td>7.3E-03</td><td>2.5E-02</td><td>BT20/MB231</td><td>MITO</td></tr>
<tr><td>>hg38_21_+_8433172_8446622_RNA45SN1_45S_50ntFlanks@4588.4614.27|CTATTTTGTTGGTTTTCGGAACTGAGG</td><td>>hg38_21_+_8433172_8446622_RNA45SN1_45S_50ntFlanks</td><td>4588</td><td>4614</td><td>27</td><td>CTATTTTGTTGGTTTTCGGAACTGAGG</td><td>675.3</td><td>-4.4</td><td>1.8</td><td>-2.4</td><td>1.4E-02</td><td>2.5E-02</td><td>BT20/MB468</td><td>MP</td></tr>
<tr><td>>trna10_GluCTC_3_-_103879524_103879595@16.42.27,trna116_GluCTC_1_-_145399233_145399304@16.42.27,trna59_GluCTC_1_+_249168447_249168518@16.42.27,trna71_GluCTC_1_-_161439189_161439260@16.42.27,trna74_GluCTC_1_-_161431809_161431880@16.42.27,trna77_GluCTC_1_-_161424398_161424469@16.42.27,trna77_GluCTC_6_+_28949976_28950047@16.42.27,trna80_GluCTC_1_-_161417018_161417089@16.42.27,trna87_GluCTC_6_-_126101393_126101464@16.42.27|TGGTTAGGATTCGGCGCTCTCACCGCC</td><td>>trna10_GluCTC_3_-_103879524_103879595</td><td>16</td><td>42</td><td>27</td><td>TGGTTAGGATTCGGCGCTCTCACCGCC</td><td>145.0</td><td>1.7</td><td>0.7</td><td>2.3</td><td>2.0E-02</td><td>2.5E-02</td><td>MB231/MB468</td><td>MP</td></tr>
<tr><td>>trnaMT_GluTTC_MT_-_14674_14742@24.67.44,trnalookalike8_GluTTC_5_-_93905172_93905240@24.67.44|ATGGTTTTTCATATCATTGGTCGTGGTTGTAGTCCGTGCGAGAA</td><td>>trnaMT_GluTTC_MT_-_14674_14742</td><td>24</td><td>67</td><td>44</td><td>ATGGTTTTTCATATCATTGGTCGTGGTTGTAGTCCGTGCGAGAA</td><td>34.3</td><td>8.4</td><td>3.1</td><td>2.8</td><td>5.9E-03</td><td>2.5E-02</td><td>BT20/MB231</td><td>MP</td></tr>
<tr><td>>hg38_21_+_8433172_8446622_RNA45SN1_45S_50ntFlanks@12988.13012.25|CAGCTCCCTCGCTGCGATCTATTGA</td><td>>hg38_21_+_8433172_8446622_RNA45SN1_45S_50ntFlanks</td><td>12988</td><td>13012</td><td>25</td><td>CAGCTCCCTCGCTGCGATCTATTGA</td><td>201.2</td><td>-2.0</td><td>0.8</td><td>-2.7</td><td>8.0E-03</td><td>2.5E-02</td><td>BT20/MB231</td><td>CYTO</td></tr>
<tr><td>>hg38_21_+_8433172_8446622_RNA45SN1_45S_50ntFlanks@3993.4011.19|GTGACTCTAGATAACCTCG</td><td>>hg38_21_+_8433172_8446622_RNA45SN1_45S_50ntFlanks</td><td>3993</td><td>4011</td><td>19</td><td>GTGACTCTAGATAACCTCG</td><td>244.2</td><td>-2.3</td><td>0.9</td><td>-2.7</td><td>8.0E-03</td><td>2.5E-02</td><td>BT20/MB231</td><td>CYTO</td></tr>
<tr><td>>hg38_21_+_8433172_8446622_RNA45SN1_45S_50ntFlanks@476.501.26|TTCGTGATCGATGTGGTGACGTCGTG</td><td>>hg38_21_+_8433172_8446622_RNA45SN1_45S_50ntFlanks</td><td>476</td><td>501</td><td>26</td><td>TTCGTGATCGATGTGGTGACGTCGTG</td><td>822.6</td><td>1.2</td><td>0.5</td><td>2.3</td><td>2.2E-02</td><td>2.5E-02</td><td>BT20/MB468</td><td>NUC</td></tr>
<tr><td>>MI0000079|hsa-mir-23a&WithFlank&19|-|13836581|13836665@15.35.21&[MIMAT0004496&hsa-miR-23a-5p&offsets|0|-1;m-447&19|-|13836630|13836651&offsets|0|-1]|GGGGTTCCTGGGGATGGGATT</td><td>>MI0000079|hsa-mir-23a&WithFlank&19|-|13836581|13836665</td><td>15</td><td>35</td><td>21</td><td>GGGGTTCCTGGGGATGGGATT</td><td>66.5</td><td>5.4</td><td>2.2</td><td>2.4</td><td>1.4E-02</td><td>2.5E-02</td><td>BT20/MB468</td><td>MP</td></tr>
<tr><td>>hg38_21_+_8433172_8446622_RNA45SN1_45S_50ntFlanks@10473.10512.40|CTCGGCCGATCGAAAGGGAGTCGGGTTCAGATCCCCGAAT</td><td>>hg38_21_+_8433172_8446622_RNA45SN1_45S_50ntFlanks</td><td>10473</td><td>10512</td><td>40</td><td>CTCGGCCGATCGAAAGGGAGTCGGGTTCAGATCCCCGAAT</td><td>240.0</td><td>-1.6</td><td>0.7</td><td>-2.3</td><td>2.2E-02</td><td>2.5E-02</td><td>MB231/MB468</td><td>CYTO</td></tr>
<tr><td>>hg38_21_+_8433172_8446622_RNA45SN1_45S_50ntFlanks@6645.6690.46|CTCGTACGACTCTTAGCGGTGGATCACTCGGCTCGTGCGTCGATGA</td><td>>hg38_21_+_8433172_8446622_RNA45SN1_45S_50ntFlanks</td><td>6645</td><td>6690</td><td>46</td><td>CTCGTACGACTCTTAGCGGTGGATCACTCGGCTCGTGCGTCGATGA</td><td>191.8</td><td>4.5</td><td>1.8</td><td>2.4</td><td>1.4E-02</td><td>2.5E-02</td><td>BT20/MB468</td><td>MP</td></tr>
<tr><td>>hg38_21_+_8433172_8446622_RNA45SN1_45S_50ntFlanks@8352.8375.24|AGTTGAAAAGAACTTTGAAGAGAG</td><td>>hg38_21_+_8433172_8446622_RNA45SN1_45S_50ntFlanks</td><td>8352</td><td>8375</td><td>24</td><td>AGTTGAAAAGAACTTTGAAGAGAG</td><td>179.0</td><td>3.6</td><td>1.6</td><td>2.3</td><td>2.0E-02</td><td>2.5E-02</td><td>MB231/MB468</td><td>MP</td></tr>
<tr><td>>hg38_21_+_8433172_8446622_RNA45SN1_45S_50ntFlanks@12626.12659.34|AATGGGGCGAAGCTACCATCTGTGGGATTATGAC</td><td>>hg38_21_+_8433172_8446622_RNA45SN1_45S_50ntFlanks</td><td>12626</td><td>12659</td><td>34</td><td>AATGGGGCGAAGCTACCATCTGTGGGATTATGAC</td><td>420.6</td><td>-1.7</td><td>0.7</td><td>-2.3</td><td>2.2E-02</td><td>2.5E-02</td><td>MB231/MB468</td><td>CYTO</td></tr>
<tr><td>>trnaMT_PheGAA_MT_+_577_647@27.60.34|AATACACTGAAAATGTTTAGACGGGCTCACATCA</td><td>>trnaMT_PheGAA_MT_+_577_647</td><td>27</td><td>60</td><td>34</td><td>AATACACTGAAAATGTTTAGACGGGCTCACATCA</td><td>34.3</td><td>8.4</td><td>3.1</td><td>2.8</td><td>6.0E-03</td><td>2.5E-02</td><td>BT20/MB231</td><td>MP</td></tr>
<tr><td>>MI0000239|hsa-mir-197&WithFlank&1|+|109598887|109598973@54.74.21&[MIMAT0000227&hsa-miR-197-3p&offsets|0|-1;m-99&1|+|109598940|109598961&offsets|0|-1]|TTCACCACCTTCTCCACCCAG</td><td>>MI0000239|hsa-mir-197&WithFlank&1|+|109598887|109598973</td><td>54</td><td>74</td><td>21</td><td>TTCACCACCTTCTCCACCCAG</td><td>362.8</td><td>2.0</td><td>0.7</td><td>2.7</td><td>7.3E-03</td><td>2.5E-02</td><td>BT20/MB231</td><td>MITO</td></tr>
<tr><td>>MI0000077|hsa-mir-21&WithFlank&17|+|59841260|59841343@14.36.23(+1G)&[MIMAT0000076&hsa-miR-21-5p&offsets|0|+1(+1G);m-1&17|+|59841273|59841294&offsets|0|+1(+1G)]|TAGCTTATCAGACTGATGTTGACG</td><td>>MI0000077|hsa-mir-21&WithFlank&17|+|59841260|59841343</td><td>14</td><td>36</td><td>23(+1G)</td><td>TAGCTTATCAGACTGATGTTGACG</td><td>16158.3</td><td>2.5</td><td>0.9</td><td>2.7</td><td>8.0E-03</td><td>2.5E-02</td><td>BT20/MB231</td><td>CYTO</td></tr>
<tr><td>>hg38_21_+_8433172_8446622_RNA45SN1_45S_50ntFlanks@11571.11594.24|GCCGACTTAGAACTGGTGCGGACC</td><td>>hg38_21_+_8433172_8446622_RNA45SN1_45S_50ntFlanks</td><td>11571</td><td>11594</td><td>24</td><td>GCCGACTTAGAACTGGTGCGGACC</td><td>334.3</td><td>-1.9</td><td>0.7</td><td>-2.7</td><td>8.0E-03</td><td>2.5E-02</td><td>BT20/MB231</td><td>CYTO</td></tr>
<tr><td>>trna15_ValAAC_5_-_180615416_180615488@1.29.29|GTTTCCGTAGTGTAGTGGTCATCACGTTC</td><td>>trna15_ValAAC_5_-_180615416_180615488</td><td>1</td><td>29</td><td>29</td><td>GTTTCCGTAGTGTAGTGGTCATCACGTTC</td><td>423.6</td><td>-3.9</td><td>1.7</td><td>-2.3</td><td>1.9E-02</td><td>2.5E-02</td><td>BT20/MB468</td><td>MITO</td></tr>
<tr><td>>am_MI0000809|hsa-mir-151a&WithFlank&8|-|140732558|140732659@53.71.19(+1A)&[MIMAT0000757&hsa-miR-151a-3p&offsets|0|-2(+1A);m-30&8|-|140732587|140732607&offsets|0|-2(+1A)]|CTAGACTGAAGCTCCTTGAA</td><td>>am_MI0000809|hsa-mir-151a&WithFlank&8|-|140732558|140732659</td><td>53</td><td>71</td><td>19(+1A)</td><td>CTAGACTGAAGCTCCTTGAA</td><td>366.4</td><td>1.8</td><td>0.6</td><td>2.7</td><td>6.5E-03</td><td>2.5E-02</td><td>BT20/MB231</td><td>NUC</td></tr>
</table>

<table>
<tr><th></th><th></th><th></th><th></th><th></th><th></th><th></th><th></th><th></th><th></th><th></th><th></th><th></th><th></th></tr>
<tr><td>>MI0000736|hsa-mir-30c-1&WithFlank&1|+|40757278|40757378@23.48.26;MI0000254|hsa-mir-30c-2&WithFlank&6|-|71376954|71377037@13.38.26&[MIMAT0000244&hsa-miR-30c-5p&offsets|0|+3;m-57&1|+|40757300|40757323&offsets|0|+2];[MIMAT0000244_1&hsa-miR-30c-5p&offsets|0|+3;m-56&6|-|71377002|71377025&offsets|0|+2]|TGTAAACATCCTACACTCTCAGCTGT</td><td>>MI0000736|hsa-mir-30c-1&WithFlank&1|+|40757278|40757378</td><td>23</td><td>48</td><td>26</td><td>TGTAAACATCCTACACTCTCAGCTGT</td><td>246.8</td><td>-3.6</td><td>1.3</td><td>-2.7</td><td>6.5E-03</td><td>2.5E-02</td><td>BT20/MB231</td><td>NUC</td></tr>
<tr><td>>hg38_21_+_8433172_8446622_RNA45SN1_45S_50ntFlanks@4006.4023.18|ACCTCGGGCCGATCGCAC</td><td>>hg38_21_+_8433172_8446622_RNA45SN1_45S_50ntFlanks</td><td>4006</td><td>4023</td><td>18</td><td>ACCTCGGGCCGATCGCAC</td><td>1163.0</td><td>2.2</td><td>1.0</td><td>2.3</td><td>2.2E-02</td><td>2.5E-02</td><td>BT20/MB468</td><td>CYTO</td></tr>
<tr><td>>hg38_21_+_8433172_8446622_RNA45SN1_45S_50ntFlanks@3813.3837.25|TTATGGTTCCTTTGGTCGCTCGCTC</td><td>>hg38_21_+_8433172_8446622_RNA45SN1_45S_50ntFlanks</td><td>3813</td><td>3837</td><td>25</td><td>TTATGGTTCCTTTGGTCGCTCGCTC</td><td>129.1</td><td>-3.3</td><td>1.3</td><td>-2.4</td><td>1.4E-02</td><td>2.5E-02</td><td>BT20/MB468</td><td>MP</td></tr>
<tr><td>>hg38_21_+_8433172_8446622_RNA45SN1_45S_50ntFlanks@12083.12106.24|CTCGCTTCTGGCGCCAAGCGCCCG</td><td>>hg38_21_+_8433172_8446622_RNA45SN1_45S_50ntFlanks</td><td>12083</td><td>12106</td><td>24</td><td>CTCGCTTCTGGCGCCAAGCGCCCG</td><td>193.0</td><td>-4.2</td><td>1.7</td><td>-2.4</td><td>1.4E-02</td><td>2.5E-02</td><td>BT20/MB468</td><td>MP</td></tr>
<tr><td>>hg38_MT_+_1621_3279_RNR2_16S_50ntFlanks@1390.1427.38|GATGGTGCAGCCGCTATTAAAGGTTCGTTTGTTCAACG</td><td>>hg38_MT_+_1621_3279_RNR2_16S_50ntFlanks</td><td>1390</td><td>1427</td><td>38</td><td>GATGGTGCAGCCGCTATTAAAGGTTCGTTTGTTCAACG</td><td>117.2</td><td>3.0</td><td>1.3</td><td>2.3</td><td>2.0E-02</td><td>2.5E-02</td><td>MB231/MB468</td><td>MP</td></tr>
<tr><td>>trna10_AlaCGC_6_+_26553731_26553802@1.19.19,trna119_AlaCGC_6_-_28641613_28641684@1.19.19,trna120_AlaAGC_6_-_28626014_28626085@1.19.19,trna13_AlaCGC_2_+_157257281_157257352@1.19.19,trna13_AlaTGC_12_-_125406301_125406372@1.19.19,trna66_AlaTGC_6_+_28611222_28611293@1.19.19,trna8_AlaTGC_12_+_125424512_125424583@1.19.19,trna8_AlaTGC_5_+_180633868_180633939@1.19.19|GGGGATGTAGCTCAGTGGT</td><td>>trna10_AlaCGC_6_+_26553731_26553802</td><td>1</td><td>19</td><td>19</td><td>GGGGATGTAGCTCAGTGGT</td><td>403.5</td><td>2.3</td><td>1.0</td><td>2.3</td><td>2.0E-02</td><td>2.5E-02</td><td>MB231/MB468</td><td>MP</td></tr>
<tr><td>>hg38_21_+_8433172_8446622_RNA45SN1_45S_50ntFlanks@10387.10424.38|TGTGAACAGCAGTTGAACATGGGTCAGTCGGTCCTGAG</td><td>>hg38_21_+_8433172_8446622_RNA45SN1_45S_50ntFlanks</td><td>10387</td><td>10424</td><td>38</td><td>TGTGAACAGCAGTTGAACATGGGTCAGTCGGTCCTGAG</td><td>485.6</td><td>1.7</td><td>0.7</td><td>2.3</td><td>2.2E-02</td><td>2.5E-02</td><td>BT20/MB468</td><td>NUC</td></tr>
<tr><td>>MI0000086|hsa-mir-28&WithFlank&3|+|188688775|188688872@60.81.22&[MIMAT0004502&hsa-miR-28-3p&offsets|0|0;m-28&3|+|188688834|188688855&offsets|0|0]|CACTAGATTGTGAGCTCCTGGA</td><td>>MI0000086|hsa-mir-28&WithFlank&3|+|188688775|188688872</td><td>60</td><td>81</td><td>22</td><td>CACTAGATTGTGAGCTCCTGGA</td><td>29609.4</td><td>2.2</td><td>0.8</td><td>2.6</td><td>8.1E-03</td><td>2.5E-02</td><td>BT20/MB231</td><td>CYTO</td></tr>
<tr><td>>trna143_LysTTT_6_-_27559593_27559665@14.33.20,trna149_LysTTT_6_-_27302769_27302841@14.33.20,trna14_LysTTT_11_-_59327808_59327880@14.33.20,trna2_LysTTT_17_+_8022473_8022545@14.33.20,trna54_LysTTT_1_+_204475655_204475727@14.33.20,trna5_LysTTT_11_+_59323902_59323974@14.33.20,trna62_LysTTT_1_-_204476158_204476230@14.33.20,trna76_LysTTT_6_+_28918806_28918878@14.33.20|AGTCGGTAGAGCATCAGACT</td><td>>trna143_LysTTT_6_-_27559593_27559665</td><td>14</td><td>33</td><td>20</td><td>AGTCGGTAGAGCATCAGACT</td><td>183.3</td><td>-4.2</td><td>1.5</td><td>-2.7</td><td>6.0E-03</td><td>2.5E-02</td><td>BT20/MB231</td><td>MP</td></tr>
<tr><td>>hg38_21_+_8433172_8446622_RNA45SN1_45S_50ntFlanks@11772.11794.23|TTAGTGACGCGCATGAATGGATG</td><td>>hg38_21_+_8433172_8446622_RNA45SN1_45S_50ntFlanks</td><td>11772</td><td>11794</td><td>23</td><td>TTAGTGACGCGCATGAATGGATG</td><td>715.1</td><td>1.6</td><td>0.7</td><td>2.3</td><td>2.2E-02</td><td>2.5E-02</td><td>MB231/MB468</td><td>CYTO</td></tr>
<tr><td>>hg38_21_+_8433172_8446622_RNA45SN1_45S_50ntFlanks@12408.12432.25|TTTGATCCTTCGATGTCGGCTCTTC</td><td>>hg38_21_+_8433172_8446622_RNA45SN1_45S_50ntFlanks</td><td>12408</td><td>12432</td><td>25</td><td>TTTGATCCTTCGATGTCGGCTCTTC</td><td>407.7</td><td>-2.2</td><td>0.9</td><td>-2.3</td><td>2.2E-02</td><td>2.5E-02</td><td>BT20/MB468</td><td>CYTO</td></tr>
<tr><td>>hg38_21_+_8433172_8446622_RNA45SN1_45S_50ntFlanks@6661.6695.35|CGGTGGATCACTCGGCTCGTGCGTCGATGAAGAAC</td><td>>hg38_21_+_8433172_8446622_RNA45SN1_45S_50ntFlanks</td><td>6661</td><td>6695</td><td>35</td><td>CGGTGGATCACTCGGCTCGTGCGTCGATGAAGAAC</td><td>452.1</td><td>1.7</td><td>0.7</td><td>2.3</td><td>2.3E-02</td><td>2.5E-02</td><td>MB231/MB468</td><td>MITO</td></tr>
<tr><td>>MI0000542|hsa-mir-320a&WithFlank&8|-|22244960|22245043@48.69.22(+1G)&[MIMAT0000510&hsa-miR-320a-3p&offsets|0|0(+1G);m-60&8|-|22244975|22244996&offsets|0|0(+1G)]|AAAAGCTGGGTTGAGAGGGCGAG</td><td>>MI0000542|hsa-mir-320a&WithFlank&8|-|22244960|22245043</td><td>48</td><td>69</td><td>22(+1G)</td><td>AAAAGCTGGGTTGAGAGGGCGAG</td><td>89.2</td><td>3.2</td><td>1.4</td><td>2.3</td><td>2.0E-02</td><td>2.5E-02</td><td>MB231/MB468</td><td>MP</td></tr>
<tr><td>>hg38_21_+_8433172_8446622_RNA45SN1_45S_50ntFlanks@6777.6806.30|TCCCGGGGCTACGCCTGTCTGAGCGTCGCT</td><td>>hg38_21_+_8433172_8446622_RNA45SN1_45S_50ntFlanks</td><td>6777</td><td>6806</td><td>30</td><td>TCCCGGGGCTACGCCTGTCTGAGCGTCGCT</td><td>154.1</td><td>1.9</td><td>0.8</td><td>2.3</td><td>2.0E-02</td><td>2.5E-02</td><td>MB231/MB468</td><td>MP</td></tr>
<tr><td>>hg38_21_+_8433172_8446622_RNA45SN1_45S_50ntFlanks@9841.9864.24|AAGCAGAACTGGCGCTGCGGGATG</td><td>>hg38_21_+_8433172_8446622_RNA45SN1_45S_50ntFlanks</td><td>9841</td><td>9864</td><td>24</td><td>AAGCAGAACTGGCGCTGCGGGATG</td><td>95.0</td><td>3.6</td><td>1.5</td><td>2.4</td><td>1.5E-02</td><td>2.5E-02</td><td>BT20/MB468</td><td>MP</td></tr>
<tr><td>>hg38_21_+_8433172_8446622_RNA45SN1_45S_50ntFlanks@4567.4587.21|TAATGGAATAGGACCGCGGTT</td><td>>hg38_21_+_8433172_8446622_RNA45SN1_45S_50ntFlanks</td><td>4567</td><td>4587</td><td>21</td><td>TAATGGAATAGGACCGCGGTT</td><td>201.4</td><td>-2.2</td><td>0.9</td><td>-2.4</td><td>1.5E-02</td><td>2.5E-02</td><td>BT20/MB468</td><td>MP</td></tr>
<tr><td>>hg38_MT_+_1621_3279_RNR2_16S_50ntFlanks@1187.1216.30|TTCGGTTGGGGCGACCTCGGAGCAGAACCC</td><td>>hg38_MT_+_1621_3279_RNR2_16S_50ntFlanks</td><td>1187</td><td>1216</td><td>30</td><td>TTCGGTTGGGGCGACCTCGGAGCAGAACCC</td><td>142.0</td><td>5.1</td><td>2.1</td><td>2.4</td><td>1.5E-02</td><td>2.5E-02</td><td>BT20/MB468</td><td>MP</td></tr>
<tr><td>>hg38_21_+_8433172_8446622_RNA45SN1_45S_50ntFlanks@4027.4045.19|CCCCGTGGCGGCGACGACC</td><td>>hg38_21_+_8433172_8446622_RNA45SN1_45S_50ntFlanks</td><td>4027</td><td>4045</td><td>19</td><td>CCCCGTGGCGGCGACGACC</td><td>113.6</td><td>-4.1</td><td>1.7</td><td>-2.4</td><td>1.5E-02</td><td>2.5E-02</td><td>BT20/MB468</td><td>MP</td></tr>
<tr><td>>MI0000299|hsa-mir-222&WithFlank&X|-|45747009|45747130@75.95.21(+1U)&[MIMAT0000279&hsa-miR-222-3p&offsets|0|0(+1U);m-140&X|-|45747033|45747056&offsets|0|-3(+1U)]|AGCTACATCTGGCTACTGGGTT</td><td>>MI0000299|hsa-mir-222&WithFlank&X|-|45747009|45747130</td><td>75</td><td>95</td><td>21(+1U)</td><td>AGCTACATCTGGCTACTGGGTT</td><td>482.5</td><td>-2.0</td><td>0.7</td><td>-2.7</td><td>7.3E-03</td><td>2.5E-02</td><td>BT20/MB231</td><td>MITO</td></tr>
<tr><td>>hg38_1_-_228634819_228635039_RNA5S12_5S_50ntFlanks@51.80.30|GTCTACGGCCATACCACCCTGAACGCGCCC</td><td>>hg38_1_-_228634819_228635039_RNA5S12_5S_50ntFlanks</td><td>51</td><td>80</td><td>30</td><td>GTCTACGGCCATACCACCCTGAACGCGCCC</td><td>1610.4</td><td>2.8</td><td>1.2</td><td>2.3</td><td>2.3E-02</td><td>2.5E-02</td><td>MB231/MB468</td><td>NUC</td></tr>
<tr><td>>trnaMT_ProTGG_MT_-_15956_16023@40.63.24|AATGGTGGAGTTAAAGACTTTTTC</td><td>>trnaMT_ProTGG_MT_-_15956_16023</td><td>40</td><td>63</td><td>24</td><td>AATGGTGGAGTTAAAGACTTTTTC</td><td>96.5</td><td>3.6</td><td>1.5</td><td>2.4</td><td>1.5E-02</td><td>2.5E-02</td><td>BT20/MB468</td><td>MP</td></tr>
<tr><td>>hg38_21_+_8433172_8446622_RNA45SN1_45S_50ntFlanks@10921.10946.26|GCTGGACGAGGCGCCGCCGCCCCCCC</td><td>>hg38_21_+_8433172_8446622_RNA45SN1_45S_50ntFlanks</td><td>10921</td><td>10946</td><td>26</td><td>GCTGGACGAGGCGCCGCCGCCCCCCC</td><td>260.0</td><td>-3.5</td><td>1.4</td><td>-2.4</td><td>1.5E-02</td><td>2.5E-02</td><td>BT20/MB468</td><td>MP</td></tr>
<tr><td>>hg38_21_+_8433172_8446622_RNA45SN1_45S_50ntFlanks@12988.13014.27|CAGCTCCCTCGCTGCGATCTATTGAAA</td><td>>hg38_21_+_8433172_8446622_RNA45SN1_45S_50ntFlanks</td><td>12988</td><td>13014</td><td>27</td><td>CAGCTCCCTCGCTGCGATCTATTGAAA</td><td>254.1</td><td>-2.1</td><td>0.8</td><td>-2.6</td><td>8.1E-03</td><td>2.5E-02</td><td>BT20/MB231</td><td>CYTO</td></tr>
<tr><td>>hg38_21_+_8433172_8446622_RNA45SN1_45S_50ntFlanks@12439.12473.35|TTGTGAAGCAGAATTCACCAAGCGTTGGATTGTTC</td><td>>hg38_21_+_8433172_8446622_RNA45SN1_45S_50ntFlanks</td><td>12439</td><td>12473</td><td>35</td><td>TTGTGAAGCAGAATTCACCAAGCGTTGGATTGTTC</td><td>490.2</td><td>1.6</td><td>0.7</td><td>2.3</td><td>2.2E-02</td><td>2.5E-02</td><td>BT20/MB468</td><td>CYTO</td></tr>
<tr><td>>MI0000082|hsa-mir-25&WithFlank&7|-|100093554|100093649@58.79.22(+1A)&[MIMAT0000081&hsa-miR-25-3p&offsets|0|0(+1A);m-23&7|-|100093571|100093592&offsets|0|0(+1A)]|CATTGCACTTGTCTCGGTCTGAA</td><td>>MI0000082|hsa-mir-25&WithFlank&7|-|100093554|100093649</td><td>58</td><td>79</td><td>22(+1A)</td><td>CATTGCACTTGTCTCGGTCTGAA</td><td>2460.3</td><td>2.0</td><td>0.8</td><td>2.6</td><td>8.1E-03</td><td>2.5E-02</td><td>BT20/MB231</td><td>CYTO</td></tr>
<tr><td>>hg38_21_+_8433172_8446622_RNA45SN1_45S_50ntFlanks@8104.8140.37|CCGCGGCGGGGCGCGGGACATGTGGCGTACGGAAGAC</td><td>>hg38_21_+_8433172_8446622_RNA45SN1_45S_50ntFlanks</td><td>8104</td><td>8140</td><td>37</td><td>CCGCGGCGGGGCGCGGGACATGTGGCGTACGGAAGAC</td><td>8242.5</td><td>-3.0</td><td>1.2</td><td>-2.4</td><td>1.5E-02</td><td>2.5E-02</td><td>BT20/MB468</td><td>MP</td></tr>
<tr><td>>hg38_21_+_8433172_8446622_RNA45SN1_45S_50ntFlanks@11707.11736.30|AAGCGCGGGTAAACGGCGGGAGTAACTATG</td><td>>hg38_21_+_8433172_8446622_RNA45SN1_45S_50ntFlanks</td><td>11707</td><td>11736</td><td>30</td><td>AAGCGCGGGTAAACGGCGGGAGTAACTATG</td><td>285.0</td><td>-1.8</td><td>0.8</td><td>-2.3</td><td>2.3E-02</td><td>2.5E-02</td><td>MB231/MB468</td><td>NUC</td></tr>
<tr><td>>hg38_21_+_8433172_8446622_RNA45SN1_45S_50ntFlanks@3837.3861.25|CCTCTCCTACTTGGATAACTGTGGT</td><td>>hg38_21_+_8433172_8446622_RNA45SN1_45S_50ntFlanks</td><td>3837</td><td>3861</td><td>25</td><td>CCTCTCCTACTTGGATAACTGTGGT</td><td>264.3</td><td>-1.1</td><td>0.5</td><td>-2.3</td><td>2.2E-02</td><td>2.5E-02</td><td>MB231/MB468</td><td>CYTO</td></tr>
<tr><td>>am_trnaMT_ArgTCG_MT_+_10405_10469@29.48.20|TTTCGACTCATTAAATTATG</td><td>>am_trnaMT_ArgTCG_MT_+_10405_10469</td><td>29</td><td>48</td><td>20</td><td>TTTCGACTCATTAAATTATG</td><td>128.3</td><td>-4.4</td><td>1.8</td><td>-2.4</td><td>1.5E-02</td><td>2.5E-02</td><td>BT20/MB468</td><td>MP</td></tr>
<tr><td>>hg38_21_+_8433172_8446622_RNA45SN1_45S_50ntFlanks@11668.11688.21|TGCCCAGTGCTCTGAATGTCA</td><td>>hg38_21_+_8433172_8446622_RNA45SN1_45S_50ntFlanks</td><td>11668</td><td>11688</td><td>21</td><td>TGCCCAGTGCTCTGAATGTCA</td><td>889.7</td><td>-2.3</td><td>0.9</td><td>-2.6</td><td>8.1E-03</td><td>2.5E-02</td><td>BT20/MB231</td><td>CYTO</td></tr>
<tr><td>>hg38_21_+_8433172_8446622_RNA45SN1_45S_50ntFlanks@475.502.28|CTTCGTGATCGATGTGGTGACGTCGTGC</td><td>>hg38_21_+_8433172_8446622_RNA45SN1_45S_50ntFlanks</td><td>475</td><td>502</td><td>28</td><td>CTTCGTGATCGATGTGGTGACGTCGTGC</td><td>257.3</td><td>2.1</td><td>0.8</td><td>2.6</td><td>8.1E-03</td><td>2.5E-02</td><td>BT20/MB231</td><td>CYTO</td></tr>
<tr><td>>hg38_21_+_8433172_8446622_RNA45SN1_45S_50ntFlanks@6640.6683.44|TCGACCTCGTACGACTCTTAGCGGTGGATCACTCGGCTCGTGCG</td><td>>hg38_21_+_8433172_8446622_RNA45SN1_45S_50ntFlanks</td><td>6640</td><td>6683</td><td>44</td><td>TCGACCTCGTACGACTCTTAGCGGTGGATCACTCGGCTCGTGCG</td><td>219.4</td><td>-3.6</td><td>1.3</td><td>-2.7</td><td>6.6E-03</td><td>2.5E-02</td><td>BT20/MB231</td><td>NUC</td></tr>
<tr><td>>hg38_21_+_8433172_8446622_RNA45SN1_45S_50ntFlanks@6652.6672.21|GACTCTTAGCGGTGGATCACT</td><td>>hg38_21_+_8433172_8446622_RNA45SN1_45S_50ntFlanks</td><td>6652</td><td>6672</td><td>21</td><td>GACTCTTAGCGGTGGATCACT</td><td>232.6</td><td>3.7</td><td>1.5</td><td>2.4</td><td>1.5E-02</td><td>2.5E-02</td><td>BT20/MB468</td><td>MP</td></tr>
<tr><td>>MI0000077|hsa-mir-21&WithFlank&17|+|59841260|59841343@13.36.24&[MIMAT0000076&hsa-miR-21-5p&offsets|-1|+1;m-1&17|+|59841273|59841294&offsets|-1|+1]|GTAGCTTATCAGACTGATGTTGAC</td><td>>MI0000077|hsa-mir-21&WithFlank&17|+|59841260|59841343</td><td>13</td><td>36</td><td>24</td><td>GTAGCTTATCAGACTGATGTTGAC</td><td>429.3</td><td>-2.0</td><td>0.8</td><td>-2.7</td><td>7.4E-03</td><td>2.5E-02</td><td>BT20/MB231</td><td>MITO</td></tr>
<tr><td>>hg38_21_+_8433172_8446622_RNA45SN1_45S_50ntFlanks@8359.8391.33|AAGAACTTTGAAGAGAGAGTTCAAGAGGGCGTG</td><td>>hg38_21_+_8433172_8446622_RNA45SN1_45S_50ntFlanks</td><td>8359</td><td>8391</td><td>33</td><td>AAGAACTTTGAAGAGAGAGTTCAAGAGGGCGTG</td><td>103.6</td><td>3.4</td><td>1.4</td><td>2.4</td><td>1.5E-02</td><td>2.5E-02</td><td>BT20/MB468</td><td>MP</td></tr>
<tr><td>>hg38_21_+_8433172_8446622_RNA45SN1_45S_50ntFlanks@10238.10267.30|CGCTACGCCGCGACGAGTAGGAGGGCCGCT</td><td>>hg38_21_+_8433172_8446622_RNA45SN1_45S_50ntFlanks</td><td>10238</td><td>10267</td><td>30</td><td>CGCTACGCCGCGACGAGTAGGAGGGCCGCT</td><td>135.3</td><td>2.3</td><td>0.9</td><td>2.4</td><td>1.5E-02</td><td>2.5E-02</td><td>BT20/MB468</td><td>MP</td></tr>
<tr><td>>MI0000810|hsa-mir-135b&WithFlank&1|-|205448296|205448404@22.44.23&[MIMAT0000758&hsa-miR-135b-5p&offsets|0|0;m-321&1|-|205448361|205448383&offsets|0|0]|TATGGCTTTTCATTCCTATGTGA</td><td>>MI0000810|hsa-mir-135b&WithFlank&1|-|205448296|205448404</td><td>22</td><td>44</td><td>23</td><td>TATGGCTTTTCATTCCTATGTGA</td><td>1706.4</td><td>1.5</td><td>0.7</td><td>2.3</td><td>2.1E-02</td><td>2.5E-02</td><td>MB231/MB468</td><td>MP</td></tr>
<tr><td>>hg38_21_+_8433172_8446622_RNA45SN1_45S_50ntFlanks@3985.4005.21|CGGCTTTGGTGACTCTAGATA</td><td>>hg38_21_+_8433172_8446622_RNA45SN1_45S_50ntFlanks</td><td>3985</td><td>4005</td><td>21</td><td>CGGCTTTGGTGACTCTAGATA</td><td>239.9</td><td>-2.5</td><td>1.1</td><td>-2.3</td><td>2.2E-02</td><td>2.5E-02</td><td>BT20/MB468</td><td>CYTO</td></tr>
<tr><td>>hg38_21_+_8433172_8446622_RNA45SN1_45S_50ntFlanks@10060.10079.20|GCAGTCGAGAGTGGACGGGA</td><td>>hg38_21_+_8433172_8446622_RNA45SN1_45S_50ntFlanks</td><td>10060</td><td>10079</td><td>20</td><td>GCAGTCGAGAGTGGACGGGA</td><td>172.3</td><td>4.5</td><td>1.6</td><td>2.7</td><td>6.0E-03</td><td>2.5E-02</td><td>BT20/MB231</td><td>MP</td></tr>
<tr><td>>trnaMT_LysTTT_MT_+_8295_8364@1.31.31,trnalookalike7_LysTTT_1_+_568844_568913@1.31.31|CACTGTAAAGCTAACTTAGCATTAACCTTTT</td><td>>trnaMT_LysTTT_MT_+_8295_8364</td><td>1</td><td>31</td><td>31</td><td>CACTGTAAAGCTAACTTAGCATTAACCTTTT</td><td>182.8</td><td>2.0</td><td>0.8</td><td>2.3</td><td>2.1E-02</td><td>2.5E-02</td><td>MB231/MB468</td><td>MP</td></tr>
<tr><td>>hg38_21_+_8433172_8446622_RNA45SN1_45S_50ntFlanks@11874.11917.44|AGAAGACCCTGTTGAGCTTGACTCTAGTCTGGCACGGTGAAGAG</td><td>>hg38_21_+_8433172_8446622_RNA45SN1_45S_50ntFlanks</td><td>11874</td><td>11917</td><td>44</td><td>AGAAGACCCTGTTGAGCTTGACTCTAGTCTGGCACGGTGAAGAG</td><td>51.2</td><td>-4.5</td><td>1.9</td><td>-2.4</td><td>1.5E-02</td><td>2.5E-02</td><td>BT20/MB468</td><td>MP</td></tr>
<tr><td>>hg38_21_+_8433172_8446622_RNA45SN1_45S_50ntFlanks@11589.11612.24|CGGACCAGGGGAATCCGACTGTTT</td><td>>hg38_21_+_8433172_8446622_RNA45SN1_45S_50ntFlanks</td><td>11589</td><td>11612</td><td>24</td><td>CGGACCAGGGGAATCCGACTGTTT</td><td>464.0</td><td>-2.2</td><td>0.9</td><td>-2.4</td><td>1.5E-02</td><td>2.5E-02</td><td>BT20/MB468</td><td>MP</td></tr>
<tr><td>>MI0000440|hsa-mir-27b&WithFlank&9|+|95085439|95085547@66.87.22&[MIMAT0000419&hsa-miR-27b-3p&offsets|-1|0;m-39&9|+|95085505|95085525&offsets|-1|0]|GTTCACAGTGGCTAAGTTCTGC</td><td>>MI0000440|hsa-mir-27b&WithFlank&9|+|95085439|95085547</td><td>66</td><td>87</td><td>22</td><td>GTTCACAGTGGCTAAGTTCTGC</td><td>88.1</td><td>3.9</td><td>1.7</td><td>2.3</td><td>2.1E-02</td><td>2.6E-02</td><td>MB231/MB468</td><td>MP</td></tr>
<tr><td>>hg38_21_+_8433172_8446622_RNA45SN1_45S_50ntFlanks@9586.9610.25|GACCTGGGTATAGGGGCGAAAGACT</td><td>>hg38_21_+_8433172_8446622_RNA45SN1_45S_50ntFlanks</td><td>9586</td><td>9610</td><td>25</td><td>GACCTGGGTATAGGGGCGAAAGACT</td><td>48.1</td><td>4.4</td><td>1.8</td><td>2.4</td><td>1.5E-02</td><td>2.6E-02</td><td>BT20/MB468</td><td>MP</td></tr>
<tr><td>>hg38_21_+_8433172_8446622_RNA45SN1_45S_50ntFlanks@7977.7996.20|CGACCTCAGATCAGACGTGG</td><td>>hg38_21_+_8433172_8446622_RNA45SN1_45S_50ntFlanks</td><td>7977</td><td>7996</td><td>20</td><td>CGACCTCAGATCAGACGTGG</td><td>143.7</td><td>-4.2</td><td>1.7</td><td>-2.4</td><td>1.5E-02</td><td>2.6E-02</td><td>BT20/MB468</td><td>MP</td></tr>
</table>

<table>
<tr><td>>trna10_AspGTC_12_-_125424193_125424264@40.72.33,trna12_AspGTC_12_-_125411891_125411962@40.72.33,trna144_AspGTC_6_-_27551236_27551307@40.72.33,trna38_AspGTC_17_-_8125556_8125627@40.72.33,trna45_AspGTC_6_+_27447453_27447524@40.72.33,trna48_AspGTC_6_+_27471523_27471594@40.72.33,trna4_AspGTC_12_+_96429799_96429870@40.72.33,trna69_AspGTC_1_-_161440205_161440276@40.72.33,trna72_AspGTC_1_-_161432824_161432895@40.72.33,trna75_AspGTC_1_-_161425414_161425485@40.72.33,trna78_AspGTC_1_-_161418033_161418104@40.72.33,trna81_AspGTC_1_-_161410615_161410686@40.72.33|CGGGAGACCGGGGTTCGATTCCCCGACGGGGAG</td><td>>trna10_AspGTC_12_-_125424193_125424264</td><td>40</td><td>72</td><td>33</td><td>CGGGAGACCGGGGTTCGATTCCCCGACGGGGAG</td><td>129.6</td><td>-4.2</td><td>1.7</td><td>-2.4</td><td>1.5E-02</td><td>2.6E-02</td><td>BT20/MB468</td><td>MP</td></tr>
<tr><td>>trna111_HisGTG_1_-_147774845_147774916@-1G.30.31,trna118_HisGTG_1_-_145396881_145396952@-1G.30.31,trna16_HisGTG_1_+_146544773_146544844@-1G.30.31,trna1_HisGTG_15_+_45493349_45493420@-1G.30.31,trna21_HisGTG_1_+_147753471_147753542@-1G.30.31,trna33_HisGTG_6_+_27125906_27125977@-1G.30.31,trna7_HisGTG_9_-_14433938_14434009@-1G.30.31,trna8_HisGTG_15_-_45492611_45492682@-1G.30.31,trna9_HisGTG_15_-_45490804_45490875@-1G.30.31|GGCCGTGATCGTATAGTGGTTAGTACTCTGC</td><td>>trna111_HisGTG_1_-_147774845_147774916</td><td>-1G</td><td>30</td><td>31</td><td>GGCCGTGATCGTATAGTGGTTAGTACTCTGC</td><td>2730.4</td><td>2.0</td><td>0.9</td><td>2.3</td><td>2.1E-02</td><td>2.6E-02</td><td>MB231/MB468</td><td>MP</td></tr>
<tr><td>>hg38_21_+_8433172_8446622_RNA45SN1_45S_50ntFlanks@9483.9506.24|CGTGTTAGGACCCGAAAGATGGTG</td><td>>hg38_21_+_8433172_8446622_RNA45SN1_45S_50ntFlanks</td><td>9483</td><td>9506</td><td>24</td><td>CGTGTTAGGACCCGAAAGATGGTG</td><td>195.8</td><td>5.6</td><td>2.1</td><td>2.7</td><td>6.0E-03</td><td>2.6E-02</td><td>BT20/MB231</td><td>MP</td></tr>
<tr><td>>hg38_21_+_8433172_8446622_RNA45SN1_45S_50ntFlanks@8177.8208.32|TTCTGATCGAGGCCCAGCCCGTGGACGGTGTG</td><td>>hg38_21_+_8433172_8446622_RNA45SN1_45S_50ntFlanks</td><td>8177</td><td>8208</td><td>32</td><td>TTCTGATCGAGGCCCAGCCCGTGGACGGTGTG</td><td>444.7</td><td>-4.5</td><td>2.0</td><td>-2.3</td><td>2.3E-02</td><td>2.6E-02</td><td>BT20/MB468</td><td>NUC</td></tr>
<tr><td>>MI0025242|hsa-mir-7706&WithFlank&15|+|85380590|85380668@50.72.23&[MIMAT0030021&hsa-miR-7706&offsets|0|-1;m-390&15|+|85380639|85380662&offsets|0|-1]|TGAAGCGCCTGTGCTCTGCCGAG</td><td>>MI0025242|hsa-mir-7706&WithFlank&15|+|85380590|85380668</td><td>50</td><td>72</td><td>23</td><td>TGAAGCGCCTGTGCTCTGCCGAG</td><td>149.1</td><td>3.4</td><td>1.4</td><td>2.4</td><td>1.5E-02</td><td>2.6E-02</td><td>BT20/MB468</td><td>MP</td></tr>
<tr><td>>hg38_21_+_8433172_8446622_RNA45SN1_45S_50ntFlanks@6721.6745.25|AATTGCAGGACACATTGATCATCGA</td><td>>hg38_21_+_8433172_8446622_RNA45SN1_45S_50ntFlanks</td><td>6721</td><td>6745</td><td>25</td><td>AATTGCAGGACACATTGATCATCGA</td><td>524.7</td><td>-4.4</td><td>1.8</td><td>-2.4</td><td>1.5E-02</td><td>2.6E-02</td><td>BT20/MB468</td><td>MP</td></tr>
<tr><td>>MI0000088|hsa-mir-30a&WithFlank&6|-|71403545|71403627@12.33.22(+1U)&[MIMAT0000087&hsa-miR-30a-5p&offsets|0|0(+1U);m-15&6|-|71403595|71403616&offsets|0|0(+1U)]|TGTAAACATCCTCGACTGGAAGT</td><td>>MI0000088|hsa-mir-30a&WithFlank&6|-|71403545|71403627</td><td>12</td><td>33</td><td>22(+1U)</td><td>TGTAAACATCCTCGACTGGAAGT</td><td>9709.9</td><td>-2.3</td><td>0.9</td><td>-2.7</td><td>7.4E-03</td><td>2.6E-02</td><td>BT20/MB231</td><td>MITO</td></tr>
<tr><td>>hg38_21_+_8433172_8446622_RNA45SN1_45S_50ntFlanks@10818.10840.23|AAGGGAAGTCGGCAAGCCGGATC</td><td>>hg38_21_+_8433172_8446622_RNA45SN1_45S_50ntFlanks</td><td>10818</td><td>10840</td><td>23</td><td>AAGGGAAGTCGGCAAGCCGGATC</td><td>191.8</td><td>-1.3</td><td>0.6</td><td>-2.3</td><td>2.2E-02</td><td>2.6E-02</td><td>MB231/MB468</td><td>CYTO</td></tr>
<tr><td>>hg38_21_+_8433172_8446622_RNA45SN1_45S_50ntFlanks@6777.6806.30|TCCCGGGGCTACGCCTGTCTGAGCGTCGCT</td><td>>hg38_21_+_8433172_8446622_RNA45SN1_45S_50ntFlanks</td><td>6777</td><td>6806</td><td>30</td><td>TCCCGGGGCTACGCCTGTCTGAGCGTCGCT</td><td>133.1</td><td>2.2</td><td>0.9</td><td>2.4</td><td>1.5E-02</td><td>2.6E-02</td><td>BT20/MB468</td><td>MP</td></tr>
<tr><td>>hg38_MT_+_1621_3279_RNR2_16S_50ntFlanks@1188.1216.29|TCGGTTGGGGCGACCTCGGAGCAGAACCC</td><td>>hg38_MT_+_1621_3279_RNR2_16S_50ntFlanks</td><td>1188</td><td>1216</td><td>29</td><td>TCGGTTGGGGCGACCTCGGAGCAGAACCC</td><td>123.0</td><td>4.3</td><td>1.8</td><td>2.4</td><td>1.5E-02</td><td>2.6E-02</td><td>BT20/MB468</td><td>MP</td></tr>
<tr><td>>hg38_21_+_8433172_8446622_RNA45SN1_45S_50ntFlanks@9532.9550.19|AGAGGAAACTCTGGTGGAG</td><td>>hg38_21_+_8433172_8446622_RNA45SN1_45S_50ntFlanks</td><td>9532</td><td>9550</td><td>19</td><td>AGAGGAAACTCTGGTGGAG</td><td>33.6</td><td>4.9</td><td>2.1</td><td>2.3</td><td>2.1E-02</td><td>2.6E-02</td><td>MB231/MB468</td><td>MP</td></tr>
<tr><td>>hg38_21_+_8433172_8446622_RNA45SN1_45S_50ntFlanks@6721.6747.27|AATTGCAGGACACATTGATCATCGACA</td><td>>hg38_21_+_8433172_8446622_RNA45SN1_45S_50ntFlanks</td><td>6721</td><td>6747</td><td>27</td><td>AATTGCAGGACACATTGATCATCGACA</td><td>1291.9</td><td>-2.8</td><td>1.1</td><td>-2.4</td><td>1.5E-02</td><td>2.6E-02</td><td>BT20/MB468</td><td>MP</td></tr>
<tr><td>>hg38_21_+_8433172_8446622_RNA45SN1_45S_50ntFlanks@12994.13016.23|CCTCGCTGCGATCTATTGAAAGT</td><td>>hg38_21_+_8433172_8446622_RNA45SN1_45S_50ntFlanks</td><td>12994</td><td>13016</td><td>23</td><td>CCTCGCTGCGATCTATTGAAAGT</td><td>275.0</td><td>-1.9</td><td>0.7</td><td>-2.6</td><td>8.2E-03</td><td>2.6E-02</td><td>BT20/MB231</td><td>CYTO</td></tr>
<tr><td>>hg38_21_+_8433172_8446622_RNA45SN1_45S_50ntFlanks@10514.10533.20|CGGAGTGGCGGAGATGGGCG</td><td>>hg38_21_+_8433172_8446622_RNA45SN1_45S_50ntFlanks</td><td>10514</td><td>10533</td><td>20</td><td>CGGAGTGGCGGAGATGGGCG</td><td>2414.0</td><td>-2.9</td><td>1.2</td><td>-2.4</td><td>1.5E-02</td><td>2.6E-02</td><td>BT20/MB468</td><td>MP</td></tr>
<tr><td>>am_trna119_LysCTT_1_-_145395522_145395594@13.32.20,trna11_LysCTT_5_-_180648979_180649051@13.32.20,trna128_LysCTT_1_-_55423542_55423614@13.32.20,trna13_LysCTT_6_+_26556774_26556846@13.32.20,trna32_LysCTT_16_-_3207406_3207478@13.32.20,trna6_LysCTT_19_-_52425396_52425468@13.32.20,trna7_LysCTT_16_+_3225692_3225764@13.32.20,trna9_LysCTT_5_+_180634755_180634827@13.32.20|CAGTCGGTAGAGCATGAGAC</td><td>>am_trna119_LysCTT_1_-_145395522_145395594</td><td>13</td><td>32</td><td>20</td><td>CAGTCGGTAGAGCATGAGAC</td><td>33.3</td><td>4.6</td><td>2.0</td><td>2.3</td><td>2.1E-02</td><td>2.6E-02</td><td>MB231/MB468</td><td>MP</td></tr>
<tr><td>>hg38_21_+_8433172_8446622_RNA45SN1_45S_50ntFlanks@3861.3899.39|TAATTCTAGAGCTAATACATGCCGACGGGCGCTGACCCC</td><td>>hg38_21_+_8433172_8446622_RNA45SN1_45S_50ntFlanks</td><td>3861</td><td>3899</td><td>39</td><td>TAATTCTAGAGCTAATACATGCCGACGGGCGCTGACCCC</td><td>128.6</td><td>-2.9</td><td>1.3</td><td>-2.3</td><td>2.2E-02</td><td>2.6E-02</td><td>BT20/MB468</td><td>CYTO</td></tr>
<tr><td>>hg38_21_+_8433172_8446622_RNA45SN1_45S_50ntFlanks@7981.8007.27|CTCAGATCAGACGTGGCGACCCGCTGA</td><td>>hg38_21_+_8433172_8446622_RNA45SN1_45S_50ntFlanks</td><td>7981</td><td>8007</td><td>27</td><td>CTCAGATCAGACGTGGCGACCCGCTGA</td><td>241.3</td><td>-2.0</td><td>0.8</td><td>-2.4</td><td>1.5E-02</td><td>2.6E-02</td><td>BT20/MB468</td><td>MP</td></tr>
<tr><td>>hg38_21_+_8433172_8446622_RNA45SN1_45S_50ntFlanks@7976.8006.31|GCGACCTCAGATCAGACGTGGCGACCCGCTG</td><td>>hg38_21_+_8433172_8446622_RNA45SN1_45S_50ntFlanks</td><td>7976</td><td>8006</td><td>31</td><td>GCGACCTCAGATCAGACGTGGCGACCCGCTG</td><td>343.7</td><td>2.9</td><td>1.1</td><td>2.7</td><td>7.5E-03</td><td>2.6E-02</td><td>BT20/MB231</td><td>MITO</td></tr>
<tr><td>>MI0000077|hsa-mir-21&WithFlank&17|+|59841260|59841343@52.73.22(+1A)&[MIMAT0004494&hsa-miR-21-3p&offsets|0|+1(+1A);m-52&17|+|59841311|59841332&offsets|0|0(+1A)]|CAACACCAGTCGATGGGCTGTCA</td><td>>MI0000077|hsa-mir-21&WithFlank&17|+|59841260|59841343</td><td>52</td><td>73</td><td>22(+1A)</td><td>CAACACCAGTCGATGGGCTGTCA</td><td>143.5</td><td>3.1</td><td>1.4</td><td>2.3</td><td>2.1E-02</td><td>2.6E-02</td><td>MB231/MB468</td><td>MP</td></tr>
<tr><td>>hg38_21_+_8433172_8446622_RNA45SN1_45S_50ntFlanks@11602.11626.25|TCCGACTGTTTAATTAAAACAAAGC</td><td>>hg38_21_+_8433172_8446622_RNA45SN1_45S_50ntFlanks</td><td>11602</td><td>11626</td><td>25</td><td>TCCGACTGTTTAATTAAAACAAAGC</td><td>94.7</td><td>3.6</td><td>1.5</td><td>2.4</td><td>1.5E-02</td><td>2.6E-02</td><td>BT20/MB468</td><td>MP</td></tr>
<tr><td>>hg38_21_+_8433172_8446622_RNA45SN1_45S_50ntFlanks@10826.10848.23|TCGGCAAGCCGGATCCGTAACTT</td><td>>hg38_21_+_8433172_8446622_RNA45SN1_45S_50ntFlanks</td><td>10826</td><td>10848</td><td>23</td><td>TCGGCAAGCCGGATCCGTAACTT</td><td>73.9</td><td>-4.1</td><td>1.7</td><td>-2.4</td><td>1.5E-02</td><td>2.6E-02</td><td>BT20/MB468</td><td>MP</td></tr>
<tr><td>>MI0000102|hsa-mir-100&WithFlank&11|-|122152223|122152314@19.38.20(+1G)&[MIMAT0000098&hsa-miR-100-5p&offsets|0|-2(+1G);m-27&11|-|122152276|122152296&offsets|0|-1(+1G)]|AACCCGTAGATCCGAACTTGG</td><td>>MI0000102|hsa-mir-100&WithFlank&11|-|122152223|122152314</td><td>19</td><td>38</td><td>20(+1G)</td><td>AACCCGTAGATCCGAACTTGG</td><td>3042.6</td><td>-3.3</td><td>1.2</td><td>-2.6</td><td>8.2E-03</td><td>2.6E-02</td><td>BT20/MB231</td><td>CYTO</td></tr>
<tr><td>>trnaMT_GluTTC_MT_-_14674_14742@38.69.32,trnalookalike8_GluTTC_5_-_93905172_93905240@38.69.32|CATTGGTCGTGGTTGTAGTCCGTGCGAGAATA</td><td>>trnaMT_GluTTC_MT_-_14674_14742</td><td>38</td><td>69</td><td>32</td><td>CATTGGTCGTGGTTGTAGTCCGTGCGAGAATA</td><td>131.6</td><td>5.4</td><td>2.2</td><td>2.4</td><td>1.5E-02</td><td>2.6E-02</td><td>BT20/MB468</td><td>MP</td></tr>
<tr><td>>hg38_MT_+_1621_3279_RNR2_16S_50ntFlanks@51.93.43|GCTAAACCTAGCCCCAAACCCACTCCACCTTACTACCAGACAA</td><td>>hg38_MT_+_1621_3279_RNR2_16S_50ntFlanks</td><td>51</td><td>93</td><td>43</td><td>GCTAAACCTAGCCCCAAACCCACTCCACCTTACTACCAGACAA</td><td>330.7</td><td>4.3</td><td>1.9</td><td>2.3</td><td>2.1E-02</td><td>2.6E-02</td><td>MB231/MB468</td><td>MP</td></tr>
<tr><td>>hg38_21_+_8433172_8446622_RNA45SN1_45S_50ntFlanks@7975.8007.33|CGCGACCTCAGATCAGACGTGGCGACCCGCTGA</td><td>>hg38_21_+_8433172_8446622_RNA45SN1_45S_50ntFlanks</td><td>7975</td><td>8007</td><td>33</td><td>CGCGACCTCAGATCAGACGTGGCGACCCGCTGA</td><td>13587.2</td><td>3.5</td><td>1.3</td><td>2.7</td><td>6.1E-03</td><td>2.6E-02</td><td>BT20/MB231</td><td>MP</td></tr>
<tr><td>>hg38_21_+_8433172_8446622_RNA45SN1_45S_50ntFlanks@12936.12969.34|AAACCATTCGTAGACGACCTGCTTCTGGGTCGGG</td><td>>hg38_21_+_8433172_8446622_RNA45SN1_45S_50ntFlanks</td><td>12936</td><td>12969</td><td>34</td><td>AAACCATTCGTAGACGACCTGCTTCTGGGTCGGG</td><td>104.6</td><td>-4.2</td><td>1.7</td><td>-2.4</td><td>1.5E-02</td><td>2.6E-02</td><td>BT20/MB468</td><td>MP</td></tr>
<tr><td>>hg38_1_-_228634819_228635039_RNA5S12_5S_50ntFlanks@129.171.43|TTGGATGGGAGACCGCCTGGGAATACCGGGTGCTGTAGGCTTT</td><td>>hg38_1_-_228634819_228635039_RNA5S12_5S_50ntFlanks</td><td>129</td><td>171</td><td>43</td><td>TTGGATGGGAGACCGCCTGGGAATACCGGGTGCTGTAGGCTTT</td><td>988.2</td><td>-2.9</td><td>1.1</td><td>-2.7</td><td>6.7E-03</td><td>2.6E-02</td><td>BT20/MB231</td><td>NUC</td></tr>
<tr><td>>MI0000471|hsa-mir-126&WithFlank&9|+|136670596|136670692@59.78.20&[MIMAT0000445&hsa-miR-126-3p&offsets|+1|-1;m-40&9|+|136670653|136670674&offsets|+1|-1]|CGTACCGTGAGTAATAATGC</td><td>>MI0000471|hsa-mir-126&WithFlank&9|+|136670596|136670692</td><td>59</td><td>78</td><td>20</td><td>CGTACCGTGAGTAATAATGC</td><td>967.1</td><td>2.6</td><td>1.0</td><td>2.6</td><td>8.3E-03</td><td>2.6E-02</td><td>BT20/MB231</td><td>CYTO</td></tr>
<tr><td>>hg38_MT_+_598_1651_RNR1_12S_50ntFlanks@424.452.29|TACGAAAGTGGCTTTAACATATCTGAACA</td><td>>hg38_MT_+_598_1651_RNR1_12S_50ntFlanks</td><td>424</td><td>452</td><td>29</td><td>TACGAAAGTGGCTTTAACATATCTGAACA</td><td>187.1</td><td>3.9</td><td>1.6</td><td>2.4</td><td>1.5E-02</td><td>2.6E-02</td><td>BT20/MB468</td><td>MP</td></tr>
<tr><td>>hg38_21_+_8433172_8446622_RNA45SN1_45S_50ntFlanks@11704.11726.23|ATGAAGCGCGGGTAAACGGCGGG</td><td>>hg38_21_+_8433172_8446622_RNA45SN1_45S_50ntFlanks</td><td>11704</td><td>11726</td><td>23</td><td>ATGAAGCGCGGGTAAACGGCGGG</td><td>155.2</td><td>-4.2</td><td>1.7</td><td>-2.4</td><td>1.5E-02</td><td>2.6E-02</td><td>BT20/MB468</td><td>MP</td></tr>
<tr><td>>MI0000082|hsa-mir-25&WithFlank&7|-|100093554|100093649@58.77.20(+1C)&[MIMAT0000081&hsa-miR-25-3p&offsets|0|-2(+1C);m-23&7|-|100093571|100093592&offsets|0|-2(+1C)]|CATTGCACTTGTCTCGGTCTC</td><td>>MI0000082|hsa-mir-25&WithFlank&7|-|100093554|100093649</td><td>58</td><td>77</td><td>20(+1C)</td><td>CATTGCACTTGTCTCGGTCTC</td><td>36.3</td><td>4.7</td><td>2.1</td><td>2.3</td><td>2.1E-02</td><td>2.6E-02</td><td>MB231/MB468</td><td>MP</td></tr>
<tr><td>>hg38_21_+_8433172_8446622_RNA45SN1_45S_50ntFlanks@6661.6693.33|CGGTGGATCACTCGGCTCGTGCGTCGATGAAGA</td><td>>hg38_21_+_8433172_8446622_RNA45SN1_45S_50ntFlanks</td><td>6661</td><td>6693</td><td>33</td><td>CGGTGGATCACTCGGCTCGTGCGTCGATGAAGA</td><td>1092.4</td><td>2.3</td><td>0.9</td><td>2.7</td><td>7.5E-03</td><td>2.6E-02</td><td>BT20/MB231</td><td>MITO</td></tr>
<tr><td>>hg38_MT_+_598_1651_RNR1_12S_50ntFlanks@916.950.35|AAGGACATTTAACTAAAACCCCTACGCATTTATAT</td><td>>hg38_MT_+_598_1651_RNR1_12S_50ntFlanks</td><td>916</td><td>950</td><td>35</td><td>AAGGACATTTAACTAAAACCCCTACGCATTTATAT</td><td>247.5</td><td>2.1</td><td>0.8</td><td>2.7</td><td>7.5E-03</td><td>2.6E-02</td><td>BT20/MB231</td><td>MITO</td></tr>
<tr><td>>trna32_LysCTT_16_-_3207406_3207478@3.33.31|CCGGCTAGCTCAGTCGGTAGAGCATGAGACC</td><td>>trna32_LysCTT_16_-_3207406_3207478</td><td>3</td><td>33</td><td>31</td><td>CCGGCTAGCTCAGTCGGTAGAGCATGAGACC</td><td>198.6</td><td>-5.6</td><td>2.1</td><td>-2.7</td><td>7.5E-03</td><td>2.6E-02</td><td>BT20/MB231</td><td>MITO</td></tr>
<tr><td>>hg38_21_+_8433172_8446622_RNA45SN1_45S_50ntFlanks@9839.9861.23|GTAAGCAGAACTGGCGCTGCGGG</td><td>>hg38_21_+_8433172_8446622_RNA45SN1_45S_50ntFlanks</td><td>9839</td><td>9861</td><td>23</td><td>GTAAGCAGAACTGGCGCTGCGGG</td><td>72.0</td><td>-4.5</td><td>1.8</td><td>-2.4</td><td>1.5E-02</td><td>2.6E-02</td><td>BT20/MB468</td><td>MP</td></tr>
<tr><td>>trnaMT_ProTGG_MT_-_15956_16023@40.61.22|AATGGTGGAGTTAAAGACTTTT</td><td>>trnaMT_ProTGG_MT_-_15956_16023</td><td>40</td><td>61</td><td>22</td><td>AATGGTGGAGTTAAAGACTTTT</td><td>52.5</td><td>3.9</td><td>1.7</td><td>2.3</td><td>2.1E-02</td><td>2.6E-02</td><td>MB231/MB468</td><td>MP</td></tr>
<tr><td>>hg38_21_+_8433172_8446622_RNA45SN1_45S_50ntFlanks@8342.8378.37|CGTAAGGGAAAGTTGAAAAGAACTTTGAAGAGAGAGT</td><td>>hg38_21_+_8433172_8446622_RNA45SN1_45S_50ntFlanks</td><td>8342</td><td>8378</td><td>37</td><td>CGTAAGGGAAAGTTGAAAAGAACTTTGAAGAGAGAGT</td><td>2890.4</td><td>-2.4</td><td>0.9</td><td>-2.6</td><td>8.3E-03</td><td>2.6E-02</td><td>BT20/MB231</td><td>CYTO</td></tr>
<tr><td>>hg38_21_+_8433172_8446622_RNA45SN1_45S_50ntFlanks@11704.11728.25|ATGAAGCGCGGGTAAACGGCGGGAG</td><td>>hg38_21_+_8433172_8446622_RNA45SN1_45S_50ntFlanks</td><td>11704</td><td>11728</td><td>25</td><td>ATGAAGCGCGGGTAAACGGCGGGAG</td><td>644.1</td><td>-3.9</td><td>1.7</td><td>-2.3</td><td>2.2E-02</td><td>2.6E-02</td><td>BT20/MB468</td><td>CYTO</td></tr>
<tr><td>>hg38_21_+_8433172_8446622_RNA45SN1_45S_50ntFlanks@13007.13039.33|TATTGAAAGTCAGCCCTCGACACAAGGGTTTGT</td><td>>hg38_21_+_8433172_8446622_RNA45SN1_45S_50ntFlanks</td><td>13007</td><td>13039</td><td>33</td><td>TATTGAAAGTCAGCCCTCGACACAAGGGTTTGT</td><td>722.6</td><td>2.5</td><td>1.0</td><td>2.4</td><td>1.5E-02</td><td>2.6E-02</td><td>BT20/MB468</td><td>MP</td></tr>
<tr><td>>hg38_21_+_8433172_8446622_RNA45SN1_45S_50ntFlanks@9572.9602.31|CAAATCGGTCGTCCGACCTGGGTATAGGGGC</td><td>>hg38_21_+_8433172_8446622_RNA45SN1_45S_50ntFlanks</td><td>9572</td><td>9602</td><td>31</td><td>CAAATCGGTCGTCCGACCTGGGTATAGGGGC</td><td>462.0</td><td>-2.3</td><td>1.0</td><td>-2.4</td><td>1.5E-02</td><td>2.6E-02</td><td>BT20/MB468</td><td>MP</td></tr>
<tr><td>>hg38_21_+_8433172_8446622_RNA45SN1_45S_50ntFlanks@11925.11948.24|AGGTGTAGAATAAGTGGGAGGCCC</td><td>>hg38_21_+_8433172_8446622_RNA45SN1_45S_50ntFlanks</td><td>11925</td><td>11948</td><td>24</td><td>AGGTGTAGAATAAGTGGGAGGCCC</td><td>120.4</td><td>5.6</td><td>2.0</td><td>2.7</td><td>6.1E-03</td><td>2.6E-02</td><td>BT20/MB231</td><td>MP</td></tr>
</table>

<table>
<tr><td>>hg38_21_+_8433172_8446622_RNA45SN1_45S_50ntFlanks@6651.6672.22|CGACTCTTAGCGGTGGATCACT</td><td>>hg38_21_+_8433172_8446622_RNA45SN1_45S_50ntFlanks</td><td>6651</td><td>6672</td><td>22</td><td>CGACTCTTAGCGGTGGATCACT</td><td>2734.5</td><td>3.4</td><td>1.2</td><td>2.7</td><td>6.1E-03</td><td>2.6E-02</td><td>BT20/MB231</td><td>MP</td></tr>
<tr><td>>MI0005566|hsa-mir-374b&WithFlank&X|-|74218541|74218624@17.37.21(+1A)&[MIMAT0004955&hsa-miR-374b-5p&offsets|0|-1(+1A);m-153&X|-|74218587|74218608&offsets|0|-1(+1A)]|ATATAATACAACCTGCTAAGTA</td><td>>MI0005566|hsa-mir-374b&WithFlank&X|-|74218541|74218624</td><td>17</td><td>37</td><td>21(+1A)</td><td>ATATAATACAACCTGCTAAGTA</td><td>259.5</td><td>-4.2</td><td>1.5</td><td>-2.7</td><td>6.1E-03</td><td>2.6E-02</td><td>BT20/MB231</td><td>MP</td></tr>
<tr><td>>hg38_MT_+_598_1651_RNR1_12S_50ntFlanks@421.455.35|GACTACGAAAGTGGCTTTAACATATCTGAACACAC</td><td>>hg38_MT_+_598_1651_RNR1_12S_50ntFlanks</td><td>421</td><td>455</td><td>35</td><td>GACTACGAAAGTGGCTTTAACATATCTGAACACAC</td><td>31.2</td><td>4.6</td><td>2.0</td><td>2.3</td><td>2.1E-02</td><td>2.6E-02</td><td>MB231/MB468</td><td>MP</td></tr>
<tr><td>>hg38_21_+_8433172_8446622_RNA45SN1_45S_50ntFlanks@12607.12626.20|TGTGCTTGGCTGAGGAGCCA</td><td>>hg38_21_+_8433172_8446622_RNA45SN1_45S_50ntFlanks</td><td>12607</td><td>12626</td><td>20</td><td>TGTGCTTGGCTGAGGAGCCA</td><td>231.5</td><td>-1.6</td><td>0.7</td><td>-2.3</td><td>2.2E-02</td><td>2.6E-02</td><td>BT20/MB468</td><td>CYTO</td></tr>
<tr><td>>hg38_21_+_8433172_8446622_RNA45SN1_45S_50ntFlanks@9503.9533.31|GGTGAACTATGCCTGGGCAGGGCGAAGCCAG</td><td>>hg38_21_+_8433172_8446622_RNA45SN1_45S_50ntFlanks</td><td>9503</td><td>9533</td><td>31</td><td>GGTGAACTATGCCTGGGCAGGGCGAAGCCAG</td><td>31.2</td><td>-8.0</td><td>3.3</td><td>-2.4</td><td>1.5E-02</td><td>2.6E-02</td><td>BT20/MB468</td><td>MP</td></tr>
<tr><td>>hg38_21_+_8433172_8446622_RNA45SN1_45S_50ntFlanks@8169.8214.46|CCAAGTCCTTCTGATCGAGGCCCAGCCCGTGGACGGTGTGAGGCCG</td><td>>hg38_21_+_8433172_8446622_RNA45SN1_45S_50ntFlanks</td><td>8169</td><td>8214</td><td>46</td><td>CCAAGTCCTTCTGATCGAGGCCCAGCCCGTGGACGGTGTGAGGCCG</td><td>225.2</td><td>-3.4</td><td>1.5</td><td>-2.3</td><td>2.2E-02</td><td>2.6E-02</td><td>BT20/MB468</td><td>CYTO</td></tr>
<tr><td>>hg38_MT_+_1621_3279_RNR2_16S_50ntFlanks@1368.1400.33|CGATGTTGGATCAGGACATCCCGATGGTGCAGC</td><td>>hg38_MT_+_1621_3279_RNR2_16S_50ntFlanks</td><td>1368</td><td>1400</td><td>33</td><td>CGATGTTGGATCAGGACATCCCGATGGTGCAGC</td><td>17.0</td><td>7.3</td><td>3.2</td><td>2.3</td><td>2.1E-02</td><td>2.6E-02</td><td>MB231/MB468</td><td>MP</td></tr>
<tr><td>>hg38_21_+_8433172_8446622_RNA45SN1_45S_50ntFlanks@10059.10082.24|GGCAGTCGAGAGTGGACGGGAGCG</td><td>>hg38_21_+_8433172_8446622_RNA45SN1_45S_50ntFlanks</td><td>10059</td><td>10082</td><td>24</td><td>GGCAGTCGAGAGTGGACGGGAGCG</td><td>85.7</td><td>5.6</td><td>2.0</td><td>2.7</td><td>6.1E-03</td><td>2.6E-02</td><td>BT20/MB231</td><td>MP</td></tr>
<tr><td>>hg38_21_+_8433172_8446622_RNA45SN1_45S_50ntFlanks@7976.8011.36|GCGACCTCAGATCAGACGTGGCGACCCGCTGAATTT</td><td>>hg38_21_+_8433172_8446622_RNA45SN1_45S_50ntFlanks</td><td>7976</td><td>8011</td><td>36</td><td>GCGACCTCAGATCAGACGTGGCGACCCGCTGAATTT</td><td>7394.1</td><td>5.1</td><td>1.9</td><td>2.7</td><td>6.1E-03</td><td>2.6E-02</td><td>BT20/MB231</td><td>MP</td></tr>
<tr><td>>MI0000072|hsa-mir-18a&WithFlank&13|+|91350745|91350827@12.33.22(+1A)&[MIMAT0000072&hsa-miR-18a-5p&offsets|0|-1(+1A);m-262&13|+|91350756|91350778&offsets|0|-1(+1A)]|TAAGGTGCATCTAGTGCAGATAA</td><td>>MI0000072|hsa-mir-18a&WithFlank&13|+|91350745|91350827</td><td>12</td><td>33</td><td>22(+1A)</td><td>TAAGGTGCATCTAGTGCAGATAA</td><td>162.6</td><td>-4.2</td><td>1.5</td><td>-2.7</td><td>6.1E-03</td><td>2.6E-02</td><td>BT20/MB231</td><td>MP</td></tr>
<tr><td>>MI0000087|hsa-mir-29a&WithFlank&7|-|130876741|130876816@48.68.21(+1G)&[MIMAT0000086&hsa-miR-29a-3p&offsets|0|-1(+1G);m-26&7|-|130876749|130876769&offsets|0|0(+1G)]|TAGCACCATCTGAAATCGGTTG</td><td>>MI0000087|hsa-mir-29a&WithFlank&7|-|130876741|130876816</td><td>48</td><td>68</td><td>21(+1G)</td><td>TAGCACCATCTGAAATCGGTTG</td><td>259.0</td><td>5.6</td><td>2.3</td><td>2.4</td><td>1.5E-02</td><td>2.6E-02</td><td>BT20/MB468</td><td>MP</td></tr>
<tr><td>>hg38_21_+_8433172_8446622_RNA45SN1_45S_50ntFlanks@10906.10926.21|TGGGCGCGCGCCGCGGCTGGA</td><td>>hg38_21_+_8433172_8446622_RNA45SN1_45S_50ntFlanks</td><td>10906</td><td>10926</td><td>21</td><td>TGGGCGCGCGCCGCGGCTGGA</td><td>57.5</td><td>-4.1</td><td>1.7</td><td>-2.4</td><td>1.5E-02</td><td>2.6E-02</td><td>BT20/MB468</td><td>MP</td></tr>
<tr><td>>trnaMT_LysTTT_MT_+_8295_8364@1.20.20,trnalookalike7_LysTTT_1_+_568844_568913@1.20.20|CACTGTAAAGCTAACTTAGC</td><td>>trnaMT_LysTTT_MT_+_8295_8364</td><td>1</td><td>20</td><td>20</td><td>CACTGTAAAGCTAACTTAGC</td><td>319.7</td><td>2.3</td><td>1.0</td><td>2.3</td><td>2.3E-02</td><td>2.6E-02</td><td>MB231/MB468</td><td>MITO</td></tr>
<tr><td>>MI0000084|hsa-mir-26b&WithFlank&2|+|218402640|218402728@18.38.21(+1C)&[MIMAT0000083&hsa-miR-26b-5p&offsets|0|0(+1C);m-53&2|+|218402657|218402678&offsets|0|-1(+1C)]|TTCAAGTAATTCAGGATAGGTC</td><td>>MI0000084|hsa-mir-26b&WithFlank&2|+|218402640|218402728</td><td>18</td><td>38</td><td>21(+1C)</td><td>TTCAAGTAATTCAGGATAGGTC</td><td>1064.2</td><td>2.4</td><td>0.9</td><td>2.6</td><td>8.3E-03</td><td>2.6E-02</td><td>BT20/MB231</td><td>CYTO</td></tr>
<tr><td>>hg38_21_+_8433172_8446622_RNA45SN1_45S_50ntFlanks@11209.11251.43|CGGCGACTCTGGACGCGAGCCGGGCCCTTCCCGTGGATCGCCC</td><td>>hg38_21_+_8433172_8446622_RNA45SN1_45S_50ntFlanks</td><td>11209</td><td>11251</td><td>43</td><td>CGGCGACTCTGGACGCGAGCCGGGCCCTTCCCGTGGATCGCCC</td><td>178.2</td><td>-3.2</td><td>1.4</td><td>-2.3</td><td>2.2E-02</td><td>2.6E-02</td><td>BT20/MB468</td><td>CYTO</td></tr>
<tr><td>>hg38_21_+_8433172_8446622_RNA45SN1_45S_50ntFlanks@7977.8017.41|CGACCTCAGATCAGACGTGGCGACCCGCTGAATTTAAGCAT</td><td>>hg38_21_+_8433172_8446622_RNA45SN1_45S_50ntFlanks</td><td>7977</td><td>8017</td><td>41</td><td>CGACCTCAGATCAGACGTGGCGACCCGCTGAATTTAAGCAT</td><td>478.4</td><td>2.4</td><td>1.1</td><td>2.3</td><td>2.3E-02</td><td>2.6E-02</td><td>MB231/MB468</td><td>MITO</td></tr>
<tr><td>>trnaMT_AsnGTT_MT_-_5657_5729@37.55.19,trnalookalike5_AsnGTT_1_-_566207_566279@37.55.19|AACTAAGTGTTTGTGGGTT</td><td>>trnaMT_AsnGTT_MT_-_5657_5729</td><td>37</td><td>55</td><td>19</td><td>AACTAAGTGTTTGTGGGTT</td><td>178.9</td><td>2.8</td><td>1.3</td><td>2.3</td><td>2.3E-02</td><td>2.6E-02</td><td>MB231/MB468</td><td>MITO</td></tr>
<tr><td>>MI0000088|hsa-mir-30a&WithFlank&6|-|71403545|71403627@12.34.23&[MIMAT0000087&hsa-miR-30a-5p&offsets|0|+1;m-15&6|-|71403595|71403616&offsets|0|+1]|TGTAAACATCCTCGACTGGAAGC</td><td>>MI0000088|hsa-mir-30a&WithFlank&6|-|71403545|71403627</td><td>12</td><td>34</td><td>23</td><td>TGTAAACATCCTCGACTGGAAGC</td><td>33577.0</td><td>-2.1</td><td>0.8</td><td>-2.7</td><td>7.6E-03</td><td>2.6E-02</td><td>BT20/MB231</td><td>MITO</td></tr>
<tr><td>>hg38_21_+_8433172_8446622_RNA45SN1_45S_50ntFlanks@10686.10708.23|GCGGCGTCCGGTGAGCTCTCGCT</td><td>>hg38_21_+_8433172_8446622_RNA45SN1_45S_50ntFlanks</td><td>10686</td><td>10708</td><td>23</td><td>GCGGCGTCCGGTGAGCTCTCGCT</td><td>3122.2</td><td>2.7</td><td>1.0</td><td>2.7</td><td>7.6E-03</td><td>2.6E-02</td><td>BT20/MB231</td><td>MITO</td></tr>
<tr><td>>hg38_21_+_8433172_8446622_RNA45SN1_45S_50ntFlanks@5069.5102.34|GGTTAATTCCGATAACGAACGAGACTCTGGCATG</td><td>>hg38_21_+_8433172_8446622_RNA45SN1_45S_50ntFlanks</td><td>5069</td><td>5102</td><td>34</td><td>GGTTAATTCCGATAACGAACGAGACTCTGGCATG</td><td>65.1</td><td>-5.1</td><td>2.2</td><td>-2.3</td><td>2.1E-02</td><td>2.6E-02</td><td>MB231/MB468</td><td>MP</td></tr>
<tr><td>>hg38_21_+_8433172_8446622_RNA45SN1_45S_50ntFlanks@5473.5499.27|CGGCCCTGGCGGAGCGCTGAGAAGACG</td><td>>hg38_21_+_8433172_8446622_RNA45SN1_45S_50ntFlanks</td><td>5473</td><td>5499</td><td>27</td><td>CGGCCCTGGCGGAGCGCTGAGAAGACG</td><td>3340.3</td><td>1.6</td><td>0.7</td><td>2.3</td><td>2.2E-02</td><td>2.6E-02</td><td>MB231/MB468</td><td>CYTO</td></tr>
<tr><td>>hg38_21_+_8433172_8446622_RNA45SN1_45S_50ntFlanks@10394.10424.31|AGCAGTTGAACATGGGTCAGTCGGTCCTGAG</td><td>>hg38_21_+_8433172_8446622_RNA45SN1_45S_50ntFlanks</td><td>10394</td><td>10424</td><td>31</td><td>AGCAGTTGAACATGGGTCAGTCGGTCCTGAG</td><td>13989.8</td><td>3.7</td><td>1.3</td><td>2.7</td><td>6.2E-03</td><td>2.6E-02</td><td>BT20/MB231</td><td>MP</td></tr>
<tr><td>>hg38_21_+_8433172_8446622_RNA45SN1_45S_50ntFlanks@10680.10706.27|GTTCCGGCGGCGTCCGGTGAGCTCTCG</td><td>>hg38_21_+_8433172_8446622_RNA45SN1_45S_50ntFlanks</td><td>10680</td><td>10706</td><td>27</td><td>GTTCCGGCGGCGTCCGGTGAGCTCTCG</td><td>306.4</td><td>6.8</td><td>2.5</td><td>2.7</td><td>6.2E-03</td><td>2.6E-02</td><td>BT20/MB231</td><td>MP</td></tr>
<tr><td>>am_MI0000786|hsa-mir-378a&WithFlank&5|+|149732819|149732896@50.70.21(+1A)&[MIMAT0000732&hsa-miR-378a-3p&offsets|+1|0(+1A);m-44&5|+|149732867|149732888&offsets|+1|0(+1A)]|CTGGACTTGGAGTCAGAAGGCA</td><td>>am_MI0000786|hsa-mir-378a&WithFlank&5|+|149732819|149732896</td><td>50</td><td>70</td><td>21(+1A)</td><td>CTGGACTTGGAGTCAGAAGGCA</td><td>1068.6</td><td>2.7</td><td>1.1</td><td>2.4</td><td>1.5E-02</td><td>2.6E-02</td><td>BT20/MB468</td><td>MP</td></tr>
<tr><td>>MI0000298|hsa-mir-221&WithFlank&X|-|45746151|45746272@71.92.22(+1A)&[MIMAT0000278&hsa-miR-221-3p&offsets|0|-1(+1A);m-91&X|-|45746181|45746202&offsets|0|0(+1A)]|AGCTACATTGTCTGCTGGGTTTA</td><td>>MI0000298|hsa-mir-221&WithFlank&X|-|45746151|45746272</td><td>71</td><td>92</td><td>22(+1A)</td><td>AGCTACATTGTCTGCTGGGTTTA</td><td>567.8</td><td>2.8</td><td>1.2</td><td>2.4</td><td>1.5E-02</td><td>2.6E-02</td><td>BT20/MB468</td><td>MP</td></tr>
<tr><td>>hg38_21_+_8433172_8446622_RNA45SN1_45S_50ntFlanks@8570.8589.20|CTCCGGCGGGTGCGGGGGTG</td><td>>hg38_21_+_8433172_8446622_RNA45SN1_45S_50ntFlanks</td><td>8570</td><td>8589</td><td>20</td><td>CTCCGGCGGGTGCGGGGGTG</td><td>182.9</td><td>-4.2</td><td>1.7</td><td>-2.4</td><td>1.5E-02</td><td>2.6E-02</td><td>BT20/MB468</td><td>MP</td></tr>
<tr><td>>hg38_21_+_8433172_8446622_RNA45SN1_45S_50ntFlanks@8381.8405.25|AAGAGGGCGTGAAACCGTTAAGAGG</td><td>>hg38_21_+_8433172_8446622_RNA45SN1_45S_50ntFlanks</td><td>8381</td><td>8405</td><td>25</td><td>AAGAGGGCGTGAAACCGTTAAGAGG</td><td>550.4</td><td>-3.1</td><td>1.3</td><td>-2.3</td><td>2.2E-02</td><td>2.6E-02</td><td>BT20/MB468</td><td>CYTO</td></tr>
<tr><td>>hg38_21_+_8433172_8446622_RNA45SN1_45S_50ntFlanks@11322.11343.22|ACGTCTCGTCGCGCGCGCGTCC</td><td>>hg38_21_+_8433172_8446622_RNA45SN1_45S_50ntFlanks</td><td>11322</td><td>11343</td><td>22</td><td>ACGTCTCGTCGCGCGCGCGTCC</td><td>172.8</td><td>-3.2</td><td>1.3</td><td>-2.4</td><td>1.5E-02</td><td>2.6E-02</td><td>BT20/MB468</td><td>MP</td></tr>
<tr><td>>hg38_21_+_8433172_8446622_RNA45SN1_45S_50ntFlanks@12003.12035.33|GCGGGCCGCCGGTGAAATACCACTACTCTGATC</td><td>>hg38_21_+_8433172_8446622_RNA45SN1_45S_50ntFlanks</td><td>12003</td><td>12035</td><td>33</td><td>GCGGGCCGCCGGTGAAATACCACTACTCTGATC</td><td>317.6</td><td>2.5</td><td>1.0</td><td>2.4</td><td>1.5E-02</td><td>2.6E-02</td><td>BT20/MB468</td><td>MP</td></tr>
<tr><td>>hg38_21_+_8433172_8446622_RNA45SN1_45S_50ntFlanks@3841.3868.28|TCCTACTTGGATAACTGTGGTAATTCTA</td><td>>hg38_21_+_8433172_8446622_RNA45SN1_45S_50ntFlanks</td><td>3841</td><td>3868</td><td>28</td><td>TCCTACTTGGATAACTGTGGTAATTCTA</td><td>648.9</td><td>1.6</td><td>0.7</td><td>2.3</td><td>2.2E-02</td><td>2.6E-02</td><td>MB231/MB468</td><td>CYTO</td></tr>
<tr><td>>hg38_MT_+_1621_3279_RNR2_16S_50ntFlanks@1090.1120.31|CAGCAAGACGAGAAGACCCTATGGAGCTTTA</td><td>>hg38_MT_+_1621_3279_RNR2_16S_50ntFlanks</td><td>1090</td><td>1120</td><td>31</td><td>CAGCAAGACGAGAAGACCCTATGGAGCTTTA</td><td>109.7</td><td>2.2</td><td>1.0</td><td>2.3</td><td>2.1E-02</td><td>2.6E-02</td><td>MB231/MB468</td><td>MP</td></tr>
<tr><td>>hg38_21_+_8433172_8446622_RNA45SN1_45S_50ntFlanks@6723.6770.48|TTGCAGGACACATTGATCATCGACACTTCGAACGCACTTGCGGCCCCG</td><td>>hg38_21_+_8433172_8446622_RNA45SN1_45S_50ntFlanks</td><td>6723</td><td>6770</td><td>48</td><td>TTGCAGGACACATTGATCATCGACACTTCGAACGCACTTGCGGCCCCG</td><td>115.2</td><td>-3.8</td><td>1.6</td><td>-2.3</td><td>2.0E-02</td><td>2.6E-02</td><td>BT20/MB468</td><td>MITO</td></tr>
<tr><td>>hg38_21_+_8433172_8446622_RNA45SN1_45S_50ntFlanks@9564.9601.38|CTGACGTGCAAATCGGTCGTCCGACCTGGGTATAGGGG</td><td>>hg38_21_+_8433172_8446622_RNA45SN1_45S_50ntFlanks</td><td>9564</td><td>9601</td><td>38</td><td>CTGACGTGCAAATCGGTCGTCCGACCTGGGTATAGGGG</td><td>78.4</td><td>-3.2</td><td>1.4</td><td>-2.3</td><td>2.1E-02</td><td>2.6E-02</td><td>MB231/MB468</td><td>MP</td></tr>
<tr><td>>hg38_21_+_8433172_8446622_RNA45SN1_45S_50ntFlanks@4975.4993.19|CCCGGACACGGACAGGATT</td><td>>hg38_21_+_8433172_8446622_RNA45SN1_45S_50ntFlanks</td><td>4975</td><td>4993</td><td>19</td><td>CCCGGACACGGACAGGATT</td><td>114.8</td><td>-3.1</td><td>1.3</td><td>-2.3</td><td>2.1E-02</td><td>2.6E-02</td><td>MB231/MB468</td><td>MP</td></tr>
<tr><td>>hg38_21_+_8433172_8446622_RNA45SN1_45S_50ntFlanks@10355.10383.29|AACTTTGAAGGCCGAAGTGGAGAAGGGTT</td><td>>hg38_21_+_8433172_8446622_RNA45SN1_45S_50ntFlanks</td><td>10355</td><td>10383</td><td>29</td><td>AACTTTGAAGGCCGAAGTGGAGAAGGGTT</td><td>100.2</td><td>-2.9</td><td>1.2</td><td>-2.3</td><td>2.0E-02</td><td>2.6E-02</td><td>BT20/MB468</td><td>MITO</td></tr>
<tr><td>>MI0000088|hsa-mir-30a&WithFlank&6|-|71403545|71403627@12.29.18(+1A)&[MIMAT0000087&hsa-miR-30a-5p&offsets|0|-4(+1A);m-15&6|-|71403595|71403616&offsets|0|-4(+1A)]|TGTAAACATCCTCGACTGA</td><td>>MI0000088|hsa-mir-30a&WithFlank&6|-|71403545|71403627</td><td>12</td><td>29</td><td>18(+1A)</td><td>TGTAAACATCCTCGACTGA</td><td>34.9</td><td>4.7</td><td>2.0</td><td>2.3</td><td>2.1E-02</td><td>2.6E-02</td><td>MB231/MB468</td><td>MP</td></tr>
<tr><td>>hg38_MT_+_598_1651_RNR1_12S_50ntFlanks@440.469.30|ACATATCTGAACACACAATAGCTAAGACCC</td><td>>hg38_MT_+_598_1651_RNR1_12S_50ntFlanks</td><td>440</td><td>469</td><td>30</td><td>ACATATCTGAACACACAATAGCTAAGACCC</td><td>130.3</td><td>4.6</td><td>1.9</td><td>2.4</td><td>1.5E-02</td><td>2.6E-02</td><td>BT20/MB468</td><td>MP</td></tr>
<tr><td>>hg38_21_+_8433172_8446622_RNA45SN1_45S_50ntFlanks@12989.13013.25|AGCTCCCTCGCTGCGATCTATTGAA</td><td>>hg38_21_+_8433172_8446622_RNA45SN1_45S_50ntFlanks</td><td>12989</td><td>13013</td><td>25</td><td>AGCTCCCTCGCTGCGATCTATTGAA</td><td>181.7</td><td>-3.7</td><td>1.4</td><td>-2.7</td><td>6.7E-03</td><td>2.6E-02</td><td>BT20/MB231</td><td>NUC</td></tr>
<tr><td>>hg38_21_+_8433172_8446622_RNA45SN1_45S_50ntFlanks@10347.10380.34|CAAACGAGAACTTTGAAGGCCGAAGTGGAGAAGG</td><td>>hg38_21_+_8433172_8446622_RNA45SN1_45S_50ntFlanks</td><td>10347</td><td>10380</td><td>34</td><td>CAAACGAGAACTTTGAAGGCCGAAGTGGAGAAGG</td><td>116.3</td><td>3.6</td><td>1.5</td><td>2.4</td><td>1.5E-02</td><td>2.6E-02</td><td>BT20/MB468</td><td>MP</td></tr>
<tr><td>>MI0000065|hsa-let-7d&WithFlank&9|+|94178828|94178926@14.36.23&[MIMAT0000065&hsa-let-7d-5p&offsets|0|+1;m-90&9|+|94178841|94178862&offsets|0|+1]|AGAGGTAGTAGGTTGCATAGTTT</td><td>>MI0000065|hsa-let-7d&WithFlank&9|+|94178828|94178926</td><td>14</td><td>36</td><td>23</td><td>AGAGGTAGTAGGTTGCATAGTTT</td><td>146.3</td><td>2.5</td><td>1.1</td><td>2.3</td><td>2.3E-02</td><td>2.6E-02</td><td>BT20/MB468</td><td>NUC</td></tr>
<tr><td>>hg38_21_+_8433172_8446622_RNA45SN1_45S_50ntFlanks@12412.12446.35|ATCCTTCGATGTCGGCTCTTCCTATCATTGTGAAG</td><td>>hg38_21_+_8433172_8446622_RNA45SN1_45S_50ntFlanks</td><td>12412</td><td>12446</td><td>35</td><td>ATCCTTCGATGTCGGCTCTTCCTATCATTGTGAAG</td><td>273.8</td><td>-3.8</td><td>1.7</td><td>-2.3</td><td>2.3E-02</td><td>2.6E-02</td><td>MB231/MB468</td><td>MITO</td></tr>
<tr><td>>MI0000088|hsa-mir-30a&WithFlank&6|-|71403545|71403627@12.35.24(+3U)&[MIMAT0000087&hsa-miR-30a-5p&offsets|0|+2(+3U);m-15&6|-|71403595|71403616&offsets|0|+2(+3U)]|TGTAAACATCCTCGACTGGAAGCTTTT</td><td>>MI0000088|hsa-mir-30a&WithFlank&6|-|71403545|71403627</td><td>12</td><td>35</td><td>24(+3U)</td><td>TGTAAACATCCTCGACTGGAAGCTTTT</td><td>141.5</td><td>-4.5</td><td>1.6</td><td>-2.7</td><td>6.8E-03</td><td>2.6E-02</td><td>BT20/MB231</td><td>NUC</td></tr>
<tr><td>>trnaMT_LysTTT_MT_+_8295_8364@1.36.36,trnalookalike7_LysTTT_1_+_568844_568913@1.36.36|CACTGTAAAGCTAACTTAGCATTAACCTTTTAAGTT</td><td>>trnaMT_LysTTT_MT_+_8295_8364</td><td>1</td><td>36</td><td>36</td><td>CACTGTAAAGCTAACTTAGCATTAACCTTTTAAGTT</td><td>266.9</td><td>3.2</td><td>1.4</td><td>2.3</td><td>2.3E-02</td><td>2.6E-02</td><td>MB231/MB468</td><td>NUC</td></tr>
<tr><td>>hg38_21_+_8433172_8446622_RNA45SN1_45S_50ntFlanks@10677.10712.36|GCGGTTCCGGCGGCGTCCGGTGAGCTCTCGCTGGCC</td><td>>hg38_21_+_8433172_8446622_RNA45SN1_45S_50ntFlanks</td><td>10677</td><td>10712</td><td>36</td><td>GCGGTTCCGGCGGCGTCCGGTGAGCTCTCGCTGGCC</td><td>1326.3</td><td>-1.6</td><td>0.7</td><td>-2.3</td><td>2.4E-02</td><td>2.6E-02</td><td>MB231/MB468</td><td>MITO</td></tr>
<tr><td>>hg38_21_+_8433172_8446622_RNA45SN1_45S_50ntFlanks@9919.9948.30|GGTGTTGGTTGATATAGACAGCAGGACGGT</td><td>>hg38_21_+_8433172_8446622_RNA45SN1_45S_50ntFlanks</td><td>9919</td><td>9948</td><td>30</td><td>GGTGTTGGTTGATATAGACAGCAGGACGGT</td><td>56.6</td><td>5.0</td><td>2.1</td><td>2.4</td><td>1.5E-02</td><td>2.6E-02</td><td>BT20/MB468</td><td>MP</td></tr>
<tr><td>>hg38_21_+_8433172_8446622_RNA45SN1_45S_50ntFlanks@11727.11744.18|AGTAACTATGACTCTCTT</td><td>>hg38_21_+_8433172_8446622_RNA45SN1_45S_50ntFlanks</td><td>11727</td><td>11744</td><td>18</td><td>AGTAACTATGACTCTCTT</td><td>886.3</td><td>1.6</td><td>0.7</td><td>2.3</td><td>2.2E-02</td><td>2.6E-02</td><td>MB231/MB468</td><td>CYTO</td></tr>
<tr><td>>MI0000100|hsa-mir-98&WithFlank&X|-|53556217|53556347@28.48.21(+1C)&[MIMAT0000096&hsa-miR-98-5p&offsets|0|-1(+1C);m-143&X|-|53556299|53556320&offsets|0|-1(+1C)]|TGAGGTAGTAAGTTGTATTGTC</td><td>>MI0000100|hsa-mir-98&WithFlank&X|-|53556217|53556347</td><td>28</td><td>48</td><td>21(+1C)</td><td>TGAGGTAGTAAGTTGTATTGTC</td><td>693.1</td><td>1.9</td><td>0.8</td><td>2.3</td><td>2.1E-02</td><td>2.6E-02</td><td>MB231/MB468</td><td>MP</td></tr>
</table>

<table>
<tr><td>>hg38_21_+_8433172_8446622_RNA45SN1_45S_50ntFlanks@12418.12447.30|CGATGTCGGCTCTTCCTATCATTGTGAAGC</td><td>>hg38_21_+_8433172_8446622_RNA45SN1_45S_50ntFlanks</td><td>12418</td><td>12447</td><td>30</td><td>CGATGTCGGCTCTTCCTATCATTGTGAAGC</td><td>541.6</td><td>-2.2</td><td>0.8</td><td>-2.6</td><td>8.4E-03</td><td>2.6E-02</td><td>BT20/MB231</td><td>CYTO</td></tr>
<tr><td>>hg38_21_+_8433172_8446622_RNA45SN1_45S_50ntFlanks@12934.12961.28|CTAAACCATTCGTAGACGACCTGCTTCT</td><td>>hg38_21_+_8433172_8446622_RNA45SN1_45S_50ntFlanks</td><td>12934</td><td>12961</td><td>28</td><td>CTAAACCATTCGTAGACGACCTGCTTCT</td><td>124.0</td><td>-4.1</td><td>1.7</td><td>-2.4</td><td>1.5E-02</td><td>2.6E-02</td><td>BT20/MB468</td><td>MP</td></tr>
<tr><td>>MI0016590|hsa-mir-3934&WithFlank&6|+|33698122|33698240@31.53.23&[MIMAT0018349&hsa-miR-3934-5p&offsets|0|+1;m-557&6|+|33698152|33698173&offsets|0|+1]|TCAGGTGTGGAAACTGAGGCAGG</td><td>>MI0016590|hsa-mir-3934&WithFlank&6|+|33698122|33698240</td><td>31</td><td>53</td><td>23</td><td>TCAGGTGTGGAAACTGAGGCAGG</td><td>42.0</td><td>4.6</td><td>2.0</td><td>2.3</td><td>2.1E-02</td><td>2.6E-02</td><td>MB231/MB468</td><td>MP</td></tr>
<tr><td>>hg38_21_+_8433172_8446622_RNA45SN1_45S_50ntFlanks@10909.10930.22|GCGCGCGCCGCGGCTGGACGAG</td><td>>hg38_21_+_8433172_8446622_RNA45SN1_45S_50ntFlanks</td><td>10909</td><td>10930</td><td>22</td><td>GCGCGCGCCGCGGCTGGACGAG</td><td>44.0</td><td>4.3</td><td>1.8</td><td>2.4</td><td>1.5E-02</td><td>2.6E-02</td><td>BT20/MB468</td><td>MP</td></tr>
<tr><td>>hg38_21_+_8433172_8446622_RNA45SN1_45S_50ntFlanks@4590.4614.25|ATTTTGTTGGTTTTCGGAACTGAGG</td><td>>hg38_21_+_8433172_8446622_RNA45SN1_45S_50ntFlanks</td><td>4590</td><td>4614</td><td>25</td><td>ATTTTGTTGGTTTTCGGAACTGAGG</td><td>192.3</td><td>-4.1</td><td>1.7</td><td>-2.4</td><td>1.5E-02</td><td>2.6E-02</td><td>BT20/MB468</td><td>MP</td></tr>
<tr><td>>hg38_21_+_8433172_8446622_RNA45SN1_45S_50ntFlanks@12640.12659.20|ACCATCTGTGGGATTATGAC</td><td>>hg38_21_+_8433172_8446622_RNA45SN1_45S_50ntFlanks</td><td>12640</td><td>12659</td><td>20</td><td>ACCATCTGTGGGATTATGAC</td><td>241.3</td><td>-1.8</td><td>0.8</td><td>-2.3</td><td>2.3E-02</td><td>2.6E-02</td><td>MB231/MB468</td><td>NUC</td></tr>
<tr><td>>MI0000073|hsa-mir-19a&WithFlank&13|+|91350885|91350978@54.76.23(+1C)&[MIMAT0000073&hsa-miR-19a-3p&offsets|-1|-1(+1C);m-201&13|+|91350939|91350961&offsets|-1|-1(+1C)]|TTGTGCAAATCTATGCAAAACTGC</td><td>>MI0000073|hsa-mir-19a&WithFlank&13|+|91350885|91350978</td><td>54</td><td>76</td><td>23(+1C)</td><td>TTGTGCAAATCTATGCAAAACTGC</td><td>47.5</td><td>3.9</td><td>1.7</td><td>2.3</td><td>2.1E-02</td><td>2.6E-02</td><td>MB231/MB468</td><td>MP</td></tr>
<tr><td>>hg38_21_+_8433172_8446622_RNA45SN1_45S_50ntFlanks@3852.3873.22|TAACTGTGGTAATTCTAGAGCT</td><td>>hg38_21_+_8433172_8446622_RNA45SN1_45S_50ntFlanks</td><td>3852</td><td>3873</td><td>22</td><td>TAACTGTGGTAATTCTAGAGCT</td><td>100.4</td><td>-4.1</td><td>1.7</td><td>-2.4</td><td>1.5E-02</td><td>2.6E-02</td><td>BT20/MB468</td><td>MP</td></tr>
<tr><td>>hg38_21_+_8433172_8446622_RNA45SN1_45S_50ntFlanks@10866.10893.28|CTAAGGGCTGGGTCGGTCGGGCTGGGGC</td><td>>hg38_21_+_8433172_8446622_RNA45SN1_45S_50ntFlanks</td><td>10866</td><td>10893</td><td>28</td><td>CTAAGGGCTGGGTCGGTCGGGCTGGGGC</td><td>101.7</td><td>-4.1</td><td>1.7</td><td>-2.4</td><td>1.5E-02</td><td>2.6E-02</td><td>BT20/MB468</td><td>MP</td></tr>
<tr><td>>hg38_21_+_8433172_8446622_RNA45SN1_45S_50ntFlanks@5445.5463.19|TCGGATCGGCCCCGCCGGG</td><td>>hg38_21_+_8433172_8446622_RNA45SN1_45S_50ntFlanks</td><td>5445</td><td>5463</td><td>19</td><td>TCGGATCGGCCCCGCCGGG</td><td>58.5</td><td>-4.2</td><td>1.7</td><td>-2.4</td><td>1.5E-02</td><td>2.6E-02</td><td>BT20/MB468</td><td>MP</td></tr>
<tr><td>>hg38_MT_+_1621_3279_RNR2_16S_50ntFlanks@1095.1122.28|AGACGAGAAGACCCTATGGAGCTTTAAT</td><td>>hg38_MT_+_1621_3279_RNR2_16S_50ntFlanks</td><td>1095</td><td>1122</td><td>28</td><td>AGACGAGAAGACCCTATGGAGCTTTAAT</td><td>54.2</td><td>4.7</td><td>2.0</td><td>2.3</td><td>2.1E-02</td><td>2.6E-02</td><td>MB231/MB468</td><td>MP</td></tr>
<tr><td>>hg38_21_+_8433172_8446622_RNA45SN1_45S_50ntFlanks@10238.10259.22|CGCTACGCCGCGACGAGTAGGA</td><td>>hg38_21_+_8433172_8446622_RNA45SN1_45S_50ntFlanks</td><td>10238</td><td>10259</td><td>22</td><td>CGCTACGCCGCGACGAGTAGGA</td><td>157.0</td><td>2.5</td><td>1.1</td><td>2.3</td><td>2.4E-02</td><td>2.6E-02</td><td>MB231/MB468</td><td>MITO</td></tr>
<tr><td>>hg38_MT_+_1621_3279_RNR2_16S_50ntFlanks@617.643.27|AAACATATAACTGAACTCCTCACACCC</td><td>>hg38_MT_+_1621_3279_RNR2_16S_50ntFlanks</td><td>617</td><td>643</td><td>27</td><td>AAACATATAACTGAACTCCTCACACCC</td><td>144.0</td><td>3.3</td><td>1.4</td><td>2.3</td><td>2.1E-02</td><td>2.6E-02</td><td>MB231/MB468</td><td>MP</td></tr>
<tr><td>>hg38_MT_+_598_1651_RNR1_12S_50ntFlanks@905.930.26|AGTATACTTCAAAGGACATTTAACTA</td><td>>hg38_MT_+_598_1651_RNR1_12S_50ntFlanks</td><td>905</td><td>930</td><td>26</td><td>AGTATACTTCAAAGGACATTTAACTA</td><td>155.8</td><td>4.1</td><td>1.8</td><td>2.3</td><td>2.1E-02</td><td>2.6E-02</td><td>MB231/MB468</td><td>MP</td></tr>
<tr><td>>hg38_21_+_8433172_8446622_RNA45SN1_45S_50ntFlanks@12416.12447.32|TTCGATGTCGGCTCTTCCTATCATTGTGAAGC</td><td>>hg38_21_+_8433172_8446622_RNA45SN1_45S_50ntFlanks</td><td>12416</td><td>12447</td><td>32</td><td>TTCGATGTCGGCTCTTCCTATCATTGTGAAGC</td><td>2670.9</td><td>-2.4</td><td>0.9</td><td>-2.6</td><td>8.4E-03</td><td>2.6E-02</td><td>BT20/MB231</td><td>CYTO</td></tr>
<tr><td>>hg38_21_+_8433172_8446622_RNA45SN1_45S_50ntFlanks@12420.12442.23|ATGTCGGCTCTTCCTATCATTGT</td><td>>hg38_21_+_8433172_8446622_RNA45SN1_45S_50ntFlanks</td><td>12420</td><td>12442</td><td>23</td><td>ATGTCGGCTCTTCCTATCATTGT</td><td>1121.8</td><td>-2.3</td><td>0.9</td><td>-2.6</td><td>8.4E-03</td><td>2.6E-02</td><td>BT20/MB231</td><td>CYTO</td></tr>
<tr><td>>hg38_21_+_8433172_8446622_RNA45SN1_45S_50ntFlanks@11573.11600.28|CGACTTAGAACTGGTGCGGACCAGGGGA</td><td>>hg38_21_+_8433172_8446622_RNA45SN1_45S_50ntFlanks</td><td>11573</td><td>11600</td><td>28</td><td>CGACTTAGAACTGGTGCGGACCAGGGGA</td><td>511.3</td><td>-1.7</td><td>0.7</td><td>-2.3</td><td>2.3E-02</td><td>2.6E-02</td><td>MB231/MB468</td><td>CYTO</td></tr>
<tr><td>>hg38_MT_+_1621_3279_RNR2_16S_50ntFlanks@1438.1477.40|TACGTGATCTGAGTTCAGACCGGAGTAATCCAGGTCGGTT</td><td>>hg38_MT_+_1621_3279_RNR2_16S_50ntFlanks</td><td>1438</td><td>1477</td><td>40</td><td>TACGTGATCTGAGTTCAGACCGGAGTAATCCAGGTCGGTT</td><td>368.2</td><td>3.8</td><td>1.6</td><td>2.4</td><td>1.5E-02</td><td>2.6E-02</td><td>BT20/MB468</td><td>MP</td></tr>
<tr><td>>hg38_21_+_8433172_8446622_RNA45SN1_45S_50ntFlanks@9381.9411.31|CGCTCGCCGGCCGAGGTGGGATCCCGAGGCC</td><td>>hg38_21_+_8433172_8446622_RNA45SN1_45S_50ntFlanks</td><td>9381</td><td>9411</td><td>31</td><td>CGCTCGCCGGCCGAGGTGGGATCCCGAGGCC</td><td>163.2</td><td>-3.9</td><td>1.7</td><td>-2.3</td><td>2.0E-02</td><td>2.6E-02</td><td>BT20/MB468</td><td>MITO</td></tr>
<tr><td>>hg38_21_+_8433172_8446622_RNA45SN1_45S_50ntFlanks@3876.3899.24|TACATGCCGACGGGCGCTGACCCC</td><td>>hg38_21_+_8433172_8446622_RNA45SN1_45S_50ntFlanks</td><td>3876</td><td>3899</td><td>24</td><td>TACATGCCGACGGGCGCTGACCCC</td><td>121.7</td><td>-2.5</td><td>1.1</td><td>-2.3</td><td>2.0E-02</td><td>2.6E-02</td><td>BT20/MB468</td><td>MITO</td></tr>
<tr><td>>MI0000812|hsa-mir-331&WithFlank&12|+|95308414|95308519@31.52.22&[MIMAT0004700&hsa-miR-331-5p&offsets|-1|-1;TJU_CMC_MD2.ID00734.5p-miR&offsets|+19|+21;m-2747&12|+|95308425|95308444&offsets|+19|+21;m-377&12|+|95308445|95308465&offsets|-1|0]|TCTAGGTATGGTCCCAGGGATC</td><td>>MI0000812|hsa-mir-331&WithFlank&12|+|95308414|95308519</td><td>31</td><td>52</td><td>22</td><td>TCTAGGTATGGTCCCAGGGATC</td><td>81.0</td><td>6.3</td><td>2.6</td><td>2.4</td><td>1.5E-02</td><td>2.6E-02</td><td>BT20/MB468</td><td>MP</td></tr>
<tr><td>>hg38_21_+_8433172_8446622_RNA45SN1_45S_50ntFlanks@12418.12453.36|CGATGTCGGCTCTTCCTATCATTGTGAAGCAGAATT</td><td>>hg38_21_+_8433172_8446622_RNA45SN1_45S_50ntFlanks</td><td>12418</td><td>12453</td><td>36</td><td>CGATGTCGGCTCTTCCTATCATTGTGAAGCAGAATT</td><td>154.4</td><td>-2.4</td><td>0.9</td><td>-2.7</td><td>6.8E-03</td><td>2.6E-02</td><td>BT20/MB231</td><td>NUC</td></tr>
<tr><td>>MI0001733|hsa-mir-452&WithFlank&X|-|151959622|151959718@23.43.21&[MIMAT0001635&hsa-miR-452-5p&offsets|+3|+2;m-188&X|-|151959677|151959699&offsets|+3|+1]|TGTTTGCAGAGGAAACTGAGA</td><td>>MI0001733|hsa-mir-452&WithFlank&X|-|151959622|151959718</td><td>23</td><td>43</td><td>21</td><td>TGTTTGCAGAGGAAACTGAGA</td><td>458.8</td><td>2.0</td><td>0.8</td><td>2.7</td><td>7.7E-03</td><td>2.6E-02</td><td>BT20/MB231</td><td>MITO</td></tr>
<tr><td>>trnaMT_SerGCT_MT_+_12207_12265@1.22.22|GAGAAAGCTCACAAGAACTGCT</td><td>>trnaMT_SerGCT_MT_+_12207_12265</td><td>1</td><td>22</td><td>22</td><td>GAGAAAGCTCACAAGAACTGCT</td><td>26795.9</td><td>3.1</td><td>1.3</td><td>2.4</td><td>1.5E-02</td><td>2.6E-02</td><td>BT20/MB468</td><td>MP</td></tr>
<tr><td>>hg38_21_+_8433172_8446622_RNA45SN1_45S_50ntFlanks@8196.8218.23|CGTGGACGGTGTGAGGCCGGTAG</td><td>>hg38_21_+_8433172_8446622_RNA45SN1_45S_50ntFlanks</td><td>8196</td><td>8218</td><td>23</td><td>CGTGGACGGTGTGAGGCCGGTAG</td><td>113.0</td><td>-3.3</td><td>1.4</td><td>-2.4</td><td>1.5E-02</td><td>2.6E-02</td><td>BT20/MB468</td><td>MP</td></tr>
<tr><td>>hg38_21_+_8433172_8446622_RNA45SN1_45S_50ntFlanks@7980.8003.24|CCTCAGATCAGACGTGGCGACCCG</td><td>>hg38_21_+_8433172_8446622_RNA45SN1_45S_50ntFlanks</td><td>7980</td><td>8003</td><td>24</td><td>CCTCAGATCAGACGTGGCGACCCG</td><td>407.2</td><td>-2.1</td><td>0.9</td><td>-2.3</td><td>2.3E-02</td><td>2.6E-02</td><td>MB231/MB468</td><td>CYTO</td></tr>
<tr><td>>hg38_21_+_8433172_8446622_RNA45SN1_45S_50ntFlanks@6647.6681.35|CGTACGACTCTTAGCGGTGGATCACTCGGCTCGTG</td><td>>hg38_21_+_8433172_8446622_RNA45SN1_45S_50ntFlanks</td><td>6647</td><td>6681</td><td>35</td><td>CGTACGACTCTTAGCGGTGGATCACTCGGCTCGTG</td><td>5284.4</td><td>-2.3</td><td>0.9</td><td>-2.4</td><td>1.5E-02</td><td>2.6E-02</td><td>BT20/MB468</td><td>MP</td></tr>
<tr><td>>hg38_21_+_8433172_8446622_RNA45SN1_45S_50ntFlanks@12204.12222.19|AAGGCGAGCTCAGGGAGGA</td><td>>hg38_21_+_8433172_8446622_RNA45SN1_45S_50ntFlanks</td><td>12204</td><td>12222</td><td>19</td><td>AAGGCGAGCTCAGGGAGGA</td><td>179.6</td><td>-2.3</td><td>1.0</td><td>-2.3</td><td>2.1E-02</td><td>2.6E-02</td><td>MB231/MB468</td><td>MP</td></tr>
<tr><td>>trnaMT_PheGAA_MT_+_577_647@38.67.30|AATGTTTAGACGGGCTCACATCACCCCATA</td><td>>trnaMT_PheGAA_MT_+_577_647</td><td>38</td><td>67</td><td>30</td><td>AATGTTTAGACGGGCTCACATCACCCCATA</td><td>24.2</td><td>4.7</td><td>2.0</td><td>2.3</td><td>2.1E-02</td><td>2.6E-02</td><td>MB231/MB468</td><td>MP</td></tr>
<tr><td>>hg38_21_+_8433172_8446622_RNA45SN1_45S_50ntFlanks@12073.12094.22|CCGAGGGGCTCTCGCTTCTGGC</td><td>>hg38_21_+_8433172_8446622_RNA45SN1_45S_50ntFlanks</td><td>12073</td><td>12094</td><td>22</td><td>CCGAGGGGCTCTCGCTTCTGGC</td><td>158.6</td><td>-2.8</td><td>1.2</td><td>-2.3</td><td>2.0E-02</td><td>2.6E-02</td><td>BT20/MB468</td><td>MITO</td></tr>
<tr><td>>hg38_21_+_8433172_8446622_RNA45SN1_45S_50ntFlanks@8357.8396.40|AAAAGAACTTTGAAGAGAGAGTTCAAGAGGGCGTGAAACC</td><td>>hg38_21_+_8433172_8446622_RNA45SN1_45S_50ntFlanks</td><td>8357</td><td>8396</td><td>40</td><td>AAAAGAACTTTGAAGAGAGAGTTCAAGAGGGCGTGAAACC</td><td>120.7</td><td>5.6</td><td>2.0</td><td>2.7</td><td>6.3E-03</td><td>2.6E-02</td><td>BT20/MB231</td><td>MP</td></tr>
<tr><td>>hg38_21_+_8433172_8446622_RNA45SN1_45S_50ntFlanks@5084.5107.24|CGAACGAGACTCTGGCATGCTAAC</td><td>>hg38_21_+_8433172_8446622_RNA45SN1_45S_50ntFlanks</td><td>5084</td><td>5107</td><td>24</td><td>CGAACGAGACTCTGGCATGCTAAC</td><td>238.4</td><td>-3.6</td><td>1.5</td><td>-2.3</td><td>2.0E-02</td><td>2.6E-02</td><td>BT20/MB468</td><td>MITO</td></tr>
<tr><td>>hg38_21_+_8433172_8446622_RNA45SN1_45S_50ntFlanks@476.497.22|TTCGTGATCGATGTGGTGACGT</td><td>>hg38_21_+_8433172_8446622_RNA45SN1_45S_50ntFlanks</td><td>476</td><td>497</td><td>22</td><td>TTCGTGATCGATGTGGTGACGT</td><td>491.7</td><td>2.6</td><td>1.2</td><td>2.3</td><td>2.3E-02</td><td>2.6E-02</td><td>BT20/MB468</td><td>NUC</td></tr>
<tr><td>>hg38_21_+_8433172_8446622_RNA45SN1_45S_50ntFlanks@12029.12058.30|TCTGATCGTTTTTTCACTGACCCGGTGAGG</td><td>>hg38_21_+_8433172_8446622_RNA45SN1_45S_50ntFlanks</td><td>12029</td><td>12058</td><td>30</td><td>TCTGATCGTTTTTTCACTGACCCGGTGAGG</td><td>188.1</td><td>5.7</td><td>2.1</td><td>2.7</td><td>6.3E-03</td><td>2.6E-02</td><td>BT20/MB231</td><td>MP</td></tr>
<tr><td>>hg38_21_+_8433172_8446622_RNA45SN1_45S_50ntFlanks@10678.10708.31|CGGTTCCGGCGGCGTCCGGTGAGCTCTCGCT</td><td>>hg38_21_+_8433172_8446622_RNA45SN1_45S_50ntFlanks</td><td>10678</td><td>10708</td><td>31</td><td>CGGTTCCGGCGGCGTCCGGTGAGCTCTCGCT</td><td>983.7</td><td>2.7</td><td>1.0</td><td>2.7</td><td>7.7E-03</td><td>2.6E-02</td><td>BT20/MB231</td><td>MITO</td></tr>
<tr><td>>MI0000097|hsa-mir-95&WithFlank&4|-|8005295|8005387@55.75.21&[MIMAT0000094&hsa-miR-95-3p&offsets|0|-1;m-328&4|-|8005313|8005333&offsets|0|0]|TTCAACGGGTATTTATTGAGC</td><td>>MI0000097|hsa-mir-95&WithFlank&4|-|8005295|8005387</td><td>55</td><td>75</td><td>21</td><td>TTCAACGGGTATTTATTGAGC</td><td>116.2</td><td>-2.0</td><td>0.9</td><td>-2.3</td><td>2.1E-02</td><td>2.6E-02</td><td>MB231/MB468</td><td>MP</td></tr>
<tr><td>>hg38_21_+_8433172_8446622_RNA45SN1_45S_50ntFlanks@11209.11229.21|CGGCGACTCTGGACGCGAGCC</td><td>>hg38_21_+_8433172_8446622_RNA45SN1_45S_50ntFlanks</td><td>11209</td><td>11229</td><td>21</td><td>CGGCGACTCTGGACGCGAGCC</td><td>789.6</td><td>-1.4</td><td>0.6</td><td>-2.3</td><td>2.3E-02</td><td>2.6E-02</td><td>BT20/MB468</td><td>CYTO</td></tr>
<tr><td>>MI0000073|hsa-mir-19a&WithFlank&13|+|91350885|91350978@54.76.23(+1U)&[MIMAT0000073&hsa-miR-19a-3p&offsets|-1|-1(+1U);m-201&13|+|91350939|91350961&offsets|-1|-1(+1U)]|TTGTGCAAATCTATGCAAAACTGT</td><td>>MI0000073|hsa-mir-19a&WithFlank&13|+|91350885|91350978</td><td>54</td><td>76</td><td>23(+1U)</td><td>TTGTGCAAATCTATGCAAAACTGT</td><td>308.9</td><td>3.2</td><td>1.4</td><td>2.3</td><td>2.1E-02</td><td>2.6E-02</td><td>MB231/MB468</td><td>MP</td></tr>
<tr><td>>hg38_21_+_8433172_8446622_RNA45SN1_45S_50ntFlanks@10535.10571.37|CGCGAGGCGTCCAGTGCGGTAACGCGACCGATCCCGG</td><td>>hg38_21_+_8433172_8446622_RNA45SN1_45S_50ntFlanks</td><td>10535</td><td>10571</td><td>37</td><td>CGCGAGGCGTCCAGTGCGGTAACGCGACCGATCCCGG</td><td>216.2</td><td>-4.2</td><td>1.7</td><td>-2.4</td><td>1.5E-02</td><td>2.6E-02</td><td>BT20/MB468</td><td>MP</td></tr>
<tr><td>>hg38_21_+_8433172_8446622_RNA45SN1_45S_50ntFlanks@12221.12248.28|GACAGAAACCTCCCGTGGAGCAGAAGGG</td><td>>hg38_21_+_8433172_8446622_RNA45SN1_45S_50ntFlanks</td><td>12221</td><td>12248</td><td>28</td><td>GACAGAAACCTCCCGTGGAGCAGAAGGG</td><td>57.5</td><td>-4.1</td><td>1.7</td><td>-2.4</td><td>1.5E-02</td><td>2.6E-02</td><td>BT20/MB468</td><td>MP</td></tr>
<tr><td>>hg38_21_+_8433172_8446622_RNA45SN1_45S_50ntFlanks@12635.12662.28|AAGCTACCATCTGTGGGATTATGACTGA</td><td>>hg38_21_+_8433172_8446622_RNA45SN1_45S_50ntFlanks</td><td>12635</td><td>12662</td><td>28</td><td>AAGCTACCATCTGTGGGATTATGACTGA</td><td>133.2</td><td>-4.1</td><td>1.7</td><td>-2.4</td><td>1.5E-02</td><td>2.6E-02</td><td>BT20/MB468</td><td>MP</td></tr>
<tr><td>>hg38_21_+_8433172_8446622_RNA45SN1_45S_50ntFlanks@3840.3859.20|CTCCTACTTGGATAACTGTG</td><td>>hg38_21_+_8433172_8446622_RNA45SN1_45S_50ntFlanks</td><td>3840</td><td>3859</td><td>20</td><td>CTCCTACTTGGATAACTGTG</td><td>735.2</td><td>1.9</td><td>0.8</td><td>2.3</td><td>2.3E-02</td><td>2.6E-02</td><td>BT20/MB468</td><td>CYTO</td></tr>
<tr><td>>MI0000289|hsa-mir-181a-1&WithFlank&1|-|198859038|198859159@30.51.22(+1G);MI0000269|hsa-mir-181a-2&WithFlank&9|+|124692436|124692557@45.66.22(+1G)&[MIMAT0000256&hsa-miR-181a-5p&offsets|0|-1(+1G);m-32&1|-|198859108|198859130&offsets|0|-1(+1G)];[MIMAT0000256_1&hsa-miR-181a-5p&offsets|0|-1(+1G);m-32&9|+|124692480|124692502&offsets|0|-1(+1G)]|AACATTCAACGCTGTCGGTGAGG</td><td>>MI0000289|hsa-mir-181a-1&WithFlank&1|-|198859038|198859159</td><td>30</td><td>51</td><td>22(+1G)</td><td>AACATTCAACGCTGTCGGTGAGG</td><td>314.7</td><td>-4.2</td><td>1.6</td><td>-2.7</td><td>6.3E-03</td><td>2.6E-02</td><td>BT20/MB231</td><td>MP</td></tr>
<tr><td>>am_trna128_GlyGCC_6_-_27870686_27870756@3.34.32,trna18_GlyGCC_16_+_70822597_70822667@3.34.32,trna19_GlyGCC_16_+_70823410_70823480@3.34.32,trna19_GlyGCC_2_-_157257659_157257729@3.34.32,trna24_GlyGCC_16_-_70812942_70813012@3.34.32,trna25_GlyGCC_16_-_70812114_70812184@3.34.32,trna5_GlyGCC_17_+_8029064_8029134@3.34.32,trna68_GlyGCC_1_-_161493637_161493707@3.34.32|ATTGGTGGTTCAGTGGTAGAATTCTCGCCTGC</td><td>>am_trna128_GlyGCC_6_-_27870686_27870756</td><td>3</td><td>34</td><td>32</td><td>ATTGGTGGTTCAGTGGTAGAATTCTCGCCTGC</td><td>149.3</td><td>-5.2</td><td>1.9</td><td>-2.7</td><td>6.3E-03</td><td>2.6E-02</td><td>BT20/MB231</td><td>MP</td></tr>
<tr><td>>hg38_21_+_8433172_8446622_RNA45SN1_45S_50ntFlanks@6652.6698.47|GACTCTTAGCGGTGGATCACTCGGCTCGTGCGTCGATGAAGAACGCA</td><td>>hg38_21_+_8433172_8446622_RNA45SN1_45S_50ntFlanks</td><td>6652</td><td>6698</td><td>47</td><td>GACTCTTAGCGGTGGATCACTCGGCTCGTGCGTCGATGAAGAACGCA</td><td>2114.4</td><td>-3.7</td><td>1.6</td><td>-2.3</td><td>2.1E-02</td><td>2.6E-02</td><td>MB231/MB468</td><td>MP</td></tr>
<tr><td>>hg38_21_+_8433172_8446622_RNA45SN1_45S_50ntFlanks@5084.5110.27|CGAACGAGACTCTGGCATGCTAACTAG</td><td>>hg38_21_+_8433172_8446622_RNA45SN1_45S_50ntFlanks</td><td>5084</td><td>5110</td><td>27</td><td>CGAACGAGACTCTGGCATGCTAACTAG</td><td>1375.7</td><td>-3.3</td><td>1.4</td><td>-2.4</td><td>1.5E-02</td><td>2.6E-02</td><td>BT20/MB468</td><td>MP</td></tr>
<tr><td>>hg38_MT_+_1621_3279_RNR2_16S_50ntFlanks@1373.1401.29|TTGGATCAGGACATCCCGATGGTGCAGCC</td><td>>hg38_MT_+_1621_3279_RNR2_16S_50ntFlanks</td><td>1373</td><td>1401</td><td>29</td><td>TTGGATCAGGACATCCCGATGGTGCAGCC</td><td>118.7</td><td>3.5</td><td>1.5</td><td>2.4</td><td>1.5E-02</td><td>2.6E-02</td><td>BT20/MB468</td><td>MP</td></tr>
<tr><td>>hg38_21_+_8433172_8446622_RNA45SN1_45S_50ntFlanks@10352.10380.29|GAGAACTTTGAAGGCCGAAGTGGAGAAGG</td><td>>hg38_21_+_8433172_8446622_RNA45SN1_45S_50ntFlanks</td><td>10352</td><td>10380</td><td>29</td><td>GAGAACTTTGAAGGCCGAAGTGGAGAAGG</td><td>87.5</td><td>3.7</td><td>1.5</td><td>2.4</td><td>1.5E-02</td><td>2.6E-02</td><td>BT20/MB468</td><td>MP</td></tr>
</table>

<table>
<tr><th></th><th></th><th></th><th></th><th></th><th></th><th></th><th></th><th></th><th></th><th></th><th></th><th></th><th></th></tr>
<tr><td>>MI0000299|hsa-mir-222&WithFlank&X|-|45747009|45747130@75.94.20(+1C)&[MIMAT0000279&hsa-miR-222-3p&offsets|0|-1(+1C);m-140&X|-|45747033|45747056&offsets|0|-4(+1C)]|AGCTACATCTGGCTACTGGGC</td><td>>MI0000299|hsa-mir-222&WithFlank&X|-|45747009|45747130</td><td>75</td><td>94</td><td>20(+1C)</td><td>AGCTACATCTGGCTACTGGGC</td><td>583.1</td><td>-2.1</td><td>0.8</td><td>-2.7</td><td>7.7E-03</td><td>2.6E-02</td><td>BT20/MB231</td><td>MITO</td></tr>
<tr><td>>MI0003137|hsa-mir-193b&WithFlank&16|+|14303961|14304055@58.78.21&[MIMAT0002819&hsa-miR-193b-3p&offsets|+1|0;m-166&16|+|14304017|14304038&offsets|+1|0]|ACTGGCCCTCAAAGTCCCGCT</td><td>>MI0003137|hsa-mir-193b&WithFlank&16|+|14303961|14304055</td><td>58</td><td>78</td><td>21</td><td>ACTGGCCCTCAAAGTCCCGCT</td><td>101.5</td><td>6.3</td><td>2.6</td><td>2.4</td><td>1.5E-02</td><td>2.6E-02</td><td>BT20/MB468</td><td>MP</td></tr>
<tr><td>>MI0000734|hsa-mir-106b&WithFlank&7|-|100093987|100094080@58.77.20(+1U)&[MIMAT0004672&hsa-miR-106b-3p&offsets|0|-2(+1U);m-87&7|-|100094003|100094023&offsets|0|-1(+1U)]|CCGCACTGTGGGTACTTGCTT</td><td>>MI0000734|hsa-mir-106b&WithFlank&7|-|100093987|100094080</td><td>58</td><td>77</td><td>20(+1U)</td><td>CCGCACTGTGGGTACTTGCTT</td><td>37.3</td><td>4.7</td><td>2.0</td><td>2.3</td><td>2.1E-02</td><td>2.6E-02</td><td>MB231/MB468</td><td>MP</td></tr>
<tr><td>>trnaMT_MetCAT_MT_+_4402_4469@41.71.31|AAATGTTGGTTATACCCTTCCCGTACTACCA</td><td>>trnaMT_MetCAT_MT_+_4402_4469</td><td>41</td><td>71</td><td>31</td><td>AAATGTTGGTTATACCCTTCCCGTACTACCA</td><td>352.2</td><td>2.8</td><td>1.2</td><td>2.4</td><td>1.5E-02</td><td>2.6E-02</td><td>BT20/MB468</td><td>MP</td></tr>
<tr><td>>hg38_21_+_8433172_8446622_RNA45SN1_45S_50ntFlanks@3852.3874.23|TAACTGTGGTAATTCTAGAGCTA</td><td>>hg38_21_+_8433172_8446622_RNA45SN1_45S_50ntFlanks</td><td>3852</td><td>3874</td><td>23</td><td>TAACTGTGGTAATTCTAGAGCTA</td><td>188.3</td><td>-1.9</td><td>0.8</td><td>-2.3</td><td>2.3E-02</td><td>2.6E-02</td><td>MB231/MB468</td><td>CYTO</td></tr>
<tr><td>>trnaMT_PheGAA_MT_+_577_647@22.62.41|AAAGCAATACACTGAAAATGTTTAGACGGGCTCACATCACC</td><td>>trnaMT_PheGAA_MT_+_577_647</td><td>22</td><td>62</td><td>41</td><td>AAAGCAATACACTGAAAATGTTTAGACGGGCTCACATCACC</td><td>33.7</td><td>8.4</td><td>3.1</td><td>2.7</td><td>6.3E-03</td><td>2.6E-02</td><td>BT20/MB231</td><td>MP</td></tr>
<tr><td>>hg38_21_+_8433172_8446622_RNA45SN1_45S_50ntFlanks@11564.11601.38|CCTAGCAGCCGACTTAGAACTGGTGCGGACCAGGGGAA</td><td>>hg38_21_+_8433172_8446622_RNA45SN1_45S_50ntFlanks</td><td>11564</td><td>11601</td><td>38</td><td>CCTAGCAGCCGACTTAGAACTGGTGCGGACCAGGGGAA</td><td>288.7</td><td>5.5</td><td>2.0</td><td>2.7</td><td>6.3E-03</td><td>2.6E-02</td><td>BT20/MB231</td><td>MP</td></tr>
<tr><td>>hg38_21_+_8433172_8446622_RNA45SN1_45S_50ntFlanks@11600.11626.27|AATCCGACTGTTTAATTAAAACAAAGC</td><td>>hg38_21_+_8433172_8446622_RNA45SN1_45S_50ntFlanks</td><td>11600</td><td>11626</td><td>27</td><td>AATCCGACTGTTTAATTAAAACAAAGC</td><td>257.9</td><td>5.5</td><td>2.0</td><td>2.7</td><td>6.3E-03</td><td>2.6E-02</td><td>BT20/MB231</td><td>MP</td></tr>
<tr><td>>hg38_21_+_8433172_8446622_RNA45SN1_45S_50ntFlanks@9794.9811.18|CGTGGAGCCGGGCGTGGA</td><td>>hg38_21_+_8433172_8446622_RNA45SN1_45S_50ntFlanks</td><td>9794</td><td>9811</td><td>18</td><td>CGTGGAGCCGGGCGTGGA</td><td>162.2</td><td>3.3</td><td>1.2</td><td>2.7</td><td>6.3E-03</td><td>2.6E-02</td><td>BT20/MB231</td><td>MP</td></tr>
<tr><td>>hg38_21_+_8433172_8446622_RNA45SN1_45S_50ntFlanks@10539.10567.29|AGGCGTCCAGTGCGGTAACGCGACCGATC</td><td>>hg38_21_+_8433172_8446622_RNA45SN1_45S_50ntFlanks</td><td>10539</td><td>10567</td><td>29</td><td>AGGCGTCCAGTGCGGTAACGCGACCGATC</td><td>134.1</td><td>-4.3</td><td>1.8</td><td>-2.4</td><td>1.5E-02</td><td>2.6E-02</td><td>BT20/MB468</td><td>MP</td></tr>
<tr><td>>hg38_21_+_8433172_8446622_RNA45SN1_45S_50ntFlanks@10350.10382.33|ACGAGAACTTTGAAGGCCGAAGTGGAGAAGGGT</td><td>>hg38_21_+_8433172_8446622_RNA45SN1_45S_50ntFlanks</td><td>10350</td><td>10382</td><td>33</td><td>ACGAGAACTTTGAAGGCCGAAGTGGAGAAGGGT</td><td>1544.4</td><td>-2.2</td><td>0.9</td><td>-2.4</td><td>1.5E-02</td><td>2.6E-02</td><td>BT20/MB468</td><td>MP</td></tr>
<tr><td>>am_trnaMT_MetCAT_MT_+_4402_4469@1.48.48|AGTAAGGTCAGCTAAATAAGCTATCGGGCCCATACCCCGAAAATGTTG</td><td>>am_trnaMT_MetCAT_MT_+_4402_4469</td><td>1</td><td>48</td><td>48</td><td>AGTAAGGTCAGCTAAATAAGCTATCGGGCCCATACCCCGAAAATGTTG</td><td>802.1</td><td>6.7</td><td>2.8</td><td>2.4</td><td>1.5E-02</td><td>2.6E-02</td><td>BT20/MB468</td><td>MP</td></tr>
<tr><td>>hg38_21_+_8433172_8446622_RNA45SN1_45S_50ntFlanks@7973.7992.20|GACGCGACCTCAGATCAGAC</td><td>>hg38_21_+_8433172_8446622_RNA45SN1_45S_50ntFlanks</td><td>7973</td><td>7992</td><td>20</td><td>GACGCGACCTCAGATCAGAC</td><td>1523.7</td><td>3.6</td><td>1.3</td><td>2.7</td><td>6.3E-03</td><td>2.6E-02</td><td>BT20/MB231</td><td>MP</td></tr>
<tr><td>>hg38_MT_+_598_1651_RNR1_12S_50ntFlanks@917.935.19|AGGACATTTAACTAAAACC</td><td>>hg38_MT_+_598_1651_RNR1_12S_50ntFlanks</td><td>917</td><td>935</td><td>19</td><td>AGGACATTTAACTAAAACC</td><td>703.7</td><td>-3.3</td><td>1.2</td><td>-2.7</td><td>6.3E-03</td><td>2.6E-02</td><td>BT20/MB231</td><td>MP</td></tr>
<tr><td>>hg38_21_+_8433172_8446622_RNA45SN1_45S_50ntFlanks@12396.12414.19|GCGACGTCGCTTTTTGATC</td><td>>hg38_21_+_8433172_8446622_RNA45SN1_45S_50ntFlanks</td><td>12396</td><td>12414</td><td>19</td><td>GCGACGTCGCTTTTTGATC</td><td>2123.7</td><td>-1.2</td><td>0.5</td><td>-2.3</td><td>2.4E-02</td><td>2.6E-02</td><td>MB231/MB468</td><td>NUC</td></tr>
<tr><td>>MI0000736|hsa-mir-30c-1&WithFlank&1|+|40757278|40757378@23.42.20;MI0000254|hsa-mir-30c-2&WithFlank&6|-|71376954|71377037@13.32.20&[MIMAT0000244&hsa-miR-30c-5p&offsets|0|-3;m-57&1|+|40757300|40757323&offsets|0|-4];[MIMAT0000244_1&hsa-miR-30c-5p&offsets|0|-3;m-56&6|-|71377002|71377025&offsets|0|-4]|TGTAAACATCCTACACTCTC</td><td>>MI0000736|hsa-mir-30c-1&WithFlank&1|+|40757278|40757378</td><td>23</td><td>42</td><td>20</td><td>TGTAAACATCCTACACTCTC</td><td>1554.3</td><td>-2.9</td><td>1.1</td><td>-2.7</td><td>6.3E-03</td><td>2.6E-02</td><td>BT20/MB231</td><td>MP</td></tr>
<tr><td>>hg38_21_+_8433172_8446622_RNA45SN1_45S_50ntFlanks@8176.8220.45|CTTCTGATCGAGGCCCAGCCCGTGGACGGTGTGAGGCCGGTAGCG</td><td>>hg38_21_+_8433172_8446622_RNA45SN1_45S_50ntFlanks</td><td>8176</td><td>8220</td><td>45</td><td>CTTCTGATCGAGGCCCAGCCCGTGGACGGTGTGAGGCCGGTAGCG</td><td>155.2</td><td>-2.8</td><td>1.1</td><td>-2.4</td><td>1.5E-02</td><td>2.6E-02</td><td>BT20/MB468</td><td>MP</td></tr>
<tr><td>>hg38_MT_+_598_1651_RNR1_12S_50ntFlanks@447.480.34|TGAACACACAATAGCTAAGACCCAAACTGGGATT</td><td>>hg38_MT_+_598_1651_RNR1_12S_50ntFlanks</td><td>447</td><td>480</td><td>34</td><td>TGAACACACAATAGCTAAGACCCAAACTGGGATT</td><td>101.2</td><td>4.6</td><td>2.0</td><td>2.3</td><td>2.1E-02</td><td>2.6E-02</td><td>MB231/MB468</td><td>MP</td></tr>
<tr><td>>MI0000086|hsa-mir-28&WithFlank&3|+|188688775|188688872@61.82.22(+1A)&[MIMAT0004502&hsa-miR-28-3p&offsets|+1|+1(+1A);m-28&3|+|188688834|188688855&offsets|+1|+1(+1A)]|ACTAGATTGTGAGCTCCTGGAGA</td><td>>MI0000086|hsa-mir-28&WithFlank&3|+|188688775|188688872</td><td>61</td><td>82</td><td>22(+1A)</td><td>ACTAGATTGTGAGCTCCTGGAGA</td><td>38.8</td><td>3.8</td><td>1.7</td><td>2.3</td><td>2.1E-02</td><td>2.6E-02</td><td>MB231/MB468</td><td>MP</td></tr>
<tr><td>>MI0000093|hsa-mir-92a-1&WithFlank&13|+|91351308|91351397@54.74.21(+1C);MI0000094|hsa-mir-92a-2&WithFlank&X|-|134169532|134169618@54.74.21(+1C)&[MIMAT0000092&hsa-miR-92a-3p&offsets|0|-1(+1C);m-9&13|+|91351361|91351382&offsets|0|-1(+1C)];[MIMAT0000092_1&hsa-miR-92a-3p&offsets|0|-1(+1C);m-11&X|-|134169544|134169565&offsets|0|-1(+1C)]|TATTGCACTTGTCCCGGCCTGC</td><td>>MI0000093|hsa-mir-92a-1&WithFlank&13|+|91351308|91351397</td><td>54</td><td>74</td><td>21(+1C)</td><td>TATTGCACTTGTCCCGGCCTGC</td><td>3262.5</td><td>2.5</td><td>1.0</td><td>2.4</td><td>1.5E-02</td><td>2.6E-02</td><td>BT20/MB468</td><td>MP</td></tr>
<tr><td>>hg38_21_+_8433172_8446622_RNA45SN1_45S_50ntFlanks@12419.12447.29|GATGTCGGCTCTTCCTATCATTGTGAAGC</td><td>>hg38_21_+_8433172_8446622_RNA45SN1_45S_50ntFlanks</td><td>12419</td><td>12447</td><td>29</td><td>GATGTCGGCTCTTCCTATCATTGTGAAGC</td><td>469.6</td><td>-2.0</td><td>0.7</td><td>-2.7</td><td>6.9E-03</td><td>2.6E-02</td><td>BT20/MB231</td><td>NUC</td></tr>
<tr><td>>hg38_MT_+_1621_3279_RNR2_16S_50ntFlanks@1370.1394.25|ATGTTGGATCAGGACATCCCGATGG</td><td>>hg38_MT_+_1621_3279_RNR2_16S_50ntFlanks</td><td>1370</td><td>1394</td><td>25</td><td>ATGTTGGATCAGGACATCCCGATGG</td><td>84.8</td><td>4.6</td><td>2.0</td><td>2.3</td><td>2.1E-02</td><td>2.6E-02</td><td>MB231/MB468</td><td>MP</td></tr>
<tr><td>>hg38_21_+_8433172_8446622_RNA45SN1_45S_50ntFlanks@11553.11591.39|CTCGGCCGGCGCCTAGCAGCCGACTTAGAACTGGTGCGG</td><td>>hg38_21_+_8433172_8446622_RNA45SN1_45S_50ntFlanks</td><td>11553</td><td>11591</td><td>39</td><td>CTCGGCCGGCGCCTAGCAGCCGACTTAGAACTGGTGCGG</td><td>106.4</td><td>-2.7</td><td>1.2</td><td>-2.3</td><td>2.3E-02</td><td>2.6E-02</td><td>BT20/MB468</td><td>CYTO</td></tr>
<tr><td>>trna10_AspGTC_12_-_125424193_125424264@35.75.41,trna12_AspGTC_12_-_125411891_125411962@35.75.41,trna144_AspGTC_6_-_27551236_27551307@35.75.41,trna38_AspGTC_17_-_8125556_8125627@35.75.41,trna45_AspGTC_6_+_27447453_27447524@35.75.41,trna48_AspGTC_6_+_27471523_27471594@35.75.41,trna4_AspGTC_12_+_96429799_96429870@35.75.41,trna69_AspGTC_1_-_161440205_161440276@35.75.41,trna72_AspGTC_1_-_161432824_161432895@35.75.41,trna75_AspGTC_1_-_161425414_161425485@35.75.41,trna78_AspGTC_1_-_161418033_161418104@35.75.41,trna81_AspGTC_1_-_161410615_161410686@35.75.41|TCACGCGGGAGACCGGGGTTCGATTCCCCGACGGGGAGCCA</td><td>>trna10_AspGTC_12_-_125424193_125424264</td><td>35</td><td>75</td><td>41</td><td>TCACGCGGGAGACCGGGGTTCGATTCCCCGACGGGGAGCCA</td><td>353.7</td><td>3.4</td><td>1.5</td><td>2.3</td><td>2.1E-02</td><td>2.6E-02</td><td>MB231/MB468</td><td>MP</td></tr>
<tr><td>>hg38_21_+_8433172_8446622_RNA45SN1_45S_50ntFlanks@11853.11886.34|CTTGGCGGAATCAGCGGGGAAAGAAGACCCTGTT</td><td>>hg38_21_+_8433172_8446622_RNA45SN1_45S_50ntFlanks</td><td>11853</td><td>11886</td><td>34</td><td>CTTGGCGGAATCAGCGGGGAAAGAAGACCCTGTT</td><td>70.9</td><td>3.4</td><td>1.4</td><td>2.4</td><td>1.5E-02</td><td>2.6E-02</td><td>BT20/MB468</td><td>MP</td></tr>
<tr><td>>hg38_21_+_8433172_8446622_RNA45SN1_45S_50ntFlanks@10457.10495.39|GATGGCCTCCGTTGCCCTCGGCCGATCGAAAGGGAGTCG</td><td>>hg38_21_+_8433172_8446622_RNA45SN1_45S_50ntFlanks</td><td>10457</td><td>10495</td><td>39</td><td>GATGGCCTCCGTTGCCCTCGGCCGATCGAAAGGGAGTCG</td><td>118.9</td><td>-3.1</td><td>1.4</td><td>-2.3</td><td>2.1E-02</td><td>2.6E-02</td><td>MB231/MB468</td><td>MP</td></tr>
<tr><td>>hg38_21_+_8433172_8446622_RNA45SN1_45S_50ntFlanks@10346.10372.27|TCAAACGAGAACTTTGAAGGCCGAAGT</td><td>>hg38_21_+_8433172_8446622_RNA45SN1_45S_50ntFlanks</td><td>10346</td><td>10372</td><td>27</td><td>TCAAACGAGAACTTTGAAGGCCGAAGT</td><td>226.5</td><td>1.2</td><td>0.5</td><td>2.3</td><td>2.3E-02</td><td>2.6E-02</td><td>MB231/MB468</td><td>CYTO</td></tr>
<tr><td>>hg38_21_+_8433172_8446622_RNA45SN1_45S_50ntFlanks@10402.10427.26|AACATGGGTCAGTCGGTCCTGAGAGA</td><td>>hg38_21_+_8433172_8446622_RNA45SN1_45S_50ntFlanks</td><td>10402</td><td>10427</td><td>26</td><td>AACATGGGTCAGTCGGTCCTGAGAGA</td><td>150.7</td><td>-3.1</td><td>1.3</td><td>-2.4</td><td>1.5E-02</td><td>2.6E-02</td><td>BT20/MB468</td><td>MP</td></tr>
<tr><td>>MI0000272|hsa-mir-182&WithFlank&7|-|129770377|129770498@29.52.24&[MIMAT0000259&hsa-miR-182-5p&offsets|0|0;m-22&7|-|129770449|129770470&offsets|0|+2]|TTTGGCAATGGTAGAACTCACACT</td><td>>MI0000272|hsa-mir-182&WithFlank&7|-|129770377|129770498</td><td>29</td><td>52</td><td>24</td><td>TTTGGCAATGGTAGAACTCACACT</td><td>23073.5</td><td>2.0</td><td>0.8</td><td>2.7</td><td>7.8E-03</td><td>2.6E-02</td><td>BT20/MB231</td><td>MITO</td></tr>
<tr><td>>trnaMT_TyrGTA_MT_-_5826_5891@24.65.42|TGGACTGTAAATCTAAAGACAGGGGTTAGGCCTCTTTTTACC</td><td>>trnaMT_TyrGTA_MT_-_5826_5891</td><td>24</td><td>65</td><td>42</td><td>TGGACTGTAAATCTAAAGACAGGGGTTAGGCCTCTTTTTACC</td><td>267.9</td><td>-2.6</td><td>1.0</td><td>-2.7</td><td>7.8E-03</td><td>2.6E-02</td><td>BT20/MB231</td><td>MITO</td></tr>
<tr><td>>hg38_1_-_228634819_228635039_RNA5S12_5S_50ntFlanks@51.82.32|GTCTACGGCCATACCACCCTGAACGCGCCCGA</td><td>>hg38_1_-_228634819_228635039_RNA5S12_5S_50ntFlanks</td><td>51</td><td>82</td><td>32</td><td>GTCTACGGCCATACCACCCTGAACGCGCCCGA</td><td>1257.7</td><td>-2.4</td><td>0.9</td><td>-2.7</td><td>7.8E-03</td><td>2.6E-02</td><td>BT20/MB231</td><td>MITO</td></tr>
<tr><td>>MI0000446|hsa-mir-125b-1&WithFlank&11|-|122099751|122099850@61.82.22&[MIMAT0004592&hsa-miR-125b-1-3p&offsets|0|0;m-485&11|-|122099769|122099790&offsets|0|0]|ACGGGTTAGGCTCTTGGGAGCT</td><td>>MI0000446|hsa-mir-125b-1&WithFlank&11|-|122099751|122099850</td><td>61</td><td>82</td><td>22</td><td>ACGGGTTAGGCTCTTGGGAGCT</td><td>530.4</td><td>-6.0</td><td>2.3</td><td>-2.7</td><td>7.8E-03</td><td>2.6E-02</td><td>BT20/MB231</td><td>MITO</td></tr>
<tr><td>>hg38_21_+_8433172_8446622_RNA45SN1_45S_50ntFlanks@5492.5510.19|AGAAGACGGTCGAACTTGA</td><td>>hg38_21_+_8433172_8446622_RNA45SN1_45S_50ntFlanks</td><td>5492</td><td>5510</td><td>19</td><td>AGAAGACGGTCGAACTTGA</td><td>636.2</td><td>-2.7</td><td>1.1</td><td>-2.4</td><td>1.5E-02</td><td>2.7E-02</td><td>BT20/MB468</td><td>MP</td></tr>
<tr><td>>trnaMT_GluTTC_MT_-_14674_14742@34.72.39,trnalookalike8_GluTTC_5_-_93905172_93905240@34.72.39|ATATCATTGGTCGTGGTTGTAGTCCGTGCGAGAATACCA</td><td>>trnaMT_GluTTC_MT_-_14674_14742</td><td>34</td><td>72</td><td>39</td><td>ATATCATTGGTCGTGGTTGTAGTCCGTGCGAGAATACCA</td><td>4320.0</td><td>3.4</td><td>1.4</td><td>2.4</td><td>1.5E-02</td><td>2.7E-02</td><td>BT20/MB468</td><td>MP</td></tr>
<tr><td>>hg38_21_+_8433172_8446622_RNA45SN1_45S_50ntFlanks@11325.11344.20|TCTCGTCGCGCGCGCGTCCG</td><td>>hg38_21_+_8433172_8446622_RNA45SN1_45S_50ntFlanks</td><td>11325</td><td>11344</td><td>20</td><td>TCTCGTCGCGCGCGCGTCCG</td><td>323.7</td><td>-4.3</td><td>1.8</td><td>-2.4</td><td>1.5E-02</td><td>2.7E-02</td><td>BT20/MB468</td><td>MP</td></tr>
<tr><td>>hg38_21_+_8433172_8446622_RNA45SN1_45S_50ntFlanks@9544.9572.29|GGTGGAGGTCCGTAGCGGTCCTGACGTGC</td><td>>hg38_21_+_8433172_8446622_RNA45SN1_45S_50ntFlanks</td><td>9544</td><td>9572</td><td>29</td><td>GGTGGAGGTCCGTAGCGGTCCTGACGTGC</td><td>131.3</td><td>-3.2</td><td>1.4</td><td>-2.3</td><td>2.1E-02</td><td>2.7E-02</td><td>MB231/MB468</td><td>MP</td></tr>
<tr><td>>hg38_21_+_8433172_8446622_RNA45SN1_45S_50ntFlanks@3860.3899.40|GTAATTCTAGAGCTAATACATGCCGACGGGCGCTGACCCC</td><td>>hg38_21_+_8433172_8446622_RNA45SN1_45S_50ntFlanks</td><td>3860</td><td>3899</td><td>40</td><td>GTAATTCTAGAGCTAATACATGCCGACGGGCGCTGACCCC</td><td>131.5</td><td>-3.0</td><td>1.3</td><td>-2.3</td><td>2.3E-02</td><td>2.7E-02</td><td>BT20/MB468</td><td>CYTO</td></tr>
<tr><td>>MI0000750|hsa-mir-26a-2&WithFlank&12|-|57824603|57824698@20.41.22(+1C);MI0000083|hsa-mir-26a-1&WithFlank&3|+|37969398|37969486@16.37.22(+1C)&[MIMAT0000082&hsa-miR-26a-5p&offsets|0|0(+1C);m-33&12|-|57824658|57824679&offsets|0|0(+1C)];[MIMAT0000082_1&hsa-miR-26a-5p&offsets|0|0(+1C);m-34&3|+|37969413|37969434&offsets|0|0(+1C)]|TTCAAGTAATCCAGGATAGGCTC</td><td>>MI0000750|hsa-mir-26a-2&WithFlank&12|-|57824603|57824698</td><td>20</td><td>41</td><td>22(+1C)</td><td>TTCAAGTAATCCAGGATAGGCTC</td><td>1469.0</td><td>-3.1</td><td>1.1</td><td>-2.7</td><td>6.4E-03</td><td>2.7E-02</td><td>BT20/MB231</td><td>MP</td></tr>
<tr><td>>hg38_21_+_8433172_8446622_RNA45SN1_45S_50ntFlanks@5494.5514.21|AAGACGGTCGAACTTGACTAT</td><td>>hg38_21_+_8433172_8446622_RNA45SN1_45S_50ntFlanks</td><td>5494</td><td>5514</td><td>21</td><td>AAGACGGTCGAACTTGACTAT</td><td>255.9</td><td>-2.1</td><td>0.9</td><td>-2.3</td><td>2.3E-02</td><td>2.7E-02</td><td>BT20/MB468</td><td>CYTO</td></tr>
<tr><td>>am_trna10_ValCAC_5_-_180649395_180649467@5.23.19,trna12_ValAAC_5_-_180645270_180645342@5.23.19,trna132_ValAAC_6_-_27721179_27721251@5.23.19,trna136_ValAAC_6_-_27648885_27648957@5.23.19,trna139_ValAAC_6_-_27618707_27618779@5.23.19,trna18_ValCAC_5_-_180529253_180529325@5.23.19,trna2_ValAAC_3_+_169490018_169490090@5.23.19,trna2_ValCAC_5_+_180524070_180524142@5.23.19,trna37_ValAAC_6_+_27203288_27203360@5.23.19,trna4_ValAAC_5_+_180591154_180591226@5.23.19,trna5_ValAAC_5_+_180596610_180596682@5.23.19,trna6_ValCAC_5_+_180600650_180600722@5.23.19,trna85_ValCAC_1_-_161369490_161369562@5.23.19,trna90_ValCAC_1_-_149684088_149684161@5.23.19,trna98_ValCAC_1_-_149298555_149298627@5.23.19,trna9_ValCAC_6_+_26538282_26538354@5.23.19|CCGTAGTGTAGTGGTTATC</td><td>>am_trna10_ValCAC_5_-_180649395_180649467</td><td>5</td><td>23</td><td>19</td><td>CCGTAGTGTAGTGGTTATC</td><td>361.2</td><td>-1.5</td><td>0.7</td><td>-2.3</td><td>2.3E-02</td><td>2.7E-02</td><td>MB231/MB468</td><td>CYTO</td></tr>
</table>

| | | | | | | | | | | | | | |
|---|---|---|---|---|---|---|---|---|---|---|---|---|---|
| >am_MI0000809\|hsa-mir-151a&WithFlank&8\|-\|140732558\|140732659@52.72.21(+1U)&[MIMAT0000757&hsa-miR-151a-3p&offsets\|-1\|-1(+1U);m-30&8\|-\|140732587\|140732607&offsets\|-1\|-1(+1U)]\|ACTAGACTGAAGCTCCTTGAGT | >am_MI0000809\|hsa-mir-151a&WithFlank&8\|-\|140732558\|140732659 | 52 | 72 | 21(+1U) | ACTAGACTGAAGCTCCTTGAGT | 47.6 | 4.6 | 2.0 | 2.3 | 2.1E-02 | 2.7E-02 | MB231/MB468 | MP |
| >trnaMT_HisGTG_MT_+_12138_12206@20.52.33\|AACATCAGATTGTGAATCTGACAACAGAGGCTT | >trnaMT_HisGTG_MT_+_12138_12206 | 20 | 52 | 33 | AACATCAGATTGTGAATCTGACAACAGAGGCTT | 41.7 | 4.5 | 2.0 | 2.3 | 2.2E-02 | 2.7E-02 | MB231/MB468 | MP |
| >hg38_21_+_8433172_8446622_RNA45SN1_45S_50ntFlanks@8184.8206.23\|CGAGGCCCAGCCCGTGGACGGTG | >hg38_21_+_8433172_8446622_RNA45SN1_45S_50ntFlanks | 8184 | 8206 | 23 | CGAGGCCCAGCCCGTGGACGGTG | 288.9 | -3.3 | 1.4 | -2.4 | 1.5E-02 | 2.7E-02 | BT20/MB468 | MP |
| >hg38_21_+_8433172_8446622_RNA45SN1_45S_50ntFlanks@10513.10535.23\|CCGGAGTGGCGGAGATGGGCGCC | >hg38_21_+_8433172_8446622_RNA45SN1_45S_50ntFlanks | 10513 | 10535 | 23 | CCGGAGTGGCGGAGATGGGCGCC | 180.1 | -3.0 | 1.3 | -2.3 | 2.2E-02 | 2.7E-02 | MB231/MB468 | MP |
| >MI0000811\|hsa-mir-148b&WithFlank&12\|+\|54337210\|54337320@30.51.22(+1U)&[MIMAT0004699&hsa-miR-148b-5p&offsets\|-1\|-1(+1U);m-371&12\|+\|54337239\|54337262&offsets\|0\|-2(+1U)]\|GAAGTTCTGTTATACACTCAGGT | >MI0000811\|hsa-mir-148b&WithFlank&12\|+\|54337210\|54337320 | 30 | 51 | 22(+1U) | GAAGTTCTGTTATACACTCAGGT | 34.8 | 4.7 | 2.0 | 2.3 | 2.2E-02 | 2.7E-02 | MB231/MB468 | MP |
| >MI0000100\|hsa-mir-98&WithFlank&X\|-\|53556217\|53556347@28.48.21(+1A)&[MIMAT0000096&hsa-miR-98-5p&offsets\|0\|-1(+1A);m-143&X\|-\|53556299\|53556320&offsets\|0\|-1(+1A)]\|TGAGGTAGTAAGTTGTATTGTA | >MI0000100\|hsa-mir-98&WithFlank&X\|-\|53556217\|53556347 | 28 | 48 | 21(+1A) | TGAGGTAGTAAGTTGTATTGTA | 235.5 | -3.2 | 1.2 | -2.7 | 6.4E-03 | 2.7E-02 | BT20/MB231 | MP |
| >hg38_21_+_8433172_8446622_RNA45SN1_45S_50ntFlanks@11573.11594.22\|CGACTTAGAACTGGTGCGGACC | >hg38_21_+_8433172_8446622_RNA45SN1_45S_50ntFlanks | 11573 | 11594 | 22 | CGACTTAGAACTGGTGCGGACC | 444.1 | -2.1 | 0.8 | -2.6 | 8.6E-03 | 2.7E-02 | BT20/MB231 | CYTO |
| >hg38_21_+_8433172_8446622_RNA45SN1_45S_50ntFlanks@4367.4406.40\|CAGTTAAAAAGCTCGTAGTTGGATCTTGGGAGCGGGCGGG | >hg38_21_+_8433172_8446622_RNA45SN1_45S_50ntFlanks | 4367 | 4406 | 40 | CAGTTAAAAAGCTCGTAGTTGGATCTTGGGAGCGGGCGGG | 117.7 | -3.6 | 1.6 | -2.3 | 2.0E-02 | 2.7E-02 | BT20/MB468 | MITO |
| >hg38_21_+_8433172_8446622_RNA45SN1_45S_50ntFlanks@6651.6692.42\|CGACTCTTAGCGGTGGATCACTCGGCTCGTGCGTCGATGAAG | >hg38_21_+_8433172_8446622_RNA45SN1_45S_50ntFlanks | 6651 | 6692 | 42 | CGACTCTTAGCGGTGGATCACTCGGCTCGTGCGTCGATGAAG | 3013.2 | -2.3 | 1.0 | -2.3 | 2.2E-02 | 2.7E-02 | MB231/MB468 | MP |
| >MI0000087\|hsa-mir-29a&WithFlank&7\|-\|130876741\|130876816@10.29.20&[MIMAT0004503&hsa-miR-29a-5p&offsets\|0\|-2;m-315&7\|-\|130876785\|130876807&offsets\|0\|-3]\|ACTGATTTCTTTTGGTGTTC | >MI0000087\|hsa-mir-29a&WithFlank&7\|-\|130876741\|130876816 | 10 | 29 | 20 | ACTGATTTCTTTTGGTGTTC | 45.9 | 4.5 | 2.0 | 2.3 | 2.2E-02 | 2.7E-02 | MB231/MB468 | MP |
| >am_trnaMT_ValTAC_MT_+_1602_1670@30.68.39\|CTTACACTTAGGAGATTTCAACTTAACTTGACCGCTCTG | >am_trnaMT_ValTAC_MT_+_1602_1670 | 30 | 68 | 39 | CTTACACTTAGGAGATTTCAACTTAACTTGACCGCTCTG | 153.3 | -4.3 | 1.9 | -2.3 | 2.3E-02 | 2.7E-02 | BT20/MB468 | CYTO |
| >MI0000088\|hsa-mir-30a&WithFlank&6\|-\|71403545\|71403627@12.33.22(+1A)&[MIMAT0000087&hsa-miR-30a-5p&offsets\|0\|0(+1A);m-15&6\|-\|71403595\|71403616&offsets\|0\|0(+1A)]\|TGTAAACATCCTCGACTGGAAGA | >MI0000088\|hsa-mir-30a&WithFlank&6\|-\|71403545\|71403627 | 12 | 33 | 22(+1A) | TGTAAACATCCTCGACTGGAAGA | 1927.9 | 3.0 | 1.2 | 2.4 | 1.5E-02 | 2.7E-02 | BT20/MB468 | MP |
| >hg38_21_+_8433172_8446622_RNA45SN1_45S_50ntFlanks@4372.4389.18\|AAAAAGCTCGTAGTTGGA | >hg38_21_+_8433172_8446622_RNA45SN1_45S_50ntFlanks | 4372 | 4389 | 18 | AAAAAGCTCGTAGTTGGA | 120.8 | -3.6 | 1.5 | -2.4 | 1.5E-02 | 2.7E-02 | BT20/MB468 | MP |
| >hg38_21_+_8433172_8446622_RNA45SN1_45S_50ntFlanks@4554.4573.20\|CCGCAGCTAGGAATAATGGA | >hg38_21_+_8433172_8446622_RNA45SN1_45S_50ntFlanks | 4554 | 4573 | 20 | CCGCAGCTAGGAATAATGGA | 175.4 | -4.2 | 1.7 | -2.4 | 1.5E-02 | 2.7E-02 | BT20/MB468 | MP |
| >trnaMT_GluTTC_MT_-_14674_14742@38.57.20,trnalookalike8_GluTTC_5_-_93905172_93905240@38.57.20\|CATTGGTCGTGGTTGTAGTC | >trnaMT_GluTTC_MT_-_14674_14742 | 38 | 57 | 20 | CATTGGTCGTGGTTGTAGTC | 68.2 | 3.0 | 1.3 | 2.3 | 2.2E-02 | 2.7E-02 | MB231/MB468 | MP |
| >hg38_21_+_8433172_8446622_RNA45SN1_45S_50ntFlanks@4046.4064.19\|CATTCGAACGTCTGCCCTA | >hg38_21_+_8433172_8446622_RNA45SN1_45S_50ntFlanks | 4046 | 4064 | 19 | CATTCGAACGTCTGCCCTA | 147.8 | -3.3 | 1.4 | -2.4 | 1.5E-02 | 2.7E-02 | BT20/MB468 | MP |
| >hg38_21_+_8433172_8446622_RNA45SN1_45S_50ntFlanks@11846.11872.27\|GAACGGGCTTGGCGGAATCAGCGGGGA | >hg38_21_+_8433172_8446622_RNA45SN1_45S_50ntFlanks | 11846 | 11872 | 27 | GAACGGGCTTGGCGGAATCAGCGGGGA | 423.5 | -4.3 | 1.8 | -2.4 | 1.5E-02 | 2.7E-02 | BT20/MB468 | MP |
| >hg38_21_+_8433172_8446622_RNA45SN1_45S_50ntFlanks@12409.12433.25\|TTGATCCTTCGATGTCGGCTCTTCC | >hg38_21_+_8433172_8446622_RNA45SN1_45S_50ntFlanks | 12409 | 12433 | 25 | TTGATCCTTCGATGTCGGCTCTTCC | 390.0 | 2.2 | 1.0 | 2.3 | 2.3E-02 | 2.7E-02 | BT20/MB468 | CYTO |
| >MI0000078\|hsa-mir-22&WithFlank&17\|-\|1713897\|1713993@59.80.22(+1A)&[MIMAT0000077&hsa-miR-22-3p&offsets\|0\|0(+1A);m-2&17\|-\|1713914\|1713935&offsets\|0\|0(+1A)]\|AAGCTGCCAGTTGAAGAACTGTA | >MI0000078\|hsa-mir-22&WithFlank&17\|-\|1713897\|1713993 | 59 | 80 | 22(+1A) | AAGCTGCCAGTTGAAGAACTGTA | 360.9 | 3.6 | 1.5 | 2.4 | 1.6E-02 | 2.7E-02 | BT20/MB468 | MP |
| >MI0000299\|hsa-mir-222&WithFlank&X\|-\|45747009\|45747130@75.96.22(+1C)&[MIMAT0000279&hsa-miR-222-3p&offsets\|0\|+1(+1C);m-140&X\|-\|45747033\|45747056&offsets\|0\|-2(+1C)]\|AGCTACATCTGGCTACTGGGTCC | >MI0000299\|hsa-mir-222&WithFlank&X\|-\|45747009\|45747130 | 75 | 96 | 22(+1C) | AGCTACATCTGGCTACTGGGTCC | 1980.0 | -2.1 | 0.8 | -2.7 | 7.9E-03 | 2.7E-02 | BT20/MB231 | MITO |
| >hg38_21_+_8433172_8446622_RNA45SN1_45S_50ntFlanks@11667.11703.37\|CTGCCCAGTGCTCTGAATGTCAAAGTGAAGAAATTCA | >hg38_21_+_8433172_8446622_RNA45SN1_45S_50ntFlanks | 11667 | 11703 | 37 | CTGCCCAGTGCTCTGAATGTCAAAGTGAAGAAATTCA | 259.0 | 2.3 | 0.9 | 2.7 | 7.9E-03 | 2.7E-02 | BT20/MB231 | MITO |
| >trnaMT_GluTTC_MT_-_14674_14742@40.72.33,trnalookalike8_GluTTC_5_-_93905172_93905240@40.72.33\|TTGGTCGTGGTTGTAGTCCGTGCGAGAATACCA | >trnaMT_GluTTC_MT_-_14674_14742 | 40 | 72 | 33 | TTGGTCGTGGTTGTAGTCCGTGCGAGAATACCA | 15840.7 | 2.3 | 1.0 | 2.3 | 2.2E-02 | 2.7E-02 | MB231/MB468 | MP |
| >hg38_21_+_8433172_8446622_RNA45SN1_45S_50ntFlanks@11588.11611.24\|GCGGACCAGGGGAATCCGACTGTT | >hg38_21_+_8433172_8446622_RNA45SN1_45S_50ntFlanks | 11588 | 11611 | 24 | GCGGACCAGGGGAATCCGACTGTT | 205.8 | -1.8 | 0.8 | -2.3 | 2.2E-02 | 2.7E-02 | MB231/MB468 | MP |
| >hg38_21_+_8433172_8446622_RNA45SN1_45S_50ntFlanks@6775.6806.32\|CCTCCCGGGGCTACGCCTGTCTGAGCGTCGCT | >hg38_21_+_8433172_8446622_RNA45SN1_45S_50ntFlanks | 6775 | 6806 | 32 | CCTCCCGGGGCTACGCCTGTCTGAGCGTCGCT | 872.9 | 1.9 | 0.8 | 2.3 | 2.4E-02 | 2.7E-02 | MB231/MB468 | MITO |
| >hg38_21_+_8433172_8446622_RNA45SN1_45S_50ntFlanks@4722.4758.37\|TCATTAATCAAGAACGAAAGTCGGAGGTTCGAAGACG | >hg38_21_+_8433172_8446622_RNA45SN1_45S_50ntFlanks | 4722 | 4758 | 37 | TCATTAATCAAGAACGAAAGTCGGAGGTTCGAAGACG | 369.2 | -2.0 | 0.8 | -2.6 | 8.6E-03 | 2.7E-02 | BT20/MB231 | CYTO |
| >MI0000809\|hsa-mir-151a&WithFlank&8\|-\|140732558\|140732659@53.74.22(+1G)&[MIMAT0000757&hsa-miR-151a-3p&offsets\|0\|+1(+1G);m-30&8\|-\|140732587\|140732607&offsets\|0\|+1(+1G)]\|CTAGACTGAAGCTCCTTGAGGAG | >MI0000809\|hsa-mir-151a&WithFlank&8\|-\|140732558\|140732659 | 53 | 74 | 22(+1G) | CTAGACTGAAGCTCCTTGAGGAG | 60.6 | 3.7 | 1.6 | 2.3 | 2.2E-02 | 2.7E-02 | MB231/MB468 | MP |
| >am_MI0000786\|hsa-mir-378a&WithFlank&5\|+\|149732819\|149732896@49.69.21(+1U)&[MIMAT0000732&hsa-miR-378a-3p&offsets\|0\|-1(+1U);m-44&5\|+\|149732867\|149732888&offsets\|0\|-1(+1U)]\|ACTGGACTTGGAGTCAGAAGGT | >am_MI0000786\|hsa-mir-378a&WithFlank&5\|+\|149732819\|149732896 | 49 | 69 | 21(+1U) | ACTGGACTTGGAGTCAGAAGGT | 15456.5 | 2.8 | 1.2 | 2.4 | 1.6E-02 | 2.7E-02 | BT20/MB468 | MP |
| >hg38_21_+_8433172_8446622_RNA45SN1_45S_50ntFlanks@7970.8006.37\|CGAGACGCGACCTCAGATCAGACGTGGCGACCCGCTG | >hg38_21_+_8433172_8446622_RNA45SN1_45S_50ntFlanks | 7970 | 8006 | 37 | CGAGACGCGACCTCAGATCAGACGTGGCGACCCGCTG | 100.6 | 1.7 | 0.8 | 2.3 | 2.4E-02 | 2.7E-02 | BT20/MB468 | NUC |
| >MI0000285\|hsa-mir-205&WithFlank&1\|+\|209432127\|209432248@40.58.19&[MIMAT0000266&hsa-miR-205-5p&offsets\|0\|-3;m-351&1\|+\|209432166\|209432187&offsets\|0\|-3]\|TCCTTCATTCCACCGGAGT | >MI0000285\|hsa-mir-205&WithFlank&1\|+\|209432127\|209432248 | 40 | 58 | 19 | TCCTTCATTCCACCGGAGT | 33.5 | 8.4 | 3.1 | 2.7 | 6.4E-03 | 2.7E-02 | BT20/MB231 | MP |
| >hg38_21_+_8433172_8446622_RNA45SN1_45S_50ntFlanks@9954.9983.30\|TGGAAGTCGGAATCCGCTAAGGAGTGTGTA | >hg38_21_+_8433172_8446622_RNA45SN1_45S_50ntFlanks | 9954 | 9983 | 30 | TGGAAGTCGGAATCCGCTAAGGAGTGTGTA | 140.8 | 5.6 | 2.0 | 2.7 | 6.4E-03 | 2.7E-02 | BT20/MB231 | MP |
| >hg38_21_+_8433172_8446622_RNA45SN1_45S_50ntFlanks@11728.11747.20\|GTAACTATGACTCTCTTAAG | >hg38_21_+_8433172_8446622_RNA45SN1_45S_50ntFlanks | 11728 | 11747 | 20 | GTAACTATGACTCTCTTAAG | 539.2 | -1.8 | 0.8 | -2.3 | 2.4E-02 | 2.7E-02 | MB231/MB468 | NUC |
| >MI0000816\|hsa-mir-335&WithFlank&7\|+\|130496105\|130496210@58.78.21(+1U)&[MIMAT0004703&hsa-miR-335-3p&offsets\|0\|-1(+1U);m-159&7\|+\|130496162\|130496183&offsets\|0\|-1(+1U)]\|TTTTTCATTATTGCTCCTGACT | >MI0000816\|hsa-mir-335&WithFlank&7\|+\|130496105\|130496210 | 58 | 78 | 21(+1U) | TTTTTCATTATTGCTCCTGACT | 140.6 | 3.7 | 1.5 | 2.4 | 1.6E-02 | 2.7E-02 | BT20/MB468 | MP |
| >trna10_LysCTT_16_+_3241501_3241573@14.33.20,trna13_LysCTT_14_-_58706613_58706685@14.33.20,trna2_LysCTT_15_+_79152904_79152976@14.33.20,trna4_LysCTT_18_-_43669271_43669343@14.33.20\|AGTCGGTAGAGCATGGGACT | >trna10_LysCTT_16_+_3241501_3241573 | 14 | 33 | 20 | AGTCGGTAGAGCATGGGACT | 63.8 | 3.8 | 1.6 | 2.3 | 2.2E-02 | 2.7E-02 | MB231/MB468 | MP |
| >hg38_21_+_8433172_8446622_RNA45SN1_45S_50ntFlanks@10186.10207.22\|TCCTCCCCTCCTCCCGCCCACG | >hg38_21_+_8433172_8446622_RNA45SN1_45S_50ntFlanks | 10186 | 10207 | 22 | TCCTCCCCTCCTCCCGCCCACG | 252.9 | -3.7 | 1.6 | -2.3 | 2.0E-02 | 2.7E-02 | BT20/MB468 | MITO |
| >trna152_ValCAC_6_-_27248049_27248121@1.29.29\|GCTTCTGTAGTGTAGTGGTTATCACGTTC | >trna152_ValCAC_6_-_27248049_27248121 | 1 | 29 | 29 | GCTTCTGTAGTGTAGTGGTTATCACGTTC | 1645.8 | -3.3 | 1.4 | -2.3 | 2.0E-02 | 2.7E-02 | BT20/MB468 | MITO |
| >trnaMT_GluTTC_MT_-_14674_14742@26.61.36,trnalookalike8_GluTTC_5_-_93905172_93905240@26.61.36\|GGTTTTTCATATCATTGGTCGTGGTTGTAGTCCGTG | >trnaMT_GluTTC_MT_-_14674_14742 | 26 | 61 | 36 | GGTTTTTCATATCATTGGTCGTGGTTGTAGTCCGTG | 466.5 | 4.7 | 2.0 | 2.4 | 1.6E-02 | 2.7E-02 | BT20/MB468 | MP |
| >hg38_21_+_8433172_8446622_RNA45SN1_45S_50ntFlanks@9569.9586.18\|GTGCAAATCGGTCGTCCG | >hg38_21_+_8433172_8446622_RNA45SN1_45S_50ntFlanks | 9569 | 9586 | 18 | GTGCAAATCGGTCGTCCG | 209.1 | -1.8 | 0.8 | -2.3 | 2.3E-02 | 2.7E-02 | BT20/MB468 | CYTO |
| >MI0000289\|hsa-mir-181a-1&WithFlank&1\|-\|198859038\|198859159@30.50.21;MI0000269\|hsa-mir-181a-2&WithFlank&9\|+\|124692436\|124692557@45.65.21&[MIMAT0000256&hsa-miR-181a-5p&offsets\|0\|-2;m-32&1\|-\|198859108\|198859130&offsets\|0\|-2];[MIMAT0000256_1&hsa-miR-181a-5p&offsets\|0\|-2;m-32&9\|+\|124692480\|124692502&offsets\|0\|-2]\|AACATTCAACGCTGTCGGTGA | >MI0000289\|hsa-mir-181a-1&WithFlank&1\|-\|198859038\|198859159 | 30 | 50 | 21 | AACATTCAACGCTGTCGGTGA | 476.3 | -1.9 | 0.7 | -2.7 | 7.0E-03 | 2.7E-02 | BT20/MB231 | NUC |
| >hg38_21_+_8433172_8446622_RNA45SN1_45S_50ntFlanks@9553.9576.24\|CCGTAGCGGTCCTGACGTGCAAAT | >hg38_21_+_8433172_8446622_RNA45SN1_45S_50ntFlanks | 9553 | 9576 | 24 | CCGTAGCGGTCCTGACGTGCAAAT | 286.4 | -2.0 | 0.8 | -2.6 | 8.7E-03 | 2.7E-02 | BT20/MB231 | CYTO |
| >hg38_21_+_8433172_8446622_RNA45SN1_45S_50ntFlanks@476.525.50\|TTCGTGATCGATGTGGTGACGTCGTGCTCTCCCGGGCCGGGTCCGAGCCG | >hg38_21_+_8433172_8446622_RNA45SN1_45S_50ntFlanks | 476 | 525 | 50 | TTCGTGATCGATGTGGTGACGTCGTGCTCTCCCGGGCCGGGTCCGAGCCG | 2103.1 | 3.4 | 1.5 | 2.3 | 2.4E-02 | 2.7E-02 | MB231/MB468 | NUC |
| >MI0001448\|hsa-mir-425&WithFlank&3\|-\|49020142\|49020240@20.41.22&[MIMAT0003393&hsa-miR-425-5p&offsets\|0\|-1;m-95&3\|-\|49020200\|49020221&offsets\|0\|0]\|AATGACACGATCACTCCCGTTG | >MI0001448\|hsa-mir-425&WithFlank&3\|-\|49020142\|49020240 | 20 | 41 | 22 | AATGACACGATCACTCCCGTTG | 336.1 | 3.9 | 1.6 | 2.4 | 1.6E-02 | 2.7E-02 | BT20/MB468 | MP |
| >am_MI0003772\|hsa-mir-151b&WithFlank&14\|-\|100109413\|100109520@66.85.20(+1C);MI0000809\|hsa-mir-151a&WithFlank&8\|-\|140732558\|140732659@17.36.20(+1C)&[MIMAT0010214&hsa-miR-151b&offsets\|0\|+2(+1C);m-179&14\|-\|100109436\|100109455&offsets\|0\|0(+1C)];[MIMAT0004697&hsa-miR-151a-5p&offsets\|0\|-1(+1C);m-75&8\|-\|140732623\|140732643&offsets\|0\|-1(+1C)]\|TCGAGGAGCTCACAGTCTAGC | >am_MI0003772\|hsa-mir-151b&WithFlank&14\|-\|100109413\|100109520 | 66 | 85 | 20(+1C) | TCGAGGAGCTCACAGTCTAGC | 386.0 | 2.5 | 1.0 | 2.6 | 8.7E-03 | 2.7E-02 | BT20/MB231 | CYTO |
| >hg38_21_+_8433172_8446622_RNA45SN1_45S_50ntFlanks@3705.3733.29\|TACCTGGTTGATCCTGCCAGTAGCATATG | >hg38_21_+_8433172_8446622_RNA45SN1_45S_50ntFlanks | 3705 | 3733 | 29 | TACCTGGTTGATCCTGCCAGTAGCATATG | 164.3 | 2.5 | 1.1 | 2.3 | 2.4E-02 | 2.7E-02 | MB231/MB468 | MITO |
| >hg38_21_+_8433172_8446622_RNA45SN1_45S_50ntFlanks@9569.9595.27\|GTGCAAATCGGTCGTCCGACCTGGGTA | >hg38_21_+_8433172_8446622_RNA45SN1_45S_50ntFlanks | 9569 | 9595 | 27 | GTGCAAATCGGTCGTCCGACCTGGGTA | 305.2 | -2.3 | 1.0 | -2.4 | 1.6E-02 | 2.7E-02 | BT20/MB468 | MP |
| >hg38_21_+_8433172_8446622_RNA45SN1_45S_50ntFlanks@476.516.41\|TTCGTGATCGATGTGGTGACGTCGTGCTCTCCCGGGCCGGG | >hg38_21_+_8433172_8446622_RNA45SN1_45S_50ntFlanks | 476 | 516 | 41 | TTCGTGATCGATGTGGTGACGTCGTGCTCTCCCGGGCCGGG | 836.6 | -2.1 | 0.8 | -2.6 | 8.7E-03 | 2.7E-02 | BT20/MB231 | CYTO |
| >hg38_MT_+_1621_3279_RNR2_16S_50ntFlanks@644.665.22\|AATTGGACCAATCTATCACCCT | >hg38_MT_+_1621_3279_RNR2_16S_50ntFlanks | 644 | 665 | 22 | AATTGGACCAATCTATCACCCT | 61.5 | 4.6 | 2.0 | 2.3 | 2.2E-02 | 2.7E-02 | MB231/MB468 | MP |

| | | | | | | | | | | | | | |
|---|---|---|---|---|---|---|---|---|---|---|---|---|---|
| >MI0000272\|hsa-mir-182&WithFlank&7\|-\|129770377\|129770498@29.47.19&[MIMAT0000259&hsa-miR-182-5p&offsets\|0\|-5;m-22&7\|-\|129770449\|129770470&offsets\|0\|-3]\|TTTGGCAATGGTAGAACTC | >MI0000272\|hsa-mir-182&WithFlank&7\|-\|129770377\|129770498 | 29 | 47 | 19 | TTTGGCAATGGTAGAACTC | 79.6 | 4.5 | 1.9 | 2.4 | 1.6E-02 | 2.7E-02 | BT20/MB468 | MP |
| >hg38_21_+_8433172_8446622_RNA45SN1_45S_50ntFlanks@6645.6681.37\|CTCGTACGACTCTTAGCGGTGGATCACTCGGCTCGTG | >hg38_21_+_8433172_8446622_RNA45SN1_45S_50ntFlanks | 6645 | 6681 | 37 | CTCGTACGACTCTTAGCGGTGGATCACTCGGCTCGTG | 1612.0 | -1.6 | 0.6 | -2.7 | 7.0E-03 | 2.7E-02 | BT20/MB231 | NUC |
| >trna10_AspGTC_12_-_125424193_125424264@1.31.31,trna12_AspGTC_12_-_125411891_125411962@1.31.31,trna38_AspGTC_17_-_8125556_8125627@1.31.31,trna45_AspGTC_6_+_27447453_27447524@1.31.31,trna48_AspGTC_6_+_27471523_27471594@1.31.31,trna4_AspGTC_12_+_96429799_96429870@1.31.31,trna69_AspGTC_1_-_161440205_161440276@1.31.31,trna72_AspGTC_1_-_161432824_161432895@1.31.31,trna75_AspGTC_1_-_161425414_161425485@1.31.31,trna78_AspGTC_1_-_161418033_161418104@1.31.31,trna81_AspGTC_1_-_161410615_161410686@1.31.31\|TCCTCGTTAGTATAGTGGTGAGTATCCCCGC | >trna10_AspGTC_12_-_125424193_125424264 | 1 | 31 | 31 | TCCTCGTTAGTATAGTGGTGAGTATCCCCGC | 196.9 | -3.7 | 1.6 | -2.3 | 2.4E-02 | 2.7E-02 | MB231/MB468 | NUC |
| >hg38_21_+_8433172_8446622_RNA45SN1_45S_50ntFlanks@6721.6742.22\|AATTGCAGGACACATTGATCAT | >hg38_21_+_8433172_8446622_RNA45SN1_45S_50ntFlanks | 6721 | 6742 | 22 | AATTGCAGGACACATTGATCAT | 170.5 | -4.1 | 1.7 | -2.4 | 1.6E-02 | 2.7E-02 | BT20/MB468 | MP |
| >hg38_1_-_228634819_228635039_RNA5S12_5S_50ntFlanks@51.75.25\|GTCTACGGCCATACCACCCTGAACG | >hg38_1_-_228634819_228635039_RNA5S12_5S_50ntFlanks | 51 | 75 | 25 | GTCTACGGCCATACCACCCTGAACG | 753.5 | 2.4 | 1.0 | 2.4 | 1.6E-02 | 2.7E-02 | BT20/MB468 | MP |
| >hg38_21_+_8433172_8446622_RNA45SN1_45S_50ntFlanks@7977.7994.18\|CGACCTCAGATCAGACGT | >hg38_21_+_8433172_8446622_RNA45SN1_45S_50ntFlanks | 7977 | 7994 | 18 | CGACCTCAGATCAGACGT | 378.7 | -1.5 | 0.6 | -2.3 | 2.3E-02 | 2.7E-02 | MB231/MB468 | CYTO |
| >MI0000067\|hsa-let-7f-1&WithFlank&9\|+\|94176341\|94176439@13.33.21(+1G);MI0000068\|hsa-let-7f-2&WithFlank&X\|-\|53557186\|53557280@14.34.21(+1G)&[MIMAT0000067&hsa-let-7f-5p&offsets\|0\|-1(+1G);m-20&9\|+\|94176353\|94176374&offsets\|0\|-1(+1G)];[MIMAT0000067_1&hsa-let-7f-5p&offsets\|0\|-1(+1G);m-19&X\|-\|53557246\|53557267&offsets\|0\|-1(+1G)]\|TGAGGTAGTAGATTGTATAGTG | >MI0000067\|hsa-let-7f-1&WithFlank&9\|+\|94176341\|94176439 | 13 | 33 | 21(+1G) | TGAGGTAGTAGATTGTATAGTG | 1828.5 | 2.2 | 0.8 | 2.6 | 8.7E-03 | 2.7E-02 | BT20/MB231 | CYTO |
| >hg38_21_+_8433172_8446622_RNA45SN1_45S_50ntFlanks@4982.5001.20\|ACGGACAGGATTGACAGATT | >hg38_21_+_8433172_8446622_RNA45SN1_45S_50ntFlanks | 4982 | 5001 | 20 | ACGGACAGGATTGACAGATT | 127.2 | -1.6 | 0.7 | -2.3 | 2.2E-02 | 2.7E-02 | MB231/MB468 | MP |
| >hg38_21_+_8433172_8446622_RNA45SN1_45S_50ntFlanks@6653.6681.29\|ACTCTTAGCGGTGGATCACTCGGCTCGTG | >hg38_21_+_8433172_8446622_RNA45SN1_45S_50ntFlanks | 6653 | 6681 | 29 | ACTCTTAGCGGTGGATCACTCGGCTCGTG | 2395.6 | -2.6 | 1.1 | -2.4 | 1.6E-02 | 2.7E-02 | BT20/MB468 | MP |
| >hg38_21_+_8433172_8446622_RNA45SN1_45S_50ntFlanks@11577.11611.35\|TTAGAACTGGTGCGGACCAGGGGAATCCGACTGTT | >hg38_21_+_8433172_8446622_RNA45SN1_45S_50ntFlanks | 11577 | 11611 | 35 | TTAGAACTGGTGCGGACCAGGGGAATCCGACTGTT | 297.4 | -2.6 | 1.1 | -2.4 | 1.6E-02 | 2.7E-02 | BT20/MB468 | MP |
| >MI0005561\|hsa-mir-877&WithFlank&6\|+\|30584326\|30584423@7.26.20&[MIMAT0004949&hsa-miR-877-5p&offsets\|0\|0;m-382&6\|+\|30584332\|30584354&offsets\|0\|-3]\|GTAGAGGAGATGGCGCAGGG | >MI0005561\|hsa-mir-877&WithFlank&6\|+\|30584326\|30584423 | 7 | 26 | 20 | GTAGAGGAGATGGCGCAGGG | 136.7 | 3.0 | 1.3 | 2.3 | 2.2E-02 | 2.7E-02 | MB231/MB468 | MP |
| >MI0003667\|hsa-mir-652&WithFlank&X\|+\|110055323\|110055432@67.87.21(+1U)&[MIMAT0003322&hsa-miR-652-3p&offsets\|0\|0(+1U);m-148&X\|+\|110055389\|110055410&offsets\|0\|-1(+1U)]\|AATGGCGCCACTAGGGTTGTGT | >MI0003667\|hsa-mir-652&WithFlank&X\|+\|110055323\|110055432 | 67 | 87 | 21(+1U) | AATGGCGCCACTAGGGTTGTGT | 455.0 | 2.1 | 0.9 | 2.3 | 2.2E-02 | 2.7E-02 | MB231/MB468 | MP |
| >hg38_21_+_8433172_8446622_RNA45SN1_45S_50ntFlanks@12994.13038.45\|CCTCGCTGCGATCTATTGAAAGTCAGCCCTCGACACAAGGGTTTG | >hg38_21_+_8433172_8446622_RNA45SN1_45S_50ntFlanks | 12994 | 13038 | 45 | CCTCGCTGCGATCTATTGAAAGTCAGCCCTCGACACAAGGGTTTG | 143.5 | 3.1 | 1.4 | 2.3 | 2.4E-02 | 2.7E-02 | MB231/MB468 | MITO |
| >trna116_GluCTC_1_-_145399233_145399304@1.39.39,trna59_GluCTC_1_+_249168447_249168518@1.39.39,trna71_GluCTC_1_-_161439189_161439260@1.39.39,trna74_GluCTC_1_-_161431809_161431880@1.39.39,trna77_GluCTC_1_-_161424398_161424469@1.39.39,trna77_GluCTC_6_+_28949976_28950047@1.39.39,trna80_GluCTC_1_-_161417018_161417089@1.39.39,trna87_GluCTC_6_-_126101393_126101464@1.39.39\|TCCCTGGTGGTCTAGTGGTTAGGATTCGGCGCTCTCACC | >trna116_GluCTC_1_-_145399233_145399304 | 1 | 39 | 39 | TCCCTGGTGGTCTAGTGGTTAGGATTCGGCGCTCTCACC | 6320.1 | 2.3 | 0.9 | 2.7 | 8.0E-03 | 2.7E-02 | BT20/MB231 | MITO |
| >hg38_21_+_8433172_8446622_RNA45SN1_45S_50ntFlanks@12991.13014.24\|CTCCCTCGCTGCGATCTATTGAAA | >hg38_21_+_8433172_8446622_RNA45SN1_45S_50ntFlanks | 12991 | 13014 | 24 | CTCCCTCGCTGCGATCTATTGAAA | 266.2 | -4.2 | 1.7 | -2.4 | 1.6E-02 | 2.7E-02 | BT20/MB468 | MP |
| >hg38_21_+_8433172_8446622_RNA45SN1_45S_50ntFlanks@12415.12454.40\|CTTCGATGTCGGCTCTTCCTATCATTGTGAAGCAGAATTC | >hg38_21_+_8433172_8446622_RNA45SN1_45S_50ntFlanks | 12415 | 12454 | 40 | CTTCGATGTCGGCTCTTCCTATCATTGTGAAGCAGAATTC | 264.9 | -1.9 | 0.8 | -2.3 | 2.4E-02 | 2.7E-02 | MB231/MB468 | MITO |
| >hg38_21_+_8433172_8446622_RNA45SN1_45S_50ntFlanks@9589.9610.22\|CTGGGTATAGGGGCGAAAGACT | >hg38_21_+_8433172_8446622_RNA45SN1_45S_50ntFlanks | 9589 | 9610 | 22 | CTGGGTATAGGGGCGAAAGACT | 41.4 | 5.0 | 2.1 | 2.4 | 1.6E-02 | 2.7E-02 | BT20/MB468 | MP |
| >hg38_MT_+_1621_3279_RNR2_16S_50ntFlanks@1368.1398.31\|CGATGTTGGATCAGGACATCCCGATGGTGCA | >hg38_MT_+_1621_3279_RNR2_16S_50ntFlanks | 1368 | 1398 | 31 | CGATGTTGGATCAGGACATCCCGATGGTGCA | 53.1 | 3.1 | 1.3 | 2.3 | 2.2E-02 | 2.7E-02 | MB231/MB468 | MP |
| >am_trnaMT_LysTTT_MT_+_8295_8364@28.52.25,trnalookalike7_LysTTT_1_+_568844_568913@28.52.25\|TTTTAAGTTAAAGATTAAGAGAACC | >am_trnaMT_LysTTT_MT_+_8295_8364 | 28 | 52 | 25 | TTTTAAGTTAAAGATTAAGAGAACC | 501.4 | 2.4 | 0.9 | 2.6 | 8.8E-03 | 2.7E-02 | BT20/MB231 | CYTO |
| >hg38_21_+_8433172_8446622_RNA45SN1_45S_50ntFlanks@12539.12572.34\|ATGATGTGTTGTTGCCATGGTAATCCTGCTCAGT | >hg38_21_+_8433172_8446622_RNA45SN1_45S_50ntFlanks | 12539 | 12572 | 34 | ATGATGTGTTGTTGCCATGGTAATCCTGCTCAGT | 121.3 | -4.1 | 1.7 | -2.4 | 1.6E-02 | 2.7E-02 | BT20/MB468 | MP |
| >hg38_21_+_8433172_8446622_RNA45SN1_45S_50ntFlanks@4922.4940.19\|CCAGGAGTGGAGCCTGCGG | >hg38_21_+_8433172_8446622_RNA45SN1_45S_50ntFlanks | 4922 | 4940 | 19 | CCAGGAGTGGAGCCTGCGG | 204.7 | -4.1 | 1.7 | -2.4 | 1.6E-02 | 2.7E-02 | BT20/MB468 | MP |
| >hg38_MT_+_1621_3279_RNR2_16S_50ntFlanks@1374.1401.28\|TGGATCAGGACATCCCGATGGTGCAGCC | >hg38_MT_+_1621_3279_RNR2_16S_50ntFlanks | 1374 | 1401 | 28 | TGGATCAGGACATCCCGATGGTGCAGCC | 82.4 | 2.9 | 1.3 | 2.3 | 2.2E-02 | 2.7E-02 | MB231/MB468 | MP |
| >hg38_21_+_8433172_8446622_RNA45SN1_45S_50ntFlanks@11567.11593.27\|AGCAGCCGACTTAGAACTGGTGCGGAC | >hg38_21_+_8433172_8446622_RNA45SN1_45S_50ntFlanks | 11567 | 11593 | 27 | AGCAGCCGACTTAGAACTGGTGCGGAC | 124.1 | -4.4 | 1.8 | -2.4 | 1.6E-02 | 2.7E-02 | BT20/MB468 | MP |
| >MI0000750\|hsa-mir-26a-2&WithFlank&12\|-\|57824603\|57824698@20.42.23(+1A);MI0000083\|hsa-mir-26a-1&WithFlank&3\|+\|37969398\|37969486@16.38.23(+1A)&[MIMAT0000082&hsa-miR-26a-5p&offsets\|0\|+1(+1A);m-33&12\|-\|57824658\|57824679&offsets\|0\|+1(+1A)];[MIMAT0000082_1&hsa-miR-26a-5p&offsets\|0\|+1(+1A);m-34&3\|+\|37969413\|37969434&offsets\|0\|+1(+1A)]\|TTCAAGTAATCCAGGATAGGCTGA | >MI0000750\|hsa-mir-26a-2&WithFlank&12\|-\|57824603\|57824698 | 20 | 42 | 23(+1A) | TTCAAGTAATCCAGGATAGGCTGA | 100.7 | 3.6 | 1.5 | 2.4 | 1.6E-02 | 2.7E-02 | BT20/MB468 | MP |
| >hg38_21_+_8433172_8446622_RNA45SN1_45S_50ntFlanks@9806.9838.33\|CGTGGAATGCGAGTGCCTAGTGGGCCACTTTTG | >hg38_21_+_8433172_8446622_RNA45SN1_45S_50ntFlanks | 9806 | 9838 | 33 | CGTGGAATGCGAGTGCCTAGTGGGCCACTTTTG | 592.5 | -4.4 | 1.8 | -2.4 | 1.6E-02 | 2.7E-02 | BT20/MB468 | MP |
| >hg38_MT_+_1621_3279_RNR2_16S_50ntFlanks@1382.1414.33\|GACATCCCGATGGTGCAGCCGCTATTAAAGGTT | >hg38_MT_+_1621_3279_RNR2_16S_50ntFlanks | 1382 | 1414 | 33 | GACATCCCGATGGTGCAGCCGCTATTAAAGGTT | 46.2 | 4.6 | 2.0 | 2.3 | 2.2E-02 | 2.7E-02 | MB231/MB468 | MP |
| >hg38_21_+_8433172_8446622_RNA45SN1_45S_50ntFlanks@7991.8011.21\|ACGTGGCGACCCGCTGAATTT | >hg38_21_+_8433172_8446622_RNA45SN1_45S_50ntFlanks | 7991 | 8011 | 21 | ACGTGGCGACCCGCTGAATTT | 867.7 | 1.6 | 0.7 | 2.3 | 2.3E-02 | 2.7E-02 | MB231/MB468 | CYTO |
| >trnaMT_ProTGG_MT_-_15956_16023@41.58.18\|ATGGTGGAGTTAAAGACT | >trnaMT_ProTGG_MT_-_15956_16023 | 41 | 58 | 18 | ATGGTGGAGTTAAAGACT | 201.8 | 3.0 | 1.3 | 2.3 | 2.2E-02 | 2.7E-02 | MB231/MB468 | MP |
| >hg38_21_+_8433172_8446622_RNA45SN1_45S_50ntFlanks@9548.9570.23\|GAGGTCCGTAGCGGTCCTGACGT | >hg38_21_+_8433172_8446622_RNA45SN1_45S_50ntFlanks | 9548 | 9570 | 23 | GAGGTCCGTAGCGGTCCTGACGT | 401.8 | 1.2 | 0.5 | 2.3 | 2.3E-02 | 2.7E-02 | MB231/MB468 | CYTO |
| >hg38_MT_+_1621_3279_RNR2_16S_50ntFlanks@617.647.31\|AAACATATAACTGAACTCCTCACACCCAATT | >hg38_MT_+_1621_3279_RNR2_16S_50ntFlanks | 617 | 647 | 31 | AAACATATAACTGAACTCCTCACACCCAATT | 16.6 | 7.3 | 3.2 | 2.3 | 2.2E-02 | 2.7E-02 | MB231/MB468 | MP |
| >hg38_21_+_8433172_8446622_RNA45SN1_45S_50ntFlanks@7991.8010.20\|ACGTGGCGACCCGCTGAATT | >hg38_21_+_8433172_8446622_RNA45SN1_45S_50ntFlanks | 7991 | 8010 | 20 | ACGTGGCGACCCGCTGAATT | 3499.9 | -2.2 | 1.0 | -2.2 | 2.4E-02 | 2.7E-02 | MB231/MB468 | MITO |
| >hg38_21_+_8433172_8446622_RNA45SN1_45S_50ntFlanks@12076.12096.21\|AGGGGCTCTCGCTTCTGGCGC | >hg38_21_+_8433172_8446622_RNA45SN1_45S_50ntFlanks | 12076 | 12096 | 21 | AGGGGCTCTCGCTTCTGGCGC | 517.0 | -2.8 | 1.2 | -2.3 | 2.3E-02 | 2.7E-02 | BT20/MB468 | CYTO |
| >hg38_21_+_8433172_8446622_RNA45SN1_45S_50ntFlanks@11999.12039.41\|CCCTGCGGGCCGCCGGTGAAATACCACTACTCTGATCGTTT | >hg38_21_+_8433172_8446622_RNA45SN1_45S_50ntFlanks | 11999 | 12039 | 41 | CCCTGCGGGCCGCCGGTGAAATACCACTACTCTGATCGTTT | 153.8 | -3.1 | 1.3 | -2.3 | 2.1E-02 | 2.7E-02 | BT20/MB468 | MITO |
| >hg38_21_+_8433172_8446622_RNA45SN1_45S_50ntFlanks@3985.4003.19\|CGGCTTTGGTGACTCTAGA | >hg38_21_+_8433172_8446622_RNA45SN1_45S_50ntFlanks | 3985 | 4003 | 19 | CGGCTTTGGTGACTCTAGA | 192.2 | -2.9 | 1.3 | -2.3 | 2.1E-02 | 2.7E-02 | BT20/MB468 | MITO |
| >hg38_21_+_8433172_8446622_RNA45SN1_45S_50ntFlanks@3838.3860.23\|CTCTCCTACTTGGATAACTGTGG | >hg38_21_+_8433172_8446622_RNA45SN1_45S_50ntFlanks | 3838 | 3860 | 23 | CTCTCCTACTTGGATAACTGTGG | 544.1 | -4.7 | 2.0 | -2.4 | 1.6E-02 | 2.7E-02 | BT20/MB468 | MP |
| >hg38_21_+_8433172_8446622_RNA45SN1_45S_50ntFlanks@6723.6743.21\|TTGCAGGACACATTGATCATC | >hg38_21_+_8433172_8446622_RNA45SN1_45S_50ntFlanks | 6723 | 6743 | 21 | TTGCAGGACACATTGATCATC | 481.3 | -2.1 | 0.9 | -2.3 | 2.3E-02 | 2.7E-02 | MB231/MB468 | CYTO |
| >MI0000440\|hsa-mir-27b&WithFlank&9\|+\|95085439\|95085547@67.88.22&[MIMAT0000419&hsa-miR-27b-3p&offsets\|0\|+1;m-39&9\|+\|95085505\|95085525&offsets\|0\|+1]\|TTCACAGTGGCTAAGTTCTGCA | >MI0000440\|hsa-mir-27b&WithFlank&9\|+\|95085439\|95085547 | 67 | 88 | 22 | TTCACAGTGGCTAAGTTCTGCA | 7750.4 | 2.2 | 0.8 | 2.6 | 8.8E-03 | 2.7E-02 | BT20/MB231 | CYTO |
| >am_trnaMT_ValTAC_MT_+_1602_1670@29.58.30\|ACTTACACTTAGGAGATTTCAACTTAACTT | >am_trnaMT_ValTAC_MT_+_1602_1670 | 29 | 58 | 30 | ACTTACACTTAGGAGATTTCAACTTAACTT | 148.2 | 5.5 | 2.3 | 2.4 | 1.6E-02 | 2.7E-02 | BT20/MB468 | MP |
| >hg38_21_+_8433172_8446622_RNA45SN1_45S_50ntFlanks@10841.10865.25\|CGTAACTTCGGGATAAGGATTGGCT | >hg38_21_+_8433172_8446622_RNA45SN1_45S_50ntFlanks | 10841 | 10865 | 25 | CGTAACTTCGGGATAAGGATTGGCT | 27811.5 | -3.6 | 1.5 | -2.4 | 1.6E-02 | 2.7E-02 | BT20/MB468 | MP |
| >hg38_21_+_8433172_8446622_RNA45SN1_45S_50ntFlanks@7990.8011.22\|GACGTGGCGACCCGCTGAATTT | >hg38_21_+_8433172_8446622_RNA45SN1_45S_50ntFlanks | 7990 | 8011 | 22 | GACGTGGCGACCCGCTGAATTT | 1259.0 | 2.3 | 0.9 | 2.4 | 1.6E-02 | 2.7E-02 | BT20/MB468 | MP |
| >am_trnaMT_SerGCT_MT_+_12207_12265@11.32.22\|ACAAGAACTGCTAACTCATGCC | >am_trnaMT_SerGCT_MT_+_12207_12265 | 11 | 32 | 22 | ACAAGAACTGCTAACTCATGCC | 81.4 | 2.9 | 1.2 | 2.4 | 1.6E-02 | 2.7E-02 | BT20/MB468 | MP |
| >trnaMT_PheGAA_MT_+_577_647@54.74.21\|CACATCACCCCATAAACACCA | >trnaMT_PheGAA_MT_+_577_647 | 54 | 74 | 21 | CACATCACCCCATAAACACCA | 830.4 | -1.8 | 0.8 | -2.2 | 2.5E-02 | 2.7E-02 | MB231/MB468 | MITO |
| >hg38_21_+_8433172_8446622_RNA45SN1_45S_50ntFlanks@7969.8011.43\|CCGAGACGCGACCTCAGATCAGACGTGGCGACCCGCTGAATTT | >hg38_21_+_8433172_8446622_RNA45SN1_45S_50ntFlanks | 7969 | 8011 | 43 | CCGAGACGCGACCTCAGATCAGACGTGGCGACCCGCTGAATTT | 170.5 | 3.9 | 1.7 | 2.3 | 2.4E-02 | 2.7E-02 | BT20/MB468 | NUC |
| >hg38_21_+_8433172_8446622_RNA45SN1_45S_50ntFlanks@12630.12654.25\|GGGCGAAGCTACCATCTGTGGGATT | >hg38_21_+_8433172_8446622_RNA45SN1_45S_50ntFlanks | 12630 | 12654 | 25 | GGGCGAAGCTACCATCTGTGGGATT | 269.3 | -4.2 | 1.7 | -2.4 | 1.6E-02 | 2.7E-02 | BT20/MB468 | MP |
| >hg38_21_+_8433172_8446622_RNA45SN1_45S_50ntFlanks@4373.4391.19\|AAAAGCTCGTAGTTGGATC | >hg38_21_+_8433172_8446622_RNA45SN1_45S_50ntFlanks | 4373 | 4391 | 19 | AAAAGCTCGTAGTTGGATC | 77.6 | -3.1 | 1.3 | -2.4 | 1.6E-02 | 2.7E-02 | BT20/MB468 | MP |
| >hg38_MT_+_1621_3279_RNR2_16S_50ntFlanks@118.139.22\|TAAAGTATAGGCGATAGAAATT | >hg38_MT_+_1621_3279_RNR2_16S_50ntFlanks | 118 | 139 | 22 | TAAAGTATAGGCGATAGAAATT | 51.0 | 6.3 | 2.6 | 2.4 | 1.6E-02 | 2.7E-02 | BT20/MB468 | MP |
| >MI0000077\|hsa-mir-21&WithFlank&17\|+\|59841260\|59841343@14.31.18&[MIMAT0000076&hsa-miR-21-5p&offsets\|0\|-4;m-1&17\|+\|59841273\|59841294&offsets\|0\|-4]\|TAGCTTATCAGACTGATG | >MI0000077\|hsa-mir-21&WithFlank&17\|+\|59841260\|59841343 | 14 | 31 | 18 | TAGCTTATCAGACTGATG | 966.8 | 2.9 | 1.2 | 2.4 | 1.6E-02 | 2.7E-02 | BT20/MB468 | MP |

<table>
<tr><td>>hg38_21_+_8433172_8446622_RNA45SN1_45S_50ntFlanks@10345.10365.21|TTCAAACGAGAACTTTGAAGG</td><td>>hg38_21_+_8433172_8446622_RNA45SN1_45S_50ntFlanks</td><td>10345</td><td>10365</td><td>21</td><td>TTCAAACGAGAACTTTGAAGG</td><td>90.3</td><td>2.0</td><td>0.8</td><td>2.4</td><td>1.6E-02</td><td>2.7E-02</td><td>BT20/MB468</td><td>MP</td></tr>
<tr><td>>MI0000067|hsa-let-7f-1&WithFlank&9|+|94176341|94176439@13.32.20(+1G);MI0000068|hsa-let-7f-2&WithFlank&X|-|53557186|53557280@14.33.20(+1G)&[MIMAT0000067&hsa-let-7f-5p&offsets|0|-2(+1G);m-20&9|+|94176353|94176374&offsets|0|-2(+1G)];[MIMAT0000067_1&hsa-let-7f-5p&offsets|0|-2(+1G);m-19&X|-|53557246|53557267&offsets|0|-2(+1G)]|TGAGGTAGTAGATTGTATAGG</td><td>>MI0000067|hsa-let-7f-1&WithFlank&9|+|94176341|94176439</td><td>13</td><td>32</td><td>20(+1G)</td><td>TGAGGTAGTAGATTGTATAGG</td><td>227.4</td><td>2.1</td><td>0.8</td><td>2.6</td><td>8.8E-03</td><td>2.7E-02</td><td>BT20/MB231</td><td>CYTO</td></tr>
<tr><td>>hg38_21_+_8433172_8446622_RNA45SN1_45S_50ntFlanks@12955.12988.34|TGCTTCTGGGTCGGGGTTTCGTACGTAGCAGAGC</td><td>>hg38_21_+_8433172_8446622_RNA45SN1_45S_50ntFlanks</td><td>12955</td><td>12988</td><td>34</td><td>TGCTTCTGGGTCGGGGTTTCGTACGTAGCAGAGC</td><td>270.4</td><td>-2.1</td><td>0.8</td><td>-2.6</td><td>8.8E-03</td><td>2.7E-02</td><td>BT20/MB231</td><td>CYTO</td></tr>
<tr><td>>hg38_21_+_8433172_8446622_RNA45SN1_45S_50ntFlanks@6772.6793.22|GTTCCTCCCGGGGCTACGCCTG</td><td>>hg38_21_+_8433172_8446622_RNA45SN1_45S_50ntFlanks</td><td>6772</td><td>6793</td><td>22</td><td>GTTCCTCCCGGGGCTACGCCTG</td><td>161.4</td><td>-3.5</td><td>1.5</td><td>-2.3</td><td>2.3E-02</td><td>2.7E-02</td><td>BT20/MB468</td><td>CYTO</td></tr>
<tr><td>>MI0000298|hsa-mir-221&WithFlank&X|-|45746151|45746272@71.93.23&[MIMAT0000278&hsa-miR-221-3p&offsets|0|0;m-91&X|-|45746181|45746202&offsets|0|+1]|AGCTACATTGTCTGCTGGGTTTC</td><td>>MI0000298|hsa-mir-221&WithFlank&X|-|45746151|45746272</td><td>71</td><td>93</td><td>23</td><td>AGCTACATTGTCTGCTGGGTTTC</td><td>18076.6</td><td>2.6</td><td>1.1</td><td>2.4</td><td>1.6E-02</td><td>2.7E-02</td><td>BT20/MB468</td><td>MP</td></tr>
<tr><td>>trna116_GluCTC_1_-_145399233_145399304@3.36.34,trna59_GluCTC_1_+_249168447_249168518@3.36.34,trna71_GluCTC_1_-_161439189_161439260@3.36.34,trna74_GluCTC_1_-_161431809_161431880@3.36.34,trna77_GluCTC_1_-_161424398_161424469@3.36.34,trna77_GluCTC_6_+_28949976_28950047@3.36.34,trna80_GluCTC_1_-_161417018_161417089@3.36.34,trna87_GluCTC_6_-_126101393_126101464@3.36.34|CCTGGTGGTCTAGTGGTTAGGATTCGGCGCTCTC</td><td>>trna116_GluCTC_1_-_145399233_145399304</td><td>3</td><td>36</td><td>34</td><td>CCTGGTGGTCTAGTGGTTAGGATTCGGCGCTCTC</td><td>276.6</td><td>-1.9</td><td>0.7</td><td>-2.6</td><td>8.9E-03</td><td>2.7E-02</td><td>BT20/MB231</td><td>CYTO</td></tr>
<tr><td>>trnaMT_GlnTTG_MT_-_4329_4400@57.75.19,trnalookalike2_GlnTTG_1_-_564879_564950@57.75.19|ATTCTCATAGTCCTAGCCA</td><td>>trnaMT_GlnTTG_MT_-_4329_4400</td><td>57</td><td>75</td><td>19</td><td>ATTCTCATAGTCCTAGCCA</td><td>152.8</td><td>3.7</td><td>1.7</td><td>2.2</td><td>2.5E-02</td><td>2.7E-02</td><td>MB231/MB468</td><td>MITO</td></tr>
<tr><td>>hg38_21_+_8433172_8446622_RNA45SN1_45S_50ntFlanks@10238.10260.23|CGCTACGCCGCGACGAGTAGGAG</td><td>>hg38_21_+_8433172_8446622_RNA45SN1_45S_50ntFlanks</td><td>10238</td><td>10260</td><td>23</td><td>CGCTACGCCGCGACGAGTAGGAG</td><td>179.0</td><td>5.3</td><td>2.0</td><td>2.7</td><td>6.5E-03</td><td>2.7E-02</td><td>BT20/MB231</td><td>MP</td></tr>
<tr><td>>MI0000090|hsa-mir-32&WithFlank&9|-|109046223|109046304@12.33.22&[MIMAT0000090&hsa-miR-32-5p&offsets|0|0;m-208&9|-|109046273|109046293&offsets|0|+1]|TATTGCACATTACTAAGTTGCA</td><td>>MI0000090|hsa-mir-32&WithFlank&9|-|109046223|109046304</td><td>12</td><td>33</td><td>22</td><td>TATTGCACATTACTAAGTTGCA</td><td>1076.4</td><td>-3.3</td><td>1.2</td><td>-2.7</td><td>6.5E-03</td><td>2.7E-02</td><td>BT20/MB231</td><td>MP</td></tr>
<tr><td>>MI0001652|hsa-mir-450a-1&WithFlank&X|-|134540335|134540437@24.43.20;MI0003187|hsa-mir-450a-2&WithFlank&X|-|134540502|134540613@29.48.20&[MIMAT0001545&hsa-miR-450a-5p&offsets|0|-2;m-288&X|-|134540393|134540414&offsets|0|-2];[MIMAT0001545_1&hsa-miR-450a-5p&offsets|0|-2;m-287&X|-|134540564|134540585&offsets|0|-2]|TTTTGCGATGTGTTCCTAAT</td><td>>MI0001652|hsa-mir-450a-1&WithFlank&X|-|134540335|134540437</td><td>24</td><td>43</td><td>20</td><td>TTTTGCGATGTGTTCCTAAT</td><td>214.0</td><td>3.0</td><td>1.1</td><td>2.6</td><td>8.9E-03</td><td>2.7E-02</td><td>BT20/MB231</td><td>CYTO</td></tr>
<tr><td>>hg38_21_+_8433172_8446622_RNA45SN1_45S_50ntFlanks@12976.12997.22|TACGTAGCAGAGCAGCTCCCTC</td><td>>hg38_21_+_8433172_8446622_RNA45SN1_45S_50ntFlanks</td><td>12976</td><td>12997</td><td>22</td><td>TACGTAGCAGAGCAGCTCCCTC</td><td>344.0</td><td>-2.1</td><td>0.8</td><td>-2.6</td><td>8.9E-03</td><td>2.7E-02</td><td>BT20/MB231</td><td>CYTO</td></tr>
<tr><td>>hg38_MT_+_598_1651_RNR1_12S_50ntFlanks@790.809.20|ACGATAGCCCTTATGAAACT</td><td>>hg38_MT_+_598_1651_RNR1_12S_50ntFlanks</td><td>790</td><td>809</td><td>20</td><td>ACGATAGCCCTTATGAAACT</td><td>156.7</td><td>2.9</td><td>1.3</td><td>2.2</td><td>2.5E-02</td><td>2.7E-02</td><td>MB231/MB468</td><td>MITO</td></tr>
<tr><td>>hg38_21_+_8433172_8446622_RNA45SN1_45S_50ntFlanks@5074.5093.20|ATTCCGATAACGAACGAGAC</td><td>>hg38_21_+_8433172_8446622_RNA45SN1_45S_50ntFlanks</td><td>5074</td><td>5093</td><td>20</td><td>ATTCCGATAACGAACGAGAC</td><td>146.5</td><td>-4.6</td><td>1.9</td><td>-2.4</td><td>1.6E-02</td><td>2.7E-02</td><td>BT20/MB468</td><td>MP</td></tr>
<tr><td>>hg38_21_+_8433172_8446622_RNA45SN1_45S_50ntFlanks@8168.8205.38|CCCAAGTCCTTCTGATCGAGGCCCAGCCCGTGGACGGT</td><td>>hg38_21_+_8433172_8446622_RNA45SN1_45S_50ntFlanks</td><td>8168</td><td>8205</td><td>38</td><td>CCCAAGTCCTTCTGATCGAGGCCCAGCCCGTGGACGGT</td><td>45.1</td><td>-4.3</td><td>1.8</td><td>-2.4</td><td>1.6E-02</td><td>2.7E-02</td><td>BT20/MB468</td><td>MP</td></tr>
<tr><td>>hg38_21_+_8433172_8446622_RNA45SN1_45S_50ntFlanks@12242.12268.27|AGAAGGGCAAAAGCTCGCTTGATCTTG</td><td>>hg38_21_+_8433172_8446622_RNA45SN1_45S_50ntFlanks</td><td>12242</td><td>12268</td><td>27</td><td>AGAAGGGCAAAAGCTCGCTTGATCTTG</td><td>263.2</td><td>-1.6</td><td>0.7</td><td>-2.3</td><td>2.1E-02</td><td>2.7E-02</td><td>BT20/MB468</td><td>MITO</td></tr>
<tr><td>>hg38_21_+_8433172_8446622_RNA45SN1_45S_50ntFlanks@10806.10827.22|AACAATGTAGGTAAGGGAAGTC</td><td>>hg38_21_+_8433172_8446622_RNA45SN1_45S_50ntFlanks</td><td>10806</td><td>10827</td><td>22</td><td>AACAATGTAGGTAAGGGAAGTC</td><td>168.4</td><td>-2.6</td><td>1.1</td><td>-2.3</td><td>2.1E-02</td><td>2.7E-02</td><td>BT20/MB468</td><td>MITO</td></tr>
<tr><td>>hg38_21_+_8433172_8446622_RNA45SN1_45S_50ntFlanks@6651.6688.38|CGACTCTTAGCGGTGGATCACTCGGCTCGTGCGTCGAT</td><td>>hg38_21_+_8433172_8446622_RNA45SN1_45S_50ntFlanks</td><td>6651</td><td>6688</td><td>38</td><td>CGACTCTTAGCGGTGGATCACTCGGCTCGTGCGTCGAT</td><td>6767.6</td><td>1.8</td><td>0.7</td><td>2.7</td><td>7.1E-03</td><td>2.7E-02</td><td>BT20/MB231</td><td>NUC</td></tr>
<tr><td>>MI0000089|hsa-mir-31&WithFlank&9|-|21512109|21512191@14.35.22(+1A)&[MIMAT0000089&hsa-miR-31-5p&offsets|0|+1(+1A);m-360&9|-|21512157|21512178&offsets|0|0(+1A)]|AGGCAAGATGCTGGCATAGCTGA</td><td>>MI0000089|hsa-mir-31&WithFlank&9|-|21512109|21512191</td><td>14</td><td>35</td><td>22(+1A)</td><td>AGGCAAGATGCTGGCATAGCTGA</td><td>365.9</td><td>2.9</td><td>1.2</td><td>2.4</td><td>1.6E-02</td><td>2.7E-02</td><td>BT20/MB468</td><td>MP</td></tr>
<tr><td>>hg38_21_+_8433172_8446622_RNA45SN1_45S_50ntFlanks@10345.10388.44|TTCAAACGAGAACTTTGAAGGCCGAAGTGGAGAAGGGTTCCATG</td><td>>hg38_21_+_8433172_8446622_RNA45SN1_45S_50ntFlanks</td><td>10345</td><td>10388</td><td>44</td><td>TTCAAACGAGAACTTTGAAGGCCGAAGTGGAGAAGGGTTCCATG</td><td>149.9</td><td>4.3</td><td>1.8</td><td>2.4</td><td>1.6E-02</td><td>2.7E-02</td><td>BT20/MB468</td><td>MP</td></tr>
<tr><td>>hg38_21_+_8433172_8446622_RNA45SN1_45S_50ntFlanks@7975.7999.25|CGCGACCTCAGATCAGACGTGGCGA</td><td>>hg38_21_+_8433172_8446622_RNA45SN1_45S_50ntFlanks</td><td>7975</td><td>7999</td><td>25</td><td>CGCGACCTCAGATCAGACGTGGCGA</td><td>210.9</td><td>2.7</td><td>1.2</td><td>2.2</td><td>2.5E-02</td><td>2.7E-02</td><td>MB231/MB468</td><td>MITO</td></tr>
<tr><td>>MI0001445|hsa-mir-423&WithFlank&17|+|30117073|30117178@58.80.23&[MIMAT0001340&hsa-miR-423-3p&offsets|-1|-1;m-100&17|+|30117131|30117153&offsets|-1|-1]|AAGCTCGGTCTGAGGCCCCTCAG</td><td>>MI0001445|hsa-mir-423&WithFlank&17|+|30117073|30117178</td><td>58</td><td>80</td><td>23</td><td>AAGCTCGGTCTGAGGCCCCTCAG</td><td>82.1</td><td>3.9</td><td>1.7</td><td>2.3</td><td>2.2E-02</td><td>2.7E-02</td><td>MB231/MB468</td><td>MP</td></tr>
<tr><td>>hg38_MT_+_598_1651_RNR1_12S_50ntFlanks@51.77.27|AATAGGTTTGGTCCTAGCCTTTCTATT</td><td>>hg38_MT_+_598_1651_RNR1_12S_50ntFlanks</td><td>51</td><td>77</td><td>27</td><td>AATAGGTTTGGTCCTAGCCTTTCTATT</td><td>1155.3</td><td>3.3</td><td>1.4</td><td>2.4</td><td>1.6E-02</td><td>2.7E-02</td><td>BT20/MB468</td><td>MP</td></tr>
<tr><td>>MI0000465|hsa-mir-191&WithFlank&3|-|49020612|49020715@22.43.22(+1C)&[MIMAT0000440&hsa-miR-191-5p&offsets|0|-1(+1C);m-48&3|-|49020672|49020694&offsets|0|-1(+1C)]|CAACGGAATCCCAAAAGCAGCTC</td><td>>MI0000465|hsa-mir-191&WithFlank&3|-|49020612|49020715</td><td>22</td><td>43</td><td>22(+1C)</td><td>CAACGGAATCCCAAAAGCAGCTC</td><td>74.2</td><td>3.6</td><td>1.5</td><td>2.4</td><td>1.6E-02</td><td>2.7E-02</td><td>BT20/MB468</td><td>MP</td></tr>
<tr><td>>hg38_21_+_8433172_8446622_RNA45SN1_45S_50ntFlanks@8104.8142.39|CCGCGGCGGGGCGCGGGACATGTGGCGTACGGAAGACCC</td><td>>hg38_21_+_8433172_8446622_RNA45SN1_45S_50ntFlanks</td><td>8104</td><td>8142</td><td>39</td><td>CCGCGGCGGGGCGCGGGACATGTGGCGTACGGAAGACCC</td><td>513.2</td><td>-3.1</td><td>1.3</td><td>-2.3</td><td>2.1E-02</td><td>2.7E-02</td><td>BT20/MB468</td><td>MITO</td></tr>
<tr><td>>MI0000264|hsa-mir-7-2&WithFlank&15|+|88611819|88611940@38.59.22;MI0000265|hsa-mir-7-3&WithFlank&19|+|4770664|4770785@37.58.22;MI0000263|hsa-mir-7-1&WithFlank&9|-|83969742|83969863@30.51.22&[MIMAT0000252&hsa-miR-7-5p&offsets|0|-2;m-400&15|+|88611856|88611879&offsets|0|-2];[MIMAT0000252_1&hsa-miR-7-5p&offsets|0|-2;m-402&19|+|4770700|4770723&offsets|0|-2];[MIMAT0000252_2&hsa-miR-7-5p&offsets|0|-2;m-401&9|-|83969811|83969834&offsets|0|-2]|TGGAAGACTAGTGATTTTGTTG</td><td>>MI0000264|hsa-mir-7-2&WithFlank&15|+|88611819|88611940</td><td>38</td><td>59</td><td>22</td><td>TGGAAGACTAGTGATTTTGTTG</td><td>938.2</td><td>2.5</td><td>0.9</td><td>2.6</td><td>8.9E-03</td><td>2.7E-02</td><td>BT20/MB231</td><td>CYTO</td></tr>
<tr><td>>hg38_21_+_8433172_8446622_RNA45SN1_45S_50ntFlanks@3840.3881.42|CTCCTACTTGGATAACTGTGGTAATTCTAGAGCTAATACATG</td><td>>hg38_21_+_8433172_8446622_RNA45SN1_45S_50ntFlanks</td><td>3840</td><td>3881</td><td>42</td><td>CTCCTACTTGGATAACTGTGGTAATTCTAGAGCTAATACATG</td><td>197.6</td><td>-2.6</td><td>1.0</td><td>-2.6</td><td>8.9E-03</td><td>2.7E-02</td><td>BT20/MB231</td><td>CYTO</td></tr>
<tr><td>>hg38_21_+_8433172_8446622_RNA45SN1_45S_50ntFlanks@12456.12479.24|CCAAGCGTTGGATTGTTCACCCAC</td><td>>hg38_21_+_8433172_8446622_RNA45SN1_45S_50ntFlanks</td><td>12456</td><td>12479</td><td>24</td><td>CCAAGCGTTGGATTGTTCACCCAC</td><td>1159.7</td><td>-4.5</td><td>1.9</td><td>-2.4</td><td>1.6E-02</td><td>2.7E-02</td><td>BT20/MB468</td><td>MP</td></tr>
<tr><td>>hg38_MT_+_598_1651_RNR1_12S_50ntFlanks@607.633.27|CTGTTCTGTAATCGATAAACCCCGATC</td><td>>hg38_MT_+_598_1651_RNR1_12S_50ntFlanks</td><td>607</td><td>633</td><td>27</td><td>CTGTTCTGTAATCGATAAACCCCGATC</td><td>118.6</td><td>3.4</td><td>1.4</td><td>2.4</td><td>1.6E-02</td><td>2.7E-02</td><td>BT20/MB468</td><td>MP</td></tr>
<tr><td>>MI0000746|hsa-mir-99b&WithFlank&19|+|51692606|51692687@51.72.22&[MIMAT0004678&hsa-miR-99b-3p&offsets|0|0;m-151&19|+|51692656|51692678&offsets|0|-1]|CAAGCTCGTGTCTGTGGGTCCG</td><td>>MI0000746|hsa-mir-99b&WithFlank&19|+|51692606|51692687</td><td>51</td><td>72</td><td>22</td><td>CAAGCTCGTGTCTGTGGGTCCG</td><td>2444.9</td><td>3.0</td><td>1.1</td><td>2.7</td><td>6.6E-03</td><td>2.7E-02</td><td>BT20/MB231</td><td>MP</td></tr>
<tr><td>>hg38_21_+_8433172_8446622_RNA45SN1_45S_50ntFlanks@9352.9376.25|CCGTGGCGCAATGAAGGTGAAGGCC</td><td>>hg38_21_+_8433172_8446622_RNA45SN1_45S_50ntFlanks</td><td>9352</td><td>9376</td><td>25</td><td>CCGTGGCGCAATGAAGGTGAAGGCC</td><td>133.0</td><td>-4.2</td><td>1.7</td><td>-2.4</td><td>1.6E-02</td><td>2.7E-02</td><td>BT20/MB468</td><td>MP</td></tr>
<tr><td>>trna119_LysCTT_1_-_145395522_145395594@4.33.30,trna11_LysCTT_5_-_180648979_180649051@4.33.30,trna13_LysCTT_6_+_26556774_26556846@4.33.30,trna7_LysCTT_16_+_3225692_3225764@4.33.30,trna9_LysCTT_5_+_180634755_180634827@4.33.30|CGGCTAGCTCAGTCGGTAGAGCATGAGACT</td><td>>trna119_LysCTT_1_-_145395522_145395594</td><td>4</td><td>33</td><td>30</td><td>CGGCTAGCTCAGTCGGTAGAGCATGAGACT</td><td>355.8</td><td>-4.6</td><td>1.8</td><td>-2.6</td><td>8.9E-03</td><td>2.7E-02</td><td>BT20/MB231</td><td>CYTO</td></tr>
<tr><td>>hg38_21_+_8433172_8446622_RNA45SN1_45S_50ntFlanks@8356.8375.20|GAAAAGAACTTTGAAGAGAG</td><td>>hg38_21_+_8433172_8446622_RNA45SN1_45S_50ntFlanks</td><td>8356</td><td>8375</td><td>20</td><td>GAAAAGAACTTTGAAGAGAG</td><td>6872.3</td><td>3.6</td><td>1.3</td><td>2.7</td><td>6.6E-03</td><td>2.7E-02</td><td>BT20/MB231</td><td>MP</td></tr>
<tr><td>>hg38_MT_+_598_1651_RNR1_12S_50ntFlanks@52.91.40|ATAGGTTTGGTCCTAGCCTTTCTATTAGCTCTTAGTAAGA</td><td>>hg38_MT_+_598_1651_RNR1_12S_50ntFlanks</td><td>52</td><td>91</td><td>40</td><td>ATAGGTTTGGTCCTAGCCTTTCTATTAGCTCTTAGTAAGA</td><td>290.2</td><td>3.1</td><td>1.4</td><td>2.2</td><td>2.5E-02</td><td>2.7E-02</td><td>MB231/MB468</td><td>MITO</td></tr>
<tr><td>>hg38_21_+_8433172_8446622_RNA45SN1_45S_50ntFlanks@12419.12458.40|GATGTCGGCTCTTCCTATCATTGTGAAGCAGAATTCACCA</td><td>>hg38_21_+_8433172_8446622_RNA45SN1_45S_50ntFlanks</td><td>12419</td><td>12458</td><td>40</td><td>GATGTCGGCTCTTCCTATCATTGTGAAGCAGAATTCACCA</td><td>125.2</td><td>3.6</td><td>1.6</td><td>2.2</td><td>2.5E-02</td><td>2.7E-02</td><td>MB231/MB468</td><td>MITO</td></tr>
<tr><td>>MI0003820|hsa-mir-454&WithFlank&17|-|59137752|59137878@70.91.22(+1C)&[MIMAT0003885&hsa-miR-454-3p&offsets|0|-1(+1C);m-272&17|-|59137787|59137809&offsets|0|-1(+1C)]|TAGTGCAATATTGCTTATAGGGC</td><td>>MI0003820|hsa-mir-454&WithFlank&17|-|59137752|59137878</td><td>70</td><td>91</td><td>22(+1C)</td><td>TAGTGCAATATTGCTTATAGGGC</td><td>73.8</td><td>3.8</td><td>1.7</td><td>2.3</td><td>2.2E-02</td><td>2.7E-02</td><td>MB231/MB468</td><td>MP</td></tr>
<tr><td>>trnaMT_LysTTT_MT_+_8295_8364@1.24.24,trnalookalike7_LysTTT_1_+_568844_568913@1.24.24|CACTGTAAAGCTAACTTAGCATTA</td><td>>trnaMT_LysTTT_MT_+_8295_8364</td><td>1</td><td>24</td><td>24</td><td>CACTGTAAAGCTAACTTAGCATTA</td><td>235.8</td><td>-2.8</td><td>1.0</td><td>-2.7</td><td>6.6E-03</td><td>2.7E-02</td><td>BT20/MB231</td><td>MP</td></tr>
<tr><td>>hg38_21_+_8433172_8446622_RNA45SN1_45S_50ntFlanks@11778.11798.21|ACGCGCATGAATGGATGAACG</td><td>>hg38_21_+_8433172_8446622_RNA45SN1_45S_50ntFlanks</td><td>11778</td><td>11798</td><td>21</td><td>ACGCGCATGAATGGATGAACG</td><td>243.3</td><td>-4.2</td><td>1.7</td><td>-2.4</td><td>1.6E-02</td><td>2.7E-02</td><td>BT20/MB468</td><td>MP</td></tr>
<tr><td>>hg38_MT_+_1621_3279_RNR2_16S_50ntFlanks@645.676.32|ATTGGACCAATCTATCACCCTATAGAAGAACT</td><td>>hg38_MT_+_1621_3279_RNR2_16S_50ntFlanks</td><td>645</td><td>676</td><td>32</td><td>ATTGGACCAATCTATCACCCTATAGAAGAACT</td><td>62.3</td><td>3.8</td><td>1.6</td><td>2.3</td><td>2.2E-02</td><td>2.7E-02</td><td>MB231/MB468</td><td>MP</td></tr>
<tr><td>>hg38_21_+_8433172_8446622_RNA45SN1_45S_50ntFlanks@6661.6691.31|CGGTGGATCACTCGGCTCGTGCGTCGATGAA</td><td>>hg38_21_+_8433172_8446622_RNA45SN1_45S_50ntFlanks</td><td>6661</td><td>6691</td><td>31</td><td>CGGTGGATCACTCGGCTCGTGCGTCGATGAA</td><td>156.1</td><td>3.3</td><td>1.2</td><td>2.7</td><td>6.6E-03</td><td>2.7E-02</td><td>BT20/MB231</td><td>MP</td></tr>
<tr><td>>hg38_21_+_8433172_8446622_RNA45SN1_45S_50ntFlanks@10346.10376.31|TCAAACGAGAACTTTGAAGGCCGAAGTGGAG</td><td>>hg38_21_+_8433172_8446622_RNA45SN1_45S_50ntFlanks</td><td>10346</td><td>10376</td><td>31</td><td>TCAAACGAGAACTTTGAAGGCCGAAGTGGAG</td><td>253.5</td><td>-2.3</td><td>1.0</td><td>-2.4</td><td>1.6E-02</td><td>2.7E-02</td><td>BT20/MB468</td><td>MP</td></tr>
<tr><td>>hg38_21_+_8433172_8446622_RNA45SN1_45S_50ntFlanks@475.495.21|CTTCGTGATCGATGTGGTGAC</td><td>>hg38_21_+_8433172_8446622_RNA45SN1_45S_50ntFlanks</td><td>475</td><td>495</td><td>21</td><td>CTTCGTGATCGATGTGGTGAC</td><td>293.2</td><td>3.9</td><td>1.7</td><td>2.3</td><td>2.2E-02</td><td>2.7E-02</td><td>MB231/MB468</td><td>MP</td></tr>
<tr><td>>hg38_21_+_8433172_8446622_RNA45SN1_45S_50ntFlanks@12907.12935.29|CACCGCACGTTCGTGGGGAACCTGGCGCT</td><td>>hg38_21_+_8433172_8446622_RNA45SN1_45S_50ntFlanks</td><td>12907</td><td>12935</td><td>29</td><td>CACCGCACGTTCGTGGGGAACCTGGCGCT</td><td>202.9</td><td>-3.6</td><td>1.6</td><td>-2.3</td><td>2.4E-02</td><td>2.7E-02</td><td>BT20/MB468</td><td>CYTO</td></tr>
<tr><td>>hg38_MT_+_1621_3279_RNR2_16S_50ntFlanks@1374.1396.23|TGGATCAGGACATCCCGATGGTG</td><td>>hg38_MT_+_1621_3279_RNR2_16S_50ntFlanks</td><td>1374</td><td>1396</td><td>23</td><td>TGGATCAGGACATCCCGATGGTG</td><td>252.4</td><td>1.8</td><td>0.8</td><td>2.3</td><td>2.2E-02</td><td>2.7E-02</td><td>MB231/MB468</td><td>MP</td></tr>
</table>

| | | | | | | | | | | | | | |
|---|---|---|---|---|---|---|---|---|---|---|---|---|---|
| >MI0003776\|hsa-mir-320b-1&WithFlank&1\|+\|116671740\|116671823@48.66.19;MI0003839\|hsa-mir-320b-2&WithFlank&1\|-\|224257034\|224257116@47.65.19;MI0003778\|hsa-mir-320c-1&WithFlank&18\|+\|21683512\|21683595@48.66.19;MI0008191\|hsa-mir-320c-2&WithFlank&18\|+\|24321669\|24321752@48.66.19;MI0000542\|hsa-mir-320a&WithFlank&8\|-\|22244960\|22245043@48.66.19&[MIMAT0005792&hsa-miR-320b&offsets\|0\|-3;m-183&1\|+\|116671787\|116671806&offsets\|0\|-1];[MIMAT0005792_1&hsa-miR-320b&offsets\|0\|-3;m-183&1\|-\|224257051\|224257070&offsets\|0\|-1];[MIMAT0005793&hsa-miR-320c&offsets\|0\|-1;m-222&18\|+\|21683559\|21683578&offsets\|0\|-1];[MIMAT0005793_1&hsa-miR-320c&offsets\|0\|-1;m-219&18\|+\|24321716\|24321735&offsets\|0\|-1];[MIMAT0000510&hsa-miR-320a-3p&offsets\|0\|-3;m-60&8\|-\|22244975\|22244996&offsets\|0\|-3]\|AAAAGCTGGGTTGAGAGGG | >MI0003776\|hsa-mir-320b-1&WithFlank&1\|+\|116671740\|116671823 | 48 | 66 | 19 | AAAAGCTGGGTTGAGAGGG | 1953.5 | 2.7 | 1.0 | 2.7 | 6.6E-03 | 2.7E-02 | BT20/MB231 | MP |
| >hg38_1_-_228634819_228635039_RNA5S12_5S_50ntFlanks@153.171.19\|ACCGGGTGCTGTAGGCTTT | >hg38_1_-_228634819_228635039_RNA5S12_5S_50ntFlanks | 153 | 171 | 19 | ACCGGGTGCTGTAGGCTTT | 21985.9 | 2.9 | 1.2 | 2.4 | 1.6E-02 | 2.7E-02 | BT20/MB468 | MP |
| >MI0000063\|hsa-let-7b&WithFlank&22\|+\|46113680\|46113774@12.29.18&[MIMAT0000063&hsa-let-7b-5p&offsets\|0\|-4;m-14&22\|+\|46113691\|46113712&offsets\|0\|-4]\|TGAGGTAGTAGGTTGTGT | >MI0000063\|hsa-let-7b&WithFlank&22\|+\|46113680\|46113774 | 12 | 29 | 18 | TGAGGTAGTAGGTTGTGT | 179.5 | 4.1 | 1.8 | 2.3 | 2.2E-02 | 2.7E-02 | MB231/MB468 | MP |
| >hg38_21_+_8433172_8446622_RNA45SN1_45S_50ntFlanks@12474.12498.25\|ACCCACTAATAGGGAACGTGAGCTG | >hg38_21_+_8433172_8446622_RNA45SN1_45S_50ntFlanks | 12474 | 12498 | 25 | ACCCACTAATAGGGAACGTGAGCTG | 309.1 | -3.7 | 1.6 | -2.3 | 2.4E-02 | 2.7E-02 | BT20/MB468 | CYTO |
| >hg38_21_+_8433172_8446622_RNA45SN1_45S_50ntFlanks@10345.10368.24\|TTCAAACGAGAACTTTGAAGGCCG | >hg38_21_+_8433172_8446622_RNA45SN1_45S_50ntFlanks | 10345 | 10368 | 24 | TTCAAACGAGAACTTTGAAGGCCG | 140.8 | -4.3 | 1.8 | -2.4 | 1.6E-02 | 2.7E-02 | BT20/MB468 | MP |
| >hg38_21_+_8433172_8446622_RNA45SN1_45S_50ntFlanks@7988.8011.24\|CAGACGTGGCGACCCGCTGAATTT | >hg38_21_+_8433172_8446622_RNA45SN1_45S_50ntFlanks | 7988 | 8011 | 24 | CAGACGTGGCGACCCGCTGAATTT | 847.4 | 4.0 | 1.5 | 2.7 | 6.6E-03 | 2.7E-02 | BT20/MB231 | MP |
| >hg38_21_+_8433172_8446622_RNA45SN1_45S_50ntFlanks@9565.9597.33\|TGACGTGCAAATCGGTCGTCCGACCTGGGTATA | >hg38_21_+_8433172_8446622_RNA45SN1_45S_50ntFlanks | 9565 | 9597 | 33 | TGACGTGCAAATCGGTCGTCCGACCTGGGTATA | 170.9 | -2.4 | 0.9 | -2.6 | 9.0E-03 | 2.7E-02 | BT20/MB231 | CYTO |
| >hg38_21_+_8433172_8446622_RNA45SN1_45S_50ntFlanks@11552.11585.34\|CCTCGGCCGGCGCCTAGCAGCCGACTTAGAACTG | >hg38_21_+_8433172_8446622_RNA45SN1_45S_50ntFlanks | 11552 | 11585 | 34 | CCTCGGCCGGCGCCTAGCAGCCGACTTAGAACTG | 74.9 | -3.2 | 1.3 | -2.4 | 1.6E-02 | 2.7E-02 | BT20/MB468 | MP |
| >trnaMT_LysTTT_MT_+_8295_8364@39.72.34,trnalookalike7_LysTTT_1_+_568844_568913@39.72.34\|AGATTAAGAGAACCAACACCTCTTTACAGTGACC | >trnaMT_LysTTT_MT_+_8295_8364 | 39 | 72 | 34 | AGATTAAGAGAACCAACACCTCTTTACAGTGACC | 78.2 | 4.5 | 2.0 | 2.3 | 2.2E-02 | 2.7E-02 | MB231/MB468 | MP |
| >hg38_MT_+_1621_3279_RNR2_16S_50ntFlanks@1369.1401.33\|GATGTTGGATCAGGACATCCCGATGGTGCAGCC | >hg38_MT_+_1621_3279_RNR2_16S_50ntFlanks | 1369 | 1401 | 33 | GATGTTGGATCAGGACATCCCGATGGTGCAGCC | 167.2 | 3.8 | 1.7 | 2.3 | 2.2E-02 | 2.7E-02 | MB231/MB468 | MP |
| >trna20_GluTTC_2_-_131094701_131094772@1.30.30,trna5_GluTTC_13_-_41634874_41634945@1.30.30\|TCCCATATGGTCTAGCGGTTAGGATTCCTG | >trna20_GluTTC_2_-_131094701_131094772 | 1 | 30 | 30 | TCCCATATGGTCTAGCGGTTAGGATTCCTG | 514.4 | -2.0 | 0.9 | -2.2 | 2.5E-02 | 2.7E-02 | MB231/MB468 | MITO |
| >trna111_HisGTG_1_-_147774845_147774916@-1T.31.32,trna118_HisGTG_1_-_145396881_145396952@-1T.31.32,trna16_HisGTG_1_+_146544773_146544844@-1T.31.32,trna1_HisGTG_15_+_45493349_45493420@-1T.31.32,trna21_HisGTG_1_+_147753471_147753542@-1T.31.32,trna33_HisGTG_6_+_27125906_27125977@-1T.31.32,trna7_HisGTG_9_-_14433938_14434009@-1T.31.32,trna8_HisGTG_15_-_45492611_45492682@-1T.31.32,trna9_HisGTG_15_-_45490804_45490875@-1T.31.32\|TGCCGTGATCGTATAGTGGTTAGTACTCTGCG | >trna111_HisGTG_1_-_147774845_147774916 | -1T | 31 | 32 | TGCCGTGATCGTATAGTGGTTAGTACTCTGCG | 27313.1 | -2.4 | 1.0 | -2.3 | 2.4E-02 | 2.7E-02 | MB231/MB468 | CYTO |
| >hg38_21_+_8433172_8446622_RNA45SN1_45S_50ntFlanks@6774.6807.34\|TCCTCCCGGGGCTACGCCTGTCTGAGCGTCGCTT | >hg38_21_+_8433172_8446622_RNA45SN1_45S_50ntFlanks | 6774 | 6807 | 34 | TCCTCCCGGGGCTACGCCTGTCTGAGCGTCGCTT | 802.0 | 1.9 | 0.7 | 2.6 | 8.1E-03 | 2.7E-02 | BT20/MB231 | MITO |
| >hg38_21_+_8433172_8446622_RNA45SN1_45S_50ntFlanks@4738.4762.25\|AAAGTCGGAGGTTCGAAGACGATCA | >hg38_21_+_8433172_8446622_RNA45SN1_45S_50ntFlanks | 4738 | 4762 | 25 | AAAGTCGGAGGTTCGAAGACGATCA | 86.9 | 3.3 | 1.4 | 2.4 | 1.6E-02 | 2.7E-02 | BT20/MB468 | MP |
| >trna10_AspGTC_12_-_125424193_125424264@39.73.35,trna12_AspGTC_12_-_125411891_125411962@39.73.35,trna144_AspGTC_6_-_27551236_27551307@39.73.35,trna38_AspGTC_17_-_8125556_8125627@39.73.35,trna45_AspGTC_6_+_27447453_27447524@39.73.35,trna48_AspGTC_6_+_27471523_27471594@39.73.35,trna4_AspGTC_12_+_96429799_96429870@39.73.35,trna69_AspGTC_1_-_161440205_161440276@39.73.35,trna72_AspGTC_1_-_161432824_161432895@39.73.35,trna75_AspGTC_1_-_161425414_161425485@39.73.35,trna78_AspGTC_1_-_161418033_161418104@39.73.35,trna81_AspGTC_1_-_161410615_161410686@39.73.35\|GCGGGAGACCGGGGTTCGATTCCCCGACGGGGAGC | >trna10_AspGTC_12_-_125424193_125424264 | 39 | 73 | 35 | GCGGGAGACCGGGGTTCGATTCCCCGACGGGGAGC | 121.3 | 3.0 | 1.3 | 2.4 | 1.6E-02 | 2.7E-02 | BT20/MB468 | MP |
| >hg38_21_+_8433172_8446622_RNA45SN1_45S_50ntFlanks@12405.12433.29\|CTTTTTGATCCTTCGATGTCGGCTCTTCC | >hg38_21_+_8433172_8446622_RNA45SN1_45S_50ntFlanks | 12405 | 12433 | 29 | CTTTTTGATCCTTCGATGTCGGCTCTTCC | 508.0 | -2.8 | 1.1 | -2.6 | 9.0E-03 | 2.7E-02 | BT20/MB231 | CYTO |
| >hg38_MT_+_1621_3279_RNR2_16S_50ntFlanks@1428.1453.26\|ATTAAAGTCCTACGTGATCTGAGTTC | >hg38_MT_+_1621_3279_RNR2_16S_50ntFlanks | 1428 | 1453 | 26 | ATTAAAGTCCTACGTGATCTGAGTTC | 441.5 | 2.9 | 1.2 | 2.4 | 1.6E-02 | 2.8E-02 | BT20/MB468 | MP |
| >MI0000811\|hsa-mir-148b&WithFlank&12\|+\|54337210\|54337320@69.89.21(+1A)&[MIMAT0000759&hsa-miR-148b-3p&offsets\|0\|-1(+1A);m-77&12\|+\|54337278\|54337299&offsets\|0\|-1(+1A)]\|TCAGTGCATCACAGAACTTTGA | >MI0000811\|hsa-mir-148b&WithFlank&12\|+\|54337210\|54337320 | 69 | 89 | 21(+1A) | TCAGTGCATCACAGAACTTTGA | 501.4 | 3.3 | 1.4 | 2.3 | 2.2E-02 | 2.8E-02 | MB231/MB468 | MP |
| >MI0000273\|hsa-mir-183&WithFlank&7\|-\|129774899\|129775020@33.56.24&[MIMAT0000261&hsa-miR-183-5p&offsets\|0\|+2;m-41&7\|-\|129774966\|129774987&offsets\|-1\|+1]\|TATGGCACTGGTAGAATTCACTGT | >MI0000273\|hsa-mir-183&WithFlank&7\|-\|129774899\|129775020 | 33 | 56 | 24 | TATGGCACTGGTAGAATTCACTGT | 152.6 | 2.3 | 1.0 | 2.4 | 1.6E-02 | 2.8E-02 | BT20/MB468 | MP |
| >hg38_21_+_8433172_8446622_RNA45SN1_45S_50ntFlanks@12474.12497.24\|ACCCACTAATAGGGAACGTGAGCT | >hg38_21_+_8433172_8446622_RNA45SN1_45S_50ntFlanks | 12474 | 12497 | 24 | ACCCACTAATAGGGAACGTGAGCT | 328.6 | 1.9 | 0.8 | 2.3 | 2.4E-02 | 2.8E-02 | BT20/MB468 | CYTO |
| >hg38_21_+_8433172_8446622_RNA45SN1_45S_50ntFlanks@6646.6695.50\|TCGTACGACTCTTAGCGGTGGATCACTCGGCTCGTGCGTCGATGAAGAAC | >hg38_21_+_8433172_8446622_RNA45SN1_45S_50ntFlanks | 6646 | 6695 | 50 | TCGTACGACTCTTAGCGGTGGATCACTCGGCTCGTGCGTCGATGAAGAAC | 1932.7 | 2.8 | 1.2 | 2.2 | 2.5E-02 | 2.8E-02 | MB231/MB468 | MITO |
| >hg38_21_+_8433172_8446622_RNA45SN1_45S_50ntFlanks@4223.4242.20\|AAAAATAACAATACAGGACT | >hg38_21_+_8433172_8446622_RNA45SN1_45S_50ntFlanks | 4223 | 4242 | 20 | AAAAATAACAATACAGGACT | 359.8 | -2.1 | 0.9 | -2.3 | 2.4E-02 | 2.8E-02 | MB231/MB468 | CYTO |
| >hg38_21_+_8433172_8446622_RNA45SN1_45S_50ntFlanks@4439.4471.33\|CCGCCCCTTGCCTCTCGGCGCCCCCTCGATGCT | >hg38_21_+_8433172_8446622_RNA45SN1_45S_50ntFlanks | 4439 | 4471 | 33 | CCGCCCCTTGCCTCTCGGCGCCCCCTCGATGCT | 47.0 | -4.2 | 1.7 | -2.4 | 1.6E-02 | 2.8E-02 | BT20/MB468 | MP |
| >hg38_MT_+_1621_3279_RNR2_16S_50ntFlanks@1368.1407.40\|CGATGTTGGATCAGGACATCCCGATGGTGCAGCCGCTATT | >hg38_MT_+_1621_3279_RNR2_16S_50ntFlanks | 1368 | 1407 | 40 | CGATGTTGGATCAGGACATCCCGATGGTGCAGCCGCTATT | 16.5 | 7.3 | 3.2 | 2.3 | 2.2E-02 | 2.8E-02 | MB231/MB468 | MP |
| >trnaMT_GluTTC_MT_-_14674_14742@45.71.27,trnalookalike8_GluTTC_5_-_93905172_93905240@45.71.27\|CGTGGTTGTAGTCCGTGCGAGAATACC | >trnaMT_GluTTC_MT_-_14674_14742 | 45 | 71 | 27 | CGTGGTTGTAGTCCGTGCGAGAATACC | 97.6 | 1.9 | 0.8 | 2.3 | 2.2E-02 | 2.8E-02 | MB231/MB468 | MP |
| >MI0033423\|hsa-mir-10399&WithFlank&7\|-\|139044094\|139044163@43.63.21(+1U)&[MIMAT0041630&hsa-miR-10399-3p&offsets\|-1\|-1(+1U);TJU_CMC_MD2.ID03061.3p-miR&offsets\|+3\|+2(+1U);m-2657&7\|-\|139044103\|139044124&offsets\|+3\|+2(+1U)]\|TCTCTCGGACAAGCTGTAGGTT | >MI0033423\|hsa-mir-10399&WithFlank&7\|-\|139044094\|139044163 | 43 | 63 | 21(+1U) | TCTCTCGGACAAGCTGTAGGTT | 232.2 | -3.3 | 1.2 | -2.7 | 7.2E-03 | 2.8E-02 | BT20/MB231 | NUC |
| >hg38_21_+_8433172_8446622_RNA45SN1_45S_50ntFlanks@5421.5444.24\|CGATTGGATGGTTTAGTGAGGCCC | >hg38_21_+_8433172_8446622_RNA45SN1_45S_50ntFlanks | 5421 | 5444 | 24 | CGATTGGATGGTTTAGTGAGGCCC | 128.0 | -4.2 | 1.7 | -2.4 | 1.6E-02 | 2.8E-02 | BT20/MB468 | MP |
| >hg38_21_+_8433172_8446622_RNA45SN1_45S_50ntFlanks@4208.4242.35\|GGGGAGGTAGTGACGAAAAATAACAATACAGGACT | >hg38_21_+_8433172_8446622_RNA45SN1_45S_50ntFlanks | 4208 | 4242 | 35 | GGGGAGGTAGTGACGAAAAATAACAATACAGGACT | 576.8 | 3.9 | 1.4 | 2.7 | 6.6E-03 | 2.8E-02 | BT20/MB231 | MP |
| >MI0000289\|hsa-mir-181a-1&WithFlank&1\|-\|198859038\|198859159@30.50.21;MI0000269\|hsa-mir-181a-2&WithFlank&9\|+\|124692436\|124692557@45.65.21&[MIMAT0000256&hsa-miR-181a-5p&offsets\|0\|-2;m-32&1\|-\|198859108\|198859130&offsets\|0\|-2];[MIMAT0000256_1&hsa-miR-181a-5p&offsets\|0\|-2;m-32&9\|+\|124692480\|124692502&offsets\|0\|-2]\|AACATTCAACGCTGTCGGTGA | >MI0000289\|hsa-mir-181a-1&WithFlank&1\|-\|198859038\|198859159 | 30 | 50 | 21 | AACATTCAACGCTGTCGGTGA | 542.8 | -2.0 | 0.7 | -2.6 | 8.2E-03 | 2.8E-02 | BT20/MB231 | MITO |
| >hg38_MT_+_1621_3279_RNR2_16S_50ntFlanks@1348.1383.36\|AACAATAGGGTTTACGACCTCGATGTTGGATCAGGA | >hg38_MT_+_1621_3279_RNR2_16S_50ntFlanks | 1348 | 1383 | 36 | AACAATAGGGTTTACGACCTCGATGTTGGATCAGGA | 71.2 | 4.5 | 2.0 | 2.3 | 2.2E-02 | 2.8E-02 | MB231/MB468 | MP |
| >MI0000478\|hsa-mir-149&WithFlank&2\|+\|240455995\|240456095@21.41.21&[MIMAT0000450&hsa-miR-149-5p&offsets\|0\|-2;m-195&2\|+\|240456015\|240456037&offsets\|0\|-2]\|TCTGGCTCCGTGTCTTCACTC | >MI0000478\|hsa-mir-149&WithFlank&2\|+\|240455995\|240456095 | 21 | 41 | 21 | TCTGGCTCCGTGTCTTCACTC | 408.7 | 1.8 | 0.7 | 2.6 | 8.2E-03 | 2.8E-02 | BT20/MB231 | MITO |
| >MI0000087\|hsa-mir-29a&WithFlank&7\|-\|130876741\|130876816@10.33.24&[MIMAT0004503&hsa-miR-29a-5p&offsets\|0\|+2;m-315&7\|-\|130876785\|130876807&offsets\|0\|+1]\|ACTGATTTCTTTTGGTGTTCAGAG | >MI0000087\|hsa-mir-29a&WithFlank&7\|-\|130876741\|130876816 | 10 | 33 | 24 | ACTGATTTCTTTTGGTGTTCAGAG | 134.0 | -5.1 | 1.9 | -2.7 | 6.7E-03 | 2.8E-02 | BT20/MB231 | MP |
| >hg38_1_-_228634819_228635039_RNA5S12_5S_50ntFlanks@128.169.42\|CTTGGATGGGAGACCGCCTGGGAATACCGGGTGCTGTAGGCT | >hg38_1_-_228634819_228635039_RNA5S12_5S_50ntFlanks | 128 | 169 | 42 | CTTGGATGGGAGACCGCCTGGGAATACCGGGTGCTGTAGGCT | 330.1 | 2.9 | 1.1 | 2.7 | 6.7E-03 | 2.8E-02 | BT20/MB231 | MP |
| >hg38_21_+_8433172_8446622_RNA45SN1_45S_50ntFlanks@6648.6674.27\|GTACGACTCTTAGCGGTGGATCACTCG | >hg38_21_+_8433172_8446622_RNA45SN1_45S_50ntFlanks | 6648 | 6674 | 27 | GTACGACTCTTAGCGGTGGATCACTCG | 433.6 | 3.6 | 1.3 | 2.7 | 6.7E-03 | 2.8E-02 | BT20/MB231 | MP |

<table>
<tr><td>>trna111_HisGTG_1_-_147774845_147774916@-1G.35.36,trna118_HisGTG_1_-_145396881_145396952@-1G.35.36,trna16_HisGTG_1_+_146544773_146544844@-1G.35.36,trna1_HisGTG_15_+_45493349_45493420@-1G.35.36,trna21_HisGTG_1_+_147753471_147753542@-1G.35.36,trna33_HisGTG_6_+_27125906_27125977@-1G.35.36,trna7_HisGTG_9_-_14433938_14434009@-1G.35.36,trna8_HisGTG_15_-_45492611_45492682@-1G.35.36,trna9_HisGTG_15_-_45490804_45490875@-1G.35.36|GGCCGTGATCGTATAGTGGTTAGTACTCTGCGTTGT</td><td>>trna111_HisGTG_1_-_147774845_147774916</td><td>-1G</td><td>35</td><td>36</td><td>GGCCGTGATCGTATAGTGGTTAGTACTCTGCGTTGT</td><td>1025.4</td><td>-3.4</td><td>1.3</td><td>-2.7</td><td>6.7E-03</td><td>2.8E-02</td><td>BT20/MB231</td><td>MP</td></tr>
<tr><td>>hg38_MT_+_598_1651_RNR1_12S_50ntFlanks@52.87.36|ATAGGTTTGGTCCTAGCCTTTCTATTAGCTCTTAGT</td><td>>hg38_MT_+_598_1651_RNR1_12S_50ntFlanks</td><td>52</td><td>87</td><td>36</td><td>ATAGGTTTGGTCCTAGCCTTTCTATTAGCTCTTAGT</td><td>1849.3</td><td>1.8</td><td>0.8</td><td>2.2</td><td>2.5E-02</td><td>2.8E-02</td><td>MB231/MB468</td><td>MITO</td></tr>
<tr><td>>MI0000683|hsa-mir-181b-2&WithFlank&9|+|124693704|124693804@22.46.25(+1U)&[MIMAT0000257_1&hsa-miR-181b-5p&offsets|0|+2(+1U);m-72&9|+|124693725|124693747&offsets|0|+2(+1U)]|AACATTCATTGCTGTCGGTGGGTTTT</td><td>>MI0000683|hsa-mir-181b-2&WithFlank&9|+|124693704|124693804</td><td>22</td><td>46</td><td>25(+1U)</td><td>AACATTCATTGCTGTCGGTGGGTTTT</td><td>264.4</td><td>-2.3</td><td>0.9</td><td>-2.7</td><td>7.2E-03</td><td>2.8E-02</td><td>BT20/MB231</td><td>NUC</td></tr>
<tr><td>>hg38_21_+_8433172_8446622_RNA45SN1_45S_50ntFlanks@12444.12476.33|AAGCAGAATTCACCAAGCGTTGGATTGTTCACC</td><td>>hg38_21_+_8433172_8446622_RNA45SN1_45S_50ntFlanks</td><td>12444</td><td>12476</td><td>33</td><td>AAGCAGAATTCACCAAGCGTTGGATTGTTCACC</td><td>283.9</td><td>-1.9</td><td>0.8</td><td>-2.3</td><td>2.1E-02</td><td>2.8E-02</td><td>BT20/MB468</td><td>MITO</td></tr>
<tr><td>>hg38_21_+_8433172_8446622_RNA45SN1_45S_50ntFlanks@10349.10369.21|AACGAGAACTTTGAAGGCCGA</td><td>>hg38_21_+_8433172_8446622_RNA45SN1_45S_50ntFlanks</td><td>10349</td><td>10369</td><td>21</td><td>AACGAGAACTTTGAAGGCCGA</td><td>389.0</td><td>1.5</td><td>0.6</td><td>2.3</td><td>2.4E-02</td><td>2.8E-02</td><td>BT20/MB468</td><td>CYTO</td></tr>
<tr><td>>MI0000746|hsa-mir-99b&WithFlank&19|+|51692606|51692687@51.73.23(+1U)&[MIMAT0004678&hsa-miR-99b-3p&offsets|0|+1(+1U);m-151&19|+|51692656|51692678&offsets|0|0(+1U)]|CAAGCTCGTGTCTGTGGGTCCGTT</td><td>>MI0000746|hsa-mir-99b&WithFlank&19|+|51692606|51692687</td><td>51</td><td>73</td><td>23(+1U)</td><td>CAAGCTCGTGTCTGTGGGTCCGTT</td><td>496.2</td><td>2.0</td><td>0.8</td><td>2.6</td><td>9.1E-03</td><td>2.8E-02</td><td>BT20/MB231</td><td>CYTO</td></tr>
<tr><td>>hg38_21_+_8433172_8446622_RNA45SN1_45S_50ntFlanks@11322.11343.22|ACGTCTCGTCGCGCGCGCGTCC</td><td>>hg38_21_+_8433172_8446622_RNA45SN1_45S_50ntFlanks</td><td>11322</td><td>11343</td><td>22</td><td>ACGTCTCGTCGCGCGCGCGTCC</td><td>3705.5</td><td>-2.3</td><td>1.0</td><td>-2.2</td><td>2.5E-02</td><td>2.8E-02</td><td>MB231/MB468</td><td>NUC</td></tr>
<tr><td>>hg38_21_+_8433172_8446622_RNA45SN1_45S_50ntFlanks@12616.12657.42|CTGAGGAGCCAATGGGGCGAAGCTACCATCTGTGGGATTATG</td><td>>hg38_21_+_8433172_8446622_RNA45SN1_45S_50ntFlanks</td><td>12616</td><td>12657</td><td>42</td><td>CTGAGGAGCCAATGGGGCGAAGCTACCATCTGTGGGATTATG</td><td>359.3</td><td>-2.0</td><td>0.9</td><td>-2.3</td><td>2.1E-02</td><td>2.8E-02</td><td>BT20/MB468</td><td>MITO</td></tr>
<tr><td>>hg38_MT_+_598_1651_RNR1_12S_50ntFlanks@904.929.26|AAGTATACTTCAAAGGACATTTAACT</td><td>>hg38_MT_+_598_1651_RNR1_12S_50ntFlanks</td><td>904</td><td>929</td><td>26</td><td>AAGTATACTTCAAAGGACATTTAACT</td><td>674.2</td><td>-3.4</td><td>1.3</td><td>-2.7</td><td>6.7E-03</td><td>2.8E-02</td><td>BT20/MB231</td><td>MP</td></tr>
<tr><td>>hg38_21_+_8433172_8446622_RNA45SN1_45S_50ntFlanks@10346.10382.37|TCAAACGAGAACTTTGAAGGCCGAAGTGGAGAAGGGT</td><td>>hg38_21_+_8433172_8446622_RNA45SN1_45S_50ntFlanks</td><td>10346</td><td>10382</td><td>37</td><td>TCAAACGAGAACTTTGAAGGCCGAAGTGGAGAAGGGT</td><td>256.6</td><td>-1.9</td><td>0.9</td><td>-2.2</td><td>2.5E-02</td><td>2.8E-02</td><td>MB231/MB468</td><td>MITO</td></tr>
<tr><td>>trnaMT_MetCAT_MT_+_4402_4469@37.71.35|CCGAAAATGTTGGTTATACCCTTCCCGTACTACCA</td><td>>trnaMT_MetCAT_MT_+_4402_4469</td><td>37</td><td>71</td><td>35</td><td>CCGAAAATGTTGGTTATACCCTTCCCGTACTACCA</td><td>856.8</td><td>-2.2</td><td>0.8</td><td>-2.6</td><td>9.1E-03</td><td>2.8E-02</td><td>BT20/MB231</td><td>CYTO</td></tr>
<tr><td>>hg38_21_+_8433172_8446622_RNA45SN1_45S_50ntFlanks@6661.6696.36|CGGTGGATCACTCGGCTCGTGCGTCGATGAAGAACG</td><td>>hg38_21_+_8433172_8446622_RNA45SN1_45S_50ntFlanks</td><td>6661</td><td>6696</td><td>36</td><td>CGGTGGATCACTCGGCTCGTGCGTCGATGAAGAACG</td><td>1324.9</td><td>2.6</td><td>1.0</td><td>2.7</td><td>6.7E-03</td><td>2.8E-02</td><td>BT20/MB231</td><td>MP</td></tr>
<tr><td>>MI0000264|hsa-mir-7-2&WithFlank&15|+|88611819|88611940@38.57.20;MI0000265|hsa-mir-7-3&WithFlank&19|+|4770664|4770785@37.56.20;MI0000263|hsa-mir-7-1&WithFlank&9|-|83969742|83969863@30.49.20&[MIMAT0000252&hsa-miR-7-5p&offsets|0|-4;m-400&15|+|88611856|88611879&offsets|0|-4];[MIMAT0000252_1&hsa-miR-7-5p&offsets|0|-4;m-402&19|+|4770700|4770723&offsets|0|-4];[MIMAT0000252_2&hsa-miR-7-5p&offsets|0|-4;m-401&9|-|83969811|83969834&offsets|0|-4]|TGGAAGACTAGTGATTTTGT</td><td>>MI0000264|hsa-mir-7-2&WithFlank&15|+|88611819|88611940</td><td>38</td><td>57</td><td>20</td><td>TGGAAGACTAGTGATTTTGT</td><td>104.6</td><td>3.7</td><td>1.5</td><td>2.4</td><td>1.6E-02</td><td>2.8E-02</td><td>BT20/MB468</td><td>MP</td></tr>
<tr><td>>hg38_MT_+_1621_3279_RNR2_16S_50ntFlanks@1203.1223.21|TCGGAGCAGAACCCAACCTCC</td><td>>hg38_MT_+_1621_3279_RNR2_16S_50ntFlanks</td><td>1203</td><td>1223</td><td>21</td><td>TCGGAGCAGAACCCAACCTCC</td><td>82.8</td><td>5.3</td><td>2.2</td><td>2.4</td><td>1.6E-02</td><td>2.8E-02</td><td>BT20/MB468</td><td>MP</td></tr>
<tr><td>>am_trnaMT_SerGCT_MT_+_12207_12265@15.40.26|GAACTGCTAACTCATGCCCCCATGTC</td><td>>am_trnaMT_SerGCT_MT_+_12207_12265</td><td>15</td><td>40</td><td>26</td><td>GAACTGCTAACTCATGCCCCCATGTC</td><td>113.3</td><td>3.9</td><td>1.6</td><td>2.4</td><td>1.6E-02</td><td>2.8E-02</td><td>BT20/MB468</td><td>MP</td></tr>
<tr><td>>MI0000063|hsa-let-7b&WithFlank&22|+|46113680|46113774@12.32.21(+1G)&[MIMAT0000063&hsa-let-7b-5p&offsets|0|-1(+1G);m-14&22|+|46113691|46113712&offsets|0|-1(+1G)]|TGAGGTAGTAGGTTGTGTGGTG</td><td>>MI0000063|hsa-let-7b&WithFlank&22|+|46113680|46113774</td><td>12</td><td>32</td><td>21(+1G)</td><td>TGAGGTAGTAGGTTGTGTGGTG</td><td>296.4</td><td>2.2</td><td>1.0</td><td>2.3</td><td>2.2E-02</td><td>2.8E-02</td><td>MB231/MB468</td><td>MP</td></tr>
<tr><td>>hg38_1_-_228634819_228635039_RNA5S12_5S_50ntFlanks@153.170.18|ACCGGGTGCTGTAGGCTT</td><td>>hg38_1_-_228634819_228635039_RNA5S12_5S_50ntFlanks</td><td>153</td><td>170</td><td>18</td><td>ACCGGGTGCTGTAGGCTT</td><td>23624.3</td><td>2.6</td><td>1.0</td><td>2.6</td><td>8.2E-03</td><td>2.8E-02</td><td>BT20/MB231</td><td>MITO</td></tr>
<tr><td>>hg38_21_+_8433172_8446622_RNA45SN1_45S_50ntFlanks@9565.9596.32|TGACGTGCAAATCGGTCGTCCGACCTGGGTAT</td><td>>hg38_21_+_8433172_8446622_RNA45SN1_45S_50ntFlanks</td><td>9565</td><td>9596</td><td>32</td><td>TGACGTGCAAATCGGTCGTCCGACCTGGGTAT</td><td>229.3</td><td>-2.4</td><td>0.9</td><td>-2.6</td><td>9.1E-03</td><td>2.8E-02</td><td>BT20/MB231</td><td>CYTO</td></tr>
<tr><td>>trna10_AspGTC_12_-_125424193_125424264@38.74.37,trna12_AspGTC_12_-_125411891_125411962@38.74.37,trna144_AspGTC_6_-_27551236_27551307@38.74.37,trna38_AspGTC_17_-_8125556_8125627@38.74.37,trna45_AspGTC_6_+_27447453_27447524@38.74.37,trna48_AspGTC_6_+_27471523_27471594@38.74.37,trna4_AspGTC_12_+_96429799_96429870@38.74.37,trna69_AspGTC_1_-_161440205_161440276@38.74.37,trna72_AspGTC_1_-_161432824_161432895@38.74.37,trna75_AspGTC_1_-_161425414_161425485@38.74.37,trna78_AspGTC_1_-_161418033_161418104@38.74.37,trna81_AspGTC_1_-_161410615_161410686@38.74.37|CGCGGGAGACCGGGGTTCGATTCCCCGACGGGGAGCC</td><td>>trna10_AspGTC_12_-_125424193_125424264</td><td>38</td><td>74</td><td>37</td><td>CGCGGGAGACCGGGGTTCGATTCCCCGACGGGGAGCC</td><td>47.3</td><td>3.1</td><td>1.3</td><td>2.3</td><td>2.2E-02</td><td>2.8E-02</td><td>MB231/MB468</td><td>MP</td></tr>
<tr><td>>MI0000298|hsa-mir-221&WithFlank&X|-|45746151|45746272@71.93.23(+2U)&[MIMAT0000278&hsa-miR-221-3p&offsets|0|0(+2U);m-91&X|-|45746181|45746202&offsets|0|+1(+2U)]|AGCTACATTGTCTGCTGGGTTTCTT</td><td>>MI0000298|hsa-mir-221&WithFlank&X|-|45746151|45746272</td><td>71</td><td>93</td><td>23(+2U)</td><td>AGCTACATTGTCTGCTGGGTTTCTT</td><td>43.7</td><td>4.6</td><td>2.0</td><td>2.3</td><td>2.2E-02</td><td>2.8E-02</td><td>MB231/MB468</td><td>MP</td></tr>
<tr><td>>hg38_21_+_8433172_8446622_RNA45SN1_45S_50ntFlanks@6651.6681.31|CGACTCTTAGCGGTGGATCACTCGGCTCGTG</td><td>>hg38_21_+_8433172_8446622_RNA45SN1_45S_50ntFlanks</td><td>6651</td><td>6681</td><td>31</td><td>CGACTCTTAGCGGTGGATCACTCGGCTCGTG</td><td>17527.8</td><td>-1.3</td><td>0.6</td><td>-2.3</td><td>2.4E-02</td><td>2.8E-02</td><td>MB231/MB468</td><td>CYTO</td></tr>
<tr><td>>hg38_21_+_8433172_8446622_RNA45SN1_45S_50ntFlanks@6651.6673.23|CGACTCTTAGCGGTGGATCACTC</td><td>>hg38_21_+_8433172_8446622_RNA45SN1_45S_50ntFlanks</td><td>6651</td><td>6673</td><td>23</td><td>CGACTCTTAGCGGTGGATCACTC</td><td>1122.7</td><td>1.6</td><td>0.7</td><td>2.2</td><td>2.5E-02</td><td>2.8E-02</td><td>BT20/MB468</td><td>NUC</td></tr>
<tr><td>>hg38_21_+_8433172_8446622_RNA45SN1_45S_50ntFlanks@6646.6664.19|TCGTACGACTCTTAGCGGT</td><td>>hg38_21_+_8433172_8446622_RNA45SN1_45S_50ntFlanks</td><td>6646</td><td>6664</td><td>19</td><td>TCGTACGACTCTTAGCGGT</td><td>52631.9</td><td>2.9</td><td>1.1</td><td>2.7</td><td>6.7E-03</td><td>2.8E-02</td><td>BT20/MB231</td><td>MP</td></tr>
<tr><td>>hg38_21_+_8433172_8446622_RNA45SN1_45S_50ntFlanks@8338.8373.36|GTACCGTAAGGGAAAGTTGAAAAGAACTTTGAAGAG</td><td>>hg38_21_+_8433172_8446622_RNA45SN1_45S_50ntFlanks</td><td>8338</td><td>8373</td><td>36</td><td>GTACCGTAAGGGAAAGTTGAAAAGAACTTTGAAGAG</td><td>590.7</td><td>4.8</td><td>1.8</td><td>2.7</td><td>6.7E-03</td><td>2.8E-02</td><td>BT20/MB231</td><td>MP</td></tr>
<tr><td>>hg38_21_+_8433172_8446622_RNA45SN1_45S_50ntFlanks@4100.4121.22|TGACCACGGGTGACGGGGAATC</td><td>>hg38_21_+_8433172_8446622_RNA45SN1_45S_50ntFlanks</td><td>4100</td><td>4121</td><td>22</td><td>TGACCACGGGTGACGGGGAATC</td><td>128.0</td><td>-4.0</td><td>1.7</td><td>-2.4</td><td>1.6E-02</td><td>2.8E-02</td><td>BT20/MB468</td><td>MP</td></tr>
<tr><td>>hg38_21_+_8433172_8446622_RNA45SN1_45S_50ntFlanks@12410.12434.25|TGATCCTTCGATGTCGGCTCTTCCT</td><td>>hg38_21_+_8433172_8446622_RNA45SN1_45S_50ntFlanks</td><td>12410</td><td>12434</td><td>25</td><td>TGATCCTTCGATGTCGGCTCTTCCT</td><td>314.8</td><td>1.5</td><td>0.7</td><td>2.2</td><td>2.5E-02</td><td>2.8E-02</td><td>MB231/MB468</td><td>NUC</td></tr>
<tr><td>>hg38_21_+_8433172_8446622_RNA45SN1_45S_50ntFlanks@10342.10362.21|ATATTCAAACGAGAACTTTGA</td><td>>hg38_21_+_8433172_8446622_RNA45SN1_45S_50ntFlanks</td><td>10342</td><td>10362</td><td>21</td><td>ATATTCAAACGAGAACTTTGA</td><td>171.9</td><td>4.4</td><td>1.6</td><td>2.7</td><td>6.7E-03</td><td>2.8E-02</td><td>BT20/MB231</td><td>MP</td></tr>
<tr><td>>hg38_21_+_8433172_8446622_RNA45SN1_45S_50ntFlanks@6699.6720.22|GCTAGCTGCGAGAATTAATGTG</td><td>>hg38_21_+_8433172_8446622_RNA45SN1_45S_50ntFlanks</td><td>6699</td><td>6720</td><td>22</td><td>GCTAGCTGCGAGAATTAATGTG</td><td>10482.9</td><td>4.6</td><td>1.7</td><td>2.7</td><td>6.7E-03</td><td>2.8E-02</td><td>BT20/MB231</td><td>MP</td></tr>
<tr><td>>hg38_21_+_8433172_8446622_RNA45SN1_45S_50ntFlanks@7975.7998.24|CGCGACCTCAGATCAGACGTGGCG</td><td>>hg38_21_+_8433172_8446622_RNA45SN1_45S_50ntFlanks</td><td>7975</td><td>7998</td><td>24</td><td>CGCGACCTCAGATCAGACGTGGCG</td><td>424.1</td><td>3.3</td><td>1.2</td><td>2.7</td><td>6.7E-03</td><td>2.8E-02</td><td>BT20/MB231</td><td>MP</td></tr>
<tr><td>>hg38_21_+_8433172_8446622_RNA45SN1_45S_50ntFlanks@9567.9605.39|ACGTGCAAATCGGTCGTCCGACCTGGGTATAGGGGCGAA</td><td>>hg38_21_+_8433172_8446622_RNA45SN1_45S_50ntFlanks</td><td>9567</td><td>9605</td><td>39</td><td>ACGTGCAAATCGGTCGTCCGACCTGGGTATAGGGGCGAA</td><td>57.5</td><td>-4.1</td><td>1.8</td><td>-2.3</td><td>2.2E-02</td><td>2.8E-02</td><td>MB231/MB468</td><td>MP</td></tr>
<tr><td>>hg38_21_+_8433172_8446622_RNA45SN1_45S_50ntFlanks@6661.6694.34|CGGTGGATCACTCGGCTCGTGCGTCGATGAAGAA</td><td>>hg38_21_+_8433172_8446622_RNA45SN1_45S_50ntFlanks</td><td>6661</td><td>6694</td><td>34</td><td>CGGTGGATCACTCGGCTCGTGCGTCGATGAAGAA</td><td>260.9</td><td>1.7</td><td>0.7</td><td>2.4</td><td>1.6E-02</td><td>2.8E-02</td><td>BT20/MB468</td><td>MP</td></tr>
<tr><td>>hg38_21_+_8433172_8446622_RNA45SN1_45S_50ntFlanks@8245.8269.25|CTTCCCGGAGTCGGGTTGCTTGGGA</td><td>>hg38_21_+_8433172_8446622_RNA45SN1_45S_50ntFlanks</td><td>8245</td><td>8269</td><td>25</td><td>CTTCCCGGAGTCGGGTTGCTTGGGA</td><td>173.4</td><td>4.2</td><td>1.5</td><td>2.7</td><td>6.7E-03</td><td>2.8E-02</td><td>BT20/MB231</td><td>MP</td></tr>
<tr><td>>hg38_21_+_8433172_8446622_RNA45SN1_45S_50ntFlanks@9454.9498.45|CCGCGCCGGGGAGGTGGAGCACGAGCGCACGTGTTAGGACCCGAA</td><td>>hg38_21_+_8433172_8446622_RNA45SN1_45S_50ntFlanks</td><td>9454</td><td>9498</td><td>45</td><td>CCGCGCCGGGGAGGTGGAGCACGAGCGCACGTGTTAGGACCCGAA</td><td>29.3</td><td>-8.0</td><td>3.3</td><td>-2.4</td><td>1.6E-02</td><td>2.8E-02</td><td>BT20/MB468</td><td>MP</td></tr>
<tr><td>>hg38_21_+_8433172_8446622_RNA45SN1_45S_50ntFlanks@9534.9571.38|AGGAAACTCTGGTGGAGGTCCGTAGCGGTCCTGACGTG</td><td>>hg38_21_+_8433172_8446622_RNA45SN1_45S_50ntFlanks</td><td>9534</td><td>9571</td><td>38</td><td>AGGAAACTCTGGTGGAGGTCCGTAGCGGTCCTGACGTG</td><td>1179.1</td><td>5.1</td><td>1.9</td><td>2.7</td><td>6.7E-03</td><td>2.8E-02</td><td>BT20/MB231</td><td>MP</td></tr>
<tr><td>>MI0000542|hsa-mir-320a&WithFlank&8|-|22244960|22245043@48.71.24(+1U)&[MIMAT0000510&hsa-miR-320a-3p&offsets|0|+2(+1U);m-60&8|-|22244975|22244996&offsets|0|+2(+1U)]|AAAAGCTGGGTTGAGAGGGCGAAAT</td><td>>MI0000542|hsa-mir-320a&WithFlank&8|-|22244960|22245043</td><td>48</td><td>71</td><td>24(+1U)</td><td>AAAAGCTGGGTTGAGAGGGCGAAAT</td><td>519.6</td><td>2.0</td><td>0.8</td><td>2.7</td><td>7.3E-03</td><td>2.8E-02</td><td>BT20/MB231</td><td>NUC</td></tr>
<tr><td>>am_trna116_GluCTC_1_-_145399233_145399304@1.28.28,trna59_GluCTC_1_+_249168447_249168518@1.28.28,trna71_GluCTC_1_-_161439189_161439260@1.28.28,trna74_GluCTC_1_-_161431809_161431880@1.28.28,trna77_GluCTC_1_-_161424398_161424469@1.28.28,trna77_GluCTC_6_+_28949976_28950047@1.28.28,trna80_GluCTC_1_-_161417018_161417089@1.28.28,trna87_GluCTC_6_-_126101393_126101464@1.28.28|TCCCTGGTGGTCTAGTGGTTAGGATTCG</td><td>>am_trna116_GluCTC_1_-_145399233_145399304</td><td>1</td><td>28</td><td>28</td><td>TCCCTGGTGGTCTAGTGGTTAGGATTCG</td><td>3795.8</td><td>4.0</td><td>1.5</td><td>2.7</td><td>6.7E-03</td><td>2.8E-02</td><td>BT20/MB231</td><td>MP</td></tr>
<tr><td>>hg38_21_+_8433172_8446622_RNA45SN1_45S_50ntFlanks@10240.10261.22|CTACGCCGCGACGAGTAGGAGG</td><td>>hg38_21_+_8433172_8446622_RNA45SN1_45S_50ntFlanks</td><td>10240</td><td>10261</td><td>22</td><td>CTACGCCGCGACGAGTAGGAGG</td><td>199.8</td><td>5.2</td><td>1.9</td><td>2.7</td><td>6.7E-03</td><td>2.8E-02</td><td>BT20/MB231</td><td>MP</td></tr>
<tr><td>>hg38_21_+_8433172_8446622_RNA45SN1_45S_50ntFlanks@8350.8375.26|AAAGTTGAAAAGAACTTTGAAGAGAG</td><td>>hg38_21_+_8433172_8446622_RNA45SN1_45S_50ntFlanks</td><td>8350</td><td>8375</td><td>26</td><td>AAAGTTGAAAAGAACTTTGAAGAGAG</td><td>5073.0</td><td>4.8</td><td>1.8</td><td>2.7</td><td>6.7E-03</td><td>2.8E-02</td><td>BT20/MB231</td><td>MP</td></tr>
<tr><td>>hg38_21_+_8433172_8446622_RNA45SN1_45S_50ntFlanks@7988.8015.28|CAGACGTGGCGACCCGCTGAATTTAAGC</td><td>>hg38_21_+_8433172_8446622_RNA45SN1_45S_50ntFlanks</td><td>7988</td><td>8015</td><td>28</td><td>CAGACGTGGCGACCCGCTGAATTTAAGC</td><td>250.0</td><td>-1.9</td><td>0.8</td><td>-2.4</td><td>1.6E-02</td><td>2.8E-02</td><td>BT20/MB468</td><td>MP</td></tr>
</table>

| | | | | | | | | | | | | | |
|---|---|---|---|---|---|---|---|---|---|---|---|---|---|
| >MI0000088\|hsa-mir-30a&WithFlank&6\|-\|71403545\|71403627@53.73.21&[MIMAT0000088&hsa-miR-30a-3p&offsets\|0\|-1;m-38&6\|-\|71403554\|71403575&offsets\|0\|-1]\|CTTTCAGTCGGATGTTTGCAG | >MI0000088\|hsa-mir-30a&WithFlank&6\|-\|71403545\|71403627 | 53 | 73 | 21 | CTTTCAGTCGGATGTTTGCAG | 427.4 | 2.3 | 0.9 | 2.4 | 1.6E-02 | 2.8E-02 | BT20/MB468 | MP |
| >hg38_MT_+_598_1651_RNR1_12S_50ntFlanks@844.868.25\|AGTAGAGTGCTTAGTTGAACAGGGC | >hg38_MT_+_598_1651_RNR1_12S_50ntFlanks | 844 | 868 | 25 | AGTAGAGTGCTTAGTTGAACAGGGC | 67.4 | 3.5 | 1.5 | 2.4 | 1.6E-02 | 2.8E-02 | BT20/MB468 | MP |
| >am_MI0000786\|hsa-mir-378a&WithFlank&5\|+\|149732819\|149732896@50.69.20(+1U)&[MIMAT0000732&hsa-miR-378a-3p&offsets\|+1\|-1(+1U);m-44&5\|+\|149732867\|149732888&offsets\|+1\|-1(+1U)]\|CTGGACTTGGAGTCAGAAGGT | >am_MI0000786\|hsa-mir-378a&WithFlank&5\|+\|149732819\|149732896 | 50 | 69 | 20(+1U) | CTGGACTTGGAGTCAGAAGGT | 373.5 | 1.5 | 0.5 | 2.7 | 7.3E-03 | 2.8E-02 | BT20/MB231 | NUC |
| >hg38_21_+_8433172_8446622_RNA45SN1_45S_50ntFlanks@10685.10709.25\|GGCGGCGTCCGGTGAGCTCTCGCTG | >hg38_21_+_8433172_8446622_RNA45SN1_45S_50ntFlanks | 10685 | 10709 | 25 | GGCGGCGTCCGGTGAGCTCTCGCTG | 540.1 | 2.7 | 1.0 | 2.6 | 8.3E-03 | 2.8E-02 | BT20/MB231 | MITO |
| >hg38_21_+_8433172_8446622_RNA45SN1_45S_50ntFlanks@12912.12935.24\|CACGTTCGTGGGGAACCTGGCGCT | >hg38_21_+_8433172_8446622_RNA45SN1_45S_50ntFlanks | 12912 | 12935 | 24 | CACGTTCGTGGGGAACCTGGCGCT | 395.1 | -3.5 | 1.5 | -2.4 | 1.6E-02 | 2.8E-02 | BT20/MB468 | MP |
| >hg38_21_+_8433172_8446622_RNA45SN1_45S_50ntFlanks@5478.5499.22\|CTGGCGGAGCGCTGAGAAGACG | >hg38_21_+_8433172_8446622_RNA45SN1_45S_50ntFlanks | 5478 | 5499 | 22 | CTGGCGGAGCGCTGAGAAGACG | 310.1 | 3.8 | 1.4 | 2.7 | 6.8E-03 | 2.8E-02 | BT20/MB231 | MP |
| >MI0000298\|hsa-mir-221&WithFlank&X\|-\|45746151\|45746272@71.90.20(+1G)&[MIMAT0000278&hsa-miR-221-3p&offsets\|0\|-3(+1G);m-91&X\|-\|45746181\|45746202&offsets\|0\|-2(+1G)]\|AGCTACATTGTCTGCTGGGTG | >MI0000298\|hsa-mir-221&WithFlank&X\|-\|45746151\|45746272 | 71 | 90 | 20(+1G) | AGCTACATTGTCTGCTGGGTG | 58.6 | 3.9 | 1.7 | 2.3 | 2.3E-02 | 2.8E-02 | MB231/MB468 | MP |
| >hg38_21_+_8433172_8446622_RNA45SN1_45S_50ntFlanks@11568.11586.19\|GCAGCCGACTTAGAACTGG | >hg38_21_+_8433172_8446622_RNA45SN1_45S_50ntFlanks | 11568 | 11586 | 19 | GCAGCCGACTTAGAACTGG | 1137.7 | -1.8 | 0.8 | -2.4 | 1.6E-02 | 2.8E-02 | BT20/MB468 | MP |
| >hg38_21_+_8433172_8446622_RNA45SN1_45S_50ntFlanks@11566.11586.21\|TAGCAGCCGACTTAGAACTGG | >hg38_21_+_8433172_8446622_RNA45SN1_45S_50ntFlanks | 11566 | 11586 | 21 | TAGCAGCCGACTTAGAACTGG | 274.9 | -2.3 | 0.9 | -2.6 | 9.2E-03 | 2.8E-02 | BT20/MB231 | CYTO |
| >hg38_21_+_8433172_8446622_RNA45SN1_45S_50ntFlanks@8195.8212.18\|CCGTGGACGGTGTGAGGC | >hg38_21_+_8433172_8446622_RNA45SN1_45S_50ntFlanks | 8195 | 8212 | 18 | CCGTGGACGGTGTGAGGC | 1322.1 | 3.7 | 1.4 | 2.7 | 6.8E-03 | 2.8E-02 | BT20/MB231 | MP |
| >hg38_21_+_8433172_8446622_RNA45SN1_45S_50ntFlanks@12072.12108.37\|CCCGAGGGGCTCTCGCTTCTGGCGCCAAGCGCCCGGC | >hg38_21_+_8433172_8446622_RNA45SN1_45S_50ntFlanks | 12072 | 12108 | 37 | CCCGAGGGGCTCTCGCTTCTGGCGCCAAGCGCCCGGC | 120.7 | -3.5 | 1.5 | -2.3 | 2.1E-02 | 2.8E-02 | BT20/MB468 | MITO |
| >hg38_21_+_8433172_8446622_RNA45SN1_45S_50ntFlanks@7975.8003.29\|CGCGACCTCAGATCAGACGTGGCGACCCG | >hg38_21_+_8433172_8446622_RNA45SN1_45S_50ntFlanks | 7975 | 8003 | 29 | CGCGACCTCAGATCAGACGTGGCGACCCG | 4374.6 | 2.0 | 0.9 | 2.3 | 2.4E-02 | 2.8E-02 | BT20/MB468 | CYTO |
| >hg38_21_+_8433172_8446622_RNA45SN1_45S_50ntFlanks@8176.8194.19\|CTTCTGATCGAGGCCCAGC | >hg38_21_+_8433172_8446622_RNA45SN1_45S_50ntFlanks | 8176 | 8194 | 19 | CTTCTGATCGAGGCCCAGC | 335.5 | 1.2 | 0.5 | 2.3 | 2.4E-02 | 2.8E-02 | BT20/MB468 | CYTO |
| >hg38_21_+_8433172_8446622_RNA45SN1_45S_50ntFlanks@9570.9603.34\|TGCAAATCGGTCGTCCGACCTGGGTATAGGGGCG | >hg38_21_+_8433172_8446622_RNA45SN1_45S_50ntFlanks | 9570 | 9603 | 34 | TGCAAATCGGTCGTCCGACCTGGGTATAGGGGCG | 454.4 | -1.7 | 0.7 | -2.2 | 2.5E-02 | 2.8E-02 | MB231/MB468 | NUC |
| >hg38_MT_+_1621_3279_RNR2_16S_50ntFlanks@1092.1120.29\|GCAAGACGAGAAGACCCTATGGAGCTTTA | >hg38_MT_+_1621_3279_RNR2_16S_50ntFlanks | 1092 | 1120 | 29 | GCAAGACGAGAAGACCCTATGGAGCTTTA | 155.6 | 2.1 | 0.9 | 2.2 | 2.5E-02 | 2.8E-02 | MB231/MB468 | MITO |
| >hg38_21_+_8433172_8446622_RNA45SN1_45S_50ntFlanks@11808.11825.18\|CTGTCCCTACCTACTATC | >hg38_21_+_8433172_8446622_RNA45SN1_45S_50ntFlanks | 11808 | 11825 | 18 | CTGTCCCTACCTACTATC | 54.9 | -4.5 | 1.9 | -2.4 | 1.6E-02 | 2.8E-02 | BT20/MB468 | MP |
| >MI0000085\|hsa-mir-27a&WithFlank&19\|-\|13836434\|13836523@57.77.21&[MIMAT0000084&hsa-miR-27a-3p&offsets\|0\|0;m-43&19\|-\|13836447\|13836467&offsets\|0\|0]\|TTCACAGTGGCTAAGTTCCGC | >MI0000085\|hsa-mir-27a&WithFlank&19\|-\|13836434\|13836523 | 57 | 77 | 21 | TTCACAGTGGCTAAGTTCCGC | 19672.0 | -3.3 | 1.2 | -2.7 | 6.8E-03 | 2.8E-02 | BT20/MB231 | MP |
| >trna14_LysTTT_11_-_59327808_59327880@1.33.33,trna2_LysTTT_17_+_8022473_8022545@1.33.33,trna54_LysTTT_1_+_204475655_204475727@1.33.33,trna5_LysTTT_11_+_59323902_59323974@1.33.33,trna62_LysTTT_1_-_204476158_204476230@1.33.33,trna76_LysTTT_6_+_28918806_28918878@1.33.33\|GCCCGGATAGCTCAGTCGGTAGAGCATCAGACT | >trna14_LysTTT_11_-_59327808_59327880 | 1 | 33 | 33 | GCCCGGATAGCTCAGTCGGTAGAGCATCAGACT | 20656.0 | -3.5 | 1.3 | -2.7 | 6.8E-03 | 2.8E-02 | BT20/MB231 | MP |
| >hg38_21_+_8433172_8446622_RNA45SN1_45S_50ntFlanks@12259.12284.26\|CTTGATCTTGATTTTCAGTACGAATA | >hg38_21_+_8433172_8446622_RNA45SN1_45S_50ntFlanks | 12259 | 12284 | 26 | CTTGATCTTGATTTTCAGTACGAATA | 262.9 | 5.6 | 2.1 | 2.7 | 6.8E-03 | 2.8E-02 | BT20/MB231 | MP |
| >hg38_21_+_8433172_8446622_RNA45SN1_45S_50ntFlanks@5493.5516.24\|GAAGACGGTCGAACTTGACTATCT | >hg38_21_+_8433172_8446622_RNA45SN1_45S_50ntFlanks | 5493 | 5516 | 24 | GAAGACGGTCGAACTTGACTATCT | 1042.6 | 3.6 | 1.3 | 2.7 | 6.8E-03 | 2.8E-02 | BT20/MB231 | MP |
| >hg38_21_+_8433172_8446622_RNA45SN1_45S_50ntFlanks@6653.6678.26\|ACTCTTAGCGGTGGATCACTCGGCTC | >hg38_21_+_8433172_8446622_RNA45SN1_45S_50ntFlanks | 6653 | 6678 | 26 | ACTCTTAGCGGTGGATCACTCGGCTC | 172.6 | 4.5 | 1.7 | 2.7 | 6.8E-03 | 2.8E-02 | BT20/MB231 | MP |
| >MI0000093\|hsa-mir-92a-1&WithFlank&13\|+\|91351308\|91351397@54.75.22(+2A);MI0000094\|hsa-mir-92a-2&WithFlank&X\|-\|134169532\|134169618@54.75.22(+2A)&[MIMAT0000092&hsa-miR-92a-3p&offsets\|0\|0(+2A);m-9&13\|+\|91351361\|91351382&offsets\|0\|0(+2A)];[MIMAT0000092_1&hsa-miR-92a-3p&offsets\|0\|0(+2A);m-11&X\|-\|134169544\|134169565&offsets\|0\|0(+2A)]\|TATTGCACTTGTCCCGGCCTGTAA | >MI0000093\|hsa-mir-92a-1&WithFlank&13\|+\|91351308\|91351397 | 54 | 75 | 22(+2A) | TATTGCACTTGTCCCGGCCTGTAA | 188.0 | -2.8 | 1.1 | -2.7 | 6.8E-03 | 2.8E-02 | BT20/MB231 | MP |
| >hg38_21_+_8433172_8446622_RNA45SN1_45S_50ntFlanks@12407.12437.31\|TTTTGATCCTTCGATGTCGGCTCTTCCTATC | >hg38_21_+_8433172_8446622_RNA45SN1_45S_50ntFlanks | 12407 | 12437 | 31 | TTTTGATCCTTCGATGTCGGCTCTTCCTATC | 760.7 | -1.4 | 0.6 | -2.3 | 2.1E-02 | 2.8E-02 | BT20/MB468 | MITO |
| >hg38_MT_+_1621_3279_RNR2_16S_50ntFlanks@1508.1535.28\|AAAGGACAAGAGAAATAAGGCCTACTTC | >hg38_MT_+_1621_3279_RNR2_16S_50ntFlanks | 1508 | 1535 | 28 | AAAGGACAAGAGAAATAAGGCCTACTTC | 16.4 | 7.3 | 3.2 | 2.3 | 2.3E-02 | 2.8E-02 | MB231/MB468 | MP |
| >hg38_MT_+_598_1651_RNR1_12S_50ntFlanks@434.469.36\|GCTTTAACATATCTGAACACACAATAGCTAAGACCC | >hg38_MT_+_598_1651_RNR1_12S_50ntFlanks | 434 | 469 | 36 | GCTTTAACATATCTGAACACACAATAGCTAAGACCC | 16.4 | 7.3 | 3.2 | 2.3 | 2.3E-02 | 2.8E-02 | MB231/MB468 | MP |
| >trnaMT_GluTTC_MT_-_14674_14742@34.72.39,trnalookalike8_GluTTC_5_-_93905172_93905240@34.72.39\|ATATCATTGGTCGTGGTTGTAGTCCGTGCGAGAATACCA | >trnaMT_GluTTC_MT_-_14674_14742 | 34 | 72 | 39 | ATATCATTGGTCGTGGTTGTAGTCCGTGCGAGAATACCA | 3951.2 | 2.7 | 1.2 | 2.3 | 2.3E-02 | 2.8E-02 | MB231/MB468 | MP |
| >hg38_21_+_8433172_8446622_RNA45SN1_45S_50ntFlanks@11206.11232.27\|CGGCGGCGACTCTGGACGCGAGCCGGG | >hg38_21_+_8433172_8446622_RNA45SN1_45S_50ntFlanks | 11206 | 11232 | 27 | CGGCGGCGACTCTGGACGCGAGCCGGG | 764.8 | -2.4 | 1.0 | -2.4 | 1.6E-02 | 2.8E-02 | BT20/MB468 | MP |
| >hg38_21_+_8433172_8446622_RNA45SN1_45S_50ntFlanks@11568.11604.37\|GCAGCCGACTTAGAACTGGTGCGGACCAGGGGAATCC | >hg38_21_+_8433172_8446622_RNA45SN1_45S_50ntFlanks | 11568 | 11604 | 37 | GCAGCCGACTTAGAACTGGTGCGGACCAGGGGAATCC | 72.3 | 3.4 | 1.4 | 2.4 | 1.6E-02 | 2.8E-02 | BT20/MB468 | MP |
| >hg38_21_+_8433172_8446622_RNA45SN1_45S_50ntFlanks@4128.4158.31\|CGATTCCGGAGAGGGAGCCTGAGAAACGGCT | >hg38_21_+_8433172_8446622_RNA45SN1_45S_50ntFlanks | 4128 | 4158 | 31 | CGATTCCGGAGAGGGAGCCTGAGAAACGGCT | 135.4 | -4.1 | 1.7 | -2.4 | 1.6E-02 | 2.8E-02 | BT20/MB468 | MP |
| >MI0000085\|hsa-mir-27a&WithFlank&19\|-\|13836434\|13836523@57.74.18;MI0000440\|hsa-mir-27b&WithFlank&9\|+\|95085439\|95085547@67.84.18&[MIMAT0000084&hsa-miR-27a-3p&offsets\|0\|-3;m-43&19\|-\|13836447\|13836467&offsets\|0\|-3];[MIMAT0000419&hsa-miR-27b-3p&offsets\|0\|-3;m-39&9\|+\|95085505\|95085525&offsets\|0\|-3]\|TTCACAGTGGCTAAGTTC | >MI0000085\|hsa-mir-27a&WithFlank&19\|-\|13836434\|13836523 | 57 | 74 | 18 | TTCACAGTGGCTAAGTTC | 569.5 | 2.7 | 1.1 | 2.4 | 1.6E-02 | 2.8E-02 | BT20/MB468 | MP |
| >trnaMT_GluTTC_MT_-_14674_14742@34.71.38,trnalookalike8_GluTTC_5_-_93905172_93905240@34.71.38\|ATATCATTGGTCGTGGTTGTAGTCCGTGCGAGAATACC | >trnaMT_GluTTC_MT_-_14674_14742 | 34 | 71 | 38 | ATATCATTGGTCGTGGTTGTAGTCCGTGCGAGAATACC | 682.2 | 3.1 | 1.4 | 2.3 | 2.3E-02 | 2.8E-02 | MB231/MB468 | MP |
| >hg38_21_+_8433172_8446622_RNA45SN1_45S_50ntFlanks@6773.6806.34\|TTCCTCCCGGGGCTACGCCTGTCTGAGCGTCGCT | >hg38_21_+_8433172_8446622_RNA45SN1_45S_50ntFlanks | 6773 | 6806 | 34 | TTCCTCCCGGGGCTACGCCTGTCTGAGCGTCGCT | 4842.1 | -1.9 | 0.9 | -2.2 | 2.5E-02 | 2.8E-02 | MB231/MB468 | MITO |
| >am_MI0000809\|hsa-mir-151a&WithFlank&8\|-\|140732558\|140732659@52.73.22(+1U)&[MIMAT0000757&hsa-miR-151a-3p&offsets\|-1\|0(+1U);m-30&8\|-\|140732587\|140732607&offsets\|-1\|0(+1U)]\|ACTAGACTGAAGCTCCTTGAGGT | >am_MI0000809\|hsa-mir-151a&WithFlank&8\|-\|140732558\|140732659 | 52 | 73 | 22(+1U) | ACTAGACTGAAGCTCCTTGAGGT | 53.4 | 3.8 | 1.7 | 2.3 | 2.3E-02 | 2.8E-02 | MB231/MB468 | MP |
| >am_trnaMT_ValTAC_MT_+_1602_1670@48.67.20\|CAACTTAACTTGACCGCTCT | >am_trnaMT_ValTAC_MT_+_1602_1670 | 48 | 67 | 20 | CAACTTAACTTGACCGCTCT | 15376.5 | 2.3 | 0.9 | 2.6 | 9.2E-03 | 2.8E-02 | BT20/MB231 | CYTO |
| >trna10_AspGTC_12_-_125424193_125424264@36.75.40,trna12_AspGTC_12_-_125411891_125411962@36.75.40,trna144_AspGTC_6_-_27551236_27551307@36.75.40,trna38_AspGTC_17_-_8125556_8125627@36.75.40,trna45_AspGTC_6_+_27447453_27447524@36.75.40,trna48_AspGTC_6_+_27471523_27471594@36.75.40,trna4_AspGTC_12_+_96429799_96429870@36.75.40,trna69_AspGTC_1_-_161440205_161440276@36.75.40,trna72_AspGTC_1_-_161432824_161432895@36.75.40,trna75_AspGTC_1_-_161425414_161425485@36.75.40,trna78_AspGTC_1_-_161418033_161418104@36.75.40,trna81_AspGTC_1_-_161410615_161410686@36.75.40\|CACGCGGGAGACCGGGGTTCGATTCCCCGACGGGGAGCCA | >trna10_AspGTC_12_-_125424193_125424264 | 36 | 75 | 40 | CACGCGGGAGACCGGGGTTCGATTCCCCGACGGGGAGCCA | 183.4 | 5.5 | 2.3 | 2.4 | 1.6E-02 | 2.8E-02 | BT20/MB468 | MP |
| >hg38_21_+_8433172_8446622_RNA45SN1_45S_50ntFlanks@10473.10494.22\|CTCGGCCGATCGAAAGGGAGTC | >hg38_21_+_8433172_8446622_RNA45SN1_45S_50ntFlanks | 10473 | 10494 | 22 | CTCGGCCGATCGAAAGGGAGTC | 192.8 | 4.5 | 1.7 | 2.7 | 6.8E-03 | 2.8E-02 | BT20/MB231 | MP |
| >hg38_21_+_8433172_8446622_RNA45SN1_45S_50ntFlanks@12942.12962.21\|TTCGTAGACGACCTGCTTCTG | >hg38_21_+_8433172_8446622_RNA45SN1_45S_50ntFlanks | 12942 | 12962 | 21 | TTCGTAGACGACCTGCTTCTG | 286.1 | -2.6 | 1.2 | -2.3 | 2.4E-02 | 2.8E-02 | BT20/MB468 | CYTO |
| >hg38_21_+_8433172_8446622_RNA45SN1_45S_50ntFlanks@10357.10376.20\|CTTTGAAGGCCGAAGTGGAG | >hg38_21_+_8433172_8446622_RNA45SN1_45S_50ntFlanks | 10357 | 10376 | 20 | CTTTGAAGGCCGAAGTGGAG | 131.6 | -2.5 | 1.1 | -2.3 | 2.4E-02 | 2.8E-02 | BT20/MB468 | CYTO |
| >hg38_21_+_8433172_8446622_RNA45SN1_45S_50ntFlanks@12894.12930.37\|CTCGCCCGTCACGCACCGCACGTTCGTGGGGAACCTG | >hg38_21_+_8433172_8446622_RNA45SN1_45S_50ntFlanks | 12894 | 12930 | 37 | CTCGCCCGTCACGCACCGCACGTTCGTGGGGAACCTG | 250.1 | -3.4 | 1.4 | -2.4 | 1.6E-02 | 2.8E-02 | BT20/MB468 | MP |
| >hg38_21_+_8433172_8446622_RNA45SN1_45S_50ntFlanks@11890.11910.21\|CTTGACTCTAGTCTGGCACGG | >hg38_21_+_8433172_8446622_RNA45SN1_45S_50ntFlanks | 11890 | 11910 | 21 | CTTGACTCTAGTCTGGCACGG | 164.1 | -3.5 | 1.5 | -2.3 | 2.4E-02 | 2.8E-02 | BT20/MB468 | CYTO |
| >hg38_21_+_8433172_8446622_RNA45SN1_45S_50ntFlanks@5467.5494.28\|GGCCCACGGCCCTGGCGGAGCGCTGAGA | >hg38_21_+_8433172_8446622_RNA45SN1_45S_50ntFlanks | 5467 | 5494 | 28 | GGCCCACGGCCCTGGCGGAGCGCTGAGA | 130.4 | -1.7 | 0.8 | -2.3 | 2.4E-02 | 2.8E-02 | MB231/MB468 | CYTO |
| >hg38_21_+_8433172_8446622_RNA45SN1_45S_50ntFlanks@10678.10707.30\|CGGTTCCGGCGGCGTCCGGTGAGCTCTCGC | >hg38_21_+_8433172_8446622_RNA45SN1_45S_50ntFlanks | 10678 | 10707 | 30 | CGGTTCCGGCGGCGTCCGGTGAGCTCTCGC | 1506.7 | -3.9 | 1.7 | -2.2 | 2.5E-02 | 2.8E-02 | BT20/MB468 | NUC |
| >MI0000748\|hsa-mir-130b&WithFlank&22\|+\|21653298\|21653391@57.77.21&[MIMAT0000691&hsa-miR-130b-3p&offsets\|0\|-1;m-205&22\|+\|21653354\|21653375&offsets\|0\|-1]\|CAGTGCAATGATGAAAGGGCA | >MI0000748\|hsa-mir-130b&WithFlank&22\|+\|21653298\|21653391 | 57 | 77 | 21 | CAGTGCAATGATGAAAGGGCA | 78.2 | 3.8 | 1.7 | 2.3 | 2.3E-02 | 2.8E-02 | MB231/MB468 | MP |
| >hg38_21_+_8433172_8446622_RNA45SN1_45S_50ntFlanks@9495.9531.37\|CGAAAGATGGTGAACTATGCCTGGGCAGGGCGAAGCC | >hg38_21_+_8433172_8446622_RNA45SN1_45S_50ntFlanks | 9495 | 9531 | 37 | CGAAAGATGGTGAACTATGCCTGGGCAGGGCGAAGCC | 314.8 | 3.8 | 1.4 | 2.7 | 6.9E-03 | 2.8E-02 | BT20/MB231 | MP |

| | | | | | | | | | | | | | |
|---|---|---|---|---|---|---|---|---|---|---|---|---|---|
| >hg38_21_+_8433172_8446622_RNA45SN1_45S_50ntFlanks@12447.12480.34\|CAGAATTCACCAAGCGTTGGATTGTTCACCCACT | >hg38_21_+_8433172_8446622_RNA45SN1_45S_50ntFlanks | 12447 | 12480 | 34 | CAGAATTCACCAAGCGTTGGATTGTTCACCCACT | 367.1 | 1.6 | 0.7 | 2.3 | 2.4E-02 | 2.8E-02 | MB231/MB468 | CYTO |
| >hg38_21_+_8433172_8446622_RNA45SN1_45S_50ntFlanks@12353.12376.24\|GTTACCACAGGGATAACTGGCTTG | >hg38_21_+_8433172_8446622_RNA45SN1_45S_50ntFlanks | 12353 | 12376 | 24 | GTTACCACAGGGATAACTGGCTTG | 66.6 | -4.0 | 1.7 | -2.4 | 1.6E-02 | 2.8E-02 | BT20/MB468 | MP |
| >hg38_21_+_8433172_8446622_RNA45SN1_45S_50ntFlanks@10393.10429.37\|CAGCAGTTGAACATGGGTCAGTCGGTCCTGAGAGATG | >hg38_21_+_8433172_8446622_RNA45SN1_45S_50ntFlanks | 10393 | 10429 | 37 | CAGCAGTTGAACATGGGTCAGTCGGTCCTGAGAGATG | 958.7 | -1.5 | 0.7 | -2.3 | 2.1E-02 | 2.8E-02 | BT20/MB468 | MITO |
| >hg38_21_+_8433172_8446622_RNA45SN1_45S_50ntFlanks@11568.11605.38\|GCAGCCGACTTAGAACTGGTGCGGACCAGGGGAATCCG | >hg38_21_+_8433172_8446622_RNA45SN1_45S_50ntFlanks | 11568 | 11605 | 38 | GCAGCCGACTTAGAACTGGTGCGGACCAGGGGAATCCG | 437.1 | -1.7 | 0.7 | -2.3 | 2.1E-02 | 2.8E-02 | BT20/MB468 | MITO |
| >hg38_21_+_8433172_8446622_RNA45SN1_45S_50ntFlanks@12935.12953.19\|TAAACCATTCGTAGACGAC | >hg38_21_+_8433172_8446622_RNA45SN1_45S_50ntFlanks | 12935 | 12953 | 19 | TAAACCATTCGTAGACGAC | 108.3 | -4.0 | 1.7 | -2.4 | 1.6E-02 | 2.8E-02 | BT20/MB468 | MP |
| >MI0003643\|hsa-mir-629&WithFlank&15\|-\|70079366\|70079474@28.48.21&[MIMAT0004810&hsa-miR-629-5p&offsets\|0\|0;m-150&15\|-\|70079427\|70079447&offsets\|0\|0]\|TGGGTTTACGTTGGGAGAACT | >MI0003643\|hsa-mir-629&WithFlank&15\|-\|70079366\|70079474 | 28 | 48 | 21 | TGGGTTTACGTTGGGAGAACT | 577.6 | 1.9 | 0.7 | 2.6 | 8.4E-03 | 2.8E-02 | BT20/MB231 | MITO |
| >hg38_MT_+_1621_3279_RNR2_16S_50ntFlanks@51.68.18\|GCTAAACCTAGCCCCAAA | >hg38_MT_+_1621_3279_RNR2_16S_50ntFlanks | 51 | 68 | 18 | GCTAAACCTAGCCCCAAA | 646.2 | 3.3 | 1.4 | 2.4 | 1.6E-02 | 2.8E-02 | BT20/MB468 | MP |
| >am_trna10_LysCTT_16_+_3241501_3241573@32.51.20,trna119_LysCTT_1_-_145395522_145395594@32.51.20,trna11_LysCTT_5_-_180648979_180649051@32.51.20,trna13_LysCTT_16_+_3246153_3246225@32.51.20,trna13_LysCTT_6_+_26556774_26556846@32.51.20,trna30_LysCTT_16_-_3230555_3230627@32.51.20,trna5_LysCTT_16_+_3214939_3215011@32.51.20,trna7_LysCTT_16_+_3225692_3225764@32.51.20,trna8_LysCTT_1_+_39970195_39970267@32.51.20,trna9_LysCTT_5_+_180634755_180634827@32.51.20\|CTCTTAATCTCAGGGTCGTG | >am_trna10_LysCTT_16_+_3241501_3241573 | 32 | 51 | 20 | CTCTTAATCTCAGGGTCGTG | 174.0 | -4.2 | 1.7 | -2.4 | 1.6E-02 | 2.8E-02 | BT20/MB468 | MP |
| >hg38_21_+_8433172_8446622_RNA45SN1_45S_50ntFlanks@4583.4608.26\|CGGTTCTATTTTGTTGGTTTTCGGAA | >hg38_21_+_8433172_8446622_RNA45SN1_45S_50ntFlanks | 4583 | 4608 | 26 | CGGTTCTATTTTGTTGGTTTTCGGAA | 64.4 | -4.1 | 1.7 | -2.4 | 1.6E-02 | 2.8E-02 | BT20/MB468 | MP |
| >hg38_21_+_8433172_8446622_RNA45SN1_45S_50ntFlanks@7988.8013.26\|CAGACGTGGCGACCCGCTGAATTTAA | >hg38_21_+_8433172_8446622_RNA45SN1_45S_50ntFlanks | 7988 | 8013 | 26 | CAGACGTGGCGACCCGCTGAATTTAA | 384.0 | -1.5 | 0.6 | -2.3 | 2.1E-02 | 2.8E-02 | BT20/MB468 | MITO |
| >hg38_1_-_228634819_228635039_RNA5S12_5S_50ntFlanks@148.170.23\|GGAATACCGGGTGCTGTAGGCTT | >hg38_1_-_228634819_228635039_RNA5S12_5S_50ntFlanks | 148 | 170 | 23 | GGAATACCGGGTGCTGTAGGCTT | 4704.7 | 1.2 | 0.5 | 2.3 | 2.1E-02 | 2.8E-02 | BT20/MB468 | MITO |
| >hg38_21_+_8433172_8446622_RNA45SN1_45S_50ntFlanks@6771.6806.36\|GGTTCCTCCCGGGGCTACGCCTGTCTGAGCGTCGCT | >hg38_21_+_8433172_8446622_RNA45SN1_45S_50ntFlanks | 6771 | 6806 | 36 | GGTTCCTCCCGGGGCTACGCCTGTCTGAGCGTCGCT | 340.5 | -1.4 | 0.6 | -2.2 | 2.5E-02 | 2.8E-02 | MB231/MB468 | NUC |
| >hg38_21_+_8433172_8446622_RNA45SN1_45S_50ntFlanks@9465.9488.24\|AGGTGGAGCACGAGCGCACGTGTT | >hg38_21_+_8433172_8446622_RNA45SN1_45S_50ntFlanks | 9465 | 9488 | 24 | AGGTGGAGCACGAGCGCACGTGTT | 251.5 | -3.0 | 1.2 | -2.4 | 1.6E-02 | 2.8E-02 | BT20/MB468 | MP |
| >MI0000063\|hsa-let-7b&WithFlank&22\|+\|46113680\|46113774@12.33.22(+1G)&[MIMAT0000063&hsa-let-7b-5p&offsets\|0\|0(+1G);m-14&22\|+\|46113691\|46113712&offsets\|0\|0(+1G)]\|TGAGGTAGTAGGTTGTGTGGTTG | >MI0000063\|hsa-let-7b&WithFlank&22\|+\|46113680\|46113774 | 12 | 33 | 22(+1G) | TGAGGTAGTAGGTTGTGTGGTTG | 157.9 | 3.0 | 1.3 | 2.3 | 2.3E-02 | 2.8E-02 | MB231/MB468 | MP |
| >hg38_21_+_8433172_8446622_RNA45SN1_45S_50ntFlanks@12996.13039.44\|TCGCTGCGATCTATTGAAAGTCAGCCCTCGACACAAGGGTTTGT | >hg38_21_+_8433172_8446622_RNA45SN1_45S_50ntFlanks | 12996 | 13039 | 44 | TCGCTGCGATCTATTGAAAGTCAGCCCTCGACACAAGGGTTTGT | 160189.2 | -3.1 | 1.1 | -2.7 | 7.4E-03 | 2.8E-02 | BT20/MB231 | NUC |
| >MI0000066\|hsa-let-7e&WithFlank&19\|+\|51692780\|51692870@14.36.23&[MIMAT0000066&hsa-let-7e-5p&offsets\|0\|+1;m-50&19\|+\|51692793\|51692814&offsets\|0\|+1]\|TGAGGTAGGAGGTTGTATAGTTG | >MI0000066\|hsa-let-7e&WithFlank&19\|+\|51692780\|51692870 | 14 | 36 | 23 | TGAGGTAGGAGGTTGTATAGTTG | 154.8 | 3.0 | 1.3 | 2.3 | 2.3E-02 | 2.8E-02 | MB231/MB468 | MP |
| >trnaMT_PheGAA_MT_+_577_647@38.55.18\|AATGTTTAGACGGGCTCA | >trnaMT_PheGAA_MT_+_577_647 | 38 | 55 | 18 | AATGTTTAGACGGGCTCA | 48.5 | 4.4 | 1.9 | 2.3 | 2.3E-02 | 2.8E-02 | MB231/MB468 | MP |
| >trna106_HisGTG_1_-_149155828_149155899@13.31.19,trna111_HisGTG_1_-_147774845_147774916@13.31.19,trna118_HisGTG_1_-_145396881_145396952@13.31.19,trna16_HisGTG_1_+_146544773_146544844@13.31.19,trna1_HisGTG_15_+_45493349_45493420@13.31.19,trna21_HisGTG_1_+_147753471_147753542@13.31.19,trna33_HisGTG_6_+_27125906_27125977@13.31.19,trna7_HisGTG_9_-_14433938_14434009@13.31.19,trna8_HisGTG_15_-_45492611_45492682@13.31.19,trna9_HisGTG_15_-_45490804_45490875@13.31.19\|TAGTGGTTAGTACTCTGCG | >trna106_HisGTG_1_-_149155828_149155899 | 13 | 31 | 19 | TAGTGGTTAGTACTCTGCG | 110.3 | -3.4 | 1.4 | -2.4 | 1.7E-02 | 2.8E-02 | BT20/MB468 | MP |
| >hg38_21_+_8433172_8446622_RNA45SN1_45S_50ntFlanks@12071.12097.27\|CCCCGAGGGGCTCTCGCTTCTGGCGCC | >hg38_21_+_8433172_8446622_RNA45SN1_45S_50ntFlanks | 12071 | 12097 | 27 | CCCCGAGGGGCTCTCGCTTCTGGCGCC | 411.9 | -3.0 | 1.3 | -2.3 | 2.4E-02 | 2.8E-02 | BT20/MB468 | CYTO |
| >MI0000081\|hsa-mir-24-2&WithFlank&19\|-\|13836281\|13836365@56.77.22(+1C);MI0000080\|hsa-mir-24-1&WithFlank&9\|+\|95086015\|95086094@50.71.22(+1C)&[MIMAT0000080&hsa-miR-24-3p&offsets\|0\|0(+1C);m-36&19\|-\|13836288\|13836310&offsets\|0\|-1(+1C)];[MIMAT0000080_1&hsa-miR-24-3p&offsets\|0\|0(+1C);m-35&9\|+\|95086064\|95086086&offsets\|0\|-1(+1C)]\|TGGCTCAGTTCAGCAGGAACAGC | >MI0000081\|hsa-mir-24-2&WithFlank&19\|-\|13836281\|13836365 | 56 | 77 | 22(+1C) | TGGCTCAGTTCAGCAGGAACAGC | 1762.1 | -2.3 | 0.9 | -2.6 | 8.4E-03 | 2.8E-02 | BT20/MB231 | MITO |
| >MI0000081\|hsa-mir-24-2&WithFlank&19\|-\|13836281\|13836365@56.74.19(+1A);MI0000080\|hsa-mir-24-1&WithFlank&9\|+\|95086015\|95086094@50.68.19(+1A)&[MIMAT0000080&hsa-miR-24-3p&offsets\|0\|-3(+1A);m-36&19\|-\|13836288\|13836310&offsets\|0\|-4(+1A)];[MIMAT0000080_1&hsa-miR-24-3p&offsets\|0\|-3(+1A);m-35&9\|+\|95086064\|95086086&offsets\|0\|-4(+1A)]\|TGGCTCAGTTCAGCAGGAAA | >MI0000081\|hsa-mir-24-2&WithFlank&19\|-\|13836281\|13836365 | 56 | 74 | 19(+1A) | TGGCTCAGTTCAGCAGGAAA | 188.8 | 2.5 | 1.0 | 2.6 | 9.3E-03 | 2.8E-02 | BT20/MB231 | CYTO |
| >hg38_21_+_8433172_8446622_RNA45SN1_45S_50ntFlanks@11585.11616.32\|GGTGCGGACCAGGGGAATCCGACTGTTTAATT | >hg38_21_+_8433172_8446622_RNA45SN1_45S_50ntFlanks | 11585 | 11616 | 32 | GGTGCGGACCAGGGGAATCCGACTGTTTAATT | 324.6 | -2.1 | 0.8 | -2.6 | 9.3E-03 | 2.8E-02 | BT20/MB231 | CYTO |
| >hg38_21_+_8433172_8446622_RNA45SN1_45S_50ntFlanks@3852.3872.21\|TAACTGTGGTAATTCTAGAGC | >hg38_21_+_8433172_8446622_RNA45SN1_45S_50ntFlanks | 3852 | 3872 | 21 | TAACTGTGGTAATTCTAGAGC | 100.9 | -4.0 | 1.7 | -2.4 | 1.7E-02 | 2.8E-02 | BT20/MB468 | MP |
| >MI0003834\|hsa-mir-769&WithFlank&19\|+\|46018926\|46019055@36.53.18&[MIMAT0003886&hsa-miR-769-5p&offsets\|0\|-4;m-216&19\|+\|46018961\|46018982&offsets\|0\|-4]\|TGAGACCTCTGGGTTCTG | >MI0003834\|hsa-mir-769&WithFlank&19\|+\|46018926\|46019055 | 36 | 53 | 18 | TGAGACCTCTGGGTTCTG | 71.0 | 3.9 | 1.7 | 2.3 | 2.3E-02 | 2.8E-02 | MB231/MB468 | MP |
| >trna2_GlyGCC_21_-_18827107_18827177@1.35.35,trna35_GlyGCC_1_+_161413094_161413164@1.35.35,trna37_GlyGCC_1_+_161420467_161420537@1.35.35,trna39_GlyGCC_1_+_161427898_161427968@1.35.35,trna41_GlyGCC_1_+_161435258_161435328@1.35.35\|GCATGGGTGGTTCAGTGGTAGAATTCTCGCCTGCC | >trna2_GlyGCC_21_-_18827107_18827177 | 1 | 35 | 35 | GCATGGGTGGTTCAGTGGTAGAATTCTCGCCTGCC | 370.5 | 3.7 | 1.4 | 2.7 | 6.9E-03 | 2.8E-02 | BT20/MB231 | MP |
| >hg38_21_+_8433172_8446622_RNA45SN1_45S_50ntFlanks@11666.11685.20\|TCTGCCCAGTGCTCTGAATG | >hg38_21_+_8433172_8446622_RNA45SN1_45S_50ntFlanks | 11666 | 11685 | 20 | TCTGCCCAGTGCTCTGAATG | 3222.0 | -3.0 | 1.3 | -2.2 | 2.5E-02 | 2.8E-02 | BT20/MB468 | NUC |
| >MI0000103\|hsa-mir-101-1&WithFlank&1\|-\|65058428\|65058514@52.73.22;MI0000739\|hsa-mir-101-2&WithFlank&9\|+\|4850291\|4850381@54.75.22&[MIMAT0000099&hsa-miR-101-3p&offsets\|-1\|0;m-17&1\|-\|65058443\|65058462&offsets\|-1\|+1];[MIMAT0000099_1&hsa-miR-101-3p&offsets\|-1\|0;m-16&9\|+\|4850345\|4850364&offsets\|-1\|+1]\|GTACAGTACTGTGATAACTGAA | >MI0000103\|hsa-mir-101-1&WithFlank&1\|-\|65058428\|65058514 | 52 | 73 | 22 | GTACAGTACTGTGATAACTGAA | 1656.9 | -2.3 | 0.9 | -2.6 | 8.4E-03 | 2.8E-02 | BT20/MB231 | MITO |
| >hg38_21_+_8433172_8446622_RNA45SN1_45S_50ntFlanks@12423.12445.23\|TCGGCTCTTCCTATCATTGTGAA | >hg38_21_+_8433172_8446622_RNA45SN1_45S_50ntFlanks | 12423 | 12445 | 23 | TCGGCTCTTCCTATCATTGTGAA | 135.9 | -2.3 | 1.0 | -2.3 | 2.4E-02 | 2.8E-02 | BT20/MB468 | CYTO |
| >hg38_21_+_8433172_8446622_RNA45SN1_45S_50ntFlanks@11321.11343.23\|CACGTCTCGTCGCGCGCGCGTCC | >hg38_21_+_8433172_8446622_RNA45SN1_45S_50ntFlanks | 11321 | 11343 | 23 | CACGTCTCGTCGCGCGCGCGTCC | 4302.8 | -4.2 | 1.9 | -2.2 | 2.5E-02 | 2.8E-02 | BT20/MB468 | NUC |
| >hg38_21_+_8433172_8446622_RNA45SN1_45S_50ntFlanks@11668.11706.39\|TGCCCAGTGCTCTGAATGTCAAAGTGAAGAAATTCAATG | >hg38_21_+_8433172_8446622_RNA45SN1_45S_50ntFlanks | 11668 | 11706 | 39 | TGCCCAGTGCTCTGAATGTCAAAGTGAAGAAATTCAATG | 50.5 | 4.9 | 2.1 | 2.4 | 1.7E-02 | 2.8E-02 | BT20/MB468 | MP |
| >MI0000273\|hsa-mir-183&WithFlank&7\|-\|129774899\|129775020@33.54.22(+1A)&[MIMAT0000261&hsa-miR-183-5p&offsets\|0\|0(+1A);m-41&7\|-\|129774966\|129774987&offsets\|-1\|-1(+1A)]\|TATGGCACTGGTAGAATTCACTA | >MI0000273\|hsa-mir-183&WithFlank&7\|-\|129774899\|129775020 | 33 | 54 | 22(+1A) | TATGGCACTGGTAGAATTCACTA | 362.9 | 1.9 | 0.7 | 2.6 | 8.4E-03 | 2.8E-02 | BT20/MB231 | MITO |
| >hg38_21_+_8433172_8446622_RNA45SN1_45S_50ntFlanks@6651.6679.29\|CGACTCTTAGCGGTGGATCACTCGGCTCG | >hg38_21_+_8433172_8446622_RNA45SN1_45S_50ntFlanks | 6651 | 6679 | 29 | CGACTCTTAGCGGTGGATCACTCGGCTCG | 88037.7 | -2.1 | 0.9 | -2.2 | 2.6E-02 | 2.8E-02 | MB231/MB468 | MITO |
| >trnaMT_GluTTC_MT_-_14674_14742@33.71.39,trnalookalike8_GluTTC_5_-_93905172_93905240@33.71.39\|CATATCATTGGTCGTGGTTGTAGTCCGTGCGAGAATACC | >trnaMT_GluTTC_MT_-_14674_14742 | 33 | 71 | 39 | CATATCATTGGTCGTGGTTGTAGTCCGTGCGAGAATACC | 125.6 | 3.8 | 1.6 | 2.4 | 1.7E-02 | 2.8E-02 | BT20/MB468 | MP |
| >MI0000749\|hsa-mir-30e&WithFlank&1\|+\|40754349\|40754452@65.86.22&[MIMAT0000693&hsa-miR-30e-3p&offsets\|0\|0;m-25&1\|+\|40754413\|40754434&offsets\|0\|0]\|CTTTCAGTCGGATGTTTACAGC | >MI0000749\|hsa-mir-30e&WithFlank&1\|+\|40754349\|40754452 | 65 | 86 | 22 | CTTTCAGTCGGATGTTTACAGC | 1917.3 | -1.8 | 0.7 | -2.6 | 8.5E-03 | 2.8E-02 | BT20/MB231 | MITO |
| >trnaMT_HisGTG_MT_+_12138_12206@43.70.28\|ACAGAGGCTTACGACCCCTTATTTACCC | >trnaMT_HisGTG_MT_+_12138_12206 | 43 | 70 | 28 | ACAGAGGCTTACGACCCCTTATTTACCC | 16.3 | 7.3 | 3.2 | 2.3 | 2.3E-02 | 2.8E-02 | MB231/MB468 | MP |
| >MI0000299\|hsa-mir-222&WithFlank&X\|-\|45747009\|45747130@75.96.22&[MIMAT0000279&hsa-miR-222-3p&offsets\|0\|+1;m-140&X\|-\|45747033\|45747056&offsets\|0\|-2]\|AGCTACATCTGGCTACTGGGTC | >MI0000299\|hsa-mir-222&WithFlank&X\|-\|45747009\|45747130 | 75 | 96 | 22 | AGCTACATCTGGCTACTGGGTC | 1876.8 | -1.4 | 0.5 | -2.7 | 7.5E-03 | 2.8E-02 | BT20/MB231 | NUC |
| >MI0005559\|hsa-mir-744&WithFlank&17\|+\|12081893\|12082002@17.36.20(+1U)&[MIMAT0004945&hsa-miR-744-5p&offsets\|0\|-2(+1U);m-172&17\|+\|12081909\|12081929&offsets\|0\|-1(+1U)]\|TGCGGGGCTAGGGCTAACAGT | >MI0005559\|hsa-mir-744&WithFlank&17\|+\|12081893\|12082002 | 17 | 36 | 20(+1U) | TGCGGGGCTAGGGCTAACAGT | 386.9 | 2.0 | 0.8 | 2.7 | 7.5E-03 | 2.8E-02 | BT20/MB231 | NUC |
| >am_MI0000809\|hsa-mir-151a&WithFlank&8\|-\|140732558\|140732659@17.37.21&[MIMAT0004697&hsa-miR-151a-5p&offsets\|0\|0;m-75&8\|-\|140732623\|140732643&offsets\|0\|0]\|TCGAGGAGCTCACAGTCTAGT | >am_MI0000809\|hsa-mir-151a&WithFlank&8\|-\|140732558\|140732659 | 17 | 37 | 21 | TCGAGGAGCTCACAGTCTAGT | 3721.5 | 1.9 | 0.7 | 2.7 | 7.5E-03 | 2.8E-02 | BT20/MB231 | NUC |
| >hg38_MT_+_598_1651_RNR1_12S_50ntFlanks@719.743.25\|TAAAGACGTTAGGTCAAGGTGTAGC | >hg38_MT_+_598_1651_RNR1_12S_50ntFlanks | 719 | 743 | 25 | TAAAGACGTTAGGTCAAGGTGTAGC | 179.1 | -2.9 | 1.2 | -2.4 | 1.7E-02 | 2.8E-02 | BT20/MB468 | MP |
| >MI0000255\|hsa-mir-30d&WithFlank&8\|-\|134804870\|134804951@12.35.24(+1A)&[MIMAT0000245&hsa-miR-30d-5p&offsets\|0\|+2(+1A);m-21&8\|-\|134804919\|134804940&offsets\|0\|+2(+1A)]\|TGTAAACATCCCCGACTGGAAGCTA | >MI0000255\|hsa-mir-30d&WithFlank&8\|-\|134804870\|134804951 | 12 | 35 | 24(+1A) | TGTAAACATCCCCGACTGGAAGCTA | 93.5 | 3.7 | 1.6 | 2.3 | 2.3E-02 | 2.8E-02 | MB231/MB468 | MP |
| >hg38_21_+_8433172_8446622_RNA45SN1_45S_50ntFlanks@7974.8015.42\|ACGCGACCTCAGATCAGACGTGGCGACCCGCTGAATTTAAGC | >hg38_21_+_8433172_8446622_RNA45SN1_45S_50ntFlanks | 7974 | 8015 | 42 | ACGCGACCTCAGATCAGACGTGGCGACCCGCTGAATTTAAGC | 280.8 | 4.3 | 1.8 | 2.4 | 1.7E-02 | 2.8E-02 | BT20/MB468 | MP |

<table>
<tr><td>>am_MI0015825|hsa-mir-378c&WithFlank&10|-|130962582|130962674@17.36.20;MI0000786|hsa-mir-378a&WithFlank&5|+|149732819|149732896@49.68.20&[MIMAT0016847&hsa-miR-378c&offsets|0|-5;m-130&10|-|130962639|130962658&offsets|0|0];[MIMAT0000732&hsa-miR-378a-3p&offsets|0|-2;m-44&5|+|149732867|149732888&offsets|0|-2]|ACTGGACTTGGAGTCAGAAG</td><td>>am_MI0015825|hsa-mir-378c&WithFlank&10|-|130962582|130962674</td><td>17</td><td>36</td><td>20</td><td>ACTGGACTTGGAGTCAGAAG</td><td>105.5</td><td>1.2</td><td>0.5</td><td>2.2</td><td>2.5E-02</td><td>2.8E-02</td><td>BT20/MB468</td><td>NUC</td></tr>
<tr><td>>MI0006444|hsa-mir-1307&WithFlank&10|-|103394247|103394407@86.108.23&[MIMAT0005951&hsa-miR-1307-3p&offsets|0|+1;m-66&10|-|103394300|103394322&offsets|0|0]|ACTCGGCGTGGCGTCGGTCGTGG</td><td>>MI0006444|hsa-mir-1307&WithFlank&10|-|103394247|103394407</td><td>86</td><td>108</td><td>23</td><td>ACTCGGCGTGGCGTCGGTCGTGG</td><td>604.6</td><td>1.3</td><td>0.6</td><td>2.2</td><td>2.5E-02</td><td>2.8E-02</td><td>BT20/MB468</td><td>NUC</td></tr>
<tr><td>>hg38_21_+_8433172_8446622_RNA45SN1_45S_50ntFlanks@9570.9603.34|TGCAAATCGGTCGTCCGACCTGGGTATAGGGGCG</td><td>>hg38_21_+_8433172_8446622_RNA45SN1_45S_50ntFlanks</td><td>9570</td><td>9603</td><td>34</td><td>TGCAAATCGGTCGTCCGACCTGGGTATAGGGGCG</td><td>268.3</td><td>-1.5</td><td>0.7</td><td>-2.2</td><td>2.5E-02</td><td>2.8E-02</td><td>BT20/MB468</td><td>CYTO</td></tr>
<tr><td>>MI0000081|hsa-mir-24-2&WithFlank&19|-|13836281|13836365@56.77.22(+1U);MI0000080|hsa-mir-24-1&WithFlank&9|+|95086015|95086094@50.71.22(+1U)&[MIMAT0000080&hsa-miR-24-3p&offsets|0|0(+1U);m-36&19|-|13836288|13836310&offsets|0|-1(+1U)];[MIMAT0000080_1&hsa-miR-24-3p&offsets|0|0(+1U);m-35&9|+|95086064|95086086&offsets|0|-1(+1U)]|TGGCTCAGTTCAGCAGGAACAGT</td><td>>MI0000081|hsa-mir-24-2&WithFlank&19|-|13836281|13836365</td><td>56</td><td>77</td><td>22(+1U)</td><td>TGGCTCAGTTCAGCAGGAACAGT</td><td>12683.7</td><td>-2.2</td><td>0.9</td><td>-2.6</td><td>8.5E-03</td><td>2.8E-02</td><td>BT20/MB231</td><td>MITO</td></tr>
<tr><td>>hg38_21_+_8433172_8446622_RNA45SN1_45S_50ntFlanks@9033.9077.45|CTCGTGGGGGGGCCGGGCCACCCCTCCCACGGCGCGACCGCTCTC</td><td>>hg38_21_+_8433172_8446622_RNA45SN1_45S_50ntFlanks</td><td>9033</td><td>9077</td><td>45</td><td>CTCGTGGGGGGGCCGGGCCACCCCTCCCACGGCGCGACCGCTCTC</td><td>28.9</td><td>-7.9</td><td>3.3</td><td>-2.4</td><td>1.7E-02</td><td>2.8E-02</td><td>BT20/MB468</td><td>MP</td></tr>
<tr><td>>hg38_21_+_8433172_8446622_RNA45SN1_45S_50ntFlanks@11212.11250.39|CGACTCTGGACGCGAGCCGGGCCCTTCCCGTGGATCGCC</td><td>>hg38_21_+_8433172_8446622_RNA45SN1_45S_50ntFlanks</td><td>11212</td><td>11250</td><td>39</td><td>CGACTCTGGACGCGAGCCGGGCCCTTCCCGTGGATCGCC</td><td>460.9</td><td>-3.0</td><td>1.4</td><td>-2.2</td><td>2.6E-02</td><td>2.8E-02</td><td>MB231/MB468</td><td>NUC</td></tr>
<tr><td>>hg38_21_+_8433172_8446622_RNA45SN1_45S_50ntFlanks@11573.11590.18|CGACTTAGAACTGGTGCG</td><td>>hg38_21_+_8433172_8446622_RNA45SN1_45S_50ntFlanks</td><td>11573</td><td>11590</td><td>18</td><td>CGACTTAGAACTGGTGCG</td><td>248.2</td><td>-1.3</td><td>0.6</td><td>-2.2</td><td>2.5E-02</td><td>2.8E-02</td><td>BT20/MB468</td><td>CYTO</td></tr>
<tr><td>>hg38_21_+_8433172_8446622_RNA45SN1_45S_50ntFlanks@12419.12439.21|GATGTCGGCTCTTCCTATCAT</td><td>>hg38_21_+_8433172_8446622_RNA45SN1_45S_50ntFlanks</td><td>12419</td><td>12439</td><td>21</td><td>GATGTCGGCTCTTCCTATCAT</td><td>3807.7</td><td>-2.1</td><td>0.8</td><td>-2.6</td><td>9.3E-03</td><td>2.8E-02</td><td>BT20/MB231</td><td>CYTO</td></tr>
<tr><td>>hg38_MT_+_1621_3279_RNR2_16S_50ntFlanks@1263.1292.30|ACTATACTCAATTGATCCAATAACTTGACC</td><td>>hg38_MT_+_1621_3279_RNR2_16S_50ntFlanks</td><td>1263</td><td>1292</td><td>30</td><td>ACTATACTCAATTGATCCAATAACTTGACC</td><td>208.2</td><td>2.3</td><td>1.0</td><td>2.3</td><td>2.3E-02</td><td>2.8E-02</td><td>MB231/MB468</td><td>MP</td></tr>
<tr><td>>hg38_21_+_8433172_8446622_RNA45SN1_45S_50ntFlanks@11786.11805.20|GAATGGATGAACGAGATTCC</td><td>>hg38_21_+_8433172_8446622_RNA45SN1_45S_50ntFlanks</td><td>11786</td><td>11805</td><td>20</td><td>GAATGGATGAACGAGATTCC</td><td>96.8</td><td>-4.0</td><td>1.7</td><td>-2.4</td><td>1.7E-02</td><td>2.8E-02</td><td>BT20/MB468</td><td>MP</td></tr>
<tr><td>>hg38_MT_+_598_1651_RNR1_12S_50ntFlanks@970.1004.35|ATGGTAAGTGTACTGGAAAGTGCACTTGGACGAAC</td><td>>hg38_MT_+_598_1651_RNR1_12S_50ntFlanks</td><td>970</td><td>1004</td><td>35</td><td>ATGGTAAGTGTACTGGAAAGTGCACTTGGACGAAC</td><td>187.2</td><td>3.8</td><td>1.7</td><td>2.3</td><td>2.3E-02</td><td>2.8E-02</td><td>MB231/MB468</td><td>MP</td></tr>
<tr><td>>hg38_21_+_8433172_8446622_RNA45SN1_45S_50ntFlanks@12420.12448.29|ATGTCGGCTCTTCCTATCATTGTGAAGCA</td><td>>hg38_21_+_8433172_8446622_RNA45SN1_45S_50ntFlanks</td><td>12420</td><td>12448</td><td>29</td><td>ATGTCGGCTCTTCCTATCATTGTGAAGCA</td><td>616.3</td><td>1.3</td><td>0.6</td><td>2.3</td><td>2.2E-02</td><td>2.8E-02</td><td>BT20/MB468</td><td>MITO</td></tr>
<tr><td>>hg38_21_+_8433172_8446622_RNA45SN1_45S_50ntFlanks@4752.4783.32|GAAGACGATCAGATACCGTCGTAGTTCCGACC</td><td>>hg38_21_+_8433172_8446622_RNA45SN1_45S_50ntFlanks</td><td>4752</td><td>4783</td><td>32</td><td>GAAGACGATCAGATACCGTCGTAGTTCCGACC</td><td>285.2</td><td>-2.4</td><td>1.1</td><td>-2.3</td><td>2.2E-02</td><td>2.8E-02</td><td>BT20/MB468</td><td>MITO</td></tr>
<tr><td>>hg38_1_-_228634819_228635039_RNA5S12_5S_50ntFlanks@128.171.44|CTTGGATGGGAGACCGCCTGGGAATACCGGGTGCTGTAGGCTTT</td><td>>hg38_1_-_228634819_228635039_RNA5S12_5S_50ntFlanks</td><td>128</td><td>171</td><td>44</td><td>CTTGGATGGGAGACCGCCTGGGAATACCGGGTGCTGTAGGCTTT</td><td>249.2</td><td>4.4</td><td>1.8</td><td>2.4</td><td>1.7E-02</td><td>2.8E-02</td><td>BT20/MB468</td><td>MP</td></tr>
<tr><td>>hg38_21_+_8433172_8446622_RNA45SN1_45S_50ntFlanks@10058.10081.24|CGGCAGTCGAGAGTGGACGGGAGC</td><td>>hg38_21_+_8433172_8446622_RNA45SN1_45S_50ntFlanks</td><td>10058</td><td>10081</td><td>24</td><td>CGGCAGTCGAGAGTGGACGGGAGC</td><td>166.0</td><td>5.5</td><td>2.0</td><td>2.7</td><td>6.9E-03</td><td>2.8E-02</td><td>BT20/MB231</td><td>MP</td></tr>
<tr><td>>hg38_21_+_8433172_8446622_RNA45SN1_45S_50ntFlanks@6652.6698.47|GACTCTTAGCGGTGGATCACTCGGCTCGTGCGTCGATGAAGAACGCA</td><td>>hg38_21_+_8433172_8446622_RNA45SN1_45S_50ntFlanks</td><td>6652</td><td>6698</td><td>47</td><td>GACTCTTAGCGGTGGATCACTCGGCTCGTGCGTCGATGAAGAACGCA</td><td>3358.7</td><td>3.5</td><td>1.3</td><td>2.7</td><td>6.9E-03</td><td>2.8E-02</td><td>BT20/MB231</td><td>MP</td></tr>
<tr><td>>MI0000299|hsa-mir-222&WithFlank&X|-|45747009|45747130@75.97.23(+1U)&[MIMAT0000279&hsa-miR-222-3p&offsets|0|+2(+1U);m-140&X|-|45747033|45747056&offsets|0|-1(+1U)]|AGCTACATCTGGCTACTGGGTCTT</td><td>>MI0000299|hsa-mir-222&WithFlank&X|-|45747009|45747130</td><td>75</td><td>97</td><td>23(+1U)</td><td>AGCTACATCTGGCTACTGGGTCTT</td><td>322.1</td><td>1.4</td><td>0.6</td><td>2.2</td><td>2.5E-02</td><td>2.8E-02</td><td>BT20/MB468</td><td>NUC</td></tr>
<tr><td>>hg38_MT_+_598_1651_RNR1_12S_50ntFlanks@910.946.37|ACTTCAAAGGACATTTAACTAAAACCCCTACGCATTT</td><td>>hg38_MT_+_598_1651_RNR1_12S_50ntFlanks</td><td>910</td><td>946</td><td>37</td><td>ACTTCAAAGGACATTTAACTAAAACCCCTACGCATTT</td><td>304.6</td><td>4.6</td><td>1.9</td><td>2.4</td><td>1.7E-02</td><td>2.8E-02</td><td>BT20/MB468</td><td>MP</td></tr>
<tr><td>>hg38_21_+_8433172_8446622_RNA45SN1_45S_50ntFlanks@12957.12974.18|CTTCTGGGTCGGGGTTTC</td><td>>hg38_21_+_8433172_8446622_RNA45SN1_45S_50ntFlanks</td><td>12957</td><td>12974</td><td>18</td><td>CTTCTGGGTCGGGGTTTC</td><td>134.5</td><td>-4.2</td><td>1.7</td><td>-2.4</td><td>1.7E-02</td><td>2.8E-02</td><td>BT20/MB468</td><td>MP</td></tr>
<tr><td>>hg38_21_+_8433172_8446622_RNA45SN1_45S_50ntFlanks@9938.9964.27|AGCAGGACGGTGGCCATGGAAGTCGGA</td><td>>hg38_21_+_8433172_8446622_RNA45SN1_45S_50ntFlanks</td><td>9938</td><td>9964</td><td>27</td><td>AGCAGGACGGTGGCCATGGAAGTCGGA</td><td>134.7</td><td>5.3</td><td>2.0</td><td>2.7</td><td>7.0E-03</td><td>2.8E-02</td><td>BT20/MB231</td><td>MP</td></tr>
<tr><td>>hg38_21_+_8433172_8446622_RNA45SN1_45S_50ntFlanks@5492.5515.24|AGAAGACGGTCGAACTTGACTATC</td><td>>hg38_21_+_8433172_8446622_RNA45SN1_45S_50ntFlanks</td><td>5492</td><td>5515</td><td>24</td><td>AGAAGACGGTCGAACTTGACTATC</td><td>292.0</td><td>-3.4</td><td>1.4</td><td>-2.4</td><td>1.7E-02</td><td>2.8E-02</td><td>BT20/MB468</td><td>MP</td></tr>
<tr><td>>trnaMT_GluTTC_MT_-_14674_14742@34.58.25,trnalookalike8_GluTTC_5_-_93905172_93905240@34.58.25|ATATCATTGGTCGTGGTTGTAGTCC</td><td>>trnaMT_GluTTC_MT_-_14674_14742</td><td>34</td><td>58</td><td>25</td><td>ATATCATTGGTCGTGGTTGTAGTCC</td><td>82.2</td><td>3.2</td><td>1.4</td><td>2.3</td><td>2.3E-02</td><td>2.8E-02</td><td>MB231/MB468</td><td>MP</td></tr>
<tr><td>>MI0006273|hsa-mir-1180&WithFlank&17|-|19344500|19344580@47.66.20&[MIMAT0005825&hsa-miR-1180-3p&offsets|0|-2;m-214&17|-|19344513|19344534&offsets|0|-2]|TTTCCGGCTCGCGTGGGTGT</td><td>>MI0006273|hsa-mir-1180&WithFlank&17|-|19344500|19344580</td><td>47</td><td>66</td><td>20</td><td>TTTCCGGCTCGCGTGGGTGT</td><td>357.4</td><td>2.1</td><td>0.8</td><td>2.6</td><td>8.5E-03</td><td>2.8E-02</td><td>BT20/MB231</td><td>MITO</td></tr>
<tr><td>>MI0000299|hsa-mir-222&WithFlank&X|-|45747009|45747130@75.96.22(+1A)&[MIMAT0000279&hsa-miR-222-3p&offsets|0|+1(+1A);m-140&X|-|45747033|45747056&offsets|0|-2(+1A)]|AGCTACATCTGGCTACTGGGTCA</td><td>>MI0000299|hsa-mir-222&WithFlank&X|-|45747009|45747130</td><td>75</td><td>96</td><td>22(+1A)</td><td>AGCTACATCTGGCTACTGGGTCA</td><td>1470.6</td><td>-2.0</td><td>0.8</td><td>-2.6</td><td>8.5E-03</td><td>2.8E-02</td><td>BT20/MB231</td><td>MITO</td></tr>
<tr><td>>hg38_21_+_8433172_8446622_RNA45SN1_45S_50ntFlanks@3841.3868.28|TCCTACTTGGATAACTGTGGTAATTCTA</td><td>>hg38_21_+_8433172_8446622_RNA45SN1_45S_50ntFlanks</td><td>3841</td><td>3868</td><td>28</td><td>TCCTACTTGGATAACTGTGGTAATTCTA</td><td>218.4</td><td>2.8</td><td>1.1</td><td>2.6</td><td>8.5E-03</td><td>2.8E-02</td><td>BT20/MB231</td><td>MITO</td></tr>
<tr><td>>hg38_21_+_8433172_8446622_RNA45SN1_45S_50ntFlanks@12416.12452.37|TTCGATGTCGGCTCTTCCTATCATTGTGAAGCAGAAT</td><td>>hg38_21_+_8433172_8446622_RNA45SN1_45S_50ntFlanks</td><td>12416</td><td>12452</td><td>37</td><td>TTCGATGTCGGCTCTTCCTATCATTGTGAAGCAGAAT</td><td>443.1</td><td>-2.1</td><td>0.8</td><td>-2.6</td><td>8.6E-03</td><td>2.8E-02</td><td>BT20/MB231</td><td>MITO</td></tr>
<tr><td>>hg38_21_+_8433172_8446622_RNA45SN1_45S_50ntFlanks@5086.5108.23|AACGAGACTCTGGCATGCTAACT</td><td>>hg38_21_+_8433172_8446622_RNA45SN1_45S_50ntFlanks</td><td>5086</td><td>5108</td><td>23</td><td>AACGAGACTCTGGCATGCTAACT</td><td>338.4</td><td>3.2</td><td>1.4</td><td>2.4</td><td>1.7E-02</td><td>2.8E-02</td><td>BT20/MB468</td><td>MP</td></tr>
<tr><td>>hg38_21_+_8433172_8446622_RNA45SN1_45S_50ntFlanks@3787.3805.19|AAACTGCGAATGGCTCATT</td><td>>hg38_21_+_8433172_8446622_RNA45SN1_45S_50ntFlanks</td><td>3787</td><td>3805</td><td>19</td><td>AAACTGCGAATGGCTCATT</td><td>180.4</td><td>-1.6</td><td>0.7</td><td>-2.2</td><td>2.6E-02</td><td>2.8E-02</td><td>MB231/MB468</td><td>NUC</td></tr>
<tr><td>>hg38_21_+_8433172_8446622_RNA45SN1_45S_50ntFlanks@5485.5526.42|AGCGCTGAGAAGACGGTCGAACTTGACTATCTAGAGGAAGTA</td><td>>hg38_21_+_8433172_8446622_RNA45SN1_45S_50ntFlanks</td><td>5485</td><td>5526</td><td>42</td><td>AGCGCTGAGAAGACGGTCGAACTTGACTATCTAGAGGAAGTA</td><td>504.1</td><td>-3.5</td><td>1.5</td><td>-2.4</td><td>1.7E-02</td><td>2.8E-02</td><td>BT20/MB468</td><td>MP</td></tr>
<tr><td>>hg38_21_+_8433172_8446622_RNA45SN1_45S_50ntFlanks@11872.11893.22|AAAGAAGACCCTGTTGAGCTTG</td><td>>hg38_21_+_8433172_8446622_RNA45SN1_45S_50ntFlanks</td><td>11872</td><td>11893</td><td>22</td><td>AAAGAAGACCCTGTTGAGCTTG</td><td>396.0</td><td>-2.1</td><td>0.9</td><td>-2.4</td><td>1.7E-02</td><td>2.8E-02</td><td>BT20/MB468</td><td>MP</td></tr>
<tr><td>>trna119_LysCTT_1_-_145395522_145395594@1.35.35,trna11_LysCTT_5_-_180648979_180649051@1.35.35,trna13_LysCTT_6_+_26556774_26556846@1.35.35,trna7_LysCTT_16_+_3225692_3225764@1.35.35,trna9_LysCTT_5_+_180634755_180634827@1.35.35|GCCCGGCTAGCTCAGTCGGTAGAGCATGAGACTCT</td><td>>trna119_LysCTT_1_-_145395522_145395594</td><td>1</td><td>35</td><td>35</td><td>GCCCGGCTAGCTCAGTCGGTAGAGCATGAGACTCT</td><td>32032.5</td><td>2.5</td><td>1.0</td><td>2.6</td><td>9.4E-03</td><td>2.8E-02</td><td>BT20/MB231</td><td>CYTO</td></tr>
<tr><td>>trnaMT_ValTAC_MT_+_1602_1670@32.71.40|TACACTTAGGAGATTTCAACTTAACTTGACCGCTCTGACC</td><td>>trnaMT_ValTAC_MT_+_1602_1670</td><td>32</td><td>71</td><td>40</td><td>TACACTTAGGAGATTTCAACTTAACTTGACCGCTCTGACC</td><td>123.2</td><td>3.9</td><td>1.7</td><td>2.3</td><td>2.3E-02</td><td>2.9E-02</td><td>MB231/MB468</td><td>MP</td></tr>
<tr><td>>MI0000483|hsa-mir-186&WithFlank&1|-|71067625|71067722@21.44.24&[MIMAT0000456&hsa-miR-186-5p&offsets|0|+2;m-84&1|-|71067681|71067702&offsets|0|+2]|CAAAGAATTCTCCTTTTGGGCTTT</td><td>>MI0000483|hsa-mir-186&WithFlank&1|-|71067625|71067722</td><td>21</td><td>44</td><td>24</td><td>CAAAGAATTCTCCTTTTGGGCTTT</td><td>2964.6</td><td>-4.2</td><td>1.6</td><td>-2.7</td><td>7.0E-03</td><td>2.9E-02</td><td>BT20/MB231</td><td>MP</td></tr>
<tr><td>>hg38_21_+_8433172_8446622_RNA45SN1_45S_50ntFlanks@8340.8375.36|ACCGTAAGGGAAAGTTGAAAAGAACTTTGAAGAGAG</td><td>>hg38_21_+_8433172_8446622_RNA45SN1_45S_50ntFlanks</td><td>8340</td><td>8375</td><td>36</td><td>ACCGTAAGGGAAAGTTGAAAAGAACTTTGAAGAGAG</td><td>1756.3</td><td>4.9</td><td>1.8</td><td>2.7</td><td>7.0E-03</td><td>2.9E-02</td><td>BT20/MB231</td><td>MP</td></tr>
<tr><td>>hg38_21_+_8433172_8446622_RNA45SN1_45S_50ntFlanks@6649.6667.19|TACGACTCTTAGCGGTGGA</td><td>>hg38_21_+_8433172_8446622_RNA45SN1_45S_50ntFlanks</td><td>6649</td><td>6667</td><td>19</td><td>TACGACTCTTAGCGGTGGA</td><td>75.6</td><td>3.3</td><td>1.4</td><td>2.4</td><td>1.7E-02</td><td>2.9E-02</td><td>BT20/MB468</td><td>MP</td></tr>
<tr><td>>MI0001445|hsa-mir-423&WithFlank&17|+|30117073|30117178@59.78.20&[MIMAT0001340&hsa-miR-423-3p&offsets|0|-3;m-100&17|+|30117131|30117153&offsets|0|-3]|AGCTCGGTCTGAGGCCCCTC</td><td>>MI0001445|hsa-mir-423&WithFlank&17|+|30117073|30117178</td><td>59</td><td>78</td><td>20</td><td>AGCTCGGTCTGAGGCCCCTC</td><td>1065.3</td><td>1.5</td><td>0.7</td><td>2.2</td><td>2.5E-02</td><td>2.9E-02</td><td>BT20/MB468</td><td>NUC</td></tr>
<tr><td>>hg38_21_+_8433172_8446622_RNA45SN1_45S_50ntFlanks@12991.13039.49|CTCCCTCGCTGCGATCTATTGAAAGTCAGCCCTCGACACAAGGGTTTGT</td><td>>hg38_21_+_8433172_8446622_RNA45SN1_45S_50ntFlanks</td><td>12991</td><td>13039</td><td>49</td><td>CTCCCTCGCTGCGATCTATTGAAAGTCAGCCCTCGACACAAGGGTTTGT</td><td>2683.9</td><td>-2.0</td><td>0.9</td><td>-2.2</td><td>2.5E-02</td><td>2.9E-02</td><td>MB231/MB468</td><td>CYTO</td></tr>
<tr><td>>hg38_21_+_8433172_8446622_RNA45SN1_45S_50ntFlanks@11671.11706.36|CCAGTGCTCTGAATGTCAAAGTGAAGAAATTCAATG</td><td>>hg38_21_+_8433172_8446622_RNA45SN1_45S_50ntFlanks</td><td>11671</td><td>11706</td><td>36</td><td>CCAGTGCTCTGAATGTCAAAGTGAAGAAATTCAATG</td><td>309.3</td><td>-2.0</td><td>0.8</td><td>-2.6</td><td>9.4E-03</td><td>2.9E-02</td><td>BT20/MB231</td><td>CYTO</td></tr>
<tr><td>>hg38_21_+_8433172_8446622_RNA45SN1_45S_50ntFlanks@12912.12951.40|CACGTTCGTGGGGAACCTGGCGCTAAACCATTCGTAGACG</td><td>>hg38_21_+_8433172_8446622_RNA45SN1_45S_50ntFlanks</td><td>12912</td><td>12951</td><td>40</td><td>CACGTTCGTGGGGAACCTGGCGCTAAACCATTCGTAGACG</td><td>319.6</td><td>-3.1</td><td>1.4</td><td>-2.2</td><td>2.6E-02</td><td>2.9E-02</td><td>MB231/MB468</td><td>MITO</td></tr>
<tr><td>>hg38_21_+_8433172_8446622_RNA45SN1_45S_50ntFlanks@12928.12972.45|CTGGCGCTAAACCATTCGTAGACGACCTGCTTCTGGGTCGGGGTT</td><td>>hg38_21_+_8433172_8446622_RNA45SN1_45S_50ntFlanks</td><td>12928</td><td>12972</td><td>45</td><td>CTGGCGCTAAACCATTCGTAGACGACCTGCTTCTGGGTCGGGGTT</td><td>213.5</td><td>-3.1</td><td>1.3</td><td>-2.3</td><td>2.2E-02</td><td>2.9E-02</td><td>BT20/MB468</td><td>MITO</td></tr>
<tr><td>>hg38_21_+_8433172_8446622_RNA45SN1_45S_50ntFlanks@6732.6753.22|ACATTGATCATCGACACTTCGA</td><td>>hg38_21_+_8433172_8446622_RNA45SN1_45S_50ntFlanks</td><td>6732</td><td>6753</td><td>22</td><td>ACATTGATCATCGACACTTCGA</td><td>279.4</td><td>-1.3</td><td>0.6</td><td>-2.2</td><td>2.5E-02</td><td>2.9E-02</td><td>MB231/MB468</td><td>CYTO</td></tr>
<tr><td>>hg38_21_+_8433172_8446622_RNA45SN1_45S_50ntFlanks@4161.4186.26|CACATCCAAGGAAGGCAGCAGGCGCG</td><td>>hg38_21_+_8433172_8446622_RNA45SN1_45S_50ntFlanks</td><td>4161</td><td>4186</td><td>26</td><td>CACATCCAAGGAAGGCAGCAGGCGCG</td><td>75.4</td><td>-4.2</td><td>1.7</td><td>-2.4</td><td>1.7E-02</td><td>2.9E-02</td><td>BT20/MB468</td><td>MP</td></tr>
<tr><td>>hg38_21_+_8433172_8446622_RNA45SN1_45S_50ntFlanks@12893.12930.38|CCTCGCCCGTCACGCACCGCACGTTCGTGGGGAACCTG</td><td>>hg38_21_+_8433172_8446622_RNA45SN1_45S_50ntFlanks</td><td>12893</td><td>12930</td><td>38</td><td>CCTCGCCCGTCACGCACCGCACGTTCGTGGGGAACCTG</td><td>206.6</td><td>-3.8</td><td>1.7</td><td>-2.2</td><td>2.6E-02</td><td>2.9E-02</td><td>MB231/MB468</td><td>NUC</td></tr>
<tr><td>>MI0000062|hsa-let-7a-3&WithFlank&22|+|46112743|46112828@58.79.22;MI0000060|hsa-let-7a-1&WithFlank&9|+|94175951|94176042@63.84.22&[MIMAT0004481&hsa-let-7a-3p&offsets|0|+1;m-180&22|+|46112800|46112821&offsets|0|0];[MIMAT0004481_1&hsa-let-7a-3p&offsets|0|+1;m-181&9|+|94176013|94176034&offsets|0|0]|CTATACAATCTACTGTCTTTCC</td><td>>MI0000062|hsa-let-7a-3&WithFlank&22|+|46112743|46112828</td><td>58</td><td>79</td><td>22</td><td>CTATACAATCTACTGTCTTTCC</td><td>499.5</td><td>-2.3</td><td>0.9</td><td>-2.6</td><td>8.6E-03</td><td>2.9E-02</td><td>BT20/MB231</td><td>MITO</td></tr>
<tr><td>>hg38_21_+_8433172_8446622_RNA45SN1_45S_50ntFlanks@12407.12444.38|TTTTGATCCTTCGATGTCGGCTCTTCCTATCATTGTGA</td><td>>hg38_21_+_8433172_8446622_RNA45SN1_45S_50ntFlanks</td><td>12407</td><td>12444</td><td>38</td><td>TTTTGATCCTTCGATGTCGGCTCTTCCTATCATTGTGA</td><td>337.7</td><td>1.9</td><td>0.9</td><td>2.2</td><td>2.5E-02</td><td>2.9E-02</td><td>MB231/MB468</td><td>CYTO</td></tr>
<tr><td>>MI0000299|hsa-mir-222&WithFlank&X|-|45747009|45747130@75.98.24(+1A)&[MIMAT0000279&hsa-miR-222-3p&offsets|0|+3(+1A);m-140&X|-|45747033|45747056&offsets|0|0(+1A)]|AGCTACATCTGGCTACTGGGTCTCA</td><td>>MI0000299|hsa-mir-222&WithFlank&X|-|45747009|45747130</td><td>75</td><td>98</td><td>24(+1A)</td><td>AGCTACATCTGGCTACTGGGTCTCA</td><td>2884.5</td><td>-1.9</td><td>0.7</td><td>-2.7</td><td>7.6E-03</td><td>2.9E-02</td><td>BT20/MB231</td><td>NUC</td></tr>
</table>

| | | | | | | | | | | | | | |
|---|---|---|---|---|---|---|---|---|---|---|---|---|---|
| >MI0000299\|hsa-mir-222&WithFlank&X\|-\|45747009\|45747130@75.99.25(+1A)&[MIMAT0000279&hsa-miR-222-3p&offsets\|0\|+4(+1A);m-140&X\|-\|45747033\|45747056&offsets\|0\|+1(+1A)]\|AGCTACATCTGGCTACTGGGTCTCTA | >MI0000299\|hsa-mir-222&WithFlank&X\|-\|45747009\|45747130 | 75 | 99 | 25(+1A) | AGCTACATCTGGCTACTGGGTCTCTA | 196.6 | -2.3 | 0.9 | -2.7 | 7.6E-03 | 2.9E-02 | BT20/MB231 | NUC |
| >hg38_MT_+_1621_3279_RNR2_16S_50ntFlanks@1369.1405.37\|GATGTTGGATCAGGACATCCCGATGGTGCAGCCGCTA | >hg38_MT_+_1621_3279_RNR2_16S_50ntFlanks | 1369 | 1405 | 37 | GATGTTGGATCAGGACATCCCGATGGTGCAGCCGCTA | 30.2 | 4.6 | 2.0 | 2.3 | 2.3E-02 | 2.9E-02 | MB231/MB468 | MP |
| >hg38_MT_+_1621_3279_RNR2_16S_50ntFlanks@51.89.39\|GCTAAACCTAGCCCCAAACCCACTCCACCTTACTACCAG | >hg38_MT_+_1621_3279_RNR2_16S_50ntFlanks | 51 | 89 | 39 | GCTAAACCTAGCCCCAAACCCACTCCACCTTACTACCAG | 556.0 | 4.3 | 1.8 | 2.4 | 1.7E-02 | 2.9E-02 | BT20/MB468 | MP |
| >hg38_21_+_8433172_8446622_RNA45SN1_45S_50ntFlanks@9534.9569.36\|AGGAAACTCTGGTGGAGGTCCGTAGCGGTCCTGACG | >hg38_21_+_8433172_8446622_RNA45SN1_45S_50ntFlanks | 9534 | 9569 | 36 | AGGAAACTCTGGTGGAGGTCCGTAGCGGTCCTGACG | 1731.0 | -2.8 | 1.2 | -2.4 | 1.7E-02 | 2.9E-02 | BT20/MB468 | MP |
| >trna10_AspGTC_12_-_125424193_125424264@39.72.34,trna12_AspGTC_12_-_125411891_125411962@39.72.34,trna144_AspGTC_6_-_27551236_27551307@39.72.34,trna38_AspGTC_17_-_8125556_8125627@39.72.34,trna45_AspGTC_6_+_27447453_27447524@39.72.34,trna48_AspGTC_6_+_27471523_27471594@39.72.34,trna4_AspGTC_12_+_96429799_96429870@39.72.34,trna69_AspGTC_1_-_161440205_161440276@39.72.34,trna72_AspGTC_1_-_161432824_161432895@39.72.34,trna75_AspGTC_1_-_161425414_161425485@39.72.34,trna78_AspGTC_1_-_161418033_161418104@39.72.34,trna81_AspGTC_1_-_161410615_161410686@39.72.34\|GCGGGAGACCGGGGTTCGATTCCCCGACGGGGAG | >trna10_AspGTC_12_-_125424193_125424264 | 39 | 72 | 34 | GCGGGAGACCGGGGTTCGATTCCCCGACGGGGAG | 118.2 | -2.0 | 0.9 | -2.3 | 2.3E-02 | 2.9E-02 | MB231/MB468 | MP |
| >hg38_MT_+_1621_3279_RNR2_16S_50ntFlanks@465.491.27\|AATTTAACTGTTAGTCCAAAGAGGAAC | >hg38_MT_+_1621_3279_RNR2_16S_50ntFlanks | 465 | 491 | 27 | AATTTAACTGTTAGTCCAAAGAGGAAC | 64.1 | 4.6 | 2.0 | 2.3 | 2.3E-02 | 2.9E-02 | MB231/MB468 | MP |
| >MI0000811\|hsa-mir-148b&WithFlank&12\|+\|54337210\|54337320@69.89.21(+1C)&[MIMAT0000759&hsa-miR-148b-3p&offsets\|0\|-1(+1C);m-77&12\|+\|54337278\|54337299&offsets\|0\|-1(+1C)]\|TCAGTGCATCACAGAACTTTGC | >MI0000811\|hsa-mir-148b&WithFlank&12\|+\|54337210\|54337320 | 69 | 89 | 21(+1C) | TCAGTGCATCACAGAACTTTGC | 1381.1 | 3.0 | 1.3 | 2.4 | 1.7E-02 | 2.9E-02 | BT20/MB468 | MP |
| >hg38_21_+_8433172_8446622_RNA45SN1_45S_50ntFlanks@6645.6669.25\|CTCGTACGACTCTTAGCGGTGGATC | >hg38_21_+_8433172_8446622_RNA45SN1_45S_50ntFlanks | 6645 | 6669 | 25 | CTCGTACGACTCTTAGCGGTGGATC | 1414.1 | -2.2 | 0.9 | -2.3 | 2.3E-02 | 2.9E-02 | MB231/MB468 | MP |
| >hg38_21_+_8433172_8446622_RNA45SN1_45S_50ntFlanks@8115.8135.21\|CGCGGGACATGTGGCGTACGG | >hg38_21_+_8433172_8446622_RNA45SN1_45S_50ntFlanks | 8115 | 8135 | 21 | CGCGGGACATGTGGCGTACGG | 323.3 | 2.6 | 1.1 | 2.4 | 1.7E-02 | 2.9E-02 | BT20/MB468 | MP |
| >hg38_21_+_8433172_8446622_RNA45SN1_45S_50ntFlanks@10798.10821.24\|CATGTTGGAACAATGTAGGTAAGG | >hg38_21_+_8433172_8446622_RNA45SN1_45S_50ntFlanks | 10798 | 10821 | 24 | CATGTTGGAACAATGTAGGTAAGG | 149.0 | -2.2 | 0.9 | -2.4 | 1.7E-02 | 2.9E-02 | BT20/MB468 | MP |
| >am_trnaMT_HisGTG_MT_+_12138_12206@26.45.20\|AGATTGTGAATCTGACAACA | >am_trnaMT_HisGTG_MT_+_12138_12206 | 26 | 45 | 20 | AGATTGTGAATCTGACAACA | 185.4 | 2.4 | 1.0 | 2.4 | 1.7E-02 | 2.9E-02 | BT20/MB468 | MP |
| >hg38_21_+_8433172_8446622_RNA45SN1_45S_50ntFlanks@7981.8007.27\|CTCAGATCAGACGTGGCGACCCGCTGA | >hg38_21_+_8433172_8446622_RNA45SN1_45S_50ntFlanks | 7981 | 8007 | 27 | CTCAGATCAGACGTGGCGACCCGCTGA | 339.2 | -1.7 | 0.8 | -2.3 | 2.2E-02 | 2.9E-02 | BT20/MB468 | MITO |
| >MI0003780\|hsa-mir-1296&WithFlank&10\|-\|63372951\|63373054@22.41.20&[MIMAT0005794&hsa-miR-1296-5p&offsets\|0\|-2;m-304&10\|-\|63373012\|63373033&offsets\|0\|-2]\|TTAGGGCCCTGGCTCCATCT | >MI0003780\|hsa-mir-1296&WithFlank&10\|-\|63372951\|63373054 | 22 | 41 | 20 | TTAGGGCCCTGGCTCCATCT | 53.6 | 4.6 | 2.0 | 2.3 | 2.3E-02 | 2.9E-02 | MB231/MB468 | MP |
| >MI0000299\|hsa-mir-222&WithFlank&X\|-\|45747009\|45747130@75.96.22&[MIMAT0000279&hsa-miR-222-3p&offsets\|0\|+1;m-140&X\|-\|45747033\|45747056&offsets\|0\|-2]\|AGCTACATCTGGCTACTGGGTC | >MI0000299\|hsa-mir-222&WithFlank&X\|-\|45747009\|45747130 | 75 | 96 | 22 | AGCTACATCTGGCTACTGGGTC | 2621.4 | -1.8 | 0.7 | -2.6 | 8.7E-03 | 2.9E-02 | BT20/MB231 | MITO |
| >hg38_1_-_228634819_228635039_RNA5S12_5S_50ntFlanks@130.171.42\|TGGATGGGAGACCGCCTGGGAATACCGGGTGCTGTAGGCTTT | >hg38_1_-_228634819_228635039_RNA5S12_5S_50ntFlanks | 130 | 171 | 42 | TGGATGGGAGACCGCCTGGGAATACCGGGTGCTGTAGGCTTT | 1275.2 | 2.3 | 0.9 | 2.6 | 8.7E-03 | 2.9E-02 | BT20/MB231 | MITO |
| >MI0000268\|hsa-mir-34a&WithFlank&1\|-\|9151662\|9151783@28.50.23&[MIMAT0000255&hsa-miR-34a-5p&offsets\|0\|+1;m-119&1\|-\|9151735\|9151756&offsets\|0\|+1]\|TGGCAGTGTCTTAGCTGGTTGTT | >MI0000268\|hsa-mir-34a&WithFlank&1\|-\|9151662\|9151783 | 28 | 50 | 23 | TGGCAGTGTCTTAGCTGGTTGTT | 702.0 | -1.9 | 0.9 | -2.3 | 2.3E-02 | 2.9E-02 | MB231/MB468 | MP |
| >hg38_21_+_8433172_8446622_RNA45SN1_45S_50ntFlanks@12992.13013.22\|TCCCTCGCTGCGATCTATTGAA | >hg38_21_+_8433172_8446622_RNA45SN1_45S_50ntFlanks | 12992 | 13013 | 22 | TCCCTCGCTGCGATCTATTGAA | 1358.9 | -1.6 | 0.7 | -2.2 | 2.5E-02 | 2.9E-02 | MB231/MB468 | CYTO |
| >hg38_21_+_8433172_8446622_RNA45SN1_45S_50ntFlanks@9861.9893.33\|GATGAACCGAACGCCGGGTTAAGGCGCCCGATG | >hg38_21_+_8433172_8446622_RNA45SN1_45S_50ntFlanks | 9861 | 9893 | 33 | GATGAACCGAACGCCGGGTTAAGGCGCCCGATG | 130.7 | 5.3 | 2.0 | 2.7 | 7.0E-03 | 2.9E-02 | BT20/MB231 | MP |
| >hg38_21_+_8433172_8446622_RNA45SN1_45S_50ntFlanks@8916.8936.21\|GAATCCCGGGGCCGAGGGAGC | >hg38_21_+_8433172_8446622_RNA45SN1_45S_50ntFlanks | 8916 | 8936 | 21 | GAATCCCGGGGCCGAGGGAGC | 31.6 | 4.8 | 2.1 | 2.3 | 2.3E-02 | 2.9E-02 | MB231/MB468 | MP |
| >trnaMT_SerGCT_MT_+_12207_12265@8.30.23\|CTCACAAGAACTGCTAACTCATG | >trnaMT_SerGCT_MT_+_12207_12265 | 8 | 30 | 23 | CTCACAAGAACTGCTAACTCATG | 118.2 | -3.5 | 1.5 | -2.4 | 1.7E-02 | 2.9E-02 | BT20/MB468 | MP |
| >trnaMT_GluTTC_MT_-_14674_14742@37.71.35,trnalookalike8_GluTTC_5_-_93905172_93905240@37.71.35\|TCATTGGTCGTGGTTGTAGTCCGTGCGAGAATACC | >trnaMT_GluTTC_MT_-_14674_14742 | 37 | 71 | 35 | TCATTGGTCGTGGTTGTAGTCCGTGCGAGAATACC | 129.4 | 2.8 | 1.2 | 2.3 | 2.3E-02 | 2.9E-02 | MB231/MB468 | MP |
| >MI0000087\|hsa-mir-29a&WithFlank&7\|-\|130876741\|130876816@10.33.24&[MIMAT0004503&hsa-miR-29a-5p&offsets\|0\|+2;m-315&7\|-\|130876785\|130876807&offsets\|0\|+1]\|ACTGATTTCTTTTGGTGTTCAGAG | >MI0000087\|hsa-mir-29a&WithFlank&7\|-\|130876741\|130876816 | 10 | 33 | 24 | ACTGATTTCTTTTGGTGTTCAGAG | 68.5 | 4.5 | 2.0 | 2.3 | 2.3E-02 | 2.9E-02 | MB231/MB468 | MP |
| >MI0003815\|hsa-mir-1301&WithFlank&2\|-\|25328634\|25328727@54.76.23&[MIMAT0005797&hsa-miR-1301-3p&offsets\|0\|-1;m-256&2\|-\|25328650\|25328674&offsets\|0\|-2]\|TTGCAGCTGCCTGGGAGTGACTT | >MI0003815\|hsa-mir-1301&WithFlank&2\|-\|25328634\|25328727 | 54 | 76 | 23 | TTGCAGCTGCCTGGGAGTGACTT | 67.9 | 5.0 | 2.1 | 2.4 | 1.7E-02 | 2.9E-02 | BT20/MB468 | MP |
| >hg38_21_+_8433172_8446622_RNA45SN1_45S_50ntFlanks@8190.8213.24\|CCAGCCCGTGGACGGTGTGAGGCC | >hg38_21_+_8433172_8446622_RNA45SN1_45S_50ntFlanks | 8190 | 8213 | 24 | CCAGCCCGTGGACGGTGTGAGGCC | 496.9 | -1.8 | 0.8 | -2.2 | 2.6E-02 | 2.9E-02 | MB231/MB468 | NUC |
| >hg38_21_+_8433172_8446622_RNA45SN1_45S_50ntFlanks@12417.12449.33\|TCGATGTCGGCTCTTCCTATCATTGTGAAGCAG | >hg38_21_+_8433172_8446622_RNA45SN1_45S_50ntFlanks | 12417 | 12449 | 33 | TCGATGTCGGCTCTTCCTATCATTGTGAAGCAG | 385.0 | 1.3 | 0.6 | 2.2 | 2.5E-02 | 2.9E-02 | MB231/MB468 | CYTO |
| >hg38_21_+_8433172_8446622_RNA45SN1_45S_50ntFlanks@12008.12033.26\|CCGCCGGTGAAATACCACTACTCTGA | >hg38_21_+_8433172_8446622_RNA45SN1_45S_50ntFlanks | 12008 | 12033 | 26 | CCGCCGGTGAAATACCACTACTCTGA | 641.2 | -3.2 | 1.4 | -2.2 | 2.5E-02 | 2.9E-02 | BT20/MB468 | CYTO |
| >hg38_21_+_8433172_8446622_RNA45SN1_45S_50ntFlanks@11999.12029.31\|CCCTGCGGGCCGCCGGTGAAATACCACTACT | >hg38_21_+_8433172_8446622_RNA45SN1_45S_50ntFlanks | 11999 | 12029 | 31 | CCCTGCGGGCCGCCGGTGAAATACCACTACT | 105.9 | -2.7 | 1.2 | -2.3 | 2.2E-02 | 2.9E-02 | BT20/MB468 | MITO |
| >hg38_21_+_8433172_8446622_RNA45SN1_45S_50ntFlanks@9940.9978.39\|CAGGACGGTGGCCATGGAAGTCGGAATCCGCTAAGGAGT | >hg38_21_+_8433172_8446622_RNA45SN1_45S_50ntFlanks | 9940 | 9978 | 39 | CAGGACGGTGGCCATGGAAGTCGGAATCCGCTAAGGAGT | 41.2 | 4.5 | 2.0 | 2.3 | 2.3E-02 | 2.9E-02 | MB231/MB468 | MP |
| >hg38_21_+_8433172_8446622_RNA45SN1_45S_50ntFlanks@10686.10706.21\|GCGGCGTCCGGTGAGCTCTCG | >hg38_21_+_8433172_8446622_RNA45SN1_45S_50ntFlanks | 10686 | 10706 | 21 | GCGGCGTCCGGTGAGCTCTCG | 792.7 | -2.8 | 1.3 | -2.2 | 2.5E-02 | 2.9E-02 | BT20/MB468 | CYTO |
| >hg38_21_+_8433172_8446622_RNA45SN1_45S_50ntFlanks@11569.11612.44\|CAGCCGACTTAGAACTGGTGCGGACCAGGGGAATCCGACTGTTT | >hg38_21_+_8433172_8446622_RNA45SN1_45S_50ntFlanks | 11569 | 11612 | 44 | CAGCCGACTTAGAACTGGTGCGGACCAGGGGAATCCGACTGTTT | 677.2 | -3.5 | 1.5 | -2.2 | 2.5E-02 | 2.9E-02 | BT20/MB468 | CYTO |
| >hg38_21_+_8433172_8446622_RNA45SN1_45S_50ntFlanks@12912.12930.19\|CACGTTCGTGGGGAACCTG | >hg38_21_+_8433172_8446622_RNA45SN1_45S_50ntFlanks | 12912 | 12930 | 19 | CACGTTCGTGGGGAACCTG | 118.9 | -3.1 | 1.4 | -2.2 | 2.5E-02 | 2.9E-02 | BT20/MB468 | CYTO |
| >hg38_MT_+_1621_3279_RNR2_16S_50ntFlanks@1425.1452.28\|ACGATTAAAGTCCTACGTGATCTGAGTT | >hg38_MT_+_1621_3279_RNR2_16S_50ntFlanks | 1425 | 1452 | 28 | ACGATTAAAGTCCTACGTGATCTGAGTT | 169.1 | 4.1 | 1.8 | 2.3 | 2.3E-02 | 2.9E-02 | MB231/MB468 | MP |
| >hg38_21_+_8433172_8446622_RNA45SN1_45S_50ntFlanks@4130.4150.21\|ATTCCGGAGAGGGAGCCTGAG | >hg38_21_+_8433172_8446622_RNA45SN1_45S_50ntFlanks | 4130 | 4150 | 21 | ATTCCGGAGAGGGAGCCTGAG | 201.5 | -2.4 | 1.0 | -2.3 | 2.2E-02 | 2.9E-02 | BT20/MB468 | MITO |
| >hg38_21_+_8433172_8446622_RNA45SN1_45S_50ntFlanks@11323.11343.21\|CGTCTCGTCGCGCGCGCGTCC | >hg38_21_+_8433172_8446622_RNA45SN1_45S_50ntFlanks | 11323 | 11343 | 21 | CGTCTCGTCGCGCGCGCGTCC | 297.5 | 2.7 | 1.1 | 2.6 | 9.5E-03 | 2.9E-02 | BT20/MB231 | CYTO |
| >hg38_MT_+_1621_3279_RNR2_16S_50ntFlanks@1367.1395.29\|TCGATGTTGGATCAGGACATCCCGATGGT | >hg38_MT_+_1621_3279_RNR2_16S_50ntFlanks | 1367 | 1395 | 29 | TCGATGTTGGATCAGGACATCCCGATGGT | 114.9 | 4.4 | 1.8 | 2.4 | 1.7E-02 | 2.9E-02 | BT20/MB468 | MP |
| >hg38_21_+_8433172_8446622_RNA45SN1_45S_50ntFlanks@7980.8010.31\|CCTCAGATCAGACGTGGCGACCCGCTGAATT | >hg38_21_+_8433172_8446622_RNA45SN1_45S_50ntFlanks | 7980 | 8010 | 31 | CCTCAGATCAGACGTGGCGACCCGCTGAATT | 374.8 | 1.7 | 0.8 | 2.3 | 2.2E-02 | 2.9E-02 | BT20/MB468 | MITO |
| >hg38_21_+_8433172_8446622_RNA45SN1_45S_50ntFlanks@4995.5024.30\|ACAGATTGATAGCTCTTTCTCGATTCCGTG | >hg38_21_+_8433172_8446622_RNA45SN1_45S_50ntFlanks | 4995 | 5024 | 30 | ACAGATTGATAGCTCTTTCTCGATTCCGTG | 1584.0 | -4.9 | 2.0 | -2.4 | 1.7E-02 | 2.9E-02 | BT20/MB468 | MP |
| >trna12_ArgCCT_16_+_3243918_3243990@35.53.19,trna18_ArgCCT_17_+_73030001_73030073@35.53.19,trna21_ArgCCT_17_-_73030526_73030598@35.53.19,trna2_ArgCCT_16_+_3202901_3202973@35.53.19,trna3_ArgCCT_7_+_139025446_139025518@35.53.19\|CTAAGCCAGGGATTGTGGG | >trna12_ArgCCT_16_+_3243918_3243990 | 35 | 53 | 19 | CTAAGCCAGGGATTGTGGG | 176.2 | -1.9 | 0.9 | -2.2 | 2.5E-02 | 2.9E-02 | MB231/MB468 | CYTO |
| >hg38_21_+_8433172_8446622_RNA45SN1_45S_50ntFlanks@9737.9765.29\|CGAAACGATCTCAACCTATTCTCAAACTT | >hg38_21_+_8433172_8446622_RNA45SN1_45S_50ntFlanks | 9737 | 9765 | 29 | CGAAACGATCTCAACCTATTCTCAAACTT | 788.0 | 1.9 | 0.9 | 2.2 | 2.6E-02 | 2.9E-02 | MB231/MB468 | NUC |
| >MI0000542\|hsa-mir-320a&WithFlank&8\|-\|22244960\|22245043@48.69.22&[MIMAT0000510&hsa-miR-320a-3p&offsets\|0\|0;m-60&8\|-\|22244975\|22244996&offsets\|0\|0]\|AAAAGCTGGGTTGAGAGGGCGA | >MI0000542\|hsa-mir-320a&WithFlank&8\|-\|22244960\|22245043 | 48 | 69 | 22 | AAAAGCTGGGTTGAGAGGGCGA | 80280.6 | 2.5 | 0.9 | 2.7 | 7.1E-03 | 2.9E-02 | BT20/MB231 | MP |
| >hg38_21_+_8433172_8446622_RNA45SN1_45S_50ntFlanks@5238.5256.19\|CTGACTGGCTCAGCGTGTG | >hg38_21_+_8433172_8446622_RNA45SN1_45S_50ntFlanks | 5238 | 5256 | 19 | CTGACTGGCTCAGCGTGTG | 287.9 | -1.9 | 0.8 | -2.3 | 2.3E-02 | 2.9E-02 | MB231/MB468 | MP |
| >trnaMT_PheGAA_MT_+_577_647@46.74.29\|GACGGGCTCACATCACCCCATAAACACCA | >trnaMT_PheGAA_MT_+_577_647 | 46 | 74 | 29 | GACGGGCTCACATCACCCCATAAACACCA | 254.4 | 2.9 | 1.2 | 2.4 | 1.7E-02 | 2.9E-02 | BT20/MB468 | MP |
| >hg38_21_+_8433172_8446622_RNA45SN1_45S_50ntFlanks@9390.9415.26\|GCCGAGGTGGGATCCCGAGGCCTCTC | >hg38_21_+_8433172_8446622_RNA45SN1_45S_50ntFlanks | 9390 | 9415 | 26 | GCCGAGGTGGGATCCCGAGGCCTCTC | 140.4 | -4.1 | 1.7 | -2.4 | 1.7E-02 | 2.9E-02 | BT20/MB468 | MP |
| >hg38_21_+_8433172_8446622_RNA45SN1_45S_50ntFlanks@10679.10709.31\|GGTTCCGGCGGCGTCCGGTGAGCTCTCGCTG | >hg38_21_+_8433172_8446622_RNA45SN1_45S_50ntFlanks | 10679 | 10709 | 31 | GGTTCCGGCGGCGTCCGGTGAGCTCTCGCTG | 3815.3 | -3.8 | 1.6 | -2.4 | 1.7E-02 | 2.9E-02 | BT20/MB468 | MP |
| >hg38_21_+_8433172_8446622_RNA45SN1_45S_50ntFlanks@6647.6665.19\|CGTACGACTCTTAGCGGTG | >hg38_21_+_8433172_8446622_RNA45SN1_45S_50ntFlanks | 6647 | 6665 | 19 | CGTACGACTCTTAGCGGTG | 17554.0 | 3.2 | 1.2 | 2.7 | 7.1E-03 | 2.9E-02 | BT20/MB231 | MP |
| >hg38_21_+_8433172_8446622_RNA45SN1_45S_50ntFlanks@6650.6681.32\|ACGACTCTTAGCGGTGGATCACTCGGCTCGTG | >hg38_21_+_8433172_8446622_RNA45SN1_45S_50ntFlanks | 6650 | 6681 | 32 | ACGACTCTTAGCGGTGGATCACTCGGCTCGTG | 1525.1 | 3.4 | 1.3 | 2.7 | 7.1E-03 | 2.9E-02 | BT20/MB231 | MP |
| >hg38_21_+_8433172_8446622_RNA45SN1_45S_50ntFlanks@6653.6681.29\|ACTCTTAGCGGTGGATCACTCGGCTCGTG | >hg38_21_+_8433172_8446622_RNA45SN1_45S_50ntFlanks | 6653 | 6681 | 29 | ACTCTTAGCGGTGGATCACTCGGCTCGTG | 430.5 | 3.2 | 1.2 | 2.7 | 7.1E-03 | 2.9E-02 | BT20/MB231 | MP |

<table>
<tr><td>>MI0000488|hsa-mir-194-1&WithFlank&1|-|220118151|220118247@21.41.21;MI0000732|hsa-mir-194-2&WithFlank&11|-|64891349|64891445@21.41.21&[MIMAT0000460&hsa-miR-194-5p&offsets|0|-1;m-131&1|-|220118206|220118227&offsets|0|-1];[MIMAT0000460_1&hsa-miR-194-5p&offsets|0|-1;m-126&11|-|64891404|64891425&offsets|0|-1]|TGTAACAGCAACTCCATGTGG</td><td>>MI0000488|hsa-mir-194-1&WithFlank&1|-|220118151|220118247</td><td>21</td><td>41</td><td>21</td><td>TGTAACAGCAACTCCATGTGG</td><td>243.7</td><td>3.0</td><td>1.3</td><td>2.4</td><td>1.7E-02</td><td>2.9E-02</td><td>BT20/MB468</td><td>MP</td></tr>
<tr><td>>hg38_21_+_8433172_8446622_RNA45SN1_45S_50ntFlanks@7990.8019.30|GACGTGGCGACCCGCTGAATTTAAGCATAT</td><td>>hg38_21_+_8433172_8446622_RNA45SN1_45S_50ntFlanks</td><td>7990</td><td>8019</td><td>30</td><td>GACGTGGCGACCCGCTGAATTTAAGCATAT</td><td>129.6</td><td>3.3</td><td>1.5</td><td>2.2</td><td>2.6E-02</td><td>2.9E-02</td><td>MB231/MB468</td><td>MITO</td></tr>
<tr><td>>hg38_21_+_8433172_8446622_RNA45SN1_45S_50ntFlanks@12407.12446.40|TTTTGATCCTTCGATGTCGGCTCTTCCTATCATTGTGAAG</td><td>>hg38_21_+_8433172_8446622_RNA45SN1_45S_50ntFlanks</td><td>12407</td><td>12446</td><td>40</td><td>TTTTGATCCTTCGATGTCGGCTCTTCCTATCATTGTGAAG</td><td>324.9</td><td>-1.6</td><td>0.7</td><td>-2.3</td><td>2.2E-02</td><td>2.9E-02</td><td>BT20/MB468</td><td>MITO</td></tr>
<tr><td>>hg38_21_+_8433172_8446622_RNA45SN1_45S_50ntFlanks@11598.11621.24|GGAATCCGACTGTTTAATTAAAAC</td><td>>hg38_21_+_8433172_8446622_RNA45SN1_45S_50ntFlanks</td><td>11598</td><td>11621</td><td>24</td><td>GGAATCCGACTGTTTAATTAAAAC</td><td>148.6</td><td>4.4</td><td>1.7</td><td>2.7</td><td>7.1E-03</td><td>2.9E-02</td><td>BT20/MB231</td><td>MP</td></tr>
<tr><td>>MI0000081|hsa-mir-24-2&WithFlank&19|-|13836281|13836365@56.75.20(+2G);MI0000080|hsa-mir-24-1&WithFlank&9|+|95086015|95086094@50.69.20(+2G)&[MIMAT0000080&hsa-miR-24-3p&offsets|0|-2(+2G);m-36&19|-|13836288|13836310&offsets|0|-3(+2G)];[MIMAT0000080_1&hsa-miR-24-3p&offsets|0|-2(+2G);m-35&9|+|95086064|95086086&offsets|0|-3(+2G)]|TGGCTCAGTTCAGCAGGAACGG</td><td>>MI0000081|hsa-mir-24-2&WithFlank&19|-|13836281|13836365</td><td>56</td><td>75</td><td>20(+2G)</td><td>TGGCTCAGTTCAGCAGGAACGG</td><td>173.2</td><td>3.6</td><td>1.5</td><td>2.4</td><td>1.7E-02</td><td>2.9E-02</td><td>BT20/MB468</td><td>MP</td></tr>
<tr><td>>MI0000095|hsa-mir-93&WithFlank&7|-|100093762|100093853@17.37.21&[MIMAT0000093&hsa-miR-93-5p&offsets|0|-2;m-29&7|-|100093815|100093837&offsets|0|-2]|CAAAGTGCTGTTCGTGCAGGT</td><td>>MI0000095|hsa-mir-93&WithFlank&7|-|100093762|100093853</td><td>17</td><td>37</td><td>21</td><td>CAAAGTGCTGTTCGTGCAGGT</td><td>2686.5</td><td>3.0</td><td>1.3</td><td>2.4</td><td>1.7E-02</td><td>2.9E-02</td><td>BT20/MB468</td><td>MP</td></tr>
<tr><td>>trnaMT_SerGCT_MT_+_12207_12265@10.38.29|CACAAGAACTGCTAACTCATGCCCCCATG</td><td>>trnaMT_SerGCT_MT_+_12207_12265</td><td>10</td><td>38</td><td>29</td><td>CACAAGAACTGCTAACTCATGCCCCCATG</td><td>245.4</td><td>-4.2</td><td>1.9</td><td>-2.2</td><td>2.5E-02</td><td>2.9E-02</td><td>BT20/MB468</td><td>CYTO</td></tr>
<tr><td>>hg38_21_+_8433172_8446622_RNA45SN1_45S_50ntFlanks@6649.6674.26|TACGACTCTTAGCGGTGGATCACTCG</td><td>>hg38_21_+_8433172_8446622_RNA45SN1_45S_50ntFlanks</td><td>6649</td><td>6674</td><td>26</td><td>TACGACTCTTAGCGGTGGATCACTCG</td><td>328.3</td><td>4.9</td><td>1.8</td><td>2.7</td><td>7.1E-03</td><td>2.9E-02</td><td>BT20/MB231</td><td>MP</td></tr>
<tr><td>>hg38_1_-_228634819_228635039_RNA5S12_5S_50ntFlanks@51.70.20|GTCTACGGCCATACCACCCT</td><td>>hg38_1_-_228634819_228635039_RNA5S12_5S_50ntFlanks</td><td>51</td><td>70</td><td>20</td><td>GTCTACGGCCATACCACCCT</td><td>724.3</td><td>3.5</td><td>1.3</td><td>2.7</td><td>7.1E-03</td><td>2.9E-02</td><td>BT20/MB231</td><td>MP</td></tr>
<tr><td>>MI0000273|hsa-mir-183&WithFlank&7|-|129774899|129775020@33.51.19&[MIMAT0000261&hsa-miR-183-5p&offsets|0|-3;m-41&7|-|129774966|129774987&offsets|-1|-4]|TATGGCACTGGTAGAATTC</td><td>>MI0000273|hsa-mir-183&WithFlank&7|-|129774899|129775020</td><td>33</td><td>51</td><td>19</td><td>TATGGCACTGGTAGAATTC</td><td>123.1</td><td>5.3</td><td>2.2</td><td>2.4</td><td>1.7E-02</td><td>2.9E-02</td><td>BT20/MB468</td><td>MP</td></tr>
<tr><td>>MI0000061|hsa-let-7a-2&WithFlank&11|-|122146516|122146599@11.32.22;MI0000062|hsa-let-7a-3&WithFlank&22|+|46112743|46112828@10.31.22;MI0000060|hsa-let-7a-1&WithFlank&9|+|94175951|94176042@12.33.22&[MIMAT0000062&hsa-let-7a-5p&offsets|0|0;m-6&11|-|122146568|122146589&offsets|0|0];[MIMAT0000062_1&hsa-let-7a-5p&offsets|0|0;m-5&22|+|46112752|46112773&offsets|0|0];[MIMAT0000062_2&hsa-let-7a-5p&offsets|0|0;m-4&9|+|94175962|94175983&offsets|0|0]|TGAGGTAGTAGGTTGTATAGTT</td><td>>MI0000061|hsa-let-7a-2&WithFlank&11|-|122146516|122146599</td><td>11</td><td>32</td><td>22</td><td>TGAGGTAGTAGGTTGTATAGTT</td><td>97653.8</td><td>-2.6</td><td>1.0</td><td>-2.7</td><td>7.1E-03</td><td>2.9E-02</td><td>BT20/MB231</td><td>MP</td></tr>
<tr><td>>trna10_AspGTC_12_-_125424193_125424264@33.69.37,trna12_AspGTC_12_-_125411891_125411962@33.69.37,trna144_AspGTC_6_-_27551236_27551307@33.69.37,trna38_AspGTC_17_-_8125556_8125627@33.69.37,trna45_AspGTC_6_+_27447453_27447524@33.69.37,trna48_AspGTC_6_+_27471523_27471594@33.69.37,trna4_AspGTC_12_+_96429799_96429870@33.69.37,trna69_AspGTC_1_-_161440205_161440276@33.69.37,trna72_AspGTC_1_-_161432824_161432895@33.69.37,trna75_AspGTC_1_-_161425414_161425485@33.69.37,trna78_AspGTC_1_-_161418033_161418104@33.69.37,trna81_AspGTC_1_-_161410615_161410686@33.69.37|TGTCACGCGGGAGACCGGGGTTCGATTCCCCGACGGG</td><td>>trna10_AspGTC_12_-_125424193_125424264</td><td>33</td><td>69</td><td>37</td><td>TGTCACGCGGGAGACCGGGGTTCGATTCCCCGACGGG</td><td>130.9</td><td>3.4</td><td>1.4</td><td>2.4</td><td>1.7E-02</td><td>2.9E-02</td><td>BT20/MB468</td><td>MP</td></tr>
<tr><td>>hg38_21_+_8433172_8446622_RNA45SN1_45S_50ntFlanks@4732.4766.35|AGAACGAAAGTCGGAGGTTCGAAGACGATCAGATA</td><td>>hg38_21_+_8433172_8446622_RNA45SN1_45S_50ntFlanks</td><td>4732</td><td>4766</td><td>35</td><td>AGAACGAAAGTCGGAGGTTCGAAGACGATCAGATA</td><td>117.6</td><td>3.3</td><td>1.4</td><td>2.4</td><td>1.7E-02</td><td>2.9E-02</td><td>BT20/MB468</td><td>MP</td></tr>
<tr><td>>hg38_21_+_8433172_8446622_RNA45SN1_45S_50ntFlanks@10347.10371.25|CAAACGAGAACTTTGAAGGCCGAAG</td><td>>hg38_21_+_8433172_8446622_RNA45SN1_45S_50ntFlanks</td><td>10347</td><td>10371</td><td>25</td><td>CAAACGAGAACTTTGAAGGCCGAAG</td><td>124.5</td><td>-1.9</td><td>0.8</td><td>-2.3</td><td>2.4E-02</td><td>2.9E-02</td><td>MB231/MB468</td><td>MP</td></tr>
<tr><td>>hg38_21_+_8433172_8446622_RNA45SN1_45S_50ntFlanks@7990.8017.28|GACGTGGCGACCCGCTGAATTTAAGCAT</td><td>>hg38_21_+_8433172_8446622_RNA45SN1_45S_50ntFlanks</td><td>7990</td><td>8017</td><td>28</td><td>GACGTGGCGACCCGCTGAATTTAAGCAT</td><td>570.9</td><td>-2.8</td><td>1.1</td><td>-2.6</td><td>9.6E-03</td><td>2.9E-02</td><td>BT20/MB231</td><td>CYTO</td></tr>
<tr><td>>hg38_21_+_8433172_8446622_RNA45SN1_45S_50ntFlanks@9621.9658.38|TCTAGTAGCTGGTTCCCTCCGAAGTTTCCCTCAGGATA</td><td>>hg38_21_+_8433172_8446622_RNA45SN1_45S_50ntFlanks</td><td>9621</td><td>9658</td><td>38</td><td>TCTAGTAGCTGGTTCCCTCCGAAGTTTCCCTCAGGATA</td><td>136.2</td><td>-4.2</td><td>1.8</td><td>-2.4</td><td>1.7E-02</td><td>2.9E-02</td><td>BT20/MB468</td><td>MP</td></tr>
<tr><td>>trna116_GluCTC_1_-_145399233_145399304@1.39.39,trna59_GluCTC_1_+_249168447_249168518@1.39.39,trna71_GluCTC_1_-_161439189_161439260@1.39.39,trna74_GluCTC_1_-_161431809_161431880@1.39.39,trna77_GluCTC_1_-_161424398_161424469@1.39.39,trna77_GluCTC_6_+_28949976_28950047@1.39.39,trna80_GluCTC_1_-_161417018_161417089@1.39.39,trna87_GluCTC_6_-_126101393_126101464@1.39.39|TCCCTGGTGGTCTAGTGGTTAGGATTCGGCGCTCTCACC</td><td>>trna116_GluCTC_1_-_145399233_145399304</td><td>1</td><td>39</td><td>39</td><td>TCCCTGGTGGTCTAGTGGTTAGGATTCGGCGCTCTCACC</td><td>36588.5</td><td>-2.1</td><td>0.8</td><td>-2.6</td><td>9.6E-03</td><td>2.9E-02</td><td>BT20/MB231</td><td>CYTO</td></tr>
<tr><td>>MI0000434|hsa-let-7i&WithFlank&12|+|62603680|62603775@12.33.22(+2A)&[MIMAT0000415&hsa-let-7i-5p&offsets|0|0(+2A);m-49&12|+|62603691|62603712&offsets|0|0(+2A)]|TGAGGTAGTAGTTTGTGCTGTTAA</td><td>>MI0000434|hsa-let-7i&WithFlank&12|+|62603680|62603775</td><td>12</td><td>33</td><td>22(+2A)</td><td>TGAGGTAGTAGTTTGTGCTGTTAA</td><td>93.2</td><td>3.4</td><td>1.4</td><td>2.4</td><td>1.7E-02</td><td>2.9E-02</td><td>BT20/MB468</td><td>MP</td></tr>
<tr><td>>hg38_MT_+_598_1651_RNR1_12S_50ntFlanks@718.745.28|GTAAAGACGTTAGGTCAAGGTGTAGCCC</td><td>>hg38_MT_+_598_1651_RNR1_12S_50ntFlanks</td><td>718</td><td>745</td><td>28</td><td>GTAAAGACGTTAGGTCAAGGTGTAGCCC</td><td>86.5</td><td>4.7</td><td>2.1</td><td>2.3</td><td>2.4E-02</td><td>2.9E-02</td><td>MB231/MB468</td><td>MP</td></tr>
<tr><td>>hg38_21_+_8433172_8446622_RNA45SN1_45S_50ntFlanks@4731.4750.20|AAGAACGAAAGTCGGAGGTT</td><td>>hg38_21_+_8433172_8446622_RNA45SN1_45S_50ntFlanks</td><td>4731</td><td>4750</td><td>20</td><td>AAGAACGAAAGTCGGAGGTT</td><td>123.5</td><td>-3.0</td><td>1.3</td><td>-2.3</td><td>2.4E-02</td><td>2.9E-02</td><td>MB231/MB468</td><td>MP</td></tr>
<tr><td>>hg38_1_-_228634819_228635039_RNA5S12_5S_50ntFlanks@146.169.24|TGGGAATACCGGGTGCTGTAGGCT</td><td>>hg38_1_-_228634819_228635039_RNA5S12_5S_50ntFlanks</td><td>146</td><td>169</td><td>24</td><td>TGGGAATACCGGGTGCTGTAGGCT</td><td>136.9</td><td>1.5</td><td>0.7</td><td>2.2</td><td>2.6E-02</td><td>2.9E-02</td><td>BT20/MB468</td><td>NUC</td></tr>
<tr><td>>hg38_21_+_8433172_8446622_RNA45SN1_45S_50ntFlanks@5237.5256.20|ACTGACTGGCTCAGCGTGTG</td><td>>hg38_21_+_8433172_8446622_RNA45SN1_45S_50ntFlanks</td><td>5237</td><td>5256</td><td>20</td><td>ACTGACTGGCTCAGCGTGTG</td><td>526.2</td><td>-2.9</td><td>1.3</td><td>-2.2</td><td>2.6E-02</td><td>2.9E-02</td><td>MB231/MB468</td><td>MITO</td></tr>
<tr><td>>hg38_21_+_8433172_8446622_RNA45SN1_45S_50ntFlanks@10508.10532.25|CGAATCCGGAGTGGCGGAGATGGGC</td><td>>hg38_21_+_8433172_8446622_RNA45SN1_45S_50ntFlanks</td><td>10508</td><td>10532</td><td>25</td><td>CGAATCCGGAGTGGCGGAGATGGGC</td><td>467.5</td><td>4.8</td><td>1.8</td><td>2.7</td><td>7.1E-03</td><td>2.9E-02</td><td>BT20/MB231</td><td>MP</td></tr>
<tr><td>>hg38_21_+_8433172_8446622_RNA45SN1_45S_50ntFlanks@7976.8010.35|GCGACCTCAGATCAGACGTGGCGACCCGCTGAATT</td><td>>hg38_21_+_8433172_8446622_RNA45SN1_45S_50ntFlanks</td><td>7976</td><td>8010</td><td>35</td><td>GCGACCTCAGATCAGACGTGGCGACCCGCTGAATT</td><td>58.6</td><td>4.6</td><td>2.1</td><td>2.3</td><td>2.4E-02</td><td>2.9E-02</td><td>MB231/MB468</td><td>MP</td></tr>
<tr><td>>hg38_21_+_8433172_8446622_RNA45SN1_45S_50ntFlanks@9533.9561.29|GAGGAAACTCTGGTGGAGGTCCGTAGCGG</td><td>>hg38_21_+_8433172_8446622_RNA45SN1_45S_50ntFlanks</td><td>9533</td><td>9561</td><td>29</td><td>GAGGAAACTCTGGTGGAGGTCCGTAGCGG</td><td>306.0</td><td>-3.2</td><td>1.3</td><td>-2.4</td><td>1.7E-02</td><td>2.9E-02</td><td>BT20/MB468</td><td>MP</td></tr>
<tr><td>>hg38_21_+_8433172_8446622_RNA45SN1_45S_50ntFlanks@12152.12179.28|TGGGGAGTTTGACTGGGGCGGTACACCT</td><td>>hg38_21_+_8433172_8446622_RNA45SN1_45S_50ntFlanks</td><td>12152</td><td>12179</td><td>28</td><td>TGGGGAGTTTGACTGGGGCGGTACACCT</td><td>172.4</td><td>-4.1</td><td>1.7</td><td>-2.4</td><td>1.7E-02</td><td>2.9E-02</td><td>BT20/MB468</td><td>MP</td></tr>
<tr><td>>hg38_21_+_8433172_8446622_RNA45SN1_45S_50ntFlanks@12405.12446.42|CTTTTTGATCCTTCGATGTCGGCTCTTCCTATCATTGTGAAG</td><td>>hg38_21_+_8433172_8446622_RNA45SN1_45S_50ntFlanks</td><td>12405</td><td>12446</td><td>42</td><td>CTTTTTGATCCTTCGATGTCGGCTCTTCCTATCATTGTGAAG</td><td>1667.5</td><td>-2.9</td><td>1.3</td><td>-2.2</td><td>2.5E-02</td><td>2.9E-02</td><td>MB231/MB468</td><td>CYTO</td></tr>
<tr><td>>hg38_21_+_8433172_8446622_RNA45SN1_45S_50ntFlanks@10679.10708.30|GGTTCCGGCGGCGTCCGGTGAGCTCTCGCT</td><td>>hg38_21_+_8433172_8446622_RNA45SN1_45S_50ntFlanks</td><td>10679</td><td>10708</td><td>30</td><td>GGTTCCGGCGGCGTCCGGTGAGCTCTCGCT</td><td>2277.7</td><td>-2.8</td><td>1.2</td><td>-2.4</td><td>1.7E-02</td><td>2.9E-02</td><td>BT20/MB468</td><td>MP</td></tr>
<tr><td>>hg38_21_+_8433172_8446622_RNA45SN1_45S_50ntFlanks@10713.10735.23|CTTGAAAATCCGGGGGAGAGGGT</td><td>>hg38_21_+_8433172_8446622_RNA45SN1_45S_50ntFlanks</td><td>10713</td><td>10735</td><td>23</td><td>CTTGAAAATCCGGGGGAGAGGGT</td><td>205.3</td><td>-2.3</td><td>1.0</td><td>-2.4</td><td>1.7E-02</td><td>2.9E-02</td><td>BT20/MB468</td><td>MP</td></tr>
<tr><td>>hg38_21_+_8433172_8446622_RNA45SN1_45S_50ntFlanks@12001.12030.30|CTGCGGGCCGCCGGTGAAATACCACTACTC</td><td>>hg38_21_+_8433172_8446622_RNA45SN1_45S_50ntFlanks</td><td>12001</td><td>12030</td><td>30</td><td>CTGCGGGCCGCCGGTGAAATACCACTACTC</td><td>194.5</td><td>-3.5</td><td>1.6</td><td>-2.2</td><td>2.5E-02</td><td>2.9E-02</td><td>BT20/MB468</td><td>CYTO</td></tr>
<tr><td>>hg38_21_+_8433172_8446622_RNA45SN1_45S_50ntFlanks@7975.8011.37|CGCGACCTCAGATCAGACGTGGCGACCCGCTGAATTT</td><td>>hg38_21_+_8433172_8446622_RNA45SN1_45S_50ntFlanks</td><td>7975</td><td>8011</td><td>37</td><td>CGCGACCTCAGATCAGACGTGGCGACCCGCTGAATTT</td><td>513127.9</td><td>2.2</td><td>0.8</td><td>2.7</td><td>7.7E-03</td><td>2.9E-02</td><td>BT20/MB231</td><td>NUC</td></tr>
<tr><td>>hg38_21_+_8433172_8446622_RNA45SN1_45S_50ntFlanks@7993.8014.22|GTGGCGACCCGCTGAATTTAAG</td><td>>hg38_21_+_8433172_8446622_RNA45SN1_45S_50ntFlanks</td><td>7993</td><td>8014</td><td>22</td><td>GTGGCGACCCGCTGAATTTAAG</td><td>175.7</td><td>-1.6</td><td>0.7</td><td>-2.3</td><td>2.2E-02</td><td>2.9E-02</td><td>BT20/MB468</td><td>MITO</td></tr>
<tr><td>>MI0000750|hsa-mir-26a-2&WithFlank&12|-|57824603|57824698@20.41.22;MI0000083|hsa-mir-26a-1&WithFlank&3|+|37969398|37969486@16.37.22&[MIMAT0000082&hsa-miR-26a-5p&offsets|0|0;m-33&12|-|57824658|57824679&offsets|0|0];[MIMAT0000082_1&hsa-miR-26a-5p&offsets|0|0;m-34&3|+|37969413|37969434&offsets|0|0]|TTCAAGTAATCCAGGATAGGCT</td><td>>MI0000750|hsa-mir-26a-2&WithFlank&12|-|57824603|57824698</td><td>20</td><td>41</td><td>22</td><td>TTCAAGTAATCCAGGATAGGCT</td><td>150821.8</td><td>-1.7</td><td>0.7</td><td>-2.6</td><td>8.8E-03</td><td>2.9E-02</td><td>BT20/MB231</td><td>MITO</td></tr>
<tr><td>>hg38_21_+_8433172_8446622_RNA45SN1_45S_50ntFlanks@11587.11616.30|TGCGGACCAGGGGAATCCGACTGTTTAATT</td><td>>hg38_21_+_8433172_8446622_RNA45SN1_45S_50ntFlanks</td><td>11587</td><td>11616</td><td>30</td><td>TGCGGACCAGGGGAATCCGACTGTTTAATT</td><td>200.0</td><td>-2.1</td><td>0.8</td><td>-2.6</td><td>9.7E-03</td><td>2.9E-02</td><td>BT20/MB231</td><td>CYTO</td></tr>
<tr><td>>hg38_21_+_8433172_8446622_RNA45SN1_45S_50ntFlanks@11454.11477.24|CCTCGGCGCGCGGCGGCGGCGGCG</td><td>>hg38_21_+_8433172_8446622_RNA45SN1_45S_50ntFlanks</td><td>11454</td><td>11477</td><td>24</td><td>CCTCGGCGCGCGGCGGCGGCGGCG</td><td>65.8</td><td>-4.0</td><td>1.7</td><td>-2.4</td><td>1.7E-02</td><td>2.9E-02</td><td>BT20/MB468</td><td>MP</td></tr>
<tr><td>>MI0000264|hsa-mir-7-2&WithFlank&15|+|88611819|88611940@38.59.22(+1A);MI0000265|hsa-mir-7-3&WithFlank&19|+|4770664|4770785@37.58.22(+1A);MI0000263|hsa-mir-7-1&WithFlank&9|-|83969742|83969863@30.51.22(+1A)&[MIMAT0000252&hsa-miR-7-5p&offsets|0|-2(+1A);m-400&15|+|88611856|88611879&offsets|0|-2(+1A)];[MIMAT0000252_1&hsa-miR-7-5p&offsets|0|-2(+1A);m-402&19|+|4770700|4770723&offsets|0|-2(+1A)];[MIMAT0000252_2&hsa-miR-7-5p&offsets|0|-2(+1A);m-401&9|-|83969811|83969834&offsets|0|-2(+1A)]|TGGAAGACTAGTGATTTTGTTGA</td><td>>MI0000264|hsa-mir-7-2&WithFlank&15|+|88611819|88611940</td><td>38</td><td>59</td><td>22(+1A)</td><td>TGGAAGACTAGTGATTTTGTTGA</td><td>224.5</td><td>3.7</td><td>1.6</td><td>2.4</td><td>1.7E-02</td><td>2.9E-02</td><td>BT20/MB468</td><td>MP</td></tr>
<tr><td>>MI0002468|hsa-mir-484&WithFlank&16|+|15643288|15643378@14.34.21&[MIMAT0002174&hsa-miR-484&offsets|0|-1;m-127&16|+|15643301|15643321&offsets|0|0]|TCAGGCTCAGTCCCCTCCCGA</td><td>>MI0002468|hsa-mir-484&WithFlank&16|+|15643288|15643378</td><td>14</td><td>34</td><td>21</td><td>TCAGGCTCAGTCCCCTCCCGA</td><td>52.4</td><td>3.0</td><td>1.3</td><td>2.3</td><td>2.4E-02</td><td>2.9E-02</td><td>MB231/MB468</td><td>MP</td></tr>
</table>

| | | | | | | | | | | | | | |
|---|---|---|---|---|---|---|---|---|---|---|---|---|---|
| >MI0000085\|hsa-mir-27a&WithFlank&19\|-\|13836434\|13836523@57.74.18;MI0000440\|hsa-mir-27b&WithFlank&9\|+\|95085439\|95085547@67.84.18&[MIMAT0000084&hsa-miR-27a-3p&offsets\|0\|-3;m-43&19\|-\|13836447\|13836467&offsets\|0\|-3];[MIMAT0000419&hsa-miR-27b-3p&offsets\|0\|-3;m-39&9\|+\|95085505\|95085525&offsets\|0\|-3]\|TTCACAGTGGCTAAGTTC | >MI0000085\|hsa-mir-27a&WithFlank&19\|-\|13836434\|13836523 | 57 | 74 | 18 | TTCACAGTGGCTAAGTTC | 2477.9 | -2.7 | 1.0 | -2.7 | 7.2E-03 | 2.9E-02 | BT20/MB231 | MP |
| >hg38_21_+_8433172_8446622_RNA45SN1_45S_50ntFlanks@10390.10424.35\|GAACAGCAGTTGAACATGGGTCAGTCGGTCCTGAG | >hg38_21_+_8433172_8446622_RNA45SN1_45S_50ntFlanks | 10390 | 10424 | 35 | GAACAGCAGTTGAACATGGGTCAGTCGGTCCTGAG | 4367.8 | 3.5 | 1.3 | 2.7 | 7.2E-03 | 2.9E-02 | BT20/MB231 | MP |
| >hg38_21_+_8433172_8446622_RNA45SN1_45S_50ntFlanks@6651.6669.19\|CGACTCTTAGCGGTGGATC | >hg38_21_+_8433172_8446622_RNA45SN1_45S_50ntFlanks | 6651 | 6669 | 19 | CGACTCTTAGCGGTGGATC | 11267.6 | 3.4 | 1.2 | 2.7 | 7.2E-03 | 2.9E-02 | BT20/MB231 | MP |
| >hg38_21_+_8433172_8446622_RNA45SN1_45S_50ntFlanks@7991.8012.22\|ACGTGGCGACCCGCTGAATTTA | >hg38_21_+_8433172_8446622_RNA45SN1_45S_50ntFlanks | 7991 | 8012 | 22 | ACGTGGCGACCCGCTGAATTTA | 354.4 | 3.7 | 1.4 | 2.7 | 7.2E-03 | 2.9E-02 | BT20/MB231 | MP |
| >hg38_21_+_8433172_8446622_RNA45SN1_45S_50ntFlanks@7970.7988.19\|CGAGACGCGACCTCAGATC | >hg38_21_+_8433172_8446622_RNA45SN1_45S_50ntFlanks | 7970 | 7988 | 19 | CGAGACGCGACCTCAGATC | 184.0 | 1.8 | 0.8 | 2.2 | 2.6E-02 | 2.9E-02 | BT20/MB468 | NUC |
| >hg38_21_+_8433172_8446622_RNA45SN1_45S_50ntFlanks@3840.3860.21\|CTCCTACTTGGATAACTGTGG | >hg38_21_+_8433172_8446622_RNA45SN1_45S_50ntFlanks | 3840 | 3860 | 21 | CTCCTACTTGGATAACTGTGG | 522.9 | 2.1 | 0.8 | 2.7 | 7.8E-03 | 2.9E-02 | BT20/MB231 | NUC |
| >hg38_MT_+_1621_3279_RNR2_16S_50ntFlanks@1352.1388.37\|ATAGGGTTTACGACCTCGATGTTGGATCAGGACATCC | >hg38_MT_+_1621_3279_RNR2_16S_50ntFlanks | 1352 | 1388 | 37 | ATAGGGTTTACGACCTCGATGTTGGATCAGGACATCC | 96.7 | 3.0 | 1.3 | 2.3 | 2.4E-02 | 2.9E-02 | MB231/MB468 | MP |
| >hg38_21_+_8433172_8446622_RNA45SN1_45S_50ntFlanks@6658.6685.28\|TAGCGGTGGATCACTCGGCTCGTGCGTC | >hg38_21_+_8433172_8446622_RNA45SN1_45S_50ntFlanks | 6658 | 6685 | 28 | TAGCGGTGGATCACTCGGCTCGTGCGTC | 102.9 | 5.3 | 2.0 | 2.7 | 7.2E-03 | 2.9E-02 | BT20/MB231 | MP |
| >MI0000433\|hsa-let-7g&WithFlank&3\|-\|52268272\|52268367@11.29.19&[MIMAT0000414&hsa-let-7g-5p&offsets\|0\|-3;m-58&3\|-\|52268336\|52268357&offsets\|0\|-3]\|TGAGGTAGTAGTTTGTACA | >MI0000433\|hsa-let-7g&WithFlank&3\|-\|52268272\|52268367 | 11 | 29 | 19 | TGAGGTAGTAGTTTGTACA | 298.4 | 2.0 | 0.8 | 2.7 | 7.8E-03 | 2.9E-02 | BT20/MB231 | NUC |
| >MI0000103\|hsa-mir-101-1&WithFlank&1\|-\|65058428\|65058514@53.74.22(+1U);MI0000739\|hsa-mir-101-2&WithFlank&9\|+\|4850291\|4850381@55.76.22(+1U)&[MIMAT0000099&hsa-miR-101-3p&offsets\|0\|+1(+1U);m-17&1\|-\|65058443\|65058462&offsets\|0\|+2(+1U)];[MIMAT0000099_1&hsa-miR-101-3p&offsets\|0\|+1(+1U);m-16&9\|+\|4850345\|4850364&offsets\|0\|+2(+1U)]\|TACAGTACTGTGATAACTGAAGT | >MI0000103\|hsa-mir-101-1&WithFlank&1\|-\|65058428\|65058514 | 53 | 74 | 22(+1U) | TACAGTACTGTGATAACTGAAGT | 821.1 | -3.1 | 1.1 | -2.7 | 7.2E-03 | 2.9E-02 | BT20/MB231 | MP |
| >hg38_21_+_8433172_8446622_RNA45SN1_45S_50ntFlanks@10107.10126.20\|CGTGTGGTGTGCGTCGGAGG | >hg38_21_+_8433172_8446622_RNA45SN1_45S_50ntFlanks | 10107 | 10126 | 20 | CGTGTGGTGTGCGTCGGAGG | 276.0 | -2.2 | 0.9 | -2.4 | 1.7E-02 | 2.9E-02 | BT20/MB468 | MP |
| >trnaMT_CysGCA_MT_-_5761_5826@48.68.21\|TTCAAACCTGCCGGGGCTTCC | >trnaMT_CysGCA_MT_-_5761_5826 | 48 | 68 | 21 | TTCAAACCTGCCGGGGCTTCC | 131.5 | 3.3 | 1.5 | 2.2 | 2.6E-02 | 2.9E-02 | MB231/MB468 | MITO |
| >MI0006273\|hsa-mir-1180&WithFlank&17\|-\|19344500\|19344580@47.67.21&[MIMAT0005825&hsa-miR-1180-3p&offsets\|0\|-1;m-214&17\|-\|19344513\|19344534&offsets\|0\|-1]\|TTTCCGGCTCGCGTGGGTGTG | >MI0006273\|hsa-mir-1180&WithFlank&17\|-\|19344500\|19344580 | 47 | 67 | 21 | TTTCCGGCTCGCGTGGGTGTG | 105.1 | 4.3 | 1.8 | 2.4 | 1.7E-02 | 2.9E-02 | BT20/MB468 | MP |
| >MI0000067\|hsa-let-7f-1&WithFlank&9\|+\|94176341\|94176439@13.33.21(+1G);MI0000068\|hsa-let-7f-2&WithFlank&X\|-\|53557186\|53557280@14.34.21(+1G)&[MIMAT0000067&hsa-let-7f-5p&offsets\|0\|-1(+1G);m-20&9\|+\|94176353\|94176374&offsets\|0\|-1(+1G)];[MIMAT0000067_1&hsa-let-7f-5p&offsets\|0\|-1(+1G);m-19&X\|-\|53557246\|53557267&offsets\|0\|-1(+1G)]\|TGAGGTAGTAGATTGTATAGTG | >MI0000067\|hsa-let-7f-1&WithFlank&9\|+\|94176341\|94176439 | 13 | 33 | 21(+1G) | TGAGGTAGTAGATTGTATAGTG | 578.3 | 2.4 | 1.1 | 2.3 | 2.4E-02 | 2.9E-02 | MB231/MB468 | MP |
| >trna12_ValAAC_5_-_180645270_180645342@1.34.34,trna132_ValAAC_6_-_27721179_27721251@1.34.34,trna136_ValAAC_6_-_27648885_27648957@1.34.34,trna139_ValAAC_6_-_27618707_27618779@1.34.34,trna2_ValAAC_3_+_169490018_169490090@1.34.34,trna4_ValAAC_5_+_180591154_180591226@1.34.34,trna5_ValAAC_5_+_180596610_180596682@1.34.34\|GTTTCCGTAGTGTAGTGGTTATCACGTTCGCCTA | >trna12_ValAAC_5_-_180645270_180645342 | 1 | 34 | 34 | GTTTCCGTAGTGTAGTGGTTATCACGTTCGCCTA | 584.0 | -3.0 | 1.2 | -2.6 | 9.7E-03 | 2.9E-02 | BT20/MB231 | CYTO |
| >MI0000088\|hsa-mir-30a&WithFlank&6\|-\|71403545\|71403627@12.33.22&[MIMAT0000087&hsa-miR-30a-5p&offsets\|0\|0;m-15&6\|-\|71403595\|71403616&offsets\|0\|0]\|TGTAAACATCCTCGACTGGAAG | >MI0000088\|hsa-mir-30a&WithFlank&6\|-\|71403545\|71403627 | 12 | 33 | 22 | TGTAAACATCCTCGACTGGAAG | 4410.9 | -2.1 | 0.8 | -2.6 | 8.8E-03 | 2.9E-02 | BT20/MB231 | MITO |
| >MI0005566\|hsa-mir-374b&WithFlank&X\|-\|74218541\|74218624@17.37.21&[MIMAT0004955&hsa-miR-374b-5p&offsets\|0\|-1;m-153&X\|-\|74218587\|74218608&offsets\|0\|-1]\|ATATAATACAACCTGCTAAGT | >MI0005566\|hsa-mir-374b&WithFlank&X\|-\|74218541\|74218624 | 17 | 37 | 21 | ATATAATACAACCTGCTAAGT | 1763.9 | -2.2 | 0.8 | -2.6 | 8.8E-03 | 2.9E-02 | BT20/MB231 | MITO |
| >am_trna128_GlyGCC_6_-_27870686_27870756@7.28.22,trna133_GlyCCC_1_-_16872434_16872504@7.28.22,trna17_GluTTC_2_-_204229131_204229201@7.28.22,trna18_GlyGCC_16_+_70822597_70822667@7.28.22,trna19_GlyGCC_16_+_70823410_70823480@7.28.22,trna19_GlyGCC_2_-_157257659_157257729@7.28.22,trna24_GlyGCC_16_-_70812942_70813012@7.28.22,trna25_GlyGCC_16_-_70812114_70812184@7.28.22,trna2_GlyGCC_21_-_18827107_18827177@7.28.22,trna35_GlyGCC_1_+_161413094_161413164@7.28.22,trna37_GlyGCC_1_+_161420467_161420537@7.28.22,trna39_GlyGCC_1_+_161427898_161427968@7.28.22,trna41_GlyGCC_1_+_161435258_161435328@7.28.22,trna4_GlyCCC_1_+_17188416_17188486@7.28.22,trna5_GlyGCC_17_+_8029064_8029134@7.28.22,trna68_GlyGCC_1_-_161493637_161493707@7.28.22,trna91_GlyCCC_1_-_149680210_149680280@7.28.22,trna99_ValCAC_1_-_149294666_149294736@7.28.22\|GTGGTTCAGTGGTAGAATTCTC | >am_trna128_GlyGCC_6_-_27870686_27870756 | 7 | 28 | 22 | GTGGTTCAGTGGTAGAATTCTC | 220.7 | -3.4 | 1.4 | -2.4 | 1.7E-02 | 2.9E-02 | BT20/MB468 | MP |
| >hg38_21_+_8433172_8446622_RNA45SN1_45S_50ntFlanks@9168.9197.30\|GGGCGGGTCGCGCCGTCGGGCCCGGGGGAG | >hg38_21_+_8433172_8446622_RNA45SN1_45S_50ntFlanks | 9168 | 9197 | 30 | GGGCGGGTCGCGCCGTCGGGCCCGGGGGAG | 123.8 | -4.4 | 1.9 | -2.4 | 1.7E-02 | 2.9E-02 | BT20/MB468 | MP |
| >hg38_21_+_8433172_8446622_RNA45SN1_45S_50ntFlanks@11584.11611.28\|TGGTGCGGACCAGGGGAATCCGACTGTT | >hg38_21_+_8433172_8446622_RNA45SN1_45S_50ntFlanks | 11584 | 11611 | 28 | TGGTGCGGACCAGGGGAATCCGACTGTT | 100.9 | -1.9 | 0.8 | -2.3 | 2.4E-02 | 2.9E-02 | MB231/MB468 | MP |
| >hg38_MT_+_1621_3279_RNR2_16S_50ntFlanks@1352.1385.34\|ATAGGGTTTACGACCTCGATGTTGGATCAGGACA | >hg38_MT_+_1621_3279_RNR2_16S_50ntFlanks | 1352 | 1385 | 34 | ATAGGGTTTACGACCTCGATGTTGGATCAGGACA | 104.0 | 2.9 | 1.3 | 2.3 | 2.4E-02 | 2.9E-02 | MB231/MB468 | MP |
| >hg38_21_+_8433172_8446622_RNA45SN1_45S_50ntFlanks@5083.5121.39\|ACGAACGAGACTCTGGCATGCTAACTAGTTACGCGACCC | >hg38_21_+_8433172_8446622_RNA45SN1_45S_50ntFlanks | 5083 | 5121 | 39 | ACGAACGAGACTCTGGCATGCTAACTAGTTACGCGACCC | 154.5 | -4.4 | 1.9 | -2.3 | 2.4E-02 | 2.9E-02 | MB231/MB468 | MP |
| >hg38_21_+_8433172_8446622_RNA45SN1_45S_50ntFlanks@11581.11609.29\|AACTGGTGCGGACCAGGGGAATCCGACTG | >hg38_21_+_8433172_8446622_RNA45SN1_45S_50ntFlanks | 11581 | 11609 | 29 | AACTGGTGCGGACCAGGGGAATCCGACTG | 381.9 | 2.1 | 0.8 | 2.6 | 8.9E-03 | 2.9E-02 | BT20/MB231 | MITO |
| >MI0000750\|hsa-mir-26a-2&WithFlank&12\|-\|57824603\|57824698@20.41.22(+1A);MI0000083\|hsa-mir-26a-1&WithFlank&3\|+\|37969398\|37969486@16.37.22(+1A)&[MIMAT0000082&hsa-miR-26a-5p&offsets\|0\|0(+1A);m-33&12\|-\|57824658\|57824679&offsets\|0\|0(+1A)];[MIMAT0000082_1&hsa-miR-26a-5p&offsets\|0\|0(+1A);m-34&3\|+\|37969413\|37969434&offsets\|0\|0(+1A)]\|TTCAAGTAATCCAGGATAGGCTA | >MI0000750\|hsa-mir-26a-2&WithFlank&12\|-\|57824603\|57824698 | 20 | 41 | 22(+1A) | TTCAAGTAATCCAGGATAGGCTA | 652.2 | 2.9 | 1.2 | 2.4 | 1.7E-02 | 2.9E-02 | BT20/MB468 | MP |
| >hg38_21_+_8433172_8446622_RNA45SN1_45S_50ntFlanks@11572.11612.41\|CCGACTTAGAACTGGTGCGGACCAGGGGAATCCGACTGTTT | >hg38_21_+_8433172_8446622_RNA45SN1_45S_50ntFlanks | 11572 | 11612 | 41 | CCGACTTAGAACTGGTGCGGACCAGGGGAATCCGACTGTTT | 1374.8 | -3.0 | 1.3 | -2.4 | 1.7E-02 | 2.9E-02 | BT20/MB468 | MP |
| >hg38_21_+_8433172_8446622_RNA45SN1_45S_50ntFlanks@11603.11627.25\|CCGACTGTTTAATTAAAACAAAGCA | >hg38_21_+_8433172_8446622_RNA45SN1_45S_50ntFlanks | 11603 | 11627 | 25 | CCGACTGTTTAATTAAAACAAAGCA | 85.1 | 3.6 | 1.5 | 2.4 | 1.7E-02 | 2.9E-02 | BT20/MB468 | MP |
| >hg38_21_+_8433172_8446622_RNA45SN1_45S_50ntFlanks@10788.10826.39\|CAGCCTCTGGCATGTTGGAACAATGTAGGTAAGGGAAGT | >hg38_21_+_8433172_8446622_RNA45SN1_45S_50ntFlanks | 10788 | 10826 | 39 | CAGCCTCTGGCATGTTGGAACAATGTAGGTAAGGGAAGT | 125.0 | -3.6 | 1.5 | -2.4 | 1.7E-02 | 2.9E-02 | BT20/MB468 | MP |
| >hg38_21_+_8433172_8446622_RNA45SN1_45S_50ntFlanks@10352.10369.18\|GAGAACTTTGAAGGCCGA | >hg38_21_+_8433172_8446622_RNA45SN1_45S_50ntFlanks | 10352 | 10369 | 18 | GAGAACTTTGAAGGCCGA | 357.4 | -2.1 | 0.8 | -2.6 | 9.7E-03 | 2.9E-02 | BT20/MB231 | CYTO |
| >hg38_21_+_8433172_8446622_RNA45SN1_45S_50ntFlanks@7992.8015.24\|CGTGGCGACCCGCTGAATTTAAGC | >hg38_21_+_8433172_8446622_RNA45SN1_45S_50ntFlanks | 7992 | 8015 | 24 | CGTGGCGACCCGCTGAATTTAAGC | 232.0 | -2.1 | 0.8 | -2.6 | 9.7E-03 | 2.9E-02 | BT20/MB231 | CYTO |
| >hg38_21_+_8433172_8446622_RNA45SN1_45S_50ntFlanks@3769.3786.18\|CGCACGGCCGGTACAGTG | >hg38_21_+_8433172_8446622_RNA45SN1_45S_50ntFlanks | 3769 | 3786 | 18 | CGCACGGCCGGTACAGTG | 194.5 | -3.3 | 1.4 | -2.4 | 1.7E-02 | 2.9E-02 | BT20/MB468 | MP |
| >hg38_21_+_8433172_8446622_RNA45SN1_45S_50ntFlanks@6699.6721.23\|GCTAGCTGCGAGAATTAATGTGA | >hg38_21_+_8433172_8446622_RNA45SN1_45S_50ntFlanks | 6699 | 6721 | 23 | GCTAGCTGCGAGAATTAATGTGA | 446.8 | -2.6 | 1.1 | -2.4 | 1.7E-02 | 2.9E-02 | BT20/MB468 | MP |
| >hg38_21_+_8433172_8446622_RNA45SN1_45S_50ntFlanks@6748.6771.24\|CTTCGAACGCACTTGCGGCCCCGG | >hg38_21_+_8433172_8446622_RNA45SN1_45S_50ntFlanks | 6748 | 6771 | 24 | CTTCGAACGCACTTGCGGCCCCGG | 638.6 | -4.3 | 1.8 | -2.4 | 1.7E-02 | 2.9E-02 | BT20/MB468 | MP |
| >MI0000298\|hsa-mir-221&WithFlank&X\|-\|45746151\|45746272@71.91.21&[MIMAT0000278&hsa-miR-221-3p&offsets\|0\|-2;m-91&X\|-\|45746181\|45746202&offsets\|0\|-1]\|AGCTACATTGTCTGCTGGGTT | >MI0000298\|hsa-mir-221&WithFlank&X\|-\|45746151\|45746272 | 71 | 91 | 21 | AGCTACATTGTCTGCTGGGTT | 10192.6 | -2.9 | 1.1 | -2.7 | 7.2E-03 | 2.9E-02 | BT20/MB231 | MP |
| >hg38_MT_+_598_1651_RNR1_12S_50ntFlanks@52.98.47\|ATAGGTTTGGTCCTAGCCTTTCTATTAGCTCTTAGTAAGATTACACA | >hg38_MT_+_598_1651_RNR1_12S_50ntFlanks | 52 | 98 | 47 | ATAGGTTTGGTCCTAGCCTTTCTATTAGCTCTTAGTAAGATTACACA | 386.9 | 6.4 | 2.7 | 2.4 | 1.7E-02 | 2.9E-02 | BT20/MB468 | MP |
| >trna27_GlyCCC_2_-_70476123_70476193@1.29.29,trna34_GlyCCC_16_-_686736_686806@1.29.29\|GCGCCGCTGGTGTAGTGGTATCATGCAAG | >trna27_GlyCCC_2_-_70476123_70476193 | 1 | 29 | 29 | GCGCCGCTGGTGTAGTGGTATCATGCAAG | 198.1 | -2.6 | 1.2 | -2.2 | 2.5E-02 | 2.9E-02 | MB231/MB468 | CYTO |
| >hg38_21_+_8433172_8446622_RNA45SN1_45S_50ntFlanks@12420.12447.28\|ATGTCGGCTCTTCCTATCATTGTGAAGC | >hg38_21_+_8433172_8446622_RNA45SN1_45S_50ntFlanks | 12420 | 12447 | 28 | ATGTCGGCTCTTCCTATCATTGTGAAGC | 712.4 | 1.7 | 0.8 | 2.2 | 2.7E-02 | 2.9E-02 | MB231/MB468 | MITO |
| >hg38_MT_+_598_1651_RNR1_12S_50ntFlanks@443.465.23\|TATCTGAACACACAATAGCTAAG | >hg38_MT_+_598_1651_RNR1_12S_50ntFlanks | 443 | 465 | 23 | TATCTGAACACACAATAGCTAAG | 299.9 | -3.2 | 1.2 | -2.7 | 7.2E-03 | 2.9E-02 | BT20/MB231 | MP |
| >hg38_21_+_8433172_8446622_RNA45SN1_45S_50ntFlanks@12474.12498.25\|ACCCACTAATAGGGAACGTGAGCTG | >hg38_21_+_8433172_8446622_RNA45SN1_45S_50ntFlanks | 12474 | 12498 | 25 | ACCCACTAATAGGGAACGTGAGCTG | 118.4 | -4.0 | 1.7 | -2.4 | 1.7E-02 | 2.9E-02 | BT20/MB468 | MP |
| >trnaMT_GlnTTG_MT_-_4329_4400@1.29.29,trnalookalike2_GlnTTG_1_-_564879_564950@1.29.29\|TAGGATGGGGTGTGATAGGTGGCACGGAG | >trnaMT_GlnTTG_MT_-_4329_4400 | 1 | 29 | 29 | TAGGATGGGGTGTGATAGGTGGCACGGAG | 1235.2 | 3.1 | 1.3 | 2.4 | 1.7E-02 | 2.9E-02 | BT20/MB468 | MP |
| >MI0000283\|hsa-mir-203a&WithFlank&14\|+\|104117399\|104117520@71.92.22(+1U)&[MIMAT0000264&hsa-miR-203a-3p&offsets\|0\|0(+1U);m-79&14\|+\|104117469\|104117490&offsets\|0\|0(+1U)]\|GTGAAATGTTTAGGACCACTAGT | >MI0000283\|hsa-mir-203a&WithFlank&14\|+\|104117399\|104117520 | 71 | 92 | 22(+1U) | GTGAAATGTTTAGGACCACTAGT | 55.3 | 4.4 | 1.9 | 2.4 | 1.7E-02 | 2.9E-02 | BT20/MB468 | MP |
| >hg38_21_+_8433172_8446622_RNA45SN1_45S_50ntFlanks@11027.11059.33\|CTCTCTCTCTCTCTCCCCCGCTCCCCGTCCTCC | >hg38_21_+_8433172_8446622_RNA45SN1_45S_50ntFlanks | 11027 | 11059 | 33 | CTCTCTCTCTCTCTCCCCCGCTCCCCGTCCTCC | 1028.9 | -4.9 | 2.2 | -2.2 | 2.6E-02 | 2.9E-02 | BT20/MB468 | NUC |

| | | | | | | | | | | | | | |
|---|---|---|---|---|---|---|---|---|---|---|---|---|---|
| >hg38_21_+_8433172_8446622_RNA45SN1_45S_50ntFlanks@10344.10383.40\|ATTCAAACGAGAACTTTGAAGGCCGAAGTGGAGAAGGGTT | >hg38_21_+_8433172_8446622_RNA45SN1_45S_50ntFlanks | 10344 | 10383 | 40 | ATTCAAACGAGAACTTTGAAGGCCGAAGTGGAGAAGGGTT | 63.6 | 4.1 | 1.7 | 2.4 | 1.7E-02 | 2.9E-02 | BT20/MB468 | MP |
| >hg38_21_+_8433172_8446622_RNA45SN1_45S_50ntFlanks@9950.9978.29\|GCCATGGAAGTCGGAATCCGCTAAGGAGT | >hg38_21_+_8433172_8446622_RNA45SN1_45S_50ntFlanks | 9950 | 9978 | 29 | GCCATGGAAGTCGGAATCCGCTAAGGAGT | 37.9 | 4.2 | 1.8 | 2.4 | 1.7E-02 | 2.9E-02 | BT20/MB468 | MP |
| >hg38_21_+_8433172_8446622_RNA45SN1_45S_50ntFlanks@10396.10426.31\|CAGTTGAACATGGGTCAGTCGGTCCTGAGAG | >hg38_21_+_8433172_8446622_RNA45SN1_45S_50ntFlanks | 10396 | 10426 | 31 | CAGTTGAACATGGGTCAGTCGGTCCTGAGAG | 110.0 | -4.1 | 1.7 | -2.4 | 1.7E-02 | 2.9E-02 | BT20/MB468 | MP |
| >hg38_21_+_8433172_8446622_RNA45SN1_45S_50ntFlanks@9535.9571.37\|GGAAACTCTGGTGGAGGTCCGTAGCGGTCCTGACGTG | >hg38_21_+_8433172_8446622_RNA45SN1_45S_50ntFlanks | 9535 | 9571 | 37 | GGAAACTCTGGTGGAGGTCCGTAGCGGTCCTGACGTG | 1391.7 | 1.6 | 0.7 | 2.2 | 2.5E-02 | 2.9E-02 | MB231/MB468 | CYTO |
| >hg38_21_+_8433172_8446622_RNA45SN1_45S_50ntFlanks@11671.11696.26\|CCAGTGCTCTGAATGTCAAAGTGAAG | >hg38_21_+_8433172_8446622_RNA45SN1_45S_50ntFlanks | 11671 | 11696 | 26 | CCAGTGCTCTGAATGTCAAAGTGAAG | 121.6 | 4.6 | 1.7 | 2.7 | 7.2E-03 | 2.9E-02 | BT20/MB231 | MP |
| >MI0033423\|hsa-mir-10399&WithFlank&7\|-\|139044094\|139044163@44.63.20(+1U)&[MIMAT0041630&hsa-miR-10399-3p&offsets\|0\|-1(+1U);TJU_CMC_MD2.ID03061.3p-miR&offsets\|+4\|+2(+1U);m-2657&7\|-\|139044103\|139044124&offsets\|+4\|+2(+1U)]\|CTCTCGGACAAGCTGTAGGTT | >MI0033423\|hsa-mir-10399&WithFlank&7\|-\|139044094\|139044163 | 44 | 63 | 20(+1U) | CTCTCGGACAAGCTGTAGGTT | 171.7 | -3.8 | 1.4 | -2.7 | 7.2E-03 | 2.9E-02 | BT20/MB231 | MP |
| >MI0003780\|hsa-mir-1296&WithFlank&10\|-\|63372951\|63373054@22.42.21(+1A)&[MIMAT0005794&hsa-miR-1296-5p&offsets\|0\|-1(+1A);m-304&10\|-\|63373012\|63373033&offsets\|0\|-1(+1A)]\|TTAGGGCCCTGGCTCCATCTCA | >MI0003780\|hsa-mir-1296&WithFlank&10\|-\|63372951\|63373054 | 22 | 42 | 21(+1A) | TTAGGGCCCTGGCTCCATCTCA | 49.3 | 6.2 | 2.6 | 2.4 | 1.7E-02 | 2.9E-02 | BT20/MB468 | MP |
| >hg38_21_+_8433172_8446622_RNA45SN1_45S_50ntFlanks@11322.11344.23\|ACGTCTCGTCGCGCGCGCGTCCG | >hg38_21_+_8433172_8446622_RNA45SN1_45S_50ntFlanks | 11322 | 11344 | 23 | ACGTCTCGTCGCGCGCGCGTCCG | 7783.8 | -2.8 | 1.2 | -2.3 | 2.2E-02 | 2.9E-02 | BT20/MB468 | MITO |
| >hg38_21_+_8433172_8446622_RNA45SN1_45S_50ntFlanks@12941.12966.26\|ATTCGTAGACGACCTGCTTCTGGGTC | >hg38_21_+_8433172_8446622_RNA45SN1_45S_50ntFlanks | 12941 | 12966 | 26 | ATTCGTAGACGACCTGCTTCTGGGTC | 162.1 | -2.0 | 0.9 | -2.3 | 2.2E-02 | 2.9E-02 | BT20/MB468 | MITO |
| >MI0000063\|hsa-let-7b&WithFlank&22\|+\|46113680\|46113774@12.34.23(+1A)&[MIMAT0000063&hsa-let-7b-5p&offsets\|0\|+1(+1A);m-14&22\|+\|46113691\|46113712&offsets\|0\|+1(+1A)]\|TGAGGTAGTAGGTTGTGTGGTTTA | >MI0000063\|hsa-let-7b&WithFlank&22\|+\|46113680\|46113774 | 12 | 34 | 23(+1A) | TGAGGTAGTAGGTTGTGTGGTTTA | 75.6 | 3.0 | 1.3 | 2.3 | 2.4E-02 | 2.9E-02 | MB231/MB468 | MP |
| >hg38_21_+_8433172_8446622_RNA45SN1_45S_50ntFlanks@10353.10384.32\|AGAACTTTGAAGGCCGAAGTGGAGAAGGGTTC | >hg38_21_+_8433172_8446622_RNA45SN1_45S_50ntFlanks | 10353 | 10384 | 32 | AGAACTTTGAAGGCCGAAGTGGAGAAGGGTTC | 56.7 | -4.4 | 1.9 | -2.3 | 2.4E-02 | 2.9E-02 | MB231/MB468 | MP |
| >hg38_21_+_8433172_8446622_RNA45SN1_45S_50ntFlanks@12948.12990.43\|GACGACCTGCTTCTGGGTCGGGGTTTCGTACGTAGCAGAGCAG | >hg38_21_+_8433172_8446622_RNA45SN1_45S_50ntFlanks | 12948 | 12990 | 43 | GACGACCTGCTTCTGGGTCGGGGTTTCGTACGTAGCAGAGCAG | 137.2 | -1.6 | 0.7 | -2.2 | 2.5E-02 | 2.9E-02 | MB231/MB468 | CYTO |
| >trnaMT_MetCAT_MT_+_4402_4469@38.71.34\|CGAAAATGTTGGTTATACCCTTCCCGTACTACCA | >trnaMT_MetCAT_MT_+_4402_4469 | 38 | 71 | 34 | CGAAAATGTTGGTTATACCCTTCCCGTACTACCA | 229.2 | -2.1 | 0.8 | -2.7 | 7.8E-03 | 2.9E-02 | BT20/MB231 | NUC |
| >hg38_21_+_8433172_8446622_RNA45SN1_45S_50ntFlanks@11315.11332.18\|CCACCCCACGTCTCGTCG | >hg38_21_+_8433172_8446622_RNA45SN1_45S_50ntFlanks | 11315 | 11332 | 18 | CCACCCCACGTCTCGTCG | 263.8 | -3.6 | 1.6 | -2.3 | 2.2E-02 | 2.9E-02 | BT20/MB468 | MITO |
| >hg38_21_+_8433172_8446622_RNA45SN1_45S_50ntFlanks@5467.5514.48\|GGCCCACGGCCCTGGCGGAGCGCTGAGAAGACGGTCGAACTTGACTAT | >hg38_21_+_8433172_8446622_RNA45SN1_45S_50ntFlanks | 5467 | 5514 | 48 | GGCCCACGGCCCTGGCGGAGCGCTGAGAAGACGGTCGAACTTGACTAT | 403.5 | -3.1 | 1.4 | -2.3 | 2.2E-02 | 2.9E-02 | BT20/MB468 | MITO |
| >hg38_21_+_8433172_8446622_RNA45SN1_45S_50ntFlanks@7984.8014.31\|AGATCAGACGTGGCGACCCGCTGAATTTAAG | >hg38_21_+_8433172_8446622_RNA45SN1_45S_50ntFlanks | 7984 | 8014 | 31 | AGATCAGACGTGGCGACCCGCTGAATTTAAG | 98.5 | -3.6 | 1.6 | -2.3 | 2.2E-02 | 2.9E-02 | BT20/MB468 | MITO |
| >hg38_21_+_8433172_8446622_RNA45SN1_45S_50ntFlanks@11873.11906.34\|AAGAAGACCCTGTTGAGCTTGACTCTAGTCTGGC | >hg38_21_+_8433172_8446622_RNA45SN1_45S_50ntFlanks | 11873 | 11906 | 34 | AAGAAGACCCTGTTGAGCTTGACTCTAGTCTGGC | 1013.2 | -4.4 | 1.9 | -2.4 | 1.7E-02 | 2.9E-02 | BT20/MB468 | MP |
| >hg38_21_+_8433172_8446622_RNA45SN1_45S_50ntFlanks@5445.5465.21\|TCGGATCGGCCCCGCCGGGGT | >hg38_21_+_8433172_8446622_RNA45SN1_45S_50ntFlanks | 5445 | 5465 | 21 | TCGGATCGGCCCCGCCGGGGT | 223.8 | -4.1 | 1.7 | -2.4 | 1.7E-02 | 2.9E-02 | BT20/MB468 | MP |
| >hg38_21_+_8433172_8446622_RNA45SN1_45S_50ntFlanks@11589.11621.33\|CGGACCAGGGGAATCCGACTGTTTAATTAAAAC | >hg38_21_+_8433172_8446622_RNA45SN1_45S_50ntFlanks | 11589 | 11621 | 33 | CGGACCAGGGGAATCCGACTGTTTAATTAAAAC | 153.3 | 4.5 | 1.7 | 2.7 | 7.3E-03 | 2.9E-02 | BT20/MB231 | MP |
| >hg38_21_+_8433172_8446622_RNA45SN1_45S_50ntFlanks@11674.11701.28\|GTGCTCTGAATGTCAAAGTGAAGAAATT | >hg38_21_+_8433172_8446622_RNA45SN1_45S_50ntFlanks | 11674 | 11701 | 28 | GTGCTCTGAATGTCAAAGTGAAGAAATT | 185.1 | 4.7 | 1.8 | 2.7 | 7.3E-03 | 2.9E-02 | BT20/MB231 | MP |
| >MI0000816\|hsa-mir-335&WithFlank&7\|+\|130496105\|130496210@22.43.22(+1C)&[MIMAT0000765&hsa-miR-335-5p&offsets\|0\|-1(+1C);m-249&7\|+\|130496126\|130496147&offsets\|0\|0(+1C)]\|TCAAGAGCAATAACGAAAAATGC | >MI0000816\|hsa-mir-335&WithFlank&7\|+\|130496105\|130496210 | 22 | 43 | 22(+1C) | TCAAGAGCAATAACGAAAAATGC | 72.4 | -5.3 | 2.0 | -2.7 | 7.3E-03 | 2.9E-02 | BT20/MB231 | MP |
| >hg38_21_+_8433172_8446622_RNA45SN1_45S_50ntFlanks@8104.8144.41\|CCGCGGCGGGGCGCGGGACATGTGGCGTACGGAAGACCCGC | >hg38_21_+_8433172_8446622_RNA45SN1_45S_50ntFlanks | 8104 | 8144 | 41 | CCGCGGCGGGGCGCGGGACATGTGGCGTACGGAAGACCCGC | 162.3 | -4.4 | 2.0 | -2.3 | 2.4E-02 | 2.9E-02 | MB231/MB468 | MP |
| >am_trna119_LysCTT_1_-_145395522_145395594@6.35.30,trna11_LysCTT_5_-_180648979_180649051@6.35.30,trna128_LysCTT_1_-_55423542_55423614@6.35.30,trna13_LysCTT_6_+_26556774_26556846@6.35.30,trna6_LysCTT_19_-_52425396_52425468@6.35.30,trna7_LysCTT_16_+_3225692_3225764@6.35.30,trna9_LysCTT_5_+_180634755_180634827@6.35.30\|GCTAGCTCAGTCGGTAGAGCATGAGACTCT | >am_trna119_LysCTT_1_-_145395522_145395594 | 6 | 35 | 30 | GCTAGCTCAGTCGGTAGAGCATGAGACTCT | 44.7 | 4.5 | 2.0 | 2.3 | 2.4E-02 | 2.9E-02 | MB231/MB468 | MP |
| >hg38_MT_+_1621_3279_RNR2_16S_50ntFlanks@51.91.41\|GCTAAACCTAGCCCCAAACCCACTCCACCTTACTACCAGAC | >hg38_MT_+_1621_3279_RNR2_16S_50ntFlanks | 51 | 91 | 41 | GCTAAACCTAGCCCCAAACCCACTCCACCTTACTACCAGAC | 512.5 | 3.3 | 1.5 | 2.3 | 2.4E-02 | 2.9E-02 | MB231/MB468 | MP |
| >MI0005756\|hsa-mir-934&WithFlank&X\|+\|136550872\|136550966@21.40.20(+1A)&[MIMAT0004977&hsa-miR-934&offsets\|0\|-2(+1A);m-943&X\|+\|136550892\|136550913&offsets\|0\|-2(+1A)]\|TGTCTACTACTGGAGACACTA | >MI0005756\|hsa-mir-934&WithFlank&X\|+\|136550872\|136550966 | 21 | 40 | 20(+1A) | TGTCTACTACTGGAGACACTA | 836.6 | 3.2 | 1.4 | 2.4 | 1.7E-02 | 2.9E-02 | BT20/MB468 | MP |
| >hg38_21_+_8433172_8446622_RNA45SN1_45S_50ntFlanks@11211.11234.24\|GCGACTCTGGACGCGAGCCGGGCC | >hg38_21_+_8433172_8446622_RNA45SN1_45S_50ntFlanks | 11211 | 11234 | 24 | GCGACTCTGGACGCGAGCCGGGCC | 97.8 | -4.0 | 1.7 | -2.4 | 1.7E-02 | 2.9E-02 | BT20/MB468 | MP |
| >hg38_21_+_8433172_8446622_RNA45SN1_45S_50ntFlanks@4364.4391.28\|CTGCAGTTAAAAAGCTCGTAGTTGGATC | >hg38_21_+_8433172_8446622_RNA45SN1_45S_50ntFlanks | 4364 | 4391 | 28 | CTGCAGTTAAAAAGCTCGTAGTTGGATC | 135.6 | -3.2 | 1.4 | -2.4 | 1.7E-02 | 2.9E-02 | BT20/MB468 | MP |
| >MI0003185\|hsa-mir-501&WithFlank&X\|+\|50009716\|50009811@57.78.22&[MIMAT0004774&hsa-miR-501-3p&offsets\|0\|0;m-168&X\|+\|50009772\|50009793&offsets\|0\|0]\|AATGCACCCGGGCAAGGATTCT | >MI0003185\|hsa-mir-501&WithFlank&X\|+\|50009716\|50009811 | 57 | 78 | 22 | AATGCACCCGGGCAAGGATTCT | 503.6 | 2.9 | 1.2 | 2.4 | 1.7E-02 | 2.9E-02 | BT20/MB468 | MP |
| >hg38_21_+_8433172_8446622_RNA45SN1_45S_50ntFlanks@6675.6696.22\|GCTCGTGCGTCGATGAAGAACG | >hg38_21_+_8433172_8446622_RNA45SN1_45S_50ntFlanks | 6675 | 6696 | 22 | GCTCGTGCGTCGATGAAGAACG | 470.3 | 4.5 | 1.7 | 2.7 | 7.3E-03 | 2.9E-02 | BT20/MB231 | MP |
| >hg38_21_+_8433172_8446622_RNA45SN1_45S_50ntFlanks@12420.12460.41\|ATGTCGGCTCTTCCTATCATTGTGAAGCAGAATTCACCAAG | >hg38_21_+_8433172_8446622_RNA45SN1_45S_50ntFlanks | 12420 | 12460 | 41 | ATGTCGGCTCTTCCTATCATTGTGAAGCAGAATTCACCAAG | 400.8 | 1.5 | 0.7 | 2.2 | 2.5E-02 | 2.9E-02 | MB231/MB468 | CYTO |
| >hg38_MT_+_598_1651_RNR1_12S_50ntFlanks@719.744.26\|TAAAGACGTTAGGTCAAGGTGTAGCC | >hg38_MT_+_598_1651_RNR1_12S_50ntFlanks | 719 | 744 | 26 | TAAAGACGTTAGGTCAAGGTGTAGCC | 516.3 | 2.5 | 1.1 | 2.3 | 2.4E-02 | 2.9E-02 | MB231/MB468 | MP |
| >MI0000093\|hsa-mir-92a-1&WithFlank&13\|+\|91351308\|91351397@54.76.23&[MIMAT0000092&hsa-miR-92a-3p&offsets\|0\|+1;m-9&13\|+\|91351361\|91351382&offsets\|0\|+1]\|TATTGCACTTGTCCCGGCCTGTT | >MI0000093\|hsa-mir-92a-1&WithFlank&13\|+\|91351308\|91351397 | 54 | 76 | 23 | TATTGCACTTGTCCCGGCCTGTT | 5108.2 | -1.7 | 0.6 | -2.7 | 7.9E-03 | 2.9E-02 | BT20/MB231 | NUC |
| >hg38_21_+_8433172_8446622_RNA45SN1_45S_50ntFlanks@12420.12437.18\|ATGTCGGCTCTTCCTATC | >hg38_21_+_8433172_8446622_RNA45SN1_45S_50ntFlanks | 12420 | 12437 | 18 | ATGTCGGCTCTTCCTATC | 9261.6 | -1.5 | 0.7 | -2.2 | 2.5E-02 | 2.9E-02 | MB231/MB468 | CYTO |
| >hg38_21_+_8433172_8446622_RNA45SN1_45S_50ntFlanks@12617.12657.41\|TGAGGAGCCAATGGGGCGAAGCTACCATCTGTGGGATTATG | >hg38_21_+_8433172_8446622_RNA45SN1_45S_50ntFlanks | 12617 | 12657 | 41 | TGAGGAGCCAATGGGGCGAAGCTACCATCTGTGGGATTATG | 120.1 | -3.7 | 1.6 | -2.3 | 2.2E-02 | 2.9E-02 | BT20/MB468 | MITO |
| >hg38_21_+_8433172_8446622_RNA45SN1_45S_50ntFlanks@12418.12438.21\|CGATGTCGGCTCTTCCTATCA | >hg38_21_+_8433172_8446622_RNA45SN1_45S_50ntFlanks | 12418 | 12438 | 21 | CGATGTCGGCTCTTCCTATCA | 2130.8 | -2.1 | 0.9 | -2.3 | 2.2E-02 | 2.9E-02 | BT20/MB468 | MITO |
| >hg38_21_+_8433172_8446622_RNA45SN1_45S_50ntFlanks@3854.3873.20\|ACTGTGGTAATTCTAGAGCT | >hg38_21_+_8433172_8446622_RNA45SN1_45S_50ntFlanks | 3854 | 3873 | 20 | ACTGTGGTAATTCTAGAGCT | 580.2 | -3.6 | 1.6 | -2.2 | 2.6E-02 | 2.9E-02 | BT20/MB468 | NUC |
| >hg38_21_+_8433172_8446622_RNA45SN1_45S_50ntFlanks@10841.10869.29\|CGTAACTTCGGGATAAGGATTGGCTCTAA | >hg38_21_+_8433172_8446622_RNA45SN1_45S_50ntFlanks | 10841 | 10869 | 29 | CGTAACTTCGGGATAAGGATTGGCTCTAA | 481.1 | 1.7 | 0.8 | 2.2 | 2.6E-02 | 2.9E-02 | BT20/MB468 | CYTO |
| >hg38_21_+_8433172_8446622_RNA45SN1_45S_50ntFlanks@12007.12031.25\|GCCGCCGGTGAAATACCACTACTCT | >hg38_21_+_8433172_8446622_RNA45SN1_45S_50ntFlanks | 12007 | 12031 | 25 | GCCGCCGGTGAAATACCACTACTCT | 600.6 | -1.4 | 0.6 | -2.2 | 2.6E-02 | 2.9E-02 | BT20/MB468 | CYTO |
| >hg38_21_+_8433172_8446622_RNA45SN1_45S_50ntFlanks@10774.10818.45\|GTCTCCAAGGTGAACAGCCTCTGGCATGTTGGAACAATGTAGGTA | >hg38_21_+_8433172_8446622_RNA45SN1_45S_50ntFlanks | 10774 | 10818 | 45 | GTCTCCAAGGTGAACAGCCTCTGGCATGTTGGAACAATGTAGGTA | 318.2 | -2.0 | 0.9 | -2.2 | 2.5E-02 | 2.9E-02 | MB231/MB468 | CYTO |
| >hg38_21_+_8433172_8446622_RNA45SN1_45S_50ntFlanks@7973.7994.22\|GACGCGACCTCAGATCAGACGT | >hg38_21_+_8433172_8446622_RNA45SN1_45S_50ntFlanks | 7973 | 7994 | 22 | GACGCGACCTCAGATCAGACGT | 480.9 | 1.5 | 0.7 | 2.2 | 2.7E-02 | 2.9E-02 | MB231/MB468 | MITO |
| >hg38_21_+_8433172_8446622_RNA45SN1_45S_50ntFlanks@11570.11590.21\|AGCCGACTTAGAACTGGTGCG | >hg38_21_+_8433172_8446622_RNA45SN1_45S_50ntFlanks | 11570 | 11590 | 21 | AGCCGACTTAGAACTGGTGCG | 134.5 | -3.3 | 1.4 | -2.4 | 1.7E-02 | 2.9E-02 | BT20/MB468 | MP |
| >hg38_21_+_8433172_8446622_RNA45SN1_45S_50ntFlanks@12202.12221.20\|CTAAGGCGAGCTCAGGGAGG | >hg38_21_+_8433172_8446622_RNA45SN1_45S_50ntFlanks | 12202 | 12221 | 20 | CTAAGGCGAGCTCAGGGAGG | 596.3 | -2.5 | 1.0 | -2.4 | 1.7E-02 | 2.9E-02 | BT20/MB468 | MP |
| >hg38_21_+_8433172_8446622_RNA45SN1_45S_50ntFlanks@12617.12654.38\|TGAGGAGCCAATGGGGCGAAGCTACCATCTGTGGGATT | >hg38_21_+_8433172_8446622_RNA45SN1_45S_50ntFlanks | 12617 | 12654 | 38 | TGAGGAGCCAATGGGGCGAAGCTACCATCTGTGGGATT | 728.3 | -2.7 | 1.1 | -2.4 | 1.7E-02 | 2.9E-02 | BT20/MB468 | MP |
| >hg38_21_+_8433172_8446622_RNA45SN1_45S_50ntFlanks@5237.5283.47\|ACTGACTGGCTCAGCGTGTGCCTACCCTACGCCGGCAGGCGCGGGTA | >hg38_21_+_8433172_8446622_RNA45SN1_45S_50ntFlanks | 5237 | 5283 | 47 | ACTGACTGGCTCAGCGTGTGCCTACCCTACGCCGGCAGGCGCGGGTA | 273.0 | 5.5 | 2.3 | 2.4 | 1.7E-02 | 2.9E-02 | BT20/MB468 | MP |
| >hg38_21_+_8433172_8446622_RNA45SN1_45S_50ntFlanks@5482.5499.18\|CGGAGCGCTGAGAAGACG | >hg38_21_+_8433172_8446622_RNA45SN1_45S_50ntFlanks | 5482 | 5499 | 18 | CGGAGCGCTGAGAAGACG | 265.0 | 4.6 | 1.7 | 2.7 | 7.3E-03 | 2.9E-02 | BT20/MB231 | MP |
| >hg38_21_+_8433172_8446622_RNA45SN1_45S_50ntFlanks@7977.8010.34\|CGACCTCAGATCAGACGTGGCGACCCGCTGAATT | >hg38_21_+_8433172_8446622_RNA45SN1_45S_50ntFlanks | 7977 | 8010 | 34 | CGACCTCAGATCAGACGTGGCGACCCGCTGAATT | 1291.3 | 4.1 | 1.5 | 2.7 | 7.3E-03 | 2.9E-02 | BT20/MB231 | MP |
| >hg38_21_+_8433172_8446622_RNA45SN1_45S_50ntFlanks@12642.12669.28\|CATCTGTGGGATTATGACTGAACGCCTC | >hg38_21_+_8433172_8446622_RNA45SN1_45S_50ntFlanks | 12642 | 12669 | 28 | CATCTGTGGGATTATGACTGAACGCCTC | 154.5 | 3.6 | 1.5 | 2.4 | 1.7E-02 | 2.9E-02 | BT20/MB468 | MP |
| >hg38_MT_+_1621_3279_RNR2_16S_50ntFlanks@840.859.20\|AAAAGTAAAAGGAACTCGGC | >hg38_MT_+_1621_3279_RNR2_16S_50ntFlanks | 840 | 859 | 20 | AAAAGTAAAAGGAACTCGGC | 107.2 | 2.7 | 1.1 | 2.4 | 1.7E-02 | 2.9E-02 | BT20/MB468 | MP |
| >MI0000456\|hsa-mir-140&WithFlank&16\|+\|69933075\|69933186@69.89.21&[MIMAT0004597&hsa-miR-140-3p&offsets\|+1\|+1;m-45&16\|+\|69933143\|69933164&offsets\|0\|-1]\|ACCACAGGGTAGAACCACGGA | >MI0000456\|hsa-mir-140&WithFlank&16\|+\|69933075\|69933186 | 69 | 89 | 21 | ACCACAGGGTAGAACCACGGA | 1050.4 | -2.1 | 0.8 | -2.6 | 8.9E-03 | 2.9E-02 | BT20/MB231 | MITO |

<table>
<tr><td>>hg38_MT_+_1621_3279_RNR2_16S_50ntFlanks@1427.1458.32|GATTAAAGTCCTACGTGATCTGAGTTCAGACC</td><td>>hg38_MT_+_1621_3279_RNR2_16S_50ntFlanks</td><td>1427</td><td>1458</td><td>32</td><td>GATTAAAGTCCTACGTGATCTGAGTTCAGACC</td><td>446.5</td><td>3.9</td><td>1.6</td><td>2.4</td><td>1.7E-02</td><td>3.0E-02</td><td>BT20/MB468</td><td>MP</td></tr>
<tr><td>>hg38_21_+_8433172_8446622_RNA45SN1_45S_50ntFlanks@11670.11691.22|CCCAGTGCTCTGAATGTCAAAG</td><td>>hg38_21_+_8433172_8446622_RNA45SN1_45S_50ntFlanks</td><td>11670</td><td>11691</td><td>22</td><td>CCCAGTGCTCTGAATGTCAAAG</td><td>370.2</td><td>4.6</td><td>1.7</td><td>2.7</td><td>7.3E-03</td><td>3.0E-02</td><td>BT20/MB231</td><td>MP</td></tr>
<tr><td>>MI0000086|hsa-mir-28&WithFlank&3|+|188688775|188688872@60.79.20(+1A)&[MIMAT0004502&hsa-miR-28-3p&offsets|0|-2(+1A);m-28&3|+|188688834|188688855&offsets|0|-2(+1A)]|CACTAGATTGTGAGCTCCTGA</td><td>>MI0000086|hsa-mir-28&WithFlank&3|+|188688775|188688872</td><td>60</td><td>79</td><td>20(+1A)</td><td>CACTAGATTGTGAGCTCCTGA</td><td>800.0</td><td>2.4</td><td>0.9</td><td>2.6</td><td>9.8E-03</td><td>3.0E-02</td><td>BT20/MB231</td><td>CYTO</td></tr>
<tr><td>>hg38_21_+_8433172_8446622_RNA45SN1_45S_50ntFlanks@12415.12453.39|CTTCGATGTCGGCTCTTCCTATCATTGTGAAGCAGAATT</td><td>>hg38_21_+_8433172_8446622_RNA45SN1_45S_50ntFlanks</td><td>12415</td><td>12453</td><td>39</td><td>CTTCGATGTCGGCTCTTCCTATCATTGTGAAGCAGAATT</td><td>206.2</td><td>-2.2</td><td>0.9</td><td>-2.6</td><td>9.8E-03</td><td>3.0E-02</td><td>BT20/MB231</td><td>CYTO</td></tr>
<tr><td>>hg38_21_+_8433172_8446622_RNA45SN1_45S_50ntFlanks@12942.12961.20|TTCGTAGACGACCTGCTTCT</td><td>>hg38_21_+_8433172_8446622_RNA45SN1_45S_50ntFlanks</td><td>12942</td><td>12961</td><td>20</td><td>TTCGTAGACGACCTGCTTCT</td><td>160.9</td><td>-2.0</td><td>0.8</td><td>-2.4</td><td>1.7E-02</td><td>3.0E-02</td><td>BT20/MB468</td><td>MP</td></tr>
<tr><td>>hg38_21_+_8433172_8446622_RNA45SN1_45S_50ntFlanks@11027.11048.22|CTCTCTCTCTCTCTCCCCCGCT</td><td>>hg38_21_+_8433172_8446622_RNA45SN1_45S_50ntFlanks</td><td>11027</td><td>11048</td><td>22</td><td>CTCTCTCTCTCTCTCCCCCGCT</td><td>1004.3</td><td>-4.9</td><td>2.2</td><td>-2.2</td><td>2.6E-02</td><td>3.0E-02</td><td>BT20/MB468</td><td>NUC</td></tr>
<tr><td>>hg38_21_+_8433172_8446622_RNA45SN1_45S_50ntFlanks@10351.10382.32|CGAGAACTTTGAAGGCCGAAGTGGAGAAGGGT</td><td>>hg38_21_+_8433172_8446622_RNA45SN1_45S_50ntFlanks</td><td>10351</td><td>10382</td><td>32</td><td>CGAGAACTTTGAAGGCCGAAGTGGAGAAGGGT</td><td>3285.1</td><td>-2.6</td><td>1.1</td><td>-2.4</td><td>1.7E-02</td><td>3.0E-02</td><td>BT20/MB468</td><td>MP</td></tr>
<tr><td>>hg38_21_+_8433172_8446622_RNA45SN1_45S_50ntFlanks@10248.10278.31|CGACGAGTAGGAGGGCCGCTGCGGTGAGCCT</td><td>>hg38_21_+_8433172_8446622_RNA45SN1_45S_50ntFlanks</td><td>10248</td><td>10278</td><td>31</td><td>CGACGAGTAGGAGGGCCGCTGCGGTGAGCCT</td><td>260.8</td><td>4.6</td><td>1.7</td><td>2.7</td><td>7.3E-03</td><td>3.0E-02</td><td>BT20/MB231</td><td>MP</td></tr>
<tr><td>>hg38_21_+_8433172_8446622_RNA45SN1_45S_50ntFlanks@11573.11602.30|CGACTTAGAACTGGTGCGGACCAGGGGAAT</td><td>>hg38_21_+_8433172_8446622_RNA45SN1_45S_50ntFlanks</td><td>11573</td><td>11602</td><td>30</td><td>CGACTTAGAACTGGTGCGGACCAGGGGAAT</td><td>459.1</td><td>4.5</td><td>1.7</td><td>2.7</td><td>7.3E-03</td><td>3.0E-02</td><td>BT20/MB231</td><td>MP</td></tr>
<tr><td>>hg38_MT_+_1621_3279_RNR2_16S_50ntFlanks@1249.1271.23|AGTCAAAGCGAACTACTATACTC</td><td>>hg38_MT_+_1621_3279_RNR2_16S_50ntFlanks</td><td>1249</td><td>1271</td><td>23</td><td>AGTCAAAGCGAACTACTATACTC</td><td>553.6</td><td>-3.2</td><td>1.2</td><td>-2.7</td><td>7.3E-03</td><td>3.0E-02</td><td>BT20/MB231</td><td>MP</td></tr>
<tr><td>>hg38_21_+_8433172_8446622_RNA45SN1_45S_50ntFlanks@11933.11956.24|AATAAGTGGGAGGCCCCCGGCGCC</td><td>>hg38_21_+_8433172_8446622_RNA45SN1_45S_50ntFlanks</td><td>11933</td><td>11956</td><td>24</td><td>AATAAGTGGGAGGCCCCCGGCGCC</td><td>104.2</td><td>-4.1</td><td>1.7</td><td>-2.4</td><td>1.7E-02</td><td>3.0E-02</td><td>BT20/MB468</td><td>MP</td></tr>
<tr><td>>hg38_21_+_8433172_8446622_RNA45SN1_45S_50ntFlanks@10350.10370.21|ACGAGAACTTTGAAGGCCGAA</td><td>>hg38_21_+_8433172_8446622_RNA45SN1_45S_50ntFlanks</td><td>10350</td><td>10370</td><td>21</td><td>ACGAGAACTTTGAAGGCCGAA</td><td>727.9</td><td>-2.5</td><td>1.0</td><td>-2.4</td><td>1.7E-02</td><td>3.0E-02</td><td>BT20/MB468</td><td>MP</td></tr>
<tr><td>>MI0000266|hsa-mir-10a&WithFlank&17|-|48579832|48579953@28.50.23(+1C)&[MIMAT0000253&hsa-miR-10a-5p&offsets|0|0(+1C);m-18&17|-|48579905|48579926&offsets|0|+1(+1C)]|TACCCTGTAGATCCGAATTTGTGC</td><td>>MI0000266|hsa-mir-10a&WithFlank&17|-|48579832|48579953</td><td>28</td><td>50</td><td>23(+1C)</td><td>TACCCTGTAGATCCGAATTTGTGC</td><td>282.8</td><td>1.9</td><td>0.8</td><td>2.6</td><td>9.8E-03</td><td>3.0E-02</td><td>BT20/MB231</td><td>CYTO</td></tr>
<tr><td>>hg38_21_+_8433172_8446622_RNA45SN1_45S_50ntFlanks@10677.10711.35|GCGGTTCCGGCGGCGTCCGGTGAGCTCTCGCTGGC</td><td>>hg38_21_+_8433172_8446622_RNA45SN1_45S_50ntFlanks</td><td>10677</td><td>10711</td><td>35</td><td>GCGGTTCCGGCGGCGTCCGGTGAGCTCTCGCTGGC</td><td>3018.8</td><td>-4.0</td><td>1.8</td><td>-2.2</td><td>2.6E-02</td><td>3.0E-02</td><td>BT20/MB468</td><td>NUC</td></tr>
<tr><td>>hg38_21_+_8433172_8446622_RNA45SN1_45S_50ntFlanks@10781.10799.19|AGGTGAACAGCCTCTGGCA</td><td>>hg38_21_+_8433172_8446622_RNA45SN1_45S_50ntFlanks</td><td>10781</td><td>10799</td><td>19</td><td>AGGTGAACAGCCTCTGGCA</td><td>242.7</td><td>-1.8</td><td>0.8</td><td>-2.2</td><td>2.6E-02</td><td>3.0E-02</td><td>MB231/MB468</td><td>CYTO</td></tr>
<tr><td>>am_MI0003581|hsa-mir-574&WithFlank&4|+|38868026|38868133@31.49.19&[MIMAT0004795&hsa-miR-574-5p&offsets|0|-4;m-213&4|+|38868056|38868078&offsets|0|-4]|TGAGTGTGTGTGTGTGAGT</td><td>>am_MI0003581|hsa-mir-574&WithFlank&4|+|38868026|38868133</td><td>31</td><td>49</td><td>19</td><td>TGAGTGTGTGTGTGTGAGT</td><td>61.8</td><td>3.8</td><td>1.7</td><td>2.3</td><td>2.4E-02</td><td>3.0E-02</td><td>MB231/MB468</td><td>MP</td></tr>
<tr><td>>hg38_21_+_8433172_8446622_RNA45SN1_45S_50ntFlanks@10689.10708.20|GCGTCCGGTGAGCTCTCGCT</td><td>>hg38_21_+_8433172_8446622_RNA45SN1_45S_50ntFlanks</td><td>10689</td><td>10708</td><td>20</td><td>GCGTCCGGTGAGCTCTCGCT</td><td>523.6</td><td>-3.5</td><td>1.5</td><td>-2.4</td><td>1.8E-02</td><td>3.0E-02</td><td>BT20/MB468</td><td>MP</td></tr>
<tr><td>>MI0000782|hsa-mir-374a&WithFlank&X|-|74287280|74287363@18.35.18&[MIMAT0000727&hsa-miR-374a-5p&offsets|0|-4;m-184&X|-|74287325|74287346&offsets|0|-4]|TTATAATACAACCTGATA</td><td>>MI0000782|hsa-mir-374a&WithFlank&X|-|74287280|74287363</td><td>18</td><td>35</td><td>18</td><td>TTATAATACAACCTGATA</td><td>247.6</td><td>-4.1</td><td>1.5</td><td>-2.7</td><td>7.4E-03</td><td>3.0E-02</td><td>BT20/MB231</td><td>MP</td></tr>
<tr><td>>hg38_21_+_8433172_8446622_RNA45SN1_45S_50ntFlanks@5059.5098.40|CGATTTGTCTGGTTAATTCCGATAACGAACGAGACTCTGG</td><td>>hg38_21_+_8433172_8446622_RNA45SN1_45S_50ntFlanks</td><td>5059</td><td>5098</td><td>40</td><td>CGATTTGTCTGGTTAATTCCGATAACGAACGAGACTCTGG</td><td>227.0</td><td>-2.9</td><td>1.3</td><td>-2.3</td><td>2.3E-02</td><td>3.0E-02</td><td>BT20/MB468</td><td>MITO</td></tr>
<tr><td>>hg38_21_+_8433172_8446622_RNA45SN1_45S_50ntFlanks@12363.12382.20|GGATAACTGGCTTGTGGCGG</td><td>>hg38_21_+_8433172_8446622_RNA45SN1_45S_50ntFlanks</td><td>12363</td><td>12382</td><td>20</td><td>GGATAACTGGCTTGTGGCGG</td><td>90.4</td><td>-4.1</td><td>1.7</td><td>-2.4</td><td>1.8E-02</td><td>3.0E-02</td><td>BT20/MB468</td><td>MP</td></tr>
<tr><td>>hg38_21_+_8433172_8446622_RNA45SN1_45S_50ntFlanks@12412.12434.23|ATCCTTCGATGTCGGCTCTTCCT</td><td>>hg38_21_+_8433172_8446622_RNA45SN1_45S_50ntFlanks</td><td>12412</td><td>12434</td><td>23</td><td>ATCCTTCGATGTCGGCTCTTCCT</td><td>1621.1</td><td>-4.1</td><td>1.9</td><td>-2.2</td><td>2.7E-02</td><td>3.0E-02</td><td>MB231/MB468</td><td>NUC</td></tr>
<tr><td>>hg38_21_+_8433172_8446622_RNA45SN1_45S_50ntFlanks@9624.9650.27|AGTAGCTGGTTCCCTCCGAAGTTTCCC</td><td>>hg38_21_+_8433172_8446622_RNA45SN1_45S_50ntFlanks</td><td>9624</td><td>9650</td><td>27</td><td>AGTAGCTGGTTCCCTCCGAAGTTTCCC</td><td>68.6</td><td>-4.1</td><td>1.7</td><td>-2.4</td><td>1.8E-02</td><td>3.0E-02</td><td>BT20/MB468</td><td>MP</td></tr>
<tr><td>>hg38_21_+_8433172_8446622_RNA45SN1_45S_50ntFlanks@4159.4182.24|ACCACATCCAAGGAAGGCAGCAGG</td><td>>hg38_21_+_8433172_8446622_RNA45SN1_45S_50ntFlanks</td><td>4159</td><td>4182</td><td>24</td><td>ACCACATCCAAGGAAGGCAGCAGG</td><td>104.4</td><td>-4.2</td><td>1.8</td><td>-2.4</td><td>1.8E-02</td><td>3.0E-02</td><td>BT20/MB468</td><td>MP</td></tr>
<tr><td>>MI0000081|hsa-mir-24-2&WithFlank&19|-|13836281|13836365@56.75.20(+1U);MI0000080|hsa-mir-24-1&WithFlank&9|+|95086015|95086094@50.69.20(+1U)&[MIMAT0000080&hsa-miR-24-3p&offsets|0|-2(+1U);m-36&19|-|13836288|13836310&offsets|0|-3(+1U)];[MIMAT0000080_1&hsa-miR-24-3p&offsets|0|-2(+1U);m-35&9|+|95086064|95086086&offsets|0|-3(+1U)]|TGGCTCAGTTCAGCAGGAACT</td><td>>MI0000081|hsa-mir-24-2&WithFlank&19|-|13836281|13836365</td><td>56</td><td>75</td><td>20(+1U)</td><td>TGGCTCAGTTCAGCAGGAACT</td><td>95.5</td><td>3.5</td><td>1.5</td><td>2.4</td><td>1.8E-02</td><td>3.0E-02</td><td>BT20/MB468</td><td>MP</td></tr>
<tr><td>>hg38_1_-_228634819_228635039_RNA5S12_5S_50ntFlanks@130.169.40|TGGATGGGAGACCGCCTGGGAATACCGGGTGCTGTAGGCT</td><td>>hg38_1_-_228634819_228635039_RNA5S12_5S_50ntFlanks</td><td>130</td><td>169</td><td>40</td><td>TGGATGGGAGACCGCCTGGGAATACCGGGTGCTGTAGGCT</td><td>1003.3</td><td>2.9</td><td>1.1</td><td>2.7</td><td>7.4E-03</td><td>3.0E-02</td><td>BT20/MB231</td><td>MP</td></tr>
<tr><td>>hg38_21_+_8433172_8446622_RNA45SN1_45S_50ntFlanks@11728.11765.38|GTAACTATGACTCTCTTAAGGTAGCCAAATGCCTCGTC</td><td>>hg38_21_+_8433172_8446622_RNA45SN1_45S_50ntFlanks</td><td>11728</td><td>11765</td><td>38</td><td>GTAACTATGACTCTCTTAAGGTAGCCAAATGCCTCGTC</td><td>296.1</td><td>-2.7</td><td>1.2</td><td>-2.2</td><td>2.6E-02</td><td>3.0E-02</td><td>MB231/MB468</td><td>CYTO</td></tr>
<tr><td>>MI0000255|hsa-mir-30d&WithFlank&8|-|134804870|134804951@52.73.22&[MIMAT0004551&hsa-miR-30d-3p&offsets|0|0;m-254&8|-|134804879|134804900&offsets|0|0]|CTTTCAGTCAGATGTTTGCTGC</td><td>>MI0000255|hsa-mir-30d&WithFlank&8|-|134804870|134804951</td><td>52</td><td>73</td><td>22</td><td>CTTTCAGTCAGATGTTTGCTGC</td><td>66.9</td><td>3.7</td><td>1.6</td><td>2.3</td><td>2.4E-02</td><td>3.0E-02</td><td>MB231/MB468</td><td>MP</td></tr>
<tr><td>>MI0000084|hsa-mir-26b&WithFlank&2|+|218402640|218402728@18.38.21&[MIMAT0000083&hsa-miR-26b-5p&offsets|0|0;m-53&2|+|218402657|218402678&offsets|0|-1]|TTCAAGTAATTCAGGATAGGT</td><td>>MI0000084|hsa-mir-26b&WithFlank&2|+|218402640|218402728</td><td>18</td><td>38</td><td>21</td><td>TTCAAGTAATTCAGGATAGGT</td><td>892.9</td><td>-2.1</td><td>0.8</td><td>-2.6</td><td>9.0E-03</td><td>3.0E-02</td><td>BT20/MB231</td><td>MITO</td></tr>
<tr><td>>hg38_MT_+_598_1651_RNR1_12S_50ntFlanks@718.744.27|GTAAAGACGTTAGGTCAAGGTGTAGCC</td><td>>hg38_MT_+_598_1651_RNR1_12S_50ntFlanks</td><td>718</td><td>744</td><td>27</td><td>GTAAAGACGTTAGGTCAAGGTGTAGCC</td><td>307.9</td><td>-2.1</td><td>0.8</td><td>-2.6</td><td>9.0E-03</td><td>3.0E-02</td><td>BT20/MB231</td><td>MITO</td></tr>
<tr><td>>hg38_21_+_8433172_8446622_RNA45SN1_45S_50ntFlanks@476.518.43|TTCGTGATCGATGTGGTGACGTCGTGCTCTCCCGGGCCGGGTC</td><td>>hg38_21_+_8433172_8446622_RNA45SN1_45S_50ntFlanks</td><td>476</td><td>518</td><td>43</td><td>TTCGTGATCGATGTGGTGACGTCGTGCTCTCCCGGGCCGGGTC</td><td>458.1</td><td>-2.4</td><td>0.9</td><td>-2.6</td><td>9.9E-03</td><td>3.0E-02</td><td>BT20/MB231</td><td>CYTO</td></tr>
<tr><td>>hg38_21_+_8433172_8446622_RNA45SN1_45S_50ntFlanks@10411.10441.31|CAGTCGGTCCTGAGAGATGGGCGAGCGCCGT</td><td>>hg38_21_+_8433172_8446622_RNA45SN1_45S_50ntFlanks</td><td>10411</td><td>10441</td><td>31</td><td>CAGTCGGTCCTGAGAGATGGGCGAGCGCCGT</td><td>332.3</td><td>-3.4</td><td>1.4</td><td>-2.4</td><td>1.8E-02</td><td>3.0E-02</td><td>BT20/MB468</td><td>MP</td></tr>
<tr><td>>MI0003602|hsa-mir-590&WithFlank&7|+|74191192|74191300@62.83.22&[MIMAT0004801&hsa-miR-590-3p&offsets|0|+1;m-332&7|+|74191253|74191273&offsets|0|+1]|TAATTTTATGTATAAGCTAGTC</td><td>>MI0003602|hsa-mir-590&WithFlank&7|+|74191192|74191300</td><td>62</td><td>83</td><td>22</td><td>TAATTTTATGTATAAGCTAGTC</td><td>37.4</td><td>4.6</td><td>2.0</td><td>2.3</td><td>2.4E-02</td><td>3.0E-02</td><td>MB231/MB468</td><td>MP</td></tr>
<tr><td>>hg38_MT_+_1621_3279_RNR2_16S_50ntFlanks@1352.1389.38|ATAGGGTTTACGACCTCGATGTTGGATCAGGACATCCC</td><td>>hg38_MT_+_1621_3279_RNR2_16S_50ntFlanks</td><td>1352</td><td>1389</td><td>38</td><td>ATAGGGTTTACGACCTCGATGTTGGATCAGGACATCCC</td><td>16.0</td><td>7.2</td><td>3.2</td><td>2.3</td><td>2.4E-02</td><td>3.0E-02</td><td>MB231/MB468</td><td>MP</td></tr>
<tr><td>>trnaMT_PheGAA_MT_+_577_647@37.62.26|AAATGTTTAGACGGGCTCACATCACC</td><td>>trnaMT_PheGAA_MT_+_577_647</td><td>37</td><td>62</td><td>26</td><td>AAATGTTTAGACGGGCTCACATCACC</td><td>125.9</td><td>5.3</td><td>2.2</td><td>2.4</td><td>1.8E-02</td><td>3.0E-02</td><td>BT20/MB468</td><td>MP</td></tr>
<tr><td>>hg38_21_+_8433172_8446622_RNA45SN1_45S_50ntFlanks@11789.11810.22|TGGATGAACGAGATTCCCACTG</td><td>>hg38_21_+_8433172_8446622_RNA45SN1_45S_50ntFlanks</td><td>11789</td><td>11810</td><td>22</td><td>TGGATGAACGAGATTCCCACTG</td><td>68.8</td><td>-5.0</td><td>2.2</td><td>-2.3</td><td>2.4E-02</td><td>3.0E-02</td><td>MB231/MB468</td><td>MP</td></tr>
<tr><td>>hg38_21_+_8433172_8446622_RNA45SN1_45S_50ntFlanks@10241.10261.21|TACGCCGCGACGAGTAGGAGG</td><td>>hg38_21_+_8433172_8446622_RNA45SN1_45S_50ntFlanks</td><td>10241</td><td>10261</td><td>21</td><td>TACGCCGCGACGAGTAGGAGG</td><td>90.2</td><td>4.2</td><td>1.6</td><td>2.7</td><td>7.4E-03</td><td>3.0E-02</td><td>BT20/MB231</td><td>MP</td></tr>
<tr><td>>hg38_21_+_8433172_8446622_RNA45SN1_45S_50ntFlanks@7990.8011.22|GACGTGGCGACCCGCTGAATTT</td><td>>hg38_21_+_8433172_8446622_RNA45SN1_45S_50ntFlanks</td><td>7990</td><td>8011</td><td>22</td><td>GACGTGGCGACCCGCTGAATTT</td><td>2479.1</td><td>3.5</td><td>1.3</td><td>2.7</td><td>7.4E-03</td><td>3.0E-02</td><td>BT20/MB231</td><td>MP</td></tr>
<tr><td>>hg38_21_+_8433172_8446622_RNA45SN1_45S_50ntFlanks@12420.12441.22|ATGTCGGCTCTTCCTATCATTG</td><td>>hg38_21_+_8433172_8446622_RNA45SN1_45S_50ntFlanks</td><td>12420</td><td>12441</td><td>22</td><td>ATGTCGGCTCTTCCTATCATTG</td><td>741.6</td><td>-2.3</td><td>0.9</td><td>-2.6</td><td>9.9E-03</td><td>3.0E-02</td><td>BT20/MB231</td><td>CYTO</td></tr>
<tr><td>>hg38_21_+_8433172_8446622_RNA45SN1_45S_50ntFlanks@6661.6679.19|CGGTGGATCACTCGGCTCG</td><td>>hg38_21_+_8433172_8446622_RNA45SN1_45S_50ntFlanks</td><td>6661</td><td>6679</td><td>19</td><td>CGGTGGATCACTCGGCTCG</td><td>831.4</td><td>2.7</td><td>1.0</td><td>2.6</td><td>9.9E-03</td><td>3.0E-02</td><td>BT20/MB231</td><td>CYTO</td></tr>
<tr><td>>hg38_21_+_8433172_8446622_RNA45SN1_45S_50ntFlanks@12259.12280.22|CTTGATCTTGATTTTCAGTACG</td><td>>hg38_21_+_8433172_8446622_RNA45SN1_45S_50ntFlanks</td><td>12259</td><td>12280</td><td>22</td><td>CTTGATCTTGATTTTCAGTACG</td><td>175.4</td><td>-3.1</td><td>1.3</td><td>-2.4</td><td>1.8E-02</td><td>3.0E-02</td><td>BT20/MB468</td><td>MP</td></tr>
<tr><td>>hg38_21_+_8433172_8446622_RNA45SN1_45S_50ntFlanks@12420.12458.39|ATGTCGGCTCTTCCTATCATTGTGAAGCAGAATTCACCA</td><td>>hg38_21_+_8433172_8446622_RNA45SN1_45S_50ntFlanks</td><td>12420</td><td>12458</td><td>39</td><td>ATGTCGGCTCTTCCTATCATTGTGAAGCAGAATTCACCA</td><td>868.0</td><td>2.3</td><td>1.0</td><td>2.2</td><td>2.7E-02</td><td>3.0E-02</td><td>MB231/MB468</td><td>MITO</td></tr>
<tr><td>>hg38_21_+_8433172_8446622_RNA45SN1_45S_50ntFlanks@6661.6690.30|CGGTGGATCACTCGGCTCGTGCGTCGATGA</td><td>>hg38_21_+_8433172_8446622_RNA45SN1_45S_50ntFlanks</td><td>6661</td><td>6690</td><td>30</td><td>CGGTGGATCACTCGGCTCGTGCGTCGATGA</td><td>574.2</td><td>2.4</td><td>0.9</td><td>2.6</td><td>9.0E-03</td><td>3.0E-02</td><td>BT20/MB231</td><td>MITO</td></tr>
<tr><td>>hg38_21_+_8433172_8446622_RNA45SN1_45S_50ntFlanks@12233.12262.30|CCGTGGAGCAGAAGGGCAAAAGCTCGCTTG</td><td>>hg38_21_+_8433172_8446622_RNA45SN1_45S_50ntFlanks</td><td>12233</td><td>12262</td><td>30</td><td>CCGTGGAGCAGAAGGGCAAAAGCTCGCTTG</td><td>110.2</td><td>-4.0</td><td>1.7</td><td>-2.4</td><td>1.8E-02</td><td>3.0E-02</td><td>BT20/MB468</td><td>MP</td></tr>
<tr><td>>MI0000085|hsa-mir-27a&WithFlank&19|-|13836434|13836523@57.76.20(+2A)&[MIMAT0000084&hsa-miR-27a-3p&offsets|0|-1(+2A);m-43&19|-|13836447|13836467&offsets|0|-1(+2A)]|TTCACAGTGGCTAAGTTCCGAA</td><td>>MI0000085|hsa-mir-27a&WithFlank&19|-|13836434|13836523</td><td>57</td><td>76</td><td>20(+2A)</td><td>TTCACAGTGGCTAAGTTCCGAA</td><td>164.6</td><td>-4.0</td><td>1.5</td><td>-2.7</td><td>7.4E-03</td><td>3.0E-02</td><td>BT20/MB231</td><td>MP</td></tr>
<tr><td>>hg38_21_+_8433172_8446622_RNA45SN1_45S_50ntFlanks@12456.12476.21|CCAAGCGTTGGATTGTTCACC</td><td>>hg38_21_+_8433172_8446622_RNA45SN1_45S_50ntFlanks</td><td>12456</td><td>12476</td><td>21</td><td>CCAAGCGTTGGATTGTTCACC</td><td>1179.3</td><td>4.5</td><td>1.7</td><td>2.7</td><td>7.4E-03</td><td>3.0E-02</td><td>BT20/MB231</td><td>MP</td></tr>
<tr><td>>hg38_21_+_8433172_8446622_RNA45SN1_45S_50ntFlanks@12646.12669.24|TGTGGGATTATGACTGAACGCCTC</td><td>>hg38_21_+_8433172_8446622_RNA45SN1_45S_50ntFlanks</td><td>12646</td><td>12669</td><td>24</td><td>TGTGGGATTATGACTGAACGCCTC</td><td>687.4</td><td>4.8</td><td>1.8</td><td>2.7</td><td>7.4E-03</td><td>3.0E-02</td><td>BT20/MB231</td><td>MP</td></tr>
<tr><td>>MI0000286|hsa-mir-210&WithFlank&11|-|568083|568204@72.93.22(+2A)&[MIMAT0000267&hsa-miR-210-3p&offsets|0|0(+2A);m-102&11|-|568112|568133&offsets|0|0(+2A)]|CTGTGCGTGTGACAGCGGCTGAAA</td><td>>MI0000286|hsa-mir-210&WithFlank&11|-|568083|568204</td><td>72</td><td>93</td><td>22(+2A)</td><td>CTGTGCGTGTGACAGCGGCTGAAA</td><td>5357.0</td><td>2.5</td><td>0.9</td><td>2.6</td><td>9.1E-03</td><td>3.0E-02</td><td>BT20/MB231</td><td>MITO</td></tr>
<tr><td>>trnaMT_PheGAA_MT_+_577_647@1.35.35|GTTTATGTAGCTTACCTCCTCAAAGCAATACACTG</td><td>>trnaMT_PheGAA_MT_+_577_647</td><td>1</td><td>35</td><td>35</td><td>GTTTATGTAGCTTACCTCCTCAAAGCAATACACTG</td><td>1246.8</td><td>-2.7</td><td>1.0</td><td>-2.6</td><td>9.1E-03</td><td>3.0E-02</td><td>BT20/MB231</td><td>MITO</td></tr>
<tr><td>>MI0000088|hsa-mir-30a&WithFlank&6|-|71403545|71403627@13.35.23(+1U)&[MIMAT0000087&hsa-miR-30a-5p&offsets|+1|+2(+1U);m-15&6|-|71403595|71403616&offsets|+1|+2(+1U)]|GTAAACATCCTCGACTGGAAGCTT</td><td>>MI0000088|hsa-mir-30a&WithFlank&6|-|71403545|71403627</td><td>13</td><td>35</td><td>23(+1U)</td><td>GTAAACATCCTCGACTGGAAGCTT</td><td>1616.2</td><td>-2.9</td><td>1.1</td><td>-2.7</td><td>7.4E-03</td><td>3.0E-02</td><td>BT20/MB231</td><td>MP</td></tr>
</table>

<table>
<tr><th></th><th></th><th></th><th></th><th></th><th></th><th></th><th></th><th></th><th></th><th></th><th></th><th></th><th></th></tr>
<tr><td>>hg38_21_+_8433172_8446622_RNA45SN1_45S_50ntFlanks@10798.10823.26|CATGTTGGAACAATGTAGGTAAGGGA</td><td>>hg38_21_+_8433172_8446622_RNA45SN1_45S_50ntFlanks</td><td>10798</td><td>10823</td><td>26</td><td>CATGTTGGAACAATGTAGGTAAGGGA</td><td>157.5</td><td>-1.5</td><td>0.7</td><td>-2.2</td><td>2.6E-02</td><td>3.0E-02</td><td>BT20/MB468</td><td>CYTO</td></tr>
<tr><td>>hg38_21_+_8433172_8446622_RNA45SN1_45S_50ntFlanks@4297.4315.19|CATTGGAGGGCAAGTCTGG</td><td>>hg38_21_+_8433172_8446622_RNA45SN1_45S_50ntFlanks</td><td>4297</td><td>4315</td><td>19</td><td>CATTGGAGGGCAAGTCTGG</td><td>411.0</td><td>2.5</td><td>1.1</td><td>2.4</td><td>1.8E-02</td><td>3.0E-02</td><td>BT20/MB468</td><td>MP</td></tr>
<tr><td>>hg38_21_+_8433172_8446622_RNA45SN1_45S_50ntFlanks@11564.11584.21|CCTAGCAGCCGACTTAGAACT</td><td>>hg38_21_+_8433172_8446622_RNA45SN1_45S_50ntFlanks</td><td>11564</td><td>11584</td><td>21</td><td>CCTAGCAGCCGACTTAGAACT</td><td>351.4</td><td>2.2</td><td>0.8</td><td>2.6</td><td>9.1E-03</td><td>3.0E-02</td><td>BT20/MB231</td><td>MITO</td></tr>
<tr><td>>trnaMT_PheGAA_MT_+_577_647@1.22.22|GTTTATGTAGCTTACCTCCTCA</td><td>>trnaMT_PheGAA_MT_+_577_647</td><td>1</td><td>22</td><td>22</td><td>GTTTATGTAGCTTACCTCCTCA</td><td>160.1</td><td>2.2</td><td>1.0</td><td>2.2</td><td>2.7E-02</td><td>3.0E-02</td><td>MB231/MB468</td><td>MITO</td></tr>
<tr><td>>trnaMT_GluTTC_MT_-_14674_14742@20.57.38,trnalookalike8_GluTTC_5_-_93905172_93905240@20.57.38|AACGATGGTTTTTCATATCATTGGTCGTGGTTGTAGTC</td><td>>trnaMT_GluTTC_MT_-_14674_14742</td><td>20</td><td>57</td><td>38</td><td>AACGATGGTTTTTCATATCATTGGTCGTGGTTGTAGTC</td><td>36.3</td><td>4.5</td><td>2.0</td><td>2.3</td><td>2.4E-02</td><td>3.0E-02</td><td>MB231/MB468</td><td>MP</td></tr>
<tr><td>>hg38_21_+_8433172_8446622_RNA45SN1_45S_50ntFlanks@8355.8392.38|TGAAAAGAACTTTGAAGAGAGAGTTCAAGAGGGCGTGA</td><td>>hg38_21_+_8433172_8446622_RNA45SN1_45S_50ntFlanks</td><td>8355</td><td>8392</td><td>38</td><td>TGAAAAGAACTTTGAAGAGAGAGTTCAAGAGGGCGTGA</td><td>108.0</td><td>4.3</td><td>1.6</td><td>2.7</td><td>7.4E-03</td><td>3.0E-02</td><td>BT20/MB231</td><td>MP</td></tr>
<tr><td>>MI0000469|hsa-mir-125a&WithFlank&19|+|51693248|51693345@21.41.21&[MIMAT0000443&hsa-miR-125a-5p&offsets|0|-3;m-63&19|+|51693268|51693289&offsets|0|-1]|TCCCTGAGACCCTTTAACCTG</td><td>>MI0000469|hsa-mir-125a&WithFlank&19|+|51693248|51693345</td><td>21</td><td>41</td><td>21</td><td>TCCCTGAGACCCTTTAACCTG</td><td>96.2</td><td>6.2</td><td>2.6</td><td>2.4</td><td>1.8E-02</td><td>3.0E-02</td><td>BT20/MB468</td><td>MP</td></tr>
<tr><td>>hg38_21_+_8433172_8446622_RNA45SN1_45S_50ntFlanks@9308.9334.27|AGGAGTCTAACACGTGCGCGAGTCGGG</td><td>>hg38_21_+_8433172_8446622_RNA45SN1_45S_50ntFlanks</td><td>9308</td><td>9334</td><td>27</td><td>AGGAGTCTAACACGTGCGCGAGTCGGG</td><td>559.2</td><td>-2.6</td><td>1.1</td><td>-2.4</td><td>1.8E-02</td><td>3.0E-02</td><td>BT20/MB468</td><td>MP</td></tr>
<tr><td>>trna14_LysTTT_11_-_59327808_59327880@1.36.36,trna2_LysTTT_17_+_8022473_8022545@1.36.36,trna54_LysTTT_1_+_204475655_204475727@1.36.36,trna5_LysTTT_11_+_59323902_59323974@1.36.36,trna62_LysTTT_1_-_204476158_204476230@1.36.36,trna76_LysTTT_6_+_28918806_28918878@1.36.36|GCCCGGATAGCTCAGTCGGTAGAGCATCAGACTTTT</td><td>>trna14_LysTTT_11_-_59327808_59327880</td><td>1</td><td>36</td><td>36</td><td>GCCCGGATAGCTCAGTCGGTAGAGCATCAGACTTTT</td><td>495.7</td><td>2.3</td><td>0.9</td><td>2.6</td><td>1.0E-02</td><td>3.0E-02</td><td>BT20/MB231</td><td>CYTO</td></tr>
<tr><td>>hg38_21_+_8433172_8446622_RNA45SN1_45S_50ntFlanks@12446.12480.35|GCAGAATTCACCAAGCGTTGGATTGTTCACCCACT</td><td>>hg38_21_+_8433172_8446622_RNA45SN1_45S_50ntFlanks</td><td>12446</td><td>12480</td><td>35</td><td>GCAGAATTCACCAAGCGTTGGATTGTTCACCCACT</td><td>530.4</td><td>-2.1</td><td>0.8</td><td>-2.6</td><td>1.0E-02</td><td>3.0E-02</td><td>BT20/MB231</td><td>CYTO</td></tr>
<tr><td>>MI0005566|hsa-mir-374b&WithFlank&X|-|74218541|74218624@17.36.20&[MIMAT0004955&hsa-miR-374b-5p&offsets|0|-2;m-153&X|-|74218587|74218608&offsets|0|-2]|ATATAATACAACCTGCTAAG</td><td>>MI0005566|hsa-mir-374b&WithFlank&X|-|74218541|74218624</td><td>17</td><td>36</td><td>20</td><td>ATATAATACAACCTGCTAAG</td><td>31.2</td><td>4.8</td><td>2.1</td><td>2.3</td><td>2.4E-02</td><td>3.0E-02</td><td>MB231/MB468</td><td>MP</td></tr>
<tr><td>>hg38_21_+_8433172_8446622_RNA45SN1_45S_50ntFlanks@3842.3859.18|CCTACTTGGATAACTGTG</td><td>>hg38_21_+_8433172_8446622_RNA45SN1_45S_50ntFlanks</td><td>3842</td><td>3859</td><td>18</td><td>CCTACTTGGATAACTGTG</td><td>195.9</td><td>-1.5</td><td>0.7</td><td>-2.3</td><td>2.3E-02</td><td>3.0E-02</td><td>BT20/MB468</td><td>MITO</td></tr>
<tr><td>>hg38_21_+_8433172_8446622_RNA45SN1_45S_50ntFlanks@10404.10430.27|CATGGGTCAGTCGGTCCTGAGAGATGG</td><td>>hg38_21_+_8433172_8446622_RNA45SN1_45S_50ntFlanks</td><td>10404</td><td>10430</td><td>27</td><td>CATGGGTCAGTCGGTCCTGAGAGATGG</td><td>331.9</td><td>-3.2</td><td>1.3</td><td>-2.4</td><td>1.8E-02</td><td>3.0E-02</td><td>BT20/MB468</td><td>MP</td></tr>
<tr><td>>am_MI0000786|hsa-mir-378a&WithFlank&5|+|149732819|149732896@50.69.20(+1U)&[MIMAT0000732&hsa-miR-378a-3p&offsets|+1|-1(+1U);m-44&5|+|149732867|149732888&offsets|+1|-1(+1U)]|CTGGACTTGGAGTCAGAAGGT</td><td>>am_MI0000786|hsa-mir-378a&WithFlank&5|+|149732819|149732896</td><td>50</td><td>69</td><td>20(+1U)</td><td>CTGGACTTGGAGTCAGAAGGT</td><td>1835.1</td><td>2.8</td><td>1.2</td><td>2.4</td><td>1.8E-02</td><td>3.0E-02</td><td>BT20/MB468</td><td>MP</td></tr>
<tr><td>>trnaMT_GluTTC_MT_-_14674_14742@40.67.28,trnalookalike8_GluTTC_5_-_93905172_93905240@40.67.28|TTGGTCGTGGTTGTAGTCCGTGCGAGAA</td><td>>trnaMT_GluTTC_MT_-_14674_14742</td><td>40</td><td>67</td><td>28</td><td>TTGGTCGTGGTTGTAGTCCGTGCGAGAA</td><td>40.1</td><td>3.7</td><td>1.7</td><td>2.3</td><td>2.4E-02</td><td>3.0E-02</td><td>MB231/MB468</td><td>MP</td></tr>
<tr><td>>hg38_21_+_8433172_8446622_RNA45SN1_45S_50ntFlanks@11668.11688.21|TGCCCAGTGCTCTGAATGTCA</td><td>>hg38_21_+_8433172_8446622_RNA45SN1_45S_50ntFlanks</td><td>11668</td><td>11688</td><td>21</td><td>TGCCCAGTGCTCTGAATGTCA</td><td>378.3</td><td>-2.6</td><td>1.0</td><td>-2.7</td><td>8.0E-03</td><td>3.0E-02</td><td>BT20/MB231</td><td>NUC</td></tr>
<tr><td>>hg38_MT_+_1621_3279_RNR2_16S_50ntFlanks@1428.1457.30|ATTAAAGTCCTACGTGATCTGAGTTCAGAC</td><td>>hg38_MT_+_1621_3279_RNR2_16S_50ntFlanks</td><td>1428</td><td>1457</td><td>30</td><td>ATTAAAGTCCTACGTGATCTGAGTTCAGAC</td><td>123.5</td><td>2.1</td><td>0.9</td><td>2.2</td><td>2.7E-02</td><td>3.0E-02</td><td>MB231/MB468</td><td>MITO</td></tr>
<tr><td>>hg38_21_+_8433172_8446622_RNA45SN1_45S_50ntFlanks@4070.4090.21|TTTCGATGGTAGTCGCCGTGC</td><td>>hg38_21_+_8433172_8446622_RNA45SN1_45S_50ntFlanks</td><td>4070</td><td>4090</td><td>21</td><td>TTTCGATGGTAGTCGCCGTGC</td><td>29.0</td><td>4.6</td><td>2.1</td><td>2.3</td><td>2.4E-02</td><td>3.0E-02</td><td>MB231/MB468</td><td>MP</td></tr>
<tr><td>>hg38_21_+_8433172_8446622_RNA45SN1_45S_50ntFlanks@12420.12440.21|ATGTCGGCTCTTCCTATCATT</td><td>>hg38_21_+_8433172_8446622_RNA45SN1_45S_50ntFlanks</td><td>12420</td><td>12440</td><td>21</td><td>ATGTCGGCTCTTCCTATCATT</td><td>1479.2</td><td>-2.3</td><td>0.9</td><td>-2.6</td><td>1.0E-02</td><td>3.0E-02</td><td>BT20/MB231</td><td>CYTO</td></tr>
<tr><td>>hg38_21_+_8433172_8446622_RNA45SN1_45S_50ntFlanks@8527.8551.25|CCTTCCCCCGCCGCCCCTCCTCCTC</td><td>>hg38_21_+_8433172_8446622_RNA45SN1_45S_50ntFlanks</td><td>8527</td><td>8551</td><td>25</td><td>CCTTCCCCCGCCGCCCCTCCTCCTC</td><td>349.8</td><td>-3.2</td><td>1.4</td><td>-2.2</td><td>2.6E-02</td><td>3.0E-02</td><td>BT20/MB468</td><td>CYTO</td></tr>
<tr><td>>hg38_21_+_8433172_8446622_RNA45SN1_45S_50ntFlanks@12072.12090.19|CCCGAGGGGCTCTCGCTTC</td><td>>hg38_21_+_8433172_8446622_RNA45SN1_45S_50ntFlanks</td><td>12072</td><td>12090</td><td>19</td><td>CCCGAGGGGCTCTCGCTTC</td><td>292.2</td><td>-3.6</td><td>1.6</td><td>-2.2</td><td>2.6E-02</td><td>3.0E-02</td><td>BT20/MB468</td><td>CYTO</td></tr>
<tr><td>>hg38_21_+_8433172_8446622_RNA45SN1_45S_50ntFlanks@7977.8011.35|CGACCTCAGATCAGACGTGGCGACCCGCTGAATTT</td><td>>hg38_21_+_8433172_8446622_RNA45SN1_45S_50ntFlanks</td><td>7977</td><td>8011</td><td>35</td><td>CGACCTCAGATCAGACGTGGCGACCCGCTGAATTT</td><td>1117.6</td><td>1.8</td><td>0.8</td><td>2.2</td><td>2.7E-02</td><td>3.0E-02</td><td>MB231/MB468</td><td>MITO</td></tr>
<tr><td>>trna15_ValAAC_5_-_180615416_180615488@1.23.23|GTTTCCGTAGTGTAGTGGTCATC</td><td>>trna15_ValAAC_5_-_180615416_180615488</td><td>1</td><td>23</td><td>23</td><td>GTTTCCGTAGTGTAGTGGTCATC</td><td>1022.7</td><td>3.2</td><td>1.4</td><td>2.4</td><td>1.8E-02</td><td>3.0E-02</td><td>BT20/MB468</td><td>MP</td></tr>
<tr><td>>hg38_21_+_8433172_8446622_RNA45SN1_45S_50ntFlanks@4980.4999.20|ACACGGACAGGATTGACAGA</td><td>>hg38_21_+_8433172_8446622_RNA45SN1_45S_50ntFlanks</td><td>4980</td><td>4999</td><td>20</td><td>ACACGGACAGGATTGACAGA</td><td>358.2</td><td>-3.1</td><td>1.3</td><td>-2.4</td><td>1.8E-02</td><td>3.0E-02</td><td>BT20/MB468</td><td>MP</td></tr>
<tr><td>>hg38_21_+_8433172_8446622_RNA45SN1_45S_50ntFlanks@12615.12657.43|GCTGAGGAGCCAATGGGGCGAAGCTACCATCTGTGGGATTATG</td><td>>hg38_21_+_8433172_8446622_RNA45SN1_45S_50ntFlanks</td><td>12615</td><td>12657</td><td>43</td><td>GCTGAGGAGCCAATGGGGCGAAGCTACCATCTGTGGGATTATG</td><td>89.1</td><td>-4.3</td><td>1.8</td><td>-2.4</td><td>1.8E-02</td><td>3.0E-02</td><td>BT20/MB468</td><td>MP</td></tr>
<tr><td>>hg38_21_+_8433172_8446622_RNA45SN1_45S_50ntFlanks@4737.4758.22|GAAAGTCGGAGGTTCGAAGACG</td><td>>hg38_21_+_8433172_8446622_RNA45SN1_45S_50ntFlanks</td><td>4737</td><td>4758</td><td>22</td><td>GAAAGTCGGAGGTTCGAAGACG</td><td>1120.2</td><td>4.1</td><td>1.5</td><td>2.7</td><td>7.5E-03</td><td>3.0E-02</td><td>BT20/MB231</td><td>MP</td></tr>
<tr><td>>hg38_21_+_8433172_8446622_RNA45SN1_45S_50ntFlanks@6648.6675.28|GTACGACTCTTAGCGGTGGATCACTCGG</td><td>>hg38_21_+_8433172_8446622_RNA45SN1_45S_50ntFlanks</td><td>6648</td><td>6675</td><td>28</td><td>GTACGACTCTTAGCGGTGGATCACTCGG</td><td>360.7</td><td>3.8</td><td>1.4</td><td>2.7</td><td>7.5E-03</td><td>3.0E-02</td><td>BT20/MB231</td><td>MP</td></tr>
<tr><td>>hg38_21_+_8433172_8446622_RNA45SN1_45S_50ntFlanks@10105.10125.21|CGCGTGTGGTGTGCGTCGGAG</td><td>>hg38_21_+_8433172_8446622_RNA45SN1_45S_50ntFlanks</td><td>10105</td><td>10125</td><td>21</td><td>CGCGTGTGGTGTGCGTCGGAG</td><td>129.0</td><td>-3.1</td><td>1.3</td><td>-2.4</td><td>1.8E-02</td><td>3.0E-02</td><td>BT20/MB468</td><td>MP</td></tr>
<tr><td>>MI0000264|hsa-mir-7-2&WithFlank&15|+|88611819|88611940@38.60.23(+1G);MI0000265|hsa-mir-7-3&WithFlank&19|+|4770664|4770785@37.59.23(+1G);MI0000263|hsa-mir-7-1&WithFlank&9|-|83969742|83969863@30.52.23(+1G)&[MIMAT0000252&hsa-miR-7-5p&offsets|0|-1(+1G);m-400&15|+|88611856|88611879&offsets|0|-1(+1G)];[MIMAT0000252_1&hsa-miR-7-5p&offsets|0|-1(+1G);m-402&19|+|4770700|4770723&offsets|0|-1(+1G)];[MIMAT0000252_2&hsa-miR-7-5p&offsets|0|-1(+1G);m-401&9|-|83969811|83969834&offsets|0|-1(+1G)]|TGGAAGACTAGTGATTTTGTTGTG</td><td>>MI0000264|hsa-mir-7-2&WithFlank&15|+|88611819|88611940</td><td>38</td><td>60</td><td>23(+1G)</td><td>TGGAAGACTAGTGATTTTGTTGTG</td><td>423.8</td><td>2.2</td><td>1.0</td><td>2.3</td><td>2.4E-02</td><td>3.0E-02</td><td>MB231/MB468</td><td>MP</td></tr>
<tr><td>>hg38_21_+_8433172_8446622_RNA45SN1_45S_50ntFlanks@11674.11691.18|GTGCTCTGAATGTCAAAG</td><td>>hg38_21_+_8433172_8446622_RNA45SN1_45S_50ntFlanks</td><td>11674</td><td>11691</td><td>18</td><td>GTGCTCTGAATGTCAAAG</td><td>86.5</td><td>3.5</td><td>1.5</td><td>2.4</td><td>1.8E-02</td><td>3.0E-02</td><td>BT20/MB468</td><td>MP</td></tr>
<tr><td>>hg38_MT_+_1621_3279_RNR2_16S_50ntFlanks@1424.1446.23|AACGATTAAAGTCCTACGTGATC</td><td>>hg38_MT_+_1621_3279_RNR2_16S_50ntFlanks</td><td>1424</td><td>1446</td><td>23</td><td>AACGATTAAAGTCCTACGTGATC</td><td>81.9</td><td>2.2</td><td>1.0</td><td>2.3</td><td>2.4E-02</td><td>3.0E-02</td><td>MB231/MB468</td><td>MP</td></tr>
<tr><td>>MI0000446|hsa-mir-125b-1&WithFlank&11|-|122099751|122099850@21.39.19;MI0000470|hsa-mir-125b-2&WithFlank&21|+|16590231|16590331@23.41.19&[MIMAT0000423&hsa-miR-125b-5p&offsets|0|-3;m-67&11|-|122099809|122099830&offsets|0|-3];[MIMAT0000423_1&hsa-miR-125b-5p&offsets|0|-3;m-68&21|+|16590253|16590274&offsets|0|-3]|TCCCTGAGACCCTAACTTG</td><td>>MI0000446|hsa-mir-125b-1&WithFlank&11|-|122099751|122099850</td><td>21</td><td>39</td><td>19</td><td>TCCCTGAGACCCTAACTTG</td><td>1373.7</td><td>-6.6</td><td>2.5</td><td>-2.6</td><td>9.2E-03</td><td>3.0E-02</td><td>BT20/MB231</td><td>MITO</td></tr>
<tr><td>>hg38_1_-_228634819_228635039_RNA5S12_5S_50ntFlanks@138.171.34|AGACCGCCTGGGAATACCGGGTGCTGTAGGCTTT</td><td>>hg38_1_-_228634819_228635039_RNA5S12_5S_50ntFlanks</td><td>138</td><td>171</td><td>34</td><td>AGACCGCCTGGGAATACCGGGTGCTGTAGGCTTT</td><td>1072.7</td><td>-2.0</td><td>0.8</td><td>-2.7</td><td>8.0E-03</td><td>3.0E-02</td><td>BT20/MB231</td><td>NUC</td></tr>
<tr><td>>hg38_21_+_8433172_8446622_RNA45SN1_45S_50ntFlanks@11580.11600.21|GAACTGGTGCGGACCAGGGGA</td><td>>hg38_21_+_8433172_8446622_RNA45SN1_45S_50ntFlanks</td><td>11580</td><td>11600</td><td>21</td><td>GAACTGGTGCGGACCAGGGGA</td><td>114.9</td><td>-1.4</td><td>0.6</td><td>-2.3</td><td>2.3E-02</td><td>3.0E-02</td><td>BT20/MB468</td><td>MITO</td></tr>
<tr><td>>MI0000264|hsa-mir-7-2&WithFlank&15|+|88611819|88611940@38.59.22;MI0000265|hsa-mir-7-3&WithFlank&19|+|4770664|4770785@37.58.22;MI0000263|hsa-mir-7-1&WithFlank&9|-|83969742|83969863@30.51.22&[MIMAT0000252&hsa-miR-7-5p&offsets|0|-2;m-400&15|+|88611856|88611879&offsets|0|-2];[MIMAT0000252_1&hsa-miR-7-5p&offsets|0|-2;m-402&19|+|4770700|4770723&offsets|0|-2];[MIMAT0000252_2&hsa-miR-7-5p&offsets|0|-2;m-401&9|-|83969811|83969834&offsets|0|-2]|TGGAAGACTAGTGATTTTGTTG</td><td>>MI0000264|hsa-mir-7-2&WithFlank&15|+|88611819|88611940</td><td>38</td><td>59</td><td>22</td><td>TGGAAGACTAGTGATTTTGTTG</td><td>3416.9</td><td>-2.4</td><td>0.9</td><td>-2.7</td><td>7.5E-03</td><td>3.0E-02</td><td>BT20/MB231</td><td>MP</td></tr>
<tr><td>>trnaMT_HisGTG_MT_+_12138_12206@32.71.40|TGAATCTGACAACAGAGGCTTACGACCCCTTATTTACCCC</td><td>>trnaMT_HisGTG_MT_+_12138_12206</td><td>32</td><td>71</td><td>40</td><td>TGAATCTGACAACAGAGGCTTACGACCCCTTATTTACCCC</td><td>1038.1</td><td>-2.6</td><td>1.0</td><td>-2.6</td><td>1.0E-02</td><td>3.0E-02</td><td>BT20/MB231</td><td>CYTO</td></tr>
<tr><td>>hg38_21_+_8433172_8446622_RNA45SN1_45S_50ntFlanks@6661.6683.23|CGGTGGATCACTCGGCTCGTGCG</td><td>>hg38_21_+_8433172_8446622_RNA45SN1_45S_50ntFlanks</td><td>6661</td><td>6683</td><td>23</td><td>CGGTGGATCACTCGGCTCGTGCG</td><td>364.0</td><td>1.7</td><td>0.7</td><td>2.4</td><td>1.8E-02</td><td>3.0E-02</td><td>BT20/MB468</td><td>MP</td></tr>
<tr><td>>trna10_AspGTC_12_-_125424193_125424264@35.70.36,trna12_AspGTC_12_-_125411891_125411962@35.70.36,trna144_AspGTC_6_-_27551236_27551307@35.70.36,trna38_AspGTC_17_-_8125556_8125627@35.70.36,trna45_AspGTC_6_+_27447453_27447524@35.70.36,trna48_AspGTC_6_+_27471523_27471594@35.70.36,trna4_AspGTC_12_+_96429799_96429870@35.70.36,trna69_AspGTC_1_-_161440205_161440276@35.70.36,trna72_AspGTC_1_-_161432824_161432895@35.70.36,trna75_AspGTC_1_-_161425414_161425485@35.70.36,trna78_AspGTC_1_-_161418033_161418104@35.70.36,trna81_AspGTC_1_-_161410615_161410686@35.70.36|TCACGCGGGAGACCGGGGTTCGATTCCCCGACGGGG</td><td>>trna10_AspGTC_12_-_125424193_125424264</td><td>35</td><td>70</td><td>36</td><td>TCACGCGGGAGACCGGGGTTCGATTCCCCGACGGGG</td><td>511.5</td><td>-2.2</td><td>1.0</td><td>-2.2</td><td>2.7E-02</td><td>3.0E-02</td><td>MB231/MB468</td><td>MITO</td></tr>
<tr><td>>hg38_21_+_8433172_8446622_RNA45SN1_45S_50ntFlanks@11918.11949.32|ACATGAGAGGTGTAGAATAAGTGGGAGGCCCC</td><td>>hg38_21_+_8433172_8446622_RNA45SN1_45S_50ntFlanks</td><td>11918</td><td>11949</td><td>32</td><td>ACATGAGAGGTGTAGAATAAGTGGGAGGCCCC</td><td>384.8</td><td>-3.3</td><td>1.4</td><td>-2.4</td><td>1.8E-02</td><td>3.0E-02</td><td>BT20/MB468</td><td>MP</td></tr>
<tr><td>>hg38_21_+_8433172_8446622_RNA45SN1_45S_50ntFlanks@8176.8223.48|CTTCTGATCGAGGCCCAGCCCGTGGACGGTGTGAGGCCGGTAGCGGCC</td><td>>hg38_21_+_8433172_8446622_RNA45SN1_45S_50ntFlanks</td><td>8176</td><td>8223</td><td>48</td><td>CTTCTGATCGAGGCCCAGCCCGTGGACGGTGTGAGGCCGGTAGCGGCC</td><td>76.7</td><td>-5.4</td><td>2.3</td><td>-2.4</td><td>1.8E-02</td><td>3.0E-02</td><td>BT20/MB468</td><td>MP</td></tr>
<tr><td>>hg38_21_+_8433172_8446622_RNA45SN1_45S_50ntFlanks@10853.10875.23|ATAAGGATTGGCTCTAAGGGCTG</td><td>>hg38_21_+_8433172_8446622_RNA45SN1_45S_50ntFlanks</td><td>10853</td><td>10875</td><td>23</td><td>ATAAGGATTGGCTCTAAGGGCTG</td><td>114.6</td><td>-3.1</td><td>1.3</td><td>-2.4</td><td>1.8E-02</td><td>3.0E-02</td><td>BT20/MB468</td><td>MP</td></tr>
<tr><td>>hg38_MT_+_1621_3279_RNR2_16S_50ntFlanks@1367.1393.27|TCGATGTTGGATCAGGACATCCCGATG</td><td>>hg38_MT_+_1621_3279_RNR2_16S_50ntFlanks</td><td>1367</td><td>1393</td><td>27</td><td>TCGATGTTGGATCAGGACATCCCGATG</td><td>115.2</td><td>3.8</td><td>1.7</td><td>2.2</td><td>2.4E-02</td><td>3.0E-02</td><td>MB231/MB468</td><td>MP</td></tr>
<tr><td>>hg38_21_+_8433172_8446622_RNA45SN1_45S_50ntFlanks@12417.12438.22|TCGATGTCGGCTCTTCCTATCA</td><td>>hg38_21_+_8433172_8446622_RNA45SN1_45S_50ntFlanks</td><td>12417</td><td>12438</td><td>22</td><td>TCGATGTCGGCTCTTCCTATCA</td><td>101.6</td><td>-4.1</td><td>1.8</td><td>-2.2</td><td>2.4E-02</td><td>3.0E-02</td><td>MB231/MB468</td><td>MP</td></tr>
</table>

<table>
<tr><th></th><th></th><th></th><th></th><th></th><th></th><th></th><th></th><th></th><th></th><th></th><th></th><th></th><th></th></tr>
<tr><td>>hg38_21_+_8433172_8446622_RNA45SN1_45S_50ntFlanks@7982.8011.30|TCAGATCAGACGTGGCGACCCGCTGAATTT</td><td>>hg38_21_+_8433172_8446622_RNA45SN1_45S_50ntFlanks</td><td>7982</td><td>8011</td><td>30</td><td>TCAGATCAGACGTGGCGACCCGCTGAATTT</td><td>82.8</td><td>-3.8</td><td>1.7</td><td>-2.2</td><td>2.4E-02</td><td>3.0E-02</td><td>MB231/MB468</td><td>MP</td></tr>
<tr><td>>hg38_21_+_8433172_8446622_RNA45SN1_45S_50ntFlanks@5236.5261.26|CACTGACTGGCTCAGCGTGTGCCTAC</td><td>>hg38_21_+_8433172_8446622_RNA45SN1_45S_50ntFlanks</td><td>5236</td><td>5261</td><td>26</td><td>CACTGACTGGCTCAGCGTGTGCCTAC</td><td>158.5</td><td>-3.4</td><td>1.5</td><td>-2.3</td><td>2.3E-02</td><td>3.0E-02</td><td>BT20/MB468</td><td>MITO</td></tr>
<tr><td>>hg38_21_+_8433172_8446622_RNA45SN1_45S_50ntFlanks@9501.9526.26|ATGGTGAACTATGCCTGGGCAGGGCG</td><td>>hg38_21_+_8433172_8446622_RNA45SN1_45S_50ntFlanks</td><td>9501</td><td>9526</td><td>26</td><td>ATGGTGAACTATGCCTGGGCAGGGCG</td><td>457.1</td><td>-3.4</td><td>1.4</td><td>-2.4</td><td>1.8E-02</td><td>3.0E-02</td><td>BT20/MB468</td><td>MP</td></tr>
<tr><td>>hg38_21_+_8433172_8446622_RNA45SN1_45S_50ntFlanks@5074.5094.21|ATTCCGATAACGAACGAGACT</td><td>>hg38_21_+_8433172_8446622_RNA45SN1_45S_50ntFlanks</td><td>5074</td><td>5094</td><td>21</td><td>ATTCCGATAACGAACGAGACT</td><td>232.6</td><td>-3.7</td><td>1.6</td><td>-2.4</td><td>1.8E-02</td><td>3.0E-02</td><td>BT20/MB468</td><td>MP</td></tr>
<tr><td>>hg38_21_+_8433172_8446622_RNA45SN1_45S_50ntFlanks@9198.9221.24|GTTCTCTCGGGGCCACGCGCGCGT</td><td>>hg38_21_+_8433172_8446622_RNA45SN1_45S_50ntFlanks</td><td>9198</td><td>9221</td><td>24</td><td>GTTCTCTCGGGGCCACGCGCGCGT</td><td>61.8</td><td>-4.0</td><td>1.7</td><td>-2.4</td><td>1.8E-02</td><td>3.0E-02</td><td>BT20/MB468</td><td>MP</td></tr>
<tr><td>>hg38_MT_+_1621_3279_RNR2_16S_50ntFlanks@1356.1388.33|GGTTTACGACCTCGATGTTGGATCAGGACATCC</td><td>>hg38_MT_+_1621_3279_RNR2_16S_50ntFlanks</td><td>1356</td><td>1388</td><td>33</td><td>GGTTTACGACCTCGATGTTGGATCAGGACATCC</td><td>187.3</td><td>3.0</td><td>1.3</td><td>2.2</td><td>2.4E-02</td><td>3.0E-02</td><td>MB231/MB468</td><td>MP</td></tr>
<tr><td>>trnaMT_CysGCA_MT_-_5761_5826@38.68.31|AAGAAGCAGCTTCAAACCTGCCGGGGCTTCC</td><td>>trnaMT_CysGCA_MT_-_5761_5826</td><td>38</td><td>68</td><td>31</td><td>AAGAAGCAGCTTCAAACCTGCCGGGGCTTCC</td><td>88.3</td><td>4.4</td><td>1.9</td><td>2.4</td><td>1.8E-02</td><td>3.0E-02</td><td>BT20/MB468</td><td>MP</td></tr>
<tr><td>>hg38_21_+_8433172_8446622_RNA45SN1_45S_50ntFlanks@12637.12664.28|GCTACCATCTGTGGGATTATGACTGAAC</td><td>>hg38_21_+_8433172_8446622_RNA45SN1_45S_50ntFlanks</td><td>12637</td><td>12664</td><td>28</td><td>GCTACCATCTGTGGGATTATGACTGAAC</td><td>132.6</td><td>5.0</td><td>2.1</td><td>2.4</td><td>1.8E-02</td><td>3.0E-02</td><td>BT20/MB468</td><td>MP</td></tr>
<tr><td>>hg38_21_+_8433172_8446622_RNA45SN1_45S_50ntFlanks@3845.3881.37|ACTTGGATAACTGTGGTAATTCTAGAGCTAATACATG</td><td>>hg38_21_+_8433172_8446622_RNA45SN1_45S_50ntFlanks</td><td>3845</td><td>3881</td><td>37</td><td>ACTTGGATAACTGTGGTAATTCTAGAGCTAATACATG</td><td>61.8</td><td>-4.3</td><td>1.8</td><td>-2.4</td><td>1.8E-02</td><td>3.0E-02</td><td>BT20/MB468</td><td>MP</td></tr>
<tr><td>>trna2_GlyGCC_21_-_18827107_18827177@1.31.31,trna35_GlyGCC_1_+_161413094_161413164@1.31.31,trna37_GlyGCC_1_+_161420467_161420537@1.31.31,trna39_GlyGCC_1_+_161427898_161427968@1.31.31,trna41_GlyGCC_1_+_161435258_161435328@1.31.31|GCATGGGTGGTTCAGTGGTAGAATTCTCGCC</td><td>>trna2_GlyGCC_21_-_18827107_18827177</td><td>1</td><td>31</td><td>31</td><td>GCATGGGTGGTTCAGTGGTAGAATTCTCGCC</td><td>1909.6</td><td>1.8</td><td>0.8</td><td>2.2</td><td>2.7E-02</td><td>3.0E-02</td><td>BT20/MB468</td><td>NUC</td></tr>
<tr><td>>hg38_21_+_8433172_8446622_RNA45SN1_45S_50ntFlanks@10795.10825.31|TGGCATGTTGGAACAATGTAGGTAAGGGAAG</td><td>>hg38_21_+_8433172_8446622_RNA45SN1_45S_50ntFlanks</td><td>10795</td><td>10825</td><td>31</td><td>TGGCATGTTGGAACAATGTAGGTAAGGGAAG</td><td>79.9</td><td>-4.0</td><td>1.7</td><td>-2.4</td><td>1.8E-02</td><td>3.0E-02</td><td>BT20/MB468</td><td>MP</td></tr>
<tr><td>>hg38_21_+_8433172_8446622_RNA45SN1_45S_50ntFlanks@12419.12450.32|GATGTCGGCTCTTCCTATCATTGTGAAGCAGA</td><td>>hg38_21_+_8433172_8446622_RNA45SN1_45S_50ntFlanks</td><td>12419</td><td>12450</td><td>32</td><td>GATGTCGGCTCTTCCTATCATTGTGAAGCAGA</td><td>120.9</td><td>2.0</td><td>0.9</td><td>2.2</td><td>2.7E-02</td><td>3.0E-02</td><td>MB231/MB468</td><td>MITO</td></tr>
<tr><td>>hg38_21_+_8433172_8446622_RNA45SN1_45S_50ntFlanks@11203.11232.30|CGGCGGCGGCGACTCTGGACGCGAGCCGGG</td><td>>hg38_21_+_8433172_8446622_RNA45SN1_45S_50ntFlanks</td><td>11203</td><td>11232</td><td>30</td><td>CGGCGGCGGCGACTCTGGACGCGAGCCGGG</td><td>90.5</td><td>5.3</td><td>2.0</td><td>2.7</td><td>7.5E-03</td><td>3.0E-02</td><td>BT20/MB231</td><td>MP</td></tr>
<tr><td>>hg38_21_+_8433172_8446622_RNA45SN1_45S_50ntFlanks@6646.6670.25|TCGTACGACTCTTAGCGGTGGATCA</td><td>>hg38_21_+_8433172_8446622_RNA45SN1_45S_50ntFlanks</td><td>6646</td><td>6670</td><td>25</td><td>TCGTACGACTCTTAGCGGTGGATCA</td><td>518.1</td><td>3.3</td><td>1.2</td><td>2.7</td><td>7.5E-03</td><td>3.0E-02</td><td>BT20/MB231</td><td>MP</td></tr>
<tr><td>>am_trnaMT_ValTAC_MT_+_1602_1670@49.67.19|AACTTAACTTGACCGCTCT</td><td>>am_trnaMT_ValTAC_MT_+_1602_1670</td><td>49</td><td>67</td><td>19</td><td>AACTTAACTTGACCGCTCT</td><td>1173.1</td><td>2.1</td><td>0.8</td><td>2.6</td><td>9.2E-03</td><td>3.0E-02</td><td>BT20/MB231</td><td>MITO</td></tr>
<tr><td>>hg38_21_+_8433172_8446622_RNA45SN1_45S_50ntFlanks@12990.13013.24|GCTCCCTCGCTGCGATCTATTGAA</td><td>>hg38_21_+_8433172_8446622_RNA45SN1_45S_50ntFlanks</td><td>12990</td><td>13013</td><td>24</td><td>GCTCCCTCGCTGCGATCTATTGAA</td><td>1186.1</td><td>-1.9</td><td>0.9</td><td>-2.2</td><td>2.6E-02</td><td>3.0E-02</td><td>BT20/MB468</td><td>CYTO</td></tr>
<tr><td>>hg38_21_+_8433172_8446622_RNA45SN1_45S_50ntFlanks@4353.4391.39|TATTAAAGTTGCTGCAGTTAAAAAGCTCGTAGTTGGATC</td><td>>hg38_21_+_8433172_8446622_RNA45SN1_45S_50ntFlanks</td><td>4353</td><td>4391</td><td>39</td><td>TATTAAAGTTGCTGCAGTTAAAAAGCTCGTAGTTGGATC</td><td>200.4</td><td>-2.1</td><td>1.0</td><td>-2.2</td><td>2.6E-02</td><td>3.0E-02</td><td>BT20/MB468</td><td>CYTO</td></tr>
<tr><td>>hg38_21_+_8433172_8446622_RNA45SN1_45S_50ntFlanks@9623.9655.33|TAGTAGCTGGTTCCCTCCGAAGTTTCCCTCAGG</td><td>>hg38_21_+_8433172_8446622_RNA45SN1_45S_50ntFlanks</td><td>9623</td><td>9655</td><td>33</td><td>TAGTAGCTGGTTCCCTCCGAAGTTTCCCTCAGG</td><td>134.7</td><td>-3.3</td><td>1.4</td><td>-2.4</td><td>1.8E-02</td><td>3.0E-02</td><td>BT20/MB468</td><td>MP</td></tr>
<tr><td>>hg38_21_+_8433172_8446622_RNA45SN1_45S_50ntFlanks@7974.7994.21|ACGCGACCTCAGATCAGACGT</td><td>>hg38_21_+_8433172_8446622_RNA45SN1_45S_50ntFlanks</td><td>7974</td><td>7994</td><td>21</td><td>ACGCGACCTCAGATCAGACGT</td><td>3700.7</td><td>2.2</td><td>0.9</td><td>2.4</td><td>1.8E-02</td><td>3.0E-02</td><td>BT20/MB468</td><td>MP</td></tr>
<tr><td>>trnaMT_SerGCT_MT_+_12207_12265@1.22.22|GAGAAAGCTCACAAGAACTGCT</td><td>>trnaMT_SerGCT_MT_+_12207_12265</td><td>1</td><td>22</td><td>22</td><td>GAGAAAGCTCACAAGAACTGCT</td><td>2860.9</td><td>2.1</td><td>0.8</td><td>2.6</td><td>8.1E-03</td><td>3.0E-02</td><td>BT20/MB231</td><td>NUC</td></tr>
<tr><td>>hg38_21_+_8433172_8446622_RNA45SN1_45S_50ntFlanks@11567.11600.34|AGCAGCCGACTTAGAACTGGTGCGGACCAGGGGA</td><td>>hg38_21_+_8433172_8446622_RNA45SN1_45S_50ntFlanks</td><td>11567</td><td>11600</td><td>34</td><td>AGCAGCCGACTTAGAACTGGTGCGGACCAGGGGA</td><td>273.5</td><td>-2.4</td><td>0.9</td><td>-2.6</td><td>1.0E-02</td><td>3.0E-02</td><td>BT20/MB231</td><td>CYTO</td></tr>
<tr><td>>hg38_21_+_8433172_8446622_RNA45SN1_45S_50ntFlanks@12415.12452.38|CTTCGATGTCGGCTCTTCCTATCATTGTGAAGCAGAAT</td><td>>hg38_21_+_8433172_8446622_RNA45SN1_45S_50ntFlanks</td><td>12415</td><td>12452</td><td>38</td><td>CTTCGATGTCGGCTCTTCCTATCATTGTGAAGCAGAAT</td><td>237.0</td><td>-2.0</td><td>0.8</td><td>-2.6</td><td>1.0E-02</td><td>3.0E-02</td><td>BT20/MB231</td><td>CYTO</td></tr>
<tr><td>>hg38_21_+_8433172_8446622_RNA45SN1_45S_50ntFlanks@7989.8013.25|AGACGTGGCGACCCGCTGAATTTAA</td><td>>hg38_21_+_8433172_8446622_RNA45SN1_45S_50ntFlanks</td><td>7989</td><td>8013</td><td>25</td><td>AGACGTGGCGACCCGCTGAATTTAA</td><td>775.7</td><td>1.9</td><td>0.8</td><td>2.3</td><td>2.3E-02</td><td>3.0E-02</td><td>BT20/MB468</td><td>MITO</td></tr>
<tr><td>>hg38_21_+_8433172_8446622_RNA45SN1_45S_50ntFlanks@12623.12657.35|GCCAATGGGGCGAAGCTACCATCTGTGGGATTATG</td><td>>hg38_21_+_8433172_8446622_RNA45SN1_45S_50ntFlanks</td><td>12623</td><td>12657</td><td>35</td><td>GCCAATGGGGCGAAGCTACCATCTGTGGGATTATG</td><td>53.5</td><td>-4.0</td><td>1.7</td><td>-2.4</td><td>1.8E-02</td><td>3.0E-02</td><td>BT20/MB468</td><td>MP</td></tr>
<tr><td>>MI0000466|hsa-mir-9-1&WithFlank&1|-|156420335|156420435@22.43.22;MI0000468|hsa-mir-9-3&WithFlank&15|+|89368011|89368112@22.43.22;MI0000467|hsa-mir-9-2&WithFlank&5|-|88666847|88666945@22.43.22&[MIMAT0000441&hsa-miR-9-5p&offsets|0|-1;m-104&1|-|156420392|156420414&offsets|0|-1];[MIMAT0000441_1&hsa-miR-9-5p&offsets|0|-1;m-105&15|+|89368032|89368054&offsets|0|-1];[MIMAT0000441_2&hsa-miR-9-5p&offsets|0|-1;m-104&5|-|88666902|88666924&offsets|0|-1]|TCTTTGGTTATCTAGCTGTATG</td><td>>MI0000466|hsa-mir-9-1&WithFlank&1|-|156420335|156420435</td><td>22</td><td>43</td><td>22</td><td>TCTTTGGTTATCTAGCTGTATG</td><td>167.4</td><td>3.5</td><td>1.5</td><td>2.4</td><td>1.8E-02</td><td>3.0E-02</td><td>BT20/MB468</td><td>MP</td></tr>
<tr><td>>MI0005561|hsa-mir-877&WithFlank&6|+|30584326|30584423@7.30.24&[MIMAT0004949&hsa-miR-877-5p&offsets|0|+4;m-382&6|+|30584332|30584354&offsets|0|+1]|GTAGAGGAGATGGCGCAGGGGACA</td><td>>MI0005561|hsa-mir-877&WithFlank&6|+|30584326|30584423</td><td>7</td><td>30</td><td>24</td><td>GTAGAGGAGATGGCGCAGGGGACA</td><td>118.2</td><td>3.4</td><td>1.4</td><td>2.4</td><td>1.8E-02</td><td>3.0E-02</td><td>BT20/MB468</td><td>MP</td></tr>
<tr><td>>hg38_21_+_8433172_8446622_RNA45SN1_45S_50ntFlanks@12043.12071.29|CACTGACCCGGTGAGGCGGGGGGGCGAGC</td><td>>hg38_21_+_8433172_8446622_RNA45SN1_45S_50ntFlanks</td><td>12043</td><td>12071</td><td>29</td><td>CACTGACCCGGTGAGGCGGGGGGGCGAGC</td><td>51.3</td><td>-4.1</td><td>1.8</td><td>-2.4</td><td>1.8E-02</td><td>3.0E-02</td><td>BT20/MB468</td><td>MP</td></tr>
<tr><td>>hg38_MT_+_1621_3279_RNR2_16S_50ntFlanks@883.906.24|AAAAACATCACCTCTAGCATCACC</td><td>>hg38_MT_+_1621_3279_RNR2_16S_50ntFlanks</td><td>883</td><td>906</td><td>24</td><td>AAAAACATCACCTCTAGCATCACC</td><td>112.2</td><td>3.7</td><td>1.7</td><td>2.2</td><td>2.7E-02</td><td>3.0E-02</td><td>MB231/MB468</td><td>MITO</td></tr>
<tr><td>>hg38_MT_+_1621_3279_RNR2_16S_50ntFlanks@1367.1398.32|TCGATGTTGGATCAGGACATCCCGATGGTGCA</td><td>>hg38_MT_+_1621_3279_RNR2_16S_50ntFlanks</td><td>1367</td><td>1398</td><td>32</td><td>TCGATGTTGGATCAGGACATCCCGATGGTGCA</td><td>83.2</td><td>4.5</td><td>2.0</td><td>2.2</td><td>2.5E-02</td><td>3.0E-02</td><td>MB231/MB468</td><td>MP</td></tr>
<tr><td>>MI0000298|hsa-mir-221&WithFlank&X|-|45746151|45746272@71.93.23(+1G)&[MIMAT0000278&hsa-miR-221-3p&offsets|0|0(+1G);m-91&X|-|45746181|45746202&offsets|0|+1(+1G)]|AGCTACATTGTCTGCTGGGTTTCG</td><td>>MI0000298|hsa-mir-221&WithFlank&X|-|45746151|45746272</td><td>71</td><td>93</td><td>23(+1G)</td><td>AGCTACATTGTCTGCTGGGTTTCG</td><td>213.0</td><td>3.0</td><td>1.3</td><td>2.2</td><td>2.5E-02</td><td>3.0E-02</td><td>MB231/MB468</td><td>MP</td></tr>
<tr><td>>hg38_21_+_8433172_8446622_RNA45SN1_45S_50ntFlanks@9534.9567.34|AGGAAACTCTGGTGGAGGTCCGTAGCGGTCCTGA</td><td>>hg38_21_+_8433172_8446622_RNA45SN1_45S_50ntFlanks</td><td>9534</td><td>9567</td><td>34</td><td>AGGAAACTCTGGTGGAGGTCCGTAGCGGTCCTGA</td><td>429.1</td><td>-4.3</td><td>1.8</td><td>-2.4</td><td>1.8E-02</td><td>3.0E-02</td><td>BT20/MB468</td><td>MP</td></tr>
<tr><td>>hg38_MT_+_598_1651_RNR1_12S_50ntFlanks@416.456.41|AAATAGACTACGAAAGTGGCTTTAACATATCTGAACACACA</td><td>>hg38_MT_+_598_1651_RNR1_12S_50ntFlanks</td><td>416</td><td>456</td><td>41</td><td>AAATAGACTACGAAAGTGGCTTTAACATATCTGAACACACA</td><td>250.7</td><td>-1.8</td><td>0.7</td><td>-2.6</td><td>9.3E-03</td><td>3.0E-02</td><td>BT20/MB231</td><td>MITO</td></tr>
<tr><td>>MI0001445|hsa-mir-423&WithFlank&17|+|30117073|30117178@58.77.20&[MIMAT0001340&hsa-miR-423-3p&offsets|-1|-4;m-100&17|+|30117131|30117153&offsets|-1|-4]|AAGCTCGGTCTGAGGCCCCT</td><td>>MI0001445|hsa-mir-423&WithFlank&17|+|30117073|30117178</td><td>58</td><td>77</td><td>20</td><td>AAGCTCGGTCTGAGGCCCCT</td><td>44.1</td><td>3.8</td><td>1.7</td><td>2.2</td><td>2.5E-02</td><td>3.0E-02</td><td>MB231/MB468</td><td>MP</td></tr>
<tr><td>>MI0000103|hsa-mir-101-1&WithFlank&1|-|65058428|65058514@53.74.22;MI0000739|hsa-mir-101-2&WithFlank&9|+|4850291|4850381@55.76.22&[MIMAT0000099&hsa-miR-101-3p&offsets|0|+1;m-17&1|-|65058443|65058462&offsets|0|+2];[MIMAT0000099_1&hsa-miR-101-3p&offsets|0|+1;m-16&9|+|4850345|4850364&offsets|0|+2]|TACAGTACTGTGATAACTGAAG</td><td>>MI0000103|hsa-mir-101-1&WithFlank&1|-|65058428|65058514</td><td>53</td><td>74</td><td>22</td><td>TACAGTACTGTGATAACTGAAG</td><td>1558.0</td><td>-2.3</td><td>0.9</td><td>-2.6</td><td>9.3E-03</td><td>3.0E-02</td><td>BT20/MB231</td><td>MITO</td></tr>
<tr><td>>hg38_21_+_8433172_8446622_RNA45SN1_45S_50ntFlanks@12447.12476.30|CAGAATTCACCAAGCGTTGGATTGTTCACC</td><td>>hg38_21_+_8433172_8446622_RNA45SN1_45S_50ntFlanks</td><td>12447</td><td>12476</td><td>30</td><td>CAGAATTCACCAAGCGTTGGATTGTTCACC</td><td>216.8</td><td>-1.7</td><td>0.8</td><td>-2.2</td><td>2.6E-02</td><td>3.0E-02</td><td>BT20/MB468</td><td>CYTO</td></tr>
<tr><td>>hg38_21_+_8433172_8446622_RNA45SN1_45S_50ntFlanks@9540.9563.24|CTCTGGTGGAGGTCCGTAGCGGTC</td><td>>hg38_21_+_8433172_8446622_RNA45SN1_45S_50ntFlanks</td><td>9540</td><td>9563</td><td>24</td><td>CTCTGGTGGAGGTCCGTAGCGGTC</td><td>65.8</td><td>-4.0</td><td>1.7</td><td>-2.4</td><td>1.8E-02</td><td>3.0E-02</td><td>BT20/MB468</td><td>MP</td></tr>
<tr><td>>hg38_21_+_8433172_8446622_RNA45SN1_45S_50ntFlanks@8349.8375.27|GAAAGTTGAAAAGAACTTTGAAGAGAG</td><td>>hg38_21_+_8433172_8446622_RNA45SN1_45S_50ntFlanks</td><td>8349</td><td>8375</td><td>27</td><td>GAAAGTTGAAAAGAACTTTGAAGAGAG</td><td>5928.3</td><td>4.8</td><td>1.8</td><td>2.7</td><td>7.6E-03</td><td>3.0E-02</td><td>BT20/MB231</td><td>MP</td></tr>
<tr><td>>hg38_21_+_8433172_8446622_RNA45SN1_45S_50ntFlanks@9954.9980.27|TGGAAGTCGGAATCCGCTAAGGAGTGT</td><td>>hg38_21_+_8433172_8446622_RNA45SN1_45S_50ntFlanks</td><td>9954</td><td>9980</td><td>27</td><td>TGGAAGTCGGAATCCGCTAAGGAGTGT</td><td>184.4</td><td>5.4</td><td>2.0</td><td>2.7</td><td>7.6E-03</td><td>3.0E-02</td><td>BT20/MB231</td><td>MP</td></tr>
<tr><td>>hg38_21_+_8433172_8446622_RNA45SN1_45S_50ntFlanks@10829.10853.25|GCAAGCCGGATCCGTAACTTCGGGA</td><td>>hg38_21_+_8433172_8446622_RNA45SN1_45S_50ntFlanks</td><td>10829</td><td>10853</td><td>25</td><td>GCAAGCCGGATCCGTAACTTCGGGA</td><td>140.7</td><td>-4.0</td><td>1.7</td><td>-2.4</td><td>1.8E-02</td><td>3.0E-02</td><td>BT20/MB468</td><td>MP</td></tr>
<tr><td>>hg38_21_+_8433172_8446622_RNA45SN1_45S_50ntFlanks@3991.4023.33|TGGTGACTCTAGATAACCTCGGGCCGATCGCAC</td><td>>hg38_21_+_8433172_8446622_RNA45SN1_45S_50ntFlanks</td><td>3991</td><td>4023</td><td>33</td><td>TGGTGACTCTAGATAACCTCGGGCCGATCGCAC</td><td>102.4</td><td>-1.8</td><td>0.8</td><td>-2.2</td><td>2.5E-02</td><td>3.0E-02</td><td>MB231/MB468</td><td>MP</td></tr>
<tr><td>>trna5_SupTTA_4_-_7325838_7325909@33.55.23|TTAGAGCCAGACTGCCTGGGTTT</td><td>>trna5_SupTTA_4_-_7325838_7325909</td><td>33</td><td>55</td><td>23</td><td>TTAGAGCCAGACTGCCTGGGTTT</td><td>154.3</td><td>3.1</td><td>1.2</td><td>2.7</td><td>7.6E-03</td><td>3.0E-02</td><td>BT20/MB231</td><td>MP</td></tr>
<tr><td>>hg38_MT_+_1621_3279_RNR2_16S_50ntFlanks@1379.1414.36|CAGGACATCCCGATGGTGCAGCCGCTATTAAAGGTT</td><td>>hg38_MT_+_1621_3279_RNR2_16S_50ntFlanks</td><td>1379</td><td>1414</td><td>36</td><td>CAGGACATCCCGATGGTGCAGCCGCTATTAAAGGTT</td><td>103.6</td><td>2.9</td><td>1.3</td><td>2.2</td><td>2.5E-02</td><td>3.0E-02</td><td>MB231/MB468</td><td>MP</td></tr>
<tr><td>>MI0003137|hsa-mir-193b&WithFlank&16|+|14303961|14304055@57.79.23&[MIMAT0002819&hsa-miR-193b-3p&offsets|0|+1;m-166&16|+|14304017|14304038&offsets|0|+1]|AACTGGCCCTCAAAGTCCCGCTT</td><td>>MI0003137|hsa-mir-193b&WithFlank&16|+|14303961|14304055</td><td>57</td><td>79</td><td>23</td><td>AACTGGCCCTCAAAGTCCCGCTT</td><td>1322.7</td><td>3.1</td><td>1.2</td><td>2.7</td><td>7.6E-03</td><td>3.0E-02</td><td>BT20/MB231</td><td>MP</td></tr>
<tr><td>>hg38_21_+_8433172_8446622_RNA45SN1_45S_50ntFlanks@4560.4589.30|CTAGGAATAATGGAATAGGACCGCGGTTCT</td><td>>hg38_21_+_8433172_8446622_RNA45SN1_45S_50ntFlanks</td><td>4560</td><td>4589</td><td>30</td><td>CTAGGAATAATGGAATAGGACCGCGGTTCT</td><td>338.9</td><td>-2.0</td><td>0.9</td><td>-2.2</td><td>2.6E-02</td><td>3.0E-02</td><td>MB231/MB468</td><td>CYTO</td></tr>
<tr><td>>hg38_21_+_8433172_8446622_RNA45SN1_45S_50ntFlanks@4734.4771.38|AACGAAAGTCGGAGGTTCGAAGACGATCAGATACCGTC</td><td>>hg38_21_+_8433172_8446622_RNA45SN1_45S_50ntFlanks</td><td>4734</td><td>4771</td><td>38</td><td>AACGAAAGTCGGAGGTTCGAAGACGATCAGATACCGTC</td><td>85.0</td><td>4.9</td><td>2.1</td><td>2.4</td><td>1.8E-02</td><td>3.0E-02</td><td>BT20/MB468</td><td>MP</td></tr>
<tr><td>>hg38_21_+_8433172_8446622_RNA45SN1_45S_50ntFlanks@5471.5510.40|CACGGCCCTGGCGGAGCGCTGAGAAGACGGTCGAACTTGA</td><td>>hg38_21_+_8433172_8446622_RNA45SN1_45S_50ntFlanks</td><td>5471</td><td>5510</td><td>40</td><td>CACGGCCCTGGCGGAGCGCTGAGAAGACGGTCGAACTTGA</td><td>50.6</td><td>-4.2</td><td>1.8</td><td>-2.4</td><td>1.8E-02</td><td>3.0E-02</td><td>BT20/MB468</td><td>MP</td></tr>
<tr><td>>hg38_21_+_8433172_8446622_RNA45SN1_45S_50ntFlanks@9308.9337.30|AGGAGTCTAACACGTGCGCGAGTCGGGGGC</td><td>>hg38_21_+_8433172_8446622_RNA45SN1_45S_50ntFlanks</td><td>9308</td><td>9337</td><td>30</td><td>AGGAGTCTAACACGTGCGCGAGTCGGGGGC</td><td>879.6</td><td>-3.6</td><td>1.5</td><td>-2.4</td><td>1.8E-02</td><td>3.0E-02</td><td>BT20/MB468</td><td>MP</td></tr>
</table>

| | | | | | | | | | | | | | |
|---|---|---|---|---|---|---|---|---|---|---|---|---|---|
| >am_trna10_ValCAC_5_-_180649395_180649467@1.18.18,trna12_ValAAC_5_-_180645270_180645342@1.18.18,trna132_ValAAC_6_-_27721179_27721251@1.18.18,trna133_ValCAC_6_-_27696327_27696399@1.18.18,trna136_ValAAC_6_-_27648885_27648957@1.18.18,trna139_ValAAC_6_-_27618707_27618779@1.18.18,trna15_ValAAC_5_-_180615416_180615488@1.18.18,trna18_ValCAC_5_-_180529253_180529325@1.18.18,trna2_ValAAC_3_+_169490018_169490090@1.18.18,trna2_ValCAC_5_+_180524070_180524142@1.18.18,trna37_ValAAC_6_+_27203288_27203360@1.18.18,trna4_ValAAC_5_+_180591154_180591226@1.18.18,trna5_ValAAC_5_+_180596610_180596682@1.18.18,trna6_ValCAC_5_+_180600650_180600722@1.18.18,trna85_ValCAC_1_-_161369490_161369562@1.18.18,trna90_ValCAC_1_-_149684088_149684161@1.18.18,trna98_ValCAC_1_-_149298555_149298627@1.18.18,trna9_ValCAC_6_+_26538282_26538354@1.18.18\|GTTTCCGTAGTGTAGTGG | >am_trna10_ValCAC_5_-_180649395_180649467 | 1 | 18 | 18 | GTTTCCGTAGTGTAGTGG | 229.9 | 2.0 | 0.9 | 2.2 | 2.8E-02 | 3.0E-02 | MB231/MB468 | MITO |
| >hg38_21_+_8433172_8446622_RNA45SN1_45S_50ntFlanks@3988.4010.23\|CTTTGGTGACTCTAGATAACCTC | >hg38_21_+_8433172_8446622_RNA45SN1_45S_50ntFlanks | 3988 | 4010 | 23 | CTTTGGTGACTCTAGATAACCTC | 273.6 | 2.1 | 0.8 | 2.6 | 1.0E-02 | 3.0E-02 | BT20/MB231 | CYTO |
| >trna116_GluCTC_1_-_145399233_145399304@37.75.39,trna134_GluTTC_1_-_16861774_16861845@37.75.39,trna71_GluCTC_1_-_161439189_161439260@37.75.39,trna74_GluCTC_1_-_161431809_161431880@37.75.39,trna77_GluCTC_1_-_161424398_161424469@37.75.39,trna77_GluCTC_6_+_28949976_28950047@37.75.39,trna80_GluCTC_1_-_161417018_161417089@37.75.39,trna84_GluTTC_1_-_161391883_161391954@37.75.39,trna87_GluCTC_6_-_126101393_126101464@37.75.39\|ACCGCCGCGGCCCGGGTTCGATTCCCGGTCAGGGAACCA | >trna116_GluCTC_1_-_145399233_145399304 | 37 | 75 | 39 | ACCGCCGCGGCCCGGGTTCGATTCCCGGTCAGGGAACCA | 4548.6 | -3.4 | 1.3 | -2.6 | 8.2E-03 | 3.0E-02 | BT20/MB231 | NUC |
| >hg38_21_+_8433172_8446622_RNA45SN1_45S_50ntFlanks@11671.11702.32\|CCAGTGCTCTGAATGTCAAAGTGAAGAAATTC | >hg38_21_+_8433172_8446622_RNA45SN1_45S_50ntFlanks | 11671 | 11702 | 32 | CCAGTGCTCTGAATGTCAAAGTGAAGAAATTC | 170.5 | -2.2 | 0.8 | -2.6 | 8.2E-03 | 3.0E-02 | BT20/MB231 | NUC |
| >hg38_21_+_8433172_8446622_RNA45SN1_45S_50ntFlanks@8383.8400.18\|GAGGGCGTGAAACCGTTA | >hg38_21_+_8433172_8446622_RNA45SN1_45S_50ntFlanks | 8383 | 8400 | 18 | GAGGGCGTGAAACCGTTA | 103.6 | -4.1 | 1.7 | -2.4 | 1.8E-02 | 3.0E-02 | BT20/MB468 | MP |
| >hg38_21_+_8433172_8446622_RNA45SN1_45S_50ntFlanks@6650.6695.46\|ACGACTCTTAGCGGTGGATCACTCGGCTCGTGCGTCGATGAAGAAC | >hg38_21_+_8433172_8446622_RNA45SN1_45S_50ntFlanks | 6650 | 6695 | 46 | ACGACTCTTAGCGGTGGATCACTCGGCTCGTGCGTCGATGAAGAAC | 268.8 | 2.5 | 1.1 | 2.2 | 2.7E-02 | 3.0E-02 | BT20/MB468 | NUC |
| >hg38_21_+_8433172_8446622_RNA45SN1_45S_50ntFlanks@12071.12096.26\|CCCCGAGGGGCTCTCGCTTCTGGCGC | >hg38_21_+_8433172_8446622_RNA45SN1_45S_50ntFlanks | 12071 | 12096 | 26 | CCCCGAGGGGCTCTCGCTTCTGGCGC | 105.9 | -4.5 | 1.9 | -2.4 | 1.8E-02 | 3.0E-02 | BT20/MB468 | MP |
| >hg38_21_+_8433172_8446622_RNA45SN1_45S_50ntFlanks@11583.11607.25\|CTGGTGCGGACCAGGGGAATCCGAC | >hg38_21_+_8433172_8446622_RNA45SN1_45S_50ntFlanks | 11583 | 11607 | 25 | CTGGTGCGGACCAGGGGAATCCGAC | 346.9 | -1.6 | 0.7 | -2.2 | 2.6E-02 | 3.0E-02 | MB231/MB468 | CYTO |
| >am_MI0000815\|hsa-mir-339&WithFlank&7\|-\|1022927\|1023032@21.40.20(+1U)&[MIMAT0000764&hsa-miR-339-5p&offsets\|0\|-3(+1U);m-163&7\|-\|1022990\|1023012&offsets\|0\|-3(+1U)]\|TCCCTGTCCTCCAGGAGCTCT | >am_MI0000815\|hsa-mir-339&WithFlank&7\|-\|1022927\|1023032 | 21 | 40 | 20(+1U) | TCCCTGTCCTCCAGGAGCTCT | 57.3 | 4.3 | 1.8 | 2.4 | 1.8E-02 | 3.0E-02 | BT20/MB468 | MP |
| >trnaMT_HisGTG_MT_+_12138_12206@26.61.36\|AGATTGTGAATCTGACAACAGAGGCTTACGACCCCT | >trnaMT_HisGTG_MT_+_12138_12206 | 26 | 61 | 36 | AGATTGTGAATCTGACAACAGAGGCTTACGACCCCT | 72.1 | 4.4 | 1.9 | 2.2 | 2.5E-02 | 3.0E-02 | MB231/MB468 | MP |
| >trnaMT_GlnTTG_MT_-_4329_4400@1.22.22,trnalookalike2_GlnTTG_1_-_564879_564950@1.22.22\|TAGGATGGGGTGTGATAGGTGG | >trnaMT_GlnTTG_MT_-_4329_4400 | 1 | 22 | 22 | TAGGATGGGGTGTGATAGGTGG | 271.6 | 4.8 | 2.0 | 2.4 | 1.8E-02 | 3.0E-02 | BT20/MB468 | MP |
| >hg38_21_+_8433172_8446622_RNA45SN1_45S_50ntFlanks@6650.6692.43\|ACGACTCTTAGCGGTGGATCACTCGGCTCGTGCGTCGATGAAG | >hg38_21_+_8433172_8446622_RNA45SN1_45S_50ntFlanks | 6650 | 6692 | 43 | ACGACTCTTAGCGGTGGATCACTCGGCTCGTGCGTCGATGAAG | 1709.1 | 1.5 | 0.7 | 2.3 | 2.3E-02 | 3.0E-02 | BT20/MB468 | MITO |
| >hg38_MT_+_1621_3279_RNR2_16S_50ntFlanks@1430.1453.24\|TAAAGTCCTACGTGATCTGAGTTC | >hg38_MT_+_1621_3279_RNR2_16S_50ntFlanks | 1430 | 1453 | 24 | TAAAGTCCTACGTGATCTGAGTTC | 115.0 | 2.8 | 1.3 | 2.2 | 2.5E-02 | 3.0E-02 | MB231/MB468 | MP |
| >hg38_21_+_8433172_8446622_RNA45SN1_45S_50ntFlanks@8182.8203.22\|ATCGAGGCCCAGCCCGTGGACG | >hg38_21_+_8433172_8446622_RNA45SN1_45S_50ntFlanks | 8182 | 8203 | 22 | ATCGAGGCCCAGCCCGTGGACG | 425.9 | -3.4 | 1.5 | -2.2 | 2.6E-02 | 3.0E-02 | MB231/MB468 | CYTO |
| >hg38_21_+_8433172_8446622_RNA45SN1_45S_50ntFlanks@6732.6752.21\|ACATTGATCATCGACACTTCG | >hg38_21_+_8433172_8446622_RNA45SN1_45S_50ntFlanks | 6732 | 6752 | 21 | ACATTGATCATCGACACTTCG | 584.6 | -1.7 | 0.8 | -2.2 | 2.8E-02 | 3.0E-02 | MB231/MB468 | MITO |
| >hg38_21_+_8433172_8446622_RNA45SN1_45S_50ntFlanks@11575.11609.35\|ACTTAGAACTGGTGCGGACCAGGGGAATCCGACTG | >hg38_21_+_8433172_8446622_RNA45SN1_45S_50ntFlanks | 11575 | 11609 | 35 | ACTTAGAACTGGTGCGGACCAGGGGAATCCGACTG | 710.5 | -2.5 | 1.1 | -2.4 | 1.8E-02 | 3.0E-02 | BT20/MB468 | MP |
| >MI0000469\|hsa-mir-125a&WithFlank&19\|+\|51693248\|51693345@21.42.22&[MIMAT0000443&hsa-miR-125a-5p&offsets\|0\|-2;m-63&19\|+\|51693268\|51693289&offsets\|0\|0]\|TCCCTGAGACCCTTTAACCTGT | >MI0000469\|hsa-mir-125a&WithFlank&19\|+\|51693248\|51693345 | 21 | 42 | 22 | TCCCTGAGACCCTTTAACCTGT | 1908.1 | 1.1 | 0.5 | 2.2 | 2.7E-02 | 3.0E-02 | BT20/MB468 | NUC |
| >am_trnaMT_ValTAC_MT_+_1602_1670@44.67.24\|ATTTCAACTTAACTTGACCGCTCT | >am_trnaMT_ValTAC_MT_+_1602_1670 | 44 | 67 | 24 | ATTTCAACTTAACTTGACCGCTCT | 417.1 | 2.3 | 0.9 | 2.6 | 1.0E-02 | 3.0E-02 | BT20/MB231 | CYTO |
| >MI0000764\|hsa-mir-363&WithFlank&X\|-\|134169372\|134169458@56.77.22&[MIMAT0000707&hsa-miR-363-3p&offsets\|0\|0;m-178&X\|-\|134169381\|134169403&offsets\|0\|-1]\|AATTGCACGGTATCCATCTGTA | >MI0000764\|hsa-mir-363&WithFlank&X\|-\|134169372\|134169458 | 56 | 77 | 22 | AATTGCACGGTATCCATCTGTA | 510.6 | 1.5 | 0.7 | 2.2 | 2.7E-02 | 3.0E-02 | BT20/MB468 | NUC |
| >MI0005559\|hsa-mir-744&WithFlank&17\|+\|12081893\|12082002@17.38.22&[MIMAT0004945&hsa-miR-744-5p&offsets\|0\|0;m-172&17\|+\|12081909\|12081929&offsets\|0\|+1]\|TGCGGGGCTAGGGCTAACAGCA | >MI0005559\|hsa-mir-744&WithFlank&17\|+\|12081893\|12082002 | 17 | 38 | 22 | TGCGGGGCTAGGGCTAACAGCA | 1169.9 | 1.8 | 0.7 | 2.6 | 9.4E-03 | 3.0E-02 | BT20/MB231 | MITO |
| >trna111_HisGTG_1_-_147774845_147774916@-1T.24.25,trna118_HisGTG_1_-_145396881_145396952@-1T.24.25,trna16_HisGTG_1_+_146544773_146544844@-1T.24.25,trna1_HisGTG_15_+_45493349_45493420@-1T.24.25,trna21_HisGTG_1_+_147753471_147753542@-1T.24.25,trna33_HisGTG_6_+_27125906_27125977@-1T.24.25,trna7_HisGTG_9_-_14433938_14434009@-1T.24.25,trna8_HisGTG_15_-_45492611_45492682@-1T.24.25,trna9_HisGTG_15_-_45490804_45490875@-1T.24.25\|TGCCGTGATCGTATAGTGGTTAGTA | >trna111_HisGTG_1_-_147774845_147774916 | -1T | 24 | 25 | TGCCGTGATCGTATAGTGGTTAGTA | 228.9 | -2.9 | 1.1 | -2.6 | 9.3E-03 | 3.0E-02 | BT20/MB231 | MITO |
| >hg38_21_+_8433172_8446622_RNA45SN1_45S_50ntFlanks@5444.5465.22\|CTCGGATCGGCCCCGCCGGGGT | >hg38_21_+_8433172_8446622_RNA45SN1_45S_50ntFlanks | 5444 | 5465 | 22 | CTCGGATCGGCCCCGCCGGGGT | 153.6 | -4.1 | 1.7 | -2.4 | 1.8E-02 | 3.0E-02 | BT20/MB468 | MP |
| >hg38_21_+_8433172_8446622_RNA45SN1_45S_50ntFlanks@7975.7993.19\|CGCGACCTCAGATCAGACG | >hg38_21_+_8433172_8446622_RNA45SN1_45S_50ntFlanks | 7975 | 7993 | 19 | CGCGACCTCAGATCAGACG | 100554.1 | 2.0 | 0.8 | 2.6 | 9.4E-03 | 3.0E-02 | BT20/MB231 | MITO |
| >hg38_21_+_8433172_8446622_RNA45SN1_45S_50ntFlanks@10128.10147.20\|CGGCGGCGGCGGCGGCGGCG | >hg38_21_+_8433172_8446622_RNA45SN1_45S_50ntFlanks | 10128 | 10147 | 20 | CGGCGGCGGCGGCGGCGGCG | 129.0 | -3.4 | 1.5 | -2.2 | 2.6E-02 | 3.0E-02 | BT20/MB468 | CYTO |
| >hg38_21_+_8433172_8446622_RNA45SN1_45S_50ntFlanks@8845.8864.20\|CGGCGGCGGCGGCGGCGGCG | >hg38_21_+_8433172_8446622_RNA45SN1_45S_50ntFlanks | 8845 | 8864 | 20 | CGGCGGCGGCGGCGGCGGCG | 129.0 | -3.4 | 1.5 | -2.2 | 2.6E-02 | 3.0E-02 | BT20/MB468 | CYTO |
| >MI0000447\|hsa-mir-128-1&WithFlank&2\|+\|135665391\|135665484@56.76.21(+1A);MI0000727\|hsa-mir-128-2&WithFlank&3\|+\|35744470\|35744565@58.78.21(+1A)&[MIMAT0000424&hsa-miR-128-3p&offsets\|0\|0(+1A);m-108&2\|+\|135665446\|135665466&offsets\|0\|0(+1A)];[MIMAT0000424_1&hsa-miR-128-3p&offsets\|0\|0(+1A);m-117&3\|+\|35744527\|35744547&offsets\|0\|0(+1A)]\|TCACAGTGAACCGGTCTCTTTA | >MI0000447\|hsa-mir-128-1&WithFlank&2\|+\|135665391\|135665484 | 56 | 76 | 21(+1A) | TCACAGTGAACCGGTCTCTTTA | 555.4 | 2.0 | 0.8 | 2.4 | 1.8E-02 | 3.0E-02 | BT20/MB468 | MP |
| >MI0000239\|hsa-mir-197&WithFlank&1\|+\|109598887\|109598973@54.74.21(+1A)&[MIMAT0000227&hsa-miR-197-3p&offsets\|0\|-1(+1A);m-99&1\|+\|109598940\|109598961&offsets\|0\|-1(+1A)]\|TTCACCACCTTCTCCACCCAGA | >MI0000239\|hsa-mir-197&WithFlank&1\|+\|109598887\|109598973 | 54 | 74 | 21(+1A) | TTCACCACCTTCTCCACCCAGA | 530.7 | 2.1 | 0.8 | 2.6 | 9.4E-03 | 3.0E-02 | BT20/MB231 | MITO |
| >hg38_21_+_8433172_8446622_RNA45SN1_45S_50ntFlanks@10348.10373.26\|AAACGAGAACTTTGAAGGCCGAAGTG | >hg38_21_+_8433172_8446622_RNA45SN1_45S_50ntFlanks | 10348 | 10373 | 26 | AAACGAGAACTTTGAAGGCCGAAGTG | 307.2 | 1.3 | 0.6 | 2.2 | 2.6E-02 | 3.0E-02 | BT20/MB468 | CYTO |
| >MI0000440\|hsa-mir-27b&WithFlank&9\|+\|95085439\|95085547@67.87.21(+1U)&[MIMAT0000419&hsa-miR-27b-3p&offsets\|0\|0(+1U);m-39&9\|+\|95085505\|95085525&offsets\|0\|0(+1U)]\|TTCACAGTGGCTAAGTTCTGCT | >MI0000440\|hsa-mir-27b&WithFlank&9\|+\|95085439\|95085547 | 67 | 87 | 21(+1U) | TTCACAGTGGCTAAGTTCTGCT | 2540.7 | -2.2 | 0.8 | -2.6 | 9.4E-03 | 3.0E-02 | BT20/MB231 | MITO |
| >hg38_21_+_8433172_8446622_RNA45SN1_45S_50ntFlanks@3840.3862.23\|CTCCTACTTGGATAACTGTGGTA | >hg38_21_+_8433172_8446622_RNA45SN1_45S_50ntFlanks | 3840 | 3862 | 23 | CTCCTACTTGGATAACTGTGGTA | 202.5 | 2.8 | 1.1 | 2.6 | 9.4E-03 | 3.0E-02 | BT20/MB231 | MITO |
| >hg38_MT_+_598_1651_RNR1_12S_50ntFlanks@439.480.42\|AACATATCTGAACACACAATAGCTAAGACCCAAACTGGGATT | >hg38_MT_+_598_1651_RNR1_12S_50ntFlanks | 439 | 480 | 42 | AACATATCTGAACACACAATAGCTAAGACCCAAACTGGGATT | 190.3 | -2.3 | 0.9 | -2.6 | 9.4E-03 | 3.0E-02 | BT20/MB231 | MITO |
| >hg38_21_+_8433172_8446622_RNA45SN1_45S_50ntFlanks@6646.6683.38\|TCGTACGACTCTTAGCGGTGGATCACTCGGCTCGTGCG | >hg38_21_+_8433172_8446622_RNA45SN1_45S_50ntFlanks | 6646 | 6683 | 38 | TCGTACGACTCTTAGCGGTGGATCACTCGGCTCGTGCG | 1388.1 | 2.1 | 0.9 | 2.4 | 1.8E-02 | 3.0E-02 | BT20/MB468 | MP |
| >hg38_21_+_8433172_8446622_RNA45SN1_45S_50ntFlanks@7977.8003.27\|CGACCTCAGATCAGACGTGGCGACCCG | >hg38_21_+_8433172_8446622_RNA45SN1_45S_50ntFlanks | 7977 | 8003 | 27 | CGACCTCAGATCAGACGTGGCGACCCG | 249.1 | -1.7 | 0.8 | -2.2 | 2.8E-02 | 3.1E-02 | MB231/MB468 | NUC |
| >hg38_21_+_8433172_8446622_RNA45SN1_45S_50ntFlanks@9549.9573.25\|AGGTCCGTAGCGGTCCTGACGTGCA | >hg38_21_+_8433172_8446622_RNA45SN1_45S_50ntFlanks | 9549 | 9573 | 25 | AGGTCCGTAGCGGTCCTGACGTGCA | 317.8 | 1.5 | 0.7 | 2.2 | 2.8E-02 | 3.1E-02 | MB231/MB468 | NUC |
| >MI0000065\|hsa-let-7d&WithFlank&9\|+\|94178828\|94178926@14.33.20&[MIMAT0000065&hsa-let-7d-5p&offsets\|0\|-2;m-90&9\|+\|94178841\|94178862&offsets\|0\|-2]\|AGAGGTAGTAGGTTGCATAG | >MI0000065\|hsa-let-7d&WithFlank&9\|+\|94178828\|94178926 | 14 | 33 | 20 | AGAGGTAGTAGGTTGCATAG | 43.6 | 3.9 | 1.7 | 2.2 | 2.5E-02 | 3.1E-02 | MB231/MB468 | MP |
| >hg38_21_+_8433172_8446622_RNA45SN1_45S_50ntFlanks@6661.6706.46\|CGGTGGATCACTCGGCTCGTGCGTCGATGAAGAACGCAGCTAGCTG | >hg38_21_+_8433172_8446622_RNA45SN1_45S_50ntFlanks | 6661 | 6706 | 46 | CGGTGGATCACTCGGCTCGTGCGTCGATGAAGAACGCAGCTAGCTG | 737.8 | 2.1 | 0.9 | 2.2 | 2.7E-02 | 3.1E-02 | BT20/MB468 | NUC |
| >hg38_21_+_8433172_8446622_RNA45SN1_45S_50ntFlanks@5472.5516.45\|ACGGCCCTGGCGGAGCGCTGAGAAGACGGTCGAACTTGACTATCT | >hg38_21_+_8433172_8446622_RNA45SN1_45S_50ntFlanks | 5472 | 5516 | 45 | ACGGCCCTGGCGGAGCGCTGAGAAGACGGTCGAACTTGACTATCT | 17329.0 | 3.0 | 1.2 | 2.6 | 9.4E-03 | 3.1E-02 | BT20/MB231 | MITO |
| >MI0005559\|hsa-mir-744&WithFlank&17\|+\|12081893\|12082002@17.38.22&[MIMAT0004945&hsa-miR-744-5p&offsets\|0\|0;m-172&17\|+\|12081909\|12081929&offsets\|0\|+1]\|TGCGGGGCTAGGGCTAACAGCA | >MI0005559\|hsa-mir-744&WithFlank&17\|+\|12081893\|12082002 | 17 | 38 | 22 | TGCGGGGCTAGGGCTAACAGCA | 716.3 | 1.7 | 0.7 | 2.6 | 8.3E-03 | 3.1E-02 | BT20/MB231 | NUC |

<table>
<thead><tr><th></th><th></th><th></th><th></th><th></th><th></th><th></th><th></th><th></th><th></th><th></th><th></th><th></th><th></th></tr></thead>
<tbody>
<tr><td>>trna111_HisGTG_1_-_147774845_147774916@-1T.31.32,trna118_HisGTG_1_-_145396881_145396952@-1T.31.32,trna16_HisGTG_1_+_146544773_146544844@-1T.31.32,trna1_HisGTG_15_+_45493349_45493420@-1T.31.32,trna21_HisGTG_1_+_147753471_147753542@-1T.31.32,trna33_HisGTG_6_+_27125906_27125977@-1T.31.32,trna7_HisGTG_9_-_14433938_14434009@-1T.31.32,trna8_HisGTG_15_-_45492611_45492682@-1T.31.32,trna9_HisGTG_15_-_45490804_45490875@-1T.31.32|TGCCGTGATCGTATAGTGGTTAGTACTCTGCG</td><td>>trna111_HisGTG_1_-_147774845_147774916</td><td>-1T</td><td>31</td><td>32</td><td>TGCCGTGATCGTATAGTGGTTAGTACTCTGCG</td><td>336.6</td><td>-2.3</td><td>0.9</td><td>-2.6</td><td>8.3E-03</td><td>3.1E-02</td><td>BT20/MB231</td><td>NUC</td></tr>
<tr><td>>hg38_21_+_8433172_8446622_RNA45SN1_45S_50ntFlanks@10792.10822.31|CTCTGGCATGTTGGAACAATGTAGGTAAGGG</td><td>>hg38_21_+_8433172_8446622_RNA45SN1_45S_50ntFlanks</td><td>10792</td><td>10822</td><td>31</td><td>CTCTGGCATGTTGGAACAATGTAGGTAAGGG</td><td>152.7</td><td>4.4</td><td>1.6</td><td>2.7</td><td>7.7E-03</td><td>3.1E-02</td><td>BT20/MB231</td><td>MP</td></tr>
<tr><td>>hg38_21_+_8433172_8446622_RNA45SN1_45S_50ntFlanks@11891.11909.19|TTGACTCTAGTCTGGCACG</td><td>>hg38_21_+_8433172_8446622_RNA45SN1_45S_50ntFlanks</td><td>11891</td><td>11909</td><td>19</td><td>TTGACTCTAGTCTGGCACG</td><td>194.6</td><td>-2.2</td><td>0.9</td><td>-2.4</td><td>1.8E-02</td><td>3.1E-02</td><td>BT20/MB468</td><td>MP</td></tr>
<tr><td>>am_trna128_GlyGCC_6_-_27870686_27870756@1.32.32,trna133_GlyCCC_1_-_16872434_16872504@1.32.32,trna18_GlyGCC_16_+_70822597_70822667@1.32.32,trna19_GlyGCC_16_+_70823410_70823480@1.32.32,trna19_GlyGCC_2_-_157257659_157257729@1.32.32,trna24_GlyGCC_16_-_70812942_70813012@1.32.32,trna25_GlyGCC_16_-_70812114_70812184@1.32.32,trna4_GlyCCC_1_+_17188416_17188486@1.32.32,trna5_GlyGCC_17_+_8029064_8029134@1.32.32,trna68_GlyGCC_1_-_161493637_161493707@1.32.32|GCATTGGTGGTTCAGTGGTAGAATTCTCGCCT</td><td>>am_trna128_GlyGCC_6_-_27870686_27870756</td><td>1</td><td>32</td><td>32</td><td>GCATTGGTGGTTCAGTGGTAGAATTCTCGCCT</td><td>32824.3</td><td>-2.4</td><td>0.9</td><td>-2.6</td><td>1.0E-02</td><td>3.1E-02</td><td>BT20/MB231</td><td>CYTO</td></tr>
<tr><td>>MI0000734|hsa-mir-106b&WithFlank&7|-|100093987|100094080@58.78.21(+1A)&[MIMAT0004672&hsa-miR-106b-3p&offsets|0|-1(+1A);m-87&7|-|100094003|100094023&offsets|0|0(+1A)]|CCGCACTGTGGGTACTTGCTGA</td><td>>MI0000734|hsa-mir-106b&WithFlank&7|-|100093987|100094080</td><td>58</td><td>78</td><td>21(+1A)</td><td>CCGCACTGTGGGTACTTGCTGA</td><td>179.2</td><td>3.9</td><td>1.7</td><td>2.2</td><td>2.5E-02</td><td>3.1E-02</td><td>MB231/MB468</td><td>MP</td></tr>
<tr><td>>hg38_21_+_8433172_8446622_RNA45SN1_45S_50ntFlanks@10546.10567.22|CAGTGCGGTAACGCGACCGATC</td><td>>hg38_21_+_8433172_8446622_RNA45SN1_45S_50ntFlanks</td><td>10546</td><td>10567</td><td>22</td><td>CAGTGCGGTAACGCGACCGATC</td><td>757.1</td><td>-2.7</td><td>1.1</td><td>-2.4</td><td>1.8E-02</td><td>3.1E-02</td><td>BT20/MB468</td><td>MP</td></tr>
<tr><td>>MI0000462|hsa-mir-152&WithFlank&17|-|48037155|48037253@60.80.21(+1U)&[MIMAT0000438&hsa-miR-152-3p&offsets|0|0(+1U);m-97&17|-|48037174|48037194&offsets|0|0(+1U)]|TCAGTGCATGACAGAACTTGGT</td><td>>MI0000462|hsa-mir-152&WithFlank&17|-|48037155|48037253</td><td>60</td><td>80</td><td>21(+1U)</td><td>TCAGTGCATGACAGAACTTGGT</td><td>1178.1</td><td>-2.8</td><td>1.1</td><td>-2.7</td><td>7.7E-03</td><td>3.1E-02</td><td>BT20/MB231</td><td>MP</td></tr>
<tr><td>>hg38_21_+_8433172_8446622_RNA45SN1_45S_50ntFlanks@10238.10267.30|CGCTACGCCGCGACGAGTAGGAGGGCCGCT</td><td>>hg38_21_+_8433172_8446622_RNA45SN1_45S_50ntFlanks</td><td>10238</td><td>10267</td><td>30</td><td>CGCTACGCCGCGACGAGTAGGAGGGCCGCT</td><td>189.3</td><td>4.4</td><td>1.7</td><td>2.7</td><td>7.7E-03</td><td>3.1E-02</td><td>BT20/MB231</td><td>MP</td></tr>
<tr><td>>hg38_21_+_8433172_8446622_RNA45SN1_45S_50ntFlanks@8354.8390.37|TTGAAAAGAACTTTGAAGAGAGAGTTCAAGAGGGCGT</td><td>>hg38_21_+_8433172_8446622_RNA45SN1_45S_50ntFlanks</td><td>8354</td><td>8390</td><td>37</td><td>TTGAAAAGAACTTTGAAGAGAGAGTTCAAGAGGGCGT</td><td>260.2</td><td>3.4</td><td>1.3</td><td>2.7</td><td>7.7E-03</td><td>3.1E-02</td><td>BT20/MB231</td><td>MP</td></tr>
<tr><td>>hg38_21_+_8433172_8446622_RNA45SN1_45S_50ntFlanks@10781.10813.33|AGGTGAACAGCCTCTGGCATGTTGGAACAATGT</td><td>>hg38_21_+_8433172_8446622_RNA45SN1_45S_50ntFlanks</td><td>10781</td><td>10813</td><td>33</td><td>AGGTGAACAGCCTCTGGCATGTTGGAACAATGT</td><td>97.7</td><td>-4.1</td><td>1.8</td><td>-2.4</td><td>1.8E-02</td><td>3.1E-02</td><td>BT20/MB468</td><td>MP</td></tr>
<tr><td>>hg38_21_+_8433172_8446622_RNA45SN1_45S_50ntFlanks@12003.12030.28|GCGGGCCGCCGGTGAAATACCACTACTC</td><td>>hg38_21_+_8433172_8446622_RNA45SN1_45S_50ntFlanks</td><td>12003</td><td>12030</td><td>28</td><td>GCGGGCCGCCGGTGAAATACCACTACTC</td><td>247.9</td><td>-1.2</td><td>0.5</td><td>-2.2</td><td>2.7E-02</td><td>3.1E-02</td><td>BT20/MB468</td><td>CYTO</td></tr>
<tr><td>>hg38_21_+_8433172_8446622_RNA45SN1_45S_50ntFlanks@9495.9515.21|CGAAAGATGGTGAACTATGCC</td><td>>hg38_21_+_8433172_8446622_RNA45SN1_45S_50ntFlanks</td><td>9495</td><td>9515</td><td>21</td><td>CGAAAGATGGTGAACTATGCC</td><td>126.9</td><td>3.3</td><td>1.4</td><td>2.4</td><td>1.8E-02</td><td>3.1E-02</td><td>BT20/MB468</td><td>MP</td></tr>
<tr><td>>trnaMT_GluTTC_MT_-_14674_14742@33.71.39,trnalookalike8_GluTTC_5_-_93905172_93905240@33.71.39|CATATCATTGGTCGTGGTTGTAGTCCGTGCGAGAATACC</td><td>>trnaMT_GluTTC_MT_-_14674_14742</td><td>33</td><td>71</td><td>39</td><td>CATATCATTGGTCGTGGTTGTAGTCCGTGCGAGAATACC</td><td>91.8</td><td>2.9</td><td>1.3</td><td>2.2</td><td>2.5E-02</td><td>3.1E-02</td><td>MB231/MB468</td><td>MP</td></tr>
<tr><td>>hg38_21_+_8433172_8446622_RNA45SN1_45S_50ntFlanks@9714.9738.25|CGAATGATTAGAGGTCTTGGGGCCG</td><td>>hg38_21_+_8433172_8446622_RNA45SN1_45S_50ntFlanks</td><td>9714</td><td>9738</td><td>25</td><td>CGAATGATTAGAGGTCTTGGGGCCG</td><td>137.0</td><td>-3.2</td><td>1.3</td><td>-2.4</td><td>1.8E-02</td><td>3.1E-02</td><td>BT20/MB468</td><td>MP</td></tr>
<tr><td>>hg38_21_+_8433172_8446622_RNA45SN1_45S_50ntFlanks@9790.9820.31|CTGGCGTGGAGCCGGGCGTGGAATGCGAGTG</td><td>>hg38_21_+_8433172_8446622_RNA45SN1_45S_50ntFlanks</td><td>9790</td><td>9820</td><td>31</td><td>CTGGCGTGGAGCCGGGCGTGGAATGCGAGTG</td><td>315.2</td><td>-1.7</td><td>0.7</td><td>-2.3</td><td>2.3E-02</td><td>3.1E-02</td><td>BT20/MB468</td><td>MITO</td></tr>
<tr><td>>MI0000089|hsa-mir-31&WithFlank&9|-|21512109|21512191@14.35.22&[MIMAT0000089&hsa-miR-31-5p&offsets|0|+1;m-360&9|-|21512157|21512178&offsets|0|0]|AGGCAAGATGCTGGCATAGCTG</td><td>>MI0000089|hsa-mir-31&WithFlank&9|-|21512109|21512191</td><td>14</td><td>35</td><td>22</td><td>AGGCAAGATGCTGGCATAGCTG</td><td>777.7</td><td>3.4</td><td>1.4</td><td>2.4</td><td>1.8E-02</td><td>3.1E-02</td><td>BT20/MB468</td><td>MP</td></tr>
<tr><td>>hg38_21_+_8433172_8446622_RNA45SN1_45S_50ntFlanks@10866.10902.37|CTAAGGGCTGGGTCGGTCGGGCTGGGGCGCGAAGCGG</td><td>>hg38_21_+_8433172_8446622_RNA45SN1_45S_50ntFlanks</td><td>10866</td><td>10902</td><td>37</td><td>CTAAGGGCTGGGTCGGTCGGGCTGGGGCGCGAAGCGG</td><td>868.2</td><td>-4.4</td><td>1.8</td><td>-2.4</td><td>1.8E-02</td><td>3.1E-02</td><td>BT20/MB468</td><td>MP</td></tr>
<tr><td>>hg38_21_+_8433172_8446622_RNA45SN1_45S_50ntFlanks@11718.11745.28|AACGGCGGGAGTAACTATGACTCTCTTA</td><td>>hg38_21_+_8433172_8446622_RNA45SN1_45S_50ntFlanks</td><td>11718</td><td>11745</td><td>28</td><td>AACGGCGGGAGTAACTATGACTCTCTTA</td><td>90.8</td><td>-2.4</td><td>1.0</td><td>-2.4</td><td>1.8E-02</td><td>3.1E-02</td><td>BT20/MB468</td><td>MP</td></tr>
<tr><td>>trna116_GluCTC_1_-_145399233_145399304@2.31.30,trna59_GluCTC_1_+_249168447_249168518@2.31.30,trna71_GluCTC_1_-_161439189_161439260@2.31.30,trna74_GluCTC_1_-_161431809_161431880@2.31.30,trna77_GluCTC_1_-_161424398_161424469@2.31.30,trna77_GluCTC_6_+_28949976_28950047@2.31.30,trna80_GluCTC_1_-_161417018_161417089@2.31.30,trna87_GluCTC_6_-_126101393_126101464@2.31.30|CCCTGGTGGTCTAGTGGTTAGGATTCGGCG</td><td>>trna116_GluCTC_1_-_145399233_145399304</td><td>2</td><td>31</td><td>30</td><td>CCCTGGTGGTCTAGTGGTTAGGATTCGGCG</td><td>753.1</td><td>-4.6</td><td>2.1</td><td>-2.2</td><td>2.7E-02</td><td>3.1E-02</td><td>BT20/MB468</td><td>NUC</td></tr>
<tr><td>>hg38_21_+_8433172_8446622_RNA45SN1_45S_50ntFlanks@9549.9574.26|AGGTCCGTAGCGGTCCTGACGTGCAA</td><td>>hg38_21_+_8433172_8446622_RNA45SN1_45S_50ntFlanks</td><td>9549</td><td>9574</td><td>26</td><td>AGGTCCGTAGCGGTCCTGACGTGCAA</td><td>108.3</td><td>5.1</td><td>1.9</td><td>2.7</td><td>7.7E-03</td><td>3.1E-02</td><td>BT20/MB231</td><td>MP</td></tr>
<tr><td>>hg38_21_+_8433172_8446622_RNA45SN1_45S_50ntFlanks@11666.11709.44|TCTGCCCAGTGCTCTGAATGTCAAAGTGAAGAAATTCAATGAAG</td><td>>hg38_21_+_8433172_8446622_RNA45SN1_45S_50ntFlanks</td><td>11666</td><td>11709</td><td>44</td><td>TCTGCCCAGTGCTCTGAATGTCAAAGTGAAGAAATTCAATGAAG</td><td>305.0</td><td>-4.6</td><td>1.8</td><td>-2.6</td><td>1.0E-02</td><td>3.1E-02</td><td>BT20/MB231</td><td>CYTO</td></tr>
<tr><td>>hg38_1_-_228634819_228635039_RNA5S12_5S_50ntFlanks@148.168.21|GGAATACCGGGTGCTGTAGGC</td><td>>hg38_1_-_228634819_228635039_RNA5S12_5S_50ntFlanks</td><td>148</td><td>168</td><td>21</td><td>GGAATACCGGGTGCTGTAGGC</td><td>1069.9</td><td>3.4</td><td>1.4</td><td>2.4</td><td>1.8E-02</td><td>3.1E-02</td><td>BT20/MB468</td><td>MP</td></tr>
<tr><td>>hg38_21_+_8433172_8446622_RNA45SN1_45S_50ntFlanks@11208.11228.21|GCGGCGACTCTGGACGCGAGC</td><td>>hg38_21_+_8433172_8446622_RNA45SN1_45S_50ntFlanks</td><td>11208</td><td>11228</td><td>21</td><td>GCGGCGACTCTGGACGCGAGC</td><td>786.6</td><td>-1.4</td><td>0.6</td><td>-2.2</td><td>2.7E-02</td><td>3.1E-02</td><td>BT20/MB468</td><td>CYTO</td></tr>
<tr><td>>hg38_21_+_8433172_8446622_RNA45SN1_45S_50ntFlanks@7991.8009.19|ACGTGGCGACCCGCTGAAT</td><td>>hg38_21_+_8433172_8446622_RNA45SN1_45S_50ntFlanks</td><td>7991</td><td>8009</td><td>19</td><td>ACGTGGCGACCCGCTGAAT</td><td>592.7</td><td>1.1</td><td>0.5</td><td>2.2</td><td>2.7E-02</td><td>3.1E-02</td><td>BT20/MB468</td><td>CYTO</td></tr>
<tr><td>>trnaMT_SerGCT_MT_+_12207_12265@14.61.48|AGAACTGCTAACTCATGCCCCCATGTCTAACAACATGGCTTTCTCACC</td><td>>trnaMT_SerGCT_MT_+_12207_12265</td><td>14</td><td>61</td><td>48</td><td>AGAACTGCTAACTCATGCCCCCATGTCTAACAACATGGCTTTCTCACC</td><td>193.0</td><td>4.5</td><td>2.0</td><td>2.2</td><td>2.8E-02</td><td>3.1E-02</td><td>MB231/MB468</td><td>MITO</td></tr>
<tr><td>>hg38_21_+_8433172_8446622_RNA45SN1_45S_50ntFlanks@10866.10892.27|CTAAGGGCTGGGTCGGTCGGGCTGGGG</td><td>>hg38_21_+_8433172_8446622_RNA45SN1_45S_50ntFlanks</td><td>10866</td><td>10892</td><td>27</td><td>CTAAGGGCTGGGTCGGTCGGGCTGGGG</td><td>493.6</td><td>-3.8</td><td>1.7</td><td>-2.2</td><td>2.7E-02</td><td>3.1E-02</td><td>BT20/MB468</td><td>CYTO</td></tr>
<tr><td>>MI0000465|hsa-mir-191&WithFlank&3|-|49020612|49020715@22.44.23(+1A)&[MIMAT0000440&hsa-miR-191-5p&offsets|0|0(+1A);m-48&3|-|49020672|49020694&offsets|0|0(+1A)]|CAACGGAATCCCAAAAGCAGCTGA</td><td>>MI0000465|hsa-mir-191&WithFlank&3|-|49020612|49020715</td><td>22</td><td>44</td><td>23(+1A)</td><td>CAACGGAATCCCAAAAGCAGCTGA</td><td>379.5</td><td>2.7</td><td>1.1</td><td>2.4</td><td>1.8E-02</td><td>3.1E-02</td><td>BT20/MB468</td><td>MP</td></tr>
<tr><td>>hg38_21_+_8433172_8446622_RNA45SN1_45S_50ntFlanks@12526.12554.29|TTTTACCCTACTGATGATGTGTTGTTGCC</td><td>>hg38_21_+_8433172_8446622_RNA45SN1_45S_50ntFlanks</td><td>12526</td><td>12554</td><td>29</td><td>TTTTACCCTACTGATGATGTGTTGTTGCC</td><td>59.9</td><td>-4.0</td><td>1.7</td><td>-2.4</td><td>1.8E-02</td><td>3.1E-02</td><td>BT20/MB468</td><td>MP</td></tr>
<tr><td>>hg38_21_+_8433172_8446622_RNA45SN1_45S_50ntFlanks@5487.5509.23|CGCTGAGAAGACGGTCGAACTTG</td><td>>hg38_21_+_8433172_8446622_RNA45SN1_45S_50ntFlanks</td><td>5487</td><td>5509</td><td>23</td><td>CGCTGAGAAGACGGTCGAACTTG</td><td>235.2</td><td>-1.9</td><td>0.8</td><td>-2.2</td><td>2.7E-02</td><td>3.1E-02</td><td>MB231/MB468</td><td>CYTO</td></tr>
<tr><td>>am_MI0003639|hsa-mir-625&WithFlank&14|+|65471096|65471192@58.80.23&[MIMAT0004808&hsa-miR-625-3p&offsets|0|+1;m-137&14|+|65471153|65471174&offsets|0|+1]|GACTATAGAACTTTCCCCCTCAT</td><td>>am_MI0003639|hsa-mir-625&WithFlank&14|+|65471096|65471192</td><td>58</td><td>80</td><td>23</td><td>GACTATAGAACTTTCCCCCTCAT</td><td>52.8</td><td>4.4</td><td>1.9</td><td>2.4</td><td>1.8E-02</td><td>3.1E-02</td><td>BT20/MB468</td><td>MP</td></tr>
<tr><td>>trnaMT_GluTTC_MT_-_14674_14742@26.71.46,trnalookalike8_GluTTC_5_-_93905172_93905240@26.71.46|GGTTTTTCATATCATTGGTCGTGGTTGTAGTCCGTGCGAGAATACC</td><td>>trnaMT_GluTTC_MT_-_14674_14742</td><td>26</td><td>71</td><td>46</td><td>GGTTTTTCATATCATTGGTCGTGGTTGTAGTCCGTGCGAGAATACC</td><td>142.5</td><td>5.4</td><td>2.3</td><td>2.4</td><td>1.8E-02</td><td>3.1E-02</td><td>BT20/MB468</td><td>MP</td></tr>
<tr><td>>hg38_21_+_8433172_8446622_RNA45SN1_45S_50ntFlanks@11571.11594.24|GCCGACTTAGAACTGGTGCGGACC</td><td>>hg38_21_+_8433172_8446622_RNA45SN1_45S_50ntFlanks</td><td>11571</td><td>11594</td><td>24</td><td>GCCGACTTAGAACTGGTGCGGACC</td><td>257.5</td><td>1.8</td><td>0.7</td><td>2.4</td><td>1.8E-02</td><td>3.1E-02</td><td>BT20/MB468</td><td>MP</td></tr>
<tr><td>>hg38_MT_+_598_1651_RNR1_12S_50ntFlanks@427.455.29|GAAAGTGGCTTTAACATATCTGAACACAC</td><td>>hg38_MT_+_598_1651_RNR1_12S_50ntFlanks</td><td>427</td><td>455</td><td>29</td><td>GAAAGTGGCTTTAACATATCTGAACACAC</td><td>148.1</td><td>3.4</td><td>1.4</td><td>2.4</td><td>1.8E-02</td><td>3.1E-02</td><td>BT20/MB468</td><td>MP</td></tr>
<tr><td>>hg38_21_+_8433172_8446622_RNA45SN1_45S_50ntFlanks@5238.5266.29|CTGACTGGCTCAGCGTGTGCCTACCCTAC</td><td>>hg38_21_+_8433172_8446622_RNA45SN1_45S_50ntFlanks</td><td>5238</td><td>5266</td><td>29</td><td>CTGACTGGCTCAGCGTGTGCCTACCCTAC</td><td>180.0</td><td>-2.1</td><td>0.8</td><td>-2.6</td><td>8.3E-03</td><td>3.1E-02</td><td>BT20/MB231</td><td>NUC</td></tr>
<tr><td>>hg38_21_+_8433172_8446622_RNA45SN1_45S_50ntFlanks@11572.11612.41|CCGACTTAGAACTGGTGCGGACCAGGGGAATCCGACTGTTT</td><td>>hg38_21_+_8433172_8446622_RNA45SN1_45S_50ntFlanks</td><td>11572</td><td>11612</td><td>41</td><td>CCGACTTAGAACTGGTGCGGACCAGGGGAATCCGACTGTTT</td><td>1117.2</td><td>-2.5</td><td>1.1</td><td>-2.2</td><td>2.7E-02</td><td>3.1E-02</td><td>BT20/MB468</td><td>CYTO</td></tr>
<tr><td>>hg38_21_+_8433172_8446622_RNA45SN1_45S_50ntFlanks@6645.6680.36|CTCGTACGACTCTTAGCGGTGGATCACTCGGCTCGT</td><td>>hg38_21_+_8433172_8446622_RNA45SN1_45S_50ntFlanks</td><td>6645</td><td>6680</td><td>36</td><td>CTCGTACGACTCTTAGCGGTGGATCACTCGGCTCGT</td><td>1248.4</td><td>3.3</td><td>1.2</td><td>2.7</td><td>7.7E-03</td><td>3.1E-02</td><td>BT20/MB231</td><td>MP</td></tr>
<tr><td>>hg38_21_+_8433172_8446622_RNA45SN1_45S_50ntFlanks@10787.10818.32|ACAGCCTCTGGCATGTTGGAACAATGTAGGTA</td><td>>hg38_21_+_8433172_8446622_RNA45SN1_45S_50ntFlanks</td><td>10787</td><td>10818</td><td>32</td><td>ACAGCCTCTGGCATGTTGGAACAATGTAGGTA</td><td>72.3</td><td>5.2</td><td>2.0</td><td>2.7</td><td>7.7E-03</td><td>3.1E-02</td><td>BT20/MB231</td><td>MP</td></tr>
<tr><td>>MI0003820|hsa-mir-454&WithFlank&17|-|59137752|59137878@70.91.22(+1C)&[MIMAT0003885&hsa-miR-454-3p&offsets|0|-1(+1C);m-272&17|-|59137787|59137809&offsets|0|-1(+1C)]|TAGTGCAATATTGCTTATAGGGC</td><td>>MI0003820|hsa-mir-454&WithFlank&17|-|59137752|59137878</td><td>70</td><td>91</td><td>22(+1C)</td><td>TAGTGCAATATTGCTTATAGGGC</td><td>127.2</td><td>-3.9</td><td>1.5</td><td>-2.7</td><td>7.7E-03</td><td>3.1E-02</td><td>BT20/MB231</td><td>MP</td></tr>
<tr><td>>MI0000114|hsa-mir-107&WithFlank&10|-|89592741|89592833@56.77.22&[MIMAT0000104&hsa-miR-107&offsets|0|-1;m-78&10|-|89592758|89592778&offsets|0|+1]|AGCAGCATTGTACAGGGCTATC</td><td>>MI0000114|hsa-mir-107&WithFlank&10|-|89592741|89592833</td><td>56</td><td>77</td><td>22</td><td>AGCAGCATTGTACAGGGCTATC</td><td>1346.0</td><td>2.2</td><td>1.0</td><td>2.2</td><td>2.5E-02</td><td>3.1E-02</td><td>MB231/MB468</td><td>MP</td></tr>
<tr><td>>trnaMT_PheGAA_MT_+_577_647@32.54.23|ACTGAAAATGTTTAGACGGGCTC</td><td>>trnaMT_PheGAA_MT_+_577_647</td><td>32</td><td>54</td><td>23</td><td>ACTGAAAATGTTTAGACGGGCTC</td><td>65.0</td><td>6.1</td><td>2.6</td><td>2.4</td><td>1.8E-02</td><td>3.1E-02</td><td>BT20/MB468</td><td>MP</td></tr>
<tr><td>>hg38_21_+_8433172_8446622_RNA45SN1_45S_50ntFlanks@11582.11603.22|ACTGGTGCGGACCAGGGGAATC</td><td>>hg38_21_+_8433172_8446622_RNA45SN1_45S_50ntFlanks</td><td>11582</td><td>11603</td><td>22</td><td>ACTGGTGCGGACCAGGGGAATC</td><td>2488.4</td><td>-2.6</td><td>1.1</td><td>-2.4</td><td>1.8E-02</td><td>3.1E-02</td><td>BT20/MB468</td><td>MP</td></tr>
<tr><td>>hg38_21_+_8433172_8446622_RNA45SN1_45S_50ntFlanks@10680.10709.30|GTTCCGGCGGCGTCCGGTGAGCTCTCGCTG</td><td>>hg38_21_+_8433172_8446622_RNA45SN1_45S_50ntFlanks</td><td>10680</td><td>10709</td><td>30</td><td>GTTCCGGCGGCGTCCGGTGAGCTCTCGCTG</td><td>2234.7</td><td>-4.5</td><td>1.9</td><td>-2.4</td><td>1.8E-02</td><td>3.1E-02</td><td>BT20/MB468</td><td>MP</td></tr>
<tr><td>>hg38_21_+_8433172_8446622_RNA45SN1_45S_50ntFlanks@4641.4665.25|CATTCGTATTGCGCCGCTAGAGGTG</td><td>>hg38_21_+_8433172_8446622_RNA45SN1_45S_50ntFlanks</td><td>4641</td><td>4665</td><td>25</td><td>CATTCGTATTGCGCCGCTAGAGGTG</td><td>235.8</td><td>-4.2</td><td>1.8</td><td>-2.4</td><td>1.8E-02</td><td>3.1E-02</td><td>BT20/MB468</td><td>MP</td></tr>
</tbody>
</table>

<table>
<tr><td>>hg38_21_+_8433172_8446622_RNA45SN1_45S_50ntFlanks@12479.12496.18|CTAATAGGGAACGTGAGC</td><td>>hg38_21_+_8433172_8446622_RNA45SN1_45S_50ntFlanks</td><td>12479</td><td>12496</td><td>18</td><td>CTAATAGGGAACGTGAGC</td><td>2144.4</td><td>1.3</td><td>0.6</td><td>2.2</td><td>2.7E-02</td><td>3.1E-02</td><td>MB231/MB468</td><td>CYTO</td></tr>
<tr><td>>MI0000749|hsa-mir-30e&WithFlank&1|+|40754349|40754452@65.85.21(+2U)&[MIMAT0000693&hsa-miR-30e-3p&offsets|0|-1(+2U);m-25&1|+|40754413|40754434&offsets|0|-1(+2U)]|CTTTCAGTCGGATGTTTACAGTT</td><td>>MI0000749|hsa-mir-30e&WithFlank&1|+|40754349|40754452</td><td>65</td><td>85</td><td>21(+2U)</td><td>CTTTCAGTCGGATGTTTACAGTT</td><td>634.2</td><td>-1.8</td><td>0.7</td><td>-2.6</td><td>9.5E-03</td><td>3.1E-02</td><td>BT20/MB231</td><td>MITO</td></tr>
<tr><td>>MI0000811|hsa-mir-148b&WithFlank&12|+|54337210|54337320@69.90.22&[MIMAT0000759&hsa-miR-148b-3p&offsets|0|0;m-77&12|+|54337278|54337299&offsets|0|0]|TCAGTGCATCACAGAACTTTGT</td><td>>MI0000811|hsa-mir-148b&WithFlank&12|+|54337210|54337320</td><td>69</td><td>90</td><td>22</td><td>TCAGTGCATCACAGAACTTTGT</td><td>9593.9</td><td>2.8</td><td>1.2</td><td>2.4</td><td>1.8E-02</td><td>3.1E-02</td><td>BT20/MB468</td><td>MP</td></tr>
<tr><td>>MI0000782|hsa-mir-374a&WithFlank&X|-|74287280|74287363@18.35.18&[MIMAT0000727&hsa-miR-374a-5p&offsets|0|-4;m-184&X|-|74287325|74287346&offsets|0|-4]|TTATAATACAACCTGATA</td><td>>MI0000782|hsa-mir-374a&WithFlank&X|-|74287280|74287363</td><td>18</td><td>35</td><td>18</td><td>TTATAATACAACCTGATA</td><td>152.4</td><td>3.2</td><td>1.4</td><td>2.2</td><td>2.5E-02</td><td>3.1E-02</td><td>MB231/MB468</td><td>MP</td></tr>
<tr><td>>MI0000234|hsa-mir-192&WithFlank&11|-|64891131|64891252@31.53.23(+1U)&[MIMAT0000222&hsa-miR-192-5p&offsets|+1|+3(+1U);m-59&11|-|64891203|64891223&offsets|+1|+3(+1U)]|TGACCTATGAATTGACAGCCAGTT</td><td>>MI0000234|hsa-mir-192&WithFlank&11|-|64891131|64891252</td><td>31</td><td>53</td><td>23(+1U)</td><td>TGACCTATGAATTGACAGCCAGTT</td><td>57.2</td><td>3.8</td><td>1.7</td><td>2.2</td><td>2.5E-02</td><td>3.1E-02</td><td>MB231/MB468</td><td>MP</td></tr>
<tr><td>>hg38_21_+_8433172_8446622_RNA45SN1_45S_50ntFlanks@12685.12711.27|CCCAGGCGGAACGATACGGCAGCGCCG</td><td>>hg38_21_+_8433172_8446622_RNA45SN1_45S_50ntFlanks</td><td>12685</td><td>12711</td><td>27</td><td>CCCAGGCGGAACGATACGGCAGCGCCG</td><td>164.5</td><td>-4.2</td><td>1.8</td><td>-2.4</td><td>1.8E-02</td><td>3.1E-02</td><td>BT20/MB468</td><td>MP</td></tr>
<tr><td>>hg38_21_+_8433172_8446622_RNA45SN1_45S_50ntFlanks@7973.8006.34|GACGCGACCTCAGATCAGACGTGGCGACCCGCTG</td><td>>hg38_21_+_8433172_8446622_RNA45SN1_45S_50ntFlanks</td><td>7973</td><td>8006</td><td>34</td><td>GACGCGACCTCAGATCAGACGTGGCGACCCGCTG</td><td>3068.0</td><td>1.6</td><td>0.7</td><td>2.3</td><td>2.4E-02</td><td>3.1E-02</td><td>BT20/MB468</td><td>MITO</td></tr>
<tr><td>>hg38_21_+_8433172_8446622_RNA45SN1_45S_50ntFlanks@3990.4023.34|TTGGTGACTCTAGATAACCTCGGGCCGATCGCAC</td><td>>hg38_21_+_8433172_8446622_RNA45SN1_45S_50ntFlanks</td><td>3990</td><td>4023</td><td>34</td><td>TTGGTGACTCTAGATAACCTCGGGCCGATCGCAC</td><td>186.7</td><td>2.2</td><td>0.9</td><td>2.4</td><td>1.8E-02</td><td>3.1E-02</td><td>BT20/MB468</td><td>MP</td></tr>
<tr><td>>hg38_21_+_8433172_8446622_RNA45SN1_45S_50ntFlanks@7989.8007.19|AGACGTGGCGACCCGCTGA</td><td>>hg38_21_+_8433172_8446622_RNA45SN1_45S_50ntFlanks</td><td>7989</td><td>8007</td><td>19</td><td>AGACGTGGCGACCCGCTGA</td><td>2249.2</td><td>-1.4</td><td>0.6</td><td>-2.3</td><td>2.4E-02</td><td>3.1E-02</td><td>BT20/MB468</td><td>MITO</td></tr>
<tr><td>>MI0000748|hsa-mir-130b&WithFlank&22|+|21653298|21653391@19.40.22&[MIMAT0004680&hsa-miR-130b-5p&offsets|0|+1;m-234&22|+|21653316|21653337&offsets|0|0]|ACTCTTTCCCTGTTGCACTACT</td><td>>MI0000748|hsa-mir-130b&WithFlank&22|+|21653298|21653391</td><td>19</td><td>40</td><td>22</td><td>ACTCTTTCCCTGTTGCACTACT</td><td>1155.0</td><td>-2.0</td><td>0.8</td><td>-2.6</td><td>9.6E-03</td><td>3.1E-02</td><td>BT20/MB231</td><td>MITO</td></tr>
<tr><td>>hg38_21_+_8433172_8446622_RNA45SN1_45S_50ntFlanks@12938.12966.29|ACCATTCGTAGACGACCTGCTTCTGGGTC</td><td>>hg38_21_+_8433172_8446622_RNA45SN1_45S_50ntFlanks</td><td>12938</td><td>12966</td><td>29</td><td>ACCATTCGTAGACGACCTGCTTCTGGGTC</td><td>92.7</td><td>-4.1</td><td>1.7</td><td>-2.4</td><td>1.8E-02</td><td>3.1E-02</td><td>BT20/MB468</td><td>MP</td></tr>
<tr><td>>hg38_21_+_8433172_8446622_RNA45SN1_45S_50ntFlanks@10840.10864.25|CCGTAACTTCGGGATAAGGATTGGC</td><td>>hg38_21_+_8433172_8446622_RNA45SN1_45S_50ntFlanks</td><td>10840</td><td>10864</td><td>25</td><td>CCGTAACTTCGGGATAAGGATTGGC</td><td>236.8</td><td>-4.2</td><td>1.8</td><td>-2.4</td><td>1.8E-02</td><td>3.1E-02</td><td>BT20/MB468</td><td>MP</td></tr>
<tr><td>>hg38_21_+_8433172_8446622_RNA45SN1_45S_50ntFlanks@12942.12964.23|TTCGTAGACGACCTGCTTCTGGG</td><td>>hg38_21_+_8433172_8446622_RNA45SN1_45S_50ntFlanks</td><td>12942</td><td>12964</td><td>23</td><td>TTCGTAGACGACCTGCTTCTGGG</td><td>142.0</td><td>-4.0</td><td>1.7</td><td>-2.4</td><td>1.8E-02</td><td>3.1E-02</td><td>BT20/MB468</td><td>MP</td></tr>
<tr><td>>hg38_21_+_8433172_8446622_RNA45SN1_45S_50ntFlanks@9858.9899.42|CGGGATGAACCGAACGCCGGGTTAAGGCGCCCGATGCCGACG</td><td>>hg38_21_+_8433172_8446622_RNA45SN1_45S_50ntFlanks</td><td>9858</td><td>9899</td><td>42</td><td>CGGGATGAACCGAACGCCGGGTTAAGGCGCCCGATGCCGACG</td><td>272.5</td><td>-3.3</td><td>1.4</td><td>-2.4</td><td>1.8E-02</td><td>3.1E-02</td><td>BT20/MB468</td><td>MP</td></tr>
<tr><td>>trnaMT_SerGCT_MT_+_12207_12265@1.45.45|GAGAAAGCTCACAAGAACTGCTAACTCATGCCCCCATGTCTAACA</td><td>>trnaMT_SerGCT_MT_+_12207_12265</td><td>1</td><td>45</td><td>45</td><td>GAGAAAGCTCACAAGAACTGCTAACTCATGCCCCCATGTCTAACA</td><td>219.7</td><td>2.7</td><td>1.2</td><td>2.2</td><td>2.8E-02</td><td>3.1E-02</td><td>MB231/MB468</td><td>MITO</td></tr>
<tr><td>>MI0000062|hsa-let-7a-3&WithFlank&22|+|46112743|46112828@58.78.21(+1U);MI0000060|hsa-let-7a-1&WithFlank&9|+|94175951|94176042@63.83.21(+1U)&[MIMAT0004481&hsa-let-7a-3p&offsets|0|0(+1U);m-180&22|+|46112800|46112821&offsets|0|-1(+1U)];[MIMAT0004481_1&hsa-let-7a-3p&offsets|0|0(+1U);m-181&9|+|94176013|94176034&offsets|0|-1(+1U)]|CTATACAATCTACTGTCTTTCT</td><td>>MI0000062|hsa-let-7a-3&WithFlank&22|+|46112743|46112828</td><td>58</td><td>78</td><td>21(+1U)</td><td>CTATACAATCTACTGTCTTTCT</td><td>331.6</td><td>1.5</td><td>0.7</td><td>2.2</td><td>2.8E-02</td><td>3.1E-02</td><td>BT20/MB468</td><td>NUC</td></tr>
<tr><td>>hg38_21_+_8433172_8446622_RNA45SN1_45S_50ntFlanks@10509.10532.24|GAATCCGGAGTGGCGGAGATGGGC</td><td>>hg38_21_+_8433172_8446622_RNA45SN1_45S_50ntFlanks</td><td>10509</td><td>10532</td><td>24</td><td>GAATCCGGAGTGGCGGAGATGGGC</td><td>142.0</td><td>-3.1</td><td>1.4</td><td>-2.2</td><td>2.5E-02</td><td>3.1E-02</td><td>MB231/MB468</td><td>MP</td></tr>
<tr><td>>hg38_21_+_8433172_8446622_RNA45SN1_45S_50ntFlanks@4730.4758.29|CAAGAACGAAAGTCGGAGGTTCGAAGACG</td><td>>hg38_21_+_8433172_8446622_RNA45SN1_45S_50ntFlanks</td><td>4730</td><td>4758</td><td>29</td><td>CAAGAACGAAAGTCGGAGGTTCGAAGACG</td><td>4863.6</td><td>-2.1</td><td>0.9</td><td>-2.4</td><td>1.8E-02</td><td>3.1E-02</td><td>BT20/MB468</td><td>MP</td></tr>
<tr><td>>MI0000440|hsa-mir-27b&WithFlank&9|+|95085439|95085547@67.86.20(+2U)&[MIMAT0000419&hsa-miR-27b-3p&offsets|0|-1(+2U);m-39&9|+|95085505|95085525&offsets|0|-1(+2U)]|TTCACAGTGGCTAAGTTCTGTT</td><td>>MI0000440|hsa-mir-27b&WithFlank&9|+|95085439|95085547</td><td>67</td><td>86</td><td>20(+2U)</td><td>TTCACAGTGGCTAAGTTCTGTT</td><td>66.5</td><td>4.5</td><td>2.0</td><td>2.2</td><td>2.5E-02</td><td>3.1E-02</td><td>MB231/MB468</td><td>MP</td></tr>
<tr><td>>hg38_21_+_8433172_8446622_RNA45SN1_45S_50ntFlanks@12416.12436.21|TTCGATGTCGGCTCTTCCTAT</td><td>>hg38_21_+_8433172_8446622_RNA45SN1_45S_50ntFlanks</td><td>12416</td><td>12436</td><td>21</td><td>TTCGATGTCGGCTCTTCCTAT</td><td>1021.7</td><td>-2.3</td><td>0.9</td><td>-2.6</td><td>1.0E-02</td><td>3.1E-02</td><td>BT20/MB231</td><td>CYTO</td></tr>
<tr><td>>hg38_21_+_8433172_8446622_RNA45SN1_45S_50ntFlanks@10061.10081.21|CAGTCGAGAGTGGACGGGAGC</td><td>>hg38_21_+_8433172_8446622_RNA45SN1_45S_50ntFlanks</td><td>10061</td><td>10081</td><td>21</td><td>CAGTCGAGAGTGGACGGGAGC</td><td>54.3</td><td>4.1</td><td>1.8</td><td>2.4</td><td>1.9E-02</td><td>3.1E-02</td><td>BT20/MB468</td><td>MP</td></tr>
<tr><td>>hg38_MT_+_1621_3279_RNR2_16S_50ntFlanks@1405.1423.19|ATTAAAGGTTCGTTTGTTC</td><td>>hg38_MT_+_1621_3279_RNR2_16S_50ntFlanks</td><td>1405</td><td>1423</td><td>19</td><td>ATTAAAGGTTCGTTTGTTC</td><td>119.7</td><td>2.1</td><td>0.9</td><td>2.2</td><td>2.5E-02</td><td>3.1E-02</td><td>MB231/MB468</td><td>MP</td></tr>
<tr><td>>hg38_21_+_8433172_8446622_RNA45SN1_45S_50ntFlanks@11570.11611.42|AGCCGACTTAGAACTGGTGCGGACCAGGGGAATCCGACTGTT</td><td>>hg38_21_+_8433172_8446622_RNA45SN1_45S_50ntFlanks</td><td>11570</td><td>11611</td><td>42</td><td>AGCCGACTTAGAACTGGTGCGGACCAGGGGAATCCGACTGTT</td><td>273.2</td><td>2.3</td><td>0.9</td><td>2.6</td><td>9.6E-03</td><td>3.1E-02</td><td>BT20/MB231</td><td>MITO</td></tr>
<tr><td>>hg38_21_+_8433172_8446622_RNA45SN1_45S_50ntFlanks@6644.6663.20|CCTCGTACGACTCTTAGCGG</td><td>>hg38_21_+_8433172_8446622_RNA45SN1_45S_50ntFlanks</td><td>6644</td><td>6663</td><td>20</td><td>CCTCGTACGACTCTTAGCGG</td><td>2444.3</td><td>-2.7</td><td>1.0</td><td>-2.6</td><td>9.6E-03</td><td>3.1E-02</td><td>BT20/MB231</td><td>MITO</td></tr>
<tr><td>>hg38_21_+_8433172_8446622_RNA45SN1_45S_50ntFlanks@11104.11133.30|AAGGGTCGGGGCGGCAGGGGCCGGCGGCGG</td><td>>hg38_21_+_8433172_8446622_RNA45SN1_45S_50ntFlanks</td><td>11104</td><td>11133</td><td>30</td><td>AAGGGTCGGGGCGGCAGGGGCCGGCGGCGG</td><td>251.8</td><td>-3.6</td><td>1.6</td><td>-2.2</td><td>2.7E-02</td><td>3.1E-02</td><td>BT20/MB468</td><td>CYTO</td></tr>
<tr><td>>hg38_21_+_8433172_8446622_RNA45SN1_45S_50ntFlanks@9956.9994.39|GAAGTCGGAATCCGCTAAGGAGTGTGTAACAACTCACCT</td><td>>hg38_21_+_8433172_8446622_RNA45SN1_45S_50ntFlanks</td><td>9956</td><td>9994</td><td>39</td><td>GAAGTCGGAATCCGCTAAGGAGTGTGTAACAACTCACCT</td><td>132.1</td><td>5.5</td><td>2.1</td><td>2.7</td><td>7.8E-03</td><td>3.1E-02</td><td>BT20/MB231</td><td>MP</td></tr>
<tr><td>>hg38_21_+_8433172_8446622_RNA45SN1_45S_50ntFlanks@12003.12033.31|GCGGGCCGCCGGTGAAATACCACTACTCTGA</td><td>>hg38_21_+_8433172_8446622_RNA45SN1_45S_50ntFlanks</td><td>12003</td><td>12033</td><td>31</td><td>GCGGGCCGCCGGTGAAATACCACTACTCTGA</td><td>158.4</td><td>-1.9</td><td>0.8</td><td>-2.2</td><td>2.5E-02</td><td>3.1E-02</td><td>MB231/MB468</td><td>MP</td></tr>
<tr><td>>trna20_GluTTC_2_-_131094701_131094772@1.30.30,trna5_GluTTC_13_-_41634874_41634945@1.30.30|TCCCATATGGTCTAGCGGTTAGGATTCCTG</td><td>>trna20_GluTTC_2_-_131094701_131094772</td><td>1</td><td>30</td><td>30</td><td>TCCCATATGGTCTAGCGGTTAGGATTCCTG</td><td>458.1</td><td>-3.5</td><td>1.5</td><td>-2.4</td><td>1.9E-02</td><td>3.1E-02</td><td>BT20/MB468</td><td>MP</td></tr>
<tr><td>>hg38_21_+_8433172_8446622_RNA45SN1_45S_50ntFlanks@10869.10904.36|AGGGCTGGGTCGGTCGGGCTGGGGCGCGAAGCGGGG</td><td>>hg38_21_+_8433172_8446622_RNA45SN1_45S_50ntFlanks</td><td>10869</td><td>10904</td><td>36</td><td>AGGGCTGGGTCGGTCGGGCTGGGGCGCGAAGCGGGG</td><td>341.1</td><td>-3.7</td><td>1.7</td><td>-2.2</td><td>2.7E-02</td><td>3.1E-02</td><td>BT20/MB468</td><td>CYTO</td></tr>
<tr><td>>hg38_21_+_8433172_8446622_RNA45SN1_45S_50ntFlanks@6676.6696.21|CTCGTGCGTCGATGAAGAACG</td><td>>hg38_21_+_8433172_8446622_RNA45SN1_45S_50ntFlanks</td><td>6676</td><td>6696</td><td>21</td><td>CTCGTGCGTCGATGAAGAACG</td><td>324.1</td><td>-2.0</td><td>0.9</td><td>-2.2</td><td>2.7E-02</td><td>3.1E-02</td><td>MB231/MB468</td><td>CYTO</td></tr>
<tr><td>>hg38_21_+_8433172_8446622_RNA45SN1_45S_50ntFlanks@11923.11940.18|AGAGGTGTAGAATAAGTG</td><td>>hg38_21_+_8433172_8446622_RNA45SN1_45S_50ntFlanks</td><td>11923</td><td>11940</td><td>18</td><td>AGAGGTGTAGAATAAGTG</td><td>117.5</td><td>4.6</td><td>1.7</td><td>2.7</td><td>7.8E-03</td><td>3.1E-02</td><td>BT20/MB231</td><td>MP</td></tr>
<tr><td>>MI0000465|hsa-mir-191&WithFlank&3|-|49020612|49020715@23.43.21&[MIMAT0000440&hsa-miR-191-5p&offsets|+1|-1;m-48&3|-|49020672|49020694&offsets|+1|-1]|AACGGAATCCCAAAAGCAGCT</td><td>>MI0000465|hsa-mir-191&WithFlank&3|-|49020612|49020715</td><td>23</td><td>43</td><td>21</td><td>AACGGAATCCCAAAAGCAGCT</td><td>234.5</td><td>-2.7</td><td>1.0</td><td>-2.7</td><td>7.8E-03</td><td>3.1E-02</td><td>BT20/MB231</td><td>MP</td></tr>
<tr><td>>MI0000286|hsa-mir-210&WithFlank&11|-|568083|568204@72.93.22(+3A)&[MIMAT0000267&hsa-miR-210-3p&offsets|0|0(+3A);m-102&11|-|568112|568133&offsets|0|0(+3A)]|CTGTGCGTGTGACAGCGGCTGAAAA</td><td>>MI0000286|hsa-mir-210&WithFlank&11|-|568083|568204</td><td>72</td><td>93</td><td>22(+3A)</td><td>CTGTGCGTGTGACAGCGGCTGAAAA</td><td>492.0</td><td>2.6</td><td>1.0</td><td>2.6</td><td>9.7E-03</td><td>3.1E-02</td><td>BT20/MB231</td><td>MITO</td></tr>
<tr><td>>MI0000786|hsa-mir-378a&WithFlank&5|+|149732819|149732896@11.32.22&[MIMAT0000731&hsa-miR-378a-5p&offsets|0|0;m-182&5|+|149732829|149732850&offsets|0|0]|CTCCTGACTCCAGGTCCTGTGT</td><td>>MI0000786|hsa-mir-378a&WithFlank&5|+|149732819|149732896</td><td>11</td><td>32</td><td>22</td><td>CTCCTGACTCCAGGTCCTGTGT</td><td>212.1</td><td>2.6</td><td>1.1</td><td>2.4</td><td>1.9E-02</td><td>3.1E-02</td><td>BT20/MB468</td><td>MP</td></tr>
<tr><td>>MI0000071|hsa-mir-17&WithFlank&13|+|91350599|91350694@20.41.22(+1A)&[MIMAT0000070&hsa-miR-17-5p&offsets|0|-1(+1A);m-88&13|+|91350618|91350640&offsets|0|-1(+1A)]|CAAAGTGCTTACAGTGCAGGTAA</td><td>>MI0000071|hsa-mir-17&WithFlank&13|+|91350599|91350694</td><td>20</td><td>41</td><td>22(+1A)</td><td>CAAAGTGCTTACAGTGCAGGTAA</td><td>190.6</td><td>1.8</td><td>0.8</td><td>2.2</td><td>2.8E-02</td><td>3.1E-02</td><td>BT20/MB468</td><td>NUC</td></tr>
<tr><td>>hg38_21_+_8433172_8446622_RNA45SN1_45S_50ntFlanks@12242.12272.31|AGAAGGGCAAAAGCTCGCTTGATCTTGATTT</td><td>>hg38_21_+_8433172_8446622_RNA45SN1_45S_50ntFlanks</td><td>12242</td><td>12272</td><td>31</td><td>AGAAGGGCAAAAGCTCGCTTGATCTTGATTT</td><td>348.4</td><td>-1.5</td><td>0.7</td><td>-2.2</td><td>2.8E-02</td><td>3.1E-02</td><td>MB231/MB468</td><td>NUC</td></tr>
<tr><td>>hg38_21_+_8433172_8446622_RNA45SN1_45S_50ntFlanks@10350.10372.23|ACGAGAACTTTGAAGGCCGAAGT</td><td>>hg38_21_+_8433172_8446622_RNA45SN1_45S_50ntFlanks</td><td>10350</td><td>10372</td><td>23</td><td>ACGAGAACTTTGAAGGCCGAAGT</td><td>684.9</td><td>-1.6</td><td>0.7</td><td>-2.2</td><td>2.8E-02</td><td>3.1E-02</td><td>MB231/MB468</td><td>MITO</td></tr>
<tr><td>>hg38_21_+_8433172_8446622_RNA45SN1_45S_50ntFlanks@12071.12097.27|CCCCGAGGGGCTCTCGCTTCTGGCGCC</td><td>>hg38_21_+_8433172_8446622_RNA45SN1_45S_50ntFlanks</td><td>12071</td><td>12097</td><td>27</td><td>CCCCGAGGGGCTCTCGCTTCTGGCGCC</td><td>494.5</td><td>-2.6</td><td>1.2</td><td>-2.2</td><td>2.7E-02</td><td>3.1E-02</td><td>MB231/MB468</td><td>CYTO</td></tr>
<tr><td>>trnaMT_ThrTGT_MT_+_15888_15953@31.52.22|TTGTAAACCGGAGATGAAAACC</td><td>>trnaMT_ThrTGT_MT_+_15888_15953</td><td>31</td><td>52</td><td>22</td><td>TTGTAAACCGGAGATGAAAACC</td><td>313.6</td><td>2.4</td><td>0.9</td><td>2.6</td><td>1.0E-02</td><td>3.1E-02</td><td>BT20/MB231</td><td>CYTO</td></tr>
<tr><td>>MI0000440|hsa-mir-27b&WithFlank&9|+|95085439|95085547@67.86.20(+1G)&[MIMAT0000419&hsa-miR-27b-3p&offsets|0|-1(+1G);m-39&9|+|95085505|95085525&offsets|0|-1(+1G)]|TTCACAGTGGCTAAGTTCTGG</td><td>>MI0000440|hsa-mir-27b&WithFlank&9|+|95085439|95085547</td><td>67</td><td>86</td><td>20(+1G)</td><td>TTCACAGTGGCTAAGTTCTGG</td><td>77.3</td><td>3.8</td><td>1.7</td><td>2.2</td><td>2.5E-02</td><td>3.1E-02</td><td>MB231/MB468</td><td>MP</td></tr>
<tr><td>>hg38_21_+_8433172_8446622_RNA45SN1_45S_50ntFlanks@10109.10127.19|TGTGGTGTGCGTCGGAGGG</td><td>>hg38_21_+_8433172_8446622_RNA45SN1_45S_50ntFlanks</td><td>10109</td><td>10127</td><td>19</td><td>TGTGGTGTGCGTCGGAGGG</td><td>198.2</td><td>-3.1</td><td>1.3</td><td>-2.4</td><td>1.9E-02</td><td>3.1E-02</td><td>BT20/MB468</td><td>MP</td></tr>
<tr><td>>hg38_21_+_8433172_8446622_RNA45SN1_45S_50ntFlanks@12457.12488.32|CAAGCGTTGGATTGTTCACCCACTAATAGGGA</td><td>>hg38_21_+_8433172_8446622_RNA45SN1_45S_50ntFlanks</td><td>12457</td><td>12488</td><td>32</td><td>CAAGCGTTGGATTGTTCACCCACTAATAGGGA</td><td>245.7</td><td>-4.1</td><td>1.8</td><td>-2.4</td><td>1.9E-02</td><td>3.1E-02</td><td>BT20/MB468</td><td>MP</td></tr>
<tr><td>>hg38_21_+_8433172_8446622_RNA45SN1_45S_50ntFlanks@6732.6752.21|ACATTGATCATCGACACTTCG</td><td>>hg38_21_+_8433172_8446622_RNA45SN1_45S_50ntFlanks</td><td>6732</td><td>6752</td><td>21</td><td>ACATTGATCATCGACACTTCG</td><td>520.1</td><td>-1.5</td><td>0.7</td><td>-2.2</td><td>2.7E-02</td><td>3.1E-02</td><td>MB231/MB468</td><td>CYTO</td></tr>
<tr><td>>hg38_21_+_8433172_8446622_RNA45SN1_45S_50ntFlanks@3771.3791.21|CACGGCCGGTACAGTGAAACT</td><td>>hg38_21_+_8433172_8446622_RNA45SN1_45S_50ntFlanks</td><td>3771</td><td>3791</td><td>21</td><td>CACGGCCGGTACAGTGAAACT</td><td>205.8</td><td>-2.0</td><td>0.9</td><td>-2.3</td><td>2.4E-02</td><td>3.1E-02</td><td>BT20/MB468</td><td>MITO</td></tr>
<tr><td>>hg38_MT_+_1621_3279_RNR2_16S_50ntFlanks@1367.1397.31|TCGATGTTGGATCAGGACATCCCGATGGTGC</td><td>>hg38_MT_+_1621_3279_RNR2_16S_50ntFlanks</td><td>1367</td><td>1397</td><td>31</td><td>TCGATGTTGGATCAGGACATCCCGATGGTGC</td><td>72.1</td><td>3.7</td><td>1.7</td><td>2.2</td><td>2.5E-02</td><td>3.1E-02</td><td>MB231/MB468</td><td>MP</td></tr>
<tr><td>>hg38_21_+_8433172_8446622_RNA45SN1_45S_50ntFlanks@6647.6686.40|CGTACGACTCTTAGCGGTGGATCACTCGGCTCGTGCGTCG</td><td>>hg38_21_+_8433172_8446622_RNA45SN1_45S_50ntFlanks</td><td>6647</td><td>6686</td><td>40</td><td>CGTACGACTCTTAGCGGTGGATCACTCGGCTCGTGCGTCG</td><td>95.8</td><td>-2.3</td><td>1.0</td><td>-2.2</td><td>2.5E-02</td><td>3.1E-02</td><td>MB231/MB468</td><td>MP</td></tr>
<tr><td>>am_trna162_MetCAT_6_-_26758550_26758622@34.54.21,trna164_MetCAT_6_-_26735574_26735646@34.54.21,trna20_MetCAT_16_+_71460396_71460468@34.54.21,trna21_MetCAT_6_+_26701712_26701784@34.54.21,trna27_MetCAT_6_+_26766444_26766516@34.54.21,trna75_MetCAT_6_+_28912352_28912424@34.54.21,trna97_MetCAT_6_-_28921042_28921114@34.54.21|CATAATCTGAAGGTCCTGAGT</td><td>>am_trna162_MetCAT_6_-_26758550_26758622</td><td>34</td><td>54</td><td>21</td><td>CATAATCTGAAGGTCCTGAGT</td><td>217.6</td><td>1.9</td><td>0.9</td><td>2.2</td><td>2.5E-02</td><td>3.1E-02</td><td>MB231/MB468</td><td>MP</td></tr>
<tr><td>>MI0000542|hsa-mir-320a&WithFlank&8|-|22244960|22245043@48.70.23&[MIMAT0000510&hsa-miR-320a-3p&offsets|0|+1;m-60&8|-|22244975|22244996&offsets|0|+1]|AAAAGCTGGGTTGAGAGGGCGAA</td><td>>MI0000542|hsa-mir-320a&WithFlank&8|-|22244960|22245043</td><td>48</td><td>70</td><td>23</td><td>AAAAGCTGGGTTGAGAGGGCGAA</td><td>7538.2</td><td>1.7</td><td>0.7</td><td>2.6</td><td>8.4E-03</td><td>3.1E-02</td><td>BT20/MB231</td><td>NUC</td></tr>
</table>

| | | | | | | | | | | | | | |
|---|---|---|---|---|---|---|---|---|---|---|---|---|---|
| >trnaMT_HisGTG_MT_+_12138_12206@28.65.38\|ATTGTGAATCTGACAACAGAGGCTTACGACCCCTTATT | >trnaMT_HisGTG_MT_+_12138_12206 | 28 | 65 | 38 | ATTGTGAATCTGACAACAGAGGCTTACGACCCCTTATT | 112.4 | 5.2 | 2.2 | 2.4 | 1.9E-02 | 3.1E-02 | BT20/MB468 | MP |
| >MI0000061\|hsa-let-7a-2&WithFlank&11\|-\|122146516\|122146599@11.30.20(+1G);MI0000062\|hsa-let-7a-3&WithFlank&22\|+\|46112743\|46112828@10.29.20(+1G);MI0000060\|hsa-let-7a-1&WithFlank&9\|+\|94175951\|94176042@12.31.20(+1G)&[MIMAT0000062&hsa-let-7a-5p&offsets\|0\|-2(+1G);m-6&11\|-\|122146568\|122146589&offsets\|0\|-2(+1G)];[MIMAT0000062_1&hsa-let-7a-5p&offsets\|0\|-2(+1G);m-5&22\|+\|46112752\|46112773&offsets\|0\|-2(+1G)];[MIMAT0000062_2&hsa-let-7a-5p&offsets\|0\|-2(+1G);m-4&9\|+\|94175962\|94175983&offsets\|0\|-2(+1G)]\|TGAGGTAGTAGGTTGTATAGG | >MI0000061\|hsa-let-7a-2&WithFlank&11\|-\|122146516\|122146599 | 11 | 30 | 20(+1G) | TGAGGTAGTAGGTTGTATAGG | 78.7 | 3.8 | 1.7 | 2.2 | 2.6E-02 | 3.1E-02 | MB231/MB468 | MP |
| >hg38_21_+_8433172_8446622_RNA45SN1_45S_50ntFlanks@9533.9569.37\|GAGGAAACTCTGGTGGAGGTCCGTAGCGGTCCTGACG | >hg38_21_+_8433172_8446622_RNA45SN1_45S_50ntFlanks | 9533 | 9569 | 37 | GAGGAAACTCTGGTGGAGGTCCGTAGCGGTCCTGACG | 359.1 | -1.9 | 0.7 | -2.6 | 1.1E-02 | 3.1E-02 | BT20/MB231 | CYTO |
| >hg38_21_+_8433172_8446622_RNA45SN1_45S_50ntFlanks@8184.8206.23\|CGAGGCCCAGCCCGTGGACGGTG | >hg38_21_+_8433172_8446622_RNA45SN1_45S_50ntFlanks | 8184 | 8206 | 23 | CGAGGCCCAGCCCGTGGACGGTG | 191.7 | -1.7 | 0.8 | -2.3 | 2.4E-02 | 3.1E-02 | BT20/MB468 | MITO |
| >MI0003185\|hsa-mir-501&WithFlank&X\|+\|50009716\|50009811@57.78.22&[MIMAT0004774&hsa-miR-501-3p&offsets\|0\|0;m-168&X\|+\|50009772\|50009793&offsets\|0\|0]\|AATGCACCCGGGCAAGGATTCT | >MI0003185\|hsa-mir-501&WithFlank&X\|+\|50009716\|50009811 | 57 | 78 | 22 | AATGCACCCGGGCAAGGATTCT | 412.7 | 1.8 | 0.8 | 2.2 | 2.6E-02 | 3.1E-02 | MB231/MB468 | MP |
| >hg38_21_+_8433172_8446622_RNA45SN1_45S_50ntFlanks@12615.12661.47\|GCTGAGGAGCCAATGGGGCGAAGCTACCATCTGTGGGATTATGACTG | >hg38_21_+_8433172_8446622_RNA45SN1_45S_50ntFlanks | 12615 | 12661 | 47 | GCTGAGGAGCCAATGGGGCGAAGCTACCATCTGTGGGATTATGACTG | 1264.4 | -3.3 | 1.4 | -2.3 | 2.4E-02 | 3.1E-02 | BT20/MB468 | MITO |
| >hg38_21_+_8433172_8446622_RNA45SN1_45S_50ntFlanks@7975.8009.35\|CGCGACCTCAGATCAGACGTGGCGACCCGCTGAAT | >hg38_21_+_8433172_8446622_RNA45SN1_45S_50ntFlanks | 7975 | 8009 | 35 | CGCGACCTCAGATCAGACGTGGCGACCCGCTGAAT | 24512.6 | 2.0 | 0.9 | 2.2 | 2.6E-02 | 3.1E-02 | MB231/MB468 | MP |
| >hg38_21_+_8433172_8446622_RNA45SN1_45S_50ntFlanks@12480.12497.18\|TAATAGGGAACGTGAGCT | >hg38_21_+_8433172_8446622_RNA45SN1_45S_50ntFlanks | 12480 | 12497 | 18 | TAATAGGGAACGTGAGCT | 331.1 | 1.6 | 0.7 | 2.2 | 2.8E-02 | 3.1E-02 | MB231/MB468 | NUC |
| >hg38_21_+_8433172_8446622_RNA45SN1_45S_50ntFlanks@9852.9899.48\|GCGCTGCGGGATGAACCGAACGCCGGGTTAAGGCGCCCGATGCCGACG | >hg38_21_+_8433172_8446622_RNA45SN1_45S_50ntFlanks | 9852 | 9899 | 48 | GCGCTGCGGGATGAACCGAACGCCGGGTTAAGGCGCCCGATGCCGACG | 199.0 | -5.1 | 2.3 | -2.3 | 2.4E-02 | 3.1E-02 | BT20/MB468 | MITO |
| >hg38_21_+_8433172_8446622_RNA45SN1_45S_50ntFlanks@12476.12497.22\|CCACTAATAGGGAACGTGAGCT | >hg38_21_+_8433172_8446622_RNA45SN1_45S_50ntFlanks | 12476 | 12497 | 22 | CCACTAATAGGGAACGTGAGCT | 378.5 | 3.4 | 1.3 | 2.7 | 7.9E-03 | 3.1E-02 | BT20/MB231 | MP |
| >hg38_21_+_8433172_8446622_RNA45SN1_45S_50ntFlanks@11672.11692.21\|CAGTGCTCTGAATGTCAAAGT | >hg38_21_+_8433172_8446622_RNA45SN1_45S_50ntFlanks | 11672 | 11692 | 21 | CAGTGCTCTGAATGTCAAAGT | 227.3 | 2.0 | 0.8 | 2.4 | 1.9E-02 | 3.1E-02 | BT20/MB468 | MP |
| >hg38_21_+_8433172_8446622_RNA45SN1_45S_50ntFlanks@9486.9506.21\|GTTAGGACCCGAAAGATGGTG | >hg38_21_+_8433172_8446622_RNA45SN1_45S_50ntFlanks | 9486 | 9506 | 21 | GTTAGGACCCGAAAGATGGTG | 61.0 | 4.1 | 1.8 | 2.4 | 1.9E-02 | 3.1E-02 | BT20/MB468 | MP |
| >hg38_21_+_8433172_8446622_RNA45SN1_45S_50ntFlanks@12004.12032.29\|CGGGCCGCCGGTGAAATACCACTACTCTG | >hg38_21_+_8433172_8446622_RNA45SN1_45S_50ntFlanks | 12004 | 12032 | 29 | CGGGCCGCCGGTGAAATACCACTACTCTG | 315.7 | -2.2 | 1.0 | -2.2 | 2.7E-02 | 3.1E-02 | MB231/MB468 | CYTO |
| >hg38_21_+_8433172_8446622_RNA45SN1_45S_50ntFlanks@10684.10705.22\|CGGCGGCGTCCGGTGAGCTCTC | >hg38_21_+_8433172_8446622_RNA45SN1_45S_50ntFlanks | 10684 | 10705 | 22 | CGGCGGCGTCCGGTGAGCTCTC | 220.9 | -1.3 | 0.6 | -2.3 | 2.4E-02 | 3.1E-02 | BT20/MB468 | MITO |
| >hg38_MT_+_1621_3279_RNR2_16S_50ntFlanks@51.71.21\|GCTAAACCTAGCCCCAAACCC | >hg38_MT_+_1621_3279_RNR2_16S_50ntFlanks | 51 | 71 | 21 | GCTAAACCTAGCCCCAAACCC | 4143.0 | 1.7 | 0.8 | 2.2 | 2.6E-02 | 3.1E-02 | MB231/MB468 | MP |
| >MI0000749\|hsa-mir-30e&WithFlank&1\|+\|40754349\|40754452@65.83.19&[MIMAT0000693&hsa-miR-30e-3p&offsets\|0\|-3;m-25&1\|+\|40754413\|40754434&offsets\|0\|-3]\|CTTTCAGTCGGATGTTTAC | >MI0000749\|hsa-mir-30e&WithFlank&1\|+\|40754349\|40754452 | 65 | 83 | 19 | CTTTCAGTCGGATGTTTAC | 549.6 | 2.1 | 0.9 | 2.2 | 2.9E-02 | 3.1E-02 | MB231/MB468 | MITO |
| >hg38_21_+_8433172_8446622_RNA45SN1_45S_50ntFlanks@7973.8007.35\|GACGCGACCTCAGATCAGACGTGGCGACCCGCTGA | >hg38_21_+_8433172_8446622_RNA45SN1_45S_50ntFlanks | 7973 | 8007 | 35 | GACGCGACCTCAGATCAGACGTGGCGACCCGCTGA | 1682.6 | 2.6 | 1.0 | 2.6 | 8.5E-03 | 3.1E-02 | BT20/MB231 | NUC |
| >hg38_21_+_8433172_8446622_RNA45SN1_45S_50ntFlanks@10350.10371.22\|ACGAGAACTTTGAAGGCCGAAG | >hg38_21_+_8433172_8446622_RNA45SN1_45S_50ntFlanks | 10350 | 10371 | 22 | ACGAGAACTTTGAAGGCCGAAG | 271.2 | -2.0 | 0.8 | -2.6 | 8.5E-03 | 3.1E-02 | BT20/MB231 | NUC |
| >MI0000103\|hsa-mir-101-1&WithFlank&1\|-\|65058428\|65058514@52.73.22(+2U);MI0000739\|hsa-mir-101-2&WithFlank&9\|+\|4850291\|4850381@54.75.22(+2U)&[MIMAT0000099&hsa-miR-101-3p&offsets\|-1\|0(+2U);m-17&1\|-\|65058443\|65058462&offsets\|-1\|+1(+2U)];[MIMAT0000099_1&hsa-miR-101-3p&offsets\|-1\|0(+2U);m-16&9\|+\|4850345\|4850364&offsets\|-1\|+1(+2U)]\|GTACAGTACTGTGATAACTGAATT | >MI0000103\|hsa-mir-101-1&WithFlank&1\|-\|65058428\|65058514 | 52 | 73 | 22(+2U) | GTACAGTACTGTGATAACTGAATT | 59.0 | 3.6 | 1.6 | 2.2 | 2.6E-02 | 3.1E-02 | MB231/MB468 | MP |
| >hg38_21_+_8433172_8446622_RNA45SN1_45S_50ntFlanks@10229.10273.45\|CCCCGCGGACGCTACGCCGCGACGAGTAGGAGGGCCGCTGCGGTG | >hg38_21_+_8433172_8446622_RNA45SN1_45S_50ntFlanks | 10229 | 10273 | 45 | CCCCGCGGACGCTACGCCGCGACGAGTAGGAGGGCCGCTGCGGTG | 31.3 | 8.3 | 3.1 | 2.7 | 7.9E-03 | 3.1E-02 | BT20/MB231 | MP |
| >hg38_MT_+_598_1651_RNR1_12S_50ntFlanks@720.744.25\|AAAGACGTTAGGTCAAGGTGTAGCC | >hg38_MT_+_598_1651_RNR1_12S_50ntFlanks | 720 | 744 | 25 | AAAGACGTTAGGTCAAGGTGTAGCC | 217.4 | -3.1 | 1.2 | -2.7 | 7.9E-03 | 3.1E-02 | BT20/MB231 | MP |
| >hg38_21_+_8433172_8446622_RNA45SN1_45S_50ntFlanks@12618.12654.37\|GAGGAGCCAATGGGGCGAAGCTACCATCTGTGGGATT | >hg38_21_+_8433172_8446622_RNA45SN1_45S_50ntFlanks | 12618 | 12654 | 37 | GAGGAGCCAATGGGGCGAAGCTACCATCTGTGGGATT | 166.2 | 4.5 | 1.7 | 2.7 | 7.9E-03 | 3.1E-02 | BT20/MB231 | MP |
| >MI0000100\|hsa-mir-98&WithFlank&X\|-\|53556217\|53556347@28.49.22&[MIMAT0000096&hsa-miR-98-5p&offsets\|0\|0;m-143&X\|-\|53556299\|53556320&offsets\|0\|0]\|TGAGGTAGTAAGTTGTATTGTT | >MI0000100\|hsa-mir-98&WithFlank&X\|-\|53556217\|53556347 | 28 | 49 | 22 | TGAGGTAGTAAGTTGTATTGTT | 4610.5 | 1.6 | 0.7 | 2.2 | 2.6E-02 | 3.1E-02 | MB231/MB468 | MP |
| >MI0000469\|hsa-mir-125a&WithFlank&19\|+\|51693248\|51693345@21.44.24&[MIMAT0000443&hsa-miR-125a-5p&offsets\|0\|0;m-63&19\|+\|51693268\|51693289&offsets\|0\|+2]\|TCCCTGAGACCCTTTAACCTGTGA | >MI0000469\|hsa-mir-125a&WithFlank&19\|+\|51693248\|51693345 | 21 | 44 | 24 | TCCCTGAGACCCTTTAACCTGTGA | 446.3 | 2.2 | 0.9 | 2.3 | 1.9E-02 | 3.1E-02 | BT20/MB468 | MP |
| >MI0000447\|hsa-mir-128-1&WithFlank&2\|+\|135665391\|135665484@56.76.21(+2A);MI0000727\|hsa-mir-128-2&WithFlank&3\|+\|35744470\|35744565@58.78.21(+2A)&[MIMAT0000424&hsa-miR-128-3p&offsets\|0\|0(+2A);m-108&2\|+\|135665446\|135665466&offsets\|0\|0(+2A)];[MIMAT0000424_1&hsa-miR-128-3p&offsets\|0\|0(+2A);m-117&3\|+\|35744527\|35744547&offsets\|0\|0(+2A)]\|TCACAGTGAACCGGTCTCTTTAA | >MI0000447\|hsa-mir-128-1&WithFlank&2\|+\|135665391\|135665484 | 56 | 76 | 21(+2A) | TCACAGTGAACCGGTCTCTTTAA | 377.3 | 2.0 | 0.9 | 2.2 | 2.9E-02 | 3.1E-02 | MB231/MB468 | MITO |
| >hg38_21_+_8433172_8446622_RNA45SN1_45S_50ntFlanks@3858.3898.41\|TGGTAATTCTAGAGCTAATACATGCCGACGGGCGCTGACCC | >hg38_21_+_8433172_8446622_RNA45SN1_45S_50ntFlanks | 3858 | 3898 | 41 | TGGTAATTCTAGAGCTAATACATGCCGACGGGCGCTGACCC | 144.5 | -2.6 | 1.2 | -2.3 | 2.4E-02 | 3.1E-02 | BT20/MB468 | MITO |
| >MI0005561\|hsa-mir-877&WithFlank&6\|+\|30584326\|30584423@7.29.23(+1U)&[MIMAT0004949&hsa-miR-877-5p&offsets\|0\|+3(+1U);m-382&6\|+\|30584332\|30584354&offsets\|0\|0(+1U)]\|GTAGAGGAGATGGCGCAGGGGACT | >MI0005561\|hsa-mir-877&WithFlank&6\|+\|30584326\|30584423 | 7 | 29 | 23(+1U) | GTAGAGGAGATGGCGCAGGGGACT | 111.2 | 2.8 | 1.3 | 2.2 | 2.6E-02 | 3.1E-02 | MB231/MB468 | MP |
| >MI0000077\|hsa-mir-21&WithFlank&17\|+\|59841260\|59841343@53.74.22&[MIMAT0004494&hsa-miR-21-3p&offsets\|+1\|+2;m-52&17\|+\|59841311\|59841332&offsets\|+1\|+1]\|AACACCAGTCGATGGGCTGTCT | >MI0000077\|hsa-mir-21&WithFlank&17\|+\|59841260\|59841343 | 53 | 74 | 22 | AACACCAGTCGATGGGCTGTCT | 153.5 | 3.7 | 1.6 | 2.2 | 2.6E-02 | 3.1E-02 | MB231/MB468 | MP |
| >MI0000749\|hsa-mir-30e&WithFlank&1\|+\|40754349\|40754452@24.46.23&[MIMAT0000692&hsa-miR-30e-5p&offsets\|+1\|+2;m-24&1\|+\|40754371\|40754394&offsets\|+1\|0]\|GTAAACATCCTTGACTGGAAGCT | >MI0000749\|hsa-mir-30e&WithFlank&1\|+\|40754349\|40754452 | 24 | 46 | 23 | GTAAACATCCTTGACTGGAAGCT | 1022.4 | 1.8 | 0.8 | 2.2 | 2.6E-02 | 3.1E-02 | MB231/MB468 | MP |
| >hg38_21_+_8433172_8446622_RNA45SN1_45S_50ntFlanks@12727.12747.21\|CCTCGGATAGCCGGTCCCCCG | >hg38_21_+_8433172_8446622_RNA45SN1_45S_50ntFlanks | 12727 | 12747 | 21 | CCTCGGATAGCCGGTCCCCCG | 67.5 | -4.2 | 1.8 | -2.3 | 1.9E-02 | 3.1E-02 | BT20/MB468 | MP |
| >MI0000065\|hsa-let-7d&WithFlank&9\|+\|94178828\|94178926@68.88.21(+1A)&[MIMAT0004484&hsa-let-7d-3p&offsets\|0\|-1(+1A);m-98&9\|+\|94178895\|94178916&offsets\|0\|-1(+1A)]\|CTATACGACCTGCTGCCTTTCA | >MI0000065\|hsa-let-7d&WithFlank&9\|+\|94178828\|94178926 | 68 | 88 | 21(+1A) | CTATACGACCTGCTGCCTTTCA | 433.9 | 1.9 | 0.7 | 2.6 | 1.1E-02 | 3.1E-02 | BT20/MB231 | CYTO |
| >hg38_21_+_8433172_8446622_RNA45SN1_45S_50ntFlanks@11669.11688.20\|GCCCAGTGCTCTGAATGTCA | >hg38_21_+_8433172_8446622_RNA45SN1_45S_50ntFlanks | 11669 | 11688 | 20 | GCCCAGTGCTCTGAATGTCA | 804.6 | -2.3 | 0.9 | -2.6 | 1.1E-02 | 3.1E-02 | BT20/MB231 | CYTO |
| >hg38_21_+_8433172_8446622_RNA45SN1_45S_50ntFlanks@12417.12437.21\|TCGATGTCGGCTCTTCCTATC | >hg38_21_+_8433172_8446622_RNA45SN1_45S_50ntFlanks | 12417 | 12437 | 21 | TCGATGTCGGCTCTTCCTATC | 2514.7 | -2.2 | 0.9 | -2.6 | 1.1E-02 | 3.2E-02 | BT20/MB231 | CYTO |
| >trnaMT_LysTTT_MT_+_8295_8364@55.73.19,trnalookalike7_LysTTT_1_+_568844_568913@55.73.19\|CACCTCTTTACAGTGACCA | >trnaMT_LysTTT_MT_+_8295_8364 | 55 | 73 | 19 | CACCTCTTTACAGTGACCA | 117.4 | 3.7 | 1.6 | 2.3 | 1.9E-02 | 3.2E-02 | BT20/MB468 | MP |
| >hg38_21_+_8433172_8446622_RNA45SN1_45S_50ntFlanks@3835.3871.37\|CTCCTCTCCTACTTGGATAACTGTGGTAATTCTAGAG | >hg38_21_+_8433172_8446622_RNA45SN1_45S_50ntFlanks | 3835 | 3871 | 37 | CTCCTCTCCTACTTGGATAACTGTGGTAATTCTAGAG | 952.1 | -2.2 | 1.0 | -2.2 | 2.7E-02 | 3.2E-02 | MB231/MB468 | CYTO |
| >hg38_21_+_8433172_8446622_RNA45SN1_45S_50ntFlanks@12435.12473.39\|ATCATTGTGAAGCAGAATTCACCAAGCGTTGGATTGTTC | >hg38_21_+_8433172_8446622_RNA45SN1_45S_50ntFlanks | 12435 | 12473 | 39 | ATCATTGTGAAGCAGAATTCACCAAGCGTTGGATTGTTC | 297.9 | -3.3 | 1.5 | -2.2 | 2.7E-02 | 3.2E-02 | MB231/MB468 | CYTO |
| >hg38_MT_+_1621_3279_RNR2_16S_50ntFlanks@1184.1205.22\|AATTTCGGTTGGGGCGACCTCG | >hg38_MT_+_1621_3279_RNR2_16S_50ntFlanks | 1184 | 1205 | 22 | AATTTCGGTTGGGGCGACCTCG | 88.4 | 6.1 | 2.6 | 2.3 | 1.9E-02 | 3.2E-02 | BT20/MB468 | MP |
| >hg38_21_+_8433172_8446622_RNA45SN1_45S_50ntFlanks@11571.11613.43\|GCCGACTTAGAACTGGTGCGGACCAGGGGAATCCGACTGTTTA | >hg38_21_+_8433172_8446622_RNA45SN1_45S_50ntFlanks | 11571 | 11613 | 43 | GCCGACTTAGAACTGGTGCGGACCAGGGGAATCCGACTGTTTA | 704.1 | 2.5 | 1.0 | 2.6 | 9.8E-03 | 3.2E-02 | BT20/MB231 | MITO |
| >hg38_21_+_8433172_8446622_RNA45SN1_45S_50ntFlanks@9915.9949.35\|AAAAGGTGTTGGTTGATATAGACAGCAGGACGGTG | >hg38_21_+_8433172_8446622_RNA45SN1_45S_50ntFlanks | 9915 | 9949 | 35 | AAAAGGTGTTGGTTGATATAGACAGCAGGACGGTG | 140.7 | 5.2 | 1.9 | 2.7 | 8.0E-03 | 3.2E-02 | BT20/MB231 | MP |
| >MI0000272\|hsa-mir-182&WithFlank&7\|-\|129770377\|129770498@29.51.23(+1C)&[MIMAT0000259&hsa-miR-182-5p&offsets\|0\|-1(+1C);m-22&7\|-\|129770449\|129770470&offsets\|0\|+1(+1C)]\|TTTGGCAATGGTAGAACTCACACC | >MI0000272\|hsa-mir-182&WithFlank&7\|-\|129770377\|129770498 | 29 | 51 | 23(+1C) | TTTGGCAATGGTAGAACTCACACC | 3316.2 | 2.0 | 0.8 | 2.6 | 9.8E-03 | 3.2E-02 | BT20/MB231 | MITO |
| >hg38_21_+_8433172_8446622_RNA45SN1_45S_50ntFlanks@6650.6668.19\|ACGACTCTTAGCGGTGGAT | >hg38_21_+_8433172_8446622_RNA45SN1_45S_50ntFlanks | 6650 | 6668 | 19 | ACGACTCTTAGCGGTGGAT | 6382.6 | 1.8 | 0.7 | 2.6 | 9.8E-03 | 3.2E-02 | BT20/MB231 | MITO |
| >hg38_21_+_8433172_8446622_RNA45SN1_45S_50ntFlanks@12417.12459.43\|TCGATGTCGGCTCTTCCTATCATTGTGAAGCAGAATTCACCAA | >hg38_21_+_8433172_8446622_RNA45SN1_45S_50ntFlanks | 12417 | 12459 | 43 | TCGATGTCGGCTCTTCCTATCATTGTGAAGCAGAATTCACCAA | 467.1 | -2.6 | 1.0 | -2.6 | 9.8E-03 | 3.2E-02 | BT20/MB231 | MITO |
| >hg38_21_+_8433172_8446622_RNA45SN1_45S_50ntFlanks@9854.9874.21\|GCTGCGGGATGAACCGAACGC | >hg38_21_+_8433172_8446622_RNA45SN1_45S_50ntFlanks | 9854 | 9874 | 21 | GCTGCGGGATGAACCGAACGC | 163.2 | 5.0 | 2.1 | 2.3 | 1.9E-02 | 3.2E-02 | BT20/MB468 | MP |
| >hg38_21_+_8433172_8446622_RNA45SN1_45S_50ntFlanks@12996.13014.19\|TCGCTGCGATCTATTGAAA | >hg38_21_+_8433172_8446622_RNA45SN1_45S_50ntFlanks | 12996 | 13014 | 19 | TCGCTGCGATCTATTGAAA | 605.1 | -2.0 | 0.8 | -2.6 | 9.9E-03 | 3.2E-02 | BT20/MB231 | MITO |
| >hg38_21_+_8433172_8446622_RNA45SN1_45S_50ntFlanks@11217.11248.32\|CTGGACGCGAGCCGGGCCCTTCCCGTGGATCG | >hg38_21_+_8433172_8446622_RNA45SN1_45S_50ntFlanks | 11217 | 11248 | 32 | CTGGACGCGAGCCGGGCCCTTCCCGTGGATCG | 543.4 | 2.4 | 1.1 | 2.3 | 2.4E-02 | 3.2E-02 | BT20/MB468 | MITO |
| >hg38_21_+_8433172_8446622_RNA45SN1_45S_50ntFlanks@3854.3874.21\|ACTGTGGTAATTCTAGAGCTA | >hg38_21_+_8433172_8446622_RNA45SN1_45S_50ntFlanks | 3854 | 3874 | 21 | ACTGTGGTAATTCTAGAGCTA | 319.5 | -1.5 | 0.7 | -2.2 | 2.7E-02 | 3.2E-02 | BT20/MB468 | CYTO |

<table>
<tr><td>>trna10_AspGTC_12_-_125424193_125424264@36.71.36,trna12_AspGTC_12_-_125411891_125411962@36.71.36,trna144_AspGTC_6_-_27551236_27551307@36.71.36,trna38_AspGTC_17_-_8125556_8125627@36.71.36,trna45_AspGTC_6_+_27447453_27447524@36.71.36,trna48_AspGTC_6_+_27471523_27471594@36.71.36,trna4_AspGTC_12_+_96429799_96429870@36.71.36,trna69_AspGTC_1_-_161440205_161440276@36.71.36,trna72_AspGTC_1_-_161432824_161432895@36.71.36,trna75_AspGTC_1_-_161425414_161425485@36.71.36,trna78_AspGTC_1_-_161418033_161418104@36.71.36,trna81_AspGTC_1_-_161410615_161410686@36.71.36|CACGCGGGAGACCGGGGTTCGATTCCCCGACGGGGA</td><td>>trna10_AspGTC_12_-_125424193_125424264</td><td>36</td><td>71</td><td>36</td><td>CACGCGGGAGACCGGGGTTCGATTCCCCGACGGGGA</td><td>332.9</td><td>-2.4</td><td>1.0</td><td>-2.6</td><td>1.1E-02</td><td>3.2E-02</td><td>BT20/MB231</td><td>CYTO</td></tr>
<tr><td>>hg38_21_+_8433172_8446622_RNA45SN1_45S_50ntFlanks@12241.12272.32|CAGAAGGGCAAAAGCTCGCTTGATCTTGATTT</td><td>>hg38_21_+_8433172_8446622_RNA45SN1_45S_50ntFlanks</td><td>12241</td><td>12272</td><td>32</td><td>CAGAAGGGCAAAAGCTCGCTTGATCTTGATTT</td><td>360.7</td><td>-1.8</td><td>0.8</td><td>-2.2</td><td>2.9E-02</td><td>3.2E-02</td><td>MB231/MB468</td><td>NUC</td></tr>
<tr><td>>hg38_21_+_8433172_8446622_RNA45SN1_45S_50ntFlanks@6677.6696.20|TCGTGCGTCGATGAAGAACG</td><td>>hg38_21_+_8433172_8446622_RNA45SN1_45S_50ntFlanks</td><td>6677</td><td>6696</td><td>20</td><td>TCGTGCGTCGATGAAGAACG</td><td>152.7</td><td>4.1</td><td>1.5</td><td>2.7</td><td>8.0E-03</td><td>3.2E-02</td><td>BT20/MB231</td><td>MP</td></tr>
<tr><td>>MI0000446|hsa-mir-125b-1&WithFlank&11|-|122099751|122099850@21.42.22(+2A);MI0000470|hsa-mir-125b-2&WithFlank&21|+|16590231|16590331@23.44.22(+2A)&[MIMAT0000423&hsa-miR-125b-5p&offsets|0|0(+2A);m-67&11|-|122099809|122099830&offsets|0|0(+2A)];[MIMAT0000423_1&hsa-miR-125b-5p&offsets|0|0(+2A);m-68&21|+|16590253|16590274&offsets|0|0(+2A)]|TCCCTGAGACCCTAACTTGTGAAA</td><td>>MI0000446|hsa-mir-125b-1&WithFlank&11|-|122099751|122099850</td><td>21</td><td>42</td><td>22(+2A)</td><td>TCCCTGAGACCCTAACTTGTGAAA</td><td>1051.8</td><td>-6.4</td><td>2.5</td><td>-2.6</td><td>9.9E-03</td><td>3.2E-02</td><td>BT20/MB231</td><td>MITO</td></tr>
<tr><td>>am_trna111_HisGTG_1_-_147774845_147774916@-1G.28.29,trna118_HisGTG_1_-_145396881_145396952@-1G.28.29,trna16_HisGTG_1_+_146544773_146544844@-1G.28.29,trna1_HisGTG_15_+_45493349_45493420@-1G.28.29,trna21_HisGTG_1_+_147753471_147753542@-1G.28.29,trna33_HisGTG_6_+_27125906_27125977@-1G.28.29,trna7_HisGTG_9_-_14433938_14434009@-1G.28.29,trna8_HisGTG_15_-_45492611_45492682@-1G.28.29,trna9_HisGTG_15_-_45490804_45490875@-1G.28.29|GGCCGTGATCGTATAGTGGTTAGTACTCT</td><td>>am_trna111_HisGTG_1_-_147774845_147774916</td><td>-1G</td><td>28</td><td>29</td><td>GGCCGTGATCGTATAGTGGTTAGTACTCT</td><td>472.3</td><td>-2.0</td><td>0.8</td><td>-2.6</td><td>9.9E-03</td><td>3.2E-02</td><td>BT20/MB231</td><td>MITO</td></tr>
<tr><td>>hg38_21_+_8433172_8446622_RNA45SN1_45S_50ntFlanks@10738.10756.19|AAATCTCGCGCCGGGCCGT</td><td>>hg38_21_+_8433172_8446622_RNA45SN1_45S_50ntFlanks</td><td>10738</td><td>10756</td><td>19</td><td>AAATCTCGCGCCGGGCCGT</td><td>598.3</td><td>-2.4</td><td>0.9</td><td>-2.6</td><td>8.6E-03</td><td>3.2E-02</td><td>BT20/MB231</td><td>NUC</td></tr>
<tr><td>>MI0003834|hsa-mir-769&WithFlank&19|+|46018926|46019055@36.56.21&[MIMAT0003886&hsa-miR-769-5p&offsets|0|-1;m-216&19|+|46018961|46018982&offsets|0|-1]|TGAGACCTCTGGGTTCTGAGC</td><td>>MI0003834|hsa-mir-769&WithFlank&19|+|46018926|46019055</td><td>36</td><td>56</td><td>21</td><td>TGAGACCTCTGGGTTCTGAGC</td><td>495.9</td><td>2.0</td><td>0.9</td><td>2.2</td><td>2.6E-02</td><td>3.2E-02</td><td>MB231/MB468</td><td>MP</td></tr>
<tr><td>>MI0000434|hsa-let-7i&WithFlank&12|+|62603680|62603775@12.33.22(+2A)&[MIMAT0000415&hsa-let-7i-5p&offsets|0|0(+2A);m-49&12|+|62603691|62603712&offsets|0|0(+2A)]|TGAGGTAGTAGTTTGTGCTGTTAA</td><td>>MI0000434|hsa-let-7i&WithFlank&12|+|62603680|62603775</td><td>12</td><td>33</td><td>22(+2A)</td><td>TGAGGTAGTAGTTTGTGCTGTTAA</td><td>108.7</td><td>3.0</td><td>1.3</td><td>2.2</td><td>2.6E-02</td><td>3.2E-02</td><td>MB231/MB468</td><td>MP</td></tr>
<tr><td>>MI0000542|hsa-mir-320a&WithFlank&8|-|22244960|22245043@48.68.21(+1U)&[MIMAT0000510&hsa-miR-320a-3p&offsets|0|-1(+1U);m-60&8|-|22244975|22244996&offsets|0|-1(+1U)]|AAAAGCTGGGTTGAGAGGGCGT</td><td>>MI0000542|hsa-mir-320a&WithFlank&8|-|22244960|22245043</td><td>48</td><td>68</td><td>21(+1U)</td><td>AAAAGCTGGGTTGAGAGGGCGT</td><td>4246.7</td><td>1.8</td><td>0.7</td><td>2.6</td><td>8.6E-03</td><td>3.2E-02</td><td>BT20/MB231</td><td>NUC</td></tr>
<tr><td>>hg38_21_+_8433172_8446622_RNA45SN1_45S_50ntFlanks@6651.6683.33|CGACTCTTAGCGGTGGATCACTCGGCTCGTGCG</td><td>>hg38_21_+_8433172_8446622_RNA45SN1_45S_50ntFlanks</td><td>6651</td><td>6683</td><td>33</td><td>CGACTCTTAGCGGTGGATCACTCGGCTCGTGCG</td><td>24107.4</td><td>-2.0</td><td>0.7</td><td>-2.6</td><td>8.6E-03</td><td>3.2E-02</td><td>BT20/MB231</td><td>NUC</td></tr>
<tr><td>>hg38_21_+_8433172_8446622_RNA45SN1_45S_50ntFlanks@10788.10805.18|CAGCCTCTGGCATGTTGG</td><td>>hg38_21_+_8433172_8446622_RNA45SN1_45S_50ntFlanks</td><td>10788</td><td>10805</td><td>18</td><td>CAGCCTCTGGCATGTTGG</td><td>75.1</td><td>-3.1</td><td>1.3</td><td>-2.3</td><td>1.9E-02</td><td>3.2E-02</td><td>BT20/MB468</td><td>MP</td></tr>
<tr><td>>MI0003643|hsa-mir-629&WithFlank&15|-|70079366|70079474@28.49.22&[MIMAT0004810&hsa-miR-629-5p&offsets|0|+1;m-150&15|-|70079427|70079447&offsets|0|+1]|TGGGTTTACGTTGGGAGAACTT</td><td>>MI0003643|hsa-mir-629&WithFlank&15|-|70079366|70079474</td><td>28</td><td>49</td><td>22</td><td>TGGGTTTACGTTGGGAGAACTT</td><td>2300.4</td><td>1.8</td><td>0.7</td><td>2.6</td><td>9.9E-03</td><td>3.2E-02</td><td>BT20/MB231</td><td>MITO</td></tr>
<tr><td>>MI0000736|hsa-mir-30c-1&WithFlank&1|+|40757278|40757378@23.46.24(+2U);MI0000254|hsa-mir-30c-2&WithFlank&6|-|71376954|71377037@13.36.24(+2U)&[MIMAT0000244&hsa-miR-30c-5p&offsets|0|+1(+2U);m-57&1|+|40757300|40757323&offsets|0|0(+2U)];[MIMAT0000244_1&hsa-miR-30c-5p&offsets|0|+1(+2U);m-56&6|-|71377002|71377025&offsets|0|0(+2U)]|TGTAAACATCCTACACTCTCAGCTTT</td><td>>MI0000736|hsa-mir-30c-1&WithFlank&1|+|40757278|40757378</td><td>23</td><td>46</td><td>24(+2U)</td><td>TGTAAACATCCTACACTCTCAGCTTT</td><td>715.3</td><td>-6.4</td><td>2.5</td><td>-2.6</td><td>9.9E-03</td><td>3.2E-02</td><td>BT20/MB231</td><td>MITO</td></tr>
<tr><td>>hg38_21_+_8433172_8446622_RNA45SN1_45S_50ntFlanks@10772.10796.25|AGGTCTCCAAGGTGAACAGCCTCTG</td><td>>hg38_21_+_8433172_8446622_RNA45SN1_45S_50ntFlanks</td><td>10772</td><td>10796</td><td>25</td><td>AGGTCTCCAAGGTGAACAGCCTCTG</td><td>312.9</td><td>-2.3</td><td>1.0</td><td>-2.3</td><td>1.9E-02</td><td>3.2E-02</td><td>BT20/MB468</td><td>MP</td></tr>
<tr><td>>hg38_MT_+_1621_3279_RNR2_16S_50ntFlanks@885.906.22|AAACATCACCTCTAGCATCACC</td><td>>hg38_MT_+_1621_3279_RNR2_16S_50ntFlanks</td><td>885</td><td>906</td><td>22</td><td>AAACATCACCTCTAGCATCACC</td><td>25.7</td><td>4.6</td><td>2.1</td><td>2.2</td><td>2.6E-02</td><td>3.2E-02</td><td>MB231/MB468</td><td>MP</td></tr>
<tr><td>>MI0000081|hsa-mir-24-2&WithFlank&19|-|13836281|13836365@18.38.21&[MIMAT0004497&hsa-miR-24-2-5p&offsets|-1|-2;m-226&19|-|13836326|13836348&offsets|0|-2]|GTGCCTACTGAGCTGAAACAC</td><td>>MI0000081|hsa-mir-24-2&WithFlank&19|-|13836281|13836365</td><td>18</td><td>38</td><td>21</td><td>GTGCCTACTGAGCTGAAACAC</td><td>100.4</td><td>3.5</td><td>1.5</td><td>2.3</td><td>1.9E-02</td><td>3.2E-02</td><td>BT20/MB468</td><td>MP</td></tr>
<tr><td>>hg38_MT_+_1621_3279_RNR2_16S_50ntFlanks@962.983.22|ACCGTGCAAAGGTAGCATAATC</td><td>>hg38_MT_+_1621_3279_RNR2_16S_50ntFlanks</td><td>962</td><td>983</td><td>22</td><td>ACCGTGCAAAGGTAGCATAATC</td><td>79.4</td><td>4.4</td><td>2.0</td><td>2.2</td><td>2.6E-02</td><td>3.2E-02</td><td>MB231/MB468</td><td>MP</td></tr>
<tr><td>>trnaMT_GluTTC_MT_-_14674_14742@38.68.31,trnalookalike8_GluTTC_5_-_93905172_93905240@38.68.31|CATTGGTCGTGGTTGTAGTCCGTGCGAGAAT</td><td>>trnaMT_GluTTC_MT_-_14674_14742</td><td>38</td><td>68</td><td>31</td><td>CATTGGTCGTGGTTGTAGTCCGTGCGAGAAT</td><td>123.4</td><td>5.3</td><td>2.3</td><td>2.3</td><td>1.9E-02</td><td>3.2E-02</td><td>BT20/MB468</td><td>MP</td></tr>
<tr><td>>hg38_21_+_8433172_8446622_RNA45SN1_45S_50ntFlanks@4735.4765.31|ACGAAAGTCGGAGGTTCGAAGACGATCAGAT</td><td>>hg38_21_+_8433172_8446622_RNA45SN1_45S_50ntFlanks</td><td>4735</td><td>4765</td><td>31</td><td>ACGAAAGTCGGAGGTTCGAAGACGATCAGAT</td><td>59.7</td><td>1.9</td><td>0.9</td><td>2.2</td><td>2.6E-02</td><td>3.2E-02</td><td>MB231/MB468</td><td>MP</td></tr>
<tr><td>>MI0000456|hsa-mir-140&WithFlank&16|+|69933075|69933186@69.89.21(+2A)&[MIMAT0004597&hsa-miR-140-3p&offsets|+1|+1(+2A);m-45&16|+|69933143|69933164&offsets|0|-1(+2A)]|ACCACAGGGTAGAACCACGGAAA</td><td>>MI0000456|hsa-mir-140&WithFlank&16|+|69933075|69933186</td><td>69</td><td>89</td><td>21(+2A)</td><td>ACCACAGGGTAGAACCACGGAAA</td><td>26.6</td><td>4.5</td><td>2.0</td><td>2.2</td><td>2.6E-02</td><td>3.2E-02</td><td>MB231/MB468</td><td>MP</td></tr>
<tr><td>>hg38_21_+_8433172_8446622_RNA45SN1_45S_50ntFlanks@11580.11607.28|GAACTGGTGCGGACCAGGGGAATCCGAC</td><td>>hg38_21_+_8433172_8446622_RNA45SN1_45S_50ntFlanks</td><td>11580</td><td>11607</td><td>28</td><td>GAACTGGTGCGGACCAGGGGAATCCGAC</td><td>392.2</td><td>-1.3</td><td>0.6</td><td>-2.2</td><td>2.8E-02</td><td>3.2E-02</td><td>BT20/MB468</td><td>CYTO</td></tr>
<tr><td>>trnaMT_ValTAC_MT_+_1602_1670@47.72.26|TCAACTTAACTTGACCGCTCTGACCA</td><td>>trnaMT_ValTAC_MT_+_1602_1670</td><td>47</td><td>72</td><td>26</td><td>TCAACTTAACTTGACCGCTCTGACCA</td><td>1077.6</td><td>-2.6</td><td>1.0</td><td>-2.6</td><td>8.7E-03</td><td>3.2E-02</td><td>BT20/MB231</td><td>NUC</td></tr>
<tr><td>>hg38_21_+_8433172_8446622_RNA45SN1_45S_50ntFlanks@6646.6682.37|TCGTACGACTCTTAGCGGTGGATCACTCGGCTCGTGC</td><td>>hg38_21_+_8433172_8446622_RNA45SN1_45S_50ntFlanks</td><td>6646</td><td>6682</td><td>37</td><td>TCGTACGACTCTTAGCGGTGGATCACTCGGCTCGTGC</td><td>148.9</td><td>2.1</td><td>1.0</td><td>2.2</td><td>2.8E-02</td><td>3.2E-02</td><td>BT20/MB468</td><td>NUC</td></tr>
<tr><td>>hg38_21_+_8433172_8446622_RNA45SN1_45S_50ntFlanks@4069.4093.25|CTTTCGATGGTAGTCGCCGTGCCTA</td><td>>hg38_21_+_8433172_8446622_RNA45SN1_45S_50ntFlanks</td><td>4069</td><td>4093</td><td>25</td><td>CTTTCGATGGTAGTCGCCGTGCCTA</td><td>148.3</td><td>-4.0</td><td>1.7</td><td>-2.3</td><td>1.9E-02</td><td>3.2E-02</td><td>BT20/MB468</td><td>MP</td></tr>
<tr><td>>hg38_21_+_8433172_8446622_RNA45SN1_45S_50ntFlanks@10879.10902.24|CGGTCGGGCTGGGGCGCGAAGCGG</td><td>>hg38_21_+_8433172_8446622_RNA45SN1_45S_50ntFlanks</td><td>10879</td><td>10902</td><td>24</td><td>CGGTCGGGCTGGGGCGCGAAGCGG</td><td>209.7</td><td>-2.5</td><td>1.1</td><td>-2.2</td><td>2.8E-02</td><td>3.2E-02</td><td>BT20/MB468</td><td>CYTO</td></tr>
<tr><td>>hg38_21_+_8433172_8446622_RNA45SN1_45S_50ntFlanks@11703.11740.38|AATGAAGCGCGGGTAAACGGCGGGAGTAACTATGACTC</td><td>>hg38_21_+_8433172_8446622_RNA45SN1_45S_50ntFlanks</td><td>11703</td><td>11740</td><td>38</td><td>AATGAAGCGCGGGTAAACGGCGGGAGTAACTATGACTC</td><td>234.0</td><td>-1.6</td><td>0.7</td><td>-2.2</td><td>2.8E-02</td><td>3.2E-02</td><td>BT20/MB468</td><td>CYTO</td></tr>
<tr><td>>MI0000088|hsa-mir-30a&WithFlank&6|-|71403545|71403627@53.73.21&[MIMAT0000088&hsa-miR-30a-3p&offsets|0|-1;m-38&6|-|71403554|71403575&offsets|0|-1]|CTTTCAGTCGGATGTTTGCAG</td><td>>MI0000088|hsa-mir-30a&WithFlank&6|-|71403545|71403627</td><td>53</td><td>73</td><td>21</td><td>CTTTCAGTCGGATGTTTGCAG</td><td>944.9</td><td>-1.8</td><td>0.7</td><td>-2.6</td><td>1.0E-02</td><td>3.2E-02</td><td>BT20/MB231</td><td>MITO</td></tr>
<tr><td>>trna116_GluCTC_1_-_145399233_145399304@37.70.34,trna134_GluTTC_1_-_16861774_16861845@37.70.34,trna71_GluCTC_1_-_161439189_161439260@37.70.34,trna74_GluCTC_1_-_161431809_161431880@37.70.34,trna77_GluCTC_1_-_161424398_161424469@37.70.34,trna77_GluCTC_6_+_28949976_28950047@37.70.34,trna80_GluCTC_1_-_161417018_161417089@37.70.34,trna84_GluTTC_1_-_161391883_161391954@37.70.34,trna87_GluCTC_6_-_126101393_126101464@37.70.34|ACCGCCGCGGCCCGGGTTCGATTCCCGGTCAGGG</td><td>>trna116_GluCTC_1_-_145399233_145399304</td><td>37</td><td>70</td><td>34</td><td>ACCGCCGCGGCCCGGGTTCGATTCCCGGTCAGGG</td><td>68.8</td><td>-3.9</td><td>1.8</td><td>-2.2</td><td>2.6E-02</td><td>3.2E-02</td><td>MB231/MB468</td><td>MP</td></tr>
<tr><td>>hg38_21_+_8433172_8446622_RNA45SN1_45S_50ntFlanks@4105.4125.21|ACGGGTGACGGGGAATCAGGG</td><td>>hg38_21_+_8433172_8446622_RNA45SN1_45S_50ntFlanks</td><td>4105</td><td>4125</td><td>21</td><td>ACGGGTGACGGGGAATCAGGG</td><td>84.8</td><td>3.7</td><td>1.6</td><td>2.3</td><td>1.9E-02</td><td>3.2E-02</td><td>BT20/MB468</td><td>MP</td></tr>
<tr><td>>hg38_21_+_8433172_8446622_RNA45SN1_45S_50ntFlanks@10682.10712.31|TCCGGCGGCGTCCGGTGAGCTCTCGCTGGCC</td><td>>hg38_21_+_8433172_8446622_RNA45SN1_45S_50ntFlanks</td><td>10682</td><td>10712</td><td>31</td><td>TCCGGCGGCGTCCGGTGAGCTCTCGCTGGCC</td><td>458.4</td><td>-1.8</td><td>0.8</td><td>-2.2</td><td>2.8E-02</td><td>3.2E-02</td><td>MB231/MB468</td><td>CYTO</td></tr>
<tr><td>>hg38_21_+_8433172_8446622_RNA45SN1_45S_50ntFlanks@12150.12181.32|GGTGGGGAGTTTGACTGGGGCGGTACACCTGT</td><td>>hg38_21_+_8433172_8446622_RNA45SN1_45S_50ntFlanks</td><td>12150</td><td>12181</td><td>32</td><td>GGTGGGGAGTTTGACTGGGGCGGTACACCTGT</td><td>51.4</td><td>-4.0</td><td>1.7</td><td>-2.3</td><td>1.9E-02</td><td>3.2E-02</td><td>BT20/MB468</td><td>MP</td></tr>
<tr><td>>trnaMT_ProTGG_MT_-_15956_16023@51.70.20|TAAAGACTTTTTCTCTGACC</td><td>>trnaMT_ProTGG_MT_-_15956_16023</td><td>51</td><td>70</td><td>20</td><td>TAAAGACTTTTTCTCTGACC</td><td>163.7</td><td>1.9</td><td>0.8</td><td>2.2</td><td>2.6E-02</td><td>3.2E-02</td><td>MB231/MB468</td><td>MP</td></tr>
<tr><td>>hg38_21_+_8433172_8446622_RNA45SN1_45S_50ntFlanks@9574.9594.21|AATCGGTCGTCCGACCTGGGT</td><td>>hg38_21_+_8433172_8446622_RNA45SN1_45S_50ntFlanks</td><td>9574</td><td>9594</td><td>21</td><td>AATCGGTCGTCCGACCTGGGT</td><td>528.4</td><td>-2.1</td><td>0.9</td><td>-2.2</td><td>2.8E-02</td><td>3.2E-02</td><td>MB231/MB468</td><td>CYTO</td></tr>
<tr><td>>hg38_21_+_8433172_8446622_RNA45SN1_45S_50ntFlanks@10344.10380.37|ATTCAAACGAGAACTTTGAAGGCCGAAGTGGAGAAGG</td><td>>hg38_21_+_8433172_8446622_RNA45SN1_45S_50ntFlanks</td><td>10344</td><td>10380</td><td>37</td><td>ATTCAAACGAGAACTTTGAAGGCCGAAGTGGAGAAGG</td><td>144.0</td><td>4.3</td><td>1.6</td><td>2.7</td><td>8.0E-03</td><td>3.2E-02</td><td>BT20/MB231</td><td>MP</td></tr>
<tr><td>>hg38_21_+_8433172_8446622_RNA45SN1_45S_50ntFlanks@12407.12433.27|TTTTGATCCTTCGATGTCGGCTCTTCC</td><td>>hg38_21_+_8433172_8446622_RNA45SN1_45S_50ntFlanks</td><td>12407</td><td>12433</td><td>27</td><td>TTTTGATCCTTCGATGTCGGCTCTTCC</td><td>1710.5</td><td>-1.2</td><td>0.5</td><td>-2.2</td><td>2.9E-02</td><td>3.2E-02</td><td>MB231/MB468</td><td>NUC</td></tr>
<tr><td>>hg38_MT_+_1621_3279_RNR2_16S_50ntFlanks@1253.1271.19|AAAGCGAACTACTATACTC</td><td>>hg38_MT_+_1621_3279_RNR2_16S_50ntFlanks</td><td>1253</td><td>1271</td><td>19</td><td>AAAGCGAACTACTATACTC</td><td>46.9</td><td>3.7</td><td>1.7</td><td>2.2</td><td>2.6E-02</td><td>3.2E-02</td><td>MB231/MB468</td><td>MP</td></tr>
<tr><td>>MI0000483|hsa-mir-186&WithFlank&1|-|71067625|71067722@21.41.21(+1C)&[MIMAT0000456&hsa-miR-186-5p&offsets|0|-1(+1C);m-84&1|-|71067681|71067702&offsets|0|-1(+1C)]|CAAAGAATTCTCCTTTTGGGCC</td><td>>MI0000483|hsa-mir-186&WithFlank&1|-|71067625|71067722</td><td>21</td><td>41</td><td>21(+1C)</td><td>CAAAGAATTCTCCTTTTGGGCC</td><td>406.6</td><td>-2.6</td><td>1.0</td><td>-2.6</td><td>8.1E-03</td><td>3.2E-02</td><td>BT20/MB231</td><td>MP</td></tr>
<tr><td>>hg38_21_+_8433172_8446622_RNA45SN1_45S_50ntFlanks@9740.9765.26|AACGATCTCAACCTATTCTCAAACTT</td><td>>hg38_21_+_8433172_8446622_RNA45SN1_45S_50ntFlanks</td><td>9740</td><td>9765</td><td>26</td><td>AACGATCTCAACCTATTCTCAAACTT</td><td>378.7</td><td>-1.5</td><td>0.7</td><td>-2.2</td><td>2.9E-02</td><td>3.2E-02</td><td>MB231/MB468</td><td>NUC</td></tr>
<tr><td>>hg38_21_+_8433172_8446622_RNA45SN1_45S_50ntFlanks@11583.11612.30|CTGGTGCGGACCAGGGGAATCCGACTGTTT</td><td>>hg38_21_+_8433172_8446622_RNA45SN1_45S_50ntFlanks</td><td>11583</td><td>11612</td><td>30</td><td>CTGGTGCGGACCAGGGGAATCCGACTGTTT</td><td>1013.0</td><td>-1.3</td><td>0.6</td><td>-2.2</td><td>2.8E-02</td><td>3.2E-02</td><td>BT20/MB468</td><td>CYTO</td></tr>
<tr><td>>hg38_21_+_8433172_8446622_RNA45SN1_45S_50ntFlanks@12417.12451.35|TCGATGTCGGCTCTTCCTATCATTGTGAAGCAGAA</td><td>>hg38_21_+_8433172_8446622_RNA45SN1_45S_50ntFlanks</td><td>12417</td><td>12451</td><td>35</td><td>TCGATGTCGGCTCTTCCTATCATTGTGAAGCAGAA</td><td>140.8</td><td>-2.4</td><td>1.1</td><td>-2.2</td><td>2.8E-02</td><td>3.2E-02</td><td>BT20/MB468</td><td>CYTO</td></tr>
</table>

<table>
<tr><td>>hg38_21_+_8433172_8446622_RNA45SN1_45S_50ntFlanks@6651.6688.38|CGACTCTTAGCGGTGGATCACTCGGCTCGTGCGTCGAT</td><td>>hg38_21_+_8433172_8446622_RNA45SN1_45S_50ntFlanks</td><td>6651</td><td>6688</td><td>38</td><td>CGACTCTTAGCGGTGGATCACTCGGCTCGTGCGTCGAT</td><td>2343.0</td><td>1.3</td><td>0.6</td><td>2.2</td><td>2.8E-02</td><td>3.2E-02</td><td>BT20/MB468</td><td>NUC</td></tr>
<tr><td>>hg38_21_+_8433172_8446622_RNA45SN1_45S_50ntFlanks@9548.9571.24|GAGGTCCGTAGCGGTCCTGACGTG</td><td>>hg38_21_+_8433172_8446622_RNA45SN1_45S_50ntFlanks</td><td>9548</td><td>9571</td><td>24</td><td>GAGGTCCGTAGCGGTCCTGACGTG</td><td>480.4</td><td>-1.8</td><td>0.8</td><td>-2.3</td><td>1.9E-02</td><td>3.2E-02</td><td>BT20/MB468</td><td>MP</td></tr>
<tr><td>>trnaMT_LysTTT_MT_+_8295_8364@1.38.38,trnalookalike7_LysTTT_1_+_568844_568913@1.38.38|CACTGTAAAGCTAACTTAGCATTAACCTTTTAAGTTAA</td><td>>trnaMT_LysTTT_MT_+_8295_8364</td><td>1</td><td>38</td><td>38</td><td>CACTGTAAAGCTAACTTAGCATTAACCTTTTAAGTTAA</td><td>1372.5</td><td>5.0</td><td>2.2</td><td>2.3</td><td>1.9E-02</td><td>3.2E-02</td><td>BT20/MB468</td><td>MP</td></tr>
<tr><td>>hg38_21_+_8433172_8446622_RNA45SN1_45S_50ntFlanks@12405.12438.34|CTTTTTGATCCTTCGATGTCGGCTCTTCCTATCA</td><td>>hg38_21_+_8433172_8446622_RNA45SN1_45S_50ntFlanks</td><td>12405</td><td>12438</td><td>34</td><td>CTTTTTGATCCTTCGATGTCGGCTCTTCCTATCA</td><td>74.0</td><td>-4.2</td><td>1.8</td><td>-2.3</td><td>1.9E-02</td><td>3.2E-02</td><td>BT20/MB468</td><td>MP</td></tr>
<tr><td>>hg38_21_+_8433172_8446622_RNA45SN1_45S_50ntFlanks@9738.9767.30|GAAACGATCTCAACCTATTCTCAAACTTTA</td><td>>hg38_21_+_8433172_8446622_RNA45SN1_45S_50ntFlanks</td><td>9738</td><td>9767</td><td>30</td><td>GAAACGATCTCAACCTATTCTCAAACTTTA</td><td>196.4</td><td>-1.8</td><td>0.8</td><td>-2.2</td><td>2.8E-02</td><td>3.2E-02</td><td>BT20/MB468</td><td>CYTO</td></tr>
<tr><td>>trnaMT_GluTTC_MT_-_14674_14742@32.72.41,trnalookalike8_GluTTC_5_-_93905172_93905240@32.72.41|TCATATCATTGGTCGTGGTTGTAGTCCGTGCGAGAATACCA</td><td>>trnaMT_GluTTC_MT_-_14674_14742</td><td>32</td><td>72</td><td>41</td><td>TCATATCATTGGTCGTGGTTGTAGTCCGTGCGAGAATACCA</td><td>473.2</td><td>-2.0</td><td>0.7</td><td>-2.6</td><td>8.7E-03</td><td>3.2E-02</td><td>BT20/MB231</td><td>NUC</td></tr>
<tr><td>>am_trnaMT_SerGCT_MT_+_12207_12265@13.38.26|AAGAACTGCTAACTCATGCCCCCATG</td><td>>am_trnaMT_SerGCT_MT_+_12207_12265</td><td>13</td><td>38</td><td>26</td><td>AAGAACTGCTAACTCATGCCCCCATG</td><td>266.4</td><td>-2.4</td><td>1.1</td><td>-2.2</td><td>2.8E-02</td><td>3.2E-02</td><td>MB231/MB468</td><td>CYTO</td></tr>
<tr><td>>hg38_MT_+_1621_3279_RNR2_16S_50ntFlanks@1431.1458.28|AAAGTCCTACGTGATCTGAGTTCAGACC</td><td>>hg38_MT_+_1621_3279_RNR2_16S_50ntFlanks</td><td>1431</td><td>1458</td><td>28</td><td>AAAGTCCTACGTGATCTGAGTTCAGACC</td><td>135.6</td><td>2.9</td><td>1.2</td><td>2.3</td><td>1.9E-02</td><td>3.2E-02</td><td>BT20/MB468</td><td>MP</td></tr>
<tr><td>>MI0000542|hsa-mir-320a&WithFlank&8|-|22244960|22245043@48.68.21(+1U)&[MIMAT0000510&hsa-miR-320a-3p&offsets|0|-1(+1U);m-60&8|-|22244975|22244996&offsets|0|-1(+1U)]|AAAAGCTGGGTTGAGAGGGCGT</td><td>>MI0000542|hsa-mir-320a&WithFlank&8|-|22244960|22245043</td><td>48</td><td>68</td><td>21(+1U)</td><td>AAAAGCTGGGTTGAGAGGGCGT</td><td>1803.3</td><td>1.8</td><td>0.7</td><td>2.6</td><td>1.0E-02</td><td>3.2E-02</td><td>BT20/MB231</td><td>MITO</td></tr>
<tr><td>>hg38_21_+_8433172_8446622_RNA45SN1_45S_50ntFlanks@9454.9488.35|CCGCGCCGGGGAGGTGGAGCACGAGCGCACGTGTT</td><td>>hg38_21_+_8433172_8446622_RNA45SN1_45S_50ntFlanks</td><td>9454</td><td>9488</td><td>35</td><td>CCGCGCCGGGGAGGTGGAGCACGAGCGCACGTGTT</td><td>760.6</td><td>-3.5</td><td>1.5</td><td>-2.3</td><td>1.9E-02</td><td>3.2E-02</td><td>BT20/MB468</td><td>MP</td></tr>
<tr><td>>hg38_21_+_8433172_8446622_RNA45SN1_45S_50ntFlanks@9303.9338.36|GACCAAGGAGTCTAACACGTGCGCGAGTCGGGGGCT</td><td>>hg38_21_+_8433172_8446622_RNA45SN1_45S_50ntFlanks</td><td>9303</td><td>9338</td><td>36</td><td>GACCAAGGAGTCTAACACGTGCGCGAGTCGGGGGCT</td><td>214.0</td><td>-4.1</td><td>1.7</td><td>-2.3</td><td>1.9E-02</td><td>3.2E-02</td><td>BT20/MB468</td><td>MP</td></tr>
<tr><td>>hg38_21_+_8433172_8446622_RNA45SN1_45S_50ntFlanks@10682.10712.31|TCCGGCGGCGTCCGGTGAGCTCTCGCTGGCC</td><td>>hg38_21_+_8433172_8446622_RNA45SN1_45S_50ntFlanks</td><td>10682</td><td>10712</td><td>31</td><td>TCCGGCGGCGTCCGGTGAGCTCTCGCTGGCC</td><td>110.7</td><td>-4.0</td><td>1.7</td><td>-2.3</td><td>1.9E-02</td><td>3.2E-02</td><td>BT20/MB468</td><td>MP</td></tr>
<tr><td>>hg38_21_+_8433172_8446622_RNA45SN1_45S_50ntFlanks@10396.10424.29|CAGTTGAACATGGGTCAGTCGGTCCTGAG</td><td>>hg38_21_+_8433172_8446622_RNA45SN1_45S_50ntFlanks</td><td>10396</td><td>10424</td><td>29</td><td>CAGTTGAACATGGGTCAGTCGGTCCTGAG</td><td>14173.8</td><td>3.1</td><td>1.2</td><td>2.6</td><td>8.1E-03</td><td>3.2E-02</td><td>BT20/MB231</td><td>MP</td></tr>
<tr><td>>hg38_21_+_8433172_8446622_RNA45SN1_45S_50ntFlanks@9956.9982.27|GAAGTCGGAATCCGCTAAGGAGTGTGT</td><td>>hg38_21_+_8433172_8446622_RNA45SN1_45S_50ntFlanks</td><td>9956</td><td>9982</td><td>27</td><td>GAAGTCGGAATCCGCTAAGGAGTGTGT</td><td>129.1</td><td>5.2</td><td>2.0</td><td>2.6</td><td>8.1E-03</td><td>3.2E-02</td><td>BT20/MB231</td><td>MP</td></tr>
<tr><td>>hg38_21_+_8433172_8446622_RNA45SN1_45S_50ntFlanks@4297.4314.18|CATTGGAGGGCAAGTCTG</td><td>>hg38_21_+_8433172_8446622_RNA45SN1_45S_50ntFlanks</td><td>4297</td><td>4314</td><td>18</td><td>CATTGGAGGGCAAGTCTG</td><td>479.2</td><td>-2.0</td><td>0.9</td><td>-2.2</td><td>2.8E-02</td><td>3.2E-02</td><td>MB231/MB468</td><td>CYTO</td></tr>
<tr><td>>hg38_21_+_8433172_8446622_RNA45SN1_45S_50ntFlanks@6775.6799.25|CCTCCCGGGGCTACGCCTGTCTGAG</td><td>>hg38_21_+_8433172_8446622_RNA45SN1_45S_50ntFlanks</td><td>6775</td><td>6799</td><td>25</td><td>CCTCCCGGGGCTACGCCTGTCTGAG</td><td>56.3</td><td>-4.0</td><td>1.7</td><td>-2.3</td><td>1.9E-02</td><td>3.2E-02</td><td>BT20/MB468</td><td>MP</td></tr>
<tr><td>>am_MI0039740|hsa-mir-12136&WithFlank&1|-|632609|632691@7.32.26(+1G)&[MIMAT0049032&hsa-miR-12136&offsets|0|+8(+1G)]|GAAAAAGTCATGGAGGCCATGGGGTTG</td><td>>am_MI0039740|hsa-mir-12136&WithFlank&1|-|632609|632691</td><td>7</td><td>32</td><td>26(+1G)</td><td>GAAAAAGTCATGGAGGCCATGGGGTTG</td><td>470.9</td><td>-1.9</td><td>0.8</td><td>-2.2</td><td>2.6E-02</td><td>3.2E-02</td><td>MB231/MB468</td><td>MP</td></tr>
<tr><td>>trnaMT_SerTGA_MT_-_7446_7514@1.27.27,trnalookalike6_SerTGA_1_-_567997_568065@1.27.27|GAAAAAGTCATGGAGGCCATGGGGTTG</td><td>>trnaMT_SerTGA_MT_-_7446_7514</td><td>1</td><td>27</td><td>27</td><td>GAAAAAGTCATGGAGGCCATGGGGTTG</td><td>470.9</td><td>-1.9</td><td>0.8</td><td>-2.2</td><td>2.6E-02</td><td>3.2E-02</td><td>MB231/MB468</td><td>MP</td></tr>
<tr><td>>hg38_21_+_8433172_8446622_RNA45SN1_45S_50ntFlanks@12939.12973.35|CCATTCGTAGACGACCTGCTTCTGGGTCGGGGTTT</td><td>>hg38_21_+_8433172_8446622_RNA45SN1_45S_50ntFlanks</td><td>12939</td><td>12973</td><td>35</td><td>CCATTCGTAGACGACCTGCTTCTGGGTCGGGGTTT</td><td>59.3</td><td>-4.1</td><td>1.7</td><td>-2.3</td><td>1.9E-02</td><td>3.2E-02</td><td>BT20/MB468</td><td>MP</td></tr>
<tr><td>>hg38_21_+_8433172_8446622_RNA45SN1_45S_50ntFlanks@9737.9754.18|CGAAACGATCTCAACCTA</td><td>>hg38_21_+_8433172_8446622_RNA45SN1_45S_50ntFlanks</td><td>9737</td><td>9754</td><td>18</td><td>CGAAACGATCTCAACCTA</td><td>180.4</td><td>-4.0</td><td>1.7</td><td>-2.3</td><td>1.9E-02</td><td>3.2E-02</td><td>BT20/MB468</td><td>MP</td></tr>
<tr><td>>hg38_1_-_228634819_228635039_RNA5S12_5S_50ntFlanks@142.171.30|CGCCTGGGAATACCGGGTGCTGTAGGCTTT</td><td>>hg38_1_-_228634819_228635039_RNA5S12_5S_50ntFlanks</td><td>142</td><td>171</td><td>30</td><td>CGCCTGGGAATACCGGGTGCTGTAGGCTTT</td><td>976.8</td><td>-1.7</td><td>0.7</td><td>-2.6</td><td>1.0E-02</td><td>3.2E-02</td><td>BT20/MB231</td><td>MITO</td></tr>
<tr><td>>MI0000815|hsa-mir-339&WithFlank&7|-|1022927|1023032@56.73.18&[MIMAT0004702&hsa-miR-339-3p&offsets|0|-5;m-235&7|-|1022957|1022977&offsets|0|-3]|TGAGCGCCTCGACGACAG</td><td>>MI0000815|hsa-mir-339&WithFlank&7|-|1022927|1023032</td><td>56</td><td>73</td><td>18</td><td>TGAGCGCCTCGACGACAG</td><td>46.4</td><td>4.5</td><td>2.0</td><td>2.2</td><td>2.6E-02</td><td>3.2E-02</td><td>MB231/MB468</td><td>MP</td></tr>
<tr><td>>hg38_MT_+_1621_3279_RNR2_16S_50ntFlanks@1367.1400.34|TCGATGTTGGATCAGGACATCCCGATGGTGCAGC</td><td>>hg38_MT_+_1621_3279_RNR2_16S_50ntFlanks</td><td>1367</td><td>1400</td><td>34</td><td>TCGATGTTGGATCAGGACATCCCGATGGTGCAGC</td><td>47.8</td><td>3.7</td><td>1.7</td><td>2.2</td><td>2.6E-02</td><td>3.2E-02</td><td>MB231/MB468</td><td>MP</td></tr>
<tr><td>>hg38_21_+_8433172_8446622_RNA45SN1_45S_50ntFlanks@9554.9595.42|CGTAGCGGTCCTGACGTGCAAATCGGTCGTCCGACCTGGGTA</td><td>>hg38_21_+_8433172_8446622_RNA45SN1_45S_50ntFlanks</td><td>9554</td><td>9595</td><td>42</td><td>CGTAGCGGTCCTGACGTGCAAATCGGTCGTCCGACCTGGGTA</td><td>56.4</td><td>-4.3</td><td>1.8</td><td>-2.3</td><td>1.9E-02</td><td>3.2E-02</td><td>BT20/MB468</td><td>MP</td></tr>
<tr><td>>hg38_21_+_8433172_8446622_RNA45SN1_45S_50ntFlanks@9175.9197.23|TCGCGCCGTCGGGCCCGGGGGAG</td><td>>hg38_21_+_8433172_8446622_RNA45SN1_45S_50ntFlanks</td><td>9175</td><td>9197</td><td>23</td><td>TCGCGCCGTCGGGCCCGGGGGAG</td><td>423.9</td><td>-2.2</td><td>1.0</td><td>-2.2</td><td>2.5E-02</td><td>3.2E-02</td><td>BT20/MB468</td><td>MITO</td></tr>
<tr><td>>hg38_MT_+_1621_3279_RNR2_16S_50ntFlanks@1349.1388.40|ACAATAGGGTTTACGACCTCGATGTTGGATCAGGACATCC</td><td>>hg38_MT_+_1621_3279_RNR2_16S_50ntFlanks</td><td>1349</td><td>1388</td><td>40</td><td>ACAATAGGGTTTACGACCTCGATGTTGGATCAGGACATCC</td><td>978.5</td><td>-1.7</td><td>0.7</td><td>-2.6</td><td>1.0E-02</td><td>3.2E-02</td><td>BT20/MB231</td><td>MITO</td></tr>
<tr><td>>hg38_21_+_8433172_8446622_RNA45SN1_45S_50ntFlanks@12999.13029.31|CTGCGATCTATTGAAAGTCAGCCCTCGACAC</td><td>>hg38_21_+_8433172_8446622_RNA45SN1_45S_50ntFlanks</td><td>12999</td><td>13029</td><td>31</td><td>CTGCGATCTATTGAAAGTCAGCCCTCGACAC</td><td>224.2</td><td>-1.6</td><td>0.8</td><td>-2.2</td><td>2.9E-02</td><td>3.2E-02</td><td>MB231/MB468</td><td>NUC</td></tr>
<tr><td>>hg38_21_+_8433172_8446622_RNA45SN1_45S_50ntFlanks@8170.8197.28|CAAGTCCTTCTGATCGAGGCCCAGCCCG</td><td>>hg38_21_+_8433172_8446622_RNA45SN1_45S_50ntFlanks</td><td>8170</td><td>8197</td><td>28</td><td>CAAGTCCTTCTGATCGAGGCCCAGCCCG</td><td>836.4</td><td>-3.9</td><td>1.8</td><td>-2.2</td><td>2.8E-02</td><td>3.2E-02</td><td>BT20/MB468</td><td>CYTO</td></tr>
<tr><td>>trna152_ValCAC_6_-_27248049_27248121@1.28.28|GCTTCTGTAGTGTAGTGGTTATCACGTT</td><td>>trna152_ValCAC_6_-_27248049_27248121</td><td>1</td><td>28</td><td>28</td><td>GCTTCTGTAGTGTAGTGGTTATCACGTT</td><td>137.6</td><td>2.3</td><td>1.0</td><td>2.3</td><td>1.9E-02</td><td>3.2E-02</td><td>BT20/MB468</td><td>MP</td></tr>
<tr><td>>hg38_21_+_8433172_8446622_RNA45SN1_45S_50ntFlanks@12934.12965.32|CTAAACCATTCGTAGACGACCTGCTTCTGGGT</td><td>>hg38_21_+_8433172_8446622_RNA45SN1_45S_50ntFlanks</td><td>12934</td><td>12965</td><td>32</td><td>CTAAACCATTCGTAGACGACCTGCTTCTGGGT</td><td>492.7</td><td>-4.3</td><td>1.9</td><td>-2.3</td><td>1.9E-02</td><td>3.2E-02</td><td>BT20/MB468</td><td>MP</td></tr>
<tr><td>>hg38_MT_+_598_1651_RNR1_12S_50ntFlanks@425.456.32|ACGAAAGTGGCTTTAACATATCTGAACACACA</td><td>>hg38_MT_+_598_1651_RNR1_12S_50ntFlanks</td><td>425</td><td>456</td><td>32</td><td>ACGAAAGTGGCTTTAACATATCTGAACACACA</td><td>103.6</td><td>3.7</td><td>1.7</td><td>2.2</td><td>2.6E-02</td><td>3.2E-02</td><td>MB231/MB468</td><td>MP</td></tr>
<tr><td>>am_trnaMT_ValTAC_MT_+_1602_1670@39.66.28|AGGAGATTTCAACTTAACTTGACCGCTC</td><td>>am_trnaMT_ValTAC_MT_+_1602_1670</td><td>39</td><td>66</td><td>28</td><td>AGGAGATTTCAACTTAACTTGACCGCTC</td><td>47.4</td><td>6.2</td><td>2.6</td><td>2.3</td><td>1.9E-02</td><td>3.2E-02</td><td>BT20/MB468</td><td>MP</td></tr>
<tr><td>>hg38_21_+_8433172_8446622_RNA45SN1_45S_50ntFlanks@10807.10827.21|ACAATGTAGGTAAGGGAAGTC</td><td>>hg38_21_+_8433172_8446622_RNA45SN1_45S_50ntFlanks</td><td>10807</td><td>10827</td><td>21</td><td>ACAATGTAGGTAAGGGAAGTC</td><td>101.3</td><td>-3.9</td><td>1.7</td><td>-2.3</td><td>1.9E-02</td><td>3.2E-02</td><td>BT20/MB468</td><td>MP</td></tr>
<tr><td>>hg38_21_+_8433172_8446622_RNA45SN1_45S_50ntFlanks@12391.12414.24|TCATAGCGACGTCGCTTTTTGATC</td><td>>hg38_21_+_8433172_8446622_RNA45SN1_45S_50ntFlanks</td><td>12391</td><td>12414</td><td>24</td><td>TCATAGCGACGTCGCTTTTTGATC</td><td>85.7</td><td>-3.9</td><td>1.7</td><td>-2.3</td><td>1.9E-02</td><td>3.2E-02</td><td>BT20/MB468</td><td>MP</td></tr>
<tr><td>>hg38_21_+_8433172_8446622_RNA45SN1_45S_50ntFlanks@6659.6689.31|AGCGGTGGATCACTCGGCTCGTGCGTCGATG</td><td>>hg38_21_+_8433172_8446622_RNA45SN1_45S_50ntFlanks</td><td>6659</td><td>6689</td><td>31</td><td>AGCGGTGGATCACTCGGCTCGTGCGTCGATG</td><td>320.3</td><td>4.2</td><td>1.6</td><td>2.6</td><td>8.2E-03</td><td>3.2E-02</td><td>BT20/MB231</td><td>MP</td></tr>
<tr><td>>hg38_21_+_8433172_8446622_RNA45SN1_45S_50ntFlanks@11575.11609.35|ACTTAGAACTGGTGCGGACCAGGGGAATCCGACTG</td><td>>hg38_21_+_8433172_8446622_RNA45SN1_45S_50ntFlanks</td><td>11575</td><td>11609</td><td>35</td><td>ACTTAGAACTGGTGCGGACCAGGGGAATCCGACTG</td><td>197.8</td><td>4.6</td><td>1.8</td><td>2.6</td><td>8.2E-03</td><td>3.2E-02</td><td>BT20/MB231</td><td>MP</td></tr>
<tr><td>>hg38_MT_+_598_1651_RNR1_12S_50ntFlanks@820.840.21|AGGTGGATTTAGCAGTAAACT</td><td>>hg38_MT_+_598_1651_RNR1_12S_50ntFlanks</td><td>820</td><td>840</td><td>21</td><td>AGGTGGATTTAGCAGTAAACT</td><td>263.5</td><td>4.6</td><td>2.0</td><td>2.3</td><td>1.9E-02</td><td>3.2E-02</td><td>BT20/MB468</td><td>MP</td></tr>
<tr><td>>hg38_21_+_8433172_8446622_RNA45SN1_45S_50ntFlanks@8017.8046.30|TATTAGTCAGCGGAGGAGAAGAAACTAACC</td><td>>hg38_21_+_8433172_8446622_RNA45SN1_45S_50ntFlanks</td><td>8017</td><td>8046</td><td>30</td><td>TATTAGTCAGCGGAGGAGAAGAAACTAACC</td><td>471.8</td><td>-4.5</td><td>1.8</td><td>-2.5</td><td>1.1E-02</td><td>3.2E-02</td><td>BT20/MB231</td><td>CYTO</td></tr>
<tr><td>>MI0006444|hsa-mir-1307&WithFlank&10|-|103394247|103394407@87.107.21&[MIMAT0005951&hsa-miR-1307-3p&offsets|+1|0;m-66&10|-|103394300|103394322&offsets|+1|-1]|CTCGGCGTGGCGTCGGTCGTG</td><td>>MI0006444|hsa-mir-1307&WithFlank&10|-|103394247|103394407</td><td>87</td><td>107</td><td>21</td><td>CTCGGCGTGGCGTCGGTCGTG</td><td>225.0</td><td>2.1</td><td>0.9</td><td>2.2</td><td>2.6E-02</td><td>3.2E-02</td><td>MB231/MB468</td><td>MP</td></tr>
<tr><td>>hg38_21_+_8433172_8446622_RNA45SN1_45S_50ntFlanks@11582.11611.30|ACTGGTGCGGACCAGGGGAATCCGACTGTT</td><td>>hg38_21_+_8433172_8446622_RNA45SN1_45S_50ntFlanks</td><td>11582</td><td>11611</td><td>30</td><td>ACTGGTGCGGACCAGGGGAATCCGACTGTT</td><td>1269.4</td><td>-1.8</td><td>0.8</td><td>-2.3</td><td>1.9E-02</td><td>3.2E-02</td><td>BT20/MB468</td><td>MP</td></tr>
<tr><td>>MI0000077|hsa-mir-21&WithFlank&17|+|59841260|59841343@13.34.22(+1U)&[MIMAT0000076&hsa-miR-21-5p&offsets|-1|-1(+1U);m-1&17|+|59841273|59841294&offsets|-1|-1(+1U)]|GTAGCTTATCAGACTGATGTTGT</td><td>>MI0000077|hsa-mir-21&WithFlank&17|+|59841260|59841343</td><td>13</td><td>34</td><td>22(+1U)</td><td>GTAGCTTATCAGACTGATGTTGT</td><td>35.9</td><td>4.5</td><td>2.0</td><td>2.2</td><td>2.6E-02</td><td>3.2E-02</td><td>MB231/MB468</td><td>MP</td></tr>
<tr><td>>hg38_21_+_8433172_8446622_RNA45SN1_45S_50ntFlanks@10385.10424.40|CATGTGAACAGCAGTTGAACATGGGTCAGTCGGTCCTGAG</td><td>>hg38_21_+_8433172_8446622_RNA45SN1_45S_50ntFlanks</td><td>10385</td><td>10424</td><td>40</td><td>CATGTGAACAGCAGTTGAACATGGGTCAGTCGGTCCTGAG</td><td>706.8</td><td>-2.6</td><td>1.0</td><td>-2.5</td><td>1.1E-02</td><td>3.2E-02</td><td>BT20/MB231</td><td>CYTO</td></tr>
<tr><td>>hg38_21_+_8433172_8446622_RNA45SN1_45S_50ntFlanks@3769.3791.23|CGCACGGCCGGTACAGTGAAACT</td><td>>hg38_21_+_8433172_8446622_RNA45SN1_45S_50ntFlanks</td><td>3769</td><td>3791</td><td>23</td><td>CGCACGGCCGGTACAGTGAAACT</td><td>294.0</td><td>-2.2</td><td>0.9</td><td>-2.5</td><td>1.1E-02</td><td>3.2E-02</td><td>BT20/MB231</td><td>CYTO</td></tr>
<tr><td>>hg38_21_+_8433172_8446622_RNA45SN1_45S_50ntFlanks@8196.8224.29|CGTGGACGGTGTGAGGCCGGTAGCGGCCC</td><td>>hg38_21_+_8433172_8446622_RNA45SN1_45S_50ntFlanks</td><td>8196</td><td>8224</td><td>29</td><td>CGTGGACGGTGTGAGGCCGGTAGCGGCCC</td><td>1733.6</td><td>-1.8</td><td>0.8</td><td>-2.2</td><td>2.8E-02</td><td>3.2E-02</td><td>MB231/MB468</td><td>CYTO</td></tr>
<tr><td>>hg38_21_+_8433172_8446622_RNA45SN1_45S_50ntFlanks@10348.10371.24|AAACGAGAACTTTGAAGGCCGAAG</td><td>>hg38_21_+_8433172_8446622_RNA45SN1_45S_50ntFlanks</td><td>10348</td><td>10371</td><td>24</td><td>AAACGAGAACTTTGAAGGCCGAAG</td><td>464.8</td><td>1.7</td><td>0.8</td><td>2.2</td><td>2.8E-02</td><td>3.2E-02</td><td>BT20/MB468</td><td>CYTO</td></tr>
<tr><td>>MI0000085|hsa-mir-27a&WithFlank&19|-|13836434|13836523@57.77.21(+2U)&[MIMAT0000084&hsa-miR-27a-3p&offsets|0|0(+2U);m-43&19|-|13836447|13836467&offsets|0|0(+2U)]|TTCACAGTGGCTAAGTTCCGCTT</td><td>>MI0000085|hsa-mir-27a&WithFlank&19|-|13836434|13836523</td><td>57</td><td>77</td><td>21(+2U)</td><td>TTCACAGTGGCTAAGTTCCGCTT</td><td>57.0</td><td>4.5</td><td>2.0</td><td>2.2</td><td>2.6E-02</td><td>3.2E-02</td><td>MB231/MB468</td><td>MP</td></tr>
<tr><td>>hg38_21_+_8433172_8446622_RNA45SN1_45S_50ntFlanks@476.495.20|TTCGTGATCGATGTGGTGAC</td><td>>hg38_21_+_8433172_8446622_RNA45SN1_45S_50ntFlanks</td><td>476</td><td>495</td><td>20</td><td>TTCGTGATCGATGTGGTGAC</td><td>5317.9</td><td>-2.1</td><td>0.8</td><td>-2.6</td><td>8.9E-03</td><td>3.2E-02</td><td>BT20/MB231</td><td>NUC</td></tr>
<tr><td>>am_trna119_LysCTT_1_-_145395522_145395594@16.33.18,trna11_LysCTT_5_-_180648979_180649051@16.33.18,trna128_LysCTT_1_-_55423542_55423614@16.33.18,trna13_LysCTT_6_+_26556774_26556846@16.33.18,trna6_LysCTT_19_-_52425396_52425468@16.33.18,trna7_LysCTT_16_+_3225692_3225764@16.33.18,trna9_LysCTT_5_+_180634755_180634827@16.33.18|TCGGTAGAGCATGAGACT</td><td>>am_trna119_LysCTT_1_-_145395522_145395594</td><td>16</td><td>33</td><td>18</td><td>TCGGTAGAGCATGAGACT</td><td>312.0</td><td>-4.9</td><td>1.9</td><td>-2.6</td><td>8.9E-03</td><td>3.2E-02</td><td>BT20/MB231</td><td>NUC</td></tr>
</table>

<table>
<tr><th></th><th></th><th></th><th></th><th></th><th></th><th></th><th></th><th></th><th></th><th></th><th></th><th></th><th></th></tr>
<tr><td>>trna10_AspGTC_12_-_125424193_125424264@37.75.39,trna12_AspGTC_12_-_125411891_125411962@37.75.39,trna144_AspGTC_6_-_27551236_27551307@37.75.39,trna38_AspGTC_17_-_8125556_8125627@37.75.39,trna45_AspGTC_6_+_27447453_27447524@37.75.39,trna48_AspGTC_6_+_27471523_27471594@37.75.39,trna4_AspGTC_12_+_96429799_96429870@37.75.39,trna69_AspGTC_1_-_161440205_161440276@37.75.39,trna72_AspGTC_1_-_161432824_161432895@37.75.39,trna75_AspGTC_1_-_161425414_161425485@37.75.39,trna78_AspGTC_1_-_161418033_161418104@37.75.39,trna81_AspGTC_1_-_161410615_161410686@37.75.39|ACGCGGGAGACCGGGGTTCGATTCCCCGACGGGGAGCCA</td><td>>trna10_AspGTC_12_-_125424193_125424264</td><td>37</td><td>75</td><td>39</td><td>ACGCGGGAGACCGGGGTTCGATTCCCCGACGGGGAGCCA</td><td>186.8</td><td>3.3</td><td>1.5</td><td>2.2</td><td>2.6E-02</td><td>3.2E-02</td><td>MB231/MB468</td><td>MP</td></tr>
<tr><td>>hg38_21_+_8433172_8446622_RNA45SN1_45S_50ntFlanks@8339.8375.37|TACCGTAAGGGAAAGTTGAAAAGAACTTTGAAGAGAG</td><td>>hg38_21_+_8433172_8446622_RNA45SN1_45S_50ntFlanks</td><td>8339</td><td>8375</td><td>37</td><td>TACCGTAAGGGAAAGTTGAAAAGAACTTTGAAGAGAG</td><td>3417.2</td><td>4.9</td><td>1.9</td><td>2.6</td><td>8.2E-03</td><td>3.2E-02</td><td>BT20/MB231</td><td>MP</td></tr>
<tr><td>>hg38_21_+_8433172_8446622_RNA45SN1_45S_50ntFlanks@10355.10377.23|AACTTTGAAGGCCGAAGTGGAGA</td><td>>hg38_21_+_8433172_8446622_RNA45SN1_45S_50ntFlanks</td><td>10355</td><td>10377</td><td>23</td><td>AACTTTGAAGGCCGAAGTGGAGA</td><td>129.9</td><td>-4.4</td><td>1.9</td><td>-2.3</td><td>1.9E-02</td><td>3.2E-02</td><td>BT20/MB468</td><td>MP</td></tr>
<tr><td>>MI0000067|hsa-let-7f-1&WithFlank&9|+|94176341|94176439@13.32.20;MI0000068|hsa-let-7f-2&WithFlank&X|-|53557186|53557280@14.33.20&[MIMAT0000067&hsa-let-7f-5p&offsets|0|-2;m-20&9|+|94176353|94176374&offsets|0|-2];[MIMAT0000067_1&hsa-let-7f-5p&offsets|0|-2;m-19&X|-|53557246|53557267&offsets|0|-2]|TGAGGTAGTAGATTGTATAG</td><td>>MI0000067|hsa-let-7f-1&WithFlank&9|+|94176341|94176439</td><td>13</td><td>32</td><td>20</td><td>TGAGGTAGTAGATTGTATAG</td><td>125.4</td><td>4.4</td><td>1.9</td><td>2.3</td><td>1.9E-02</td><td>3.2E-02</td><td>BT20/MB468</td><td>MP</td></tr>
<tr><td>>am_MI0039740|hsa-mir-12136&WithFlank&1|-|632609|632691@7.30.24(+1C)&[MIMAT0049032&hsa-miR-12136&offsets|0|+6(+1C)]|GAAAAAGTCATGGAGGCCATGGGGC</td><td>>am_MI0039740|hsa-mir-12136&WithFlank&1|-|632609|632691</td><td>7</td><td>30</td><td>24(+1C)</td><td>GAAAAAGTCATGGAGGCCATGGGGC</td><td>440.2</td><td>4.2</td><td>1.8</td><td>2.3</td><td>1.9E-02</td><td>3.2E-02</td><td>BT20/MB468</td><td>MP</td></tr>
<tr><td>>hg38_21_+_8433172_8446622_RNA45SN1_45S_50ntFlanks@12073.12094.22|CCGAGGGGCTCTCGCTTCTGGC</td><td>>hg38_21_+_8433172_8446622_RNA45SN1_45S_50ntFlanks</td><td>12073</td><td>12094</td><td>22</td><td>CCGAGGGGCTCTCGCTTCTGGC</td><td>354.0</td><td>-4.2</td><td>1.8</td><td>-2.3</td><td>1.9E-02</td><td>3.2E-02</td><td>BT20/MB468</td><td>MP</td></tr>
<tr><td>>hg38_21_+_8433172_8446622_RNA45SN1_45S_50ntFlanks@10682.10712.31|TCCGGCGGCGTCCGGTGAGCTCTCGCTGGCC</td><td>>hg38_21_+_8433172_8446622_RNA45SN1_45S_50ntFlanks</td><td>10682</td><td>10712</td><td>31</td><td>TCCGGCGGCGTCCGGTGAGCTCTCGCTGGCC</td><td>309.8</td><td>-2.8</td><td>1.2</td><td>-2.2</td><td>2.5E-02</td><td>3.2E-02</td><td>BT20/MB468</td><td>MITO</td></tr>
<tr><td>>hg38_21_+_8433172_8446622_RNA45SN1_45S_50ntFlanks@12893.12932.40|CCTCGCCCGTCACGCACCGCACGTTCGTGGGGAACCTGGC</td><td>>hg38_21_+_8433172_8446622_RNA45SN1_45S_50ntFlanks</td><td>12893</td><td>12932</td><td>40</td><td>CCTCGCCCGTCACGCACCGCACGTTCGTGGGGAACCTGGC</td><td>193.9</td><td>-1.5</td><td>0.7</td><td>-2.2</td><td>2.8E-02</td><td>3.2E-02</td><td>BT20/MB468</td><td>CYTO</td></tr>
<tr><td>>hg38_21_+_8433172_8446622_RNA45SN1_45S_50ntFlanks@9564.9603.40|CTGACGTGCAAATCGGTCGTCCGACCTGGGTATAGGGGCG</td><td>>hg38_21_+_8433172_8446622_RNA45SN1_45S_50ntFlanks</td><td>9564</td><td>9603</td><td>40</td><td>CTGACGTGCAAATCGGTCGTCCGACCTGGGTATAGGGGCG</td><td>141.2</td><td>-3.1</td><td>1.4</td><td>-2.2</td><td>2.7E-02</td><td>3.2E-02</td><td>MB231/MB468</td><td>MP</td></tr>
<tr><td>>hg38_MT_+_598_1651_RNR1_12S_50ntFlanks@606.633.28|CCTGTTCTGTAATCGATAAACCCCGATC</td><td>>hg38_MT_+_598_1651_RNR1_12S_50ntFlanks</td><td>606</td><td>633</td><td>28</td><td>CCTGTTCTGTAATCGATAAACCCCGATC</td><td>215.7</td><td>2.2</td><td>1.0</td><td>2.2</td><td>2.9E-02</td><td>3.2E-02</td><td>MB231/MB468</td><td>MITO</td></tr>
<tr><td>>MI0000542|hsa-mir-320a&WithFlank&8|-|22244960|22245043@50.70.21(+1U)&[MIMAT0000510&hsa-miR-320a-3p&offsets|+2|+1(+1U);m-60&8|-|22244975|22244996&offsets|+2|+1(+1U)]|AAGCTGGGTTGAGAGGGCGAAT</td><td>>MI0000542|hsa-mir-320a&WithFlank&8|-|22244960|22245043</td><td>50</td><td>70</td><td>21(+1U)</td><td>AAGCTGGGTTGAGAGGGCGAAT</td><td>109.8</td><td>3.0</td><td>1.4</td><td>2.2</td><td>2.7E-02</td><td>3.2E-02</td><td>MB231/MB468</td><td>MP</td></tr>
<tr><td>>hg38_21_+_8433172_8446622_RNA45SN1_45S_50ntFlanks@10353.10388.36|AGAACTTTGAAGGCCGAAGTGGAGAAGGGTTCCATG</td><td>>hg38_21_+_8433172_8446622_RNA45SN1_45S_50ntFlanks</td><td>10353</td><td>10388</td><td>36</td><td>AGAACTTTGAAGGCCGAAGTGGAGAAGGGTTCCATG</td><td>455.6</td><td>-2.1</td><td>1.0</td><td>-2.2</td><td>2.8E-02</td><td>3.2E-02</td><td>MB231/MB468</td><td>CYTO</td></tr>
<tr><td>>hg38_21_+_8433172_8446622_RNA45SN1_45S_50ntFlanks@9534.9571.38|AGGAAACTCTGGTGGAGGTCCGTAGCGGTCCTGACGTG</td><td>>hg38_21_+_8433172_8446622_RNA45SN1_45S_50ntFlanks</td><td>9534</td><td>9571</td><td>38</td><td>AGGAAACTCTGGTGGAGGTCCGTAGCGGTCCTGACGTG</td><td>326.0</td><td>2.4</td><td>0.9</td><td>2.6</td><td>1.0E-02</td><td>3.2E-02</td><td>BT20/MB231</td><td>MITO</td></tr>
<tr><td>>MI0000254|hsa-mir-30c-2&WithFlank&6|-|71376954|71377037@53.73.21(+1A)&[MIMAT0004550&hsa-miR-30c-2-3p&offsets|0|-1(+1A);m-206&6|-|71376964|71376985&offsets|0|-1(+1A)]|CTGGGAGAAGGCTGTTTACTCA</td><td>>MI0000254|hsa-mir-30c-2&WithFlank&6|-|71376954|71377037</td><td>53</td><td>73</td><td>21(+1A)</td><td>CTGGGAGAAGGCTGTTTACTCA</td><td>36.1</td><td>4.5</td><td>2.0</td><td>2.2</td><td>2.7E-02</td><td>3.2E-02</td><td>MB231/MB468</td><td>MP</td></tr>
<tr><td>>hg38_21_+_8433172_8446622_RNA45SN1_45S_50ntFlanks@10128.10145.18|CGGCGGCGGCGGCGGCGG</td><td>>hg38_21_+_8433172_8446622_RNA45SN1_45S_50ntFlanks</td><td>10128</td><td>10145</td><td>18</td><td>CGGCGGCGGCGGCGGCGG</td><td>568.2</td><td>-3.8</td><td>1.7</td><td>-2.2</td><td>2.5E-02</td><td>3.2E-02</td><td>BT20/MB468</td><td>MITO</td></tr>
<tr><td>>hg38_21_+_8433172_8446622_RNA45SN1_45S_50ntFlanks@10131.10148.18|CGGCGGCGGCGGCGGCGG</td><td>>hg38_21_+_8433172_8446622_RNA45SN1_45S_50ntFlanks</td><td>10131</td><td>10148</td><td>18</td><td>CGGCGGCGGCGGCGGCGG</td><td>568.2</td><td>-3.8</td><td>1.7</td><td>-2.2</td><td>2.5E-02</td><td>3.2E-02</td><td>BT20/MB468</td><td>MITO</td></tr>
<tr><td>>hg38_21_+_8433172_8446622_RNA45SN1_45S_50ntFlanks@8824.8841.18|CGGCGGCGGCGGCGGCGG</td><td>>hg38_21_+_8433172_8446622_RNA45SN1_45S_50ntFlanks</td><td>8824</td><td>8841</td><td>18</td><td>CGGCGGCGGCGGCGGCGG</td><td>568.2</td><td>-3.8</td><td>1.7</td><td>-2.2</td><td>2.5E-02</td><td>3.2E-02</td><td>BT20/MB468</td><td>MITO</td></tr>
<tr><td>>hg38_21_+_8433172_8446622_RNA45SN1_45S_50ntFlanks@8845.8862.18|CGGCGGCGGCGGCGGCGG</td><td>>hg38_21_+_8433172_8446622_RNA45SN1_45S_50ntFlanks</td><td>8845</td><td>8862</td><td>18</td><td>CGGCGGCGGCGGCGGCGG</td><td>568.2</td><td>-3.8</td><td>1.7</td><td>-2.2</td><td>2.5E-02</td><td>3.2E-02</td><td>BT20/MB468</td><td>MITO</td></tr>
<tr><td>>hg38_21_+_8433172_8446622_RNA45SN1_45S_50ntFlanks@8848.8865.18|CGGCGGCGGCGGCGGCGG</td><td>>hg38_21_+_8433172_8446622_RNA45SN1_45S_50ntFlanks</td><td>8848</td><td>8865</td><td>18</td><td>CGGCGGCGGCGGCGGCGG</td><td>568.2</td><td>-3.8</td><td>1.7</td><td>-2.2</td><td>2.5E-02</td><td>3.2E-02</td><td>BT20/MB468</td><td>MITO</td></tr>
<tr><td>>hg38_21_+_8433172_8446622_RNA45SN1_45S_50ntFlanks@12997.13038.42|CGCTGCGATCTATTGAAAGTCAGCCCTCGACACAAGGGTTTG</td><td>>hg38_21_+_8433172_8446622_RNA45SN1_45S_50ntFlanks</td><td>12997</td><td>13038</td><td>42</td><td>CGCTGCGATCTATTGAAAGTCAGCCCTCGACACAAGGGTTTG</td><td>5207.1</td><td>-2.8</td><td>1.1</td><td>-2.5</td><td>1.1E-02</td><td>3.2E-02</td><td>BT20/MB231</td><td>CYTO</td></tr>
<tr><td>>hg38_21_+_8433172_8446622_RNA45SN1_45S_50ntFlanks@12419.12436.18|GATGTCGGCTCTTCCTAT</td><td>>hg38_21_+_8433172_8446622_RNA45SN1_45S_50ntFlanks</td><td>12419</td><td>12436</td><td>18</td><td>GATGTCGGCTCTTCCTAT</td><td>2970.5</td><td>-1.3</td><td>0.6</td><td>-2.2</td><td>2.8E-02</td><td>3.2E-02</td><td>MB231/MB468</td><td>CYTO</td></tr>
<tr><td>>trnaMT_ValTAC_MT_+_1602_1670@39.72.34|AGGAGATTTCAACTTAACTTGACCGCTCTGACCA</td><td>>trnaMT_ValTAC_MT_+_1602_1670</td><td>39</td><td>72</td><td>34</td><td>AGGAGATTTCAACTTAACTTGACCGCTCTGACCA</td><td>177.6</td><td>2.2</td><td>1.0</td><td>2.2</td><td>2.7E-02</td><td>3.2E-02</td><td>MB231/MB468</td><td>MP</td></tr>
<tr><td>>hg38_21_+_8433172_8446622_RNA45SN1_45S_50ntFlanks@9788.9820.33|CGCTGGCGTGGAGCCGGGCGTGGAATGCGAGTG</td><td>>hg38_21_+_8433172_8446622_RNA45SN1_45S_50ntFlanks</td><td>9788</td><td>9820</td><td>33</td><td>CGCTGGCGTGGAGCCGGGCGTGGAATGCGAGTG</td><td>337.2</td><td>-4.3</td><td>2.0</td><td>-2.2</td><td>2.9E-02</td><td>3.3E-02</td><td>BT20/MB468</td><td>NUC</td></tr>
<tr><td>>hg38_21_+_8433172_8446622_RNA45SN1_45S_50ntFlanks@6655.6681.27|TCTTAGCGGTGGATCACTCGGCTCGTG</td><td>>hg38_21_+_8433172_8446622_RNA45SN1_45S_50ntFlanks</td><td>6655</td><td>6681</td><td>27</td><td>TCTTAGCGGTGGATCACTCGGCTCGTG</td><td>1737.6</td><td>-2.3</td><td>1.0</td><td>-2.3</td><td>2.0E-02</td><td>3.3E-02</td><td>BT20/MB468</td><td>MP</td></tr>
<tr><td>>am_trnaMT_ValTAC_MT_+_1602_1670@48.65.18|CAACTTAACTTGACCGCT</td><td>>am_trnaMT_ValTAC_MT_+_1602_1670</td><td>48</td><td>65</td><td>18</td><td>CAACTTAACTTGACCGCT</td><td>52.3</td><td>6.1</td><td>2.6</td><td>2.3</td><td>2.0E-02</td><td>3.3E-02</td><td>BT20/MB468</td><td>MP</td></tr>
<tr><td>>hg38_21_+_8433172_8446622_RNA45SN1_45S_50ntFlanks@9939.9960.22|GCAGGACGGTGGCCATGGAAGT</td><td>>hg38_21_+_8433172_8446622_RNA45SN1_45S_50ntFlanks</td><td>9939</td><td>9960</td><td>22</td><td>GCAGGACGGTGGCCATGGAAGT</td><td>480.3</td><td>2.8</td><td>1.1</td><td>2.6</td><td>8.3E-03</td><td>3.3E-02</td><td>BT20/MB231</td><td>MP</td></tr>
<tr><td>>MI0003776|hsa-mir-320b-1&WithFlank&1|+|116671740|116671823@48.66.19(+2A);MI0003839|hsa-mir-320b-2&WithFlank&1|-|224257034|224257116@47.65.19(+2A);MI0003778|hsa-mir-320c-1&WithFlank&18|+|21683512|21683595@48.66.19(+2A);MI0008191|hsa-mir-320c-2&WithFlank&18|+|24321669|24321752@48.66.19(+2A);MI0000542|hsa-mir-320a&WithFlank&8|-|22244960|22245043@48.66.19(+2A)&[MIMAT0005792&hsa-miR-320b&offsets|0|-3(+2A);m-183&1|+|116671787|116671806&offsets|0|-1(+2A)];[MIMAT0005792_1&hsa-miR-320b&offsets|0|-3(+2A);m-183&1|-|224257051|224257070&offsets|0|-1(+2A)];[MIMAT0005793&hsa-miR-320c&offsets|0|-1(+2A);m-222&18|+|21683559|21683578&offsets|0|-1(+2A)];[MIMAT0005793_1&hsa-miR-320c&offsets|0|-1(+2A);m-219&18|+|24321716|24321735&offsets|0|-1(+2A)];[MIMAT0000510&hsa-miR-320a-3p&offsets|0|-3(+2A);m-60&8|-|22244975|22244996&offsets|0|-3(+2A)]|AAAAGCTGGGTTGAGAGGGAA</td><td>>MI0003776|hsa-mir-320b-1&WithFlank&1|+|116671740|116671823</td><td>48</td><td>66</td><td>19(+2A)</td><td>AAAAGCTGGGTTGAGAGGGAA</td><td>137.5</td><td>3.0</td><td>1.1</td><td>2.6</td><td>8.3E-03</td><td>3.3E-02</td><td>BT20/MB231</td><td>MP</td></tr>
<tr><td>>hg38_21_+_8433172_8446622_RNA45SN1_45S_50ntFlanks@12894.12930.37|CTCGCCCGTCACGCACCGCACGTTCGTGGGGAACCTG</td><td>>hg38_21_+_8433172_8446622_RNA45SN1_45S_50ntFlanks</td><td>12894</td><td>12930</td><td>37</td><td>CTCGCCCGTCACGCACCGCACGTTCGTGGGGAACCTG</td><td>237.3</td><td>-3.6</td><td>1.6</td><td>-2.2</td><td>2.5E-02</td><td>3.3E-02</td><td>BT20/MB468</td><td>MITO</td></tr>
<tr><td>>MI0000285|hsa-mir-205&WithFlank&1|+|209432127|209432248@40.60.21&[MIMAT0000266&hsa-miR-205-5p&offsets|0|-1;m-351&1|+|209432166|209432187&offsets|0|-1]|TCCTTCATTCCACCGGAGTCT</td><td>>MI0000285|hsa-mir-205&WithFlank&1|+|209432127|209432248</td><td>40</td><td>60</td><td>21</td><td>TCCTTCATTCCACCGGAGTCT</td><td>6481.7</td><td>2.9</td><td>1.3</td><td>2.3</td><td>2.0E-02</td><td>3.3E-02</td><td>BT20/MB468</td><td>MP</td></tr>
<tr><td>>hg38_MT_+_598_1651_RNR1_12S_50ntFlanks@905.929.25|AGTATACTTCAAAGGACATTTAACT</td><td>>hg38_MT_+_598_1651_RNR1_12S_50ntFlanks</td><td>905</td><td>929</td><td>25</td><td>AGTATACTTCAAAGGACATTTAACT</td><td>673.5</td><td>2.9</td><td>1.3</td><td>2.3</td><td>2.0E-02</td><td>3.3E-02</td><td>BT20/MB468</td><td>MP</td></tr>
<tr><td>>MI0000061|hsa-let-7a-2&WithFlank&11|-|122146516|122146599@11.33.23;MI0000062|hsa-let-7a-3&WithFlank&22|+|46112743|46112828@10.32.23;MI0000060|hsa-let-7a-1&WithFlank&9|+|94175951|94176042@12.34.23&[MIMAT0000062&hsa-let-7a-5p&offsets|0|+1;m-6&11|-|122146568|122146589&offsets|0|+1];[MIMAT0000062_1&hsa-let-7a-5p&offsets|0|+1;m-5&22|+|46112752|46112773&offsets|0|+1];[MIMAT0000062_2&hsa-let-7a-5p&offsets|0|+1;m-4&9|+|94175962|94175983&offsets|0|+1]|TGAGGTAGTAGGTTGTATAGTTT</td><td>>MI0000061|hsa-let-7a-2&WithFlank&11|-|122146516|122146599</td><td>11</td><td>33</td><td>23</td><td>TGAGGTAGTAGGTTGTATAGTTT</td><td>3911.7</td><td>-1.8</td><td>0.7</td><td>-2.6</td><td>1.0E-02</td><td>3.3E-02</td><td>BT20/MB231</td><td>MITO</td></tr>
<tr><td>>hg38_21_+_8433172_8446622_RNA45SN1_45S_50ntFlanks@9906.9928.23|AGACCCCAGAAAAGGTGTTGGTT</td><td>>hg38_21_+_8433172_8446622_RNA45SN1_45S_50ntFlanks</td><td>9906</td><td>9928</td><td>23</td><td>AGACCCCAGAAAAGGTGTTGGTT</td><td>370.3</td><td>5.9</td><td>2.3</td><td>2.6</td><td>1.0E-02</td><td>3.3E-02</td><td>BT20/MB231</td><td>MITO</td></tr>
<tr><td>>trna27_GlyCCC_2_-_70476123_70476193@4.34.31,trna34_GlyCCC_16_-_686736_686806@4.34.31|CCGCTGGTGTAGTGGTATCATGCAAGATTCC</td><td>>trna27_GlyCCC_2_-_70476123_70476193</td><td>4</td><td>34</td><td>31</td><td>CCGCTGGTGTAGTGGTATCATGCAAGATTCC</td><td>218.3</td><td>-1.8</td><td>0.7</td><td>-2.6</td><td>1.0E-02</td><td>3.3E-02</td><td>BT20/MB231</td><td>MITO</td></tr>
<tr><td>>MI0000255|hsa-mir-30d&WithFlank&8|-|134804870|134804951@12.34.23(+1G)&[MIMAT0000245&hsa-miR-30d-5p&offsets|0|+1(+1G);m-21&8|-|134804919|134804940&offsets|0|+1(+1G)]|TGTAAACATCCCCGACTGGAAGCG</td><td>>MI0000255|hsa-mir-30d&WithFlank&8|-|134804870|134804951</td><td>12</td><td>34</td><td>23(+1G)</td><td>TGTAAACATCCCCGACTGGAAGCG</td><td>2670.9</td><td>2.4</td><td>0.9</td><td>2.5</td><td>1.1E-02</td><td>3.3E-02</td><td>BT20/MB231</td><td>CYTO</td></tr>
<tr><td>>trnaMT_HisGTG_MT_+_12138_12206@28.62.35|ATTGTGAATCTGACAACAGAGGCTTACGACCCCTT</td><td>>trnaMT_HisGTG_MT_+_12138_12206</td><td>28</td><td>62</td><td>35</td><td>ATTGTGAATCTGACAACAGAGGCTTACGACCCCTT</td><td>124.2</td><td>-4.4</td><td>2.0</td><td>-2.2</td><td>2.8E-02</td><td>3.3E-02</td><td>MB231/MB468</td><td>CYTO</td></tr>
<tr><td>>hg38_21_+_8433172_8446622_RNA45SN1_45S_50ntFlanks@8351.8373.23|AAGTTGAAAAGAACTTTGAAGAG</td><td>>hg38_21_+_8433172_8446622_RNA45SN1_45S_50ntFlanks</td><td>8351</td><td>8373</td><td>23</td><td>AAGTTGAAAAGAACTTTGAAGAG</td><td>2573.2</td><td>4.6</td><td>1.7</td><td>2.6</td><td>8.3E-03</td><td>3.3E-02</td><td>BT20/MB231</td><td>MP</td></tr>
<tr><td>>hg38_21_+_8433172_8446622_RNA45SN1_45S_50ntFlanks@10770.10796.27|GCAGGTCTCCAAGGTGAACAGCCTCTG</td><td>>hg38_21_+_8433172_8446622_RNA45SN1_45S_50ntFlanks</td><td>10770</td><td>10796</td><td>27</td><td>GCAGGTCTCCAAGGTGAACAGCCTCTG</td><td>384.5</td><td>-1.2</td><td>0.5</td><td>-2.2</td><td>2.5E-02</td><td>3.3E-02</td><td>BT20/MB468</td><td>MITO</td></tr>
<tr><td>>hg38_21_+_8433172_8446622_RNA45SN1_45S_50ntFlanks@10873.10904.32|CTGGGTCGGTCGGGCTGGGGCGCGAAGCGGGG</td><td>>hg38_21_+_8433172_8446622_RNA45SN1_45S_50ntFlanks</td><td>10873</td><td>10904</td><td>32</td><td>CTGGGTCGGTCGGGCTGGGGCGCGAAGCGGGG</td><td>165.8</td><td>-4.0</td><td>1.7</td><td>-2.3</td><td>2.0E-02</td><td>3.3E-02</td><td>BT20/MB468</td><td>MP</td></tr>
</table>

<table>
<tr><th></th><th></th><th></th><th></th><th></th><th></th><th></th><th></th><th></th><th></th><th></th><th></th><th></th><th></th></tr>
<tr><td>>am_trna10_ValCAC_5_-_180649395_180649467@5.23.19,trna12_ValAAC_5_-_180645270_180645342@5.23.19,trna132_ValAAC_6_-_27721179_27721251@5.23.19,trna136_ValAAC_6_-_27648885_27648957@5.23.19,trna139_ValAAC_6_-_27618707_27618779@5.23.19,trna18_ValCAC_5_-_180529253_180529325@5.23.19,trna2_ValAAC_3_+_169490018_169490090@5.23.19,trna2_ValCAC_5_+_180524070_180524142@5.23.19,trna37_ValAAC_6_+_27203288_27203360@5.23.19,trna4_ValAAC_5_+_180591154_180591226@5.23.19,trna5_ValAAC_5_+_180596610_180596682@5.23.19,trna6_ValCAC_5_+_180600650_180600722@5.23.19,trna85_ValCAC_1_-_161369490_161369562@5.23.19,trna90_ValCAC_1_-_149684088_149684161@5.23.19,trna98_ValCAC_1_-_149298555_149298627@5.23.19,trna9_ValCAC_6_+_26538282_26538354@5.23.19|CCGTAGTGTAGTGGTTATC</td><td>>am_trna10_ValCAC_5_-_180649395_180649467</td><td>5</td><td>23</td><td>19</td><td>CCGTAGTGTAGTGGTTATC</td><td>159.0</td><td>1.7</td><td>0.8</td><td>2.2</td><td>3.0E-02</td><td>3.3E-02</td><td>MB231/MB468</td><td>MITO</td></tr>
<tr><td>>hg38_21_+_8433172_8446622_RNA45SN1_45S_50ntFlanks@8342.8377.36|CGTAAGGGAAAGTTGAAAAGAACTTTGAAGAGAGAG</td><td>>hg38_21_+_8433172_8446622_RNA45SN1_45S_50ntFlanks</td><td>8342</td><td>8377</td><td>36</td><td>CGTAAGGGAAAGTTGAAAAGAACTTTGAAGAGAGAG</td><td>639.8</td><td>-2.2</td><td>0.9</td><td>-2.5</td><td>1.1E-02</td><td>3.3E-02</td><td>BT20/MB231</td><td>CYTO</td></tr>
<tr><td>>MI0000093|hsa-mir-92a-1&WithFlank&13|+|91351308|91351397@54.74.21(+1G);MI0000094|hsa-mir-92a-2&WithFlank&X|-|134169532|134169618@54.74.21(+1G)&[MIMAT0000092&hsa-miR-92a-3p&offsets|0|-1(+1G);m-9&13|+|91351361|91351382&offsets|0|-1(+1G)];[MIMAT0000092_1&hsa-miR-92a-3p&offsets|0|-1(+1G);m-11&X|-|134169544|134169565&offsets|0|-1(+1G)]|TATTGCACTTGTCCCGGCCTGG</td><td>>MI0000093|hsa-mir-92a-1&WithFlank&13|+|91351308|91351397</td><td>54</td><td>74</td><td>21(+1G)</td><td>TATTGCACTTGTCCCGGCCTGG</td><td>807.4</td><td>2.2</td><td>1.0</td><td>2.2</td><td>2.7E-02</td><td>3.3E-02</td><td>MB231/MB468</td><td>MP</td></tr>
<tr><td>>am_MI0003581|hsa-mir-574&WithFlank&4|+|38868026|38868133@31.50.20&[MIMAT0004795&hsa-miR-574-5p&offsets|0|-3;m-213&4|+|38868056|38868078&offsets|0|-3]|TGAGTGTGTGTGTGTGAGTG</td><td>>am_MI0003581|hsa-mir-574&WithFlank&4|+|38868026|38868133</td><td>31</td><td>50</td><td>20</td><td>TGAGTGTGTGTGTGTGAGTG</td><td>32.2</td><td>4.6</td><td>2.1</td><td>2.2</td><td>2.7E-02</td><td>3.3E-02</td><td>MB231/MB468</td><td>MP</td></tr>
<tr><td>>hg38_21_+_8433172_8446622_RNA45SN1_45S_50ntFlanks@11571.11601.31|GCCGACTTAGAACTGGTGCGGACCAGGGGAA</td><td>>hg38_21_+_8433172_8446622_RNA45SN1_45S_50ntFlanks</td><td>11571</td><td>11601</td><td>31</td><td>GCCGACTTAGAACTGGTGCGGACCAGGGGAA</td><td>510.6</td><td>3.6</td><td>1.4</td><td>2.6</td><td>8.3E-03</td><td>3.3E-02</td><td>BT20/MB231</td><td>MP</td></tr>
<tr><td>>hg38_21_+_8433172_8446622_RNA45SN1_45S_50ntFlanks@5086.5124.39|AACGAGACTCTGGCATGCTAACTAGTTACGCGACCCCCG</td><td>>hg38_21_+_8433172_8446622_RNA45SN1_45S_50ntFlanks</td><td>5086</td><td>5124</td><td>39</td><td>AACGAGACTCTGGCATGCTAACTAGTTACGCGACCCCCG</td><td>159.7</td><td>4.4</td><td>1.7</td><td>2.6</td><td>8.3E-03</td><td>3.3E-02</td><td>BT20/MB231</td><td>MP</td></tr>
<tr><td>>hg38_21_+_8433172_8446622_RNA45SN1_45S_50ntFlanks@11567.11588.22|AGCAGCCGACTTAGAACTGGTG</td><td>>hg38_21_+_8433172_8446622_RNA45SN1_45S_50ntFlanks</td><td>11567</td><td>11588</td><td>22</td><td>AGCAGCCGACTTAGAACTGGTG</td><td>326.4</td><td>1.2</td><td>0.5</td><td>2.2</td><td>2.5E-02</td><td>3.3E-02</td><td>BT20/MB468</td><td>MITO</td></tr>
<tr><td>>hg38_MT_+_598_1651_RNR1_12S_50ntFlanks@420.462.43|AGACTACGAAAGTGGCTTTAACATATCTGAACACACAATAGCT</td><td>>hg38_MT_+_598_1651_RNR1_12S_50ntFlanks</td><td>420</td><td>462</td><td>43</td><td>AGACTACGAAAGTGGCTTTAACATATCTGAACACACAATAGCT</td><td>107.5</td><td>5.2</td><td>2.2</td><td>2.3</td><td>2.0E-02</td><td>3.3E-02</td><td>BT20/MB468</td><td>MP</td></tr>
<tr><td>>hg38_21_+_8433172_8446622_RNA45SN1_45S_50ntFlanks@12475.12497.23|CCCACTAATAGGGAACGTGAGCT</td><td>>hg38_21_+_8433172_8446622_RNA45SN1_45S_50ntFlanks</td><td>12475</td><td>12497</td><td>23</td><td>CCCACTAATAGGGAACGTGAGCT</td><td>138.0</td><td>-2.8</td><td>1.3</td><td>-2.2</td><td>2.7E-02</td><td>3.3E-02</td><td>MB231/MB468</td><td>MP</td></tr>
<tr><td>>hg38_21_+_8433172_8446622_RNA45SN1_45S_50ntFlanks@9534.9568.35|AGGAAACTCTGGTGGAGGTCCGTAGCGGTCCTGAC</td><td>>hg38_21_+_8433172_8446622_RNA45SN1_45S_50ntFlanks</td><td>9534</td><td>9568</td><td>35</td><td>AGGAAACTCTGGTGGAGGTCCGTAGCGGTCCTGAC</td><td>684.6</td><td>-1.9</td><td>0.9</td><td>-2.2</td><td>2.8E-02</td><td>3.3E-02</td><td>MB231/MB468</td><td>CYTO</td></tr>
<tr><td>>hg38_21_+_8433172_8446622_RNA45SN1_45S_50ntFlanks@12457.12488.32|CAAGCGTTGGATTGTTCACCCACTAATAGGGA</td><td>>hg38_21_+_8433172_8446622_RNA45SN1_45S_50ntFlanks</td><td>12457</td><td>12488</td><td>32</td><td>CAAGCGTTGGATTGTTCACCCACTAATAGGGA</td><td>190.2</td><td>-2.6</td><td>1.2</td><td>-2.2</td><td>2.5E-02</td><td>3.3E-02</td><td>BT20/MB468</td><td>MITO</td></tr>
<tr><td>>MI0000298|hsa-mir-221&WithFlank&X|-|45746151|45746272@71.93.23(+2U)&[MIMAT0000278&hsa-miR-221-3p&offsets|0|0(+2U);m-91&X|-|45746181|45746202&offsets|0|+1(+2U)]|AGCTACATTGTCTGCTGGGTTTCTT</td><td>>MI0000298|hsa-mir-221&WithFlank&X|-|45746151|45746272</td><td>71</td><td>93</td><td>23(+2U)</td><td>AGCTACATTGTCTGCTGGGTTTCTT</td><td>217.6</td><td>-1.9</td><td>0.7</td><td>-2.6</td><td>1.0E-02</td><td>3.3E-02</td><td>BT20/MB231</td><td>MITO</td></tr>
<tr><td>>hg38_21_+_8433172_8446622_RNA45SN1_45S_50ntFlanks@8348.8379.32|GGAAAGTTGAAAAGAACTTTGAAGAGAGAGTT</td><td>>hg38_21_+_8433172_8446622_RNA45SN1_45S_50ntFlanks</td><td>8348</td><td>8379</td><td>32</td><td>GGAAAGTTGAAAAGAACTTTGAAGAGAGAGTT</td><td>83.1</td><td>3.2</td><td>1.4</td><td>2.3</td><td>2.0E-02</td><td>3.3E-02</td><td>BT20/MB468</td><td>MP</td></tr>
<tr><td>>hg38_MT_+_598_1651_RNR1_12S_50ntFlanks@428.456.29|AAAGTGGCTTTAACATATCTGAACACACA</td><td>>hg38_MT_+_598_1651_RNR1_12S_50ntFlanks</td><td>428</td><td>456</td><td>29</td><td>AAAGTGGCTTTAACATATCTGAACACACA</td><td>306.5</td><td>3.7</td><td>1.6</td><td>2.3</td><td>2.0E-02</td><td>3.3E-02</td><td>BT20/MB468</td><td>MP</td></tr>
<tr><td>>hg38_21_+_8433172_8446622_RNA45SN1_45S_50ntFlanks@6653.6677.25|ACTCTTAGCGGTGGATCACTCGGCT</td><td>>hg38_21_+_8433172_8446622_RNA45SN1_45S_50ntFlanks</td><td>6653</td><td>6677</td><td>25</td><td>ACTCTTAGCGGTGGATCACTCGGCT</td><td>430.1</td><td>3.1</td><td>1.2</td><td>2.6</td><td>8.3E-03</td><td>3.3E-02</td><td>BT20/MB231</td><td>MP</td></tr>
<tr><td>>hg38_21_+_8433172_8446622_RNA45SN1_45S_50ntFlanks@8347.8378.32|GGGAAAGTTGAAAAGAACTTTGAAGAGAGAGT</td><td>>hg38_21_+_8433172_8446622_RNA45SN1_45S_50ntFlanks</td><td>8347</td><td>8378</td><td>32</td><td>GGGAAAGTTGAAAAGAACTTTGAAGAGAGAGT</td><td>325.9</td><td>2.9</td><td>1.1</td><td>2.6</td><td>8.3E-03</td><td>3.3E-02</td><td>BT20/MB231</td><td>MP</td></tr>
<tr><td>>hg38_21_+_8433172_8446622_RNA45SN1_45S_50ntFlanks@9951.9981.31|CCATGGAAGTCGGAATCCGCTAAGGAGTGTG</td><td>>hg38_21_+_8433172_8446622_RNA45SN1_45S_50ntFlanks</td><td>9951</td><td>9981</td><td>31</td><td>CCATGGAAGTCGGAATCCGCTAAGGAGTGTG</td><td>115.8</td><td>5.3</td><td>2.0</td><td>2.6</td><td>8.3E-03</td><td>3.3E-02</td><td>BT20/MB231</td><td>MP</td></tr>
<tr><td>>hg38_21_+_8433172_8446622_RNA45SN1_45S_50ntFlanks@9457.9488.32|CGCCGGGGAGGTGGAGCACGAGCGCACGTGTT</td><td>>hg38_21_+_8433172_8446622_RNA45SN1_45S_50ntFlanks</td><td>9457</td><td>9488</td><td>32</td><td>CGCCGGGGAGGTGGAGCACGAGCGCACGTGTT</td><td>199.8</td><td>-3.2</td><td>1.4</td><td>-2.3</td><td>2.0E-02</td><td>3.3E-02</td><td>BT20/MB468</td><td>MP</td></tr>
<tr><td>>hg38_21_+_8433172_8446622_RNA45SN1_45S_50ntFlanks@12415.12440.26|CTTCGATGTCGGCTCTTCCTATCATT</td><td>>hg38_21_+_8433172_8446622_RNA45SN1_45S_50ntFlanks</td><td>12415</td><td>12440</td><td>26</td><td>CTTCGATGTCGGCTCTTCCTATCATT</td><td>563.8</td><td>1.6</td><td>0.7</td><td>2.2</td><td>2.5E-02</td><td>3.3E-02</td><td>BT20/MB468</td><td>MITO</td></tr>
<tr><td>>hg38_21_+_8433172_8446622_RNA45SN1_45S_50ntFlanks@12926.12956.31|ACCTGGCGCTAAACCATTCGTAGACGACCTG</td><td>>hg38_21_+_8433172_8446622_RNA45SN1_45S_50ntFlanks</td><td>12926</td><td>12956</td><td>31</td><td>ACCTGGCGCTAAACCATTCGTAGACGACCTG</td><td>167.1</td><td>-4.0</td><td>1.7</td><td>-2.3</td><td>2.0E-02</td><td>3.3E-02</td><td>BT20/MB468</td><td>MP</td></tr>
<tr><td>>hg38_21_+_8433172_8446622_RNA45SN1_45S_50ntFlanks@7992.8016.25|CGTGGCGACCCGCTGAATTTAAGCA</td><td>>hg38_21_+_8433172_8446622_RNA45SN1_45S_50ntFlanks</td><td>7992</td><td>8016</td><td>25</td><td>CGTGGCGACCCGCTGAATTTAAGCA</td><td>763.3</td><td>-1.5</td><td>0.7</td><td>-2.2</td><td>3.0E-02</td><td>3.3E-02</td><td>MB231/MB468</td><td>MITO</td></tr>
<tr><td>>trnaMT_GlnTTG_MT_-_4329_4400@1.21.21,trnalookalike2_GlnTTG_1_-_564879_564950@1.21.21|TAGGATGGGGTGTGATAGGTG</td><td>>trnaMT_GlnTTG_MT_-_4329_4400</td><td>1</td><td>21</td><td>21</td><td>TAGGATGGGGTGTGATAGGTG</td><td>129.2</td><td>1.7</td><td>0.8</td><td>2.2</td><td>2.7E-02</td><td>3.3E-02</td><td>MB231/MB468</td><td>MP</td></tr>
<tr><td>>hg38_21_+_8433172_8446622_RNA45SN1_45S_50ntFlanks@7972.8010.39|AGACGCGACCTCAGATCAGACGTGGCGACCCGCTGAATT</td><td>>hg38_21_+_8433172_8446622_RNA45SN1_45S_50ntFlanks</td><td>7972</td><td>8010</td><td>39</td><td>AGACGCGACCTCAGATCAGACGTGGCGACCCGCTGAATT</td><td>1532.4</td><td>2.7</td><td>1.0</td><td>2.6</td><td>9.0E-03</td><td>3.3E-02</td><td>BT20/MB231</td><td>NUC</td></tr>
<tr><td>>hg38_21_+_8433172_8446622_RNA45SN1_45S_50ntFlanks@11572.11611.40|CCGACTTAGAACTGGTGCGGACCAGGGGAATCCGACTGTT</td><td>>hg38_21_+_8433172_8446622_RNA45SN1_45S_50ntFlanks</td><td>11572</td><td>11611</td><td>40</td><td>CCGACTTAGAACTGGTGCGGACCAGGGGAATCCGACTGTT</td><td>840.5</td><td>-2.6</td><td>1.1</td><td>-2.3</td><td>2.0E-02</td><td>3.3E-02</td><td>BT20/MB468</td><td>MP</td></tr>
<tr><td>>hg38_21_+_8433172_8446622_RNA45SN1_45S_50ntFlanks@10921.10939.19|GCTGGACGAGGCGCCGCCG</td><td>>hg38_21_+_8433172_8446622_RNA45SN1_45S_50ntFlanks</td><td>10921</td><td>10939</td><td>19</td><td>GCTGGACGAGGCGCCGCCG</td><td>221.0</td><td>-2.6</td><td>1.2</td><td>-2.2</td><td>2.9E-02</td><td>3.3E-02</td><td>BT20/MB468</td><td>CYTO</td></tr>
<tr><td>>hg38_21_+_8433172_8446622_RNA45SN1_45S_50ntFlanks@3853.3884.32|AACTGTGGTAATTCTAGAGCTAATACATGCCG</td><td>>hg38_21_+_8433172_8446622_RNA45SN1_45S_50ntFlanks</td><td>3853</td><td>3884</td><td>32</td><td>AACTGTGGTAATTCTAGAGCTAATACATGCCG</td><td>759.2</td><td>2.2</td><td>1.0</td><td>2.2</td><td>2.8E-02</td><td>3.3E-02</td><td>MB231/MB468</td><td>CYTO</td></tr>
<tr><td>>hg38_21_+_8433172_8446622_RNA45SN1_45S_50ntFlanks@9951.9983.33|CCATGGAAGTCGGAATCCGCTAAGGAGTGTGTA</td><td>>hg38_21_+_8433172_8446622_RNA45SN1_45S_50ntFlanks</td><td>9951</td><td>9983</td><td>33</td><td>CCATGGAAGTCGGAATCCGCTAAGGAGTGTGTA</td><td>45.5</td><td>3.4</td><td>1.5</td><td>2.3</td><td>2.0E-02</td><td>3.3E-02</td><td>BT20/MB468</td><td>MP</td></tr>
<tr><td>>trna10_AspGTC_12_-_125424193_125424264@32.70.39,trna12_AspGTC_12_-_125411891_125411962@32.70.39,trna144_AspGTC_6_-_27551236_27551307@32.70.39,trna38_AspGTC_17_-_8125556_8125627@32.70.39,trna45_AspGTC_6_+_27447453_27447524@32.70.39,trna48_AspGTC_6_+_27471523_27471594@32.70.39,trna4_AspGTC_12_+_96429799_96429870@32.70.39,trna69_AspGTC_1_-_161440205_161440276@32.70.39,trna72_AspGTC_1_-_161432824_161432895@32.70.39,trna75_AspGTC_1_-_161425414_161425485@32.70.39,trna78_AspGTC_1_-_161418033_161418104@32.70.39,trna81_AspGTC_1_-_161410615_161410686@32.70.39|CTGTCACGCGGGAGACCGGGGTTCGATTCCCCGACGGGG</td><td>>trna10_AspGTC_12_-_125424193_125424264</td><td>32</td><td>70</td><td>39</td><td>CTGTCACGCGGGAGACCGGGGTTCGATTCCCCGACGGGG</td><td>317.9</td><td>-3.4</td><td>1.5</td><td>-2.3</td><td>2.0E-02</td><td>3.3E-02</td><td>BT20/MB468</td><td>MP</td></tr>
<tr><td>>trnaMT_ValTAC_MT_+_1602_1670@32.72.41|TACACTTAGGAGATTTCAACTTAACTTGACCGCTCTGACCA</td><td>>trnaMT_ValTAC_MT_+_1602_1670</td><td>32</td><td>72</td><td>41</td><td>TACACTTAGGAGATTTCAACTTAACTTGACCGCTCTGACCA</td><td>188.8</td><td>3.3</td><td>1.5</td><td>2.2</td><td>2.7E-02</td><td>3.3E-02</td><td>MB231/MB468</td><td>MP</td></tr>
<tr><td>>hg38_21_+_8433172_8446622_RNA45SN1_45S_50ntFlanks@5496.5525.30|GACGGTCGAACTTGACTATCTAGAGGAAGT</td><td>>hg38_21_+_8433172_8446622_RNA45SN1_45S_50ntFlanks</td><td>5496</td><td>5525</td><td>30</td><td>GACGGTCGAACTTGACTATCTAGAGGAAGT</td><td>129.2</td><td>3.7</td><td>1.6</td><td>2.3</td><td>2.0E-02</td><td>3.3E-02</td><td>BT20/MB468</td><td>MP</td></tr>
<tr><td>>hg38_21_+_8433172_8446622_RNA45SN1_45S_50ntFlanks@4730.4751.22|CAAGAACGAAAGTCGGAGGTTC</td><td>>hg38_21_+_8433172_8446622_RNA45SN1_45S_50ntFlanks</td><td>4730</td><td>4751</td><td>22</td><td>CAAGAACGAAAGTCGGAGGTTC</td><td>460.1</td><td>-1.5</td><td>0.7</td><td>-2.2</td><td>2.8E-02</td><td>3.3E-02</td><td>MB231/MB468</td><td>CYTO</td></tr>
<tr><td>>hg38_21_+_8433172_8446622_RNA45SN1_45S_50ntFlanks@11784.11813.30|ATGAATGGATGAACGAGATTCCCACTGTCC</td><td>>hg38_21_+_8433172_8446622_RNA45SN1_45S_50ntFlanks</td><td>11784</td><td>11813</td><td>30</td><td>ATGAATGGATGAACGAGATTCCCACTGTCC</td><td>66.7</td><td>2.3</td><td>1.0</td><td>2.3</td><td>2.0E-02</td><td>3.3E-02</td><td>BT20/MB468</td><td>MP</td></tr>
<tr><td>>hg38_21_+_8433172_8446622_RNA45SN1_45S_50ntFlanks@8115.8141.27|CGCGGGACATGTGGCGTACGGAAGACC</td><td>>hg38_21_+_8433172_8446622_RNA45SN1_45S_50ntFlanks</td><td>8115</td><td>8141</td><td>27</td><td>CGCGGGACATGTGGCGTACGGAAGACC</td><td>3285.2</td><td>-2.3</td><td>1.0</td><td>-2.3</td><td>2.0E-02</td><td>3.3E-02</td><td>BT20/MB468</td><td>MP</td></tr>
<tr><td>>trna27_GlyCCC_2_-_70476123_70476193@1.29.29,trna34_GlyCCC_16_-_686736_686806@1.29.29|GCGCCGCTGGTGTAGTGGTATCATGCAAG</td><td>>trna27_GlyCCC_2_-_70476123_70476193</td><td>1</td><td>29</td><td>29</td><td>GCGCCGCTGGTGTAGTGGTATCATGCAAG</td><td>163.9</td><td>-2.5</td><td>1.2</td><td>-2.2</td><td>2.9E-02</td><td>3.3E-02</td><td>BT20/MB468</td><td>CYTO</td></tr>
<tr><td>>MI0003205|hsa-mir-532&WithFlank&X|+|50003142|50003244@26.47.22&[MIMAT0002888&hsa-miR-532-5p&offsets|0|0;m-65&X|+|50003167|50003188&offsets|0|0]|CATGCCTTGAGTGTAGGACCGT</td><td>>MI0003205|hsa-mir-532&WithFlank&X|+|50003142|50003244</td><td>26</td><td>47</td><td>22</td><td>CATGCCTTGAGTGTAGGACCGT</td><td>3744.3</td><td>1.7</td><td>0.7</td><td>2.2</td><td>2.7E-02</td><td>3.3E-02</td><td>MB231/MB468</td><td>MP</td></tr>
<tr><td>>MI0001445|hsa-mir-423&WithFlank&17|+|30117073|30117178@59.80.22(+1G)&[MIMAT0001340&hsa-miR-423-3p&offsets|0|-1(+1G);m-100&17|+|30117131|30117153&offsets|0|-1(+1G)]|AGCTCGGTCTGAGGCCCCTCAGG</td><td>>MI0001445|hsa-mir-423&WithFlank&17|+|30117073|30117178</td><td>59</td><td>80</td><td>22(+1G)</td><td>AGCTCGGTCTGAGGCCCCTCAGG</td><td>92.9</td><td>3.8</td><td>1.7</td><td>2.2</td><td>2.7E-02</td><td>3.3E-02</td><td>MB231/MB468</td><td>MP</td></tr>
<tr><td>>MI0000103|hsa-mir-101-1&WithFlank&1|-|65058428|65058514@53.72.20;MI0000739|hsa-mir-101-2&WithFlank&9|+|4850291|4850381@55.74.20&[MIMAT0000099&hsa-miR-101-3p&offsets|0|-1;m-17&1|-|65058443|65058462&offsets|0|0];[MIMAT0000099_1&hsa-miR-101-3p&offsets|0|-1;m-16&9|+|4850345|4850364&offsets|0|0]|TACAGTACTGTGATAACTGA</td><td>>MI0000103|hsa-mir-101-1&WithFlank&1|-|65058428|65058514</td><td>53</td><td>72</td><td>20</td><td>TACAGTACTGTGATAACTGA</td><td>1536.3</td><td>2.7</td><td>1.2</td><td>2.3</td><td>2.0E-02</td><td>3.3E-02</td><td>BT20/MB468</td><td>MP</td></tr>
<tr><td>>am_MI0000786|hsa-mir-378a&WithFlank&5|+|149732819|149732896@49.70.22(+2A)&[MIMAT0000732&hsa-miR-378a-3p&offsets|0|0(+2A);m-44&5|+|149732867|149732888&offsets|0|0(+2A)]|ACTGGACTTGGAGTCAGAAGGCAA</td><td>>am_MI0000786|hsa-mir-378a&WithFlank&5|+|149732819|149732896</td><td>49</td><td>70</td><td>22(+2A)</td><td>ACTGGACTTGGAGTCAGAAGGCAA</td><td>196.8</td><td>1.9</td><td>0.8</td><td>2.5</td><td>1.1E-02</td><td>3.3E-02</td><td>BT20/MB231</td><td>CYTO</td></tr>
</table>

<table>
<tr><th></th><th></th><th></th><th></th><th></th><th></th><th></th><th></th><th></th><th></th><th></th><th></th><th></th><th></th></tr>
<tr><td>>hg38_21_+_8433172_8446622_RNA45SN1_45S_50ntFlanks@12417.12446.30|TCGATGTCGGCTCTTCCTATCATTGTGAAG</td><td>>hg38_21_+_8433172_8446622_RNA45SN1_45S_50ntFlanks</td><td>12417</td><td>12446</td><td>30</td><td>TCGATGTCGGCTCTTCCTATCATTGTGAAG</td><td>926.0</td><td>-2.4</td><td>1.0</td><td>-2.5</td><td>1.1E-02</td><td>3.3E-02</td><td>BT20/MB231</td><td>CYTO</td></tr>
<tr><td>>hg38_21_+_8433172_8446622_RNA45SN1_45S_50ntFlanks@4684.4722.39|AAGACGGACCAGAGCGAAAGCATTTGCCAAGAATGTTTT</td><td>>hg38_21_+_8433172_8446622_RNA45SN1_45S_50ntFlanks</td><td>4684</td><td>4722</td><td>39</td><td>AAGACGGACCAGAGCGAAAGCATTTGCCAAGAATGTTTT</td><td>60.9</td><td>-4.8</td><td>2.2</td><td>-2.2</td><td>2.7E-02</td><td>3.3E-02</td><td>MB231/MB468</td><td>MP</td></tr>
<tr><td>>hg38_21_+_8433172_8446622_RNA45SN1_45S_50ntFlanks@12410.12434.25|TGATCCTTCGATGTCGGCTCTTCCT</td><td>>hg38_21_+_8433172_8446622_RNA45SN1_45S_50ntFlanks</td><td>12410</td><td>12434</td><td>25</td><td>TGATCCTTCGATGTCGGCTCTTCCT</td><td>1872.7</td><td>3.5</td><td>1.3</td><td>2.6</td><td>9.1E-03</td><td>3.3E-02</td><td>BT20/MB231</td><td>NUC</td></tr>
<tr><td>>MI0000298|hsa-mir-221&WithFlank&X|-|45746151|45746272@31.53.23&[MIMAT0004568&hsa-miR-221-5p&offsets|0|+1;m-405&X|-|45746220|45746242&offsets|0|0]|ACCTGGCATACAATGTAGATTTC</td><td>>MI0000298|hsa-mir-221&WithFlank&X|-|45746151|45746272</td><td>31</td><td>53</td><td>23</td><td>ACCTGGCATACAATGTAGATTTC</td><td>554.7</td><td>-1.5</td><td>0.6</td><td>-2.6</td><td>9.1E-03</td><td>3.3E-02</td><td>BT20/MB231</td><td>NUC</td></tr>
<tr><td>>hg38_21_+_8433172_8446622_RNA45SN1_45S_50ntFlanks@11672.11707.36|CAGTGCTCTGAATGTCAAAGTGAAGAAATTCAATGA</td><td>>hg38_21_+_8433172_8446622_RNA45SN1_45S_50ntFlanks</td><td>11672</td><td>11707</td><td>36</td><td>CAGTGCTCTGAATGTCAAAGTGAAGAAATTCAATGA</td><td>713.1</td><td>-2.2</td><td>0.9</td><td>-2.5</td><td>1.1E-02</td><td>3.3E-02</td><td>BT20/MB231</td><td>CYTO</td></tr>
<tr><td>>hg38_1_-_228634819_228635039_RNA5S12_5S_50ntFlanks@152.171.20|TACCGGGTGCTGTAGGCTTT</td><td>>hg38_1_-_228634819_228635039_RNA5S12_5S_50ntFlanks</td><td>152</td><td>171</td><td>20</td><td>TACCGGGTGCTGTAGGCTTT</td><td>1570.7</td><td>-1.5</td><td>0.7</td><td>-2.2</td><td>3.0E-02</td><td>3.3E-02</td><td>MB231/MB468</td><td>MITO</td></tr>
<tr><td>>hg38_21_+_8433172_8446622_RNA45SN1_45S_50ntFlanks@12479.12496.18|CTAATAGGGAACGTGAGC</td><td>>hg38_21_+_8433172_8446622_RNA45SN1_45S_50ntFlanks</td><td>12479</td><td>12496</td><td>18</td><td>CTAATAGGGAACGTGAGC</td><td>285.8</td><td>-1.7</td><td>0.8</td><td>-2.2</td><td>3.0E-02</td><td>3.3E-02</td><td>MB231/MB468</td><td>MITO</td></tr>
<tr><td>>hg38_1_-_228634819_228635039_RNA5S12_5S_50ntFlanks@148.171.24|GGAATACCGGGTGCTGTAGGCTTT</td><td>>hg38_1_-_228634819_228635039_RNA5S12_5S_50ntFlanks</td><td>148</td><td>171</td><td>24</td><td>GGAATACCGGGTGCTGTAGGCTTT</td><td>1530.0</td><td>1.9</td><td>0.7</td><td>2.6</td><td>1.0E-02</td><td>3.3E-02</td><td>BT20/MB231</td><td>MITO</td></tr>
<tr><td>>hg38_21_+_8433172_8446622_RNA45SN1_45S_50ntFlanks@9266.9285.20|CGACGTCGGCTACCCACCCG</td><td>>hg38_21_+_8433172_8446622_RNA45SN1_45S_50ntFlanks</td><td>9266</td><td>9285</td><td>20</td><td>CGACGTCGGCTACCCACCCG</td><td>408.6</td><td>-2.6</td><td>1.2</td><td>-2.2</td><td>2.9E-02</td><td>3.3E-02</td><td>BT20/MB468</td><td>CYTO</td></tr>
<tr><td>>hg38_21_+_8433172_8446622_RNA45SN1_45S_50ntFlanks@4067.4101.35|AACTTTCGATGGTAGTCGCCGTGCCTACCATGGTG</td><td>>hg38_21_+_8433172_8446622_RNA45SN1_45S_50ntFlanks</td><td>4067</td><td>4101</td><td>35</td><td>AACTTTCGATGGTAGTCGCCGTGCCTACCATGGTG</td><td>182.0</td><td>1.5</td><td>0.7</td><td>2.2</td><td>2.7E-02</td><td>3.3E-02</td><td>MB231/MB468</td><td>MP</td></tr>
<tr><td>>hg38_21_+_8433172_8446622_RNA45SN1_45S_50ntFlanks@10820.10838.19|GGGAAGTCGGCAAGCCGGA</td><td>>hg38_21_+_8433172_8446622_RNA45SN1_45S_50ntFlanks</td><td>10820</td><td>10838</td><td>19</td><td>GGGAAGTCGGCAAGCCGGA</td><td>202.5</td><td>-2.0</td><td>0.9</td><td>-2.3</td><td>2.0E-02</td><td>3.3E-02</td><td>BT20/MB468</td><td>MP</td></tr>
<tr><td>>hg38_21_+_8433172_8446622_RNA45SN1_45S_50ntFlanks@12420.12455.36|ATGTCGGCTCTTCCTATCATTGTGAAGCAGAATTCA</td><td>>hg38_21_+_8433172_8446622_RNA45SN1_45S_50ntFlanks</td><td>12420</td><td>12455</td><td>36</td><td>ATGTCGGCTCTTCCTATCATTGTGAAGCAGAATTCA</td><td>836.5</td><td>-4.4</td><td>1.9</td><td>-2.3</td><td>2.0E-02</td><td>3.3E-02</td><td>BT20/MB468</td><td>MP</td></tr>
<tr><td>>hg38_21_+_8433172_8446622_RNA45SN1_45S_50ntFlanks@9454.9494.41|CCGCGCCGGGGAGGTGGAGCACGAGCGCACGTGTTAGGACC</td><td>>hg38_21_+_8433172_8446622_RNA45SN1_45S_50ntFlanks</td><td>9454</td><td>9494</td><td>41</td><td>CCGCGCCGGGGAGGTGGAGCACGAGCGCACGTGTTAGGACC</td><td>358.7</td><td>-4.3</td><td>1.8</td><td>-2.3</td><td>2.0E-02</td><td>3.3E-02</td><td>BT20/MB468</td><td>MP</td></tr>
<tr><td>>hg38_21_+_8433172_8446622_RNA45SN1_45S_50ntFlanks@9860.9893.34|GGATGAACCGAACGCCGGGTTAAGGCGCCCGATG</td><td>>hg38_21_+_8433172_8446622_RNA45SN1_45S_50ntFlanks</td><td>9860</td><td>9893</td><td>34</td><td>GGATGAACCGAACGCCGGGTTAAGGCGCCCGATG</td><td>134.9</td><td>-2.6</td><td>1.1</td><td>-2.3</td><td>2.0E-02</td><td>3.3E-02</td><td>BT20/MB468</td><td>MP</td></tr>
<tr><td>>hg38_MT_+_598_1651_RNR1_12S_50ntFlanks@981.1001.21|ACTGGAAAGTGCACTTGGACG</td><td>>hg38_MT_+_598_1651_RNR1_12S_50ntFlanks</td><td>981</td><td>1001</td><td>21</td><td>ACTGGAAAGTGCACTTGGACG</td><td>99.3</td><td>4.2</td><td>1.8</td><td>2.3</td><td>2.0E-02</td><td>3.3E-02</td><td>BT20/MB468</td><td>MP</td></tr>
<tr><td>>hg38_21_+_8433172_8446622_RNA45SN1_45S_50ntFlanks@11575.11611.37|ACTTAGAACTGGTGCGGACCAGGGGAATCCGACTGTT</td><td>>hg38_21_+_8433172_8446622_RNA45SN1_45S_50ntFlanks</td><td>11575</td><td>11611</td><td>37</td><td>ACTTAGAACTGGTGCGGACCAGGGGAATCCGACTGTT</td><td>1427.2</td><td>-1.6</td><td>0.7</td><td>-2.2</td><td>2.5E-02</td><td>3.3E-02</td><td>BT20/MB468</td><td>MITO</td></tr>
<tr><td>>hg38_21_+_8433172_8446622_RNA45SN1_45S_50ntFlanks@11572.11599.28|CCGACTTAGAACTGGTGCGGACCAGGGG</td><td>>hg38_21_+_8433172_8446622_RNA45SN1_45S_50ntFlanks</td><td>11572</td><td>11599</td><td>28</td><td>CCGACTTAGAACTGGTGCGGACCAGGGG</td><td>733.0</td><td>-1.8</td><td>0.8</td><td>-2.3</td><td>2.0E-02</td><td>3.3E-02</td><td>BT20/MB468</td><td>MP</td></tr>
<tr><td>>hg38_21_+_8433172_8446622_RNA45SN1_45S_50ntFlanks@8358.8392.35|AAAGAACTTTGAAGAGAGAGTTCAAGAGGGCGTGA</td><td>>hg38_21_+_8433172_8446622_RNA45SN1_45S_50ntFlanks</td><td>8358</td><td>8392</td><td>35</td><td>AAAGAACTTTGAAGAGAGAGTTCAAGAGGGCGTGA</td><td>114.8</td><td>5.2</td><td>2.0</td><td>2.6</td><td>8.4E-03</td><td>3.3E-02</td><td>BT20/MB231</td><td>MP</td></tr>
<tr><td>>hg38_21_+_8433172_8446622_RNA45SN1_45S_50ntFlanks@8117.8138.22|CGGGACATGTGGCGTACGGAAG</td><td>>hg38_21_+_8433172_8446622_RNA45SN1_45S_50ntFlanks</td><td>8117</td><td>8138</td><td>22</td><td>CGGGACATGTGGCGTACGGAAG</td><td>856.7</td><td>-2.1</td><td>0.9</td><td>-2.3</td><td>2.0E-02</td><td>3.3E-02</td><td>BT20/MB468</td><td>MP</td></tr>
<tr><td>>hg38_MT_+_598_1651_RNR1_12S_50ntFlanks@719.744.26|TAAAGACGTTAGGTCAAGGTGTAGCC</td><td>>hg38_MT_+_598_1651_RNR1_12S_50ntFlanks</td><td>719</td><td>744</td><td>26</td><td>TAAAGACGTTAGGTCAAGGTGTAGCC</td><td>385.6</td><td>2.1</td><td>1.0</td><td>2.2</td><td>3.0E-02</td><td>3.3E-02</td><td>MB231/MB468</td><td>MITO</td></tr>
<tr><td>>MI0000088|hsa-mir-30a&WithFlank&6|-|71403545|71403627@12.33.22(+2A)&[MIMAT0000087&hsa-miR-30a-5p&offsets|0|0(+2A);m-15&6|-|71403595|71403616&offsets|0|0(+2A)]|TGTAAACATCCTCGACTGGAAGAA</td><td>>MI0000088|hsa-mir-30a&WithFlank&6|-|71403545|71403627</td><td>12</td><td>33</td><td>22(+2A)</td><td>TGTAAACATCCTCGACTGGAAGAA</td><td>137.9</td><td>-4.9</td><td>1.9</td><td>-2.6</td><td>8.4E-03</td><td>3.3E-02</td><td>BT20/MB231</td><td>MP</td></tr>
<tr><td>>hg38_21_+_8433172_8446622_RNA45SN1_45S_50ntFlanks@4734.4761.28|AACGAAAGTCGGAGGTTCGAAGACGATC</td><td>>hg38_21_+_8433172_8446622_RNA45SN1_45S_50ntFlanks</td><td>4734</td><td>4761</td><td>28</td><td>AACGAAAGTCGGAGGTTCGAAGACGATC</td><td>2669.8</td><td>3.6</td><td>1.4</td><td>2.6</td><td>8.4E-03</td><td>3.3E-02</td><td>BT20/MB231</td><td>MP</td></tr>
<tr><td>>MI0000115|hsa-mir-16-2&WithFlank&3|+|160404739|160404831@58.79.22(+1U)&[MIMAT0004518&hsa-miR-16-2-3p&offsets|-1|-1(+1U);m-252&3|+|160404796|160404816&offsets|0|+1(+1U)]|ACCAATATTACTGTGCTGCTTTT</td><td>>MI0000115|hsa-mir-16-2&WithFlank&3|+|160404739|160404831</td><td>58</td><td>79</td><td>22(+1U)</td><td>ACCAATATTACTGTGCTGCTTTT</td><td>204.1</td><td>-2.1</td><td>0.8</td><td>-2.6</td><td>1.0E-02</td><td>3.3E-02</td><td>BT20/MB231</td><td>MITO</td></tr>
<tr><td>>hg38_21_+_8433172_8446622_RNA45SN1_45S_50ntFlanks@11208.11235.28|GCGGCGACTCTGGACGCGAGCCGGGCCC</td><td>>hg38_21_+_8433172_8446622_RNA45SN1_45S_50ntFlanks</td><td>11208</td><td>11235</td><td>28</td><td>GCGGCGACTCTGGACGCGAGCCGGGCCC</td><td>280.0</td><td>-1.7</td><td>0.8</td><td>-2.2</td><td>2.9E-02</td><td>3.3E-02</td><td>BT20/MB468</td><td>CYTO</td></tr>
<tr><td>>hg38_21_+_8433172_8446622_RNA45SN1_45S_50ntFlanks@8836.8866.31|CGGCGGTGGCGGCGGCGGCGGCGGCGGCGGG</td><td>>hg38_21_+_8433172_8446622_RNA45SN1_45S_50ntFlanks</td><td>8836</td><td>8866</td><td>31</td><td>CGGCGGTGGCGGCGGCGGCGGCGGCGGCGGG</td><td>59.8</td><td>-5.2</td><td>2.2</td><td>-2.3</td><td>2.0E-02</td><td>3.3E-02</td><td>BT20/MB468</td><td>MP</td></tr>
<tr><td>>hg38_21_+_8433172_8446622_RNA45SN1_45S_50ntFlanks@9573.9603.31|AAATCGGTCGTCCGACCTGGGTATAGGGGCG</td><td>>hg38_21_+_8433172_8446622_RNA45SN1_45S_50ntFlanks</td><td>9573</td><td>9603</td><td>31</td><td>AAATCGGTCGTCCGACCTGGGTATAGGGGCG</td><td>504.5</td><td>-1.2</td><td>0.6</td><td>-2.2</td><td>2.9E-02</td><td>3.3E-02</td><td>BT20/MB468</td><td>CYTO</td></tr>
<tr><td>>MI0000736|hsa-mir-30c-1&WithFlank&1|+|40757278|40757378@23.45.23;MI0000254|hsa-mir-30c-2&WithFlank&6|-|71376954|71377037@13.35.23&[MIMAT0000244&hsa-miR-30c-5p&offsets|0|0;m-57&1|+|40757300|40757323&offsets|0|-1];[MIMAT0000244_1&hsa-miR-30c-5p&offsets|0|0;m-56&6|-|71377002|71377025&offsets|0|-1]|TGTAAACATCCTACACTCTCAGC</td><td>>MI0000736|hsa-mir-30c-1&WithFlank&1|+|40757278|40757378</td><td>23</td><td>45</td><td>23</td><td>TGTAAACATCCTACACTCTCAGC</td><td>1778.9</td><td>2.2</td><td>0.9</td><td>2.3</td><td>2.0E-02</td><td>3.3E-02</td><td>BT20/MB468</td><td>MP</td></tr>
<tr><td>>hg38_21_+_8433172_8446622_RNA45SN1_45S_50ntFlanks@6651.6675.25|CGACTCTTAGCGGTGGATCACTCGG</td><td>>hg38_21_+_8433172_8446622_RNA45SN1_45S_50ntFlanks</td><td>6651</td><td>6675</td><td>25</td><td>CGACTCTTAGCGGTGGATCACTCGG</td><td>5511.4</td><td>2.4</td><td>1.1</td><td>2.2</td><td>2.9E-02</td><td>3.3E-02</td><td>BT20/MB468</td><td>CYTO</td></tr>
<tr><td>>hg38_MT_+_1621_3279_RNR2_16S_50ntFlanks@1428.1452.25|ATTAAAGTCCTACGTGATCTGAGTT</td><td>>hg38_MT_+_1621_3279_RNR2_16S_50ntFlanks</td><td>1428</td><td>1452</td><td>25</td><td>ATTAAAGTCCTACGTGATCTGAGTT</td><td>335.3</td><td>2.3</td><td>1.1</td><td>2.2</td><td>2.7E-02</td><td>3.3E-02</td><td>MB231/MB468</td><td>MP</td></tr>
<tr><td>>hg38_MT_+_598_1651_RNR1_12S_50ntFlanks@558.580.23|GCTTAAAACTCAAAGGACCTGGC</td><td>>hg38_MT_+_598_1651_RNR1_12S_50ntFlanks</td><td>558</td><td>580</td><td>23</td><td>GCTTAAAACTCAAAGGACCTGGC</td><td>61.2</td><td>3.9</td><td>1.8</td><td>2.2</td><td>2.7E-02</td><td>3.3E-02</td><td>MB231/MB468</td><td>MP</td></tr>
<tr><td>>hg38_21_+_8433172_8446622_RNA45SN1_45S_50ntFlanks@10796.10817.22|GGCATGTTGGAACAATGTAGGT</td><td>>hg38_21_+_8433172_8446622_RNA45SN1_45S_50ntFlanks</td><td>10796</td><td>10817</td><td>22</td><td>GGCATGTTGGAACAATGTAGGT</td><td>193.8</td><td>-2.0</td><td>0.8</td><td>-2.3</td><td>2.0E-02</td><td>3.3E-02</td><td>BT20/MB468</td><td>MP</td></tr>
<tr><td>>hg38_21_+_8433172_8446622_RNA45SN1_45S_50ntFlanks@3781.3801.21|ACAGTGAAACTGCGAATGGCT</td><td>>hg38_21_+_8433172_8446622_RNA45SN1_45S_50ntFlanks</td><td>3781</td><td>3801</td><td>21</td><td>ACAGTGAAACTGCGAATGGCT</td><td>268.9</td><td>-2.9</td><td>1.3</td><td>-2.2</td><td>2.5E-02</td><td>3.3E-02</td><td>BT20/MB468</td><td>MITO</td></tr>
<tr><td>>hg38_21_+_8433172_8446622_RNA45SN1_45S_50ntFlanks@10677.10706.30|GCGGTTCCGGCGGCGTCCGGTGAGCTCTCG</td><td>>hg38_21_+_8433172_8446622_RNA45SN1_45S_50ntFlanks</td><td>10677</td><td>10706</td><td>30</td><td>GCGGTTCCGGCGGCGTCCGGTGAGCTCTCG</td><td>217.0</td><td>-2.7</td><td>1.3</td><td>-2.2</td><td>3.0E-02</td><td>3.3E-02</td><td>MB231/MB468</td><td>NUC</td></tr>
<tr><td>>hg38_21_+_8433172_8446622_RNA45SN1_45S_50ntFlanks@9485.9506.22|TGTTAGGACCCGAAAGATGGTG</td><td>>hg38_21_+_8433172_8446622_RNA45SN1_45S_50ntFlanks</td><td>9485</td><td>9506</td><td>22</td><td>TGTTAGGACCCGAAAGATGGTG</td><td>122.4</td><td>5.2</td><td>2.0</td><td>2.6</td><td>8.5E-03</td><td>3.3E-02</td><td>BT20/MB231</td><td>MP</td></tr>
<tr><td>>hg38_21_+_8433172_8446622_RNA45SN1_45S_50ntFlanks@10772.10805.34|AGGTCTCCAAGGTGAACAGCCTCTGGCATGTTGG</td><td>>hg38_21_+_8433172_8446622_RNA45SN1_45S_50ntFlanks</td><td>10772</td><td>10805</td><td>34</td><td>AGGTCTCCAAGGTGAACAGCCTCTGGCATGTTGG</td><td>143.2</td><td>-1.6</td><td>0.7</td><td>-2.2</td><td>2.5E-02</td><td>3.3E-02</td><td>BT20/MB468</td><td>MITO</td></tr>
<tr><td>>hg38_21_+_8433172_8446622_RNA45SN1_45S_50ntFlanks@4067.4095.29|AACTTTCGATGGTAGTCGCCGTGCCTACC</td><td>>hg38_21_+_8433172_8446622_RNA45SN1_45S_50ntFlanks</td><td>4067</td><td>4095</td><td>29</td><td>AACTTTCGATGGTAGTCGCCGTGCCTACC</td><td>236.8</td><td>-4.1</td><td>1.8</td><td>-2.3</td><td>2.0E-02</td><td>3.3E-02</td><td>BT20/MB468</td><td>MP</td></tr>
<tr><td>>hg38_21_+_8433172_8446622_RNA45SN1_45S_50ntFlanks@6661.6695.35|CGGTGGATCACTCGGCTCGTGCGTCGATGAAGAAC</td><td>>hg38_21_+_8433172_8446622_RNA45SN1_45S_50ntFlanks</td><td>6661</td><td>6695</td><td>35</td><td>CGGTGGATCACTCGGCTCGTGCGTCGATGAAGAAC</td><td>3199.2</td><td>2.8</td><td>1.1</td><td>2.6</td><td>8.5E-03</td><td>3.3E-02</td><td>BT20/MB231</td><td>MP</td></tr>
<tr><td>>MI0000077|hsa-mir-21&WithFlank&17|+|59841260|59841343@14.35.22(+2A)&[MIMAT0000076&hsa-miR-21-5p&offsets|0|0(+2A);m-1&17|+|59841273|59841294&offsets|0|0(+2A)]|TAGCTTATCAGACTGATGTTGAAA</td><td>>MI0000077|hsa-mir-21&WithFlank&17|+|59841260|59841343</td><td>14</td><td>35</td><td>22(+2A)</td><td>TAGCTTATCAGACTGATGTTGAAA</td><td>134.7</td><td>2.4</td><td>1.0</td><td>2.3</td><td>2.0E-02</td><td>3.3E-02</td><td>BT20/MB468</td><td>MP</td></tr>
<tr><td>>trnaMT_LeuTAA_MT_+_3230_3304@13.30.18|GAGCCCGGTAATCGCATA</td><td>>trnaMT_LeuTAA_MT_+_3230_3304</td><td>13</td><td>30</td><td>18</td><td>GAGCCCGGTAATCGCATA</td><td>72.1</td><td>6.1</td><td>2.6</td><td>2.3</td><td>2.0E-02</td><td>3.3E-02</td><td>BT20/MB468</td><td>MP</td></tr>
<tr><td>>hg38_21_+_8433172_8446622_RNA45SN1_45S_50ntFlanks@11767.11803.37|TCTAATTAGTGACGCGCATGAATGGATGAACGAGATT</td><td>>hg38_21_+_8433172_8446622_RNA45SN1_45S_50ntFlanks</td><td>11767</td><td>11803</td><td>37</td><td>TCTAATTAGTGACGCGCATGAATGGATGAACGAGATT</td><td>154.6</td><td>5.4</td><td>2.1</td><td>2.6</td><td>8.5E-03</td><td>3.3E-02</td><td>BT20/MB231</td><td>MP</td></tr>
<tr><td>>hg38_21_+_8433172_8446622_RNA45SN1_45S_50ntFlanks@4738.4758.21|AAAGTCGGAGGTTCGAAGACG</td><td>>hg38_21_+_8433172_8446622_RNA45SN1_45S_50ntFlanks</td><td>4738</td><td>4758</td><td>21</td><td>AAAGTCGGAGGTTCGAAGACG</td><td>953.4</td><td>-1.8</td><td>0.7</td><td>-2.6</td><td>1.1E-02</td><td>3.3E-02</td><td>BT20/MB231</td><td>MITO</td></tr>
<tr><td>>hg38_21_+_8433172_8446622_RNA45SN1_45S_50ntFlanks@6647.6684.38|CGTACGACTCTTAGCGGTGGATCACTCGGCTCGTGCGT</td><td>>hg38_21_+_8433172_8446622_RNA45SN1_45S_50ntFlanks</td><td>6647</td><td>6684</td><td>38</td><td>CGTACGACTCTTAGCGGTGGATCACTCGGCTCGTGCGT</td><td>51.8</td><td>4.2</td><td>1.8</td><td>2.3</td><td>2.0E-02</td><td>3.3E-02</td><td>BT20/MB468</td><td>MP</td></tr>
<tr><td>>hg38_21_+_8433172_8446622_RNA45SN1_45S_50ntFlanks@5527.5553.27|AAAGTCGTAACAAGGTTTCCGTAGGTG</td><td>>hg38_21_+_8433172_8446622_RNA45SN1_45S_50ntFlanks</td><td>5527</td><td>5553</td><td>27</td><td>AAAGTCGTAACAAGGTTTCCGTAGGTG</td><td>73.1</td><td>-3.9</td><td>1.7</td><td>-2.3</td><td>2.0E-02</td><td>3.3E-02</td><td>BT20/MB468</td><td>MP</td></tr>
<tr><td>>MI0000471|hsa-mir-126&WithFlank&9|+|136670596|136670692@58.77.20(+1A)&[MIMAT0000445&hsa-miR-126-3p&offsets|0|-2(+1A);m-40&9|+|136670653|136670674&offsets|0|-2(+1A)]|TCGTACCGTGAGTAATAATGA</td><td>>MI0000471|hsa-mir-126&WithFlank&9|+|136670596|136670692</td><td>58</td><td>77</td><td>20(+1A)</td><td>TCGTACCGTGAGTAATAATGA</td><td>619.1</td><td>-2.8</td><td>1.1</td><td>-2.6</td><td>8.5E-03</td><td>3.3E-02</td><td>BT20/MB231</td><td>MP</td></tr>
<tr><td>>hg38_21_+_8433172_8446622_RNA45SN1_45S_50ntFlanks@9481.9505.25|CACGTGTTAGGACCCGAAAGATGGT</td><td>>hg38_21_+_8433172_8446622_RNA45SN1_45S_50ntFlanks</td><td>9481</td><td>9505</td><td>25</td><td>CACGTGTTAGGACCCGAAAGATGGT</td><td>64.6</td><td>6.0</td><td>2.6</td><td>2.3</td><td>2.0E-02</td><td>3.3E-02</td><td>BT20/MB468</td><td>MP</td></tr>
<tr><td>>hg38_21_+_8433172_8446622_RNA45SN1_45S_50ntFlanks@6652.6673.22|GACTCTTAGCGGTGGATCACTC</td><td>>hg38_21_+_8433172_8446622_RNA45SN1_45S_50ntFlanks</td><td>6652</td><td>6673</td><td>22</td><td>GACTCTTAGCGGTGGATCACTC</td><td>623.8</td><td>2.9</td><td>1.1</td><td>2.6</td><td>8.5E-03</td><td>3.3E-02</td><td>BT20/MB231</td><td>MP</td></tr>
<tr><td>>hg38_21_+_8433172_8446622_RNA45SN1_45S_50ntFlanks@9943.9978.36|GACGGTGGCCATGGAAGTCGGAATCCGCTAAGGAGT</td><td>>hg38_21_+_8433172_8446622_RNA45SN1_45S_50ntFlanks</td><td>9943</td><td>9978</td><td>36</td><td>GACGGTGGCCATGGAAGTCGGAATCCGCTAAGGAGT</td><td>145.4</td><td>5.4</td><td>2.0</td><td>2.6</td><td>8.5E-03</td><td>3.3E-02</td><td>BT20/MB231</td><td>MP</td></tr>
<tr><td>>hg38_21_+_8433172_8446622_RNA45SN1_45S_50ntFlanks@9953.9982.30|ATGGAAGTCGGAATCCGCTAAGGAGTGTGT</td><td>>hg38_21_+_8433172_8446622_RNA45SN1_45S_50ntFlanks</td><td>9953</td><td>9982</td><td>30</td><td>ATGGAAGTCGGAATCCGCTAAGGAGTGTGT</td><td>105.8</td><td>5.4</td><td>2.0</td><td>2.6</td><td>8.5E-03</td><td>3.3E-02</td><td>BT20/MB231</td><td>MP</td></tr>
</table>

| | | | | | | | | | | | | | |
|---|---|---|---|---|---|---|---|---|---|---|---|---|---|
| >MI0000264\|hsa-mir-7-2&WithFlank&15\|+\|88611819\|88611940@38.57.20(+1C);MI0000265\|hsa-mir-7-3&WithFlank&19\|+\|4770664\|4770785@37.56.20(+1C);MI0000263\|hsa-mir-7-1&WithFlank&9\|-\|83969742\|83969863@30.49.20(+1C)&[MIMAT0000252&hsa-miR-7-5p&offsets\|0\|-4(+1C);m-400&15\|+\|88611856\|88611879&offsets\|0\|-4(+1C)];[MIMAT0000252_1&hsa-miR-7-5p&offsets\|0\|-4(+1C);m-402&19\|+\|4770700\|4770723&offsets\|0\|-4(+1C)];[MIMAT0000252_2&hsa-miR-7-5p&offsets\|0\|-4(+1C);m-401&9\|-\|83969811\|83969834&offsets\|0\|-4(+1C)]\|TGGAAGACTAGTGATTTTGTC | >MI0000264\|hsa-mir-7-2&WithFlank&15\|+\|88611819\|88611940 | 38 | 57 | 20(+1C) | TGGAAGACTAGTGATTTTGTC | 543.1 | -2.9 | 1.1 | -2.6 | 8.5E-03 | 3.3E-02 | BT20/MB231 | MP |
| >hg38_21_+_8433172_8446622_RNA45SN1_45S_50ntFlanks@12928.12951.24\|CTGGCGCTAAACCATTCGTAGACG | >hg38_21_+_8433172_8446622_RNA45SN1_45S_50ntFlanks | 12928 | 12951 | 24 | CTGGCGCTAAACCATTCGTAGACG | 148.8 | -3.2 | 1.4 | -2.3 | 2.0E-02 | 3.3E-02 | BT20/MB468 | MP |
| >hg38_21_+_8433172_8446622_RNA45SN1_45S_50ntFlanks@4738.4755.18\|AAAGTCGGAGGTTCGAAG | >hg38_21_+_8433172_8446622_RNA45SN1_45S_50ntFlanks | 4738 | 4755 | 18 | AAAGTCGGAGGTTCGAAG | 263.3 | -2.1 | 0.9 | -2.3 | 2.0E-02 | 3.3E-02 | BT20/MB468 | MP |
| >hg38_21_+_8433172_8446622_RNA45SN1_45S_50ntFlanks@10820.10840.21\|GGGAAGTCGGCAAGCCGGATC | >hg38_21_+_8433172_8446622_RNA45SN1_45S_50ntFlanks | 10820 | 10840 | 21 | GGGAAGTCGGCAAGCCGGATC | 1904.0 | -2.6 | 1.1 | -2.3 | 2.0E-02 | 3.3E-02 | BT20/MB468 | MP |
| >hg38_21_+_8433172_8446622_RNA45SN1_45S_50ntFlanks@11566.11584.19\|TAGCAGCCGACTTAGAACT | >hg38_21_+_8433172_8446622_RNA45SN1_45S_50ntFlanks | 11566 | 11584 | 19 | TAGCAGCCGACTTAGAACT | 76.6 | 3.0 | 1.4 | 2.2 | 2.7E-02 | 3.3E-02 | MB231/MB468 | MP |
| >hg38_21_+_8433172_8446622_RNA45SN1_45S_50ntFlanks@5077.5102.26\|CCGATAACGAACGAGACTCTGGCATG | >hg38_21_+_8433172_8446622_RNA45SN1_45S_50ntFlanks | 5077 | 5102 | 26 | CCGATAACGAACGAGACTCTGGCATG | 202.2 | -4.1 | 1.8 | -2.3 | 2.0E-02 | 3.3E-02 | BT20/MB468 | MP |
| >hg38_21_+_8433172_8446622_RNA45SN1_45S_50ntFlanks@12934.12955.22\|CTAAACCATTCGTAGACGACCT | >hg38_21_+_8433172_8446622_RNA45SN1_45S_50ntFlanks | 12934 | 12955 | 22 | CTAAACCATTCGTAGACGACCT | 150.0 | -3.9 | 1.7 | -2.3 | 2.0E-02 | 3.3E-02 | BT20/MB468 | MP |
| >hg38_21_+_8433172_8446622_RNA45SN1_45S_50ntFlanks@6652.6689.38\|GACTCTTAGCGGTGGATCACTCGGCTCGTGCGTCGATG | >hg38_21_+_8433172_8446622_RNA45SN1_45S_50ntFlanks | 6652 | 6689 | 38 | GACTCTTAGCGGTGGATCACTCGGCTCGTGCGTCGATG | 14326.3 | 2.1 | 0.8 | 2.6 | 1.1E-02 | 3.3E-02 | BT20/MB231 | MITO |
| >hg38_1_-_228634819_228635039_RNA5S12_5S_50ntFlanks@135.171.37\|GGGAGACCGCCTGGGAATACCGGGTGCTGTAGGCTTT | >hg38_1_-_228634819_228635039_RNA5S12_5S_50ntFlanks | 135 | 171 | 37 | GGGAGACCGCCTGGGAATACCGGGTGCTGTAGGCTTT | 3455.1 | 1.8 | 0.8 | 2.2 | 2.7E-02 | 3.3E-02 | MB231/MB468 | MP |
| >hg38_MT_+_598_1651_RNR1_12S_50ntFlanks@916.950.35\|AAGGACATTTAACTAAAACCCCTACGCATTTATAT | >hg38_MT_+_598_1651_RNR1_12S_50ntFlanks | 916 | 950 | 35 | AAGGACATTTAACTAAAACCCCTACGCATTTATAT | 92.0 | 3.7 | 1.7 | 2.2 | 2.7E-02 | 3.3E-02 | MB231/MB468 | MP |
| >MI0000261\|hsa-mir-139&WithFlank&11\|-\|72615057\|72615136@13.34.22&[MIMAT0000250&hsa-miR-139-5p&offsets\|0\|-1;m-186&11\|-\|72615102\|72615124&offsets\|0\|-1]\|TCTACAGTGCACGTGTCTCCAG | >MI0000261\|hsa-mir-139&WithFlank&11\|-\|72615057\|72615136 | 13 | 34 | 22 | TCTACAGTGCACGTGTCTCCAG | 148.8 | -2.9 | 1.1 | -2.6 | 8.5E-03 | 3.3E-02 | BT20/MB231 | MP |
| >hg38_21_+_8433172_8446622_RNA45SN1_45S_50ntFlanks@12415.12434.20\|CTTCGATGTCGGCTCTTCCT | >hg38_21_+_8433172_8446622_RNA45SN1_45S_50ntFlanks | 12415 | 12434 | 20 | CTTCGATGTCGGCTCTTCCT | 12485.9 | -2.4 | 1.1 | -2.2 | 3.0E-02 | 3.3E-02 | BT20/MB468 | NUC |
| >hg38_MT_+_1621_3279_RNR2_16S_50ntFlanks@1423.1451.29\|CAACGATTAAAGTCCTACGTGATCTGAGT | >hg38_MT_+_1621_3279_RNR2_16S_50ntFlanks | 1423 | 1451 | 29 | CAACGATTAAAGTCCTACGTGATCTGAGT | 24.8 | 4.5 | 2.1 | 2.2 | 2.7E-02 | 3.3E-02 | MB231/MB468 | MP |
| >MI0000542\|hsa-mir-320a&WithFlank&8\|-\|22244960\|22245043@50.68.19(+1U)&[MIMAT0000510&hsa-miR-320a-3p&offsets\|+2\|-1(+1U);m-60&8\|-\|22244975\|22244996&offsets\|+2\|-1(+1U)]\|AAGCTGGGTTGAGAGGGCGT | >MI0000542\|hsa-mir-320a&WithFlank&8\|-\|22244960\|22245043 | 50 | 68 | 19(+1U) | AAGCTGGGTTGAGAGGGCGT | 42.7 | 4.6 | 2.1 | 2.2 | 2.7E-02 | 3.3E-02 | MB231/MB468 | MP |
| >trnaMT_GluTTC_MT_-_14674_14742@44.71.28,trnalookalike8_GluTTC_5_-_93905172_93905240@44.71.28\|TCGTGGTTGTAGTCCGTGCGAGAATACC | >trnaMT_GluTTC_MT_-_14674_14742 | 44 | 71 | 28 | TCGTGGTTGTAGTCCGTGCGAGAATACC | 223.6 | 2.4 | 1.1 | 2.2 | 2.7E-02 | 3.3E-02 | MB231/MB468 | MP |
| >trna16_TyrGTA_14_-_21131351_21131444@59.76.18,trna17_TyrGTA_14_-_21128117_21128210@59.76.18,trna19_TyrGTA_14_-_21121258_21121351@59.76.18,trna2_TyrGTA_2_+_27273650_27273738@59.76.18,trna4_TyrGTA_8_+_67025602_67025694@59.76.18,trna5_TyrGTA_14_+_21151432_21151520@59.76.18,trna5_TyrGTA_8_+_67026223_67026311@59.76.18\|TTCCGGCTCGAAGGACCA | >trna16_TyrGTA_14_-_21131351_21131444 | 59 | 76 | 18 | TTCCGGCTCGAAGGACCA | 3212.4 | 2.0 | 0.8 | 2.5 | 1.1E-02 | 3.3E-02 | BT20/MB231 | CYTO |
| >hg38_21_+_8433172_8446622_RNA45SN1_45S_50ntFlanks@12414.12442.29\|CCTTCGATGTCGGCTCTTCCTATCATTGT | >hg38_21_+_8433172_8446622_RNA45SN1_45S_50ntFlanks | 12414 | 12442 | 29 | CCTTCGATGTCGGCTCTTCCTATCATTGT | 471.2 | -2.2 | 0.9 | -2.5 | 1.1E-02 | 3.3E-02 | BT20/MB231 | CYTO |
| >hg38_21_+_8433172_8446622_RNA45SN1_45S_50ntFlanks@5557.5580.24\|CTGCGGAAGGATCATTAACGGAGC | >hg38_21_+_8433172_8446622_RNA45SN1_45S_50ntFlanks | 5557 | 5580 | 24 | CTGCGGAAGGATCATTAACGGAGC | 244.6 | -2.0 | 0.8 | -2.5 | 1.1E-02 | 3.3E-02 | BT20/MB231 | CYTO |
| >MI0003666\|hsa-mir-651&WithFlank&X\|+\|8126959\|8127067@22.42.21&[MIMAT0003321&hsa-miR-651-5p&offsets\|0\|-1;m-399&X\|+\|8126980\|8127001&offsets\|0\|-1]\|TTTAGGATAAGCTTGACTTTT | >MI0003666\|hsa-mir-651&WithFlank&X\|+\|8126959\|8127067 | 22 | 42 | 21 | TTTAGGATAAGCTTGACTTTT | 34.8 | 3.8 | 1.7 | 2.2 | 2.7E-02 | 3.3E-02 | MB231/MB468 | MP |
| >MI0003190\|hsa-mir-505&WithFlank&X\|-\|139924142\|139924237@57.80.24&[MIMAT0002876&hsa-miR-505-3p&offsets\|0\|+2;m-196&X\|-\|139924160\|139924181&offsets\|0\|+2]\|CGTCAACACTTGCTGGTTTCCTCT | >MI0003190\|hsa-mir-505&WithFlank&X\|-\|139924142\|139924237 | 57 | 80 | 24 | CGTCAACACTTGCTGGTTTCCTCT | 236.7 | 1.7 | 0.7 | 2.6 | 9.2E-03 | 3.3E-02 | BT20/MB231 | NUC |
| >trnaMT_LysTTT_MT_+_8295_8364@37.73.37,trnalookalike7_LysTTT_1_+_568844_568913@37.73.37\|AAAGATTAAGAGAACCAACACCTCTTTACAGTGACCA | >trnaMT_LysTTT_MT_+_8295_8364 | 37 | 73 | 37 | AAAGATTAAGAGAACCAACACCTCTTTACAGTGACCA | 407.7 | 4.2 | 1.8 | 2.3 | 2.0E-02 | 3.3E-02 | BT20/MB468 | MP |
| >hg38_21_+_8433172_8446622_RNA45SN1_45S_50ntFlanks@4002.4026.25\|GATAACCTCGGGCCGATCGCACGCC | >hg38_21_+_8433172_8446622_RNA45SN1_45S_50ntFlanks | 4002 | 4026 | 25 | GATAACCTCGGGCCGATCGCACGCC | 240.8 | -4.0 | 1.7 | -2.3 | 2.0E-02 | 3.3E-02 | BT20/MB468 | MP |
| >hg38_21_+_8433172_8446622_RNA45SN1_45S_50ntFlanks@11768.11806.39\|CTAATTAGTGACGCGCATGAATGGATGAACGAGATTCCC | >hg38_21_+_8433172_8446622_RNA45SN1_45S_50ntFlanks | 11768 | 11806 | 39 | CTAATTAGTGACGCGCATGAATGGATGAACGAGATTCCC | 118.7 | -4.1 | 1.8 | -2.3 | 2.0E-02 | 3.3E-02 | BT20/MB468 | MP |
| >hg38_21_+_8433172_8446622_RNA45SN1_45S_50ntFlanks@9459.9495.37\|CCGGGGAGGTGGAGCACGAGCGCACGTGTTAGGACCC | >hg38_21_+_8433172_8446622_RNA45SN1_45S_50ntFlanks | 9459 | 9495 | 37 | CCGGGGAGGTGGAGCACGAGCGCACGTGTTAGGACCC | 71.8 | -4.0 | 1.7 | -2.3 | 2.0E-02 | 3.3E-02 | BT20/MB468 | MP |
| >hg38_21_+_8433172_8446622_RNA45SN1_45S_50ntFlanks@5421.5466.46\|CGATTGGATGGTTTAGTGAGGCCCTCGGATCGGCCCCGCCGGGGTC | >hg38_21_+_8433172_8446622_RNA45SN1_45S_50ntFlanks | 5421 | 5466 | 46 | CGATTGGATGGTTTAGTGAGGCCCTCGGATCGGCCCCGCCGGGGTC | 83.4 | -3.6 | 1.6 | -2.2 | 2.6E-02 | 3.3E-02 | BT20/MB468 | MITO |
| >hg38_21_+_8433172_8446622_RNA45SN1_45S_50ntFlanks@475.496.22\|CTTCGTGATCGATGTGGTGACG | >hg38_21_+_8433172_8446622_RNA45SN1_45S_50ntFlanks | 475 | 496 | 22 | CTTCGTGATCGATGTGGTGACG | 1323.3 | 4.2 | 1.8 | 2.3 | 2.0E-02 | 3.3E-02 | BT20/MB468 | MP |
| >hg38_21_+_8433172_8446622_RNA45SN1_45S_50ntFlanks@3705.3743.39\|TACCTGGTTGATCCTGCCAGTAGCATATGCTTGTCTCAA | >hg38_21_+_8433172_8446622_RNA45SN1_45S_50ntFlanks | 3705 | 3743 | 39 | TACCTGGTTGATCCTGCCAGTAGCATATGCTTGTCTCAA | 817.9 | 1.8 | 0.8 | 2.2 | 3.0E-02 | 3.3E-02 | MB231/MB468 | MITO |
| >hg38_21_+_8433172_8446622_RNA45SN1_45S_50ntFlanks@11873.11901.29\|AAGAAGACCCTGTTGAGCTTGACTCTAGT | >hg38_21_+_8433172_8446622_RNA45SN1_45S_50ntFlanks | 11873 | 11901 | 29 | AAGAAGACCCTGTTGAGCTTGACTCTAGT | 176.5 | -1.0 | 0.5 | -2.2 | 2.9E-02 | 3.3E-02 | BT20/MB468 | CYTO |
| >hg38_MT_+_598_1651_RNR1_12S_50ntFlanks@908.935.28\|ATACTTCAAAGGACATTTAACTAAAACC | >hg38_MT_+_598_1651_RNR1_12S_50ntFlanks | 908 | 935 | 28 | ATACTTCAAAGGACATTTAACTAAAACC | 74.2 | 3.8 | 1.7 | 2.2 | 2.7E-02 | 3.3E-02 | MB231/MB468 | MP |
| >hg38_21_+_8433172_8446622_RNA45SN1_45S_50ntFlanks@12937.12955.19\|AACCATTCGTAGACGACCT | >hg38_21_+_8433172_8446622_RNA45SN1_45S_50ntFlanks | 12937 | 12955 | 19 | AACCATTCGTAGACGACCT | 140.3 | -1.4 | 0.6 | -2.2 | 2.6E-02 | 3.3E-02 | BT20/MB468 | MITO |
| >am_trnaMT_PheGAA_MT_+_577_647@45.63.19\|AGACGGGCTCACATCACCC | >am_trnaMT_PheGAA_MT_+_577_647 | 45 | 63 | 19 | AGACGGGCTCACATCACCC | 99.6 | -2.8 | 1.3 | -2.2 | 2.7E-02 | 3.3E-02 | MB231/MB468 | MP |
| >trnaMT_GluTTC_MT_-_14674_14742@37.71.35,trnalookalike8_GluTTC_5_-_93905172_93905240@37.71.35\|TCATTGGTCGTGGTTGTAGTCCGTGCGAGAATACC | >trnaMT_GluTTC_MT_-_14674_14742 | 37 | 71 | 35 | TCATTGGTCGTGGTTGTAGTCCGTGCGAGAATACC | 131.0 | 3.4 | 1.5 | 2.3 | 2.0E-02 | 3.3E-02 | BT20/MB468 | MP |
| >MI0000299\|hsa-mir-222&WithFlank&X\|-\|45747009\|45747130@75.94.20&[MIMAT0000279&hsa-miR-222-3p&offsets\|0\|-1;m-140&X\|-\|45747033\|45747056&offsets\|0\|-4]\|AGCTACATCTGGCTACTGGG | >MI0000299\|hsa-mir-222&WithFlank&X\|-\|45747009\|45747130 | 75 | 94 | 20 | AGCTACATCTGGCTACTGGG | 259.9 | -1.8 | 0.7 | -2.6 | 1.1E-02 | 3.3E-02 | BT20/MB231 | MITO |
| >trna111_HisGTG_1_-_147774845_147774916@1.30.30,trna118_HisGTG_1_-_145396881_145396952@1.30.30,trna16_HisGTG_1_+_146544773_146544844@1.30.30,trna1_HisGTG_15_+_45493349_45493420@1.30.30,trna21_HisGTG_1_+_147753471_147753542@1.30.30,trna33_HisGTG_6_+_27125906_27125977@1.30.30,trna7_HisGTG_9_-_14433938_14434009@1.30.30,trna8_HisGTG_15_-_45492611_45492682@1.30.30,trna9_HisGTG_15_-_45490804_45490875@1.30.30\|GCCGTGATCGTATAGTGGTTAGTACTCTGC | >trna111_HisGTG_1_-_147774845_147774916 | 1 | 30 | 30 | GCCGTGATCGTATAGTGGTTAGTACTCTGC | 1072.5 | -2.1 | 0.8 | -2.6 | 1.1E-02 | 3.3E-02 | BT20/MB231 | MITO |
| >hg38_21_+_8433172_8446622_RNA45SN1_45S_50ntFlanks@4669.4691.23\|TTCTTGGACCGGCGCAAGACGGA | >hg38_21_+_8433172_8446622_RNA45SN1_45S_50ntFlanks | 4669 | 4691 | 23 | TTCTTGGACCGGCGCAAGACGGA | 138.6 | 4.5 | 1.9 | 2.3 | 2.0E-02 | 3.3E-02 | BT20/MB468 | MP |
| >hg38_21_+_8433172_8446622_RNA45SN1_45S_50ntFlanks@9540.9571.32\|CTCTGGTGGAGGTCCGTAGCGGTCCTGACGTG | >hg38_21_+_8433172_8446622_RNA45SN1_45S_50ntFlanks | 9540 | 9571 | 32 | CTCTGGTGGAGGTCCGTAGCGGTCCTGACGTG | 134.0 | 5.3 | 2.0 | 2.6 | 8.6E-03 | 3.3E-02 | BT20/MB231 | MP |
| >hg38_21_+_8433172_8446622_RNA45SN1_45S_50ntFlanks@8175.8213.39\|CCTTCTGATCGAGGCCCAGCCCGTGGACGGTGTGAGGCC | >hg38_21_+_8433172_8446622_RNA45SN1_45S_50ntFlanks | 8175 | 8213 | 39 | CCTTCTGATCGAGGCCCAGCCCGTGGACGGTGTGAGGCC | 409.1 | 4.6 | 1.8 | 2.6 | 8.6E-03 | 3.3E-02 | BT20/MB231 | MP |
| >hg38_21_+_8433172_8446622_RNA45SN1_45S_50ntFlanks@9496.9520.25\|GAAAGATGGTGAACTATGCCTGGGC | >hg38_21_+_8433172_8446622_RNA45SN1_45S_50ntFlanks | 9496 | 9520 | 25 | GAAAGATGGTGAACTATGCCTGGGC | 296.2 | 3.5 | 1.3 | 2.6 | 8.6E-03 | 3.3E-02 | BT20/MB231 | MP |
| >MI0000084\|hsa-mir-26b&WithFlank&2\|+\|218402640\|218402728@18.38.21(+1C)&[MIMAT0000083&hsa-miR-26b-5p&offsets\|0\|0(+1C);m-53&2\|+\|218402657\|218402678&offsets\|0\|-1(+1C)]\|TTCAAGTAATTCAGGATAGGTC | >MI0000084\|hsa-mir-26b&WithFlank&2\|+\|218402640\|218402728 | 18 | 38 | 21(+1C) | TTCAAGTAATTCAGGATAGGTC | 1332.2 | 1.6 | 0.6 | 2.6 | 9.3E-03 | 3.3E-02 | BT20/MB231 | NUC |
| >trnaMT_SerGCT_MT_+_12207_12265@1.23.23\|GAGAAAGCTCACAAGAACTGCTA | >trnaMT_SerGCT_MT_+_12207_12265 | 1 | 23 | 23 | GAGAAAGCTCACAAGAACTGCTA | 3296.6 | 3.3 | 1.4 | 2.3 | 2.0E-02 | 3.4E-02 | BT20/MB468 | MP |
| >hg38_21_+_8433172_8446622_RNA45SN1_45S_50ntFlanks@12850.12868.19\|CTTCGTCCTGGGAAACGGG | >hg38_21_+_8433172_8446622_RNA45SN1_45S_50ntFlanks | 12850 | 12868 | 19 | CTTCGTCCTGGGAAACGGG | 133.2 | -4.2 | 1.8 | -2.3 | 2.0E-02 | 3.4E-02 | BT20/MB468 | MP |
| >hg38_21_+_8433172_8446622_RNA45SN1_45S_50ntFlanks@12420.12448.29\|ATGTCGGCTCTTCCTATCATTGTGAAGCA | >hg38_21_+_8433172_8446622_RNA45SN1_45S_50ntFlanks | 12420 | 12448 | 29 | ATGTCGGCTCTTCCTATCATTGTGAAGCA | 2754.4 | -2.3 | 0.9 | -2.5 | 1.1E-02 | 3.4E-02 | BT20/MB231 | CYTO |
| >hg38_21_+_8433172_8446622_RNA45SN1_45S_50ntFlanks@12007.12026.20\|GCCGCCGGTGAAATACCACT | >hg38_21_+_8433172_8446622_RNA45SN1_45S_50ntFlanks | 12007 | 12026 | 20 | GCCGCCGGTGAAATACCACT | 362.9 | 4.8 | 2.1 | 2.3 | 2.0E-02 | 3.4E-02 | BT20/MB468 | MP |
| >hg38_21_+_8433172_8446622_RNA45SN1_45S_50ntFlanks@10398.10424.27\|GTTGAACATGGGTCAGTCGGTCCTGAG | >hg38_21_+_8433172_8446622_RNA45SN1_45S_50ntFlanks | 10398 | 10424 | 27 | GTTGAACATGGGTCAGTCGGTCCTGAG | 10471.9 | 3.3 | 1.3 | 2.6 | 8.6E-03 | 3.4E-02 | BT20/MB231 | MP |
| >hg38_21_+_8433172_8446622_RNA45SN1_45S_50ntFlanks@9534.9569.36\|AGGAAACTCTGGTGGAGGTCCGTAGCGGTCCTGACG | >hg38_21_+_8433172_8446622_RNA45SN1_45S_50ntFlanks | 9534 | 9569 | 36 | AGGAAACTCTGGTGGAGGTCCGTAGCGGTCCTGACG | 470.4 | 5.6 | 2.2 | 2.6 | 8.6E-03 | 3.4E-02 | BT20/MB231 | MP |

<table>
<tr><th></th><th></th><th></th><th></th><th></th><th></th><th></th><th></th><th></th><th></th><th></th><th></th><th></th><th></th></tr>
<tr><td>>MI0000061|hsa-let-7a-2&WithFlank&11|-|122146516|122146599@11.32.22(+1C);MI0000062|hsa-let-7a-3&WithFlank&22|+|46112743|46112828@10.31.22(+1C);MI0000060|hsa-let-7a-1&WithFlank&9|+|94175951|94176042@12.33.22(+1C)&[MIMAT0000062&hsa-let-7a-5p&offsets|0|0(+1C);m-6&11|-|122146568|122146589&offsets|0|0(+1C)];[MIMAT0000062_1&hsa-let-7a-5p&offsets|0|0(+1C);m-5&22|+|46112752|46112773&offsets|0|0(+1C)];[MIMAT0000062_2&hsa-let-7a-5p&offsets|0|0(+1C);m-4&9|+|94175962|94175983&offsets|0|0(+1C)]|TGAGGTAGTAGGTTGTATAGTTC</td><td>>MI0000061|hsa-let-7a-2&WithFlank&11|-|122146516|122146599</td><td>11</td><td>32</td><td>22(+1C)</td><td>TGAGGTAGTAGGTTGTATAGTTC</td><td>1314.4</td><td>-2.7</td><td>1.0</td><td>-2.6</td><td>8.6E-03</td><td>3.4E-02</td><td>BT20/MB231</td><td>MP</td></tr>
<tr><td>>hg38_21_+_8433172_8446622_RNA45SN1_45S_50ntFlanks@11207.11229.23|GGCGGCGACTCTGGACGCGAGCC</td><td>>hg38_21_+_8433172_8446622_RNA45SN1_45S_50ntFlanks</td><td>11207</td><td>11229</td><td>23</td><td>GGCGGCGACTCTGGACGCGAGCC</td><td>247.6</td><td>4.6</td><td>1.8</td><td>2.6</td><td>8.6E-03</td><td>3.4E-02</td><td>BT20/MB231</td><td>MP</td></tr>
<tr><td>>hg38_21_+_8433172_8446622_RNA45SN1_45S_50ntFlanks@8192.8213.22|AGCCCGTGGACGGTGTGAGGCC</td><td>>hg38_21_+_8433172_8446622_RNA45SN1_45S_50ntFlanks</td><td>8192</td><td>8213</td><td>22</td><td>AGCCCGTGGACGGTGTGAGGCC</td><td>455.4</td><td>4.4</td><td>1.7</td><td>2.6</td><td>8.6E-03</td><td>3.4E-02</td><td>BT20/MB231</td><td>MP</td></tr>
<tr><td>>hg38_21_+_8433172_8446622_RNA45SN1_45S_50ntFlanks@9921.9949.29|TGTTGGTTGATATAGACAGCAGGACGGTG</td><td>>hg38_21_+_8433172_8446622_RNA45SN1_45S_50ntFlanks</td><td>9921</td><td>9949</td><td>29</td><td>TGTTGGTTGATATAGACAGCAGGACGGTG</td><td>97.4</td><td>5.3</td><td>2.0</td><td>2.6</td><td>8.6E-03</td><td>3.4E-02</td><td>BT20/MB231</td><td>MP</td></tr>
<tr><td>>hg38_21_+_8433172_8446622_RNA45SN1_45S_50ntFlanks@11729.11746.18|TAACTATGACTCTCTTAA</td><td>>hg38_21_+_8433172_8446622_RNA45SN1_45S_50ntFlanks</td><td>11729</td><td>11746</td><td>18</td><td>TAACTATGACTCTCTTAA</td><td>48.8</td><td>-4.1</td><td>1.8</td><td>-2.3</td><td>2.0E-02</td><td>3.4E-02</td><td>BT20/MB468</td><td>MP</td></tr>
<tr><td>>hg38_21_+_8433172_8446622_RNA45SN1_45S_50ntFlanks@12973.12991.19|TCGTACGTAGCAGAGCAGC</td><td>>hg38_21_+_8433172_8446622_RNA45SN1_45S_50ntFlanks</td><td>12973</td><td>12991</td><td>19</td><td>TCGTACGTAGCAGAGCAGC</td><td>1609.8</td><td>-3.7</td><td>1.6</td><td>-2.3</td><td>2.0E-02</td><td>3.4E-02</td><td>BT20/MB468</td><td>MP</td></tr>
<tr><td>>hg38_21_+_8433172_8446622_RNA45SN1_45S_50ntFlanks@12444.12481.38|AAGCAGAATTCACCAAGCGTTGGATTGTTCACCCACTA</td><td>>hg38_21_+_8433172_8446622_RNA45SN1_45S_50ntFlanks</td><td>12444</td><td>12481</td><td>38</td><td>AAGCAGAATTCACCAAGCGTTGGATTGTTCACCCACTA</td><td>90.3</td><td>-2.9</td><td>1.3</td><td>-2.2</td><td>2.6E-02</td><td>3.4E-02</td><td>BT20/MB468</td><td>MITO</td></tr>
<tr><td>>hg38_21_+_8433172_8446622_RNA45SN1_45S_50ntFlanks@12418.12445.28|CGATGTCGGCTCTTCCTATCATTGTGAA</td><td>>hg38_21_+_8433172_8446622_RNA45SN1_45S_50ntFlanks</td><td>12418</td><td>12445</td><td>28</td><td>CGATGTCGGCTCTTCCTATCATTGTGAA</td><td>1241.7</td><td>-1.3</td><td>0.6</td><td>-2.2</td><td>2.9E-02</td><td>3.4E-02</td><td>MB231/MB468</td><td>CYTO</td></tr>
<tr><td>>trnaMT_GluTTC_MT_-_14674_14742@35.72.38,trnalookalike8_GluTTC_5_-_93905172_93905240@35.72.38|TATCATTGGTCGTGGTTGTAGTCCGTGCGAGAATACCA</td><td>>trnaMT_GluTTC_MT_-_14674_14742</td><td>35</td><td>72</td><td>38</td><td>TATCATTGGTCGTGGTTGTAGTCCGTGCGAGAATACCA</td><td>849.3</td><td>-2.2</td><td>1.0</td><td>-2.2</td><td>3.0E-02</td><td>3.4E-02</td><td>BT20/MB468</td><td>NUC</td></tr>
<tr><td>>hg38_1_-_228634819_228635039_RNA5S12_5S_50ntFlanks@127.171.45|ACTTGGATGGGAGACCGCCTGGGAATACCGGGTGCTGTAGGCTTT</td><td>>hg38_1_-_228634819_228635039_RNA5S12_5S_50ntFlanks</td><td>127</td><td>171</td><td>45</td><td>ACTTGGATGGGAGACCGCCTGGGAATACCGGGTGCTGTAGGCTTT</td><td>667.3</td><td>-3.2</td><td>1.2</td><td>-2.6</td><td>9.3E-03</td><td>3.4E-02</td><td>BT20/MB231</td><td>NUC</td></tr>
<tr><td>>hg38_21_+_8433172_8446622_RNA45SN1_45S_50ntFlanks@10473.10495.23|CTCGGCCGATCGAAAGGGAGTCG</td><td>>hg38_21_+_8433172_8446622_RNA45SN1_45S_50ntFlanks</td><td>10473</td><td>10495</td><td>23</td><td>CTCGGCCGATCGAAAGGGAGTCG</td><td>140.0</td><td>-3.4</td><td>1.5</td><td>-2.3</td><td>2.0E-02</td><td>3.4E-02</td><td>BT20/MB468</td><td>MP</td></tr>
<tr><td>>MI0003591|hsa-mir-584&WithFlank&5|-|149062307|149062415@22.43.22(+1A)&[MIMAT0003249&hsa-miR-584-5p&offsets|0|0(+1A);m-167&5|-|149062374|149062394&offsets|0|+1(+1A)]|TTATGGTTTGCCTGGGACTGAGA</td><td>>MI0003591|hsa-mir-584&WithFlank&5|-|149062307|149062415</td><td>22</td><td>43</td><td>22(+1A)</td><td>TTATGGTTTGCCTGGGACTGAGA</td><td>96.4</td><td>3.6</td><td>1.6</td><td>2.2</td><td>2.8E-02</td><td>3.4E-02</td><td>MB231/MB468</td><td>MP</td></tr>
<tr><td>>hg38_21_+_8433172_8446622_RNA45SN1_45S_50ntFlanks@5515.5545.31|CTAGAGGAAGTAAAAGTCGTAACAAGGTTTC</td><td>>hg38_21_+_8433172_8446622_RNA45SN1_45S_50ntFlanks</td><td>5515</td><td>5545</td><td>31</td><td>CTAGAGGAAGTAAAAGTCGTAACAAGGTTTC</td><td>112.8</td><td>-4.0</td><td>1.7</td><td>-2.3</td><td>2.0E-02</td><td>3.4E-02</td><td>BT20/MB468</td><td>MP</td></tr>
<tr><td>>MI0000095|hsa-mir-93&WithFlank&7|-|100093762|100093853@18.39.22&[MIMAT0000093&hsa-miR-93-5p&offsets|+1|0;m-29&7|-|100093815|100093837&offsets|+1|0]|AAAGTGCTGTTCGTGCAGGTAG</td><td>>MI0000095|hsa-mir-93&WithFlank&7|-|100093762|100093853</td><td>18</td><td>39</td><td>22</td><td>AAAGTGCTGTTCGTGCAGGTAG</td><td>88.4</td><td>4.5</td><td>1.9</td><td>2.3</td><td>2.0E-02</td><td>3.4E-02</td><td>BT20/MB468</td><td>MP</td></tr>
<tr><td>>hg38_21_+_8433172_8446622_RNA45SN1_45S_50ntFlanks@10345.10369.25|TTCAAACGAGAACTTTGAAGGCCGA</td><td>>hg38_21_+_8433172_8446622_RNA45SN1_45S_50ntFlanks</td><td>10345</td><td>10369</td><td>25</td><td>TTCAAACGAGAACTTTGAAGGCCGA</td><td>112.2</td><td>2.3</td><td>1.0</td><td>2.3</td><td>2.0E-02</td><td>3.4E-02</td><td>BT20/MB468</td><td>MP</td></tr>
<tr><td>>hg38_MT_+_598_1651_RNR1_12S_50ntFlanks@425.455.31|ACGAAAGTGGCTTTAACATATCTGAACACAC</td><td>>hg38_MT_+_598_1651_RNR1_12S_50ntFlanks</td><td>425</td><td>455</td><td>31</td><td>ACGAAAGTGGCTTTAACATATCTGAACACAC</td><td>160.1</td><td>2.8</td><td>1.3</td><td>2.2</td><td>2.8E-02</td><td>3.4E-02</td><td>MB231/MB468</td><td>MP</td></tr>
<tr><td>>MI0000070|hsa-mir-16-1&WithFlank&13|-|50048967|50049067@20.40.21;MI0000115|hsa-mir-16-2&WithFlank&3|+|160404739|160404831@16.36.21&[MIMAT0000069&hsa-miR-16-5p&offsets|0|-1;m-62&13|-|50049027|50049048&offsets|0|-1];[MIMAT0000069_1&hsa-miR-16-5p&offsets|0|-1;m-61&3|+|160404754|160404775&offsets|0|-1]|TAGCAGCACGTAAATATTGGC</td><td>>MI0000070|hsa-mir-16-1&WithFlank&13|-|50048967|50049067</td><td>20</td><td>40</td><td>21</td><td>TAGCAGCACGTAAATATTGGC</td><td>1274.0</td><td>-2.0</td><td>0.8</td><td>-2.6</td><td>1.1E-02</td><td>3.4E-02</td><td>BT20/MB231</td><td>MITO</td></tr>
<tr><td>>MI0000077|hsa-mir-21&WithFlank&17|+|59841260|59841343@14.34.21(+1G)&[MIMAT0000076&hsa-miR-21-5p&offsets|0|-1(+1G);m-1&17|+|59841273|59841294&offsets|0|-1(+1G)]|TAGCTTATCAGACTGATGTTGG</td><td>>MI0000077|hsa-mir-21&WithFlank&17|+|59841260|59841343</td><td>14</td><td>34</td><td>21(+1G)</td><td>TAGCTTATCAGACTGATGTTGG</td><td>497.6</td><td>2.6</td><td>1.2</td><td>2.2</td><td>2.9E-02</td><td>3.4E-02</td><td>MB231/MB468</td><td>CYTO</td></tr>
<tr><td>>hg38_21_+_8433172_8446622_RNA45SN1_45S_50ntFlanks@6654.6688.35|CTCTTAGCGGTGGATCACTCGGCTCGTGCGTCGAT</td><td>>hg38_21_+_8433172_8446622_RNA45SN1_45S_50ntFlanks</td><td>6654</td><td>6688</td><td>35</td><td>CTCTTAGCGGTGGATCACTCGGCTCGTGCGTCGAT</td><td>447.0</td><td>1.5</td><td>0.7</td><td>2.2</td><td>2.6E-02</td><td>3.4E-02</td><td>BT20/MB468</td><td>MITO</td></tr>
<tr><td>>hg38_21_+_8433172_8446622_RNA45SN1_45S_50ntFlanks@4595.4616.22|GTTGGTTTTCGGAACTGAGGCC</td><td>>hg38_21_+_8433172_8446622_RNA45SN1_45S_50ntFlanks</td><td>4595</td><td>4616</td><td>22</td><td>GTTGGTTTTCGGAACTGAGGCC</td><td>170.8</td><td>4.3</td><td>1.6</td><td>2.6</td><td>8.7E-03</td><td>3.4E-02</td><td>BT20/MB231</td><td>MP</td></tr>
<tr><td>>hg38_21_+_8433172_8446622_RNA45SN1_45S_50ntFlanks@6722.6760.39|ATTGCAGGACACATTGATCATCGACACTTCGAACGCACT</td><td>>hg38_21_+_8433172_8446622_RNA45SN1_45S_50ntFlanks</td><td>6722</td><td>6760</td><td>39</td><td>ATTGCAGGACACATTGATCATCGACACTTCGAACGCACT</td><td>173.7</td><td>-2.0</td><td>0.8</td><td>-2.5</td><td>1.2E-02</td><td>3.4E-02</td><td>BT20/MB231</td><td>CYTO</td></tr>
<tr><td>>trnaMT_PheGAA_MT_+_577_647@37.57.21|AAATGTTTAGACGGGCTCACA</td><td>>trnaMT_PheGAA_MT_+_577_647</td><td>37</td><td>57</td><td>21</td><td>AAATGTTTAGACGGGCTCACA</td><td>32.8</td><td>3.7</td><td>1.7</td><td>2.2</td><td>2.8E-02</td><td>3.4E-02</td><td>MB231/MB468</td><td>MP</td></tr>
<tr><td>>hg38_MT_+_1621_3279_RNR2_16S_50ntFlanks@1425.1450.26|ACGATTAAAGTCCTACGTGATCTGAG</td><td>>hg38_MT_+_1621_3279_RNR2_16S_50ntFlanks</td><td>1425</td><td>1450</td><td>26</td><td>ACGATTAAAGTCCTACGTGATCTGAG</td><td>53.1</td><td>4.2</td><td>1.8</td><td>2.3</td><td>2.0E-02</td><td>3.4E-02</td><td>BT20/MB468</td><td>MP</td></tr>
<tr><td>>hg38_21_+_8433172_8446622_RNA45SN1_45S_50ntFlanks@8154.8181.28|CCGCTCGTGGGGGGCCCAAGTCCTTCTG</td><td>>hg38_21_+_8433172_8446622_RNA45SN1_45S_50ntFlanks</td><td>8154</td><td>8181</td><td>28</td><td>CCGCTCGTGGGGGGCCCAAGTCCTTCTG</td><td>242.8</td><td>-2.6</td><td>1.2</td><td>-2.2</td><td>2.9E-02</td><td>3.4E-02</td><td>BT20/MB468</td><td>CYTO</td></tr>
<tr><td>>hg38_21_+_8433172_8446622_RNA45SN1_45S_50ntFlanks@8123.8142.20|ATGTGGCGTACGGAAGACCC</td><td>>hg38_21_+_8433172_8446622_RNA45SN1_45S_50ntFlanks</td><td>8123</td><td>8142</td><td>20</td><td>ATGTGGCGTACGGAAGACCC</td><td>100.3</td><td>5.0</td><td>2.2</td><td>2.3</td><td>2.0E-02</td><td>3.4E-02</td><td>BT20/MB468</td><td>MP</td></tr>
<tr><td>>hg38_21_+_8433172_8446622_RNA45SN1_45S_50ntFlanks@11571.11612.42|GCCGACTTAGAACTGGTGCGGACCAGGGGAATCCGACTGTTT</td><td>>hg38_21_+_8433172_8446622_RNA45SN1_45S_50ntFlanks</td><td>11571</td><td>11612</td><td>42</td><td>GCCGACTTAGAACTGGTGCGGACCAGGGGAATCCGACTGTTT</td><td>508.8</td><td>1.9</td><td>0.9</td><td>2.2</td><td>2.6E-02</td><td>3.4E-02</td><td>BT20/MB468</td><td>MITO</td></tr>
<tr><td>>hg38_21_+_8433172_8446622_RNA45SN1_45S_50ntFlanks@12416.12439.24|TTCGATGTCGGCTCTTCCTATCAT</td><td>>hg38_21_+_8433172_8446622_RNA45SN1_45S_50ntFlanks</td><td>12416</td><td>12439</td><td>24</td><td>TTCGATGTCGGCTCTTCCTATCAT</td><td>3200.4</td><td>-2.2</td><td>0.9</td><td>-2.5</td><td>1.2E-02</td><td>3.4E-02</td><td>BT20/MB231</td><td>CYTO</td></tr>
<tr><td>>hg38_21_+_8433172_8446622_RNA45SN1_45S_50ntFlanks@11542.11584.43|CGGCGCCTCGCCTCGGCCGGCGCCTAGCAGCCGACTTAGAACT</td><td>>hg38_21_+_8433172_8446622_RNA45SN1_45S_50ntFlanks</td><td>11542</td><td>11584</td><td>43</td><td>CGGCGCCTCGCCTCGGCCGGCGCCTAGCAGCCGACTTAGAACT</td><td>411.7</td><td>-2.0</td><td>0.8</td><td>-2.5</td><td>1.2E-02</td><td>3.4E-02</td><td>BT20/MB231</td><td>CYTO</td></tr>
<tr><td>>hg38_21_+_8433172_8446622_RNA45SN1_45S_50ntFlanks@7985.8018.34|GATCAGACGTGGCGACCCGCTGAATTTAAGCATA</td><td>>hg38_21_+_8433172_8446622_RNA45SN1_45S_50ntFlanks</td><td>7985</td><td>8018</td><td>34</td><td>GATCAGACGTGGCGACCCGCTGAATTTAAGCATA</td><td>356.7</td><td>-3.7</td><td>1.7</td><td>-2.2</td><td>3.1E-02</td><td>3.4E-02</td><td>MB231/MB468</td><td>MITO</td></tr>
<tr><td>>trnaMT_GluTTC_MT_-_14674_14742@39.70.32,trnalookalike8_GluTTC_5_-_93905172_93905240@39.70.32|ATTGGTCGTGGTTGTAGTCCGTGCGAGAATAC</td><td>>trnaMT_GluTTC_MT_-_14674_14742</td><td>39</td><td>70</td><td>32</td><td>ATTGGTCGTGGTTGTAGTCCGTGCGAGAATAC</td><td>75.6</td><td>4.4</td><td>2.0</td><td>2.2</td><td>2.8E-02</td><td>3.4E-02</td><td>MB231/MB468</td><td>MP</td></tr>
<tr><td>>MI0000253|hsa-mir-148a&WithFlank&7|-|25949913|25949992@50.70.21(+1C)&[MIMAT0000243&hsa-miR-148a-3p&offsets|0|-1(+1C);m-7&7|-|25949922|25949943&offsets|0|-1(+1C)]|TCAGTGCACTACAGAACTTTGC</td><td>>MI0000253|hsa-mir-148a&WithFlank&7|-|25949913|25949992</td><td>50</td><td>70</td><td>21(+1C)</td><td>TCAGTGCACTACAGAACTTTGC</td><td>1672.6</td><td>2.3</td><td>1.1</td><td>2.2</td><td>3.1E-02</td><td>3.4E-02</td><td>MB231/MB468</td><td>MITO</td></tr>
<tr><td>>MI0000077|hsa-mir-21&WithFlank&17|+|59841260|59841343@14.35.22(+1U)&[MIMAT0000076&hsa-miR-21-5p&offsets|0|0(+1U);m-1&17|+|59841273|59841294&offsets|0|0(+1U)]|TAGCTTATCAGACTGATGTTGAT</td><td>>MI0000077|hsa-mir-21&WithFlank&17|+|59841260|59841343</td><td>14</td><td>35</td><td>22(+1U)</td><td>TAGCTTATCAGACTGATGTTGAT</td><td>177082.2</td><td>2.5</td><td>1.0</td><td>2.5</td><td>1.2E-02</td><td>3.4E-02</td><td>BT20/MB231</td><td>CYTO</td></tr>
<tr><td>>hg38_1_-_228634819_228635039_RNA5S12_5S_50ntFlanks@51.81.31|GTCTACGGCCATACCACCCTGAACGCGCCCG</td><td>>hg38_1_-_228634819_228635039_RNA5S12_5S_50ntFlanks</td><td>51</td><td>81</td><td>31</td><td>GTCTACGGCCATACCACCCTGAACGCGCCCG</td><td>655.7</td><td>2.5</td><td>1.0</td><td>2.6</td><td>8.7E-03</td><td>3.4E-02</td><td>BT20/MB231</td><td>MP</td></tr>
<tr><td>>hg38_21_+_8433172_8446622_RNA45SN1_45S_50ntFlanks@11598.11626.29|GGAATCCGACTGTTTAATTAAAACAAAGC</td><td>>hg38_21_+_8433172_8446622_RNA45SN1_45S_50ntFlanks</td><td>11598</td><td>11626</td><td>29</td><td>GGAATCCGACTGTTTAATTAAAACAAAGC</td><td>158.6</td><td>5.3</td><td>2.0</td><td>2.6</td><td>8.7E-03</td><td>3.4E-02</td><td>BT20/MB231</td><td>MP</td></tr>
<tr><td>>trnaMT_SerGCT_MT_+_12207_12265@27.62.36|CATGCCCCCATGTCTAACAACATGGCTTTCTCACCA</td><td>>trnaMT_SerGCT_MT_+_12207_12265</td><td>27</td><td>62</td><td>36</td><td>CATGCCCCCATGTCTAACAACATGGCTTTCTCACCA</td><td>156.7</td><td>3.8</td><td>1.6</td><td>2.3</td><td>2.0E-02</td><td>3.4E-02</td><td>BT20/MB468</td><td>MP</td></tr>
<tr><td>>hg38_21_+_8433172_8446622_RNA45SN1_45S_50ntFlanks@5473.5500.28|CGGCCCTGGCGGAGCGCTGAGAAGACGG</td><td>>hg38_21_+_8433172_8446622_RNA45SN1_45S_50ntFlanks</td><td>5473</td><td>5500</td><td>28</td><td>CGGCCCTGGCGGAGCGCTGAGAAGACGG</td><td>208.0</td><td>3.5</td><td>1.3</td><td>2.6</td><td>8.7E-03</td><td>3.4E-02</td><td>BT20/MB231</td><td>MP</td></tr>
<tr><td>>trnaMT_HisGTG_MT_+_12138_12206@28.63.36|ATTGTGAATCTGACAACAGAGGCTTACGACCCCTTA</td><td>>trnaMT_HisGTG_MT_+_12138_12206</td><td>28</td><td>63</td><td>36</td><td>ATTGTGAATCTGACAACAGAGGCTTACGACCCCTTA</td><td>303.5</td><td>3.1</td><td>1.3</td><td>2.3</td><td>2.0E-02</td><td>3.4E-02</td><td>BT20/MB468</td><td>MP</td></tr>
<tr><td>>trnaMT_ValTAC_MT_+_1602_1670@46.72.27|TTCAACTTAACTTGACCGCTCTGACCA</td><td>>trnaMT_ValTAC_MT_+_1602_1670</td><td>46</td><td>72</td><td>27</td><td>TTCAACTTAACTTGACCGCTCTGACCA</td><td>241.2</td><td>2.5</td><td>1.1</td><td>2.3</td><td>2.0E-02</td><td>3.4E-02</td><td>BT20/MB468</td><td>MP</td></tr>
<tr><td>>hg38_21_+_8433172_8446622_RNA45SN1_45S_50ntFlanks@9931.9950.20|TATAGACAGCAGGACGGTGG</td><td>>hg38_21_+_8433172_8446622_RNA45SN1_45S_50ntFlanks</td><td>9931</td><td>9950</td><td>20</td><td>TATAGACAGCAGGACGGTGG</td><td>114.8</td><td>4.3</td><td>1.9</td><td>2.3</td><td>2.0E-02</td><td>3.4E-02</td><td>BT20/MB468</td><td>MP</td></tr>
<tr><td>>MI0000767|hsa-mir-365a&WithFlank&16|+|14309279|14309377@22.45.24&[MIMAT0009199&hsa-miR-365a-5p&offsets|0|+1;m-595&16|+|14309300|14309322&offsets|0|+1]|AGGGACTTTTGGGGGCAGATGTGT</td><td>>MI0000767|hsa-mir-365a&WithFlank&16|+|14309279|14309377</td><td>22</td><td>45</td><td>24</td><td>AGGGACTTTTGGGGGCAGATGTGT</td><td>234.8</td><td>2.8</td><td>1.1</td><td>2.6</td><td>8.7E-03</td><td>3.4E-02</td><td>BT20/MB231</td><td>MP</td></tr>
<tr><td>>MI0000299|hsa-mir-222&WithFlank&X|-|45747009|45747130@75.99.25(+1C)&[MIMAT0000279&hsa-miR-222-3p&offsets|0|+4(+1C);m-140&X|-|45747033|45747056&offsets|0|+1(+1C)]|AGCTACATCTGGCTACTGGGTCTCTC</td><td>>MI0000299|hsa-mir-222&WithFlank&X|-|45747009|45747130</td><td>75</td><td>99</td><td>25(+1C)</td><td>AGCTACATCTGGCTACTGGGTCTCTC</td><td>183.5</td><td>-3.1</td><td>1.2</td><td>-2.6</td><td>8.7E-03</td><td>3.4E-02</td><td>BT20/MB231</td><td>MP</td></tr>
<tr><td>>MI0000811|hsa-mir-148b&WithFlank&12|+|54337210|54337320@69.88.20(+1A)&[MIMAT0000759&hsa-miR-148b-3p&offsets|0|-2(+1A);m-77&12|+|54337278|54337299&offsets|0|-2(+1A)]|TCAGTGCATCACAGAACTTTA</td><td>>MI0000811|hsa-mir-148b&WithFlank&12|+|54337210|54337320</td><td>69</td><td>88</td><td>20(+1A)</td><td>TCAGTGCATCACAGAACTTTA</td><td>69.5</td><td>4.5</td><td>2.0</td><td>2.2</td><td>2.8E-02</td><td>3.4E-02</td><td>MB231/MB468</td><td>MP</td></tr>
<tr><td>>MI0000746|hsa-mir-99b&WithFlank&19|+|51692606|51692687@51.71.21(+1A)&[MIMAT0004678&hsa-miR-99b-3p&offsets|0|-1(+1A);m-151&19|+|51692656|51692678&offsets|0|-2(+1A)]|CAAGCTCGTGTCTGTGGGTCCA</td><td>>MI0000746|hsa-mir-99b&WithFlank&19|+|51692606|51692687</td><td>51</td><td>71</td><td>21(+1A)</td><td>CAAGCTCGTGTCTGTGGGTCCA</td><td>404.9</td><td>1.9</td><td>0.8</td><td>2.5</td><td>1.1E-02</td><td>3.4E-02</td><td>BT20/MB231</td><td>MITO</td></tr>
<tr><td>>hg38_21_+_8433172_8446622_RNA45SN1_45S_50ntFlanks@9837.9854.18|TGGTAAGCAGAACTGGCG</td><td>>hg38_21_+_8433172_8446622_RNA45SN1_45S_50ntFlanks</td><td>9837</td><td>9854</td><td>18</td><td>TGGTAAGCAGAACTGGCG</td><td>58.6</td><td>6.0</td><td>2.6</td><td>2.3</td><td>2.0E-02</td><td>3.4E-02</td><td>BT20/MB468</td><td>MP</td></tr>
<tr><td>>trna16_GlnTTG_17_+_47269890_47269961@1.29.29,trna64_GlnTTG_6_+_28557156_28557227@1.29.29,trna84_GlnTTG_6_+_145503859_145503930@1.29.29|GGTCCCATGGTGTAATGGTTAGCACTCTG</td><td>>trna16_GlnTTG_17_+_47269890_47269961</td><td>1</td><td>29</td><td>29</td><td>GGTCCCATGGTGTAATGGTTAGCACTCTG</td><td>893.4</td><td>-3.5</td><td>1.5</td><td>-2.3</td><td>2.0E-02</td><td>3.4E-02</td><td>BT20/MB468</td><td>MP</td></tr>
<tr><td>>hg38_21_+_8433172_8446622_RNA45SN1_45S_50ntFlanks@9940.9959.20|CAGGACGGTGGCCATGGAAG</td><td>>hg38_21_+_8433172_8446622_RNA45SN1_45S_50ntFlanks</td><td>9940</td><td>9959</td><td>20</td><td>CAGGACGGTGGCCATGGAAG</td><td>193.4</td><td>4.2</td><td>1.8</td><td>2.3</td><td>2.0E-02</td><td>3.4E-02</td><td>BT20/MB468</td><td>MP</td></tr>
<tr><td>>hg38_21_+_8433172_8446622_RNA45SN1_45S_50ntFlanks@9953.9979.27|ATGGAAGTCGGAATCCGCTAAGGAGTG</td><td>>hg38_21_+_8433172_8446622_RNA45SN1_45S_50ntFlanks</td><td>9953</td><td>9979</td><td>27</td><td>ATGGAAGTCGGAATCCGCTAAGGAGTG</td><td>206.5</td><td>2.5</td><td>1.1</td><td>2.3</td><td>2.0E-02</td><td>3.4E-02</td><td>BT20/MB468</td><td>MP</td></tr>
</table>

<table>
<tr><th></th><th></th><th></th><th></th><th></th><th></th><th></th><th></th><th></th><th></th><th></th><th></th><th></th><th></th></tr>
<tr><td>>hg38_21_+_8433172_8446622_RNA45SN1_45S_50ntFlanks@12474.12499.26|ACCCACTAATAGGGAACGTGAGCTGG</td><td>>hg38_21_+_8433172_8446622_RNA45SN1_45S_50ntFlanks</td><td>12474</td><td>12499</td><td>26</td><td>ACCCACTAATAGGGAACGTGAGCTGG</td><td>167.0</td><td>-2.0</td><td>0.9</td><td>-2.2</td><td>2.9E-02</td><td>3.4E-02</td><td>MB231/MB468</td><td>CYTO</td></tr>
<tr><td>>MI0000465|hsa-mir-191&WithFlank&3|-|49020612|49020715@22.43.22&[MIMAT0000440&hsa-miR-191-5p&offsets|0|-1;m-48&3|-|49020672|49020694&offsets|0|-1]|CAACGGAATCCCAAAAGCAGCT</td><td>>MI0000465|hsa-mir-191&WithFlank&3|-|49020612|49020715</td><td>22</td><td>43</td><td>22</td><td>CAACGGAATCCCAAAAGCAGCT</td><td>4444.9</td><td>2.6</td><td>1.1</td><td>2.3</td><td>2.0E-02</td><td>3.4E-02</td><td>BT20/MB468</td><td>MP</td></tr>
<tr><td>>hg38_21_+_8433172_8446622_RNA45SN1_45S_50ntFlanks@4884.4904.21|CAAAGCTGAAACTTAAAGGAA</td><td>>hg38_21_+_8433172_8446622_RNA45SN1_45S_50ntFlanks</td><td>4884</td><td>4904</td><td>21</td><td>CAAAGCTGAAACTTAAAGGAA</td><td>108.0</td><td>-3.2</td><td>1.4</td><td>-2.3</td><td>2.0E-02</td><td>3.4E-02</td><td>BT20/MB468</td><td>MP</td></tr>
<tr><td>>hg38_21_+_8433172_8446622_RNA45SN1_45S_50ntFlanks@6645.6666.22|CTCGTACGACTCTTAGCGGTGG</td><td>>hg38_21_+_8433172_8446622_RNA45SN1_45S_50ntFlanks</td><td>6645</td><td>6666</td><td>22</td><td>CTCGTACGACTCTTAGCGGTGG</td><td>2852.7</td><td>1.8</td><td>0.8</td><td>2.2</td><td>3.1E-02</td><td>3.4E-02</td><td>MB231/MB468</td><td>NUC</td></tr>
<tr><td>>hg38_21_+_8433172_8446622_RNA45SN1_45S_50ntFlanks@11667.11702.36|CTGCCCAGTGCTCTGAATGTCAAAGTGAAGAAATTC</td><td>>hg38_21_+_8433172_8446622_RNA45SN1_45S_50ntFlanks</td><td>11667</td><td>11702</td><td>36</td><td>CTGCCCAGTGCTCTGAATGTCAAAGTGAAGAAATTC</td><td>113.1</td><td>4.1</td><td>1.8</td><td>2.3</td><td>2.0E-02</td><td>3.4E-02</td><td>BT20/MB468</td><td>MP</td></tr>
<tr><td>>hg38_21_+_8433172_8446622_RNA45SN1_45S_50ntFlanks@9546.9571.26|TGGAGGTCCGTAGCGGTCCTGACGTG</td><td>>hg38_21_+_8433172_8446622_RNA45SN1_45S_50ntFlanks</td><td>9546</td><td>9571</td><td>26</td><td>TGGAGGTCCGTAGCGGTCCTGACGTG</td><td>1819.3</td><td>-2.1</td><td>0.8</td><td>-2.5</td><td>1.2E-02</td><td>3.4E-02</td><td>BT20/MB231</td><td>CYTO</td></tr>
<tr><td>>hg38_21_+_8433172_8446622_RNA45SN1_45S_50ntFlanks@9939.9956.18|GCAGGACGGTGGCCATGG</td><td>>hg38_21_+_8433172_8446622_RNA45SN1_45S_50ntFlanks</td><td>9939</td><td>9956</td><td>18</td><td>GCAGGACGGTGGCCATGG</td><td>451.3</td><td>3.1</td><td>1.2</td><td>2.6</td><td>8.7E-03</td><td>3.4E-02</td><td>BT20/MB231</td><td>MP</td></tr>
<tr><td>>hg38_1_-_228634819_228635039_RNA5S12_5S_50ntFlanks@134.172.39|TGGGAGACCGCCTGGGAATACCGGGTGCTGTAGGCTTTT</td><td>>hg38_1_-_228634819_228635039_RNA5S12_5S_50ntFlanks</td><td>134</td><td>172</td><td>39</td><td>TGGGAGACCGCCTGGGAATACCGGGTGCTGTAGGCTTTT</td><td>835.4</td><td>-4.8</td><td>1.8</td><td>-2.6</td><td>8.7E-03</td><td>3.4E-02</td><td>BT20/MB231</td><td>MP</td></tr>
<tr><td>>trnaMT_ValTAC_MT_+_1602_1670@52.72.21|TTAACTTGACCGCTCTGACCA</td><td>>trnaMT_ValTAC_MT_+_1602_1670</td><td>52</td><td>72</td><td>21</td><td>TTAACTTGACCGCTCTGACCA</td><td>809.8</td><td>2.0</td><td>0.9</td><td>2.2</td><td>3.1E-02</td><td>3.4E-02</td><td>MB231/MB468</td><td>MITO</td></tr>
<tr><td>>hg38_21_+_8433172_8446622_RNA45SN1_45S_50ntFlanks@10386.10424.39|ATGTGAACAGCAGTTGAACATGGGTCAGTCGGTCCTGAG</td><td>>hg38_21_+_8433172_8446622_RNA45SN1_45S_50ntFlanks</td><td>10386</td><td>10424</td><td>39</td><td>ATGTGAACAGCAGTTGAACATGGGTCAGTCGGTCCTGAG</td><td>326.6</td><td>3.9</td><td>1.8</td><td>2.2</td><td>2.8E-02</td><td>3.4E-02</td><td>MB231/MB468</td><td>MP</td></tr>
<tr><td>>hg38_21_+_8433172_8446622_RNA45SN1_45S_50ntFlanks@11579.11607.29|AGAACTGGTGCGGACCAGGGGAATCCGAC</td><td>>hg38_21_+_8433172_8446622_RNA45SN1_45S_50ntFlanks</td><td>11579</td><td>11607</td><td>29</td><td>AGAACTGGTGCGGACCAGGGGAATCCGAC</td><td>570.9</td><td>-2.4</td><td>1.1</td><td>-2.3</td><td>2.1E-02</td><td>3.4E-02</td><td>BT20/MB468</td><td>MP</td></tr>
<tr><td>>trna119_LysCTT_1_-_145395522_145395594@1.36.36,trna11_LysCTT_5_-_180648979_180649051@1.36.36,trna13_LysCTT_6_+_26556774_26556846@1.36.36,trna7_LysCTT_16_+_3225692_3225764@1.36.36,trna9_LysCTT_5_+_180634755_180634827@1.36.36|GCCCGGCTAGCTCAGTCGGTAGAGCATGAGACTCTT</td><td>>trna119_LysCTT_1_-_145395522_145395594</td><td>1</td><td>36</td><td>36</td><td>GCCCGGCTAGCTCAGTCGGTAGAGCATGAGACTCTT</td><td>4666.5</td><td>2.5</td><td>1.0</td><td>2.5</td><td>1.2E-02</td><td>3.4E-02</td><td>BT20/MB231</td><td>CYTO</td></tr>
<tr><td>>hg38_1_-_228634819_228635039_RNA5S12_5S_50ntFlanks@150.170.21|AATACCGGGTGCTGTAGGCTT</td><td>>hg38_1_-_228634819_228635039_RNA5S12_5S_50ntFlanks</td><td>150</td><td>170</td><td>21</td><td>AATACCGGGTGCTGTAGGCTT</td><td>1625.9</td><td>2.3</td><td>1.1</td><td>2.2</td><td>3.1E-02</td><td>3.4E-02</td><td>MB231/MB468</td><td>NUC</td></tr>
<tr><td>>hg38_21_+_8433172_8446622_RNA45SN1_45S_50ntFlanks@10795.10822.28|TGGCATGTTGGAACAATGTAGGTAAGGG</td><td>>hg38_21_+_8433172_8446622_RNA45SN1_45S_50ntFlanks</td><td>10795</td><td>10822</td><td>28</td><td>TGGCATGTTGGAACAATGTAGGTAAGGG</td><td>154.5</td><td>-1.9</td><td>0.8</td><td>-2.3</td><td>2.1E-02</td><td>3.4E-02</td><td>BT20/MB468</td><td>MP</td></tr>
<tr><td>>trnaMT_PheGAA_MT_+_577_647@42.73.32|TTTAGACGGGCTCACATCACCCCATAAACACC</td><td>>trnaMT_PheGAA_MT_+_577_647</td><td>42</td><td>73</td><td>32</td><td>TTTAGACGGGCTCACATCACCCCATAAACACC</td><td>92.4</td><td>2.0</td><td>0.9</td><td>2.2</td><td>2.8E-02</td><td>3.4E-02</td><td>MB231/MB468</td><td>MP</td></tr>
<tr><td>>hg38_21_+_8433172_8446622_RNA45SN1_45S_50ntFlanks@8168.8187.20|CCCAAGTCCTTCTGATCGAG</td><td>>hg38_21_+_8433172_8446622_RNA45SN1_45S_50ntFlanks</td><td>8168</td><td>8187</td><td>20</td><td>CCCAAGTCCTTCTGATCGAG</td><td>351.4</td><td>-3.8</td><td>1.7</td><td>-2.2</td><td>3.0E-02</td><td>3.4E-02</td><td>BT20/MB468</td><td>CYTO</td></tr>
<tr><td>>hg38_21_+_8433172_8446622_RNA45SN1_45S_50ntFlanks@7974.8003.30|ACGCGACCTCAGATCAGACGTGGCGACCCG</td><td>>hg38_21_+_8433172_8446622_RNA45SN1_45S_50ntFlanks</td><td>7974</td><td>8003</td><td>30</td><td>ACGCGACCTCAGATCAGACGTGGCGACCCG</td><td>677.6</td><td>2.1</td><td>1.0</td><td>2.2</td><td>3.0E-02</td><td>3.4E-02</td><td>BT20/MB468</td><td>CYTO</td></tr>
<tr><td>>hg38_21_+_8433172_8446622_RNA45SN1_45S_50ntFlanks@10873.10902.30|CTGGGTCGGTCGGGCTGGGGCGCGAAGCGG</td><td>>hg38_21_+_8433172_8446622_RNA45SN1_45S_50ntFlanks</td><td>10873</td><td>10902</td><td>30</td><td>CTGGGTCGGTCGGGCTGGGGCGCGAAGCGG</td><td>336.2</td><td>-2.6</td><td>1.2</td><td>-2.2</td><td>2.6E-02</td><td>3.4E-02</td><td>BT20/MB468</td><td>MITO</td></tr>
<tr><td>>hg38_21_+_8433172_8446622_RNA45SN1_45S_50ntFlanks@12417.12450.34|TCGATGTCGGCTCTTCCTATCATTGTGAAGCAGA</td><td>>hg38_21_+_8433172_8446622_RNA45SN1_45S_50ntFlanks</td><td>12417</td><td>12450</td><td>34</td><td>TCGATGTCGGCTCTTCCTATCATTGTGAAGCAGA</td><td>194.3</td><td>-2.0</td><td>0.8</td><td>-2.5</td><td>1.2E-02</td><td>3.4E-02</td><td>BT20/MB231</td><td>CYTO</td></tr>
<tr><td>>hg38_21_+_8433172_8446622_RNA45SN1_45S_50ntFlanks@9534.9573.40|AGGAAACTCTGGTGGAGGTCCGTAGCGGTCCTGACGTGCA</td><td>>hg38_21_+_8433172_8446622_RNA45SN1_45S_50ntFlanks</td><td>9534</td><td>9573</td><td>40</td><td>AGGAAACTCTGGTGGAGGTCCGTAGCGGTCCTGACGTGCA</td><td>433.5</td><td>-2.1</td><td>0.8</td><td>-2.5</td><td>1.2E-02</td><td>3.4E-02</td><td>BT20/MB231</td><td>CYTO</td></tr>
<tr><td>>hg38_21_+_8433172_8446622_RNA45SN1_45S_50ntFlanks@10391.10433.43|AACAGCAGTTGAACATGGGTCAGTCGGTCCTGAGAGATGGGCG</td><td>>hg38_21_+_8433172_8446622_RNA45SN1_45S_50ntFlanks</td><td>10391</td><td>10433</td><td>43</td><td>AACAGCAGTTGAACATGGGTCAGTCGGTCCTGAGAGATGGGCG</td><td>112.4</td><td>-3.6</td><td>1.5</td><td>-2.3</td><td>2.1E-02</td><td>3.4E-02</td><td>BT20/MB468</td><td>MP</td></tr>
<tr><td>>trnaMT_SerGCT_MT_+_12207_12265@10.31.22|CACAAGAACTGCTAACTCATGC</td><td>>trnaMT_SerGCT_MT_+_12207_12265</td><td>10</td><td>31</td><td>22</td><td>CACAAGAACTGCTAACTCATGC</td><td>152.1</td><td>-2.5</td><td>1.2</td><td>-2.2</td><td>3.0E-02</td><td>3.4E-02</td><td>MB231/MB468</td><td>CYTO</td></tr>
<tr><td>>MI0000286|hsa-mir-210&WithFlank&11|-|568083|568204@72.92.21&[MIMAT0000267&hsa-miR-210-3p&offsets|0|-1;m-102&11|-|568112|568133&offsets|0|-1]|CTGTGCGTGTGACAGCGGCTG</td><td>>MI0000286|hsa-mir-210&WithFlank&11|-|568083|568204</td><td>72</td><td>92</td><td>21</td><td>CTGTGCGTGTGACAGCGGCTG</td><td>78.6</td><td>3.1</td><td>1.4</td><td>2.2</td><td>2.8E-02</td><td>3.4E-02</td><td>MB231/MB468</td><td>MP</td></tr>
<tr><td>>trnaMT_LysTTT_MT_+_8295_8364@53.73.21,trnalookalike7_LysTTT_1_+_568844_568913@53.73.21|AACACCTCTTTACAGTGACCA</td><td>>trnaMT_LysTTT_MT_+_8295_8364</td><td>53</td><td>73</td><td>21</td><td>AACACCTCTTTACAGTGACCA</td><td>755.2</td><td>-3.1</td><td>1.2</td><td>-2.6</td><td>8.8E-03</td><td>3.4E-02</td><td>BT20/MB231</td><td>MP</td></tr>
<tr><td>>hg38_21_+_8433172_8446622_RNA45SN1_45S_50ntFlanks@6723.6746.24|TTGCAGGACACATTGATCATCGAC</td><td>>hg38_21_+_8433172_8446622_RNA45SN1_45S_50ntFlanks</td><td>6723</td><td>6746</td><td>24</td><td>TTGCAGGACACATTGATCATCGAC</td><td>105.8</td><td>-3.9</td><td>1.7</td><td>-2.3</td><td>2.1E-02</td><td>3.4E-02</td><td>BT20/MB468</td><td>MP</td></tr>
<tr><td>>hg38_21_+_8433172_8446622_RNA45SN1_45S_50ntFlanks@6651.6668.18|CGACTCTTAGCGGTGGAT</td><td>>hg38_21_+_8433172_8446622_RNA45SN1_45S_50ntFlanks</td><td>6651</td><td>6668</td><td>18</td><td>CGACTCTTAGCGGTGGAT</td><td>9424.5</td><td>3.0</td><td>1.1</td><td>2.6</td><td>8.8E-03</td><td>3.4E-02</td><td>BT20/MB231</td><td>MP</td></tr>
<tr><td>>MI0000103|hsa-mir-101-1&WithFlank&1|-|65058428|65058514@53.73.21(+1U);MI0000739|hsa-mir-101-2&WithFlank&9|+|4850291|4850381@55.75.21(+1U)&[MIMAT0000099&hsa-miR-101-3p&offsets|0|0(+1U);m-17&1|-|65058443|65058462&offsets|0|+1(+1U)];[MIMAT0000099_1&hsa-miR-101-3p&offsets|0|0(+1U);m-16&9|+|4850345|4850364&offsets|0|+1(+1U)]|TACAGTACTGTGATAACTGAAT</td><td>>MI0000103|hsa-mir-101-1&WithFlank&1|-|65058428|65058514</td><td>53</td><td>73</td><td>21(+1U)</td><td>TACAGTACTGTGATAACTGAAT</td><td>776.9</td><td>-4.0</td><td>1.5</td><td>-2.6</td><td>8.8E-03</td><td>3.4E-02</td><td>BT20/MB231</td><td>MP</td></tr>
<tr><td>>hg38_21_+_8433172_8446622_RNA45SN1_45S_50ntFlanks@9574.9610.37|AATCGGTCGTCCGACCTGGGTATAGGGGCGAAAGACT</td><td>>hg38_21_+_8433172_8446622_RNA45SN1_45S_50ntFlanks</td><td>9574</td><td>9610</td><td>37</td><td>AATCGGTCGTCCGACCTGGGTATAGGGGCGAAAGACT</td><td>156.4</td><td>2.1</td><td>0.9</td><td>2.2</td><td>2.8E-02</td><td>3.4E-02</td><td>MB231/MB468</td><td>MP</td></tr>
<tr><td>>hg38_21_+_8433172_8446622_RNA45SN1_45S_50ntFlanks@10387.10407.21|TGTGAACAGCAGTTGAACATG</td><td>>hg38_21_+_8433172_8446622_RNA45SN1_45S_50ntFlanks</td><td>10387</td><td>10407</td><td>21</td><td>TGTGAACAGCAGTTGAACATG</td><td>163.3</td><td>-1.7</td><td>0.8</td><td>-2.2</td><td>2.6E-02</td><td>3.4E-02</td><td>BT20/MB468</td><td>MITO</td></tr>
<tr><td>>hg38_21_+_8433172_8446622_RNA45SN1_45S_50ntFlanks@9937.9955.19|CAGCAGGACGGTGGCCATG</td><td>>hg38_21_+_8433172_8446622_RNA45SN1_45S_50ntFlanks</td><td>9937</td><td>9955</td><td>19</td><td>CAGCAGGACGGTGGCCATG</td><td>310.5</td><td>4.1</td><td>1.6</td><td>2.6</td><td>8.8E-03</td><td>3.4E-02</td><td>BT20/MB231</td><td>MP</td></tr>
<tr><td>>MI0001448|hsa-mir-425&WithFlank&3|-|49020142|49020240@20.43.24(+1C)&[MIMAT0003393&hsa-miR-425-5p&offsets|0|+1(+1C);m-95&3|-|49020200|49020221&offsets|0|+2(+1C)]|AATGACACGATCACTCCCGTTGAGC</td><td>>MI0001448|hsa-mir-425&WithFlank&3|-|49020142|49020240</td><td>20</td><td>43</td><td>24(+1C)</td><td>AATGACACGATCACTCCCGTTGAGC</td><td>179.3</td><td>3.0</td><td>1.4</td><td>2.2</td><td>2.8E-02</td><td>3.4E-02</td><td>MB231/MB468</td><td>MP</td></tr>
<tr><td>>hg38_21_+_8433172_8446622_RNA45SN1_45S_50ntFlanks@11568.11586.19|GCAGCCGACTTAGAACTGG</td><td>>hg38_21_+_8433172_8446622_RNA45SN1_45S_50ntFlanks</td><td>11568</td><td>11586</td><td>19</td><td>GCAGCCGACTTAGAACTGG</td><td>554.6</td><td>-2.2</td><td>0.9</td><td>-2.5</td><td>1.2E-02</td><td>3.4E-02</td><td>BT20/MB231</td><td>CYTO</td></tr>
<tr><td>>trnaMT_PheGAA_MT_+_577_647@1.34.34|GTTTATGTAGCTTACCTCCTCAAAGCAATACACT</td><td>>trnaMT_PheGAA_MT_+_577_647</td><td>1</td><td>34</td><td>34</td><td>GTTTATGTAGCTTACCTCCTCAAAGCAATACACT</td><td>578.2</td><td>3.5</td><td>1.5</td><td>2.3</td><td>2.1E-02</td><td>3.4E-02</td><td>BT20/MB468</td><td>MP</td></tr>
<tr><td>>trnaMT_PheGAA_MT_+_577_647@39.60.22|ATGTTTAGACGGGCTCACATCA</td><td>>trnaMT_PheGAA_MT_+_577_647</td><td>39</td><td>60</td><td>22</td><td>ATGTTTAGACGGGCTCACATCA</td><td>62.0</td><td>2.0</td><td>0.9</td><td>2.2</td><td>2.8E-02</td><td>3.4E-02</td><td>MB231/MB468</td><td>MP</td></tr>
<tr><td>>hg38_21_+_8433172_8446622_RNA45SN1_45S_50ntFlanks@9794.9819.26|CGTGGAGCCGGGCGTGGAATGCGAGT</td><td>>hg38_21_+_8433172_8446622_RNA45SN1_45S_50ntFlanks</td><td>9794</td><td>9819</td><td>26</td><td>CGTGGAGCCGGGCGTGGAATGCGAGT</td><td>107.0</td><td>3.3</td><td>1.4</td><td>2.3</td><td>2.1E-02</td><td>3.4E-02</td><td>BT20/MB468</td><td>MP</td></tr>
<tr><td>>hg38_21_+_8433172_8446622_RNA45SN1_45S_50ntFlanks@7975.8007.33|CGCGACCTCAGATCAGACGTGGCGACCCGCTGA</td><td>>hg38_21_+_8433172_8446622_RNA45SN1_45S_50ntFlanks</td><td>7975</td><td>8007</td><td>33</td><td>CGCGACCTCAGATCAGACGTGGCGACCCGCTGA</td><td>7890.8</td><td>2.0</td><td>0.9</td><td>2.3</td><td>2.1E-02</td><td>3.4E-02</td><td>BT20/MB468</td><td>MP</td></tr>
<tr><td>>am_trnaMT_PheGAA_MT_+_577_647@40.64.25|TGTTTAGACGGGCTCACATCACCCC</td><td>>am_trnaMT_PheGAA_MT_+_577_647</td><td>40</td><td>64</td><td>25</td><td>TGTTTAGACGGGCTCACATCACCCC</td><td>24.4</td><td>4.5</td><td>2.1</td><td>2.2</td><td>2.8E-02</td><td>3.4E-02</td><td>MB231/MB468</td><td>MP</td></tr>
<tr><td>>MI0000746|hsa-mir-99b&WithFlank&19|+|51692606|51692687@13.33.21(+1A)&[MIMAT0000689&hsa-miR-99b-5p&offsets|0|-1(+1A);m-10&19|+|51692618|51692639&offsets|0|-1(+1A)]|CACCCGTAGAACCGACCTTGCA</td><td>>MI0000746|hsa-mir-99b&WithFlank&19|+|51692606|51692687</td><td>13</td><td>33</td><td>21(+1A)</td><td>CACCCGTAGAACCGACCTTGCA</td><td>25565.3</td><td>2.3</td><td>0.9</td><td>2.5</td><td>1.2E-02</td><td>3.4E-02</td><td>BT20/MB231</td><td>CYTO</td></tr>
<tr><td>>hg38_21_+_8433172_8446622_RNA45SN1_45S_50ntFlanks@11573.11600.28|CGACTTAGAACTGGTGCGGACCAGGGGA</td><td>>hg38_21_+_8433172_8446622_RNA45SN1_45S_50ntFlanks</td><td>11573</td><td>11600</td><td>28</td><td>CGACTTAGAACTGGTGCGGACCAGGGGA</td><td>325.9</td><td>1.3</td><td>0.6</td><td>2.2</td><td>2.6E-02</td><td>3.4E-02</td><td>BT20/MB468</td><td>MITO</td></tr>
<tr><td>>hg38_21_+_8433172_8446622_RNA45SN1_45S_50ntFlanks@7983.8016.34|CAGATCAGACGTGGCGACCCGCTGAATTTAAGCA</td><td>>hg38_21_+_8433172_8446622_RNA45SN1_45S_50ntFlanks</td><td>7983</td><td>8016</td><td>34</td><td>CAGATCAGACGTGGCGACCCGCTGAATTTAAGCA</td><td>112.6</td><td>-3.3</td><td>1.4</td><td>-2.3</td><td>2.1E-02</td><td>3.4E-02</td><td>BT20/MB468</td><td>MP</td></tr>
<tr><td>>hg38_21_+_8433172_8446622_RNA45SN1_45S_50ntFlanks@10679.10705.27|GGTTCCGGCGGCGTCCGGTGAGCTCTC</td><td>>hg38_21_+_8433172_8446622_RNA45SN1_45S_50ntFlanks</td><td>10679</td><td>10705</td><td>27</td><td>GGTTCCGGCGGCGTCCGGTGAGCTCTC</td><td>237.4</td><td>-3.6</td><td>1.6</td><td>-2.2</td><td>2.6E-02</td><td>3.4E-02</td><td>BT20/MB468</td><td>MITO</td></tr>
<tr><td>>hg38_21_+_8433172_8446622_RNA45SN1_45S_50ntFlanks@10513.10530.18|CCGGAGTGGCGGAGATGG</td><td>>hg38_21_+_8433172_8446622_RNA45SN1_45S_50ntFlanks</td><td>10513</td><td>10530</td><td>18</td><td>CCGGAGTGGCGGAGATGG</td><td>253.0</td><td>2.5</td><td>1.1</td><td>2.3</td><td>2.1E-02</td><td>3.4E-02</td><td>BT20/MB468</td><td>MP</td></tr>
<tr><td>>hg38_21_+_8433172_8446622_RNA45SN1_45S_50ntFlanks@10827.10848.22|CGGCAAGCCGGATCCGTAACTT</td><td>>hg38_21_+_8433172_8446622_RNA45SN1_45S_50ntFlanks</td><td>10827</td><td>10848</td><td>22</td><td>CGGCAAGCCGGATCCGTAACTT</td><td>67.9</td><td>-3.9</td><td>1.7</td><td>-2.3</td><td>2.1E-02</td><td>3.4E-02</td><td>BT20/MB468</td><td>MP</td></tr>
<tr><td>>hg38_21_+_8433172_8446622_RNA45SN1_45S_50ntFlanks@7980.8003.24|CCTCAGATCAGACGTGGCGACCCG</td><td>>hg38_21_+_8433172_8446622_RNA45SN1_45S_50ntFlanks</td><td>7980</td><td>8003</td><td>24</td><td>CCTCAGATCAGACGTGGCGACCCG</td><td>423.7</td><td>-2.2</td><td>1.0</td><td>-2.3</td><td>2.1E-02</td><td>3.4E-02</td><td>BT20/MB468</td><td>MP</td></tr>
<tr><td>>MI0000471|hsa-mir-126&WithFlank&9|+|136670596|136670692@58.78.21&[MIMAT0000445&hsa-miR-126-3p&offsets|0|-1;m-40&9|+|136670653|136670674&offsets|0|-1]|TCGTACCGTGAGTAATAATGC</td><td>>MI0000471|hsa-mir-126&WithFlank&9|+|136670596|136670692</td><td>58</td><td>78</td><td>21</td><td>TCGTACCGTGAGTAATAATGC</td><td>7938.5</td><td>-2.3</td><td>0.9</td><td>-2.6</td><td>8.8E-03</td><td>3.4E-02</td><td>BT20/MB231</td><td>MP</td></tr>
<tr><td>>hg38_21_+_8433172_8446622_RNA45SN1_45S_50ntFlanks@8104.8141.38|CCGCGGCGGGGCGCGGGACATGTGGCGTACGGAAGACC</td><td>>hg38_21_+_8433172_8446622_RNA45SN1_45S_50ntFlanks</td><td>8104</td><td>8141</td><td>38</td><td>CCGCGGCGGGGCGCGGGACATGTGGCGTACGGAAGACC</td><td>450.5</td><td>4.2</td><td>1.6</td><td>2.6</td><td>8.8E-03</td><td>3.4E-02</td><td>BT20/MB231</td><td>MP</td></tr>
<tr><td>>MI0000268|hsa-mir-34a&WithFlank&1|-|9151662|9151783@28.48.21(+1C)&[MIMAT0000255&hsa-miR-34a-5p&offsets|0|-1(+1C);m-119&1|-|9151735|9151756&offsets|0|-1(+1C)]|TGGCAGTGTCTTAGCTGGTTGC</td><td>>MI0000268|hsa-mir-34a&WithFlank&1|-|9151662|9151783</td><td>28</td><td>48</td><td>21(+1C)</td><td>TGGCAGTGTCTTAGCTGGTTGC</td><td>131.3</td><td>-2.1</td><td>1.0</td><td>-2.2</td><td>2.8E-02</td><td>3.4E-02</td><td>MB231/MB468</td><td>MP</td></tr>
<tr><td>>hg38_21_+_8433172_8446622_RNA45SN1_45S_50ntFlanks@12458.12487.30|AAGCGTTGGATTGTTCACCCACTAATAGGG</td><td>>hg38_21_+_8433172_8446622_RNA45SN1_45S_50ntFlanks</td><td>12458</td><td>12487</td><td>30</td><td>AAGCGTTGGATTGTTCACCCACTAATAGGG</td><td>155.5</td><td>-3.3</td><td>1.4</td><td>-2.3</td><td>2.1E-02</td><td>3.4E-02</td><td>BT20/MB468</td><td>MP</td></tr>
<tr><td>>hg38_21_+_8433172_8446622_RNA45SN1_45S_50ntFlanks@3842.3869.28|CCTACTTGGATAACTGTGGTAATTCTAG</td><td>>hg38_21_+_8433172_8446622_RNA45SN1_45S_50ntFlanks</td><td>3842</td><td>3869</td><td>28</td><td>CCTACTTGGATAACTGTGGTAATTCTAG</td><td>73.6</td><td>-4.1</td><td>1.8</td><td>-2.3</td><td>2.1E-02</td><td>3.4E-02</td><td>BT20/MB468</td><td>MP</td></tr>
<tr><td>>hg38_21_+_8433172_8446622_RNA45SN1_45S_50ntFlanks@12479.12499.21|CTAATAGGGAACGTGAGCTGG</td><td>>hg38_21_+_8433172_8446622_RNA45SN1_45S_50ntFlanks</td><td>12479</td><td>12499</td><td>21</td><td>CTAATAGGGAACGTGAGCTGG</td><td>673.0</td><td>-1.4</td><td>0.7</td><td>-2.2</td><td>3.0E-02</td><td>3.4E-02</td><td>MB231/MB468</td><td>CYTO</td></tr>
<tr><td>>hg38_21_+_8433172_8446622_RNA45SN1_45S_50ntFlanks@12636.12664.29|AGCTACCATCTGTGGGATTATGACTGAAC</td><td>>hg38_21_+_8433172_8446622_RNA45SN1_45S_50ntFlanks</td><td>12636</td><td>12664</td><td>29</td><td>AGCTACCATCTGTGGGATTATGACTGAAC</td><td>111.8</td><td>5.2</td><td>2.0</td><td>2.6</td><td>8.8E-03</td><td>3.4E-02</td><td>BT20/MB231</td><td>MP</td></tr>
</table>

| | | | | | | | | | | | | | |
|---|---|---|---|---|---|---|---|---|---|---|---|---|---|
| >hg38_21_+_8433172_8446622_RNA45SN1_45S_50ntFlanks@5473.5501.29\|CGGCCCTGGCGGAGCGCTGAGAAGACGGT | >hg38_21_+_8433172_8446622_RNA45SN1_45S_50ntFlanks | 5473 | 5501 | 29 | CGGCCCTGGCGGAGCGCTGAGAAGACGGT | 252.3 | 4.1 | 1.6 | 2.6 | 8.8E-03 | 3.4E-02 | BT20/MB231 | MP |
| >MI0000078\|hsa-mir-22&WithFlank&17\|-\|1713897\|1713993@21.42.22&[MIMAT0004495&hsa-miR-22-5p&offsets\|0\|0;m-217&17\|-\|1713953\|1713973&offsets\|0\|+1]\|AGTTCTTCAGTGGCAAGCTTTA | >MI0000078\|hsa-mir-22&WithFlank&17\|-\|1713897\|1713993 | 21 | 42 | 22 | AGTTCTTCAGTGGCAAGCTTTA | 1326.1 | -2.8 | 1.1 | -2.6 | 8.9E-03 | 3.4E-02 | BT20/MB231 | MP |
| >MI0000270\|hsa-mir-181b-1&WithFlank&1\|-\|198858867\|198858988@42.65.24(+1A);MI0000683\|hsa-mir-181b-2&WithFlank&9\|+\|124693704\|124693804@22.45.24(+1A)&[MIMAT0000257&hsa-miR-181b-5p&offsets\|0\|+1(+1A);m-73&1\|-\|198858925\|198858947&offsets\|0\|+1(+1A)];[MIMAT0000257_1&hsa-miR-181b-5p&offsets\|0\|+1(+1A);m-72&9\|+\|124693725\|124693747&offsets\|0\|+1(+1A)]\|AACATTCATTGCTGTCGGTGGGTTA | >MI0000270\|hsa-mir-181b-1&WithFlank&1\|-\|198858867\|198858988 | 42 | 65 | 24(+1A) | AACATTCATTGCTGTCGGTGGGTTA | 111.7 | -5.0 | 1.9 | -2.6 | 8.9E-03 | 3.4E-02 | BT20/MB231 | MP |
| >hg38_21_+_8433172_8446622_RNA45SN1_45S_50ntFlanks@6652.6692.41\|GACTCTTAGCGGTGGATCACTCGGCTCGTGCGTCGATGAAG | >hg38_21_+_8433172_8446622_RNA45SN1_45S_50ntFlanks | 6652 | 6692 | 41 | GACTCTTAGCGGTGGATCACTCGGCTCGTGCGTCGATGAAG | 1476.4 | 2.9 | 1.1 | 2.6 | 8.9E-03 | 3.4E-02 | BT20/MB231 | MP |
| >hg38_21_+_8433172_8446622_RNA45SN1_45S_50ntFlanks@11920.11948.29\|ATGAGAGGTGTAGAATAAGTGGGAGGCCC | >hg38_21_+_8433172_8446622_RNA45SN1_45S_50ntFlanks | 11920 | 11948 | 29 | ATGAGAGGTGTAGAATAAGTGGGAGGCCC | 398.4 | -2.4 | 1.1 | -2.2 | 3.0E-02 | 3.4E-02 | MB231/MB468 | CYTO |
| >hg38_21_+_8433172_8446622_RNA45SN1_45S_50ntFlanks@3784.3801.18\|GTGAAACTGCGAATGGCT | >hg38_21_+_8433172_8446622_RNA45SN1_45S_50ntFlanks | 3784 | 3801 | 18 | GTGAAACTGCGAATGGCT | 213.2 | -1.2 | 0.6 | -2.2 | 3.0E-02 | 3.4E-02 | MB231/MB468 | CYTO |
| >MI0006273\|hsa-mir-1180&WithFlank&17\|-\|19344500\|19344580@47.68.22&[MIMAT0005825&hsa-miR-1180-3p&offsets\|0\|0;m-214&17\|-\|19344513\|19344534&offsets\|0\|0]\|TTTCCGGCTCGCGTGGGTGTGT | >MI0006273\|hsa-mir-1180&WithFlank&17\|-\|19344500\|19344580 | 47 | 68 | 22 | TTTCCGGCTCGCGTGGGTGTGT | 511.5 | 1.7 | 0.8 | 2.2 | 2.8E-02 | 3.4E-02 | MB231/MB468 | MP |
| >hg38_21_+_8433172_8446622_RNA45SN1_45S_50ntFlanks@6734.6772.39\|ATTGATCATCGACACTTCGAACGCACTTGCGGCCCCGGG | >hg38_21_+_8433172_8446622_RNA45SN1_45S_50ntFlanks | 6734 | 6772 | 39 | ATTGATCATCGACACTTCGAACGCACTTGCGGCCCCGGG | 746.8 | 1.3 | 0.6 | 2.2 | 2.6E-02 | 3.4E-02 | BT20/MB468 | MITO |
| >hg38_21_+_8433172_8446622_RNA45SN1_45S_50ntFlanks@475.492.18\|CTTCGTGATCGATGTGGT | >hg38_21_+_8433172_8446622_RNA45SN1_45S_50ntFlanks | 475 | 492 | 18 | CTTCGTGATCGATGTGGT | 2535.8 | 2.0 | 0.8 | 2.6 | 9.5E-03 | 3.4E-02 | BT20/MB231 | NUC |
| >hg38_21_+_8433172_8446622_RNA45SN1_45S_50ntFlanks@10798.10823.26\|CATGTTGGAACAATGTAGGTAAGGGA | >hg38_21_+_8433172_8446622_RNA45SN1_45S_50ntFlanks | 10798 | 10823 | 26 | CATGTTGGAACAATGTAGGTAAGGGA | 480.1 | -1.8 | 0.8 | -2.3 | 2.1E-02 | 3.4E-02 | BT20/MB468 | MP |
| >hg38_21_+_8433172_8446622_RNA45SN1_45S_50ntFlanks@8081.8100.20\|GAGCCCAGCGCCGAATCCCC | >hg38_21_+_8433172_8446622_RNA45SN1_45S_50ntFlanks | 8081 | 8100 | 20 | GAGCCCAGCGCCGAATCCCC | 299.8 | 6.2 | 2.7 | 2.3 | 2.1E-02 | 3.4E-02 | BT20/MB468 | MP |
| >hg38_21_+_8433172_8446622_RNA45SN1_45S_50ntFlanks@9927.9950.24\|TTGATATAGACAGCAGGACGGTGG | >hg38_21_+_8433172_8446622_RNA45SN1_45S_50ntFlanks | 9927 | 9950 | 24 | TTGATATAGACAGCAGGACGGTGG | 122.0 | 5.0 | 1.9 | 2.6 | 8.9E-03 | 3.4E-02 | BT20/MB231 | MP |
| >hg38_21_+_8433172_8446622_RNA45SN1_45S_50ntFlanks@10348.10365.18\|AAACGAGAACTTTGAAGG | >hg38_21_+_8433172_8446622_RNA45SN1_45S_50ntFlanks | 10348 | 10365 | 18 | AAACGAGAACTTTGAAGG | 248.5 | -1.5 | 0.7 | -2.2 | 3.0E-02 | 3.4E-02 | MB231/MB468 | CYTO |
| >hg38_1_-_228634819_228635039_RNA5S12_5S_50ntFlanks@132.171.40\|GATGGGAGACCGCCTGGGAATACCGGGTGCTGTAGGCTTT | >hg38_1_-_228634819_228635039_RNA5S12_5S_50ntFlanks | 132 | 171 | 40 | GATGGGAGACCGCCTGGGAATACCGGGTGCTGTAGGCTTT | 6198.4 | 1.8 | 0.8 | 2.2 | 3.1E-02 | 3.4E-02 | MB231/MB468 | NUC |
| >MI0000100\|hsa-mir-98&WithFlank&X\|-\|53556217\|53556347@28.48.21(+1C)&[MIMAT0000096&hsa-miR-98-5p&offsets\|0\|-1(+1C);m-143&X\|-\|53556299\|53556320&offsets\|0\|-1(+1C)]\|TGAGGTAGTAAGTTGTATTGTC | >MI0000100\|hsa-mir-98&WithFlank&X\|-\|53556217\|53556347 | 28 | 48 | 21(+1C) | TGAGGTAGTAAGTTGTATTGTC | 672.6 | 2.3 | 0.9 | 2.5 | 1.2E-02 | 3.4E-02 | BT20/MB231 | CYTO |
| >hg38_21_+_8433172_8446622_RNA45SN1_45S_50ntFlanks@8179.8206.28\|CTGATCGAGGCCCAGCCCGTGGACGGTG | >hg38_21_+_8433172_8446622_RNA45SN1_45S_50ntFlanks | 8179 | 8206 | 28 | CTGATCGAGGCCCAGCCCGTGGACGGTG | 648.8 | -4.5 | 2.1 | -2.2 | 3.1E-02 | 3.4E-02 | BT20/MB468 | NUC |
| >am_trna116_GluCTC_1_-_145399233_145399304@1.29.29,trna59_GluCTC_1_+_249168447_249168518@1.29.29,trna71_GluCTC_1_-_161439189_161439260@1.29.29,trna74_GluCTC_1_-_161431809_161431880@1.29.29,trna77_GluCTC_1_-_161424398_161424469@1.29.29,trna77_GluCTC_6_+_28949976_28950047@1.29.29,trna80_GluCTC_1_-_161417018_161417089@1.29.29,trna87_GluCTC_6_-_126101393_126101464@1.29.29\|TCCCTGGTGGTCTAGTGGTTAGGATTCGG | >am_trna116_GluCTC_1_-_145399233_145399304 | 1 | 29 | 29 | TCCCTGGTGGTCTAGTGGTTAGGATTCGG | 12961.1 | -2.7 | 1.2 | -2.3 | 2.1E-02 | 3.4E-02 | BT20/MB468 | MP |
| >hg38_21_+_8433172_8446622_RNA45SN1_45S_50ntFlanks@5473.5512.40\|CGGCCCTGGCGGAGCGCTGAGAAGACGGTCGAACTTGACT | >hg38_21_+_8433172_8446622_RNA45SN1_45S_50ntFlanks | 5473 | 5512 | 40 | CGGCCCTGGCGGAGCGCTGAGAAGACGGTCGAACTTGACT | 108.8 | 2.0 | 0.9 | 2.2 | 3.1E-02 | 3.4E-02 | MB231/MB468 | MITO |
| >hg38_21_+_8433172_8446622_RNA45SN1_45S_50ntFlanks@6732.6758.27\|ACATTGATCATCGACACTTCGAACGCA | >hg38_21_+_8433172_8446622_RNA45SN1_45S_50ntFlanks | 6732 | 6758 | 27 | ACATTGATCATCGACACTTCGAACGCA | 251.4 | -1.1 | 0.5 | -2.2 | 3.1E-02 | 3.4E-02 | MB231/MB468 | NUC |
| >hg38_21_+_8433172_8446622_RNA45SN1_45S_50ntFlanks@6772.6802.31\|GTTCCTCCCGGGGCTACGCCTGTCTGAGCGT | >hg38_21_+_8433172_8446622_RNA45SN1_45S_50ntFlanks | 6772 | 6802 | 31 | GTTCCTCCCGGGGCTACGCCTGTCTGAGCGT | 1068.3 | 4.3 | 1.9 | 2.3 | 2.1E-02 | 3.4E-02 | BT20/MB468 | MP |
| >hg38_21_+_8433172_8446622_RNA45SN1_45S_50ntFlanks@5082.5108.27\|AACGAACGAGACTCTGGCATGCTAACT | >hg38_21_+_8433172_8446622_RNA45SN1_45S_50ntFlanks | 5082 | 5108 | 27 | AACGAACGAGACTCTGGCATGCTAACT | 368.9 | 3.0 | 1.3 | 2.3 | 2.1E-02 | 3.4E-02 | BT20/MB468 | MP |
| >hg38_21_+_8433172_8446622_RNA45SN1_45S_50ntFlanks@8380.8405.26\|CAAGAGGGCGTGAAACCGTTAAGAGG | >hg38_21_+_8433172_8446622_RNA45SN1_45S_50ntFlanks | 8380 | 8405 | 26 | CAAGAGGGCGTGAAACCGTTAAGAGG | 148.6 | 3.5 | 1.3 | 2.6 | 8.9E-03 | 3.4E-02 | BT20/MB231 | MP |
| >MI0000734\|hsa-mir-106b&WithFlank&7\|-\|100093987\|100094080@18.38.21(+1U)&[MIMAT0000680&hsa-miR-106b-5p&offsets\|0\|0(+1U);m-103&7\|-\|100094043\|100094063&offsets\|0\|0(+1U)]\|TAAAGTGCTGACAGTGCAGATT | >MI0000734\|hsa-mir-106b&WithFlank&7\|-\|100093987\|100094080 | 18 | 38 | 21(+1U) | TAAAGTGCTGACAGTGCAGATT | 251.9 | -3.9 | 1.5 | -2.6 | 8.9E-03 | 3.4E-02 | BT20/MB231 | MP |
| >MI0003137\|hsa-mir-193b&WithFlank&16\|+\|14303961\|14304055@20.40.21(+1U)&[MIMAT0004767&hsa-miR-193b-5p&offsets\|0\|-1(+1U);m-343&16\|+\|14303980\|14304001&offsets\|0\|-1(+1U)]\|CGGGGTTTTGAGGGCGAGATGT | >MI0003137\|hsa-mir-193b&WithFlank&16\|+\|14303961\|14304055 | 20 | 40 | 21(+1U) | CGGGGTTTTGAGGGCGAGATGT | 157.3 | 4.0 | 1.5 | 2.6 | 8.9E-03 | 3.4E-02 | BT20/MB231 | MP |
| >hg38_21_+_8433172_8446622_RNA45SN1_45S_50ntFlanks@8179.8207.29\|CTGATCGAGGCCCAGCCCGTGGACGGTGT | >hg38_21_+_8433172_8446622_RNA45SN1_45S_50ntFlanks | 8179 | 8207 | 29 | CTGATCGAGGCCCAGCCCGTGGACGGTGT | 249.2 | 3.8 | 1.7 | 2.3 | 2.1E-02 | 3.4E-02 | BT20/MB468 | MP |
| >trna20_GluTTC_2_-_131094701_131094772@1.30.30,trna5_GluTTC_13_-_41634874_41634945@1.30.30\|TCCCATATGGTCTAGCGGTTAGGATTCCTG | >trna20_GluTTC_2_-_131094701_131094772 | 1 | 30 | 30 | TCCCATATGGTCTAGCGGTTAGGATTCCTG | 778.7 | -2.2 | 1.0 | -2.2 | 2.8E-02 | 3.4E-02 | MB231/MB468 | MP |
| >hg38_21_+_8433172_8446622_RNA45SN1_45S_50ntFlanks@5473.5491.19\|CGGCCCTGGCGGAGCGCTG | >hg38_21_+_8433172_8446622_RNA45SN1_45S_50ntFlanks | 5473 | 5491 | 19 | CGGCCCTGGCGGAGCGCTG | 181.4 | -1.9 | 0.8 | -2.3 | 2.1E-02 | 3.4E-02 | BT20/MB468 | MP |
| >hg38_21_+_8433172_8446622_RNA45SN1_45S_50ntFlanks@9566.9596.31\|GACGTGCAAATCGGTCGTCCGACCTGGGTAT | >hg38_21_+_8433172_8446622_RNA45SN1_45S_50ntFlanks | 9566 | 9596 | 31 | GACGTGCAAATCGGTCGTCCGACCTGGGTAT | 1128.3 | -3.6 | 1.5 | -2.3 | 2.1E-02 | 3.4E-02 | BT20/MB468 | MP |
| >hg38_21_+_8433172_8446622_RNA45SN1_45S_50ntFlanks@9252.9272.21\|CACGGGGTCGGCGGCGACGTC | >hg38_21_+_8433172_8446622_RNA45SN1_45S_50ntFlanks | 9252 | 9272 | 21 | CACGGGGTCGGCGGCGACGTC | 237.7 | -2.1 | 0.9 | -2.3 | 2.1E-02 | 3.4E-02 | BT20/MB468 | MP |
| >am_trnaMT_ValTAC_MT_+_1602_1670@33.57.25\|ACACTTAGGAGATTTCAACTTAACT | >am_trnaMT_ValTAC_MT_+_1602_1670 | 33 | 57 | 25 | ACACTTAGGAGATTTCAACTTAACT | 77.5 | 6.0 | 2.6 | 2.3 | 2.1E-02 | 3.4E-02 | BT20/MB468 | MP |
| >hg38_MT_+_1621_3279_RNR2_16S_50ntFlanks@1434.1458.25\|GTCCTACGTGATCTGAGTTCAGACC | >hg38_MT_+_1621_3279_RNR2_16S_50ntFlanks | 1434 | 1458 | 25 | GTCCTACGTGATCTGAGTTCAGACC | 57.5 | 3.7 | 1.7 | 2.2 | 2.8E-02 | 3.4E-02 | MB231/MB468 | MP |
| >hg38_MT_+_1621_3279_RNR2_16S_50ntFlanks@1430.1451.22\|TAAAGTCCTACGTGATCTGAGT | >hg38_MT_+_1621_3279_RNR2_16S_50ntFlanks | 1430 | 1451 | 22 | TAAAGTCCTACGTGATCTGAGT | 44.3 | 4.5 | 2.0 | 2.2 | 2.8E-02 | 3.4E-02 | MB231/MB468 | MP |
| >MI0000089\|hsa-mir-31&WithFlank&9\|-\|21512109\|21512191@14.35.22(+1C)&[MIMAT0000089&hsa-miR-31-5p&offsets\|0\|+1(+1C);m-360&9\|-\|21512157\|21512178&offsets\|0\|0(+1C)]\|AGGCAAGATGCTGGCATAGCTGC | >MI0000089\|hsa-mir-31&WithFlank&9\|-\|21512109\|21512191 | 14 | 35 | 22(+1C) | AGGCAAGATGCTGGCATAGCTGC | 233.3 | 3.8 | 1.6 | 2.3 | 2.1E-02 | 3.4E-02 | BT20/MB468 | MP |
| >hg38_MT_+_1621_3279_RNR2_16S_50ntFlanks@293.311.19\|GACGAGCTACCTAAGAACA | >hg38_MT_+_1621_3279_RNR2_16S_50ntFlanks | 293 | 311 | 19 | GACGAGCTACCTAAGAACA | 1081.3 | 3.6 | 1.6 | 2.3 | 2.1E-02 | 3.4E-02 | BT20/MB468 | MP |
| >hg38_21_+_8433172_8446622_RNA45SN1_45S_50ntFlanks@9485.9509.25\|TGTTAGGACCCGAAAGATGGTGAAC | >hg38_21_+_8433172_8446622_RNA45SN1_45S_50ntFlanks | 9485 | 9509 | 25 | TGTTAGGACCCGAAAGATGGTGAAC | 157.7 | 5.3 | 2.0 | 2.6 | 8.9E-03 | 3.4E-02 | BT20/MB231 | MP |
| >hg38_MT_+_1621_3279_RNR2_16S_50ntFlanks@842.864.23\|AAGTAAAAGGAACTCGGCAAATC | >hg38_MT_+_1621_3279_RNR2_16S_50ntFlanks | 842 | 864 | 23 | AAGTAAAAGGAACTCGGCAAATC | 61.5 | 3.7 | 1.7 | 2.2 | 2.8E-02 | 3.4E-02 | MB231/MB468 | MP |
| >trnaMT_ValTAC_MT_+_1602_1670@52.72.21\|TTAACTTGACCGCTCTGACCA | >trnaMT_ValTAC_MT_+_1602_1670 | 52 | 72 | 21 | TTAACTTGACCGCTCTGACCA | 546.3 | 2.1 | 0.8 | 2.5 | 1.2E-02 | 3.4E-02 | BT20/MB231 | CYTO |
| >hg38_21_+_8433172_8446622_RNA45SN1_45S_50ntFlanks@12416.12433.18\|TTCGATGTCGGCTCTTCC | >hg38_21_+_8433172_8446622_RNA45SN1_45S_50ntFlanks | 12416 | 12433 | 18 | TTCGATGTCGGCTCTTCC | 11228.4 | -2.5 | 1.0 | -2.5 | 1.2E-02 | 3.4E-02 | BT20/MB231 | CYTO |
| >hg38_21_+_8433172_8446622_RNA45SN1_45S_50ntFlanks@12416.12442.27\|TTCGATGTCGGCTCTTCCTATCATTGT | >hg38_21_+_8433172_8446622_RNA45SN1_45S_50ntFlanks | 12416 | 12442 | 27 | TTCGATGTCGGCTCTTCCTATCATTGT | 3322.5 | -2.2 | 0.9 | -2.5 | 1.2E-02 | 3.4E-02 | BT20/MB231 | CYTO |
| >hg38_21_+_8433172_8446622_RNA45SN1_45S_50ntFlanks@11577.11594.18\|TTAGAACTGGTGCGGACC | >hg38_21_+_8433172_8446622_RNA45SN1_45S_50ntFlanks | 11577 | 11594 | 18 | TTAGAACTGGTGCGGACC | 208.1 | -1.9 | 0.8 | -2.5 | 1.2E-02 | 3.4E-02 | BT20/MB231 | CYTO |
| >hg38_21_+_8433172_8446622_RNA45SN1_45S_50ntFlanks@8020.8046.27\|TAGTCAGCGGAGGAGAAGAAACTAACC | >hg38_21_+_8433172_8446622_RNA45SN1_45S_50ntFlanks | 8020 | 8046 | 27 | TAGTCAGCGGAGGAGAAGAAACTAACC | 613.6 | -4.5 | 1.8 | -2.5 | 1.2E-02 | 3.4E-02 | BT20/MB231 | CYTO |
| >hg38_21_+_8433172_8446622_RNA45SN1_45S_50ntFlanks@13003.13027.25\|GATCTATTGAAAGTCAGCCCTCGAC | >hg38_21_+_8433172_8446622_RNA45SN1_45S_50ntFlanks | 13003 | 13027 | 25 | GATCTATTGAAAGTCAGCCCTCGAC | 218.3 | -3.1 | 1.4 | -2.2 | 3.0E-02 | 3.4E-02 | MB231/MB468 | CYTO |
| >hg38_21_+_8433172_8446622_RNA45SN1_45S_50ntFlanks@6646.6668.23\|TCGTACGACTCTTAGCGGTGGAT | >hg38_21_+_8433172_8446622_RNA45SN1_45S_50ntFlanks | 6646 | 6668 | 23 | TCGTACGACTCTTAGCGGTGGAT | 11321.1 | 1.9 | 0.7 | 2.6 | 9.5E-03 | 3.4E-02 | BT20/MB231 | NUC |
| >hg38_MT_+_598_1651_RNR1_12S_50ntFlanks@443.480.38\|TATCTGAACACACAATAGCTAAGACCCAAACTGGGATT | >hg38_MT_+_598_1651_RNR1_12S_50ntFlanks | 443 | 480 | 38 | TATCTGAACACACAATAGCTAAGACCCAAACTGGGATT | 325.7 | 3.1 | 1.4 | 2.2 | 2.8E-02 | 3.4E-02 | MB231/MB468 | MP |
| >MI0000102\|hsa-mir-100&WithFlank&11\|-\|122152223\|122152314@22.39.18&[MIMAT0000098&hsa-miR-100-5p&offsets\|+3\|-1;m-27&11\|-\|122152276\|122152296&offsets\|+3\|0]\|CCGTAGATCCGAACTTGT | >MI0000102\|hsa-mir-100&WithFlank&11\|-\|122152223\|122152314 | 22 | 39 | 18 | CCGTAGATCCGAACTTGT | 131.8 | -10.1 | 3.9 | -2.6 | 9.6E-03 | 3.4E-02 | BT20/MB231 | NUC |
| >trnaMT_GluTTC_MT_-_14674_14742@26.66.41,trnalookalike8_GluTTC_5_-_93905172_93905240@26.66.41\|GGTTTTTCATATCATTGGTCGTGGTTGTAGTCCGTGCGAGA | >trnaMT_GluTTC_MT_-_14674_14742 | 26 | 66 | 41 | GGTTTTTCATATCATTGGTCGTGGTTGTAGTCCGTGCGAGA | 81.0 | 4.4 | 1.9 | 2.3 | 2.1E-02 | 3.4E-02 | BT20/MB468 | MP |
| >hg38_21_+_8433172_8446622_RNA45SN1_45S_50ntFlanks@11999.12037.39\|CCCTGCGGGCCGCCGGTGAAATACCACTACTCTGATCGT | >hg38_21_+_8433172_8446622_RNA45SN1_45S_50ntFlanks | 11999 | 12037 | 39 | CCCTGCGGGCCGCCGGTGAAATACCACTACTCTGATCGT | 159.2 | -1.6 | 0.7 | -2.2 | 2.7E-02 | 3.4E-02 | BT20/MB468 | MITO |
| >MI0000471\|hsa-mir-126&WithFlank&9\|+\|136670596\|136670692@21.41.21(+1U)&[MIMAT0000444&hsa-miR-126-5p&offsets\|0\|0(+1U);m-114&9\|+\|136670616\|136670636&offsets\|0\|0(+1U)]\|CATTATTACTTTTGGTACGCGT | >MI0000471\|hsa-mir-126&WithFlank&9\|+\|136670596\|136670692 | 21 | 41 | 21(+1U) | CATTATTACTTTTGGTACGCGT | 224.3 | -5.1 | 1.9 | -2.6 | 9.0E-03 | 3.4E-02 | BT20/MB231 | MP |
| >hg38_21_+_8433172_8446622_RNA45SN1_45S_50ntFlanks@11694.11728.35\|AAGAAATTCAATGAAGCGCGGGTAAACGGCGGGAG | >hg38_21_+_8433172_8446622_RNA45SN1_45S_50ntFlanks | 11694 | 11728 | 35 | AAGAAATTCAATGAAGCGCGGGTAAACGGCGGGAG | 131.2 | -4.0 | 1.7 | -2.3 | 2.1E-02 | 3.4E-02 | BT20/MB468 | MP |
| >hg38_MT_+_1621_3279_RNR2_16S_50ntFlanks@1367.1395.29\|TCGATGTTGGATCAGGACATCCCGATGGT | >hg38_MT_+_1621_3279_RNR2_16S_50ntFlanks | 1367 | 1395 | 29 | TCGATGTTGGATCAGGACATCCCGATGGT | 115.9 | 3.7 | 1.7 | 2.2 | 2.8E-02 | 3.4E-02 | MB231/MB468 | MP |

<table>
<tr><th></th><th></th><th></th><th></th><th></th><th></th><th></th><th></th><th></th><th></th><th></th><th></th><th></th><th></th></tr>
<tr><td>>hg38_21_+_8433172_8446622_RNA45SN1_45S_50ntFlanks@4102.4129.28|ACCACGGGTGACGGGGAATCAGGGTTCG</td><td>>hg38_21_+_8433172_8446622_RNA45SN1_45S_50ntFlanks</td><td>4102</td><td>4129</td><td>28</td><td>ACCACGGGTGACGGGGAATCAGGGTTCG</td><td>103.7</td><td>-3.2</td><td>1.4</td><td>-2.3</td><td>2.1E-02</td><td>3.4E-02</td><td>BT20/MB468</td><td>MP</td></tr>
<tr><td>>MI0000061|hsa-let-7a-2&WithFlank&11|-|122146516|122146599@11.31.21(+1A);MI0000062|hsa-let-7a-3&WithFlank&22|+|46112743|46112828@10.30.21(+1A);MI0000060|hsa-let-7a-1&WithFlank&9|+|94175951|94176042@12.32.21(+1A)&[MIMAT0000062&hsa-let-7a-5p&offsets|0|-1(+1A);m-6&11|-|122146568|122146589&offsets|0|-1(+1A)];[MIMAT0000062_1&hsa-let-7a-5p&offsets|0|-1(+1A);m-5&22|+|46112752|46112773&offsets|0|-1(+1A)];[MIMAT0000062_2&hsa-let-7a-5p&offsets|0|-1(+1A);m-4&9|+|94175962|94175983&offsets|0|-1(+1A)]|TGAGGTAGTAGGTTGTATAGTA</td><td>>MI0000061|hsa-let-7a-2&WithFlank&11|-|122146516|122146599</td><td>11</td><td>31</td><td>21(+1A)</td><td>TGAGGTAGTAGGTTGTATAGTA</td><td>365.7</td><td>2.5</td><td>1.1</td><td>2.3</td><td>2.1E-02</td><td>3.4E-02</td><td>BT20/MB468</td><td>MP</td></tr>
<tr><td>>hg38_21_+_8433172_8446622_RNA45SN1_45S_50ntFlanks@8104.8138.35|CCGCGGCGGGGCGCGGGACATGTGGCGTACGGAAG</td><td>>hg38_21_+_8433172_8446622_RNA45SN1_45S_50ntFlanks</td><td>8104</td><td>8138</td><td>35</td><td>CCGCGGCGGGGCGCGGGACATGTGGCGTACGGAAG</td><td>655.4</td><td>-4.4</td><td>2.1</td><td>-2.2</td><td>3.1E-02</td><td>3.4E-02</td><td>BT20/MB468</td><td>NUC</td></tr>
<tr><td>>hg38_21_+_8433172_8446622_RNA45SN1_45S_50ntFlanks@11562.11603.42|CGCCTAGCAGCCGACTTAGAACTGGTGCGGACCAGGGGAATC</td><td>>hg38_21_+_8433172_8446622_RNA45SN1_45S_50ntFlanks</td><td>11562</td><td>11603</td><td>42</td><td>CGCCTAGCAGCCGACTTAGAACTGGTGCGGACCAGGGGAATC</td><td>616.1</td><td>-2.0</td><td>0.9</td><td>-2.2</td><td>2.7E-02</td><td>3.4E-02</td><td>BT20/MB468</td><td>MITO</td></tr>
<tr><td>>hg38_21_+_8433172_8446622_RNA45SN1_45S_50ntFlanks@4881.4903.23|TTGCAAAGCTGAAACTTAAAGGA</td><td>>hg38_21_+_8433172_8446622_RNA45SN1_45S_50ntFlanks</td><td>4881</td><td>4903</td><td>23</td><td>TTGCAAAGCTGAAACTTAAAGGA</td><td>87.1</td><td>2.3</td><td>1.0</td><td>2.3</td><td>2.1E-02</td><td>3.4E-02</td><td>BT20/MB468</td><td>MP</td></tr>
<tr><td>>hg38_21_+_8433172_8446622_RNA45SN1_45S_50ntFlanks@9548.9570.23|GAGGTCCGTAGCGGTCCTGACGT</td><td>>hg38_21_+_8433172_8446622_RNA45SN1_45S_50ntFlanks</td><td>9548</td><td>9570</td><td>23</td><td>GAGGTCCGTAGCGGTCCTGACGT</td><td>368.3</td><td>1.4</td><td>0.6</td><td>2.2</td><td>3.0E-02</td><td>3.4E-02</td><td>BT20/MB468</td><td>CYTO</td></tr>
<tr><td>>hg38_21_+_8433172_8446622_RNA45SN1_45S_50ntFlanks@4069.4094.26|CTTTCGATGGTAGTCGCCGTGCCTAC</td><td>>hg38_21_+_8433172_8446622_RNA45SN1_45S_50ntFlanks</td><td>4069</td><td>4094</td><td>26</td><td>CTTTCGATGGTAGTCGCCGTGCCTAC</td><td>276.4</td><td>-2.1</td><td>1.0</td><td>-2.2</td><td>3.0E-02</td><td>3.4E-02</td><td>MB231/MB468</td><td>CYTO</td></tr>
<tr><td>>hg38_21_+_8433172_8446622_RNA45SN1_45S_50ntFlanks@9552.9571.20|TCCGTAGCGGTCCTGACGTG</td><td>>hg38_21_+_8433172_8446622_RNA45SN1_45S_50ntFlanks</td><td>9552</td><td>9571</td><td>20</td><td>TCCGTAGCGGTCCTGACGTG</td><td>289.4</td><td>-1.3</td><td>0.6</td><td>-2.2</td><td>2.7E-02</td><td>3.4E-02</td><td>BT20/MB468</td><td>MITO</td></tr>
<tr><td>>hg38_21_+_8433172_8446622_RNA45SN1_45S_50ntFlanks@6647.6669.23|CGTACGACTCTTAGCGGTGGATC</td><td>>hg38_21_+_8433172_8446622_RNA45SN1_45S_50ntFlanks</td><td>6647</td><td>6669</td><td>23</td><td>CGTACGACTCTTAGCGGTGGATC</td><td>1260.0</td><td>1.2</td><td>0.6</td><td>2.2</td><td>3.1E-02</td><td>3.4E-02</td><td>MB231/MB468</td><td>NUC</td></tr>
<tr><td>>hg38_21_+_8433172_8446622_RNA45SN1_45S_50ntFlanks@12181.12220.40|TCAAACGGTAACGCAGGTGTCCTAAGGCGAGCTCAGGGAG</td><td>>hg38_21_+_8433172_8446622_RNA45SN1_45S_50ntFlanks</td><td>12181</td><td>12220</td><td>40</td><td>TCAAACGGTAACGCAGGTGTCCTAAGGCGAGCTCAGGGAG</td><td>97.7</td><td>-2.6</td><td>1.2</td><td>-2.2</td><td>2.7E-02</td><td>3.5E-02</td><td>BT20/MB468</td><td>MITO</td></tr>
<tr><td>>MI0000471|hsa-mir-126&WithFlank&9|+|136670596|136670692@21.41.21&[MIMAT0000444&hsa-miR-126-5p&offsets|0|0;m-114&9|+|136670616|136670636&offsets|0|0]|CATTATTACTTTTGGTACGCG</td><td>>MI0000471|hsa-mir-126&WithFlank&9|+|136670596|136670692</td><td>21</td><td>41</td><td>21</td><td>CATTATTACTTTTGGTACGCG</td><td>524.4</td><td>-2.8</td><td>1.1</td><td>-2.6</td><td>9.0E-03</td><td>3.5E-02</td><td>BT20/MB231</td><td>MP</td></tr>
<tr><td>>hg38_21_+_8433172_8446622_RNA45SN1_45S_50ntFlanks@11565.11584.20|CTAGCAGCCGACTTAGAACT</td><td>>hg38_21_+_8433172_8446622_RNA45SN1_45S_50ntFlanks</td><td>11565</td><td>11584</td><td>20</td><td>CTAGCAGCCGACTTAGAACT</td><td>263.4</td><td>4.4</td><td>1.7</td><td>2.6</td><td>9.0E-03</td><td>3.5E-02</td><td>BT20/MB231</td><td>MP</td></tr>
<tr><td>>hg38_21_+_8433172_8446622_RNA45SN1_45S_50ntFlanks@10685.10709.25|GGCGGCGTCCGGTGAGCTCTCGCTG</td><td>>hg38_21_+_8433172_8446622_RNA45SN1_45S_50ntFlanks</td><td>10685</td><td>10709</td><td>25</td><td>GGCGGCGTCCGGTGAGCTCTCGCTG</td><td>167.5</td><td>-1.4</td><td>0.7</td><td>-2.2</td><td>3.0E-02</td><td>3.5E-02</td><td>MB231/MB468</td><td>CYTO</td></tr>
<tr><td>>hg38_21_+_8433172_8446622_RNA45SN1_45S_50ntFlanks@12628.12657.30|TGGGGCGAAGCTACCATCTGTGGGATTATG</td><td>>hg38_21_+_8433172_8446622_RNA45SN1_45S_50ntFlanks</td><td>12628</td><td>12657</td><td>30</td><td>TGGGGCGAAGCTACCATCTGTGGGATTATG</td><td>248.0</td><td>-4.1</td><td>1.8</td><td>-2.3</td><td>2.1E-02</td><td>3.5E-02</td><td>BT20/MB468</td><td>MP</td></tr>
<tr><td>>hg38_21_+_8433172_8446622_RNA45SN1_45S_50ntFlanks@12201.12220.20|CCTAAGGCGAGCTCAGGGAG</td><td>>hg38_21_+_8433172_8446622_RNA45SN1_45S_50ntFlanks</td><td>12201</td><td>12220</td><td>20</td><td>CCTAAGGCGAGCTCAGGGAG</td><td>116.6</td><td>-3.9</td><td>1.7</td><td>-2.3</td><td>2.1E-02</td><td>3.5E-02</td><td>BT20/MB468</td><td>MP</td></tr>
<tr><td>>MI0000746|hsa-mir-99b&WithFlank&19|+|51692606|51692687@13.32.20&[MIMAT0000689&hsa-miR-99b-5p&offsets|0|-2;m-10&19|+|51692618|51692639&offsets|0|-2]|CACCCGTAGAACCGACCTTG</td><td>>MI0000746|hsa-mir-99b&WithFlank&19|+|51692606|51692687</td><td>13</td><td>32</td><td>20</td><td>CACCCGTAGAACCGACCTTG</td><td>4170.5</td><td>2.2</td><td>0.9</td><td>2.5</td><td>1.2E-02</td><td>3.5E-02</td><td>BT20/MB231</td><td>CYTO</td></tr>
<tr><td>>MI0003760|hsa-mir-671&WithFlank&7|+|151238415|151238544@35.59.25(+4U)&[MIMAT0003880&hsa-miR-671-5p&offsets|0|+2(+4U);m-316&7|+|151238449|151238472&offsets|0|+1(+4U)]|AGGAAGCCCTGGAGGGGCTGGAGGTTTTT</td><td>>MI0003760|hsa-mir-671&WithFlank&7|+|151238415|151238544</td><td>35</td><td>59</td><td>25(+4U)</td><td>AGGAAGCCCTGGAGGGGCTGGAGGTTTTT</td><td>39.5</td><td>4.5</td><td>2.0</td><td>2.2</td><td>2.8E-02</td><td>3.5E-02</td><td>MB231/MB468</td><td>MP</td></tr>
<tr><td>>trnaMT_GluTTC_MT_-_14674_14742@40.68.29,trnalookalike8_GluTTC_5_-_93905172_93905240@40.68.29|TTGGTCGTGGTTGTAGTCCGTGCGAGAAT</td><td>>trnaMT_GluTTC_MT_-_14674_14742</td><td>40</td><td>68</td><td>29</td><td>TTGGTCGTGGTTGTAGTCCGTGCGAGAAT</td><td>164.9</td><td>3.8</td><td>1.7</td><td>2.3</td><td>2.1E-02</td><td>3.5E-02</td><td>BT20/MB468</td><td>MP</td></tr>
<tr><td>>hg38_21_+_8433172_8446622_RNA45SN1_45S_50ntFlanks@10349.10383.35|AACGAGAACTTTGAAGGCCGAAGTGGAGAAGGGTT</td><td>>hg38_21_+_8433172_8446622_RNA45SN1_45S_50ntFlanks</td><td>10349</td><td>10383</td><td>35</td><td>AACGAGAACTTTGAAGGCCGAAGTGGAGAAGGGTT</td><td>2443.6</td><td>-1.6</td><td>0.7</td><td>-2.2</td><td>2.7E-02</td><td>3.5E-02</td><td>BT20/MB468</td><td>MITO</td></tr>
<tr><td>>MI0000815|hsa-mir-339&WithFlank&7|-|1022927|1023032@56.76.21&[MIMAT0004702&hsa-miR-339-3p&offsets|0|-2;m-235&7|-|1022957|1022977&offsets|0|0]|TGAGCGCCTCGACGACAGAGC</td><td>>MI0000815|hsa-mir-339&WithFlank&7|-|1022927|1023032</td><td>56</td><td>76</td><td>21</td><td>TGAGCGCCTCGACGACAGAGC</td><td>407.8</td><td>2.0</td><td>0.8</td><td>2.5</td><td>1.2E-02</td><td>3.5E-02</td><td>BT20/MB231</td><td>CYTO</td></tr>
<tr><td>>hg38_21_+_8433172_8446622_RNA45SN1_45S_50ntFlanks@11662.11684.23|GATTTCTGCCCAGTGCTCTGAAT</td><td>>hg38_21_+_8433172_8446622_RNA45SN1_45S_50ntFlanks</td><td>11662</td><td>11684</td><td>23</td><td>GATTTCTGCCCAGTGCTCTGAAT</td><td>51.4</td><td>3.3</td><td>1.4</td><td>2.3</td><td>2.1E-02</td><td>3.5E-02</td><td>BT20/MB468</td><td>MP</td></tr>
<tr><td>>hg38_21_+_8433172_8446622_RNA45SN1_45S_50ntFlanks@4685.4723.39|AGACGGACCAGAGCGAAAGCATTTGCCAAGAATGTTTTC</td><td>>hg38_21_+_8433172_8446622_RNA45SN1_45S_50ntFlanks</td><td>4685</td><td>4723</td><td>39</td><td>AGACGGACCAGAGCGAAAGCATTTGCCAAGAATGTTTTC</td><td>468.6</td><td>2.1</td><td>1.0</td><td>2.2</td><td>3.0E-02</td><td>3.5E-02</td><td>MB231/MB468</td><td>CYTO</td></tr>
<tr><td>>hg38_21_+_8433172_8446622_RNA45SN1_45S_50ntFlanks@4129.4159.31|GATTCCGGAGAGGGAGCCTGAGAAACGGCTA</td><td>>hg38_21_+_8433172_8446622_RNA45SN1_45S_50ntFlanks</td><td>4129</td><td>4159</td><td>31</td><td>GATTCCGGAGAGGGAGCCTGAGAAACGGCTA</td><td>38.0</td><td>-4.1</td><td>1.8</td><td>-2.3</td><td>2.1E-02</td><td>3.5E-02</td><td>BT20/MB468</td><td>MP</td></tr>
<tr><td>>hg38_21_+_8433172_8446622_RNA45SN1_45S_50ntFlanks@5487.5513.27|CGCTGAGAAGACGGTCGAACTTGACTA</td><td>>hg38_21_+_8433172_8446622_RNA45SN1_45S_50ntFlanks</td><td>5487</td><td>5513</td><td>27</td><td>CGCTGAGAAGACGGTCGAACTTGACTA</td><td>193.9</td><td>-1.6</td><td>0.8</td><td>-2.2</td><td>3.0E-02</td><td>3.5E-02</td><td>MB231/MB468</td><td>CYTO</td></tr>
<tr><td>>MI0000270|hsa-mir-181b-1&WithFlank&1|-|198858867|198858988@42.62.21;MI0000683|hsa-mir-181b-2&WithFlank&9|+|124693704|124693804@22.42.21&[MIMAT0000257&hsa-miR-181b-5p&offsets|0|-2;m-73&1|-|198858925|198858947&offsets|0|-2];[MIMAT0000257_1&hsa-miR-181b-5p&offsets|0|-2;m-72&9|+|124693725|124693747&offsets|0|-2]|AACATTCATTGCTGTCGGTGG</td><td>>MI0000270|hsa-mir-181b-1&WithFlank&1|-|198858867|198858988</td><td>42</td><td>62</td><td>21</td><td>AACATTCATTGCTGTCGGTGG</td><td>542.2</td><td>-2.7</td><td>1.0</td><td>-2.6</td><td>9.0E-03</td><td>3.5E-02</td><td>BT20/MB231</td><td>MP</td></tr>
<tr><td>>hg38_21_+_8433172_8446622_RNA45SN1_45S_50ntFlanks@8179.8207.29|CTGATCGAGGCCCAGCCCGTGGACGGTGT</td><td>>hg38_21_+_8433172_8446622_RNA45SN1_45S_50ntFlanks</td><td>8179</td><td>8207</td><td>29</td><td>CTGATCGAGGCCCAGCCCGTGGACGGTGT</td><td>227.6</td><td>6.3</td><td>2.4</td><td>2.6</td><td>9.0E-03</td><td>3.5E-02</td><td>BT20/MB231</td><td>MP</td></tr>
<tr><td>>hg38_21_+_8433172_8446622_RNA45SN1_45S_50ntFlanks@3840.3861.22|CTCCTACTTGGATAACTGTGGT</td><td>>hg38_21_+_8433172_8446622_RNA45SN1_45S_50ntFlanks</td><td>3840</td><td>3861</td><td>22</td><td>CTCCTACTTGGATAACTGTGGT</td><td>1102.7</td><td>1.8</td><td>0.7</td><td>2.5</td><td>1.2E-02</td><td>3.5E-02</td><td>BT20/MB231</td><td>CYTO</td></tr>
<tr><td>>MI0000108|hsa-mir-103a-2&WithFlank&20|+|3917488|3917577@56.76.21;MI0000109|hsa-mir-103a-1&WithFlank&5|-|168560890|168560979@56.76.21&[MIMAT0000101&hsa-miR-103a-3p&offsets|+2|0;m-12&20|+|3917541|3917563&offsets|+2|0];[MIMAT0000101_1&hsa-miR-103a-3p&offsets|+2|0;m-13&5|-|168560904|168560926&offsets|+2|0;m-44220&5|-|168560908|168560926&offsets|+2|+4]|CAGCATTGTACAGGGCTATGA</td><td>>MI0000108|hsa-mir-103a-2&WithFlank&20|+|3917488|3917577</td><td>56</td><td>76</td><td>21</td><td>CAGCATTGTACAGGGCTATGA</td><td>479.7</td><td>-2.6</td><td>1.0</td><td>-2.6</td><td>9.0E-03</td><td>3.5E-02</td><td>BT20/MB231</td><td>MP</td></tr>
<tr><td>>MI0000289|hsa-mir-181a-1&WithFlank&1|-|198859038|198859159@30.50.21;MI0000269|hsa-mir-181a-2&WithFlank&9|+|124692436|124692557@45.65.21&[MIMAT0000256&hsa-miR-181a-5p&offsets|0|-2;m-32&1|-|198859108|198859130&offsets|0|-2];[MIMAT0000256_1&hsa-miR-181a-5p&offsets|0|-2;m-32&9|+|124692480|124692502&offsets|0|-2]|AACATTCAACGCTGTCGGTGA</td><td>>MI0000289|hsa-mir-181a-1&WithFlank&1|-|198859038|198859159</td><td>30</td><td>50</td><td>21</td><td>AACATTCAACGCTGTCGGTGA</td><td>1005.0</td><td>-2.6</td><td>1.0</td><td>-2.6</td><td>9.0E-03</td><td>3.5E-02</td><td>BT20/MB231</td><td>MP</td></tr>
<tr><td>>hg38_21_+_8433172_8446622_RNA45SN1_45S_50ntFlanks@12637.12664.28|GCTACCATCTGTGGGATTATGACTGAAC</td><td>>hg38_21_+_8433172_8446622_RNA45SN1_45S_50ntFlanks</td><td>12637</td><td>12664</td><td>28</td><td>GCTACCATCTGTGGGATTATGACTGAAC</td><td>196.0</td><td>5.0</td><td>1.9</td><td>2.6</td><td>9.0E-03</td><td>3.5E-02</td><td>BT20/MB231</td><td>MP</td></tr>
<tr><td>>trnaMT_SerGCT_MT_+_12207_12265@1.25.25|GAGAAAGCTCACAAGAACTGCTAAC</td><td>>trnaMT_SerGCT_MT_+_12207_12265</td><td>1</td><td>25</td><td>25</td><td>GAGAAAGCTCACAAGAACTGCTAAC</td><td>76954.4</td><td>-2.5</td><td>1.0</td><td>-2.6</td><td>9.1E-03</td><td>3.5E-02</td><td>BT20/MB231</td><td>MP</td></tr>
<tr><td>>hg38_21_+_8433172_8446622_RNA45SN1_45S_50ntFlanks@12941.12965.25|ATTCGTAGACGACCTGCTTCTGGGT</td><td>>hg38_21_+_8433172_8446622_RNA45SN1_45S_50ntFlanks</td><td>12941</td><td>12965</td><td>25</td><td>ATTCGTAGACGACCTGCTTCTGGGT</td><td>206.8</td><td>-2.1</td><td>0.9</td><td>-2.3</td><td>2.1E-02</td><td>3.5E-02</td><td>BT20/MB468</td><td>MP</td></tr>
<tr><td>>hg38_21_+_8433172_8446622_RNA45SN1_45S_50ntFlanks@7975.7994.20|CGCGACCTCAGATCAGACGT</td><td>>hg38_21_+_8433172_8446622_RNA45SN1_45S_50ntFlanks</td><td>7975</td><td>7994</td><td>20</td><td>CGCGACCTCAGATCAGACGT</td><td>233010.5</td><td>3.2</td><td>1.2</td><td>2.6</td><td>9.1E-03</td><td>3.5E-02</td><td>BT20/MB231</td><td>MP</td></tr>
<tr><td>>hg38_21_+_8433172_8446622_RNA45SN1_45S_50ntFlanks@11672.11692.21|CAGTGCTCTGAATGTCAAAGT</td><td>>hg38_21_+_8433172_8446622_RNA45SN1_45S_50ntFlanks</td><td>11672</td><td>11692</td><td>21</td><td>CAGTGCTCTGAATGTCAAAGT</td><td>109.5</td><td>-1.4</td><td>0.7</td><td>-2.2</td><td>2.9E-02</td><td>3.5E-02</td><td>MB231/MB468</td><td>MP</td></tr>
<tr><td>>hg38_21_+_8433172_8446622_RNA45SN1_45S_50ntFlanks@12450.12476.27|AATTCACCAAGCGTTGGATTGTTCACC</td><td>>hg38_21_+_8433172_8446622_RNA45SN1_45S_50ntFlanks</td><td>12450</td><td>12476</td><td>27</td><td>AATTCACCAAGCGTTGGATTGTTCACC</td><td>243.6</td><td>-2.1</td><td>0.8</td><td>-2.5</td><td>1.2E-02</td><td>3.5E-02</td><td>BT20/MB231</td><td>CYTO</td></tr>
<tr><td>>hg38_21_+_8433172_8446622_RNA45SN1_45S_50ntFlanks@11775.11794.20|GTGACGCGCATGAATGGATG</td><td>>hg38_21_+_8433172_8446622_RNA45SN1_45S_50ntFlanks</td><td>11775</td><td>11794</td><td>20</td><td>GTGACGCGCATGAATGGATG</td><td>205.9</td><td>-2.1</td><td>1.0</td><td>-2.2</td><td>2.9E-02</td><td>3.5E-02</td><td>MB231/MB468</td><td>MP</td></tr>
<tr><td>>MI0000289|hsa-mir-181a-1&WithFlank&1|-|198859038|198859159@70.90.21(+1U)&[MIMAT0000270&hsa-miR-181a-3p&offsets|0|-1(+1U);m-125&1|-|198859069|198859090&offsets|0|-1(+1U)]|ACCATCGACCGTTGATTGTACT</td><td>>MI0000289|hsa-mir-181a-1&WithFlank&1|-|198859038|198859159</td><td>70</td><td>90</td><td>21(+1U)</td><td>ACCATCGACCGTTGATTGTACT</td><td>648.9</td><td>-2.8</td><td>1.1</td><td>-2.5</td><td>1.2E-02</td><td>3.5E-02</td><td>BT20/MB231</td><td>CYTO</td></tr>
<tr><td>>hg38_21_+_8433172_8446622_RNA45SN1_45S_50ntFlanks@9551.9568.18|GTCCGTAGCGGTCCTGAC</td><td>>hg38_21_+_8433172_8446622_RNA45SN1_45S_50ntFlanks</td><td>9551</td><td>9568</td><td>18</td><td>GTCCGTAGCGGTCCTGAC</td><td>403.5</td><td>-2.1</td><td>0.8</td><td>-2.5</td><td>1.2E-02</td><td>3.5E-02</td><td>BT20/MB231</td><td>CYTO</td></tr>
<tr><td>>hg38_MT_+_1621_3279_RNR2_16S_50ntFlanks@1370.1402.33|ATGTTGGATCAGGACATCCCGATGGTGCAGCCG</td><td>>hg38_MT_+_1621_3279_RNR2_16S_50ntFlanks</td><td>1370</td><td>1402</td><td>33</td><td>ATGTTGGATCAGGACATCCCGATGGTGCAGCCG</td><td>184.6</td><td>2.9</td><td>1.3</td><td>2.3</td><td>2.1E-02</td><td>3.5E-02</td><td>BT20/MB468</td><td>MP</td></tr>
<tr><td>>hg38_21_+_8433172_8446622_RNA45SN1_45S_50ntFlanks@6647.6665.19|CGTACGACTCTTAGCGGTG</td><td>>hg38_21_+_8433172_8446622_RNA45SN1_45S_50ntFlanks</td><td>6647</td><td>6665</td><td>19</td><td>CGTACGACTCTTAGCGGTG</td><td>8664.3</td><td>1.9</td><td>0.7</td><td>2.6</td><td>9.7E-03</td><td>3.5E-02</td><td>BT20/MB231</td><td>NUC</td></tr>
<tr><td>>hg38_21_+_8433172_8446622_RNA45SN1_45S_50ntFlanks@6648.6683.36|GTACGACTCTTAGCGGTGGATCACTCGGCTCGTGCG</td><td>>hg38_21_+_8433172_8446622_RNA45SN1_45S_50ntFlanks</td><td>6648</td><td>6683</td><td>36</td><td>GTACGACTCTTAGCGGTGGATCACTCGGCTCGTGCG</td><td>260.4</td><td>-2.5</td><td>1.0</td><td>-2.6</td><td>9.7E-03</td><td>3.5E-02</td><td>BT20/MB231</td><td>NUC</td></tr>
<tr><td>>MI0000446|hsa-mir-125b-1&WithFlank&11|-|122099751|122099850@21.40.20;MI0000470|hsa-mir-125b-2&WithFlank&21|+|16590231|16590331@23.42.20&[MIMAT0000423&hsa-miR-125b-5p&offsets|0|-2;m-67&11|-|122099809|122099830&offsets|0|-2];[MIMAT0000423_1&hsa-miR-125b-5p&offsets|0|-2;m-68&21|+|16590253|16590274&offsets|0|-2]|TCCCTGAGACCCTAACTTGT</td><td>>MI0000446|hsa-mir-125b-1&WithFlank&11|-|122099751|122099850</td><td>21</td><td>40</td><td>20</td><td>TCCCTGAGACCCTAACTTGT</td><td>2275.3</td><td>-6.6</td><td>2.6</td><td>-2.5</td><td>1.1E-02</td><td>3.5E-02</td><td>BT20/MB231</td><td>MITO</td></tr>
<tr><td>>hg38_21_+_8433172_8446622_RNA45SN1_45S_50ntFlanks@12481.12499.19|AATAGGGAACGTGAGCTGG</td><td>>hg38_21_+_8433172_8446622_RNA45SN1_45S_50ntFlanks</td><td>12481</td><td>12499</td><td>19</td><td>AATAGGGAACGTGAGCTGG</td><td>418.0</td><td>4.7</td><td>1.8</td><td>2.6</td><td>9.1E-03</td><td>3.5E-02</td><td>BT20/MB231</td><td>MP</td></tr>
<tr><td>>hg38_21_+_8433172_8446622_RNA45SN1_45S_50ntFlanks@9918.9946.29|AGGTGTTGGTTGATATAGACAGCAGGACG</td><td>>hg38_21_+_8433172_8446622_RNA45SN1_45S_50ntFlanks</td><td>9918</td><td>9946</td><td>29</td><td>AGGTGTTGGTTGATATAGACAGCAGGACG</td><td>60.8</td><td>4.8</td><td>2.1</td><td>2.3</td><td>2.1E-02</td><td>3.5E-02</td><td>BT20/MB468</td><td>MP</td></tr>
<tr><td>>trnaMT_ValTAC_MT_+_1602_1670@29.71.43|ACTTACACTTAGGAGATTTCAACTTAACTTGACCGCTCTGACC</td><td>>trnaMT_ValTAC_MT_+_1602_1670</td><td>29</td><td>71</td><td>43</td><td>ACTTACACTTAGGAGATTTCAACTTAACTTGACCGCTCTGACC</td><td>390.1</td><td>-2.2</td><td>0.9</td><td>-2.5</td><td>1.1E-02</td><td>3.5E-02</td><td>BT20/MB231</td><td>MITO</td></tr>
</table>

<table>
<tr><td></td><td></td><td></td><td></td><td></td><td></td><td></td><td></td><td></td><td></td><td></td><td></td><td></td><td></td></tr>
<tr><td>>am_trna117_GlyTCC_1_-_145397864_145397935@1.29.29,trna45_GlyTCC_1_+_161500903_161500974@1.29.29,trna70_GlyTCC_1_-_161439547_161439618@1.29.29,trna73_GlyTCC_1_-_161432166_161432237@1.29.29,trna76_GlyTCC_1_-_161424756_161424827@1.29.29,trna79_GlyTCC_1_-_161417375_161417446@1.29.29|GCGTTGGTGGTATAGTGGTGAGCATAGCT</td><td>>am_trna117_GlyTCC_1_-_145397864_145397935</td><td>1</td><td>29</td><td>29</td><td>GCGTTGGTGGTATAGTGGTGAGCATAGCT</td><td>1190.9</td><td>-3.4</td><td>1.5</td><td>-2.3</td><td>2.1E-02</td><td>3.5E-02</td><td>BT20/MB468</td><td>MP</td></tr>
<tr><td>>hg38_21_+_8433172_8446622_RNA45SN1_45S_50ntFlanks@10404.10429.26|CATGGGTCAGTCGGTCCTGAGAGATG</td><td>>hg38_21_+_8433172_8446622_RNA45SN1_45S_50ntFlanks</td><td>10404</td><td>10429</td><td>26</td><td>CATGGGTCAGTCGGTCCTGAGAGATG</td><td>1456.6</td><td>-1.8</td><td>0.8</td><td>-2.3</td><td>2.1E-02</td><td>3.5E-02</td><td>BT20/MB468</td><td>MP</td></tr>
<tr><td>>MI0000289|hsa-mir-181a-1&WithFlank&1|-|198859038|198859159@70.90.21(+1A)&[MIMAT0000270&hsa-miR-181a-3p&offsets|0|-1(+1A);m-125&1|-|198859069|198859090&offsets|0|-1(+1A)]|ACCATCGACCGTTGATTGTACA</td><td>>MI0000289|hsa-mir-181a-1&WithFlank&1|-|198859038|198859159</td><td>70</td><td>90</td><td>21(+1A)</td><td>ACCATCGACCGTTGATTGTACA</td><td>305.5</td><td>-2.8</td><td>1.1</td><td>-2.5</td><td>1.2E-02</td><td>3.5E-02</td><td>BT20/MB231</td><td>CYTO</td></tr>
<tr><td>>hg38_21_+_8433172_8446622_RNA45SN1_45S_50ntFlanks@12418.12460.43|CGATGTCGGCTCTTCCTATCATTGTGAAGCAGAATTCACCAAG</td><td>>hg38_21_+_8433172_8446622_RNA45SN1_45S_50ntFlanks</td><td>12418</td><td>12460</td><td>43</td><td>CGATGTCGGCTCTTCCTATCATTGTGAAGCAGAATTCACCAAG</td><td>587.8</td><td>-2.8</td><td>1.3</td><td>-2.2</td><td>3.1E-02</td><td>3.5E-02</td><td>MB231/MB468</td><td>NUC</td></tr>
<tr><td>>MI0000465|hsa-mir-191&WithFlank&3|-|49020612|49020715@22.41.20(+1U)&[MIMAT0000440&hsa-miR-191-5p&offsets|0|-3(+1U);m-48&3|-|49020672|49020694&offsets|0|-3(+1U)]|CAACGGAATCCCAAAAGCAGT</td><td>>MI0000465|hsa-mir-191&WithFlank&3|-|49020612|49020715</td><td>22</td><td>41</td><td>20(+1U)</td><td>CAACGGAATCCCAAAAGCAGT</td><td>123.7</td><td>3.7</td><td>1.6</td><td>2.3</td><td>2.1E-02</td><td>3.5E-02</td><td>BT20/MB468</td><td>MP</td></tr>
<tr><td>>hg38_21_+_8433172_8446622_RNA45SN1_45S_50ntFlanks@10319.10341.23|GCAGATCTTGGTGGTAGTAGCAA</td><td>>hg38_21_+_8433172_8446622_RNA45SN1_45S_50ntFlanks</td><td>10319</td><td>10341</td><td>23</td><td>GCAGATCTTGGTGGTAGTAGCAA</td><td>85.5</td><td>3.3</td><td>1.4</td><td>2.3</td><td>2.1E-02</td><td>3.5E-02</td><td>BT20/MB468</td><td>MP</td></tr>
<tr><td>>hg38_21_+_8433172_8446622_RNA45SN1_45S_50ntFlanks@10101.10126.26|CGCGCGCGTGTGGTGTGCGTCGGAGG</td><td>>hg38_21_+_8433172_8446622_RNA45SN1_45S_50ntFlanks</td><td>10101</td><td>10126</td><td>26</td><td>CGCGCGCGTGTGGTGTGCGTCGGAGG</td><td>605.3</td><td>-3.8</td><td>1.7</td><td>-2.2</td><td>2.7E-02</td><td>3.5E-02</td><td>BT20/MB468</td><td>MITO</td></tr>
<tr><td>>hg38_1_-_228634819_228635039_RNA5S12_5S_50ntFlanks@132.171.40|GATGGGAGACCGCCTGGGAATACCGGGTGCTGTAGGCTTT</td><td>>hg38_1_-_228634819_228635039_RNA5S12_5S_50ntFlanks</td><td>132</td><td>171</td><td>40</td><td>GATGGGAGACCGCCTGGGAATACCGGGTGCTGTAGGCTTT</td><td>10410.7</td><td>2.3</td><td>0.9</td><td>2.5</td><td>1.1E-02</td><td>3.5E-02</td><td>BT20/MB231</td><td>MITO</td></tr>
<tr><td>>hg38_21_+_8433172_8446622_RNA45SN1_45S_50ntFlanks@9921.9948.28|TGTTGGTTGATATAGACAGCAGGACGGT</td><td>>hg38_21_+_8433172_8446622_RNA45SN1_45S_50ntFlanks</td><td>9921</td><td>9948</td><td>28</td><td>TGTTGGTTGATATAGACAGCAGGACGGT</td><td>91.2</td><td>4.3</td><td>1.7</td><td>2.6</td><td>9.1E-03</td><td>3.5E-02</td><td>BT20/MB231</td><td>MP</td></tr>
<tr><td>>hg38_21_+_8433172_8446622_RNA45SN1_45S_50ntFlanks@6650.6697.48|ACGACTCTTAGCGGTGGATCACTCGGCTCGTGCGTCGATGAAGAACGC</td><td>>hg38_21_+_8433172_8446622_RNA45SN1_45S_50ntFlanks</td><td>6650</td><td>6697</td><td>48</td><td>ACGACTCTTAGCGGTGGATCACTCGGCTCGTGCGTCGATGAAGAACGC</td><td>1017.4</td><td>2.1</td><td>1.0</td><td>2.2</td><td>2.7E-02</td><td>3.5E-02</td><td>BT20/MB468</td><td>MITO</td></tr>
<tr><td>>trnaMT_SerGCT_MT_+_12207_12265@8.31.24|CTCACAAGAACTGCTAACTCATGC</td><td>>trnaMT_SerGCT_MT_+_12207_12265</td><td>8</td><td>31</td><td>24</td><td>CTCACAAGAACTGCTAACTCATGC</td><td>182.4</td><td>-2.9</td><td>1.3</td><td>-2.2</td><td>3.0E-02</td><td>3.5E-02</td><td>BT20/MB468</td><td>CYTO</td></tr>
<tr><td>>hg38_21_+_8433172_8446622_RNA45SN1_45S_50ntFlanks@12416.12455.40|TTCGATGTCGGCTCTTCCTATCATTGTGAAGCAGAATTCA</td><td>>hg38_21_+_8433172_8446622_RNA45SN1_45S_50ntFlanks</td><td>12416</td><td>12455</td><td>40</td><td>TTCGATGTCGGCTCTTCCTATCATTGTGAAGCAGAATTCA</td><td>872.8</td><td>-2.2</td><td>1.0</td><td>-2.1</td><td>3.2E-02</td><td>3.5E-02</td><td>MB231/MB468</td><td>MITO</td></tr>
<tr><td>>hg38_21_+_8433172_8446622_RNA45SN1_45S_50ntFlanks@4735.4759.25|ACGAAAGTCGGAGGTTCGAAGACGA</td><td>>hg38_21_+_8433172_8446622_RNA45SN1_45S_50ntFlanks</td><td>4735</td><td>4759</td><td>25</td><td>ACGAAAGTCGGAGGTTCGAAGACGA</td><td>648.4</td><td>1.8</td><td>0.8</td><td>2.2</td><td>3.0E-02</td><td>3.5E-02</td><td>MB231/MB468</td><td>CYTO</td></tr>
<tr><td>>hg38_21_+_8433172_8446622_RNA45SN1_45S_50ntFlanks@5489.5509.21|CTGAGAAGACGGTCGAACTTG</td><td>>hg38_21_+_8433172_8446622_RNA45SN1_45S_50ntFlanks</td><td>5489</td><td>5509</td><td>21</td><td>CTGAGAAGACGGTCGAACTTG</td><td>1155.8</td><td>-2.7</td><td>1.2</td><td>-2.3</td><td>2.1E-02</td><td>3.5E-02</td><td>BT20/MB468</td><td>MP</td></tr>
<tr><td>>hg38_MT_+_598_1651_RNR1_12S_50ntFlanks@973.1004.32|GTAAGTGTACTGGAAAGTGCACTTGGACGAAC</td><td>>hg38_MT_+_598_1651_RNR1_12S_50ntFlanks</td><td>973</td><td>1004</td><td>32</td><td>GTAAGTGTACTGGAAAGTGCACTTGGACGAAC</td><td>137.0</td><td>3.0</td><td>1.4</td><td>2.2</td><td>2.9E-02</td><td>3.5E-02</td><td>MB231/MB468</td><td>MP</td></tr>
<tr><td>>trnaMT_GluTTC_MT_-_14674_14742@27.61.35,trnalookalike8_GluTTC_5_-_93905172_93905240@27.61.35|GTTTTTCATATCATTGGTCGTGGTTGTAGTCCGTG</td><td>>trnaMT_GluTTC_MT_-_14674_14742</td><td>27</td><td>61</td><td>35</td><td>GTTTTTCATATCATTGGTCGTGGTTGTAGTCCGTG</td><td>178.3</td><td>-2.2</td><td>1.0</td><td>-2.2</td><td>3.0E-02</td><td>3.5E-02</td><td>MB231/MB468</td><td>CYTO</td></tr>
<tr><td>>MI0005559|hsa-mir-744&WithFlank&17|+|12081893|12082002@17.38.22&[MIMAT0004945&hsa-miR-744-5p&offsets|0|0;m-172&17|+|12081909|12081929&offsets|0|+1]|TGCGGGGCTAGGGCTAACAGCA</td><td>>MI0005559|hsa-mir-744&WithFlank&17|+|12081893|12082002</td><td>17</td><td>38</td><td>22</td><td>TGCGGGGCTAGGGCTAACAGCA</td><td>1011.1</td><td>2.0</td><td>0.8</td><td>2.5</td><td>1.2E-02</td><td>3.5E-02</td><td>BT20/MB231</td><td>CYTO</td></tr>
<tr><td>>MI0000067|hsa-let-7f-1&WithFlank&9|+|94176341|94176439@13.35.23&[MIMAT0000067&hsa-let-7f-5p&offsets|0|+1;m-20&9|+|94176353|94176374&offsets|0|+1]|TGAGGTAGTAGATTGTATAGTTG</td><td>>MI0000067|hsa-let-7f-1&WithFlank&9|+|94176341|94176439</td><td>13</td><td>35</td><td>23</td><td>TGAGGTAGTAGATTGTATAGTTG</td><td>361.4</td><td>1.6</td><td>0.6</td><td>2.6</td><td>9.7E-03</td><td>3.5E-02</td><td>BT20/MB231</td><td>NUC</td></tr>
<tr><td>>hg38_21_+_8433172_8446622_RNA45SN1_45S_50ntFlanks@11104.11129.26|AAGGGTCGGGGCGGCAGGGGCCGGCG</td><td>>hg38_21_+_8433172_8446622_RNA45SN1_45S_50ntFlanks</td><td>11104</td><td>11129</td><td>26</td><td>AAGGGTCGGGGCGGCAGGGGCCGGCG</td><td>124.1</td><td>-3.1</td><td>1.3</td><td>-2.3</td><td>2.1E-02</td><td>3.5E-02</td><td>BT20/MB468</td><td>MP</td></tr>
<tr><td>>hg38_21_+_8433172_8446622_RNA45SN1_45S_50ntFlanks@4752.4786.35|GAAGACGATCAGATACCGTCGTAGTTCCGACCATA</td><td>>hg38_21_+_8433172_8446622_RNA45SN1_45S_50ntFlanks</td><td>4752</td><td>4786</td><td>35</td><td>GAAGACGATCAGATACCGTCGTAGTTCCGACCATA</td><td>235.5</td><td>1.6</td><td>0.8</td><td>2.2</td><td>3.0E-02</td><td>3.5E-02</td><td>MB231/MB468</td><td>CYTO</td></tr>
<tr><td>>hg38_21_+_8433172_8446622_RNA45SN1_45S_50ntFlanks@11919.11940.22|CATGAGAGGTGTAGAATAAGTG</td><td>>hg38_21_+_8433172_8446622_RNA45SN1_45S_50ntFlanks</td><td>11919</td><td>11940</td><td>22</td><td>CATGAGAGGTGTAGAATAAGTG</td><td>290.9</td><td>4.6</td><td>1.8</td><td>2.6</td><td>9.1E-03</td><td>3.5E-02</td><td>BT20/MB231</td><td>MP</td></tr>
<tr><td>>hg38_21_+_8433172_8446622_RNA45SN1_45S_50ntFlanks@6663.6680.18|GTGGATCACTCGGCTCGT</td><td>>hg38_21_+_8433172_8446622_RNA45SN1_45S_50ntFlanks</td><td>6663</td><td>6680</td><td>18</td><td>GTGGATCACTCGGCTCGT</td><td>287.7</td><td>-3.3</td><td>1.4</td><td>-2.3</td><td>2.1E-02</td><td>3.5E-02</td><td>BT20/MB468</td><td>MP</td></tr>
<tr><td>>hg38_21_+_8433172_8446622_RNA45SN1_45S_50ntFlanks@9622.9652.31|CTAGTAGCTGGTTCCCTCCGAAGTTTCCCTC</td><td>>hg38_21_+_8433172_8446622_RNA45SN1_45S_50ntFlanks</td><td>9622</td><td>9652</td><td>31</td><td>CTAGTAGCTGGTTCCCTCCGAAGTTTCCCTC</td><td>245.2</td><td>-4.0</td><td>1.7</td><td>-2.3</td><td>2.1E-02</td><td>3.5E-02</td><td>BT20/MB468</td><td>MP</td></tr>
<tr><td>>MI0000103|hsa-mir-101-1&WithFlank&1|-|65058428|65058514@53.73.21;MI0000739|hsa-mir-101-2&WithFlank&9|+|4850291|4850381@55.75.21&[MIMAT0000099&hsa-miR-101-3p&offsets|0|0;m-17&1|-|65058443|65058462&offsets|0|+1];[MIMAT0000099_1&hsa-miR-101-3p&offsets|0|0;m-16&9|+|4850345|4850364&offsets|0|+1]|TACAGTACTGTGATAACTGAA</td><td>>MI0000103|hsa-mir-101-1&WithFlank&1|-|65058428|65058514</td><td>53</td><td>73</td><td>21</td><td>TACAGTACTGTGATAACTGAA</td><td>153.7</td><td>2.6</td><td>1.2</td><td>2.2</td><td>3.1E-02</td><td>3.5E-02</td><td>BT20/MB468</td><td>NUC</td></tr>
<tr><td>>hg38_21_+_8433172_8446622_RNA45SN1_45S_50ntFlanks@10391.10422.32|AACAGCAGTTGAACATGGGTCAGTCGGTCCTG</td><td>>hg38_21_+_8433172_8446622_RNA45SN1_45S_50ntFlanks</td><td>10391</td><td>10422</td><td>32</td><td>AACAGCAGTTGAACATGGGTCAGTCGGTCCTG</td><td>239.3</td><td>-2.0</td><td>0.9</td><td>-2.3</td><td>2.1E-02</td><td>3.5E-02</td><td>BT20/MB468</td><td>MP</td></tr>
<tr><td>>MI0000487|hsa-mir-193a&WithFlank&17|+|31559990|31560089@27.46.20&[MIMAT0004614&hsa-miR-193a-5p&offsets|0|-2;m-123&17|+|31560016|31560037&offsets|0|-2]|TGGGTCTTTGCGGGCGAGAT</td><td>>MI0000487|hsa-mir-193a&WithFlank&17|+|31559990|31560089</td><td>27</td><td>46</td><td>20</td><td>TGGGTCTTTGCGGGCGAGAT</td><td>90.4</td><td>5.1</td><td>1.9</td><td>2.6</td><td>9.2E-03</td><td>3.5E-02</td><td>BT20/MB231</td><td>MP</td></tr>
<tr><td>>hg38_21_+_8433172_8446622_RNA45SN1_45S_50ntFlanks@12481.12498.18|AATAGGGAACGTGAGCTG</td><td>>hg38_21_+_8433172_8446622_RNA45SN1_45S_50ntFlanks</td><td>12481</td><td>12498</td><td>18</td><td>AATAGGGAACGTGAGCTG</td><td>580.2</td><td>5.0</td><td>1.9</td><td>2.6</td><td>9.1E-03</td><td>3.5E-02</td><td>BT20/MB231</td><td>MP</td></tr>
<tr><td>>hg38_21_+_8433172_8446622_RNA45SN1_45S_50ntFlanks@11918.11958.41|ACATGAGAGGTGTAGAATAAGTGGGAGGCCCCCGGCGCCCC</td><td>>hg38_21_+_8433172_8446622_RNA45SN1_45S_50ntFlanks</td><td>11918</td><td>11958</td><td>41</td><td>ACATGAGAGGTGTAGAATAAGTGGGAGGCCCCCGGCGCCCC</td><td>61.0</td><td>-4.2</td><td>1.8</td><td>-2.3</td><td>2.1E-02</td><td>3.5E-02</td><td>BT20/MB468</td><td>MP</td></tr>
<tr><td>>hg38_21_+_8433172_8446622_RNA45SN1_45S_50ntFlanks@6739.6771.33|TCATCGACACTTCGAACGCACTTGCGGCCCCGG</td><td>>hg38_21_+_8433172_8446622_RNA45SN1_45S_50ntFlanks</td><td>6739</td><td>6771</td><td>33</td><td>TCATCGACACTTCGAACGCACTTGCGGCCCCGG</td><td>368.1</td><td>-4.2</td><td>1.8</td><td>-2.3</td><td>2.1E-02</td><td>3.5E-02</td><td>BT20/MB468</td><td>MP</td></tr>
<tr><td>>am_trnaMT_PheGAA_MT_+_577_647@45.64.20|AGACGGGCTCACATCACCCC</td><td>>am_trnaMT_PheGAA_MT_+_577_647</td><td>45</td><td>64</td><td>20</td><td>AGACGGGCTCACATCACCCC</td><td>84.8</td><td>-2.8</td><td>1.3</td><td>-2.2</td><td>2.9E-02</td><td>3.5E-02</td><td>MB231/MB468</td><td>MP</td></tr>
<tr><td>>am_trnaMT_PheGAA_MT_+_577_647@41.59.19|GTTTAGACGGGCTCACATC</td><td>>am_trnaMT_PheGAA_MT_+_577_647</td><td>41</td><td>59</td><td>19</td><td>GTTTAGACGGGCTCACATC</td><td>148.7</td><td>1.6</td><td>0.7</td><td>2.2</td><td>2.9E-02</td><td>3.5E-02</td><td>MB231/MB468</td><td>MP</td></tr>
<tr><td>>hg38_21_+_8433172_8446622_RNA45SN1_45S_50ntFlanks@9571.9595.25|GCAAATCGGTCGTCCGACCTGGGTA</td><td>>hg38_21_+_8433172_8446622_RNA45SN1_45S_50ntFlanks</td><td>9571</td><td>9595</td><td>25</td><td>GCAAATCGGTCGTCCGACCTGGGTA</td><td>221.5</td><td>-2.3</td><td>1.0</td><td>-2.3</td><td>2.1E-02</td><td>3.5E-02</td><td>BT20/MB468</td><td>MP</td></tr>
<tr><td>>hg38_21_+_8433172_8446622_RNA45SN1_45S_50ntFlanks@9170.9197.28|GCGGGTCGCGCCGTCGGGCCCGGGGGAG</td><td>>hg38_21_+_8433172_8446622_RNA45SN1_45S_50ntFlanks</td><td>9170</td><td>9197</td><td>28</td><td>GCGGGTCGCGCCGTCGGGCCCGGGGGAG</td><td>160.9</td><td>-4.4</td><td>1.9</td><td>-2.3</td><td>2.1E-02</td><td>3.5E-02</td><td>BT20/MB468</td><td>MP</td></tr>
<tr><td>>hg38_21_+_8433172_8446622_RNA45SN1_45S_50ntFlanks@8354.8373.20|TTGAAAAGAACTTTGAAGAG</td><td>>hg38_21_+_8433172_8446622_RNA45SN1_45S_50ntFlanks</td><td>8354</td><td>8373</td><td>20</td><td>TTGAAAAGAACTTTGAAGAG</td><td>1198.9</td><td>4.5</td><td>1.7</td><td>2.6</td><td>9.2E-03</td><td>3.5E-02</td><td>BT20/MB231</td><td>MP</td></tr>
<tr><td>>hg38_21_+_8433172_8446622_RNA45SN1_45S_50ntFlanks@11581.11612.32|AACTGGTGCGGACCAGGGGAATCCGACTGTTT</td><td>>hg38_21_+_8433172_8446622_RNA45SN1_45S_50ntFlanks</td><td>11581</td><td>11612</td><td>32</td><td>AACTGGTGCGGACCAGGGGAATCCGACTGTTT</td><td>2712.9</td><td>-1.6</td><td>0.7</td><td>-2.2</td><td>2.7E-02</td><td>3.5E-02</td><td>BT20/MB468</td><td>MITO</td></tr>
<tr><td>>hg38_21_+_8433172_8446622_RNA45SN1_45S_50ntFlanks@9502.9525.24|TGGTGAACTATGCCTGGGCAGGGC</td><td>>hg38_21_+_8433172_8446622_RNA45SN1_45S_50ntFlanks</td><td>9502</td><td>9525</td><td>24</td><td>TGGTGAACTATGCCTGGGCAGGGC</td><td>152.7</td><td>6.4</td><td>2.5</td><td>2.6</td><td>9.2E-03</td><td>3.5E-02</td><td>BT20/MB231</td><td>MP</td></tr>
<tr><td>>hg38_21_+_8433172_8446622_RNA45SN1_45S_50ntFlanks@12418.12451.34|CGATGTCGGCTCTTCCTATCATTGTGAAGCAGAA</td><td>>hg38_21_+_8433172_8446622_RNA45SN1_45S_50ntFlanks</td><td>12418</td><td>12451</td><td>34</td><td>CGATGTCGGCTCTTCCTATCATTGTGAAGCAGAA</td><td>290.3</td><td>-1.3</td><td>0.6</td><td>-2.2</td><td>3.1E-02</td><td>3.5E-02</td><td>MB231/MB468</td><td>CYTO</td></tr>
<tr><td>>hg38_21_+_8433172_8446622_RNA45SN1_45S_50ntFlanks@11668.11687.20|TGCCCAGTGCTCTGAATGTC</td><td>>hg38_21_+_8433172_8446622_RNA45SN1_45S_50ntFlanks</td><td>11668</td><td>11687</td><td>20</td><td>TGCCCAGTGCTCTGAATGTC</td><td>2867.9</td><td>-2.1</td><td>0.8</td><td>-2.5</td><td>1.2E-02</td><td>3.5E-02</td><td>BT20/MB231</td><td>CYTO</td></tr>
<tr><td>>hg38_21_+_8433172_8446622_RNA45SN1_45S_50ntFlanks@5485.5512.28|AGCGCTGAGAAGACGGTCGAACTTGACT</td><td>>hg38_21_+_8433172_8446622_RNA45SN1_45S_50ntFlanks</td><td>5485</td><td>5512</td><td>28</td><td>AGCGCTGAGAAGACGGTCGAACTTGACT</td><td>624.6</td><td>2.1</td><td>1.0</td><td>2.2</td><td>2.7E-02</td><td>3.5E-02</td><td>BT20/MB468</td><td>MITO</td></tr>
<tr><td>>hg38_21_+_8433172_8446622_RNA45SN1_45S_50ntFlanks@10771.10798.28|CAGGTCTCCAAGGTGAACAGCCTCTGGC</td><td>>hg38_21_+_8433172_8446622_RNA45SN1_45S_50ntFlanks</td><td>10771</td><td>10798</td><td>28</td><td>CAGGTCTCCAAGGTGAACAGCCTCTGGC</td><td>64.9</td><td>-3.9</td><td>1.7</td><td>-2.3</td><td>2.1E-02</td><td>3.5E-02</td><td>BT20/MB468</td><td>MP</td></tr>
<tr><td>>hg38_21_+_8433172_8446622_RNA45SN1_45S_50ntFlanks@11027.11055.29|CTCTCTCTCTCTCTCCCCCGCTCCCCGTC</td><td>>hg38_21_+_8433172_8446622_RNA45SN1_45S_50ntFlanks</td><td>11027</td><td>11055</td><td>29</td><td>CTCTCTCTCTCTCTCCCCCGCTCCCCGTC</td><td>1639.2</td><td>3.9</td><td>1.8</td><td>2.2</td><td>2.7E-02</td><td>3.5E-02</td><td>BT20/MB468</td><td>MITO</td></tr>
<tr><td>>hg38_MT_+_1621_3279_RNR2_16S_50ntFlanks@115.136.22|AAATAAAGTATAGGCGATAGAA</td><td>>hg38_MT_+_1621_3279_RNR2_16S_50ntFlanks</td><td>115</td><td>136</td><td>22</td><td>AAATAAAGTATAGGCGATAGAA</td><td>144.7</td><td>3.0</td><td>1.4</td><td>2.2</td><td>2.9E-02</td><td>3.5E-02</td><td>MB231/MB468</td><td>MP</td></tr>
<tr><td>>hg38_MT_+_1621_3279_RNR2_16S_50ntFlanks@1389.1418.30|CGATGGTGCAGCCGCTATTAAAGGTTCGTT</td><td>>hg38_MT_+_1621_3279_RNR2_16S_50ntFlanks</td><td>1389</td><td>1418</td><td>30</td><td>CGATGGTGCAGCCGCTATTAAAGGTTCGTT</td><td>40.5</td><td>4.5</td><td>2.0</td><td>2.2</td><td>2.9E-02</td><td>3.5E-02</td><td>MB231/MB468</td><td>MP</td></tr>
<tr><td>>hg38_21_+_8433172_8446622_RNA45SN1_45S_50ntFlanks@12638.12664.27|CTACCATCTGTGGGATTATGACTGAAC</td><td>>hg38_21_+_8433172_8446622_RNA45SN1_45S_50ntFlanks</td><td>12638</td><td>12664</td><td>27</td><td>CTACCATCTGTGGGATTATGACTGAAC</td><td>91.3</td><td>2.2</td><td>1.0</td><td>2.3</td><td>2.1E-02</td><td>3.5E-02</td><td>BT20/MB468</td><td>MP</td></tr>
<tr><td>>hg38_21_+_8433172_8446622_RNA45SN1_45S_50ntFlanks@11666.11688.23|TCTGCCCAGTGCTCTGAATGTCA</td><td>>hg38_21_+_8433172_8446622_RNA45SN1_45S_50ntFlanks</td><td>11666</td><td>11688</td><td>23</td><td>TCTGCCCAGTGCTCTGAATGTCA</td><td>218.3</td><td>1.6</td><td>0.7</td><td>2.1</td><td>3.2E-02</td><td>3.5E-02</td><td>MB231/MB468</td><td>NUC</td></tr>
<tr><td>>hg38_MT_+_1621_3279_RNR2_16S_50ntFlanks@1389.1423.35|CGATGGTGCAGCCGCTATTAAAGGTTCGTTTGTTC</td><td>>hg38_MT_+_1621_3279_RNR2_16S_50ntFlanks</td><td>1389</td><td>1423</td><td>35</td><td>CGATGGTGCAGCCGCTATTAAAGGTTCGTTTGTTC</td><td>67.4</td><td>3.7</td><td>1.7</td><td>2.2</td><td>2.9E-02</td><td>3.5E-02</td><td>MB231/MB468</td><td>MP</td></tr>
<tr><td>>trna119_LysCTT_1_-_145395522_145395594@1.31.31,trna11_LysCTT_5_-_180648979_180649051@1.31.31,trna13_LysCTT_6_+_26556774_26556846@1.31.31,trna32_LysCTT_16_-_3207406_3207478@1.31.31,trna7_LysCTT_16_+_3225692_3225764@1.31.31,trna9_LysCTT_5_+_180634755_180634827@1.31.31|GCCCGGCTAGCTCAGTCGGTAGAGCATGAGA</td><td>>trna119_LysCTT_1_-_145395522_145395594</td><td>1</td><td>31</td><td>31</td><td>GCCCGGCTAGCTCAGTCGGTAGAGCATGAGA</td><td>1115.8</td><td>-3.0</td><td>1.4</td><td>-2.2</td><td>2.7E-02</td><td>3.5E-02</td><td>BT20/MB468</td><td>MITO</td></tr>
<tr><td>>hg38_21_+_8433172_8446622_RNA45SN1_45S_50ntFlanks@11712.11736.25|CGGGTAAACGGCGGGAGTAACTATG</td><td>>hg38_21_+_8433172_8446622_RNA45SN1_45S_50ntFlanks</td><td>11712</td><td>11736</td><td>25</td><td>CGGGTAAACGGCGGGAGTAACTATG</td><td>125.7</td><td>-2.1</td><td>0.9</td><td>-2.3</td><td>2.1E-02</td><td>3.5E-02</td><td>BT20/MB468</td><td>MP</td></tr>
</table>

<table>
<tr><td></td><td></td><td></td><td></td><td></td><td></td><td></td><td></td><td></td><td></td><td></td><td></td><td></td><td></td></tr>
<tr><td>>hg38_21_+_8433172_8446622_RNA45SN1_45S_50ntFlanks@10820.10840.21|GGGAAGTCGGCAAGCCGGATC</td><td>>hg38_21_+_8433172_8446622_RNA45SN1_45S_50ntFlanks</td><td>10820</td><td>10840</td><td>21</td><td>GGGAAGTCGGCAAGCCGGATC</td><td>579.7</td><td>4.8</td><td>1.8</td><td>2.6</td><td>9.2E-03</td><td>3.5E-02</td><td>BT20/MB231</td><td>MP</td></tr>
<tr><td>>hg38_21_+_8433172_8446622_RNA45SN1_45S_50ntFlanks@4884.4917.34|CAAAGCTGAAACTTAAAGGAATTGACGGAAGGGC</td><td>>hg38_21_+_8433172_8446622_RNA45SN1_45S_50ntFlanks</td><td>4884</td><td>4917</td><td>34</td><td>CAAAGCTGAAACTTAAAGGAATTGACGGAAGGGC</td><td>284.3</td><td>-3.1</td><td>1.4</td><td>-2.3</td><td>2.1E-02</td><td>3.5E-02</td><td>BT20/MB468</td><td>MP</td></tr>
<tr><td>>MI0000085|hsa-mir-27a&WithFlank&19|-|13836434|13836523@57.75.19(+1U)&[MIMAT0000084&hsa-miR-27a-3p&offsets|0|-2(+1U);m-43&19|-|13836447|13836467&offsets|0|-2(+1U)]|TTCACAGTGGCTAAGTTCCT</td><td>>MI0000085|hsa-mir-27a&WithFlank&19|-|13836434|13836523</td><td>57</td><td>75</td><td>19(+1U)</td><td>TTCACAGTGGCTAAGTTCCT</td><td>931.0</td><td>-3.2</td><td>1.2</td><td>-2.6</td><td>9.2E-03</td><td>3.5E-02</td><td>BT20/MB231</td><td>MP</td></tr>
<tr><td>>MI0000088|hsa-mir-30a&WithFlank&6|-|71403545|71403627@54.71.18&[MIMAT0000088&hsa-miR-30a-3p&offsets|+1|-3;m-38&6|-|71403554|71403575&offsets|+1|-3]|TTTCAGTCGGATGTTTGC</td><td>>MI0000088|hsa-mir-30a&WithFlank&6|-|71403545|71403627</td><td>54</td><td>71</td><td>18</td><td>TTTCAGTCGGATGTTTGC</td><td>750.5</td><td>-1.7</td><td>0.7</td><td>-2.5</td><td>1.1E-02</td><td>3.5E-02</td><td>BT20/MB231</td><td>MITO</td></tr>
<tr><td>>MI0000542|hsa-mir-320a&WithFlank&8|-|22244960|22245043@49.69.21&[MIMAT0000510&hsa-miR-320a-3p&offsets|+1|0;m-60&8|-|22244975|22244996&offsets|+1|0]|AAAGCTGGGTTGAGAGGGCGA</td><td>>MI0000542|hsa-mir-320a&WithFlank&8|-|22244960|22245043</td><td>49</td><td>69</td><td>21</td><td>AAAGCTGGGTTGAGAGGGCGA</td><td>3497.9</td><td>2.4</td><td>0.9</td><td>2.6</td><td>9.2E-03</td><td>3.5E-02</td><td>BT20/MB231</td><td>MP</td></tr>
<tr><td>>hg38_21_+_8433172_8446622_RNA45SN1_45S_50ntFlanks@12625.12654.30|CAATGGGGCGAAGCTACCATCTGTGGGATT</td><td>>hg38_21_+_8433172_8446622_RNA45SN1_45S_50ntFlanks</td><td>12625</td><td>12654</td><td>30</td><td>CAATGGGGCGAAGCTACCATCTGTGGGATT</td><td>262.6</td><td>4.7</td><td>1.8</td><td>2.6</td><td>9.2E-03</td><td>3.5E-02</td><td>BT20/MB231</td><td>MP</td></tr>
<tr><td>>hg38_MT_+_598_1651_RNR1_12S_50ntFlanks@982.1004.23|CTGGAAAGTGCACTTGGACGAAC</td><td>>hg38_MT_+_598_1651_RNR1_12S_50ntFlanks</td><td>982</td><td>1004</td><td>23</td><td>CTGGAAAGTGCACTTGGACGAAC</td><td>114.1</td><td>2.2</td><td>1.0</td><td>2.1</td><td>3.2E-02</td><td>3.5E-02</td><td>MB231/MB468</td><td>MITO</td></tr>
<tr><td>>hg38_21_+_8433172_8446622_RNA45SN1_45S_50ntFlanks@11582.11602.21|ACTGGTGCGGACCAGGGGAAT</td><td>>hg38_21_+_8433172_8446622_RNA45SN1_45S_50ntFlanks</td><td>11582</td><td>11602</td><td>21</td><td>ACTGGTGCGGACCAGGGGAAT</td><td>458.4</td><td>-2.3</td><td>1.0</td><td>-2.3</td><td>2.1E-02</td><td>3.5E-02</td><td>BT20/MB468</td><td>MP</td></tr>
<tr><td>>hg38_21_+_8433172_8446622_RNA45SN1_45S_50ntFlanks@12418.12441.24|CGATGTCGGCTCTTCCTATCATTG</td><td>>hg38_21_+_8433172_8446622_RNA45SN1_45S_50ntFlanks</td><td>12418</td><td>12441</td><td>24</td><td>CGATGTCGGCTCTTCCTATCATTG</td><td>299.3</td><td>-4.1</td><td>1.8</td><td>-2.3</td><td>2.1E-02</td><td>3.5E-02</td><td>BT20/MB468</td><td>MP</td></tr>
<tr><td>>am_MI0000809|hsa-mir-151a&WithFlank&8|-|140732558|140732659@17.38.22&[MIMAT0004697&hsa-miR-151a-5p&offsets|0|+1;m-75&8|-|140732623|140732643&offsets|0|+1]|TCGAGGAGCTCACAGTCTAGTA</td><td>>am_MI0000809|hsa-mir-151a&WithFlank&8|-|140732558|140732659</td><td>17</td><td>38</td><td>22</td><td>TCGAGGAGCTCACAGTCTAGTA</td><td>1396.3</td><td>1.8</td><td>0.7</td><td>2.6</td><td>9.9E-03</td><td>3.5E-02</td><td>BT20/MB231</td><td>NUC</td></tr>
<tr><td>>hg38_21_+_8433172_8446622_RNA45SN1_45S_50ntFlanks@10350.10383.34|ACGAGAACTTTGAAGGCCGAAGTGGAGAAGGGTT</td><td>>hg38_21_+_8433172_8446622_RNA45SN1_45S_50ntFlanks</td><td>10350</td><td>10383</td><td>34</td><td>ACGAGAACTTTGAAGGCCGAAGTGGAGAAGGGTT</td><td>1257.4</td><td>3.9</td><td>1.5</td><td>2.6</td><td>9.3E-03</td><td>3.5E-02</td><td>BT20/MB231</td><td>MP</td></tr>
<tr><td>>hg38_21_+_8433172_8446622_RNA45SN1_45S_50ntFlanks@8319.8356.38|CGAGACCGATAGTCAACAAGTACCGTAAGGGAAAGTTG</td><td>>hg38_21_+_8433172_8446622_RNA45SN1_45S_50ntFlanks</td><td>8319</td><td>8356</td><td>38</td><td>CGAGACCGATAGTCAACAAGTACCGTAAGGGAAAGTTG</td><td>278.3</td><td>-2.4</td><td>1.0</td><td>-2.3</td><td>2.2E-02</td><td>3.5E-02</td><td>BT20/MB468</td><td>MP</td></tr>
<tr><td>>hg38_21_+_8433172_8446622_RNA45SN1_45S_50ntFlanks@12632.12664.33|GCGAAGCTACCATCTGTGGGATTATGACTGAAC</td><td>>hg38_21_+_8433172_8446622_RNA45SN1_45S_50ntFlanks</td><td>12632</td><td>12664</td><td>33</td><td>GCGAAGCTACCATCTGTGGGATTATGACTGAAC</td><td>302.5</td><td>3.6</td><td>1.4</td><td>2.6</td><td>9.3E-03</td><td>3.5E-02</td><td>BT20/MB231</td><td>MP</td></tr>
<tr><td>>hg38_21_+_8433172_8446622_RNA45SN1_45S_50ntFlanks@6647.6679.33|CGTACGACTCTTAGCGGTGGATCACTCGGCTCG</td><td>>hg38_21_+_8433172_8446622_RNA45SN1_45S_50ntFlanks</td><td>6647</td><td>6679</td><td>33</td><td>CGTACGACTCTTAGCGGTGGATCACTCGGCTCG</td><td>302.7</td><td>1.7</td><td>0.8</td><td>2.2</td><td>3.1E-02</td><td>3.5E-02</td><td>BT20/MB468</td><td>CYTO</td></tr>
<tr><td>>hg38_21_+_8433172_8446622_RNA45SN1_45S_50ntFlanks@9300.9322.23|ACGGACCAAGGAGTCTAACACGT</td><td>>hg38_21_+_8433172_8446622_RNA45SN1_45S_50ntFlanks</td><td>9300</td><td>9322</td><td>23</td><td>ACGGACCAAGGAGTCTAACACGT</td><td>86.5</td><td>4.2</td><td>1.6</td><td>2.6</td><td>9.3E-03</td><td>3.5E-02</td><td>BT20/MB231</td><td>MP</td></tr>
<tr><td>>hg38_21_+_8433172_8446622_RNA45SN1_45S_50ntFlanks@9955.9982.28|GGAAGTCGGAATCCGCTAAGGAGTGTGT</td><td>>hg38_21_+_8433172_8446622_RNA45SN1_45S_50ntFlanks</td><td>9955</td><td>9982</td><td>28</td><td>GGAAGTCGGAATCCGCTAAGGAGTGTGT</td><td>152.9</td><td>4.3</td><td>1.6</td><td>2.6</td><td>9.3E-03</td><td>3.5E-02</td><td>BT20/MB231</td><td>MP</td></tr>
<tr><td>>hg38_21_+_8433172_8446622_RNA45SN1_45S_50ntFlanks@12420.12445.26|ATGTCGGCTCTTCCTATCATTGTGAA</td><td>>hg38_21_+_8433172_8446622_RNA45SN1_45S_50ntFlanks</td><td>12420</td><td>12445</td><td>26</td><td>ATGTCGGCTCTTCCTATCATTGTGAA</td><td>1544.8</td><td>-1.2</td><td>0.6</td><td>-2.2</td><td>3.1E-02</td><td>3.5E-02</td><td>MB231/MB468</td><td>CYTO</td></tr>
<tr><td>>trnaMT_GlnTTG_MT_-_4329_4400@1.26.26,trnalookalike2_GlnTTG_1_-_564879_564950@1.26.26|TAGGATGGGGTGTGATAGGTGGCACG</td><td>>trnaMT_GlnTTG_MT_-_4329_4400</td><td>1</td><td>26</td><td>26</td><td>TAGGATGGGGTGTGATAGGTGGCACG</td><td>253.0</td><td>2.2</td><td>1.0</td><td>2.2</td><td>2.9E-02</td><td>3.5E-02</td><td>MB231/MB468</td><td>MP</td></tr>
<tr><td>>MI0003763|hsa-mir-767&WithFlank&X|-|152393415|152393535@33.56.24&[MIMAT0003882&hsa-miR-767-5p&offsets|0|+1;m-788&X|-|152393481|152393503&offsets|0|+1]|TGCACCATGGTTGTCTGAGCATGC</td><td>>MI0003763|hsa-mir-767&WithFlank&X|-|152393415|152393535</td><td>33</td><td>56</td><td>24</td><td>TGCACCATGGTTGTCTGAGCATGC</td><td>119.2</td><td>2.4</td><td>1.0</td><td>2.3</td><td>2.2E-02</td><td>3.5E-02</td><td>BT20/MB468</td><td>MP</td></tr>
<tr><td>>MI0000078|hsa-mir-22&WithFlank&17|-|1713897|1713993@59.80.22(+1C)&[MIMAT0000077&hsa-miR-22-3p&offsets|0|0(+1C);m-2&17|-|1713914|1713935&offsets|0|0(+1C)]|AAGCTGCCAGTTGAAGAACTGTC</td><td>>MI0000078|hsa-mir-22&WithFlank&17|-|1713897|1713993</td><td>59</td><td>80</td><td>22(+1C)</td><td>AAGCTGCCAGTTGAAGAACTGTC</td><td>275.4</td><td>-4.1</td><td>1.6</td><td>-2.6</td><td>9.3E-03</td><td>3.5E-02</td><td>BT20/MB231</td><td>MP</td></tr>
<tr><td>>hg38_21_+_8433172_8446622_RNA45SN1_45S_50ntFlanks@12007.12039.33|GCCGCCGGTGAAATACCACTACTCTGATCGTTT</td><td>>hg38_21_+_8433172_8446622_RNA45SN1_45S_50ntFlanks</td><td>12007</td><td>12039</td><td>33</td><td>GCCGCCGGTGAAATACCACTACTCTGATCGTTT</td><td>490.4</td><td>2.0</td><td>0.9</td><td>2.2</td><td>2.7E-02</td><td>3.5E-02</td><td>BT20/MB468</td><td>MITO</td></tr>
<tr><td>>hg38_21_+_8433172_8446622_RNA45SN1_45S_50ntFlanks@3990.4024.35|TTGGTGACTCTAGATAACCTCGGGCCGATCGCACG</td><td>>hg38_21_+_8433172_8446622_RNA45SN1_45S_50ntFlanks</td><td>3990</td><td>4024</td><td>35</td><td>TTGGTGACTCTAGATAACCTCGGGCCGATCGCACG</td><td>564.2</td><td>1.7</td><td>0.8</td><td>2.2</td><td>3.1E-02</td><td>3.5E-02</td><td>MB231/MB468</td><td>CYTO</td></tr>
<tr><td>>MI0000750|hsa-mir-26a-2&WithFlank&12|-|57824603|57824698@20.37.18;MI0000083|hsa-mir-26a-1&WithFlank&3|+|37969398|37969486@16.33.18&[MIMAT0000082&hsa-miR-26a-5p&offsets|0|-4;m-33&12|-|57824658|57824679&offsets|0|-4];[MIMAT0000082_1&hsa-miR-26a-5p&offsets|0|-4;m-34&3|+|37969413|37969434&offsets|0|-4]|TTCAAGTAATCCAGGATA</td><td>>MI0000750|hsa-mir-26a-2&WithFlank&12|-|57824603|57824698</td><td>20</td><td>37</td><td>18</td><td>TTCAAGTAATCCAGGATA</td><td>98.0</td><td>3.8</td><td>1.7</td><td>2.2</td><td>2.9E-02</td><td>3.5E-02</td><td>MB231/MB468</td><td>MP</td></tr>
<tr><td>>hg38_21_+_8433172_8446622_RNA45SN1_45S_50ntFlanks@4734.4760.27|AACGAAAGTCGGAGGTTCGAAGACGAT</td><td>>hg38_21_+_8433172_8446622_RNA45SN1_45S_50ntFlanks</td><td>4734</td><td>4760</td><td>27</td><td>AACGAAAGTCGGAGGTTCGAAGACGAT</td><td>205.4</td><td>4.5</td><td>1.7</td><td>2.6</td><td>9.3E-03</td><td>3.5E-02</td><td>BT20/MB231</td><td>MP</td></tr>
<tr><td>>hg38_21_+_8433172_8446622_RNA45SN1_45S_50ntFlanks@12417.12450.34|TCGATGTCGGCTCTTCCTATCATTGTGAAGCAGA</td><td>>hg38_21_+_8433172_8446622_RNA45SN1_45S_50ntFlanks</td><td>12417</td><td>12450</td><td>34</td><td>TCGATGTCGGCTCTTCCTATCATTGTGAAGCAGA</td><td>260.3</td><td>1.2</td><td>0.6</td><td>2.2</td><td>3.1E-02</td><td>3.5E-02</td><td>MB231/MB468</td><td>CYTO</td></tr>
<tr><td>>trnaMT_PheGAA_MT_+_577_647@1.44.44|GTTTATGTAGCTTACCTCCTCAAAGCAATACACTGAAAATGTTT</td><td>>trnaMT_PheGAA_MT_+_577_647</td><td>1</td><td>44</td><td>44</td><td>GTTTATGTAGCTTACCTCCTCAAAGCAATACACTGAAAATGTTT</td><td>468.3</td><td>2.3</td><td>0.9</td><td>2.5</td><td>1.1E-02</td><td>3.5E-02</td><td>BT20/MB231</td><td>MITO</td></tr>
<tr><td>>hg38_21_+_8433172_8446622_RNA45SN1_45S_50ntFlanks@4069.4101.33|CTTTCGATGGTAGTCGCCGTGCCTACCATGGTG</td><td>>hg38_21_+_8433172_8446622_RNA45SN1_45S_50ntFlanks</td><td>4069</td><td>4101</td><td>33</td><td>CTTTCGATGGTAGTCGCCGTGCCTACCATGGTG</td><td>354.0</td><td>-2.0</td><td>0.9</td><td>-2.3</td><td>2.2E-02</td><td>3.5E-02</td><td>BT20/MB468</td><td>MP</td></tr>
<tr><td>>hg38_21_+_8433172_8446622_RNA45SN1_45S_50ntFlanks@10682.10711.30|TCCGGCGGCGTCCGGTGAGCTCTCGCTGGC</td><td>>hg38_21_+_8433172_8446622_RNA45SN1_45S_50ntFlanks</td><td>10682</td><td>10711</td><td>30</td><td>TCCGGCGGCGTCCGGTGAGCTCTCGCTGGC</td><td>159.1</td><td>-2.6</td><td>1.2</td><td>-2.2</td><td>2.7E-02</td><td>3.5E-02</td><td>BT20/MB468</td><td>MITO</td></tr>
<tr><td>>MI0000081|hsa-mir-24-2&WithFlank&19|-|13836281|13836365@56.75.20(+1U);MI0000080|hsa-mir-24-1&WithFlank&9|+|95086015|95086094@50.69.20(+1U)&[MIMAT0000080&hsa-miR-24-3p&offsets|0|-2(+1U);m-36&19|-|13836288|13836310&offsets|0|-3(+1U)];[MIMAT0000080_1&hsa-miR-24-3p&offsets|0|-2(+1U);m-35&9|+|95086064|95086086&offsets|0|-3(+1U)]|TGGCTCAGTTCAGCAGGAACT</td><td>>MI0000081|hsa-mir-24-2&WithFlank&19|-|13836281|13836365</td><td>56</td><td>75</td><td>20(+1U)</td><td>TGGCTCAGTTCAGCAGGAACT</td><td>108.8</td><td>3.0</td><td>1.4</td><td>2.2</td><td>2.9E-02</td><td>3.5E-02</td><td>MB231/MB468</td><td>MP</td></tr>
<tr><td>>trnaMT_ProTGG_MT_-_15956_16023@40.63.24|AATGGTGGAGTTAAAGACTTTTTC</td><td>>trnaMT_ProTGG_MT_-_15956_16023</td><td>40</td><td>63</td><td>24</td><td>AATGGTGGAGTTAAAGACTTTTTC</td><td>78.0</td><td>2.7</td><td>1.2</td><td>2.2</td><td>2.9E-02</td><td>3.5E-02</td><td>MB231/MB468</td><td>MP</td></tr>
<tr><td>>hg38_21_+_8433172_8446622_RNA45SN1_45S_50ntFlanks@5472.5512.41|ACGGCCCTGGCGGAGCGCTGAGAAGACGGTCGAACTTGACT</td><td>>hg38_21_+_8433172_8446622_RNA45SN1_45S_50ntFlanks</td><td>5472</td><td>5512</td><td>41</td><td>ACGGCCCTGGCGGAGCGCTGAGAAGACGGTCGAACTTGACT</td><td>345.5</td><td>1.6</td><td>0.8</td><td>2.1</td><td>3.2E-02</td><td>3.5E-02</td><td>BT20/MB468</td><td>NUC</td></tr>
<tr><td>>hg38_21_+_8433172_8446622_RNA45SN1_45S_50ntFlanks@480.497.18|TGATCGATGTGGTGACGT</td><td>>hg38_21_+_8433172_8446622_RNA45SN1_45S_50ntFlanks</td><td>480</td><td>497</td><td>18</td><td>TGATCGATGTGGTGACGT</td><td>439.9</td><td>-2.3</td><td>0.9</td><td>-2.6</td><td>9.9E-03</td><td>3.5E-02</td><td>BT20/MB231</td><td>NUC</td></tr>
<tr><td>>MI0000111|hsa-mir-105-1&WithFlank&X|-|152392213|152392305@21.41.21(+1A);MI0000112|hsa-mir-105-2&WithFlank&X|-|152394406|152394498@21.41.21(+1A)&[MIMAT0000102&hsa-miR-105-5p&offsets|+2|0(+1A);m-735&X|-|152392266|152392287&offsets|+2|+1(+1A)];[MIMAT0000102_1&hsa-miR-105-5p&offsets|+2|0(+1A);m-735&X|-|152394459|152394480&offsets|+2|+1(+1A)]|AAATGCTCAGACTCCTGTGGTA</td><td>>MI0000111|hsa-mir-105-1&WithFlank&X|-|152392213|152392305</td><td>21</td><td>41</td><td>21(+1A)</td><td>AAATGCTCAGACTCCTGTGGTA</td><td>29.7</td><td>8.2</td><td>3.2</td><td>2.6</td><td>9.4E-03</td><td>3.5E-02</td><td>BT20/MB231</td><td>MP</td></tr>
<tr><td>>hg38_21_+_8433172_8446622_RNA45SN1_45S_50ntFlanks@11538.11584.47|CGCGCGGCGCCTCGCCTCGGCCGGCGCCTAGCAGCCGACTTAGAACT</td><td>>hg38_21_+_8433172_8446622_RNA45SN1_45S_50ntFlanks</td><td>11538</td><td>11584</td><td>47</td><td>CGCGCGGCGCCTCGCCTCGGCCGGCGCCTAGCAGCCGACTTAGAACT</td><td>122.7</td><td>3.5</td><td>1.6</td><td>2.1</td><td>3.2E-02</td><td>3.6E-02</td><td>MB231/MB468</td><td>MITO</td></tr>
<tr><td>>trnaMT_GlnTTG_MT_-_4329_4400@1.36.36,trnalookalike2_GlnTTG_1_-_564879_564950@1.36.36|TAGGATGGGGTGTGATAGGTGGCACGGAGAATTTTG</td><td>>trnaMT_GlnTTG_MT_-_4329_4400</td><td>1</td><td>36</td><td>36</td><td>TAGGATGGGGTGTGATAGGTGGCACGGAGAATTTTG</td><td>153.8</td><td>2.7</td><td>1.2</td><td>2.2</td><td>2.9E-02</td><td>3.6E-02</td><td>MB231/MB468</td><td>MP</td></tr>
<tr><td>>hg38_21_+_8433172_8446622_RNA45SN1_45S_50ntFlanks@12475.12498.24|CCCACTAATAGGGAACGTGAGCTG</td><td>>hg38_21_+_8433172_8446622_RNA45SN1_45S_50ntFlanks</td><td>12475</td><td>12498</td><td>24</td><td>CCCACTAATAGGGAACGTGAGCTG</td><td>196.4</td><td>-4.0</td><td>1.7</td><td>-2.3</td><td>2.2E-02</td><td>3.6E-02</td><td>BT20/MB468</td><td>MP</td></tr>
<tr><td>>trna111_HisGTG_1_-_147774845_147774916@-1T.34.35,trna118_HisGTG_1_-_145396881_145396952@-1T.34.35,trna16_HisGTG_1_+_146544773_146544844@-1T.34.35,trna1_HisGTG_15_+_45493349_45493420@-1T.34.35,trna21_HisGTG_1_+_147753471_147753542@-1T.34.35,trna33_HisGTG_6_+_27125906_27125977@-1T.34.35,trna7_HisGTG_9_-_14433938_14434009@-1T.34.35,trna8_HisGTG_15_-_45492611_45492682@-1T.34.35,trna9_HisGTG_15_-_45490804_45490875@-1T.34.35|TGCCGTGATCGTATAGTGGTTAGTACTCTGCGTTG</td><td>>trna111_HisGTG_1_-_147774845_147774916</td><td>-1T</td><td>34</td><td>35</td><td>TGCCGTGATCGTATAGTGGTTAGTACTCTGCGTTG</td><td>5542.8</td><td>-2.5</td><td>1.0</td><td>-2.5</td><td>1.2E-02</td><td>3.6E-02</td><td>BT20/MB231</td><td>CYTO</td></tr>
<tr><td>>hg38_21_+_8433172_8446622_RNA45SN1_45S_50ntFlanks@11727.11748.22|AGTAACTATGACTCTCTTAAGG</td><td>>hg38_21_+_8433172_8446622_RNA45SN1_45S_50ntFlanks</td><td>11727</td><td>11748</td><td>22</td><td>AGTAACTATGACTCTCTTAAGG</td><td>467.0</td><td>-2.0</td><td>0.8</td><td>-2.5</td><td>1.2E-02</td><td>3.6E-02</td><td>BT20/MB231</td><td>CYTO</td></tr>
<tr><td>>hg38_21_+_8433172_8446622_RNA45SN1_45S_50ntFlanks@12420.12454.35|ATGTCGGCTCTTCCTATCATTGTGAAGCAGAATTC</td><td>>hg38_21_+_8433172_8446622_RNA45SN1_45S_50ntFlanks</td><td>12420</td><td>12454</td><td>35</td><td>ATGTCGGCTCTTCCTATCATTGTGAAGCAGAATTC</td><td>120.5</td><td>-2.1</td><td>0.9</td><td>-2.3</td><td>2.2E-02</td><td>3.6E-02</td><td>BT20/MB468</td><td>MP</td></tr>
<tr><td>>hg38_21_+_8433172_8446622_RNA45SN1_45S_50ntFlanks@5471.5488.18|CACGGCCCTGGCGGAGCG</td><td>>hg38_21_+_8433172_8446622_RNA45SN1_45S_50ntFlanks</td><td>5471</td><td>5488</td><td>18</td><td>CACGGCCCTGGCGGAGCG</td><td>92.5</td><td>-4.1</td><td>1.8</td><td>-2.3</td><td>2.2E-02</td><td>3.6E-02</td><td>BT20/MB468</td><td>MP</td></tr>
<tr><td>>MI0000270|hsa-mir-181b-1&WithFlank&1|-|198858867|198858988@42.61.20;MI0000683|hsa-mir-181b-2&WithFlank&9|+|124693704|124693804@22.41.20&[MIMAT0000257&hsa-miR-181b-5p&offsets|0|-3;m-73&1|-|198858925|198858947&offsets|0|-3];[MIMAT0000257_1&hsa-miR-181b-5p&offsets|0|-3;m-72&9|+|124693725|124693747&offsets|0|-3]|AACATTCATTGCTGTCGGTG</td><td>>MI0000270|hsa-mir-181b-1&WithFlank&1|-|198858867|198858988</td><td>42</td><td>61</td><td>20</td><td>AACATTCATTGCTGTCGGTG</td><td>609.8</td><td>-2.4</td><td>0.9</td><td>-2.6</td><td>9.4E-03</td><td>3.6E-02</td><td>BT20/MB231</td><td>MP</td></tr>
<tr><td>>hg38_1_-_228634819_228635039_RNA5S12_5S_50ntFlanks@130.171.42|TGGATGGGAGACCGCCTGGGAATACCGGGTGCTGTAGGCTTT</td><td>>hg38_1_-_228634819_228635039_RNA5S12_5S_50ntFlanks</td><td>130</td><td>171</td><td>42</td><td>TGGATGGGAGACCGCCTGGGAATACCGGGTGCTGTAGGCTTT</td><td>170.2</td><td>3.5</td><td>1.6</td><td>2.1</td><td>3.2E-02</td><td>3.6E-02</td><td>MB231/MB468</td><td>MITO</td></tr>
<tr><td>>hg38_21_+_8433172_8446622_RNA45SN1_45S_50ntFlanks@6647.6664.18|CGTACGACTCTTAGCGGT</td><td>>hg38_21_+_8433172_8446622_RNA45SN1_45S_50ntFlanks</td><td>6647</td><td>6664</td><td>18</td><td>CGTACGACTCTTAGCGGT</td><td>1388.2</td><td>1.3</td><td>0.6</td><td>2.2</td><td>2.9E-02</td><td>3.6E-02</td><td>MB231/MB468</td><td>MP</td></tr>
<tr><td>>hg38_21_+_8433172_8446622_RNA45SN1_45S_50ntFlanks@11568.11597.30|GCAGCCGACTTAGAACTGGTGCGGACCAGG</td><td>>hg38_21_+_8433172_8446622_RNA45SN1_45S_50ntFlanks</td><td>11568</td><td>11597</td><td>30</td><td>GCAGCCGACTTAGAACTGGTGCGGACCAGG</td><td>143.0</td><td>1.8</td><td>0.8</td><td>2.3</td><td>2.2E-02</td><td>3.6E-02</td><td>BT20/MB468</td><td>MP</td></tr>
</table>

<table>
<tr><th></th><th></th><th></th><th></th><th></th><th></th><th></th><th></th><th></th><th></th><th></th><th></th><th></th><th></th></tr>
<tr><td>>hg38_21_+_8433172_8446622_RNA45SN1_45S_50ntFlanks@10685.10706.22|GGCGGCGTCCGGTGAGCTCTCG</td><td>>hg38_21_+_8433172_8446622_RNA45SN1_45S_50ntFlanks</td><td>10685</td><td>10706</td><td>22</td><td>GGCGGCGTCCGGTGAGCTCTCG</td><td>1039.9</td><td>-4.6</td><td>2.1</td><td>-2.1</td><td>3.2E-02</td><td>3.6E-02</td><td>BT20/MB468</td><td>NUC</td></tr>
<tr><td>>MI0003513|hsa-mir-455&WithFlank&9|+|114209428|114209535@22.43.22(+1A)&[MIMAT0003150&hsa-miR-455-5p&offsets|0|0(+1A);m-232&9|+|114209449|114209470&offsets|0|0(+1A)]|TATGTGCCTTTGGACTACATCGA</td><td>>MI0003513|hsa-mir-455&WithFlank&9|+|114209428|114209535</td><td>22</td><td>43</td><td>22(+1A)</td><td>TATGTGCCTTTGGACTACATCGA</td><td>106.8</td><td>-4.9</td><td>1.9</td><td>-2.6</td><td>9.4E-03</td><td>3.6E-02</td><td>BT20/MB231</td><td>MP</td></tr>
<tr><td>>hg38_MT_+_1621_3279_RNR2_16S_50ntFlanks@1370.1407.38|ATGTTGGATCAGGACATCCCGATGGTGCAGCCGCTATT</td><td>>hg38_MT_+_1621_3279_RNR2_16S_50ntFlanks</td><td>1370</td><td>1407</td><td>38</td><td>ATGTTGGATCAGGACATCCCGATGGTGCAGCCGCTATT</td><td>87.4</td><td>3.5</td><td>1.5</td><td>2.3</td><td>2.2E-02</td><td>3.6E-02</td><td>BT20/MB468</td><td>MP</td></tr>
<tr><td>>MI0003643|hsa-mir-629&WithFlank&15|-|70079366|70079474@28.48.21&[MIMAT0004810&hsa-miR-629-5p&offsets|0|0;m-150&15|-|70079427|70079447&offsets|0|0]|TGGGTTTACGTTGGGAGAACT</td><td>>MI0003643|hsa-mir-629&WithFlank&15|-|70079366|70079474</td><td>28</td><td>48</td><td>21</td><td>TGGGTTTACGTTGGGAGAACT</td><td>889.3</td><td>1.9</td><td>0.8</td><td>2.5</td><td>1.2E-02</td><td>3.6E-02</td><td>BT20/MB231</td><td>CYTO</td></tr>
<tr><td>>am_trnaMT_ValTAC_MT_+_1602_1670@48.66.19|CAACTTAACTTGACCGCTC</td><td>>am_trnaMT_ValTAC_MT_+_1602_1670</td><td>48</td><td>66</td><td>19</td><td>CAACTTAACTTGACCGCTC</td><td>960.6</td><td>2.2</td><td>0.9</td><td>2.5</td><td>1.2E-02</td><td>3.6E-02</td><td>BT20/MB231</td><td>CYTO</td></tr>
<tr><td>>MI0000811|hsa-mir-148b&WithFlank&12|+|54337210|54337320@69.91.23&[MIMAT0000759&hsa-miR-148b-3p&offsets|0|+1;m-77&12|+|54337278|54337299&offsets|0|+1]|TCAGTGCATCACAGAACTTTGTC</td><td>>MI0000811|hsa-mir-148b&WithFlank&12|+|54337210|54337320</td><td>69</td><td>91</td><td>23</td><td>TCAGTGCATCACAGAACTTTGTC</td><td>696.2</td><td>-3.1</td><td>1.2</td><td>-2.6</td><td>9.4E-03</td><td>3.6E-02</td><td>BT20/MB231</td><td>MP</td></tr>
<tr><td>>hg38_21_+_8433172_8446622_RNA45SN1_45S_50ntFlanks@12415.12436.22|CTTCGATGTCGGCTCTTCCTAT</td><td>>hg38_21_+_8433172_8446622_RNA45SN1_45S_50ntFlanks</td><td>12415</td><td>12436</td><td>22</td><td>CTTCGATGTCGGCTCTTCCTAT</td><td>497.4</td><td>-4.1</td><td>1.8</td><td>-2.3</td><td>2.2E-02</td><td>3.6E-02</td><td>BT20/MB468</td><td>MP</td></tr>
<tr><td>>hg38_21_+_8433172_8446622_RNA45SN1_45S_50ntFlanks@11594.11613.20|CAGGGGAATCCGACTGTTTA</td><td>>hg38_21_+_8433172_8446622_RNA45SN1_45S_50ntFlanks</td><td>11594</td><td>11613</td><td>20</td><td>CAGGGGAATCCGACTGTTTA</td><td>235.6</td><td>-3.1</td><td>1.3</td><td>-2.3</td><td>2.2E-02</td><td>3.6E-02</td><td>BT20/MB468</td><td>MP</td></tr>
<tr><td>>hg38_21_+_8433172_8446622_RNA45SN1_45S_50ntFlanks@6647.6668.22|CGTACGACTCTTAGCGGTGGAT</td><td>>hg38_21_+_8433172_8446622_RNA45SN1_45S_50ntFlanks</td><td>6647</td><td>6668</td><td>22</td><td>CGTACGACTCTTAGCGGTGGAT</td><td>1302.0</td><td>1.5</td><td>0.7</td><td>2.2</td><td>2.9E-02</td><td>3.6E-02</td><td>MB231/MB468</td><td>MP</td></tr>
<tr><td>>hg38_21_+_8433172_8446622_RNA45SN1_45S_50ntFlanks@7975.8011.37|CGCGACCTCAGATCAGACGTGGCGACCCGCTGAATTT</td><td>>hg38_21_+_8433172_8446622_RNA45SN1_45S_50ntFlanks</td><td>7975</td><td>8011</td><td>37</td><td>CGCGACCTCAGATCAGACGTGGCGACCCGCTGAATTT</td><td>576587.3</td><td>1.6</td><td>0.6</td><td>2.5</td><td>1.1E-02</td><td>3.6E-02</td><td>BT20/MB231</td><td>MITO</td></tr>
<tr><td>>MI0000440|hsa-mir-27b&WithFlank&9|+|95085439|95085547@67.87.21(+1U)&[MIMAT0000419&hsa-miR-27b-3p&offsets|0|0(+1U);m-39&9|+|95085505|95085525&offsets|0|0(+1U)]|TTCACAGTGGCTAAGTTCTGCT</td><td>>MI0000440|hsa-mir-27b&WithFlank&9|+|95085439|95085547</td><td>67</td><td>87</td><td>21(+1U)</td><td>TTCACAGTGGCTAAGTTCTGCT</td><td>416.8</td><td>2.5</td><td>1.1</td><td>2.3</td><td>2.2E-02</td><td>3.6E-02</td><td>BT20/MB468</td><td>MP</td></tr>
<tr><td>>hg38_21_+_8433172_8446622_RNA45SN1_45S_50ntFlanks@12396.12418.23|GCGACGTCGCTTTTTGATCCTTC</td><td>>hg38_21_+_8433172_8446622_RNA45SN1_45S_50ntFlanks</td><td>12396</td><td>12418</td><td>23</td><td>GCGACGTCGCTTTTTGATCCTTC</td><td>359.7</td><td>1.6</td><td>0.7</td><td>2.1</td><td>3.2E-02</td><td>3.6E-02</td><td>MB231/MB468</td><td>NUC</td></tr>
<tr><td>>hg38_21_+_8433172_8446622_RNA45SN1_45S_50ntFlanks@5236.5282.47|CACTGACTGGCTCAGCGTGTGCCTACCCTACGCCGGCAGGCGCGGGT</td><td>>hg38_21_+_8433172_8446622_RNA45SN1_45S_50ntFlanks</td><td>5236</td><td>5282</td><td>47</td><td>CACTGACTGGCTCAGCGTGTGCCTACCCTACGCCGGCAGGCGCGGGT</td><td>291.0</td><td>5.3</td><td>2.3</td><td>2.3</td><td>2.2E-02</td><td>3.6E-02</td><td>BT20/MB468</td><td>MP</td></tr>
<tr><td>>hg38_21_+_8433172_8446622_RNA45SN1_45S_50ntFlanks@4296.4314.19|CCATTGGAGGGCAAGTCTG</td><td>>hg38_21_+_8433172_8446622_RNA45SN1_45S_50ntFlanks</td><td>4296</td><td>4314</td><td>19</td><td>CCATTGGAGGGCAAGTCTG</td><td>88.9</td><td>-3.9</td><td>1.7</td><td>-2.3</td><td>2.2E-02</td><td>3.6E-02</td><td>BT20/MB468</td><td>MP</td></tr>
<tr><td>>trnaMT_ThrTGT_MT_+_15888_15953@39.69.31|CGGAGATGAAAACCTTTTTCCAAGGACACCA</td><td>>trnaMT_ThrTGT_MT_+_15888_15953</td><td>39</td><td>69</td><td>31</td><td>CGGAGATGAAAACCTTTTTCCAAGGACACCA</td><td>131.4</td><td>2.4</td><td>1.1</td><td>2.2</td><td>2.9E-02</td><td>3.6E-02</td><td>MB231/MB468</td><td>MP</td></tr>
<tr><td>>hg38_21_+_8433172_8446622_RNA45SN1_45S_50ntFlanks@10349.10380.32|AACGAGAACTTTGAAGGCCGAAGTGGAGAAGG</td><td>>hg38_21_+_8433172_8446622_RNA45SN1_45S_50ntFlanks</td><td>10349</td><td>10380</td><td>32</td><td>AACGAGAACTTTGAAGGCCGAAGTGGAGAAGG</td><td>290.5</td><td>-2.0</td><td>0.9</td><td>-2.2</td><td>2.9E-02</td><td>3.6E-02</td><td>MB231/MB468</td><td>MP</td></tr>
<tr><td>>MI0000734|hsa-mir-106b&WithFlank&7|-|100093987|100094080@58.79.22(+1A)&[MIMAT0004672&hsa-miR-106b-3p&offsets|0|0(+1A);m-87&7|-|100094003|100094023&offsets|0|+1(+1A)]|CCGCACTGTGGGTACTTGCTGCA</td><td>>MI0000734|hsa-mir-106b&WithFlank&7|-|100093987|100094080</td><td>58</td><td>79</td><td>22(+1A)</td><td>CCGCACTGTGGGTACTTGCTGCA</td><td>61.7</td><td>4.1</td><td>1.8</td><td>2.3</td><td>2.2E-02</td><td>3.6E-02</td><td>BT20/MB468</td><td>MP</td></tr>
<tr><td>>trnaMT_GluTTC_MT_-_14674_14742@39.59.21,trnalookalike8_GluTTC_5_-_93905172_93905240@39.59.21|ATTGGTCGTGGTTGTAGTCCG</td><td>>trnaMT_GluTTC_MT_-_14674_14742</td><td>39</td><td>59</td><td>21</td><td>ATTGGTCGTGGTTGTAGTCCG</td><td>88.3</td><td>3.0</td><td>1.4</td><td>2.2</td><td>2.9E-02</td><td>3.6E-02</td><td>MB231/MB468</td><td>MP</td></tr>
<tr><td>>hg38_21_+_8433172_8446622_RNA45SN1_45S_50ntFlanks@9595.9612.18|ATAGGGGCGAAAGACTAA</td><td>>hg38_21_+_8433172_8446622_RNA45SN1_45S_50ntFlanks</td><td>9595</td><td>9612</td><td>18</td><td>ATAGGGGCGAAAGACTAA</td><td>47.5</td><td>3.4</td><td>1.5</td><td>2.3</td><td>2.2E-02</td><td>3.6E-02</td><td>BT20/MB468</td><td>MP</td></tr>
<tr><td>>hg38_21_+_8433172_8446622_RNA45SN1_45S_50ntFlanks@7992.8009.18|CGTGGCGACCCGCTGAAT</td><td>>hg38_21_+_8433172_8446622_RNA45SN1_45S_50ntFlanks</td><td>7992</td><td>8009</td><td>18</td><td>CGTGGCGACCCGCTGAAT</td><td>943.8</td><td>-1.2</td><td>0.5</td><td>-2.2</td><td>2.8E-02</td><td>3.6E-02</td><td>BT20/MB468</td><td>MITO</td></tr>
<tr><td>>MI0000764|hsa-mir-363&WithFlank&X|-|134169372|134169458@56.78.23&[MIMAT0000707&hsa-miR-363-3p&offsets|0|+1;m-178&X|-|134169381|134169403&offsets|0|0]|AATTGCACGGTATCCATCTGTAA</td><td>>MI0000764|hsa-mir-363&WithFlank&X|-|134169372|134169458</td><td>56</td><td>78</td><td>23</td><td>AATTGCACGGTATCCATCTGTAA</td><td>147.5</td><td>2.3</td><td>1.0</td><td>2.3</td><td>2.2E-02</td><td>3.6E-02</td><td>BT20/MB468</td><td>MP</td></tr>
<tr><td>>hg38_21_+_8433172_8446622_RNA45SN1_45S_50ntFlanks@11535.11560.26|TTCCGCGCGGCGCCTCGCCTCGGCCG</td><td>>hg38_21_+_8433172_8446622_RNA45SN1_45S_50ntFlanks</td><td>11535</td><td>11560</td><td>26</td><td>TTCCGCGCGGCGCCTCGCCTCGGCCG</td><td>55.8</td><td>-4.0</td><td>1.7</td><td>-2.3</td><td>2.2E-02</td><td>3.6E-02</td><td>BT20/MB468</td><td>MP</td></tr>
<tr><td>>MI0000746|hsa-mir-99b&WithFlank&19|+|51692606|51692687@13.34.22(+1A)&[MIMAT0000689&hsa-miR-99b-5p&offsets|0|0(+1A);m-10&19|+|51692618|51692639&offsets|0|0(+1A)]|CACCCGTAGAACCGACCTTGCGA</td><td>>MI0000746|hsa-mir-99b&WithFlank&19|+|51692606|51692687</td><td>13</td><td>34</td><td>22(+1A)</td><td>CACCCGTAGAACCGACCTTGCGA</td><td>293.5</td><td>1.7</td><td>0.8</td><td>2.2</td><td>2.9E-02</td><td>3.6E-02</td><td>MB231/MB468</td><td>MP</td></tr>
<tr><td>>hg38_21_+_8433172_8446622_RNA45SN1_45S_50ntFlanks@6646.6669.24|TCGTACGACTCTTAGCGGTGGATC</td><td>>hg38_21_+_8433172_8446622_RNA45SN1_45S_50ntFlanks</td><td>6646</td><td>6669</td><td>24</td><td>TCGTACGACTCTTAGCGGTGGATC</td><td>6758.3</td><td>1.6</td><td>0.8</td><td>2.2</td><td>3.1E-02</td><td>3.6E-02</td><td>MB231/MB468</td><td>CYTO</td></tr>
<tr><td>>hg38_21_+_8433172_8446622_RNA45SN1_45S_50ntFlanks@12642.12661.20|CATCTGTGGGATTATGACTG</td><td>>hg38_21_+_8433172_8446622_RNA45SN1_45S_50ntFlanks</td><td>12642</td><td>12661</td><td>20</td><td>CATCTGTGGGATTATGACTG</td><td>850.2</td><td>-1.8</td><td>0.8</td><td>-2.2</td><td>2.8E-02</td><td>3.6E-02</td><td>BT20/MB468</td><td>MITO</td></tr>
<tr><td>>MI0000811|hsa-mir-148b&WithFlank&12|+|54337210|54337320@69.90.22&[MIMAT0000759&hsa-miR-148b-3p&offsets|0|0;m-77&12|+|54337278|54337299&offsets|0|0]|TCAGTGCATCACAGAACTTTGT</td><td>>MI0000811|hsa-mir-148b&WithFlank&12|+|54337210|54337320</td><td>69</td><td>90</td><td>22</td><td>TCAGTGCATCACAGAACTTTGT</td><td>7200.2</td><td>-2.2</td><td>0.9</td><td>-2.5</td><td>1.2E-02</td><td>3.6E-02</td><td>BT20/MB231</td><td>MITO</td></tr>
<tr><td>>hg38_21_+_8433172_8446622_RNA45SN1_45S_50ntFlanks@6660.6685.26|GCGGTGGATCACTCGGCTCGTGCGTC</td><td>>hg38_21_+_8433172_8446622_RNA45SN1_45S_50ntFlanks</td><td>6660</td><td>6685</td><td>26</td><td>GCGGTGGATCACTCGGCTCGTGCGTC</td><td>272.7</td><td>2.8</td><td>1.1</td><td>2.5</td><td>1.2E-02</td><td>3.6E-02</td><td>BT20/MB231</td><td>MITO</td></tr>
<tr><td>>hg38_21_+_8433172_8446622_RNA45SN1_45S_50ntFlanks@12420.12450.31|ATGTCGGCTCTTCCTATCATTGTGAAGCAGA</td><td>>hg38_21_+_8433172_8446622_RNA45SN1_45S_50ntFlanks</td><td>12420</td><td>12450</td><td>31</td><td>ATGTCGGCTCTTCCTATCATTGTGAAGCAGA</td><td>200.0</td><td>-1.9</td><td>0.8</td><td>-2.5</td><td>1.3E-02</td><td>3.6E-02</td><td>BT20/MB231</td><td>CYTO</td></tr>
<tr><td>>MI0000542|hsa-mir-320a&WithFlank&8|-|22244960|22245043@48.70.23(+1U)&[MIMAT0000510&hsa-miR-320a-3p&offsets|0|+1(+1U);m-60&8|-|22244975|22244996&offsets|0|+1(+1U)]|AAAAGCTGGGTTGAGAGGGCGAAT</td><td>>MI0000542|hsa-mir-320a&WithFlank&8|-|22244960|22245043</td><td>48</td><td>70</td><td>23(+1U)</td><td>AAAAGCTGGGTTGAGAGGGCGAAT</td><td>3182.6</td><td>1.8</td><td>0.7</td><td>2.5</td><td>1.2E-02</td><td>3.6E-02</td><td>BT20/MB231</td><td>MITO</td></tr>
<tr><td>>hg38_21_+_8433172_8446622_RNA45SN1_45S_50ntFlanks@12894.12932.39|CTCGCCCGTCACGCACCGCACGTTCGTGGGGAACCTGGC</td><td>>hg38_21_+_8433172_8446622_RNA45SN1_45S_50ntFlanks</td><td>12894</td><td>12932</td><td>39</td><td>CTCGCCCGTCACGCACCGCACGTTCGTGGGGAACCTGGC</td><td>370.8</td><td>-1.3</td><td>0.6</td><td>-2.2</td><td>2.8E-02</td><td>3.6E-02</td><td>BT20/MB468</td><td>MITO</td></tr>
<tr><td>>hg38_1_-_228634819_228635039_RNA5S12_5S_50ntFlanks@133.167.35|ATGGGAGACCGCCTGGGAATACCGGGTGCTGTAGG</td><td>>hg38_1_-_228634819_228635039_RNA5S12_5S_50ntFlanks</td><td>133</td><td>167</td><td>35</td><td>ATGGGAGACCGCCTGGGAATACCGGGTGCTGTAGG</td><td>161.1</td><td>5.4</td><td>2.1</td><td>2.6</td><td>9.5E-03</td><td>3.6E-02</td><td>BT20/MB231</td><td>MP</td></tr>
<tr><td>>hg38_21_+_8433172_8446622_RNA45SN1_45S_50ntFlanks@12477.12496.20|CACTAATAGGGAACGTGAGC</td><td>>hg38_21_+_8433172_8446622_RNA45SN1_45S_50ntFlanks</td><td>12477</td><td>12496</td><td>20</td><td>CACTAATAGGGAACGTGAGC</td><td>491.4</td><td>-1.6</td><td>0.8</td><td>-2.2</td><td>3.0E-02</td><td>3.6E-02</td><td>MB231/MB468</td><td>MP</td></tr>
<tr><td>>MI0000299|hsa-mir-222&WithFlank&X|-|45747009|45747130@75.97.23&[MIMAT0000279&hsa-miR-222-3p&offsets|0|+2;m-140&X|-|45747033|45747056&offsets|0|-1]|AGCTACATCTGGCTACTGGGTCT</td><td>>MI0000299|hsa-mir-222&WithFlank&X|-|45747009|45747130</td><td>75</td><td>97</td><td>23</td><td>AGCTACATCTGGCTACTGGGTCT</td><td>11742.4</td><td>-1.9</td><td>0.8</td><td>-2.5</td><td>1.2E-02</td><td>3.6E-02</td><td>BT20/MB231</td><td>MITO</td></tr>
<tr><td>>hg38_21_+_8433172_8446622_RNA45SN1_45S_50ntFlanks@9495.9520.26|CGAAAGATGGTGAACTATGCCTGGGC</td><td>>hg38_21_+_8433172_8446622_RNA45SN1_45S_50ntFlanks</td><td>9495</td><td>9520</td><td>26</td><td>CGAAAGATGGTGAACTATGCCTGGGC</td><td>171.8</td><td>4.5</td><td>1.7</td><td>2.6</td><td>9.5E-03</td><td>3.6E-02</td><td>BT20/MB231</td><td>MP</td></tr>
<tr><td>>MI0000093|hsa-mir-92a-1&WithFlank&13|+|91351308|91351397@54.74.21(+1C);MI0000094|hsa-mir-92a-2&WithFlank&X|-|134169532|134169618@54.74.21(+1C)&[MIMAT0000092&hsa-miR-92a-3p&offsets|0|-1(+1C);m-9&13|+|91351361|91351382&offsets|0|-1(+1C)];[MIMAT0000092_1&hsa-miR-92a-3p&offsets|0|-1(+1C);m-11&X|-|134169544|134169565&offsets|0|-1(+1C)]|TATTGCACTTGTCCCGGCCTGC</td><td>>MI0000093|hsa-mir-92a-1&WithFlank&13|+|91351308|91351397</td><td>54</td><td>74</td><td>21(+1C)</td><td>TATTGCACTTGTCCCGGCCTGC</td><td>14039.5</td><td>-2.0</td><td>0.8</td><td>-2.5</td><td>1.2E-02</td><td>3.6E-02</td><td>BT20/MB231</td><td>MITO</td></tr>
<tr><td>>hg38_21_+_8433172_8446622_RNA45SN1_45S_50ntFlanks@6646.6668.23|TCGTACGACTCTTAGCGGTGGAT</td><td>>hg38_21_+_8433172_8446622_RNA45SN1_45S_50ntFlanks</td><td>6646</td><td>6668</td><td>23</td><td>TCGTACGACTCTTAGCGGTGGAT</td><td>37540.9</td><td>1.8</td><td>0.7</td><td>2.5</td><td>1.2E-02</td><td>3.6E-02</td><td>BT20/MB231</td><td>MITO</td></tr>
<tr><td>>MI0000108|hsa-mir-103a-2&WithFlank&20|+|3917488|3917577@54.76.23;MI0000109|hsa-mir-103a-1&WithFlank&5|-|168560890|168560979@54.76.23&[MIMAT0000101&hsa-miR-103a-3p&offsets|0|0;m-12&20|+|3917541|3917563&offsets|0|0];[MIMAT0000101_1&hsa-miR-103a-3p&offsets|0|0;m-13&5|-|168560904|168560926&offsets|0|0;m-44220&5|-|168560908|168560926&offsets|0|+4]|AGCAGCATTGTACAGGGCTATGA</td><td>>MI0000108|hsa-mir-103a-2&WithFlank&20|+|3917488|3917577</td><td>54</td><td>76</td><td>23</td><td>AGCAGCATTGTACAGGGCTATGA</td><td>30756.9</td><td>2.0</td><td>0.9</td><td>2.2</td><td>3.0E-02</td><td>3.6E-02</td><td>MB231/MB468</td><td>MP</td></tr>
<tr><td>>hg38_21_+_8433172_8446622_RNA45SN1_45S_50ntFlanks@4995.5023.29|ACAGATTGATAGCTCTTTCTCGATTCCGT</td><td>>hg38_21_+_8433172_8446622_RNA45SN1_45S_50ntFlanks</td><td>4995</td><td>5023</td><td>29</td><td>ACAGATTGATAGCTCTTTCTCGATTCCGT</td><td>171.9</td><td>-4.3</td><td>1.9</td><td>-2.3</td><td>2.2E-02</td><td>3.6E-02</td><td>BT20/MB468</td><td>MP</td></tr>
<tr><td>>hg38_21_+_8433172_8446622_RNA45SN1_45S_50ntFlanks@7991.8008.18|ACGTGGCGACCCGCTGAA</td><td>>hg38_21_+_8433172_8446622_RNA45SN1_45S_50ntFlanks</td><td>7991</td><td>8008</td><td>18</td><td>ACGTGGCGACCCGCTGAA</td><td>1215.2</td><td>-1.6</td><td>0.8</td><td>-2.2</td><td>3.1E-02</td><td>3.6E-02</td><td>BT20/MB468</td><td>CYTO</td></tr>
<tr><td>>MI0000542|hsa-mir-320a&WithFlank&8|-|22244960|22245043@49.70.22(+1U)&[MIMAT0000510&hsa-miR-320a-3p&offsets|+1|+1(+1U);m-60&8|-|22244975|22244996&offsets|+1|+1(+1U)]|AAAGCTGGGTTGAGAGGGCGAAT</td><td>>MI0000542|hsa-mir-320a&WithFlank&8|-|22244960|22245043</td><td>49</td><td>70</td><td>22(+1U)</td><td>AAAGCTGGGTTGAGAGGGCGAAT</td><td>385.2</td><td>1.9</td><td>0.7</td><td>2.6</td><td>1.0E-02</td><td>3.6E-02</td><td>BT20/MB231</td><td>NUC</td></tr>
<tr><td>>hg38_21_+_8433172_8446622_RNA45SN1_45S_50ntFlanks@12995.13012.18|CTCGCTGCGATCTATTGA</td><td>>hg38_21_+_8433172_8446622_RNA45SN1_45S_50ntFlanks</td><td>12995</td><td>13012</td><td>18</td><td>CTCGCTGCGATCTATTGA</td><td>792.9</td><td>1.1</td><td>0.5</td><td>2.1</td><td>3.3E-02</td><td>3.6E-02</td><td>MB231/MB468</td><td>NUC</td></tr>
<tr><td>>MI0000542|hsa-mir-320a&WithFlank&8|-|22244960|22245043@49.68.20&[MIMAT0000510&hsa-miR-320a-3p&offsets|+1|-1;m-60&8|-|22244975|22244996&offsets|+1|-1]|AAAGCTGGGTTGAGAGGGCG</td><td>>MI0000542|hsa-mir-320a&WithFlank&8|-|22244960|22245043</td><td>49</td><td>68</td><td>20</td><td>AAAGCTGGGTTGAGAGGGCG</td><td>1032.2</td><td>2.5</td><td>1.0</td><td>2.6</td><td>9.5E-03</td><td>3.6E-02</td><td>BT20/MB231</td><td>MP</td></tr>
<tr><td>>hg38_21_+_8433172_8446622_RNA45SN1_45S_50ntFlanks@9926.9947.22|GTTGATATAGACAGCAGGACGG</td><td>>hg38_21_+_8433172_8446622_RNA45SN1_45S_50ntFlanks</td><td>9926</td><td>9947</td><td>22</td><td>GTTGATATAGACAGCAGGACGG</td><td>71.8</td><td>4.8</td><td>2.1</td><td>2.3</td><td>2.2E-02</td><td>3.6E-02</td><td>BT20/MB468</td><td>MP</td></tr>
<tr><td>>hg38_21_+_8433172_8446622_RNA45SN1_45S_50ntFlanks@11668.11685.18|TGCCCAGTGCTCTGAATG</td><td>>hg38_21_+_8433172_8446622_RNA45SN1_45S_50ntFlanks</td><td>11668</td><td>11685</td><td>18</td><td>TGCCCAGTGCTCTGAATG</td><td>496.4</td><td>-2.1</td><td>0.8</td><td>-2.5</td><td>1.3E-02</td><td>3.6E-02</td><td>BT20/MB231</td><td>CYTO</td></tr>
<tr><td>>MI0001445|hsa-mir-423&WithFlank&17|+|30117073|30117178@58.78.21&[MIMAT0001340&hsa-miR-423-3p&offsets|-1|-3;m-100&17|+|30117131|30117153&offsets|-1|-3]|AAGCTCGGTCTGAGGCCCCTC</td><td>>MI0001445|hsa-mir-423&WithFlank&17|+|30117073|30117178</td><td>58</td><td>78</td><td>21</td><td>AAGCTCGGTCTGAGGCCCCTC</td><td>205.5</td><td>3.0</td><td>1.4</td><td>2.2</td><td>3.0E-02</td><td>3.6E-02</td><td>MB231/MB468</td><td>MP</td></tr>
<tr><td>>hg38_21_+_8433172_8446622_RNA45SN1_45S_50ntFlanks@7976.7994.19|GCGACCTCAGATCAGACGT</td><td>>hg38_21_+_8433172_8446622_RNA45SN1_45S_50ntFlanks</td><td>7976</td><td>7994</td><td>19</td><td>GCGACCTCAGATCAGACGT</td><td>300.5</td><td>1.9</td><td>0.7</td><td>2.6</td><td>1.0E-02</td><td>3.6E-02</td><td>BT20/MB231</td><td>NUC</td></tr>
<tr><td>>hg38_21_+_8433172_8446622_RNA45SN1_45S_50ntFlanks@6701.6720.20|TAGCTGCGAGAATTAATGTG</td><td>>hg38_21_+_8433172_8446622_RNA45SN1_45S_50ntFlanks</td><td>6701</td><td>6720</td><td>20</td><td>TAGCTGCGAGAATTAATGTG</td><td>200.7</td><td>-2.9</td><td>1.3</td><td>-2.2</td><td>3.0E-02</td><td>3.6E-02</td><td>MB231/MB468</td><td>MP</td></tr>
<tr><td>>hg38_21_+_8433172_8446622_RNA45SN1_45S_50ntFlanks@10500.10524.25|CAGATCCCCGAATCCGGAGTGGCGG</td><td>>hg38_21_+_8433172_8446622_RNA45SN1_45S_50ntFlanks</td><td>10500</td><td>10524</td><td>25</td><td>CAGATCCCCGAATCCGGAGTGGCGG</td><td>72.4</td><td>-3.9</td><td>1.7</td><td>-2.3</td><td>2.2E-02</td><td>3.6E-02</td><td>BT20/MB468</td><td>MP</td></tr>
<tr><td>>hg38_21_+_8433172_8446622_RNA45SN1_45S_50ntFlanks@13008.13025.18|ATTGAAAGTCAGCCCTCG</td><td>>hg38_21_+_8433172_8446622_RNA45SN1_45S_50ntFlanks</td><td>13008</td><td>13025</td><td>18</td><td>ATTGAAAGTCAGCCCTCG</td><td>308.0</td><td>-4.1</td><td>1.8</td><td>-2.3</td><td>2.2E-02</td><td>3.6E-02</td><td>BT20/MB468</td><td>MP</td></tr>
</table>

| | | | | | | | | | | | | | |
|---|---|---|---|---|---|---|---|---|---|---|---|---|---|
| >hg38_21_+_8433172_8446622_RNA45SN1_45S_50ntFlanks@12942.12965.24\|TTCGTAGACGACCTGCTTCTGGGT | >hg38_21_+_8433172_8446622_RNA45SN1_45S_50ntFlanks | 12942 | 12965 | 24 | TTCGTAGACGACCTGCTTCTGGGT | 257.4 | -4.0 | 1.8 | -2.3 | 2.2E-02 | 3.6E-02 | BT20/MB468 | MP |
| >MI0001445\|hsa-mir-423&WithFlank&17\|+\|30117073\|30117178@59.81.23(+1A)&[MIMAT0001340&hsa-miR-423-3p&offsets\|0\|0(+1A);m-100&17\|+\|30117131\|30117153&offsets\|0\|0(+1A)]\|AGCTCGGTCTGAGGCCCCTCAGTA | >MI0001445\|hsa-mir-423&WithFlank&17\|+\|30117073\|30117178 | 59 | 81 | 23(+1A) | AGCTCGGTCTGAGGCCCCTCAGTA | 63.7 | 3.7 | 1.7 | 2.2 | 3.0E-02 | 3.6E-02 | MB231/MB468 | MP |
| >hg38_21_+_8433172_8446622_RNA45SN1_45S_50ntFlanks@11663.11697.35\|ATTTCTGCCCAGTGCTCTGAATGTCAAAGTGAAGA | >hg38_21_+_8433172_8446622_RNA45SN1_45S_50ntFlanks | 11663 | 11697 | 35 | ATTTCTGCCCAGTGCTCTGAATGTCAAAGTGAAGA | 177.5 | 5.4 | 2.1 | 2.6 | 9.5E-03 | 3.6E-02 | BT20/MB231 | MP |
| >hg38_21_+_8433172_8446622_RNA45SN1_45S_50ntFlanks@10388.10424.37\|GTGAACAGCAGTTGAACATGGGTCAGTCGGTCCTGAG | >hg38_21_+_8433172_8446622_RNA45SN1_45S_50ntFlanks | 10388 | 10424 | 37 | GTGAACAGCAGTTGAACATGGGTCAGTCGGTCCTGAG | 540.0 | 2.7 | 1.2 | 2.2 | 3.0E-02 | 3.6E-02 | MB231/MB468 | MP |
| >hg38_21_+_8433172_8446622_RNA45SN1_45S_50ntFlanks@9927.9951.25\|TTGATATAGACAGCAGGACGGTGGC | >hg38_21_+_8433172_8446622_RNA45SN1_45S_50ntFlanks | 9927 | 9951 | 25 | TTGATATAGACAGCAGGACGGTGGC | 138.4 | 3.9 | 1.5 | 2.6 | 9.5E-03 | 3.6E-02 | BT20/MB231 | MP |
| >hg38_21_+_8433172_8446622_RNA45SN1_45S_50ntFlanks@10349.10373.25\|AACGAGAACTTTGAAGGCCGAAGTG | >hg38_21_+_8433172_8446622_RNA45SN1_45S_50ntFlanks | 10349 | 10373 | 25 | AACGAGAACTTTGAAGGCCGAAGTG | 862.3 | -3.2 | 1.5 | -2.2 | 3.0E-02 | 3.6E-02 | MB231/MB468 | MP |
| >MI0000736\|hsa-mir-30c-1&WithFlank&1\|+\|40757278\|40757378@23.46.24(+1U);MI0000254\|hsa-mir-30c-2&WithFlank&6\|-\|71376954\|71377037@13.36.24(+1U)&[MIMAT0000244&hsa-miR-30c-5p&offsets\|0\|+1(+1U);m-57&1\|+\|40757300\|40757323&offsets\|0\|0(+1U)];[MIMAT0000244_1&hsa-miR-30c-5p&offsets\|0\|+1(+1U);m-56&6\|-\|71377002\|71377025&offsets\|0\|0(+1U)]\|TGTAAACATCCTACACTCTCAGCTT | >MI0000736\|hsa-mir-30c-1&WithFlank&1\|+\|40757278\|40757378 | 23 | 46 | 24(+1U) | TGTAAACATCCTACACTCTCAGCTT | 1974.1 | -4.9 | 1.9 | -2.5 | 1.2E-02 | 3.6E-02 | BT20/MB231 | MITO |
| >hg38_21_+_8433172_8446622_RNA45SN1_45S_50ntFlanks@12417.12456.40\|TCGATGTCGGCTCTTCCTATCATTGTGAAGCAGAATTCAC | >hg38_21_+_8433172_8446622_RNA45SN1_45S_50ntFlanks | 12417 | 12456 | 40 | TCGATGTCGGCTCTTCCTATCATTGTGAAGCAGAATTCAC | 242.5 | -2.2 | 0.9 | -2.5 | 1.2E-02 | 3.6E-02 | BT20/MB231 | MITO |
| >hg38_21_+_8433172_8446622_RNA45SN1_45S_50ntFlanks@10912.10931.20\|CGCGCCGCGGCTGGACGAGG | >hg38_21_+_8433172_8446622_RNA45SN1_45S_50ntFlanks | 10912 | 10931 | 20 | CGCGCCGCGGCTGGACGAGG | 205.8 | -2.7 | 1.2 | -2.2 | 3.2E-02 | 3.6E-02 | BT20/MB468 | CYTO |
| >am_trna128_GlyGCC_6_-_27870686_27870756@1.33.33,trna18_GlyGCC_16_+_70822597_70822667@1.33.33,trna19_GlyGCC_16_+_70823410_70823480@1.33.33,trna19_GlyGCC_2_-_157257659_157257729@1.33.33,trna24_GlyGCC_16_-_70812942_70813012@1.33.33,trna25_GlyGCC_16_-_70812114_70812184@1.33.33,trna5_GlyGCC_17_+_8029064_8029134@1.33.33,trna68_GlyGCC_1_-_161493637_161493707@1.33.33\|GCATTGGTGGTTCAGTGGTAGAATTCTCGCCTG | >am_trna128_GlyGCC_6_-_27870686_27870756 | 1 | 33 | 33 | GCATTGGTGGTTCAGTGGTAGAATTCTCGCCTG | 4957.0 | -2.0 | 0.8 | -2.5 | 1.2E-02 | 3.6E-02 | BT20/MB231 | MITO |
| >hg38_21_+_8433172_8446622_RNA45SN1_45S_50ntFlanks@11596.11621.26\|GGGGAATCCGACTGTTTAATTAAAAC | >hg38_21_+_8433172_8446622_RNA45SN1_45S_50ntFlanks | 11596 | 11621 | 26 | GGGGAATCCGACTGTTTAATTAAAAC | 132.1 | 5.4 | 2.1 | 2.6 | 9.6E-03 | 3.6E-02 | BT20/MB231 | MP |
| >hg38_21_+_8433172_8446622_RNA45SN1_45S_50ntFlanks@5495.5516.22\|AGACGGTCGAACTTGACTATCT | >hg38_21_+_8433172_8446622_RNA45SN1_45S_50ntFlanks | 5495 | 5516 | 22 | AGACGGTCGAACTTGACTATCT | 615.9 | 3.4 | 1.3 | 2.6 | 9.6E-03 | 3.6E-02 | BT20/MB231 | MP |
| >trna119_LysCTT_1_-_145395522_145395594@1.32.32,trna11_LysCTT_5_-_180648979_180649051@1.32.32,trna13_LysCTT_6_+_26556774_26556846@1.32.32,trna32_LysCTT_16_-_3207406_3207478@1.32.32,trna7_LysCTT_16_+_3225692_3225764@1.32.32,trna9_LysCTT_5_+_180634755_180634827@1.32.32\|GCCCGGCTAGCTCAGTCGGTAGAGCATGAGAC | >trna119_LysCTT_1_-_145395522_145395594 | 1 | 32 | 32 | GCCCGGCTAGCTCAGTCGGTAGAGCATGAGAC | 4996.8 | -2.5 | 1.0 | -2.5 | 1.3E-02 | 3.6E-02 | BT20/MB231 | CYTO |
| >hg38_21_+_8433172_8446622_RNA45SN1_45S_50ntFlanks@4722.4761.40\|TCATTAATCAAGAACGAAAGTCGGAGGTTCGAAGACGATC | >hg38_21_+_8433172_8446622_RNA45SN1_45S_50ntFlanks | 4722 | 4761 | 40 | TCATTAATCAAGAACGAAAGTCGGAGGTTCGAAGACGATC | 207.4 | -2.1 | 0.8 | -2.5 | 1.3E-02 | 3.6E-02 | BT20/MB231 | CYTO |
| >hg38_21_+_8433172_8446622_RNA45SN1_45S_50ntFlanks@11906.11931.26\|CACGGTGAAGAGACATGAGAGGTGTA | >hg38_21_+_8433172_8446622_RNA45SN1_45S_50ntFlanks | 11906 | 11931 | 26 | CACGGTGAAGAGACATGAGAGGTGTA | 125.9 | 3.3 | 1.4 | 2.3 | 2.2E-02 | 3.6E-02 | BT20/MB468 | MP |
| >hg38_21_+_8433172_8446622_RNA45SN1_45S_50ntFlanks@10128.10146.19\|CGGCGGCGGCGGCGGCGGC | >hg38_21_+_8433172_8446622_RNA45SN1_45S_50ntFlanks | 10128 | 10146 | 19 | CGGCGGCGGCGGCGGCGGC | 69.0 | -4.0 | 1.7 | -2.3 | 2.2E-02 | 3.6E-02 | BT20/MB468 | MP |
| >hg38_21_+_8433172_8446622_RNA45SN1_45S_50ntFlanks@8845.8863.19\|CGGCGGCGGCGGCGGCGGC | >hg38_21_+_8433172_8446622_RNA45SN1_45S_50ntFlanks | 8845 | 8863 | 19 | CGGCGGCGGCGGCGGCGGC | 69.0 | -4.0 | 1.7 | -2.3 | 2.2E-02 | 3.6E-02 | BT20/MB468 | MP |
| >hg38_21_+_8433172_8446622_RNA45SN1_45S_50ntFlanks@6647.6696.50\|CGTACGACTCTTAGCGGTGGATCACTCGGCTCGTGCGTCGATGAAGAACG | >hg38_21_+_8433172_8446622_RNA45SN1_45S_50ntFlanks | 6647 | 6696 | 50 | CGTACGACTCTTAGCGGTGGATCACTCGGCTCGTGCGTCGATGAAGAACG | 952.0 | 2.2 | 1.0 | 2.2 | 2.8E-02 | 3.6E-02 | BT20/MB468 | MITO |
| >trna116_GluCTC_1_-_145399233_145399304@3.35.33,trna59_GluCTC_1_+_249168447_249168518@3.35.33,trna71_GluCTC_1_-_161439189_161439260@3.35.33,trna74_GluCTC_1_-_161431809_161431880@3.35.33,trna77_GluCTC_1_-_161424398_161424469@3.35.33,trna77_GluCTC_6_+_28949976_28950047@3.35.33,trna80_GluCTC_1_-_161417018_161417089@3.35.33,trna87_GluCTC_6_-_126101393_126101464@3.35.33\|CCTGGTGGTCTAGTGGTTAGGATTCGGCGCTCT | >trna116_GluCTC_1_-_145399233_145399304 | 3 | 35 | 33 | CCTGGTGGTCTAGTGGTTAGGATTCGGCGCTCT | 65.1 | 2.6 | 1.1 | 2.3 | 2.2E-02 | 3.6E-02 | BT20/MB468 | MP |
| >hg38_1_-_228634819_228635039_RNA5S12_5S_50ntFlanks@51.75.25\|GTCTACGGCCATACCACCCTGAACG | >hg38_1_-_228634819_228635039_RNA5S12_5S_50ntFlanks | 51 | 75 | 25 | GTCTACGGCCATACCACCCTGAACG | 738.2 | 2.5 | 1.0 | 2.6 | 9.6E-03 | 3.6E-02 | BT20/MB231 | MP |
| >hg38_21_+_8433172_8446622_RNA45SN1_45S_50ntFlanks@11581.11608.28\|AACTGGTGCGGACCAGGGGAATCCGACT | >hg38_21_+_8433172_8446622_RNA45SN1_45S_50ntFlanks | 11581 | 11608 | 28 | AACTGGTGCGGACCAGGGGAATCCGACT | 142.9 | -3.0 | 1.3 | -2.3 | 2.2E-02 | 3.6E-02 | BT20/MB468 | MP |
| >hg38_MT_+_1621_3279_RNR2_16S_50ntFlanks@1152.1169.18\|CACAGGTCCTAAACTACC | >hg38_MT_+_1621_3279_RNR2_16S_50ntFlanks | 1152 | 1169 | 18 | CACAGGTCCTAAACTACC | 84.0 | 4.4 | 2.0 | 2.2 | 3.0E-02 | 3.6E-02 | MB231/MB468 | MP |
| >hg38_21_+_8433172_8446622_RNA45SN1_45S_50ntFlanks@12416.12450.35\|TTCGATGTCGGCTCTTCCTATCATTGTGAAGCAGA | >hg38_21_+_8433172_8446622_RNA45SN1_45S_50ntFlanks | 12416 | 12450 | 35 | TTCGATGTCGGCTCTTCCTATCATTGTGAAGCAGA | 334.9 | -1.9 | 0.8 | -2.5 | 1.2E-02 | 3.6E-02 | BT20/MB231 | MITO |
| >hg38_21_+_8433172_8446622_RNA45SN1_45S_50ntFlanks@8381.8406.26\|AAGAGGGCGTGAAACCGTTAAGAGGT | >hg38_21_+_8433172_8446622_RNA45SN1_45S_50ntFlanks | 8381 | 8406 | 26 | AAGAGGGCGTGAAACCGTTAAGAGGT | 1848.2 | -2.0 | 0.9 | -2.2 | 3.0E-02 | 3.6E-02 | MB231/MB468 | MP |
| >MI0000542\|hsa-mir-320a&WithFlank&8\|-\|22244960\|22245043@50.72.23&[MIMAT0000510&hsa-miR-320a-3p&offsets\|+2\|+3;m-60&8\|-\|22244975\|22244996&offsets\|+2\|+3]\|AAGCTGGGTTGAGAGGGCGAAAA | >MI0000542\|hsa-mir-320a&WithFlank&8\|-\|22244960\|22245043 | 50 | 72 | 23 | AAGCTGGGTTGAGAGGGCGAAAA | 75.9 | 4.9 | 1.9 | 2.6 | 9.6E-03 | 3.6E-02 | BT20/MB231 | MP |
| >hg38_21_+_8433172_8446622_RNA45SN1_45S_50ntFlanks@10349.10375.27\|AACGAGAACTTTGAAGGCCGAAGTGGA | >hg38_21_+_8433172_8446622_RNA45SN1_45S_50ntFlanks | 10349 | 10375 | 27 | AACGAGAACTTTGAAGGCCGAAGTGGA | 137.6 | -3.1 | 1.4 | -2.2 | 3.0E-02 | 3.6E-02 | MB231/MB468 | MP |
| >hg38_21_+_8433172_8446622_RNA45SN1_45S_50ntFlanks@10677.10707.31\|GCGGTTCCGGCGGCGTCCGGTGAGCTCTCGC | >hg38_21_+_8433172_8446622_RNA45SN1_45S_50ntFlanks | 10677 | 10707 | 31 | GCGGTTCCGGCGGCGTCCGGTGAGCTCTCGC | 412.7 | 2.9 | 1.1 | 2.6 | 9.6E-03 | 3.6E-02 | BT20/MB231 | MP |
| >trnaMT_SerGCT_MT_+_12207_12265@1.34.34\|GAGAAAGCTCACAAGAACTGCTAACTCATGCCCC | >trnaMT_SerGCT_MT_+_12207_12265 | 1 | 34 | 34 | GAGAAAGCTCACAAGAACTGCTAACTCATGCCCC | 239.3 | 4.0 | 1.7 | 2.3 | 2.2E-02 | 3.6E-02 | BT20/MB468 | MP |
| >hg38_21_+_8433172_8446622_RNA45SN1_45S_50ntFlanks@4358.4391.34\|AAGTTGCTGCAGTTAAAAAGCTCGTAGTTGGATC | >hg38_21_+_8433172_8446622_RNA45SN1_45S_50ntFlanks | 4358 | 4391 | 34 | AAGTTGCTGCAGTTAAAAAGCTCGTAGTTGGATC | 113.3 | -3.2 | 1.4 | -2.3 | 2.2E-02 | 3.6E-02 | BT20/MB468 | MP |
| >hg38_21_+_8433172_8446622_RNA45SN1_45S_50ntFlanks@11561.11601.41\|GCGCCTAGCAGCCGACTTAGAACTGGTGCGGACCAGGGGAA | >hg38_21_+_8433172_8446622_RNA45SN1_45S_50ntFlanks | 11561 | 11601 | 41 | GCGCCTAGCAGCCGACTTAGAACTGGTGCGGACCAGGGGAA | 682.5 | -3.9 | 1.8 | -2.1 | 3.2E-02 | 3.6E-02 | BT20/MB468 | CYTO |
| >MI0006444\|hsa-mir-1307&WithFlank&10\|-\|103394247\|103394407@87.108.22&[MIMAT0005951&hsa-miR-1307-3p&offsets\|+1\|+1;m-66&10\|-\|103394300\|103394322&offsets\|+1\|0]\|CTCGGCGTGGCGTCGGTCGTGG | >MI0006444\|hsa-mir-1307&WithFlank&10\|-\|103394247\|103394407 | 87 | 108 | 22 | CTCGGCGTGGCGTCGGTCGTGG | 647.0 | 1.9 | 0.9 | 2.2 | 3.0E-02 | 3.6E-02 | MB231/MB468 | MP |
| >MI0000746\|hsa-mir-99b&WithFlank&19\|+\|51692606\|51692687@13.34.22(+1A)&[MIMAT0000689&hsa-miR-99b-5p&offsets\|0\|0(+1A);m-10&19\|+\|51692618\|51692639&offsets\|0\|0(+1A)]\|CACCCGTAGAACCGACCTTGCGA | >MI0000746\|hsa-mir-99b&WithFlank&19\|+\|51692606\|51692687 | 13 | 34 | 22(+1A) | CACCCGTAGAACCGACCTTGCGA | 4012.2 | 2.0 | 0.8 | 2.5 | 1.3E-02 | 3.6E-02 | BT20/MB231 | CYTO |
| >hg38_21_+_8433172_8446622_RNA45SN1_45S_50ntFlanks@3850.3883.34\|GATAACTGTGGTAATTCTAGAGCTAATACATGCC | >hg38_21_+_8433172_8446622_RNA45SN1_45S_50ntFlanks | 3850 | 3883 | 34 | GATAACTGTGGTAATTCTAGAGCTAATACATGCC | 424.3 | -2.0 | 0.8 | -2.5 | 1.3E-02 | 3.6E-02 | BT20/MB231 | CYTO |
| >hg38_21_+_8433172_8446622_RNA45SN1_45S_50ntFlanks@10361.10382.22\|GAAGGCCGAAGTGGAGAAGGGT | >hg38_21_+_8433172_8446622_RNA45SN1_45S_50ntFlanks | 10361 | 10382 | 22 | GAAGGCCGAAGTGGAGAAGGGT | 146.2 | -2.9 | 1.3 | -2.2 | 3.0E-02 | 3.6E-02 | MB231/MB468 | MP |
| >hg38_21_+_8433172_8446622_RNA45SN1_45S_50ntFlanks@8176.8217.42\|CTTCTGATCGAGGCCCAGCCCGTGGACGGTGTGAGGCCGGTA | >hg38_21_+_8433172_8446622_RNA45SN1_45S_50ntFlanks | 8176 | 8217 | 42 | CTTCTGATCGAGGCCCAGCCCGTGGACGGTGTGAGGCCGGTA | 134.3 | -2.1 | 1.0 | -2.2 | 3.0E-02 | 3.6E-02 | MB231/MB468 | MP |
| >hg38_21_+_8433172_8446622_RNA45SN1_45S_50ntFlanks@4767.4786.20\|CCGTCGTAGTTCCGACCATA | >hg38_21_+_8433172_8446622_RNA45SN1_45S_50ntFlanks | 4767 | 4786 | 20 | CCGTCGTAGTTCCGACCATA | 67.9 | -2.1 | 1.0 | -2.2 | 3.0E-02 | 3.6E-02 | MB231/MB468 | MP |
| >hg38_21_+_8433172_8446622_RNA45SN1_45S_50ntFlanks@12397.12420.24\|CGACGTCGCTTTTTGATCCTTCGA | >hg38_21_+_8433172_8446622_RNA45SN1_45S_50ntFlanks | 12397 | 12420 | 24 | CGACGTCGCTTTTTGATCCTTCGA | 59.7 | -3.9 | 1.8 | -2.2 | 3.0E-02 | 3.6E-02 | MB231/MB468 | MP |
| >MI0005756\|hsa-mir-934&WithFlank&X\|+\|136550872\|136550966@21.41.21&[MIMAT0004977&hsa-miR-934&offsets\|0\|-1;m-943&X\|+\|136550892\|136550913&offsets\|0\|-1]\|TGTCTACTACTGGAGACACTG | >MI0005756\|hsa-mir-934&WithFlank&X\|+\|136550872\|136550966 | 21 | 41 | 21 | TGTCTACTACTGGAGACACTG | 5852.8 | 2.9 | 1.3 | 2.3 | 2.2E-02 | 3.6E-02 | BT20/MB468 | MP |
| >hg38_21_+_8433172_8446622_RNA45SN1_45S_50ntFlanks@8196.8214.19\|CGTGGACGGTGTGAGGCCG | >hg38_21_+_8433172_8446622_RNA45SN1_45S_50ntFlanks | 8196 | 8214 | 19 | CGTGGACGGTGTGAGGCCG | 3182.8 | -1.9 | 0.9 | -2.1 | 3.2E-02 | 3.6E-02 | MB231/MB468 | CYTO |
| >hg38_21_+_8433172_8446622_RNA45SN1_45S_50ntFlanks@9597.9616.20\|AGGGGCGAAAGACTAATCGA | >hg38_21_+_8433172_8446622_RNA45SN1_45S_50ntFlanks | 9597 | 9616 | 20 | AGGGGCGAAAGACTAATCGA | 126.8 | -3.0 | 1.3 | -2.3 | 2.2E-02 | 3.6E-02 | BT20/MB468 | MP |
| >hg38_21_+_8433172_8446622_RNA45SN1_45S_50ntFlanks@11604.11621.18\|CGACTGTTTAATTAAAAC | >hg38_21_+_8433172_8446622_RNA45SN1_45S_50ntFlanks | 11604 | 11621 | 18 | CGACTGTTTAATTAAAAC | 325.4 | 4.4 | 1.7 | 2.6 | 9.7E-03 | 3.6E-02 | BT20/MB231 | MP |
| >hg38_MT_+_1621_3279_RNR2_16S_50ntFlanks@1424.1458.35\|AACGATTAAAGTCCTACGTGATCTGAGTTCAGACC | >hg38_MT_+_1621_3279_RNR2_16S_50ntFlanks | 1424 | 1458 | 35 | AACGATTAAAGTCCTACGTGATCTGAGTTCAGACC | 41.3 | 3.6 | 1.7 | 2.2 | 3.0E-02 | 3.6E-02 | MB231/MB468 | MP |
| >hg38_21_+_8433172_8446622_RNA45SN1_45S_50ntFlanks@7975.8016.42\|CGCGACCTCAGATCAGACGTGGCGACCCGCTGAATTTAAGCA | >hg38_21_+_8433172_8446622_RNA45SN1_45S_50ntFlanks | 7975 | 8016 | 42 | CGCGACCTCAGATCAGACGTGGCGACCCGCTGAATTTAAGCA | 5667.9 | 2.4 | 1.1 | 2.1 | 3.3E-02 | 3.6E-02 | BT20/MB468 | NUC |
| >hg38_21_+_8433172_8446622_RNA45SN1_45S_50ntFlanks@12992.13011.20\|TCCCTCGCTGCGATCTATTG | >hg38_21_+_8433172_8446622_RNA45SN1_45S_50ntFlanks | 12992 | 13011 | 20 | TCCCTCGCTGCGATCTATTG | 316.5 | -2.3 | 1.1 | -2.1 | 3.2E-02 | 3.6E-02 | MB231/MB468 | CYTO |

<table>
<tr><td></td><td></td><td></td><td></td><td></td><td></td><td></td><td></td><td></td><td></td><td></td><td></td><td></td><td></td></tr>
<tr><td>>hg38_21_+_8433172_8446622_RNA45SN1_45S_50ntFlanks@10840.10865.26|CCGTAACTTCGGGATAAGGATTGGCT</td><td>>hg38_21_+_8433172_8446622_RNA45SN1_45S_50ntFlanks</td><td>10840</td><td>10865</td><td>26</td><td>CCGTAACTTCGGGATAAGGATTGGCT</td><td>1975.4</td><td>-3.6</td><td>1.6</td><td>-2.3</td><td>2.2E-02</td><td>3.6E-02</td><td>BT20/MB468</td><td>MP</td></tr>
<tr><td>>hg38_21_+_8433172_8446622_RNA45SN1_45S_50ntFlanks@8144.8163.20|CTCCCCGGCGCCGCTCGTGG</td><td>>hg38_21_+_8433172_8446622_RNA45SN1_45S_50ntFlanks</td><td>8144</td><td>8163</td><td>20</td><td>CTCCCCGGCGCCGCTCGTGG</td><td>57.3</td><td>-3.9</td><td>1.7</td><td>-2.3</td><td>2.2E-02</td><td>3.7E-02</td><td>BT20/MB468</td><td>MP</td></tr>
<tr><td>>hg38_21_+_8433172_8446622_RNA45SN1_45S_50ntFlanks@9604.9630.27|AAAGACTAATCGAACCATCTAGTAGCT</td><td>>hg38_21_+_8433172_8446622_RNA45SN1_45S_50ntFlanks</td><td>9604</td><td>9630</td><td>27</td><td>AAAGACTAATCGAACCATCTAGTAGCT</td><td>194.6</td><td>-1.6</td><td>0.6</td><td>-2.6</td><td>1.0E-02</td><td>3.7E-02</td><td>BT20/MB231</td><td>NUC</td></tr>
<tr><td>>hg38_21_+_8433172_8446622_RNA45SN1_45S_50ntFlanks@12233.12258.26|CCGTGGAGCAGAAGGGCAAAAGCTCG</td><td>>hg38_21_+_8433172_8446622_RNA45SN1_45S_50ntFlanks</td><td>12233</td><td>12258</td><td>26</td><td>CCGTGGAGCAGAAGGGCAAAAGCTCG</td><td>156.9</td><td>-3.9</td><td>1.7</td><td>-2.3</td><td>2.2E-02</td><td>3.7E-02</td><td>BT20/MB468</td><td>MP</td></tr>
<tr><td>>MI0000447|hsa-mir-128-1&WithFlank&2|+|135665391|135665484@56.77.22&[MIMAT0000424&hsa-miR-128-3p&offsets|0|+1;m-108&2|+|135665446|135665466&offsets|0|+1]|TCACAGTGAACCGGTCTCTTTT</td><td>>MI0000447|hsa-mir-128-1&WithFlank&2|+|135665391|135665484</td><td>56</td><td>77</td><td>22</td><td>TCACAGTGAACCGGTCTCTTTT</td><td>1150.5</td><td>2.0</td><td>0.9</td><td>2.3</td><td>2.2E-02</td><td>3.7E-02</td><td>BT20/MB468</td><td>MP</td></tr>
<tr><td>>hg38_21_+_8433172_8446622_RNA45SN1_45S_50ntFlanks@10798.10823.26|CATGTTGGAACAATGTAGGTAAGGGA</td><td>>hg38_21_+_8433172_8446622_RNA45SN1_45S_50ntFlanks</td><td>10798</td><td>10823</td><td>26</td><td>CATGTTGGAACAATGTAGGTAAGGGA</td><td>118.5</td><td>5.1</td><td>2.0</td><td>2.6</td><td>9.7E-03</td><td>3.7E-02</td><td>BT20/MB231</td><td>MP</td></tr>
<tr><td>>hg38_21_+_8433172_8446622_RNA45SN1_45S_50ntFlanks@11217.11243.27|CTGGACGCGAGCCGGGCCCTTCCCGTG</td><td>>hg38_21_+_8433172_8446622_RNA45SN1_45S_50ntFlanks</td><td>11217</td><td>11243</td><td>27</td><td>CTGGACGCGAGCCGGGCCCTTCCCGTG</td><td>73.9</td><td>5.2</td><td>2.0</td><td>2.6</td><td>9.7E-03</td><td>3.7E-02</td><td>BT20/MB231</td><td>MP</td></tr>
<tr><td>>hg38_21_+_8433172_8446622_RNA45SN1_45S_50ntFlanks@6652.6689.38|GACTCTTAGCGGTGGATCACTCGGCTCGTGCGTCGATG</td><td>>hg38_21_+_8433172_8446622_RNA45SN1_45S_50ntFlanks</td><td>6652</td><td>6689</td><td>38</td><td>GACTCTTAGCGGTGGATCACTCGGCTCGTGCGTCGATG</td><td>8796.1</td><td>3.2</td><td>1.3</td><td>2.6</td><td>9.7E-03</td><td>3.7E-02</td><td>BT20/MB231</td><td>MP</td></tr>
<tr><td>>hg38_MT_+_598_1651_RNR1_12S_50ntFlanks@822.840.19|GTGGATTTAGCAGTAAACT</td><td>>hg38_MT_+_598_1651_RNR1_12S_50ntFlanks</td><td>822</td><td>840</td><td>19</td><td>GTGGATTTAGCAGTAAACT</td><td>459.1</td><td>2.7</td><td>1.0</td><td>2.6</td><td>9.7E-03</td><td>3.7E-02</td><td>BT20/MB231</td><td>MP</td></tr>
<tr><td>>MI0000077|hsa-mir-21&WithFlank&17|+|59841260|59841343@14.35.22&[MIMAT0000076&hsa-miR-21-5p&offsets|0|0;m-1&17|+|59841273|59841294&offsets|0|0]|TAGCTTATCAGACTGATGTTGA</td><td>>MI0000077|hsa-mir-21&WithFlank&17|+|59841260|59841343</td><td>14</td><td>35</td><td>22</td><td>TAGCTTATCAGACTGATGTTGA</td><td>124201.7</td><td>-1.9</td><td>0.8</td><td>-2.5</td><td>1.2E-02</td><td>3.7E-02</td><td>BT20/MB231</td><td>MITO</td></tr>
<tr><td>>hg38_21_+_8433172_8446622_RNA45SN1_45S_50ntFlanks@12455.12487.33|ACCAAGCGTTGGATTGTTCACCCACTAATAGGG</td><td>>hg38_21_+_8433172_8446622_RNA45SN1_45S_50ntFlanks</td><td>12455</td><td>12487</td><td>33</td><td>ACCAAGCGTTGGATTGTTCACCCACTAATAGGG</td><td>100.0</td><td>-3.3</td><td>1.4</td><td>-2.3</td><td>2.2E-02</td><td>3.7E-02</td><td>BT20/MB468</td><td>MP</td></tr>
<tr><td>>hg38_21_+_8433172_8446622_RNA45SN1_45S_50ntFlanks@6651.6681.31|CGACTCTTAGCGGTGGATCACTCGGCTCGTG</td><td>>hg38_21_+_8433172_8446622_RNA45SN1_45S_50ntFlanks</td><td>6651</td><td>6681</td><td>31</td><td>CGACTCTTAGCGGTGGATCACTCGGCTCGTG</td><td>52717.6</td><td>-2.3</td><td>1.0</td><td>-2.3</td><td>2.2E-02</td><td>3.7E-02</td><td>BT20/MB468</td><td>MP</td></tr>
<tr><td>>hg38_1_-_228634819_228635039_RNA5S12_5S_50ntFlanks@130.170.41|TGGATGGGAGACCGCCTGGGAATACCGGGTGCTGTAGGCTT</td><td>>hg38_1_-_228634819_228635039_RNA5S12_5S_50ntFlanks</td><td>130</td><td>170</td><td>41</td><td>TGGATGGGAGACCGCCTGGGAATACCGGGTGCTGTAGGCTT</td><td>6128.1</td><td>2.7</td><td>1.0</td><td>2.6</td><td>9.7E-03</td><td>3.7E-02</td><td>BT20/MB231</td><td>MP</td></tr>
<tr><td>>hg38_MT_+_598_1651_RNR1_12S_50ntFlanks@326.352.27|AATAGAAGCCGGCGTAAAGAGTGTTTT</td><td>>hg38_MT_+_598_1651_RNR1_12S_50ntFlanks</td><td>326</td><td>352</td><td>27</td><td>AATAGAAGCCGGCGTAAAGAGTGTTTT</td><td>389.9</td><td>4.5</td><td>2.0</td><td>2.3</td><td>2.2E-02</td><td>3.7E-02</td><td>BT20/MB468</td><td>MP</td></tr>
<tr><td>>hg38_21_+_8433172_8446622_RNA45SN1_45S_50ntFlanks@11568.11600.33|GCAGCCGACTTAGAACTGGTGCGGACCAGGGGA</td><td>>hg38_21_+_8433172_8446622_RNA45SN1_45S_50ntFlanks</td><td>11568</td><td>11600</td><td>33</td><td>GCAGCCGACTTAGAACTGGTGCGGACCAGGGGA</td><td>410.9</td><td>1.3</td><td>0.6</td><td>2.2</td><td>2.8E-02</td><td>3.7E-02</td><td>BT20/MB468</td><td>MITO</td></tr>
<tr><td>>am_trnaMT_PheGAA_MT_+_577_647@41.67.27|GTTTAGACGGGCTCACATCACCCCATA</td><td>>am_trnaMT_PheGAA_MT_+_577_647</td><td>41</td><td>67</td><td>27</td><td>GTTTAGACGGGCTCACATCACCCCATA</td><td>78.4</td><td>2.6</td><td>1.2</td><td>2.2</td><td>3.0E-02</td><td>3.7E-02</td><td>MB231/MB468</td><td>MP</td></tr>
<tr><td>>hg38_MT_+_1621_3279_RNR2_16S_50ntFlanks@51.89.39|GCTAAACCTAGCCCCAAACCCACTCCACCTTACTACCAG</td><td>>hg38_MT_+_1621_3279_RNR2_16S_50ntFlanks</td><td>51</td><td>89</td><td>39</td><td>GCTAAACCTAGCCCCAAACCCACTCCACCTTACTACCAG</td><td>315.0</td><td>3.1</td><td>1.4</td><td>2.2</td><td>3.0E-02</td><td>3.7E-02</td><td>MB231/MB468</td><td>MP</td></tr>
<tr><td>>am_trnaMT_LysTTT_MT_+_8295_8364@27.52.26,trnalookalike7_LysTTT_1_+_568844_568913@27.52.26|CTTTTAAGTTAAAGATTAAGAGAACC</td><td>>am_trnaMT_LysTTT_MT_+_8295_8364</td><td>27</td><td>52</td><td>26</td><td>CTTTTAAGTTAAAGATTAAGAGAACC</td><td>161.3</td><td>3.1</td><td>1.4</td><td>2.2</td><td>3.0E-02</td><td>3.7E-02</td><td>MB231/MB468</td><td>MP</td></tr>
<tr><td>>trnaMT_MetCAT_MT_+_4402_4469@37.71.35|CCGAAAATGTTGGTTATACCCTTCCCGTACTACCA</td><td>>trnaMT_MetCAT_MT_+_4402_4469</td><td>37</td><td>71</td><td>35</td><td>CCGAAAATGTTGGTTATACCCTTCCCGTACTACCA</td><td>248.5</td><td>2.9</td><td>1.4</td><td>2.2</td><td>3.0E-02</td><td>3.7E-02</td><td>MB231/MB468</td><td>MP</td></tr>
<tr><td>>MI0000825|hsa-mir-345&WithFlank&14|+|100307853|100307962@24.44.21(+1U)&[MIMAT0000772&hsa-miR-345-5p&offsets|0|-1(+1U);m-193&14|+|100307876|100307896&offsets|0|0(+1U)]|GCTGACTCCTAGTCCAGGGCTT</td><td>>MI0000825|hsa-mir-345&WithFlank&14|+|100307853|100307962</td><td>24</td><td>44</td><td>21(+1U)</td><td>GCTGACTCCTAGTCCAGGGCTT</td><td>81.7</td><td>4.3</td><td>1.9</td><td>2.3</td><td>2.2E-02</td><td>3.7E-02</td><td>BT20/MB468</td><td>MP</td></tr>
<tr><td>>hg38_21_+_8433172_8446622_RNA45SN1_45S_50ntFlanks@11170.11189.20|CCGCGAGGGGGGCCCGGGCA</td><td>>hg38_21_+_8433172_8446622_RNA45SN1_45S_50ntFlanks</td><td>11170</td><td>11189</td><td>20</td><td>CCGCGAGGGGGGCCCGGGCA</td><td>435.5</td><td>-2.4</td><td>1.0</td><td>-2.5</td><td>1.3E-02</td><td>3.7E-02</td><td>BT20/MB231</td><td>CYTO</td></tr>
<tr><td>>hg38_21_+_8433172_8446622_RNA45SN1_45S_50ntFlanks@12896.12932.37|CGCCCGTCACGCACCGCACGTTCGTGGGGAACCTGGC</td><td>>hg38_21_+_8433172_8446622_RNA45SN1_45S_50ntFlanks</td><td>12896</td><td>12932</td><td>37</td><td>CGCCCGTCACGCACCGCACGTTCGTGGGGAACCTGGC</td><td>72.5</td><td>-3.9</td><td>1.7</td><td>-2.3</td><td>2.2E-02</td><td>3.7E-02</td><td>BT20/MB468</td><td>MP</td></tr>
<tr><td>>MI0000299|hsa-mir-222&WithFlank&X|-|45747009|45747130@75.96.22(+1A)&[MIMAT0000279&hsa-miR-222-3p&offsets|0|+1(+1A);m-140&X|-|45747033|45747056&offsets|0|-2(+1A)]|AGCTACATCTGGCTACTGGGTCA</td><td>>MI0000299|hsa-mir-222&WithFlank&X|-|45747009|45747130</td><td>75</td><td>96</td><td>22(+1A)</td><td>AGCTACATCTGGCTACTGGGTCA</td><td>260.1</td><td>3.5</td><td>1.6</td><td>2.3</td><td>2.2E-02</td><td>3.7E-02</td><td>BT20/MB468</td><td>MP</td></tr>
<tr><td>>hg38_21_+_8433172_8446622_RNA45SN1_45S_50ntFlanks@12635.12669.35|AAGCTACCATCTGTGGGATTATGACTGAACGCCTC</td><td>>hg38_21_+_8433172_8446622_RNA45SN1_45S_50ntFlanks</td><td>12635</td><td>12669</td><td>35</td><td>AAGCTACCATCTGTGGGATTATGACTGAACGCCTC</td><td>297.1</td><td>3.5</td><td>1.4</td><td>2.6</td><td>9.8E-03</td><td>3.7E-02</td><td>BT20/MB231</td><td>MP</td></tr>
<tr><td>>am_trnaMT_ValTAC_MT_+_1602_1670@50.68.19|ACTTAACTTGACCGCTCTG</td><td>>am_trnaMT_ValTAC_MT_+_1602_1670</td><td>50</td><td>68</td><td>19</td><td>ACTTAACTTGACCGCTCTG</td><td>293.3</td><td>4.0</td><td>1.5</td><td>2.6</td><td>9.8E-03</td><td>3.7E-02</td><td>BT20/MB231</td><td>MP</td></tr>
<tr><td>>hg38_21_+_8433172_8446622_RNA45SN1_45S_50ntFlanks@12249.12269.21|CAAAAGCTCGCTTGATCTTGA</td><td>>hg38_21_+_8433172_8446622_RNA45SN1_45S_50ntFlanks</td><td>12249</td><td>12269</td><td>21</td><td>CAAAAGCTCGCTTGATCTTGA</td><td>81.4</td><td>-3.9</td><td>1.7</td><td>-2.3</td><td>2.3E-02</td><td>3.7E-02</td><td>BT20/MB468</td><td>MP</td></tr>
<tr><td>>MI0000088|hsa-mir-30a&WithFlank&6|-|71403545|71403627@53.74.22(+1A)&[MIMAT0000088&hsa-miR-30a-3p&offsets|0|0(+1A);m-38&6|-|71403554|71403575&offsets|0|0(+1A)]|CTTTCAGTCGGATGTTTGCAGCA</td><td>>MI0000088|hsa-mir-30a&WithFlank&6|-|71403545|71403627</td><td>53</td><td>74</td><td>22(+1A)</td><td>CTTTCAGTCGGATGTTTGCAGCA</td><td>105.6</td><td>2.9</td><td>1.3</td><td>2.2</td><td>3.0E-02</td><td>3.7E-02</td><td>MB231/MB468</td><td>MP</td></tr>
<tr><td>>hg38_21_+_8433172_8446622_RNA45SN1_45S_50ntFlanks@9305.9330.26|CCAAGGAGTCTAACACGTGCGCGAGT</td><td>>hg38_21_+_8433172_8446622_RNA45SN1_45S_50ntFlanks</td><td>9305</td><td>9330</td><td>26</td><td>CCAAGGAGTCTAACACGTGCGCGAGT</td><td>59.4</td><td>5.0</td><td>1.9</td><td>2.6</td><td>9.8E-03</td><td>3.7E-02</td><td>BT20/MB231</td><td>MP</td></tr>
<tr><td>>hg38_21_+_8433172_8446622_RNA45SN1_45S_50ntFlanks@5073.5092.20|AATTCCGATAACGAACGAGA</td><td>>hg38_21_+_8433172_8446622_RNA45SN1_45S_50ntFlanks</td><td>5073</td><td>5092</td><td>20</td><td>AATTCCGATAACGAACGAGA</td><td>137.6</td><td>-4.2</td><td>1.8</td><td>-2.3</td><td>2.3E-02</td><td>3.7E-02</td><td>BT20/MB468</td><td>MP</td></tr>
<tr><td>>hg38_21_+_8433172_8446622_RNA45SN1_45S_50ntFlanks@4472.4494.23|CTTAGCTGAGTGTCCCGCGGGGC</td><td>>hg38_21_+_8433172_8446622_RNA45SN1_45S_50ntFlanks</td><td>4472</td><td>4494</td><td>23</td><td>CTTAGCTGAGTGTCCCGCGGGGC</td><td>750.9</td><td>-4.2</td><td>1.9</td><td>-2.3</td><td>2.3E-02</td><td>3.7E-02</td><td>BT20/MB468</td><td>MP</td></tr>
<tr><td>>hg38_21_+_8433172_8446622_RNA45SN1_45S_50ntFlanks@12247.12273.27|GGCAAAAGCTCGCTTGATCTTGATTTT</td><td>>hg38_21_+_8433172_8446622_RNA45SN1_45S_50ntFlanks</td><td>12247</td><td>12273</td><td>27</td><td>GGCAAAAGCTCGCTTGATCTTGATTTT</td><td>135.0</td><td>-3.6</td><td>1.7</td><td>-2.2</td><td>2.8E-02</td><td>3.7E-02</td><td>BT20/MB468</td><td>MITO</td></tr>
<tr><td>>hg38_21_+_8433172_8446622_RNA45SN1_45S_50ntFlanks@6664.6689.26|TGGATCACTCGGCTCGTGCGTCGATG</td><td>>hg38_21_+_8433172_8446622_RNA45SN1_45S_50ntFlanks</td><td>6664</td><td>6689</td><td>26</td><td>TGGATCACTCGGCTCGTGCGTCGATG</td><td>585.1</td><td>-1.5</td><td>0.7</td><td>-2.2</td><td>2.8E-02</td><td>3.7E-02</td><td>BT20/MB468</td><td>MITO</td></tr>
<tr><td>>hg38_21_+_8433172_8446622_RNA45SN1_45S_50ntFlanks@9303.9336.34|GACCAAGGAGTCTAACACGTGCGCGAGTCGGGGG</td><td>>hg38_21_+_8433172_8446622_RNA45SN1_45S_50ntFlanks</td><td>9303</td><td>9336</td><td>34</td><td>GACCAAGGAGTCTAACACGTGCGCGAGTCGGGGG</td><td>100.4</td><td>-3.9</td><td>1.7</td><td>-2.3</td><td>2.3E-02</td><td>3.7E-02</td><td>BT20/MB468</td><td>MP</td></tr>
<tr><td>>hg38_21_+_8433172_8446622_RNA45SN1_45S_50ntFlanks@6651.6679.29|CGACTCTTAGCGGTGGATCACTCGGCTCG</td><td>>hg38_21_+_8433172_8446622_RNA45SN1_45S_50ntFlanks</td><td>6651</td><td>6679</td><td>29</td><td>CGACTCTTAGCGGTGGATCACTCGGCTCG</td><td>10150.7</td><td>3.9</td><td>1.5</td><td>2.6</td><td>9.8E-03</td><td>3.7E-02</td><td>BT20/MB231</td><td>MP</td></tr>
<tr><td>>hg38_21_+_8433172_8446622_RNA45SN1_45S_50ntFlanks@11661.11679.19|TGATTTCTGCCCAGTGCTC</td><td>>hg38_21_+_8433172_8446622_RNA45SN1_45S_50ntFlanks</td><td>11661</td><td>11679</td><td>19</td><td>TGATTTCTGCCCAGTGCTC</td><td>192.6</td><td>-4.0</td><td>1.7</td><td>-2.3</td><td>2.3E-02</td><td>3.7E-02</td><td>BT20/MB468</td><td>MP</td></tr>
<tr><td>>hg38_21_+_8433172_8446622_RNA45SN1_45S_50ntFlanks@5071.5097.27|TTAATTCCGATAACGAACGAGACTCTG</td><td>>hg38_21_+_8433172_8446622_RNA45SN1_45S_50ntFlanks</td><td>5071</td><td>5097</td><td>27</td><td>TTAATTCCGATAACGAACGAGACTCTG</td><td>227.9</td><td>-2.2</td><td>1.0</td><td>-2.2</td><td>2.8E-02</td><td>3.7E-02</td><td>BT20/MB468</td><td>MITO</td></tr>
<tr><td>>hg38_MT_+_1621_3279_RNR2_16S_50ntFlanks@1000.1017.18|GGGACCTGTATGAATGGC</td><td>>hg38_MT_+_1621_3279_RNR2_16S_50ntFlanks</td><td>1000</td><td>1017</td><td>18</td><td>GGGACCTGTATGAATGGC</td><td>271.7</td><td>2.9</td><td>1.4</td><td>2.2</td><td>3.0E-02</td><td>3.7E-02</td><td>MB231/MB468</td><td>MP</td></tr>
<tr><td>>hg38_21_+_8433172_8446622_RNA45SN1_45S_50ntFlanks@4130.4148.19|ATTCCGGAGAGGGAGCCTG</td><td>>hg38_21_+_8433172_8446622_RNA45SN1_45S_50ntFlanks</td><td>4130</td><td>4148</td><td>19</td><td>ATTCCGGAGAGGGAGCCTG</td><td>188.8</td><td>-2.0</td><td>0.9</td><td>-2.3</td><td>2.3E-02</td><td>3.7E-02</td><td>BT20/MB468</td><td>MP</td></tr>
<tr><td>>trnaMT_ValTAC_MT_+_1602_1670@25.72.48|CCCAACTTACACTTAGGAGATTTCAACTTAACTTGACCGCTCTGACCA</td><td>>trnaMT_ValTAC_MT_+_1602_1670</td><td>25</td><td>72</td><td>48</td><td>CCCAACTTACACTTAGGAGATTTCAACTTAACTTGACCGCTCTGACCA</td><td>165.7</td><td>3.9</td><td>1.8</td><td>2.1</td><td>3.4E-02</td><td>3.7E-02</td><td>MB231/MB468</td><td>MITO</td></tr>
<tr><td>>hg38_21_+_8433172_8446622_RNA45SN1_45S_50ntFlanks@3850.3873.24|GATAACTGTGGTAATTCTAGAGCT</td><td>>hg38_21_+_8433172_8446622_RNA45SN1_45S_50ntFlanks</td><td>3850</td><td>3873</td><td>24</td><td>GATAACTGTGGTAATTCTAGAGCT</td><td>481.9</td><td>-2.1</td><td>0.9</td><td>-2.3</td><td>2.3E-02</td><td>3.7E-02</td><td>BT20/MB468</td><td>MP</td></tr>
<tr><td>>hg38_21_+_8433172_8446622_RNA45SN1_45S_50ntFlanks@12417.12445.29|TCGATGTCGGCTCTTCCTATCATTGTGAA</td><td>>hg38_21_+_8433172_8446622_RNA45SN1_45S_50ntFlanks</td><td>12417</td><td>12445</td><td>29</td><td>TCGATGTCGGCTCTTCCTATCATTGTGAA</td><td>933.1</td><td>-1.3</td><td>0.6</td><td>-2.1</td><td>3.2E-02</td><td>3.7E-02</td><td>BT20/MB468</td><td>CYTO</td></tr>
<tr><td>>hg38_21_+_8433172_8446622_RNA45SN1_45S_50ntFlanks@12444.12486.43|AAGCAGAATTCACCAAGCGTTGGATTGTTCACCCACTAATAGG</td><td>>hg38_21_+_8433172_8446622_RNA45SN1_45S_50ntFlanks</td><td>12444</td><td>12486</td><td>43</td><td>AAGCAGAATTCACCAAGCGTTGGATTGTTCACCCACTAATAGG</td><td>233.3</td><td>-2.9</td><td>1.4</td><td>-2.1</td><td>3.2E-02</td><td>3.7E-02</td><td>BT20/MB468</td><td>CYTO</td></tr>
<tr><td>>trnaMT_ProTGG_MT_-_15956_16023@46.70.25|GGAGTTAAAGACTTTTTCTCTGACC</td><td>>trnaMT_ProTGG_MT_-_15956_16023</td><td>46</td><td>70</td><td>25</td><td>GGAGTTAAAGACTTTTTCTCTGACC</td><td>126.0</td><td>2.8</td><td>1.2</td><td>2.3</td><td>2.3E-02</td><td>3.7E-02</td><td>BT20/MB468</td><td>MP</td></tr>
<tr><td>>MI0000746|hsa-mir-99b&WithFlank&19|+|51692606|51692687@51.72.22(+1A)&[MIMAT0004678&hsa-miR-99b-3p&offsets|0|0(+1A);m-151&19|+|51692656|51692678&offsets|0|-1(+1A)]|CAAGCTCGTGTCTGTGGGTCCGA</td><td>>MI0000746|hsa-mir-99b&WithFlank&19|+|51692606|51692687</td><td>51</td><td>72</td><td>22(+1A)</td><td>CAAGCTCGTGTCTGTGGGTCCGA</td><td>439.4</td><td>1.9</td><td>0.8</td><td>2.5</td><td>1.2E-02</td><td>3.7E-02</td><td>BT20/MB231</td><td>MITO</td></tr>
<tr><td>>trnaMT_GluTTC_MT_-_14674_14742@39.66.28,trnalookalike8_GluTTC_5_-_93905172_93905240@39.66.28|ATTGGTCGTGGTTGTAGTCCGTGCGAGA</td><td>>trnaMT_GluTTC_MT_-_14674_14742</td><td>39</td><td>66</td><td>28</td><td>ATTGGTCGTGGTTGTAGTCCGTGCGAGA</td><td>1091.0</td><td>1.8</td><td>0.7</td><td>2.5</td><td>1.2E-02</td><td>3.7E-02</td><td>BT20/MB231</td><td>MITO</td></tr>
<tr><td>>hg38_21_+_8433172_8446622_RNA45SN1_45S_50ntFlanks@6652.6669.18|GACTCTTAGCGGTGGATC</td><td>>hg38_21_+_8433172_8446622_RNA45SN1_45S_50ntFlanks</td><td>6652</td><td>6669</td><td>18</td><td>GACTCTTAGCGGTGGATC</td><td>1140.9</td><td>1.9</td><td>0.8</td><td>2.5</td><td>1.2E-02</td><td>3.7E-02</td><td>BT20/MB231</td><td>MITO</td></tr>
<tr><td>>MI0000270|hsa-mir-181b-1&WithFlank&1|-|198858867|198858988@42.61.20;MI0000683|hsa-mir-181b-2&WithFlank&9|+|124693704|124693804@22.41.20&[MIMAT0000257&hsa-miR-181b-5p&offsets|0|-3;m-73&1|-|198858925|198858947&offsets|0|-3];[MIMAT0000257_1&hsa-miR-181b-5p&offsets|0|-3;m-72&9|+|124693725|124693747&offsets|0|-3]|AACATTCATTGCTGTCGGTG</td><td>>MI0000270|hsa-mir-181b-1&WithFlank&1|-|198858867|198858988</td><td>42</td><td>61</td><td>20</td><td>AACATTCATTGCTGTCGGTG</td><td>224.0</td><td>-1.9</td><td>0.7</td><td>-2.5</td><td>1.2E-02</td><td>3.7E-02</td><td>BT20/MB231</td><td>MITO</td></tr>
<tr><td>>hg38_MT_+_598_1651_RNR1_12S_50ntFlanks@967.1004.38|AACATGGTAAGTGTACTGGAAAGTGCACTTGGACGAAC</td><td>>hg38_MT_+_598_1651_RNR1_12S_50ntFlanks</td><td>967</td><td>1004</td><td>38</td><td>AACATGGTAAGTGTACTGGAAAGTGCACTTGGACGAAC</td><td>355.6</td><td>-2.0</td><td>0.8</td><td>-2.5</td><td>1.2E-02</td><td>3.7E-02</td><td>BT20/MB231</td><td>MITO</td></tr>
<tr><td>>hg38_21_+_8433172_8446622_RNA45SN1_45S_50ntFlanks@6649.6696.48|TACGACTCTTAGCGGTGGATCACTCGGCTCGTGCGTCGATGAAGAACG</td><td>>hg38_21_+_8433172_8446622_RNA45SN1_45S_50ntFlanks</td><td>6649</td><td>6696</td><td>48</td><td>TACGACTCTTAGCGGTGGATCACTCGGCTCGTGCGTCGATGAAGAACG</td><td>597.4</td><td>2.2</td><td>1.0</td><td>2.2</td><td>2.9E-02</td><td>3.7E-02</td><td>BT20/MB468</td><td>MITO</td></tr>
<tr><td>>hg38_21_+_8433172_8446622_RNA45SN1_45S_50ntFlanks@12242.12272.31|AGAAGGGCAAAAGCTCGCTTGATCTTGATTT</td><td>>hg38_21_+_8433172_8446622_RNA45SN1_45S_50ntFlanks</td><td>12242</td><td>12272</td><td>31</td><td>AGAAGGGCAAAAGCTCGCTTGATCTTGATTT</td><td>495.7</td><td>-1.8</td><td>0.8</td><td>-2.2</td><td>2.9E-02</td><td>3.7E-02</td><td>BT20/MB468</td><td>MITO</td></tr>
<tr><td>>hg38_21_+_8433172_8446622_RNA45SN1_45S_50ntFlanks@11568.11590.23|GCAGCCGACTTAGAACTGGTGCG</td><td>>hg38_21_+_8433172_8446622_RNA45SN1_45S_50ntFlanks</td><td>11568</td><td>11590</td><td>23</td><td>GCAGCCGACTTAGAACTGGTGCG</td><td>175.6</td><td>-3.4</td><td>1.5</td><td>-2.3</td><td>2.3E-02</td><td>3.7E-02</td><td>BT20/MB468</td><td>MP</td></tr>
<tr><td>>hg38_21_+_8433172_8446622_RNA45SN1_45S_50ntFlanks@6646.6673.28|TCGTACGACTCTTAGCGGTGGATCACTC</td><td>>hg38_21_+_8433172_8446622_RNA45SN1_45S_50ntFlanks</td><td>6646</td><td>6673</td><td>28</td><td>TCGTACGACTCTTAGCGGTGGATCACTC</td><td>662.7</td><td>1.4</td><td>0.7</td><td>2.1</td><td>3.3E-02</td><td>3.7E-02</td><td>BT20/MB468</td><td>NUC</td></tr>
</table>

<table>
<tr><td>>am_MI0000809|hsa-mir-151a&WithFlank&8|-|140732558|140732659@53.72.20(+1U)&[MIMAT0000757&hsa-miR-151a-3p&offsets|0|-1(+1U);m-30&8|-|140732587|140732607&offsets|0|-1(+1U)]|CTAGACTGAAGCTCCTTGAGT</td><td>>am_MI0000809|hsa-mir-151a&WithFlank&8|-|140732558|140732659</td><td>53</td><td>72</td><td>20(+1U)</td><td>CTAGACTGAAGCTCCTTGAGT</td><td>6165.6</td><td>1.9</td><td>0.7</td><td>2.5</td><td>1.3E-02</td><td>3.7E-02</td><td>BT20/MB231</td><td>CYTO</td></tr>
<tr><td>>hg38_21_+_8433172_8446622_RNA45SN1_45S_50ntFlanks@9842.9864.23|AGCAGAACTGGCGCTGCGGGATG</td><td>>hg38_21_+_8433172_8446622_RNA45SN1_45S_50ntFlanks</td><td>9842</td><td>9864</td><td>23</td><td>AGCAGAACTGGCGCTGCGGGATG</td><td>92.0</td><td>-2.3</td><td>1.1</td><td>-2.2</td><td>3.0E-02</td><td>3.7E-02</td><td>MB231/MB468</td><td>MP</td></tr>
<tr><td>>MI0000286|hsa-mir-210&WithFlank&11|-|568083|568204@72.94.23&[MIMAT0000267&hsa-miR-210-3p&offsets|0|+1;m-102&11|-|568112|568133&offsets|0|+1]|CTGTGCGTGTGACAGCGGCTGAT</td><td>>MI0000286|hsa-mir-210&WithFlank&11|-|568083|568204</td><td>72</td><td>94</td><td>23</td><td>CTGTGCGTGTGACAGCGGCTGAT</td><td>420.2</td><td>2.3</td><td>0.9</td><td>2.6</td><td>1.0E-02</td><td>3.7E-02</td><td>BT20/MB231</td><td>NUC</td></tr>
<tr><td>>hg38_21_+_8433172_8446622_RNA45SN1_45S_50ntFlanks@10346.10373.28|TCAAACGAGAACTTTGAAGGCCGAAGTG</td><td>>hg38_21_+_8433172_8446622_RNA45SN1_45S_50ntFlanks</td><td>10346</td><td>10373</td><td>28</td><td>TCAAACGAGAACTTTGAAGGCCGAAGTG</td><td>579.3</td><td>-2.2</td><td>0.9</td><td>-2.5</td><td>1.3E-02</td><td>3.7E-02</td><td>BT20/MB231</td><td>CYTO</td></tr>
<tr><td>>hg38_21_+_8433172_8446622_RNA45SN1_45S_50ntFlanks@5491.5522.32|GAGAAGACGGTCGAACTTGACTATCTAGAGGA</td><td>>hg38_21_+_8433172_8446622_RNA45SN1_45S_50ntFlanks</td><td>5491</td><td>5522</td><td>32</td><td>GAGAAGACGGTCGAACTTGACTATCTAGAGGA</td><td>58.6</td><td>3.9</td><td>1.7</td><td>2.3</td><td>2.3E-02</td><td>3.7E-02</td><td>BT20/MB468</td><td>MP</td></tr>
<tr><td>>hg38_21_+_8433172_8446622_RNA45SN1_45S_50ntFlanks@10350.10369.20|ACGAGAACTTTGAAGGCCGA</td><td>>hg38_21_+_8433172_8446622_RNA45SN1_45S_50ntFlanks</td><td>10350</td><td>10369</td><td>20</td><td>ACGAGAACTTTGAAGGCCGA</td><td>355.5</td><td>-2.1</td><td>0.8</td><td>-2.5</td><td>1.3E-02</td><td>3.7E-02</td><td>BT20/MB231</td><td>CYTO</td></tr>
<tr><td>>hg38_21_+_8433172_8446622_RNA45SN1_45S_50ntFlanks@4987.5016.30|CAGGATTGACAGATTGATAGCTCTTTCTCG</td><td>>hg38_21_+_8433172_8446622_RNA45SN1_45S_50ntFlanks</td><td>4987</td><td>5016</td><td>30</td><td>CAGGATTGACAGATTGATAGCTCTTTCTCG</td><td>93.5</td><td>3.6</td><td>1.6</td><td>2.3</td><td>2.3E-02</td><td>3.7E-02</td><td>BT20/MB468</td><td>MP</td></tr>
<tr><td>>hg38_21_+_8433172_8446622_RNA45SN1_45S_50ntFlanks@3860.3899.40|GTAATTCTAGAGCTAATACATGCCGACGGGCGCTGACCCC</td><td>>hg38_21_+_8433172_8446622_RNA45SN1_45S_50ntFlanks</td><td>3860</td><td>3899</td><td>40</td><td>GTAATTCTAGAGCTAATACATGCCGACGGGCGCTGACCCC</td><td>172.5</td><td>-1.8</td><td>0.8</td><td>-2.1</td><td>3.2E-02</td><td>3.7E-02</td><td>MB231/MB468</td><td>CYTO</td></tr>
<tr><td>>MI0003591|hsa-mir-584&WithFlank&5|-|149062307|149062415@22.41.20(+1G)&[MIMAT0003249&hsa-miR-584-5p&offsets|0|-2(+1G);m-167&5|-|149062374|149062394&offsets|0|-1(+1G)]|TTATGGTTTGCCTGGGACTGG</td><td>>MI0003591|hsa-mir-584&WithFlank&5|-|149062307|149062415</td><td>22</td><td>41</td><td>20(+1G)</td><td>TTATGGTTTGCCTGGGACTGG</td><td>56.3</td><td>3.6</td><td>1.7</td><td>2.2</td><td>3.1E-02</td><td>3.7E-02</td><td>MB231/MB468</td><td>MP</td></tr>
<tr><td>>hg38_21_+_8433172_8446622_RNA45SN1_45S_50ntFlanks@12632.12657.26|GCGAAGCTACCATCTGTGGGATTATG</td><td>>hg38_21_+_8433172_8446622_RNA45SN1_45S_50ntFlanks</td><td>12632</td><td>12657</td><td>26</td><td>GCGAAGCTACCATCTGTGGGATTATG</td><td>405.5</td><td>-2.5</td><td>1.1</td><td>-2.3</td><td>2.3E-02</td><td>3.7E-02</td><td>BT20/MB468</td><td>MP</td></tr>
<tr><td>>hg38_21_+_8433172_8446622_RNA45SN1_45S_50ntFlanks@9494.9515.22|CCGAAAGATGGTGAACTATGCC</td><td>>hg38_21_+_8433172_8446622_RNA45SN1_45S_50ntFlanks</td><td>9494</td><td>9515</td><td>22</td><td>CCGAAAGATGGTGAACTATGCC</td><td>205.1</td><td>4.3</td><td>1.7</td><td>2.6</td><td>9.9E-03</td><td>3.7E-02</td><td>BT20/MB231</td><td>MP</td></tr>
<tr><td>>MI0000441|hsa-mir-30b&WithFlank&8|-|134800514|134800613@23.43.21(+1C)&[MIMAT0000420&hsa-miR-30b-5p&offsets|0|-1(+1C);m-76&8|-|134800570|134800591&offsets|0|-1(+1C)]|TGTAAACATCCTACACTCAGCC</td><td>>MI0000441|hsa-mir-30b&WithFlank&8|-|134800514|134800613</td><td>23</td><td>43</td><td>21(+1C)</td><td>TGTAAACATCCTACACTCAGCC</td><td>485.4</td><td>-3.0</td><td>1.2</td><td>-2.6</td><td>9.9E-03</td><td>3.7E-02</td><td>BT20/MB231</td><td>MP</td></tr>
<tr><td>>hg38_21_+_8433172_8446622_RNA45SN1_45S_50ntFlanks@11784.11802.19|ATGAATGGATGAACGAGAT</td><td>>hg38_21_+_8433172_8446622_RNA45SN1_45S_50ntFlanks</td><td>11784</td><td>11802</td><td>19</td><td>ATGAATGGATGAACGAGAT</td><td>253.4</td><td>-4.1</td><td>1.8</td><td>-2.3</td><td>2.3E-02</td><td>3.7E-02</td><td>BT20/MB468</td><td>MP</td></tr>
<tr><td>>hg38_21_+_8433172_8446622_RNA45SN1_45S_50ntFlanks@11670.11702.33|CCCAGTGCTCTGAATGTCAAAGTGAAGAAATTC</td><td>>hg38_21_+_8433172_8446622_RNA45SN1_45S_50ntFlanks</td><td>11670</td><td>11702</td><td>33</td><td>CCCAGTGCTCTGAATGTCAAAGTGAAGAAATTC</td><td>927.1</td><td>-2.5</td><td>1.1</td><td>-2.3</td><td>2.3E-02</td><td>3.7E-02</td><td>BT20/MB468</td><td>MP</td></tr>
<tr><td>>hg38_21_+_8433172_8446622_RNA45SN1_45S_50ntFlanks@10383.10424.42|TCCATGTGAACAGCAGTTGAACATGGGTCAGTCGGTCCTGAG</td><td>>hg38_21_+_8433172_8446622_RNA45SN1_45S_50ntFlanks</td><td>10383</td><td>10424</td><td>42</td><td>TCCATGTGAACAGCAGTTGAACATGGGTCAGTCGGTCCTGAG</td><td>3924.4</td><td>-2.3</td><td>0.9</td><td>-2.5</td><td>1.3E-02</td><td>3.7E-02</td><td>BT20/MB231</td><td>CYTO</td></tr>
<tr><td>>MI0000071|hsa-mir-17&WithFlank&13|+|91350599|91350694@20.41.22&[MIMAT0000070&hsa-miR-17-5p&offsets|0|-1;m-88&13|+|91350618|91350640&offsets|0|-1]|CAAAGTGCTTACAGTGCAGGTA</td><td>>MI0000071|hsa-mir-17&WithFlank&13|+|91350599|91350694</td><td>20</td><td>41</td><td>22</td><td>CAAAGTGCTTACAGTGCAGGTA</td><td>1033.3</td><td>1.9</td><td>0.9</td><td>2.2</td><td>3.1E-02</td><td>3.7E-02</td><td>MB231/MB468</td><td>MP</td></tr>
<tr><td>>hg38_21_+_8433172_8446622_RNA45SN1_45S_50ntFlanks@3705.3747.43|TACCTGGTTGATCCTGCCAGTAGCATATGCTTGTCTCAAAGAT</td><td>>hg38_21_+_8433172_8446622_RNA45SN1_45S_50ntFlanks</td><td>3705</td><td>3747</td><td>43</td><td>TACCTGGTTGATCCTGCCAGTAGCATATGCTTGTCTCAAAGAT</td><td>292.3</td><td>-1.9</td><td>0.8</td><td>-2.5</td><td>1.2E-02</td><td>3.7E-02</td><td>BT20/MB231</td><td>MITO</td></tr>
<tr><td>>hg38_21_+_8433172_8446622_RNA45SN1_45S_50ntFlanks@12476.12509.34|CCACTAATAGGGAACGTGAGCTGGGTTTAGACCG</td><td>>hg38_21_+_8433172_8446622_RNA45SN1_45S_50ntFlanks</td><td>12476</td><td>12509</td><td>34</td><td>CCACTAATAGGGAACGTGAGCTGGGTTTAGACCG</td><td>102.7</td><td>-3.3</td><td>1.5</td><td>-2.1</td><td>3.2E-02</td><td>3.7E-02</td><td>BT20/MB468</td><td>CYTO</td></tr>
<tr><td>>hg38_MT_+_1621_3279_RNR2_16S_50ntFlanks@1349.1383.35|ACAATAGGGTTTACGACCTCGATGTTGGATCAGGA</td><td>>hg38_MT_+_1621_3279_RNR2_16S_50ntFlanks</td><td>1349</td><td>1383</td><td>35</td><td>ACAATAGGGTTTACGACCTCGATGTTGGATCAGGA</td><td>68.1</td><td>3.7</td><td>1.7</td><td>2.2</td><td>3.1E-02</td><td>3.7E-02</td><td>MB231/MB468</td><td>MP</td></tr>
<tr><td>>hg38_21_+_8433172_8446622_RNA45SN1_45S_50ntFlanks@476.497.22|TTCGTGATCGATGTGGTGACGT</td><td>>hg38_21_+_8433172_8446622_RNA45SN1_45S_50ntFlanks</td><td>476</td><td>497</td><td>22</td><td>TTCGTGATCGATGTGGTGACGT</td><td>405.9</td><td>-4.0</td><td>1.6</td><td>-2.6</td><td>9.9E-03</td><td>3.7E-02</td><td>BT20/MB231</td><td>MP</td></tr>
<tr><td>>hg38_21_+_8433172_8446622_RNA45SN1_45S_50ntFlanks@4924.4944.21|AGGAGTGGAGCCTGCGGCTTA</td><td>>hg38_21_+_8433172_8446622_RNA45SN1_45S_50ntFlanks</td><td>4924</td><td>4944</td><td>21</td><td>AGGAGTGGAGCCTGCGGCTTA</td><td>453.7</td><td>3.9</td><td>1.7</td><td>2.3</td><td>2.3E-02</td><td>3.7E-02</td><td>BT20/MB468</td><td>MP</td></tr>
<tr><td>>trna10_AspGTC_12_-_125424193_125424264@36.74.39,trna12_AspGTC_12_-_125411891_125411962@36.74.39,trna144_AspGTC_6_-_27551236_27551307@36.74.39,trna38_AspGTC_17_-_8125556_8125627@36.74.39,trna45_AspGTC_6_+_27447453_27447524@36.74.39,trna48_AspGTC_6_+_27471523_27471594@36.74.39,trna4_AspGTC_12_+_96429799_96429870@36.74.39,trna69_AspGTC_1_-_161440205_161440276@36.74.39,trna72_AspGTC_1_-_161432824_161432895@36.74.39,trna75_AspGTC_1_-_161425414_161425485@36.74.39,trna78_AspGTC_1_-_161418033_161418104@36.74.39,trna81_AspGTC_1_-_161410615_161410686@36.74.39|CACGCGGGAGACCGGGGTTCGATTCCCCGACGGGGAGCC</td><td>>trna10_AspGTC_12_-_125424193_125424264</td><td>36</td><td>74</td><td>39</td><td>CACGCGGGAGACCGGGGTTCGATTCCCCGACGGGGAGCC</td><td>182.8</td><td>-2.0</td><td>0.8</td><td>-2.5</td><td>1.2E-02</td><td>3.7E-02</td><td>BT20/MB231</td><td>MITO</td></tr>
<tr><td>>trnaMT_SerGCT_MT_+_12207_12265@1.24.24|GAGAAAGCTCACAAGAACTGCTAA</td><td>>trnaMT_SerGCT_MT_+_12207_12265</td><td>1</td><td>24</td><td>24</td><td>GAGAAAGCTCACAAGAACTGCTAA</td><td>2479.5</td><td>3.4</td><td>1.5</td><td>2.3</td><td>2.3E-02</td><td>3.7E-02</td><td>BT20/MB468</td><td>MP</td></tr>
<tr><td>>hg38_21_+_8433172_8446622_RNA45SN1_45S_50ntFlanks@11670.11702.33|CCCAGTGCTCTGAATGTCAAAGTGAAGAAATTC</td><td>>hg38_21_+_8433172_8446622_RNA45SN1_45S_50ntFlanks</td><td>11670</td><td>11702</td><td>33</td><td>CCCAGTGCTCTGAATGTCAAAGTGAAGAAATTC</td><td>2258.7</td><td>-1.8</td><td>0.8</td><td>-2.1</td><td>3.2E-02</td><td>3.7E-02</td><td>MB231/MB468</td><td>CYTO</td></tr>
<tr><td>>hg38_21_+_8433172_8446622_RNA45SN1_45S_50ntFlanks@8185.8208.24|GAGGCCCAGCCCGTGGACGGTGTG</td><td>>hg38_21_+_8433172_8446622_RNA45SN1_45S_50ntFlanks</td><td>8185</td><td>8208</td><td>24</td><td>GAGGCCCAGCCCGTGGACGGTGTG</td><td>216.5</td><td>3.4</td><td>1.5</td><td>2.3</td><td>2.3E-02</td><td>3.7E-02</td><td>BT20/MB468</td><td>MP</td></tr>
<tr><td>>hg38_21_+_8433172_8446622_RNA45SN1_45S_50ntFlanks@7971.8011.41|GAGACGCGACCTCAGATCAGACGTGGCGACCCGCTGAATTT</td><td>>hg38_21_+_8433172_8446622_RNA45SN1_45S_50ntFlanks</td><td>7971</td><td>8011</td><td>41</td><td>GAGACGCGACCTCAGATCAGACGTGGCGACCCGCTGAATTT</td><td>429.2</td><td>2.3</td><td>0.9</td><td>2.6</td><td>1.0E-02</td><td>3.7E-02</td><td>BT20/MB231</td><td>NUC</td></tr>
<tr><td>>hg38_21_+_8433172_8446622_RNA45SN1_45S_50ntFlanks@7976.8009.34|GCGACCTCAGATCAGACGTGGCGACCCGCTGAAT</td><td>>hg38_21_+_8433172_8446622_RNA45SN1_45S_50ntFlanks</td><td>7976</td><td>8009</td><td>34</td><td>GCGACCTCAGATCAGACGTGGCGACCCGCTGAAT</td><td>369.0</td><td>1.8</td><td>0.8</td><td>2.1</td><td>3.4E-02</td><td>3.7E-02</td><td>MB231/MB468</td><td>MITO</td></tr>
<tr><td>>hg38_MT_+_1621_3279_RNR2_16S_50ntFlanks@842.859.18|AAGTAAAAGGAACTCGGC</td><td>>hg38_MT_+_1621_3279_RNR2_16S_50ntFlanks</td><td>842</td><td>859</td><td>18</td><td>AAGTAAAAGGAACTCGGC</td><td>75.1</td><td>2.6</td><td>1.1</td><td>2.3</td><td>2.3E-02</td><td>3.7E-02</td><td>BT20/MB468</td><td>MP</td></tr>
<tr><td>>am_MI0003581|hsa-mir-574&WithFlank&4|+|38868026|38868133@31.53.23(+1A)&[MIMAT0004795&hsa-miR-574-5p&offsets|0|0(+1A);m-213&4|+|38868056|38868078&offsets|0|0(+1A)]|TGAGTGTGTGTGTGTGAGTGTGTA</td><td>>am_MI0003581|hsa-mir-574&WithFlank&4|+|38868026|38868133</td><td>31</td><td>53</td><td>23(+1A)</td><td>TGAGTGTGTGTGTGTGAGTGTGTA</td><td>84.6</td><td>3.7</td><td>1.7</td><td>2.2</td><td>3.1E-02</td><td>3.7E-02</td><td>MB231/MB468</td><td>MP</td></tr>
<tr><td>>trna111_HisGTG_1_-_147774845_147774916@3.30.28,trna118_HisGTG_1_-_145396881_145396952@3.30.28,trna16_HisGTG_1_+_146544773_146544844@3.30.28,trna1_HisGTG_15_+_45493349_45493420@3.30.28,trna21_HisGTG_1_+_147753471_147753542@3.30.28,trna33_HisGTG_6_+_27125906_27125977@3.30.28,trna7_HisGTG_9_-_14433938_14434009@3.30.28,trna8_HisGTG_15_-_45492611_45492682@3.30.28,trna9_HisGTG_15_-_45490804_45490875@3.30.28|CGTGATCGTATAGTGGTTAGTACTCTGC</td><td>>trna111_HisGTG_1_-_147774845_147774916</td><td>3</td><td>30</td><td>28</td><td>CGTGATCGTATAGTGGTTAGTACTCTGC</td><td>675.1</td><td>1.9</td><td>0.8</td><td>2.5</td><td>1.3E-02</td><td>3.7E-02</td><td>BT20/MB231</td><td>CYTO</td></tr>
<tr><td>>hg38_21_+_8433172_8446622_RNA45SN1_45S_50ntFlanks@6646.6684.39|TCGTACGACTCTTAGCGGTGGATCACTCGGCTCGTGCGT</td><td>>hg38_21_+_8433172_8446622_RNA45SN1_45S_50ntFlanks</td><td>6646</td><td>6684</td><td>39</td><td>TCGTACGACTCTTAGCGGTGGATCACTCGGCTCGTGCGT</td><td>98.7</td><td>2.2</td><td>1.0</td><td>2.2</td><td>3.1E-02</td><td>3.7E-02</td><td>MB231/MB468</td><td>MP</td></tr>
<tr><td>>hg38_21_+_8433172_8446622_RNA45SN1_45S_50ntFlanks@12457.12486.30|CAAGCGTTGGATTGTTCACCCACTAATAGG</td><td>>hg38_21_+_8433172_8446622_RNA45SN1_45S_50ntFlanks</td><td>12457</td><td>12486</td><td>30</td><td>CAAGCGTTGGATTGTTCACCCACTAATAGG</td><td>227.5</td><td>-3.5</td><td>1.7</td><td>-2.1</td><td>3.2E-02</td><td>3.7E-02</td><td>BT20/MB468</td><td>CYTO</td></tr>
<tr><td>>hg38_21_+_8433172_8446622_RNA45SN1_45S_50ntFlanks@4761.4782.22|CAGATACCGTCGTAGTTCCGAC</td><td>>hg38_21_+_8433172_8446622_RNA45SN1_45S_50ntFlanks</td><td>4761</td><td>4782</td><td>22</td><td>CAGATACCGTCGTAGTTCCGAC</td><td>70.6</td><td>-3.8</td><td>1.7</td><td>-2.3</td><td>2.3E-02</td><td>3.7E-02</td><td>BT20/MB468</td><td>MP</td></tr>
<tr><td>>hg38_21_+_8433172_8446622_RNA45SN1_45S_50ntFlanks@8176.8194.19|CTTCTGATCGAGGCCCAGC</td><td>>hg38_21_+_8433172_8446622_RNA45SN1_45S_50ntFlanks</td><td>8176</td><td>8194</td><td>19</td><td>CTTCTGATCGAGGCCCAGC</td><td>733.0</td><td>-2.2</td><td>0.9</td><td>-2.5</td><td>1.3E-02</td><td>3.7E-02</td><td>BT20/MB231</td><td>CYTO</td></tr>
<tr><td>>hg38_21_+_8433172_8446622_RNA45SN1_45S_50ntFlanks@11582.11615.34|ACTGGTGCGGACCAGGGGAATCCGACTGTTTAAT</td><td>>hg38_21_+_8433172_8446622_RNA45SN1_45S_50ntFlanks</td><td>11582</td><td>11615</td><td>34</td><td>ACTGGTGCGGACCAGGGGAATCCGACTGTTTAAT</td><td>68.3</td><td>4.8</td><td>2.1</td><td>2.3</td><td>2.3E-02</td><td>3.7E-02</td><td>BT20/MB468</td><td>MP</td></tr>
<tr><td>>hg38_21_+_8433172_8446622_RNA45SN1_45S_50ntFlanks@10781.10825.45|AGGTGAACAGCCTCTGGCATGTTGGAACAATGTAGGTAAGGGAAG</td><td>>hg38_21_+_8433172_8446622_RNA45SN1_45S_50ntFlanks</td><td>10781</td><td>10825</td><td>45</td><td>AGGTGAACAGCCTCTGGCATGTTGGAACAATGTAGGTAAGGGAAG</td><td>167.3</td><td>-2.8</td><td>1.3</td><td>-2.2</td><td>2.9E-02</td><td>3.7E-02</td><td>BT20/MB468</td><td>MITO</td></tr>
<tr><td>>MI0000469|hsa-mir-125a&WithFlank&19|+|51693248|51693345@21.43.23&[MIMAT0000443&hsa-miR-125a-5p&offsets|0|-1;m-63&19|+|51693268|51693289&offsets|0|+1]|TCCCTGAGACCCTTTAACCTGTG</td><td>>MI0000469|hsa-mir-125a&WithFlank&19|+|51693248|51693345</td><td>21</td><td>43</td><td>23</td><td>TCCCTGAGACCCTTTAACCTGTG</td><td>37268.0</td><td>2.3</td><td>0.9</td><td>2.5</td><td>1.3E-02</td><td>3.7E-02</td><td>BT20/MB231</td><td>CYTO</td></tr>
<tr><td>>hg38_21_+_8433172_8446622_RNA45SN1_45S_50ntFlanks@10789.10818.30|AGCCTCTGGCATGTTGGAACAATGTAGGTA</td><td>>hg38_21_+_8433172_8446622_RNA45SN1_45S_50ntFlanks</td><td>10789</td><td>10818</td><td>30</td><td>AGCCTCTGGCATGTTGGAACAATGTAGGTA</td><td>645.8</td><td>-1.8</td><td>0.8</td><td>-2.3</td><td>2.3E-02</td><td>3.7E-02</td><td>BT20/MB468</td><td>MP</td></tr>
<tr><td>>hg38_21_+_8433172_8446622_RNA45SN1_45S_50ntFlanks@10514.10532.19|CGGAGTGGCGGAGATGGGC</td><td>>hg38_21_+_8433172_8446622_RNA45SN1_45S_50ntFlanks</td><td>10514</td><td>10532</td><td>19</td><td>CGGAGTGGCGGAGATGGGC</td><td>160.4</td><td>-3.8</td><td>1.7</td><td>-2.2</td><td>3.1E-02</td><td>3.7E-02</td><td>MB231/MB468</td><td>MP</td></tr>
<tr><td>>hg38_21_+_8433172_8446622_RNA45SN1_45S_50ntFlanks@11846.11867.22|GAACGGGCTTGGCGGAATCAGC</td><td>>hg38_21_+_8433172_8446622_RNA45SN1_45S_50ntFlanks</td><td>11846</td><td>11867</td><td>22</td><td>GAACGGGCTTGGCGGAATCAGC</td><td>180.7</td><td>-2.1</td><td>0.9</td><td>-2.5</td><td>1.3E-02</td><td>3.7E-02</td><td>BT20/MB231</td><td>CYTO</td></tr>
<tr><td>>hg38_21_+_8433172_8446622_RNA45SN1_45S_50ntFlanks@9951.9982.32|CCATGGAAGTCGGAATCCGCTAAGGAGTGTGT</td><td>>hg38_21_+_8433172_8446622_RNA45SN1_45S_50ntFlanks</td><td>9951</td><td>9982</td><td>32</td><td>CCATGGAAGTCGGAATCCGCTAAGGAGTGTGT</td><td>99.2</td><td>5.3</td><td>2.0</td><td>2.6</td><td>9.9E-03</td><td>3.7E-02</td><td>BT20/MB231</td><td>MP</td></tr>
<tr><td>>hg38_MT_+_1621_3279_RNR2_16S_50ntFlanks@1433.1451.19|AGTCCTACGTGATCTGAGT</td><td>>hg38_MT_+_1621_3279_RNR2_16S_50ntFlanks</td><td>1433</td><td>1451</td><td>19</td><td>AGTCCTACGTGATCTGAGT</td><td>38.6</td><td>3.7</td><td>1.7</td><td>2.2</td><td>3.1E-02</td><td>3.7E-02</td><td>MB231/MB468</td><td>MP</td></tr>
<tr><td>>hg38_21_+_8433172_8446622_RNA45SN1_45S_50ntFlanks@8914.8936.23|CCGAATCCCGGGGCCGAGGGAGC</td><td>>hg38_21_+_8433172_8446622_RNA45SN1_45S_50ntFlanks</td><td>8914</td><td>8936</td><td>23</td><td>CCGAATCCCGGGGCCGAGGGAGC</td><td>87.3</td><td>5.0</td><td>1.9</td><td>2.6</td><td>1.0E-02</td><td>3.7E-02</td><td>BT20/MB231</td><td>MP</td></tr>
<tr><td>>hg38_21_+_8433172_8446622_RNA45SN1_45S_50ntFlanks@11602.11621.20|TCCGACTGTTTAATTAAAAC</td><td>>hg38_21_+_8433172_8446622_RNA45SN1_45S_50ntFlanks</td><td>11602</td><td>11621</td><td>20</td><td>TCCGACTGTTTAATTAAAAC</td><td>335.9</td><td>-1.3</td><td>0.6</td><td>-2.1</td><td>3.4E-02</td><td>3.7E-02</td><td>MB231/MB468</td><td>NUC</td></tr>
</table>

<table>
<tr><td>>MI0000811|hsa-mir-148b&WithFlank&12|+|54337210|54337320@69.89.21(+1C)&[MIMAT0000759&hsa-miR-148b-3p&offsets|0|-1(+1C);m-77&12|+|54337278|54337299&offsets|0|-1(+1C)]|TCAGTGCATCACAGAACTTTGC</td><td>>MI0000811|hsa-mir-148b&WithFlank&12|+|54337210|54337320</td><td>69</td><td>89</td><td>21(+1C)</td><td>TCAGTGCATCACAGAACTTTGC</td><td>978.4</td><td>-2.3</td><td>0.9</td><td>-2.5</td><td>1.2E-02</td><td>3.7E-02</td><td>BT20/MB231</td><td>MITO</td></tr>
<tr><td>>hg38_21_+_8433172_8446622_RNA45SN1_45S_50ntFlanks@10236.10259.24|GACGCTACGCCGCGACGAGTAGGA</td><td>>hg38_21_+_8433172_8446622_RNA45SN1_45S_50ntFlanks</td><td>10236</td><td>10259</td><td>24</td><td>GACGCTACGCCGCGACGAGTAGGA</td><td>149.8</td><td>3.4</td><td>1.5</td><td>2.3</td><td>2.3E-02</td><td>3.7E-02</td><td>BT20/MB468</td><td>MP</td></tr>
<tr><td>>am_trnaMT_GluTTC_MT_-_14674_14742@27.55.29,trnalookalike8_GluTTC_5_-_93905172_93905240@27.55.29|GTTTTTCATATCATTGGTCGTGGTTGTAG</td><td>>am_trnaMT_GluTTC_MT_-_14674_14742</td><td>27</td><td>55</td><td>29</td><td>GTTTTTCATATCATTGGTCGTGGTTGTAG</td><td>146.4</td><td>-4.1</td><td>1.8</td><td>-2.3</td><td>2.3E-02</td><td>3.7E-02</td><td>BT20/MB468</td><td>MP</td></tr>
<tr><td>>hg38_21_+_8433172_8446622_RNA45SN1_45S_50ntFlanks@11579.11604.26|AGAACTGGTGCGGACCAGGGGAATCC</td><td>>hg38_21_+_8433172_8446622_RNA45SN1_45S_50ntFlanks</td><td>11579</td><td>11604</td><td>26</td><td>AGAACTGGTGCGGACCAGGGGAATCC</td><td>142.6</td><td>-1.1</td><td>0.5</td><td>-2.1</td><td>3.3E-02</td><td>3.7E-02</td><td>BT20/MB468</td><td>CYTO</td></tr>
<tr><td>>hg38_21_+_8433172_8446622_RNA45SN1_45S_50ntFlanks@12417.12439.23|TCGATGTCGGCTCTTCCTATCAT</td><td>>hg38_21_+_8433172_8446622_RNA45SN1_45S_50ntFlanks</td><td>12417</td><td>12439</td><td>23</td><td>TCGATGTCGGCTCTTCCTATCAT</td><td>750.4</td><td>1.4</td><td>0.7</td><td>2.1</td><td>3.3E-02</td><td>3.7E-02</td><td>BT20/MB468</td><td>CYTO</td></tr>
<tr><td>>MI0000471|hsa-mir-126&WithFlank&9|+|136670596|136670692@58.77.20(+1U)&[MIMAT0000445&hsa-miR-126-3p&offsets|0|-2(+1U);m-40&9|+|136670653|136670674&offsets|0|-2(+1U)]|TCGTACCGTGAGTAATAATGT</td><td>>MI0000471|hsa-mir-126&WithFlank&9|+|136670596|136670692</td><td>58</td><td>77</td><td>20(+1U)</td><td>TCGTACCGTGAGTAATAATGT</td><td>2743.2</td><td>-2.7</td><td>1.0</td><td>-2.6</td><td>1.0E-02</td><td>3.7E-02</td><td>BT20/MB231</td><td>MP</td></tr>
<tr><td>>hg38_21_+_8433172_8446622_RNA45SN1_45S_50ntFlanks@7975.7993.19|CGCGACCTCAGATCAGACG</td><td>>hg38_21_+_8433172_8446622_RNA45SN1_45S_50ntFlanks</td><td>7975</td><td>7993</td><td>19</td><td>CGCGACCTCAGATCAGACG</td><td>46309.1</td><td>2.8</td><td>1.1</td><td>2.6</td><td>1.0E-02</td><td>3.7E-02</td><td>BT20/MB231</td><td>MP</td></tr>
<tr><td>>MI0000446|hsa-mir-125b-1&WithFlank&11|-|122099751|122099850@61.78.18&[MIMAT0004592&hsa-miR-125b-1-3p&offsets|0|-4;m-485&11|-|122099769|122099790&offsets|0|-4]|ACGGGTTAGGCTCTTGGG</td><td>>MI0000446|hsa-mir-125b-1&WithFlank&11|-|122099751|122099850</td><td>61</td><td>78</td><td>18</td><td>ACGGGTTAGGCTCTTGGG</td><td>79.6</td><td>-5.0</td><td>2.0</td><td>-2.6</td><td>1.0E-02</td><td>3.7E-02</td><td>BT20/MB231</td><td>MP</td></tr>
<tr><td>>MI0000269|hsa-mir-181a-2&WithFlank&9|+|124692436|124692557@83.103.21(+1A)&[MIMAT0004558&hsa-miR-181a-2-3p&offsets|0|-1(+1A);m-71&9|+|124692518|124692539&offsets|0|-1(+1A)]|ACCACTGACCGTTGACTGTACA</td><td>>MI0000269|hsa-mir-181a-2&WithFlank&9|+|124692436|124692557</td><td>83</td><td>103</td><td>21(+1A)</td><td>ACCACTGACCGTTGACTGTACA</td><td>149.0</td><td>-2.8</td><td>1.1</td><td>-2.6</td><td>1.0E-02</td><td>3.7E-02</td><td>BT20/MB231</td><td>MP</td></tr>
<tr><td>>hg38_21_+_8433172_8446622_RNA45SN1_45S_50ntFlanks@4975.5001.27|CCCGGACACGGACAGGATTGACAGATT</td><td>>hg38_21_+_8433172_8446622_RNA45SN1_45S_50ntFlanks</td><td>4975</td><td>5001</td><td>27</td><td>CCCGGACACGGACAGGATTGACAGATT</td><td>197.6</td><td>1.1</td><td>0.5</td><td>2.1</td><td>3.4E-02</td><td>3.7E-02</td><td>BT20/MB468</td><td>NUC</td></tr>
<tr><td>>hg38_21_+_8433172_8446622_RNA45SN1_45S_50ntFlanks@9225.9249.25|CCGAAGAGGGGGACGGCGGAGCGAG</td><td>>hg38_21_+_8433172_8446622_RNA45SN1_45S_50ntFlanks</td><td>9225</td><td>9249</td><td>25</td><td>CCGAAGAGGGGGACGGCGGAGCGAG</td><td>76.4</td><td>-3.9</td><td>1.7</td><td>-2.3</td><td>2.3E-02</td><td>3.7E-02</td><td>BT20/MB468</td><td>MP</td></tr>
<tr><td>>MI0000542|hsa-mir-320a&WithFlank&8|-|22244960|22245043@48.69.22(+3U)&[MIMAT0000510&hsa-miR-320a-3p&offsets|0|0(+3U);m-60&8|-|22244975|22244996&offsets|0|0(+3U)]|AAAAGCTGGGTTGAGAGGGCGATTT</td><td>>MI0000542|hsa-mir-320a&WithFlank&8|-|22244960|22245043</td><td>48</td><td>69</td><td>22(+3U)</td><td>AAAAGCTGGGTTGAGAGGGCGATTT</td><td>71.6</td><td>3.8</td><td>1.8</td><td>2.2</td><td>3.1E-02</td><td>3.7E-02</td><td>MB231/MB468</td><td>MP</td></tr>
<tr><td>>hg38_21_+_8433172_8446622_RNA45SN1_45S_50ntFlanks@10869.10892.24|AGGGCTGGGTCGGTCGGGCTGGGG</td><td>>hg38_21_+_8433172_8446622_RNA45SN1_45S_50ntFlanks</td><td>10869</td><td>10892</td><td>24</td><td>AGGGCTGGGTCGGTCGGGCTGGGG</td><td>355.9</td><td>-3.0</td><td>1.4</td><td>-2.2</td><td>2.9E-02</td><td>3.7E-02</td><td>BT20/MB468</td><td>MITO</td></tr>
<tr><td>>hg38_21_+_8433172_8446622_RNA45SN1_45S_50ntFlanks@10907.10929.23|GGGCGCGCGCCGCGGCTGGACGA</td><td>>hg38_21_+_8433172_8446622_RNA45SN1_45S_50ntFlanks</td><td>10907</td><td>10929</td><td>23</td><td>GGGCGCGCGCCGCGGCTGGACGA</td><td>159.2</td><td>5.2</td><td>2.0</td><td>2.6</td><td>1.0E-02</td><td>3.7E-02</td><td>BT20/MB231</td><td>MP</td></tr>
<tr><td>>hg38_21_+_8433172_8446622_RNA45SN1_45S_50ntFlanks@3984.4005.22|GCGGCTTTGGTGACTCTAGATA</td><td>>hg38_21_+_8433172_8446622_RNA45SN1_45S_50ntFlanks</td><td>3984</td><td>4005</td><td>22</td><td>GCGGCTTTGGTGACTCTAGATA</td><td>430.8</td><td>-4.1</td><td>1.8</td><td>-2.3</td><td>2.3E-02</td><td>3.7E-02</td><td>BT20/MB468</td><td>MP</td></tr>
<tr><td>>hg38_21_+_8433172_8446622_RNA45SN1_45S_50ntFlanks@4128.4150.23|CGATTCCGGAGAGGGAGCCTGAG</td><td>>hg38_21_+_8433172_8446622_RNA45SN1_45S_50ntFlanks</td><td>4128</td><td>4150</td><td>23</td><td>CGATTCCGGAGAGGGAGCCTGAG</td><td>108.1</td><td>-1.9</td><td>0.8</td><td>-2.3</td><td>2.3E-02</td><td>3.7E-02</td><td>BT20/MB468</td><td>MP</td></tr>
<tr><td>>MI0000077|hsa-mir-21&WithFlank&17|+|59841260|59841343@52.72.21(+1A)&[MIMAT0004494&hsa-miR-21-3p&offsets|0|0(+1A);m-52&17|+|59841311|59841332&offsets|0|-1(+1A)]|CAACACCAGTCGATGGGCTGTA</td><td>>MI0000077|hsa-mir-21&WithFlank&17|+|59841260|59841343</td><td>52</td><td>72</td><td>21(+1A)</td><td>CAACACCAGTCGATGGGCTGTA</td><td>351.7</td><td>5.3</td><td>2.3</td><td>2.3</td><td>2.3E-02</td><td>3.7E-02</td><td>BT20/MB468</td><td>MP</td></tr>
<tr><td>>hg38_MT_+_1621_3279_RNR2_16S_50ntFlanks@51.85.35|GCTAAACCTAGCCCCAAACCCACTCCACCTTACTA</td><td>>hg38_MT_+_1621_3279_RNR2_16S_50ntFlanks</td><td>51</td><td>85</td><td>35</td><td>GCTAAACCTAGCCCCAAACCCACTCCACCTTACTA</td><td>475.4</td><td>2.3</td><td>1.1</td><td>2.2</td><td>3.1E-02</td><td>3.7E-02</td><td>MB231/MB468</td><td>MP</td></tr>
<tr><td>>MI0000812|hsa-mir-331&WithFlank&12|+|95308414|95308519@67.87.21(+1U)&[MIMAT0000760&hsa-miR-331-3p&offsets|0|0(+1U);m-192&12|+|95308480|95308501&offsets|0|-1(+1U)]|GCCCCTGGGCCTATCCTAGAAT</td><td>>MI0000812|hsa-mir-331&WithFlank&12|+|95308414|95308519</td><td>67</td><td>87</td><td>21(+1U)</td><td>GCCCCTGGGCCTATCCTAGAAT</td><td>43.3</td><td>3.7</td><td>1.7</td><td>2.2</td><td>3.1E-02</td><td>3.7E-02</td><td>MB231/MB468</td><td>MP</td></tr>
<tr><td>>hg38_21_+_8433172_8446622_RNA45SN1_45S_50ntFlanks@6663.6696.34|GTGGATCACTCGGCTCGTGCGTCGATGAAGAACG</td><td>>hg38_21_+_8433172_8446622_RNA45SN1_45S_50ntFlanks</td><td>6663</td><td>6696</td><td>34</td><td>GTGGATCACTCGGCTCGTGCGTCGATGAAGAACG</td><td>261.9</td><td>2.7</td><td>1.1</td><td>2.5</td><td>1.2E-02</td><td>3.7E-02</td><td>BT20/MB231</td><td>MITO</td></tr>
<tr><td>>hg38_21_+_8433172_8446622_RNA45SN1_45S_50ntFlanks@7977.8019.43|CGACCTCAGATCAGACGTGGCGACCCGCTGAATTTAAGCATAT</td><td>>hg38_21_+_8433172_8446622_RNA45SN1_45S_50ntFlanks</td><td>7977</td><td>8019</td><td>43</td><td>CGACCTCAGATCAGACGTGGCGACCCGCTGAATTTAAGCATAT</td><td>720.4</td><td>-2.8</td><td>1.1</td><td>-2.5</td><td>1.2E-02</td><td>3.8E-02</td><td>BT20/MB231</td><td>MITO</td></tr>
<tr><td>>hg38_21_+_8433172_8446622_RNA45SN1_45S_50ntFlanks@5475.5516.42|GCCCTGGCGGAGCGCTGAGAAGACGGTCGAACTTGACTATCT</td><td>>hg38_21_+_8433172_8446622_RNA45SN1_45S_50ntFlanks</td><td>5475</td><td>5516</td><td>42</td><td>GCCCTGGCGGAGCGCTGAGAAGACGGTCGAACTTGACTATCT</td><td>1373.1</td><td>-1.8</td><td>0.7</td><td>-2.5</td><td>1.3E-02</td><td>3.8E-02</td><td>BT20/MB231</td><td>CYTO</td></tr>
<tr><td>>hg38_21_+_8433172_8446622_RNA45SN1_45S_50ntFlanks@7983.8017.35|CAGATCAGACGTGGCGACCCGCTGAATTTAAGCAT</td><td>>hg38_21_+_8433172_8446622_RNA45SN1_45S_50ntFlanks</td><td>7983</td><td>8017</td><td>35</td><td>CAGATCAGACGTGGCGACCCGCTGAATTTAAGCAT</td><td>108.8</td><td>-3.3</td><td>1.4</td><td>-2.3</td><td>2.3E-02</td><td>3.8E-02</td><td>BT20/MB468</td><td>MP</td></tr>
<tr><td>>hg38_21_+_8433172_8446622_RNA45SN1_45S_50ntFlanks@4657.4686.30|CTAGAGGTGAAATTCTTGGACCGGCGCAAG</td><td>>hg38_21_+_8433172_8446622_RNA45SN1_45S_50ntFlanks</td><td>4657</td><td>4686</td><td>30</td><td>CTAGAGGTGAAATTCTTGGACCGGCGCAAG</td><td>335.1</td><td>-4.1</td><td>1.8</td><td>-2.3</td><td>2.3E-02</td><td>3.8E-02</td><td>BT20/MB468</td><td>MP</td></tr>
<tr><td>>MI0000084|hsa-mir-26b&WithFlank&2|+|218402640|218402728@18.37.20(+1A)&[MIMAT0000083&hsa-miR-26b-5p&offsets|0|-1(+1A);m-53&2|+|218402657|218402678&offsets|0|-2(+1A)]|TTCAAGTAATTCAGGATAGGA</td><td>>MI0000084|hsa-mir-26b&WithFlank&2|+|218402640|218402728</td><td>18</td><td>37</td><td>20(+1A)</td><td>TTCAAGTAATTCAGGATAGGA</td><td>61.2</td><td>3.2</td><td>1.5</td><td>2.2</td><td>3.1E-02</td><td>3.8E-02</td><td>MB231/MB468</td><td>MP</td></tr>
<tr><td>>hg38_21_+_8433172_8446622_RNA45SN1_45S_50ntFlanks@4369.4389.21|GTTAAAAAGCTCGTAGTTGGA</td><td>>hg38_21_+_8433172_8446622_RNA45SN1_45S_50ntFlanks</td><td>4369</td><td>4389</td><td>21</td><td>GTTAAAAAGCTCGTAGTTGGA</td><td>87.1</td><td>-4.0</td><td>1.7</td><td>-2.3</td><td>2.3E-02</td><td>3.8E-02</td><td>BT20/MB468</td><td>MP</td></tr>
<tr><td>>MI0000749|hsa-mir-30e&WithFlank&1|+|40754349|40754452@23.46.24&[MIMAT0000692&hsa-miR-30e-5p&offsets|0|+2;m-24&1|+|40754371|40754394&offsets|0|0]|TGTAAACATCCTTGACTGGAAGCT</td><td>>MI0000749|hsa-mir-30e&WithFlank&1|+|40754349|40754452</td><td>23</td><td>46</td><td>24</td><td>TGTAAACATCCTTGACTGGAAGCT</td><td>1550.5</td><td>2.2</td><td>1.1</td><td>2.1</td><td>3.4E-02</td><td>3.8E-02</td><td>BT20/MB468</td><td>NUC</td></tr>
<tr><td>>am_MI0003581|hsa-mir-574&WithFlank&4|+|38868026|38868133@31.54.24(+1C)&[MIMAT0004795&hsa-miR-574-5p&offsets|0|+1(+1C);m-213&4|+|38868056|38868078&offsets|0|+1(+1C)]|TGAGTGTGTGTGTGTGAGTGTGTGC</td><td>>am_MI0003581|hsa-mir-574&WithFlank&4|+|38868026|38868133</td><td>31</td><td>54</td><td>24(+1C)</td><td>TGAGTGTGTGTGTGTGAGTGTGTGC</td><td>204.4</td><td>-4.8</td><td>1.9</td><td>-2.6</td><td>1.0E-02</td><td>3.8E-02</td><td>BT20/MB231</td><td>MP</td></tr>
<tr><td>>hg38_21_+_8433172_8446622_RNA45SN1_45S_50ntFlanks@3843.3861.19|CTACTTGGATAACTGTGGT</td><td>>hg38_21_+_8433172_8446622_RNA45SN1_45S_50ntFlanks</td><td>3843</td><td>3861</td><td>19</td><td>CTACTTGGATAACTGTGGT</td><td>593.9</td><td>-1.9</td><td>0.9</td><td>-2.2</td><td>2.9E-02</td><td>3.8E-02</td><td>BT20/MB468</td><td>MITO</td></tr>
<tr><td>>hg38_21_+_8433172_8446622_RNA45SN1_45S_50ntFlanks@6653.6679.27|ACTCTTAGCGGTGGATCACTCGGCTCG</td><td>>hg38_21_+_8433172_8446622_RNA45SN1_45S_50ntFlanks</td><td>6653</td><td>6679</td><td>27</td><td>ACTCTTAGCGGTGGATCACTCGGCTCG</td><td>648.7</td><td>1.5</td><td>0.7</td><td>2.2</td><td>2.9E-02</td><td>3.8E-02</td><td>BT20/MB468</td><td>MITO</td></tr>
<tr><td>>hg38_21_+_8433172_8446622_RNA45SN1_45S_50ntFlanks@12449.12480.32|GAATTCACCAAGCGTTGGATTGTTCACCCACT</td><td>>hg38_21_+_8433172_8446622_RNA45SN1_45S_50ntFlanks</td><td>12449</td><td>12480</td><td>32</td><td>GAATTCACCAAGCGTTGGATTGTTCACCCACT</td><td>63.2</td><td>2.9</td><td>1.3</td><td>2.2</td><td>3.1E-02</td><td>3.8E-02</td><td>MB231/MB468</td><td>MP</td></tr>
<tr><td>>hg38_21_+_8433172_8446622_RNA45SN1_45S_50ntFlanks@3903.3937.35|CGCGGGGGGGATGCGTGCATTTATCAGATCAAAAC</td><td>>hg38_21_+_8433172_8446622_RNA45SN1_45S_50ntFlanks</td><td>3903</td><td>3937</td><td>35</td><td>CGCGGGGGGGATGCGTGCATTTATCAGATCAAAAC</td><td>74.0</td><td>-5.1</td><td>2.2</td><td>-2.3</td><td>2.3E-02</td><td>3.8E-02</td><td>BT20/MB468</td><td>MP</td></tr>
<tr><td>>hg38_21_+_8433172_8446622_RNA45SN1_45S_50ntFlanks@12458.12478.21|AAGCGTTGGATTGTTCACCCA</td><td>>hg38_21_+_8433172_8446622_RNA45SN1_45S_50ntFlanks</td><td>12458</td><td>12478</td><td>21</td><td>AAGCGTTGGATTGTTCACCCA</td><td>403.0</td><td>-1.3</td><td>0.6</td><td>-2.2</td><td>2.9E-02</td><td>3.8E-02</td><td>BT20/MB468</td><td>MITO</td></tr>
<tr><td>>hg38_21_+_8433172_8446622_RNA45SN1_45S_50ntFlanks@11874.11909.36|AGAAGACCCTGTTGAGCTTGACTCTAGTCTGGCACG</td><td>>hg38_21_+_8433172_8446622_RNA45SN1_45S_50ntFlanks</td><td>11874</td><td>11909</td><td>36</td><td>AGAAGACCCTGTTGAGCTTGACTCTAGTCTGGCACG</td><td>165.1</td><td>-4.0</td><td>1.8</td><td>-2.3</td><td>2.3E-02</td><td>3.8E-02</td><td>BT20/MB468</td><td>MP</td></tr>
<tr><td>>MI0000084|hsa-mir-26b&WithFlank&2|+|218402640|218402728@18.39.22(+1C)&[MIMAT0000083&hsa-miR-26b-5p&offsets|0|+1(+1C);m-53&2|+|218402657|218402678&offsets|0|0(+1C)]|TTCAAGTAATTCAGGATAGGTTC</td><td>>MI0000084|hsa-mir-26b&WithFlank&2|+|218402640|218402728</td><td>18</td><td>39</td><td>22(+1C)</td><td>TTCAAGTAATTCAGGATAGGTTC</td><td>246.9</td><td>-3.9</td><td>1.5</td><td>-2.6</td><td>1.0E-02</td><td>3.8E-02</td><td>BT20/MB231</td><td>MP</td></tr>
<tr><td>>trnaMT_GluTTC_MT_-_14674_14742@45.70.26,trnalookalike8_GluTTC_5_-_93905172_93905240@45.70.26|CGTGGTTGTAGTCCGTGCGAGAATAC</td><td>>trnaMT_GluTTC_MT_-_14674_14742</td><td>45</td><td>70</td><td>26</td><td>CGTGGTTGTAGTCCGTGCGAGAATAC</td><td>95.2</td><td>3.6</td><td>1.7</td><td>2.2</td><td>3.1E-02</td><td>3.8E-02</td><td>MB231/MB468</td><td>MP</td></tr>
<tr><td>>trna111_HisGTG_1_-_147774845_147774916@-1T.35.36,trna118_HisGTG_1_-_145396881_145396952@-1T.35.36,trna16_HisGTG_1_+_146544773_146544844@-1T.35.36,trna1_HisGTG_15_+_45493349_45493420@-1T.35.36,trna21_HisGTG_1_+_147753471_147753542@-1T.35.36,trna33_HisGTG_6_+_27125906_27125977@-1T.35.36,trna7_HisGTG_9_-_14433938_14434009@-1T.35.36,trna8_HisGTG_15_-_45492611_45492682@-1T.35.36,trna9_HisGTG_15_-_45490804_45490875@-1T.35.36|TGCCGTGATCGTATAGTGGTTAGTACTCTGCGTTGT</td><td>>trna111_HisGTG_1_-_147774845_147774916</td><td>-1T</td><td>35</td><td>36</td><td>TGCCGTGATCGTATAGTGGTTAGTACTCTGCGTTGT</td><td>252.5</td><td>-2.8</td><td>1.1</td><td>-2.5</td><td>1.2E-02</td><td>3.8E-02</td><td>BT20/MB231</td><td>MITO</td></tr>
<tr><td>>am_MI0000786|hsa-mir-378a&WithFlank&5|+|149732819|149732896@50.69.20(+1A)&[MIMAT0000732&hsa-miR-378a-3p&offsets|+1|-1(+1A);m-44&5|+|149732867|149732888&offsets|+1|-1(+1A)]|CTGGACTTGGAGTCAGAAGGA</td><td>>am_MI0000786|hsa-mir-378a&WithFlank&5|+|149732819|149732896</td><td>50</td><td>69</td><td>20(+1A)</td><td>CTGGACTTGGAGTCAGAAGGA</td><td>297.0</td><td>2.2</td><td>0.9</td><td>2.5</td><td>1.3E-02</td><td>3.8E-02</td><td>BT20/MB231</td><td>CYTO</td></tr>
<tr><td>>MI0000115|hsa-mir-16-2&WithFlank&3|+|160404739|160404831@59.79.21&[MIMAT0004518&hsa-miR-16-2-3p&offsets|0|-1;m-252&3|+|160404796|160404816&offsets|+1|+1]|CCAATATTACTGTGCTGCTTT</td><td>>MI0000115|hsa-mir-16-2&WithFlank&3|+|160404739|160404831</td><td>59</td><td>79</td><td>21</td><td>CCAATATTACTGTGCTGCTTT</td><td>122.6</td><td>3.8</td><td>1.8</td><td>2.2</td><td>3.1E-02</td><td>3.8E-02</td><td>MB231/MB468</td><td>MP</td></tr>
<tr><td>>hg38_21_+_8433172_8446622_RNA45SN1_45S_50ntFlanks@8196.8224.29|CGTGGACGGTGTGAGGCCGGTAGCGGCCC</td><td>>hg38_21_+_8433172_8446622_RNA45SN1_45S_50ntFlanks</td><td>8196</td><td>8224</td><td>29</td><td>CGTGGACGGTGTGAGGCCGGTAGCGGCCC</td><td>358.2</td><td>-2.5</td><td>1.0</td><td>-2.5</td><td>1.3E-02</td><td>3.8E-02</td><td>BT20/MB231</td><td>CYTO</td></tr>
</table>

<table>
<tr><td>>trna10_AspGTC_12_-_125424193_125424264@1.34.34,trna12_AspGTC_12_-_125411891_125411962@1.34.34,trna38_AspGTC_17_-_8125556_8125627@1.34.34,trna45_AspGTC_6_+_27447453_27447524@1.34.34,trna48_AspGTC_6_+_27471523_27471594@1.34.34,trna4_AspGTC_12_+_96429799_96429870@1.34.34,trna69_AspGTC_1_-_161440205_161440276@1.34.34,trna72_AspGTC_1_-_161432824_161432895@1.34.34,trna75_AspGTC_1_-_161425414_161425485@1.34.34,trna78_AspGTC_1_-_161418033_161418104@1.34.34,trna81_AspGTC_1_-_161410615_161410686@1.34.34|TCCTCGTTAGTATAGTGGTGAGTATCCCCGCCTG</td><td>>trna10_AspGTC_12_-_125424193_125424264</td><td>1</td><td>34</td><td>34</td><td>TCCTCGTTAGTATAGTGGTGAGTATCCCCGCCTG</td><td>1011.6</td><td>-3.0</td><td>1.2</td><td>-2.5</td><td>1.3E-02</td><td>3.8E-02</td><td>BT20/MB231</td><td>CYTO</td></tr>
<tr><td>>hg38_21_+_8433172_8446622_RNA45SN1_45S_50ntFlanks@4163.4183.21|CATCCAAGGAAGGCAGCAGGC</td><td>>hg38_21_+_8433172_8446622_RNA45SN1_45S_50ntFlanks</td><td>4163</td><td>4183</td><td>21</td><td>CATCCAAGGAAGGCAGCAGGC</td><td>85.7</td><td>-3.9</td><td>1.7</td><td>-2.3</td><td>2.3E-02</td><td>3.8E-02</td><td>BT20/MB468</td><td>MP</td></tr>
<tr><td>>hg38_21_+_8433172_8446622_RNA45SN1_45S_50ntFlanks@9381.9409.29|CGCTCGCCGGCCGAGGTGGGATCCCGAGG</td><td>>hg38_21_+_8433172_8446622_RNA45SN1_45S_50ntFlanks</td><td>9381</td><td>9409</td><td>29</td><td>CGCTCGCCGGCCGAGGTGGGATCCCGAGG</td><td>2497.7</td><td>-2.7</td><td>1.2</td><td>-2.2</td><td>2.9E-02</td><td>3.8E-02</td><td>BT20/MB468</td><td>MITO</td></tr>
<tr><td>>MI0000483|hsa-mir-186&WithFlank&1|-|71067625|71067722@21.44.24&[MIMAT0000456&hsa-miR-186-5p&offsets|0|+2;m-84&1|-|71067681|71067702&offsets|0|+2]|CAAAGAATTCTCCTTTTGGGCTTT</td><td>>MI0000483|hsa-mir-186&WithFlank&1|-|71067625|71067722</td><td>21</td><td>44</td><td>24</td><td>CAAAGAATTCTCCTTTTGGGCTTT</td><td>1820.3</td><td>2.4</td><td>1.1</td><td>2.2</td><td>3.1E-02</td><td>3.8E-02</td><td>MB231/MB468</td><td>MP</td></tr>
<tr><td>>hg38_21_+_8433172_8446622_RNA45SN1_45S_50ntFlanks@10362.10380.19|AAGGCCGAAGTGGAGAAGG</td><td>>hg38_21_+_8433172_8446622_RNA45SN1_45S_50ntFlanks</td><td>10362</td><td>10380</td><td>19</td><td>AAGGCCGAAGTGGAGAAGG</td><td>142.5</td><td>-2.0</td><td>0.9</td><td>-2.3</td><td>2.3E-02</td><td>3.8E-02</td><td>BT20/MB468</td><td>MP</td></tr>
<tr><td>>hg38_21_+_8433172_8446622_RNA45SN1_45S_50ntFlanks@6649.6675.27|TACGACTCTTAGCGGTGGATCACTCGG</td><td>>hg38_21_+_8433172_8446622_RNA45SN1_45S_50ntFlanks</td><td>6649</td><td>6675</td><td>27</td><td>TACGACTCTTAGCGGTGGATCACTCGG</td><td>277.5</td><td>2.5</td><td>1.1</td><td>2.3</td><td>2.3E-02</td><td>3.8E-02</td><td>BT20/MB468</td><td>MP</td></tr>
<tr><td>>MI0000093|hsa-mir-92a-1&WithFlank&13|+|91351308|91351397@54.74.21(+1A);MI0000094|hsa-mir-92a-2&WithFlank&X|-|134169532|134169618@54.74.21(+1A)&[MIMAT0000092&hsa-miR-92a-3p&offsets|0|-1(+1A);m-9&13|+|91351361|91351382&offsets|0|-1(+1A)];[MIMAT0000092_1&hsa-miR-92a-3p&offsets|0|-1(+1A);m-11&X|-|134169544|134169565&offsets|0|-1(+1A)]|TATTGCACTTGTCCCGGCCTGA</td><td>>MI0000093|hsa-mir-92a-1&WithFlank&13|+|91351308|91351397</td><td>54</td><td>74</td><td>21(+1A)</td><td>TATTGCACTTGTCCCGGCCTGA</td><td>857.1</td><td>2.6</td><td>1.1</td><td>2.3</td><td>2.3E-02</td><td>3.8E-02</td><td>BT20/MB468</td><td>MP</td></tr>
<tr><td>>hg38_21_+_8433172_8446622_RNA45SN1_45S_50ntFlanks@10348.10367.20|AAACGAGAACTTTGAAGGCC</td><td>>hg38_21_+_8433172_8446622_RNA45SN1_45S_50ntFlanks</td><td>10348</td><td>10367</td><td>20</td><td>AAACGAGAACTTTGAAGGCC</td><td>674.6</td><td>-2.4</td><td>1.1</td><td>-2.2</td><td>3.1E-02</td><td>3.8E-02</td><td>MB231/MB468</td><td>MP</td></tr>
<tr><td>>hg38_21_+_8433172_8446622_RNA45SN1_45S_50ntFlanks@11568.11602.35|GCAGCCGACTTAGAACTGGTGCGGACCAGGGGAAT</td><td>>hg38_21_+_8433172_8446622_RNA45SN1_45S_50ntFlanks</td><td>11568</td><td>11602</td><td>35</td><td>GCAGCCGACTTAGAACTGGTGCGGACCAGGGGAAT</td><td>395.8</td><td>3.2</td><td>1.3</td><td>2.6</td><td>1.0E-02</td><td>3.8E-02</td><td>BT20/MB231</td><td>MP</td></tr>
<tr><td>>hg38_21_+_8433172_8446622_RNA45SN1_45S_50ntFlanks@7976.8010.35|GCGACCTCAGATCAGACGTGGCGACCCGCTGAATT</td><td>>hg38_21_+_8433172_8446622_RNA45SN1_45S_50ntFlanks</td><td>7976</td><td>8010</td><td>35</td><td>GCGACCTCAGATCAGACGTGGCGACCCGCTGAATT</td><td>1175.0</td><td>-2.0</td><td>0.8</td><td>-2.5</td><td>1.3E-02</td><td>3.8E-02</td><td>BT20/MB231</td><td>CYTO</td></tr>
<tr><td>>hg38_21_+_8433172_8446622_RNA45SN1_45S_50ntFlanks@11566.11585.20|TAGCAGCCGACTTAGAACTG</td><td>>hg38_21_+_8433172_8446622_RNA45SN1_45S_50ntFlanks</td><td>11566</td><td>11585</td><td>20</td><td>TAGCAGCCGACTTAGAACTG</td><td>362.7</td><td>-2.0</td><td>0.8</td><td>-2.5</td><td>1.3E-02</td><td>3.8E-02</td><td>BT20/MB231</td><td>CYTO</td></tr>
<tr><td>>hg38_21_+_8433172_8446622_RNA45SN1_45S_50ntFlanks@5086.5121.36|AACGAGACTCTGGCATGCTAACTAGTTACGCGACCC</td><td>>hg38_21_+_8433172_8446622_RNA45SN1_45S_50ntFlanks</td><td>5086</td><td>5121</td><td>36</td><td>AACGAGACTCTGGCATGCTAACTAGTTACGCGACCC</td><td>87.7</td><td>-4.0</td><td>1.9</td><td>-2.2</td><td>3.1E-02</td><td>3.8E-02</td><td>MB231/MB468</td><td>MP</td></tr>
<tr><td>>hg38_21_+_8433172_8446622_RNA45SN1_45S_50ntFlanks@10107.10131.25|CGTGTGGTGTGCGTCGGAGGGCGGC</td><td>>hg38_21_+_8433172_8446622_RNA45SN1_45S_50ntFlanks</td><td>10107</td><td>10131</td><td>25</td><td>CGTGTGGTGTGCGTCGGAGGGCGGC</td><td>141.1</td><td>3.4</td><td>1.5</td><td>2.3</td><td>2.3E-02</td><td>3.8E-02</td><td>BT20/MB468</td><td>MP</td></tr>
<tr><td>>hg38_21_+_8433172_8446622_RNA45SN1_45S_50ntFlanks@6721.6750.30|AATTGCAGGACACATTGATCATCGACACTT</td><td>>hg38_21_+_8433172_8446622_RNA45SN1_45S_50ntFlanks</td><td>6721</td><td>6750</td><td>30</td><td>AATTGCAGGACACATTGATCATCGACACTT</td><td>203.4</td><td>-3.4</td><td>1.6</td><td>-2.1</td><td>3.3E-02</td><td>3.8E-02</td><td>BT20/MB468</td><td>CYTO</td></tr>
<tr><td>>hg38_21_+_8433172_8446622_RNA45SN1_45S_50ntFlanks@7981.8005.25|CTCAGATCAGACGTGGCGACCCGCT</td><td>>hg38_21_+_8433172_8446622_RNA45SN1_45S_50ntFlanks</td><td>7981</td><td>8005</td><td>25</td><td>CTCAGATCAGACGTGGCGACCCGCT</td><td>119.9</td><td>-2.3</td><td>1.1</td><td>-2.1</td><td>3.3E-02</td><td>3.8E-02</td><td>BT20/MB468</td><td>CYTO</td></tr>
<tr><td>>hg38_21_+_8433172_8446622_RNA45SN1_45S_50ntFlanks@4243.4271.29|CTTTCGAGGCCCTGTAATTGGAATGAGTC</td><td>>hg38_21_+_8433172_8446622_RNA45SN1_45S_50ntFlanks</td><td>4243</td><td>4271</td><td>29</td><td>CTTTCGAGGCCCTGTAATTGGAATGAGTC</td><td>2211.0</td><td>4.5</td><td>2.0</td><td>2.3</td><td>2.3E-02</td><td>3.8E-02</td><td>BT20/MB468</td><td>MP</td></tr>
<tr><td>>MI0000077|hsa-mir-21&WithFlank&17|+|59841260|59841343@14.35.22&[MIMAT0000076&hsa-miR-21-5p&offsets|0|0;m-1&17|+|59841273|59841294&offsets|0|0]|TAGCTTATCAGACTGATGTTGA</td><td>>MI0000077|hsa-mir-21&WithFlank&17|+|59841260|59841343</td><td>14</td><td>35</td><td>22</td><td>TAGCTTATCAGACTGATGTTGA</td><td>338971.2</td><td>2.9</td><td>1.3</td><td>2.3</td><td>2.3E-02</td><td>3.8E-02</td><td>BT20/MB468</td><td>MP</td></tr>
<tr><td>>hg38_MT_+_1621_3279_RNR2_16S_50ntFlanks@1370.1392.23|ATGTTGGATCAGGACATCCCGAT</td><td>>hg38_MT_+_1621_3279_RNR2_16S_50ntFlanks</td><td>1370</td><td>1392</td><td>23</td><td>ATGTTGGATCAGGACATCCCGAT</td><td>115.2</td><td>2.9</td><td>1.4</td><td>2.2</td><td>3.1E-02</td><td>3.8E-02</td><td>MB231/MB468</td><td>MP</td></tr>
<tr><td>>hg38_1_-_228634819_228635039_RNA5S12_5S_50ntFlanks@51.91.41|GTCTACGGCCATACCACCCTGAACGCGCCCGATCTCGTCTG</td><td>>hg38_1_-_228634819_228635039_RNA5S12_5S_50ntFlanks</td><td>51</td><td>91</td><td>41</td><td>GTCTACGGCCATACCACCCTGAACGCGCCCGATCTCGTCTG</td><td>3461.8</td><td>2.6</td><td>1.0</td><td>2.6</td><td>1.0E-02</td><td>3.8E-02</td><td>BT20/MB231</td><td>MP</td></tr>
<tr><td>>MI0000465|hsa-mir-191&WithFlank&3|-|49020612|49020715@22.42.21(+1C)&[MIMAT0000440&hsa-miR-191-5p&offsets|0|-2(+1C);m-48&3|-|49020672|49020694&offsets|0|-2(+1C)]|CAACGGAATCCCAAAAGCAGCC</td><td>>MI0000465|hsa-mir-191&WithFlank&3|-|49020612|49020715</td><td>22</td><td>42</td><td>21(+1C)</td><td>CAACGGAATCCCAAAAGCAGCC</td><td>663.2</td><td>2.6</td><td>1.1</td><td>2.3</td><td>2.3E-02</td><td>3.8E-02</td><td>BT20/MB468</td><td>MP</td></tr>
<tr><td>>hg38_21_+_8433172_8446622_RNA45SN1_45S_50ntFlanks@3776.3798.23|CCGGTACAGTGAAACTGCGAATG</td><td>>hg38_21_+_8433172_8446622_RNA45SN1_45S_50ntFlanks</td><td>3776</td><td>3798</td><td>23</td><td>CCGGTACAGTGAAACTGCGAATG</td><td>106.7</td><td>-3.9</td><td>1.7</td><td>-2.3</td><td>2.3E-02</td><td>3.8E-02</td><td>BT20/MB468</td><td>MP</td></tr>
<tr><td>>hg38_21_+_8433172_8446622_RNA45SN1_45S_50ntFlanks@6707.6734.28|CGAGAATTAATGTGAATTGCAGGACACA</td><td>>hg38_21_+_8433172_8446622_RNA45SN1_45S_50ntFlanks</td><td>6707</td><td>6734</td><td>28</td><td>CGAGAATTAATGTGAATTGCAGGACACA</td><td>64.9</td><td>-3.9</td><td>1.7</td><td>-2.3</td><td>2.3E-02</td><td>3.8E-02</td><td>BT20/MB468</td><td>MP</td></tr>
<tr><td>>hg38_21_+_8433172_8446622_RNA45SN1_45S_50ntFlanks@11770.11794.25|AATTAGTGACGCGCATGAATGGATG</td><td>>hg38_21_+_8433172_8446622_RNA45SN1_45S_50ntFlanks</td><td>11770</td><td>11794</td><td>25</td><td>AATTAGTGACGCGCATGAATGGATG</td><td>633.2</td><td>-1.3</td><td>0.6</td><td>-2.1</td><td>3.4E-02</td><td>3.8E-02</td><td>MB231/MB468</td><td>NUC</td></tr>
<tr><td>>MI0000088|hsa-mir-30a&WithFlank&6|-|71403545|71403627@12.33.22&[MIMAT0000087&hsa-miR-30a-5p&offsets|0|0;m-15&6|-|71403595|71403616&offsets|0|0]|TGTAAACATCCTCGACTGGAAG</td><td>>MI0000088|hsa-mir-30a&WithFlank&6|-|71403545|71403627</td><td>12</td><td>33</td><td>22</td><td>TGTAAACATCCTCGACTGGAAG</td><td>18730.3</td><td>-2.5</td><td>1.0</td><td>-2.6</td><td>1.0E-02</td><td>3.8E-02</td><td>BT20/MB231</td><td>MP</td></tr>
<tr><td>>hg38_21_+_8433172_8446622_RNA45SN1_45S_50ntFlanks@11572.11599.28|CCGACTTAGAACTGGTGCGGACCAGGGG</td><td>>hg38_21_+_8433172_8446622_RNA45SN1_45S_50ntFlanks</td><td>11572</td><td>11599</td><td>28</td><td>CCGACTTAGAACTGGTGCGGACCAGGGG</td><td>232.8</td><td>4.3</td><td>1.7</td><td>2.6</td><td>1.0E-02</td><td>3.8E-02</td><td>BT20/MB231</td><td>MP</td></tr>
<tr><td>>hg38_21_+_8433172_8446622_RNA45SN1_45S_50ntFlanks@6660.6680.21|GCGGTGGATCACTCGGCTCGT</td><td>>hg38_21_+_8433172_8446622_RNA45SN1_45S_50ntFlanks</td><td>6660</td><td>6680</td><td>21</td><td>GCGGTGGATCACTCGGCTCGT</td><td>84.7</td><td>4.2</td><td>1.6</td><td>2.6</td><td>1.0E-02</td><td>3.8E-02</td><td>BT20/MB231</td><td>MP</td></tr>
<tr><td>>hg38_21_+_8433172_8446622_RNA45SN1_45S_50ntFlanks@8336.8375.40|AAGTACCGTAAGGGAAAGTTGAAAAGAACTTTGAAGAGAG</td><td>>hg38_21_+_8433172_8446622_RNA45SN1_45S_50ntFlanks</td><td>8336</td><td>8375</td><td>40</td><td>AAGTACCGTAAGGGAAAGTTGAAAAGAACTTTGAAGAGAG</td><td>1335.7</td><td>4.6</td><td>1.8</td><td>2.6</td><td>1.0E-02</td><td>3.8E-02</td><td>BT20/MB231</td><td>MP</td></tr>
<tr><td>>hg38_21_+_8433172_8446622_RNA45SN1_45S_50ntFlanks@9954.9982.29|TGGAAGTCGGAATCCGCTAAGGAGTGTGT</td><td>>hg38_21_+_8433172_8446622_RNA45SN1_45S_50ntFlanks</td><td>9954</td><td>9982</td><td>29</td><td>TGGAAGTCGGAATCCGCTAAGGAGTGTGT</td><td>118.7</td><td>4.4</td><td>1.7</td><td>2.6</td><td>1.0E-02</td><td>3.8E-02</td><td>BT20/MB231</td><td>MP</td></tr>
<tr><td>>MI0003581|hsa-mir-574&WithFlank&4|+|38868026|38868133@67.86.20(+1U)&[MIMAT0003239&hsa-miR-574-3p&offsets|0|-2(+1U);m-149&4|+|38868092|38868113&offsets|0|-2(+1U)]|CACGCTCATGCACACACCCAT</td><td>>MI0003581|hsa-mir-574&WithFlank&4|+|38868026|38868133</td><td>67</td><td>86</td><td>20(+1U)</td><td>CACGCTCATGCACACACCCAT</td><td>158.3</td><td>-3.8</td><td>1.5</td><td>-2.6</td><td>1.0E-02</td><td>3.8E-02</td><td>BT20/MB231</td><td>MP</td></tr>
<tr><td>>hg38_21_+_8433172_8446622_RNA45SN1_45S_50ntFlanks@476.504.29|TTCGTGATCGATGTGGTGACGTCGTGCTC</td><td>>hg38_21_+_8433172_8446622_RNA45SN1_45S_50ntFlanks</td><td>476</td><td>504</td><td>29</td><td>TTCGTGATCGATGTGGTGACGTCGTGCTC</td><td>134.5</td><td>3.6</td><td>1.7</td><td>2.2</td><td>3.1E-02</td><td>3.8E-02</td><td>MB231/MB468</td><td>MP</td></tr>
<tr><td>>hg38_MT_+_1621_3279_RNR2_16S_50ntFlanks@1188.1215.28|TCGGTTGGGGCGACCTCGGAGCAGAACC</td><td>>hg38_MT_+_1621_3279_RNR2_16S_50ntFlanks</td><td>1188</td><td>1215</td><td>28</td><td>TCGGTTGGGGCGACCTCGGAGCAGAACC</td><td>67.7</td><td>2.7</td><td>1.3</td><td>2.2</td><td>3.1E-02</td><td>3.8E-02</td><td>MB231/MB468</td><td>MP</td></tr>
<tr><td>>hg38_21_+_8433172_8446622_RNA45SN1_45S_50ntFlanks@4980.5001.22|ACACGGACAGGATTGACAGATT</td><td>>hg38_21_+_8433172_8446622_RNA45SN1_45S_50ntFlanks</td><td>4980</td><td>5001</td><td>22</td><td>ACACGGACAGGATTGACAGATT</td><td>204.4</td><td>3.2</td><td>1.2</td><td>2.6</td><td>1.0E-02</td><td>3.8E-02</td><td>BT20/MB231</td><td>MP</td></tr>
<tr><td>>trnaMT_SerGCT_MT_+_12207_12265@14.47.34|AGAACTGCTAACTCATGCCCCCATGTCTAACAAC</td><td>>trnaMT_SerGCT_MT_+_12207_12265</td><td>14</td><td>47</td><td>34</td><td>AGAACTGCTAACTCATGCCCCCATGTCTAACAAC</td><td>81.4</td><td>2.8</td><td>1.3</td><td>2.2</td><td>3.1E-02</td><td>3.8E-02</td><td>MB231/MB468</td><td>MP</td></tr>
<tr><td>>hg38_21_+_8433172_8446622_RNA45SN1_45S_50ntFlanks@10808.10844.37|CAATGTAGGTAAGGGAAGTCGGCAAGCCGGATCCGTA</td><td>>hg38_21_+_8433172_8446622_RNA45SN1_45S_50ntFlanks</td><td>10808</td><td>10844</td><td>37</td><td>CAATGTAGGTAAGGGAAGTCGGCAAGCCGGATCCGTA</td><td>85.0</td><td>-3.3</td><td>1.5</td><td>-2.3</td><td>2.3E-02</td><td>3.8E-02</td><td>BT20/MB468</td><td>MP</td></tr>
<tr><td>>trnaMT_LysTTT_MT_+_8295_8364@37.73.37,trnalookalike7_LysTTT_1_+_568844_568913@37.73.37|AAAGATTAAGAGAACCAACACCTCTTTACAGTGACCA</td><td>>trnaMT_LysTTT_MT_+_8295_8364</td><td>37</td><td>73</td><td>37</td><td>AAAGATTAAGAGAACCAACACCTCTTTACAGTGACCA</td><td>151.7</td><td>2.5</td><td>1.2</td><td>2.1</td><td>3.5E-02</td><td>3.8E-02</td><td>MB231/MB468</td><td>MITO</td></tr>
<tr><td>>hg38_21_+_8433172_8446622_RNA45SN1_45S_50ntFlanks@6651.6680.30|CGACTCTTAGCGGTGGATCACTCGGCTCGT</td><td>>hg38_21_+_8433172_8446622_RNA45SN1_45S_50ntFlanks</td><td>6651</td><td>6680</td><td>30</td><td>CGACTCTTAGCGGTGGATCACTCGGCTCGT</td><td>4553.6</td><td>3.3</td><td>1.3</td><td>2.6</td><td>1.0E-02</td><td>3.8E-02</td><td>BT20/MB231</td><td>MP</td></tr>
<tr><td>>MI0000084|hsa-mir-26b&WithFlank&2|+|218402640|218402728@18.38.21(+1A)&[MIMAT0000083&hsa-miR-26b-5p&offsets|0|0(+1A);m-53&2|+|218402657|218402678&offsets|0|-1(+1A)]|TTCAAGTAATTCAGGATAGGTA</td><td>>MI0000084|hsa-mir-26b&WithFlank&2|+|218402640|218402728</td><td>18</td><td>38</td><td>21(+1A)</td><td>TTCAAGTAATTCAGGATAGGTA</td><td>209.3</td><td>3.2</td><td>1.5</td><td>2.2</td><td>3.1E-02</td><td>3.8E-02</td><td>MB231/MB468</td><td>MP</td></tr>
<tr><td>>MI0000081|hsa-mir-24-2&WithFlank&19|-|13836281|13836365@56.77.22;MI0000080|hsa-mir-24-1&WithFlank&9|+|95086015|95086094@50.71.22&[MIMAT0000080&hsa-miR-24-3p&offsets|0|0;m-36&19|-|13836288|13836310&offsets|0|-1];[MIMAT0000080_1&hsa-miR-24-3p&offsets|0|0;m-35&9|+|95086064|95086086&offsets|0|-1]|TGGCTCAGTTCAGCAGGAACAG</td><td>>MI0000081|hsa-mir-24-2&WithFlank&19|-|13836281|13836365</td><td>56</td><td>77</td><td>22</td><td>TGGCTCAGTTCAGCAGGAACAG</td><td>3909.9</td><td>1.0</td><td>0.5</td><td>2.1</td><td>3.4E-02</td><td>3.8E-02</td><td>BT20/MB468</td><td>NUC</td></tr>
<tr><td>>hg38_21_+_8433172_8446622_RNA45SN1_45S_50ntFlanks@12936.12956.21|AAACCATTCGTAGACGACCTG</td><td>>hg38_21_+_8433172_8446622_RNA45SN1_45S_50ntFlanks</td><td>12936</td><td>12956</td><td>21</td><td>AAACCATTCGTAGACGACCTG</td><td>3102.2</td><td>-3.7</td><td>1.6</td><td>-2.3</td><td>2.3E-02</td><td>3.8E-02</td><td>BT20/MB468</td><td>MP</td></tr>
<tr><td>>hg38_MT_+_598_1651_RNR1_12S_50ntFlanks@428.468.41|AAAGTGGCTTTAACATATCTGAACACACAATAGCTAAGACC</td><td>>hg38_MT_+_598_1651_RNR1_12S_50ntFlanks</td><td>428</td><td>468</td><td>41</td><td>AAAGTGGCTTTAACATATCTGAACACACAATAGCTAAGACC</td><td>257.8</td><td>-4.2</td><td>1.7</td><td>-2.5</td><td>1.4E-02</td><td>3.8E-02</td><td>BT20/MB231</td><td>CYTO</td></tr>
<tr><td>>hg38_MT_+_1621_3279_RNR2_16S_50ntFlanks@999.1026.28|AGGGACCTGTATGAATGGCTCCACGAGG</td><td>>hg38_MT_+_1621_3279_RNR2_16S_50ntFlanks</td><td>999</td><td>1026</td><td>28</td><td>AGGGACCTGTATGAATGGCTCCACGAGG</td><td>52.4</td><td>3.6</td><td>1.7</td><td>2.2</td><td>3.1E-02</td><td>3.8E-02</td><td>MB231/MB468</td><td>MP</td></tr>
<tr><td>>MI0000815|hsa-mir-339&WithFlank&7|-|1022927|1023032@56.77.22&[MIMAT0004702&hsa-miR-339-3p&offsets|0|-1;m-235&7|-|1022957|1022977&offsets|0|+1]|TGAGCGCCTCGACGACAGAGCC</td><td>>MI0000815|hsa-mir-339&WithFlank&7|-|1022927|1023032</td><td>56</td><td>77</td><td>22</td><td>TGAGCGCCTCGACGACAGAGCC</td><td>112.2</td><td>3.7</td><td>1.7</td><td>2.2</td><td>3.2E-02</td><td>3.8E-02</td><td>MB231/MB468</td><td>MP</td></tr>
<tr><td>>hg38_21_+_8433172_8446622_RNA45SN1_45S_50ntFlanks@12406.12433.28|TTTTTGATCCTTCGATGTCGGCTCTTCC</td><td>>hg38_21_+_8433172_8446622_RNA45SN1_45S_50ntFlanks</td><td>12406</td><td>12433</td><td>28</td><td>TTTTTGATCCTTCGATGTCGGCTCTTCC</td><td>618.5</td><td>-2.6</td><td>1.1</td><td>-2.5</td><td>1.4E-02</td><td>3.8E-02</td><td>BT20/MB231</td><td>CYTO</td></tr>
<tr><td>>hg38_1_-_228634819_228635039_RNA5S12_5S_50ntFlanks@135.169.35|GGGAGACCGCCTGGGAATACCGGGTGCTGTAGGCT</td><td>>hg38_1_-_228634819_228635039_RNA5S12_5S_50ntFlanks</td><td>135</td><td>169</td><td>35</td><td>GGGAGACCGCCTGGGAATACCGGGTGCTGTAGGCT</td><td>1601.5</td><td>2.3</td><td>0.9</td><td>2.5</td><td>1.3E-02</td><td>3.8E-02</td><td>BT20/MB231</td><td>MITO</td></tr>
</table>

<table>
<tr><td>>MI0000067|hsa-let-7f-1&WithFlank&9|+|94176341|94176439@13.33.21;MI0000068|hsa-let-7f-2&WithFlank&X|-|53557186|53557280@14.34.21&[MIMAT0000067&hsa-let-7f-5p&offsets|0|-1;m-20&9|+|94176353|94176374&offsets|0|-1];[MIMAT0000067_1&hsa-let-7f-5p&offsets|0|-1;m-19&X|-|53557246|53557267&offsets|0|-1]|TGAGGTAGTAGATTGTATAGT</td><td>>MI0000067|hsa-let-7f-1&WithFlank&9|+|94176341|94176439</td><td>13</td><td>33</td><td>21</td><td>TGAGGTAGTAGATTGTATAGT</td><td>1636.8</td><td>2.1</td><td>0.9</td><td>2.3</td><td>2.4E-02</td><td>3.8E-02</td><td>BT20/MB468</td><td>MP</td></tr>
<tr><td>>hg38_21_+_8433172_8446622_RNA45SN1_45S_50ntFlanks@11921.11940.20|TGAGAGGTGTAGAATAAGTG</td><td>>hg38_21_+_8433172_8446622_RNA45SN1_45S_50ntFlanks</td><td>11921</td><td>11940</td><td>20</td><td>TGAGAGGTGTAGAATAAGTG</td><td>56.5</td><td>4.0</td><td>1.8</td><td>2.3</td><td>2.4E-02</td><td>3.8E-02</td><td>BT20/MB468</td><td>MP</td></tr>
<tr><td>>am_trna119_LysCTT_1_-_145395522_145395594@14.34.21,trna11_LysCTT_5_-_180648979_180649051@14.34.21,trna128_LysCTT_1_-_55423542_55423614@14.34.21,trna13_LysCTT_6_+_26556774_26556846@14.34.21,trna6_LysCTT_19_-_52425396_52425468@14.34.21,trna7_LysCTT_16_+_3225692_3225764@14.34.21,trna9_LysCTT_5_+_180634755_180634827@14.34.21|AGTCGGTAGAGCATGAGACTC</td><td>>am_trna119_LysCTT_1_-_145395522_145395594</td><td>14</td><td>34</td><td>21</td><td>AGTCGGTAGAGCATGAGACTC</td><td>218.5</td><td>-4.0</td><td>1.6</td><td>-2.6</td><td>1.0E-02</td><td>3.8E-02</td><td>BT20/MB231</td><td>MP</td></tr>
<tr><td>>hg38_21_+_8433172_8446622_RNA45SN1_45S_50ntFlanks@10794.10825.32|CTGGCATGTTGGAACAATGTAGGTAAGGGAAG</td><td>>hg38_21_+_8433172_8446622_RNA45SN1_45S_50ntFlanks</td><td>10794</td><td>10825</td><td>32</td><td>CTGGCATGTTGGAACAATGTAGGTAAGGGAAG</td><td>299.8</td><td>-2.9</td><td>1.3</td><td>-2.1</td><td>3.3E-02</td><td>3.8E-02</td><td>BT20/MB468</td><td>CYTO</td></tr>
<tr><td>>hg38_21_+_8433172_8446622_RNA45SN1_45S_50ntFlanks@8552.8579.28|CTCCCCGGAGGGGGCGGGCTCCGGCGGG</td><td>>hg38_21_+_8433172_8446622_RNA45SN1_45S_50ntFlanks</td><td>8552</td><td>8579</td><td>28</td><td>CTCCCCGGAGGGGGCGGGCTCCGGCGGG</td><td>155.6</td><td>-3.3</td><td>1.5</td><td>-2.1</td><td>3.3E-02</td><td>3.8E-02</td><td>BT20/MB468</td><td>CYTO</td></tr>
<tr><td>>MI0000650|hsa-mir-200c&WithFlank&12|+|6963693|6963772@11.32.22&[MIMAT0004657&hsa-miR-200c-5p&offsets|0|0;m-483&12|+|6963703|6963724&offsets|0|0]|CGTCTTACCCAGCAGTGTTTGG</td><td>>MI0000650|hsa-mir-200c&WithFlank&12|+|6963693|6963772</td><td>11</td><td>32</td><td>22</td><td>CGTCTTACCCAGCAGTGTTTGG</td><td>58.4</td><td>5.1</td><td>2.2</td><td>2.3</td><td>2.4E-02</td><td>3.8E-02</td><td>BT20/MB468</td><td>MP</td></tr>
<tr><td>>hg38_21_+_8433172_8446622_RNA45SN1_45S_50ntFlanks@11561.11601.41|GCGCCTAGCAGCCGACTTAGAACTGGTGCGGACCAGGGGAA</td><td>>hg38_21_+_8433172_8446622_RNA45SN1_45S_50ntFlanks</td><td>11561</td><td>11601</td><td>41</td><td>GCGCCTAGCAGCCGACTTAGAACTGGTGCGGACCAGGGGAA</td><td>1252.3</td><td>-4.7</td><td>2.1</td><td>-2.3</td><td>2.4E-02</td><td>3.8E-02</td><td>BT20/MB468</td><td>MP</td></tr>
<tr><td>>hg38_21_+_8433172_8446622_RNA45SN1_45S_50ntFlanks@475.495.21|CTTCGTGATCGATGTGGTGAC</td><td>>hg38_21_+_8433172_8446622_RNA45SN1_45S_50ntFlanks</td><td>475</td><td>495</td><td>21</td><td>CTTCGTGATCGATGTGGTGAC</td><td>262.6</td><td>4.2</td><td>1.9</td><td>2.3</td><td>2.4E-02</td><td>3.8E-02</td><td>BT20/MB468</td><td>MP</td></tr>
<tr><td>>hg38_MT_+_598_1651_RNR1_12S_50ntFlanks@821.840.20|GGTGGATTTAGCAGTAAACT</td><td>>hg38_MT_+_598_1651_RNR1_12S_50ntFlanks</td><td>821</td><td>840</td><td>20</td><td>GGTGGATTTAGCAGTAAACT</td><td>96.7</td><td>5.9</td><td>2.6</td><td>2.3</td><td>2.4E-02</td><td>3.8E-02</td><td>BT20/MB468</td><td>MP</td></tr>
<tr><td>>MI0000542|hsa-mir-320a&WithFlank&8|-|22244960|22245043@48.70.23(+1G)&[MIMAT0000510&hsa-miR-320a-3p&offsets|0|+1(+1G);m-60&8|-|22244975|22244996&offsets|0|+1(+1G)]|AAAAGCTGGGTTGAGAGGGCGAAG</td><td>>MI0000542|hsa-mir-320a&WithFlank&8|-|22244960|22245043</td><td>48</td><td>70</td><td>23(+1G)</td><td>AAAAGCTGGGTTGAGAGGGCGAAG</td><td>1111.4</td><td>2.5</td><td>1.0</td><td>2.6</td><td>1.0E-02</td><td>3.8E-02</td><td>BT20/MB231</td><td>MP</td></tr>
<tr><td>>hg38_21_+_8433172_8446622_RNA45SN1_45S_50ntFlanks@12488.12524.37|AACGTGAGCTGGGTTTAGACCGTCGTGAGACAGGTTA</td><td>>hg38_21_+_8433172_8446622_RNA45SN1_45S_50ntFlanks</td><td>12488</td><td>12524</td><td>37</td><td>AACGTGAGCTGGGTTTAGACCGTCGTGAGACAGGTTA</td><td>170.7</td><td>-3.3</td><td>1.6</td><td>-2.1</td><td>3.4E-02</td><td>3.8E-02</td><td>BT20/MB468</td><td>CYTO</td></tr>
<tr><td>>hg38_21_+_8433172_8446622_RNA45SN1_45S_50ntFlanks@5045.5063.19|CTTAGTTGGTGGAGCGATT</td><td>>hg38_21_+_8433172_8446622_RNA45SN1_45S_50ntFlanks</td><td>5045</td><td>5063</td><td>19</td><td>CTTAGTTGGTGGAGCGATT</td><td>320.9</td><td>3.5</td><td>1.4</td><td>2.6</td><td>1.0E-02</td><td>3.8E-02</td><td>BT20/MB231</td><td>MP</td></tr>
<tr><td>>MI0000077|hsa-mir-21&WithFlank&17|+|59841260|59841343@13.35.23&[MIMAT0000076&hsa-miR-21-5p&offsets|-1|0;m-1&17|+|59841273|59841294&offsets|-1|0]|GTAGCTTATCAGACTGATGTTGA</td><td>>MI0000077|hsa-mir-21&WithFlank&17|+|59841260|59841343</td><td>13</td><td>35</td><td>23</td><td>GTAGCTTATCAGACTGATGTTGA</td><td>829.3</td><td>2.4</td><td>1.0</td><td>2.5</td><td>1.4E-02</td><td>3.8E-02</td><td>BT20/MB231</td><td>CYTO</td></tr>
<tr><td>>hg38_21_+_8433172_8446622_RNA45SN1_45S_50ntFlanks@12259.12282.24|CTTGATCTTGATTTTCAGTACGAA</td><td>>hg38_21_+_8433172_8446622_RNA45SN1_45S_50ntFlanks</td><td>12259</td><td>12282</td><td>24</td><td>CTTGATCTTGATTTTCAGTACGAA</td><td>687.5</td><td>-3.5</td><td>1.5</td><td>-2.3</td><td>2.4E-02</td><td>3.8E-02</td><td>BT20/MB468</td><td>MP</td></tr>
<tr><td>>trna10_AlaCGC_6_+_26553731_26553802@1.18.18,trna119_AlaCGC_6_-_28641613_28641684@1.18.18,trna120_AlaAGC_6_-_28626014_28626085@1.18.18,trna13_AlaCGC_2_+_157257281_157257352@1.18.18,trna13_AlaTGC_12_-_125406301_125406372@1.18.18,trna66_AlaTGC_6_+_28611222_28611293@1.18.18,trna8_AlaTGC_12_+_125424512_125424583@1.18.18,trna8_AlaTGC_5_+_180633868_180633939@1.18.18|GGGGATGTAGCTCAGTGG</td><td>>trna10_AlaCGC_6_+_26553731_26553802</td><td>1</td><td>18</td><td>18</td><td>GGGGATGTAGCTCAGTGG</td><td>172.6</td><td>2.9</td><td>1.4</td><td>2.1</td><td>3.5E-02</td><td>3.8E-02</td><td>MB231/MB468</td><td>MITO</td></tr>
<tr><td>>hg38_21_+_8433172_8446622_RNA45SN1_45S_50ntFlanks@4686.4707.22|GACGGACCAGAGCGAAAGCATT</td><td>>hg38_21_+_8433172_8446622_RNA45SN1_45S_50ntFlanks</td><td>4686</td><td>4707</td><td>22</td><td>GACGGACCAGAGCGAAAGCATT</td><td>779.3</td><td>4.2</td><td>1.9</td><td>2.3</td><td>2.4E-02</td><td>3.8E-02</td><td>BT20/MB468</td><td>MP</td></tr>
<tr><td>>MI0000542|hsa-mir-320a&WithFlank&8|-|22244960|22245043@48.71.24(+1U)&[MIMAT0000510&hsa-miR-320a-3p&offsets|0|+2(+1U);m-60&8|-|22244975|22244996&offsets|0|+2(+1U)]|AAAAGCTGGGTTGAGAGGGCGAAAT</td><td>>MI0000542|hsa-mir-320a&WithFlank&8|-|22244960|22245043</td><td>48</td><td>71</td><td>24(+1U)</td><td>AAAAGCTGGGTTGAGAGGGCGAAAT</td><td>319.3</td><td>1.9</td><td>0.8</td><td>2.5</td><td>1.3E-02</td><td>3.8E-02</td><td>BT20/MB231</td><td>MITO</td></tr>
<tr><td>>hg38_21_+_8433172_8446622_RNA45SN1_45S_50ntFlanks@11667.11703.37|CTGCCCAGTGCTCTGAATGTCAAAGTGAAGAAATTCA</td><td>>hg38_21_+_8433172_8446622_RNA45SN1_45S_50ntFlanks</td><td>11667</td><td>11703</td><td>37</td><td>CTGCCCAGTGCTCTGAATGTCAAAGTGAAGAAATTCA</td><td>54.3</td><td>4.7</td><td>2.1</td><td>2.3</td><td>2.4E-02</td><td>3.8E-02</td><td>BT20/MB468</td><td>MP</td></tr>
<tr><td>>hg38_21_+_8433172_8446622_RNA45SN1_45S_50ntFlanks@11665.11685.21|TTCTGCCCAGTGCTCTGAATG</td><td>>hg38_21_+_8433172_8446622_RNA45SN1_45S_50ntFlanks</td><td>11665</td><td>11685</td><td>21</td><td>TTCTGCCCAGTGCTCTGAATG</td><td>596.7</td><td>-2.6</td><td>1.2</td><td>-2.1</td><td>3.4E-02</td><td>3.8E-02</td><td>BT20/MB468</td><td>NUC</td></tr>
<tr><td>>hg38_21_+_8433172_8446622_RNA45SN1_45S_50ntFlanks@6660.6696.37|GCGGTGGATCACTCGGCTCGTGCGTCGATGAAGAACG</td><td>>hg38_21_+_8433172_8446622_RNA45SN1_45S_50ntFlanks</td><td>6660</td><td>6696</td><td>37</td><td>GCGGTGGATCACTCGGCTCGTGCGTCGATGAAGAACG</td><td>689.9</td><td>2.4</td><td>1.1</td><td>2.3</td><td>2.4E-02</td><td>3.8E-02</td><td>BT20/MB468</td><td>MP</td></tr>
<tr><td>>hg38_21_+_8433172_8446622_RNA45SN1_45S_50ntFlanks@9794.9816.23|CGTGGAGCCGGGCGTGGAATGCG</td><td>>hg38_21_+_8433172_8446622_RNA45SN1_45S_50ntFlanks</td><td>9794</td><td>9816</td><td>23</td><td>CGTGGAGCCGGGCGTGGAATGCG</td><td>219.1</td><td>2.8</td><td>1.2</td><td>2.3</td><td>2.4E-02</td><td>3.8E-02</td><td>BT20/MB468</td><td>MP</td></tr>
<tr><td>>hg38_21_+_8433172_8446622_RNA45SN1_45S_50ntFlanks@11906.11933.28|CACGGTGAAGAGACATGAGAGGTGTAGA</td><td>>hg38_21_+_8433172_8446622_RNA45SN1_45S_50ntFlanks</td><td>11906</td><td>11933</td><td>28</td><td>CACGGTGAAGAGACATGAGAGGTGTAGA</td><td>42.7</td><td>4.9</td><td>2.1</td><td>2.3</td><td>2.4E-02</td><td>3.8E-02</td><td>BT20/MB468</td><td>MP</td></tr>
<tr><td>>am_trna128_GlyGCC_6_-_27870686_27870756@7.35.29,trna18_GlyGCC_16_+_70822597_70822667@7.35.29,trna19_GlyGCC_16_+_70823410_70823480@7.35.29,trna19_GlyGCC_2_-_157257659_157257729@7.35.29,trna24_GlyGCC_16_-_70812942_70813012@7.35.29,trna25_GlyGCC_16_-_70812114_70812184@7.35.29,trna2_GlyGCC_21_-_18827107_18827177@7.35.29,trna35_GlyGCC_1_+_161413094_161413164@7.35.29,trna37_GlyGCC_1_+_161420467_161420537@7.35.29,trna39_GlyGCC_1_+_161427898_161427968@7.35.29,trna41_GlyGCC_1_+_161435258_161435328@7.35.29,trna5_GlyGCC_17_+_8029064_8029134@7.35.29,trna68_GlyGCC_1_-_161493637_161493707@7.35.29|GTGGTTCAGTGGTAGAATTCTCGCCTGCC</td><td>>am_trna128_GlyGCC_6_-_27870686_27870756</td><td>7</td><td>35</td><td>29</td><td>GTGGTTCAGTGGTAGAATTCTCGCCTGCC</td><td>79.0</td><td>3.6</td><td>1.7</td><td>2.1</td><td>3.2E-02</td><td>3.8E-02</td><td>MB231/MB468</td><td>MP</td></tr>
<tr><td>>hg38_21_+_8433172_8446622_RNA45SN1_45S_50ntFlanks@6650.6670.21|ACGACTCTTAGCGGTGGATCA</td><td>>hg38_21_+_8433172_8446622_RNA45SN1_45S_50ntFlanks</td><td>6650</td><td>6670</td><td>21</td><td>ACGACTCTTAGCGGTGGATCA</td><td>116.6</td><td>-3.2</td><td>1.4</td><td>-2.3</td><td>2.4E-02</td><td>3.8E-02</td><td>BT20/MB468</td><td>MP</td></tr>
<tr><td>>MI0000103|hsa-mir-101-1&WithFlank&1|-|65058428|65058514@53.72.20(+2C);MI0000739|hsa-mir-101-2&WithFlank&9|+|4850291|4850381@55.74.20(+2C)&[MIMAT0000099&hsa-miR-101-3p&offsets|0|-1(+2C);m-17&1|-|65058443|65058462&offsets|0|0(+2C)];[MIMAT0000099_1&hsa-miR-101-3p&offsets|0|-1(+2C);m-16&9|+|4850345|4850364&offsets|0|0(+2C)]|TACAGTACTGTGATAACTGACC</td><td>>MI0000103|hsa-mir-101-1&WithFlank&1|-|65058428|65058514</td><td>53</td><td>72</td><td>20(+2C)</td><td>TACAGTACTGTGATAACTGACC</td><td>41.1</td><td>4.4</td><td>2.0</td><td>2.1</td><td>3.2E-02</td><td>3.8E-02</td><td>MB231/MB468</td><td>MP</td></tr>
<tr><td>>MI0000077|hsa-mir-21&WithFlank&17|+|59841260|59841343@14.34.21(+2U)&[MIMAT0000076&hsa-miR-21-5p&offsets|0|-1(+2U);m-1&17|+|59841273|59841294&offsets|0|-1(+2U)]|TAGCTTATCAGACTGATGTTGTT</td><td>>MI0000077|hsa-mir-21&WithFlank&17|+|59841260|59841343</td><td>14</td><td>34</td><td>21(+2U)</td><td>TAGCTTATCAGACTGATGTTGTT</td><td>74.5</td><td>3.6</td><td>1.7</td><td>2.1</td><td>3.2E-02</td><td>3.8E-02</td><td>MB231/MB468</td><td>MP</td></tr>
<tr><td>>hg38_21_+_8433172_8446622_RNA45SN1_45S_50ntFlanks@6652.6697.46|GACTCTTAGCGGTGGATCACTCGGCTCGTGCGTCGATGAAGAACGC</td><td>>hg38_21_+_8433172_8446622_RNA45SN1_45S_50ntFlanks</td><td>6652</td><td>6697</td><td>46</td><td>GACTCTTAGCGGTGGATCACTCGGCTCGTGCGTCGATGAAGAACGC</td><td>698.2</td><td>-2.3</td><td>0.9</td><td>-2.5</td><td>1.4E-02</td><td>3.8E-02</td><td>BT20/MB231</td><td>CYTO</td></tr>
<tr><td>>hg38_21_+_8433172_8446622_RNA45SN1_45S_50ntFlanks@11703.11726.24|AATGAAGCGCGGGTAAACGGCGGG</td><td>>hg38_21_+_8433172_8446622_RNA45SN1_45S_50ntFlanks</td><td>11703</td><td>11726</td><td>24</td><td>AATGAAGCGCGGGTAAACGGCGGG</td><td>160.8</td><td>-2.5</td><td>1.1</td><td>-2.2</td><td>3.0E-02</td><td>3.8E-02</td><td>BT20/MB468</td><td>MITO</td></tr>
<tr><td>>hg38_21_+_8433172_8446622_RNA45SN1_45S_50ntFlanks@7993.8011.19|GTGGCGACCCGCTGAATTT</td><td>>hg38_21_+_8433172_8446622_RNA45SN1_45S_50ntFlanks</td><td>7993</td><td>8011</td><td>19</td><td>GTGGCGACCCGCTGAATTT</td><td>281.3</td><td>1.6</td><td>0.6</td><td>2.5</td><td>1.1E-02</td><td>3.8E-02</td><td>BT20/MB231</td><td>NUC</td></tr>
<tr><td>>MI0000285|hsa-mir-205&WithFlank&1|+|209432127|209432248@40.59.20&[MIMAT0000266&hsa-miR-205-5p&offsets|0|-2;m-351&1|+|209432166|209432187&offsets|0|-2]|TCCTTCATTCCACCGGAGTC</td><td>>MI0000285|hsa-mir-205&WithFlank&1|+|209432127|209432248</td><td>40</td><td>59</td><td>20</td><td>TCCTTCATTCCACCGGAGTC</td><td>624.9</td><td>3.2</td><td>1.4</td><td>2.3</td><td>2.4E-02</td><td>3.8E-02</td><td>BT20/MB468</td><td>MP</td></tr>
<tr><td>>hg38_MT_+_598_1651_RNR1_12S_50ntFlanks@427.453.27|GAAAGTGGCTTTAACATATCTGAACAC</td><td>>hg38_MT_+_598_1651_RNR1_12S_50ntFlanks</td><td>427</td><td>453</td><td>27</td><td>GAAAGTGGCTTTAACATATCTGAACAC</td><td>79.7</td><td>2.9</td><td>1.3</td><td>2.1</td><td>3.2E-02</td><td>3.9E-02</td><td>MB231/MB468</td><td>MP</td></tr>
<tr><td>>hg38_21_+_8433172_8446622_RNA45SN1_45S_50ntFlanks@12913.12933.21|ACGTTCGTGGGGAACCTGGCG</td><td>>hg38_21_+_8433172_8446622_RNA45SN1_45S_50ntFlanks</td><td>12913</td><td>12933</td><td>21</td><td>ACGTTCGTGGGGAACCTGGCG</td><td>177.6</td><td>-1.9</td><td>0.9</td><td>-2.2</td><td>3.0E-02</td><td>3.9E-02</td><td>BT20/MB468</td><td>MITO</td></tr>
<tr><td>>hg38_21_+_8433172_8446622_RNA45SN1_45S_50ntFlanks@7982.8012.31|TCAGATCAGACGTGGCGACCCGCTGAATTTA</td><td>>hg38_21_+_8433172_8446622_RNA45SN1_45S_50ntFlanks</td><td>7982</td><td>8012</td><td>31</td><td>TCAGATCAGACGTGGCGACCCGCTGAATTTA</td><td>284.2</td><td>-2.8</td><td>1.3</td><td>-2.2</td><td>3.0E-02</td><td>3.9E-02</td><td>BT20/MB468</td><td>MITO</td></tr>
<tr><td>>hg38_21_+_8433172_8446622_RNA45SN1_45S_50ntFlanks@10819.10839.21|AGGGAAGTCGGCAAGCCGGAT</td><td>>hg38_21_+_8433172_8446622_RNA45SN1_45S_50ntFlanks</td><td>10819</td><td>10839</td><td>21</td><td>AGGGAAGTCGGCAAGCCGGAT</td><td>102.4</td><td>4.4</td><td>1.7</td><td>2.6</td><td>1.0E-02</td><td>3.9E-02</td><td>BT20/MB231</td><td>MP</td></tr>
<tr><td>>hg38_21_+_8433172_8446622_RNA45SN1_45S_50ntFlanks@5475.5516.42|GCCCTGGCGGAGCGCTGAGAAGACGGTCGAACTTGACTATCT</td><td>>hg38_21_+_8433172_8446622_RNA45SN1_45S_50ntFlanks</td><td>5475</td><td>5516</td><td>42</td><td>GCCCTGGCGGAGCGCTGAGAAGACGGTCGAACTTGACTATCT</td><td>290.0</td><td>4.3</td><td>1.7</td><td>2.6</td><td>1.0E-02</td><td>3.9E-02</td><td>BT20/MB231</td><td>MP</td></tr>
<tr><td>>MI0000088|hsa-mir-30a&WithFlank&6|-|71403545|71403627@12.32.21(+1A)&[MIMAT0000087&hsa-miR-30a-5p&offsets|0|-1(+1A);m-15&6|-|71403595|71403616&offsets|0|-1(+1A)]|TGTAAACATCCTCGACTGGAAA</td><td>>MI0000088|hsa-mir-30a&WithFlank&6|-|71403545|71403627</td><td>12</td><td>32</td><td>21(+1A)</td><td>TGTAAACATCCTCGACTGGAAA</td><td>1531.2</td><td>-2.6</td><td>1.0</td><td>-2.6</td><td>1.0E-02</td><td>3.9E-02</td><td>BT20/MB231</td><td>MP</td></tr>
<tr><td>>hg38_21_+_8433172_8446622_RNA45SN1_45S_50ntFlanks@12399.12437.39|ACGTCGCTTTTTGATCCTTCGATGTCGGCTCTTCCTATC</td><td>>hg38_21_+_8433172_8446622_RNA45SN1_45S_50ntFlanks</td><td>12399</td><td>12437</td><td>39</td><td>ACGTCGCTTTTTGATCCTTCGATGTCGGCTCTTCCTATC</td><td>195.5</td><td>-1.5</td><td>0.7</td><td>-2.2</td><td>3.0E-02</td><td>3.9E-02</td><td>BT20/MB468</td><td>MITO</td></tr>
<tr><td>>MI0000440|hsa-mir-27b&WithFlank&9|+|95085439|95085547@67.85.19(+1A)&[MIMAT0000419&hsa-miR-27b-3p&offsets|0|-2(+1A);m-39&9|+|95085505|95085525&offsets|0|-2(+1A)]|TTCACAGTGGCTAAGTTCTA</td><td>>MI0000440|hsa-mir-27b&WithFlank&9|+|95085439|95085547</td><td>67</td><td>85</td><td>19(+1A)</td><td>TTCACAGTGGCTAAGTTCTA</td><td>2039.4</td><td>2.4</td><td>1.0</td><td>2.5</td><td>1.4E-02</td><td>3.9E-02</td><td>BT20/MB231</td><td>CYTO</td></tr>
<tr><td>>hg38_21_+_8433172_8446622_RNA45SN1_45S_50ntFlanks@12686.12703.18|CCAGGCGGAACGATACGG</td><td>>hg38_21_+_8433172_8446622_RNA45SN1_45S_50ntFlanks</td><td>12686</td><td>12703</td><td>18</td><td>CCAGGCGGAACGATACGG</td><td>198.1</td><td>-4.0</td><td>1.8</td><td>-2.3</td><td>2.4E-02</td><td>3.9E-02</td><td>BT20/MB468</td><td>MP</td></tr>
<tr><td>>hg38_21_+_8433172_8446622_RNA45SN1_45S_50ntFlanks@6646.6689.44|TCGTACGACTCTTAGCGGTGGATCACTCGGCTCGTGCGTCGATG</td><td>>hg38_21_+_8433172_8446622_RNA45SN1_45S_50ntFlanks</td><td>6646</td><td>6689</td><td>44</td><td>TCGTACGACTCTTAGCGGTGGATCACTCGGCTCGTGCGTCGATG</td><td>1681.6</td><td>2.2</td><td>0.9</td><td>2.5</td><td>1.4E-02</td><td>3.9E-02</td><td>BT20/MB231</td><td>CYTO</td></tr>
</table>

| | | | | | | | | | | | | | |
|---|---|---|---|---|---|---|---|---|---|---|---|---|---|
| >am_trna119_LysCTT_1_-_145395522_145395594@16.33.18,trna11_LysCTT_5_-_180648979_180649051@16.33.18,trna128_LysCTT_1_-_55423542_55423614@16.33.18,trna13_LysCTT_6_+_26556774_26556846@16.33.18,trna6_LysCTT_19_-_52425396_52425468@16.33.18,trna7_LysCTT_16_+_3225692_3225764@16.33.18,trna9_LysCTT_5_+_180634755_180634827@16.33.18\|TCGGTAGAGCATGAGACT | >am_trna119_LysCTT_1_-_145395522_145395594 | 16 | 33 | 18 | TCGGTAGAGCATGAGACT | 291.6 | -2.6 | 1.0 | -2.5 | 1.4E-02 | 3.9E-02 | BT20/MB231 | CYTO |
| >hg38_21_+_8433172_8446622_RNA45SN1_45S_50ntFlanks@6741.6771.31\|ATCGACACTTCGAACGCACTTGCGGCCCCGG | >hg38_21_+_8433172_8446622_RNA45SN1_45S_50ntFlanks | 6741 | 6771 | 31 | ATCGACACTTCGAACGCACTTGCGGCCCCGG | 1391.2 | -4.3 | 1.9 | -2.3 | 2.4E-02 | 3.9E-02 | BT20/MB468 | MP |
| >hg38_21_+_8433172_8446622_RNA45SN1_45S_50ntFlanks@12395.12421.27\|AGCGACGTCGCTTTTTGATCCTTCGAT | >hg38_21_+_8433172_8446622_RNA45SN1_45S_50ntFlanks | 12395 | 12421 | 27 | AGCGACGTCGCTTTTTGATCCTTCGAT | 174.8 | -1.8 | 0.7 | -2.5 | 1.1E-02 | 3.9E-02 | BT20/MB231 | NUC |
| >trnaMT_TyrGTA_MT_-_5826_5891@33.69.37\|AATCTAAAGACAGGGGTTAGGCCTCTTTTTACCACCA | >trnaMT_TyrGTA_MT_-_5826_5891 | 33 | 69 | 37 | AATCTAAAGACAGGGGTTAGGCCTCTTTTTACCACCA | 178.6 | 4.6 | 2.2 | 2.1 | 3.2E-02 | 3.9E-02 | MB231/MB468 | MP |
| >hg38_21_+_8433172_8446622_RNA45SN1_45S_50ntFlanks@11571.11600.30\|GCCGACTTAGAACTGGTGCGGACCAGGGGA | >hg38_21_+_8433172_8446622_RNA45SN1_45S_50ntFlanks | 11571 | 11600 | 30 | GCCGACTTAGAACTGGTGCGGACCAGGGGA | 855.9 | 3.4 | 1.3 | 2.6 | 1.0E-02 | 3.9E-02 | BT20/MB231 | MP |
| >hg38_21_+_8433172_8446622_RNA45SN1_45S_50ntFlanks@10116.10136.21\|TGCGTCGGAGGGCGGCGGCGG | >hg38_21_+_8433172_8446622_RNA45SN1_45S_50ntFlanks | 10116 | 10136 | 21 | TGCGTCGGAGGGCGGCGGCGG | 475.6 | -3.6 | 1.7 | -2.2 | 3.0E-02 | 3.9E-02 | BT20/MB468 | MITO |
| >hg38_21_+_8433172_8446622_RNA45SN1_45S_50ntFlanks@4297.4317.21\|CATTGGAGGGCAAGTCTGGTG | >hg38_21_+_8433172_8446622_RNA45SN1_45S_50ntFlanks | 4297 | 4317 | 21 | CATTGGAGGGCAAGTCTGGTG | 564.0 | 4.4 | 1.7 | 2.6 | 1.0E-02 | 3.9E-02 | BT20/MB231 | MP |
| >hg38_21_+_8433172_8446622_RNA45SN1_45S_50ntFlanks@9593.9610.18\|GTATAGGGGCGAAAGACT | >hg38_21_+_8433172_8446622_RNA45SN1_45S_50ntFlanks | 9593 | 9610 | 18 | GTATAGGGGCGAAAGACT | 161.1 | 4.3 | 1.7 | 2.6 | 1.0E-02 | 3.9E-02 | BT20/MB231 | MP |
| >hg38_21_+_8433172_8446622_RNA45SN1_45S_50ntFlanks@12456.12496.41\|CCAAGCGTTGGATTGTTCACCCACTAATAGGGAACGTGAGC | >hg38_21_+_8433172_8446622_RNA45SN1_45S_50ntFlanks | 12456 | 12496 | 41 | CCAAGCGTTGGATTGTTCACCCACTAATAGGGAACGTGAGC | 273.8 | -1.8 | 0.7 | -2.5 | 1.4E-02 | 3.9E-02 | BT20/MB231 | CYTO |
| >hg38_MT_+_1621_3279_RNR2_16S_50ntFlanks@883.906.24\|AAAAACATCACCTCTAGCATCACC | >hg38_MT_+_1621_3279_RNR2_16S_50ntFlanks | 883 | 906 | 24 | AAAAACATCACCTCTAGCATCACC | 106.4 | 3.1 | 1.4 | 2.3 | 2.4E-02 | 3.9E-02 | BT20/MB468 | MP |
| >MI0000447\|hsa-mir-128-1&WithFlank&2\|+\|135665391\|135665484@56.74.19;MI0000727\|hsa-mir-128-2&WithFlank&3\|+\|35744470\|35744565@58.76.19&[MIMAT0000424&hsa-miR-128-3p&offsets\|0\|-2;m-108&2\|+\|135665446\|135665466&offsets\|0\|-2];[MIMAT0000424_1&hsa-miR-128-3p&offsets\|0\|-2;m-117&3\|+\|35744527\|35744547&offsets\|0\|-2]\|TCACAGTGAACCGGTCTCT | >MI0000447\|hsa-mir-128-1&WithFlank&2\|+\|135665391\|135665484 | 56 | 74 | 19 | TCACAGTGAACCGGTCTCT | 60.5 | 3.3 | 1.4 | 2.3 | 2.4E-02 | 3.9E-02 | BT20/MB468 | MP |
| >hg38_21_+_8433172_8446622_RNA45SN1_45S_50ntFlanks@12407.12443.37\|TTTTGATCCTTCGATGTCGGCTCTTCCTATCATTGTG | >hg38_21_+_8433172_8446622_RNA45SN1_45S_50ntFlanks | 12407 | 12443 | 37 | TTTTGATCCTTCGATGTCGGCTCTTCCTATCATTGTG | 657.8 | -2.7 | 1.1 | -2.5 | 1.4E-02 | 3.9E-02 | BT20/MB231 | CYTO |
| >hg38_21_+_8433172_8446622_RNA45SN1_45S_50ntFlanks@4723.4761.39\|CATTAATCAAGAACGAAAGTCGGAGGTTCGAAGACGATC | >hg38_21_+_8433172_8446622_RNA45SN1_45S_50ntFlanks | 4723 | 4761 | 39 | CATTAATCAAGAACGAAAGTCGGAGGTTCGAAGACGATC | 353.1 | -1.9 | 0.8 | -2.5 | 1.4E-02 | 3.9E-02 | BT20/MB231 | CYTO |
| >hg38_21_+_8433172_8446622_RNA45SN1_45S_50ntFlanks@12420.12444.25\|ATGTCGGCTCTTCCTATCATTGTGA | >hg38_21_+_8433172_8446622_RNA45SN1_45S_50ntFlanks | 12420 | 12444 | 25 | ATGTCGGCTCTTCCTATCATTGTGA | 1012.7 | -2.1 | 0.9 | -2.5 | 1.4E-02 | 3.9E-02 | BT20/MB231 | CYTO |
| >MI0001448\|hsa-mir-425&WithFlank&3\|-\|49020142\|49020240@20.41.22&[MIMAT0003393&hsa-miR-425-5p&offsets\|0\|-1;m-95&3\|-\|49020200\|49020221&offsets\|0\|0]\|AATGACACGATCACTCCCGTTG | >MI0001448\|hsa-mir-425&WithFlank&3\|-\|49020142\|49020240 | 20 | 41 | 22 | AATGACACGATCACTCCCGTTG | 535.8 | 2.3 | 0.9 | 2.5 | 1.4E-02 | 3.9E-02 | BT20/MB231 | CYTO |
| >MI0000264\|hsa-mir-7-2&WithFlank&15\|+\|88611819\|88611940@38.60.23(+1A);MI0000265\|hsa-mir-7-3&WithFlank&19\|+\|4770664\|4770785@37.59.23(+1A);MI0000263\|hsa-mir-7-1&WithFlank&9\|-\|83969742\|83969863@30.52.23(+1A)&[MIMAT0000252&hsa-miR-7-5p&offsets\|0\|-1(+1A);m-400&15\|+\|88611856\|88611879&offsets\|0\|-1(+1A)];[MIMAT0000252_1&hsa-miR-7-5p&offsets\|0\|-1(+1A);m-402&19\|+\|4770700\|4770723&offsets\|0\|-1(+1A)];[MIMAT0000252_2&hsa-miR-7-5p&offsets\|0\|-1(+1A);m-401&9\|-\|83969811\|83969834&offsets\|0\|-1(+1A)]\|TGGAAGACTAGTGATTTTGTTGTA | >MI0000264\|hsa-mir-7-2&WithFlank&15\|+\|88611819\|88611940 | 38 | 60 | 23(+1A) | TGGAAGACTAGTGATTTTGTTGTA | 424.5 | 2.4 | 1.0 | 2.3 | 2.4E-02 | 3.9E-02 | BT20/MB468 | MP |
| >hg38_21_+_8433172_8446622_RNA45SN1_45S_50ntFlanks@7988.8009.22\|CAGACGTGGCGACCCGCTGAAT | >hg38_21_+_8433172_8446622_RNA45SN1_45S_50ntFlanks | 7988 | 8009 | 22 | CAGACGTGGCGACCCGCTGAAT | 784.5 | 3.4 | 1.3 | 2.6 | 1.0E-02 | 3.9E-02 | BT20/MB231 | MP |
| >hg38_MT_+_1621_3279_RNR2_16S_50ntFlanks@1425.1448.24\|ACGATTAAAGTCCTACGTGATCTG | >hg38_MT_+_1621_3279_RNR2_16S_50ntFlanks | 1425 | 1448 | 24 | ACGATTAAAGTCCTACGTGATCTG | 463.6 | -3.2 | 1.2 | -2.6 | 1.0E-02 | 3.9E-02 | BT20/MB231 | MP |
| >trna111_HisGTG_1_-_147774845_147774916@-1G.34.35,trna118_HisGTG_1_-_145396881_145396952@-1G.34.35,trna16_HisGTG_1_+_146544773_146544844@-1G.34.35,trna1_HisGTG_15_+_45493349_45493420@-1G.34.35,trna21_HisGTG_1_+_147753471_147753542@-1G.34.35,trna33_HisGTG_6_+_27125906_27125977@-1G.34.35,trna7_HisGTG_9_-_14433938_14434009@-1G.34.35,trna8_HisGTG_15_-_45492611_45492682@-1G.34.35,trna9_HisGTG_15_-_45490804_45490875@-1G.34.35\|GGCCGTGATCGTATAGTGGTTAGTACTCTGCGTTG | >trna111_HisGTG_1_-_147774845_147774916 | -1G | 34 | 35 | GGCCGTGATCGTATAGTGGTTAGTACTCTGCGTTG | 564.5 | -2.5 | 1.2 | -2.1 | 3.5E-02 | 3.9E-02 | BT20/MB468 | NUC |
| >hg38_MT_+_598_1651_RNR1_12S_50ntFlanks@439.480.42\|AACATATCTGAACACACAATAGCTAAGACCCAAACTGGGATT | >hg38_MT_+_598_1651_RNR1_12S_50ntFlanks | 439 | 480 | 42 | AACATATCTGAACACACAATAGCTAAGACCCAAACTGGGATT | 148.9 | 2.4 | 1.2 | 2.1 | 3.5E-02 | 3.9E-02 | MB231/MB468 | MITO |
| >am_trnaMT_HisGTG_MT_+_12138_12206@26.44.19\|AGATTGTGAATCTGACAAC | >am_trnaMT_HisGTG_MT_+_12138_12206 | 26 | 44 | 19 | AGATTGTGAATCTGACAAC | 943.9 | 2.4 | 1.1 | 2.1 | 3.5E-02 | 3.9E-02 | MB231/MB468 | MITO |
| >hg38_21_+_8433172_8446622_RNA45SN1_45S_50ntFlanks@10515.10532.18\|GGAGTGGCGGAGATGGGC | >hg38_21_+_8433172_8446622_RNA45SN1_45S_50ntFlanks | 10515 | 10532 | 18 | GGAGTGGCGGAGATGGGC | 147.2 | 4.2 | 1.9 | 2.3 | 2.4E-02 | 3.9E-02 | BT20/MB468 | MP |
| >hg38_MT_+_1621_3279_RNR2_16S_50ntFlanks@625.656.32\|AACTGAACTCCTCACACCCAATTGGACCAATC | >hg38_MT_+_1621_3279_RNR2_16S_50ntFlanks | 625 | 656 | 32 | AACTGAACTCCTCACACCCAATTGGACCAATC | 64.1 | 4.1 | 1.8 | 2.3 | 2.4E-02 | 3.9E-02 | BT20/MB468 | MP |
| >hg38_21_+_8433172_8446622_RNA45SN1_45S_50ntFlanks@10349.10370.22\|AACGAGAACTTTGAAGGCCGAA | >hg38_21_+_8433172_8446622_RNA45SN1_45S_50ntFlanks | 10349 | 10370 | 22 | AACGAGAACTTTGAAGGCCGAA | 494.0 | -2.1 | 1.0 | -2.1 | 3.2E-02 | 3.9E-02 | MB231/MB468 | MP |
| >hg38_21_+_8433172_8446622_RNA45SN1_45S_50ntFlanks@9484.9503.20\|GTGTTAGGACCCGAAAGATG | >hg38_21_+_8433172_8446622_RNA45SN1_45S_50ntFlanks | 9484 | 9503 | 20 | GTGTTAGGACCCGAAAGATG | 229.7 | 5.2 | 2.0 | 2.6 | 1.0E-02 | 3.9E-02 | BT20/MB231 | MP |
| >hg38_21_+_8433172_8446622_RNA45SN1_45S_50ntFlanks@11876.11900.25\|AAGACCCTGTTGAGCTTGACTCTAG | >hg38_21_+_8433172_8446622_RNA45SN1_45S_50ntFlanks | 11876 | 11900 | 25 | AAGACCCTGTTGAGCTTGACTCTAG | 235.5 | -3.9 | 1.7 | -2.3 | 2.4E-02 | 3.9E-02 | BT20/MB468 | MP |
| >MI0000542\|hsa-mir-320a&WithFlank&8\|-\|22244960\|22245043@49.70.22&[MIMAT0000510&hsa-miR-320a-3p&offsets\|+1\|+1;m-60&8\|-\|22244975\|22244996&offsets\|+1\|+1]\|AAAGCTGGGTTGAGAGGGCGAA | >MI0000542\|hsa-mir-320a&WithFlank&8\|-\|22244960\|22245043 | 49 | 70 | 22 | AAAGCTGGGTTGAGAGGGCGAA | 142.4 | 3.0 | 1.4 | 2.1 | 3.2E-02 | 3.9E-02 | MB231/MB468 | MP |
| >hg38_21_+_8433172_8446622_RNA45SN1_45S_50ntFlanks@12420.12452.33\|ATGTCGGCTCTTCCTATCATTGTGAAGCAGAAT | >hg38_21_+_8433172_8446622_RNA45SN1_45S_50ntFlanks | 12420 | 12452 | 33 | ATGTCGGCTCTTCCTATCATTGTGAAGCAGAAT | 246.7 | 1.7 | 0.8 | 2.1 | 3.5E-02 | 3.9E-02 | MB231/MB468 | NUC |
| >hg38_21_+_8433172_8446622_RNA45SN1_45S_50ntFlanks@9533.9560.28\|GAGGAAACTCTGGTGGAGGTCCGTAGCG | >hg38_21_+_8433172_8446622_RNA45SN1_45S_50ntFlanks | 9533 | 9560 | 28 | GAGGAAACTCTGGTGGAGGTCCGTAGCG | 189.6 | -2.1 | 1.0 | -2.1 | 3.2E-02 | 3.9E-02 | MB231/MB468 | MP |
| >hg38_MT_+_598_1651_RNR1_12S_50ntFlanks@52.91.40\|ATAGGTTTGGTCCTAGCCTTTCTATTAGCTCTTAGTAAGA | >hg38_MT_+_598_1651_RNR1_12S_50ntFlanks | 52 | 91 | 40 | ATAGGTTTGGTCCTAGCCTTTCTATTAGCTCTTAGTAAGA | 1013.1 | 3.5 | 1.6 | 2.3 | 2.4E-02 | 3.9E-02 | BT20/MB468 | MP |
| >hg38_21_+_8433172_8446622_RNA45SN1_45S_50ntFlanks@7976.7994.19\|GCGACCTCAGATCAGACGT | >hg38_21_+_8433172_8446622_RNA45SN1_45S_50ntFlanks | 7976 | 7994 | 19 | GCGACCTCAGATCAGACGT | 456.9 | 3.1 | 1.2 | 2.6 | 1.1E-02 | 3.9E-02 | BT20/MB231 | MP |
| >MI0000465\|hsa-mir-191&WithFlank&3\|-\|49020612\|49020715@22.45.24&[MIMAT0000440&hsa-miR-191-5p&offsets\|0\|+1;m-48&3\|-\|49020672\|49020694&offsets\|0\|+1]\|CAACGGAATCCCAAAAGCAGCTGT | >MI0000465\|hsa-mir-191&WithFlank&3\|-\|49020612\|49020715 | 22 | 45 | 24 | CAACGGAATCCCAAAAGCAGCTGT | 6843.0 | -2.3 | 0.9 | -2.5 | 1.3E-02 | 3.9E-02 | BT20/MB231 | MITO |
| >am_trna119_LysCTT_1_-_145395522_145395594@14.33.20,trna11_LysCTT_5_-_180648979_180649051@14.33.20,trna128_LysCTT_1_-_55423542_55423614@14.33.20,trna13_LysCTT_6_+_26556774_26556846@14.33.20,trna6_LysCTT_19_-_52425396_52425468@14.33.20,trna7_LysCTT_16_+_3225692_3225764@14.33.20,trna9_LysCTT_5_+_180634755_180634827@14.33.20\|AGTCGGTAGAGCATGAGACT | >am_trna119_LysCTT_1_-_145395522_145395594 | 14 | 33 | 20 | AGTCGGTAGAGCATGAGACT | 110.7 | 2.8 | 1.3 | 2.1 | 3.2E-02 | 3.9E-02 | MB231/MB468 | MP |
| >hg38_21_+_8433172_8446622_RNA45SN1_45S_50ntFlanks@10241.10259.19\|TACGCCGCGACGAGTAGGA | >hg38_21_+_8433172_8446622_RNA45SN1_45S_50ntFlanks | 10241 | 10259 | 19 | TACGCCGCGACGAGTAGGA | 158.2 | 5.2 | 2.0 | 2.6 | 1.1E-02 | 3.9E-02 | BT20/MB231 | MP |
| >hg38_21_+_8433172_8446622_RNA45SN1_45S_50ntFlanks@9862.9902.41\|ATGAACCGAACGCCGGGTTAAGGCGCCCGATGCCGACGCTC | >hg38_21_+_8433172_8446622_RNA45SN1_45S_50ntFlanks | 9862 | 9902 | 41 | ATGAACCGAACGCCGGGTTAAGGCGCCCGATGCCGACGCTC | 133.6 | -3.8 | 1.7 | -2.2 | 3.0E-02 | 3.9E-02 | BT20/MB468 | MITO |
| >hg38_MT_+_598_1651_RNR1_12S_50ntFlanks@915.946.32\|AAAGGACATTTAACTAAAACCCCTACGCATTT | >hg38_MT_+_598_1651_RNR1_12S_50ntFlanks | 915 | 946 | 32 | AAAGGACATTTAACTAAAACCCCTACGCATTT | 255.9 | 2.8 | 1.3 | 2.1 | 3.2E-02 | 3.9E-02 | MB231/MB468 | MP |
| >MI0000448\|hsa-mir-130a&WithFlank&11\|+\|57641192\|57641292@21.42.22&[MIMAT0004593&hsa-miR-130a-5p&offsets\|0\|0;m-622&11\|+\|57641212\|57641233&offsets\|0\|0]\|GCTCTTTTCACATTGTGCTACT | >MI0000448\|hsa-mir-130a&WithFlank&11\|+\|57641192\|57641292 | 21 | 42 | 22 | GCTCTTTTCACATTGTGCTACT | 100.0 | 4.3 | 2.0 | 2.1 | 3.2E-02 | 3.9E-02 | MB231/MB468 | MP |
| >hg38_21_+_8433172_8446622_RNA45SN1_45S_50ntFlanks@8186.8208.23\|AGGCCCAGCCCGTGGACGGTGTG | >hg38_21_+_8433172_8446622_RNA45SN1_45S_50ntFlanks | 8186 | 8208 | 23 | AGGCCCAGCCCGTGGACGGTGTG | 138.7 | -3.1 | 1.4 | -2.1 | 3.2E-02 | 3.9E-02 | MB231/MB468 | MP |
| >hg38_21_+_8433172_8446622_RNA45SN1_45S_50ntFlanks@12957.12975.19\|CTTCTGGGTCGGGGTTTCG | >hg38_21_+_8433172_8446622_RNA45SN1_45S_50ntFlanks | 12957 | 12975 | 19 | CTTCTGGGTCGGGGTTTCG | 144.8 | -2.5 | 1.2 | -2.2 | 3.0E-02 | 3.9E-02 | BT20/MB468 | MITO |
| >hg38_21_+_8433172_8446622_RNA45SN1_45S_50ntFlanks@12999.13039.41\|CTGCGATCTATTGAAAGTCAGCCCTCGACACAAGGGTTTGT | >hg38_21_+_8433172_8446622_RNA45SN1_45S_50ntFlanks | 12999 | 13039 | 41 | CTGCGATCTATTGAAAGTCAGCCCTCGACACAAGGGTTTGT | 500.0 | 2.3 | 1.1 | 2.1 | 3.2E-02 | 3.9E-02 | MB231/MB468 | MP |
| >hg38_21_+_8433172_8446622_RNA45SN1_45S_50ntFlanks@9533.9570.38\|GAGGAAACTCTGGTGGAGGTCCGTAGCGGTCCTGACGT | >hg38_21_+_8433172_8446622_RNA45SN1_45S_50ntFlanks | 9533 | 9570 | 38 | GAGGAAACTCTGGTGGAGGTCCGTAGCGGTCCTGACGT | 813.1 | 1.4 | 0.7 | 2.1 | 3.4E-02 | 3.9E-02 | MB231/MB468 | CYTO |
| >hg38_21_+_8433172_8446622_RNA45SN1_45S_50ntFlanks@12073.12097.25\|CCGAGGGGCTCTCGCTTCTGGCGCC | >hg38_21_+_8433172_8446622_RNA45SN1_45S_50ntFlanks | 12073 | 12097 | 25 | CCGAGGGGCTCTCGCTTCTGGCGCC | 516.6 | -4.1 | 1.8 | -2.3 | 2.4E-02 | 3.9E-02 | BT20/MB468 | MP |
| >MI0000077\|hsa-mir-21&WithFlank&17\|+\|59841260\|59841343@15.36.22&[MIMAT0000076&hsa-miR-21-5p&offsets\|+1\|+1;m-1&17\|+\|59841273\|59841294&offsets\|+1\|+1]\|AGCTTATCAGACTGATGTTGAC | >MI0000077\|hsa-mir-21&WithFlank&17\|+\|59841260\|59841343 | 15 | 36 | 22 | AGCTTATCAGACTGATGTTGAC | 989.8 | 2.1 | 1.0 | 2.1 | 3.2E-02 | 3.9E-02 | MB231/MB468 | MP |
| >hg38_21_+_8433172_8446622_RNA45SN1_45S_50ntFlanks@10291.10328.38\|CGCGGGCCCGGGTGGAGCCGCCGCAGGTGCAGATCTTG | >hg38_21_+_8433172_8446622_RNA45SN1_45S_50ntFlanks | 10291 | 10328 | 38 | CGCGGGCCCGGGTGGAGCCGCCGCAGGTGCAGATCTTG | 49.6 | -3.9 | 1.7 | -2.3 | 2.4E-02 | 3.9E-02 | BT20/MB468 | MP |
| >hg38_MT_+_1621_3279_RNR2_16S_50ntFlanks@841.859.19\|AAAGTAAAAGGAACTCGGC | >hg38_MT_+_1621_3279_RNR2_16S_50ntFlanks | 841 | 859 | 19 | AAAGTAAAAGGAACTCGGC | 92.9 | 2.5 | 1.1 | 2.3 | 2.4E-02 | 3.9E-02 | BT20/MB468 | MP |

| | | | | | | | | | | | | | |
|---|---|---|---|---|---|---|---|---|---|---|---|---|---|
| >hg38_21_+_8433172_8446622_RNA45SN1_45S_50ntFlanks@12360.12384.25\|CAGGGATAACTGGCTTGTGGCGGCC | >hg38_21_+_8433172_8446622_RNA45SN1_45S_50ntFlanks | 12360 | 12384 | 25 | CAGGGATAACTGGCTTGTGGCGGCC | 177.9 | -3.9 | 1.7 | -2.3 | 2.4E-02 | 3.9E-02 | BT20/MB468 | MP |
| >MI0017294\|hsa-mir-4664&WithFlank&8\|-\|143733077\|143733159@50.71.22&[MIMAT0019738&hsa-miR-4664-3p&offsets\|0\|0;m-1025&8\|-\|143733094\|143733110&offsets\|0\|+5]\|CTTCCGGTCTGTGAGCCCCGTC | >MI0017294\|hsa-mir-4664&WithFlank&8\|-\|143733077\|143733159 | 50 | 71 | 22 | CTTCCGGTCTGTGAGCCCCGTC | 59.1 | 4.3 | 1.9 | 2.3 | 2.4E-02 | 3.9E-02 | BT20/MB468 | MP |
| >hg38_21_+_8433172_8446622_RNA45SN1_45S_50ntFlanks@11970.11998.29\|CGCGAGGGGCCCGGGGCGGGGTCCGCCGG | >hg38_21_+_8433172_8446622_RNA45SN1_45S_50ntFlanks | 11970 | 11998 | 29 | CGCGAGGGGCCCGGGGCGGGGTCCGCCGG | 166.7 | -4.1 | 1.8 | -2.3 | 2.4E-02 | 3.9E-02 | BT20/MB468 | MP |
| >MI0003137\|hsa-mir-193b&WithFlank&16\|+\|14303961\|14304055@57.77.21&[MIMAT0002819&hsa-miR-193b-3p&offsets\|0\|-1;m-166&16\|+\|14304017\|14304038&offsets\|0\|-1]\|AACTGGCCCTCAAAGTCCCGC | >MI0003137\|hsa-mir-193b&WithFlank&16\|+\|14303961\|14304055 | 57 | 77 | 21 | AACTGGCCCTCAAAGTCCCGC | 1046.8 | 2.6 | 1.0 | 2.6 | 1.1E-02 | 3.9E-02 | BT20/MB231 | MP |
| >MI0000085\|hsa-mir-27a&WithFlank&19\|-\|13836434\|13836523@57.76.20&[MIMAT0000084&hsa-miR-27a-3p&offsets\|0\|-1;m-43&19\|-\|13836447\|13836467&offsets\|0\|-1]\|TTCACAGTGGCTAAGTTCCG | >MI0000085\|hsa-mir-27a&WithFlank&19\|-\|13836434\|13836523 | 57 | 76 | 20 | TTCACAGTGGCTAAGTTCCG | 27888.4 | -2.7 | 1.1 | -2.6 | 1.1E-02 | 3.9E-02 | BT20/MB231 | MP |
| >hg38_21_+_8433172_8446622_RNA45SN1_45S_50ntFlanks@4752.4786.35\|GAAGACGATCAGATACCGTCGTAGTTCCGACCATA | >hg38_21_+_8433172_8446622_RNA45SN1_45S_50ntFlanks | 4752 | 4786 | 35 | GAAGACGATCAGATACCGTCGTAGTTCCGACCATA | 187.1 | -2.7 | 1.3 | -2.2 | 3.0E-02 | 3.9E-02 | BT20/MB468 | MITO |
| >hg38_MT_+_598_1651_RNR1_12S_50ntFlanks@817.840.24\|CGAAGGTGGATTTAGCAGTAAACT | >hg38_MT_+_598_1651_RNR1_12S_50ntFlanks | 817 | 840 | 24 | CGAAGGTGGATTTAGCAGTAAACT | 102.6 | 4.0 | 1.6 | 2.6 | 1.1E-02 | 3.9E-02 | BT20/MB231 | MP |
| >hg38_21_+_8433172_8446622_RNA45SN1_45S_50ntFlanks@9534.9563.30\|AGGAAACTCTGGTGGAGGTCCGTAGCGGTC | >hg38_21_+_8433172_8446622_RNA45SN1_45S_50ntFlanks | 9534 | 9563 | 30 | AGGAAACTCTGGTGGAGGTCCGTAGCGGTC | 485.6 | 4.0 | 1.6 | 2.6 | 1.1E-02 | 3.9E-02 | BT20/MB231 | MP |
| >am_trnaMT_SerGCT_MT_+_12207_12265@14.35.22\|AGAACTGCTAACTCATGCCCCC | >am_trnaMT_SerGCT_MT_+_12207_12265 | 14 | 35 | 22 | AGAACTGCTAACTCATGCCCCC | 78.6 | 3.8 | 1.8 | 2.1 | 3.2E-02 | 3.9E-02 | MB231/MB468 | MP |
| >hg38_21_+_8433172_8446622_RNA45SN1_45S_50ntFlanks@6652.6675.24\|GACTCTTAGCGGTGGATCACTCGG | >hg38_21_+_8433172_8446622_RNA45SN1_45S_50ntFlanks | 6652 | 6675 | 24 | GACTCTTAGCGGTGGATCACTCGG | 1159.7 | 2.1 | 1.0 | 2.2 | 3.0E-02 | 3.9E-02 | BT20/MB468 | MITO |
| >trnaMT_GlnTTG_MT_-_4329_4400@1.36.36,trnalookalike2_GlnTTG_1_-_564879_564950@1.36.36\|TAGGATGGGGTGTGATAGGTGGCACGGAGAATTTTG | >trnaMT_GlnTTG_MT_-_4329_4400 | 1 | 36 | 36 | TAGGATGGGGTGTGATAGGTGGCACGGAGAATTTTG | 509.2 | 2.1 | 0.8 | 2.5 | 1.3E-02 | 3.9E-02 | BT20/MB231 | MITO |
| >hg38_21_+_8433172_8446622_RNA45SN1_45S_50ntFlanks@4982.5020.39\|ACGGACAGGATTGACAGATTGATAGCTCTTTCTCGATTC | >hg38_21_+_8433172_8446622_RNA45SN1_45S_50ntFlanks | 4982 | 5020 | 39 | ACGGACAGGATTGACAGATTGATAGCTCTTTCTCGATTC | 84.0 | -4.7 | 2.2 | -2.1 | 3.2E-02 | 3.9E-02 | MB231/MB468 | MP |
| >hg38_21_+_8433172_8446622_RNA45SN1_45S_50ntFlanks@5059.5098.40\|CGATTTGTCTGGTTAATTCCGATAACGAACGAGACTCTGG | >hg38_21_+_8433172_8446622_RNA45SN1_45S_50ntFlanks | 5059 | 5098 | 40 | CGATTTGTCTGGTTAATTCCGATAACGAACGAGACTCTGG | 780.3 | -4.3 | 1.9 | -2.3 | 2.4E-02 | 3.9E-02 | BT20/MB468 | MP |
| >hg38_1_-_228634819_228635039_RNA5S12_5S_50ntFlanks@51.83.33\|GTCTACGGCCATACCACCCTGAACGCGCCCGAT | >hg38_1_-_228634819_228635039_RNA5S12_5S_50ntFlanks | 51 | 83 | 33 | GTCTACGGCCATACCACCCTGAACGCGCCCGAT | 2772.0 | 3.3 | 1.3 | 2.6 | 1.1E-02 | 3.9E-02 | BT20/MB231 | MP |
| >hg38_21_+_8433172_8446622_RNA45SN1_45S_50ntFlanks@11671.11697.27\|CCAGTGCTCTGAATGTCAAAGTGAAGA | >hg38_21_+_8433172_8446622_RNA45SN1_45S_50ntFlanks | 11671 | 11697 | 27 | CCAGTGCTCTGAATGTCAAAGTGAAGA | 244.4 | 4.4 | 1.7 | 2.6 | 1.1E-02 | 3.9E-02 | BT20/MB231 | MP |
| >hg38_21_+_8433172_8446622_RNA45SN1_45S_50ntFlanks@12420.12444.25\|ATGTCGGCTCTTCCTATCATTGTGA | >hg38_21_+_8433172_8446622_RNA45SN1_45S_50ntFlanks | 12420 | 12444 | 25 | ATGTCGGCTCTTCCTATCATTGTGA | 617.7 | -1.0 | 0.5 | -2.2 | 3.0E-02 | 3.9E-02 | BT20/MB468 | MITO |
| >hg38_21_+_8433172_8446622_RNA45SN1_45S_50ntFlanks@4922.4941.20\|CCAGGAGTGGAGCCTGCGGC | >hg38_21_+_8433172_8446622_RNA45SN1_45S_50ntFlanks | 4922 | 4941 | 20 | CCAGGAGTGGAGCCTGCGGC | 304.4 | -2.3 | 1.0 | -2.2 | 3.0E-02 | 3.9E-02 | BT20/MB468 | MITO |
| >MI0000071\|hsa-mir-17&WithFlank&13\|+\|91350599\|91350694@20.41.22(+1C)&[MIMAT0000070&hsa-miR-17-5p&offsets\|0\|-1(+1C);m-88&13\|+\|91350618\|91350640&offsets\|0\|-1(+1C)]\|CAAAGTGCTTACAGTGCAGGTAC | >MI0000071\|hsa-mir-17&WithFlank&13\|+\|91350599\|91350694 | 20 | 41 | 22(+1C) | CAAAGTGCTTACAGTGCAGGTAC | 113.2 | -3.9 | 1.5 | -2.6 | 1.1E-02 | 3.9E-02 | BT20/MB231 | MP |
| >MI0000270\|hsa-mir-181b-1&WithFlank&1\|-\|198858867\|198858988@42.63.22;MI0000683\|hsa-mir-181b-2&WithFlank&9\|+\|124693704\|124693804@22.43.22&[MIMAT0000257&hsa-miR-181b-5p&offsets\|0\|-1;m-73&1\|-\|198858925\|198858947&offsets\|0\|-1];[MIMAT0000257_1&hsa-miR-181b-5p&offsets\|0\|-1;m-72&9\|+\|124693725\|124693747&offsets\|0\|-1]\|AACATTCATTGCTGTCGGTGGG | >MI0000270\|hsa-mir-181b-1&WithFlank&1\|-\|198858867\|198858988 | 42 | 63 | 22 | AACATTCATTGCTGTCGGTGGG | 200.2 | 3.0 | 1.4 | 2.1 | 3.2E-02 | 3.9E-02 | MB231/MB468 | MP |
| >hg38_21_+_8433172_8446622_RNA45SN1_45S_50ntFlanks@10049.10081.33\|GGCCGTCGCCGGCAGTCGAGAGTGGACGGGAGC | >hg38_21_+_8433172_8446622_RNA45SN1_45S_50ntFlanks | 10049 | 10081 | 33 | GGCCGTCGCCGGCAGTCGAGAGTGGACGGGAGC | 80.8 | 5.1 | 2.0 | 2.6 | 1.1E-02 | 3.9E-02 | BT20/MB231 | MP |
| >hg38_21_+_8433172_8446622_RNA45SN1_45S_50ntFlanks@11578.11613.36\|TAGAACTGGTGCGGACCAGGGGAATCCGACTGTTTA | >hg38_21_+_8433172_8446622_RNA45SN1_45S_50ntFlanks | 11578 | 11613 | 36 | TAGAACTGGTGCGGACCAGGGGAATCCGACTGTTTA | 145.5 | 4.1 | 1.6 | 2.6 | 1.1E-02 | 3.9E-02 | BT20/MB231 | MP |
| >MI0000085\|hsa-mir-27a&WithFlank&19\|-\|13836434\|13836523@57.76.20(+1G)&[MIMAT0000084&hsa-miR-27a-3p&offsets\|0\|-1(+1G);m-43&19\|-\|13836447\|13836467&offsets\|0\|-1(+1G)]\|TTCACAGTGGCTAAGTTCCGG | >MI0000085\|hsa-mir-27a&WithFlank&19\|-\|13836434\|13836523 | 57 | 76 | 20(+1G) | TTCACAGTGGCTAAGTTCCGG | 273.0 | 1.9 | 0.9 | 2.1 | 3.2E-02 | 3.9E-02 | MB231/MB468 | MP |
| >hg38_MT_+_598_1651_RNR1_12S_50ntFlanks@916.941.26\|AAGGACATTTAACTAAAACCCCTACG | >hg38_MT_+_598_1651_RNR1_12S_50ntFlanks | 916 | 941 | 26 | AAGGACATTTAACTAAAACCCCTACG | 167.9 | 3.8 | 1.8 | 2.1 | 3.2E-02 | 3.9E-02 | MB231/MB468 | MP |
| >hg38_21_+_8433172_8446622_RNA45SN1_45S_50ntFlanks@8104.8138.35\|CCGCGGCGGGGCGCGGGACATGTGGCGTACGGAAG | >hg38_21_+_8433172_8446622_RNA45SN1_45S_50ntFlanks | 8104 | 8138 | 35 | CCGCGGCGGGGCGCGGGACATGTGGCGTACGGAAG | 283.5 | 4.2 | 1.6 | 2.6 | 1.1E-02 | 3.9E-02 | BT20/MB231 | MP |
| >MI0000816\|hsa-mir-335&WithFlank&7\|+\|130496105\|130496210@58.78.21(+1A)&[MIMAT0004703&hsa-miR-335-3p&offsets\|0\|-1(+1A);m-159&7\|+\|130496162\|130496183&offsets\|0\|-1(+1A)]\|TTTTTCATTATTGCTCCTGACA | >MI0000816\|hsa-mir-335&WithFlank&7\|+\|130496105\|130496210 | 58 | 78 | 21(+1A) | TTTTTCATTATTGCTCCTGACA | 257.1 | -3.0 | 1.2 | -2.6 | 1.1E-02 | 3.9E-02 | BT20/MB231 | MP |
| >hg38_MT_+_598_1651_RNR1_12S_50ntFlanks@428.454.27\|AAAGTGGCTTTAACATATCTGAACACA | >hg38_MT_+_598_1651_RNR1_12S_50ntFlanks | 428 | 454 | 27 | AAAGTGGCTTTAACATATCTGAACACA | 136.1 | 3.4 | 1.5 | 2.3 | 2.4E-02 | 3.9E-02 | BT20/MB468 | MP |
| >hg38_21_+_8433172_8446622_RNA45SN1_45S_50ntFlanks@6780.6806.27\|CGGGGCTACGCCTGTCTGAGCGTCGCT | >hg38_21_+_8433172_8446622_RNA45SN1_45S_50ntFlanks | 6780 | 6806 | 27 | CGGGGCTACGCCTGTCTGAGCGTCGCT | 543.3 | 1.7 | 0.8 | 2.1 | 3.6E-02 | 3.9E-02 | MB231/MB468 | MITO |
| >MI0002470\|hsa-mir-486-1&WithFlank&8\|-\|41660435\|41660514@10.31.22(+1U);MI0023622\|hsa-mir-486-2&WithFlank&8\|+\|41660438\|41660513@7.28.22(+1U)&[MIMAT0002177&hsa-miR-486-5p&offsets\|0\|0(+1U);m-107&8\|-\|41660484\|41660505&offsets\|0\|0(+1U)];[MIMAT0002177_1&hsa-miR-486-5p&offsets\|0\|0(+1U);TJU_CMC_MD2.ID03138.5p-miR&offsets\|0\|-2(+1U);m-106&8\|+\|41660444\|41660465&offsets\|0\|0(+1U)]\|TCCTGTACTGAGCTGCCCCGAGT | >MI0002470\|hsa-mir-486-1&WithFlank&8\|-\|41660435\|41660514 | 10 | 31 | 22(+1U) | TCCTGTACTGAGCTGCCCCGAGT | 455.4 | -4.6 | 1.9 | -2.5 | 1.4E-02 | 3.9E-02 | BT20/MB231 | CYTO |
| >am_trnaMT_ValTAC_MT_+_1602_1670@48.68.21\|CAACTTAACTTGACCGCTCTG | >am_trnaMT_ValTAC_MT_+_1602_1670 | 48 | 68 | 21 | CAACTTAACTTGACCGCTCTG | 823.2 | 1.9 | 0.8 | 2.5 | 1.4E-02 | 3.9E-02 | BT20/MB231 | CYTO |
| >hg38_21_+_8433172_8446622_RNA45SN1_45S_50ntFlanks@12995.13012.18\|CTCGCTGCGATCTATTGA | >hg38_21_+_8433172_8446622_RNA45SN1_45S_50ntFlanks | 12995 | 13012 | 18 | CTCGCTGCGATCTATTGA | 936.2 | -1.7 | 0.7 | -2.5 | 1.4E-02 | 3.9E-02 | BT20/MB231 | CYTO |
| >hg38_21_+_8433172_8446622_RNA45SN1_45S_50ntFlanks@12918.12940.23\|CGTGGGGAACCTGGCGCTAAACC | >hg38_21_+_8433172_8446622_RNA45SN1_45S_50ntFlanks | 12918 | 12940 | 23 | CGTGGGGAACCTGGCGCTAAACC | 213.2 | -2.0 | 0.8 | -2.5 | 1.4E-02 | 3.9E-02 | BT20/MB231 | CYTO |
| >am_trnaMT_ValTAC_MT_+_1602_1670@38.57.20\|TAGGAGATTTCAACTTAACT | >am_trnaMT_ValTAC_MT_+_1602_1670 | 38 | 57 | 20 | TAGGAGATTTCAACTTAACT | 356.1 | 4.0 | 1.8 | 2.3 | 2.4E-02 | 3.9E-02 | BT20/MB468 | MP |
| >hg38_21_+_8433172_8446622_RNA45SN1_45S_50ntFlanks@11569.11600.32\|CAGCCGACTTAGAACTGGTGCGGACCAGGGGA | >hg38_21_+_8433172_8446622_RNA45SN1_45S_50ntFlanks | 11569 | 11600 | 32 | CAGCCGACTTAGAACTGGTGCGGACCAGGGGA | 382.3 | 4.3 | 1.7 | 2.6 | 1.1E-02 | 3.9E-02 | BT20/MB231 | MP |
| >hg38_21_+_8433172_8446622_RNA45SN1_45S_50ntFlanks@6647.6673.27\|CGTACGACTCTTAGCGGTGGATCACTC | >hg38_21_+_8433172_8446622_RNA45SN1_45S_50ntFlanks | 6647 | 6673 | 27 | CGTACGACTCTTAGCGGTGGATCACTC | 108.5 | 1.2 | 0.6 | 2.1 | 3.5E-02 | 3.9E-02 | BT20/MB468 | NUC |
| >hg38_21_+_8433172_8446622_RNA45SN1_45S_50ntFlanks@4045.4066.22\|CCATTCGAACGTCTGCCCTATC | >hg38_21_+_8433172_8446622_RNA45SN1_45S_50ntFlanks | 4045 | 4066 | 22 | CCATTCGAACGTCTGCCCTATC | 615.0 | -2.3 | 0.9 | -2.5 | 1.4E-02 | 3.9E-02 | BT20/MB231 | CYTO |
| >trnaMT_ValTAC_MT_+_1602_1670@38.71.34\|TAGGAGATTTCAACTTAACTTGACCGCTCTGACC | >trnaMT_ValTAC_MT_+_1602_1670 | 38 | 71 | 34 | TAGGAGATTTCAACTTAACTTGACCGCTCTGACC | 115.7 | 4.3 | 2.0 | 2.1 | 3.3E-02 | 3.9E-02 | MB231/MB468 | MP |
| >hg38_21_+_8433172_8446622_RNA45SN1_45S_50ntFlanks@5498.5532.35\|CGGTCGAACTTGACTATCTAGAGGAAGTAAAAGTC | >hg38_21_+_8433172_8446622_RNA45SN1_45S_50ntFlanks | 5498 | 5532 | 35 | CGGTCGAACTTGACTATCTAGAGGAAGTAAAAGTC | 63.8 | 3.1 | 1.4 | 2.3 | 2.4E-02 | 3.9E-02 | BT20/MB468 | MP |
| >hg38_21_+_8433172_8446622_RNA45SN1_45S_50ntFlanks@10793.10817.25\|TCTGGCATGTTGGAACAATGTAGGT | >hg38_21_+_8433172_8446622_RNA45SN1_45S_50ntFlanks | 10793 | 10817 | 25 | TCTGGCATGTTGGAACAATGTAGGT | 91.6 | -2.9 | 1.3 | -2.3 | 2.4E-02 | 3.9E-02 | BT20/MB468 | MP |
| >MI0006444\|hsa-mir-1307&WithFlank&10\|-\|103394247\|103394407@86.107.22&[MIMAT0005951&hsa-miR-1307-3p&offsets\|0\|0;m-66&10\|-\|103394300\|103394322&offsets\|0\|-1]\|ACTCGGCGTGGCGTCGGTCGTG | >MI0006444\|hsa-mir-1307&WithFlank&10\|-\|103394247\|103394407 | 86 | 107 | 22 | ACTCGGCGTGGCGTCGGTCGTG | 136.7 | 1.1 | 0.5 | 2.1 | 3.5E-02 | 3.9E-02 | BT20/MB468 | NUC |
| >MI0000440\|hsa-mir-27b&WithFlank&9\|+\|95085439\|95085547@67.87.21(+1C)&[MIMAT0000419&hsa-miR-27b-3p&offsets\|0\|0(+1C);m-39&9\|+\|95085505\|95085525&offsets\|0\|0(+1C)]\|TTCACAGTGGCTAAGTTCTGCC | >MI0000440\|hsa-mir-27b&WithFlank&9\|+\|95085439\|95085547 | 67 | 87 | 21(+1C) | TTCACAGTGGCTAAGTTCTGCC | 411.9 | -2.2 | 0.9 | -2.5 | 1.3E-02 | 3.9E-02 | BT20/MB231 | MITO |
| >hg38_21_+_8433172_8446622_RNA45SN1_45S_50ntFlanks@4738.4761.24\|AAAGTCGGAGGTTCGAAGACGATC | >hg38_21_+_8433172_8446622_RNA45SN1_45S_50ntFlanks | 4738 | 4761 | 24 | AAAGTCGGAGGTTCGAAGACGATC | 730.2 | -1.5 | 0.7 | -2.1 | 3.3E-02 | 3.9E-02 | MB231/MB468 | MP |
| >MI0000750\|hsa-mir-26a-2&WithFlank&12\|-\|57824603\|57824698@20.40.21(+1G);MI0000083\|hsa-mir-26a-1&WithFlank&3\|+\|37969398\|37969486@16.36.21(+1G)&[MIMAT0000082&hsa-miR-26a-5p&offsets\|0\|-1(+1G);m-33&12\|-\|57824658\|57824679&offsets\|0\|-1(+1G)];[MIMAT0000082_1&hsa-miR-26a-5p&offsets\|0\|-1(+1G);m-34&3\|+\|37969413\|37969434&offsets\|0\|-1(+1G)]\|TTCAAGTAATCCAGGATAGGCG | >MI0000750\|hsa-mir-26a-2&WithFlank&12\|-\|57824603\|57824698 | 20 | 40 | 21(+1G) | TTCAAGTAATCCAGGATAGGCG | 2760.9 | -2.0 | 0.8 | -2.5 | 1.3E-02 | 3.9E-02 | BT20/MB231 | MITO |
| >hg38_21_+_8433172_8446622_RNA45SN1_45S_50ntFlanks@11541.11584.44\|GCGGCGCCTCGCCTCGGCCGGCGCCTAGCAGCCGACTTAGAACT | >hg38_21_+_8433172_8446622_RNA45SN1_45S_50ntFlanks | 11541 | 11584 | 44 | GCGGCGCCTCGCCTCGGCCGGCGCCTAGCAGCCGACTTAGAACT | 263.2 | 3.8 | 1.8 | 2.1 | 3.6E-02 | 3.9E-02 | MB231/MB468 | NUC |
| >hg38_21_+_8433172_8446622_RNA45SN1_45S_50ntFlanks@4735.4759.25\|ACGAAAGTCGGAGGTTCGAAGACGA | >hg38_21_+_8433172_8446622_RNA45SN1_45S_50ntFlanks | 4735 | 4759 | 25 | ACGAAAGTCGGAGGTTCGAAGACGA | 509.7 | 1.7 | 0.8 | 2.1 | 3.4E-02 | 3.9E-02 | BT20/MB468 | CYTO |
| >trna116_GluCTC_1_-_145399233_145399304@1.40.40,trna59_GluCTC_1_+_249168447_249168518@1.40.40,trna71_GluCTC_1_-_161439189_161439260@1.40.40,trna74_GluCTC_1_-_161431809_161431880@1.40.40,trna77_GluCTC_1_-_161424398_161424469@1.40.40,trna77_GluCTC_6_+_28949976_28950047@1.40.40,trna80_GluCTC_1_-_161417018_161417089@1.40.40,trna87_GluCTC_6_-_126101393_126101464@1.40.40\|TCCCTGGTGGTCTAGTGGTTAGGATTCGGCGCTCTCACCG | >trna116_GluCTC_1_-_145399233_145399304 | 1 | 40 | 40 | TCCCTGGTGGTCTAGTGGTTAGGATTCGGCGCTCTCACCG | 510.4 | 2.8 | 1.1 | 2.5 | 1.3E-02 | 3.9E-02 | BT20/MB231 | MITO |

| | | | | | | | | | | | | | |
|---|---|---|---|---|---|---|---|---|---|---|---|---|---|
| >MI0000108\|hsa-mir-103a-2&WithFlank&20\|+\|3917488\|3917577@54.76.23(+1C);MI0000109\|hsa-mir-103a-1&WithFlank&5\|-\|168560890\|168560979@54.76.23(+1C)&[MIMAT0000101&hsa-miR-103a-3p&offsets\|0\|0(+1C);m-12&20\|+\|3917541\|3917563&offsets\|0\|0(+1C)];[MIMAT0000101_1&hsa-miR-103a-3p&offsets\|0\|0(+1C);m-13&5\|-\|168560904\|168560926&offsets\|0\|0(+1C);m-44220&5\|-\|168560908\|168560926&offsets\|0\|+4(+1C)]\|AGCAGCATTGTACAGGGCTATGAC | >MI0000108\|hsa-mir-103a-2&WithFlank&20\|+\|3917488\|3917577 | 54 | 76 | 23(+1C) | AGCAGCATTGTACAGGGCTATGAC | 108.4 | 2.9 | 1.4 | 2.1 | 3.3E-02 | 3.9E-02 | MB231/MB468 | MP |
| >hg38_21_+_8433172_8446622_RNA45SN1_45S_50ntFlanks@10346.10382.37\|TCAAACGAGAACTTTGAAGGCCGAAGTGGAGAAGGGT | >hg38_21_+_8433172_8446622_RNA45SN1_45S_50ntFlanks | 10346 | 10382 | 37 | TCAAACGAGAACTTTGAAGGCCGAAGTGGAGAAGGGT | 172.8 | -1.8 | 0.9 | -2.1 | 3.3E-02 | 3.9E-02 | MB231/MB468 | MP |
| >hg38_21_+_8433172_8446622_RNA45SN1_45S_50ntFlanks@5472.5512.41\|ACGGCCCTGGCGGAGCGCTGAGAAGACGGTCGAACTTGACT | >hg38_21_+_8433172_8446622_RNA45SN1_45S_50ntFlanks | 5472 | 5512 | 41 | ACGGCCCTGGCGGAGCGCTGAGAAGACGGTCGAACTTGACT | 169.7 | 2.8 | 1.3 | 2.1 | 3.3E-02 | 3.9E-02 | MB231/MB468 | MP |
| >trnaMT_GlnTTG_MT_-_4329_4400@1.30.30,trnalookalike2_GlnTTG_1_-_564879_564950@1.30.30\|TAGGATGGGGTGTGATAGGTGGCACGGAGA | >trnaMT_GlnTTG_MT_-_4329_4400 | 1 | 30 | 30 | TAGGATGGGGTGTGATAGGTGGCACGGAGA | 2651.2 | 3.1 | 1.4 | 2.3 | 2.4E-02 | 3.9E-02 | BT20/MB468 | MP |
| >hg38_21_+_8433172_8446622_RNA45SN1_45S_50ntFlanks@12328.12346.19\|GTTTTAAGCAGGAGGTGTC | >hg38_21_+_8433172_8446622_RNA45SN1_45S_50ntFlanks | 12328 | 12346 | 19 | GTTTTAAGCAGGAGGTGTC | 128.1 | -4.0 | 1.8 | -2.3 | 2.4E-02 | 3.9E-02 | BT20/MB468 | MP |
| >trna111_HisGTG_1_-_147774845_147774916@-1T.22.23,trna118_HisGTG_1_-_145396881_145396952@-1T.22.23,trna16_HisGTG_1_+_146544773_146544844@-1T.22.23,trna1_HisGTG_15_+_45493349_45493420@-1T.22.23,trna21_HisGTG_1_+_147753471_147753542@-1T.22.23,trna33_HisGTG_6_+_27125906_27125977@-1T.22.23,trna7_HisGTG_9_-_14433938_14434009@-1T.22.23,trna8_HisGTG_15_-_45492611_45492682@-1T.22.23,trna9_HisGTG_15_-_45490804_45490875@-1T.22.23\|TGCCGTGATCGTATAGTGGTTAG | >trna111_HisGTG_1_-_147774845_147774916 | -1T | 22 | 23 | TGCCGTGATCGTATAGTGGTTAG | 478.6 | -3.9 | 1.5 | -2.6 | 1.1E-02 | 3.9E-02 | BT20/MB231 | MP |
| >hg38_21_+_8433172_8446622_RNA45SN1_45S_50ntFlanks@8336.8373.38\|AAGTACCGTAAGGGAAAGTTGAAAAGAACTTTGAAGAG | >hg38_21_+_8433172_8446622_RNA45SN1_45S_50ntFlanks | 8336 | 8373 | 38 | AAGTACCGTAAGGGAAAGTTGAAAAGAACTTTGAAGAG | 740.9 | 4.7 | 1.8 | 2.6 | 1.1E-02 | 3.9E-02 | BT20/MB231 | MP |
| >hg38_21_+_8433172_8446622_RNA45SN1_45S_50ntFlanks@12944.12972.29\|CGTAGACGACCTGCTTCTGGGTCGGGGTT | >hg38_21_+_8433172_8446622_RNA45SN1_45S_50ntFlanks | 12944 | 12972 | 29 | CGTAGACGACCTGCTTCTGGGTCGGGGTT | 120.2 | -3.4 | 1.6 | -2.2 | 3.1E-02 | 3.9E-02 | BT20/MB468 | MITO |
| >hg38_21_+_8433172_8446622_RNA45SN1_45S_50ntFlanks@8159.8178.20\|CGTGGGGGGCCCAAGTCCTT | >hg38_21_+_8433172_8446622_RNA45SN1_45S_50ntFlanks | 8159 | 8178 | 20 | CGTGGGGGGCCCAAGTCCTT | 339.2 | -3.2 | 1.5 | -2.2 | 3.1E-02 | 3.9E-02 | BT20/MB468 | MITO |
| >hg38_21_+_8433172_8446622_RNA45SN1_45S_50ntFlanks@9198.9220.23\|GTTCTCTCGGGGCCACGCGCGCG | >hg38_21_+_8433172_8446622_RNA45SN1_45S_50ntFlanks | 9198 | 9220 | 23 | GTTCTCTCGGGGCCACGCGCGCG | 892.0 | -3.9 | 1.8 | -2.2 | 3.1E-02 | 3.9E-02 | BT20/MB468 | MITO |
| >MI0000077\|hsa-mir-21&WithFlank&17\|+\|59841260\|59841343@18.36.19&[MIMAT0000076&hsa-miR-21-5p&offsets\|+4\|+1;m-1&17\|+\|59841273\|59841294&offsets\|+4\|+1]\|TTATCAGACTGATGTTGAC | >MI0000077\|hsa-mir-21&WithFlank&17\|+\|59841260\|59841343 | 18 | 36 | 19 | TTATCAGACTGATGTTGAC | 40.5 | 4.4 | 2.0 | 2.1 | 3.3E-02 | 3.9E-02 | MB231/MB468 | MP |
| >MI0003760\|hsa-mir-671&WithFlank&7\|+\|151238415\|151238544@35.58.24&[MIMAT0003880&hsa-miR-671-5p&offsets\|0\|+1;m-316&7\|+\|151238449\|151238472&offsets\|0\|0]\|AGGAAGCCCTGGAGGGGCTGGAGG | >MI0003760\|hsa-mir-671&WithFlank&7\|+\|151238415\|151238544 | 35 | 58 | 24 | AGGAAGCCCTGGAGGGGCTGGAGG | 294.9 | 2.3 | 0.9 | 2.5 | 1.4E-02 | 3.9E-02 | BT20/MB231 | CYTO |
| >hg38_21_+_8433172_8446622_RNA45SN1_45S_50ntFlanks@10663.10706.44\|CCTTGGAAAGCGTCGCGGTTCCGGCGGCGTCCGGTGAGCTCTCG | >hg38_21_+_8433172_8446622_RNA45SN1_45S_50ntFlanks | 10663 | 10706 | 44 | CCTTGGAAAGCGTCGCGGTTCCGGCGGCGTCCGGTGAGCTCTCG | 281.0 | -2.8 | 1.3 | -2.2 | 3.1E-02 | 3.9E-02 | BT20/MB468 | MITO |
| >hg38_21_+_8433172_8446622_RNA45SN1_45S_50ntFlanks@10349.10377.29\|AACGAGAACTTTGAAGGCCGAAGTGGAGA | >hg38_21_+_8433172_8446622_RNA45SN1_45S_50ntFlanks | 10349 | 10377 | 29 | AACGAGAACTTTGAAGGCCGAAGTGGAGA | 979.5 | -1.4 | 0.7 | -2.1 | 3.4E-02 | 3.9E-02 | MB231/MB468 | CYTO |
| >hg38_21_+_8433172_8446622_RNA45SN1_45S_50ntFlanks@12418.12449.32\|CGATGTCGGCTCTTCCTATCATTGTGAAGCAG | >hg38_21_+_8433172_8446622_RNA45SN1_45S_50ntFlanks | 12418 | 12449 | 32 | CGATGTCGGCTCTTCCTATCATTGTGAAGCAG | 598.9 | -1.2 | 0.6 | -2.1 | 3.6E-02 | 3.9E-02 | MB231/MB468 | NUC |
| >hg38_21_+_8433172_8446622_RNA45SN1_45S_50ntFlanks@6656.6686.31\|CTTAGCGGTGGATCACTCGGCTCGTGCGTCG | >hg38_21_+_8433172_8446622_RNA45SN1_45S_50ntFlanks | 6656 | 6686 | 31 | CTTAGCGGTGGATCACTCGGCTCGTGCGTCG | 280.9 | -1.3 | 0.6 | -2.2 | 3.1E-02 | 3.9E-02 | BT20/MB468 | MITO |
| >hg38_21_+_8433172_8446622_RNA45SN1_45S_50ntFlanks@10411.10445.35\|CAGTCGGTCCTGAGAGATGGGCGAGCGCCGTTCCG | >hg38_21_+_8433172_8446622_RNA45SN1_45S_50ntFlanks | 10411 | 10445 | 35 | CAGTCGGTCCTGAGAGATGGGCGAGCGCCGTTCCG | 129.2 | -3.9 | 1.7 | -2.3 | 2.4E-02 | 3.9E-02 | BT20/MB468 | MP |
| >hg38_21_+_8433172_8446622_RNA45SN1_45S_50ntFlanks@10770.10818.49\|GCAGGTCTCCAAGGTGAACAGCCTCTGGCATGTTGGAACAATGTAGGTA | >hg38_21_+_8433172_8446622_RNA45SN1_45S_50ntFlanks | 10770 | 10818 | 49 | GCAGGTCTCCAAGGTGAACAGCCTCTGGCATGTTGGAACAATGTAGGTA | 139.5 | -3.9 | 1.8 | -2.1 | 3.5E-02 | 3.9E-02 | BT20/MB468 | CYTO |
| >MI0000065\|hsa-let-7d&WithFlank&9\|+\|94178828\|94178926@14.34.21(+1C)&[MIMAT0000065&hsa-let-7d-5p&offsets\|0\|-1(+1C);m-90&9\|+\|94178841\|94178862&offsets\|0\|-1(+1C)]\|AGAGGTAGTAGGTTGCATAGTC | >MI0000065\|hsa-let-7d&WithFlank&9\|+\|94178828\|94178926 | 14 | 34 | 21(+1C) | AGAGGTAGTAGGTTGCATAGTC | 2142.1 | 2.2 | 0.9 | 2.5 | 1.4E-02 | 3.9E-02 | BT20/MB231 | CYTO |
| >am_MI0000809\|hsa-mir-151a&WithFlank&8\|-\|140732558\|140732659@53.70.18&[MIMAT0000757&hsa-miR-151a-3p&offsets\|0\|-3;m-30&8\|-\|140732587\|140732607&offsets\|0\|-3]\|CTAGACTGAAGCTCCTTG | >am_MI0000809\|hsa-mir-151a&WithFlank&8\|-\|140732558\|140732659 | 53 | 70 | 18 | CTAGACTGAAGCTCCTTG | 1945.7 | 1.9 | 0.8 | 2.5 | 1.4E-02 | 3.9E-02 | BT20/MB231 | CYTO |
| >hg38_21_+_8433172_8446622_RNA45SN1_45S_50ntFlanks@6697.6720.24\|CAGCTAGCTGCGAGAATTAATGTG | >hg38_21_+_8433172_8446622_RNA45SN1_45S_50ntFlanks | 6697 | 6720 | 24 | CAGCTAGCTGCGAGAATTAATGTG | 1472.3 | -2.1 | 0.9 | -2.5 | 1.4E-02 | 3.9E-02 | BT20/MB231 | CYTO |
| >hg38_21_+_8433172_8446622_RNA45SN1_45S_50ntFlanks@10443.10460.18\|CCGAAGGGACGGGCGATG | >hg38_21_+_8433172_8446622_RNA45SN1_45S_50ntFlanks | 10443 | 10460 | 18 | CCGAAGGGACGGGCGATG | 131.0 | 4.4 | 1.7 | 2.6 | 1.1E-02 | 3.9E-02 | BT20/MB231 | MP |
| >hg38_21_+_8433172_8446622_RNA45SN1_45S_50ntFlanks@5237.5264.28\|ACTGACTGGCTCAGCGTGTGCCTACCCT | >hg38_21_+_8433172_8446622_RNA45SN1_45S_50ntFlanks | 5237 | 5264 | 28 | ACTGACTGGCTCAGCGTGTGCCTACCCT | 222.6 | 4.2 | 1.7 | 2.6 | 1.1E-02 | 3.9E-02 | BT20/MB231 | MP |
| >hg38_21_+_8433172_8446622_RNA45SN1_45S_50ntFlanks@9482.9505.24\|ACGTGTTAGGACCCGAAAGATGGT | >hg38_21_+_8433172_8446622_RNA45SN1_45S_50ntFlanks | 9482 | 9505 | 24 | ACGTGTTAGGACCCGAAAGATGGT | 254.3 | 6.4 | 2.5 | 2.6 | 1.1E-02 | 3.9E-02 | BT20/MB231 | MP |
| >trnaMT_SerGCT_MT_+_12207_12265@29.62.34\|TGCCCCCATGTCTAACAACATGGCTTTCTCACCA | >trnaMT_SerGCT_MT_+_12207_12265 | 29 | 62 | 34 | TGCCCCCATGTCTAACAACATGGCTTTCTCACCA | 369.1 | -3.1 | 1.2 | -2.6 | 1.1E-02 | 3.9E-02 | BT20/MB231 | MP |
| >hg38_21_+_8433172_8446622_RNA45SN1_45S_50ntFlanks@6651.6678.28\|CGACTCTTAGCGGTGGATCACTCGGCTC | >hg38_21_+_8433172_8446622_RNA45SN1_45S_50ntFlanks | 6651 | 6678 | 28 | CGACTCTTAGCGGTGGATCACTCGGCTC | 1366.8 | -1.7 | 0.8 | -2.1 | 3.3E-02 | 3.9E-02 | MB231/MB468 | MP |
| >am_trnaMT_ProTGG_MT_-_15956_16023@10.32.23\|GTTTAAATTAGAATCTTAGCTTT | >am_trnaMT_ProTGG_MT_-_15956_16023 | 10 | 32 | 23 | GTTTAAATTAGAATCTTAGCTTT | 48.3 | 2.9 | 1.3 | 2.1 | 3.3E-02 | 3.9E-02 | MB231/MB468 | MP |
| >trnaMT_CysGCA_MT_-_5761_5826@47.69.23\|CTTCAAACCTGCCGGGGCTTCCA | >trnaMT_CysGCA_MT_-_5761_5826 | 47 | 69 | 23 | CTTCAAACCTGCCGGGGCTTCCA | 189.8 | -2.1 | 1.0 | -2.1 | 3.4E-02 | 3.9E-02 | MB231/MB468 | CYTO |
| >hg38_21_+_8433172_8446622_RNA45SN1_45S_50ntFlanks@11914.11943.30\|AGAGACATGAGAGGTGTAGAATAAGTGGGA | >hg38_21_+_8433172_8446622_RNA45SN1_45S_50ntFlanks | 11914 | 11943 | 30 | AGAGACATGAGAGGTGTAGAATAAGTGGGA | 95.5 | 5.2 | 2.0 | 2.6 | 1.1E-02 | 3.9E-02 | BT20/MB231 | MP |
| >MI0000253\|hsa-mir-148a&WithFlank&7\|-\|25949913\|25949992@50.71.22&[MIMAT0000243&hsa-miR-148a-3p&offsets\|0\|0;m-7&7\|-\|25949922\|25949943&offsets\|0\|0]\|TCAGTGCACTACAGAACTTTGT | >MI0000253\|hsa-mir-148a&WithFlank&7\|-\|25949913\|25949992 | 50 | 71 | 22 | TCAGTGCACTACAGAACTTTGT | 12303.4 | 2.1 | 1.0 | 2.1 | 3.6E-02 | 3.9E-02 | MB231/MB468 | MITO |
| >MI0000081\|hsa-mir-24-2&WithFlank&19\|-\|13836281\|13836365@56.73.18;MI0000080\|hsa-mir-24-1&WithFlank&9\|+\|95086015\|95086094@50.67.18&[MIMAT0000080&hsa-miR-24-3p&offsets\|0\|-4;m-36&19\|-\|13836288\|13836310&offsets\|0\|-5];[MIMAT0000080_1&hsa-miR-24-3p&offsets\|0\|-4;m-35&9\|+\|95086064\|95086086&offsets\|0\|-5]\|TGGCTCAGTTCAGCAGGA | >MI0000081\|hsa-mir-24-2&WithFlank&19\|-\|13836281\|13836365 | 56 | 73 | 18 | TGGCTCAGTTCAGCAGGA | 113.8 | 3.8 | 1.8 | 2.1 | 3.3E-02 | 3.9E-02 | MB231/MB468 | MP |
| >trna111_HisGTG_1_-_147774845_147774916@-1T.26.27,trna118_HisGTG_1_-_145396881_145396952@-1T.26.27,trna16_HisGTG_1_+_146544773_146544844@-1T.26.27,trna1_HisGTG_15_+_45493349_45493420@-1T.26.27,trna21_HisGTG_1_+_147753471_147753542@-1T.26.27,trna33_HisGTG_6_+_27125906_27125977@-1T.26.27,trna7_HisGTG_9_-_14433938_14434009@-1T.26.27,trna8_HisGTG_15_-_45492611_45492682@-1T.26.27,trna9_HisGTG_15_-_45490804_45490875@-1T.26.27\|TGCCGTGATCGTATAGTGGTTAGTACT | >trna111_HisGTG_1_-_147774845_147774916 | -1T | 26 | 27 | TGCCGTGATCGTATAGTGGTTAGTACT | 3580.8 | -2.6 | 1.0 | -2.5 | 1.1E-02 | 3.9E-02 | BT20/MB231 | NUC |
| >hg38_21_+_8433172_8446622_RNA45SN1_45S_50ntFlanks@10247.10284.38\|GCGACGAGTAGGAGGGCCGCTGCGGTGAGCCTTGAAGC | >hg38_21_+_8433172_8446622_RNA45SN1_45S_50ntFlanks | 10247 | 10284 | 38 | GCGACGAGTAGGAGGGCCGCTGCGGTGAGCCTTGAAGC | 151.9 | 5.1 | 2.0 | 2.5 | 1.1E-02 | 4.0E-02 | BT20/MB231 | MP |
| >am_trna10_ValCAC_5_-_180649395_180649467@1.36.36,trna18_ValCAC_5_-_180529253_180529325@1.36.36,trna2_ValCAC_5_+_180524070_180524142@1.36.36,trna6_ValCAC_5_+_180600650_180600722@1.36.36,trna85_ValCAC_1_-_161369490_161369562@1.36.36,trna90_ValCAC_1_-_149684088_149684161@1.36.36,trna98_ValCAC_1_-_149298555_149298627@1.36.36,trna9_ValCAC_6_+_26538282_26538354@1.36.36\|GTTTCCGTAGTGTAGTGGTTATCACGTTCGCCTCAC | >am_trna10_ValCAC_5_-_180649395_180649467 | 1 | 36 | 36 | GTTTCCGTAGTGTAGTGGTTATCACGTTCGCCTCAC | 628.3 | -2.1 | 0.9 | -2.5 | 1.3E-02 | 4.0E-02 | BT20/MB231 | MITO |
| >MI0000471\|hsa-mir-126&WithFlank&9\|+\|136670596\|136670692@58.77.20(+1U)&[MIMAT0000445&hsa-miR-126-3p&offsets\|0\|-2(+1U);m-40&9\|+\|136670653\|136670674&offsets\|0\|-2(+1U)]\|TCGTACCGTGAGTAATAATGT | >MI0000471\|hsa-mir-126&WithFlank&9\|+\|136670596\|136670692 | 58 | 77 | 20(+1U) | TCGTACCGTGAGTAATAATGT | 770.7 | 2.3 | 0.9 | 2.4 | 1.4E-02 | 4.0E-02 | BT20/MB231 | CYTO |
| >trna111_HisGTG_1_-_147774845_147774916@-1T.32.33,trna118_HisGTG_1_-_145396881_145396952@-1T.32.33,trna16_HisGTG_1_+_146544773_146544844@-1T.32.33,trna1_HisGTG_15_+_45493349_45493420@-1T.32.33,trna21_HisGTG_1_+_147753471_147753542@-1T.32.33,trna33_HisGTG_6_+_27125906_27125977@-1T.32.33,trna7_HisGTG_9_-_14433938_14434009@-1T.32.33,trna8_HisGTG_15_-_45492611_45492682@-1T.32.33,trna9_HisGTG_15_-_45490804_45490875@-1T.32.33\|TGCCGTGATCGTATAGTGGTTAGTACTCTGCGT | >trna111_HisGTG_1_-_147774845_147774916 | -1T | 32 | 33 | TGCCGTGATCGTATAGTGGTTAGTACTCTGCGT | 723.9 | -2.3 | 0.9 | -2.4 | 1.4E-02 | 4.0E-02 | BT20/MB231 | CYTO |
| >hg38_21_+_8433172_8446622_RNA45SN1_45S_50ntFlanks@9544.9576.33\|GGTGGAGGTCCGTAGCGGTCCTGACGTGCAAAT | >hg38_21_+_8433172_8446622_RNA45SN1_45S_50ntFlanks | 9544 | 9576 | 33 | GGTGGAGGTCCGTAGCGGTCCTGACGTGCAAAT | 224.3 | -2.1 | 0.9 | -2.4 | 1.4E-02 | 4.0E-02 | BT20/MB231 | CYTO |

<table>
<tr><th></th><th></th><th></th><th></th><th></th><th></th><th></th><th></th><th></th><th></th><th></th><th></th><th></th><th></th></tr>
<tr><td>>MI0000107|hsa-mir-29b-2&WithFlank&1|-|207802437|207802529@57.77.21(+1U);MI0000105|hsa-mir-29b-1&WithFlank&7|-|130877453|130877545@56.76.21(+1U)&[MIMAT0000100&hsa-miR-29b-3p&offsets|-1|-3(+1U);m-81&1|-|207802450|207802472&offsets|-1|-3(+1U)];[MIMAT0000100_1&hsa-miR-29b-3p&offsets|-1|-3(+1U);m-82&7|-|130877467|130877489&offsets|-1|-3(+1U)]|CTAGCACCATTTGAAATCAGTT</td><td>>MI0000107|hsa-mir-29b-2&WithFlank&1|-|207802437|207802529</td><td>57</td><td>77</td><td>21(+1U)</td><td>CTAGCACCATTTGAAATCAGTT</td><td>38.3</td><td>4.4</td><td>2.1</td><td>2.1</td><td>3.3E-02</td><td>4.0E-02</td><td>MB231/MB468</td><td>MP</td></tr>
<tr><td>>MI0000085|hsa-mir-27a&WithFlank&19|-|13836434|13836523@57.75.19(+1A)&[MIMAT0000084&hsa-miR-27a-3p&offsets|0|-2(+1A);m-43&19|-|13836447|13836467&offsets|0|-2(+1A)]|TTCACAGTGGCTAAGTTCCA</td><td>>MI0000085|hsa-mir-27a&WithFlank&19|-|13836434|13836523</td><td>57</td><td>75</td><td>19(+1A)</td><td>TTCACAGTGGCTAAGTTCCA</td><td>1175.1</td><td>2.4</td><td>1.1</td><td>2.2</td><td>2.4E-02</td><td>4.0E-02</td><td>BT20/MB468</td><td>MP</td></tr>
<tr><td>>hg38_21_+_8433172_8446622_RNA45SN1_45S_50ntFlanks@9307.9331.25|AAGGAGTCTAACACGTGCGCGAGTC</td><td>>hg38_21_+_8433172_8446622_RNA45SN1_45S_50ntFlanks</td><td>9307</td><td>9331</td><td>25</td><td>AAGGAGTCTAACACGTGCGCGAGTC</td><td>126.8</td><td>3.0</td><td>1.2</td><td>2.5</td><td>1.1E-02</td><td>4.0E-02</td><td>BT20/MB231</td><td>MP</td></tr>
<tr><td>>hg38_21_+_8433172_8446622_RNA45SN1_45S_50ntFlanks@12418.12455.38|CGATGTCGGCTCTTCCTATCATTGTGAAGCAGAATTCA</td><td>>hg38_21_+_8433172_8446622_RNA45SN1_45S_50ntFlanks</td><td>12418</td><td>12455</td><td>38</td><td>CGATGTCGGCTCTTCCTATCATTGTGAAGCAGAATTCA</td><td>641.2</td><td>-3.7</td><td>1.6</td><td>-2.2</td><td>2.5E-02</td><td>4.0E-02</td><td>BT20/MB468</td><td>MP</td></tr>
<tr><td>>hg38_21_+_8433172_8446622_RNA45SN1_45S_50ntFlanks@4736.4758.23|CGAAAGTCGGAGGTTCGAAGACG</td><td>>hg38_21_+_8433172_8446622_RNA45SN1_45S_50ntFlanks</td><td>4736</td><td>4758</td><td>23</td><td>CGAAAGTCGGAGGTTCGAAGACG</td><td>742.0</td><td>-2.3</td><td>1.0</td><td>-2.2</td><td>2.5E-02</td><td>4.0E-02</td><td>BT20/MB468</td><td>MP</td></tr>
<tr><td>>hg38_21_+_8433172_8446622_RNA45SN1_45S_50ntFlanks@12076.12096.21|AGGGGCTCTCGCTTCTGGCGC</td><td>>hg38_21_+_8433172_8446622_RNA45SN1_45S_50ntFlanks</td><td>12076</td><td>12096</td><td>21</td><td>AGGGGCTCTCGCTTCTGGCGC</td><td>81.7</td><td>-4.0</td><td>1.8</td><td>-2.2</td><td>2.5E-02</td><td>4.0E-02</td><td>BT20/MB468</td><td>MP</td></tr>
<tr><td>>hg38_21_+_8433172_8446622_RNA45SN1_45S_50ntFlanks@10906.10926.21|TGGGCGCGCGCCGCGGCTGGA</td><td>>hg38_21_+_8433172_8446622_RNA45SN1_45S_50ntFlanks</td><td>10906</td><td>10926</td><td>21</td><td>TGGGCGCGCGCCGCGGCTGGA</td><td>232.2</td><td>-2.1</td><td>1.0</td><td>-2.1</td><td>3.5E-02</td><td>4.0E-02</td><td>MB231/MB468</td><td>CYTO</td></tr>
<tr><td>>hg38_MT_+_1621_3279_RNR2_16S_50ntFlanks@203.230.28|ATAGCAAGGACTAACCCCTATACCTTCT</td><td>>hg38_MT_+_1621_3279_RNR2_16S_50ntFlanks</td><td>203</td><td>230</td><td>28</td><td>ATAGCAAGGACTAACCCCTATACCTTCT</td><td>25.1</td><td>4.4</td><td>2.1</td><td>2.1</td><td>3.3E-02</td><td>4.0E-02</td><td>MB231/MB468</td><td>MP</td></tr>
<tr><td>>hg38_21_+_8433172_8446622_RNA45SN1_45S_50ntFlanks@8159.8188.30|CGTGGGGGGCCCAAGTCCTTCTGATCGAGG</td><td>>hg38_21_+_8433172_8446622_RNA45SN1_45S_50ntFlanks</td><td>8159</td><td>8188</td><td>30</td><td>CGTGGGGGGCCCAAGTCCTTCTGATCGAGG</td><td>84.5</td><td>-2.2</td><td>1.0</td><td>-2.2</td><td>2.5E-02</td><td>4.0E-02</td><td>BT20/MB468</td><td>MP</td></tr>
<tr><td>>hg38_21_+_8433172_8446622_RNA45SN1_45S_50ntFlanks@12539.12567.29|ATGATGTGTTGTTGCCATGGTAATCCTGC</td><td>>hg38_21_+_8433172_8446622_RNA45SN1_45S_50ntFlanks</td><td>12539</td><td>12567</td><td>29</td><td>ATGATGTGTTGTTGCCATGGTAATCCTGC</td><td>822.7</td><td>-4.2</td><td>1.9</td><td>-2.2</td><td>2.5E-02</td><td>4.0E-02</td><td>BT20/MB468</td><td>MP</td></tr>
<tr><td>>trna111_HisGTG_1_-_147774845_147774916@1.30.30,trna118_HisGTG_1_-_145396881_145396952@1.30.30,trna16_HisGTG_1_+_146544773_146544844@1.30.30,trna1_HisGTG_15_+_45493349_45493420@1.30.30,trna21_HisGTG_1_+_147753471_147753542@1.30.30,trna33_HisGTG_6_+_27125906_27125977@1.30.30,trna7_HisGTG_9_-_14433938_14434009@1.30.30,trna8_HisGTG_15_-_45492611_45492682@1.30.30,trna9_HisGTG_15_-_45490804_45490875@1.30.30|GCCGTGATCGTATAGTGGTTAGTACTCTGC</td><td>>trna111_HisGTG_1_-_147774845_147774916</td><td>1</td><td>30</td><td>30</td><td>GCCGTGATCGTATAGTGGTTAGTACTCTGC</td><td>1182.7</td><td>1.8</td><td>0.8</td><td>2.1</td><td>3.3E-02</td><td>4.0E-02</td><td>MB231/MB468</td><td>MP</td></tr>
<tr><td>>MI0000433|hsa-let-7g&WithFlank&3|-|52268272|52268367@11.32.22(+1A)&[MIMAT0000414&hsa-let-7g-5p&offsets|0|0(+1A);m-58&3|-|52268336|52268357&offsets|0|0(+1A)]|TGAGGTAGTAGTTTGTACAGTTA</td><td>>MI0000433|hsa-let-7g&WithFlank&3|-|52268272|52268367</td><td>11</td><td>32</td><td>22(+1A)</td><td>TGAGGTAGTAGTTTGTACAGTTA</td><td>1899.9</td><td>2.4</td><td>1.0</td><td>2.4</td><td>1.4E-02</td><td>4.0E-02</td><td>BT20/MB231</td><td>CYTO</td></tr>
<tr><td>>hg38_21_+_8433172_8446622_RNA45SN1_45S_50ntFlanks@6661.6707.47|CGGTGGATCACTCGGCTCGTGCGTCGATGAAGAACGCAGCTAGCTGC</td><td>>hg38_21_+_8433172_8446622_RNA45SN1_45S_50ntFlanks</td><td>6661</td><td>6707</td><td>47</td><td>CGGTGGATCACTCGGCTCGTGCGTCGATGAAGAACGCAGCTAGCTGC</td><td>1657.6</td><td>3.3</td><td>1.3</td><td>2.5</td><td>1.1E-02</td><td>4.0E-02</td><td>BT20/MB231</td><td>NUC</td></tr>
<tr><td>>hg38_21_+_8433172_8446622_RNA45SN1_45S_50ntFlanks@12274.12295.22|CAGTACGAATACAGACCGTGAA</td><td>>hg38_21_+_8433172_8446622_RNA45SN1_45S_50ntFlanks</td><td>12274</td><td>12295</td><td>22</td><td>CAGTACGAATACAGACCGTGAA</td><td>81.0</td><td>-3.8</td><td>1.7</td><td>-2.2</td><td>2.5E-02</td><td>4.0E-02</td><td>BT20/MB468</td><td>MP</td></tr>
<tr><td>>hg38_21_+_8433172_8446622_RNA45SN1_45S_50ntFlanks@4390.4423.34|TCTTGGGAGCGGGCGGGCGGTCCGCCGCGAGGCG</td><td>>hg38_21_+_8433172_8446622_RNA45SN1_45S_50ntFlanks</td><td>4390</td><td>4423</td><td>34</td><td>TCTTGGGAGCGGGCGGGCGGTCCGCCGCGAGGCG</td><td>95.8</td><td>-3.9</td><td>1.7</td><td>-2.2</td><td>2.5E-02</td><td>4.0E-02</td><td>BT20/MB468</td><td>MP</td></tr>
<tr><td>>trna152_ValCAC_6_-_27248049_27248121@1.29.29|GCTTCTGTAGTGTAGTGGTTATCACGTTC</td><td>>trna152_ValCAC_6_-_27248049_27248121</td><td>1</td><td>29</td><td>29</td><td>GCTTCTGTAGTGTAGTGGTTATCACGTTC</td><td>7371.9</td><td>-2.5</td><td>1.1</td><td>-2.2</td><td>2.5E-02</td><td>4.0E-02</td><td>BT20/MB468</td><td>MP</td></tr>
<tr><td>>MI0000489|hsa-mir-195&WithFlank&17|-|7017609|7017707@21.41.21&[MIMAT0000461&hsa-miR-195-5p&offsets|0|0;m-191&17|-|7017666|7017687&offsets|0|-1]|TAGCAGCACAGAAATATTGGC</td><td>>MI0000489|hsa-mir-195&WithFlank&17|-|7017609|7017707</td><td>21</td><td>41</td><td>21</td><td>TAGCAGCACAGAAATATTGGC</td><td>94.4</td><td>4.4</td><td>2.0</td><td>2.2</td><td>2.5E-02</td><td>4.0E-02</td><td>BT20/MB468</td><td>MP</td></tr>
<tr><td>>hg38_MT_+_598_1651_RNR1_12S_50ntFlanks@971.1004.34|TGGTAAGTGTACTGGAAAGTGCACTTGGACGAAC</td><td>>hg38_MT_+_598_1651_RNR1_12S_50ntFlanks</td><td>971</td><td>1004</td><td>34</td><td>TGGTAAGTGTACTGGAAAGTGCACTTGGACGAAC</td><td>375.1</td><td>-3.0</td><td>1.2</td><td>-2.5</td><td>1.1E-02</td><td>4.0E-02</td><td>BT20/MB231</td><td>MP</td></tr>
<tr><td>>MI0000298|hsa-mir-221&WithFlank&X|-|45746151|45746272@71.90.20(+1A)&[MIMAT0000278&hsa-miR-221-3p&offsets|0|-3(+1A);m-91&X|-|45746181|45746202&offsets|0|-2(+1A)]|AGCTACATTGTCTGCTGGGTA</td><td>>MI0000298|hsa-mir-221&WithFlank&X|-|45746151|45746272</td><td>71</td><td>90</td><td>20(+1A)</td><td>AGCTACATTGTCTGCTGGGTA</td><td>83.8</td><td>4.4</td><td>2.0</td><td>2.2</td><td>2.5E-02</td><td>4.0E-02</td><td>BT20/MB468</td><td>MP</td></tr>
<tr><td>>hg38_21_+_8433172_8446622_RNA45SN1_45S_50ntFlanks@12474.12496.23|ACCCACTAATAGGGAACGTGAGC</td><td>>hg38_21_+_8433172_8446622_RNA45SN1_45S_50ntFlanks</td><td>12474</td><td>12496</td><td>23</td><td>ACCCACTAATAGGGAACGTGAGC</td><td>876.8</td><td>1.9</td><td>0.9</td><td>2.1</td><td>3.6E-02</td><td>4.0E-02</td><td>MB231/MB468</td><td>NUC</td></tr>
<tr><td>>hg38_21_+_8433172_8446622_RNA45SN1_45S_50ntFlanks@12396.12418.23|GCGACGTCGCTTTTTGATCCTTC</td><td>>hg38_21_+_8433172_8446622_RNA45SN1_45S_50ntFlanks</td><td>12396</td><td>12418</td><td>23</td><td>GCGACGTCGCTTTTTGATCCTTC</td><td>778.2</td><td>-2.6</td><td>1.1</td><td>-2.4</td><td>1.4E-02</td><td>4.0E-02</td><td>BT20/MB231</td><td>CYTO</td></tr>
<tr><td>>hg38_21_+_8433172_8446622_RNA45SN1_45S_50ntFlanks@13008.13039.32|ATTGAAAGTCAGCCCTCGACACAAGGGTTTGT</td><td>>hg38_21_+_8433172_8446622_RNA45SN1_45S_50ntFlanks</td><td>13008</td><td>13039</td><td>32</td><td>ATTGAAAGTCAGCCCTCGACACAAGGGTTTGT</td><td>1802.3</td><td>2.3</td><td>1.0</td><td>2.2</td><td>2.5E-02</td><td>4.0E-02</td><td>BT20/MB468</td><td>MP</td></tr>
<tr><td>>hg38_21_+_8433172_8446622_RNA45SN1_45S_50ntFlanks@5485.5518.34|AGCGCTGAGAAGACGGTCGAACTTGACTATCTAG</td><td>>hg38_21_+_8433172_8446622_RNA45SN1_45S_50ntFlanks</td><td>5485</td><td>5518</td><td>34</td><td>AGCGCTGAGAAGACGGTCGAACTTGACTATCTAG</td><td>698.6</td><td>-3.7</td><td>1.7</td><td>-2.2</td><td>3.1E-02</td><td>4.0E-02</td><td>BT20/MB468</td><td>MITO</td></tr>
<tr><td>>MI0000272|hsa-mir-182&WithFlank&7|-|129770377|129770498@29.51.23(+1A)&[MIMAT0000259&hsa-miR-182-5p&offsets|0|-1(+1A);m-22&7|-|129770449|129770470&offsets|0|+1(+1A)]|TTTGGCAATGGTAGAACTCACACA</td><td>>MI0000272|hsa-mir-182&WithFlank&7|-|129770377|129770498</td><td>29</td><td>51</td><td>23(+1A)</td><td>TTTGGCAATGGTAGAACTCACACA</td><td>1424.7</td><td>1.9</td><td>0.8</td><td>2.5</td><td>1.3E-02</td><td>4.0E-02</td><td>BT20/MB231</td><td>MITO</td></tr>
<tr><td>>hg38_21_+_8433172_8446622_RNA45SN1_45S_50ntFlanks@4130.4150.21|ATTCCGGAGAGGGAGCCTGAG</td><td>>hg38_21_+_8433172_8446622_RNA45SN1_45S_50ntFlanks</td><td>4130</td><td>4150</td><td>21</td><td>ATTCCGGAGAGGGAGCCTGAG</td><td>421.7</td><td>-4.2</td><td>2.0</td><td>-2.1</td><td>3.6E-02</td><td>4.0E-02</td><td>BT20/MB468</td><td>NUC</td></tr>
<tr><td>>hg38_21_+_8433172_8446622_RNA45SN1_45S_50ntFlanks@12249.12272.24|CAAAAGCTCGCTTGATCTTGATTT</td><td>>hg38_21_+_8433172_8446622_RNA45SN1_45S_50ntFlanks</td><td>12249</td><td>12272</td><td>24</td><td>CAAAAGCTCGCTTGATCTTGATTT</td><td>251.9</td><td>-2.1</td><td>1.0</td><td>-2.1</td><td>3.5E-02</td><td>4.0E-02</td><td>MB231/MB468</td><td>CYTO</td></tr>
<tr><td>>trna116_GluCTC_1_-_145399233_145399304@32.73.42,trna71_GluCTC_1_-_161439189_161439260@32.73.42,trna74_GluCTC_1_-_161431809_161431880@32.73.42,trna77_GluCTC_1_-_161424398_161424469@32.73.42,trna77_GluCTC_6_+_28949976_28950047@32.73.42,trna80_GluCTC_1_-_161417018_161417089@32.73.42,trna87_GluCTC_6_-_126101393_126101464@32.73.42|CTCTCACCGCCGCGGCCCGGGTTCGATTCCCGGTCAGGGAAC</td><td>>trna116_GluCTC_1_-_145399233_145399304</td><td>32</td><td>73</td><td>42</td><td>CTCTCACCGCCGCGGCCCGGGTTCGATTCCCGGTCAGGGAAC</td><td>152.2</td><td>-3.7</td><td>1.7</td><td>-2.2</td><td>3.1E-02</td><td>4.0E-02</td><td>BT20/MB468</td><td>MITO</td></tr>
<tr><td>>hg38_21_+_8433172_8446622_RNA45SN1_45S_50ntFlanks@12958.12993.36|TTCTGGGTCGGGGTTTCGTACGTAGCAGAGCAGCTC</td><td>>hg38_21_+_8433172_8446622_RNA45SN1_45S_50ntFlanks</td><td>12958</td><td>12993</td><td>36</td><td>TTCTGGGTCGGGGTTTCGTACGTAGCAGAGCAGCTC</td><td>1398.3</td><td>-2.4</td><td>1.0</td><td>-2.4</td><td>1.4E-02</td><td>4.0E-02</td><td>BT20/MB231</td><td>CYTO</td></tr>
<tr><td>>hg38_1_-_228634819_228635039_RNA5S12_5S_50ntFlanks@130.170.41|TGGATGGGAGACCGCCTGGGAATACCGGGTGCTGTAGGCTT</td><td>>hg38_1_-_228634819_228635039_RNA5S12_5S_50ntFlanks</td><td>130</td><td>170</td><td>41</td><td>TGGATGGGAGACCGCCTGGGAATACCGGGTGCTGTAGGCTT</td><td>184.0</td><td>2.7</td><td>1.3</td><td>2.1</td><td>3.6E-02</td><td>4.0E-02</td><td>MB231/MB468</td><td>MITO</td></tr>
<tr><td>>trna116_GluCTC_1_-_145399233_145399304@39.73.35,trna134_GluTTC_1_-_16861774_16861845@39.73.35,trna71_GluCTC_1_-_161439189_161439260@39.73.35,trna74_GluCTC_1_-_161431809_161431880@39.73.35,trna77_GluCTC_1_-_161424398_161424469@39.73.35,trna77_GluCTC_6_+_28949976_28950047@39.73.35,trna80_GluCTC_1_-_161417018_161417089@39.73.35,trna84_GluTTC_1_-_161391883_161391954@39.73.35,trna87_GluCTC_6_-_126101393_126101464@39.73.35|CGCCGCGGCCCGGGTTCGATTCCCGGTCAGGGAAC</td><td>>trna116_GluCTC_1_-_145399233_145399304</td><td>39</td><td>73</td><td>35</td><td>CGCCGCGGCCCGGGTTCGATTCCCGGTCAGGGAAC</td><td>418.7</td><td>-2.9</td><td>1.2</td><td>-2.4</td><td>1.5E-02</td><td>4.0E-02</td><td>BT20/MB231</td><td>CYTO</td></tr>
<tr><td>>MI0000483|hsa-mir-186&WithFlank&1|-|71067625|71067722@21.42.22&[MIMAT0000456&hsa-miR-186-5p&offsets|0|0;m-84&1|-|71067681|71067702&offsets|0|0]|CAAAGAATTCTCCTTTTGGGCT</td><td>>MI0000483|hsa-mir-186&WithFlank&1|-|71067625|71067722</td><td>21</td><td>42</td><td>22</td><td>CAAAGAATTCTCCTTTTGGGCT</td><td>2613.1</td><td>-2.2</td><td>0.9</td><td>-2.5</td><td>1.1E-02</td><td>4.0E-02</td><td>BT20/MB231</td><td>MP</td></tr>
<tr><td>>hg38_21_+_8433172_8446622_RNA45SN1_45S_50ntFlanks@9853.9874.22|CGCTGCGGGATGAACCGAACGC</td><td>>hg38_21_+_8433172_8446622_RNA45SN1_45S_50ntFlanks</td><td>9853</td><td>9874</td><td>22</td><td>CGCTGCGGGATGAACCGAACGC</td><td>145.1</td><td>4.3</td><td>1.7</td><td>2.5</td><td>1.1E-02</td><td>4.0E-02</td><td>BT20/MB231</td><td>MP</td></tr>
<tr><td>>MI0000077|hsa-mir-21&WithFlank&17|+|59841260|59841343@14.34.21(+1U)&[MIMAT0000076&hsa-miR-21-5p&offsets|0|-1(+1U);m-1&17|+|59841273|59841294&offsets|0|-1(+1U)]|TAGCTTATCAGACTGATGTTGT</td><td>>MI0000077|hsa-mir-21&WithFlank&17|+|59841260|59841343</td><td>14</td><td>34</td><td>21(+1U)</td><td>TAGCTTATCAGACTGATGTTGT</td><td>5383.7</td><td>3.2</td><td>1.4</td><td>2.2</td><td>2.5E-02</td><td>4.0E-02</td><td>BT20/MB468</td><td>MP</td></tr>
<tr><td>>hg38_21_+_8433172_8446622_RNA45SN1_45S_50ntFlanks@12414.12448.35|CCTTCGATGTCGGCTCTTCCTATCATTGTGAAGCA</td><td>>hg38_21_+_8433172_8446622_RNA45SN1_45S_50ntFlanks</td><td>12414</td><td>12448</td><td>35</td><td>CCTTCGATGTCGGCTCTTCCTATCATTGTGAAGCA</td><td>1034.9</td><td>-2.1</td><td>0.9</td><td>-2.4</td><td>1.5E-02</td><td>4.0E-02</td><td>BT20/MB231</td><td>CYTO</td></tr>
<tr><td>>hg38_21_+_8433172_8446622_RNA45SN1_45S_50ntFlanks@4047.4066.20|ATTCGAACGTCTGCCCTATC</td><td>>hg38_21_+_8433172_8446622_RNA45SN1_45S_50ntFlanks</td><td>4047</td><td>4066</td><td>20</td><td>ATTCGAACGTCTGCCCTATC</td><td>846.7</td><td>-2.1</td><td>0.9</td><td>-2.4</td><td>1.5E-02</td><td>4.0E-02</td><td>BT20/MB231</td><td>CYTO</td></tr>
<tr><td>>hg38_21_+_8433172_8446622_RNA45SN1_45S_50ntFlanks@9932.9955.24|ATAGACAGCAGGACGGTGGCCATG</td><td>>hg38_21_+_8433172_8446622_RNA45SN1_45S_50ntFlanks</td><td>9932</td><td>9955</td><td>24</td><td>ATAGACAGCAGGACGGTGGCCATG</td><td>214.7</td><td>5.0</td><td>1.9</td><td>2.5</td><td>1.1E-02</td><td>4.0E-02</td><td>BT20/MB231</td><td>MP</td></tr>
<tr><td>>MI0000070|hsa-mir-16-1&WithFlank&13|-|50048967|50049067@20.40.21(+1A);MI0000115|hsa-mir-16-2&WithFlank&3|+|160404739|160404831@16.36.21(+1A)&[MIMAT0000069&hsa-miR-16-5p&offsets|0|-1(+1A);m-62&13|-|50049027|50049048&offsets|0|-1(+1A)];[MIMAT0000069_1&hsa-miR-16-5p&offsets|0|-1(+1A);m-61&3|+|160404754|160404775&offsets|0|-1(+1A)]|TAGCAGCACGTAAATATTGGCA</td><td>>MI0000070|hsa-mir-16-1&WithFlank&13|-|50048967|50049067</td><td>20</td><td>40</td><td>21(+1A)</td><td>TAGCAGCACGTAAATATTGGCA</td><td>2106.1</td><td>2.0</td><td>1.0</td><td>2.1</td><td>3.3E-02</td><td>4.0E-02</td><td>MB231/MB468</td><td>MP</td></tr>
<tr><td>>hg38_21_+_8433172_8446622_RNA45SN1_45S_50ntFlanks@9937.9979.43|CAGCAGGACGGTGGCCATGGAAGTCGGAATCCGCTAAGGAGTG</td><td>>hg38_21_+_8433172_8446622_RNA45SN1_45S_50ntFlanks</td><td>9937</td><td>9979</td><td>43</td><td>CAGCAGGACGGTGGCCATGGAAGTCGGAATCCGCTAAGGAGTG</td><td>39.2</td><td>4.8</td><td>2.1</td><td>2.2</td><td>2.5E-02</td><td>4.0E-02</td><td>BT20/MB468</td><td>MP</td></tr>
</table>

<table>
<tr><th></th><th></th><th></th><th></th><th></th><th></th><th></th><th></th><th></th><th></th><th></th><th></th><th></th><th></th></tr>
<tr><td>>trna111_HisGTG_1_-_147774845_147774916@1.35.35,trna118_HisGTG_1_-_145396881_145396952@1.35.35,trna16_HisGTG_1_+_146544773_146544844@1.35.35,trna1_HisGTG_15_+_45493349_45493420@1.35.35,trna21_HisGTG_1_+_147753471_147753542@1.35.35,trna33_HisGTG_6_+_27125906_27125977@1.35.35,trna7_HisGTG_9_-_14433938_14434009@1.35.35,trna8_HisGTG_15_-_45492611_45492682@1.35.35,trna9_HisGTG_15_-_45490804_45490875@1.35.35|GCCGTGATCGTATAGTGGTTAGTACTCTGCGTTGT</td><td>>trna111_HisGTG_1_-_147774845_147774916</td><td>1</td><td>35</td><td>35</td><td>GCCGTGATCGTATAGTGGTTAGTACTCTGCGTTGT</td><td>3294.1</td><td>2.9</td><td>1.2</td><td>2.4</td><td>1.5E-02</td><td>4.0E-02</td><td>BT20/MB231</td><td>CYTO</td></tr>
<tr><td>>hg38_21_+_8433172_8446622_RNA45SN1_45S_50ntFlanks@5073.5104.32|AATTCCGATAACGAACGAGACTCTGGCATGCT</td><td>>hg38_21_+_8433172_8446622_RNA45SN1_45S_50ntFlanks</td><td>5073</td><td>5104</td><td>32</td><td>AATTCCGATAACGAACGAGACTCTGGCATGCT</td><td>177.7</td><td>-2.3</td><td>1.1</td><td>-2.2</td><td>3.1E-02</td><td>4.0E-02</td><td>BT20/MB468</td><td>MITO</td></tr>
<tr><td>>hg38_21_+_8433172_8446622_RNA45SN1_45S_50ntFlanks@4130.4158.29|ATTCCGGAGAGGGAGCCTGAGAAACGGCT</td><td>>hg38_21_+_8433172_8446622_RNA45SN1_45S_50ntFlanks</td><td>4130</td><td>4158</td><td>29</td><td>ATTCCGGAGAGGGAGCCTGAGAAACGGCT</td><td>83.9</td><td>-3.9</td><td>1.7</td><td>-2.2</td><td>2.5E-02</td><td>4.0E-02</td><td>BT20/MB468</td><td>MP</td></tr>
<tr><td>>trnaMT_AsnGTT_MT_-_5657_5729@38.76.39,trnalookalike5_AsnGTT_1_-_566207_566279@38.76.39|ACTAAGTGTTTGTGGGTTTAAGTCCCATTGGTCTAGCCA</td><td>>trnaMT_AsnGTT_MT_-_5657_5729</td><td>38</td><td>76</td><td>39</td><td>ACTAAGTGTTTGTGGGTTTAAGTCCCATTGGTCTAGCCA</td><td>379.0</td><td>3.2</td><td>1.5</td><td>2.1</td><td>3.7E-02</td><td>4.0E-02</td><td>MB231/MB468</td><td>MITO</td></tr>
<tr><td>>hg38_1_-_228634819_228635039_RNA5S12_5S_50ntFlanks@136.170.35|GGAGACCGCCTGGGAATACCGGGTGCTGTAGGCTT</td><td>>hg38_1_-_228634819_228635039_RNA5S12_5S_50ntFlanks</td><td>136</td><td>170</td><td>35</td><td>GGAGACCGCCTGGGAATACCGGGTGCTGTAGGCTT</td><td>1438.5</td><td>-1.3</td><td>0.6</td><td>-2.1</td><td>3.3E-02</td><td>4.0E-02</td><td>MB231/MB468</td><td>MP</td></tr>
<tr><td>>MI0000469|hsa-mir-125a&WithFlank&19|+|51693248|51693345@59.79.21&[MIMAT0004602&hsa-miR-125a-3p&offsets|0|-1;m-333&19|+|51693306|51693326&offsets|0|0]|ACAGGTGAGGTTCTTGGGAGC</td><td>>MI0000469|hsa-mir-125a&WithFlank&19|+|51693248|51693345</td><td>59</td><td>79</td><td>21</td><td>ACAGGTGAGGTTCTTGGGAGC</td><td>98.8</td><td>2.9</td><td>1.4</td><td>2.1</td><td>3.3E-02</td><td>4.0E-02</td><td>MB231/MB468</td><td>MP</td></tr>
<tr><td>>hg38_21_+_8433172_8446622_RNA45SN1_45S_50ntFlanks@12715.12747.33|AGCCTCGGTTGGCCTCGGATAGCCGGTCCCCCG</td><td>>hg38_21_+_8433172_8446622_RNA45SN1_45S_50ntFlanks</td><td>12715</td><td>12747</td><td>33</td><td>AGCCTCGGTTGGCCTCGGATAGCCGGTCCCCCG</td><td>430.2</td><td>-4.2</td><td>1.9</td><td>-2.2</td><td>2.5E-02</td><td>4.0E-02</td><td>BT20/MB468</td><td>MP</td></tr>
<tr><td>>hg38_21_+_8433172_8446622_RNA45SN1_45S_50ntFlanks@5490.5516.27|TGAGAAGACGGTCGAACTTGACTATCT</td><td>>hg38_21_+_8433172_8446622_RNA45SN1_45S_50ntFlanks</td><td>5490</td><td>5516</td><td>27</td><td>TGAGAAGACGGTCGAACTTGACTATCT</td><td>2654.3</td><td>3.2</td><td>1.3</td><td>2.5</td><td>1.1E-02</td><td>4.0E-02</td><td>BT20/MB231</td><td>NUC</td></tr>
<tr><td>>hg38_21_+_8433172_8446622_RNA45SN1_45S_50ntFlanks@4722.4759.38|TCATTAATCAAGAACGAAAGTCGGAGGTTCGAAGACGA</td><td>>hg38_21_+_8433172_8446622_RNA45SN1_45S_50ntFlanks</td><td>4722</td><td>4759</td><td>38</td><td>TCATTAATCAAGAACGAAAGTCGGAGGTTCGAAGACGA</td><td>1735.9</td><td>-2.9</td><td>1.3</td><td>-2.2</td><td>2.5E-02</td><td>4.0E-02</td><td>BT20/MB468</td><td>MP</td></tr>
<tr><td>>hg38_21_+_8433172_8446622_RNA45SN1_45S_50ntFlanks@7977.8013.37|CGACCTCAGATCAGACGTGGCGACCCGCTGAATTTAA</td><td>>hg38_21_+_8433172_8446622_RNA45SN1_45S_50ntFlanks</td><td>7977</td><td>8013</td><td>37</td><td>CGACCTCAGATCAGACGTGGCGACCCGCTGAATTTAA</td><td>177.1</td><td>2.1</td><td>1.0</td><td>2.1</td><td>3.7E-02</td><td>4.0E-02</td><td>MB231/MB468</td><td>MITO</td></tr>
<tr><td>>am_trna119_LysCTT_1_-_145395522_145395594@7.35.29,trna11_LysCTT_5_-_180648979_180649051@7.35.29,trna128_LysCTT_1_-_55423542_55423614@7.35.29,trna13_LysCTT_6_+_26556774_26556846@7.35.29,trna6_LysCTT_19_-_52425396_52425468@7.35.29,trna7_LysCTT_16_+_3225692_3225764@7.35.29,trna9_LysCTT_5_+_180634755_180634827@7.35.29|CTAGCTCAGTCGGTAGAGCATGAGACTCT</td><td>>am_trna119_LysCTT_1_-_145395522_145395594</td><td>7</td><td>35</td><td>29</td><td>CTAGCTCAGTCGGTAGAGCATGAGACTCT</td><td>173.7</td><td>-3.8</td><td>1.5</td><td>-2.5</td><td>1.1E-02</td><td>4.0E-02</td><td>BT20/MB231</td><td>MP</td></tr>
<tr><td>>hg38_MT_+_1621_3279_RNR2_16S_50ntFlanks@1439.1457.19|ACGTGATCTGAGTTCAGAC</td><td>>hg38_MT_+_1621_3279_RNR2_16S_50ntFlanks</td><td>1439</td><td>1457</td><td>19</td><td>ACGTGATCTGAGTTCAGAC</td><td>528.0</td><td>-2.7</td><td>1.1</td><td>-2.5</td><td>1.1E-02</td><td>4.0E-02</td><td>BT20/MB231</td><td>MP</td></tr>
<tr><td>>hg38_21_+_8433172_8446622_RNA45SN1_45S_50ntFlanks@6719.6756.38|TGAATTGCAGGACACATTGATCATCGACACTTCGAACG</td><td>>hg38_21_+_8433172_8446622_RNA45SN1_45S_50ntFlanks</td><td>6719</td><td>6756</td><td>38</td><td>TGAATTGCAGGACACATTGATCATCGACACTTCGAACG</td><td>226.6</td><td>-1.3</td><td>0.6</td><td>-2.1</td><td>3.5E-02</td><td>4.0E-02</td><td>MB231/MB468</td><td>CYTO</td></tr>
<tr><td>>hg38_21_+_8433172_8446622_RNA45SN1_45S_50ntFlanks@7975.8012.38|CGCGACCTCAGATCAGACGTGGCGACCCGCTGAATTTA</td><td>>hg38_21_+_8433172_8446622_RNA45SN1_45S_50ntFlanks</td><td>7975</td><td>8012</td><td>38</td><td>CGCGACCTCAGATCAGACGTGGCGACCCGCTGAATTTA</td><td>75640.1</td><td>-1.7</td><td>0.8</td><td>-2.1</td><td>3.7E-02</td><td>4.0E-02</td><td>MB231/MB468</td><td>MITO</td></tr>
<tr><td>>MI0000465|hsa-mir-191&WithFlank&3|-|49020612|49020715@22.43.22(+1A)&[MIMAT0000440&hsa-miR-191-5p&offsets|0|-1(+1A);m-48&3|-|49020672|49020694&offsets|0|-1(+1A)]|CAACGGAATCCCAAAAGCAGCTA</td><td>>MI0000465|hsa-mir-191&WithFlank&3|-|49020612|49020715</td><td>22</td><td>43</td><td>22(+1A)</td><td>CAACGGAATCCCAAAAGCAGCTA</td><td>1960.0</td><td>1.4</td><td>0.6</td><td>2.5</td><td>1.1E-02</td><td>4.0E-02</td><td>BT20/MB231</td><td>NUC</td></tr>
<tr><td>>hg38_21_+_8433172_8446622_RNA45SN1_45S_50ntFlanks@476.503.28|TTCGTGATCGATGTGGTGACGTCGTGCT</td><td>>hg38_21_+_8433172_8446622_RNA45SN1_45S_50ntFlanks</td><td>476</td><td>503</td><td>28</td><td>TTCGTGATCGATGTGGTGACGTCGTGCT</td><td>107.1</td><td>4.0</td><td>1.8</td><td>2.2</td><td>2.5E-02</td><td>4.0E-02</td><td>BT20/MB468</td><td>MP</td></tr>
<tr><td>>hg38_21_+_8433172_8446622_RNA45SN1_45S_50ntFlanks@5469.5499.31|CCCACGGCCCTGGCGGAGCGCTGAGAAGACG</td><td>>hg38_21_+_8433172_8446622_RNA45SN1_45S_50ntFlanks</td><td>5469</td><td>5499</td><td>31</td><td>CCCACGGCCCTGGCGGAGCGCTGAGAAGACG</td><td>439.7</td><td>-2.7</td><td>1.3</td><td>-2.1</td><td>3.5E-02</td><td>4.0E-02</td><td>MB231/MB468</td><td>CYTO</td></tr>
<tr><td>>hg38_21_+_8433172_8446622_RNA45SN1_45S_50ntFlanks@8360.8378.19|AGAACTTTGAAGAGAGAGT</td><td>>hg38_21_+_8433172_8446622_RNA45SN1_45S_50ntFlanks</td><td>8360</td><td>8378</td><td>19</td><td>AGAACTTTGAAGAGAGAGT</td><td>84.0</td><td>-4.1</td><td>1.8</td><td>-2.2</td><td>2.5E-02</td><td>4.0E-02</td><td>BT20/MB468</td><td>MP</td></tr>
<tr><td>>trna27_GlyCCC_2_-_70476123_70476193@4.35.32,trna34_GlyCCC_16_-_686736_686806@4.35.32|CCGCTGGTGTAGTGGTATCATGCAAGATTCCC</td><td>>trna27_GlyCCC_2_-_70476123_70476193</td><td>4</td><td>35</td><td>32</td><td>CCGCTGGTGTAGTGGTATCATGCAAGATTCCC</td><td>176.9</td><td>-2.6</td><td>1.0</td><td>-2.5</td><td>1.3E-02</td><td>4.0E-02</td><td>BT20/MB231</td><td>MITO</td></tr>
<tr><td>>hg38_21_+_8433172_8446622_RNA45SN1_45S_50ntFlanks@11910.11933.24|GTGAAGAGACATGAGAGGTGTAGA</td><td>>hg38_21_+_8433172_8446622_RNA45SN1_45S_50ntFlanks</td><td>11910</td><td>11933</td><td>24</td><td>GTGAAGAGACATGAGAGGTGTAGA</td><td>33.9</td><td>5.9</td><td>2.6</td><td>2.2</td><td>2.5E-02</td><td>4.0E-02</td><td>BT20/MB468</td><td>MP</td></tr>
<tr><td>>hg38_21_+_8433172_8446622_RNA45SN1_45S_50ntFlanks@13008.13039.32|ATTGAAAGTCAGCCCTCGACACAAGGGTTTGT</td><td>>hg38_21_+_8433172_8446622_RNA45SN1_45S_50ntFlanks</td><td>13008</td><td>13039</td><td>32</td><td>ATTGAAAGTCAGCCCTCGACACAAGGGTTTGT</td><td>1682.5</td><td>2.1</td><td>1.0</td><td>2.1</td><td>3.6E-02</td><td>4.0E-02</td><td>MB231/MB468</td><td>NUC</td></tr>
<tr><td>>hg38_1_-_228634819_228635039_RNA5S12_5S_50ntFlanks@133.169.37|ATGGGAGACCGCCTGGGAATACCGGGTGCTGTAGGCT</td><td>>hg38_1_-_228634819_228635039_RNA5S12_5S_50ntFlanks</td><td>133</td><td>169</td><td>37</td><td>ATGGGAGACCGCCTGGGAATACCGGGTGCTGTAGGCT</td><td>595.5</td><td>-2.1</td><td>1.0</td><td>-2.1</td><td>3.7E-02</td><td>4.0E-02</td><td>MB231/MB468</td><td>MITO</td></tr>
<tr><td>>hg38_MT_+_1621_3279_RNR2_16S_50ntFlanks@1191.1215.25|GTTGGGGCGACCTCGGAGCAGAACC</td><td>>hg38_MT_+_1621_3279_RNR2_16S_50ntFlanks</td><td>1191</td><td>1215</td><td>25</td><td>GTTGGGGCGACCTCGGAGCAGAACC</td><td>110.0</td><td>4.8</td><td>2.2</td><td>2.2</td><td>2.5E-02</td><td>4.0E-02</td><td>BT20/MB468</td><td>MP</td></tr>
<tr><td>>hg38_21_+_8433172_8446622_RNA45SN1_45S_50ntFlanks@12658.12685.28|ACTGAACGCCTCTAAGTCAGAATCCCGC</td><td>>hg38_21_+_8433172_8446622_RNA45SN1_45S_50ntFlanks</td><td>12658</td><td>12685</td><td>28</td><td>ACTGAACGCCTCTAAGTCAGAATCCCGC</td><td>70.7</td><td>-3.8</td><td>1.7</td><td>-2.2</td><td>2.5E-02</td><td>4.0E-02</td><td>BT20/MB468</td><td>MP</td></tr>
<tr><td>>hg38_21_+_8433172_8446622_RNA45SN1_45S_50ntFlanks@9457.9494.38|CGCCGGGGAGGTGGAGCACGAGCGCACGTGTTAGGACC</td><td>>hg38_21_+_8433172_8446622_RNA45SN1_45S_50ntFlanks</td><td>9457</td><td>9494</td><td>38</td><td>CGCCGGGGAGGTGGAGCACGAGCGCACGTGTTAGGACC</td><td>96.8</td><td>-4.0</td><td>1.8</td><td>-2.2</td><td>2.5E-02</td><td>4.0E-02</td><td>BT20/MB468</td><td>MP</td></tr>
<tr><td>>hg38_21_+_8433172_8446622_RNA45SN1_45S_50ntFlanks@475.498.24|CTTCGTGATCGATGTGGTGACGTC</td><td>>hg38_21_+_8433172_8446622_RNA45SN1_45S_50ntFlanks</td><td>475</td><td>498</td><td>24</td><td>CTTCGTGATCGATGTGGTGACGTC</td><td>7728.0</td><td>1.6</td><td>0.8</td><td>2.1</td><td>3.6E-02</td><td>4.0E-02</td><td>BT20/MB468</td><td>NUC</td></tr>
<tr><td>>am_trnaMT_ProTGG_MT_-_15956_16023@53.70.18|AAGACTTTTTCTCTGACC</td><td>>am_trnaMT_ProTGG_MT_-_15956_16023</td><td>53</td><td>70</td><td>18</td><td>AAGACTTTTTCTCTGACC</td><td>2291.9</td><td>-2.6</td><td>1.0</td><td>-2.5</td><td>1.1E-02</td><td>4.0E-02</td><td>BT20/MB231</td><td>MP</td></tr>
<tr><td>>MI0000072|hsa-mir-18a&WithFlank&13|+|91350745|91350827@12.32.21&[MIMAT0000072&hsa-miR-18a-5p&offsets|0|-2;m-262&13|+|91350756|91350778&offsets|0|-2]|TAAGGTGCATCTAGTGCAGAT</td><td>>MI0000072|hsa-mir-18a&WithFlank&13|+|91350745|91350827</td><td>12</td><td>32</td><td>21</td><td>TAAGGTGCATCTAGTGCAGAT</td><td>155.4</td><td>-4.8</td><td>1.9</td><td>-2.5</td><td>1.1E-02</td><td>4.0E-02</td><td>BT20/MB231</td><td>MP</td></tr>
<tr><td>>hg38_21_+_8433172_8446622_RNA45SN1_45S_50ntFlanks@9554.9577.24|CGTAGCGGTCCTGACGTGCAAATC</td><td>>hg38_21_+_8433172_8446622_RNA45SN1_45S_50ntFlanks</td><td>9554</td><td>9577</td><td>24</td><td>CGTAGCGGTCCTGACGTGCAAATC</td><td>923.8</td><td>-2.2</td><td>0.9</td><td>-2.5</td><td>1.1E-02</td><td>4.0E-02</td><td>BT20/MB231</td><td>NUC</td></tr>
<tr><td>>hg38_21_+_8433172_8446622_RNA45SN1_45S_50ntFlanks@9307.9334.28|AAGGAGTCTAACACGTGCGCGAGTCGGG</td><td>>hg38_21_+_8433172_8446622_RNA45SN1_45S_50ntFlanks</td><td>9307</td><td>9334</td><td>28</td><td>AAGGAGTCTAACACGTGCGCGAGTCGGG</td><td>377.1</td><td>4.3</td><td>1.7</td><td>2.5</td><td>1.1E-02</td><td>4.0E-02</td><td>BT20/MB231</td><td>MP</td></tr>
<tr><td>>hg38_21_+_8433172_8446622_RNA45SN1_45S_50ntFlanks@8189.8213.25|CCCAGCCCGTGGACGGTGTGAGGCC</td><td>>hg38_21_+_8433172_8446622_RNA45SN1_45S_50ntFlanks</td><td>8189</td><td>8213</td><td>25</td><td>CCCAGCCCGTGGACGGTGTGAGGCC</td><td>157.0</td><td>-1.6</td><td>0.7</td><td>-2.2</td><td>3.1E-02</td><td>4.0E-02</td><td>BT20/MB468</td><td>MITO</td></tr>
<tr><td>>MI0000434|hsa-let-7i&WithFlank&12|+|62603680|62603775@12.33.22(+2U)&[MIMAT0000415&hsa-let-7i-5p&offsets|0|0(+2U);m-49&12|+|62603691|62603712&offsets|0|0(+2U)]|TGAGGTAGTAGTTTGTGCTGTTTT</td><td>>MI0000434|hsa-let-7i&WithFlank&12|+|62603680|62603775</td><td>12</td><td>33</td><td>22(+2U)</td><td>TGAGGTAGTAGTTTGTGCTGTTTT</td><td>524.7</td><td>-3.8</td><td>1.5</td><td>-2.5</td><td>1.1E-02</td><td>4.0E-02</td><td>BT20/MB231</td><td>MP</td></tr>
<tr><td>>hg38_21_+_8433172_8446622_RNA45SN1_45S_50ntFlanks@11571.11602.32|GCCGACTTAGAACTGGTGCGGACCAGGGGAAT</td><td>>hg38_21_+_8433172_8446622_RNA45SN1_45S_50ntFlanks</td><td>11571</td><td>11602</td><td>32</td><td>GCCGACTTAGAACTGGTGCGGACCAGGGGAAT</td><td>669.2</td><td>3.3</td><td>1.3</td><td>2.5</td><td>1.1E-02</td><td>4.0E-02</td><td>BT20/MB231</td><td>MP</td></tr>
<tr><td>>hg38_21_+_8433172_8446622_RNA45SN1_45S_50ntFlanks@11321.11342.22|CACGTCTCGTCGCGCGCGCGTC</td><td>>hg38_21_+_8433172_8446622_RNA45SN1_45S_50ntFlanks</td><td>11321</td><td>11342</td><td>22</td><td>CACGTCTCGTCGCGCGCGCGTC</td><td>4951.9</td><td>-4.9</td><td>2.3</td><td>-2.1</td><td>3.6E-02</td><td>4.0E-02</td><td>BT20/MB468</td><td>NUC</td></tr>
<tr><td>>trnaMT_GluTTC_MT_-_14674_14742@31.71.41,trnalookalike8_GluTTC_5_-_93905172_93905240@31.71.41|TTCATATCATTGGTCGTGGTTGTAGTCCGTGCGAGAATACC</td><td>>trnaMT_GluTTC_MT_-_14674_14742</td><td>31</td><td>71</td><td>41</td><td>TTCATATCATTGGTCGTGGTTGTAGTCCGTGCGAGAATACC</td><td>94.3</td><td>4.4</td><td>2.1</td><td>2.1</td><td>3.3E-02</td><td>4.0E-02</td><td>MB231/MB468</td><td>MP</td></tr>
<tr><td>>MI0000088|hsa-mir-30a&WithFlank&6|-|71403545|71403627@12.33.22(+1A)&[MIMAT0000087&hsa-miR-30a-5p&offsets|0|0(+1A);m-15&6|-|71403595|71403616&offsets|0|0(+1A)]|TGTAAACATCCTCGACTGGAAGA</td><td>>MI0000088|hsa-mir-30a&WithFlank&6|-|71403545|71403627</td><td>12</td><td>33</td><td>22(+1A)</td><td>TGTAAACATCCTCGACTGGAAGA</td><td>2096.2</td><td>-2.0</td><td>0.8</td><td>-2.5</td><td>1.3E-02</td><td>4.0E-02</td><td>BT20/MB231</td><td>MITO</td></tr>
<tr><td>>hg38_21_+_8433172_8446622_RNA45SN1_45S_50ntFlanks@6649.6669.21|TACGACTCTTAGCGGTGGATC</td><td>>hg38_21_+_8433172_8446622_RNA45SN1_45S_50ntFlanks</td><td>6649</td><td>6669</td><td>21</td><td>TACGACTCTTAGCGGTGGATC</td><td>254.7</td><td>1.2</td><td>0.6</td><td>2.1</td><td>3.7E-02</td><td>4.0E-02</td><td>MB231/MB468</td><td>NUC</td></tr>
<tr><td>>hg38_21_+_8433172_8446622_RNA45SN1_45S_50ntFlanks@5237.5267.31|ACTGACTGGCTCAGCGTGTGCCTACCCTACG</td><td>>hg38_21_+_8433172_8446622_RNA45SN1_45S_50ntFlanks</td><td>5237</td><td>5267</td><td>31</td><td>ACTGACTGGCTCAGCGTGTGCCTACCCTACG</td><td>137.0</td><td>-2.9</td><td>1.3</td><td>-2.2</td><td>3.1E-02</td><td>4.0E-02</td><td>BT20/MB468</td><td>MITO</td></tr>
<tr><td>>hg38_21_+_8433172_8446622_RNA45SN1_45S_50ntFlanks@6646.6681.36|TCGTACGACTCTTAGCGGTGGATCACTCGGCTCGTG</td><td>>hg38_21_+_8433172_8446622_RNA45SN1_45S_50ntFlanks</td><td>6646</td><td>6681</td><td>36</td><td>TCGTACGACTCTTAGCGGTGGATCACTCGGCTCGTG</td><td>25174.1</td><td>-2.3</td><td>1.0</td><td>-2.2</td><td>2.5E-02</td><td>4.0E-02</td><td>BT20/MB468</td><td>MP</td></tr>
<tr><td>>hg38_21_+_8433172_8446622_RNA45SN1_45S_50ntFlanks@12630.12667.38|GGGCGAAGCTACCATCTGTGGGATTATGACTGAACGCC</td><td>>hg38_21_+_8433172_8446622_RNA45SN1_45S_50ntFlanks</td><td>12630</td><td>12667</td><td>38</td><td>GGGCGAAGCTACCATCTGTGGGATTATGACTGAACGCC</td><td>93.8</td><td>3.5</td><td>1.7</td><td>2.1</td><td>3.7E-02</td><td>4.0E-02</td><td>MB231/MB468</td><td>MITO</td></tr>
<tr><td>>trnaMT_SerGCT_MT_+_12207_12265@1.23.23|GAGAAAGCTCACAAGAACTGCTA</td><td>>trnaMT_SerGCT_MT_+_12207_12265</td><td>1</td><td>23</td><td>23</td><td>GAGAAAGCTCACAAGAACTGCTA</td><td>1771.9</td><td>1.6</td><td>0.8</td><td>2.1</td><td>3.4E-02</td><td>4.0E-02</td><td>MB231/MB468</td><td>MP</td></tr>
<tr><td>>hg38_21_+_8433172_8446622_RNA45SN1_45S_50ntFlanks@6648.6665.18|GTACGACTCTTAGCGGTG</td><td>>hg38_21_+_8433172_8446622_RNA45SN1_45S_50ntFlanks</td><td>6648</td><td>6665</td><td>18</td><td>GTACGACTCTTAGCGGTG</td><td>1852.6</td><td>3.0</td><td>1.2</td><td>2.5</td><td>1.1E-02</td><td>4.0E-02</td><td>BT20/MB231</td><td>MP</td></tr>
<tr><td>>hg38_21_+_8433172_8446622_RNA45SN1_45S_50ntFlanks@10571.10608.38|GAGAAGCCGGCGGGAGCCCCGGGGAGAGTTCTCTTTTC</td><td>>hg38_21_+_8433172_8446622_RNA45SN1_45S_50ntFlanks</td><td>10571</td><td>10608</td><td>38</td><td>GAGAAGCCGGCGGGAGCCCCGGGGAGAGTTCTCTTTTC</td><td>100.1</td><td>-3.8</td><td>1.7</td><td>-2.2</td><td>2.5E-02</td><td>4.0E-02</td><td>BT20/MB468</td><td>MP</td></tr>
<tr><td>>hg38_21_+_8433172_8446622_RNA45SN1_45S_50ntFlanks@12942.12959.18|TTCGTAGACGACCTGCTT</td><td>>hg38_21_+_8433172_8446622_RNA45SN1_45S_50ntFlanks</td><td>12942</td><td>12959</td><td>18</td><td>TTCGTAGACGACCTGCTT</td><td>963.4</td><td>-1.3</td><td>0.6</td><td>-2.1</td><td>3.5E-02</td><td>4.0E-02</td><td>BT20/MB468</td><td>CYTO</td></tr>
<tr><td>>hg38_21_+_8433172_8446622_RNA45SN1_45S_50ntFlanks@11570.11599.30|AGCCGACTTAGAACTGGTGCGGACCAGGGG</td><td>>hg38_21_+_8433172_8446622_RNA45SN1_45S_50ntFlanks</td><td>11570</td><td>11599</td><td>30</td><td>AGCCGACTTAGAACTGGTGCGGACCAGGGG</td><td>239.0</td><td>1.8</td><td>0.8</td><td>2.2</td><td>2.5E-02</td><td>4.0E-02</td><td>BT20/MB468</td><td>MP</td></tr>
<tr><td>>MI0000077|hsa-mir-21&WithFlank&17|+|59841260|59841343@14.34.21(+1G)&[MIMAT0000076&hsa-miR-21-5p&offsets|0|-1(+1G);m-1&17|+|59841273|59841294&offsets|0|-1(+1G)]|TAGCTTATCAGACTGATGTTGG</td><td>>MI0000077|hsa-mir-21&WithFlank&17|+|59841260|59841343</td><td>14</td><td>34</td><td>21(+1G)</td><td>TAGCTTATCAGACTGATGTTGG</td><td>1676.4</td><td>-2.1</td><td>0.8</td><td>-2.5</td><td>1.3E-02</td><td>4.0E-02</td><td>BT20/MB231</td><td>MITO</td></tr>
<tr><td>>hg38_1_-_228634819_228635039_RNA5S12_5S_50ntFlanks@152.169.18|TACCGGGTGCTGTAGGCT</td><td>>hg38_1_-_228634819_228635039_RNA5S12_5S_50ntFlanks</td><td>152</td><td>169</td><td>18</td><td>TACCGGGTGCTGTAGGCT</td><td>296.6</td><td>1.8</td><td>0.7</td><td>2.5</td><td>1.3E-02</td><td>4.0E-02</td><td>BT20/MB231</td><td>MITO</td></tr>
<tr><td>>hg38_21_+_8433172_8446622_RNA45SN1_45S_50ntFlanks@3841.3873.33|TCCTACTTGGATAACTGTGGTAATTCTAGAGCT</td><td>>hg38_21_+_8433172_8446622_RNA45SN1_45S_50ntFlanks</td><td>3841</td><td>3873</td><td>33</td><td>TCCTACTTGGATAACTGTGGTAATTCTAGAGCT</td><td>400.3</td><td>2.1</td><td>0.8</td><td>2.4</td><td>1.5E-02</td><td>4.0E-02</td><td>BT20/MB231</td><td>CYTO</td></tr>
</table>

<table>
<tr><td>>hg38_21_+_8433172_8446622_RNA45SN1_45S_50ntFlanks@4926.4947.22|GAGTGGAGCCTGCGGCTTAATT</td><td>>hg38_21_+_8433172_8446622_RNA45SN1_45S_50ntFlanks</td><td>4926</td><td>4947</td><td>22</td><td>GAGTGGAGCCTGCGGCTTAATT</td><td>71.3</td><td>4.7</td><td>2.1</td><td>2.2</td><td>2.5E-02</td><td>4.0E-02</td><td>BT20/MB468</td><td>MP</td></tr>
<tr><td>>hg38_MT_+_598_1651_RNR1_12S_50ntFlanks@638.657.20|TCACCACCTCTTGCTCAGCC</td><td>>hg38_MT_+_598_1651_RNR1_12S_50ntFlanks</td><td>638</td><td>657</td><td>20</td><td>TCACCACCTCTTGCTCAGCC</td><td>37.8</td><td>4.3</td><td>2.0</td><td>2.1</td><td>3.4E-02</td><td>4.0E-02</td><td>MB231/MB468</td><td>MP</td></tr>
<tr><td>>hg38_MT_+_1621_3279_RNR2_16S_50ntFlanks@840.864.25|AAAAGTAAAAGGAACTCGGCAAATC</td><td>>hg38_MT_+_1621_3279_RNR2_16S_50ntFlanks</td><td>840</td><td>864</td><td>25</td><td>AAAAGTAAAAGGAACTCGGCAAATC</td><td>59.8</td><td>4.3</td><td>2.0</td><td>2.1</td><td>3.4E-02</td><td>4.0E-02</td><td>MB231/MB468</td><td>MP</td></tr>
<tr><td>>hg38_21_+_8433172_8446622_RNA45SN1_45S_50ntFlanks@6645.6681.37|CTCGTACGACTCTTAGCGGTGGATCACTCGGCTCGTG</td><td>>hg38_21_+_8433172_8446622_RNA45SN1_45S_50ntFlanks</td><td>6645</td><td>6681</td><td>37</td><td>CTCGTACGACTCTTAGCGGTGGATCACTCGGCTCGTG</td><td>3683.5</td><td>-2.2</td><td>1.0</td><td>-2.2</td><td>2.5E-02</td><td>4.1E-02</td><td>BT20/MB468</td><td>MP</td></tr>
<tr><td>>hg38_21_+_8433172_8446622_RNA45SN1_45S_50ntFlanks@11920.11950.31|ATGAGAGGTGTAGAATAAGTGGGAGGCCCCC</td><td>>hg38_21_+_8433172_8446622_RNA45SN1_45S_50ntFlanks</td><td>11920</td><td>11950</td><td>31</td><td>ATGAGAGGTGTAGAATAAGTGGGAGGCCCCC</td><td>471.0</td><td>4.9</td><td>1.9</td><td>2.5</td><td>1.1E-02</td><td>4.1E-02</td><td>BT20/MB231</td><td>MP</td></tr>
<tr><td>>hg38_21_+_8433172_8446622_RNA45SN1_45S_50ntFlanks@12936.12959.24|AAACCATTCGTAGACGACCTGCTT</td><td>>hg38_21_+_8433172_8446622_RNA45SN1_45S_50ntFlanks</td><td>12936</td><td>12959</td><td>24</td><td>AAACCATTCGTAGACGACCTGCTT</td><td>286.9</td><td>-1.0</td><td>0.5</td><td>-2.1</td><td>3.6E-02</td><td>4.1E-02</td><td>BT20/MB468</td><td>CYTO</td></tr>
<tr><td>>hg38_21_+_8433172_8446622_RNA45SN1_45S_50ntFlanks@10248.10270.23|CGACGAGTAGGAGGGCCGCTGCG</td><td>>hg38_21_+_8433172_8446622_RNA45SN1_45S_50ntFlanks</td><td>10248</td><td>10270</td><td>23</td><td>CGACGAGTAGGAGGGCCGCTGCG</td><td>129.5</td><td>3.3</td><td>1.5</td><td>2.2</td><td>2.5E-02</td><td>4.1E-02</td><td>BT20/MB468</td><td>MP</td></tr>
<tr><td>>hg38_21_+_8433172_8446622_RNA45SN1_45S_50ntFlanks@3777.3794.18|CGGTACAGTGAAACTGCG</td><td>>hg38_21_+_8433172_8446622_RNA45SN1_45S_50ntFlanks</td><td>3777</td><td>3794</td><td>18</td><td>CGGTACAGTGAAACTGCG</td><td>115.0</td><td>-3.2</td><td>1.5</td><td>-2.1</td><td>3.6E-02</td><td>4.1E-02</td><td>BT20/MB468</td><td>CYTO</td></tr>
<tr><td>>MI0000084|hsa-mir-26b&WithFlank&2|+|218402640|218402728@18.37.20(+1C)&[MIMAT0000083&hsa-miR-26b-5p&offsets|0|-1(+1C);m-53&2|+|218402657|218402678&offsets|0|-2(+1C)]|TTCAAGTAATTCAGGATAGGC</td><td>>MI0000084|hsa-mir-26b&WithFlank&2|+|218402640|218402728</td><td>18</td><td>37</td><td>20(+1C)</td><td>TTCAAGTAATTCAGGATAGGC</td><td>263.1</td><td>3.1</td><td>1.5</td><td>2.1</td><td>3.4E-02</td><td>4.1E-02</td><td>MB231/MB468</td><td>MP</td></tr>
<tr><td>>MI0000085|hsa-mir-27a&WithFlank&19|-|13836434|13836523@16.36.21&[MIMAT0004501&hsa-miR-27a-5p&offsets|0|-1;m-270&19|-|13836487|13836508&offsets|0|-1]|AGGGCTTAGCTGCTTGTGAGC</td><td>>MI0000085|hsa-mir-27a&WithFlank&19|-|13836434|13836523</td><td>16</td><td>36</td><td>21</td><td>AGGGCTTAGCTGCTTGTGAGC</td><td>514.2</td><td>-2.5</td><td>1.0</td><td>-2.5</td><td>1.1E-02</td><td>4.1E-02</td><td>BT20/MB231</td><td>MP</td></tr>
<tr><td>>trnaMT_SerGCT_MT_+_12207_12265@30.62.33|GCCCCCATGTCTAACAACATGGCTTTCTCACCA</td><td>>trnaMT_SerGCT_MT_+_12207_12265</td><td>30</td><td>62</td><td>33</td><td>GCCCCCATGTCTAACAACATGGCTTTCTCACCA</td><td>210.1</td><td>-4.2</td><td>1.7</td><td>-2.5</td><td>1.1E-02</td><td>4.1E-02</td><td>BT20/MB231</td><td>MP</td></tr>
<tr><td>>hg38_21_+_8433172_8446622_RNA45SN1_45S_50ntFlanks@10856.10879.24|AGGATTGGCTCTAAGGGCTGGGTC</td><td>>hg38_21_+_8433172_8446622_RNA45SN1_45S_50ntFlanks</td><td>10856</td><td>10879</td><td>24</td><td>AGGATTGGCTCTAAGGGCTGGGTC</td><td>101.8</td><td>4.3</td><td>1.7</td><td>2.5</td><td>1.1E-02</td><td>4.1E-02</td><td>BT20/MB231</td><td>MP</td></tr>
<tr><td>>MI0000078|hsa-mir-22&WithFlank&17|-|1713897|1713993@59.80.22&[MIMAT0000077&hsa-miR-22-3p&offsets|0|0;m-2&17|-|1713914|1713935&offsets|0|0]|AAGCTGCCAGTTGAAGAACTGT</td><td>>MI0000078|hsa-mir-22&WithFlank&17|-|1713897|1713993</td><td>59</td><td>80</td><td>22</td><td>AAGCTGCCAGTTGAAGAACTGT</td><td>14393.0</td><td>2.3</td><td>1.0</td><td>2.4</td><td>1.5E-02</td><td>4.1E-02</td><td>BT20/MB231</td><td>CYTO</td></tr>
<tr><td>>hg38_21_+_8433172_8446622_RNA45SN1_45S_50ntFlanks@11552.11585.34|CCTCGGCCGGCGCCTAGCAGCCGACTTAGAACTG</td><td>>hg38_21_+_8433172_8446622_RNA45SN1_45S_50ntFlanks</td><td>11552</td><td>11585</td><td>34</td><td>CCTCGGCCGGCGCCTAGCAGCCGACTTAGAACTG</td><td>186.5</td><td>-1.2</td><td>0.5</td><td>-2.1</td><td>3.2E-02</td><td>4.1E-02</td><td>BT20/MB468</td><td>MITO</td></tr>
<tr><td>>hg38_21_+_8433172_8446622_RNA45SN1_45S_50ntFlanks@11583.11611.29|CTGGTGCGGACCAGGGGAATCCGACTGTT</td><td>>hg38_21_+_8433172_8446622_RNA45SN1_45S_50ntFlanks</td><td>11583</td><td>11611</td><td>29</td><td>CTGGTGCGGACCAGGGGAATCCGACTGTT</td><td>391.6</td><td>2.9</td><td>1.2</td><td>2.5</td><td>1.1E-02</td><td>4.1E-02</td><td>BT20/MB231</td><td>MP</td></tr>
<tr><td>>hg38_21_+_8433172_8446622_RNA45SN1_45S_50ntFlanks@6648.6675.28|GTACGACTCTTAGCGGTGGATCACTCGG</td><td>>hg38_21_+_8433172_8446622_RNA45SN1_45S_50ntFlanks</td><td>6648</td><td>6675</td><td>28</td><td>GTACGACTCTTAGCGGTGGATCACTCGG</td><td>532.5</td><td>-1.9</td><td>0.8</td><td>-2.5</td><td>1.2E-02</td><td>4.1E-02</td><td>BT20/MB231</td><td>NUC</td></tr>
<tr><td>>hg38_21_+_8433172_8446622_RNA45SN1_45S_50ntFlanks@10879.10905.27|CGGTCGGGCTGGGGCGCGAAGCGGGGC</td><td>>hg38_21_+_8433172_8446622_RNA45SN1_45S_50ntFlanks</td><td>10879</td><td>10905</td><td>27</td><td>CGGTCGGGCTGGGGCGCGAAGCGGGGC</td><td>43.3</td><td>4.0</td><td>1.8</td><td>2.2</td><td>2.5E-02</td><td>4.1E-02</td><td>BT20/MB468</td><td>MP</td></tr>
<tr><td>>hg38_21_+_8433172_8446622_RNA45SN1_45S_50ntFlanks@11920.11952.33|ATGAGAGGTGTAGAATAAGTGGGAGGCCCCCGG</td><td>>hg38_21_+_8433172_8446622_RNA45SN1_45S_50ntFlanks</td><td>11920</td><td>11952</td><td>33</td><td>ATGAGAGGTGTAGAATAAGTGGGAGGCCCCCGG</td><td>95.0</td><td>-3.8</td><td>1.7</td><td>-2.2</td><td>2.5E-02</td><td>4.1E-02</td><td>BT20/MB468</td><td>MP</td></tr>
<tr><td>>trnaMT_HisGTG_MT_+_12138_12206@19.53.35|AAACATCAGATTGTGAATCTGACAACAGAGGCTTA</td><td>>trnaMT_HisGTG_MT_+_12138_12206</td><td>19</td><td>53</td><td>35</td><td>AAACATCAGATTGTGAATCTGACAACAGAGGCTTA</td><td>167.9</td><td>5.1</td><td>2.3</td><td>2.2</td><td>2.5E-02</td><td>4.1E-02</td><td>BT20/MB468</td><td>MP</td></tr>
<tr><td>>hg38_21_+_8433172_8446622_RNA45SN1_45S_50ntFlanks@10822.10865.44|GAAGTCGGCAAGCCGGATCCGTAACTTCGGGATAAGGATTGGCT</td><td>>hg38_21_+_8433172_8446622_RNA45SN1_45S_50ntFlanks</td><td>10822</td><td>10865</td><td>44</td><td>GAAGTCGGCAAGCCGGATCCGTAACTTCGGGATAAGGATTGGCT</td><td>184.0</td><td>-2.7</td><td>1.2</td><td>-2.1</td><td>3.2E-02</td><td>4.1E-02</td><td>BT20/MB468</td><td>MITO</td></tr>
<tr><td>>trnaMT_ValTAC_MT_+_1602_1670@55.72.18|ACTTGACCGCTCTGACCA</td><td>>trnaMT_ValTAC_MT_+_1602_1670</td><td>55</td><td>72</td><td>18</td><td>ACTTGACCGCTCTGACCA</td><td>1257.8</td><td>1.9</td><td>0.8</td><td>2.2</td><td>2.5E-02</td><td>4.1E-02</td><td>BT20/MB468</td><td>MP</td></tr>
<tr><td>>hg38_21_+_8433172_8446622_RNA45SN1_45S_50ntFlanks@11671.11688.18|CCAGTGCTCTGAATGTCA</td><td>>hg38_21_+_8433172_8446622_RNA45SN1_45S_50ntFlanks</td><td>11671</td><td>11688</td><td>18</td><td>CCAGTGCTCTGAATGTCA</td><td>194.3</td><td>-3.0</td><td>1.3</td><td>-2.2</td><td>2.5E-02</td><td>4.1E-02</td><td>BT20/MB468</td><td>MP</td></tr>
<tr><td>>hg38_21_+_8433172_8446622_RNA45SN1_45S_50ntFlanks@12419.12452.34|GATGTCGGCTCTTCCTATCATTGTGAAGCAGAAT</td><td>>hg38_21_+_8433172_8446622_RNA45SN1_45S_50ntFlanks</td><td>12419</td><td>12452</td><td>34</td><td>GATGTCGGCTCTTCCTATCATTGTGAAGCAGAAT</td><td>54.0</td><td>4.3</td><td>2.0</td><td>2.1</td><td>3.4E-02</td><td>4.1E-02</td><td>MB231/MB468</td><td>MP</td></tr>
<tr><td>>hg38_21_+_8433172_8446622_RNA45SN1_45S_50ntFlanks@8911.8936.26|TCGCCGAATCCCGGGGCCGAGGGAGC</td><td>>hg38_21_+_8433172_8446622_RNA45SN1_45S_50ntFlanks</td><td>8911</td><td>8936</td><td>26</td><td>TCGCCGAATCCCGGGGCCGAGGGAGC</td><td>89.5</td><td>4.9</td><td>1.9</td><td>2.5</td><td>1.1E-02</td><td>4.1E-02</td><td>BT20/MB231</td><td>MP</td></tr>
<tr><td>>MI0000108|hsa-mir-103a-2&WithFlank&20|+|3917488|3917577@54.76.23(+2U);MI0000109|hsa-mir-103a-1&WithFlank&5|-|168560890|168560979@54.76.23(+2U)&[MIMAT0000101&hsa-miR-103a-3p&offsets|0|0(+2U);m-12&20|+|3917541|3917563&offsets|0|0(+2U)];[MIMAT0000101_1&hsa-miR-103a-3p&offsets|0|0(+2U);m-13&5|-|168560904|168560926&offsets|0|0(+2U);m-44220&5|-|168560908|168560926&offsets|0|+4(+2U)]|AGCAGCATTGTACAGGGCTATGATT</td><td>>MI0000108|hsa-mir-103a-2&WithFlank&20|+|3917488|3917577</td><td>54</td><td>76</td><td>23(+2U)</td><td>AGCAGCATTGTACAGGGCTATGATT</td><td>739.2</td><td>-1.6</td><td>0.6</td><td>-2.5</td><td>1.2E-02</td><td>4.1E-02</td><td>BT20/MB231</td><td>NUC</td></tr>
<tr><td>>trnaMT_PheGAA_MT_+_577_647@1.20.20|GTTTATGTAGCTTACCTCCT</td><td>>trnaMT_PheGAA_MT_+_577_647</td><td>1</td><td>20</td><td>20</td><td>GTTTATGTAGCTTACCTCCT</td><td>279.7</td><td>-2.1</td><td>0.8</td><td>-2.5</td><td>1.2E-02</td><td>4.1E-02</td><td>BT20/MB231</td><td>NUC</td></tr>
<tr><td>>hg38_21_+_8433172_8446622_RNA45SN1_45S_50ntFlanks@10401.10439.39|GAACATGGGTCAGTCGGTCCTGAGAGATGGGCGAGCGCC</td><td>>hg38_21_+_8433172_8446622_RNA45SN1_45S_50ntFlanks</td><td>10401</td><td>10439</td><td>39</td><td>GAACATGGGTCAGTCGGTCCTGAGAGATGGGCGAGCGCC</td><td>281.6</td><td>-2.5</td><td>1.0</td><td>-2.5</td><td>1.2E-02</td><td>4.1E-02</td><td>BT20/MB231</td><td>NUC</td></tr>
<tr><td>>hg38_21_+_8433172_8446622_RNA45SN1_45S_50ntFlanks@11669.11707.39|GCCCAGTGCTCTGAATGTCAAAGTGAAGAAATTCAATGA</td><td>>hg38_21_+_8433172_8446622_RNA45SN1_45S_50ntFlanks</td><td>11669</td><td>11707</td><td>39</td><td>GCCCAGTGCTCTGAATGTCAAAGTGAAGAAATTCAATGA</td><td>262.9</td><td>-2.5</td><td>1.0</td><td>-2.5</td><td>1.2E-02</td><td>4.1E-02</td><td>BT20/MB231</td><td>NUC</td></tr>
<tr><td>>hg38_21_+_8433172_8446622_RNA45SN1_45S_50ntFlanks@11884.11903.20|GTTGAGCTTGACTCTAGTCT</td><td>>hg38_21_+_8433172_8446622_RNA45SN1_45S_50ntFlanks</td><td>11884</td><td>11903</td><td>20</td><td>GTTGAGCTTGACTCTAGTCT</td><td>1751.3</td><td>4.6</td><td>1.8</td><td>2.5</td><td>1.1E-02</td><td>4.1E-02</td><td>BT20/MB231</td><td>MP</td></tr>
<tr><td>>hg38_21_+_8433172_8446622_RNA45SN1_45S_50ntFlanks@11456.11473.18|TCGGCGCGCGGCGGCGGC</td><td>>hg38_21_+_8433172_8446622_RNA45SN1_45S_50ntFlanks</td><td>11456</td><td>11473</td><td>18</td><td>TCGGCGCGCGGCGGCGGC</td><td>378.8</td><td>-4.6</td><td>1.9</td><td>-2.4</td><td>1.5E-02</td><td>4.1E-02</td><td>BT20/MB231</td><td>CYTO</td></tr>
<tr><td>>hg38_21_+_8433172_8446622_RNA45SN1_45S_50ntFlanks@12409.12446.38|TTGATCCTTCGATGTCGGCTCTTCCTATCATTGTGAAG</td><td>>hg38_21_+_8433172_8446622_RNA45SN1_45S_50ntFlanks</td><td>12409</td><td>12446</td><td>38</td><td>TTGATCCTTCGATGTCGGCTCTTCCTATCATTGTGAAG</td><td>210.0</td><td>-2.5</td><td>1.0</td><td>-2.4</td><td>1.5E-02</td><td>4.1E-02</td><td>BT20/MB231</td><td>CYTO</td></tr>
<tr><td>>hg38_21_+_8433172_8446622_RNA45SN1_45S_50ntFlanks@7976.8006.31|GCGACCTCAGATCAGACGTGGCGACCCGCTG</td><td>>hg38_21_+_8433172_8446622_RNA45SN1_45S_50ntFlanks</td><td>7976</td><td>8006</td><td>31</td><td>GCGACCTCAGATCAGACGTGGCGACCCGCTG</td><td>804.6</td><td>1.5</td><td>0.7</td><td>2.1</td><td>3.2E-02</td><td>4.1E-02</td><td>BT20/MB468</td><td>MITO</td></tr>
<tr><td>>hg38_21_+_8433172_8446622_RNA45SN1_45S_50ntFlanks@6719.6741.23|TGAATTGCAGGACACATTGATCA</td><td>>hg38_21_+_8433172_8446622_RNA45SN1_45S_50ntFlanks</td><td>6719</td><td>6741</td><td>23</td><td>TGAATTGCAGGACACATTGATCA</td><td>182.2</td><td>-1.4</td><td>0.7</td><td>-2.1</td><td>3.2E-02</td><td>4.1E-02</td><td>BT20/MB468</td><td>MITO</td></tr>
<tr><td>>hg38_21_+_8433172_8446622_RNA45SN1_45S_50ntFlanks@10808.10836.29|CAATGTAGGTAAGGGAAGTCGGCAAGCCG</td><td>>hg38_21_+_8433172_8446622_RNA45SN1_45S_50ntFlanks</td><td>10808</td><td>10836</td><td>29</td><td>CAATGTAGGTAAGGGAAGTCGGCAAGCCG</td><td>161.7</td><td>-3.4</td><td>1.6</td><td>-2.1</td><td>3.2E-02</td><td>4.1E-02</td><td>BT20/MB468</td><td>MITO</td></tr>
<tr><td>>hg38_21_+_8433172_8446622_RNA45SN1_45S_50ntFlanks@12456.12493.38|CCAAGCGTTGGATTGTTCACCCACTAATAGGGAACGTG</td><td>>hg38_21_+_8433172_8446622_RNA45SN1_45S_50ntFlanks</td><td>12456</td><td>12493</td><td>38</td><td>CCAAGCGTTGGATTGTTCACCCACTAATAGGGAACGTG</td><td>486.9</td><td>-1.6</td><td>0.7</td><td>-2.1</td><td>3.2E-02</td><td>4.1E-02</td><td>BT20/MB468</td><td>MITO</td></tr>
<tr><td>>hg38_21_+_8433172_8446622_RNA45SN1_45S_50ntFlanks@10234.10255.22|CGGACGCTACGCCGCGACGAGT</td><td>>hg38_21_+_8433172_8446622_RNA45SN1_45S_50ntFlanks</td><td>10234</td><td>10255</td><td>22</td><td>CGGACGCTACGCCGCGACGAGT</td><td>83.9</td><td>3.9</td><td>1.5</td><td>2.5</td><td>1.1E-02</td><td>4.1E-02</td><td>BT20/MB231</td><td>MP</td></tr>
<tr><td>>hg38_21_+_8433172_8446622_RNA45SN1_45S_50ntFlanks@10792.10816.25|CTCTGGCATGTTGGAACAATGTAGG</td><td>>hg38_21_+_8433172_8446622_RNA45SN1_45S_50ntFlanks</td><td>10792</td><td>10816</td><td>25</td><td>CTCTGGCATGTTGGAACAATGTAGG</td><td>355.1</td><td>-2.0</td><td>0.9</td><td>-2.2</td><td>2.5E-02</td><td>4.1E-02</td><td>BT20/MB468</td><td>MP</td></tr>
<tr><td>>hg38_21_+_8433172_8446622_RNA45SN1_45S_50ntFlanks@5086.5117.32|AACGAGACTCTGGCATGCTAACTAGTTACGCG</td><td>>hg38_21_+_8433172_8446622_RNA45SN1_45S_50ntFlanks</td><td>5086</td><td>5117</td><td>32</td><td>AACGAGACTCTGGCATGCTAACTAGTTACGCG</td><td>177.5</td><td>-2.5</td><td>1.2</td><td>-2.1</td><td>3.4E-02</td><td>4.1E-02</td><td>MB231/MB468</td><td>MP</td></tr>
<tr><td>>MI0000285|hsa-mir-205&WithFlank&1|+|209432127|209432248@40.59.20(+1C)&[MIMAT0000266&hsa-miR-205-5p&offsets|0|-2(+1C);m-351&1|+|209432166|209432187&offsets|0|-2(+1C)]|TCCTTCATTCCACCGGAGTCC</td><td>>MI0000285|hsa-mir-205&WithFlank&1|+|209432127|209432248</td><td>40</td><td>59</td><td>20(+1C)</td><td>TCCTTCATTCCACCGGAGTCC</td><td>935.5</td><td>2.8</td><td>1.3</td><td>2.2</td><td>2.5E-02</td><td>4.1E-02</td><td>BT20/MB468</td><td>MP</td></tr>
<tr><td>>hg38_21_+_8433172_8446622_RNA45SN1_45S_50ntFlanks@10057.10081.25|CCGGCAGTCGAGAGTGGACGGGAGC</td><td>>hg38_21_+_8433172_8446622_RNA45SN1_45S_50ntFlanks</td><td>10057</td><td>10081</td><td>25</td><td>CCGGCAGTCGAGAGTGGACGGGAGC</td><td>37.6</td><td>4.8</td><td>2.1</td><td>2.2</td><td>2.5E-02</td><td>4.1E-02</td><td>BT20/MB468</td><td>MP</td></tr>
<tr><td>>hg38_21_+_8433172_8446622_RNA45SN1_45S_50ntFlanks@12393.12419.27|ATAGCGACGTCGCTTTTTGATCCTTCG</td><td>>hg38_21_+_8433172_8446622_RNA45SN1_45S_50ntFlanks</td><td>12393</td><td>12419</td><td>27</td><td>ATAGCGACGTCGCTTTTTGATCCTTCG</td><td>39.6</td><td>-3.9</td><td>1.8</td><td>-2.2</td><td>2.5E-02</td><td>4.1E-02</td><td>BT20/MB468</td><td>MP</td></tr>
<tr><td>>MI0000101|hsa-mir-99a&WithFlank&21|+|16539083|16539175@19.40.22&[MIMAT0000097&hsa-miR-99a-5p&offsets|0|0;m-64&21|+|16539101|16539121&offsets|0|+1]|AACCCGTAGATCCGATCTTGTG</td><td>>MI0000101|hsa-mir-99a&WithFlank&21|+|16539083|16539175</td><td>19</td><td>40</td><td>22</td><td>AACCCGTAGATCCGATCTTGTG</td><td>842.8</td><td>-1.1</td><td>0.5</td><td>-2.1</td><td>3.6E-02</td><td>4.1E-02</td><td>BT20/MB468</td><td>CYTO</td></tr>
<tr><td>>hg38_21_+_8433172_8446622_RNA45SN1_45S_50ntFlanks@6775.6803.29|CCTCCCGGGGCTACGCCTGTCTGAGCGTC</td><td>>hg38_21_+_8433172_8446622_RNA45SN1_45S_50ntFlanks</td><td>6775</td><td>6803</td><td>29</td><td>CCTCCCGGGGCTACGCCTGTCTGAGCGTC</td><td>224.6</td><td>-1.1</td><td>0.5</td><td>-2.1</td><td>3.6E-02</td><td>4.1E-02</td><td>BT20/MB468</td><td>CYTO</td></tr>
<tr><td>>MI0000061|hsa-let-7a-2&WithFlank&11|-|122146516|122146599@11.31.21;MI0000062|hsa-let-7a-3&WithFlank&22|+|46112743|46112828@10.30.21;MI0000060|hsa-let-7a-1&WithFlank&9|+|94175951|94176042@12.32.21&[MIMAT0000062&hsa-let-7a-5p&offsets|0|-1;m-6&11|-|122146568|122146589&offsets|0|-1];[MIMAT0000062_1&hsa-let-7a-5p&offsets|0|-1;m-5&22|+|46112752|46112773&offsets|0|-1];[MIMAT0000062_2&hsa-let-7a-5p&offsets|0|-1;m-4&9|+|94175962|94175983&offsets|0|-1]|TGAGGTAGTAGGTTGTATAGT</td><td>>MI0000061|hsa-let-7a-2&WithFlank&11|-|122146516|122146599</td><td>11</td><td>31</td><td>21</td><td>TGAGGTAGTAGGTTGTATAGT</td><td>11412.6</td><td>-2.5</td><td>1.0</td><td>-2.5</td><td>1.1E-02</td><td>4.1E-02</td><td>BT20/MB231</td><td>MP</td></tr>
<tr><td>>MI0000440|hsa-mir-27b&WithFlank&9|+|95085439|95085547@67.86.20(+1U)&[MIMAT0000419&hsa-miR-27b-3p&offsets|0|-1(+1U);m-39&9|+|95085505|95085525&offsets|0|-1(+1U)]|TTCACAGTGGCTAAGTTCTGT</td><td>>MI0000440|hsa-mir-27b&WithFlank&9|+|95085439|95085547</td><td>67</td><td>86</td><td>20(+1U)</td><td>TTCACAGTGGCTAAGTTCTGT</td><td>12946.2</td><td>1.4</td><td>0.5</td><td>2.5</td><td>1.2E-02</td><td>4.1E-02</td><td>BT20/MB231</td><td>NUC</td></tr>
<tr><td>>hg38_1_-_228634819_228635039_RNA5S12_5S_50ntFlanks@129.169.41|TTGGATGGGAGACCGCCTGGGAATACCGGGTGCTGTAGGCT</td><td>>hg38_1_-_228634819_228635039_RNA5S12_5S_50ntFlanks</td><td>129</td><td>169</td><td>41</td><td>TTGGATGGGAGACCGCCTGGGAATACCGGGTGCTGTAGGCT</td><td>684.2</td><td>2.8</td><td>1.1</td><td>2.5</td><td>1.1E-02</td><td>4.1E-02</td><td>BT20/MB231</td><td>MP</td></tr>
<tr><td>>hg38_21_+_8433172_8446622_RNA45SN1_45S_50ntFlanks@8115.8138.24|CGCGGGACATGTGGCGTACGGAAG</td><td>>hg38_21_+_8433172_8446622_RNA45SN1_45S_50ntFlanks</td><td>8115</td><td>8138</td><td>24</td><td>CGCGGGACATGTGGCGTACGGAAG</td><td>983.0</td><td>-1.8</td><td>0.8</td><td>-2.1</td><td>3.2E-02</td><td>4.1E-02</td><td>BT20/MB468</td><td>MITO</td></tr>
<tr><td>>hg38_21_+_8433172_8446622_RNA45SN1_45S_50ntFlanks@12644.12669.26|TCTGTGGGATTATGACTGAACGCCTC</td><td>>hg38_21_+_8433172_8446622_RNA45SN1_45S_50ntFlanks</td><td>12644</td><td>12669</td><td>26</td><td>TCTGTGGGATTATGACTGAACGCCTC</td><td>346.9</td><td>-2.0</td><td>0.8</td><td>-2.4</td><td>1.5E-02</td><td>4.1E-02</td><td>BT20/MB231</td><td>CYTO</td></tr>
</table>

<table>
<tr><td>>MI0000088|hsa-mir-30a&WithFlank&6|-|71403545|71403627@12.34.23(+1G)&[MIMAT0000087&hsa-miR-30a-5p&offsets|0|+1(+1G);m-15&6|-|71403595|71403616&offsets|0|+1(+1G)]|TGTAAACATCCTCGACTGGAAGCG</td><td>>MI0000088|hsa-mir-30a&WithFlank&6|-|71403545|71403627</td><td>12</td><td>34</td><td>23(+1G)</td><td>TGTAAACATCCTCGACTGGAAGCG</td><td>10172.4</td><td>-2.8</td><td>1.1</td><td>-2.5</td><td>1.1E-02</td><td>4.1E-02</td><td>BT20/MB231</td><td>MP</td></tr>
<tr><td>>hg38_21_+_8433172_8446622_RNA45SN1_45S_50ntFlanks@8120.8141.22|GACATGTGGCGTACGGAAGACC</td><td>>hg38_21_+_8433172_8446622_RNA45SN1_45S_50ntFlanks</td><td>8120</td><td>8141</td><td>22</td><td>GACATGTGGCGTACGGAAGACC</td><td>97.7</td><td>4.9</td><td>2.0</td><td>2.5</td><td>1.1E-02</td><td>4.1E-02</td><td>BT20/MB231</td><td>MP</td></tr>
<tr><td>>hg38_21_+_8433172_8446622_RNA45SN1_45S_50ntFlanks@9792.9813.22|GGCGTGGAGCCGGGCGTGGAAT</td><td>>hg38_21_+_8433172_8446622_RNA45SN1_45S_50ntFlanks</td><td>9792</td><td>9813</td><td>22</td><td>GGCGTGGAGCCGGGCGTGGAAT</td><td>108.1</td><td>5.1</td><td>2.0</td><td>2.5</td><td>1.1E-02</td><td>4.1E-02</td><td>BT20/MB231</td><td>MP</td></tr>
<tr><td>>MI0005561|hsa-mir-877&WithFlank&6|+|30584326|30584423@7.28.22&[MIMAT0004949&hsa-miR-877-5p&offsets|0|+2;m-382&6|+|30584332|30584354&offsets|0|-1]|GTAGAGGAGATGGCGCAGGGGA</td><td>>MI0005561|hsa-mir-877&WithFlank&6|+|30584326|30584423</td><td>7</td><td>28</td><td>22</td><td>GTAGAGGAGATGGCGCAGGGGA</td><td>98.3</td><td>2.8</td><td>1.3</td><td>2.1</td><td>3.4E-02</td><td>4.1E-02</td><td>MB231/MB468</td><td>MP</td></tr>
<tr><td>>hg38_21_+_8433172_8446622_RNA45SN1_45S_50ntFlanks@7973.8006.34|GACGCGACCTCAGATCAGACGTGGCGACCCGCTG</td><td>>hg38_21_+_8433172_8446622_RNA45SN1_45S_50ntFlanks</td><td>7973</td><td>8006</td><td>34</td><td>GACGCGACCTCAGATCAGACGTGGCGACCCGCTG</td><td>618.3</td><td>-1.4</td><td>0.7</td><td>-2.1</td><td>3.7E-02</td><td>4.1E-02</td><td>MB231/MB468</td><td>NUC</td></tr>
<tr><td>>hg38_21_+_8433172_8446622_RNA45SN1_45S_50ntFlanks@10059.10081.23|GGCAGTCGAGAGTGGACGGGAGC</td><td>>hg38_21_+_8433172_8446622_RNA45SN1_45S_50ntFlanks</td><td>10059</td><td>10081</td><td>23</td><td>GGCAGTCGAGAGTGGACGGGAGC</td><td>220.0</td><td>3.9</td><td>1.5</td><td>2.5</td><td>1.1E-02</td><td>4.1E-02</td><td>BT20/MB231</td><td>MP</td></tr>
<tr><td>>hg38_21_+_8433172_8446622_RNA45SN1_45S_50ntFlanks@5473.5507.35|CGGCCCTGGCGGAGCGCTGAGAAGACGGTCGAACT</td><td>>hg38_21_+_8433172_8446622_RNA45SN1_45S_50ntFlanks</td><td>5473</td><td>5507</td><td>35</td><td>CGGCCCTGGCGGAGCGCTGAGAAGACGGTCGAACT</td><td>1295.6</td><td>2.5</td><td>1.0</td><td>2.4</td><td>1.5E-02</td><td>4.1E-02</td><td>BT20/MB231</td><td>CYTO</td></tr>
<tr><td>>hg38_21_+_8433172_8446622_RNA45SN1_45S_50ntFlanks@11668.11701.34|TGCCCAGTGCTCTGAATGTCAAAGTGAAGAAATT</td><td>>hg38_21_+_8433172_8446622_RNA45SN1_45S_50ntFlanks</td><td>11668</td><td>11701</td><td>34</td><td>TGCCCAGTGCTCTGAATGTCAAAGTGAAGAAATT</td><td>588.7</td><td>-2.2</td><td>0.9</td><td>-2.4</td><td>1.5E-02</td><td>4.1E-02</td><td>BT20/MB231</td><td>CYTO</td></tr>
<tr><td>>hg38_MT_+_1621_3279_RNR2_16S_50ntFlanks@488.518.31|GAACAGCTCTTTGGACACTAGGAAAAAACCT</td><td>>hg38_MT_+_1621_3279_RNR2_16S_50ntFlanks</td><td>488</td><td>518</td><td>31</td><td>GAACAGCTCTTTGGACACTAGGAAAAAACCT</td><td>101.5</td><td>2.7</td><td>1.3</td><td>2.1</td><td>3.4E-02</td><td>4.1E-02</td><td>MB231/MB468</td><td>MP</td></tr>
<tr><td>>MI0000067|hsa-let-7f-1&WithFlank&9|+|94176341|94176439@13.32.20(+1A);MI0000068|hsa-let-7f-2&WithFlank&X|-|53557186|53557280@14.33.20(+1A)&[MIMAT0000067&hsa-let-7f-5p&offsets|0|-2(+1A);m-20&9|+|94176353|94176374&offsets|0|-2(+1A)];[MIMAT0000067_1&hsa-let-7f-5p&offsets|0|-2(+1A);m-19&X|-|53557246|53557267&offsets|0|-2(+1A)]|TGAGGTAGTAGATTGTATAGA</td><td>>MI0000067|hsa-let-7f-1&WithFlank&9|+|94176341|94176439</td><td>13</td><td>32</td><td>20(+1A)</td><td>TGAGGTAGTAGATTGTATAGA</td><td>257.6</td><td>-3.1</td><td>1.2</td><td>-2.5</td><td>1.1E-02</td><td>4.1E-02</td><td>BT20/MB231</td><td>MP</td></tr>
<tr><td>>MI0005561|hsa-mir-877&WithFlank&6|+|30584326|30584423@7.29.23&[MIMAT0004949&hsa-miR-877-5p&offsets|0|+3;m-382&6|+|30584332|30584354&offsets|0|0]|GTAGAGGAGATGGCGCAGGGGAC</td><td>>MI0005561|hsa-mir-877&WithFlank&6|+|30584326|30584423</td><td>7</td><td>29</td><td>23</td><td>GTAGAGGAGATGGCGCAGGGGAC</td><td>99.8</td><td>2.7</td><td>1.3</td><td>2.1</td><td>3.4E-02</td><td>4.1E-02</td><td>MB231/MB468</td><td>MP</td></tr>
<tr><td>>MI0000253|hsa-mir-148a&WithFlank&7|-|25949913|25949992@50.71.22&[MIMAT0000243&hsa-miR-148a-3p&offsets|0|0;m-7&7|-|25949922|25949943&offsets|0|0]|TCAGTGCACTACAGAACTTTGT</td><td>>MI0000253|hsa-mir-148a&WithFlank&7|-|25949913|25949992</td><td>50</td><td>71</td><td>22</td><td>TCAGTGCACTACAGAACTTTGT</td><td>8691.6</td><td>1.8</td><td>0.9</td><td>2.1</td><td>3.6E-02</td><td>4.1E-02</td><td>MB231/MB468</td><td>CYTO</td></tr>
<tr><td>>hg38_1_-_228634819_228635039_RNA5S12_5S_50ntFlanks@154.171.18|CCGGGTGCTGTAGGCTTT</td><td>>hg38_1_-_228634819_228635039_RNA5S12_5S_50ntFlanks</td><td>154</td><td>171</td><td>18</td><td>CCGGGTGCTGTAGGCTTT</td><td>10114.1</td><td>3.6</td><td>1.4</td><td>2.5</td><td>1.1E-02</td><td>4.1E-02</td><td>BT20/MB231</td><td>MP</td></tr>
<tr><td>>hg38_21_+_8433172_8446622_RNA45SN1_45S_50ntFlanks@10821.10865.45|GGAAGTCGGCAAGCCGGATCCGTAACTTCGGGATAAGGATTGGCT</td><td>>hg38_21_+_8433172_8446622_RNA45SN1_45S_50ntFlanks</td><td>10821</td><td>10865</td><td>45</td><td>GGAAGTCGGCAAGCCGGATCCGTAACTTCGGGATAAGGATTGGCT</td><td>282.2</td><td>-2.8</td><td>1.3</td><td>-2.1</td><td>3.2E-02</td><td>4.1E-02</td><td>BT20/MB468</td><td>MITO</td></tr>
<tr><td>>hg38_MT_+_1621_3279_RNR2_16S_50ntFlanks@1430.1453.24|TAAAGTCCTACGTGATCTGAGTTC</td><td>>hg38_MT_+_1621_3279_RNR2_16S_50ntFlanks</td><td>1430</td><td>1453</td><td>24</td><td>TAAAGTCCTACGTGATCTGAGTTC</td><td>123.5</td><td>3.5</td><td>1.6</td><td>2.2</td><td>2.6E-02</td><td>4.1E-02</td><td>BT20/MB468</td><td>MP</td></tr>
<tr><td>>hg38_21_+_8433172_8446622_RNA45SN1_45S_50ntFlanks@6653.6686.34|ACTCTTAGCGGTGGATCACTCGGCTCGTGCGTCG</td><td>>hg38_21_+_8433172_8446622_RNA45SN1_45S_50ntFlanks</td><td>6653</td><td>6686</td><td>34</td><td>ACTCTTAGCGGTGGATCACTCGGCTCGTGCGTCG</td><td>924.0</td><td>1.4</td><td>0.7</td><td>2.1</td><td>3.7E-02</td><td>4.1E-02</td><td>MB231/MB468</td><td>NUC</td></tr>
<tr><td>>hg38_21_+_8433172_8446622_RNA45SN1_45S_50ntFlanks@11582.11611.30|ACTGGTGCGGACCAGGGGAATCCGACTGTT</td><td>>hg38_21_+_8433172_8446622_RNA45SN1_45S_50ntFlanks</td><td>11582</td><td>11611</td><td>30</td><td>ACTGGTGCGGACCAGGGGAATCCGACTGTT</td><td>352.8</td><td>2.7</td><td>1.1</td><td>2.5</td><td>1.1E-02</td><td>4.1E-02</td><td>BT20/MB231</td><td>MP</td></tr>
<tr><td>>am_trna116_GluCTC_1_-_145399233_145399304@7.29.23,trna59_GluCTC_1_+_249168447_249168518@7.29.23,trna71_GluCTC_1_-_161439189_161439260@7.29.23,trna74_GluCTC_1_-_161431809_161431880@7.29.23,trna77_GluCTC_1_-_161424398_161424469@7.29.23,trna77_GluCTC_6_+_28949976_28950047@7.29.23,trna80_GluCTC_1_-_161417018_161417089@7.29.23,trna87_GluCTC_6_-_126101393_126101464@7.29.23|GTGGTCTAGTGGTTAGGATTCGG</td><td>>am_trna116_GluCTC_1_-_145399233_145399304</td><td>7</td><td>29</td><td>23</td><td>GTGGTCTAGTGGTTAGGATTCGG</td><td>99.4</td><td>-3.8</td><td>1.7</td><td>-2.2</td><td>2.6E-02</td><td>4.1E-02</td><td>BT20/MB468</td><td>MP</td></tr>
<tr><td>>hg38_21_+_8433172_8446622_RNA45SN1_45S_50ntFlanks@9621.9647.27|TCTAGTAGCTGGTTCCCTCCGAAGTTT</td><td>>hg38_21_+_8433172_8446622_RNA45SN1_45S_50ntFlanks</td><td>9621</td><td>9647</td><td>27</td><td>TCTAGTAGCTGGTTCCCTCCGAAGTTT</td><td>119.0</td><td>-1.6</td><td>0.7</td><td>-2.1</td><td>3.2E-02</td><td>4.1E-02</td><td>BT20/MB468</td><td>MITO</td></tr>
<tr><td>>hg38_21_+_8433172_8446622_RNA45SN1_45S_50ntFlanks@10425.10453.29|AGATGGGCGAGCGCCGTTCCGAAGGGACG</td><td>>hg38_21_+_8433172_8446622_RNA45SN1_45S_50ntFlanks</td><td>10425</td><td>10453</td><td>29</td><td>AGATGGGCGAGCGCCGTTCCGAAGGGACG</td><td>774.2</td><td>-1.7</td><td>0.8</td><td>-2.1</td><td>3.6E-02</td><td>4.1E-02</td><td>MB231/MB468</td><td>CYTO</td></tr>
<tr><td>>hg38_21_+_8433172_8446622_RNA45SN1_45S_50ntFlanks@10239.10267.29|GCTACGCCGCGACGAGTAGGAGGGCCGCT</td><td>>hg38_21_+_8433172_8446622_RNA45SN1_45S_50ntFlanks</td><td>10239</td><td>10267</td><td>29</td><td>GCTACGCCGCGACGAGTAGGAGGGCCGCT</td><td>156.6</td><td>5.2</td><td>2.0</td><td>2.5</td><td>1.1E-02</td><td>4.1E-02</td><td>BT20/MB231</td><td>MP</td></tr>
<tr><td>>MI0000077|hsa-mir-21&WithFlank&17|+|59841260|59841343@14.34.21&[MIMAT0000076&hsa-miR-21-5p&offsets|0|-1;m-1&17|+|59841273|59841294&offsets|0|-1]|TAGCTTATCAGACTGATGTTG</td><td>>MI0000077|hsa-mir-21&WithFlank&17|+|59841260|59841343</td><td>14</td><td>34</td><td>21</td><td>TAGCTTATCAGACTGATGTTG</td><td>18421.9</td><td>2.6</td><td>1.1</td><td>2.4</td><td>1.5E-02</td><td>4.1E-02</td><td>BT20/MB231</td><td>CYTO</td></tr>
<tr><td>>hg38_21_+_8433172_8446622_RNA45SN1_45S_50ntFlanks@6700.6720.21|CTAGCTGCGAGAATTAATGTG</td><td>>hg38_21_+_8433172_8446622_RNA45SN1_45S_50ntFlanks</td><td>6700</td><td>6720</td><td>21</td><td>CTAGCTGCGAGAATTAATGTG</td><td>1184.8</td><td>-2.1</td><td>0.9</td><td>-2.4</td><td>1.5E-02</td><td>4.1E-02</td><td>BT20/MB231</td><td>CYTO</td></tr>
<tr><td>>trna20_GluTTC_2_-_131094701_131094772@1.32.32,trna5_GluTTC_13_-_41634874_41634945@1.32.32|TCCCATATGGTCTAGCGGTTAGGATTCCTGGT</td><td>>trna20_GluTTC_2_-_131094701_131094772</td><td>1</td><td>32</td><td>32</td><td>TCCCATATGGTCTAGCGGTTAGGATTCCTGGT</td><td>496.4</td><td>-3.9</td><td>1.5</td><td>-2.5</td><td>1.1E-02</td><td>4.1E-02</td><td>BT20/MB231</td><td>MP</td></tr>
<tr><td>>hg38_21_+_8433172_8446622_RNA45SN1_45S_50ntFlanks@5487.5515.29|CGCTGAGAAGACGGTCGAACTTGACTATC</td><td>>hg38_21_+_8433172_8446622_RNA45SN1_45S_50ntFlanks</td><td>5487</td><td>5515</td><td>29</td><td>CGCTGAGAAGACGGTCGAACTTGACTATC</td><td>520.0</td><td>1.2</td><td>0.6</td><td>2.1</td><td>3.6E-02</td><td>4.1E-02</td><td>BT20/MB468</td><td>CYTO</td></tr>
<tr><td>>MI0000084|hsa-mir-26b&WithFlank&2|+|218402640|218402728@18.38.21&[MIMAT0000083&hsa-miR-26b-5p&offsets|0|0;m-53&2|+|218402657|218402678&offsets|0|-1]|TTCAAGTAATTCAGGATAGGT</td><td>>MI0000084|hsa-mir-26b&WithFlank&2|+|218402640|218402728</td><td>18</td><td>38</td><td>21</td><td>TTCAAGTAATTCAGGATAGGT</td><td>2663.6</td><td>-2.9</td><td>1.1</td><td>-2.5</td><td>1.1E-02</td><td>4.1E-02</td><td>BT20/MB231</td><td>MP</td></tr>
<tr><td>>hg38_MT_+_1621_3279_RNR2_16S_50ntFlanks@51.79.29|GCTAAACCTAGCCCCAAACCCACTCCACC</td><td>>hg38_MT_+_1621_3279_RNR2_16S_50ntFlanks</td><td>51</td><td>79</td><td>29</td><td>GCTAAACCTAGCCCCAAACCCACTCCACC</td><td>266.5</td><td>1.0</td><td>0.5</td><td>2.1</td><td>3.7E-02</td><td>4.1E-02</td><td>BT20/MB468</td><td>NUC</td></tr>
<tr><td>>hg38_21_+_8433172_8446622_RNA45SN1_45S_50ntFlanks@4685.4722.38|AGACGGACCAGAGCGAAAGCATTTGCCAAGAATGTTTT</td><td>>hg38_21_+_8433172_8446622_RNA45SN1_45S_50ntFlanks</td><td>4685</td><td>4722</td><td>38</td><td>AGACGGACCAGAGCGAAAGCATTTGCCAAGAATGTTTT</td><td>333.2</td><td>-1.8</td><td>0.9</td><td>-2.1</td><td>3.2E-02</td><td>4.1E-02</td><td>BT20/MB468</td><td>MITO</td></tr>
<tr><td>>hg38_21_+_8433172_8446622_RNA45SN1_45S_50ntFlanks@10809.10826.18|AATGTAGGTAAGGGAAGT</td><td>>hg38_21_+_8433172_8446622_RNA45SN1_45S_50ntFlanks</td><td>10809</td><td>10826</td><td>18</td><td>AATGTAGGTAAGGGAAGT</td><td>903.0</td><td>2.3</td><td>1.1</td><td>2.1</td><td>3.6E-02</td><td>4.1E-02</td><td>MB231/MB468</td><td>CYTO</td></tr>
<tr><td>>hg38_MT_+_1621_3279_RNR2_16S_50ntFlanks@1390.1426.37|GATGGTGCAGCCGCTATTAAAGGTTCGTTTGTTCAAC</td><td>>hg38_MT_+_1621_3279_RNR2_16S_50ntFlanks</td><td>1390</td><td>1426</td><td>37</td><td>GATGGTGCAGCCGCTATTAAAGGTTCGTTTGTTCAAC</td><td>55.0</td><td>4.2</td><td>2.0</td><td>2.1</td><td>3.4E-02</td><td>4.1E-02</td><td>MB231/MB468</td><td>MP</td></tr>
<tr><td>>MI0000803|hsa-mir-330&WithFlank&19|-|45638988|45639093@64.86.23&[MIMAT0000751&hsa-miR-330-3p&offsets|+1|+1;m-347&19|-|45639009|45639030&offsets|0|+1]|CAAAGCACACGGCCTGCAGAGAG</td><td>>MI0000803|hsa-mir-330&WithFlank&19|-|45638988|45639093</td><td>64</td><td>86</td><td>23</td><td>CAAAGCACACGGCCTGCAGAGAG</td><td>40.5</td><td>4.4</td><td>2.1</td><td>2.1</td><td>3.4E-02</td><td>4.1E-02</td><td>MB231/MB468</td><td>MP</td></tr>
<tr><td>>hg38_21_+_8433172_8446622_RNA45SN1_45S_50ntFlanks@11667.11693.27|CTGCCCAGTGCTCTGAATGTCAAAGTG</td><td>>hg38_21_+_8433172_8446622_RNA45SN1_45S_50ntFlanks</td><td>11667</td><td>11693</td><td>27</td><td>CTGCCCAGTGCTCTGAATGTCAAAGTG</td><td>53.2</td><td>4.0</td><td>1.8</td><td>2.2</td><td>2.6E-02</td><td>4.1E-02</td><td>BT20/MB468</td><td>MP</td></tr>
<tr><td>>hg38_MT_+_1621_3279_RNR2_16S_50ntFlanks@1281.1301.21|AATAACTTGACCAACGGAACA</td><td>>hg38_MT_+_1621_3279_RNR2_16S_50ntFlanks</td><td>1281</td><td>1301</td><td>21</td><td>AATAACTTGACCAACGGAACA</td><td>56.3</td><td>2.6</td><td>1.2</td><td>2.1</td><td>3.4E-02</td><td>4.1E-02</td><td>MB231/MB468</td><td>MP</td></tr>
<tr><td>>am_trnaMT_SerGCT_MT_+_12207_12265@15.48.34|GAACTGCTAACTCATGCCCCCATGTCTAACAACA</td><td>>am_trnaMT_SerGCT_MT_+_12207_12265</td><td>15</td><td>48</td><td>34</td><td>GAACTGCTAACTCATGCCCCCATGTCTAACAACA</td><td>65.7</td><td>4.3</td><td>2.0</td><td>2.1</td><td>3.4E-02</td><td>4.1E-02</td><td>MB231/MB468</td><td>MP</td></tr>
<tr><td>>hg38_21_+_8433172_8446622_RNA45SN1_45S_50ntFlanks@10458.10478.21|ATGGCCTCCGTTGCCCTCGGC</td><td>>hg38_21_+_8433172_8446622_RNA45SN1_45S_50ntFlanks</td><td>10458</td><td>10478</td><td>21</td><td>ATGGCCTCCGTTGCCCTCGGC</td><td>289.1</td><td>-3.9</td><td>1.8</td><td>-2.2</td><td>2.6E-02</td><td>4.1E-02</td><td>BT20/MB468</td><td>MP</td></tr>
<tr><td>>am_MI0000809|hsa-mir-151a&WithFlank&8|-|140732558|140732659@53.70.18(+3G)&[MIMAT0000757&hsa-miR-151a-3p&offsets|0|-3(+3G);m-30&8|-|140732587|140732607&offsets|0|-3(+3G)]|CTAGACTGAAGCTCCTTGGGG</td><td>>am_MI0000809|hsa-mir-151a&WithFlank&8|-|140732558|140732659</td><td>53</td><td>70</td><td>18(+3G)</td><td>CTAGACTGAAGCTCCTTGGGG</td><td>603.2</td><td>1.8</td><td>0.8</td><td>2.4</td><td>1.5E-02</td><td>4.1E-02</td><td>BT20/MB231</td><td>CYTO</td></tr>
<tr><td>>MI0003760|hsa-mir-671&WithFlank&7|+|151238415|151238544@35.58.24&[MIMAT0003880&hsa-miR-671-5p&offsets|0|+1;m-316&7|+|151238449|151238472&offsets|0|0]|AGGAAGCCCTGGAGGGGCTGGAGG</td><td>>MI0003760|hsa-mir-671&WithFlank&7|+|151238415|151238544</td><td>35</td><td>58</td><td>24</td><td>AGGAAGCCCTGGAGGGGCTGGAGG</td><td>453.4</td><td>3.9</td><td>1.7</td><td>2.2</td><td>2.6E-02</td><td>4.1E-02</td><td>BT20/MB468</td><td>MP</td></tr>
<tr><td>>am_trnaMT_SerGCT_MT_+_12207_12265@12.31.20|CAAGAACTGCTAACTCATGC</td><td>>am_trnaMT_SerGCT_MT_+_12207_12265</td><td>12</td><td>31</td><td>20</td><td>CAAGAACTGCTAACTCATGC</td><td>380.5</td><td>2.2</td><td>1.0</td><td>2.1</td><td>3.6E-02</td><td>4.1E-02</td><td>BT20/MB468</td><td>CYTO</td></tr>
<tr><td>>hg38_21_+_8433172_8446622_RNA45SN1_45S_50ntFlanks@5482.5516.35|CGGAGCGCTGAGAAGACGGTCGAACTTGACTATCT</td><td>>hg38_21_+_8433172_8446622_RNA45SN1_45S_50ntFlanks</td><td>5482</td><td>5516</td><td>35</td><td>CGGAGCGCTGAGAAGACGGTCGAACTTGACTATCT</td><td>350.7</td><td>4.4</td><td>1.8</td><td>2.5</td><td>1.1E-02</td><td>4.1E-02</td><td>BT20/MB231</td><td>MP</td></tr>
<tr><td>>hg38_21_+_8433172_8446622_RNA45SN1_45S_50ntFlanks@10798.10825.28|CATGTTGGAACAATGTAGGTAAGGGAAG</td><td>>hg38_21_+_8433172_8446622_RNA45SN1_45S_50ntFlanks</td><td>10798</td><td>10825</td><td>28</td><td>CATGTTGGAACAATGTAGGTAAGGGAAG</td><td>241.9</td><td>-2.5</td><td>1.2</td><td>-2.1</td><td>3.2E-02</td><td>4.1E-02</td><td>BT20/MB468</td><td>MITO</td></tr>
<tr><td>>hg38_21_+_8433172_8446622_RNA45SN1_45S_50ntFlanks@11874.11906.33|AGAAGACCCTGTTGAGCTTGACTCTAGTCTGGC</td><td>>hg38_21_+_8433172_8446622_RNA45SN1_45S_50ntFlanks</td><td>11874</td><td>11906</td><td>33</td><td>AGAAGACCCTGTTGAGCTTGACTCTAGTCTGGC</td><td>1071.0</td><td>-4.2</td><td>1.9</td><td>-2.2</td><td>2.6E-02</td><td>4.1E-02</td><td>BT20/MB468</td><td>MP</td></tr>
<tr><td>>MI0000254|hsa-mir-30c-2&WithFlank&6|-|71376954|71377037@54.74.21&[MIMAT0004550&hsa-miR-30c-2-3p&offsets|+1|0;m-206&6|-|71376964|71376985&offsets|+1|0]|TGGGAGAAGGCTGTTTACTCT</td><td>>MI0000254|hsa-mir-30c-2&WithFlank&6|-|71376954|71377037</td><td>54</td><td>74</td><td>21</td><td>TGGGAGAAGGCTGTTTACTCT</td><td>104.7</td><td>-3.7</td><td>1.5</td><td>-2.5</td><td>1.1E-02</td><td>4.1E-02</td><td>BT20/MB231</td><td>MP</td></tr>
<tr><td>>hg38_21_+_8433172_8446622_RNA45SN1_45S_50ntFlanks@12639.12659.21|TACCATCTGTGGGATTATGAC</td><td>>hg38_21_+_8433172_8446622_RNA45SN1_45S_50ntFlanks</td><td>12639</td><td>12659</td><td>21</td><td>TACCATCTGTGGGATTATGAC</td><td>196.2</td><td>-3.1</td><td>1.4</td><td>-2.2</td><td>2.6E-02</td><td>4.1E-02</td><td>BT20/MB468</td><td>MP</td></tr>
<tr><td>>hg38_21_+_8433172_8446622_RNA45SN1_45S_50ntFlanks@10391.10429.39|AACAGCAGTTGAACATGGGTCAGTCGGTCCTGAGAGATG</td><td>>hg38_21_+_8433172_8446622_RNA45SN1_45S_50ntFlanks</td><td>10391</td><td>10429</td><td>39</td><td>AACAGCAGTTGAACATGGGTCAGTCGGTCCTGAGAGATG</td><td>826.0</td><td>-2.3</td><td>1.0</td><td>-2.2</td><td>2.6E-02</td><td>4.1E-02</td><td>BT20/MB468</td><td>MP</td></tr>
<tr><td>>trnaMT_LeuTAA_MT_+_3230_3304@22.44.23|AATCGCATAAAACTTAAAACTTT</td><td>>trnaMT_LeuTAA_MT_+_3230_3304</td><td>22</td><td>44</td><td>23</td><td>AATCGCATAAAACTTAAAACTTT</td><td>59.4</td><td>-4.0</td><td>1.8</td><td>-2.2</td><td>2.6E-02</td><td>4.1E-02</td><td>BT20/MB468</td><td>MP</td></tr>
<tr><td>>hg38_21_+_8433172_8446622_RNA45SN1_45S_50ntFlanks@9552.9569.18|TCCGTAGCGGTCCTGACG</td><td>>hg38_21_+_8433172_8446622_RNA45SN1_45S_50ntFlanks</td><td>9552</td><td>9569</td><td>18</td><td>TCCGTAGCGGTCCTGACG</td><td>456.4</td><td>-1.8</td><td>0.7</td><td>-2.4</td><td>1.5E-02</td><td>4.1E-02</td><td>BT20/MB231</td><td>CYTO</td></tr>
<tr><td>>trnaMT_ThrTGT_MT_+_15888_15953@37.68.32|ACCGGAGATGAAAACCTTTTTCCAAGGACACC</td><td>>trnaMT_ThrTGT_MT_+_15888_15953</td><td>37</td><td>68</td><td>32</td><td>ACCGGAGATGAAAACCTTTTTCCAAGGACACC</td><td>107.1</td><td>5.1</td><td>2.3</td><td>2.2</td><td>2.6E-02</td><td>4.1E-02</td><td>BT20/MB468</td><td>MP</td></tr>
<tr><td>>hg38_21_+_8433172_8446622_RNA45SN1_45S_50ntFlanks@12407.12436.30|TTTTGATCCTTCGATGTCGGCTCTTCCTAT</td><td>>hg38_21_+_8433172_8446622_RNA45SN1_45S_50ntFlanks</td><td>12407</td><td>12436</td><td>30</td><td>TTTTGATCCTTCGATGTCGGCTCTTCCTAT</td><td>170.3</td><td>-3.9</td><td>1.7</td><td>-2.2</td><td>2.6E-02</td><td>4.1E-02</td><td>BT20/MB468</td><td>MP</td></tr>
<tr><td>>hg38_21_+_8433172_8446622_RNA45SN1_45S_50ntFlanks@9793.9819.27|GCGTGGAGCCGGGCGTGGAATGCGAGT</td><td>>hg38_21_+_8433172_8446622_RNA45SN1_45S_50ntFlanks</td><td>9793</td><td>9819</td><td>27</td><td>GCGTGGAGCCGGGCGTGGAATGCGAGT</td><td>56.7</td><td>3.1</td><td>1.4</td><td>2.2</td><td>2.6E-02</td><td>4.1E-02</td><td>BT20/MB468</td><td>MP</td></tr>
<tr><td>>hg38_21_+_8433172_8446622_RNA45SN1_45S_50ntFlanks@12444.12480.37|AAGCAGAATTCACCAAGCGTTGGATTGTTCACCCACT</td><td>>hg38_21_+_8433172_8446622_RNA45SN1_45S_50ntFlanks</td><td>12444</td><td>12480</td><td>37</td><td>AAGCAGAATTCACCAAGCGTTGGATTGTTCACCCACT</td><td>500.4</td><td>-2.0</td><td>0.8</td><td>-2.4</td><td>1.5E-02</td><td>4.1E-02</td><td>BT20/MB231</td><td>CYTO</td></tr>
</table>

<table>
<tr><td>>am_trnaMT_ValTAC_MT_+_1602_1670@27.48.22|CAACTTACACTTAGGAGATTTC</td><td>>am_trnaMT_ValTAC_MT_+_1602_1670</td><td>27</td><td>48</td><td>22</td><td>CAACTTACACTTAGGAGATTTC</td><td>70.5</td><td>5.9</td><td>2.6</td><td>2.2</td><td>2.6E-02</td><td>4.1E-02</td><td>BT20/MB468</td><td>MP</td></tr>
<tr><td>>hg38_21_+_8433172_8446622_RNA45SN1_45S_50ntFlanks@5422.5450.29|GATTGGATGGTTTAGTGAGGCCCTCGGAT</td><td>>hg38_21_+_8433172_8446622_RNA45SN1_45S_50ntFlanks</td><td>5422</td><td>5450</td><td>29</td><td>GATTGGATGGTTTAGTGAGGCCCTCGGAT</td><td>1990.7</td><td>3.9</td><td>1.5</td><td>2.5</td><td>1.1E-02</td><td>4.1E-02</td><td>BT20/MB231</td><td>MP</td></tr>
<tr><td>>hg38_21_+_8433172_8446622_RNA45SN1_45S_50ntFlanks@8068.8103.36|AGTGAACAGGGAAGAGCCCAGCGCCGAATCCCCGCC</td><td>>hg38_21_+_8433172_8446622_RNA45SN1_45S_50ntFlanks</td><td>8068</td><td>8103</td><td>36</td><td>AGTGAACAGGGAAGAGCCCAGCGCCGAATCCCCGCC</td><td>138.0</td><td>-2.8</td><td>1.3</td><td>-2.1</td><td>3.4E-02</td><td>4.1E-02</td><td>MB231/MB468</td><td>MP</td></tr>
<tr><td>>hg38_21_+_8433172_8446622_RNA45SN1_45S_50ntFlanks@9533.9577.45|GAGGAAACTCTGGTGGAGGTCCGTAGCGGTCCTGACGTGCAAATC</td><td>>hg38_21_+_8433172_8446622_RNA45SN1_45S_50ntFlanks</td><td>9533</td><td>9577</td><td>45</td><td>GAGGAAACTCTGGTGGAGGTCCGTAGCGGTCCTGACGTGCAAATC</td><td>296.8</td><td>-4.4</td><td>1.8</td><td>-2.4</td><td>1.5E-02</td><td>4.1E-02</td><td>BT20/MB231</td><td>CYTO</td></tr>
<tr><td>>hg38_21_+_8433172_8446622_RNA45SN1_45S_50ntFlanks@10240.10263.24|CTACGCCGCGACGAGTAGGAGGGC</td><td>>hg38_21_+_8433172_8446622_RNA45SN1_45S_50ntFlanks</td><td>10240</td><td>10263</td><td>24</td><td>CTACGCCGCGACGAGTAGGAGGGC</td><td>233.4</td><td>2.2</td><td>1.0</td><td>2.2</td><td>2.6E-02</td><td>4.1E-02</td><td>BT20/MB468</td><td>MP</td></tr>
<tr><td>>MI0000077|hsa-mir-21&WithFlank&17|+|59841260|59841343@14.36.23(+1C)&[MIMAT0000076&hsa-miR-21-5p&offsets|0|+1(+1C);m-1&17|+|59841273|59841294&offsets|0|+1(+1C)]|TAGCTTATCAGACTGATGTTGACC</td><td>>MI0000077|hsa-mir-21&WithFlank&17|+|59841260|59841343</td><td>14</td><td>36</td><td>23(+1C)</td><td>TAGCTTATCAGACTGATGTTGACC</td><td>49953.4</td><td>-1.8</td><td>0.7</td><td>-2.5</td><td>1.4E-02</td><td>4.1E-02</td><td>BT20/MB231</td><td>MITO</td></tr>
<tr><td>>hg38_21_+_8433172_8446622_RNA45SN1_45S_50ntFlanks@5186.5205.20|CAATAACAGGTCTGTGATGC</td><td>>hg38_21_+_8433172_8446622_RNA45SN1_45S_50ntFlanks</td><td>5186</td><td>5205</td><td>20</td><td>CAATAACAGGTCTGTGATGC</td><td>358.4</td><td>-2.2</td><td>1.0</td><td>-2.1</td><td>3.2E-02</td><td>4.1E-02</td><td>BT20/MB468</td><td>MITO</td></tr>
<tr><td>>hg38_21_+_8433172_8446622_RNA45SN1_45S_50ntFlanks@6661.6690.30|CGGTGGATCACTCGGCTCGTGCGTCGATGA</td><td>>hg38_21_+_8433172_8446622_RNA45SN1_45S_50ntFlanks</td><td>6661</td><td>6690</td><td>30</td><td>CGGTGGATCACTCGGCTCGTGCGTCGATGA</td><td>297.9</td><td>1.5</td><td>0.6</td><td>2.5</td><td>1.2E-02</td><td>4.1E-02</td><td>BT20/MB231</td><td>NUC</td></tr>
<tr><td>>hg38_21_+_8433172_8446622_RNA45SN1_45S_50ntFlanks@6651.6687.37|CGACTCTTAGCGGTGGATCACTCGGCTCGTGCGTCGA</td><td>>hg38_21_+_8433172_8446622_RNA45SN1_45S_50ntFlanks</td><td>6651</td><td>6687</td><td>37</td><td>CGACTCTTAGCGGTGGATCACTCGGCTCGTGCGTCGA</td><td>819.2</td><td>1.3</td><td>0.6</td><td>2.1</td><td>3.7E-02</td><td>4.1E-02</td><td>BT20/MB468</td><td>NUC</td></tr>
<tr><td>>hg38_21_+_8433172_8446622_RNA45SN1_45S_50ntFlanks@10409.10432.24|GTCAGTCGGTCCTGAGAGATGGGC</td><td>>hg38_21_+_8433172_8446622_RNA45SN1_45S_50ntFlanks</td><td>10409</td><td>10432</td><td>24</td><td>GTCAGTCGGTCCTGAGAGATGGGC</td><td>233.0</td><td>-1.1</td><td>0.5</td><td>-2.1</td><td>3.6E-02</td><td>4.1E-02</td><td>BT20/MB468</td><td>CYTO</td></tr>
<tr><td>>MI0000087|hsa-mir-29a&WithFlank&7|-|130876741|130876816@48.67.20(+1G)&[MIMAT0000086&hsa-miR-29a-3p&offsets|0|-2(+1G);m-26&7|-|130876749|130876769&offsets|0|-1(+1G)]|TAGCACCATCTGAAATCGGTG</td><td>>MI0000087|hsa-mir-29a&WithFlank&7|-|130876741|130876816</td><td>48</td><td>67</td><td>20(+1G)</td><td>TAGCACCATCTGAAATCGGTG</td><td>214.2</td><td>5.2</td><td>2.3</td><td>2.2</td><td>2.6E-02</td><td>4.1E-02</td><td>BT20/MB468</td><td>MP</td></tr>
<tr><td>>hg38_21_+_8433172_8446622_RNA45SN1_45S_50ntFlanks@12488.12519.32|AACGTGAGCTGGGTTTAGACCGTCGTGAGACA</td><td>>hg38_21_+_8433172_8446622_RNA45SN1_45S_50ntFlanks</td><td>12488</td><td>12519</td><td>32</td><td>AACGTGAGCTGGGTTTAGACCGTCGTGAGACA</td><td>131.3</td><td>-2.5</td><td>1.2</td><td>-2.1</td><td>3.2E-02</td><td>4.1E-02</td><td>BT20/MB468</td><td>MITO</td></tr>
<tr><td>>hg38_1_-_228634819_228635039_RNA5S12_5S_50ntFlanks@140.170.31|ACCGCCTGGGAATACCGGGTGCTGTAGGCTT</td><td>>hg38_1_-_228634819_228635039_RNA5S12_5S_50ntFlanks</td><td>140</td><td>170</td><td>31</td><td>ACCGCCTGGGAATACCGGGTGCTGTAGGCTT</td><td>12153.4</td><td>1.5</td><td>0.7</td><td>2.1</td><td>3.2E-02</td><td>4.2E-02</td><td>BT20/MB468</td><td>MITO</td></tr>
<tr><td>>am_trna128_GlyGCC_6_-_27870686_27870756@3.35.33,trna18_GlyGCC_16_+_70822597_70822667@3.35.33,trna19_GlyGCC_16_+_70823410_70823480@3.35.33,trna19_GlyGCC_2_-_157257659_157257729@3.35.33,trna24_GlyGCC_16_-_70812942_70813012@3.35.33,trna25_GlyGCC_16_-_70812114_70812184@3.35.33,trna5_GlyGCC_17_+_8029064_8029134@3.35.33,trna68_GlyGCC_1_-_161493637_161493707@3.35.33|ATTGGTGGTTCAGTGGTAGAATTCTCGCCTGCC</td><td>>am_trna128_GlyGCC_6_-_27870686_27870756</td><td>3</td><td>35</td><td>33</td><td>ATTGGTGGTTCAGTGGTAGAATTCTCGCCTGCC</td><td>84.0</td><td>3.4</td><td>1.5</td><td>2.2</td><td>2.6E-02</td><td>4.2E-02</td><td>BT20/MB468</td><td>MP</td></tr>
<tr><td>>hg38_21_+_8433172_8446622_RNA45SN1_45S_50ntFlanks@10401.10432.32|GAACATGGGTCAGTCGGTCCTGAGAGATGGGC</td><td>>hg38_21_+_8433172_8446622_RNA45SN1_45S_50ntFlanks</td><td>10401</td><td>10432</td><td>32</td><td>GAACATGGGTCAGTCGGTCCTGAGAGATGGGC</td><td>330.1</td><td>4.4</td><td>1.7</td><td>2.5</td><td>1.2E-02</td><td>4.2E-02</td><td>BT20/MB231</td><td>MP</td></tr>
<tr><td>>hg38_21_+_8433172_8446622_RNA45SN1_45S_50ntFlanks@7975.7994.20|CGCGACCTCAGATCAGACGT</td><td>>hg38_21_+_8433172_8446622_RNA45SN1_45S_50ntFlanks</td><td>7975</td><td>7994</td><td>20</td><td>CGCGACCTCAGATCAGACGT</td><td>57669.2</td><td>1.4</td><td>0.7</td><td>2.1</td><td>3.8E-02</td><td>4.2E-02</td><td>MB231/MB468</td><td>NUC</td></tr>
<tr><td>>hg38_21_+_8433172_8446622_RNA45SN1_45S_50ntFlanks@3822.3852.31|CTTTGGTCGCTCGCTCCTCTCCTACTTGGAT</td><td>>hg38_21_+_8433172_8446622_RNA45SN1_45S_50ntFlanks</td><td>3822</td><td>3852</td><td>31</td><td>CTTTGGTCGCTCGCTCCTCTCCTACTTGGAT</td><td>157.4</td><td>-2.8</td><td>1.3</td><td>-2.1</td><td>3.2E-02</td><td>4.2E-02</td><td>BT20/MB468</td><td>MITO</td></tr>
<tr><td>>hg38_21_+_8433172_8446622_RNA45SN1_45S_50ntFlanks@9924.9948.25|TGGTTGATATAGACAGCAGGACGGT</td><td>>hg38_21_+_8433172_8446622_RNA45SN1_45S_50ntFlanks</td><td>9924</td><td>9948</td><td>25</td><td>TGGTTGATATAGACAGCAGGACGGT</td><td>91.8</td><td>4.1</td><td>1.6</td><td>2.5</td><td>1.2E-02</td><td>4.2E-02</td><td>BT20/MB231</td><td>MP</td></tr>
<tr><td>>am_MI0000815|hsa-mir-339&WithFlank&7|-|1022927|1023032@21.42.22&[MIMAT0000764&hsa-miR-339-5p&offsets|0|-1;m-163&7|-|1022990|1023012&offsets|0|-1]|TCCCTGTCCTCCAGGAGCTCAC</td><td>>am_MI0000815|hsa-mir-339&WithFlank&7|-|1022927|1023032</td><td>21</td><td>42</td><td>22</td><td>TCCCTGTCCTCCAGGAGCTCAC</td><td>55.1</td><td>4.2</td><td>1.9</td><td>2.2</td><td>2.6E-02</td><td>4.2E-02</td><td>BT20/MB468</td><td>MP</td></tr>
<tr><td>>hg38_21_+_8433172_8446622_RNA45SN1_45S_50ntFlanks@12633.12655.23|CGAAGCTACCATCTGTGGGATTA</td><td>>hg38_21_+_8433172_8446622_RNA45SN1_45S_50ntFlanks</td><td>12633</td><td>12655</td><td>23</td><td>CGAAGCTACCATCTGTGGGATTA</td><td>51.0</td><td>-3.8</td><td>1.7</td><td>-2.2</td><td>2.6E-02</td><td>4.2E-02</td><td>BT20/MB468</td><td>MP</td></tr>
<tr><td>>MI0000746|hsa-mir-99b&WithFlank&19|+|51692606|51692687@13.33.21(+1C)&[MIMAT0000689&hsa-miR-99b-5p&offsets|0|-1(+1C);m-10&19|+|51692618|51692639&offsets|0|-1(+1C)]|CACCCGTAGAACCGACCTTGCC</td><td>>MI0000746|hsa-mir-99b&WithFlank&19|+|51692606|51692687</td><td>13</td><td>33</td><td>21(+1C)</td><td>CACCCGTAGAACCGACCTTGCC</td><td>945.5</td><td>2.2</td><td>0.9</td><td>2.4</td><td>1.5E-02</td><td>4.2E-02</td><td>BT20/MB231</td><td>CYTO</td></tr>
<tr><td>>hg38_21_+_8433172_8446622_RNA45SN1_45S_50ntFlanks@8836.8867.32|CGGCGGTGGCGGCGGCGGCGGCGGCGGCGGGA</td><td>>hg38_21_+_8433172_8446622_RNA45SN1_45S_50ntFlanks</td><td>8836</td><td>8867</td><td>32</td><td>CGGCGGTGGCGGCGGCGGCGGCGGCGGCGGGA</td><td>53.0</td><td>-5.0</td><td>2.2</td><td>-2.2</td><td>2.6E-02</td><td>4.2E-02</td><td>BT20/MB468</td><td>MP</td></tr>
<tr><td>>MI0000085|hsa-mir-27a&WithFlank&19|-|13836434|13836523@57.76.20(+1A)&[MIMAT0000084&hsa-miR-27a-3p&offsets|0|-1(+1A);m-43&19|-|13836447|13836467&offsets|0|-1(+1A)]|TTCACAGTGGCTAAGTTCCGA</td><td>>MI0000085|hsa-mir-27a&WithFlank&19|-|13836434|13836523</td><td>57</td><td>76</td><td>20(+1A)</td><td>TTCACAGTGGCTAAGTTCCGA</td><td>8662.6</td><td>-2.6</td><td>1.0</td><td>-2.5</td><td>1.2E-02</td><td>4.2E-02</td><td>BT20/MB231</td><td>MP</td></tr>
<tr><td>>hg38_21_+_8433172_8446622_RNA45SN1_45S_50ntFlanks@5473.5516.44|CGGCCCTGGCGGAGCGCTGAGAAGACGGTCGAACTTGACTATCT</td><td>>hg38_21_+_8433172_8446622_RNA45SN1_45S_50ntFlanks</td><td>5473</td><td>5516</td><td>44</td><td>CGGCCCTGGCGGAGCGCTGAGAAGACGGTCGAACTTGACTATCT</td><td>233.0</td><td>2.0</td><td>1.0</td><td>2.1</td><td>3.7E-02</td><td>4.2E-02</td><td>BT20/MB468</td><td>NUC</td></tr>
<tr><td>>hg38_21_+_8433172_8446622_RNA45SN1_45S_50ntFlanks@10871.10888.18|GGCTGGGTCGGTCGGGCT</td><td>>hg38_21_+_8433172_8446622_RNA45SN1_45S_50ntFlanks</td><td>10871</td><td>10888</td><td>18</td><td>GGCTGGGTCGGTCGGGCT</td><td>95.0</td><td>3.3</td><td>1.5</td><td>2.2</td><td>2.6E-02</td><td>4.2E-02</td><td>BT20/MB468</td><td>MP</td></tr>
<tr><td>>hg38_21_+_8433172_8446622_RNA45SN1_45S_50ntFlanks@9566.9597.32|GACGTGCAAATCGGTCGTCCGACCTGGGTATA</td><td>>hg38_21_+_8433172_8446622_RNA45SN1_45S_50ntFlanks</td><td>9566</td><td>9597</td><td>32</td><td>GACGTGCAAATCGGTCGTCCGACCTGGGTATA</td><td>553.8</td><td>-2.4</td><td>1.1</td><td>-2.2</td><td>2.6E-02</td><td>4.2E-02</td><td>BT20/MB468</td><td>MP</td></tr>
<tr><td>>am_trnaMT_SerGCT_MT_+_12207_12265@11.38.28|ACAAGAACTGCTAACTCATGCCCCCATG</td><td>>am_trnaMT_SerGCT_MT_+_12207_12265</td><td>11</td><td>38</td><td>28</td><td>ACAAGAACTGCTAACTCATGCCCCCATG</td><td>113.1</td><td>-2.7</td><td>1.3</td><td>-2.1</td><td>3.5E-02</td><td>4.2E-02</td><td>MB231/MB468</td><td>MP</td></tr>
<tr><td>>MI0000736|hsa-mir-30c-1&WithFlank&1|+|40757278|40757378@23.45.23;MI0000254|hsa-mir-30c-2&WithFlank&6|-|71376954|71377037@13.35.23&[MIMAT0000244&hsa-miR-30c-5p&offsets|0|0;m-57&1|+|40757300|40757323&offsets|0|-1];[MIMAT0000244_1&hsa-miR-30c-5p&offsets|0|0;m-56&6|-|71377002|71377025&offsets|0|-1]|TGTAAACATCCTACACTCTCAGC</td><td>>MI0000736|hsa-mir-30c-1&WithFlank&1|+|40757278|40757378</td><td>23</td><td>45</td><td>23</td><td>TGTAAACATCCTACACTCTCAGC</td><td>32018.2</td><td>-1.6</td><td>0.6</td><td>-2.5</td><td>1.2E-02</td><td>4.2E-02</td><td>BT20/MB231</td><td>NUC</td></tr>
<tr><td>>MI0000441|hsa-mir-30b&WithFlank&8|-|134800514|134800613@61.82.22&[MIMAT0004589&hsa-miR-30b-3p&offsets|0|0;m-362&8|-|134800532|134800553&offsets|0|0]|CTGGGAGGTGGATGTTTACTTC</td><td>>MI0000441|hsa-mir-30b&WithFlank&8|-|134800514|134800613</td><td>61</td><td>82</td><td>22</td><td>CTGGGAGGTGGATGTTTACTTC</td><td>51.3</td><td>3.7</td><td>1.7</td><td>2.1</td><td>3.5E-02</td><td>4.2E-02</td><td>MB231/MB468</td><td>MP</td></tr>
<tr><td>>hg38_21_+_8433172_8446622_RNA45SN1_45S_50ntFlanks@4498.4524.27|AAGCGTTTACTTTGAAAAAATTAGAGT</td><td>>hg38_21_+_8433172_8446622_RNA45SN1_45S_50ntFlanks</td><td>4498</td><td>4524</td><td>27</td><td>AAGCGTTTACTTTGAAAAAATTAGAGT</td><td>325.7</td><td>4.5</td><td>1.9</td><td>2.4</td><td>1.5E-02</td><td>4.2E-02</td><td>BT20/MB231</td><td>CYTO</td></tr>
<tr><td>>hg38_21_+_8433172_8446622_RNA45SN1_45S_50ntFlanks@3842.3871.30|CCTACTTGGATAACTGTGGTAATTCTAGAG</td><td>>hg38_21_+_8433172_8446622_RNA45SN1_45S_50ntFlanks</td><td>3842</td><td>3871</td><td>30</td><td>CCTACTTGGATAACTGTGGTAATTCTAGAG</td><td>81.6</td><td>-3.8</td><td>1.7</td><td>-2.2</td><td>2.6E-02</td><td>4.2E-02</td><td>BT20/MB468</td><td>MP</td></tr>
<tr><td>>am_trna10_ValCAC_5_-_180649395_180649467@1.30.30,trna12_ValAAC_5_-_180645270_180645342@1.30.30,trna132_ValAAC_6_-_27721179_27721251@1.30.30,trna136_ValAAC_6_-_27648885_27648957@1.30.30,trna139_ValAAC_6_-_27618707_27618779@1.30.30,trna18_ValCAC_5_-_180529253_180529325@1.30.30,trna2_ValAAC_3_+_169490018_169490090@1.30.30,trna2_ValCAC_5_+_180524070_180524142@1.30.30,trna4_ValAAC_5_+_180591154_180591226@1.30.30,trna5_ValAAC_5_+_180596610_180596682@1.30.30,trna6_ValCAC_5_+_180600650_180600722@1.30.30,trna85_ValCAC_1_-_161369490_161369562@1.30.30,trna90_ValCAC_1_-_149684088_149684161@1.30.30,trna98_ValCAC_1_-_149298555_149298627@1.30.30,trna9_ValCAC_6_+_26538282_26538354@1.30.30|GTTTCCGTAGTGTAGTGGTTATCACGTTCG</td><td>>am_trna10_ValCAC_5_-_180649395_180649467</td><td>1</td><td>30</td><td>30</td><td>GTTTCCGTAGTGTAGTGGTTATCACGTTCG</td><td>234.5</td><td>-3.4</td><td>1.6</td><td>-2.1</td><td>3.7E-02</td><td>4.2E-02</td><td>BT20/MB468</td><td>CYTO</td></tr>
<tr><td>>hg38_21_+_8433172_8446622_RNA45SN1_45S_50ntFlanks@11208.11254.47|GCGGCGACTCTGGACGCGAGCCGGGCCCTTCCCGTGGATCGCCCCAG</td><td>>hg38_21_+_8433172_8446622_RNA45SN1_45S_50ntFlanks</td><td>11208</td><td>11254</td><td>47</td><td>GCGGCGACTCTGGACGCGAGCCGGGCCCTTCCCGTGGATCGCCCCAG</td><td>655.7</td><td>-3.7</td><td>1.7</td><td>-2.2</td><td>2.6E-02</td><td>4.2E-02</td><td>BT20/MB468</td><td>MP</td></tr>
<tr><td>>hg38_21_+_8433172_8446622_RNA45SN1_45S_50ntFlanks@13007.13027.21|TATTGAAAGTCAGCCCTCGAC</td><td>>hg38_21_+_8433172_8446622_RNA45SN1_45S_50ntFlanks</td><td>13007</td><td>13027</td><td>21</td><td>TATTGAAAGTCAGCCCTCGAC</td><td>290.7</td><td>-2.6</td><td>1.3</td><td>-2.1</td><td>3.8E-02</td><td>4.2E-02</td><td>MB231/MB468</td><td>NUC</td></tr>
<tr><td>>hg38_21_+_8433172_8446622_RNA45SN1_45S_50ntFlanks@6646.6664.19|TCGTACGACTCTTAGCGGT</td><td>>hg38_21_+_8433172_8446622_RNA45SN1_45S_50ntFlanks</td><td>6646</td><td>6664</td><td>19</td><td>TCGTACGACTCTTAGCGGT</td><td>40538.2</td><td>1.5</td><td>0.6</td><td>2.5</td><td>1.2E-02</td><td>4.2E-02</td><td>BT20/MB231</td><td>NUC</td></tr>
<tr><td>>hg38_21_+_8433172_8446622_RNA45SN1_45S_50ntFlanks@9534.9570.37|AGGAAACTCTGGTGGAGGTCCGTAGCGGTCCTGACGT</td><td>>hg38_21_+_8433172_8446622_RNA45SN1_45S_50ntFlanks</td><td>9534</td><td>9570</td><td>37</td><td>AGGAAACTCTGGTGGAGGTCCGTAGCGGTCCTGACGT</td><td>587.1</td><td>-2.0</td><td>0.8</td><td>-2.4</td><td>1.5E-02</td><td>4.2E-02</td><td>BT20/MB231</td><td>CYTO</td></tr>
<tr><td>>hg38_MT_+_598_1651_RNR1_12S_50ntFlanks@904.922.19|AAGTATACTTCAAAGGACA</td><td>>hg38_MT_+_598_1651_RNR1_12S_50ntFlanks</td><td>904</td><td>922</td><td>19</td><td>AAGTATACTTCAAAGGACA</td><td>75.6</td><td>3.0</td><td>1.4</td><td>2.1</td><td>3.5E-02</td><td>4.2E-02</td><td>MB231/MB468</td><td>MP</td></tr>
<tr><td>>MI0000095|hsa-mir-93&WithFlank&7|-|100093762|100093853@56.78.23&[MIMAT0004509&hsa-miR-93-3p&offsets|0|+1;m-250&7|-|100093776|100093798&offsets|0|0]|ACTGCTGAGCTAGCACTTCCCGA</td><td>>MI0000095|hsa-mir-93&WithFlank&7|-|100093762|100093853</td><td>56</td><td>78</td><td>23</td><td>ACTGCTGAGCTAGCACTTCCCGA</td><td>240.2</td><td>2.1</td><td>0.9</td><td>2.4</td><td>1.5E-02</td><td>4.2E-02</td><td>BT20/MB231</td><td>CYTO</td></tr>
<tr><td>>MI0000749|hsa-mir-30e&WithFlank&1|+|40754349|40754452@65.85.21(+1U)&[MIMAT0000693&hsa-miR-30e-3p&offsets|0|-1(+1U);m-25&1|+|40754413|40754434&offsets|0|-1(+1U)]|CTTTCAGTCGGATGTTTACAGT</td><td>>MI0000749|hsa-mir-30e&WithFlank&1|+|40754349|40754452</td><td>65</td><td>85</td><td>21(+1U)</td><td>CTTTCAGTCGGATGTTTACAGT</td><td>2719.6</td><td>-1.8</td><td>0.7</td><td>-2.5</td><td>1.4E-02</td><td>4.2E-02</td><td>BT20/MB231</td><td>MITO</td></tr>
<tr><td>>MI0000767|hsa-mir-365a&WithFlank&16|+|14309279|14309377@62.81.20;MI0000769|hsa-mir-365b&WithFlank&17|+|31575405|31575527@74.93.20&[MIMAT0000710&hsa-miR-365a-3p&offsets|0|-2;m-176&16|+|14309340|14309361&offsets|0|-2];[MIMAT0022834&hsa-miR-365b-3p&offsets|0|-2;m-175&17|+|31575478|31575499&offsets|0|-2]|TAATGCCCCTAAAAATCCTT</td><td>>MI0000767|hsa-mir-365a&WithFlank&16|+|14309279|14309377</td><td>62</td><td>81</td><td>20</td><td>TAATGCCCCTAAAAATCCTT</td><td>308.0</td><td>2.0</td><td>0.8</td><td>2.5</td><td>1.4E-02</td><td>4.2E-02</td><td>BT20/MB231</td><td>MITO</td></tr>
</table>

<table>
<tr><td>>am_trna10_ValCAC_5_-_180649395_180649467@1.32.32,trna12_ValAAC_5_-_180645270_180645342@1.32.32,trna132_ValAAC_6_-_27721179_27721251@1.32.32,trna136_ValAAC_6_-_27648885_27648957@1.32.32,trna139_ValAAC_6_-_27618707_27618779@1.32.32,trna18_ValCAC_5_-_180529253_180529325@1.32.32,trna2_ValAAC_3_+_169490018_169490090@1.32.32,trna2_ValCAC_5_+_180524070_180524142@1.32.32,trna4_ValAAC_5_+_180591154_180591226@1.32.32,trna5_ValAAC_5_+_180596610_180596682@1.32.32,trna6_ValCAC_5_+_180600650_180600722@1.32.32,trna85_ValCAC_1_-_161369490_161369562@1.32.32,trna90_ValCAC_1_-_149684088_149684161@1.32.32,trna98_ValCAC_1_-_149298555_149298627@1.32.32,trna9_ValCAC_6_+_26538282_26538354@1.32.32|GTTTCCGTAGTGTAGTGGTTATCACGTTCGCC</td><td>>am_trna10_ValCAC_5_-_180649395_180649467</td><td>1</td><td>32</td><td>32</td><td>GTTTCCGTAGTGTAGTGGTTATCACGTTCGCC</td><td>21468.6</td><td>-2.6</td><td>1.1</td><td>-2.4</td><td>1.5E-02</td><td>4.2E-02</td><td>BT20/MB231</td><td>CYTO</td></tr>
<tr><td>>hg38_21_+_8433172_8446622_RNA45SN1_45S_50ntFlanks@9947.9978.32|GTGGCCATGGAAGTCGGAATCCGCTAAGGAGT</td><td>>hg38_21_+_8433172_8446622_RNA45SN1_45S_50ntFlanks</td><td>9947</td><td>9978</td><td>32</td><td>GTGGCCATGGAAGTCGGAATCCGCTAAGGAGT</td><td>104.3</td><td>5.0</td><td>2.0</td><td>2.5</td><td>1.2E-02</td><td>4.2E-02</td><td>BT20/MB231</td><td>MP</td></tr>
<tr><td>>MI0000782|hsa-mir-374a&WithFlank&X|-|74287280|74287363@18.37.20(+1C)&[MIMAT0000727&hsa-miR-374a-5p&offsets|0|-2(+1C);m-184&X|-|74287325|74287346&offsets|0|-2(+1C)]|TTATAATACAACCTGATAAGC</td><td>>MI0000782|hsa-mir-374a&WithFlank&X|-|74287280|74287363</td><td>18</td><td>37</td><td>20(+1C)</td><td>TTATAATACAACCTGATAAGC</td><td>108.1</td><td>-5.0</td><td>2.0</td><td>-2.5</td><td>1.2E-02</td><td>4.2E-02</td><td>BT20/MB231</td><td>MP</td></tr>
<tr><td>>MI0000299|hsa-mir-222&WithFlank&X|-|45747009|45747130@75.96.22(+1C)&[MIMAT0000279&hsa-miR-222-3p&offsets|0|+1(+1C);m-140&X|-|45747033|45747056&offsets|0|-2(+1C)]|AGCTACATCTGGCTACTGGGTCC</td><td>>MI0000299|hsa-mir-222&WithFlank&X|-|45747009|45747130</td><td>75</td><td>96</td><td>22(+1C)</td><td>AGCTACATCTGGCTACTGGGTCC</td><td>142.0</td><td>1.3</td><td>0.6</td><td>2.1</td><td>3.8E-02</td><td>4.2E-02</td><td>BT20/MB468</td><td>NUC</td></tr>
<tr><td>>hg38_MT_+_1621_3279_RNR2_16S_50ntFlanks@1368.1385.18|CGATGTTGGATCAGGACA</td><td>>hg38_MT_+_1621_3279_RNR2_16S_50ntFlanks</td><td>1368</td><td>1385</td><td>18</td><td>CGATGTTGGATCAGGACA</td><td>138.3</td><td>1.8</td><td>0.8</td><td>2.1</td><td>3.5E-02</td><td>4.2E-02</td><td>MB231/MB468</td><td>MP</td></tr>
<tr><td>>hg38_21_+_8433172_8446622_RNA45SN1_45S_50ntFlanks@10353.10376.24|AGAACTTTGAAGGCCGAAGTGGAG</td><td>>hg38_21_+_8433172_8446622_RNA45SN1_45S_50ntFlanks</td><td>10353</td><td>10376</td><td>24</td><td>AGAACTTTGAAGGCCGAAGTGGAG</td><td>67.0</td><td>-4.0</td><td>1.8</td><td>-2.2</td><td>2.6E-02</td><td>4.2E-02</td><td>BT20/MB468</td><td>MP</td></tr>
<tr><td>>hg38_21_+_8433172_8446622_RNA45SN1_45S_50ntFlanks@12949.12990.42|ACGACCTGCTTCTGGGTCGGGGTTTCGTACGTAGCAGAGCAG</td><td>>hg38_21_+_8433172_8446622_RNA45SN1_45S_50ntFlanks</td><td>12949</td><td>12990</td><td>42</td><td>ACGACCTGCTTCTGGGTCGGGGTTTCGTACGTAGCAGAGCAG</td><td>133.6</td><td>-3.4</td><td>1.6</td><td>-2.1</td><td>3.3E-02</td><td>4.2E-02</td><td>BT20/MB468</td><td>MITO</td></tr>
<tr><td>>hg38_21_+_8433172_8446622_RNA45SN1_45S_50ntFlanks@4669.4691.23|TTCTTGGACCGGCGCAAGACGGA</td><td>>hg38_21_+_8433172_8446622_RNA45SN1_45S_50ntFlanks</td><td>4669</td><td>4691</td><td>23</td><td>TTCTTGGACCGGCGCAAGACGGA</td><td>68.8</td><td>2.7</td><td>1.3</td><td>2.1</td><td>3.5E-02</td><td>4.2E-02</td><td>MB231/MB468</td><td>MP</td></tr>
<tr><td>>hg38_21_+_8433172_8446622_RNA45SN1_45S_50ntFlanks@7977.8009.33|CGACCTCAGATCAGACGTGGCGACCCGCTGAAT</td><td>>hg38_21_+_8433172_8446622_RNA45SN1_45S_50ntFlanks</td><td>7977</td><td>8009</td><td>33</td><td>CGACCTCAGATCAGACGTGGCGACCCGCTGAAT</td><td>1867.5</td><td>3.7</td><td>1.5</td><td>2.5</td><td>1.2E-02</td><td>4.2E-02</td><td>BT20/MB231</td><td>MP</td></tr>
<tr><td>>MI0000088|hsa-mir-30a&WithFlank&6|-|71403545|71403627@12.30.19&[MIMAT0000087&hsa-miR-30a-5p&offsets|0|-3;m-15&6|-|71403595|71403616&offsets|0|-3]|TGTAAACATCCTCGACTGG</td><td>>MI0000088|hsa-mir-30a&WithFlank&6|-|71403545|71403627</td><td>12</td><td>30</td><td>19</td><td>TGTAAACATCCTCGACTGG</td><td>166.2</td><td>3.4</td><td>1.5</td><td>2.2</td><td>2.6E-02</td><td>4.2E-02</td><td>BT20/MB468</td><td>MP</td></tr>
<tr><td>>MI0003820|hsa-mir-454&WithFlank&17|-|59137752|59137878@70.92.23&[MIMAT0003885&hsa-miR-454-3p&offsets|0|0;m-272&17|-|59137787|59137809&offsets|0|0]|TAGTGCAATATTGCTTATAGGGT</td><td>>MI0003820|hsa-mir-454&WithFlank&17|-|59137752|59137878</td><td>70</td><td>92</td><td>23</td><td>TAGTGCAATATTGCTTATAGGGT</td><td>804.2</td><td>-3.8</td><td>1.5</td><td>-2.5</td><td>1.2E-02</td><td>4.2E-02</td><td>BT20/MB231</td><td>MP</td></tr>
<tr><td>>hg38_21_+_8433172_8446622_RNA45SN1_45S_50ntFlanks@8240.8261.22|CGGGTCTTCCCGGAGTCGGGTT</td><td>>hg38_21_+_8433172_8446622_RNA45SN1_45S_50ntFlanks</td><td>8240</td><td>8261</td><td>22</td><td>CGGGTCTTCCCGGAGTCGGGTT</td><td>89.4</td><td>4.9</td><td>1.9</td><td>2.5</td><td>1.2E-02</td><td>4.2E-02</td><td>BT20/MB231</td><td>MP</td></tr>
<tr><td>>hg38_21_+_8433172_8446622_RNA45SN1_45S_50ntFlanks@6646.6684.39|TCGTACGACTCTTAGCGGTGGATCACTCGGCTCGTGCGT</td><td>>hg38_21_+_8433172_8446622_RNA45SN1_45S_50ntFlanks</td><td>6646</td><td>6684</td><td>39</td><td>TCGTACGACTCTTAGCGGTGGATCACTCGGCTCGTGCGT</td><td>246.1</td><td>1.9</td><td>0.7</td><td>2.5</td><td>1.2E-02</td><td>4.2E-02</td><td>BT20/MB231</td><td>NUC</td></tr>
<tr><td>>hg38_21_+_8433172_8446622_RNA45SN1_45S_50ntFlanks@10409.10436.28|GTCAGTCGGTCCTGAGAGATGGGCGAGC</td><td>>hg38_21_+_8433172_8446622_RNA45SN1_45S_50ntFlanks</td><td>10409</td><td>10436</td><td>28</td><td>GTCAGTCGGTCCTGAGAGATGGGCGAGC</td><td>71.1</td><td>3.3</td><td>1.5</td><td>2.2</td><td>2.6E-02</td><td>4.2E-02</td><td>BT20/MB468</td><td>MP</td></tr>
<tr><td>>hg38_21_+_8433172_8446622_RNA45SN1_45S_50ntFlanks@11584.11611.28|TGGTGCGGACCAGGGGAATCCGACTGTT</td><td>>hg38_21_+_8433172_8446622_RNA45SN1_45S_50ntFlanks</td><td>11584</td><td>11611</td><td>28</td><td>TGGTGCGGACCAGGGGAATCCGACTGTT</td><td>253.7</td><td>2.1</td><td>0.9</td><td>2.2</td><td>2.6E-02</td><td>4.2E-02</td><td>BT20/MB468</td><td>MP</td></tr>
<tr><td>>hg38_21_+_8433172_8446622_RNA45SN1_45S_50ntFlanks@5491.5516.26|GAGAAGACGGTCGAACTTGACTATCT</td><td>>hg38_21_+_8433172_8446622_RNA45SN1_45S_50ntFlanks</td><td>5491</td><td>5516</td><td>26</td><td>GAGAAGACGGTCGAACTTGACTATCT</td><td>5346.3</td><td>3.0</td><td>1.2</td><td>2.5</td><td>1.2E-02</td><td>4.2E-02</td><td>BT20/MB231</td><td>MP</td></tr>
<tr><td>>hg38_21_+_8433172_8446622_RNA45SN1_45S_50ntFlanks@6702.6719.18|AGCTGCGAGAATTAATGT</td><td>>hg38_21_+_8433172_8446622_RNA45SN1_45S_50ntFlanks</td><td>6702</td><td>6719</td><td>18</td><td>AGCTGCGAGAATTAATGT</td><td>79.2</td><td>-3.6</td><td>1.7</td><td>-2.1</td><td>3.5E-02</td><td>4.2E-02</td><td>MB231/MB468</td><td>MP</td></tr>
<tr><td>>hg38_21_+_8433172_8446622_RNA45SN1_45S_50ntFlanks@7988.8010.23|CAGACGTGGCGACCCGCTGAATT</td><td>>hg38_21_+_8433172_8446622_RNA45SN1_45S_50ntFlanks</td><td>7988</td><td>8010</td><td>23</td><td>CAGACGTGGCGACCCGCTGAATT</td><td>858.0</td><td>-2.1</td><td>1.0</td><td>-2.2</td><td>2.6E-02</td><td>4.2E-02</td><td>BT20/MB468</td><td>MP</td></tr>
<tr><td>>hg38_MT_+_598_1651_RNR1_12S_50ntFlanks@994.1012.19|CTTGGACGAACCAGAGTGT</td><td>>hg38_MT_+_598_1651_RNR1_12S_50ntFlanks</td><td>994</td><td>1012</td><td>19</td><td>CTTGGACGAACCAGAGTGT</td><td>38.5</td><td>3.6</td><td>1.7</td><td>2.1</td><td>3.5E-02</td><td>4.2E-02</td><td>MB231/MB468</td><td>MP</td></tr>
<tr><td>>hg38_MT_+_598_1651_RNR1_12S_50ntFlanks@983.1001.19|TGGAAAGTGCACTTGGACG</td><td>>hg38_MT_+_598_1651_RNR1_12S_50ntFlanks</td><td>983</td><td>1001</td><td>19</td><td>TGGAAAGTGCACTTGGACG</td><td>94.8</td><td>2.4</td><td>1.1</td><td>2.2</td><td>2.6E-02</td><td>4.2E-02</td><td>BT20/MB468</td><td>MP</td></tr>
<tr><td>>hg38_21_+_8433172_8446622_RNA45SN1_45S_50ntFlanks@12423.12462.40|TCGGCTCTTCCTATCATTGTGAAGCAGAATTCACCAAGCG</td><td>>hg38_21_+_8433172_8446622_RNA45SN1_45S_50ntFlanks</td><td>12423</td><td>12462</td><td>40</td><td>TCGGCTCTTCCTATCATTGTGAAGCAGAATTCACCAAGCG</td><td>148.8</td><td>-5.3</td><td>2.2</td><td>-2.4</td><td>1.5E-02</td><td>4.2E-02</td><td>BT20/MB231</td><td>CYTO</td></tr>
<tr><td>>hg38_21_+_8433172_8446622_RNA45SN1_45S_50ntFlanks@10677.10710.34|GCGGTTCCGGCGGCGTCCGGTGAGCTCTCGCTGG</td><td>>hg38_21_+_8433172_8446622_RNA45SN1_45S_50ntFlanks</td><td>10677</td><td>10710</td><td>34</td><td>GCGGTTCCGGCGGCGTCCGGTGAGCTCTCGCTGG</td><td>264.4</td><td>3.3</td><td>1.3</td><td>2.5</td><td>1.2E-02</td><td>4.2E-02</td><td>BT20/MB231</td><td>MP</td></tr>
<tr><td>>hg38_21_+_8433172_8446622_RNA45SN1_45S_50ntFlanks@11847.11887.41|AACGGGCTTGGCGGAATCAGCGGGGAAAGAAGACCCTGTTG</td><td>>hg38_21_+_8433172_8446622_RNA45SN1_45S_50ntFlanks</td><td>11847</td><td>11887</td><td>41</td><td>AACGGGCTTGGCGGAATCAGCGGGGAAAGAAGACCCTGTTG</td><td>84.9</td><td>3.0</td><td>1.4</td><td>2.2</td><td>2.6E-02</td><td>4.2E-02</td><td>BT20/MB468</td><td>MP</td></tr>
<tr><td>>hg38_21_+_8433172_8446622_RNA45SN1_45S_50ntFlanks@5473.5514.42|CGGCCCTGGCGGAGCGCTGAGAAGACGGTCGAACTTGACTAT</td><td>>hg38_21_+_8433172_8446622_RNA45SN1_45S_50ntFlanks</td><td>5473</td><td>5514</td><td>42</td><td>CGGCCCTGGCGGAGCGCTGAGAAGACGGTCGAACTTGACTAT</td><td>579.5</td><td>-2.9</td><td>1.3</td><td>-2.2</td><td>2.6E-02</td><td>4.2E-02</td><td>BT20/MB468</td><td>MP</td></tr>
<tr><td>>hg38_21_+_8433172_8446622_RNA45SN1_45S_50ntFlanks@4661.4689.29|AGGTGAAATTCTTGGACCGGCGCAAGACG</td><td>>hg38_21_+_8433172_8446622_RNA45SN1_45S_50ntFlanks</td><td>4661</td><td>4689</td><td>29</td><td>AGGTGAAATTCTTGGACCGGCGCAAGACG</td><td>224.9</td><td>-1.9</td><td>0.9</td><td>-2.1</td><td>3.7E-02</td><td>4.2E-02</td><td>BT20/MB468</td><td>CYTO</td></tr>
<tr><td>>hg38_21_+_8433172_8446622_RNA45SN1_45S_50ntFlanks@6658.6689.32|TAGCGGTGGATCACTCGGCTCGTGCGTCGATG</td><td>>hg38_21_+_8433172_8446622_RNA45SN1_45S_50ntFlanks</td><td>6658</td><td>6689</td><td>32</td><td>TAGCGGTGGATCACTCGGCTCGTGCGTCGATG</td><td>215.7</td><td>-1.3</td><td>0.6</td><td>-2.1</td><td>3.8E-02</td><td>4.2E-02</td><td>MB231/MB468</td><td>NUC</td></tr>
<tr><td>>am_MI0000786|hsa-mir-378a&WithFlank&5|+|149732819|149732896@49.70.22(+1A)&[MIMAT0000732&hsa-miR-378a-3p&offsets|0|0(+1A);m-44&5|+|149732867|149732888&offsets|0|0(+1A)]|ACTGGACTTGGAGTCAGAAGGCA</td><td>>am_MI0000786|hsa-mir-378a&WithFlank&5|+|149732819|149732896</td><td>49</td><td>70</td><td>22(+1A)</td><td>ACTGGACTTGGAGTCAGAAGGCA</td><td>6858.2</td><td>2.4</td><td>1.1</td><td>2.2</td><td>2.6E-02</td><td>4.2E-02</td><td>BT20/MB468</td><td>MP</td></tr>
<tr><td>>trna119_LysCTT_1_-_145395522_145395594@1.29.29,trna11_LysCTT_5_-_180648979_180649051@1.29.29,trna13_LysCTT_6_+_26556774_26556846@1.29.29,trna32_LysCTT_16_-_3207406_3207478@1.29.29,trna7_LysCTT_16_+_3225692_3225764@1.29.29,trna9_LysCTT_5_+_180634755_180634827@1.29.29|GCCCGGCTAGCTCAGTCGGTAGAGCATGA</td><td>>trna119_LysCTT_1_-_145395522_145395594</td><td>1</td><td>29</td><td>29</td><td>GCCCGGCTAGCTCAGTCGGTAGAGCATGA</td><td>479.1</td><td>-2.9</td><td>1.4</td><td>-2.1</td><td>3.3E-02</td><td>4.2E-02</td><td>BT20/MB468</td><td>MITO</td></tr>
<tr><td>>hg38_21_+_8433172_8446622_RNA45SN1_45S_50ntFlanks@4772.4793.22|GTAGTTCCGACCATAAACGATG</td><td>>hg38_21_+_8433172_8446622_RNA45SN1_45S_50ntFlanks</td><td>4772</td><td>4793</td><td>22</td><td>GTAGTTCCGACCATAAACGATG</td><td>146.9</td><td>-1.2</td><td>0.5</td><td>-2.1</td><td>3.3E-02</td><td>4.2E-02</td><td>BT20/MB468</td><td>MITO</td></tr>
<tr><td>>hg38_21_+_8433172_8446622_RNA45SN1_45S_50ntFlanks@5492.5525.34|AGAAGACGGTCGAACTTGACTATCTAGAGGAAGT</td><td>>hg38_21_+_8433172_8446622_RNA45SN1_45S_50ntFlanks</td><td>5492</td><td>5525</td><td>34</td><td>AGAAGACGGTCGAACTTGACTATCTAGAGGAAGT</td><td>1019.8</td><td>2.4</td><td>1.0</td><td>2.4</td><td>1.6E-02</td><td>4.2E-02</td><td>BT20/MB231</td><td>CYTO</td></tr>
<tr><td>>hg38_21_+_8433172_8446622_RNA45SN1_45S_50ntFlanks@12642.12669.28|CATCTGTGGGATTATGACTGAACGCCTC</td><td>>hg38_21_+_8433172_8446622_RNA45SN1_45S_50ntFlanks</td><td>12642</td><td>12669</td><td>28</td><td>CATCTGTGGGATTATGACTGAACGCCTC</td><td>314.6</td><td>-2.3</td><td>1.0</td><td>-2.4</td><td>1.6E-02</td><td>4.2E-02</td><td>BT20/MB231</td><td>CYTO</td></tr>
<tr><td>>hg38_21_+_8433172_8446622_RNA45SN1_45S_50ntFlanks@12996.13037.42|TCGCTGCGATCTATTGAAAGTCAGCCCTCGACACAAGGGTTT</td><td>>hg38_21_+_8433172_8446622_RNA45SN1_45S_50ntFlanks</td><td>12996</td><td>13037</td><td>42</td><td>TCGCTGCGATCTATTGAAAGTCAGCCCTCGACACAAGGGTTT</td><td>1248.5</td><td>-2.7</td><td>1.1</td><td>-2.4</td><td>1.6E-02</td><td>4.2E-02</td><td>BT20/MB231</td><td>CYTO</td></tr>
<tr><td>>am_trna2_LeuTAA_X_-_55207755_55207829@1.18.18,trna4_LeuTAA_20_-_55933428_55933502@1.18.18,trna60_LeuTAA_1_-_238106956_238107030@1.18.18,trna9_LeuTAA_2_+_117781417_117781491@1.18.18,trnaMT_LeuTAA_MT_+_3230_3304@1.18.18|GTTAAGATGGCAGAGCCC</td><td>>am_trna2_LeuTAA_X_-_55207755_55207829</td><td>1</td><td>18</td><td>18</td><td>GTTAAGATGGCAGAGCCC</td><td>249.2</td><td>3.4</td><td>1.4</td><td>2.5</td><td>1.2E-02</td><td>4.2E-02</td><td>BT20/MB231</td><td>NUC</td></tr>
<tr><td>>MI0000088|hsa-mir-30a&WithFlank&6|-|71403545|71403627@53.71.19&[MIMAT0000088&hsa-miR-30a-3p&offsets|0|-3;m-38&6|-|71403554|71403575&offsets|0|-3]|CTTTCAGTCGGATGTTTGC</td><td>>MI0000088|hsa-mir-30a&WithFlank&6|-|71403545|71403627</td><td>53</td><td>71</td><td>19</td><td>CTTTCAGTCGGATGTTTGC</td><td>434.9</td><td>2.0</td><td>0.9</td><td>2.2</td><td>2.6E-02</td><td>4.2E-02</td><td>BT20/MB468</td><td>MP</td></tr>
<tr><td>>hg38_21_+_8433172_8446622_RNA45SN1_45S_50ntFlanks@3846.3868.23|CTTGGATAACTGTGGTAATTCTA</td><td>>hg38_21_+_8433172_8446622_RNA45SN1_45S_50ntFlanks</td><td>3846</td><td>3868</td><td>23</td><td>CTTGGATAACTGTGGTAATTCTA</td><td>677.2</td><td>-2.4</td><td>1.1</td><td>-2.2</td><td>2.6E-02</td><td>4.2E-02</td><td>BT20/MB468</td><td>MP</td></tr>
<tr><td>>hg38_21_+_8433172_8446622_RNA45SN1_45S_50ntFlanks@12007.12029.23|GCCGCCGGTGAAATACCACTACT</td><td>>hg38_21_+_8433172_8446622_RNA45SN1_45S_50ntFlanks</td><td>12007</td><td>12029</td><td>23</td><td>GCCGCCGGTGAAATACCACTACT</td><td>552.9</td><td>-3.4</td><td>1.5</td><td>-2.2</td><td>2.6E-02</td><td>4.2E-02</td><td>BT20/MB468</td><td>MP</td></tr>
<tr><td>>MI0000483|hsa-mir-186&WithFlank&1|-|71067625|71067722@21.43.23&[MIMAT0000456&hsa-miR-186-5p&offsets|0|+1;m-84&1|-|71067681|71067702&offsets|0|+1]|CAAAGAATTCTCCTTTTGGGCTT</td><td>>MI0000483|hsa-mir-186&WithFlank&1|-|71067625|71067722</td><td>21</td><td>43</td><td>23</td><td>CAAAGAATTCTCCTTTTGGGCTT</td><td>1379.6</td><td>1.9</td><td>0.8</td><td>2.5</td><td>1.2E-02</td><td>4.2E-02</td><td>BT20/MB231</td><td>NUC</td></tr>
<tr><td>>hg38_1_-_228634819_228635039_RNA5S12_5S_50ntFlanks@133.171.39|ATGGGAGACCGCCTGGGAATACCGGGTGCTGTAGGCTTT</td><td>>hg38_1_-_228634819_228635039_RNA5S12_5S_50ntFlanks</td><td>133</td><td>171</td><td>39</td><td>ATGGGAGACCGCCTGGGAATACCGGGTGCTGTAGGCTTT</td><td>12075.6</td><td>3.6</td><td>1.4</td><td>2.5</td><td>1.2E-02</td><td>4.2E-02</td><td>BT20/MB231</td><td>MP</td></tr>
<tr><td>>hg38_21_+_8433172_8446622_RNA45SN1_45S_50ntFlanks@6661.6693.33|CGGTGGATCACTCGGCTCGTGCGTCGATGAAGA</td><td>>hg38_21_+_8433172_8446622_RNA45SN1_45S_50ntFlanks</td><td>6661</td><td>6693</td><td>33</td><td>CGGTGGATCACTCGGCTCGTGCGTCGATGAAGA</td><td>189.8</td><td>-1.6</td><td>0.7</td><td>-2.1</td><td>3.5E-02</td><td>4.2E-02</td><td>MB231/MB468</td><td>MP</td></tr>
<tr><td>>hg38_MT_+_1621_3279_RNR2_16S_50ntFlanks@1390.1423.34|GATGGTGCAGCCGCTATTAAAGGTTCGTTTGTTC</td><td>>hg38_MT_+_1621_3279_RNR2_16S_50ntFlanks</td><td>1390</td><td>1423</td><td>34</td><td>GATGGTGCAGCCGCTATTAAAGGTTCGTTTGTTC</td><td>35.5</td><td>4.3</td><td>2.0</td><td>2.1</td><td>3.5E-02</td><td>4.2E-02</td><td>MB231/MB468</td><td>MP</td></tr>
<tr><td>>trna116_GluCTC_1_-_145399233_145399304@3.36.34,trna59_GluCTC_1_+_249168447_249168518@3.36.34,trna71_GluCTC_1_-_161439189_161439260@3.36.34,trna74_GluCTC_1_-_161431809_161431880@3.36.34,trna77_GluCTC_1_-_161424398_161424469@3.36.34,trna77_GluCTC_6_+_28949976_28950047@3.36.34,trna80_GluCTC_1_-_161417018_161417089@3.36.34,trna87_GluCTC_6_-_126101393_126101464@3.36.34|CCTGGTGGTCTAGTGGTTAGGATTCGGCGCTCTC</td><td>>trna116_GluCTC_1_-_145399233_145399304</td><td>3</td><td>36</td><td>34</td><td>CCTGGTGGTCTAGTGGTTAGGATTCGGCGCTCTC</td><td>1052.5</td><td>-1.7</td><td>0.7</td><td>-2.5</td><td>1.4E-02</td><td>4.2E-02</td><td>BT20/MB231</td><td>MITO</td></tr>
</table>

<table>
<tr><td>>hg38_21_+_8433172_8446622_RNA45SN1_45S_50ntFlanks@12945.12982.38|GTAGACGACCTGCTTCTGGGTCGGGGTTTCGTACGTAG</td><td>>hg38_21_+_8433172_8446622_RNA45SN1_45S_50ntFlanks</td><td>12945</td><td>12982</td><td>38</td><td>GTAGACGACCTGCTTCTGGGTCGGGGTTTCGTACGTAG</td><td>165.3</td><td>-1.5</td><td>0.7</td><td>-2.1</td><td>3.7E-02</td><td>4.2E-02</td><td>MB231/MB468</td><td>CYTO</td></tr>
<tr><td>>hg38_MT_+_1621_3279_RNR2_16S_50ntFlanks@1432.1452.21|AAGTCCTACGTGATCTGAGTT</td><td>>hg38_MT_+_1621_3279_RNR2_16S_50ntFlanks</td><td>1432</td><td>1452</td><td>21</td><td>AAGTCCTACGTGATCTGAGTT</td><td>98.7</td><td>2.8</td><td>1.3</td><td>2.1</td><td>3.5E-02</td><td>4.2E-02</td><td>MB231/MB468</td><td>MP</td></tr>
<tr><td>>hg38_21_+_8433172_8446622_RNA45SN1_45S_50ntFlanks@11999.12032.34|CCCTGCGGGCCGCCGGTGAAATACCACTACTCTG</td><td>>hg38_21_+_8433172_8446622_RNA45SN1_45S_50ntFlanks</td><td>11999</td><td>12032</td><td>34</td><td>CCCTGCGGGCCGCCGGTGAAATACCACTACTCTG</td><td>11436.1</td><td>-4.4</td><td>2.1</td><td>-2.1</td><td>3.7E-02</td><td>4.2E-02</td><td>BT20/MB468</td><td>CYTO</td></tr>
<tr><td>>hg38_21_+_8433172_8446622_RNA45SN1_45S_50ntFlanks@6731.6753.23|CACATTGATCATCGACACTTCGA</td><td>>hg38_21_+_8433172_8446622_RNA45SN1_45S_50ntFlanks</td><td>6731</td><td>6753</td><td>23</td><td>CACATTGATCATCGACACTTCGA</td><td>328.6</td><td>-2.5</td><td>1.2</td><td>-2.1</td><td>3.7E-02</td><td>4.2E-02</td><td>BT20/MB468</td><td>CYTO</td></tr>
<tr><td>>hg38_MT_+_1621_3279_RNR2_16S_50ntFlanks@1424.1451.28|AACGATTAAAGTCCTACGTGATCTGAGT</td><td>>hg38_MT_+_1621_3279_RNR2_16S_50ntFlanks</td><td>1424</td><td>1451</td><td>28</td><td>AACGATTAAAGTCCTACGTGATCTGAGT</td><td>66.2</td><td>3.5</td><td>1.7</td><td>2.1</td><td>3.5E-02</td><td>4.2E-02</td><td>MB231/MB468</td><td>MP</td></tr>
<tr><td>>MI0000077|hsa-mir-21&WithFlank&17|+|59841260|59841343@14.36.23(+1A)&[MIMAT0000076&hsa-miR-21-5p&offsets|0|+1(+1A);m-1&17|+|59841273|59841294&offsets|0|+1(+1A)]|TAGCTTATCAGACTGATGTTGACA</td><td>>MI0000077|hsa-mir-21&WithFlank&17|+|59841260|59841343</td><td>14</td><td>36</td><td>23(+1A)</td><td>TAGCTTATCAGACTGATGTTGACA</td><td>189895.3</td><td>2.2</td><td>0.9</td><td>2.4</td><td>1.6E-02</td><td>4.2E-02</td><td>BT20/MB231</td><td>CYTO</td></tr>
<tr><td>>am_MI0000809|hsa-mir-151a&WithFlank&8|-|140732558|140732659@53.73.21(+1C)&[MIMAT0000757&hsa-miR-151a-3p&offsets|0|0(+1C);m-30&8|-|140732587|140732607&offsets|0|0(+1C)]|CTAGACTGAAGCTCCTTGAGGC</td><td>>am_MI0000809|hsa-mir-151a&WithFlank&8|-|140732558|140732659</td><td>53</td><td>73</td><td>21(+1C)</td><td>CTAGACTGAAGCTCCTTGAGGC</td><td>2353.4</td><td>1.8</td><td>0.7</td><td>2.4</td><td>1.6E-02</td><td>4.2E-02</td><td>BT20/MB231</td><td>CYTO</td></tr>
<tr><td>>hg38_21_+_8433172_8446622_RNA45SN1_45S_50ntFlanks@10383.10429.47|TCCATGTGAACAGCAGTTGAACATGGGTCAGTCGGTCCTGAGAGATG</td><td>>hg38_21_+_8433172_8446622_RNA45SN1_45S_50ntFlanks</td><td>10383</td><td>10429</td><td>47</td><td>TCCATGTGAACAGCAGTTGAACATGGGTCAGTCGGTCCTGAGAGATG</td><td>400.5</td><td>-2.6</td><td>1.1</td><td>-2.4</td><td>1.6E-02</td><td>4.2E-02</td><td>BT20/MB231</td><td>CYTO</td></tr>
<tr><td>>hg38_21_+_8433172_8446622_RNA45SN1_45S_50ntFlanks@12414.12446.33|CCTTCGATGTCGGCTCTTCCTATCATTGTGAAG</td><td>>hg38_21_+_8433172_8446622_RNA45SN1_45S_50ntFlanks</td><td>12414</td><td>12446</td><td>33</td><td>CCTTCGATGTCGGCTCTTCCTATCATTGTGAAG</td><td>391.9</td><td>-1.9</td><td>0.8</td><td>-2.4</td><td>1.6E-02</td><td>4.2E-02</td><td>BT20/MB231</td><td>CYTO</td></tr>
<tr><td>>hg38_21_+_8433172_8446622_RNA45SN1_45S_50ntFlanks@12420.12454.35|ATGTCGGCTCTTCCTATCATTGTGAAGCAGAATTC</td><td>>hg38_21_+_8433172_8446622_RNA45SN1_45S_50ntFlanks</td><td>12420</td><td>12454</td><td>35</td><td>ATGTCGGCTCTTCCTATCATTGTGAAGCAGAATTC</td><td>200.3</td><td>-2.0</td><td>0.8</td><td>-2.4</td><td>1.6E-02</td><td>4.2E-02</td><td>BT20/MB231</td><td>CYTO</td></tr>
<tr><td>>MI0000087|hsa-mir-29a&WithFlank&7|-|130876741|130876816@48.69.22(+1A)&[MIMAT0000086&hsa-miR-29a-3p&offsets|0|0(+1A);m-26&7|-|130876749|130876769&offsets|0|+1(+1A)]|TAGCACCATCTGAAATCGGTTAA</td><td>>MI0000087|hsa-mir-29a&WithFlank&7|-|130876741|130876816</td><td>48</td><td>69</td><td>22(+1A)</td><td>TAGCACCATCTGAAATCGGTTAA</td><td>383.2</td><td>-2.3</td><td>0.9</td><td>-2.5</td><td>1.4E-02</td><td>4.2E-02</td><td>BT20/MB231</td><td>MITO</td></tr>
<tr><td>>trnaMT_MetCAT_MT_+_4402_4469@43.70.28|ATGTTGGTTATACCCTTCCCGTACTACC</td><td>>trnaMT_MetCAT_MT_+_4402_4469</td><td>43</td><td>70</td><td>28</td><td>ATGTTGGTTATACCCTTCCCGTACTACC</td><td>59.2</td><td>3.5</td><td>1.6</td><td>2.1</td><td>3.5E-02</td><td>4.2E-02</td><td>MB231/MB468</td><td>MP</td></tr>
<tr><td>>hg38_21_+_8433172_8446622_RNA45SN1_45S_50ntFlanks@7975.8013.39|CGCGACCTCAGATCAGACGTGGCGACCCGCTGAATTTAA</td><td>>hg38_21_+_8433172_8446622_RNA45SN1_45S_50ntFlanks</td><td>7975</td><td>8013</td><td>39</td><td>CGCGACCTCAGATCAGACGTGGCGACCCGCTGAATTTAA</td><td>13974.2</td><td>-2.3</td><td>0.9</td><td>-2.4</td><td>1.6E-02</td><td>4.2E-02</td><td>BT20/MB231</td><td>CYTO</td></tr>
<tr><td>>trna143_LysTTT_6_-_27559593_27559665@11.28.18,trna149_LysTTT_6_-_27302769_27302841@11.28.18,trna14_LysTTT_11_-_59327808_59327880@11.28.18,trna2_LysTTT_17_+_8022473_8022545@11.28.18,trna54_LysTTT_1_+_204475655_204475727@11.28.18,trna5_LysTTT_11_+_59323902_59323974@11.28.18,trna62_LysTTT_1_-_204476158_204476230@11.28.18,trna76_LysTTT_6_+_28918806_28918878@11.28.18|CTCAGTCGGTAGAGCATC</td><td>>trna143_LysTTT_6_-_27559593_27559665</td><td>11</td><td>28</td><td>18</td><td>CTCAGTCGGTAGAGCATC</td><td>56.4</td><td>-3.9</td><td>1.9</td><td>-2.1</td><td>3.5E-02</td><td>4.2E-02</td><td>MB231/MB468</td><td>MP</td></tr>
<tr><td>>hg38_21_+_8433172_8446622_RNA45SN1_45S_50ntFlanks@3846.3868.23|CTTGGATAACTGTGGTAATTCTA</td><td>>hg38_21_+_8433172_8446622_RNA45SN1_45S_50ntFlanks</td><td>3846</td><td>3868</td><td>23</td><td>CTTGGATAACTGTGGTAATTCTA</td><td>210.9</td><td>4.5</td><td>1.8</td><td>2.5</td><td>1.2E-02</td><td>4.2E-02</td><td>BT20/MB231</td><td>MP</td></tr>
<tr><td>>hg38_21_+_8433172_8446622_RNA45SN1_45S_50ntFlanks@10792.10815.24|CTCTGGCATGTTGGAACAATGTAG</td><td>>hg38_21_+_8433172_8446622_RNA45SN1_45S_50ntFlanks</td><td>10792</td><td>10815</td><td>24</td><td>CTCTGGCATGTTGGAACAATGTAG</td><td>197.8</td><td>-1.8</td><td>0.9</td><td>-2.1</td><td>3.7E-02</td><td>4.2E-02</td><td>BT20/MB468</td><td>CYTO</td></tr>
<tr><td>>hg38_21_+_8433172_8446622_RNA45SN1_45S_50ntFlanks@12473.12496.24|CACCCACTAATAGGGAACGTGAGC</td><td>>hg38_21_+_8433172_8446622_RNA45SN1_45S_50ntFlanks</td><td>12473</td><td>12496</td><td>24</td><td>CACCCACTAATAGGGAACGTGAGC</td><td>537.3</td><td>-2.1</td><td>0.9</td><td>-2.4</td><td>1.6E-02</td><td>4.2E-02</td><td>BT20/MB231</td><td>CYTO</td></tr>
<tr><td>>trna116_GluCTC_1_-_145399233_145399304@39.74.36,trna134_GluTTC_1_-_16861774_16861845@39.74.36,trna71_GluCTC_1_-_161439189_161439260@39.74.36,trna74_GluCTC_1_-_161431809_161431880@39.74.36,trna77_GluCTC_1_-_161424398_161424469@39.74.36,trna77_GluCTC_6_+_28949976_28950047@39.74.36,trna80_GluCTC_1_-_161417018_161417089@39.74.36,trna84_GluTTC_1_-_161391883_161391954@39.74.36,trna87_GluCTC_6_-_126101393_126101464@39.74.36|CGCCGCGGCCCGGGTTCGATTCCCGGTCAGGGAACC</td><td>>trna116_GluCTC_1_-_145399233_145399304</td><td>39</td><td>74</td><td>36</td><td>CGCCGCGGCCCGGGTTCGATTCCCGGTCAGGGAACC</td><td>1071.9</td><td>-2.5</td><td>1.0</td><td>-2.4</td><td>1.6E-02</td><td>4.2E-02</td><td>BT20/MB231</td><td>CYTO</td></tr>
<tr><td>>hg38_21_+_8433172_8446622_RNA45SN1_45S_50ntFlanks@11571.11588.18|GCCGACTTAGAACTGGTG</td><td>>hg38_21_+_8433172_8446622_RNA45SN1_45S_50ntFlanks</td><td>11571</td><td>11588</td><td>18</td><td>GCCGACTTAGAACTGGTG</td><td>239.8</td><td>-2.7</td><td>1.3</td><td>-2.1</td><td>3.5E-02</td><td>4.2E-02</td><td>MB231/MB468</td><td>MP</td></tr>
<tr><td>>MI0000448|hsa-mir-130a&WithFlank&11|+|57641192|57641292@61.79.19(+1U)&[MIMAT0000425&hsa-miR-130a-3p&offsets|0|-3(+1U);m-139&11|+|57641252|57641273&offsets|0|-3(+1U)]|CAGTGCAATGTTAAAAGGGT</td><td>>MI0000448|hsa-mir-130a&WithFlank&11|+|57641192|57641292</td><td>61</td><td>79</td><td>19(+1U)</td><td>CAGTGCAATGTTAAAAGGGT</td><td>72.0</td><td>4.4</td><td>2.1</td><td>2.1</td><td>3.5E-02</td><td>4.2E-02</td><td>MB231/MB468</td><td>MP</td></tr>
<tr><td>>hg38_21_+_8433172_8446622_RNA45SN1_45S_50ntFlanks@12995.13038.44|CTCGCTGCGATCTATTGAAAGTCAGCCCTCGACACAAGGGTTTG</td><td>>hg38_21_+_8433172_8446622_RNA45SN1_45S_50ntFlanks</td><td>12995</td><td>13038</td><td>44</td><td>CTCGCTGCGATCTATTGAAAGTCAGCCCTCGACACAAGGGTTTG</td><td>972.9</td><td>-2.3</td><td>0.9</td><td>-2.5</td><td>1.2E-02</td><td>4.2E-02</td><td>BT20/MB231</td><td>NUC</td></tr>
<tr><td>>MI0000234|hsa-mir-192&WithFlank&11|-|64891131|64891252@30.51.22(+2U)&[MIMAT0000222&hsa-miR-192-5p&offsets|0|+1(+2U);m-59&11|-|64891203|64891223&offsets|0|+1(+2U)]|CTGACCTATGAATTGACAGCCATT</td><td>>MI0000234|hsa-mir-192&WithFlank&11|-|64891131|64891252</td><td>30</td><td>51</td><td>22(+2U)</td><td>CTGACCTATGAATTGACAGCCATT</td><td>113.7</td><td>2.8</td><td>1.3</td><td>2.1</td><td>3.5E-02</td><td>4.2E-02</td><td>MB231/MB468</td><td>MP</td></tr>
<tr><td>>hg38_21_+_8433172_8446622_RNA45SN1_45S_50ntFlanks@8358.8376.19|AAAGAACTTTGAAGAGAGA</td><td>>hg38_21_+_8433172_8446622_RNA45SN1_45S_50ntFlanks</td><td>8358</td><td>8376</td><td>19</td><td>AAAGAACTTTGAAGAGAGA</td><td>105.0</td><td>-3.0</td><td>1.4</td><td>-2.1</td><td>3.5E-02</td><td>4.2E-02</td><td>MB231/MB468</td><td>MP</td></tr>
<tr><td>>hg38_21_+_8433172_8446622_RNA45SN1_45S_50ntFlanks@9554.9576.23|CGTAGCGGTCCTGACGTGCAAAT</td><td>>hg38_21_+_8433172_8446622_RNA45SN1_45S_50ntFlanks</td><td>9554</td><td>9576</td><td>23</td><td>CGTAGCGGTCCTGACGTGCAAAT</td><td>257.1</td><td>1.2</td><td>0.6</td><td>2.1</td><td>3.3E-02</td><td>4.2E-02</td><td>BT20/MB468</td><td>MITO</td></tr>
<tr><td>>hg38_MT_+_598_1651_RNR1_12S_50ntFlanks@52.74.23|ATAGGTTTGGTCCTAGCCTTTCT</td><td>>hg38_MT_+_598_1651_RNR1_12S_50ntFlanks</td><td>52</td><td>74</td><td>23</td><td>ATAGGTTTGGTCCTAGCCTTTCT</td><td>367.0</td><td>2.4</td><td>1.1</td><td>2.1</td><td>3.9E-02</td><td>4.2E-02</td><td>MB231/MB468</td><td>MITO</td></tr>
<tr><td>>hg38_1_-_228634819_228635039_RNA5S12_5S_50ntFlanks@51.90.40|GTCTACGGCCATACCACCCTGAACGCGCCCGATCTCGTCT</td><td>>hg38_1_-_228634819_228635039_RNA5S12_5S_50ntFlanks</td><td>51</td><td>90</td><td>40</td><td>GTCTACGGCCATACCACCCTGAACGCGCCCGATCTCGTCT</td><td>510.3</td><td>2.0</td><td>1.0</td><td>2.1</td><td>3.8E-02</td><td>4.2E-02</td><td>BT20/MB468</td><td>NUC</td></tr>
<tr><td>>hg38_21_+_8433172_8446622_RNA45SN1_45S_50ntFlanks@10571.10599.29|GAGAAGCCGGCGGGAGCCCCGGGGAGAGT</td><td>>hg38_21_+_8433172_8446622_RNA45SN1_45S_50ntFlanks</td><td>10571</td><td>10599</td><td>29</td><td>GAGAAGCCGGCGGGAGCCCCGGGGAGAGT</td><td>101.2</td><td>-3.0</td><td>1.4</td><td>-2.1</td><td>3.5E-02</td><td>4.2E-02</td><td>MB231/MB468</td><td>MP</td></tr>
<tr><td>>hg38_21_+_8433172_8446622_RNA45SN1_45S_50ntFlanks@8358.8391.34|AAAGAACTTTGAAGAGAGAGTTCAAGAGGGCGTG</td><td>>hg38_21_+_8433172_8446622_RNA45SN1_45S_50ntFlanks</td><td>8358</td><td>8391</td><td>34</td><td>AAAGAACTTTGAAGAGAGAGTTCAAGAGGGCGTG</td><td>244.0</td><td>3.3</td><td>1.3</td><td>2.5</td><td>1.2E-02</td><td>4.2E-02</td><td>BT20/MB231</td><td>MP</td></tr>
<tr><td>>hg38_21_+_8433172_8446622_RNA45SN1_45S_50ntFlanks@9565.9597.33|TGACGTGCAAATCGGTCGTCCGACCTGGGTATA</td><td>>hg38_21_+_8433172_8446622_RNA45SN1_45S_50ntFlanks</td><td>9565</td><td>9597</td><td>33</td><td>TGACGTGCAAATCGGTCGTCCGACCTGGGTATA</td><td>1065.9</td><td>-3.4</td><td>1.5</td><td>-2.2</td><td>2.6E-02</td><td>4.2E-02</td><td>BT20/MB468</td><td>MP</td></tr>
<tr><td>>hg38_21_+_8433172_8446622_RNA45SN1_45S_50ntFlanks@11207.11234.28|GGCGGCGACTCTGGACGCGAGCCGGGCC</td><td>>hg38_21_+_8433172_8446622_RNA45SN1_45S_50ntFlanks</td><td>11207</td><td>11234</td><td>28</td><td>GGCGGCGACTCTGGACGCGAGCCGGGCC</td><td>179.2</td><td>-3.9</td><td>1.8</td><td>-2.2</td><td>2.6E-02</td><td>4.2E-02</td><td>BT20/MB468</td><td>MP</td></tr>
<tr><td>>trnaMT_SerGCT_MT_+_12207_12265@8.38.31|CTCACAAGAACTGCTAACTCATGCCCCCATG</td><td>>trnaMT_SerGCT_MT_+_12207_12265</td><td>8</td><td>38</td><td>31</td><td>CTCACAAGAACTGCTAACTCATGCCCCCATG</td><td>136.1</td><td>-3.3</td><td>1.5</td><td>-2.2</td><td>2.6E-02</td><td>4.2E-02</td><td>BT20/MB468</td><td>MP</td></tr>
<tr><td>>hg38_21_+_8433172_8446622_RNA45SN1_45S_50ntFlanks@11667.11703.37|CTGCCCAGTGCTCTGAATGTCAAAGTGAAGAAATTCA</td><td>>hg38_21_+_8433172_8446622_RNA45SN1_45S_50ntFlanks</td><td>11667</td><td>11703</td><td>37</td><td>CTGCCCAGTGCTCTGAATGTCAAAGTGAAGAAATTCA</td><td>632.4</td><td>-2.0</td><td>0.8</td><td>-2.4</td><td>1.6E-02</td><td>4.2E-02</td><td>BT20/MB231</td><td>CYTO</td></tr>
<tr><td>>hg38_MT_+_598_1651_RNR1_12S_50ntFlanks@410.427.18|GACACAAAATAGACTACG</td><td>>hg38_MT_+_598_1651_RNR1_12S_50ntFlanks</td><td>410</td><td>427</td><td>18</td><td>GACACAAAATAGACTACG</td><td>44.8</td><td>4.4</td><td>2.1</td><td>2.1</td><td>3.5E-02</td><td>4.2E-02</td><td>MB231/MB468</td><td>MP</td></tr>
<tr><td>>MI0000071|hsa-mir-17&WithFlank&13|+|91350599|91350694@20.42.23(+1C)&[MIMAT0000070&hsa-miR-17-5p&offsets|0|0(+1C);m-88&13|+|91350618|91350640&offsets|0|0(+1C)]|CAAAGTGCTTACAGTGCAGGTAGC</td><td>>MI0000071|hsa-mir-17&WithFlank&13|+|91350599|91350694</td><td>20</td><td>42</td><td>23(+1C)</td><td>CAAAGTGCTTACAGTGCAGGTAGC</td><td>172.0</td><td>3.0</td><td>1.4</td><td>2.1</td><td>3.5E-02</td><td>4.2E-02</td><td>MB231/MB468</td><td>MP</td></tr>
<tr><td>>hg38_21_+_8433172_8446622_RNA45SN1_45S_50ntFlanks@11727.11749.23|AGTAACTATGACTCTCTTAAGGT</td><td>>hg38_21_+_8433172_8446622_RNA45SN1_45S_50ntFlanks</td><td>11727</td><td>11749</td><td>23</td><td>AGTAACTATGACTCTCTTAAGGT</td><td>216.5</td><td>-1.2</td><td>0.6</td><td>-2.1</td><td>3.9E-02</td><td>4.2E-02</td><td>MB231/MB468</td><td>NUC</td></tr>
<tr><td>>hg38_21_+_8433172_8446622_RNA45SN1_45S_50ntFlanks@3842.3862.21|CCTACTTGGATAACTGTGGTA</td><td>>hg38_21_+_8433172_8446622_RNA45SN1_45S_50ntFlanks</td><td>3842</td><td>3862</td><td>21</td><td>CCTACTTGGATAACTGTGGTA</td><td>96.4</td><td>-2.2</td><td>1.0</td><td>-2.2</td><td>2.7E-02</td><td>4.2E-02</td><td>BT20/MB468</td><td>MP</td></tr>
<tr><td>>MI0000465|hsa-mir-191&WithFlank&3|-|49020612|49020715@23.45.23&[MIMAT0000440&hsa-miR-191-5p&offsets|+1|+1;m-48&3|-|49020672|49020694&offsets|+1|+1]|AACGGAATCCCAAAAGCAGCTGT</td><td>>MI0000465|hsa-mir-191&WithFlank&3|-|49020612|49020715</td><td>23</td><td>45</td><td>23</td><td>AACGGAATCCCAAAAGCAGCTGT</td><td>92.4</td><td>5.9</td><td>2.6</td><td>2.2</td><td>2.7E-02</td><td>4.2E-02</td><td>BT20/MB468</td><td>MP</td></tr>
<tr><td>>hg38_21_+_8433172_8446622_RNA45SN1_45S_50ntFlanks@12407.12434.28|TTTTGATCCTTCGATGTCGGCTCTTCCT</td><td>>hg38_21_+_8433172_8446622_RNA45SN1_45S_50ntFlanks</td><td>12407</td><td>12434</td><td>28</td><td>TTTTGATCCTTCGATGTCGGCTCTTCCT</td><td>1654.9</td><td>-1.4</td><td>0.7</td><td>-2.1</td><td>3.7E-02</td><td>4.3E-02</td><td>BT20/MB468</td><td>CYTO</td></tr>
<tr><td>>hg38_21_+_8433172_8446622_RNA45SN1_45S_50ntFlanks@3833.3860.28|CGCTCCTCTCCTACTTGGATAACTGTGG</td><td>>hg38_21_+_8433172_8446622_RNA45SN1_45S_50ntFlanks</td><td>3833</td><td>3860</td><td>28</td><td>CGCTCCTCTCCTACTTGGATAACTGTGG</td><td>258.2</td><td>-1.9</td><td>0.8</td><td>-2.4</td><td>1.6E-02</td><td>4.3E-02</td><td>BT20/MB231</td><td>CYTO</td></tr>
<tr><td>>hg38_21_+_8433172_8446622_RNA45SN1_45S_50ntFlanks@9653.9679.27|AGGATAGCTGGCGCTCTCGCAGACCCG</td><td>>hg38_21_+_8433172_8446622_RNA45SN1_45S_50ntFlanks</td><td>9653</td><td>9679</td><td>27</td><td>AGGATAGCTGGCGCTCTCGCAGACCCG</td><td>420.1</td><td>-1.6</td><td>0.7</td><td>-2.1</td><td>3.3E-02</td><td>4.3E-02</td><td>BT20/MB468</td><td>MITO</td></tr>
<tr><td>>hg38_21_+_8433172_8446622_RNA45SN1_45S_50ntFlanks@5479.5501.23|TGGCGGAGCGCTGAGAAGACGGT</td><td>>hg38_21_+_8433172_8446622_RNA45SN1_45S_50ntFlanks</td><td>5479</td><td>5501</td><td>23</td><td>TGGCGGAGCGCTGAGAAGACGGT</td><td>89.6</td><td>5.1</td><td>2.0</td><td>2.5</td><td>1.2E-02</td><td>4.3E-02</td><td>BT20/MB231</td><td>MP</td></tr>
<tr><td>>MI0000077|hsa-mir-21&WithFlank&17|+|59841260|59841343@14.39.26(+1U)&[MIMAT0000076&hsa-miR-21-5p&offsets|0|+4(+1U);m-1&17|+|59841273|59841294&offsets|0|+4(+1U)]|TAGCTTATCAGACTGATGTTGACTGTT</td><td>>MI0000077|hsa-mir-21&WithFlank&17|+|59841260|59841343</td><td>14</td><td>39</td><td>26(+1U)</td><td>TAGCTTATCAGACTGATGTTGACTGTT</td><td>305.4</td><td>1.9</td><td>0.8</td><td>2.4</td><td>1.6E-02</td><td>4.3E-02</td><td>BT20/MB231</td><td>CYTO</td></tr>
<tr><td>>hg38_21_+_8433172_8446622_RNA45SN1_45S_50ntFlanks@12999.13028.30|CTGCGATCTATTGAAAGTCAGCCCTCGACA</td><td>>hg38_21_+_8433172_8446622_RNA45SN1_45S_50ntFlanks</td><td>12999</td><td>13028</td><td>30</td><td>CTGCGATCTATTGAAAGTCAGCCCTCGACA</td><td>65.9</td><td>-3.8</td><td>1.7</td><td>-2.2</td><td>2.7E-02</td><td>4.3E-02</td><td>BT20/MB468</td><td>MP</td></tr>
<tr><td>>trnaMT_PheGAA_MT_+_577_647@39.67.29|ATGTTTAGACGGGCTCACATCACCCCATA</td><td>>trnaMT_PheGAA_MT_+_577_647</td><td>39</td><td>67</td><td>29</td><td>ATGTTTAGACGGGCTCACATCACCCCATA</td><td>170.9</td><td>3.8</td><td>1.7</td><td>2.2</td><td>2.7E-02</td><td>4.3E-02</td><td>BT20/MB468</td><td>MP</td></tr>
<tr><td>>hg38_21_+_8433172_8446622_RNA45SN1_45S_50ntFlanks@4047.4073.27|ATTCGAACGTCTGCCCTATCAACTTTC</td><td>>hg38_21_+_8433172_8446622_RNA45SN1_45S_50ntFlanks</td><td>4047</td><td>4073</td><td>27</td><td>ATTCGAACGTCTGCCCTATCAACTTTC</td><td>124.9</td><td>-3.8</td><td>1.7</td><td>-2.2</td><td>2.7E-02</td><td>4.3E-02</td><td>BT20/MB468</td><td>MP</td></tr>
<tr><td>>trnaMT_LysTTT_MT_+_8295_8364@42.72.31,trnalookalike7_LysTTT_1_+_568844_568913@42.72.31|TTAAGAGAACCAACACCTCTTTACAGTGACC</td><td>>trnaMT_LysTTT_MT_+_8295_8364</td><td>42</td><td>72</td><td>31</td><td>TTAAGAGAACCAACACCTCTTTACAGTGACC</td><td>39.6</td><td>4.2</td><td>2.0</td><td>2.1</td><td>3.5E-02</td><td>4.3E-02</td><td>MB231/MB468</td><td>MP</td></tr>
</table>

| | | | | | | | | | | | | | |
|---|---|---|---|---|---|---|---|---|---|---|---|---|---|
| >MI0000465\|hsa-mir-191&WithFlank&3\|-\|49020612\|49020715@22.44.23(+1C)&[MIMAT0000440&hsa-miR-191-5p&offsets\|0\|0(+1C);m-48&3\|-\|49020672\|49020694&offsets\|0\|0(+1C)]\|CAACGGAATCCCAAAAGCAGCTGC | >MI0000465\|hsa-mir-191&WithFlank&3\|-\|49020612\|49020715 | 22 | 44 | 23(+1C) | CAACGGAATCCCAAAAGCAGCTGC | 2066.1 | -3.0 | 1.2 | -2.5 | 1.2E-02 | 4.3E-02 | BT20/MB231 | MP |
| >hg38_21_+_8433172_8446622_RNA45SN1_45S_50ntFlanks@10411.10442.32\|CAGTCGGTCCTGAGAGATGGGCGAGCGCCGTT | >hg38_21_+_8433172_8446622_RNA45SN1_45S_50ntFlanks | 10411 | 10442 | 32 | CAGTCGGTCCTGAGAGATGGGCGAGCGCCGTT | 118.1 | 3.0 | 1.2 | 2.5 | 1.2E-02 | 4.3E-02 | BT20/MB231 | MP |
| >hg38_MT_+_1621_3279_RNR2_16S_50ntFlanks@486.521.36\|AGGAACAGCTCTTTGGACACTAGGAAAAAACCTTGT | >hg38_MT_+_1621_3279_RNR2_16S_50ntFlanks | 486 | 521 | 36 | AGGAACAGCTCTTTGGACACTAGGAAAAAACCTTGT | 80.0 | 4.7 | 2.1 | 2.2 | 2.7E-02 | 4.3E-02 | BT20/MB468 | MP |
| >hg38_21_+_8433172_8446622_RNA45SN1_45S_50ntFlanks@3843.3871.29\|CTACTTGGATAACTGTGGTAATTCTAGAG | >hg38_21_+_8433172_8446622_RNA45SN1_45S_50ntFlanks | 3843 | 3871 | 29 | CTACTTGGATAACTGTGGTAATTCTAGAG | 590.3 | -1.8 | 0.9 | -2.1 | 3.7E-02 | 4.3E-02 | BT20/MB468 | CYTO |
| >hg38_1_-_228634819_228635039_RNA5S12_5S_50ntFlanks@132.170.39\|GATGGGAGACCGCCTGGGAATACCGGGTGCTGTAGGCTT | >hg38_1_-_228634819_228635039_RNA5S12_5S_50ntFlanks | 132 | 170 | 39 | GATGGGAGACCGCCTGGGAATACCGGGTGCTGTAGGCTT | 49065.1 | 1.9 | 0.9 | 2.1 | 3.7E-02 | 4.3E-02 | MB231/MB468 | CYTO |
| >hg38_21_+_8433172_8446622_RNA45SN1_45S_50ntFlanks@4742.4761.20\|TCGGAGGTTCGAAGACGATC | >hg38_21_+_8433172_8446622_RNA45SN1_45S_50ntFlanks | 4742 | 4761 | 20 | TCGGAGGTTCGAAGACGATC | 106.9 | 2.2 | 1.0 | 2.2 | 2.7E-02 | 4.3E-02 | BT20/MB468 | MP |
| >trna3_ArgTCT_11_+_59318767_59318852@1.29.29,trna4_ArgTCT_17_+_8024243_8024330@1.29.29,trna4_ArgTCT_9_-_131102355_131102445@1.29.29,trna52_ArgTCT_6_+_27529963_27530049@1.29.29\|GGCTCTGTGGCGCAATGGATAGCGCATTG | >trna3_ArgTCT_11_+_59318767_59318852 | 1 | 29 | 29 | GGCTCTGTGGCGCAATGGATAGCGCATTG | 89.5 | -4.1 | 1.9 | -2.2 | 2.7E-02 | 4.3E-02 | BT20/MB468 | MP |
| >hg38_21_+_8433172_8446622_RNA45SN1_45S_50ntFlanks@6655.6681.27\|TCTTAGCGGTGGATCACTCGGCTCGTG | >hg38_21_+_8433172_8446622_RNA45SN1_45S_50ntFlanks | 6655 | 6681 | 27 | TCTTAGCGGTGGATCACTCGGCTCGTG | 2042.3 | -1.3 | 0.6 | -2.1 | 3.9E-02 | 4.3E-02 | MB231/MB468 | NUC |
| >hg38_21_+_8433172_8446622_RNA45SN1_45S_50ntFlanks@5237.5256.20\|ACTGACTGGCTCAGCGTGTG | >hg38_21_+_8433172_8446622_RNA45SN1_45S_50ntFlanks | 5237 | 5256 | 20 | ACTGACTGGCTCAGCGTGTG | 410.9 | -1.8 | 0.8 | -2.1 | 3.6E-02 | 4.3E-02 | MB231/MB468 | MP |
| >hg38_21_+_8433172_8446622_RNA45SN1_45S_50ntFlanks@4380.4408.29\|CGTAGTTGGATCTTGGGAGCGGGCGGGCG | >hg38_21_+_8433172_8446622_RNA45SN1_45S_50ntFlanks | 4380 | 4408 | 29 | CGTAGTTGGATCTTGGGAGCGGGCGGGCG | 126.1 | -2.4 | 1.1 | -2.1 | 3.7E-02 | 4.3E-02 | BT20/MB468 | CYTO |
| >hg38_MT_+_598_1651_RNR1_12S_50ntFlanks@427.455.29\|GAAAGTGGCTTTAACATATCTGAACACAC | >hg38_MT_+_598_1651_RNR1_12S_50ntFlanks | 427 | 455 | 29 | GAAAGTGGCTTTAACATATCTGAACACAC | 137.8 | 2.7 | 1.3 | 2.1 | 3.6E-02 | 4.3E-02 | MB231/MB468 | MP |
| >hg38_21_+_8433172_8446622_RNA45SN1_45S_50ntFlanks@4498.4517.20\|AAGCGTTTACTTTGAAAAAA | >hg38_21_+_8433172_8446622_RNA45SN1_45S_50ntFlanks | 4498 | 4517 | 20 | AAGCGTTTACTTTGAAAAAA | 376.5 | -2.4 | 1.1 | -2.1 | 3.3E-02 | 4.3E-02 | BT20/MB468 | MITO |
| >hg38_MT_+_1621_3279_RNR2_16S_50ntFlanks@1415.1446.32\|CGTTTGTTCAACGATTAAAGTCCTACGTGATC | >hg38_MT_+_1621_3279_RNR2_16S_50ntFlanks | 1415 | 1446 | 32 | CGTTTGTTCAACGATTAAAGTCCTACGTGATC | 71.1 | 2.6 | 1.2 | 2.1 | 3.6E-02 | 4.3E-02 | MB231/MB468 | MP |
| >hg38_21_+_8433172_8446622_RNA45SN1_45S_50ntFlanks@10683.10708.26\|CCGGCGGCGTCCGGTGAGCTCTCGCT | >hg38_21_+_8433172_8446622_RNA45SN1_45S_50ntFlanks | 10683 | 10708 | 26 | CCGGCGGCGTCCGGTGAGCTCTCGCT | 192.3 | -3.0 | 1.4 | -2.2 | 2.7E-02 | 4.3E-02 | BT20/MB468 | MP |
| >am_trnaMT_MetCAT_MT_+_4402_4469@1.47.47\|AGTAAGGTCAGCTAAATAAGCTATCGGGCCCATACCCCGAAAATGTT | >am_trnaMT_MetCAT_MT_+_4402_4469 | 1 | 47 | 47 | AGTAAGGTCAGCTAAATAAGCTATCGGGCCCATACCCCGAAAATGTT | 1626.9 | 6.4 | 2.9 | 2.2 | 2.7E-02 | 4.3E-02 | BT20/MB468 | MP |
| >hg38_21_+_8433172_8446622_RNA45SN1_45S_50ntFlanks@12251.12284.34\|AAAGCTCGCTTGATCTTGATTTTCAGTACGAATA | >hg38_21_+_8433172_8446622_RNA45SN1_45S_50ntFlanks | 12251 | 12284 | 34 | AAAGCTCGCTTGATCTTGATTTTCAGTACGAATA | 430.0 | -1.8 | 0.9 | -2.1 | 3.3E-02 | 4.3E-02 | BT20/MB468 | MITO |
| >trna111_HisGTG_1_-_147774845_147774916@-1A.31.32,trna118_HisGTG_1_-_145396881_145396952@-1A.31.32,trna16_HisGTG_1_+_146544773_146544844@-1A.31.32,trna1_HisGTG_15_+_45493349_45493420@-1A.31.32,trna21_HisGTG_1_+_147753471_147753542@-1A.31.32,trna33_HisGTG_6_+_27125906_27125977@-1A.31.32,trna7_HisGTG_9_-_14433938_14434009@-1A.31.32,trna8_HisGTG_15_-_45492611_45492682@-1A.31.32,trna9_HisGTG_15_-_45490804_45490875@-1A.31.32\|AGCCGTGATCGTATAGTGGTTAGTACTCTGCG | >trna111_HisGTG_1_-_147774845_147774916 | -1A | 31 | 32 | AGCCGTGATCGTATAGTGGTTAGTACTCTGCG | 3902.6 | -2.2 | 1.1 | -2.1 | 3.7E-02 | 4.3E-02 | MB231/MB468 | CYTO |
| >hg38_21_+_8433172_8446622_RNA45SN1_45S_50ntFlanks@11587.11611.25\|TGCGGACCAGGGGAATCCGACTGTT | >hg38_21_+_8433172_8446622_RNA45SN1_45S_50ntFlanks | 11587 | 11611 | 25 | TGCGGACCAGGGGAATCCGACTGTT | 661.0 | -1.2 | 0.6 | -2.1 | 3.7E-02 | 4.3E-02 | MB231/MB468 | CYTO |
| >hg38_21_+_8433172_8446622_RNA45SN1_45S_50ntFlanks@11589.11611.23\|CGGACCAGGGGAATCCGACTGTT | >hg38_21_+_8433172_8446622_RNA45SN1_45S_50ntFlanks | 11589 | 11611 | 23 | CGGACCAGGGGAATCCGACTGTT | 343.0 | 1.8 | 0.7 | 2.5 | 1.2E-02 | 4.3E-02 | BT20/MB231 | NUC |
| >hg38_21_+_8433172_8446622_RNA45SN1_45S_50ntFlanks@12451.12476.26\|ATTCACCAAGCGTTGGATTGTTCACC | >hg38_21_+_8433172_8446622_RNA45SN1_45S_50ntFlanks | 12451 | 12476 | 26 | ATTCACCAAGCGTTGGATTGTTCACC | 234.2 | -2.2 | 0.9 | -2.5 | 1.2E-02 | 4.3E-02 | BT20/MB231 | NUC |
| >hg38_MT_+_1621_3279_RNR2_16S_50ntFlanks@1425.1453.29\|ACGATTAAAGTCCTACGTGATCTGAGTTC | >hg38_MT_+_1621_3279_RNR2_16S_50ntFlanks | 1425 | 1453 | 29 | ACGATTAAAGTCCTACGTGATCTGAGTTC | 133.5 | 3.7 | 1.7 | 2.2 | 2.7E-02 | 4.3E-02 | BT20/MB468 | MP |
| >hg38_21_+_8433172_8446622_RNA45SN1_45S_50ntFlanks@5059.5081.23\|CGATTTGTCTGGTTAATTCCGAT | >hg38_21_+_8433172_8446622_RNA45SN1_45S_50ntFlanks | 5059 | 5081 | 23 | CGATTTGTCTGGTTAATTCCGAT | 204.6 | -2.6 | 1.2 | -2.2 | 2.7E-02 | 4.3E-02 | BT20/MB468 | MP |
| >hg38_21_+_8433172_8446622_RNA45SN1_45S_50ntFlanks@10784.10801.18\|TGAACAGCCTCTGGCATG | >hg38_21_+_8433172_8446622_RNA45SN1_45S_50ntFlanks | 10784 | 10801 | 18 | TGAACAGCCTCTGGCATG | 76.9 | -4.9 | 2.2 | -2.2 | 2.7E-02 | 4.3E-02 | BT20/MB468 | MP |
| >hg38_21_+_8433172_8446622_RNA45SN1_45S_50ntFlanks@7988.8012.25\|CAGACGTGGCGACCCGCTGAATTTA | >hg38_21_+_8433172_8446622_RNA45SN1_45S_50ntFlanks | 7988 | 8012 | 25 | CAGACGTGGCGACCCGCTGAATTTA | 351.3 | -1.3 | 0.6 | -2.1 | 3.8E-02 | 4.3E-02 | BT20/MB468 | CYTO |
| >hg38_21_+_8433172_8446622_RNA45SN1_45S_50ntFlanks@10421.10456.36\|TGAGAGATGGGCGAGCGCCGTTCCGAAGGGACGGGC | >hg38_21_+_8433172_8446622_RNA45SN1_45S_50ntFlanks | 10421 | 10456 | 36 | TGAGAGATGGGCGAGCGCCGTTCCGAAGGGACGGGC | 2849.2 | -1.8 | 0.9 | -2.1 | 3.7E-02 | 4.3E-02 | MB231/MB468 | CYTO |
| >MI0000266\|hsa-mir-10a&WithFlank&17\|-\|48579832\|48579953@29.49.21&[MIMAT0000253&hsa-miR-10a-5p&offsets\|+1\|-1;m-18&17\|-\|48579905\|48579926&offsets\|+1\|0]\|ACCCTGTAGATCCGAATTTGT | >MI0000266\|hsa-mir-10a&WithFlank&17\|-\|48579832\|48579953 | 29 | 49 | 21 | ACCCTGTAGATCCGAATTTGT | 4488.2 | 1.9 | 0.8 | 2.4 | 1.6E-02 | 4.3E-02 | BT20/MB231 | CYTO |
| >hg38_21_+_8433172_8446622_RNA45SN1_45S_50ntFlanks@10809.10826.18\|AATGTAGGTAAGGGAAGT | >hg38_21_+_8433172_8446622_RNA45SN1_45S_50ntFlanks | 10809 | 10826 | 18 | AATGTAGGTAAGGGAAGT | 509.4 | -2.5 | 1.0 | -2.4 | 1.6E-02 | 4.3E-02 | BT20/MB231 | CYTO |
| >hg38_21_+_8433172_8446622_RNA45SN1_45S_50ntFlanks@12247.12272.26\|GGCAAAAGCTCGCTTGATCTTGATTT | >hg38_21_+_8433172_8446622_RNA45SN1_45S_50ntFlanks | 12247 | 12272 | 26 | GGCAAAAGCTCGCTTGATCTTGATTT | 148.1 | -2.0 | 0.9 | -2.1 | 3.3E-02 | 4.3E-02 | BT20/MB468 | MITO |
| >hg38_21_+_8433172_8446622_RNA45SN1_45S_50ntFlanks@6732.6752.21\|ACATTGATCATCGACACTTCG | >hg38_21_+_8433172_8446622_RNA45SN1_45S_50ntFlanks | 6732 | 6752 | 21 | ACATTGATCATCGACACTTCG | 450.0 | -1.4 | 0.7 | -2.1 | 3.8E-02 | 4.3E-02 | BT20/MB468 | CYTO |
| >hg38_21_+_8433172_8446622_RNA45SN1_45S_50ntFlanks@12417.12444.28\|TCGATGTCGGCTCTTCCTATCATTGTGA | >hg38_21_+_8433172_8446622_RNA45SN1_45S_50ntFlanks | 12417 | 12444 | 28 | TCGATGTCGGCTCTTCCTATCATTGTGA | 1325.3 | -2.1 | 0.9 | -2.4 | 1.6E-02 | 4.3E-02 | BT20/MB231 | CYTO |
| >hg38_21_+_8433172_8446622_RNA45SN1_45S_50ntFlanks@4207.4242.36\|CGGGGAGGTAGTGACGAAAAATAACAATACAGGACT | >hg38_21_+_8433172_8446622_RNA45SN1_45S_50ntFlanks | 4207 | 4242 | 36 | CGGGGAGGTAGTGACGAAAAATAACAATACAGGACT | 303.2 | -2.1 | 0.9 | -2.4 | 1.6E-02 | 4.3E-02 | BT20/MB231 | CYTO |
| >hg38_21_+_8433172_8446622_RNA45SN1_45S_50ntFlanks@10346.10371.26\|TCAAACGAGAACTTTGAAGGCCGAAG | >hg38_21_+_8433172_8446622_RNA45SN1_45S_50ntFlanks | 10346 | 10371 | 26 | TCAAACGAGAACTTTGAAGGCCGAAG | 286.4 | 3.5 | 1.4 | 2.5 | 1.2E-02 | 4.3E-02 | BT20/MB231 | MP |
| >hg38_21_+_8433172_8446622_RNA45SN1_45S_50ntFlanks@11567.11600.34\|AGCAGCCGACTTAGAACTGGTGCGGACCAGGGGA | >hg38_21_+_8433172_8446622_RNA45SN1_45S_50ntFlanks | 11567 | 11600 | 34 | AGCAGCCGACTTAGAACTGGTGCGGACCAGGGGA | 428.7 | 3.4 | 1.4 | 2.5 | 1.2E-02 | 4.3E-02 | BT20/MB231 | MP |
| >am_MI0000786\|hsa-mir-378a&WithFlank&5\|+\|149732819\|149732896@49.69.21(+1U)&[MIMAT0000732&hsa-miR-378a-3p&offsets\|0\|-1(+1U);m-44&5\|+\|149732867\|149732888&offsets\|0\|-1(+1U)]\|ACTGGACTTGGAGTCAGAAGGT | >am_MI0000786\|hsa-mir-378a&WithFlank&5\|+\|149732819\|149732896 | 49 | 69 | 21(+1U) | ACTGGACTTGGAGTCAGAAGGT | 11740.8 | 1.6 | 0.8 | 2.1 | 3.6E-02 | 4.3E-02 | MB231/MB468 | MP |
| >hg38_1_-_228634819_228635039_RNA5S12_5S_50ntFlanks@149.171.23\|GAATACCGGGTGCTGTAGGCTTT | >hg38_1_-_228634819_228635039_RNA5S12_5S_50ntFlanks | 149 | 171 | 23 | GAATACCGGGTGCTGTAGGCTTT | 5967.1 | 3.6 | 1.4 | 2.5 | 1.2E-02 | 4.3E-02 | BT20/MB231 | MP |
| >MI0000270\|hsa-mir-181b-1&WithFlank&1\|-\|198858867\|198858988@42.63.22(+1A);MI0000683\|hsa-mir-181b-2&WithFlank&9\|+\|124693704\|124693804@22.43.22(+1A)&[MIMAT0000257&hsa-miR-181b-5p&offsets\|0\|-1(+1A);m-73&1\|-\|198858925\|198858947&offsets\|0\|-1(+1A)];[MIMAT0000257_1&hsa-miR-181b-5p&offsets\|0\|-1(+1A);m-72&9\|+\|124693725\|124693747&offsets\|0\|-1(+1A)]\|AACATTCATTGCTGTCGGTGGGA | >MI0000270\|hsa-mir-181b-1&WithFlank&1\|-\|198858867\|198858988 | 42 | 63 | 22(+1A) | AACATTCATTGCTGTCGGTGGGA | 198.8 | 2.9 | 1.4 | 2.1 | 3.6E-02 | 4.3E-02 | MB231/MB468 | MP |
| >hg38_21_+_8433172_8446622_RNA45SN1_45S_50ntFlanks@11189.11225.37\|ACCCGGGGGGCCGGCGGCGGCGGCGACTCTGGACGCG | >hg38_21_+_8433172_8446622_RNA45SN1_45S_50ntFlanks | 11189 | 11225 | 37 | ACCCGGGGGGCCGGCGGCGGCGGCGACTCTGGACGCG | 138.9 | -4.0 | 1.8 | -2.2 | 2.7E-02 | 4.3E-02 | BT20/MB468 | MP |
| >trna119_LysCTT_1_-_145395522_145395594@1.32.32,trna11_LysCTT_5_-_180648979_180649051@1.32.32,trna13_LysCTT_6_+_26556774_26556846@1.32.32,trna32_LysCTT_16_-_3207406_3207478@1.32.32,trna7_LysCTT_16_+_3225692_3225764@1.32.32,trna9_LysCTT_5_+_180634755_180634827@1.32.32\|GCCCGGCTAGCTCAGTCGGTAGAGCATGAGAC | >trna119_LysCTT_1_-_145395522_145395594 | 1 | 32 | 32 | GCCCGGCTAGCTCAGTCGGTAGAGCATGAGAC | 813.5 | 1.4 | 0.7 | 2.1 | 3.6E-02 | 4.3E-02 | MB231/MB468 | MP |
| >hg38_MT_+_1621_3279_RNR2_16S_50ntFlanks@468.491.24\|TTAACTGTTAGTCCAAAGAGGAAC | >hg38_MT_+_1621_3279_RNR2_16S_50ntFlanks | 468 | 491 | 24 | TTAACTGTTAGTCCAAAGAGGAAC | 37.0 | 3.6 | 1.7 | 2.1 | 3.6E-02 | 4.3E-02 | MB231/MB468 | MP |
| >hg38_21_+_8433172_8446622_RNA45SN1_45S_50ntFlanks@7981.8011.31\|CTCAGATCAGACGTGGCGACCCGCTGAATTT | >hg38_21_+_8433172_8446622_RNA45SN1_45S_50ntFlanks | 7981 | 8011 | 31 | CTCAGATCAGACGTGGCGACCCGCTGAATTT | 392.7 | 1.5 | 0.7 | 2.1 | 3.4E-02 | 4.3E-02 | BT20/MB468 | MITO |
| >MI0000760\|hsa-mir-361&WithFlank&X\|-\|85903630\|85903713@51.73.23&[MIMAT0004682&hsa-miR-361-3p&offsets\|0\|0;m-124&X\|-\|85903642\|85903663&offsets\|0\|+1]\|TCCCCCAGGTGTGATTCTGATTT | >MI0000760\|hsa-mir-361&WithFlank&X\|-\|85903630\|85903713 | 51 | 73 | 23 | TCCCCCAGGTGTGATTCTGATTT | 93.8 | 2.2 | 1.0 | 2.2 | 2.7E-02 | 4.3E-02 | BT20/MB468 | MP |
| >hg38_21_+_8433172_8446622_RNA45SN1_45S_50ntFlanks@8105.8144.40\|CGCGGCGGGGCGCGGGACATGTGGCGTACGGAAGACCCGC | >hg38_21_+_8433172_8446622_RNA45SN1_45S_50ntFlanks | 8105 | 8144 | 40 | CGCGGCGGGGCGCGGGACATGTGGCGTACGGAAGACCCGC | 463.9 | -3.7 | 1.8 | -2.1 | 3.6E-02 | 4.3E-02 | MB231/MB468 | MP |
| >hg38_21_+_8433172_8446622_RNA45SN1_45S_50ntFlanks@6699.6719.21\|GCTAGCTGCGAGAATTAATGT | >hg38_21_+_8433172_8446622_RNA45SN1_45S_50ntFlanks | 6699 | 6719 | 21 | GCTAGCTGCGAGAATTAATGT | 2129.3 | -3.7 | 1.7 | -2.2 | 2.7E-02 | 4.3E-02 | BT20/MB468 | MP |
| >hg38_21_+_8433172_8446622_RNA45SN1_45S_50ntFlanks@11217.11239.23\|CTGGACGCGAGCCGGGCCCTTCC | >hg38_21_+_8433172_8446622_RNA45SN1_45S_50ntFlanks | 11217 | 11239 | 23 | CTGGACGCGAGCCGGGCCCTTCC | 289.7 | -2.5 | 1.1 | -2.2 | 2.7E-02 | 4.3E-02 | BT20/MB468 | MP |
| >MI0000076\|hsa-mir-20a&WithFlank&13\|+\|91351059\|91351141@14.35.22&[MIMAT0000075&hsa-miR-20a-5p&offsets\|0\|-1;m-96&13\|+\|91351072\|91351094&offsets\|0\|-1]\|TAAAGTGCTTATAGTGCAGGTA | >MI0000076\|hsa-mir-20a&WithFlank&13\|+\|91351059\|91351141 | 14 | 35 | 22 | TAAAGTGCTTATAGTGCAGGTA | 551.6 | -6.1 | 2.5 | -2.4 | 1.4E-02 | 4.3E-02 | BT20/MB231 | MITO |
| >hg38_21_+_8433172_8446622_RNA45SN1_45S_50ntFlanks@11321.11340.20\|CACGTCTCGTCGCGCGCGCG | >hg38_21_+_8433172_8446622_RNA45SN1_45S_50ntFlanks | 11321 | 11340 | 20 | CACGTCTCGTCGCGCGCGCG | 108.7 | -3.8 | 1.7 | -2.2 | 2.7E-02 | 4.3E-02 | BT20/MB468 | MP |
| >MI0003780\|hsa-mir-1296&WithFlank&10\|-\|63372951\|63373054@22.41.20&[MIMAT0005794&hsa-miR-1296-5p&offsets\|0\|-2;m-304&10\|-\|63373012\|63373033&offsets\|0\|-2]\|TTAGGGCCCTGGCTCCATCT | >MI0003780\|hsa-mir-1296&WithFlank&10\|-\|63372951\|63373054 | 22 | 41 | 20 | TTAGGGCCCTGGCTCCATCT | 83.2 | 5.8 | 2.6 | 2.2 | 2.7E-02 | 4.3E-02 | BT20/MB468 | MP |

| | | | | | | | | | | | | | |
|---|---|---|---|---|---|---|---|---|---|---|---|---|---|
| >hg38_21_+_8433172_8446622_RNA45SN1_45S_50ntFlanks@3773.3791.19\|CGGCCGGTACAGTGAAACT | >hg38_21_+_8433172_8446622_RNA45SN1_45S_50ntFlanks | 3773 | 3791 | 19 | CGGCCGGTACAGTGAAACT | 265.8 | -2.2 | 0.9 | -2.4 | 1.6E-02 | 4.3E-02 | BT20/MB231 | CYTO |
| >hg38_21_+_8433172_8446622_RNA45SN1_45S_50ntFlanks@10427.10456.30\|ATGGGCGAGCGCCGTTCCGAAGGGACGGGC | >hg38_21_+_8433172_8446622_RNA45SN1_45S_50ntFlanks | 10427 | 10456 | 30 | ATGGGCGAGCGCCGTTCCGAAGGGACGGGC | 199.0 | 4.3 | 1.7 | 2.5 | 1.2E-02 | 4.3E-02 | BT20/MB231 | MP |
| >MI0003188\|hsa-mir-503&WithFlank&X\|-\|134546322\|134546404@12.31.20(+1A)&[MIMAT0002874&hsa-miR-503-5p&offsets\|0\|-3(+1A);m-297&X\|-\|134546371\|134546393&offsets\|0\|-3(+1A)]\|TAGCAGCGGGAACAGTTCTGA | >MI0003188\|hsa-mir-503&WithFlank&X\|-\|134546322\|134546404 | 12 | 31 | 20(+1A) | TAGCAGCGGGAACAGTTCTGA | 325.1 | 2.1 | 0.9 | 2.4 | 1.6E-02 | 4.3E-02 | BT20/MB231 | CYTO |
| >hg38_21_+_8433172_8446622_RNA45SN1_45S_50ntFlanks@9554.9577.24\|CGTAGCGGTCCTGACGTGCAAATC | >hg38_21_+_8433172_8446622_RNA45SN1_45S_50ntFlanks | 9554 | 9577 | 24 | CGTAGCGGTCCTGACGTGCAAATC | 3007.5 | -2.0 | 0.8 | -2.4 | 1.6E-02 | 4.3E-02 | BT20/MB231 | CYTO |
| >hg38_21_+_8433172_8446622_RNA45SN1_45S_50ntFlanks@4051.4070.20\|GAACGTCTGCCCTATCAACT | >hg38_21_+_8433172_8446622_RNA45SN1_45S_50ntFlanks | 4051 | 4070 | 20 | GAACGTCTGCCCTATCAACT | 341.1 | -2.0 | 0.9 | -2.4 | 1.6E-02 | 4.3E-02 | BT20/MB231 | CYTO |
| >hg38_21_+_8433172_8446622_RNA45SN1_45S_50ntFlanks@4824.4844.21\|TGACCCGCCGGGCAGCTTCCG | >hg38_21_+_8433172_8446622_RNA45SN1_45S_50ntFlanks | 4824 | 4844 | 21 | TGACCCGCCGGGCAGCTTCCG | 919.5 | -2.2 | 0.9 | -2.4 | 1.6E-02 | 4.3E-02 | BT20/MB231 | CYTO |
| >hg38_21_+_8433172_8446622_RNA45SN1_45S_50ntFlanks@10664.10709.46\|CTTGGAAAGCGTCGCGGTTCCGGCGGCGTCCGGTGAGCTCTCGCTG | >hg38_21_+_8433172_8446622_RNA45SN1_45S_50ntFlanks | 10664 | 10709 | 46 | CTTGGAAAGCGTCGCGGTTCCGGCGGCGTCCGGTGAGCTCTCGCTG | 142.7 | -3.6 | 1.7 | -2.1 | 3.4E-02 | 4.3E-02 | BT20/MB468 | MITO |
| >hg38_21_+_8433172_8446622_RNA45SN1_45S_50ntFlanks@12406.12433.28\|TTTTTGATCCTTCGATGTCGGCTCTTCC | >hg38_21_+_8433172_8446622_RNA45SN1_45S_50ntFlanks | 12406 | 12433 | 28 | TTTTTGATCCTTCGATGTCGGCTCTTCC | 721.6 | 2.0 | 1.0 | 2.1 | 3.8E-02 | 4.3E-02 | MB231/MB468 | CYTO |
| >hg38_21_+_8433172_8446622_RNA45SN1_45S_50ntFlanks@6645.6665.21\|CTCGTACGACTCTTAGCGGTG | >hg38_21_+_8433172_8446622_RNA45SN1_45S_50ntFlanks | 6645 | 6665 | 21 | CTCGTACGACTCTTAGCGGTG | 21475.8 | 3.1 | 1.2 | 2.5 | 1.2E-02 | 4.3E-02 | BT20/MB231 | MP |
| >hg38_21_+_8433172_8446622_RNA45SN1_45S_50ntFlanks@12458.12480.23\|AAGCGTTGGATTGTTCACCCACT | >hg38_21_+_8433172_8446622_RNA45SN1_45S_50ntFlanks | 12458 | 12480 | 23 | AAGCGTTGGATTGTTCACCCACT | 127.0 | -2.8 | 1.4 | -2.1 | 3.6E-02 | 4.3E-02 | MB231/MB468 | MP |
| >hg38_21_+_8433172_8446622_RNA45SN1_45S_50ntFlanks@7975.8012.38\|CGCGACCTCAGATCAGACGTGGCGACCCGCTGAATTTA | >hg38_21_+_8433172_8446622_RNA45SN1_45S_50ntFlanks | 7975 | 8012 | 38 | CGCGACCTCAGATCAGACGTGGCGACCCGCTGAATTTA | 33494.8 | 1.3 | 0.6 | 2.1 | 3.9E-02 | 4.3E-02 | MB231/MB468 | NUC |
| >trnaMT_PheGAA_MT_+_577_647@37.55.19\|AAATGTTTAGACGGGCTCA | >trnaMT_PheGAA_MT_+_577_647 | 37 | 55 | 19 | AAATGTTTAGACGGGCTCA | 44.1 | 3.4 | 1.6 | 2.1 | 3.6E-02 | 4.3E-02 | MB231/MB468 | MP |
| >hg38_21_+_8433172_8446622_RNA45SN1_45S_50ntFlanks@4876.4903.28\|TATGGTTGCAAAGCTGAAACTTAAAGGA | >hg38_21_+_8433172_8446622_RNA45SN1_45S_50ntFlanks | 4876 | 4903 | 28 | TATGGTTGCAAAGCTGAAACTTAAAGGA | 83.0 | 2.3 | 1.0 | 2.2 | 2.7E-02 | 4.3E-02 | BT20/MB468 | MP |
| >hg38_MT_+_1621_3279_RNR2_16S_50ntFlanks@617.643.27\|AAACATATAACTGAACTCCTCACACCC | >hg38_MT_+_1621_3279_RNR2_16S_50ntFlanks | 617 | 643 | 27 | AAACATATAACTGAACTCCTCACACCC | 230.1 | 4.6 | 2.1 | 2.2 | 2.7E-02 | 4.3E-02 | BT20/MB468 | MP |
| >hg38_21_+_8433172_8446622_RNA45SN1_45S_50ntFlanks@9419.9452.34\|TCCGCCGAGGGCGCACCACCGGCCCGTCTCGCCC | >hg38_21_+_8433172_8446622_RNA45SN1_45S_50ntFlanks | 9419 | 9452 | 34 | TCCGCCGAGGGCGCACCACCGGCCCGTCTCGCCC | 79.8 | -3.8 | 1.7 | -2.2 | 2.7E-02 | 4.3E-02 | BT20/MB468 | MP |
| >MI0000063\|hsa-let-7b&WithFlank&22\|+\|46113680\|46113774@66.87.22&[MIMAT0004482&hsa-let-7b-3p&offsets\|0\|0;m-171&22\|+\|46113745\|46113766&offsets\|0\|0]\|CTATACAACCTACTGCCTTCCC | >MI0000063\|hsa-let-7b&WithFlank&22\|+\|46113680\|46113774 | 66 | 87 | 22 | CTATACAACCTACTGCCTTCCC | 29.2 | 4.5 | 2.2 | 2.1 | 3.6E-02 | 4.3E-02 | MB231/MB468 | MP |
| >MI0000749\|hsa-mir-30e&WithFlank&1\|+\|40754349\|40754452@23.45.23(+1C)&[MIMAT0000692&hsa-miR-30e-5p&offsets\|0\|+1(+1C);m-24&1\|+\|40754371\|40754394&offsets\|0\|-1(+1C)]\|TGTAAACATCCTTGACTGGAAGCC | >MI0000749\|hsa-mir-30e&WithFlank&1\|+\|40754349\|40754452 | 23 | 45 | 23(+1C) | TGTAAACATCCTTGACTGGAAGCC | 3699.6 | 1.7 | 0.8 | 2.1 | 3.6E-02 | 4.3E-02 | MB231/MB468 | MP |
| >hg38_21_+_8433172_8446622_RNA45SN1_45S_50ntFlanks@10392.10424.33\|ACAGCAGTTGAACATGGGTCAGTCGGTCCTGAG | >hg38_21_+_8433172_8446622_RNA45SN1_45S_50ntFlanks | 10392 | 10424 | 33 | ACAGCAGTTGAACATGGGTCAGTCGGTCCTGAG | 2097.7 | -2.3 | 1.1 | -2.1 | 3.6E-02 | 4.3E-02 | MB231/MB468 | MP |
| >hg38_21_+_8433172_8446622_RNA45SN1_45S_50ntFlanks@10352.10376.25\|GAGAACTTTGAAGGCCGAAGTGGAG | >hg38_21_+_8433172_8446622_RNA45SN1_45S_50ntFlanks | 10352 | 10376 | 25 | GAGAACTTTGAAGGCCGAAGTGGAG | 59.5 | -4.7 | 2.2 | -2.1 | 3.6E-02 | 4.3E-02 | MB231/MB468 | MP |
| >hg38_MT_+_1621_3279_RNR2_16S_50ntFlanks@1369.1402.34\|GATGTTGGATCAGGACATCCCGATGGTGCAGCCG | >hg38_MT_+_1621_3279_RNR2_16S_50ntFlanks | 1369 | 1402 | 34 | GATGTTGGATCAGGACATCCCGATGGTGCAGCCG | 54.6 | 3.5 | 1.7 | 2.1 | 3.6E-02 | 4.3E-02 | MB231/MB468 | MP |
| >hg38_21_+_8433172_8446622_RNA45SN1_45S_50ntFlanks@3840.3865.26\|CTCCTACTTGGATAACTGTGGTAATT | >hg38_21_+_8433172_8446622_RNA45SN1_45S_50ntFlanks | 3840 | 3865 | 26 | CTCCTACTTGGATAACTGTGGTAATT | 791.2 | 1.8 | 0.7 | 2.4 | 1.6E-02 | 4.3E-02 | BT20/MB231 | CYTO |
| >hg38_21_+_8433172_8446622_RNA45SN1_45S_50ntFlanks@11672.11706.35\|CAGTGCTCTGAATGTCAAAGTGAAGAAATTCAATG | >hg38_21_+_8433172_8446622_RNA45SN1_45S_50ntFlanks | 11672 | 11706 | 35 | CAGTGCTCTGAATGTCAAAGTGAAGAAATTCAATG | 672.8 | -2.1 | 0.9 | -2.4 | 1.6E-02 | 4.3E-02 | BT20/MB231 | CYTO |
| >hg38_21_+_8433172_8446622_RNA45SN1_45S_50ntFlanks@4051.4071.21\|GAACGTCTGCCCTATCAACTT | >hg38_21_+_8433172_8446622_RNA45SN1_45S_50ntFlanks | 4051 | 4071 | 21 | GAACGTCTGCCCTATCAACTT | 172.7 | -2.1 | 0.9 | -2.4 | 1.6E-02 | 4.3E-02 | BT20/MB231 | CYTO |
| >hg38_21_+_8433172_8446622_RNA45SN1_45S_50ntFlanks@8381.8406.26\|AAGAGGGCGTGAAACCGTTAAGAGGT | >hg38_21_+_8433172_8446622_RNA45SN1_45S_50ntFlanks | 8381 | 8406 | 26 | AAGAGGGCGTGAAACCGTTAAGAGGT | 1247.9 | -1.7 | 0.7 | -2.2 | 2.7E-02 | 4.3E-02 | BT20/MB468 | MP |
| >hg38_21_+_8433172_8446622_RNA45SN1_45S_50ntFlanks@10347.10380.34\|CAAACGAGAACTTTGAAGGCCGAAGTGGAGAAGG | >hg38_21_+_8433172_8446622_RNA45SN1_45S_50ntFlanks | 10347 | 10380 | 34 | CAAACGAGAACTTTGAAGGCCGAAGTGGAGAAGG | 190.2 | 4.1 | 1.6 | 2.5 | 1.2E-02 | 4.3E-02 | BT20/MB231 | MP |
| >hg38_21_+_8433172_8446622_RNA45SN1_45S_50ntFlanks@5082.5104.23\|AACGAACGAGACTCTGGCATGCT | >hg38_21_+_8433172_8446622_RNA45SN1_45S_50ntFlanks | 5082 | 5104 | 23 | AACGAACGAGACTCTGGCATGCT | 1071.4 | 3.2 | 1.3 | 2.5 | 1.2E-02 | 4.3E-02 | BT20/MB231 | MP |
| >hg38_21_+_8433172_8446622_RNA45SN1_45S_50ntFlanks@6645.6663.19\|CTCGTACGACTCTTAGCGG | >hg38_21_+_8433172_8446622_RNA45SN1_45S_50ntFlanks | 6645 | 6663 | 19 | CTCGTACGACTCTTAGCGG | 1452.2 | -1.3 | 0.6 | -2.1 | 3.6E-02 | 4.3E-02 | MB231/MB468 | MP |
| >hg38_MT_+_598_1651_RNR1_12S_50ntFlanks@905.935.31\|AGTATACTTCAAAGGACATTTAACTAAAACC | >hg38_MT_+_598_1651_RNR1_12S_50ntFlanks | 905 | 935 | 31 | AGTATACTTCAAAGGACATTTAACTAAAACC | 217.9 | 3.9 | 1.7 | 2.2 | 2.7E-02 | 4.3E-02 | BT20/MB468 | MP |
| >MI0000440\|hsa-mir-27b&WithFlank&9\|+\|95085439\|95085547@67.85.19(+2U)&[MIMAT0000419&hsa-miR-27b-3p&offsets\|0\|-2(+2U);m-39&9\|+\|95085505\|95085525&offsets\|0\|-2(+2U)]\|TTCACAGTGGCTAAGTTCTTT | >MI0000440\|hsa-mir-27b&WithFlank&9\|+\|95085439\|95085547 | 67 | 85 | 19(+2U) | TTCACAGTGGCTAAGTTCTTT | 86.7 | 1.8 | 0.9 | 2.1 | 3.6E-02 | 4.3E-02 | MB231/MB468 | MP |
| >MI0016003\|hsa-mir-3613&WithFlank&13\|-\|49996409\|49996507@22.43.22&[MIMAT0017990&hsa-miR-3613-5p&offsets\|0\|0;m-259&13\|-\|49996465\|49996486&offsets\|0\|0]\|TGTTGTACTTTTTTTTTTGTTC | >MI0016003\|hsa-mir-3613&WithFlank&13\|-\|49996409\|49996507 | 22 | 43 | 22 | TGTTGTACTTTTTTTTTTGTTC | 457.2 | 2.0 | 0.8 | 2.5 | 1.3E-02 | 4.3E-02 | BT20/MB231 | NUC |
| >hg38_21_+_8433172_8446622_RNA45SN1_45S_50ntFlanks@12072.12110.39\|CCCGAGGGGCTCTCGCTTCTGGCGCCAAGCGCCCGGCCG | >hg38_21_+_8433172_8446622_RNA45SN1_45S_50ntFlanks | 12072 | 12110 | 39 | CCCGAGGGGCTCTCGCTTCTGGCGCCAAGCGCCCGGCCG | 42.3 | -4.0 | 1.8 | -2.2 | 2.7E-02 | 4.3E-02 | BT20/MB468 | MP |
| >hg38_21_+_8433172_8446622_RNA45SN1_45S_50ntFlanks@12160.12180.21\|TTGACTGGGGCGGTACACCTG | >hg38_21_+_8433172_8446622_RNA45SN1_45S_50ntFlanks | 12160 | 12180 | 21 | TTGACTGGGGCGGTACACCTG | 136.7 | -2.6 | 1.2 | -2.1 | 3.4E-02 | 4.3E-02 | BT20/MB468 | MITO |
| >am_trna119_LysCTT_1_-_145395522_145395594@14.34.21,trna11_LysCTT_5_-_180648979_180649051@14.34.21,trna128_LysCTT_1_-_55423542_55423614@14.34.21,trna13_LysCTT_6_+_26556774_26556846@14.34.21,trna6_LysCTT_19_-_52425396_52425468@14.34.21,trna7_LysCTT_16_+_3225692_3225764@14.34.21,trna9_LysCTT_5_+_180634755_180634827@14.34.21\|AGTCGGTAGAGCATGAGACTC | >am_trna119_LysCTT_1_-_145395522_145395594 | 14 | 34 | 21 | AGTCGGTAGAGCATGAGACTC | 122.6 | 2.7 | 1.3 | 2.1 | 3.6E-02 | 4.3E-02 | MB231/MB468 | MP |
| >hg38_21_+_8433172_8446622_RNA45SN1_45S_50ntFlanks@4683.4722.40\|CAAGACGGACCAGAGCGAAAGCATTTGCCAAGAATGTTTT | >hg38_21_+_8433172_8446622_RNA45SN1_45S_50ntFlanks | 4683 | 4722 | 40 | CAAGACGGACCAGAGCGAAAGCATTTGCCAAGAATGTTTT | 442.6 | -2.7 | 1.3 | -2.1 | 3.4E-02 | 4.3E-02 | BT20/MB468 | MITO |
| >MI0000102\|hsa-mir-100&WithFlank&11\|-\|122152223\|122152314@20.40.21&[MIMAT0000098&hsa-miR-100-5p&offsets\|+1\|0;m-27&11\|-\|122152276\|122152296&offsets\|+1\|+1]\|ACCCGTAGATCCGAACTTGTG | >MI0000102\|hsa-mir-100&WithFlank&11\|-\|122152223\|122152314 | 20 | 40 | 21 | ACCCGTAGATCCGAACTTGTG | 1509.4 | -4.7 | 1.9 | -2.4 | 1.6E-02 | 4.3E-02 | BT20/MB231 | CYTO |
| >hg38_21_+_8433172_8446622_RNA45SN1_45S_50ntFlanks@11200.11221.22\|CGGCGGCGGCGGCGACTCTGGA | >hg38_21_+_8433172_8446622_RNA45SN1_45S_50ntFlanks | 11200 | 11221 | 22 | CGGCGGCGGCGGCGACTCTGGA | 48.0 | -4.0 | 1.8 | -2.2 | 2.7E-02 | 4.3E-02 | BT20/MB468 | MP |
| >hg38_21_+_8433172_8446622_RNA45SN1_45S_50ntFlanks@11208.11238.31\|GCGGCGACTCTGGACGCGAGCCGGGCCCTTC | >hg38_21_+_8433172_8446622_RNA45SN1_45S_50ntFlanks | 11208 | 11238 | 31 | GCGGCGACTCTGGACGCGAGCCGGGCCCTTC | 433.2 | -1.7 | 0.8 | -2.1 | 3.8E-02 | 4.3E-02 | BT20/MB468 | CYTO |
| >hg38_21_+_8433172_8446622_RNA45SN1_45S_50ntFlanks@10821.10865.45\|GGAAGTCGGCAAGCCGGATCCGTAACTTCGGGATAAGGATTGGCT | >hg38_21_+_8433172_8446622_RNA45SN1_45S_50ntFlanks | 10821 | 10865 | 45 | GGAAGTCGGCAAGCCGGATCCGTAACTTCGGGATAAGGATTGGCT | 99.2 | -3.5 | 1.6 | -2.2 | 2.7E-02 | 4.3E-02 | BT20/MB468 | MP |
| >hg38_21_+_8433172_8446622_RNA45SN1_45S_50ntFlanks@9353.9370.18\|CGTGGCGCAATGAAGGTG | >hg38_21_+_8433172_8446622_RNA45SN1_45S_50ntFlanks | 9353 | 9370 | 18 | CGTGGCGCAATGAAGGTG | 1063.6 | -4.3 | 1.9 | -2.2 | 2.7E-02 | 4.3E-02 | BT20/MB468 | MP |
| >hg38_21_+_8433172_8446622_RNA45SN1_45S_50ntFlanks@11568.11612.45\|GCAGCCGACTTAGAACTGGTGCGGACCAGGGGAATCCGACTGTTT | >hg38_21_+_8433172_8446622_RNA45SN1_45S_50ntFlanks | 11568 | 11612 | 45 | GCAGCCGACTTAGAACTGGTGCGGACCAGGGGAATCCGACTGTTT | 378.4 | 2.6 | 1.2 | 2.1 | 3.4E-02 | 4.3E-02 | BT20/MB468 | MITO |
| >trna10_AspGTC_12_-_125424193_125424264@32.70.39,trna12_AspGTC_12_-_125411891_125411962@32.70.39,trna144_AspGTC_6_-_27551236_27551307@32.70.39,trna38_AspGTC_17_-_8125556_8125627@32.70.39,trna45_AspGTC_6_+_27447453_27447524@32.70.39,trna48_AspGTC_6_+_27471523_27471594@32.70.39,trna4_AspGTC_12_+_96429799_96429870@32.70.39,trna69_AspGTC_1_-_161440205_161440276@32.70.39,trna72_AspGTC_1_-_161432824_161432895@32.70.39,trna75_AspGTC_1_-_161425414_161425485@32.70.39,trna78_AspGTC_1_-_161418033_161418104@32.70.39,trna81_AspGTC_1_-_161410615_161410686@32.70.39\|CTGTCACGCGGGAGACCGGGGTTCGATTCCCCGACGGGG | >trna10_AspGTC_12_-_125424193_125424264 | 32 | 70 | 39 | CTGTCACGCGGGAGACCGGGGTTCGATTCCCCGACGGGG | 117.9 | -2.9 | 1.4 | -2.1 | 3.4E-02 | 4.3E-02 | BT20/MB468 | MITO |
| >hg38_21_+_8433172_8446622_RNA45SN1_45S_50ntFlanks@12407.12431.25\|TTTTGATCCTTCGATGTCGGCTCTT | >hg38_21_+_8433172_8446622_RNA45SN1_45S_50ntFlanks | 12407 | 12431 | 25 | TTTTGATCCTTCGATGTCGGCTCTT | 359.1 | -1.4 | 0.7 | -2.1 | 3.4E-02 | 4.3E-02 | BT20/MB468 | MITO |
| >hg38_21_+_8433172_8446622_RNA45SN1_45S_50ntFlanks@4004.4042.39\|TAACCTCGGGCCGATCGCACGCCCCCCGTGGCGGCGACG | >hg38_21_+_8433172_8446622_RNA45SN1_45S_50ntFlanks | 4004 | 4042 | 39 | TAACCTCGGGCCGATCGCACGCCCCCCGTGGCGGCGACG | 130.6 | -1.8 | 0.9 | -2.1 | 3.8E-02 | 4.3E-02 | BT20/MB468 | CYTO |
| >MI0000064\|hsa-let-7c&WithFlank&21\|+\|16539822\|16539917@17.38.22&[MIMAT0000064&hsa-let-7c-5p&offsets\|0\|0;m-37&21\|+\|16539838\|16539859&offsets\|0\|0]\|TGAGGTAGTAGGTTGTATGGTT | >MI0000064\|hsa-let-7c&WithFlank&21\|+\|16539822\|16539917 | 17 | 38 | 22 | TGAGGTAGTAGGTTGTATGGTT | 447.3 | -1.8 | 0.7 | -2.4 | 1.4E-02 | 4.3E-02 | BT20/MB231 | MITO |
| >hg38_21_+_8433172_8446622_RNA45SN1_45S_50ntFlanks@4731.4758.28\|AAGAACGAAAGTCGGAGGTTCGAAGACG | >hg38_21_+_8433172_8446622_RNA45SN1_45S_50ntFlanks | 4731 | 4758 | 28 | AAGAACGAAAGTCGGAGGTTCGAAGACG | 14407.9 | -2.2 | 1.0 | -2.2 | 2.7E-02 | 4.3E-02 | BT20/MB468 | MP |
| >trnaMT_LysTTT_MT_+_8295_8364@1.43.43,trnalookalike7_LysTTT_1_+_568844_568913@1.43.43\|CACTGTAAAGCTAACTTAGCATTAACCTTTTAAGTTAAAGATT | >trnaMT_LysTTT_MT_+_8295_8364 | 1 | 43 | 43 | CACTGTAAAGCTAACTTAGCATTAACCTTTTAAGTTAAAGATT | 331.7 | 2.2 | 1.1 | 2.1 | 4.0E-02 | 4.3E-02 | MB231/MB468 | MITO |
| >hg38_21_+_8433172_8446622_RNA45SN1_45S_50ntFlanks@12409.12443.35\|TTGATCCTTCGATGTCGGCTCTTCCTATCATTGTG | >hg38_21_+_8433172_8446622_RNA45SN1_45S_50ntFlanks | 12409 | 12443 | 35 | TTGATCCTTCGATGTCGGCTCTTCCTATCATTGTG | 595.6 | -1.1 | 0.5 | -2.1 | 3.9E-02 | 4.3E-02 | MB231/MB468 | NUC |
| >hg38_21_+_8433172_8446622_RNA45SN1_45S_50ntFlanks@9597.9614.18\|AGGGGCGAAAGACTAATC | >hg38_21_+_8433172_8446622_RNA45SN1_45S_50ntFlanks | 9597 | 9614 | 18 | AGGGGCGAAAGACTAATC | 300.8 | -3.0 | 1.4 | -2.2 | 2.7E-02 | 4.3E-02 | BT20/MB468 | MP |

<table>
<tr><th></th><th></th><th></th><th></th><th></th><th></th><th></th><th></th><th></th><th></th><th></th><th></th><th></th><th></th></tr>
<tr><td>>MI0000085|hsa-mir-27a&WithFlank&19|-|13836434|13836523@57.75.19(+1U)&[MIMAT0000084&hsa-miR-27a-3p&offsets|0|-2(+1U);m-43&19|-|13836447|13836467&offsets|0|-2(+1U)]|TTCACAGTGGCTAAGTTCCT</td><td>>MI0000085|hsa-mir-27a&WithFlank&19|-|13836434|13836523</td><td>57</td><td>75</td><td>19(+1U)</td><td>TTCACAGTGGCTAAGTTCCT</td><td>168.7</td><td>2.5</td><td>1.1</td><td>2.2</td><td>2.7E-02</td><td>4.3E-02</td><td>BT20/MB468</td><td>MP</td></tr>
<tr><td>>hg38_21_+_8433172_8446622_RNA45SN1_45S_50ntFlanks@8182.8209.28|ATCGAGGCCCAGCCCGTGGACGGTGTGA</td><td>>hg38_21_+_8433172_8446622_RNA45SN1_45S_50ntFlanks</td><td>8182</td><td>8209</td><td>28</td><td>ATCGAGGCCCAGCCCGTGGACGGTGTGA</td><td>403.2</td><td>-3.6</td><td>1.8</td><td>-2.1</td><td>3.9E-02</td><td>4.3E-02</td><td>MB231/MB468</td><td>NUC</td></tr>
<tr><td>>hg38_21_+_8433172_8446622_RNA45SN1_45S_50ntFlanks@10420.10456.37|CTGAGAGATGGGCGAGCGCCGTTCCGAAGGGACGGGC</td><td>>hg38_21_+_8433172_8446622_RNA45SN1_45S_50ntFlanks</td><td>10420</td><td>10456</td><td>37</td><td>CTGAGAGATGGGCGAGCGCCGTTCCGAAGGGACGGGC</td><td>5001.1</td><td>-1.8</td><td>0.9</td><td>-2.1</td><td>3.8E-02</td><td>4.3E-02</td><td>MB231/MB468</td><td>CYTO</td></tr>
<tr><td>>hg38_21_+_8433172_8446622_RNA45SN1_45S_50ntFlanks@5485.5516.32|AGCGCTGAGAAGACGGTCGAACTTGACTATCT</td><td>>hg38_21_+_8433172_8446622_RNA45SN1_45S_50ntFlanks</td><td>5485</td><td>5516</td><td>32</td><td>AGCGCTGAGAAGACGGTCGAACTTGACTATCT</td><td>2999.0</td><td>1.4</td><td>0.7</td><td>2.1</td><td>3.8E-02</td><td>4.3E-02</td><td>MB231/MB468</td><td>CYTO</td></tr>
<tr><td>>hg38_21_+_8433172_8446622_RNA45SN1_45S_50ntFlanks@10807.10829.23|ACAATGTAGGTAAGGGAAGTCGG</td><td>>hg38_21_+_8433172_8446622_RNA45SN1_45S_50ntFlanks</td><td>10807</td><td>10829</td><td>23</td><td>ACAATGTAGGTAAGGGAAGTCGG</td><td>151.7</td><td>-2.3</td><td>1.1</td><td>-2.1</td><td>3.8E-02</td><td>4.3E-02</td><td>MB231/MB468</td><td>CYTO</td></tr>
<tr><td>>hg38_21_+_8433172_8446622_RNA45SN1_45S_50ntFlanks@5491.5509.19|GAGAAGACGGTCGAACTTG</td><td>>hg38_21_+_8433172_8446622_RNA45SN1_45S_50ntFlanks</td><td>5491</td><td>5509</td><td>19</td><td>GAGAAGACGGTCGAACTTG</td><td>181.6</td><td>3.3</td><td>1.3</td><td>2.5</td><td>1.2E-02</td><td>4.3E-02</td><td>BT20/MB231</td><td>MP</td></tr>
<tr><td>>hg38_21_+_8433172_8446622_RNA45SN1_45S_50ntFlanks@8105.8142.38|CGCGGCGGGGCGCGGGACATGTGGCGTACGGAAGACCC</td><td>>hg38_21_+_8433172_8446622_RNA45SN1_45S_50ntFlanks</td><td>8105</td><td>8142</td><td>38</td><td>CGCGGCGGGGCGCGGGACATGTGGCGTACGGAAGACCC</td><td>295.7</td><td>5.3</td><td>2.1</td><td>2.5</td><td>1.2E-02</td><td>4.3E-02</td><td>BT20/MB231</td><td>MP</td></tr>
<tr><td>>hg38_21_+_8433172_8446622_RNA45SN1_45S_50ntFlanks@12934.12960.27|CTAAACCATTCGTAGACGACCTGCTTC</td><td>>hg38_21_+_8433172_8446622_RNA45SN1_45S_50ntFlanks</td><td>12934</td><td>12960</td><td>27</td><td>CTAAACCATTCGTAGACGACCTGCTTC</td><td>311.8</td><td>-1.2</td><td>0.6</td><td>-2.1</td><td>3.4E-02</td><td>4.3E-02</td><td>BT20/MB468</td><td>MITO</td></tr>
<tr><td>>hg38_21_+_8433172_8446622_RNA45SN1_45S_50ntFlanks@11664.11685.22|TTTCTGCCCAGTGCTCTGAATG</td><td>>hg38_21_+_8433172_8446622_RNA45SN1_45S_50ntFlanks</td><td>11664</td><td>11685</td><td>22</td><td>TTTCTGCCCAGTGCTCTGAATG</td><td>334.6</td><td>-2.2</td><td>1.0</td><td>-2.2</td><td>2.7E-02</td><td>4.3E-02</td><td>BT20/MB468</td><td>MP</td></tr>
<tr><td>>hg38_21_+_8433172_8446622_RNA45SN1_45S_50ntFlanks@11667.11691.25|CTGCCCAGTGCTCTGAATGTCAAAG</td><td>>hg38_21_+_8433172_8446622_RNA45SN1_45S_50ntFlanks</td><td>11667</td><td>11691</td><td>25</td><td>CTGCCCAGTGCTCTGAATGTCAAAG</td><td>198.7</td><td>4.0</td><td>1.8</td><td>2.2</td><td>2.7E-02</td><td>4.3E-02</td><td>BT20/MB468</td><td>MP</td></tr>
<tr><td>>hg38_21_+_8433172_8446622_RNA45SN1_45S_50ntFlanks@7978.8012.35|GACCTCAGATCAGACGTGGCGACCCGCTGAATTTA</td><td>>hg38_21_+_8433172_8446622_RNA45SN1_45S_50ntFlanks</td><td>7978</td><td>8012</td><td>35</td><td>GACCTCAGATCAGACGTGGCGACCCGCTGAATTTA</td><td>206.0</td><td>1.7</td><td>0.8</td><td>2.1</td><td>3.9E-02</td><td>4.3E-02</td><td>MB231/MB468</td><td>NUC</td></tr>
<tr><td>>trnaMT_HisGTG_MT_+_12138_12206@42.71.30|AACAGAGGCTTACGACCCCTTATTTACCCC</td><td>>trnaMT_HisGTG_MT_+_12138_12206</td><td>42</td><td>71</td><td>30</td><td>AACAGAGGCTTACGACCCCTTATTTACCCC</td><td>166.3</td><td>4.5</td><td>2.0</td><td>2.2</td><td>2.7E-02</td><td>4.3E-02</td><td>BT20/MB468</td><td>MP</td></tr>
<tr><td>>hg38_21_+_8433172_8446622_RNA45SN1_45S_50ntFlanks@483.506.24|TCGATGTGGTGACGTCGTGCTCTC</td><td>>hg38_21_+_8433172_8446622_RNA45SN1_45S_50ntFlanks</td><td>483</td><td>506</td><td>24</td><td>TCGATGTGGTGACGTCGTGCTCTC</td><td>226.7</td><td>1.7</td><td>0.8</td><td>2.1</td><td>3.9E-02</td><td>4.3E-02</td><td>BT20/MB468</td><td>NUC</td></tr>
<tr><td>>hg38_21_+_8433172_8446622_RNA45SN1_45S_50ntFlanks@5492.5524.33|AGAAGACGGTCGAACTTGACTATCTAGAGGAAG</td><td>>hg38_21_+_8433172_8446622_RNA45SN1_45S_50ntFlanks</td><td>5492</td><td>5524</td><td>33</td><td>AGAAGACGGTCGAACTTGACTATCTAGAGGAAG</td><td>221.9</td><td>-3.0</td><td>1.4</td><td>-2.2</td><td>2.7E-02</td><td>4.3E-02</td><td>BT20/MB468</td><td>MP</td></tr>
<tr><td>>hg38_21_+_8433172_8446622_RNA45SN1_45S_50ntFlanks@11540.11562.23|CGCGGCGCCTCGCCTCGGCCGGC</td><td>>hg38_21_+_8433172_8446622_RNA45SN1_45S_50ntFlanks</td><td>11540</td><td>11562</td><td>23</td><td>CGCGGCGCCTCGCCTCGGCCGGC</td><td>199.3</td><td>-4.3</td><td>1.8</td><td>-2.4</td><td>1.6E-02</td><td>4.3E-02</td><td>BT20/MB231</td><td>CYTO</td></tr>
<tr><td>>MI0000746|hsa-mir-99b&WithFlank&19|+|51692606|51692687@51.70.20&[MIMAT0004678&hsa-miR-99b-3p&offsets|0|-2;m-151&19|+|51692656|51692678&offsets|0|-3]|CAAGCTCGTGTCTGTGGGTC</td><td>>MI0000746|hsa-mir-99b&WithFlank&19|+|51692606|51692687</td><td>51</td><td>70</td><td>20</td><td>CAAGCTCGTGTCTGTGGGTC</td><td>93.1</td><td>1.9</td><td>0.9</td><td>2.1</td><td>3.6E-02</td><td>4.3E-02</td><td>MB231/MB468</td><td>MP</td></tr>
<tr><td>>hg38_21_+_8433172_8446622_RNA45SN1_45S_50ntFlanks@12247.12268.22|GGCAAAAGCTCGCTTGATCTTG</td><td>>hg38_21_+_8433172_8446622_RNA45SN1_45S_50ntFlanks</td><td>12247</td><td>12268</td><td>22</td><td>GGCAAAAGCTCGCTTGATCTTG</td><td>63.9</td><td>-3.8</td><td>1.7</td><td>-2.2</td><td>2.7E-02</td><td>4.3E-02</td><td>BT20/MB468</td><td>MP</td></tr>
<tr><td>>hg38_MT_+_1621_3279_RNR2_16S_50ntFlanks@114.136.23|CAAATAAAGTATAGGCGATAGAA</td><td>>hg38_MT_+_1621_3279_RNR2_16S_50ntFlanks</td><td>114</td><td>136</td><td>23</td><td>CAAATAAAGTATAGGCGATAGAA</td><td>68.8</td><td>3.7</td><td>1.8</td><td>2.1</td><td>3.6E-02</td><td>4.3E-02</td><td>MB231/MB468</td><td>MP</td></tr>
<tr><td>>hg38_21_+_8433172_8446622_RNA45SN1_45S_50ntFlanks@11702.11728.27|CAATGAAGCGCGGGTAAACGGCGGGAG</td><td>>hg38_21_+_8433172_8446622_RNA45SN1_45S_50ntFlanks</td><td>11702</td><td>11728</td><td>27</td><td>CAATGAAGCGCGGGTAAACGGCGGGAG</td><td>177.4</td><td>-3.9</td><td>1.8</td><td>-2.2</td><td>2.7E-02</td><td>4.3E-02</td><td>BT20/MB468</td><td>MP</td></tr>
<tr><td>>hg38_21_+_8433172_8446622_RNA45SN1_45S_50ntFlanks@6746.6771.26|CACTTCGAACGCACTTGCGGCCCCGG</td><td>>hg38_21_+_8433172_8446622_RNA45SN1_45S_50ntFlanks</td><td>6746</td><td>6771</td><td>26</td><td>CACTTCGAACGCACTTGCGGCCCCGG</td><td>1316.3</td><td>-4.2</td><td>1.9</td><td>-2.2</td><td>2.7E-02</td><td>4.3E-02</td><td>BT20/MB468</td><td>MP</td></tr>
<tr><td>>hg38_21_+_8433172_8446622_RNA45SN1_45S_50ntFlanks@6648.6695.48|GTACGACTCTTAGCGGTGGATCACTCGGCTCGTGCGTCGATGAAGAAC</td><td>>hg38_21_+_8433172_8446622_RNA45SN1_45S_50ntFlanks</td><td>6648</td><td>6695</td><td>48</td><td>GTACGACTCTTAGCGGTGGATCACTCGGCTCGTGCGTCGATGAAGAAC</td><td>227.6</td><td>4.1</td><td>1.8</td><td>2.2</td><td>2.7E-02</td><td>4.4E-02</td><td>BT20/MB468</td><td>MP</td></tr>
<tr><td>>hg38_21_+_8433172_8446622_RNA45SN1_45S_50ntFlanks@3843.3879.37|CTACTTGGATAACTGTGGTAATTCTAGAGCTAATACA</td><td>>hg38_21_+_8433172_8446622_RNA45SN1_45S_50ntFlanks</td><td>3843</td><td>3879</td><td>37</td><td>CTACTTGGATAACTGTGGTAATTCTAGAGCTAATACA</td><td>62.8</td><td>-4.7</td><td>2.2</td><td>-2.1</td><td>3.6E-02</td><td>4.4E-02</td><td>MB231/MB468</td><td>MP</td></tr>
<tr><td>>hg38_21_+_8433172_8446622_RNA45SN1_45S_50ntFlanks@11922.11940.19|GAGAGGTGTAGAATAAGTG</td><td>>hg38_21_+_8433172_8446622_RNA45SN1_45S_50ntFlanks</td><td>11922</td><td>11940</td><td>19</td><td>GAGAGGTGTAGAATAAGTG</td><td>118.0</td><td>4.3</td><td>1.7</td><td>2.5</td><td>1.2E-02</td><td>4.4E-02</td><td>BT20/MB231</td><td>MP</td></tr>
<tr><td>>MI0003834|hsa-mir-769&WithFlank&19|+|46018926|46019055@36.56.21(+1A)&[MIMAT0003886&hsa-miR-769-5p&offsets|0|-1(+1A);m-216&19|+|46018961|46018982&offsets|0|-1(+1A)]|TGAGACCTCTGGGTTCTGAGCA</td><td>>MI0003834|hsa-mir-769&WithFlank&19|+|46018926|46019055</td><td>36</td><td>56</td><td>21(+1A)</td><td>TGAGACCTCTGGGTTCTGAGCA</td><td>174.4</td><td>2.8</td><td>1.3</td><td>2.1</td><td>3.6E-02</td><td>4.4E-02</td><td>MB231/MB468</td><td>MP</td></tr>
<tr><td>>hg38_21_+_8433172_8446622_RNA45SN1_45S_50ntFlanks@9546.9572.27|TGGAGGTCCGTAGCGGTCCTGACGTGC</td><td>>hg38_21_+_8433172_8446622_RNA45SN1_45S_50ntFlanks</td><td>9546</td><td>9572</td><td>27</td><td>TGGAGGTCCGTAGCGGTCCTGACGTGC</td><td>627.6</td><td>3.7</td><td>1.5</td><td>2.5</td><td>1.2E-02</td><td>4.4E-02</td><td>BT20/MB231</td><td>MP</td></tr>
<tr><td>>trnaMT_GluTTC_MT_-_14674_14742@38.61.24,trnalookalike8_GluTTC_5_-_93905172_93905240@38.61.24|CATTGGTCGTGGTTGTAGTCCGTG</td><td>>trnaMT_GluTTC_MT_-_14674_14742</td><td>38</td><td>61</td><td>24</td><td>CATTGGTCGTGGTTGTAGTCCGTG</td><td>108.5</td><td>1.6</td><td>0.8</td><td>2.1</td><td>3.6E-02</td><td>4.4E-02</td><td>MB231/MB468</td><td>MP</td></tr>
<tr><td>>hg38_21_+_8433172_8446622_RNA45SN1_45S_50ntFlanks@4555.4572.18|CGCAGCTAGGAATAATGG</td><td>>hg38_21_+_8433172_8446622_RNA45SN1_45S_50ntFlanks</td><td>4555</td><td>4572</td><td>18</td><td>CGCAGCTAGGAATAATGG</td><td>457.4</td><td>-1.5</td><td>0.7</td><td>-2.1</td><td>3.8E-02</td><td>4.4E-02</td><td>BT20/MB468</td><td>CYTO</td></tr>
<tr><td>>hg38_21_+_8433172_8446622_RNA45SN1_45S_50ntFlanks@10921.10946.26|GCTGGACGAGGCGCCGCCGCCCCCCC</td><td>>hg38_21_+_8433172_8446622_RNA45SN1_45S_50ntFlanks</td><td>10921</td><td>10946</td><td>26</td><td>GCTGGACGAGGCGCCGCCGCCCCCCC</td><td>601.6</td><td>-4.3</td><td>2.1</td><td>-2.1</td><td>3.9E-02</td><td>4.4E-02</td><td>BT20/MB468</td><td>NUC</td></tr>
<tr><td>>hg38_21_+_8433172_8446622_RNA45SN1_45S_50ntFlanks@11322.11340.19|ACGTCTCGTCGCGCGCGCG</td><td>>hg38_21_+_8433172_8446622_RNA45SN1_45S_50ntFlanks</td><td>11322</td><td>11340</td><td>19</td><td>ACGTCTCGTCGCGCGCGCG</td><td>537.8</td><td>-2.9</td><td>1.4</td><td>-2.1</td><td>4.0E-02</td><td>4.4E-02</td><td>MB231/MB468</td><td>NUC</td></tr>
<tr><td>>MI0000100|hsa-mir-98&WithFlank&X|-|53556217|53556347@28.49.22(+1U)&[MIMAT0000096&hsa-miR-98-5p&offsets|0|0(+1U);m-143&X|-|53556299|53556320&offsets|0|0(+1U)]|TGAGGTAGTAAGTTGTATTGTTT</td><td>>MI0000100|hsa-mir-98&WithFlank&X|-|53556217|53556347</td><td>28</td><td>49</td><td>22(+1U)</td><td>TGAGGTAGTAAGTTGTATTGTTT</td><td>140.9</td><td>2.7</td><td>1.3</td><td>2.1</td><td>3.6E-02</td><td>4.4E-02</td><td>MB231/MB468</td><td>MP</td></tr>
<tr><td>>hg38_21_+_8433172_8446622_RNA45SN1_45S_50ntFlanks@5082.5111.30|AACGAACGAGACTCTGGCATGCTAACTAGT</td><td>>hg38_21_+_8433172_8446622_RNA45SN1_45S_50ntFlanks</td><td>5082</td><td>5111</td><td>30</td><td>AACGAACGAGACTCTGGCATGCTAACTAGT</td><td>155.0</td><td>3.9</td><td>1.6</td><td>2.5</td><td>1.2E-02</td><td>4.4E-02</td><td>BT20/MB231</td><td>MP</td></tr>
<tr><td>>hg38_21_+_8433172_8446622_RNA45SN1_45S_50ntFlanks@12418.12438.21|CGATGTCGGCTCTTCCTATCA</td><td>>hg38_21_+_8433172_8446622_RNA45SN1_45S_50ntFlanks</td><td>12418</td><td>12438</td><td>21</td><td>CGATGTCGGCTCTTCCTATCA</td><td>1032.2</td><td>1.4</td><td>0.7</td><td>2.1</td><td>3.8E-02</td><td>4.4E-02</td><td>MB231/MB468</td><td>CYTO</td></tr>
<tr><td>>hg38_21_+_8433172_8446622_RNA45SN1_45S_50ntFlanks@6644.6664.21|CCTCGTACGACTCTTAGCGGT</td><td>>hg38_21_+_8433172_8446622_RNA45SN1_45S_50ntFlanks</td><td>6644</td><td>6664</td><td>21</td><td>CCTCGTACGACTCTTAGCGGT</td><td>51.1</td><td>1.7</td><td>0.8</td><td>2.1</td><td>3.6E-02</td><td>4.4E-02</td><td>MB231/MB468</td><td>MP</td></tr>
<tr><td>>hg38_21_+_8433172_8446622_RNA45SN1_45S_50ntFlanks@11561.11589.29|GCGCCTAGCAGCCGACTTAGAACTGGTGC</td><td>>hg38_21_+_8433172_8446622_RNA45SN1_45S_50ntFlanks</td><td>11561</td><td>11589</td><td>29</td><td>GCGCCTAGCAGCCGACTTAGAACTGGTGC</td><td>232.2</td><td>-3.1</td><td>1.4</td><td>-2.2</td><td>2.7E-02</td><td>4.4E-02</td><td>BT20/MB468</td><td>MP</td></tr>
<tr><td>>hg38_21_+_8433172_8446622_RNA45SN1_45S_50ntFlanks@12420.12439.20|ATGTCGGCTCTTCCTATCAT</td><td>>hg38_21_+_8433172_8446622_RNA45SN1_45S_50ntFlanks</td><td>12420</td><td>12439</td><td>20</td><td>ATGTCGGCTCTTCCTATCAT</td><td>148.8</td><td>-3.7</td><td>1.8</td><td>-2.1</td><td>3.6E-02</td><td>4.4E-02</td><td>MB231/MB468</td><td>MP</td></tr>
<tr><td>>MI0000483|hsa-mir-186&WithFlank&1|-|71067625|71067722@21.41.21(+1A)&[MIMAT0000456&hsa-miR-186-5p&offsets|0|-1(+1A);m-84&1|-|71067681|71067702&offsets|0|-1(+1A)]|CAAAGAATTCTCCTTTTGGGCA</td><td>>MI0000483|hsa-mir-186&WithFlank&1|-|71067625|71067722</td><td>21</td><td>41</td><td>21(+1A)</td><td>CAAAGAATTCTCCTTTTGGGCA</td><td>348.3</td><td>-2.4</td><td>1.0</td><td>-2.5</td><td>1.2E-02</td><td>4.4E-02</td><td>BT20/MB231</td><td>MP</td></tr>
<tr><td>>MI0000811|hsa-mir-148b&WithFlank&12|+|54337210|54337320@30.51.22(+1U)&[MIMAT0004699&hsa-miR-148b-5p&offsets|-1|-1(+1U);m-371&12|+|54337239|54337262&offsets|0|-2(+1U)]|GAAGTTCTGTTATACACTCAGGT</td><td>>MI0000811|hsa-mir-148b&WithFlank&12|+|54337210|54337320</td><td>30</td><td>51</td><td>22(+1U)</td><td>GAAGTTCTGTTATACACTCAGGT</td><td>61.2</td><td>-5.0</td><td>2.0</td><td>-2.5</td><td>1.2E-02</td><td>4.4E-02</td><td>BT20/MB231</td><td>MP</td></tr>
<tr><td>>MI0000063|hsa-let-7b&WithFlank&22|+|46113680|46113774@12.33.22(+2A)&[MIMAT0000063&hsa-let-7b-5p&offsets|0|0(+2A);m-14&22|+|46113691|46113712&offsets|0|0(+2A)]|TGAGGTAGTAGGTTGTGTGGTTAA</td><td>>MI0000063|hsa-let-7b&WithFlank&22|+|46113680|46113774</td><td>12</td><td>33</td><td>22(+2A)</td><td>TGAGGTAGTAGGTTGTGTGGTTAA</td><td>130.0</td><td>2.8</td><td>1.3</td><td>2.1</td><td>3.6E-02</td><td>4.4E-02</td><td>MB231/MB468</td><td>MP</td></tr>
<tr><td>>hg38_21_+_8433172_8446622_RNA45SN1_45S_50ntFlanks@6650.6683.34|ACGACTCTTAGCGGTGGATCACTCGGCTCGTGCG</td><td>>hg38_21_+_8433172_8446622_RNA45SN1_45S_50ntFlanks</td><td>6650</td><td>6683</td><td>34</td><td>ACGACTCTTAGCGGTGGATCACTCGGCTCGTGCG</td><td>6135.6</td><td>-1.9</td><td>0.8</td><td>-2.5</td><td>1.3E-02</td><td>4.4E-02</td><td>BT20/MB231</td><td>NUC</td></tr>
<tr><td>>hg38_MT_+_1621_3279_RNR2_16S_50ntFlanks@51.88.38|GCTAAACCTAGCCCCAAACCCACTCCACCTTACTACCA</td><td>>hg38_MT_+_1621_3279_RNR2_16S_50ntFlanks</td><td>51</td><td>88</td><td>38</td><td>GCTAAACCTAGCCCCAAACCCACTCCACCTTACTACCA</td><td>11747.6</td><td>-2.2</td><td>0.9</td><td>-2.4</td><td>1.5E-02</td><td>4.4E-02</td><td>BT20/MB231</td><td>MITO</td></tr>
<tr><td>>hg38_21_+_8433172_8446622_RNA45SN1_45S_50ntFlanks@6646.6681.36|TCGTACGACTCTTAGCGGTGGATCACTCGGCTCGTG</td><td>>hg38_21_+_8433172_8446622_RNA45SN1_45S_50ntFlanks</td><td>6646</td><td>6681</td><td>36</td><td>TCGTACGACTCTTAGCGGTGGATCACTCGGCTCGTG</td><td>7644.5</td><td>3.2</td><td>1.3</td><td>2.5</td><td>1.2E-02</td><td>4.4E-02</td><td>BT20/MB231</td><td>MP</td></tr>
<tr><td>>hg38_21_+_8433172_8446622_RNA45SN1_45S_50ntFlanks@4225.4242.18|AAATAACAATACAGGACT</td><td>>hg38_21_+_8433172_8446622_RNA45SN1_45S_50ntFlanks</td><td>4225</td><td>4242</td><td>18</td><td>AAATAACAATACAGGACT</td><td>1145.6</td><td>-3.3</td><td>1.5</td><td>-2.2</td><td>2.7E-02</td><td>4.4E-02</td><td>BT20/MB468</td><td>MP</td></tr>
<tr><td>>MI0000088|hsa-mir-30a&WithFlank&6|-|71403545|71403627@13.34.22(+1G)&[MIMAT0000087&hsa-miR-30a-5p&offsets|+1|+1(+1G);m-15&6|-|71403595|71403616&offsets|+1|+1(+1G)]|GTAAACATCCTCGACTGGAAGCG</td><td>>MI0000088|hsa-mir-30a&WithFlank&6|-|71403545|71403627</td><td>13</td><td>34</td><td>22(+1G)</td><td>GTAAACATCCTCGACTGGAAGCG</td><td>67.9</td><td>3.5</td><td>1.7</td><td>2.1</td><td>3.7E-02</td><td>4.4E-02</td><td>MB231/MB468</td><td>MP</td></tr>
<tr><td>>hg38_21_+_8433172_8446622_RNA45SN1_45S_50ntFlanks@11567.11585.19|AGCAGCCGACTTAGAACTG</td><td>>hg38_21_+_8433172_8446622_RNA45SN1_45S_50ntFlanks</td><td>11567</td><td>11585</td><td>19</td><td>AGCAGCCGACTTAGAACTG</td><td>1260.9</td><td>-2.2</td><td>0.9</td><td>-2.5</td><td>1.3E-02</td><td>4.4E-02</td><td>BT20/MB231</td><td>NUC</td></tr>
<tr><td>>hg38_21_+_8433172_8446622_RNA45SN1_45S_50ntFlanks@8176.8215.40|CTTCTGATCGAGGCCCAGCCCGTGGACGGTGTGAGGCCGG</td><td>>hg38_21_+_8433172_8446622_RNA45SN1_45S_50ntFlanks</td><td>8176</td><td>8215</td><td>40</td><td>CTTCTGATCGAGGCCCAGCCCGTGGACGGTGTGAGGCCGG</td><td>332.9</td><td>-1.4</td><td>0.7</td><td>-2.1</td><td>3.4E-02</td><td>4.4E-02</td><td>BT20/MB468</td><td>MITO</td></tr>
<tr><td>>hg38_21_+_8433172_8446622_RNA45SN1_45S_50ntFlanks@4297.4321.25|CATTGGAGGGCAAGTCTGGTGCCAG</td><td>>hg38_21_+_8433172_8446622_RNA45SN1_45S_50ntFlanks</td><td>4297</td><td>4321</td><td>25</td><td>CATTGGAGGGCAAGTCTGGTGCCAG</td><td>238.8</td><td>-3.9</td><td>1.8</td><td>-2.2</td><td>2.7E-02</td><td>4.4E-02</td><td>BT20/MB468</td><td>MP</td></tr>
<tr><td>>hg38_MT_+_598_1651_RNR1_12S_50ntFlanks@968.1004.37|ACATGGTAAGTGTACTGGAAAGTGCACTTGGACGAAC</td><td>>hg38_MT_+_598_1651_RNR1_12S_50ntFlanks</td><td>968</td><td>1004</td><td>37</td><td>ACATGGTAAGTGTACTGGAAAGTGCACTTGGACGAAC</td><td>129.8</td><td>2.9</td><td>1.4</td><td>2.1</td><td>3.7E-02</td><td>4.4E-02</td><td>MB231/MB468</td><td>MP</td></tr>
<tr><td>>MI0000465|hsa-mir-191&WithFlank&3|-|49020612|49020715@22.43.22(+2U)&[MIMAT0000440&hsa-miR-191-5p&offsets|0|-1(+2U);m-48&3|-|49020672|49020694&offsets|0|-1(+2U)]|CAACGGAATCCCAAAAGCAGCTTT</td><td>>MI0000465|hsa-mir-191&WithFlank&3|-|49020612|49020715</td><td>22</td><td>43</td><td>22(+2U)</td><td>CAACGGAATCCCAAAAGCAGCTTT</td><td>228.4</td><td>-2.5</td><td>1.0</td><td>-2.5</td><td>1.2E-02</td><td>4.4E-02</td><td>BT20/MB231</td><td>MP</td></tr>
<tr><td>>trnaMT_MetCAT_MT_+_4402_4469@33.71.39|TACCCCGAAAATGTTGGTTATACCCTTCCCGTACTACCA</td><td>>trnaMT_MetCAT_MT_+_4402_4469</td><td>33</td><td>71</td><td>39</td><td>TACCCCGAAAATGTTGGTTATACCCTTCCCGTACTACCA</td><td>762.3</td><td>-2.2</td><td>0.9</td><td>-2.4</td><td>1.5E-02</td><td>4.4E-02</td><td>BT20/MB231</td><td>MITO</td></tr>
<tr><td>>am_MI0003639|hsa-mir-625&WithFlank&14|+|65471096|65471192@58.78.21&[MIMAT0004808&hsa-miR-625-3p&offsets|0|-1;m-137&14|+|65471153|65471174&offsets|0|-1]|GACTATAGAACTTTCCCCCTC</td><td>>am_MI0003639|hsa-mir-625&WithFlank&14|+|65471096|65471192</td><td>58</td><td>78</td><td>21</td><td>GACTATAGAACTTTCCCCCTC</td><td>172.7</td><td>-2.7</td><td>1.1</td><td>-2.5</td><td>1.2E-02</td><td>4.4E-02</td><td>BT20/MB231</td><td>MP</td></tr>
</table>

| | | | | | | | | | | | | | |
|---|---|---|---|---|---|---|---|---|---|---|---|---|---|
| >MI0000101\|hsa-mir-99a&WithFlank&21\|+\|16539083\|16539175@19.38.20(+1C)&[MIMAT0000097&hsa-miR-99a-5p&offsets\|0\|-2(+1C);m-64&21\|+\|16539101\|16539121&offsets\|0\|-1(+1C)]\|AACCCGTAGATCCGATCTTGC | >MI0000101\|hsa-mir-99a&WithFlank&21\|+\|16539083\|16539175 | 19 | 38 | 20(+1C) | AACCCGTAGATCCGATCTTGC | 712.8 | -1.7 | 0.8 | -2.1 | 3.7E-02 | 4.4E-02 | MB231/MB468 | MP |
| >hg38_21_+_8433172_8446622_RNA45SN1_45S_50ntFlanks@3845.3872.28\|ACTTGGATAACTGTGGTAATTCTAGAGC | >hg38_21_+_8433172_8446622_RNA45SN1_45S_50ntFlanks | 3845 | 3872 | 28 | ACTTGGATAACTGTGGTAATTCTAGAGC | 57.7 | -3.8 | 1.7 | -2.2 | 2.8E-02 | 4.4E-02 | BT20/MB468 | MP |
| >hg38_MT_+_598_1651_RNR1_12S_50ntFlanks@908.946.39\|ATACTTCAAAGGACATTTAACTAAAACCCCTACGCATTT | >hg38_MT_+_598_1651_RNR1_12S_50ntFlanks | 908 | 946 | 39 | ATACTTCAAAGGACATTTAACTAAAACCCCTACGCATTT | 65.8 | 3.5 | 1.7 | 2.1 | 3.7E-02 | 4.4E-02 | MB231/MB468 | MP |
| >hg38_21_+_8433172_8446622_RNA45SN1_45S_50ntFlanks@9929.9960.32\|GATATAGACAGCAGGACGGTGGCCATGGAAGT | >hg38_21_+_8433172_8446622_RNA45SN1_45S_50ntFlanks | 9929 | 9960 | 32 | GATATAGACAGCAGGACGGTGGCCATGGAAGT | 172.2 | 3.0 | 1.2 | 2.5 | 1.2E-02 | 4.4E-02 | BT20/MB231 | MP |
| >hg38_21_+_8433172_8446622_RNA45SN1_45S_50ntFlanks@12028.12055.28\|CTCTGATCGTTTTTTCACTGACCCGGTG | >hg38_21_+_8433172_8446622_RNA45SN1_45S_50ntFlanks | 12028 | 12055 | 28 | CTCTGATCGTTTTTTCACTGACCCGGTG | 310.5 | 4.3 | 1.7 | 2.5 | 1.2E-02 | 4.4E-02 | BT20/MB231 | MP |
| >hg38_21_+_8433172_8446622_RNA45SN1_45S_50ntFlanks@11582.11613.32\|ACTGGTGCGGACCAGGGGAATCCGACTGTTTA | >hg38_21_+_8433172_8446622_RNA45SN1_45S_50ntFlanks | 11582 | 11613 | 32 | ACTGGTGCGGACCAGGGGAATCCGACTGTTTA | 214.0 | 2.6 | 1.0 | 2.5 | 1.2E-02 | 4.4E-02 | BT20/MB231 | MP |
| >hg38_21_+_8433172_8446622_RNA45SN1_45S_50ntFlanks@9300.9335.36\|ACGGACCAAGGAGTCTAACACGTGCGCGAGTCGGGG | >hg38_21_+_8433172_8446622_RNA45SN1_45S_50ntFlanks | 9300 | 9335 | 36 | ACGGACCAAGGAGTCTAACACGTGCGCGAGTCGGGG | 56.9 | 2.1 | 0.9 | 2.2 | 2.8E-02 | 4.4E-02 | BT20/MB468 | MP |
| >hg38_21_+_8433172_8446622_RNA45SN1_45S_50ntFlanks@5446.5464.19\|CGGATCGGCCCCGCCGGGG | >hg38_21_+_8433172_8446622_RNA45SN1_45S_50ntFlanks | 5446 | 5464 | 19 | CGGATCGGCCCCGCCGGGG | 138.2 | -3.8 | 1.7 | -2.2 | 2.8E-02 | 4.4E-02 | BT20/MB468 | MP |
| >hg38_MT_+_1621_3279_RNR2_16S_50ntFlanks@1217.1248.32\|AACCTCCGAGCAGTACATGCTAAGACTTCACC | >hg38_MT_+_1621_3279_RNR2_16S_50ntFlanks | 1217 | 1248 | 32 | AACCTCCGAGCAGTACATGCTAAGACTTCACC | 126.1 | 2.9 | 1.4 | 2.1 | 3.7E-02 | 4.4E-02 | MB231/MB468 | MP |
| >hg38_21_+_8433172_8446622_RNA45SN1_45S_50ntFlanks@9602.9630.29\|CGAAAGACTAATCGAACCATCTAGTAGCT | >hg38_21_+_8433172_8446622_RNA45SN1_45S_50ntFlanks | 9602 | 9630 | 29 | CGAAAGACTAATCGAACCATCTAGTAGCT | 135.8 | 2.2 | 1.0 | 2.2 | 2.8E-02 | 4.4E-02 | BT20/MB468 | MP |
| >hg38_21_+_8433172_8446622_RNA45SN1_45S_50ntFlanks@4997.5024.28\|AGATTGATAGCTCTTTCTCGATTCCGTG | >hg38_21_+_8433172_8446622_RNA45SN1_45S_50ntFlanks | 4997 | 5024 | 28 | AGATTGATAGCTCTTTCTCGATTCCGTG | 66.2 | -4.1 | 1.9 | -2.2 | 2.8E-02 | 4.4E-02 | BT20/MB468 | MP |
| >hg38_21_+_8433172_8446622_RNA45SN1_45S_50ntFlanks@10685.10707.23\|GGCGGCGTCCGGTGAGCTCTCGC | >hg38_21_+_8433172_8446622_RNA45SN1_45S_50ntFlanks | 10685 | 10707 | 23 | GGCGGCGTCCGGTGAGCTCTCGC | 715.7 | 5.5 | 2.2 | 2.5 | 1.2E-02 | 4.4E-02 | BT20/MB231 | MP |
| >hg38_21_+_8433172_8446622_RNA45SN1_45S_50ntFlanks@10866.10904.39\|CTAAGGGCTGGGTCGGTCGGGCTGGGGCGCGAAGCGGGG | >hg38_21_+_8433172_8446622_RNA45SN1_45S_50ntFlanks | 10866 | 10904 | 39 | CTAAGGGCTGGGTCGGTCGGGCTGGGGCGCGAAGCGGGG | 312.6 | -3.5 | 1.7 | -2.1 | 3.9E-02 | 4.4E-02 | BT20/MB468 | CYTO |
| >hg38_21_+_8433172_8446622_RNA45SN1_45S_50ntFlanks@12907.12934.28\|CACCGCACGTTCGTGGGGAACCTGGCGC | >hg38_21_+_8433172_8446622_RNA45SN1_45S_50ntFlanks | 12907 | 12934 | 28 | CACCGCACGTTCGTGGGGAACCTGGCGC | 53.7 | -3.8 | 1.7 | -2.2 | 2.8E-02 | 4.4E-02 | BT20/MB468 | MP |
| >hg38_21_+_8433172_8446622_RNA45SN1_45S_50ntFlanks@5492.5523.32\|AGAAGACGGTCGAACTTGACTATCTAGAGGAA | >hg38_21_+_8433172_8446622_RNA45SN1_45S_50ntFlanks | 5492 | 5523 | 32 | AGAAGACGGTCGAACTTGACTATCTAGAGGAA | 115.0 | -4.0 | 1.8 | -2.2 | 2.8E-02 | 4.4E-02 | BT20/MB468 | MP |
| >hg38_21_+_8433172_8446622_RNA45SN1_45S_50ntFlanks@12242.12268.27\|AGAAGGGCAAAAGCTCGCTTGATCTTG | >hg38_21_+_8433172_8446622_RNA45SN1_45S_50ntFlanks | 12242 | 12268 | 27 | AGAAGGGCAAAAGCTCGCTTGATCTTG | 201.6 | -2.6 | 1.3 | -2.1 | 4.0E-02 | 4.4E-02 | MB231/MB468 | NUC |
| >hg38_21_+_8433172_8446622_RNA45SN1_45S_50ntFlanks@3996.4024.29\|ACTCTAGATAACCTCGGGCCGATCGCACG | >hg38_21_+_8433172_8446622_RNA45SN1_45S_50ntFlanks | 3996 | 4024 | 29 | ACTCTAGATAACCTCGGGCCGATCGCACG | 142.7 | -3.8 | 1.7 | -2.2 | 2.8E-02 | 4.4E-02 | BT20/MB468 | MP |
| >hg38_21_+_8433172_8446622_RNA45SN1_45S_50ntFlanks@10663.10706.44\|CCTTGGAAAGCGTCGCGGTTCCGGCGGCGTCCGGTGAGCTCTCG | >hg38_21_+_8433172_8446622_RNA45SN1_45S_50ntFlanks | 10663 | 10706 | 44 | CCTTGGAAAGCGTCGCGGTTCCGGCGGCGTCCGGTGAGCTCTCG | 67.3 | -4.2 | 1.9 | -2.2 | 2.8E-02 | 4.4E-02 | BT20/MB468 | MP |
| >hg38_21_+_8433172_8446622_RNA45SN1_45S_50ntFlanks@12942.12959.18\|TTCGTAGACGACCTGCTT | >hg38_21_+_8433172_8446622_RNA45SN1_45S_50ntFlanks | 12942 | 12959 | 18 | TTCGTAGACGACCTGCTT | 416.2 | -1.2 | 0.6 | -2.1 | 4.0E-02 | 4.4E-02 | MB231/MB468 | NUC |
| >hg38_21_+_8433172_8446622_RNA45SN1_45S_50ntFlanks@11217.11235.19\|CTGGACGCGAGCCGGGCCC | >hg38_21_+_8433172_8446622_RNA45SN1_45S_50ntFlanks | 11217 | 11235 | 19 | CTGGACGCGAGCCGGGCCC | 87.7 | -3.7 | 1.8 | -2.1 | 3.7E-02 | 4.4E-02 | MB231/MB468 | MP |
| >hg38_21_+_8433172_8446622_RNA45SN1_45S_50ntFlanks@10351.10370.20\|CGAGAACTTTGAAGGCCGAA | >hg38_21_+_8433172_8446622_RNA45SN1_45S_50ntFlanks | 10351 | 10370 | 20 | CGAGAACTTTGAAGGCCGAA | 203.9 | -1.3 | 0.6 | -2.1 | 3.5E-02 | 4.4E-02 | BT20/MB468 | MITO |
| >hg38_21_+_8433172_8446622_RNA45SN1_45S_50ntFlanks@9604.9623.20\|AAAGACTAATCGAACCATCT | >hg38_21_+_8433172_8446622_RNA45SN1_45S_50ntFlanks | 9604 | 9623 | 20 | AAAGACTAATCGAACCATCT | 171.6 | -1.3 | 0.6 | -2.1 | 3.9E-02 | 4.4E-02 | MB231/MB468 | CYTO |
| >trnaMT_GlnTTG_MT_-_4329_4400@1.30.30,trnalookalike2_GlnTTG_1_-_564879_564950@1.30.30\|TAGGATGGGGTGTGATAGGTGGCACGGAGA | >trnaMT_GlnTTG_MT_-_4329_4400 | 1 | 30 | 30 | TAGGATGGGGTGTGATAGGTGGCACGGAGA | 398.4 | 3.0 | 1.5 | 2.1 | 3.9E-02 | 4.4E-02 | BT20/MB468 | CYTO |
| >hg38_21_+_8433172_8446622_RNA45SN1_45S_50ntFlanks@11561.11586.26\|GCGCCTAGCAGCCGACTTAGAACTGG | >hg38_21_+_8433172_8446622_RNA45SN1_45S_50ntFlanks | 11561 | 11586 | 26 | GCGCCTAGCAGCCGACTTAGAACTGG | 1074.1 | -3.4 | 1.6 | -2.2 | 2.8E-02 | 4.4E-02 | BT20/MB468 | MP |
| >hg38_21_+_8433172_8446622_RNA45SN1_45S_50ntFlanks@12413.12436.24\|TCCTTCGATGTCGGCTCTTCCTAT | >hg38_21_+_8433172_8446622_RNA45SN1_45S_50ntFlanks | 12413 | 12436 | 24 | TCCTTCGATGTCGGCTCTTCCTAT | 210.8 | -2.6 | 1.2 | -2.1 | 3.5E-02 | 4.4E-02 | BT20/MB468 | MITO |
| >hg38_21_+_8433172_8446622_RNA45SN1_45S_50ntFlanks@13022.13039.18\|CTCGACACAAGGGTTTGT | >hg38_21_+_8433172_8446622_RNA45SN1_45S_50ntFlanks | 13022 | 13039 | 18 | CTCGACACAAGGGTTTGT | 280.7 | 1.6 | 0.7 | 2.2 | 2.8E-02 | 4.4E-02 | BT20/MB468 | MP |
| >hg38_21_+_8433172_8446622_RNA45SN1_45S_50ntFlanks@5436.5466.31\|GTGAGGCCCTCGGATCGGCCCCGCCGGGGTC | >hg38_21_+_8433172_8446622_RNA45SN1_45S_50ntFlanks | 5436 | 5466 | 31 | GTGAGGCCCTCGGATCGGCCCCGCCGGGGTC | 203.4 | -3.4 | 1.6 | -2.1 | 3.5E-02 | 4.4E-02 | BT20/MB468 | MITO |
| >hg38_21_+_8433172_8446622_RNA45SN1_45S_50ntFlanks@5237.5257.21\|ACTGACTGGCTCAGCGTGTGC | >hg38_21_+_8433172_8446622_RNA45SN1_45S_50ntFlanks | 5237 | 5257 | 21 | ACTGACTGGCTCAGCGTGTGC | 215.7 | 3.5 | 1.4 | 2.5 | 1.2E-02 | 4.4E-02 | BT20/MB231 | MP |
| >hg38_21_+_8433172_8446622_RNA45SN1_45S_50ntFlanks@12456.12496.41\|CCAAGCGTTGGATTGTTCACCCACTAATAGGGAACGTGAGC | >hg38_21_+_8433172_8446622_RNA45SN1_45S_50ntFlanks | 12456 | 12496 | 41 | CCAAGCGTTGGATTGTTCACCCACTAATAGGGAACGTGAGC | 449.9 | -1.7 | 0.8 | -2.1 | 3.5E-02 | 4.4E-02 | BT20/MB468 | MITO |
| >hg38_21_+_8433172_8446622_RNA45SN1_45S_50ntFlanks@7989.8017.29\|AGACGTGGCGACCCGCTGAATTTAAGCAT | >hg38_21_+_8433172_8446622_RNA45SN1_45S_50ntFlanks | 7989 | 8017 | 29 | AGACGTGGCGACCCGCTGAATTTAAGCAT | 667.5 | -1.5 | 0.7 | -2.1 | 4.0E-02 | 4.4E-02 | MB231/MB468 | MITO |
| >hg38_MT_+_1621_3279_RNR2_16S_50ntFlanks@1458.1477.20\|CGGAGTAATCCAGGTCGGTT | >hg38_MT_+_1621_3279_RNR2_16S_50ntFlanks | 1458 | 1477 | 20 | CGGAGTAATCCAGGTCGGTT | 104.3 | 3.5 | 1.7 | 2.1 | 3.7E-02 | 4.4E-02 | MB231/MB468 | MP |
| >hg38_MT_+_1621_3279_RNR2_16S_50ntFlanks@488.517.30\|GAACAGCTCTTTGGACACTAGGAAAAAACC | >hg38_MT_+_1621_3279_RNR2_16S_50ntFlanks | 488 | 517 | 30 | GAACAGCTCTTTGGACACTAGGAAAAAACC | 72.2 | 2.6 | 1.3 | 2.1 | 3.7E-02 | 4.4E-02 | MB231/MB468 | MP |
| >hg38_21_+_8433172_8446622_RNA45SN1_45S_50ntFlanks@9391.9412.22\|CCGAGGTGGGATCCCGAGGCCT | >hg38_21_+_8433172_8446622_RNA45SN1_45S_50ntFlanks | 9391 | 9412 | 22 | CCGAGGTGGGATCCCGAGGCCT | 97.7 | -3.8 | 1.7 | -2.2 | 2.8E-02 | 4.4E-02 | BT20/MB468 | MP |
| >MI0000749\|hsa-mir-30e&WithFlank&1\|+\|40754349\|40754452@24.46.23(+1U)&[MIMAT0000692&hsa-miR-30e-5p&offsets\|+1\|+2(+1U);m-24&1\|+\|40754371\|40754394&offsets\|+1\|0(+1U)]\|GTAAACATCCTTGACTGGAAGCTT | >MI0000749\|hsa-mir-30e&WithFlank&1\|+\|40754349\|40754452 | 24 | 46 | 23(+1U) | GTAAACATCCTTGACTGGAAGCTT | 172.6 | 3.6 | 1.6 | 2.2 | 2.8E-02 | 4.4E-02 | BT20/MB468 | MP |
| >hg38_21_+_8433172_8446622_RNA45SN1_45S_50ntFlanks@6779.6802.24\|CCGGGGCTACGCCTGTCTGAGCGT | >hg38_21_+_8433172_8446622_RNA45SN1_45S_50ntFlanks | 6779 | 6802 | 24 | CCGGGGCTACGCCTGTCTGAGCGT | 796.7 | 3.7 | 1.5 | 2.5 | 1.2E-02 | 4.4E-02 | BT20/MB231 | MP |
| >trnaMT_LysTTT_MT_+_8295_8364@36.72.37,trnalookalike7_LysTTT_1_+_568844_568913@36.72.37\|TAAAGATTAAGAGAACCAACACCTCTTTACAGTGACC | >trnaMT_LysTTT_MT_+_8295_8364 | 36 | 72 | 37 | TAAAGATTAAGAGAACCAACACCTCTTTACAGTGACC | 615.6 | -3.2 | 1.3 | -2.5 | 1.2E-02 | 4.4E-02 | BT20/MB231 | MP |
| >hg38_21_+_8433172_8446622_RNA45SN1_45S_50ntFlanks@6652.6688.37\|GACTCTTAGCGGTGGATCACTCGGCTCGTGCGTCGAT | >hg38_21_+_8433172_8446622_RNA45SN1_45S_50ntFlanks | 6652 | 6688 | 37 | GACTCTTAGCGGTGGATCACTCGGCTCGTGCGTCGAT | 2149.6 | 2.4 | 1.1 | 2.2 | 2.8E-02 | 4.4E-02 | BT20/MB468 | MP |
| >hg38_21_+_8433172_8446622_RNA45SN1_45S_50ntFlanks@5471.5516.46\|CACGGCCCTGGCGGAGCGCTGAGAAGACGGTCGAACTTGACTATCT | >hg38_21_+_8433172_8446622_RNA45SN1_45S_50ntFlanks | 5471 | 5516 | 46 | CACGGCCCTGGCGGAGCGCTGAGAAGACGGTCGAACTTGACTATCT | 2020.7 | 2.9 | 1.2 | 2.4 | 1.5E-02 | 4.4E-02 | BT20/MB231 | MITO |
| >MI0000253\|hsa-mir-148a&WithFlank&7\|-\|25949913\|25949992@50.70.21&[MIMAT0000243&hsa-miR-148a-3p&offsets\|0\|-1;m-7&7\|-\|25949922\|25949943&offsets\|0\|-1]\|TCAGTGCACTACAGAACTTTG | >MI0000253\|hsa-mir-148a&WithFlank&7\|-\|25949913\|25949992 | 50 | 70 | 21 | TCAGTGCACTACAGAACTTTG | 380.1 | 3.0 | 1.2 | 2.5 | 1.3E-02 | 4.4E-02 | BT20/MB231 | MP |
| >hg38_21_+_8433172_8446622_RNA45SN1_45S_50ntFlanks@7973.8004.32\|GACGCGACCTCAGATCAGACGTGGCGACCCGC | >hg38_21_+_8433172_8446622_RNA45SN1_45S_50ntFlanks | 7973 | 8004 | 32 | GACGCGACCTCAGATCAGACGTGGCGACCCGC | 199.8 | 1.6 | 0.8 | 2.0 | 4.0E-02 | 4.4E-02 | MB231/MB468 | MITO |
| >hg38_21_+_8433172_8446622_RNA45SN1_45S_50ntFlanks@9927.9946.20\|TTGATATAGACAGCAGGACG | >hg38_21_+_8433172_8446622_RNA45SN1_45S_50ntFlanks | 9927 | 9946 | 20 | TTGATATAGACAGCAGGACG | 85.7 | 3.9 | 1.6 | 2.5 | 1.3E-02 | 4.4E-02 | BT20/MB231 | MP |
| >hg38_MT_+_1621_3279_RNR2_16S_50ntFlanks@1390.1415.26\|GATGGTGCAGCCGCTATTAAAGGTTC | >hg38_MT_+_1621_3279_RNR2_16S_50ntFlanks | 1390 | 1415 | 26 | GATGGTGCAGCCGCTATTAAAGGTTC | 256.4 | -3.9 | 1.6 | -2.5 | 1.3E-02 | 4.4E-02 | BT20/MB231 | MP |
| >MI0000465\|hsa-mir-191&WithFlank&3\|-\|49020612\|49020715@22.42.21(+1A)&[MIMAT0000440&hsa-miR-191-5p&offsets\|0\|-2(+1A);m-48&3\|-\|49020672\|49020694&offsets\|0\|-2(+1A)]\|CAACGGAATCCCAAAAGCAGCA | >MI0000465\|hsa-mir-191&WithFlank&3\|-\|49020612\|49020715 | 22 | 42 | 21(+1A) | CAACGGAATCCCAAAAGCAGCA | 1314.7 | 2.2 | 0.9 | 2.4 | 1.7E-02 | 4.4E-02 | BT20/MB231 | CYTO |
| >MI0003560\|hsa-mir-92b&WithFlank&1\|+\|155195171\|155195278@67.88.22(+1A)&[MIMAT0003218&hsa-miR-92b-3p&offsets\|0\|0(+1A);m-115&1\|+\|155195237\|155195258&offsets\|0\|0(+1A)]\|TATTGCACTCGTCCCGGCCTCCA | >MI0003560\|hsa-mir-92b&WithFlank&1\|+\|155195171\|155195278 | 67 | 88 | 22(+1A) | TATTGCACTCGTCCCGGCCTCCA | 702.1 | 1.8 | 0.8 | 2.4 | 1.7E-02 | 4.4E-02 | BT20/MB231 | CYTO |
| >hg38_21_+_8433172_8446622_RNA45SN1_45S_50ntFlanks@12416.12444.29\|TTCGATGTCGGCTCTTCCTATCATTGTGA | >hg38_21_+_8433172_8446622_RNA45SN1_45S_50ntFlanks | 12416 | 12444 | 29 | TTCGATGTCGGCTCTTCCTATCATTGTGA | 3900.8 | -2.1 | 0.9 | -2.4 | 1.7E-02 | 4.4E-02 | BT20/MB231 | CYTO |
| >hg38_21_+_8433172_8446622_RNA45SN1_45S_50ntFlanks@7990.8019.30\|GACGTGGCGACCCGCTGAATTTAAGCATAT | >hg38_21_+_8433172_8446622_RNA45SN1_45S_50ntFlanks | 7990 | 8019 | 30 | GACGTGGCGACCCGCTGAATTTAAGCATAT | 970.8 | -2.6 | 1.1 | -2.4 | 1.7E-02 | 4.4E-02 | BT20/MB231 | CYTO |
| >hg38_21_+_8433172_8446622_RNA45SN1_45S_50ntFlanks@12640.12669.30\|ACCATCTGTGGGATTATGACTGAACGCCTC | >hg38_21_+_8433172_8446622_RNA45SN1_45S_50ntFlanks | 12640 | 12669 | 30 | ACCATCTGTGGGATTATGACTGAACGCCTC | 190.3 | 2.2 | 1.0 | 2.2 | 2.8E-02 | 4.4E-02 | BT20/MB468 | MP |
| >hg38_21_+_8433172_8446622_RNA45SN1_45S_50ntFlanks@6647.6690.44\|CGTACGACTCTTAGCGGTGGATCACTCGGCTCGTGCGTCGATGA | >hg38_21_+_8433172_8446622_RNA45SN1_45S_50ntFlanks | 6647 | 6690 | 44 | CGTACGACTCTTAGCGGTGGATCACTCGGCTCGTGCGTCGATGA | 199.2 | 3.4 | 1.5 | 2.2 | 2.8E-02 | 4.4E-02 | BT20/MB468 | MP |
| >hg38_21_+_8433172_8446622_RNA45SN1_45S_50ntFlanks@6650.6669.20\|ACGACTCTTAGCGGTGGATC | >hg38_21_+_8433172_8446622_RNA45SN1_45S_50ntFlanks | 6650 | 6669 | 20 | ACGACTCTTAGCGGTGGATC | 1908.6 | 3.3 | 1.3 | 2.5 | 1.3E-02 | 4.4E-02 | BT20/MB231 | MP |
| >hg38_MT_+_598_1651_RNR1_12S_50ntFlanks@427.456.30\|GAAAGTGGCTTTAACATATCTGAACACACA | >hg38_MT_+_598_1651_RNR1_12S_50ntFlanks | 427 | 456 | 30 | GAAAGTGGCTTTAACATATCTGAACACACA | 77.3 | 4.8 | 2.2 | 2.2 | 2.8E-02 | 4.4E-02 | BT20/MB468 | MP |
| >MI0001448\|hsa-mir-425&WithFlank&3\|-\|49020142\|49020240@20.43.24(+1A)&[MIMAT0003393&hsa-miR-425-5p&offsets\|0\|+1(+1A);m-95&3\|-\|49020200\|49020221&offsets\|0\|+2(+1A)]\|AATGACACGATCACTCCCGTTGAGA | >MI0001448\|hsa-mir-425&WithFlank&3\|-\|49020142\|49020240 | 20 | 43 | 24(+1A) | AATGACACGATCACTCCCGTTGAGA | 396.3 | 2.1 | 0.9 | 2.4 | 1.7E-02 | 4.4E-02 | BT20/MB231 | CYTO |

<table>
<tr><td>>am_trna16_TyrGTA_14_-_21131351_21131444@1.25.25,trna17_TyrGTA_14_-_21128117_21128210@1.25.25,trna17_TyrGTA_6_+_26595102_26595190@1.25.25,trna18_TyrGTA_14_-_21125623_21125716@1.25.25,trna19_TyrGTA_14_-_21121258_21121351@1.25.25,trna4_TyrGTA_8_+_67025602_67025694@1.25.25,trna5_TyrGTA_14_+_21151432_21151520@1.25.25,trna5_TyrGTA_8_+_67026223_67026311@1.25.25|CCTTCGATAGCTCAGCTGGTAGAGC</td><td>>am_trna16_TyrGTA_14_-_21131351_21131444</td><td>1</td><td>25</td><td>25</td><td>CCTTCGATAGCTCAGCTGGTAGAGC</td><td>920.5</td><td>2.9</td><td>1.2</td><td>2.4</td><td>1.7E-02</td><td>4.4E-02</td><td>BT20/MB231</td><td>CYTO</td></tr>
<tr><td>>trnaMT_LysTTT_MT_+_8295_8364@55.73.19,trnalookalike7_LysTTT_1_+_568844_568913@55.73.19|CACCTCTTTACAGTGACCA</td><td>>trnaMT_LysTTT_MT_+_8295_8364</td><td>55</td><td>73</td><td>19</td><td>CACCTCTTTACAGTGACCA</td><td>99.7</td><td>2.8</td><td>1.3</td><td>2.1</td><td>3.7E-02</td><td>4.4E-02</td><td>MB231/MB468</td><td>MP</td></tr>
<tr><td>>hg38_MT_+_1621_3279_RNR2_16S_50ntFlanks@1185.1215.31|ATTTCGGTTGGGGCGACCTCGGAGCAGAACC</td><td>>hg38_MT_+_1621_3279_RNR2_16S_50ntFlanks</td><td>1185</td><td>1215</td><td>31</td><td>ATTTCGGTTGGGGCGACCTCGGAGCAGAACC</td><td>124.3</td><td>2.6</td><td>1.2</td><td>2.1</td><td>3.7E-02</td><td>4.4E-02</td><td>MB231/MB468</td><td>MP</td></tr>
<tr><td>>hg38_21_+_8433172_8446622_RNA45SN1_45S_50ntFlanks@483.532.50|TCGATGTGGTGACGTCGTGCTCTCCCGGGCCGGGTCCGAGCCGCGACGGG</td><td>>hg38_21_+_8433172_8446622_RNA45SN1_45S_50ntFlanks</td><td>483</td><td>532</td><td>50</td><td>TCGATGTGGTGACGTCGTGCTCTCCCGGGCCGGGTCCGAGCCGCGACGGG</td><td>4662.1</td><td>3.2</td><td>1.5</td><td>2.1</td><td>4.0E-02</td><td>4.4E-02</td><td>MB231/MB468</td><td>NUC</td></tr>
<tr><td>>am_trnaMT_ValTAC_MT_+_1602_1670@49.68.20|AACTTAACTTGACCGCTCTG</td><td>>am_trnaMT_ValTAC_MT_+_1602_1670</td><td>49</td><td>68</td><td>20</td><td>AACTTAACTTGACCGCTCTG</td><td>172.7</td><td>3.8</td><td>1.5</td><td>2.5</td><td>1.3E-02</td><td>4.4E-02</td><td>BT20/MB231</td><td>MP</td></tr>
<tr><td>>hg38_21_+_8433172_8446622_RNA45SN1_45S_50ntFlanks@5077.5117.41|CCGATAACGAACGAGACTCTGGCATGCTAACTAGTTACGCG</td><td>>hg38_21_+_8433172_8446622_RNA45SN1_45S_50ntFlanks</td><td>5077</td><td>5117</td><td>41</td><td>CCGATAACGAACGAGACTCTGGCATGCTAACTAGTTACGCG</td><td>91.4</td><td>-4.1</td><td>1.9</td><td>-2.2</td><td>2.8E-02</td><td>4.4E-02</td><td>BT20/MB468</td><td>MP</td></tr>
<tr><td>>hg38_21_+_8433172_8446622_RNA45SN1_45S_50ntFlanks@9925.9947.23|GGTTGATATAGACAGCAGGACGG</td><td>>hg38_21_+_8433172_8446622_RNA45SN1_45S_50ntFlanks</td><td>9925</td><td>9947</td><td>23</td><td>GGTTGATATAGACAGCAGGACGG</td><td>50.5</td><td>4.7</td><td>2.1</td><td>2.2</td><td>2.8E-02</td><td>4.4E-02</td><td>BT20/MB468</td><td>MP</td></tr>
<tr><td>>hg38_21_+_8433172_8446622_RNA45SN1_45S_50ntFlanks@11666.11690.25|TCTGCCCAGTGCTCTGAATGTCAAA</td><td>>hg38_21_+_8433172_8446622_RNA45SN1_45S_50ntFlanks</td><td>11666</td><td>11690</td><td>25</td><td>TCTGCCCAGTGCTCTGAATGTCAAA</td><td>84.4</td><td>5.1</td><td>2.0</td><td>2.5</td><td>1.3E-02</td><td>4.4E-02</td><td>BT20/MB231</td><td>MP</td></tr>
<tr><td>>MI0000816|hsa-mir-335&WithFlank&7|+|130496105|130496210@58.78.21(+1U)&[MIMAT0004703&hsa-miR-335-3p&offsets|0|-1(+1U);m-159&7|+|130496162|130496183&offsets|0|-1(+1U)]|TTTTTCATTATTGCTCCTGACT</td><td>>MI0000816|hsa-mir-335&WithFlank&7|+|130496105|130496210</td><td>58</td><td>78</td><td>21(+1U)</td><td>TTTTTCATTATTGCTCCTGACT</td><td>647.5</td><td>-2.8</td><td>1.1</td><td>-2.5</td><td>1.3E-02</td><td>4.4E-02</td><td>BT20/MB231</td><td>MP</td></tr>
<tr><td>>hg38_21_+_8433172_8446622_RNA45SN1_45S_50ntFlanks@11918.11943.26|ACATGAGAGGTGTAGAATAAGTGGGA</td><td>>hg38_21_+_8433172_8446622_RNA45SN1_45S_50ntFlanks</td><td>11918</td><td>11943</td><td>26</td><td>ACATGAGAGGTGTAGAATAAGTGGGA</td><td>44.9</td><td>5.0</td><td>2.3</td><td>2.2</td><td>2.8E-02</td><td>4.4E-02</td><td>BT20/MB468</td><td>MP</td></tr>
<tr><td>>hg38_21_+_8433172_8446622_RNA45SN1_45S_50ntFlanks@12616.12659.44|CTGAGGAGCCAATGGGGCGAAGCTACCATCTGTGGGATTATGAC</td><td>>hg38_21_+_8433172_8446622_RNA45SN1_45S_50ntFlanks</td><td>12616</td><td>12659</td><td>44</td><td>CTGAGGAGCCAATGGGGCGAAGCTACCATCTGTGGGATTATGAC</td><td>148.1</td><td>-2.7</td><td>1.3</td><td>-2.1</td><td>3.5E-02</td><td>4.4E-02</td><td>BT20/MB468</td><td>MITO</td></tr>
<tr><td>>hg38_21_+_8433172_8446622_RNA45SN1_45S_50ntFlanks@11668.11685.18|TGCCCAGTGCTCTGAATG</td><td>>hg38_21_+_8433172_8446622_RNA45SN1_45S_50ntFlanks</td><td>11668</td><td>11685</td><td>18</td><td>TGCCCAGTGCTCTGAATG</td><td>479.2</td><td>-2.4</td><td>1.0</td><td>-2.5</td><td>1.3E-02</td><td>4.4E-02</td><td>BT20/MB231</td><td>NUC</td></tr>
<tr><td>>MI0000746|hsa-mir-99b&WithFlank&19|+|51692606|51692687@13.32.20(+1U)&[MIMAT0000689&hsa-miR-99b-5p&offsets|0|-2(+1U);m-10&19|+|51692618|51692639&offsets|0|-2(+1U)]|CACCCGTAGAACCGACCTTGT</td><td>>MI0000746|hsa-mir-99b&WithFlank&19|+|51692606|51692687</td><td>13</td><td>32</td><td>20(+1U)</td><td>CACCCGTAGAACCGACCTTGT</td><td>5405.0</td><td>1.7</td><td>0.7</td><td>2.4</td><td>1.5E-02</td><td>4.4E-02</td><td>BT20/MB231</td><td>MITO</td></tr>
<tr><td>>hg38_21_+_8433172_8446622_RNA45SN1_45S_50ntFlanks@12998.13017.20|GCTGCGATCTATTGAAAGTC</td><td>>hg38_21_+_8433172_8446622_RNA45SN1_45S_50ntFlanks</td><td>12998</td><td>13017</td><td>20</td><td>GCTGCGATCTATTGAAAGTC</td><td>254.1</td><td>-2.1</td><td>0.9</td><td>-2.5</td><td>1.3E-02</td><td>4.4E-02</td><td>BT20/MB231</td><td>NUC</td></tr>
<tr><td>>hg38_21_+_8433172_8446622_RNA45SN1_45S_50ntFlanks@11317.11338.22|ACCCCACGTCTCGTCGCGCGCG</td><td>>hg38_21_+_8433172_8446622_RNA45SN1_45S_50ntFlanks</td><td>11317</td><td>11338</td><td>22</td><td>ACCCCACGTCTCGTCGCGCGCG</td><td>1635.4</td><td>-3.2</td><td>1.5</td><td>-2.1</td><td>3.5E-02</td><td>4.4E-02</td><td>BT20/MB468</td><td>MITO</td></tr>
<tr><td>>hg38_1_-_228634819_228635039_RNA5S12_5S_50ntFlanks@141.170.30|CCGCCTGGGAATACCGGGTGCTGTAGGCTT</td><td>>hg38_1_-_228634819_228635039_RNA5S12_5S_50ntFlanks</td><td>141</td><td>170</td><td>30</td><td>CCGCCTGGGAATACCGGGTGCTGTAGGCTT</td><td>1328.6</td><td>1.8</td><td>0.8</td><td>2.1</td><td>3.5E-02</td><td>4.4E-02</td><td>BT20/MB468</td><td>MITO</td></tr>
<tr><td>>MI0000072|hsa-mir-18a&WithFlank&13|+|91350745|91350827@12.33.22(+1A)&[MIMAT0000072&hsa-miR-18a-5p&offsets|0|-1(+1A);m-262&13|+|91350756|91350778&offsets|0|-1(+1A)]|TAAGGTGCATCTAGTGCAGATAA</td><td>>MI0000072|hsa-mir-18a&WithFlank&13|+|91350745|91350827</td><td>12</td><td>33</td><td>22(+1A)</td><td>TAAGGTGCATCTAGTGCAGATAA</td><td>97.3</td><td>3.6</td><td>1.7</td><td>2.1</td><td>3.7E-02</td><td>4.4E-02</td><td>MB231/MB468</td><td>MP</td></tr>
<tr><td>>hg38_21_+_8433172_8446622_RNA45SN1_45S_50ntFlanks@6648.6695.48|GTACGACTCTTAGCGGTGGATCACTCGGCTCGTGCGTCGATGAAGAAC</td><td>>hg38_21_+_8433172_8446622_RNA45SN1_45S_50ntFlanks</td><td>6648</td><td>6695</td><td>48</td><td>GTACGACTCTTAGCGGTGGATCACTCGGCTCGTGCGTCGATGAAGAAC</td><td>462.1</td><td>-2.3</td><td>1.0</td><td>-2.4</td><td>1.7E-02</td><td>4.4E-02</td><td>BT20/MB231</td><td>CYTO</td></tr>
<tr><td>>hg38_21_+_8433172_8446622_RNA45SN1_45S_50ntFlanks@10404.10424.21|CATGGGTCAGTCGGTCCTGAG</td><td>>hg38_21_+_8433172_8446622_RNA45SN1_45S_50ntFlanks</td><td>10404</td><td>10424</td><td>21</td><td>CATGGGTCAGTCGGTCCTGAG</td><td>2067.2</td><td>1.3</td><td>0.7</td><td>2.0</td><td>4.0E-02</td><td>4.4E-02</td><td>MB231/MB468</td><td>NUC</td></tr>
<tr><td>>trnaMT_ValTAC_MT_+_1602_1670@38.72.35|TAGGAGATTTCAACTTAACTTGACCGCTCTGACCA</td><td>>trnaMT_ValTAC_MT_+_1602_1670</td><td>38</td><td>72</td><td>35</td><td>TAGGAGATTTCAACTTAACTTGACCGCTCTGACCA</td><td>207.3</td><td>5.8</td><td>2.7</td><td>2.2</td><td>2.8E-02</td><td>4.5E-02</td><td>BT20/MB468</td><td>MP</td></tr>
<tr><td>>hg38_21_+_8433172_8446622_RNA45SN1_45S_50ntFlanks@10318.10340.23|TGCAGATCTTGGTGGTAGTAGCA</td><td>>hg38_21_+_8433172_8446622_RNA45SN1_45S_50ntFlanks</td><td>10318</td><td>10340</td><td>23</td><td>TGCAGATCTTGGTGGTAGTAGCA</td><td>169.0</td><td>2.3</td><td>1.0</td><td>2.2</td><td>2.8E-02</td><td>4.5E-02</td><td>BT20/MB468</td><td>MP</td></tr>
<tr><td>>hg38_21_+_8433172_8446622_RNA45SN1_45S_50ntFlanks@3845.3874.30|ACTTGGATAACTGTGGTAATTCTAGAGCTA</td><td>>hg38_21_+_8433172_8446622_RNA45SN1_45S_50ntFlanks</td><td>3845</td><td>3874</td><td>30</td><td>ACTTGGATAACTGTGGTAATTCTAGAGCTA</td><td>110.2</td><td>-2.2</td><td>1.0</td><td>-2.2</td><td>2.8E-02</td><td>4.5E-02</td><td>BT20/MB468</td><td>MP</td></tr>
<tr><td>>hg38_21_+_8433172_8446622_RNA45SN1_45S_50ntFlanks@5446.5463.18|CGGATCGGCCCCGCCGGG</td><td>>hg38_21_+_8433172_8446622_RNA45SN1_45S_50ntFlanks</td><td>5446</td><td>5463</td><td>18</td><td>CGGATCGGCCCCGCCGGG</td><td>185.6</td><td>-4.0</td><td>1.8</td><td>-2.2</td><td>2.8E-02</td><td>4.5E-02</td><td>BT20/MB468</td><td>MP</td></tr>
<tr><td>>MI0000095|hsa-mir-93&WithFlank&7|-|100093762|100093853@17.36.20(+1C)&[MIMAT0000093&hsa-miR-93-5p&offsets|0|-3(+1C);m-29&7|-|100093815|100093837&offsets|0|-3(+1C)]|CAAAGTGCTGTTCGTGCAGGC</td><td>>MI0000095|hsa-mir-93&WithFlank&7|-|100093762|100093853</td><td>17</td><td>36</td><td>20(+1C)</td><td>CAAAGTGCTGTTCGTGCAGGC</td><td>372.0</td><td>3.0</td><td>1.3</td><td>2.2</td><td>2.8E-02</td><td>4.5E-02</td><td>BT20/MB468</td><td>MP</td></tr>
<tr><td>>hg38_21_+_8433172_8446622_RNA45SN1_45S_50ntFlanks@11868.11893.26|GGGGAAAGAAGACCCTGTTGAGCTTG</td><td>>hg38_21_+_8433172_8446622_RNA45SN1_45S_50ntFlanks</td><td>11868</td><td>11893</td><td>26</td><td>GGGGAAAGAAGACCCTGTTGAGCTTG</td><td>56.3</td><td>3.9</td><td>1.8</td><td>2.2</td><td>2.8E-02</td><td>4.5E-02</td><td>BT20/MB468</td><td>MP</td></tr>
<tr><td>>hg38_MT_+_598_1651_RNR1_12S_50ntFlanks@52.77.26|ATAGGTTTGGTCCTAGCCTTTCTATT</td><td>>hg38_MT_+_598_1651_RNR1_12S_50ntFlanks</td><td>52</td><td>77</td><td>26</td><td>ATAGGTTTGGTCCTAGCCTTTCTATT</td><td>108.8</td><td>1.6</td><td>0.8</td><td>2.1</td><td>4.0E-02</td><td>4.5E-02</td><td>BT20/MB468</td><td>NUC</td></tr>
<tr><td>>hg38_21_+_8433172_8446622_RNA45SN1_45S_50ntFlanks@3993.4024.32|GTGACTCTAGATAACCTCGGGCCGATCGCACG</td><td>>hg38_21_+_8433172_8446622_RNA45SN1_45S_50ntFlanks</td><td>3993</td><td>4024</td><td>32</td><td>GTGACTCTAGATAACCTCGGGCCGATCGCACG</td><td>1108.0</td><td>-1.5</td><td>0.8</td><td>-2.0</td><td>4.0E-02</td><td>4.5E-02</td><td>MB231/MB468</td><td>NUC</td></tr>
<tr><td>>MI0000066|hsa-let-7e&WithFlank&19|+|51692780|51692870@14.34.21(+1G)&[MIMAT0000066&hsa-let-7e-5p&offsets|0|-1(+1G);m-50&19|+|51692793|51692814&offsets|0|-1(+1G)]|TGAGGTAGGAGGTTGTATAGTG</td><td>>MI0000066|hsa-let-7e&WithFlank&19|+|51692780|51692870</td><td>14</td><td>34</td><td>21(+1G)</td><td>TGAGGTAGGAGGTTGTATAGTG</td><td>507.3</td><td>2.4</td><td>1.0</td><td>2.4</td><td>1.7E-02</td><td>4.5E-02</td><td>BT20/MB231</td><td>CYTO</td></tr>
<tr><td>>MI0000469|hsa-mir-125a&WithFlank&19|+|51693248|51693345@21.41.21(+1C)&[MIMAT0000443&hsa-miR-125a-5p&offsets|0|-3(+1C);m-63&19|+|51693268|51693289&offsets|0|-1(+1C)]|TCCCTGAGACCCTTTAACCTGC</td><td>>MI0000469|hsa-mir-125a&WithFlank&19|+|51693248|51693345</td><td>21</td><td>41</td><td>21(+1C)</td><td>TCCCTGAGACCCTTTAACCTGC</td><td>983.1</td><td>2.1</td><td>0.9</td><td>2.4</td><td>1.7E-02</td><td>4.5E-02</td><td>BT20/MB231</td><td>CYTO</td></tr>
<tr><td>>MI0001448|hsa-mir-425&WithFlank&3|-|49020142|49020240@20.42.23&[MIMAT0003393&hsa-miR-425-5p&offsets|0|0;m-95&3|-|49020200|49020221&offsets|0|+1]|AATGACACGATCACTCCCGTTGA</td><td>>MI0001448|hsa-mir-425&WithFlank&3|-|49020142|49020240</td><td>20</td><td>42</td><td>23</td><td>AATGACACGATCACTCCCGTTGA</td><td>549.8</td><td>2.3</td><td>1.0</td><td>2.4</td><td>1.7E-02</td><td>4.5E-02</td><td>BT20/MB231</td><td>CYTO</td></tr>
<tr><td>>trna20_GluTTC_2_-_131094701_131094772@1.33.33,trna5_GluTTC_13_-_41634874_41634945@1.33.33|TCCCATATGGTCTAGCGGTTAGGATTCCTGGTT</td><td>>trna20_GluTTC_2_-_131094701_131094772</td><td>1</td><td>33</td><td>33</td><td>TCCCATATGGTCTAGCGGTTAGGATTCCTGGTT</td><td>3242.1</td><td>-1.9</td><td>0.8</td><td>-2.4</td><td>1.7E-02</td><td>4.5E-02</td><td>BT20/MB231</td><td>CYTO</td></tr>
<tr><td>>hg38_21_+_8433172_8446622_RNA45SN1_45S_50ntFlanks@10678.10711.34|CGGTTCCGGCGGCGTCCGGTGAGCTCTCGCTGGC</td><td>>hg38_21_+_8433172_8446622_RNA45SN1_45S_50ntFlanks</td><td>10678</td><td>10711</td><td>34</td><td>CGGTTCCGGCGGCGTCCGGTGAGCTCTCGCTGGC</td><td>839.5</td><td>2.8</td><td>1.2</td><td>2.4</td><td>1.7E-02</td><td>4.5E-02</td><td>BT20/MB231</td><td>CYTO</td></tr>
<tr><td>>hg38_21_+_8433172_8446622_RNA45SN1_45S_50ntFlanks@5078.5117.40|CGATAACGAACGAGACTCTGGCATGCTAACTAGTTACGCG</td><td>>hg38_21_+_8433172_8446622_RNA45SN1_45S_50ntFlanks</td><td>5078</td><td>5117</td><td>40</td><td>CGATAACGAACGAGACTCTGGCATGCTAACTAGTTACGCG</td><td>239.5</td><td>-1.8</td><td>0.8</td><td>-2.4</td><td>1.7E-02</td><td>4.5E-02</td><td>BT20/MB231</td><td>CYTO</td></tr>
<tr><td>>hg38_21_+_8433172_8446622_RNA45SN1_45S_50ntFlanks@11212.11238.27|CGACTCTGGACGCGAGCCGGGCCCTTC</td><td>>hg38_21_+_8433172_8446622_RNA45SN1_45S_50ntFlanks</td><td>11212</td><td>11238</td><td>27</td><td>CGACTCTGGACGCGAGCCGGGCCCTTC</td><td>116.8</td><td>-2.4</td><td>1.1</td><td>-2.2</td><td>2.8E-02</td><td>4.5E-02</td><td>BT20/MB468</td><td>MP</td></tr>
<tr><td>>hg38_21_+_8433172_8446622_RNA45SN1_45S_50ntFlanks@11553.11589.37|CTCGGCCGGCGCCTAGCAGCCGACTTAGAACTGGTGC</td><td>>hg38_21_+_8433172_8446622_RNA45SN1_45S_50ntFlanks</td><td>11553</td><td>11589</td><td>37</td><td>CTCGGCCGGCGCCTAGCAGCCGACTTAGAACTGGTGC</td><td>142.6</td><td>4.3</td><td>1.7</td><td>2.5</td><td>1.3E-02</td><td>4.5E-02</td><td>BT20/MB231</td><td>MP</td></tr>
<tr><td>>hg38_21_+_8433172_8446622_RNA45SN1_45S_50ntFlanks@9497.9525.29|AAAGATGGTGAACTATGCCTGGGCAGGGC</td><td>>hg38_21_+_8433172_8446622_RNA45SN1_45S_50ntFlanks</td><td>9497</td><td>9525</td><td>29</td><td>AAAGATGGTGAACTATGCCTGGGCAGGGC</td><td>128.8</td><td>5.1</td><td>2.0</td><td>2.5</td><td>1.3E-02</td><td>4.5E-02</td><td>BT20/MB231</td><td>MP</td></tr>
<tr><td>>hg38_21_+_8433172_8446622_RNA45SN1_45S_50ntFlanks@6649.6676.28|TACGACTCTTAGCGGTGGATCACTCGGC</td><td>>hg38_21_+_8433172_8446622_RNA45SN1_45S_50ntFlanks</td><td>6649</td><td>6676</td><td>28</td><td>TACGACTCTTAGCGGTGGATCACTCGGC</td><td>567.2</td><td>3.6</td><td>1.4</td><td>2.5</td><td>1.3E-02</td><td>4.5E-02</td><td>BT20/MB231</td><td>MP</td></tr>
<tr><td>>hg38_21_+_8433172_8446622_RNA45SN1_45S_50ntFlanks@6648.6668.21|GTACGACTCTTAGCGGTGGAT</td><td>>hg38_21_+_8433172_8446622_RNA45SN1_45S_50ntFlanks</td><td>6648</td><td>6668</td><td>21</td><td>GTACGACTCTTAGCGGTGGAT</td><td>758.0</td><td>1.9</td><td>0.8</td><td>2.4</td><td>1.7E-02</td><td>4.5E-02</td><td>BT20/MB231</td><td>CYTO</td></tr>
<tr><td>>hg38_21_+_8433172_8446622_RNA45SN1_45S_50ntFlanks@12420.12443.24|ATGTCGGCTCTTCCTATCATTGTG</td><td>>hg38_21_+_8433172_8446622_RNA45SN1_45S_50ntFlanks</td><td>12420</td><td>12443</td><td>24</td><td>ATGTCGGCTCTTCCTATCATTGTG</td><td>1020.1</td><td>-2.2</td><td>0.9</td><td>-2.4</td><td>1.7E-02</td><td>4.5E-02</td><td>BT20/MB231</td><td>CYTO</td></tr>
<tr><td>>hg38_21_+_8433172_8446622_RNA45SN1_45S_50ntFlanks@5473.5509.37|CGGCCCTGGCGGAGCGCTGAGAAGACGGTCGAACTTG</td><td>>hg38_21_+_8433172_8446622_RNA45SN1_45S_50ntFlanks</td><td>5473</td><td>5509</td><td>37</td><td>CGGCCCTGGCGGAGCGCTGAGAAGACGGTCGAACTTG</td><td>1530.6</td><td>2.2</td><td>0.9</td><td>2.4</td><td>1.7E-02</td><td>4.5E-02</td><td>BT20/MB231</td><td>CYTO</td></tr>
<tr><td>>hg38_21_+_8433172_8446622_RNA45SN1_45S_50ntFlanks@7974.8012.39|ACGCGACCTCAGATCAGACGTGGCGACCCGCTGAATTTA</td><td>>hg38_21_+_8433172_8446622_RNA45SN1_45S_50ntFlanks</td><td>7974</td><td>8012</td><td>39</td><td>ACGCGACCTCAGATCAGACGTGGCGACCCGCTGAATTTA</td><td>2755.0</td><td>-2.2</td><td>0.9</td><td>-2.4</td><td>1.7E-02</td><td>4.5E-02</td><td>BT20/MB231</td><td>CYTO</td></tr>
<tr><td>>hg38_21_+_8433172_8446622_RNA45SN1_45S_50ntFlanks@6660.6678.19|GCGGTGGATCACTCGGCTC</td><td>>hg38_21_+_8433172_8446622_RNA45SN1_45S_50ntFlanks</td><td>6660</td><td>6678</td><td>19</td><td>GCGGTGGATCACTCGGCTC</td><td>176.4</td><td>4.8</td><td>1.9</td><td>2.5</td><td>1.3E-02</td><td>4.5E-02</td><td>BT20/MB231</td><td>MP</td></tr>
<tr><td>>hg38_MT_+_1621_3279_RNR2_16S_50ntFlanks@1438.1457.20|TACGTGATCTGAGTTCAGAC</td><td>>hg38_MT_+_1621_3279_RNR2_16S_50ntFlanks</td><td>1438</td><td>1457</td><td>20</td><td>TACGTGATCTGAGTTCAGAC</td><td>447.2</td><td>2.2</td><td>1.1</td><td>2.1</td><td>3.7E-02</td><td>4.5E-02</td><td>MB231/MB468</td><td>MP</td></tr>
<tr><td>>hg38_21_+_8433172_8446622_RNA45SN1_45S_50ntFlanks@10403.10424.22|ACATGGGTCAGTCGGTCCTGAG</td><td>>hg38_21_+_8433172_8446622_RNA45SN1_45S_50ntFlanks</td><td>10403</td><td>10424</td><td>22</td><td>ACATGGGTCAGTCGGTCCTGAG</td><td>779.3</td><td>1.5</td><td>0.7</td><td>2.1</td><td>3.7E-02</td><td>4.5E-02</td><td>MB231/MB468</td><td>MP</td></tr>
<tr><td>>hg38_21_+_8433172_8446622_RNA45SN1_45S_50ntFlanks@9923.9955.33|TTGGTTGATATAGACAGCAGGACGGTGGCCATG</td><td>>hg38_21_+_8433172_8446622_RNA45SN1_45S_50ntFlanks</td><td>9923</td><td>9955</td><td>33</td><td>TTGGTTGATATAGACAGCAGGACGGTGGCCATG</td><td>90.9</td><td>5.1</td><td>2.1</td><td>2.5</td><td>1.3E-02</td><td>4.5E-02</td><td>BT20/MB231</td><td>MP</td></tr>
<tr><td>>hg38_21_+_8433172_8446622_RNA45SN1_45S_50ntFlanks@6718.6743.26|GTGAATTGCAGGACACATTGATCATC</td><td>>hg38_21_+_8433172_8446622_RNA45SN1_45S_50ntFlanks</td><td>6718</td><td>6743</td><td>26</td><td>GTGAATTGCAGGACACATTGATCATC</td><td>313.9</td><td>2.1</td><td>0.9</td><td>2.4</td><td>1.5E-02</td><td>4.5E-02</td><td>BT20/MB231</td><td>MITO</td></tr>
<tr><td>>hg38_21_+_8433172_8446622_RNA45SN1_45S_50ntFlanks@11672.11706.35|CAGTGCTCTGAATGTCAAAGTGAAGAAATTCAATG</td><td>>hg38_21_+_8433172_8446622_RNA45SN1_45S_50ntFlanks</td><td>11672</td><td>11706</td><td>35</td><td>CAGTGCTCTGAATGTCAAAGTGAAGAAATTCAATG</td><td>283.6</td><td>2.0</td><td>0.8</td><td>2.4</td><td>1.5E-02</td><td>4.5E-02</td><td>BT20/MB231</td><td>MITO</td></tr>
<tr><td>>trnaMT_GluTTC_MT_-_14674_14742@38.71.34,trnalookalike8_GluTTC_5_-_93905172_93905240@38.71.34|CATTGGTCGTGGTTGTAGTCCGTGCGAGAATACC</td><td>>trnaMT_GluTTC_MT_-_14674_14742</td><td>38</td><td>71</td><td>34</td><td>CATTGGTCGTGGTTGTAGTCCGTGCGAGAATACC</td><td>372.7</td><td>-1.9</td><td>0.8</td><td>-2.5</td><td>1.3E-02</td><td>4.5E-02</td><td>BT20/MB231</td><td>NUC</td></tr>
</table>

| | | | | | | | | | | | | | |
|---|---|---|---|---|---|---|---|---|---|---|---|---|---|
| >MI0000086\|hsa-mir-28&WithFlank&3\|+\|188688775\|188688872@62.82.21(+1U)&[MIMAT0004502&hsa-miR-28-3p&offsets\|+2\|+1(+1U);m-28&3\|+\|188688834\|188688855&offsets\|+2\|+1(+1U)]\|CTAGATTGTGAGCTCCTGGAGT | >MI0000086\|hsa-mir-28&WithFlank&3\|+\|188688775\|188688872 | 62 | 82 | 21(+1U) | CTAGATTGTGAGCTCCTGGAGT | 580.5 | 1.9 | 0.8 | 2.4 | 1.7E-02 | 4.5E-02 | BT20/MB231 | CYTO |
| >hg38_21_+_8433172_8446622_RNA45SN1_45S_50ntFlanks@12418.12437.20\|CGATGTCGGCTCTTCCTATC | >hg38_21_+_8433172_8446622_RNA45SN1_45S_50ntFlanks | 12418 | 12437 | 20 | CGATGTCGGCTCTTCCTATC | 1699.0 | -1.9 | 0.8 | -2.4 | 1.7E-02 | 4.5E-02 | BT20/MB231 | CYTO |
| >hg38_21_+_8433172_8446622_RNA45SN1_45S_50ntFlanks@4727.4768.42\|AATCAAGAACGAAAGTCGGAGGTTCGAAGACGATCAGATACC | >hg38_21_+_8433172_8446622_RNA45SN1_45S_50ntFlanks | 4727 | 4768 | 42 | AATCAAGAACGAAAGTCGGAGGTTCGAAGACGATCAGATACC | 3427.1 | -2.4 | 1.0 | -2.4 | 1.7E-02 | 4.5E-02 | BT20/MB231 | CYTO |
| >hg38_21_+_8433172_8446622_RNA45SN1_45S_50ntFlanks@6649.6668.20\|TACGACTCTTAGCGGTGGAT | >hg38_21_+_8433172_8446622_RNA45SN1_45S_50ntFlanks | 6649 | 6668 | 20 | TACGACTCTTAGCGGTGGAT | 345.7 | 1.7 | 0.7 | 2.4 | 1.7E-02 | 4.5E-02 | BT20/MB231 | CYTO |
| >hg38_21_+_8433172_8446622_RNA45SN1_45S_50ntFlanks@10351.10369.19\|CGAGAACTTTGAAGGCCGA | >hg38_21_+_8433172_8446622_RNA45SN1_45S_50ntFlanks | 10351 | 10369 | 19 | CGAGAACTTTGAAGGCCGA | 212.9 | -2.0 | 0.8 | -2.4 | 1.7E-02 | 4.5E-02 | BT20/MB231 | CYTO |
| >hg38_21_+_8433172_8446622_RNA45SN1_45S_50ntFlanks@8194.8213.20\|CCCGTGGACGGTGTGAGGCC | >hg38_21_+_8433172_8446622_RNA45SN1_45S_50ntFlanks | 8194 | 8213 | 20 | CCCGTGGACGGTGTGAGGCC | 348.6 | -2.1 | 0.9 | -2.4 | 1.7E-02 | 4.5E-02 | BT20/MB231 | CYTO |
| >hg38_21_+_8433172_8446622_RNA45SN1_45S_50ntFlanks@5478.5500.23\|CTGGCGGAGCGCTGAGAAGACGG | >hg38_21_+_8433172_8446622_RNA45SN1_45S_50ntFlanks | 5478 | 5500 | 23 | CTGGCGGAGCGCTGAGAAGACGG | 521.9 | 3.6 | 1.4 | 2.5 | 1.3E-02 | 4.5E-02 | BT20/MB231 | MP |
| >hg38_21_+_8433172_8446622_RNA45SN1_45S_50ntFlanks@11564.11605.42\|CCTAGCAGCCGACTTAGAACTGGTGCGGACCAGGGGAATCCG | >hg38_21_+_8433172_8446622_RNA45SN1_45S_50ntFlanks | 11564 | 11605 | 42 | CCTAGCAGCCGACTTAGAACTGGTGCGGACCAGGGGAATCCG | 172.6 | -4.3 | 1.9 | -2.2 | 2.8E-02 | 4.5E-02 | BT20/MB468 | MP |
| >hg38_21_+_8433172_8446622_RNA45SN1_45S_50ntFlanks@12606.12624.19\|ATGTGCTTGGCTGAGGAGC | >hg38_21_+_8433172_8446622_RNA45SN1_45S_50ntFlanks | 12606 | 12624 | 19 | ATGTGCTTGGCTGAGGAGC | 288.0 | 1.3 | 0.6 | 2.1 | 3.9E-02 | 4.5E-02 | MB231/MB468 | CYTO |
| >MI0000100\|hsa-mir-98&WithFlank&X\|-\|53556217\|53556347@28.48.21&[MIMAT0000096&hsa-miR-98-5p&offsets\|0\|-1;m-143&X\|-\|53556299\|53556320&offsets\|0\|-1]\|TGAGGTAGTAAGTTGTATTGT | >MI0000100\|hsa-mir-98&WithFlank&X\|-\|53556217\|53556347 | 28 | 48 | 21 | TGAGGTAGTAAGTTGTATTGT | 681.8 | -2.7 | 1.1 | -2.5 | 1.3E-02 | 4.5E-02 | BT20/MB231 | MP |
| >hg38_21_+_8433172_8446622_RNA45SN1_45S_50ntFlanks@10057.10079.23\|CCGGCAGTCGAGAGTGGACGGGA | >hg38_21_+_8433172_8446622_RNA45SN1_45S_50ntFlanks | 10057 | 10079 | 23 | CCGGCAGTCGAGAGTGGACGGGA | 91.9 | 4.0 | 1.6 | 2.5 | 1.3E-02 | 4.5E-02 | BT20/MB231 | MP |
| >hg38_21_+_8433172_8446622_RNA45SN1_45S_50ntFlanks@4729.4759.31\|TCAAGAACGAAAGTCGGAGGTTCGAAGACGA | >hg38_21_+_8433172_8446622_RNA45SN1_45S_50ntFlanks | 4729 | 4759 | 31 | TCAAGAACGAAAGTCGGAGGTTCGAAGACGA | 369.9 | 4.4 | 1.8 | 2.5 | 1.3E-02 | 4.5E-02 | BT20/MB231 | MP |
| >hg38_21_+_8433172_8446622_RNA45SN1_45S_50ntFlanks@4733.4758.26\|GAACGAAAGTCGGAGGTTCGAAGACG | >hg38_21_+_8433172_8446622_RNA45SN1_45S_50ntFlanks | 4733 | 4758 | 26 | GAACGAAAGTCGGAGGTTCGAAGACG | 4433.7 | 3.7 | 1.5 | 2.5 | 1.3E-02 | 4.5E-02 | BT20/MB231 | MP |
| >trnaMT_ProTGG_MT_-_15956_16023@1.24.24\|CAGAGAATAGTTTAAATTAGAATC | >trnaMT_ProTGG_MT_-_15956_16023 | 1 | 24 | 24 | CAGAGAATAGTTTAAATTAGAATC | 237.0 | 1.8 | 0.9 | 2.1 | 3.7E-02 | 4.5E-02 | MB231/MB468 | MP |
| >hg38_21_+_8433172_8446622_RNA45SN1_45S_50ntFlanks@4738.4756.19\|AAAGTCGGAGGTTCGAAGA | >hg38_21_+_8433172_8446622_RNA45SN1_45S_50ntFlanks | 4738 | 4756 | 19 | AAAGTCGGAGGTTCGAAGA | 167.3 | 2.3 | 1.0 | 2.2 | 2.8E-02 | 4.5E-02 | BT20/MB468 | MP |
| >hg38_21_+_8433172_8446622_RNA45SN1_45S_50ntFlanks@9934.9960.27\|AGACAGCAGGACGGTGGCCATGGAAGT | >hg38_21_+_8433172_8446622_RNA45SN1_45S_50ntFlanks | 9934 | 9960 | 27 | AGACAGCAGGACGGTGGCCATGGAAGT | 115.3 | 5.1 | 2.0 | 2.5 | 1.3E-02 | 4.5E-02 | BT20/MB231 | MP |
| >hg38_21_+_8433172_8446622_RNA45SN1_45S_50ntFlanks@7973.8005.33\|GACGCGACCTCAGATCAGACGTGGCGACCCGCT | >hg38_21_+_8433172_8446622_RNA45SN1_45S_50ntFlanks | 7973 | 8005 | 33 | GACGCGACCTCAGATCAGACGTGGCGACCCGCT | 5508.3 | 1.9 | 0.8 | 2.4 | 1.5E-02 | 4.5E-02 | BT20/MB231 | MITO |
| >trna10_LysCTT_16_+_3241501_3241573@11.35.25,trna13_LysCTT_14_-_58706613_58706685@11.35.25,trna2_LysCTT_15_+_79152904_79152976@11.35.25,trna4_LysCTT_18_-_43669271_43669343@11.35.25\|CTCAGTCGGTAGAGCATGGGACTCT | >trna10_LysCTT_16_+_3241501_3241573 | 11 | 35 | 25 | CTCAGTCGGTAGAGCATGGGACTCT | 293.8 | -5.5 | 2.3 | -2.4 | 1.5E-02 | 4.5E-02 | BT20/MB231 | MITO |
| >hg38_21_+_8433172_8446622_RNA45SN1_45S_50ntFlanks@9738.9758.21\|GAAACGATCTCAACCTATTCT | >hg38_21_+_8433172_8446622_RNA45SN1_45S_50ntFlanks | 9738 | 9758 | 21 | GAAACGATCTCAACCTATTCT | 1922.8 | -4.3 | 1.9 | -2.2 | 2.8E-02 | 4.5E-02 | BT20/MB468 | MP |
| >hg38_21_+_8433172_8446622_RNA45SN1_45S_50ntFlanks@8338.8375.38\|GTACCGTAAGGGAAAGTTGAAAAGAACTTTGAAGAGAG | >hg38_21_+_8433172_8446622_RNA45SN1_45S_50ntFlanks | 8338 | 8375 | 38 | GTACCGTAAGGGAAAGTTGAAAAGAACTTTGAAGAGAG | 1490.4 | 4.5 | 1.8 | 2.5 | 1.3E-02 | 4.5E-02 | BT20/MB231 | MP |
| >hg38_21_+_8433172_8446622_RNA45SN1_45S_50ntFlanks@10715.10736.22\|TGAAAATCCGGGGGAGAGGGTG | >hg38_21_+_8433172_8446622_RNA45SN1_45S_50ntFlanks | 10715 | 10736 | 22 | TGAAAATCCGGGGGAGAGGGTG | 138.3 | -2.8 | 1.3 | -2.2 | 2.8E-02 | 4.5E-02 | BT20/MB468 | MP |
| >hg38_21_+_8433172_8446622_RNA45SN1_45S_50ntFlanks@12634.12657.24\|GAAGCTACCATCTGTGGGATTATG | >hg38_21_+_8433172_8446622_RNA45SN1_45S_50ntFlanks | 12634 | 12657 | 24 | GAAGCTACCATCTGTGGGATTATG | 162.9 | -3.0 | 1.4 | -2.2 | 2.8E-02 | 4.5E-02 | BT20/MB468 | MP |
| >MI0003656\|hsa-mir-641&WithFlank&19\|-\|40282537\|40282647@72.93.22&[TJU_CMC_MD2.ID01715.3p-miR&offsets\|0\|0;m-3103&19\|-\|40282555\|40282576&offsets\|0\|0]\|TGACTGTCCTATGTCTTTCCTT | >MI0003656\|hsa-mir-641&WithFlank&19\|-\|40282537\|40282647 | 72 | 93 | 22 | TGACTGTCCTATGTCTTTCCTT | 35.9 | 4.3 | 2.1 | 2.1 | 3.7E-02 | 4.5E-02 | MB231/MB468 | MP |
| >trna116_GluCTC_1_-_145399233_145399304@3.33.31,trna59_GluCTC_1_+_249168447_249168518@3.33.31,trna71_GluCTC_1_-_161439189_161439260@3.33.31,trna74_GluCTC_1_-_161431809_161431880@3.33.31,trna77_GluCTC_1_-_161424398_161424469@3.33.31,trna77_GluCTC_6_+_28949976_28950047@3.33.31,trna80_GluCTC_1_-_161417018_161417089@3.33.31,trna87_GluCTC_6_-_126101393_126101464@3.33.31\|CCTGGTGGTCTAGTGGTTAGGATTCGGCGCT | >trna116_GluCTC_1_-_145399233_145399304 | 3 | 33 | 31 | CCTGGTGGTCTAGTGGTTAGGATTCGGCGCT | 125.7 | 3.6 | 1.7 | 2.1 | 3.7E-02 | 4.5E-02 | MB231/MB468 | MP |
| >MI0001448\|hsa-mir-425&WithFlank&3\|-\|49020142\|49020240@20.42.23(+1A)&[MIMAT0003393&hsa-miR-425-5p&offsets\|0\|0(+1A);m-95&3\|-\|49020200\|49020221&offsets\|0\|+1(+1A)]\|AATGACACGATCACTCCCGTTGAA | >MI0001448\|hsa-mir-425&WithFlank&3\|-\|49020142\|49020240 | 20 | 42 | 23(+1A) | AATGACACGATCACTCCCGTTGAA | 111.6 | -4.8 | 1.9 | -2.5 | 1.3E-02 | 4.5E-02 | BT20/MB231 | MP |
| >hg38_21_+_8433172_8446622_RNA45SN1_45S_50ntFlanks@6644.6681.38\|CCTCGTACGACTCTTAGCGGTGGATCACTCGGCTCGTG | >hg38_21_+_8433172_8446622_RNA45SN1_45S_50ntFlanks | 6644 | 6681 | 38 | CCTCGTACGACTCTTAGCGGTGGATCACTCGGCTCGTG | 149.6 | -2.4 | 1.1 | -2.2 | 2.8E-02 | 4.5E-02 | BT20/MB468 | MP |
| >trna119_LysCTT_1_-_145395522_145395594@3.35.33,trna11_LysCTT_5_-_180648979_180649051@3.35.33,trna13_LysCTT_6_+_26556774_26556846@3.35.33,trna7_LysCTT_16_+_3225692_3225764@3.35.33,trna9_LysCTT_5_+_180634755_180634827@3.35.33\|CCGGCTAGCTCAGTCGGTAGAGCATGAGACTCT | >trna119_LysCTT_1_-_145395522_145395594 | 3 | 35 | 33 | CCGGCTAGCTCAGTCGGTAGAGCATGAGACTCT | 335.0 | -3.8 | 1.5 | -2.5 | 1.3E-02 | 4.5E-02 | BT20/MB231 | MP |
| >hg38_21_+_8433172_8446622_RNA45SN1_45S_50ntFlanks@8192.8212.21\|AGCCCGTGGACGGTGTGAGGC | >hg38_21_+_8433172_8446622_RNA45SN1_45S_50ntFlanks | 8192 | 8212 | 21 | AGCCCGTGGACGGTGTGAGGC | 420.7 | 3.1 | 1.3 | 2.5 | 1.3E-02 | 4.5E-02 | BT20/MB231 | MP |
| >hg38_21_+_8433172_8446622_RNA45SN1_45S_50ntFlanks@9928.9947.20\|TGATATAGACAGCAGGACGG | >hg38_21_+_8433172_8446622_RNA45SN1_45S_50ntFlanks | 9928 | 9947 | 20 | TGATATAGACAGCAGGACGG | 164.0 | 4.8 | 2.2 | 2.2 | 2.8E-02 | 4.5E-02 | BT20/MB468 | MP |
| >hg38_21_+_8433172_8446622_RNA45SN1_45S_50ntFlanks@10788.10818.31\|CAGCCTCTGGCATGTTGGAACAATGTAGGTA | >hg38_21_+_8433172_8446622_RNA45SN1_45S_50ntFlanks | 10788 | 10818 | 31 | CAGCCTCTGGCATGTTGGAACAATGTAGGTA | 136.0 | 4.0 | 1.6 | 2.5 | 1.3E-02 | 4.5E-02 | BT20/MB231 | MP |
| >hg38_21_+_8433172_8446622_RNA45SN1_45S_50ntFlanks@10378.10406.29\|AGGGTTCCATGTGAACAGCAGTTGAACAT | >hg38_21_+_8433172_8446622_RNA45SN1_45S_50ntFlanks | 10378 | 10406 | 29 | AGGGTTCCATGTGAACAGCAGTTGAACAT | 59.6 | -3.7 | 1.7 | -2.2 | 2.8E-02 | 4.5E-02 | BT20/MB468 | MP |
| >hg38_21_+_8433172_8446622_RNA45SN1_45S_50ntFlanks@4727.4765.39\|AATCAAGAACGAAAGTCGGAGGTTCGAAGACGATCAGAT | >hg38_21_+_8433172_8446622_RNA45SN1_45S_50ntFlanks | 4727 | 4765 | 39 | AATCAAGAACGAAAGTCGGAGGTTCGAAGACGATCAGAT | 136.3 | 2.0 | 1.0 | 2.0 | 4.1E-02 | 4.5E-02 | MB231/MB468 | MITO |
| >am_trna119_LysCTT_1_-_145395522_145395594@8.35.28,trna11_LysCTT_5_-_180648979_180649051@8.35.28,trna128_LysCTT_1_-_55423542_55423614@8.35.28,trna13_LysCTT_6_+_26556774_26556846@8.35.28,trna6_LysCTT_19_-_52425396_52425468@8.35.28,trna7_LysCTT_16_+_3225692_3225764@8.35.28,trna9_LysCTT_5_+_180634755_180634827@8.35.28\|TAGCTCAGTCGGTAGAGCATGAGACTCT | >am_trna119_LysCTT_1_-_145395522_145395594 | 8 | 35 | 28 | TAGCTCAGTCGGTAGAGCATGAGACTCT | 36.4 | 4.2 | 2.0 | 2.1 | 3.8E-02 | 4.5E-02 | MB231/MB468 | MP |
| >trnaMT_LeuTAA_MT_+_3230_3304@18.44.27\|CGGTAATCGCATAAAACTTAAAACTTT | >trnaMT_LeuTAA_MT_+_3230_3304 | 18 | 44 | 27 | CGGTAATCGCATAAAACTTAAAACTTT | 472.3 | -1.9 | 0.9 | -2.1 | 3.9E-02 | 4.5E-02 | MB231/MB468 | CYTO |
| >trnaMT_SerGCT_MT_+_12207_12265@23.62.40\|AACTCATGCCCCCATGTCTAACAACATGGCTTTCTCACCA | >trnaMT_SerGCT_MT_+_12207_12265 | 23 | 62 | 40 | AACTCATGCCCCCATGTCTAACAACATGGCTTTCTCACCA | 105.7 | 5.0 | 2.3 | 2.2 | 2.8E-02 | 4.5E-02 | BT20/MB468 | MP |
| >hg38_21_+_8433172_8446622_RNA45SN1_45S_50ntFlanks@10686.10708.23\|GCGGCGTCCGGTGAGCTCTCGCT | >hg38_21_+_8433172_8446622_RNA45SN1_45S_50ntFlanks | 10686 | 10708 | 23 | GCGGCGTCCGGTGAGCTCTCGCT | 279.7 | -3.1 | 1.5 | -2.1 | 3.8E-02 | 4.5E-02 | MB231/MB468 | MP |
| >MI0000286\|hsa-mir-210&WithFlank&11\|-\|568083\|568204@72.92.21(+1U)&[MIMAT0000267&hsa-miR-210-3p&offsets\|0\|-1(+1U);m-102&11\|-\|568112\|568133&offsets\|0\|-1(+1U)]\|CTGTGCGTGTGACAGCGGCTGT | >MI0000286\|hsa-mir-210&WithFlank&11\|-\|568083\|568204 | 72 | 92 | 21(+1U) | CTGTGCGTGTGACAGCGGCTGT | 229.0 | 4.4 | 2.1 | 2.1 | 3.8E-02 | 4.5E-02 | MB231/MB468 | MP |
| >hg38_21_+_8433172_8446622_RNA45SN1_45S_50ntFlanks@11781.11805.25\|CGCATGAATGGATGAACGAGATTCC | >hg38_21_+_8433172_8446622_RNA45SN1_45S_50ntFlanks | 11781 | 11805 | 25 | CGCATGAATGGATGAACGAGATTCC | 492.2 | -2.2 | 1.0 | -2.2 | 2.8E-02 | 4.5E-02 | BT20/MB468 | MP |
| >hg38_21_+_8433172_8446622_RNA45SN1_45S_50ntFlanks@6661.6680.20\|CGGTGGATCACTCGGCTCGT | >hg38_21_+_8433172_8446622_RNA45SN1_45S_50ntFlanks | 6661 | 6680 | 20 | CGGTGGATCACTCGGCTCGT | 456.7 | 1.6 | 0.6 | 2.5 | 1.3E-02 | 4.5E-02 | BT20/MB231 | NUC |
| >hg38_21_+_8433172_8446622_RNA45SN1_45S_50ntFlanks@11568.11586.19\|GCAGCCGACTTAGAACTGG | >hg38_21_+_8433172_8446622_RNA45SN1_45S_50ntFlanks | 11568 | 11586 | 19 | GCAGCCGACTTAGAACTGG | 639.6 | -2.2 | 0.9 | -2.5 | 1.3E-02 | 4.5E-02 | BT20/MB231 | NUC |
| >hg38_21_+_8433172_8446622_RNA45SN1_45S_50ntFlanks@8345.8375.31\|AAGGGAAAGTTGAAAAGAACTTTGAAGAGAG | >hg38_21_+_8433172_8446622_RNA45SN1_45S_50ntFlanks | 8345 | 8375 | 31 | AAGGGAAAGTTGAAAAGAACTTTGAAGAGAG | 4367.6 | 3.8 | 1.5 | 2.5 | 1.3E-02 | 4.5E-02 | BT20/MB231 | MP |
| >hg38_MT_+_1621_3279_RNR2_16S_50ntFlanks@1383.1414.32\|ACATCCCGATGGTGCAGCCGCTATTAAAGGTT | >hg38_MT_+_1621_3279_RNR2_16S_50ntFlanks | 1383 | 1414 | 32 | ACATCCCGATGGTGCAGCCGCTATTAAAGGTT | 125.9 | -4.8 | 1.9 | -2.5 | 1.3E-02 | 4.5E-02 | BT20/MB231 | MP |
| >hg38_21_+_8433172_8446622_RNA45SN1_45S_50ntFlanks@9858.9880.23\|CGGGATGAACCGAACGCCGGGTT | >hg38_21_+_8433172_8446622_RNA45SN1_45S_50ntFlanks | 9858 | 9880 | 23 | CGGGATGAACCGAACGCCGGGTT | 315.0 | -3.4 | 1.6 | -2.2 | 2.8E-02 | 4.5E-02 | BT20/MB468 | MP |
| >MI0000285\|hsa-mir-205&WithFlank&1\|+\|209432127\|209432248@40.60.21(+1C)&[MIMAT0000266&hsa-miR-205-5p&offsets\|0\|-1(+1C);m-351&1\|+\|209432166\|209432187&offsets\|0\|-1(+1C)]\|TCCTTCATTCCACCGGAGTCTC | >MI0000285\|hsa-mir-205&WithFlank&1\|+\|209432127\|209432248 | 40 | 60 | 21(+1C) | TCCTTCATTCCACCGGAGTCTC | 236.4 | 2.8 | 1.3 | 2.2 | 2.8E-02 | 4.5E-02 | BT20/MB468 | MP |
| >am_MI0000786\|hsa-mir-378a&WithFlank&5\|+\|149732819\|149732896@50.69.20(+1U)&[MIMAT0000732&hsa-miR-378a-3p&offsets\|+1\|-1(+1U);m-44&5\|+\|149732867\|149732888&offsets\|+1\|-1(+1U)]\|CTGGACTTGGAGTCAGAAGGT | >am_MI0000786\|hsa-mir-378a&WithFlank&5\|+\|149732819\|149732896 | 50 | 69 | 20(+1U) | CTGGACTTGGAGTCAGAAGGT | 857.2 | 1.9 | 0.8 | 2.4 | 1.7E-02 | 4.5E-02 | BT20/MB231 | CYTO |
| >hg38_21_+_8433172_8446622_RNA45SN1_45S_50ntFlanks@6722.6744.23\|ATTGCAGGACACATTGATCATCG | >hg38_21_+_8433172_8446622_RNA45SN1_45S_50ntFlanks | 6722 | 6744 | 23 | ATTGCAGGACACATTGATCATCG | 470.9 | -4.0 | 1.8 | -2.2 | 2.8E-02 | 4.5E-02 | BT20/MB468 | MP |
| >hg38_21_+_8433172_8446622_RNA45SN1_45S_50ntFlanks@7993.8012.20\|GTGGCGACCCGCTGAATTTA | >hg38_21_+_8433172_8446622_RNA45SN1_45S_50ntFlanks | 7993 | 8012 | 20 | GTGGCGACCCGCTGAATTTA | 124.4 | -3.8 | 1.7 | -2.2 | 2.8E-02 | 4.5E-02 | BT20/MB468 | MP |

<table>
<tr><td></td><td></td><td></td><td></td><td></td><td></td><td></td><td></td><td></td><td></td><td></td><td></td><td></td><td></td></tr>
<tr><td>>hg38_21_+_8433172_8446622_RNA45SN1_45S_50ntFlanks@4732.4752.21|AGAACGAAAGTCGGAGGTTCG</td><td>>hg38_21_+_8433172_8446622_RNA45SN1_45S_50ntFlanks</td><td>4732</td><td>4752</td><td>21</td><td>AGAACGAAAGTCGGAGGTTCG</td><td>257.3</td><td>-1.2</td><td>0.6</td><td>-2.1</td><td>3.9E-02</td><td>4.5E-02</td><td>MB231/MB468</td><td>CYTO</td></tr>
<tr><td>>hg38_21_+_8433172_8446622_RNA45SN1_45S_50ntFlanks@3705.3741.37|TACCTGGTTGATCCTGCCAGTAGCATATGCTTGTCTC</td><td>>hg38_21_+_8433172_8446622_RNA45SN1_45S_50ntFlanks</td><td>3705</td><td>3741</td><td>37</td><td>TACCTGGTTGATCCTGCCAGTAGCATATGCTTGTCTC</td><td>430.4</td><td>1.1</td><td>0.5</td><td>2.0</td><td>4.1E-02</td><td>4.5E-02</td><td>MB231/MB468</td><td>NUC</td></tr>
<tr><td>>MI0000542|hsa-mir-320a&WithFlank&8|-|22244960|22245043@49.68.20(+1C)&[MIMAT0000510&hsa-miR-320a-3p&offsets|+1|-1(+1C);m-60&8|-|22244975|22244996&offsets|+1|-1(+1C)]|AAAGCTGGGTTGAGAGGGCGC</td><td>>MI0000542|hsa-mir-320a&WithFlank&8|-|22244960|22245043</td><td>49</td><td>68</td><td>20(+1C)</td><td>AAAGCTGGGTTGAGAGGGCGC</td><td>28.5</td><td>4.5</td><td>2.2</td><td>2.1</td><td>3.8E-02</td><td>4.5E-02</td><td>MB231/MB468</td><td>MP</td></tr>
<tr><td>>hg38_21_+_8433172_8446622_RNA45SN1_45S_50ntFlanks@11583.11612.30|CTGGTGCGGACCAGGGGAATCCGACTGTTT</td><td>>hg38_21_+_8433172_8446622_RNA45SN1_45S_50ntFlanks</td><td>11583</td><td>11612</td><td>30</td><td>CTGGTGCGGACCAGGGGAATCCGACTGTTT</td><td>2239.5</td><td>-2.1</td><td>1.0</td><td>-2.0</td><td>4.1E-02</td><td>4.5E-02</td><td>BT20/MB468</td><td>NUC</td></tr>
<tr><td>>hg38_21_+_8433172_8446622_RNA45SN1_45S_50ntFlanks@11567.11586.20|AGCAGCCGACTTAGAACTGG</td><td>>hg38_21_+_8433172_8446622_RNA45SN1_45S_50ntFlanks</td><td>11567</td><td>11586</td><td>20</td><td>AGCAGCCGACTTAGAACTGG</td><td>845.1</td><td>-2.1</td><td>0.9</td><td>-2.4</td><td>1.7E-02</td><td>4.5E-02</td><td>BT20/MB231</td><td>CYTO</td></tr>
<tr><td>>hg38_21_+_8433172_8446622_RNA45SN1_45S_50ntFlanks@4050.4079.30|CGAACGTCTGCCCTATCAACTTTCGATGGT</td><td>>hg38_21_+_8433172_8446622_RNA45SN1_45S_50ntFlanks</td><td>4050</td><td>4079</td><td>30</td><td>CGAACGTCTGCCCTATCAACTTTCGATGGT</td><td>117.2</td><td>2.3</td><td>1.0</td><td>2.2</td><td>2.8E-02</td><td>4.5E-02</td><td>BT20/MB468</td><td>MP</td></tr>
<tr><td>>hg38_21_+_8433172_8446622_RNA45SN1_45S_50ntFlanks@6731.6772.42|CACATTGATCATCGACACTTCGAACGCACTTGCGGCCCCGGG</td><td>>hg38_21_+_8433172_8446622_RNA45SN1_45S_50ntFlanks</td><td>6731</td><td>6772</td><td>42</td><td>CACATTGATCATCGACACTTCGAACGCACTTGCGGCCCCGGG</td><td>104.4</td><td>-5.1</td><td>2.3</td><td>-2.2</td><td>2.8E-02</td><td>4.5E-02</td><td>BT20/MB468</td><td>MP</td></tr>
<tr><td>>hg38_21_+_8433172_8446622_RNA45SN1_45S_50ntFlanks@8912.8936.25|CGCCGAATCCCGGGGCCGAGGGAGC</td><td>>hg38_21_+_8433172_8446622_RNA45SN1_45S_50ntFlanks</td><td>8912</td><td>8936</td><td>25</td><td>CGCCGAATCCCGGGGCCGAGGGAGC</td><td>41.5</td><td>3.6</td><td>1.8</td><td>2.1</td><td>3.8E-02</td><td>4.5E-02</td><td>MB231/MB468</td><td>MP</td></tr>
<tr><td>>hg38_21_+_8433172_8446622_RNA45SN1_45S_50ntFlanks@10362.10379.18|AAGGCCGAAGTGGAGAAG</td><td>>hg38_21_+_8433172_8446622_RNA45SN1_45S_50ntFlanks</td><td>10362</td><td>10379</td><td>18</td><td>AAGGCCGAAGTGGAGAAG</td><td>197.4</td><td>-3.8</td><td>1.7</td><td>-2.2</td><td>2.8E-02</td><td>4.5E-02</td><td>BT20/MB468</td><td>MP</td></tr>
<tr><td>>trna152_ValCAC_6_-_27248049_27248121@1.32.32|GCTTCTGTAGTGTAGTGGTTATCACGTTCGCC</td><td>>trna152_ValCAC_6_-_27248049_27248121</td><td>1</td><td>32</td><td>32</td><td>GCTTCTGTAGTGTAGTGGTTATCACGTTCGCC</td><td>2339.5</td><td>-2.3</td><td>0.9</td><td>-2.4</td><td>1.5E-02</td><td>4.5E-02</td><td>BT20/MB231</td><td>MITO</td></tr>
<tr><td>>hg38_21_+_8433172_8446622_RNA45SN1_45S_50ntFlanks@9545.9563.19|GTGGAGGTCCGTAGCGGTC</td><td>>hg38_21_+_8433172_8446622_RNA45SN1_45S_50ntFlanks</td><td>9545</td><td>9563</td><td>19</td><td>GTGGAGGTCCGTAGCGGTC</td><td>1995.4</td><td>-3.6</td><td>1.6</td><td>-2.2</td><td>2.8E-02</td><td>4.5E-02</td><td>BT20/MB468</td><td>MP</td></tr>
<tr><td>>hg38_1_-_228634819_228635039_RNA5S12_5S_50ntFlanks@151.170.20|ATACCGGGTGCTGTAGGCTT</td><td>>hg38_1_-_228634819_228635039_RNA5S12_5S_50ntFlanks</td><td>151</td><td>170</td><td>20</td><td>ATACCGGGTGCTGTAGGCTT</td><td>2291.8</td><td>1.1</td><td>0.5</td><td>2.1</td><td>3.5E-02</td><td>4.5E-02</td><td>BT20/MB468</td><td>MITO</td></tr>
<tr><td>>hg38_21_+_8433172_8446622_RNA45SN1_45S_50ntFlanks@4919.4937.19|CCACCAGGAGTGGAGCCTG</td><td>>hg38_21_+_8433172_8446622_RNA45SN1_45S_50ntFlanks</td><td>4919</td><td>4937</td><td>19</td><td>CCACCAGGAGTGGAGCCTG</td><td>583.7</td><td>-2.4</td><td>1.1</td><td>-2.1</td><td>3.5E-02</td><td>4.5E-02</td><td>BT20/MB468</td><td>MITO</td></tr>
<tr><td>>MI0006444|hsa-mir-1307&WithFlank&10|-|103394247|103394407@86.109.24&[MIMAT0005951&hsa-miR-1307-3p&offsets|0|+2;m-66&10|-|103394300|103394322&offsets|0|+1]|ACTCGGCGTGGCGTCGGTCGTGGT</td><td>>MI0006444|hsa-mir-1307&WithFlank&10|-|103394247|103394407</td><td>86</td><td>109</td><td>24</td><td>ACTCGGCGTGGCGTCGGTCGTGGT</td><td>65.0</td><td>3.6</td><td>1.8</td><td>2.1</td><td>3.8E-02</td><td>4.5E-02</td><td>MB231/MB468</td><td>MP</td></tr>
<tr><td>>hg38_MT_+_1621_3279_RNR2_16S_50ntFlanks@115.145.31|AAATAAAGTATAGGCGATAGAAATTGAAACC</td><td>>hg38_MT_+_1621_3279_RNR2_16S_50ntFlanks</td><td>115</td><td>145</td><td>31</td><td>AAATAAAGTATAGGCGATAGAAATTGAAACC</td><td>104.6</td><td>5.8</td><td>2.6</td><td>2.2</td><td>2.8E-02</td><td>4.5E-02</td><td>BT20/MB468</td><td>MP</td></tr>
<tr><td>>hg38_21_+_8433172_8446622_RNA45SN1_45S_50ntFlanks@11771.11803.33|ATTAGTGACGCGCATGAATGGATGAACGAGATT</td><td>>hg38_21_+_8433172_8446622_RNA45SN1_45S_50ntFlanks</td><td>11771</td><td>11803</td><td>33</td><td>ATTAGTGACGCGCATGAATGGATGAACGAGATT</td><td>84.5</td><td>-3.6</td><td>1.7</td><td>-2.1</td><td>3.8E-02</td><td>4.5E-02</td><td>MB231/MB468</td><td>MP</td></tr>
<tr><td>>hg38_21_+_8433172_8446622_RNA45SN1_45S_50ntFlanks@11568.11601.34|GCAGCCGACTTAGAACTGGTGCGGACCAGGGGAA</td><td>>hg38_21_+_8433172_8446622_RNA45SN1_45S_50ntFlanks</td><td>11568</td><td>11601</td><td>34</td><td>GCAGCCGACTTAGAACTGGTGCGGACCAGGGGAA</td><td>331.7</td><td>3.3</td><td>1.3</td><td>2.5</td><td>1.3E-02</td><td>4.5E-02</td><td>BT20/MB231</td><td>MP</td></tr>
<tr><td>>trnaMT_ValTAC_MT_+_1602_1670@31.72.42|TTACACTTAGGAGATTTCAACTTAACTTGACCGCTCTGACCA</td><td>>trnaMT_ValTAC_MT_+_1602_1670</td><td>31</td><td>72</td><td>42</td><td>TTACACTTAGGAGATTTCAACTTAACTTGACCGCTCTGACCA</td><td>290.9</td><td>4.5</td><td>2.1</td><td>2.2</td><td>2.8E-02</td><td>4.5E-02</td><td>BT20/MB468</td><td>MP</td></tr>
<tr><td>>hg38_21_+_8433172_8446622_RNA45SN1_45S_50ntFlanks@11703.11730.28|AATGAAGCGCGGGTAAACGGCGGGAGTA</td><td>>hg38_21_+_8433172_8446622_RNA45SN1_45S_50ntFlanks</td><td>11703</td><td>11730</td><td>28</td><td>AATGAAGCGCGGGTAAACGGCGGGAGTA</td><td>65.1</td><td>-3.0</td><td>1.4</td><td>-2.2</td><td>2.8E-02</td><td>4.5E-02</td><td>BT20/MB468</td><td>MP</td></tr>
<tr><td>>hg38_21_+_8433172_8446622_RNA45SN1_45S_50ntFlanks@12417.12452.36|TCGATGTCGGCTCTTCCTATCATTGTGAAGCAGAAT</td><td>>hg38_21_+_8433172_8446622_RNA45SN1_45S_50ntFlanks</td><td>12417</td><td>12452</td><td>36</td><td>TCGATGTCGGCTCTTCCTATCATTGTGAAGCAGAAT</td><td>155.1</td><td>-2.2</td><td>0.9</td><td>-2.5</td><td>1.3E-02</td><td>4.5E-02</td><td>BT20/MB231</td><td>NUC</td></tr>
<tr><td>>hg38_21_+_8433172_8446622_RNA45SN1_45S_50ntFlanks@7974.8004.31|ACGCGACCTCAGATCAGACGTGGCGACCCGC</td><td>>hg38_21_+_8433172_8446622_RNA45SN1_45S_50ntFlanks</td><td>7974</td><td>8004</td><td>31</td><td>ACGCGACCTCAGATCAGACGTGGCGACCCGC</td><td>3430.9</td><td>2.7</td><td>1.1</td><td>2.4</td><td>1.7E-02</td><td>4.5E-02</td><td>BT20/MB231</td><td>CYTO</td></tr>
<tr><td>>hg38_21_+_8433172_8446622_RNA45SN1_45S_50ntFlanks@10690.10709.20|CGTCCGGTGAGCTCTCGCTG</td><td>>hg38_21_+_8433172_8446622_RNA45SN1_45S_50ntFlanks</td><td>10690</td><td>10709</td><td>20</td><td>CGTCCGGTGAGCTCTCGCTG</td><td>842.9</td><td>-4.1</td><td>1.9</td><td>-2.2</td><td>2.9E-02</td><td>4.5E-02</td><td>BT20/MB468</td><td>MP</td></tr>
<tr><td>>hg38_21_+_8433172_8446622_RNA45SN1_45S_50ntFlanks@8174.8216.43|TCCTTCTGATCGAGGCCCAGCCCGTGGACGGTGTGAGGCCGGT</td><td>>hg38_21_+_8433172_8446622_RNA45SN1_45S_50ntFlanks</td><td>8174</td><td>8216</td><td>43</td><td>TCCTTCTGATCGAGGCCCAGCCCGTGGACGGTGTGAGGCCGGT</td><td>352.2</td><td>1.9</td><td>0.8</td><td>2.4</td><td>1.5E-02</td><td>4.5E-02</td><td>BT20/MB231</td><td>MITO</td></tr>
<tr><td>>hg38_21_+_8433172_8446622_RNA45SN1_45S_50ntFlanks@6649.6681.33|TACGACTCTTAGCGGTGGATCACTCGGCTCGTG</td><td>>hg38_21_+_8433172_8446622_RNA45SN1_45S_50ntFlanks</td><td>6649</td><td>6681</td><td>33</td><td>TACGACTCTTAGCGGTGGATCACTCGGCTCGTG</td><td>985.8</td><td>1.5</td><td>0.7</td><td>2.0</td><td>4.1E-02</td><td>4.5E-02</td><td>MB231/MB468</td><td>NUC</td></tr>
<tr><td>>hg38_MT_+_598_1651_RNR1_12S_50ntFlanks@984.1004.21|GGAAAGTGCACTTGGACGAAC</td><td>>hg38_MT_+_598_1651_RNR1_12S_50ntFlanks</td><td>984</td><td>1004</td><td>21</td><td>GGAAAGTGCACTTGGACGAAC</td><td>76.5</td><td>2.7</td><td>1.3</td><td>2.1</td><td>3.8E-02</td><td>4.5E-02</td><td>MB231/MB468</td><td>MP</td></tr>
<tr><td>>MI0000085|hsa-mir-27a&WithFlank&19|-|13836434|13836523@57.76.20(+1A)&[MIMAT0000084&hsa-miR-27a-3p&offsets|0|-1(+1A);m-43&19|-|13836447|13836467&offsets|0|-1(+1A)]|TTCACAGTGGCTAAGTTCCGA</td><td>>MI0000085|hsa-mir-27a&WithFlank&19|-|13836434|13836523</td><td>57</td><td>76</td><td>20(+1A)</td><td>TTCACAGTGGCTAAGTTCCGA</td><td>492.8</td><td>1.3</td><td>0.6</td><td>2.0</td><td>4.1E-02</td><td>4.5E-02</td><td>BT20/MB468</td><td>NUC</td></tr>
<tr><td>>hg38_21_+_8433172_8446622_RNA45SN1_45S_50ntFlanks@12003.12031.29|GCGGGCCGCCGGTGAAATACCACTACTCT</td><td>>hg38_21_+_8433172_8446622_RNA45SN1_45S_50ntFlanks</td><td>12003</td><td>12031</td><td>29</td><td>GCGGGCCGCCGGTGAAATACCACTACTCT</td><td>535.1</td><td>-1.5</td><td>0.7</td><td>-2.1</td><td>4.0E-02</td><td>4.5E-02</td><td>BT20/MB468</td><td>CYTO</td></tr>
<tr><td>>hg38_21_+_8433172_8446622_RNA45SN1_45S_50ntFlanks@10352.10383.32|GAGAACTTTGAAGGCCGAAGTGGAGAAGGGTT</td><td>>hg38_21_+_8433172_8446622_RNA45SN1_45S_50ntFlanks</td><td>10352</td><td>10383</td><td>32</td><td>GAGAACTTTGAAGGCCGAAGTGGAGAAGGGTT</td><td>599.3</td><td>3.8</td><td>1.5</td><td>2.5</td><td>1.3E-02</td><td>4.5E-02</td><td>BT20/MB231</td><td>MP</td></tr>
<tr><td>>hg38_21_+_8433172_8446622_RNA45SN1_45S_50ntFlanks@12657.12674.18|GACTGAACGCCTCTAAGT</td><td>>hg38_21_+_8433172_8446622_RNA45SN1_45S_50ntFlanks</td><td>12657</td><td>12674</td><td>18</td><td>GACTGAACGCCTCTAAGT</td><td>172.7</td><td>-1.1</td><td>0.5</td><td>-2.1</td><td>3.6E-02</td><td>4.5E-02</td><td>BT20/MB468</td><td>MITO</td></tr>
<tr><td>>hg38_21_+_8433172_8446622_RNA45SN1_45S_50ntFlanks@11582.11612.31|ACTGGTGCGGACCAGGGGAATCCGACTGTTT</td><td>>hg38_21_+_8433172_8446622_RNA45SN1_45S_50ntFlanks</td><td>11582</td><td>11612</td><td>31</td><td>ACTGGTGCGGACCAGGGGAATCCGACTGTTT</td><td>273.0</td><td>1.6</td><td>0.7</td><td>2.4</td><td>1.5E-02</td><td>4.5E-02</td><td>BT20/MB231</td><td>MITO</td></tr>
<tr><td>>hg38_21_+_8433172_8446622_RNA45SN1_45S_50ntFlanks@4122.4145.24|AGGGTTCGATTCCGGAGAGGGAGC</td><td>>hg38_21_+_8433172_8446622_RNA45SN1_45S_50ntFlanks</td><td>4122</td><td>4145</td><td>24</td><td>AGGGTTCGATTCCGGAGAGGGAGC</td><td>99.2</td><td>-3.2</td><td>1.5</td><td>-2.1</td><td>4.0E-02</td><td>4.5E-02</td><td>BT20/MB468</td><td>CYTO</td></tr>
<tr><td>>MI0000239|hsa-mir-197&WithFlank&1|+|109598887|109598973@54.74.21(+1U)&[MIMAT0000227&hsa-miR-197-3p&offsets|0|-1(+1U);m-99&1|+|109598940|109598961&offsets|0|-1(+1U)]|TTCACCACCTTCTCCACCCAGT</td><td>>MI0000239|hsa-mir-197&WithFlank&1|+|109598887|109598973</td><td>54</td><td>74</td><td>21(+1U)</td><td>TTCACCACCTTCTCCACCCAGT</td><td>2216.9</td><td>1.9</td><td>0.8</td><td>2.4</td><td>1.5E-02</td><td>4.5E-02</td><td>BT20/MB231</td><td>MITO</td></tr>
<tr><td>>hg38_21_+_8433172_8446622_RNA45SN1_45S_50ntFlanks@10878.10902.25|TCGGTCGGGCTGGGGCGCGAAGCGG</td><td>>hg38_21_+_8433172_8446622_RNA45SN1_45S_50ntFlanks</td><td>10878</td><td>10902</td><td>25</td><td>TCGGTCGGGCTGGGGCGCGAAGCGG</td><td>146.3</td><td>-1.4</td><td>0.7</td><td>-2.1</td><td>3.6E-02</td><td>4.5E-02</td><td>BT20/MB468</td><td>MITO</td></tr>
<tr><td>>hg38_21_+_8433172_8446622_RNA45SN1_45S_50ntFlanks@10348.10367.20|AAACGAGAACTTTGAAGGCC</td><td>>hg38_21_+_8433172_8446622_RNA45SN1_45S_50ntFlanks</td><td>10348</td><td>10367</td><td>20</td><td>AAACGAGAACTTTGAAGGCC</td><td>794.4</td><td>1.3</td><td>0.6</td><td>2.1</td><td>4.0E-02</td><td>4.5E-02</td><td>MB231/MB468</td><td>CYTO</td></tr>
<tr><td>>hg38_21_+_8433172_8446622_RNA45SN1_45S_50ntFlanks@7991.8010.20|ACGTGGCGACCCGCTGAATT</td><td>>hg38_21_+_8433172_8446622_RNA45SN1_45S_50ntFlanks</td><td>7991</td><td>8010</td><td>20</td><td>ACGTGGCGACCCGCTGAATT</td><td>565.3</td><td>1.8</td><td>0.9</td><td>2.0</td><td>4.1E-02</td><td>4.5E-02</td><td>MB231/MB468</td><td>NUC</td></tr>
<tr><td>>MI0000737|hsa-mir-200a&WithFlank&1|+|1167857|1167958@60.82.23&[MIMAT0000682&hsa-miR-200a-3p&offsets|0|+1;m-142&1|+|1167916|1167938&offsets|0|0]|TAACACTGTCTGGTAACGATGTT</td><td>>MI0000737|hsa-mir-200a&WithFlank&1|+|1167857|1167958</td><td>60</td><td>82</td><td>23</td><td>TAACACTGTCTGGTAACGATGTT</td><td>277.0</td><td>-2.5</td><td>1.2</td><td>-2.0</td><td>4.1E-02</td><td>4.5E-02</td><td>MB231/MB468</td><td>NUC</td></tr>
<tr><td>>hg38_21_+_8433172_8446622_RNA45SN1_45S_50ntFlanks@10351.10379.29|CGAGAACTTTGAAGGCCGAAGTGGAGAAG</td><td>>hg38_21_+_8433172_8446622_RNA45SN1_45S_50ntFlanks</td><td>10351</td><td>10379</td><td>29</td><td>CGAGAACTTTGAAGGCCGAAGTGGAGAAG</td><td>255.8</td><td>4.5</td><td>1.8</td><td>2.5</td><td>1.3E-02</td><td>4.5E-02</td><td>BT20/MB231</td><td>MP</td></tr>
<tr><td>>MI0000088|hsa-mir-30a&WithFlank&6|-|71403545|71403627@13.33.21&[MIMAT0000087&hsa-miR-30a-5p&offsets|+1|0;m-15&6|-|71403595|71403616&offsets|+1|0]|GTAAACATCCTCGACTGGAAG</td><td>>MI0000088|hsa-mir-30a&WithFlank&6|-|71403545|71403627</td><td>13</td><td>33</td><td>21</td><td>GTAAACATCCTCGACTGGAAG</td><td>50.0</td><td>3.4</td><td>1.6</td><td>2.1</td><td>3.8E-02</td><td>4.5E-02</td><td>MB231/MB468</td><td>MP</td></tr>
<tr><td>>hg38_21_+_8433172_8446622_RNA45SN1_45S_50ntFlanks@9344.9370.27|GAAAGCCGCCGTGGCGCAATGAAGGTG</td><td>>hg38_21_+_8433172_8446622_RNA45SN1_45S_50ntFlanks</td><td>9344</td><td>9370</td><td>27</td><td>GAAAGCCGCCGTGGCGCAATGAAGGTG</td><td>86.2</td><td>-3.8</td><td>1.7</td><td>-2.2</td><td>2.9E-02</td><td>4.5E-02</td><td>BT20/MB468</td><td>MP</td></tr>
<tr><td>>hg38_21_+_8433172_8446622_RNA45SN1_45S_50ntFlanks@4738.4761.24|AAAGTCGGAGGTTCGAAGACGATC</td><td>>hg38_21_+_8433172_8446622_RNA45SN1_45S_50ntFlanks</td><td>4738</td><td>4761</td><td>24</td><td>AAAGTCGGAGGTTCGAAGACGATC</td><td>501.9</td><td>1.2</td><td>0.6</td><td>2.0</td><td>4.1E-02</td><td>4.5E-02</td><td>MB231/MB468</td><td>NUC</td></tr>
<tr><td>>hg38_21_+_8433172_8446622_RNA45SN1_45S_50ntFlanks@4724.4767.44|ATTAATCAAGAACGAAAGTCGGAGGTTCGAAGACGATCAGATAC</td><td>>hg38_21_+_8433172_8446622_RNA45SN1_45S_50ntFlanks</td><td>4724</td><td>4767</td><td>44</td><td>ATTAATCAAGAACGAAAGTCGGAGGTTCGAAGACGATCAGATAC</td><td>616.5</td><td>-2.5</td><td>1.0</td><td>-2.4</td><td>1.7E-02</td><td>4.5E-02</td><td>BT20/MB231</td><td>CYTO</td></tr>
<tr><td>>MI0000298|hsa-mir-221&WithFlank&X|-|45746151|45746272@71.91.21(+1G)&[MIMAT0000278&hsa-miR-221-3p&offsets|0|-2(+1G);m-91&X|-|45746181|45746202&offsets|0|-1(+1G)]|AGCTACATTGTCTGCTGGGTTG</td><td>>MI0000298|hsa-mir-221&WithFlank&X|-|45746151|45746272</td><td>71</td><td>91</td><td>21(+1G)</td><td>AGCTACATTGTCTGCTGGGTTG</td><td>668.3</td><td>-2.8</td><td>1.1</td><td>-2.5</td><td>1.3E-02</td><td>4.5E-02</td><td>BT20/MB231</td><td>MP</td></tr>
<tr><td>>hg38_21_+_8433172_8446622_RNA45SN1_45S_50ntFlanks@10069.10090.22|AGTGGACGGGAGCGGCGGGGGC</td><td>>hg38_21_+_8433172_8446622_RNA45SN1_45S_50ntFlanks</td><td>10069</td><td>10090</td><td>22</td><td>AGTGGACGGGAGCGGCGGGGGC</td><td>134.7</td><td>4.8</td><td>1.9</td><td>2.5</td><td>1.3E-02</td><td>4.5E-02</td><td>BT20/MB231</td><td>MP</td></tr>
<tr><td>>hg38_21_+_8433172_8446622_RNA45SN1_45S_50ntFlanks@5472.5513.42|ACGGCCCTGGCGGAGCGCTGAGAAGACGGTCGAACTTGACTA</td><td>>hg38_21_+_8433172_8446622_RNA45SN1_45S_50ntFlanks</td><td>5472</td><td>5513</td><td>42</td><td>ACGGCCCTGGCGGAGCGCTGAGAAGACGGTCGAACTTGACTA</td><td>1604.4</td><td>-2.1</td><td>0.9</td><td>-2.4</td><td>1.7E-02</td><td>4.5E-02</td><td>BT20/MB231</td><td>CYTO</td></tr>
<tr><td>>hg38_21_+_8433172_8446622_RNA45SN1_45S_50ntFlanks@10821.10838.18|GGAAGTCGGCAAGCCGGA</td><td>>hg38_21_+_8433172_8446622_RNA45SN1_45S_50ntFlanks</td><td>10821</td><td>10838</td><td>18</td><td>GGAAGTCGGCAAGCCGGA</td><td>147.0</td><td>4.1</td><td>1.7</td><td>2.5</td><td>1.3E-02</td><td>4.5E-02</td><td>BT20/MB231</td><td>MP</td></tr>
<tr><td>>hg38_21_+_8433172_8446622_RNA45SN1_45S_50ntFlanks@9475.9503.29|CGAGCGCACGTGTTAGGACCCGAAAGATG</td><td>>hg38_21_+_8433172_8446622_RNA45SN1_45S_50ntFlanks</td><td>9475</td><td>9503</td><td>29</td><td>CGAGCGCACGTGTTAGGACCCGAAAGATG</td><td>576.1</td><td>-1.9</td><td>0.9</td><td>-2.1</td><td>3.6E-02</td><td>4.5E-02</td><td>BT20/MB468</td><td>MITO</td></tr>
<tr><td>>MI0001445|hsa-mir-423&WithFlank&17|+|30117073|30117178@23.41.19&[MIMAT0004748&hsa-miR-423-5p&offsets|0|-4;m-141&17|+|30117095|30117117&offsets|0|-4]|TGAGGGGCAGAGAGCGAGA</td><td>>MI0001445|hsa-mir-423&WithFlank&17|+|30117073|30117178</td><td>23</td><td>41</td><td>19</td><td>TGAGGGGCAGAGAGCGAGA</td><td>53.0</td><td>4.2</td><td>2.0</td><td>2.1</td><td>3.8E-02</td><td>4.5E-02</td><td>MB231/MB468</td><td>MP</td></tr>
<tr><td>>trna116_GluCTC_1_-_145399233_145399304@3.31.29,trna59_GluCTC_1_+_249168447_249168518@3.31.29,trna71_GluCTC_1_-_161439189_161439260@3.31.29,trna74_GluCTC_1_-_161431809_161431880@3.31.29,trna77_GluCTC_1_-_161424398_161424469@3.31.29,trna77_GluCTC_6_+_28949976_28950047@3.31.29,trna80_GluCTC_1_-_161417018_161417089@3.31.29,trna87_GluCTC_6_-_126101393_126101464@3.31.29|CCTGGTGGTCTAGTGGTTAGGATTCGGCG</td><td>>trna116_GluCTC_1_-_145399233_145399304</td><td>3</td><td>31</td><td>29</td><td>CCTGGTGGTCTAGTGGTTAGGATTCGGCG</td><td>91.7</td><td>-1.9</td><td>0.9</td><td>-2.1</td><td>3.8E-02</td><td>4.5E-02</td><td>MB231/MB468</td><td>MP</td></tr>
<tr><td>>hg38_21_+_8433172_8446622_RNA45SN1_45S_50ntFlanks@9860.9899.40|GGATGAACCGAACGCCGGGTTAAGGCGCCCGATGCCGACG</td><td>>hg38_21_+_8433172_8446622_RNA45SN1_45S_50ntFlanks</td><td>9860</td><td>9899</td><td>40</td><td>GGATGAACCGAACGCCGGGTTAAGGCGCCCGATGCCGACG</td><td>92.1</td><td>-3.4</td><td>1.5</td><td>-2.2</td><td>2.9E-02</td><td>4.5E-02</td><td>BT20/MB468</td><td>MP</td></tr>
</table>

| | | | | | | | | | | | | | |
|---|---|---|---|---|---|---|---|---|---|---|---|---|---|
| >hg38_21_+_8433172_8446622_RNA45SN1_45S_50ntFlanks@11322.11343.22\|ACGTCTCGTCGCGCGCGCGTCC | >hg38_21_+_8433172_8446622_RNA45SN1_45S_50ntFlanks | 11322 | 11343 | 22 | ACGTCTCGTCGCGCGCGCGTCC | 1329.9 | -1.4 | 0.7 | -2.1 | 4.0E-02 | 4.5E-02 | BT20/MB468 | CYTO |
| >hg38_21_+_8433172_8446622_RNA45SN1_45S_50ntFlanks@9551.9569.19\|GTCCGTAGCGGTCCTGACG | >hg38_21_+_8433172_8446622_RNA45SN1_45S_50ntFlanks | 9551 | 9569 | 19 | GTCCGTAGCGGTCCTGACG | 856.6 | -2.0 | 0.8 | -2.4 | 1.8E-02 | 4.5E-02 | BT20/MB231 | CYTO |
| >hg38_21_+_8433172_8446622_RNA45SN1_45S_50ntFlanks@11710.11740.31\|CGCGGGTAAACGGCGGGAGTAACTATGACTC | >hg38_21_+_8433172_8446622_RNA45SN1_45S_50ntFlanks | 11710 | 11740 | 31 | CGCGGGTAAACGGCGGGAGTAACTATGACTC | 455.9 | -1.9 | 0.9 | -2.1 | 4.0E-02 | 4.5E-02 | MB231/MB468 | CYTO |
| >hg38_21_+_8433172_8446622_RNA45SN1_45S_50ntFlanks@8123.8140.18\|ATGTGGCGTACGGAAGAC | >hg38_21_+_8433172_8446622_RNA45SN1_45S_50ntFlanks | 8123 | 8140 | 18 | ATGTGGCGTACGGAAGAC | 86.2 | 2.3 | 1.1 | 2.2 | 2.9E-02 | 4.6E-02 | BT20/MB468 | MP |
| >am_MI0003639\|hsa-mir-625&WithFlank&14\|+\|65471096\|65471192@58.79.22&[MIMAT0004808&hsa-miR-625-3p&offsets\|0\|0;m-137&14\|+\|65471153\|65471174&offsets\|0\|0]\|GACTATAGAACTTTCCCCCTCA | >am_MI0003639\|hsa-mir-625&WithFlank&14\|+\|65471096\|65471192 | 58 | 79 | 22 | GACTATAGAACTTTCCCCCTCA | 395.9 | -2.5 | 1.0 | -2.5 | 1.3E-02 | 4.6E-02 | BT20/MB231 | MP |
| >hg38_MT_+_598_1651_RNR1_12S_50ntFlanks@905.922.18\|AGTATACTTCAAAGGACA | >hg38_MT_+_598_1651_RNR1_12S_50ntFlanks | 905 | 922 | 18 | AGTATACTTCAAAGGACA | 243.8 | -3.8 | 1.5 | -2.5 | 1.3E-02 | 4.6E-02 | BT20/MB231 | MP |
| >hg38_21_+_8433172_8446622_RNA45SN1_45S_50ntFlanks@9485.9506.22\|TGTTAGGACCCGAAAGATGGTG | >hg38_21_+_8433172_8446622_RNA45SN1_45S_50ntFlanks | 9485 | 9506 | 22 | TGTTAGGACCCGAAAGATGGTG | 87.6 | 3.9 | 1.8 | 2.2 | 2.9E-02 | 4.6E-02 | BT20/MB468 | MP |
| >MI0000803\|hsa-mir-330&WithFlank&19\|-\|45638988\|45639093@63.85.23&[MIMAT0000751&hsa-miR-330-3p&offsets\|0\|0;m-347&19\|-\|45639009\|45639030&offsets\|-1\|0]\|GCAAAGCACACGGCCTGCAGAGA | >MI0000803\|hsa-mir-330&WithFlank&19\|-\|45638988\|45639093 | 63 | 85 | 23 | GCAAAGCACACGGCCTGCAGAGA | 378.0 | 2.0 | 0.8 | 2.4 | 1.8E-02 | 4.6E-02 | BT20/MB231 | CYTO |
| >hg38_21_+_8433172_8446622_RNA45SN1_45S_50ntFlanks@10498.10524.27\|TTCAGATCCCCGAATCCGGAGTGGCGG | >hg38_21_+_8433172_8446622_RNA45SN1_45S_50ntFlanks | 10498 | 10524 | 27 | TTCAGATCCCCGAATCCGGAGTGGCGG | 81.9 | -3.7 | 1.7 | -2.2 | 2.9E-02 | 4.6E-02 | BT20/MB468 | MP |
| >hg38_21_+_8433172_8446622_RNA45SN1_45S_50ntFlanks@11784.11818.35\|ATGAATGGATGAACGAGATTCCCACTGTCCCTACC | >hg38_21_+_8433172_8446622_RNA45SN1_45S_50ntFlanks | 11784 | 11818 | 35 | ATGAATGGATGAACGAGATTCCCACTGTCCCTACC | 77.5 | 2.2 | 1.0 | 2.2 | 2.9E-02 | 4.6E-02 | BT20/MB468 | MP |
| >hg38_21_+_8433172_8446622_RNA45SN1_45S_50ntFlanks@4212.4242.31\|AGGTAGTGACGAAAAATAACAATACAGGACT | >hg38_21_+_8433172_8446622_RNA45SN1_45S_50ntFlanks | 4212 | 4242 | 31 | AGGTAGTGACGAAAAATAACAATACAGGACT | 667.3 | 3.2 | 1.4 | 2.2 | 2.9E-02 | 4.6E-02 | BT20/MB468 | MP |
| >hg38_21_+_8433172_8446622_RNA45SN1_45S_50ntFlanks@11671.11704.34\|CCAGTGCTCTGAATGTCAAAGTGAAGAAATTCAA | >hg38_21_+_8433172_8446622_RNA45SN1_45S_50ntFlanks | 11671 | 11704 | 34 | CCAGTGCTCTGAATGTCAAAGTGAAGAAATTCAA | 427.4 | -1.8 | 0.9 | -2.1 | 4.0E-02 | 4.6E-02 | MB231/MB468 | CYTO |
| >hg38_21_+_8433172_8446622_RNA45SN1_45S_50ntFlanks@7989.8019.31\|AGACGTGGCGACCCGCTGAATTTAAGCATAT | >hg38_21_+_8433172_8446622_RNA45SN1_45S_50ntFlanks | 7989 | 8019 | 31 | AGACGTGGCGACCCGCTGAATTTAAGCATAT | 1048.7 | -2.4 | 1.0 | -2.4 | 1.8E-02 | 4.6E-02 | BT20/MB231 | CYTO |
| >hg38_21_+_8433172_8446622_RNA45SN1_45S_50ntFlanks@4732.4761.30\|AGAACGAAAGTCGGAGGTTCGAAGACGATC | >hg38_21_+_8433172_8446622_RNA45SN1_45S_50ntFlanks | 4732 | 4761 | 30 | AGAACGAAAGTCGGAGGTTCGAAGACGATC | 4424.2 | 3.8 | 1.5 | 2.5 | 1.3E-02 | 4.6E-02 | BT20/MB231 | MP |
| >hg38_MT_+_1621_3279_RNR2_16S_50ntFlanks@1383.1401.19\|ACATCCCGATGGTGCAGCC | >hg38_MT_+_1621_3279_RNR2_16S_50ntFlanks | 1383 | 1401 | 19 | ACATCCCGATGGTGCAGCC | 69.6 | -4.9 | 2.0 | -2.5 | 1.3E-02 | 4.6E-02 | BT20/MB231 | MP |
| >hg38_21_+_8433172_8446622_RNA45SN1_45S_50ntFlanks@11642.11660.19\|GGCGGGTGTTGACGCGATG | >hg38_21_+_8433172_8446622_RNA45SN1_45S_50ntFlanks | 11642 | 11660 | 19 | GGCGGGTGTTGACGCGATG | 1124.8 | -2.8 | 1.3 | -2.2 | 2.9E-02 | 4.6E-02 | BT20/MB468 | MP |
| >hg38_21_+_8433172_8446622_RNA45SN1_45S_50ntFlanks@6653.6676.24\|ACTCTTAGCGGTGGATCACTCGGC | >hg38_21_+_8433172_8446622_RNA45SN1_45S_50ntFlanks | 6653 | 6676 | 24 | ACTCTTAGCGGTGGATCACTCGGC | 359.5 | 3.1 | 1.2 | 2.5 | 1.3E-02 | 4.6E-02 | BT20/MB231 | MP |
| >hg38_21_+_8433172_8446622_RNA45SN1_45S_50ntFlanks@12455.12477.23\|ACCAAGCGTTGGATTGTTCACCC | >hg38_21_+_8433172_8446622_RNA45SN1_45S_50ntFlanks | 12455 | 12477 | 23 | ACCAAGCGTTGGATTGTTCACCC | 297.0 | -1.9 | 0.8 | -2.4 | 1.8E-02 | 4.6E-02 | BT20/MB231 | CYTO |
| >hg38_21_+_8433172_8446622_RNA45SN1_45S_50ntFlanks@10781.10823.43\|AGGTGAACAGCCTCTGGCATGTTGGAACAATGTAGGTAAGGGA | >hg38_21_+_8433172_8446622_RNA45SN1_45S_50ntFlanks | 10781 | 10823 | 43 | AGGTGAACAGCCTCTGGCATGTTGGAACAATGTAGGTAAGGGA | 180.6 | -2.7 | 1.3 | -2.1 | 3.6E-02 | 4.6E-02 | BT20/MB468 | MITO |
| >trnaMT_ProTGG_MT_-_15956_16023@50.70.21\|TTAAAGACTTTTTCTCTGACC | >trnaMT_ProTGG_MT_-_15956_16023 | 50 | 70 | 21 | TTAAAGACTTTTTCTCTGACC | 351.5 | -2.6 | 1.0 | -2.5 | 1.3E-02 | 4.6E-02 | BT20/MB231 | MP |
| >hg38_21_+_8433172_8446622_RNA45SN1_45S_50ntFlanks@6731.6753.23\|CACATTGATCATCGACACTTCGA | >hg38_21_+_8433172_8446622_RNA45SN1_45S_50ntFlanks | 6731 | 6753 | 23 | CACATTGATCATCGACACTTCGA | 203.6 | -3.8 | 1.8 | -2.2 | 2.9E-02 | 4.6E-02 | BT20/MB468 | MP |
| >hg38_MT_+_1621_3279_RNR2_16S_50ntFlanks@1369.1393.25\|GATGTTGGATCAGGACATCCCGATG | >hg38_MT_+_1621_3279_RNR2_16S_50ntFlanks | 1369 | 1393 | 25 | GATGTTGGATCAGGACATCCCGATG | 38.4 | 3.5 | 1.7 | 2.1 | 3.8E-02 | 4.6E-02 | MB231/MB468 | MP |
| >hg38_21_+_8433172_8446622_RNA45SN1_45S_50ntFlanks@4588.4607.20\|CTATTTTGTTGGTTTTCGGA | >hg38_21_+_8433172_8446622_RNA45SN1_45S_50ntFlanks | 4588 | 4607 | 20 | CTATTTTGTTGGTTTTCGGA | 155.1 | -1.7 | 0.8 | -2.1 | 3.8E-02 | 4.6E-02 | MB231/MB468 | MP |
| >hg38_21_+_8433172_8446622_RNA45SN1_45S_50ntFlanks@4732.4757.26\|AGAACGAAAGTCGGAGGTTCGAAGAC | >hg38_21_+_8433172_8446622_RNA45SN1_45S_50ntFlanks | 4732 | 4757 | 26 | AGAACGAAAGTCGGAGGTTCGAAGAC | 104.5 | -2.8 | 1.3 | -2.1 | 3.8E-02 | 4.6E-02 | MB231/MB468 | MP |
| >MI0000809\|hsa-mir-151a&WithFlank&8\|-\|140732558\|140732659@51.73.23(+1U)&[MIMAT0000757&hsa-miR-151a-3p&offsets\|-2\|0(+1U);m-30&8\|-\|140732587\|140732607&offsets\|-2\|0(+1U)]\|TACTAGACTGAAGCTCCTTGAGGT | >MI0000809\|hsa-mir-151a&WithFlank&8\|-\|140732558\|140732659 | 51 | 73 | 23(+1U) | TACTAGACTGAAGCTCCTTGAGGT | 322.0 | -2.3 | 0.9 | -2.5 | 1.3E-02 | 4.6E-02 | BT20/MB231 | MP |
| >hg38_21_+_8433172_8446622_RNA45SN1_45S_50ntFlanks@6647.6677.31\|CGTACGACTCTTAGCGGTGGATCACTCGGCT | >hg38_21_+_8433172_8446622_RNA45SN1_45S_50ntFlanks | 6647 | 6677 | 31 | CGTACGACTCTTAGCGGTGGATCACTCGGCT | 1227.6 | 3.0 | 1.2 | 2.5 | 1.3E-02 | 4.6E-02 | BT20/MB231 | MP |
| >MI0000270\|hsa-mir-181b-1&WithFlank&1\|-\|198858867\|198858988@42.60.19(+1A);MI0000683\|hsa-mir-181b-2&WithFlank&9\|+\|124693704\|124693804@22.40.19(+1A)&[MIMAT0000257&hsa-miR-181b-5p&offsets\|0\|-4(+1A);m-73&1\|-\|198858925\|198858947&offsets\|0\|-4(+1A)];[MIMAT0000257_1&hsa-miR-181b-5p&offsets\|0\|-4(+1A);m-72&9\|+\|124693725\|124693747&offsets\|0\|-4(+1A)]\|AACATTCATTGCTGTCGGTA | >MI0000270\|hsa-mir-181b-1&WithFlank&1\|-\|198858867\|198858988 | 42 | 60 | 19(+1A) | AACATTCATTGCTGTCGGTA | 108.3 | -3.0 | 1.2 | -2.5 | 1.3E-02 | 4.6E-02 | BT20/MB231 | MP |
| >hg38_MT_+_1621_3279_RNR2_16S_50ntFlanks@931.959.29\|GTGACACATGTTTAACGGCCGCGGTACCC | >hg38_MT_+_1621_3279_RNR2_16S_50ntFlanks | 931 | 959 | 29 | GTGACACATGTTTAACGGCCGCGGTACCC | 48.0 | 4.2 | 2.0 | 2.1 | 3.8E-02 | 4.6E-02 | MB231/MB468 | MP |
| >MI0000469\|hsa-mir-125a&WithFlank&19\|+\|51693248\|51693345@21.43.23&[MIMAT0000443&hsa-miR-125a-5p&offsets\|0\|-1;m-63&19\|+\|51693268\|51693289&offsets\|0\|+1]\|TCCCTGAGACCCTTTAACCTGTG | >MI0000469\|hsa-mir-125a&WithFlank&19\|+\|51693248\|51693345 | 21 | 43 | 23 | TCCCTGAGACCCTTTAACCTGTG | 23976.3 | 1.4 | 0.6 | 2.5 | 1.3E-02 | 4.6E-02 | BT20/MB231 | NUC |
| >MI0003137\|hsa-mir-193b&WithFlank&16\|+\|14303961\|14304055@57.76.20&[MIMAT0002819&hsa-miR-193b-3p&offsets\|0\|-2;m-166&16\|+\|14304017\|14304038&offsets\|0\|-2]\|AACTGGCCCTCAAAGTCCCG | >MI0003137\|hsa-mir-193b&WithFlank&16\|+\|14303961\|14304055 | 57 | 76 | 20 | AACTGGCCCTCAAAGTCCCG | 218.4 | 3.1 | 1.3 | 2.5 | 1.3E-02 | 4.6E-02 | BT20/MB231 | MP |
| >hg38_21_+_8433172_8446622_RNA45SN1_45S_50ntFlanks@10484.10512.29\|GAAAGGGAGTCGGGTTCAGATCCCCGAAT | >hg38_21_+_8433172_8446622_RNA45SN1_45S_50ntFlanks | 10484 | 10512 | 29 | GAAAGGGAGTCGGGTTCAGATCCCCGAAT | 71.9 | 3.2 | 1.5 | 2.2 | 2.9E-02 | 4.6E-02 | BT20/MB468 | MP |
| >trna111_HisGTG_1_-_147774845_147774916@-1C.31.32,trna118_HisGTG_1_-_145396881_145396952@-1C.31.32,trna16_HisGTG_1_+_146544773_146544844@-1C.31.32,trna1_HisGTG_15_+_45493349_45493420@-1C.31.32,trna21_HisGTG_1_+_147753471_147753542@-1C.31.32,trna33_HisGTG_6_+_27125906_27125977@-1C.31.32,trna7_HisGTG_9_-_14433938_14434009@-1C.31.32,trna8_HisGTG_15_-_45492611_45492682@-1C.31.32,trna9_HisGTG_15_-_45490804_45490875@-1C.31.32\|CGCCGTGATCGTATAGTGGTTAGTACTCTGCG | >trna111_HisGTG_1_-_147774845_147774916 | -1C | 31 | 32 | CGCCGTGATCGTATAGTGGTTAGTACTCTGCG | 97.8 | -2.8 | 1.3 | -2.1 | 3.8E-02 | 4.6E-02 | MB231/MB468 | MP |
| >hg38_21_+_8433172_8446622_RNA45SN1_45S_50ntFlanks@11573.11612.40\|CGACTTAGAACTGGTGCGGACCAGGGGAATCCGACTGTTT | >hg38_21_+_8433172_8446622_RNA45SN1_45S_50ntFlanks | 11573 | 11612 | 40 | CGACTTAGAACTGGTGCGGACCAGGGGAATCCGACTGTTT | 125.3 | -2.5 | 1.2 | -2.1 | 3.8E-02 | 4.6E-02 | MB231/MB468 | MP |
| >hg38_21_+_8433172_8446622_RNA45SN1_45S_50ntFlanks@7978.8009.32\|GACCTCAGATCAGACGTGGCGACCCGCTGAAT | >hg38_21_+_8433172_8446622_RNA45SN1_45S_50ntFlanks | 7978 | 8009 | 32 | GACCTCAGATCAGACGTGGCGACCCGCTGAAT | 543.9 | -1.9 | 0.8 | -2.4 | 1.8E-02 | 4.6E-02 | BT20/MB231 | CYTO |
| >hg38_21_+_8433172_8446622_RNA45SN1_45S_50ntFlanks@12416.12453.38\|TTCGATGTCGGCTCTTCCTATCATTGTGAAGCAGAATT | >hg38_21_+_8433172_8446622_RNA45SN1_45S_50ntFlanks | 12416 | 12453 | 38 | TTCGATGTCGGCTCTTCCTATCATTGTGAAGCAGAATT | 254.5 | -2.1 | 0.9 | -2.5 | 1.3E-02 | 4.6E-02 | BT20/MB231 | NUC |
| >MI0000342\|hsa-mir-200b&WithFlank&1\|+\|1167098\|1167204@63.84.22(+1U)&[MIMAT0000318&hsa-miR-200b-3p&offsets\|0\|0(+1U);m-93&1\|+\|1167160\|1167182&offsets\|0\|-1(+1U)]\|TAATACTGCCTGGTAATGATGAT | >MI0000342\|hsa-mir-200b&WithFlank&1\|+\|1167098\|1167204 | 63 | 84 | 22(+1U) | TAATACTGCCTGGTAATGATGAT | 279.1 | -2.9 | 1.4 | -2.0 | 4.2E-02 | 4.6E-02 | MB231/MB468 | MITO |
| >trnaMT_SerGCT_MT_+_12207_12265@14.40.27\|AGAACTGCTAACTCATGCCCCCATGTC | >trnaMT_SerGCT_MT_+_12207_12265 | 14 | 40 | 27 | AGAACTGCTAACTCATGCCCCCATGTC | 127.7 | 2.8 | 1.4 | 2.0 | 4.2E-02 | 4.6E-02 | MB231/MB468 | MITO |
| >hg38_21_+_8433172_8446622_RNA45SN1_45S_50ntFlanks@12990.13013.24\|GCTCCCTCGCTGCGATCTATTGAA | >hg38_21_+_8433172_8446622_RNA45SN1_45S_50ntFlanks | 12990 | 13013 | 24 | GCTCCCTCGCTGCGATCTATTGAA | 289.1 | -3.7 | 1.5 | -2.5 | 1.4E-02 | 4.6E-02 | BT20/MB231 | NUC |
| >hg38_21_+_8433172_8446622_RNA45SN1_45S_50ntFlanks@9544.9573.30\|GGTGGAGGTCCGTAGCGGTCCTGACGTGCA | >hg38_21_+_8433172_8446622_RNA45SN1_45S_50ntFlanks | 9544 | 9573 | 30 | GGTGGAGGTCCGTAGCGGTCCTGACGTGCA | 623.5 | -1.6 | 0.8 | -2.0 | 4.2E-02 | 4.6E-02 | MB231/MB468 | NUC |
| >hg38_21_+_8433172_8446622_RNA45SN1_45S_50ntFlanks@12252.12284.33\|AAGCTCGCTTGATCTTGATTTTCAGTACGAATA | >hg38_21_+_8433172_8446622_RNA45SN1_45S_50ntFlanks | 12252 | 12284 | 33 | AAGCTCGCTTGATCTTGATTTTCAGTACGAATA | 347.4 | -1.9 | 0.9 | -2.1 | 3.6E-02 | 4.6E-02 | BT20/MB468 | MITO |
| >hg38_MT_+_1621_3279_RNR2_16S_50ntFlanks@460.485.26\|TTGTAAATTTAACTGTTAGTCCAAAG | >hg38_MT_+_1621_3279_RNR2_16S_50ntFlanks | 460 | 485 | 26 | TTGTAAATTTAACTGTTAGTCCAAAG | 68.7 | 3.1 | 1.5 | 2.1 | 3.8E-02 | 4.6E-02 | MB231/MB468 | MP |
| >hg38_21_+_8433172_8446622_RNA45SN1_45S_50ntFlanks@11561.11594.34\|GCGCCTAGCAGCCGACTTAGAACTGGTGCGGACC | >hg38_21_+_8433172_8446622_RNA45SN1_45S_50ntFlanks | 11561 | 11594 | 34 | GCGCCTAGCAGCCGACTTAGAACTGGTGCGGACC | 245.1 | -2.1 | 1.0 | -2.2 | 2.9E-02 | 4.6E-02 | BT20/MB468 | MP |
| >hg38_21_+_8433172_8446622_RNA45SN1_45S_50ntFlanks@12938.12957.20\|ACCATTCGTAGACGACCTGC | >hg38_21_+_8433172_8446622_RNA45SN1_45S_50ntFlanks | 12938 | 12957 | 20 | ACCATTCGTAGACGACCTGC | 392.1 | -2.0 | 0.8 | -2.4 | 1.8E-02 | 4.6E-02 | BT20/MB231 | CYTO |
| >hg38_21_+_8433172_8446622_RNA45SN1_45S_50ntFlanks@10130.10149.20\|GCGGCGGCGGCGGCGGCGGG | >hg38_21_+_8433172_8446622_RNA45SN1_45S_50ntFlanks | 10130 | 10149 | 20 | GCGGCGGCGGCGGCGGCGGG | 206.0 | -3.6 | 1.7 | -2.1 | 3.6E-02 | 4.6E-02 | BT20/MB468 | MITO |
| >hg38_21_+_8433172_8446622_RNA45SN1_45S_50ntFlanks@8847.8866.20\|GCGGCGGCGGCGGCGGCGGG | >hg38_21_+_8433172_8446622_RNA45SN1_45S_50ntFlanks | 8847 | 8866 | 20 | GCGGCGGCGGCGGCGGCGGG | 206.0 | -3.6 | 1.7 | -2.1 | 3.6E-02 | 4.6E-02 | BT20/MB468 | MITO |
| >hg38_21_+_8433172_8446622_RNA45SN1_45S_50ntFlanks@4008.4025.18\|CTCGGGCCGATCGCACGC | >hg38_21_+_8433172_8446622_RNA45SN1_45S_50ntFlanks | 4008 | 4025 | 18 | CTCGGGCCGATCGCACGC | 257.7 | -2.4 | 1.2 | -2.0 | 4.0E-02 | 4.6E-02 | BT20/MB468 | CYTO |
| >hg38_21_+_8433172_8446622_RNA45SN1_45S_50ntFlanks@9604.9629.26\|AAAGACTAATCGAACCATCTAGTAGC | >hg38_21_+_8433172_8446622_RNA45SN1_45S_50ntFlanks | 9604 | 9629 | 26 | AAAGACTAATCGAACCATCTAGTAGC | 195.1 | 5.1 | 2.1 | 2.5 | 1.3E-02 | 4.6E-02 | BT20/MB231 | MP |
| >MI0000299\|hsa-mir-222&WithFlank&X\|-\|45747009\|45747130@75.96.22(+1G)&[MIMAT0000279&hsa-miR-222-3p&offsets\|0\|+1(+1G);m-140&X\|-\|45747033\|45747056&offsets\|0\|-2(+1G)]\|AGCTACATCTGGCTACTGGGTCG | >MI0000299\|hsa-mir-222&WithFlank&X\|-\|45747009\|45747130 | 75 | 96 | 22(+1G) | AGCTACATCTGGCTACTGGGTCG | 312.0 | -3.0 | 1.2 | -2.5 | 1.3E-02 | 4.6E-02 | BT20/MB231 | MP |
| >hg38_21_+_8433172_8446622_RNA45SN1_45S_50ntFlanks@11874.11899.26\|AGAAGACCCTGTTGAGCTTGACTCTA | >hg38_21_+_8433172_8446622_RNA45SN1_45S_50ntFlanks | 11874 | 11899 | 26 | AGAAGACCCTGTTGAGCTTGACTCTA | 438.9 | -1.3 | 0.6 | -2.1 | 3.6E-02 | 4.6E-02 | BT20/MB468 | MITO |

| | | | | | | | | | | | | | |
|---|---|---|---|---|---|---|---|---|---|---|---|---|---|
| >MI0000061\|hsa-let-7a-2&WithFlank&11\|-\|122146516\|122146599@11.30.20(+1C);MI0000062\|hsa-let-7a-3&WithFlank&22\|+\|46112743\|46112828@10.29.20(+1C);MI0000060\|hsa-let-7a-1&WithFlank&9\|+\|94175951\|94176042@12.31.20(+1C)&[MIMAT0000062&hsa-let-7a-5p&offsets\|0\|-2(+1C);m-6&11\|-\|122146568\|122146589&offsets\|0\|-2(+1C)];[MIMAT0000062_1&hsa-let-7a-5p&offsets\|0\|-2(+1C);m-5&22\|+\|46112752\|46112773&offsets\|0\|-2(+1C)];[MIMAT0000062_2&hsa-let-7a-5p&offsets\|0\|-2(+1C);m-4&9\|+\|94175962\|94175983&offsets\|0\|-2(+1C)]\|TGAGGTAGTAGGTTGTATAGC | >MI0000061\|hsa-let-7a-2&WithFlank&11\|-\|122146516\|122146599 | 11 | 30 | 20(+1C) | TGAGGTAGTAGGTTGTATAGC | 1619.5 | -2.6 | 1.1 | -2.5 | 1.3E-02 | 4.6E-02 | BT20/MB231 | MP |
| >hg38_21_+_8433172_8446622_RNA45SN1_45S_50ntFlanks@9035.9056.22\|CGTGGGGGGGCCGGGCCACCCC | >hg38_21_+_8433172_8446622_RNA45SN1_45S_50ntFlanks | 9035 | 9056 | 22 | CGTGGGGGGGCCGGGCCACCCC | 159.4 | -3.5 | 1.7 | -2.1 | 3.6E-02 | 4.6E-02 | BT20/MB468 | MITO |
| >trna116_GluCTC_1_-_145399233_145399304@37.75.39,trna134_GluTTC_1_-_16861774_16861845@37.75.39,trna71_GluCTC_1_-_161439189_161439260@37.75.39,trna74_GluCTC_1_-_161431809_161431880@37.75.39,trna77_GluCTC_1_-_161424398_161424469@37.75.39,trna77_GluCTC_6_+_28949976_28950047@37.75.39,trna80_GluCTC_1_-_161417018_161417089@37.75.39,trna84_GluTTC_1_-_161391883_161391954@37.75.39,trna87_GluCTC_6_-_126101393_126101464@37.75.39\|ACCGCCGCGGCCCGGGTTCGATTCCCGGTCAGGGAACCA | >trna116_GluCTC_1_-_145399233_145399304 | 37 | 75 | 39 | ACCGCCGCGGCCCGGGTTCGATTCCCGGTCAGGGAACCA | 600.8 | -2.2 | 0.9 | -2.4 | 1.6E-02 | 4.6E-02 | BT20/MB231 | MITO |
| >hg38_MT_+_1621_3279_RNR2_16S_50ntFlanks@460.492.33\|TTGTAAATTTAACTGTTAGTCCAAAGAGGAACA | >hg38_MT_+_1621_3279_RNR2_16S_50ntFlanks | 460 | 492 | 33 | TTGTAAATTTAACTGTTAGTCCAAAGAGGAACA | 476.4 | -1.7 | 0.7 | -2.4 | 1.6E-02 | 4.6E-02 | BT20/MB231 | MITO |
| >trna116_GluCTC_1_-_145399233_145399304@41.75.35,trna134_GluTTC_1_-_16861774_16861845@41.75.35,trna71_GluCTC_1_-_161439189_161439260@41.75.35,trna74_GluCTC_1_-_161431809_161431880@41.75.35,trna77_GluCTC_1_-_161424398_161424469@41.75.35,trna77_GluCTC_6_+_28949976_28950047@41.75.35,trna80_GluCTC_1_-_161417018_161417089@41.75.35,trna84_GluTTC_1_-_161391883_161391954@41.75.35,trna87_GluCTC_6_-_126101393_126101464@41.75.35\|CCGCGGCCCGGGTTCGATTCCCGGTCAGGGAACCA | >trna116_GluCTC_1_-_145399233_145399304 | 41 | 75 | 35 | CCGCGGCCCGGGTTCGATTCCCGGTCAGGGAACCA | 60.6 | 2.3 | 1.1 | 2.1 | 3.9E-02 | 4.6E-02 | MB231/MB468 | MP |
| >trna111_HisGTG_1_-_147774845_147774916@-1G.31.32,trna118_HisGTG_1_-_145396881_145396952@-1G.31.32,trna16_HisGTG_1_+_146544773_146544844@-1G.31.32,trna1_HisGTG_15_+_45493349_45493420@-1G.31.32,trna21_HisGTG_1_+_147753471_147753542@-1G.31.32,trna33_HisGTG_6_+_27125906_27125977@-1G.31.32,trna7_HisGTG_9_-_14433938_14434009@-1G.31.32,trna8_HisGTG_15_-_45492611_45492682@-1G.31.32,trna9_HisGTG_15_-_45490804_45490875@-1G.31.32\|GGCCGTGATCGTATAGTGGTTAGTACTCTGCG | >trna111_HisGTG_1_-_147774845_147774916 | -1G | 31 | 32 | GGCCGTGATCGTATAGTGGTTAGTACTCTGCG | 30895.4 | -1.9 | 1.0 | -2.0 | 4.2E-02 | 4.6E-02 | MB231/MB468 | NUC |
| >hg38_21_+_8433172_8446622_RNA45SN1_45S_50ntFlanks@10866.10890.25\|CTAAGGGCTGGGTCGGTCGGGCTGG | >hg38_21_+_8433172_8446622_RNA45SN1_45S_50ntFlanks | 10866 | 10890 | 25 | CTAAGGGCTGGGTCGGTCGGGCTGG | 608.7 | -2.5 | 1.1 | -2.2 | 2.9E-02 | 4.6E-02 | BT20/MB468 | MP |
| >hg38_21_+_8433172_8446622_RNA45SN1_45S_50ntFlanks@4297.4316.20\|CATTGGAGGGCAAGTCTGGT | >hg38_21_+_8433172_8446622_RNA45SN1_45S_50ntFlanks | 4297 | 4316 | 20 | CATTGGAGGGCAAGTCTGGT | 305.3 | -1.7 | 0.8 | -2.0 | 4.0E-02 | 4.6E-02 | MB231/MB468 | CYTO |
| >hg38_21_+_8433172_8446622_RNA45SN1_45S_50ntFlanks@3991.4020.30\|TGGTGACTCTAGATAACCTCGGGCCGATCG | >hg38_21_+_8433172_8446622_RNA45SN1_45S_50ntFlanks | 3991 | 4020 | 30 | TGGTGACTCTAGATAACCTCGGGCCGATCG | 50.4 | -3.7 | 1.7 | -2.2 | 2.9E-02 | 4.6E-02 | BT20/MB468 | MP |
| >hg38_21_+_8433172_8446622_RNA45SN1_45S_50ntFlanks@8109.8138.30\|GCGGGGCGCGGGACATGTGGCGTACGGAAG | >hg38_21_+_8433172_8446622_RNA45SN1_45S_50ntFlanks | 8109 | 8138 | 30 | GCGGGGCGCGGGACATGTGGCGTACGGAAG | 323.6 | -2.5 | 1.1 | -2.2 | 2.9E-02 | 4.6E-02 | BT20/MB468 | MP |
| >hg38_21_+_8433172_8446622_RNA45SN1_45S_50ntFlanks@12994.13012.19\|CCTCGCTGCGATCTATTGA | >hg38_21_+_8433172_8446622_RNA45SN1_45S_50ntFlanks | 12994 | 13012 | 19 | CCTCGCTGCGATCTATTGA | 469.6 | -1.3 | 0.6 | -2.0 | 4.0E-02 | 4.6E-02 | MB231/MB468 | CYTO |
| >MI0000542\|hsa-mir-320a&WithFlank&8\|-\|22244960\|22245043@50.70.21&[MIMAT0000510&hsa-miR-320a-3p&offsets\|+2\|+1;m-60&8\|-\|22244975\|22244996&offsets\|+2\|+1]\|AAGCTGGGTTGAGAGGGCGAA | >MI0000542\|hsa-mir-320a&WithFlank&8\|-\|22244960\|22245043 | 50 | 70 | 21 | AAGCTGGGTTGAGAGGGCGAA | 63.3 | 3.6 | 1.7 | 2.1 | 3.9E-02 | 4.6E-02 | MB231/MB468 | MP |
| >hg38_21_+_8433172_8446622_RNA45SN1_45S_50ntFlanks@4738.4759.22\|AAAGTCGGAGGTTCGAAGACGA | >hg38_21_+_8433172_8446622_RNA45SN1_45S_50ntFlanks | 4738 | 4759 | 22 | AAAGTCGGAGGTTCGAAGACGA | 392.2 | -1.3 | 0.6 | -2.0 | 4.1E-02 | 4.6E-02 | BT20/MB468 | CYTO |
| >hg38_MT_+_1621_3279_RNR2_16S_50ntFlanks@842.859.18\|AAGTAAAAGGAACTCGGC | >hg38_MT_+_1621_3279_RNR2_16S_50ntFlanks | 842 | 859 | 18 | AAGTAAAAGGAACTCGGC | 67.0 | 1.8 | 0.9 | 2.1 | 3.9E-02 | 4.6E-02 | MB231/MB468 | MP |
| >trna5_AspGTC_12_+_98897281_98897352@36.73.38\|CACGCGGGAGACCGGGGTTCAATTCCCCGACGGGGAGC | >trna5_AspGTC_12_+_98897281_98897352 | 36 | 73 | 38 | CACGCGGGAGACCGGGGTTCAATTCCCCGACGGGGAGC | 710.9 | -4.6 | 1.9 | -2.4 | 1.8E-02 | 4.6E-02 | BT20/MB231 | CYTO |
| >hg38_21_+_8433172_8446622_RNA45SN1_45S_50ntFlanks@9300.9323.24\|ACGGACCAAGGAGTCTAACACGTG | >hg38_21_+_8433172_8446622_RNA45SN1_45S_50ntFlanks | 9300 | 9323 | 24 | ACGGACCAAGGAGTCTAACACGTG | 130.5 | 2.1 | 0.9 | 2.2 | 2.9E-02 | 4.6E-02 | BT20/MB468 | MP |
| >MI0000301\|hsa-mir-224&WithFlank&X\|-\|151958572\|151958664@14.37.24&[MIMAT0000281&hsa-miR-224-5p&offsets\|+1\|0;m-238&X\|-\|151958630\|151958651&offsets\|0\|+2]\|CAAGTCACTAGTGGTTCCGTTTAG | >MI0000301\|hsa-mir-224&WithFlank&X\|-\|151958572\|151958664 | 14 | 37 | 24 | CAAGTCACTAGTGGTTCCGTTTAG | 4587.6 | 2.0 | 0.9 | 2.4 | 1.8E-02 | 4.6E-02 | BT20/MB231 | CYTO |
| >am_MI0000809\|hsa-mir-151a&WithFlank&8\|-\|140732558\|140732659@52.73.22&[MIMAT0000757&hsa-miR-151a-3p&offsets\|-1\|0;m-30&8\|-\|140732587\|140732607&offsets\|-1\|0]\|ACTAGACTGAAGCTCCTTGAGG | >am_MI0000809\|hsa-mir-151a&WithFlank&8\|-\|140732558\|140732659 | 52 | 73 | 22 | ACTAGACTGAAGCTCCTTGAGG | 2908.7 | 1.8 | 0.7 | 2.4 | 1.8E-02 | 4.6E-02 | BT20/MB231 | CYTO |
| >MI0000299\|hsa-mir-222&WithFlank&X\|-\|45747009\|45747130@75.93.19&[MIMAT0000279&hsa-miR-222-3p&offsets\|0\|-2;m-140&X\|-\|45747033\|45747056&offsets\|0\|-5]\|AGCTACATCTGGCTACTGG | >MI0000299\|hsa-mir-222&WithFlank&X\|-\|45747009\|45747130 | 75 | 93 | 19 | AGCTACATCTGGCTACTGG | 64.3 | 4.2 | 2.0 | 2.1 | 3.9E-02 | 4.6E-02 | MB231/MB468 | MP |
| >hg38_21_+_8433172_8446622_RNA45SN1_45S_50ntFlanks@10349.10373.25\|AACGAGAACTTTGAAGGCCGAAGTG | >hg38_21_+_8433172_8446622_RNA45SN1_45S_50ntFlanks | 10349 | 10373 | 25 | AACGAGAACTTTGAAGGCCGAAGTG | 634.3 | -2.4 | 1.1 | -2.2 | 2.9E-02 | 4.6E-02 | BT20/MB468 | MP |
| >MI0000298\|hsa-mir-221&WithFlank&X\|-\|45746151\|45746272@71.91.21(+2C)&[MIMAT0000278&hsa-miR-221-3p&offsets\|0\|-2(+2C);m-91&X\|-\|45746181\|45746202&offsets\|0\|-1(+2C)]\|AGCTACATTGTCTGCTGGGTTCC | >MI0000298\|hsa-mir-221&WithFlank&X\|-\|45746151\|45746272 | 71 | 91 | 21(+2C) | AGCTACATTGTCTGCTGGGTTCC | 144.3 | 2.8 | 1.3 | 2.1 | 3.9E-02 | 4.6E-02 | MB231/MB468 | MP |
| >hg38_21_+_8433172_8446622_RNA45SN1_45S_50ntFlanks@11602.11623.22\|TCCGACTGTTTAATTAAAACAA | >hg38_21_+_8433172_8446622_RNA45SN1_45S_50ntFlanks | 11602 | 11623 | 22 | TCCGACTGTTTAATTAAAACAA | 323.2 | 4.3 | 1.7 | 2.5 | 1.3E-02 | 4.6E-02 | BT20/MB231 | MP |
| >hg38_MT_+_1621_3279_RNR2_16S_50ntFlanks@1363.1390.28\|GACCTCGATGTTGGATCAGGACATCCCG | >hg38_MT_+_1621_3279_RNR2_16S_50ntFlanks | 1363 | 1390 | 28 | GACCTCGATGTTGGATCAGGACATCCCG | 110.8 | 2.2 | 1.1 | 2.1 | 3.9E-02 | 4.6E-02 | MB231/MB468 | MP |
| >hg38_21_+_8433172_8446622_RNA45SN1_45S_50ntFlanks@6650.6698.49\|ACGACTCTTAGCGGTGGATCACTCGGCTCGTGCGTCGATGAAGAACGCA | >hg38_21_+_8433172_8446622_RNA45SN1_45S_50ntFlanks | 6650 | 6698 | 49 | ACGACTCTTAGCGGTGGATCACTCGGCTCGTGCGTCGATGAAGAACGCA | 572.4 | -4.4 | 2.1 | -2.1 | 3.9E-02 | 4.6E-02 | MB231/MB468 | MP |
| >hg38_21_+_8433172_8446622_RNA45SN1_45S_50ntFlanks@6647.6666.20\|CGTACGACTCTTAGCGGTGG | >hg38_21_+_8433172_8446622_RNA45SN1_45S_50ntFlanks | 6647 | 6666 | 20 | CGTACGACTCTTAGCGGTGG | 3411.4 | 2.5 | 1.0 | 2.5 | 1.3E-02 | 4.6E-02 | BT20/MB231 | MP |
| >MI0000465\|hsa-mir-191&WithFlank&3\|-\|49020612\|49020715@22.45.24(+1A)&[MIMAT0000440&hsa-miR-191-5p&offsets\|0\|+1(+1A);m-48&3\|-\|49020672\|49020694&offsets\|0\|+1(+1A)]\|CAACGGAATCCCAAAAGCAGCTGTA | >MI0000465\|hsa-mir-191&WithFlank&3\|-\|49020612\|49020715 | 22 | 45 | 24(+1A) | CAACGGAATCCCAAAAGCAGCTGTA | 47.9 | 3.6 | 1.7 | 2.1 | 3.9E-02 | 4.6E-02 | MB231/MB468 | MP |
| >hg38_21_+_8433172_8446622_RNA45SN1_45S_50ntFlanks@12420.12454.35\|ATGTCGGCTCTTCCTATCATTGTGAAGCAGAATTC | >hg38_21_+_8433172_8446622_RNA45SN1_45S_50ntFlanks | 12420 | 12454 | 35 | ATGTCGGCTCTTCCTATCATTGTGAAGCAGAATTC | 158.9 | -2.7 | 1.3 | -2.1 | 3.9E-02 | 4.6E-02 | MB231/MB468 | MP |
| >trnaMT_HisGTG_MT_+_12138_12206@32.70.39\|TGAATCTGACAACAGAGGCTTACGACCCCTTATTTACCC | >trnaMT_HisGTG_MT_+_12138_12206 | 32 | 70 | 39 | TGAATCTGACAACAGAGGCTTACGACCCCTTATTTACCC | 9444.1 | 2.0 | 1.0 | 2.0 | 4.2E-02 | 4.6E-02 | MB231/MB468 | MITO |
| >hg38_21_+_8433172_8446622_RNA45SN1_45S_50ntFlanks@9305.9337.33\|CCAAGGAGTCTAACACGTGCGCGAGTCGGGGGC | >hg38_21_+_8433172_8446622_RNA45SN1_45S_50ntFlanks | 9305 | 9337 | 33 | CCAAGGAGTCTAACACGTGCGCGAGTCGGGGGC | 604.0 | -4.0 | 1.8 | -2.2 | 2.9E-02 | 4.6E-02 | BT20/MB468 | MP |
| >MI0000542\|hsa-mir-320a&WithFlank&8\|-\|22244960\|22245043@49.69.21&[MIMAT0000510&hsa-miR-320a-3p&offsets\|+1\|0;m-60&8\|-\|22244975\|22244996&offsets\|+1\|0]\|AAAGCTGGGTTGAGAGGGCGA | >MI0000542\|hsa-mir-320a&WithFlank&8\|-\|22244960\|22245043 | 49 | 69 | 21 | AAAGCTGGGTTGAGAGGGCGA | 364.7 | 2.9 | 1.4 | 2.1 | 3.9E-02 | 4.6E-02 | MB231/MB468 | MP |
| >hg38_21_+_8433172_8446622_RNA45SN1_45S_50ntFlanks@10806.10837.32\|AACAATGTAGGTAAGGGAAGTCGGCAAGCCGG | >hg38_21_+_8433172_8446622_RNA45SN1_45S_50ntFlanks | 10806 | 10837 | 32 | AACAATGTAGGTAAGGGAAGTCGGCAAGCCGG | 97.6 | -2.8 | 1.3 | -2.2 | 2.9E-02 | 4.6E-02 | BT20/MB468 | MP |
| >hg38_21_+_8433172_8446622_RNA45SN1_45S_50ntFlanks@9625.9658.34\|GTAGCTGGTTCCCTCCGAAGTTTCCCTCAGGATA | >hg38_21_+_8433172_8446622_RNA45SN1_45S_50ntFlanks | 9625 | 9658 | 34 | GTAGCTGGTTCCCTCCGAAGTTTCCCTCAGGATA | 211.1 | -1.1 | 0.5 | -2.1 | 3.6E-02 | 4.6E-02 | BT20/MB468 | MITO |
| >hg38_21_+_8433172_8446622_RNA45SN1_45S_50ntFlanks@3853.3883.31\|AACTGTGGTAATTCTAGAGCTAATACATGCC | >hg38_21_+_8433172_8446622_RNA45SN1_45S_50ntFlanks | 3853 | 3883 | 31 | AACTGTGGTAATTCTAGAGCTAATACATGCC | 218.0 | -1.1 | 0.6 | -2.0 | 4.2E-02 | 4.6E-02 | MB231/MB468 | NUC |
| >hg38_MT_+_1621_3279_RNR2_16S_50ntFlanks@1436.1457.22\|CCTACGTGATCTGAGTTCAGAC | >hg38_MT_+_1621_3279_RNR2_16S_50ntFlanks | 1436 | 1457 | 22 | CCTACGTGATCTGAGTTCAGAC | 90.9 | 2.6 | 1.3 | 2.1 | 3.9E-02 | 4.6E-02 | MB231/MB468 | MP |
| >MI0005566\|hsa-mir-374b&WithFlank&X\|-\|74218541\|74218624@17.36.20(+1C)&[MIMAT0004955&hsa-miR-374b-5p&offsets\|0\|-2(+1C);m-153&X\|-\|74218587\|74218608&offsets\|0\|-2(+1C)]\|ATATAATACAACCTGCTAAGC | >MI0005566\|hsa-mir-374b&WithFlank&X\|-\|74218541\|74218624 | 17 | 36 | 20(+1C) | ATATAATACAACCTGCTAAGC | 71.6 | 3.7 | 1.8 | 2.1 | 3.9E-02 | 4.6E-02 | MB231/MB468 | MP |
| >hg38_21_+_8433172_8446622_RNA45SN1_45S_50ntFlanks@10678.10703.26\|CGGTTCCGGCGGCGTCCGGTGAGCTC | >hg38_21_+_8433172_8446622_RNA45SN1_45S_50ntFlanks | 10678 | 10703 | 26 | CGGTTCCGGCGGCGTCCGGTGAGCTC | 103.1 | -2.8 | 1.3 | -2.2 | 2.9E-02 | 4.6E-02 | BT20/MB468 | MP |
| >hg38_21_+_8433172_8446622_RNA45SN1_45S_50ntFlanks@4040.4059.20\|ACGACCCATTCGAACGTCTG | >hg38_21_+_8433172_8446622_RNA45SN1_45S_50ntFlanks | 4040 | 4059 | 20 | ACGACCCATTCGAACGTCTG | 171.2 | -3.8 | 1.7 | -2.2 | 2.9E-02 | 4.6E-02 | BT20/MB468 | MP |
| >trnaMT_LysTTT_MT_+_8295_8364@36.73.38,trnalookalike7_LysTTT_1_+_568844_568913@36.73.38\|TAAAGATTAAGAGAACCAACACCTCTTTACAGTGACCA | >trnaMT_LysTTT_MT_+_8295_8364 | 36 | 73 | 38 | TAAAGATTAAGAGAACCAACACCTCTTTACAGTGACCA | 138.4 | 3.5 | 1.7 | 2.0 | 4.2E-02 | 4.6E-02 | MB231/MB468 | MITO |
| >hg38_21_+_8433172_8446622_RNA45SN1_45S_50ntFlanks@6671.6690.20\|CTCGGCTCGTGCGTCGATGA | >hg38_21_+_8433172_8446622_RNA45SN1_45S_50ntFlanks | 6671 | 6690 | 20 | CTCGGCTCGTGCGTCGATGA | 106.0 | -3.7 | 1.7 | -2.2 | 2.9E-02 | 4.6E-02 | BT20/MB468 | MP |
| >MI0000076\|hsa-mir-20a&WithFlank&13\|+\|91351059\|91351141@14.35.22(+1U)&[MIMAT0000075&hsa-miR-20a-5p&offsets\|0\|-1(+1U);m-96&13\|+\|91351072\|91351094&offsets\|0\|-1(+1U)]\|TAAAGTGCTTATAGTGCAGGTAT | >MI0000076\|hsa-mir-20a&WithFlank&13\|+\|91351059\|91351141 | 14 | 35 | 22(+1U) | TAAAGTGCTTATAGTGCAGGTAT | 289.7 | 2.9 | 1.4 | 2.1 | 3.9E-02 | 4.6E-02 | MB231/MB468 | MP |
| >hg38_21_+_8433172_8446622_RNA45SN1_45S_50ntFlanks@6649.6695.47\|TACGACTCTTAGCGGTGGATCACTCGGCTCGTGCGTCGATGAAGAAC | >hg38_21_+_8433172_8446622_RNA45SN1_45S_50ntFlanks | 6649 | 6695 | 47 | TACGACTCTTAGCGGTGGATCACTCGGCTCGTGCGTCGATGAAGAAC | 77.9 | 3.9 | 1.8 | 2.2 | 2.9E-02 | 4.6E-02 | BT20/MB468 | MP |

| | | | | | | | | | | | | | |
|---|---|---|---|---|---|---|---|---|---|---|---|---|---|
| >hg38_21_+_8433172_8446622_RNA45SN1_45S_50ntFlanks@12418.12444.27\|CGATGTCGGCTCTTCCTATCATTGTGA | >hg38_21_+_8433172_8446622_RNA45SN1_45S_50ntFlanks | 12418 | 12444 | 27 | CGATGTCGGCTCTTCCTATCATTGTGA | 782.6 | -2.0 | 0.8 | -2.4 | 1.8E-02 | 4.6E-02 | BT20/MB231 | CYTO |
| >hg38_MT_+_1621_3279_RNR2_16S_50ntFlanks@486.504.19\|AGGAACAGCTCTTTGGACA | >hg38_MT_+_1621_3279_RNR2_16S_50ntFlanks | 486 | 504 | 19 | AGGAACAGCTCTTTGGACA | 149.2 | -2.6 | 1.3 | -2.1 | 3.9E-02 | 4.6E-02 | MB231/MB468 | MP |
| >trna119_LysCTT_1_-_145395522_145395594@1.34.34,trna11_LysCTT_5_-_180648979_180649051@1.34.34,trna13_LysCTT_6_+_26556774_26556846@1.34.34,trna7_LysCTT_16_+_3225692_3225764@1.34.34,trna9_LysCTT_5_+_180634755_180634827@1.34.34\|GCCCGGCTAGCTCAGTCGGTAGAGCATGAGACTC | >trna119_LysCTT_1_-_145395522_145395594 | 1 | 34 | 34 | GCCCGGCTAGCTCAGTCGGTAGAGCATGAGACTC | 14985.9 | -2.5 | 1.1 | -2.4 | 1.8E-02 | 4.6E-02 | BT20/MB231 | CYTO |
| >hg38_21_+_8433172_8446622_RNA45SN1_45S_50ntFlanks@11322.11340.19\|ACGTCTCGTCGCGCGCGCG | >hg38_21_+_8433172_8446622_RNA45SN1_45S_50ntFlanks | 11322 | 11340 | 19 | ACGTCTCGTCGCGCGCGCG | 160.2 | -2.8 | 1.3 | -2.2 | 2.9E-02 | 4.6E-02 | BT20/MB468 | MP |
| >hg38_21_+_8433172_8446622_RNA45SN1_45S_50ntFlanks@11566.11600.35\|TAGCAGCCGACTTAGAACTGGTGCGGACCAGGGGA | >hg38_21_+_8433172_8446622_RNA45SN1_45S_50ntFlanks | 11566 | 11600 | 35 | TAGCAGCCGACTTAGAACTGGTGCGGACCAGGGGA | 129.5 | 2.2 | 1.0 | 2.2 | 2.9E-02 | 4.6E-02 | BT20/MB468 | MP |
| >hg38_1_-_228634819_228635039_RNA5S12_5S_50ntFlanks@133.167.35\|ATGGGAGACCGCCTGGGAATACCGGGTGCTGTAGG | >hg38_1_-_228634819_228635039_RNA5S12_5S_50ntFlanks | 133 | 167 | 35 | ATGGGAGACCGCCTGGGAATACCGGGTGCTGTAGG | 81.4 | 3.2 | 1.5 | 2.2 | 2.9E-02 | 4.6E-02 | BT20/MB468 | MP |
| >am_trnaMT_SerGCT_MT_+_12207_12265@14.34.21\|AGAACTGCTAACTCATGCCCC | >am_trnaMT_SerGCT_MT_+_12207_12265 | 14 | 34 | 21 | AGAACTGCTAACTCATGCCCC | 65.5 | 2.9 | 1.4 | 2.1 | 3.9E-02 | 4.6E-02 | MB231/MB468 | MP |
| >hg38_21_+_8433172_8446622_RNA45SN1_45S_50ntFlanks@475.492.18\|CTTCGTGATCGATGTGGT | >hg38_21_+_8433172_8446622_RNA45SN1_45S_50ntFlanks | 475 | 492 | 18 | CTTCGTGATCGATGTGGT | 398.1 | 2.1 | 0.9 | 2.4 | 1.6E-02 | 4.6E-02 | BT20/MB231 | MITO |
| >hg38_21_+_8433172_8446622_RNA45SN1_45S_50ntFlanks@12244.12273.30\|AAGGGCAAAAGCTCGCTTGATCTTGATTTT | >hg38_21_+_8433172_8446622_RNA45SN1_45S_50ntFlanks | 12244 | 12273 | 30 | AAGGGCAAAAGCTCGCTTGATCTTGATTTT | 419.3 | -1.7 | 0.8 | -2.1 | 3.6E-02 | 4.6E-02 | BT20/MB468 | MITO |
| >hg38_21_+_8433172_8446622_RNA45SN1_45S_50ntFlanks@9554.9592.39\|CGTAGCGGTCCTGACGTGCAAATCGGTCGTCCGACCTGG | >hg38_21_+_8433172_8446622_RNA45SN1_45S_50ntFlanks | 9554 | 9592 | 39 | CGTAGCGGTCCTGACGTGCAAATCGGTCGTCCGACCTGG | 61.1 | -4.1 | 1.9 | -2.2 | 2.9E-02 | 4.6E-02 | BT20/MB468 | MP |
| >hg38_21_+_8433172_8446622_RNA45SN1_45S_50ntFlanks@11539.11583.45\|GCGCGGCGCCTCGCCTCGGCCGGCGCCTAGCAGCCGACTTAGAAC | >hg38_21_+_8433172_8446622_RNA45SN1_45S_50ntFlanks | 11539 | 11583 | 45 | GCGCGGCGCCTCGCCTCGGCCGGCGCCTAGCAGCCGACTTAGAAC | 246.3 | 3.6 | 1.8 | 2.0 | 4.2E-02 | 4.6E-02 | MB231/MB468 | NUC |
| >hg38_21_+_8433172_8446622_RNA45SN1_45S_50ntFlanks@10907.10930.24\|GGGCGCGCGCCGCGGCTGGACGAG | >hg38_21_+_8433172_8446622_RNA45SN1_45S_50ntFlanks | 10907 | 10930 | 24 | GGGCGCGCGCCGCGGCTGGACGAG | 265.0 | 3.1 | 1.3 | 2.5 | 1.3E-02 | 4.6E-02 | BT20/MB231 | MP |
| >hg38_21_+_8433172_8446622_RNA45SN1_45S_50ntFlanks@5486.5509.24\|GCGCTGAGAAGACGGTCGAACTTG | >hg38_21_+_8433172_8446622_RNA45SN1_45S_50ntFlanks | 5486 | 5509 | 24 | GCGCTGAGAAGACGGTCGAACTTG | 247.9 | 5.1 | 2.1 | 2.5 | 1.3E-02 | 4.6E-02 | BT20/MB231 | MP |
| >am_MI0000786\|hsa-mir-378a&WithFlank&5\|+\|149732819\|149732896@49.71.23&[MIMAT0000732&hsa-miR-378a-3p&offsets\|0\|+1;m-44&5\|+\|149732867\|149732888&offsets\|0\|+1]\|ACTGGACTTGGAGTCAGAAGGCC | >am_MI0000786\|hsa-mir-378a&WithFlank&5\|+\|149732819\|149732896 | 49 | 71 | 23 | ACTGGACTTGGAGTCAGAAGGCC | 897.1 | 1.5 | 0.7 | 2.1 | 3.9E-02 | 4.6E-02 | MB231/MB468 | MP |
| >hg38_21_+_8433172_8446622_RNA45SN1_45S_50ntFlanks@9936.9956.21\|ACAGCAGGACGGTGGCCATGG | >hg38_21_+_8433172_8446622_RNA45SN1_45S_50ntFlanks | 9936 | 9956 | 21 | ACAGCAGGACGGTGGCCATGG | 66.3 | 5.0 | 2.0 | 2.5 | 1.3E-02 | 4.6E-02 | BT20/MB231 | MP |
| >hg38_21_+_8433172_8446622_RNA45SN1_45S_50ntFlanks@9462.9494.33\|GGGAGGTGGAGCACGAGCGCACGTGTTAGGACC | >hg38_21_+_8433172_8446622_RNA45SN1_45S_50ntFlanks | 9462 | 9494 | 33 | GGGAGGTGGAGCACGAGCGCACGTGTTAGGACC | 70.2 | -3.9 | 1.8 | -2.2 | 2.9E-02 | 4.6E-02 | BT20/MB468 | MP |
| >MI0000082\|hsa-mir-25&WithFlank&7\|-\|100093554\|100093649@58.78.21(+1G)&[MIMAT0000081&hsa-miR-25-3p&offsets\|0\|-1(+1G);m-23&7\|-\|100093571\|100093592&offsets\|0\|-1(+1G)]\|CATTGCACTTGTCTCGGTCTGG | >MI0000082\|hsa-mir-25&WithFlank&7\|-\|100093554\|100093649 | 58 | 78 | 21(+1G) | CATTGCACTTGTCTCGGTCTGG | 392.0 | 1.3 | 0.7 | 2.0 | 4.2E-02 | 4.6E-02 | BT20/MB468 | NUC |
| >hg38_21_+_8433172_8446622_RNA45SN1_45S_50ntFlanks@12542.12573.32\|ATGTGTTGTTGCCATGGTAATCCTGCTCAGTA | >hg38_21_+_8433172_8446622_RNA45SN1_45S_50ntFlanks | 12542 | 12573 | 32 | ATGTGTTGTTGCCATGGTAATCCTGCTCAGTA | 65.0 | -3.7 | 1.7 | -2.2 | 2.9E-02 | 4.6E-02 | BT20/MB468 | MP |
| >hg38_21_+_8433172_8446622_RNA45SN1_45S_50ntFlanks@10362.10382.21\|AAGGCCGAAGTGGAGAAGGGT | >hg38_21_+_8433172_8446622_RNA45SN1_45S_50ntFlanks | 10362 | 10382 | 21 | AAGGCCGAAGTGGAGAAGGGT | 325.8 | -1.6 | 0.8 | -2.0 | 4.1E-02 | 4.6E-02 | MB231/MB468 | CYTO |
| >hg38_21_+_8433172_8446622_RNA45SN1_45S_50ntFlanks@3782.3800.19\|CAGTGAAACTGCGAATGGC | >hg38_21_+_8433172_8446622_RNA45SN1_45S_50ntFlanks | 3782 | 3800 | 19 | CAGTGAAACTGCGAATGGC | 932.7 | -2.8 | 1.3 | -2.2 | 2.9E-02 | 4.6E-02 | BT20/MB468 | MP |
| >hg38_MT_+_598_1651_RNR1_12S_50ntFlanks@52.93.42\|ATAGGTTTGGTCCTAGCCTTTCTATTAGCTCTTAGTAAGATT | >hg38_MT_+_598_1651_RNR1_12S_50ntFlanks | 52 | 93 | 42 | ATAGGTTTGGTCCTAGCCTTTCTATTAGCTCTTAGTAAGATT | 63.3 | 2.8 | 1.4 | 2.0 | 4.2E-02 | 4.6E-02 | BT20/MB468 | NUC |
| >hg38_21_+_8433172_8446622_RNA45SN1_45S_50ntFlanks@12604.12624.21\|GTATGTGCTTGGCTGAGGAGC | >hg38_21_+_8433172_8446622_RNA45SN1_45S_50ntFlanks | 12604 | 12624 | 21 | GTATGTGCTTGGCTGAGGAGC | 360.6 | -2.2 | 0.9 | -2.4 | 1.8E-02 | 4.6E-02 | BT20/MB231 | CYTO |
| >hg38_21_+_8433172_8446622_RNA45SN1_45S_50ntFlanks@6645.6663.19\|CTCGTACGACTCTTAGCGG | >hg38_21_+_8433172_8446622_RNA45SN1_45S_50ntFlanks | 6645 | 6663 | 19 | CTCGTACGACTCTTAGCGG | 4626.8 | 2.5 | 1.0 | 2.5 | 1.3E-02 | 4.6E-02 | BT20/MB231 | MP |
| >hg38_21_+_8433172_8446622_RNA45SN1_45S_50ntFlanks@10477.10495.19\|GCCGATCGAAAGGGAGTCG | >hg38_21_+_8433172_8446622_RNA45SN1_45S_50ntFlanks | 10477 | 10495 | 19 | GCCGATCGAAAGGGAGTCG | 225.0 | -3.9 | 1.8 | -2.2 | 2.9E-02 | 4.6E-02 | BT20/MB468 | MP |
| >hg38_21_+_8433172_8446622_RNA45SN1_45S_50ntFlanks@12457.12487.31\|CAAGCGTTGGATTGTTCACCCACTAATAGGG | >hg38_21_+_8433172_8446622_RNA45SN1_45S_50ntFlanks | 12457 | 12487 | 31 | CAAGCGTTGGATTGTTCACCCACTAATAGGG | 553.4 | -4.0 | 1.9 | -2.2 | 2.9E-02 | 4.6E-02 | BT20/MB468 | MP |
| >hg38_21_+_8433172_8446622_RNA45SN1_45S_50ntFlanks@4241.4275.35\|CTCTTTCGAGGCCCTGTAATTGGAATGAGTCCACT | >hg38_21_+_8433172_8446622_RNA45SN1_45S_50ntFlanks | 4241 | 4275 | 35 | CTCTTTCGAGGCCCTGTAATTGGAATGAGTCCACT | 155.2 | -2.7 | 1.3 | -2.1 | 3.7E-02 | 4.6E-02 | BT20/MB468 | MITO |
| >hg38_21_+_8433172_8446622_RNA45SN1_45S_50ntFlanks@6646.6671.26\|TCGTACGACTCTTAGCGGTGGATCAC | >hg38_21_+_8433172_8446622_RNA45SN1_45S_50ntFlanks | 6646 | 6671 | 26 | TCGTACGACTCTTAGCGGTGGATCAC | 182.7 | 1.8 | 0.8 | 2.2 | 2.9E-02 | 4.7E-02 | BT20/MB468 | MP |
| >hg38_MT_+_1621_3279_RNR2_16S_50ntFlanks@1432.1453.22\|AAGTCCTACGTGATCTGAGTTC | >hg38_MT_+_1621_3279_RNR2_16S_50ntFlanks | 1432 | 1453 | 22 | AAGTCCTACGTGATCTGAGTTC | 194.4 | 2.7 | 1.3 | 2.1 | 3.9E-02 | 4.7E-02 | MB231/MB468 | MP |
| >hg38_21_+_8433172_8446622_RNA45SN1_45S_50ntFlanks@11567.11590.24\|AGCAGCCGACTTAGAACTGGTGCG | >hg38_21_+_8433172_8446622_RNA45SN1_45S_50ntFlanks | 11567 | 11590 | 24 | AGCAGCCGACTTAGAACTGGTGCG | 180.4 | -2.4 | 1.1 | -2.2 | 2.9E-02 | 4.7E-02 | BT20/MB468 | MP |
| >hg38_MT_+_598_1651_RNR1_12S_50ntFlanks@911.950.40\|CTTCAAAGGACATTTAACTAAAACCCCTACGCATTTATAT | >hg38_MT_+_598_1651_RNR1_12S_50ntFlanks | 911 | 950 | 40 | CTTCAAAGGACATTTAACTAAAACCCCTACGCATTTATAT | 94.6 | 4.9 | 2.3 | 2.2 | 2.9E-02 | 4.7E-02 | BT20/MB468 | MP |
| >hg38_21_+_8433172_8446622_RNA45SN1_45S_50ntFlanks@10387.10424.38\|TGTGAACAGCAGTTGAACATGGGTCAGTCGGTCCTGAG | >hg38_21_+_8433172_8446622_RNA45SN1_45S_50ntFlanks | 10387 | 10424 | 38 | TGTGAACAGCAGTTGAACATGGGTCAGTCGGTCCTGAG | 267.9 | 3.6 | 1.8 | 2.1 | 3.9E-02 | 4.7E-02 | MB231/MB468 | MP |
| >MI0000263\|hsa-mir-7-1&WithFlank&9\|-\|83969742\|83969863@30.54.25&[MIMAT0000252_2&hsa-miR-7-5p&offsets\|0\|+1;m-401&9\|-\|83969811\|83969834&offsets\|0\|+1]\|TGGAAGACTAGTGATTTTGTTGTTT | >MI0000263\|hsa-mir-7-1&WithFlank&9\|-\|83969742\|83969863 | 30 | 54 | 25 | TGGAAGACTAGTGATTTTGTTGTTT | 2383.1 | -1.9 | 0.8 | -2.4 | 1.6E-02 | 4.7E-02 | BT20/MB231 | MITO |
| >MI0003820\|hsa-mir-454&WithFlank&17\|-\|59137752\|59137878@70.92.23&[MIMAT0003885&hsa-miR-454-3p&offsets\|0\|0;m-272&17\|-\|59137787\|59137809&offsets\|0\|0]\|TAGTGCAATATTGCTTATAGGGT | >MI0003820\|hsa-mir-454&WithFlank&17\|-\|59137752\|59137878 | 70 | 92 | 23 | TAGTGCAATATTGCTTATAGGGT | 360.3 | 2.2 | 0.9 | 2.4 | 1.8E-02 | 4.7E-02 | BT20/MB231 | CYTO |
| >hg38_21_+_8433172_8446622_RNA45SN1_45S_50ntFlanks@5473.5491.19\|CGGCCCTGGCGGAGCGCTG | >hg38_21_+_8433172_8446622_RNA45SN1_45S_50ntFlanks | 5473 | 5491 | 19 | CGGCCCTGGCGGAGCGCTG | 967.2 | 2.5 | 1.1 | 2.4 | 1.8E-02 | 4.7E-02 | BT20/MB231 | CYTO |
| >hg38_21_+_8433172_8446622_RNA45SN1_45S_50ntFlanks@10353.10371.19\|AGAACTTTGAAGGCCGAAG | >hg38_21_+_8433172_8446622_RNA45SN1_45S_50ntFlanks | 10353 | 10371 | 19 | AGAACTTTGAAGGCCGAAG | 44.6 | -3.9 | 1.8 | -2.2 | 3.0E-02 | 4.7E-02 | BT20/MB468 | MP |
| >hg38_21_+_8433172_8446622_RNA45SN1_45S_50ntFlanks@9740.9757.18\|AACGATCTCAACCTATTC | >hg38_21_+_8433172_8446622_RNA45SN1_45S_50ntFlanks | 9740 | 9757 | 18 | AACGATCTCAACCTATTC | 203.4 | -3.8 | 1.8 | -2.2 | 3.0E-02 | 4.7E-02 | BT20/MB468 | MP |
| >hg38_21_+_8433172_8446622_RNA45SN1_45S_50ntFlanks@7988.8019.32\|CAGACGTGGCGACCCGCTGAATTTAAGCATAT | >hg38_21_+_8433172_8446622_RNA45SN1_45S_50ntFlanks | 7988 | 8019 | 32 | CAGACGTGGCGACCCGCTGAATTTAAGCATAT | 70.2 | -3.7 | 1.8 | -2.1 | 3.9E-02 | 4.7E-02 | MB231/MB468 | MP |
| >trnaMT_GluTTC_MT_-_14674_14742@47.66.20,trnalookalike8_GluTTC_5_-_93905172_93905240@47.66.20\|TGGTTGTAGTCCGTGCGAGA | >trnaMT_GluTTC_MT_-_14674_14742 | 47 | 66 | 20 | TGGTTGTAGTCCGTGCGAGA | 171.6 | 1.3 | 0.6 | 2.1 | 3.9E-02 | 4.7E-02 | MB231/MB468 | MP |
| >hg38_21_+_8433172_8446622_RNA45SN1_45S_50ntFlanks@8196.8214.19\|CGTGGACGGTGTGAGGCCG | >hg38_21_+_8433172_8446622_RNA45SN1_45S_50ntFlanks | 8196 | 8214 | 19 | CGTGGACGGTGTGAGGCCG | 617.6 | -2.4 | 1.0 | -2.4 | 1.8E-02 | 4.7E-02 | BT20/MB231 | CYTO |
| >hg38_21_+_8433172_8446622_RNA45SN1_45S_50ntFlanks@3846.3874.29\|CTTGGATAACTGTGGTAATTCTAGAGCTA | >hg38_21_+_8433172_8446622_RNA45SN1_45S_50ntFlanks | 3846 | 3874 | 29 | CTTGGATAACTGTGGTAATTCTAGAGCTA | 201.3 | -1.1 | 0.5 | -2.0 | 4.1E-02 | 4.7E-02 | MB231/MB468 | CYTO |
| >hg38_21_+_8433172_8446622_RNA45SN1_45S_50ntFlanks@4202.4219.18\|CGACCCGGGGAGGTAGTG | >hg38_21_+_8433172_8446622_RNA45SN1_45S_50ntFlanks | 4202 | 4219 | 18 | CGACCCGGGGAGGTAGTG | 557.7 | -4.0 | 1.8 | -2.2 | 3.0E-02 | 4.7E-02 | BT20/MB468 | MP |
| >hg38_21_+_8433172_8446622_RNA45SN1_45S_50ntFlanks@6646.6670.25\|TCGTACGACTCTTAGCGGTGGATCA | >hg38_21_+_8433172_8446622_RNA45SN1_45S_50ntFlanks | 6646 | 6670 | 25 | TCGTACGACTCTTAGCGGTGGATCA | 1301.4 | 1.6 | 0.7 | 2.4 | 1.8E-02 | 4.7E-02 | BT20/MB231 | CYTO |
| >MI0000234\|hsa-mir-192&WithFlank&11\|-\|64891131\|64891252@31.51.21(+1C)&[MIMAT0000222&hsa-miR-192-5p&offsets\|+1\|+1(+1C);m-59&11\|-\|64891203\|64891223&offsets\|+1\|+1(+1C)]\|TGACCTATGAATTGACAGCCAC | >MI0000234\|hsa-mir-192&WithFlank&11\|-\|64891131\|64891252 | 31 | 51 | 21(+1C) | TGACCTATGAATTGACAGCCAC | 55.8 | 2.7 | 1.3 | 2.1 | 3.9E-02 | 4.7E-02 | MB231/MB468 | MP |
| >hg38_1_-_228634819_228635039_RNA5S12_5S_50ntFlanks@127.169.43\|ACTTGGATGGGAGACCGCCTGGGAATACCGGGTGCTGTAGGCT | >hg38_1_-_228634819_228635039_RNA5S12_5S_50ntFlanks | 127 | 169 | 43 | ACTTGGATGGGAGACCGCCTGGGAATACCGGGTGCTGTAGGCT | 289.9 | 2.9 | 1.2 | 2.5 | 1.3E-02 | 4.7E-02 | BT20/MB231 | MP |
| >hg38_MT_+_1621_3279_RNR2_16S_50ntFlanks@292.312.21\|AGACGAGCTACCTAAGAACAG | >hg38_MT_+_1621_3279_RNR2_16S_50ntFlanks | 292 | 312 | 21 | AGACGAGCTACCTAAGAACAG | 121.9 | 4.2 | 1.9 | 2.2 | 3.0E-02 | 4.7E-02 | BT20/MB468 | MP |
| >MI0000070\|hsa-mir-16-1&WithFlank&13\|-\|50048967\|50049067@20.41.22;MI0000115\|hsa-mir-16-2&WithFlank&3\|+\|160404739\|160404831@16.37.22&[MIMAT0000069&hsa-miR-16-5p&offsets\|0\|0;m-62&13\|-\|50049027\|50049048&offsets\|0\|0];[MIMAT0000069_1&hsa-miR-16-5p&offsets\|0\|0;m-61&3\|+\|160404754\|160404775&offsets\|0\|0]\|TAGCAGCACGTAAATATTGGCG | >MI0000070\|hsa-mir-16-1&WithFlank&13\|-\|50048967\|50049067 | 20 | 41 | 22 | TAGCAGCACGTAAATATTGGCG | 11411.8 | 2.8 | 1.3 | 2.2 | 3.0E-02 | 4.7E-02 | BT20/MB468 | MP |
| >MI0001448\|hsa-mir-425&WithFlank&3\|-\|49020142\|49020240@20.39.20&[MIMAT0003393&hsa-miR-425-5p&offsets\|0\|-3;m-95&3\|-\|49020200\|49020221&offsets\|0\|-2]\|AATGACACGATCACTCCCGT | >MI0001448\|hsa-mir-425&WithFlank&3\|-\|49020142\|49020240 | 20 | 39 | 20 | AATGACACGATCACTCCCGT | 59.8 | 4.4 | 2.1 | 2.1 | 3.9E-02 | 4.7E-02 | MB231/MB468 | MP |
| >hg38_21_+_8433172_8446622_RNA45SN1_45S_50ntFlanks@6649.6668.20\|TACGACTCTTAGCGGTGGAT | >hg38_21_+_8433172_8446622_RNA45SN1_45S_50ntFlanks | 6649 | 6668 | 20 | TACGACTCTTAGCGGTGGAT | 746.4 | 1.6 | 0.8 | 2.0 | 4.3E-02 | 4.7E-02 | MB231/MB468 | MITO |
| >hg38_21_+_8433172_8446622_RNA45SN1_45S_50ntFlanks@10408.10453.46\|GGTCAGTCGGTCCTGAGAGATGGGCGAGCGCCGTTCCGAAGGGACG | >hg38_21_+_8433172_8446622_RNA45SN1_45S_50ntFlanks | 10408 | 10453 | 46 | GGTCAGTCGGTCCTGAGAGATGGGCGAGCGCCGTTCCGAAGGGACG | 274.2 | -2.2 | 1.1 | -2.0 | 4.1E-02 | 4.7E-02 | MB231/MB468 | CYTO |
| >trna10_LysCTT_16_+_3241501_3241573@13.33.21,trna13_LysCTT_14_-_58706613_58706685@13.33.21,trna2_LysCTT_15_+_79152904_79152976@13.33.21,trna4_LysCTT_18_-_43669271_43669343@13.33.21\|CAGTCGGTAGAGCATGGGACT | >trna10_LysCTT_16_+_3241501_3241573 | 13 | 33 | 21 | CAGTCGGTAGAGCATGGGACT | 209.3 | -2.7 | 1.1 | -2.4 | 1.6E-02 | 4.7E-02 | BT20/MB231 | MITO |
| >hg38_MT_+_598_1651_RNR1_12S_50ntFlanks@969.1004.36\|CATGGTAAGTGTACTGGAAAGTGCACTTGGACGAAC | >hg38_MT_+_598_1651_RNR1_12S_50ntFlanks | 969 | 1004 | 36 | CATGGTAAGTGTACTGGAAAGTGCACTTGGACGAAC | 272.7 | -2.1 | 0.9 | -2.4 | 1.6E-02 | 4.7E-02 | BT20/MB231 | MITO |
| >hg38_21_+_8433172_8446622_RNA45SN1_45S_50ntFlanks@6646.6666.21\|TCGTACGACTCTTAGCGGTGG | >hg38_21_+_8433172_8446622_RNA45SN1_45S_50ntFlanks | 6646 | 6666 | 21 | TCGTACGACTCTTAGCGGTGG | 8145.3 | 1.7 | 0.8 | 2.0 | 4.3E-02 | 4.7E-02 | MB231/MB468 | NUC |

| | | | | | | | | | | | | | |
|---|---|---|---|---|---|---|---|---|---|---|---|---|---|
| >MI0000111\|hsa-mir-105-1&WithFlank&X\|-\|152392213\|152392305@21.41.21;MI0000112\|hsa-mir-105-2&WithFlank&X\|-\|152394406\|152394498@21.41.21&[MIMAT0000102&hsa-miR-105-5p&offsets\|+2\|0;m-735&X\|-\|152392266\|152392287&offsets\|+2\|+1];[MIMAT0000102_1&hsa-miR-105-5p&offsets\|+2\|0;m-735&X\|-\|152394459\|152394480&offsets\|+2\|+1]\|AAATGCTCAGACTCCTGTGGT | >MI0000111\|hsa-mir-105-1&WithFlank&X\|-\|152392213\|152392305 | 21 | 41 | 21 | AAATGCTCAGACTCCTGTGGT | 26.2 | 8.0 | 3.3 | 2.5 | 1.4E-02 | 4.7E-02 | BT20/MB231 | MP |
| >MI0016005\|hsa-mir-3615&WithFlank&17\|+\|74748607\|74748705@57.78.22&[MIMAT0017994&hsa-miR-3615&offsets\|0\|+1;m-271&17\|+\|74748663\|74748684&offsets\|0\|0]\|TCTCTCGGCTCCTCGCGGCTCG | >MI0016005\|hsa-mir-3615&WithFlank&17\|+\|74748607\|74748705 | 57 | 78 | 22 | TCTCTCGGCTCCTCGCGGCTCG | 614.2 | -1.7 | 0.7 | -2.4 | 1.6E-02 | 4.7E-02 | BT20/MB231 | MITO |
| >MI0000299\|hsa-mir-222&WithFlank&X\|-\|45747009\|45747130@75.95.21&[MIMAT0000279&hsa-miR-222-3p&offsets\|0\|0;m-140&X\|-\|45747033\|45747056&offsets\|0\|-3]\|AGCTACATCTGGCTACTGGGT | >MI0000299\|hsa-mir-222&WithFlank&X\|-\|45747009\|45747130 | 75 | 95 | 21 | AGCTACATCTGGCTACTGGGT | 431.5 | 1.2 | 0.6 | 2.0 | 4.2E-02 | 4.7E-02 | BT20/MB468 | NUC |
| >hg38_1_-_228634819_228635039_RNA5S12_5S_50ntFlanks@144.171.28\|CCTGGGAATACCGGGTGCTGTAGGCTTT | >hg38_1_-_228634819_228635039_RNA5S12_5S_50ntFlanks | 144 | 171 | 28 | CCTGGGAATACCGGGTGCTGTAGGCTTT | 4662.1 | -1.9 | 0.8 | -2.5 | 1.4E-02 | 4.7E-02 | BT20/MB231 | NUC |
| >MI0000084\|hsa-mir-26b&WithFlank&2\|+\|218402640\|218402728@18.39.22&[MIMAT0000083&hsa-miR-26b-5p&offsets\|0\|+1;m-53&2\|+\|218402657\|218402678&offsets\|0\|0]\|TTCAAGTAATTCAGGATAGGTT | >MI0000084\|hsa-mir-26b&WithFlank&2\|+\|218402640\|218402728 | 18 | 39 | 22 | TTCAAGTAATTCAGGATAGGTT | 8055.8 | 2.1 | 0.9 | 2.4 | 1.8E-02 | 4.7E-02 | BT20/MB231 | CYTO |
| >hg38_21_+_8433172_8446622_RNA45SN1_45S_50ntFlanks@476.502.27\|TTCGTGATCGATGTGGTGACGTCGTGC | >hg38_21_+_8433172_8446622_RNA45SN1_45S_50ntFlanks | 476 | 502 | 27 | TTCGTGATCGATGTGGTGACGTCGTGC | 81.6 | 3.4 | 1.7 | 2.1 | 3.9E-02 | 4.7E-02 | MB231/MB468 | MP |
| >hg38_21_+_8433172_8446622_RNA45SN1_45S_50ntFlanks@12456.12478.23\|CCAAGCGTTGGATTGTTCACCCA | >hg38_21_+_8433172_8446622_RNA45SN1_45S_50ntFlanks | 12456 | 12478 | 23 | CCAAGCGTTGGATTGTTCACCCA | 6132.8 | -4.5 | 2.1 | -2.2 | 3.0E-02 | 4.7E-02 | BT20/MB468 | MP |
| >hg38_21_+_8433172_8446622_RNA45SN1_45S_50ntFlanks@11703.11736.34\|AATGAAGCGCGGGTAAACGGCGGGAGTAACTATG | >hg38_21_+_8433172_8446622_RNA45SN1_45S_50ntFlanks | 11703 | 11736 | 34 | AATGAAGCGCGGGTAAACGGCGGGAGTAACTATG | 661.8 | 1.6 | 0.8 | 2.0 | 4.1E-02 | 4.7E-02 | MB231/MB468 | CYTO |
| >MI0000093\|hsa-mir-92a-1&WithFlank&13\|+\|91351308\|91351397@54.75.22(+2A);MI0000094\|hsa-mir-92a-2&WithFlank&X\|-\|134169532\|134169618@54.75.22(+2A)&[MIMAT0000092&hsa-miR-92a-3p&offsets\|0\|0(+2A);m-9&13\|+\|91351361\|91351382&offsets\|0\|0(+2A)];[MIMAT0000092_1&hsa-miR-92a-3p&offsets\|0\|0(+2A);m-11&X\|-\|134169544\|134169565&offsets\|0\|0(+2A)]\|TATTGCACTTGTCCCGGCCTGTAA | >MI0000093\|hsa-mir-92a-1&WithFlank&13\|+\|91351308\|91351397 | 54 | 75 | 22(+2A) | TATTGCACTTGTCCCGGCCTGTAA | 381.8 | -1.2 | 0.6 | -2.0 | 4.2E-02 | 4.7E-02 | BT20/MB468 | NUC |
| >MI0000066\|hsa-let-7e&WithFlank&19\|+\|51692780\|51692870@14.34.21(+1C)&[MIMAT0000066&hsa-let-7e-5p&offsets\|0\|-1(+1C);m-50&19\|+\|51692793\|51692814&offsets\|0\|-1(+1C)]\|TGAGGTAGGAGGTTGTATAGTC | >MI0000066\|hsa-let-7e&WithFlank&19\|+\|51692780\|51692870 | 14 | 34 | 21(+1C) | TGAGGTAGGAGGTTGTATAGTC | 298.9 | 2.4 | 1.1 | 2.2 | 3.0E-02 | 4.7E-02 | BT20/MB468 | MP |
| >hg38_21_+_8433172_8446622_RNA45SN1_45S_50ntFlanks@6780.6802.23\|CGGGGCTACGCCTGTCTGAGCGT | >hg38_21_+_8433172_8446622_RNA45SN1_45S_50ntFlanks | 6780 | 6802 | 23 | CGGGGCTACGCCTGTCTGAGCGT | 734.0 | -1.9 | 0.8 | -2.4 | 1.8E-02 | 4.7E-02 | BT20/MB231 | CYTO |
| >hg38_21_+_8433172_8446622_RNA45SN1_45S_50ntFlanks@11572.11612.41\|CCGACTTAGAACTGGTGCGGACCAGGGGAATCCGACTGTTT | >hg38_21_+_8433172_8446622_RNA45SN1_45S_50ntFlanks | 11572 | 11612 | 41 | CCGACTTAGAACTGGTGCGGACCAGGGGAATCCGACTGTTT | 397.3 | -2.4 | 1.0 | -2.4 | 1.8E-02 | 4.7E-02 | BT20/MB231 | CYTO |
| >hg38_21_+_8433172_8446622_RNA45SN1_45S_50ntFlanks@6661.6689.29\|CGGTGGATCACTCGGCTCGTGCGTCGATG | >hg38_21_+_8433172_8446622_RNA45SN1_45S_50ntFlanks | 6661 | 6689 | 29 | CGGTGGATCACTCGGCTCGTGCGTCGATG | 1489.9 | 2.1 | 0.9 | 2.4 | 1.8E-02 | 4.7E-02 | BT20/MB231 | CYTO |
| >hg38_21_+_8433172_8446622_RNA45SN1_45S_50ntFlanks@10408.10435.28\|GGTCAGTCGGTCCTGAGAGATGGGCGAG | >hg38_21_+_8433172_8446622_RNA45SN1_45S_50ntFlanks | 10408 | 10435 | 28 | GGTCAGTCGGTCCTGAGAGATGGGCGAG | 1417.6 | -4.2 | 1.9 | -2.2 | 3.0E-02 | 4.7E-02 | BT20/MB468 | MP |
| >hg38_21_+_8433172_8446622_RNA45SN1_45S_50ntFlanks@10349.10384.36\|AACGAGAACTTTGAAGGCCGAAGTGGAGAAGGGTTC | >hg38_21_+_8433172_8446622_RNA45SN1_45S_50ntFlanks | 10349 | 10384 | 36 | AACGAGAACTTTGAAGGCCGAAGTGGAGAAGGGTTC | 895.2 | -1.3 | 0.6 | -2.0 | 4.1E-02 | 4.7E-02 | MB231/MB468 | CYTO |
| >hg38_21_+_8433172_8446622_RNA45SN1_45S_50ntFlanks@4730.4761.32\|CAAGAACGAAAGTCGGAGGTTCGAAGACGATC | >hg38_21_+_8433172_8446622_RNA45SN1_45S_50ntFlanks | 4730 | 4761 | 32 | CAAGAACGAAAGTCGGAGGTTCGAAGACGATC | 315.2 | 2.2 | 1.0 | 2.2 | 3.0E-02 | 4.7E-02 | BT20/MB468 | MP |
| >hg38_21_+_8433172_8446622_RNA45SN1_45S_50ntFlanks@6648.6692.45\|GTACGACTCTTAGCGGTGGATCACTCGGCTCGTGCGTCGATGAAG | >hg38_21_+_8433172_8446622_RNA45SN1_45S_50ntFlanks | 6648 | 6692 | 45 | GTACGACTCTTAGCGGTGGATCACTCGGCTCGTGCGTCGATGAAG | 1292.9 | 1.3 | 0.6 | 2.1 | 3.7E-02 | 4.7E-02 | BT20/MB468 | MITO |
| >am_trna11_ProAGG_16_+_3241989_3242060@1.29.29,trna12_ProAGG_6_+_26555498_26555569@1.29.29,trna12_ProTGG_11_-_75946869_75946940@1.29.29,trna14_ProTGG_5_-_180615854_180615925@1.29.29,trna22_ProAGG_14_-_21081560_21081631@1.29.29,trna23_ProAGG_14_-_21077495_21077566@1.29.29,trna28_ProTGG_16_-_3234133_3234204@1.29.29,trna29_ProAGG_16_-_3232635_3232706@1.29.29,trna2_ProAGG_7_+_128423504_128423575@1.29.29,trna30_ProCGG_6_+_27059521_27059592@1.29.29,trna37_ProCGG_17_-_8126151_8126222@1.29.29,trna3_ProTGG_16_+_3208923_3208994@1.29.29,trna52_ProCGG_1_+_167683962_167684033@1.29.29,trna65_ProAGG_1_-_167684725_167684796@1.29.29,trna6_ProCGG_16_+_3222049_3222120@1.29.29,trna6_ProTGG_14_+_21152175_21152246@1.29.29,trna8_ProTGG_16_+_3238094_3238165@1.29.29,trna9_ProAGG_11_+_75946557_75946628@1.29.29,trna9_ProAGG_16_+_3239634_3239705@1.29.29\|GGCTCGTTGGTCTAGGGGTATGATTCTCG | >am_trna11_ProAGG_16_+_3241989_3242060 | 1 | 29 | 29 | GGCTCGTTGGTCTAGGGGTATGATTCTCG | 247.1 | -3.3 | 1.5 | -2.2 | 3.0E-02 | 4.7E-02 | BT20/MB468 | MP |
| >hg38_21_+_8433172_8446622_RNA45SN1_45S_50ntFlanks@12417.12448.32\|TCGATGTCGGCTCTTCCTATCATTGTGAAGCA | >hg38_21_+_8433172_8446622_RNA45SN1_45S_50ntFlanks | 12417 | 12448 | 32 | TCGATGTCGGCTCTTCCTATCATTGTGAAGCA | 2996.0 | -2.2 | 0.9 | -2.4 | 1.8E-02 | 4.7E-02 | BT20/MB231 | CYTO |
| >hg38_21_+_8433172_8446622_RNA45SN1_45S_50ntFlanks@3878.3898.21\|CATGCCGACGGGCGCTGACCC | >hg38_21_+_8433172_8446622_RNA45SN1_45S_50ntFlanks | 3878 | 3898 | 21 | CATGCCGACGGGCGCTGACCC | 432.4 | -1.4 | 0.7 | -2.0 | 4.2E-02 | 4.7E-02 | BT20/MB468 | CYTO |
| >hg38_21_+_8433172_8446622_RNA45SN1_45S_50ntFlanks@4003.4024.22\|ATAACCTCGGGCCGATCGCACG | >hg38_21_+_8433172_8446622_RNA45SN1_45S_50ntFlanks | 4003 | 4024 | 22 | ATAACCTCGGGCCGATCGCACG | 1313.1 | -4.1 | 1.9 | -2.2 | 3.0E-02 | 4.7E-02 | BT20/MB468 | MP |
| >hg38_21_+_8433172_8446622_RNA45SN1_45S_50ntFlanks@11581.11609.29\|AACTGGTGCGGACCAGGGGAATCCGACTG | >hg38_21_+_8433172_8446622_RNA45SN1_45S_50ntFlanks | 11581 | 11609 | 29 | AACTGGTGCGGACCAGGGGAATCCGACTG | 338.3 | 3.1 | 1.2 | 2.5 | 1.4E-02 | 4.7E-02 | BT20/MB231 | MP |
| >hg38_21_+_8433172_8446622_RNA45SN1_45S_50ntFlanks@11561.11584.24\|GCGCCTAGCAGCCGACTTAGAACT | >hg38_21_+_8433172_8446622_RNA45SN1_45S_50ntFlanks | 11561 | 11584 | 24 | GCGCCTAGCAGCCGACTTAGAACT | 303.4 | -1.2 | 0.6 | -2.0 | 4.3E-02 | 4.7E-02 | MB231/MB468 | NUC |
| >MI0000440\|hsa-mir-27b&WithFlank&9\|+\|95085439\|95085547@67.87.21(+2U)&[MIMAT0000419&hsa-miR-27b-3p&offsets\|0\|0(+2U);m-39&9\|+\|95085505\|95085525&offsets\|0\|0(+2U)]\|TTCACAGTGGCTAAGTTCTGCTT | >MI0000440\|hsa-mir-27b&WithFlank&9\|+\|95085439\|95085547 | 67 | 87 | 21(+2U) | TTCACAGTGGCTAAGTTCTGCTT | 939.9 | -2.4 | 1.0 | -2.5 | 1.4E-02 | 4.7E-02 | BT20/MB231 | MP |
| >hg38_21_+_8433172_8446622_RNA45SN1_45S_50ntFlanks@8358.8390.33\|AAAGAACTTTGAAGAGAGAGTTCAAGAGGGCGT | >hg38_21_+_8433172_8446622_RNA45SN1_45S_50ntFlanks | 8358 | 8390 | 33 | AAAGAACTTTGAAGAGAGAGTTCAAGAGGGCGT | 127.6 | 3.2 | 1.3 | 2.5 | 1.4E-02 | 4.7E-02 | BT20/MB231 | MP |
| >hg38_21_+_8433172_8446622_RNA45SN1_45S_50ntFlanks@9930.9955.26\|ATATAGACAGCAGGACGGTGGCCATG | >hg38_21_+_8433172_8446622_RNA45SN1_45S_50ntFlanks | 9930 | 9955 | 26 | ATATAGACAGCAGGACGGTGGCCATG | 127.4 | 4.8 | 1.9 | 2.5 | 1.4E-02 | 4.7E-02 | BT20/MB231 | MP |
| >hg38_21_+_8433172_8446622_RNA45SN1_45S_50ntFlanks@6698.6720.23\|AGCTAGCTGCGAGAATTAATGTG | >hg38_21_+_8433172_8446622_RNA45SN1_45S_50ntFlanks | 6698 | 6720 | 23 | AGCTAGCTGCGAGAATTAATGTG | 2190.8 | -2.1 | 0.9 | -2.4 | 1.8E-02 | 4.7E-02 | BT20/MB231 | CYTO |
| >am_trna111_HisGTG_1_-_147774845_147774916@-1G.28.29,trna118_HisGTG_1_-_145396881_145396952@-1G.28.29,trna16_HisGTG_1_+_146544773_146544844@-1G.28.29,trna1_HisGTG_15_+_45493349_45493420@-1G.28.29,trna21_HisGTG_1_+_147753471_147753542@-1G.28.29,trna33_HisGTG_6_+_27125906_27125977@-1G.28.29,trna7_HisGTG_9_-_14433938_14434009@-1G.28.29,trna8_HisGTG_15_-_45492611_45492682@-1G.28.29,trna9_HisGTG_15_-_45490804_45490875@-1G.28.29\|GGCCGTGATCGTATAGTGGTTAGTACTCT | >am_trna111_HisGTG_1_-_147774845_147774916 | -1G | 28 | 29 | GGCCGTGATCGTATAGTGGTTAGTACTCT | 682.8 | 2.2 | 1.0 | 2.2 | 3.0E-02 | 4.7E-02 | BT20/MB468 | MP |
| >trnaMT_PheGAA_MT_+_577_647@47.74.28\|ACGGGCTCACATCACCCCATAAACACCA | >trnaMT_PheGAA_MT_+_577_647 | 47 | 74 | 28 | ACGGGCTCACATCACCCCATAAACACCA | 89.0 | 2.6 | 1.2 | 2.2 | 3.0E-02 | 4.7E-02 | BT20/MB468 | MP |
| >hg38_21_+_8433172_8446622_RNA45SN1_45S_50ntFlanks@11569.11595.27\|CAGCCGACTTAGAACTGGTGCGGACCA | >hg38_21_+_8433172_8446622_RNA45SN1_45S_50ntFlanks | 11569 | 11595 | 27 | CAGCCGACTTAGAACTGGTGCGGACCA | 97.4 | -3.0 | 1.4 | -2.2 | 3.0E-02 | 4.7E-02 | BT20/MB468 | MP |
| >hg38_21_+_8433172_8446622_RNA45SN1_45S_50ntFlanks@12407.12433.27\|TTTTGATCCTTCGATGTCGGCTCTTCC | >hg38_21_+_8433172_8446622_RNA45SN1_45S_50ntFlanks | 12407 | 12433 | 27 | TTTTGATCCTTCGATGTCGGCTCTTCC | 2444.5 | 2.0 | 1.0 | 2.0 | 4.2E-02 | 4.7E-02 | MB231/MB468 | CYTO |
| >hg38_21_+_8433172_8446622_RNA45SN1_45S_50ntFlanks@11603.11623.21\|CCGACTGTTTAATTAAAACAA | >hg38_21_+_8433172_8446622_RNA45SN1_45S_50ntFlanks | 11603 | 11623 | 21 | CCGACTGTTTAATTAAAACAA | 314.9 | 3.5 | 1.6 | 2.2 | 3.0E-02 | 4.7E-02 | BT20/MB468 | MP |
| >trnaMT_PheGAA_MT_+_577_647@1.35.35\|GTTTATGTAGCTTACCTCCTCAAAGCAATACACTG | >trnaMT_PheGAA_MT_+_577_647 | 1 | 35 | 35 | GTTTATGTAGCTTACCTCCTCAAAGCAATACACTG | 481.2 | -4.8 | 1.9 | -2.5 | 1.4E-02 | 4.7E-02 | BT20/MB231 | MP |
| >hg38_21_+_8433172_8446622_RNA45SN1_45S_50ntFlanks@6661.6698.38\|CGGTGGATCACTCGGCTCGTGCGTCGATGAAGAACGCA | >hg38_21_+_8433172_8446622_RNA45SN1_45S_50ntFlanks | 6661 | 6698 | 38 | CGGTGGATCACTCGGCTCGTGCGTCGATGAAGAACGCA | 291.7 | 1.6 | 0.7 | 2.4 | 1.8E-02 | 4.7E-02 | BT20/MB231 | CYTO |
| >hg38_21_+_8433172_8446622_RNA45SN1_45S_50ntFlanks@3766.3794.29\|GTACGCACGGCCGGTACAGTGAAACTGCG | >hg38_21_+_8433172_8446622_RNA45SN1_45S_50ntFlanks | 3766 | 3794 | 29 | GTACGCACGGCCGGTACAGTGAAACTGCG | 369.9 | -2.3 | 1.0 | -2.4 | 1.8E-02 | 4.7E-02 | BT20/MB231 | CYTO |
| >hg38_21_+_8433172_8446622_RNA45SN1_45S_50ntFlanks@12444.12483.40\|AAGCAGAATTCACCAAGCGTTGGATTGTTCACCCACTAAT | >hg38_21_+_8433172_8446622_RNA45SN1_45S_50ntFlanks | 12444 | 12483 | 40 | AAGCAGAATTCACCAAGCGTTGGATTGTTCACCCACTAAT | 118.5 | -2.0 | 0.9 | -2.1 | 3.7E-02 | 4.7E-02 | BT20/MB468 | MITO |
| >hg38_21_+_8433172_8446622_RNA45SN1_45S_50ntFlanks@9736.9759.24\|CCGAAACGATCTCAACCTATTCTC | >hg38_21_+_8433172_8446622_RNA45SN1_45S_50ntFlanks | 9736 | 9759 | 24 | CCGAAACGATCTCAACCTATTCTC | 91.6 | -3.7 | 1.7 | -2.2 | 3.0E-02 | 4.7E-02 | BT20/MB468 | MP |
| >hg38_21_+_8433172_8446622_RNA45SN1_45S_50ntFlanks@4555.4573.19\|CGCAGCTAGGAATAATGGA | >hg38_21_+_8433172_8446622_RNA45SN1_45S_50ntFlanks | 4555 | 4573 | 19 | CGCAGCTAGGAATAATGGA | 108.5 | -2.2 | 1.1 | -2.0 | 4.2E-02 | 4.7E-02 | BT20/MB468 | CYTO |
| >hg38_21_+_8433172_8446622_RNA45SN1_45S_50ntFlanks@12656.12675.20\|TGACTGAACGCCTCTAAGTC | >hg38_21_+_8433172_8446622_RNA45SN1_45S_50ntFlanks | 12656 | 12675 | 20 | TGACTGAACGCCTCTAAGTC | 162.5 | -1.6 | 0.8 | -2.1 | 3.7E-02 | 4.7E-02 | BT20/MB468 | MITO |
| >hg38_21_+_8433172_8446622_RNA45SN1_45S_50ntFlanks@12907.12935.29\|CACCGCACGTTCGTGGGGAACCTGGCGCT | >hg38_21_+_8433172_8446622_RNA45SN1_45S_50ntFlanks | 12907 | 12935 | 29 | CACCGCACGTTCGTGGGGAACCTGGCGCT | 259.5 | -2.9 | 1.4 | -2.0 | 4.2E-02 | 4.7E-02 | MB231/MB468 | CYTO |

<table>
<tr><td>>hg38_21_+_8433172_8446622_RNA45SN1_45S_50ntFlanks@475.502.28|CTTCGTGATCGATGTGGTGACGTCGTGC</td><td>>hg38_21_+_8433172_8446622_RNA45SN1_45S_50ntFlanks</td><td>475</td><td>502</td><td>28</td><td>CTTCGTGATCGATGTGGTGACGTCGTGC</td><td>250.6</td><td>4.1</td><td>1.9</td><td>2.2</td><td>3.0E-02</td><td>4.7E-02</td><td>BT20/MB468</td><td>MP</td></tr>
<tr><td>>hg38_21_+_8433172_8446622_RNA45SN1_45S_50ntFlanks@4128.4146.19|CGATTCCGGAGAGGGAGCC</td><td>>hg38_21_+_8433172_8446622_RNA45SN1_45S_50ntFlanks</td><td>4128</td><td>4146</td><td>19</td><td>CGATTCCGGAGAGGGAGCC</td><td>134.7</td><td>-3.7</td><td>1.7</td><td>-2.2</td><td>3.0E-02</td><td>4.7E-02</td><td>BT20/MB468</td><td>MP</td></tr>
<tr><td>>hg38_21_+_8433172_8446622_RNA45SN1_45S_50ntFlanks@12416.12449.34|TTCGATGTCGGCTCTTCCTATCATTGTGAAGCAG</td><td>>hg38_21_+_8433172_8446622_RNA45SN1_45S_50ntFlanks</td><td>12416</td><td>12449</td><td>34</td><td>TTCGATGTCGGCTCTTCCTATCATTGTGAAGCAG</td><td>490.0</td><td>-2.5</td><td>1.0</td><td>-2.5</td><td>1.4E-02</td><td>4.7E-02</td><td>BT20/MB231</td><td>NUC</td></tr>
<tr><td>>hg38_21_+_8433172_8446622_RNA45SN1_45S_50ntFlanks@4732.4758.27|AGAACGAAAGTCGGAGGTTCGAAGACG</td><td>>hg38_21_+_8433172_8446622_RNA45SN1_45S_50ntFlanks</td><td>4732</td><td>4758</td><td>27</td><td>AGAACGAAAGTCGGAGGTTCGAAGACG</td><td>2929.0</td><td>3.7</td><td>1.5</td><td>2.5</td><td>1.4E-02</td><td>4.7E-02</td><td>BT20/MB231</td><td>MP</td></tr>
<tr><td>>hg38_21_+_8433172_8446622_RNA45SN1_45S_50ntFlanks@8113.8142.30|GGCGCGGGACATGTGGCGTACGGAAGACCC</td><td>>hg38_21_+_8433172_8446622_RNA45SN1_45S_50ntFlanks</td><td>8113</td><td>8142</td><td>30</td><td>GGCGCGGGACATGTGGCGTACGGAAGACCC</td><td>233.6</td><td>-3.8</td><td>1.8</td><td>-2.2</td><td>3.0E-02</td><td>4.7E-02</td><td>BT20/MB468</td><td>MP</td></tr>
<tr><td>>hg38_21_+_8433172_8446622_RNA45SN1_45S_50ntFlanks@4726.4765.40|TAATCAAGAACGAAAGTCGGAGGTTCGAAGACGATCAGAT</td><td>>hg38_21_+_8433172_8446622_RNA45SN1_45S_50ntFlanks</td><td>4726</td><td>4765</td><td>40</td><td>TAATCAAGAACGAAAGTCGGAGGTTCGAAGACGATCAGAT</td><td>54.5</td><td>4.6</td><td>2.1</td><td>2.2</td><td>3.0E-02</td><td>4.7E-02</td><td>BT20/MB468</td><td>MP</td></tr>
<tr><td>>hg38_21_+_8433172_8446622_RNA45SN1_45S_50ntFlanks@7990.8011.22|GACGTGGCGACCCGCTGAATTT</td><td>>hg38_21_+_8433172_8446622_RNA45SN1_45S_50ntFlanks</td><td>7990</td><td>8011</td><td>22</td><td>GACGTGGCGACCCGCTGAATTT</td><td>500.2</td><td>-1.4</td><td>0.7</td><td>-2.1</td><td>4.0E-02</td><td>4.7E-02</td><td>MB231/MB468</td><td>MP</td></tr>
<tr><td>>am_MI0003583|hsa-mir-576&WithFlank&4|+|109488692|109488801@61.82.22(+1A)&[MIMAT0004796&hsa-miR-576-3p&offsets|0|0(+1A);m-524&4|+|109488752|109488772&offsets|0|+1(+1A)]|AAGATGTGGAAAAATTGGAATCA</td><td>>am_MI0003583|hsa-mir-576&WithFlank&4|+|109488692|109488801</td><td>61</td><td>82</td><td>22(+1A)</td><td>AAGATGTGGAAAAATTGGAATCA</td><td>80.4</td><td>4.2</td><td>2.0</td><td>2.1</td><td>4.0E-02</td><td>4.7E-02</td><td>MB231/MB468</td><td>MP</td></tr>
<tr><td>>trnaMT_ValTAC_MT_+_1602_1670@47.72.26|TCAACTTAACTTGACCGCTCTGACCA</td><td>>trnaMT_ValTAC_MT_+_1602_1670</td><td>47</td><td>72</td><td>26</td><td>TCAACTTAACTTGACCGCTCTGACCA</td><td>221.0</td><td>2.1</td><td>1.0</td><td>2.2</td><td>3.0E-02</td><td>4.7E-02</td><td>BT20/MB468</td><td>MP</td></tr>
<tr><td>>hg38_21_+_8433172_8446622_RNA45SN1_45S_50ntFlanks@11326.11344.19|CTCGTCGCGCGCGCGTCCG</td><td>>hg38_21_+_8433172_8446622_RNA45SN1_45S_50ntFlanks</td><td>11326</td><td>11344</td><td>19</td><td>CTCGTCGCGCGCGCGTCCG</td><td>548.3</td><td>2.1</td><td>1.0</td><td>2.0</td><td>4.3E-02</td><td>4.8E-02</td><td>MB231/MB468</td><td>NUC</td></tr>
<tr><td>>MI0000299|hsa-mir-222&WithFlank&X|-|45747009|45747130@75.95.21(+1A)&[MIMAT0000279&hsa-miR-222-3p&offsets|0|0(+1A);m-140&X|-|45747033|45747056&offsets|0|-3(+1A)]|AGCTACATCTGGCTACTGGGTA</td><td>>MI0000299|hsa-mir-222&WithFlank&X|-|45747009|45747130</td><td>75</td><td>95</td><td>21(+1A)</td><td>AGCTACATCTGGCTACTGGGTA</td><td>364.1</td><td>-2.7</td><td>1.1</td><td>-2.5</td><td>1.4E-02</td><td>4.8E-02</td><td>BT20/MB231</td><td>MP</td></tr>
<tr><td>>hg38_21_+_8433172_8446622_RNA45SN1_45S_50ntFlanks@11579.11607.29|AGAACTGGTGCGGACCAGGGGAATCCGAC</td><td>>hg38_21_+_8433172_8446622_RNA45SN1_45S_50ntFlanks</td><td>11579</td><td>11607</td><td>29</td><td>AGAACTGGTGCGGACCAGGGGAATCCGAC</td><td>523.7</td><td>-1.4</td><td>0.7</td><td>-2.0</td><td>4.2E-02</td><td>4.8E-02</td><td>MB231/MB468</td><td>CYTO</td></tr>
<tr><td>>hg38_21_+_8433172_8446622_RNA45SN1_45S_50ntFlanks@10829.10858.30|GCAAGCCGGATCCGTAACTTCGGGATAAGG</td><td>>hg38_21_+_8433172_8446622_RNA45SN1_45S_50ntFlanks</td><td>10829</td><td>10858</td><td>30</td><td>GCAAGCCGGATCCGTAACTTCGGGATAAGG</td><td>133.6</td><td>-2.7</td><td>1.3</td><td>-2.1</td><td>3.7E-02</td><td>4.8E-02</td><td>BT20/MB468</td><td>MITO</td></tr>
<tr><td>>MI0000085|hsa-mir-27a&WithFlank&19|-|13836434|13836523@57.76.20(+3A)&[MIMAT0000084&hsa-miR-27a-3p&offsets|0|-1(+3A);m-43&19|-|13836447|13836467&offsets|0|-1(+3A)]|TTCACAGTGGCTAAGTTCCGAAA</td><td>>MI0000085|hsa-mir-27a&WithFlank&19|-|13836434|13836523</td><td>57</td><td>76</td><td>20(+3A)</td><td>TTCACAGTGGCTAAGTTCCGAAA</td><td>35.4</td><td>4.3</td><td>2.1</td><td>2.1</td><td>4.0E-02</td><td>4.8E-02</td><td>MB231/MB468</td><td>MP</td></tr>
<tr><td>>hg38_MT_+_598_1651_RNR1_12S_50ntFlanks@905.925.21|AGTATACTTCAAAGGACATTT</td><td>>hg38_MT_+_598_1651_RNR1_12S_50ntFlanks</td><td>905</td><td>925</td><td>21</td><td>AGTATACTTCAAAGGACATTT</td><td>52.2</td><td>3.6</td><td>1.7</td><td>2.1</td><td>4.0E-02</td><td>4.8E-02</td><td>MB231/MB468</td><td>MP</td></tr>
<tr><td>>hg38_21_+_8433172_8446622_RNA45SN1_45S_50ntFlanks@10342.10361.20|ATATTCAAACGAGAACTTTG</td><td>>hg38_21_+_8433172_8446622_RNA45SN1_45S_50ntFlanks</td><td>10342</td><td>10361</td><td>20</td><td>ATATTCAAACGAGAACTTTG</td><td>280.9</td><td>-1.5</td><td>0.8</td><td>-2.0</td><td>4.3E-02</td><td>4.8E-02</td><td>MB231/MB468</td><td>NUC</td></tr>
<tr><td>>hg38_21_+_8433172_8446622_RNA45SN1_45S_50ntFlanks@9485.9503.19|TGTTAGGACCCGAAAGATG</td><td>>hg38_21_+_8433172_8446622_RNA45SN1_45S_50ntFlanks</td><td>9485</td><td>9503</td><td>19</td><td>TGTTAGGACCCGAAAGATG</td><td>210.3</td><td>4.9</td><td>2.0</td><td>2.5</td><td>1.4E-02</td><td>4.8E-02</td><td>BT20/MB231</td><td>MP</td></tr>
<tr><td>>hg38_21_+_8433172_8446622_RNA45SN1_45S_50ntFlanks@10849.10867.19|CGGGATAAGGATTGGCTCT</td><td>>hg38_21_+_8433172_8446622_RNA45SN1_45S_50ntFlanks</td><td>10849</td><td>10867</td><td>19</td><td>CGGGATAAGGATTGGCTCT</td><td>462.3</td><td>-1.7</td><td>0.8</td><td>-2.0</td><td>4.2E-02</td><td>4.8E-02</td><td>MB231/MB468</td><td>CYTO</td></tr>
<tr><td>>MI0000063|hsa-let-7b&WithFlank&22|+|46113680|46113774@12.30.19(+1A)&[MIMAT0000063&hsa-let-7b-5p&offsets|0|-3(+1A);m-14&22|+|46113691|46113712&offsets|0|-3(+1A)]|TGAGGTAGTAGGTTGTGTGA</td><td>>MI0000063|hsa-let-7b&WithFlank&22|+|46113680|46113774</td><td>12</td><td>30</td><td>19(+1A)</td><td>TGAGGTAGTAGGTTGTGTGA</td><td>64.2</td><td>2.9</td><td>1.4</td><td>2.1</td><td>4.0E-02</td><td>4.8E-02</td><td>MB231/MB468</td><td>MP</td></tr>
<tr><td>>hg38_21_+_8433172_8446622_RNA45SN1_45S_50ntFlanks@11568.11588.21|GCAGCCGACTTAGAACTGGTG</td><td>>hg38_21_+_8433172_8446622_RNA45SN1_45S_50ntFlanks</td><td>11568</td><td>11588</td><td>21</td><td>GCAGCCGACTTAGAACTGGTG</td><td>1153.8</td><td>-2.1</td><td>1.0</td><td>-2.2</td><td>3.0E-02</td><td>4.8E-02</td><td>BT20/MB468</td><td>MP</td></tr>
<tr><td>>hg38_MT_+_1621_3279_RNR2_16S_50ntFlanks@1249.1275.27|AGTCAAAGCGAACTACTATACTCAATT</td><td>>hg38_MT_+_1621_3279_RNR2_16S_50ntFlanks</td><td>1249</td><td>1275</td><td>27</td><td>AGTCAAAGCGAACTACTATACTCAATT</td><td>172.9</td><td>3.6</td><td>1.7</td><td>2.2</td><td>3.0E-02</td><td>4.8E-02</td><td>BT20/MB468</td><td>MP</td></tr>
<tr><td>>hg38_21_+_8433172_8446622_RNA45SN1_45S_50ntFlanks@5078.5102.25|CGATAACGAACGAGACTCTGGCATG</td><td>>hg38_21_+_8433172_8446622_RNA45SN1_45S_50ntFlanks</td><td>5078</td><td>5102</td><td>25</td><td>CGATAACGAACGAGACTCTGGCATG</td><td>1046.8</td><td>-2.9</td><td>1.3</td><td>-2.2</td><td>3.0E-02</td><td>4.8E-02</td><td>BT20/MB468</td><td>MP</td></tr>
<tr><td>>hg38_21_+_8433172_8446622_RNA45SN1_45S_50ntFlanks@12323.12346.24|TTTGGGTTTTAAGCAGGAGGTGTC</td><td>>hg38_21_+_8433172_8446622_RNA45SN1_45S_50ntFlanks</td><td>12323</td><td>12346</td><td>24</td><td>TTTGGGTTTTAAGCAGGAGGTGTC</td><td>138.3</td><td>-3.7</td><td>1.7</td><td>-2.2</td><td>3.0E-02</td><td>4.8E-02</td><td>BT20/MB468</td><td>MP</td></tr>
<tr><td>>MI0000082|hsa-mir-25&WithFlank&7|-|100093554|100093649@58.79.22(+1A)&[MIMAT0000081&hsa-miR-25-3p&offsets|0|0(+1A);m-23&7|-|100093571|100093592&offsets|0|0(+1A)]|CATTGCACTTGTCTCGGTCTGAA</td><td>>MI0000082|hsa-mir-25&WithFlank&7|-|100093554|100093649</td><td>58</td><td>79</td><td>22(+1A)</td><td>CATTGCACTTGTCTCGGTCTGAA</td><td>297.1</td><td>2.8</td><td>1.4</td><td>2.1</td><td>4.0E-02</td><td>4.8E-02</td><td>MB231/MB468</td><td>MP</td></tr>
<tr><td>>MI0000081|hsa-mir-24-2&WithFlank&19|-|13836281|13836365@56.74.19;MI0000080|hsa-mir-24-1&WithFlank&9|+|95086015|95086094@50.68.19&[MIMAT0000080&hsa-miR-24-3p&offsets|0|-3;m-36&19|-|13836288|13836310&offsets|0|-4];[MIMAT0000080_1&hsa-miR-24-3p&offsets|0|-3;m-35&9|+|95086064|95086086&offsets|0|-4]|TGGCTCAGTTCAGCAGGAA</td><td>>MI0000081|hsa-mir-24-2&WithFlank&19|-|13836281|13836365</td><td>56</td><td>74</td><td>19</td><td>TGGCTCAGTTCAGCAGGAA</td><td>69.9</td><td>2.8</td><td>1.4</td><td>2.1</td><td>4.0E-02</td><td>4.8E-02</td><td>MB231/MB468</td><td>MP</td></tr>
<tr><td>>MI0000077|hsa-mir-21&WithFlank&17|+|59841260|59841343@16.36.21&[MIMAT0000076&hsa-miR-21-5p&offsets|+2|+1;m-1&17|+|59841273|59841294&offsets|+2|+1]|GCTTATCAGACTGATGTTGAC</td><td>>MI0000077|hsa-mir-21&WithFlank&17|+|59841260|59841343</td><td>16</td><td>36</td><td>21</td><td>GCTTATCAGACTGATGTTGAC</td><td>507.6</td><td>2.1</td><td>0.9</td><td>2.4</td><td>1.9E-02</td><td>4.8E-02</td><td>BT20/MB231</td><td>CYTO</td></tr>
<tr><td>>hg38_21_+_8433172_8446622_RNA45SN1_45S_50ntFlanks@3711.3729.19|GTTGATCCTGCCAGTAGCA</td><td>>hg38_21_+_8433172_8446622_RNA45SN1_45S_50ntFlanks</td><td>3711</td><td>3729</td><td>19</td><td>GTTGATCCTGCCAGTAGCA</td><td>676.4</td><td>-1.9</td><td>0.9</td><td>-2.0</td><td>4.4E-02</td><td>4.8E-02</td><td>MB231/MB468</td><td>MITO</td></tr>
<tr><td>>hg38_21_+_8433172_8446622_RNA45SN1_45S_50ntFlanks@9568.9601.34|CGTGCAAATCGGTCGTCCGACCTGGGTATAGGGG</td><td>>hg38_21_+_8433172_8446622_RNA45SN1_45S_50ntFlanks</td><td>9568</td><td>9601</td><td>34</td><td>CGTGCAAATCGGTCGTCCGACCTGGGTATAGGGG</td><td>231.1</td><td>-1.4</td><td>0.7</td><td>-2.1</td><td>3.8E-02</td><td>4.8E-02</td><td>BT20/MB468</td><td>MITO</td></tr>
<tr><td>>MI0000465|hsa-mir-191&WithFlank&3|-|49020612|49020715@22.42.21(+1A)&[MIMAT0000440&hsa-miR-191-5p&offsets|0|-2(+1A);m-48&3|-|49020672|49020694&offsets|0|-2(+1A)]|CAACGGAATCCCAAAAGCAGCA</td><td>>MI0000465|hsa-mir-191&WithFlank&3|-|49020612|49020715</td><td>22</td><td>42</td><td>21(+1A)</td><td>CAACGGAATCCCAAAAGCAGCA</td><td>1071.4</td><td>1.4</td><td>0.6</td><td>2.5</td><td>1.4E-02</td><td>4.8E-02</td><td>BT20/MB231</td><td>NUC</td></tr>
<tr><td>>hg38_21_+_8433172_8446622_RNA45SN1_45S_50ntFlanks@6646.6686.41|TCGTACGACTCTTAGCGGTGGATCACTCGGCTCGTGCGTCG</td><td>>hg38_21_+_8433172_8446622_RNA45SN1_45S_50ntFlanks</td><td>6646</td><td>6686</td><td>41</td><td>TCGTACGACTCTTAGCGGTGGATCACTCGGCTCGTGCGTCG</td><td>309.8</td><td>1.8</td><td>0.7</td><td>2.5</td><td>1.4E-02</td><td>4.8E-02</td><td>BT20/MB231</td><td>NUC</td></tr>
<tr><td>>hg38_21_+_8433172_8446622_RNA45SN1_45S_50ntFlanks@9806.9835.30|CGTGGAATGCGAGTGCCTAGTGGGCCACTT</td><td>>hg38_21_+_8433172_8446622_RNA45SN1_45S_50ntFlanks</td><td>9806</td><td>9835</td><td>30</td><td>CGTGGAATGCGAGTGCCTAGTGGGCCACTT</td><td>77.1</td><td>-3.3</td><td>1.5</td><td>-2.2</td><td>3.0E-02</td><td>4.8E-02</td><td>BT20/MB468</td><td>MP</td></tr>
<tr><td>>hg38_21_+_8433172_8446622_RNA45SN1_45S_50ntFlanks@12907.12931.25|CACCGCACGTTCGTGGGGAACCTGG</td><td>>hg38_21_+_8433172_8446622_RNA45SN1_45S_50ntFlanks</td><td>12907</td><td>12931</td><td>25</td><td>CACCGCACGTTCGTGGGGAACCTGG</td><td>162.3</td><td>-3.1</td><td>1.5</td><td>-2.0</td><td>4.2E-02</td><td>4.8E-02</td><td>BT20/MB468</td><td>CYTO</td></tr>
<tr><td>>hg38_21_+_8433172_8446622_RNA45SN1_45S_50ntFlanks@8356.8395.40|GAAAAGAACTTTGAAGAGAGAGTTCAAGAGGGCGTGAAAC</td><td>>hg38_21_+_8433172_8446622_RNA45SN1_45S_50ntFlanks</td><td>8356</td><td>8395</td><td>40</td><td>GAAAAGAACTTTGAAGAGAGAGTTCAAGAGGGCGTGAAAC</td><td>52.1</td><td>4.6</td><td>2.1</td><td>2.2</td><td>3.0E-02</td><td>4.8E-02</td><td>BT20/MB468</td><td>MP</td></tr>
<tr><td>>hg38_21_+_8433172_8446622_RNA45SN1_45S_50ntFlanks@10769.10796.28|AGCAGGTCTCCAAGGTGAACAGCCTCTG</td><td>>hg38_21_+_8433172_8446622_RNA45SN1_45S_50ntFlanks</td><td>10769</td><td>10796</td><td>28</td><td>AGCAGGTCTCCAAGGTGAACAGCCTCTG</td><td>272.3</td><td>-2.8</td><td>1.4</td><td>-2.1</td><td>4.0E-02</td><td>4.8E-02</td><td>MB231/MB468</td><td>MP</td></tr>
<tr><td>>am_MI0000786|hsa-mir-378a&WithFlank&5|+|149732819|149732896@49.70.22&[MIMAT0000732&hsa-miR-378a-3p&offsets|0|0;m-44&5|+|149732867|149732888&offsets|0|0]|ACTGGACTTGGAGTCAGAAGGC</td><td>>am_MI0000786|hsa-mir-378a&WithFlank&5|+|149732819|149732896</td><td>49</td><td>70</td><td>22</td><td>ACTGGACTTGGAGTCAGAAGGC</td><td>56820.5</td><td>2.5</td><td>1.1</td><td>2.2</td><td>3.0E-02</td><td>4.8E-02</td><td>BT20/MB468</td><td>MP</td></tr>
<tr><td>>trnaMT_SerGCT_MT_+_12207_12265@1.19.19|GAGAAAGCTCACAAGAACT</td><td>>trnaMT_SerGCT_MT_+_12207_12265</td><td>1</td><td>19</td><td>19</td><td>GAGAAAGCTCACAAGAACT</td><td>1104.0</td><td>1.7</td><td>0.7</td><td>2.4</td><td>1.9E-02</td><td>4.8E-02</td><td>BT20/MB231</td><td>CYTO</td></tr>
<tr><td>>hg38_21_+_8433172_8446622_RNA45SN1_45S_50ntFlanks@4731.4761.31|AAGAACGAAAGTCGGAGGTTCGAAGACGATC</td><td>>hg38_21_+_8433172_8446622_RNA45SN1_45S_50ntFlanks</td><td>4731</td><td>4761</td><td>31</td><td>AAGAACGAAAGTCGGAGGTTCGAAGACGATC</td><td>4769.3</td><td>3.6</td><td>1.5</td><td>2.5</td><td>1.4E-02</td><td>4.8E-02</td><td>BT20/MB231</td><td>MP</td></tr>
<tr><td>>hg38_21_+_8433172_8446622_RNA45SN1_45S_50ntFlanks@6652.6676.25|GACTCTTAGCGGTGGATCACTCGGC</td><td>>hg38_21_+_8433172_8446622_RNA45SN1_45S_50ntFlanks</td><td>6652</td><td>6676</td><td>25</td><td>GACTCTTAGCGGTGGATCACTCGGC</td><td>8596.1</td><td>2.9</td><td>1.2</td><td>2.4</td><td>1.9E-02</td><td>4.8E-02</td><td>BT20/MB231</td><td>CYTO</td></tr>
<tr><td>>hg38_21_+_8433172_8446622_RNA45SN1_45S_50ntFlanks@10869.10892.24|AGGGCTGGGTCGGTCGGGCTGGGG</td><td>>hg38_21_+_8433172_8446622_RNA45SN1_45S_50ntFlanks</td><td>10869</td><td>10892</td><td>24</td><td>AGGGCTGGGTCGGTCGGGCTGGGG</td><td>630.9</td><td>-3.8</td><td>1.9</td><td>-2.0</td><td>4.2E-02</td><td>4.8E-02</td><td>BT20/MB468</td><td>CYTO</td></tr>
<tr><td>>MI0000760|hsa-mir-361&WithFlank&X|-|85903630|85903713@12.32.21&[MIMAT0000703&hsa-miR-361-5p&offsets|0|-1;m-89&X|-|85903681|85903702&offsets|0|-1]|TTATCAGAATCTCCAGGGGTA</td><td>>MI0000760|hsa-mir-361&WithFlank&X|-|85903630|85903713</td><td>12</td><td>32</td><td>21</td><td>TTATCAGAATCTCCAGGGGTA</td><td>86.9</td><td>4.3</td><td>2.0</td><td>2.2</td><td>3.0E-02</td><td>4.8E-02</td><td>BT20/MB468</td><td>MP</td></tr>
<tr><td>>hg38_MT_+_598_1651_RNR1_12S_50ntFlanks@441.480.40|CATATCTGAACACACAATAGCTAAGACCCAAACTGGGATT</td><td>>hg38_MT_+_598_1651_RNR1_12S_50ntFlanks</td><td>441</td><td>480</td><td>40</td><td>CATATCTGAACACACAATAGCTAAGACCCAAACTGGGATT</td><td>331.0</td><td>-2.1</td><td>0.9</td><td>-2.4</td><td>1.6E-02</td><td>4.8E-02</td><td>BT20/MB231</td><td>MITO</td></tr>
<tr><td>>hg38_21_+_8433172_8446622_RNA45SN1_45S_50ntFlanks@8170.8213.44|CAAGTCCTTCTGATCGAGGCCCAGCCCGTGGACGGTGTGAGGCC</td><td>>hg38_21_+_8433172_8446622_RNA45SN1_45S_50ntFlanks</td><td>8170</td><td>8213</td><td>44</td><td>CAAGTCCTTCTGATCGAGGCCCAGCCCGTGGACGGTGTGAGGCC</td><td>361.6</td><td>-2.8</td><td>1.4</td><td>-2.0</td><td>4.2E-02</td><td>4.8E-02</td><td>BT20/MB468</td><td>CYTO</td></tr>
<tr><td>>hg38_21_+_8433172_8446622_RNA45SN1_45S_50ntFlanks@12054.12071.18|TGAGGCGGGGGGGCGAGC</td><td>>hg38_21_+_8433172_8446622_RNA45SN1_45S_50ntFlanks</td><td>12054</td><td>12071</td><td>18</td><td>TGAGGCGGGGGGGCGAGC</td><td>141.4</td><td>-3.3</td><td>1.6</td><td>-2.0</td><td>4.2E-02</td><td>4.8E-02</td><td>BT20/MB468</td><td>CYTO</td></tr>
<tr><td>>hg38_21_+_8433172_8446622_RNA45SN1_45S_50ntFlanks@12419.12437.19|GATGTCGGCTCTTCCTATC</td><td>>hg38_21_+_8433172_8446622_RNA45SN1_45S_50ntFlanks</td><td>12419</td><td>12437</td><td>19</td><td>GATGTCGGCTCTTCCTATC</td><td>4142.6</td><td>1.2</td><td>0.6</td><td>2.0</td><td>4.4E-02</td><td>4.8E-02</td><td>MB231/MB468</td><td>NUC</td></tr>
<tr><td>>hg38_21_+_8433172_8446622_RNA45SN1_45S_50ntFlanks@11582.11611.30|ACTGGTGCGGACCAGGGGAATCCGACTGTT</td><td>>hg38_21_+_8433172_8446622_RNA45SN1_45S_50ntFlanks</td><td>11582</td><td>11611</td><td>30</td><td>ACTGGTGCGGACCAGGGGAATCCGACTGTT</td><td>517.9</td><td>1.6</td><td>0.7</td><td>2.4</td><td>1.6E-02</td><td>4.8E-02</td><td>BT20/MB231</td><td>MITO</td></tr>
<tr><td>>hg38_21_+_8433172_8446622_RNA45SN1_45S_50ntFlanks@12646.12669.24|TGTGGGATTATGACTGAACGCCTC</td><td>>hg38_21_+_8433172_8446622_RNA45SN1_45S_50ntFlanks</td><td>12646</td><td>12669</td><td>24</td><td>TGTGGGATTATGACTGAACGCCTC</td><td>809.1</td><td>2.1</td><td>0.9</td><td>2.4</td><td>1.6E-02</td><td>4.8E-02</td><td>BT20/MB231</td><td>MITO</td></tr>
<tr><td>>hg38_1_-_228634819_228635039_RNA5S12_5S_50ntFlanks@146.170.25|TGGGAATACCGGGTGCTGTAGGCTT</td><td>>hg38_1_-_228634819_228635039_RNA5S12_5S_50ntFlanks</td><td>146</td><td>170</td><td>25</td><td>TGGGAATACCGGGTGCTGTAGGCTT</td><td>526.4</td><td>1.4</td><td>0.7</td><td>2.0</td><td>4.3E-02</td><td>4.8E-02</td><td>BT20/MB468</td><td>NUC</td></tr>
<tr><td>>hg38_21_+_8433172_8446622_RNA45SN1_45S_50ntFlanks@12274.12317.44|CAGTACGAATACAGACCGTGAAAGCGGGGCCTCACGATCCTTCT</td><td>>hg38_21_+_8433172_8446622_RNA45SN1_45S_50ntFlanks</td><td>12274</td><td>12317</td><td>44</td><td>CAGTACGAATACAGACCGTGAAAGCGGGGCCTCACGATCCTTCT</td><td>109.6</td><td>-2.7</td><td>1.3</td><td>-2.1</td><td>3.8E-02</td><td>4.8E-02</td><td>BT20/MB468</td><td>MITO</td></tr>
<tr><td>>hg38_21_+_8433172_8446622_RNA45SN1_45S_50ntFlanks@12073.12090.18|CCGAGGGGCTCTCGCTTC</td><td>>hg38_21_+_8433172_8446622_RNA45SN1_45S_50ntFlanks</td><td>12073</td><td>12090</td><td>18</td><td>CCGAGGGGCTCTCGCTTC</td><td>608.8</td><td>-2.9</td><td>1.4</td><td>-2.0</td><td>4.2E-02</td><td>4.8E-02</td><td>BT20/MB468</td><td>CYTO</td></tr>
<tr><td>>hg38_21_+_8433172_8446622_RNA45SN1_45S_50ntFlanks@4733.4751.19|GAACGAAAGTCGGAGGTTC</td><td>>hg38_21_+_8433172_8446622_RNA45SN1_45S_50ntFlanks</td><td>4733</td><td>4751</td><td>19</td><td>GAACGAAAGTCGGAGGTTC</td><td>398.3</td><td>1.2</td><td>0.6</td><td>2.0</td><td>4.2E-02</td><td>4.8E-02</td><td>BT20/MB468</td><td>CYTO</td></tr>
<tr><td>>MI0000446|hsa-mir-125b-1&WithFlank&11|-|122099751|122099850@61.80.20(+1U)&[MIMAT0004592&hsa-miR-125b-1-3p&offsets|0|-2(+1U);m-485&11|-|122099769|122099790&offsets|0|-2(+1U)]|ACGGGTTAGGCTCTTGGGAGT</td><td>>MI0000446|hsa-mir-125b-1&WithFlank&11|-|122099751|122099850</td><td>61</td><td>80</td><td>20(+1U)</td><td>ACGGGTTAGGCTCTTGGGAGT</td><td>165.7</td><td>-5.0</td><td>2.1</td><td>-2.4</td><td>1.6E-02</td><td>4.8E-02</td><td>BT20/MB231</td><td>MITO</td></tr>
</table>

<table>
<tr><th></th><th></th><th></th><th></th><th></th><th></th><th></th><th></th><th></th><th></th><th></th><th></th><th></th><th></th></tr>
<tr><td>>trna10_LysCTT_16_+_3241501_3241573@9.35.27,trna13_LysCTT_14_-_58706613_58706685@9.35.27,trna2_LysCTT_15_+_79152904_79152976@9.35.27,trna4_LysCTT_18_-_43669271_43669343@9.35.27|AGCTCAGTCGGTAGAGCATGGGACTCT</td><td>>trna10_LysCTT_16_+_3241501_3241573</td><td>9</td><td>35</td><td>27</td><td>AGCTCAGTCGGTAGAGCATGGGACTCT</td><td>289.9</td><td>-5.4</td><td>2.2</td><td>-2.4</td><td>1.6E-02</td><td>4.8E-02</td><td>BT20/MB231</td><td>MITO</td></tr>
<tr><td>>trna119_LysCTT_1_-_145395522_145395594@1.32.32,trna11_LysCTT_5_-_180648979_180649051@1.32.32,trna13_LysCTT_6_+_26556774_26556846@1.32.32,trna32_LysCTT_16_-_3207406_3207478@1.32.32,trna7_LysCTT_16_+_3225692_3225764@1.32.32,trna9_LysCTT_5_+_180634755_180634827@1.32.32|GCCCGGCTAGCTCAGTCGGTAGAGCATGAGAC</td><td>>trna119_LysCTT_1_-_145395522_145395594</td><td>1</td><td>32</td><td>32</td><td>GCCCGGCTAGCTCAGTCGGTAGAGCATGAGAC</td><td>3221.0</td><td>-2.5</td><td>1.1</td><td>-2.4</td><td>1.6E-02</td><td>4.8E-02</td><td>BT20/MB231</td><td>MITO</td></tr>
<tr><td>>trnaMT_ValTAC_MT_+_1602_1670@27.71.45|CAACTTACACTTAGGAGATTTCAACTTAACTTGACCGCTCTGACC</td><td>>trnaMT_ValTAC_MT_+_1602_1670</td><td>27</td><td>71</td><td>45</td><td>CAACTTACACTTAGGAGATTTCAACTTAACTTGACCGCTCTGACC</td><td>255.7</td><td>-2.1</td><td>0.9</td><td>-2.4</td><td>1.6E-02</td><td>4.8E-02</td><td>BT20/MB231</td><td>MITO</td></tr>
<tr><td>>hg38_21_+_8433172_8446622_RNA45SN1_45S_50ntFlanks@12416.12455.40|TTCGATGTCGGCTCTTCCTATCATTGTGAAGCAGAATTCA</td><td>>hg38_21_+_8433172_8446622_RNA45SN1_45S_50ntFlanks</td><td>12416</td><td>12455</td><td>40</td><td>TTCGATGTCGGCTCTTCCTATCATTGTGAAGCAGAATTCA</td><td>269.0</td><td>-2.2</td><td>0.9</td><td>-2.4</td><td>1.6E-02</td><td>4.8E-02</td><td>BT20/MB231</td><td>MITO</td></tr>
<tr><td>>hg38_MT_+_598_1651_RNR1_12S_50ntFlanks@440.480.41|ACATATCTGAACACACAATAGCTAAGACCCAAACTGGGATT</td><td>>hg38_MT_+_598_1651_RNR1_12S_50ntFlanks</td><td>440</td><td>480</td><td>41</td><td>ACATATCTGAACACACAATAGCTAAGACCCAAACTGGGATT</td><td>237.0</td><td>-2.2</td><td>0.9</td><td>-2.4</td><td>1.6E-02</td><td>4.8E-02</td><td>BT20/MB231</td><td>MITO</td></tr>
<tr><td>>MI0000103|hsa-mir-101-1&WithFlank&1|-|65058428|65058514@52.72.21;MI0000739|hsa-mir-101-2&WithFlank&9|+|4850291|4850381@54.74.21&[MIMAT0000099&hsa-miR-101-3p&offsets|-1|-1;m-17&1|-|65058443|65058462&offsets|-1|0];[MIMAT0000099_1&hsa-miR-101-3p&offsets|-1|-1;m-16&9|+|4850345|4850364&offsets|-1|0]|GTACAGTACTGTGATAACTGA</td><td>>MI0000103|hsa-mir-101-1&WithFlank&1|-|65058428|65058514</td><td>52</td><td>72</td><td>21</td><td>GTACAGTACTGTGATAACTGA</td><td>625.3</td><td>-2.2</td><td>0.9</td><td>-2.4</td><td>1.7E-02</td><td>4.8E-02</td><td>BT20/MB231</td><td>MITO</td></tr>
<tr><td>>hg38_MT_+_1621_3279_RNR2_16S_50ntFlanks@1369.1400.32|GATGTTGGATCAGGACATCCCGATGGTGCAGC</td><td>>hg38_MT_+_1621_3279_RNR2_16S_50ntFlanks</td><td>1369</td><td>1400</td><td>32</td><td>GATGTTGGATCAGGACATCCCGATGGTGCAGC</td><td>47.8</td><td>4.2</td><td>2.0</td><td>2.1</td><td>4.0E-02</td><td>4.8E-02</td><td>MB231/MB468</td><td>MP</td></tr>
<tr><td>>MI0000070|hsa-mir-16-1&WithFlank&13|-|50048967|50049067@20.41.22;MI0000115|hsa-mir-16-2&WithFlank&3|+|160404739|160404831@16.37.22&[MIMAT0000069&hsa-miR-16-5p&offsets|0|0;m-62&13|-|50049027|50049048&offsets|0|0];[MIMAT0000069_1&hsa-miR-16-5p&offsets|0|0;m-61&3|+|160404754|160404775&offsets|0|0]|TAGCAGCACGTAAATATTGGCG</td><td>>MI0000070|hsa-mir-16-1&WithFlank&13|-|50048967|50049067</td><td>20</td><td>41</td><td>22</td><td>TAGCAGCACGTAAATATTGGCG</td><td>4937.7</td><td>2.3</td><td>1.0</td><td>2.3</td><td>1.9E-02</td><td>4.8E-02</td><td>BT20/MB231</td><td>CYTO</td></tr>
<tr><td>>trnaMT_ValTAC_MT_+_1602_1670@55.72.18|ACTTGACCGCTCTGACCA</td><td>>trnaMT_ValTAC_MT_+_1602_1670</td><td>55</td><td>72</td><td>18</td><td>ACTTGACCGCTCTGACCA</td><td>655.1</td><td>-1.1</td><td>0.5</td><td>-2.1</td><td>4.0E-02</td><td>4.8E-02</td><td>MB231/MB468</td><td>MP</td></tr>
<tr><td>>hg38_21_+_8433172_8446622_RNA45SN1_45S_50ntFlanks@3811.3834.24|AGTTATGGTTCCTTTGGTCGCTCG</td><td>>hg38_21_+_8433172_8446622_RNA45SN1_45S_50ntFlanks</td><td>3811</td><td>3834</td><td>24</td><td>AGTTATGGTTCCTTTGGTCGCTCG</td><td>133.1</td><td>-2.6</td><td>1.2</td><td>-2.1</td><td>3.8E-02</td><td>4.8E-02</td><td>BT20/MB468</td><td>MITO</td></tr>
<tr><td>>MI0000070|hsa-mir-16-1&WithFlank&13|-|50048967|50049067@20.43.24&[MIMAT0000069&hsa-miR-16-5p&offsets|0|+2;m-62&13|-|50049027|50049048&offsets|0|+2]|TAGCAGCACGTAAATATTGGCGTT</td><td>>MI0000070|hsa-mir-16-1&WithFlank&13|-|50048967|50049067</td><td>20</td><td>43</td><td>24</td><td>TAGCAGCACGTAAATATTGGCGTT</td><td>177.1</td><td>2.9</td><td>1.4</td><td>2.1</td><td>4.0E-02</td><td>4.8E-02</td><td>MB231/MB468</td><td>MP</td></tr>
<tr><td>>MI0000102|hsa-mir-100&WithFlank&11|-|122152223|122152314@19.38.20(+1A)&[MIMAT0000098&hsa-miR-100-5p&offsets|0|-2(+1A);m-27&11|-|122152276|122152296&offsets|0|-1(+1A)]|AACCCGTAGATCCGAACTTGA</td><td>>MI0000102|hsa-mir-100&WithFlank&11|-|122152223|122152314</td><td>19</td><td>38</td><td>20(+1A)</td><td>AACCCGTAGATCCGAACTTGA</td><td>5263.7</td><td>-2.9</td><td>1.3</td><td>-2.3</td><td>1.9E-02</td><td>4.8E-02</td><td>BT20/MB231</td><td>CYTO</td></tr>
<tr><td>>hg38_MT_+_1621_3279_RNR2_16S_50ntFlanks@1370.1404.35|ATGTTGGATCAGGACATCCCGATGGTGCAGCCGCT</td><td>>hg38_MT_+_1621_3279_RNR2_16S_50ntFlanks</td><td>1370</td><td>1404</td><td>35</td><td>ATGTTGGATCAGGACATCCCGATGGTGCAGCCGCT</td><td>210.0</td><td>4.2</td><td>2.0</td><td>2.2</td><td>3.1E-02</td><td>4.8E-02</td><td>BT20/MB468</td><td>MP</td></tr>
<tr><td>>hg38_21_+_8433172_8446622_RNA45SN1_45S_50ntFlanks@11568.11602.35|GCAGCCGACTTAGAACTGGTGCGGACCAGGGGAAT</td><td>>hg38_21_+_8433172_8446622_RNA45SN1_45S_50ntFlanks</td><td>11568</td><td>11602</td><td>35</td><td>GCAGCCGACTTAGAACTGGTGCGGACCAGGGGAAT</td><td>302.0</td><td>2.1</td><td>0.9</td><td>2.4</td><td>1.7E-02</td><td>4.8E-02</td><td>BT20/MB231</td><td>MITO</td></tr>
<tr><td>>hg38_21_+_8433172_8446622_RNA45SN1_45S_50ntFlanks@6646.6685.40|TCGTACGACTCTTAGCGGTGGATCACTCGGCTCGTGCGTC</td><td>>hg38_21_+_8433172_8446622_RNA45SN1_45S_50ntFlanks</td><td>6646</td><td>6685</td><td>40</td><td>TCGTACGACTCTTAGCGGTGGATCACTCGGCTCGTGCGTC</td><td>20307.2</td><td>2.2</td><td>0.9</td><td>2.4</td><td>1.7E-02</td><td>4.8E-02</td><td>BT20/MB231</td><td>MITO</td></tr>
<tr><td>>MI0003139|hsa-mir-181d&WithFlank&19|+|13874869|13875017@42.65.24&[MIMAT0002821&hsa-miR-181d-5p&offsets|0|+1;m-194&19|+|13874910|13874932&offsets|0|+1]|AACATTCATTGTTGTCGGTGGGTT</td><td>>MI0003139|hsa-mir-181d&WithFlank&19|+|13874869|13875017</td><td>42</td><td>65</td><td>24</td><td>AACATTCATTGTTGTCGGTGGGTT</td><td>140.7</td><td>2.6</td><td>1.2</td><td>2.2</td><td>3.1E-02</td><td>4.8E-02</td><td>BT20/MB468</td><td>MP</td></tr>
<tr><td>>hg38_21_+_8433172_8446622_RNA45SN1_45S_50ntFlanks@12081.12106.26|CTCTCGCTTCTGGCGCCAAGCGCCCG</td><td>>hg38_21_+_8433172_8446622_RNA45SN1_45S_50ntFlanks</td><td>12081</td><td>12106</td><td>26</td><td>CTCTCGCTTCTGGCGCCAAGCGCCCG</td><td>81.2</td><td>-3.8</td><td>1.7</td><td>-2.2</td><td>3.1E-02</td><td>4.8E-02</td><td>BT20/MB468</td><td>MP</td></tr>
<tr><td>>trnaMT_SerGCT_MT_+_12207_12265@12.48.37|CAAGAACTGCTAACTCATGCCCCCATGTCTAACAACA</td><td>>trnaMT_SerGCT_MT_+_12207_12265</td><td>12</td><td>48</td><td>37</td><td>CAAGAACTGCTAACTCATGCCCCCATGTCTAACAACA</td><td>69.6</td><td>2.6</td><td>1.3</td><td>2.0</td><td>4.0E-02</td><td>4.8E-02</td><td>MB231/MB468</td><td>MP</td></tr>
<tr><td>>hg38_21_+_8433172_8446622_RNA45SN1_45S_50ntFlanks@12452.12476.25|TTCACCAAGCGTTGGATTGTTCACC</td><td>>hg38_21_+_8433172_8446622_RNA45SN1_45S_50ntFlanks</td><td>12452</td><td>12476</td><td>25</td><td>TTCACCAAGCGTTGGATTGTTCACC</td><td>97.6</td><td>-2.8</td><td>1.4</td><td>-2.0</td><td>4.0E-02</td><td>4.8E-02</td><td>MB231/MB468</td><td>MP</td></tr>
<tr><td>>hg38_21_+_8433172_8446622_RNA45SN1_45S_50ntFlanks@11569.11612.44|CAGCCGACTTAGAACTGGTGCGGACCAGGGGAATCCGACTGTTT</td><td>>hg38_21_+_8433172_8446622_RNA45SN1_45S_50ntFlanks</td><td>11569</td><td>11612</td><td>44</td><td>CAGCCGACTTAGAACTGGTGCGGACCAGGGGAATCCGACTGTTT</td><td>580.0</td><td>2.4</td><td>1.0</td><td>2.4</td><td>1.7E-02</td><td>4.8E-02</td><td>BT20/MB231</td><td>MITO</td></tr>
<tr><td>>hg38_21_+_8433172_8446622_RNA45SN1_45S_50ntFlanks@12418.12460.43|CGATGTCGGCTCTTCCTATCATTGTGAAGCAGAATTCACCAAG</td><td>>hg38_21_+_8433172_8446622_RNA45SN1_45S_50ntFlanks</td><td>12418</td><td>12460</td><td>43</td><td>CGATGTCGGCTCTTCCTATCATTGTGAAGCAGAATTCACCAAG</td><td>383.3</td><td>-2.4</td><td>1.0</td><td>-2.4</td><td>1.7E-02</td><td>4.8E-02</td><td>BT20/MB231</td><td>MITO</td></tr>
<tr><td>>MI0000108|hsa-mir-103a-2&WithFlank&20|+|3917488|3917577@54.75.22(+2U);MI0000109|hsa-mir-103a-1&WithFlank&5|-|168560890|168560979@54.75.22(+2U)&[MIMAT0000101&hsa-miR-103a-3p&offsets|0|-1(+2U);m-12&20|+|3917541|3917563&offsets|0|-1(+2U)];[MIMAT0000101_1&hsa-miR-103a-3p&offsets|0|-1(+2U);m-13&5|-|168560904|168560926&offsets|0|-1(+2U);m-44220&5|-|168560908|168560926&offsets|0|+3(+2U)]|AGCAGCATTGTACAGGGCTATGTT</td><td>>MI0000108|hsa-mir-103a-2&WithFlank&20|+|3917488|3917577</td><td>54</td><td>75</td><td>22(+2U)</td><td>AGCAGCATTGTACAGGGCTATGTT</td><td>312.5</td><td>-1.8</td><td>0.8</td><td>-2.4</td><td>1.7E-02</td><td>4.8E-02</td><td>BT20/MB231</td><td>MITO</td></tr>
<tr><td>>hg38_21_+_8433172_8446622_RNA45SN1_45S_50ntFlanks@10350.10373.24|ACGAGAACTTTGAAGGCCGAAGTG</td><td>>hg38_21_+_8433172_8446622_RNA45SN1_45S_50ntFlanks</td><td>10350</td><td>10373</td><td>24</td><td>ACGAGAACTTTGAAGGCCGAAGTG</td><td>606.6</td><td>-1.5</td><td>0.7</td><td>-2.0</td><td>4.4E-02</td><td>4.8E-02</td><td>MB231/MB468</td><td>NUC</td></tr>
<tr><td>>hg38_21_+_8433172_8446622_RNA45SN1_45S_50ntFlanks@6733.6753.21|CATTGATCATCGACACTTCGA</td><td>>hg38_21_+_8433172_8446622_RNA45SN1_45S_50ntFlanks</td><td>6733</td><td>6753</td><td>21</td><td>CATTGATCATCGACACTTCGA</td><td>673.6</td><td>1.6</td><td>0.8</td><td>2.1</td><td>3.8E-02</td><td>4.8E-02</td><td>BT20/MB468</td><td>MITO</td></tr>
<tr><td>>trnaMT_GluTTC_MT_-_14674_14742@41.72.32,trnalookalike8_GluTTC_5_-_93905172_93905240@41.72.32|TGGTCGTGGTTGTAGTCCGTGCGAGAATACCA</td><td>>trnaMT_GluTTC_MT_-_14674_14742</td><td>41</td><td>72</td><td>32</td><td>TGGTCGTGGTTGTAGTCCGTGCGAGAATACCA</td><td>9284.4</td><td>2.6</td><td>1.2</td><td>2.2</td><td>3.1E-02</td><td>4.8E-02</td><td>BT20/MB468</td><td>MP</td></tr>
<tr><td>>hg38_21_+_8433172_8446622_RNA45SN1_45S_50ntFlanks@6651.6687.37|CGACTCTTAGCGGTGGATCACTCGGCTCGTGCGTCGA</td><td>>hg38_21_+_8433172_8446622_RNA45SN1_45S_50ntFlanks</td><td>6651</td><td>6687</td><td>37</td><td>CGACTCTTAGCGGTGGATCACTCGGCTCGTGCGTCGA</td><td>3992.7</td><td>1.7</td><td>0.9</td><td>2.0</td><td>4.4E-02</td><td>4.8E-02</td><td>MB231/MB468</td><td>MITO</td></tr>
<tr><td>>MI0000063|hsa-let-7b&WithFlank&22|+|46113680|46113774@12.34.23(+1U)&[MIMAT0000063&hsa-let-7b-5p&offsets|0|+1(+1U);m-14&22|+|46113691|46113712&offsets|0|+1(+1U)]|TGAGGTAGTAGGTTGTGTGGTTTT</td><td>>MI0000063|hsa-let-7b&WithFlank&22|+|46113680|46113774</td><td>12</td><td>34</td><td>23(+1U)</td><td>TGAGGTAGTAGGTTGTGTGGTTTT</td><td>1103.1</td><td>1.8</td><td>0.8</td><td>2.3</td><td>1.9E-02</td><td>4.8E-02</td><td>BT20/MB231</td><td>CYTO</td></tr>
<tr><td>>MI0000746|hsa-mir-99b&WithFlank&19|+|51692606|51692687@13.32.20(+1U)&[MIMAT0000689&hsa-miR-99b-5p&offsets|0|-2(+1U);m-10&19|+|51692618|51692639&offsets|0|-2(+1U)]|CACCCGTAGAACCGACCTTGT</td><td>>MI0000746|hsa-mir-99b&WithFlank&19|+|51692606|51692687</td><td>13</td><td>32</td><td>20(+1U)</td><td>CACCCGTAGAACCGACCTTGT</td><td>7663.6</td><td>2.2</td><td>0.9</td><td>2.5</td><td>1.4E-02</td><td>4.8E-02</td><td>BT20/MB231</td><td>MP</td></tr>
<tr><td>>hg38_21_+_8433172_8446622_RNA45SN1_45S_50ntFlanks@6652.6688.37|GACTCTTAGCGGTGGATCACTCGGCTCGTGCGTCGAT</td><td>>hg38_21_+_8433172_8446622_RNA45SN1_45S_50ntFlanks</td><td>6652</td><td>6688</td><td>37</td><td>GACTCTTAGCGGTGGATCACTCGGCTCGTGCGTCGAT</td><td>2192.2</td><td>2.4</td><td>1.0</td><td>2.5</td><td>1.4E-02</td><td>4.8E-02</td><td>BT20/MB231</td><td>MP</td></tr>
<tr><td>>trnaMT_PheGAA_MT_+_577_647@1.18.18|GTTTATGTAGCTTACCTC</td><td>>trnaMT_PheGAA_MT_+_577_647</td><td>1</td><td>18</td><td>18</td><td>GTTTATGTAGCTTACCTC</td><td>1115.5</td><td>-3.0</td><td>1.2</td><td>-2.5</td><td>1.4E-02</td><td>4.8E-02</td><td>BT20/MB231</td><td>MP</td></tr>
<tr><td>>hg38_21_+_8433172_8446622_RNA45SN1_45S_50ntFlanks@11808.11825.18|CTGTCCCTACCTACTATC</td><td>>hg38_21_+_8433172_8446622_RNA45SN1_45S_50ntFlanks</td><td>11808</td><td>11825</td><td>18</td><td>CTGTCCCTACCTACTATC</td><td>81.8</td><td>-3.1</td><td>1.5</td><td>-2.0</td><td>4.0E-02</td><td>4.8E-02</td><td>MB231/MB468</td><td>MP</td></tr>
<tr><td>>hg38_21_+_8433172_8446622_RNA45SN1_45S_50ntFlanks@9573.9610.38|AAATCGGTCGTCCGACCTGGGTATAGGGGCGAAAGACT</td><td>>hg38_21_+_8433172_8446622_RNA45SN1_45S_50ntFlanks</td><td>9573</td><td>9610</td><td>38</td><td>AAATCGGTCGTCCGACCTGGGTATAGGGGCGAAAGACT</td><td>188.4</td><td>2.1</td><td>1.0</td><td>2.0</td><td>4.4E-02</td><td>4.8E-02</td><td>MB231/MB468</td><td>MITO</td></tr>
<tr><td>>hg38_21_+_8433172_8446622_RNA45SN1_45S_50ntFlanks@8243.8269.27|GTCTTCCCGGAGTCGGGTTGCTTGGGA</td><td>>hg38_21_+_8433172_8446622_RNA45SN1_45S_50ntFlanks</td><td>8243</td><td>8269</td><td>27</td><td>GTCTTCCCGGAGTCGGGTTGCTTGGGA</td><td>201.7</td><td>2.2</td><td>1.0</td><td>2.2</td><td>3.1E-02</td><td>4.8E-02</td><td>BT20/MB468</td><td>MP</td></tr>
<tr><td>>MI0000456|hsa-mir-140&WithFlank&16|+|69933075|69933186@29.48.20&[MIMAT0000431&hsa-miR-140-5p&offsets|0|-2;m-173&16|+|69933103|69933124&offsets|0|-2]|CAGTGGTTTTACCCTATGGT</td><td>>MI0000456|hsa-mir-140&WithFlank&16|+|69933075|69933186</td><td>29</td><td>48</td><td>20</td><td>CAGTGGTTTTACCCTATGGT</td><td>667.1</td><td>-3.1</td><td>1.3</td><td>-2.5</td><td>1.4E-02</td><td>4.8E-02</td><td>BT20/MB231</td><td>MP</td></tr>
<tr><td>>hg38_21_+_8433172_8446622_RNA45SN1_45S_50ntFlanks@11566.11600.35|TAGCAGCCGACTTAGAACTGGTGCGGACCAGGGGA</td><td>>hg38_21_+_8433172_8446622_RNA45SN1_45S_50ntFlanks</td><td>11566</td><td>11600</td><td>35</td><td>TAGCAGCCGACTTAGAACTGGTGCGGACCAGGGGA</td><td>216.9</td><td>4.1</td><td>1.7</td><td>2.5</td><td>1.4E-02</td><td>4.8E-02</td><td>BT20/MB231</td><td>MP</td></tr>
<tr><td>>hg38_21_+_8433172_8446622_RNA45SN1_45S_50ntFlanks@8356.8391.36|GAAAAGAACTTTGAAGAGAGAGTTCAAGAGGGCGTG</td><td>>hg38_21_+_8433172_8446622_RNA45SN1_45S_50ntFlanks</td><td>8356</td><td>8391</td><td>36</td><td>GAAAAGAACTTTGAAGAGAGAGTTCAAGAGGGCGTG</td><td>198.4</td><td>3.2</td><td>1.3</td><td>2.5</td><td>1.4E-02</td><td>4.8E-02</td><td>BT20/MB231</td><td>MP</td></tr>
<tr><td>>hg38_21_+_8433172_8446622_RNA45SN1_45S_50ntFlanks@12419.12449.31|GATGTCGGCTCTTCCTATCATTGTGAAGCAG</td><td>>hg38_21_+_8433172_8446622_RNA45SN1_45S_50ntFlanks</td><td>12419</td><td>12449</td><td>31</td><td>GATGTCGGCTCTTCCTATCATTGTGAAGCAG</td><td>449.7</td><td>-2.3</td><td>0.9</td><td>-2.4</td><td>1.4E-02</td><td>4.8E-02</td><td>BT20/MB231</td><td>NUC</td></tr>
<tr><td>>trna111_HisGTG_1_-_147774845_147774916@-1A.34.35,trna118_HisGTG_1_-_145396881_145396952@-1A.34.35,trna16_HisGTG_1_+_146544773_146544844@-1A.34.35,trna1_HisGTG_15_+_45493349_45493420@-1A.34.35,trna21_HisGTG_1_+_147753471_147753542@-1A.34.35,trna33_HisGTG_6_+_27125906_27125977@-1A.34.35,trna7_HisGTG_9_-_14433938_14434009@-1A.34.35,trna8_HisGTG_15_-_45492611_45492682@-1A.34.35,trna9_HisGTG_15_-_45490804_45490875@-1A.34.35|AGCCGTGATCGTATAGTGGTTAGTACTCTGCGTTG</td><td>>trna111_HisGTG_1_-_147774845_147774916</td><td>-1A</td><td>34</td><td>35</td><td>AGCCGTGATCGTATAGTGGTTAGTACTCTGCGTTG</td><td>1134.7</td><td>-2.5</td><td>1.1</td><td>-2.3</td><td>1.9E-02</td><td>4.8E-02</td><td>BT20/MB231</td><td>CYTO</td></tr>
<tr><td>>hg38_21_+_8433172_8446622_RNA45SN1_45S_50ntFlanks@6733.6753.21|CATTGATCATCGACACTTCGA</td><td>>hg38_21_+_8433172_8446622_RNA45SN1_45S_50ntFlanks</td><td>6733</td><td>6753</td><td>21</td><td>CATTGATCATCGACACTTCGA</td><td>343.2</td><td>-1.3</td><td>0.7</td><td>-2.0</td><td>4.3E-02</td><td>4.8E-02</td><td>BT20/MB468</td><td>CYTO</td></tr>
<tr><td>>am_trnaMT_SerGCT_MT_+_12207_12265@12.33.22|CAAGAACTGCTAACTCATGCCC</td><td>>am_trnaMT_SerGCT_MT_+_12207_12265</td><td>12</td><td>33</td><td>22</td><td>CAAGAACTGCTAACTCATGCCC</td><td>76.4</td><td>4.1</td><td>1.9</td><td>2.2</td><td>3.1E-02</td><td>4.8E-02</td><td>BT20/MB468</td><td>MP</td></tr>
<tr><td>>trnaMT_GluTTC_MT_-_14674_14742@39.68.30,trnalookalike8_GluTTC_5_-_93905172_93905240@39.68.30|ATTGGTCGTGGTTGTAGTCCGTGCGAGAAT</td><td>>trnaMT_GluTTC_MT_-_14674_14742</td><td>39</td><td>68</td><td>30</td><td>ATTGGTCGTGGTTGTAGTCCGTGCGAGAAT</td><td>141.5</td><td>1.7</td><td>0.8</td><td>2.0</td><td>4.1E-02</td><td>4.8E-02</td><td>MB231/MB468</td><td>MP</td></tr>
<tr><td>>hg38_21_+_8433172_8446622_RNA45SN1_45S_50ntFlanks@10677.10712.36|GCGGTTCCGGCGGCGTCCGGTGAGCTCTCGCTGGCC</td><td>>hg38_21_+_8433172_8446622_RNA45SN1_45S_50ntFlanks</td><td>10677</td><td>10712</td><td>36</td><td>GCGGTTCCGGCGGCGTCCGGTGAGCTCTCGCTGGCC</td><td>2018.8</td><td>2.2</td><td>1.0</td><td>2.1</td><td>3.8E-02</td><td>4.8E-02</td><td>BT20/MB468</td><td>MITO</td></tr>
<tr><td>>hg38_21_+_8433172_8446622_RNA45SN1_45S_50ntFlanks@6661.6694.34|CGGTGGATCACTCGGCTCGTGCGTCGATGAAGAA</td><td>>hg38_21_+_8433172_8446622_RNA45SN1_45S_50ntFlanks</td><td>6661</td><td>6694</td><td>34</td><td>CGGTGGATCACTCGGCTCGTGCGTCGATGAAGAA</td><td>571.2</td><td>2.1</td><td>0.9</td><td>2.4</td><td>1.7E-02</td><td>4.8E-02</td><td>BT20/MB231</td><td>MITO</td></tr>
<tr><td>>hg38_21_+_8433172_8446622_RNA45SN1_45S_50ntFlanks@11665.11685.21|TTCTGCCCAGTGCTCTGAATG</td><td>>hg38_21_+_8433172_8446622_RNA45SN1_45S_50ntFlanks</td><td>11665</td><td>11685</td><td>21</td><td>TTCTGCCCAGTGCTCTGAATG</td><td>337.4</td><td>-1.2</td><td>0.6</td><td>-2.0</td><td>4.4E-02</td><td>4.8E-02</td><td>MB231/MB468</td><td>NUC</td></tr>
</table>

<table>
<tr><td>>hg38_21_+_8433172_8446622_RNA45SN1_45S_50ntFlanks@476.498.23|TTCGTGATCGATGTGGTGACGTC</td><td>>hg38_21_+_8433172_8446622_RNA45SN1_45S_50ntFlanks</td><td>476</td><td>498</td><td>23</td><td>TTCGTGATCGATGTGGTGACGTC</td><td>841.7</td><td>-2.7</td><td>1.1</td><td>-2.5</td><td>1.4E-02</td><td>4.8E-02</td><td>BT20/MB231</td><td>MP</td></tr>
<tr><td>>hg38_21_+_8433172_8446622_RNA45SN1_45S_50ntFlanks@6662.6685.24|GGTGGATCACTCGGCTCGTGCGTC</td><td>>hg38_21_+_8433172_8446622_RNA45SN1_45S_50ntFlanks</td><td>6662</td><td>6685</td><td>24</td><td>GGTGGATCACTCGGCTCGTGCGTC</td><td>251.9</td><td>4.8</td><td>2.0</td><td>2.5</td><td>1.4E-02</td><td>4.8E-02</td><td>BT20/MB231</td><td>MP</td></tr>
<tr><td>>hg38_21_+_8433172_8446622_RNA45SN1_45S_50ntFlanks@6647.6683.37|CGTACGACTCTTAGCGGTGGATCACTCGGCTCGTGCG</td><td>>hg38_21_+_8433172_8446622_RNA45SN1_45S_50ntFlanks</td><td>6647</td><td>6683</td><td>37</td><td>CGTACGACTCTTAGCGGTGGATCACTCGGCTCGTGCG</td><td>149.1</td><td>2.1</td><td>1.0</td><td>2.2</td><td>3.1E-02</td><td>4.8E-02</td><td>BT20/MB468</td><td>MP</td></tr>
<tr><td>>hg38_21_+_8433172_8446622_RNA45SN1_45S_50ntFlanks@9935.9957.23|GACAGCAGGACGGTGGCCATGGA</td><td>>hg38_21_+_8433172_8446622_RNA45SN1_45S_50ntFlanks</td><td>9935</td><td>9957</td><td>23</td><td>GACAGCAGGACGGTGGCCATGGA</td><td>59.1</td><td>4.6</td><td>2.1</td><td>2.2</td><td>3.1E-02</td><td>4.8E-02</td><td>BT20/MB468</td><td>MP</td></tr>
<tr><td>>hg38_21_+_8433172_8446622_RNA45SN1_45S_50ntFlanks@4560.4589.30|CTAGGAATAATGGAATAGGACCGCGGTTCT</td><td>>hg38_21_+_8433172_8446622_RNA45SN1_45S_50ntFlanks</td><td>4560</td><td>4589</td><td>30</td><td>CTAGGAATAATGGAATAGGACCGCGGTTCT</td><td>181.9</td><td>-3.0</td><td>1.4</td><td>-2.2</td><td>3.1E-02</td><td>4.8E-02</td><td>BT20/MB468</td><td>MP</td></tr>
<tr><td>>MI0000076|hsa-mir-20a&WithFlank&13|+|91351059|91351141@14.36.23(+1C)&[MIMAT0000075&hsa-miR-20a-5p&offsets|0|0(+1C);m-96&13|+|91351072|91351094&offsets|0|0(+1C)]|TAAAGTGCTTATAGTGCAGGTAGC</td><td>>MI0000076|hsa-mir-20a&WithFlank&13|+|91351059|91351141</td><td>14</td><td>36</td><td>23(+1C)</td><td>TAAAGTGCTTATAGTGCAGGTAGC</td><td>75.8</td><td>3.5</td><td>1.7</td><td>2.0</td><td>4.1E-02</td><td>4.8E-02</td><td>MB231/MB468</td><td>MP</td></tr>
<tr><td>>MI0000234|hsa-mir-192&WithFlank&11|-|64891131|64891252@31.51.21(+1U)&[MIMAT0000222&hsa-miR-192-5p&offsets|+1|+1(+1U);m-59&11|-|64891203|64891223&offsets|+1|+1(+1U)]|TGACCTATGAATTGACAGCCAT</td><td>>MI0000234|hsa-mir-192&WithFlank&11|-|64891131|64891252</td><td>31</td><td>51</td><td>21(+1U)</td><td>TGACCTATGAATTGACAGCCAT</td><td>260.3</td><td>-1.8</td><td>0.8</td><td>-2.4</td><td>1.7E-02</td><td>4.8E-02</td><td>BT20/MB231</td><td>MITO</td></tr>
<tr><td>>hg38_21_+_8433172_8446622_RNA45SN1_45S_50ntFlanks@12419.12438.20|GATGTCGGCTCTTCCTATCA</td><td>>hg38_21_+_8433172_8446622_RNA45SN1_45S_50ntFlanks</td><td>12419</td><td>12438</td><td>20</td><td>GATGTCGGCTCTTCCTATCA</td><td>566.6</td><td>1.8</td><td>0.9</td><td>2.0</td><td>4.4E-02</td><td>4.8E-02</td><td>MB231/MB468</td><td>MITO</td></tr>
<tr><td>>trnaMT_ValTAC_MT_+_1602_1670@55.72.18|ACTTGACCGCTCTGACCA</td><td>>trnaMT_ValTAC_MT_+_1602_1670</td><td>55</td><td>72</td><td>18</td><td>ACTTGACCGCTCTGACCA</td><td>11731.1</td><td>2.2</td><td>1.1</td><td>2.1</td><td>3.8E-02</td><td>4.8E-02</td><td>BT20/MB468</td><td>MITO</td></tr>
<tr><td>>hg38_21_+_8433172_8446622_RNA45SN1_45S_50ntFlanks@12007.12030.24|GCCGCCGGTGAAATACCACTACTC</td><td>>hg38_21_+_8433172_8446622_RNA45SN1_45S_50ntFlanks</td><td>12007</td><td>12030</td><td>24</td><td>GCCGCCGGTGAAATACCACTACTC</td><td>1871.7</td><td>-2.8</td><td>1.4</td><td>-2.1</td><td>3.8E-02</td><td>4.8E-02</td><td>BT20/MB468</td><td>MITO</td></tr>
<tr><td>>MI0005561|hsa-mir-877&WithFlank&6|+|30584326|30584423@7.25.19(+1A)&[MIMAT0004949&hsa-miR-877-5p&offsets|0|-1(+1A);m-382&6|+|30584332|30584354&offsets|0|-4(+1A)]|GTAGAGGAGATGGCGCAGGA</td><td>>MI0005561|hsa-mir-877&WithFlank&6|+|30584326|30584423</td><td>7</td><td>25</td><td>19(+1A)</td><td>GTAGAGGAGATGGCGCAGGA</td><td>37.6</td><td>3.6</td><td>1.8</td><td>2.0</td><td>4.1E-02</td><td>4.8E-02</td><td>MB231/MB468</td><td>MP</td></tr>
<tr><td>>hg38_21_+_8433172_8446622_RNA45SN1_45S_50ntFlanks@8168.8195.28|CCCAAGTCCTTCTGATCGAGGCCCAGCC</td><td>>hg38_21_+_8433172_8446622_RNA45SN1_45S_50ntFlanks</td><td>8168</td><td>8195</td><td>28</td><td>CCCAAGTCCTTCTGATCGAGGCCCAGCC</td><td>206.8</td><td>-1.5</td><td>0.7</td><td>-2.0</td><td>4.3E-02</td><td>4.8E-02</td><td>BT20/MB468</td><td>CYTO</td></tr>
<tr><td>>MI0000101|hsa-mir-99a&WithFlank&21|+|16539083|16539175@19.38.20(+1G)&[MIMAT0000097&hsa-miR-99a-5p&offsets|0|-2(+1G);m-64&21|+|16539101|16539121&offsets|0|-1(+1G)]|AACCCGTAGATCCGATCTTGG</td><td>>MI0000101|hsa-mir-99a&WithFlank&21|+|16539083|16539175</td><td>19</td><td>38</td><td>20(+1G)</td><td>AACCCGTAGATCCGATCTTGG</td><td>110.5</td><td>-2.7</td><td>1.3</td><td>-2.0</td><td>4.1E-02</td><td>4.8E-02</td><td>MB231/MB468</td><td>MP</td></tr>
<tr><td>>hg38_21_+_8433172_8446622_RNA45SN1_45S_50ntFlanks@10787.10826.40|ACAGCCTCTGGCATGTTGGAACAATGTAGGTAAGGGAAGT</td><td>>hg38_21_+_8433172_8446622_RNA45SN1_45S_50ntFlanks</td><td>10787</td><td>10826</td><td>40</td><td>ACAGCCTCTGGCATGTTGGAACAATGTAGGTAAGGGAAGT</td><td>400.5</td><td>-1.5</td><td>0.7</td><td>-2.1</td><td>3.8E-02</td><td>4.8E-02</td><td>BT20/MB468</td><td>MITO</td></tr>
<tr><td>>hg38_21_+_8433172_8446622_RNA45SN1_45S_50ntFlanks@6648.6680.33|GTACGACTCTTAGCGGTGGATCACTCGGCTCGT</td><td>>hg38_21_+_8433172_8446622_RNA45SN1_45S_50ntFlanks</td><td>6648</td><td>6680</td><td>33</td><td>GTACGACTCTTAGCGGTGGATCACTCGGCTCGT</td><td>868.6</td><td>3.1</td><td>1.3</td><td>2.5</td><td>1.4E-02</td><td>4.8E-02</td><td>BT20/MB231</td><td>MP</td></tr>
<tr><td>>trnaMT_GluTTC_MT_-_14674_14742@31.72.42,trnalookalike8_GluTTC_5_-_93905172_93905240@31.72.42|TTCATATCATTGGTCGTGGTTGTAGTCCGTGCGAGAATACCA</td><td>>trnaMT_GluTTC_MT_-_14674_14742</td><td>31</td><td>72</td><td>42</td><td>TTCATATCATTGGTCGTGGTTGTAGTCCGTGCGAGAATACCA</td><td>2385.5</td><td>-1.9</td><td>0.8</td><td>-2.4</td><td>1.7E-02</td><td>4.8E-02</td><td>BT20/MB231</td><td>MITO</td></tr>
<tr><td>>hg38_21_+_8433172_8446622_RNA45SN1_45S_50ntFlanks@12644.12669.26|TCTGTGGGATTATGACTGAACGCCTC</td><td>>hg38_21_+_8433172_8446622_RNA45SN1_45S_50ntFlanks</td><td>12644</td><td>12669</td><td>26</td><td>TCTGTGGGATTATGACTGAACGCCTC</td><td>394.3</td><td>2.1</td><td>0.9</td><td>2.4</td><td>1.7E-02</td><td>4.8E-02</td><td>BT20/MB231</td><td>MITO</td></tr>
<tr><td>>trnaMT_PheGAA_MT_+_577_647@38.62.25|AATGTTTAGACGGGCTCACATCACC</td><td>>trnaMT_PheGAA_MT_+_577_647</td><td>38</td><td>62</td><td>25</td><td>AATGTTTAGACGGGCTCACATCACC</td><td>92.5</td><td>1.9</td><td>0.9</td><td>2.0</td><td>4.1E-02</td><td>4.8E-02</td><td>MB231/MB468</td><td>MP</td></tr>
<tr><td>>hg38_21_+_8433172_8446622_RNA45SN1_45S_50ntFlanks@12709.12730.22|CCGCGGAGCCTCGGTTGGCCTC</td><td>>hg38_21_+_8433172_8446622_RNA45SN1_45S_50ntFlanks</td><td>12709</td><td>12730</td><td>22</td><td>CCGCGGAGCCTCGGTTGGCCTC</td><td>126.9</td><td>-4.0</td><td>1.8</td><td>-2.2</td><td>3.1E-02</td><td>4.9E-02</td><td>BT20/MB468</td><td>MP</td></tr>
<tr><td>>hg38_21_+_8433172_8446622_RNA45SN1_45S_50ntFlanks@6648.6676.29|GTACGACTCTTAGCGGTGGATCACTCGGC</td><td>>hg38_21_+_8433172_8446622_RNA45SN1_45S_50ntFlanks</td><td>6648</td><td>6676</td><td>29</td><td>GTACGACTCTTAGCGGTGGATCACTCGGC</td><td>2274.6</td><td>3.3</td><td>1.4</td><td>2.5</td><td>1.4E-02</td><td>4.9E-02</td><td>BT20/MB231</td><td>MP</td></tr>
<tr><td>>hg38_21_+_8433172_8446622_RNA45SN1_45S_50ntFlanks@12475.12496.22|CCCACTAATAGGGAACGTGAGC</td><td>>hg38_21_+_8433172_8446622_RNA45SN1_45S_50ntFlanks</td><td>12475</td><td>12496</td><td>22</td><td>CCCACTAATAGGGAACGTGAGC</td><td>892.0</td><td>-2.0</td><td>0.9</td><td>-2.3</td><td>1.9E-02</td><td>4.9E-02</td><td>BT20/MB231</td><td>CYTO</td></tr>
<tr><td>>hg38_21_+_8433172_8446622_RNA45SN1_45S_50ntFlanks@6652.6679.28|GACTCTTAGCGGTGGATCACTCGGCTCG</td><td>>hg38_21_+_8433172_8446622_RNA45SN1_45S_50ntFlanks</td><td>6652</td><td>6679</td><td>28</td><td>GACTCTTAGCGGTGGATCACTCGGCTCG</td><td>3778.8</td><td>2.8</td><td>1.2</td><td>2.3</td><td>1.9E-02</td><td>4.9E-02</td><td>BT20/MB231</td><td>CYTO</td></tr>
<tr><td>>hg38_21_+_8433172_8446622_RNA45SN1_45S_50ntFlanks@4734.4758.25|AACGAAAGTCGGAGGTTCGAAGACG</td><td>>hg38_21_+_8433172_8446622_RNA45SN1_45S_50ntFlanks</td><td>4734</td><td>4758</td><td>25</td><td>AACGAAAGTCGGAGGTTCGAAGACG</td><td>4943.4</td><td>-1.5</td><td>0.7</td><td>-2.0</td><td>4.1E-02</td><td>4.9E-02</td><td>MB231/MB468</td><td>MP</td></tr>
<tr><td>>trna116_GluCTC_1_-_145399233_145399304@37.74.38,trna134_GluTTC_1_-_16861774_16861845@37.74.38,trna71_GluCTC_1_-_161439189_161439260@37.74.38,trna74_GluCTC_1_-_161431809_161431880@37.74.38,trna77_GluCTC_1_-_161424398_161424469@37.74.38,trna77_GluCTC_6_+_28949976_28950047@37.74.38,trna80_GluCTC_1_-_161417018_161417089@37.74.38,trna84_GluTTC_1_-_161391883_161391954@37.74.38,trna87_GluCTC_6_-_126101393_126101464@37.74.38|ACCGCCGCGGCCCGGGTTCGATTCCCGGTCAGGGAACC</td><td>>trna116_GluCTC_1_-_145399233_145399304</td><td>37</td><td>74</td><td>38</td><td>ACCGCCGCGGCCCGGGTTCGATTCCCGGTCAGGGAACC</td><td>44.0</td><td>5.4</td><td>2.6</td><td>2.0</td><td>4.1E-02</td><td>4.9E-02</td><td>MB231/MB468</td><td>MP</td></tr>
<tr><td>>MI0000066|hsa-let-7e&WithFlank&19|+|51692780|51692870@59.80.22&[MIMAT0004485&hsa-let-7e-3p&offsets|0|0;m-260&19|+|51692838|51692859&offsets|0|0]|CTATACGGCCTCCTAGCTTTCC</td><td>>MI0000066|hsa-let-7e&WithFlank&19|+|51692780|51692870</td><td>59</td><td>80</td><td>22</td><td>CTATACGGCCTCCTAGCTTTCC</td><td>215.9</td><td>1.8</td><td>0.8</td><td>2.3</td><td>1.9E-02</td><td>4.9E-02</td><td>BT20/MB231</td><td>CYTO</td></tr>
<tr><td>>hg38_21_+_8433172_8446622_RNA45SN1_45S_50ntFlanks@11641.11685.45|CGGCGGGTGTTGACGCGATGTGATTTCTGCCCAGTGCTCTGAATG</td><td>>hg38_21_+_8433172_8446622_RNA45SN1_45S_50ntFlanks</td><td>11641</td><td>11685</td><td>45</td><td>CGGCGGGTGTTGACGCGATGTGATTTCTGCCCAGTGCTCTGAATG</td><td>609.7</td><td>-3.7</td><td>1.7</td><td>-2.2</td><td>3.1E-02</td><td>4.9E-02</td><td>BT20/MB468</td><td>MP</td></tr>
<tr><td>>MI0000264|hsa-mir-7-2&WithFlank&15|+|88611819|88611940@38.58.21;MI0000265|hsa-mir-7-3&WithFlank&19|+|4770664|4770785@37.57.21;MI0000263|hsa-mir-7-1&WithFlank&9|-|83969742|83969863@30.50.21&[MIMAT0000252&hsa-miR-7-5p&offsets|0|-3;m-400&15|+|88611856|88611879&offsets|0|-3];[MIMAT0000252_1&hsa-miR-7-5p&offsets|0|-3;m-402&19|+|4770700|4770723&offsets|0|-3];[MIMAT0000252_2&hsa-miR-7-5p&offsets|0|-3;m-401&9|-|83969811|83969834&offsets|0|-3]|TGGAAGACTAGTGATTTTGTT</td><td>>MI0000264|hsa-mir-7-2&WithFlank&15|+|88611819|88611940</td><td>38</td><td>58</td><td>21</td><td>TGGAAGACTAGTGATTTTGTT</td><td>1047.5</td><td>2.0</td><td>0.9</td><td>2.3</td><td>1.9E-02</td><td>4.9E-02</td><td>BT20/MB231</td><td>CYTO</td></tr>
<tr><td>>hg38_21_+_8433172_8446622_RNA45SN1_45S_50ntFlanks@10866.10894.29|CTAAGGGCTGGGTCGGTCGGGCTGGGGCG</td><td>>hg38_21_+_8433172_8446622_RNA45SN1_45S_50ntFlanks</td><td>10866</td><td>10894</td><td>29</td><td>CTAAGGGCTGGGTCGGTCGGGCTGGGGCG</td><td>365.5</td><td>-3.4</td><td>1.7</td><td>-2.0</td><td>4.3E-02</td><td>4.9E-02</td><td>BT20/MB468</td><td>CYTO</td></tr>
<tr><td>>hg38_MT_+_1621_3279_RNR2_16S_50ntFlanks@1370.1404.35|ATGTTGGATCAGGACATCCCGATGGTGCAGCCGCT</td><td>>hg38_MT_+_1621_3279_RNR2_16S_50ntFlanks</td><td>1370</td><td>1404</td><td>35</td><td>ATGTTGGATCAGGACATCCCGATGGTGCAGCCGCT</td><td>163.3</td><td>3.4</td><td>1.7</td><td>2.0</td><td>4.1E-02</td><td>4.9E-02</td><td>MB231/MB468</td><td>MP</td></tr>
<tr><td>>trnaMT_PheGAA_MT_+_577_647@1.20.20|GTTTATGTAGCTTACCTCCT</td><td>>trnaMT_PheGAA_MT_+_577_647</td><td>1</td><td>20</td><td>20</td><td>GTTTATGTAGCTTACCTCCT</td><td>424.3</td><td>2.0</td><td>1.0</td><td>2.0</td><td>4.5E-02</td><td>4.9E-02</td><td>MB231/MB468</td><td>MITO</td></tr>
<tr><td>>MI0000107|hsa-mir-29b-2&WithFlank&1|-|207802437|207802529@58.80.23;MI0000105|hsa-mir-29b-1&WithFlank&7|-|130877453|130877545@57.79.23&[MIMAT0000100&hsa-miR-29b-3p&offsets|0|0;m-81&1|-|207802450|207802472&offsets|0|0];[MIMAT0000100_1&hsa-miR-29b-3p&offsets|0|0;m-82&7|-|130877467|130877489&offsets|0|0]|TAGCACCATTTGAAATCAGTGTT</td><td>>MI0000107|hsa-mir-29b-2&WithFlank&1|-|207802437|207802529</td><td>58</td><td>80</td><td>23</td><td>TAGCACCATTTGAAATCAGTGTT</td><td>1229.3</td><td>2.5</td><td>1.2</td><td>2.0</td><td>4.1E-02</td><td>4.9E-02</td><td>MB231/MB468</td><td>MP</td></tr>
<tr><td>>hg38_21_+_8433172_8446622_RNA45SN1_45S_50ntFlanks@6779.6807.29|CCGGGGCTACGCCTGTCTGAGCGTCGCTT</td><td>>hg38_21_+_8433172_8446622_RNA45SN1_45S_50ntFlanks</td><td>6779</td><td>6807</td><td>29</td><td>CCGGGGCTACGCCTGTCTGAGCGTCGCTT</td><td>115.9</td><td>-1.5</td><td>0.7</td><td>-2.0</td><td>4.1E-02</td><td>4.9E-02</td><td>MB231/MB468</td><td>MP</td></tr>
<tr><td>>hg38_21_+_8433172_8446622_RNA45SN1_45S_50ntFlanks@10808.10833.26|CAATGTAGGTAAGGGAAGTCGGCAAG</td><td>>hg38_21_+_8433172_8446622_RNA45SN1_45S_50ntFlanks</td><td>10808</td><td>10833</td><td>26</td><td>CAATGTAGGTAAGGGAAGTCGGCAAG</td><td>72.6</td><td>-2.8</td><td>1.3</td><td>-2.2</td><td>3.1E-02</td><td>4.9E-02</td><td>BT20/MB468</td><td>MP</td></tr>
<tr><td>>hg38_21_+_8433172_8446622_RNA45SN1_45S_50ntFlanks@11572.11607.36|CCGACTTAGAACTGGTGCGGACCAGGGGAATCCGAC</td><td>>hg38_21_+_8433172_8446622_RNA45SN1_45S_50ntFlanks</td><td>11572</td><td>11607</td><td>36</td><td>CCGACTTAGAACTGGTGCGGACCAGGGGAATCCGAC</td><td>281.1</td><td>-1.4</td><td>0.7</td><td>-2.0</td><td>4.3E-02</td><td>4.9E-02</td><td>BT20/MB468</td><td>CYTO</td></tr>
<tr><td>>hg38_21_+_8433172_8446622_RNA45SN1_45S_50ntFlanks@10402.10443.42|AACATGGGTCAGTCGGTCCTGAGAGATGGGCGAGCGCCGTTC</td><td>>hg38_21_+_8433172_8446622_RNA45SN1_45S_50ntFlanks</td><td>10402</td><td>10443</td><td>42</td><td>AACATGGGTCAGTCGGTCCTGAGAGATGGGCGAGCGCCGTTC</td><td>126.2</td><td>-4.0</td><td>1.9</td><td>-2.1</td><td>3.8E-02</td><td>4.9E-02</td><td>BT20/MB468</td><td>MITO</td></tr>
<tr><td>>hg38_MT_+_1621_3279_RNR2_16S_50ntFlanks@1367.1402.36|TCGATGTTGGATCAGGACATCCCGATGGTGCAGCCG</td><td>>hg38_MT_+_1621_3279_RNR2_16S_50ntFlanks</td><td>1367</td><td>1402</td><td>36</td><td>TCGATGTTGGATCAGGACATCCCGATGGTGCAGCCG</td><td>43.5</td><td>4.2</td><td>2.0</td><td>2.0</td><td>4.1E-02</td><td>4.9E-02</td><td>MB231/MB468</td><td>MP</td></tr>
<tr><td>>hg38_21_+_8433172_8446622_RNA45SN1_45S_50ntFlanks@12415.12455.41|CTTCGATGTCGGCTCTTCCTATCATTGTGAAGCAGAATTCA</td><td>>hg38_21_+_8433172_8446622_RNA45SN1_45S_50ntFlanks</td><td>12415</td><td>12455</td><td>41</td><td>CTTCGATGTCGGCTCTTCCTATCATTGTGAAGCAGAATTCA</td><td>577.7</td><td>-2.0</td><td>1.0</td><td>-2.0</td><td>4.3E-02</td><td>4.9E-02</td><td>MB231/MB468</td><td>CYTO</td></tr>
<tr><td>>hg38_21_+_8433172_8446622_RNA45SN1_45S_50ntFlanks@7976.8008.33|GCGACCTCAGATCAGACGTGGCGACCCGCTGAA</td><td>>hg38_21_+_8433172_8446622_RNA45SN1_45S_50ntFlanks</td><td>7976</td><td>8008</td><td>33</td><td>GCGACCTCAGATCAGACGTGGCGACCCGCTGAA</td><td>334.2</td><td>-1.1</td><td>0.6</td><td>-2.0</td><td>4.4E-02</td><td>4.9E-02</td><td>BT20/MB468</td><td>NUC</td></tr>
<tr><td>>hg38_MT_+_1621_3279_RNR2_16S_50ntFlanks@1373.1414.42|TTGGATCAGGACATCCCGATGGTGCAGCCGCTATTAAAGGTT</td><td>>hg38_MT_+_1621_3279_RNR2_16S_50ntFlanks</td><td>1373</td><td>1414</td><td>42</td><td>TTGGATCAGGACATCCCGATGGTGCAGCCGCTATTAAAGGTT</td><td>101.0</td><td>4.2</td><td>1.9</td><td>2.2</td><td>3.1E-02</td><td>4.9E-02</td><td>BT20/MB468</td><td>MP</td></tr>
<tr><td>>hg38_21_+_8433172_8446622_RNA45SN1_45S_50ntFlanks@6660.6677.18|GCGGTGGATCACTCGGCT</td><td>>hg38_21_+_8433172_8446622_RNA45SN1_45S_50ntFlanks</td><td>6660</td><td>6677</td><td>18</td><td>GCGGTGGATCACTCGGCT</td><td>76.4</td><td>2.1</td><td>1.0</td><td>2.2</td><td>3.1E-02</td><td>4.9E-02</td><td>BT20/MB468</td><td>MP</td></tr>
<tr><td>>MI0000077|hsa-mir-21&WithFlank&17|+|59841260|59841343@53.74.22&[MIMAT0004494&hsa-miR-21-3p&offsets|+1|+2;m-52&17|+|59841311|59841332&offsets|+1|+1]|AACACCAGTCGATGGGCTGTCT</td><td>>MI0000077|hsa-mir-21&WithFlank&17|+|59841260|59841343</td><td>53</td><td>74</td><td>22</td><td>AACACCAGTCGATGGGCTGTCT</td><td>258.7</td><td>5.0</td><td>2.3</td><td>2.2</td><td>3.1E-02</td><td>4.9E-02</td><td>BT20/MB468</td><td>MP</td></tr>
<tr><td>>hg38_21_+_8433172_8446622_RNA45SN1_45S_50ntFlanks@12419.12440.22|GATGTCGGCTCTTCCTATCATT</td><td>>hg38_21_+_8433172_8446622_RNA45SN1_45S_50ntFlanks</td><td>12419</td><td>12440</td><td>22</td><td>GATGTCGGCTCTTCCTATCATT</td><td>2078.8</td><td>-2.0</td><td>0.9</td><td>-2.3</td><td>1.9E-02</td><td>4.9E-02</td><td>BT20/MB231</td><td>CYTO</td></tr>
<tr><td>>hg38_21_+_8433172_8446622_RNA45SN1_45S_50ntFlanks@3840.3858.19|CTCCTACTTGGATAACTGT</td><td>>hg38_21_+_8433172_8446622_RNA45SN1_45S_50ntFlanks</td><td>3840</td><td>3858</td><td>19</td><td>CTCCTACTTGGATAACTGT</td><td>235.4</td><td>1.7</td><td>0.7</td><td>2.3</td><td>1.9E-02</td><td>4.9E-02</td><td>BT20/MB231</td><td>CYTO</td></tr>
<tr><td>>hg38_21_+_8433172_8446622_RNA45SN1_45S_50ntFlanks@8686.8706.21|CCGGCTCCGGGACGGCTGGGA</td><td>>hg38_21_+_8433172_8446622_RNA45SN1_45S_50ntFlanks</td><td>8686</td><td>8706</td><td>21</td><td>CCGGCTCCGGGACGGCTGGGA</td><td>1460.6</td><td>-2.0</td><td>0.9</td><td>-2.3</td><td>1.9E-02</td><td>4.9E-02</td><td>BT20/MB231</td><td>CYTO</td></tr>
<tr><td>>MI0000734|hsa-mir-106b&WithFlank&7|-|100093987|100094080@58.80.23&[MIMAT0004672&hsa-miR-106b-3p&offsets|0|+1;m-87&7|-|100094003|100094023&offsets|0|+2]|CCGCACTGTGGGTACTTGCTGCT</td><td>>MI0000734|hsa-mir-106b&WithFlank&7|-|100093987|100094080</td><td>58</td><td>80</td><td>23</td><td>CCGCACTGTGGGTACTTGCTGCT</td><td>94.2</td><td>2.9</td><td>1.4</td><td>2.0</td><td>4.1E-02</td><td>4.9E-02</td><td>MB231/MB468</td><td>MP</td></tr>
</table>

<table>
<tr><td>>MI0000286|hsa-mir-210&WithFlank&11|-|568083|568204@72.93.22&[MIMAT0000267&hsa-miR-210-3p&offsets|0|0;m-102&11|-|568112|568133&offsets|0|0]|CTGTGCGTGTGACAGCGGCTGA</td><td>>MI0000286|hsa-mir-210&WithFlank&11|-|568083|568204</td><td>72</td><td>93</td><td>22</td><td>CTGTGCGTGTGACAGCGGCTGA</td><td>910.6</td><td>3.1</td><td>1.5</td><td>2.0</td><td>4.1E-02</td><td>4.9E-02</td><td>MB231/MB468</td><td>MP</td></tr>
<tr><td>>hg38_21_+_8433172_8446622_RNA45SN1_45S_50ntFlanks@7973.7995.23|GACGCGACCTCAGATCAGACGTG</td><td>>hg38_21_+_8433172_8446622_RNA45SN1_45S_50ntFlanks</td><td>7973</td><td>7995</td><td>23</td><td>GACGCGACCTCAGATCAGACGTG</td><td>331.7</td><td>-1.5</td><td>0.8</td><td>-2.0</td><td>4.1E-02</td><td>4.9E-02</td><td>MB231/MB468</td><td>MP</td></tr>
<tr><td>>trnaMT_PheGAA_MT_+_577_647@1.21.21|GTTTATGTAGCTTACCTCCTC</td><td>>trnaMT_PheGAA_MT_+_577_647</td><td>1</td><td>21</td><td>21</td><td>GTTTATGTAGCTTACCTCCTC</td><td>256.6</td><td>1.8</td><td>0.9</td><td>2.0</td><td>4.5E-02</td><td>4.9E-02</td><td>MB231/MB468</td><td>MITO</td></tr>
<tr><td>>hg38_21_+_8433172_8446622_RNA45SN1_45S_50ntFlanks@8195.8223.29|CCGTGGACGGTGTGAGGCCGGTAGCGGCC</td><td>>hg38_21_+_8433172_8446622_RNA45SN1_45S_50ntFlanks</td><td>8195</td><td>8223</td><td>29</td><td>CCGTGGACGGTGTGAGGCCGGTAGCGGCC</td><td>71.6</td><td>-3.7</td><td>1.7</td><td>-2.2</td><td>3.1E-02</td><td>4.9E-02</td><td>BT20/MB468</td><td>MP</td></tr>
<tr><td>>hg38_21_+_8433172_8446622_RNA45SN1_45S_50ntFlanks@9573.9602.30|AAATCGGTCGTCCGACCTGGGTATAGGGGC</td><td>>hg38_21_+_8433172_8446622_RNA45SN1_45S_50ntFlanks</td><td>9573</td><td>9602</td><td>30</td><td>AAATCGGTCGTCCGACCTGGGTATAGGGGC</td><td>304.3</td><td>-1.9</td><td>0.9</td><td>-2.0</td><td>4.1E-02</td><td>4.9E-02</td><td>MB231/MB468</td><td>MP</td></tr>
<tr><td>>am_trnaMT_ValTAC_MT_+_1602_1670@49.67.19|AACTTAACTTGACCGCTCT</td><td>>am_trnaMT_ValTAC_MT_+_1602_1670</td><td>49</td><td>67</td><td>19</td><td>AACTTAACTTGACCGCTCT</td><td>1580.6</td><td>2.2</td><td>0.9</td><td>2.4</td><td>1.5E-02</td><td>4.9E-02</td><td>BT20/MB231</td><td>NUC</td></tr>
<tr><td>>hg38_21_+_8433172_8446622_RNA45SN1_45S_50ntFlanks@6659.6683.25|AGCGGTGGATCACTCGGCTCGTGCG</td><td>>hg38_21_+_8433172_8446622_RNA45SN1_45S_50ntFlanks</td><td>6659</td><td>6683</td><td>25</td><td>AGCGGTGGATCACTCGGCTCGTGCG</td><td>281.0</td><td>-2.0</td><td>0.8</td><td>-2.4</td><td>1.5E-02</td><td>4.9E-02</td><td>BT20/MB231</td><td>NUC</td></tr>
<tr><td>>hg38_21_+_8433172_8446622_RNA45SN1_45S_50ntFlanks@9546.9573.28|TGGAGGTCCGTAGCGGTCCTGACGTGCA</td><td>>hg38_21_+_8433172_8446622_RNA45SN1_45S_50ntFlanks</td><td>9546</td><td>9573</td><td>28</td><td>TGGAGGTCCGTAGCGGTCCTGACGTGCA</td><td>318.7</td><td>-2.5</td><td>1.0</td><td>-2.4</td><td>1.5E-02</td><td>4.9E-02</td><td>BT20/MB231</td><td>NUC</td></tr>
<tr><td>>hg38_21_+_8433172_8446622_RNA45SN1_45S_50ntFlanks@6646.6690.45|TCGTACGACTCTTAGCGGTGGATCACTCGGCTCGTGCGTCGATGA</td><td>>hg38_21_+_8433172_8446622_RNA45SN1_45S_50ntFlanks</td><td>6646</td><td>6690</td><td>45</td><td>TCGTACGACTCTTAGCGGTGGATCACTCGGCTCGTGCGTCGATGA</td><td>606.2</td><td>2.2</td><td>0.9</td><td>2.4</td><td>1.5E-02</td><td>4.9E-02</td><td>BT20/MB231</td><td>NUC</td></tr>
<tr><td>>hg38_21_+_8433172_8446622_RNA45SN1_45S_50ntFlanks@11541.11584.44|GCGGCGCCTCGCCTCGGCCGGCGCCTAGCAGCCGACTTAGAACT</td><td>>hg38_21_+_8433172_8446622_RNA45SN1_45S_50ntFlanks</td><td>11541</td><td>11584</td><td>44</td><td>GCGGCGCCTCGCCTCGGCCGGCGCCTAGCAGCCGACTTAGAACT</td><td>243.8</td><td>-2.1</td><td>0.9</td><td>-2.4</td><td>1.5E-02</td><td>4.9E-02</td><td>BT20/MB231</td><td>NUC</td></tr>
<tr><td>>hg38_21_+_8433172_8446622_RNA45SN1_45S_50ntFlanks@12420.12459.40|ATGTCGGCTCTTCCTATCATTGTGAAGCAGAATTCACCAA</td><td>>hg38_21_+_8433172_8446622_RNA45SN1_45S_50ntFlanks</td><td>12420</td><td>12459</td><td>40</td><td>ATGTCGGCTCTTCCTATCATTGTGAAGCAGAATTCACCAA</td><td>214.7</td><td>-4.4</td><td>1.8</td><td>-2.4</td><td>1.5E-02</td><td>4.9E-02</td><td>BT20/MB231</td><td>NUC</td></tr>
<tr><td>>hg38_MT_+_1621_3279_RNR2_16S_50ntFlanks@459.492.34|CTTGTAAATTTAACTGTTAGTCCAAAGAGGAACA</td><td>>hg38_MT_+_1621_3279_RNR2_16S_50ntFlanks</td><td>459</td><td>492</td><td>34</td><td>CTTGTAAATTTAACTGTTAGTCCAAAGAGGAACA</td><td>250.5</td><td>-2.2</td><td>0.9</td><td>-2.4</td><td>1.5E-02</td><td>4.9E-02</td><td>BT20/MB231</td><td>NUC</td></tr>
<tr><td>>MI0000088|hsa-mir-30a&WithFlank&6|-|71403545|71403627@53.73.21&[MIMAT0000088&hsa-miR-30a-3p&offsets|0|-1;m-38&6|-|71403554|71403575&offsets|0|-1]|CTTTCAGTCGGATGTTTGCAG</td><td>>MI0000088|hsa-mir-30a&WithFlank&6|-|71403545|71403627</td><td>53</td><td>73</td><td>21</td><td>CTTTCAGTCGGATGTTTGCAG</td><td>1467.7</td><td>-1.4</td><td>0.6</td><td>-2.4</td><td>1.5E-02</td><td>4.9E-02</td><td>BT20/MB231</td><td>NUC</td></tr>
<tr><td>>hg38_21_+_8433172_8446622_RNA45SN1_45S_50ntFlanks@10866.10889.24|CTAAGGGCTGGGTCGGTCGGGCTG</td><td>>hg38_21_+_8433172_8446622_RNA45SN1_45S_50ntFlanks</td><td>10866</td><td>10889</td><td>24</td><td>CTAAGGGCTGGGTCGGTCGGGCTG</td><td>456.0</td><td>-3.5</td><td>1.7</td><td>-2.0</td><td>4.3E-02</td><td>4.9E-02</td><td>BT20/MB468</td><td>CYTO</td></tr>
<tr><td>>hg38_21_+_8433172_8446622_RNA45SN1_45S_50ntFlanks@4547.4573.27|CTGGATACCGCAGCTAGGAATAATGGA</td><td>>hg38_21_+_8433172_8446622_RNA45SN1_45S_50ntFlanks</td><td>4547</td><td>4573</td><td>27</td><td>CTGGATACCGCAGCTAGGAATAATGGA</td><td>128.2</td><td>-3.9</td><td>1.8</td><td>-2.2</td><td>3.1E-02</td><td>4.9E-02</td><td>BT20/MB468</td><td>MP</td></tr>
<tr><td>>hg38_21_+_8433172_8446622_RNA45SN1_45S_50ntFlanks@4380.4407.28|CGTAGTTGGATCTTGGGAGCGGGCGGGC</td><td>>hg38_21_+_8433172_8446622_RNA45SN1_45S_50ntFlanks</td><td>4380</td><td>4407</td><td>28</td><td>CGTAGTTGGATCTTGGGAGCGGGCGGGC</td><td>1617.7</td><td>4.9</td><td>2.3</td><td>2.2</td><td>3.1E-02</td><td>4.9E-02</td><td>BT20/MB468</td><td>MP</td></tr>
<tr><td>>hg38_MT_+_1621_3279_RNR2_16S_50ntFlanks@1492.1518.27|AATTCCTCCCTGTACGAAAGGACAAGA</td><td>>hg38_MT_+_1621_3279_RNR2_16S_50ntFlanks</td><td>1492</td><td>1518</td><td>27</td><td>AATTCCTCCCTGTACGAAAGGACAAGA</td><td>45.8</td><td>3.6</td><td>1.7</td><td>2.0</td><td>4.1E-02</td><td>4.9E-02</td><td>MB231/MB468</td><td>MP</td></tr>
<tr><td>>MI0003205|hsa-mir-532&WithFlank&X|+|50003142|50003244@26.47.22(+1A)&[MIMAT0002888&hsa-miR-532-5p&offsets|0|0(+1A);m-65&X|+|50003167|50003188&offsets|0|0(+1A)]|CATGCCTTGAGTGTAGGACCGTA</td><td>>MI0003205|hsa-mir-532&WithFlank&X|+|50003142|50003244</td><td>26</td><td>47</td><td>22(+1A)</td><td>CATGCCTTGAGTGTAGGACCGTA</td><td>184.9</td><td>1.9</td><td>0.8</td><td>2.3</td><td>1.9E-02</td><td>4.9E-02</td><td>BT20/MB231</td><td>CYTO</td></tr>
<tr><td>>hg38_21_+_8433172_8446622_RNA45SN1_45S_50ntFlanks@11578.11601.24|TAGAACTGGTGCGGACCAGGGGAA</td><td>>hg38_21_+_8433172_8446622_RNA45SN1_45S_50ntFlanks</td><td>11578</td><td>11601</td><td>24</td><td>TAGAACTGGTGCGGACCAGGGGAA</td><td>97.4</td><td>4.9</td><td>2.0</td><td>2.5</td><td>1.4E-02</td><td>4.9E-02</td><td>BT20/MB231</td><td>MP</td></tr>
<tr><td>>hg38_21_+_8433172_8446622_RNA45SN1_45S_50ntFlanks@6653.6683.31|ACTCTTAGCGGTGGATCACTCGGCTCGTGCG</td><td>>hg38_21_+_8433172_8446622_RNA45SN1_45S_50ntFlanks</td><td>6653</td><td>6683</td><td>31</td><td>ACTCTTAGCGGTGGATCACTCGGCTCGTGCG</td><td>3655.5</td><td>-1.8</td><td>0.7</td><td>-2.4</td><td>1.5E-02</td><td>4.9E-02</td><td>BT20/MB231</td><td>NUC</td></tr>
<tr><td>>hg38_21_+_8433172_8446622_RNA45SN1_45S_50ntFlanks@9533.9562.30|GAGGAAACTCTGGTGGAGGTCCGTAGCGGT</td><td>>hg38_21_+_8433172_8446622_RNA45SN1_45S_50ntFlanks</td><td>9533</td><td>9562</td><td>30</td><td>GAGGAAACTCTGGTGGAGGTCCGTAGCGGT</td><td>431.3</td><td>-1.5</td><td>0.7</td><td>-2.0</td><td>4.3E-02</td><td>4.9E-02</td><td>MB231/MB468</td><td>CYTO</td></tr>
<tr><td>>hg38_21_+_8433172_8446622_RNA45SN1_45S_50ntFlanks@11587.11612.26|TGCGGACCAGGGGAATCCGACTGTTT</td><td>>hg38_21_+_8433172_8446622_RNA45SN1_45S_50ntFlanks</td><td>11587</td><td>11612</td><td>26</td><td>TGCGGACCAGGGGAATCCGACTGTTT</td><td>216.6</td><td>-1.4</td><td>0.7</td><td>-2.0</td><td>4.5E-02</td><td>4.9E-02</td><td>MB231/MB468</td><td>NUC</td></tr>
<tr><td>>hg38_21_+_8433172_8446622_RNA45SN1_45S_50ntFlanks@11190.11216.27|CCCGGGGGGCCGGCGGCGGCGGCGACT</td><td>>hg38_21_+_8433172_8446622_RNA45SN1_45S_50ntFlanks</td><td>11190</td><td>11216</td><td>27</td><td>CCCGGGGGGCCGGCGGCGGCGGCGACT</td><td>76.3</td><td>-3.2</td><td>1.5</td><td>-2.2</td><td>3.1E-02</td><td>4.9E-02</td><td>BT20/MB468</td><td>MP</td></tr>
<tr><td>>trna10_LysCTT_16_+_3241501_3241573@11.35.25,trna13_LysCTT_14_-_58706613_58706685@11.35.25,trna2_LysCTT_15_+_79152904_79152976@11.35.25,trna4_LysCTT_18_-_43669271_43669343@11.35.25|CTCAGTCGGTAGAGCATGGGACTCT</td><td>>trna10_LysCTT_16_+_3241501_3241573</td><td>11</td><td>35</td><td>25</td><td>CTCAGTCGGTAGAGCATGGGACTCT</td><td>80.3</td><td>-3.9</td><td>1.6</td><td>-2.5</td><td>1.4E-02</td><td>4.9E-02</td><td>BT20/MB231</td><td>MP</td></tr>
<tr><td>>hg38_21_+_8433172_8446622_RNA45SN1_45S_50ntFlanks@475.517.43|CTTCGTGATCGATGTGGTGACGTCGTGCTCTCCCGGGCCGGGT</td><td>>hg38_21_+_8433172_8446622_RNA45SN1_45S_50ntFlanks</td><td>475</td><td>517</td><td>43</td><td>CTTCGTGATCGATGTGGTGACGTCGTGCTCTCCCGGGCCGGGT</td><td>1177.9</td><td>-2.3</td><td>1.0</td><td>-2.3</td><td>1.9E-02</td><td>4.9E-02</td><td>BT20/MB231</td><td>CYTO</td></tr>
<tr><td>>hg38_21_+_8433172_8446622_RNA45SN1_45S_50ntFlanks@11583.11609.27|CTGGTGCGGACCAGGGGAATCCGACTG</td><td>>hg38_21_+_8433172_8446622_RNA45SN1_45S_50ntFlanks</td><td>11583</td><td>11609</td><td>27</td><td>CTGGTGCGGACCAGGGGAATCCGACTG</td><td>715.2</td><td>-1.5</td><td>0.7</td><td>-2.0</td><td>4.3E-02</td><td>4.9E-02</td><td>BT20/MB468</td><td>CYTO</td></tr>
<tr><td>>hg38_21_+_8433172_8446622_RNA45SN1_45S_50ntFlanks@11572.11599.28|CCGACTTAGAACTGGTGCGGACCAGGGG</td><td>>hg38_21_+_8433172_8446622_RNA45SN1_45S_50ntFlanks</td><td>11572</td><td>11599</td><td>28</td><td>CCGACTTAGAACTGGTGCGGACCAGGGG</td><td>365.6</td><td>1.1</td><td>0.6</td><td>2.1</td><td>3.9E-02</td><td>4.9E-02</td><td>BT20/MB468</td><td>MITO</td></tr>
<tr><td>>am_MI0003639|hsa-mir-625&WithFlank&14|+|65471096|65471192@58.78.21(+1U)&[MIMAT0004808&hsa-miR-625-3p&offsets|0|-1(+1U);m-137&14|+|65471153|65471174&offsets|0|-1(+1U)]|GACTATAGAACTTTCCCCCTCT</td><td>>am_MI0003639|hsa-mir-625&WithFlank&14|+|65471096|65471192</td><td>58</td><td>78</td><td>21(+1U)</td><td>GACTATAGAACTTTCCCCCTCT</td><td>384.4</td><td>-2.4</td><td>1.0</td><td>-2.5</td><td>1.4E-02</td><td>4.9E-02</td><td>BT20/MB231</td><td>MP</td></tr>
<tr><td>>am_trna119_LysCTT_1_-_145395522_145395594@8.35.28,trna11_LysCTT_5_-_180648979_180649051@8.35.28,trna128_LysCTT_1_-_55423542_55423614@8.35.28,trna13_LysCTT_6_+_26556774_26556846@8.35.28,trna6_LysCTT_19_-_52425396_52425468@8.35.28,trna7_LysCTT_16_+_3225692_3225764@8.35.28,trna9_LysCTT_5_+_180634755_180634827@8.35.28|TAGCTCAGTCGGTAGAGCATGAGACTCT</td><td>>am_trna119_LysCTT_1_-_145395522_145395594</td><td>8</td><td>35</td><td>28</td><td>TAGCTCAGTCGGTAGAGCATGAGACTCT</td><td>69.0</td><td>-4.8</td><td>2.0</td><td>-2.5</td><td>1.4E-02</td><td>4.9E-02</td><td>BT20/MB231</td><td>MP</td></tr>
<tr><td>>hg38_21_+_8433172_8446622_RNA45SN1_45S_50ntFlanks@3705.3722.18|TACCTGGTTGATCCTGCC</td><td>>hg38_21_+_8433172_8446622_RNA45SN1_45S_50ntFlanks</td><td>3705</td><td>3722</td><td>18</td><td>TACCTGGTTGATCCTGCC</td><td>4465.5</td><td>3.3</td><td>1.4</td><td>2.5</td><td>1.4E-02</td><td>4.9E-02</td><td>BT20/MB231</td><td>MP</td></tr>
<tr><td>>trnaMT_GluTTC_MT_-_14674_14742@34.65.32,trnalookalike8_GluTTC_5_-_93905172_93905240@34.65.32|ATATCATTGGTCGTGGTTGTAGTCCGTGCGAG</td><td>>trnaMT_GluTTC_MT_-_14674_14742</td><td>34</td><td>65</td><td>32</td><td>ATATCATTGGTCGTGGTTGTAGTCCGTGCGAG</td><td>90.8</td><td>2.0</td><td>1.0</td><td>2.0</td><td>4.1E-02</td><td>4.9E-02</td><td>MB231/MB468</td><td>MP</td></tr>
<tr><td>>hg38_21_+_8433172_8446622_RNA45SN1_45S_50ntFlanks@7980.8005.26|CCTCAGATCAGACGTGGCGACCCGCT</td><td>>hg38_21_+_8433172_8446622_RNA45SN1_45S_50ntFlanks</td><td>7980</td><td>8005</td><td>26</td><td>CCTCAGATCAGACGTGGCGACCCGCT</td><td>146.3</td><td>-2.3</td><td>1.1</td><td>-2.0</td><td>4.3E-02</td><td>4.9E-02</td><td>BT20/MB468</td><td>CYTO</td></tr>
<tr><td>>hg38_21_+_8433172_8446622_RNA45SN1_45S_50ntFlanks@11668.11691.24|TGCCCAGTGCTCTGAATGTCAAAG</td><td>>hg38_21_+_8433172_8446622_RNA45SN1_45S_50ntFlanks</td><td>11668</td><td>11691</td><td>24</td><td>TGCCCAGTGCTCTGAATGTCAAAG</td><td>687.5</td><td>-2.0</td><td>0.8</td><td>-2.3</td><td>1.9E-02</td><td>4.9E-02</td><td>BT20/MB231</td><td>CYTO</td></tr>
<tr><td>>trna10_AspGTC_12_-_125424193_125424264@1.36.36,trna12_AspGTC_12_-_125411891_125411962@1.36.36,trna38_AspGTC_17_-_8125556_8125627@1.36.36,trna45_AspGTC_6_+_27447453_27447524@1.36.36,trna48_AspGTC_6_+_27471523_27471594@1.36.36,trna4_AspGTC_12_+_96429799_96429870@1.36.36,trna69_AspGTC_1_-_161440205_161440276@1.36.36,trna72_AspGTC_1_-_161432824_161432895@1.36.36,trna75_AspGTC_1_-_161425414_161425485@1.36.36,trna78_AspGTC_1_-_161418033_161418104@1.36.36,trna81_AspGTC_1_-_161410615_161410686@1.36.36|TCCTCGTTAGTATAGTGGTGAGTATCCCCGCCTGTC</td><td>>trna10_AspGTC_12_-_125424193_125424264</td><td>1</td><td>36</td><td>36</td><td>TCCTCGTTAGTATAGTGGTGAGTATCCCCGCCTGTC</td><td>924.4</td><td>-4.9</td><td>2.1</td><td>-2.3</td><td>1.9E-02</td><td>4.9E-02</td><td>BT20/MB231</td><td>CYTO</td></tr>
<tr><td>>hg38_21_+_8433172_8446622_RNA45SN1_45S_50ntFlanks@11584.11623.40|TGGTGCGGACCAGGGGAATCCGACTGTTTAATTAAAACAA</td><td>>hg38_21_+_8433172_8446622_RNA45SN1_45S_50ntFlanks</td><td>11584</td><td>11623</td><td>40</td><td>TGGTGCGGACCAGGGGAATCCGACTGTTTAATTAAAACAA</td><td>96.2</td><td>5.3</td><td>2.6</td><td>2.0</td><td>4.1E-02</td><td>4.9E-02</td><td>MB231/MB468</td><td>MP</td></tr>
<tr><td>>hg38_21_+_8433172_8446622_RNA45SN1_45S_50ntFlanks@4380.4423.44|CGTAGTTGGATCTTGGGAGCGGGCGGGCGGTCCGCCGCGAGGCG</td><td>>hg38_21_+_8433172_8446622_RNA45SN1_45S_50ntFlanks</td><td>4380</td><td>4423</td><td>44</td><td>CGTAGTTGGATCTTGGGAGCGGGCGGGCGGTCCGCCGCGAGGCG</td><td>161.3</td><td>-2.8</td><td>1.4</td><td>-2.0</td><td>4.3E-02</td><td>4.9E-02</td><td>BT20/MB468</td><td>CYTO</td></tr>
<tr><td>>hg38_21_+_8433172_8446622_RNA45SN1_45S_50ntFlanks@6731.6752.22|CACATTGATCATCGACACTTCG</td><td>>hg38_21_+_8433172_8446622_RNA45SN1_45S_50ntFlanks</td><td>6731</td><td>6752</td><td>22</td><td>CACATTGATCATCGACACTTCG</td><td>257.1</td><td>-3.8</td><td>1.8</td><td>-2.2</td><td>3.1E-02</td><td>4.9E-02</td><td>BT20/MB468</td><td>MP</td></tr>
<tr><td>>hg38_21_+_8433172_8446622_RNA45SN1_45S_50ntFlanks@10677.10712.36|GCGGTTCCGGCGGCGTCCGGTGAGCTCTCGCTGGCC</td><td>>hg38_21_+_8433172_8446622_RNA45SN1_45S_50ntFlanks</td><td>10677</td><td>10712</td><td>36</td><td>GCGGTTCCGGCGGCGTCCGGTGAGCTCTCGCTGGCC</td><td>224.5</td><td>-1.7</td><td>0.8</td><td>-2.0</td><td>4.1E-02</td><td>4.9E-02</td><td>MB231/MB468</td><td>MP</td></tr>
<tr><td>>trnaMT_ValTAC_MT_+_1602_1670@28.71.44|AACTTACACTTAGGAGATTTCAACTTAACTTGACCGCTCTGACC</td><td>>trnaMT_ValTAC_MT_+_1602_1670</td><td>28</td><td>71</td><td>44</td><td>AACTTACACTTAGGAGATTTCAACTTAACTTGACCGCTCTGACC</td><td>334.5</td><td>5.1</td><td>2.4</td><td>2.2</td><td>3.1E-02</td><td>4.9E-02</td><td>BT20/MB468</td><td>MP</td></tr>
<tr><td>>trnaMT_GluTTC_MT_-_14674_14742@39.62.24,trnalookalike8_GluTTC_5_-_93905172_93905240@39.62.24|ATTGGTCGTGGTTGTAGTCCGTGC</td><td>>trnaMT_GluTTC_MT_-_14674_14742</td><td>39</td><td>62</td><td>24</td><td>ATTGGTCGTGGTTGTAGTCCGTGC</td><td>76.3</td><td>2.7</td><td>1.3</td><td>2.0</td><td>4.1E-02</td><td>4.9E-02</td><td>MB231/MB468</td><td>MP</td></tr>
<tr><td>>hg38_21_+_8433172_8446622_RNA45SN1_45S_50ntFlanks@11324.11344.21|GTCTCGTCGCGCGCGCGTCCG</td><td>>hg38_21_+_8433172_8446622_RNA45SN1_45S_50ntFlanks</td><td>11324</td><td>11344</td><td>21</td><td>GTCTCGTCGCGCGCGCGTCCG</td><td>116.3</td><td>-2.8</td><td>1.3</td><td>-2.2</td><td>3.1E-02</td><td>4.9E-02</td><td>BT20/MB468</td><td>MP</td></tr>
<tr><td>>trna111_HisGTG_1_-_147774845_147774916@1.31.31,trna118_HisGTG_1_-_145396881_145396952@1.31.31,trna16_HisGTG_1_+_146544773_146544844@1.31.31,trna1_HisGTG_15_+_45493349_45493420@1.31.31,trna21_HisGTG_1_+_147753471_147753542@1.31.31,trna33_HisGTG_6_+_27125906_27125977@1.31.31,trna7_HisGTG_9_-_14433938_14434009@1.31.31,trna8_HisGTG_15_-_45492611_45492682@1.31.31,trna9_HisGTG_15_-_45490804_45490875@1.31.31|GCCGTGATCGTATAGTGGTTAGTACTCTGCG</td><td>>trna111_HisGTG_1_-_147774845_147774916</td><td>1</td><td>31</td><td>31</td><td>GCCGTGATCGTATAGTGGTTAGTACTCTGCG</td><td>10032.6</td><td>-2.1</td><td>0.9</td><td>-2.3</td><td>2.0E-02</td><td>4.9E-02</td><td>BT20/MB231</td><td>CYTO</td></tr>
<tr><td>>hg38_21_+_8433172_8446622_RNA45SN1_45S_50ntFlanks@12474.12496.23|ACCCACTAATAGGGAACGTGAGC</td><td>>hg38_21_+_8433172_8446622_RNA45SN1_45S_50ntFlanks</td><td>12474</td><td>12496</td><td>23</td><td>ACCCACTAATAGGGAACGTGAGC</td><td>1554.4</td><td>-2.0</td><td>0.9</td><td>-2.3</td><td>2.0E-02</td><td>4.9E-02</td><td>BT20/MB231</td><td>CYTO</td></tr>
</table>

<table>
<tr><td></td><td></td><td></td><td></td><td></td><td></td><td></td><td></td><td></td><td></td><td></td><td></td><td></td><td></td></tr>
<tr><td>>hg38_MT_+_1621_3279_RNR2_16S_50ntFlanks@1439.1477.39|ACGTGATCTGAGTTCAGACCGGAGTAATCCAGGTCGGTT</td><td>>hg38_MT_+_1621_3279_RNR2_16S_50ntFlanks</td><td>1439</td><td>1477</td><td>39</td><td>ACGTGATCTGAGTTCAGACCGGAGTAATCCAGGTCGGTT</td><td>245.8</td><td>3.2</td><td>1.5</td><td>2.2</td><td>3.1E-02</td><td>4.9E-02</td><td>BT20/MB468</td><td>MP</td></tr>
<tr><td>>hg38_21_+_8433172_8446622_RNA45SN1_45S_50ntFlanks@4734.4766.33|AACGAAAGTCGGAGGTTCGAAGACGATCAGATA</td><td>>hg38_21_+_8433172_8446622_RNA45SN1_45S_50ntFlanks</td><td>4734</td><td>4766</td><td>33</td><td>AACGAAAGTCGGAGGTTCGAAGACGATCAGATA</td><td>83.1</td><td>3.2</td><td>1.5</td><td>2.2</td><td>3.1E-02</td><td>4.9E-02</td><td>BT20/MB468</td><td>MP</td></tr>
<tr><td>>hg38_MT_+_1621_3279_RNR2_16S_50ntFlanks@1370.1405.36|ATGTTGGATCAGGACATCCCGATGGTGCAGCCGCTA</td><td>>hg38_MT_+_1621_3279_RNR2_16S_50ntFlanks</td><td>1370</td><td>1405</td><td>36</td><td>ATGTTGGATCAGGACATCCCGATGGTGCAGCCGCTA</td><td>86.2</td><td>4.2</td><td>2.1</td><td>2.0</td><td>4.1E-02</td><td>4.9E-02</td><td>MB231/MB468</td><td>MP</td></tr>
<tr><td>>hg38_21_+_8433172_8446622_RNA45SN1_45S_50ntFlanks@3837.3871.35|CCTCTCCTACTTGGATAACTGTGGTAATTCTAGAG</td><td>>hg38_21_+_8433172_8446622_RNA45SN1_45S_50ntFlanks</td><td>3837</td><td>3871</td><td>35</td><td>CCTCTCCTACTTGGATAACTGTGGTAATTCTAGAG</td><td>392.2</td><td>-1.7</td><td>0.9</td><td>-2.0</td><td>4.3E-02</td><td>4.9E-02</td><td>MB231/MB468</td><td>CYTO</td></tr>
<tr><td>>MI0000082|hsa-mir-25&WithFlank&7|-|100093554|100093649@58.78.21(+1G)&[MIMAT0000081&hsa-miR-25-3p&offsets|0|-1(+1G);m-23&7|-|100093571|100093592&offsets|0|-1(+1G)]|CATTGCACTTGTCTCGGTCTGG</td><td>>MI0000082|hsa-mir-25&WithFlank&7|-|100093554|100093649</td><td>58</td><td>78</td><td>21(+1G)</td><td>CATTGCACTTGTCTCGGTCTGG</td><td>1646.2</td><td>1.9</td><td>0.8</td><td>2.3</td><td>2.0E-02</td><td>4.9E-02</td><td>BT20/MB231</td><td>CYTO</td></tr>
<tr><td>>hg38_21_+_8433172_8446622_RNA45SN1_45S_50ntFlanks@10349.10379.31|AACGAGAACTTTGAAGGCCGAAGTGGAGAAG</td><td>>hg38_21_+_8433172_8446622_RNA45SN1_45S_50ntFlanks</td><td>10349</td><td>10379</td><td>31</td><td>AACGAGAACTTTGAAGGCCGAAGTGGAGAAG</td><td>449.0</td><td>1.5</td><td>0.7</td><td>2.0</td><td>4.3E-02</td><td>4.9E-02</td><td>MB231/MB468</td><td>CYTO</td></tr>
<tr><td>>hg38_21_+_8433172_8446622_RNA45SN1_45S_50ntFlanks@7983.8017.35|CAGATCAGACGTGGCGACCCGCTGAATTTAAGCAT</td><td>>hg38_21_+_8433172_8446622_RNA45SN1_45S_50ntFlanks</td><td>7983</td><td>8017</td><td>35</td><td>CAGATCAGACGTGGCGACCCGCTGAATTTAAGCAT</td><td>480.0</td><td>-1.9</td><td>0.9</td><td>-2.0</td><td>4.5E-02</td><td>4.9E-02</td><td>MB231/MB468</td><td>MITO</td></tr>
<tr><td>>hg38_21_+_8433172_8446622_RNA45SN1_45S_50ntFlanks@12748.12769.22|CCTGTCCCCGCCGGCGGGCCGC</td><td>>hg38_21_+_8433172_8446622_RNA45SN1_45S_50ntFlanks</td><td>12748</td><td>12769</td><td>22</td><td>CCTGTCCCCGCCGGCGGGCCGC</td><td>210.2</td><td>-3.4</td><td>1.6</td><td>-2.1</td><td>3.9E-02</td><td>4.9E-02</td><td>BT20/MB468</td><td>MITO</td></tr>
<tr><td>>hg38_MT_+_1621_3279_RNR2_16S_50ntFlanks@1363.1387.25|GACCTCGATGTTGGATCAGGACATC</td><td>>hg38_MT_+_1621_3279_RNR2_16S_50ntFlanks</td><td>1363</td><td>1387</td><td>25</td><td>GACCTCGATGTTGGATCAGGACATC</td><td>58.1</td><td>3.6</td><td>1.8</td><td>2.0</td><td>4.2E-02</td><td>4.9E-02</td><td>MB231/MB468</td><td>MP</td></tr>
<tr><td>>hg38_21_+_8433172_8446622_RNA45SN1_45S_50ntFlanks@4099.4120.22|GTGACCACGGGTGACGGGGAAT</td><td>>hg38_21_+_8433172_8446622_RNA45SN1_45S_50ntFlanks</td><td>4099</td><td>4120</td><td>22</td><td>GTGACCACGGGTGACGGGGAAT</td><td>65.9</td><td>-3.7</td><td>1.7</td><td>-2.2</td><td>3.1E-02</td><td>4.9E-02</td><td>BT20/MB468</td><td>MP</td></tr>
<tr><td>>hg38_21_+_8433172_8446622_RNA45SN1_45S_50ntFlanks@3785.3802.18|TGAAACTGCGAATGGCTC</td><td>>hg38_21_+_8433172_8446622_RNA45SN1_45S_50ntFlanks</td><td>3785</td><td>3802</td><td>18</td><td>TGAAACTGCGAATGGCTC</td><td>231.9</td><td>-3.2</td><td>1.5</td><td>-2.2</td><td>3.1E-02</td><td>4.9E-02</td><td>BT20/MB468</td><td>MP</td></tr>
<tr><td>>hg38_21_+_8433172_8446622_RNA45SN1_45S_50ntFlanks@4102.4125.24|ACCACGGGTGACGGGGAATCAGGG</td><td>>hg38_21_+_8433172_8446622_RNA45SN1_45S_50ntFlanks</td><td>4102</td><td>4125</td><td>24</td><td>ACCACGGGTGACGGGGAATCAGGG</td><td>281.4</td><td>2.4</td><td>1.1</td><td>2.2</td><td>3.1E-02</td><td>4.9E-02</td><td>BT20/MB468</td><td>MP</td></tr>
<tr><td>>hg38_21_+_8433172_8446622_RNA45SN1_45S_50ntFlanks@10099.10126.28|CGCGCGCGCGTGTGGTGTGCGTCGGAGG</td><td>>hg38_21_+_8433172_8446622_RNA45SN1_45S_50ntFlanks</td><td>10099</td><td>10126</td><td>28</td><td>CGCGCGCGCGTGTGGTGTGCGTCGGAGG</td><td>114.5</td><td>-3.9</td><td>1.8</td><td>-2.2</td><td>3.1E-02</td><td>4.9E-02</td><td>BT20/MB468</td><td>MP</td></tr>
<tr><td>>hg38_21_+_8433172_8446622_RNA45SN1_45S_50ntFlanks@5485.5506.22|AGCGCTGAGAAGACGGTCGAAC</td><td>>hg38_21_+_8433172_8446622_RNA45SN1_45S_50ntFlanks</td><td>5485</td><td>5506</td><td>22</td><td>AGCGCTGAGAAGACGGTCGAAC</td><td>1661.7</td><td>-2.1</td><td>1.0</td><td>-2.2</td><td>3.1E-02</td><td>4.9E-02</td><td>BT20/MB468</td><td>MP</td></tr>
<tr><td>>hg38_21_+_8433172_8446622_RNA45SN1_45S_50ntFlanks@7976.8006.31|GCGACCTCAGATCAGACGTGGCGACCCGCTG</td><td>>hg38_21_+_8433172_8446622_RNA45SN1_45S_50ntFlanks</td><td>7976</td><td>8006</td><td>31</td><td>GCGACCTCAGATCAGACGTGGCGACCCGCTG</td><td>431.3</td><td>2.5</td><td>1.2</td><td>2.2</td><td>3.1E-02</td><td>4.9E-02</td><td>BT20/MB468</td><td>MP</td></tr>
<tr><td>>trna111_HisGTG_1_-_147774845_147774916@3.30.28,trna118_HisGTG_1_-_145396881_145396952@3.30.28,trna16_HisGTG_1_+_146544773_146544844@3.30.28,trna1_HisGTG_15_+_45493349_45493420@3.30.28,trna21_HisGTG_1_+_147753471_147753542@3.30.28,trna33_HisGTG_6_+_27125906_27125977@3.30.28,trna7_HisGTG_9_-_14433938_14434009@3.30.28,trna8_HisGTG_15_-_45492611_45492682@3.30.28,trna9_HisGTG_15_-_45490804_45490875@3.30.28|CGTGATCGTATAGTGGTTAGTACTCTGC</td><td>>trna111_HisGTG_1_-_147774845_147774916</td><td>3</td><td>30</td><td>28</td><td>CGTGATCGTATAGTGGTTAGTACTCTGC</td><td>61.5</td><td>1.7</td><td>0.8</td><td>2.0</td><td>4.2E-02</td><td>4.9E-02</td><td>MB231/MB468</td><td>MP</td></tr>
<tr><td>>hg38_21_+_8433172_8446622_RNA45SN1_45S_50ntFlanks@9590.9610.21|TGGGTATAGGGGCGAAAGACT</td><td>>hg38_21_+_8433172_8446622_RNA45SN1_45S_50ntFlanks</td><td>9590</td><td>9610</td><td>21</td><td>TGGGTATAGGGGCGAAAGACT</td><td>381.3</td><td>-1.9</td><td>0.8</td><td>-2.3</td><td>2.0E-02</td><td>4.9E-02</td><td>BT20/MB231</td><td>CYTO</td></tr>
<tr><td>>hg38_21_+_8433172_8446622_RNA45SN1_45S_50ntFlanks@12405.12429.25|CTTTTTGATCCTTCGATGTCGGCTC</td><td>>hg38_21_+_8433172_8446622_RNA45SN1_45S_50ntFlanks</td><td>12405</td><td>12429</td><td>25</td><td>CTTTTTGATCCTTCGATGTCGGCTC</td><td>1079.9</td><td>-4.1</td><td>1.9</td><td>-2.2</td><td>3.2E-02</td><td>4.9E-02</td><td>BT20/MB468</td><td>MP</td></tr>
<tr><td>>hg38_21_+_8433172_8446622_RNA45SN1_45S_50ntFlanks@11920.11958.39|ATGAGAGGTGTAGAATAAGTGGGAGGCCCCCGGCGCCCC</td><td>>hg38_21_+_8433172_8446622_RNA45SN1_45S_50ntFlanks</td><td>11920</td><td>11958</td><td>39</td><td>ATGAGAGGTGTAGAATAAGTGGGAGGCCCCCGGCGCCCC</td><td>85.2</td><td>-3.9</td><td>1.8</td><td>-2.2</td><td>3.2E-02</td><td>4.9E-02</td><td>BT20/MB468</td><td>MP</td></tr>
<tr><td>>hg38_MT_+_598_1651_RNR1_12S_50ntFlanks@429.452.24|AAGTGGCTTTAACATATCTGAACA</td><td>>hg38_MT_+_598_1651_RNR1_12S_50ntFlanks</td><td>429</td><td>452</td><td>24</td><td>AAGTGGCTTTAACATATCTGAACA</td><td>113.1</td><td>3.9</td><td>1.8</td><td>2.2</td><td>3.2E-02</td><td>4.9E-02</td><td>BT20/MB468</td><td>MP</td></tr>
<tr><td>>hg38_21_+_8433172_8446622_RNA45SN1_45S_50ntFlanks@8379.8399.21|TCAAGAGGGCGTGAAACCGTT</td><td>>hg38_21_+_8433172_8446622_RNA45SN1_45S_50ntFlanks</td><td>8379</td><td>8399</td><td>21</td><td>TCAAGAGGGCGTGAAACCGTT</td><td>151.8</td><td>-2.1</td><td>1.0</td><td>-2.2</td><td>3.2E-02</td><td>4.9E-02</td><td>BT20/MB468</td><td>MP</td></tr>
<tr><td>>am_trnaMT_SerGCT_MT_+_12207_12265@12.38.27|CAAGAACTGCTAACTCATGCCCCCATG</td><td>>am_trnaMT_SerGCT_MT_+_12207_12265</td><td>12</td><td>38</td><td>27</td><td>CAAGAACTGCTAACTCATGCCCCCATG</td><td>273.2</td><td>-2.3</td><td>1.1</td><td>-2.0</td><td>4.4E-02</td><td>4.9E-02</td><td>MB231/MB468</td><td>CYTO</td></tr>
<tr><td>>hg38_21_+_8433172_8446622_RNA45SN1_45S_50ntFlanks@11104.11131.28|AAGGGTCGGGGCGGCAGGGGCCGGCGGC</td><td>>hg38_21_+_8433172_8446622_RNA45SN1_45S_50ntFlanks</td><td>11104</td><td>11131</td><td>28</td><td>AAGGGTCGGGGCGGCAGGGGCCGGCGGC</td><td>172.4</td><td>-3.0</td><td>1.4</td><td>-2.1</td><td>3.2E-02</td><td>4.9E-02</td><td>BT20/MB468</td><td>MP</td></tr>
<tr><td>>hg38_1_-_228634819_228635039_RNA5S12_5S_50ntFlanks@51.87.37|GTCTACGGCCATACCACCCTGAACGCGCCCGATCTCG</td><td>>hg38_1_-_228634819_228635039_RNA5S12_5S_50ntFlanks</td><td>51</td><td>87</td><td>37</td><td>GTCTACGGCCATACCACCCTGAACGCGCCCGATCTCG</td><td>899.6</td><td>2.5</td><td>1.2</td><td>2.0</td><td>4.5E-02</td><td>5.0E-02</td><td>BT20/MB468</td><td>NUC</td></tr>
<tr><td>>hg38_21_+_8433172_8446622_RNA45SN1_45S_50ntFlanks@4983.5020.38|CGGACAGGATTGACAGATTGATAGCTCTTTCTCGATTC</td><td>>hg38_21_+_8433172_8446622_RNA45SN1_45S_50ntFlanks</td><td>4983</td><td>5020</td><td>38</td><td>CGGACAGGATTGACAGATTGATAGCTCTTTCTCGATTC</td><td>129.6</td><td>4.4</td><td>1.8</td><td>2.4</td><td>1.4E-02</td><td>5.0E-02</td><td>BT20/MB231</td><td>MP</td></tr>
<tr><td>>MI0000736|hsa-mir-30c-1&WithFlank&1|+|40757278|40757378@23.46.24(+2U);MI0000254|hsa-mir-30c-2&WithFlank&6|-|71376954|71377037@13.36.24(+2U)&[MIMAT0000244&hsa-miR-30c-5p&offsets|0|+1(+2U);m-57&1|+|40757300|40757323&offsets|0|0(+2U)];[MIMAT0000244_1&hsa-miR-30c-5p&offsets|0|+1(+2U);m-56&6|-|71377002|71377025&offsets|0|0(+2U)]|TGTAAACATCCTACACTCTCAGCTTT</td><td>>MI0000736|hsa-mir-30c-1&WithFlank&1|+|40757278|40757378</td><td>23</td><td>46</td><td>24(+2U)</td><td>TGTAAACATCCTACACTCTCAGCTTT</td><td>291.2</td><td>-3.4</td><td>1.4</td><td>-2.4</td><td>1.5E-02</td><td>5.0E-02</td><td>BT20/MB231</td><td>NUC</td></tr>
<tr><td>>hg38_MT_+_1621_3279_RNR2_16S_50ntFlanks@1391.1422.32|ATGGTGCAGCCGCTATTAAAGGTTCGTTTGTT</td><td>>hg38_MT_+_1621_3279_RNR2_16S_50ntFlanks</td><td>1391</td><td>1422</td><td>32</td><td>ATGGTGCAGCCGCTATTAAAGGTTCGTTTGTT</td><td>39.4</td><td>3.5</td><td>1.7</td><td>2.0</td><td>4.2E-02</td><td>5.0E-02</td><td>MB231/MB468</td><td>MP</td></tr>
<tr><td>>hg38_21_+_8433172_8446622_RNA45SN1_45S_50ntFlanks@12202.12232.31|CTAAGGCGAGCTCAGGGAGGACAGAAACCTC</td><td>>hg38_21_+_8433172_8446622_RNA45SN1_45S_50ntFlanks</td><td>12202</td><td>12232</td><td>31</td><td>CTAAGGCGAGCTCAGGGAGGACAGAAACCTC</td><td>225.9</td><td>-3.2</td><td>1.5</td><td>-2.1</td><td>3.2E-02</td><td>5.0E-02</td><td>BT20/MB468</td><td>MP</td></tr>
<tr><td>>hg38_21_+_8433172_8446622_RNA45SN1_45S_50ntFlanks@5086.5110.25|AACGAGACTCTGGCATGCTAACTAG</td><td>>hg38_21_+_8433172_8446622_RNA45SN1_45S_50ntFlanks</td><td>5086</td><td>5110</td><td>25</td><td>AACGAGACTCTGGCATGCTAACTAG</td><td>368.7</td><td>3.3</td><td>1.4</td><td>2.4</td><td>1.4E-02</td><td>5.0E-02</td><td>BT20/MB231</td><td>MP</td></tr>
<tr><td>>am_trnaMT_HisGTG_MT_+_12138_12206@27.44.18|GATTGTGAATCTGACAAC</td><td>>am_trnaMT_HisGTG_MT_+_12138_12206</td><td>27</td><td>44</td><td>18</td><td>GATTGTGAATCTGACAAC</td><td>629.5</td><td>2.2</td><td>1.1</td><td>2.0</td><td>4.5E-02</td><td>5.0E-02</td><td>MB231/MB468</td><td>MITO</td></tr>
<tr><td>>hg38_21_+_8433172_8446622_RNA45SN1_45S_50ntFlanks@12416.12449.34|TTCGATGTCGGCTCTTCCTATCATTGTGAAGCAG</td><td>>hg38_21_+_8433172_8446622_RNA45SN1_45S_50ntFlanks</td><td>12416</td><td>12449</td><td>34</td><td>TTCGATGTCGGCTCTTCCTATCATTGTGAAGCAG</td><td>64.2</td><td>1.9</td><td>0.9</td><td>2.0</td><td>4.2E-02</td><td>5.0E-02</td><td>MB231/MB468</td><td>MP</td></tr>
<tr><td>>hg38_21_+_8433172_8446622_RNA45SN1_45S_50ntFlanks@12306.12323.18|CACGATCCTTCTGACCTT</td><td>>hg38_21_+_8433172_8446622_RNA45SN1_45S_50ntFlanks</td><td>12306</td><td>12323</td><td>18</td><td>CACGATCCTTCTGACCTT</td><td>173.8</td><td>-3.8</td><td>1.8</td><td>-2.1</td><td>3.2E-02</td><td>5.0E-02</td><td>BT20/MB468</td><td>MP</td></tr>
<tr><td>>hg38_21_+_8433172_8446622_RNA45SN1_45S_50ntFlanks@475.503.29|CTTCGTGATCGATGTGGTGACGTCGTGCT</td><td>>hg38_21_+_8433172_8446622_RNA45SN1_45S_50ntFlanks</td><td>475</td><td>503</td><td>29</td><td>CTTCGTGATCGATGTGGTGACGTCGTGCT</td><td>930.4</td><td>1.8</td><td>0.8</td><td>2.3</td><td>2.0E-02</td><td>5.0E-02</td><td>BT20/MB231</td><td>CYTO</td></tr>
<tr><td>>hg38_21_+_8433172_8446622_RNA45SN1_45S_50ntFlanks@11027.11059.33|CTCTCTCTCTCTCTCCCCCGCTCCCCGTCCTCC</td><td>>hg38_21_+_8433172_8446622_RNA45SN1_45S_50ntFlanks</td><td>11027</td><td>11059</td><td>33</td><td>CTCTCTCTCTCTCTCCCCCGCTCCCCGTCCTCC</td><td>914.4</td><td>3.5</td><td>1.7</td><td>2.1</td><td>3.9E-02</td><td>5.0E-02</td><td>BT20/MB468</td><td>MITO</td></tr>
<tr><td>>hg38_21_+_8433172_8446622_RNA45SN1_45S_50ntFlanks@10348.10373.26|AAACGAGAACTTTGAAGGCCGAAGTG</td><td>>hg38_21_+_8433172_8446622_RNA45SN1_45S_50ntFlanks</td><td>10348</td><td>10373</td><td>26</td><td>AAACGAGAACTTTGAAGGCCGAAGTG</td><td>389.1</td><td>1.5</td><td>0.7</td><td>2.0</td><td>4.4E-02</td><td>5.0E-02</td><td>MB231/MB468</td><td>CYTO</td></tr>
<tr><td>>am_trna111_HisGTG_1_-_147774845_147774916@-1G.28.29,trna118_HisGTG_1_-_145396881_145396952@-1G.28.29,trna16_HisGTG_1_+_146544773_146544844@-1G.28.29,trna1_HisGTG_15_+_45493349_45493420@-1G.28.29,trna21_HisGTG_1_+_147753471_147753542@-1G.28.29,trna33_HisGTG_6_+_27125906_27125977@-1G.28.29,trna7_HisGTG_9_-_14433938_14434009@-1G.28.29,trna8_HisGTG_15_-_45492611_45492682@-1G.28.29,trna9_HisGTG_15_-_45490804_45490875@-1G.28.29|GGCCGTGATCGTATAGTGGTTAGTACTCT</td><td>>am_trna111_HisGTG_1_-_147774845_147774916</td><td>-1G</td><td>28</td><td>29</td><td>GGCCGTGATCGTATAGTGGTTAGTACTCT</td><td>3276.8</td><td>-2.7</td><td>1.2</td><td>-2.3</td><td>2.0E-02</td><td>5.0E-02</td><td>BT20/MB231</td><td>CYTO</td></tr>
<tr><td>>MI0000825|hsa-mir-345&WithFlank&14|+|100307853|100307962@24.44.21&[MIMAT0000772&hsa-miR-345-5p&offsets|0|-1;m-193&14|+|100307876|100307896&offsets|0|0]|GCTGACTCCTAGTCCAGGGCT</td><td>>MI0000825|hsa-mir-345&WithFlank&14|+|100307853|100307962</td><td>24</td><td>44</td><td>21</td><td>GCTGACTCCTAGTCCAGGGCT</td><td>158.6</td><td>3.5</td><td>1.6</td><td>2.1</td><td>3.2E-02</td><td>5.0E-02</td><td>BT20/MB468</td><td>MP</td></tr>
<tr><td>>hg38_21_+_8433172_8446622_RNA45SN1_45S_50ntFlanks@12416.12448.33|TTCGATGTCGGCTCTTCCTATCATTGTGAAGCA</td><td>>hg38_21_+_8433172_8446622_RNA45SN1_45S_50ntFlanks</td><td>12416</td><td>12448</td><td>33</td><td>TTCGATGTCGGCTCTTCCTATCATTGTGAAGCA</td><td>75.0</td><td>3.8</td><td>1.8</td><td>2.1</td><td>3.2E-02</td><td>5.0E-02</td><td>BT20/MB468</td><td>MP</td></tr>
<tr><td>>hg38_21_+_8433172_8446622_RNA45SN1_45S_50ntFlanks@4753.4782.30|AAGACGATCAGATACCGTCGTAGTTCCGAC</td><td>>hg38_21_+_8433172_8446622_RNA45SN1_45S_50ntFlanks</td><td>4753</td><td>4782</td><td>30</td><td>AAGACGATCAGATACCGTCGTAGTTCCGAC</td><td>376.5</td><td>-4.0</td><td>1.9</td><td>-2.1</td><td>3.2E-02</td><td>5.0E-02</td><td>BT20/MB468</td><td>MP</td></tr>
<tr><td>>trnaMT_GluTTC_MT_-_14674_14742@41.71.31,trnalookalike8_GluTTC_5_-_93905172_93905240@41.71.31|TGGTCGTGGTTGTAGTCCGTGCGAGAATACC</td><td>>trnaMT_GluTTC_MT_-_14674_14742</td><td>41</td><td>71</td><td>31</td><td>TGGTCGTGGTTGTAGTCCGTGCGAGAATACC</td><td>248.2</td><td>2.0</td><td>1.0</td><td>2.0</td><td>4.6E-02</td><td>5.0E-02</td><td>MB231/MB468</td><td>MITO</td></tr>
<tr><td>>MI0000067|hsa-let-7f-1&WithFlank&9|+|94176341|94176439@13.31.19;MI0000068|hsa-let-7f-2&WithFlank&X|-|53557186|53557280@14.32.19&[MIMAT0000067&hsa-let-7f-5p&offsets|0|-3;m-20&9|+|94176353|94176374&offsets|0|-3];[MIMAT0000067_1&hsa-let-7f-5p&offsets|0|-3;m-19&X|-|53557246|53557267&offsets|0|-3]|TGAGGTAGTAGATTGTATA</td><td>>MI0000067|hsa-let-7f-1&WithFlank&9|+|94176341|94176439</td><td>13</td><td>31</td><td>19</td><td>TGAGGTAGTAGATTGTATA</td><td>459.9</td><td>1.7</td><td>0.9</td><td>2.0</td><td>4.5E-02</td><td>5.0E-02</td><td>BT20/MB468</td><td>NUC</td></tr>
<tr><td>>hg38_21_+_8433172_8446622_RNA45SN1_45S_50ntFlanks@11209.11229.21|CGGCGACTCTGGACGCGAGCC</td><td>>hg38_21_+_8433172_8446622_RNA45SN1_45S_50ntFlanks</td><td>11209</td><td>11229</td><td>21</td><td>CGGCGACTCTGGACGCGAGCC</td><td>1187.5</td><td>-3.4</td><td>1.7</td><td>-2.0</td><td>4.5E-02</td><td>5.0E-02</td><td>BT20/MB468</td><td>NUC</td></tr>
<tr><td>>MI0000085|hsa-mir-27a&WithFlank&19|-|13836434|13836523@16.36.21&[MIMAT0004501&hsa-miR-27a-5p&offsets|0|-1;m-270&19|-|13836487|13836508&offsets|0|-1]|AGGGCTTAGCTGCTTGTGAGC</td><td>>MI0000085|hsa-mir-27a&WithFlank&19|-|13836434|13836523</td><td>16</td><td>36</td><td>21</td><td>AGGGCTTAGCTGCTTGTGAGC</td><td>119.5</td><td>4.0</td><td>1.9</td><td>2.1</td><td>3.2E-02</td><td>5.0E-02</td><td>BT20/MB468</td><td>MP</td></tr>
<tr><td>>hg38_21_+_8433172_8446622_RNA45SN1_45S_50ntFlanks@6661.6701.41|CGGTGGATCACTCGGCTCGTGCGTCGATGAAGAACGCAGCT</td><td>>hg38_21_+_8433172_8446622_RNA45SN1_45S_50ntFlanks</td><td>6661</td><td>6701</td><td>41</td><td>CGGTGGATCACTCGGCTCGTGCGTCGATGAAGAACGCAGCT</td><td>5147.1</td><td>1.9</td><td>0.8</td><td>2.4</td><td>1.7E-02</td><td>5.0E-02</td><td>BT20/MB231</td><td>MITO</td></tr>
<tr><td>>hg38_21_+_8433172_8446622_RNA45SN1_45S_50ntFlanks@11719.11742.24|ACGGCGGGAGTAACTATGACTCTC</td><td>>hg38_21_+_8433172_8446622_RNA45SN1_45S_50ntFlanks</td><td>11719</td><td>11742</td><td>24</td><td>ACGGCGGGAGTAACTATGACTCTC</td><td>102.2</td><td>-3.0</td><td>1.4</td><td>-2.1</td><td>3.2E-02</td><td>5.0E-02</td><td>BT20/MB468</td><td>MP</td></tr>
<tr><td>>am_trnaMT_MetCAT_MT_+_4402_4469@1.28.28|AGTAAGGTCAGCTAAATAAGCTATCGGG</td><td>>am_trnaMT_MetCAT_MT_+_4402_4469</td><td>1</td><td>28</td><td>28</td><td>AGTAAGGTCAGCTAAATAAGCTATCGGG</td><td>62.0</td><td>-4.5</td><td>2.2</td><td>-2.0</td><td>4.2E-02</td><td>5.0E-02</td><td>MB231/MB468</td><td>MP</td></tr>
</table>

<table>
<tr><td></td><td></td><td></td><td></td><td></td><td></td><td></td><td></td><td></td><td></td><td></td><td></td><td></td><td></td></tr>
<tr><td>>hg38_21_+_8433172_8446622_RNA45SN1_45S_50ntFlanks@11587.11611.25|TGCGGACCAGGGGAATCCGACTGTT</td><td>>hg38_21_+_8433172_8446622_RNA45SN1_45S_50ntFlanks</td><td>11587</td><td>11611</td><td>25</td><td>TGCGGACCAGGGGAATCCGACTGTT</td><td>561.2</td><td>-1.2</td><td>0.6</td><td>-2.0</td><td>4.4E-02</td><td>5.0E-02</td><td>BT20/MB468</td><td>CYTO</td></tr>
<tr><td>>hg38_21_+_8433172_8446622_RNA45SN1_45S_50ntFlanks@7975.8008.34|CGCGACCTCAGATCAGACGTGGCGACCCGCTGAA</td><td>>hg38_21_+_8433172_8446622_RNA45SN1_45S_50ntFlanks</td><td>7975</td><td>8008</td><td>34</td><td>CGCGACCTCAGATCAGACGTGGCGACCCGCTGAA</td><td>171494.9</td><td>4.0</td><td>1.6</td><td>2.4</td><td>1.5E-02</td><td>5.0E-02</td><td>BT20/MB231</td><td>MP</td></tr>
<tr><td>>hg38_21_+_8433172_8446622_RNA45SN1_45S_50ntFlanks@12007.12033.27|GCCGCCGGTGAAATACCACTACTCTGA</td><td>>hg38_21_+_8433172_8446622_RNA45SN1_45S_50ntFlanks</td><td>12007</td><td>12033</td><td>27</td><td>GCCGCCGGTGAAATACCACTACTCTGA</td><td>619.1</td><td>2.1</td><td>0.9</td><td>2.4</td><td>1.7E-02</td><td>5.0E-02</td><td>BT20/MB231</td><td>MITO</td></tr>
<tr><td>>MI0000078|hsa-mir-22&WithFlank&17|-|1713897|1713993@59.81.23&[MIMAT0000077&hsa-miR-22-3p&offsets|0|+1;m-2&17|-|1713914|1713935&offsets|0|+1]|AAGCTGCCAGTTGAAGAACTGTT</td><td>>MI0000078|hsa-mir-22&WithFlank&17|-|1713897|1713993</td><td>59</td><td>81</td><td>23</td><td>AAGCTGCCAGTTGAAGAACTGTT</td><td>652.4</td><td>-2.0</td><td>0.8</td><td>-2.4</td><td>1.7E-02</td><td>5.0E-02</td><td>BT20/MB231</td><td>MITO</td></tr>
<tr><td>>hg38_21_+_8433172_8446622_RNA45SN1_45S_50ntFlanks@11565.11584.20|CTAGCAGCCGACTTAGAACT</td><td>>hg38_21_+_8433172_8446622_RNA45SN1_45S_50ntFlanks</td><td>11565</td><td>11584</td><td>20</td><td>CTAGCAGCCGACTTAGAACT</td><td>323.4</td><td>1.9</td><td>0.8</td><td>2.4</td><td>1.7E-02</td><td>5.0E-02</td><td>BT20/MB231</td><td>MITO</td></tr>
<tr><td>>trna10_LysCTT_16_+_3241501_3241573@1.29.29,trna13_LysCTT_14_-_58706613_58706685@1.29.29,trna2_LysCTT_15_+_79152904_79152976@1.29.29|GCCCGGCTAGCTCAGTCGGTAGAGCATGG</td><td>>trna10_LysCTT_16_+_3241501_3241573</td><td>1</td><td>29</td><td>29</td><td>GCCCGGCTAGCTCAGTCGGTAGAGCATGG</td><td>162.4</td><td>-3.2</td><td>1.6</td><td>-2.1</td><td>3.9E-02</td><td>5.0E-02</td><td>BT20/MB468</td><td>MITO</td></tr>
<tr><td>>hg38_21_+_8433172_8446622_RNA45SN1_45S_50ntFlanks@6661.6686.26|CGGTGGATCACTCGGCTCGTGCGTCG</td><td>>hg38_21_+_8433172_8446622_RNA45SN1_45S_50ntFlanks</td><td>6661</td><td>6686</td><td>26</td><td>CGGTGGATCACTCGGCTCGTGCGTCG</td><td>2056.8</td><td>2.0</td><td>0.8</td><td>2.4</td><td>1.7E-02</td><td>5.0E-02</td><td>BT20/MB231</td><td>MITO</td></tr>
<tr><td>>MI0000813|hsa-mir-324&WithFlank&17|-|7223291|7223385@22.43.22&[MIMAT0000761&hsa-miR-324-5p&offsets|0|0;m-212&17|-|7223342|7223364&offsets|0|-1]|CGCATCCCCTAGGGCATTGGTG</td><td>>MI0000813|hsa-mir-324&WithFlank&17|-|7223291|7223385</td><td>22</td><td>43</td><td>22</td><td>CGCATCCCCTAGGGCATTGGTG</td><td>126.4</td><td>4.3</td><td>2.0</td><td>2.1</td><td>3.2E-02</td><td>5.0E-02</td><td>BT20/MB468</td><td>MP</td></tr>
<tr><td>>hg38_21_+_8433172_8446622_RNA45SN1_45S_50ntFlanks@10389.10430.42|TGAACAGCAGTTGAACATGGGTCAGTCGGTCCTGAGAGATGG</td><td>>hg38_21_+_8433172_8446622_RNA45SN1_45S_50ntFlanks</td><td>10389</td><td>10430</td><td>42</td><td>TGAACAGCAGTTGAACATGGGTCAGTCGGTCCTGAGAGATGG</td><td>101.5</td><td>-3.2</td><td>1.5</td><td>-2.1</td><td>3.2E-02</td><td>5.0E-02</td><td>BT20/MB468</td><td>MP</td></tr>
<tr><td>>MI0000255|hsa-mir-30d&WithFlank&8|-|134804870|134804951@12.34.23(+1C)&[MIMAT0000245&hsa-miR-30d-5p&offsets|0|+1(+1C);m-21&8|-|134804919|134804940&offsets|0|+1(+1C)]|TGTAAACATCCCCGACTGGAAGCC</td><td>>MI0000255|hsa-mir-30d&WithFlank&8|-|134804870|134804951</td><td>12</td><td>34</td><td>23(+1C)</td><td>TGTAAACATCCCCGACTGGAAGCC</td><td>10982.9</td><td>1.9</td><td>0.8</td><td>2.3</td><td>2.0E-02</td><td>5.0E-02</td><td>BT20/MB231</td><td>CYTO</td></tr>
<tr><td>>trna116_GluCTC_1_-_145399233_145399304@3.35.33,trna59_GluCTC_1_+_249168447_249168518@3.35.33,trna71_GluCTC_1_-_161439189_161439260@3.35.33,trna74_GluCTC_1_-_161431809_161431880@3.35.33,trna77_GluCTC_1_-_161424398_161424469@3.35.33,trna77_GluCTC_6_+_28949976_28950047@3.35.33,trna80_GluCTC_1_-_161417018_161417089@3.35.33,trna87_GluCTC_6_-_126101393_126101464@3.35.33|CCTGGTGGTCTAGTGGTTAGGATTCGGCGCTCT</td><td>>trna116_GluCTC_1_-_145399233_145399304</td><td>3</td><td>35</td><td>33</td><td>CCTGGTGGTCTAGTGGTTAGGATTCGGCGCTCT</td><td>275.8</td><td>-1.7</td><td>0.7</td><td>-2.3</td><td>2.0E-02</td><td>5.0E-02</td><td>BT20/MB231</td><td>CYTO</td></tr>
<tr><td>>hg38_1_-_228634819_228635039_RNA5S12_5S_50ntFlanks@147.171.25|GGGAATACCGGGTGCTGTAGGCTTT</td><td>>hg38_1_-_228634819_228635039_RNA5S12_5S_50ntFlanks</td><td>147</td><td>171</td><td>25</td><td>GGGAATACCGGGTGCTGTAGGCTTT</td><td>10039.9</td><td>2.7</td><td>1.3</td><td>2.1</td><td>3.2E-02</td><td>5.0E-02</td><td>BT20/MB468</td><td>MP</td></tr>
<tr><td>>hg38_21_+_8433172_8446622_RNA45SN1_45S_50ntFlanks@6647.6669.23|CGTACGACTCTTAGCGGTGGATC</td><td>>hg38_21_+_8433172_8446622_RNA45SN1_45S_50ntFlanks</td><td>6647</td><td>6669</td><td>23</td><td>CGTACGACTCTTAGCGGTGGATC</td><td>3409.3</td><td>1.5</td><td>0.7</td><td>2.1</td><td>3.2E-02</td><td>5.0E-02</td><td>BT20/MB468</td><td>MP</td></tr>
<tr><td>>hg38_21_+_8433172_8446622_RNA45SN1_45S_50ntFlanks@6661.6679.19|CGGTGGATCACTCGGCTCG</td><td>>hg38_21_+_8433172_8446622_RNA45SN1_45S_50ntFlanks</td><td>6661</td><td>6679</td><td>19</td><td>CGGTGGATCACTCGGCTCG</td><td>456.7</td><td>1.8</td><td>0.8</td><td>2.1</td><td>3.2E-02</td><td>5.0E-02</td><td>BT20/MB468</td><td>MP</td></tr>
<tr><td>>hg38_21_+_8433172_8446622_RNA45SN1_45S_50ntFlanks@10684.10708.25|CGGCGGCGTCCGGTGAGCTCTCGCT</td><td>>hg38_21_+_8433172_8446622_RNA45SN1_45S_50ntFlanks</td><td>10684</td><td>10708</td><td>25</td><td>CGGCGGCGTCCGGTGAGCTCTCGCT</td><td>413.2</td><td>-1.9</td><td>0.9</td><td>-2.1</td><td>3.2E-02</td><td>5.0E-02</td><td>BT20/MB468</td><td>MP</td></tr>
<tr><td>>hg38_21_+_8433172_8446622_RNA45SN1_45S_50ntFlanks@12991.13039.49|CTCCCTCGCTGCGATCTATTGAAAGTCAGCCCTCGACACAAGGGTTTGT</td><td>>hg38_21_+_8433172_8446622_RNA45SN1_45S_50ntFlanks</td><td>12991</td><td>13039</td><td>49</td><td>CTCCCTCGCTGCGATCTATTGAAAGTCAGCCCTCGACACAAGGGTTTGT</td><td>296.0</td><td>-4.0</td><td>1.9</td><td>-2.1</td><td>3.2E-02</td><td>5.0E-02</td><td>BT20/MB468</td><td>MP</td></tr>
<tr><td>>MI0000266|hsa-mir-10a&WithFlank&17|-|48579832|48579953@28.48.21&[MIMAT0000253&hsa-miR-10a-5p&offsets|0|-2;m-18&17|-|48579905|48579926&offsets|0|-1]|TACCCTGTAGATCCGAATTTG</td><td>>MI0000266|hsa-mir-10a&WithFlank&17|-|48579832|48579953</td><td>28</td><td>48</td><td>21</td><td>TACCCTGTAGATCCGAATTTG</td><td>975.5</td><td>2.1</td><td>0.9</td><td>2.3</td><td>2.0E-02</td><td>5.0E-02</td><td>BT20/MB231</td><td>CYTO</td></tr>
<tr><td>>am_trna10_ValCAC_5_-_180649395_180649467@1.33.33,trna12_ValAAC_5_-_180645270_180645342@1.33.33,trna132_ValAAC_6_-_27721179_27721251@1.33.33,trna136_ValAAC_6_-_27648885_27648957@1.33.33,trna139_ValAAC_6_-_27618707_27618779@1.33.33,trna18_ValCAC_5_-_180529253_180529325@1.33.33,trna2_ValAAC_3_+_169490018_169490090@1.33.33,trna2_ValCAC_5_+_180524070_180524142@1.33.33,trna4_ValAAC_5_+_180591154_180591226@1.33.33,trna5_ValAAC_5_+_180596610_180596682@1.33.33,trna6_ValCAC_5_+_180600650_180600722@1.33.33,trna85_ValCAC_1_-_161369490_161369562@1.33.33,trna90_ValCAC_1_-_149684088_149684161@1.33.33,trna98_ValCAC_1_-_149298555_149298627@1.33.33,trna9_ValCAC_6_+_26538282_26538354@1.33.33|GTTTCCGTAGTGTAGTGGTTATCACGTTCGCCT</td><td>>am_trna10_ValCAC_5_-_180649395_180649467</td><td>1</td><td>33</td><td>33</td><td>GTTTCCGTAGTGTAGTGGTTATCACGTTCGCCT</td><td>8689.4</td><td>-2.8</td><td>1.2</td><td>-2.3</td><td>2.0E-02</td><td>5.0E-02</td><td>BT20/MB231</td><td>CYTO</td></tr>
<tr><td>>hg38_21_+_8433172_8446622_RNA45SN1_45S_50ntFlanks@11572.11613.42|CCGACTTAGAACTGGTGCGGACCAGGGGAATCCGACTGTTTA</td><td>>hg38_21_+_8433172_8446622_RNA45SN1_45S_50ntFlanks</td><td>11572</td><td>11613</td><td>42</td><td>CCGACTTAGAACTGGTGCGGACCAGGGGAATCCGACTGTTTA</td><td>549.1</td><td>1.7</td><td>0.9</td><td>2.0</td><td>4.4E-02</td><td>5.0E-02</td><td>MB231/MB468</td><td>CYTO</td></tr>
<tr><td>>hg38_MT_+_598_1651_RNR1_12S_50ntFlanks@980.1001.22|TACTGGAAAGTGCACTTGGACG</td><td>>hg38_MT_+_598_1651_RNR1_12S_50ntFlanks</td><td>980</td><td>1001</td><td>22</td><td>TACTGGAAAGTGCACTTGGACG</td><td>103.3</td><td>3.6</td><td>1.8</td><td>2.0</td><td>4.2E-02</td><td>5.0E-02</td><td>MB231/MB468</td><td>MP</td></tr>
<tr><td>>MI0000465|hsa-mir-191&WithFlank&3|-|49020612|49020715@22.45.24&[MIMAT0000440&hsa-miR-191-5p&offsets|0|+1;m-48&3|-|49020672|49020694&offsets|0|+1]|CAACGGAATCCCAAAAGCAGCTGT</td><td>>MI0000465|hsa-mir-191&WithFlank&3|-|49020612|49020715</td><td>22</td><td>45</td><td>24</td><td>CAACGGAATCCCAAAAGCAGCTGT</td><td>12682.8</td><td>-2.8</td><td>1.2</td><td>-2.4</td><td>1.5E-02</td><td>5.0E-02</td><td>BT20/MB231</td><td>MP</td></tr>
<tr><td>>hg38_1_-_228634819_228635039_RNA5S12_5S_50ntFlanks@146.171.26|TGGGAATACCGGGTGCTGTAGGCTTT</td><td>>hg38_1_-_228634819_228635039_RNA5S12_5S_50ntFlanks</td><td>146</td><td>171</td><td>26</td><td>TGGGAATACCGGGTGCTGTAGGCTTT</td><td>4113.0</td><td>3.4</td><td>1.4</td><td>2.4</td><td>1.5E-02</td><td>5.0E-02</td><td>BT20/MB231</td><td>MP</td></tr>
<tr><td>>hg38_21_+_8433172_8446622_RNA45SN1_45S_50ntFlanks@12419.12453.35|GATGTCGGCTCTTCCTATCATTGTGAAGCAGAATT</td><td>>hg38_21_+_8433172_8446622_RNA45SN1_45S_50ntFlanks</td><td>12419</td><td>12453</td><td>35</td><td>GATGTCGGCTCTTCCTATCATTGTGAAGCAGAATT</td><td>167.7</td><td>3.3</td><td>1.7</td><td>2.0</td><td>4.6E-02</td><td>5.0E-02</td><td>MB231/MB468</td><td>MITO</td></tr>
<tr><td>>hg38_21_+_8433172_8446622_RNA45SN1_45S_50ntFlanks@4727.4758.32|AATCAAGAACGAAAGTCGGAGGTTCGAAGACG</td><td>>hg38_21_+_8433172_8446622_RNA45SN1_45S_50ntFlanks</td><td>4727</td><td>4758</td><td>32</td><td>AATCAAGAACGAAAGTCGGAGGTTCGAAGACG</td><td>1019.1</td><td>1.4</td><td>0.7</td><td>2.0</td><td>4.4E-02</td><td>5.0E-02</td><td>MB231/MB468</td><td>CYTO</td></tr>
<tr><td>>hg38_21_+_8433172_8446622_RNA45SN1_45S_50ntFlanks@11873.11897.25|AAGAAGACCCTGTTGAGCTTGACTC</td><td>>hg38_21_+_8433172_8446622_RNA45SN1_45S_50ntFlanks</td><td>11873</td><td>11897</td><td>25</td><td>AAGAAGACCCTGTTGAGCTTGACTC</td><td>526.2</td><td>-2.2</td><td>1.0</td><td>-2.1</td><td>3.2E-02</td><td>5.0E-02</td><td>BT20/MB468</td><td>MP</td></tr>
<tr><td>>hg38_21_+_8433172_8446622_RNA45SN1_45S_50ntFlanks@12993.13013.21|CCCTCGCTGCGATCTATTGAA</td><td>>hg38_21_+_8433172_8446622_RNA45SN1_45S_50ntFlanks</td><td>12993</td><td>13013</td><td>21</td><td>CCCTCGCTGCGATCTATTGAA</td><td>86.5</td><td>-3.0</td><td>1.4</td><td>-2.1</td><td>3.2E-02</td><td>5.0E-02</td><td>BT20/MB468</td><td>MP</td></tr>
<tr><td>>MI0000070|hsa-mir-16-1&WithFlank&13|-|50048967|50049067@20.38.19(+1A);MI0000115|hsa-mir-16-2&WithFlank&3|+|160404739|160404831@16.34.19(+1A)&[MIMAT0000069&hsa-miR-16-5p&offsets|0|-3(+1A);m-62&13|-|50049027|50049048&offsets|0|-3(+1A)];[MIMAT0000069_1&hsa-miR-16-5p&offsets|0|-3(+1A);m-61&3|+|160404754|160404775&offsets|0|-3(+1A)]|TAGCAGCACGTAAATATTGA</td><td>>MI0000070|hsa-mir-16-1&WithFlank&13|-|50048967|50049067</td><td>20</td><td>38</td><td>19(+1A)</td><td>TAGCAGCACGTAAATATTGA</td><td>75.6</td><td>4.9</td><td>2.3</td><td>2.1</td><td>3.2E-02</td><td>5.0E-02</td><td>BT20/MB468</td><td>MP</td></tr>
<tr><td>>hg38_21_+_8433172_8446622_RNA45SN1_45S_50ntFlanks@8378.8405.28|TTCAAGAGGGCGTGAAACCGTTAAGAGG</td><td>>hg38_21_+_8433172_8446622_RNA45SN1_45S_50ntFlanks</td><td>8378</td><td>8405</td><td>28</td><td>TTCAAGAGGGCGTGAAACCGTTAAGAGG</td><td>104.9</td><td>-1.8</td><td>0.9</td><td>-2.1</td><td>3.2E-02</td><td>5.0E-02</td><td>BT20/MB468</td><td>MP</td></tr>
<tr><td>>MI0000272|hsa-mir-182&WithFlank&7|-|129770377|129770498@29.53.25(+1A)&[MIMAT0000259&hsa-miR-182-5p&offsets|0|+1(+1A);m-22&7|-|129770449|129770470&offsets|0|+3(+1A)]|TTTGGCAATGGTAGAACTCACACTGA</td><td>>MI0000272|hsa-mir-182&WithFlank&7|-|129770377|129770498</td><td>29</td><td>53</td><td>25(+1A)</td><td>TTTGGCAATGGTAGAACTCACACTGA</td><td>157.5</td><td>3.2</td><td>1.5</td><td>2.1</td><td>3.2E-02</td><td>5.0E-02</td><td>BT20/MB468</td><td>MP</td></tr>
<tr><td>>hg38_21_+_8433172_8446622_RNA45SN1_45S_50ntFlanks@11573.11593.21|CGACTTAGAACTGGTGCGGAC</td><td>>hg38_21_+_8433172_8446622_RNA45SN1_45S_50ntFlanks</td><td>11573</td><td>11593</td><td>21</td><td>CGACTTAGAACTGGTGCGGAC</td><td>457.7</td><td>-2.1</td><td>1.0</td><td>-2.1</td><td>3.2E-02</td><td>5.0E-02</td><td>BT20/MB468</td><td>MP</td></tr>
<tr><td>>hg38_21_+_8433172_8446622_RNA45SN1_45S_50ntFlanks@10466.10483.18|CGTTGCCCTCGGCCGATC</td><td>>hg38_21_+_8433172_8446622_RNA45SN1_45S_50ntFlanks</td><td>10466</td><td>10483</td><td>18</td><td>CGTTGCCCTCGGCCGATC</td><td>56.2</td><td>-3.7</td><td>1.7</td><td>-2.1</td><td>3.2E-02</td><td>5.0E-02</td><td>BT20/MB468</td><td>MP</td></tr>
<tr><td>>hg38_21_+_8433172_8446622_RNA45SN1_45S_50ntFlanks@12413.12434.22|TCCTTCGATGTCGGCTCTTCCT</td><td>>hg38_21_+_8433172_8446622_RNA45SN1_45S_50ntFlanks</td><td>12413</td><td>12434</td><td>22</td><td>TCCTTCGATGTCGGCTCTTCCT</td><td>413.4</td><td>-1.5</td><td>0.7</td><td>-2.0</td><td>4.4E-02</td><td>5.0E-02</td><td>MB231/MB468</td><td>CYTO</td></tr>
<tr><td>>hg38_21_+_8433172_8446622_RNA45SN1_45S_50ntFlanks@5086.5112.27|AACGAGACTCTGGCATGCTAACTAGTT</td><td>>hg38_21_+_8433172_8446622_RNA45SN1_45S_50ntFlanks</td><td>5086</td><td>5112</td><td>27</td><td>AACGAGACTCTGGCATGCTAACTAGTT</td><td>710.4</td><td>3.2</td><td>1.3</td><td>2.4</td><td>1.5E-02</td><td>5.0E-02</td><td>BT20/MB231</td><td>MP</td></tr>
<tr><td>>MI0000465|hsa-mir-191&WithFlank&3|-|49020612|49020715@22.42.21(+1A)&[MIMAT0000440&hsa-miR-191-5p&offsets|0|-2(+1A);m-48&3|-|49020672|49020694&offsets|0|-2(+1A)]|CAACGGAATCCCAAAAGCAGCA</td><td>>MI0000465|hsa-mir-191&WithFlank&3|-|49020612|49020715</td><td>22</td><td>42</td><td>21(+1A)</td><td>CAACGGAATCCCAAAAGCAGCA</td><td>410.4</td><td>2.5</td><td>1.2</td><td>2.1</td><td>3.2E-02</td><td>5.0E-02</td><td>BT20/MB468</td><td>MP</td></tr>
<tr><td>>hg38_21_+_8433172_8446622_RNA45SN1_45S_50ntFlanks@10375.10406.32|AGAAGGGTTCCATGTGAACAGCAGTTGAACAT</td><td>>hg38_21_+_8433172_8446622_RNA45SN1_45S_50ntFlanks</td><td>10375</td><td>10406</td><td>32</td><td>AGAAGGGTTCCATGTGAACAGCAGTTGAACAT</td><td>151.7</td><td>-3.8</td><td>1.8</td><td>-2.1</td><td>3.2E-02</td><td>5.0E-02</td><td>BT20/MB468</td><td>MP</td></tr>
<tr><td>>hg38_21_+_8433172_8446622_RNA45SN1_45S_50ntFlanks@4566.4587.22|ATAATGGAATAGGACCGCGGTT</td><td>>hg38_21_+_8433172_8446622_RNA45SN1_45S_50ntFlanks</td><td>4566</td><td>4587</td><td>22</td><td>ATAATGGAATAGGACCGCGGTT</td><td>127.9</td><td>-2.1</td><td>1.0</td><td>-2.1</td><td>3.2E-02</td><td>5.0E-02</td><td>BT20/MB468</td><td>MP</td></tr>
<tr><td>>hg38_MT_+_1621_3279_RNR2_16S_50ntFlanks@1366.1401.36|CTCGATGTTGGATCAGGACATCCCGATGGTGCAGCC</td><td>>hg38_MT_+_1621_3279_RNR2_16S_50ntFlanks</td><td>1366</td><td>1401</td><td>36</td><td>CTCGATGTTGGATCAGGACATCCCGATGGTGCAGCC</td><td>31.1</td><td>4.2</td><td>2.1</td><td>2.0</td><td>4.2E-02</td><td>5.0E-02</td><td>MB231/MB468</td><td>MP</td></tr>
</table>
